## Supplemental Information for "A Murine Database of Structural Variants Enables the Genetic Architecture of a Spontaneous Murine Lymphoma to be Characterized"

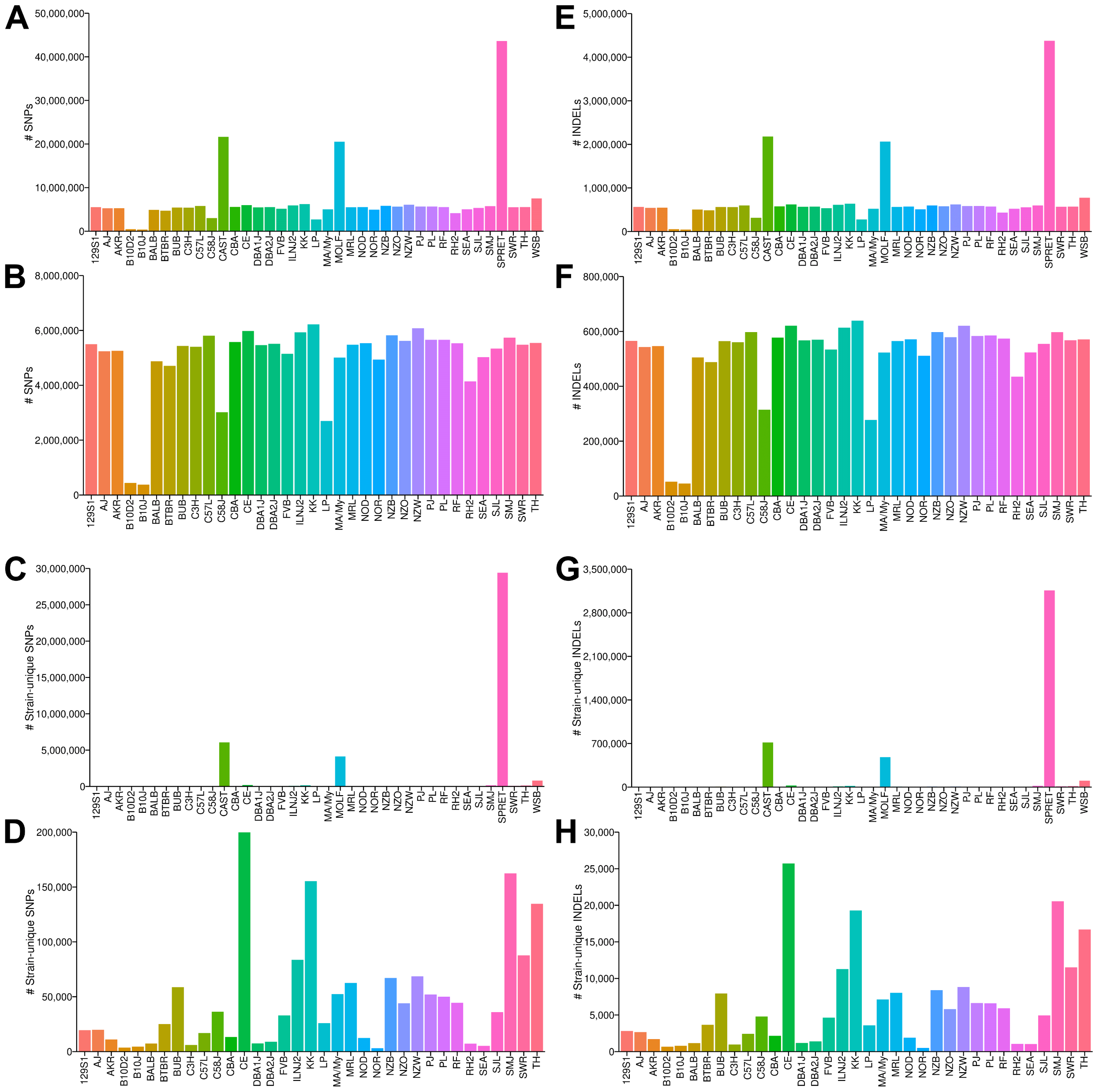

**Figure S1. SNP and INDEL discovery in all 39 strains and 35 classical inbred mouse strains, Related to Figure 1.**

(A-B) The number of SNP sites in (A) all 39 inbred strains or (B) in the 35 classical inbred strains against the C57BL/6 reference.

(C-D) The number of sites with strain-unique SNP alleles in (C) all 39 inbred strains or (D) in the 35 classical inbred strains. Most of the strain-unique SNPs are present in the four wild derived strains (CAST, SPRET, MOLF, WSB).

(E-F) The number of INDEL sites identified in (E) all 39 inbred strains or (F) in the 35 classical inbred strains against the C57BL/6 reference.

(G-H) The number of sites with strain-unique INDEL alleles identified in (G) all 39 inbred strains or (H) in the 35 classical inbred strains. Most strain-unique INDELs are present in the four wild derived strains (CAST, SPRET, MOLF, WSB).

**A**

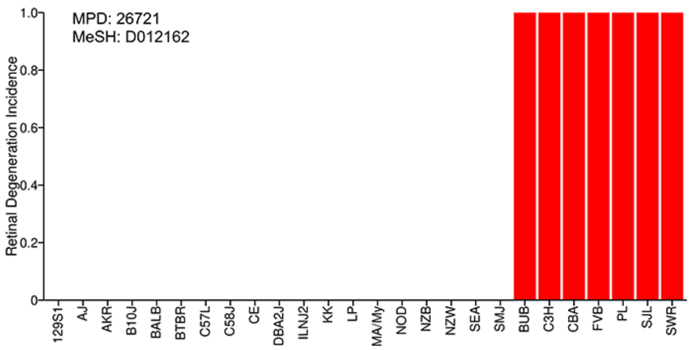

| # | Gene | SNP Flag | Haplotype | EffectSize | Pvalue | FDR | Position | LitScore | PubMed |
|---|---|---|---|---|---|---|---|---|---|
| 0 | Btbd8 | 2 | | 1.000 | 0 | 0 | 5:107582693-107585921 | 0.398 | Indirect |
| 1 | Pcgf3 | 2 | | 1.000 | 0 | 0 | 5:108609082-108616075 | 0.793 | Indirect |
| 2 | 1700028K03Rik | 2 | | 1.000 | 0 | 0 | 5:107677220-107681660 | 0.927 | Indirect |
| 3 | Pde6b | 2 | | 1.000 | 0 | 0 | 5:108537951-108543167 | 0.968 | 27203441,235932... |
| 4 | Tmem175 | 2 | | 1.000 | 0 | 0 | 5:108788563-108791304 | 0.805 | Indirect |
| 5 | Pole | 2 | | 1.000 | 0 | 0 | 5:110485520-110503524 | 0.939 | Indirect |

**B**

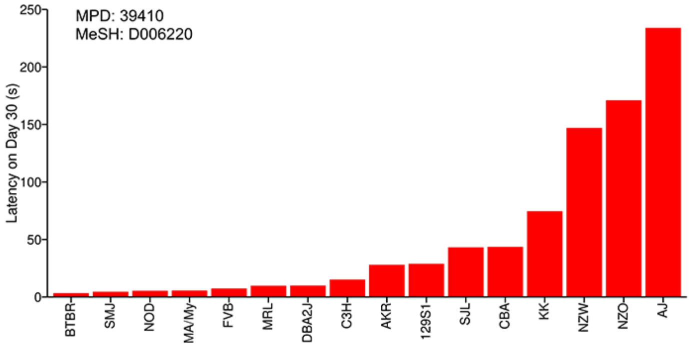

| # | Gene | SNP Flag | Haplotype | EffectSize | Pvalue | FDR | Position | LitScore | PubMed |
|---|---|---|---|---|---|---|---|---|---|
| 0 | Ctnnbl1 | 1 | | 0.955 | 4.25283e-9 | 0.0000143434 | 2:157591691-157608294 | 0.921 | Indirect |
| 1 | Vstm2l | 0 | | 0.955 | 4.25283e-9 | 0.0000143434 | 2:157759007-157762152 | 0.080 | Indirect |
| 2 | Sp8 | 1 | | 0.951 | 7.54079e-9 | 0.0000837927 | 12:118793977-118812596 | 0.940 | Indirect |
| 3 | Itgb8 | 1 | | 0.957 | 2.37158e-8 | 0.0000837927 | 12:119163326-119202835 | 0.883 | Indirect |
| 4 | Tti1 | 1 | | 0.955 | 3.04391e-8 | 0.000047382 | 2:157850645-157865836 | 0.497 | Indirect |
| 5 | Rprd1b | 0 | | 0.955 | 3.04391e-8 | 0.000047382 | 2:157893077-157926550 | 0.651 | Indirect |
| 6 | Ankrd27 | 1 | | 0.936 | 3.66389e-8 | 0.000179774 | 7:35319572-35322079 | 0.718 | Indirect |
| 7 | Pdcd5 | 1 | | 0.934 | 4.51109e-8 | 0.000179774 | 7:35345690-35345943 | 0.847 | Indirect |
| 8 | D630003M21Rik | 1 | | 0.951 | 5.05249e-8 | 0.0000681615 | 2:158011353-158037054 | 0.177 | Indirect |
| 9 | Kcns3 | 1 | | 0.904 | 6.51218e-8 | 0.0000837927 | 12:11167045-11168921 | 0.729 | Indirect |
| 10 | Abcb5 | 1 | | 0.904 | 6.51218e-8 | 0.0000837927 | 12:118822004-118834711 | 0.874 | 25647612 |
| 11 | Macc1 | 2 | | 0.904 | 6.51218e-8 | 0.0000837927 | 12:119313294-119316265 | 0.845 | Indirect |
| 12 | Slc7a10 | 1 | | 0.904 | 6.51218e-8 | 0.000179774 | 7:34889472-34889817 | 0.861 | Indirect |
| 13 | Wdr88 | 1 | | 0.904 | 6.51218e-8 | 0.000179774 | 7:34942968-34943134 | 0.000 | Indirect |
| 14 | Gpatch1 | 1 | | 0.904 | 6.51218e-8 | 0.000179774 | 7:34978249-34978731 | 0.258 | Indirect |
| 15 | Il1rapl2 | 1 | | 0.904 | 6.51218e-8 | 0.000102437 | X:136524539-136537114 | 0.598 | Indirect |

**Figure S2. Using the SNP database developed here, the AI pipeline correctly identifies the causative genetic factors for retinal degeneration (A) and resistance to the extrapyramidal effects of haloperidol (B), Related to Figure 1.** The Mouse Phenome Database (MPD) has datasets examining the incidence of retinal degeneration in 26 inbred strains and the latency to move after 30 days of haloperidol treatment in 16 inbred strains, which are shown in the top graph of each panel. The AI pipeline outputs the gene symbols for that the genes it identifies as having the strongest genetic association with the phenotypic response pattern upon analysis of the published literature. The output includes the calculated p-value for the genetic association, the genetic effect size, and the chromosome with the starting and ending position of each haplotype. The color of each square within the haplotype diagram indicates the haplotype of that strain. The literature score shows the strength of the association of that gene with the measured trait, which was performed using the indicated MeSH term. Of the genes associated with retinal degeneration, *Pde6b* has a perfect genetic association score, the highest literature association score, and it is the only gene with a direct association with retinal degeneration as indicated by the indicated PubMed identification number. Of the genes associated with susceptibility to haloperidol toxicity, *Abcb5* has a strong genetic association (genetic effect size 0.9) and is the only gene with a PubMed paper that is directly linked with the trait. *Pde6b* and *Abcb5* were previously shown to contain the causative murine genetic factors for retinal degeneration and susceptibility to haloperidol toxicity, respectively.

A

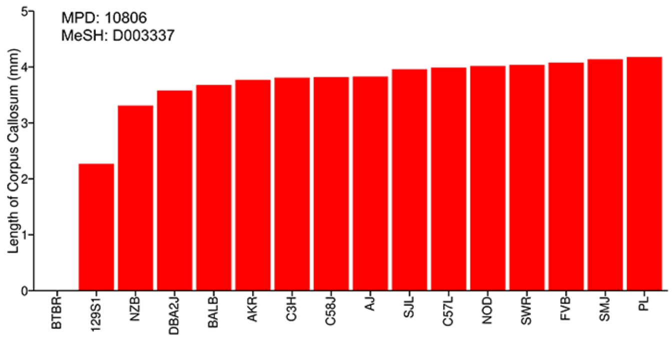

| # | Gene | SNP Flag | Haplotype | EffectSize | Pvalue | FDR | Position | LitScore | PubMed |
|---|---|---|---|---|---|---|---|---|---|
| 0 | Cd180 | 2 | | 0.785 | 0.00000211382 | 0.000659641 | 13:102850747-102859716 | 0.912 | Indirect |
| 1 | Parp10 | 2 | | 0.785 | 0.00000211382 | 0.000236118 | 15:76127135-76127135 | 0.852 | Indirect |
| 2 | Draxin | 2 | | 0.785 | 0.00000211382 | 0.000342675 | 4:148200038-148200038 | 0.832 | 19150847,23206892 |
| 3 | Hcfc1r1 | 2 | | 0.771 | 0.0000750136 | 0.00511472 | 17:23890264-23903432 | 0.871 | Indirect |
| 4 | Slc22a16 | 2 | | 0.415 | 0.00345635 | 0.0706927 | 10:40353401-40503418 | 0.961 | Indirect |
| 5 | Galntl5 | 2 | | 0.337 | 0.042536 | 0.47682 | 5:25377931-25513025 | 0.424 | Indirect |

B

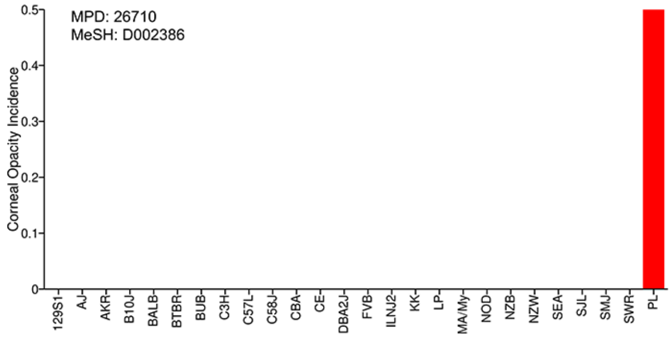

| # | Gene | SNP Flag | Haplotype | EffectSize | Pvalue | FDR | Position | LitScore | PubMed |
|---|---|---|---|---|---|---|---|---|---|
| 0 | Nid1 | 2 | | 1.000 | 0 | 0 | 13:13640077-13640077 | 0.813 | 31877171,25347398 |
| 1 | Ephx3 | 2 | | 1.000 | 0 | 0 | 17:32408705-32413853 | 0.678 | Indirect |
| 2 | Dthd1 | 2 | | 1.000 | 0 | 0 | 5:63039664-63039664 | 0.128 | Indirect |
| 3 | Mtss2 | 2 | | 1.000 | 0 | 0 | 8:111448935-111455804 | 0.476 | Indirect |
| 4 | Gabarap | 2 | | 0.425 | 0.000542049 | 0.0394737 | 11:69883098-69883526 | 0.908 | Indirect |
| 5 | Ankrd55 | 2 | | 0.182 | 0.0156415 | 0.877158 | 13:112496559-112499085 | 0.516 | Indirect |
| 6 | Hcfc1r1 | 2 | | 0.206 | 0.0414612 | 0.373036 | 17:23890264-23903432 | 0.828 | Indirect |

**Figure S3. Using the INDEL database developed here, the AI pipeline correctly identifies the causative genetic factors for agenesis of the corpus callosum (A) and cataract formation (B), Related to Figure 1.** The Mouse Phenome Database (MPD) has datasets measuring the length of the corpus callosum in 16 inbred strains and the incidence of corneal opacity appearing in 26 inbred strains, which are shown in the top graph of each panel. The AI pipeline outputs the gene symbols for the genes it identifies as having the strongest genetic association with the phenotypic response pattern based upon analysis of the published literature. The output includes the calculated p-value for the genetic association, the genetic effect size, and the chromosome with the starting and ending position of each haplotype block. The color of each square within the haplotype diagram indicates the haplotype of that strain. The literature score shows the strength of the association of that gene with the measured trait, which was performed using the indicated MeSH term. Of the genes associated with agenesis of the corpus callosum, *Draxin* has a high impact INDEL, and it is the only gene with a direct association with the corpus callosum as indicated by the indicated PubMed identification number. Of the genes associated with corneal opacity, *Nid1* has a strong genetic association (genetic effect size 1.0) and is the only gene with a PubMed paper that is directly linked with the trait. *Draxin* and *Nid1* were previously shown to contain INDELs that are the murine causative genetic factors for agenesis of the corpus collosum and cataract, respectively.

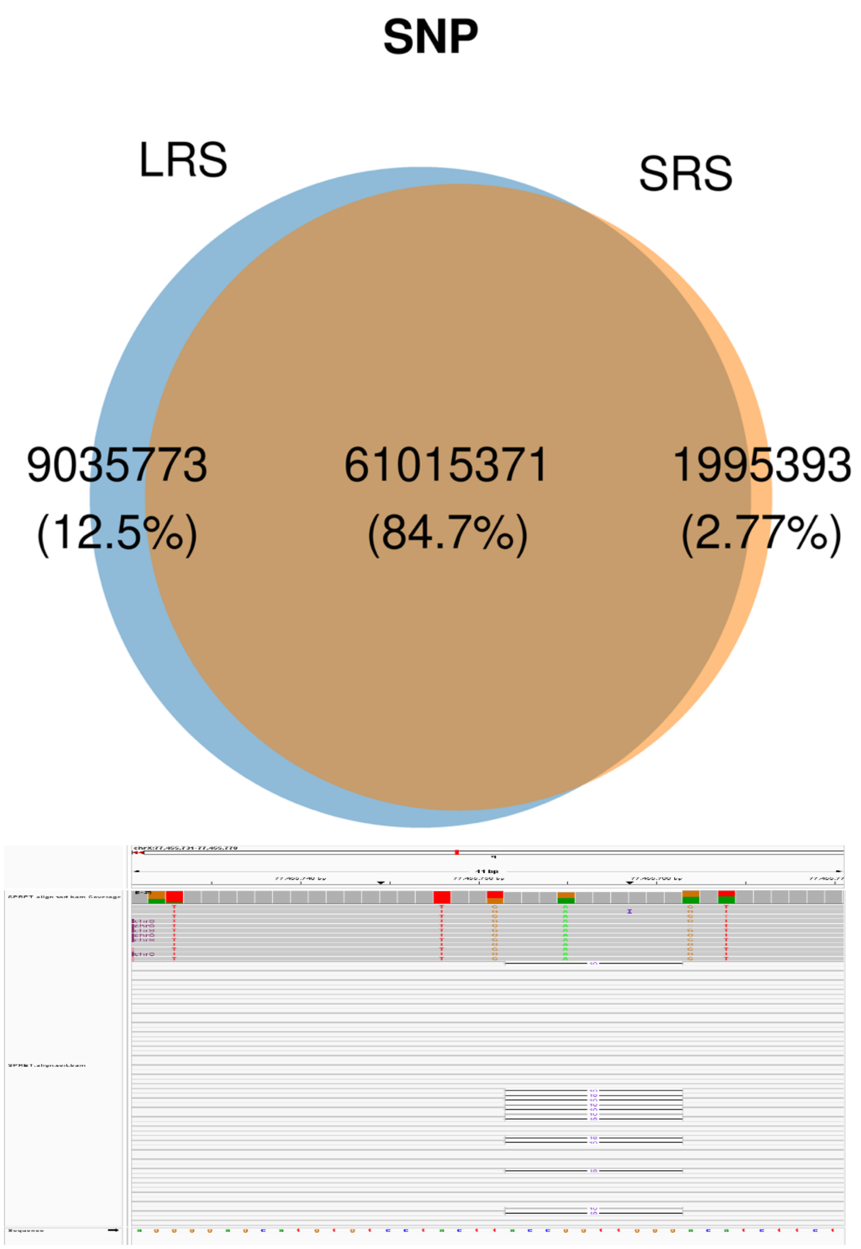

**Figure S4. Comparison of SNPs identified by LRS and SRS and visualization of a misclassified SPRET-unique SNP, Related to Figure 1.** *Top:* A Venn diagram comparing the SNPs identified in this study (LRS), which analyzed 39 inbred strains, with those identified in a previous study using short range sequencing (SRS). The number of SNPs (and the percentages are shown in parenthesis) that are uniquely found in one dataset or are common to both datasets are shown. *Bottom:* An IGV image a SPRET-unique SNP at *chrX:77455751* whose genotype was misclassified as homozygous when it should have been labelled as heterozygous.

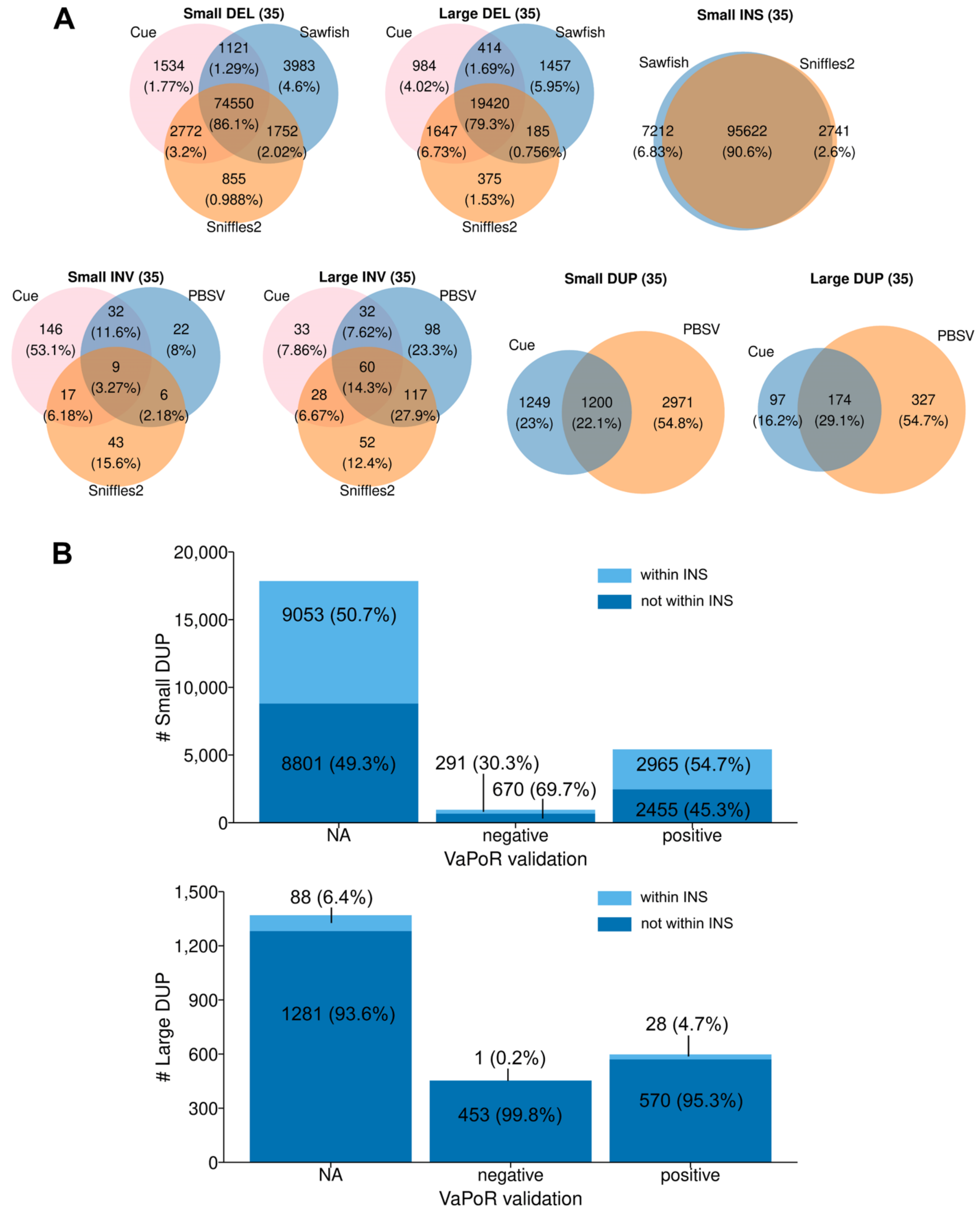

**Figure S5. Assessing the performance of different SV identification programs in 35 classical inbred strains, Related to Figures 1-3.** (**A**) SV identification programs have an acceptable level of agreement for identification of small (50 – 1000 bp) and large (1 kb – 500 kb) DELs, and for small INSs. The results obtained for the 35 classical inbred strains are shown in separate Venn diagrams. The number of each type of SV identified by each individual or combination of programs are shown within the circles (and percentages are shown within parenthesis). However, there was substantial divergence in the small and large INVs identified by Cue, PBSV and Sniffles2. Only the INVs that were validated by visual inspection (using the IGV) are counted. (**B**) Small and large DUPs identified by Cue and PBSV for the 35 inbred strains, and the VaPoR validation results are shown. Most DUPs could not be assessed (NA) by VaPoR. The assessable DUPs that passed (positive) or failed (negative) VaPoR validation are also shown. Also, 55% of the small DUPs were located within an INS, while most large DUPs were not within an INS.

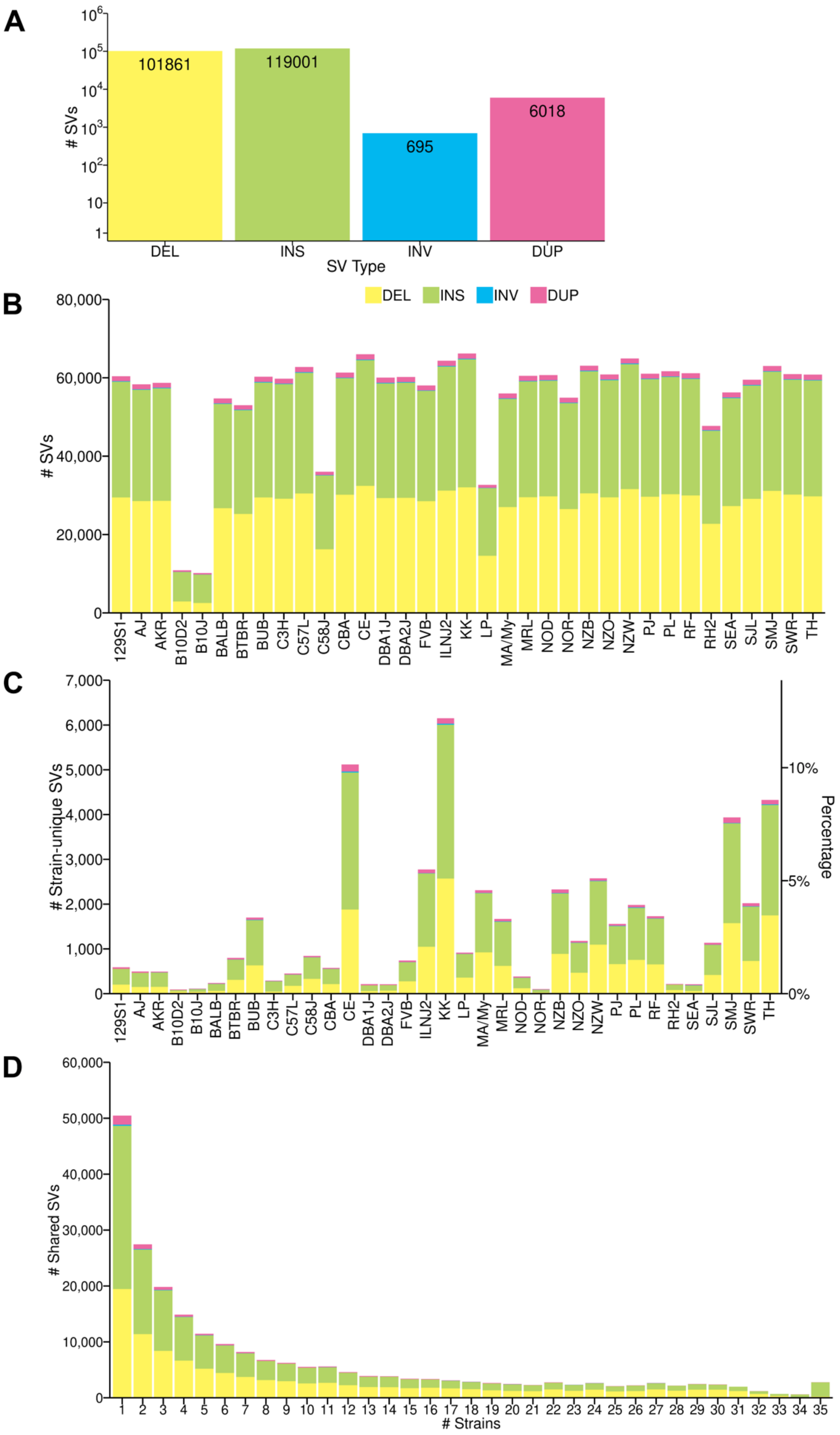

A
### SVs
101861
119001
695
6018
DEL
INS
INV
DUP
SV Type
B
DEL INS INV DUP
### SVs
129S1 AJ AKR B10D2 B10J BALB BTBR BUB C3H C57L C58J CBA CE DBA1J DBA2J FVB ILNJ2 KK LP MA/My MRL NOD NOR NZB NZO NZW PJ PL RF RH2 SEA SJL SMJ SWR TH
C
### Strain-unique SVs
Percentage
D
### Shared SVs
### Strains

**Figure S6. Characteristics or the SV alleles among the 35 classical inbred strains, Related to Figures 1-3.** (**A**) The number of different types of SVs [deletion (DEL), insertion (INS), inversion (INV) or duplication (DUP)] detected in the genome of the 35 classical inbred strains. The Y-axis is log-scale. (**B**) The total number of SVs detected in each strain in 35 classical inbred strains. The type of SV is indicated by the color within the bar (as shown in A). (**C**) The number (left y-axis) and percentage (right y-axis) of strain-unique SVs detected in the 35 classical inbred strains. (**D**) The number of strains with a shared SV allele of the indicated type are shown for the 35 classical inbred strains. Most of the minor SV alleles are shared by 1-3 strains.

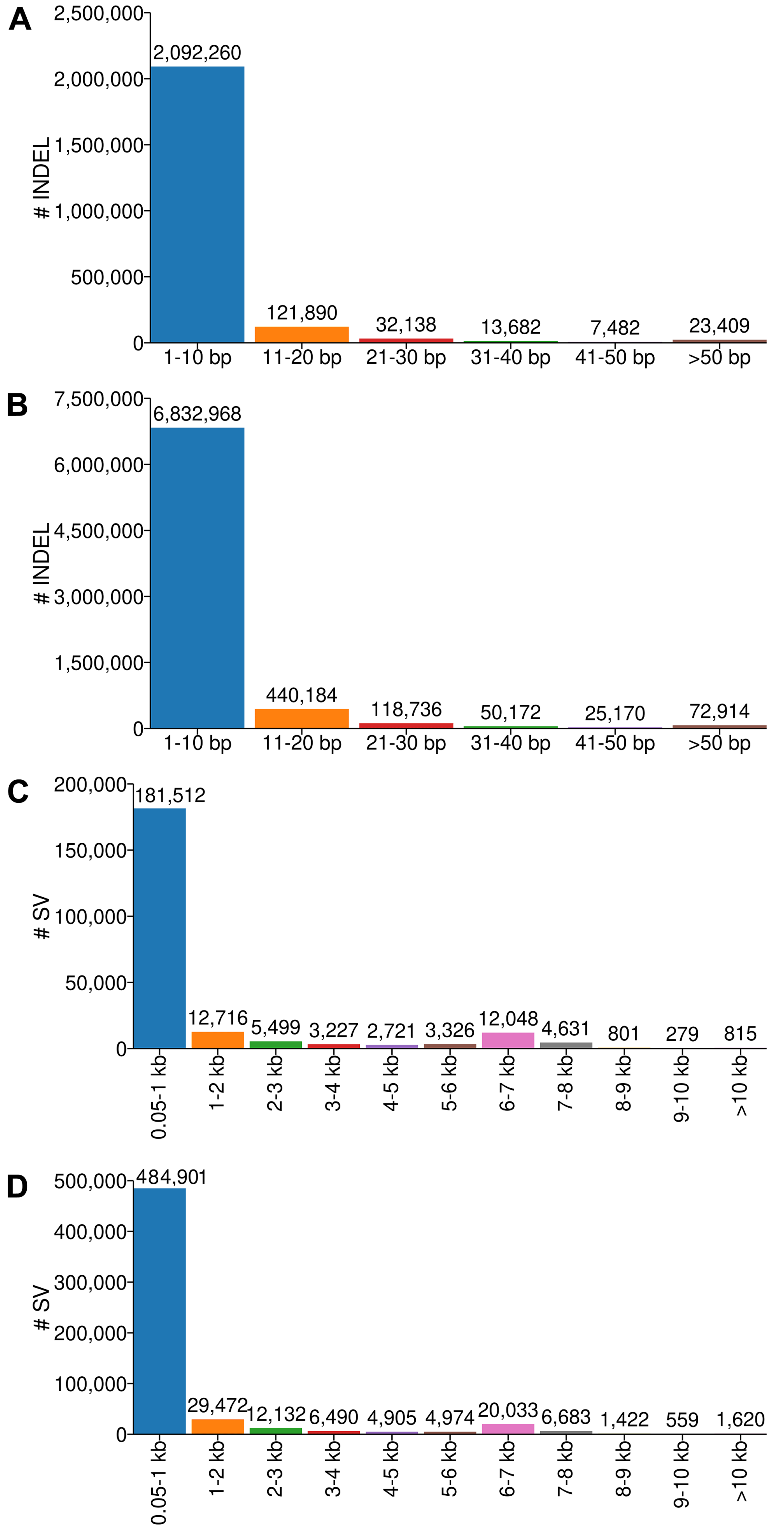

**Figure S7. Length distributions of INDELs and SVs identified in 35 classical and 39 inbred mouse strains, Related to Figures 1-3.** (A, B) The length distribution of the INDELs identified in the 35 classical inbred strains (A) or in all 39 inbred strains (B). (C, D) The length distribution of SVs identified in the 35 classical inbred strains (C) or in all 39 inbred strains (D).

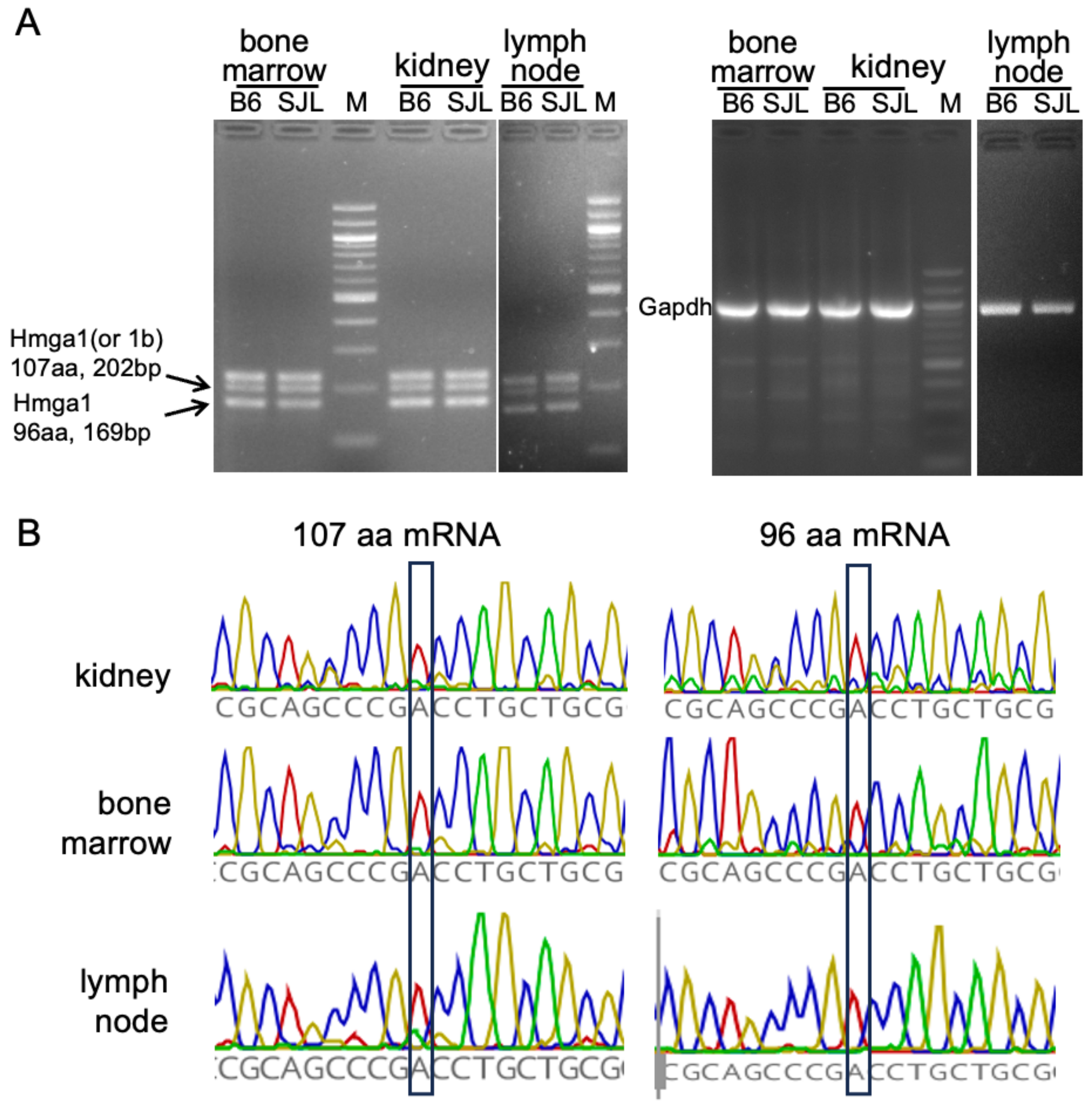

**Figure S8. *Hmga1* encodes the mRNAs expressed in C57BL/6 and SJL bone marrow, kidney and lymph nodes, Related to Figure 6.** (**A**) RT-PCR was performed on hind limb bone marrow (femur, tibia), kidney and inguinal lymph node tissue obtained from C57BL/6 (B6) and SJL mice. The amplicons encoding the Hmga1 (96 amino acids) and Hmga1/Hmga1b 107 amino acid proteins are indicated by arrows (as in Figure 6A). The patterns of amplicon expression in kidney, bone marrow and lymph nodes (left image) are identical to that in liver and spleen obtained from B6 and SJL mice (see Figure 6B). RT-PCR of *GAPDH* amplicons are shown as a control (right image). (**B**) RNA was prepared from kidney, bone marrow and inguinal lymph nodes obtained from C57BL/6 mice, and RT-PCR amplicons from *Hmga1* and *Hmga1b* mRNAs were generated and sequenced. The 3' UTR of *Hmga1* mRNA of C57BL/6 mice has an A, while *Hmga1b* mRNA has a G at the boxed position (as shown in Figure 6). The sequencing results indicate that the mRNAs in C57BL/6 kidney, bone marrow and lymph nodes are encoded by *Hmga1*. This differs from the mRNAs in thymus, which are encoded by *Hmga1b* (Figure 6).

**Table S1. Characteristics of the LRS data obtained from inbred 40 strains, Related to STAR METHODS.** The strain name, Jackson Lab #, sequence amount in gigabases (GB); and the mean, median and number of reads (in millions) for each strain are shown. The N50 read-length distribution statistic assesses read-length quality: it represents the length of the shortest read in the group of the longest sequences that together represent >50% of the nucleotides in that sequence set. Sequencing was performed with a PacBio Revio instrument using the HiFi system.

| Strain | JAX Strain # | GB | # Reads (M) | Mean Read Length | Median Read Length | N50 Read length (bp) |
|---|---|---|---|---|---|---|
| A/J | #000646 | 85.12 | 4.69 | 18132 | 18061 | 20255 |
| C57BL/6J | #000664 | 87.65 | 5.05 | 17359 | 17127 | 18965 |
| C57BL/10J | #000665 | 92.54 | 5.76 | 16071 | 15825 | 17362 |
| C57L/J | #000668 | 92.20 | 5.92 | 15583 | 15347 | 16863 |
| CBA/J | #000656 | 88.19 | 5.12 | 17230 | 17062 | 19088 |
| DBA/1J | #000670 | 109.58 | 6.58 | 16644 | 16300 | 17989 |
| DBA/2J | #000671 | 106.62 | 6.03 | 17694 | 17341 | 19069 |
| FVB/NJ | #001800 | 97.39 | 6.07 | 16039 | 15682 | 17481 |
| KK.Cg-Ay/J | #002468 | 88.64 | 5.40 | 16426 | 16085 | 17674 |
| LP/J | #000676 | 89.05 | 5.49 | 16210 | 16089 | 17873 |
| NOD/ShiLtJ | #001976 | 87.21 | 4.92 | 17720 | 17496 | 19586 |
| NOR/LtJ | #002050 | 93.00 | 5.77 | 16114 | 15819 | 17722 |
| NZB/BlNJ | #000684 | 87.23 | 5.27 | 16550 | 16390 | 18197 |
| NZW/LacJ | #001058 | 90.83 | 6.10 | 14884 | 14629 | 15914 |
| TALLYHO/JngJ | #005314 | 85.01 | 5.59 | 15211 | 14949 | 16694 |
| AKR/J | #000648 | 94.11 | 5.63 | 16725 | 16477 | 18181 |
| B10.D2-Hc1 H2d H2-T18c/nSnJ | #000463 | 98.45 | 5.89 | 16722 | 16546 | 18087 |
| C3H/HeJ | #000659 | 87.20 | 5.53 | 15754 | 15717 | 17701 |
| RF/J | #000682 | 92.80 | 5.49 | 16893 | 16648 | 18797 |
| SJL/J | #000686 | 101.17 | 6.45 | 15679 | 15487 | 17057 |
| MRL/MpJ | #000486 | 96.00 | 6.17 | 15566 | 15309 | 16825 |
| NZO/HlLtJ | #002105 | 88.20 | 5.03 | 17538 | 17314 | 19153 |
| WSB/EiJ | #001145 | 97.26 | 6.03 | 16135 | 15869 | 17391 |
| CAST/EiJ | #000928 | 79.22 | 5.94 | 13344 | 12413 | 15657 |
| MOLF/EiJ | #000550 | 84.45 | 5.50 | 15345 | 14993 | 16945 |
| SPRET/EiJ | #001146 | 82.31 | 4.86 | 16936 | 16748 | 18858 |
| SWR/J | #000689 | 85.84 | 7.04 | 12194 | 10771 | 14418 |
| 129S1/SvImJ | #002448 | 84.49 | 5.16 | 16387 | 16181 | 17795 |
| BALB/cJ | #000651 | 108.88 | 6.67 | 16333 | 16056 | 18050 |
| BTBR T+ Itpr3tf/J | #002282 | 82.31 | 4.72 | 17433 | 17224 | 18906 |
| BUB/BnJ | #000653 | 81.32 | 6.38 | 12741 | 12688 | 14612 |
| MA/MyJ | #000677 | 87.84 | 6.22 | 14119 | 14041 | 15684 |
| C58/J | #000669 | 79.26 | 7.21 | 10997 | 10550 | 13356 |
| SM/J | #000687 | 46.31 | 4.70 | 9864 | 9138 | 11960 |
| CE/J | #000657 | 74.83 | 6.10 | 12266 | 12004 | 13752 |
| PL/J | #000680 | 70.32 | 5.96 | 11803 | 11505 | 13551 |
| SEA/GnJ | #000644 | 90.93 | 7.14 | 12737 | 12637 | 14857 |
| I/LnJ | #000674 | 91.93 | 6.65 | 13829 | 13785 | 15595 |
| RHJ/LeJ | #001591 | 81.86 | 7.67 | 10668 | 10493 | 12523 |
| P/J | #000679 | 92.88 | 5.80 | 16150 | 15778 | 17645 |
| **Total** | | **3540.45** | **233.69** | **15301** | | |
| **Mean** | | **88.51** | **5.84** | | | |

**Table S2. Summary of simulation study results performed to assess the effect of three distinct parameter configurations on simulation results, Related to Table 1.** (**A**) The Default (D), Stringent (S) or Relaxed (R) parameters used for the parameter effect studies. For an explanation of the parameters, please refer to the GitHub page: https://github.com/acenglish/truvari/wiki/bench. In brief, the three distinct parameter configurations in Truvari varied in three criteria: **refdist** (the maximum allowable distance between the base call's start/end and the comparison call), **pctsize** (the ratio of the minimum to maximum size between the base and comparison calls), and **pctovl** (the ratio of overlapping bases between the two calls relative to the longest span).

| **Parameters** | **Default** | **Stringent** | **Relaxed** |
|---|---|---|---|
| **refdist** | **500** | **100** | **1000** |
| pctseq | 0.0 | 0.0 | 0.0 |
| minhaplen | 50 | 50 | 50 |
| **pctsize** | **0.7** | **0.9** | **0.5** |
| **pctovl** | **0.7** | **0.9** | **0.5** |
| typeignore | false | false | false |
| chunksize | 1000 | 1000 | 1000 |
| dup_to_ins | false | false | false |
| sizemin | 50 | 50 | 50 |
| sizefilt | 50 | 50 | 50 |
| sizemax | 100000000 | 100000000 | 100000000 |
| passonly | true | true | true |

**Table S3. The results of simulations assessing the ability of Cue, Sniffles2, Sawfish, PBSV, and Dipcall to identify the indicated types of SVs using the D, S, and R parameters, Related to Table 1.** For each simulation, 1000 SVs of a single type (DEL, INS, INV, DUP) within a defined size [small 50–1000 bp; or large 1–100 kb] were artificially inserted at random positions into the GRCm39 reference sequence. Truvari was used to evaluate precision (P), recall (R), and F1 statistic (F1) for different SV types/sizes using the three D, S, R parameter configurations, which were calculated based on the ability of each program to detect the inserted SVs. Cue, Sniffles2, Sawfish, and PBSV analyzed HiFi long-read sequencing data (30× coverage) that was aligned using minimap2. The simulations were performed using PBSIM3. The genomic sequences analyzed by Dipcall were assembled using HiFiasm. NA: not assessed. W* and $W^{\$}$: withdrawn for the reasons discussed in the manuscript. Only small differences were obtained with different parameters (which were mostly seen for small DUPs). These results indicates that parameter differences would not alter our choice of SV calling program.

| | | Cue | | | Sniffles2 | | | Sawfish | | | PBSV | | | Dipcall | | |
|---|---|---|---|---|---|---|---|---|---|---|---|---|---|---|---|---|
| | | D | S | R | D | S | R | D | S | R | D | S | R | D | S | R |
| Small DEL | P | 1 | 1 | 1 | 1 | 1 | 1 | 1 | 1 | 1 | 1 | 1 | 1 | 0.99 | 0.99 | 0.99 |
| | R | 0.994 | 0.994 | 0.994 | 0.999 | 0.999 | 0.999 | 1 | 1 | 1 | 1 | 1 | 1 | 0.702 | 0.702 | 0.702 |
| | F1 | 0.996 | 0.996 | 0.996 | 0.9995 | 0.9995 | 0.9995 | 1 | 1 | 1 | 1 | 1 | 1 | 0.821 | 0.821 | 0.821 |
| Large DEL | P | 1 | 1 | 1 | 0.991 | 0.991 | 0.991 | 0.989 | 0.989 | 0.989 | 0.997 | 0.997 | 0.997 | 1 | 1 | 1 |
| | R | 0.998 | 0.998 | 0.998 | 0.548 | 0.548 | 0.548 | 0.995 | 0.995 | 0.995 | 0.648 | 0.648 | 0.648 | 0.494 | 0.494 | 0.494 |
| | F1 | 0.999 | 0.999 | 0.999 | 0.706 | 0.706 | 0.706 | 0.992 | 0.992 | 0.992 | 0.785 | 0.785 | 0.785 | 0.661 | 0.661 | 0.661 |
| Small INS | P | NA | | | 0.998 | 0.998 | 0.998 | 1 | 1 | 1 | 1 | 1 | 1 | 0.987 | 0.985 | 0.987 |
| | R | NA | | | 0.978 | 0.978 | 0.978 | 0.982 | 0.982 | 0.982 | 0.979 | 0.979 | 0.979 | 0.811 | 0.81 | 0.811 |
| | F1 | NA | | | 0.988 | 0.988 | 0.988 | 0.991 | 0.991 | 0.991 | 0.989 | 0.989 | 0.989 | 0.89 | 0.889 | 0.89 |
| Large INS | P | NA | | | 1 | 1 | 1 | 1 | 1 | 1 | 1 | 1 | 1 | 1 | 1 | 1 |
| | R | NA | | | 0.182 | 0.182 | 0.182 | 0.358 | 0.358 | 0.358 | 0.13 | 0.13 | 0.13 | 0.936 | 0.936 | 0.936 |
| | F1 | NA | | | 0.308 | 0.308 | 0.308 | 0.527 | 0.527 | 0.527 | 0.23 | 0.23 | 0.23 | 0.967 | 0.967 | 0.967 |
| Small INV | P | 0.999 | 0.974 | 1.0 | 0.908 | 0.906 | 0.929 | W* | | | 0.967 | 0.967 | 0.967 | NA | | |
| | R | 0.909 | 0.886 | 0.91 | 0.434 | 0.433 | 0.444 | W* | | | 0.758 | 0.758 | 0.758 | NA | | |
| | F1 | 0.952 | 0.928 | 0953 | 0.587 | 0.586 | 0.601 | W* | | | 0.85 | 0.85 | 0.85 | NA | | |
| Large INV | P | 0.999 | 0.999 | 0.999 | 0.994 | 0.994 | 0.994 | W* | | | 0.994 | 0.994 | 0.994 | NA | | |
| | R | 0.999 | 0.999 | 0.999 | 0.998 | 0.998 | 0.998 | W* | | | 0.901 | 0.901 | 0.901 | NA | | |
| | F1 | 0.999 | 0.999 | 0.999 | 0.996 | 0.996 | 0.996 | W* | | | 0.9454 | 0.9454 | 0.9454 | NA | | |
| Small DUP | P | 0.997 | 0.942 | 1.0 | 0 | | | $W^{\$}$ | | | 0.792 | 0.777 | 0.792 | NA | | |
| | R | 0.976 | 0.921 | 0.978 | 0 | | | $W^{\$}$ | | | 0.795 | 0.78 | 0.795 | NA | | |
| | F1 | 0.987 | 0.931 | 0.989 | 0 | | | $W^{\$}$ | | | 0.793 | 0.778 | 0.793 | NA | | |
| Large DUP | P | 0.999 | 0.999 | 0.999 | 0.92 | 0.92 | 0.934 | 0.956 | 0.956 | 0.96 | 0.986 | 0.984 | 0.986 | NA | | |
| | R | 0.993 | 0.993 | 0.993 | 0.916 | 0.916 | 0.93 | 0.762 | 0.762 | 0.765 | 0.982 | 0.98 | 0.982 | NA | | |
| | F1 | 0.996 | 0.996 | 0.996 | 0.918 | 0.918 | 0.932 | 0.848 | 0.848 | 0.851 | 0.984 | 0.982 | 0.984 | NA | | |

**Table S5. The functional consequences of SVs that were predicted by VEP, Related to Figures 1-3.** Their estimated severity (HIGH, MODERATE, LOW, MODIFIER) and the type of effect (consequences) was determined for SVs in all 39 strains or those in the 35 classical inbred strains.

| Impact | Consequences | #SV 39 strains | #SV 35 strains |
|---|---|---|---|
| HIGH | transcript_ablation | 2068 | 1079 |
| | splice_acceptor_variant | 33 | 14 |
| | splice_donor_variant | 44 | 24 |
| | stop_gained | 29 | 10 |
| | frameshift_variant | 102 | 55 |
| | stop_lost | 171 | 92 |
| | start_lost | 4 | 1 |
| | transcript_amplification | 191 | 130 |
| | feature_elongation | 5823 | 2538 |
| | feature_truncation | 5555 | 2388 |
| MODERATE | inframe_insertion | 39 | 24 |
| | inframe_deletion | 183 | 88 |
| | protein_altering_variant | 5 | 2 |
| LOW | splice_donor_5th_base_variant | 44 | 24 |
| | splice_region_variant | 19 | 9 |
| | splice_donor_region_variant | 7 | 3 |
| | splice_polypyrimidine_tract_variant | 275 | 98 |
| | start_retained_variant | 4 | 1 |
| | stop_retained_variant | 19 | 12 |
| MODIFIER | coding_sequence_variant | 2389 | 1398 |
| | mature_miRNA_variant | 1 | 0 |
| | 5_prime_UTR_variant | 1171 | 636 |
| | 3_prime_UTR_variant | 4100 | 1808 |
| | non_coding_transcript_exon_variant | 7140 | 3161 |
| | intron_variant | 244001 | 97134 |
| | non_coding_transcript_variant | 41051 | 17103 |
| | coding_transcript_variant | 108 | 68 |
| | upstream_gene_variant | 11946 | 4955 |
| | downstream_gene_variant | 14283 | 5984 |
| | regulatory_region_ablation | 7130 | 3537 |
| | regulatory_region_amplification | 358 | 253 |
| | regulatory_region_variant | 66411 | 29290 |
| | intergenic_variant | 290358 | 117312 |

**Table S6. The number of insertion and deletion SVs identified by VEP analysis as having a high impact (protein coding) in all 39 strains or in the 35 classical inbred strains are shown, Related to Figures 1-3.**

| | 39 strains | | 35 strains | |
|---|---|---|---|---|
| | # SV | # Unique Genes | # SV | # Unique Genes |
| Small INS | 2082 | 1899 | 801 | 765 |
| Large INS | 822 | 740 | 518 | 487 |
| Small DEL | 1688 | 1535 | 705 | 654 |
| Large DEL | 590 | 536 | 293 | 270 |

**Table S12. Summary of the SNPs, INDELs, and SVs identified when our C57BL/6 LRS data was compared with the GRCm39 reference sequence for C57Bl/6, Related to Figure 1.**

| Variant Type | Subtype | Count |
|---|---|---|
| SNP | - | 11751 |
| INDEL | - | 9851 |
| Deletion | Small DEL | 603 |
| | Large DEL | 114 |
| | **Total DEL** | **717** |
| Insertion | Small INS | 4040 |
| | Large INS | 228 |
| | **Total INS** | **4268** |
| Inversion | Small INV | 5 |
| | Large INV | 9 |
| | **Total INV** | **14** |
| Duplication | Small DUP | 269 |
| | Large DUP | 23 |
| | **Total DUP** | **292** |

**Table S4. Lists of SVs identified in our study and those identified in a prior study.** The data are organized into four categories: deletions (DEL), insertions (INS), inversions (INV), and duplications (DUP). The lists include common variants shared between the studies and unique variants that are specific to each dataset.

| Identified_By_Us | Identified_By_JAX |
|---|---|
| chr1-3101582-DEL-357 | chr1-3101575-DEL-358 |
| chr1-3108754-DEL-66 | chr1-3108755-DEL-66 |
| chr1-3123169-DEL-6969 | chr1-3123168-DEL-6969 |
| chr1-3153418-DEL-64 | chr1-3153414-DEL-68 |
| chr1-3161192-DEL-716 | chr1-3161193-DEL-716 |
| chr1-3206789-DEL-3069 | chr1-3206790-DEL-3069 |
| chr1-3231001-DEL-3515 | chr1-3231002-DEL-3515 |
| chr1-3250730-DEL-64 | chr1-3250731-DEL-64 |
| chr1-3263166-DEL-54 | chr1-3263167-DEL-54 |
| chr1-3263200-DEL-650 | chr1-3263222-DEL-643 |
| chr1-3263864-DEL-661 | chr1-3263890-DEL-637 |
| chr1-3376545-DEL-167 | chr1-3376546-DEL-167 |
| chr1-3378160-DEL-1012 | chr1-3378160-DEL-1008 |
| chr1-3387796-DEL-1580 | chr1-3387795-DEL-1581 |
| chr1-3422257-DEL-406 | chr1-3422258-DEL-406 |
| chr1-3455462-DEL-64 | chr1-3455463-DEL-64 |
| chr1-3478634-DEL-84 | chr1-3478624-DEL-78 |
| chr1-3487447-DEL-67 | chr1-3487453-DEL-62 |
| chr1-3542338-DEL-64 | chr1-3542306-DEL-66 |
| chr1-3559078-DEL-68 | chr1-3559078-DEL-68 |
| chr1-3569711-DEL-2811 | chr1-3569712-DEL-2811 |
| chr1-3577082-DEL-60 | chr1-3577082-DEL-60 |
| chr1-3610990-DEL-186 | chr1-3610991-DEL-186 |
| chr1-3705933-DEL-1644 | chr1-3705934-DEL-1644 |
| chr1-3756748-DEL-52 | chr1-3756749-DEL-52 |
| chr1-3770696-DEL-191 | chr1-3770697-DEL-191 |
| chr1-3771229-DEL-179 | chr1-3771230-DEL-179 |
| chr1-3779826-DEL-112 | chr1-3779828-DEL-111 |
| chr1-3794652-DEL-53 | chr1-3794653-DEL-53 |
| chr1-3805385-DEL-975 | chr1-3805385-DEL-975 |
| chr1-3816507-DEL-63 | chr1-3816508-DEL-63 |
| chr1-3826327-DEL-203 | chr1-3826327-DEL-203 |
| chr1-3847957-DEL-1268 | chr1-3847958-DEL-1268 |
| chr1-3850113-DEL-66 | chr1-3850114-DEL-78 |
| chr1-3851558-DEL-57 | chr1-3851559-DEL-57 |
| chr1-3856844-DEL-3714 | chr1-3856845-DEL-3714 |
| chr1-3875140-DEL-304 | chr1-3875138-DEL-305 |
| chr1-3880934-DEL-1108 | chr1-3880935-DEL-1108 |
| chr1-3894054-DEL-852 | chr1-3894055-DEL-852 |
| chr1-3917075-DEL-1911 | chr1-3917076-DEL-1911 |
| chr1-3923103-DEL-199 | chr1-3923103-DEL-199 |
| chr1-3932741-DEL-161 | chr1-3932742-DEL-161 |
| chr1-3940751-DEL-54 | chr1-3940752-DEL-54 |
| chr1-3956980-DEL-52 | chr1-3956981-DEL-52 |
| chr1-3966422-DEL-2182 | chr1-3966407-DEL-2183 |
| chr1-3991709-DEL-3042 | chr1-3991710-DEL-3042 |
| chr1-3994970-DEL-951 | chr1-3994975-DEL-938 |
| chr1-4027581-DEL-483 | chr1-4027582-DEL-483 |
| chr1-4068054-DEL-329 | chr1-4068054-DEL-330 |
| chr1-4149505-DEL-6561 | chr1-4149511-DEL-6561 |
| chr1-4194032-DEL-52 | chr1-4194029-DEL-56 |
| chr1-4202396-DEL-343 | chr1-4202397-DEL-343 |
| chr1-4212720-DEL-60 | chr1-4212721-DEL-60 |
| chr1-4245284-DEL-74 | chr1-4245240-DEL-102 |
| chr1-4268856-DEL-4107 | chr1-4268857-DEL-4107 |
| chr1-4282895-DEL-1261 | chr1-4282896-DEL-1261 |
| chr1-4288915-DEL-337 | chr1-4288916-DEL-337 |
| chr1-4315520-DEL-154 | chr1-4315521-DEL-154 |
| chr1-4318205-DEL-383 | chr1-4318205-DEL-383 |
| chr1-4405456-DEL-204 | chr1-4405457-DEL-204 |
| chr1-4450916-DEL-121 | chr1-4450917-DEL-121 |
| chr1-4510705-DEL-133 | chr1-4510706-DEL-133 |
| chr1-4525104-DEL-56 | chr1-4525093-DEL-63 |
| chr1-4554076-DEL-3145 | chr1-4554077-DEL-3145 |
| chr1-4559763-DEL-106 | chr1-4559764-DEL-106 |
| chr1-4591168-DEL-1149 | chr1-4591169-DEL-1149 |
| chr1-4597343-DEL-86 | chr1-4597361-DEL-60 |
| chr1-4600779-DEL-4764 | chr1-4600780-DEL-4764 |
| chr1-4617032-DEL-53 | chr1-4617033-DEL-53 |
| chr1-4650900-DEL-197 | chr1-4650901-DEL-197 |
| chr1-4671436-DEL-58 | chr1-4671436-DEL-58 |
| chr1-4782637-DEL-497 | chr1-4782638-DEL-496 |
| chr1-4784878-DEL-181 | chr1-4784879-DEL-181 |
| chr1-4792182-DEL-218 | chr1-4792183-DEL-218 |
| chr1-4797326-DEL-581 | chr1-4797327-DEL-581 |
| chr1-4827562-DEL-62 | chr1-4827569-DEL-56 |
| chr1-4828304-DEL-255 | chr1-4828305-DEL-255 |

| | |
|---|---|
| chr1-4849601-DEL-244 | chr1-4849602-DEL-244 |
| chr1-4851850-DEL-68 | chr1-4851851-DEL-68 |
| chr1-4859389-DEL-191 | chr1-4859390-DEL-191 |
| chr1-4870268-DEL-599 | chr1-4870269-DEL-599 |
| chr1-4881262-DEL-81 | chr1-4881262-DEL-81 |
| chr1-4882390-DEL-181 | chr1-4882391-DEL-181 |
| chr1-4886136-DEL-169 | chr1-4886137-DEL-169 |
| chr1-4893826-DEL-175 | chr1-4893827-DEL-175 |
| chr1-4896717-DEL-171 | chr1-4896718-DEL-171 |
| chr1-4903687-DEL-181 | chr1-4903688-DEL-181 |
| chr1-4904195-DEL-98 | chr1-4904189-DEL-100 |
| chr1-4955480-DEL-78 | chr1-4955469-DEL-70 |
| chr1-4964240-DEL-86 | chr1-4964238-DEL-62 |
| chr1-4965267-DEL-52 | chr1-4965268-DEL-52 |
| chr1-4991203-DEL-206 | chr1-4991204-DEL-206 |
| chr1-5017343-DEL-6361 | chr1-5017343-DEL-6361 |
| chr1-5028540-DEL-77 | chr1-5028550-DEL-68 |
| chr1-5055585-DEL-2663 | chr1-5055586-DEL-2663 |
| chr1-5062224-DEL-105 | chr1-5062225-DEL-105 |
| chr1-5103121-DEL-72 | chr1-5103122-DEL-72 |
| chr1-5122474-DEL-101 | chr1-5122474-DEL-101 |
| chr1-5172971-DEL-196 | chr1-5172972-DEL-196 |
| chr1-5179802-DEL-1529 | chr1-5179802-DEL-1529 |
| chr1-5259365-DEL-6581 | chr1-5259366-DEL-6581 |
| chr1-5259584-DEL-214 | chr1-5259598-DEL-214 |
| chr1-5267512-DEL-343 | chr1-5267516-DEL-343 |
| chr1-5269257-DEL-3224 | chr1-5269258-DEL-3224 |
| chr1-5347112-DEL-304 | chr1-5347113-DEL-304 |
| chr1-5366407-DEL-839 | chr1-5366408-DEL-839 |
| chr1-5378128-DEL-2124 | chr1-5378128-DEL-2124 |
| chr1-5420602-DEL-6885 | chr1-5420603-DEL-6885 |
| chr1-5447679-DEL-83 | chr1-5447676-DEL-84 |
| chr1-5557298-DEL-79 | chr1-5557302-DEL-76 |
| chr1-5573007-DEL-86 | chr1-5572985-DEL-90 |
| chr1-5588237-DEL-164 | chr1-5588237-DEL-164 |
| chr1-5621698-DEL-2466 | chr1-5621692-DEL-2468 |
| chr1-5627573-DEL-1209 | chr1-5627574-DEL-1209 |
| chr1-5637425-DEL-58 | chr1-5637426-DEL-58 |
| chr1-5697538-DEL-1166 | chr1-5697539-DEL-1166 |
| chr1-5770785-DEL-1889 | chr1-5770786-DEL-1888 |
| chr1-5804588-DEL-135 | chr1-5804589-DEL-135 |
| chr1-5858257-DEL-237 | chr1-5858258-DEL-237 |
| chr1-5898159-DEL-341 | chr1-5898160-DEL-341 |
| chr1-5910742-DEL-940 | chr1-5910743-DEL-940 |
| chr1-5920398-DEL-178 | chr1-5920398-DEL-178 |
| chr1-5928255-DEL-302 | chr1-5928256-DEL-302 |
| chr1-5930047-DEL-60 | chr1-5930051-DEL-57 |
| chr1-5932023-DEL-1812 | chr1-5932024-DEL-1812 |
| chr1-5945133-DEL-204 | chr1-5945134-DEL-204 |
| chr1-5997847-DEL-54 | chr1-5997848-DEL-54 |
| chr1-6049271-DEL-430 | chr1-6049272-DEL-430 |
| chr1-6063304-DEL-6271 | chr1-6063305-DEL-6270 |
| chr1-6070548-DEL-70 | chr1-6070547-DEL-72 |
| chr1-6072954-DEL-90 | chr1-6072955-DEL-85 |
| chr1-6075838-DEL-70 | chr1-6075839-DEL-60 |
| chr1-6079243-DEL-4405 | chr1-6079244-DEL-4405 |
| chr1-6105424-DEL-59 | chr1-6105418-DEL-59 |
| chr1-6131401-DEL-112 | chr1-6131406-DEL-108 |
| chr1-6154411-DEL-206 | chr1-6154412-DEL-206 |
| chr1-6160355-DEL-88 | chr1-6160356-DEL-88 |
| chr1-6162102-DEL-175 | chr1-6162103-DEL-175 |
| chr1-6176031-DEL-68 | chr1-6176031-DEL-60 |
| chr1-6177806-DEL-1354 | chr1-6177803-DEL-1355 |
| chr1-6180297-DEL-53 | chr1-6180297-DEL-53 |
| chr1-6185743-DEL-453 | chr1-6185743-DEL-442 |
| chr1-6187004-DEL-178 | chr1-6187005-DEL-178 |
| chr1-6193337-DEL-613 | chr1-6193338-DEL-613 |
| chr1-6194385-DEL-189 | chr1-6194386-DEL-189 |
| chr1-6203867-DEL-55 | chr1-6203861-DEL-50 |
| chr1-6208010-DEL-138 | chr1-6208011-DEL-138 |
| chr1-6238326-DEL-4710 | chr1-6238327-DEL-4710 |
| chr1-6343746-DEL-658 | chr1-6343747-DEL-658 |
| chr1-6375372-DEL-52 | chr1-6375375-DEL-50 |
| chr1-6438536-DEL-214 | chr1-6438537-DEL-214 |
| chr1-6455861-DEL-630 | chr1-6455861-DEL-630 |
| chr1-6497374-DEL-204 | chr1-6497375-DEL-204 |
| chr1-6500598-DEL-7111 | chr1-6500599-DEL-7111 |
| chr1-6527006-DEL-1171 | chr1-6527007-DEL-1171 |
| chr1-6533280-DEL-2603 | chr1-6533281-DEL-2603 |
| chr1-6534526-DEL-257 | chr1-6534527-DEL-257 |

| | |
|---|---|
| chr1-6568481-DEL-1069 | chr1-6568482-DEL-1069 |
| chr1-6571574-DEL-51 | chr1-6571575-DEL-51 |
| chr1-6574639-DEL-3712 | chr1-6574640-DEL-3712 |
| chr1-6616443-DEL-562 | chr1-6616444-DEL-562 |
| chr1-6627435-DEL-55 | chr1-6627432-DEL-56 |
| chr1-6628609-DEL-510 | chr1-6628597-DEL-509 |
| chr1-6629217-DEL-346 | chr1-6629218-DEL-346 |
| chr1-6633047-DEL-6195 | chr1-6633042-DEL-6194 |
| chr1-6637419-DEL-72 | chr1-6637420-DEL-72 |
| chr1-6644714-DEL-240 | chr1-6644719-DEL-243 |
| chr1-6660897-DEL-63 | chr1-6660898-DEL-63 |
| chr1-6662067-DEL-54 | chr1-6662067-DEL-54 |
| chr1-6671151-DEL-54 | chr1-6671152-DEL-54 |
| chr1-6696649-DEL-5647 | chr1-6696650-DEL-5647 |
| chr1-6706244-DEL-1512 | chr1-6706245-DEL-1512 |
| chr1-6715673-DEL-72 | chr1-6715674-DEL-72 |
| chr1-6734938-DEL-68 | chr1-6734939-DEL-68 |
| chr1-6735641-DEL-570 | chr1-6735642-DEL-570 |
| chr1-6747127-DEL-770 | chr1-6747128-DEL-770 |
| chr1-6749998-DEL-54 | chr1-6749999-DEL-54 |
| chr1-6760851-DEL-3165 | chr1-6760852-DEL-3165 |
| chr1-6769351-DEL-1047 | chr1-6769352-DEL-1047 |
| chr1-6785752-DEL-1142 | chr1-6785753-DEL-1142 |
| chr1-6811666-DEL-84 | chr1-6811667-DEL-84 |
| chr1-6821405-DEL-72 | chr1-6821406-DEL-72 |
| chr1-6835227-DEL-120 | chr1-6835228-DEL-120 |
| chr1-6857344-DEL-70 | chr1-6857345-DEL-70 |
| chr1-6889838-DEL-72 | chr1-6889839-DEL-72 |
| chr1-6935993-DEL-193 | chr1-6935994-DEL-193 |
| chr1-6984442-DEL-109 | chr1-6984443-DEL-109 |
| chr1-7004291-DEL-85 | chr1-7004292-DEL-85 |
| chr1-7014198-DEL-291 | chr1-7014199-DEL-291 |
| chr1-7024679-DEL-87 | chr1-7024662-DEL-103 |
| chr1-7026498-DEL-231 | chr1-7026499-DEL-231 |
| chr1-7078921-DEL-54 | chr1-7078922-DEL-54 |
| chr1-7088577-DEL-1201 | chr1-7088578-DEL-1201 |
| chr1-7099957-DEL-1111 | chr1-7099959-DEL-1112 |
| chr1-7109727-DEL-85 | chr1-7109721-DEL-60 |
| chr1-7116708-DEL-222 | chr1-7116709-DEL-222 |
| chr1-7143514-DEL-2019 | chr1-7143516-DEL-2018 |
| chr1-7174343-DEL-651 | chr1-7174340-DEL-653 |
| chr1-7194398-DEL-302 | chr1-7194399-DEL-302 |
| chr1-7227444-DEL-57 | chr1-7227445-DEL-57 |
| chr1-7247898-DEL-2117 | chr1-7247899-DEL-2122 |
| chr1-7260437-DEL-6649 | chr1-7260438-DEL-6649 |
| chr1-7301039-DEL-6407 | chr1-7301040-DEL-6407 |
| chr1-7342187-DEL-117 | chr1-7342188-DEL-117 |
| chr1-7398871-DEL-7093 | chr1-7398872-DEL-7093 |
| chr1-7423033-DEL-1250 | chr1-7423034-DEL-1250 |
| chr1-7429335-DEL-156 | chr1-7429336-DEL-156 |
| chr1-7443386-DEL-961 | chr1-7443386-DEL-961 |
| chr1-7481210-DEL-67 | chr1-7481211-DEL-67 |
| chr1-7515869-DEL-138 | chr1-7515869-DEL-138 |
| chr1-7522899-DEL-1494 | chr1-7522899-DEL-1494 |
| chr1-7528266-DEL-5284 | chr1-7528265-DEL-5284 |
| chr1-7546766-DEL-12060 | chr1-7546751-DEL-12075 |
| chr1-7546807-DEL-5514 | chr1-7546808-DEL-5514 |
| chr1-7568501-DEL-538 | chr1-7568502-DEL-538 |
| chr1-7570832-DEL-431 | chr1-7570833-DEL-431 |
| chr1-7596402-DEL-216 | chr1-7596402-DEL-216 |
| chr1-7600326-DEL-67 | chr1-7600326-DEL-67 |
| chr1-7605011-DEL-63 | chr1-7605012-DEL-63 |
| chr1-7614612-DEL-79 | chr1-7614602-DEL-117 |
| chr1-7628369-DEL-988 | chr1-7628377-DEL-988 |
| chr1-7675429-DEL-1889 | chr1-7675430-DEL-1889 |
| chr1-7678414-DEL-3983 | chr1-7678415-DEL-3983 |
| chr1-7681913-DEL-86 | chr1-7681914-DEL-86 |
| chr1-7682566-DEL-428 | chr1-7682557-DEL-463 |
| chr1-7685787-DEL-5463 | chr1-7685788-DEL-5463 |
| chr1-7701813-DEL-1499 | chr1-7701814-DEL-1499 |
| chr1-7713692-DEL-681 | chr1-7713693-DEL-681 |
| chr1-7745975-DEL-51 | chr1-7745971-DEL-55 |
| chr1-7766130-DEL-212 | chr1-7766131-DEL-212 |
| chr1-7783125-DEL-218 | chr1-7783126-DEL-218 |
| chr1-7788550-DEL-2688 | chr1-7788551-DEL-2688 |
| chr1-7792016-DEL-5853 | chr1-7792017-DEL-5853 |
| chr1-7800288-DEL-118 | chr1-7800294-DEL-118 |
| chr1-7809351-DEL-3820 | chr1-7809352-DEL-3820 |
| chr1-7815563-DEL-180 | chr1-7815564-DEL-180 |
| chr1-7842083-DEL-497 | chr1-7842084-DEL-497 |

| | |
|---|---|
| chr1-7844813-DEL-53 | chr1-7844812-DEL-54 |
| chr1-7855738-DEL-431 | chr1-7855738-DEL-431 |
| chr1-7873886-DEL-1109 | chr1-7873886-DEL-1109 |
| chr1-7908211-DEL-1544 | chr1-7908212-DEL-1544 |
| chr1-7910787-DEL-1320 | chr1-7910788-DEL-1320 |
| chr1-7913507-DEL-6553 | chr1-7913508-DEL-6553 |
| chr1-7969237-DEL-60 | chr1-7969223-DEL-61 |
| chr1-7969941-DEL-459 | chr1-7969942-DEL-455 |
| chr1-7972706-DEL-68 | chr1-7972715-DEL-52 |
| chr1-7979553-DEL-51 | chr1-7979549-DEL-55 |
| chr1-7984443-DEL-390 | chr1-7984444-DEL-390 |
| chr1-7986815-DEL-114 | chr1-7986816-DEL-114 |
| chr1-7992067-DEL-326 | chr1-7992067-DEL-326 |
| chr1-8001056-DEL-869 | chr1-8001056-DEL-869 |
| chr1-8012883-DEL-7461 | chr1-8012884-DEL-7461 |
| chr1-8063924-DEL-7111 | chr1-8063925-DEL-7111 |
| chr1-8073108-DEL-81 | chr1-8073149-DEL-79 |
| chr1-8080499-DEL-58 | chr1-8080500-DEL-58 |
| chr1-8097198-DEL-3574 | chr1-8097199-DEL-3574 |
| chr1-8169280-DEL-474 | chr1-8169280-DEL-474 |
| chr1-8174976-DEL-82 | chr1-8174977-DEL-82 |
| chr1-8181638-DEL-6699 | chr1-8181649-DEL-6688 |
| chr1-8193204-DEL-198 | chr1-8193205-DEL-198 |
| chr1-8197206-DEL-2237 | chr1-8197207-DEL-2237 |
| chr1-8200217-DEL-248 | chr1-8200224-DEL-249 |
| chr1-8204370-DEL-2214 | chr1-8204368-DEL-2215 |
| chr1-8204944-DEL-126 | chr1-8204945-DEL-126 |
| chr1-8248565-DEL-2684 | chr1-8248566-DEL-2684 |
| chr1-8282686-DEL-6590 | chr1-8282676-DEL-6600 |
| chr1-8305778-DEL-249 | chr1-8305787-DEL-245 |
| chr1-8313294-DEL-1385 | chr1-8313275-DEL-1399 |
| chr1-8319164-DEL-6985 | chr1-8319165-DEL-6985 |
| chr1-8331549-DEL-328 | chr1-8331550-DEL-328 |
| chr1-8334407-DEL-109 | chr1-8334408-DEL-108 |
| chr1-8347139-DEL-645 | chr1-8347140-DEL-645 |
| chr1-8411504-DEL-567 | chr1-8411505-DEL-567 |
| chr1-8422435-DEL-114 | chr1-8422474-DEL-82 |
| chr1-8423864-DEL-51 | chr1-8423855-DEL-50 |
| chr1-8480905-DEL-468 | chr1-8480906-DEL-468 |
| chr1-8522208-DEL-730 | chr1-8522210-DEL-726 |
| chr1-8526542-DEL-162 | chr1-8526566-DEL-170 |
| chr1-8553621-DEL-519 | chr1-8553621-DEL-519 |
| chr1-8560260-DEL-209 | chr1-8560260-DEL-209 |
| chr1-8604799-DEL-1284 | chr1-8604800-DEL-1284 |
| chr1-8642219-DEL-314 | chr1-8642220-DEL-314 |
| chr1-8655990-DEL-204 | chr1-8655991-DEL-204 |
| chr1-8669051-DEL-2620 | chr1-8669042-DEL-2617 |
| chr1-8698691-DEL-6604 | chr1-8698691-DEL-6604 |
| chr1-8703556-DEL-83 | chr1-8703557-DEL-83 |
| chr1-8711257-DEL-214 | chr1-8711258-DEL-214 |
| chr1-8712999-DEL-108 | chr1-8713000-DEL-108 |
| chr1-8725754-DEL-68 | chr1-8725752-DEL-69 |
| chr1-8728572-DEL-4181 | chr1-8728573-DEL-4181 |
| chr1-8734317-DEL-6220 | chr1-8734318-DEL-6220 |
| chr1-8817538-DEL-188 | chr1-8817538-DEL-188 |
| chr1-8820711-DEL-74 | chr1-8820722-DEL-54 |
| chr1-8827633-DEL-213 | chr1-8827633-DEL-213 |
| chr1-8847508-DEL-431 | chr1-8847509-DEL-431 |
| chr1-8850199-DEL-740 | chr1-8850200-DEL-740 |
| chr1-8857502-DEL-80 | chr1-8857490-DEL-81 |
| chr1-8861865-DEL-590 | chr1-8861865-DEL-590 |
| chr1-8867821-DEL-1891 | chr1-8867822-DEL-1891 |
| chr1-8893133-DEL-416 | chr1-8893132-DEL-416 |
| chr1-8907769-DEL-212 | chr1-8907770-DEL-212 |
| chr1-8908986-DEL-548 | chr1-8908987-DEL-548 |
| chr1-8915727-DEL-573 | chr1-8915728-DEL-573 |
| chr1-8934044-DEL-98 | chr1-8934045-DEL-98 |
| chr1-8973966-DEL-219 | chr1-8973967-DEL-219 |
| chr1-8986875-DEL-433 | chr1-8986876-DEL-433 |
| chr1-8994959-DEL-52 | chr1-8994960-DEL-52 |
| chr1-9000885-DEL-840 | chr1-9000886-DEL-840 |
| chr1-9024533-DEL-209 | chr1-9024534-DEL-209 |
| chr1-9030822-DEL-51 | chr1-9030831-DEL-61 |
| chr1-9039520-DEL-223 | chr1-9039518-DEL-211 |
| chr1-9048896-DEL-723 | chr1-9048897-DEL-723 |
| chr1-9081013-DEL-56 | chr1-9081014-DEL-56 |
| chr1-9098803-DEL-692 | chr1-9098804-DEL-692 |
| chr1-9102858-DEL-77 | chr1-9102859-DEL-77 |
| chr1-9103386-DEL-2077 | chr1-9103387-DEL-2077 |
| chr1-9109923-DEL-3810 | chr1-9109922-DEL-3809 |

| | |
|---|---|
| chr1-9115055-DEL-6376 | chr1-9115042-DEL-6377 |
| chr1-9163939-DEL-197 | chr1-9163940-DEL-197 |
| chr1-9190974-DEL-4560 | chr1-9190975-DEL-4560 |
| chr1-9205461-DEL-1227 | chr1-9205461-DEL-1228 |
| chr1-9210548-DEL-656 | chr1-9210549-DEL-656 |
| chr1-9219089-DEL-60 | chr1-9219092-DEL-69 |
| chr1-9236546-DEL-168 | chr1-9236547-DEL-168 |
| chr1-9245610-DEL-6590 | chr1-9245611-DEL-6590 |
| chr1-9260572-DEL-384 | chr1-9260573-DEL-384 |
| chr1-9263297-DEL-51 | chr1-9263298-DEL-51 |
| chr1-9305250-DEL-5363 | chr1-9305251-DEL-5363 |
| chr1-9310789-DEL-214 | chr1-9310790-DEL-214 |
| chr1-9321206-DEL-201 | chr1-9321207-DEL-201 |
| chr1-9347104-DEL-445 | chr1-9347105-DEL-445 |
| chr1-9373801-DEL-1215 | chr1-9373801-DEL-1215 |
| chr1-9414127-DEL-535 | chr1-9414116-DEL-549 |
| chr1-9418431-DEL-206 | chr1-9418432-DEL-206 |
| chr1-9450446-DEL-498 | chr1-9450445-DEL-499 |
| chr1-9456810-DEL-210 | chr1-9456811-DEL-210 |
| chr1-9471984-DEL-197 | chr1-9471950-DEL-199 |
| chr1-9500861-DEL-2266 | chr1-9500862-DEL-2266 |
| chr1-9506903-DEL-634 | chr1-9506903-DEL-634 |
| chr1-9523404-DEL-4667 | chr1-9523407-DEL-4664 |
| chr1-9523783-DEL-739 | chr1-9523784-DEL-739 |
| chr1-9524722-DEL-346 | chr1-9524722-DEL-340 |
| chr1-9550021-DEL-775 | chr1-9550022-DEL-775 |
| chr1-9557019-DEL-78 | chr1-9557017-DEL-79 |
| chr1-9563717-DEL-325 | chr1-9563717-DEL-325 |
| chr1-9573699-DEL-394 | chr1-9573700-DEL-394 |
| chr1-9576686-DEL-356 | chr1-9576687-DEL-356 |
| chr1-9584569-DEL-92 | chr1-9584570-DEL-92 |
| chr1-9604744-DEL-438 | chr1-9604745-DEL-438 |
| chr1-9608088-DEL-251 | chr1-9608089-DEL-251 |
| chr1-9611152-DEL-81 | chr1-9611152-DEL-82 |
| chr1-9611657-DEL-56 | chr1-9611658-DEL-56 |
| chr1-9644746-DEL-1536 | chr1-9644747-DEL-1536 |
| chr1-9651968-DEL-96 | chr1-9651971-DEL-103 |
| chr1-9661686-DEL-54 | chr1-9661687-DEL-54 |
| chr1-9675833-DEL-144 | chr1-9675842-DEL-135 |
| chr1-9685072-DEL-124 | chr1-9685110-DEL-93 |
| chr1-9698661-DEL-199 | chr1-9698662-DEL-199 |
| chr1-9705912-DEL-120 | chr1-9705913-DEL-120 |
| chr1-9798024-DEL-137 | chr1-9798025-DEL-137 |
| chr1-9804635-DEL-53 | chr1-9804636-DEL-53 |
| chr1-9821627-DEL-164 | chr1-9821628-DEL-164 |
| chr1-9822894-DEL-52 | chr1-9822895-DEL-52 |
| chr1-9826713-DEL-118 | chr1-9826712-DEL-118 |
| chr1-9832532-DEL-215 | chr1-9832533-DEL-215 |
| chr1-9844154-DEL-207 | chr1-9844155-DEL-207 |
| chr1-9847849-DEL-318 | chr1-9847850-DEL-318 |
| chr1-9848624-DEL-486 | chr1-9848625-DEL-486 |
| chr1-9874737-DEL-395 | chr1-9874738-DEL-395 |
| chr1-9877726-DEL-1081 | chr1-9877727-DEL-1081 |
| chr1-9888194-DEL-920 | chr1-9888195-DEL-920 |
| chr1-9896043-DEL-98 | chr1-9896058-DEL-70 |
| chr1-9902077-DEL-100 | chr1-9902077-DEL-100 |
| chr1-9960772-DEL-494 | chr1-9960757-DEL-492 |
| chr1-9963760-DEL-69 | chr1-9963763-DEL-67 |
| chr1-10004408-DEL-209 | chr1-10004409-DEL-209 |
| chr1-10019107-DEL-91 | chr1-10019108-DEL-91 |
| chr1-10042934-DEL-64 | chr1-10042935-DEL-64 |
| chr1-10050326-DEL-1365 | chr1-10050327-DEL-1365 |
| chr1-10065656-DEL-514 | chr1-10065657-DEL-514 |
| chr1-10066467-DEL-195 | chr1-10066468-DEL-195 |
| chr1-10135751-DEL-182 | chr1-10135752-DEL-182 |
| chr1-10147662-DEL-145 | chr1-10147674-DEL-98 |
| chr1-10151520-DEL-64 | chr1-10151521-DEL-62 |
| chr1-10187564-DEL-81 | chr1-10187563-DEL-50 |
| chr1-10205011-DEL-231 | chr1-10205012-DEL-231 |
| chr1-10256887-DEL-117 | chr1-10256928-DEL-81 |
| chr1-10309002-DEL-173 | chr1-10309012-DEL-177 |
| chr1-10314970-DEL-96 | chr1-10314941-DEL-96 |
| chr1-10352706-DEL-246 | chr1-10352707-DEL-246 |
| chr1-10356272-DEL-55 | chr1-10356294-DEL-56 |
| chr1-10361094-DEL-96 | chr1-10361093-DEL-108 |
| chr1-10364867-DEL-305 | chr1-10364868-DEL-305 |
| chr1-10378329-DEL-149 | chr1-10378330-DEL-149 |
| chr1-10383303-DEL-175 | chr1-10383304-DEL-175 |
| chr1-10415561-DEL-173 | chr1-10415562-DEL-173 |
| chr1-10421354-DEL-85 | chr1-10421352-DEL-56 |

| | |
|---|---|
| chr1-10445252-DEL-158 | chr1-10445253-DEL-158 |
| chr1-10536404-DEL-3028 | chr1-10536405-DEL-3028 |
| chr1-10540401-DEL-478 | chr1-10540402-DEL-478 |
| chr1-10549159-DEL-259 | chr1-10549160-DEL-259 |
| chr1-10549705-DEL-464 | chr1-10549706-DEL-464 |
| chr1-10565648-DEL-1061 | chr1-10565649-DEL-1061 |
| chr1-10581500-DEL-280 | chr1-10581501-DEL-280 |
| chr1-10599169-DEL-6423 | chr1-10599162-DEL-6424 |
| chr1-10610647-DEL-3331 | chr1-10610650-DEL-3332 |
| chr1-10619367-DEL-373 | chr1-10619367-DEL-373 |
| chr1-10677545-DEL-309 | chr1-10677546-DEL-309 |
| chr1-10680941-DEL-307 | chr1-10680942-DEL-307 |
| chr1-10716942-DEL-2017 | chr1-10716941-DEL-2018 |
| chr1-10753983-DEL-98 | chr1-10753991-DEL-88 |
| chr1-10757762-DEL-66 | chr1-10757763-DEL-66 |
| chr1-10781195-DEL-83 | chr1-10781188-DEL-60 |
| chr1-10784389-DEL-59 | chr1-10784390-DEL-51 |
| chr1-10800096-DEL-7126 | chr1-10800097-DEL-7126 |
| chr1-10825785-DEL-65 | chr1-10825779-DEL-71 |
| chr1-10850315-DEL-97 | chr1-10850310-DEL-103 |
| chr1-10855082-DEL-1174 | chr1-10855083-DEL-1175 |
| chr1-10857065-DEL-6162 | chr1-10857066-DEL-6162 |
| chr1-10875486-DEL-2431 | chr1-10875487-DEL-2431 |
| chr1-10940517-DEL-938 | chr1-10940517-DEL-938 |
| chr1-10946128-DEL-63 | chr1-10946129-DEL-63 |
| chr1-10952488-DEL-6429 | chr1-10952489-DEL-6429 |
| chr1-10959854-DEL-6440 | chr1-10959855-DEL-6440 |
| chr1-10976785-DEL-114 | chr1-10976785-DEL-114 |
| chr1-10992594-DEL-3269 | chr1-10992595-DEL-3269 |
| chr1-10997592-DEL-59 | chr1-10997592-DEL-53 |
| chr1-11006776-DEL-434 | chr1-11006777-DEL-434 |
| chr1-11030884-DEL-57 | chr1-11030890-DEL-55 |
| chr1-11040872-DEL-13296 | chr1-11040873-DEL-13296 |
| chr1-11051881-DEL-445 | chr1-11051882-DEL-445 |
| chr1-11058144-DEL-593 | chr1-11058146-DEL-592 |
| chr1-11058954-DEL-408 | chr1-11058955-DEL-408 |
| chr1-11085132-DEL-392 | chr1-11085135-DEL-390 |
| chr1-11093932-DEL-78 | chr1-11093933-DEL-78 |
| chr1-11099785-DEL-3320 | chr1-11099786-DEL-3320 |
| chr1-11110018-DEL-62 | chr1-11110019-DEL-62 |
| chr1-11132367-DEL-51 | chr1-11132353-DEL-58 |
| chr1-11147521-DEL-61 | chr1-11147522-DEL-61 |
| chr1-11170500-DEL-173 | chr1-11170522-DEL-157 |
| chr1-11191414-DEL-139 | chr1-11191415-DEL-139 |
| chr1-11224566-DEL-133 | chr1-11224567-DEL-135 |
| chr1-11244629-DEL-54 | chr1-11244630-DEL-54 |
| chr1-11254095-DEL-369 | chr1-11254096-DEL-369 |
| chr1-11270396-DEL-503 | chr1-11270397-DEL-503 |
| chr1-11271222-DEL-1726 | chr1-11271223-DEL-1726 |
| chr1-11284526-DEL-706 | chr1-11284526-DEL-703 |
| chr1-11299789-DEL-58 | chr1-11299790-DEL-58 |
| chr1-11336757-DEL-106 | chr1-11336758-DEL-106 |
| chr1-11354499-DEL-105 | chr1-11354502-DEL-107 |
| chr1-11361912-DEL-81 | chr1-11361913-DEL-81 |
| chr1-11362963-DEL-69 | chr1-11362964-DEL-69 |
| chr1-11406584-DEL-134 | chr1-11406585-DEL-134 |
| chr1-11413678-DEL-333 | chr1-11413676-DEL-334 |
| chr1-11437708-DEL-2726 | chr1-11437709-DEL-2726 |
| chr1-11456078-DEL-88 | chr1-11456079-DEL-88 |
| chr1-11489636-DEL-433 | chr1-11489640-DEL-430 |
| chr1-11503380-DEL-1392 | chr1-11503381-DEL-1392 |
| chr1-11509533-DEL-202 | chr1-11509534-DEL-202 |
| chr1-11512000-DEL-206 | chr1-11512001-DEL-206 |
| chr1-11514893-DEL-101 | chr1-11514904-DEL-86 |
| chr1-11533331-DEL-292 | chr1-11533332-DEL-292 |
| chr1-11537738-DEL-137 | chr1-11537739-DEL-137 |
| chr1-11540289-DEL-896 | chr1-11540285-DEL-897 |
| chr1-11542800-DEL-1283 | chr1-11542801-DEL-1283 |
| chr1-11553515-DEL-121 | chr1-11553515-DEL-121 |
| chr1-11575379-DEL-140 | chr1-11575380-DEL-140 |
| chr1-11587729-DEL-54 | chr1-11587734-DEL-50 |
| chr1-11591203-DEL-287 | chr1-11591204-DEL-287 |
| chr1-11604666-DEL-326 | chr1-11604667-DEL-326 |
| chr1-11638045-DEL-995 | chr1-11638046-DEL-995 |
| chr1-11652512-DEL-54 | chr1-11652513-DEL-54 |
| chr1-11685084-DEL-12156 | chr1-11685085-DEL-12156 |
| chr1-11707804-DEL-670 | chr1-11707805-DEL-670 |
| chr1-11744177-DEL-6406 | chr1-11744178-DEL-6406 |
| chr1-11757502-DEL-338 | chr1-11757503-DEL-338 |
| chr1-11776690-DEL-1887 | chr1-11776691-DEL-1887 |

| | |
|---|---|
| chr1-11780091-DEL-1124 | chr1-11780092-DEL-1124 |
| chr1-11793444-DEL-6636 | chr1-11793445-DEL-6636 |
| chr1-11823376-DEL-104 | chr1-11823385-DEL-144 |
| chr1-11838029-DEL-60 | chr1-11838028-DEL-54 |
| chr1-11846266-DEL-327 | chr1-11846267-DEL-327 |
| chr1-11847014-DEL-6980 | chr1-11847015-DEL-6980 |
| chr1-11856383-DEL-176 | chr1-11856384-DEL-176 |
| chr1-11892833-DEL-461 | chr1-11892834-DEL-461 |
| chr1-11922521-DEL-401 | chr1-11922521-DEL-401 |
| chr1-11923203-DEL-214 | chr1-11923203-DEL-213 |
| chr1-11923931-DEL-6158 | chr1-11923917-DEL-6172 |
| chr1-11957332-DEL-4568 | chr1-11957333-DEL-4568 |
| chr1-11961956-DEL-2678 | chr1-11961956-DEL-2678 |
| chr1-11985803-DEL-69 | chr1-11985804-DEL-69 |
| chr1-12011254-DEL-95 | chr1-12011270-DEL-85 |
| chr1-12016865-DEL-4268 | chr1-12016865-DEL-4268 |
| chr1-12020399-DEL-75 | chr1-12020400-DEL-75 |
| chr1-12024577-DEL-57 | chr1-12024554-DEL-80 |
| chr1-12035394-DEL-561 | chr1-12035395-DEL-561 |
| chr1-12108709-DEL-3076 | chr1-12108716-DEL-3046 |
| chr1-12123916-DEL-109 | chr1-12123907-DEL-110 |
| chr1-12165559-DEL-325 | chr1-12165560-DEL-325 |
| chr1-12176599-DEL-217 | chr1-12176592-DEL-219 |
| chr1-12202378-DEL-1615 | chr1-12202379-DEL-1615 |
| chr1-12287419-DEL-6304 | chr1-12287418-DEL-6304 |
| chr1-12310886-DEL-51 | chr1-12310887-DEL-50 |
| chr1-12341626-DEL-63 | chr1-12341627-DEL-63 |
| chr1-12342645-DEL-193 | chr1-12342646-DEL-193 |
| chr1-12353558-DEL-366 | chr1-12353559-DEL-366 |
| chr1-12382738-DEL-59 | chr1-12382738-DEL-59 |
| chr1-12401712-DEL-1894 | chr1-12401713-DEL-1894 |
| chr1-12427184-DEL-1382 | chr1-12427185-DEL-1382 |
| chr1-12436223-DEL-427 | chr1-12436224-DEL-427 |
| chr1-12441066-DEL-54 | chr1-12441092-DEL-50 |
| chr1-12445145-DEL-5400 | chr1-12445146-DEL-5400 |
| chr1-12497013-DEL-71 | chr1-12497014-DEL-71 |
| chr1-12505164-DEL-52 | chr1-12505165-DEL-52 |
| chr1-12507361-DEL-56 | chr1-12507362-DEL-52 |
| chr1-12541289-DEL-128 | chr1-12541289-DEL-127 |
| chr1-12542734-DEL-173 | chr1-12542735-DEL-173 |
| chr1-12552605-DEL-93 | chr1-12552606-DEL-93 |
| chr1-12557978-DEL-65 | chr1-12557979-DEL-65 |
| chr1-12564304-DEL-120 | chr1-12564305-DEL-120 |
| chr1-12612979-DEL-121 | chr1-12613014-DEL-117 |
| chr1-12613687-DEL-116 | chr1-12613685-DEL-85 |
| chr1-12615868-DEL-331 | chr1-12615869-DEL-331 |
| chr1-12620266-DEL-796 | chr1-12620267-DEL-795 |
| chr1-12700342-DEL-65 | chr1-12700343-DEL-65 |
| chr1-12705409-DEL-215 | chr1-12705410-DEL-215 |
| chr1-12715999-DEL-133 | chr1-12716000-DEL-133 |
| chr1-12722511-DEL-56 | chr1-12722475-DEL-80 |
| chr1-12723417-DEL-368 | chr1-12723448-DEL-351 |
| chr1-12729865-DEL-60 | chr1-12729866-DEL-60 |
| chr1-12733061-DEL-5695 | chr1-12733062-DEL-5695 |
| chr1-12770770-DEL-91 | chr1-12770771-DEL-91 |
| chr1-12776315-DEL-201 | chr1-12776344-DEL-200 |
| chr1-12778596-DEL-230 | chr1-12778574-DEL-223 |
| chr1-12816294-DEL-209 | chr1-12816295-DEL-209 |
| chr1-12833118-DEL-210 | chr1-12833119-DEL-210 |
| chr1-12842803-DEL-90 | chr1-12842804-DEL-90 |
| chr1-12873964-DEL-59 | chr1-12873966-DEL-58 |
| chr1-12935455-DEL-76 | chr1-12935475-DEL-57 |
| chr1-12956527-DEL-92 | chr1-12956528-DEL-92 |
| chr1-12962135-DEL-63 | chr1-12962136-DEL-58 |
| chr1-12972746-DEL-496 | chr1-12972747-DEL-496 |
| chr1-12987410-DEL-214 | chr1-12987411-DEL-214 |
| chr1-13032688-DEL-56 | chr1-13032688-DEL-56 |
| chr1-13054313-DEL-75 | chr1-13054305-DEL-84 |
| chr1-13066434-DEL-145 | chr1-13066435-DEL-145 |
| chr1-13070920-DEL-206 | chr1-13070921-DEL-206 |
| chr1-13071796-DEL-204 | chr1-13071799-DEL-205 |
| chr1-13072952-DEL-113 | chr1-13072957-DEL-107 |
| chr1-13101000-DEL-207 | chr1-13101001-DEL-207 |
| chr1-13105015-DEL-389 | chr1-13105015-DEL-389 |
| chr1-13111148-DEL-215 | chr1-13111149-DEL-215 |
| chr1-13123152-DEL-96 | chr1-13123170-DEL-86 |
| chr1-13124461-DEL-206 | chr1-13124461-DEL-206 |
| chr1-13145763-DEL-1038 | chr1-13145746-DEL-1053 |
| chr1-13159517-DEL-208 | chr1-13159504-DEL-207 |
| chr1-13162997-DEL-411 | chr1-13162998-DEL-411 |

| | |
|---|---|
| chr1-13182678-DEL-596 | chr1-13182679-DEL-596 |
| chr1-13188427-DEL-195 | chr1-13188428-DEL-195 |
| chr1-13193389-DEL-71 | chr1-13193408-DEL-53 |
| chr1-13200371-DEL-110 | chr1-13200404-DEL-78 |
| chr1-13208187-DEL-98 | chr1-13208188-DEL-98 |
| chr1-13240138-DEL-171 | chr1-13240139-DEL-171 |
| chr1-13248627-DEL-216 | chr1-13248628-DEL-216 |
| chr1-13273578-DEL-175 | chr1-13273587-DEL-174 |
| chr1-13277781-DEL-6439 | chr1-13277782-DEL-6440 |
| chr1-13292252-DEL-2158 | chr1-13292253-DEL-2158 |
| chr1-13330179-DEL-70 | chr1-13330180-DEL-70 |
| chr1-13338632-DEL-128 | chr1-13338633-DEL-128 |
| chr1-13343221-DEL-117 | chr1-13343240-DEL-102 |
| chr1-13359577-DEL-122 | chr1-13359578-DEL-122 |
| chr1-13394498-DEL-224 | chr1-13394499-DEL-224 |
| chr1-13405058-DEL-206 | chr1-13405059-DEL-205 |
| chr1-13408924-DEL-121 | chr1-13408917-DEL-121 |
| chr1-13414932-DEL-91 | chr1-13414932-DEL-64 |
| chr1-13438422-DEL-59 | chr1-13438395-DEL-60 |
| chr1-13447640-DEL-198 | chr1-13447641-DEL-197 |
| chr1-13448668-DEL-1787 | chr1-13448669-DEL-1787 |
| chr1-13451041-DEL-52 | chr1-13451042-DEL-52 |
| chr1-13469385-DEL-278 | chr1-13469386-DEL-279 |
| chr1-13483626-DEL-290 | chr1-13483628-DEL-286 |
| chr1-13484993-DEL-343 | chr1-13484994-DEL-343 |
| chr1-13489526-DEL-595 | chr1-13489527-DEL-595 |
| chr1-13504639-DEL-79 | chr1-13504605-DEL-74 |
| chr1-13505631-DEL-363 | chr1-13505632-DEL-365 |
| chr1-13508859-DEL-52 | chr1-13508871-DEL-59 |
| chr1-13515523-DEL-204 | chr1-13515524-DEL-204 |
| chr1-13531671-DEL-74 | chr1-13531664-DEL-82 |
| chr1-13546703-DEL-7312 | chr1-13546702-DEL-7313 |
| chr1-13583713-DEL-394 | chr1-13583714-DEL-394 |
| chr1-13586253-DEL-210 | chr1-13586254-DEL-210 |
| chr1-13593577-DEL-57 | chr1-13593578-DEL-57 |
| chr1-13619249-DEL-9668 | chr1-13619230-DEL-9698 |
| chr1-13662634-DEL-118 | chr1-13662634-DEL-118 |
| chr1-13663991-DEL-73 | chr1-13663983-DEL-74 |
| chr1-13721097-DEL-68 | chr1-13721098-DEL-68 |
| chr1-13736012-DEL-252 | chr1-13736004-DEL-253 |
| chr1-13779210-DEL-75 | chr1-13779211-DEL-75 |
| chr1-13788348-DEL-90 | chr1-13788384-DEL-68 |
| chr1-13801103-DEL-6450 | chr1-13801112-DEL-6451 |
| chr1-13809586-DEL-104 | chr1-13809587-DEL-104 |
| chr1-13832786-DEL-235 | chr1-13832792-DEL-238 |
| chr1-13833882-DEL-6164 | chr1-13833872-DEL-6174 |
| chr1-13874228-DEL-61 | chr1-13874231-DEL-58 |
| chr1-13875660-DEL-210 | chr1-13875663-DEL-208 |
| chr1-13915286-DEL-75 | chr1-13915286-DEL-75 |
| chr1-13932513-DEL-66 | chr1-13932514-DEL-68 |
| chr1-13936617-DEL-58 | chr1-13936617-DEL-58 |
| chr1-13967340-DEL-213 | chr1-13967341-DEL-213 |
| chr1-14009512-DEL-155 | chr1-14009513-DEL-155 |
| chr1-14022454-DEL-495 | chr1-14022442-DEL-507 |
| chr1-14030931-DEL-2744 | chr1-14030931-DEL-2745 |
| chr1-14033850-DEL-468 | chr1-14033851-DEL-468 |
| chr1-14068791-DEL-344 | chr1-14068792-DEL-344 |
| chr1-14122910-DEL-51 | chr1-14122911-DEL-51 |
| chr1-14131807-DEL-175 | chr1-14131808-DEL-175 |
| chr1-14155076-DEL-6880 | chr1-14155077-DEL-6880 |
| chr1-14164783-DEL-496 | chr1-14164784-DEL-496 |
| chr1-14183448-DEL-6854 | chr1-14183449-DEL-6854 |
| chr1-14183628-DEL-212 | chr1-14183629-DEL-212 |
| chr1-14226226-DEL-344 | chr1-14226227-DEL-344 |
| chr1-14228973-DEL-993 | chr1-14228974-DEL-993 |
| chr1-14238140-DEL-278 | chr1-14238141-DEL-278 |
| chr1-14280474-DEL-624 | chr1-14280475-DEL-624 |
| chr1-14316261-DEL-68 | chr1-14316262-DEL-68 |
| chr1-14333607-DEL-66 | chr1-14333608-DEL-62 |
| chr1-14336352-DEL-172 | chr1-14336387-DEL-144 |
| chr1-14355102-DEL-60 | chr1-14355103-DEL-60 |
| chr1-14435403-DEL-2114 | chr1-14435396-DEL-2109 |
| chr1-14437784-DEL-468 | chr1-14437785-DEL-468 |
| chr1-14465634-DEL-807 | chr1-14465635-DEL-807 |
| chr1-14467842-DEL-118 | chr1-14467843-DEL-118 |
| chr1-14468375-DEL-323 | chr1-14468387-DEL-314 |
| chr1-14472327-DEL-121 | chr1-14472328-DEL-121 |
| chr1-14498159-DEL-57 | chr1-14498164-DEL-53 |
| chr1-14502060-DEL-76 | chr1-14502061-DEL-76 |
| chr1-14511435-DEL-203 | chr1-14511436-DEL-203 |

| | |
|---|---|
| chr1-14516509-DEL-318 | chr1-14516509-DEL-319 |
| chr1-14517718-DEL-6228 | chr1-14517718-DEL-6228 |
| chr1-14525652-DEL-245 | chr1-14525653-DEL-245 |
| chr1-14546419-DEL-6284 | chr1-14546420-DEL-6284 |
| chr1-14563972-DEL-626 | chr1-14563973-DEL-623 |
| chr1-14568256-DEL-120 | chr1-14568287-DEL-90 |
| chr1-14614216-DEL-96 | chr1-14614217-DEL-96 |
| chr1-14702057-DEL-1168 | chr1-14702058-DEL-1168 |
| chr1-14728703-DEL-185 | chr1-14728707-DEL-182 |
| chr1-14734874-DEL-208 | chr1-14734875-DEL-208 |
| chr1-14753990-DEL-335 | chr1-14753991-DEL-335 |
| chr1-14800943-DEL-108 | chr1-14800944-DEL-108 |
| chr1-14807201-DEL-417 | chr1-14807202-DEL-417 |
| chr1-14813608-DEL-983 | chr1-14813609-DEL-983 |
| chr1-14815330-DEL-1496 | chr1-14815331-DEL-1496 |
| chr1-14822485-DEL-189 | chr1-14822486-DEL-189 |
| chr1-14840514-DEL-61 | chr1-14840514-DEL-62 |
| chr1-14854059-DEL-778 | chr1-14854061-DEL-777 |
| chr1-14901858-DEL-569 | chr1-14901859-DEL-569 |
| chr1-14909462-DEL-1024 | chr1-14909463-DEL-1024 |
| chr1-14931727-DEL-353 | chr1-14931726-DEL-352 |
| chr1-14933114-DEL-643 | chr1-14933115-DEL-643 |
| chr1-14935226-DEL-1058 | chr1-14935227-DEL-1058 |
| chr1-14958178-DEL-863 | chr1-14958177-DEL-864 |
| chr1-14999082-DEL-99 | chr1-14999075-DEL-129 |
| chr1-15012794-DEL-207 | chr1-15012795-DEL-207 |
| chr1-15029570-DEL-1297 | chr1-15029571-DEL-1297 |
| chr1-15083912-DEL-1429 | chr1-15083913-DEL-1429 |
| chr1-15090512-DEL-6194 | chr1-15090513-DEL-6194 |
| chr1-15103592-DEL-647 | chr1-15103593-DEL-647 |
| chr1-15121506-DEL-828 | chr1-15121507-DEL-828 |
| chr1-15122413-DEL-166 | chr1-15122414-DEL-166 |
| chr1-15148243-DEL-14958 | chr1-15148244-DEL-14958 |
| chr1-15150422-DEL-3050 | chr1-15150423-DEL-3050 |
| chr1-15153719-DEL-7130 | chr1-15153720-DEL-7130 |
| chr1-15169421-DEL-550 | chr1-15169421-DEL-550 |
| chr1-15170621-DEL-1259 | chr1-15170618-DEL-1260 |
| chr1-15171676-DEL-61 | chr1-15171677-DEL-61 |
| chr1-15209262-DEL-935 | chr1-15209273-DEL-927 |
| chr1-15242746-DEL-98 | chr1-15242747-DEL-98 |
| chr1-15247856-DEL-994 | chr1-15247848-DEL-1002 |
| chr1-15305281-DEL-76 | chr1-15305283-DEL-75 |
| chr1-15328099-DEL-6485 | chr1-15328100-DEL-6485 |
| chr1-15347005-DEL-51 | chr1-15346994-DEL-54 |
| chr1-15363832-DEL-1514 | chr1-15363833-DEL-1514 |
| chr1-15414838-DEL-80 | chr1-15414863-DEL-84 |
| chr1-15421091-DEL-54 | chr1-15421092-DEL-54 |
| chr1-15437788-DEL-340 | chr1-15437789-DEL-340 |
| chr1-15439159-DEL-689 | chr1-15439160-DEL-689 |
| chr1-15442389-DEL-2470 | chr1-15442389-DEL-2470 |
| chr1-15443328-DEL-96 | chr1-15443329-DEL-96 |
| chr1-15466156-DEL-703 | chr1-15466157-DEL-703 |
| chr1-15491841-DEL-72 | chr1-15491841-DEL-72 |
| chr1-15492406-DEL-703 | chr1-15492407-DEL-703 |
| chr1-15514439-DEL-219 | chr1-15514440-DEL-219 |
| chr1-15532231-DEL-176 | chr1-15532232-DEL-176 |
| chr1-15534180-DEL-528 | chr1-15534181-DEL-529 |
| chr1-15554464-DEL-88 | chr1-15554478-DEL-72 |
| chr1-15561618-DEL-6301 | chr1-15561619-DEL-6301 |
| chr1-15589547-DEL-241 | chr1-15589549-DEL-240 |
| chr1-15595784-DEL-1669 | chr1-15595785-DEL-1669 |
| chr1-15607115-DEL-77 | chr1-15607113-DEL-79 |
| chr1-15626542-DEL-601 | chr1-15626543-DEL-601 |
| chr1-15637379-DEL-91 | chr1-15637381-DEL-53 |
| chr1-15639982-DEL-60 | chr1-15639983-DEL-60 |
| chr1-15656145-DEL-515 | chr1-15656146-DEL-515 |
| chr1-15667087-DEL-183 | chr1-15667088-DEL-183 |
| chr1-15688992-DEL-623 | chr1-15688989-DEL-624 |
| chr1-15691549-DEL-170 | chr1-15691550-DEL-170 |
| chr1-15715633-DEL-52 | chr1-15715634-DEL-52 |
| chr1-15734826-DEL-59 | chr1-15734813-DEL-69 |
| chr1-15759853-DEL-97 | chr1-15759854-DEL-97 |
| chr1-15771043-DEL-66 | chr1-15771047-DEL-104 |
| chr1-15776512-DEL-155 | chr1-15776494-DEL-136 |
| chr1-15777919-DEL-210 | chr1-15777920-DEL-210 |
| chr1-15806782-DEL-7025 | chr1-15806783-DEL-7025 |
| chr1-15843225-DEL-108 | chr1-15843205-DEL-95 |
| chr1-15856113-DEL-62 | chr1-15856113-DEL-62 |
| chr1-15874666-DEL-549 | chr1-15874666-DEL-549 |
| chr1-15915015-DEL-184 | chr1-15915016-DEL-184 |

| | |
|---|---|
| chr1-15932590-DEL-5511 | chr1-15932591-DEL-5511 |
| chr1-15940148-DEL-80 | chr1-15940149-DEL-80 |
| chr1-15971708-DEL-173 | chr1-15971715-DEL-172 |
| chr1-15998091-DEL-6819 | chr1-15998092-DEL-6819 |
| chr1-16018106-DEL-396 | chr1-16018107-DEL-396 |
| chr1-16020248-DEL-6215 | chr1-16020249-DEL-6215 |
| chr1-16031524-DEL-208 | chr1-16031525-DEL-208 |
| chr1-16032217-DEL-132 | chr1-16032218-DEL-132 |
| chr1-16035190-DEL-54 | chr1-16035190-DEL-54 |
| chr1-16047425-DEL-51 | chr1-16047426-DEL-51 |
| chr1-16059805-DEL-266 | chr1-16059806-DEL-266 |
| chr1-16100642-DEL-6165 | chr1-16100643-DEL-6165 |
| chr1-16108463-DEL-205 | chr1-16108464-DEL-205 |
| chr1-16114474-DEL-77 | chr1-16114475-DEL-77 |
| chr1-16130555-DEL-118 | chr1-16130554-DEL-119 |
| chr1-16157230-DEL-516 | chr1-16157231-DEL-516 |
| chr1-16159999-DEL-393 | chr1-16160008-DEL-385 |
| chr1-16204627-DEL-65 | chr1-16204628-DEL-65 |
| chr1-16218206-DEL-76 | chr1-16218207-DEL-76 |
| chr1-16260064-DEL-702 | chr1-16260065-DEL-702 |
| chr1-16275735-DEL-345 | chr1-16275736-DEL-345 |
| chr1-16280641-DEL-80 | chr1-16280642-DEL-80 |
| chr1-16321323-DEL-117 | chr1-16321324-DEL-117 |
| chr1-16357562-DEL-52 | chr1-16357541-DEL-76 |
| chr1-16360228-DEL-53 | chr1-16360229-DEL-53 |
| chr1-16386121-DEL-110 | chr1-16386122-DEL-110 |
| chr1-16451374-DEL-1887 | chr1-16451375-DEL-1887 |
| chr1-16523103-DEL-5040 | chr1-16523103-DEL-5040 |
| chr1-16528709-DEL-401 | chr1-16528710-DEL-401 |
| chr1-16537781-DEL-785 | chr1-16537782-DEL-785 |
| chr1-16548214-DEL-515 | chr1-16548215-DEL-515 |
| chr1-16573919-DEL-57 | chr1-16573920-DEL-57 |
| chr1-16577624-DEL-58 | chr1-16577625-DEL-58 |
| chr1-16581615-DEL-4252 | chr1-16581616-DEL-4252 |
| chr1-16582754-DEL-59 | chr1-16582755-DEL-59 |
| chr1-16583029-DEL-353 | chr1-16583027-DEL-354 |
| chr1-16586544-DEL-380 | chr1-16586526-DEL-374 |
| chr1-16598396-DEL-1456 | chr1-16598397-DEL-1456 |
| chr1-16602493-DEL-64 | chr1-16602494-DEL-64 |
| chr1-16624679-DEL-65 | chr1-16624680-DEL-65 |
| chr1-16680335-DEL-180 | chr1-16680336-DEL-180 |
| chr1-16692156-DEL-4149 | chr1-16692157-DEL-4149 |
| chr1-16716263-DEL-203 | chr1-16716264-DEL-203 |
| chr1-16743758-DEL-125 | chr1-16743759-DEL-125 |
| chr1-16749431-DEL-247 | chr1-16749432-DEL-247 |
| chr1-16750684-DEL-513 | chr1-16750685-DEL-513 |
| chr1-16756073-DEL-67 | chr1-16756070-DEL-68 |
| chr1-16772302-DEL-522 | chr1-16772303-DEL-522 |
| chr1-16781161-DEL-3158 | chr1-16781156-DEL-3177 |
| chr1-16787191-DEL-201 | chr1-16787192-DEL-201 |
| chr1-16788916-DEL-1187 | chr1-16788917-DEL-1187 |
| chr1-16792282-DEL-185 | chr1-16792283-DEL-185 |
| chr1-16808491-DEL-7057 | chr1-16808492-DEL-7057 |
| chr1-16821274-DEL-89 | chr1-16821314-DEL-50 |
| chr1-16832381-DEL-427 | chr1-16832382-DEL-427 |
| chr1-16846077-DEL-327 | chr1-16846077-DEL-325 |
| chr1-16868545-DEL-85 | chr1-16868513-DEL-86 |
| chr1-16898208-DEL-7457 | chr1-16898209-DEL-7457 |
| chr1-16918353-DEL-358 | chr1-16918354-DEL-358 |
| chr1-16958577-DEL-176 | chr1-16958578-DEL-176 |
| chr1-16959076-DEL-788 | chr1-16959077-DEL-788 |
| chr1-16962116-DEL-6401 | chr1-16962117-DEL-6401 |
| chr1-16971627-DEL-1488 | chr1-16971628-DEL-1488 |
| chr1-16974352-DEL-376 | chr1-16974353-DEL-376 |
| chr1-16983082-DEL-2708 | chr1-16983083-DEL-2708 |
| chr1-17017234-DEL-6906 | chr1-17017234-DEL-6906 |
| chr1-17041134-DEL-1114 | chr1-17041135-DEL-1114 |
| chr1-17051897-DEL-422 | chr1-17051916-DEL-397 |
| chr1-17061901-DEL-87 | chr1-17061901-DEL-87 |
| chr1-17066740-DEL-51 | chr1-17066740-DEL-51 |
| chr1-17071969-DEL-77 | chr1-17071958-DEL-77 |
| chr1-17143826-DEL-1659 | chr1-17143826-DEL-1659 |
| chr1-17189387-DEL-73 | chr1-17189345-DEL-100 |
| chr1-17235633-DEL-202 | chr1-17235634-DEL-202 |
| chr1-17263146-DEL-204 | chr1-17263139-DEL-205 |
| chr1-17272426-DEL-107 | chr1-17272427-DEL-107 |
| chr1-17304806-DEL-219 | chr1-17304795-DEL-220 |
| chr1-17331969-DEL-1879 | chr1-17331963-DEL-1880 |
| chr1-17346078-DEL-816 | chr1-17346079-DEL-816 |
| chr1-17352933-DEL-67 | chr1-17352934-DEL-67 |

| | |
|---|---|
| chr1-17381601-DEL-402 | chr1-17381602-DEL-403 |
| chr1-17389912-DEL-1067 | chr1-17389913-DEL-1067 |
| chr1-17399123-DEL-6171 | chr1-17399124-DEL-6171 |
| chr1-17422260-DEL-69 | chr1-17422261-DEL-69 |
| chr1-17428480-DEL-2437 | chr1-17428481-DEL-2435 |
| chr1-17446933-DEL-321 | chr1-17446934-DEL-321 |
| chr1-17453498-DEL-6886 | chr1-17453499-DEL-6886 |
| chr1-17465836-DEL-1621 | chr1-17465837-DEL-1621 |
| chr1-17471936-DEL-3139 | chr1-17471937-DEL-3139 |
| chr1-17473752-DEL-5502 | chr1-17473753-DEL-5502 |
| chr1-17492798-DEL-326 | chr1-17492799-DEL-326 |
| chr1-17525197-DEL-968 | chr1-17525198-DEL-968 |
| chr1-17530015-DEL-206 | chr1-17530016-DEL-206 |
| chr1-17537064-DEL-6685 | chr1-17537065-DEL-6685 |
| chr1-17562474-DEL-55 | chr1-17562475-DEL-55 |
| chr1-17562892-DEL-190 | chr1-17562893-DEL-190 |
| chr1-17578170-DEL-129 | chr1-17578171-DEL-129 |
| chr1-17582170-DEL-1272 | chr1-17582171-DEL-1272 |
| chr1-17615925-DEL-57 | chr1-17615926-DEL-57 |
| chr1-17635787-DEL-87 | chr1-17635788-DEL-87 |
| chr1-17663191-DEL-53 | chr1-17663186-DEL-55 |
| chr1-17669172-DEL-819 | chr1-17669173-DEL-819 |
| chr1-17704772-DEL-59 | chr1-17704801-DEL-50 |
| chr1-17731835-DEL-403 | chr1-17731836-DEL-403 |
| chr1-17733113-DEL-396 | chr1-17733114-DEL-396 |
| chr1-17750057-DEL-6367 | chr1-17750058-DEL-6367 |
| chr1-17769137-DEL-6619 | chr1-17769138-DEL-6619 |
| chr1-17775257-DEL-416 | chr1-17775258-DEL-416 |
| chr1-17783514-DEL-269 | chr1-17783494-DEL-286 |
| chr1-17858546-DEL-193 | chr1-17858547-DEL-193 |
| chr1-17863187-DEL-500 | chr1-17863188-DEL-500 |
| chr1-17886129-DEL-1993 | chr1-17886129-DEL-1994 |
| chr1-17903841-DEL-312 | chr1-17903842-DEL-305 |
| chr1-17912209-DEL-171 | chr1-17912210-DEL-171 |
| chr1-17974081-DEL-215 | chr1-17974082-DEL-215 |
| chr1-17986343-DEL-6469 | chr1-17986344-DEL-6469 |
| chr1-18001310-DEL-54 | chr1-18001311-DEL-54 |
| chr1-18008547-DEL-53 | chr1-18008548-DEL-53 |
| chr1-18021033-DEL-231 | chr1-18021033-DEL-231 |
| chr1-18075248-DEL-552 | chr1-18075253-DEL-553 |
| chr1-18108709-DEL-1370 | chr1-18108710-DEL-1370 |
| chr1-18122031-DEL-54 | chr1-18122032-DEL-54 |
| chr1-18124933-DEL-216 | chr1-18124934-DEL-216 |
| chr1-18148174-DEL-9628 | chr1-18148161-DEL-9617 |
| chr1-18153162-DEL-2305 | chr1-18153161-DEL-2305 |
| chr1-18183008-DEL-379 | chr1-18183009-DEL-379 |
| chr1-18209202-DEL-1970 | chr1-18209203-DEL-1970 |
| chr1-18226357-DEL-1889 | chr1-18226358-DEL-1889 |
| chr1-18234360-DEL-314 | chr1-18234361-DEL-314 |
| chr1-18236811-DEL-65 | chr1-18236830-DEL-50 |
| chr1-18259899-DEL-113 | chr1-18259892-DEL-121 |
| chr1-18264329-DEL-83 | chr1-18264330-DEL-83 |
| chr1-18277851-DEL-1519 | chr1-18277852-DEL-1519 |
| chr1-18299296-DEL-62 | chr1-18299297-DEL-62 |
| chr1-18322698-DEL-2668 | chr1-18322699-DEL-2668 |
| chr1-18354173-DEL-86 | chr1-18354174-DEL-86 |
| chr1-18363993-DEL-849 | chr1-18363989-DEL-850 |
| chr1-18369136-DEL-996 | chr1-18369137-DEL-996 |
| chr1-18372894-DEL-1519 | chr1-18372895-DEL-1519 |
| chr1-18388888-DEL-52 | chr1-18388889-DEL-52 |
| chr1-18390906-DEL-79 | chr1-18390907-DEL-79 |
| chr1-18394450-DEL-336 | chr1-18394451-DEL-336 |
| chr1-18427772-DEL-205 | chr1-18427773-DEL-205 |
| chr1-18429392-DEL-604 | chr1-18429390-DEL-605 |
| chr1-18450590-DEL-148 | chr1-18450601-DEL-124 |
| chr1-18462133-DEL-258 | chr1-18462134-DEL-258 |
| chr1-18474334-DEL-103 | chr1-18474315-DEL-106 |
| chr1-18491199-DEL-72 | chr1-18491200-DEL-72 |
| chr1-18502594-DEL-132 | chr1-18502595-DEL-132 |
| chr1-18504610-DEL-86 | chr1-18504611-DEL-86 |
| chr1-18506591-DEL-125 | chr1-18506596-DEL-119 |
| chr1-18507642-DEL-501 | chr1-18507643-DEL-501 |
| chr1-18515123-DEL-1508 | chr1-18515124-DEL-1508 |
| chr1-18519261-DEL-3659 | chr1-18519262-DEL-3659 |
| chr1-18542898-DEL-6543 | chr1-18542901-DEL-6553 |
| chr1-18560986-DEL-66 | chr1-18560975-DEL-59 |
| chr1-18587057-DEL-4379 | chr1-18587058-DEL-4379 |
| chr1-18607236-DEL-7530 | chr1-18607237-DEL-7530 |
| chr1-18608057-DEL-212 | chr1-18608058-DEL-212 |
| chr1-18615100-DEL-613 | chr1-18615101-DEL-613 |

| | |
|---|---|
| chr1-18616592-DEL-1031 | chr1-18616593-DEL-1032 |
| chr1-18661670-DEL-64 | chr1-18661671-DEL-64 |
| chr1-18685786-DEL-146 | chr1-18685796-DEL-128 |
| chr1-18696746-DEL-705 | chr1-18696747-DEL-705 |
| chr1-18698764-DEL-432 | chr1-18698765-DEL-432 |
| chr1-18700207-DEL-160 | chr1-18700208-DEL-160 |
| chr1-18706495-DEL-1294 | chr1-18706493-DEL-1296 |
| chr1-18711242-DEL-1652 | chr1-18711243-DEL-1652 |
| chr1-18712011-DEL-62 | chr1-18712012-DEL-62 |
| chr1-18722923-DEL-54 | chr1-18722924-DEL-54 |
| chr1-18728835-DEL-6661 | chr1-18728836-DEL-6661 |
| chr1-18755586-DEL-1209 | chr1-18755587-DEL-1209 |
| chr1-18798968-DEL-52 | chr1-18798960-DEL-72 |
| chr1-18818360-DEL-53 | chr1-18818356-DEL-52 |
| chr1-18830843-DEL-2773 | chr1-18830844-DEL-2773 |
| chr1-18869876-DEL-843 | chr1-18869877-DEL-843 |
| chr1-18873494-DEL-2076 | chr1-18873494-DEL-2076 |
| chr1-18876281-DEL-266 | chr1-18876281-DEL-267 |
| chr1-18885174-DEL-6171 | chr1-18885174-DEL-6171 |
| chr1-18931651-DEL-329 | chr1-18931652-DEL-329 |
| chr1-18966851-DEL-1174 | chr1-18966852-DEL-1174 |
| chr1-18970301-DEL-1239 | chr1-18970302-DEL-1239 |
| chr1-19025771-DEL-76 | chr1-19025790-DEL-58 |
| chr1-19084202-DEL-1580 | chr1-19084186-DEL-1578 |
| chr1-19086586-DEL-108 | chr1-19086587-DEL-108 |
| chr1-19148418-DEL-1147 | chr1-19148419-DEL-1147 |
| chr1-19153814-DEL-388 | chr1-19153816-DEL-387 |
| chr1-19165887-DEL-59 | chr1-19165888-DEL-59 |
| chr1-19183926-DEL-77 | chr1-19183927-DEL-78 |
| chr1-19195937-DEL-6560 | chr1-19195938-DEL-6560 |
| chr1-19217252-DEL-64 | chr1-19217253-DEL-64 |
| chr1-19228317-DEL-457 | chr1-19228318-DEL-457 |
| chr1-19246828-DEL-218 | chr1-19246829-DEL-218 |
| chr1-19300188-DEL-62 | chr1-19300181-DEL-80 |
| chr1-19327744-DEL-238 | chr1-19327745-DEL-237 |
| chr1-19329014-DEL-61 | chr1-19329015-DEL-61 |
| chr1-19335711-DEL-210 | chr1-19335712-DEL-210 |
| chr1-19353448-DEL-3093 | chr1-19353449-DEL-3093 |
| chr1-19418860-DEL-73 | chr1-19418854-DEL-74 |
| chr1-19442495-DEL-6593 | chr1-19442496-DEL-6593 |
| chr1-19442527-DEL-212 | chr1-19442527-DEL-212 |
| chr1-19451442-DEL-1210 | chr1-19451443-DEL-1210 |
| chr1-19463913-DEL-385 | chr1-19463914-DEL-385 |
| chr1-19469010-DEL-1814 | chr1-19469010-DEL-1814 |
| chr1-19472063-DEL-3209 | chr1-19472064-DEL-3209 |
| chr1-19540868-DEL-1380 | chr1-19540869-DEL-1380 |
| chr1-19553568-DEL-73 | chr1-19553569-DEL-73 |
| chr1-19558627-DEL-92 | chr1-19558628-DEL-92 |
| chr1-19574410-DEL-167 | chr1-19574411-DEL-167 |
| chr1-19588421-DEL-397 | chr1-19588422-DEL-395 |
| chr1-19608383-DEL-54 | chr1-19608384-DEL-51 |
| chr1-19612400-DEL-58 | chr1-19612401-DEL-54 |
| chr1-19654729-DEL-78 | chr1-19654700-DEL-100 |
| chr1-19671636-DEL-3333 | chr1-19671637-DEL-3333 |
| chr1-19692572-DEL-7223 | chr1-19692573-DEL-7223 |
| chr1-19707242-DEL-77 | chr1-19707243-DEL-77 |
| chr1-19736906-DEL-1245 | chr1-19736907-DEL-1245 |
| chr1-19757376-DEL-6399 | chr1-19757377-DEL-6399 |
| chr1-19777109-DEL-6353 | chr1-19777110-DEL-6353 |
| chr1-19777222-DEL-211 | chr1-19777223-DEL-211 |
| chr1-19809924-DEL-168 | chr1-19809925-DEL-168 |
| chr1-19869550-DEL-6792 | chr1-19869545-DEL-6793 |
| chr1-19877806-DEL-52 | chr1-19877809-DEL-88 |
| chr1-19899983-DEL-51 | chr1-19899984-DEL-51 |
| chr1-19907138-DEL-1205 | chr1-19907139-DEL-1205 |
| chr1-19909089-DEL-6843 | chr1-19909090-DEL-6843 |
| chr1-19909473-DEL-206 | chr1-19909474-DEL-206 |
| chr1-19960327-DEL-58 | chr1-19960328-DEL-58 |
| chr1-19977248-DEL-219 | chr1-19977254-DEL-218 |
| chr1-20016898-DEL-70 | chr1-20016899-DEL-70 |
| chr1-20064492-DEL-5400 | chr1-20064493-DEL-5400 |
| chr1-20072167-DEL-85 | chr1-20072167-DEL-85 |
| chr1-20073752-DEL-70 | chr1-20073753-DEL-70 |
| chr1-20074731-DEL-504 | chr1-20074732-DEL-504 |
| chr1-20080043-DEL-134 | chr1-20080044-DEL-129 |
| chr1-20110369-DEL-5018 | chr1-20110363-DEL-5018 |
| chr1-20115088-DEL-193 | chr1-20115089-DEL-193 |
| chr1-20124910-DEL-79 | chr1-20124911-DEL-79 |
| chr1-20125448-DEL-146 | chr1-20125449-DEL-142 |
| chr1-20131649-DEL-79 | chr1-20131648-DEL-80 |

| | |
|---|---|
| chr1-20137279-DEL-420 | chr1-20137280-DEL-420 |
| chr1-20140963-DEL-288 | chr1-20140963-DEL-288 |
| chr1-20174469-DEL-53 | chr1-20174471-DEL-52 |
| chr1-20190484-DEL-57 | chr1-20190485-DEL-57 |
| chr1-20191324-DEL-104 | chr1-20191325-DEL-104 |
| chr1-20196848-DEL-1740 | chr1-20196849-DEL-1740 |
| chr1-20273997-DEL-2966 | chr1-20273998-DEL-2966 |
| chr1-20280541-DEL-85 | chr1-20280542-DEL-85 |
| chr1-20292264-DEL-82 | chr1-20292287-DEL-58 |
| chr1-20299183-DEL-6134 | chr1-20299184-DEL-6134 |
| chr1-20319340-DEL-2065 | chr1-20319341-DEL-2065 |
| chr1-20327648-DEL-364 | chr1-20327651-DEL-362 |
| chr1-20332319-DEL-806 | chr1-20332320-DEL-806 |
| chr1-20340963-DEL-543 | chr1-20340964-DEL-543 |
| chr1-20348153-DEL-6873 | chr1-20348149-DEL-6872 |
| chr1-20385573-DEL-6318 | chr1-20385574-DEL-6318 |
| chr1-20394055-DEL-399 | chr1-20394056-DEL-399 |
| chr1-20418651-DEL-1031 | chr1-20418652-DEL-1031 |
| chr1-20421131-DEL-197 | chr1-20421132-DEL-197 |
| chr1-20463037-DEL-1474 | chr1-20463038-DEL-1474 |
| chr1-20502433-DEL-7415 | chr1-20502425-DEL-7416 |
| chr1-20521379-DEL-67 | chr1-20521380-DEL-67 |
| chr1-20554891-DEL-80 | chr1-20554893-DEL-79 |
| chr1-20556602-DEL-7238 | chr1-20556603-DEL-7238 |
| chr1-20579467-DEL-54 | chr1-20579463-DEL-58 |
| chr1-20584430-DEL-118 | chr1-20584424-DEL-119 |
| chr1-20586217-DEL-51 | chr1-20586218-DEL-51 |
| chr1-20594891-DEL-886 | chr1-20594892-DEL-886 |
| chr1-20601968-DEL-254 | chr1-20601965-DEL-250 |
| chr1-20619337-DEL-148 | chr1-20619338-DEL-148 |
| chr1-20630856-DEL-461 | chr1-20630857-DEL-461 |
| chr1-20642679-DEL-1558 | chr1-20642680-DEL-1558 |
| chr1-20644704-DEL-6582 | chr1-20644705-DEL-6582 |
| chr1-20664872-DEL-1325 | chr1-20664873-DEL-1325 |
| chr1-20702972-DEL-489 | chr1-20702973-DEL-489 |
| chr1-20707837-DEL-2277 | chr1-20707838-DEL-2278 |
| chr1-20712571-DEL-6178 | chr1-20712572-DEL-6177 |
| chr1-20744066-DEL-82 | chr1-20744066-DEL-84 |
| chr1-20745910-DEL-110 | chr1-20745911-DEL-110 |
| chr1-20775625-DEL-7087 | chr1-20775626-DEL-7087 |
| chr1-20779695-DEL-897 | chr1-20779682-DEL-898 |
| chr1-20814151-DEL-238 | chr1-20814135-DEL-221 |
| chr1-20858238-DEL-97 | chr1-20858239-DEL-97 |
| chr1-20869600-DEL-124 | chr1-20869601-DEL-123 |
| chr1-20890168-DEL-62 | chr1-20890171-DEL-62 |
| chr1-20892702-DEL-1394 | chr1-20892716-DEL-1385 |
| chr1-20900620-DEL-216 | chr1-20900621-DEL-201 |
| chr1-20901705-DEL-81 | chr1-20901706-DEL-81 |
| chr1-20913260-DEL-60 | chr1-20913262-DEL-61 |
| chr1-20944917-DEL-54 | chr1-20944918-DEL-54 |
| chr1-20968150-DEL-55 | chr1-20968151-DEL-55 |
| chr1-20973320-DEL-58 | chr1-20973321-DEL-58 |
| chr1-20984767-DEL-255 | chr1-20984785-DEL-255 |
| chr1-20988956-DEL-617 | chr1-20988957-DEL-617 |
| chr1-21000092-DEL-749 | chr1-21000093-DEL-749 |
| chr1-21001878-DEL-211 | chr1-21001879-DEL-211 |
| chr1-21014074-DEL-60 | chr1-21014075-DEL-60 |
| chr1-21015074-DEL-92 | chr1-21015073-DEL-96 |
| chr1-21022403-DEL-241 | chr1-21022404-DEL-241 |
| chr1-21053876-DEL-655 | chr1-21053877-DEL-656 |
| chr1-21055464-DEL-54 | chr1-21055465-DEL-52 |
| chr1-21064135-DEL-719 | chr1-21064136-DEL-719 |
| chr1-21095892-DEL-55 | chr1-21095892-DEL-61 |
| chr1-21107881-DEL-137 | chr1-21107882-DEL-137 |
| chr1-21118199-DEL-56 | chr1-21118200-DEL-56 |
| chr1-21168739-DEL-736 | chr1-21168740-DEL-736 |
| chr1-21185077-DEL-7116 | chr1-21185078-DEL-7116 |
| chr1-21204972-DEL-3400 | chr1-21204973-DEL-3400 |
| chr1-21210333-DEL-60 | chr1-21210334-DEL-57 |
| chr1-21224450-DEL-52 | chr1-21224451-DEL-52 |
| chr1-21228438-DEL-128 | chr1-21228439-DEL-128 |
| chr1-21233026-DEL-142 | chr1-21233027-DEL-142 |
| chr1-21235146-DEL-81 | chr1-21235147-DEL-81 |
| chr1-21239871-DEL-158 | chr1-21239871-DEL-158 |
| chr1-21241272-DEL-203 | chr1-21241272-DEL-203 |
| chr1-21265558-DEL-6879 | chr1-21265559-DEL-6879 |
| chr1-21281730-DEL-111 | chr1-21281731-DEL-111 |
| chr1-21342533-DEL-815 | chr1-21342523-DEL-816 |
| chr1-21348037-DEL-185 | chr1-21348038-DEL-185 |
| chr1-21349594-DEL-97 | chr1-21349596-DEL-96 |

| | |
|---|---|
| chr1-21354263-DEL-1014 | chr1-21354264-DEL-1014 |
| chr1-21360049-DEL-1006 | chr1-21360050-DEL-1006 |
| chr1-21366079-DEL-59 | chr1-21366080-DEL-59 |
| chr1-21384913-DEL-193 | chr1-21384914-DEL-193 |
| chr1-21389372-DEL-816 | chr1-21389373-DEL-816 |
| chr1-21405195-DEL-205 | chr1-21405196-DEL-205 |
| chr1-21419699-DEL-110 | chr1-21419699-DEL-110 |
| chr1-21423383-DEL-186 | chr1-21423382-DEL-186 |
| chr1-21430598-DEL-3943 | chr1-21430597-DEL-3943 |
| chr1-21444795-DEL-177 | chr1-21444794-DEL-177 |
| chr1-21459652-DEL-363 | chr1-21459653-DEL-361 |
| chr1-21462964-DEL-296 | chr1-21462965-DEL-296 |
| chr1-21490265-DEL-465 | chr1-21490266-DEL-465 |
| chr1-21493204-DEL-54 | chr1-21493205-DEL-54 |
| chr1-21497083-DEL-51 | chr1-21497087-DEL-53 |
| chr1-21510611-DEL-59 | chr1-21510612-DEL-59 |
| chr1-21519117-DEL-135 | chr1-21519125-DEL-103 |
| chr1-21542523-DEL-85 | chr1-21542524-DEL-85 |
| chr1-21551722-DEL-219 | chr1-21551723-DEL-219 |
| chr1-21570637-DEL-132 | chr1-21570638-DEL-132 |
| chr1-21571834-DEL-87 | chr1-21571821-DEL-84 |
| chr1-21578795-DEL-88 | chr1-21578835-DEL-56 |
| chr1-21596872-DEL-439 | chr1-21596877-DEL-436 |
| chr1-21611597-DEL-207 | chr1-21611598-DEL-207 |
| chr1-21619000-DEL-501 | chr1-21619001-DEL-501 |
| chr1-21632414-DEL-54 | chr1-21632414-DEL-50 |
| chr1-21651845-DEL-3424 | chr1-21651842-DEL-3424 |
| chr1-21667305-DEL-63 | chr1-21667306-DEL-63 |
| chr1-21702957-DEL-240 | chr1-21702958-DEL-240 |
| chr1-21717823-DEL-650 | chr1-21717824-DEL-650 |
| chr1-21727670-DEL-510 | chr1-21727671-DEL-510 |
| chr1-21738358-DEL-1763 | chr1-21738359-DEL-1763 |
| chr1-21740519-DEL-1320 | chr1-21740520-DEL-1320 |
| chr1-21762688-DEL-1005 | chr1-21762689-DEL-1005 |
| chr1-21800777-DEL-60 | chr1-21800778-DEL-60 |
| chr1-21813173-DEL-116 | chr1-21813174-DEL-116 |
| chr1-21849720-DEL-106 | chr1-21849758-DEL-68 |
| chr1-21863392-DEL-115 | chr1-21863396-DEL-112 |
| chr1-21866998-DEL-69 | chr1-21866999-DEL-69 |
| chr1-21879924-DEL-70 | chr1-21879925-DEL-70 |
| chr1-21943402-DEL-773 | chr1-21943403-DEL-773 |
| chr1-21996618-DEL-738 | chr1-21996619-DEL-738 |
| chr1-22016204-DEL-65 | chr1-22016205-DEL-65 |
| chr1-22065474-DEL-6868 | chr1-22065474-DEL-6868 |
| chr1-22076812-DEL-212 | chr1-22076813-DEL-212 |
| chr1-22124715-DEL-7026 | chr1-22124712-DEL-7027 |
| chr1-22148623-DEL-203 | chr1-22148652-DEL-204 |
| chr1-22150481-DEL-391 | chr1-22150482-DEL-391 |
| chr1-22162501-DEL-6221 | chr1-22162502-DEL-6221 |
| chr1-22177586-DEL-318 | chr1-22177583-DEL-319 |
| chr1-22181084-DEL-53 | chr1-22181085-DEL-53 |
| chr1-22199635-DEL-967 | chr1-22199636-DEL-967 |
| chr1-22223022-DEL-10663 | chr1-22223023-DEL-10663 |
| chr1-22256421-DEL-2034 | chr1-22256422-DEL-2034 |
| chr1-22288398-DEL-204 | chr1-22288399-DEL-204 |
| chr1-22317839-DEL-366 | chr1-22317840-DEL-366 |
| chr1-22319399-DEL-51 | chr1-22319377-DEL-52 |
| chr1-22340676-DEL-911 | chr1-22340677-DEL-911 |
| chr1-22352197-DEL-483 | chr1-22352198-DEL-483 |
| chr1-22353241-DEL-434 | chr1-22353242-DEL-434 |
| chr1-22374066-DEL-261 | chr1-22374067-DEL-261 |
| chr1-22382462-DEL-54 | chr1-22382464-DEL-52 |
| chr1-22391605-DEL-80 | chr1-22391606-DEL-80 |
| chr1-22398969-DEL-905 | chr1-22398970-DEL-905 |
| chr1-22406132-DEL-175 | chr1-22406133-DEL-175 |
| chr1-22431716-DEL-54 | chr1-22431717-DEL-54 |
| chr1-22444138-DEL-332 | chr1-22444139-DEL-332 |
| chr1-22446140-DEL-258 | chr1-22446141-DEL-258 |
| chr1-22537352-DEL-1366 | chr1-22537342-DEL-1366 |
| chr1-22564141-DEL-140 | chr1-22564141-DEL-128 |
| chr1-22593058-DEL-74 | chr1-22593059-DEL-74 |
| chr1-22606167-DEL-532 | chr1-22606168-DEL-531 |
| chr1-22606879-DEL-762 | chr1-22606880-DEL-762 |
| chr1-22608929-DEL-85 | chr1-22608930-DEL-85 |
| chr1-22645529-DEL-75 | chr1-22645519-DEL-82 |
| chr1-22672355-DEL-80 | chr1-22672372-DEL-64 |
| chr1-22676073-DEL-1245 | chr1-22676074-DEL-1245 |
| chr1-22685737-DEL-342 | chr1-22685738-DEL-342 |
| chr1-22696077-DEL-72 | chr1-22696078-DEL-72 |
| chr1-22735407-DEL-1957 | chr1-22735408-DEL-1957 |

| | |
|---|---|
| chr1-22737906-DEL-2071 | chr1-22737908-DEL-2070 |
| chr1-22740553-DEL-552 | chr1-22740554-DEL-552 |
| chr1-22741345-DEL-73 | chr1-22741352-DEL-68 |
| chr1-22752808-DEL-86 | chr1-22752809-DEL-86 |
| chr1-22805122-DEL-837 | chr1-22805123-DEL-837 |
| chr1-22836543-DEL-51 | chr1-22836544-DEL-51 |
| chr1-22853976-DEL-90 | chr1-22853975-DEL-87 |
| chr1-22877245-DEL-504 | chr1-22877246-DEL-504 |
| chr1-22907018-DEL-6633 | chr1-22907019-DEL-6633 |
| chr1-22923656-DEL-164 | chr1-22923659-DEL-148 |
| chr1-22947559-DEL-248 | chr1-22947560-DEL-248 |
| chr1-22955280-DEL-6383 | chr1-22955281-DEL-6383 |
| chr1-22986203-DEL-640 | chr1-22986204-DEL-640 |
| chr1-22987940-DEL-492 | chr1-22987941-DEL-492 |
| chr1-22988870-DEL-1882 | chr1-22988871-DEL-1882 |
| chr1-22991131-DEL-369 | chr1-22991132-DEL-369 |
| chr1-23020828-DEL-1541 | chr1-23020829-DEL-1541 |
| chr1-23022691-DEL-1678 | chr1-23022692-DEL-1678 |
| chr1-23039711-DEL-1069 | chr1-23039712-DEL-1069 |
| chr1-23075306-DEL-192 | chr1-23075293-DEL-193 |
| chr1-23094116-DEL-624 | chr1-23094133-DEL-620 |
| chr1-23146134-DEL-404 | chr1-23146135-DEL-404 |
| chr1-23185926-DEL-3482 | chr1-23185927-DEL-3482 |
| chr1-23189812-DEL-68 | chr1-23189813-DEL-63 |
| chr1-23205143-DEL-56 | chr1-23205134-DEL-57 |
| chr1-23259596-DEL-70 | chr1-23259599-DEL-68 |
| chr1-23267283-DEL-92 | chr1-23267284-DEL-92 |
| chr1-23275441-DEL-399 | chr1-23275442-DEL-399 |
| chr1-23334792-DEL-11705 | chr1-23334793-DEL-11705 |
| chr1-23340191-DEL-5246 | chr1-23340191-DEL-5246 |
| chr1-23354217-DEL-1267 | chr1-23354218-DEL-1266 |
| chr1-23357291-DEL-961 | chr1-23357292-DEL-961 |
| chr1-23377210-DEL-157 | chr1-23377211-DEL-157 |
| chr1-23399306-DEL-120 | chr1-23399307-DEL-120 |
| chr1-23404185-DEL-747 | chr1-23404184-DEL-748 |
| chr1-23449932-DEL-64 | chr1-23449933-DEL-64 |
| chr1-23481320-DEL-90 | chr1-23481331-DEL-80 |
| chr1-23508794-DEL-2033 | chr1-23508795-DEL-2033 |
| chr1-23525509-DEL-86 | chr1-23525510-DEL-86 |
| chr1-23539513-DEL-81 | chr1-23539514-DEL-81 |
| chr1-23547710-DEL-72 | chr1-23547713-DEL-83 |
| chr1-23595522-DEL-147 | chr1-23595523-DEL-147 |
| chr1-23617767-DEL-59 | chr1-23617796-DEL-59 |
| chr1-23621182-DEL-232 | chr1-23621185-DEL-231 |
| chr1-23626206-DEL-104 | chr1-23626209-DEL-102 |
| chr1-23683803-DEL-70 | chr1-23683805-DEL-69 |
| chr1-23685967-DEL-94 | chr1-23685968-DEL-94 |
| chr1-23713750-DEL-197 | chr1-23713751-DEL-197 |
| chr1-23721034-DEL-360 | chr1-23721035-DEL-360 |
| chr1-23723181-DEL-223 | chr1-23723182-DEL-223 |
| chr1-23740565-DEL-339 | chr1-23740566-DEL-339 |
| chr1-23751452-DEL-58 | chr1-23751453-DEL-58 |
| chr1-23755326-DEL-57 | chr1-23755304-DEL-57 |
| chr1-23775091-DEL-6334 | chr1-23775096-DEL-6333 |
| chr1-23788461-DEL-81 | chr1-23788462-DEL-81 |
| chr1-23805914-DEL-51 | chr1-23805901-DEL-52 |
| chr1-23817815-DEL-2190 | chr1-23817816-DEL-2190 |
| chr1-23821397-DEL-4172 | chr1-23821398-DEL-4172 |
| chr1-23829627-DEL-135 | chr1-23829628-DEL-135 |
| chr1-23835044-DEL-133 | chr1-23835034-DEL-144 |
| chr1-23862793-DEL-51 | chr1-23862794-DEL-51 |
| chr1-23879948-DEL-104 | chr1-23879955-DEL-85 |
| chr1-23895166-DEL-176 | chr1-23895167-DEL-174 |
| chr1-23912103-DEL-191 | chr1-23912104-DEL-191 |
| chr1-23917085-DEL-791 | chr1-23917086-DEL-791 |
| chr1-23921310-DEL-66 | chr1-23921311-DEL-66 |
| chr1-23923285-DEL-122 | chr1-23923286-DEL-122 |
| chr1-23941032-DEL-165 | chr1-23941034-DEL-163 |
| chr1-23952880-DEL-91 | chr1-23952876-DEL-91 |
| chr1-23974737-DEL-58 | chr1-23974738-DEL-58 |
| chr1-23984345-DEL-63 | chr1-23984346-DEL-63 |
| chr1-23990324-DEL-58 | chr1-23990325-DEL-58 |
| chr1-23999828-DEL-73 | chr1-23999829-DEL-73 |
| chr1-24000911-DEL-91 | chr1-24000912-DEL-91 |
| chr1-24002220-DEL-126 | chr1-24002221-DEL-126 |
| chr1-24010544-DEL-325 | chr1-24010544-DEL-325 |
| chr1-24037187-DEL-594 | chr1-24037188-DEL-594 |
| chr1-24055406-DEL-87 | chr1-24055407-DEL-87 |
| chr1-24075379-DEL-739 | chr1-24075380-DEL-739 |
| chr1-24107702-DEL-279 | chr1-24107703-DEL-279 |

| | |
|---|---|
| chr1-24114265-DEL-6460 | chr1-24114266-DEL-6460 |
| chr1-24121052-DEL-122 | chr1-24121053-DEL-122 |
| chr1-24127462-DEL-1053 | chr1-24127463-DEL-1026 |
| chr1-24150373-DEL-52 | chr1-24150375-DEL-56 |
| chr1-24156197-DEL-757 | chr1-24156198-DEL-757 |
| chr1-24185403-DEL-326 | chr1-24185404-DEL-321 |
| chr1-24249548-DEL-93 | chr1-24249519-DEL-93 |
| chr1-24254557-DEL-412 | chr1-24254557-DEL-412 |
| chr1-24312470-DEL-121 | chr1-24312471-DEL-121 |
| chr1-24344469-DEL-762 | chr1-24344470-DEL-762 |
| chr1-24355348-DEL-203 | chr1-24355349-DEL-203 |
| chr1-24370338-DEL-524 | chr1-24370341-DEL-506 |
| chr1-24384660-DEL-1145 | chr1-24384661-DEL-1145 |
| chr1-24414358-DEL-55 | chr1-24414358-DEL-55 |
| chr1-24454309-DEL-589 | chr1-24454310-DEL-589 |
| chr1-24492639-DEL-62 | chr1-24492668-DEL-74 |
| chr1-24501623-DEL-1859 | chr1-24501624-DEL-1859 |
| chr1-24522411-DEL-282 | chr1-24522412-DEL-282 |
| chr1-24527586-DEL-70 | chr1-24527587-DEL-70 |
| chr1-24532886-DEL-160 | chr1-24532887-DEL-160 |
| chr1-24535410-DEL-6583 | chr1-24535411-DEL-6583 |
| chr1-24572667-DEL-109 | chr1-24572673-DEL-118 |
| chr1-24574557-DEL-207 | chr1-24574558-DEL-207 |
| chr1-24584362-DEL-1343 | chr1-24584363-DEL-1343 |
| chr1-24599069-DEL-69 | chr1-24599069-DEL-73 |
| chr1-24627499-DEL-100 | chr1-24627500-DEL-100 |
| chr1-24633558-DEL-914 | chr1-24633559-DEL-914 |
| chr1-24650299-DEL-5605 | chr1-24650300-DEL-5605 |
| chr1-24664180-DEL-64 | chr1-24664180-DEL-60 |
| chr1-24667271-DEL-144 | chr1-24667272-DEL-144 |
| chr1-24670117-DEL-136 | chr1-24670118-DEL-136 |
| chr1-24678806-DEL-6456 | chr1-24678807-DEL-6456 |
| chr1-24711408-DEL-53 | chr1-24711409-DEL-53 |
| chr1-24714557-DEL-54 | chr1-24714558-DEL-54 |
| chr1-24728766-DEL-314 | chr1-24728762-DEL-313 |
| chr1-24749817-DEL-70 | chr1-24749818-DEL-70 |
| chr1-24755832-DEL-52 | chr1-24755823-DEL-60 |
| chr1-24760380-DEL-6538 | chr1-24760381-DEL-6538 |
| chr1-24768349-DEL-65 | chr1-24768361-DEL-66 |
| chr1-24832567-DEL-63 | chr1-24832568-DEL-63 |
| chr1-24840799-DEL-10600 | chr1-24840784-DEL-10615 |
| chr1-24871873-DEL-58 | chr1-24871874-DEL-58 |
| chr1-24876433-DEL-5987 | chr1-24876426-DEL-5988 |
| chr1-24892323-DEL-485 | chr1-24892322-DEL-486 |
| chr1-24908496-DEL-75 | chr1-24908496-DEL-77 |
| chr1-24911293-DEL-67 | chr1-24911288-DEL-72 |
| chr1-24929686-DEL-280 | chr1-24929686-DEL-280 |
| chr1-24936395-DEL-311 | chr1-24936402-DEL-305 |
| chr1-24947590-DEL-798 | chr1-24947591-DEL-798 |
| chr1-24949299-DEL-52 | chr1-24949300-DEL-52 |
| chr1-24968614-DEL-962 | chr1-24968615-DEL-962 |
| chr1-24980724-DEL-85 | chr1-24980740-DEL-70 |
| chr1-25023600-DEL-341 | chr1-25023601-DEL-341 |
| chr1-25046104-DEL-143 | chr1-25046105-DEL-143 |
| chr1-25053899-DEL-4924 | chr1-25053900-DEL-4924 |
| chr1-25059325-DEL-750 | chr1-25059326-DEL-749 |
| chr1-25078819-DEL-613 | chr1-25078820-DEL-608 |
| chr1-25088548-DEL-833 | chr1-25088549-DEL-833 |
| chr1-25098952-DEL-277 | chr1-25098953-DEL-277 |
| chr1-25144124-DEL-161 | chr1-25144125-DEL-162 |
| chr1-25158848-DEL-6716 | chr1-25158849-DEL-6716 |
| chr1-25202008-DEL-57 | chr1-25202003-DEL-63 |
| chr1-25227112-DEL-332 | chr1-25227113-DEL-332 |
| chr1-25238232-DEL-53 | chr1-25238195-DEL-53 |
| chr1-25241479-DEL-5120 | chr1-25241480-DEL-5120 |
| chr1-25299163-DEL-185 | chr1-25299164-DEL-185 |
| chr1-25313760-DEL-1308 | chr1-25313761-DEL-1308 |
| chr1-25317283-DEL-88 | chr1-25317283-DEL-86 |
| chr1-25321590-DEL-645 | chr1-25321591-DEL-645 |
| chr1-25324515-DEL-1188 | chr1-25324506-DEL-1189 |
| chr1-25325820-DEL-499 | chr1-25325821-DEL-499 |
| chr1-25360274-DEL-185 | chr1-25360275-DEL-185 |
| chr1-25412709-DEL-78 | chr1-25412710-DEL-78 |
| chr1-25462293-DEL-130 | chr1-25462293-DEL-94 |
| chr1-25506458-DEL-70 | chr1-25506438-DEL-90 |
| chr1-25507386-DEL-69 | chr1-25507377-DEL-93 |
| chr1-25510913-DEL-701 | chr1-25510914-DEL-701 |
| chr1-25517487-DEL-82 | chr1-25517480-DEL-82 |
| chr1-25526819-DEL-86 | chr1-25526819-DEL-86 |
| chr1-25559409-DEL-6216 | chr1-25559410-DEL-6216 |

| | |
|---|---|
| chr1-25611852-DEL-2430 | chr1-25611853-DEL-2430 |
| chr1-25629899-DEL-6490 | chr1-25629899-DEL-6490 |
| chr1-25652304-DEL-621 | chr1-25652305-DEL-621 |
| chr1-25668631-DEL-121 | chr1-25668631-DEL-121 |
| chr1-25701120-DEL-7262 | chr1-25701121-DEL-7262 |
| chr1-25733208-DEL-63 | chr1-25733208-DEL-57 |
| chr1-25743271-DEL-291 | chr1-25743272-DEL-291 |
| chr1-25745359-DEL-60 | chr1-25745338-DEL-50 |
| chr1-25750056-DEL-54 | chr1-25750057-DEL-54 |
| chr1-25760332-DEL-231 | chr1-25760332-DEL-230 |
| chr1-25772584-DEL-69 | chr1-25772593-DEL-61 |
| chr1-25779516-DEL-207 | chr1-25779517-DEL-207 |
| chr1-25795169-DEL-315 | chr1-25795171-DEL-335 |
| chr1-25847881-DEL-100 | chr1-25847882-DEL-100 |
| chr1-25879876-DEL-262 | chr1-25879877-DEL-262 |
| chr1-25881626-DEL-78 | chr1-25881615-DEL-78 |
| chr1-25954788-DEL-3389 | chr1-25954789-DEL-3389 |
| chr1-25971651-DEL-53 | chr1-25971652-DEL-53 |
| chr1-25973093-DEL-601 | chr1-25973094-DEL-601 |
| chr1-25974495-DEL-82 | chr1-25974496-DEL-82 |
| chr1-25990659-DEL-56 | chr1-25990660-DEL-56 |
| chr1-25996759-DEL-64 | chr1-25996760-DEL-64 |
| chr1-26009747-DEL-60 | chr1-26009748-DEL-59 |
| chr1-26014968-DEL-200 | chr1-26014969-DEL-200 |
| chr1-26019378-DEL-60 | chr1-26019379-DEL-60 |
| chr1-26031567-DEL-490 | chr1-26031567-DEL-489 |
| chr1-26089269-DEL-372 | chr1-26089270-DEL-372 |
| chr1-26120332-DEL-129 | chr1-26120333-DEL-129 |
| chr1-26150976-DEL-106 | chr1-26150975-DEL-108 |
| chr1-26156953-DEL-1457 | chr1-26156953-DEL-1457 |
| chr1-26181381-DEL-78 | chr1-26181382-DEL-78 |
| chr1-26211580-DEL-2346 | chr1-26211581-DEL-2346 |
| chr1-26230874-DEL-58 | chr1-26230872-DEL-67 |
| chr1-26245031-DEL-721 | chr1-26245032-DEL-721 |
| chr1-26249453-DEL-677 | chr1-26249454-DEL-677 |
| chr1-26250271-DEL-700 | chr1-26250272-DEL-701 |
| chr1-26252218-DEL-103 | chr1-26252219-DEL-103 |
| chr1-26257458-DEL-910 | chr1-26257459-DEL-910 |
| chr1-26260306-DEL-80 | chr1-26260307-DEL-80 |
| chr1-26271523-DEL-372 | chr1-26271524-DEL-372 |
| chr1-26278141-DEL-456 | chr1-26278142-DEL-452 |
| chr1-26285375-DEL-83 | chr1-26285373-DEL-84 |
| chr1-26287364-DEL-4675 | chr1-26287365-DEL-4675 |
| chr1-26297991-DEL-2221 | chr1-26297992-DEL-2221 |
| chr1-26308576-DEL-1701 | chr1-26308577-DEL-1705 |
| chr1-26324446-DEL-2203 | chr1-26324447-DEL-2203 |
| chr1-26325841-DEL-68 | chr1-26325842-DEL-68 |
| chr1-26342748-DEL-1155 | chr1-26342749-DEL-1155 |
| chr1-26361251-DEL-64 | chr1-26361252-DEL-64 |
| chr1-26362709-DEL-296 | chr1-26362710-DEL-296 |
| chr1-26388259-DEL-5195 | chr1-26388260-DEL-5195 |
| chr1-26453079-DEL-7084 | chr1-26453080-DEL-7084 |
| chr1-26467443-DEL-4399 | chr1-26467444-DEL-4399 |
| chr1-26473345-DEL-4373 | chr1-26473346-DEL-4373 |
| chr1-26480007-DEL-6194 | chr1-26480008-DEL-6194 |
| chr1-26493773-DEL-116 | chr1-26493750-DEL-140 |
| chr1-26509438-DEL-5868 | chr1-26509439-DEL-5868 |
| chr1-26541097-DEL-937 | chr1-26541098-DEL-937 |
| chr1-26563340-DEL-322 | chr1-26563341-DEL-318 |
| chr1-26579317-DEL-425 | chr1-26579318-DEL-425 |
| chr1-26584006-DEL-196 | chr1-26584007-DEL-196 |
| chr1-26587378-DEL-366 | chr1-26587379-DEL-365 |
| chr1-26588844-DEL-54 | chr1-26588845-DEL-54 |
| chr1-26599222-DEL-116 | chr1-26599223-DEL-116 |
| chr1-26604920-DEL-400 | chr1-26604921-DEL-400 |
| chr1-26606426-DEL-101 | chr1-26606427-DEL-101 |
| chr1-26625282-DEL-6395 | chr1-26625283-DEL-6394 |
| chr1-26660693-DEL-3454 | chr1-26660694-DEL-3454 |
| chr1-26674802-DEL-359 | chr1-26674803-DEL-359 |
| chr1-26692584-DEL-2367 | chr1-26692585-DEL-2367 |
| chr1-26696902-DEL-1040 | chr1-26696903-DEL-1040 |
| chr1-26700615-DEL-71 | chr1-26700616-DEL-70 |
| chr1-26706698-DEL-2290 | chr1-26706699-DEL-2290 |
| chr1-26738026-DEL-843 | chr1-26738027-DEL-843 |
| chr1-26749712-DEL-547 | chr1-26749713-DEL-547 |
| chr1-26761296-DEL-56 | chr1-26761295-DEL-70 |
| chr1-26773967-DEL-148 | chr1-26773968-DEL-148 |
| chr1-26784078-DEL-175 | chr1-26784079-DEL-175 |
| chr1-26791023-DEL-182 | chr1-26791024-DEL-182 |
| chr1-26806628-DEL-67 | chr1-26806629-DEL-67 |

| | |
|---|---|
| chr1-26808645-DEL-75 | chr1-26808649-DEL-66 |
| chr1-26812073-DEL-58 | chr1-26812074-DEL-54 |
| chr1-26822443-DEL-59 | chr1-26822444-DEL-59 |
| chr1-26825755-DEL-212 | chr1-26825774-DEL-194 |
| chr1-26828038-DEL-54 | chr1-26828061-DEL-74 |
| chr1-26840410-DEL-2673 | chr1-26840411-DEL-2673 |
| chr1-26845473-DEL-244 | chr1-26845474-DEL-244 |
| chr1-26875720-DEL-1178 | chr1-26875709-DEL-1177 |
| chr1-26925308-DEL-62 | chr1-26925282-DEL-63 |
| chr1-26978007-DEL-2552 | chr1-26978008-DEL-2552 |
| chr1-26983682-DEL-1859 | chr1-26983683-DEL-1858 |
| chr1-27024636-DEL-61 | chr1-27024641-DEL-60 |
| chr1-27028242-DEL-126 | chr1-27028240-DEL-125 |
| chr1-27060842-DEL-824 | chr1-27060843-DEL-824 |
| chr1-27072104-DEL-1663 | chr1-27072105-DEL-1663 |
| chr1-27084752-DEL-79 | chr1-27084781-DEL-52 |
| chr1-27095840-DEL-77 | chr1-27095841-DEL-77 |
| chr1-27096538-DEL-86 | chr1-27096539-DEL-86 |
| chr1-27128255-DEL-476 | chr1-27128256-DEL-476 |
| chr1-27131836-DEL-6201 | chr1-27131831-DEL-6202 |
| chr1-27147968-DEL-60 | chr1-27147969-DEL-90 |
| chr1-27167278-DEL-58 | chr1-27167277-DEL-60 |
| chr1-27175587-DEL-2213 | chr1-27175588-DEL-2213 |
| chr1-27178799-DEL-5954 | chr1-27178800-DEL-5954 |
| chr1-27186627-DEL-133 | chr1-27186628-DEL-133 |
| chr1-27198489-DEL-6816 | chr1-27198473-DEL-6817 |
| chr1-27209924-DEL-242 | chr1-27209925-DEL-242 |
| chr1-27219211-DEL-7130 | chr1-27219212-DEL-7130 |
| chr1-27246487-DEL-52 | chr1-27246488-DEL-52 |
| chr1-27259159-DEL-67 | chr1-27259160-DEL-67 |
| chr1-27309348-DEL-6472 | chr1-27309349-DEL-6472 |
| chr1-27315347-DEL-199 | chr1-27315348-DEL-199 |
| chr1-27326273-DEL-559 | chr1-27326274-DEL-560 |
| chr1-27332626-DEL-161 | chr1-27332627-DEL-185 |
| chr1-27342090-DEL-67 | chr1-27342091-DEL-67 |
| chr1-27346682-DEL-445 | chr1-27346683-DEL-445 |
| chr1-27358143-DEL-203 | chr1-27358144-DEL-203 |
| chr1-27370774-DEL-74 | chr1-27370773-DEL-75 |
| chr1-27406064-DEL-432 | chr1-27406065-DEL-432 |
| chr1-27408808-DEL-633 | chr1-27408811-DEL-631 |
| chr1-27544695-DEL-571 | chr1-27544696-DEL-571 |
| chr1-27729885-DEL-104 | chr1-27729886-DEL-104 |
| chr1-27730714-DEL-1133 | chr1-27730713-DEL-1134 |
| chr1-27794159-DEL-6244 | chr1-27794160-DEL-6244 |
| chr1-27812939-DEL-1109 | chr1-27812940-DEL-1109 |
| chr1-27871125-DEL-920 | chr1-27871126-DEL-920 |
| chr1-27895248-DEL-2142 | chr1-27895248-DEL-2142 |
| chr1-27933783-DEL-51 | chr1-27933783-DEL-51 |
| chr1-28006132-DEL-91 | chr1-28006133-DEL-91 |
| chr1-28060051-DEL-58 | chr1-28060052-DEL-52 |
| chr1-28066386-DEL-38758 | chr1-28066386-DEL-38758 |
| chr1-28067380-DEL-2082 | chr1-28067381-DEL-2082 |
| chr1-28073763-DEL-448 | chr1-28073764-DEL-448 |
| chr1-28115439-DEL-1138 | chr1-28115440-DEL-1138 |
| chr1-28117055-DEL-677 | chr1-28117056-DEL-677 |
| chr1-28160877-DEL-52 | chr1-28160880-DEL-50 |
| chr1-28171918-DEL-539 | chr1-28171918-DEL-539 |
| chr1-28225128-DEL-6848 | chr1-28225129-DEL-6848 |
| chr1-28287657-DEL-3100 | chr1-28287657-DEL-3101 |
| chr1-28295594-DEL-64 | chr1-28295603-DEL-56 |
| chr1-28326666-DEL-7267 | chr1-28326650-DEL-7281 |
| chr1-28327689-DEL-212 | chr1-28327690-DEL-212 |
| chr1-28359323-DEL-342 | chr1-28359324-DEL-342 |
| chr1-28360987-DEL-70 | chr1-28360993-DEL-64 |
| chr1-28415066-DEL-205 | chr1-28415066-DEL-205 |
| chr1-28488023-DEL-227 | chr1-28488024-DEL-227 |
| chr1-28495210-DEL-3014 | chr1-28495210-DEL-3014 |
| chr1-28516729-DEL-70 | chr1-28516737-DEL-62 |
| chr1-28537982-DEL-399 | chr1-28537982-DEL-399 |
| chr1-28546296-DEL-52 | chr1-28546270-DEL-54 |
| chr1-28549546-DEL-1018 | chr1-28549537-DEL-1017 |
| chr1-28572020-DEL-449 | chr1-28572020-DEL-449 |
| chr1-28615571-DEL-76 | chr1-28615569-DEL-74 |
| chr1-28619348-DEL-517 | chr1-28619349-DEL-517 |
| chr1-28639938-DEL-159 | chr1-28639940-DEL-159 |
| chr1-28643230-DEL-790 | chr1-28643231-DEL-790 |
| chr1-28657569-DEL-2483 | chr1-28657570-DEL-2483 |
| chr1-28671527-DEL-902 | chr1-28671528-DEL-902 |
| chr1-28673984-DEL-139 | chr1-28673985-DEL-139 |
| chr1-28703524-DEL-313 | chr1-28703524-DEL-313 |

| | |
|---|---|
| chr1-28706832-DEL-1207 | chr1-28706832-DEL-1207 |
| chr1-28709011-DEL-397 | chr1-28709012-DEL-397 |
| chr1-28711368-DEL-210 | chr1-28711369-DEL-210 |
| chr1-28826739-DEL-71 | chr1-28826736-DEL-74 |
| chr1-28828023-DEL-101 | chr1-28828024-DEL-101 |
| chr1-28852559-DEL-52 | chr1-28852560-DEL-52 |
| chr1-28853058-DEL-68 | chr1-28853059-DEL-68 |
| chr1-28889347-DEL-108 | chr1-28889347-DEL-107 |
| chr1-28890151-DEL-92 | chr1-28890152-DEL-92 |
| chr1-28914082-DEL-54 | chr1-28914076-DEL-55 |
| chr1-28928087-DEL-2818 | chr1-28928067-DEL-2819 |
| chr1-28945643-DEL-92 | chr1-28945639-DEL-103 |
| chr1-28961035-DEL-78 | chr1-28961048-DEL-66 |
| chr1-28975418-DEL-92 | chr1-28975419-DEL-98 |
| chr1-28981907-DEL-86 | chr1-28981908-DEL-86 |
| chr1-29024032-DEL-58 | chr1-29024033-DEL-58 |
| chr1-29064455-DEL-89 | chr1-29064456-DEL-89 |
| chr1-29072462-DEL-170 | chr1-29072466-DEL-158 |
| chr1-29099667-DEL-332 | chr1-29099668-DEL-332 |
| chr1-29104547-DEL-401 | chr1-29104547-DEL-401 |
| chr1-29125231-DEL-402 | chr1-29125232-DEL-402 |
| chr1-29140061-DEL-79 | chr1-29140095-DEL-52 |
| chr1-29148379-DEL-890 | chr1-29148380-DEL-890 |
| chr1-29155595-DEL-73 | chr1-29155596-DEL-73 |
| chr1-29165596-DEL-530 | chr1-29165597-DEL-530 |
| chr1-29171305-DEL-581 | chr1-29171306-DEL-581 |
| chr1-29227663-DEL-597 | chr1-29227664-DEL-597 |
| chr1-29257375-DEL-301 | chr1-29257378-DEL-298 |
| chr1-29282301-DEL-78 | chr1-29282326-DEL-54 |
| chr1-29293140-DEL-399 | chr1-29293141-DEL-399 |
| chr1-29318178-DEL-307 | chr1-29318179-DEL-306 |
| chr1-29346154-DEL-56 | chr1-29346157-DEL-54 |
| chr1-29356790-DEL-92 | chr1-29356791-DEL-92 |
| chr1-29407295-DEL-6612 | chr1-29407296-DEL-6612 |
| chr1-29420602-DEL-97 | chr1-29420622-DEL-56 |
| chr1-29442414-DEL-454 | chr1-29442415-DEL-454 |
| chr1-29445517-DEL-111 | chr1-29445517-DEL-105 |
| chr1-29446060-DEL-6507 | chr1-29446060-DEL-6507 |
| chr1-29475261-DEL-4697 | chr1-29475260-DEL-4697 |
| chr1-29530127-DEL-257 | chr1-29530128-DEL-257 |
| chr1-29550178-DEL-74 | chr1-29550179-DEL-74 |
| chr1-29584513-DEL-152 | chr1-29584518-DEL-152 |
| chr1-29605784-DEL-278 | chr1-29605785-DEL-278 |
| chr1-29619284-DEL-6405 | chr1-29619285-DEL-6405 |
| chr1-29625340-DEL-208 | chr1-29625341-DEL-208 |
| chr1-29635769-DEL-485 | chr1-29635769-DEL-485 |
| chr1-29645500-DEL-1950 | chr1-29645501-DEL-1950 |
| chr1-29652380-DEL-66 | chr1-29652381-DEL-66 |
| chr1-29682341-DEL-6440 | chr1-29682342-DEL-6440 |
| chr1-29694250-DEL-81 | chr1-29694251-DEL-81 |
| chr1-29713721-DEL-568 | chr1-29713722-DEL-568 |
| chr1-29717030-DEL-567 | chr1-29717030-DEL-567 |
| chr1-29737096-DEL-2885 | chr1-29737097-DEL-2885 |
| chr1-29742491-DEL-6468 | chr1-29742492-DEL-6468 |
| chr1-29776269-DEL-4184 | chr1-29776260-DEL-4185 |
| chr1-29901414-DEL-785 | chr1-29901415-DEL-785 |
| chr1-29917645-DEL-136 | chr1-29917669-DEL-115 |
| chr1-29927978-DEL-6435 | chr1-29927974-DEL-6435 |
| chr1-29964927-DEL-129 | chr1-29964928-DEL-129 |
| chr1-29981821-DEL-6249 | chr1-29981821-DEL-6249 |
| chr1-30005428-DEL-192 | chr1-30005429-DEL-192 |
| chr1-30017400-DEL-828 | chr1-30017394-DEL-837 |
| chr1-30039816-DEL-56 | chr1-30039813-DEL-60 |
| chr1-30051526-DEL-63 | chr1-30051530-DEL-68 |
| chr1-30058438-DEL-831 | chr1-30058439-DEL-831 |
| chr1-30067641-DEL-765 | chr1-30067642-DEL-765 |
| chr1-30075207-DEL-7339 | chr1-30075208-DEL-7339 |
| chr1-30084255-DEL-3453 | chr1-30084255-DEL-3453 |
| chr1-30105796-DEL-6719 | chr1-30105797-DEL-6719 |
| chr1-30137968-DEL-990 | chr1-30137969-DEL-990 |
| chr1-30141165-DEL-401 | chr1-30141166-DEL-400 |
| chr1-30147150-DEL-70 | chr1-30147153-DEL-68 |
| chr1-30169568-DEL-776 | chr1-30169569-DEL-776 |
| chr1-30189392-DEL-118 | chr1-30189387-DEL-124 |
| chr1-30244945-DEL-78 | chr1-30244955-DEL-69 |
| chr1-30288923-DEL-76 | chr1-30288922-DEL-62 |
| chr1-30293716-DEL-1513 | chr1-30293716-DEL-1514 |
| chr1-30330542-DEL-207 | chr1-30330543-DEL-207 |
| chr1-30343124-DEL-9753 | chr1-30343125-DEL-9753 |
| chr1-30349916-DEL-208 | chr1-30349917-DEL-208 |

| | |
|---|---|
| chr1-30365303-DEL-376 | chr1-30365290-DEL-378 |
| chr1-30372192-DEL-6392 | chr1-30372176-DEL-6406 |
| chr1-30404620-DEL-66 | chr1-30404597-DEL-90 |
| chr1-30458249-DEL-178 | chr1-30458250-DEL-178 |
| chr1-30473407-DEL-79 | chr1-30473408-DEL-79 |
| chr1-30479039-DEL-221 | chr1-30479040-DEL-221 |
| chr1-30501224-DEL-1164 | chr1-30501225-DEL-1164 |
| chr1-30504487-DEL-60 | chr1-30504488-DEL-60 |
| chr1-30507627-DEL-3319 | chr1-30507628-DEL-3319 |
| chr1-30533173-DEL-88 | chr1-30533172-DEL-86 |
| chr1-30565696-DEL-75 | chr1-30565697-DEL-75 |
| chr1-30577730-DEL-612 | chr1-30577731-DEL-612 |
| chr1-30588416-DEL-180 | chr1-30588417-DEL-180 |
| chr1-30623283-DEL-213 | chr1-30623284-DEL-213 |
| chr1-30655512-DEL-499 | chr1-30655513-DEL-499 |
| chr1-30670391-DEL-204 | chr1-30670388-DEL-205 |
| chr1-30674967-DEL-128 | chr1-30674968-DEL-124 |
| chr1-30691437-DEL-272 | chr1-30691438-DEL-272 |
| chr1-30694325-DEL-188 | chr1-30694326-DEL-188 |
| chr1-30699637-DEL-6498 | chr1-30699638-DEL-6498 |
| chr1-30726801-DEL-2669 | chr1-30726802-DEL-2669 |
| chr1-30738907-DEL-88 | chr1-30738908-DEL-88 |
| chr1-30744822-DEL-425 | chr1-30744822-DEL-425 |
| chr1-30747312-DEL-1020 | chr1-30747313-DEL-1020 |
| chr1-30756447-DEL-61 | chr1-30756448-DEL-61 |
| chr1-30760406-DEL-66 | chr1-30760407-DEL-66 |
| chr1-30769637-DEL-214 | chr1-30769637-DEL-214 |
| chr1-30780581-DEL-84 | chr1-30780582-DEL-84 |
| chr1-30892742-DEL-6282 | chr1-30892743-DEL-6282 |
| chr1-30903274-DEL-319 | chr1-30903275-DEL-319 |
| chr1-30922475-DEL-2395 | chr1-30922476-DEL-2395 |
| chr1-30932666-DEL-1889 | chr1-30932667-DEL-1889 |
| chr1-30940158-DEL-125 | chr1-30940159-DEL-125 |
| chr1-30948934-DEL-58 | chr1-30948935-DEL-58 |
| chr1-30952077-DEL-484 | chr1-30952078-DEL-484 |
| chr1-30952917-DEL-417 | chr1-30952918-DEL-417 |
| chr1-31033476-DEL-284 | chr1-31033477-DEL-284 |
| chr1-31069167-DEL-1045 | chr1-31069168-DEL-1045 |
| chr1-31081135-DEL-425 | chr1-31081136-DEL-425 |
| chr1-31110745-DEL-648 | chr1-31110746-DEL-648 |
| chr1-31114020-DEL-7173 | chr1-31114020-DEL-7171 |
| chr1-31146872-DEL-97 | chr1-31146873-DEL-95 |
| chr1-31152067-DEL-68 | chr1-31152066-DEL-70 |
| chr1-31157827-DEL-60 | chr1-31157828-DEL-50 |
| chr1-31168566-DEL-696 | chr1-31168565-DEL-696 |
| chr1-31208606-DEL-1875 | chr1-31208607-DEL-1875 |
| chr1-31213239-DEL-196 | chr1-31213240-DEL-196 |
| chr1-31250009-DEL-1003 | chr1-31250010-DEL-1003 |
| chr1-31256959-DEL-715 | chr1-31256960-DEL-715 |
| chr1-31296152-DEL-62 | chr1-31296153-DEL-62 |
| chr1-31316478-DEL-408 | chr1-31316479-DEL-408 |
| chr1-31324649-DEL-65 | chr1-31324650-DEL-65 |
| chr1-31334175-DEL-6244 | chr1-31334176-DEL-6244 |
| chr1-31360182-DEL-99 | chr1-31360183-DEL-99 |
| chr1-31365367-DEL-185 | chr1-31365370-DEL-183 |
| chr1-31384898-DEL-509 | chr1-31384899-DEL-509 |
| chr1-31388890-DEL-207 | chr1-31388891-DEL-207 |
| chr1-31389777-DEL-6459 | chr1-31389778-DEL-6459 |
| chr1-31398782-DEL-60 | chr1-31398780-DEL-50 |
| chr1-31430185-DEL-59 | chr1-31430172-DEL-52 |
| chr1-31449434-DEL-6359 | chr1-31449435-DEL-6359 |
| chr1-31473906-DEL-55 | chr1-31473904-DEL-59 |
| chr1-31525379-DEL-6392 | chr1-31525380-DEL-6392 |
| chr1-31557423-DEL-64 | chr1-31557424-DEL-62 |
| chr1-31591218-DEL-74 | chr1-31591219-DEL-74 |
| chr1-31623596-DEL-2910 | chr1-31623597-DEL-2910 |
| chr1-31654363-DEL-1589 | chr1-31654347-DEL-1590 |
| chr1-31691864-DEL-72 | chr1-31691866-DEL-70 |
| chr1-31709714-DEL-54 | chr1-31709715-DEL-54 |
| chr1-31720858-DEL-770 | chr1-31720859-DEL-770 |
| chr1-31727613-DEL-328 | chr1-31727614-DEL-328 |
| chr1-31733881-DEL-6682 | chr1-31733882-DEL-6682 |
| chr1-31756400-DEL-1736 | chr1-31756401-DEL-1735 |
| chr1-31765498-DEL-328 | chr1-31765499-DEL-328 |
| chr1-31769616-DEL-4145 | chr1-31769617-DEL-4145 |
| chr1-31771021-DEL-491 | chr1-31771022-DEL-491 |
| chr1-31831491-DEL-787 | chr1-31831492-DEL-787 |
| chr1-31847990-DEL-55 | chr1-31847990-DEL-55 |
| chr1-31853316-DEL-6208 | chr1-31853317-DEL-6208 |
| chr1-31883712-DEL-428 | chr1-31883713-DEL-428 |

| | |
|---|---|
| chr1-31889531-DEL-2324 | chr1-31889532-DEL-2324 |
| chr1-31913695-DEL-65 | chr1-31913696-DEL-65 |
| chr1-31969461-DEL-515 | chr1-31969462-DEL-515 |
| chr1-31975873-DEL-457 | chr1-31975874-DEL-457 |
| chr1-32053660-DEL-121 | chr1-32053655-DEL-123 |
| chr1-32085198-DEL-227 | chr1-32085199-DEL-227 |
| chr1-32088743-DEL-105 | chr1-32088755-DEL-80 |
| chr1-32112531-DEL-424 | chr1-32112532-DEL-424 |
| chr1-32120512-DEL-762 | chr1-32120513-DEL-762 |
| chr1-32201576-DEL-1982 | chr1-32201577-DEL-1982 |
| chr1-32218511-DEL-422 | chr1-32218512-DEL-422 |
| chr1-32270677-DEL-60 | chr1-32270678-DEL-60 |
| chr1-32298619-DEL-6564 | chr1-32298620-DEL-6564 |
| chr1-32309756-DEL-282 | chr1-32309757-DEL-282 |
| chr1-32313556-DEL-79 | chr1-32313557-DEL-79 |
| chr1-32317692-DEL-1357 | chr1-32317691-DEL-1360 |
| chr1-32362363-DEL-951 | chr1-32362364-DEL-951 |
| chr1-32367275-DEL-139 | chr1-32367276-DEL-139 |
| chr1-32384706-DEL-55 | chr1-32384704-DEL-55 |
| chr1-32400242-DEL-54 | chr1-32400243-DEL-54 |
| chr1-32402298-DEL-25742 | chr1-32402298-DEL-25742 |
| chr1-32411438-DEL-6367 | chr1-32411439-DEL-6367 |
| chr1-32421172-DEL-236 | chr1-32421173-DEL-235 |
| chr1-32426050-DEL-74 | chr1-32426057-DEL-68 |
| chr1-32440746-DEL-950 | chr1-32440747-DEL-950 |
| chr1-32454441-DEL-490 | chr1-32454442-DEL-490 |
| chr1-32475525-DEL-201 | chr1-32475526-DEL-201 |
| chr1-32520711-DEL-5735 | chr1-32520712-DEL-5735 |
| chr1-32524189-DEL-1888 | chr1-32524190-DEL-1888 |
| chr1-32577646-DEL-475 | chr1-32577647-DEL-475 |
| chr1-32627317-DEL-818 | chr1-32627318-DEL-818 |
| chr1-32633407-DEL-168 | chr1-32633408-DEL-168 |
| chr1-32650564-DEL-214 | chr1-32650565-DEL-214 |
| chr1-32661887-DEL-333 | chr1-32661888-DEL-333 |
| chr1-32663883-DEL-201 | chr1-32663884-DEL-201 |
| chr1-32684344-DEL-52 | chr1-32684345-DEL-52 |
| chr1-32704726-DEL-6435 | chr1-32704727-DEL-6435 |
| chr1-32712829-DEL-59 | chr1-32712830-DEL-59 |
| chr1-32729249-DEL-398 | chr1-32729242-DEL-399 |
| chr1-32730636-DEL-256 | chr1-32730637-DEL-256 |
| chr1-32741280-DEL-327 | chr1-32741278-DEL-328 |
| chr1-32767209-DEL-57 | chr1-32767222-DEL-66 |
| chr1-32767509-DEL-401 | chr1-32767510-DEL-401 |
| chr1-32793790-DEL-4285 | chr1-32793791-DEL-4285 |
| chr1-32818402-DEL-2226 | chr1-32818403-DEL-2226 |
| chr1-32821863-DEL-264 | chr1-32821864-DEL-264 |
| chr1-32833528-DEL-457 | chr1-32833529-DEL-457 |
| chr1-32850114-DEL-1965 | chr1-32850135-DEL-1946 |
| chr1-32852175-DEL-870 | chr1-32852175-DEL-870 |
| chr1-32852794-DEL-1303 | chr1-32852795-DEL-1303 |
| chr1-32860589-DEL-52 | chr1-32860596-DEL-58 |
| chr1-32888428-DEL-3598 | chr1-32888429-DEL-3636 |
| chr1-32908550-DEL-1174 | chr1-32908551-DEL-1174 |
| chr1-32915517-DEL-92 | chr1-32915516-DEL-92 |
| chr1-32918133-DEL-416 | chr1-32918134-DEL-416 |
| chr1-32948112-DEL-237 | chr1-32948113-DEL-237 |
| chr1-32968964-DEL-470 | chr1-32968965-DEL-470 |
| chr1-32970318-DEL-80 | chr1-32970319-DEL-80 |
| chr1-32989090-DEL-75 | chr1-32989091-DEL-75 |
| chr1-32992414-DEL-60 | chr1-32992415-DEL-60 |
| chr1-32997401-DEL-52 | chr1-32997402-DEL-52 |
| chr1-33014388-DEL-4452 | chr1-33014390-DEL-4451 |
| chr1-33036011-DEL-114 | chr1-33036012-DEL-114 |
| chr1-33052419-DEL-6902 | chr1-33052420-DEL-6902 |
| chr1-33067898-DEL-371 | chr1-33067901-DEL-370 |
| chr1-33073113-DEL-690 | chr1-33073114-DEL-686 |
| chr1-33076131-DEL-875 | chr1-33076128-DEL-876 |
| chr1-33085044-DEL-6451 | chr1-33085030-DEL-6452 |
| chr1-33103739-DEL-1895 | chr1-33103740-DEL-1895 |
| chr1-33125128-DEL-55 | chr1-33125119-DEL-50 |
| chr1-33135897-DEL-53 | chr1-33135896-DEL-55 |
| chr1-33154343-DEL-3229 | chr1-33154344-DEL-3229 |
| chr1-33160803-DEL-630 | chr1-33160804-DEL-630 |
| chr1-33184855-DEL-797 | chr1-33184856-DEL-797 |
| chr1-33186589-DEL-54 | chr1-33186594-DEL-52 |
| chr1-33192521-DEL-56 | chr1-33192520-DEL-58 |
| chr1-33201608-DEL-8233 | chr1-33201609-DEL-8233 |
| chr1-33207831-DEL-1882 | chr1-33207831-DEL-1882 |
| chr1-33210796-DEL-56 | chr1-33210797-DEL-61 |
| chr1-33220124-DEL-194 | chr1-33220125-DEL-194 |

| | |
|---|---|
| chr1-33229187-DEL-129 | chr1-33229188-DEL-129 |
| chr1-33234053-DEL-202 | chr1-33234054-DEL-202 |
| chr1-33234672-DEL-128 | chr1-33234665-DEL-136 |
| chr1-33238941-DEL-64 | chr1-33238940-DEL-68 |
| chr1-33257329-DEL-57 | chr1-33257330-DEL-69 |
| chr1-33275380-DEL-307 | chr1-33275381-DEL-307 |
| chr1-33311217-DEL-2684 | chr1-33311218-DEL-2684 |
| chr1-33338667-DEL-6380 | chr1-33338653-DEL-6394 |
| chr1-33372821-DEL-200 | chr1-33372822-DEL-200 |
| chr1-33379501-DEL-3598 | chr1-33379502-DEL-3598 |
| chr1-33414351-DEL-60 | chr1-33414352-DEL-60 |
| chr1-33442948-DEL-581 | chr1-33442949-DEL-581 |
| chr1-33446488-DEL-7462 | chr1-33446489-DEL-7462 |
| chr1-33446813-DEL-834 | chr1-33446814-DEL-834 |
| chr1-33472063-DEL-576 | chr1-33472063-DEL-576 |
| chr1-33474042-DEL-209 | chr1-33474036-DEL-209 |
| chr1-33501007-DEL-108 | chr1-33501008-DEL-108 |
| chr1-33511019-DEL-56 | chr1-33511020-DEL-56 |
| chr1-33525770-DEL-190 | chr1-33525806-DEL-198 |
| chr1-33538988-DEL-740 | chr1-33538989-DEL-740 |
| chr1-33577534-DEL-1714 | chr1-33577535-DEL-1714 |
| chr1-33580231-DEL-119 | chr1-33580232-DEL-119 |
| chr1-33589759-DEL-3603 | chr1-33589760-DEL-3603 |
| chr1-33602198-DEL-52 | chr1-33602199-DEL-52 |
| chr1-33605577-DEL-327 | chr1-33605578-DEL-327 |
| chr1-33614735-DEL-1036 | chr1-33614736-DEL-1036 |
| chr1-33639446-DEL-115 | chr1-33639447-DEL-115 |
| chr1-33655416-DEL-11504 | chr1-33655417-DEL-11504 |
| chr1-33655791-DEL-905 | chr1-33655781-DEL-899 |
| chr1-33659019-DEL-1726 | chr1-33659020-DEL-1726 |
| chr1-33675734-DEL-1732 | chr1-33675735-DEL-1732 |
| chr1-33719561-DEL-584 | chr1-33719562-DEL-584 |
| chr1-33729733-DEL-60 | chr1-33729754-DEL-86 |
| chr1-33745443-DEL-365 | chr1-33745443-DEL-365 |
| chr1-33754727-DEL-58 | chr1-33754728-DEL-58 |
| chr1-33757006-DEL-212 | chr1-33757007-DEL-212 |
| chr1-33761838-DEL-209 | chr1-33761839-DEL-209 |
| chr1-33775586-DEL-94 | chr1-33775586-DEL-94 |
| chr1-33783871-DEL-189 | chr1-33783871-DEL-189 |
| chr1-33789038-DEL-201 | chr1-33789039-DEL-201 |
| chr1-33790903-DEL-181 | chr1-33790896-DEL-182 |
| chr1-33792154-DEL-62 | chr1-33792155-DEL-62 |
| chr1-33824626-DEL-334 | chr1-33824627-DEL-334 |
| chr1-33863623-DEL-415 | chr1-33863624-DEL-415 |
| chr1-33870471-DEL-54 | chr1-33870472-DEL-54 |
| chr1-33885069-DEL-594 | chr1-33885070-DEL-593 |
| chr1-33888096-DEL-1518 | chr1-33888097-DEL-1518 |
| chr1-33911601-DEL-203 | chr1-33911602-DEL-203 |
| chr1-33924386-DEL-202 | chr1-33924387-DEL-202 |
| chr1-33952347-DEL-310 | chr1-33952348-DEL-310 |
| chr1-33956504-DEL-143 | chr1-33956505-DEL-143 |
| chr1-33981519-DEL-181 | chr1-33981520-DEL-181 |
| chr1-33989608-DEL-1525 | chr1-33989607-DEL-1526 |
| chr1-33990338-DEL-69 | chr1-33990339-DEL-69 |
| chr1-33998491-DEL-351 | chr1-33998492-DEL-351 |
| chr1-34018460-DEL-54 | chr1-34018461-DEL-54 |
| chr1-34030808-DEL-254 | chr1-34030809-DEL-253 |
| chr1-34041759-DEL-203 | chr1-34041760-DEL-203 |
| chr1-34068792-DEL-214 | chr1-34068793-DEL-214 |
| chr1-34071537-DEL-232 | chr1-34071538-DEL-232 |
| chr1-34076135-DEL-57 | chr1-34076136-DEL-57 |
| chr1-34088733-DEL-68 | chr1-34088734-DEL-68 |
| chr1-34099087-DEL-168 | chr1-34099088-DEL-168 |
| chr1-34117433-DEL-6402 | chr1-34117434-DEL-6402 |
| chr1-34127555-DEL-180 | chr1-34127543-DEL-181 |
| chr1-34129544-DEL-190 | chr1-34129543-DEL-192 |
| chr1-34136729-DEL-618 | chr1-34136730-DEL-618 |
| chr1-34138548-DEL-272 | chr1-34138565-DEL-280 |
| chr1-34151658-DEL-175 | chr1-34151659-DEL-175 |
| chr1-34175228-DEL-318 | chr1-34175229-DEL-318 |
| chr1-34283247-DEL-80 | chr1-34283244-DEL-80 |
| chr1-34297770-DEL-510 | chr1-34297777-DEL-508 |
| chr1-34359758-DEL-201 | chr1-34359759-DEL-201 |
| chr1-34366257-DEL-72 | chr1-34366258-DEL-72 |
| chr1-34373757-DEL-522 | chr1-34373758-DEL-522 |
| chr1-34379406-DEL-204 | chr1-34379407-DEL-204 |
| chr1-34383939-DEL-66 | chr1-34383940-DEL-66 |
| chr1-34386531-DEL-3919 | chr1-34386532-DEL-3919 |
| chr1-34391229-DEL-105 | chr1-34391230-DEL-105 |
| chr1-34397207-DEL-504 | chr1-34397208-DEL-504 |

| | |
|---|---|
| chr1-34411238-DEL-124 | chr1-34411239-DEL-124 |
| chr1-34412087-DEL-52 | chr1-34412088-DEL-52 |
| chr1-34423557-DEL-66 | chr1-34423534-DEL-110 |
| chr1-34442974-DEL-150 | chr1-34443005-DEL-120 |
| chr1-34449760-DEL-64 | chr1-34449755-DEL-68 |
| chr1-34450328-DEL-430 | chr1-34450329-DEL-430 |
| chr1-34456638-DEL-6783 | chr1-34456639-DEL-6783 |
| chr1-34480312-DEL-292 | chr1-34480313-DEL-292 |
| chr1-34487946-DEL-83 | chr1-34487947-DEL-83 |
| chr1-34489542-DEL-83 | chr1-34489543-DEL-83 |
| chr1-34508048-DEL-205 | chr1-34508038-DEL-194 |
| chr1-34526785-DEL-51 | chr1-34526782-DEL-52 |
| chr1-34558883-DEL-1612 | chr1-34558884-DEL-1612 |
| chr1-34559294-DEL-73 | chr1-34559295-DEL-73 |
| chr1-34571709-DEL-115 | chr1-34571710-DEL-115 |
| chr1-34603077-DEL-208 | chr1-34603079-DEL-206 |
| chr1-34607763-DEL-118 | chr1-34607764-DEL-118 |
| chr1-34609428-DEL-628 | chr1-34609431-DEL-628 |
| chr1-34616134-DEL-78 | chr1-34616135-DEL-78 |
| chr1-34641775-DEL-3033 | chr1-34641776-DEL-3033 |
| chr1-34659156-DEL-364 | chr1-34659142-DEL-370 |
| chr1-34659720-DEL-999 | chr1-34659721-DEL-999 |
| chr1-34661428-DEL-516 | chr1-34661422-DEL-517 |
| chr1-34670330-DEL-82 | chr1-34670331-DEL-82 |
| chr1-34677177-DEL-93 | chr1-34677178-DEL-93 |
| chr1-34683728-DEL-6401 | chr1-34683729-DEL-6401 |
| chr1-34695051-DEL-135 | chr1-34695052-DEL-135 |
| chr1-34695581-DEL-151 | chr1-34695564-DEL-152 |
| chr1-34697712-DEL-3870 | chr1-34697713-DEL-3870 |
| chr1-34698017-DEL-54 | chr1-34698018-DEL-54 |
| chr1-34699713-DEL-229 | chr1-34699714-DEL-229 |
| chr1-34741675-DEL-449 | chr1-34741676-DEL-450 |
| chr1-34803847-DEL-54 | chr1-34803848-DEL-54 |
| chr1-34815500-DEL-2686 | chr1-34815502-DEL-2685 |
| chr1-34822149-DEL-54 | chr1-34822150-DEL-54 |
| chr1-34834316-DEL-204 | chr1-34834317-DEL-204 |
| chr1-34842997-DEL-52 | chr1-34842998-DEL-52 |
| chr1-34886962-DEL-72 | chr1-34886963-DEL-72 |
| chr1-34919134-DEL-194 | chr1-34919122-DEL-195 |
| chr1-34954152-DEL-4106 | chr1-34954153-DEL-4106 |
| chr1-34969525-DEL-3974 | chr1-34969527-DEL-3973 |
| chr1-34971668-DEL-67 | chr1-34971665-DEL-72 |
| chr1-34977430-DEL-66 | chr1-34977426-DEL-70 |
| chr1-34984107-DEL-172 | chr1-34984108-DEL-172 |
| chr1-34985321-DEL-191 | chr1-34985322-DEL-191 |
| chr1-35026365-DEL-228 | chr1-35026364-DEL-226 |
| chr1-35051132-DEL-432 | chr1-35051133-DEL-432 |
| chr1-35102767-DEL-5772 | chr1-35102767-DEL-5772 |
| chr1-35113203-DEL-238 | chr1-35113204-DEL-238 |
| chr1-35120086-DEL-2941 | chr1-35120087-DEL-2941 |
| chr1-35123793-DEL-52 | chr1-35123794-DEL-52 |
| chr1-35125092-DEL-859 | chr1-35125093-DEL-859 |
| chr1-35125171-DEL-111 | chr1-35125178-DEL-99 |
| chr1-35141899-DEL-1621 | chr1-35141921-DEL-1622 |
| chr1-35145809-DEL-80 | chr1-35145792-DEL-98 |
| chr1-35155368-DEL-126 | chr1-35155369-DEL-126 |
| chr1-35163539-DEL-344 | chr1-35163540-DEL-344 |
| chr1-35174018-DEL-238 | chr1-35174019-DEL-238 |
| chr1-35176668-DEL-206 | chr1-35176669-DEL-206 |
| chr1-35186687-DEL-63 | chr1-35186687-DEL-81 |
| chr1-35193850-DEL-4855 | chr1-35193851-DEL-4855 |
| chr1-35214342-DEL-234 | chr1-35214348-DEL-232 |
| chr1-35215018-DEL-400 | chr1-35215019-DEL-400 |
| chr1-35249759-DEL-99 | chr1-35249760-DEL-99 |
| chr1-35292371-DEL-74 | chr1-35292372-DEL-74 |
| chr1-35295510-DEL-87 | chr1-35295505-DEL-87 |
| chr1-35298125-DEL-134 | chr1-35298145-DEL-139 |
| chr1-35308464-DEL-97 | chr1-35308451-DEL-88 |
| chr1-35310045-DEL-3697 | chr1-35310046-DEL-3697 |
| chr1-35340130-DEL-2472 | chr1-35340131-DEL-2472 |
| chr1-35346156-DEL-400 | chr1-35346157-DEL-400 |
| chr1-35355389-DEL-2842 | chr1-35355390-DEL-2842 |
| chr1-35363199-DEL-460 | chr1-35363182-DEL-461 |
| chr1-35376410-DEL-182 | chr1-35376398-DEL-183 |
| chr1-35378563-DEL-182 | chr1-35378564-DEL-182 |
| chr1-35397018-DEL-118 | chr1-35397024-DEL-110 |
| chr1-35407810-DEL-59 | chr1-35407804-DEL-60 |
| chr1-35447279-DEL-6338 | chr1-35447280-DEL-6338 |
| chr1-35504997-DEL-70 | chr1-35504998-DEL-70 |
| chr1-35516413-DEL-1035 | chr1-35516414-DEL-1035 |

| | |
|---|---|
| chr1-35528664-DEL-1716 | chr1-35528665-DEL-1716 |
| chr1-35530609-DEL-493 | chr1-35530610-DEL-493 |
| chr1-35542337-DEL-101 | chr1-35542342-DEL-97 |
| chr1-35548028-DEL-506 | chr1-35548029-DEL-506 |
| chr1-35571116-DEL-615 | chr1-35571117-DEL-615 |
| chr1-35603852-DEL-54 | chr1-35603853-DEL-54 |
| chr1-35634817-DEL-233 | chr1-35634815-DEL-234 |
| chr1-35638467-DEL-370 | chr1-35638462-DEL-371 |
| chr1-35639783-DEL-73 | chr1-35639784-DEL-73 |
| chr1-35655922-DEL-69 | chr1-35655923-DEL-69 |
| chr1-35708260-DEL-277 | chr1-35708261-DEL-277 |
| chr1-35735604-DEL-58 | chr1-35735605-DEL-58 |
| chr1-35743048-DEL-204 | chr1-35743049-DEL-204 |
| chr1-35744496-DEL-170 | chr1-35744537-DEL-132 |
| chr1-35746625-DEL-915 | chr1-35746626-DEL-915 |
| chr1-35765719-DEL-109 | chr1-35765720-DEL-109 |
| chr1-35771368-DEL-1647 | chr1-35771369-DEL-1647 |
| chr1-35793195-DEL-139 | chr1-35793196-DEL-139 |
| chr1-35794172-DEL-342 | chr1-35794173-DEL-342 |
| chr1-35821089-DEL-77 | chr1-35821072-DEL-54 |
| chr1-35827579-DEL-7083 | chr1-35827580-DEL-7083 |
| chr1-35879596-DEL-98 | chr1-35879596-DEL-56 |
| chr1-35897064-DEL-66 | chr1-35897062-DEL-65 |
| chr1-35903832-DEL-70 | chr1-35903841-DEL-73 |
| chr1-35906484-DEL-1890 | chr1-35906485-DEL-1890 |
| chr1-35925754-DEL-75 | chr1-35925755-DEL-75 |
| chr1-35935193-DEL-3309 | chr1-35935195-DEL-3308 |
| chr1-35942587-DEL-307 | chr1-35942588-DEL-306 |
| chr1-35953541-DEL-128 | chr1-35953524-DEL-129 |
| chr1-35976295-DEL-70 | chr1-35976282-DEL-68 |
| chr1-36004193-DEL-67 | chr1-36004194-DEL-67 |
| chr1-36005108-DEL-207 | chr1-36005109-DEL-207 |
| chr1-36015919-DEL-56 | chr1-36015920-DEL-104 |
| chr1-36019575-DEL-212 | chr1-36019575-DEL-212 |
| chr1-36024164-DEL-130 | chr1-36024165-DEL-130 |
| chr1-36027838-DEL-216 | chr1-36027839-DEL-216 |
| chr1-36029433-DEL-213 | chr1-36029432-DEL-214 |
| chr1-36070551-DEL-6442 | chr1-36070552-DEL-6442 |
| chr1-36086675-DEL-440 | chr1-36086676-DEL-440 |
| chr1-36090763-DEL-219 | chr1-36090764-DEL-219 |
| chr1-36092823-DEL-635 | chr1-36092823-DEL-631 |
| chr1-36147097-DEL-67 | chr1-36147146-DEL-65 |
| chr1-36174005-DEL-59 | chr1-36174022-DEL-78 |
| chr1-36198879-DEL-86 | chr1-36198880-DEL-86 |
| chr1-36202833-DEL-206 | chr1-36202834-DEL-206 |
| chr1-36206473-DEL-315 | chr1-36206474-DEL-314 |
| chr1-36238014-DEL-95 | chr1-36238015-DEL-95 |
| chr1-36245957-DEL-242 | chr1-36245958-DEL-242 |
| chr1-36261998-DEL-88 | chr1-36261983-DEL-62 |
| chr1-36317457-DEL-490 | chr1-36317458-DEL-490 |
| chr1-36325326-DEL-498 | chr1-36325327-DEL-498 |
| chr1-36417182-DEL-216 | chr1-36417183-DEL-216 |
| chr1-36434083-DEL-124 | chr1-36434087-DEL-121 |
| chr1-36505684-DEL-62 | chr1-36505661-DEL-61 |
| chr1-36676862-DEL-72 | chr1-36676864-DEL-80 |
| chr1-36678350-DEL-1089 | chr1-36678351-DEL-1089 |
| chr1-36689160-DEL-135 | chr1-36689161-DEL-135 |
| chr1-36736335-DEL-51 | chr1-36736336-DEL-51 |
| chr1-36780822-DEL-51 | chr1-36780823-DEL-51 |
| chr1-36789078-DEL-1896 | chr1-36789079-DEL-1896 |
| chr1-36794551-DEL-60 | chr1-36794552-DEL-60 |
| chr1-36798915-DEL-53 | chr1-36798916-DEL-53 |
| chr1-36803259-DEL-76 | chr1-36803260-DEL-76 |
| chr1-36841705-DEL-400 | chr1-36841706-DEL-400 |
| chr1-36858706-DEL-198 | chr1-36858707-DEL-198 |
| chr1-36871325-DEL-82 | chr1-36871296-DEL-91 |
| chr1-36886530-DEL-1071 | chr1-36886531-DEL-1071 |
| chr1-36892232-DEL-200 | chr1-36892233-DEL-200 |
| chr1-36949319-DEL-64 | chr1-36949320-DEL-80 |
| chr1-36982002-DEL-56 | chr1-36982003-DEL-56 |
| chr1-36990254-DEL-6295 | chr1-36990255-DEL-6295 |
| chr1-37004027-DEL-94 | chr1-37004028-DEL-94 |
| chr1-37072498-DEL-363 | chr1-37072493-DEL-364 |
| chr1-37082066-DEL-1779 | chr1-37082067-DEL-1779 |
| chr1-37085827-DEL-124 | chr1-37085828-DEL-124 |
| chr1-37086715-DEL-1146 | chr1-37086716-DEL-1146 |
| chr1-37106976-DEL-107 | chr1-37106977-DEL-107 |
| chr1-37123703-DEL-117 | chr1-37123704-DEL-117 |
| chr1-37130677-DEL-169 | chr1-37130666-DEL-145 |
| chr1-37157664-DEL-200 | chr1-37157664-DEL-200 |

| | |
|---|---|
| chr1-37165914-DEL-126 | chr1-37165915-DEL-126 |
| chr1-37185340-DEL-364 | chr1-37185352-DEL-354 |
| chr1-37197589-DEL-1070 | chr1-37197590-DEL-1070 |
| chr1-37206004-DEL-64 | chr1-37206017-DEL-60 |
| chr1-37219654-DEL-1576 | chr1-37219655-DEL-1576 |
| chr1-37234881-DEL-52 | chr1-37234881-DEL-52 |
| chr1-37239637-DEL-311 | chr1-37239638-DEL-311 |
| chr1-37248210-DEL-408 | chr1-37248211-DEL-408 |
| chr1-37262213-DEL-287 | chr1-37262214-DEL-287 |
| chr1-37275426-DEL-207 | chr1-37275426-DEL-208 |
| chr1-37282702-DEL-531 | chr1-37282703-DEL-531 |
| chr1-37296311-DEL-104 | chr1-37296312-DEL-100 |
| chr1-37317881-DEL-4829 | chr1-37317882-DEL-4829 |
| chr1-37358392-DEL-166 | chr1-37358426-DEL-154 |
| chr1-37366095-DEL-115 | chr1-37366096-DEL-115 |
| chr1-37433916-DEL-90 | chr1-37433917-DEL-90 |
| chr1-37512772-DEL-58 | chr1-37512775-DEL-50 |
| chr1-37521354-DEL-204 | chr1-37521355-DEL-204 |
| chr1-37523533-DEL-248 | chr1-37523534-DEL-248 |
| chr1-37542575-DEL-61 | chr1-37542576-DEL-61 |
| chr1-37545113-DEL-54 | chr1-37545112-DEL-56 |
| chr1-37547396-DEL-188 | chr1-37547397-DEL-188 |
| chr1-37559954-DEL-66 | chr1-37559955-DEL-66 |
| chr1-37562963-DEL-104 | chr1-37562985-DEL-130 |
| chr1-37569427-DEL-1650 | chr1-37569428-DEL-1650 |
| chr1-37589543-DEL-80 | chr1-37589544-DEL-80 |
| chr1-37592501-DEL-412 | chr1-37592505-DEL-409 |
| chr1-37595012-DEL-897 | chr1-37595013-DEL-897 |
| chr1-37606538-DEL-144 | chr1-37606531-DEL-152 |
| chr1-37643037-DEL-1893 | chr1-37643038-DEL-1893 |
| chr1-37677207-DEL-403 | chr1-37677208-DEL-403 |
| chr1-37691808-DEL-76 | chr1-37691809-DEL-76 |
| chr1-37693938-DEL-208 | chr1-37693939-DEL-208 |
| chr1-37701041-DEL-70 | chr1-37701049-DEL-63 |
| chr1-37714156-DEL-82 | chr1-37714157-DEL-82 |
| chr1-37719011-DEL-284 | chr1-37719012-DEL-284 |
| chr1-37783396-DEL-54 | chr1-37783397-DEL-54 |
| chr1-37812965-DEL-4726 | chr1-37812966-DEL-4726 |
| chr1-37833980-DEL-84 | chr1-37833980-DEL-66 |
| chr1-37842302-DEL-92 | chr1-37842303-DEL-92 |
| chr1-37847474-DEL-67 | chr1-37847472-DEL-65 |
| chr1-37969534-DEL-147 | chr1-37969535-DEL-147 |
| chr1-37970026-DEL-722 | chr1-37970027-DEL-722 |
| chr1-37973061-DEL-180 | chr1-37973062-DEL-180 |
| chr1-37983110-DEL-596 | chr1-37983111-DEL-596 |
| chr1-38000130-DEL-59 | chr1-38000131-DEL-64 |
| chr1-38006794-DEL-618 | chr1-38006791-DEL-619 |
| chr1-38039631-DEL-159 | chr1-38039632-DEL-159 |
| chr1-38053068-DEL-198 | chr1-38053069-DEL-198 |
| chr1-38055811-DEL-226 | chr1-38055812-DEL-226 |
| chr1-38140311-DEL-110 | chr1-38140312-DEL-110 |
| chr1-38207463-DEL-687 | chr1-38207464-DEL-659 |
| chr1-38229825-DEL-216 | chr1-38229826-DEL-216 |
| chr1-38251955-DEL-98 | chr1-38251956-DEL-98 |
| chr1-38264809-DEL-340 | chr1-38264810-DEL-340 |
| chr1-38275886-DEL-367 | chr1-38275887-DEL-367 |
| chr1-38277597-DEL-712 | chr1-38277598-DEL-712 |
| chr1-38296068-DEL-51 | chr1-38296069-DEL-51 |
| chr1-38298415-DEL-52 | chr1-38298415-DEL-52 |
| chr1-38315213-DEL-75 | chr1-38315204-DEL-76 |
| chr1-38322159-DEL-53 | chr1-38322160-DEL-53 |
| chr1-38338383-DEL-3755 | chr1-38338384-DEL-3755 |
| chr1-38354330-DEL-234 | chr1-38354376-DEL-195 |
| chr1-38369319-DEL-315 | chr1-38369320-DEL-315 |
| chr1-38370634-DEL-1335 | chr1-38370635-DEL-1335 |
| chr1-38394103-DEL-2844 | chr1-38394104-DEL-2844 |
| chr1-38404809-DEL-83 | chr1-38404833-DEL-59 |
| chr1-38425944-DEL-120 | chr1-38425907-DEL-120 |
| chr1-38427563-DEL-52 | chr1-38427564-DEL-52 |
| chr1-38443373-DEL-95 | chr1-38443346-DEL-96 |
| chr1-38451024-DEL-622 | chr1-38451025-DEL-622 |
| chr1-38463623-DEL-114 | chr1-38463668-DEL-70 |
| chr1-38493214-DEL-508 | chr1-38493215-DEL-507 |
| chr1-38511168-DEL-161 | chr1-38511192-DEL-138 |
| chr1-38530387-DEL-1515 | chr1-38530388-DEL-1515 |
| chr1-38582857-DEL-100 | chr1-38582858-DEL-100 |
| chr1-38587102-DEL-117 | chr1-38587103-DEL-117 |
| chr1-38603771-DEL-244 | chr1-38603772-DEL-244 |
| chr1-38643891-DEL-116 | chr1-38643892-DEL-116 |
| chr1-38644657-DEL-124 | chr1-38644658-DEL-124 |

| | |
|---|---|
| chr1-38653176-DEL-63 | chr1-38653177-DEL-63 |
| chr1-38692992-DEL-531 | chr1-38692993-DEL-531 |
| chr1-38726060-DEL-58 | chr1-38726059-DEL-60 |
| chr1-38743874-DEL-4219 | chr1-38743875-DEL-4219 |
| chr1-38772993-DEL-1404 | chr1-38772993-DEL-1404 |
| chr1-38798826-DEL-69 | chr1-38798776-DEL-80 |
| chr1-38822535-DEL-105 | chr1-38822545-DEL-114 |
| chr1-38833128-DEL-60 | chr1-38833129-DEL-60 |
| chr1-38847925-DEL-1749 | chr1-38847926-DEL-1749 |
| chr1-38848745-DEL-659 | chr1-38848740-DEL-660 |
| chr1-38858887-DEL-58 | chr1-38858888-DEL-58 |
| chr1-38890395-DEL-57 | chr1-38890396-DEL-57 |
| chr1-38893371-DEL-126 | chr1-38893372-DEL-126 |
| chr1-38906384-DEL-195 | chr1-38906385-DEL-195 |
| chr1-38926268-DEL-77 | chr1-38926268-DEL-77 |
| chr1-38931243-DEL-199 | chr1-38931244-DEL-199 |
| chr1-38941548-DEL-71 | chr1-38941547-DEL-73 |
| chr1-38949869-DEL-63 | chr1-38949870-DEL-63 |
| chr1-38958921-DEL-52 | chr1-38958922-DEL-52 |
| chr1-38974890-DEL-208 | chr1-38974891-DEL-208 |
| chr1-38999682-DEL-70 | chr1-38999678-DEL-75 |
| chr1-39027929-DEL-172 | chr1-39027930-DEL-172 |
| chr1-39029206-DEL-289 | chr1-39029207-DEL-289 |
| chr1-39038312-DEL-159 | chr1-39038317-DEL-160 |
| chr1-39038731-DEL-527 | chr1-39038732-DEL-527 |
| chr1-39069987-DEL-76 | chr1-39069988-DEL-76 |
| chr1-39071472-DEL-76 | chr1-39071473-DEL-76 |
| chr1-39081785-DEL-57 | chr1-39081786-DEL-57 |
| chr1-39097028-DEL-2825 | chr1-39097029-DEL-2825 |
| chr1-39118772-DEL-1111 | chr1-39118773-DEL-1111 |
| chr1-39120259-DEL-2636 | chr1-39120260-DEL-2636 |
| chr1-39131421-DEL-94 | chr1-39131422-DEL-94 |
| chr1-39226540-DEL-60 | chr1-39226541-DEL-60 |
| chr1-39285683-DEL-205 | chr1-39285684-DEL-205 |
| chr1-39303819-DEL-543 | chr1-39303820-DEL-543 |
| chr1-39331553-DEL-76 | chr1-39331554-DEL-76 |
| chr1-39341803-DEL-340 | chr1-39341804-DEL-340 |
| chr1-39391988-DEL-443 | chr1-39391988-DEL-431 |
| chr1-39420898-DEL-794 | chr1-39420902-DEL-792 |
| chr1-39454437-DEL-82 | chr1-39454438-DEL-82 |
| chr1-39471660-DEL-475 | chr1-39471661-DEL-475 |
| chr1-39477256-DEL-78 | chr1-39477257-DEL-78 |
| chr1-39493045-DEL-140 | chr1-39493046-DEL-140 |
| chr1-39523490-DEL-94 | chr1-39523491-DEL-94 |
| chr1-39567984-DEL-56 | chr1-39567985-DEL-56 |
| chr1-39620556-DEL-68 | chr1-39620557-DEL-68 |
| chr1-39621147-DEL-153 | chr1-39621151-DEL-146 |
| chr1-39634481-DEL-90 | chr1-39634473-DEL-82 |
| chr1-39635788-DEL-126 | chr1-39635790-DEL-127 |
| chr1-39659555-DEL-128 | chr1-39659594-DEL-90 |
| chr1-39666892-DEL-62 | chr1-39666892-DEL-62 |
| chr1-39683264-DEL-207 | chr1-39683265-DEL-207 |
| chr1-39684948-DEL-84 | chr1-39684971-DEL-52 |
| chr1-39712355-DEL-80 | chr1-39712370-DEL-80 |
| chr1-39724118-DEL-104 | chr1-39724096-DEL-87 |
| chr1-39732773-DEL-90 | chr1-39732774-DEL-90 |
| chr1-39739269-DEL-745 | chr1-39739270-DEL-743 |
| chr1-39740577-DEL-1441 | chr1-39740578-DEL-1441 |
| chr1-39756922-DEL-386 | chr1-39756923-DEL-386 |
| chr1-39761119-DEL-6227 | chr1-39761127-DEL-6220 |
| chr1-39785141-DEL-62 | chr1-39785142-DEL-62 |
| chr1-39786285-DEL-50 | chr1-39786253-DEL-83 |
| chr1-39787070-DEL-196 | chr1-39787071-DEL-196 |
| chr1-39794913-DEL-85 | chr1-39794920-DEL-93 |
| chr1-39794977-DEL-1015 | chr1-39794955-DEL-1000 |
| chr1-39804278-DEL-61 | chr1-39804278-DEL-61 |
| chr1-39808455-DEL-206 | chr1-39808456-DEL-206 |
| chr1-39816692-DEL-60 | chr1-39816693-DEL-60 |
| chr1-39839535-DEL-126 | chr1-39839527-DEL-137 |
| chr1-39841385-DEL-774 | chr1-39841386-DEL-774 |
| chr1-39849595-DEL-573 | chr1-39849596-DEL-573 |
| chr1-39867176-DEL-1937 | chr1-39867177-DEL-1937 |
| chr1-39906775-DEL-96 | chr1-39906768-DEL-114 |
| chr1-39925649-DEL-58 | chr1-39925650-DEL-58 |
| chr1-39929598-DEL-198 | chr1-39929599-DEL-198 |
| chr1-39932431-DEL-60 | chr1-39932432-DEL-60 |
| chr1-39936545-DEL-82 | chr1-39936546-DEL-82 |
| chr1-39954831-DEL-90 | chr1-39954832-DEL-90 |
| chr1-39956146-DEL-59 | chr1-39956153-DEL-55 |
| chr1-39965415-DEL-92 | chr1-39965411-DEL-93 |

| | |
|---|---|
| chr1-40021707-DEL-56 | chr1-40021707-DEL-56 |
| chr1-40025417-DEL-56 | chr1-40025418-DEL-56 |
| chr1-40045209-DEL-391 | chr1-40045210-DEL-391 |
| chr1-40057944-DEL-71 | chr1-40057945-DEL-71 |
| chr1-40086103-DEL-1929 | chr1-40086107-DEL-1927 |
| chr1-40102414-DEL-550 | chr1-40102415-DEL-550 |
| chr1-40141344-DEL-66 | chr1-40141345-DEL-60 |
| chr1-40171671-DEL-108 | chr1-40171680-DEL-100 |
| chr1-40174674-DEL-52 | chr1-40174675-DEL-52 |
| chr1-40180907-DEL-52 | chr1-40180907-DEL-82 |
| chr1-40181382-DEL-6035 | chr1-40181383-DEL-6035 |
| chr1-40207240-DEL-894 | chr1-40207241-DEL-894 |
| chr1-40234244-DEL-6518 | chr1-40234245-DEL-6518 |
| chr1-40244566-DEL-66 | chr1-40244567-DEL-66 |
| chr1-40250825-DEL-284 | chr1-40250826-DEL-284 |
| chr1-40278068-DEL-241 | chr1-40278069-DEL-241 |
| chr1-40290193-DEL-92 | chr1-40290202-DEL-84 |
| chr1-40297730-DEL-80 | chr1-40297710-DEL-81 |
| chr1-40308288-DEL-338 | chr1-40308296-DEL-334 |
| chr1-40333785-DEL-72 | chr1-40333785-DEL-72 |
| chr1-40346029-DEL-2871 | chr1-40346030-DEL-2871 |
| chr1-40362147-DEL-54 | chr1-40362148-DEL-54 |
| chr1-40375186-DEL-76 | chr1-40375211-DEL-52 |
| chr1-40376983-DEL-1574 | chr1-40376984-DEL-1574 |
| chr1-40426534-DEL-57 | chr1-40426535-DEL-57 |
| chr1-40434016-DEL-66 | chr1-40434015-DEL-50 |
| chr1-40452596-DEL-66 | chr1-40452597-DEL-66 |
| chr1-40460433-DEL-56 | chr1-40460399-DEL-88 |
| chr1-40461372-DEL-406 | chr1-40461373-DEL-408 |
| chr1-40462737-DEL-72 | chr1-40462698-DEL-76 |
| chr1-40464917-DEL-116 | chr1-40464918-DEL-116 |
| chr1-40486214-DEL-2466 | chr1-40486213-DEL-2462 |
| chr1-40491505-DEL-6229 | chr1-40491506-DEL-6229 |
| chr1-40524675-DEL-228 | chr1-40524658-DEL-263 |
| chr1-40527966-DEL-102 | chr1-40527957-DEL-93 |
| chr1-40558756-DEL-2029 | chr1-40558757-DEL-2030 |
| chr1-40575887-DEL-224 | chr1-40575888-DEL-224 |
| chr1-40584160-DEL-880 | chr1-40584161-DEL-880 |
| chr1-40603895-DEL-199 | chr1-40603890-DEL-198 |
| chr1-40604576-DEL-80 | chr1-40604577-DEL-80 |
| chr1-40706884-DEL-73 | chr1-40706884-DEL-73 |
| chr1-40710247-DEL-322 | chr1-40710289-DEL-323 |
| chr1-40716798-DEL-114 | chr1-40716799-DEL-114 |
| chr1-40737150-DEL-4462 | chr1-40737151-DEL-4462 |
| chr1-40750196-DEL-406 | chr1-40750197-DEL-406 |
| chr1-40777424-DEL-288 | chr1-40777425-DEL-288 |
| chr1-40798178-DEL-315 | chr1-40798179-DEL-315 |
| chr1-40821358-DEL-153 | chr1-40821357-DEL-169 |
| chr1-40831513-DEL-52 | chr1-40831514-DEL-52 |
| chr1-40856833-DEL-74 | chr1-40856821-DEL-77 |
| chr1-40869378-DEL-595 | chr1-40869419-DEL-551 |
| chr1-40909097-DEL-212 | chr1-40909098-DEL-212 |
| chr1-40919171-DEL-118 | chr1-40919203-DEL-70 |
| chr1-40920209-DEL-940 | chr1-40920210-DEL-940 |
| chr1-40954353-DEL-1891 | chr1-40954354-DEL-1891 |
| chr1-40966715-DEL-100 | chr1-40966716-DEL-100 |
| chr1-40985873-DEL-56 | chr1-40985875-DEL-55 |
| chr1-40995061-DEL-62 | chr1-40995062-DEL-64 |
| chr1-40996764-DEL-52 | chr1-40996765-DEL-58 |
| chr1-41056973-DEL-52 | chr1-41056974-DEL-52 |
| chr1-41062049-DEL-535 | chr1-41062030-DEL-542 |
| chr1-41077949-DEL-80 | chr1-41077963-DEL-82 |
| chr1-41096342-DEL-83 | chr1-41096346-DEL-53 |
| chr1-41106167-DEL-55 | chr1-41106168-DEL-55 |
| chr1-41114879-DEL-2395 | chr1-41114880-DEL-2395 |
| chr1-41117522-DEL-900 | chr1-41117522-DEL-900 |
| chr1-41119911-DEL-57 | chr1-41119911-DEL-58 |
| chr1-41121980-DEL-177 | chr1-41121980-DEL-176 |
| chr1-41123451-DEL-56 | chr1-41123452-DEL-56 |
| chr1-41124050-DEL-208 | chr1-41124050-DEL-208 |
| chr1-41127286-DEL-3060 | chr1-41127287-DEL-3060 |
| chr1-41150370-DEL-106 | chr1-41150371-DEL-106 |
| chr1-41156311-DEL-75 | chr1-41156312-DEL-75 |
| chr1-41182966-DEL-694 | chr1-41182967-DEL-694 |
| chr1-41185305-DEL-61 | chr1-41185308-DEL-59 |
| chr1-41216060-DEL-971 | chr1-41216061-DEL-971 |
| chr1-41221627-DEL-802 | chr1-41221621-DEL-803 |
| chr1-41293540-DEL-91 | chr1-41293545-DEL-92 |
| chr1-41327998-DEL-90 | chr1-41327999-DEL-90 |
| chr1-41348829-DEL-200 | chr1-41348779-DEL-201 |

| | |
|---|---|
| chr1-41357787-DEL-706 | chr1-41357787-DEL-706 |
| chr1-41358885-DEL-413 | chr1-41358885-DEL-413 |
| chr1-41369157-DEL-2555 | chr1-41369158-DEL-2555 |
| chr1-41369466-DEL-602 | chr1-41369461-DEL-603 |
| chr1-41409364-DEL-347 | chr1-41409364-DEL-347 |
| chr1-41437603-DEL-223 | chr1-41437583-DEL-236 |
| chr1-41491051-DEL-92 | chr1-41491020-DEL-121 |
| chr1-41494767-DEL-558 | chr1-41494768-DEL-558 |
| chr1-41506947-DEL-6223 | chr1-41506948-DEL-6223 |
| chr1-41532054-DEL-436 | chr1-41532055-DEL-436 |
| chr1-41546234-DEL-100 | chr1-41546235-DEL-94 |
| chr1-41596915-DEL-520 | chr1-41596915-DEL-520 |
| chr1-41600299-DEL-143 | chr1-41600299-DEL-143 |
| chr1-41616516-DEL-62 | chr1-41616517-DEL-62 |
| chr1-41675996-DEL-148 | chr1-41675994-DEL-160 |
| chr1-41682937-DEL-2338 | chr1-41682939-DEL-2336 |
| chr1-41703331-DEL-74 | chr1-41703332-DEL-74 |
| chr1-41703482-DEL-807 | chr1-41703483-DEL-807 |
| chr1-41717308-DEL-2994 | chr1-41717309-DEL-2994 |
| chr1-41757154-DEL-53 | chr1-41757155-DEL-53 |
| chr1-41776683-DEL-528 | chr1-41776683-DEL-528 |
| chr1-41804710-DEL-74 | chr1-41804711-DEL-74 |
| chr1-41809564-DEL-214 | chr1-41809565-DEL-214 |
| chr1-41831873-DEL-140 | chr1-41831882-DEL-132 |
| chr1-41887706-DEL-603 | chr1-41887710-DEL-602 |
| chr1-41898460-DEL-526 | chr1-41898461-DEL-526 |
| chr1-41906831-DEL-71 | chr1-41906814-DEL-61 |
| chr1-41960919-DEL-762 | chr1-41960918-DEL-762 |
| chr1-41979717-DEL-959 | chr1-41979733-DEL-950 |
| chr1-41997601-DEL-105 | chr1-41997598-DEL-108 |
| chr1-42025472-DEL-57 | chr1-42025472-DEL-58 |
| chr1-42037984-DEL-78 | chr1-42037981-DEL-62 |
| chr1-42053200-DEL-2228 | chr1-42053201-DEL-2228 |
| chr1-42060408-DEL-199 | chr1-42060397-DEL-211 |
| chr1-42069809-DEL-887 | chr1-42069810-DEL-886 |
| chr1-42077700-DEL-124 | chr1-42077701-DEL-124 |
| chr1-42133120-DEL-71 | chr1-42133121-DEL-71 |
| chr1-42176571-DEL-6387 | chr1-42176572-DEL-6387 |
| chr1-42219260-DEL-222 | chr1-42219261-DEL-222 |
| chr1-42220202-DEL-117 | chr1-42220203-DEL-117 |
| chr1-42229233-DEL-58 | chr1-42229234-DEL-51 |
| chr1-42237823-DEL-158 | chr1-42237824-DEL-155 |
| chr1-42249226-DEL-114 | chr1-42249227-DEL-114 |
| chr1-42276468-DEL-852 | chr1-42276469-DEL-852 |
| chr1-42319914-DEL-54 | chr1-42319915-DEL-54 |
| chr1-42373717-DEL-6639 | chr1-42373718-DEL-6639 |
| chr1-42404561-DEL-54 | chr1-42404562-DEL-60 |
| chr1-42417508-DEL-1283 | chr1-42417509-DEL-1283 |
| chr1-42454387-DEL-57 | chr1-42454387-DEL-57 |
| chr1-42481671-DEL-65 | chr1-42481671-DEL-87 |
| chr1-42522379-DEL-69 | chr1-42522380-DEL-69 |
| chr1-42584872-DEL-70 | chr1-42584878-DEL-61 |
| chr1-42607747-DEL-54 | chr1-42607748-DEL-60 |
| chr1-42646847-DEL-243 | chr1-42646848-DEL-225 |
| chr1-42650083-DEL-61 | chr1-42650076-DEL-63 |
| chr1-42678873-DEL-118 | chr1-42678878-DEL-122 |
| chr1-42685257-DEL-1891 | chr1-42685258-DEL-1891 |
| chr1-42714798-DEL-55 | chr1-42714799-DEL-55 |
| chr1-42825103-DEL-110 | chr1-42825104-DEL-110 |
| chr1-42828404-DEL-218 | chr1-42828413-DEL-209 |
| chr1-42836483-DEL-70 | chr1-42836484-DEL-70 |
| chr1-42857979-DEL-84 | chr1-42857980-DEL-94 |
| chr1-42869289-DEL-272 | chr1-42869302-DEL-260 |
| chr1-42902328-DEL-119 | chr1-42902329-DEL-119 |
| chr1-42922622-DEL-225 | chr1-42922623-DEL-225 |
| chr1-42957636-DEL-3732 | chr1-42957637-DEL-3732 |
| chr1-42973634-DEL-65 | chr1-42973637-DEL-66 |
| chr1-42982396-DEL-76 | chr1-42982395-DEL-76 |
| chr1-43011418-DEL-64 | chr1-43011419-DEL-54 |
| chr1-43012153-DEL-4696 | chr1-43012154-DEL-4696 |
| chr1-43067443-DEL-58 | chr1-43067444-DEL-58 |
| chr1-43069955-DEL-400 | chr1-43069956-DEL-400 |
| chr1-43077782-DEL-52 | chr1-43077783-DEL-52 |
| chr1-43093073-DEL-229 | chr1-43093071-DEL-227 |
| chr1-43121763-DEL-52 | chr1-43121764-DEL-50 |
| chr1-43160561-DEL-241 | chr1-43160562-DEL-241 |
| chr1-43175827-DEL-185 | chr1-43175852-DEL-165 |
| chr1-43188018-DEL-716 | chr1-43188019-DEL-716 |
| chr1-43225058-DEL-102 | chr1-43225065-DEL-66 |
| chr1-43228198-DEL-733 | chr1-43228199-DEL-733 |

| | |
|---|---|
| chr1-43239560-DEL-207 | chr1-43239529-DEL-206 |
| chr1-43277374-DEL-56 | chr1-43277375-DEL-66 |
| chr1-43306365-DEL-269 | chr1-43306366-DEL-268 |
| chr1-43383050-DEL-123 | chr1-43383051-DEL-123 |
| chr1-43392241-DEL-51 | chr1-43392242-DEL-51 |
| chr1-43408367-DEL-1312 | chr1-43408368-DEL-1312 |
| chr1-43415368-DEL-504 | chr1-43415369-DEL-504 |
| chr1-43421582-DEL-2037 | chr1-43421583-DEL-2037 |
| chr1-43446375-DEL-62 | chr1-43446376-DEL-62 |
| chr1-43453281-DEL-56 | chr1-43453282-DEL-56 |
| chr1-43471511-DEL-76 | chr1-43471520-DEL-68 |
| chr1-43552394-DEL-481 | chr1-43552392-DEL-481 |
| chr1-43605243-DEL-232 | chr1-43605244-DEL-232 |
| chr1-43605937-DEL-55 | chr1-43605938-DEL-55 |
| chr1-43610514-DEL-470 | chr1-43610515-DEL-470 |
| chr1-43626625-DEL-91 | chr1-43626630-DEL-87 |
| chr1-43629416-DEL-94 | chr1-43629402-DEL-98 |
| chr1-43641598-DEL-52 | chr1-43641588-DEL-57 |
| chr1-43650107-DEL-643 | chr1-43650108-DEL-643 |
| chr1-43657731-DEL-343 | chr1-43657732-DEL-343 |
| chr1-43658974-DEL-2994 | chr1-43658975-DEL-2994 |
| chr1-43667721-DEL-1904 | chr1-43667719-DEL-1904 |
| chr1-43669906-DEL-826 | chr1-43669905-DEL-827 |
| chr1-43697519-DEL-214 | chr1-43697525-DEL-194 |
| chr1-43717029-DEL-417 | chr1-43717040-DEL-420 |
| chr1-43720772-DEL-311 | chr1-43720773-DEL-311 |
| chr1-43727135-DEL-3993 | chr1-43727136-DEL-3993 |
| chr1-43736702-DEL-214 | chr1-43736703-DEL-214 |
| chr1-43739983-DEL-126 | chr1-43740032-DEL-88 |
| chr1-43741906-DEL-447 | chr1-43741907-DEL-447 |
| chr1-43770446-DEL-51 | chr1-43770447-DEL-51 |
| chr1-43772674-DEL-186 | chr1-43772675-DEL-186 |
| chr1-43833287-DEL-131 | chr1-43833288-DEL-131 |
| chr1-43872825-DEL-1068 | chr1-43872824-DEL-1070 |
| chr1-43880929-DEL-204 | chr1-43880929-DEL-199 |
| chr1-43887119-DEL-7307 | chr1-43887120-DEL-7307 |
| chr1-43887456-DEL-889 | chr1-43887458-DEL-887 |
| chr1-43894057-DEL-208 | chr1-43894058-DEL-208 |
| chr1-43910178-DEL-65 | chr1-43910179-DEL-55 |
| chr1-43922888-DEL-59 | chr1-43922894-DEL-55 |
| chr1-43932151-DEL-205 | chr1-43932152-DEL-205 |
| chr1-43943615-DEL-6030 | chr1-43943614-DEL-6031 |
| chr1-43945756-DEL-112 | chr1-43945757-DEL-112 |
| chr1-43970560-DEL-174 | chr1-43970561-DEL-174 |
| chr1-44015181-DEL-192 | chr1-44015182-DEL-192 |
| chr1-44057356-DEL-461 | chr1-44057357-DEL-461 |
| chr1-44132394-DEL-76 | chr1-44132395-DEL-76 |
| chr1-44141976-DEL-59 | chr1-44142015-DEL-67 |
| chr1-44173489-DEL-557 | chr1-44173490-DEL-557 |
| chr1-44175627-DEL-60 | chr1-44175627-DEL-58 |
| chr1-44190143-DEL-1213 | chr1-44190144-DEL-1213 |
| chr1-44192744-DEL-495 | chr1-44192745-DEL-495 |
| chr1-44214663-DEL-51 | chr1-44214664-DEL-51 |
| chr1-44226215-DEL-60 | chr1-44226216-DEL-60 |
| chr1-44242123-DEL-323 | chr1-44242124-DEL-323 |
| chr1-44255129-DEL-108 | chr1-44255130-DEL-108 |
| chr1-44284224-DEL-6201 | chr1-44284225-DEL-6201 |
| chr1-44296749-DEL-389 | chr1-44296749-DEL-389 |
| chr1-44318413-DEL-6383 | chr1-44318414-DEL-6383 |
| chr1-44380141-DEL-2637 | chr1-44380137-DEL-2638 |
| chr1-44407386-DEL-7417 | chr1-44407387-DEL-7417 |
| chr1-44463649-DEL-505 | chr1-44463652-DEL-505 |
| chr1-44470587-DEL-431 | chr1-44470588-DEL-432 |
| chr1-44472165-DEL-911 | chr1-44472166-DEL-911 |
| chr1-44494937-DEL-60 | chr1-44494938-DEL-60 |
| chr1-44500210-DEL-334 | chr1-44500211-DEL-334 |
| chr1-44506994-DEL-62 | chr1-44506995-DEL-62 |
| chr1-44511573-DEL-1268 | chr1-44511574-DEL-1266 |
| chr1-44514023-DEL-16941 | chr1-44514024-DEL-16941 |
| chr1-44536592-DEL-203 | chr1-44536593-DEL-203 |
| chr1-44554432-DEL-768 | chr1-44554433-DEL-768 |
| chr1-44557954-DEL-75 | chr1-44557927-DEL-103 |
| chr1-44558661-DEL-2793 | chr1-44558662-DEL-2793 |
| chr1-44566877-DEL-602 | chr1-44566878-DEL-602 |
| chr1-44568462-DEL-213 | chr1-44568463-DEL-211 |
| chr1-44584256-DEL-173 | chr1-44584257-DEL-173 |
| chr1-44607135-DEL-52 | chr1-44607136-DEL-52 |
| chr1-44630775-DEL-64 | chr1-44630776-DEL-64 |
| chr1-44641247-DEL-7168 | chr1-44641248-DEL-7168 |
| chr1-44665760-DEL-631 | chr1-44665761-DEL-631 |

| | |
|---|---|
| chr1-44667162-DEL-469 | chr1-44667165-DEL-471 |
| chr1-44690869-DEL-2162 | chr1-44690873-DEL-2163 |
| chr1-44704452-DEL-58 | chr1-44704451-DEL-59 |
| chr1-44722108-DEL-192 | chr1-44722120-DEL-193 |
| chr1-44731724-DEL-6776 | chr1-44731712-DEL-6777 |
| chr1-44739998-DEL-52 | chr1-44739999-DEL-52 |
| chr1-44749306-DEL-693 | chr1-44749325-DEL-675 |
| chr1-44761527-DEL-7591 | chr1-44761528-DEL-7591 |
| chr1-44768017-DEL-1101 | chr1-44768018-DEL-1101 |
| chr1-44788948-DEL-187 | chr1-44788948-DEL-187 |
| chr1-44821822-DEL-1077 | chr1-44821823-DEL-1077 |
| chr1-44837841-DEL-93 | chr1-44837842-DEL-93 |
| chr1-44851227-DEL-397 | chr1-44851228-DEL-397 |
| chr1-44859949-DEL-5542 | chr1-44859950-DEL-5542 |
| chr1-44867880-DEL-1959 | chr1-44867833-DEL-2004 |
| chr1-44880072-DEL-827 | chr1-44880073-DEL-827 |
| chr1-44883731-DEL-1466 | chr1-44883732-DEL-1466 |
| chr1-44916123-DEL-1157 | chr1-44916124-DEL-1157 |
| chr1-44925440-DEL-205 | chr1-44925441-DEL-205 |
| chr1-44936193-DEL-6168 | chr1-44936194-DEL-6168 |
| chr1-44942453-DEL-6050 | chr1-44942453-DEL-6050 |
| chr1-44971102-DEL-595 | chr1-44971103-DEL-595 |
| chr1-44979456-DEL-82 | chr1-44979457-DEL-82 |
| chr1-44980659-DEL-136 | chr1-44980660-DEL-136 |
| chr1-44998405-DEL-7321 | chr1-44998406-DEL-7321 |
| chr1-45040746-DEL-1281 | chr1-45040747-DEL-1281 |
| chr1-45083454-DEL-62 | chr1-45083455-DEL-62 |
| chr1-45086609-DEL-5554 | chr1-45086599-DEL-5555 |
| chr1-45092320-DEL-77 | chr1-45092319-DEL-78 |
| chr1-45092738-DEL-1895 | chr1-45092739-DEL-1895 |
| chr1-45095118-DEL-72 | chr1-45095121-DEL-68 |
| chr1-45125046-DEL-1899 | chr1-45125046-DEL-1899 |
| chr1-45126687-DEL-59 | chr1-45126688-DEL-54 |
| chr1-45204558-DEL-6845 | chr1-45204559-DEL-6845 |
| chr1-45204600-DEL-215 | chr1-45204601-DEL-215 |
| chr1-45228154-DEL-1007 | chr1-45228155-DEL-1007 |
| chr1-45238160-DEL-7019 | chr1-45238161-DEL-7019 |
| chr1-45266269-DEL-117 | chr1-45266270-DEL-118 |
| chr1-45270111-DEL-6254 | chr1-45270112-DEL-6254 |
| chr1-45277656-DEL-768 | chr1-45277657-DEL-768 |
| chr1-45368695-DEL-134 | chr1-45368698-DEL-132 |
| chr1-45370516-DEL-117 | chr1-45370517-DEL-117 |
| chr1-45376715-DEL-198 | chr1-45376716-DEL-192 |
| chr1-45399201-DEL-1596 | chr1-45399202-DEL-1596 |
| chr1-45401006-DEL-198 | chr1-45401007-DEL-198 |
| chr1-45403956-DEL-8283 | chr1-45403957-DEL-8283 |
| chr1-45438580-DEL-62 | chr1-45438581-DEL-62 |
| chr1-45440775-DEL-51 | chr1-45440776-DEL-51 |
| chr1-45446549-DEL-74 | chr1-45446550-DEL-82 |
| chr1-45447292-DEL-97 | chr1-45447331-DEL-63 |
| chr1-45470542-DEL-205 | chr1-45470543-DEL-205 |
| chr1-45472818-DEL-51 | chr1-45472797-DEL-72 |
| chr1-45479749-DEL-627 | chr1-45479750-DEL-627 |
| chr1-45489831-DEL-100 | chr1-45489832-DEL-98 |
| chr1-45504734-DEL-247 | chr1-45504735-DEL-248 |
| chr1-45514047-DEL-460 | chr1-45514047-DEL-461 |
| chr1-45533026-DEL-1166 | chr1-45533027-DEL-1166 |
| chr1-45549759-DEL-3668 | chr1-45549759-DEL-3668 |
| chr1-45560576-DEL-2801 | chr1-45560577-DEL-2801 |
| chr1-45578928-DEL-527 | chr1-45578929-DEL-527 |
| chr1-45590760-DEL-77 | chr1-45590764-DEL-76 |
| chr1-45596308-DEL-7412 | chr1-45596301-DEL-7417 |
| chr1-45608057-DEL-1345 | chr1-45608058-DEL-1336 |
| chr1-45629068-DEL-119 | chr1-45629069-DEL-119 |
| chr1-45649763-DEL-2825 | chr1-45649759-DEL-2826 |
| chr1-45670025-DEL-1184 | chr1-45670025-DEL-1184 |
| chr1-45676377-DEL-122 | chr1-45676378-DEL-122 |
| chr1-45677599-DEL-827 | chr1-45677600-DEL-827 |
| chr1-45689342-DEL-185 | chr1-45689343-DEL-185 |
| chr1-45693375-DEL-6276 | chr1-45693367-DEL-6277 |
| chr1-45699246-DEL-87 | chr1-45699247-DEL-87 |
| chr1-45700460-DEL-400 | chr1-45700461-DEL-400 |
| chr1-45722407-DEL-180 | chr1-45722408-DEL-180 |
| chr1-45773229-DEL-5283 | chr1-45773230-DEL-5283 |
| chr1-45796424-DEL-796 | chr1-45796425-DEL-796 |
| chr1-45819512-DEL-3296 | chr1-45819513-DEL-3296 |
| chr1-45825146-DEL-201 | chr1-45825147-DEL-201 |
| chr1-45827841-DEL-73 | chr1-45827842-DEL-74 |
| chr1-45864269-DEL-168 | chr1-45864270-DEL-168 |
| chr1-45896319-DEL-671 | chr1-45896320-DEL-671 |

| | |
|---|---|
| chr1-45938709-DEL-1377 | chr1-45938709-DEL-1377 |
| chr1-45983606-DEL-2141 | chr1-45983607-DEL-2141 |
| chr1-46001025-DEL-163 | chr1-46001026-DEL-163 |
| chr1-46010285-DEL-11495 | chr1-46010286-DEL-11495 |
| chr1-46072709-DEL-709 | chr1-46072710-DEL-709 |
| chr1-46088466-DEL-2762 | chr1-46088467-DEL-2762 |
| chr1-46088662-DEL-11923 | chr1-46088663-DEL-11923 |
| chr1-46103565-DEL-61 | chr1-46103566-DEL-61 |
| chr1-46220035-DEL-361 | chr1-46220036-DEL-361 |
| chr1-46310392-DEL-63 | chr1-46310393-DEL-63 |
| chr1-46323132-DEL-82 | chr1-46323133-DEL-82 |
| chr1-46323595-DEL-186 | chr1-46323596-DEL-186 |
| chr1-46367881-DEL-55 | chr1-46367871-DEL-65 |
| chr1-46384824-DEL-6259 | chr1-46384825-DEL-6259 |
| chr1-46403586-DEL-465 | chr1-46403585-DEL-424 |
| chr1-46414388-DEL-677 | chr1-46414389-DEL-677 |
| chr1-46422265-DEL-6058 | chr1-46422267-DEL-6056 |
| chr1-46422501-DEL-128 | chr1-46422502-DEL-128 |
| chr1-46482118-DEL-7101 | chr1-46482119-DEL-7101 |
| chr1-46495866-DEL-70 | chr1-46495867-DEL-70 |
| chr1-46522409-DEL-63 | chr1-46522396-DEL-77 |
| chr1-46531265-DEL-58 | chr1-46531254-DEL-70 |
| chr1-46599027-DEL-208 | chr1-46599028-DEL-208 |
| chr1-46601961-DEL-6380 | chr1-46601962-DEL-6380 |
| chr1-46622504-DEL-6257 | chr1-46622505-DEL-6257 |
| chr1-46740655-DEL-1626 | chr1-46740655-DEL-1626 |
| chr1-46756394-DEL-73 | chr1-46756397-DEL-75 |
| chr1-46788256-DEL-93 | chr1-46788257-DEL-93 |
| chr1-46790642-DEL-199 | chr1-46790643-DEL-199 |
| chr1-46825065-DEL-231 | chr1-46825066-DEL-231 |
| chr1-46834284-DEL-113 | chr1-46834285-DEL-113 |
| chr1-46839499-DEL-495 | chr1-46839499-DEL-495 |
| chr1-46853709-DEL-322 | chr1-46853710-DEL-322 |
| chr1-46868885-DEL-116 | chr1-46868886-DEL-116 |
| chr1-46875700-DEL-255 | chr1-46875701-DEL-255 |
| chr1-46887089-DEL-99 | chr1-46887090-DEL-99 |
| chr1-46894388-DEL-52 | chr1-46894389-DEL-52 |
| chr1-46921997-DEL-213 | chr1-46921998-DEL-213 |
| chr1-46925878-DEL-51 | chr1-46925879-DEL-51 |
| chr1-46989933-DEL-117 | chr1-46989934-DEL-117 |
| chr1-46990302-DEL-6225 | chr1-46990303-DEL-6225 |
| chr1-47030443-DEL-6824 | chr1-47030444-DEL-6824 |
| chr1-47071837-DEL-55 | chr1-47071800-DEL-80 |
| chr1-47100556-DEL-341 | chr1-47100557-DEL-340 |
| chr1-47139367-DEL-187 | chr1-47139368-DEL-187 |
| chr1-47144973-DEL-6444 | chr1-47144974-DEL-6444 |
| chr1-47161548-DEL-177 | chr1-47161549-DEL-177 |
| chr1-47172821-DEL-1869 | chr1-47172822-DEL-1869 |
| chr1-47182099-DEL-6748 | chr1-47182100-DEL-6748 |
| chr1-47192290-DEL-499 | chr1-47192291-DEL-499 |
| chr1-47207434-DEL-220 | chr1-47207435-DEL-220 |
| chr1-47214150-DEL-6623 | chr1-47214151-DEL-6623 |
| chr1-47233266-DEL-397 | chr1-47233267-DEL-397 |
| chr1-47245023-DEL-195 | chr1-47245024-DEL-195 |
| chr1-47249079-DEL-1770 | chr1-47249080-DEL-1770 |
| chr1-47262608-DEL-4508 | chr1-47262609-DEL-4508 |
| chr1-47290202-DEL-81 | chr1-47290204-DEL-79 |
| chr1-47375262-DEL-96 | chr1-47375290-DEL-69 |
| chr1-47418885-DEL-1433 | chr1-47418886-DEL-1433 |
| chr1-47420793-DEL-723 | chr1-47420794-DEL-723 |
| chr1-47469173-DEL-930 | chr1-47469174-DEL-930 |
| chr1-47481155-DEL-541 | chr1-47481156-DEL-541 |
| chr1-47491343-DEL-1087 | chr1-47491344-DEL-1087 |
| chr1-47495553-DEL-390 | chr1-47495554-DEL-390 |
| chr1-47506654-DEL-351 | chr1-47506662-DEL-350 |
| chr1-47525970-DEL-219 | chr1-47525971-DEL-219 |
| chr1-47557076-DEL-53 | chr1-47557077-DEL-53 |
| chr1-47557490-DEL-6181 | chr1-47557491-DEL-6181 |
| chr1-47579759-DEL-644 | chr1-47579760-DEL-644 |
| chr1-47589025-DEL-61 | chr1-47589026-DEL-61 |
| chr1-47626836-DEL-6254 | chr1-47626837-DEL-6254 |
| chr1-47654509-DEL-1236 | chr1-47654510-DEL-1236 |
| chr1-47668098-DEL-8867 | chr1-47668099-DEL-8867 |
| chr1-47696709-DEL-78 | chr1-47696711-DEL-75 |
| chr1-47704050-DEL-89 | chr1-47704019-DEL-121 |
| chr1-47714381-DEL-6203 | chr1-47714382-DEL-6203 |
| chr1-47724753-DEL-4831 | chr1-47724754-DEL-4831 |
| chr1-47753953-DEL-6717 | chr1-47753965-DEL-6717 |
| chr1-47813238-DEL-212 | chr1-47813239-DEL-212 |
| chr1-47826464-DEL-77 | chr1-47826441-DEL-78 |

| | |
|---|---|
| chr1-47871604-DEL-78 | chr1-47871611-DEL-72 |
| chr1-47881315-DEL-322 | chr1-47881315-DEL-322 |
| chr1-47882268-DEL-718 | chr1-47882269-DEL-718 |
| chr1-47912346-DEL-89 | chr1-47912347-DEL-68 |
| chr1-47917546-DEL-912 | chr1-47917546-DEL-912 |
| chr1-47918595-DEL-6399 | chr1-47918596-DEL-6399 |
| chr1-47975751-DEL-1912 | chr1-47975752-DEL-1912 |
| chr1-47994972-DEL-2641 | chr1-47994973-DEL-2641 |
| chr1-47999671-DEL-317 | chr1-47999672-DEL-317 |
| chr1-48025168-DEL-92 | chr1-48025169-DEL-92 |
| chr1-48032325-DEL-199 | chr1-48032326-DEL-199 |
| chr1-48038713-DEL-153 | chr1-48038714-DEL-153 |
| chr1-48043654-DEL-52 | chr1-48043655-DEL-52 |
| chr1-48101864-DEL-200 | chr1-48101865-DEL-200 |
| chr1-48110620-DEL-129 | chr1-48110621-DEL-129 |
| chr1-48113974-DEL-788 | chr1-48113975-DEL-788 |
| chr1-48128152-DEL-5295 | chr1-48128153-DEL-5295 |
| chr1-48188659-DEL-80 | chr1-48188660-DEL-80 |
| chr1-48241799-DEL-798 | chr1-48241800-DEL-798 |
| chr1-48247210-DEL-182 | chr1-48247211-DEL-182 |
| chr1-48263007-DEL-424 | chr1-48263008-DEL-424 |
| chr1-48264359-DEL-6689 | chr1-48264360-DEL-6689 |
| chr1-48285686-DEL-1922 | chr1-48285687-DEL-1921 |
| chr1-48289218-DEL-52 | chr1-48289219-DEL-52 |
| chr1-48291424-DEL-503 | chr1-48291425-DEL-503 |
| chr1-48301071-DEL-136 | chr1-48301072-DEL-136 |
| chr1-48301577-DEL-6539 | chr1-48301578-DEL-6539 |
| chr1-48303153-DEL-2308 | chr1-48303154-DEL-2308 |
| chr1-48311697-DEL-428 | chr1-48311698-DEL-428 |
| chr1-48323351-DEL-3178 | chr1-48323353-DEL-3167 |
| chr1-48327132-DEL-480 | chr1-48327133-DEL-480 |
| chr1-48333294-DEL-76 | chr1-48333322-DEL-56 |
| chr1-48338379-DEL-1428 | chr1-48338380-DEL-1428 |
| chr1-48378594-DEL-1176 | chr1-48378594-DEL-1176 |
| chr1-48378949-DEL-245 | chr1-48378950-DEL-245 |
| chr1-48393503-DEL-8967 | chr1-48393504-DEL-8967 |
| chr1-48424125-DEL-82 | chr1-48424146-DEL-52 |
| chr1-48452571-DEL-3753 | chr1-48452572-DEL-3753 |
| chr1-48456947-DEL-6562 | chr1-48456952-DEL-6560 |
| chr1-48467932-DEL-490 | chr1-48467927-DEL-492 |
| chr1-48469419-DEL-530 | chr1-48469419-DEL-530 |
| chr1-48473058-DEL-87 | chr1-48473059-DEL-87 |
| chr1-48508708-DEL-82 | chr1-48508709-DEL-82 |
| chr1-48532991-DEL-66 | chr1-48532992-DEL-66 |
| chr1-48546430-DEL-498 | chr1-48546431-DEL-498 |
| chr1-48565229-DEL-108 | chr1-48565230-DEL-108 |
| chr1-48567258-DEL-448 | chr1-48567259-DEL-448 |
| chr1-48584691-DEL-7709 | chr1-48584692-DEL-7709 |
| chr1-48595841-DEL-66 | chr1-48595842-DEL-52 |
| chr1-48596849-DEL-82 | chr1-48596839-DEL-112 |
| chr1-48617137-DEL-704 | chr1-48617138-DEL-704 |
| chr1-48691506-DEL-121 | chr1-48691507-DEL-120 |
| chr1-48698043-DEL-60 | chr1-48698044-DEL-62 |
| chr1-48720704-DEL-109 | chr1-48720705-DEL-109 |
| chr1-48722348-DEL-288 | chr1-48722335-DEL-302 |
| chr1-48769556-DEL-86 | chr1-48769557-DEL-86 |
| chr1-48776956-DEL-4172 | chr1-48776957-DEL-4172 |
| chr1-48840072-DEL-162 | chr1-48840073-DEL-162 |
| chr1-48840715-DEL-2174 | chr1-48840716-DEL-2174 |
| chr1-48854133-DEL-6481 | chr1-48854137-DEL-6482 |
| chr1-48861863-DEL-6223 | chr1-48861864-DEL-6223 |
| chr1-48922439-DEL-59 | chr1-48922441-DEL-64 |
| chr1-48938730-DEL-740 | chr1-48938731-DEL-740 |
| chr1-48944141-DEL-198 | chr1-48944142-DEL-198 |
| chr1-48954188-DEL-938 | chr1-48954189-DEL-938 |
| chr1-48960860-DEL-6567 | chr1-48960861-DEL-6567 |
| chr1-48990052-DEL-66 | chr1-48990053-DEL-66 |
| chr1-49005912-DEL-58 | chr1-49005931-DEL-50 |
| chr1-49023534-DEL-246 | chr1-49023535-DEL-246 |
| chr1-49036406-DEL-457 | chr1-49036406-DEL-457 |
| chr1-49044484-DEL-143 | chr1-49044484-DEL-143 |
| chr1-49056438-DEL-62 | chr1-49056439-DEL-62 |
| chr1-49057444-DEL-113 | chr1-49057476-DEL-84 |
| chr1-49074157-DEL-53 | chr1-49074157-DEL-55 |
| chr1-49076117-DEL-101 | chr1-49076118-DEL-101 |
| chr1-49081400-DEL-210 | chr1-49081401-DEL-210 |
| chr1-49095297-DEL-442 | chr1-49095298-DEL-442 |
| chr1-49109227-DEL-3284 | chr1-49109228-DEL-3284 |
| chr1-49145696-DEL-56 | chr1-49145699-DEL-68 |
| chr1-49170328-DEL-589 | chr1-49170340-DEL-577 |

| | |
|---|---|
| chr1-49220837-DEL-861 | chr1-49220838-DEL-861 |
| chr1-49226817-DEL-106 | chr1-49226818-DEL-106 |
| chr1-49231112-DEL-883 | chr1-49231113-DEL-883 |
| chr1-49254192-DEL-92 | chr1-49254199-DEL-64 |
| chr1-49279425-DEL-66 | chr1-49279421-DEL-72 |
| chr1-49289893-DEL-6199 | chr1-49289894-DEL-6199 |
| chr1-49334969-DEL-1862 | chr1-49334970-DEL-1863 |
| chr1-49341782-DEL-3127 | chr1-49341783-DEL-3127 |
| chr1-49346914-DEL-474 | chr1-49346915-DEL-474 |
| chr1-49360189-DEL-8372 | chr1-49360190-DEL-8372 |
| chr1-49371791-DEL-52 | chr1-49371775-DEL-50 |
| chr1-49383238-DEL-2600 | chr1-49383239-DEL-2600 |
| chr1-49387338-DEL-61 | chr1-49387340-DEL-60 |
| chr1-49416513-DEL-248 | chr1-49416514-DEL-248 |
| chr1-49437353-DEL-1090 | chr1-49437354-DEL-1090 |
| chr1-49496338-DEL-4181 | chr1-49496339-DEL-4181 |
| chr1-49503392-DEL-160 | chr1-49503393-DEL-160 |
| chr1-49504507-DEL-62 | chr1-49504508-DEL-62 |
| chr1-49571120-DEL-381 | chr1-49571120-DEL-381 |
| chr1-49576163-DEL-434 | chr1-49576164-DEL-434 |
| chr1-49588507-DEL-1466 | chr1-49588508-DEL-1466 |
| chr1-49594548-DEL-594 | chr1-49594597-DEL-545 |
| chr1-49619201-DEL-68 | chr1-49619202-DEL-68 |
| chr1-49622262-DEL-1953 | chr1-49622263-DEL-1953 |
| chr1-49637178-DEL-641 | chr1-49637179-DEL-641 |
| chr1-49752172-DEL-71 | chr1-49752171-DEL-72 |
| chr1-49756029-DEL-56 | chr1-49756030-DEL-56 |
| chr1-49781484-DEL-57 | chr1-49781485-DEL-57 |
| chr1-49801545-DEL-428 | chr1-49801546-DEL-428 |
| chr1-49835507-DEL-152 | chr1-49835508-DEL-152 |
| chr1-49904136-DEL-740 | chr1-49904136-DEL-740 |
| chr1-49907317-DEL-694 | chr1-49907318-DEL-694 |
| chr1-49911885-DEL-368 | chr1-49911886-DEL-368 |
| chr1-49917770-DEL-500 | chr1-49917771-DEL-500 |
| chr1-49921819-DEL-58 | chr1-49921819-DEL-90 |
| chr1-49924033-DEL-74 | chr1-49924034-DEL-74 |
| chr1-49925200-DEL-2758 | chr1-49925201-DEL-2758 |
| chr1-49926263-DEL-232 | chr1-49926264-DEL-232 |
| chr1-49945651-DEL-70 | chr1-49945692-DEL-60 |
| chr1-49952718-DEL-371 | chr1-49952719-DEL-371 |
| chr1-49954901-DEL-205 | chr1-49954902-DEL-205 |
| chr1-49966270-DEL-312 | chr1-49966266-DEL-313 |
| chr1-49973779-DEL-5772 | chr1-49973780-DEL-5772 |
| chr1-50011013-DEL-1884 | chr1-50011013-DEL-1887 |
| chr1-50017839-DEL-50 | chr1-50017838-DEL-52 |
| chr1-50037027-DEL-95 | chr1-50037055-DEL-89 |
| chr1-50093834-DEL-61 | chr1-50093835-DEL-61 |
| chr1-50117882-DEL-84 | chr1-50117879-DEL-85 |
| chr1-50131716-DEL-1061 | chr1-50131718-DEL-1062 |
| chr1-50157341-DEL-524 | chr1-50157342-DEL-524 |
| chr1-50198618-DEL-3123 | chr1-50198619-DEL-3123 |
| chr1-50199124-DEL-60 | chr1-50199125-DEL-60 |
| chr1-50206077-DEL-62 | chr1-50206078-DEL-62 |
| chr1-50215669-DEL-556 | chr1-50215670-DEL-556 |
| chr1-50216416-DEL-52 | chr1-50216424-DEL-71 |
| chr1-50234007-DEL-630 | chr1-50234008-DEL-630 |
| chr1-50278666-DEL-54 | chr1-50278667-DEL-54 |
| chr1-50300462-DEL-982 | chr1-50300462-DEL-982 |
| chr1-50303875-DEL-2765 | chr1-50303876-DEL-2765 |
| chr1-50315850-DEL-746 | chr1-50315850-DEL-746 |
| chr1-50321937-DEL-445 | chr1-50321937-DEL-445 |
| chr1-50327006-DEL-70 | chr1-50327023-DEL-52 |
| chr1-50328453-DEL-58 | chr1-50328454-DEL-58 |
| chr1-50329767-DEL-56 | chr1-50329748-DEL-68 |
| chr1-50330264-DEL-1354 | chr1-50330265-DEL-1354 |
| chr1-50331750-DEL-697 | chr1-50331751-DEL-697 |
| chr1-50332804-DEL-3904 | chr1-50332806-DEL-3904 |
| chr1-50343434-DEL-105 | chr1-50343439-DEL-100 |
| chr1-50359547-DEL-58 | chr1-50359560-DEL-68 |
| chr1-50369116-DEL-6442 | chr1-50369117-DEL-6442 |
| chr1-50389359-DEL-51 | chr1-50389358-DEL-53 |
| chr1-50402039-DEL-1764 | chr1-50402040-DEL-1764 |
| chr1-50448297-DEL-54 | chr1-50448289-DEL-68 |
| chr1-50455832-DEL-2508 | chr1-50455833-DEL-2508 |
| chr1-50475081-DEL-2911 | chr1-50475082-DEL-2911 |
| chr1-50478516-DEL-57 | chr1-50478517-DEL-57 |
| chr1-50491305-DEL-61 | chr1-50491306-DEL-61 |
| chr1-50497541-DEL-63 | chr1-50497543-DEL-62 |
| chr1-50499213-DEL-1885 | chr1-50499214-DEL-1885 |
| chr1-50525788-DEL-2563 | chr1-50525789-DEL-2563 |

| | |
|---|---|
| chr1-50569270-DEL-698 | chr1-50569271-DEL-698 |
| chr1-50575703-DEL-215 | chr1-50575704-DEL-215 |
| chr1-50604551-DEL-1249 | chr1-50604552-DEL-1249 |
| chr1-50606669-DEL-1077 | chr1-50606670-DEL-1077 |
| chr1-50616366-DEL-97 | chr1-50616367-DEL-97 |
| chr1-50631882-DEL-2643 | chr1-50631883-DEL-2643 |
| chr1-50649941-DEL-953 | chr1-50649942-DEL-953 |
| chr1-50669757-DEL-90 | chr1-50669757-DEL-91 |
| chr1-50738941-DEL-438 | chr1-50738942-DEL-438 |
| chr1-50782893-DEL-108 | chr1-50782894-DEL-102 |
| chr1-50853750-DEL-565 | chr1-50853751-DEL-565 |
| chr1-50866284-DEL-62 | chr1-50866287-DEL-60 |
| chr1-50887062-DEL-1055 | chr1-50887063-DEL-1055 |
| chr1-50887487-DEL-127 | chr1-50887488-DEL-127 |
| chr1-50896038-DEL-1549 | chr1-50896039-DEL-1549 |
| chr1-50914877-DEL-6217 | chr1-50914878-DEL-6217 |
| chr1-50928603-DEL-462 | chr1-50928604-DEL-462 |
| chr1-50941989-DEL-2593 | chr1-50941990-DEL-2593 |
| chr1-51081424-DEL-97 | chr1-51081425-DEL-97 |
| chr1-51141674-DEL-1768 | chr1-51141675-DEL-1768 |
| chr1-51168122-DEL-137 | chr1-51168133-DEL-124 |
| chr1-51181542-DEL-180 | chr1-51181543-DEL-180 |
| chr1-51205070-DEL-202 | chr1-51205071-DEL-202 |
| chr1-51207803-DEL-706 | chr1-51207817-DEL-699 |
| chr1-51216080-DEL-2544 | chr1-51216081-DEL-2544 |
| chr1-51269282-DEL-152 | chr1-51269283-DEL-152 |
| chr1-51274642-DEL-77 | chr1-51274643-DEL-77 |
| chr1-51290920-DEL-5710 | chr1-51290921-DEL-5710 |
| chr1-51317786-DEL-565 | chr1-51317787-DEL-565 |
| chr1-51322447-DEL-182 | chr1-51322488-DEL-138 |
| chr1-51351698-DEL-80 | chr1-51351712-DEL-81 |
| chr1-51355738-DEL-58 | chr1-51355739-DEL-58 |
| chr1-51409415-DEL-496 | chr1-51409455-DEL-466 |
| chr1-51415183-DEL-433 | chr1-51415184-DEL-433 |
| chr1-51424064-DEL-635 | chr1-51424065-DEL-635 |
| chr1-51447581-DEL-873 | chr1-51447582-DEL-873 |
| chr1-51457097-DEL-52 | chr1-51457073-DEL-82 |
| chr1-51472574-DEL-6551 | chr1-51472558-DEL-6552 |
| chr1-51495213-DEL-546 | chr1-51495214-DEL-540 |
| chr1-51501031-DEL-181 | chr1-51501032-DEL-181 |
| chr1-51519629-DEL-52 | chr1-51519630-DEL-52 |
| chr1-51543680-DEL-181 | chr1-51543681-DEL-181 |
| chr1-51582313-DEL-4562 | chr1-51582314-DEL-4562 |
| chr1-51584056-DEL-1854 | chr1-51584057-DEL-1854 |
| chr1-51624851-DEL-193 | chr1-51624852-DEL-193 |
| chr1-51676808-DEL-188 | chr1-51676809-DEL-188 |
| chr1-51685150-DEL-83 | chr1-51685151-DEL-83 |
| chr1-51691086-DEL-2727 | chr1-51691087-DEL-2727 |
| chr1-51697678-DEL-60 | chr1-51697679-DEL-60 |
| chr1-51707268-DEL-212 | chr1-51707269-DEL-212 |
| chr1-51722537-DEL-70 | chr1-51722538-DEL-70 |
| chr1-51733018-DEL-52 | chr1-51733019-DEL-52 |
| chr1-51743753-DEL-908 | chr1-51743754-DEL-908 |
| chr1-51750391-DEL-6442 | chr1-51750392-DEL-6442 |
| chr1-51790904-DEL-51 | chr1-51790913-DEL-52 |
| chr1-51811505-DEL-71 | chr1-51811506-DEL-71 |
| chr1-51851641-DEL-152 | chr1-51851642-DEL-152 |
| chr1-51880763-DEL-61 | chr1-51880764-DEL-61 |
| chr1-51906549-DEL-2826 | chr1-51906550-DEL-2826 |
| chr1-51978019-DEL-65 | chr1-51978022-DEL-66 |
| chr1-51978698-DEL-145 | chr1-51978699-DEL-145 |
| chr1-51992380-DEL-72 | chr1-51992336-DEL-68 |
| chr1-52020699-DEL-436 | chr1-52020700-DEL-436 |
| chr1-52026870-DEL-628 | chr1-52026871-DEL-628 |
| chr1-52045340-DEL-54 | chr1-52045305-DEL-90 |
| chr1-52054421-DEL-1215 | chr1-52054407-DEL-1218 |
| chr1-52076660-DEL-206 | chr1-52076661-DEL-206 |
| chr1-52118306-DEL-64 | chr1-52118307-DEL-64 |
| chr1-52131295-DEL-230 | chr1-52131296-DEL-230 |
| chr1-52162646-DEL-100 | chr1-52162647-DEL-100 |
| chr1-52166942-DEL-889 | chr1-52166943-DEL-889 |
| chr1-52209886-DEL-66 | chr1-52209887-DEL-66 |
| chr1-52213345-DEL-229 | chr1-52213346-DEL-229 |
| chr1-52279297-DEL-586 | chr1-52279298-DEL-587 |
| chr1-52284149-DEL-88 | chr1-52284150-DEL-88 |
| chr1-52304026-DEL-70 | chr1-52304027-DEL-70 |
| chr1-52343806-DEL-168 | chr1-52343807-DEL-168 |
| chr1-52348874-DEL-75 | chr1-52348875-DEL-75 |
| chr1-52351861-DEL-8430 | chr1-52351862-DEL-8430 |
| chr1-52366251-DEL-734 | chr1-52366252-DEL-734 |

| | |
|---|---|
| chr1-52379292-DEL-1564 | chr1-52379293-DEL-1564 |
| chr1-52395400-DEL-912 | chr1-52395400-DEL-912 |
| chr1-52409702-DEL-1893 | chr1-52409703-DEL-1893 |
| chr1-52420551-DEL-193 | chr1-52420532-DEL-179 |
| chr1-52430921-DEL-93 | chr1-52430922-DEL-65 |
| chr1-52439159-DEL-636 | chr1-52439160-DEL-636 |
| chr1-52439756-DEL-212 | chr1-52439755-DEL-212 |
| chr1-52499844-DEL-72 | chr1-52499845-DEL-72 |
| chr1-52581816-DEL-135 | chr1-52581817-DEL-135 |
| chr1-52621206-DEL-193 | chr1-52621207-DEL-193 |
| chr1-52623592-DEL-1269 | chr1-52623582-DEL-1270 |
| chr1-52629018-DEL-4401 | chr1-52629024-DEL-4402 |
| chr1-52694563-DEL-64 | chr1-52694564-DEL-64 |
| chr1-52730551-DEL-214 | chr1-52730552-DEL-214 |
| chr1-52776086-DEL-782 | chr1-52776087-DEL-782 |
| chr1-52776858-DEL-293 | chr1-52776858-DEL-293 |
| chr1-52780941-DEL-3453 | chr1-52780942-DEL-3453 |
| chr1-52783120-DEL-956 | chr1-52783121-DEL-956 |
| chr1-52784505-DEL-80 | chr1-52784506-DEL-80 |
| chr1-52862446-DEL-328 | chr1-52862446-DEL-328 |
| chr1-52876349-DEL-1480 | chr1-52876350-DEL-1480 |
| chr1-52880836-DEL-1910 | chr1-52880837-DEL-1910 |
| chr1-52897168-DEL-1838 | chr1-52897169-DEL-1838 |
| chr1-52900681-DEL-545 | chr1-52900682-DEL-545 |
| chr1-52919614-DEL-134 | chr1-52919614-DEL-135 |
| chr1-52921746-DEL-170 | chr1-52921747-DEL-170 |
| chr1-52926877-DEL-6401 | chr1-52926878-DEL-6401 |
| chr1-52932966-DEL-201 | chr1-52932967-DEL-201 |
| chr1-52942639-DEL-56 | chr1-52942640-DEL-56 |
| chr1-52949339-DEL-211 | chr1-52949340-DEL-211 |
| chr1-52958569-DEL-67 | chr1-52958596-DEL-65 |
| chr1-52978653-DEL-6907 | chr1-52978654-DEL-6906 |
| chr1-52978947-DEL-211 | chr1-52978948-DEL-211 |
| chr1-52987461-DEL-312 | chr1-52987462-DEL-312 |
| chr1-53015716-DEL-6424 | chr1-53015717-DEL-6424 |
| chr1-53047989-DEL-185 | chr1-53047990-DEL-185 |
| chr1-53055119-DEL-2007 | chr1-53055120-DEL-2007 |
| chr1-53064325-DEL-498 | chr1-53064326-DEL-498 |
| chr1-53098048-DEL-63 | chr1-53098050-DEL-67 |
| chr1-53120268-DEL-111 | chr1-53120277-DEL-103 |
| chr1-53142215-DEL-62 | chr1-53142249-DEL-58 |
| chr1-53143146-DEL-80 | chr1-53143135-DEL-92 |
| chr1-53168739-DEL-5225 | chr1-53168740-DEL-5225 |
| chr1-53206450-DEL-191 | chr1-53206451-DEL-194 |
| chr1-53210328-DEL-351 | chr1-53210329-DEL-351 |
| chr1-53219373-DEL-118 | chr1-53219337-DEL-154 |
| chr1-53267974-DEL-358 | chr1-53267975-DEL-358 |
| chr1-53280713-DEL-533 | chr1-53280713-DEL-533 |
| chr1-53283516-DEL-1748 | chr1-53283515-DEL-1749 |
| chr1-53304010-DEL-99 | chr1-53303994-DEL-99 |
| chr1-53315539-DEL-149 | chr1-53315540-DEL-148 |
| chr1-53393618-DEL-1639 | chr1-53393619-DEL-1639 |
| chr1-53403767-DEL-82 | chr1-53403768-DEL-62 |
| chr1-53424607-DEL-252 | chr1-53424608-DEL-252 |
| chr1-53446702-DEL-1898 | chr1-53446703-DEL-1898 |
| chr1-53543851-DEL-3958 | chr1-53543852-DEL-3958 |
| chr1-53558332-DEL-1261 | chr1-53558333-DEL-1262 |
| chr1-53600767-DEL-96 | chr1-53600772-DEL-96 |
| chr1-53637596-DEL-52 | chr1-53637584-DEL-56 |
| chr1-53731936-DEL-3890 | chr1-53731937-DEL-3890 |
| chr1-53771375-DEL-179 | chr1-53771376-DEL-179 |
| chr1-53789750-DEL-65 | chr1-53789751-DEL-65 |
| chr1-53852829-DEL-134 | chr1-53852829-DEL-135 |
| chr1-53872056-DEL-98 | chr1-53872056-DEL-98 |
| chr1-53874844-DEL-1049 | chr1-53874845-DEL-1049 |
| chr1-53919165-DEL-175 | chr1-53919198-DEL-167 |
| chr1-53955503-DEL-51 | chr1-53955536-DEL-66 |
| chr1-53987552-DEL-54 | chr1-53987554-DEL-65 |
| chr1-53994630-DEL-6086 | chr1-53994631-DEL-6086 |
| chr1-54027028-DEL-587 | chr1-54027029-DEL-587 |
| chr1-54046869-DEL-184 | chr1-54046870-DEL-184 |
| chr1-54065346-DEL-6231 | chr1-54065347-DEL-6231 |
| chr1-54080904-DEL-203 | chr1-54080905-DEL-203 |
| chr1-54090766-DEL-186 | chr1-54090767-DEL-186 |
| chr1-54137294-DEL-1414 | chr1-54137295-DEL-1414 |
| chr1-54154911-DEL-78 | chr1-54154927-DEL-54 |
| chr1-54207751-DEL-53 | chr1-54207738-DEL-57 |
| chr1-54237643-DEL-1259 | chr1-54237644-DEL-1259 |
| chr1-54256549-DEL-66 | chr1-54256550-DEL-66 |
| chr1-54283330-DEL-5037 | chr1-54283331-DEL-5037 |

| | |
|---|---|
| chr1-54304617-DEL-6640 | chr1-54304618-DEL-6640 |
| chr1-54328649-DEL-92 | chr1-54328659-DEL-56 |
| chr1-54355756-DEL-436 | chr1-54355757-DEL-436 |
| chr1-54373225-DEL-114 | chr1-54373227-DEL-112 |
| chr1-54375904-DEL-433 | chr1-54375905-DEL-433 |
| chr1-54387451-DEL-89 | chr1-54387452-DEL-89 |
| chr1-54389904-DEL-76 | chr1-54389905-DEL-76 |
| chr1-54396597-DEL-54 | chr1-54396584-DEL-68 |
| chr1-54400499-DEL-82 | chr1-54400500-DEL-82 |
| chr1-54411751-DEL-54 | chr1-54411752-DEL-54 |
| chr1-54412843-DEL-51 | chr1-54412844-DEL-51 |
| chr1-54426321-DEL-6587 | chr1-54426322-DEL-6587 |
| chr1-54435781-DEL-346 | chr1-54435782-DEL-346 |
| chr1-54456166-DEL-232 | chr1-54456167-DEL-232 |
| chr1-54473567-DEL-60 | chr1-54473553-DEL-80 |
| chr1-54501508-DEL-453 | chr1-54501509-DEL-453 |
| chr1-54552368-DEL-96 | chr1-54552369-DEL-96 |
| chr1-54564336-DEL-70 | chr1-54564337-DEL-70 |
| chr1-54573772-DEL-51 | chr1-54573803-DEL-52 |
| chr1-54623443-DEL-55 | chr1-54623458-DEL-60 |
| chr1-54648178-DEL-2126 | chr1-54648179-DEL-2126 |
| chr1-54658112-DEL-403 | chr1-54658113-DEL-403 |
| chr1-54661483-DEL-102 | chr1-54661480-DEL-101 |
| chr1-54669786-DEL-638 | chr1-54669787-DEL-638 |
| chr1-54748469-DEL-110 | chr1-54748446-DEL-111 |
| chr1-54763343-DEL-1153 | chr1-54763344-DEL-1153 |
| chr1-54820504-DEL-2242 | chr1-54820505-DEL-2242 |
| chr1-54826914-DEL-67 | chr1-54826916-DEL-73 |
| chr1-54841193-DEL-55 | chr1-54841194-DEL-55 |
| chr1-54844246-DEL-272 | chr1-54844247-DEL-272 |
| chr1-54863117-DEL-160 | chr1-54863117-DEL-158 |
| chr1-54870120-DEL-395 | chr1-54870121-DEL-395 |
| chr1-54915269-DEL-207 | chr1-54915270-DEL-207 |
| chr1-54926318-DEL-219 | chr1-54926319-DEL-219 |
| chr1-54928970-DEL-6401 | chr1-54928971-DEL-6401 |
| chr1-54942393-DEL-179 | chr1-54942394-DEL-179 |
| chr1-54985275-DEL-51 | chr1-54985276-DEL-51 |
| chr1-54987074-DEL-206 | chr1-54987076-DEL-205 |
| chr1-54995188-DEL-103 | chr1-54995189-DEL-103 |
| chr1-55009969-DEL-84 | chr1-55009970-DEL-60 |
| chr1-55011537-DEL-58 | chr1-55011538-DEL-58 |
| chr1-55047897-DEL-58 | chr1-55047904-DEL-52 |
| chr1-55077439-DEL-2721 | chr1-55077441-DEL-2720 |
| chr1-55083599-DEL-401 | chr1-55083600-DEL-401 |
| chr1-55088713-DEL-104 | chr1-55088714-DEL-100 |
| chr1-55108430-DEL-200 | chr1-55108430-DEL-200 |
| chr1-55144235-DEL-216 | chr1-55144236-DEL-216 |
| chr1-55145657-DEL-199 | chr1-55145656-DEL-199 |
| chr1-55154621-DEL-412 | chr1-55154622-DEL-412 |
| chr1-55173830-DEL-57 | chr1-55173831-DEL-57 |
| chr1-55173863-DEL-600 | chr1-55173864-DEL-600 |
| chr1-55194310-DEL-52 | chr1-55194311-DEL-52 |
| chr1-55211427-DEL-102 | chr1-55211388-DEL-136 |
| chr1-55291838-DEL-151 | chr1-55291838-DEL-151 |
| chr1-55313978-DEL-772 | chr1-55313979-DEL-772 |
| chr1-55335880-DEL-58 | chr1-55335881-DEL-58 |
| chr1-55368540-DEL-51 | chr1-55368541-DEL-51 |
| chr1-55405639-DEL-2472 | chr1-55405640-DEL-2472 |
| chr1-55480424-DEL-746 | chr1-55480431-DEL-704 |
| chr1-55482839-DEL-1382 | chr1-55482840-DEL-1382 |
| chr1-55504544-DEL-1830 | chr1-55504545-DEL-1830 |
| chr1-55509537-DEL-1782 | chr1-55509538-DEL-1782 |
| chr1-55515410-DEL-6478 | chr1-55515411-DEL-6478 |
| chr1-55526777-DEL-55 | chr1-55526777-DEL-55 |
| chr1-55530607-DEL-183 | chr1-55530608-DEL-183 |
| chr1-55549224-DEL-53 | chr1-55549225-DEL-53 |
| chr1-55589694-DEL-153 | chr1-55589695-DEL-153 |
| chr1-55603471-DEL-78 | chr1-55603499-DEL-81 |
| chr1-55621830-DEL-53 | chr1-55621831-DEL-53 |
| chr1-55623861-DEL-54 | chr1-55623862-DEL-54 |
| chr1-55654222-DEL-1594 | chr1-55654223-DEL-1594 |
| chr1-55677752-DEL-360 | chr1-55677752-DEL-360 |
| chr1-55682637-DEL-120 | chr1-55682638-DEL-120 |
| chr1-55701239-DEL-54 | chr1-55701258-DEL-60 |
| chr1-55742479-DEL-353 | chr1-55742480-DEL-353 |
| chr1-55755161-DEL-50 | chr1-55755130-DEL-83 |
| chr1-55817514-DEL-4925 | chr1-55817515-DEL-4925 |
| chr1-55830064-DEL-851 | chr1-55830065-DEL-851 |
| chr1-55832418-DEL-96 | chr1-55832443-DEL-72 |
| chr1-55861769-DEL-101 | chr1-55861770-DEL-96 |

| | |
|---|---|
| chr1-55866972-DEL-99 | chr1-55867006-DEL-66 |
| chr1-55880110-DEL-352 | chr1-55880111-DEL-352 |
| chr1-55892524-DEL-66 | chr1-55892525-DEL-66 |
| chr1-55896063-DEL-83 | chr1-55896064-DEL-83 |
| chr1-55905976-DEL-157 | chr1-55905975-DEL-158 |
| chr1-55911732-DEL-85 | chr1-55911766-DEL-52 |
| chr1-55962783-DEL-6440 | chr1-55962784-DEL-6440 |
| chr1-56020280-DEL-60 | chr1-56020301-DEL-56 |
| chr1-56047598-DEL-259 | chr1-56047598-DEL-240 |
| chr1-56065625-DEL-431 | chr1-56065626-DEL-431 |
| chr1-56069372-DEL-1890 | chr1-56069373-DEL-1890 |
| chr1-56114580-DEL-262 | chr1-56114581-DEL-262 |
| chr1-56121008-DEL-55 | chr1-56121009-DEL-55 |
| chr1-56130380-DEL-498 | chr1-56130381-DEL-498 |
| chr1-56131290-DEL-7135 | chr1-56131291-DEL-7135 |
| chr1-56146915-DEL-1297 | chr1-56146913-DEL-1300 |
| chr1-56176173-DEL-6383 | chr1-56176174-DEL-6383 |
| chr1-56247498-DEL-153 | chr1-56247456-DEL-183 |
| chr1-56256337-DEL-122 | chr1-56256338-DEL-122 |
| chr1-56313420-DEL-58 | chr1-56313420-DEL-59 |
| chr1-56313532-DEL-3128 | chr1-56313533-DEL-3128 |
| chr1-56319897-DEL-2137 | chr1-56319898-DEL-2137 |
| chr1-56331733-DEL-1015 | chr1-56331734-DEL-1015 |
| chr1-56333118-DEL-58 | chr1-56333119-DEL-58 |
| chr1-56334778-DEL-321 | chr1-56334779-DEL-321 |
| chr1-56337771-DEL-785 | chr1-56337772-DEL-785 |
| chr1-56355633-DEL-110 | chr1-56355678-DEL-71 |
| chr1-56375097-DEL-135 | chr1-56375076-DEL-156 |
| chr1-56419480-DEL-6446 | chr1-56419481-DEL-6446 |
| chr1-56419495-DEL-212 | chr1-56419496-DEL-212 |
| chr1-56431737-DEL-120 | chr1-56431738-DEL-120 |
| chr1-56452363-DEL-997 | chr1-56452364-DEL-997 |
| chr1-56456996-DEL-230 | chr1-56456996-DEL-230 |
| chr1-56473381-DEL-322 | chr1-56473382-DEL-322 |
| chr1-56490512-DEL-909 | chr1-56490513-DEL-909 |
| chr1-56497573-DEL-66 | chr1-56497602-DEL-66 |
| chr1-56497799-DEL-6206 | chr1-56497776-DEL-6227 |
| chr1-56535630-DEL-520 | chr1-56535644-DEL-495 |
| chr1-56540605-DEL-58 | chr1-56540604-DEL-58 |
| chr1-56551078-DEL-70 | chr1-56551083-DEL-52 |
| chr1-56567865-DEL-531 | chr1-56567866-DEL-531 |
| chr1-56605948-DEL-88 | chr1-56605949-DEL-88 |
| chr1-56619363-DEL-63 | chr1-56619364-DEL-63 |
| chr1-56634017-DEL-389 | chr1-56634018-DEL-389 |
| chr1-56654462-DEL-219 | chr1-56654469-DEL-219 |
| chr1-56663726-DEL-233 | chr1-56663727-DEL-232 |
| chr1-56671771-DEL-414 | chr1-56671772-DEL-398 |
| chr1-56706759-DEL-287 | chr1-56706760-DEL-287 |
| chr1-56721457-DEL-163 | chr1-56721458-DEL-163 |
| chr1-56728899-DEL-52 | chr1-56728905-DEL-53 |
| chr1-56749901-DEL-1217 | chr1-56749902-DEL-1217 |
| chr1-56753522-DEL-6819 | chr1-56753523-DEL-6819 |
| chr1-56753809-DEL-212 | chr1-56753809-DEL-212 |
| chr1-56810073-DEL-470 | chr1-56810073-DEL-470 |
| chr1-56811401-DEL-7046 | chr1-56811402-DEL-7046 |
| chr1-56825303-DEL-54 | chr1-56825308-DEL-50 |
| chr1-56830318-DEL-70 | chr1-56830319-DEL-70 |
| chr1-56907811-DEL-144 | chr1-56907812-DEL-144 |
| chr1-56914401-DEL-85 | chr1-56914414-DEL-72 |
| chr1-56919990-DEL-68 | chr1-56919994-DEL-65 |
| chr1-57035754-DEL-401 | chr1-57035745-DEL-403 |
| chr1-57051425-DEL-76 | chr1-57051426-DEL-76 |
| chr1-57074422-DEL-57 | chr1-57074402-DEL-78 |
| chr1-57078907-DEL-65 | chr1-57078908-DEL-50 |
| chr1-57103445-DEL-6579 | chr1-57103446-DEL-6579 |
| chr1-57111584-DEL-62 | chr1-57111585-DEL-62 |
| chr1-57148152-DEL-1512 | chr1-57148153-DEL-1512 |
| chr1-57171634-DEL-710 | chr1-57171635-DEL-710 |
| chr1-57286143-DEL-71 | chr1-57286144-DEL-71 |
| chr1-57319511-DEL-1236 | chr1-57319512-DEL-1236 |
| chr1-57343067-DEL-74 | chr1-57343068-DEL-74 |
| chr1-57347497-DEL-3012 | chr1-57347498-DEL-3012 |
| chr1-57371515-DEL-455 | chr1-57371516-DEL-455 |
| chr1-57419894-DEL-96 | chr1-57419895-DEL-96 |
| chr1-57486860-DEL-75 | chr1-57486877-DEL-59 |
| chr1-57498161-DEL-84 | chr1-57498166-DEL-97 |
| chr1-57515352-DEL-110 | chr1-57515353-DEL-110 |
| chr1-57519311-DEL-114 | chr1-57519312-DEL-114 |
| chr1-57524746-DEL-272 | chr1-57524747-DEL-272 |
| chr1-57549566-DEL-58 | chr1-57549567-DEL-55 |

| | |
|---|---|
| chr1-57560479-DEL-830 | chr1-57560480-DEL-830 |
| chr1-57574117-DEL-53 | chr1-57574118-DEL-53 |
| chr1-57581346-DEL-486 | chr1-57581324-DEL-482 |
| chr1-57605995-DEL-66 | chr1-57605996-DEL-66 |
| chr1-57624252-DEL-3723 | chr1-57624253-DEL-3723 |
| chr1-57749210-DEL-60 | chr1-57749211-DEL-60 |
| chr1-57761619-DEL-1071 | chr1-57761620-DEL-1071 |
| chr1-57770497-DEL-125 | chr1-57770498-DEL-125 |
| chr1-57792232-DEL-92 | chr1-57792233-DEL-92 |
| chr1-57894194-DEL-58 | chr1-57894195-DEL-58 |
| chr1-57898287-DEL-401 | chr1-57898288-DEL-401 |
| chr1-57953691-DEL-2570 | chr1-57953692-DEL-2571 |
| chr1-57959204-DEL-78 | chr1-57959205-DEL-78 |
| chr1-58013123-DEL-2633 | chr1-58013124-DEL-2633 |
| chr1-58016356-DEL-1200 | chr1-58016357-DEL-1200 |
| chr1-58051014-DEL-69 | chr1-58051015-DEL-69 |
| chr1-58059758-DEL-92 | chr1-58059756-DEL-93 |
| chr1-58062219-DEL-70 | chr1-58062220-DEL-70 |
| chr1-58121502-DEL-768 | chr1-58121503-DEL-768 |
| chr1-58122924-DEL-83 | chr1-58122928-DEL-80 |
| chr1-58126495-DEL-1363 | chr1-58126496-DEL-1363 |
| chr1-58134474-DEL-58 | chr1-58134457-DEL-74 |
| chr1-58150820-DEL-84 | chr1-58150819-DEL-86 |
| chr1-58169220-DEL-277 | chr1-58169221-DEL-277 |
| chr1-58170334-DEL-526 | chr1-58170335-DEL-526 |
| chr1-58181182-DEL-96 | chr1-58181182-DEL-96 |
| chr1-58189216-DEL-200 | chr1-58189217-DEL-200 |
| chr1-58194236-DEL-707 | chr1-58194237-DEL-707 |
| chr1-58202018-DEL-62 | chr1-58202019-DEL-64 |
| chr1-58266380-DEL-811 | chr1-58266381-DEL-811 |
| chr1-58275392-DEL-72 | chr1-58275391-DEL-73 |
| chr1-58297361-DEL-142 | chr1-58297362-DEL-142 |
| chr1-58309663-DEL-76 | chr1-58309662-DEL-78 |
| chr1-58310347-DEL-66 | chr1-58310358-DEL-56 |
| chr1-58319678-DEL-81 | chr1-58319679-DEL-81 |
| chr1-58341009-DEL-68 | chr1-58341006-DEL-72 |
| chr1-58355598-DEL-6131 | chr1-58355599-DEL-6131 |
| chr1-58362499-DEL-7129 | chr1-58362500-DEL-7129 |
| chr1-58392606-DEL-119 | chr1-58392607-DEL-119 |
| chr1-58397148-DEL-51 | chr1-58397105-DEL-50 |
| chr1-58403302-DEL-57 | chr1-58403303-DEL-57 |
| chr1-58423289-DEL-223 | chr1-58423290-DEL-223 |
| chr1-58429872-DEL-211 | chr1-58429874-DEL-207 |
| chr1-58431551-DEL-569 | chr1-58431552-DEL-569 |
| chr1-58473884-DEL-202 | chr1-58473885-DEL-202 |
| chr1-58475573-DEL-206 | chr1-58475574-DEL-206 |
| chr1-58480158-DEL-115 | chr1-58480159-DEL-115 |
| chr1-58491709-DEL-204 | chr1-58491710-DEL-204 |
| chr1-58510407-DEL-223 | chr1-58510408-DEL-223 |
| chr1-58524239-DEL-3875 | chr1-58524240-DEL-3875 |
| chr1-58528357-DEL-1015 | chr1-58528358-DEL-1015 |
| chr1-58536059-DEL-218 | chr1-58536060-DEL-218 |
| chr1-58540213-DEL-203 | chr1-58540212-DEL-204 |
| chr1-58614402-DEL-213 | chr1-58614403-DEL-213 |
| chr1-58633151-DEL-310 | chr1-58633152-DEL-310 |
| chr1-58646001-DEL-567 | chr1-58646002-DEL-567 |
| chr1-58662589-DEL-68 | chr1-58662590-DEL-68 |
| chr1-58685739-DEL-1750 | chr1-58685740-DEL-1750 |
| chr1-58726395-DEL-2960 | chr1-58726396-DEL-2960 |
| chr1-58732709-DEL-681 | chr1-58732710-DEL-681 |
| chr1-58733266-DEL-226 | chr1-58733257-DEL-227 |
| chr1-58737185-DEL-813 | chr1-58737185-DEL-813 |
| chr1-58769501-DEL-400 | chr1-58769502-DEL-400 |
| chr1-58784604-DEL-51 | chr1-58784602-DEL-52 |
| chr1-58788669-DEL-2543 | chr1-58788684-DEL-2539 |
| chr1-58806314-DEL-498 | chr1-58806315-DEL-498 |
| chr1-58835660-DEL-82 | chr1-58835661-DEL-52 |
| chr1-58843707-DEL-65 | chr1-58843713-DEL-60 |
| chr1-58850235-DEL-8533 | chr1-58850236-DEL-8533 |
| chr1-58870538-DEL-66 | chr1-58870543-DEL-56 |
| chr1-58888707-DEL-81 | chr1-58888704-DEL-79 |
| chr1-58910730-DEL-731 | chr1-58910731-DEL-731 |
| chr1-58911496-DEL-404 | chr1-58911497-DEL-404 |
| chr1-58943658-DEL-62 | chr1-58943652-DEL-60 |
| chr1-58969903-DEL-4465 | chr1-58969904-DEL-4465 |
| chr1-58998379-DEL-69 | chr1-58998389-DEL-68 |
| chr1-59013231-DEL-53 | chr1-59013229-DEL-56 |
| chr1-59070599-DEL-794 | chr1-59070600-DEL-794 |
| chr1-59117655-DEL-85 | chr1-59117656-DEL-85 |
| chr1-59119854-DEL-67 | chr1-59119855-DEL-67 |

| | |
|---|---|
| chr1-59137117-DEL-79 | chr1-59137117-DEL-80 |
| chr1-59150707-DEL-74 | chr1-59150708-DEL-74 |
| chr1-59161513-DEL-62 | chr1-59161508-DEL-64 |
| chr1-59196964-DEL-301 | chr1-59196965-DEL-301 |
| chr1-59215582-DEL-156 | chr1-59215583-DEL-156 |
| chr1-59216317-DEL-140 | chr1-59216318-DEL-140 |
| chr1-59217563-DEL-86 | chr1-59217564-DEL-86 |
| chr1-59225452-DEL-59 | chr1-59225453-DEL-59 |
| chr1-59238885-DEL-126 | chr1-59238886-DEL-126 |
| chr1-59264199-DEL-55 | chr1-59264200-DEL-55 |
| chr1-59265456-DEL-198 | chr1-59265448-DEL-207 |
| chr1-59271285-DEL-206 | chr1-59271286-DEL-206 |
| chr1-59272700-DEL-597 | chr1-59272701-DEL-597 |
| chr1-59281859-DEL-435 | chr1-59281860-DEL-435 |
| chr1-59286135-DEL-202 | chr1-59286136-DEL-202 |
| chr1-59297325-DEL-110 | chr1-59297343-DEL-99 |
| chr1-59304015-DEL-227 | chr1-59304016-DEL-227 |
| chr1-59338331-DEL-323 | chr1-59338330-DEL-323 |
| chr1-59355010-DEL-60 | chr1-59355011-DEL-56 |
| chr1-59363923-DEL-328 | chr1-59363924-DEL-328 |
| chr1-59384299-DEL-5039 | chr1-59384300-DEL-5039 |
| chr1-59427668-DEL-60 | chr1-59427669-DEL-60 |
| chr1-59428160-DEL-91 | chr1-59428161-DEL-91 |
| chr1-59447034-DEL-204 | chr1-59447035-DEL-204 |
| chr1-59456445-DEL-169 | chr1-59456446-DEL-169 |
| chr1-59481414-DEL-136 | chr1-59481415-DEL-135 |
| chr1-59484175-DEL-103 | chr1-59484176-DEL-103 |
| chr1-59537971-DEL-57 | chr1-59537972-DEL-57 |
| chr1-59557162-DEL-433 | chr1-59557163-DEL-433 |
| chr1-59560247-DEL-383 | chr1-59560248-DEL-383 |
| chr1-59576398-DEL-653 | chr1-59576399-DEL-653 |
| chr1-59598198-DEL-1858 | chr1-59598199-DEL-1858 |
| chr1-59601069-DEL-140 | chr1-59601069-DEL-168 |
| chr1-59625057-DEL-64 | chr1-59625067-DEL-74 |
| chr1-59635542-DEL-70 | chr1-59635537-DEL-75 |
| chr1-59645890-DEL-754 | chr1-59645891-DEL-754 |
| chr1-59646986-DEL-138 | chr1-59646987-DEL-138 |
| chr1-59682187-DEL-105 | chr1-59682188-DEL-105 |
| chr1-59685793-DEL-63 | chr1-59685794-DEL-63 |
| chr1-59691343-DEL-143 | chr1-59691344-DEL-143 |
| chr1-59698047-DEL-216 | chr1-59698048-DEL-216 |
| chr1-59706889-DEL-557 | chr1-59706890-DEL-557 |
| chr1-59714283-DEL-51 | chr1-59714260-DEL-66 |
| chr1-59715017-DEL-199 | chr1-59715018-DEL-199 |
| chr1-59716310-DEL-164 | chr1-59716311-DEL-164 |
| chr1-59734460-DEL-208 | chr1-59734461-DEL-208 |
| chr1-59740967-DEL-93 | chr1-59740968-DEL-93 |
| chr1-59756065-DEL-103 | chr1-59756063-DEL-106 |
| chr1-59766556-DEL-112 | chr1-59766557-DEL-109 |
| chr1-59776504-DEL-206 | chr1-59776505-DEL-206 |
| chr1-59779893-DEL-215 | chr1-59779894-DEL-215 |
| chr1-59780603-DEL-78 | chr1-59780604-DEL-78 |
| chr1-59786896-DEL-198 | chr1-59786893-DEL-199 |
| chr1-59795789-DEL-192 | chr1-59795790-DEL-192 |
| chr1-59798334-DEL-202 | chr1-59798335-DEL-202 |
| chr1-59818123-DEL-320 | chr1-59818124-DEL-320 |
| chr1-59823348-DEL-202 | chr1-59823349-DEL-202 |
| chr1-59825444-DEL-56 | chr1-59825423-DEL-52 |
| chr1-59829959-DEL-93 | chr1-59829960-DEL-93 |
| chr1-59846770-DEL-1163 | chr1-59846771-DEL-1163 |
| chr1-59849826-DEL-400 | chr1-59849827-DEL-400 |
| chr1-59860690-DEL-843 | chr1-59860691-DEL-843 |
| chr1-59928185-DEL-63 | chr1-59928193-DEL-56 |
| chr1-59938228-DEL-83 | chr1-59938229-DEL-83 |
| chr1-59942852-DEL-168 | chr1-59942853-DEL-168 |
| chr1-59955721-DEL-215 | chr1-59955722-DEL-215 |
| chr1-59993330-DEL-476 | chr1-59993331-DEL-476 |
| chr1-60023657-DEL-52 | chr1-60023658-DEL-52 |
| chr1-60043223-DEL-147 | chr1-60043216-DEL-128 |
| chr1-60051628-DEL-81 | chr1-60051629-DEL-81 |
| chr1-60059830-DEL-865 | chr1-60059831-DEL-865 |
| chr1-60077761-DEL-497 | chr1-60077762-DEL-497 |
| chr1-60087899-DEL-358 | chr1-60087900-DEL-358 |
| chr1-60101391-DEL-183 | chr1-60101392-DEL-182 |
| chr1-60105481-DEL-202 | chr1-60105482-DEL-202 |
| chr1-60113778-DEL-198 | chr1-60113779-DEL-197 |
| chr1-60140361-DEL-95 | chr1-60140377-DEL-101 |
| chr1-60146600-DEL-70 | chr1-60146601-DEL-70 |
| chr1-60171826-DEL-174 | chr1-60171827-DEL-174 |
| chr1-60173432-DEL-187 | chr1-60173424-DEL-188 |

| | |
|---|---|
| chr1-60182364-DEL-58 | chr1-60182365-DEL-58 |
| chr1-60196947-DEL-4511 | chr1-60196952-DEL-4507 |
| chr1-60200106-DEL-213 | chr1-60200107-DEL-213 |
| chr1-60206503-DEL-224 | chr1-60206504-DEL-224 |
| chr1-60222876-DEL-52 | chr1-60222877-DEL-52 |
| chr1-60228146-DEL-211 | chr1-60228147-DEL-211 |
| chr1-60278770-DEL-205 | chr1-60278771-DEL-205 |
| chr1-60284986-DEL-206 | chr1-60284987-DEL-206 |
| chr1-60336465-DEL-51 | chr1-60336466-DEL-51 |
| chr1-60359261-DEL-7576 | chr1-60359262-DEL-7576 |
| chr1-60367806-DEL-121 | chr1-60367797-DEL-129 |
| chr1-60380138-DEL-212 | chr1-60380139-DEL-212 |
| chr1-60396097-DEL-51 | chr1-60396078-DEL-80 |
| chr1-60399430-DEL-196 | chr1-60399431-DEL-196 |
| chr1-60414661-DEL-331 | chr1-60414662-DEL-331 |
| chr1-60433873-DEL-62 | chr1-60433874-DEL-62 |
| chr1-60437225-DEL-602 | chr1-60437228-DEL-602 |
| chr1-60513739-DEL-210 | chr1-60513740-DEL-210 |
| chr1-60561785-DEL-461 | chr1-60561786-DEL-461 |
| chr1-60581676-DEL-70 | chr1-60581677-DEL-70 |
| chr1-60585742-DEL-94 | chr1-60585742-DEL-94 |
| chr1-60613745-DEL-69 | chr1-60613745-DEL-69 |
| chr1-60630470-DEL-1433 | chr1-60630470-DEL-1433 |
| chr1-60637356-DEL-148 | chr1-60637357-DEL-148 |
| chr1-60664066-DEL-175 | chr1-60664023-DEL-201 |
| chr1-60669745-DEL-6221 | chr1-60669746-DEL-6221 |
| chr1-60721071-DEL-168 | chr1-60721072-DEL-168 |
| chr1-60726972-DEL-6248 | chr1-60726979-DEL-6228 |
| chr1-60742692-DEL-141 | chr1-60742693-DEL-141 |
| chr1-60743057-DEL-1808 | chr1-60743075-DEL-1803 |
| chr1-60759234-DEL-60 | chr1-60759234-DEL-60 |
| chr1-60764016-DEL-7088 | chr1-60764017-DEL-7088 |
| chr1-60790005-DEL-262 | chr1-60790006-DEL-262 |
| chr1-60825474-DEL-5497 | chr1-60825475-DEL-5497 |
| chr1-60854579-DEL-169 | chr1-60854580-DEL-169 |
| chr1-60865885-DEL-68 | chr1-60865886-DEL-68 |
| chr1-60875930-DEL-213 | chr1-60875931-DEL-213 |
| chr1-60985765-DEL-70 | chr1-60985766-DEL-70 |
| chr1-61022034-DEL-2334 | chr1-61022035-DEL-2334 |
| chr1-61114120-DEL-271 | chr1-61114121-DEL-271 |
| chr1-61147700-DEL-56 | chr1-61147701-DEL-56 |
| chr1-61201665-DEL-629 | chr1-61201666-DEL-629 |
| chr1-61205754-DEL-252 | chr1-61205755-DEL-252 |
| chr1-61237479-DEL-53 | chr1-61237480-DEL-53 |
| chr1-61248707-DEL-6377 | chr1-61248708-DEL-6377 |
| chr1-61315889-DEL-112 | chr1-61315888-DEL-114 |
| chr1-61391914-DEL-86 | chr1-61391915-DEL-86 |
| chr1-61400633-DEL-7174 | chr1-61400634-DEL-7174 |
| chr1-61470871-DEL-1153 | chr1-61470872-DEL-1153 |
| chr1-61485257-DEL-3248 | chr1-61485258-DEL-3248 |
| chr1-61503088-DEL-521 | chr1-61503089-DEL-521 |
| chr1-61506540-DEL-124 | chr1-61506550-DEL-115 |
| chr1-61608402-DEL-4185 | chr1-61608403-DEL-4185 |
| chr1-61621451-DEL-5001 | chr1-61621452-DEL-5001 |
| chr1-61638500-DEL-70 | chr1-61638501-DEL-70 |
| chr1-61656070-DEL-576 | chr1-61656072-DEL-575 |
| chr1-61668332-DEL-74 | chr1-61668327-DEL-67 |
| chr1-61683403-DEL-159 | chr1-61683402-DEL-161 |
| chr1-61709879-DEL-6604 | chr1-61709880-DEL-6604 |
| chr1-61774242-DEL-181 | chr1-61774243-DEL-181 |
| chr1-61787688-DEL-982 | chr1-61787680-DEL-983 |
| chr1-61792486-DEL-2039 | chr1-61792487-DEL-2039 |
| chr1-61830728-DEL-58 | chr1-61830729-DEL-58 |
| chr1-61839107-DEL-348 | chr1-61839108-DEL-348 |
| chr1-61866784-DEL-96 | chr1-61866791-DEL-90 |
| chr1-61884417-DEL-80 | chr1-61884418-DEL-80 |
| chr1-61931916-DEL-2875 | chr1-61931917-DEL-2875 |
| chr1-61991912-DEL-51 | chr1-61991904-DEL-52 |
| chr1-62004836-DEL-70 | chr1-62004837-DEL-70 |
| chr1-62025624-DEL-391 | chr1-62025625-DEL-387 |
| chr1-62127582-DEL-54 | chr1-62127583-DEL-54 |
| chr1-62139095-DEL-5751 | chr1-62139085-DEL-5750 |
| chr1-62178105-DEL-132 | chr1-62178082-DEL-128 |
| chr1-62183493-DEL-72 | chr1-62183494-DEL-72 |
| chr1-62188799-DEL-127 | chr1-62188800-DEL-127 |
| chr1-62245821-DEL-219 | chr1-62245822-DEL-219 |
| chr1-62259353-DEL-6199 | chr1-62259354-DEL-6199 |
| chr1-62267150-DEL-63 | chr1-62267163-DEL-63 |
| chr1-62291845-DEL-224 | chr1-62291846-DEL-224 |
| chr1-62334539-DEL-6957 | chr1-62334540-DEL-6957 |

| | |
|---|---|
| chr1-62342475-DEL-80 | chr1-62342465-DEL-62 |
| chr1-62347610-DEL-200 | chr1-62347595-DEL-216 |
| chr1-62351156-DEL-93 | chr1-62351157-DEL-93 |
| chr1-62438856-DEL-64 | chr1-62438831-DEL-89 |
| chr1-62441328-DEL-593 | chr1-62441329-DEL-593 |
| chr1-62462610-DEL-211 | chr1-62462611-DEL-211 |
| chr1-62496773-DEL-941 | chr1-62496774-DEL-941 |
| chr1-62573142-DEL-7311 | chr1-62573182-DEL-7268 |
| chr1-62623103-DEL-158 | chr1-62623130-DEL-118 |
| chr1-62643925-DEL-68 | chr1-62643925-DEL-68 |
| chr1-62671550-DEL-100 | chr1-62671551-DEL-100 |
| chr1-62682757-DEL-333 | chr1-62682758-DEL-333 |
| chr1-62710300-DEL-6421 | chr1-62710301-DEL-6421 |
| chr1-62795096-DEL-64 | chr1-62795097-DEL-58 |
| chr1-62832971-DEL-60 | chr1-62832972-DEL-60 |
| chr1-62847066-DEL-56 | chr1-62847061-DEL-56 |
| chr1-62884042-DEL-70 | chr1-62884043-DEL-70 |
| chr1-62884971-DEL-60 | chr1-62884972-DEL-60 |
| chr1-62904055-DEL-6075 | chr1-62904056-DEL-6075 |
| chr1-62926722-DEL-57 | chr1-62926725-DEL-54 |
| chr1-63000054-DEL-638 | chr1-63000055-DEL-638 |
| chr1-63001375-DEL-365 | chr1-63001376-DEL-365 |
| chr1-63025445-DEL-95 | chr1-63025430-DEL-86 |
| chr1-63042334-DEL-584 | chr1-63042335-DEL-584 |
| chr1-63052695-DEL-124 | chr1-63052696-DEL-124 |
| chr1-63053797-DEL-212 | chr1-63053798-DEL-212 |
| chr1-63073643-DEL-400 | chr1-63073644-DEL-400 |
| chr1-63077022-DEL-79 | chr1-63077026-DEL-80 |
| chr1-63080508-DEL-185 | chr1-63080509-DEL-185 |
| chr1-63085186-DEL-206 | chr1-63085187-DEL-206 |
| chr1-63128196-DEL-53 | chr1-63128197-DEL-55 |
| chr1-63173238-DEL-2089 | chr1-63173237-DEL-2089 |
| chr1-63179683-DEL-85 | chr1-63179684-DEL-85 |
| chr1-63223736-DEL-746 | chr1-63223737-DEL-746 |
| chr1-63259763-DEL-104 | chr1-63259751-DEL-118 |
| chr1-63261231-DEL-99 | chr1-63261217-DEL-118 |
| chr1-63283890-DEL-188 | chr1-63283891-DEL-188 |
| chr1-63293295-DEL-218 | chr1-63293296-DEL-218 |
| chr1-63305741-DEL-82 | chr1-63305741-DEL-83 |
| chr1-63337786-DEL-158 | chr1-63337787-DEL-158 |
| chr1-63385571-DEL-205 | chr1-63385572-DEL-205 |
| chr1-63411163-DEL-146 | chr1-63411164-DEL-146 |
| chr1-63419040-DEL-51 | chr1-63419027-DEL-66 |
| chr1-63437655-DEL-1267 | chr1-63437656-DEL-1267 |
| chr1-63458716-DEL-52 | chr1-63458717-DEL-54 |
| chr1-63459776-DEL-499 | chr1-63459777-DEL-499 |
| chr1-63465728-DEL-573 | chr1-63465729-DEL-573 |
| chr1-63466585-DEL-191 | chr1-63466586-DEL-191 |
| chr1-63482489-DEL-52 | chr1-63482490-DEL-52 |
| chr1-63496555-DEL-138 | chr1-63496565-DEL-115 |
| chr1-63536662-DEL-366 | chr1-63536663-DEL-366 |
| chr1-63563531-DEL-54 | chr1-63563530-DEL-56 |
| chr1-63580936-DEL-271 | chr1-63580937-DEL-271 |
| chr1-63638287-DEL-927 | chr1-63638287-DEL-927 |
| chr1-63650938-DEL-211 | chr1-63650939-DEL-211 |
| chr1-63656396-DEL-270 | chr1-63656397-DEL-270 |
| chr1-63671875-DEL-92 | chr1-63671878-DEL-88 |
| chr1-63708361-DEL-73 | chr1-63708354-DEL-82 |
| chr1-63713029-DEL-53 | chr1-63713029-DEL-53 |
| chr1-63732239-DEL-1891 | chr1-63732240-DEL-1891 |
| chr1-63740244-DEL-57 | chr1-63740245-DEL-57 |
| chr1-63776133-DEL-543 | chr1-63776133-DEL-543 |
| chr1-63778978-DEL-1472 | chr1-63778979-DEL-1472 |
| chr1-63798325-DEL-416 | chr1-63798326-DEL-416 |
| chr1-63811565-DEL-1901 | chr1-63811566-DEL-1901 |
| chr1-63816864-DEL-227 | chr1-63816865-DEL-227 |
| chr1-63820721-DEL-1818 | chr1-63820722-DEL-1818 |
| chr1-63834547-DEL-80 | chr1-63834548-DEL-80 |
| chr1-63869095-DEL-6597 | chr1-63869096-DEL-6597 |
| chr1-63891587-DEL-143 | chr1-63891587-DEL-143 |
| chr1-63898273-DEL-6412 | chr1-63898274-DEL-6412 |
| chr1-63914699-DEL-71 | chr1-63914697-DEL-68 |
| chr1-63925357-DEL-7130 | chr1-63925358-DEL-7130 |
| chr1-63965809-DEL-51 | chr1-63965779-DEL-60 |
| chr1-63999883-DEL-64 | chr1-63999884-DEL-64 |
| chr1-64056004-DEL-66 | chr1-64055997-DEL-74 |
| chr1-64141471-DEL-58 | chr1-64141472-DEL-58 |
| chr1-64190450-DEL-1895 | chr1-64190451-DEL-1895 |
| chr1-64217353-DEL-7093 | chr1-64217354-DEL-7093 |
| chr1-64244852-DEL-66 | chr1-64244873-DEL-73 |

| | |
|---|---|
| chr1-64246046-DEL-65 | chr1-64246047-DEL-65 |
| chr1-64275749-DEL-467 | chr1-64275750-DEL-467 |
| chr1-64292399-DEL-150 | chr1-64292421-DEL-144 |
| chr1-64298947-DEL-203 | chr1-64298983-DEL-202 |
| chr1-64352788-DEL-2810 | chr1-64352789-DEL-2810 |
| chr1-64365391-DEL-2048 | chr1-64365392-DEL-2048 |
| chr1-64389063-DEL-64 | chr1-64389047-DEL-84 |
| chr1-64411662-DEL-355 | chr1-64411657-DEL-357 |
| chr1-64415808-DEL-204 | chr1-64415809-DEL-204 |
| chr1-64422239-DEL-7111 | chr1-64422240-DEL-7111 |
| chr1-64432315-DEL-124 | chr1-64432345-DEL-102 |
| chr1-64435832-DEL-674 | chr1-64435830-DEL-672 |
| chr1-64444728-DEL-1065 | chr1-64444729-DEL-1065 |
| chr1-64456045-DEL-68 | chr1-64456054-DEL-50 |
| chr1-64464045-DEL-113 | chr1-64464046-DEL-113 |
| chr1-64464863-DEL-910 | chr1-64464864-DEL-910 |
| chr1-64465989-DEL-1187 | chr1-64465990-DEL-1187 |
| chr1-64475666-DEL-514 | chr1-64475667-DEL-514 |
| chr1-64510677-DEL-2282 | chr1-64510677-DEL-2282 |
| chr1-64528372-DEL-61 | chr1-64528373-DEL-60 |
| chr1-64530371-DEL-293 | chr1-64530372-DEL-295 |
| chr1-64549766-DEL-401 | chr1-64549767-DEL-401 |
| chr1-64553157-DEL-52 | chr1-64553158-DEL-52 |
| chr1-64575922-DEL-326 | chr1-64575923-DEL-326 |
| chr1-64597702-DEL-381 | chr1-64597703-DEL-381 |
| chr1-64619849-DEL-69 | chr1-64619850-DEL-69 |
| chr1-64629472-DEL-181 | chr1-64629473-DEL-181 |
| chr1-64652446-DEL-1090 | chr1-64652447-DEL-1090 |
| chr1-64661017-DEL-86 | chr1-64661021-DEL-87 |
| chr1-64675932-DEL-204 | chr1-64675932-DEL-205 |
| chr1-64680831-DEL-197 | chr1-64680832-DEL-197 |
| chr1-64681932-DEL-355 | chr1-64681933-DEL-355 |
| chr1-64709511-DEL-452 | chr1-64709512-DEL-452 |
| chr1-64752719-DEL-51 | chr1-64752720-DEL-51 |
| chr1-64786279-DEL-386 | chr1-64786280-DEL-386 |
| chr1-64796354-DEL-81 | chr1-64796355-DEL-81 |
| chr1-64998287-DEL-62 | chr1-64998288-DEL-62 |
| chr1-64999464-DEL-7083 | chr1-64999465-DEL-7083 |
| chr1-65033020-DEL-772 | chr1-65033020-DEL-773 |
| chr1-65066851-DEL-176 | chr1-65066866-DEL-162 |
| chr1-65073313-DEL-56 | chr1-65073314-DEL-56 |
| chr1-65076031-DEL-440 | chr1-65076032-DEL-439 |
| chr1-65095389-DEL-194 | chr1-65095389-DEL-190 |
| chr1-65110420-DEL-65 | chr1-65110418-DEL-64 |
| chr1-65132381-DEL-2549 | chr1-65132401-DEL-2548 |
| chr1-65140137-DEL-601 | chr1-65140138-DEL-601 |
| chr1-65143093-DEL-233 | chr1-65143094-DEL-233 |
| chr1-65143944-DEL-52 | chr1-65143945-DEL-52 |
| chr1-65153349-DEL-163 | chr1-65153356-DEL-157 |
| chr1-65153915-DEL-186 | chr1-65153916-DEL-186 |
| chr1-65162034-DEL-52 | chr1-65162035-DEL-52 |
| chr1-65164322-DEL-2897 | chr1-65164323-DEL-2897 |
| chr1-65176462-DEL-249 | chr1-65176463-DEL-249 |
| chr1-65188690-DEL-52 | chr1-65188691-DEL-52 |
| chr1-65189870-DEL-76 | chr1-65189871-DEL-76 |
| chr1-65213469-DEL-127 | chr1-65213470-DEL-128 |
| chr1-65267291-DEL-76 | chr1-65267292-DEL-76 |
| chr1-65272723-DEL-85 | chr1-65272724-DEL-85 |
| chr1-65274734-DEL-698 | chr1-65274735-DEL-698 |
| chr1-65286902-DEL-61 | chr1-65286912-DEL-63 |
| chr1-65341033-DEL-966 | chr1-65341034-DEL-966 |
| chr1-65365638-DEL-111 | chr1-65365637-DEL-112 |
| chr1-65376432-DEL-364 | chr1-65376433-DEL-364 |
| chr1-65398059-DEL-2417 | chr1-65398060-DEL-2417 |
| chr1-65413900-DEL-176 | chr1-65413901-DEL-176 |
| chr1-65476912-DEL-63 | chr1-65476913-DEL-63 |
| chr1-65481956-DEL-66 | chr1-65481957-DEL-66 |
| chr1-65487775-DEL-2371 | chr1-65487776-DEL-2371 |
| chr1-65505182-DEL-265 | chr1-65505182-DEL-265 |
| chr1-65509767-DEL-1282 | chr1-65509725-DEL-1300 |
| chr1-65523332-DEL-1484 | chr1-65523333-DEL-1484 |
| chr1-65529556-DEL-171 | chr1-65529557-DEL-171 |
| chr1-65532519-DEL-1294 | chr1-65532520-DEL-1292 |
| chr1-65538232-DEL-138 | chr1-65538232-DEL-138 |
| chr1-65539012-DEL-2616 | chr1-65539010-DEL-2616 |
| chr1-65546695-DEL-516 | chr1-65546696-DEL-516 |
| chr1-65578680-DEL-3126 | chr1-65578681-DEL-3126 |
| chr1-65591992-DEL-459 | chr1-65591993-DEL-459 |
| chr1-65594609-DEL-58 | chr1-65594610-DEL-58 |
| chr1-65666836-DEL-52 | chr1-65666830-DEL-54 |

| | |
|---|---|
| chr1-65673348-DEL-54 | chr1-65673348-DEL-50 |
| chr1-65707825-DEL-505 | chr1-65707826-DEL-505 |
| chr1-65729729-DEL-61 | chr1-65729726-DEL-65 |
| chr1-65730583-DEL-198 | chr1-65730584-DEL-199 |
| chr1-65737964-DEL-914 | chr1-65737964-DEL-916 |
| chr1-65747274-DEL-2197 | chr1-65747275-DEL-2197 |
| chr1-65754046-DEL-378 | chr1-65754047-DEL-378 |
| chr1-65766218-DEL-851 | chr1-65766220-DEL-852 |
| chr1-65770037-DEL-1148 | chr1-65770038-DEL-1148 |
| chr1-65776301-DEL-53 | chr1-65776299-DEL-51 |
| chr1-65788168-DEL-6199 | chr1-65788169-DEL-6199 |
| chr1-65819065-DEL-201 | chr1-65819066-DEL-201 |
| chr1-65822387-DEL-333 | chr1-65822388-DEL-333 |
| chr1-65854212-DEL-2652 | chr1-65854213-DEL-2651 |
| chr1-65869033-DEL-72 | chr1-65869034-DEL-58 |
| chr1-65888848-DEL-64 | chr1-65888849-DEL-64 |
| chr1-65897635-DEL-96 | chr1-65897636-DEL-92 |
| chr1-65911674-DEL-4635 | chr1-65911675-DEL-4635 |
| chr1-65923360-DEL-57 | chr1-65923361-DEL-57 |
| chr1-65925414-DEL-190 | chr1-65925427-DEL-186 |
| chr1-65928749-DEL-2590 | chr1-65928750-DEL-2590 |
| chr1-65949209-DEL-52 | chr1-65949210-DEL-54 |
| chr1-65951271-DEL-124 | chr1-65951272-DEL-124 |
| chr1-65961757-DEL-532 | chr1-65961758-DEL-532 |
| chr1-65968405-DEL-65 | chr1-65968410-DEL-61 |
| chr1-65981120-DEL-110 | chr1-65981121-DEL-110 |
| chr1-65987106-DEL-2248 | chr1-65987107-DEL-2248 |
| chr1-66012597-DEL-6715 | chr1-66012598-DEL-6715 |
| chr1-66087683-DEL-6166 | chr1-66087684-DEL-6166 |
| chr1-66114594-DEL-442 | chr1-66114581-DEL-450 |
| chr1-66129601-DEL-1065 | chr1-66129602-DEL-1065 |
| chr1-66143943-DEL-552 | chr1-66143954-DEL-551 |
| chr1-66156563-DEL-52 | chr1-66156564-DEL-52 |
| chr1-66171393-DEL-63 | chr1-66171394-DEL-63 |
| chr1-66216929-DEL-52 | chr1-66216930-DEL-52 |
| chr1-66222363-DEL-56 | chr1-66222364-DEL-56 |
| chr1-66227980-DEL-320 | chr1-66227981-DEL-320 |
| chr1-66231136-DEL-74 | chr1-66231163-DEL-55 |
| chr1-66233268-DEL-189 | chr1-66233269-DEL-190 |
| chr1-66269851-DEL-1042 | chr1-66269852-DEL-1042 |
| chr1-66279220-DEL-53 | chr1-66279221-DEL-53 |
| chr1-66281296-DEL-57 | chr1-66281297-DEL-57 |
| chr1-66298406-DEL-1988 | chr1-66298407-DEL-1988 |
| chr1-66310218-DEL-6681 | chr1-66310219-DEL-6681 |
| chr1-66372346-DEL-74 | chr1-66372347-DEL-74 |
| chr1-66380932-DEL-93 | chr1-66380933-DEL-93 |
| chr1-66397486-DEL-116 | chr1-66397493-DEL-110 |
| chr1-66474678-DEL-83 | chr1-66474683-DEL-79 |
| chr1-66486980-DEL-1161 | chr1-66486981-DEL-1161 |
| chr1-66505365-DEL-196 | chr1-66505366-DEL-196 |
| chr1-66535744-DEL-394 | chr1-66535745-DEL-394 |
| chr1-66554170-DEL-6202 | chr1-66554171-DEL-6202 |
| chr1-66582053-DEL-6332 | chr1-66582054-DEL-6332 |
| chr1-66614533-DEL-6205 | chr1-66614534-DEL-6205 |
| chr1-66628958-DEL-56 | chr1-66628965-DEL-54 |
| chr1-66638289-DEL-800 | chr1-66638290-DEL-800 |
| chr1-66644069-DEL-387 | chr1-66644070-DEL-387 |
| chr1-66654185-DEL-2083 | chr1-66654192-DEL-2077 |
| chr1-66671210-DEL-98 | chr1-66671207-DEL-98 |
| chr1-66673693-DEL-64 | chr1-66673693-DEL-64 |
| chr1-66721267-DEL-420 | chr1-66721268-DEL-420 |
| chr1-66744983-DEL-201 | chr1-66744991-DEL-202 |
| chr1-66754503-DEL-304 | chr1-66754504-DEL-304 |
| chr1-66775219-DEL-111 | chr1-66775220-DEL-111 |
| chr1-66796886-DEL-177 | chr1-66796887-DEL-173 |
| chr1-66848453-DEL-61 | chr1-66848427-DEL-52 |
| chr1-66874475-DEL-70 | chr1-66874476-DEL-70 |
| chr1-66876002-DEL-62 | chr1-66876007-DEL-58 |
| chr1-66882880-DEL-1008 | chr1-66882881-DEL-1008 |
| chr1-66888189-DEL-662 | chr1-66888187-DEL-663 |
| chr1-66908840-DEL-452 | chr1-66908841-DEL-452 |
| chr1-66929500-DEL-197 | chr1-66929501-DEL-197 |
| chr1-66934107-DEL-78 | chr1-66934108-DEL-78 |
| chr1-66943877-DEL-91 | chr1-66943873-DEL-84 |
| chr1-66959365-DEL-389 | chr1-66959366-DEL-389 |
| chr1-66962737-DEL-76 | chr1-66962738-DEL-76 |
| chr1-66987651-DEL-498 | chr1-66987652-DEL-498 |
| chr1-66988570-DEL-1447 | chr1-66988571-DEL-1447 |
| chr1-66999186-DEL-114 | chr1-66999187-DEL-114 |
| chr1-67017917-DEL-4307 | chr1-67017918-DEL-4307 |

| | |
|---|---|
| chr1-67073184-DEL-57 | chr1-67073176-DEL-58 |
| chr1-67076802-DEL-59 | chr1-67076803-DEL-59 |
| chr1-67112341-DEL-97 | chr1-67112342-DEL-97 |
| chr1-67130096-DEL-154 | chr1-67130093-DEL-154 |
| chr1-67141847-DEL-123 | chr1-67141848-DEL-123 |
| chr1-67145137-DEL-2747 | chr1-67145138-DEL-2746 |
| chr1-67148222-DEL-51 | chr1-67148223-DEL-51 |
| chr1-67163430-DEL-98 | chr1-67163431-DEL-98 |
| chr1-67168844-DEL-54 | chr1-67168845-DEL-54 |
| chr1-67171801-DEL-59 | chr1-67171802-DEL-60 |
| chr1-67195108-DEL-54 | chr1-67195109-DEL-53 |
| chr1-67198822-DEL-68 | chr1-67198815-DEL-76 |
| chr1-67206324-DEL-52 | chr1-67206315-DEL-56 |
| chr1-67207287-DEL-50 | chr1-67207283-DEL-54 |
| chr1-67279283-DEL-199 | chr1-67279244-DEL-199 |
| chr1-67298476-DEL-365 | chr1-67298477-DEL-365 |
| chr1-67305707-DEL-200 | chr1-67305708-DEL-200 |
| chr1-67316270-DEL-6197 | chr1-67316271-DEL-6197 |
| chr1-67327156-DEL-56 | chr1-67327156-DEL-56 |
| chr1-67357741-DEL-176 | chr1-67357743-DEL-175 |
| chr1-67358952-DEL-152 | chr1-67358953-DEL-152 |
| chr1-67373548-DEL-140 | chr1-67373549-DEL-140 |
| chr1-67383971-DEL-109 | chr1-67383972-DEL-151 |
| chr1-67425702-DEL-90 | chr1-67425703-DEL-90 |
| chr1-67481196-DEL-391 | chr1-67481197-DEL-391 |
| chr1-67532400-DEL-342 | chr1-67532401-DEL-342 |
| chr1-67537529-DEL-246 | chr1-67537537-DEL-239 |
| chr1-67550344-DEL-50 | chr1-67550331-DEL-64 |
| chr1-67558738-DEL-169 | chr1-67558739-DEL-169 |
| chr1-67568028-DEL-156 | chr1-67568029-DEL-156 |
| chr1-67585385-DEL-52 | chr1-67585386-DEL-52 |
| chr1-67606035-DEL-430 | chr1-67606043-DEL-429 |
| chr1-67616794-DEL-165 | chr1-67616795-DEL-165 |
| chr1-67620456-DEL-161 | chr1-67620473-DEL-137 |
| chr1-67620689-DEL-1891 | chr1-67620690-DEL-1891 |
| chr1-67626567-DEL-6208 | chr1-67626568-DEL-6208 |
| chr1-67636579-DEL-1357 | chr1-67636580-DEL-1357 |
| chr1-67642949-DEL-757 | chr1-67642950-DEL-757 |
| chr1-67647115-DEL-195 | chr1-67647111-DEL-205 |
| chr1-67647993-DEL-358 | chr1-67647994-DEL-358 |
| chr1-67658796-DEL-6223 | chr1-67658797-DEL-6223 |
| chr1-67665702-DEL-373 | chr1-67665703-DEL-373 |
| chr1-67678210-DEL-195 | chr1-67678211-DEL-195 |
| chr1-67721661-DEL-6256 | chr1-67721662-DEL-6256 |
| chr1-67746525-DEL-55 | chr1-67746526-DEL-55 |
| chr1-67756428-DEL-2290 | chr1-67756426-DEL-2291 |
| chr1-67782357-DEL-604 | chr1-67782358-DEL-604 |
| chr1-67790597-DEL-7098 | chr1-67790598-DEL-7098 |
| chr1-67824309-DEL-123 | chr1-67824333-DEL-106 |
| chr1-67830077-DEL-88 | chr1-67830078-DEL-88 |
| chr1-67842932-DEL-93 | chr1-67842933-DEL-89 |
| chr1-67869073-DEL-481 | chr1-67869074-DEL-481 |
| chr1-67905301-DEL-52 | chr1-67905302-DEL-52 |
| chr1-67946230-DEL-78 | chr1-67946231-DEL-78 |
| chr1-67955022-DEL-57 | chr1-67955023-DEL-56 |
| chr1-68018695-DEL-62 | chr1-68018696-DEL-62 |
| chr1-68020825-DEL-52 | chr1-68020826-DEL-52 |
| chr1-68036361-DEL-1574 | chr1-68036362-DEL-1574 |
| chr1-68103339-DEL-152 | chr1-68103343-DEL-148 |
| chr1-68116671-DEL-540 | chr1-68116672-DEL-540 |
| chr1-68131686-DEL-109 | chr1-68131687-DEL-109 |
| chr1-68141736-DEL-120 | chr1-68141735-DEL-121 |
| chr1-68144486-DEL-68 | chr1-68144446-DEL-100 |
| chr1-68148972-DEL-54 | chr1-68148973-DEL-56 |
| chr1-68165645-DEL-473 | chr1-68165646-DEL-473 |
| chr1-68166928-DEL-3523 | chr1-68166929-DEL-3522 |
| chr1-68179634-DEL-222 | chr1-68179635-DEL-222 |
| chr1-68187740-DEL-2110 | chr1-68187742-DEL-2109 |
| chr1-68190589-DEL-1377 | chr1-68190590-DEL-1377 |
| chr1-68198413-DEL-612 | chr1-68198413-DEL-612 |
| chr1-68227035-DEL-72 | chr1-68227036-DEL-72 |
| chr1-68248501-DEL-661 | chr1-68248502-DEL-661 |
| chr1-68279523-DEL-126 | chr1-68279524-DEL-126 |
| chr1-68283532-DEL-181 | chr1-68283533-DEL-181 |
| chr1-68286943-DEL-2092 | chr1-68286944-DEL-2092 |
| chr1-68307273-DEL-6377 | chr1-68307274-DEL-6377 |
| chr1-68317175-DEL-74 | chr1-68317176-DEL-74 |
| chr1-68341705-DEL-457 | chr1-68341706-DEL-457 |
| chr1-68380109-DEL-439 | chr1-68380108-DEL-430 |
| chr1-68388897-DEL-371 | chr1-68388898-DEL-371 |

| | |
|---|---|
| chr1-68406791-DEL-642 | chr1-68406792-DEL-642 |
| chr1-68424845-DEL-69 | chr1-68424841-DEL-62 |
| chr1-68444994-DEL-6225 | chr1-68444995-DEL-6225 |
| chr1-68464334-DEL-806 | chr1-68464335-DEL-806 |
| chr1-68469062-DEL-207 | chr1-68469063-DEL-207 |
| chr1-68473903-DEL-58 | chr1-68473905-DEL-54 |
| chr1-68508103-DEL-6610 | chr1-68508104-DEL-6610 |
| chr1-68530411-DEL-80 | chr1-68530412-DEL-80 |
| chr1-68533592-DEL-661 | chr1-68533587-DEL-655 |
| chr1-68543653-DEL-114 | chr1-68543654-DEL-114 |
| chr1-68558136-DEL-1204 | chr1-68558137-DEL-1204 |
| chr1-68610432-DEL-176 | chr1-68610433-DEL-176 |
| chr1-68617163-DEL-92 | chr1-68617164-DEL-92 |
| chr1-68622166-DEL-6683 | chr1-68622167-DEL-6683 |
| chr1-68649868-DEL-2886 | chr1-68649869-DEL-2886 |
| chr1-68664057-DEL-6609 | chr1-68664058-DEL-6609 |
| chr1-68686249-DEL-425 | chr1-68686250-DEL-425 |
| chr1-68741531-DEL-90 | chr1-68741532-DEL-90 |
| chr1-68746751-DEL-54 | chr1-68746752-DEL-54 |
| chr1-68779078-DEL-65 | chr1-68779079-DEL-65 |
| chr1-68805473-DEL-175 | chr1-68805474-DEL-175 |
| chr1-68860007-DEL-74 | chr1-68860030-DEL-58 |
| chr1-68921493-DEL-277 | chr1-68921494-DEL-277 |
| chr1-68922019-DEL-892 | chr1-68922020-DEL-892 |
| chr1-68924597-DEL-62 | chr1-68924586-DEL-86 |
| chr1-68932725-DEL-181 | chr1-68932726-DEL-181 |
| chr1-68940720-DEL-217 | chr1-68940721-DEL-217 |
| chr1-68942246-DEL-2420 | chr1-68942247-DEL-2420 |
| chr1-68956432-DEL-52 | chr1-68956433-DEL-52 |
| chr1-68962445-DEL-188 | chr1-68962446-DEL-188 |
| chr1-68970027-DEL-56 | chr1-68970028-DEL-56 |
| chr1-68982143-DEL-94 | chr1-68982144-DEL-99 |
| chr1-69066022-DEL-133 | chr1-69066022-DEL-133 |
| chr1-69068598-DEL-63 | chr1-69068599-DEL-63 |
| chr1-69121583-DEL-425 | chr1-69121583-DEL-426 |
| chr1-69160760-DEL-135 | chr1-69160761-DEL-135 |
| chr1-69165205-DEL-974 | chr1-69165206-DEL-974 |
| chr1-69180314-DEL-64 | chr1-69180315-DEL-64 |
| chr1-69184994-DEL-72 | chr1-69184965-DEL-108 |
| chr1-69207038-DEL-1443 | chr1-69207039-DEL-1443 |
| chr1-69230003-DEL-571 | chr1-69230004-DEL-571 |
| chr1-69230656-DEL-6424 | chr1-69230657-DEL-6424 |
| chr1-69270581-DEL-7105 | chr1-69270582-DEL-7105 |
| chr1-69280590-DEL-88 | chr1-69280591-DEL-88 |
| chr1-69286019-DEL-184 | chr1-69286020-DEL-184 |
| chr1-69297454-DEL-64 | chr1-69297455-DEL-64 |
| chr1-69299095-DEL-2960 | chr1-69299094-DEL-2961 |
| chr1-69321852-DEL-451 | chr1-69321866-DEL-449 |
| chr1-69352422-DEL-108 | chr1-69352423-DEL-108 |
| chr1-69392037-DEL-687 | chr1-69392038-DEL-687 |
| chr1-69412204-DEL-56 | chr1-69412205-DEL-56 |
| chr1-69413273-DEL-184 | chr1-69413274-DEL-184 |
| chr1-69466196-DEL-440 | chr1-69466197-DEL-440 |
| chr1-69526482-DEL-272 | chr1-69526483-DEL-272 |
| chr1-69567628-DEL-1616 | chr1-69567629-DEL-1616 |
| chr1-69580382-DEL-56 | chr1-69580388-DEL-58 |
| chr1-69618113-DEL-1451 | chr1-69618114-DEL-1451 |
| chr1-69626370-DEL-386 | chr1-69626371-DEL-386 |
| chr1-69639823-DEL-297 | chr1-69639824-DEL-295 |
| chr1-69737154-DEL-5463 | chr1-69737155-DEL-5463 |
| chr1-69771542-DEL-76 | chr1-69771516-DEL-88 |
| chr1-69801386-DEL-59 | chr1-69801386-DEL-60 |
| chr1-69889016-DEL-2287 | chr1-69889017-DEL-2287 |
| chr1-69914434-DEL-830 | chr1-69914443-DEL-832 |
| chr1-69916091-DEL-91 | chr1-69916092-DEL-91 |
| chr1-69922131-DEL-549 | chr1-69922132-DEL-549 |
| chr1-69937581-DEL-55 | chr1-69937582-DEL-55 |
| chr1-69946251-DEL-56 | chr1-69946242-DEL-53 |
| chr1-69955089-DEL-205 | chr1-69955089-DEL-205 |
| chr1-69956333-DEL-67 | chr1-69956334-DEL-60 |
| chr1-69979479-DEL-6380 | chr1-69979480-DEL-6380 |
| chr1-70001718-DEL-141 | chr1-70001719-DEL-139 |
| chr1-70078467-DEL-855 | chr1-70078463-DEL-850 |
| chr1-70081755-DEL-400 | chr1-70081756-DEL-400 |
| chr1-70087783-DEL-791 | chr1-70087782-DEL-790 |
| chr1-70100616-DEL-3009 | chr1-70100617-DEL-3009 |
| chr1-70104277-DEL-51 | chr1-70104278-DEL-51 |
| chr1-70104517-DEL-429 | chr1-70104518-DEL-429 |
| chr1-70114564-DEL-181 | chr1-70114565-DEL-181 |
| chr1-70152463-DEL-157 | chr1-70152464-DEL-157 |

| | |
|---|---|
| chr1-70158360-DEL-1360 | chr1-70158367-DEL-1360 |
| chr1-70169087-DEL-1705 | chr1-70169088-DEL-1704 |
| chr1-70237153-DEL-1383 | chr1-70237154-DEL-1383 |
| chr1-70246165-DEL-262 | chr1-70246164-DEL-263 |
| chr1-70247858-DEL-6459 | chr1-70247859-DEL-6459 |
| chr1-70273677-DEL-498 | chr1-70273678-DEL-498 |
| chr1-70276233-DEL-4536 | chr1-70276234-DEL-4536 |
| chr1-70297133-DEL-82 | chr1-70297150-DEL-66 |
| chr1-70303912-DEL-51 | chr1-70303913-DEL-51 |
| chr1-70316801-DEL-75 | chr1-70316808-DEL-84 |
| chr1-70350106-DEL-152 | chr1-70350107-DEL-152 |
| chr1-70368278-DEL-78 | chr1-70368290-DEL-50 |
| chr1-70376724-DEL-65 | chr1-70376725-DEL-64 |
| chr1-70383695-DEL-125 | chr1-70383696-DEL-125 |
| chr1-70403366-DEL-383 | chr1-70403364-DEL-384 |
| chr1-70406442-DEL-108 | chr1-70406443-DEL-108 |
| chr1-70408610-DEL-676 | chr1-70408611-DEL-676 |
| chr1-70456266-DEL-84 | chr1-70456278-DEL-61 |
| chr1-70490350-DEL-1896 | chr1-70490351-DEL-1896 |
| chr1-70496028-DEL-1515 | chr1-70496028-DEL-1507 |
| chr1-70528195-DEL-67 | chr1-70528198-DEL-65 |
| chr1-70551917-DEL-74 | chr1-70551922-DEL-70 |
| chr1-70598905-DEL-97 | chr1-70598908-DEL-94 |
| chr1-70599893-DEL-92 | chr1-70599906-DEL-75 |
| chr1-70616793-DEL-1095 | chr1-70616794-DEL-1095 |
| chr1-70626439-DEL-55 | chr1-70626439-DEL-55 |
| chr1-70654031-DEL-840 | chr1-70654033-DEL-841 |
| chr1-70668087-DEL-6635 | chr1-70668087-DEL-6635 |
| chr1-70705393-DEL-1125 | chr1-70705394-DEL-1125 |
| chr1-70709834-DEL-897 | chr1-70709836-DEL-896 |
| chr1-70723047-DEL-63 | chr1-70723048-DEL-65 |
| chr1-70726286-DEL-3727 | chr1-70726287-DEL-3727 |
| chr1-70796603-DEL-172 | chr1-70796604-DEL-172 |
| chr1-70809446-DEL-62 | chr1-70809447-DEL-62 |
| chr1-70813771-DEL-644 | chr1-70813772-DEL-644 |
| chr1-70854951-DEL-948 | chr1-70854952-DEL-948 |
| chr1-70883958-DEL-50 | chr1-70883974-DEL-60 |
| chr1-70911996-DEL-4520 | chr1-70911997-DEL-4520 |
| chr1-70948629-DEL-313 | chr1-70948630-DEL-313 |
| chr1-70958622-DEL-97 | chr1-70958626-DEL-94 |
| chr1-70975244-DEL-556 | chr1-70975245-DEL-555 |
| chr1-70982564-DEL-849 | chr1-70982565-DEL-849 |
| chr1-70991849-DEL-1477 | chr1-70991850-DEL-1477 |
| chr1-71013561-DEL-52 | chr1-71013528-DEL-90 |
| chr1-71019230-DEL-6801 | chr1-71019231-DEL-6801 |
| chr1-71027685-DEL-1492 | chr1-71027673-DEL-1509 |
| chr1-71044991-DEL-96 | chr1-71044960-DEL-111 |
| chr1-71065810-DEL-76 | chr1-71065811-DEL-70 |
| chr1-71077536-DEL-69 | chr1-71077540-DEL-59 |
| chr1-71081932-DEL-214 | chr1-71081938-DEL-214 |
| chr1-71104952-DEL-95 | chr1-71104953-DEL-95 |
| chr1-71118740-DEL-6842 | chr1-71118741-DEL-6842 |
| chr1-71171854-DEL-88 | chr1-71171855-DEL-88 |
| chr1-71195869-DEL-72 | chr1-71195870-DEL-72 |
| chr1-71210823-DEL-96 | chr1-71210794-DEL-84 |
| chr1-71226805-DEL-843 | chr1-71226806-DEL-843 |
| chr1-71265225-DEL-576 | chr1-71265226-DEL-576 |
| chr1-71289701-DEL-6221 | chr1-71289702-DEL-6221 |
| chr1-71340364-DEL-76 | chr1-71340365-DEL-76 |
| chr1-71377665-DEL-54 | chr1-71377666-DEL-54 |
| chr1-71475157-DEL-173 | chr1-71475158-DEL-173 |
| chr1-71489041-DEL-52 | chr1-71489042-DEL-52 |
| chr1-71489915-DEL-1702 | chr1-71489916-DEL-1702 |
| chr1-71503588-DEL-132 | chr1-71503562-DEL-132 |
| chr1-71505099-DEL-880 | chr1-71505099-DEL-880 |
| chr1-71507626-DEL-75 | chr1-71507627-DEL-75 |
| chr1-71556059-DEL-64 | chr1-71556060-DEL-64 |
| chr1-71558333-DEL-70 | chr1-71558338-DEL-66 |
| chr1-71571199-DEL-4085 | chr1-71571200-DEL-4085 |
| chr1-71575443-DEL-72 | chr1-71575452-DEL-73 |
| chr1-71599562-DEL-189 | chr1-71599563-DEL-189 |
| chr1-71609384-DEL-51 | chr1-71609385-DEL-51 |
| chr1-71621185-DEL-206 | chr1-71621186-DEL-206 |
| chr1-71634446-DEL-199 | chr1-71634447-DEL-199 |
| chr1-71664510-DEL-163 | chr1-71664511-DEL-163 |
| chr1-71683612-DEL-186 | chr1-71683613-DEL-186 |
| chr1-71703695-DEL-72 | chr1-71703696-DEL-72 |
| chr1-71717377-DEL-116 | chr1-71717378-DEL-116 |
| chr1-71724409-DEL-57 | chr1-71724410-DEL-57 |
| chr1-71733279-DEL-165 | chr1-71733280-DEL-165 |

| | |
|---|---|
| chr1-71744785-DEL-214 | chr1-71744782-DEL-209 |
| chr1-71756970-DEL-56 | chr1-71756971-DEL-56 |
| chr1-71758017-DEL-178 | chr1-71758018-DEL-178 |
| chr1-71765644-DEL-52 | chr1-71765672-DEL-53 |
| chr1-71777362-DEL-243 | chr1-71777363-DEL-243 |
| chr1-71806665-DEL-55 | chr1-71806671-DEL-52 |
| chr1-71826310-DEL-2233 | chr1-71826311-DEL-2233 |
| chr1-71856517-DEL-104 | chr1-71856519-DEL-103 |
| chr1-71869819-DEL-65 | chr1-71869818-DEL-66 |
| chr1-71895342-DEL-356 | chr1-71895343-DEL-356 |
| chr1-71896815-DEL-313 | chr1-71896816-DEL-313 |
| chr1-71899098-DEL-288 | chr1-71899090-DEL-289 |
| chr1-71908847-DEL-918 | chr1-71908848-DEL-918 |
| chr1-71945235-DEL-55 | chr1-71945235-DEL-55 |
| chr1-71947904-DEL-199 | chr1-71947905-DEL-199 |
| chr1-71950349-DEL-68 | chr1-71950331-DEL-51 |
| chr1-71989106-DEL-229 | chr1-71989107-DEL-229 |
| chr1-72014971-DEL-120 | chr1-72014972-DEL-120 |
| chr1-72019688-DEL-3064 | chr1-72019689-DEL-3064 |
| chr1-72049419-DEL-207 | chr1-72049420-DEL-207 |
| chr1-72049793-DEL-1117 | chr1-72049826-DEL-1085 |
| chr1-72073444-DEL-85 | chr1-72073445-DEL-85 |
| chr1-72081061-DEL-105 | chr1-72081081-DEL-68 |
| chr1-72116341-DEL-131 | chr1-72116319-DEL-136 |
| chr1-72117116-DEL-263 | chr1-72117117-DEL-263 |
| chr1-72126974-DEL-738 | chr1-72126975-DEL-738 |
| chr1-72138458-DEL-118 | chr1-72138459-DEL-118 |
| chr1-72143083-DEL-102 | chr1-72143084-DEL-102 |
| chr1-72156018-DEL-752 | chr1-72156019-DEL-752 |
| chr1-72164708-DEL-7193 | chr1-72164709-DEL-7193 |
| chr1-72181987-DEL-62 | chr1-72181989-DEL-56 |
| chr1-72183350-DEL-53 | chr1-72183351-DEL-51 |
| chr1-72189372-DEL-96 | chr1-72189373-DEL-96 |
| chr1-72200771-DEL-88 | chr1-72200768-DEL-92 |
| chr1-72264379-DEL-55 | chr1-72264374-DEL-56 |
| chr1-72266607-DEL-118 | chr1-72266608-DEL-118 |
| chr1-72272618-DEL-836 | chr1-72272619-DEL-836 |
| chr1-72273258-DEL-158 | chr1-72273249-DEL-128 |
| chr1-72278480-DEL-196 | chr1-72278480-DEL-197 |
| chr1-72284240-DEL-53 | chr1-72284241-DEL-53 |
| chr1-72285233-DEL-4499 | chr1-72285234-DEL-4499 |
| chr1-72286344-DEL-212 | chr1-72286343-DEL-207 |
| chr1-72288788-DEL-407 | chr1-72288789-DEL-407 |
| chr1-72309781-DEL-214 | chr1-72309782-DEL-214 |
| chr1-72316967-DEL-61 | chr1-72316964-DEL-50 |
| chr1-72322279-DEL-58 | chr1-72322280-DEL-58 |
| chr1-72336185-DEL-197 | chr1-72336185-DEL-197 |
| chr1-72337645-DEL-370 | chr1-72337646-DEL-370 |
| chr1-72350188-DEL-1048 | chr1-72350189-DEL-1048 |
| chr1-72352224-DEL-67 | chr1-72352193-DEL-85 |
| chr1-72434585-DEL-751 | chr1-72434586-DEL-751 |
| chr1-72436245-DEL-465 | chr1-72436246-DEL-465 |
| chr1-72437254-DEL-338 | chr1-72437255-DEL-338 |
| chr1-72447813-DEL-52 | chr1-72447814-DEL-50 |
| chr1-72485362-DEL-210 | chr1-72485363-DEL-210 |
| chr1-72541328-DEL-117 | chr1-72541329-DEL-117 |
| chr1-72546506-DEL-518 | chr1-72546507-DEL-518 |
| chr1-72581239-DEL-6344 | chr1-72581240-DEL-6344 |
| chr1-72592445-DEL-144 | chr1-72592446-DEL-144 |
| chr1-72615717-DEL-249 | chr1-72615718-DEL-249 |
| chr1-72641116-DEL-66 | chr1-72641114-DEL-80 |
| chr1-72658038-DEL-625 | chr1-72658039-DEL-625 |
| chr1-72711568-DEL-89 | chr1-72711543-DEL-90 |
| chr1-72713534-DEL-613 | chr1-72713535-DEL-613 |
| chr1-72717462-DEL-1003 | chr1-72717463-DEL-1003 |
| chr1-72720564-DEL-57 | chr1-72720563-DEL-60 |
| chr1-72747126-DEL-819 | chr1-72747129-DEL-817 |
| chr1-72748699-DEL-511 | chr1-72748700-DEL-509 |
| chr1-72765414-DEL-63 | chr1-72765425-DEL-54 |
| chr1-72811304-DEL-58 | chr1-72811304-DEL-58 |
| chr1-72814689-DEL-567 | chr1-72814690-DEL-567 |
| chr1-72857195-DEL-760 | chr1-72857196-DEL-760 |
| chr1-72869999-DEL-295 | chr1-72870000-DEL-295 |
| chr1-72880781-DEL-437 | chr1-72880790-DEL-432 |
| chr1-72889436-DEL-403 | chr1-72889437-DEL-403 |
| chr1-72935556-DEL-474 | chr1-72935557-DEL-474 |
| chr1-72971912-DEL-127 | chr1-72971913-DEL-127 |
| chr1-73025584-DEL-198 | chr1-73025608-DEL-197 |
| chr1-73037149-DEL-2568 | chr1-73037150-DEL-2568 |
| chr1-73039308-DEL-457 | chr1-73039310-DEL-457 |

| | |
|---|---|
| chr1-73050446-DEL-281 | chr1-73050447-DEL-281 |
| chr1-73051512-DEL-54 | chr1-73051511-DEL-91 |
| chr1-73060707-DEL-745 | chr1-73060708-DEL-745 |
| chr1-73104169-DEL-108 | chr1-73104170-DEL-108 |
| chr1-73115555-DEL-70 | chr1-73115541-DEL-84 |
| chr1-73144074-DEL-64 | chr1-73144075-DEL-64 |
| chr1-73146732-DEL-142 | chr1-73146735-DEL-143 |
| chr1-73149731-DEL-344 | chr1-73149724-DEL-345 |
| chr1-73173999-DEL-1156 | chr1-73174000-DEL-1156 |
| chr1-73240102-DEL-447 | chr1-73240103-DEL-447 |
| chr1-73254964-DEL-59 | chr1-73254965-DEL-57 |
| chr1-73304085-DEL-52 | chr1-73304086-DEL-52 |
| chr1-73331057-DEL-6220 | chr1-73331058-DEL-6220 |
| chr1-73350400-DEL-106 | chr1-73350401-DEL-106 |
| chr1-73354570-DEL-464 | chr1-73354571-DEL-464 |
| chr1-73356949-DEL-6744 | chr1-73356949-DEL-6744 |
| chr1-73363098-DEL-205 | chr1-73363099-DEL-204 |
| chr1-73377104-DEL-324 | chr1-73377105-DEL-324 |
| chr1-73392758-DEL-61 | chr1-73392759-DEL-61 |
| chr1-73423774-DEL-785 | chr1-73423786-DEL-769 |
| chr1-73441883-DEL-1996 | chr1-73441884-DEL-1996 |
| chr1-73447411-DEL-533 | chr1-73447412-DEL-533 |
| chr1-73486674-DEL-120 | chr1-73486675-DEL-120 |
| chr1-73506035-DEL-3357 | chr1-73506036-DEL-3357 |
| chr1-73511848-DEL-6187 | chr1-73511849-DEL-6187 |
| chr1-73517827-DEL-64 | chr1-73517828-DEL-64 |
| chr1-73587724-DEL-6796 | chr1-73587725-DEL-6796 |
| chr1-73662778-DEL-1554 | chr1-73662779-DEL-1554 |
| chr1-73686087-DEL-6389 | chr1-73686088-DEL-6389 |
| chr1-73697403-DEL-76 | chr1-73697403-DEL-76 |
| chr1-73705227-DEL-102 | chr1-73705228-DEL-102 |
| chr1-73706986-DEL-157 | chr1-73706999-DEL-155 |
| chr1-73758784-DEL-64 | chr1-73758784-DEL-63 |
| chr1-73774241-DEL-67 | chr1-73774237-DEL-68 |
| chr1-73840118-DEL-99 | chr1-73840119-DEL-55 |
| chr1-73891103-DEL-354 | chr1-73891104-DEL-354 |
| chr1-73897573-DEL-110 | chr1-73897574-DEL-110 |
| chr1-73907011-DEL-6398 | chr1-73907012-DEL-6398 |
| chr1-73924411-DEL-212 | chr1-73924412-DEL-212 |
| chr1-73934373-DEL-204 | chr1-73934374-DEL-204 |
| chr1-73945393-DEL-1868 | chr1-73945394-DEL-1868 |
| chr1-73946455-DEL-97 | chr1-73946456-DEL-97 |
| chr1-74016504-DEL-193 | chr1-74016505-DEL-193 |
| chr1-74026151-DEL-76 | chr1-74026180-DEL-52 |
| chr1-74091666-DEL-72 | chr1-74091669-DEL-70 |
| chr1-74103714-DEL-615 | chr1-74103715-DEL-615 |
| chr1-74117120-DEL-322 | chr1-74117121-DEL-322 |
| chr1-74149708-DEL-62 | chr1-74149709-DEL-62 |
| chr1-74154073-DEL-2393 | chr1-74154070-DEL-2393 |
| chr1-74164829-DEL-180 | chr1-74164830-DEL-180 |
| chr1-74166506-DEL-84 | chr1-74166507-DEL-90 |
| chr1-74199028-DEL-193 | chr1-74199029-DEL-193 |
| chr1-74208745-DEL-118 | chr1-74208716-DEL-123 |
| chr1-74212990-DEL-209 | chr1-74212991-DEL-207 |
| chr1-74215497-DEL-68 | chr1-74215469-DEL-66 |
| chr1-74221928-DEL-175 | chr1-74221929-DEL-175 |
| chr1-74225488-DEL-217 | chr1-74225489-DEL-217 |
| chr1-74238671-DEL-60 | chr1-74238672-DEL-60 |
| chr1-74251880-DEL-1737 | chr1-74251881-DEL-1737 |
| chr1-74254175-DEL-417 | chr1-74254176-DEL-417 |
| chr1-74265197-DEL-203 | chr1-74265198-DEL-203 |
| chr1-74267972-DEL-627 | chr1-74267973-DEL-627 |
| chr1-74269445-DEL-213 | chr1-74269446-DEL-213 |
| chr1-74270465-DEL-109 | chr1-74270466-DEL-109 |
| chr1-74282902-DEL-211 | chr1-74282882-DEL-232 |
| chr1-74296914-DEL-228 | chr1-74296915-DEL-228 |
| chr1-74322803-DEL-62 | chr1-74322804-DEL-62 |
| chr1-74366091-DEL-241 | chr1-74366113-DEL-218 |
| chr1-74369785-DEL-65 | chr1-74369793-DEL-56 |
| chr1-74389488-DEL-56 | chr1-74389518-DEL-56 |
| chr1-74404598-DEL-208 | chr1-74404591-DEL-226 |
| chr1-74511874-DEL-221 | chr1-74511875-DEL-221 |
| chr1-74574857-DEL-202 | chr1-74574858-DEL-202 |
| chr1-74578530-DEL-77 | chr1-74578531-DEL-77 |
| chr1-74632341-DEL-209 | chr1-74632342-DEL-209 |
| chr1-74635338-DEL-206 | chr1-74635339-DEL-206 |
| chr1-74645209-DEL-2844 | chr1-74645210-DEL-2844 |
| chr1-74648464-DEL-158 | chr1-74648465-DEL-158 |
| chr1-74652400-DEL-202 | chr1-74652400-DEL-203 |
| chr1-74680435-DEL-52 | chr1-74680436-DEL-52 |

| | |
|---|---|
| chr1-74692573-DEL-7185 | chr1-74692574-DEL-7185 |
| chr1-74722425-DEL-240 | chr1-74722426-DEL-240 |
| chr1-74746489-DEL-91 | chr1-74746518-DEL-63 |
| chr1-74749444-DEL-216 | chr1-74749445-DEL-216 |
| chr1-74766765-DEL-832 | chr1-74766766-DEL-832 |
| chr1-74824750-DEL-55 | chr1-74824753-DEL-53 |
| chr1-74858234-DEL-243 | chr1-74858235-DEL-243 |
| chr1-74867073-DEL-199 | chr1-74867074-DEL-199 |
| chr1-74872038-DEL-51 | chr1-74872007-DEL-54 |
| chr1-74878963-DEL-54 | chr1-74878964-DEL-54 |
| chr1-74886259-DEL-55 | chr1-74886260-DEL-55 |
| chr1-74915462-DEL-93 | chr1-74915463-DEL-93 |
| chr1-74919579-DEL-208 | chr1-74919580-DEL-208 |
| chr1-74928079-DEL-78 | chr1-74928084-DEL-96 |
| chr1-74973023-DEL-51 | chr1-74973021-DEL-52 |
| chr1-75026759-DEL-152 | chr1-75026760-DEL-152 |
| chr1-75086888-DEL-203 | chr1-75086889-DEL-203 |
| chr1-75132920-DEL-98 | chr1-75132921-DEL-98 |
| chr1-75205516-DEL-62 | chr1-75205520-DEL-59 |
| chr1-75219230-DEL-83 | chr1-75219237-DEL-81 |
| chr1-75225541-DEL-129 | chr1-75225528-DEL-135 |
| chr1-75235818-DEL-56 | chr1-75235819-DEL-56 |
| chr1-75269230-DEL-160 | chr1-75269231-DEL-160 |
| chr1-75272818-DEL-72 | chr1-75272819-DEL-72 |
| chr1-75300259-DEL-208 | chr1-75300260-DEL-208 |
| chr1-75310740-DEL-51 | chr1-75310741-DEL-51 |
| chr1-75373170-DEL-63 | chr1-75373172-DEL-62 |
| chr1-75389709-DEL-93 | chr1-75389710-DEL-93 |
| chr1-75402111-DEL-204 | chr1-75402112-DEL-204 |
| chr1-75479319-DEL-82 | chr1-75479329-DEL-73 |
| chr1-75561517-DEL-206 | chr1-75561518-DEL-206 |
| chr1-75567481-DEL-186 | chr1-75567467-DEL-187 |
| chr1-75570306-DEL-79 | chr1-75570293-DEL-69 |
| chr1-75581338-DEL-77 | chr1-75581339-DEL-77 |
| chr1-75591709-DEL-162 | chr1-75591721-DEL-165 |
| chr1-75601455-DEL-96 | chr1-75601456-DEL-96 |
| chr1-75602660-DEL-51 | chr1-75602661-DEL-51 |
| chr1-75610389-DEL-139 | chr1-75610390-DEL-138 |
| chr1-75620009-DEL-497 | chr1-75620010-DEL-497 |
| chr1-75663582-DEL-1889 | chr1-75663583-DEL-1889 |
| chr1-75675425-DEL-1208 | chr1-75675425-DEL-1210 |
| chr1-75684401-DEL-7734 | chr1-75684402-DEL-7734 |
| chr1-75719482-DEL-95 | chr1-75719483-DEL-95 |
| chr1-75751141-DEL-315 | chr1-75751142-DEL-315 |
| chr1-75767211-DEL-1255 | chr1-75767212-DEL-1255 |
| chr1-75768612-DEL-871 | chr1-75768613-DEL-871 |
| chr1-75840968-DEL-113 | chr1-75840969-DEL-113 |
| chr1-75848490-DEL-701 | chr1-75848491-DEL-701 |
| chr1-75905293-DEL-530 | chr1-75905294-DEL-530 |
| chr1-75974371-DEL-165 | chr1-75974408-DEL-129 |
| chr1-75979545-DEL-2933 | chr1-75979546-DEL-2933 |
| chr1-76010204-DEL-59 | chr1-76010182-DEL-50 |
| chr1-76032416-DEL-4435 | chr1-76032417-DEL-4435 |
| chr1-76058096-DEL-84 | chr1-76058097-DEL-84 |
| chr1-76063789-DEL-55 | chr1-76063790-DEL-55 |
| chr1-76107290-DEL-1761 | chr1-76107291-DEL-1761 |
| chr1-76115237-DEL-122 | chr1-76115204-DEL-127 |
| chr1-76180625-DEL-316 | chr1-76180626-DEL-314 |
| chr1-76268652-DEL-205 | chr1-76268653-DEL-205 |
| chr1-76288581-DEL-6419 | chr1-76288582-DEL-6420 |
| chr1-76327949-DEL-6430 | chr1-76327950-DEL-6430 |
| chr1-76406330-DEL-120 | chr1-76406331-DEL-120 |
| chr1-76436616-DEL-6448 | chr1-76436617-DEL-6448 |
| chr1-76455617-DEL-83 | chr1-76455618-DEL-83 |
| chr1-76478762-DEL-66 | chr1-76478763-DEL-66 |
| chr1-76490833-DEL-196 | chr1-76490833-DEL-194 |
| chr1-76545571-DEL-51 | chr1-76545571-DEL-51 |
| chr1-76565092-DEL-2264 | chr1-76565092-DEL-2264 |
| chr1-76567454-DEL-1806 | chr1-76567454-DEL-1806 |
| chr1-76591424-DEL-4442 | chr1-76591425-DEL-4442 |
| chr1-76690646-DEL-54 | chr1-76690647-DEL-54 |
| chr1-76697448-DEL-685 | chr1-76697449-DEL-685 |
| chr1-76703146-DEL-212 | chr1-76703147-DEL-213 |
| chr1-76706747-DEL-58 | chr1-76706748-DEL-106 |
| chr1-76715017-DEL-64 | chr1-76715018-DEL-64 |
| chr1-76716254-DEL-3166 | chr1-76716255-DEL-3166 |
| chr1-76726805-DEL-214 | chr1-76726806-DEL-214 |
| chr1-76738193-DEL-4976 | chr1-76738194-DEL-4976 |
| chr1-76743356-DEL-402 | chr1-76743357-DEL-402 |
| chr1-76749972-DEL-4676 | chr1-76749973-DEL-4676 |

| | |
|---|---|
| chr1-76778669-DEL-1355 | chr1-76778670-DEL-1355 |
| chr1-76825726-DEL-96 | chr1-76825727-DEL-96 |
| chr1-76853757-DEL-66 | chr1-76853758-DEL-66 |
| chr1-76900096-DEL-54 | chr1-76900097-DEL-54 |
| chr1-76929537-DEL-291 | chr1-76929538-DEL-291 |
| chr1-76941974-DEL-67 | chr1-76941976-DEL-66 |
| chr1-76971634-DEL-1133 | chr1-76971635-DEL-1133 |
| chr1-76973865-DEL-56 | chr1-76973855-DEL-92 |
| chr1-77054810-DEL-152 | chr1-77054811-DEL-152 |
| chr1-77069383-DEL-54 | chr1-77069377-DEL-60 |
| chr1-77149805-DEL-454 | chr1-77149806-DEL-454 |
| chr1-77197986-DEL-59 | chr1-77197987-DEL-63 |
| chr1-77207810-DEL-1463 | chr1-77207811-DEL-1463 |
| chr1-77211222-DEL-100 | chr1-77211228-DEL-95 |
| chr1-77276675-DEL-233 | chr1-77276676-DEL-233 |
| chr1-77328549-DEL-54 | chr1-77328549-DEL-54 |
| chr1-77331496-DEL-56 | chr1-77331497-DEL-56 |
| chr1-77334537-DEL-57 | chr1-77334537-DEL-57 |
| chr1-77335442-DEL-145 | chr1-77335436-DEL-153 |
| chr1-77392049-DEL-282 | chr1-77392050-DEL-282 |
| chr1-77425023-DEL-198 | chr1-77425024-DEL-198 |
| chr1-77454148-DEL-6415 | chr1-77454167-DEL-6395 |
| chr1-77481500-DEL-83 | chr1-77481501-DEL-83 |
| chr1-77495076-DEL-56 | chr1-77495082-DEL-56 |
| chr1-77512216-DEL-118 | chr1-77512217-DEL-112 |
| chr1-77517553-DEL-60 | chr1-77517554-DEL-56 |
| chr1-77548878-DEL-2374 | chr1-77548878-DEL-2371 |
| chr1-77553721-DEL-62 | chr1-77553722-DEL-62 |
| chr1-77575450-DEL-52 | chr1-77575450-DEL-52 |
| chr1-77644274-DEL-4246 | chr1-77644275-DEL-4246 |
| chr1-77677760-DEL-62 | chr1-77677749-DEL-74 |
| chr1-77738517-DEL-575 | chr1-77738518-DEL-575 |
| chr1-77740101-DEL-69 | chr1-77740100-DEL-71 |
| chr1-77754531-DEL-59 | chr1-77754532-DEL-59 |
| chr1-77760112-DEL-6245 | chr1-77760113-DEL-6246 |
| chr1-77773603-DEL-806 | chr1-77773604-DEL-806 |
| chr1-77779274-DEL-70 | chr1-77779265-DEL-72 |
| chr1-77781125-DEL-122 | chr1-77781126-DEL-122 |
| chr1-77791529-DEL-152 | chr1-77791530-DEL-152 |
| chr1-77832605-DEL-365 | chr1-77832606-DEL-365 |
| chr1-77838239-DEL-541 | chr1-77838234-DEL-509 |
| chr1-77891213-DEL-158 | chr1-77891204-DEL-124 |
| chr1-77904042-DEL-143 | chr1-77904069-DEL-114 |
| chr1-77913259-DEL-5957 | chr1-77913256-DEL-5958 |
| chr1-77932702-DEL-107 | chr1-77932703-DEL-107 |
| chr1-77938596-DEL-1371 | chr1-77938597-DEL-1371 |
| chr1-77960861-DEL-70 | chr1-77960841-DEL-60 |
| chr1-77964655-DEL-119 | chr1-77964656-DEL-119 |
| chr1-77966067-DEL-1901 | chr1-77966067-DEL-1901 |
| chr1-77975480-DEL-945 | chr1-77975481-DEL-945 |
| chr1-77982395-DEL-76 | chr1-77982396-DEL-78 |
| chr1-77997171-DEL-52 | chr1-77997172-DEL-52 |
| chr1-78022804-DEL-496 | chr1-78022805-DEL-496 |
| chr1-78023567-DEL-1253 | chr1-78023568-DEL-1253 |
| chr1-78049381-DEL-57 | chr1-78049385-DEL-56 |
| chr1-78054402-DEL-82 | chr1-78054428-DEL-57 |
| chr1-78061301-DEL-648 | chr1-78061302-DEL-648 |
| chr1-78063669-DEL-86 | chr1-78063668-DEL-86 |
| chr1-78106882-DEL-56 | chr1-78106879-DEL-68 |
| chr1-78112016-DEL-130 | chr1-78112017-DEL-130 |
| chr1-78179005-DEL-52 | chr1-78179006-DEL-54 |
| chr1-78208259-DEL-111 | chr1-78208261-DEL-72 |
| chr1-78212539-DEL-924 | chr1-78212538-DEL-925 |
| chr1-78214046-DEL-152 | chr1-78214046-DEL-152 |
| chr1-78216492-DEL-7451 | chr1-78216493-DEL-7451 |
| chr1-78224389-DEL-58 | chr1-78224374-DEL-68 |
| chr1-78243827-DEL-173 | chr1-78243828-DEL-173 |
| chr1-78249429-DEL-67 | chr1-78249430-DEL-67 |
| chr1-78259318-DEL-86 | chr1-78259319-DEL-86 |
| chr1-78277866-DEL-400 | chr1-78277867-DEL-400 |
| chr1-78292769-DEL-219 | chr1-78292770-DEL-219 |
| chr1-78332685-DEL-124 | chr1-78332686-DEL-124 |
| chr1-78333713-DEL-93 | chr1-78333714-DEL-93 |
| chr1-78337659-DEL-128 | chr1-78337679-DEL-109 |
| chr1-78342077-DEL-99 | chr1-78342113-DEL-99 |
| chr1-78364932-DEL-328 | chr1-78364933-DEL-328 |
| chr1-78366155-DEL-121 | chr1-78366151-DEL-126 |
| chr1-78392245-DEL-223 | chr1-78392246-DEL-223 |
| chr1-78409495-DEL-56 | chr1-78409496-DEL-58 |
| chr1-78486778-DEL-84 | chr1-78486779-DEL-108 |

| | |
|---|---|
| chr1-78512306-DEL-54 | chr1-78512306-DEL-54 |
| chr1-78520103-DEL-210 | chr1-78520104-DEL-210 |
| chr1-78548563-DEL-200 | chr1-78548576-DEL-186 |
| chr1-78550327-DEL-916 | chr1-78550328-DEL-916 |
| chr1-78578597-DEL-98 | chr1-78578598-DEL-98 |
| chr1-78584093-DEL-60 | chr1-78584094-DEL-60 |
| chr1-78585520-DEL-1390 | chr1-78585521-DEL-1390 |
| chr1-78589988-DEL-89 | chr1-78589995-DEL-95 |
| chr1-78619828-DEL-83 | chr1-78619827-DEL-81 |
| chr1-78624212-DEL-88 | chr1-78624213-DEL-86 |
| chr1-78648417-DEL-62 | chr1-78648426-DEL-54 |
| chr1-78657964-DEL-554 | chr1-78657965-DEL-554 |
| chr1-78696637-DEL-93 | chr1-78696624-DEL-91 |
| chr1-78697546-DEL-66 | chr1-78697547-DEL-66 |
| chr1-78728755-DEL-120 | chr1-78728756-DEL-120 |
| chr1-78735138-DEL-131 | chr1-78735113-DEL-112 |
| chr1-78748783-DEL-417 | chr1-78748784-DEL-418 |
| chr1-78749173-DEL-1218 | chr1-78749174-DEL-1218 |
| chr1-78755463-DEL-378 | chr1-78755464-DEL-375 |
| chr1-78763067-DEL-62 | chr1-78763068-DEL-62 |
| chr1-78765048-DEL-57 | chr1-78765046-DEL-56 |
| chr1-78768430-DEL-2526 | chr1-78768431-DEL-2526 |
| chr1-78777754-DEL-7129 | chr1-78777755-DEL-7139 |
| chr1-78785298-DEL-743 | chr1-78785282-DEL-755 |
| chr1-78805544-DEL-3208 | chr1-78805545-DEL-3208 |
| chr1-78812800-DEL-912 | chr1-78812801-DEL-912 |
| chr1-78820534-DEL-77 | chr1-78820535-DEL-77 |
| chr1-78828088-DEL-373 | chr1-78828089-DEL-373 |
| chr1-78836199-DEL-486 | chr1-78836201-DEL-487 |
| chr1-78852032-DEL-68 | chr1-78852020-DEL-59 |
| chr1-78858606-DEL-120 | chr1-78858607-DEL-120 |
| chr1-78886257-DEL-561 | chr1-78886247-DEL-567 |
| chr1-78889992-DEL-58 | chr1-78889993-DEL-58 |
| chr1-78908221-DEL-788 | chr1-78908222-DEL-788 |
| chr1-78926807-DEL-6362 | chr1-78926808-DEL-6362 |
| chr1-78961520-DEL-6619 | chr1-78961521-DEL-6619 |
| chr1-78972500-DEL-3392 | chr1-78972506-DEL-3389 |
| chr1-78996528-DEL-4118 | chr1-78996529-DEL-4118 |
| chr1-79047981-DEL-3351 | chr1-79047981-DEL-3351 |
| chr1-79051421-DEL-3285 | chr1-79051422-DEL-3284 |
| chr1-79076114-DEL-6576 | chr1-79076115-DEL-6576 |
| chr1-79079810-DEL-7071 | chr1-79079811-DEL-7071 |
| chr1-79083326-DEL-54 | chr1-79083327-DEL-54 |
| chr1-79097925-DEL-6526 | chr1-79097916-DEL-6527 |
| chr1-79111678-DEL-143 | chr1-79111677-DEL-144 |
| chr1-79117649-DEL-240 | chr1-79117650-DEL-240 |
| chr1-79120244-DEL-54 | chr1-79120245-DEL-54 |
| chr1-79122670-DEL-111 | chr1-79122671-DEL-111 |
| chr1-79127342-DEL-222 | chr1-79127340-DEL-223 |
| chr1-79128707-DEL-200 | chr1-79128708-DEL-200 |
| chr1-79145148-DEL-79 | chr1-79145143-DEL-75 |
| chr1-79162932-DEL-64 | chr1-79162906-DEL-54 |
| chr1-79176596-DEL-90 | chr1-79176597-DEL-90 |
| chr1-79197437-DEL-69 | chr1-79197438-DEL-69 |
| chr1-79225815-DEL-9542 | chr1-79225816-DEL-9542 |
| chr1-79256846-DEL-401 | chr1-79256845-DEL-401 |
| chr1-79317976-DEL-1892 | chr1-79317977-DEL-1892 |
| chr1-79336302-DEL-1454 | chr1-79336303-DEL-1454 |
| chr1-79338941-DEL-1238 | chr1-79338942-DEL-1238 |
| chr1-79341011-DEL-259 | chr1-79341012-DEL-258 |
| chr1-79349532-DEL-303 | chr1-79349533-DEL-303 |
| chr1-79353844-DEL-52 | chr1-79353839-DEL-52 |
| chr1-79354594-DEL-670 | chr1-79354595-DEL-670 |
| chr1-79358538-DEL-6707 | chr1-79358539-DEL-6707 |
| chr1-79373197-DEL-95 | chr1-79373197-DEL-103 |
| chr1-79376866-DEL-400 | chr1-79376867-DEL-400 |
| chr1-79391541-DEL-501 | chr1-79391542-DEL-501 |
| chr1-79435087-DEL-4911 | chr1-79435088-DEL-4911 |
| chr1-79455253-DEL-525 | chr1-79455254-DEL-522 |
| chr1-79466066-DEL-6177 | chr1-79466067-DEL-6177 |
| chr1-79472790-DEL-868 | chr1-79472790-DEL-868 |
| chr1-79486271-DEL-1891 | chr1-79486272-DEL-1891 |
| chr1-79506949-DEL-266 | chr1-79506950-DEL-256 |
| chr1-79524008-DEL-1631 | chr1-79524009-DEL-1631 |
| chr1-79525341-DEL-153 | chr1-79525342-DEL-153 |
| chr1-79527791-DEL-6288 | chr1-79527792-DEL-6288 |
| chr1-79543558-DEL-112 | chr1-79543565-DEL-112 |
| chr1-79562864-DEL-179 | chr1-79562866-DEL-165 |
| chr1-79584843-DEL-317 | chr1-79584844-DEL-317 |
| chr1-79612389-DEL-56 | chr1-79612390-DEL-56 |

| | |
|---|---|
| chr1-79630196-DEL-2494 | chr1-79630197-DEL-2494 |
| chr1-79647247-DEL-60 | chr1-79647248-DEL-60 |
| chr1-79666429-DEL-1745 | chr1-79666430-DEL-1745 |
| chr1-79676187-DEL-498 | chr1-79676188-DEL-498 |
| chr1-79710102-DEL-122 | chr1-79710099-DEL-119 |
| chr1-79722754-DEL-400 | chr1-79722755-DEL-400 |
| chr1-79725785-DEL-158 | chr1-79725786-DEL-158 |
| chr1-79727814-DEL-175 | chr1-79727815-DEL-125 |
| chr1-79729171-DEL-116 | chr1-79729172-DEL-116 |
| chr1-79803874-DEL-1984 | chr1-79803875-DEL-1984 |
| chr1-79831672-DEL-80 | chr1-79831660-DEL-98 |
| chr1-79841653-DEL-530 | chr1-79841657-DEL-529 |
| chr1-79843865-DEL-214 | chr1-79843866-DEL-214 |
| chr1-79844976-DEL-239 | chr1-79844976-DEL-239 |
| chr1-79857516-DEL-78 | chr1-79857517-DEL-78 |
| chr1-79885904-DEL-59 | chr1-79885905-DEL-59 |
| chr1-79888799-DEL-371 | chr1-79888800-DEL-333 |
| chr1-79914788-DEL-7088 | chr1-79914789-DEL-7088 |
| chr1-79925220-DEL-1180 | chr1-79925221-DEL-1180 |
| chr1-79926132-DEL-54 | chr1-79926133-DEL-54 |
| chr1-79939472-DEL-6401 | chr1-79939473-DEL-6401 |
| chr1-79967258-DEL-52 | chr1-79967259-DEL-52 |
| chr1-80049109-DEL-62 | chr1-80049110-DEL-62 |
| chr1-80062209-DEL-2356 | chr1-80062210-DEL-2356 |
| chr1-80062744-DEL-540 | chr1-80062745-DEL-541 |
| chr1-80084851-DEL-89 | chr1-80084852-DEL-88 |
| chr1-80091713-DEL-59 | chr1-80091710-DEL-50 |
| chr1-80101134-DEL-66 | chr1-80101135-DEL-66 |
| chr1-80102218-DEL-3218 | chr1-80102243-DEL-3202 |
| chr1-80113166-DEL-4987 | chr1-80113167-DEL-4987 |
| chr1-80119499-DEL-5052 | chr1-80119500-DEL-5052 |
| chr1-80125331-DEL-83 | chr1-80125332-DEL-83 |
| chr1-80153474-DEL-996 | chr1-80153471-DEL-997 |
| chr1-80160033-DEL-52 | chr1-80160034-DEL-50 |
| chr1-80182689-DEL-154 | chr1-80182681-DEL-154 |
| chr1-80204297-DEL-60 | chr1-80204307-DEL-64 |
| chr1-80209672-DEL-84 | chr1-80209673-DEL-84 |
| chr1-80239198-DEL-213 | chr1-80239199-DEL-213 |
| chr1-80276648-DEL-276 | chr1-80276649-DEL-276 |
| chr1-80279230-DEL-66 | chr1-80279231-DEL-66 |
| chr1-80294555-DEL-220 | chr1-80294556-DEL-220 |
| chr1-80319294-DEL-364 | chr1-80319295-DEL-363 |
| chr1-80324848-DEL-324 | chr1-80324848-DEL-324 |
| chr1-80355895-DEL-104 | chr1-80355893-DEL-60 |
| chr1-80366161-DEL-53 | chr1-80366162-DEL-53 |
| chr1-80376386-DEL-99 | chr1-80376420-DEL-60 |
| chr1-80384115-DEL-172 | chr1-80384117-DEL-161 |
| chr1-80397165-DEL-75 | chr1-80397166-DEL-75 |
| chr1-80413432-DEL-205 | chr1-80413433-DEL-205 |
| chr1-80420605-DEL-1020 | chr1-80420605-DEL-1020 |
| chr1-80435783-DEL-141 | chr1-80435784-DEL-141 |
| chr1-80441303-DEL-145 | chr1-80441272-DEL-148 |
| chr1-80457911-DEL-1056 | chr1-80457912-DEL-1056 |
| chr1-80475823-DEL-82 | chr1-80475774-DEL-90 |
| chr1-80484563-DEL-129 | chr1-80484566-DEL-127 |
| chr1-80493085-DEL-70 | chr1-80493085-DEL-70 |
| chr1-80504480-DEL-284 | chr1-80504481-DEL-284 |
| chr1-80522303-DEL-166 | chr1-80522304-DEL-166 |
| chr1-80532467-DEL-52 | chr1-80532468-DEL-52 |
| chr1-80533488-DEL-92 | chr1-80533455-DEL-91 |
| chr1-80548744-DEL-507 | chr1-80548745-DEL-507 |
| chr1-80553215-DEL-77 | chr1-80553216-DEL-77 |
| chr1-80576411-DEL-69 | chr1-80576412-DEL-69 |
| chr1-80592806-DEL-109 | chr1-80592807-DEL-109 |
| chr1-80632755-DEL-64 | chr1-80632754-DEL-66 |
| chr1-80656763-DEL-438 | chr1-80656780-DEL-422 |
| chr1-80664371-DEL-8206 | chr1-80664371-DEL-8206 |
| chr1-80672936-DEL-177 | chr1-80672937-DEL-177 |
| chr1-80674351-DEL-1122 | chr1-80674352-DEL-1122 |
| chr1-80715772-DEL-94 | chr1-80715773-DEL-94 |
| chr1-80718696-DEL-133 | chr1-80718697-DEL-133 |
| chr1-80723128-DEL-120 | chr1-80723129-DEL-120 |
| chr1-80763486-DEL-141 | chr1-80763487-DEL-141 |
| chr1-80773710-DEL-338 | chr1-80773711-DEL-338 |
| chr1-80776730-DEL-69 | chr1-80776776-DEL-51 |
| chr1-80798173-DEL-77 | chr1-80798176-DEL-64 |
| chr1-80840829-DEL-1184 | chr1-80840830-DEL-1184 |
| chr1-80855455-DEL-155 | chr1-80855456-DEL-155 |
| chr1-80861919-DEL-493 | chr1-80861920-DEL-493 |
| chr1-80866120-DEL-225 | chr1-80866117-DEL-229 |

| | |
|---|---|
| chr1-80921195-DEL-105 | chr1-80921196-DEL-105 |
| chr1-80923772-DEL-82 | chr1-80923775-DEL-80 |
| chr1-80958612-DEL-1886 | chr1-80958613-DEL-1886 |
| chr1-80961482-DEL-6628 | chr1-80961483-DEL-6628 |
| chr1-80993941-DEL-81 | chr1-80993941-DEL-81 |
| chr1-81000519-DEL-223 | chr1-81000520-DEL-223 |
| chr1-81014600-DEL-121 | chr1-81014599-DEL-75 |
| chr1-81015180-DEL-136 | chr1-81015185-DEL-132 |
| chr1-81015577-DEL-941 | chr1-81015578-DEL-940 |
| chr1-81019643-DEL-177 | chr1-81019640-DEL-203 |
| chr1-81022740-DEL-689 | chr1-81022741-DEL-689 |
| chr1-81026286-DEL-50 | chr1-81026257-DEL-50 |
| chr1-81031672-DEL-1121 | chr1-81031673-DEL-1121 |
| chr1-81033719-DEL-426 | chr1-81033720-DEL-426 |
| chr1-81153288-DEL-87 | chr1-81153289-DEL-87 |
| chr1-81184485-DEL-457 | chr1-81184498-DEL-441 |
| chr1-81261159-DEL-298 | chr1-81261160-DEL-296 |
| chr1-81267983-DEL-56 | chr1-81267992-DEL-56 |
| chr1-81298615-DEL-191 | chr1-81298580-DEL-208 |
| chr1-81307927-DEL-254 | chr1-81307928-DEL-254 |
| chr1-81328632-DEL-250 | chr1-81328643-DEL-252 |
| chr1-81329504-DEL-54 | chr1-81329509-DEL-50 |
| chr1-81339066-DEL-395 | chr1-81339067-DEL-395 |
| chr1-81384366-DEL-169 | chr1-81384367-DEL-169 |
| chr1-81397367-DEL-7007 | chr1-81397368-DEL-7007 |
| chr1-81438755-DEL-2125 | chr1-81438756-DEL-2125 |
| chr1-81476176-DEL-6486 | chr1-81476177-DEL-6486 |
| chr1-81489347-DEL-1252 | chr1-81489357-DEL-1243 |
| chr1-81541876-DEL-7087 | chr1-81541877-DEL-7087 |
| chr1-81625584-DEL-175 | chr1-81625585-DEL-175 |
| chr1-81646225-DEL-655 | chr1-81646243-DEL-633 |
| chr1-81669403-DEL-1759 | chr1-81669404-DEL-1759 |
| chr1-81670133-DEL-387 | chr1-81670134-DEL-387 |
| chr1-81699526-DEL-1947 | chr1-81699532-DEL-1948 |
| chr1-81729384-DEL-582 | chr1-81729385-DEL-582 |
| chr1-81767139-DEL-53 | chr1-81767145-DEL-60 |
| chr1-81790888-DEL-1278 | chr1-81790887-DEL-1280 |
| chr1-81845922-DEL-196 | chr1-81845923-DEL-186 |
| chr1-81850600-DEL-278 | chr1-81850601-DEL-278 |
| chr1-81897077-DEL-529 | chr1-81897065-DEL-529 |
| chr1-81901261-DEL-1009 | chr1-81901262-DEL-1009 |
| chr1-81903707-DEL-139 | chr1-81903708-DEL-137 |
| chr1-81906860-DEL-76 | chr1-81906872-DEL-62 |
| chr1-81938768-DEL-115 | chr1-81938769-DEL-115 |
| chr1-81952692-DEL-56 | chr1-81952711-DEL-58 |
| chr1-81960271-DEL-2671 | chr1-81960272-DEL-2671 |
| chr1-82027382-DEL-305 | chr1-82027383-DEL-305 |
| chr1-82029772-DEL-120 | chr1-82029776-DEL-118 |
| chr1-82034532-DEL-60 | chr1-82034533-DEL-60 |
| chr1-82035311-DEL-173 | chr1-82035338-DEL-149 |
| chr1-82036762-DEL-358 | chr1-82036763-DEL-358 |
| chr1-82039272-DEL-1848 | chr1-82039273-DEL-1848 |
| chr1-82059244-DEL-58 | chr1-82059241-DEL-58 |
| chr1-82142866-DEL-82 | chr1-82142855-DEL-52 |
| chr1-82151340-DEL-6478 | chr1-82151325-DEL-6461 |
| chr1-82151428-DEL-214 | chr1-82151429-DEL-214 |
| chr1-82207101-DEL-447 | chr1-82207102-DEL-447 |
| chr1-82245972-DEL-62 | chr1-82245973-DEL-62 |
| chr1-82315084-DEL-257 | chr1-82315085-DEL-257 |
| chr1-82324662-DEL-398 | chr1-82324663-DEL-398 |
| chr1-82356473-DEL-424 | chr1-82356474-DEL-424 |
| chr1-82374297-DEL-6467 | chr1-82374298-DEL-6467 |
| chr1-82398831-DEL-6387 | chr1-82398832-DEL-6387 |
| chr1-82455450-DEL-282 | chr1-82455483-DEL-274 |
| chr1-82478753-DEL-4957 | chr1-82478754-DEL-4957 |
| chr1-82483921-DEL-153 | chr1-82483936-DEL-165 |
| chr1-82498161-DEL-60 | chr1-82498147-DEL-60 |
| chr1-82515163-DEL-335 | chr1-82515164-DEL-335 |
| chr1-82524809-DEL-304 | chr1-82524810-DEL-304 |
| chr1-82525415-DEL-4900 | chr1-82525416-DEL-4900 |
| chr1-82545486-DEL-56 | chr1-82545487-DEL-56 |
| chr1-82572273-DEL-232 | chr1-82572274-DEL-232 |
| chr1-82577662-DEL-79 | chr1-82577663-DEL-79 |
| chr1-82588586-DEL-54 | chr1-82588587-DEL-54 |
| chr1-82596944-DEL-186 | chr1-82596932-DEL-197 |
| chr1-82606650-DEL-86 | chr1-82606651-DEL-86 |
| chr1-82622926-DEL-575 | chr1-82622927-DEL-575 |
| chr1-82658642-DEL-194 | chr1-82658643-DEL-194 |
| chr1-82668812-DEL-468 | chr1-82668812-DEL-468 |
| chr1-82671632-DEL-55 | chr1-82671633-DEL-53 |

| | |
|---|---|
| chr1-82680840-DEL-201 | chr1-82680841-DEL-201 |
| chr1-82772748-DEL-78 | chr1-82772749-DEL-77 |
| chr1-82777534-DEL-60 | chr1-82777529-DEL-51 |
| chr1-82865346-DEL-73 | chr1-82865350-DEL-58 |
| chr1-82885499-DEL-422 | chr1-82885500-DEL-422 |
| chr1-82886288-DEL-81 | chr1-82886273-DEL-101 |
| chr1-82887678-DEL-617 | chr1-82887679-DEL-617 |
| chr1-82908668-DEL-11989 | chr1-82908667-DEL-11989 |
| chr1-82934704-DEL-1817 | chr1-82934703-DEL-1817 |
| chr1-82938076-DEL-63 | chr1-82938050-DEL-90 |
| chr1-82975990-DEL-628 | chr1-82975991-DEL-628 |
| chr1-82994886-DEL-1037 | chr1-82994887-DEL-1037 |
| chr1-83000916-DEL-420 | chr1-83000915-DEL-420 |
| chr1-83009030-DEL-53 | chr1-83009031-DEL-53 |
| chr1-83014583-DEL-416 | chr1-83014584-DEL-416 |
| chr1-83038001-DEL-64 | chr1-83038005-DEL-62 |
| chr1-83094079-DEL-56 | chr1-83094080-DEL-56 |
| chr1-83116707-DEL-1028 | chr1-83116708-DEL-1028 |
| chr1-83123483-DEL-394 | chr1-83123484-DEL-393 |
| chr1-83133552-DEL-84 | chr1-83133537-DEL-54 |
| chr1-83152403-DEL-167 | chr1-83152402-DEL-168 |
| chr1-83201574-DEL-498 | chr1-83201575-DEL-498 |
| chr1-83206555-DEL-620 | chr1-83206555-DEL-620 |
| chr1-83223028-DEL-1371 | chr1-83223025-DEL-1375 |
| chr1-83282100-DEL-60 | chr1-83282101-DEL-60 |
| chr1-83296102-DEL-132 | chr1-83296133-DEL-145 |
| chr1-83301473-DEL-127 | chr1-83301474-DEL-127 |
| chr1-83308260-DEL-277 | chr1-83308263-DEL-275 |
| chr1-83343450-DEL-458 | chr1-83343450-DEL-459 |
| chr1-83347406-DEL-6234 | chr1-83347407-DEL-6234 |
| chr1-83382744-DEL-152 | chr1-83382745-DEL-151 |
| chr1-83383413-DEL-145 | chr1-83383414-DEL-145 |
| chr1-83396870-DEL-86 | chr1-83396871-DEL-85 |
| chr1-83412249-DEL-794 | chr1-83412250-DEL-788 |
| chr1-83434512-DEL-6617 | chr1-83434513-DEL-6617 |
| chr1-83455587-DEL-53 | chr1-83455588-DEL-53 |
| chr1-83488605-DEL-191 | chr1-83488606-DEL-193 |
| chr1-83488765-DEL-6464 | chr1-83488766-DEL-6464 |
| chr1-83500668-DEL-200 | chr1-83500669-DEL-200 |
| chr1-83504329-DEL-111 | chr1-83504330-DEL-111 |
| chr1-83505923-DEL-159 | chr1-83505923-DEL-154 |
| chr1-83517194-DEL-68 | chr1-83517195-DEL-68 |
| chr1-83533397-DEL-1347 | chr1-83533399-DEL-1346 |
| chr1-83575481-DEL-54 | chr1-83575482-DEL-54 |
| chr1-83580641-DEL-402 | chr1-83580642-DEL-402 |
| chr1-83593012-DEL-6210 | chr1-83593013-DEL-6210 |
| chr1-83600376-DEL-71 | chr1-83600373-DEL-72 |
| chr1-83608038-DEL-224 | chr1-83608039-DEL-223 |
| chr1-83681988-DEL-475 | chr1-83681964-DEL-500 |
| chr1-83715282-DEL-56 | chr1-83715283-DEL-56 |
| chr1-83715898-DEL-136 | chr1-83715877-DEL-108 |
| chr1-83732077-DEL-118 | chr1-83732078-DEL-118 |
| chr1-83744107-DEL-75 | chr1-83744108-DEL-69 |
| chr1-83766401-DEL-71 | chr1-83766402-DEL-70 |
| chr1-83767023-DEL-187 | chr1-83767024-DEL-187 |
| chr1-83768926-DEL-54 | chr1-83768927-DEL-54 |
| chr1-83797330-DEL-8316 | chr1-83797320-DEL-8317 |
| chr1-83822085-DEL-551 | chr1-83822086-DEL-551 |
| chr1-83823085-DEL-60 | chr1-83823086-DEL-56 |
| chr1-83827673-DEL-511 | chr1-83827674-DEL-511 |
| chr1-83863516-DEL-558 | chr1-83863517-DEL-558 |
| chr1-83883908-DEL-120 | chr1-83883891-DEL-100 |
| chr1-83888489-DEL-198 | chr1-83888490-DEL-198 |
| chr1-83958824-DEL-8419 | chr1-83958825-DEL-8419 |
| chr1-84022311-DEL-87 | chr1-84022311-DEL-65 |
| chr1-84099413-DEL-6461 | chr1-84099391-DEL-6466 |
| chr1-84107398-DEL-82 | chr1-84107401-DEL-72 |
| chr1-84111820-DEL-52 | chr1-84111825-DEL-50 |
| chr1-84139048-DEL-6392 | chr1-84139049-DEL-6392 |
| chr1-84150078-DEL-69 | chr1-84150109-DEL-88 |
| chr1-84180806-DEL-243 | chr1-84180807-DEL-243 |
| chr1-84239768-DEL-947 | chr1-84239769-DEL-947 |
| chr1-84241205-DEL-7231 | chr1-84241206-DEL-7228 |
| chr1-84255365-DEL-81 | chr1-84255382-DEL-77 |
| chr1-84284067-DEL-288 | chr1-84284068-DEL-288 |
| chr1-84293827-DEL-232 | chr1-84293828-DEL-232 |
| chr1-84309866-DEL-549 | chr1-84309867-DEL-549 |
| chr1-84336829-DEL-328 | chr1-84336830-DEL-328 |
| chr1-84351104-DEL-6290 | chr1-84351105-DEL-6290 |
| chr1-84394714-DEL-318 | chr1-84394706-DEL-315 |

| | |
|---|---|
| chr1-84404975-DEL-465 | chr1-84404976-DEL-465 |
| chr1-84406224-DEL-373 | chr1-84406225-DEL-373 |
| chr1-84420708-DEL-91 | chr1-84420709-DEL-91 |
| chr1-84423444-DEL-1442 | chr1-84423445-DEL-1442 |
| chr1-84429386-DEL-6536 | chr1-84429387-DEL-6536 |
| chr1-84453750-DEL-61 | chr1-84453751-DEL-61 |
| chr1-84474843-DEL-118 | chr1-84474886-DEL-76 |
| chr1-84520139-DEL-6371 | chr1-84520140-DEL-6371 |
| chr1-84543567-DEL-4414 | chr1-84543568-DEL-4414 |
| chr1-84549448-DEL-4798 | chr1-84549449-DEL-4798 |
| chr1-84609773-DEL-114 | chr1-84609774-DEL-114 |
| chr1-84653589-DEL-55 | chr1-84653590-DEL-55 |
| chr1-84665726-DEL-520 | chr1-84665690-DEL-554 |
| chr1-84678311-DEL-6603 | chr1-84678312-DEL-6603 |
| chr1-84678328-DEL-208 | chr1-84678329-DEL-208 |
| chr1-84690923-DEL-185 | chr1-84690924-DEL-185 |
| chr1-84695039-DEL-80 | chr1-84695017-DEL-102 |
| chr1-84790205-DEL-58 | chr1-84790206-DEL-58 |
| chr1-84796176-DEL-209 | chr1-84796177-DEL-209 |
| chr1-84803672-DEL-64 | chr1-84803673-DEL-60 |
| chr1-84843781-DEL-215 | chr1-84843770-DEL-216 |
| chr1-84851926-DEL-1705 | chr1-84851927-DEL-1705 |
| chr1-84856629-DEL-109 | chr1-84856630-DEL-109 |
| chr1-84869382-DEL-182 | chr1-84869383-DEL-182 |
| chr1-84880028-DEL-70 | chr1-84880044-DEL-52 |
| chr1-84894805-DEL-216 | chr1-84894806-DEL-216 |
| chr1-84897408-DEL-220 | chr1-84897409-DEL-220 |
| chr1-84899751-DEL-209 | chr1-84899752-DEL-209 |
| chr1-84945383-DEL-256 | chr1-84945383-DEL-256 |
| chr1-84949337-DEL-227 | chr1-84949337-DEL-227 |
| chr1-84973617-DEL-4664 | chr1-84973597-DEL-4704 |
| chr1-84983604-DEL-209 | chr1-84983605-DEL-209 |
| chr1-84998275-DEL-208 | chr1-84998276-DEL-208 |
| chr1-85084162-DEL-54 | chr1-85084163-DEL-53 |
| chr1-85088591-DEL-176 | chr1-85088593-DEL-148 |
| chr1-85109719-DEL-191 | chr1-85109720-DEL-191 |
| chr1-85191260-DEL-250 | chr1-85191260-DEL-250 |
| chr1-85194795-DEL-203 | chr1-85194796-DEL-203 |
| chr1-85300681-DEL-54 | chr1-85300692-DEL-50 |
| chr1-85469858-DEL-4189 | chr1-85469858-DEL-4189 |
| chr1-85488305-DEL-204 | chr1-85488309-DEL-205 |
| chr1-85517753-DEL-56 | chr1-85517754-DEL-56 |
| chr1-85527275-DEL-116 | chr1-85527282-DEL-108 |
| chr1-85565867-DEL-700 | chr1-85565866-DEL-709 |
| chr1-85601802-DEL-89 | chr1-85601818-DEL-60 |
| chr1-85603652-DEL-210 | chr1-85603653-DEL-210 |
| chr1-85619249-DEL-90 | chr1-85619237-DEL-125 |
| chr1-85620439-DEL-61 | chr1-85620440-DEL-61 |
| chr1-85654544-DEL-54 | chr1-85654542-DEL-57 |
| chr1-85654766-DEL-4445 | chr1-85654767-DEL-4445 |
| chr1-85682209-DEL-66 | chr1-85682210-DEL-66 |
| chr1-85687337-DEL-77 | chr1-85687340-DEL-95 |
| chr1-85707620-DEL-69 | chr1-85707621-DEL-69 |
| chr1-85741774-DEL-307 | chr1-85741775-DEL-307 |
| chr1-85790395-DEL-165 | chr1-85790396-DEL-165 |
| chr1-85813297-DEL-212 | chr1-85813298-DEL-212 |
| chr1-85840011-DEL-592 | chr1-85840006-DEL-598 |
| chr1-85855110-DEL-65 | chr1-85855089-DEL-51 |
| chr1-85869892-DEL-66 | chr1-85869893-DEL-66 |
| chr1-85903311-DEL-81 | chr1-85903324-DEL-80 |
| chr1-85916116-DEL-78 | chr1-85916120-DEL-79 |
| chr1-85931755-DEL-488 | chr1-85931756-DEL-488 |
| chr1-85977219-DEL-211 | chr1-85977220-DEL-211 |
| chr1-85979676-DEL-119 | chr1-85979677-DEL-127 |
| chr1-86017448-DEL-142 | chr1-86017449-DEL-143 |
| chr1-86088685-DEL-82 | chr1-86088654-DEL-82 |
| chr1-86158239-DEL-120 | chr1-86158240-DEL-120 |
| chr1-86193874-DEL-112 | chr1-86193845-DEL-138 |
| chr1-86210401-DEL-214 | chr1-86210402-DEL-214 |
| chr1-86211634-DEL-199 | chr1-86211635-DEL-199 |
| chr1-86245209-DEL-3443 | chr1-86245210-DEL-3443 |
| chr1-86256022-DEL-62 | chr1-86256023-DEL-62 |
| chr1-86260439-DEL-201 | chr1-86260440-DEL-201 |
| chr1-86302025-DEL-84 | chr1-86302026-DEL-78 |
| chr1-86307761-DEL-170 | chr1-86307762-DEL-170 |
| chr1-86329801-DEL-194 | chr1-86329802-DEL-194 |
| chr1-86333928-DEL-136 | chr1-86333929-DEL-135 |
| chr1-86347020-DEL-52 | chr1-86347021-DEL-52 |
| chr1-86354517-DEL-90 | chr1-86354518-DEL-90 |
| chr1-86372717-DEL-165 | chr1-86372718-DEL-165 |

| | |
|---|---|
| chr1-86388607-DEL-511 | chr1-86388615-DEL-504 |
| chr1-86395563-DEL-216 | chr1-86395564-DEL-216 |
| chr1-86402707-DEL-71 | chr1-86402708-DEL-71 |
| chr1-86406009-DEL-368 | chr1-86406010-DEL-368 |
| chr1-86445290-DEL-160 | chr1-86445291-DEL-160 |
| chr1-86466843-DEL-202 | chr1-86466844-DEL-202 |
| chr1-86505750-DEL-262 | chr1-86505751-DEL-262 |
| chr1-86521531-DEL-1216 | chr1-86521532-DEL-1216 |
| chr1-86563562-DEL-563 | chr1-86563563-DEL-563 |
| chr1-86578627-DEL-221 | chr1-86578628-DEL-221 |
| chr1-86579761-DEL-176 | chr1-86579762-DEL-176 |
| chr1-86582463-DEL-212 | chr1-86582464-DEL-212 |
| chr1-86614118-DEL-140 | chr1-86614119-DEL-140 |
| chr1-86622954-DEL-450 | chr1-86622955-DEL-450 |
| chr1-86669467-DEL-87 | chr1-86669468-DEL-87 |
| chr1-86741513-DEL-68 | chr1-86741496-DEL-60 |
| chr1-86795079-DEL-644 | chr1-86795080-DEL-644 |
| chr1-86805567-DEL-310 | chr1-86805568-DEL-310 |
| chr1-86812682-DEL-470 | chr1-86812683-DEL-470 |
| chr1-86831362-DEL-276 | chr1-86831363-DEL-276 |
| chr1-86834509-DEL-1050 | chr1-86834509-DEL-1049 |
| chr1-86868071-DEL-2437 | chr1-86868072-DEL-2437 |
| chr1-86877453-DEL-186 | chr1-86877405-DEL-186 |
| chr1-86914397-DEL-1666 | chr1-86914398-DEL-1666 |
| chr1-86917702-DEL-59 | chr1-86917703-DEL-59 |
| chr1-86934399-DEL-89 | chr1-86934400-DEL-89 |
| chr1-86940809-DEL-498 | chr1-86940810-DEL-498 |
| chr1-87003111-DEL-5401 | chr1-87003112-DEL-5401 |
| chr1-87033637-DEL-56 | chr1-87033638-DEL-56 |
| chr1-87061634-DEL-6936 | chr1-87061635-DEL-6936 |
| chr1-87087529-DEL-150 | chr1-87087534-DEL-146 |
| chr1-87096323-DEL-58 | chr1-87096330-DEL-68 |
| chr1-87168502-DEL-208 | chr1-87168503-DEL-208 |
| chr1-87173478-DEL-227 | chr1-87173479-DEL-227 |
| chr1-87223198-DEL-173 | chr1-87223199-DEL-173 |
| chr1-87230898-DEL-198 | chr1-87230899-DEL-198 |
| chr1-87365218-DEL-68 | chr1-87365207-DEL-87 |
| chr1-87387846-DEL-7135 | chr1-87387847-DEL-7135 |
| chr1-87411182-DEL-120 | chr1-87411183-DEL-120 |
| chr1-87427899-DEL-67 | chr1-87427903-DEL-64 |
| chr1-87441920-DEL-595 | chr1-87441930-DEL-593 |
| chr1-87450257-DEL-580 | chr1-87450258-DEL-576 |
| chr1-87457194-DEL-2228 | chr1-87457195-DEL-2228 |
| chr1-87459515-DEL-209 | chr1-87459516-DEL-209 |
| chr1-87493428-DEL-63 | chr1-87493424-DEL-88 |
| chr1-87518863-DEL-66 | chr1-87518864-DEL-66 |
| chr1-87528897-DEL-58 | chr1-87528902-DEL-54 |
| chr1-87537948-DEL-55 | chr1-87537949-DEL-55 |
| chr1-87544868-DEL-1888 | chr1-87544869-DEL-1888 |
| chr1-87566911-DEL-209 | chr1-87566912-DEL-209 |
| chr1-87632855-DEL-225 | chr1-87632856-DEL-225 |
| chr1-87659184-DEL-1083 | chr1-87659185-DEL-1083 |
| chr1-87669230-DEL-6250 | chr1-87669231-DEL-6250 |
| chr1-87746527-DEL-1198 | chr1-87746528-DEL-1198 |
| chr1-87752555-DEL-210 | chr1-87752556-DEL-210 |
| chr1-87758246-DEL-181 | chr1-87758233-DEL-179 |
| chr1-87760608-DEL-437 | chr1-87760609-DEL-437 |
| chr1-87766111-DEL-70 | chr1-87766112-DEL-70 |
| chr1-87770141-DEL-213 | chr1-87770142-DEL-213 |
| chr1-87774926-DEL-91 | chr1-87774925-DEL-93 |
| chr1-87829669-DEL-232 | chr1-87829670-DEL-232 |
| chr1-87946987-DEL-609 | chr1-87946988-DEL-609 |
| chr1-87956343-DEL-6806 | chr1-87956344-DEL-6806 |
| chr1-87993335-DEL-71 | chr1-87993335-DEL-71 |
| chr1-88039473-DEL-948 | chr1-88039474-DEL-948 |
| chr1-88057187-DEL-78 | chr1-88057184-DEL-82 |
| chr1-88103495-DEL-80 | chr1-88103496-DEL-80 |
| chr1-88123126-DEL-78 | chr1-88123081-DEL-92 |
| chr1-88194002-DEL-51 | chr1-88194001-DEL-51 |
| chr1-88198039-DEL-1126 | chr1-88198038-DEL-1126 |
| chr1-88199974-DEL-240 | chr1-88199974-DEL-240 |
| chr1-88220344-DEL-1689 | chr1-88220344-DEL-1689 |
| chr1-88256116-DEL-324 | chr1-88256117-DEL-324 |
| chr1-88258711-DEL-718 | chr1-88258712-DEL-718 |
| chr1-88296644-DEL-60 | chr1-88296645-DEL-60 |
| chr1-88326670-DEL-88 | chr1-88326671-DEL-80 |
| chr1-88352085-DEL-117 | chr1-88352086-DEL-96 |
| chr1-88419099-DEL-57 | chr1-88419101-DEL-77 |
| chr1-88440531-DEL-100 | chr1-88440528-DEL-104 |
| chr1-88476856-DEL-144 | chr1-88476836-DEL-117 |

| | |
|---|---|
| chr1-88478932-DEL-350 | chr1-88478933-DEL-350 |
| chr1-88500964-DEL-250 | chr1-88500965-DEL-248 |
| chr1-88516731-DEL-187 | chr1-88516758-DEL-163 |
| chr1-88519544-DEL-58 | chr1-88519545-DEL-58 |
| chr1-88523300-DEL-192 | chr1-88523301-DEL-192 |
| chr1-88525240-DEL-2454 | chr1-88525241-DEL-2453 |
| chr1-88531121-DEL-192 | chr1-88531119-DEL-195 |
| chr1-88539296-DEL-350 | chr1-88539297-DEL-350 |
| chr1-88541719-DEL-1864 | chr1-88541720-DEL-1864 |
| chr1-88553629-DEL-63 | chr1-88553630-DEL-63 |
| chr1-88554448-DEL-149 | chr1-88554449-DEL-149 |
| chr1-88559484-DEL-422 | chr1-88559485-DEL-422 |
| chr1-88624677-DEL-209 | chr1-88624678-DEL-209 |
| chr1-88643320-DEL-144 | chr1-88643321-DEL-144 |
| chr1-88657709-DEL-84 | chr1-88657710-DEL-84 |
| chr1-88667720-DEL-108 | chr1-88667733-DEL-106 |
| chr1-88712397-DEL-298 | chr1-88712395-DEL-300 |
| chr1-88718388-DEL-435 | chr1-88718389-DEL-435 |
| chr1-88721476-DEL-68 | chr1-88721477-DEL-68 |
| chr1-88723186-DEL-214 | chr1-88723187-DEL-214 |
| chr1-88724108-DEL-105 | chr1-88724109-DEL-105 |
| chr1-88745657-DEL-731 | chr1-88745658-DEL-722 |
| chr1-88757443-DEL-6442 | chr1-88757444-DEL-6442 |
| chr1-88773720-DEL-104 | chr1-88773721-DEL-102 |
| chr1-88794209-DEL-69 | chr1-88794206-DEL-69 |
| chr1-88824123-DEL-199 | chr1-88824135-DEL-237 |
| chr1-88833700-DEL-76 | chr1-88833707-DEL-56 |
| chr1-88839963-DEL-76 | chr1-88839964-DEL-76 |
| chr1-88847539-DEL-72 | chr1-88847548-DEL-54 |
| chr1-88848562-DEL-161 | chr1-88848563-DEL-161 |
| chr1-88869706-DEL-52 | chr1-88869709-DEL-50 |
| chr1-88878738-DEL-1134 | chr1-88878739-DEL-1134 |
| chr1-88880321-DEL-76 | chr1-88880322-DEL-76 |
| chr1-88900922-DEL-64 | chr1-88900923-DEL-64 |
| chr1-88905130-DEL-477 | chr1-88905131-DEL-477 |
| chr1-88970534-DEL-251 | chr1-88970535-DEL-251 |
| chr1-88975518-DEL-64 | chr1-88975520-DEL-57 |
| chr1-88979118-DEL-242 | chr1-88979119-DEL-242 |
| chr1-88994590-DEL-2483 | chr1-88994591-DEL-2483 |
| chr1-89024480-DEL-5556 | chr1-89024481-DEL-5556 |
| chr1-89041060-DEL-51 | chr1-89041061-DEL-51 |
| chr1-89046275-DEL-105 | chr1-89046276-DEL-105 |
| chr1-89055224-DEL-100 | chr1-89055225-DEL-101 |
| chr1-89055736-DEL-95 | chr1-89055737-DEL-95 |
| chr1-89087900-DEL-114 | chr1-89087901-DEL-114 |
| chr1-89088800-DEL-605 | chr1-89088800-DEL-603 |
| chr1-89100312-DEL-397 | chr1-89100318-DEL-392 |
| chr1-89106109-DEL-108 | chr1-89106110-DEL-108 |
| chr1-89126789-DEL-4192 | chr1-89126790-DEL-4192 |
| chr1-89134118-DEL-54 | chr1-89134115-DEL-58 |
| chr1-89143291-DEL-8417 | chr1-89143286-DEL-8418 |
| chr1-89153975-DEL-88 | chr1-89153976-DEL-88 |
| chr1-89180433-DEL-84 | chr1-89180433-DEL-84 |
| chr1-89180604-DEL-1063 | chr1-89180606-DEL-1060 |
| chr1-89213390-DEL-68 | chr1-89213390-DEL-68 |
| chr1-89215442-DEL-68 | chr1-89215443-DEL-68 |
| chr1-89230618-DEL-135 | chr1-89230619-DEL-135 |
| chr1-89235173-DEL-63 | chr1-89235173-DEL-63 |
| chr1-89252936-DEL-889 | chr1-89252933-DEL-890 |
| chr1-89327323-DEL-64 | chr1-89327324-DEL-64 |
| chr1-89354622-DEL-60 | chr1-89354623-DEL-60 |
| chr1-89360571-DEL-56 | chr1-89360572-DEL-56 |
| chr1-89365639-DEL-106 | chr1-89365640-DEL-106 |
| chr1-89387082-DEL-98 | chr1-89387083-DEL-98 |
| chr1-89429681-DEL-65 | chr1-89429682-DEL-65 |
| chr1-89451241-DEL-57 | chr1-89451242-DEL-57 |
| chr1-89473649-DEL-89 | chr1-89473623-DEL-90 |
| chr1-89518164-DEL-202 | chr1-89518165-DEL-202 |
| chr1-89532898-DEL-61 | chr1-89532903-DEL-68 |
| chr1-89546191-DEL-554 | chr1-89546192-DEL-554 |
| chr1-89556138-DEL-95 | chr1-89556139-DEL-95 |
| chr1-89570660-DEL-101 | chr1-89570661-DEL-101 |
| chr1-89655471-DEL-58 | chr1-89655474-DEL-52 |
| chr1-89681857-DEL-209 | chr1-89681858-DEL-209 |
| chr1-89682923-DEL-108 | chr1-89682924-DEL-108 |
| chr1-89686352-DEL-1043 | chr1-89686353-DEL-1043 |
| chr1-89698585-DEL-85 | chr1-89698586-DEL-85 |
| chr1-89706955-DEL-392 | chr1-89706956-DEL-392 |
| chr1-89735474-DEL-160 | chr1-89735475-DEL-160 |
| chr1-89767045-DEL-391 | chr1-89767046-DEL-391 |

| | |
|---|---|
| chr1-89783338-DEL-70 | chr1-89783339-DEL-70 |
| chr1-89785048-DEL-202 | chr1-89785049-DEL-202 |
| chr1-89843842-DEL-195 | chr1-89843843-DEL-195 |
| chr1-89878530-DEL-482 | chr1-89878533-DEL-483 |
| chr1-89889737-DEL-55 | chr1-89889735-DEL-55 |
| chr1-89918527-DEL-770 | chr1-89918528-DEL-770 |
| chr1-89974653-DEL-85 | chr1-89974654-DEL-85 |
| chr1-90033773-DEL-2721 | chr1-90033774-DEL-2721 |
| chr1-90052103-DEL-69 | chr1-90052104-DEL-69 |
| chr1-90115395-DEL-390 | chr1-90115396-DEL-390 |
| chr1-90125032-DEL-2047 | chr1-90125033-DEL-2047 |
| chr1-90169324-DEL-55 | chr1-90169325-DEL-55 |
| chr1-90214914-DEL-57 | chr1-90214914-DEL-75 |
| chr1-90248852-DEL-132 | chr1-90248858-DEL-131 |
| chr1-90296013-DEL-54 | chr1-90296014-DEL-54 |
| chr1-90320840-DEL-56 | chr1-90320841-DEL-56 |
| chr1-90326801-DEL-630 | chr1-90326802-DEL-630 |
| chr1-90375256-DEL-793 | chr1-90375256-DEL-793 |
| chr1-90429413-DEL-1208 | chr1-90429414-DEL-1208 |
| chr1-90442709-DEL-61 | chr1-90442710-DEL-61 |
| chr1-90478240-DEL-494 | chr1-90478241-DEL-494 |
| chr1-90483241-DEL-89 | chr1-90483242-DEL-81 |
| chr1-90506586-DEL-74 | chr1-90506571-DEL-74 |
| chr1-90513754-DEL-54 | chr1-90513751-DEL-50 |
| chr1-90522156-DEL-158 | chr1-90522157-DEL-158 |
| chr1-90546548-DEL-52 | chr1-90546549-DEL-52 |
| chr1-90550724-DEL-5626 | chr1-90550725-DEL-5626 |
| chr1-90612159-DEL-2246 | chr1-90612160-DEL-2222 |
| chr1-90642228-DEL-78 | chr1-90642235-DEL-72 |
| chr1-90662499-DEL-82 | chr1-90662500-DEL-80 |
| chr1-90755367-DEL-73 | chr1-90755368-DEL-73 |
| chr1-90777325-DEL-94 | chr1-90777326-DEL-94 |
| chr1-90802428-DEL-63 | chr1-90802435-DEL-57 |
| chr1-90808578-DEL-436 | chr1-90808579-DEL-436 |
| chr1-90812012-DEL-106 | chr1-90812013-DEL-106 |
| chr1-90821768-DEL-112 | chr1-90821769-DEL-112 |
| chr1-90839680-DEL-109 | chr1-90839681-DEL-87 |
| chr1-90840223-DEL-52 | chr1-90840224-DEL-52 |
| chr1-90882296-DEL-75 | chr1-90882317-DEL-54 |
| chr1-90883226-DEL-66 | chr1-90883227-DEL-66 |
| chr1-90892309-DEL-2988 | chr1-90892310-DEL-2986 |
| chr1-90893170-DEL-112 | chr1-90893171-DEL-112 |
| chr1-90896837-DEL-57 | chr1-90896805-DEL-59 |
| chr1-90916444-DEL-7119 | chr1-90916449-DEL-7125 |
| chr1-90930259-DEL-64 | chr1-90930260-DEL-64 |
| chr1-90933216-DEL-129 | chr1-90933216-DEL-113 |
| chr1-90942514-DEL-96 | chr1-90942515-DEL-128 |
| chr1-90953179-DEL-129 | chr1-90953180-DEL-129 |
| chr1-90963152-DEL-94 | chr1-90963153-DEL-94 |
| chr1-90994203-DEL-167 | chr1-90994204-DEL-167 |
| chr1-91058267-DEL-5495 | chr1-91058268-DEL-5495 |
| chr1-91093481-DEL-62 | chr1-91093482-DEL-62 |
| chr1-91171684-DEL-66 | chr1-91171685-DEL-66 |
| chr1-91197945-DEL-340 | chr1-91197946-DEL-340 |
| chr1-91234473-DEL-177 | chr1-91234474-DEL-177 |
| chr1-91259679-DEL-56 | chr1-91259680-DEL-56 |
| chr1-91291456-DEL-56 | chr1-91291457-DEL-56 |
| chr1-91331333-DEL-163 | chr1-91331334-DEL-163 |
| chr1-91346237-DEL-78 | chr1-91346238-DEL-78 |
| chr1-91371348-DEL-864 | chr1-91371348-DEL-864 |
| chr1-91398949-DEL-7086 | chr1-91398950-DEL-7086 |
| chr1-91443061-DEL-53 | chr1-91443062-DEL-53 |
| chr1-91452313-DEL-200 | chr1-91452314-DEL-200 |
| chr1-91463721-DEL-75 | chr1-91463722-DEL-75 |
| chr1-91501129-DEL-53 | chr1-91501129-DEL-84 |
| chr1-91504121-DEL-60 | chr1-91504122-DEL-60 |
| chr1-91540712-DEL-103 | chr1-91540713-DEL-103 |
| chr1-91549454-DEL-1069 | chr1-91549454-DEL-1069 |
| chr1-91567381-DEL-793 | chr1-91567380-DEL-791 |
| chr1-91567984-DEL-59 | chr1-91567989-DEL-56 |
| chr1-91581828-DEL-920 | chr1-91581829-DEL-920 |
| chr1-91592946-DEL-1551 | chr1-91592949-DEL-1551 |
| chr1-91647727-DEL-225 | chr1-91647729-DEL-223 |
| chr1-91653696-DEL-612 | chr1-91653697-DEL-611 |
| chr1-91662057-DEL-116 | chr1-91662102-DEL-102 |
| chr1-91680362-DEL-98 | chr1-91680373-DEL-88 |
| chr1-91724958-DEL-84 | chr1-91724958-DEL-60 |
| chr1-91764815-DEL-193 | chr1-91764836-DEL-191 |
| chr1-91795118-DEL-407 | chr1-91795119-DEL-407 |
| chr1-91801173-DEL-67 | chr1-91801174-DEL-67 |

| | |
|---|---|
| chr1-91845388-DEL-1781 | chr1-91845388-DEL-1781 |
| chr1-92000910-DEL-61 | chr1-92000909-DEL-60 |
| chr1-92002851-DEL-263 | chr1-92002852-DEL-263 |
| chr1-92088306-DEL-63 | chr1-92088307-DEL-63 |
| chr1-92117622-DEL-53 | chr1-92117640-DEL-73 |
| chr1-92148303-DEL-413 | chr1-92148304-DEL-413 |
| chr1-92173223-DEL-400 | chr1-92173224-DEL-400 |
| chr1-92201220-DEL-60 | chr1-92201220-DEL-60 |
| chr1-92201241-DEL-2423 | chr1-92201242-DEL-2423 |
| chr1-92218761-DEL-85 | chr1-92218776-DEL-71 |
| chr1-92229561-DEL-323 | chr1-92229562-DEL-323 |
| chr1-92234893-DEL-65 | chr1-92234893-DEL-65 |
| chr1-92260065-DEL-74 | chr1-92260066-DEL-74 |
| chr1-92279919-DEL-1887 | chr1-92279920-DEL-1887 |
| chr1-92313255-DEL-211 | chr1-92313256-DEL-211 |
| chr1-92331458-DEL-485 | chr1-92331462-DEL-484 |
| chr1-92339927-DEL-6389 | chr1-92339928-DEL-6389 |
| chr1-92361496-DEL-54 | chr1-92361522-DEL-53 |
| chr1-92385525-DEL-64 | chr1-92385526-DEL-64 |
| chr1-92414944-DEL-2225 | chr1-92414955-DEL-2216 |
| chr1-92454302-DEL-2998 | chr1-92454303-DEL-2998 |
| chr1-92467396-DEL-116 | chr1-92467397-DEL-116 |
| chr1-92478952-DEL-4031 | chr1-92478953-DEL-4031 |
| chr1-92479881-DEL-328 | chr1-92479882-DEL-328 |
| chr1-92481711-DEL-66 | chr1-92481711-DEL-66 |
| chr1-92492386-DEL-3595 | chr1-92492387-DEL-3595 |
| chr1-92503950-DEL-66 | chr1-92503951-DEL-66 |
| chr1-92504661-DEL-210 | chr1-92504662-DEL-210 |
| chr1-92562238-DEL-58 | chr1-92562239-DEL-58 |
| chr1-92564188-DEL-78 | chr1-92564189-DEL-78 |
| chr1-92566407-DEL-55 | chr1-92566408-DEL-55 |
| chr1-92571272-DEL-72 | chr1-92571240-DEL-72 |
| chr1-92605225-DEL-402 | chr1-92605226-DEL-402 |
| chr1-92650257-DEL-515 | chr1-92650258-DEL-515 |
| chr1-92668745-DEL-54 | chr1-92668752-DEL-58 |
| chr1-92687954-DEL-54 | chr1-92687925-DEL-94 |
| chr1-92728093-DEL-454 | chr1-92728094-DEL-454 |
| chr1-92818779-DEL-144 | chr1-92818780-DEL-144 |
| chr1-92820975-DEL-98 | chr1-92820976-DEL-98 |
| chr1-93015157-DEL-80 | chr1-93015158-DEL-80 |
| chr1-93016826-DEL-64 | chr1-93016827-DEL-64 |
| chr1-93035084-DEL-4359 | chr1-93035085-DEL-4359 |
| chr1-93066773-DEL-56 | chr1-93066800-DEL-56 |
| chr1-93074630-DEL-82 | chr1-93074655-DEL-58 |
| chr1-93108191-DEL-199 | chr1-93108194-DEL-198 |
| chr1-93123664-DEL-325 | chr1-93123666-DEL-324 |
| chr1-93193713-DEL-218 | chr1-93193714-DEL-218 |
| chr1-93217503-DEL-80 | chr1-93217504-DEL-100 |
| chr1-93268710-DEL-54 | chr1-93268709-DEL-55 |
| chr1-93384992-DEL-60 | chr1-93384993-DEL-60 |
| chr1-93414254-DEL-204 | chr1-93414255-DEL-204 |
| chr1-93422734-DEL-472 | chr1-93422735-DEL-472 |
| chr1-93470443-DEL-1599 | chr1-93470442-DEL-1599 |
| chr1-93503107-DEL-231 | chr1-93503108-DEL-231 |
| chr1-93513368-DEL-194 | chr1-93513357-DEL-195 |
| chr1-93532346-DEL-170 | chr1-93532347-DEL-170 |
| chr1-93557465-DEL-338 | chr1-93557505-DEL-338 |
| chr1-93559927-DEL-175 | chr1-93559928-DEL-175 |
| chr1-93589707-DEL-150 | chr1-93589707-DEL-142 |
| chr1-93594961-DEL-51 | chr1-93594962-DEL-51 |
| chr1-93601984-DEL-422 | chr1-93601996-DEL-414 |
| chr1-93638378-DEL-180 | chr1-93638379-DEL-180 |
| chr1-93643293-DEL-384 | chr1-93643294-DEL-384 |
| chr1-93655248-DEL-457 | chr1-93655249-DEL-457 |
| chr1-93662173-DEL-54 | chr1-93662174-DEL-54 |
| chr1-93663153-DEL-226 | chr1-93663154-DEL-226 |
| chr1-93671285-DEL-69 | chr1-93671287-DEL-68 |
| chr1-93688353-DEL-243 | chr1-93688354-DEL-243 |
| chr1-93758251-DEL-440 | chr1-93758256-DEL-436 |
| chr1-93791929-DEL-251 | chr1-93791922-DEL-259 |
| chr1-93805836-DEL-128 | chr1-93805837-DEL-128 |
| chr1-93813806-DEL-215 | chr1-93813807-DEL-215 |
| chr1-93823639-DEL-261 | chr1-93823640-DEL-261 |
| chr1-93841322-DEL-98 | chr1-93841322-DEL-98 |
| chr1-93910363-DEL-60 | chr1-93910363-DEL-60 |
| chr1-93917816-DEL-470 | chr1-93917816-DEL-470 |
| chr1-93920922-DEL-222 | chr1-93920923-DEL-222 |
| chr1-93977795-DEL-54 | chr1-93977796-DEL-54 |
| chr1-93987910-DEL-507 | chr1-93987903-DEL-510 |
| chr1-93996337-DEL-1092 | chr1-93996338-DEL-1092 |

| | |
|---|---|
| chr1-94017981-DEL-197 | chr1-94017982-DEL-197 |
| chr1-94027285-DEL-80 | chr1-94027285-DEL-80 |
| chr1-94032890-DEL-2218 | chr1-94032891-DEL-2218 |
| chr1-94042188-DEL-619 | chr1-94042189-DEL-619 |
| chr1-94069470-DEL-1266 | chr1-94069471-DEL-1266 |
| chr1-94078552-DEL-6739 | chr1-94078553-DEL-6739 |
| chr1-94087928-DEL-1198 | chr1-94087929-DEL-1198 |
| chr1-94112653-DEL-619 | chr1-94112654-DEL-619 |
| chr1-94121729-DEL-5029 | chr1-94121730-DEL-5029 |
| chr1-94127681-DEL-1140 | chr1-94127682-DEL-1140 |
| chr1-94129187-DEL-210 | chr1-94129188-DEL-210 |
| chr1-94133073-DEL-80 | chr1-94133074-DEL-80 |
| chr1-94141348-DEL-57 | chr1-94141355-DEL-58 |
| chr1-94151053-DEL-56 | chr1-94151054-DEL-56 |
| chr1-94182209-DEL-92 | chr1-94182214-DEL-88 |
| chr1-94194616-DEL-58 | chr1-94194617-DEL-58 |
| chr1-94211473-DEL-351 | chr1-94211474-DEL-351 |
| chr1-94216104-DEL-565 | chr1-94216104-DEL-565 |
| chr1-94235829-DEL-61 | chr1-94235830-DEL-61 |
| chr1-94237081-DEL-61 | chr1-94237049-DEL-81 |
| chr1-94266088-DEL-2747 | chr1-94266089-DEL-2747 |
| chr1-94268689-DEL-3953 | chr1-94268690-DEL-3953 |
| chr1-94280306-DEL-102 | chr1-94280307-DEL-102 |
| chr1-94285401-DEL-234 | chr1-94285402-DEL-234 |
| chr1-94319090-DEL-1669 | chr1-94319091-DEL-1669 |
| chr1-94394735-DEL-3179 | chr1-94394736-DEL-3179 |
| chr1-94402902-DEL-7047 | chr1-94402903-DEL-7047 |
| chr1-94432990-DEL-1245 | chr1-94432991-DEL-1245 |
| chr1-94456206-DEL-171 | chr1-94456205-DEL-172 |
| chr1-94456988-DEL-552 | chr1-94456989-DEL-552 |
| chr1-94476709-DEL-324 | chr1-94476710-DEL-313 |
| chr1-94499275-DEL-1925 | chr1-94499276-DEL-1925 |
| chr1-94524780-DEL-1863 | chr1-94524781-DEL-1863 |
| chr1-94529814-DEL-78 | chr1-94529815-DEL-78 |
| chr1-94543100-DEL-110 | chr1-94543108-DEL-76 |
| chr1-94573039-DEL-3212 | chr1-94573039-DEL-3212 |
| chr1-94580953-DEL-1146 | chr1-94580954-DEL-1146 |
| chr1-94605186-DEL-57 | chr1-94605178-DEL-57 |
| chr1-94620835-DEL-102 | chr1-94620836-DEL-102 |
| chr1-94650608-DEL-6858 | chr1-94650609-DEL-6858 |
| chr1-94668338-DEL-70 | chr1-94668362-DEL-60 |
| chr1-94677732-DEL-66 | chr1-94677720-DEL-78 |
| chr1-94678016-DEL-684 | chr1-94677993-DEL-679 |
| chr1-94682047-DEL-494 | chr1-94682048-DEL-494 |
| chr1-94696492-DEL-123 | chr1-94696442-DEL-126 |
| chr1-94698238-DEL-3106 | chr1-94698237-DEL-3105 |
| chr1-94731138-DEL-196 | chr1-94731139-DEL-196 |
| chr1-94759057-DEL-377 | chr1-94759057-DEL-377 |
| chr1-94763453-DEL-92 | chr1-94763454-DEL-92 |
| chr1-94766025-DEL-458 | chr1-94766026-DEL-458 |
| chr1-94768919-DEL-266 | chr1-94768920-DEL-266 |
| chr1-94778899-DEL-380 | chr1-94778901-DEL-379 |
| chr1-94781003-DEL-2955 | chr1-94780991-DEL-2956 |
| chr1-94813495-DEL-223 | chr1-94813496-DEL-223 |
| chr1-94819274-DEL-603 | chr1-94819275-DEL-603 |
| chr1-94849181-DEL-6335 | chr1-94849182-DEL-6335 |
| chr1-94867301-DEL-2576 | chr1-94867302-DEL-2576 |
| chr1-94873112-DEL-1225 | chr1-94873112-DEL-1225 |
| chr1-94916003-DEL-53 | chr1-94916006-DEL-51 |
| chr1-94918827-DEL-225 | chr1-94918828-DEL-225 |
| chr1-94992328-DEL-326 | chr1-94992329-DEL-326 |
| chr1-95041980-DEL-649 | chr1-95041981-DEL-649 |
| chr1-95042783-DEL-312 | chr1-95042784-DEL-304 |
| chr1-95058397-DEL-63 | chr1-95058394-DEL-59 |
| chr1-95072652-DEL-1163 | chr1-95072653-DEL-1163 |
| chr1-95074987-DEL-55 | chr1-95074989-DEL-68 |
| chr1-95130630-DEL-191 | chr1-95130631-DEL-191 |
| chr1-95153197-DEL-4201 | chr1-95153198-DEL-4201 |
| chr1-95160112-DEL-6984 | chr1-95160100-DEL-7004 |
| chr1-95177438-DEL-51 | chr1-95177439-DEL-51 |
| chr1-95178922-DEL-78 | chr1-95178947-DEL-54 |
| chr1-95188108-DEL-76 | chr1-95188117-DEL-52 |
| chr1-95192282-DEL-109 | chr1-95192241-DEL-124 |
| chr1-95198466-DEL-2123 | chr1-95198467-DEL-2123 |
| chr1-95220754-DEL-222 | chr1-95220755-DEL-222 |
| chr1-95248935-DEL-146 | chr1-95248936-DEL-146 |
| chr1-95304148-DEL-365 | chr1-95304149-DEL-365 |
| chr1-95314174-DEL-1681 | chr1-95314175-DEL-1681 |
| chr1-95379646-DEL-348 | chr1-95379649-DEL-346 |
| chr1-95398644-DEL-201 | chr1-95398645-DEL-201 |

| | |
|---|---|
| chr1-95426293-DEL-640 | chr1-95426293-DEL-640 |
| chr1-95439123-DEL-6716 | chr1-95439124-DEL-6716 |
| chr1-95446335-DEL-74 | chr1-95446343-DEL-66 |
| chr1-95480014-DEL-56 | chr1-95480015-DEL-56 |
| chr1-95493915-DEL-66 | chr1-95493916-DEL-66 |
| chr1-95524001-DEL-220 | chr1-95524002-DEL-220 |
| chr1-95562927-DEL-1454 | chr1-95562928-DEL-1454 |
| chr1-95622874-DEL-4488 | chr1-95622875-DEL-4488 |
| chr1-95637371-DEL-54 | chr1-95637372-DEL-54 |
| chr1-95651051-DEL-177 | chr1-95651052-DEL-177 |
| chr1-95665597-DEL-1944 | chr1-95665598-DEL-1944 |
| chr1-95688854-DEL-277 | chr1-95688866-DEL-272 |
| chr1-95711061-DEL-6409 | chr1-95711061-DEL-6409 |
| chr1-95731516-DEL-74 | chr1-95731516-DEL-74 |
| chr1-95737433-DEL-4890 | chr1-95737434-DEL-4890 |
| chr1-95748302-DEL-516 | chr1-95748303-DEL-516 |
| chr1-95756109-DEL-439 | chr1-95756110-DEL-439 |
| chr1-95758437-DEL-1523 | chr1-95758438-DEL-1523 |
| chr1-95791481-DEL-197 | chr1-95791482-DEL-197 |
| chr1-95834485-DEL-206 | chr1-95834486-DEL-206 |
| chr1-95880862-DEL-603 | chr1-95880863-DEL-603 |
| chr1-95886878-DEL-198 | chr1-95886879-DEL-198 |
| chr1-95902449-DEL-125 | chr1-95902426-DEL-123 |
| chr1-95908056-DEL-63 | chr1-95908058-DEL-62 |
| chr1-95914238-DEL-1206 | chr1-95914239-DEL-1206 |
| chr1-95963234-DEL-3144 | chr1-95963236-DEL-3144 |
| chr1-95978632-DEL-96 | chr1-95978609-DEL-74 |
| chr1-95997444-DEL-6129 | chr1-95997445-DEL-6129 |
| chr1-96006793-DEL-690 | chr1-96006794-DEL-690 |
| chr1-96052026-DEL-168 | chr1-96052027-DEL-168 |
| chr1-96068354-DEL-10300 | chr1-96068355-DEL-10300 |
| chr1-96076667-DEL-342 | chr1-96076668-DEL-342 |
| chr1-96077414-DEL-2681 | chr1-96077415-DEL-2681 |
| chr1-96119001-DEL-6358 | chr1-96119002-DEL-6358 |
| chr1-96139766-DEL-54 | chr1-96139766-DEL-50 |
| chr1-96146492-DEL-64 | chr1-96146493-DEL-64 |
| chr1-96151558-DEL-700 | chr1-96151558-DEL-700 |
| chr1-96152764-DEL-2629 | chr1-96152765-DEL-2629 |
| chr1-96168540-DEL-6324 | chr1-96168541-DEL-6324 |
| chr1-96177422-DEL-1358 | chr1-96177423-DEL-1358 |
| chr1-96200131-DEL-166 | chr1-96200132-DEL-166 |
| chr1-96205079-DEL-276 | chr1-96205080-DEL-276 |
| chr1-96251987-DEL-66 | chr1-96252006-DEL-54 |
| chr1-96261440-DEL-689 | chr1-96261441-DEL-689 |
| chr1-96284032-DEL-117 | chr1-96284033-DEL-117 |
| chr1-96305483-DEL-56 | chr1-96305484-DEL-56 |
| chr1-96324022-DEL-558 | chr1-96324023-DEL-558 |
| chr1-96371365-DEL-306 | chr1-96371366-DEL-306 |
| chr1-96378947-DEL-60 | chr1-96378948-DEL-60 |
| chr1-96382891-DEL-408 | chr1-96382892-DEL-408 |
| chr1-96414892-DEL-436 | chr1-96414892-DEL-436 |
| chr1-96422359-DEL-1604 | chr1-96422361-DEL-1603 |
| chr1-96425209-DEL-97 | chr1-96425210-DEL-97 |
| chr1-96435028-DEL-1065 | chr1-96435029-DEL-1065 |
| chr1-96452786-DEL-141 | chr1-96452787-DEL-141 |
| chr1-96461071-DEL-5497 | chr1-96461071-DEL-5497 |
| chr1-96481544-DEL-53 | chr1-96481552-DEL-50 |
| chr1-96493309-DEL-780 | chr1-96493312-DEL-777 |
| chr1-96500135-DEL-6416 | chr1-96500135-DEL-6416 |
| chr1-96519411-DEL-2785 | chr1-96519412-DEL-2785 |
| chr1-96520452-DEL-82 | chr1-96520453-DEL-82 |
| chr1-96557532-DEL-439 | chr1-96557533-DEL-439 |
| chr1-96565832-DEL-58 | chr1-96565833-DEL-58 |
| chr1-96568471-DEL-337 | chr1-96568472-DEL-337 |
| chr1-96570202-DEL-3642 | chr1-96570203-DEL-3642 |
| chr1-96576619-DEL-2767 | chr1-96576619-DEL-2767 |
| chr1-96577301-DEL-626 | chr1-96577308-DEL-620 |
| chr1-96607087-DEL-2157 | chr1-96607087-DEL-2157 |
| chr1-96622466-DEL-174 | chr1-96622467-DEL-174 |
| chr1-96632924-DEL-1796 | chr1-96632923-DEL-1796 |
| chr1-96645689-DEL-1140 | chr1-96645690-DEL-1140 |
| chr1-96651654-DEL-65 | chr1-96651654-DEL-66 |
| chr1-96660579-DEL-73 | chr1-96660574-DEL-78 |
| chr1-96668732-DEL-3798 | chr1-96668735-DEL-3802 |
| chr1-96695081-DEL-56 | chr1-96695082-DEL-62 |
| chr1-96697987-DEL-78 | chr1-96697985-DEL-86 |
| chr1-96711820-DEL-6046 | chr1-96711821-DEL-6046 |
| chr1-96748075-DEL-51 | chr1-96748037-DEL-92 |
| chr1-96748540-DEL-98 | chr1-96748542-DEL-94 |
| chr1-96755167-DEL-866 | chr1-96755168-DEL-866 |

| | |
|---|---|
| chr1-96775686-DEL-59 | chr1-96775690-DEL-55 |
| chr1-96786589-DEL-114 | chr1-96786590-DEL-114 |
| chr1-96830223-DEL-202 | chr1-96830224-DEL-201 |
| chr1-96870082-DEL-2947 | chr1-96870083-DEL-2947 |
| chr1-96870683-DEL-54 | chr1-96870722-DEL-61 |
| chr1-96896635-DEL-81 | chr1-96896636-DEL-81 |
| chr1-96931016-DEL-52 | chr1-96931017-DEL-52 |
| chr1-96949576-DEL-55 | chr1-96949555-DEL-88 |
| chr1-97018482-DEL-172 | chr1-97018483-DEL-172 |
| chr1-97024863-DEL-788 | chr1-97024864-DEL-788 |
| chr1-97106225-DEL-1886 | chr1-97106226-DEL-1886 |
| chr1-97119029-DEL-1864 | chr1-97119030-DEL-1864 |
| chr1-97207260-DEL-199 | chr1-97207261-DEL-199 |
| chr1-97245716-DEL-94 | chr1-97245715-DEL-95 |
| chr1-97308138-DEL-213 | chr1-97308135-DEL-213 |
| chr1-97344783-DEL-6478 | chr1-97344784-DEL-6478 |
| chr1-97373591-DEL-3960 | chr1-97373592-DEL-3959 |
| chr1-97381655-DEL-2724 | chr1-97381655-DEL-2724 |
| chr1-97384577-DEL-66 | chr1-97384578-DEL-66 |
| chr1-97385360-DEL-53 | chr1-97385354-DEL-72 |
| chr1-97426947-DEL-181 | chr1-97426950-DEL-182 |
| chr1-97440758-DEL-199 | chr1-97440759-DEL-199 |
| chr1-97458840-DEL-2143 | chr1-97458841-DEL-2143 |
| chr1-97496782-DEL-52 | chr1-97496783-DEL-52 |
| chr1-97506661-DEL-1370 | chr1-97506662-DEL-1370 |
| chr1-97513162-DEL-1697 | chr1-97513165-DEL-1695 |
| chr1-97519318-DEL-113 | chr1-97519333-DEL-99 |
| chr1-97519891-DEL-272 | chr1-97519892-DEL-272 |
| chr1-97552793-DEL-134 | chr1-97552764-DEL-142 |
| chr1-97570709-DEL-61 | chr1-97570705-DEL-63 |
| chr1-97574367-DEL-104 | chr1-97574368-DEL-104 |
| chr1-97578335-DEL-52 | chr1-97578338-DEL-50 |
| chr1-97593171-DEL-1021 | chr1-97593172-DEL-1021 |
| chr1-97608247-DEL-2570 | chr1-97608246-DEL-2570 |
| chr1-97609505-DEL-237 | chr1-97609506-DEL-237 |
| chr1-97619963-DEL-108 | chr1-97620009-DEL-63 |
| chr1-97651807-DEL-1119 | chr1-97651808-DEL-1119 |
| chr1-97655559-DEL-323 | chr1-97655560-DEL-323 |
| chr1-97706306-DEL-3108 | chr1-97706307-DEL-3108 |
| chr1-97717834-DEL-59 | chr1-97717811-DEL-68 |
| chr1-97719792-DEL-177 | chr1-97719792-DEL-177 |
| chr1-97739629-DEL-325 | chr1-97739629-DEL-325 |
| chr1-97740746-DEL-247 | chr1-97740747-DEL-247 |
| chr1-97793405-DEL-6400 | chr1-97793406-DEL-6400 |
| chr1-97826151-DEL-56 | chr1-97826152-DEL-56 |
| chr1-97838203-DEL-60 | chr1-97838198-DEL-57 |
| chr1-97856515-DEL-68 | chr1-97856516-DEL-68 |
| chr1-97863031-DEL-171 | chr1-97863069-DEL-155 |
| chr1-97876353-DEL-875 | chr1-97876354-DEL-875 |
| chr1-97900650-DEL-66 | chr1-97900651-DEL-65 |
| chr1-97926943-DEL-166 | chr1-97926944-DEL-166 |
| chr1-97929998-DEL-51 | chr1-97929984-DEL-69 |
| chr1-98014019-DEL-52 | chr1-98014019-DEL-52 |
| chr1-98037949-DEL-6199 | chr1-98037950-DEL-6199 |
| chr1-98045041-DEL-53 | chr1-98045041-DEL-53 |
| chr1-98089527-DEL-123 | chr1-98089528-DEL-123 |
| chr1-98110659-DEL-696 | chr1-98110660-DEL-696 |
| chr1-98122818-DEL-104 | chr1-98122861-DEL-102 |
| chr1-98128861-DEL-6398 | chr1-98128862-DEL-6398 |
| chr1-98141297-DEL-373 | chr1-98141298-DEL-373 |
| chr1-98214567-DEL-8633 | chr1-98214568-DEL-8633 |
| chr1-98241570-DEL-547 | chr1-98241576-DEL-548 |
| chr1-98245583-DEL-700 | chr1-98245584-DEL-700 |
| chr1-98294147-DEL-6653 | chr1-98294148-DEL-6653 |
| chr1-98294165-DEL-424 | chr1-98294165-DEL-424 |
| chr1-98385429-DEL-2289 | chr1-98385429-DEL-2289 |
| chr1-98438579-DEL-959 | chr1-98438580-DEL-959 |
| chr1-98442741-DEL-66 | chr1-98442742-DEL-66 |
| chr1-98446534-DEL-197 | chr1-98446535-DEL-197 |
| chr1-98455395-DEL-859 | chr1-98455396-DEL-859 |
| chr1-98457194-DEL-391 | chr1-98457204-DEL-384 |
| chr1-98461413-DEL-180 | chr1-98461414-DEL-180 |
| chr1-98464956-DEL-358 | chr1-98464957-DEL-358 |
| chr1-98528118-DEL-3344 | chr1-98528112-DEL-3350 |
| chr1-98541323-DEL-762 | chr1-98541324-DEL-762 |
| chr1-98542325-DEL-6394 | chr1-98542325-DEL-6394 |
| chr1-98555610-DEL-444 | chr1-98555611-DEL-444 |
| chr1-98558248-DEL-297 | chr1-98558249-DEL-297 |
| chr1-98569895-DEL-1891 | chr1-98569896-DEL-1891 |
| chr1-98572693-DEL-6412 | chr1-98572694-DEL-6412 |

| | |
|---|---|
| chr1-98587598-DEL-170 | chr1-98587599-DEL-170 |
| chr1-98592201-DEL-362 | chr1-98592202-DEL-362 |
| chr1-98593900-DEL-75 | chr1-98593886-DEL-61 |
| chr1-98594925-DEL-68 | chr1-98594889-DEL-101 |
| chr1-98637670-DEL-92 | chr1-98637671-DEL-92 |
| chr1-98641659-DEL-272 | chr1-98641660-DEL-272 |
| chr1-98652544-DEL-401 | chr1-98652545-DEL-401 |
| chr1-98671722-DEL-366 | chr1-98671717-DEL-367 |
| chr1-98686862-DEL-165 | chr1-98686863-DEL-165 |
| chr1-98710452-DEL-82 | chr1-98710453-DEL-72 |
| chr1-98715036-DEL-404 | chr1-98715037-DEL-404 |
| chr1-98739500-DEL-822 | chr1-98739500-DEL-822 |
| chr1-98740540-DEL-1057 | chr1-98740541-DEL-1057 |
| chr1-98744473-DEL-809 | chr1-98744474-DEL-809 |
| chr1-98745533-DEL-872 | chr1-98745534-DEL-873 |
| chr1-98767857-DEL-6212 | chr1-98767858-DEL-6212 |
| chr1-98793920-DEL-806 | chr1-98793921-DEL-806 |
| chr1-98805098-DEL-897 | chr1-98805099-DEL-897 |
| chr1-98855006-DEL-1233 | chr1-98855017-DEL-1223 |
| chr1-98856973-DEL-317 | chr1-98856974-DEL-317 |
| chr1-98866622-DEL-77 | chr1-98866609-DEL-67 |
| chr1-98872825-DEL-719 | chr1-98872826-DEL-720 |
| chr1-98874496-DEL-6186 | chr1-98874497-DEL-6186 |
| chr1-98896290-DEL-1655 | chr1-98896291-DEL-1655 |
| chr1-98907355-DEL-1601 | chr1-98907356-DEL-1601 |
| chr1-98911601-DEL-139 | chr1-98911602-DEL-139 |
| chr1-98912539-DEL-262 | chr1-98912540-DEL-259 |
| chr1-98913873-DEL-100 | chr1-98913885-DEL-77 |
| chr1-98917237-DEL-90 | chr1-98917238-DEL-90 |
| chr1-98918961-DEL-52 | chr1-98918962-DEL-52 |
| chr1-98920156-DEL-62 | chr1-98920157-DEL-62 |
| chr1-98927678-DEL-2747 | chr1-98927679-DEL-2742 |
| chr1-98950737-DEL-786 | chr1-98950738-DEL-786 |
| chr1-98966459-DEL-1545 | chr1-98966460-DEL-1545 |
| chr1-98969148-DEL-4434 | chr1-98969147-DEL-4435 |
| chr1-98977603-DEL-58 | chr1-98977604-DEL-58 |
| chr1-99007667-DEL-1252 | chr1-99007668-DEL-1252 |
| chr1-99010157-DEL-497 | chr1-99010157-DEL-497 |
| chr1-99027397-DEL-1877 | chr1-99027397-DEL-1877 |
| chr1-99030938-DEL-442 | chr1-99030939-DEL-442 |
| chr1-99031676-DEL-545 | chr1-99031677-DEL-545 |
| chr1-99034477-DEL-1487 | chr1-99034478-DEL-1487 |
| chr1-99044878-DEL-109 | chr1-99044879-DEL-95 |
| chr1-99056929-DEL-3022 | chr1-99056930-DEL-3022 |
| chr1-99070508-DEL-1687 | chr1-99070509-DEL-1687 |
| chr1-99077383-DEL-59 | chr1-99077384-DEL-63 |
| chr1-99093230-DEL-95 | chr1-99093227-DEL-99 |
| chr1-99104360-DEL-316 | chr1-99104361-DEL-316 |
| chr1-99110879-DEL-7213 | chr1-99110880-DEL-7213 |
| chr1-99130770-DEL-6396 | chr1-99130771-DEL-6396 |
| chr1-99138842-DEL-52 | chr1-99138843-DEL-52 |
| chr1-99150985-DEL-1282 | chr1-99150986-DEL-1282 |
| chr1-99156879-DEL-203 | chr1-99156880-DEL-203 |
| chr1-99190290-DEL-4100 | chr1-99190281-DEL-4099 |
| chr1-99195380-DEL-60 | chr1-99195381-DEL-60 |
| chr1-99196986-DEL-2254 | chr1-99196989-DEL-2255 |
| chr1-99204305-DEL-410 | chr1-99204306-DEL-410 |
| chr1-99210287-DEL-4547 | chr1-99210288-DEL-4547 |
| chr1-99214983-DEL-584 | chr1-99214984-DEL-572 |
| chr1-99219600-DEL-1570 | chr1-99219601-DEL-1570 |
| chr1-99222983-DEL-1183 | chr1-99222995-DEL-1189 |
| chr1-99234858-DEL-128 | chr1-99234862-DEL-108 |
| chr1-99238980-DEL-3158 | chr1-99238975-DEL-3159 |
| chr1-99242740-DEL-50 | chr1-99242741-DEL-51 |
| chr1-99273096-DEL-6587 | chr1-99273097-DEL-6587 |
| chr1-99283115-DEL-84 | chr1-99283116-DEL-78 |
| chr1-99285603-DEL-59 | chr1-99285605-DEL-57 |
| chr1-99296286-DEL-412 | chr1-99296287-DEL-414 |
| chr1-99299222-DEL-401 | chr1-99299222-DEL-401 |
| chr1-99309633-DEL-54 | chr1-99309618-DEL-75 |
| chr1-99324089-DEL-1962 | chr1-99324090-DEL-1962 |
| chr1-99353916-DEL-114 | chr1-99353916-DEL-114 |
| chr1-99358647-DEL-108 | chr1-99358648-DEL-108 |
| chr1-99362914-DEL-201 | chr1-99362914-DEL-201 |
| chr1-99365278-DEL-1806 | chr1-99365280-DEL-1807 |
| chr1-99370085-DEL-76 | chr1-99370088-DEL-59 |
| chr1-99379068-DEL-245 | chr1-99379068-DEL-245 |
| chr1-99385801-DEL-1470 | chr1-99385801-DEL-1470 |
| chr1-99435304-DEL-87 | chr1-99435308-DEL-96 |
| chr1-99446571-DEL-61 | chr1-99446572-DEL-61 |

| | |
|---|---|
| chr1-99447809-DEL-101 | chr1-99447830-DEL-101 |
| chr1-99454524-DEL-232 | chr1-99454525-DEL-232 |
| chr1-99455775-DEL-274 | chr1-99455776-DEL-273 |
| chr1-99462286-DEL-895 | chr1-99462286-DEL-895 |
| chr1-99470103-DEL-4176 | chr1-99470101-DEL-4176 |
| chr1-99479945-DEL-984 | chr1-99479946-DEL-984 |
| chr1-99481281-DEL-98 | chr1-99481292-DEL-73 |
| chr1-99488957-DEL-872 | chr1-99488965-DEL-874 |
| chr1-99490082-DEL-268 | chr1-99490083-DEL-268 |
| chr1-99491649-DEL-1469 | chr1-99491650-DEL-1469 |
| chr1-99515893-DEL-53 | chr1-99515894-DEL-53 |
| chr1-99541166-DEL-1935 | chr1-99541165-DEL-1936 |
| chr1-99558814-DEL-300 | chr1-99558808-DEL-323 |
| chr1-99560586-DEL-62 | chr1-99560587-DEL-62 |
| chr1-99562758-DEL-441 | chr1-99562738-DEL-441 |
| chr1-99575891-DEL-2186 | chr1-99575892-DEL-2186 |
| chr1-99582107-DEL-64 | chr1-99582120-DEL-52 |
| chr1-99586893-DEL-1066 | chr1-99586893-DEL-1066 |
| chr1-99590800-DEL-51 | chr1-99590798-DEL-54 |
| chr1-99629360-DEL-1348 | chr1-99629361-DEL-1348 |
| chr1-99635673-DEL-2789 | chr1-99635674-DEL-2789 |
| chr1-99658039-DEL-1167 | chr1-99658039-DEL-1169 |
| chr1-99662567-DEL-193 | chr1-99662568-DEL-193 |
| chr1-99664703-DEL-52 | chr1-99664704-DEL-54 |
| chr1-99685438-DEL-210 | chr1-99685439-DEL-210 |
| chr1-99706219-DEL-394 | chr1-99706220-DEL-393 |
| chr1-99718672-DEL-2728 | chr1-99718673-DEL-2728 |
| chr1-99722945-DEL-83 | chr1-99722920-DEL-127 |
| chr1-99724014-DEL-548 | chr1-99724015-DEL-548 |
| chr1-99745629-DEL-6398 | chr1-99745630-DEL-6398 |
| chr1-99761903-DEL-86 | chr1-99761884-DEL-122 |
| chr1-99786793-DEL-62 | chr1-99786796-DEL-54 |
| chr1-99789059-DEL-243 | chr1-99789060-DEL-243 |
| chr1-99819638-DEL-344 | chr1-99819639-DEL-344 |
| chr1-99885928-DEL-180 | chr1-99885914-DEL-198 |
| chr1-99901106-DEL-167 | chr1-99901107-DEL-167 |
| chr1-99919814-DEL-56 | chr1-99919830-DEL-73 |
| chr1-99938678-DEL-1044 | chr1-99938676-DEL-1045 |
| chr1-99951565-DEL-399 | chr1-99951566-DEL-399 |
| chr1-99953429-DEL-1563 | chr1-99953430-DEL-1563 |
| chr1-99956941-DEL-76 | chr1-99956942-DEL-76 |
| chr1-99990312-DEL-402 | chr1-99990313-DEL-402 |
| chr1-100009893-DEL-248 | chr1-100009899-DEL-245 |
| chr1-100021712-DEL-129 | chr1-100021712-DEL-129 |
| chr1-100046943-DEL-124 | chr1-100046944-DEL-124 |
| chr1-100050693-DEL-1608 | chr1-100050693-DEL-1608 |
| chr1-100059916-DEL-296 | chr1-100059917-DEL-296 |
| chr1-100063108-DEL-636 | chr1-100063109-DEL-636 |
| chr1-100066082-DEL-191 | chr1-100066083-DEL-188 |
| chr1-100110851-DEL-921 | chr1-100110852-DEL-921 |
| chr1-100147304-DEL-72 | chr1-100147325-DEL-62 |
| chr1-100151541-DEL-8324 | chr1-100151542-DEL-8324 |
| chr1-100160654-DEL-58 | chr1-100160655-DEL-58 |
| chr1-100168206-DEL-75 | chr1-100168226-DEL-62 |
| chr1-100198341-DEL-76 | chr1-100198342-DEL-76 |
| chr1-100234373-DEL-1285 | chr1-100234373-DEL-1284 |
| chr1-100240230-DEL-52 | chr1-100240231-DEL-52 |
| chr1-100242008-DEL-6451 | chr1-100242009-DEL-6451 |
| chr1-100248894-DEL-88 | chr1-100248910-DEL-67 |
| chr1-100249679-DEL-138 | chr1-100249680-DEL-138 |
| chr1-100260307-DEL-62 | chr1-100260308-DEL-68 |
| chr1-100278636-DEL-1765 | chr1-100278637-DEL-1765 |
| chr1-100285060-DEL-831 | chr1-100285061-DEL-831 |
| chr1-100303506-DEL-2009 | chr1-100303507-DEL-2009 |
| chr1-100307159-DEL-1093 | chr1-100307160-DEL-1093 |
| chr1-100321208-DEL-1001 | chr1-100321209-DEL-1001 |
| chr1-100335002-DEL-3080 | chr1-100335003-DEL-3080 |
| chr1-100376527-DEL-188 | chr1-100376528-DEL-188 |
| chr1-100377882-DEL-100 | chr1-100377917-DEL-103 |
| chr1-100378591-DEL-100 | chr1-100378592-DEL-100 |
| chr1-100430007-DEL-1470 | chr1-100430008-DEL-1470 |
| chr1-100436909-DEL-1367 | chr1-100436910-DEL-1365 |
| chr1-100451656-DEL-277 | chr1-100451657-DEL-277 |
| chr1-100458838-DEL-78 | chr1-100458839-DEL-78 |
| chr1-100459731-DEL-70 | chr1-100459737-DEL-51 |
| chr1-100572476-DEL-52 | chr1-100572467-DEL-50 |
| chr1-100574716-DEL-248 | chr1-100574717-DEL-248 |
| chr1-100613703-DEL-240 | chr1-100613704-DEL-240 |
| chr1-100617066-DEL-645 | chr1-100617067-DEL-645 |
| chr1-100641838-DEL-421 | chr1-100641839-DEL-415 |

| | |
|---|---|
| chr1-100649249-DEL-138 | chr1-100649274-DEL-113 |
| chr1-100665686-DEL-452 | chr1-100665687-DEL-452 |
| chr1-100682014-DEL-7029 | chr1-100682015-DEL-7029 |
| chr1-100682621-DEL-208 | chr1-100682622-DEL-208 |
| chr1-100690501-DEL-608 | chr1-100690502-DEL-610 |
| chr1-100694103-DEL-1129 | chr1-100694100-DEL-1130 |
| chr1-100712236-DEL-441 | chr1-100712237-DEL-441 |
| chr1-100731206-DEL-63 | chr1-100731192-DEL-66 |
| chr1-100736728-DEL-116 | chr1-100736728-DEL-114 |
| chr1-100756752-DEL-1919 | chr1-100756753-DEL-1919 |
| chr1-100776469-DEL-619 | chr1-100776470-DEL-619 |
| chr1-100787978-DEL-9565 | chr1-100787979-DEL-9565 |
| chr1-100801834-DEL-826 | chr1-100801835-DEL-826 |
| chr1-100802924-DEL-175 | chr1-100802925-DEL-175 |
| chr1-100834145-DEL-429 | chr1-100834146-DEL-429 |
| chr1-100835757-DEL-64 | chr1-100835752-DEL-70 |
| chr1-100867849-DEL-120 | chr1-100867851-DEL-119 |
| chr1-100871988-DEL-1387 | chr1-100872002-DEL-1388 |
| chr1-100882589-DEL-336 | chr1-100882590-DEL-336 |
| chr1-100914215-DEL-208 | chr1-100914216-DEL-208 |
| chr1-100929906-DEL-1654 | chr1-100929907-DEL-1654 |
| chr1-100944423-DEL-86 | chr1-100944424-DEL-86 |
| chr1-100967816-DEL-6171 | chr1-100967817-DEL-6171 |
| chr1-100991676-DEL-233 | chr1-100991677-DEL-233 |
| chr1-101023578-DEL-620 | chr1-101023579-DEL-620 |
| chr1-101049235-DEL-222 | chr1-101049236-DEL-222 |
| chr1-101055657-DEL-348 | chr1-101055658-DEL-348 |
| chr1-101059502-DEL-1495 | chr1-101059510-DEL-1487 |
| chr1-101061495-DEL-84 | chr1-101061494-DEL-74 |
| chr1-101073794-DEL-308 | chr1-101073795-DEL-303 |
| chr1-101075696-DEL-2252 | chr1-101075697-DEL-2252 |
| chr1-101078670-DEL-211 | chr1-101078671-DEL-211 |
| chr1-101121477-DEL-5847 | chr1-101121458-DEL-5867 |
| chr1-101133415-DEL-52 | chr1-101133416-DEL-52 |
| chr1-101160908-DEL-51 | chr1-101160909-DEL-51 |
| chr1-101172807-DEL-3094 | chr1-101172808-DEL-3094 |
| chr1-101190937-DEL-6184 | chr1-101190937-DEL-6185 |
| chr1-101230127-DEL-6466 | chr1-101230128-DEL-6466 |
| chr1-101243078-DEL-1334 | chr1-101243074-DEL-1335 |
| chr1-101258325-DEL-6334 | chr1-101258326-DEL-6334 |
| chr1-101294560-DEL-109 | chr1-101294561-DEL-109 |
| chr1-101302030-DEL-198 | chr1-101302031-DEL-198 |
| chr1-101322013-DEL-161 | chr1-101322014-DEL-161 |
| chr1-101325980-DEL-56 | chr1-101325981-DEL-56 |
| chr1-101332628-DEL-1767 | chr1-101332629-DEL-1767 |
| chr1-101345342-DEL-3430 | chr1-101345343-DEL-3430 |
| chr1-101360880-DEL-64 | chr1-101360881-DEL-68 |
| chr1-101383340-DEL-6171 | chr1-101383341-DEL-6171 |
| chr1-101390643-DEL-76 | chr1-101390649-DEL-64 |
| chr1-101396486-DEL-52 | chr1-101396487-DEL-52 |
| chr1-101403504-DEL-336 | chr1-101403507-DEL-335 |
| chr1-101454002-DEL-344 | chr1-101454003-DEL-344 |
| chr1-101463652-DEL-375 | chr1-101463653-DEL-375 |
| chr1-101498161-DEL-123 | chr1-101498162-DEL-123 |
| chr1-101506881-DEL-206 | chr1-101506882-DEL-206 |
| chr1-101513857-DEL-62 | chr1-101513858-DEL-58 |
| chr1-101524396-DEL-435 | chr1-101524397-DEL-435 |
| chr1-101528487-DEL-6600 | chr1-101528488-DEL-6600 |
| chr1-101562925-DEL-95 | chr1-101562935-DEL-95 |
| chr1-101593221-DEL-2069 | chr1-101593221-DEL-2069 |
| chr1-101668410-DEL-57 | chr1-101668411-DEL-57 |
| chr1-101671478-DEL-286 | chr1-101671479-DEL-286 |
| chr1-101672913-DEL-58 | chr1-101672914-DEL-58 |
| chr1-101680555-DEL-91 | chr1-101680522-DEL-87 |
| chr1-101684711-DEL-163 | chr1-101684712-DEL-163 |
| chr1-101686077-DEL-6421 | chr1-101686078-DEL-6421 |
| chr1-101739324-DEL-95 | chr1-101739366-DEL-62 |
| chr1-101767065-DEL-2095 | chr1-101767065-DEL-2096 |
| chr1-101783645-DEL-64 | chr1-101783646-DEL-64 |
| chr1-101787031-DEL-5711 | chr1-101787032-DEL-5711 |
| chr1-101803210-DEL-101 | chr1-101803236-DEL-52 |
| chr1-101814705-DEL-82 | chr1-101814706-DEL-82 |
| chr1-101846288-DEL-62 | chr1-101846289-DEL-64 |
| chr1-101848377-DEL-195 | chr1-101848378-DEL-195 |
| chr1-101855420-DEL-1053 | chr1-101855419-DEL-1053 |
| chr1-101856209-DEL-6783 | chr1-101856210-DEL-6783 |
| chr1-101905887-DEL-7134 | chr1-101905888-DEL-7134 |
| chr1-101917291-DEL-2550 | chr1-101917292-DEL-2554 |
| chr1-101922424-DEL-498 | chr1-101922424-DEL-498 |
| chr1-101925075-DEL-348 | chr1-101925076-DEL-348 |

| | |
|---|---|
| chr1-101948375-DEL-52 | chr1-101948376-DEL-52 |
| chr1-101958644-DEL-182 | chr1-101958683-DEL-142 |
| chr1-101961047-DEL-292 | chr1-101961048-DEL-292 |
| chr1-101962364-DEL-59 | chr1-101962362-DEL-61 |
| chr1-101971905-DEL-303 | chr1-101971905-DEL-303 |
| chr1-101972762-DEL-758 | chr1-101972762-DEL-758 |
| chr1-102020958-DEL-201 | chr1-102020959-DEL-201 |
| chr1-102075984-DEL-216 | chr1-102075984-DEL-216 |
| chr1-102077327-DEL-2868 | chr1-102077323-DEL-2869 |
| chr1-102086672-DEL-61 | chr1-102086676-DEL-64 |
| chr1-102089712-DEL-871 | chr1-102089713-DEL-871 |
| chr1-102122190-DEL-1893 | chr1-102122191-DEL-1893 |
| chr1-102129137-DEL-399 | chr1-102129138-DEL-399 |
| chr1-102132552-DEL-1531 | chr1-102132551-DEL-1532 |
| chr1-102144550-DEL-54 | chr1-102144547-DEL-54 |
| chr1-102169982-DEL-1063 | chr1-102169978-DEL-1067 |
| chr1-102171593-DEL-12735 | chr1-102171591-DEL-12736 |
| chr1-102177241-DEL-6132 | chr1-102177242-DEL-6132 |
| chr1-102193711-DEL-242 | chr1-102193712-DEL-242 |
| chr1-102260540-DEL-118 | chr1-102260541-DEL-118 |
| chr1-102282565-DEL-52 | chr1-102282566-DEL-52 |
| chr1-102316398-DEL-6181 | chr1-102316399-DEL-6181 |
| chr1-102323630-DEL-177 | chr1-102323631-DEL-176 |
| chr1-102347244-DEL-67 | chr1-102347267-DEL-69 |
| chr1-102362937-DEL-274 | chr1-102362937-DEL-274 |
| chr1-102387438-DEL-562 | chr1-102387440-DEL-563 |
| chr1-102389657-DEL-2998 | chr1-102389658-DEL-2998 |
| chr1-102404912-DEL-4161 | chr1-102404913-DEL-4161 |
| chr1-102419108-DEL-73 | chr1-102419108-DEL-73 |
| chr1-102436331-DEL-72 | chr1-102436331-DEL-72 |
| chr1-102456213-DEL-1524 | chr1-102456212-DEL-1526 |
| chr1-102531542-DEL-1989 | chr1-102531543-DEL-1989 |
| chr1-102535060-DEL-214 | chr1-102535061-DEL-214 |
| chr1-102539947-DEL-366 | chr1-102539947-DEL-366 |
| chr1-102557424-DEL-977 | chr1-102557425-DEL-977 |
| chr1-102569930-DEL-2121 | chr1-102569931-DEL-2121 |
| chr1-102613558-DEL-11753 | chr1-102613559-DEL-11753 |
| chr1-102623824-DEL-93 | chr1-102623825-DEL-93 |
| chr1-102631146-DEL-108 | chr1-102631147-DEL-108 |
| chr1-102654531-DEL-54 | chr1-102654534-DEL-91 |
| chr1-102659763-DEL-769 | chr1-102659764-DEL-769 |
| chr1-102670692-DEL-2064 | chr1-102670693-DEL-2064 |
| chr1-102684149-DEL-624 | chr1-102684150-DEL-624 |
| chr1-102690225-DEL-384 | chr1-102690226-DEL-384 |
| chr1-102692566-DEL-565 | chr1-102692567-DEL-565 |
| chr1-102697505-DEL-2098 | chr1-102697505-DEL-2097 |
| chr1-102707651-DEL-289 | chr1-102707652-DEL-289 |
| chr1-102723823-DEL-368 | chr1-102723823-DEL-368 |
| chr1-102738306-DEL-52 | chr1-102738306-DEL-52 |
| chr1-102745742-DEL-468 | chr1-102745736-DEL-474 |
| chr1-102749962-DEL-2444 | chr1-102749955-DEL-2448 |
| chr1-102757389-DEL-452 | chr1-102757389-DEL-452 |
| chr1-102759926-DEL-1443 | chr1-102759926-DEL-1443 |
| chr1-102761923-DEL-691 | chr1-102761923-DEL-691 |
| chr1-102765416-DEL-142 | chr1-102765416-DEL-144 |
| chr1-102777482-DEL-92 | chr1-102777479-DEL-91 |
| chr1-102787834-DEL-87 | chr1-102787835-DEL-87 |
| chr1-102788541-DEL-358 | chr1-102788542-DEL-358 |
| chr1-102789890-DEL-135 | chr1-102789891-DEL-135 |
| chr1-102794438-DEL-853 | chr1-102794439-DEL-853 |
| chr1-102818238-DEL-58 | chr1-102818239-DEL-58 |
| chr1-102825317-DEL-113 | chr1-102825317-DEL-111 |
| chr1-102836931-DEL-491 | chr1-102836932-DEL-491 |
| chr1-102859056-DEL-916 | chr1-102859056-DEL-916 |
| chr1-102869542-DEL-88 | chr1-102869543-DEL-86 |
| chr1-102870753-DEL-54 | chr1-102870772-DEL-56 |
| chr1-102891018-DEL-199 | chr1-102891019-DEL-199 |
| chr1-102897812-DEL-88 | chr1-102897802-DEL-56 |
| chr1-102912974-DEL-115 | chr1-102912968-DEL-104 |
| chr1-102941338-DEL-71 | chr1-102941354-DEL-62 |
| chr1-102953653-DEL-859 | chr1-102953654-DEL-859 |
| chr1-102977011-DEL-54 | chr1-102977012-DEL-54 |
| chr1-102978574-DEL-168 | chr1-102978575-DEL-168 |
| chr1-102990609-DEL-1441 | chr1-102990610-DEL-1441 |
| chr1-102994690-DEL-846 | chr1-102994691-DEL-846 |
| chr1-103006682-DEL-79 | chr1-103006663-DEL-105 |
| chr1-103031878-DEL-69 | chr1-103031888-DEL-54 |
| chr1-103037404-DEL-9102 | chr1-103037404-DEL-9102 |
| chr1-103049720-DEL-1926 | chr1-103049720-DEL-1926 |
| chr1-103062936-DEL-153 | chr1-103062937-DEL-153 |

| | |
|---|---|
| chr1-103075486-DEL-6658 | chr1-103075487-DEL-6658 |
| chr1-103082642-DEL-1881 | chr1-103082635-DEL-1894 |
| chr1-103085006-DEL-1780 | chr1-103085007-DEL-1780 |
| chr1-103095234-DEL-283 | chr1-103095235-DEL-283 |
| chr1-103096587-DEL-74 | chr1-103096590-DEL-72 |
| chr1-103122147-DEL-221 | chr1-103122148-DEL-216 |
| chr1-103132884-DEL-7131 | chr1-103132885-DEL-7131 |
| chr1-103144734-DEL-1726 | chr1-103144734-DEL-1726 |
| chr1-103166702-DEL-98 | chr1-103166715-DEL-86 |
| chr1-103174303-DEL-2375 | chr1-103174304-DEL-2375 |
| chr1-103183376-DEL-1696 | chr1-103183377-DEL-1696 |
| chr1-103188576-DEL-65 | chr1-103188577-DEL-65 |
| chr1-103210411-DEL-214 | chr1-103210411-DEL-214 |
| chr1-103233440-DEL-134 | chr1-103233441-DEL-134 |
| chr1-103235433-DEL-508 | chr1-103235434-DEL-508 |
| chr1-103289293-DEL-78 | chr1-103289294-DEL-78 |
| chr1-103303423-DEL-449 | chr1-103303423-DEL-449 |
| chr1-103307162-DEL-687 | chr1-103307153-DEL-701 |
| chr1-103308758-DEL-193 | chr1-103308758-DEL-193 |
| chr1-103319734-DEL-55 | chr1-103319735-DEL-55 |
| chr1-103383307-DEL-2226 | chr1-103383308-DEL-2227 |
| chr1-103406546-DEL-2640 | chr1-103406547-DEL-2640 |
| chr1-103427275-DEL-118 | chr1-103427275-DEL-118 |
| chr1-103443257-DEL-68 | chr1-103443257-DEL-68 |
| chr1-103452090-DEL-73 | chr1-103452096-DEL-67 |
| chr1-103482731-DEL-411 | chr1-103482761-DEL-382 |
| chr1-103497862-DEL-229 | chr1-103497863-DEL-229 |
| chr1-103505050-DEL-54 | chr1-103505051-DEL-54 |
| chr1-103512092-DEL-69 | chr1-103512096-DEL-57 |
| chr1-103512417-DEL-11323 | chr1-103512418-DEL-11323 |
| chr1-103516432-DEL-5360 | chr1-103516433-DEL-5360 |
| chr1-103538940-DEL-1508 | chr1-103538941-DEL-1508 |
| chr1-103540875-DEL-336 | chr1-103540876-DEL-336 |
| chr1-103541900-DEL-1607 | chr1-103541901-DEL-1607 |
| chr1-103542624-DEL-550 | chr1-103542624-DEL-550 |
| chr1-103580056-DEL-1907 | chr1-103580057-DEL-1907 |
| chr1-103594095-DEL-60 | chr1-103594096-DEL-60 |
| chr1-103596886-DEL-184 | chr1-103596886-DEL-184 |
| chr1-103612605-DEL-92 | chr1-103612592-DEL-106 |
| chr1-103614515-DEL-401 | chr1-103614516-DEL-401 |
| chr1-103621971-DEL-889 | chr1-103621972-DEL-889 |
| chr1-103630384-DEL-2053 | chr1-103630385-DEL-2053 |
| chr1-103669627-DEL-4505 | chr1-103669627-DEL-4505 |
| chr1-103674709-DEL-182 | chr1-103674710-DEL-182 |
| chr1-103677663-DEL-72 | chr1-103677676-DEL-60 |
| chr1-103677978-DEL-612 | chr1-103677978-DEL-613 |
| chr1-103680616-DEL-430 | chr1-103680599-DEL-448 |
| chr1-103684435-DEL-86 | chr1-103684436-DEL-86 |
| chr1-103720395-DEL-480 | chr1-103720396-DEL-480 |
| chr1-103739328-DEL-447 | chr1-103739329-DEL-447 |
| chr1-103744790-DEL-56 | chr1-103744742-DEL-66 |
| chr1-103775069-DEL-293 | chr1-103775070-DEL-293 |
| chr1-103782699-DEL-122 | chr1-103782689-DEL-127 |
| chr1-103833783-DEL-287 | chr1-103833784-DEL-287 |
| chr1-103836648-DEL-135 | chr1-103836649-DEL-135 |
| chr1-103837599-DEL-270 | chr1-103837600-DEL-270 |
| chr1-103841238-DEL-497 | chr1-103841239-DEL-497 |
| chr1-103847614-DEL-6620 | chr1-103847615-DEL-6620 |
| chr1-103855530-DEL-52 | chr1-103855503-DEL-55 |
| chr1-103859758-DEL-920 | chr1-103859751-DEL-917 |
| chr1-103865179-DEL-790 | chr1-103865180-DEL-790 |
| chr1-103897444-DEL-65 | chr1-103897431-DEL-66 |
| chr1-103910956-DEL-61 | chr1-103910963-DEL-51 |
| chr1-103927117-DEL-200 | chr1-103927118-DEL-200 |
| chr1-103964341-DEL-393 | chr1-103964341-DEL-393 |
| chr1-103990377-DEL-61 | chr1-103990378-DEL-61 |
| chr1-104086787-DEL-64 | chr1-104086788-DEL-64 |
| chr1-104090460-DEL-576 | chr1-104090461-DEL-576 |
| chr1-104132638-DEL-197 | chr1-104132639-DEL-197 |
| chr1-104133663-DEL-1687 | chr1-104133660-DEL-1705 |
| chr1-104144509-DEL-69 | chr1-104144510-DEL-69 |
| chr1-104145374-DEL-7031 | chr1-104145375-DEL-7031 |
| chr1-104177676-DEL-179 | chr1-104177677-DEL-179 |
| chr1-104217296-DEL-5244 | chr1-104217297-DEL-5244 |
| chr1-104236968-DEL-235 | chr1-104236969-DEL-235 |
| chr1-104238912-DEL-53 | chr1-104238913-DEL-53 |
| chr1-104247038-DEL-261 | chr1-104247071-DEL-236 |
| chr1-104292079-DEL-2085 | chr1-104292080-DEL-2085 |
| chr1-104296381-DEL-204 | chr1-104296382-DEL-204 |
| chr1-104370399-DEL-152 | chr1-104370400-DEL-152 |

| | |
|---|---|
| chr1-104374310-DEL-100 | chr1-104374328-DEL-65 |
| chr1-104508881-DEL-3032 | chr1-104508882-DEL-3032 |
| chr1-104525974-DEL-86 | chr1-104525977-DEL-70 |
| chr1-104554237-DEL-184 | chr1-104554229-DEL-185 |
| chr1-104565347-DEL-2157 | chr1-104565348-DEL-2157 |
| chr1-104570090-DEL-977 | chr1-104570092-DEL-974 |
| chr1-104597737-DEL-58 | chr1-104597698-DEL-88 |
| chr1-104598791-DEL-519 | chr1-104598792-DEL-519 |
| chr1-104601731-DEL-382 | chr1-104601731-DEL-379 |
| chr1-104672160-DEL-63 | chr1-104672161-DEL-63 |
| chr1-104689216-DEL-1081 | chr1-104689217-DEL-1081 |
| chr1-104701842-DEL-109 | chr1-104701842-DEL-110 |
| chr1-104759745-DEL-83 | chr1-104759746-DEL-83 |
| chr1-104802089-DEL-201 | chr1-104802088-DEL-202 |
| chr1-104821701-DEL-361 | chr1-104821702-DEL-361 |
| chr1-104853109-DEL-1255 | chr1-104853110-DEL-1255 |
| chr1-104890332-DEL-7001 | chr1-104890335-DEL-7003 |
| chr1-104904726-DEL-69 | chr1-104904727-DEL-69 |
| chr1-104907384-DEL-77 | chr1-104907377-DEL-78 |
| chr1-104910545-DEL-184 | chr1-104910549-DEL-180 |
| chr1-104941408-DEL-7342 | chr1-104941409-DEL-7342 |
| chr1-104947812-DEL-212 | chr1-104947813-DEL-212 |
| chr1-104954651-DEL-206 | chr1-104954652-DEL-206 |
| chr1-104957571-DEL-376 | chr1-104957572-DEL-376 |
| chr1-104965366-DEL-132 | chr1-104965367-DEL-122 |
| chr1-104970200-DEL-180 | chr1-104970199-DEL-180 |
| chr1-104977531-DEL-159 | chr1-104977524-DEL-167 |
| chr1-105107545-DEL-56 | chr1-105107555-DEL-63 |
| chr1-105129258-DEL-2175 | chr1-105129259-DEL-2175 |
| chr1-105131703-DEL-237 | chr1-105131704-DEL-237 |
| chr1-105132245-DEL-67 | chr1-105132243-DEL-67 |
| chr1-105132557-DEL-483 | chr1-105132558-DEL-481 |
| chr1-105143670-DEL-241 | chr1-105143671-DEL-241 |
| chr1-105146796-DEL-443 | chr1-105146797-DEL-443 |
| chr1-105154449-DEL-3984 | chr1-105154441-DEL-3997 |
| chr1-105180073-DEL-56 | chr1-105180078-DEL-52 |
| chr1-105220079-DEL-61 | chr1-105220082-DEL-59 |
| chr1-105241781-DEL-83 | chr1-105241796-DEL-69 |
| chr1-105251498-DEL-52 | chr1-105251499-DEL-52 |
| chr1-105296812-DEL-100 | chr1-105296813-DEL-100 |
| chr1-105311533-DEL-118 | chr1-105311500-DEL-110 |
| chr1-105316497-DEL-1890 | chr1-105316498-DEL-1890 |
| chr1-105334019-DEL-63 | chr1-105334020-DEL-63 |
| chr1-105341650-DEL-2545 | chr1-105341651-DEL-2545 |
| chr1-105369200-DEL-1942 | chr1-105369201-DEL-1942 |
| chr1-105369708-DEL-58 | chr1-105369709-DEL-58 |
| chr1-105383651-DEL-2981 | chr1-105383652-DEL-2981 |
| chr1-105388485-DEL-108 | chr1-105388486-DEL-108 |
| chr1-105405111-DEL-203 | chr1-105405112-DEL-203 |
| chr1-105457051-DEL-581 | chr1-105457052-DEL-581 |
| chr1-105460178-DEL-943 | chr1-105460179-DEL-943 |
| chr1-105495742-DEL-6409 | chr1-105495743-DEL-6409 |
| chr1-105521759-DEL-169 | chr1-105521760-DEL-169 |
| chr1-105537519-DEL-5940 | chr1-105537519-DEL-5941 |
| chr1-105573038-DEL-252 | chr1-105573039-DEL-252 |
| chr1-105597182-DEL-211 | chr1-105597183-DEL-211 |
| chr1-105633063-DEL-771 | chr1-105633064-DEL-771 |
| chr1-105646749-DEL-208 | chr1-105646750-DEL-208 |
| chr1-105660188-DEL-216 | chr1-105660189-DEL-216 |
| chr1-105711815-DEL-86 | chr1-105711798-DEL-84 |
| chr1-105753989-DEL-220 | chr1-105753990-DEL-220 |
| chr1-105757207-DEL-106 | chr1-105757208-DEL-106 |
| chr1-105761069-DEL-6275 | chr1-105761070-DEL-6275 |
| chr1-105769778-DEL-401 | chr1-105769779-DEL-401 |
| chr1-105787985-DEL-1112 | chr1-105787986-DEL-1112 |
| chr1-105795257-DEL-286 | chr1-105795244-DEL-296 |
| chr1-105834575-DEL-83 | chr1-105834566-DEL-81 |
| chr1-105846447-DEL-95 | chr1-105846448-DEL-93 |
| chr1-105857034-DEL-64 | chr1-105857035-DEL-62 |
| chr1-105865881-DEL-338 | chr1-105865879-DEL-339 |
| chr1-105869460-DEL-74 | chr1-105869461-DEL-74 |
| chr1-105870089-DEL-85 | chr1-105870090-DEL-85 |
| chr1-105873518-DEL-215 | chr1-105873519-DEL-215 |
| chr1-105884702-DEL-155 | chr1-105884703-DEL-155 |
| chr1-105888778-DEL-76 | chr1-105888779-DEL-76 |
| chr1-105889855-DEL-476 | chr1-105889856-DEL-476 |
| chr1-105971963-DEL-200 | chr1-105971964-DEL-200 |
| chr1-105975210-DEL-61 | chr1-105975211-DEL-61 |
| chr1-105996190-DEL-377 | chr1-105996191-DEL-377 |
| chr1-106007003-DEL-1570 | chr1-106007004-DEL-1570 |

| | |
|---|---|
| chr1-106015907-DEL-400 | chr1-106015908-DEL-400 |
| chr1-106032855-DEL-355 | chr1-106032856-DEL-355 |
| chr1-106040701-DEL-6689 | chr1-106040702-DEL-6689 |
| chr1-106077688-DEL-122 | chr1-106077689-DEL-122 |
| chr1-106109404-DEL-203 | chr1-106109405-DEL-203 |
| chr1-106179310-DEL-974 | chr1-106179311-DEL-974 |
| chr1-106221441-DEL-7023 | chr1-106221442-DEL-7023 |
| chr1-106251363-DEL-642 | chr1-106251364-DEL-642 |
| chr1-106298278-DEL-58 | chr1-106298279-DEL-58 |
| chr1-106316328-DEL-64 | chr1-106316329-DEL-64 |
| chr1-106316762-DEL-95 | chr1-106316767-DEL-106 |
| chr1-106349201-DEL-62 | chr1-106349202-DEL-62 |
| chr1-106371937-DEL-837 | chr1-106371938-DEL-837 |
| chr1-106391199-DEL-1032 | chr1-106391200-DEL-1032 |
| chr1-106397106-DEL-682 | chr1-106397106-DEL-712 |
| chr1-106400726-DEL-240 | chr1-106400727-DEL-240 |
| chr1-106447507-DEL-116 | chr1-106447548-DEL-94 |
| chr1-106459870-DEL-55 | chr1-106459871-DEL-55 |
| chr1-106542698-DEL-62 | chr1-106542698-DEL-66 |
| chr1-106569680-DEL-276 | chr1-106569681-DEL-276 |
| chr1-106609984-DEL-60 | chr1-106609987-DEL-58 |
| chr1-106620065-DEL-322 | chr1-106620066-DEL-322 |
| chr1-106627975-DEL-184 | chr1-106627960-DEL-200 |
| chr1-106650933-DEL-63 | chr1-106650934-DEL-63 |
| chr1-106658434-DEL-2881 | chr1-106658435-DEL-2881 |
| chr1-106665114-DEL-214 | chr1-106665114-DEL-214 |
| chr1-106673251-DEL-62 | chr1-106673252-DEL-62 |
| chr1-106682155-DEL-77 | chr1-106682156-DEL-77 |
| chr1-106712070-DEL-519 | chr1-106712071-DEL-519 |
| chr1-106717515-DEL-92 | chr1-106717516-DEL-92 |
| chr1-106729447-DEL-5715 | chr1-106729448-DEL-5715 |
| chr1-106744818-DEL-244 | chr1-106744820-DEL-243 |
| chr1-106751763-DEL-408 | chr1-106751765-DEL-408 |
| chr1-106765734-DEL-122 | chr1-106765735-DEL-122 |
| chr1-106776531-DEL-6694 | chr1-106776565-DEL-6665 |
| chr1-106785128-DEL-54 | chr1-106785129-DEL-54 |
| chr1-106814303-DEL-71 | chr1-106814304-DEL-71 |
| chr1-106832100-DEL-100 | chr1-106832100-DEL-101 |
| chr1-106837154-DEL-108 | chr1-106837149-DEL-114 |
| chr1-106846526-DEL-281 | chr1-106846527-DEL-281 |
| chr1-106849363-DEL-2865 | chr1-106849364-DEL-2865 |
| chr1-106862958-DEL-62 | chr1-106862959-DEL-62 |
| chr1-106867550-DEL-341 | chr1-106867551-DEL-341 |
| chr1-106873769-DEL-62 | chr1-106873770-DEL-62 |
| chr1-106892848-DEL-69 | chr1-106892849-DEL-69 |
| chr1-106965364-DEL-1123 | chr1-106965365-DEL-1123 |
| chr1-107111184-DEL-392 | chr1-107111185-DEL-392 |
| chr1-107121043-DEL-67 | chr1-107121043-DEL-67 |
| chr1-107130416-DEL-6874 | chr1-107130417-DEL-6874 |
| chr1-107161155-DEL-202 | chr1-107161140-DEL-202 |
| chr1-107196015-DEL-486 | chr1-107196015-DEL-486 |
| chr1-107266255-DEL-140 | chr1-107266275-DEL-105 |
| chr1-107283841-DEL-376 | chr1-107283845-DEL-373 |
| chr1-107333919-DEL-124 | chr1-107333921-DEL-109 |
| chr1-107346305-DEL-72 | chr1-107346306-DEL-72 |
| chr1-107365313-DEL-1150 | chr1-107365314-DEL-1150 |
| chr1-107371504-DEL-55 | chr1-107371502-DEL-56 |
| chr1-107388268-DEL-229 | chr1-107388277-DEL-225 |
| chr1-107389331-DEL-995 | chr1-107389289-DEL-996 |
| chr1-107391741-DEL-489 | chr1-107391742-DEL-489 |
| chr1-107397974-DEL-71 | chr1-107397977-DEL-72 |
| chr1-107402247-DEL-2030 | chr1-107402248-DEL-2030 |
| chr1-107436905-DEL-60 | chr1-107436912-DEL-58 |
| chr1-107454442-DEL-348 | chr1-107454451-DEL-345 |
| chr1-107488413-DEL-198 | chr1-107488414-DEL-198 |
| chr1-107515832-DEL-82 | chr1-107515855-DEL-60 |
| chr1-107523754-DEL-60 | chr1-107523755-DEL-60 |
| chr1-107553904-DEL-61 | chr1-107553905-DEL-61 |
| chr1-107555980-DEL-4351 | chr1-107555981-DEL-4351 |
| chr1-107560712-DEL-401 | chr1-107560718-DEL-399 |
| chr1-107577290-DEL-75 | chr1-107577291-DEL-75 |
| chr1-107578686-DEL-54 | chr1-107578687-DEL-54 |
| chr1-107615197-DEL-186 | chr1-107615198-DEL-186 |
| chr1-107635641-DEL-60 | chr1-107635642-DEL-58 |
| chr1-107672376-DEL-155 | chr1-107672346-DEL-136 |
| chr1-107679489-DEL-4949 | chr1-107679490-DEL-4949 |
| chr1-107718719-DEL-924 | chr1-107718720-DEL-924 |
| chr1-107721583-DEL-6487 | chr1-107721584-DEL-6487 |
| chr1-107734037-DEL-80 | chr1-107733994-DEL-124 |
| chr1-107735274-DEL-74 | chr1-107735270-DEL-74 |

| | |
|---|---|
| chr1-107750506-DEL-7218 | chr1-107750507-DEL-7218 |
| chr1-107767905-DEL-123 | chr1-107767906-DEL-123 |
| chr1-107775938-DEL-70 | chr1-107775943-DEL-66 |
| chr1-107809044-DEL-9183 | chr1-107809057-DEL-9184 |
| chr1-107809219-DEL-416 | chr1-107809220-DEL-416 |
| chr1-107823379-DEL-315 | chr1-107823380-DEL-314 |
| chr1-107824667-DEL-81 | chr1-107824678-DEL-94 |
| chr1-107847681-DEL-6490 | chr1-107847682-DEL-6490 |
| chr1-107858780-DEL-919 | chr1-107858781-DEL-919 |
| chr1-107900978-DEL-683 | chr1-107900979-DEL-683 |
| chr1-107921908-DEL-1337 | chr1-107921909-DEL-1337 |
| chr1-107923668-DEL-401 | chr1-107923669-DEL-401 |
| chr1-107931113-DEL-69 | chr1-107931103-DEL-66 |
| chr1-107947263-DEL-64 | chr1-107947264-DEL-64 |
| chr1-107966525-DEL-2094 | chr1-107966533-DEL-2092 |
| chr1-107985480-DEL-215 | chr1-107985481-DEL-215 |
| chr1-107987902-DEL-82 | chr1-107987892-DEL-95 |
| chr1-107991046-DEL-518 | chr1-107991047-DEL-518 |
| chr1-108018717-DEL-66 | chr1-108018703-DEL-81 |
| chr1-108033608-DEL-357 | chr1-108033609-DEL-356 |
| chr1-108036554-DEL-491 | chr1-108036552-DEL-490 |
| chr1-108064919-DEL-2707 | chr1-108064920-DEL-2707 |
| chr1-108074827-DEL-80 | chr1-108074828-DEL-80 |
| chr1-108095863-DEL-950 | chr1-108095864-DEL-950 |
| chr1-108107704-DEL-190 | chr1-108107705-DEL-190 |
| chr1-108132388-DEL-1409 | chr1-108132389-DEL-1409 |
| chr1-108156046-DEL-262 | chr1-108156047-DEL-262 |
| chr1-108173586-DEL-64 | chr1-108173587-DEL-66 |
| chr1-108194470-DEL-52 | chr1-108194467-DEL-56 |
| chr1-108215106-DEL-6243 | chr1-108215107-DEL-6243 |
| chr1-108223733-DEL-280 | chr1-108223734-DEL-280 |
| chr1-108249122-DEL-7158 | chr1-108249123-DEL-7158 |
| chr1-108278750-DEL-6147 | chr1-108278751-DEL-6147 |
| chr1-108325826-DEL-204 | chr1-108325827-DEL-204 |
| chr1-108332780-DEL-11837 | chr1-108332766-DEL-11842 |
| chr1-108347051-DEL-1718 | chr1-108347052-DEL-1718 |
| chr1-108348307-DEL-1385 | chr1-108348308-DEL-1387 |
| chr1-108373986-DEL-501 | chr1-108373984-DEL-501 |
| chr1-108378706-DEL-54 | chr1-108378707-DEL-70 |
| chr1-108384975-DEL-56 | chr1-108384972-DEL-58 |
| chr1-108387139-DEL-58 | chr1-108387139-DEL-58 |
| chr1-108415886-DEL-51 | chr1-108415856-DEL-52 |
| chr1-108431232-DEL-839 | chr1-108431234-DEL-838 |
| chr1-108445637-DEL-6228 | chr1-108445638-DEL-6228 |
| chr1-108488123-DEL-55 | chr1-108488124-DEL-55 |
| chr1-108538107-DEL-64 | chr1-108538108-DEL-68 |
| chr1-108552381-DEL-203 | chr1-108552382-DEL-203 |
| chr1-108571870-DEL-60 | chr1-108571851-DEL-56 |
| chr1-108590245-DEL-390 | chr1-108590246-DEL-390 |
| chr1-108592497-DEL-108 | chr1-108592498-DEL-108 |
| chr1-108612348-DEL-750 | chr1-108612349-DEL-750 |
| chr1-108616156-DEL-2515 | chr1-108616171-DEL-2506 |
| chr1-108630753-DEL-214 | chr1-108630754-DEL-214 |
| chr1-108633348-DEL-51 | chr1-108633349-DEL-51 |
| chr1-108640660-DEL-835 | chr1-108640665-DEL-831 |
| chr1-108660966-DEL-6460 | chr1-108660967-DEL-6460 |
| chr1-108669826-DEL-1061 | chr1-108669827-DEL-1061 |
| chr1-108676125-DEL-236 | chr1-108676126-DEL-236 |
| chr1-108693065-DEL-93 | chr1-108693046-DEL-85 |
| chr1-108708387-DEL-4435 | chr1-108708387-DEL-4435 |
| chr1-108720727-DEL-470 | chr1-108720728-DEL-470 |
| chr1-108773505-DEL-1236 | chr1-108773505-DEL-1226 |
| chr1-108777852-DEL-233 | chr1-108777853-DEL-233 |
| chr1-108785831-DEL-7926 | chr1-108785832-DEL-7926 |
| chr1-108798798-DEL-90 | chr1-108798799-DEL-90 |
| chr1-108803349-DEL-99 | chr1-108803350-DEL-99 |
| chr1-108806784-DEL-1026 | chr1-108806789-DEL-1022 |
| chr1-108808004-DEL-68 | chr1-108808005-DEL-68 |
| chr1-108838299-DEL-66 | chr1-108838270-DEL-58 |
| chr1-108839388-DEL-370 | chr1-108839389-DEL-370 |
| chr1-108848714-DEL-63 | chr1-108848714-DEL-63 |
| chr1-108865180-DEL-500 | chr1-108865181-DEL-500 |
| chr1-108867776-DEL-629 | chr1-108867776-DEL-627 |
| chr1-108869163-DEL-112 | chr1-108869164-DEL-112 |
| chr1-108872861-DEL-3968 | chr1-108872862-DEL-3968 |
| chr1-108895693-DEL-55 | chr1-108895694-DEL-55 |
| chr1-108896300-DEL-64 | chr1-108896300-DEL-64 |
| chr1-108914243-DEL-2826 | chr1-108914243-DEL-2826 |
| chr1-108925969-DEL-2166 | chr1-108925970-DEL-2166 |
| chr1-108928593-DEL-71 | chr1-108928570-DEL-72 |

| | |
|---|---|
| chr1-108943721-DEL-866 | chr1-108943722-DEL-866 |
| chr1-108962378-DEL-6850 | chr1-108962364-DEL-6875 |
| chr1-108962751-DEL-424 | chr1-108962752-DEL-424 |
| chr1-108977875-DEL-2188 | chr1-108977876-DEL-2188 |
| chr1-108985313-DEL-62 | chr1-108985347-DEL-56 |
| chr1-108992629-DEL-700 | chr1-108992630-DEL-700 |
| chr1-108994278-DEL-6473 | chr1-108994279-DEL-6473 |
| chr1-109028488-DEL-1120 | chr1-109028489-DEL-1120 |
| chr1-109031472-DEL-64 | chr1-109031473-DEL-64 |
| chr1-109056399-DEL-5262 | chr1-109056400-DEL-5262 |
| chr1-109071682-DEL-542 | chr1-109071684-DEL-528 |
| chr1-109085579-DEL-819 | chr1-109085580-DEL-819 |
| chr1-109095014-DEL-434 | chr1-109095015-DEL-434 |
| chr1-109097773-DEL-88 | chr1-109097762-DEL-101 |
| chr1-109113439-DEL-202 | chr1-109113440-DEL-202 |
| chr1-109115333-DEL-58 | chr1-109115334-DEL-58 |
| chr1-109127665-DEL-6188 | chr1-109127666-DEL-6188 |
| chr1-109185015-DEL-1628 | chr1-109185016-DEL-1628 |
| chr1-109219065-DEL-2688 | chr1-109219066-DEL-2688 |
| chr1-109249622-DEL-63 | chr1-109249623-DEL-63 |
| chr1-109262167-DEL-1316 | chr1-109262167-DEL-1316 |
| chr1-109263533-DEL-1004 | chr1-109263533-DEL-1004 |
| chr1-109265794-DEL-93 | chr1-109265795-DEL-93 |
| chr1-109271179-DEL-6450 | chr1-109271180-DEL-6450 |
| chr1-109277325-DEL-214 | chr1-109277326-DEL-214 |
| chr1-109286135-DEL-59 | chr1-109286135-DEL-59 |
| chr1-109298555-DEL-369 | chr1-109298556-DEL-369 |
| chr1-109303700-DEL-568 | chr1-109303679-DEL-575 |
| chr1-109317583-DEL-965 | chr1-109317584-DEL-965 |
| chr1-109329723-DEL-64 | chr1-109329721-DEL-60 |
| chr1-109338971-DEL-199 | chr1-109338972-DEL-199 |
| chr1-109351807-DEL-179 | chr1-109351808-DEL-179 |
| chr1-109354610-DEL-229 | chr1-109354611-DEL-229 |
| chr1-109368420-DEL-3602 | chr1-109368421-DEL-3602 |
| chr1-109374516-DEL-17345 | chr1-109374517-DEL-17345 |
| chr1-109386531-DEL-1260 | chr1-109386532-DEL-1260 |
| chr1-109393938-DEL-99 | chr1-109393938-DEL-99 |
| chr1-109410855-DEL-142 | chr1-109410856-DEL-142 |
| chr1-109429379-DEL-359 | chr1-109429379-DEL-359 |
| chr1-109429778-DEL-4716 | chr1-109429779-DEL-4716 |
| chr1-109438977-DEL-158 | chr1-109438978-DEL-158 |
| chr1-109447952-DEL-90 | chr1-109447955-DEL-88 |
| chr1-109453518-DEL-248 | chr1-109453519-DEL-249 |
| chr1-109469501-DEL-1977 | chr1-109469502-DEL-1977 |
| chr1-109491263-DEL-52 | chr1-109491280-DEL-52 |
| chr1-109496155-DEL-73 | chr1-109496156-DEL-73 |
| chr1-109504062-DEL-68 | chr1-109504071-DEL-60 |
| chr1-109518699-DEL-75 | chr1-109518700-DEL-75 |
| chr1-109532318-DEL-414 | chr1-109532319-DEL-414 |
| chr1-109539841-DEL-272 | chr1-109539842-DEL-272 |
| chr1-109541116-DEL-7146 | chr1-109541117-DEL-7146 |
| chr1-109557521-DEL-619 | chr1-109557522-DEL-619 |
| chr1-109608589-DEL-1280 | chr1-109608590-DEL-1280 |
| chr1-109644178-DEL-569 | chr1-109644179-DEL-569 |
| chr1-109678497-DEL-2560 | chr1-109678498-DEL-2560 |
| chr1-109704394-DEL-568 | chr1-109704395-DEL-568 |
| chr1-109708299-DEL-122 | chr1-109708300-DEL-122 |
| chr1-109719890-DEL-365 | chr1-109719891-DEL-365 |
| chr1-109747976-DEL-59 | chr1-109747976-DEL-63 |
| chr1-109753453-DEL-997 | chr1-109753454-DEL-996 |
| chr1-109771300-DEL-1804 | chr1-109771297-DEL-1806 |
| chr1-109799253-DEL-6636 | chr1-109799254-DEL-6636 |
| chr1-109808883-DEL-56 | chr1-109808884-DEL-56 |
| chr1-109820654-DEL-289 | chr1-109820655-DEL-289 |
| chr1-109836545-DEL-70 | chr1-109836546-DEL-70 |
| chr1-109935025-DEL-201 | chr1-109935026-DEL-201 |
| chr1-109952161-DEL-63 | chr1-109952162-DEL-63 |
| chr1-110019369-DEL-2514 | chr1-110019370-DEL-2514 |
| chr1-110069475-DEL-60 | chr1-110069476-DEL-60 |
| chr1-110080592-DEL-111 | chr1-110080602-DEL-102 |
| chr1-110098395-DEL-7743 | chr1-110098396-DEL-7743 |
| chr1-110108971-DEL-1241 | chr1-110108972-DEL-1241 |
| chr1-110112941-DEL-1728 | chr1-110112942-DEL-1728 |
| chr1-110131527-DEL-54 | chr1-110131525-DEL-52 |
| chr1-110158930-DEL-92 | chr1-110158934-DEL-88 |
| chr1-110189179-DEL-82 | chr1-110189198-DEL-64 |
| chr1-110192437-DEL-1054 | chr1-110192438-DEL-1054 |
| chr1-110212250-DEL-4415 | chr1-110212251-DEL-4415 |
| chr1-110232962-DEL-1701 | chr1-110232966-DEL-1698 |
| chr1-110234473-DEL-91 | chr1-110234474-DEL-91 |

| | |
|---|---|
| chr1-110256839-DEL-5151 | chr1-110256840-DEL-5151 |
| chr1-110315681-DEL-852 | chr1-110315682-DEL-852 |
| chr1-110328048-DEL-680 | chr1-110328049-DEL-680 |
| chr1-110349689-DEL-65 | chr1-110349690-DEL-65 |
| chr1-110355731-DEL-197 | chr1-110355728-DEL-195 |
| chr1-110360224-DEL-424 | chr1-110360225-DEL-423 |
| chr1-110403023-DEL-2383 | chr1-110403010-DEL-2371 |
| chr1-110415648-DEL-552 | chr1-110415653-DEL-550 |
| chr1-110438472-DEL-991 | chr1-110438480-DEL-995 |
| chr1-110460413-DEL-535 | chr1-110460414-DEL-535 |
| chr1-110498184-DEL-357 | chr1-110498185-DEL-357 |
| chr1-110501448-DEL-72 | chr1-110501449-DEL-72 |
| chr1-110502996-DEL-669 | chr1-110502997-DEL-669 |
| chr1-110517653-DEL-124 | chr1-110517654-DEL-124 |
| chr1-110536554-DEL-62 | chr1-110536555-DEL-60 |
| chr1-110540719-DEL-463 | chr1-110540720-DEL-463 |
| chr1-110541798-DEL-69 | chr1-110541801-DEL-67 |
| chr1-110571449-DEL-316 | chr1-110571450-DEL-316 |
| chr1-110573861-DEL-1672 | chr1-110573862-DEL-1672 |
| chr1-110579090-DEL-536 | chr1-110579091-DEL-537 |
| chr1-110603081-DEL-1469 | chr1-110603082-DEL-1469 |
| chr1-110605051-DEL-4258 | chr1-110605052-DEL-4258 |
| chr1-110617473-DEL-413 | chr1-110617474-DEL-413 |
| chr1-110636239-DEL-105 | chr1-110636240-DEL-105 |
| chr1-110652228-DEL-92 | chr1-110652229-DEL-92 |
| chr1-110656231-DEL-116 | chr1-110656240-DEL-108 |
| chr1-110667178-DEL-123 | chr1-110667186-DEL-105 |
| chr1-110738732-DEL-1242 | chr1-110738733-DEL-1242 |
| chr1-110766752-DEL-90 | chr1-110766753-DEL-90 |
| chr1-110776342-DEL-16518 | chr1-110776343-DEL-16518 |
| chr1-110807303-DEL-56 | chr1-110807304-DEL-56 |
| chr1-110825600-DEL-52 | chr1-110825601-DEL-52 |
| chr1-110827522-DEL-106 | chr1-110827524-DEL-75 |
| chr1-110856041-DEL-208 | chr1-110856042-DEL-208 |
| chr1-110865332-DEL-4837 | chr1-110865333-DEL-4837 |
| chr1-110898032-DEL-56 | chr1-110898033-DEL-56 |
| chr1-110916951-DEL-137 | chr1-110916952-DEL-137 |
| chr1-110928531-DEL-561 | chr1-110928532-DEL-561 |
| chr1-110930527-DEL-55 | chr1-110930528-DEL-55 |
| chr1-110950954-DEL-354 | chr1-110950955-DEL-354 |
| chr1-111035241-DEL-1586 | chr1-111035241-DEL-1585 |
| chr1-111046586-DEL-94 | chr1-111046589-DEL-92 |
| chr1-111079617-DEL-73 | chr1-111079618-DEL-73 |
| chr1-111102072-DEL-1314 | chr1-111102073-DEL-1314 |
| chr1-111113900-DEL-203 | chr1-111113901-DEL-203 |
| chr1-111181343-DEL-7106 | chr1-111181344-DEL-7106 |
| chr1-111233956-DEL-1519 | chr1-111233957-DEL-1519 |
| chr1-111355610-DEL-113 | chr1-111355610-DEL-113 |
| chr1-111388419-DEL-6284 | chr1-111388439-DEL-6255 |
| chr1-111397653-DEL-733 | chr1-111397654-DEL-733 |
| chr1-111433199-DEL-555 | chr1-111433200-DEL-555 |
| chr1-111478203-DEL-206 | chr1-111478204-DEL-206 |
| chr1-111501379-DEL-7827 | chr1-111501380-DEL-7827 |
| chr1-111508680-DEL-204 | chr1-111508681-DEL-204 |
| chr1-111541176-DEL-530 | chr1-111541157-DEL-549 |
| chr1-111543929-DEL-1753 | chr1-111543926-DEL-1756 |
| chr1-111572913-DEL-1897 | chr1-111572914-DEL-1897 |
| chr1-111581714-DEL-183 | chr1-111581715-DEL-183 |
| chr1-111585893-DEL-385 | chr1-111585894-DEL-385 |
| chr1-111657654-DEL-56 | chr1-111657655-DEL-62 |
| chr1-111666567-DEL-56 | chr1-111666568-DEL-56 |
| chr1-111675703-DEL-640 | chr1-111675704-DEL-640 |
| chr1-111682176-DEL-105 | chr1-111682177-DEL-105 |
| chr1-111686968-DEL-665 | chr1-111686969-DEL-673 |
| chr1-111697975-DEL-92 | chr1-111697976-DEL-92 |
| chr1-111700331-DEL-82 | chr1-111700335-DEL-52 |
| chr1-111700344-DEL-3872 | chr1-111700336-DEL-3880 |
| chr1-111713592-DEL-447 | chr1-111713593-DEL-447 |
| chr1-111718148-DEL-73 | chr1-111718172-DEL-83 |
| chr1-111729068-DEL-57 | chr1-111729070-DEL-56 |
| chr1-111738581-DEL-1615 | chr1-111738582-DEL-1615 |
| chr1-111741025-DEL-1913 | chr1-111741025-DEL-1914 |
| chr1-111741114-DEL-291 | chr1-111741121-DEL-285 |
| chr1-111753622-DEL-3164 | chr1-111753622-DEL-3165 |
| chr1-111765897-DEL-61 | chr1-111765900-DEL-76 |
| chr1-111779202-DEL-72 | chr1-111779200-DEL-120 |
| chr1-111801427-DEL-132 | chr1-111801421-DEL-133 |
| chr1-111808165-DEL-273 | chr1-111808166-DEL-273 |
| chr1-111812908-DEL-6337 | chr1-111812909-DEL-6336 |
| chr1-111822677-DEL-380 | chr1-111822678-DEL-380 |

| | |
|---|---|
| chr1-111824481-DEL-213 | chr1-111824468-DEL-211 |
| chr1-111827435-DEL-56 | chr1-111827436-DEL-56 |
| chr1-111834952-DEL-262 | chr1-111834953-DEL-262 |
| chr1-111850502-DEL-4139 | chr1-111850503-DEL-4139 |
| chr1-111854999-DEL-103 | chr1-111855000-DEL-103 |
| chr1-111866690-DEL-107 | chr1-111866691-DEL-107 |
| chr1-111868732-DEL-68 | chr1-111868733-DEL-68 |
| chr1-111878117-DEL-2061 | chr1-111878118-DEL-2060 |
| chr1-111885060-DEL-6281 | chr1-111885057-DEL-6280 |
| chr1-111906623-DEL-106 | chr1-111906624-DEL-106 |
| chr1-111908240-DEL-4451 | chr1-111908241-DEL-4451 |
| chr1-111922737-DEL-1013 | chr1-111922738-DEL-1013 |
| chr1-111937026-DEL-4188 | chr1-111937027-DEL-4188 |
| chr1-111942344-DEL-59 | chr1-111942345-DEL-59 |
| chr1-111969369-DEL-5189 | chr1-111969370-DEL-5190 |
| chr1-111979419-DEL-741 | chr1-111979420-DEL-741 |
| chr1-111989831-DEL-72 | chr1-111989803-DEL-84 |
| chr1-112008372-DEL-2238 | chr1-112008391-DEL-2220 |
| chr1-112015258-DEL-2306 | chr1-112015259-DEL-2306 |
| chr1-112048354-DEL-157 | chr1-112048355-DEL-157 |
| chr1-112053536-DEL-2283 | chr1-112053545-DEL-2280 |
| chr1-112061638-DEL-85 | chr1-112061639-DEL-85 |
| chr1-112064992-DEL-488 | chr1-112064989-DEL-489 |
| chr1-112075544-DEL-51 | chr1-112075544-DEL-62 |
| chr1-112121568-DEL-79 | chr1-112121576-DEL-75 |
| chr1-112139808-DEL-55 | chr1-112139809-DEL-55 |
| chr1-112141810-DEL-6658 | chr1-112141811-DEL-6658 |
| chr1-112148153-DEL-212 | chr1-112148154-DEL-212 |
| chr1-112181067-DEL-57 | chr1-112181067-DEL-58 |
| chr1-112181701-DEL-190 | chr1-112181700-DEL-186 |
| chr1-112184583-DEL-334 | chr1-112184584-DEL-334 |
| chr1-112188750-DEL-107 | chr1-112188751-DEL-107 |
| chr1-112215231-DEL-314 | chr1-112215231-DEL-314 |
| chr1-112243201-DEL-76 | chr1-112243194-DEL-68 |
| chr1-112257309-DEL-1534 | chr1-112257310-DEL-1534 |
| chr1-112262658-DEL-422 | chr1-112262658-DEL-422 |
| chr1-112267690-DEL-99 | chr1-112267691-DEL-99 |
| chr1-112303058-DEL-311 | chr1-112303055-DEL-312 |
| chr1-112311207-DEL-156 | chr1-112311208-DEL-156 |
| chr1-112328833-DEL-83 | chr1-112328834-DEL-83 |
| chr1-112329772-DEL-919 | chr1-112329773-DEL-919 |
| chr1-112337603-DEL-55 | chr1-112337604-DEL-55 |
| chr1-112342960-DEL-75 | chr1-112342959-DEL-108 |
| chr1-112346978-DEL-52 | chr1-112346979-DEL-52 |
| chr1-112347656-DEL-65 | chr1-112347657-DEL-65 |
| chr1-112355710-DEL-379 | chr1-112355711-DEL-379 |
| chr1-112359562-DEL-278 | chr1-112359563-DEL-278 |
| chr1-112364200-DEL-54 | chr1-112364199-DEL-51 |
| chr1-112386664-DEL-146 | chr1-112386687-DEL-152 |
| chr1-112392207-DEL-65 | chr1-112392203-DEL-66 |
| chr1-112412264-DEL-56 | chr1-112412265-DEL-56 |
| chr1-112434340-DEL-70 | chr1-112434341-DEL-70 |
| chr1-112435277-DEL-1268 | chr1-112435281-DEL-1240 |
| chr1-112439827-DEL-488 | chr1-112439828-DEL-488 |
| chr1-112441510-DEL-139 | chr1-112441510-DEL-139 |
| chr1-112459556-DEL-5555 | chr1-112459556-DEL-5555 |
| chr1-112493505-DEL-1888 | chr1-112493506-DEL-1888 |
| chr1-112506489-DEL-1157 | chr1-112506489-DEL-1157 |
| chr1-112520224-DEL-8271 | chr1-112520224-DEL-8271 |
| chr1-112527839-DEL-75 | chr1-112527840-DEL-75 |
| chr1-112552980-DEL-745 | chr1-112552981-DEL-745 |
| chr1-112560022-DEL-146 | chr1-112560015-DEL-163 |
| chr1-112571988-DEL-189 | chr1-112571989-DEL-153 |
| chr1-112606713-DEL-83 | chr1-112606753-DEL-81 |
| chr1-112613519-DEL-74 | chr1-112613520-DEL-50 |
| chr1-112618250-DEL-91 | chr1-112618251-DEL-91 |
| chr1-112626853-DEL-1701 | chr1-112626854-DEL-1701 |
| chr1-112648947-DEL-53 | chr1-112648948-DEL-53 |
| chr1-112656291-DEL-87 | chr1-112656295-DEL-86 |
| chr1-112659769-DEL-538 | chr1-112659770-DEL-538 |
| chr1-112677488-DEL-186 | chr1-112677489-DEL-186 |
| chr1-112680274-DEL-90 | chr1-112680275-DEL-80 |
| chr1-112692070-DEL-1700 | chr1-112692071-DEL-1700 |
| chr1-112692561-DEL-206 | chr1-112692562-DEL-206 |
| chr1-112707210-DEL-116 | chr1-112707210-DEL-116 |
| chr1-112713886-DEL-84 | chr1-112713887-DEL-84 |
| chr1-112714466-DEL-51 | chr1-112714469-DEL-51 |
| chr1-112717993-DEL-1142 | chr1-112717994-DEL-1142 |
| chr1-112738325-DEL-57 | chr1-112738326-DEL-57 |
| chr1-112743915-DEL-6284 | chr1-112743916-DEL-6284 |

| | |
|---|---|
| chr1-112765833-DEL-6924 | chr1-112765834-DEL-6924 |
| chr1-112800834-DEL-61 | chr1-112800835-DEL-56 |
| chr1-112820275-DEL-79 | chr1-112820272-DEL-56 |
| chr1-112857119-DEL-6062 | chr1-112857120-DEL-6062 |
| chr1-112868546-DEL-301 | chr1-112868547-DEL-301 |
| chr1-112874310-DEL-129 | chr1-112874311-DEL-129 |
| chr1-112894419-DEL-1005 | chr1-112894420-DEL-1005 |
| chr1-112943798-DEL-7128 | chr1-112943798-DEL-7128 |
| chr1-112953350-DEL-64 | chr1-112953351-DEL-64 |
| chr1-112968984-DEL-56 | chr1-112968983-DEL-60 |
| chr1-112972483-DEL-63 | chr1-112972484-DEL-63 |
| chr1-112978783-DEL-6608 | chr1-112978784-DEL-6608 |
| chr1-112999019-DEL-110 | chr1-112999020-DEL-110 |
| chr1-113026437-DEL-638 | chr1-113026438-DEL-638 |
| chr1-113031154-DEL-80 | chr1-113031155-DEL-80 |
| chr1-113053553-DEL-52 | chr1-113053554-DEL-52 |
| chr1-113057794-DEL-398 | chr1-113057795-DEL-398 |
| chr1-113072898-DEL-394 | chr1-113072899-DEL-394 |
| chr1-113075651-DEL-138 | chr1-113075651-DEL-138 |
| chr1-113077152-DEL-95 | chr1-113077150-DEL-95 |
| chr1-113079880-DEL-149 | chr1-113079881-DEL-149 |
| chr1-113095565-DEL-1553 | chr1-113095566-DEL-1553 |
| chr1-113099940-DEL-1077 | chr1-113099941-DEL-1077 |
| chr1-113103489-DEL-64 | chr1-113103489-DEL-64 |
| chr1-113111414-DEL-6328 | chr1-113111415-DEL-6328 |
| chr1-113121538-DEL-63 | chr1-113121502-DEL-52 |
| chr1-113130065-DEL-1538 | chr1-113130063-DEL-1537 |
| chr1-113162153-DEL-1413 | chr1-113162153-DEL-1413 |
| chr1-113169983-DEL-809 | chr1-113169983-DEL-809 |
| chr1-113174834-DEL-858 | chr1-113174834-DEL-858 |
| chr1-113199580-DEL-337 | chr1-113199581-DEL-337 |
| chr1-113203012-DEL-1164 | chr1-113203013-DEL-1164 |
| chr1-113215876-DEL-102 | chr1-113215877-DEL-82 |
| chr1-113222263-DEL-271 | chr1-113222264-DEL-271 |
| chr1-113235161-DEL-72 | chr1-113235161-DEL-72 |
| chr1-113239727-DEL-417 | chr1-113239720-DEL-417 |
| chr1-113264513-DEL-6486 | chr1-113264514-DEL-6486 |
| chr1-113295966-DEL-485 | chr1-113295966-DEL-485 |
| chr1-113330795-DEL-829 | chr1-113330796-DEL-829 |
| chr1-113356986-DEL-1082 | chr1-113356987-DEL-1082 |
| chr1-113358450-DEL-1386 | chr1-113358452-DEL-1385 |
| chr1-113371837-DEL-90 | chr1-113371831-DEL-98 |
| chr1-113374233-DEL-207 | chr1-113374237-DEL-166 |
| chr1-113399786-DEL-60 | chr1-113399781-DEL-58 |
| chr1-113402530-DEL-102 | chr1-113402531-DEL-99 |
| chr1-113405407-DEL-807 | chr1-113405408-DEL-807 |
| chr1-113408034-DEL-1253 | chr1-113408035-DEL-1253 |
| chr1-113411017-DEL-1548 | chr1-113411018-DEL-1548 |
| chr1-113421386-DEL-141 | chr1-113421401-DEL-120 |
| chr1-113451348-DEL-187 | chr1-113451348-DEL-187 |
| chr1-113465819-DEL-1381 | chr1-113465819-DEL-1381 |
| chr1-113491616-DEL-1062 | chr1-113491608-DEL-1070 |
| chr1-113492980-DEL-156 | chr1-113492980-DEL-156 |
| chr1-113507554-DEL-363 | chr1-113507555-DEL-359 |
| chr1-113515078-DEL-1297 | chr1-113515079-DEL-1297 |
| chr1-113530380-DEL-2214 | chr1-113530381-DEL-2214 |
| chr1-113546101-DEL-91 | chr1-113546093-DEL-89 |
| chr1-113548498-DEL-8503 | chr1-113548492-DEL-8509 |
| chr1-113553525-DEL-224 | chr1-113553526-DEL-224 |
| chr1-113572259-DEL-2604 | chr1-113572259-DEL-2604 |
| chr1-113574659-DEL-66 | chr1-113574660-DEL-66 |
| chr1-113577286-DEL-956 | chr1-113577286-DEL-956 |
| chr1-113593329-DEL-638 | chr1-113593330-DEL-638 |
| chr1-113606038-DEL-74 | chr1-113606049-DEL-62 |
| chr1-113607672-DEL-2724 | chr1-113607673-DEL-2724 |
| chr1-113614222-DEL-206 | chr1-113614226-DEL-205 |
| chr1-113624134-DEL-1963 | chr1-113624137-DEL-1963 |
| chr1-113628176-DEL-124 | chr1-113628177-DEL-124 |
| chr1-113635129-DEL-270 | chr1-113635128-DEL-270 |
| chr1-113641959-DEL-3585 | chr1-113641960-DEL-3585 |
| chr1-113649676-DEL-201 | chr1-113649678-DEL-197 |
| chr1-113654747-DEL-169 | chr1-113654748-DEL-169 |
| chr1-113666713-DEL-1892 | chr1-113666717-DEL-1889 |
| chr1-113678238-DEL-201 | chr1-113678238-DEL-201 |
| chr1-113698130-DEL-2133 | chr1-113698131-DEL-2133 |
| chr1-113711038-DEL-52 | chr1-113711045-DEL-50 |
| chr1-113723248-DEL-96 | chr1-113723249-DEL-96 |
| chr1-113731488-DEL-2499 | chr1-113731489-DEL-2499 |
| chr1-113749609-DEL-399 | chr1-113749610-DEL-399 |
| chr1-113752378-DEL-60 | chr1-113752383-DEL-56 |

| | |
|---|---|
| chr1-113754345-DEL-873 | chr1-113754345-DEL-873 |
| chr1-113771934-DEL-554 | chr1-113771934-DEL-554 |
| chr1-113780890-DEL-181 | chr1-113780926-DEL-154 |
| chr1-113782337-DEL-237 | chr1-113782332-DEL-243 |
| chr1-113799403-DEL-80 | chr1-113799415-DEL-50 |
| chr1-113804423-DEL-520 | chr1-113804424-DEL-519 |
| chr1-113813731-DEL-60 | chr1-113813732-DEL-60 |
| chr1-113821543-DEL-189 | chr1-113821544-DEL-189 |
| chr1-113823258-DEL-2533 | chr1-113823259-DEL-2533 |
| chr1-113849045-DEL-57 | chr1-113849045-DEL-56 |
| chr1-113852713-DEL-301 | chr1-113852710-DEL-301 |
| chr1-113858830-DEL-635 | chr1-113858831-DEL-635 |
| chr1-113859776-DEL-76 | chr1-113859778-DEL-74 |
| chr1-113868767-DEL-1362 | chr1-113868768-DEL-1362 |
| chr1-113902606-DEL-2353 | chr1-113902608-DEL-2354 |
| chr1-113906011-DEL-78 | chr1-113906012-DEL-78 |
| chr1-113923020-DEL-62 | chr1-113923009-DEL-58 |
| chr1-113929154-DEL-56 | chr1-113929163-DEL-50 |
| chr1-113957230-DEL-6618 | chr1-113957231-DEL-6618 |
| chr1-113974111-DEL-58 | chr1-113974115-DEL-56 |
| chr1-113981078-DEL-2189 | chr1-113981079-DEL-2189 |
| chr1-113996887-DEL-10161 | chr1-113996888-DEL-10161 |
| chr1-114010632-DEL-446 | chr1-114010633-DEL-446 |
| chr1-114019657-DEL-112 | chr1-114019658-DEL-112 |
| chr1-114025552-DEL-767 | chr1-114025553-DEL-767 |
| chr1-114039024-DEL-3894 | chr1-114039005-DEL-3899 |
| chr1-114044088-DEL-76 | chr1-114044088-DEL-76 |
| chr1-114055992-DEL-6626 | chr1-114055991-DEL-6627 |
| chr1-114068304-DEL-84 | chr1-114068300-DEL-82 |
| chr1-114068748-DEL-1155 | chr1-114068749-DEL-1155 |
| chr1-114104349-DEL-7337 | chr1-114104350-DEL-7337 |
| chr1-114110688-DEL-212 | chr1-114110689-DEL-212 |
| chr1-114112943-DEL-5343 | chr1-114112944-DEL-5343 |
| chr1-114130903-DEL-4290 | chr1-114130913-DEL-4298 |
| chr1-114139103-DEL-4143 | chr1-114139104-DEL-4143 |
| chr1-114151502-DEL-6573 | chr1-114151503-DEL-6573 |
| chr1-114186624-DEL-80 | chr1-114186625-DEL-80 |
| chr1-114198463-DEL-6086 | chr1-114198452-DEL-6097 |
| chr1-114233977-DEL-437 | chr1-114233978-DEL-437 |
| chr1-114284057-DEL-51 | chr1-114284058-DEL-51 |
| chr1-114289759-DEL-462 | chr1-114289750-DEL-461 |
| chr1-114296274-DEL-580 | chr1-114296273-DEL-581 |
| chr1-114335661-DEL-222 | chr1-114335661-DEL-222 |
| chr1-114338526-DEL-4794 | chr1-114338527-DEL-4794 |
| chr1-114348005-DEL-162 | chr1-114348003-DEL-162 |
| chr1-114348864-DEL-1231 | chr1-114348866-DEL-1230 |
| chr1-114353451-DEL-292 | chr1-114353452-DEL-292 |
| chr1-114358507-DEL-56 | chr1-114358508-DEL-64 |
| chr1-114376177-DEL-78 | chr1-114376192-DEL-66 |
| chr1-114379169-DEL-57 | chr1-114379170-DEL-57 |
| chr1-114397745-DEL-58 | chr1-114397751-DEL-70 |
| chr1-114409187-DEL-53 | chr1-114409188-DEL-53 |
| chr1-114409895-DEL-1043 | chr1-114409896-DEL-1043 |
| chr1-114423599-DEL-775 | chr1-114423600-DEL-775 |
| chr1-114461141-DEL-86 | chr1-114461140-DEL-88 |
| chr1-114479820-DEL-56 | chr1-114479821-DEL-56 |
| chr1-114485020-DEL-7133 | chr1-114485021-DEL-7133 |
| chr1-114485481-DEL-424 | chr1-114485482-DEL-424 |
| chr1-114505788-DEL-1084 | chr1-114505788-DEL-1084 |
| chr1-114507740-DEL-1578 | chr1-114507741-DEL-1578 |
| chr1-114510414-DEL-3363 | chr1-114510415-DEL-3363 |
| chr1-114518573-DEL-336 | chr1-114518574-DEL-336 |
| chr1-114523189-DEL-74 | chr1-114523188-DEL-75 |
| chr1-114529869-DEL-218 | chr1-114529870-DEL-218 |
| chr1-114531199-DEL-6489 | chr1-114531199-DEL-6482 |
| chr1-114565454-DEL-60 | chr1-114565455-DEL-60 |
| chr1-114575094-DEL-64 | chr1-114575095-DEL-64 |
| chr1-114584791-DEL-5978 | chr1-114584779-DEL-5988 |
| chr1-114588479-DEL-3713 | chr1-114588480-DEL-3713 |
| chr1-114604236-DEL-1227 | chr1-114604236-DEL-1227 |
| chr1-114613334-DEL-783 | chr1-114613335-DEL-783 |
| chr1-114627361-DEL-197 | chr1-114627362-DEL-197 |
| chr1-114628180-DEL-4184 | chr1-114628181-DEL-4184 |
| chr1-114659383-DEL-8151 | chr1-114659381-DEL-8151 |
| chr1-114688110-DEL-1992 | chr1-114688110-DEL-1992 |
| chr1-114692234-DEL-6666 | chr1-114692237-DEL-6664 |
| chr1-114719381-DEL-523 | chr1-114719381-DEL-523 |
| chr1-114723405-DEL-491 | chr1-114723405-DEL-491 |
| chr1-114730047-DEL-7082 | chr1-114730048-DEL-7082 |
| chr1-114783326-DEL-1956 | chr1-114783327-DEL-1956 |

| | |
|---|---|
| chr1-114784366-DEL-117 | chr1-114784371-DEL-101 |
| chr1-114794781-DEL-76 | chr1-114794782-DEL-76 |
| chr1-114795614-DEL-1187 | chr1-114795615-DEL-1187 |
| chr1-114802568-DEL-1004 | chr1-114802569-DEL-1001 |
| chr1-114832457-DEL-63 | chr1-114832457-DEL-63 |
| chr1-114848875-DEL-11390 | chr1-114848876-DEL-11390 |
| chr1-114854027-DEL-192 | chr1-114854028-DEL-192 |
| chr1-114862090-DEL-585 | chr1-114862086-DEL-589 |
| chr1-114867876-DEL-710 | chr1-114867889-DEL-704 |
| chr1-114871479-DEL-1837 | chr1-114871480-DEL-1837 |
| chr1-114909023-DEL-68 | chr1-114909032-DEL-60 |
| chr1-114912213-DEL-1981 | chr1-114912214-DEL-1981 |
| chr1-114918344-DEL-878 | chr1-114918345-DEL-878 |
| chr1-114920764-DEL-2789 | chr1-114920765-DEL-2789 |
| chr1-114923642-DEL-62 | chr1-114923645-DEL-60 |
| chr1-114924667-DEL-56 | chr1-114924668-DEL-56 |
| chr1-114930703-DEL-614 | chr1-114930699-DEL-618 |
| chr1-114941586-DEL-142 | chr1-114941587-DEL-142 |
| chr1-114959830-DEL-275 | chr1-114959831-DEL-275 |
| chr1-114961190-DEL-319 | chr1-114961191-DEL-319 |
| chr1-114975331-DEL-4020 | chr1-114975332-DEL-4020 |
| chr1-114987584-DEL-411 | chr1-114987585-DEL-411 |
| chr1-115023888-DEL-62 | chr1-115023889-DEL-62 |
| chr1-115036281-DEL-426 | chr1-115036282-DEL-426 |
| chr1-115039992-DEL-51 | chr1-115040007-DEL-56 |
| chr1-115040380-DEL-1465 | chr1-115040381-DEL-1465 |
| chr1-115073449-DEL-492 | chr1-115073450-DEL-492 |
| chr1-115086830-DEL-56 | chr1-115086835-DEL-52 |
| chr1-115163035-DEL-58 | chr1-115163044-DEL-50 |
| chr1-115187113-DEL-331 | chr1-115187114-DEL-331 |
| chr1-115206327-DEL-111 | chr1-115206327-DEL-112 |
| chr1-115206433-DEL-823 | chr1-115206434-DEL-823 |
| chr1-115251819-DEL-315 | chr1-115251819-DEL-315 |
| chr1-115268589-DEL-117 | chr1-115268553-DEL-108 |
| chr1-115285821-DEL-1828 | chr1-115285821-DEL-1828 |
| chr1-115292011-DEL-657 | chr1-115292011-DEL-657 |
| chr1-115297077-DEL-205 | chr1-115297078-DEL-205 |
| chr1-115297723-DEL-58 | chr1-115297722-DEL-70 |
| chr1-115302853-DEL-498 | chr1-115302852-DEL-498 |
| chr1-115339462-DEL-414 | chr1-115339463-DEL-414 |
| chr1-115344985-DEL-126 | chr1-115344986-DEL-126 |
| chr1-115352503-DEL-90 | chr1-115352504-DEL-90 |
| chr1-115355432-DEL-2319 | chr1-115355433-DEL-2319 |
| chr1-115374961-DEL-6850 | chr1-115374962-DEL-6850 |
| chr1-115387825-DEL-132 | chr1-115387826-DEL-132 |
| chr1-115390630-DEL-716 | chr1-115390631-DEL-716 |
| chr1-115397922-DEL-483 | chr1-115397923-DEL-483 |
| chr1-115398940-DEL-529 | chr1-115398941-DEL-529 |
| chr1-115404398-DEL-6211 | chr1-115404399-DEL-6211 |
| chr1-115431779-DEL-388 | chr1-115431780-DEL-388 |
| chr1-115457959-DEL-3142 | chr1-115457960-DEL-3142 |
| chr1-115462512-DEL-52 | chr1-115462512-DEL-70 |
| chr1-115486331-DEL-527 | chr1-115486336-DEL-522 |
| chr1-115492562-DEL-735 | chr1-115492563-DEL-735 |
| chr1-115502056-DEL-70 | chr1-115502062-DEL-66 |
| chr1-115507018-DEL-137 | chr1-115507016-DEL-138 |
| chr1-115520316-DEL-5734 | chr1-115520317-DEL-5733 |
| chr1-115534543-DEL-70 | chr1-115534539-DEL-56 |
| chr1-115543787-DEL-85 | chr1-115543788-DEL-85 |
| chr1-115569068-DEL-1052 | chr1-115569069-DEL-1052 |
| chr1-115572372-DEL-108 | chr1-115572373-DEL-108 |
| chr1-115573852-DEL-179 | chr1-115573852-DEL-179 |
| chr1-115639632-DEL-52 | chr1-115639653-DEL-60 |
| chr1-115678005-DEL-747 | chr1-115678006-DEL-747 |
| chr1-115710926-DEL-75 | chr1-115710944-DEL-50 |
| chr1-115727149-DEL-7185 | chr1-115727150-DEL-7185 |
| chr1-115742946-DEL-84 | chr1-115742947-DEL-84 |
| chr1-115752700-DEL-96 | chr1-115752709-DEL-78 |
| chr1-115801334-DEL-101 | chr1-115801337-DEL-93 |
| chr1-115803909-DEL-71 | chr1-115803917-DEL-64 |
| chr1-115812151-DEL-69 | chr1-115812152-DEL-69 |
| chr1-115812981-DEL-5453 | chr1-115812982-DEL-5453 |
| chr1-115830930-DEL-53 | chr1-115830931-DEL-53 |
| chr1-115867192-DEL-62 | chr1-115867193-DEL-62 |
| chr1-115891634-DEL-403 | chr1-115891622-DEL-416 |
| chr1-115925490-DEL-554 | chr1-115925491-DEL-554 |
| chr1-115933725-DEL-181 | chr1-115933726-DEL-181 |
| chr1-115940445-DEL-96 | chr1-115940483-DEL-52 |
| chr1-115971114-DEL-54 | chr1-115971115-DEL-54 |
| chr1-115978682-DEL-204 | chr1-115978683-DEL-204 |

| | |
|---|---|
| chr1-115982438-DEL-195 | chr1-115982437-DEL-198 |
| chr1-116050588-DEL-312 | chr1-116050589-DEL-311 |
| chr1-116087769-DEL-63 | chr1-116087814-DEL-62 |
| chr1-116150950-DEL-540 | chr1-116150951-DEL-540 |
| chr1-116171418-DEL-169 | chr1-116171419-DEL-169 |
| chr1-116178473-DEL-53 | chr1-116178474-DEL-53 |
| chr1-116203687-DEL-296 | chr1-116203688-DEL-296 |
| chr1-116214666-DEL-83 | chr1-116214663-DEL-77 |
| chr1-116249710-DEL-52 | chr1-116249703-DEL-60 |
| chr1-116271098-DEL-1175 | chr1-116271099-DEL-1175 |
| chr1-116284409-DEL-6907 | chr1-116284410-DEL-6907 |
| chr1-116306616-DEL-78 | chr1-116306599-DEL-76 |
| chr1-116311117-DEL-1048 | chr1-116311117-DEL-1045 |
| chr1-116330639-DEL-62 | chr1-116330638-DEL-64 |
| chr1-116340069-DEL-83 | chr1-116340070-DEL-65 |
| chr1-116380997-DEL-97 | chr1-116380998-DEL-97 |
| chr1-116394271-DEL-69 | chr1-116394272-DEL-69 |
| chr1-116417957-DEL-491 | chr1-116417956-DEL-489 |
| chr1-116419533-DEL-2412 | chr1-116419519-DEL-2426 |
| chr1-116442748-DEL-77 | chr1-116442749-DEL-77 |
| chr1-116461490-DEL-1696 | chr1-116461485-DEL-1704 |
| chr1-116473932-DEL-66 | chr1-116473933-DEL-66 |
| chr1-116475310-DEL-421 | chr1-116475310-DEL-421 |
| chr1-116477071-DEL-132 | chr1-116477072-DEL-132 |
| chr1-116502118-DEL-935 | chr1-116502115-DEL-934 |
| chr1-116516231-DEL-858 | chr1-116516232-DEL-858 |
| chr1-116540350-DEL-97 | chr1-116540358-DEL-108 |
| chr1-116550584-DEL-3639 | chr1-116550585-DEL-3639 |
| chr1-116567575-DEL-61 | chr1-116567578-DEL-59 |
| chr1-116575989-DEL-7133 | chr1-116575990-DEL-7133 |
| chr1-116610824-DEL-4614 | chr1-116610825-DEL-4614 |
| chr1-116620240-DEL-566 | chr1-116620241-DEL-566 |
| chr1-116629191-DEL-2458 | chr1-116629182-DEL-2467 |
| chr1-116643994-DEL-555 | chr1-116643995-DEL-555 |
| chr1-116644508-DEL-168 | chr1-116644509-DEL-168 |
| chr1-116649763-DEL-67 | chr1-116649757-DEL-65 |
| chr1-116659541-DEL-399 | chr1-116659541-DEL-399 |
| chr1-116698648-DEL-565 | chr1-116698649-DEL-565 |
| chr1-116703972-DEL-95 | chr1-116703973-DEL-95 |
| chr1-116721403-DEL-480 | chr1-116721408-DEL-479 |
| chr1-116733864-DEL-4795 | chr1-116733865-DEL-4794 |
| chr1-116757376-DEL-1974 | chr1-116757372-DEL-1976 |
| chr1-116768274-DEL-281 | chr1-116768275-DEL-281 |
| chr1-116775721-DEL-2340 | chr1-116775722-DEL-2340 |
| chr1-116783199-DEL-7084 | chr1-116783200-DEL-7084 |
| chr1-116789341-DEL-212 | chr1-116789342-DEL-212 |
| chr1-116831102-DEL-648 | chr1-116831103-DEL-648 |
| chr1-116870401-DEL-1846 | chr1-116870402-DEL-1846 |
| chr1-116913990-DEL-76 | chr1-116913991-DEL-76 |
| chr1-116949501-DEL-52 | chr1-116949501-DEL-52 |
| chr1-116956581-DEL-100 | chr1-116956582-DEL-100 |
| chr1-116978606-DEL-91 | chr1-116978607-DEL-91 |
| chr1-117007848-DEL-1090 | chr1-117007849-DEL-1090 |
| chr1-117009947-DEL-182 | chr1-117009948-DEL-182 |
| chr1-117016757-DEL-2885 | chr1-117016758-DEL-2885 |
| chr1-117037196-DEL-61 | chr1-117037197-DEL-61 |
| chr1-117144970-DEL-88 | chr1-117144980-DEL-85 |
| chr1-117146939-DEL-75 | chr1-117146940-DEL-75 |
| chr1-117159370-DEL-7870 | chr1-117159371-DEL-7870 |
| chr1-117181296-DEL-7394 | chr1-117181297-DEL-7394 |
| chr1-117204275-DEL-4163 | chr1-117204276-DEL-4163 |
| chr1-117221914-DEL-3013 | chr1-117221923-DEL-3005 |
| chr1-117226433-DEL-458 | chr1-117226434-DEL-458 |
| chr1-117243715-DEL-149 | chr1-117243712-DEL-149 |
| chr1-117249045-DEL-372 | chr1-117249045-DEL-372 |
| chr1-117252539-DEL-333 | chr1-117252539-DEL-333 |
| chr1-117253676-DEL-366 | chr1-117253677-DEL-366 |
| chr1-117264745-DEL-51 | chr1-117264745-DEL-52 |
| chr1-117266485-DEL-3858 | chr1-117266486-DEL-3858 |
| chr1-117278930-DEL-107 | chr1-117278932-DEL-106 |
| chr1-117289872-DEL-68 | chr1-117289889-DEL-58 |
| chr1-117297490-DEL-6870 | chr1-117297491-DEL-6870 |
| chr1-117316699-DEL-109 | chr1-117316703-DEL-148 |
| chr1-117328613-DEL-6288 | chr1-117328613-DEL-6288 |
| chr1-117347353-DEL-418 | chr1-117347354-DEL-418 |
| chr1-117355690-DEL-6482 | chr1-117355689-DEL-6482 |
| chr1-117363386-DEL-795 | chr1-117363387-DEL-795 |
| chr1-117363991-DEL-1942 | chr1-117363992-DEL-1942 |
| chr1-117368425-DEL-80 | chr1-117368427-DEL-79 |
| chr1-117392919-DEL-197 | chr1-117392905-DEL-195 |

| | |
|---|---|
| chr1-117399272-DEL-3000 | chr1-117399273-DEL-3000 |
| chr1-117403871-DEL-51 | chr1-117403872-DEL-51 |
| chr1-117407348-DEL-6886 | chr1-117407349-DEL-6886 |
| chr1-117416189-DEL-206 | chr1-117416188-DEL-206 |
| chr1-117416451-DEL-526 | chr1-117416452-DEL-526 |
| chr1-117417276-DEL-189 | chr1-117417277-DEL-189 |
| chr1-117424489-DEL-200 | chr1-117424490-DEL-200 |
| chr1-117427747-DEL-1843 | chr1-117427744-DEL-1843 |
| chr1-117460044-DEL-198 | chr1-117460045-DEL-198 |
| chr1-117481897-DEL-2269 | chr1-117481898-DEL-2269 |
| chr1-117488744-DEL-525 | chr1-117488745-DEL-525 |
| chr1-117511656-DEL-984 | chr1-117511657-DEL-984 |
| chr1-117515212-DEL-223 | chr1-117515213-DEL-223 |
| chr1-117540746-DEL-3340 | chr1-117540747-DEL-3340 |
| chr1-117604656-DEL-84 | chr1-117604657-DEL-84 |
| chr1-117625300-DEL-206 | chr1-117625301-DEL-205 |
| chr1-117629660-DEL-7083 | chr1-117629661-DEL-7083 |
| chr1-117725059-DEL-431 | chr1-117725060-DEL-431 |
| chr1-117874152-DEL-205 | chr1-117874155-DEL-206 |
| chr1-117918028-DEL-2397 | chr1-117918041-DEL-2374 |
| chr1-118018994-DEL-886 | chr1-118018995-DEL-886 |
| chr1-118019934-DEL-438 | chr1-118019934-DEL-438 |
| chr1-118049823-DEL-6760 | chr1-118049824-DEL-6760 |
| chr1-118127436-DEL-84 | chr1-118127437-DEL-84 |
| chr1-118159264-DEL-73 | chr1-118159259-DEL-79 |
| chr1-118216220-DEL-2785 | chr1-118216221-DEL-2785 |
| chr1-118252575-DEL-92 | chr1-118252576-DEL-92 |
| chr1-118264976-DEL-185 | chr1-118264977-DEL-185 |
| chr1-118268149-DEL-76 | chr1-118268150-DEL-72 |
| chr1-118280891-DEL-63 | chr1-118280892-DEL-63 |
| chr1-118322832-DEL-75 | chr1-118322833-DEL-75 |
| chr1-118329132-DEL-80 | chr1-118329133-DEL-80 |
| chr1-118339507-DEL-187 | chr1-118339508-DEL-187 |
| chr1-118349683-DEL-66 | chr1-118349640-DEL-62 |
| chr1-118385442-DEL-143 | chr1-118385443-DEL-138 |
| chr1-118417326-DEL-1385 | chr1-118417325-DEL-1385 |
| chr1-118509506-DEL-6800 | chr1-118509507-DEL-6800 |
| chr1-118515494-DEL-208 | chr1-118515494-DEL-208 |
| chr1-118548771-DEL-141 | chr1-118548772-DEL-141 |
| chr1-118551506-DEL-61 | chr1-118551507-DEL-61 |
| chr1-118560315-DEL-195 | chr1-118560316-DEL-195 |
| chr1-118565111-DEL-3574 | chr1-118565112-DEL-3575 |
| chr1-118575364-DEL-93 | chr1-118575365-DEL-93 |
| chr1-118579753-DEL-53 | chr1-118579751-DEL-54 |
| chr1-118580404-DEL-171 | chr1-118580405-DEL-171 |
| chr1-118597978-DEL-386 | chr1-118597979-DEL-386 |
| chr1-118598744-DEL-118 | chr1-118598745-DEL-118 |
| chr1-118600361-DEL-216 | chr1-118600368-DEL-210 |
| chr1-118652923-DEL-202 | chr1-118652924-DEL-202 |
| chr1-118667289-DEL-332 | chr1-118667290-DEL-332 |
| chr1-118683073-DEL-144 | chr1-118683080-DEL-108 |
| chr1-118689941-DEL-64 | chr1-118689939-DEL-62 |
| chr1-118690700-DEL-1054 | chr1-118690701-DEL-1054 |
| chr1-118707145-DEL-59 | chr1-118707138-DEL-66 |
| chr1-118719520-DEL-1688 | chr1-118719521-DEL-1688 |
| chr1-118734553-DEL-119 | chr1-118734554-DEL-119 |
| chr1-118749750-DEL-70 | chr1-118749746-DEL-80 |
| chr1-118751091-DEL-150 | chr1-118751094-DEL-148 |
| chr1-118803977-DEL-406 | chr1-118803978-DEL-406 |
| chr1-118833544-DEL-216 | chr1-118833545-DEL-216 |
| chr1-118919801-DEL-126 | chr1-118919802-DEL-132 |
| chr1-118936940-DEL-78 | chr1-118936939-DEL-79 |
| chr1-118941021-DEL-62 | chr1-118941032-DEL-52 |
| chr1-118957385-DEL-129 | chr1-118957388-DEL-127 |
| chr1-119003557-DEL-63 | chr1-119003558-DEL-63 |
| chr1-119045693-DEL-285 | chr1-119045694-DEL-285 |
| chr1-119052959-DEL-145 | chr1-119052960-DEL-145 |
| chr1-119070918-DEL-77 | chr1-119070919-DEL-77 |
| chr1-119082444-DEL-1053 | chr1-119082445-DEL-1053 |
| chr1-119088437-DEL-54 | chr1-119088438-DEL-54 |
| chr1-119094666-DEL-56 | chr1-119094667-DEL-56 |
| chr1-119099846-DEL-72 | chr1-119099847-DEL-72 |
| chr1-119106363-DEL-324 | chr1-119106364-DEL-324 |
| chr1-119113290-DEL-126 | chr1-119113240-DEL-136 |
| chr1-119172952-DEL-88 | chr1-119172953-DEL-84 |
| chr1-119223741-DEL-58 | chr1-119223742-DEL-58 |
| chr1-119254633-DEL-84 | chr1-119254634-DEL-84 |
| chr1-119290544-DEL-1713 | chr1-119290544-DEL-1714 |
| chr1-119357508-DEL-100 | chr1-119357509-DEL-100 |
| chr1-119361665-DEL-202 | chr1-119361666-DEL-202 |

| | |
|---|---|
| chr1-119403021-DEL-86 | chr1-119403022-DEL-129 |
| chr1-119439831-DEL-93 | chr1-119439832-DEL-93 |
| chr1-119457867-DEL-68 | chr1-119457868-DEL-68 |
| chr1-119502053-DEL-2849 | chr1-119502054-DEL-2849 |
| chr1-119518144-DEL-165 | chr1-119518145-DEL-165 |
| chr1-119545512-DEL-883 | chr1-119545513-DEL-883 |
| chr1-119566748-DEL-131 | chr1-119566749-DEL-129 |
| chr1-119617178-DEL-140 | chr1-119617179-DEL-140 |
| chr1-119623144-DEL-51 | chr1-119623145-DEL-51 |
| chr1-119672272-DEL-214 | chr1-119672273-DEL-214 |
| chr1-119714046-DEL-881 | chr1-119714047-DEL-881 |
| chr1-119749460-DEL-89 | chr1-119749450-DEL-100 |
| chr1-119786059-DEL-113 | chr1-119786107-DEL-80 |
| chr1-119831977-DEL-68 | chr1-119831986-DEL-64 |
| chr1-119851873-DEL-150 | chr1-119851874-DEL-150 |
| chr1-119953369-DEL-56 | chr1-119953370-DEL-56 |
| chr1-120021565-DEL-222 | chr1-120021566-DEL-222 |
| chr1-120026477-DEL-55 | chr1-120026478-DEL-55 |
| chr1-120035150-DEL-656 | chr1-120035151-DEL-656 |
| chr1-120063683-DEL-259 | chr1-120063684-DEL-259 |
| chr1-120068331-DEL-64 | chr1-120068332-DEL-60 |
| chr1-120071763-DEL-79 | chr1-120071764-DEL-79 |
| chr1-120127397-DEL-82 | chr1-120127398-DEL-82 |
| chr1-120161836-DEL-59 | chr1-120161837-DEL-59 |
| chr1-120226633-DEL-212 | chr1-120226633-DEL-213 |
| chr1-120229032-DEL-134 | chr1-120229033-DEL-132 |
| chr1-120247471-DEL-212 | chr1-120247472-DEL-212 |
| chr1-120253181-DEL-98 | chr1-120253182-DEL-98 |
| chr1-120286262-DEL-94 | chr1-120286269-DEL-74 |
| chr1-120288918-DEL-52 | chr1-120288919-DEL-52 |
| chr1-120360549-DEL-739 | chr1-120360550-DEL-739 |
| chr1-120389136-DEL-262 | chr1-120389137-DEL-262 |
| chr1-120420558-DEL-100 | chr1-120420559-DEL-100 |
| chr1-120422950-DEL-412 | chr1-120422958-DEL-404 |
| chr1-120434160-DEL-407 | chr1-120434161-DEL-407 |
| chr1-120458805-DEL-195 | chr1-120458794-DEL-241 |
| chr1-120470704-DEL-67 | chr1-120470685-DEL-68 |
| chr1-120478302-DEL-79 | chr1-120478303-DEL-79 |
| chr1-120488623-DEL-79 | chr1-120488624-DEL-79 |
| chr1-120489241-DEL-437 | chr1-120489242-DEL-431 |
| chr1-120494087-DEL-118 | chr1-120494096-DEL-110 |
| chr1-120498593-DEL-5143 | chr1-120498594-DEL-5143 |
| chr1-120519492-DEL-62 | chr1-120519493-DEL-66 |
| chr1-120556790-DEL-78 | chr1-120556791-DEL-78 |
| chr1-120589903-DEL-52 | chr1-120589904-DEL-52 |
| chr1-120594262-DEL-344 | chr1-120594263-DEL-344 |
| chr1-120616592-DEL-183 | chr1-120616593-DEL-183 |
| chr1-120650670-DEL-116 | chr1-120650666-DEL-116 |
| chr1-120751874-DEL-197 | chr1-120751875-DEL-197 |
| chr1-120763269-DEL-62 | chr1-120763270-DEL-62 |
| chr1-120779433-DEL-106 | chr1-120779434-DEL-106 |
| chr1-120816103-DEL-132 | chr1-120816104-DEL-132 |
| chr1-120865983-DEL-259 | chr1-120865978-DEL-260 |
| chr1-120867541-DEL-64 | chr1-120867542-DEL-64 |
| chr1-120882246-DEL-663 | chr1-120882247-DEL-663 |
| chr1-120892655-DEL-422 | chr1-120892654-DEL-423 |
| chr1-120912175-DEL-96 | chr1-120912176-DEL-96 |
| chr1-120921703-DEL-6461 | chr1-120921704-DEL-6461 |
| chr1-120929008-DEL-62 | chr1-120929041-DEL-54 |
| chr1-120949339-DEL-85 | chr1-120949341-DEL-84 |
| chr1-120953176-DEL-108 | chr1-120953177-DEL-108 |
| chr1-120956549-DEL-739 | chr1-120956550-DEL-739 |
| chr1-120963369-DEL-6328 | chr1-120963370-DEL-6328 |
| chr1-121065449-DEL-425 | chr1-121065450-DEL-425 |
| chr1-121077515-DEL-72 | chr1-121077482-DEL-78 |
| chr1-121080934-DEL-66 | chr1-121080926-DEL-75 |
| chr1-121089147-DEL-1330 | chr1-121089148-DEL-1330 |
| chr1-121124262-DEL-1649 | chr1-121124263-DEL-1649 |
| chr1-121130314-DEL-8476 | chr1-121130315-DEL-8476 |
| chr1-121140920-DEL-1241 | chr1-121140921-DEL-1241 |
| chr1-121172367-DEL-484 | chr1-121172368-DEL-484 |
| chr1-121209591-DEL-93 | chr1-121209592-DEL-93 |
| chr1-121210642-DEL-60 | chr1-121210643-DEL-60 |
| chr1-121217384-DEL-60 | chr1-121217387-DEL-70 |
| chr1-121221868-DEL-60 | chr1-121221869-DEL-60 |
| chr1-121224193-DEL-68 | chr1-121224194-DEL-68 |
| chr1-121226631-DEL-126 | chr1-121226632-DEL-126 |
| chr1-121235488-DEL-51 | chr1-121235486-DEL-52 |
| chr1-121245679-DEL-1581 | chr1-121245680-DEL-1581 |
| chr1-121263969-DEL-111 | chr1-121263970-DEL-111 |

| | |
|---|---|
| chr1-121267310-DEL-504 | chr1-121267311-DEL-504 |
| chr1-121302367-DEL-51 | chr1-121302368-DEL-52 |
| chr1-121308894-DEL-127 | chr1-121308891-DEL-123 |
| chr1-121369750-DEL-74 | chr1-121369751-DEL-74 |
| chr1-121381955-DEL-225 | chr1-121381956-DEL-225 |
| chr1-121382572-DEL-127 | chr1-121382573-DEL-99 |
| chr1-121385375-DEL-824 | chr1-121385376-DEL-824 |
| chr1-121411166-DEL-1040 | chr1-121411193-DEL-1014 |
| chr1-121411754-DEL-128 | chr1-121411704-DEL-130 |
| chr1-121414127-DEL-109 | chr1-121414128-DEL-109 |
| chr1-121432868-DEL-167 | chr1-121432868-DEL-165 |
| chr1-121434715-DEL-465 | chr1-121434716-DEL-465 |
| chr1-121440737-DEL-67 | chr1-121440738-DEL-67 |
| chr1-121443147-DEL-1421 | chr1-121443148-DEL-1421 |
| chr1-121466985-DEL-72 | chr1-121466986-DEL-72 |
| chr1-121518346-DEL-266 | chr1-121518347-DEL-266 |
| chr1-121533941-DEL-103 | chr1-121533943-DEL-101 |
| chr1-121537805-DEL-61 | chr1-121537812-DEL-54 |
| chr1-121549596-DEL-6705 | chr1-121549597-DEL-6705 |
| chr1-121549602-DEL-498 | chr1-121549603-DEL-498 |
| chr1-121565945-DEL-304 | chr1-121565946-DEL-304 |
| chr1-121566354-DEL-823 | chr1-121566355-DEL-824 |
| chr1-121584610-DEL-72 | chr1-121584611-DEL-51 |
| chr1-121585741-DEL-470 | chr1-121585742-DEL-470 |
| chr1-121591303-DEL-70 | chr1-121591308-DEL-66 |
| chr1-121591559-DEL-6950 | chr1-121591560-DEL-6950 |
| chr1-121592485-DEL-122 | chr1-121592486-DEL-122 |
| chr1-121600477-DEL-3104 | chr1-121600478-DEL-3104 |
| chr1-121620321-DEL-772 | chr1-121620322-DEL-772 |
| chr1-121624632-DEL-70 | chr1-121624633-DEL-70 |
| chr1-121665595-DEL-6370 | chr1-121665596-DEL-6370 |
| chr1-121704969-DEL-4466 | chr1-121704970-DEL-4450 |
| chr1-121714478-DEL-330 | chr1-121714478-DEL-330 |
| chr1-121719813-DEL-56 | chr1-121719825-DEL-52 |
| chr1-121720810-DEL-108 | chr1-121720806-DEL-108 |
| chr1-121726507-DEL-90 | chr1-121726495-DEL-98 |
| chr1-121748938-DEL-1076 | chr1-121748939-DEL-1076 |
| chr1-121765668-DEL-1286 | chr1-121765669-DEL-1286 |
| chr1-121771177-DEL-159 | chr1-121771186-DEL-151 |
| chr1-121785786-DEL-342 | chr1-121785787-DEL-342 |
| chr1-121813923-DEL-60 | chr1-121813916-DEL-64 |
| chr1-121818969-DEL-449 | chr1-121818970-DEL-449 |
| chr1-121828497-DEL-305 | chr1-121828498-DEL-305 |
| chr1-121831640-DEL-60 | chr1-121831641-DEL-60 |
| chr1-121911652-DEL-53 | chr1-121911649-DEL-52 |
| chr1-121917405-DEL-377 | chr1-121917409-DEL-368 |
| chr1-121930354-DEL-752 | chr1-121930355-DEL-752 |
| chr1-121933536-DEL-73 | chr1-121933536-DEL-73 |
| chr1-121936117-DEL-73 | chr1-121936118-DEL-73 |
| chr1-121965400-DEL-199 | chr1-121965400-DEL-199 |
| chr1-122008592-DEL-145 | chr1-122008593-DEL-145 |
| chr1-122014133-DEL-7060 | chr1-122014134-DEL-7060 |
| chr1-122054242-DEL-312 | chr1-122054243-DEL-312 |
| chr1-122063294-DEL-263 | chr1-122063295-DEL-263 |
| chr1-122064778-DEL-2166 | chr1-122064779-DEL-2166 |
| chr1-122186838-DEL-65 | chr1-122186839-DEL-65 |
| chr1-122187691-DEL-78 | chr1-122187692-DEL-78 |
| chr1-122188432-DEL-55 | chr1-122188433-DEL-51 |
| chr1-122254681-DEL-170 | chr1-122254688-DEL-164 |
| chr1-122263049-DEL-77 | chr1-122263055-DEL-72 |
| chr1-122268973-DEL-504 | chr1-122268974-DEL-504 |
| chr1-122284851-DEL-223 | chr1-122284852-DEL-223 |
| chr1-122286814-DEL-156 | chr1-122286813-DEL-152 |
| chr1-122311385-DEL-106 | chr1-122311398-DEL-101 |
| chr1-122312855-DEL-899 | chr1-122312856-DEL-899 |
| chr1-122317132-DEL-60 | chr1-122317137-DEL-54 |
| chr1-122320604-DEL-121 | chr1-122320605-DEL-121 |
| chr1-122380572-DEL-134 | chr1-122380573-DEL-134 |
| chr1-122384828-DEL-158 | chr1-122384830-DEL-157 |
| chr1-122448494-DEL-54 | chr1-122448484-DEL-60 |
| chr1-122450736-DEL-1846 | chr1-122450737-DEL-1846 |
| chr1-122494448-DEL-173 | chr1-122494449-DEL-173 |
| chr1-122505539-DEL-209 | chr1-122505540-DEL-209 |
| chr1-122506159-DEL-54 | chr1-122506160-DEL-54 |
| chr1-122506866-DEL-353 | chr1-122506867-DEL-353 |
| chr1-122508815-DEL-644 | chr1-122508816-DEL-643 |
| chr1-122533401-DEL-1546 | chr1-122533402-DEL-1546 |
| chr1-122554473-DEL-65 | chr1-122554474-DEL-50 |
| chr1-122563562-DEL-497 | chr1-122563563-DEL-497 |
| chr1-122572755-DEL-6207 | chr1-122572756-DEL-6207 |

| | |
|---|---|
| chr1-122579184-DEL-130 | chr1-122579165-DEL-150 |
| chr1-122579738-DEL-6210 | chr1-122579739-DEL-6210 |
| chr1-122616624-DEL-361 | chr1-122616630-DEL-329 |
| chr1-122623624-DEL-1451 | chr1-122623625-DEL-1451 |
| chr1-122625644-DEL-61 | chr1-122625598-DEL-66 |
| chr1-122627873-DEL-64 | chr1-122627874-DEL-64 |
| chr1-122639179-DEL-204 | chr1-122639180-DEL-204 |
| chr1-122643005-DEL-28567 | chr1-122643000-DEL-28572 |
| chr1-122670735-DEL-68 | chr1-122670719-DEL-72 |
| chr1-122674919-DEL-120 | chr1-122674934-DEL-80 |
| chr1-122693840-DEL-7640 | chr1-122693842-DEL-7639 |
| chr1-122700897-DEL-208 | chr1-122700897-DEL-208 |
| chr1-122704656-DEL-780 | chr1-122704657-DEL-780 |
| chr1-122709861-DEL-110 | chr1-122709862-DEL-103 |
| chr1-122714342-DEL-1325 | chr1-122714343-DEL-1325 |
| chr1-122720206-DEL-68 | chr1-122720207-DEL-68 |
| chr1-122721275-DEL-230 | chr1-122721276-DEL-230 |
| chr1-122737531-DEL-248 | chr1-122737532-DEL-248 |
| chr1-122754971-DEL-71 | chr1-122754941-DEL-96 |
| chr1-122759650-DEL-414 | chr1-122759651-DEL-414 |
| chr1-122774531-DEL-1075 | chr1-122774532-DEL-1075 |
| chr1-122775379-DEL-77 | chr1-122775380-DEL-77 |
| chr1-122786371-DEL-93 | chr1-122786371-DEL-92 |
| chr1-122801326-DEL-2403 | chr1-122801327-DEL-2403 |
| chr1-122819382-DEL-3207 | chr1-122819383-DEL-3207 |
| chr1-122819748-DEL-507 | chr1-122819749-DEL-507 |
| chr1-122820579-DEL-1589 | chr1-122820580-DEL-1589 |
| chr1-122830864-DEL-54 | chr1-122830864-DEL-54 |
| chr1-122837771-DEL-1948 | chr1-122837772-DEL-1948 |
| chr1-122880597-DEL-6389 | chr1-122880597-DEL-6389 |
| chr1-122880647-DEL-201 | chr1-122880648-DEL-201 |
| chr1-122892579-DEL-161 | chr1-122892580-DEL-161 |
| chr1-122914161-DEL-69 | chr1-122914162-DEL-69 |
| chr1-122937752-DEL-115 | chr1-122937753-DEL-115 |
| chr1-122976779-DEL-424 | chr1-122976780-DEL-424 |
| chr1-123000324-DEL-908 | chr1-123000327-DEL-907 |
| chr1-123001949-DEL-166 | chr1-123001938-DEL-169 |
| chr1-123045899-DEL-60 | chr1-123045900-DEL-60 |
| chr1-123073232-DEL-2570 | chr1-123073231-DEL-2571 |
| chr1-123096892-DEL-166 | chr1-123096892-DEL-167 |
| chr1-123099607-DEL-205 | chr1-123099608-DEL-205 |
| chr1-123111719-DEL-1593 | chr1-123111721-DEL-1592 |
| chr1-123124374-DEL-6149 | chr1-123124375-DEL-6149 |
| chr1-123131094-DEL-166 | chr1-123131095-DEL-166 |
| chr1-123137853-DEL-1999 | chr1-123137853-DEL-1999 |
| chr1-123139897-DEL-3338 | chr1-123139898-DEL-3338 |
| chr1-123157860-DEL-66 | chr1-123157865-DEL-65 |
| chr1-123168956-DEL-91 | chr1-123168957-DEL-91 |
| chr1-123171975-DEL-58 | chr1-123171948-DEL-86 |
| chr1-123178313-DEL-100 | chr1-123178314-DEL-99 |
| chr1-123231538-DEL-115 | chr1-123231539-DEL-115 |
| chr1-123246613-DEL-576 | chr1-123246614-DEL-576 |
| chr1-123267727-DEL-239 | chr1-123267726-DEL-256 |
| chr1-123298383-DEL-63 | chr1-123298403-DEL-52 |
| chr1-123311000-DEL-916 | chr1-123311001-DEL-916 |
| chr1-123326412-DEL-756 | chr1-123326413-DEL-747 |
| chr1-123332146-DEL-1165 | chr1-123332147-DEL-1165 |
| chr1-123334359-DEL-2232 | chr1-123334360-DEL-2232 |
| chr1-123335578-DEL-1533 | chr1-123335579-DEL-1533 |
| chr1-123346442-DEL-154 | chr1-123346443-DEL-154 |
| chr1-123366164-DEL-110 | chr1-123366167-DEL-108 |
| chr1-123369565-DEL-52 | chr1-123369558-DEL-60 |
| chr1-123380522-DEL-51 | chr1-123380526-DEL-52 |
| chr1-123392957-DEL-60 | chr1-123392958-DEL-60 |
| chr1-123435822-DEL-79 | chr1-123435823-DEL-79 |
| chr1-123436822-DEL-60 | chr1-123436837-DEL-50 |
| chr1-123464240-DEL-120 | chr1-123464255-DEL-104 |
| chr1-123471379-DEL-60 | chr1-123471380-DEL-54 |
| chr1-123477071-DEL-217 | chr1-123477072-DEL-217 |
| chr1-123495243-DEL-1208 | chr1-123495244-DEL-1206 |
| chr1-123559382-DEL-80 | chr1-123559395-DEL-58 |
| chr1-123561610-DEL-684 | chr1-123561611-DEL-684 |
| chr1-123577150-DEL-63 | chr1-123577166-DEL-74 |
| chr1-123578929-DEL-1688 | chr1-123578930-DEL-1688 |
| chr1-123590275-DEL-530 | chr1-123590272-DEL-524 |
| chr1-123596958-DEL-62 | chr1-123596930-DEL-68 |
| chr1-123613085-DEL-706 | chr1-123613086-DEL-706 |
| chr1-123616858-DEL-80 | chr1-123616853-DEL-85 |
| chr1-123632340-DEL-906 | chr1-123632348-DEL-899 |
| chr1-123709339-DEL-176 | chr1-123709340-DEL-170 |

| | |
|---|---|
| chr1-123738199-DEL-184 | chr1-123738200-DEL-184 |
| chr1-123741220-DEL-52 | chr1-123741221-DEL-52 |
| chr1-123747239-DEL-1338 | chr1-123747224-DEL-1332 |
| chr1-123784754-DEL-6465 | chr1-123784755-DEL-6465 |
| chr1-123811834-DEL-58 | chr1-123811842-DEL-58 |
| chr1-123828429-DEL-68 | chr1-123828422-DEL-69 |
| chr1-123837174-DEL-221 | chr1-123837175-DEL-216 |
| chr1-123845278-DEL-128 | chr1-123845297-DEL-123 |
| chr1-123852463-DEL-68 | chr1-123852464-DEL-64 |
| chr1-123875261-DEL-62 | chr1-123875261-DEL-68 |
| chr1-123934750-DEL-1451 | chr1-123934751-DEL-1451 |
| chr1-124013272-DEL-6240 | chr1-124013273-DEL-6240 |
| chr1-124026693-DEL-491 | chr1-124026694-DEL-491 |
| chr1-124035654-DEL-678 | chr1-124035655-DEL-678 |
| chr1-124045962-DEL-945 | chr1-124045963-DEL-945 |
| chr1-124050740-DEL-63 | chr1-124050741-DEL-63 |
| chr1-124064714-DEL-251 | chr1-124064715-DEL-251 |
| chr1-124069067-DEL-6461 | chr1-124069068-DEL-6461 |
| chr1-124077270-DEL-54 | chr1-124077271-DEL-54 |
| chr1-124092014-DEL-1848 | chr1-124092010-DEL-1846 |
| chr1-124101251-DEL-68 | chr1-124101255-DEL-50 |
| chr1-124104605-DEL-592 | chr1-124104640-DEL-589 |
| chr1-124109757-DEL-51 | chr1-124109758-DEL-51 |
| chr1-124120128-DEL-113 | chr1-124120164-DEL-74 |
| chr1-124123773-DEL-6133 | chr1-124123774-DEL-6133 |
| chr1-124184232-DEL-153 | chr1-124184233-DEL-153 |
| chr1-124216670-DEL-84 | chr1-124216683-DEL-74 |
| chr1-124236354-DEL-571 | chr1-124236355-DEL-571 |
| chr1-124237599-DEL-567 | chr1-124237600-DEL-567 |
| chr1-124272350-DEL-354 | chr1-124272350-DEL-354 |
| chr1-124334718-DEL-130 | chr1-124334709-DEL-136 |
| chr1-124348083-DEL-75 | chr1-124348080-DEL-85 |
| chr1-124366108-DEL-24390 | chr1-124366108-DEL-24390 |
| chr1-124374502-DEL-410 | chr1-124374502-DEL-410 |
| chr1-124404267-DEL-1186 | chr1-124404268-DEL-1186 |
| chr1-124412730-DEL-52 | chr1-124412731-DEL-52 |
| chr1-124421570-DEL-209 | chr1-124421572-DEL-207 |
| chr1-124435241-DEL-54 | chr1-124435245-DEL-51 |
| chr1-124447561-DEL-974 | chr1-124447561-DEL-974 |
| chr1-124449522-DEL-51 | chr1-124449540-DEL-50 |
| chr1-124452605-DEL-78 | chr1-124452605-DEL-74 |
| chr1-124464192-DEL-65 | chr1-124464192-DEL-65 |
| chr1-124465956-DEL-1338 | chr1-124465957-DEL-1338 |
| chr1-124471141-DEL-78 | chr1-124471142-DEL-78 |
| chr1-124490041-DEL-62 | chr1-124490042-DEL-56 |
| chr1-124505769-DEL-58 | chr1-124505770-DEL-58 |
| chr1-124541822-DEL-533 | chr1-124541823-DEL-532 |
| chr1-124560108-DEL-756 | chr1-124560091-DEL-773 |
| chr1-124569455-DEL-3061 | chr1-124569452-DEL-3062 |
| chr1-124575502-DEL-56 | chr1-124575503-DEL-56 |
| chr1-124588290-DEL-535 | chr1-124588291-DEL-535 |
| chr1-124602660-DEL-52 | chr1-124602652-DEL-66 |
| chr1-124629711-DEL-80 | chr1-124629696-DEL-74 |
| chr1-124643198-DEL-142 | chr1-124643198-DEL-142 |
| chr1-124649786-DEL-98 | chr1-124649793-DEL-92 |
| chr1-124658839-DEL-65 | chr1-124658840-DEL-65 |
| chr1-124688899-DEL-56 | chr1-124688900-DEL-56 |
| chr1-124712464-DEL-98 | chr1-124712423-DEL-98 |
| chr1-124778405-DEL-107 | chr1-124778406-DEL-107 |
| chr1-124807124-DEL-110 | chr1-124807125-DEL-110 |
| chr1-124808488-DEL-392 | chr1-124808489-DEL-392 |
| chr1-124826687-DEL-126 | chr1-124826656-DEL-170 |
| chr1-124840422-DEL-576 | chr1-124840427-DEL-575 |
| chr1-124849759-DEL-214 | chr1-124849765-DEL-214 |
| chr1-124850450-DEL-556 | chr1-124850451-DEL-556 |
| chr1-124869999-DEL-90 | chr1-124870000-DEL-90 |
| chr1-124889917-DEL-1244 | chr1-124889918-DEL-1244 |
| chr1-124893448-DEL-6684 | chr1-124893449-DEL-6684 |
| chr1-124901419-DEL-70 | chr1-124901420-DEL-70 |
| chr1-124905122-DEL-63 | chr1-124905123-DEL-63 |
| chr1-124928243-DEL-401 | chr1-124928243-DEL-400 |
| chr1-124935174-DEL-579 | chr1-124935175-DEL-579 |
| chr1-124941105-DEL-727 | chr1-124941066-DEL-729 |
| chr1-124952831-DEL-327 | chr1-124952832-DEL-327 |
| chr1-124961884-DEL-56 | chr1-124961870-DEL-70 |
| chr1-124978945-DEL-2195 | chr1-124978946-DEL-2196 |
| chr1-124989722-DEL-249 | chr1-124989723-DEL-249 |
| chr1-125018685-DEL-6367 | chr1-125018686-DEL-6367 |
| chr1-125041448-DEL-1135 | chr1-125041449-DEL-1135 |
| chr1-125059970-DEL-12609 | chr1-125059971-DEL-12609 |

| | |
|---|---|
| chr1-125084682-DEL-244 | chr1-125084684-DEL-237 |
| chr1-125089028-DEL-1585 | chr1-125089023-DEL-1586 |
| chr1-125091237-DEL-3309 | chr1-125091238-DEL-3309 |
| chr1-125100721-DEL-79 | chr1-125100716-DEL-84 |
| chr1-125120456-DEL-1306 | chr1-125120457-DEL-1306 |
| chr1-125129774-DEL-83 | chr1-125129771-DEL-87 |
| chr1-125165379-DEL-54 | chr1-125165380-DEL-60 |
| chr1-125171968-DEL-3062 | chr1-125171969-DEL-3062 |
| chr1-125185006-DEL-2539 | chr1-125185007-DEL-2539 |
| chr1-125185971-DEL-400 | chr1-125185972-DEL-400 |
| chr1-125203876-DEL-2508 | chr1-125203874-DEL-2511 |
| chr1-125208528-DEL-63 | chr1-125208529-DEL-63 |
| chr1-125210710-DEL-92 | chr1-125210701-DEL-110 |
| chr1-125244494-DEL-7215 | chr1-125244495-DEL-7215 |
| chr1-125268934-DEL-726 | chr1-125268935-DEL-726 |
| chr1-125273868-DEL-4195 | chr1-125273869-DEL-4195 |
| chr1-125282050-DEL-66 | chr1-125282071-DEL-58 |
| chr1-125283214-DEL-7311 | chr1-125283234-DEL-7293 |
| chr1-125289555-DEL-825 | chr1-125289556-DEL-825 |
| chr1-125317588-DEL-382 | chr1-125317589-DEL-382 |
| chr1-125317879-DEL-173 | chr1-125317879-DEL-173 |
| chr1-125404422-DEL-86 | chr1-125404423-DEL-84 |
| chr1-125406045-DEL-107 | chr1-125406026-DEL-108 |
| chr1-125412105-DEL-2891 | chr1-125412106-DEL-2891 |
| chr1-125423663-DEL-216 | chr1-125423664-DEL-216 |
| chr1-125438560-DEL-2765 | chr1-125438561-DEL-2765 |
| chr1-125443093-DEL-52 | chr1-125443093-DEL-52 |
| chr1-125446356-DEL-56 | chr1-125446357-DEL-56 |
| chr1-125480407-DEL-3184 | chr1-125480407-DEL-3184 |
| chr1-125511431-DEL-70 | chr1-125511402-DEL-86 |
| chr1-125538415-DEL-102 | chr1-125538416-DEL-102 |
| chr1-125539434-DEL-70 | chr1-125539435-DEL-70 |
| chr1-125541894-DEL-101 | chr1-125541904-DEL-98 |
| chr1-125556960-DEL-53 | chr1-125556957-DEL-57 |
| chr1-125615061-DEL-102 | chr1-125615052-DEL-100 |
| chr1-125617377-DEL-52 | chr1-125617377-DEL-52 |
| chr1-125674616-DEL-57 | chr1-125674617-DEL-57 |
| chr1-125692480-DEL-69 | chr1-125692443-DEL-92 |
| chr1-125710188-DEL-1391 | chr1-125710189-DEL-1391 |
| chr1-125719564-DEL-57 | chr1-125719563-DEL-50 |
| chr1-125753270-DEL-57 | chr1-125753282-DEL-56 |
| chr1-125759383-DEL-58 | chr1-125759384-DEL-57 |
| chr1-125775275-DEL-72 | chr1-125775276-DEL-72 |
| chr1-125791170-DEL-93 | chr1-125791171-DEL-93 |
| chr1-125819878-DEL-58 | chr1-125819879-DEL-58 |
| chr1-125865076-DEL-51 | chr1-125865077-DEL-51 |
| chr1-125867139-DEL-495 | chr1-125867140-DEL-495 |
| chr1-125901842-DEL-66 | chr1-125901841-DEL-68 |
| chr1-125940908-DEL-79 | chr1-125940909-DEL-79 |
| chr1-125969725-DEL-215 | chr1-125969726-DEL-215 |
| chr1-125993597-DEL-66 | chr1-125993608-DEL-66 |
| chr1-126006459-DEL-60 | chr1-126006468-DEL-50 |
| chr1-126011767-DEL-140 | chr1-126011768-DEL-139 |
| chr1-126023226-DEL-58 | chr1-126023239-DEL-54 |
| chr1-126031372-DEL-606 | chr1-126031373-DEL-606 |
| chr1-126039684-DEL-195 | chr1-126039685-DEL-189 |
| chr1-126040620-DEL-70 | chr1-126040621-DEL-70 |
| chr1-126048853-DEL-74 | chr1-126048854-DEL-74 |
| chr1-126077603-DEL-132 | chr1-126077604-DEL-132 |
| chr1-126078361-DEL-77 | chr1-126078362-DEL-77 |
| chr1-126091765-DEL-1254 | chr1-126091766-DEL-1254 |
| chr1-126150767-DEL-166 | chr1-126150768-DEL-166 |
| chr1-126165386-DEL-68 | chr1-126165388-DEL-59 |
| chr1-126186960-DEL-101 | chr1-126186961-DEL-101 |
| chr1-126214535-DEL-53 | chr1-126214534-DEL-53 |
| chr1-126217973-DEL-80 | chr1-126217974-DEL-80 |
| chr1-126242311-DEL-473 | chr1-126242312-DEL-473 |
| chr1-126247044-DEL-180 | chr1-126247045-DEL-180 |
| chr1-126256118-DEL-60 | chr1-126256119-DEL-56 |
| chr1-126266183-DEL-128 | chr1-126266184-DEL-128 |
| chr1-126270375-DEL-178 | chr1-126270376-DEL-178 |
| chr1-126286079-DEL-54 | chr1-126286079-DEL-54 |
| chr1-126309192-DEL-54 | chr1-126309193-DEL-54 |
| chr1-126339416-DEL-113 | chr1-126339417-DEL-113 |
| chr1-126343955-DEL-1358 | chr1-126343956-DEL-1358 |
| chr1-126351755-DEL-5676 | chr1-126351756-DEL-5676 |
| chr1-126400162-DEL-5168 | chr1-126400163-DEL-5168 |
| chr1-126401988-DEL-118 | chr1-126401966-DEL-146 |
| chr1-126406538-DEL-422 | chr1-126406539-DEL-422 |
| chr1-126413089-DEL-66 | chr1-126413090-DEL-66 |

| | |
|---|---|
| chr1-126425944-DEL-93 | chr1-126425963-DEL-68 |
| chr1-126428965-DEL-94 | chr1-126428966-DEL-94 |
| chr1-126485695-DEL-3448 | chr1-126485696-DEL-3448 |
| chr1-126513239-DEL-418 | chr1-126513240-DEL-418 |
| chr1-126527727-DEL-123 | chr1-126527728-DEL-123 |
| chr1-126538350-DEL-138 | chr1-126538364-DEL-126 |
| chr1-126547732-DEL-399 | chr1-126547731-DEL-399 |
| chr1-126555148-DEL-2913 | chr1-126555149-DEL-2913 |
| chr1-126560675-DEL-1080 | chr1-126560676-DEL-1080 |
| chr1-126563506-DEL-806 | chr1-126563502-DEL-803 |
| chr1-126565116-DEL-76 | chr1-126565114-DEL-76 |
| chr1-126589946-DEL-384 | chr1-126589947-DEL-384 |
| chr1-126591403-DEL-6489 | chr1-126591404-DEL-6489 |
| chr1-126610222-DEL-65 | chr1-126610223-DEL-65 |
| chr1-126648413-DEL-51 | chr1-126648414-DEL-51 |
| chr1-126652772-DEL-68 | chr1-126652773-DEL-52 |
| chr1-126662396-DEL-461 | chr1-126662397-DEL-461 |
| chr1-126674081-DEL-146 | chr1-126674105-DEL-110 |
| chr1-126684738-DEL-62 | chr1-126684739-DEL-64 |
| chr1-126688095-DEL-54 | chr1-126688096-DEL-54 |
| chr1-126709804-DEL-733 | chr1-126709805-DEL-733 |
| chr1-126719931-DEL-6163 | chr1-126719932-DEL-6163 |
| chr1-126735773-DEL-100 | chr1-126735795-DEL-110 |
| chr1-126804632-DEL-54 | chr1-126804633-DEL-54 |
| chr1-126812018-DEL-169 | chr1-126812024-DEL-163 |
| chr1-126832626-DEL-317 | chr1-126832631-DEL-313 |
| chr1-126860739-DEL-108 | chr1-126860740-DEL-108 |
| chr1-126866902-DEL-5217 | chr1-126866903-DEL-5217 |
| chr1-126897894-DEL-796 | chr1-126897895-DEL-796 |
| chr1-126926617-DEL-51 | chr1-126926618-DEL-51 |
| chr1-126935899-DEL-85 | chr1-126935900-DEL-96 |
| chr1-126941401-DEL-127 | chr1-126941402-DEL-127 |
| chr1-126949686-DEL-54 | chr1-126949686-DEL-54 |
| chr1-126952907-DEL-58 | chr1-126952918-DEL-52 |
| chr1-126955140-DEL-518 | chr1-126955141-DEL-518 |
| chr1-127016724-DEL-1123 | chr1-127016725-DEL-1123 |
| chr1-127050863-DEL-88 | chr1-127050864-DEL-88 |
| chr1-127057787-DEL-480 | chr1-127057790-DEL-481 |
| chr1-127061476-DEL-182 | chr1-127061477-DEL-182 |
| chr1-127087292-DEL-84 | chr1-127087284-DEL-75 |
| chr1-127124893-DEL-76 | chr1-127124892-DEL-50 |
| chr1-127141784-DEL-446 | chr1-127141785-DEL-446 |
| chr1-127164785-DEL-708 | chr1-127164783-DEL-702 |
| chr1-127209229-DEL-2409 | chr1-127209230-DEL-2409 |
| chr1-127213977-DEL-473 | chr1-127213975-DEL-473 |
| chr1-127216108-DEL-87 | chr1-127216109-DEL-87 |
| chr1-127260667-DEL-171 | chr1-127260668-DEL-171 |
| chr1-127290100-DEL-145 | chr1-127290101-DEL-145 |
| chr1-127316907-DEL-1020 | chr1-127316905-DEL-1021 |
| chr1-127319124-DEL-141 | chr1-127319125-DEL-141 |
| chr1-127329704-DEL-52 | chr1-127329705-DEL-52 |
| chr1-127363870-DEL-90 | chr1-127363871-DEL-90 |
| chr1-127382647-DEL-439 | chr1-127382648-DEL-439 |
| chr1-127391344-DEL-53 | chr1-127391344-DEL-54 |
| chr1-127407756-DEL-2411 | chr1-127407757-DEL-2411 |
| chr1-127432040-DEL-116 | chr1-127432053-DEL-84 |
| chr1-127447818-DEL-64 | chr1-127447819-DEL-64 |
| chr1-127453787-DEL-647 | chr1-127453788-DEL-647 |
| chr1-127454751-DEL-2151 | chr1-127454752-DEL-2151 |
| chr1-127460357-DEL-108 | chr1-127460400-DEL-97 |
| chr1-127462623-DEL-75 | chr1-127462624-DEL-75 |
| chr1-127463945-DEL-129 | chr1-127463946-DEL-129 |
| chr1-127483475-DEL-51 | chr1-127483476-DEL-51 |
| chr1-127485987-DEL-7233 | chr1-127485988-DEL-7233 |
| chr1-127497562-DEL-922 | chr1-127497560-DEL-919 |
| chr1-127499038-DEL-247 | chr1-127499035-DEL-285 |
| chr1-127518494-DEL-3137 | chr1-127518495-DEL-3137 |
| chr1-127538099-DEL-56 | chr1-127538077-DEL-104 |
| chr1-127542493-DEL-879 | chr1-127542494-DEL-879 |
| chr1-127581122-DEL-132 | chr1-127581124-DEL-130 |
| chr1-127611613-DEL-2459 | chr1-127611614-DEL-2459 |
| chr1-127618418-DEL-144 | chr1-127618425-DEL-138 |
| chr1-127624658-DEL-68 | chr1-127624613-DEL-100 |
| chr1-127628902-DEL-340 | chr1-127628903-DEL-340 |
| chr1-127630586-DEL-56 | chr1-127630587-DEL-50 |
| chr1-127647026-DEL-743 | chr1-127647027-DEL-743 |
| chr1-127653044-DEL-64 | chr1-127653031-DEL-75 |
| chr1-127763294-DEL-71 | chr1-127763295-DEL-71 |
| chr1-127772498-DEL-542 | chr1-127772499-DEL-542 |
| chr1-127773686-DEL-226 | chr1-127773698-DEL-220 |

| | |
|---|---|
| chr1-127785383-DEL-117 | chr1-127785410-DEL-73 |
| chr1-127795413-DEL-102 | chr1-127795414-DEL-102 |
| chr1-127832715-DEL-1554 | chr1-127832716-DEL-1554 |
| chr1-127834541-DEL-206 | chr1-127834540-DEL-206 |
| chr1-127836341-DEL-189 | chr1-127836342-DEL-189 |
| chr1-127856301-DEL-58 | chr1-127856316-DEL-56 |
| chr1-127857835-DEL-214 | chr1-127857836-DEL-214 |
| chr1-127899993-DEL-63 | chr1-127900000-DEL-75 |
| chr1-127961682-DEL-84 | chr1-127961683-DEL-84 |
| chr1-127966243-DEL-537 | chr1-127966244-DEL-537 |
| chr1-127992824-DEL-1797 | chr1-127992805-DEL-1798 |
| chr1-127995977-DEL-62 | chr1-127995984-DEL-56 |
| chr1-128049734-DEL-203 | chr1-128049735-DEL-203 |
| chr1-128117447-DEL-60 | chr1-128117448-DEL-60 |
| chr1-128136588-DEL-213 | chr1-128136589-DEL-213 |
| chr1-128156095-DEL-1939 | chr1-128156096-DEL-1939 |
| chr1-128161024-DEL-639 | chr1-128161037-DEL-627 |
| chr1-128177890-DEL-150 | chr1-128177891-DEL-150 |
| chr1-128190155-DEL-224 | chr1-128190156-DEL-224 |
| chr1-128209881-DEL-208 | chr1-128209882-DEL-208 |
| chr1-128210499-DEL-82 | chr1-128210500-DEL-82 |
| chr1-128243408-DEL-1862 | chr1-128243409-DEL-1862 |
| chr1-128273175-DEL-148 | chr1-128273176-DEL-148 |
| chr1-128275941-DEL-84 | chr1-128275942-DEL-84 |
| chr1-128292889-DEL-213 | chr1-128292889-DEL-213 |
| chr1-128385930-DEL-97 | chr1-128385931-DEL-97 |
| chr1-128393631-DEL-220 | chr1-128393611-DEL-264 |
| chr1-128397393-DEL-51 | chr1-128397372-DEL-66 |
| chr1-128420235-DEL-245 | chr1-128420236-DEL-245 |
| chr1-128428600-DEL-3115 | chr1-128428601-DEL-3115 |
| chr1-128480131-DEL-82 | chr1-128480132-DEL-80 |
| chr1-128482026-DEL-80 | chr1-128482027-DEL-80 |
| chr1-128487321-DEL-56 | chr1-128487321-DEL-56 |
| chr1-128505409-DEL-60 | chr1-128505410-DEL-76 |
| chr1-128518197-DEL-60 | chr1-128518206-DEL-52 |
| chr1-128528889-DEL-91 | chr1-128528890-DEL-91 |
| chr1-128529692-DEL-60 | chr1-128529685-DEL-68 |
| chr1-128537426-DEL-52 | chr1-128537427-DEL-52 |
| chr1-128548327-DEL-472 | chr1-128548328-DEL-472 |
| chr1-128618314-DEL-74 | chr1-128618315-DEL-74 |
| chr1-128621113-DEL-129 | chr1-128621114-DEL-129 |
| chr1-128676755-DEL-52 | chr1-128676756-DEL-52 |
| chr1-128678075-DEL-61 | chr1-128678078-DEL-59 |
| chr1-128683906-DEL-104 | chr1-128683903-DEL-105 |
| chr1-128692843-DEL-66 | chr1-128692867-DEL-65 |
| chr1-128700918-DEL-6517 | chr1-128700919-DEL-6517 |
| chr1-128711799-DEL-103 | chr1-128711800-DEL-103 |
| chr1-128719314-DEL-56 | chr1-128719314-DEL-56 |
| chr1-128738613-DEL-51 | chr1-128738622-DEL-50 |
| chr1-128774488-DEL-6597 | chr1-128774488-DEL-6597 |
| chr1-128820478-DEL-107 | chr1-128820459-DEL-102 |
| chr1-128828199-DEL-59 | chr1-128828198-DEL-59 |
| chr1-128847068-DEL-825 | chr1-128847069-DEL-825 |
| chr1-128853107-DEL-80 | chr1-128853108-DEL-80 |
| chr1-128890001-DEL-1794 | chr1-128890002-DEL-1794 |
| chr1-128892540-DEL-99 | chr1-128892541-DEL-99 |
| chr1-128914643-DEL-795 | chr1-128914641-DEL-796 |
| chr1-128948717-DEL-324 | chr1-128948718-DEL-324 |
| chr1-128976621-DEL-98 | chr1-128976622-DEL-98 |
| chr1-128979519-DEL-78 | chr1-128979520-DEL-78 |
| chr1-129001167-DEL-54 | chr1-129001168-DEL-54 |
| chr1-129059944-DEL-1429 | chr1-129059945-DEL-1429 |
| chr1-129069892-DEL-232 | chr1-129069891-DEL-233 |
| chr1-129110297-DEL-1666 | chr1-129110312-DEL-1651 |
| chr1-129126752-DEL-56 | chr1-129126753-DEL-56 |
| chr1-129150581-DEL-67 | chr1-129150581-DEL-67 |
| chr1-129160833-DEL-495 | chr1-129160834-DEL-495 |
| chr1-129183454-DEL-91 | chr1-129183455-DEL-91 |
| chr1-129214429-DEL-60 | chr1-129214429-DEL-54 |
| chr1-129257436-DEL-7705 | chr1-129257437-DEL-7705 |
| chr1-129258607-DEL-1278 | chr1-129258607-DEL-1278 |
| chr1-129286168-DEL-93 | chr1-129286169-DEL-93 |
| chr1-129290205-DEL-52 | chr1-129290200-DEL-53 |
| chr1-129305475-DEL-92 | chr1-129305476-DEL-90 |
| chr1-129321799-DEL-6333 | chr1-129321796-DEL-6337 |
| chr1-129333024-DEL-1913 | chr1-129333024-DEL-1913 |
| chr1-129347233-DEL-148 | chr1-129347232-DEL-150 |
| chr1-129364013-DEL-6677 | chr1-129364014-DEL-6677 |
| chr1-129412989-DEL-210 | chr1-129412990-DEL-210 |
| chr1-129460493-DEL-337 | chr1-129460494-DEL-337 |

| | |
|---|---|
| chr1-129633025-DEL-969 | chr1-129633026-DEL-969 |
| chr1-129634930-DEL-57 | chr1-129634882-DEL-106 |
| chr1-129646959-DEL-196 | chr1-129646960-DEL-196 |
| chr1-129695599-DEL-6222 | chr1-129695600-DEL-6222 |
| chr1-129699818-DEL-133 | chr1-129699821-DEL-131 |
| chr1-129711392-DEL-789 | chr1-129711393-DEL-789 |
| chr1-129713212-DEL-52 | chr1-129713213-DEL-52 |
| chr1-129726672-DEL-56 | chr1-129726669-DEL-78 |
| chr1-129740396-DEL-922 | chr1-129740391-DEL-923 |
| chr1-129815107-DEL-50 | chr1-129815097-DEL-60 |
| chr1-129838633-DEL-60 | chr1-129838634-DEL-60 |
| chr1-129839190-DEL-1968 | chr1-129839191-DEL-1968 |
| chr1-129841496-DEL-607 | chr1-129841488-DEL-610 |
| chr1-129855588-DEL-61 | chr1-129855589-DEL-61 |
| chr1-129925481-DEL-225 | chr1-129925483-DEL-223 |
| chr1-129934643-DEL-202 | chr1-129934602-DEL-203 |
| chr1-129959527-DEL-6610 | chr1-129959528-DEL-6610 |
| chr1-130067221-DEL-78 | chr1-130067222-DEL-78 |
| chr1-130107369-DEL-52 | chr1-130107370-DEL-52 |
| chr1-130115828-DEL-193 | chr1-130115824-DEL-193 |
| chr1-130119010-DEL-111 | chr1-130119011-DEL-111 |
| chr1-130127374-DEL-6235 | chr1-130127375-DEL-6235 |
| chr1-130140153-DEL-338 | chr1-130140154-DEL-338 |
| chr1-130157844-DEL-1494 | chr1-130157844-DEL-1494 |
| chr1-130170777-DEL-101 | chr1-130170768-DEL-108 |
| chr1-130202366-DEL-2338 | chr1-130202367-DEL-2338 |
| chr1-130221393-DEL-541 | chr1-130221394-DEL-541 |
| chr1-130239875-DEL-806 | chr1-130239876-DEL-806 |
| chr1-130253969-DEL-86 | chr1-130253970-DEL-82 |
| chr1-130258085-DEL-450 | chr1-130258085-DEL-450 |
| chr1-130270098-DEL-78 | chr1-130270100-DEL-81 |
| chr1-130282259-DEL-57 | chr1-130282258-DEL-58 |
| chr1-130331238-DEL-164 | chr1-130331239-DEL-164 |
| chr1-130334668-DEL-866 | chr1-130334669-DEL-867 |
| chr1-130348663-DEL-91 | chr1-130348645-DEL-64 |
| chr1-130351980-DEL-66 | chr1-130351992-DEL-66 |
| chr1-130414505-DEL-302 | chr1-130414506-DEL-302 |
| chr1-130418822-DEL-6649 | chr1-130418823-DEL-6649 |
| chr1-130419183-DEL-212 | chr1-130419184-DEL-212 |
| chr1-130430371-DEL-66 | chr1-130430372-DEL-66 |
| chr1-130447645-DEL-935 | chr1-130447646-DEL-935 |
| chr1-130454398-DEL-413 | chr1-130454397-DEL-414 |
| chr1-130462035-DEL-4926 | chr1-130462036-DEL-4926 |
| chr1-130496573-DEL-52 | chr1-130496574-DEL-52 |
| chr1-130499087-DEL-62 | chr1-130499093-DEL-63 |
| chr1-130505032-DEL-106 | chr1-130505068-DEL-64 |
| chr1-130511506-DEL-78 | chr1-130511506-DEL-78 |
| chr1-130521057-DEL-955 | chr1-130521058-DEL-955 |
| chr1-130527629-DEL-7523 | chr1-130527628-DEL-7528 |
| chr1-130538758-DEL-59 | chr1-130538759-DEL-59 |
| chr1-130545501-DEL-71 | chr1-130545501-DEL-71 |
| chr1-130561328-DEL-171 | chr1-130561331-DEL-169 |
| chr1-130596336-DEL-93 | chr1-130596336-DEL-75 |
| chr1-130600674-DEL-70 | chr1-130600674-DEL-71 |
| chr1-130638009-DEL-54 | chr1-130638000-DEL-66 |
| chr1-130645927-DEL-60 | chr1-130645922-DEL-66 |
| chr1-130674152-DEL-171 | chr1-130674153-DEL-171 |
| chr1-130675987-DEL-452 | chr1-130675988-DEL-452 |
| chr1-130682177-DEL-1715 | chr1-130682178-DEL-1715 |
| chr1-130684121-DEL-305 | chr1-130684122-DEL-305 |
| chr1-130686023-DEL-1543 | chr1-130686024-DEL-1542 |
| chr1-130744004-DEL-52 | chr1-130744005-DEL-52 |
| chr1-130780527-DEL-72 | chr1-130780534-DEL-66 |
| chr1-130796171-DEL-309 | chr1-130796172-DEL-309 |
| chr1-130806547-DEL-54 | chr1-130806554-DEL-66 |
| chr1-130822678-DEL-55 | chr1-130822648-DEL-62 |
| chr1-130823685-DEL-111 | chr1-130823686-DEL-111 |
| chr1-130877378-DEL-54 | chr1-130877381-DEL-55 |
| chr1-130881211-DEL-53 | chr1-130881212-DEL-53 |
| chr1-130882127-DEL-5357 | chr1-130882128-DEL-5357 |
| chr1-130898264-DEL-515 | chr1-130898265-DEL-515 |
| chr1-130909447-DEL-1117 | chr1-130909448-DEL-1117 |
| chr1-130962010-DEL-94 | chr1-130962011-DEL-94 |
| chr1-130994981-DEL-54 | chr1-130994982-DEL-54 |
| chr1-131016356-DEL-78 | chr1-131016357-DEL-78 |
| chr1-131043946-DEL-218 | chr1-131043947-DEL-218 |
| chr1-131045484-DEL-117 | chr1-131045485-DEL-117 |
| chr1-131084349-DEL-964 | chr1-131084357-DEL-957 |
| chr1-131085566-DEL-198 | chr1-131085567-DEL-198 |
| chr1-131087727-DEL-140 | chr1-131087756-DEL-112 |

| | |
|---|---|
| chr1-131106720-DEL-76 | chr1-131106721-DEL-76 |
| chr1-131119569-DEL-1527 | chr1-131119570-DEL-1527 |
| chr1-131123692-DEL-114 | chr1-131123692-DEL-115 |
| chr1-131145479-DEL-80 | chr1-131145480-DEL-80 |
| chr1-131153657-DEL-1139 | chr1-131153648-DEL-1112 |
| chr1-131159480-DEL-59 | chr1-131159481-DEL-59 |
| chr1-131176807-DEL-387 | chr1-131176808-DEL-386 |
| chr1-131190554-DEL-69 | chr1-131190557-DEL-67 |
| chr1-131209109-DEL-112 | chr1-131209062-DEL-110 |
| chr1-131229823-DEL-6851 | chr1-131229824-DEL-6851 |
| chr1-131242698-DEL-213 | chr1-131242697-DEL-214 |
| chr1-131256489-DEL-466 | chr1-131256490-DEL-466 |
| chr1-131288883-DEL-105 | chr1-131288884-DEL-105 |
| chr1-131294455-DEL-453 | chr1-131294458-DEL-454 |
| chr1-131305582-DEL-201 | chr1-131305583-DEL-201 |
| chr1-131363205-DEL-87 | chr1-131363206-DEL-87 |
| chr1-131367219-DEL-202 | chr1-131367220-DEL-202 |
| chr1-131402426-DEL-158 | chr1-131402427-DEL-158 |
| chr1-131416501-DEL-88 | chr1-131416502-DEL-88 |
| chr1-131442421-DEL-55 | chr1-131442414-DEL-53 |
| chr1-131492125-DEL-204 | chr1-131492126-DEL-204 |
| chr1-131501514-DEL-309 | chr1-131501515-DEL-309 |
| chr1-131568319-DEL-57 | chr1-131568320-DEL-57 |
| chr1-131570298-DEL-9058 | chr1-131570317-DEL-9054 |
| chr1-131579952-DEL-64 | chr1-131579943-DEL-74 |
| chr1-131601146-DEL-1084 | chr1-131601147-DEL-1084 |
| chr1-131604399-DEL-895 | chr1-131604400-DEL-895 |
| chr1-131608266-DEL-6463 | chr1-131608267-DEL-6463 |
| chr1-131619649-DEL-64 | chr1-131619650-DEL-64 |
| chr1-131653394-DEL-52 | chr1-131653395-DEL-52 |
| chr1-131656575-DEL-80 | chr1-131656576-DEL-80 |
| chr1-131662251-DEL-60 | chr1-131662252-DEL-66 |
| chr1-131728573-DEL-53 | chr1-131728574-DEL-53 |
| chr1-131779645-DEL-159 | chr1-131779646-DEL-159 |
| chr1-131799066-DEL-204 | chr1-131799067-DEL-204 |
| chr1-131812645-DEL-95 | chr1-131812646-DEL-95 |
| chr1-131825519-DEL-198 | chr1-131825520-DEL-198 |
| chr1-131827799-DEL-53 | chr1-131827800-DEL-53 |
| chr1-131830171-DEL-195 | chr1-131830172-DEL-195 |
| chr1-131878713-DEL-83 | chr1-131878714-DEL-83 |
| chr1-131907656-DEL-198 | chr1-131907657-DEL-198 |
| chr1-131912733-DEL-218 | chr1-131912740-DEL-212 |
| chr1-131926321-DEL-365 | chr1-131926322-DEL-365 |
| chr1-131927366-DEL-208 | chr1-131927354-DEL-209 |
| chr1-131938028-DEL-207 | chr1-131938029-DEL-207 |
| chr1-132001466-DEL-53 | chr1-132001464-DEL-54 |
| chr1-132022115-DEL-123 | chr1-132022116-DEL-123 |
| chr1-132023580-DEL-845 | chr1-132023581-DEL-845 |
| chr1-132032051-DEL-67 | chr1-132032046-DEL-67 |
| chr1-132074855-DEL-57 | chr1-132074858-DEL-50 |
| chr1-132100843-DEL-218 | chr1-132100844-DEL-218 |
| chr1-132102501-DEL-2884 | chr1-132102502-DEL-2884 |
| chr1-132142160-DEL-66 | chr1-132142173-DEL-66 |
| chr1-132151337-DEL-1094 | chr1-132151338-DEL-1094 |
| chr1-132152903-DEL-196 | chr1-132152904-DEL-196 |
| chr1-132159410-DEL-102 | chr1-132159386-DEL-103 |
| chr1-132191121-DEL-581 | chr1-132191118-DEL-582 |
| chr1-132201714-DEL-65 | chr1-132201713-DEL-67 |
| chr1-132211756-DEL-1496 | chr1-132211757-DEL-1496 |
| chr1-132256961-DEL-227 | chr1-132256975-DEL-232 |
| chr1-132262607-DEL-71 | chr1-132262608-DEL-71 |
| chr1-132268530-DEL-58 | chr1-132268505-DEL-58 |
| chr1-132271674-DEL-156 | chr1-132271713-DEL-150 |
| chr1-132276999-DEL-299 | chr1-132277000-DEL-322 |
| chr1-132278018-DEL-91 | chr1-132278019-DEL-91 |
| chr1-132289921-DEL-1005 | chr1-132289920-DEL-1006 |
| chr1-132297139-DEL-84 | chr1-132297152-DEL-72 |
| chr1-132326757-DEL-54 | chr1-132326758-DEL-54 |
| chr1-132337843-DEL-366 | chr1-132337844-DEL-366 |
| chr1-132349223-DEL-231 | chr1-132349205-DEL-203 |
| chr1-132372070-DEL-205 | chr1-132372078-DEL-198 |
| chr1-132419520-DEL-200 | chr1-132419521-DEL-200 |
| chr1-132464000-DEL-1335 | chr1-132464001-DEL-1335 |
| chr1-132466893-DEL-72 | chr1-132466894-DEL-72 |
| chr1-132474286-DEL-692 | chr1-132474285-DEL-694 |
| chr1-132481382-DEL-385 | chr1-132481383-DEL-385 |
| chr1-132535567-DEL-66 | chr1-132535570-DEL-64 |
| chr1-132560797-DEL-99 | chr1-132560798-DEL-99 |
| chr1-132589951-DEL-163 | chr1-132589966-DEL-151 |
| chr1-132595105-DEL-173 | chr1-132595106-DEL-172 |

| | |
|---|---|
| chr1-132596498-DEL-116 | chr1-132596496-DEL-117 |
| chr1-132616627-DEL-217 | chr1-132616640-DEL-204 |
| chr1-132672080-DEL-765 | chr1-132672081-DEL-765 |
| chr1-132680802-DEL-369 | chr1-132680798-DEL-367 |
| chr1-132689389-DEL-196 | chr1-132689390-DEL-196 |
| chr1-132700431-DEL-94 | chr1-132700425-DEL-100 |
| chr1-132701469-DEL-53 | chr1-132701470-DEL-53 |
| chr1-132752505-DEL-219 | chr1-132752506-DEL-219 |
| chr1-132753412-DEL-78 | chr1-132753413-DEL-54 |
| chr1-132769288-DEL-302 | chr1-132769289-DEL-301 |
| chr1-132777494-DEL-625 | chr1-132777495-DEL-625 |
| chr1-132795608-DEL-249 | chr1-132795609-DEL-249 |
| chr1-132802324-DEL-54 | chr1-132802325-DEL-66 |
| chr1-132845865-DEL-78 | chr1-132845866-DEL-78 |
| chr1-132857797-DEL-74 | chr1-132857798-DEL-60 |
| chr1-132861251-DEL-65 | chr1-132861251-DEL-65 |
| chr1-132893795-DEL-213 | chr1-132893796-DEL-213 |
| chr1-132894107-DEL-1035 | chr1-132894116-DEL-1027 |
| chr1-132898862-DEL-72 | chr1-132898863-DEL-72 |
| chr1-132906069-DEL-127 | chr1-132906054-DEL-128 |
| chr1-132908912-DEL-604 | chr1-132908913-DEL-604 |
| chr1-132939948-DEL-177 | chr1-132939947-DEL-178 |
| chr1-132955365-DEL-254 | chr1-132955366-DEL-254 |
| chr1-132956818-DEL-184 | chr1-132956819-DEL-184 |
| chr1-132971506-DEL-205 | chr1-132971507-DEL-205 |
| chr1-132989276-DEL-245 | chr1-132989276-DEL-221 |
| chr1-133014376-DEL-71 | chr1-133014377-DEL-71 |
| chr1-133040310-DEL-1012 | chr1-133040311-DEL-1012 |
| chr1-133042300-DEL-179 | chr1-133042301-DEL-179 |
| chr1-133051684-DEL-93 | chr1-133051685-DEL-93 |
| chr1-133053336-DEL-52 | chr1-133053337-DEL-52 |
| chr1-133070860-DEL-173 | chr1-133070861-DEL-173 |
| chr1-133087551-DEL-125 | chr1-133087552-DEL-125 |
| chr1-133134787-DEL-449 | chr1-133134788-DEL-449 |
| chr1-133138995-DEL-228 | chr1-133138986-DEL-229 |
| chr1-133152044-DEL-67 | chr1-133152039-DEL-59 |
| chr1-133233569-DEL-506 | chr1-133233570-DEL-506 |
| chr1-133235831-DEL-60 | chr1-133235832-DEL-60 |
| chr1-133242610-DEL-84 | chr1-133242611-DEL-84 |
| chr1-133261448-DEL-205 | chr1-133261449-DEL-205 |
| chr1-133302823-DEL-73 | chr1-133302824-DEL-73 |
| chr1-133320093-DEL-98 | chr1-133320094-DEL-96 |
| chr1-133323719-DEL-607 | chr1-133323720-DEL-607 |
| chr1-133333676-DEL-56 | chr1-133333677-DEL-56 |
| chr1-133366796-DEL-250 | chr1-133366797-DEL-250 |
| chr1-133374791-DEL-94 | chr1-133374810-DEL-80 |
| chr1-133383619-DEL-53 | chr1-133383615-DEL-66 |
| chr1-133400297-DEL-86 | chr1-133400297-DEL-81 |
| chr1-133406383-DEL-66 | chr1-133406384-DEL-67 |
| chr1-133459500-DEL-238 | chr1-133459503-DEL-235 |
| chr1-133481210-DEL-208 | chr1-133481211-DEL-208 |
| chr1-133502389-DEL-186 | chr1-133502378-DEL-186 |
| chr1-133521040-DEL-203 | chr1-133521040-DEL-204 |
| chr1-133600435-DEL-208 | chr1-133600436-DEL-208 |
| chr1-133604999-DEL-239 | chr1-133605000-DEL-239 |
| chr1-133606725-DEL-74 | chr1-133606726-DEL-74 |
| chr1-133614218-DEL-186 | chr1-133614219-DEL-186 |
| chr1-133617397-DEL-56 | chr1-133617398-DEL-54 |
| chr1-133618464-DEL-76 | chr1-133618465-DEL-60 |
| chr1-133659037-DEL-211 | chr1-133659038-DEL-211 |
| chr1-133668452-DEL-61 | chr1-133668453-DEL-61 |
| chr1-133738808-DEL-128 | chr1-133738805-DEL-129 |
| chr1-133747861-DEL-52 | chr1-133747862-DEL-52 |
| chr1-133767572-DEL-71 | chr1-133767573-DEL-71 |
| chr1-133806130-DEL-278 | chr1-133806131-DEL-278 |
| chr1-133867126-DEL-371 | chr1-133867127-DEL-371 |
| chr1-133888937-DEL-123 | chr1-133888930-DEL-131 |
| chr1-133894562-DEL-225 | chr1-133894563-DEL-225 |
| chr1-133900073-DEL-163 | chr1-133900074-DEL-163 |
| chr1-133904335-DEL-52 | chr1-133904316-DEL-60 |
| chr1-133922277-DEL-372 | chr1-133922277-DEL-372 |
| chr1-133926904-DEL-3694 | chr1-133926905-DEL-3694 |
| chr1-133928445-DEL-826 | chr1-133928446-DEL-826 |
| chr1-133969008-DEL-61 | chr1-133969009-DEL-61 |
| chr1-133979384-DEL-190 | chr1-133979385-DEL-190 |
| chr1-134029933-DEL-640 | chr1-134029934-DEL-640 |
| chr1-134044291-DEL-70 | chr1-134044292-DEL-70 |
| chr1-134047205-DEL-75 | chr1-134047206-DEL-75 |
| chr1-134052533-DEL-5148 | chr1-134052534-DEL-5148 |
| chr1-134069721-DEL-234 | chr1-134069722-DEL-234 |

| | |
|---|---|
| chr1-134102898-DEL-62 | chr1-134102899-DEL-62 |
| chr1-134148752-DEL-126 | chr1-134148727-DEL-124 |
| chr1-134177438-DEL-79 | chr1-134177395-DEL-80 |
| chr1-134184519-DEL-1873 | chr1-134184520-DEL-1873 |
| chr1-134187360-DEL-283 | chr1-134187360-DEL-283 |
| chr1-134198847-DEL-357 | chr1-134198848-DEL-357 |
| chr1-134224284-DEL-188 | chr1-134224279-DEL-227 |
| chr1-134276318-DEL-498 | chr1-134276319-DEL-498 |
| chr1-134284694-DEL-204 | chr1-134284695-DEL-204 |
| chr1-134294856-DEL-202 | chr1-134294857-DEL-202 |
| chr1-134295650-DEL-202 | chr1-134295651-DEL-202 |
| chr1-134327807-DEL-5522 | chr1-134327808-DEL-5522 |
| chr1-134338504-DEL-64 | chr1-134338505-DEL-64 |
| chr1-134376225-DEL-208 | chr1-134376226-DEL-208 |
| chr1-134408757-DEL-1550 | chr1-134408756-DEL-1551 |
| chr1-134411775-DEL-96 | chr1-134411776-DEL-96 |
| chr1-134436504-DEL-201 | chr1-134436505-DEL-201 |
| chr1-134438747-DEL-70 | chr1-134438768-DEL-50 |
| chr1-134452508-DEL-199 | chr1-134452509-DEL-199 |
| chr1-134459579-DEL-54 | chr1-134459580-DEL-54 |
| chr1-134475527-DEL-52 | chr1-134475484-DEL-89 |
| chr1-134605939-DEL-51 | chr1-134605940-DEL-57 |
| chr1-134618656-DEL-83 | chr1-134618644-DEL-86 |
| chr1-134623521-DEL-197 | chr1-134623522-DEL-197 |
| chr1-134640098-DEL-206 | chr1-134640099-DEL-206 |
| chr1-134670163-DEL-432 | chr1-134670164-DEL-432 |
| chr1-134722694-DEL-65 | chr1-134722695-DEL-65 |
| chr1-134780786-DEL-188 | chr1-134780781-DEL-188 |
| chr1-134793011-DEL-147 | chr1-134793012-DEL-147 |
| chr1-134818115-DEL-68 | chr1-134818111-DEL-88 |
| chr1-134847346-DEL-85 | chr1-134847347-DEL-85 |
| chr1-134865164-DEL-210 | chr1-134865165-DEL-210 |
| chr1-134867248-DEL-101 | chr1-134867238-DEL-74 |
| chr1-134867739-DEL-117 | chr1-134867740-DEL-117 |
| chr1-134877091-DEL-148 | chr1-134877096-DEL-131 |
| chr1-134921067-DEL-324 | chr1-134921068-DEL-324 |
| chr1-134923046-DEL-53 | chr1-134923047-DEL-52 |
| chr1-134974784-DEL-192 | chr1-134974785-DEL-192 |
| chr1-135021296-DEL-55 | chr1-135021296-DEL-55 |
| chr1-135030353-DEL-52 | chr1-135030354-DEL-52 |
| chr1-135039271-DEL-212 | chr1-135039272-DEL-212 |
| chr1-135043522-DEL-202 | chr1-135043523-DEL-202 |
| chr1-135058417-DEL-68 | chr1-135058418-DEL-68 |
| chr1-135065762-DEL-53 | chr1-135065763-DEL-53 |
| chr1-135077165-DEL-188 | chr1-135077166-DEL-188 |
| chr1-135102083-DEL-56 | chr1-135102084-DEL-56 |
| chr1-135115075-DEL-84 | chr1-135115075-DEL-84 |
| chr1-135116311-DEL-212 | chr1-135116312-DEL-212 |
| chr1-135117539-DEL-434 | chr1-135117540-DEL-434 |
| chr1-135129411-DEL-68 | chr1-135129412-DEL-68 |
| chr1-135134650-DEL-80 | chr1-135134651-DEL-80 |
| chr1-135139663-DEL-186 | chr1-135139664-DEL-186 |
| chr1-135146665-DEL-77 | chr1-135146659-DEL-60 |
| chr1-135156736-DEL-213 | chr1-135156737-DEL-213 |
| chr1-135159715-DEL-1431 | chr1-135159716-DEL-1431 |
| chr1-135164022-DEL-280 | chr1-135164023-DEL-280 |
| chr1-135172619-DEL-56 | chr1-135172660-DEL-50 |
| chr1-135198735-DEL-191 | chr1-135198736-DEL-191 |
| chr1-135201160-DEL-70 | chr1-135201157-DEL-67 |
| chr1-135203532-DEL-54 | chr1-135203541-DEL-50 |
| chr1-135226311-DEL-1279 | chr1-135226312-DEL-1279 |
| chr1-135234697-DEL-122 | chr1-135234698-DEL-122 |
| chr1-135249281-DEL-55 | chr1-135249272-DEL-64 |
| chr1-135320087-DEL-63 | chr1-135320099-DEL-60 |
| chr1-135334873-DEL-207 | chr1-135334874-DEL-207 |
| chr1-135377051-DEL-197 | chr1-135377051-DEL-197 |
| chr1-135412216-DEL-215 | chr1-135412217-DEL-215 |
| chr1-135431114-DEL-114 | chr1-135431120-DEL-114 |
| chr1-135440582-DEL-56 | chr1-135440583-DEL-56 |
| chr1-135447860-DEL-202 | chr1-135447862-DEL-201 |
| chr1-135454234-DEL-223 | chr1-135454238-DEL-230 |
| chr1-135469579-DEL-217 | chr1-135469565-DEL-196 |
| chr1-135528837-DEL-93 | chr1-135528836-DEL-94 |
| chr1-135536080-DEL-60 | chr1-135536069-DEL-100 |
| chr1-135541399-DEL-79 | chr1-135541400-DEL-97 |
| chr1-135548313-DEL-76 | chr1-135548326-DEL-64 |
| chr1-135562779-DEL-401 | chr1-135562780-DEL-401 |
| chr1-135599532-DEL-77 | chr1-135599533-DEL-77 |
| chr1-135605076-DEL-77 | chr1-135605077-DEL-79 |
| chr1-135624085-DEL-247 | chr1-135624086-DEL-247 |

| | |
|---|---|
| chr1-135628077-DEL-218 | chr1-135628078-DEL-218 |
| chr1-135635877-DEL-483 | chr1-135635878-DEL-483 |
| chr1-135651078-DEL-92 | chr1-135651079-DEL-92 |
| chr1-135652158-DEL-759 | chr1-135652159-DEL-759 |
| chr1-135669489-DEL-62 | chr1-135669490-DEL-62 |
| chr1-135669709-DEL-446 | chr1-135669710-DEL-446 |
| chr1-135708910-DEL-94 | chr1-135708911-DEL-94 |
| chr1-135714102-DEL-52 | chr1-135714102-DEL-52 |
| chr1-135729685-DEL-51 | chr1-135729686-DEL-51 |
| chr1-135751060-DEL-103 | chr1-135751061-DEL-103 |
| chr1-135797517-DEL-544 | chr1-135797518-DEL-544 |
| chr1-135815253-DEL-69 | chr1-135815227-DEL-98 |
| chr1-135861162-DEL-58 | chr1-135861163-DEL-58 |
| chr1-135871165-DEL-80 | chr1-135871166-DEL-80 |
| chr1-135942084-DEL-254 | chr1-135942084-DEL-255 |
| chr1-135952470-DEL-220 | chr1-135952471-DEL-220 |
| chr1-135974236-DEL-63 | chr1-135974234-DEL-66 |
| chr1-136009401-DEL-206 | chr1-136009405-DEL-207 |
| chr1-136014123-DEL-65 | chr1-136014124-DEL-65 |
| chr1-136021587-DEL-51 | chr1-136021588-DEL-51 |
| chr1-136120503-DEL-104 | chr1-136120505-DEL-83 |
| chr1-136172206-DEL-115 | chr1-136172208-DEL-93 |
| chr1-136172888-DEL-108 | chr1-136172895-DEL-102 |
| chr1-136233726-DEL-134 | chr1-136233727-DEL-134 |
| chr1-136258144-DEL-52 | chr1-136258145-DEL-52 |
| chr1-136260216-DEL-204 | chr1-136260217-DEL-204 |
| chr1-136264487-DEL-87 | chr1-136264487-DEL-87 |
| chr1-136303307-DEL-315 | chr1-136303306-DEL-313 |
| chr1-136314565-DEL-56 | chr1-136314578-DEL-93 |
| chr1-136325191-DEL-76 | chr1-136325194-DEL-72 |
| chr1-136333626-DEL-170 | chr1-136333627-DEL-170 |
| chr1-136354309-DEL-84 | chr1-136354288-DEL-63 |
| chr1-136376016-DEL-206 | chr1-136376014-DEL-204 |
| chr1-136411638-DEL-940 | chr1-136411639-DEL-940 |
| chr1-136413105-DEL-183 | chr1-136413106-DEL-181 |
| chr1-136470572-DEL-83 | chr1-136470573-DEL-83 |
| chr1-136477319-DEL-1090 | chr1-136477320-DEL-1090 |
| chr1-136479925-DEL-98 | chr1-136479892-DEL-133 |
| chr1-136485031-DEL-203 | chr1-136485032-DEL-203 |
| chr1-136486229-DEL-228 | chr1-136486230-DEL-228 |
| chr1-136487984-DEL-191 | chr1-136487973-DEL-192 |
| chr1-136507508-DEL-498 | chr1-136507509-DEL-498 |
| chr1-136509441-DEL-214 | chr1-136509442-DEL-214 |
| chr1-136512603-DEL-188 | chr1-136512604-DEL-188 |
| chr1-136518999-DEL-98 | chr1-136519000-DEL-98 |
| chr1-136532471-DEL-66 | chr1-136532448-DEL-90 |
| chr1-136562594-DEL-203 | chr1-136562595-DEL-203 |
| chr1-136574781-DEL-85 | chr1-136574782-DEL-85 |
| chr1-136577589-DEL-195 | chr1-136577590-DEL-195 |
| chr1-136591978-DEL-227 | chr1-136591979-DEL-227 |
| chr1-136617169-DEL-212 | chr1-136617170-DEL-212 |
| chr1-136622035-DEL-227 | chr1-136622065-DEL-198 |
| chr1-136629945-DEL-217 | chr1-136629946-DEL-217 |
| chr1-136630459-DEL-534 | chr1-136630460-DEL-534 |
| chr1-136640493-DEL-765 | chr1-136640500-DEL-767 |
| chr1-136648971-DEL-1325 | chr1-136648972-DEL-1325 |
| chr1-136675816-DEL-95 | chr1-136675817-DEL-95 |
| chr1-136678645-DEL-117 | chr1-136678644-DEL-106 |
| chr1-136682969-DEL-229 | chr1-136682984-DEL-205 |
| chr1-136716747-DEL-833 | chr1-136716748-DEL-833 |
| chr1-136719692-DEL-203 | chr1-136719693-DEL-203 |
| chr1-136720747-DEL-845 | chr1-136720748-DEL-845 |
| chr1-136725247-DEL-80 | chr1-136725204-DEL-101 |
| chr1-136729020-DEL-94 | chr1-136729021-DEL-94 |
| chr1-136735718-DEL-2733 | chr1-136735719-DEL-2733 |
| chr1-136760904-DEL-550 | chr1-136760905-DEL-550 |
| chr1-136784666-DEL-354 | chr1-136784663-DEL-358 |
| chr1-136792037-DEL-961 | chr1-136792038-DEL-961 |
| chr1-136801552-DEL-100 | chr1-136801553-DEL-100 |
| chr1-136829012-DEL-54 | chr1-136829013-DEL-54 |
| chr1-136850990-DEL-229 | chr1-136850991-DEL-229 |
| chr1-136889700-DEL-109 | chr1-136889701-DEL-109 |
| chr1-136920417-DEL-207 | chr1-136920418-DEL-207 |
| chr1-136946552-DEL-204 | chr1-136946542-DEL-203 |
| chr1-136948306-DEL-52 | chr1-136948307-DEL-52 |
| chr1-136949041-DEL-168 | chr1-136949041-DEL-169 |
| chr1-136975898-DEL-99 | chr1-136975899-DEL-99 |
| chr1-136978374-DEL-57 | chr1-136978375-DEL-57 |
| chr1-136987580-DEL-603 | chr1-136987581-DEL-603 |
| chr1-136990170-DEL-6415 | chr1-136990171-DEL-6415 |

| | |
|---|---|
| chr1-137008411-DEL-577 | chr1-137008412-DEL-577 |
| chr1-137009293-DEL-54 | chr1-137009294-DEL-64 |
| chr1-137055166-DEL-122 | chr1-137055167-DEL-122 |
| chr1-137070874-DEL-120 | chr1-137070875-DEL-120 |
| chr1-137077356-DEL-6818 | chr1-137077357-DEL-6818 |
| chr1-137096108-DEL-363 | chr1-137096109-DEL-363 |
| chr1-137126311-DEL-58 | chr1-137126312-DEL-58 |
| chr1-137136622-DEL-246 | chr1-137136616-DEL-275 |
| chr1-137145769-DEL-200 | chr1-137145770-DEL-200 |
| chr1-137193200-DEL-86 | chr1-137193205-DEL-62 |
| chr1-137201672-DEL-4335 | chr1-137201673-DEL-4335 |
| chr1-137216805-DEL-435 | chr1-137216812-DEL-408 |
| chr1-137231752-DEL-275 | chr1-137231753-DEL-275 |
| chr1-137239913-DEL-69 | chr1-137239909-DEL-56 |
| chr1-137250062-DEL-93 | chr1-137250061-DEL-90 |
| chr1-137265151-DEL-13159 | chr1-137265158-DEL-13156 |
| chr1-137290074-DEL-76 | chr1-137290046-DEL-77 |
| chr1-137398119-DEL-93 | chr1-137398120-DEL-93 |
| chr1-137454809-DEL-64 | chr1-137454810-DEL-64 |
| chr1-137507024-DEL-799 | chr1-137507025-DEL-799 |
| chr1-137511624-DEL-171 | chr1-137511625-DEL-171 |
| chr1-137513851-DEL-331 | chr1-137513852-DEL-332 |
| chr1-137515813-DEL-62 | chr1-137515818-DEL-62 |
| chr1-137518922-DEL-97 | chr1-137518923-DEL-97 |
| chr1-137538944-DEL-852 | chr1-137538943-DEL-851 |
| chr1-137579392-DEL-84 | chr1-137579385-DEL-87 |
| chr1-137605254-DEL-245 | chr1-137605255-DEL-245 |
| chr1-137608577-DEL-935 | chr1-137608578-DEL-935 |
| chr1-137637696-DEL-6649 | chr1-137637697-DEL-6649 |
| chr1-137651213-DEL-1135 | chr1-137651218-DEL-1130 |
| chr1-137655834-DEL-536 | chr1-137655835-DEL-536 |
| chr1-137680853-DEL-475 | chr1-137680854-DEL-475 |
| chr1-137682261-DEL-3036 | chr1-137682262-DEL-3036 |
| chr1-137711955-DEL-103 | chr1-137711956-DEL-103 |
| chr1-137712507-DEL-218 | chr1-137712508-DEL-218 |
| chr1-137730895-DEL-3232 | chr1-137730896-DEL-3232 |
| chr1-137777632-DEL-323 | chr1-137777633-DEL-323 |
| chr1-137811282-DEL-54 | chr1-137811283-DEL-54 |
| chr1-137869696-DEL-58 | chr1-137869697-DEL-58 |
| chr1-137882371-DEL-68 | chr1-137882372-DEL-68 |
| chr1-137893218-DEL-53 | chr1-137893219-DEL-53 |
| chr1-137897721-DEL-65 | chr1-137897696-DEL-83 |
| chr1-137902962-DEL-68 | chr1-137902963-DEL-68 |
| chr1-137906297-DEL-66 | chr1-137906300-DEL-64 |
| chr1-137907324-DEL-4559 | chr1-137907325-DEL-4559 |
| chr1-137937824-DEL-81 | chr1-137937825-DEL-81 |
| chr1-137945896-DEL-54 | chr1-137945869-DEL-53 |
| chr1-137954353-DEL-128 | chr1-137954357-DEL-125 |
| chr1-137955097-DEL-463 | chr1-137955098-DEL-463 |
| chr1-137969586-DEL-96 | chr1-137969587-DEL-96 |
| chr1-138011024-DEL-70 | chr1-138011025-DEL-70 |
| chr1-138022511-DEL-56 | chr1-138022512-DEL-64 |
| chr1-138045976-DEL-1179 | chr1-138045977-DEL-1179 |
| chr1-138054394-DEL-59 | chr1-138054396-DEL-71 |
| chr1-138129317-DEL-464 | chr1-138129316-DEL-465 |
| chr1-138148245-DEL-70 | chr1-138148246-DEL-70 |
| chr1-138168820-DEL-186 | chr1-138168821-DEL-186 |
| chr1-138169328-DEL-2283 | chr1-138169331-DEL-2283 |
| chr1-138193507-DEL-66 | chr1-138193508-DEL-54 |
| chr1-138228435-DEL-56 | chr1-138228436-DEL-56 |
| chr1-138233150-DEL-6941 | chr1-138233151-DEL-6941 |
| chr1-138241732-DEL-357 | chr1-138241733-DEL-357 |
| chr1-138267066-DEL-1209 | chr1-138267067-DEL-1206 |
| chr1-138268368-DEL-89 | chr1-138268383-DEL-75 |
| chr1-138268710-DEL-343 | chr1-138268711-DEL-343 |
| chr1-138326766-DEL-670 | chr1-138326767-DEL-670 |
| chr1-138329433-DEL-123 | chr1-138329434-DEL-123 |
| chr1-138352859-DEL-60 | chr1-138352860-DEL-60 |
| chr1-138356626-DEL-839 | chr1-138356627-DEL-839 |
| chr1-138379714-DEL-6203 | chr1-138379715-DEL-6203 |
| chr1-138453858-DEL-2503 | chr1-138453859-DEL-2503 |
| chr1-138506771-DEL-59 | chr1-138506760-DEL-58 |
| chr1-138526690-DEL-121 | chr1-138526690-DEL-122 |
| chr1-138529330-DEL-623 | chr1-138529331-DEL-623 |
| chr1-138537499-DEL-1894 | chr1-138537504-DEL-1893 |
| chr1-138557203-DEL-66 | chr1-138557222-DEL-52 |
| chr1-138578762-DEL-69 | chr1-138578763-DEL-69 |
| chr1-138628604-DEL-126 | chr1-138628605-DEL-126 |
| chr1-138671225-DEL-2170 | chr1-138671226-DEL-2167 |
| chr1-138743108-DEL-186 | chr1-138743109-DEL-186 |

| | |
|---|---|
| chr1-138777800-DEL-210 | chr1-138777801-DEL-210 |
| chr1-138785359-DEL-196 | chr1-138785361-DEL-194 |
| chr1-138799702-DEL-210 | chr1-138799703-DEL-210 |
| chr1-138830791-DEL-184 | chr1-138830792-DEL-184 |
| chr1-138874035-DEL-219 | chr1-138874036-DEL-219 |
| chr1-138874745-DEL-3606 | chr1-138874746-DEL-3606 |
| chr1-138874978-DEL-70 | chr1-138874979-DEL-70 |
| chr1-138877915-DEL-78 | chr1-138877916-DEL-78 |
| chr1-138907001-DEL-1195 | chr1-138907002-DEL-1195 |
| chr1-138910541-DEL-92 | chr1-138910542-DEL-92 |
| chr1-138913524-DEL-121 | chr1-138913525-DEL-121 |
| chr1-138925187-DEL-76 | chr1-138925183-DEL-77 |
| chr1-139001442-DEL-53 | chr1-139001443-DEL-53 |
| chr1-139044226-DEL-6289 | chr1-139044227-DEL-6289 |
| chr1-139065066-DEL-783 | chr1-139065067-DEL-783 |
| chr1-139067720-DEL-171 | chr1-139067721-DEL-171 |
| chr1-139082511-DEL-299 | chr1-139082512-DEL-299 |
| chr1-139123485-DEL-432 | chr1-139123486-DEL-432 |
| chr1-139151230-DEL-95 | chr1-139151231-DEL-95 |
| chr1-139154477-DEL-302 | chr1-139154478-DEL-302 |
| chr1-139180693-DEL-1065 | chr1-139180694-DEL-1065 |
| chr1-139213347-DEL-63 | chr1-139213348-DEL-68 |
| chr1-139234307-DEL-69 | chr1-139234332-DEL-63 |
| chr1-139248063-DEL-101 | chr1-139248064-DEL-101 |
| chr1-139283853-DEL-109 | chr1-139283893-DEL-70 |
| chr1-139297091-DEL-1135 | chr1-139297092-DEL-1135 |
| chr1-139334991-DEL-84 | chr1-139334994-DEL-80 |
| chr1-139338828-DEL-210 | chr1-139338829-DEL-210 |
| chr1-139455426-DEL-2418 | chr1-139455383-DEL-2418 |
| chr1-139457447-DEL-353 | chr1-139457448-DEL-353 |
| chr1-139468066-DEL-6596 | chr1-139468067-DEL-6596 |
| chr1-139486718-DEL-72 | chr1-139486719-DEL-72 |
| chr1-139529919-DEL-102 | chr1-139529920-DEL-102 |
| chr1-139531547-DEL-1043 | chr1-139531548-DEL-1043 |
| chr1-139541138-DEL-7484 | chr1-139541139-DEL-7484 |
| chr1-139552458-DEL-146 | chr1-139552458-DEL-116 |
| chr1-139620581-DEL-394 | chr1-139620582-DEL-394 |
| chr1-139687881-DEL-772 | chr1-139687882-DEL-772 |
| chr1-139697900-DEL-873 | chr1-139697901-DEL-873 |
| chr1-139720664-DEL-3325 | chr1-139720665-DEL-3325 |
| chr1-139752062-DEL-1097 | chr1-139752063-DEL-1097 |
| chr1-139754913-DEL-97 | chr1-139754915-DEL-96 |
| chr1-139793141-DEL-359 | chr1-139793142-DEL-359 |
| chr1-139835064-DEL-6734 | chr1-139835065-DEL-6734 |
| chr1-139852181-DEL-503 | chr1-139852182-DEL-503 |
| chr1-139880202-DEL-6413 | chr1-139880203-DEL-6413 |
| chr1-139919038-DEL-139 | chr1-139919039-DEL-139 |
| chr1-139965217-DEL-349 | chr1-139965218-DEL-349 |
| chr1-139970911-DEL-5106 | chr1-139970911-DEL-5106 |
| chr1-140036698-DEL-2758 | chr1-140036699-DEL-2758 |
| chr1-140143130-DEL-1592 | chr1-140143131-DEL-1592 |
| chr1-140279838-DEL-458 | chr1-140279839-DEL-458 |
| chr1-140338746-DEL-52 | chr1-140338747-DEL-52 |
| chr1-140364206-DEL-64 | chr1-140364203-DEL-66 |
| chr1-140402435-DEL-170 | chr1-140402436-DEL-167 |
| chr1-140424592-DEL-6660 | chr1-140424593-DEL-6660 |
| chr1-140442382-DEL-3674 | chr1-140442383-DEL-3674 |
| chr1-140469130-DEL-59 | chr1-140469131-DEL-59 |
| chr1-140492119-DEL-302 | chr1-140492120-DEL-302 |
| chr1-140561295-DEL-654 | chr1-140561296-DEL-654 |
| chr1-140569106-DEL-80 | chr1-140569107-DEL-80 |
| chr1-140570537-DEL-6257 | chr1-140570538-DEL-6257 |
| chr1-140579211-DEL-52 | chr1-140579212-DEL-52 |
| chr1-140584091-DEL-340 | chr1-140584092-DEL-340 |
| chr1-140592023-DEL-1328 | chr1-140592024-DEL-1328 |
| chr1-140595190-DEL-1553 | chr1-140595191-DEL-1553 |
| chr1-140605200-DEL-6769 | chr1-140605201-DEL-6769 |
| chr1-140618396-DEL-2187 | chr1-140618397-DEL-2187 |
| chr1-140669906-DEL-3017 | chr1-140669907-DEL-3017 |
| chr1-140674843-DEL-144 | chr1-140674844-DEL-144 |
| chr1-140678415-DEL-117 | chr1-140678412-DEL-116 |
| chr1-140687120-DEL-264 | chr1-140687121-DEL-264 |
| chr1-140690414-DEL-1253 | chr1-140690415-DEL-1253 |
| chr1-140709270-DEL-1817 | chr1-140709271-DEL-1817 |
| chr1-140714482-DEL-4789 | chr1-140714483-DEL-4789 |
| chr1-140716659-DEL-1068 | chr1-140716657-DEL-1059 |
| chr1-140722156-DEL-3579 | chr1-140722157-DEL-3579 |
| chr1-140744492-DEL-7215 | chr1-140744493-DEL-7215 |
| chr1-140762715-DEL-1354 | chr1-140762716-DEL-1354 |
| chr1-140785126-DEL-54 | chr1-140785127-DEL-54 |

| | |
|---|---|
| chr1-140789343-DEL-80 | chr1-140789362-DEL-86 |
| chr1-140805602-DEL-485 | chr1-140805604-DEL-485 |
| chr1-140823011-DEL-1002 | chr1-140823011-DEL-1002 |
| chr1-140824683-DEL-82 | chr1-140824686-DEL-80 |
| chr1-140835359-DEL-2622 | chr1-140835360-DEL-2622 |
| chr1-140846341-DEL-200 | chr1-140846342-DEL-200 |
| chr1-140851657-DEL-212 | chr1-140851658-DEL-212 |
| chr1-140875759-DEL-262 | chr1-140875760-DEL-262 |
| chr1-140884804-DEL-51 | chr1-140884805-DEL-51 |
| chr1-140891682-DEL-8683 | chr1-140891683-DEL-8683 |
| chr1-140906515-DEL-55 | chr1-140906514-DEL-56 |
| chr1-140933638-DEL-399 | chr1-140933638-DEL-399 |
| chr1-140963097-DEL-7143 | chr1-140963098-DEL-7143 |
| chr1-140986061-DEL-626 | chr1-140986062-DEL-626 |
| chr1-140997095-DEL-1036 | chr1-140997096-DEL-1036 |
| chr1-141001179-DEL-247 | chr1-141001180-DEL-247 |
| chr1-141014137-DEL-101 | chr1-141014138-DEL-101 |
| chr1-141015957-DEL-101 | chr1-141015958-DEL-101 |
| chr1-141021624-DEL-615 | chr1-141021625-DEL-615 |
| chr1-141033659-DEL-51 | chr1-141033677-DEL-52 |
| chr1-141063208-DEL-231 | chr1-141063209-DEL-231 |
| chr1-141063969-DEL-98 | chr1-141063970-DEL-98 |
| chr1-141088916-DEL-61 | chr1-141088917-DEL-61 |
| chr1-141089267-DEL-702 | chr1-141089268-DEL-702 |
| chr1-141093962-DEL-296 | chr1-141093963-DEL-296 |
| chr1-141096656-DEL-365 | chr1-141096657-DEL-365 |
| chr1-141114525-DEL-201 | chr1-141114526-DEL-201 |
| chr1-141117820-DEL-424 | chr1-141117821-DEL-424 |
| chr1-141148395-DEL-448 | chr1-141148396-DEL-448 |
| chr1-141154175-DEL-99 | chr1-141154175-DEL-100 |
| chr1-141157050-DEL-85 | chr1-141157051-DEL-85 |
| chr1-141158354-DEL-120 | chr1-141158355-DEL-120 |
| chr1-141161988-DEL-496 | chr1-141161989-DEL-496 |
| chr1-141189538-DEL-6594 | chr1-141189539-DEL-6594 |
| chr1-141222658-DEL-72 | chr1-141222687-DEL-50 |
| chr1-141228183-DEL-397 | chr1-141228184-DEL-397 |
| chr1-141229393-DEL-506 | chr1-141229394-DEL-506 |
| chr1-141232787-DEL-89 | chr1-141232788-DEL-89 |
| chr1-141277483-DEL-2410 | chr1-141277484-DEL-2410 |
| chr1-141295827-DEL-55 | chr1-141295828-DEL-55 |
| chr1-141299750-DEL-2031 | chr1-141299751-DEL-2031 |
| chr1-141312022-DEL-326 | chr1-141312023-DEL-326 |
| chr1-141360974-DEL-661 | chr1-141360975-DEL-661 |
| chr1-141364206-DEL-247 | chr1-141364207-DEL-247 |
| chr1-141371840-DEL-964 | chr1-141371841-DEL-964 |
| chr1-141387184-DEL-799 | chr1-141387185-DEL-799 |
| chr1-141389693-DEL-1002 | chr1-141389694-DEL-1002 |
| chr1-141442557-DEL-460 | chr1-141442577-DEL-451 |
| chr1-141480482-DEL-6275 | chr1-141480491-DEL-6277 |
| chr1-141505669-DEL-64 | chr1-141505669-DEL-62 |
| chr1-141516022-DEL-382 | chr1-141516023-DEL-382 |
| chr1-141528198-DEL-389 | chr1-141528199-DEL-386 |
| chr1-141554804-DEL-56 | chr1-141554806-DEL-54 |
| chr1-141569112-DEL-206 | chr1-141569113-DEL-206 |
| chr1-141577287-DEL-1861 | chr1-141577288-DEL-1861 |
| chr1-141602770-DEL-56 | chr1-141602769-DEL-57 |
| chr1-141604477-DEL-350 | chr1-141604478-DEL-350 |
| chr1-141625773-DEL-6198 | chr1-141625774-DEL-6198 |
| chr1-141632406-DEL-224 | chr1-141632406-DEL-224 |
| chr1-141638470-DEL-1552 | chr1-141638471-DEL-1552 |
| chr1-141645815-DEL-1476 | chr1-141645816-DEL-1476 |
| chr1-141657020-DEL-189 | chr1-141657021-DEL-189 |
| chr1-141659051-DEL-207 | chr1-141659052-DEL-207 |
| chr1-141666522-DEL-61 | chr1-141666523-DEL-61 |
| chr1-141679150-DEL-1894 | chr1-141679151-DEL-1894 |
| chr1-141683692-DEL-296 | chr1-141683691-DEL-295 |
| chr1-141708306-DEL-253 | chr1-141708303-DEL-249 |
| chr1-141713733-DEL-69 | chr1-141713733-DEL-69 |
| chr1-141719990-DEL-93 | chr1-141719991-DEL-94 |
| chr1-141725503-DEL-839 | chr1-141725504-DEL-839 |
| chr1-141780456-DEL-625 | chr1-141780457-DEL-625 |
| chr1-141837428-DEL-655 | chr1-141837429-DEL-655 |
| chr1-141840232-DEL-51 | chr1-141840263-DEL-62 |
| chr1-141852712-DEL-1270 | chr1-141852713-DEL-1270 |
| chr1-141857120-DEL-64 | chr1-141857121-DEL-64 |
| chr1-141858191-DEL-1896 | chr1-141858177-DEL-1901 |
| chr1-141865487-DEL-65 | chr1-141865494-DEL-55 |
| chr1-141872854-DEL-5133 | chr1-141872855-DEL-5133 |
| chr1-141897083-DEL-56 | chr1-141897084-DEL-55 |
| chr1-141917528-DEL-78 | chr1-141917541-DEL-72 |

| | |
|---|---|
| chr1-141922301-DEL-690 | chr1-141922302-DEL-690 |
| chr1-141924952-DEL-89 | chr1-141924953-DEL-89 |
| chr1-141996219-DEL-204 | chr1-141996220-DEL-204 |
| chr1-142035005-DEL-52 | chr1-142035006-DEL-52 |
| chr1-142052479-DEL-3075 | chr1-142052480-DEL-3075 |
| chr1-142061500-DEL-123 | chr1-142061501-DEL-117 |
| chr1-142072747-DEL-372 | chr1-142072748-DEL-371 |
| chr1-142131803-DEL-215 | chr1-142131804-DEL-215 |
| chr1-142132877-DEL-225 | chr1-142132886-DEL-222 |
| chr1-142144803-DEL-52 | chr1-142144828-DEL-52 |
| chr1-142145784-DEL-440 | chr1-142145785-DEL-440 |
| chr1-142162531-DEL-58 | chr1-142162532-DEL-58 |
| chr1-142216544-DEL-362 | chr1-142216545-DEL-362 |
| chr1-142217378-DEL-60 | chr1-142217379-DEL-60 |
| chr1-142270020-DEL-107 | chr1-142270021-DEL-107 |
| chr1-142270496-DEL-433 | chr1-142270497-DEL-433 |
| chr1-142278480-DEL-1665 | chr1-142278481-DEL-1665 |
| chr1-142283707-DEL-8005 | chr1-142283708-DEL-8005 |
| chr1-142327222-DEL-58 | chr1-142327223-DEL-52 |
| chr1-142346428-DEL-154 | chr1-142346429-DEL-154 |
| chr1-142370298-DEL-60 | chr1-142370299-DEL-60 |
| chr1-142400947-DEL-75 | chr1-142400948-DEL-75 |
| chr1-142421950-DEL-6556 | chr1-142421951-DEL-6556 |
| chr1-142444728-DEL-633 | chr1-142444729-DEL-633 |
| chr1-142452848-DEL-64 | chr1-142452867-DEL-55 |
| chr1-142474377-DEL-426 | chr1-142474378-DEL-426 |
| chr1-142498423-DEL-1629 | chr1-142498424-DEL-1629 |
| chr1-142501092-DEL-2010 | chr1-142501093-DEL-2010 |
| chr1-142510745-DEL-62 | chr1-142510746-DEL-62 |
| chr1-142512622-DEL-5265 | chr1-142512623-DEL-5265 |
| chr1-142574193-DEL-158 | chr1-142574194-DEL-158 |
| chr1-142574256-DEL-1530 | chr1-142574257-DEL-1530 |
| chr1-142610549-DEL-177 | chr1-142610550-DEL-177 |
| chr1-142621265-DEL-60 | chr1-142621270-DEL-70 |
| chr1-142651053-DEL-6218 | chr1-142651054-DEL-6218 |
| chr1-142651226-DEL-64 | chr1-142651227-DEL-64 |
| chr1-142698147-DEL-1128 | chr1-142698148-DEL-1128 |
| chr1-142700852-DEL-938 | chr1-142700853-DEL-938 |
| chr1-142724128-DEL-267 | chr1-142724162-DEL-266 |
| chr1-142763371-DEL-57 | chr1-142763372-DEL-57 |
| chr1-142825759-DEL-148 | chr1-142825760-DEL-148 |
| chr1-142846560-DEL-896 | chr1-142846561-DEL-896 |
| chr1-142872686-DEL-58 | chr1-142872686-DEL-58 |
| chr1-142891312-DEL-377 | chr1-142891313-DEL-377 |
| chr1-142907046-DEL-1281 | chr1-142907047-DEL-1281 |
| chr1-142948858-DEL-6227 | chr1-142948857-DEL-6227 |
| chr1-142960725-DEL-1771 | chr1-142960725-DEL-1768 |
| chr1-142964845-DEL-1274 | chr1-142964846-DEL-1274 |
| chr1-143005309-DEL-141 | chr1-143005312-DEL-139 |
| chr1-143070961-DEL-667 | chr1-143070962-DEL-667 |
| chr1-143088282-DEL-8505 | chr1-143088283-DEL-8505 |
| chr1-143103303-DEL-683 | chr1-143103304-DEL-683 |
| chr1-143108258-DEL-114 | chr1-143108275-DEL-90 |
| chr1-143117691-DEL-330 | chr1-143117692-DEL-330 |
| chr1-143168720-DEL-3006 | chr1-143168720-DEL-3006 |
| chr1-143175256-DEL-1086 | chr1-143175212-DEL-1087 |
| chr1-143180041-DEL-1036 | chr1-143180042-DEL-1036 |
| chr1-143193943-DEL-135 | chr1-143193941-DEL-136 |
| chr1-143253541-DEL-6683 | chr1-143253542-DEL-6683 |
| chr1-143296902-DEL-63 | chr1-143296896-DEL-97 |
| chr1-143298475-DEL-809 | chr1-143298476-DEL-809 |
| chr1-143376590-DEL-64 | chr1-143376593-DEL-58 |
| chr1-143387636-DEL-214 | chr1-143387637-DEL-214 |
| chr1-143405163-DEL-212 | chr1-143405164-DEL-212 |
| chr1-143406604-DEL-58 | chr1-143406605-DEL-58 |
| chr1-143419230-DEL-62 | chr1-143419231-DEL-62 |
| chr1-143456825-DEL-302 | chr1-143456826-DEL-302 |
| chr1-143463528-DEL-498 | chr1-143463529-DEL-498 |
| chr1-143488535-DEL-329 | chr1-143488540-DEL-325 |
| chr1-143506312-DEL-164 | chr1-143506313-DEL-164 |
| chr1-143517913-DEL-166 | chr1-143517914-DEL-166 |
| chr1-143552506-DEL-184 | chr1-143552507-DEL-184 |
| chr1-143555791-DEL-214 | chr1-143555792-DEL-214 |
| chr1-143585636-DEL-81 | chr1-143585637-DEL-81 |
| chr1-143586778-DEL-6148 | chr1-143586779-DEL-6148 |
| chr1-143672632-DEL-206 | chr1-143672633-DEL-206 |
| chr1-143711871-DEL-210 | chr1-143711872-DEL-210 |
| chr1-143724596-DEL-1176 | chr1-143724609-DEL-1185 |
| chr1-143737826-DEL-6665 | chr1-143737827-DEL-6665 |
| chr1-143738199-DEL-212 | chr1-143738200-DEL-212 |

| | |
|---|---|
| chr1-143745994-DEL-637 | chr1-143745995-DEL-637 |
| chr1-143766162-DEL-6201 | chr1-143766163-DEL-6201 |
| chr1-143785405-DEL-54 | chr1-143785403-DEL-56 |
| chr1-143794451-DEL-238 | chr1-143794452-DEL-237 |
| chr1-143840743-DEL-6632 | chr1-143840744-DEL-6632 |
| chr1-143846873-DEL-200 | chr1-143846874-DEL-200 |
| chr1-143859377-DEL-176 | chr1-143859378-DEL-176 |
| chr1-143872923-DEL-157 | chr1-143872905-DEL-162 |
| chr1-143909829-DEL-110 | chr1-143909828-DEL-80 |
| chr1-143932208-DEL-238 | chr1-143932209-DEL-238 |
| chr1-143940981-DEL-6390 | chr1-143940982-DEL-6390 |
| chr1-143981931-DEL-123 | chr1-143981932-DEL-123 |
| chr1-144031161-DEL-1584 | chr1-144031162-DEL-1584 |
| chr1-144041257-DEL-196 | chr1-144041258-DEL-196 |
| chr1-144044095-DEL-507 | chr1-144044096-DEL-506 |
| chr1-144056892-DEL-51 | chr1-144056893-DEL-51 |
| chr1-144077929-DEL-66 | chr1-144077930-DEL-66 |
| chr1-144091068-DEL-66 | chr1-144091069-DEL-66 |
| chr1-144099993-DEL-2134 | chr1-144099994-DEL-2134 |
| chr1-144162768-DEL-54 | chr1-144162769-DEL-54 |
| chr1-144170091-DEL-261 | chr1-144170092-DEL-261 |
| chr1-144174892-DEL-1105 | chr1-144174893-DEL-1105 |
| chr1-144196632-DEL-6463 | chr1-144196633-DEL-6462 |
| chr1-144196789-DEL-214 | chr1-144196790-DEL-214 |
| chr1-144258686-DEL-314 | chr1-144258683-DEL-315 |
| chr1-144259781-DEL-7320 | chr1-144259782-DEL-7320 |
| chr1-144270433-DEL-3029 | chr1-144270434-DEL-3029 |
| chr1-144277859-DEL-1845 | chr1-144277860-DEL-1845 |
| chr1-144293514-DEL-86 | chr1-144293515-DEL-86 |
| chr1-144296428-DEL-196 | chr1-144296429-DEL-196 |
| chr1-144324916-DEL-72 | chr1-144324917-DEL-68 |
| chr1-144333963-DEL-6154 | chr1-144333964-DEL-6154 |
| chr1-144373475-DEL-152 | chr1-144373482-DEL-145 |
| chr1-144399675-DEL-114 | chr1-144399672-DEL-118 |
| chr1-144494756-DEL-51 | chr1-144494749-DEL-59 |
| chr1-144496324-DEL-484 | chr1-144496325-DEL-484 |
| chr1-144550223-DEL-5943 | chr1-144550222-DEL-5943 |
| chr1-144572138-DEL-74 | chr1-144572139-DEL-74 |
| chr1-144602200-DEL-234 | chr1-144602201-DEL-234 |
| chr1-144614911-DEL-6274 | chr1-144614912-DEL-6274 |
| chr1-144614948-DEL-64 | chr1-144614949-DEL-64 |
| chr1-144623297-DEL-4036 | chr1-144623298-DEL-4036 |
| chr1-144659847-DEL-81 | chr1-144659848-DEL-81 |
| chr1-144662690-DEL-66 | chr1-144662691-DEL-66 |
| chr1-144663748-DEL-56 | chr1-144663749-DEL-56 |
| chr1-144681107-DEL-603 | chr1-144681108-DEL-603 |
| chr1-144682647-DEL-6363 | chr1-144682648-DEL-6363 |
| chr1-144682793-DEL-208 | chr1-144682794-DEL-208 |
| chr1-144690938-DEL-179 | chr1-144690939-DEL-179 |
| chr1-144699106-DEL-966 | chr1-144699107-DEL-966 |
| chr1-144730406-DEL-7126 | chr1-144730407-DEL-7126 |
| chr1-144757545-DEL-1124 | chr1-144757546-DEL-1124 |
| chr1-144875838-DEL-565 | chr1-144875839-DEL-565 |
| chr1-144877141-DEL-81 | chr1-144877144-DEL-82 |
| chr1-144953100-DEL-67 | chr1-144953093-DEL-75 |
| chr1-144954215-DEL-53 | chr1-144954216-DEL-53 |
| chr1-144962157-DEL-1034 | chr1-144962158-DEL-1034 |
| chr1-144976084-DEL-108 | chr1-144976085-DEL-102 |
| chr1-144976775-DEL-4043 | chr1-144976776-DEL-4043 |
| chr1-144986632-DEL-52 | chr1-144986633-DEL-56 |
| chr1-144999076-DEL-608 | chr1-144999077-DEL-608 |
| chr1-145035422-DEL-59 | chr1-145035430-DEL-75 |
| chr1-145035943-DEL-1177 | chr1-145035942-DEL-1177 |
| chr1-145064900-DEL-58 | chr1-145064901-DEL-58 |
| chr1-145124749-DEL-6490 | chr1-145124750-DEL-6490 |
| chr1-145130896-DEL-212 | chr1-145130897-DEL-212 |
| chr1-145152543-DEL-1360 | chr1-145152544-DEL-1360 |
| chr1-145171829-DEL-60 | chr1-145171800-DEL-90 |
| chr1-145181570-DEL-2537 | chr1-145181566-DEL-2537 |
| chr1-145185037-DEL-226 | chr1-145185038-DEL-226 |
| chr1-145230980-DEL-713 | chr1-145230981-DEL-713 |
| chr1-145320190-DEL-6028 | chr1-145320191-DEL-6028 |
| chr1-145326354-DEL-933 | chr1-145326355-DEL-933 |
| chr1-145345605-DEL-6303 | chr1-145345606-DEL-6303 |
| chr1-145403074-DEL-8527 | chr1-145403070-DEL-8528 |
| chr1-145413038-DEL-124 | chr1-145413039-DEL-124 |
| chr1-145421871-DEL-2294 | chr1-145421869-DEL-2295 |
| chr1-145429111-DEL-549 | chr1-145429113-DEL-548 |
| chr1-145436283-DEL-75 | chr1-145436280-DEL-73 |
| chr1-145437545-DEL-522 | chr1-145437546-DEL-522 |

| | |
|---|---|
| chr1-145439160-DEL-449 | chr1-145439159-DEL-448 |
| chr1-145445014-DEL-2138 | chr1-145445016-DEL-2137 |
| chr1-145457832-DEL-508 | chr1-145457833-DEL-508 |
| chr1-145465667-DEL-1306 | chr1-145465668-DEL-1306 |
| chr1-145497721-DEL-2536 | chr1-145497718-DEL-2537 |
| chr1-145503043-DEL-154 | chr1-145503043-DEL-154 |
| chr1-145515488-DEL-1827 | chr1-145515489-DEL-1827 |
| chr1-145528544-DEL-119 | chr1-145528545-DEL-119 |
| chr1-145540157-DEL-1978 | chr1-145540158-DEL-1978 |
| chr1-145608325-DEL-1794 | chr1-145608315-DEL-1795 |
| chr1-145625408-DEL-368 | chr1-145625441-DEL-346 |
| chr1-145633705-DEL-370 | chr1-145633706-DEL-370 |
| chr1-145657832-DEL-2408 | chr1-145657833-DEL-2408 |
| chr1-145669579-DEL-5871 | chr1-145669580-DEL-5871 |
| chr1-145676849-DEL-782 | chr1-145676850-DEL-782 |
| chr1-145693065-DEL-1778 | chr1-145693066-DEL-1777 |
| chr1-145694812-DEL-6486 | chr1-145694813-DEL-6486 |
| chr1-145719670-DEL-1813 | chr1-145719671-DEL-1812 |
| chr1-145739993-DEL-183 | chr1-145739994-DEL-183 |
| chr1-145748626-DEL-92 | chr1-145748627-DEL-92 |
| chr1-145751896-DEL-205 | chr1-145751897-DEL-205 |
| chr1-145776523-DEL-1648 | chr1-145776524-DEL-1648 |
| chr1-145780321-DEL-60 | chr1-145780321-DEL-60 |
| chr1-145791524-DEL-1120 | chr1-145791525-DEL-1120 |
| chr1-145813655-DEL-390 | chr1-145813656-DEL-390 |
| chr1-145824031-DEL-73 | chr1-145824032-DEL-73 |
| chr1-145841450-DEL-781 | chr1-145841451-DEL-781 |
| chr1-145841499-DEL-208 | chr1-145841500-DEL-208 |
| chr1-145844067-DEL-59 | chr1-145844068-DEL-59 |
| chr1-145844385-DEL-538 | chr1-145844385-DEL-539 |
| chr1-145850750-DEL-854 | chr1-145850748-DEL-846 |
| chr1-145880082-DEL-6959 | chr1-145880069-DEL-6960 |
| chr1-145908368-DEL-432 | chr1-145908369-DEL-432 |
| chr1-145916998-DEL-381 | chr1-145916999-DEL-381 |
| chr1-145918769-DEL-79 | chr1-145918770-DEL-79 |
| chr1-145926592-DEL-96 | chr1-145926624-DEL-76 |
| chr1-145961533-DEL-171 | chr1-145961534-DEL-171 |
| chr1-145966199-DEL-3964 | chr1-145966200-DEL-3964 |
| chr1-145985900-DEL-65 | chr1-145985901-DEL-65 |
| chr1-145985986-DEL-1519 | chr1-145985985-DEL-1520 |
| chr1-145988637-DEL-198 | chr1-145988638-DEL-198 |
| chr1-145990883-DEL-53 | chr1-145990884-DEL-53 |
| chr1-145991206-DEL-1657 | chr1-145991207-DEL-1657 |
| chr1-145996715-DEL-7070 | chr1-145996716-DEL-7070 |
| chr1-146013679-DEL-57 | chr1-146013680-DEL-61 |
| chr1-146025382-DEL-3153 | chr1-146025383-DEL-3153 |
| chr1-146034119-DEL-365 | chr1-146034120-DEL-365 |
| chr1-146046540-DEL-1467 | chr1-146046541-DEL-1467 |
| chr1-146049742-DEL-414 | chr1-146049743-DEL-414 |
| chr1-146067708-DEL-1815 | chr1-146067710-DEL-1814 |
| chr1-146087038-DEL-57 | chr1-146087039-DEL-57 |
| chr1-146102577-DEL-61 | chr1-146102578-DEL-61 |
| chr1-146118841-DEL-1134 | chr1-146118842-DEL-1134 |
| chr1-146154155-DEL-231 | chr1-146154156-DEL-231 |
| chr1-146157813-DEL-7112 | chr1-146157814-DEL-7112 |
| chr1-146175658-DEL-2528 | chr1-146175659-DEL-2528 |
| chr1-146179786-DEL-311 | chr1-146179787-DEL-311 |
| chr1-146183887-DEL-377 | chr1-146183888-DEL-377 |
| chr1-146196930-DEL-64 | chr1-146196930-DEL-65 |
| chr1-146218007-DEL-125 | chr1-146218008-DEL-125 |
| chr1-146226252-DEL-76 | chr1-146226253-DEL-76 |
| chr1-146257022-DEL-1343 | chr1-146257023-DEL-1343 |
| chr1-146263120-DEL-606 | chr1-146263122-DEL-606 |
| chr1-146272937-DEL-7089 | chr1-146272938-DEL-7089 |
| chr1-146298501-DEL-55 | chr1-146298501-DEL-60 |
| chr1-146312241-DEL-52 | chr1-146312242-DEL-50 |
| chr1-146320996-DEL-6592 | chr1-146320997-DEL-6592 |
| chr1-146337766-DEL-275 | chr1-146337767-DEL-275 |
| chr1-146394575-DEL-54 | chr1-146394576-DEL-54 |
| chr1-146409237-DEL-186 | chr1-146409238-DEL-186 |
| chr1-146437129-DEL-343 | chr1-146437130-DEL-343 |
| chr1-146476247-DEL-87 | chr1-146476231-DEL-81 |
| chr1-146479785-DEL-6291 | chr1-146479786-DEL-6291 |
| chr1-146487745-DEL-82 | chr1-146487743-DEL-81 |
| chr1-146497900-DEL-66 | chr1-146497901-DEL-66 |
| chr1-146511480-DEL-55 | chr1-146511481-DEL-55 |
| chr1-146553508-DEL-360 | chr1-146553509-DEL-360 |
| chr1-146572301-DEL-144 | chr1-146572310-DEL-96 |
| chr1-146600623-DEL-4609 | chr1-146600624-DEL-4609 |
| chr1-146644382-DEL-128 | chr1-146644383-DEL-128 |

| | |
|---|---|
| chr1-146681523-DEL-464 | chr1-146681524-DEL-464 |
| chr1-146698122-DEL-117 | chr1-146698123-DEL-117 |
| chr1-146699426-DEL-66 | chr1-146699427-DEL-54 |
| chr1-146712758-DEL-682 | chr1-146712759-DEL-682 |
| chr1-146737904-DEL-236 | chr1-146737905-DEL-236 |
| chr1-146739077-DEL-6169 | chr1-146739078-DEL-6169 |
| chr1-146781734-DEL-767 | chr1-146781735-DEL-767 |
| chr1-146788985-DEL-72 | chr1-146788986-DEL-73 |
| chr1-146809296-DEL-100 | chr1-146809296-DEL-100 |
| chr1-146811364-DEL-806 | chr1-146811365-DEL-806 |
| chr1-146814561-DEL-157 | chr1-146814562-DEL-157 |
| chr1-146865031-DEL-1103 | chr1-146865032-DEL-1103 |
| chr1-146869619-DEL-1725 | chr1-146869620-DEL-1725 |
| chr1-146895568-DEL-148 | chr1-146895569-DEL-148 |
| chr1-146938048-DEL-435 | chr1-146938049-DEL-435 |
| chr1-146956866-DEL-1001 | chr1-146956867-DEL-1001 |
| chr1-146985294-DEL-2948 | chr1-146985295-DEL-2948 |
| chr1-147065610-DEL-586 | chr1-147065613-DEL-585 |
| chr1-147073522-DEL-400 | chr1-147073523-DEL-400 |
| chr1-147112706-DEL-328 | chr1-147112707-DEL-328 |
| chr1-147116216-DEL-70 | chr1-147116217-DEL-70 |
| chr1-147126147-DEL-56 | chr1-147126148-DEL-56 |
| chr1-147127747-DEL-60 | chr1-147127748-DEL-60 |
| chr1-147131707-DEL-94 | chr1-147131708-DEL-94 |
| chr1-147132652-DEL-56 | chr1-147132653-DEL-56 |
| chr1-147150825-DEL-528 | chr1-147150829-DEL-523 |
| chr1-147158108-DEL-6396 | chr1-147158109-DEL-6396 |
| chr1-147169860-DEL-56 | chr1-147169861-DEL-58 |
| chr1-147173804-DEL-60 | chr1-147173805-DEL-60 |
| chr1-147174779-DEL-463 | chr1-147174776-DEL-460 |
| chr1-147229606-DEL-66 | chr1-147229607-DEL-66 |
| chr1-147235358-DEL-397 | chr1-147235359-DEL-397 |
| chr1-147259769-DEL-730 | chr1-147259770-DEL-730 |
| chr1-147267564-DEL-246 | chr1-147267565-DEL-246 |
| chr1-147291840-DEL-2744 | chr1-147291841-DEL-2744 |
| chr1-147306224-DEL-2752 | chr1-147306225-DEL-2752 |
| chr1-147333378-DEL-60 | chr1-147333379-DEL-60 |
| chr1-147406973-DEL-56 | chr1-147407005-DEL-66 |
| chr1-147426063-DEL-112 | chr1-147426064-DEL-112 |
| chr1-147427887-DEL-545 | chr1-147427888-DEL-545 |
| chr1-147434895-DEL-1878 | chr1-147434896-DEL-1878 |
| chr1-147443415-DEL-60 | chr1-147443416-DEL-60 |
| chr1-147447280-DEL-94 | chr1-147447281-DEL-94 |
| chr1-147454835-DEL-264 | chr1-147454836-DEL-264 |
| chr1-147467641-DEL-6388 | chr1-147467641-DEL-6388 |
| chr1-147479794-DEL-90 | chr1-147479795-DEL-90 |
| chr1-147520427-DEL-69 | chr1-147520387-DEL-109 |
| chr1-147546983-DEL-6485 | chr1-147547002-DEL-6474 |
| chr1-147576189-DEL-50 | chr1-147576206-DEL-52 |
| chr1-147580744-DEL-62 | chr1-147580751-DEL-56 |
| chr1-147593976-DEL-133 | chr1-147593978-DEL-124 |
| chr1-147602949-DEL-354 | chr1-147602950-DEL-354 |
| chr1-147605555-DEL-320 | chr1-147605555-DEL-320 |
| chr1-147608824-DEL-1253 | chr1-147608825-DEL-1253 |
| chr1-147617751-DEL-1751 | chr1-147617752-DEL-1751 |
| chr1-147626007-DEL-425 | chr1-147625999-DEL-426 |
| chr1-147630820-DEL-661 | chr1-147630821-DEL-659 |
| chr1-147632500-DEL-90 | chr1-147632501-DEL-90 |
| chr1-147639548-DEL-477 | chr1-147639546-DEL-467 |
| chr1-147647053-DEL-1788 | chr1-147647054-DEL-1788 |
| chr1-147672098-DEL-10528 | chr1-147672099-DEL-10528 |
| chr1-147789200-DEL-990 | chr1-147789186-DEL-990 |
| chr1-147840406-DEL-325 | chr1-147840407-DEL-325 |
| chr1-147860775-DEL-4568 | chr1-147860776-DEL-4568 |
| chr1-147872933-DEL-1382 | chr1-147872934-DEL-1381 |
| chr1-147921521-DEL-273 | chr1-147921522-DEL-273 |
| chr1-147934629-DEL-127 | chr1-147934630-DEL-127 |
| chr1-147942911-DEL-58 | chr1-147942876-DEL-70 |
| chr1-147945030-DEL-3874 | chr1-147945031-DEL-3874 |
| chr1-147950657-DEL-7217 | chr1-147950658-DEL-7217 |
| chr1-147956855-DEL-208 | chr1-147956851-DEL-201 |
| chr1-147977623-DEL-59 | chr1-147977623-DEL-59 |
| chr1-147986232-DEL-971 | chr1-147986233-DEL-971 |
| chr1-148026588-DEL-1068 | chr1-148026589-DEL-1068 |
| chr1-148044345-DEL-5771 | chr1-148044346-DEL-5771 |
| chr1-148078141-DEL-402 | chr1-148078142-DEL-402 |
| chr1-148088038-DEL-6259 | chr1-148088039-DEL-6259 |
| chr1-148096032-DEL-1688 | chr1-148096033-DEL-1688 |
| chr1-148131010-DEL-85 | chr1-148131011-DEL-85 |
| chr1-148149834-DEL-5511 | chr1-148149835-DEL-5511 |

| | |
|---|---|
| chr1-148163967-DEL-191 | chr1-148163968-DEL-191 |
| chr1-148171748-DEL-6342 | chr1-148171749-DEL-6342 |
| chr1-148179651-DEL-72 | chr1-148179652-DEL-72 |
| chr1-148213761-DEL-728 | chr1-148213762-DEL-727 |
| chr1-148239368-DEL-6381 | chr1-148239369-DEL-6381 |
| chr1-148254577-DEL-208 | chr1-148254577-DEL-208 |
| chr1-148269254-DEL-1383 | chr1-148269255-DEL-1381 |
| chr1-148295743-DEL-59 | chr1-148295742-DEL-56 |
| chr1-148312293-DEL-529 | chr1-148312294-DEL-529 |
| chr1-148332494-DEL-7158 | chr1-148332495-DEL-7158 |
| chr1-148344056-DEL-286 | chr1-148344052-DEL-286 |
| chr1-148361190-DEL-5272 | chr1-148361191-DEL-5272 |
| chr1-148372294-DEL-136 | chr1-148372295-DEL-136 |
| chr1-148388380-DEL-52 | chr1-148388385-DEL-80 |
| chr1-148391447-DEL-6658 | chr1-148391448-DEL-6658 |
| chr1-148417840-DEL-6224 | chr1-148417841-DEL-6224 |
| chr1-148435499-DEL-82 | chr1-148435517-DEL-65 |
| chr1-148447682-DEL-133 | chr1-148447683-DEL-133 |
| chr1-148453952-DEL-831 | chr1-148453953-DEL-831 |
| chr1-148457919-DEL-359 | chr1-148457920-DEL-359 |
| chr1-148460567-DEL-1072 | chr1-148460568-DEL-1072 |
| chr1-148476833-DEL-616 | chr1-148476834-DEL-616 |
| chr1-148491270-DEL-52 | chr1-148491271-DEL-52 |
| chr1-148495854-DEL-147 | chr1-148495855-DEL-147 |
| chr1-148512229-DEL-1360 | chr1-148512216-DEL-1373 |
| chr1-148547447-DEL-54 | chr1-148547448-DEL-54 |
| chr1-148555448-DEL-156 | chr1-148555449-DEL-156 |
| chr1-148577608-DEL-609 | chr1-148577609-DEL-609 |
| chr1-148579572-DEL-68 | chr1-148579573-DEL-66 |
| chr1-148610459-DEL-452 | chr1-148610460-DEL-452 |
| chr1-148614144-DEL-60 | chr1-148614145-DEL-60 |
| chr1-148635882-DEL-1890 | chr1-148635883-DEL-1891 |
| chr1-148647580-DEL-575 | chr1-148647580-DEL-575 |
| chr1-148688299-DEL-69 | chr1-148688300-DEL-69 |
| chr1-148760175-DEL-192 | chr1-148760176-DEL-192 |
| chr1-148761865-DEL-56 | chr1-148761866-DEL-56 |
| chr1-148776168-DEL-74 | chr1-148776160-DEL-79 |
| chr1-148802198-DEL-237 | chr1-148802206-DEL-235 |
| chr1-148804386-DEL-205 | chr1-148804387-DEL-205 |
| chr1-148825072-DEL-75 | chr1-148825076-DEL-70 |
| chr1-148841961-DEL-1476 | chr1-148841961-DEL-1476 |
| chr1-148888567-DEL-1208 | chr1-148888568-DEL-1208 |
| chr1-148899029-DEL-2863 | chr1-148899025-DEL-2864 |
| chr1-148902205-DEL-354 | chr1-148902204-DEL-354 |
| chr1-149017436-DEL-473 | chr1-149017437-DEL-473 |
| chr1-149052587-DEL-3315 | chr1-149052588-DEL-3315 |
| chr1-149059313-DEL-1481 | chr1-149059314-DEL-1481 |
| chr1-149069095-DEL-56 | chr1-149069096-DEL-56 |
| chr1-149086032-DEL-67 | chr1-149086035-DEL-65 |
| chr1-149092527-DEL-70 | chr1-149092528-DEL-70 |
| chr1-149095791-DEL-5474 | chr1-149095792-DEL-5474 |
| chr1-149125527-DEL-58 | chr1-149125527-DEL-58 |
| chr1-149157970-DEL-751 | chr1-149157971-DEL-751 |
| chr1-149162336-DEL-1926 | chr1-149162337-DEL-1926 |
| chr1-149164904-DEL-201 | chr1-149164905-DEL-201 |
| chr1-149172180-DEL-6780 | chr1-149172181-DEL-6780 |
| chr1-149190445-DEL-1341 | chr1-149190446-DEL-1341 |
| chr1-149269024-DEL-7082 | chr1-149269025-DEL-7082 |
| chr1-149292630-DEL-1239 | chr1-149292631-DEL-1233 |
| chr1-149293834-DEL-54 | chr1-149293835-DEL-54 |
| chr1-149309130-DEL-93 | chr1-149309125-DEL-99 |
| chr1-149326768-DEL-6296 | chr1-149326769-DEL-6296 |
| chr1-149353514-DEL-580 | chr1-149353515-DEL-580 |
| chr1-149374086-DEL-68 | chr1-149374087-DEL-68 |
| chr1-149376929-DEL-60 | chr1-149376930-DEL-60 |
| chr1-149382035-DEL-7169 | chr1-149382036-DEL-7169 |
| chr1-149399005-DEL-117 | chr1-149399019-DEL-119 |
| chr1-149401845-DEL-101 | chr1-149401846-DEL-101 |
| chr1-149435974-DEL-184 | chr1-149435975-DEL-184 |
| chr1-149491103-DEL-70 | chr1-149491105-DEL-68 |
| chr1-149507202-DEL-402 | chr1-149507203-DEL-402 |
| chr1-149555556-DEL-66 | chr1-149555557-DEL-66 |
| chr1-149595469-DEL-52 | chr1-149595469-DEL-52 |
| chr1-149611349-DEL-1491 | chr1-149611350-DEL-1491 |
| chr1-149616379-DEL-205 | chr1-149616380-DEL-205 |
| chr1-149652664-DEL-167 | chr1-149652665-DEL-167 |
| chr1-149683305-DEL-56 | chr1-149683304-DEL-88 |
| chr1-149713006-DEL-171 | chr1-149712987-DEL-191 |
| chr1-149718370-DEL-811 | chr1-149718371-DEL-811 |
| chr1-149754565-DEL-142 | chr1-149754566-DEL-142 |

| | |
|---|---|
| chr1-149764082-DEL-90 | chr1-149764088-DEL-84 |
| chr1-149765366-DEL-517 | chr1-149765367-DEL-517 |
| chr1-149789625-DEL-647 | chr1-149789626-DEL-647 |
| chr1-149806241-DEL-56 | chr1-149806246-DEL-52 |
| chr1-149809546-DEL-135 | chr1-149809551-DEL-131 |
| chr1-149811772-DEL-1049 | chr1-149811773-DEL-1049 |
| chr1-149814443-DEL-59 | chr1-149814440-DEL-61 |
| chr1-149829316-DEL-56 | chr1-149829316-DEL-80 |
| chr1-149850165-DEL-55 | chr1-149850166-DEL-51 |
| chr1-149852391-DEL-59 | chr1-149852392-DEL-67 |
| chr1-149857286-DEL-323 | chr1-149857287-DEL-323 |
| chr1-149867307-DEL-172 | chr1-149867307-DEL-172 |
| chr1-149867880-DEL-352 | chr1-149867881-DEL-352 |
| chr1-149872947-DEL-6199 | chr1-149872946-DEL-6199 |
| chr1-149880294-DEL-1082 | chr1-149880295-DEL-1082 |
| chr1-149885960-DEL-110 | chr1-149885961-DEL-110 |
| chr1-149904289-DEL-415 | chr1-149904278-DEL-416 |
| chr1-149921480-DEL-64 | chr1-149921481-DEL-64 |
| chr1-149943192-DEL-160 | chr1-149943193-DEL-159 |
| chr1-149957628-DEL-559 | chr1-149957629-DEL-559 |
| chr1-149999748-DEL-78 | chr1-149999749-DEL-78 |
| chr1-150004879-DEL-66 | chr1-150004890-DEL-68 |
| chr1-150023720-DEL-62 | chr1-150023721-DEL-56 |
| chr1-150025737-DEL-54 | chr1-150025738-DEL-54 |
| chr1-150029265-DEL-55 | chr1-150029266-DEL-55 |
| chr1-150044631-DEL-705 | chr1-150044632-DEL-705 |
| chr1-150079567-DEL-670 | chr1-150079568-DEL-670 |
| chr1-150097162-DEL-2609 | chr1-150097163-DEL-2609 |
| chr1-150105429-DEL-1783 | chr1-150105430-DEL-1783 |
| chr1-150108168-DEL-7243 | chr1-150108169-DEL-7243 |
| chr1-150140643-DEL-1181 | chr1-150140644-DEL-1181 |
| chr1-150148738-DEL-8064 | chr1-150148738-DEL-8064 |
| chr1-150169402-DEL-73 | chr1-150169403-DEL-73 |
| chr1-150199128-DEL-94 | chr1-150199129-DEL-94 |
| chr1-150213602-DEL-6390 | chr1-150213603-DEL-6390 |
| chr1-150220907-DEL-722 | chr1-150220901-DEL-723 |
| chr1-150285399-DEL-155 | chr1-150285400-DEL-155 |
| chr1-150287529-DEL-57 | chr1-150287514-DEL-89 |
| chr1-150330659-DEL-216 | chr1-150330660-DEL-216 |
| chr1-150359523-DEL-53 | chr1-150359524-DEL-53 |
| chr1-150388571-DEL-4204 | chr1-150388572-DEL-4204 |
| chr1-150397035-DEL-62 | chr1-150397068-DEL-62 |
| chr1-150399462-DEL-3948 | chr1-150399463-DEL-3948 |
| chr1-150441321-DEL-628 | chr1-150441322-DEL-628 |
| chr1-150501166-DEL-439 | chr1-150501167-DEL-439 |
| chr1-150505453-DEL-467 | chr1-150505454-DEL-467 |
| chr1-150552094-DEL-52 | chr1-150552095-DEL-53 |
| chr1-150590784-DEL-330 | chr1-150590794-DEL-326 |
| chr1-150663954-DEL-379 | chr1-150663955-DEL-377 |
| chr1-150668945-DEL-140 | chr1-150668954-DEL-132 |
| chr1-150714562-DEL-1258 | chr1-150714573-DEL-1243 |
| chr1-150716134-DEL-1132 | chr1-150716132-DEL-1133 |
| chr1-150727283-DEL-1530 | chr1-150727283-DEL-1530 |
| chr1-150729661-DEL-132 | chr1-150729662-DEL-132 |
| chr1-150744641-DEL-221 | chr1-150744642-DEL-221 |
| chr1-150788182-DEL-732 | chr1-150788183-DEL-732 |
| chr1-150795233-DEL-67 | chr1-150795218-DEL-63 |
| chr1-150890072-DEL-130 | chr1-150890074-DEL-131 |
| chr1-150899395-DEL-423 | chr1-150899394-DEL-424 |
| chr1-150923874-DEL-90 | chr1-150923872-DEL-68 |
| chr1-150936260-DEL-107 | chr1-150936261-DEL-107 |
| chr1-150938860-DEL-58 | chr1-150938861-DEL-58 |
| chr1-150959316-DEL-224 | chr1-150959316-DEL-225 |
| chr1-150960033-DEL-73 | chr1-150960000-DEL-61 |
| chr1-150992786-DEL-85 | chr1-150992772-DEL-81 |
| chr1-150997066-DEL-72 | chr1-150997067-DEL-72 |
| chr1-150997766-DEL-1969 | chr1-150997767-DEL-1969 |
| chr1-151017354-DEL-57 | chr1-151017355-DEL-57 |
| chr1-151052062-DEL-263 | chr1-151052063-DEL-263 |
| chr1-151063434-DEL-3799 | chr1-151063435-DEL-3799 |
| chr1-151088593-DEL-70 | chr1-151088594-DEL-70 |
| chr1-151117513-DEL-77 | chr1-151117513-DEL-77 |
| chr1-151120745-DEL-85 | chr1-151120746-DEL-85 |
| chr1-151122687-DEL-198 | chr1-151122688-DEL-198 |
| chr1-151127977-DEL-373 | chr1-151127978-DEL-373 |
| chr1-151136613-DEL-52 | chr1-151136614-DEL-52 |
| chr1-151188545-DEL-92 | chr1-151188545-DEL-95 |
| chr1-151192138-DEL-215 | chr1-151192139-DEL-214 |
| chr1-151193328-DEL-201 | chr1-151193329-DEL-201 |
| chr1-151202603-DEL-745 | chr1-151202604-DEL-745 |

| | |
|---|---|
| chr1-151207822-DEL-70 | chr1-151207823-DEL-71 |
| chr1-151223990-DEL-197 | chr1-151223991-DEL-197 |
| chr1-151247329-DEL-478 | chr1-151247330-DEL-478 |
| chr1-151249662-DEL-79 | chr1-151249671-DEL-63 |
| chr1-151250274-DEL-100 | chr1-151250274-DEL-100 |
| chr1-151251332-DEL-61 | chr1-151251336-DEL-64 |
| chr1-151260369-DEL-159 | chr1-151260374-DEL-155 |
| chr1-151262295-DEL-94 | chr1-151262296-DEL-94 |
| chr1-151271518-DEL-212 | chr1-151271519-DEL-212 |
| chr1-151272439-DEL-72 | chr1-151272407-DEL-57 |
| chr1-151279248-DEL-77 | chr1-151279249-DEL-77 |
| chr1-151301622-DEL-710 | chr1-151301623-DEL-710 |
| chr1-151334144-DEL-6125 | chr1-151334145-DEL-6125 |
| chr1-151334612-DEL-63 | chr1-151334613-DEL-61 |
| chr1-151347840-DEL-234 | chr1-151347841-DEL-234 |
| chr1-151383056-DEL-68 | chr1-151383057-DEL-68 |
| chr1-151385321-DEL-55 | chr1-151385321-DEL-55 |
| chr1-151393559-DEL-789 | chr1-151393559-DEL-789 |
| chr1-151408924-DEL-627 | chr1-151408925-DEL-627 |
| chr1-151415335-DEL-207 | chr1-151415336-DEL-207 |
| chr1-151422640-DEL-8144 | chr1-151422640-DEL-8142 |
| chr1-151429200-DEL-213 | chr1-151429201-DEL-213 |
| chr1-151441798-DEL-129 | chr1-151441836-DEL-108 |
| chr1-151465125-DEL-1990 | chr1-151465126-DEL-1990 |
| chr1-151469810-DEL-64 | chr1-151469811-DEL-64 |
| chr1-151482266-DEL-112 | chr1-151482267-DEL-112 |
| chr1-151494380-DEL-64 | chr1-151494381-DEL-64 |
| chr1-151517348-DEL-591 | chr1-151517349-DEL-591 |
| chr1-151518752-DEL-212 | chr1-151518753-DEL-212 |
| chr1-151527780-DEL-62 | chr1-151527780-DEL-62 |
| chr1-151568468-DEL-111 | chr1-151568469-DEL-108 |
| chr1-151605188-DEL-3215 | chr1-151605178-DEL-3180 |
| chr1-151618571-DEL-1870 | chr1-151618571-DEL-1870 |
| chr1-151640880-DEL-54 | chr1-151640881-DEL-54 |
| chr1-151647913-DEL-181 | chr1-151647914-DEL-181 |
| chr1-151715358-DEL-167 | chr1-151715359-DEL-167 |
| chr1-151729355-DEL-52 | chr1-151729354-DEL-54 |
| chr1-151735187-DEL-52 | chr1-151735187-DEL-52 |
| chr1-151741642-DEL-6397 | chr1-151741636-DEL-6401 |
| chr1-151753115-DEL-70 | chr1-151753161-DEL-74 |
| chr1-151773628-DEL-123 | chr1-151773629-DEL-126 |
| chr1-151782430-DEL-54 | chr1-151782431-DEL-54 |
| chr1-151792894-DEL-944 | chr1-151792895-DEL-944 |
| chr1-151794260-DEL-1183 | chr1-151794261-DEL-1183 |
| chr1-151801890-DEL-80 | chr1-151801891-DEL-80 |
| chr1-151831294-DEL-170 | chr1-151831296-DEL-171 |
| chr1-151845264-DEL-1038 | chr1-151845270-DEL-1041 |
| chr1-151884075-DEL-53 | chr1-151884068-DEL-60 |
| chr1-151900314-DEL-66 | chr1-151900315-DEL-66 |
| chr1-151919619-DEL-206 | chr1-151919620-DEL-205 |
| chr1-151938834-DEL-96 | chr1-151938835-DEL-96 |
| chr1-151972345-DEL-126 | chr1-151972346-DEL-84 |
| chr1-151975613-DEL-162 | chr1-151975614-DEL-162 |
| chr1-152004472-DEL-118 | chr1-152004473-DEL-118 |
| chr1-152008412-DEL-92 | chr1-152008413-DEL-92 |
| chr1-152014316-DEL-120 | chr1-152014317-DEL-120 |
| chr1-152066030-DEL-2011 | chr1-152066029-DEL-2011 |
| chr1-152076728-DEL-227 | chr1-152076729-DEL-227 |
| chr1-152080477-DEL-1848 | chr1-152080478-DEL-1848 |
| chr1-152084330-DEL-66 | chr1-152084331-DEL-66 |
| chr1-152115918-DEL-52 | chr1-152115919-DEL-52 |
| chr1-152117162-DEL-219 | chr1-152117163-DEL-219 |
| chr1-152117808-DEL-6156 | chr1-152117809-DEL-6156 |
| chr1-152117894-DEL-214 | chr1-152117895-DEL-214 |
| chr1-152132958-DEL-181 | chr1-152132958-DEL-182 |
| chr1-152138131-DEL-71 | chr1-152138144-DEL-68 |
| chr1-152139387-DEL-51 | chr1-152139388-DEL-51 |
| chr1-152156999-DEL-144 | chr1-152157000-DEL-144 |
| chr1-152160497-DEL-70 | chr1-152160489-DEL-70 |
| chr1-152170560-DEL-1326 | chr1-152170561-DEL-1326 |
| chr1-152186567-DEL-206 | chr1-152186568-DEL-206 |
| chr1-152194140-DEL-2020 | chr1-152194141-DEL-2020 |
| chr1-152215442-DEL-3453 | chr1-152215443-DEL-3453 |
| chr1-152222502-DEL-86 | chr1-152222525-DEL-50 |
| chr1-152241088-DEL-62 | chr1-152241091-DEL-60 |
| chr1-152242728-DEL-52 | chr1-152242728-DEL-52 |
| chr1-152267440-DEL-5291 | chr1-152267441-DEL-5291 |
| chr1-152268033-DEL-156 | chr1-152268033-DEL-156 |
| chr1-152275099-DEL-53 | chr1-152275100-DEL-58 |
| chr1-152278920-DEL-52 | chr1-152278879-DEL-52 |

| | |
|---|---|
| chr1-152289303-DEL-332 | chr1-152289304-DEL-333 |
| chr1-152305234-DEL-51 | chr1-152305259-DEL-51 |
| chr1-152311356-DEL-2291 | chr1-152311356-DEL-2291 |
| chr1-152318395-DEL-166 | chr1-152318396-DEL-166 |
| chr1-152327970-DEL-108 | chr1-152327971-DEL-108 |
| chr1-152328344-DEL-402 | chr1-152328345-DEL-402 |
| chr1-152338517-DEL-2108 | chr1-152338518-DEL-2108 |
| chr1-152353277-DEL-6475 | chr1-152353276-DEL-6476 |
| chr1-152380902-DEL-1124 | chr1-152380903-DEL-1124 |
| chr1-152413125-DEL-73 | chr1-152413127-DEL-70 |
| chr1-152437872-DEL-120 | chr1-152437893-DEL-100 |
| chr1-152446337-DEL-68 | chr1-152446338-DEL-68 |
| chr1-152530114-DEL-63 | chr1-152530115-DEL-63 |
| chr1-152569666-DEL-62 | chr1-152569672-DEL-56 |
| chr1-152587462-DEL-1206 | chr1-152587463-DEL-1206 |
| chr1-152600429-DEL-86 | chr1-152600440-DEL-62 |
| chr1-152603924-DEL-58 | chr1-152603914-DEL-64 |
| chr1-152610853-DEL-276 | chr1-152610828-DEL-279 |
| chr1-152630414-DEL-186 | chr1-152630409-DEL-187 |
| chr1-152669153-DEL-108 | chr1-152669154-DEL-108 |
| chr1-152691373-DEL-106 | chr1-152691374-DEL-104 |
| chr1-152704698-DEL-52 | chr1-152704699-DEL-50 |
| chr1-152732608-DEL-52 | chr1-152732609-DEL-52 |
| chr1-152763769-DEL-214 | chr1-152763770-DEL-214 |
| chr1-152801286-DEL-207 | chr1-152801287-DEL-207 |
| chr1-152834892-DEL-59 | chr1-152834893-DEL-59 |
| chr1-152875496-DEL-65 | chr1-152875499-DEL-61 |
| chr1-152896429-DEL-180 | chr1-152896430-DEL-180 |
| chr1-152902899-DEL-56 | chr1-152902897-DEL-57 |
| chr1-152906444-DEL-216 | chr1-152906445-DEL-216 |
| chr1-152908179-DEL-61 | chr1-152908180-DEL-60 |
| chr1-152919451-DEL-192 | chr1-152919452-DEL-192 |
| chr1-152934781-DEL-331 | chr1-152934782-DEL-331 |
| chr1-152937850-DEL-194 | chr1-152937851-DEL-194 |
| chr1-152943182-DEL-76 | chr1-152943199-DEL-109 |
| chr1-152955242-DEL-1440 | chr1-152955243-DEL-1440 |
| chr1-152964330-DEL-52 | chr1-152964311-DEL-72 |
| chr1-152965060-DEL-216 | chr1-152965061-DEL-216 |
| chr1-152978853-DEL-178 | chr1-152978854-DEL-178 |
| chr1-152984115-DEL-206 | chr1-152984110-DEL-207 |
| chr1-153015152-DEL-80 | chr1-153015153-DEL-80 |
| chr1-153018224-DEL-488 | chr1-153018225-DEL-488 |
| chr1-153048667-DEL-75 | chr1-153048668-DEL-75 |
| chr1-153058050-DEL-123 | chr1-153058037-DEL-145 |
| chr1-153059845-DEL-78 | chr1-153059846-DEL-65 |
| chr1-153060328-DEL-94 | chr1-153060328-DEL-94 |
| chr1-153065959-DEL-678 | chr1-153065966-DEL-664 |
| chr1-153074460-DEL-400 | chr1-153074461-DEL-400 |
| chr1-153085520-DEL-56 | chr1-153085520-DEL-56 |
| chr1-153109747-DEL-87 | chr1-153109748-DEL-87 |
| chr1-153145894-DEL-67 | chr1-153145895-DEL-67 |
| chr1-153177517-DEL-483 | chr1-153177518-DEL-483 |
| chr1-153182018-DEL-95 | chr1-153182019-DEL-95 |
| chr1-153185316-DEL-82 | chr1-153185317-DEL-82 |
| chr1-153213931-DEL-71 | chr1-153213936-DEL-70 |
| chr1-153214455-DEL-801 | chr1-153214456-DEL-801 |
| chr1-153235533-DEL-54 | chr1-153235534-DEL-54 |
| chr1-153247089-DEL-86 | chr1-153247090-DEL-86 |
| chr1-153251930-DEL-102 | chr1-153251935-DEL-98 |
| chr1-153257763-DEL-57 | chr1-153257763-DEL-57 |
| chr1-153258080-DEL-1911 | chr1-153258098-DEL-1891 |
| chr1-153261487-DEL-106 | chr1-153261486-DEL-107 |
| chr1-153287078-DEL-159 | chr1-153287079-DEL-159 |
| chr1-153370118-DEL-181 | chr1-153370127-DEL-180 |
| chr1-153372778-DEL-2804 | chr1-153372779-DEL-2804 |
| chr1-153376612-DEL-199 | chr1-153376613-DEL-199 |
| chr1-153382325-DEL-616 | chr1-153382298-DEL-612 |
| chr1-153384254-DEL-64 | chr1-153384255-DEL-64 |
| chr1-153385261-DEL-74 | chr1-153385262-DEL-74 |
| chr1-153393650-DEL-182 | chr1-153393666-DEL-197 |
| chr1-153401791-DEL-90 | chr1-153401792-DEL-90 |
| chr1-153410176-DEL-570 | chr1-153410177-DEL-570 |
| chr1-153412586-DEL-60 | chr1-153412555-DEL-53 |
| chr1-153414516-DEL-184 | chr1-153414517-DEL-184 |
| chr1-153416404-DEL-96 | chr1-153416415-DEL-86 |
| chr1-153435316-DEL-66 | chr1-153435310-DEL-68 |
| chr1-153441021-DEL-102 | chr1-153441019-DEL-150 |
| chr1-153456460-DEL-65 | chr1-153456461-DEL-65 |
| chr1-153468452-DEL-209 | chr1-153468453-DEL-209 |
| chr1-153478845-DEL-88 | chr1-153478846-DEL-88 |

| | |
|---|---|
| chr1-153495197-DEL-1277 | chr1-153495198-DEL-1277 |
| chr1-153539252-DEL-161 | chr1-153539253-DEL-161 |
| chr1-153539876-DEL-124 | chr1-153539881-DEL-92 |
| chr1-153559620-DEL-52 | chr1-153559621-DEL-52 |
| chr1-153584283-DEL-51 | chr1-153584281-DEL-54 |
| chr1-153601131-DEL-183 | chr1-153601132-DEL-183 |
| chr1-153631594-DEL-56 | chr1-153631599-DEL-54 |
| chr1-153653232-DEL-935 | chr1-153653233-DEL-935 |
| chr1-153657029-DEL-66 | chr1-153657054-DEL-82 |
| chr1-153658185-DEL-1230 | chr1-153658186-DEL-1231 |
| chr1-153690169-DEL-68 | chr1-153690170-DEL-68 |
| chr1-153710716-DEL-60 | chr1-153710715-DEL-62 |
| chr1-153738193-DEL-104 | chr1-153738194-DEL-104 |
| chr1-153741387-DEL-224 | chr1-153741388-DEL-224 |
| chr1-153752071-DEL-203 | chr1-153752072-DEL-203 |
| chr1-153756202-DEL-495 | chr1-153756203-DEL-495 |
| chr1-153758663-DEL-257 | chr1-153758663-DEL-257 |
| chr1-153759496-DEL-460 | chr1-153759497-DEL-460 |
| chr1-153773797-DEL-129 | chr1-153773799-DEL-130 |
| chr1-153779751-DEL-288 | chr1-153779752-DEL-288 |
| chr1-153799790-DEL-241 | chr1-153799791-DEL-241 |
| chr1-153803581-DEL-73 | chr1-153803576-DEL-91 |
| chr1-153804217-DEL-54 | chr1-153804218-DEL-52 |
| chr1-153832007-DEL-87 | chr1-153831999-DEL-115 |
| chr1-153840210-DEL-72 | chr1-153840211-DEL-72 |
| chr1-153852243-DEL-72 | chr1-153852269-DEL-53 |
| chr1-153881005-DEL-51 | chr1-153881006-DEL-51 |
| chr1-153883834-DEL-204 | chr1-153883832-DEL-205 |
| chr1-153888513-DEL-1840 | chr1-153888514-DEL-1840 |
| chr1-153897338-DEL-848 | chr1-153897335-DEL-855 |
| chr1-153927059-DEL-93 | chr1-153927060-DEL-94 |
| chr1-153927862-DEL-112 | chr1-153927863-DEL-111 |
| chr1-153935162-DEL-89 | chr1-153935163-DEL-87 |
| chr1-153937104-DEL-176 | chr1-153937105-DEL-176 |
| chr1-153963657-DEL-87 | chr1-153963650-DEL-95 |
| chr1-153967689-DEL-109 | chr1-153967690-DEL-109 |
| chr1-153990649-DEL-377 | chr1-153990650-DEL-377 |
| chr1-153998904-DEL-56 | chr1-153998911-DEL-50 |
| chr1-154007709-DEL-6213 | chr1-154007710-DEL-6213 |
| chr1-154020139-DEL-63 | chr1-154020140-DEL-63 |
| chr1-154025538-DEL-52 | chr1-154025543-DEL-64 |
| chr1-154062726-DEL-430 | chr1-154062727-DEL-430 |
| chr1-154067654-DEL-337 | chr1-154067655-DEL-337 |
| chr1-154092733-DEL-68 | chr1-154092734-DEL-62 |
| chr1-154101373-DEL-122 | chr1-154101394-DEL-98 |
| chr1-154118633-DEL-6338 | chr1-154118634-DEL-6338 |
| chr1-154131168-DEL-132 | chr1-154131173-DEL-100 |
| chr1-154156269-DEL-72 | chr1-154156286-DEL-52 |
| chr1-154176498-DEL-51 | chr1-154176499-DEL-51 |
| chr1-154195685-DEL-586 | chr1-154195686-DEL-586 |
| chr1-154203865-DEL-1221 | chr1-154203866-DEL-1224 |
| chr1-154223671-DEL-75 | chr1-154223672-DEL-75 |
| chr1-154247411-DEL-1261 | chr1-154247412-DEL-1261 |
| chr1-154330022-DEL-2946 | chr1-154330023-DEL-2946 |
| chr1-154418002-DEL-82 | chr1-154418010-DEL-74 |
| chr1-154436581-DEL-62 | chr1-154436569-DEL-60 |
| chr1-154529051-DEL-52 | chr1-154529052-DEL-52 |
| chr1-154565572-DEL-2658 | chr1-154565573-DEL-2658 |
| chr1-154575974-DEL-64 | chr1-154575975-DEL-64 |
| chr1-154630093-DEL-52 | chr1-154630094-DEL-52 |
| chr1-154634758-DEL-431 | chr1-154634742-DEL-432 |
| chr1-154669881-DEL-52 | chr1-154669882-DEL-52 |
| chr1-154709459-DEL-116 | chr1-154709460-DEL-116 |
| chr1-154724837-DEL-606 | chr1-154724838-DEL-606 |
| chr1-154774712-DEL-7095 | chr1-154774713-DEL-7095 |
| chr1-154787535-DEL-64 | chr1-154787512-DEL-64 |
| chr1-154793001-DEL-76 | chr1-154793007-DEL-68 |
| chr1-154810039-DEL-1577 | chr1-154810039-DEL-1577 |
| chr1-154825464-DEL-64 | chr1-154825465-DEL-64 |
| chr1-154835631-DEL-3132 | chr1-154835632-DEL-3132 |
| chr1-154842377-DEL-1860 | chr1-154842378-DEL-1860 |
| chr1-154855394-DEL-58 | chr1-154855395-DEL-64 |
| chr1-154862507-DEL-935 | chr1-154862508-DEL-935 |
| chr1-154864292-DEL-96 | chr1-154864294-DEL-71 |
| chr1-154869489-DEL-99 | chr1-154869496-DEL-88 |
| chr1-154896895-DEL-104 | chr1-154896912-DEL-120 |
| chr1-154932319-DEL-262 | chr1-154932320-DEL-262 |
| chr1-154935732-DEL-139 | chr1-154935733-DEL-153 |
| chr1-154952498-DEL-83 | chr1-154952472-DEL-80 |
| chr1-154985081-DEL-52 | chr1-154985081-DEL-50 |

| | |
|---|---|
| chr1-155000616-DEL-209 | chr1-155000617-DEL-209 |
| chr1-155024373-DEL-406 | chr1-155024374-DEL-406 |
| chr1-155031781-DEL-252 | chr1-155031782-DEL-252 |
| chr1-155049260-DEL-116 | chr1-155049261-DEL-116 |
| chr1-155094238-DEL-400 | chr1-155094228-DEL-401 |
| chr1-155110372-DEL-265 | chr1-155110374-DEL-278 |
| chr1-155210655-DEL-62 | chr1-155210671-DEL-62 |
| chr1-155234228-DEL-60 | chr1-155234222-DEL-66 |
| chr1-155255482-DEL-4085 | chr1-155255483-DEL-4085 |
| chr1-155261540-DEL-643 | chr1-155261545-DEL-639 |
| chr1-155271527-DEL-727 | chr1-155271528-DEL-727 |
| chr1-155320630-DEL-145 | chr1-155320629-DEL-127 |
| chr1-155323799-DEL-55 | chr1-155323800-DEL-55 |
| chr1-155351214-DEL-65 | chr1-155351215-DEL-65 |
| chr1-155366821-DEL-61 | chr1-155366806-DEL-56 |
| chr1-155390607-DEL-203 | chr1-155390608-DEL-203 |
| chr1-155414335-DEL-6444 | chr1-155414337-DEL-6442 |
| chr1-155418937-DEL-179 | chr1-155418938-DEL-179 |
| chr1-155419796-DEL-567 | chr1-155419797-DEL-567 |
| chr1-155424706-DEL-182 | chr1-155424708-DEL-177 |
| chr1-155445752-DEL-220 | chr1-155445753-DEL-220 |
| chr1-155464005-DEL-345 | chr1-155464007-DEL-339 |
| chr1-155479371-DEL-87 | chr1-155479372-DEL-87 |
| chr1-155511136-DEL-114 | chr1-155511139-DEL-127 |
| chr1-155552038-DEL-223 | chr1-155552039-DEL-223 |
| chr1-155598410-DEL-71 | chr1-155598411-DEL-71 |
| chr1-155631103-DEL-1789 | chr1-155631104-DEL-1789 |
| chr1-155640469-DEL-56 | chr1-155640470-DEL-54 |
| chr1-155656593-DEL-218 | chr1-155656594-DEL-218 |
| chr1-155694114-DEL-64 | chr1-155694125-DEL-62 |
| chr1-155694496-DEL-282 | chr1-155694497-DEL-235 |
| chr1-155761492-DEL-62 | chr1-155761474-DEL-62 |
| chr1-155777543-DEL-245 | chr1-155777544-DEL-245 |
| chr1-155792234-DEL-65 | chr1-155792235-DEL-65 |
| chr1-155798998-DEL-126 | chr1-155799001-DEL-124 |
| chr1-155813994-DEL-626 | chr1-155813992-DEL-627 |
| chr1-155830306-DEL-707 | chr1-155830307-DEL-707 |
| chr1-155863855-DEL-307 | chr1-155863856-DEL-307 |
| chr1-155871700-DEL-79 | chr1-155871703-DEL-80 |
| chr1-155960372-DEL-85 | chr1-155960383-DEL-86 |
| chr1-155969421-DEL-248 | chr1-155969422-DEL-245 |
| chr1-155976103-DEL-465 | chr1-155976104-DEL-465 |
| chr1-156015425-DEL-240 | chr1-156015426-DEL-240 |
| chr1-156016529-DEL-977 | chr1-156016530-DEL-977 |
| chr1-156042807-DEL-62 | chr1-156042808-DEL-62 |
| chr1-156046661-DEL-429 | chr1-156046662-DEL-429 |
| chr1-156056099-DEL-74 | chr1-156056100-DEL-74 |
| chr1-156065887-DEL-1298 | chr1-156065889-DEL-1298 |
| chr1-156101670-DEL-251 | chr1-156101671-DEL-251 |
| chr1-156109891-DEL-225 | chr1-156109892-DEL-225 |
| chr1-156116708-DEL-946 | chr1-156116709-DEL-946 |
| chr1-156120104-DEL-166 | chr1-156120105-DEL-166 |
| chr1-156146532-DEL-377 | chr1-156146533-DEL-377 |
| chr1-156148157-DEL-194 | chr1-156148158-DEL-194 |
| chr1-156160914-DEL-762 | chr1-156160915-DEL-762 |
| chr1-156183039-DEL-64 | chr1-156183040-DEL-64 |
| chr1-156190498-DEL-196 | chr1-156190499-DEL-196 |
| chr1-156193593-DEL-507 | chr1-156193594-DEL-507 |
| chr1-156206276-DEL-216 | chr1-156206277-DEL-216 |
| chr1-156223313-DEL-199 | chr1-156223314-DEL-199 |
| chr1-156277445-DEL-90 | chr1-156277439-DEL-91 |
| chr1-156279490-DEL-221 | chr1-156279491-DEL-221 |
| chr1-156291248-DEL-399 | chr1-156291249-DEL-399 |
| chr1-156295062-DEL-120 | chr1-156295060-DEL-110 |
| chr1-156310811-DEL-2273 | chr1-156310812-DEL-2273 |
| chr1-156313324-DEL-160 | chr1-156313325-DEL-160 |
| chr1-156327747-DEL-254 | chr1-156327748-DEL-254 |
| chr1-156344115-DEL-195 | chr1-156344116-DEL-195 |
| chr1-156345256-DEL-170 | chr1-156345281-DEL-146 |
| chr1-156350301-DEL-235 | chr1-156350302-DEL-235 |
| chr1-156351525-DEL-2725 | chr1-156351526-DEL-2725 |
| chr1-156357991-DEL-136 | chr1-156357992-DEL-136 |
| chr1-156361342-DEL-203 | chr1-156361343-DEL-203 |
| chr1-156362279-DEL-109 | chr1-156362291-DEL-116 |
| chr1-156373845-DEL-57 | chr1-156373845-DEL-57 |
| chr1-156379498-DEL-250 | chr1-156379498-DEL-251 |
| chr1-156383076-DEL-51 | chr1-156383077-DEL-51 |
| chr1-156427066-DEL-68 | chr1-156427067-DEL-68 |
| chr1-156437471-DEL-59 | chr1-156437472-DEL-59 |
| chr1-156454677-DEL-114 | chr1-156454678-DEL-114 |

| | |
|---|---|
| chr1-156463796-DEL-60 | chr1-156463796-DEL-60 |
| chr1-156488173-DEL-336 | chr1-156488174-DEL-336 |
| chr1-156492043-DEL-123 | chr1-156492044-DEL-123 |
| chr1-156510393-DEL-217 | chr1-156510391-DEL-216 |
| chr1-156512724-DEL-146 | chr1-156512725-DEL-146 |
| chr1-156523561-DEL-133 | chr1-156523562-DEL-133 |
| chr1-156540039-DEL-201 | chr1-156540040-DEL-201 |
| chr1-156605353-DEL-75 | chr1-156605352-DEL-68 |
| chr1-156621024-DEL-918 | chr1-156621025-DEL-918 |
| chr1-156628672-DEL-184 | chr1-156628668-DEL-185 |
| chr1-156638850-DEL-86 | chr1-156638846-DEL-99 |
| chr1-156663504-DEL-112 | chr1-156663505-DEL-112 |
| chr1-156676421-DEL-1975 | chr1-156676422-DEL-1974 |
| chr1-156682830-DEL-199 | chr1-156682831-DEL-199 |
| chr1-156702209-DEL-123 | chr1-156702210-DEL-123 |
| chr1-156706649-DEL-206 | chr1-156706650-DEL-206 |
| chr1-156739581-DEL-80 | chr1-156739582-DEL-80 |
| chr1-156761936-DEL-139 | chr1-156761904-DEL-140 |
| chr1-156762324-DEL-436 | chr1-156762325-DEL-436 |
| chr1-156763502-DEL-69 | chr1-156763480-DEL-70 |
| chr1-156770376-DEL-216 | chr1-156770377-DEL-216 |
| chr1-156774010-DEL-58 | chr1-156774011-DEL-58 |
| chr1-156782143-DEL-916 | chr1-156782144-DEL-916 |
| chr1-156783121-DEL-77 | chr1-156783122-DEL-77 |
| chr1-156796210-DEL-1248 | chr1-156796211-DEL-1248 |
| chr1-156803194-DEL-305 | chr1-156803195-DEL-305 |
| chr1-156880739-DEL-223 | chr1-156880740-DEL-223 |
| chr1-156894945-DEL-363 | chr1-156894946-DEL-363 |
| chr1-156904096-DEL-163 | chr1-156904115-DEL-153 |
| chr1-156918051-DEL-60 | chr1-156918052-DEL-57 |
| chr1-156920930-DEL-221 | chr1-156920931-DEL-221 |
| chr1-156966251-DEL-101 | chr1-156966252-DEL-101 |
| chr1-156970048-DEL-177 | chr1-156970047-DEL-152 |
| chr1-157019204-DEL-66 | chr1-157019205-DEL-66 |
| chr1-157023319-DEL-297 | chr1-157023320-DEL-297 |
| chr1-157052848-DEL-211 | chr1-157052849-DEL-211 |
| chr1-157068034-DEL-217 | chr1-157068035-DEL-217 |
| chr1-157087554-DEL-1123 | chr1-157087554-DEL-1123 |
| chr1-157147376-DEL-102 | chr1-157147374-DEL-103 |
| chr1-157163948-DEL-210 | chr1-157163949-DEL-210 |
| chr1-157178352-DEL-25219 | chr1-157178335-DEL-25236 |
| chr1-157216696-DEL-5238 | chr1-157216697-DEL-5238 |
| chr1-157231293-DEL-126 | chr1-157231294-DEL-126 |
| chr1-157244731-DEL-62 | chr1-157244732-DEL-62 |
| chr1-157248472-DEL-177 | chr1-157248473-DEL-177 |
| chr1-157251192-DEL-148 | chr1-157251197-DEL-149 |
| chr1-157261372-DEL-353 | chr1-157261373-DEL-353 |
| chr1-157272880-DEL-227 | chr1-157272881-DEL-227 |
| chr1-157279568-DEL-215 | chr1-157279569-DEL-215 |
| chr1-157291513-DEL-87 | chr1-157291521-DEL-80 |
| chr1-157301840-DEL-73 | chr1-157301840-DEL-74 |
| chr1-157310043-DEL-218 | chr1-157310043-DEL-218 |
| chr1-157326533-DEL-60 | chr1-157326513-DEL-61 |
| chr1-157328345-DEL-134 | chr1-157328346-DEL-134 |
| chr1-157332244-DEL-1064 | chr1-157332245-DEL-1064 |
| chr1-157367930-DEL-512 | chr1-157367931-DEL-512 |
| chr1-157375573-DEL-59 | chr1-157375574-DEL-59 |
| chr1-157379860-DEL-223 | chr1-157379861-DEL-223 |
| chr1-157426656-DEL-503 | chr1-157426657-DEL-503 |
| chr1-157434322-DEL-6652 | chr1-157434323-DEL-6652 |
| chr1-157445299-DEL-129 | chr1-157445300-DEL-129 |
| chr1-157455062-DEL-71 | chr1-157455063-DEL-71 |
| chr1-157505938-DEL-132 | chr1-157505939-DEL-132 |
| chr1-157527563-DEL-58 | chr1-157527564-DEL-58 |
| chr1-157535168-DEL-440 | chr1-157535169-DEL-440 |
| chr1-157537204-DEL-70 | chr1-157537205-DEL-70 |
| chr1-157552783-DEL-363 | chr1-157552784-DEL-357 |
| chr1-157575482-DEL-57 | chr1-157575493-DEL-93 |
| chr1-157640903-DEL-210 | chr1-157640904-DEL-210 |
| chr1-157704792-DEL-6809 | chr1-157704793-DEL-6809 |
| chr1-157731022-DEL-58 | chr1-157731023-DEL-52 |
| chr1-157761114-DEL-1264 | chr1-157761115-DEL-1264 |
| chr1-157813585-DEL-60 | chr1-157813586-DEL-60 |
| chr1-157840249-DEL-354 | chr1-157840250-DEL-354 |
| chr1-157857010-DEL-196 | chr1-157857011-DEL-196 |
| chr1-157945313-DEL-345 | chr1-157945312-DEL-338 |
| chr1-157948208-DEL-69 | chr1-157948209-DEL-69 |
| chr1-157971436-DEL-78 | chr1-157971440-DEL-53 |
| chr1-157976690-DEL-3537 | chr1-157976691-DEL-3537 |
| chr1-158091755-DEL-177 | chr1-158091756-DEL-177 |

| | |
|---|---|
| chr1-158115172-DEL-53 | chr1-158115162-DEL-65 |
| chr1-158118571-DEL-269 | chr1-158118572-DEL-269 |
| chr1-158136201-DEL-159 | chr1-158136202-DEL-159 |
| chr1-158159077-DEL-7089 | chr1-158159078-DEL-7089 |
| chr1-158159528-DEL-212 | chr1-158159529-DEL-212 |
| chr1-158187362-DEL-88 | chr1-158187363-DEL-88 |
| chr1-158204758-DEL-202 | chr1-158204761-DEL-200 |
| chr1-158380451-DEL-316 | chr1-158380450-DEL-315 |
| chr1-158470098-DEL-6272 | chr1-158470099-DEL-6272 |
| chr1-158531254-DEL-52 | chr1-158531255-DEL-52 |
| chr1-158531272-DEL-619 | chr1-158531273-DEL-619 |
| chr1-158584633-DEL-322 | chr1-158584633-DEL-318 |
| chr1-158606645-DEL-1991 | chr1-158606646-DEL-1991 |
| chr1-158633279-DEL-1890 | chr1-158633280-DEL-1890 |
| chr1-158637284-DEL-75 | chr1-158637310-DEL-50 |
| chr1-158637721-DEL-1204 | chr1-158637722-DEL-1204 |
| chr1-158666607-DEL-1976 | chr1-158666608-DEL-1976 |
| chr1-158707131-DEL-6747 | chr1-158707132-DEL-6747 |
| chr1-158729725-DEL-156 | chr1-158729720-DEL-168 |
| chr1-158766378-DEL-52 | chr1-158766379-DEL-52 |
| chr1-158839299-DEL-112 | chr1-158839299-DEL-114 |
| chr1-158849530-DEL-427 | chr1-158849523-DEL-428 |
| chr1-158867202-DEL-64 | chr1-158867165-DEL-108 |
| chr1-158875116-DEL-349 | chr1-158875117-DEL-349 |
| chr1-158893449-DEL-414 | chr1-158893450-DEL-414 |
| chr1-158915267-DEL-143 | chr1-158915268-DEL-143 |
| chr1-158944358-DEL-3314 | chr1-158944359-DEL-3314 |
| chr1-158967583-DEL-139 | chr1-158967583-DEL-139 |
| chr1-159038249-DEL-109 | chr1-159038206-DEL-152 |
| chr1-159081887-DEL-633 | chr1-159081890-DEL-628 |
| chr1-159110696-DEL-248 | chr1-159110697-DEL-248 |
| chr1-159111245-DEL-286 | chr1-159111246-DEL-286 |
| chr1-159121378-DEL-392 | chr1-159121378-DEL-392 |
| chr1-159123237-DEL-365 | chr1-159123238-DEL-365 |
| chr1-159171787-DEL-96 | chr1-159171788-DEL-96 |
| chr1-159207303-DEL-64 | chr1-159207305-DEL-100 |
| chr1-159209663-DEL-58 | chr1-159209664-DEL-58 |
| chr1-159211456-DEL-172 | chr1-159211457-DEL-172 |
| chr1-159219830-DEL-64 | chr1-159219806-DEL-97 |
| chr1-159226926-DEL-623 | chr1-159226927-DEL-623 |
| chr1-159230262-DEL-506 | chr1-159230263-DEL-506 |
| chr1-159271454-DEL-52 | chr1-159271451-DEL-62 |
| chr1-159401242-DEL-70 | chr1-159401243-DEL-70 |
| chr1-159489603-DEL-58 | chr1-159489604-DEL-58 |
| chr1-159535700-DEL-76 | chr1-159535701-DEL-76 |
| chr1-159549752-DEL-77 | chr1-159549753-DEL-77 |
| chr1-159560394-DEL-52 | chr1-159560383-DEL-64 |
| chr1-159570859-DEL-160 | chr1-159570892-DEL-120 |
| chr1-159571443-DEL-851 | chr1-159571444-DEL-851 |
| chr1-159580776-DEL-65 | chr1-159580776-DEL-68 |
| chr1-159585218-DEL-82 | chr1-159585191-DEL-84 |
| chr1-159603786-DEL-93 | chr1-159603787-DEL-93 |
| chr1-159615526-DEL-116 | chr1-159615525-DEL-115 |
| chr1-159670946-DEL-71 | chr1-159670947-DEL-71 |
| chr1-159726087-DEL-106 | chr1-159726088-DEL-106 |
| chr1-159736307-DEL-63 | chr1-159736302-DEL-63 |
| chr1-159753852-DEL-67 | chr1-159753853-DEL-69 |
| chr1-159761617-DEL-784 | chr1-159761618-DEL-784 |
| chr1-159773857-DEL-313 | chr1-159773858-DEL-313 |
| chr1-159788728-DEL-99 | chr1-159788729-DEL-99 |
| chr1-159797333-DEL-7087 | chr1-159797334-DEL-7087 |
| chr1-159877746-DEL-100 | chr1-159877747-DEL-100 |
| chr1-159879266-DEL-167 | chr1-159879243-DEL-168 |
| chr1-159885754-DEL-56 | chr1-159885755-DEL-56 |
| chr1-159957831-DEL-62 | chr1-159957832-DEL-70 |
| chr1-159974579-DEL-195 | chr1-159974580-DEL-195 |
| chr1-159992521-DEL-190 | chr1-159992522-DEL-190 |
| chr1-160017698-DEL-378 | chr1-160017699-DEL-378 |
| chr1-160044616-DEL-53 | chr1-160044617-DEL-53 |
| chr1-160077233-DEL-78 | chr1-160077234-DEL-72 |
| chr1-160081640-DEL-443 | chr1-160081640-DEL-443 |
| chr1-160107701-DEL-96 | chr1-160107702-DEL-96 |
| chr1-160148811-DEL-413 | chr1-160148831-DEL-391 |
| chr1-160156839-DEL-251 | chr1-160156840-DEL-251 |
| chr1-160237380-DEL-141 | chr1-160237380-DEL-141 |
| chr1-160240285-DEL-234 | chr1-160240286-DEL-234 |
| chr1-160248876-DEL-7010 | chr1-160248877-DEL-7010 |
| chr1-160249099-DEL-624 | chr1-160249098-DEL-624 |
| chr1-160288584-DEL-72 | chr1-160288589-DEL-68 |
| chr1-160370110-DEL-455 | chr1-160370111-DEL-455 |

| | |
|---|---|
| chr1-160416307-DEL-62 | chr1-160416308-DEL-62 |
| chr1-160462667-DEL-175 | chr1-160462666-DEL-177 |
| chr1-160465481-DEL-5173 | chr1-160465482-DEL-5173 |
| chr1-160529444-DEL-54 | chr1-160529445-DEL-54 |
| chr1-160561667-DEL-74 | chr1-160561668-DEL-74 |
| chr1-160591733-DEL-1321 | chr1-160591734-DEL-1322 |
| chr1-160643461-DEL-5135 | chr1-160643462-DEL-5135 |
| chr1-160669168-DEL-809 | chr1-160669169-DEL-809 |
| chr1-160671573-DEL-177 | chr1-160671574-DEL-177 |
| chr1-160703686-DEL-58 | chr1-160703687-DEL-61 |
| chr1-160704287-DEL-263 | chr1-160704288-DEL-263 |
| chr1-160740102-DEL-104 | chr1-160740103-DEL-104 |
| chr1-160822639-DEL-76 | chr1-160822632-DEL-84 |
| chr1-160825953-DEL-90 | chr1-160825952-DEL-80 |
| chr1-160841517-DEL-1799 | chr1-160841518-DEL-1800 |
| chr1-160849701-DEL-243 | chr1-160849702-DEL-243 |
| chr1-160851405-DEL-180 | chr1-160851406-DEL-180 |
| chr1-160854786-DEL-70 | chr1-160854787-DEL-70 |
| chr1-160866404-DEL-160 | chr1-160866402-DEL-160 |
| chr1-160871868-DEL-173 | chr1-160871869-DEL-173 |
| chr1-160875313-DEL-52 | chr1-160875314-DEL-52 |
| chr1-160875853-DEL-206 | chr1-160875854-DEL-206 |
| chr1-160891780-DEL-76 | chr1-160891781-DEL-76 |
| chr1-160892177-DEL-185 | chr1-160892178-DEL-185 |
| chr1-160900731-DEL-127 | chr1-160900732-DEL-127 |
| chr1-160945705-DEL-1776 | chr1-160945706-DEL-1776 |
| chr1-160946030-DEL-401 | chr1-160946031-DEL-401 |
| chr1-160966827-DEL-105 | chr1-160966828-DEL-105 |
| chr1-160974975-DEL-332 | chr1-160974973-DEL-330 |
| chr1-160992624-DEL-204 | chr1-160992625-DEL-204 |
| chr1-160995601-DEL-121 | chr1-160995626-DEL-73 |
| chr1-161013604-DEL-479 | chr1-161013604-DEL-480 |
| chr1-161016987-DEL-978 | chr1-161016988-DEL-978 |
| chr1-161059774-DEL-207 | chr1-161059775-DEL-207 |
| chr1-161084229-DEL-61 | chr1-161084230-DEL-61 |
| chr1-161091533-DEL-104 | chr1-161091534-DEL-104 |
| chr1-161105995-DEL-217 | chr1-161105996-DEL-217 |
| chr1-161144577-DEL-59 | chr1-161144578-DEL-59 |
| chr1-161152253-DEL-78 | chr1-161152254-DEL-78 |
| chr1-161228593-DEL-142 | chr1-161228594-DEL-142 |
| chr1-161240748-DEL-111 | chr1-161240745-DEL-110 |
| chr1-161256697-DEL-2366 | chr1-161256698-DEL-2366 |
| chr1-161275880-DEL-220 | chr1-161275878-DEL-220 |
| chr1-161289597-DEL-188 | chr1-161289598-DEL-188 |
| chr1-161310545-DEL-56 | chr1-161310546-DEL-56 |
| chr1-161348546-DEL-354 | chr1-161348547-DEL-354 |
| chr1-161430373-DEL-222 | chr1-161430374-DEL-222 |
| chr1-161446157-DEL-187 | chr1-161446158-DEL-187 |
| chr1-161453464-DEL-61 | chr1-161453465-DEL-61 |
| chr1-161500998-DEL-62 | chr1-161500999-DEL-50 |
| chr1-161539452-DEL-68 | chr1-161539451-DEL-68 |
| chr1-161543102-DEL-101 | chr1-161543130-DEL-71 |
| chr1-161553074-DEL-2187 | chr1-161553075-DEL-2187 |
| chr1-161587479-DEL-359 | chr1-161587471-DEL-360 |
| chr1-161605942-DEL-62 | chr1-161605943-DEL-52 |
| chr1-161668381-DEL-51 | chr1-161668382-DEL-51 |
| chr1-161688265-DEL-96 | chr1-161688266-DEL-96 |
| chr1-161709407-DEL-1347 | chr1-161709408-DEL-1347 |
| chr1-161717025-DEL-222 | chr1-161717026-DEL-222 |
| chr1-161744808-DEL-63 | chr1-161744809-DEL-63 |
| chr1-161745903-DEL-4075 | chr1-161745904-DEL-4075 |
| chr1-161760388-DEL-6340 | chr1-161760389-DEL-6340 |
| chr1-161773633-DEL-1354 | chr1-161773634-DEL-1354 |
| chr1-161850176-DEL-82 | chr1-161850177-DEL-82 |
| chr1-161891139-DEL-6601 | chr1-161891140-DEL-6601 |
| chr1-161898842-DEL-856 | chr1-161898843-DEL-856 |
| chr1-161908743-DEL-203 | chr1-161908745-DEL-202 |
| chr1-161914852-DEL-1244 | chr1-161914853-DEL-1244 |
| chr1-161930470-DEL-419 | chr1-161930471-DEL-419 |
| chr1-161941807-DEL-6306 | chr1-161941808-DEL-6306 |
| chr1-162026555-DEL-138 | chr1-162026556-DEL-138 |
| chr1-162072675-DEL-807 | chr1-162072676-DEL-807 |
| chr1-162076051-DEL-58 | chr1-162076052-DEL-56 |
| chr1-162089542-DEL-90 | chr1-162089543-DEL-90 |
| chr1-162091589-DEL-6602 | chr1-162091590-DEL-6602 |
| chr1-162115300-DEL-66 | chr1-162115301-DEL-66 |
| chr1-162132737-DEL-57 | chr1-162132737-DEL-55 |
| chr1-162144523-DEL-83 | chr1-162144524-DEL-84 |
| chr1-162161782-DEL-78 | chr1-162161783-DEL-78 |
| chr1-162163248-DEL-168 | chr1-162163215-DEL-179 |

| | |
|---|---|
| chr1-162166048-DEL-6902 | chr1-162166049-DEL-6902 |
| chr1-162182191-DEL-58 | chr1-162182192-DEL-58 |
| chr1-162186596-DEL-922 | chr1-162186597-DEL-922 |
| chr1-162199261-DEL-53 | chr1-162199261-DEL-53 |
| chr1-162215691-DEL-133 | chr1-162215738-DEL-116 |
| chr1-162247245-DEL-243 | chr1-162247246-DEL-243 |
| chr1-162251354-DEL-2172 | chr1-162251355-DEL-2172 |
| chr1-162252704-DEL-330 | chr1-162252705-DEL-330 |
| chr1-162259226-DEL-107 | chr1-162259227-DEL-107 |
| chr1-162272001-DEL-70 | chr1-162272006-DEL-64 |
| chr1-162278730-DEL-76 | chr1-162278731-DEL-76 |
| chr1-162304516-DEL-160 | chr1-162304517-DEL-160 |
| chr1-162316227-DEL-79 | chr1-162316262-DEL-92 |
| chr1-162317095-DEL-6511 | chr1-162317096-DEL-6511 |
| chr1-162351431-DEL-133 | chr1-162351431-DEL-130 |
| chr1-162352471-DEL-932 | chr1-162352472-DEL-932 |
| chr1-162383543-DEL-120 | chr1-162383542-DEL-122 |
| chr1-162395363-DEL-59 | chr1-162395364-DEL-59 |
| chr1-162432251-DEL-180 | chr1-162432252-DEL-180 |
| chr1-162459386-DEL-745 | chr1-162459387-DEL-745 |
| chr1-162463073-DEL-119 | chr1-162463074-DEL-70 |
| chr1-162492368-DEL-54 | chr1-162492369-DEL-52 |
| chr1-162590863-DEL-81 | chr1-162590863-DEL-81 |
| chr1-162593438-DEL-81 | chr1-162593439-DEL-81 |
| chr1-162595927-DEL-66 | chr1-162595928-DEL-66 |
| chr1-162610549-DEL-199 | chr1-162610550-DEL-199 |
| chr1-162627979-DEL-2160 | chr1-162627980-DEL-2160 |
| chr1-162642088-DEL-3890 | chr1-162642089-DEL-3890 |
| chr1-162647447-DEL-6456 | chr1-162647448-DEL-6456 |
| chr1-162668100-DEL-7111 | chr1-162668101-DEL-7111 |
| chr1-162706166-DEL-70 | chr1-162706167-DEL-70 |
| chr1-162734100-DEL-85 | chr1-162734101-DEL-85 |
| chr1-162744879-DEL-72 | chr1-162744857-DEL-74 |
| chr1-162776330-DEL-152 | chr1-162776331-DEL-152 |
| chr1-162777660-DEL-1890 | chr1-162777661-DEL-1890 |
| chr1-162786737-DEL-64 | chr1-162786737-DEL-64 |
| chr1-162807888-DEL-1076 | chr1-162807882-DEL-1123 |
| chr1-162809016-DEL-904 | chr1-162809039-DEL-881 |
| chr1-162822832-DEL-1413 | chr1-162822833-DEL-1413 |
| chr1-162825252-DEL-2044 | chr1-162825252-DEL-2043 |
| chr1-162839280-DEL-7134 | chr1-162839271-DEL-7135 |
| chr1-162890406-DEL-131 | chr1-162890407-DEL-131 |
| chr1-162890539-DEL-1459 | chr1-162890542-DEL-1457 |
| chr1-162894620-DEL-2120 | chr1-162894624-DEL-2112 |
| chr1-162924583-DEL-238 | chr1-162924584-DEL-238 |
| chr1-162934357-DEL-65 | chr1-162934358-DEL-65 |
| chr1-162935061-DEL-56 | chr1-162935064-DEL-54 |
| chr1-162951690-DEL-761 | chr1-162951691-DEL-761 |
| chr1-162988379-DEL-865 | chr1-162988380-DEL-865 |
| chr1-163029926-DEL-674 | chr1-163029927-DEL-672 |
| chr1-163050896-DEL-1448 | chr1-163050897-DEL-1448 |
| chr1-163068990-DEL-2388 | chr1-163068991-DEL-2388 |
| chr1-163069797-DEL-739 | chr1-163069798-DEL-739 |
| chr1-163090477-DEL-77 | chr1-163090478-DEL-77 |
| chr1-163103899-DEL-1273 | chr1-163103900-DEL-1273 |
| chr1-163105993-DEL-1227 | chr1-163105994-DEL-1227 |
| chr1-163124525-DEL-584 | chr1-163124526-DEL-584 |
| chr1-163155861-DEL-922 | chr1-163155862-DEL-922 |
| chr1-163159529-DEL-1624 | chr1-163159530-DEL-1624 |
| chr1-163162887-DEL-195 | chr1-163162888-DEL-195 |
| chr1-163170183-DEL-178 | chr1-163170186-DEL-176 |
| chr1-163198449-DEL-67 | chr1-163198450-DEL-67 |
| chr1-163199992-DEL-128 | chr1-163200007-DEL-114 |
| chr1-163210307-DEL-180 | chr1-163210308-DEL-180 |
| chr1-163296080-DEL-4127 | chr1-163296081-DEL-4127 |
| chr1-163393377-DEL-1514 | chr1-163393371-DEL-1521 |
| chr1-163461029-DEL-952 | chr1-163461030-DEL-952 |
| chr1-163475540-DEL-80 | chr1-163475543-DEL-80 |
| chr1-163503001-DEL-3100 | chr1-163503002-DEL-3100 |
| chr1-163508236-DEL-183 | chr1-163508237-DEL-183 |
| chr1-163517576-DEL-486 | chr1-163517577-DEL-486 |
| chr1-163520530-DEL-375 | chr1-163520531-DEL-375 |
| chr1-163531166-DEL-66 | chr1-163531141-DEL-69 |
| chr1-163537637-DEL-6533 | chr1-163537637-DEL-6533 |
| chr1-163547473-DEL-204 | chr1-163547474-DEL-204 |
| chr1-163568654-DEL-51 | chr1-163568651-DEL-55 |
| chr1-163609749-DEL-52 | chr1-163609733-DEL-76 |
| chr1-163611387-DEL-6659 | chr1-163611388-DEL-6659 |
| chr1-163616212-DEL-3261 | chr1-163616213-DEL-3261 |
| chr1-163619375-DEL-73 | chr1-163619376-DEL-72 |

| | |
|---|---|
| chr1-163634333-DEL-6178 | chr1-163634334-DEL-6178 |
| chr1-163679689-DEL-89 | chr1-163679690-DEL-89 |
| chr1-163700716-DEL-101 | chr1-163700681-DEL-102 |
| chr1-163703950-DEL-606 | chr1-163703951-DEL-606 |
| chr1-163716795-DEL-62 | chr1-163716817-DEL-54 |
| chr1-163730614-DEL-128 | chr1-163730615-DEL-128 |
| chr1-163802547-DEL-134 | chr1-163802536-DEL-150 |
| chr1-163828102-DEL-96 | chr1-163828103-DEL-96 |
| chr1-163831127-DEL-60 | chr1-163831128-DEL-60 |
| chr1-163841619-DEL-330 | chr1-163841617-DEL-330 |
| chr1-163846831-DEL-204 | chr1-163846832-DEL-204 |
| chr1-163848193-DEL-66 | chr1-163848192-DEL-66 |
| chr1-163852437-DEL-214 | chr1-163852438-DEL-214 |
| chr1-163858811-DEL-4544 | chr1-163858812-DEL-4544 |
| chr1-163917989-DEL-440 | chr1-163917990-DEL-440 |
| chr1-163922813-DEL-209 | chr1-163922814-DEL-209 |
| chr1-163935591-DEL-171 | chr1-163935592-DEL-176 |
| chr1-163955299-DEL-198 | chr1-163955300-DEL-198 |
| chr1-163968435-DEL-401 | chr1-163968436-DEL-401 |
| chr1-163973137-DEL-69 | chr1-163973143-DEL-68 |
| chr1-163974188-DEL-180 | chr1-163974189-DEL-180 |
| chr1-163976789-DEL-76 | chr1-163976790-DEL-76 |
| chr1-164024505-DEL-249 | chr1-164024506-DEL-249 |
| chr1-164049172-DEL-816 | chr1-164049173-DEL-816 |
| chr1-164051371-DEL-5750 | chr1-164051372-DEL-5750 |
| chr1-164059300-DEL-541 | chr1-164059301-DEL-541 |
| chr1-164069721-DEL-76 | chr1-164069722-DEL-76 |
| chr1-164075226-DEL-451 | chr1-164075227-DEL-451 |
| chr1-164095612-DEL-2445 | chr1-164095613-DEL-2445 |
| chr1-164137506-DEL-66 | chr1-164137491-DEL-79 |
| chr1-164153136-DEL-1194 | chr1-164153137-DEL-1194 |
| chr1-164156736-DEL-57 | chr1-164156734-DEL-58 |
| chr1-164157652-DEL-340 | chr1-164157651-DEL-340 |
| chr1-164175682-DEL-197 | chr1-164175683-DEL-197 |
| chr1-164179018-DEL-626 | chr1-164179019-DEL-626 |
| chr1-164215259-DEL-150 | chr1-164215243-DEL-138 |
| chr1-164221604-DEL-276 | chr1-164221605-DEL-276 |
| chr1-164236641-DEL-398 | chr1-164236642-DEL-398 |
| chr1-164257988-DEL-205 | chr1-164257989-DEL-205 |
| chr1-164291659-DEL-55 | chr1-164291665-DEL-62 |
| chr1-164311807-DEL-4448 | chr1-164311808-DEL-4448 |
| chr1-164323498-DEL-1368 | chr1-164323498-DEL-1326 |
| chr1-164326947-DEL-996 | chr1-164326948-DEL-996 |
| chr1-164337172-DEL-1335 | chr1-164337173-DEL-1335 |
| chr1-164352342-DEL-147 | chr1-164352335-DEL-148 |
| chr1-164390433-DEL-71 | chr1-164390434-DEL-71 |
| chr1-164435256-DEL-141 | chr1-164435256-DEL-136 |
| chr1-164454607-DEL-870 | chr1-164454608-DEL-869 |
| chr1-164468230-DEL-60 | chr1-164468223-DEL-68 |
| chr1-164479811-DEL-90 | chr1-164479812-DEL-90 |
| chr1-164484412-DEL-3113 | chr1-164484413-DEL-3113 |
| chr1-164493785-DEL-506 | chr1-164493786-DEL-506 |
| chr1-164495858-DEL-52 | chr1-164495859-DEL-52 |
| chr1-164521166-DEL-197 | chr1-164521167-DEL-197 |
| chr1-164551358-DEL-6385 | chr1-164551359-DEL-6385 |
| chr1-164564131-DEL-6484 | chr1-164564132-DEL-6484 |
| chr1-164570299-DEL-212 | chr1-164570300-DEL-212 |
| chr1-164573894-DEL-86 | chr1-164573855-DEL-98 |
| chr1-164575086-DEL-57 | chr1-164575085-DEL-69 |
| chr1-164605345-DEL-359 | chr1-164605346-DEL-359 |
| chr1-164606645-DEL-58 | chr1-164606605-DEL-58 |
| chr1-164639381-DEL-880 | chr1-164639382-DEL-880 |
| chr1-164664251-DEL-178 | chr1-164664252-DEL-178 |
| chr1-164684578-DEL-356 | chr1-164684579-DEL-356 |
| chr1-164691479-DEL-74 | chr1-164691480-DEL-74 |
| chr1-164705599-DEL-110 | chr1-164705603-DEL-112 |
| chr1-164706779-DEL-205 | chr1-164706780-DEL-205 |
| chr1-164712754-DEL-62 | chr1-164712763-DEL-54 |
| chr1-164718408-DEL-780 | chr1-164718409-DEL-780 |
| chr1-164732299-DEL-207 | chr1-164732295-DEL-208 |
| chr1-164741717-DEL-178 | chr1-164741731-DEL-190 |
| chr1-164744073-DEL-6249 | chr1-164744067-DEL-6250 |
| chr1-164744112-DEL-198 | chr1-164744097-DEL-199 |
| chr1-164803887-DEL-174 | chr1-164803889-DEL-173 |
| chr1-164818257-DEL-685 | chr1-164818258-DEL-685 |
| chr1-164851908-DEL-176 | chr1-164851909-DEL-176 |
| chr1-164853498-DEL-6393 | chr1-164853499-DEL-6393 |
| chr1-164914261-DEL-419 | chr1-164914262-DEL-419 |
| chr1-164914782-DEL-1268 | chr1-164914783-DEL-1268 |
| chr1-164924145-DEL-502 | chr1-164924140-DEL-503 |

| | |
|---|---|
| chr1-164925262-DEL-327 | chr1-164925263-DEL-327 |
| chr1-164931394-DEL-57 | chr1-164931385-DEL-58 |
| chr1-164933939-DEL-357 | chr1-164933940-DEL-355 |
| chr1-164935088-DEL-62 | chr1-164935089-DEL-62 |
| chr1-164937742-DEL-68 | chr1-164937701-DEL-102 |
| chr1-164992331-DEL-68 | chr1-164992316-DEL-69 |
| chr1-164999148-DEL-2639 | chr1-164999149-DEL-2613 |
| chr1-165004860-DEL-60 | chr1-165004861-DEL-60 |
| chr1-165007956-DEL-52 | chr1-165007953-DEL-53 |
| chr1-165023768-DEL-274 | chr1-165023770-DEL-273 |
| chr1-165060739-DEL-52 | chr1-165060725-DEL-72 |
| chr1-165093102-DEL-251 | chr1-165093103-DEL-251 |
| chr1-165094819-DEL-6440 | chr1-165094820-DEL-6440 |
| chr1-165116553-DEL-202 | chr1-165116554-DEL-202 |
| chr1-165146623-DEL-62 | chr1-165146624-DEL-62 |
| chr1-165205203-DEL-287 | chr1-165205207-DEL-284 |
| chr1-165207611-DEL-4093 | chr1-165207611-DEL-4093 |
| chr1-165232178-DEL-6743 | chr1-165232179-DEL-6743 |
| chr1-165240868-DEL-834 | chr1-165240869-DEL-834 |
| chr1-165295820-DEL-56 | chr1-165295821-DEL-56 |
| chr1-165335962-DEL-64 | chr1-165335965-DEL-62 |
| chr1-165338503-DEL-690 | chr1-165338504-DEL-690 |
| chr1-165354168-DEL-7361 | chr1-165354169-DEL-7361 |
| chr1-165367048-DEL-68 | chr1-165367049-DEL-68 |
| chr1-165377052-DEL-202 | chr1-165377053-DEL-202 |
| chr1-165408144-DEL-455 | chr1-165408145-DEL-455 |
| chr1-165427387-DEL-910 | chr1-165427388-DEL-910 |
| chr1-165458491-DEL-70 | chr1-165458476-DEL-56 |
| chr1-165472191-DEL-57 | chr1-165472190-DEL-58 |
| chr1-165481741-DEL-68 | chr1-165481742-DEL-68 |
| chr1-165498310-DEL-94 | chr1-165498293-DEL-106 |
| chr1-165525028-DEL-88 | chr1-165525050-DEL-78 |
| chr1-165539960-DEL-528 | chr1-165539960-DEL-528 |
| chr1-165543470-DEL-52 | chr1-165543503-DEL-54 |
| chr1-165549017-DEL-113 | chr1-165549018-DEL-113 |
| chr1-165559058-DEL-53 | chr1-165559057-DEL-54 |
| chr1-165571955-DEL-251 | chr1-165571956-DEL-251 |
| chr1-165586867-DEL-580 | chr1-165586868-DEL-580 |
| chr1-165599725-DEL-60 | chr1-165599726-DEL-60 |
| chr1-165610910-DEL-71 | chr1-165610911-DEL-71 |
| chr1-165626301-DEL-100 | chr1-165626302-DEL-100 |
| chr1-165637068-DEL-439 | chr1-165637061-DEL-438 |
| chr1-165665767-DEL-873 | chr1-165665768-DEL-873 |
| chr1-165682330-DEL-137 | chr1-165682331-DEL-137 |
| chr1-165690962-DEL-864 | chr1-165690963-DEL-864 |
| chr1-165797413-DEL-1183 | chr1-165797414-DEL-1183 |
| chr1-165811276-DEL-139 | chr1-165811284-DEL-134 |
| chr1-165822769-DEL-216 | chr1-165822770-DEL-216 |
| chr1-165833559-DEL-401 | chr1-165833560-DEL-401 |
| chr1-165848434-DEL-203 | chr1-165848435-DEL-203 |
| chr1-165904397-DEL-920 | chr1-165904398-DEL-920 |
| chr1-165907109-DEL-7160 | chr1-165907110-DEL-7160 |
| chr1-165919776-DEL-189 | chr1-165919777-DEL-189 |
| chr1-165942249-DEL-214 | chr1-165942250-DEL-214 |
| chr1-165946727-DEL-88 | chr1-165946728-DEL-88 |
| chr1-165953284-DEL-355 | chr1-165953273-DEL-356 |
| chr1-165969439-DEL-176 | chr1-165969439-DEL-176 |
| chr1-165996100-DEL-227 | chr1-165996101-DEL-227 |
| chr1-165997141-DEL-58 | chr1-165997165-DEL-69 |
| chr1-166002413-DEL-5667 | chr1-166002414-DEL-5667 |
| chr1-166094270-DEL-241 | chr1-166094257-DEL-242 |
| chr1-166101539-DEL-2038 | chr1-166101540-DEL-2038 |
| chr1-166162048-DEL-91 | chr1-166162049-DEL-91 |
| chr1-166164253-DEL-126 | chr1-166164254-DEL-126 |
| chr1-166167862-DEL-80 | chr1-166167863-DEL-80 |
| chr1-166176306-DEL-77 | chr1-166176307-DEL-77 |
| chr1-166184274-DEL-2381 | chr1-166184275-DEL-2381 |
| chr1-166246061-DEL-6477 | chr1-166246062-DEL-6477 |
| chr1-166259065-DEL-398 | chr1-166259066-DEL-398 |
| chr1-166291400-DEL-62 | chr1-166291401-DEL-62 |
| chr1-166296721-DEL-6349 | chr1-166296722-DEL-6349 |
| chr1-166353398-DEL-78 | chr1-166353399-DEL-78 |
| chr1-166362948-DEL-281 | chr1-166362949-DEL-281 |
| chr1-166386243-DEL-54 | chr1-166386244-DEL-54 |
| chr1-166394205-DEL-499 | chr1-166394206-DEL-498 |
| chr1-166408695-DEL-3458 | chr1-166408696-DEL-3458 |
| chr1-166474127-DEL-54 | chr1-166474128-DEL-54 |
| chr1-166489286-DEL-138 | chr1-166489287-DEL-138 |
| chr1-166513529-DEL-508 | chr1-166513530-DEL-508 |
| chr1-166522025-DEL-550 | chr1-166522026-DEL-549 |

| | |
|---|---|
| chr1-166531072-DEL-53 | chr1-166531073-DEL-51 |
| chr1-166549752-DEL-1676 | chr1-166549753-DEL-1676 |
| chr1-166566926-DEL-1232 | chr1-166566931-DEL-1232 |
| chr1-166590700-DEL-6746 | chr1-166590701-DEL-6746 |
| chr1-166606658-DEL-220 | chr1-166606659-DEL-220 |
| chr1-166617308-DEL-117 | chr1-166617307-DEL-118 |
| chr1-166686461-DEL-82 | chr1-166686462-DEL-82 |
| chr1-166699880-DEL-3629 | chr1-166699881-DEL-3629 |
| chr1-166740018-DEL-6439 | chr1-166740019-DEL-6439 |
| chr1-166767575-DEL-474 | chr1-166767576-DEL-474 |
| chr1-166773097-DEL-889 | chr1-166773098-DEL-889 |
| chr1-166784809-DEL-78 | chr1-166784804-DEL-72 |
| chr1-166788240-DEL-205 | chr1-166788241-DEL-205 |
| chr1-166798672-DEL-1183 | chr1-166798673-DEL-1184 |
| chr1-166813568-DEL-158 | chr1-166813569-DEL-158 |
| chr1-166815294-DEL-6350 | chr1-166815295-DEL-6350 |
| chr1-166842496-DEL-126 | chr1-166842497-DEL-126 |
| chr1-166850013-DEL-61 | chr1-166850010-DEL-75 |
| chr1-166875153-DEL-56 | chr1-166875148-DEL-66 |
| chr1-166899515-DEL-721 | chr1-166899516-DEL-721 |
| chr1-166903807-DEL-303 | chr1-166903816-DEL-291 |
| chr1-166925899-DEL-252 | chr1-166925899-DEL-252 |
| chr1-166953690-DEL-1553 | chr1-166953691-DEL-1553 |
| chr1-166968689-DEL-2142 | chr1-166968690-DEL-2142 |
| chr1-166974971-DEL-262 | chr1-166974972-DEL-262 |
| chr1-167002847-DEL-139 | chr1-167002840-DEL-139 |
| chr1-167024493-DEL-6611 | chr1-167024494-DEL-6611 |
| chr1-167024572-DEL-214 | chr1-167024572-DEL-214 |
| chr1-167040289-DEL-67 | chr1-167040290-DEL-66 |
| chr1-167041620-DEL-128 | chr1-167041619-DEL-126 |
| chr1-167042821-DEL-62 | chr1-167042824-DEL-60 |
| chr1-167045164-DEL-1274 | chr1-167045165-DEL-1274 |
| chr1-167047805-DEL-66 | chr1-167047806-DEL-57 |
| chr1-167058302-DEL-225 | chr1-167058303-DEL-225 |
| chr1-167085684-DEL-240 | chr1-167085685-DEL-240 |
| chr1-167086461-DEL-61 | chr1-167086460-DEL-61 |
| chr1-167094031-DEL-52 | chr1-167094032-DEL-52 |
| chr1-167124588-DEL-206 | chr1-167124586-DEL-206 |
| chr1-167137321-DEL-54 | chr1-167137322-DEL-54 |
| chr1-167140267-DEL-84 | chr1-167140261-DEL-84 |
| chr1-167143386-DEL-220 | chr1-167143387-DEL-220 |
| chr1-167144569-DEL-83 | chr1-167144570-DEL-83 |
| chr1-167166711-DEL-4201 | chr1-167166712-DEL-4201 |
| chr1-167171371-DEL-405 | chr1-167171372-DEL-405 |
| chr1-167172727-DEL-646 | chr1-167172728-DEL-646 |
| chr1-167225512-DEL-3366 | chr1-167225513-DEL-3366 |
| chr1-167260192-DEL-133 | chr1-167260162-DEL-175 |
| chr1-167277240-DEL-127 | chr1-167277241-DEL-127 |
| chr1-167302351-DEL-6229 | chr1-167302352-DEL-6229 |
| chr1-167334644-DEL-53 | chr1-167334644-DEL-53 |
| chr1-167364405-DEL-71 | chr1-167364407-DEL-72 |
| chr1-167384698-DEL-1976 | chr1-167384699-DEL-1976 |
| chr1-167390048-DEL-63 | chr1-167390081-DEL-50 |
| chr1-167431160-DEL-183 | chr1-167431161-DEL-183 |
| chr1-167442057-DEL-245 | chr1-167442058-DEL-245 |
| chr1-167445407-DEL-83 | chr1-167445408-DEL-83 |
| chr1-167449459-DEL-645 | chr1-167449460-DEL-645 |
| chr1-167498096-DEL-160 | chr1-167498097-DEL-160 |
| chr1-167498764-DEL-77 | chr1-167498765-DEL-84 |
| chr1-167535985-DEL-52 | chr1-167535958-DEL-90 |
| chr1-167548294-DEL-65 | chr1-167548295-DEL-61 |
| chr1-167568843-DEL-61 | chr1-167568844-DEL-61 |
| chr1-167594734-DEL-757 | chr1-167594735-DEL-757 |
| chr1-167607619-DEL-52 | chr1-167607616-DEL-54 |
| chr1-167612372-DEL-710 | chr1-167612364-DEL-711 |
| chr1-167618004-DEL-92 | chr1-167617957-DEL-110 |
| chr1-167677787-DEL-75 | chr1-167677796-DEL-65 |
| chr1-167680478-DEL-423 | chr1-167680481-DEL-421 |
| chr1-167691079-DEL-123 | chr1-167691080-DEL-123 |
| chr1-167709045-DEL-218 | chr1-167709046-DEL-218 |
| chr1-167717654-DEL-1116 | chr1-167717660-DEL-1110 |
| chr1-167726199-DEL-122 | chr1-167726200-DEL-122 |
| chr1-167742810-DEL-123 | chr1-167742811-DEL-123 |
| chr1-167783152-DEL-240 | chr1-167783153-DEL-240 |
| chr1-167871982-DEL-66 | chr1-167871987-DEL-62 |
| chr1-167875746-DEL-67 | chr1-167875747-DEL-67 |
| chr1-167907902-DEL-112 | chr1-167907903-DEL-112 |
| chr1-167938108-DEL-66 | chr1-167938103-DEL-72 |
| chr1-167943739-DEL-1415 | chr1-167943740-DEL-1415 |
| chr1-167996318-DEL-133 | chr1-167996319-DEL-133 |

| | |
|---|---|
| chr1-168072830-DEL-53 | chr1-168072830-DEL-54 |
| chr1-168122396-DEL-68 | chr1-168122413-DEL-67 |
| chr1-168141317-DEL-494 | chr1-168141318-DEL-494 |
| chr1-168143368-DEL-345 | chr1-168143369-DEL-345 |
| chr1-168153747-DEL-83 | chr1-168153748-DEL-83 |
| chr1-168161515-DEL-123 | chr1-168161516-DEL-123 |
| chr1-168199920-DEL-63 | chr1-168199921-DEL-63 |
| chr1-168223902-DEL-216 | chr1-168223903-DEL-216 |
| chr1-168229064-DEL-192 | chr1-168229065-DEL-192 |
| chr1-168285489-DEL-157 | chr1-168285495-DEL-152 |
| chr1-168320553-DEL-1148 | chr1-168320554-DEL-1148 |
| chr1-168334375-DEL-55 | chr1-168334376-DEL-55 |
| chr1-168393927-DEL-216 | chr1-168393927-DEL-216 |
| chr1-168394392-DEL-478 | chr1-168394393-DEL-478 |
| chr1-168400666-DEL-91 | chr1-168400659-DEL-98 |
| chr1-168407941-DEL-155 | chr1-168407942-DEL-143 |
| chr1-168414189-DEL-68 | chr1-168414190-DEL-68 |
| chr1-168424113-DEL-210 | chr1-168424114-DEL-210 |
| chr1-168428614-DEL-502 | chr1-168428615-DEL-502 |
| chr1-168429835-DEL-149 | chr1-168429836-DEL-124 |
| chr1-168450776-DEL-6631 | chr1-168450777-DEL-6631 |
| chr1-168471109-DEL-6580 | chr1-168471110-DEL-6580 |
| chr1-168485586-DEL-200 | chr1-168485587-DEL-200 |
| chr1-168518204-DEL-52 | chr1-168518205-DEL-52 |
| chr1-168528098-DEL-124 | chr1-168528099-DEL-124 |
| chr1-168538660-DEL-88 | chr1-168538661-DEL-86 |
| chr1-168541180-DEL-88 | chr1-168541189-DEL-80 |
| chr1-168547899-DEL-6475 | chr1-168547900-DEL-6475 |
| chr1-168587273-DEL-53 | chr1-168587273-DEL-53 |
| chr1-168607121-DEL-280 | chr1-168607114-DEL-288 |
| chr1-168630599-DEL-2468 | chr1-168630600-DEL-2468 |
| chr1-168654221-DEL-67 | chr1-168654222-DEL-75 |
| chr1-168705076-DEL-51 | chr1-168705077-DEL-51 |
| chr1-168754423-DEL-2536 | chr1-168754424-DEL-2536 |
| chr1-168754704-DEL-84 | chr1-168754705-DEL-84 |
| chr1-168757190-DEL-1106 | chr1-168757191-DEL-1106 |
| chr1-168810964-DEL-6354 | chr1-168810965-DEL-6354 |
| chr1-168830152-DEL-59 | chr1-168830135-DEL-60 |
| chr1-168840911-DEL-66 | chr1-168840888-DEL-80 |
| chr1-168869665-DEL-70 | chr1-168869666-DEL-70 |
| chr1-168878498-DEL-54 | chr1-168878499-DEL-54 |
| chr1-168883194-DEL-60 | chr1-168883193-DEL-50 |
| chr1-168919572-DEL-52 | chr1-168919573-DEL-52 |
| chr1-168930215-DEL-58 | chr1-168930216-DEL-58 |
| chr1-168934774-DEL-6198 | chr1-168934775-DEL-6198 |
| chr1-168968456-DEL-1391 | chr1-168968457-DEL-1391 |
| chr1-168981182-DEL-741 | chr1-168981183-DEL-741 |
| chr1-168982065-DEL-148 | chr1-168982066-DEL-148 |
| chr1-168988049-DEL-52 | chr1-168988050-DEL-52 |
| chr1-168992297-DEL-85 | chr1-168992298-DEL-61 |
| chr1-168996579-DEL-279 | chr1-168996580-DEL-279 |
| chr1-168997236-DEL-587 | chr1-168997237-DEL-587 |
| chr1-169002280-DEL-6664 | chr1-169002281-DEL-6664 |
| chr1-169017881-DEL-4500 | chr1-169017879-DEL-4493 |
| chr1-169030804-DEL-478 | chr1-169030805-DEL-478 |
| chr1-169047825-DEL-1252 | chr1-169047826-DEL-1252 |
| chr1-169060328-DEL-60 | chr1-169060329-DEL-58 |
| chr1-169083171-DEL-88 | chr1-169083147-DEL-81 |
| chr1-169112352-DEL-6383 | chr1-169112353-DEL-6383 |
| chr1-169123538-DEL-64 | chr1-169123539-DEL-64 |
| chr1-169130565-DEL-327 | chr1-169130566-DEL-327 |
| chr1-169142199-DEL-84 | chr1-169142200-DEL-84 |
| chr1-169148304-DEL-645 | chr1-169148305-DEL-645 |
| chr1-169150656-DEL-1297 | chr1-169150657-DEL-1297 |
| chr1-169166892-DEL-1407 | chr1-169166893-DEL-1407 |
| chr1-169173613-DEL-80 | chr1-169173613-DEL-80 |
| chr1-169205760-DEL-68 | chr1-169205761-DEL-68 |
| chr1-169208838-DEL-725 | chr1-169208839-DEL-725 |
| chr1-169215053-DEL-57 | chr1-169215054-DEL-57 |
| chr1-169216887-DEL-281 | chr1-169216880-DEL-314 |
| chr1-169249647-DEL-82 | chr1-169249684-DEL-82 |
| chr1-169253435-DEL-6276 | chr1-169253437-DEL-6274 |
| chr1-169284865-DEL-70 | chr1-169284858-DEL-75 |
| chr1-169341227-DEL-73 | chr1-169341193-DEL-117 |
| chr1-169352730-DEL-61 | chr1-169352731-DEL-61 |
| chr1-169369982-DEL-6477 | chr1-169369983-DEL-6477 |
| chr1-169376671-DEL-401 | chr1-169376672-DEL-401 |
| chr1-169378275-DEL-60 | chr1-169378276-DEL-60 |
| chr1-169387051-DEL-62 | chr1-169387060-DEL-61 |
| chr1-169390358-DEL-216 | chr1-169390359-DEL-198 |

| | |
|---|---|
| chr1-169431831-DEL-62 | chr1-169431825-DEL-74 |
| chr1-169446081-DEL-319 | chr1-169446082-DEL-319 |
| chr1-169457701-DEL-154 | chr1-169457724-DEL-139 |
| chr1-169472780-DEL-1076 | chr1-169472780-DEL-1076 |
| chr1-169475520-DEL-145 | chr1-169475521-DEL-145 |
| chr1-169481589-DEL-54 | chr1-169481598-DEL-52 |
| chr1-169498174-DEL-62 | chr1-169498175-DEL-62 |
| chr1-169529021-DEL-775 | chr1-169529017-DEL-776 |
| chr1-169531745-DEL-150 | chr1-169531746-DEL-150 |
| chr1-169540363-DEL-199 | chr1-169540366-DEL-196 |
| chr1-169567032-DEL-491 | chr1-169567033-DEL-487 |
| chr1-169586154-DEL-71 | chr1-169586155-DEL-71 |
| chr1-169620224-DEL-77 | chr1-169620225-DEL-77 |
| chr1-169632971-DEL-1158 | chr1-169632972-DEL-1158 |
| chr1-169648145-DEL-109 | chr1-169648146-DEL-109 |
| chr1-169653665-DEL-2301 | chr1-169653664-DEL-2300 |
| chr1-169672488-DEL-59 | chr1-169672486-DEL-58 |
| chr1-169690203-DEL-1317 | chr1-169690204-DEL-1317 |
| chr1-169707302-DEL-120 | chr1-169707294-DEL-100 |
| chr1-169733551-DEL-73 | chr1-169733552-DEL-73 |
| chr1-169734381-DEL-182 | chr1-169734340-DEL-224 |
| chr1-169753299-DEL-52 | chr1-169753298-DEL-60 |
| chr1-169798805-DEL-162 | chr1-169798803-DEL-160 |
| chr1-169864957-DEL-70 | chr1-169864963-DEL-65 |
| chr1-169874264-DEL-62 | chr1-169874264-DEL-50 |
| chr1-169893335-DEL-196 | chr1-169893336-DEL-196 |
| chr1-169894357-DEL-1316 | chr1-169894358-DEL-1316 |
| chr1-169897192-DEL-971 | chr1-169897193-DEL-971 |
| chr1-169900347-DEL-200 | chr1-169900348-DEL-200 |
| chr1-169909772-DEL-281 | chr1-169909773-DEL-281 |
| chr1-169928074-DEL-213 | chr1-169928074-DEL-213 |
| chr1-169933641-DEL-241 | chr1-169933642-DEL-241 |
| chr1-169943360-DEL-7090 | chr1-169943361-DEL-7090 |
| chr1-169962837-DEL-74 | chr1-169962838-DEL-74 |
| chr1-169969183-DEL-54 | chr1-169969184-DEL-54 |
| chr1-169972707-DEL-55 | chr1-169972708-DEL-55 |
| chr1-169984133-DEL-212 | chr1-169984133-DEL-212 |
| chr1-170056540-DEL-3184 | chr1-170056541-DEL-3184 |
| chr1-170065947-DEL-74 | chr1-170065964-DEL-50 |
| chr1-170081519-DEL-95 | chr1-170081520-DEL-95 |
| chr1-170083727-DEL-6208 | chr1-170083728-DEL-6208 |
| chr1-170103879-DEL-181 | chr1-170103888-DEL-173 |
| chr1-170124334-DEL-142 | chr1-170124331-DEL-163 |
| chr1-170140559-DEL-58 | chr1-170140556-DEL-59 |
| chr1-170142132-DEL-662 | chr1-170142139-DEL-656 |
| chr1-170156258-DEL-195 | chr1-170156259-DEL-195 |
| chr1-170160670-DEL-212 | chr1-170160671-DEL-212 |
| chr1-170189892-DEL-90 | chr1-170189896-DEL-97 |
| chr1-170204051-DEL-158 | chr1-170204054-DEL-150 |
| chr1-170204712-DEL-55 | chr1-170204713-DEL-55 |
| chr1-170209570-DEL-87 | chr1-170209595-DEL-57 |
| chr1-170209595-DEL-1197 | chr1-170209596-DEL-1197 |
| chr1-170212329-DEL-60 | chr1-170212330-DEL-60 |
| chr1-170311177-DEL-124 | chr1-170311177-DEL-93 |
| chr1-170314432-DEL-56 | chr1-170314437-DEL-52 |
| chr1-170324724-DEL-240 | chr1-170324725-DEL-240 |
| chr1-170418023-DEL-55 | chr1-170418024-DEL-55 |
| chr1-170437145-DEL-84 | chr1-170437146-DEL-88 |
| chr1-170456054-DEL-61 | chr1-170456054-DEL-61 |
| chr1-170461099-DEL-163 | chr1-170461100-DEL-163 |
| chr1-170462502-DEL-109 | chr1-170462499-DEL-102 |
| chr1-170470191-DEL-169 | chr1-170470192-DEL-169 |
| chr1-170515571-DEL-1833 | chr1-170515572-DEL-1833 |
| chr1-170518756-DEL-70 | chr1-170518761-DEL-66 |
| chr1-170539122-DEL-129 | chr1-170539123-DEL-129 |
| chr1-170541092-DEL-51 | chr1-170541093-DEL-51 |
| chr1-170564593-DEL-99 | chr1-170564594-DEL-103 |
| chr1-170636145-DEL-206 | chr1-170636146-DEL-206 |
| chr1-170644166-DEL-64 | chr1-170644167-DEL-64 |
| chr1-170654072-DEL-75 | chr1-170654085-DEL-91 |
| chr1-170661962-DEL-84 | chr1-170661963-DEL-84 |
| chr1-170671183-DEL-66 | chr1-170671140-DEL-99 |
| chr1-170674772-DEL-62 | chr1-170674763-DEL-60 |
| chr1-170686351-DEL-209 | chr1-170686352-DEL-209 |
| chr1-170700381-DEL-400 | chr1-170700382-DEL-400 |
| chr1-170705120-DEL-625 | chr1-170705131-DEL-625 |
| chr1-170715089-DEL-61 | chr1-170715086-DEL-63 |
| chr1-170721483-DEL-51 | chr1-170721484-DEL-51 |
| chr1-170747381-DEL-565 | chr1-170747382-DEL-565 |
| chr1-170757166-DEL-210 | chr1-170757167-DEL-210 |

| | |
|---|---|
| chr1-170769084-DEL-8661 | chr1-170769085-DEL-8661 |
| chr1-170787235-DEL-73 | chr1-170787254-DEL-51 |
| chr1-170814990-DEL-64 | chr1-170814993-DEL-68 |
| chr1-170836086-DEL-171 | chr1-170836087-DEL-171 |
| chr1-170838080-DEL-703 | chr1-170838081-DEL-703 |
| chr1-170912419-DEL-51 | chr1-170912420-DEL-51 |
| chr1-170914999-DEL-1246 | chr1-170915000-DEL-1246 |
| chr1-170947604-DEL-62 | chr1-170947605-DEL-62 |
| chr1-170970781-DEL-219 | chr1-170970782-DEL-219 |
| chr1-170990178-DEL-54 | chr1-170990179-DEL-54 |
| chr1-171008272-DEL-72 | chr1-171008273-DEL-72 |
| chr1-171035437-DEL-210 | chr1-171035438-DEL-210 |
| chr1-171058945-DEL-118 | chr1-171058946-DEL-106 |
| chr1-171082373-DEL-65 | chr1-171082352-DEL-64 |
| chr1-171090694-DEL-52 | chr1-171090672-DEL-61 |
| chr1-171092141-DEL-362 | chr1-171092154-DEL-359 |
| chr1-171093330-DEL-68 | chr1-171093305-DEL-81 |
| chr1-171095761-DEL-208 | chr1-171095762-DEL-208 |
| chr1-171127512-DEL-61 | chr1-171127513-DEL-61 |
| chr1-171128257-DEL-193 | chr1-171128252-DEL-195 |
| chr1-171130886-DEL-86 | chr1-171130887-DEL-86 |
| chr1-171132623-DEL-503 | chr1-171132624-DEL-503 |
| chr1-171139023-DEL-5063 | chr1-171139023-DEL-5063 |
| chr1-171139990-DEL-2359 | chr1-171139991-DEL-2355 |
| chr1-171142583-DEL-543 | chr1-171142584-DEL-543 |
| chr1-171145199-DEL-176 | chr1-171145200-DEL-176 |
| chr1-171154396-DEL-205 | chr1-171154397-DEL-204 |
| chr1-171177482-DEL-333 | chr1-171177477-DEL-334 |
| chr1-171180294-DEL-59 | chr1-171180295-DEL-59 |
| chr1-171183752-DEL-70 | chr1-171183753-DEL-70 |
| chr1-171255203-DEL-80 | chr1-171255204-DEL-80 |
| chr1-171259404-DEL-202 | chr1-171259392-DEL-203 |
| chr1-171260234-DEL-50 | chr1-171260235-DEL-50 |
| chr1-171279286-DEL-209 | chr1-171279287-DEL-209 |
| chr1-171283677-DEL-208 | chr1-171283678-DEL-208 |
| chr1-171285990-DEL-74 | chr1-171285990-DEL-74 |
| chr1-171296029-DEL-224 | chr1-171296074-DEL-177 |
| chr1-171308709-DEL-8674 | chr1-171308710-DEL-8674 |
| chr1-171318041-DEL-282 | chr1-171318029-DEL-280 |
| chr1-171318797-DEL-481 | chr1-171318807-DEL-471 |
| chr1-171320573-DEL-67 | chr1-171320574-DEL-63 |
| chr1-171321373-DEL-185 | chr1-171321365-DEL-186 |
| chr1-171323942-DEL-204 | chr1-171323942-DEL-205 |
| chr1-171326996-DEL-67 | chr1-171326997-DEL-67 |
| chr1-171329643-DEL-203 | chr1-171329644-DEL-203 |
| chr1-171330498-DEL-62 | chr1-171330498-DEL-62 |
| chr1-171362475-DEL-206 | chr1-171362476-DEL-205 |
| chr1-171364260-DEL-825 | chr1-171364260-DEL-825 |
| chr1-171374780-DEL-63 | chr1-171374781-DEL-63 |
| chr1-171397779-DEL-476 | chr1-171397774-DEL-477 |
| chr1-171402818-DEL-63 | chr1-171402822-DEL-62 |
| chr1-171451972-DEL-2799 | chr1-171451973-DEL-2799 |
| chr1-171538629-DEL-487 | chr1-171538630-DEL-487 |
| chr1-171559007-DEL-141 | chr1-171559008-DEL-141 |
| chr1-171588259-DEL-54 | chr1-171588259-DEL-54 |
| chr1-171657272-DEL-3027 | chr1-171657273-DEL-3027 |
| chr1-171663134-DEL-508 | chr1-171663135-DEL-512 |
| chr1-171684719-DEL-7247 | chr1-171684720-DEL-7247 |
| chr1-171701233-DEL-226 | chr1-171701234-DEL-226 |
| chr1-171723802-DEL-53 | chr1-171723803-DEL-53 |
| chr1-171728037-DEL-59 | chr1-171728033-DEL-64 |
| chr1-171730464-DEL-545 | chr1-171730465-DEL-545 |
| chr1-171740994-DEL-196 | chr1-171740995-DEL-196 |
| chr1-171768946-DEL-83 | chr1-171768947-DEL-83 |
| chr1-171831308-DEL-116 | chr1-171831327-DEL-126 |
| chr1-171848941-DEL-52 | chr1-171848942-DEL-52 |
| chr1-171859494-DEL-206 | chr1-171859495-DEL-206 |
| chr1-171860738-DEL-2347 | chr1-171860739-DEL-2347 |
| chr1-171866622-DEL-58 | chr1-171866622-DEL-56 |
| chr1-171866780-DEL-772 | chr1-171866781-DEL-772 |
| chr1-171869925-DEL-794 | chr1-171869926-DEL-794 |
| chr1-171875244-DEL-76 | chr1-171875239-DEL-76 |
| chr1-171915456-DEL-198 | chr1-171915457-DEL-198 |
| chr1-171927823-DEL-225 | chr1-171927824-DEL-225 |
| chr1-171938522-DEL-198 | chr1-171938523-DEL-198 |
| chr1-171959036-DEL-219 | chr1-171959037-DEL-219 |
| chr1-171971449-DEL-1083 | chr1-171971450-DEL-1083 |
| chr1-171982743-DEL-203 | chr1-171982744-DEL-203 |
| chr1-172005871-DEL-192 | chr1-172005872-DEL-191 |
| chr1-172013608-DEL-60 | chr1-172013609-DEL-60 |

| | |
|---|---|
| chr1-172020608-DEL-576 | chr1-172020619-DEL-571 |
| chr1-172098446-DEL-206 | chr1-172098447-DEL-206 |
| chr1-172110201-DEL-123 | chr1-172110202-DEL-123 |
| chr1-172152599-DEL-159 | chr1-172152600-DEL-159 |
| chr1-172167485-DEL-68 | chr1-172167486-DEL-68 |
| chr1-172198490-DEL-56 | chr1-172198493-DEL-54 |
| chr1-172247064-DEL-68 | chr1-172247065-DEL-68 |
| chr1-172266006-DEL-116 | chr1-172266007-DEL-116 |
| chr1-172274103-DEL-194 | chr1-172274104-DEL-194 |
| chr1-172293375-DEL-206 | chr1-172293364-DEL-207 |
| chr1-172296765-DEL-205 | chr1-172296766-DEL-205 |
| chr1-172355884-DEL-66 | chr1-172355885-DEL-66 |
| chr1-172370658-DEL-105 | chr1-172370659-DEL-105 |
| chr1-172374289-DEL-120 | chr1-172374286-DEL-124 |
| chr1-172427106-DEL-109 | chr1-172427073-DEL-143 |
| chr1-172486283-DEL-98 | chr1-172486285-DEL-97 |
| chr1-172486808-DEL-145 | chr1-172486809-DEL-145 |
| chr1-172505322-DEL-52 | chr1-172505323-DEL-52 |
| chr1-172527105-DEL-60 | chr1-172527106-DEL-60 |
| chr1-172578422-DEL-63 | chr1-172578423-DEL-63 |
| chr1-172579416-DEL-177 | chr1-172579409-DEL-178 |
| chr1-172599641-DEL-251 | chr1-172599642-DEL-251 |
| chr1-172601753-DEL-155 | chr1-172601744-DEL-160 |
| chr1-172628521-DEL-498 | chr1-172628522-DEL-498 |
| chr1-172640462-DEL-60 | chr1-172640463-DEL-60 |
| chr1-172664502-DEL-80 | chr1-172664503-DEL-80 |
| chr1-172666122-DEL-117 | chr1-172666086-DEL-118 |
| chr1-172679518-DEL-1479 | chr1-172679511-DEL-1480 |
| chr1-172706161-DEL-6548 | chr1-172706164-DEL-6551 |
| chr1-172721093-DEL-62 | chr1-172721082-DEL-74 |
| chr1-172741352-DEL-178 | chr1-172741349-DEL-180 |
| chr1-172743539-DEL-134 | chr1-172743540-DEL-134 |
| chr1-172770009-DEL-346 | chr1-172770010-DEL-346 |
| chr1-172786073-DEL-101 | chr1-172786074-DEL-101 |
| chr1-172795143-DEL-54 | chr1-172795130-DEL-55 |
| chr1-172798307-DEL-200 | chr1-172798308-DEL-200 |
| chr1-172799955-DEL-2499 | chr1-172799956-DEL-2499 |
| chr1-172804694-DEL-4146 | chr1-172804692-DEL-4131 |
| chr1-172828643-DEL-2619 | chr1-172828644-DEL-2619 |
| chr1-172861384-DEL-280 | chr1-172861385-DEL-280 |
| chr1-172899348-DEL-66 | chr1-172899349-DEL-64 |
| chr1-172916609-DEL-62 | chr1-172916610-DEL-62 |
| chr1-172929788-DEL-65 | chr1-172929789-DEL-65 |
| chr1-172944618-DEL-76 | chr1-172944619-DEL-76 |
| chr1-172983793-DEL-6215 | chr1-172983794-DEL-6215 |
| chr1-172996460-DEL-351 | chr1-172996462-DEL-350 |
| chr1-173003298-DEL-4796 | chr1-173003299-DEL-4796 |
| chr1-173017081-DEL-56 | chr1-173017037-DEL-54 |
| chr1-173069479-DEL-66 | chr1-173069480-DEL-66 |
| chr1-173070574-DEL-92 | chr1-173070575-DEL-92 |
| chr1-173081968-DEL-6809 | chr1-173081969-DEL-6809 |
| chr1-173094497-DEL-56 | chr1-173094498-DEL-56 |
| chr1-173180171-DEL-76 | chr1-173180172-DEL-76 |
| chr1-173185138-DEL-89 | chr1-173185139-DEL-89 |
| chr1-173211326-DEL-211 | chr1-173211320-DEL-212 |
| chr1-173227981-DEL-785 | chr1-173227982-DEL-784 |
| chr1-173236114-DEL-200 | chr1-173236115-DEL-200 |
| chr1-173302473-DEL-90 | chr1-173302474-DEL-82 |
| chr1-173318629-DEL-58 | chr1-173318630-DEL-56 |
| chr1-173320523-DEL-175 | chr1-173320524-DEL-175 |
| chr1-173324771-DEL-79 | chr1-173324774-DEL-118 |
| chr1-173406141-DEL-3289 | chr1-173406141-DEL-3289 |
| chr1-173428053-DEL-84 | chr1-173428054-DEL-84 |
| chr1-173436912-DEL-267 | chr1-173436896-DEL-291 |
| chr1-173454844-DEL-56 | chr1-173454845-DEL-56 |
| chr1-173488589-DEL-51 | chr1-173488606-DEL-78 |
| chr1-173508398-DEL-159 | chr1-173508398-DEL-159 |
| chr1-173539597-DEL-56 | chr1-173539564-DEL-56 |
| chr1-173542182-DEL-57 | chr1-173542156-DEL-56 |
| chr1-173562681-DEL-63 | chr1-173562682-DEL-63 |
| chr1-173578860-DEL-113 | chr1-173578857-DEL-114 |
| chr1-173660497-DEL-6492 | chr1-173660496-DEL-6492 |
| chr1-173726330-DEL-92 | chr1-173726331-DEL-92 |
| chr1-173747497-DEL-116 | chr1-173747482-DEL-84 |
| chr1-173749075-DEL-54 | chr1-173749075-DEL-54 |
| chr1-173784572-DEL-2788 | chr1-173784573-DEL-2788 |
| chr1-173795461-DEL-2893 | chr1-173795461-DEL-2893 |
| chr1-173800107-DEL-90 | chr1-173800108-DEL-90 |
| chr1-173805010-DEL-60 | chr1-173805010-DEL-61 |
| chr1-173815799-DEL-6132 | chr1-173815799-DEL-6131 |

| | |
|---|---|
| chr1-173845462-DEL-6345 | chr1-173845463-DEL-6345 |
| chr1-173852804-DEL-374 | chr1-173852805-DEL-374 |
| chr1-173866661-DEL-591 | chr1-173866662-DEL-591 |
| chr1-173870790-DEL-86 | chr1-173870781-DEL-87 |
| chr1-173875650-DEL-53 | chr1-173875651-DEL-53 |
| chr1-173884941-DEL-180 | chr1-173884942-DEL-180 |
| chr1-173886811-DEL-2685 | chr1-173886812-DEL-2685 |
| chr1-173900597-DEL-1417 | chr1-173900591-DEL-1416 |
| chr1-173921501-DEL-203 | chr1-173921502-DEL-203 |
| chr1-173924806-DEL-75 | chr1-173924807-DEL-75 |
| chr1-173933596-DEL-95 | chr1-173933607-DEL-85 |
| chr1-173959994-DEL-3230 | chr1-173959997-DEL-3227 |
| chr1-173988262-DEL-292 | chr1-173988263-DEL-292 |
| chr1-173995591-DEL-646 | chr1-173995592-DEL-646 |
| chr1-173999899-DEL-100 | chr1-173999920-DEL-84 |
| chr1-174006734-DEL-190 | chr1-174006734-DEL-190 |
| chr1-174010419-DEL-804 | chr1-174010420-DEL-804 |
| chr1-174034026-DEL-89 | chr1-174034027-DEL-89 |
| chr1-174041802-DEL-61 | chr1-174041803-DEL-61 |
| chr1-174043746-DEL-56 | chr1-174043747-DEL-55 |
| chr1-174048267-DEL-970 | chr1-174048268-DEL-970 |
| chr1-174059579-DEL-58 | chr1-174059567-DEL-54 |
| chr1-174059905-DEL-3141 | chr1-174059906-DEL-3141 |
| chr1-174090477-DEL-80 | chr1-174090446-DEL-72 |
| chr1-174134255-DEL-6185 | chr1-174134255-DEL-6185 |
| chr1-174147029-DEL-3049 | chr1-174147029-DEL-3049 |
| chr1-174188213-DEL-1058 | chr1-174188214-DEL-1058 |
| chr1-174225765-DEL-212 | chr1-174225766-DEL-212 |
| chr1-174239495-DEL-434 | chr1-174239496-DEL-434 |
| chr1-174249357-DEL-937 | chr1-174249358-DEL-937 |
| chr1-174263183-DEL-180 | chr1-174263184-DEL-208 |
| chr1-174272514-DEL-74 | chr1-174272517-DEL-72 |
| chr1-174275153-DEL-219 | chr1-174275154-DEL-219 |
| chr1-174284150-DEL-7115 | chr1-174284151-DEL-7115 |
| chr1-174300993-DEL-100 | chr1-174300994-DEL-100 |
| chr1-174303268-DEL-221 | chr1-174303269-DEL-221 |
| chr1-174314297-DEL-51 | chr1-174314296-DEL-53 |
| chr1-174318735-DEL-60 | chr1-174318736-DEL-60 |
| chr1-174344976-DEL-51 | chr1-174344993-DEL-51 |
| chr1-174358655-DEL-86 | chr1-174358658-DEL-84 |
| chr1-174362563-DEL-795 | chr1-174362564-DEL-794 |
| chr1-174371382-DEL-51 | chr1-174371379-DEL-51 |
| chr1-174373981-DEL-186 | chr1-174373981-DEL-187 |
| chr1-174376580-DEL-69 | chr1-174376581-DEL-69 |
| chr1-174396006-DEL-62 | chr1-174396007-DEL-62 |
| chr1-174406989-DEL-60 | chr1-174406990-DEL-58 |
| chr1-174413140-DEL-62 | chr1-174413141-DEL-62 |
| chr1-174423922-DEL-100 | chr1-174423923-DEL-100 |
| chr1-174431355-DEL-138 | chr1-174431356-DEL-138 |
| chr1-174445989-DEL-64 | chr1-174445989-DEL-64 |
| chr1-174448073-DEL-95 | chr1-174448054-DEL-119 |
| chr1-174523716-DEL-100 | chr1-174523723-DEL-88 |
| chr1-174526673-DEL-52 | chr1-174526674-DEL-52 |
| chr1-174535402-DEL-6201 | chr1-174535403-DEL-6201 |
| chr1-174546158-DEL-52 | chr1-174546158-DEL-52 |
| chr1-174547943-DEL-857 | chr1-174547944-DEL-857 |
| chr1-174555327-DEL-108 | chr1-174555328-DEL-108 |
| chr1-174567568-DEL-1139 | chr1-174567569-DEL-1139 |
| chr1-174569242-DEL-87 | chr1-174569270-DEL-54 |
| chr1-174596008-DEL-114 | chr1-174596009-DEL-114 |
| chr1-174612051-DEL-3294 | chr1-174612052-DEL-3294 |
| chr1-174615122-DEL-201 | chr1-174615123-DEL-201 |
| chr1-174630434-DEL-118 | chr1-174630435-DEL-118 |
| chr1-174645539-DEL-61 | chr1-174645536-DEL-53 |
| chr1-174650557-DEL-99 | chr1-174650555-DEL-100 |
| chr1-174659523-DEL-686 | chr1-174659524-DEL-686 |
| chr1-174669022-DEL-88 | chr1-174669023-DEL-88 |
| chr1-174687433-DEL-56 | chr1-174687420-DEL-70 |
| chr1-174699619-DEL-143 | chr1-174699589-DEL-125 |
| chr1-174713215-DEL-67 | chr1-174713216-DEL-67 |
| chr1-174751883-DEL-56 | chr1-174751884-DEL-52 |
| chr1-174770536-DEL-6473 | chr1-174770537-DEL-6473 |
| chr1-174770699-DEL-214 | chr1-174770700-DEL-214 |
| chr1-174805081-DEL-1056 | chr1-174805082-DEL-1056 |
| chr1-174819121-DEL-55 | chr1-174819093-DEL-60 |
| chr1-174835765-DEL-7205 | chr1-174835766-DEL-7205 |
| chr1-174858610-DEL-144 | chr1-174858610-DEL-144 |
| chr1-174867551-DEL-54 | chr1-174867516-DEL-90 |
| chr1-174869561-DEL-6291 | chr1-174869562-DEL-6291 |
| chr1-174902344-DEL-104 | chr1-174902345-DEL-104 |

| | |
|---|---|
| chr1-174904070-DEL-294 | chr1-174904071-DEL-294 |
| chr1-174906075-DEL-55 | chr1-174906076-DEL-55 |
| chr1-174914195-DEL-64 | chr1-174914197-DEL-63 |
| chr1-174914758-DEL-1857 | chr1-174914759-DEL-1857 |
| chr1-174928595-DEL-52 | chr1-174928598-DEL-62 |
| chr1-174940798-DEL-52 | chr1-174940799-DEL-52 |
| chr1-174944476-DEL-1892 | chr1-174944477-DEL-1892 |
| chr1-174963043-DEL-171 | chr1-174963044-DEL-171 |
| chr1-174966754-DEL-2514 | chr1-174966755-DEL-2514 |
| chr1-174996034-DEL-70 | chr1-174996049-DEL-56 |
| chr1-174999128-DEL-189 | chr1-174999129-DEL-189 |
| chr1-175008618-DEL-83 | chr1-175008619-DEL-83 |
| chr1-175010708-DEL-80 | chr1-175010709-DEL-80 |
| chr1-175019663-DEL-1385 | chr1-175019664-DEL-1381 |
| chr1-175048057-DEL-80 | chr1-175048058-DEL-80 |
| chr1-175050938-DEL-1793 | chr1-175050939-DEL-1793 |
| chr1-175059662-DEL-179 | chr1-175059663-DEL-180 |
| chr1-175073832-DEL-583 | chr1-175073836-DEL-578 |
| chr1-175077472-DEL-1426 | chr1-175077473-DEL-1426 |
| chr1-175079424-DEL-2764 | chr1-175079425-DEL-2764 |
| chr1-175083589-DEL-52 | chr1-175083590-DEL-52 |
| chr1-175084132-DEL-76 | chr1-175084138-DEL-75 |
| chr1-175094064-DEL-247 | chr1-175094033-DEL-241 |
| chr1-175101980-DEL-959 | chr1-175101981-DEL-959 |
| chr1-175107955-DEL-5031 | chr1-175107947-DEL-5032 |
| chr1-175136545-DEL-55 | chr1-175136546-DEL-50 |
| chr1-175166566-DEL-6504 | chr1-175166567-DEL-6504 |
| chr1-175215586-DEL-109 | chr1-175215588-DEL-110 |
| chr1-175239663-DEL-193 | chr1-175239664-DEL-193 |
| chr1-175241023-DEL-201 | chr1-175241024-DEL-201 |
| chr1-175249824-DEL-59 | chr1-175249825-DEL-59 |
| chr1-175279279-DEL-893 | chr1-175279280-DEL-897 |
| chr1-175294353-DEL-58 | chr1-175294354-DEL-62 |
| chr1-175307816-DEL-64 | chr1-175307817-DEL-64 |
| chr1-175316423-DEL-81 | chr1-175316394-DEL-80 |
| chr1-175318420-DEL-110 | chr1-175318413-DEL-92 |
| chr1-175339912-DEL-84 | chr1-175339913-DEL-84 |
| chr1-175340568-DEL-235 | chr1-175340569-DEL-235 |
| chr1-175365498-DEL-72 | chr1-175365499-DEL-72 |
| chr1-175366465-DEL-401 | chr1-175366466-DEL-409 |
| chr1-175379240-DEL-103 | chr1-175379265-DEL-93 |
| chr1-175381424-DEL-56 | chr1-175381430-DEL-52 |
| chr1-175461789-DEL-147 | chr1-175461790-DEL-147 |
| chr1-175473598-DEL-837 | chr1-175473599-DEL-837 |
| chr1-175484477-DEL-78 | chr1-175484478-DEL-78 |
| chr1-175488631-DEL-68 | chr1-175488632-DEL-68 |
| chr1-175550033-DEL-59 | chr1-175549989-DEL-104 |
| chr1-175576957-DEL-75 | chr1-175576950-DEL-72 |
| chr1-175581863-DEL-169 | chr1-175581864-DEL-169 |
| chr1-175585617-DEL-445 | chr1-175585618-DEL-445 |
| chr1-175604667-DEL-85 | chr1-175604664-DEL-72 |
| chr1-175623342-DEL-3140 | chr1-175623342-DEL-3141 |
| chr1-175644411-DEL-6233 | chr1-175644412-DEL-6233 |
| chr1-175664736-DEL-60 | chr1-175664713-DEL-80 |
| chr1-175670108-DEL-6413 | chr1-175670109-DEL-6413 |
| chr1-175705237-DEL-188 | chr1-175705237-DEL-187 |
| chr1-175713797-DEL-116 | chr1-175713795-DEL-121 |
| chr1-175717272-DEL-1865 | chr1-175717273-DEL-1865 |
| chr1-175744764-DEL-244 | chr1-175744765-DEL-244 |
| chr1-175750943-DEL-366 | chr1-175750944-DEL-366 |
| chr1-175764842-DEL-68 | chr1-175764825-DEL-52 |
| chr1-175795029-DEL-93 | chr1-175795030-DEL-93 |
| chr1-175797322-DEL-224 | chr1-175797323-DEL-224 |
| chr1-175812819-DEL-89 | chr1-175812820-DEL-89 |
| chr1-175828806-DEL-200 | chr1-175828807-DEL-200 |
| chr1-175845049-DEL-53 | chr1-175845042-DEL-60 |
| chr1-175853997-DEL-99 | chr1-175853998-DEL-99 |
| chr1-175881891-DEL-1262 | chr1-175881892-DEL-1262 |
| chr1-175900905-DEL-274 | chr1-175900896-DEL-275 |
| chr1-175944208-DEL-141 | chr1-175944209-DEL-141 |
| chr1-175952883-DEL-101 | chr1-175952929-DEL-52 |
| chr1-175987327-DEL-223 | chr1-175987328-DEL-223 |
| chr1-175992383-DEL-62 | chr1-175992384-DEL-62 |
| chr1-176004167-DEL-708 | chr1-176004167-DEL-708 |
| chr1-176028819-DEL-50 | chr1-176028827-DEL-56 |
| chr1-176039801-DEL-430 | chr1-176039802-DEL-430 |
| chr1-176048915-DEL-69 | chr1-176048913-DEL-70 |
| chr1-176054279-DEL-130 | chr1-176054292-DEL-122 |
| chr1-176058158-DEL-56 | chr1-176058136-DEL-58 |
| chr1-176096469-DEL-665 | chr1-176096470-DEL-665 |

| | |
|---|---|
| chr1-176107540-DEL-440 | chr1-176107541-DEL-440 |
| chr1-176109654-DEL-6796 | chr1-176109655-DEL-6796 |
| chr1-176139972-DEL-244 | chr1-176139973-DEL-244 |
| chr1-176140840-DEL-54 | chr1-176140841-DEL-54 |
| chr1-176211205-DEL-5735 | chr1-176211206-DEL-5735 |
| chr1-176238288-DEL-584 | chr1-176238289-DEL-583 |
| chr1-176242576-DEL-201 | chr1-176242577-DEL-201 |
| chr1-176268175-DEL-242 | chr1-176268176-DEL-242 |
| chr1-176288087-DEL-130 | chr1-176288088-DEL-130 |
| chr1-176293297-DEL-78 | chr1-176293303-DEL-72 |
| chr1-176305357-DEL-1798 | chr1-176305357-DEL-1799 |
| chr1-176329479-DEL-2329 | chr1-176329480-DEL-2329 |
| chr1-176332223-DEL-57 | chr1-176332224-DEL-57 |
| chr1-176368674-DEL-13522 | chr1-176368675-DEL-13522 |
| chr1-176452083-DEL-68 | chr1-176452084-DEL-68 |
| chr1-176477371-DEL-1866 | chr1-176477372-DEL-1866 |
| chr1-176499898-DEL-356 | chr1-176499897-DEL-357 |
| chr1-176539556-DEL-232 | chr1-176539555-DEL-233 |
| chr1-176546350-DEL-61 | chr1-176546348-DEL-61 |
| chr1-176546729-DEL-494 | chr1-176546730-DEL-494 |
| chr1-176552684-DEL-78 | chr1-176552685-DEL-78 |
| chr1-176576508-DEL-61 | chr1-176576509-DEL-61 |
| chr1-176680004-DEL-60 | chr1-176680005-DEL-60 |
| chr1-176702477-DEL-52 | chr1-176702478-DEL-52 |
| chr1-176720872-DEL-6593 | chr1-176720873-DEL-6593 |
| chr1-176747584-DEL-1695 | chr1-176747585-DEL-1695 |
| chr1-176773074-DEL-80 | chr1-176773074-DEL-80 |
| chr1-176802002-DEL-135 | chr1-176801956-DEL-170 |
| chr1-176828031-DEL-84 | chr1-176828032-DEL-84 |
| chr1-176986250-DEL-132 | chr1-176986249-DEL-133 |
| chr1-177040598-DEL-6438 | chr1-177040599-DEL-6438 |
| chr1-177134274-DEL-56 | chr1-177134275-DEL-52 |
| chr1-177139365-DEL-60 | chr1-177139365-DEL-63 |
| chr1-177156094-DEL-134 | chr1-177156107-DEL-122 |
| chr1-177156804-DEL-409 | chr1-177156805-DEL-409 |
| chr1-177166934-DEL-193 | chr1-177166935-DEL-193 |
| chr1-177184575-DEL-160 | chr1-177184570-DEL-166 |
| chr1-177191105-DEL-62 | chr1-177191109-DEL-60 |
| chr1-177202165-DEL-52 | chr1-177202166-DEL-52 |
| chr1-177225541-DEL-118 | chr1-177225542-DEL-118 |
| chr1-177387387-DEL-187 | chr1-177387388-DEL-187 |
| chr1-177451802-DEL-100 | chr1-177451803-DEL-100 |
| chr1-177453001-DEL-149 | chr1-177453000-DEL-148 |
| chr1-177456448-DEL-59 | chr1-177456441-DEL-60 |
| chr1-177480801-DEL-7113 | chr1-177480802-DEL-7113 |
| chr1-177513324-DEL-2134 | chr1-177513325-DEL-2134 |
| chr1-177515350-DEL-199 | chr1-177515351-DEL-199 |
| chr1-177565661-DEL-121 | chr1-177565686-DEL-97 |
| chr1-177567596-DEL-201 | chr1-177567586-DEL-198 |
| chr1-177585669-DEL-2921 | chr1-177585670-DEL-2921 |
| chr1-177625277-DEL-1986 | chr1-177625280-DEL-1983 |
| chr1-177686702-DEL-457 | chr1-177686703-DEL-457 |
| chr1-177692474-DEL-398 | chr1-177692475-DEL-398 |
| chr1-177714903-DEL-78 | chr1-177714904-DEL-78 |
| chr1-177731799-DEL-1107 | chr1-177731800-DEL-1111 |
| chr1-177751077-DEL-2850 | chr1-177751078-DEL-2850 |
| chr1-177756903-DEL-6456 | chr1-177756904-DEL-6456 |
| chr1-177790613-DEL-6998 | chr1-177790612-DEL-6998 |
| chr1-177832038-DEL-6445 | chr1-177832039-DEL-6445 |
| chr1-177849719-DEL-3566 | chr1-177849720-DEL-3566 |
| chr1-177855396-DEL-308 | chr1-177855397-DEL-308 |
| chr1-177870622-DEL-59 | chr1-177870623-DEL-59 |
| chr1-177898561-DEL-621 | chr1-177898562-DEL-621 |
| chr1-177907362-DEL-6515 | chr1-177907363-DEL-6515 |
| chr1-177949747-DEL-56 | chr1-177949748-DEL-56 |
| chr1-177962589-DEL-203 | chr1-177962590-DEL-203 |
| chr1-178001673-DEL-7090 | chr1-178001674-DEL-7090 |
| chr1-178041183-DEL-158 | chr1-178041184-DEL-158 |
| chr1-178070434-DEL-220 | chr1-178070435-DEL-220 |
| chr1-178075813-DEL-133 | chr1-178075812-DEL-135 |
| chr1-178112728-DEL-167 | chr1-178112729-DEL-167 |
| chr1-178131607-DEL-57 | chr1-178131608-DEL-60 |
| chr1-178136273-DEL-214 | chr1-178136272-DEL-214 |
| chr1-178184400-DEL-619 | chr1-178184401-DEL-619 |
| chr1-178198767-DEL-203 | chr1-178198768-DEL-203 |
| chr1-178203943-DEL-2612 | chr1-178203944-DEL-2612 |
| chr1-178229193-DEL-816 | chr1-178229194-DEL-816 |
| chr1-178241993-DEL-406 | chr1-178241994-DEL-406 |
| chr1-178247655-DEL-62 | chr1-178247655-DEL-62 |
| chr1-178251778-DEL-86 | chr1-178251779-DEL-86 |

| | |
|---|---|
| chr1-178270806-DEL-144 | chr1-178270807-DEL-144 |
| chr1-178275187-DEL-72 | chr1-178275179-DEL-75 |
| chr1-178285002-DEL-91 | chr1-178285003-DEL-91 |
| chr1-178299063-DEL-1888 | chr1-178299064-DEL-1888 |
| chr1-178351764-DEL-72 | chr1-178351765-DEL-72 |
| chr1-178352450-DEL-190 | chr1-178352440-DEL-191 |
| chr1-178426937-DEL-121 | chr1-178426938-DEL-121 |
| chr1-178444530-DEL-82 | chr1-178444528-DEL-84 |
| chr1-178534045-DEL-1260 | chr1-178534046-DEL-1260 |
| chr1-178536634-DEL-486 | chr1-178536635-DEL-486 |
| chr1-178568551-DEL-51 | chr1-178568546-DEL-51 |
| chr1-178590210-DEL-126 | chr1-178590221-DEL-108 |
| chr1-178607526-DEL-70 | chr1-178607522-DEL-71 |
| chr1-178672037-DEL-442 | chr1-178672039-DEL-441 |
| chr1-178736266-DEL-132 | chr1-178736262-DEL-99 |
| chr1-178793022-DEL-61 | chr1-178793023-DEL-61 |
| chr1-178823167-DEL-134 | chr1-178823170-DEL-132 |
| chr1-178867286-DEL-54 | chr1-178867287-DEL-54 |
| chr1-178885204-DEL-279 | chr1-178885209-DEL-275 |
| chr1-178890570-DEL-109 | chr1-178890571-DEL-109 |
| chr1-178934782-DEL-56 | chr1-178934783-DEL-56 |
| chr1-178996581-DEL-119 | chr1-178996582-DEL-119 |
| chr1-179007265-DEL-500 | chr1-179007266-DEL-500 |
| chr1-179019252-DEL-221 | chr1-179019253-DEL-221 |
| chr1-179026655-DEL-184 | chr1-179026656-DEL-184 |
| chr1-179046723-DEL-200 | chr1-179046724-DEL-200 |
| chr1-179054864-DEL-60 | chr1-179054865-DEL-60 |
| chr1-179087612-DEL-299 | chr1-179087613-DEL-299 |
| chr1-179098291-DEL-1128 | chr1-179098292-DEL-1128 |
| chr1-179113194-DEL-55 | chr1-179113194-DEL-67 |
| chr1-179143052-DEL-78 | chr1-179143032-DEL-56 |
| chr1-179167405-DEL-139 | chr1-179167448-DEL-139 |
| chr1-179204180-DEL-241 | chr1-179204181-DEL-241 |
| chr1-179204867-DEL-78 | chr1-179204868-DEL-78 |
| chr1-179206166-DEL-147 | chr1-179206165-DEL-147 |
| chr1-179209021-DEL-64 | chr1-179209022-DEL-64 |
| chr1-179216529-DEL-486 | chr1-179216530-DEL-486 |
| chr1-179218289-DEL-2965 | chr1-179218290-DEL-2965 |
| chr1-179222346-DEL-180 | chr1-179222347-DEL-180 |
| chr1-179242237-DEL-201 | chr1-179242238-DEL-201 |
| chr1-179287692-DEL-812 | chr1-179287693-DEL-812 |
| chr1-179289026-DEL-93 | chr1-179289073-DEL-54 |
| chr1-179292588-DEL-1659 | chr1-179292589-DEL-1659 |
| chr1-179294426-DEL-56 | chr1-179294399-DEL-84 |
| chr1-179296876-DEL-201 | chr1-179296877-DEL-201 |
| chr1-179297780-DEL-315 | chr1-179297781-DEL-315 |
| chr1-179321998-DEL-177 | chr1-179321999-DEL-177 |
| chr1-179333294-DEL-236 | chr1-179333295-DEL-236 |
| chr1-179335899-DEL-60 | chr1-179335900-DEL-60 |
| chr1-179384006-DEL-51 | chr1-179384007-DEL-51 |
| chr1-179413268-DEL-180 | chr1-179413233-DEL-180 |
| chr1-179430288-DEL-180 | chr1-179430289-DEL-180 |
| chr1-179430291-DEL-720 | chr1-179430291-DEL-720 |
| chr1-179445654-DEL-1898 | chr1-179445654-DEL-1898 |
| chr1-179463120-DEL-848 | chr1-179463120-DEL-848 |
| chr1-179476184-DEL-225 | chr1-179476185-DEL-225 |
| chr1-179479099-DEL-549 | chr1-179479100-DEL-549 |
| chr1-179481597-DEL-7066 | chr1-179481598-DEL-7066 |
| chr1-179510244-DEL-221 | chr1-179510245-DEL-221 |
| chr1-179517536-DEL-3646 | chr1-179517537-DEL-3646 |
| chr1-179522506-DEL-77 | chr1-179522507-DEL-77 |
| chr1-179528628-DEL-200 | chr1-179528629-DEL-200 |
| chr1-179542163-DEL-209 | chr1-179542166-DEL-209 |
| chr1-179544845-DEL-514 | chr1-179544846-DEL-514 |
| chr1-179570691-DEL-140 | chr1-179570692-DEL-140 |
| chr1-179610271-DEL-207 | chr1-179610272-DEL-207 |
| chr1-179636730-DEL-68 | chr1-179636714-DEL-50 |
| chr1-179645929-DEL-58 | chr1-179645930-DEL-58 |
| chr1-179668941-DEL-370 | chr1-179668942-DEL-370 |
| chr1-179677379-DEL-111 | chr1-179677380-DEL-111 |
| chr1-179682487-DEL-204 | chr1-179682487-DEL-204 |
| chr1-179683841-DEL-813 | chr1-179683842-DEL-813 |
| chr1-179710695-DEL-61 | chr1-179710696-DEL-61 |
| chr1-179713792-DEL-238 | chr1-179713793-DEL-238 |
| chr1-179721458-DEL-7275 | chr1-179721459-DEL-7275 |
| chr1-179753959-DEL-59 | chr1-179753960-DEL-59 |
| chr1-179782802-DEL-130 | chr1-179782801-DEL-139 |
| chr1-179798855-DEL-51 | chr1-179798856-DEL-51 |
| chr1-179811193-DEL-52 | chr1-179811187-DEL-53 |
| chr1-179838246-DEL-2499 | chr1-179838247-DEL-2499 |

| | |
|---|---|
| chr1-179857487-DEL-117 | chr1-179857488-DEL-117 |
| chr1-179862462-DEL-510 | chr1-179862463-DEL-510 |
| chr1-179869932-DEL-75 | chr1-179869894-DEL-94 |
| chr1-180026175-DEL-95 | chr1-180026176-DEL-95 |
| chr1-180033429-DEL-68 | chr1-180033430-DEL-68 |
| chr1-180033808-DEL-323 | chr1-180033809-DEL-323 |
| chr1-180050731-DEL-329 | chr1-180050732-DEL-329 |
| chr1-180110857-DEL-87 | chr1-180110859-DEL-86 |
| chr1-180170308-DEL-51 | chr1-180170309-DEL-57 |
| chr1-180200236-DEL-77 | chr1-180200236-DEL-63 |
| chr1-180227007-DEL-57 | chr1-180227010-DEL-55 |
| chr1-180281219-DEL-54 | chr1-180281220-DEL-54 |
| chr1-180297925-DEL-455 | chr1-180297926-DEL-455 |
| chr1-180325675-DEL-1351 | chr1-180325676-DEL-1351 |
| chr1-180337150-DEL-98 | chr1-180337151-DEL-105 |
| chr1-180339335-DEL-400 | chr1-180339336-DEL-400 |
| chr1-180345623-DEL-66 | chr1-180345624-DEL-66 |
| chr1-180378976-DEL-209 | chr1-180378977-DEL-209 |
| chr1-180403080-DEL-213 | chr1-180403081-DEL-213 |
| chr1-180420708-DEL-209 | chr1-180420709-DEL-209 |
| chr1-180437161-DEL-248 | chr1-180437162-DEL-248 |
| chr1-180445589-DEL-205 | chr1-180445590-DEL-205 |
| chr1-180462353-DEL-776 | chr1-180462348-DEL-777 |
| chr1-180482884-DEL-169 | chr1-180482880-DEL-170 |
| chr1-180494498-DEL-202 | chr1-180494499-DEL-202 |
| chr1-180543313-DEL-128 | chr1-180543338-DEL-132 |
| chr1-180568273-DEL-354 | chr1-180568274-DEL-354 |
| chr1-180587974-DEL-61 | chr1-180587975-DEL-53 |
| chr1-180598017-DEL-90 | chr1-180598033-DEL-81 |
| chr1-180604625-DEL-51 | chr1-180604626-DEL-51 |
| chr1-180610150-DEL-213 | chr1-180610151-DEL-213 |
| chr1-180618028-DEL-945 | chr1-180618024-DEL-947 |
| chr1-180627208-DEL-212 | chr1-180627209-DEL-212 |
| chr1-180634707-DEL-99 | chr1-180634708-DEL-98 |
| chr1-180663635-DEL-183 | chr1-180663636-DEL-183 |
| chr1-180667374-DEL-100 | chr1-180667412-DEL-84 |
| chr1-180679467-DEL-3845 | chr1-180679468-DEL-3845 |
| chr1-180710501-DEL-59 | chr1-180710502-DEL-59 |
| chr1-180713073-DEL-107 | chr1-180713074-DEL-107 |
| chr1-180729320-DEL-1053 | chr1-180729321-DEL-1041 |
| chr1-180747654-DEL-112 | chr1-180747652-DEL-119 |
| chr1-180753103-DEL-217 | chr1-180753104-DEL-217 |
| chr1-180768231-DEL-68 | chr1-180768185-DEL-98 |
| chr1-180785833-DEL-115 | chr1-180785834-DEL-115 |
| chr1-180787197-DEL-75 | chr1-180787194-DEL-110 |
| chr1-180792898-DEL-141 | chr1-180792899-DEL-141 |
| chr1-180806955-DEL-122 | chr1-180806956-DEL-122 |
| chr1-180836422-DEL-83 | chr1-180836423-DEL-83 |
| chr1-180846656-DEL-91 | chr1-180846659-DEL-90 |
| chr1-180861586-DEL-79 | chr1-180861636-DEL-66 |
| chr1-180864517-DEL-220 | chr1-180864518-DEL-220 |
| chr1-180893692-DEL-209 | chr1-180893693-DEL-209 |
| chr1-180898013-DEL-60 | chr1-180897982-DEL-84 |
| chr1-180902491-DEL-951 | chr1-180902515-DEL-928 |
| chr1-180932170-DEL-207 | chr1-180932172-DEL-206 |
| chr1-180942420-DEL-54 | chr1-180942419-DEL-55 |
| chr1-180952127-DEL-668 | chr1-180952128-DEL-668 |
| chr1-180958182-DEL-78 | chr1-180958183-DEL-78 |
| chr1-180959235-DEL-117 | chr1-180959236-DEL-117 |
| chr1-180963433-DEL-201 | chr1-180963434-DEL-201 |
| chr1-180964277-DEL-68 | chr1-180964270-DEL-50 |
| chr1-181018836-DEL-218 | chr1-181018837-DEL-218 |
| chr1-181051279-DEL-208 | chr1-181051280-DEL-208 |
| chr1-181069467-DEL-481 | chr1-181069468-DEL-481 |
| chr1-181070470-DEL-89 | chr1-181070463-DEL-90 |
| chr1-181091084-DEL-7087 | chr1-181091085-DEL-7087 |
| chr1-181098504-DEL-815 | chr1-181098543-DEL-799 |
| chr1-181100010-DEL-73 | chr1-181099979-DEL-70 |
| chr1-181122408-DEL-251 | chr1-181122409-DEL-251 |
| chr1-181136571-DEL-83 | chr1-181136572-DEL-83 |
| chr1-181137627-DEL-230 | chr1-181137628-DEL-230 |
| chr1-181147984-DEL-60 | chr1-181147985-DEL-60 |
| chr1-181156193-DEL-1423 | chr1-181156194-DEL-1423 |
| chr1-181182587-DEL-65 | chr1-181182588-DEL-65 |
| chr1-181187592-DEL-90 | chr1-181187579-DEL-135 |
| chr1-181223486-DEL-2669 | chr1-181223487-DEL-2670 |
| chr1-181227835-DEL-93 | chr1-181227816-DEL-99 |
| chr1-181239768-DEL-86 | chr1-181239769-DEL-86 |
| chr1-181250081-DEL-61 | chr1-181250082-DEL-61 |
| chr1-181272535-DEL-52 | chr1-181272536-DEL-52 |

| | |
|---|---|
| chr1-181305926-DEL-195 | chr1-181305927-DEL-195 |
| chr1-181317898-DEL-222 | chr1-181317899-DEL-222 |
| chr1-181320468-DEL-6578 | chr1-181320469-DEL-6578 |
| chr1-181351719-DEL-1789 | chr1-181351720-DEL-1789 |
| chr1-181381591-DEL-67 | chr1-181381606-DEL-66 |
| chr1-181382487-DEL-145 | chr1-181382488-DEL-145 |
| chr1-181399121-DEL-796 | chr1-181399134-DEL-775 |
| chr1-181445328-DEL-67 | chr1-181445329-DEL-67 |
| chr1-181502058-DEL-250 | chr1-181502059-DEL-250 |
| chr1-181531571-DEL-73 | chr1-181531570-DEL-67 |
| chr1-181603479-DEL-104 | chr1-181603480-DEL-104 |
| chr1-181636061-DEL-204 | chr1-181636054-DEL-205 |
| chr1-181655158-DEL-54 | chr1-181655158-DEL-54 |
| chr1-181681514-DEL-74 | chr1-181681512-DEL-66 |
| chr1-181681920-DEL-165 | chr1-181681923-DEL-162 |
| chr1-181682700-DEL-208 | chr1-181682701-DEL-208 |
| chr1-181695728-DEL-3899 | chr1-181695729-DEL-3899 |
| chr1-181703041-DEL-208 | chr1-181703041-DEL-208 |
| chr1-181707459-DEL-88 | chr1-181707460-DEL-88 |
| chr1-181708133-DEL-173 | chr1-181708134-DEL-173 |
| chr1-181717001-DEL-197 | chr1-181717002-DEL-197 |
| chr1-181729865-DEL-84 | chr1-181729876-DEL-66 |
| chr1-181767316-DEL-211 | chr1-181767317-DEL-211 |
| chr1-181770102-DEL-55 | chr1-181770103-DEL-55 |
| chr1-181787286-DEL-74 | chr1-181787287-DEL-74 |
| chr1-181809310-DEL-190 | chr1-181809311-DEL-190 |
| chr1-181861294-DEL-90 | chr1-181861252-DEL-100 |
| chr1-181866613-DEL-180 | chr1-181866606-DEL-181 |
| chr1-181868812-DEL-1469 | chr1-181868813-DEL-1469 |
| chr1-181902548-DEL-200 | chr1-181902549-DEL-200 |
| chr1-181910413-DEL-207 | chr1-181910414-DEL-207 |
| chr1-181911350-DEL-233 | chr1-181911351-DEL-233 |
| chr1-181930676-DEL-210 | chr1-181930677-DEL-210 |
| chr1-181937709-DEL-72 | chr1-181937710-DEL-72 |
| chr1-181950329-DEL-223 | chr1-181950322-DEL-224 |
| chr1-181958234-DEL-73 | chr1-181958235-DEL-72 |
| chr1-181960754-DEL-389 | chr1-181960755-DEL-389 |
| chr1-181995625-DEL-103 | chr1-181995622-DEL-101 |
| chr1-182019026-DEL-207 | chr1-182019027-DEL-207 |
| chr1-182020483-DEL-214 | chr1-182020484-DEL-214 |
| chr1-182028452-DEL-8425 | chr1-182028453-DEL-8425 |
| chr1-182046310-DEL-102 | chr1-182046335-DEL-78 |
| chr1-182047272-DEL-324 | chr1-182047273-DEL-324 |
| chr1-182084680-DEL-8985 | chr1-182084681-DEL-8985 |
| chr1-182102730-DEL-150 | chr1-182102731-DEL-150 |
| chr1-182154267-DEL-62 | chr1-182154268-DEL-62 |
| chr1-182159836-DEL-127 | chr1-182159837-DEL-127 |
| chr1-182163043-DEL-216 | chr1-182163044-DEL-216 |
| chr1-182202862-DEL-91 | chr1-182202863-DEL-91 |
| chr1-182277951-DEL-971 | chr1-182277952-DEL-971 |
| chr1-182387368-DEL-441 | chr1-182387369-DEL-441 |
| chr1-182404588-DEL-105 | chr1-182404608-DEL-84 |
| chr1-182442935-DEL-111 | chr1-182442923-DEL-86 |
| chr1-182471940-DEL-53 | chr1-182471941-DEL-53 |
| chr1-182473257-DEL-60 | chr1-182473258-DEL-60 |
| chr1-182496914-DEL-64 | chr1-182496921-DEL-58 |
| chr1-182505723-DEL-65 | chr1-182505724-DEL-59 |
| chr1-182526828-DEL-68 | chr1-182526847-DEL-60 |
| chr1-182559517-DEL-207 | chr1-182559518-DEL-207 |
| chr1-182563592-DEL-261 | chr1-182563593-DEL-261 |
| chr1-182588879-DEL-472 | chr1-182588880-DEL-472 |
| chr1-182615749-DEL-177 | chr1-182615750-DEL-177 |
| chr1-182639378-DEL-101 | chr1-182639391-DEL-103 |
| chr1-182705082-DEL-69 | chr1-182705082-DEL-69 |
| chr1-182711913-DEL-210 | chr1-182711914-DEL-180 |
| chr1-182720221-DEL-1728 | chr1-182720247-DEL-1686 |
| chr1-182720479-DEL-84 | chr1-182720480-DEL-84 |
| chr1-182738502-DEL-60 | chr1-182738501-DEL-61 |
| chr1-182777760-DEL-68 | chr1-182777759-DEL-63 |
| chr1-182782185-DEL-51 | chr1-182782186-DEL-51 |
| chr1-182854596-DEL-1899 | chr1-182854597-DEL-1899 |
| chr1-182857825-DEL-70 | chr1-182857826-DEL-70 |
| chr1-182860332-DEL-83 | chr1-182860333-DEL-83 |
| chr1-182862371-DEL-99 | chr1-182862372-DEL-99 |
| chr1-182876353-DEL-56 | chr1-182876380-DEL-58 |
| chr1-182885144-DEL-184 | chr1-182885145-DEL-184 |
| chr1-182892278-DEL-917 | chr1-182892279-DEL-915 |
| chr1-182895640-DEL-53 | chr1-182895641-DEL-53 |
| chr1-182984029-DEL-258 | chr1-182983996-DEL-294 |
| chr1-183016010-DEL-514 | chr1-183016011-DEL-514 |

| | |
|---|---|
| chr1-183038168-DEL-163 | chr1-183038182-DEL-153 |
| chr1-183072121-DEL-57 | chr1-183072122-DEL-57 |
| chr1-183080929-DEL-190 | chr1-183080930-DEL-190 |
| chr1-183090749-DEL-88 | chr1-183090750-DEL-88 |
| chr1-183118756-DEL-102 | chr1-183118757-DEL-102 |
| chr1-183145186-DEL-66 | chr1-183145187-DEL-66 |
| chr1-183159175-DEL-176 | chr1-183159176-DEL-176 |
| chr1-183188967-DEL-284 | chr1-183188968-DEL-284 |
| chr1-183214997-DEL-203 | chr1-183214998-DEL-203 |
| chr1-183235026-DEL-94 | chr1-183235027-DEL-94 |
| chr1-183271550-DEL-6442 | chr1-183271551-DEL-6442 |
| chr1-183282204-DEL-104 | chr1-183282205-DEL-104 |
| chr1-183296654-DEL-6783 | chr1-183296655-DEL-6783 |
| chr1-183316500-DEL-301 | chr1-183316501-DEL-265 |
| chr1-183323834-DEL-59 | chr1-183323835-DEL-59 |
| chr1-183329243-DEL-62 | chr1-183329227-DEL-51 |
| chr1-183330249-DEL-1042 | chr1-183330250-DEL-1042 |
| chr1-183349918-DEL-164 | chr1-183349891-DEL-150 |
| chr1-183353161-DEL-498 | chr1-183353162-DEL-498 |
| chr1-183353667-DEL-102 | chr1-183353668-DEL-102 |
| chr1-183377603-DEL-60 | chr1-183377618-DEL-60 |
| chr1-183411014-DEL-359 | chr1-183411014-DEL-360 |
| chr1-183413457-DEL-228 | chr1-183413458-DEL-228 |
| chr1-183420943-DEL-363 | chr1-183420944-DEL-363 |
| chr1-183498421-DEL-62 | chr1-183498421-DEL-62 |
| chr1-183509268-DEL-85 | chr1-183509269-DEL-85 |
| chr1-183548985-DEL-184 | chr1-183548986-DEL-184 |
| chr1-183569478-DEL-65 | chr1-183569479-DEL-65 |
| chr1-183635342-DEL-70 | chr1-183635340-DEL-67 |
| chr1-183643809-DEL-90 | chr1-183643810-DEL-90 |
| chr1-183647327-DEL-58 | chr1-183647328-DEL-58 |
| chr1-183742539-DEL-64 | chr1-183742552-DEL-52 |
| chr1-183744504-DEL-226 | chr1-183744505-DEL-226 |
| chr1-183809573-DEL-122 | chr1-183809573-DEL-122 |
| chr1-183817526-DEL-61 | chr1-183817525-DEL-51 |
| chr1-183866562-DEL-617 | chr1-183866563-DEL-617 |
| chr1-183877630-DEL-87 | chr1-183877631-DEL-87 |
| chr1-183880164-DEL-87 | chr1-183880165-DEL-87 |
| chr1-183883494-DEL-4267 | chr1-183883493-DEL-4267 |
| chr1-183956679-DEL-266 | chr1-183956685-DEL-263 |
| chr1-183962165-DEL-70 | chr1-183962166-DEL-60 |
| chr1-183997614-DEL-433 | chr1-183997609-DEL-438 |
| chr1-184004075-DEL-82 | chr1-184004069-DEL-77 |
| chr1-184071682-DEL-63 | chr1-184071636-DEL-71 |
| chr1-184083858-DEL-83 | chr1-184083859-DEL-83 |
| chr1-184138037-DEL-69 | chr1-184138038-DEL-69 |
| chr1-184193135-DEL-63 | chr1-184193137-DEL-61 |
| chr1-184196572-DEL-358 | chr1-184196573-DEL-358 |
| chr1-184205416-DEL-684 | chr1-184205417-DEL-684 |
| chr1-184272690-DEL-1222 | chr1-184272709-DEL-1223 |
| chr1-184383734-DEL-134 | chr1-184383737-DEL-129 |
| chr1-184405828-DEL-94 | chr1-184405831-DEL-50 |
| chr1-184410671-DEL-100 | chr1-184410697-DEL-75 |
| chr1-184429368-DEL-73 | chr1-184429361-DEL-119 |
| chr1-184430019-DEL-82 | chr1-184430016-DEL-80 |
| chr1-184441627-DEL-83 | chr1-184441628-DEL-83 |
| chr1-184473132-DEL-84 | chr1-184473136-DEL-81 |
| chr1-184489908-DEL-330 | chr1-184489909-DEL-330 |
| chr1-184536653-DEL-364 | chr1-184536654-DEL-364 |
| chr1-184537322-DEL-58 | chr1-184537323-DEL-58 |
| chr1-184539537-DEL-128 | chr1-184539538-DEL-128 |
| chr1-184552270-DEL-190 | chr1-184552271-DEL-190 |
| chr1-184555735-DEL-54 | chr1-184555734-DEL-56 |
| chr1-184560411-DEL-98 | chr1-184560382-DEL-99 |
| chr1-184566879-DEL-180 | chr1-184566927-DEL-132 |
| chr1-184573149-DEL-64 | chr1-184573150-DEL-64 |
| chr1-184590843-DEL-54 | chr1-184590844-DEL-54 |
| chr1-184622619-DEL-438 | chr1-184622618-DEL-439 |
| chr1-184643526-DEL-63 | chr1-184643528-DEL-62 |
| chr1-184671262-DEL-91 | chr1-184671265-DEL-89 |
| chr1-184686028-DEL-654 | chr1-184686029-DEL-654 |
| chr1-184691795-DEL-148 | chr1-184691796-DEL-148 |
| chr1-184722672-DEL-115 | chr1-184722673-DEL-115 |
| chr1-184728909-DEL-72 | chr1-184728910-DEL-72 |
| chr1-184763054-DEL-652 | chr1-184763068-DEL-628 |
| chr1-184769833-DEL-52 | chr1-184769834-DEL-52 |
| chr1-184788401-DEL-293 | chr1-184788431-DEL-260 |
| chr1-184812436-DEL-3038 | chr1-184812437-DEL-3038 |
| chr1-184871137-DEL-77 | chr1-184871138-DEL-77 |
| chr1-184879321-DEL-372 | chr1-184879309-DEL-361 |

| | |
|---|---|
| chr1-184887728-DEL-6485 | chr1-184887729-DEL-6485 |
| chr1-184904125-DEL-514 | chr1-184904126-DEL-514 |
| chr1-184910944-DEL-95 | chr1-184910945-DEL-95 |
| chr1-184927784-DEL-79 | chr1-184927785-DEL-78 |
| chr1-184932269-DEL-52 | chr1-184932270-DEL-52 |
| chr1-184945846-DEL-265 | chr1-184945847-DEL-265 |
| chr1-184946863-DEL-180 | chr1-184946909-DEL-135 |
| chr1-184953049-DEL-363 | chr1-184953050-DEL-363 |
| chr1-184969036-DEL-405 | chr1-184969037-DEL-405 |
| chr1-184986880-DEL-129 | chr1-184986885-DEL-121 |
| chr1-184996090-DEL-154 | chr1-184996090-DEL-154 |
| chr1-185000792-DEL-244 | chr1-185000793-DEL-244 |
| chr1-185035689-DEL-3017 | chr1-185035690-DEL-3017 |
| chr1-185042332-DEL-400 | chr1-185042333-DEL-400 |
| chr1-185072856-DEL-66 | chr1-185072834-DEL-64 |
| chr1-185080233-DEL-69 | chr1-185080234-DEL-69 |
| chr1-185103405-DEL-232 | chr1-185103406-DEL-232 |
| chr1-185115370-DEL-197 | chr1-185115371-DEL-197 |
| chr1-185122912-DEL-271 | chr1-185122913-DEL-271 |
| chr1-185131954-DEL-201 | chr1-185131954-DEL-201 |
| chr1-185149138-DEL-182 | chr1-185149139-DEL-182 |
| chr1-185177576-DEL-208 | chr1-185177577-DEL-208 |
| chr1-185182423-DEL-56 | chr1-185182426-DEL-54 |
| chr1-185213520-DEL-216 | chr1-185213521-DEL-216 |
| chr1-185252990-DEL-203 | chr1-185252991-DEL-203 |
| chr1-185254047-DEL-128 | chr1-185254048-DEL-128 |
| chr1-185261517-DEL-54 | chr1-185261518-DEL-54 |
| chr1-185282492-DEL-100 | chr1-185282493-DEL-100 |
| chr1-185295823-DEL-6443 | chr1-185295824-DEL-6443 |
| chr1-185337265-DEL-286 | chr1-185337266-DEL-285 |
| chr1-185353399-DEL-6442 | chr1-185353400-DEL-6442 |
| chr1-185361649-DEL-98 | chr1-185361650-DEL-98 |
| chr1-185380832-DEL-55 | chr1-185380829-DEL-67 |
| chr1-185415237-DEL-80 | chr1-185415234-DEL-52 |
| chr1-185422739-DEL-596 | chr1-185422740-DEL-596 |
| chr1-185435838-DEL-68 | chr1-185435835-DEL-55 |
| chr1-185462929-DEL-81 | chr1-185462953-DEL-62 |
| chr1-185485120-DEL-125 | chr1-185485121-DEL-125 |
| chr1-185517316-DEL-1574 | chr1-185517317-DEL-1574 |
| chr1-185520645-DEL-110 | chr1-185520646-DEL-110 |
| chr1-185522140-DEL-78 | chr1-185522141-DEL-78 |
| chr1-185545924-DEL-90 | chr1-185545921-DEL-87 |
| chr1-185552321-DEL-53 | chr1-185552322-DEL-53 |
| chr1-185584370-DEL-86 | chr1-185584371-DEL-86 |
| chr1-185587816-DEL-97 | chr1-185587818-DEL-95 |
| chr1-185592906-DEL-82 | chr1-185592905-DEL-84 |
| chr1-185605530-DEL-1298 | chr1-185605531-DEL-1298 |
| chr1-185628607-DEL-69 | chr1-185628608-DEL-69 |
| chr1-185653639-DEL-134 | chr1-185653640-DEL-134 |
| chr1-185703098-DEL-210 | chr1-185703129-DEL-180 |
| chr1-185733482-DEL-351 | chr1-185733483-DEL-351 |
| chr1-185739440-DEL-88 | chr1-185739445-DEL-84 |
| chr1-185749818-DEL-98 | chr1-185749819-DEL-98 |
| chr1-185752058-DEL-3985 | chr1-185752059-DEL-3985 |
| chr1-185782262-DEL-104 | chr1-185782263-DEL-104 |
| chr1-185784887-DEL-437 | chr1-185784888-DEL-437 |
| chr1-185830243-DEL-66 | chr1-185830244-DEL-66 |
| chr1-185836354-DEL-6407 | chr1-185836355-DEL-6407 |
| chr1-185864801-DEL-98 | chr1-185864802-DEL-98 |
| chr1-185874359-DEL-69 | chr1-185874353-DEL-70 |
| chr1-185879661-DEL-52 | chr1-185879661-DEL-53 |
| chr1-185886307-DEL-174 | chr1-185886308-DEL-174 |
| chr1-185886800-DEL-1380 | chr1-185886801-DEL-1380 |
| chr1-185891422-DEL-61 | chr1-185891423-DEL-61 |
| chr1-185916052-DEL-68 | chr1-185916025-DEL-68 |
| chr1-186018854-DEL-70 | chr1-186018855-DEL-70 |
| chr1-186069236-DEL-464 | chr1-186069234-DEL-462 |
| chr1-186110800-DEL-58 | chr1-186110801-DEL-58 |
| chr1-186213703-DEL-59 | chr1-186213704-DEL-59 |
| chr1-186216412-DEL-86 | chr1-186216429-DEL-107 |
| chr1-186219208-DEL-69 | chr1-186219166-DEL-98 |
| chr1-186253619-DEL-55 | chr1-186253599-DEL-56 |
| chr1-186298206-DEL-506 | chr1-186298207-DEL-506 |
| chr1-186324112-DEL-6228 | chr1-186324125-DEL-6229 |
| chr1-186351666-DEL-423 | chr1-186351667-DEL-423 |
| chr1-186430556-DEL-96 | chr1-186430557-DEL-96 |
| chr1-186441943-DEL-262 | chr1-186441945-DEL-218 |
| chr1-186457878-DEL-9295 | chr1-186457879-DEL-9295 |
| chr1-186498913-DEL-793 | chr1-186498914-DEL-793 |
| chr1-186601703-DEL-6454 | chr1-186601704-DEL-6454 |

| | |
|---|---|
| chr1-186611073-DEL-162 | chr1-186611076-DEL-160 |
| chr1-186634419-DEL-351 | chr1-186634420-DEL-351 |
| chr1-186656008-DEL-115 | chr1-186656007-DEL-116 |
| chr1-186680206-DEL-347 | chr1-186680207-DEL-347 |
| chr1-186692524-DEL-61 | chr1-186692525-DEL-61 |
| chr1-186695459-DEL-568 | chr1-186695460-DEL-568 |
| chr1-186707884-DEL-222 | chr1-186707885-DEL-222 |
| chr1-186723379-DEL-73 | chr1-186723380-DEL-73 |
| chr1-186794833-DEL-6679 | chr1-186794834-DEL-6679 |
| chr1-186844362-DEL-146 | chr1-186844363-DEL-146 |
| chr1-186888716-DEL-467 | chr1-186888713-DEL-468 |
| chr1-186891253-DEL-73 | chr1-186891267-DEL-60 |
| chr1-186923613-DEL-159 | chr1-186923614-DEL-159 |
| chr1-186945613-DEL-165 | chr1-186945614-DEL-165 |
| chr1-186948700-DEL-225 | chr1-186948701-DEL-225 |
| chr1-186951061-DEL-85 | chr1-186951057-DEL-86 |
| chr1-186953922-DEL-60 | chr1-186953923-DEL-60 |
| chr1-186971328-DEL-197 | chr1-186971329-DEL-197 |
| chr1-186985111-DEL-61 | chr1-186985087-DEL-62 |
| chr1-186995982-DEL-57 | chr1-186995983-DEL-57 |
| chr1-187014370-DEL-95 | chr1-187014346-DEL-72 |
| chr1-187046729-DEL-59 | chr1-187046715-DEL-50 |
| chr1-187085931-DEL-744 | chr1-187085932-DEL-744 |
| chr1-187105622-DEL-95 | chr1-187105614-DEL-89 |
| chr1-187124927-DEL-7235 | chr1-187124928-DEL-7235 |
| chr1-187162579-DEL-968 | chr1-187162580-DEL-968 |
| chr1-187209196-DEL-51 | chr1-187209197-DEL-51 |
| chr1-187225774-DEL-233 | chr1-187225772-DEL-234 |
| chr1-187225830-DEL-717 | chr1-187225831-DEL-717 |
| chr1-187255642-DEL-1340 | chr1-187255643-DEL-1340 |
| chr1-187273541-DEL-99 | chr1-187273542-DEL-99 |
| chr1-187331601-DEL-58 | chr1-187331603-DEL-58 |
| chr1-187351620-DEL-600 | chr1-187351605-DEL-575 |
| chr1-187410803-DEL-86 | chr1-187410804-DEL-86 |
| chr1-187428865-DEL-51 | chr1-187428866-DEL-51 |
| chr1-187498132-DEL-56 | chr1-187498133-DEL-56 |
| chr1-187568244-DEL-60 | chr1-187568245-DEL-60 |
| chr1-187660653-DEL-126 | chr1-187660654-DEL-126 |
| chr1-187720654-DEL-419 | chr1-187720649-DEL-421 |
| chr1-187872512-DEL-543 | chr1-187872513-DEL-543 |
| chr1-187914436-DEL-239 | chr1-187914437-DEL-239 |
| chr1-187963007-DEL-367 | chr1-187963008-DEL-367 |
| chr1-188007115-DEL-103 | chr1-188007116-DEL-102 |
| chr1-188041622-DEL-310 | chr1-188041623-DEL-310 |
| chr1-188054480-DEL-1231 | chr1-188054481-DEL-1231 |
| chr1-188056473-DEL-78 | chr1-188056474-DEL-78 |
| chr1-188056494-DEL-1178 | chr1-188056495-DEL-1178 |
| chr1-188067937-DEL-6955 | chr1-188067938-DEL-6955 |
| chr1-188106346-DEL-1524 | chr1-188106347-DEL-1524 |
| chr1-188157573-DEL-72 | chr1-188157574-DEL-72 |
| chr1-188185282-DEL-313 | chr1-188185283-DEL-313 |
| chr1-188197973-DEL-108 | chr1-188198017-DEL-81 |
| chr1-188202747-DEL-252 | chr1-188202748-DEL-252 |
| chr1-188225051-DEL-1218 | chr1-188225052-DEL-1218 |
| chr1-188267531-DEL-126 | chr1-188267532-DEL-126 |
| chr1-188275315-DEL-359 | chr1-188275316-DEL-359 |
| chr1-188339068-DEL-64 | chr1-188339068-DEL-64 |
| chr1-188340239-DEL-6763 | chr1-188340238-DEL-6763 |
| chr1-188360560-DEL-62 | chr1-188360561-DEL-62 |
| chr1-188382155-DEL-58 | chr1-188382156-DEL-58 |
| chr1-188406184-DEL-384 | chr1-188406185-DEL-384 |
| chr1-188425103-DEL-58 | chr1-188425104-DEL-58 |
| chr1-188451573-DEL-84 | chr1-188451574-DEL-84 |
| chr1-188463605-DEL-125 | chr1-188463606-DEL-125 |
| chr1-188483794-DEL-428 | chr1-188483795-DEL-428 |
| chr1-188484829-DEL-405 | chr1-188484830-DEL-405 |
| chr1-188523654-DEL-118 | chr1-188523655-DEL-118 |
| chr1-188525192-DEL-99 | chr1-188525194-DEL-99 |
| chr1-188525618-DEL-710 | chr1-188525605-DEL-758 |
| chr1-188539014-DEL-68 | chr1-188539015-DEL-68 |
| chr1-188540131-DEL-133 | chr1-188540132-DEL-133 |
| chr1-188555039-DEL-150 | chr1-188555039-DEL-147 |
| chr1-188577745-DEL-81 | chr1-188577746-DEL-81 |
| chr1-188619817-DEL-108 | chr1-188619807-DEL-79 |
| chr1-188622993-DEL-342 | chr1-188622991-DEL-378 |
| chr1-188655601-DEL-6473 | chr1-188655602-DEL-6473 |
| chr1-188676101-DEL-74 | chr1-188676102-DEL-82 |
| chr1-188690473-DEL-1287 | chr1-188690474-DEL-1287 |
| chr1-188697970-DEL-386 | chr1-188697971-DEL-386 |
| chr1-188714851-DEL-102 | chr1-188714851-DEL-103 |

| | |
|---|---|
| chr1-188722561-DEL-205 | chr1-188722562-DEL-205 |
| chr1-188729700-DEL-215 | chr1-188729701-DEL-215 |
| chr1-188754182-DEL-468 | chr1-188754183-DEL-468 |
| chr1-188788846-DEL-6136 | chr1-188788847-DEL-6136 |
| chr1-188804448-DEL-60 | chr1-188804449-DEL-60 |
| chr1-188834912-DEL-98 | chr1-188834912-DEL-111 |
| chr1-188851380-DEL-88 | chr1-188851381-DEL-88 |
| chr1-188891636-DEL-435 | chr1-188891637-DEL-435 |
| chr1-188926742-DEL-539 | chr1-188926743-DEL-539 |
| chr1-188964879-DEL-57 | chr1-188964880-DEL-52 |
| chr1-188971935-DEL-1890 | chr1-188971937-DEL-1889 |
| chr1-188980571-DEL-64 | chr1-188980575-DEL-57 |
| chr1-189001170-DEL-1445 | chr1-189001171-DEL-1445 |
| chr1-189009496-DEL-544 | chr1-189009505-DEL-545 |
| chr1-189052354-DEL-78 | chr1-189052355-DEL-74 |
| chr1-189061540-DEL-72 | chr1-189061547-DEL-66 |
| chr1-189113705-DEL-132 | chr1-189113706-DEL-132 |
| chr1-189122675-DEL-6755 | chr1-189122676-DEL-6755 |
| chr1-189177912-DEL-9377 | chr1-189177929-DEL-9365 |
| chr1-189183980-DEL-3127 | chr1-189183981-DEL-3127 |
| chr1-189186073-DEL-73 | chr1-189186074-DEL-73 |
| chr1-189188357-DEL-265 | chr1-189188322-DEL-301 |
| chr1-189198913-DEL-91 | chr1-189198914-DEL-91 |
| chr1-189214587-DEL-341 | chr1-189214620-DEL-357 |
| chr1-189224834-DEL-892 | chr1-189224835-DEL-892 |
| chr1-189249201-DEL-54 | chr1-189249183-DEL-72 |
| chr1-189255676-DEL-51 | chr1-189255677-DEL-51 |
| chr1-189265985-DEL-2416 | chr1-189265986-DEL-2416 |
| chr1-189269365-DEL-179 | chr1-189269366-DEL-179 |
| chr1-189282489-DEL-220 | chr1-189282490-DEL-220 |
| chr1-189298162-DEL-240 | chr1-189298162-DEL-239 |
| chr1-189304229-DEL-62 | chr1-189304230-DEL-62 |
| chr1-189330871-DEL-221 | chr1-189330872-DEL-221 |
| chr1-189339958-DEL-81 | chr1-189339959-DEL-81 |
| chr1-189342124-DEL-136 | chr1-189342159-DEL-96 |
| chr1-189350405-DEL-95 | chr1-189350398-DEL-96 |
| chr1-189353115-DEL-65 | chr1-189353114-DEL-66 |
| chr1-189356270-DEL-299 | chr1-189356254-DEL-300 |
| chr1-189430211-DEL-117 | chr1-189430212-DEL-117 |
| chr1-189432961-DEL-148 | chr1-189432962-DEL-148 |
| chr1-189450033-DEL-203 | chr1-189450034-DEL-203 |
| chr1-189454215-DEL-218 | chr1-189454216-DEL-218 |
| chr1-189457134-DEL-108 | chr1-189457135-DEL-108 |
| chr1-189466258-DEL-719 | chr1-189466262-DEL-720 |
| chr1-189468268-DEL-123 | chr1-189468267-DEL-123 |
| chr1-189481331-DEL-172 | chr1-189481339-DEL-165 |
| chr1-189490825-DEL-95 | chr1-189490826-DEL-114 |
| chr1-189505544-DEL-94 | chr1-189505545-DEL-94 |
| chr1-189526100-DEL-72 | chr1-189526101-DEL-72 |
| chr1-189547481-DEL-72 | chr1-189547482-DEL-72 |
| chr1-189570150-DEL-69 | chr1-189570150-DEL-69 |
| chr1-189591590-DEL-202 | chr1-189591591-DEL-202 |
| chr1-189611895-DEL-112 | chr1-189611918-DEL-75 |
| chr1-189625335-DEL-68 | chr1-189625336-DEL-70 |
| chr1-189627366-DEL-91 | chr1-189627323-DEL-89 |
| chr1-189672692-DEL-213 | chr1-189672693-DEL-215 |
| chr1-189673470-DEL-51 | chr1-189673464-DEL-51 |
| chr1-189675804-DEL-184 | chr1-189675817-DEL-172 |
| chr1-189680632-DEL-78 | chr1-189680633-DEL-78 |
| chr1-189683226-DEL-83 | chr1-189683192-DEL-84 |
| chr1-189720806-DEL-152 | chr1-189720807-DEL-152 |
| chr1-189729687-DEL-54 | chr1-189729690-DEL-50 |
| chr1-189763906-DEL-813 | chr1-189763907-DEL-813 |
| chr1-189805859-DEL-109 | chr1-189805860-DEL-109 |
| chr1-189858571-DEL-92 | chr1-189858572-DEL-92 |
| chr1-189928833-DEL-52 | chr1-189928834-DEL-52 |
| chr1-189943848-DEL-67 | chr1-189943838-DEL-60 |
| chr1-189982678-DEL-76 | chr1-189982679-DEL-76 |
| chr1-190006449-DEL-99 | chr1-190006450-DEL-99 |
| chr1-190018480-DEL-77 | chr1-190018481-DEL-77 |
| chr1-190057297-DEL-498 | chr1-190057298-DEL-498 |
| chr1-190083159-DEL-1053 | chr1-190083160-DEL-1053 |
| chr1-190136891-DEL-56 | chr1-190136891-DEL-56 |
| chr1-190141927-DEL-136 | chr1-190141927-DEL-132 |
| chr1-190165305-DEL-64 | chr1-190165306-DEL-64 |
| chr1-190169908-DEL-164 | chr1-190169909-DEL-164 |
| chr1-190176070-DEL-145 | chr1-190176071-DEL-145 |
| chr1-190187381-DEL-1044 | chr1-190187382-DEL-1044 |
| chr1-190188478-DEL-119 | chr1-190188479-DEL-119 |
| chr1-190191482-DEL-690 | chr1-190191483-DEL-690 |

| | |
|---|---|
| chr1-190209212-DEL-211 | chr1-190209213-DEL-211 |
| chr1-190250097-DEL-59 | chr1-190250098-DEL-59 |
| chr1-190262093-DEL-57 | chr1-190262093-DEL-57 |
| chr1-190327780-DEL-52 | chr1-190327781-DEL-52 |
| chr1-190342528-DEL-126 | chr1-190342529-DEL-118 |
| chr1-190346447-DEL-2961 | chr1-190346448-DEL-2961 |
| chr1-190355153-DEL-138 | chr1-190355154-DEL-138 |
| chr1-190399461-DEL-71 | chr1-190399462-DEL-71 |
| chr1-190525506-DEL-195 | chr1-190525507-DEL-194 |
| chr1-190529744-DEL-121 | chr1-190529745-DEL-121 |
| chr1-190537142-DEL-135 | chr1-190537143-DEL-135 |
| chr1-190549270-DEL-62 | chr1-190549271-DEL-62 |
| chr1-190560148-DEL-87 | chr1-190560139-DEL-88 |
| chr1-190603205-DEL-55 | chr1-190603205-DEL-55 |
| chr1-190614521-DEL-777 | chr1-190614522-DEL-777 |
| chr1-190619595-DEL-70 | chr1-190619584-DEL-77 |
| chr1-190648712-DEL-95 | chr1-190648713-DEL-95 |
| chr1-190649457-DEL-187 | chr1-190649451-DEL-187 |
| chr1-190693250-DEL-151 | chr1-190693250-DEL-151 |
| chr1-190698635-DEL-262 | chr1-190698636-DEL-262 |
| chr1-190703497-DEL-161 | chr1-190703497-DEL-160 |
| chr1-190714942-DEL-2847 | chr1-190714943-DEL-2847 |
| chr1-190743024-DEL-210 | chr1-190743018-DEL-211 |
| chr1-190755100-DEL-160 | chr1-190755099-DEL-208 |
| chr1-190756212-DEL-200 | chr1-190756213-DEL-200 |
| chr1-190769923-DEL-56 | chr1-190769917-DEL-66 |
| chr1-190773245-DEL-213 | chr1-190773246-DEL-213 |
| chr1-190779188-DEL-204 | chr1-190779189-DEL-204 |
| chr1-190791581-DEL-167 | chr1-190791582-DEL-167 |
| chr1-190815833-DEL-53 | chr1-190815834-DEL-53 |
| chr1-190835517-DEL-70 | chr1-190835518-DEL-70 |
| chr1-190848785-DEL-233 | chr1-190848800-DEL-229 |
| chr1-190878911-DEL-1892 | chr1-190878912-DEL-1892 |
| chr1-191016516-DEL-75 | chr1-191016517-DEL-75 |
| chr1-191023264-DEL-203 | chr1-191023265-DEL-203 |
| chr1-191025960-DEL-62 | chr1-191025961-DEL-62 |
| chr1-191038069-DEL-87 | chr1-191038070-DEL-87 |
| chr1-191053690-DEL-463 | chr1-191053689-DEL-464 |
| chr1-191061523-DEL-244 | chr1-191061524-DEL-244 |
| chr1-191076912-DEL-134 | chr1-191076914-DEL-133 |
| chr1-191079462-DEL-200 | chr1-191079463-DEL-200 |
| chr1-191113103-DEL-104 | chr1-191113104-DEL-104 |
| chr1-191117550-DEL-3031 | chr1-191117550-DEL-3031 |
| chr1-191133962-DEL-203 | chr1-191133963-DEL-203 |
| chr1-191138316-DEL-210 | chr1-191138317-DEL-210 |
| chr1-191144295-DEL-200 | chr1-191144296-DEL-200 |
| chr1-191188883-DEL-265 | chr1-191188884-DEL-227 |
| chr1-191200892-DEL-105 | chr1-191200893-DEL-105 |
| chr1-191233693-DEL-2987 | chr1-191233693-DEL-2987 |
| chr1-191239856-DEL-60 | chr1-191239857-DEL-60 |
| chr1-191240279-DEL-230 | chr1-191240280-DEL-230 |
| chr1-191267692-DEL-149 | chr1-191267693-DEL-149 |
| chr1-191285352-DEL-229 | chr1-191285386-DEL-196 |
| chr1-191286380-DEL-211 | chr1-191286381-DEL-211 |
| chr1-191294273-DEL-204 | chr1-191294274-DEL-204 |
| chr1-191316732-DEL-964 | chr1-191316733-DEL-964 |
| chr1-191343313-DEL-215 | chr1-191343314-DEL-215 |
| chr1-191344479-DEL-105 | chr1-191344480-DEL-105 |
| chr1-191397099-DEL-184 | chr1-191397094-DEL-185 |
| chr1-191417252-DEL-716 | chr1-191417253-DEL-716 |
| chr1-191418098-DEL-650 | chr1-191418099-DEL-650 |
| chr1-191426882-DEL-59 | chr1-191426883-DEL-59 |
| chr1-191433969-DEL-70 | chr1-191433970-DEL-70 |
| chr1-191479025-DEL-625 | chr1-191479036-DEL-615 |
| chr1-191487316-DEL-176 | chr1-191487317-DEL-176 |
| chr1-191546090-DEL-92 | chr1-191546074-DEL-93 |
| chr1-191586249-DEL-106 | chr1-191586250-DEL-106 |
| chr1-191601447-DEL-92 | chr1-191601448-DEL-92 |
| chr1-191602272-DEL-1891 | chr1-191602273-DEL-1891 |
| chr1-191654557-DEL-422 | chr1-191654558-DEL-421 |
| chr1-191665720-DEL-8985 | chr1-191665721-DEL-8985 |
| chr1-191682139-DEL-60 | chr1-191682139-DEL-60 |
| chr1-191698510-DEL-123 | chr1-191698510-DEL-123 |
| chr1-191703926-DEL-200 | chr1-191703927-DEL-200 |
| chr1-191758022-DEL-804 | chr1-191758023-DEL-804 |
| chr1-191764893-DEL-51 | chr1-191764894-DEL-51 |
| chr1-191839717-DEL-4814 | chr1-191839718-DEL-4814 |
| chr1-191862760-DEL-292 | chr1-191862757-DEL-295 |
| chr1-191881449-DEL-811 | chr1-191881450-DEL-811 |
| chr1-191888193-DEL-58 | chr1-191888202-DEL-50 |

| | |
|---|---|
| chr1-191889497-DEL-50 | chr1-191889530-DEL-52 |
| chr1-191892407-DEL-87 | chr1-191892411-DEL-88 |
| chr1-191901827-DEL-187 | chr1-191901828-DEL-187 |
| chr1-191906606-DEL-61 | chr1-191906607-DEL-61 |
| chr1-191912222-DEL-1680 | chr1-191912222-DEL-1680 |
| chr1-191912507-DEL-384 | chr1-191912508-DEL-384 |
| chr1-191925815-DEL-221 | chr1-191925816-DEL-221 |
| chr1-191934117-DEL-204 | chr1-191934118-DEL-204 |
| chr1-191937661-DEL-112 | chr1-191937662-DEL-112 |
| chr1-191938893-DEL-936 | chr1-191938894-DEL-936 |
| chr1-191990828-DEL-726 | chr1-191990829-DEL-726 |
| chr1-191996373-DEL-118 | chr1-191996374-DEL-118 |
| chr1-192013266-DEL-209 | chr1-192013267-DEL-209 |
| chr1-192049369-DEL-892 | chr1-192049370-DEL-892 |
| chr1-192057508-DEL-96 | chr1-192057508-DEL-96 |
| chr1-192091448-DEL-212 | chr1-192091450-DEL-213 |
| chr1-192123375-DEL-80 | chr1-192123376-DEL-80 |
| chr1-192124404-DEL-406 | chr1-192124405-DEL-406 |
| chr1-192132900-DEL-80 | chr1-192132899-DEL-50 |
| chr1-192175272-DEL-52 | chr1-192175273-DEL-52 |
| chr1-192195948-DEL-64 | chr1-192195949-DEL-64 |
| chr1-192235914-DEL-656 | chr1-192235915-DEL-656 |
| chr1-192251874-DEL-400 | chr1-192251869-DEL-401 |
| chr1-192253891-DEL-6536 | chr1-192253892-DEL-6536 |
| chr1-192308502-DEL-7505 | chr1-192308503-DEL-7505 |
| chr1-192340144-DEL-58 | chr1-192340156-DEL-59 |
| chr1-192341780-DEL-245 | chr1-192341781-DEL-245 |
| chr1-192348068-DEL-262 | chr1-192348069-DEL-262 |
| chr1-192368278-DEL-96 | chr1-192368278-DEL-94 |
| chr1-192384311-DEL-340 | chr1-192384312-DEL-340 |
| chr1-192398248-DEL-72 | chr1-192398227-DEL-122 |
| chr1-192410954-DEL-118 | chr1-192410955-DEL-118 |
| chr1-192473385-DEL-168 | chr1-192473386-DEL-168 |
| chr1-192492729-DEL-79 | chr1-192492726-DEL-55 |
| chr1-192600704-DEL-168 | chr1-192600705-DEL-168 |
| chr1-192603565-DEL-80 | chr1-192603573-DEL-80 |
| chr1-192621688-DEL-75 | chr1-192621689-DEL-75 |
| chr1-192638419-DEL-2379 | chr1-192638420-DEL-2379 |
| chr1-192653171-DEL-89 | chr1-192653172-DEL-84 |
| chr1-192711137-DEL-237 | chr1-192711138-DEL-237 |
| chr1-192752915-DEL-7088 | chr1-192752916-DEL-7088 |
| chr1-192806270-DEL-143 | chr1-192806271-DEL-143 |
| chr1-192821386-DEL-87 | chr1-192821387-DEL-87 |
| chr1-192890521-DEL-2110 | chr1-192890522-DEL-2110 |
| chr1-192892379-DEL-198 | chr1-192892380-DEL-198 |
| chr1-192910118-DEL-107 | chr1-192910117-DEL-108 |
| chr1-192915948-DEL-76 | chr1-192915949-DEL-76 |
| chr1-192929491-DEL-76 | chr1-192929496-DEL-72 |
| chr1-192945090-DEL-103 | chr1-192945091-DEL-103 |
| chr1-192962336-DEL-70 | chr1-192962337-DEL-70 |
| chr1-192974411-DEL-186 | chr1-192974412-DEL-186 |
| chr1-192975912-DEL-66 | chr1-192975913-DEL-66 |
| chr1-192985083-DEL-60 | chr1-192985083-DEL-60 |
| chr1-193056004-DEL-60 | chr1-193056005-DEL-60 |
| chr1-193070621-DEL-65 | chr1-193070622-DEL-65 |
| chr1-193089284-DEL-110 | chr1-193089285-DEL-110 |
| chr1-193129484-DEL-229 | chr1-193129485-DEL-229 |
| chr1-193141244-DEL-69 | chr1-193141242-DEL-103 |
| chr1-193152221-DEL-1909 | chr1-193152222-DEL-1909 |
| chr1-193177304-DEL-52 | chr1-193177313-DEL-51 |
| chr1-193183607-DEL-234 | chr1-193183627-DEL-260 |
| chr1-193215426-DEL-218 | chr1-193215427-DEL-218 |
| chr1-193235086-DEL-123 | chr1-193235077-DEL-133 |
| chr1-193351102-DEL-980 | chr1-193351102-DEL-980 |
| chr1-193421154-DEL-75 | chr1-193421155-DEL-75 |
| chr1-193433848-DEL-56 | chr1-193433849-DEL-56 |
| chr1-193486192-DEL-6247 | chr1-193486193-DEL-6247 |
| chr1-193552834-DEL-194 | chr1-193552835-DEL-194 |
| chr1-193577771-DEL-57 | chr1-193577772-DEL-57 |
| chr1-193584558-DEL-87 | chr1-193584559-DEL-87 |
| chr1-193703442-DEL-62 | chr1-193703440-DEL-52 |
| chr1-193753695-DEL-4448 | chr1-193753696-DEL-4448 |
| chr1-193762675-DEL-6556 | chr1-193762676-DEL-6556 |
| chr1-193806670-DEL-352 | chr1-193806671-DEL-352 |
| chr1-193810242-DEL-559 | chr1-193810241-DEL-561 |
| chr1-193819740-DEL-143 | chr1-193819741-DEL-143 |
| chr1-193827177-DEL-581 | chr1-193827177-DEL-581 |
| chr1-194007487-DEL-85 | chr1-194007503-DEL-50 |
| chr1-194099935-DEL-62 | chr1-194099928-DEL-58 |
| chr1-194109549-DEL-431 | chr1-194109550-DEL-431 |

| | |
|---|---|
| chr1-194111022-DEL-73 | chr1-194111022-DEL-73 |
| chr1-194225059-DEL-245 | chr1-194225060-DEL-245 |
| chr1-194285389-DEL-52 | chr1-194285386-DEL-60 |
| chr1-194336123-DEL-3082 | chr1-194336124-DEL-3082 |
| chr1-194373568-DEL-55 | chr1-194373569-DEL-55 |
| chr1-194552210-DEL-160 | chr1-194552204-DEL-148 |
| chr1-194628276-DEL-67 | chr1-194628277-DEL-67 |
| chr1-194720239-DEL-67 | chr1-194720240-DEL-67 |
| chr1-194731998-DEL-6997 | chr1-194731999-DEL-6997 |
| chr1-194777982-DEL-415 | chr1-194777983-DEL-415 |
| chr1-194821673-DEL-51 | chr1-194821674-DEL-51 |
| chr1-194887455-DEL-6822 | chr1-194887456-DEL-6822 |
| chr1-194963954-DEL-406 | chr1-194963955-DEL-406 |
| chr1-195002589-DEL-7165 | chr1-195002590-DEL-7165 |
| chr1-195018272-DEL-11627 | chr1-195018271-DEL-11627 |
| chr1-195027700-DEL-201 | chr1-195027701-DEL-201 |
| chr1-195041562-DEL-64 | chr1-195041558-DEL-68 |
| chr10-3191820-DEL-13194 | chr10-3191851-DEL-13169 |
| chr10-3225227-DEL-1321 | chr10-3225228-DEL-1321 |
| chr10-3289781-DEL-64 | chr10-3289784-DEL-62 |
| chr10-3337698-DEL-156 | chr10-3337699-DEL-156 |
| chr10-3375089-DEL-84 | chr10-3375091-DEL-89 |
| chr10-3390538-DEL-88 | chr10-3390538-DEL-132 |
| chr10-3427163-DEL-108 | chr10-3427164-DEL-108 |
| chr10-3450241-DEL-238 | chr10-3450242-DEL-238 |
| chr10-3477083-DEL-205 | chr10-3477084-DEL-205 |
| chr10-3485637-DEL-593 | chr10-3485635-DEL-594 |
| chr10-3486461-DEL-516 | chr10-3486462-DEL-516 |
| chr10-3521725-DEL-1322 | chr10-3521726-DEL-1322 |
| chr10-3526829-DEL-6264 | chr10-3526830-DEL-6264 |
| chr10-3550937-DEL-7089 | chr10-3550938-DEL-7089 |
| chr10-3583576-DEL-71 | chr10-3583577-DEL-71 |
| chr10-3619351-DEL-182 | chr10-3619352-DEL-182 |
| chr10-3635065-DEL-240 | chr10-3635066-DEL-216 |
| chr10-3644491-DEL-179 | chr10-3644492-DEL-179 |
| chr10-3652596-DEL-61 | chr10-3652597-DEL-61 |
| chr10-3658839-DEL-164 | chr10-3658818-DEL-189 |
| chr10-3710970-DEL-108 | chr10-3710942-DEL-88 |
| chr10-3713587-DEL-74 | chr10-3713588-DEL-58 |
| chr10-3790952-DEL-212 | chr10-3790953-DEL-212 |
| chr10-3820347-DEL-74 | chr10-3820325-DEL-50 |
| chr10-3830891-DEL-51 | chr10-3830884-DEL-59 |
| chr10-3884838-DEL-56 | chr10-3884839-DEL-56 |
| chr10-3885356-DEL-203 | chr10-3885357-DEL-203 |
| chr10-3889204-DEL-76 | chr10-3889205-DEL-65 |
| chr10-3912340-DEL-90 | chr10-3912341-DEL-92 |
| chr10-3950804-DEL-53 | chr10-3950805-DEL-53 |
| chr10-3953785-DEL-88 | chr10-3953810-DEL-63 |
| chr10-3955918-DEL-72 | chr10-3955919-DEL-72 |
| chr10-3976225-DEL-73 | chr10-3976226-DEL-73 |
| chr10-3985865-DEL-184 | chr10-3985864-DEL-186 |
| chr10-4003145-DEL-83 | chr10-4003134-DEL-92 |
| chr10-4005397-DEL-180 | chr10-4005398-DEL-180 |
| chr10-4006329-DEL-175 | chr10-4006330-DEL-175 |
| chr10-4026320-DEL-68 | chr10-4026321-DEL-68 |
| chr10-4026504-DEL-915 | chr10-4026505-DEL-915 |
| chr10-4042210-DEL-172 | chr10-4042211-DEL-172 |
| chr10-4065351-DEL-583 | chr10-4065354-DEL-581 |
| chr10-4067440-DEL-60 | chr10-4067441-DEL-60 |
| chr10-4081820-DEL-65 | chr10-4081827-DEL-58 |
| chr10-4097065-DEL-224 | chr10-4097065-DEL-224 |
| chr10-4111873-DEL-138 | chr10-4111874-DEL-156 |
| chr10-4141094-DEL-6441 | chr10-4141095-DEL-6441 |
| chr10-4150254-DEL-56 | chr10-4150255-DEL-56 |
| chr10-4151001-DEL-197 | chr10-4151002-DEL-197 |
| chr10-4152232-DEL-208 | chr10-4152233-DEL-208 |
| chr10-4153799-DEL-336 | chr10-4153800-DEL-336 |
| chr10-4164780-DEL-563 | chr10-4164781-DEL-563 |
| chr10-4165951-DEL-265 | chr10-4165952-DEL-265 |
| chr10-4168753-DEL-87 | chr10-4168754-DEL-87 |
| chr10-4170297-DEL-259 | chr10-4170298-DEL-259 |
| chr10-4174453-DEL-170 | chr10-4174456-DEL-168 |
| chr10-4178417-DEL-56 | chr10-4178418-DEL-56 |
| chr10-4180211-DEL-174 | chr10-4180212-DEL-174 |
| chr10-4202872-DEL-186 | chr10-4202873-DEL-186 |
| chr10-4204050-DEL-147 | chr10-4204071-DEL-140 |
| chr10-4225286-DEL-213 | chr10-4225287-DEL-213 |
| chr10-4230897-DEL-56 | chr10-4230897-DEL-60 |
| chr10-4237873-DEL-165 | chr10-4237874-DEL-165 |
| chr10-4249520-DEL-53 | chr10-4249521-DEL-53 |

| | |
|---|---|
| chr10-4255375-DEL-212 | chr10-4255379-DEL-213 |
| chr10-4264882-DEL-137 | chr10-4264883-DEL-137 |
| chr10-4270097-DEL-354 | chr10-4270098-DEL-354 |
| chr10-4273343-DEL-58 | chr10-4273344-DEL-58 |
| chr10-4280880-DEL-200 | chr10-4280881-DEL-200 |
| chr10-4289852-DEL-52 | chr10-4289851-DEL-51 |
| chr10-4295080-DEL-173 | chr10-4295081-DEL-173 |
| chr10-4301194-DEL-211 | chr10-4301195-DEL-211 |
| chr10-4332397-DEL-102 | chr10-4332398-DEL-102 |
| chr10-4349574-DEL-1340 | chr10-4349575-DEL-1340 |
| chr10-4351343-DEL-205 | chr10-4351344-DEL-205 |
| chr10-4354831-DEL-159 | chr10-4354832-DEL-159 |
| chr10-4384407-DEL-71 | chr10-4384424-DEL-71 |
| chr10-4393129-DEL-3579 | chr10-4393130-DEL-3579 |
| chr10-4402353-DEL-67 | chr10-4402365-DEL-54 |
| chr10-4412089-DEL-477 | chr10-4412090-DEL-477 |
| chr10-4416582-DEL-55 | chr10-4416547-DEL-55 |
| chr10-4439101-DEL-202 | chr10-4439102-DEL-202 |
| chr10-4440501-DEL-868 | chr10-4440502-DEL-868 |
| chr10-4445968-DEL-91 | chr10-4445969-DEL-95 |
| chr10-4450054-DEL-217 | chr10-4450054-DEL-217 |
| chr10-4452809-DEL-649 | chr10-4452810-DEL-649 |
| chr10-4457314-DEL-78 | chr10-4457339-DEL-54 |
| chr10-4477189-DEL-63 | chr10-4477202-DEL-70 |
| chr10-4482059-DEL-62 | chr10-4482060-DEL-62 |
| chr10-4545660-DEL-1550 | chr10-4545661-DEL-1550 |
| chr10-4547926-DEL-176 | chr10-4547927-DEL-176 |
| chr10-4552672-DEL-66 | chr10-4552673-DEL-66 |
| chr10-4567482-DEL-124 | chr10-4567483-DEL-124 |
| chr10-4574064-DEL-95 | chr10-4574070-DEL-77 |
| chr10-4593049-DEL-474 | chr10-4593050-DEL-474 |
| chr10-4600971-DEL-52 | chr10-4600972-DEL-52 |
| chr10-4622649-DEL-256 | chr10-4622604-DEL-257 |
| chr10-4624506-DEL-51 | chr10-4624507-DEL-51 |
| chr10-4627112-DEL-92 | chr10-4627110-DEL-92 |
| chr10-4651553-DEL-162 | chr10-4651554-DEL-162 |
| chr10-4654300-DEL-62 | chr10-4654301-DEL-62 |
| chr10-4680218-DEL-78 | chr10-4680222-DEL-75 |
| chr10-4711727-DEL-1290 | chr10-4711727-DEL-1290 |
| chr10-4712530-DEL-2093 | chr10-4712531-DEL-2093 |
| chr10-4729348-DEL-108 | chr10-4729349-DEL-108 |
| chr10-4761177-DEL-52 | chr10-4761176-DEL-54 |
| chr10-4780067-DEL-205 | chr10-4780068-DEL-205 |
| chr10-4781511-DEL-6419 | chr10-4781512-DEL-6419 |
| chr10-4822917-DEL-56 | chr10-4822934-DEL-52 |
| chr10-4861268-DEL-7283 | chr10-4861247-DEL-7306 |
| chr10-4861857-DEL-636 | chr10-4861858-DEL-636 |
| chr10-4885206-DEL-2836 | chr10-4885214-DEL-2820 |
| chr10-4913892-DEL-61 | chr10-4913893-DEL-61 |
| chr10-4965434-DEL-68 | chr10-4965434-DEL-66 |
| chr10-5008564-DEL-134 | chr10-5008573-DEL-114 |
| chr10-5011041-DEL-76 | chr10-5011042-DEL-76 |
| chr10-5039768-DEL-6879 | chr10-5039769-DEL-6879 |
| chr10-5039828-DEL-423 | chr10-5039816-DEL-424 |
| chr10-5063709-DEL-217 | chr10-5063710-DEL-217 |
| chr10-5072116-DEL-53 | chr10-5072117-DEL-53 |
| chr10-5116504-DEL-119 | chr10-5116510-DEL-109 |
| chr10-5130805-DEL-223 | chr10-5130806-DEL-217 |
| chr10-5159075-DEL-54 | chr10-5159072-DEL-58 |
| chr10-5162532-DEL-105 | chr10-5162533-DEL-56 |
| chr10-5168975-DEL-54 | chr10-5168989-DEL-52 |
| chr10-5180656-DEL-130 | chr10-5180691-DEL-138 |
| chr10-5192187-DEL-432 | chr10-5192188-DEL-432 |
| chr10-5256366-DEL-207 | chr10-5256363-DEL-207 |
| chr10-5298373-DEL-115 | chr10-5298375-DEL-114 |
| chr10-5320969-DEL-197 | chr10-5320970-DEL-197 |
| chr10-5336407-DEL-2189 | chr10-5336407-DEL-2189 |
| chr10-5338656-DEL-6808 | chr10-5338657-DEL-6808 |
| chr10-5432631-DEL-52 | chr10-5432652-DEL-69 |
| chr10-5487606-DEL-830 | chr10-5487607-DEL-830 |
| chr10-5489471-DEL-162 | chr10-5489472-DEL-162 |
| chr10-5530173-DEL-6401 | chr10-5530174-DEL-6401 |
| chr10-5546755-DEL-175 | chr10-5546756-DEL-174 |
| chr10-5550897-DEL-387 | chr10-5550915-DEL-389 |
| chr10-5557234-DEL-740 | chr10-5557235-DEL-738 |
| chr10-5574196-DEL-220 | chr10-5574197-DEL-220 |
| chr10-5615379-DEL-206 | chr10-5615380-DEL-206 |
| chr10-5636385-DEL-582 | chr10-5636386-DEL-582 |
| chr10-5644972-DEL-61 | chr10-5644973-DEL-61 |
| chr10-5647433-DEL-63 | chr10-5647434-DEL-63 |

| | |
|---|---|
| chr10-5662388-DEL-66 | chr10-5662389-DEL-63 |
| chr10-5664480-DEL-109 | chr10-5664481-DEL-84 |
| chr10-5665927-DEL-349 | chr10-5665928-DEL-349 |
| chr10-5667712-DEL-9043 | chr10-5667713-DEL-9043 |
| chr10-5677346-DEL-92 | chr10-5677347-DEL-92 |
| chr10-5696613-DEL-403 | chr10-5696614-DEL-403 |
| chr10-5717526-DEL-486 | chr10-5717527-DEL-486 |
| chr10-5724397-DEL-175 | chr10-5724398-DEL-175 |
| chr10-5749076-DEL-60 | chr10-5749068-DEL-56 |
| chr10-5766375-DEL-209 | chr10-5766369-DEL-210 |
| chr10-5801484-DEL-615 | chr10-5801485-DEL-615 |
| chr10-5819339-DEL-75 | chr10-5819340-DEL-75 |
| chr10-5842203-DEL-1607 | chr10-5842204-DEL-1607 |
| chr10-5860990-DEL-502 | chr10-5860991-DEL-502 |
| chr10-5877767-DEL-402 | chr10-5877768-DEL-402 |
| chr10-5880216-DEL-1589 | chr10-5880216-DEL-1589 |
| chr10-5891462-DEL-52 | chr10-5891479-DEL-56 |
| chr10-5971414-DEL-95 | chr10-5971415-DEL-95 |
| chr10-6030153-DEL-8813 | chr10-6030154-DEL-8813 |
| chr10-6036235-DEL-815 | chr10-6036235-DEL-815 |
| chr10-6049980-DEL-531 | chr10-6049981-DEL-531 |
| chr10-6054426-DEL-1431 | chr10-6054427-DEL-1431 |
| chr10-6056901-DEL-486 | chr10-6056902-DEL-486 |
| chr10-6071219-DEL-189 | chr10-6071220-DEL-189 |
| chr10-6088001-DEL-179 | chr10-6088002-DEL-179 |
| chr10-6101127-DEL-164 | chr10-6101125-DEL-165 |
| chr10-6119591-DEL-6360 | chr10-6119592-DEL-6360 |
| chr10-6142306-DEL-987 | chr10-6142307-DEL-987 |
| chr10-6147425-DEL-416 | chr10-6147426-DEL-416 |
| chr10-6167784-DEL-54 | chr10-6167785-DEL-54 |
| chr10-6171236-DEL-509 | chr10-6171237-DEL-509 |
| chr10-6179152-DEL-707 | chr10-6179153-DEL-707 |
| chr10-6256784-DEL-64 | chr10-6256785-DEL-64 |
| chr10-6265181-DEL-317 | chr10-6265182-DEL-317 |
| chr10-6276015-DEL-52 | chr10-6275984-DEL-76 |
| chr10-6288281-DEL-97 | chr10-6288232-DEL-127 |
| chr10-6293827-DEL-8330 | chr10-6293828-DEL-8330 |
| chr10-6314659-DEL-65 | chr10-6314660-DEL-65 |
| chr10-6323861-DEL-96 | chr10-6323862-DEL-90 |
| chr10-6325076-DEL-61 | chr10-6325071-DEL-61 |
| chr10-6334071-DEL-85 | chr10-6334072-DEL-94 |
| chr10-6364239-DEL-382 | chr10-6364240-DEL-382 |
| chr10-6367259-DEL-837 | chr10-6367260-DEL-837 |
| chr10-6393024-DEL-263 | chr10-6393027-DEL-260 |
| chr10-6393964-DEL-131 | chr10-6393965-DEL-131 |
| chr10-6395355-DEL-158 | chr10-6395360-DEL-123 |
| chr10-6399464-DEL-5969 | chr10-6399465-DEL-5969 |
| chr10-6411642-DEL-229 | chr10-6411643-DEL-229 |
| chr10-6429779-DEL-273 | chr10-6429804-DEL-249 |
| chr10-6449925-DEL-92 | chr10-6449926-DEL-92 |
| chr10-6464482-DEL-51 | chr10-6464482-DEL-51 |
| chr10-6473761-DEL-929 | chr10-6473762-DEL-929 |
| chr10-6481035-DEL-390 | chr10-6481036-DEL-390 |
| chr10-6485031-DEL-1666 | chr10-6485030-DEL-1666 |
| chr10-6494504-DEL-670 | chr10-6494505-DEL-670 |
| chr10-6499639-DEL-328 | chr10-6499640-DEL-328 |
| chr10-6510805-DEL-116 | chr10-6510806-DEL-116 |
| chr10-6516697-DEL-4895 | chr10-6516698-DEL-4895 |
| chr10-6528071-DEL-434 | chr10-6528072-DEL-434 |
| chr10-6549390-DEL-131 | chr10-6549401-DEL-122 |
| chr10-6596957-DEL-81 | chr10-6596946-DEL-90 |
| chr10-6604594-DEL-6541 | chr10-6604595-DEL-6541 |
| chr10-6632243-DEL-749 | chr10-6632245-DEL-746 |
| chr10-6648092-DEL-82 | chr10-6648126-DEL-85 |
| chr10-6691611-DEL-1729 | chr10-6691596-DEL-1730 |
| chr10-6698439-DEL-56 | chr10-6698447-DEL-59 |
| chr10-6755097-DEL-153 | chr10-6755084-DEL-167 |
| chr10-6764402-DEL-71 | chr10-6764403-DEL-71 |
| chr10-6775410-DEL-52 | chr10-6775411-DEL-52 |
| chr10-6783528-DEL-81 | chr10-6783508-DEL-66 |
| chr10-6809384-DEL-54 | chr10-6809388-DEL-50 |
| chr10-6824356-DEL-53 | chr10-6824352-DEL-54 |
| chr10-6846251-DEL-201 | chr10-6846215-DEL-203 |
| chr10-6849629-DEL-58 | chr10-6849630-DEL-58 |
| chr10-6875449-DEL-226 | chr10-6875450-DEL-226 |
| chr10-6889313-DEL-225 | chr10-6889314-DEL-225 |
| chr10-6951975-DEL-583 | chr10-6951973-DEL-583 |
| chr10-6954070-DEL-105 | chr10-6954071-DEL-105 |
| chr10-6954691-DEL-74 | chr10-6954723-DEL-58 |
| chr10-6964225-DEL-80 | chr10-6964226-DEL-80 |

| | |
|---|---|
| chr10-6964811-DEL-146 | chr10-6964810-DEL-147 |
| chr10-6988654-DEL-187 | chr10-6988655-DEL-187 |
| chr10-7073450-DEL-84 | chr10-7073467-DEL-68 |
| chr10-7091850-DEL-81 | chr10-7091851-DEL-81 |
| chr10-7150213-DEL-157 | chr10-7150214-DEL-157 |
| chr10-7166962-DEL-271 | chr10-7166963-DEL-271 |
| chr10-7182633-DEL-66 | chr10-7182621-DEL-51 |
| chr10-7191282-DEL-2697 | chr10-7191288-DEL-2699 |
| chr10-7235422-DEL-73 | chr10-7235406-DEL-74 |
| chr10-7236856-DEL-2991 | chr10-7236858-DEL-2992 |
| chr10-7239306-DEL-124 | chr10-7239307-DEL-124 |
| chr10-7253716-DEL-64 | chr10-7253703-DEL-78 |
| chr10-7277852-DEL-457 | chr10-7277853-DEL-457 |
| chr10-7278885-DEL-2664 | chr10-7278874-DEL-2665 |
| chr10-7284731-DEL-54 | chr10-7284732-DEL-54 |
| chr10-7293910-DEL-102 | chr10-7293911-DEL-102 |
| chr10-7294812-DEL-540 | chr10-7294814-DEL-542 |
| chr10-7303568-DEL-93 | chr10-7303583-DEL-79 |
| chr10-7315584-DEL-75 | chr10-7315585-DEL-75 |
| chr10-7339423-DEL-597 | chr10-7339423-DEL-597 |
| chr10-7344550-DEL-280 | chr10-7344550-DEL-280 |
| chr10-7345956-DEL-199 | chr10-7345957-DEL-199 |
| chr10-7373408-DEL-6401 | chr10-7373409-DEL-6401 |
| chr10-7396499-DEL-64 | chr10-7396513-DEL-66 |
| chr10-7408303-DEL-637 | chr10-7408304-DEL-637 |
| chr10-7424434-DEL-8455 | chr10-7424435-DEL-8455 |
| chr10-7436660-DEL-268 | chr10-7436654-DEL-275 |
| chr10-7445543-DEL-1721 | chr10-7445543-DEL-1721 |
| chr10-7447953-DEL-1627 | chr10-7447955-DEL-1627 |
| chr10-7450943-DEL-199 | chr10-7450944-DEL-199 |
| chr10-7459399-DEL-2111 | chr10-7459400-DEL-2111 |
| chr10-7461202-DEL-76 | chr10-7461205-DEL-74 |
| chr10-7477621-DEL-210 | chr10-7477625-DEL-211 |
| chr10-7483505-DEL-163 | chr10-7483506-DEL-163 |
| chr10-7484501-DEL-107 | chr10-7484500-DEL-143 |
| chr10-7496608-DEL-128 | chr10-7496609-DEL-128 |
| chr10-7509645-DEL-240 | chr10-7509646-DEL-240 |
| chr10-7511824-DEL-300 | chr10-7511825-DEL-300 |
| chr10-7544762-DEL-200 | chr10-7544763-DEL-202 |
| chr10-7545979-DEL-76 | chr10-7545980-DEL-76 |
| chr10-7573055-DEL-99 | chr10-7573056-DEL-95 |
| chr10-7573790-DEL-61 | chr10-7573791-DEL-61 |
| chr10-7609016-DEL-190 | chr10-7609017-DEL-190 |
| chr10-7621206-DEL-215 | chr10-7621207-DEL-215 |
| chr10-7625473-DEL-1398 | chr10-7625474-DEL-1398 |
| chr10-7671702-DEL-445 | chr10-7671703-DEL-445 |
| chr10-7765887-DEL-7252 | chr10-7765888-DEL-7252 |
| chr10-7784454-DEL-89 | chr10-7784455-DEL-89 |
| chr10-7817612-DEL-221 | chr10-7817613-DEL-221 |
| chr10-7846010-DEL-449 | chr10-7846011-DEL-449 |
| chr10-7849750-DEL-333 | chr10-7849751-DEL-333 |
| chr10-7912638-DEL-52 | chr10-7912635-DEL-62 |
| chr10-7913117-DEL-216 | chr10-7913118-DEL-216 |
| chr10-7944304-DEL-1194 | chr10-7944305-DEL-1194 |
| chr10-7952114-DEL-448 | chr10-7952115-DEL-448 |
| chr10-7956290-DEL-1623 | chr10-7956291-DEL-1623 |
| chr10-7981700-DEL-95 | chr10-7981703-DEL-125 |
| chr10-8013296-DEL-237 | chr10-8013297-DEL-237 |
| chr10-8021801-DEL-1013 | chr10-8021802-DEL-1013 |
| chr10-8031181-DEL-1586 | chr10-8031182-DEL-1586 |
| chr10-8037724-DEL-9284 | chr10-8037725-DEL-9284 |
| chr10-8046896-DEL-60 | chr10-8046897-DEL-60 |
| chr10-8067885-DEL-110 | chr10-8067848-DEL-98 |
| chr10-8127431-DEL-109 | chr10-8127433-DEL-108 |
| chr10-8139135-DEL-6143 | chr10-8139123-DEL-6147 |
| chr10-8147738-DEL-1231 | chr10-8147739-DEL-1231 |
| chr10-8184841-DEL-80 | chr10-8184842-DEL-80 |
| chr10-8186746-DEL-9381 | chr10-8186747-DEL-9381 |
| chr10-8188084-DEL-2548 | chr10-8188085-DEL-2548 |
| chr10-8242028-DEL-92 | chr10-8242028-DEL-107 |
| chr10-8245125-DEL-219 | chr10-8245126-DEL-219 |
| chr10-8249928-DEL-136 | chr10-8249939-DEL-129 |
| chr10-8258142-DEL-1292 | chr10-8258143-DEL-1292 |
| chr10-8272923-DEL-7163 | chr10-8272924-DEL-7163 |
| chr10-8273470-DEL-78 | chr10-8273471-DEL-78 |
| chr10-8296830-DEL-66 | chr10-8296831-DEL-66 |
| chr10-8311521-DEL-64 | chr10-8311522-DEL-64 |
| chr10-8321119-DEL-56 | chr10-8321120-DEL-56 |
| chr10-8327756-DEL-81 | chr10-8327748-DEL-82 |
| chr10-8352938-DEL-148 | chr10-8352939-DEL-148 |

| | |
|---|---|
| chr10-8397631-DEL-82 | chr10-8397631-DEL-70 |
| chr10-8401346-DEL-153 | chr10-8401347-DEL-153 |
| chr10-8402182-DEL-82 | chr10-8402183-DEL-82 |
| chr10-8403667-DEL-208 | chr10-8403668-DEL-208 |
| chr10-8407163-DEL-78 | chr10-8407140-DEL-78 |
| chr10-8419894-DEL-7138 | chr10-8419895-DEL-7138 |
| chr10-8428820-DEL-328 | chr10-8428821-DEL-328 |
| chr10-8433919-DEL-432 | chr10-8433920-DEL-432 |
| chr10-8474980-DEL-7166 | chr10-8474979-DEL-7166 |
| chr10-8511012-DEL-2593 | chr10-8511013-DEL-2593 |
| chr10-8519004-DEL-147 | chr10-8519005-DEL-147 |
| chr10-8531465-DEL-52 | chr10-8531430-DEL-82 |
| chr10-8545641-DEL-133 | chr10-8545642-DEL-133 |
| chr10-8553004-DEL-292 | chr10-8553005-DEL-292 |
| chr10-8554082-DEL-99 | chr10-8554083-DEL-99 |
| chr10-8576408-DEL-6587 | chr10-8576409-DEL-6587 |
| chr10-8576764-DEL-197 | chr10-8576799-DEL-199 |
| chr10-8587914-DEL-180 | chr10-8587915-DEL-180 |
| chr10-8614941-DEL-88 | chr10-8614938-DEL-62 |
| chr10-8654307-DEL-103 | chr10-8654308-DEL-103 |
| chr10-8666474-DEL-470 | chr10-8666475-DEL-470 |
| chr10-8705586-DEL-64 | chr10-8705574-DEL-52 |
| chr10-8724261-DEL-227 | chr10-8724256-DEL-228 |
| chr10-8744130-DEL-211 | chr10-8744131-DEL-211 |
| chr10-8759117-DEL-194 | chr10-8759114-DEL-197 |
| chr10-8816653-DEL-52 | chr10-8816641-DEL-75 |
| chr10-8851562-DEL-1435 | chr10-8851563-DEL-1435 |
| chr10-8854822-DEL-76 | chr10-8854823-DEL-76 |
| chr10-8877889-DEL-125 | chr10-8877926-DEL-90 |
| chr10-8889212-DEL-76 | chr10-8889212-DEL-74 |
| chr10-8911445-DEL-2047 | chr10-8911466-DEL-2059 |
| chr10-8943952-DEL-96 | chr10-8943953-DEL-96 |
| chr10-8948076-DEL-56 | chr10-8948119-DEL-56 |
| chr10-8956459-DEL-352 | chr10-8956460-DEL-352 |
| chr10-8999218-DEL-395 | chr10-8999219-DEL-395 |
| chr10-9015149-DEL-53 | chr10-9015121-DEL-55 |
| chr10-9030943-DEL-150 | chr10-9030944-DEL-150 |
| chr10-9071070-DEL-73 | chr10-9071119-DEL-66 |
| chr10-9072838-DEL-206 | chr10-9072839-DEL-206 |
| chr10-9079403-DEL-55 | chr10-9079404-DEL-53 |
| chr10-9098011-DEL-66 | chr10-9098003-DEL-76 |
| chr10-9128281-DEL-330 | chr10-9128282-DEL-330 |
| chr10-9134743-DEL-57 | chr10-9134744-DEL-57 |
| chr10-9154826-DEL-120 | chr10-9154827-DEL-120 |
| chr10-9169221-DEL-91 | chr10-9169235-DEL-52 |
| chr10-9174457-DEL-51 | chr10-9174458-DEL-51 |
| chr10-9181282-DEL-62 | chr10-9181289-DEL-80 |
| chr10-9183823-DEL-444 | chr10-9183824-DEL-444 |
| chr10-9203626-DEL-100 | chr10-9203627-DEL-100 |
| chr10-9212949-DEL-431 | chr10-9212950-DEL-431 |
| chr10-9268583-DEL-277 | chr10-9268584-DEL-277 |
| chr10-9294410-DEL-87 | chr10-9294411-DEL-87 |
| chr10-9316511-DEL-2515 | chr10-9316512-DEL-2515 |
| chr10-9336250-DEL-53 | chr10-9336251-DEL-53 |
| chr10-9364482-DEL-124 | chr10-9364483-DEL-124 |
| chr10-9381807-DEL-246 | chr10-9381808-DEL-246 |
| chr10-9396642-DEL-60 | chr10-9396649-DEL-54 |
| chr10-9438677-DEL-56 | chr10-9438678-DEL-54 |
| chr10-9447176-DEL-74 | chr10-9447172-DEL-68 |
| chr10-9453264-DEL-1528 | chr10-9453265-DEL-1528 |
| chr10-9470033-DEL-59 | chr10-9470034-DEL-59 |
| chr10-9472825-DEL-312 | chr10-9472826-DEL-332 |
| chr10-9489815-DEL-82 | chr10-9489814-DEL-96 |
| chr10-9545798-DEL-131 | chr10-9545799-DEL-131 |
| chr10-9565092-DEL-217 | chr10-9565093-DEL-217 |
| chr10-9605561-DEL-1993 | chr10-9605562-DEL-1993 |
| chr10-9607907-DEL-59 | chr10-9607913-DEL-58 |
| chr10-9620005-DEL-3759 | chr10-9620006-DEL-3759 |
| chr10-9627476-DEL-194 | chr10-9627479-DEL-192 |
| chr10-9671452-DEL-56 | chr10-9671428-DEL-62 |
| chr10-9688465-DEL-392 | chr10-9688466-DEL-392 |
| chr10-9703961-DEL-173 | chr10-9703964-DEL-161 |
| chr10-9709511-DEL-882 | chr10-9709512-DEL-882 |
| chr10-9722769-DEL-4761 | chr10-9722770-DEL-4761 |
| chr10-9741168-DEL-92 | chr10-9741169-DEL-92 |
| chr10-9766239-DEL-147 | chr10-9766211-DEL-185 |
| chr10-9790748-DEL-51 | chr10-9790749-DEL-51 |
| chr10-9792915-DEL-241 | chr10-9792916-DEL-241 |
| chr10-9810154-DEL-78 | chr10-9810155-DEL-78 |
| chr10-9831577-DEL-100 | chr10-9831586-DEL-95 |

| | |
|---|---|
| chr10-9844707-DEL-885 | chr10-9844706-DEL-852 |
| chr10-9875133-DEL-114 | chr10-9875108-DEL-111 |
| chr10-9893707-DEL-115 | chr10-9893708-DEL-115 |
| chr10-9902463-DEL-6668 | chr10-9902464-DEL-6668 |
| chr10-9915961-DEL-697 | chr10-9915962-DEL-697 |
| chr10-9927253-DEL-135 | chr10-9927254-DEL-135 |
| chr10-9954704-DEL-6218 | chr10-9954694-DEL-6229 |
| chr10-9994152-DEL-109 | chr10-9994153-DEL-109 |
| chr10-10005166-DEL-278 | chr10-10005167-DEL-278 |
| chr10-10029611-DEL-76 | chr10-10029588-DEL-80 |
| chr10-10064857-DEL-2760 | chr10-10064858-DEL-2760 |
| chr10-10139455-DEL-498 | chr10-10139456-DEL-498 |
| chr10-10149285-DEL-133 | chr10-10149253-DEL-163 |
| chr10-10221585-DEL-78 | chr10-10221586-DEL-78 |
| chr10-10238664-DEL-129 | chr10-10238664-DEL-126 |
| chr10-10295752-DEL-776 | chr10-10295750-DEL-777 |
| chr10-10330825-DEL-839 | chr10-10330826-DEL-839 |
| chr10-10333841-DEL-84 | chr10-10333863-DEL-84 |
| chr10-10356477-DEL-125 | chr10-10356477-DEL-125 |
| chr10-10393505-DEL-5198 | chr10-10393504-DEL-5199 |
| chr10-10425114-DEL-54 | chr10-10425115-DEL-54 |
| chr10-10442636-DEL-6478 | chr10-10442637-DEL-6478 |
| chr10-10460166-DEL-6795 | chr10-10460169-DEL-6792 |
| chr10-10467659-DEL-2988 | chr10-10467660-DEL-2988 |
| chr10-10471177-DEL-1382 | chr10-10471178-DEL-1382 |
| chr10-10483940-DEL-80 | chr10-10483941-DEL-80 |
| chr10-10496416-DEL-82 | chr10-10496421-DEL-80 |
| chr10-10526621-DEL-1157 | chr10-10526622-DEL-1157 |
| chr10-10542849-DEL-6541 | chr10-10542850-DEL-6541 |
| chr10-10587486-DEL-2700 | chr10-10587487-DEL-2700 |
| chr10-10609802-DEL-104 | chr10-10609815-DEL-62 |
| chr10-10666394-DEL-54 | chr10-10666424-DEL-54 |
| chr10-10699464-DEL-979 | chr10-10699465-DEL-979 |
| chr10-10706100-DEL-488 | chr10-10706101-DEL-488 |
| chr10-10727470-DEL-1622 | chr10-10727471-DEL-1622 |
| chr10-10734953-DEL-241 | chr10-10734954-DEL-241 |
| chr10-10738207-DEL-148 | chr10-10738208-DEL-148 |
| chr10-10753640-DEL-114 | chr10-10753641-DEL-114 |
| chr10-10779918-DEL-51 | chr10-10779919-DEL-51 |
| chr10-10809397-DEL-70 | chr10-10809398-DEL-70 |
| chr10-10823062-DEL-878 | chr10-10823063-DEL-878 |
| chr10-10831533-DEL-6404 | chr10-10831520-DEL-6409 |
| chr10-10846806-DEL-82 | chr10-10846807-DEL-82 |
| chr10-10849333-DEL-56 | chr10-10849334-DEL-56 |
| chr10-10851102-DEL-6516 | chr10-10851103-DEL-6516 |
| chr10-10861817-DEL-111 | chr10-10861815-DEL-114 |
| chr10-10865915-DEL-343 | chr10-10865930-DEL-336 |
| chr10-10875718-DEL-183 | chr10-10875719-DEL-183 |
| chr10-10891899-DEL-82 | chr10-10891925-DEL-56 |
| chr10-10894740-DEL-86 | chr10-10894765-DEL-52 |
| chr10-10921173-DEL-124 | chr10-10921174-DEL-124 |
| chr10-10987297-DEL-179 | chr10-10987299-DEL-178 |
| chr10-11011805-DEL-834 | chr10-11011806-DEL-834 |
| chr10-11016242-DEL-192 | chr10-11016243-DEL-192 |
| chr10-11019855-DEL-56 | chr10-11019840-DEL-72 |
| chr10-11094144-DEL-6401 | chr10-11094145-DEL-6401 |
| chr10-11110646-DEL-203 | chr10-11110647-DEL-203 |
| chr10-11119367-DEL-306 | chr10-11119368-DEL-306 |
| chr10-11201838-DEL-54 | chr10-11201839-DEL-54 |
| chr10-11202511-DEL-78 | chr10-11202503-DEL-79 |
| chr10-11226385-DEL-211 | chr10-11226388-DEL-209 |
| chr10-11228389-DEL-600 | chr10-11228390-DEL-600 |
| chr10-11230103-DEL-377 | chr10-11230104-DEL-377 |
| chr10-11236749-DEL-492 | chr10-11236750-DEL-492 |
| chr10-11258111-DEL-215 | chr10-11258112-DEL-215 |
| chr10-11293830-DEL-60 | chr10-11293833-DEL-58 |
| chr10-11302201-DEL-86 | chr10-11302204-DEL-74 |
| chr10-11347266-DEL-182 | chr10-11347267-DEL-182 |
| chr10-11362943-DEL-5813 | chr10-11362943-DEL-5813 |
| chr10-11371279-DEL-81 | chr10-11371276-DEL-55 |
| chr10-11374963-DEL-202 | chr10-11374963-DEL-203 |
| chr10-11400792-DEL-1430 | chr10-11400792-DEL-1430 |
| chr10-11408403-DEL-358 | chr10-11408404-DEL-358 |
| chr10-11429023-DEL-2127 | chr10-11429024-DEL-2127 |
| chr10-11448091-DEL-62 | chr10-11448092-DEL-62 |
| chr10-11455801-DEL-6210 | chr10-11455802-DEL-6210 |
| chr10-11484951-DEL-1272 | chr10-11484952-DEL-1272 |
| chr10-11540290-DEL-76 | chr10-11540290-DEL-80 |
| chr10-11556179-DEL-222 | chr10-11556180-DEL-222 |
| chr10-11590614-DEL-1151 | chr10-11590615-DEL-1151 |

| | |
|---|---|
| chr10-11599612-DEL-103 | chr10-11599608-DEL-100 |
| chr10-11715205-DEL-105 | chr10-11715206-DEL-105 |
| chr10-11743470-DEL-157 | chr10-11743471-DEL-125 |
| chr10-11776330-DEL-3225 | chr10-11776331-DEL-3225 |
| chr10-11786258-DEL-1000 | chr10-11786259-DEL-1000 |
| chr10-11794757-DEL-60 | chr10-11794758-DEL-60 |
| chr10-11816690-DEL-81 | chr10-11816682-DEL-82 |
| chr10-11925681-DEL-1786 | chr10-11925667-DEL-1784 |
| chr10-11935994-DEL-98 | chr10-11935995-DEL-64 |
| chr10-11971173-DEL-557 | chr10-11971174-DEL-557 |
| chr10-11986470-DEL-237 | chr10-11986471-DEL-237 |
| chr10-12001763-DEL-87 | chr10-12001764-DEL-87 |
| chr10-12092448-DEL-52 | chr10-12092455-DEL-52 |
| chr10-12099993-DEL-558 | chr10-12099994-DEL-558 |
| chr10-12110998-DEL-5515 | chr10-12110999-DEL-5511 |
| chr10-12119737-DEL-265 | chr10-12119760-DEL-253 |
| chr10-12120919-DEL-64 | chr10-12120921-DEL-62 |
| chr10-12134879-DEL-377 | chr10-12134879-DEL-377 |
| chr10-12142190-DEL-6589 | chr10-12142191-DEL-6589 |
| chr10-12163597-DEL-85 | chr10-12163598-DEL-85 |
| chr10-12195081-DEL-65 | chr10-12195082-DEL-50 |
| chr10-12231118-DEL-6183 | chr10-12231119-DEL-6183 |
| chr10-12244073-DEL-195 | chr10-12244074-DEL-195 |
| chr10-12261611-DEL-10163 | chr10-12261612-DEL-10163 |
| chr10-12293690-DEL-56 | chr10-12293691-DEL-56 |
| chr10-12305290-DEL-224 | chr10-12305291-DEL-224 |
| chr10-12313184-DEL-54 | chr10-12313185-DEL-81 |
| chr10-12342675-DEL-60 | chr10-12342674-DEL-54 |
| chr10-12375054-DEL-51 | chr10-12375054-DEL-51 |
| chr10-12376801-DEL-67 | chr10-12376803-DEL-66 |
| chr10-12414474-DEL-100 | chr10-12414475-DEL-100 |
| chr10-12450750-DEL-71 | chr10-12450751-DEL-71 |
| chr10-12521413-DEL-3290 | chr10-12521414-DEL-3290 |
| chr10-12554504-DEL-73 | chr10-12554505-DEL-73 |
| chr10-12573516-DEL-64 | chr10-12573517-DEL-64 |
| chr10-12605115-DEL-163 | chr10-12605116-DEL-163 |
| chr10-12608848-DEL-230 | chr10-12608848-DEL-224 |
| chr10-12614513-DEL-72 | chr10-12614514-DEL-72 |
| chr10-12650546-DEL-192 | chr10-12650536-DEL-193 |
| chr10-12658729-DEL-208 | chr10-12658730-DEL-208 |
| chr10-12669651-DEL-67 | chr10-12669652-DEL-67 |
| chr10-12710446-DEL-53 | chr10-12710446-DEL-53 |
| chr10-12748989-DEL-58 | chr10-12748990-DEL-58 |
| chr10-12751200-DEL-68 | chr10-12751201-DEL-68 |
| chr10-12760767-DEL-71 | chr10-12760768-DEL-71 |
| chr10-12781053-DEL-105 | chr10-12781054-DEL-105 |
| chr10-12787091-DEL-68 | chr10-12787092-DEL-68 |
| chr10-12790383-DEL-59 | chr10-12790384-DEL-59 |
| chr10-12812499-DEL-80 | chr10-12812512-DEL-68 |
| chr10-12813012-DEL-53 | chr10-12813013-DEL-84 |
| chr10-12819047-DEL-68 | chr10-12819051-DEL-64 |
| chr10-12827915-DEL-64 | chr10-12827916-DEL-60 |
| chr10-12835207-DEL-108 | chr10-12835208-DEL-108 |
| chr10-12837321-DEL-141 | chr10-12837322-DEL-141 |
| chr10-12854328-DEL-201 | chr10-12854329-DEL-201 |
| chr10-12857934-DEL-1729 | chr10-12857935-DEL-1730 |
| chr10-12858582-DEL-166 | chr10-12858583-DEL-166 |
| chr10-12866102-DEL-56 | chr10-12866103-DEL-56 |
| chr10-12873093-DEL-6541 | chr10-12873094-DEL-6541 |
| chr10-12887231-DEL-193 | chr10-12887237-DEL-188 |
| chr10-12896716-DEL-63 | chr10-12896718-DEL-63 |
| chr10-12900875-DEL-82 | chr10-12900908-DEL-92 |
| chr10-12920727-DEL-94 | chr10-12920728-DEL-94 |
| chr10-12944066-DEL-72 | chr10-12944058-DEL-75 |
| chr10-12955588-DEL-73 | chr10-12955589-DEL-74 |
| chr10-13036428-DEL-69 | chr10-13036423-DEL-75 |
| chr10-13066778-DEL-164 | chr10-13066779-DEL-164 |
| chr10-13081518-DEL-1082 | chr10-13081519-DEL-1082 |
| chr10-13099930-DEL-54 | chr10-13099931-DEL-64 |
| chr10-13183127-DEL-70 | chr10-13183127-DEL-69 |
| chr10-13201290-DEL-86 | chr10-13201291-DEL-86 |
| chr10-13224776-DEL-180 | chr10-13224777-DEL-180 |
| chr10-13247395-DEL-60 | chr10-13247398-DEL-58 |
| chr10-13249000-DEL-165 | chr10-13249001-DEL-165 |
| chr10-13277588-DEL-1643 | chr10-13277585-DEL-1644 |
| chr10-13284773-DEL-53 | chr10-13284772-DEL-53 |
| chr10-13326863-DEL-58 | chr10-13326863-DEL-58 |
| chr10-13352775-DEL-66 | chr10-13352776-DEL-66 |
| chr10-13357257-DEL-1562 | chr10-13357258-DEL-1562 |
| chr10-13379705-DEL-302 | chr10-13379706-DEL-302 |

| | |
|---|---|
| chr10-13404352-DEL-178 | chr10-13404353-DEL-178 |
| chr10-13408808-DEL-51 | chr10-13408809-DEL-51 |
| chr10-13442699-DEL-89 | chr10-13442683-DEL-96 |
| chr10-13470680-DEL-74 | chr10-13470681-DEL-74 |
| chr10-13481197-DEL-6680 | chr10-13481198-DEL-6680 |
| chr10-13482169-DEL-204 | chr10-13482170-DEL-204 |
| chr10-13494820-DEL-54 | chr10-13494821-DEL-54 |
| chr10-13509873-DEL-63 | chr10-13509874-DEL-63 |
| chr10-13537965-DEL-881 | chr10-13537966-DEL-881 |
| chr10-13544822-DEL-443 | chr10-13544823-DEL-443 |
| chr10-13579071-DEL-68 | chr10-13579072-DEL-70 |
| chr10-13625725-DEL-673 | chr10-13625723-DEL-671 |
| chr10-13652418-DEL-287 | chr10-13652419-DEL-287 |
| chr10-13659408-DEL-351 | chr10-13659409-DEL-351 |
| chr10-13678443-DEL-785 | chr10-13678444-DEL-785 |
| chr10-13711093-DEL-51 | chr10-13711092-DEL-52 |
| chr10-13724810-DEL-68 | chr10-13724811-DEL-68 |
| chr10-13734091-DEL-64 | chr10-13734092-DEL-64 |
| chr10-13768749-DEL-8240 | chr10-13768750-DEL-8240 |
| chr10-13769194-DEL-1369 | chr10-13769195-DEL-1369 |
| chr10-13777238-DEL-308 | chr10-13777239-DEL-308 |
| chr10-13781151-DEL-56 | chr10-13781152-DEL-56 |
| chr10-13788116-DEL-1076 | chr10-13788117-DEL-1072 |
| chr10-13816522-DEL-100 | chr10-13816517-DEL-94 |
| chr10-13818236-DEL-214 | chr10-13818237-DEL-214 |
| chr10-13833352-DEL-51 | chr10-13833353-DEL-51 |
| chr10-13875985-DEL-52 | chr10-13875954-DEL-52 |
| chr10-13886176-DEL-77 | chr10-13886177-DEL-77 |
| chr10-13888568-DEL-67 | chr10-13888571-DEL-67 |
| chr10-13903369-DEL-126 | chr10-13903370-DEL-126 |
| chr10-13927745-DEL-155 | chr10-13927766-DEL-120 |
| chr10-13942629-DEL-862 | chr10-13942630-DEL-862 |
| chr10-13942720-DEL-250 | chr10-13942721-DEL-250 |
| chr10-13951817-DEL-53 | chr10-13951818-DEL-53 |
| chr10-13992062-DEL-462 | chr10-13992065-DEL-443 |
| chr10-13995535-DEL-82 | chr10-13995548-DEL-60 |
| chr10-14057811-DEL-594 | chr10-14057812-DEL-594 |
| chr10-14097277-DEL-2942 | chr10-14097278-DEL-2942 |
| chr10-14117523-DEL-769 | chr10-14117522-DEL-772 |
| chr10-14137787-DEL-87 | chr10-14137788-DEL-87 |
| chr10-14143931-DEL-1337 | chr10-14143932-DEL-1337 |
| chr10-14149600-DEL-552 | chr10-14149601-DEL-552 |
| chr10-14164722-DEL-60 | chr10-14164723-DEL-60 |
| chr10-14165032-DEL-6864 | chr10-14165033-DEL-6864 |
| chr10-14172589-DEL-64 | chr10-14172590-DEL-64 |
| chr10-14184201-DEL-2327 | chr10-14184202-DEL-2327 |
| chr10-14194968-DEL-367 | chr10-14194968-DEL-366 |
| chr10-14217932-DEL-124 | chr10-14217929-DEL-125 |
| chr10-14223962-DEL-61 | chr10-14223963-DEL-61 |
| chr10-14270497-DEL-855 | chr10-14270468-DEL-860 |
| chr10-14272756-DEL-92 | chr10-14272773-DEL-82 |
| chr10-14325749-DEL-185 | chr10-14325750-DEL-185 |
| chr10-14359354-DEL-5597 | chr10-14359355-DEL-5597 |
| chr10-14370904-DEL-57 | chr10-14370905-DEL-57 |
| chr10-14375043-DEL-689 | chr10-14375044-DEL-689 |
| chr10-14398786-DEL-555 | chr10-14398788-DEL-550 |
| chr10-14430561-DEL-62 | chr10-14430554-DEL-63 |
| chr10-14434977-DEL-2954 | chr10-14434978-DEL-2955 |
| chr10-14442337-DEL-111 | chr10-14442338-DEL-111 |
| chr10-14447587-DEL-103 | chr10-14447619-DEL-96 |
| chr10-14459630-DEL-556 | chr10-14459631-DEL-556 |
| chr10-14468555-DEL-199 | chr10-14468556-DEL-199 |
| chr10-14478950-DEL-375 | chr10-14478951-DEL-375 |
| chr10-14480173-DEL-192 | chr10-14480174-DEL-192 |
| chr10-14485091-DEL-4420 | chr10-14485092-DEL-4420 |
| chr10-14566212-DEL-1682 | chr10-14566213-DEL-1682 |
| chr10-14585014-DEL-3531 | chr10-14585015-DEL-3531 |
| chr10-14596826-DEL-61 | chr10-14596827-DEL-73 |
| chr10-14706214-DEL-4605 | chr10-14706215-DEL-4605 |
| chr10-14727148-DEL-3135 | chr10-14727145-DEL-3131 |
| chr10-14735819-DEL-119 | chr10-14735820-DEL-117 |
| chr10-14741373-DEL-873 | chr10-14741374-DEL-873 |
| chr10-14751696-DEL-80 | chr10-14751701-DEL-76 |
| chr10-14778260-DEL-883 | chr10-14778261-DEL-883 |
| chr10-14798802-DEL-214 | chr10-14798803-DEL-214 |
| chr10-14844231-DEL-64 | chr10-14844217-DEL-70 |
| chr10-14852148-DEL-87 | chr10-14852149-DEL-88 |
| chr10-14870022-DEL-61 | chr10-14870018-DEL-72 |
| chr10-14897146-DEL-1995 | chr10-14897147-DEL-1995 |
| chr10-14907333-DEL-1148 | chr10-14907334-DEL-1148 |

| | |
|---|---|
| chr10-14939161-DEL-52 | chr10-14939161-DEL-52 |
| chr10-14939539-DEL-6910 | chr10-14939540-DEL-6910 |
| chr10-14939956-DEL-214 | chr10-14939957-DEL-214 |
| chr10-14961127-DEL-66 | chr10-14961128-DEL-66 |
| chr10-14973604-DEL-60 | chr10-14973605-DEL-60 |
| chr10-14973629-DEL-3399 | chr10-14973647-DEL-3387 |
| chr10-15007133-DEL-1148 | chr10-15007134-DEL-1148 |
| chr10-15021557-DEL-50 | chr10-15021555-DEL-52 |
| chr10-15055614-DEL-3835 | chr10-15055615-DEL-3835 |
| chr10-15068519-DEL-11334 | chr10-15068520-DEL-11334 |
| chr10-15099218-DEL-66 | chr10-15099224-DEL-56 |
| chr10-15127262-DEL-76 | chr10-15127219-DEL-124 |
| chr10-15138027-DEL-76 | chr10-15138026-DEL-71 |
| chr10-15153937-DEL-209 | chr10-15153938-DEL-209 |
| chr10-15177519-DEL-256 | chr10-15177520-DEL-256 |
| chr10-15184131-DEL-3681 | chr10-15184132-DEL-3681 |
| chr10-15193448-DEL-52 | chr10-15193437-DEL-64 |
| chr10-15256892-DEL-6325 | chr10-15256893-DEL-6323 |
| chr10-15284659-DEL-52 | chr10-15284659-DEL-52 |
| chr10-15293280-DEL-424 | chr10-15293281-DEL-424 |
| chr10-15303546-DEL-447 | chr10-15303547-DEL-446 |
| chr10-15305083-DEL-60 | chr10-15305083-DEL-60 |
| chr10-15307016-DEL-42093 | chr10-15307016-DEL-42093 |
| chr10-15313034-DEL-896 | chr10-15313035-DEL-896 |
| chr10-15327942-DEL-58 | chr10-15327945-DEL-56 |
| chr10-15332534-DEL-526 | chr10-15332535-DEL-526 |
| chr10-15336142-DEL-100 | chr10-15336143-DEL-100 |
| chr10-15340050-DEL-6396 | chr10-15340051-DEL-6396 |
| chr10-15352360-DEL-412 | chr10-15352361-DEL-412 |
| chr10-15358019-DEL-2988 | chr10-15358015-DEL-2985 |
| chr10-15358200-DEL-54 | chr10-15358201-DEL-54 |
| chr10-15366004-DEL-699 | chr10-15366005-DEL-699 |
| chr10-15376212-DEL-455 | chr10-15376213-DEL-455 |
| chr10-15383730-DEL-535 | chr10-15383731-DEL-535 |
| chr10-15393580-DEL-1079 | chr10-15393581-DEL-1079 |
| chr10-15420409-DEL-62 | chr10-15420447-DEL-70 |
| chr10-15470738-DEL-6394 | chr10-15470739-DEL-6394 |
| chr10-15495927-DEL-71 | chr10-15495928-DEL-71 |
| chr10-15501602-DEL-416 | chr10-15501603-DEL-416 |
| chr10-15511152-DEL-212 | chr10-15511151-DEL-212 |
| chr10-15527218-DEL-95 | chr10-15527219-DEL-95 |
| chr10-15527268-DEL-1600 | chr10-15527269-DEL-1597 |
| chr10-15545931-DEL-51 | chr10-15545932-DEL-51 |
| chr10-15553557-DEL-398 | chr10-15553558-DEL-398 |
| chr10-15586069-DEL-273 | chr10-15586070-DEL-273 |
| chr10-15601769-DEL-108 | chr10-15601811-DEL-65 |
| chr10-15615146-DEL-400 | chr10-15615157-DEL-390 |
| chr10-15616760-DEL-161 | chr10-15616761-DEL-161 |
| chr10-15649258-DEL-866 | chr10-15649259-DEL-866 |
| chr10-15650326-DEL-6564 | chr10-15650327-DEL-6564 |
| chr10-15664963-DEL-121 | chr10-15664964-DEL-121 |
| chr10-15671921-DEL-57 | chr10-15671922-DEL-57 |
| chr10-15675374-DEL-113 | chr10-15675375-DEL-113 |
| chr10-15721815-DEL-150 | chr10-15721816-DEL-150 |
| chr10-15735036-DEL-415 | chr10-15735037-DEL-415 |
| chr10-15781275-DEL-90 | chr10-15781276-DEL-90 |
| chr10-15810333-DEL-246 | chr10-15810290-DEL-283 |
| chr10-15811015-DEL-2682 | chr10-15811016-DEL-2682 |
| chr10-15825468-DEL-52 | chr10-15825469-DEL-52 |
| chr10-15825898-DEL-422 | chr10-15825899-DEL-422 |
| chr10-15826820-DEL-1056 | chr10-15826810-DEL-1057 |
| chr10-15833654-DEL-563 | chr10-15833643-DEL-574 |
| chr10-15842792-DEL-491 | chr10-15842792-DEL-491 |
| chr10-15896782-DEL-175 | chr10-15896783-DEL-175 |
| chr10-15897486-DEL-904 | chr10-15897488-DEL-905 |
| chr10-15904381-DEL-54 | chr10-15904382-DEL-54 |
| chr10-15910533-DEL-2004 | chr10-15910534-DEL-2004 |
| chr10-15912680-DEL-7009 | chr10-15912681-DEL-7009 |
| chr10-15929447-DEL-80 | chr10-15929459-DEL-68 |
| chr10-15985096-DEL-425 | chr10-15985097-DEL-425 |
| chr10-15991031-DEL-68 | chr10-15991034-DEL-66 |
| chr10-16018668-DEL-158 | chr10-16018669-DEL-158 |
| chr10-16036212-DEL-84 | chr10-16036196-DEL-85 |
| chr10-16063272-DEL-164 | chr10-16063273-DEL-164 |
| chr10-16066391-DEL-1549 | chr10-16066392-DEL-1549 |
| chr10-16068909-DEL-60 | chr10-16068910-DEL-60 |
| chr10-16071159-DEL-173 | chr10-16071160-DEL-173 |
| chr10-16113675-DEL-4270 | chr10-16113676-DEL-4270 |
| chr10-16146313-DEL-56 | chr10-16146312-DEL-53 |
| chr10-16185604-DEL-1884 | chr10-16185605-DEL-1884 |

| | |
|---|---|
| chr10-16204111-DEL-68 | chr10-16204113-DEL-56 |
| chr10-16206495-DEL-589 | chr10-16206495-DEL-589 |
| chr10-16284684-DEL-55 | chr10-16284685-DEL-55 |
| chr10-16293988-DEL-945 | chr10-16293989-DEL-945 |
| chr10-16312997-DEL-850 | chr10-16312993-DEL-850 |
| chr10-16388351-DEL-100 | chr10-16388343-DEL-92 |
| chr10-16393272-DEL-84 | chr10-16393273-DEL-84 |
| chr10-16417289-DEL-2028 | chr10-16417290-DEL-2028 |
| chr10-16455638-DEL-117 | chr10-16455642-DEL-113 |
| chr10-16486989-DEL-72 | chr10-16486990-DEL-72 |
| chr10-16535969-DEL-1087 | chr10-16535970-DEL-1087 |
| chr10-16547183-DEL-58 | chr10-16547184-DEL-58 |
| chr10-16567846-DEL-6532 | chr10-16567847-DEL-6532 |
| chr10-16623704-DEL-2261 | chr10-16623705-DEL-2261 |
| chr10-16634042-DEL-506 | chr10-16634043-DEL-506 |
| chr10-16647072-DEL-6715 | chr10-16647073-DEL-6715 |
| chr10-16656670-DEL-133 | chr10-16656646-DEL-134 |
| chr10-16690252-DEL-60 | chr10-16690257-DEL-54 |
| chr10-16696072-DEL-6642 | chr10-16696073-DEL-6641 |
| chr10-16706080-DEL-372 | chr10-16706081-DEL-372 |
| chr10-16777151-DEL-348 | chr10-16777152-DEL-348 |
| chr10-16781247-DEL-855 | chr10-16781248-DEL-852 |
| chr10-16800608-DEL-330 | chr10-16800609-DEL-330 |
| chr10-16841853-DEL-512 | chr10-16841852-DEL-512 |
| chr10-16877790-DEL-698 | chr10-16877791-DEL-698 |
| chr10-16937449-DEL-60 | chr10-16937463-DEL-90 |
| chr10-16986414-DEL-55 | chr10-16986415-DEL-55 |
| chr10-16987563-DEL-4091 | chr10-16987564-DEL-4091 |
| chr10-16992345-DEL-64 | chr10-16992352-DEL-58 |
| chr10-17019276-DEL-6266 | chr10-17019276-DEL-6266 |
| chr10-17192805-DEL-7606 | chr10-17192806-DEL-7606 |
| chr10-17234965-DEL-840 | chr10-17234964-DEL-841 |
| chr10-17242480-DEL-58 | chr10-17242502-DEL-75 |
| chr10-17256635-DEL-52 | chr10-17256636-DEL-52 |
| chr10-17259552-DEL-167 | chr10-17259553-DEL-167 |
| chr10-17319947-DEL-404 | chr10-17319952-DEL-400 |
| chr10-17323875-DEL-78 | chr10-17323876-DEL-78 |
| chr10-17341484-DEL-61 | chr10-17341483-DEL-61 |
| chr10-17345045-DEL-63 | chr10-17345046-DEL-63 |
| chr10-17375421-DEL-1897 | chr10-17375422-DEL-1897 |
| chr10-17447755-DEL-68 | chr10-17447706-DEL-68 |
| chr10-17497324-DEL-90 | chr10-17497333-DEL-82 |
| chr10-17508161-DEL-57 | chr10-17508146-DEL-55 |
| chr10-17555032-DEL-70 | chr10-17555017-DEL-71 |
| chr10-17564954-DEL-60 | chr10-17564906-DEL-97 |
| chr10-17568465-DEL-115 | chr10-17568433-DEL-100 |
| chr10-17609838-DEL-353 | chr10-17609836-DEL-351 |
| chr10-17610782-DEL-399 | chr10-17610783-DEL-399 |
| chr10-17621851-DEL-236 | chr10-17621852-DEL-236 |
| chr10-17647485-DEL-229 | chr10-17647486-DEL-229 |
| chr10-17650534-DEL-196 | chr10-17650535-DEL-196 |
| chr10-17654011-DEL-120 | chr10-17654012-DEL-120 |
| chr10-17658382-DEL-209 | chr10-17658383-DEL-209 |
| chr10-17680874-DEL-58 | chr10-17680875-DEL-58 |
| chr10-17686699-DEL-2509 | chr10-17686700-DEL-2509 |
| chr10-17711646-DEL-85 | chr10-17711655-DEL-79 |
| chr10-17726105-DEL-52 | chr10-17726106-DEL-52 |
| chr10-17728496-DEL-1889 | chr10-17728497-DEL-1889 |
| chr10-17735481-DEL-488 | chr10-17735482-DEL-488 |
| chr10-17767692-DEL-127 | chr10-17767693-DEL-127 |
| chr10-17789721-DEL-68 | chr10-17789736-DEL-52 |
| chr10-17885495-DEL-60 | chr10-17885496-DEL-60 |
| chr10-17891471-DEL-400 | chr10-17891472-DEL-400 |
| chr10-17900389-DEL-343 | chr10-17900390-DEL-343 |
| chr10-17907871-DEL-51 | chr10-17907872-DEL-51 |
| chr10-17963402-DEL-189 | chr10-17963403-DEL-189 |
| chr10-18006606-DEL-413 | chr10-18006607-DEL-413 |
| chr10-18014862-DEL-483 | chr10-18014863-DEL-483 |
| chr10-18022263-DEL-80 | chr10-18022264-DEL-80 |
| chr10-18028010-DEL-401 | chr10-18028018-DEL-401 |
| chr10-18046425-DEL-1294 | chr10-18046426-DEL-1294 |
| chr10-18075132-DEL-516 | chr10-18075133-DEL-516 |
| chr10-18094600-DEL-80 | chr10-18094601-DEL-80 |
| chr10-18099746-DEL-64 | chr10-18099747-DEL-64 |
| chr10-18104269-DEL-1891 | chr10-18104270-DEL-1891 |
| chr10-18114851-DEL-2672 | chr10-18114852-DEL-2672 |
| chr10-18129098-DEL-54 | chr10-18129099-DEL-54 |
| chr10-18132065-DEL-1227 | chr10-18132066-DEL-1227 |
| chr10-18154492-DEL-2888 | chr10-18154493-DEL-2888 |
| chr10-18168088-DEL-3524 | chr10-18168089-DEL-3524 |

| | |
|---|---|
| chr10-18174831-DEL-80 | chr10-18174832-DEL-80 |
| chr10-18198604-DEL-227 | chr10-18198605-DEL-227 |
| chr10-18204674-DEL-67 | chr10-18204675-DEL-67 |
| chr10-18243680-DEL-4695 | chr10-18243681-DEL-4695 |
| chr10-18249450-DEL-63 | chr10-18249470-DEL-51 |
| chr10-18252544-DEL-54 | chr10-18252545-DEL-50 |
| chr10-18254232-DEL-752 | chr10-18254233-DEL-752 |
| chr10-18269104-DEL-68 | chr10-18269133-DEL-50 |
| chr10-18295704-DEL-86 | chr10-18295705-DEL-86 |
| chr10-18309491-DEL-974 | chr10-18309492-DEL-974 |
| chr10-18337444-DEL-278 | chr10-18337441-DEL-279 |
| chr10-18351971-DEL-69 | chr10-18351971-DEL-69 |
| chr10-18361029-DEL-211 | chr10-18361029-DEL-211 |
| chr10-18384911-DEL-58 | chr10-18384912-DEL-58 |
| chr10-18395835-DEL-196 | chr10-18395836-DEL-196 |
| chr10-18431198-DEL-3009 | chr10-18431199-DEL-3009 |
| chr10-18441070-DEL-54 | chr10-18441059-DEL-66 |
| chr10-18446966-DEL-61 | chr10-18446967-DEL-61 |
| chr10-18448136-DEL-164 | chr10-18448137-DEL-164 |
| chr10-18489357-DEL-77 | chr10-18489363-DEL-52 |
| chr10-18495974-DEL-786 | chr10-18495975-DEL-786 |
| chr10-18498365-DEL-197 | chr10-18498366-DEL-197 |
| chr10-18513658-DEL-78 | chr10-18513659-DEL-61 |
| chr10-18518928-DEL-120 | chr10-18518929-DEL-118 |
| chr10-18528829-DEL-374 | chr10-18528812-DEL-352 |
| chr10-18564531-DEL-434 | chr10-18564532-DEL-434 |
| chr10-18581608-DEL-211 | chr10-18581606-DEL-212 |
| chr10-18592241-DEL-347 | chr10-18592238-DEL-348 |
| chr10-18635351-DEL-215 | chr10-18635352-DEL-215 |
| chr10-18689536-DEL-552 | chr10-18689537-DEL-552 |
| chr10-18710487-DEL-64 | chr10-18710493-DEL-56 |
| chr10-18761418-DEL-352 | chr10-18761419-DEL-352 |
| chr10-18763609-DEL-322 | chr10-18763610-DEL-322 |
| chr10-18776963-DEL-60 | chr10-18776960-DEL-61 |
| chr10-18785401-DEL-937 | chr10-18785402-DEL-937 |
| chr10-18797883-DEL-109 | chr10-18797884-DEL-109 |
| chr10-18814089-DEL-59 | chr10-18814090-DEL-59 |
| chr10-18815528-DEL-752 | chr10-18815529-DEL-752 |
| chr10-18822347-DEL-126 | chr10-18822348-DEL-126 |
| chr10-18835747-DEL-177 | chr10-18835748-DEL-176 |
| chr10-18866328-DEL-71 | chr10-18866308-DEL-89 |
| chr10-18924382-DEL-282 | chr10-18924379-DEL-284 |
| chr10-18959145-DEL-53 | chr10-18959146-DEL-53 |
| chr10-18963155-DEL-98 | chr10-18963149-DEL-99 |
| chr10-18971630-DEL-115 | chr10-18971631-DEL-115 |
| chr10-18978444-DEL-357 | chr10-18978445-DEL-357 |
| chr10-18999377-DEL-53 | chr10-18999370-DEL-61 |
| chr10-19004187-DEL-828 | chr10-19004188-DEL-828 |
| chr10-19011208-DEL-2190 | chr10-19011209-DEL-2190 |
| chr10-19022109-DEL-56 | chr10-19022110-DEL-60 |
| chr10-19034967-DEL-80 | chr10-19034968-DEL-80 |
| chr10-19036210-DEL-70 | chr10-19036211-DEL-70 |
| chr10-19051923-DEL-72 | chr10-19051901-DEL-56 |
| chr10-19064069-DEL-85 | chr10-19064070-DEL-85 |
| chr10-19094219-DEL-54 | chr10-19094220-DEL-54 |
| chr10-19097914-DEL-1790 | chr10-19097906-DEL-1788 |
| chr10-19112496-DEL-195 | chr10-19112497-DEL-195 |
| chr10-19115026-DEL-216 | chr10-19115027-DEL-216 |
| chr10-19145922-DEL-3331 | chr10-19145912-DEL-3332 |
| chr10-19163312-DEL-147 | chr10-19163313-DEL-147 |
| chr10-19169759-DEL-341 | chr10-19169761-DEL-339 |
| chr10-19177398-DEL-102 | chr10-19177391-DEL-62 |
| chr10-19181677-DEL-184 | chr10-19181678-DEL-184 |
| chr10-19197228-DEL-1279 | chr10-19197229-DEL-1279 |
| chr10-19223594-DEL-65 | chr10-19223595-DEL-65 |
| chr10-19255873-DEL-831 | chr10-19255874-DEL-831 |
| chr10-19260312-DEL-1748 | chr10-19260313-DEL-1748 |
| chr10-19298922-DEL-124 | chr10-19298923-DEL-123 |
| chr10-19304316-DEL-185 | chr10-19304317-DEL-185 |
| chr10-19321134-DEL-57 | chr10-19321134-DEL-57 |
| chr10-19325493-DEL-269 | chr10-19325494-DEL-269 |
| chr10-19358468-DEL-133 | chr10-19358469-DEL-133 |
| chr10-19368085-DEL-811 | chr10-19368086-DEL-811 |
| chr10-19374850-DEL-342 | chr10-19374849-DEL-343 |
| chr10-19379454-DEL-91 | chr10-19379497-DEL-73 |
| chr10-19414650-DEL-7104 | chr10-19414651-DEL-7104 |
| chr10-19443594-DEL-183 | chr10-19443595-DEL-182 |
| chr10-19477272-DEL-81 | chr10-19477275-DEL-79 |
| chr10-19505224-DEL-470 | chr10-19505225-DEL-470 |
| chr10-19528406-DEL-60 | chr10-19528401-DEL-56 |

| | |
|---|---|
| chr10-19540556-DEL-78 | chr10-19540557-DEL-78 |
| chr10-19572815-DEL-1093 | chr10-19572815-DEL-1093 |
| chr10-19591506-DEL-713 | chr10-19591507-DEL-713 |
| chr10-19605287-DEL-6476 | chr10-19605288-DEL-6476 |
| chr10-19647892-DEL-6184 | chr10-19647893-DEL-6184 |
| chr10-19658587-DEL-4631 | chr10-19658588-DEL-4631 |
| chr10-19673021-DEL-4240 | chr10-19673022-DEL-4240 |
| chr10-19677457-DEL-297 | chr10-19677458-DEL-297 |
| chr10-19748300-DEL-64 | chr10-19748301-DEL-64 |
| chr10-19768530-DEL-832 | chr10-19768512-DEL-828 |
| chr10-19775974-DEL-52 | chr10-19775975-DEL-52 |
| chr10-19787246-DEL-2741 | chr10-19787247-DEL-2741 |
| chr10-19795861-DEL-98 | chr10-19795848-DEL-118 |
| chr10-19847178-DEL-176 | chr10-19847179-DEL-176 |
| chr10-19854913-DEL-6462 | chr10-19854914-DEL-6462 |
| chr10-19872258-DEL-56 | chr10-19872263-DEL-52 |
| chr10-19883077-DEL-198 | chr10-19883092-DEL-193 |
| chr10-19886374-DEL-7159 | chr10-19886375-DEL-7159 |
| chr10-19909851-DEL-526 | chr10-19909852-DEL-527 |
| chr10-19954364-DEL-203 | chr10-19954365-DEL-203 |
| chr10-19961294-DEL-218 | chr10-19961295-DEL-218 |
| chr10-20007231-DEL-220 | chr10-20007232-DEL-220 |
| chr10-20009819-DEL-70 | chr10-20009820-DEL-70 |
| chr10-20011228-DEL-77 | chr10-20011229-DEL-76 |
| chr10-20063032-DEL-55 | chr10-20063033-DEL-55 |
| chr10-20079651-DEL-243 | chr10-20079652-DEL-243 |
| chr10-20086891-DEL-78 | chr10-20086888-DEL-57 |
| chr10-20108904-DEL-57 | chr10-20108904-DEL-57 |
| chr10-20112094-DEL-69 | chr10-20112094-DEL-69 |
| chr10-20191673-DEL-203 | chr10-20191674-DEL-203 |
| chr10-20224367-DEL-143 | chr10-20224368-DEL-143 |
| chr10-20224943-DEL-261 | chr10-20224944-DEL-261 |
| chr10-20247046-DEL-193 | chr10-20247045-DEL-193 |
| chr10-20280796-DEL-56 | chr10-20280795-DEL-68 |
| chr10-20305472-DEL-66 | chr10-20305472-DEL-50 |
| chr10-20306823-DEL-111 | chr10-20306824-DEL-111 |
| chr10-20322857-DEL-62 | chr10-20322852-DEL-68 |
| chr10-20335577-DEL-104 | chr10-20335548-DEL-130 |
| chr10-20341334-DEL-112 | chr10-20341342-DEL-114 |
| chr10-20354945-DEL-55 | chr10-20354945-DEL-56 |
| chr10-20393927-DEL-413 | chr10-20393928-DEL-413 |
| chr10-20411134-DEL-71 | chr10-20411135-DEL-71 |
| chr10-20452021-DEL-72 | chr10-20452050-DEL-50 |
| chr10-20518731-DEL-76 | chr10-20518732-DEL-58 |
| chr10-20519023-DEL-6174 | chr10-20519024-DEL-6174 |
| chr10-20536365-DEL-976 | chr10-20536365-DEL-975 |
| chr10-20583997-DEL-68 | chr10-20583998-DEL-66 |
| chr10-20621491-DEL-64 | chr10-20621483-DEL-78 |
| chr10-20623289-DEL-517 | chr10-20623288-DEL-512 |
| chr10-20628974-DEL-645 | chr10-20628975-DEL-645 |
| chr10-20631198-DEL-202 | chr10-20631198-DEL-204 |
| chr10-20637313-DEL-101 | chr10-20637314-DEL-101 |
| chr10-20647714-DEL-3721 | chr10-20647715-DEL-3721 |
| chr10-20662774-DEL-988 | chr10-20662775-DEL-988 |
| chr10-20664467-DEL-498 | chr10-20664468-DEL-498 |
| chr10-20666104-DEL-51 | chr10-20666100-DEL-62 |
| chr10-20686730-DEL-398 | chr10-20686731-DEL-398 |
| chr10-20689044-DEL-67 | chr10-20689043-DEL-68 |
| chr10-20748334-DEL-83 | chr10-20748335-DEL-83 |
| chr10-20753498-DEL-403 | chr10-20753499-DEL-403 |
| chr10-20826631-DEL-56 | chr10-20826636-DEL-52 |
| chr10-20830905-DEL-110 | chr10-20830904-DEL-112 |
| chr10-20836100-DEL-96 | chr10-20836101-DEL-96 |
| chr10-20842102-DEL-64 | chr10-20842103-DEL-64 |
| chr10-20886118-DEL-65 | chr10-20886118-DEL-64 |
| chr10-20925572-DEL-116 | chr10-20925584-DEL-104 |
| chr10-20936340-DEL-73 | chr10-20936341-DEL-73 |
| chr10-20938588-DEL-72 | chr10-20938587-DEL-70 |
| chr10-20940219-DEL-65 | chr10-20940220-DEL-65 |
| chr10-20973115-DEL-185 | chr10-20973116-DEL-185 |
| chr10-21016649-DEL-472 | chr10-21016650-DEL-472 |
| chr10-21025892-DEL-194 | chr10-21025927-DEL-148 |
| chr10-21060610-DEL-86 | chr10-21060611-DEL-86 |
| chr10-21084271-DEL-1569 | chr10-21084272-DEL-1569 |
| chr10-21087098-DEL-5357 | chr10-21087099-DEL-5357 |
| chr10-21101683-DEL-89 | chr10-21101703-DEL-85 |
| chr10-21107232-DEL-58 | chr10-21107235-DEL-50 |
| chr10-21154325-DEL-5881 | chr10-21154312-DEL-5883 |
| chr10-21182632-DEL-412 | chr10-21182633-DEL-411 |
| chr10-21200163-DEL-74 | chr10-21200164-DEL-74 |

| | |
|---|---|
| chr10-21283592-DEL-2687 | chr10-21283593-DEL-2687 |
| chr10-21286401-DEL-1014 | chr10-21286402-DEL-1014 |
| chr10-21366055-DEL-399 | chr10-21366056-DEL-399 |
| chr10-21397316-DEL-323 | chr10-21397317-DEL-323 |
| chr10-21406976-DEL-454 | chr10-21406977-DEL-454 |
| chr10-21427499-DEL-238 | chr10-21427500-DEL-238 |
| chr10-21452797-DEL-4126 | chr10-21452798-DEL-4126 |
| chr10-21494083-DEL-206 | chr10-21494084-DEL-206 |
| chr10-21500810-DEL-944 | chr10-21500811-DEL-944 |
| chr10-21518178-DEL-7241 | chr10-21518179-DEL-7241 |
| chr10-21536243-DEL-52 | chr10-21536245-DEL-51 |
| chr10-21537112-DEL-491 | chr10-21537113-DEL-491 |
| chr10-21571870-DEL-167 | chr10-21571863-DEL-178 |
| chr10-21581423-DEL-764 | chr10-21581424-DEL-764 |
| chr10-21590052-DEL-226 | chr10-21590053-DEL-226 |
| chr10-21591776-DEL-327 | chr10-21591777-DEL-327 |
| chr10-21602233-DEL-309 | chr10-21602239-DEL-289 |
| chr10-21603789-DEL-8456 | chr10-21603790-DEL-8456 |
| chr10-21678078-DEL-198 | chr10-21678080-DEL-197 |
| chr10-21679482-DEL-69 | chr10-21679480-DEL-64 |
| chr10-21751261-DEL-58 | chr10-21751254-DEL-50 |
| chr10-21751860-DEL-106 | chr10-21751844-DEL-107 |
| chr10-21829470-DEL-100 | chr10-21829471-DEL-100 |
| chr10-21873366-DEL-234 | chr10-21873367-DEL-234 |
| chr10-21891761-DEL-180 | chr10-21891762-DEL-180 |
| chr10-21914649-DEL-1738 | chr10-21914649-DEL-1738 |
| chr10-21920561-DEL-2371 | chr10-21920561-DEL-2371 |
| chr10-21932777-DEL-204 | chr10-21932777-DEL-204 |
| chr10-22028972-DEL-202 | chr10-22028973-DEL-202 |
| chr10-22030355-DEL-103 | chr10-22030356-DEL-103 |
| chr10-22094042-DEL-87 | chr10-22094076-DEL-54 |
| chr10-22096603-DEL-438 | chr10-22096604-DEL-438 |
| chr10-22098779-DEL-52 | chr10-22098779-DEL-52 |
| chr10-22099891-DEL-56 | chr10-22099890-DEL-56 |
| chr10-22133983-DEL-53 | chr10-22133984-DEL-53 |
| chr10-22234025-DEL-303 | chr10-22234026-DEL-303 |
| chr10-22244777-DEL-1854 | chr10-22244778-DEL-1854 |
| chr10-22248528-DEL-996 | chr10-22248528-DEL-996 |
| chr10-22252140-DEL-96 | chr10-22252117-DEL-120 |
| chr10-22255755-DEL-52 | chr10-22255758-DEL-50 |
| chr10-22284763-DEL-54 | chr10-22284764-DEL-54 |
| chr10-22336129-DEL-216 | chr10-22336128-DEL-218 |
| chr10-22337754-DEL-238 | chr10-22337755-DEL-238 |
| chr10-22352383-DEL-550 | chr10-22352384-DEL-550 |
| chr10-22359646-DEL-159 | chr10-22359646-DEL-159 |
| chr10-22360310-DEL-989 | chr10-22360310-DEL-989 |
| chr10-22362249-DEL-124 | chr10-22362249-DEL-147 |
| chr10-22365244-DEL-66 | chr10-22365244-DEL-67 |
| chr10-22375265-DEL-1389 | chr10-22375265-DEL-1389 |
| chr10-22376204-DEL-440 | chr10-22376205-DEL-440 |
| chr10-22377849-DEL-90 | chr10-22377849-DEL-90 |
| chr10-22441536-DEL-122 | chr10-22441535-DEL-102 |
| chr10-22445034-DEL-63 | chr10-22445035-DEL-63 |
| chr10-22455990-DEL-181 | chr10-22455990-DEL-181 |
| chr10-22465563-DEL-66 | chr10-22465564-DEL-66 |
| chr10-22497398-DEL-6481 | chr10-22497399-DEL-6481 |
| chr10-22499620-DEL-128 | chr10-22499620-DEL-128 |
| chr10-22511016-DEL-248 | chr10-22511017-DEL-248 |
| chr10-22531822-DEL-69 | chr10-22531822-DEL-69 |
| chr10-22545242-DEL-66 | chr10-22545219-DEL-78 |
| chr10-22545618-DEL-194 | chr10-22545619-DEL-194 |
| chr10-22559113-DEL-7330 | chr10-22559114-DEL-7330 |
| chr10-22610173-DEL-9049 | chr10-22610174-DEL-9049 |
| chr10-22643627-DEL-97 | chr10-22643666-DEL-67 |
| chr10-22644710-DEL-848 | chr10-22644711-DEL-848 |
| chr10-22648254-DEL-84 | chr10-22648243-DEL-84 |
| chr10-22648983-DEL-121 | chr10-22648982-DEL-122 |
| chr10-22662824-DEL-178 | chr10-22662825-DEL-178 |
| chr10-22673155-DEL-6459 | chr10-22673139-DEL-6475 |
| chr10-22687056-DEL-79 | chr10-22687049-DEL-87 |
| chr10-22702456-DEL-109 | chr10-22702457-DEL-109 |
| chr10-22725560-DEL-124 | chr10-22725599-DEL-124 |
| chr10-22735204-DEL-6249 | chr10-22735205-DEL-6249 |
| chr10-22748072-DEL-90 | chr10-22748073-DEL-90 |
| chr10-22771523-DEL-101 | chr10-22771524-DEL-101 |
| chr10-22773256-DEL-328 | chr10-22773257-DEL-328 |
| chr10-22804939-DEL-416 | chr10-22804942-DEL-414 |
| chr10-22842415-DEL-6465 | chr10-22842416-DEL-6465 |
| chr10-22848634-DEL-201 | chr10-22848634-DEL-201 |
| chr10-22859725-DEL-52 | chr10-22859698-DEL-64 |

| | |
|---|---|
| chr10-22894361-DEL-74 | chr10-22894370-DEL-64 |
| chr10-22931092-DEL-510 | chr10-22931093-DEL-510 |
| chr10-22938481-DEL-304 | chr10-22938482-DEL-304 |
| chr10-22954851-DEL-75 | chr10-22954852-DEL-75 |
| chr10-22961502-DEL-581 | chr10-22961503-DEL-581 |
| chr10-22973615-DEL-1330 | chr10-22973616-DEL-1330 |
| chr10-23032498-DEL-76 | chr10-23032497-DEL-78 |
| chr10-23082890-DEL-249 | chr10-23082891-DEL-249 |
| chr10-23087770-DEL-72 | chr10-23087765-DEL-70 |
| chr10-23111359-DEL-93 | chr10-23111360-DEL-93 |
| chr10-23125819-DEL-66 | chr10-23125815-DEL-56 |
| chr10-23150475-DEL-529 | chr10-23150476-DEL-529 |
| chr10-23153987-DEL-1200 | chr10-23153988-DEL-1200 |
| chr10-23173601-DEL-1329 | chr10-23173602-DEL-1329 |
| chr10-23178489-DEL-65 | chr10-23178490-DEL-65 |
| chr10-23179366-DEL-553 | chr10-23179367-DEL-551 |
| chr10-23201880-DEL-390 | chr10-23201881-DEL-390 |
| chr10-23218910-DEL-51 | chr10-23218911-DEL-51 |
| chr10-23244878-DEL-69 | chr10-23244844-DEL-54 |
| chr10-23246630-DEL-167 | chr10-23246631-DEL-167 |
| chr10-23261297-DEL-93 | chr10-23261301-DEL-90 |
| chr10-23264532-DEL-263 | chr10-23264536-DEL-260 |
| chr10-23280448-DEL-2741 | chr10-23280448-DEL-2742 |
| chr10-23282306-DEL-83 | chr10-23282307-DEL-83 |
| chr10-23292605-DEL-54 | chr10-23292606-DEL-54 |
| chr10-23303917-DEL-1238 | chr10-23303918-DEL-1238 |
| chr10-23335180-DEL-4124 | chr10-23335181-DEL-4124 |
| chr10-23376900-DEL-6504 | chr10-23376901-DEL-6504 |
| chr10-23403636-DEL-2035 | chr10-23403637-DEL-2035 |
| chr10-23406512-DEL-56 | chr10-23406513-DEL-56 |
| chr10-23431307-DEL-294 | chr10-23431308-DEL-294 |
| chr10-23441922-DEL-159 | chr10-23441923-DEL-159 |
| chr10-23444841-DEL-57 | chr10-23444845-DEL-54 |
| chr10-23449403-DEL-651 | chr10-23449404-DEL-651 |
| chr10-23466389-DEL-52 | chr10-23466386-DEL-56 |
| chr10-23471042-DEL-512 | chr10-23471043-DEL-512 |
| chr10-23484048-DEL-56 | chr10-23484049-DEL-56 |
| chr10-23497196-DEL-260 | chr10-23497181-DEL-258 |
| chr10-23498940-DEL-2361 | chr10-23498940-DEL-2362 |
| chr10-23504622-DEL-67 | chr10-23504628-DEL-62 |
| chr10-23507546-DEL-6432 | chr10-23507547-DEL-6432 |
| chr10-23525879-DEL-222 | chr10-23525880-DEL-222 |
| chr10-23567898-DEL-400 | chr10-23567899-DEL-400 |
| chr10-23602232-DEL-479 | chr10-23602233-DEL-479 |
| chr10-23647691-DEL-217 | chr10-23647693-DEL-219 |
| chr10-23652760-DEL-295 | chr10-23652765-DEL-291 |
| chr10-23654576-DEL-157 | chr10-23654577-DEL-157 |
| chr10-23655088-DEL-1836 | chr10-23655089-DEL-1836 |
| chr10-23670010-DEL-211 | chr10-23670011-DEL-211 |
| chr10-23673915-DEL-66 | chr10-23673916-DEL-66 |
| chr10-23675182-DEL-52 | chr10-23675183-DEL-52 |
| chr10-23682853-DEL-60 | chr10-23682854-DEL-60 |
| chr10-23690341-DEL-189 | chr10-23690342-DEL-189 |
| chr10-23706315-DEL-1894 | chr10-23706316-DEL-1894 |
| chr10-23754790-DEL-236 | chr10-23754791-DEL-230 |
| chr10-23758506-DEL-124 | chr10-23758507-DEL-124 |
| chr10-23762520-DEL-401 | chr10-23762521-DEL-401 |
| chr10-23766161-DEL-201 | chr10-23766162-DEL-201 |
| chr10-23782869-DEL-3530 | chr10-23782869-DEL-3530 |
| chr10-23809538-DEL-53 | chr10-23809538-DEL-53 |
| chr10-23812420-DEL-222 | chr10-23812421-DEL-222 |
| chr10-23819770-DEL-222 | chr10-23819771-DEL-222 |
| chr10-23856383-DEL-2232 | chr10-23856384-DEL-2232 |
| chr10-23858239-DEL-414 | chr10-23858240-DEL-414 |
| chr10-23859004-DEL-59 | chr10-23859005-DEL-59 |
| chr10-23867399-DEL-242 | chr10-23867400-DEL-242 |
| chr10-23922158-DEL-90 | chr10-23922159-DEL-90 |
| chr10-23933273-DEL-878 | chr10-23933274-DEL-878 |
| chr10-23958559-DEL-64 | chr10-23958560-DEL-64 |
| chr10-23960580-DEL-210 | chr10-23960581-DEL-210 |
| chr10-23964171-DEL-113 | chr10-23964192-DEL-87 |
| chr10-24003916-DEL-55 | chr10-24003906-DEL-52 |
| chr10-24009012-DEL-88 | chr10-24009013-DEL-88 |
| chr10-24021428-DEL-199 | chr10-24021429-DEL-199 |
| chr10-24071967-DEL-204 | chr10-24071964-DEL-205 |
| chr10-24074219-DEL-108 | chr10-24074220-DEL-108 |
| chr10-24092788-DEL-240 | chr10-24092789-DEL-240 |
| chr10-24102322-DEL-58 | chr10-24102323-DEL-58 |
| chr10-24140667-DEL-211 | chr10-24140668-DEL-211 |
| chr10-24144990-DEL-110 | chr10-24145006-DEL-94 |

| | |
|---|---|
| chr10-24174176-DEL-191 | chr10-24174177-DEL-191 |
| chr10-24188007-DEL-53 | chr10-24187964-DEL-90 |
| chr10-24201837-DEL-5503 | chr10-24201838-DEL-5503 |
| chr10-24231062-DEL-335 | chr10-24231057-DEL-338 |
| chr10-24263654-DEL-96 | chr10-24263677-DEL-74 |
| chr10-24266808-DEL-67 | chr10-24266808-DEL-68 |
| chr10-24269140-DEL-106 | chr10-24269141-DEL-106 |
| chr10-24296637-DEL-501 | chr10-24296638-DEL-501 |
| chr10-24311123-DEL-559 | chr10-24311124-DEL-559 |
| chr10-24314451-DEL-508 | chr10-24314452-DEL-508 |
| chr10-24326716-DEL-160 | chr10-24326717-DEL-160 |
| chr10-24372700-DEL-523 | chr10-24372701-DEL-522 |
| chr10-24388505-DEL-66 | chr10-24388506-DEL-66 |
| chr10-24400644-DEL-156 | chr10-24400645-DEL-156 |
| chr10-24426221-DEL-64 | chr10-24426222-DEL-64 |
| chr10-24427278-DEL-57 | chr10-24427278-DEL-57 |
| chr10-24429437-DEL-89 | chr10-24429438-DEL-89 |
| chr10-24437197-DEL-2406 | chr10-24437198-DEL-2406 |
| chr10-24443176-DEL-536 | chr10-24443177-DEL-536 |
| chr10-24492089-DEL-446 | chr10-24492069-DEL-434 |
| chr10-24495863-DEL-51 | chr10-24495864-DEL-51 |
| chr10-24508509-DEL-181 | chr10-24508510-DEL-180 |
| chr10-24536770-DEL-388 | chr10-24536771-DEL-388 |
| chr10-24550896-DEL-215 | chr10-24550897-DEL-215 |
| chr10-24562322-DEL-5190 | chr10-24562323-DEL-5190 |
| chr10-24580710-DEL-131 | chr10-24580711-DEL-131 |
| chr10-24604646-DEL-87 | chr10-24604647-DEL-87 |
| chr10-24610103-DEL-184 | chr10-24610104-DEL-184 |
| chr10-24629312-DEL-84 | chr10-24629327-DEL-70 |
| chr10-24668166-DEL-75 | chr10-24668167-DEL-75 |
| chr10-24690585-DEL-132 | chr10-24690586-DEL-126 |
| chr10-24692122-DEL-1216 | chr10-24692123-DEL-1216 |
| chr10-24694796-DEL-154 | chr10-24694805-DEL-139 |
| chr10-24705877-DEL-56 | chr10-24705878-DEL-52 |
| chr10-24715870-DEL-227 | chr10-24715871-DEL-227 |
| chr10-24724193-DEL-381 | chr10-24724190-DEL-385 |
| chr10-24725977-DEL-67 | chr10-24725983-DEL-66 |
| chr10-24730406-DEL-5607 | chr10-24730407-DEL-5607 |
| chr10-24799904-DEL-112 | chr10-24799905-DEL-112 |
| chr10-24816936-DEL-54 | chr10-24816965-DEL-50 |
| chr10-24817609-DEL-57 | chr10-24817610-DEL-57 |
| chr10-24820067-DEL-212 | chr10-24820068-DEL-212 |
| chr10-24837581-DEL-52 | chr10-24837584-DEL-52 |
| chr10-24845403-DEL-53 | chr10-24845404-DEL-53 |
| chr10-24865278-DEL-76 | chr10-24865279-DEL-76 |
| chr10-24881794-DEL-1584 | chr10-24881795-DEL-1583 |
| chr10-24914357-DEL-202 | chr10-24914362-DEL-198 |
| chr10-24916380-DEL-134 | chr10-24916385-DEL-128 |
| chr10-24918265-DEL-64 | chr10-24918264-DEL-65 |
| chr10-24918904-DEL-75 | chr10-24918905-DEL-75 |
| chr10-24921711-DEL-58 | chr10-24921718-DEL-50 |
| chr10-24966912-DEL-288 | chr10-24966913-DEL-288 |
| chr10-24977147-DEL-6236 | chr10-24977122-DEL-6246 |
| chr10-24985520-DEL-1637 | chr10-24985521-DEL-1637 |
| chr10-25006888-DEL-879 | chr10-25006889-DEL-879 |
| chr10-25039903-DEL-143 | chr10-25039904-DEL-143 |
| chr10-25042352-DEL-208 | chr10-25042351-DEL-208 |
| chr10-25043593-DEL-235 | chr10-25043594-DEL-235 |
| chr10-25080739-DEL-817 | chr10-25080740-DEL-817 |
| chr10-25084790-DEL-57 | chr10-25084797-DEL-51 |
| chr10-25094784-DEL-87 | chr10-25094785-DEL-87 |
| chr10-25098287-DEL-69 | chr10-25098288-DEL-69 |
| chr10-25121189-DEL-147 | chr10-25121190-DEL-147 |
| chr10-25129099-DEL-1773 | chr10-25129100-DEL-1773 |
| chr10-25140508-DEL-169 | chr10-25140509-DEL-169 |
| chr10-25177217-DEL-208 | chr10-25177218-DEL-208 |
| chr10-25196177-DEL-172 | chr10-25196178-DEL-172 |
| chr10-25204030-DEL-477 | chr10-25204031-DEL-477 |
| chr10-25209978-DEL-57 | chr10-25209979-DEL-57 |
| chr10-25216014-DEL-438 | chr10-25216015-DEL-438 |
| chr10-25224004-DEL-210 | chr10-25224005-DEL-210 |
| chr10-25258876-DEL-136 | chr10-25258877-DEL-136 |
| chr10-25294817-DEL-6657 | chr10-25294818-DEL-6657 |
| chr10-25304441-DEL-122 | chr10-25304442-DEL-122 |
| chr10-25305944-DEL-122 | chr10-25305955-DEL-86 |
| chr10-25318295-DEL-78 | chr10-25318296-DEL-78 |
| chr10-25322316-DEL-105 | chr10-25322317-DEL-105 |
| chr10-25327471-DEL-72 | chr10-25327472-DEL-66 |
| chr10-25328490-DEL-392 | chr10-25328491-DEL-392 |
| chr10-25349844-DEL-1544 | chr10-25349845-DEL-1544 |

| | |
|---|---|
| chr10-25361602-DEL-201 | chr10-25361603-DEL-201 |
| chr10-25364287-DEL-69 | chr10-25364291-DEL-104 |
| chr10-25372986-DEL-118 | chr10-25372987-DEL-118 |
| chr10-25393720-DEL-425 | chr10-25393721-DEL-425 |
| chr10-25400798-DEL-77 | chr10-25400799-DEL-76 |
| chr10-25400982-DEL-454 | chr10-25400983-DEL-454 |
| chr10-25416289-DEL-55 | chr10-25416294-DEL-51 |
| chr10-25457211-DEL-69 | chr10-25457212-DEL-60 |
| chr10-25463119-DEL-513 | chr10-25463120-DEL-513 |
| chr10-25475624-DEL-1655 | chr10-25475625-DEL-1655 |
| chr10-25478577-DEL-1272 | chr10-25478578-DEL-1254 |
| chr10-25498867-DEL-6440 | chr10-25498868-DEL-6440 |
| chr10-25533699-DEL-181 | chr10-25533700-DEL-181 |
| chr10-25539776-DEL-6177 | chr10-25539777-DEL-6177 |
| chr10-25556813-DEL-314 | chr10-25556826-DEL-310 |
| chr10-25559327-DEL-59 | chr10-25559328-DEL-59 |
| chr10-25586276-DEL-98 | chr10-25586278-DEL-89 |
| chr10-25611356-DEL-2479 | chr10-25611357-DEL-2479 |
| chr10-25621721-DEL-177 | chr10-25621722-DEL-177 |
| chr10-25668543-DEL-1796 | chr10-25668544-DEL-1796 |
| chr10-25690655-DEL-213 | chr10-25690656-DEL-213 |
| chr10-25696943-DEL-84 | chr10-25696944-DEL-84 |
| chr10-25711495-DEL-57 | chr10-25711512-DEL-56 |
| chr10-25740455-DEL-74 | chr10-25740456-DEL-68 |
| chr10-25763543-DEL-549 | chr10-25763544-DEL-549 |
| chr10-25770068-DEL-529 | chr10-25770069-DEL-529 |
| chr10-25771504-DEL-8419 | chr10-25771505-DEL-8419 |
| chr10-25812169-DEL-6833 | chr10-25812170-DEL-6833 |
| chr10-25844311-DEL-4872 | chr10-25844312-DEL-4872 |
| chr10-25849498-DEL-750 | chr10-25849499-DEL-749 |
| chr10-25854220-DEL-312 | chr10-25854222-DEL-310 |
| chr10-25870497-DEL-92 | chr10-25870464-DEL-123 |
| chr10-25893963-DEL-54 | chr10-25893965-DEL-55 |
| chr10-25949368-DEL-58 | chr10-25949369-DEL-54 |
| chr10-25967956-DEL-79 | chr10-25967947-DEL-88 |
| chr10-25983632-DEL-1015 | chr10-25983633-DEL-1015 |
| chr10-26002818-DEL-6003 | chr10-26002818-DEL-6003 |
| chr10-26029593-DEL-237 | chr10-26029594-DEL-237 |
| chr10-26066222-DEL-62 | chr10-26066223-DEL-60 |
| chr10-26071750-DEL-404 | chr10-26071751-DEL-404 |
| chr10-26076094-DEL-993 | chr10-26076095-DEL-993 |
| chr10-26106287-DEL-83 | chr10-26106288-DEL-83 |
| chr10-26126701-DEL-563 | chr10-26126702-DEL-563 |
| chr10-26141900-DEL-88 | chr10-26141896-DEL-71 |
| chr10-26169988-DEL-898 | chr10-26169989-DEL-898 |
| chr10-26185938-DEL-418 | chr10-26185939-DEL-418 |
| chr10-26209642-DEL-169 | chr10-26209642-DEL-169 |
| chr10-26241783-DEL-86 | chr10-26241782-DEL-129 |
| chr10-26262526-DEL-68 | chr10-26262527-DEL-68 |
| chr10-26271466-DEL-209 | chr10-26271467-DEL-209 |
| chr10-26273490-DEL-245 | chr10-26273487-DEL-245 |
| chr10-26278544-DEL-66 | chr10-26278534-DEL-67 |
| chr10-26279588-DEL-4174 | chr10-26279589-DEL-4174 |
| chr10-26284981-DEL-182 | chr10-26284982-DEL-182 |
| chr10-26287212-DEL-76 | chr10-26287213-DEL-76 |
| chr10-26325391-DEL-216 | chr10-26325392-DEL-216 |
| chr10-26423110-DEL-96 | chr10-26423130-DEL-103 |
| chr10-26425482-DEL-1460 | chr10-26425483-DEL-1460 |
| chr10-26450237-DEL-84 | chr10-26450238-DEL-78 |
| chr10-26488411-DEL-1591 | chr10-26488412-DEL-1591 |
| chr10-26516936-DEL-3579 | chr10-26516937-DEL-3579 |
| chr10-26527791-DEL-60 | chr10-26527792-DEL-60 |
| chr10-26532686-DEL-6637 | chr10-26532687-DEL-6637 |
| chr10-26538920-DEL-400 | chr10-26538929-DEL-395 |
| chr10-26539997-DEL-287 | chr10-26539998-DEL-287 |
| chr10-26553498-DEL-476 | chr10-26553499-DEL-476 |
| chr10-26557200-DEL-70 | chr10-26557201-DEL-70 |
| chr10-26557775-DEL-181 | chr10-26557776-DEL-181 |
| chr10-26559407-DEL-812 | chr10-26559407-DEL-812 |
| chr10-26563342-DEL-69 | chr10-26563343-DEL-69 |
| chr10-26606258-DEL-504 | chr10-26606259-DEL-496 |
| chr10-26606872-DEL-3551 | chr10-26606873-DEL-3551 |
| chr10-26629736-DEL-124 | chr10-26629737-DEL-124 |
| chr10-26630421-DEL-100 | chr10-26630422-DEL-100 |
| chr10-26657584-DEL-1545 | chr10-26657585-DEL-1546 |
| chr10-26671819-DEL-68 | chr10-26671820-DEL-68 |
| chr10-26688971-DEL-182 | chr10-26688972-DEL-182 |
| chr10-26738696-DEL-78 | chr10-26738697-DEL-78 |
| chr10-26757361-DEL-491 | chr10-26757362-DEL-491 |
| chr10-26769665-DEL-257 | chr10-26769664-DEL-258 |

| | |
|---|---|
| chr10-26782896-DEL-215 | chr10-26782897-DEL-215 |
| chr10-26802037-DEL-478 | chr10-26802037-DEL-478 |
| chr10-26802963-DEL-8411 | chr10-26802964-DEL-8411 |
| chr10-26814189-DEL-549 | chr10-26814190-DEL-549 |
| chr10-26841181-DEL-193 | chr10-26841182-DEL-193 |
| chr10-26842423-DEL-52 | chr10-26842424-DEL-52 |
| chr10-26874406-DEL-207 | chr10-26874407-DEL-207 |
| chr10-26878587-DEL-144 | chr10-26878629-DEL-144 |
| chr10-26888050-DEL-1315 | chr10-26888051-DEL-1315 |
| chr10-26891992-DEL-58 | chr10-26891993-DEL-58 |
| chr10-26892450-DEL-1238 | chr10-26892447-DEL-1239 |
| chr10-26946445-DEL-511 | chr10-26946446-DEL-511 |
| chr10-26952280-DEL-625 | chr10-26952281-DEL-625 |
| chr10-26963671-DEL-62 | chr10-26963672-DEL-60 |
| chr10-26986135-DEL-93 | chr10-26986117-DEL-94 |
| chr10-26988261-DEL-108 | chr10-26988261-DEL-109 |
| chr10-27005877-DEL-116 | chr10-27005878-DEL-116 |
| chr10-27006651-DEL-280 | chr10-27006652-DEL-280 |
| chr10-27022664-DEL-177 | chr10-27022665-DEL-177 |
| chr10-27032790-DEL-599 | chr10-27032791-DEL-599 |
| chr10-27048781-DEL-52 | chr10-27048782-DEL-51 |
| chr10-27130664-DEL-9616 | chr10-27130665-DEL-9616 |
| chr10-27188143-DEL-103 | chr10-27188142-DEL-104 |
| chr10-27229408-DEL-79 | chr10-27229409-DEL-75 |
| chr10-27229926-DEL-6584 | chr10-27229927-DEL-6584 |
| chr10-27277264-DEL-311 | chr10-27277270-DEL-301 |
| chr10-27279085-DEL-5024 | chr10-27279085-DEL-5022 |
| chr10-27306541-DEL-54 | chr10-27306542-DEL-54 |
| chr10-27307653-DEL-55 | chr10-27307654-DEL-55 |
| chr10-27410074-DEL-311 | chr10-27410075-DEL-310 |
| chr10-27416509-DEL-289 | chr10-27416510-DEL-289 |
| chr10-27422593-DEL-68 | chr10-27422594-DEL-68 |
| chr10-27432745-DEL-681 | chr10-27432746-DEL-681 |
| chr10-27469936-DEL-496 | chr10-27469922-DEL-492 |
| chr10-27477155-DEL-265 | chr10-27477157-DEL-262 |
| chr10-27479939-DEL-354 | chr10-27479940-DEL-354 |
| chr10-27480831-DEL-260 | chr10-27480832-DEL-260 |
| chr10-27515897-DEL-3560 | chr10-27515898-DEL-3560 |
| chr10-27538468-DEL-84 | chr10-27538502-DEL-63 |
| chr10-27538690-DEL-395 | chr10-27538685-DEL-401 |
| chr10-27557770-DEL-372 | chr10-27557771-DEL-372 |
| chr10-27617742-DEL-1254 | chr10-27617740-DEL-1255 |
| chr10-27626526-DEL-4618 | chr10-27626527-DEL-4618 |
| chr10-27665983-DEL-7915 | chr10-27665984-DEL-7915 |
| chr10-27688510-DEL-13252 | chr10-27688511-DEL-13252 |
| chr10-27728376-DEL-52 | chr10-27728377-DEL-52 |
| chr10-27734811-DEL-662 | chr10-27734812-DEL-662 |
| chr10-27743424-DEL-560 | chr10-27743425-DEL-560 |
| chr10-27747555-DEL-2014 | chr10-27747556-DEL-2014 |
| chr10-27760882-DEL-497 | chr10-27760883-DEL-497 |
| chr10-27770420-DEL-1212 | chr10-27770421-DEL-1212 |
| chr10-27779910-DEL-5864 | chr10-27779911-DEL-5864 |
| chr10-27787713-DEL-3429 | chr10-27787714-DEL-3429 |
| chr10-27791362-DEL-78 | chr10-27791363-DEL-78 |
| chr10-27812322-DEL-736 | chr10-27812323-DEL-736 |
| chr10-27814100-DEL-73 | chr10-27814101-DEL-73 |
| chr10-27862016-DEL-151 | chr10-27862017-DEL-151 |
| chr10-27893851-DEL-56 | chr10-27893852-DEL-56 |
| chr10-27896275-DEL-228 | chr10-27896276-DEL-228 |
| chr10-27897563-DEL-105 | chr10-27897564-DEL-105 |
| chr10-27916972-DEL-6240 | chr10-27916973-DEL-6240 |
| chr10-27928604-DEL-54 | chr10-27928603-DEL-84 |
| chr10-27930087-DEL-515 | chr10-27930089-DEL-514 |
| chr10-27998731-DEL-108 | chr10-27998731-DEL-106 |
| chr10-28012175-DEL-136 | chr10-28012176-DEL-132 |
| chr10-28023285-DEL-238 | chr10-28023286-DEL-238 |
| chr10-28071456-DEL-1600 | chr10-28071457-DEL-1600 |
| chr10-28077995-DEL-55 | chr10-28077992-DEL-59 |
| chr10-28092422-DEL-426 | chr10-28092423-DEL-427 |
| chr10-28113448-DEL-203 | chr10-28113449-DEL-203 |
| chr10-28134491-DEL-1199 | chr10-28134492-DEL-1199 |
| chr10-28141384-DEL-65 | chr10-28141385-DEL-65 |
| chr10-28164307-DEL-66 | chr10-28164306-DEL-70 |
| chr10-28172395-DEL-1156 | chr10-28172395-DEL-1156 |
| chr10-28217593-DEL-1888 | chr10-28217594-DEL-1888 |
| chr10-28244653-DEL-230 | chr10-28244654-DEL-230 |
| chr10-28253308-DEL-587 | chr10-28253299-DEL-588 |
| chr10-28303148-DEL-352 | chr10-28303149-DEL-349 |
| chr10-28345020-DEL-51 | chr10-28344994-DEL-52 |
| chr10-28353606-DEL-5154 | chr10-28353607-DEL-5154 |

| | |
|---|---|
| chr10-28402794-DEL-6539 | chr10-28402795-DEL-6539 |
| chr10-28452701-DEL-460 | chr10-28452659-DEL-461 |
| chr10-28486533-DEL-584 | chr10-28486534-DEL-584 |
| chr10-28495381-DEL-61 | chr10-28495380-DEL-63 |
| chr10-28498551-DEL-87 | chr10-28498552-DEL-87 |
| chr10-28515453-DEL-127 | chr10-28515451-DEL-133 |
| chr10-28537642-DEL-66 | chr10-28537643-DEL-66 |
| chr10-28583230-DEL-4537 | chr10-28583231-DEL-4537 |
| chr10-28590370-DEL-2384 | chr10-28590365-DEL-2383 |
| chr10-28617855-DEL-610 | chr10-28617856-DEL-610 |
| chr10-28660031-DEL-65 | chr10-28660031-DEL-65 |
| chr10-28677012-DEL-108 | chr10-28677040-DEL-60 |
| chr10-28685836-DEL-1636 | chr10-28685837-DEL-1636 |
| chr10-28700858-DEL-59 | chr10-28700857-DEL-64 |
| chr10-28704267-DEL-74 | chr10-28704268-DEL-74 |
| chr10-28724360-DEL-1191 | chr10-28724359-DEL-1191 |
| chr10-28751412-DEL-58 | chr10-28751413-DEL-58 |
| chr10-28781516-DEL-71 | chr10-28781518-DEL-68 |
| chr10-28781663-DEL-699 | chr10-28781664-DEL-699 |
| chr10-28783588-DEL-12373 | chr10-28783585-DEL-12367 |
| chr10-28798288-DEL-440 | chr10-28798289-DEL-440 |
| chr10-28804672-DEL-403 | chr10-28804673-DEL-403 |
| chr10-28810087-DEL-6596 | chr10-28810075-DEL-6597 |
| chr10-28825557-DEL-312 | chr10-28825556-DEL-313 |
| chr10-28826579-DEL-867 | chr10-28826580-DEL-867 |
| chr10-28827984-DEL-62 | chr10-28828003-DEL-62 |
| chr10-28886278-DEL-3988 | chr10-28886279-DEL-3988 |
| chr10-28886385-DEL-80 | chr10-28886386-DEL-80 |
| chr10-28906054-DEL-1249 | chr10-28906055-DEL-1249 |
| chr10-28918815-DEL-201 | chr10-28918815-DEL-201 |
| chr10-28927610-DEL-91 | chr10-28927610-DEL-91 |
| chr10-28929129-DEL-59 | chr10-28929112-DEL-78 |
| chr10-28937240-DEL-55 | chr10-28937240-DEL-55 |
| chr10-29021031-DEL-62 | chr10-29021025-DEL-50 |
| chr10-29043055-DEL-72 | chr10-29043072-DEL-74 |
| chr10-29060390-DEL-2902 | chr10-29060391-DEL-2902 |
| chr10-29077868-DEL-1151 | chr10-29077869-DEL-1151 |
| chr10-29083174-DEL-156 | chr10-29083175-DEL-156 |
| chr10-29091266-DEL-673 | chr10-29091267-DEL-673 |
| chr10-29106245-DEL-58 | chr10-29106238-DEL-66 |
| chr10-29117978-DEL-451 | chr10-29117979-DEL-451 |
| chr10-29119024-DEL-6609 | chr10-29119008-DEL-6609 |
| chr10-29119369-DEL-204 | chr10-29119370-DEL-204 |
| chr10-29163446-DEL-2011 | chr10-29163454-DEL-2009 |
| chr10-29165898-DEL-55 | chr10-29165876-DEL-68 |
| chr10-29168781-DEL-511 | chr10-29168782-DEL-511 |
| chr10-29182506-DEL-396 | chr10-29182507-DEL-396 |
| chr10-29205947-DEL-79 | chr10-29205948-DEL-79 |
| chr10-29252651-DEL-737 | chr10-29252652-DEL-737 |
| chr10-29263810-DEL-88 | chr10-29263809-DEL-92 |
| chr10-29270753-DEL-168 | chr10-29270754-DEL-168 |
| chr10-29271147-DEL-475 | chr10-29271148-DEL-474 |
| chr10-29282747-DEL-202 | chr10-29282748-DEL-202 |
| chr10-29284389-DEL-1022 | chr10-29284390-DEL-1022 |
| chr10-29350784-DEL-198 | chr10-29350785-DEL-198 |
| chr10-29450965-DEL-550 | chr10-29450966-DEL-550 |
| chr10-29467179-DEL-68 | chr10-29467180-DEL-66 |
| chr10-29481943-DEL-452 | chr10-29481936-DEL-448 |
| chr10-29495615-DEL-6514 | chr10-29495616-DEL-6514 |
| chr10-29532689-DEL-1395 | chr10-29532691-DEL-1396 |
| chr10-29550847-DEL-62 | chr10-29550866-DEL-59 |
| chr10-29553247-DEL-1609 | chr10-29553248-DEL-1609 |
| chr10-29599750-DEL-471 | chr10-29599751-DEL-471 |
| chr10-29636952-DEL-3162 | chr10-29636954-DEL-3157 |
| chr10-29643838-DEL-6266 | chr10-29643839-DEL-6266 |
| chr10-29673104-DEL-1891 | chr10-29673105-DEL-1891 |
| chr10-29747821-DEL-3594 | chr10-29747822-DEL-3594 |
| chr10-29766631-DEL-441 | chr10-29766632-DEL-441 |
| chr10-29772690-DEL-213 | chr10-29772691-DEL-213 |
| chr10-29800272-DEL-6396 | chr10-29800273-DEL-6396 |
| chr10-29807112-DEL-65 | chr10-29807113-DEL-70 |
| chr10-29822875-DEL-166 | chr10-29822918-DEL-120 |
| chr10-29840335-DEL-1342 | chr10-29840336-DEL-1342 |
| chr10-29846952-DEL-82 | chr10-29846955-DEL-54 |
| chr10-29848674-DEL-99 | chr10-29848671-DEL-90 |
| chr10-29898134-DEL-62 | chr10-29898135-DEL-62 |
| chr10-29962982-DEL-185 | chr10-29962983-DEL-185 |
| chr10-29989554-DEL-452 | chr10-29989555-DEL-452 |
| chr10-29999401-DEL-75 | chr10-29999402-DEL-75 |
| chr10-30009350-DEL-431 | chr10-30009351-DEL-431 |

| | |
|---|---|
| chr10-30018056-DEL-381 | chr10-30018057-DEL-381 |
| chr10-30021460-DEL-417 | chr10-30021461-DEL-417 |
| chr10-30023794-DEL-409 | chr10-30023795-DEL-409 |
| chr10-30032796-DEL-769 | chr10-30032797-DEL-769 |
| chr10-30034782-DEL-201 | chr10-30034783-DEL-201 |
| chr10-30041880-DEL-1091 | chr10-30041880-DEL-1091 |
| chr10-30091994-DEL-7108 | chr10-30091995-DEL-7108 |
| chr10-30142246-DEL-62 | chr10-30142233-DEL-62 |
| chr10-30157090-DEL-58 | chr10-30157091-DEL-58 |
| chr10-30172111-DEL-5507 | chr10-30172112-DEL-5507 |
| chr10-30183983-DEL-481 | chr10-30183980-DEL-482 |
| chr10-30189687-DEL-586 | chr10-30189688-DEL-586 |
| chr10-30190437-DEL-349 | chr10-30190438-DEL-349 |
| chr10-30201409-DEL-1077 | chr10-30201408-DEL-1075 |
| chr10-30209556-DEL-54 | chr10-30209557-DEL-54 |
| chr10-30223588-DEL-147 | chr10-30223586-DEL-147 |
| chr10-30268676-DEL-92 | chr10-30268679-DEL-75 |
| chr10-30268953-DEL-2324 | chr10-30268954-DEL-2324 |
| chr10-30285527-DEL-60 | chr10-30285529-DEL-54 |
| chr10-30292008-DEL-68 | chr10-30292002-DEL-78 |
| chr10-30299498-DEL-54 | chr10-30299499-DEL-54 |
| chr10-30304808-DEL-1499 | chr10-30304809-DEL-1497 |
| chr10-30344302-DEL-1996 | chr10-30344303-DEL-1996 |
| chr10-30357230-DEL-4476 | chr10-30357231-DEL-4476 |
| chr10-30399173-DEL-2279 | chr10-30399158-DEL-2280 |
| chr10-30425562-DEL-83 | chr10-30425516-DEL-82 |
| chr10-30428949-DEL-1837 | chr10-30428950-DEL-1837 |
| chr10-30431040-DEL-498 | chr10-30431041-DEL-498 |
| chr10-30446707-DEL-6457 | chr10-30446708-DEL-6457 |
| chr10-30478492-DEL-201 | chr10-30478493-DEL-201 |
| chr10-30490472-DEL-221 | chr10-30490473-DEL-221 |
| chr10-30535292-DEL-513 | chr10-30535293-DEL-513 |
| chr10-30544466-DEL-90 | chr10-30544467-DEL-90 |
| chr10-30562744-DEL-62 | chr10-30562745-DEL-62 |
| chr10-30582573-DEL-52 | chr10-30582574-DEL-52 |
| chr10-30588900-DEL-879 | chr10-30588901-DEL-879 |
| chr10-30608600-DEL-150 | chr10-30608601-DEL-150 |
| chr10-30615911-DEL-81 | chr10-30615912-DEL-81 |
| chr10-30652737-DEL-1020 | chr10-30652738-DEL-1020 |
| chr10-30660896-DEL-83 | chr10-30660897-DEL-83 |
| chr10-30689983-DEL-7096 | chr10-30689984-DEL-7096 |
| chr10-30699674-DEL-52 | chr10-30699673-DEL-53 |
| chr10-30738680-DEL-129 | chr10-30738681-DEL-129 |
| chr10-30801055-DEL-58 | chr10-30801058-DEL-56 |
| chr10-30818791-DEL-87 | chr10-30818792-DEL-87 |
| chr10-30841384-DEL-52 | chr10-30841385-DEL-52 |
| chr10-30842100-DEL-658 | chr10-30842101-DEL-658 |
| chr10-30866864-DEL-105 | chr10-30866893-DEL-77 |
| chr10-30876450-DEL-788 | chr10-30876451-DEL-788 |
| chr10-30896444-DEL-166 | chr10-30896444-DEL-166 |
| chr10-30930864-DEL-6657 | chr10-30930865-DEL-6657 |
| chr10-30938237-DEL-122 | chr10-30938238-DEL-76 |
| chr10-30942055-DEL-99 | chr10-30942055-DEL-99 |
| chr10-30944259-DEL-74 | chr10-30944258-DEL-70 |
| chr10-30946993-DEL-227 | chr10-30946994-DEL-227 |
| chr10-30956750-DEL-6252 | chr10-30956751-DEL-6252 |
| chr10-30964558-DEL-7263 | chr10-30964559-DEL-7263 |
| chr10-30991990-DEL-60 | chr10-30991991-DEL-60 |
| chr10-30998556-DEL-1243 | chr10-30998556-DEL-1244 |
| chr10-31037342-DEL-227 | chr10-31037355-DEL-251 |
| chr10-31061562-DEL-2932 | chr10-31061563-DEL-2932 |
| chr10-31091962-DEL-113 | chr10-31091990-DEL-133 |
| chr10-31109148-DEL-765 | chr10-31109149-DEL-765 |
| chr10-31125487-DEL-63 | chr10-31125488-DEL-63 |
| chr10-31135077-DEL-90 | chr10-31135076-DEL-91 |
| chr10-31143070-DEL-2036 | chr10-31143071-DEL-2036 |
| chr10-31152342-DEL-691 | chr10-31152343-DEL-689 |
| chr10-31156006-DEL-458 | chr10-31156007-DEL-458 |
| chr10-31189211-DEL-105 | chr10-31189217-DEL-126 |
| chr10-31192519-DEL-63 | chr10-31192520-DEL-63 |
| chr10-31209230-DEL-575 | chr10-31209231-DEL-575 |
| chr10-31229698-DEL-96 | chr10-31229685-DEL-104 |
| chr10-31236984-DEL-112 | chr10-31236985-DEL-112 |
| chr10-31239492-DEL-219 | chr10-31239493-DEL-219 |
| chr10-31246075-DEL-332 | chr10-31246076-DEL-332 |
| chr10-31248128-DEL-58 | chr10-31248129-DEL-58 |
| chr10-31249113-DEL-92 | chr10-31249114-DEL-92 |
| chr10-31276291-DEL-168 | chr10-31276292-DEL-168 |
| chr10-31281169-DEL-6443 | chr10-31281170-DEL-6443 |
| chr10-31289464-DEL-733 | chr10-31289465-DEL-733 |

| | |
|---|---|
| chr10-31312510-DEL-115 | chr10-31312511-DEL-115 |
| chr10-31327991-DEL-423 | chr10-31327992-DEL-423 |
| chr10-31334199-DEL-214 | chr10-31334200-DEL-214 |
| chr10-31353054-DEL-386 | chr10-31353055-DEL-386 |
| chr10-31358620-DEL-7090 | chr10-31358621-DEL-7090 |
| chr10-31385042-DEL-107 | chr10-31385043-DEL-107 |
| chr10-31425770-DEL-206 | chr10-31425771-DEL-206 |
| chr10-31442245-DEL-206 | chr10-31442246-DEL-206 |
| chr10-31444312-DEL-64 | chr10-31444313-DEL-66 |
| chr10-31451699-DEL-88 | chr10-31451700-DEL-88 |
| chr10-31487505-DEL-52 | chr10-31487492-DEL-66 |
| chr10-31520983-DEL-61 | chr10-31520968-DEL-77 |
| chr10-31574605-DEL-63 | chr10-31574606-DEL-63 |
| chr10-31623013-DEL-55 | chr10-31623014-DEL-55 |
| chr10-31624842-DEL-64 | chr10-31624843-DEL-64 |
| chr10-31626111-DEL-4775 | chr10-31626112-DEL-4775 |
| chr10-31657677-DEL-1571 | chr10-31657678-DEL-1571 |
| chr10-31682791-DEL-96 | chr10-31682792-DEL-96 |
| chr10-31742016-DEL-414 | chr10-31742014-DEL-417 |
| chr10-31761691-DEL-214 | chr10-31761691-DEL-214 |
| chr10-31763137-DEL-88 | chr10-31763165-DEL-72 |
| chr10-31765287-DEL-2247 | chr10-31765288-DEL-2247 |
| chr10-31774776-DEL-271 | chr10-31774776-DEL-271 |
| chr10-31855988-DEL-6499 | chr10-31855989-DEL-6499 |
| chr10-31873179-DEL-3335 | chr10-31873193-DEL-3329 |
| chr10-31879767-DEL-121 | chr10-31879768-DEL-121 |
| chr10-31900239-DEL-283 | chr10-31900240-DEL-283 |
| chr10-31915507-DEL-6478 | chr10-31915508-DEL-6478 |
| chr10-31946832-DEL-814 | chr10-31946833-DEL-814 |
| chr10-31958467-DEL-644 | chr10-31958478-DEL-636 |
| chr10-31987783-DEL-73 | chr10-31987786-DEL-71 |
| chr10-32037631-DEL-64 | chr10-32037632-DEL-64 |
| chr10-32039482-DEL-68 | chr10-32039481-DEL-62 |
| chr10-32058147-DEL-64 | chr10-32058140-DEL-64 |
| chr10-32085629-DEL-6038 | chr10-32085629-DEL-6039 |
| chr10-32113925-DEL-57 | chr10-32113886-DEL-67 |
| chr10-32114473-DEL-1679 | chr10-32114474-DEL-1679 |
| chr10-32116541-DEL-63 | chr10-32116510-DEL-95 |
| chr10-32164027-DEL-61 | chr10-32164028-DEL-60 |
| chr10-32168220-DEL-292 | chr10-32168221-DEL-292 |
| chr10-32175680-DEL-94 | chr10-32175681-DEL-94 |
| chr10-32212556-DEL-1988 | chr10-32212557-DEL-1988 |
| chr10-32226388-DEL-8830 | chr10-32226389-DEL-8830 |
| chr10-32248741-DEL-101 | chr10-32248742-DEL-101 |
| chr10-32255626-DEL-59 | chr10-32255619-DEL-60 |
| chr10-32258170-DEL-59 | chr10-32258162-DEL-94 |
| chr10-32280833-DEL-427 | chr10-32280834-DEL-427 |
| chr10-32321202-DEL-260 | chr10-32321200-DEL-255 |
| chr10-32324288-DEL-479 | chr10-32324289-DEL-479 |
| chr10-32325468-DEL-52 | chr10-32325469-DEL-52 |
| chr10-32326254-DEL-927 | chr10-32326255-DEL-927 |
| chr10-32375130-DEL-6655 | chr10-32375131-DEL-6655 |
| chr10-32410494-DEL-213 | chr10-32410495-DEL-213 |
| chr10-32441751-DEL-422 | chr10-32441752-DEL-422 |
| chr10-32480158-DEL-56 | chr10-32480159-DEL-56 |
| chr10-32483598-DEL-3139 | chr10-32483599-DEL-3139 |
| chr10-32488081-DEL-114 | chr10-32488082-DEL-114 |
| chr10-32488541-DEL-2015 | chr10-32488542-DEL-2015 |
| chr10-32511445-DEL-3665 | chr10-32511446-DEL-3665 |
| chr10-32564500-DEL-1071 | chr10-32564499-DEL-1072 |
| chr10-32583204-DEL-66 | chr10-32583205-DEL-66 |
| chr10-32584839-DEL-2065 | chr10-32584840-DEL-2065 |
| chr10-32605583-DEL-110 | chr10-32605584-DEL-110 |
| chr10-32630656-DEL-174 | chr10-32630654-DEL-130 |
| chr10-32634314-DEL-355 | chr10-32634315-DEL-355 |
| chr10-32634992-DEL-101 | chr10-32634993-DEL-76 |
| chr10-32686218-DEL-56 | chr10-32686219-DEL-56 |
| chr10-32692596-DEL-7044 | chr10-32692597-DEL-7044 |
| chr10-32714469-DEL-2257 | chr10-32714470-DEL-2257 |
| chr10-32718886-DEL-496 | chr10-32718887-DEL-496 |
| chr10-32734944-DEL-263 | chr10-32734945-DEL-263 |
| chr10-32744465-DEL-53 | chr10-32744466-DEL-53 |
| chr10-32774367-DEL-1201 | chr10-32774368-DEL-1201 |
| chr10-32785214-DEL-112 | chr10-32785175-DEL-140 |
| chr10-32787707-DEL-1963 | chr10-32787707-DEL-1964 |
| chr10-32812604-DEL-2678 | chr10-32812604-DEL-2678 |
| chr10-32829955-DEL-59 | chr10-32829955-DEL-59 |
| chr10-32836207-DEL-1089 | chr10-32836208-DEL-1089 |
| chr10-32841954-DEL-6747 | chr10-32841955-DEL-6747 |
| chr10-32893770-DEL-579 | chr10-32893770-DEL-579 |

| | |
|---|---|
| chr10-32900097-DEL-143 | chr10-32900097-DEL-143 |
| chr10-32939792-DEL-339 | chr10-32939793-DEL-339 |
| chr10-32987535-DEL-69 | chr10-32987533-DEL-65 |
| chr10-32988444-DEL-60 | chr10-32988446-DEL-59 |
| chr10-33008711-DEL-118 | chr10-33008717-DEL-113 |
| chr10-33009642-DEL-453 | chr10-33009643-DEL-453 |
| chr10-33025889-DEL-75 | chr10-33025890-DEL-75 |
| chr10-33027482-DEL-188 | chr10-33027483-DEL-188 |
| chr10-33032224-DEL-85 | chr10-33032225-DEL-85 |
| chr10-33070653-DEL-78 | chr10-33070653-DEL-78 |
| chr10-33073921-DEL-101 | chr10-33073918-DEL-105 |
| chr10-33111287-DEL-700 | chr10-33111288-DEL-700 |
| chr10-33119786-DEL-80 | chr10-33119786-DEL-80 |
| chr10-33120117-DEL-3912 | chr10-33120108-DEL-3913 |
| chr10-33138795-DEL-6227 | chr10-33138796-DEL-6227 |
| chr10-33145443-DEL-402 | chr10-33145444-DEL-402 |
| chr10-33148142-DEL-680 | chr10-33148143-DEL-680 |
| chr10-33188347-DEL-453 | chr10-33188348-DEL-453 |
| chr10-33189748-DEL-6106 | chr10-33189749-DEL-6106 |
| chr10-33234978-DEL-211 | chr10-33234979-DEL-211 |
| chr10-33236340-DEL-224 | chr10-33236341-DEL-224 |
| chr10-33254302-DEL-261 | chr10-33254303-DEL-261 |
| chr10-33276775-DEL-64 | chr10-33276776-DEL-64 |
| chr10-33298422-DEL-68 | chr10-33298420-DEL-69 |
| chr10-33332075-DEL-289 | chr10-33332075-DEL-289 |
| chr10-33339529-DEL-338 | chr10-33339529-DEL-338 |
| chr10-33347291-DEL-156 | chr10-33347292-DEL-156 |
| chr10-33378777-DEL-6554 | chr10-33378777-DEL-6554 |
| chr10-33390347-DEL-230 | chr10-33390347-DEL-230 |
| chr10-33401602-DEL-244 | chr10-33401603-DEL-244 |
| chr10-33448262-DEL-6364 | chr10-33448263-DEL-6364 |
| chr10-33454735-DEL-111 | chr10-33454735-DEL-111 |
| chr10-33470532-DEL-83 | chr10-33470525-DEL-83 |
| chr10-33477863-DEL-108 | chr10-33477866-DEL-106 |
| chr10-33481762-DEL-6353 | chr10-33481749-DEL-6362 |
| chr10-33506305-DEL-83 | chr10-33506306-DEL-83 |
| chr10-33512854-DEL-6390 | chr10-33512847-DEL-6386 |
| chr10-33567498-DEL-4275 | chr10-33567494-DEL-4275 |
| chr10-33576585-DEL-351 | chr10-33576563-DEL-362 |
| chr10-33580791-DEL-631 | chr10-33580792-DEL-631 |
| chr10-33597462-DEL-197 | chr10-33597463-DEL-197 |
| chr10-33697496-DEL-6459 | chr10-33697496-DEL-6459 |
| chr10-33743207-DEL-970 | chr10-33743208-DEL-970 |
| chr10-33762775-DEL-4314 | chr10-33762776-DEL-4314 |
| chr10-33794661-DEL-210 | chr10-33794662-DEL-210 |
| chr10-33823304-DEL-135 | chr10-33823305-DEL-135 |
| chr10-33833158-DEL-1730 | chr10-33833159-DEL-1730 |
| chr10-33833871-DEL-187 | chr10-33833872-DEL-187 |
| chr10-33850306-DEL-415 | chr10-33850307-DEL-415 |
| chr10-33869045-DEL-212 | chr10-33869046-DEL-212 |
| chr10-33870897-DEL-231 | chr10-33870898-DEL-231 |
| chr10-33891641-DEL-152 | chr10-33891681-DEL-113 |
| chr10-33899848-DEL-8163 | chr10-33899850-DEL-8162 |
| chr10-33910576-DEL-57 | chr10-33910579-DEL-55 |
| chr10-33913760-DEL-956 | chr10-33913761-DEL-956 |
| chr10-33948677-DEL-749 | chr10-33948678-DEL-749 |
| chr10-33953466-DEL-200 | chr10-33953465-DEL-202 |
| chr10-33957082-DEL-178 | chr10-33957083-DEL-178 |
| chr10-33960990-DEL-81 | chr10-33960991-DEL-81 |
| chr10-34012719-DEL-2119 | chr10-34012720-DEL-2119 |
| chr10-34022289-DEL-285 | chr10-34022290-DEL-285 |
| chr10-34044400-DEL-211 | chr10-34044401-DEL-211 |
| chr10-34054173-DEL-56 | chr10-34054171-DEL-61 |
| chr10-34058603-DEL-54 | chr10-34058604-DEL-54 |
| chr10-34081436-DEL-385 | chr10-34081435-DEL-387 |
| chr10-34095854-DEL-176 | chr10-34095856-DEL-175 |
| chr10-34097770-DEL-223 | chr10-34097771-DEL-224 |
| chr10-34116105-DEL-184 | chr10-34116106-DEL-184 |
| chr10-34124728-DEL-61 | chr10-34124729-DEL-61 |
| chr10-34141058-DEL-213 | chr10-34141059-DEL-213 |
| chr10-34143080-DEL-1999 | chr10-34143085-DEL-1996 |
| chr10-34151793-DEL-51 | chr10-34151794-DEL-51 |
| chr10-34157727-DEL-98 | chr10-34157728-DEL-98 |
| chr10-34165045-DEL-454 | chr10-34165046-DEL-454 |
| chr10-34169200-DEL-243 | chr10-34169201-DEL-243 |
| chr10-34181888-DEL-201 | chr10-34181889-DEL-201 |
| chr10-34190905-DEL-53 | chr10-34190906-DEL-53 |
| chr10-34192169-DEL-88 | chr10-34192175-DEL-88 |
| chr10-34204736-DEL-71 | chr10-34204737-DEL-70 |
| chr10-34221834-DEL-65 | chr10-34221835-DEL-65 |

| | |
|---|---|
| chr10-34224124-DEL-3988 | chr10-34224125-DEL-3988 |
| chr10-34250727-DEL-1402 | chr10-34250728-DEL-1402 |
| chr10-34288957-DEL-81 | chr10-34288966-DEL-52 |
| chr10-34310407-DEL-7190 | chr10-34310408-DEL-7190 |
| chr10-34344869-DEL-75 | chr10-34344874-DEL-71 |
| chr10-34361910-DEL-4280 | chr10-34361911-DEL-4280 |
| chr10-34427568-DEL-57 | chr10-34427521-DEL-74 |
| chr10-34435555-DEL-1051 | chr10-34435556-DEL-1051 |
| chr10-34437758-DEL-72 | chr10-34437759-DEL-72 |
| chr10-34441581-DEL-629 | chr10-34441582-DEL-629 |
| chr10-34471075-DEL-1156 | chr10-34471077-DEL-1155 |
| chr10-34476211-DEL-52 | chr10-34476212-DEL-52 |
| chr10-34506569-DEL-328 | chr10-34506570-DEL-328 |
| chr10-34521709-DEL-195 | chr10-34521710-DEL-195 |
| chr10-34569719-DEL-332 | chr10-34569720-DEL-332 |
| chr10-34590724-DEL-125 | chr10-34590725-DEL-125 |
| chr10-34601870-DEL-3248 | chr10-34601871-DEL-3248 |
| chr10-34614930-DEL-10801 | chr10-34614930-DEL-10801 |
| chr10-34648981-DEL-3115 | chr10-34648982-DEL-3115 |
| chr10-34664158-DEL-74 | chr10-34664159-DEL-74 |
| chr10-34681681-DEL-89 | chr10-34681682-DEL-89 |
| chr10-34686603-DEL-327 | chr10-34686604-DEL-327 |
| chr10-34694106-DEL-172 | chr10-34694105-DEL-173 |
| chr10-34695851-DEL-919 | chr10-34695852-DEL-919 |
| chr10-34701811-DEL-212 | chr10-34701812-DEL-212 |
| chr10-34746280-DEL-60 | chr10-34746289-DEL-52 |
| chr10-34764000-DEL-522 | chr10-34764001-DEL-522 |
| chr10-34854301-DEL-384 | chr10-34854302-DEL-384 |
| chr10-34949535-DEL-6520 | chr10-34949536-DEL-6520 |
| chr10-34950994-DEL-332 | chr10-34950995-DEL-332 |
| chr10-34959591-DEL-3182 | chr10-34959592-DEL-3182 |
| chr10-34987782-DEL-60 | chr10-34987783-DEL-60 |
| chr10-34989045-DEL-777 | chr10-34989046-DEL-777 |
| chr10-35020522-DEL-852 | chr10-35020523-DEL-852 |
| chr10-35022421-DEL-98 | chr10-35022462-DEL-50 |
| chr10-35034228-DEL-1503 | chr10-35034229-DEL-1503 |
| chr10-35047144-DEL-244 | chr10-35047145-DEL-244 |
| chr10-35060887-DEL-336 | chr10-35060888-DEL-336 |
| chr10-35064201-DEL-479 | chr10-35064202-DEL-479 |
| chr10-35081352-DEL-160 | chr10-35081353-DEL-160 |
| chr10-35100653-DEL-666 | chr10-35100654-DEL-666 |
| chr10-35113156-DEL-59 | chr10-35113157-DEL-59 |
| chr10-35160128-DEL-184 | chr10-35160128-DEL-184 |
| chr10-35165557-DEL-327 | chr10-35165557-DEL-327 |
| chr10-35168380-DEL-60 | chr10-35168385-DEL-56 |
| chr10-35186069-DEL-7015 | chr10-35186070-DEL-7015 |
| chr10-35208864-DEL-192 | chr10-35208865-DEL-192 |
| chr10-35209243-DEL-433 | chr10-35209244-DEL-416 |
| chr10-35227063-DEL-6605 | chr10-35227064-DEL-6605 |
| chr10-35244637-DEL-72 | chr10-35244638-DEL-72 |
| chr10-35293611-DEL-431 | chr10-35293611-DEL-431 |
| chr10-35294446-DEL-54 | chr10-35294447-DEL-54 |
| chr10-35304687-DEL-327 | chr10-35304688-DEL-327 |
| chr10-35326883-DEL-5461 | chr10-35326884-DEL-5461 |
| chr10-35359407-DEL-76 | chr10-35359408-DEL-76 |
| chr10-35381464-DEL-495 | chr10-35381465-DEL-495 |
| chr10-35391273-DEL-6657 | chr10-35391274-DEL-6657 |
| chr10-35409538-DEL-51 | chr10-35409539-DEL-51 |
| chr10-35417362-DEL-2308 | chr10-35417363-DEL-2308 |
| chr10-35424061-DEL-441 | chr10-35424064-DEL-438 |
| chr10-35434818-DEL-326 | chr10-35434819-DEL-326 |
| chr10-35441733-DEL-1061 | chr10-35441734-DEL-1059 |
| chr10-35448125-DEL-58 | chr10-35448126-DEL-62 |
| chr10-35510443-DEL-6348 | chr10-35510444-DEL-6348 |
| chr10-35527706-DEL-53 | chr10-35527697-DEL-54 |
| chr10-35537174-DEL-133 | chr10-35537175-DEL-133 |
| chr10-35544778-DEL-6378 | chr10-35544779-DEL-6378 |
| chr10-35552473-DEL-7825 | chr10-35552474-DEL-7825 |
| chr10-35565150-DEL-209 | chr10-35565151-DEL-209 |
| chr10-35584603-DEL-4373 | chr10-35584604-DEL-4373 |
| chr10-35589251-DEL-353 | chr10-35589252-DEL-350 |
| chr10-35598627-DEL-2918 | chr10-35598628-DEL-2918 |
| chr10-35600744-DEL-75 | chr10-35600745-DEL-75 |
| chr10-35609776-DEL-64 | chr10-35609777-DEL-64 |
| chr10-35614597-DEL-243 | chr10-35614598-DEL-243 |
| chr10-35628197-DEL-140 | chr10-35628198-DEL-152 |
| chr10-35669566-DEL-1263 | chr10-35669567-DEL-1263 |
| chr10-35672289-DEL-92 | chr10-35672294-DEL-94 |
| chr10-35728981-DEL-976 | chr10-35728982-DEL-976 |
| chr10-35730712-DEL-251 | chr10-35730726-DEL-243 |

| | |
|---|---|
| chr10-35736758-DEL-2875 | chr10-35736759-DEL-2875 |
| chr10-35744151-DEL-152 | chr10-35744145-DEL-159 |
| chr10-35840370-DEL-400 | chr10-35840371-DEL-400 |
| chr10-35848219-DEL-581 | chr10-35848214-DEL-571 |
| chr10-35862194-DEL-162 | chr10-35862195-DEL-162 |
| chr10-35868583-DEL-200 | chr10-35868584-DEL-200 |
| chr10-35943271-DEL-281 | chr10-35943272-DEL-281 |
| chr10-35971657-DEL-67 | chr10-35971677-DEL-52 |
| chr10-35999361-DEL-84 | chr10-35999361-DEL-84 |
| chr10-36036664-DEL-6197 | chr10-36036665-DEL-6197 |
| chr10-36046305-DEL-892 | chr10-36046307-DEL-893 |
| chr10-36062936-DEL-649 | chr10-36062937-DEL-649 |
| chr10-36101367-DEL-214 | chr10-36101368-DEL-214 |
| chr10-36103196-DEL-52 | chr10-36103171-DEL-84 |
| chr10-36110868-DEL-207 | chr10-36110869-DEL-207 |
| chr10-36140077-DEL-247 | chr10-36140078-DEL-247 |
| chr10-36172445-DEL-1904 | chr10-36172446-DEL-1904 |
| chr10-36176860-DEL-189 | chr10-36176861-DEL-177 |
| chr10-36181862-DEL-253 | chr10-36181863-DEL-253 |
| chr10-36202650-DEL-60 | chr10-36202651-DEL-70 |
| chr10-36295949-DEL-2845 | chr10-36295950-DEL-2845 |
| chr10-36300837-DEL-382 | chr10-36300882-DEL-332 |
| chr10-36387672-DEL-506 | chr10-36387673-DEL-506 |
| chr10-36414969-DEL-95 | chr10-36414972-DEL-96 |
| chr10-36438127-DEL-1206 | chr10-36438128-DEL-1206 |
| chr10-36470997-DEL-1893 | chr10-36470998-DEL-1893 |
| chr10-36512467-DEL-172 | chr10-36512468-DEL-170 |
| chr10-36513751-DEL-7198 | chr10-36513752-DEL-7198 |
| chr10-36544638-DEL-1315 | chr10-36544639-DEL-1315 |
| chr10-36613610-DEL-1863 | chr10-36613611-DEL-1863 |
| chr10-36621692-DEL-940 | chr10-36621693-DEL-940 |
| chr10-36632748-DEL-855 | chr10-36632746-DEL-856 |
| chr10-36632971-DEL-2080 | chr10-36632967-DEL-2081 |
| chr10-36646275-DEL-1331 | chr10-36646276-DEL-1331 |
| chr10-36653219-DEL-4151 | chr10-36653220-DEL-4151 |
| chr10-36662085-DEL-165 | chr10-36662086-DEL-165 |
| chr10-36683469-DEL-54 | chr10-36683462-DEL-62 |
| chr10-36697862-DEL-1452 | chr10-36697863-DEL-1452 |
| chr10-36724919-DEL-161 | chr10-36724911-DEL-132 |
| chr10-36757450-DEL-539 | chr10-36757451-DEL-539 |
| chr10-36758420-DEL-1364 | chr10-36758424-DEL-1367 |
| chr10-36770228-DEL-94 | chr10-36770229-DEL-94 |
| chr10-36778845-DEL-74 | chr10-36778846-DEL-74 |
| chr10-36800664-DEL-60 | chr10-36800662-DEL-51 |
| chr10-36811538-DEL-55 | chr10-36811526-DEL-54 |
| chr10-36818308-DEL-6477 | chr10-36818309-DEL-6477 |
| chr10-36830074-DEL-228 | chr10-36830076-DEL-227 |
| chr10-36844148-DEL-73 | chr10-36844115-DEL-73 |
| chr10-36844742-DEL-57 | chr10-36844743-DEL-69 |
| chr10-36878980-DEL-137 | chr10-36878981-DEL-137 |
| chr10-36885005-DEL-66 | chr10-36885006-DEL-66 |
| chr10-36885496-DEL-445 | chr10-36885497-DEL-445 |
| chr10-36886919-DEL-660 | chr10-36886920-DEL-660 |
| chr10-36892777-DEL-208 | chr10-36892778-DEL-208 |
| chr10-36927369-DEL-2085 | chr10-36927371-DEL-2084 |
| chr10-36960297-DEL-79 | chr10-36960283-DEL-76 |
| chr10-36999807-DEL-77 | chr10-36999808-DEL-77 |
| chr10-37026215-DEL-61 | chr10-37026216-DEL-61 |
| chr10-37049743-DEL-246 | chr10-37049748-DEL-238 |
| chr10-37082002-DEL-158 | chr10-37081983-DEL-159 |
| chr10-37087788-DEL-124 | chr10-37087789-DEL-124 |
| chr10-37093505-DEL-811 | chr10-37093506-DEL-811 |
| chr10-37139920-DEL-608 | chr10-37139921-DEL-608 |
| chr10-37151057-DEL-62 | chr10-37151058-DEL-62 |
| chr10-37172646-DEL-164 | chr10-37172647-DEL-164 |
| chr10-37217256-DEL-61 | chr10-37217257-DEL-61 |
| chr10-37293075-DEL-68 | chr10-37293076-DEL-68 |
| chr10-37297560-DEL-62 | chr10-37297570-DEL-56 |
| chr10-37308635-DEL-58 | chr10-37308636-DEL-58 |
| chr10-37352172-DEL-340 | chr10-37352173-DEL-339 |
| chr10-37358604-DEL-105 | chr10-37358605-DEL-105 |
| chr10-37372604-DEL-448 | chr10-37372605-DEL-448 |
| chr10-37385563-DEL-3268 | chr10-37385564-DEL-3268 |
| chr10-37398595-DEL-2393 | chr10-37398589-DEL-2394 |
| chr10-37411291-DEL-1593 | chr10-37411292-DEL-1593 |
| chr10-37483915-DEL-2209 | chr10-37483916-DEL-2209 |
| chr10-37484546-DEL-586 | chr10-37484547-DEL-586 |
| chr10-37501211-DEL-70 | chr10-37501209-DEL-71 |
| chr10-37528563-DEL-1021 | chr10-37528564-DEL-1021 |
| chr10-37577715-DEL-6887 | chr10-37577716-DEL-6887 |

| | |
|---|---|
| chr10-37601231-DEL-463 | chr10-37601232-DEL-463 |
| chr10-37606061-DEL-6348 | chr10-37606062-DEL-6348 |
| chr10-37622520-DEL-1059 | chr10-37622521-DEL-1059 |
| chr10-37634000-DEL-469 | chr10-37634001-DEL-454 |
| chr10-37659192-DEL-423 | chr10-37659193-DEL-423 |
| chr10-37676031-DEL-1480 | chr10-37676031-DEL-1480 |
| chr10-37696472-DEL-59 | chr10-37696467-DEL-66 |
| chr10-37715420-DEL-99 | chr10-37715421-DEL-99 |
| chr10-37727458-DEL-2231 | chr10-37727433-DEL-2231 |
| chr10-37760096-DEL-148 | chr10-37760097-DEL-123 |
| chr10-37760758-DEL-526 | chr10-37760759-DEL-526 |
| chr10-37765248-DEL-68 | chr10-37765252-DEL-68 |
| chr10-37799465-DEL-407 | chr10-37799466-DEL-407 |
| chr10-37811271-DEL-6363 | chr10-37811272-DEL-6363 |
| chr10-37823314-DEL-2347 | chr10-37823315-DEL-2347 |
| chr10-37837582-DEL-402 | chr10-37837583-DEL-402 |
| chr10-37840523-DEL-6614 | chr10-37840533-DEL-6601 |
| chr10-37875597-DEL-3251 | chr10-37875598-DEL-3251 |
| chr10-37889441-DEL-51 | chr10-37889441-DEL-57 |
| chr10-37919733-DEL-1437 | chr10-37919732-DEL-1438 |
| chr10-37938172-DEL-255 | chr10-37938173-DEL-255 |
| chr10-37941382-DEL-54 | chr10-37941370-DEL-70 |
| chr10-37953267-DEL-54 | chr10-37953268-DEL-54 |
| chr10-37997694-DEL-88 | chr10-37997695-DEL-84 |
| chr10-38008350-DEL-72 | chr10-38008351-DEL-72 |
| chr10-38020208-DEL-2617 | chr10-38020208-DEL-2618 |
| chr10-38020879-DEL-70 | chr10-38020880-DEL-70 |
| chr10-38027581-DEL-207 | chr10-38027582-DEL-207 |
| chr10-38030892-DEL-351 | chr10-38030893-DEL-351 |
| chr10-38042772-DEL-6309 | chr10-38042773-DEL-6309 |
| chr10-38049162-DEL-3105 | chr10-38049163-DEL-3105 |
| chr10-38086080-DEL-202 | chr10-38086082-DEL-199 |
| chr10-38094401-DEL-6448 | chr10-38094402-DEL-6448 |
| chr10-38101553-DEL-71 | chr10-38101554-DEL-71 |
| chr10-38102899-DEL-332 | chr10-38102900-DEL-332 |
| chr10-38128127-DEL-1913 | chr10-38128127-DEL-1913 |
| chr10-38134106-DEL-6468 | chr10-38134107-DEL-6468 |
| chr10-38145567-DEL-3787 | chr10-38145568-DEL-3787 |
| chr10-38153817-DEL-80 | chr10-38153818-DEL-80 |
| chr10-38177574-DEL-80 | chr10-38177575-DEL-80 |
| chr10-38258839-DEL-96 | chr10-38258856-DEL-82 |
| chr10-38261089-DEL-1722 | chr10-38261090-DEL-1722 |
| chr10-38278503-DEL-97 | chr10-38278503-DEL-97 |
| chr10-38280442-DEL-1138 | chr10-38280443-DEL-1138 |
| chr10-38287731-DEL-177 | chr10-38287722-DEL-186 |
| chr10-38293459-DEL-1769 | chr10-38293460-DEL-1769 |
| chr10-38311489-DEL-63 | chr10-38311490-DEL-63 |
| chr10-38336482-DEL-77 | chr10-38336483-DEL-77 |
| chr10-38375174-DEL-499 | chr10-38375175-DEL-498 |
| chr10-38375996-DEL-54 | chr10-38375997-DEL-52 |
| chr10-38376778-DEL-2376 | chr10-38376779-DEL-2376 |
| chr10-38377434-DEL-859 | chr10-38377435-DEL-859 |
| chr10-38381937-DEL-55 | chr10-38381938-DEL-65 |
| chr10-38400058-DEL-464 | chr10-38400059-DEL-464 |
| chr10-38423125-DEL-504 | chr10-38423126-DEL-504 |
| chr10-38424247-DEL-68 | chr10-38424245-DEL-64 |
| chr10-38443828-DEL-78 | chr10-38443827-DEL-82 |
| chr10-38451601-DEL-208 | chr10-38451645-DEL-164 |
| chr10-38454309-DEL-66 | chr10-38454310-DEL-66 |
| chr10-38456259-DEL-66 | chr10-38456260-DEL-66 |
| chr10-38466080-DEL-3817 | chr10-38466090-DEL-3800 |
| chr10-38477503-DEL-1924 | chr10-38477504-DEL-1924 |
| chr10-38489071-DEL-122 | chr10-38489072-DEL-122 |
| chr10-38512448-DEL-56 | chr10-38512449-DEL-56 |
| chr10-38513835-DEL-6414 | chr10-38513836-DEL-6414 |
| chr10-38523771-DEL-1548 | chr10-38523772-DEL-1549 |
| chr10-38533627-DEL-63 | chr10-38533625-DEL-62 |
| chr10-38554467-DEL-298 | chr10-38554468-DEL-298 |
| chr10-38583026-DEL-4757 | chr10-38583028-DEL-4754 |
| chr10-38609855-DEL-818 | chr10-38609856-DEL-818 |
| chr10-38618812-DEL-200 | chr10-38618792-DEL-201 |
| chr10-38726801-DEL-196 | chr10-38726802-DEL-196 |
| chr10-38733106-DEL-214 | chr10-38733107-DEL-214 |
| chr10-38775596-DEL-1885 | chr10-38775598-DEL-1885 |
| chr10-38782229-DEL-61 | chr10-38782230-DEL-61 |
| chr10-38805515-DEL-62 | chr10-38805516-DEL-62 |
| chr10-38808616-DEL-7154 | chr10-38808617-DEL-7154 |
| chr10-38827060-DEL-6362 | chr10-38827061-DEL-6362 |
| chr10-38839910-DEL-458 | chr10-38839896-DEL-456 |
| chr10-38847740-DEL-221 | chr10-38847741-DEL-214 |

| | |
|---|---|
| chr10-38875026-DEL-100 | chr10-38874998-DEL-98 |
| chr10-38880540-DEL-99 | chr10-38880500-DEL-100 |
| chr10-38939555-DEL-62 | chr10-38939556-DEL-62 |
| chr10-38965279-DEL-53 | chr10-38965289-DEL-54 |
| chr10-39000301-DEL-400 | chr10-39000302-DEL-400 |
| chr10-39000918-DEL-113 | chr10-39000927-DEL-105 |
| chr10-39006538-DEL-650 | chr10-39006539-DEL-650 |
| chr10-39027901-DEL-76 | chr10-39027902-DEL-74 |
| chr10-39037965-DEL-87 | chr10-39037974-DEL-79 |
| chr10-39075602-DEL-217 | chr10-39075603-DEL-207 |
| chr10-39077069-DEL-1319 | chr10-39077070-DEL-1318 |
| chr10-39082684-DEL-195 | chr10-39082685-DEL-195 |
| chr10-39084653-DEL-74 | chr10-39084654-DEL-74 |
| chr10-39124280-DEL-136 | chr10-39124280-DEL-135 |
| chr10-39160486-DEL-1200 | chr10-39160487-DEL-1200 |
| chr10-39170355-DEL-406 | chr10-39170356-DEL-406 |
| chr10-39227599-DEL-97 | chr10-39227600-DEL-94 |
| chr10-39230437-DEL-86 | chr10-39230402-DEL-118 |
| chr10-39237399-DEL-54 | chr10-39237401-DEL-56 |
| chr10-39281328-DEL-56 | chr10-39281332-DEL-55 |
| chr10-39289946-DEL-52 | chr10-39289948-DEL-51 |
| chr10-39345910-DEL-147 | chr10-39345911-DEL-146 |
| chr10-39351548-DEL-97 | chr10-39351534-DEL-92 |
| chr10-39419906-DEL-112 | chr10-39419892-DEL-110 |
| chr10-39460370-DEL-191 | chr10-39460351-DEL-189 |
| chr10-39481574-DEL-401 | chr10-39481575-DEL-401 |
| chr10-39520889-DEL-216 | chr10-39520887-DEL-220 |
| chr10-39525607-DEL-110 | chr10-39525608-DEL-110 |
| chr10-39636766-DEL-3976 | chr10-39636767-DEL-3976 |
| chr10-39644844-DEL-245 | chr10-39644845-DEL-245 |
| chr10-39646653-DEL-232 | chr10-39646654-DEL-232 |
| chr10-39777068-DEL-204 | chr10-39777065-DEL-203 |
| chr10-39781055-DEL-54 | chr10-39781038-DEL-55 |
| chr10-39791846-DEL-75 | chr10-39791847-DEL-75 |
| chr10-39795031-DEL-53 | chr10-39795032-DEL-53 |
| chr10-39817934-DEL-102 | chr10-39817929-DEL-104 |
| chr10-39838211-DEL-649 | chr10-39838212-DEL-649 |
| chr10-39848696-DEL-61 | chr10-39848697-DEL-54 |
| chr10-39860224-DEL-54 | chr10-39860225-DEL-54 |
| chr10-39864023-DEL-67 | chr10-39864001-DEL-91 |
| chr10-39884469-DEL-72 | chr10-39884501-DEL-54 |
| chr10-39905572-DEL-54 | chr10-39905573-DEL-54 |
| chr10-39957594-DEL-235 | chr10-39957595-DEL-235 |
| chr10-39959665-DEL-68 | chr10-39959666-DEL-68 |
| chr10-40008658-DEL-222 | chr10-40008659-DEL-222 |
| chr10-40010176-DEL-54 | chr10-40010140-DEL-51 |
| chr10-40030654-DEL-116 | chr10-40030655-DEL-116 |
| chr10-40050209-DEL-64 | chr10-40050210-DEL-64 |
| chr10-40061222-DEL-1789 | chr10-40061223-DEL-1789 |
| chr10-40076213-DEL-1146 | chr10-40076214-DEL-1146 |
| chr10-40104446-DEL-205 | chr10-40104447-DEL-205 |
| chr10-40104842-DEL-3226 | chr10-40104843-DEL-3226 |
| chr10-40110540-DEL-55 | chr10-40110540-DEL-54 |
| chr10-40130021-DEL-220 | chr10-40130022-DEL-220 |
| chr10-40130690-DEL-208 | chr10-40130683-DEL-209 |
| chr10-40140430-DEL-55 | chr10-40140431-DEL-55 |
| chr10-40145037-DEL-202 | chr10-40145038-DEL-202 |
| chr10-40181693-DEL-57 | chr10-40181694-DEL-57 |
| chr10-40195365-DEL-196 | chr10-40195366-DEL-196 |
| chr10-40197798-DEL-3969 | chr10-40197799-DEL-3969 |
| chr10-40237024-DEL-641 | chr10-40237067-DEL-645 |
| chr10-40267045-DEL-191 | chr10-40267046-DEL-191 |
| chr10-40300367-DEL-202 | chr10-40300368-DEL-202 |
| chr10-40305920-DEL-209 | chr10-40305927-DEL-208 |
| chr10-40331179-DEL-200 | chr10-40331180-DEL-200 |
| chr10-40341968-DEL-126 | chr10-40341968-DEL-88 |
| chr10-40361363-DEL-70 | chr10-40361387-DEL-70 |
| chr10-40366173-DEL-138 | chr10-40366174-DEL-138 |
| chr10-40414095-DEL-1564 | chr10-40414096-DEL-1564 |
| chr10-40431047-DEL-201 | chr10-40431048-DEL-201 |
| chr10-40431751-DEL-519 | chr10-40431752-DEL-519 |
| chr10-40463240-DEL-121 | chr10-40463241-DEL-121 |
| chr10-40485053-DEL-163 | chr10-40485053-DEL-163 |
| chr10-40493944-DEL-51 | chr10-40493930-DEL-52 |
| chr10-40531658-DEL-63 | chr10-40531645-DEL-50 |
| chr10-40544598-DEL-108 | chr10-40544599-DEL-108 |
| chr10-40563785-DEL-123 | chr10-40563785-DEL-123 |
| chr10-40587823-DEL-52 | chr10-40587811-DEL-60 |
| chr10-40589540-DEL-51 | chr10-40589541-DEL-51 |
| chr10-40611786-DEL-69 | chr10-40611794-DEL-56 |

| | |
|---|---|
| chr10-40635664-DEL-373 | chr10-40635644-DEL-384 |
| chr10-40642862-DEL-1913 | chr10-40642863-DEL-1913 |
| chr10-40649669-DEL-227 | chr10-40649670-DEL-222 |
| chr10-40658978-DEL-61 | chr10-40658979-DEL-60 |
| chr10-40664611-DEL-59 | chr10-40664610-DEL-60 |
| chr10-40670292-DEL-52 | chr10-40670295-DEL-50 |
| chr10-40688367-DEL-2835 | chr10-40688368-DEL-2835 |
| chr10-40693715-DEL-823 | chr10-40693716-DEL-823 |
| chr10-40699969-DEL-64 | chr10-40699981-DEL-62 |
| chr10-40705672-DEL-492 | chr10-40705673-DEL-492 |
| chr10-40735769-DEL-956 | chr10-40735804-DEL-924 |
| chr10-40762606-DEL-812 | chr10-40762607-DEL-812 |
| chr10-40789435-DEL-435 | chr10-40789436-DEL-435 |
| chr10-40831923-DEL-56 | chr10-40831924-DEL-56 |
| chr10-40840909-DEL-436 | chr10-40840910-DEL-436 |
| chr10-40894239-DEL-208 | chr10-40894227-DEL-205 |
| chr10-40905327-DEL-865 | chr10-40905328-DEL-865 |
| chr10-40916722-DEL-64 | chr10-40916731-DEL-90 |
| chr10-40920714-DEL-54 | chr10-40920715-DEL-54 |
| chr10-40998357-DEL-52 | chr10-40998357-DEL-54 |
| chr10-41003128-DEL-6749 | chr10-41003128-DEL-6747 |
| chr10-41025991-DEL-115 | chr10-41025981-DEL-123 |
| chr10-41064100-DEL-53 | chr10-41064100-DEL-54 |
| chr10-41105066-DEL-116 | chr10-41105067-DEL-116 |
| chr10-41107130-DEL-62 | chr10-41107130-DEL-62 |
| chr10-41129035-DEL-427 | chr10-41129036-DEL-427 |
| chr10-41139373-DEL-340 | chr10-41139374-DEL-340 |
| chr10-41160406-DEL-83 | chr10-41160407-DEL-83 |
| chr10-41185229-DEL-212 | chr10-41185230-DEL-212 |
| chr10-41193875-DEL-199 | chr10-41193876-DEL-199 |
| chr10-41197149-DEL-246 | chr10-41197150-DEL-246 |
| chr10-41209149-DEL-1697 | chr10-41209150-DEL-1697 |
| chr10-41298547-DEL-51 | chr10-41298548-DEL-51 |
| chr10-41310039-DEL-219 | chr10-41310040-DEL-219 |
| chr10-41312639-DEL-8985 | chr10-41312640-DEL-8985 |
| chr10-41324415-DEL-123 | chr10-41324416-DEL-105 |
| chr10-41368237-DEL-215 | chr10-41368238-DEL-215 |
| chr10-41383917-DEL-140 | chr10-41383918-DEL-140 |
| chr10-41480356-DEL-251 | chr10-41480349-DEL-259 |
| chr10-41485993-DEL-103 | chr10-41485994-DEL-103 |
| chr10-41547512-DEL-100 | chr10-41547513-DEL-100 |
| chr10-41555702-DEL-254 | chr10-41555701-DEL-255 |
| chr10-41561447-DEL-686 | chr10-41561448-DEL-686 |
| chr10-41571411-DEL-203 | chr10-41571412-DEL-209 |
| chr10-41574623-DEL-105 | chr10-41574624-DEL-105 |
| chr10-41588430-DEL-212 | chr10-41588431-DEL-212 |
| chr10-41596198-DEL-100 | chr10-41596199-DEL-100 |
| chr10-41609010-DEL-160 | chr10-41609011-DEL-160 |
| chr10-41699522-DEL-188 | chr10-41699523-DEL-188 |
| chr10-41706774-DEL-269 | chr10-41706775-DEL-269 |
| chr10-41716521-DEL-7110 | chr10-41716522-DEL-7110 |
| chr10-41737996-DEL-1282 | chr10-41737997-DEL-1282 |
| chr10-41768038-DEL-194 | chr10-41768019-DEL-214 |
| chr10-41789973-DEL-52 | chr10-41789983-DEL-50 |
| chr10-41812500-DEL-76 | chr10-41812507-DEL-58 |
| chr10-41828561-DEL-436 | chr10-41828562-DEL-436 |
| chr10-41846204-DEL-225 | chr10-41846205-DEL-225 |
| chr10-41851764-DEL-81 | chr10-41851765-DEL-81 |
| chr10-41869687-DEL-68 | chr10-41869686-DEL-68 |
| chr10-41910734-DEL-851 | chr10-41910725-DEL-852 |
| chr10-41928524-DEL-2947 | chr10-41928525-DEL-2947 |
| chr10-41936819-DEL-189 | chr10-41936814-DEL-187 |
| chr10-41942917-DEL-328 | chr10-41942918-DEL-328 |
| chr10-41949664-DEL-242 | chr10-41949665-DEL-242 |
| chr10-41981153-DEL-95 | chr10-41981168-DEL-108 |
| chr10-41983132-DEL-80 | chr10-41983121-DEL-92 |
| chr10-41989729-DEL-107 | chr10-41989711-DEL-108 |
| chr10-41998625-DEL-68 | chr10-41998618-DEL-64 |
| chr10-42018605-DEL-957 | chr10-42018606-DEL-957 |
| chr10-42029390-DEL-113 | chr10-42029374-DEL-114 |
| chr10-42092301-DEL-2420 | chr10-42092298-DEL-2421 |
| chr10-42096422-DEL-80 | chr10-42096423-DEL-80 |
| chr10-42103248-DEL-68 | chr10-42103249-DEL-68 |
| chr10-42112188-DEL-62 | chr10-42112173-DEL-60 |
| chr10-42131935-DEL-56 | chr10-42131907-DEL-58 |
| chr10-42160529-DEL-193 | chr10-42160530-DEL-193 |
| chr10-42210231-DEL-54 | chr10-42210232-DEL-54 |
| chr10-42236593-DEL-56 | chr10-42236600-DEL-50 |
| chr10-42250748-DEL-7248 | chr10-42250749-DEL-7248 |
| chr10-42270517-DEL-169 | chr10-42270519-DEL-168 |

| | |
|---|---|
| chr10-42282128-DEL-190 | chr10-42282129-DEL-190 |
| chr10-42297340-DEL-58 | chr10-42297341-DEL-58 |
| chr10-42298422-DEL-116 | chr10-42298423-DEL-116 |
| chr10-42302193-DEL-57 | chr10-42302194-DEL-57 |
| chr10-42305946-DEL-401 | chr10-42305947-DEL-401 |
| chr10-42306771-DEL-208 | chr10-42306767-DEL-209 |
| chr10-42308105-DEL-195 | chr10-42308106-DEL-195 |
| chr10-42315357-DEL-55 | chr10-42315349-DEL-56 |
| chr10-42318470-DEL-205 | chr10-42318508-DEL-178 |
| chr10-42335450-DEL-214 | chr10-42335451-DEL-214 |
| chr10-42338988-DEL-279 | chr10-42338951-DEL-281 |
| chr10-42341749-DEL-109 | chr10-42341750-DEL-109 |
| chr10-42358508-DEL-191 | chr10-42358509-DEL-191 |
| chr10-42397494-DEL-368 | chr10-42397495-DEL-368 |
| chr10-42408016-DEL-750 | chr10-42408020-DEL-751 |
| chr10-42426787-DEL-129 | chr10-42426788-DEL-129 |
| chr10-42430052-DEL-109 | chr10-42430045-DEL-85 |
| chr10-42451017-DEL-53 | chr10-42451018-DEL-53 |
| chr10-42452279-DEL-213 | chr10-42452280-DEL-213 |
| chr10-42472889-DEL-86 | chr10-42472871-DEL-77 |
| chr10-42476443-DEL-216 | chr10-42476444-DEL-214 |
| chr10-42479724-DEL-78 | chr10-42479725-DEL-78 |
| chr10-42496922-DEL-682 | chr10-42496923-DEL-682 |
| chr10-42503597-DEL-102 | chr10-42503610-DEL-106 |
| chr10-42517877-DEL-62 | chr10-42517878-DEL-62 |
| chr10-42523886-DEL-192 | chr10-42523887-DEL-192 |
| chr10-42528679-DEL-192 | chr10-42528699-DEL-179 |
| chr10-42530847-DEL-124 | chr10-42530847-DEL-131 |
| chr10-42536354-DEL-197 | chr10-42536355-DEL-197 |
| chr10-42583007-DEL-77 | chr10-42583004-DEL-81 |
| chr10-42630997-DEL-121 | chr10-42630998-DEL-121 |
| chr10-42693861-DEL-1278 | chr10-42693862-DEL-1278 |
| chr10-42697932-DEL-144 | chr10-42697929-DEL-148 |
| chr10-42725555-DEL-124 | chr10-42725557-DEL-123 |
| chr10-42733480-DEL-82 | chr10-42733470-DEL-78 |
| chr10-42773982-DEL-194 | chr10-42773983-DEL-194 |
| chr10-42779322-DEL-510 | chr10-42779325-DEL-508 |
| chr10-42782634-DEL-943 | chr10-42782635-DEL-943 |
| chr10-42803190-DEL-239 | chr10-42803191-DEL-239 |
| chr10-42808550-DEL-204 | chr10-42808551-DEL-204 |
| chr10-42815550-DEL-71 | chr10-42815552-DEL-69 |
| chr10-42830139-DEL-68 | chr10-42830152-DEL-88 |
| chr10-42858345-DEL-99 | chr10-42858327-DEL-70 |
| chr10-42860128-DEL-277 | chr10-42860127-DEL-278 |
| chr10-42876857-DEL-51 | chr10-42876857-DEL-51 |
| chr10-42895850-DEL-84 | chr10-42895851-DEL-84 |
| chr10-42937676-DEL-60 | chr10-42937649-DEL-60 |
| chr10-42955036-DEL-61 | chr10-42955037-DEL-61 |
| chr10-42957698-DEL-709 | chr10-42957698-DEL-709 |
| chr10-42985795-DEL-61 | chr10-42985796-DEL-61 |
| chr10-43098353-DEL-152 | chr10-43098354-DEL-152 |
| chr10-43114748-DEL-230 | chr10-43114749-DEL-230 |
| chr10-43131601-DEL-390 | chr10-43131602-DEL-390 |
| chr10-43134869-DEL-3632 | chr10-43134871-DEL-3631 |
| chr10-43183618-DEL-52 | chr10-43183576-DEL-51 |
| chr10-43213351-DEL-3156 | chr10-43213352-DEL-3156 |
| chr10-43219029-DEL-52 | chr10-43219030-DEL-56 |
| chr10-43231179-DEL-1878 | chr10-43231180-DEL-1878 |
| chr10-43252449-DEL-414 | chr10-43252450-DEL-414 |
| chr10-43287238-DEL-87 | chr10-43287239-DEL-88 |
| chr10-43318173-DEL-185 | chr10-43318174-DEL-185 |
| chr10-43324390-DEL-159 | chr10-43324391-DEL-159 |
| chr10-43326166-DEL-532 | chr10-43326167-DEL-532 |
| chr10-43337063-DEL-70 | chr10-43337092-DEL-68 |
| chr10-43380380-DEL-81 | chr10-43380381-DEL-81 |
| chr10-43399778-DEL-235 | chr10-43399779-DEL-235 |
| chr10-43400068-DEL-570 | chr10-43400069-DEL-570 |
| chr10-43410028-DEL-56 | chr10-43410029-DEL-52 |
| chr10-43422790-DEL-432 | chr10-43422791-DEL-432 |
| chr10-43433626-DEL-144 | chr10-43433628-DEL-142 |
| chr10-43438174-DEL-208 | chr10-43438175-DEL-208 |
| chr10-43439454-DEL-193 | chr10-43439455-DEL-193 |
| chr10-43440981-DEL-116 | chr10-43441006-DEL-92 |
| chr10-43448745-DEL-206 | chr10-43448737-DEL-207 |
| chr10-43574810-DEL-56 | chr10-43574783-DEL-83 |
| chr10-43645474-DEL-4672 | chr10-43645428-DEL-4673 |
| chr10-43684798-DEL-99 | chr10-43684798-DEL-99 |
| chr10-43684956-DEL-3881 | chr10-43684940-DEL-3882 |
| chr10-43709622-DEL-3260 | chr10-43709623-DEL-3260 |
| chr10-43728500-DEL-3023 | chr10-43728501-DEL-3023 |

| | |
|---|---|
| chr10-43753075-DEL-2972 | chr10-43753076-DEL-2972 |
| chr10-43801722-DEL-171 | chr10-43801732-DEL-170 |
| chr10-43809671-DEL-69 | chr10-43809667-DEL-115 |
| chr10-43825552-DEL-240 | chr10-43825553-DEL-240 |
| chr10-43853062-DEL-64 | chr10-43853090-DEL-50 |
| chr10-43857006-DEL-400 | chr10-43857007-DEL-400 |
| chr10-43858053-DEL-148 | chr10-43858056-DEL-146 |
| chr10-43888146-DEL-84 | chr10-43888147-DEL-84 |
| chr10-43902377-DEL-500 | chr10-43902378-DEL-500 |
| chr10-43908428-DEL-109 | chr10-43908429-DEL-109 |
| chr10-43920589-DEL-58 | chr10-43920590-DEL-58 |
| chr10-43923826-DEL-54 | chr10-43923827-DEL-54 |
| chr10-43949973-DEL-54 | chr10-43949974-DEL-50 |
| chr10-43958354-DEL-160 | chr10-43958355-DEL-160 |
| chr10-43966598-DEL-137 | chr10-43966598-DEL-137 |
| chr10-43968124-DEL-54 | chr10-43968125-DEL-54 |
| chr10-43985502-DEL-80 | chr10-43985503-DEL-80 |
| chr10-43997767-DEL-109 | chr10-43997767-DEL-110 |
| chr10-44003349-DEL-133 | chr10-44003350-DEL-133 |
| chr10-44018587-DEL-139 | chr10-44018595-DEL-108 |
| chr10-44033224-DEL-82 | chr10-44033226-DEL-83 |
| chr10-44033807-DEL-76 | chr10-44033808-DEL-76 |
| chr10-44088138-DEL-1449 | chr10-44088138-DEL-1450 |
| chr10-44095345-DEL-70 | chr10-44095346-DEL-70 |
| chr10-44104858-DEL-598 | chr10-44104859-DEL-598 |
| chr10-44196020-DEL-1060 | chr10-44196021-DEL-1060 |
| chr10-44262703-DEL-64 | chr10-44262704-DEL-64 |
| chr10-44298647-DEL-140 | chr10-44298661-DEL-121 |
| chr10-44344371-DEL-69 | chr10-44344374-DEL-67 |
| chr10-44387831-DEL-100 | chr10-44387850-DEL-80 |
| chr10-44466545-DEL-1880 | chr10-44466546-DEL-1880 |
| chr10-44487088-DEL-107 | chr10-44487119-DEL-58 |
| chr10-44509554-DEL-66 | chr10-44509555-DEL-68 |
| chr10-44516913-DEL-341 | chr10-44516914-DEL-341 |
| chr10-44528624-DEL-65 | chr10-44528624-DEL-70 |
| chr10-44611379-DEL-2344 | chr10-44611380-DEL-2344 |
| chr10-44664290-DEL-627 | chr10-44664286-DEL-621 |
| chr10-44668151-DEL-194 | chr10-44668152-DEL-194 |
| chr10-44678285-DEL-63 | chr10-44678289-DEL-64 |
| chr10-44691269-DEL-1137 | chr10-44691270-DEL-1137 |
| chr10-44705304-DEL-872 | chr10-44705305-DEL-873 |
| chr10-44744885-DEL-89 | chr10-44744886-DEL-89 |
| chr10-44779902-DEL-55 | chr10-44779903-DEL-55 |
| chr10-44781745-DEL-6823 | chr10-44781746-DEL-6823 |
| chr10-44781959-DEL-416 | chr10-44781960-DEL-416 |
| chr10-44796211-DEL-347 | chr10-44796220-DEL-320 |
| chr10-44839988-DEL-183 | chr10-44839989-DEL-183 |
| chr10-44893038-DEL-81 | chr10-44893039-DEL-91 |
| chr10-44910396-DEL-64 | chr10-44910397-DEL-64 |
| chr10-44927904-DEL-198 | chr10-44927905-DEL-198 |
| chr10-44939744-DEL-358 | chr10-44939745-DEL-358 |
| chr10-44953017-DEL-62 | chr10-44953018-DEL-62 |
| chr10-44955507-DEL-140 | chr10-44955508-DEL-140 |
| chr10-44992031-DEL-573 | chr10-44992009-DEL-560 |
| chr10-44998356-DEL-81 | chr10-44998357-DEL-81 |
| chr10-45043239-DEL-495 | chr10-45043240-DEL-495 |
| chr10-45053816-DEL-550 | chr10-45053817-DEL-550 |
| chr10-45062529-DEL-72 | chr10-45062542-DEL-50 |
| chr10-45066193-DEL-116 | chr10-45066223-DEL-116 |
| chr10-45078752-DEL-1186 | chr10-45078752-DEL-1186 |
| chr10-45093678-DEL-919 | chr10-45093679-DEL-919 |
| chr10-45094990-DEL-836 | chr10-45094985-DEL-837 |
| chr10-45102333-DEL-52 | chr10-45102320-DEL-72 |
| chr10-45104351-DEL-231 | chr10-45104374-DEL-222 |
| chr10-45111681-DEL-1648 | chr10-45111682-DEL-1648 |
| chr10-45125139-DEL-65 | chr10-45125137-DEL-68 |
| chr10-45128397-DEL-505 | chr10-45128398-DEL-505 |
| chr10-45167367-DEL-77 | chr10-45167368-DEL-73 |
| chr10-45172827-DEL-207 | chr10-45172828-DEL-207 |
| chr10-45176310-DEL-816 | chr10-45176311-DEL-816 |
| chr10-45200822-DEL-242 | chr10-45200823-DEL-242 |
| chr10-45229371-DEL-97 | chr10-45229377-DEL-102 |
| chr10-45237286-DEL-2185 | chr10-45237286-DEL-2185 |
| chr10-45243683-DEL-524 | chr10-45243684-DEL-524 |
| chr10-45250744-DEL-90 | chr10-45250745-DEL-90 |
| chr10-45271663-DEL-632 | chr10-45271652-DEL-644 |
| chr10-45299803-DEL-53 | chr10-45299804-DEL-53 |
| chr10-45358064-DEL-222 | chr10-45358065-DEL-222 |
| chr10-45358579-DEL-60 | chr10-45358575-DEL-59 |
| chr10-45430899-DEL-5477 | chr10-45430900-DEL-5477 |

| | |
|---|---|
| chr10-45443422-DEL-1888 | chr10-45443423-DEL-1888 |
| chr10-45461526-DEL-2638 | chr10-45461527-DEL-2638 |
| chr10-45482609-DEL-60 | chr10-45482610-DEL-60 |
| chr10-45501208-DEL-654 | chr10-45501209-DEL-654 |
| chr10-45513421-DEL-177 | chr10-45513425-DEL-216 |
| chr10-45533314-DEL-286 | chr10-45533315-DEL-286 |
| chr10-45564023-DEL-6202 | chr10-45564024-DEL-6202 |
| chr10-45574140-DEL-561 | chr10-45574141-DEL-561 |
| chr10-45597381-DEL-67 | chr10-45597366-DEL-64 |
| chr10-45600117-DEL-7579 | chr10-45600118-DEL-7579 |
| chr10-45613468-DEL-2054 | chr10-45613469-DEL-2054 |
| chr10-45623320-DEL-277 | chr10-45623321-DEL-277 |
| chr10-45628263-DEL-349 | chr10-45628264-DEL-349 |
| chr10-45646478-DEL-64 | chr10-45646479-DEL-64 |
| chr10-45668103-DEL-56 | chr10-45668104-DEL-56 |
| chr10-45715048-DEL-6463 | chr10-45715042-DEL-6470 |
| chr10-45733430-DEL-333 | chr10-45733431-DEL-333 |
| chr10-45751371-DEL-814 | chr10-45751372-DEL-814 |
| chr10-45757340-DEL-748 | chr10-45757340-DEL-748 |
| chr10-45768035-DEL-171 | chr10-45768034-DEL-173 |
| chr10-45781671-DEL-290 | chr10-45781648-DEL-324 |
| chr10-45785679-DEL-1263 | chr10-45785681-DEL-1263 |
| chr10-45787825-DEL-151 | chr10-45787826-DEL-151 |
| chr10-45809020-DEL-781 | chr10-45809021-DEL-781 |
| chr10-45817980-DEL-84 | chr10-45817981-DEL-102 |
| chr10-45819932-DEL-172 | chr10-45819933-DEL-172 |
| chr10-45839211-DEL-507 | chr10-45839212-DEL-507 |
| chr10-45851858-DEL-350 | chr10-45851859-DEL-350 |
| chr10-45935867-DEL-139 | chr10-45935882-DEL-119 |
| chr10-46004013-DEL-2470 | chr10-46004013-DEL-2465 |
| chr10-46012591-DEL-2828 | chr10-46012592-DEL-2828 |
| chr10-46042893-DEL-68 | chr10-46042894-DEL-68 |
| chr10-46066906-DEL-172 | chr10-46066907-DEL-172 |
| chr10-46121522-DEL-699 | chr10-46121523-DEL-699 |
| chr10-46127651-DEL-76 | chr10-46127655-DEL-122 |
| chr10-46145463-DEL-82 | chr10-46145472-DEL-74 |
| chr10-46196985-DEL-80 | chr10-46196986-DEL-80 |
| chr10-46199512-DEL-356 | chr10-46199512-DEL-357 |
| chr10-46204627-DEL-56 | chr10-46204632-DEL-52 |
| chr10-46214919-DEL-319 | chr10-46214919-DEL-317 |
| chr10-46235330-DEL-1611 | chr10-46235331-DEL-1611 |
| chr10-46237013-DEL-1938 | chr10-46237014-DEL-1938 |
| chr10-46240539-DEL-6493 | chr10-46240542-DEL-6492 |
| chr10-46256489-DEL-201 | chr10-46256474-DEL-201 |
| chr10-46281318-DEL-294 | chr10-46281319-DEL-293 |
| chr10-46326435-DEL-154 | chr10-46326451-DEL-126 |
| chr10-46367298-DEL-327 | chr10-46367299-DEL-327 |
| chr10-46394112-DEL-257 | chr10-46394113-DEL-257 |
| chr10-46399233-DEL-791 | chr10-46399233-DEL-791 |
| chr10-46412975-DEL-1887 | chr10-46412976-DEL-1887 |
| chr10-46429294-DEL-587 | chr10-46429295-DEL-587 |
| chr10-46445658-DEL-1605 | chr10-46445659-DEL-1605 |
| chr10-46456093-DEL-7851 | chr10-46456094-DEL-7851 |
| chr10-46471984-DEL-526 | chr10-46471985-DEL-526 |
| chr10-46475373-DEL-1551 | chr10-46475374-DEL-1551 |
| chr10-46486497-DEL-759 | chr10-46486498-DEL-759 |
| chr10-46499536-DEL-576 | chr10-46499537-DEL-576 |
| chr10-46519607-DEL-60 | chr10-46519592-DEL-70 |
| chr10-46522658-DEL-3546 | chr10-46522659-DEL-3546 |
| chr10-46551725-DEL-63 | chr10-46551692-DEL-64 |
| chr10-46578863-DEL-174 | chr10-46578864-DEL-174 |
| chr10-46597987-DEL-475 | chr10-46597987-DEL-475 |
| chr10-46617456-DEL-3194 | chr10-46617457-DEL-3194 |
| chr10-46627982-DEL-7009 | chr10-46627983-DEL-7009 |
| chr10-46664166-DEL-2558 | chr10-46664166-DEL-2558 |
| chr10-46667968-DEL-54 | chr10-46667967-DEL-56 |
| chr10-46673619-DEL-333 | chr10-46673620-DEL-333 |
| chr10-46676278-DEL-361 | chr10-46676279-DEL-361 |
| chr10-46682180-DEL-6417 | chr10-46682181-DEL-6417 |
| chr10-46721420-DEL-51 | chr10-46721421-DEL-51 |
| chr10-46724533-DEL-94 | chr10-46724534-DEL-94 |
| chr10-46738504-DEL-175 | chr10-46738505-DEL-175 |
| chr10-46765708-DEL-238 | chr10-46765709-DEL-230 |
| chr10-46789029-DEL-6415 | chr10-46789030-DEL-6415 |
| chr10-46799822-DEL-62 | chr10-46799823-DEL-62 |
| chr10-46842776-DEL-7999 | chr10-46842777-DEL-7999 |
| chr10-46875594-DEL-67 | chr10-46875595-DEL-67 |
| chr10-46876298-DEL-6189 | chr10-46876302-DEL-6184 |
| chr10-46883806-DEL-3163 | chr10-46883807-DEL-3163 |
| chr10-46898979-DEL-126 | chr10-46899001-DEL-120 |

| | |
|---|---|
| chr10-46944334-DEL-6399 | chr10-46944335-DEL-6399 |
| chr10-46987461-DEL-56 | chr10-46987461-DEL-71 |
| chr10-46988608-DEL-1102 | chr10-46988608-DEL-1102 |
| chr10-47004657-DEL-87 | chr10-47004635-DEL-105 |
| chr10-47050967-DEL-174 | chr10-47050972-DEL-173 |
| chr10-47064044-DEL-827 | chr10-47064045-DEL-827 |
| chr10-47071872-DEL-3446 | chr10-47071873-DEL-3446 |
| chr10-47082400-DEL-70 | chr10-47082407-DEL-74 |
| chr10-47085569-DEL-162 | chr10-47085570-DEL-162 |
| chr10-47092275-DEL-222 | chr10-47092276-DEL-222 |
| chr10-47095913-DEL-326 | chr10-47095913-DEL-326 |
| chr10-47097463-DEL-6431 | chr10-47097463-DEL-6431 |
| chr10-47104741-DEL-384 | chr10-47104742-DEL-384 |
| chr10-47113665-DEL-77 | chr10-47113666-DEL-71 |
| chr10-47201963-DEL-3615 | chr10-47201956-DEL-3614 |
| chr10-47206879-DEL-6429 | chr10-47206880-DEL-6429 |
| chr10-47214076-DEL-72 | chr10-47214077-DEL-72 |
| chr10-47215090-DEL-60 | chr10-47215090-DEL-60 |
| chr10-47235063-DEL-170 | chr10-47235062-DEL-171 |
| chr10-47240607-DEL-56 | chr10-47240608-DEL-56 |
| chr10-47306002-DEL-72 | chr10-47306003-DEL-50 |
| chr10-47314369-DEL-6860 | chr10-47314369-DEL-6860 |
| chr10-47328203-DEL-4612 | chr10-47328204-DEL-4612 |
| chr10-47348519-DEL-84 | chr10-47348520-DEL-84 |
| chr10-47415628-DEL-349 | chr10-47415629-DEL-349 |
| chr10-47416566-DEL-51 | chr10-47416567-DEL-51 |
| chr10-47450308-DEL-6193 | chr10-47450309-DEL-6193 |
| chr10-47466901-DEL-328 | chr10-47466902-DEL-328 |
| chr10-47468214-DEL-231 | chr10-47468215-DEL-231 |
| chr10-47468806-DEL-1125 | chr10-47468807-DEL-1125 |
| chr10-47486466-DEL-2820 | chr10-47486467-DEL-2820 |
| chr10-47495932-DEL-1520 | chr10-47495932-DEL-1520 |
| chr10-47507291-DEL-129 | chr10-47507292-DEL-129 |
| chr10-47512071-DEL-2719 | chr10-47512072-DEL-2718 |
| chr10-47516149-DEL-558 | chr10-47516150-DEL-558 |
| chr10-47516934-DEL-1470 | chr10-47516935-DEL-1470 |
| chr10-47518710-DEL-1175 | chr10-47518711-DEL-1175 |
| chr10-47520552-DEL-57 | chr10-47520553-DEL-57 |
| chr10-47526180-DEL-92 | chr10-47526181-DEL-92 |
| chr10-47539657-DEL-6391 | chr10-47539658-DEL-6391 |
| chr10-47546596-DEL-4107 | chr10-47546597-DEL-4107 |
| chr10-47556693-DEL-2905 | chr10-47556695-DEL-2906 |
| chr10-47596404-DEL-2494 | chr10-47596405-DEL-2494 |
| chr10-47601289-DEL-197 | chr10-47601289-DEL-197 |
| chr10-47609049-DEL-346 | chr10-47609049-DEL-346 |
| chr10-47613195-DEL-962 | chr10-47613196-DEL-962 |
| chr10-47619615-DEL-15378 | chr10-47619616-DEL-15378 |
| chr10-47647385-DEL-55 | chr10-47647385-DEL-55 |
| chr10-47657234-DEL-695 | chr10-47657235-DEL-695 |
| chr10-47658775-DEL-234 | chr10-47658775-DEL-234 |
| chr10-47675470-DEL-6468 | chr10-47675471-DEL-6468 |
| chr10-47727786-DEL-1026 | chr10-47727786-DEL-1027 |
| chr10-47737322-DEL-174 | chr10-47737323-DEL-174 |
| chr10-47785713-DEL-1613 | chr10-47785708-DEL-1612 |
| chr10-47802889-DEL-81 | chr10-47802907-DEL-63 |
| chr10-47832411-DEL-57 | chr10-47832417-DEL-62 |
| chr10-47834077-DEL-97 | chr10-47834077-DEL-94 |
| chr10-47840925-DEL-68 | chr10-47840926-DEL-66 |
| chr10-47862634-DEL-306 | chr10-47862634-DEL-306 |
| chr10-47869112-DEL-52 | chr10-47869111-DEL-52 |
| chr10-47891819-DEL-69 | chr10-47891813-DEL-90 |
| chr10-47912740-DEL-114 | chr10-47912741-DEL-114 |
| chr10-47918291-DEL-4100 | chr10-47918279-DEL-4092 |
| chr10-47924542-DEL-1836 | chr10-47924541-DEL-1836 |
| chr10-47928664-DEL-1397 | chr10-47928693-DEL-1369 |
| chr10-47932150-DEL-2772 | chr10-47932151-DEL-2772 |
| chr10-47946124-DEL-1422 | chr10-47946119-DEL-1427 |
| chr10-47948717-DEL-93 | chr10-47948718-DEL-93 |
| chr10-47951921-DEL-168 | chr10-47951922-DEL-168 |
| chr10-47960218-DEL-102 | chr10-47960219-DEL-102 |
| chr10-47963505-DEL-52 | chr10-47963506-DEL-52 |
| chr10-47996441-DEL-77 | chr10-47996442-DEL-77 |
| chr10-48010369-DEL-105 | chr10-48010370-DEL-78 |
| chr10-48075590-DEL-6225 | chr10-48075599-DEL-6225 |
| chr10-48082471-DEL-309 | chr10-48082472-DEL-309 |
| chr10-48086557-DEL-408 | chr10-48086558-DEL-408 |
| chr10-48089931-DEL-2480 | chr10-48089932-DEL-2480 |
| chr10-48091072-DEL-209 | chr10-48091073-DEL-209 |
| chr10-48095658-DEL-67 | chr10-48095652-DEL-86 |
| chr10-48105169-DEL-12790 | chr10-48105161-DEL-12796 |

| | |
|---|---|
| chr10-48107919-DEL-212 | chr10-48107920-DEL-212 |
| chr10-48119219-DEL-61 | chr10-48119220-DEL-59 |
| chr10-48119595-DEL-4492 | chr10-48119596-DEL-4492 |
| chr10-48125001-DEL-182 | chr10-48125036-DEL-172 |
| chr10-48130758-DEL-899 | chr10-48130755-DEL-911 |
| chr10-48140040-DEL-111 | chr10-48140032-DEL-112 |
| chr10-48141672-DEL-1290 | chr10-48141673-DEL-1290 |
| chr10-48155895-DEL-202 | chr10-48155896-DEL-202 |
| chr10-48180488-DEL-68 | chr10-48180481-DEL-76 |
| chr10-48188241-DEL-75 | chr10-48188238-DEL-78 |
| chr10-48191641-DEL-189 | chr10-48191642-DEL-189 |
| chr10-48200327-DEL-87 | chr10-48200328-DEL-86 |
| chr10-48201877-DEL-1154 | chr10-48201878-DEL-1154 |
| chr10-48210885-DEL-633 | chr10-48210885-DEL-633 |
| chr10-48213369-DEL-613 | chr10-48213370-DEL-613 |
| chr10-48222845-DEL-66 | chr10-48222846-DEL-66 |
| chr10-48228184-DEL-93 | chr10-48228185-DEL-93 |
| chr10-48249920-DEL-360 | chr10-48249921-DEL-360 |
| chr10-48254667-DEL-56 | chr10-48254668-DEL-56 |
| chr10-48258431-DEL-6484 | chr10-48258432-DEL-6484 |
| chr10-48274181-DEL-588 | chr10-48274157-DEL-584 |
| chr10-48297243-DEL-1483 | chr10-48297244-DEL-1483 |
| chr10-48315122-DEL-1690 | chr10-48315107-DEL-1703 |
| chr10-48321769-DEL-52 | chr10-48321776-DEL-54 |
| chr10-48335778-DEL-1170 | chr10-48335779-DEL-1170 |
| chr10-48340519-DEL-70 | chr10-48340520-DEL-70 |
| chr10-48389129-DEL-671 | chr10-48389130-DEL-671 |
| chr10-48409822-DEL-3219 | chr10-48409802-DEL-3195 |
| chr10-48431771-DEL-440 | chr10-48431771-DEL-440 |
| chr10-48439097-DEL-72 | chr10-48439102-DEL-67 |
| chr10-48450515-DEL-4137 | chr10-48450517-DEL-4135 |
| chr10-48492338-DEL-15794 | chr10-48492339-DEL-15794 |
| chr10-48524997-DEL-3549 | chr10-48524998-DEL-3549 |
| chr10-48533639-DEL-52 | chr10-48533637-DEL-50 |
| chr10-48541955-DEL-1078 | chr10-48541962-DEL-1078 |
| chr10-48553245-DEL-6597 | chr10-48553246-DEL-6597 |
| chr10-48608796-DEL-343 | chr10-48608797-DEL-343 |
| chr10-48613973-DEL-755 | chr10-48613974-DEL-755 |
| chr10-48630051-DEL-75 | chr10-48630052-DEL-75 |
| chr10-48701569-DEL-6775 | chr10-48701570-DEL-6775 |
| chr10-48714844-DEL-68 | chr10-48714845-DEL-68 |
| chr10-48736131-DEL-80 | chr10-48736132-DEL-80 |
| chr10-48786017-DEL-106 | chr10-48786018-DEL-106 |
| chr10-48803738-DEL-66 | chr10-48803739-DEL-66 |
| chr10-48821140-DEL-1489 | chr10-48821140-DEL-1489 |
| chr10-48823710-DEL-359 | chr10-48823711-DEL-359 |
| chr10-48842357-DEL-351 | chr10-48842356-DEL-352 |
| chr10-48844875-DEL-3889 | chr10-48844876-DEL-3889 |
| chr10-48855234-DEL-219 | chr10-48855235-DEL-219 |
| chr10-48859660-DEL-306 | chr10-48859661-DEL-306 |
| chr10-48862807-DEL-665 | chr10-48862808-DEL-665 |
| chr10-48863881-DEL-88 | chr10-48863882-DEL-88 |
| chr10-48865509-DEL-55 | chr10-48865510-DEL-55 |
| chr10-48868046-DEL-4049 | chr10-48868047-DEL-4049 |
| chr10-48871184-DEL-150 | chr10-48871185-DEL-150 |
| chr10-48876053-DEL-1683 | chr10-48876054-DEL-1683 |
| chr10-48920719-DEL-527 | chr10-48920720-DEL-527 |
| chr10-48959053-DEL-2559 | chr10-48959054-DEL-2559 |
| chr10-48992230-DEL-51 | chr10-48992231-DEL-51 |
| chr10-48996943-DEL-111 | chr10-48996944-DEL-111 |
| chr10-49058563-DEL-60 | chr10-49058560-DEL-64 |
| chr10-49080282-DEL-89 | chr10-49080283-DEL-89 |
| chr10-49084848-DEL-219 | chr10-49084849-DEL-219 |
| chr10-49101118-DEL-6286 | chr10-49101119-DEL-6286 |
| chr10-49153418-DEL-3574 | chr10-49153419-DEL-3574 |
| chr10-49171199-DEL-1868 | chr10-49171200-DEL-1868 |
| chr10-49202013-DEL-52 | chr10-49202014-DEL-52 |
| chr10-49223132-DEL-268 | chr10-49223133-DEL-268 |
| chr10-49250649-DEL-1216 | chr10-49250650-DEL-1216 |
| chr10-49284701-DEL-62 | chr10-49284705-DEL-58 |
| chr10-49290507-DEL-937 | chr10-49290507-DEL-920 |
| chr10-49300821-DEL-3308 | chr10-49300822-DEL-3308 |
| chr10-49309993-DEL-218 | chr10-49309992-DEL-216 |
| chr10-49318282-DEL-1002 | chr10-49318282-DEL-1002 |
| chr10-49365193-DEL-576 | chr10-49365200-DEL-574 |
| chr10-49379881-DEL-1762 | chr10-49379881-DEL-1762 |
| chr10-49388662-DEL-2506 | chr10-49388663-DEL-2506 |
| chr10-49404985-DEL-616 | chr10-49404986-DEL-616 |
| chr10-49414071-DEL-52 | chr10-49414072-DEL-52 |
| chr10-49443962-DEL-254 | chr10-49443963-DEL-254 |

| | |
|---|---|
| chr10-49451522-DEL-58 | chr10-49451508-DEL-87 |
| chr10-49453843-DEL-222 | chr10-49453818-DEL-252 |
| chr10-49454669-DEL-212 | chr10-49454668-DEL-213 |
| chr10-49466826-DEL-1496 | chr10-49466827-DEL-1496 |
| chr10-49488983-DEL-3999 | chr10-49488984-DEL-3999 |
| chr10-49500891-DEL-3149 | chr10-49500891-DEL-3163 |
| chr10-49509511-DEL-52 | chr10-49509512-DEL-52 |
| chr10-49516685-DEL-55 | chr10-49516686-DEL-55 |
| chr10-49532225-DEL-74 | chr10-49532223-DEL-76 |
| chr10-49544563-DEL-57 | chr10-49544564-DEL-57 |
| chr10-49561838-DEL-58 | chr10-49561853-DEL-87 |
| chr10-49578243-DEL-350 | chr10-49578244-DEL-347 |
| chr10-49579806-DEL-413 | chr10-49579807-DEL-413 |
| chr10-49582736-DEL-308 | chr10-49582737-DEL-307 |
| chr10-49595588-DEL-1030 | chr10-49595597-DEL-1023 |
| chr10-49601461-DEL-232 | chr10-49601462-DEL-232 |
| chr10-49614628-DEL-112 | chr10-49614673-DEL-78 |
| chr10-49619751-DEL-2451 | chr10-49619752-DEL-2451 |
| chr10-49630557-DEL-72 | chr10-49630558-DEL-84 |
| chr10-49639231-DEL-1001 | chr10-49639232-DEL-1001 |
| chr10-49674988-DEL-386 | chr10-49674989-DEL-386 |
| chr10-49679069-DEL-53 | chr10-49679070-DEL-74 |
| chr10-49699578-DEL-7342 | chr10-49699579-DEL-7342 |
| chr10-49724075-DEL-492 | chr10-49724076-DEL-491 |
| chr10-49729526-DEL-3838 | chr10-49729527-DEL-3838 |
| chr10-49738840-DEL-200 | chr10-49738841-DEL-200 |
| chr10-49745590-DEL-217 | chr10-49745591-DEL-219 |
| chr10-49773366-DEL-557 | chr10-49773367-DEL-557 |
| chr10-49776345-DEL-647 | chr10-49776346-DEL-647 |
| chr10-49779343-DEL-4784 | chr10-49779344-DEL-4784 |
| chr10-49795873-DEL-67 | chr10-49795873-DEL-67 |
| chr10-49818206-DEL-1791 | chr10-49818207-DEL-1790 |
| chr10-49823559-DEL-1844 | chr10-49823560-DEL-1844 |
| chr10-49828039-DEL-61 | chr10-49828040-DEL-61 |
| chr10-49843386-DEL-6046 | chr10-49843387-DEL-6045 |
| chr10-49893635-DEL-740 | chr10-49893636-DEL-740 |
| chr10-49905270-DEL-6272 | chr10-49905271-DEL-6272 |
| chr10-49911997-DEL-591 | chr10-49911998-DEL-591 |
| chr10-49914941-DEL-423 | chr10-49914942-DEL-423 |
| chr10-49924631-DEL-52 | chr10-49924632-DEL-52 |
| chr10-49940018-DEL-932 | chr10-49940019-DEL-932 |
| chr10-49981022-DEL-118 | chr10-49981023-DEL-118 |
| chr10-49985200-DEL-6436 | chr10-49985201-DEL-6436 |
| chr10-50011127-DEL-78 | chr10-50011150-DEL-54 |
| chr10-50054548-DEL-6430 | chr10-50054549-DEL-6429 |
| chr10-50084806-DEL-146 | chr10-50084807-DEL-146 |
| chr10-50087109-DEL-313 | chr10-50087110-DEL-313 |
| chr10-50103781-DEL-3693 | chr10-50103782-DEL-3693 |
| chr10-50117796-DEL-119 | chr10-50117797-DEL-119 |
| chr10-50130037-DEL-199 | chr10-50130038-DEL-199 |
| chr10-50166089-DEL-6440 | chr10-50166090-DEL-6440 |
| chr10-50193292-DEL-3031 | chr10-50193293-DEL-3031 |
| chr10-50207693-DEL-148 | chr10-50207712-DEL-123 |
| chr10-50209557-DEL-1874 | chr10-50209558-DEL-1874 |
| chr10-50213975-DEL-644 | chr10-50213976-DEL-644 |
| chr10-50221992-DEL-329 | chr10-50221989-DEL-327 |
| chr10-50238995-DEL-6191 | chr10-50238993-DEL-6202 |
| chr10-50239008-DEL-200 | chr10-50239009-DEL-200 |
| chr10-50258990-DEL-62 | chr10-50258991-DEL-62 |
| chr10-50262685-DEL-6329 | chr10-50262686-DEL-6329 |
| chr10-50271199-DEL-639 | chr10-50271200-DEL-639 |
| chr10-50272179-DEL-202 | chr10-50272180-DEL-202 |
| chr10-50295640-DEL-8031 | chr10-50295641-DEL-8031 |
| chr10-50323166-DEL-358 | chr10-50323167-DEL-358 |
| chr10-50333237-DEL-5147 | chr10-50333238-DEL-5147 |
| chr10-50391109-DEL-106 | chr10-50391110-DEL-106 |
| chr10-50399884-DEL-4337 | chr10-50399885-DEL-4337 |
| chr10-50418706-DEL-2287 | chr10-50418702-DEL-2286 |
| chr10-50549165-DEL-6455 | chr10-50549166-DEL-6455 |
| chr10-50566744-DEL-120 | chr10-50566725-DEL-127 |
| chr10-50567784-DEL-202 | chr10-50567785-DEL-202 |
| chr10-50570772-DEL-83 | chr10-50570774-DEL-81 |
| chr10-50627055-DEL-518 | chr10-50627056-DEL-518 |
| chr10-50679982-DEL-2313 | chr10-50679975-DEL-2314 |
| chr10-50728472-DEL-336 | chr10-50728473-DEL-336 |
| chr10-50754977-DEL-58 | chr10-50754979-DEL-57 |
| chr10-50807990-DEL-713 | chr10-50807991-DEL-713 |
| chr10-50903878-DEL-212 | chr10-50903878-DEL-209 |
| chr10-50910381-DEL-6806 | chr10-50910382-DEL-6806 |
| chr10-50923993-DEL-69 | chr10-50923996-DEL-67 |

| | |
|---|---|
| chr10-50924678-DEL-3935 | chr10-50924671-DEL-3936 |
| chr10-50928922-DEL-109 | chr10-50928942-DEL-89 |
| chr10-50955632-DEL-324 | chr10-50955633-DEL-324 |
| chr10-50975711-DEL-163 | chr10-50975712-DEL-163 |
| chr10-50981366-DEL-66 | chr10-50981367-DEL-66 |
| chr10-50983339-DEL-277 | chr10-50983339-DEL-275 |
| chr10-50985732-DEL-928 | chr10-50985733-DEL-928 |
| chr10-51002110-DEL-6983 | chr10-51002110-DEL-6983 |
| chr10-51006271-DEL-172 | chr10-51006272-DEL-172 |
| chr10-51008605-DEL-399 | chr10-51008606-DEL-399 |
| chr10-51035915-DEL-505 | chr10-51035916-DEL-505 |
| chr10-51045112-DEL-196 | chr10-51045113-DEL-196 |
| chr10-51060096-DEL-690 | chr10-51060097-DEL-690 |
| chr10-51069033-DEL-52 | chr10-51069029-DEL-52 |
| chr10-51097977-DEL-7371 | chr10-51097981-DEL-7378 |
| chr10-51117852-DEL-54 | chr10-51117853-DEL-54 |
| chr10-51151342-DEL-98 | chr10-51151343-DEL-98 |
| chr10-51206386-DEL-736 | chr10-51206381-DEL-742 |
| chr10-51266036-DEL-53 | chr10-51266037-DEL-53 |
| chr10-51311153-DEL-387 | chr10-51311154-DEL-386 |
| chr10-51312721-DEL-6448 | chr10-51312722-DEL-6448 |
| chr10-51341210-DEL-53 | chr10-51341205-DEL-55 |
| chr10-51342864-DEL-56 | chr10-51342865-DEL-56 |
| chr10-51346221-DEL-6116 | chr10-51346210-DEL-6117 |
| chr10-51365191-DEL-195 | chr10-51365192-DEL-195 |
| chr10-51370722-DEL-60 | chr10-51370723-DEL-60 |
| chr10-51377321-DEL-12172 | chr10-51377322-DEL-12172 |
| chr10-51381351-DEL-62 | chr10-51381352-DEL-62 |
| chr10-51404888-DEL-1497 | chr10-51404888-DEL-1500 |
| chr10-51438257-DEL-98 | chr10-51438256-DEL-95 |
| chr10-51452473-DEL-6135 | chr10-51452481-DEL-6128 |
| chr10-51472033-DEL-100 | chr10-51472034-DEL-100 |
| chr10-51477329-DEL-2795 | chr10-51477330-DEL-2795 |
| chr10-51481640-DEL-174 | chr10-51481641-DEL-174 |
| chr10-51529984-DEL-579 | chr10-51529985-DEL-579 |
| chr10-51535739-DEL-133 | chr10-51535740-DEL-133 |
| chr10-51549537-DEL-196 | chr10-51549538-DEL-196 |
| chr10-51600147-DEL-80 | chr10-51600153-DEL-83 |
| chr10-51618317-DEL-201 | chr10-51618314-DEL-199 |
| chr10-51620200-DEL-204 | chr10-51620201-DEL-204 |
| chr10-51667042-DEL-1093 | chr10-51667043-DEL-1093 |
| chr10-51677462-DEL-2276 | chr10-51677463-DEL-2276 |
| chr10-51686098-DEL-224 | chr10-51686099-DEL-224 |
| chr10-51686821-DEL-199 | chr10-51686822-DEL-199 |
| chr10-51709448-DEL-1313 | chr10-51709449-DEL-1313 |
| chr10-51723474-DEL-64 | chr10-51723475-DEL-64 |
| chr10-51749942-DEL-56 | chr10-51749956-DEL-76 |
| chr10-51759117-DEL-4886 | chr10-51759118-DEL-4886 |
| chr10-51776498-DEL-325 | chr10-51776499-DEL-325 |
| chr10-51848064-DEL-793 | chr10-51848058-DEL-794 |
| chr10-51860619-DEL-157 | chr10-51860620-DEL-157 |
| chr10-51861125-DEL-6474 | chr10-51861126-DEL-6474 |
| chr10-51868817-DEL-6624 | chr10-51868818-DEL-6624 |
| chr10-51885484-DEL-230 | chr10-51885485-DEL-230 |
| chr10-51895402-DEL-171 | chr10-51895403-DEL-163 |
| chr10-51928617-DEL-6401 | chr10-51928618-DEL-6401 |
| chr10-51946493-DEL-505 | chr10-51946494-DEL-505 |
| chr10-51965657-DEL-1281 | chr10-51965658-DEL-1281 |
| chr10-51981790-DEL-553 | chr10-51981788-DEL-554 |
| chr10-51984970-DEL-6396 | chr10-51984971-DEL-6396 |
| chr10-51993819-DEL-52 | chr10-51993820-DEL-52 |
| chr10-51999772-DEL-54 | chr10-51999738-DEL-59 |
| chr10-52010649-DEL-6621 | chr10-52010650-DEL-6621 |
| chr10-52016801-DEL-208 | chr10-52016802-DEL-208 |
| chr10-52018988-DEL-57 | chr10-52018974-DEL-103 |
| chr10-52051577-DEL-171 | chr10-52051588-DEL-159 |
| chr10-52054074-DEL-52 | chr10-52054075-DEL-52 |
| chr10-52066857-DEL-962 | chr10-52066857-DEL-962 |
| chr10-52090020-DEL-194 | chr10-52090021-DEL-194 |
| chr10-52090976-DEL-206 | chr10-52090977-DEL-207 |
| chr10-52095358-DEL-98 | chr10-52095359-DEL-98 |
| chr10-52101979-DEL-117 | chr10-52101980-DEL-117 |
| chr10-52115584-DEL-168 | chr10-52115581-DEL-168 |
| chr10-52119870-DEL-110 | chr10-52119871-DEL-110 |
| chr10-52121056-DEL-186 | chr10-52121057-DEL-186 |
| chr10-52125380-DEL-209 | chr10-52125381-DEL-209 |
| chr10-52143949-DEL-220 | chr10-52143942-DEL-221 |
| chr10-52144245-DEL-530 | chr10-52144246-DEL-530 |
| chr10-52167205-DEL-150 | chr10-52167206-DEL-150 |
| chr10-52174649-DEL-75 | chr10-52174650-DEL-80 |

| | |
|---|---|
| chr10-52177391-DEL-187 | chr10-52177392-DEL-187 |
| chr10-52203879-DEL-745 | chr10-52203880-DEL-745 |
| chr10-52205913-DEL-60 | chr10-52205914-DEL-60 |
| chr10-52210356-DEL-165 | chr10-52210357-DEL-165 |
| chr10-52216377-DEL-5306 | chr10-52216378-DEL-5306 |
| chr10-52219076-DEL-428 | chr10-52219077-DEL-430 |
| chr10-52244814-DEL-324 | chr10-52244815-DEL-324 |
| chr10-52245371-DEL-372 | chr10-52245416-DEL-360 |
| chr10-52268214-DEL-210 | chr10-52268215-DEL-210 |
| chr10-52271938-DEL-54 | chr10-52271937-DEL-56 |
| chr10-52346875-DEL-70 | chr10-52346864-DEL-58 |
| chr10-52364450-DEL-723 | chr10-52364451-DEL-723 |
| chr10-52371132-DEL-3650 | chr10-52371132-DEL-3650 |
| chr10-52381042-DEL-137 | chr10-52381043-DEL-137 |
| chr10-52447349-DEL-64 | chr10-52447376-DEL-64 |
| chr10-52524941-DEL-396 | chr10-52524942-DEL-396 |
| chr10-52538115-DEL-399 | chr10-52538116-DEL-399 |
| chr10-52541225-DEL-52 | chr10-52541226-DEL-54 |
| chr10-52543701-DEL-550 | chr10-52543702-DEL-550 |
| chr10-52553988-DEL-170 | chr10-52553988-DEL-170 |
| chr10-52584203-DEL-322 | chr10-52584204-DEL-322 |
| chr10-52616196-DEL-115 | chr10-52616193-DEL-117 |
| chr10-52636974-DEL-6323 | chr10-52636974-DEL-6324 |
| chr10-52647935-DEL-875 | chr10-52647935-DEL-868 |
| chr10-52659226-DEL-3399 | chr10-52659227-DEL-3399 |
| chr10-52668139-DEL-816 | chr10-52668138-DEL-816 |
| chr10-52678638-DEL-65 | chr10-52678641-DEL-59 |
| chr10-52820929-DEL-6554 | chr10-52820930-DEL-6554 |
| chr10-52863408-DEL-1012 | chr10-52863409-DEL-1012 |
| chr10-52867163-DEL-405 | chr10-52867164-DEL-404 |
| chr10-52891727-DEL-240 | chr10-52891728-DEL-240 |
| chr10-52893075-DEL-105 | chr10-52893070-DEL-111 |
| chr10-52913738-DEL-98 | chr10-52913710-DEL-87 |
| chr10-52919147-DEL-132 | chr10-52919148-DEL-128 |
| chr10-52924286-DEL-162 | chr10-52924287-DEL-162 |
| chr10-52928411-DEL-54 | chr10-52928411-DEL-54 |
| chr10-52949887-DEL-1963 | chr10-52949888-DEL-1963 |
| chr10-53005102-DEL-201 | chr10-53005103-DEL-201 |
| chr10-53032114-DEL-60 | chr10-53032115-DEL-60 |
| chr10-53044736-DEL-440 | chr10-53044737-DEL-440 |
| chr10-53050722-DEL-56 | chr10-53050723-DEL-56 |
| chr10-53052278-DEL-128 | chr10-53052279-DEL-128 |
| chr10-53121421-DEL-6352 | chr10-53121414-DEL-6350 |
| chr10-53131867-DEL-57 | chr10-53131875-DEL-53 |
| chr10-53189688-DEL-81 | chr10-53189689-DEL-81 |
| chr10-53244394-DEL-55 | chr10-53244395-DEL-55 |
| chr10-53247894-DEL-73 | chr10-53247895-DEL-73 |
| chr10-53259645-DEL-55 | chr10-53259645-DEL-55 |
| chr10-53267732-DEL-195 | chr10-53267733-DEL-195 |
| chr10-53294848-DEL-151 | chr10-53294848-DEL-175 |
| chr10-53316081-DEL-65 | chr10-53316054-DEL-50 |
| chr10-53318998-DEL-54 | chr10-53318963-DEL-96 |
| chr10-53320114-DEL-80 | chr10-53320136-DEL-89 |
| chr10-53341152-DEL-104 | chr10-53341157-DEL-100 |
| chr10-53348529-DEL-392 | chr10-53348530-DEL-392 |
| chr10-53350633-DEL-3584 | chr10-53350634-DEL-3584 |
| chr10-53380730-DEL-309 | chr10-53380731-DEL-309 |
| chr10-53390826-DEL-209 | chr10-53390827-DEL-209 |
| chr10-53410215-DEL-82 | chr10-53410216-DEL-70 |
| chr10-53445711-DEL-6440 | chr10-53445712-DEL-6440 |
| chr10-53496547-DEL-87 | chr10-53496550-DEL-115 |
| chr10-53531702-DEL-120 | chr10-53531703-DEL-120 |
| chr10-53540865-DEL-389 | chr10-53540864-DEL-390 |
| chr10-53542400-DEL-6054 | chr10-53542401-DEL-6054 |
| chr10-53552275-DEL-142 | chr10-53552276-DEL-142 |
| chr10-53554883-DEL-405 | chr10-53554883-DEL-404 |
| chr10-53610541-DEL-402 | chr10-53610542-DEL-402 |
| chr10-53624540-DEL-72 | chr10-53624519-DEL-72 |
| chr10-53644981-DEL-1343 | chr10-53644982-DEL-1343 |
| chr10-53657674-DEL-1137 | chr10-53657697-DEL-1131 |
| chr10-53659058-DEL-56 | chr10-53659058-DEL-54 |
| chr10-53720362-DEL-53 | chr10-53720347-DEL-69 |
| chr10-53758741-DEL-1281 | chr10-53758742-DEL-1281 |
| chr10-53761868-DEL-520 | chr10-53761869-DEL-520 |
| chr10-53769714-DEL-6747 | chr10-53769715-DEL-6747 |
| chr10-53813025-DEL-59 | chr10-53813026-DEL-59 |
| chr10-53842234-DEL-995 | chr10-53842235-DEL-995 |
| chr10-53845849-DEL-222 | chr10-53845840-DEL-223 |
| chr10-53940673-DEL-234 | chr10-53940674-DEL-234 |
| chr10-53959270-DEL-965 | chr10-53959271-DEL-965 |

| | |
|---|---|
| chr10-53980151-DEL-263 | chr10-53980152-DEL-263 |
| chr10-53986819-DEL-305 | chr10-53986820-DEL-306 |
| chr10-54001404-DEL-1733 | chr10-54001405-DEL-1733 |
| chr10-54013614-DEL-401 | chr10-54013615-DEL-401 |
| chr10-54018241-DEL-6815 | chr10-54018242-DEL-6815 |
| chr10-54025400-DEL-99 | chr10-54025398-DEL-99 |
| chr10-54029245-DEL-1819 | chr10-54029245-DEL-1819 |
| chr10-54060737-DEL-398 | chr10-54060738-DEL-398 |
| chr10-54073456-DEL-62 | chr10-54073476-DEL-85 |
| chr10-54087390-DEL-12532 | chr10-54087391-DEL-12532 |
| chr10-54162166-DEL-886 | chr10-54162167-DEL-886 |
| chr10-54186631-DEL-400 | chr10-54186632-DEL-400 |
| chr10-54193836-DEL-237 | chr10-54193836-DEL-237 |
| chr10-54221891-DEL-338 | chr10-54221892-DEL-338 |
| chr10-54271502-DEL-683 | chr10-54271502-DEL-683 |
| chr10-54274611-DEL-465 | chr10-54274611-DEL-465 |
| chr10-54293338-DEL-379 | chr10-54293339-DEL-379 |
| chr10-54330165-DEL-301 | chr10-54330166-DEL-301 |
| chr10-54354377-DEL-656 | chr10-54354378-DEL-656 |
| chr10-54356720-DEL-538 | chr10-54356721-DEL-538 |
| chr10-54359979-DEL-1890 | chr10-54359980-DEL-1890 |
| chr10-54360984-DEL-1134 | chr10-54360985-DEL-1134 |
| chr10-54367847-DEL-786 | chr10-54367848-DEL-786 |
| chr10-54380074-DEL-75 | chr10-54380075-DEL-75 |
| chr10-54390242-DEL-87 | chr10-54390248-DEL-82 |
| chr10-54431466-DEL-94 | chr10-54431467-DEL-94 |
| chr10-54463748-DEL-76 | chr10-54463751-DEL-68 |
| chr10-54464864-DEL-4740 | chr10-54464865-DEL-4740 |
| chr10-54485318-DEL-130 | chr10-54485319-DEL-130 |
| chr10-54489423-DEL-6594 | chr10-54489424-DEL-6594 |
| chr10-54504542-DEL-686 | chr10-54504543-DEL-686 |
| chr10-54506751-DEL-155 | chr10-54506756-DEL-151 |
| chr10-54517082-DEL-186 | chr10-54517083-DEL-186 |
| chr10-54593002-DEL-439 | chr10-54593003-DEL-439 |
| chr10-54622681-DEL-57 | chr10-54622652-DEL-84 |
| chr10-54682394-DEL-2509 | chr10-54682395-DEL-2509 |
| chr10-54713090-DEL-2036 | chr10-54713091-DEL-2037 |
| chr10-54744211-DEL-74 | chr10-54744216-DEL-75 |
| chr10-54746494-DEL-79 | chr10-54746495-DEL-79 |
| chr10-54757082-DEL-112 | chr10-54757083-DEL-112 |
| chr10-54762828-DEL-4321 | chr10-54762829-DEL-4321 |
| chr10-54777806-DEL-6478 | chr10-54777807-DEL-6478 |
| chr10-54785348-DEL-62 | chr10-54785348-DEL-58 |
| chr10-54808613-DEL-58 | chr10-54808614-DEL-58 |
| chr10-54812563-DEL-2048 | chr10-54812564-DEL-2046 |
| chr10-54816201-DEL-61 | chr10-54816202-DEL-61 |
| chr10-54827875-DEL-239 | chr10-54827876-DEL-239 |
| chr10-54864603-DEL-59 | chr10-54864604-DEL-59 |
| chr10-54920913-DEL-87 | chr10-54920914-DEL-87 |
| chr10-54927652-DEL-946 | chr10-54927653-DEL-946 |
| chr10-54944056-DEL-2040 | chr10-54944057-DEL-2040 |
| chr10-54955941-DEL-2940 | chr10-54955942-DEL-2940 |
| chr10-54973286-DEL-56 | chr10-54973283-DEL-60 |
| chr10-54995507-DEL-897 | chr10-54995507-DEL-897 |
| chr10-55005284-DEL-54 | chr10-55005281-DEL-62 |
| chr10-55035988-DEL-155 | chr10-55036013-DEL-131 |
| chr10-55037214-DEL-2851 | chr10-55037215-DEL-2851 |
| chr10-55177269-DEL-231 | chr10-55177270-DEL-231 |
| chr10-55177965-DEL-1849 | chr10-55177962-DEL-1847 |
| chr10-55202356-DEL-65 | chr10-55202339-DEL-62 |
| chr10-55258659-DEL-399 | chr10-55258660-DEL-399 |
| chr10-55263235-DEL-113 | chr10-55263236-DEL-87 |
| chr10-55267730-DEL-66 | chr10-55267737-DEL-60 |
| chr10-55282738-DEL-70 | chr10-55282737-DEL-71 |
| chr10-55288541-DEL-53 | chr10-55288542-DEL-54 |
| chr10-55293774-DEL-52 | chr10-55293775-DEL-54 |
| chr10-55301898-DEL-1038 | chr10-55301899-DEL-1038 |
| chr10-55310356-DEL-653 | chr10-55310357-DEL-653 |
| chr10-55314098-DEL-60 | chr10-55314089-DEL-63 |
| chr10-55315836-DEL-69 | chr10-55315837-DEL-69 |
| chr10-55322519-DEL-2161 | chr10-55322520-DEL-2161 |
| chr10-55329972-DEL-882 | chr10-55329973-DEL-882 |
| chr10-55331145-DEL-3062 | chr10-55331146-DEL-3062 |
| chr10-55359923-DEL-1603 | chr10-55359924-DEL-1603 |
| chr10-55364501-DEL-340 | chr10-55364502-DEL-325 |
| chr10-55374099-DEL-3669 | chr10-55374099-DEL-3669 |
| chr10-55382321-DEL-130 | chr10-55382322-DEL-130 |
| chr10-55417711-DEL-2016 | chr10-55417712-DEL-2016 |
| chr10-55445513-DEL-277 | chr10-55445513-DEL-277 |
| chr10-55446076-DEL-102 | chr10-55446081-DEL-98 |

| | |
|---|---|
| chr10-55450915-DEL-6327 | chr10-55450916-DEL-6327 |
| chr10-55450966-DEL-204 | chr10-55450967-DEL-204 |
| chr10-55477608-DEL-68 | chr10-55477613-DEL-60 |
| chr10-55503861-DEL-8842 | chr10-55503882-DEL-8833 |
| chr10-55529271-DEL-199 | chr10-55529272-DEL-199 |
| chr10-55539152-DEL-65 | chr10-55539153-DEL-65 |
| chr10-55540177-DEL-6287 | chr10-55540178-DEL-6287 |
| chr10-55553416-DEL-147 | chr10-55553417-DEL-143 |
| chr10-55563867-DEL-60 | chr10-55563870-DEL-58 |
| chr10-55565880-DEL-138 | chr10-55565881-DEL-138 |
| chr10-55591624-DEL-5307 | chr10-55591624-DEL-5307 |
| chr10-55601395-DEL-73 | chr10-55601396-DEL-73 |
| chr10-55621790-DEL-4280 | chr10-55621791-DEL-4280 |
| chr10-55627480-DEL-95 | chr10-55627481-DEL-90 |
| chr10-55661539-DEL-347 | chr10-55661552-DEL-320 |
| chr10-55675612-DEL-1415 | chr10-55675613-DEL-1415 |
| chr10-55680422-DEL-68 | chr10-55680423-DEL-68 |
| chr10-55681159-DEL-808 | chr10-55681160-DEL-808 |
| chr10-55719493-DEL-460 | chr10-55719494-DEL-460 |
| chr10-55781876-DEL-667 | chr10-55781877-DEL-667 |
| chr10-55784498-DEL-1444 | chr10-55784499-DEL-1443 |
| chr10-55817185-DEL-132 | chr10-55817186-DEL-132 |
| chr10-55820762-DEL-1438 | chr10-55820762-DEL-1438 |
| chr10-55837986-DEL-66 | chr10-55837985-DEL-68 |
| chr10-55866037-DEL-91 | chr10-55866011-DEL-118 |
| chr10-55907167-DEL-6485 | chr10-55907168-DEL-6485 |
| chr10-55967796-DEL-58 | chr10-55967797-DEL-58 |
| chr10-55973524-DEL-1022 | chr10-55973525-DEL-1022 |
| chr10-55997820-DEL-56 | chr10-55997820-DEL-63 |
| chr10-56005763-DEL-52 | chr10-56005764-DEL-52 |
| chr10-56008879-DEL-3501 | chr10-56008874-DEL-3502 |
| chr10-56042070-DEL-83 | chr10-56042071-DEL-83 |
| chr10-56044643-DEL-96 | chr10-56044644-DEL-96 |
| chr10-56051576-DEL-681 | chr10-56051577-DEL-681 |
| chr10-56058617-DEL-74 | chr10-56058615-DEL-52 |
| chr10-56059913-DEL-52 | chr10-56059906-DEL-56 |
| chr10-56062779-DEL-93 | chr10-56062780-DEL-93 |
| chr10-56088001-DEL-1793 | chr10-56088002-DEL-1793 |
| chr10-56097024-DEL-57 | chr10-56097025-DEL-57 |
| chr10-56099615-DEL-227 | chr10-56099616-DEL-227 |
| chr10-56126444-DEL-6764 | chr10-56126445-DEL-6764 |
| chr10-56139092-DEL-680 | chr10-56139093-DEL-680 |
| chr10-56163328-DEL-6708 | chr10-56163329-DEL-6708 |
| chr10-56185194-DEL-194 | chr10-56185195-DEL-194 |
| chr10-56204401-DEL-435 | chr10-56204402-DEL-435 |
| chr10-56230496-DEL-716 | chr10-56230470-DEL-725 |
| chr10-56239093-DEL-196 | chr10-56239094-DEL-196 |
| chr10-56241895-DEL-1095 | chr10-56241896-DEL-1095 |
| chr10-56258439-DEL-347 | chr10-56258440-DEL-346 |
| chr10-56306474-DEL-192 | chr10-56306475-DEL-192 |
| chr10-56314065-DEL-55 | chr10-56314062-DEL-50 |
| chr10-56351513-DEL-76 | chr10-56351514-DEL-76 |
| chr10-56367138-DEL-459 | chr10-56367139-DEL-459 |
| chr10-56370699-DEL-56 | chr10-56370660-DEL-52 |
| chr10-56385249-DEL-129 | chr10-56385250-DEL-129 |
| chr10-56386026-DEL-204 | chr10-56385998-DEL-205 |
| chr10-56400332-DEL-52 | chr10-56400333-DEL-52 |
| chr10-56404867-DEL-114 | chr10-56404868-DEL-114 |
| chr10-56452099-DEL-4768 | chr10-56452100-DEL-4768 |
| chr10-56463008-DEL-326 | chr10-56463005-DEL-322 |
| chr10-56511390-DEL-63 | chr10-56511391-DEL-63 |
| chr10-56513777-DEL-308 | chr10-56513778-DEL-308 |
| chr10-56518406-DEL-81 | chr10-56518407-DEL-81 |
| chr10-56521856-DEL-6329 | chr10-56521857-DEL-6329 |
| chr10-56531987-DEL-1193 | chr10-56531988-DEL-1192 |
| chr10-56540149-DEL-52 | chr10-56540150-DEL-52 |
| chr10-56543118-DEL-60 | chr10-56543119-DEL-58 |
| chr10-56543773-DEL-791 | chr10-56543774-DEL-791 |
| chr10-56577060-DEL-7200 | chr10-56577061-DEL-7200 |
| chr10-56596839-DEL-7129 | chr10-56596840-DEL-7129 |
| chr10-56617022-DEL-122 | chr10-56617023-DEL-122 |
| chr10-56628711-DEL-75 | chr10-56628701-DEL-63 |
| chr10-56657521-DEL-436 | chr10-56657519-DEL-437 |
| chr10-56670613-DEL-735 | chr10-56670614-DEL-735 |
| chr10-56674286-DEL-62 | chr10-56674272-DEL-72 |
| chr10-56680554-DEL-54 | chr10-56680552-DEL-57 |
| chr10-56692597-DEL-2178 | chr10-56692598-DEL-2178 |
| chr10-56747174-DEL-325 | chr10-56747175-DEL-325 |
| chr10-56788591-DEL-201 | chr10-56788592-DEL-201 |
| chr10-56804943-DEL-1089 | chr10-56804944-DEL-1089 |

| | |
|---|---|
| chr10-56829800-DEL-78 | chr10-56829801-DEL-78 |
| chr10-56852453-DEL-1140 | chr10-56852454-DEL-1140 |
| chr10-56891872-DEL-334 | chr10-56891873-DEL-334 |
| chr10-56898887-DEL-449 | chr10-56898888-DEL-449 |
| chr10-56899530-DEL-6425 | chr10-56899531-DEL-6425 |
| chr10-56926590-DEL-71 | chr10-56926593-DEL-72 |
| chr10-56927805-DEL-1868 | chr10-56927806-DEL-1868 |
| chr10-56931348-DEL-73 | chr10-56931349-DEL-69 |
| chr10-56944910-DEL-151 | chr10-56944911-DEL-151 |
| chr10-56949087-DEL-4164 | chr10-56949088-DEL-4164 |
| chr10-56961139-DEL-83 | chr10-56961140-DEL-83 |
| chr10-56994132-DEL-91 | chr10-56994133-DEL-59 |
| chr10-57013791-DEL-7084 | chr10-57013792-DEL-7084 |
| chr10-57031152-DEL-541 | chr10-57031153-DEL-541 |
| chr10-57039038-DEL-63 | chr10-57039039-DEL-63 |
| chr10-57063743-DEL-77 | chr10-57063744-DEL-77 |
| chr10-57070311-DEL-187 | chr10-57070312-DEL-187 |
| chr10-57081753-DEL-1659 | chr10-57081754-DEL-1659 |
| chr10-57085135-DEL-431 | chr10-57085136-DEL-431 |
| chr10-57087147-DEL-66 | chr10-57087148-DEL-64 |
| chr10-57094563-DEL-696 | chr10-57094564-DEL-696 |
| chr10-57110698-DEL-90 | chr10-57110699-DEL-90 |
| chr10-57115143-DEL-6420 | chr10-57115144-DEL-6420 |
| chr10-57129723-DEL-63 | chr10-57129724-DEL-63 |
| chr10-57145805-DEL-177 | chr10-57145804-DEL-177 |
| chr10-57155484-DEL-51 | chr10-57155485-DEL-51 |
| chr10-57161140-DEL-344 | chr10-57161140-DEL-344 |
| chr10-57179680-DEL-51 | chr10-57179690-DEL-63 |
| chr10-57202952-DEL-7141 | chr10-57202953-DEL-7141 |
| chr10-57232184-DEL-80 | chr10-57232184-DEL-94 |
| chr10-57264065-DEL-3030 | chr10-57264066-DEL-3030 |
| chr10-57276410-DEL-1464 | chr10-57276411-DEL-1464 |
| chr10-57293580-DEL-803 | chr10-57293581-DEL-803 |
| chr10-57295748-DEL-338 | chr10-57295749-DEL-342 |
| chr10-57342487-DEL-219 | chr10-57342488-DEL-219 |
| chr10-57351661-DEL-59 | chr10-57351651-DEL-52 |
| chr10-57371376-DEL-61 | chr10-57371359-DEL-60 |
| chr10-57381494-DEL-195 | chr10-57381487-DEL-196 |
| chr10-57411509-DEL-963 | chr10-57411510-DEL-963 |
| chr10-57419171-DEL-137 | chr10-57419169-DEL-138 |
| chr10-57432294-DEL-905 | chr10-57432294-DEL-904 |
| chr10-57441191-DEL-329 | chr10-57441210-DEL-303 |
| chr10-57493871-DEL-51 | chr10-57493870-DEL-55 |
| chr10-57502202-DEL-1209 | chr10-57502203-DEL-1209 |
| chr10-57511575-DEL-100 | chr10-57511592-DEL-60 |
| chr10-57539254-DEL-94 | chr10-57539254-DEL-94 |
| chr10-57551775-DEL-74 | chr10-57551749-DEL-111 |
| chr10-57554598-DEL-6360 | chr10-57554599-DEL-6360 |
| chr10-57591624-DEL-78 | chr10-57591601-DEL-100 |
| chr10-57599590-DEL-551 | chr10-57599591-DEL-551 |
| chr10-57642738-DEL-6668 | chr10-57642724-DEL-6680 |
| chr10-57653516-DEL-62 | chr10-57653517-DEL-62 |
| chr10-57658115-DEL-209 | chr10-57658116-DEL-209 |
| chr10-57660006-DEL-80 | chr10-57660006-DEL-80 |
| chr10-57746526-DEL-52 | chr10-57746526-DEL-52 |
| chr10-58006364-DEL-280 | chr10-58006364-DEL-280 |
| chr10-58021191-DEL-70 | chr10-58021191-DEL-69 |
| chr10-58042284-DEL-67 | chr10-58042285-DEL-67 |
| chr10-58099327-DEL-206 | chr10-58099328-DEL-206 |
| chr10-58154934-DEL-227 | chr10-58154935-DEL-227 |
| chr10-58158018-DEL-56 | chr10-58158019-DEL-56 |
| chr10-58184757-DEL-201 | chr10-58184763-DEL-195 |
| chr10-58233222-DEL-198 | chr10-58233223-DEL-198 |
| chr10-58271166-DEL-190 | chr10-58271167-DEL-190 |
| chr10-58321929-DEL-592 | chr10-58321930-DEL-592 |
| chr10-58351193-DEL-399 | chr10-58351194-DEL-399 |
| chr10-58388653-DEL-3235 | chr10-58388654-DEL-3235 |
| chr10-58394403-DEL-217 | chr10-58394404-DEL-217 |
| chr10-58456347-DEL-53 | chr10-58456348-DEL-53 |
| chr10-58468715-DEL-110 | chr10-58468716-DEL-110 |
| chr10-58480346-DEL-65 | chr10-58480346-DEL-63 |
| chr10-58531634-DEL-194 | chr10-58531634-DEL-192 |
| chr10-58559998-DEL-1012 | chr10-58559999-DEL-1012 |
| chr10-58578686-DEL-70 | chr10-58578692-DEL-67 |
| chr10-58581827-DEL-52 | chr10-58581866-DEL-56 |
| chr10-58590021-DEL-87 | chr10-58590022-DEL-87 |
| chr10-58596360-DEL-6990 | chr10-58596360-DEL-6990 |
| chr10-58650995-DEL-725 | chr10-58650996-DEL-725 |
| chr10-58652820-DEL-265 | chr10-58652821-DEL-265 |
| chr10-58734432-DEL-63 | chr10-58734433-DEL-63 |

| | |
|---|---|
| chr10-58742193-DEL-318 | chr10-58742194-DEL-318 |
| chr10-58784859-DEL-486 | chr10-58784860-DEL-486 |
| chr10-58798586-DEL-2364 | chr10-58798587-DEL-2364 |
| chr10-58825301-DEL-57 | chr10-58825302-DEL-57 |
| chr10-58863047-DEL-7799 | chr10-58863048-DEL-7799 |
| chr10-58893131-DEL-319 | chr10-58893132-DEL-319 |
| chr10-58896613-DEL-78 | chr10-58896614-DEL-78 |
| chr10-58961636-DEL-228 | chr10-58961637-DEL-228 |
| chr10-58972898-DEL-75 | chr10-58972891-DEL-76 |
| chr10-58974951-DEL-52 | chr10-58974952-DEL-52 |
| chr10-58978204-DEL-114 | chr10-58978205-DEL-114 |
| chr10-58981592-DEL-208 | chr10-58981593-DEL-208 |
| chr10-58992762-DEL-338 | chr10-58992763-DEL-338 |
| chr10-58995178-DEL-52 | chr10-58995179-DEL-52 |
| chr10-59013582-DEL-66 | chr10-59013581-DEL-70 |
| chr10-59080790-DEL-101 | chr10-59080811-DEL-81 |
| chr10-59165276-DEL-200 | chr10-59165292-DEL-200 |
| chr10-59174429-DEL-201 | chr10-59174430-DEL-201 |
| chr10-59217614-DEL-5268 | chr10-59217615-DEL-5268 |
| chr10-59231424-DEL-1158 | chr10-59231425-DEL-1158 |
| chr10-59235829-DEL-383 | chr10-59235789-DEL-373 |
| chr10-59262753-DEL-128 | chr10-59262754-DEL-128 |
| chr10-59267960-DEL-133 | chr10-59267963-DEL-131 |
| chr10-59294610-DEL-146 | chr10-59294595-DEL-116 |
| chr10-59300023-DEL-190 | chr10-59300037-DEL-191 |
| chr10-59309879-DEL-316 | chr10-59309880-DEL-316 |
| chr10-59353663-DEL-132 | chr10-59353681-DEL-134 |
| chr10-59410815-DEL-70 | chr10-59410806-DEL-68 |
| chr10-59415415-DEL-84 | chr10-59415416-DEL-84 |
| chr10-59416043-DEL-204 | chr10-59416043-DEL-204 |
| chr10-59425208-DEL-107 | chr10-59425214-DEL-102 |
| chr10-59480253-DEL-86 | chr10-59480253-DEL-80 |
| chr10-59487118-DEL-233 | chr10-59487119-DEL-233 |
| chr10-59489256-DEL-226 | chr10-59489279-DEL-226 |
| chr10-59521255-DEL-116 | chr10-59521224-DEL-102 |
| chr10-59522760-DEL-149 | chr10-59522761-DEL-149 |
| chr10-59535426-DEL-807 | chr10-59535427-DEL-807 |
| chr10-59548686-DEL-230 | chr10-59548687-DEL-230 |
| chr10-59550840-DEL-54 | chr10-59550825-DEL-54 |
| chr10-59573249-DEL-401 | chr10-59573250-DEL-401 |
| chr10-59585713-DEL-250 | chr10-59585713-DEL-250 |
| chr10-59590461-DEL-264 | chr10-59590462-DEL-264 |
| chr10-59635820-DEL-80 | chr10-59635827-DEL-96 |
| chr10-59636819-DEL-67 | chr10-59636820-DEL-67 |
| chr10-59642318-DEL-1525 | chr10-59642319-DEL-1525 |
| chr10-59646398-DEL-62 | chr10-59646399-DEL-62 |
| chr10-59651046-DEL-207 | chr10-59651047-DEL-207 |
| chr10-59666605-DEL-105 | chr10-59666652-DEL-105 |
| chr10-59679446-DEL-76 | chr10-59679405-DEL-77 |
| chr10-59686334-DEL-6413 | chr10-59686335-DEL-6413 |
| chr10-59713178-DEL-119 | chr10-59713134-DEL-120 |
| chr10-59740893-DEL-179 | chr10-59740894-DEL-179 |
| chr10-59746016-DEL-54 | chr10-59746017-DEL-54 |
| chr10-59771678-DEL-63 | chr10-59771679-DEL-67 |
| chr10-59796057-DEL-214 | chr10-59796058-DEL-214 |
| chr10-59829858-DEL-221 | chr10-59829859-DEL-221 |
| chr10-59835115-DEL-228 | chr10-59835116-DEL-228 |
| chr10-59849711-DEL-210 | chr10-59849705-DEL-211 |
| chr10-59858974-DEL-202 | chr10-59858975-DEL-202 |
| chr10-59872185-DEL-548 | chr10-59872186-DEL-548 |
| chr10-59908741-DEL-644 | chr10-59908742-DEL-644 |
| chr10-59909507-DEL-665 | chr10-59909508-DEL-665 |
| chr10-59922343-DEL-208 | chr10-59922310-DEL-211 |
| chr10-59927422-DEL-199 | chr10-59927423-DEL-199 |
| chr10-59937085-DEL-204 | chr10-59937087-DEL-203 |
| chr10-59983250-DEL-435 | chr10-59983251-DEL-435 |
| chr10-59984287-DEL-108 | chr10-59984288-DEL-108 |
| chr10-59988568-DEL-942 | chr10-59988569-DEL-942 |
| chr10-59996137-DEL-558 | chr10-59996138-DEL-558 |
| chr10-60009569-DEL-516 | chr10-60009570-DEL-516 |
| chr10-60034479-DEL-266 | chr10-60034480-DEL-266 |
| chr10-60043476-DEL-203 | chr10-60043477-DEL-203 |
| chr10-60044885-DEL-63 | chr10-60044883-DEL-54 |
| chr10-60047033-DEL-64 | chr10-60047031-DEL-56 |
| chr10-60076722-DEL-184 | chr10-60076723-DEL-184 |
| chr10-60087280-DEL-190 | chr10-60087281-DEL-190 |
| chr10-60104592-DEL-88 | chr10-60104593-DEL-88 |
| chr10-60106701-DEL-56 | chr10-60106700-DEL-58 |
| chr10-60147603-DEL-76 | chr10-60147598-DEL-76 |
| chr10-60188280-DEL-72 | chr10-60188281-DEL-72 |

| | |
|---|---|
| chr10-60210745-DEL-99 | chr10-60210748-DEL-97 |
| chr10-60346399-DEL-80 | chr10-60346400-DEL-80 |
| chr10-60354449-DEL-294 | chr10-60354451-DEL-293 |
| chr10-60382377-DEL-6452 | chr10-60382378-DEL-6452 |
| chr10-60404709-DEL-431 | chr10-60404710-DEL-431 |
| chr10-60408764-DEL-59 | chr10-60408765-DEL-59 |
| chr10-60430077-DEL-56 | chr10-60430078-DEL-56 |
| chr10-60467416-DEL-63 | chr10-60467417-DEL-63 |
| chr10-60605081-DEL-189 | chr10-60605082-DEL-189 |
| chr10-60614323-DEL-52 | chr10-60614324-DEL-52 |
| chr10-60646930-DEL-219 | chr10-60646931-DEL-219 |
| chr10-60683217-DEL-53 | chr10-60683218-DEL-53 |
| chr10-60702476-DEL-1647 | chr10-60702477-DEL-1647 |
| chr10-60718263-DEL-1347 | chr10-60718264-DEL-1347 |
| chr10-60728251-DEL-203 | chr10-60728252-DEL-203 |
| chr10-60734528-DEL-56 | chr10-60734543-DEL-64 |
| chr10-60757330-DEL-94 | chr10-60757331-DEL-94 |
| chr10-60773955-DEL-205 | chr10-60773956-DEL-205 |
| chr10-60787993-DEL-68 | chr10-60787994-DEL-68 |
| chr10-60806894-DEL-60 | chr10-60806895-DEL-56 |
| chr10-60818455-DEL-88 | chr10-60818452-DEL-88 |
| chr10-60932230-DEL-215 | chr10-60932237-DEL-210 |
| chr10-60947493-DEL-197 | chr10-60947494-DEL-197 |
| chr10-60963385-DEL-809 | chr10-60963386-DEL-809 |
| chr10-60972239-DEL-67 | chr10-60972240-DEL-64 |
| chr10-60982995-DEL-68 | chr10-60982947-DEL-84 |
| chr10-60986730-DEL-1734 | chr10-60986747-DEL-1718 |
| chr10-61004542-DEL-52 | chr10-61004538-DEL-53 |
| chr10-61013056-DEL-84 | chr10-61013067-DEL-71 |
| chr10-61017730-DEL-65 | chr10-61017726-DEL-50 |
| chr10-61029821-DEL-175 | chr10-61029822-DEL-175 |
| chr10-61046537-DEL-210 | chr10-61046538-DEL-210 |
| chr10-61055873-DEL-205 | chr10-61055873-DEL-205 |
| chr10-61071604-DEL-69 | chr10-61071605-DEL-69 |
| chr10-61079508-DEL-819 | chr10-61079509-DEL-819 |
| chr10-61083172-DEL-516 | chr10-61083173-DEL-516 |
| chr10-61118993-DEL-385 | chr10-61118994-DEL-385 |
| chr10-61129293-DEL-188 | chr10-61129295-DEL-201 |
| chr10-61154729-DEL-221 | chr10-61154730-DEL-221 |
| chr10-61159748-DEL-137 | chr10-61159749-DEL-137 |
| chr10-61171728-DEL-116 | chr10-61171778-DEL-93 |
| chr10-61173999-DEL-163 | chr10-61174044-DEL-147 |
| chr10-61199519-DEL-210 | chr10-61199518-DEL-211 |
| chr10-61221760-DEL-68 | chr10-61221761-DEL-68 |
| chr10-61232832-DEL-209 | chr10-61232833-DEL-209 |
| chr10-61239508-DEL-492 | chr10-61239509-DEL-492 |
| chr10-61249530-DEL-134 | chr10-61249539-DEL-126 |
| chr10-61253241-DEL-200 | chr10-61253242-DEL-200 |
| chr10-61257811-DEL-56 | chr10-61257837-DEL-66 |
| chr10-61271347-DEL-386 | chr10-61271348-DEL-386 |
| chr10-61272455-DEL-685 | chr10-61272456-DEL-685 |
| chr10-61288868-DEL-137 | chr10-61288866-DEL-117 |
| chr10-61298088-DEL-7703 | chr10-61298089-DEL-7703 |
| chr10-61307798-DEL-185 | chr10-61307799-DEL-185 |
| chr10-61308619-DEL-98 | chr10-61308620-DEL-98 |
| chr10-61348960-DEL-75 | chr10-61348961-DEL-75 |
| chr10-61349439-DEL-1987 | chr10-61349437-DEL-1985 |
| chr10-61373122-DEL-491 | chr10-61373123-DEL-491 |
| chr10-61380341-DEL-144 | chr10-61380342-DEL-144 |
| chr10-61387569-DEL-212 | chr10-61387571-DEL-211 |
| chr10-61445124-DEL-161 | chr10-61445114-DEL-168 |
| chr10-61451681-DEL-79 | chr10-61451638-DEL-80 |
| chr10-61471272-DEL-87 | chr10-61471273-DEL-87 |
| chr10-61483581-DEL-96 | chr10-61483582-DEL-96 |
| chr10-61486183-DEL-203 | chr10-61486184-DEL-203 |
| chr10-61511628-DEL-78 | chr10-61511631-DEL-76 |
| chr10-61548783-DEL-51 | chr10-61548781-DEL-60 |
| chr10-61592451-DEL-68 | chr10-61592452-DEL-68 |
| chr10-61629457-DEL-237 | chr10-61629458-DEL-237 |
| chr10-61652138-DEL-52 | chr10-61652139-DEL-52 |
| chr10-61753234-DEL-161 | chr10-61753235-DEL-161 |
| chr10-61770099-DEL-52 | chr10-61770099-DEL-52 |
| chr10-61802330-DEL-359 | chr10-61802292-DEL-394 |
| chr10-61806988-DEL-58 | chr10-61806989-DEL-58 |
| chr10-61817918-DEL-85 | chr10-61817919-DEL-85 |
| chr10-61854727-DEL-57 | chr10-61854728-DEL-58 |
| chr10-61870430-DEL-88 | chr10-61870431-DEL-88 |
| chr10-61878716-DEL-448 | chr10-61878717-DEL-448 |
| chr10-61883455-DEL-200 | chr10-61883456-DEL-200 |
| chr10-61912602-DEL-76 | chr10-61912572-DEL-120 |

| | |
|---|---|
| chr10-61927141-DEL-429 | chr10-61927142-DEL-429 |
| chr10-61931987-DEL-281 | chr10-61931988-DEL-281 |
| chr10-62025483-DEL-68 | chr10-62025484-DEL-68 |
| chr10-62035797-DEL-113 | chr10-62035798-DEL-113 |
| chr10-62059693-DEL-64 | chr10-62059693-DEL-64 |
| chr10-62075740-DEL-78 | chr10-62075744-DEL-78 |
| chr10-62082269-DEL-59 | chr10-62082269-DEL-59 |
| chr10-62108332-DEL-91 | chr10-62108332-DEL-92 |
| chr10-62124437-DEL-305 | chr10-62124438-DEL-305 |
| chr10-62126736-DEL-131 | chr10-62126737-DEL-131 |
| chr10-62201534-DEL-69 | chr10-62201543-DEL-72 |
| chr10-62203197-DEL-3605 | chr10-62203198-DEL-3605 |
| chr10-62255998-DEL-51 | chr10-62255963-DEL-52 |
| chr10-62258638-DEL-181 | chr10-62258639-DEL-182 |
| chr10-62263196-DEL-214 | chr10-62263197-DEL-214 |
| chr10-62275259-DEL-67 | chr10-62275264-DEL-66 |
| chr10-62308556-DEL-449 | chr10-62308552-DEL-447 |
| chr10-62314439-DEL-195 | chr10-62314440-DEL-195 |
| chr10-62321063-DEL-180 | chr10-62321064-DEL-180 |
| chr10-62328119-DEL-187 | chr10-62328132-DEL-145 |
| chr10-62407702-DEL-211 | chr10-62407702-DEL-214 |
| chr10-62444661-DEL-4447 | chr10-62444661-DEL-4447 |
| chr10-62525683-DEL-220 | chr10-62525684-DEL-220 |
| chr10-62528462-DEL-68 | chr10-62528471-DEL-58 |
| chr10-62556608-DEL-203 | chr10-62556609-DEL-203 |
| chr10-62563643-DEL-169 | chr10-62563629-DEL-184 |
| chr10-62569201-DEL-2936 | chr10-62569202-DEL-2936 |
| chr10-62634731-DEL-211 | chr10-62634732-DEL-211 |
| chr10-62664947-DEL-210 | chr10-62664947-DEL-218 |
| chr10-62682216-DEL-259 | chr10-62682254-DEL-243 |
| chr10-62685737-DEL-68 | chr10-62685726-DEL-80 |
| chr10-62688476-DEL-210 | chr10-62688477-DEL-210 |
| chr10-62706485-DEL-211 | chr10-62706488-DEL-209 |
| chr10-62725870-DEL-716 | chr10-62725871-DEL-716 |
| chr10-62733553-DEL-207 | chr10-62733553-DEL-207 |
| chr10-62735857-DEL-91 | chr10-62735858-DEL-91 |
| chr10-62740676-DEL-277 | chr10-62740690-DEL-262 |
| chr10-62745097-DEL-200 | chr10-62745098-DEL-200 |
| chr10-62753888-DEL-176 | chr10-62753884-DEL-181 |
| chr10-62760574-DEL-94 | chr10-62760575-DEL-94 |
| chr10-62761851-DEL-410 | chr10-62761850-DEL-409 |
| chr10-62762919-DEL-211 | chr10-62762920-DEL-211 |
| chr10-62769317-DEL-196 | chr10-62769318-DEL-196 |
| chr10-62772206-DEL-208 | chr10-62772213-DEL-201 |
| chr10-62783265-DEL-62 | chr10-62783266-DEL-60 |
| chr10-62813371-DEL-51 | chr10-62813372-DEL-51 |
| chr10-62818686-DEL-202 | chr10-62818687-DEL-202 |
| chr10-62831859-DEL-227 | chr10-62831860-DEL-227 |
| chr10-62834515-DEL-180 | chr10-62834516-DEL-180 |
| chr10-62849147-DEL-66 | chr10-62849152-DEL-62 |
| chr10-62871079-DEL-254 | chr10-62871080-DEL-254 |
| chr10-62872115-DEL-178 | chr10-62872116-DEL-178 |
| chr10-62872943-DEL-307 | chr10-62872946-DEL-310 |
| chr10-62884843-DEL-154 | chr10-62884844-DEL-154 |
| chr10-62909118-DEL-781 | chr10-62909127-DEL-749 |
| chr10-62911088-DEL-135 | chr10-62911089-DEL-135 |
| chr10-62913116-DEL-206 | chr10-62913117-DEL-206 |
| chr10-62913937-DEL-433 | chr10-62913938-DEL-433 |
| chr10-62930755-DEL-1318 | chr10-62930756-DEL-1318 |
| chr10-62939715-DEL-7142 | chr10-62939716-DEL-7142 |
| chr10-62986233-DEL-69 | chr10-62986234-DEL-69 |
| chr10-63012702-DEL-81 | chr10-63012703-DEL-81 |
| chr10-63036456-DEL-86 | chr10-63036443-DEL-84 |
| chr10-63083814-DEL-64 | chr10-63083815-DEL-64 |
| chr10-63129101-DEL-59 | chr10-63129102-DEL-59 |
| chr10-63160996-DEL-206 | chr10-63160997-DEL-206 |
| chr10-63185172-DEL-112 | chr10-63185181-DEL-104 |
| chr10-63196231-DEL-73 | chr10-63196232-DEL-73 |
| chr10-63204477-DEL-62 | chr10-63204478-DEL-62 |
| chr10-63208979-DEL-108 | chr10-63208974-DEL-111 |
| chr10-63222147-DEL-66 | chr10-63222148-DEL-66 |
| chr10-63226010-DEL-5065 | chr10-63226052-DEL-5023 |
| chr10-63241085-DEL-203 | chr10-63241086-DEL-203 |
| chr10-63269677-DEL-127 | chr10-63269675-DEL-128 |
| chr10-63298460-DEL-188 | chr10-63298461-DEL-187 |
| chr10-63310851-DEL-107 | chr10-63310852-DEL-107 |
| chr10-63319343-DEL-87 | chr10-63319344-DEL-87 |
| chr10-63344563-DEL-2541 | chr10-63344564-DEL-2541 |
| chr10-63348427-DEL-6467 | chr10-63348424-DEL-6468 |
| chr10-63381486-DEL-324 | chr10-63381487-DEL-324 |

| | |
|---|---|
| chr10-63386058-DEL-317 | chr10-63386059-DEL-317 |
| chr10-63414183-DEL-66 | chr10-63414147-DEL-58 |
| chr10-63428896-DEL-451 | chr10-63428883-DEL-431 |
| chr10-63440844-DEL-919 | chr10-63440847-DEL-917 |
| chr10-63454945-DEL-756 | chr10-63454948-DEL-750 |
| chr10-63483995-DEL-54 | chr10-63484000-DEL-50 |
| chr10-63490040-DEL-70 | chr10-63490039-DEL-74 |
| chr10-63491408-DEL-72 | chr10-63491417-DEL-72 |
| chr10-63498973-DEL-6543 | chr10-63498974-DEL-6543 |
| chr10-63534878-DEL-90 | chr10-63534851-DEL-90 |
| chr10-63539603-DEL-54 | chr10-63539604-DEL-54 |
| chr10-63559277-DEL-6947 | chr10-63559278-DEL-6947 |
| chr10-63609760-DEL-6433 | chr10-63609761-DEL-6433 |
| chr10-63623568-DEL-610 | chr10-63623569-DEL-610 |
| chr10-63624500-DEL-1019 | chr10-63624501-DEL-1019 |
| chr10-63658355-DEL-126 | chr10-63658356-DEL-132 |
| chr10-63669554-DEL-1193 | chr10-63669555-DEL-1191 |
| chr10-63695196-DEL-58 | chr10-63695197-DEL-58 |
| chr10-63732122-DEL-477 | chr10-63732121-DEL-474 |
| chr10-63752162-DEL-319 | chr10-63752163-DEL-319 |
| chr10-63803971-DEL-398 | chr10-63803972-DEL-398 |
| chr10-63810008-DEL-638 | chr10-63810023-DEL-624 |
| chr10-63811127-DEL-555 | chr10-63811127-DEL-555 |
| chr10-63831872-DEL-2365 | chr10-63831873-DEL-2365 |
| chr10-63909109-DEL-164 | chr10-63909110-DEL-164 |
| chr10-63956707-DEL-574 | chr10-63956708-DEL-574 |
| chr10-63959308-DEL-98 | chr10-63959303-DEL-86 |
| chr10-63963805-DEL-52 | chr10-63963806-DEL-52 |
| chr10-64029409-DEL-328 | chr10-64029410-DEL-328 |
| chr10-64037048-DEL-6840 | chr10-64037049-DEL-6840 |
| chr10-64092926-DEL-187 | chr10-64092927-DEL-187 |
| chr10-64182225-DEL-130 | chr10-64182226-DEL-128 |
| chr10-64182857-DEL-62 | chr10-64182858-DEL-74 |
| chr10-64227037-DEL-229 | chr10-64227033-DEL-230 |
| chr10-64229637-DEL-2400 | chr10-64229638-DEL-2400 |
| chr10-64234352-DEL-57 | chr10-64234357-DEL-53 |
| chr10-64249391-DEL-68 | chr10-64249391-DEL-106 |
| chr10-64277062-DEL-78 | chr10-64277062-DEL-78 |
| chr10-64300881-DEL-105 | chr10-64300888-DEL-99 |
| chr10-64302020-DEL-67 | chr10-64302002-DEL-85 |
| chr10-64315074-DEL-161 | chr10-64315075-DEL-161 |
| chr10-64317211-DEL-313 | chr10-64317218-DEL-307 |
| chr10-64328909-DEL-56 | chr10-64328910-DEL-56 |
| chr10-64335713-DEL-436 | chr10-64335712-DEL-436 |
| chr10-64341009-DEL-1097 | chr10-64341010-DEL-1097 |
| chr10-64375297-DEL-1885 | chr10-64375290-DEL-1888 |
| chr10-64433296-DEL-1451 | chr10-64433297-DEL-1451 |
| chr10-64457461-DEL-2780 | chr10-64457462-DEL-2780 |
| chr10-64463454-DEL-207 | chr10-64463455-DEL-207 |
| chr10-64485221-DEL-67 | chr10-64485242-DEL-52 |
| chr10-64488203-DEL-134 | chr10-64488203-DEL-108 |
| chr10-64505141-DEL-1553 | chr10-64505142-DEL-1552 |
| chr10-64509743-DEL-63 | chr10-64509748-DEL-78 |
| chr10-64544423-DEL-213 | chr10-64544424-DEL-213 |
| chr10-64549801-DEL-90 | chr10-64549808-DEL-128 |
| chr10-64587797-DEL-242 | chr10-64587798-DEL-242 |
| chr10-64588888-DEL-326 | chr10-64588889-DEL-326 |
| chr10-64652166-DEL-809 | chr10-64652167-DEL-809 |
| chr10-64656222-DEL-2406 | chr10-64656222-DEL-2404 |
| chr10-64658847-DEL-586 | chr10-64658848-DEL-586 |
| chr10-64662732-DEL-56 | chr10-64662733-DEL-56 |
| chr10-64669214-DEL-137 | chr10-64669215-DEL-137 |
| chr10-64696591-DEL-543 | chr10-64696588-DEL-547 |
| chr10-64714242-DEL-329 | chr10-64714245-DEL-328 |
| chr10-64764678-DEL-6091 | chr10-64764679-DEL-6091 |
| chr10-64773583-DEL-1939 | chr10-64773584-DEL-1939 |
| chr10-64776144-DEL-482 | chr10-64776144-DEL-482 |
| chr10-64784594-DEL-204 | chr10-64784595-DEL-204 |
| chr10-64786050-DEL-191 | chr10-64786051-DEL-191 |
| chr10-64859246-DEL-209 | chr10-64859247-DEL-209 |
| chr10-64929993-DEL-6193 | chr10-64929994-DEL-6193 |
| chr10-64970067-DEL-3123 | chr10-64970068-DEL-3123 |
| chr10-64976543-DEL-108 | chr10-64976542-DEL-113 |
| chr10-65001628-DEL-6781 | chr10-65001629-DEL-6781 |
| chr10-65021615-DEL-68 | chr10-65021613-DEL-70 |
| chr10-65042459-DEL-58 | chr10-65042462-DEL-56 |
| chr10-65049776-DEL-85 | chr10-65049783-DEL-64 |
| chr10-65088565-DEL-86 | chr10-65088566-DEL-86 |
| chr10-65136653-DEL-83 | chr10-65136654-DEL-83 |
| chr10-65158606-DEL-581 | chr10-65158608-DEL-580 |

| | |
|---|---|
| chr10-65168766-DEL-51 | chr10-65168767-DEL-55 |
| chr10-65218984-DEL-6664 | chr10-65218985-DEL-6664 |
| chr10-65251717-DEL-108 | chr10-65251718-DEL-108 |
| chr10-65280452-DEL-149 | chr10-65280453-DEL-149 |
| chr10-65282862-DEL-194 | chr10-65282867-DEL-200 |
| chr10-65311664-DEL-1889 | chr10-65311665-DEL-1889 |
| chr10-65356252-DEL-1027 | chr10-65356252-DEL-1027 |
| chr10-65367984-DEL-1914 | chr10-65367985-DEL-1914 |
| chr10-65382033-DEL-130 | chr10-65382034-DEL-130 |
| chr10-65390306-DEL-157 | chr10-65390307-DEL-157 |
| chr10-65429194-DEL-2377 | chr10-65429195-DEL-2377 |
| chr10-65474707-DEL-6010 | chr10-65474708-DEL-6010 |
| chr10-65492763-DEL-6665 | chr10-65492764-DEL-6665 |
| chr10-65510277-DEL-254 | chr10-65510285-DEL-254 |
| chr10-65514650-DEL-131 | chr10-65514651-DEL-131 |
| chr10-65563228-DEL-56 | chr10-65563225-DEL-65 |
| chr10-65573990-DEL-56 | chr10-65573991-DEL-56 |
| chr10-65580214-DEL-458 | chr10-65580215-DEL-458 |
| chr10-65585722-DEL-215 | chr10-65585723-DEL-215 |
| chr10-65597736-DEL-4020 | chr10-65597737-DEL-4020 |
| chr10-65598852-DEL-54 | chr10-65598852-DEL-54 |
| chr10-65602682-DEL-255 | chr10-65602710-DEL-228 |
| chr10-65606302-DEL-68 | chr10-65606303-DEL-68 |
| chr10-65629146-DEL-810 | chr10-65629147-DEL-810 |
| chr10-65652538-DEL-17342 | chr10-65652522-DEL-17343 |
| chr10-65674773-DEL-5975 | chr10-65674774-DEL-5975 |
| chr10-65698638-DEL-1613 | chr10-65698639-DEL-1613 |
| chr10-65730821-DEL-328 | chr10-65730822-DEL-328 |
| chr10-65733383-DEL-62 | chr10-65733376-DEL-70 |
| chr10-65736638-DEL-6611 | chr10-65736639-DEL-6611 |
| chr10-65736695-DEL-402 | chr10-65736651-DEL-402 |
| chr10-65745847-DEL-1590 | chr10-65745846-DEL-1590 |
| chr10-65750066-DEL-51 | chr10-65750067-DEL-51 |
| chr10-65813998-DEL-69 | chr10-65813999-DEL-69 |
| chr10-65832584-DEL-6135 | chr10-65832585-DEL-6135 |
| chr10-65870664-DEL-92 | chr10-65870665-DEL-92 |
| chr10-65880317-DEL-568 | chr10-65880318-DEL-568 |
| chr10-65902392-DEL-1887 | chr10-65902393-DEL-1887 |
| chr10-65908396-DEL-6680 | chr10-65908397-DEL-6680 |
| chr10-65928781-DEL-890 | chr10-65928782-DEL-890 |
| chr10-65940311-DEL-60 | chr10-65940304-DEL-68 |
| chr10-65987543-DEL-57 | chr10-65987544-DEL-57 |
| chr10-65992425-DEL-1131 | chr10-65992426-DEL-1131 |
| chr10-66005118-DEL-1166 | chr10-66005119-DEL-1166 |
| chr10-66035205-DEL-432 | chr10-66035206-DEL-432 |
| chr10-66042928-DEL-2090 | chr10-66042929-DEL-2090 |
| chr10-66071492-DEL-347 | chr10-66071493-DEL-347 |
| chr10-66072100-DEL-116 | chr10-66072093-DEL-96 |
| chr10-66102525-DEL-198 | chr10-66102526-DEL-198 |
| chr10-66125775-DEL-80 | chr10-66125776-DEL-80 |
| chr10-66143035-DEL-3076 | chr10-66143036-DEL-3076 |
| chr10-66158365-DEL-162 | chr10-66158348-DEL-179 |
| chr10-66213377-DEL-580 | chr10-66213377-DEL-580 |
| chr10-66214911-DEL-399 | chr10-66214912-DEL-399 |
| chr10-66230906-DEL-112 | chr10-66230907-DEL-112 |
| chr10-66251019-DEL-53 | chr10-66251025-DEL-52 |
| chr10-66252001-DEL-67 | chr10-66251995-DEL-67 |
| chr10-66257345-DEL-10235 | chr10-66257337-DEL-10229 |
| chr10-66257347-DEL-201 | chr10-66257370-DEL-201 |
| chr10-66289319-DEL-3480 | chr10-66289320-DEL-3480 |
| chr10-66296603-DEL-73 | chr10-66296603-DEL-75 |
| chr10-66298202-DEL-6595 | chr10-66298203-DEL-6595 |
| chr10-66327548-DEL-247 | chr10-66327549-DEL-247 |
| chr10-66330742-DEL-252 | chr10-66330723-DEL-248 |
| chr10-66332077-DEL-52 | chr10-66332078-DEL-52 |
| chr10-66336606-DEL-56 | chr10-66336607-DEL-56 |
| chr10-66346321-DEL-134 | chr10-66346322-DEL-134 |
| chr10-66367808-DEL-270 | chr10-66367809-DEL-270 |
| chr10-66370435-DEL-60 | chr10-66370391-DEL-55 |
| chr10-66389126-DEL-122 | chr10-66389125-DEL-123 |
| chr10-66389357-DEL-802 | chr10-66389358-DEL-802 |
| chr10-66402303-DEL-384 | chr10-66402304-DEL-384 |
| chr10-66417975-DEL-6871 | chr10-66417975-DEL-6871 |
| chr10-66462107-DEL-77 | chr10-66462072-DEL-64 |
| chr10-66469021-DEL-527 | chr10-66469029-DEL-500 |
| chr10-66470441-DEL-90 | chr10-66470438-DEL-94 |
| chr10-66475095-DEL-61 | chr10-66475096-DEL-61 |
| chr10-66477925-DEL-433 | chr10-66477926-DEL-433 |
| chr10-66507715-DEL-199 | chr10-66507716-DEL-199 |
| chr10-66535076-DEL-183 | chr10-66535077-DEL-183 |

| | |
|---|---|
| chr10-66536488-DEL-197 | chr10-66536489-DEL-197 |
| chr10-66558256-DEL-7125 | chr10-66558257-DEL-7125 |
| chr10-66581977-DEL-325 | chr10-66581978-DEL-325 |
| chr10-66590873-DEL-241 | chr10-66590874-DEL-241 |
| chr10-66645015-DEL-799 | chr10-66645013-DEL-798 |
| chr10-66656008-DEL-153 | chr10-66656014-DEL-145 |
| chr10-66657750-DEL-55 | chr10-66657751-DEL-55 |
| chr10-66659526-DEL-59 | chr10-66659527-DEL-60 |
| chr10-66669448-DEL-197 | chr10-66669437-DEL-198 |
| chr10-66672861-DEL-395 | chr10-66672862-DEL-395 |
| chr10-66684065-DEL-83 | chr10-66684066-DEL-83 |
| chr10-66686462-DEL-52 | chr10-66686442-DEL-72 |
| chr10-66689816-DEL-79 | chr10-66689813-DEL-80 |
| chr10-66702610-DEL-54 | chr10-66702611-DEL-54 |
| chr10-66708690-DEL-200 | chr10-66708691-DEL-200 |
| chr10-66722312-DEL-214 | chr10-66722313-DEL-214 |
| chr10-66740092-DEL-207 | chr10-66740093-DEL-207 |
| chr10-66796742-DEL-2136 | chr10-66796744-DEL-2135 |
| chr10-66799491-DEL-98 | chr10-66799492-DEL-98 |
| chr10-66817390-DEL-531 | chr10-66817379-DEL-534 |
| chr10-66819089-DEL-159 | chr10-66819090-DEL-159 |
| chr10-66826188-DEL-988 | chr10-66826189-DEL-988 |
| chr10-66843784-DEL-61 | chr10-66843785-DEL-58 |
| chr10-66885983-DEL-72 | chr10-66885982-DEL-64 |
| chr10-66896535-DEL-127 | chr10-66896536-DEL-127 |
| chr10-66906460-DEL-100 | chr10-66906461-DEL-100 |
| chr10-66907924-DEL-556 | chr10-66907925-DEL-556 |
| chr10-66915375-DEL-222 | chr10-66915376-DEL-222 |
| chr10-66916514-DEL-85 | chr10-66916516-DEL-76 |
| chr10-66918836-DEL-215 | chr10-66918837-DEL-215 |
| chr10-66998205-DEL-199 | chr10-66998206-DEL-199 |
| chr10-67011979-DEL-85 | chr10-67011980-DEL-85 |
| chr10-67035273-DEL-221 | chr10-67035274-DEL-221 |
| chr10-67048082-DEL-203 | chr10-67048083-DEL-203 |
| chr10-67121796-DEL-216 | chr10-67121797-DEL-216 |
| chr10-67146803-DEL-66 | chr10-67146806-DEL-64 |
| chr10-67150490-DEL-253 | chr10-67150491-DEL-253 |
| chr10-67181704-DEL-157 | chr10-67181705-DEL-157 |
| chr10-67182836-DEL-90 | chr10-67182837-DEL-90 |
| chr10-67187818-DEL-211 | chr10-67187819-DEL-211 |
| chr10-67205423-DEL-99 | chr10-67205424-DEL-95 |
| chr10-67241928-DEL-123 | chr10-67241929-DEL-123 |
| chr10-67249224-DEL-57 | chr10-67249207-DEL-87 |
| chr10-67275514-DEL-120 | chr10-67275515-DEL-120 |
| chr10-67281912-DEL-119 | chr10-67281913-DEL-119 |
| chr10-67331068-DEL-55 | chr10-67331023-DEL-81 |
| chr10-67389960-DEL-81 | chr10-67389948-DEL-94 |
| chr10-67391839-DEL-244 | chr10-67391840-DEL-244 |
| chr10-67403359-DEL-72 | chr10-67403360-DEL-72 |
| chr10-67413137-DEL-272 | chr10-67413138-DEL-272 |
| chr10-67415036-DEL-650 | chr10-67415036-DEL-650 |
| chr10-67420357-DEL-92 | chr10-67420358-DEL-92 |
| chr10-67474756-DEL-82 | chr10-67474757-DEL-82 |
| chr10-67495984-DEL-1240 | chr10-67495985-DEL-1240 |
| chr10-67502618-DEL-65 | chr10-67502619-DEL-65 |
| chr10-67507408-DEL-497 | chr10-67507409-DEL-481 |
| chr10-67510500-DEL-440 | chr10-67510501-DEL-440 |
| chr10-67512521-DEL-2081 | chr10-67512522-DEL-2081 |
| chr10-67517229-DEL-96 | chr10-67517230-DEL-96 |
| chr10-67539794-DEL-60 | chr10-67539795-DEL-60 |
| chr10-67543091-DEL-52 | chr10-67543111-DEL-70 |
| chr10-67544292-DEL-106 | chr10-67544293-DEL-106 |
| chr10-67592349-DEL-104 | chr10-67592344-DEL-110 |
| chr10-67593310-DEL-82 | chr10-67593272-DEL-116 |
| chr10-67595649-DEL-66 | chr10-67595636-DEL-82 |
| chr10-67631156-DEL-76 | chr10-67631135-DEL-74 |
| chr10-67632821-DEL-50 | chr10-67632808-DEL-64 |
| chr10-67644179-DEL-1274 | chr10-67644180-DEL-1274 |
| chr10-67748956-DEL-59 | chr10-67748957-DEL-59 |
| chr10-67751323-DEL-86 | chr10-67751318-DEL-84 |
| chr10-67761358-DEL-1880 | chr10-67761359-DEL-1880 |
| chr10-67779773-DEL-841 | chr10-67779774-DEL-841 |
| chr10-67794133-DEL-348 | chr10-67794134-DEL-348 |
| chr10-67805022-DEL-741 | chr10-67805023-DEL-741 |
| chr10-67824601-DEL-2645 | chr10-67824602-DEL-2645 |
| chr10-67845628-DEL-78 | chr10-67845629-DEL-78 |
| chr10-67854225-DEL-150 | chr10-67854226-DEL-150 |
| chr10-67858969-DEL-60 | chr10-67858936-DEL-59 |
| chr10-67904931-DEL-59 | chr10-67904932-DEL-59 |
| chr10-67909046-DEL-57 | chr10-67909036-DEL-58 |

| | |
|---|---|
| chr10-67914439-DEL-205 | chr10-67914440-DEL-205 |
| chr10-67918516-DEL-184 | chr10-67918515-DEL-190 |
| chr10-67947671-DEL-61 | chr10-67947655-DEL-58 |
| chr10-68012521-DEL-68 | chr10-68012522-DEL-62 |
| chr10-68016892-DEL-140 | chr10-68016893-DEL-140 |
| chr10-68060813-DEL-660 | chr10-68060821-DEL-676 |
| chr10-68079180-DEL-3100 | chr10-68079181-DEL-3100 |
| chr10-68123291-DEL-56 | chr10-68123284-DEL-66 |
| chr10-68123475-DEL-733 | chr10-68123476-DEL-733 |
| chr10-68145416-DEL-207 | chr10-68145411-DEL-205 |
| chr10-68161470-DEL-2417 | chr10-68161471-DEL-2417 |
| chr10-68172012-DEL-52 | chr10-68172013-DEL-52 |
| chr10-68211079-DEL-857 | chr10-68211077-DEL-840 |
| chr10-68242800-DEL-972 | chr10-68242801-DEL-972 |
| chr10-68246796-DEL-3117 | chr10-68246797-DEL-3117 |
| chr10-68250074-DEL-69 | chr10-68250072-DEL-73 |
| chr10-68267169-DEL-51 | chr10-68267167-DEL-53 |
| chr10-68336683-DEL-438 | chr10-68336684-DEL-438 |
| chr10-68343057-DEL-59 | chr10-68343058-DEL-59 |
| chr10-68351593-DEL-91 | chr10-68351580-DEL-104 |
| chr10-68402799-DEL-89 | chr10-68402800-DEL-89 |
| chr10-68407831-DEL-139 | chr10-68407832-DEL-139 |
| chr10-68419782-DEL-54 | chr10-68419783-DEL-54 |
| chr10-68445592-DEL-66 | chr10-68445591-DEL-64 |
| chr10-68446319-DEL-64 | chr10-68446318-DEL-62 |
| chr10-68478972-DEL-54 | chr10-68478968-DEL-70 |
| chr10-68489905-DEL-70 | chr10-68489920-DEL-50 |
| chr10-68516885-DEL-52 | chr10-68516886-DEL-52 |
| chr10-68630781-DEL-113 | chr10-68630785-DEL-120 |
| chr10-68642756-DEL-55 | chr10-68642757-DEL-55 |
| chr10-68681469-DEL-744 | chr10-68681470-DEL-744 |
| chr10-68706544-DEL-517 | chr10-68706557-DEL-513 |
| chr10-68744373-DEL-56 | chr10-68744376-DEL-54 |
| chr10-68753463-DEL-70 | chr10-68753463-DEL-76 |
| chr10-68756603-DEL-490 | chr10-68756604-DEL-490 |
| chr10-68759940-DEL-171 | chr10-68759941-DEL-171 |
| chr10-68778400-DEL-115 | chr10-68778401-DEL-115 |
| chr10-68795412-DEL-99 | chr10-68795412-DEL-99 |
| chr10-68829294-DEL-56 | chr10-68829295-DEL-56 |
| chr10-68844594-DEL-84 | chr10-68844595-DEL-84 |
| chr10-68848690-DEL-805 | chr10-68848692-DEL-804 |
| chr10-68854998-DEL-720 | chr10-68854999-DEL-720 |
| chr10-68860807-DEL-53 | chr10-68860768-DEL-54 |
| chr10-68870755-DEL-2654 | chr10-68870756-DEL-2654 |
| chr10-68895797-DEL-6238 | chr10-68895799-DEL-6237 |
| chr10-68898722-DEL-13724 | chr10-68898723-DEL-13724 |
| chr10-68916970-DEL-54 | chr10-68916975-DEL-98 |
| chr10-68918233-DEL-130 | chr10-68918248-DEL-134 |
| chr10-68922699-DEL-427 | chr10-68922700-DEL-427 |
| chr10-68939231-DEL-61 | chr10-68939216-DEL-68 |
| chr10-68948068-DEL-75 | chr10-68948069-DEL-75 |
| chr10-68998716-DEL-60 | chr10-68998715-DEL-62 |
| chr10-69012826-DEL-88 | chr10-69012796-DEL-114 |
| chr10-69032440-DEL-6436 | chr10-69032441-DEL-6436 |
| chr10-69040062-DEL-63 | chr10-69040029-DEL-55 |
| chr10-69042829-DEL-159 | chr10-69042830-DEL-159 |
| chr10-69057782-DEL-76 | chr10-69057766-DEL-56 |
| chr10-69064579-DEL-60 | chr10-69064588-DEL-52 |
| chr10-69065257-DEL-6667 | chr10-69065258-DEL-6667 |
| chr10-69083399-DEL-283 | chr10-69083400-DEL-283 |
| chr10-69100311-DEL-100 | chr10-69100312-DEL-100 |
| chr10-69105344-DEL-72 | chr10-69105345-DEL-72 |
| chr10-69113275-DEL-71 | chr10-69113276-DEL-71 |
| chr10-69116397-DEL-64 | chr10-69116398-DEL-64 |
| chr10-69136594-DEL-166 | chr10-69136595-DEL-166 |
| chr10-69155115-DEL-58 | chr10-69155116-DEL-58 |
| chr10-69159645-DEL-186 | chr10-69159646-DEL-186 |
| chr10-69166487-DEL-200 | chr10-69166488-DEL-200 |
| chr10-69181423-DEL-68 | chr10-69181424-DEL-68 |
| chr10-69238022-DEL-399 | chr10-69238023-DEL-399 |
| chr10-69244054-DEL-360 | chr10-69244055-DEL-360 |
| chr10-69255303-DEL-262 | chr10-69255304-DEL-262 |
| chr10-69261190-DEL-58 | chr10-69261191-DEL-58 |
| chr10-69265909-DEL-52 | chr10-69265864-DEL-98 |
| chr10-69268426-DEL-151 | chr10-69268428-DEL-149 |
| chr10-69276815-DEL-1449 | chr10-69276803-DEL-1469 |
| chr10-69305631-DEL-60 | chr10-69305627-DEL-70 |
| chr10-69309555-DEL-2820 | chr10-69309556-DEL-2820 |
| chr10-69318770-DEL-199 | chr10-69318771-DEL-199 |
| chr10-69326777-DEL-124 | chr10-69326802-DEL-106 |

| | |
|---|---|
| chr10-69329489-DEL-909 | chr10-69329490-DEL-909 |
| chr10-69359872-DEL-93 | chr10-69359878-DEL-68 |
| chr10-69373211-DEL-142 | chr10-69373212-DEL-142 |
| chr10-69391429-DEL-78 | chr10-69391430-DEL-78 |
| chr10-69403929-DEL-72 | chr10-69403930-DEL-72 |
| chr10-69435046-DEL-130 | chr10-69435047-DEL-120 |
| chr10-69445839-DEL-51 | chr10-69445839-DEL-51 |
| chr10-69446508-DEL-504 | chr10-69446509-DEL-504 |
| chr10-69478611-DEL-69 | chr10-69478608-DEL-66 |
| chr10-69496376-DEL-216 | chr10-69496377-DEL-216 |
| chr10-69515546-DEL-178 | chr10-69515547-DEL-178 |
| chr10-69549522-DEL-89 | chr10-69549523-DEL-59 |
| chr10-69559228-DEL-99 | chr10-69559229-DEL-99 |
| chr10-69589645-DEL-198 | chr10-69589645-DEL-195 |
| chr10-69593615-DEL-74 | chr10-69593612-DEL-74 |
| chr10-69596227-DEL-72 | chr10-69596228-DEL-72 |
| chr10-69597761-DEL-52 | chr10-69597762-DEL-52 |
| chr10-69599438-DEL-56 | chr10-69599439-DEL-56 |
| chr10-69600912-DEL-52 | chr10-69600911-DEL-52 |
| chr10-69636263-DEL-92 | chr10-69636264-DEL-92 |
| chr10-69638670-DEL-119 | chr10-69638669-DEL-99 |
| chr10-69655426-DEL-1102 | chr10-69655427-DEL-1102 |
| chr10-69693830-DEL-69 | chr10-69693829-DEL-69 |
| chr10-69700142-DEL-66 | chr10-69700145-DEL-64 |
| chr10-69743378-DEL-180 | chr10-69743379-DEL-180 |
| chr10-69746398-DEL-276 | chr10-69746399-DEL-276 |
| chr10-69783970-DEL-58 | chr10-69783966-DEL-62 |
| chr10-69786389-DEL-189 | chr10-69786391-DEL-188 |
| chr10-69802508-DEL-1087 | chr10-69802509-DEL-1087 |
| chr10-69845235-DEL-141 | chr10-69845247-DEL-135 |
| chr10-69873885-DEL-58 | chr10-69873886-DEL-58 |
| chr10-69883848-DEL-216 | chr10-69883857-DEL-227 |
| chr10-69897190-DEL-175 | chr10-69897191-DEL-175 |
| chr10-69898550-DEL-533 | chr10-69898551-DEL-533 |
| chr10-69918324-DEL-2255 | chr10-69918325-DEL-2255 |
| chr10-69918346-DEL-239 | chr10-69918347-DEL-239 |
| chr10-69923379-DEL-133 | chr10-69923380-DEL-133 |
| chr10-69952662-DEL-325 | chr10-69952651-DEL-323 |
| chr10-69958374-DEL-89 | chr10-69958375-DEL-89 |
| chr10-69959705-DEL-203 | chr10-69959706-DEL-203 |
| chr10-69987781-DEL-169 | chr10-69987782-DEL-169 |
| chr10-69992831-DEL-51 | chr10-69992832-DEL-51 |
| chr10-70047121-DEL-1572 | chr10-70047122-DEL-1572 |
| chr10-70053860-DEL-736 | chr10-70053861-DEL-736 |
| chr10-70058186-DEL-175 | chr10-70058186-DEL-175 |
| chr10-70142429-DEL-56 | chr10-70142430-DEL-56 |
| chr10-70160258-DEL-424 | chr10-70160259-DEL-424 |
| chr10-70162478-DEL-94 | chr10-70162496-DEL-114 |
| chr10-70164294-DEL-508 | chr10-70164295-DEL-508 |
| chr10-70169438-DEL-486 | chr10-70169439-DEL-486 |
| chr10-70173606-DEL-397 | chr10-70173607-DEL-397 |
| chr10-70206647-DEL-129 | chr10-70206648-DEL-129 |
| chr10-70241902-DEL-75 | chr10-70241903-DEL-75 |
| chr10-70243577-DEL-433 | chr10-70243578-DEL-433 |
| chr10-70295727-DEL-1289 | chr10-70295728-DEL-1289 |
| chr10-70316282-DEL-64 | chr10-70316239-DEL-60 |
| chr10-70320118-DEL-187 | chr10-70320119-DEL-187 |
| chr10-70351200-DEL-54 | chr10-70351201-DEL-54 |
| chr10-70373402-DEL-135 | chr10-70373376-DEL-136 |
| chr10-70384313-DEL-121 | chr10-70384313-DEL-121 |
| chr10-70386176-DEL-149 | chr10-70386179-DEL-147 |
| chr10-70440269-DEL-153 | chr10-70440270-DEL-153 |
| chr10-70470433-DEL-89 | chr10-70470447-DEL-72 |
| chr10-70477970-DEL-51 | chr10-70477971-DEL-51 |
| chr10-70480174-DEL-101 | chr10-70480198-DEL-77 |
| chr10-70483440-DEL-104 | chr10-70483443-DEL-100 |
| chr10-70493236-DEL-3119 | chr10-70493237-DEL-3119 |
| chr10-70498263-DEL-6217 | chr10-70498290-DEL-6218 |
| chr10-70507511-DEL-66 | chr10-70507522-DEL-58 |
| chr10-70516829-DEL-336 | chr10-70516830-DEL-336 |
| chr10-70538369-DEL-52 | chr10-70538370-DEL-52 |
| chr10-70550950-DEL-54 | chr10-70550951-DEL-54 |
| chr10-70608967-DEL-128 | chr10-70608968-DEL-128 |
| chr10-70615306-DEL-1011 | chr10-70615307-DEL-1011 |
| chr10-70620668-DEL-3334 | chr10-70620669-DEL-3334 |
| chr10-70627207-DEL-7161 | chr10-70627208-DEL-7161 |
| chr10-70662485-DEL-106 | chr10-70662486-DEL-106 |
| chr10-70663776-DEL-78 | chr10-70663777-DEL-78 |
| chr10-70688966-DEL-129 | chr10-70688967-DEL-129 |
| chr10-70715126-DEL-11478 | chr10-70715124-DEL-11479 |

| | |
|---|---|
| chr10-70717920-DEL-5008 | chr10-70717921-DEL-5008 |
| chr10-70726095-DEL-128 | chr10-70726096-DEL-128 |
| chr10-70729125-DEL-2238 | chr10-70729126-DEL-2238 |
| chr10-70736585-DEL-94 | chr10-70736576-DEL-103 |
| chr10-70747716-DEL-1806 | chr10-70747717-DEL-1806 |
| chr10-70750581-DEL-391 | chr10-70750582-DEL-391 |
| chr10-70750840-DEL-1055 | chr10-70750841-DEL-1055 |
| chr10-70768515-DEL-215 | chr10-70768542-DEL-172 |
| chr10-70782088-DEL-156 | chr10-70782089-DEL-156 |
| chr10-70787635-DEL-471 | chr10-70787636-DEL-471 |
| chr10-70796319-DEL-293 | chr10-70796303-DEL-294 |
| chr10-70806746-DEL-162 | chr10-70806747-DEL-162 |
| chr10-70809175-DEL-200 | chr10-70809176-DEL-200 |
| chr10-70820587-DEL-1697 | chr10-70820588-DEL-1697 |
| chr10-70822388-DEL-90 | chr10-70822389-DEL-90 |
| chr10-70832272-DEL-94 | chr10-70832291-DEL-58 |
| chr10-70837746-DEL-72 | chr10-70837735-DEL-84 |
| chr10-70841540-DEL-74 | chr10-70841531-DEL-104 |
| chr10-70848556-DEL-6442 | chr10-70848557-DEL-6442 |
| chr10-70865644-DEL-96 | chr10-70865671-DEL-64 |
| chr10-70878507-DEL-156 | chr10-70878508-DEL-156 |
| chr10-70902610-DEL-360 | chr10-70902611-DEL-360 |
| chr10-70909467-DEL-53 | chr10-70909468-DEL-53 |
| chr10-70909638-DEL-2670 | chr10-70909639-DEL-2670 |
| chr10-70912706-DEL-139 | chr10-70912707-DEL-139 |
| chr10-70913323-DEL-98 | chr10-70913324-DEL-98 |
| chr10-70918441-DEL-174 | chr10-70918442-DEL-174 |
| chr10-70933337-DEL-1124 | chr10-70933338-DEL-1124 |
| chr10-70947005-DEL-3032 | chr10-70947006-DEL-3032 |
| chr10-70969090-DEL-235 | chr10-70969091-DEL-235 |
| chr10-70981248-DEL-88 | chr10-70981249-DEL-88 |
| chr10-70993807-DEL-65 | chr10-70993808-DEL-65 |
| chr10-71023146-DEL-61 | chr10-71023147-DEL-61 |
| chr10-71053951-DEL-275 | chr10-71053952-DEL-275 |
| chr10-71065731-DEL-130 | chr10-71065732-DEL-130 |
| chr10-71069485-DEL-133 | chr10-71069483-DEL-133 |
| chr10-71080842-DEL-504 | chr10-71080843-DEL-504 |
| chr10-71088688-DEL-205 | chr10-71088689-DEL-205 |
| chr10-71128058-DEL-53 | chr10-71128059-DEL-54 |
| chr10-71131206-DEL-1851 | chr10-71131205-DEL-1851 |
| chr10-71145059-DEL-78 | chr10-71145057-DEL-60 |
| chr10-71147469-DEL-178 | chr10-71147491-DEL-188 |
| chr10-71152738-DEL-197 | chr10-71152739-DEL-197 |
| chr10-71159271-DEL-211 | chr10-71159272-DEL-211 |
| chr10-71160533-DEL-72 | chr10-71160534-DEL-60 |
| chr10-71210313-DEL-275 | chr10-71210314-DEL-275 |
| chr10-71218544-DEL-98 | chr10-71218545-DEL-98 |
| chr10-71224982-DEL-1166 | chr10-71224981-DEL-1167 |
| chr10-71232655-DEL-460 | chr10-71232656-DEL-460 |
| chr10-71253857-DEL-128 | chr10-71253858-DEL-128 |
| chr10-71254895-DEL-1549 | chr10-71254896-DEL-1549 |
| chr10-71274735-DEL-583 | chr10-71274723-DEL-579 |
| chr10-71277241-DEL-176 | chr10-71277242-DEL-176 |
| chr10-71283533-DEL-137 | chr10-71283490-DEL-135 |
| chr10-71287915-DEL-1889 | chr10-71287916-DEL-1889 |
| chr10-71291131-DEL-78 | chr10-71291150-DEL-72 |
| chr10-71316043-DEL-713 | chr10-71316044-DEL-713 |
| chr10-71328207-DEL-60 | chr10-71328208-DEL-60 |
| chr10-71386799-DEL-2737 | chr10-71386800-DEL-2737 |
| chr10-71443211-DEL-1566 | chr10-71443212-DEL-1566 |
| chr10-71498846-DEL-59 | chr10-71498813-DEL-75 |
| chr10-71500071-DEL-142 | chr10-71500072-DEL-130 |
| chr10-71505368-DEL-1709 | chr10-71505369-DEL-1709 |
| chr10-71527228-DEL-7117 | chr10-71527229-DEL-7117 |
| chr10-71540102-DEL-3610 | chr10-71540103-DEL-3610 |
| chr10-71541877-DEL-429 | chr10-71541878-DEL-429 |
| chr10-71586576-DEL-140 | chr10-71586573-DEL-152 |
| chr10-71594916-DEL-151 | chr10-71594917-DEL-151 |
| chr10-71623340-DEL-339 | chr10-71623343-DEL-337 |
| chr10-71634430-DEL-93 | chr10-71634431-DEL-93 |
| chr10-71654198-DEL-6587 | chr10-71654199-DEL-6587 |
| chr10-71685398-DEL-6261 | chr10-71685399-DEL-6261 |
| chr10-71718301-DEL-205 | chr10-71718257-DEL-204 |
| chr10-71728888-DEL-65 | chr10-71728888-DEL-65 |
| chr10-71759493-DEL-305 | chr10-71759494-DEL-305 |
| chr10-71760037-DEL-88 | chr10-71760028-DEL-105 |
| chr10-71812069-DEL-251 | chr10-71812070-DEL-251 |
| chr10-71820464-DEL-242 | chr10-71820476-DEL-240 |
| chr10-71843492-DEL-6983 | chr10-71843493-DEL-6983 |
| chr10-71895973-DEL-394 | chr10-71895974-DEL-394 |

| | |
|---|---|
| chr10-72001680-DEL-70 | chr10-72001680-DEL-70 |
| chr10-72074397-DEL-2107 | chr10-72074398-DEL-2107 |
| chr10-72085788-DEL-6235 | chr10-72085803-DEL-6219 |
| chr10-72095724-DEL-2675 | chr10-72095725-DEL-2675 |
| chr10-72098338-DEL-62 | chr10-72098336-DEL-52 |
| chr10-72129388-DEL-69 | chr10-72129389-DEL-69 |
| chr10-72133006-DEL-1254 | chr10-72133007-DEL-1254 |
| chr10-72145012-DEL-5019 | chr10-72145014-DEL-5018 |
| chr10-72169860-DEL-1704 | chr10-72169861-DEL-1704 |
| chr10-72190945-DEL-989 | chr10-72190946-DEL-989 |
| chr10-72211691-DEL-268 | chr10-72211692-DEL-268 |
| chr10-72240315-DEL-980 | chr10-72240316-DEL-980 |
| chr10-72244618-DEL-773 | chr10-72244619-DEL-773 |
| chr10-72265299-DEL-1142 | chr10-72265300-DEL-1142 |
| chr10-72290980-DEL-139 | chr10-72290981-DEL-137 |
| chr10-72315478-DEL-199 | chr10-72315479-DEL-199 |
| chr10-72327116-DEL-55 | chr10-72327115-DEL-63 |
| chr10-72328644-DEL-6447 | chr10-72328643-DEL-6448 |
| chr10-72356537-DEL-79 | chr10-72356538-DEL-81 |
| chr10-72367628-DEL-87 | chr10-72367629-DEL-87 |
| chr10-72379935-DEL-5131 | chr10-72379935-DEL-5130 |
| chr10-72411454-DEL-51 | chr10-72411455-DEL-51 |
| chr10-72411848-DEL-726 | chr10-72411849-DEL-726 |
| chr10-72425501-DEL-2153 | chr10-72425501-DEL-2153 |
| chr10-72486483-DEL-434 | chr10-72486484-DEL-434 |
| chr10-72507336-DEL-58 | chr10-72507335-DEL-58 |
| chr10-72510827-DEL-176 | chr10-72510828-DEL-176 |
| chr10-72522748-DEL-261 | chr10-72522749-DEL-261 |
| chr10-72559839-DEL-229 | chr10-72559840-DEL-229 |
| chr10-72574910-DEL-244 | chr10-72574911-DEL-244 |
| chr10-72622921-DEL-314 | chr10-72622922-DEL-314 |
| chr10-72632852-DEL-60 | chr10-72632823-DEL-76 |
| chr10-72643514-DEL-518 | chr10-72643519-DEL-513 |
| chr10-72655022-DEL-490 | chr10-72655022-DEL-488 |
| chr10-72719467-DEL-111 | chr10-72719468-DEL-111 |
| chr10-72722785-DEL-253 | chr10-72722786-DEL-253 |
| chr10-72767688-DEL-6399 | chr10-72767689-DEL-6399 |
| chr10-72778816-DEL-126 | chr10-72778817-DEL-126 |
| chr10-72789595-DEL-80 | chr10-72789594-DEL-82 |
| chr10-72806980-DEL-70 | chr10-72806981-DEL-70 |
| chr10-72816086-DEL-58 | chr10-72816087-DEL-58 |
| chr10-72863157-DEL-1067 | chr10-72863158-DEL-1067 |
| chr10-72885214-DEL-755 | chr10-72885215-DEL-755 |
| chr10-72976403-DEL-244 | chr10-72976388-DEL-244 |
| chr10-72984568-DEL-63 | chr10-72984569-DEL-63 |
| chr10-72988466-DEL-169 | chr10-72988465-DEL-170 |
| chr10-72991538-DEL-88 | chr10-72991531-DEL-98 |
| chr10-72992706-DEL-443 | chr10-72992707-DEL-443 |
| chr10-73027305-DEL-445 | chr10-73027306-DEL-445 |
| chr10-73029047-DEL-2125 | chr10-73029048-DEL-2125 |
| chr10-73032530-DEL-265 | chr10-73032531-DEL-265 |
| chr10-73080718-DEL-52 | chr10-73080719-DEL-52 |
| chr10-73088781-DEL-1453 | chr10-73088782-DEL-1453 |
| chr10-73091177-DEL-212 | chr10-73091179-DEL-211 |
| chr10-73124213-DEL-6543 | chr10-73124214-DEL-6543 |
| chr10-73159882-DEL-79 | chr10-73159884-DEL-76 |
| chr10-73162915-DEL-55 | chr10-73162871-DEL-56 |
| chr10-73229530-DEL-130 | chr10-73229531-DEL-130 |
| chr10-73273338-DEL-6848 | chr10-73273339-DEL-6848 |
| chr10-73273643-DEL-424 | chr10-73273644-DEL-424 |
| chr10-73298973-DEL-69 | chr10-73298979-DEL-55 |
| chr10-73331632-DEL-408 | chr10-73331633-DEL-408 |
| chr10-73344176-DEL-51 | chr10-73344176-DEL-51 |
| chr10-73346266-DEL-76 | chr10-73346266-DEL-76 |
| chr10-73354770-DEL-66 | chr10-73354771-DEL-64 |
| chr10-73362026-DEL-1933 | chr10-73362028-DEL-1931 |
| chr10-73373914-DEL-52 | chr10-73373915-DEL-52 |
| chr10-73377143-DEL-120 | chr10-73377152-DEL-123 |
| chr10-73378887-DEL-191 | chr10-73378888-DEL-191 |
| chr10-73383852-DEL-3205 | chr10-73383857-DEL-3198 |
| chr10-73412110-DEL-6828 | chr10-73412111-DEL-6828 |
| chr10-73421561-DEL-91 | chr10-73421577-DEL-84 |
| chr10-73472520-DEL-499 | chr10-73472521-DEL-499 |
| chr10-73481940-DEL-276 | chr10-73481941-DEL-276 |
| chr10-73485651-DEL-57 | chr10-73485635-DEL-54 |
| chr10-73514034-DEL-1552 | chr10-73514034-DEL-1552 |
| chr10-73541045-DEL-126 | chr10-73541044-DEL-127 |
| chr10-73565671-DEL-66 | chr10-73565666-DEL-68 |
| chr10-73572891-DEL-457 | chr10-73572859-DEL-458 |
| chr10-73573514-DEL-1356 | chr10-73573515-DEL-1356 |

| | |
|---|---|
| chr10-73581357-DEL-102 | chr10-73581336-DEL-83 |
| chr10-73590946-DEL-67 | chr10-73590953-DEL-69 |
| chr10-73600664-DEL-6751 | chr10-73600657-DEL-6752 |
| chr10-73641518-DEL-4212 | chr10-73641517-DEL-4211 |
| chr10-73690549-DEL-2734 | chr10-73690550-DEL-2734 |
| chr10-73696522-DEL-122 | chr10-73696523-DEL-122 |
| chr10-73716782-DEL-52 | chr10-73716799-DEL-53 |
| chr10-73738669-DEL-72 | chr10-73738636-DEL-96 |
| chr10-73798554-DEL-272 | chr10-73798555-DEL-272 |
| chr10-73806175-DEL-57 | chr10-73806173-DEL-52 |
| chr10-73830490-DEL-447 | chr10-73830491-DEL-447 |
| chr10-73847595-DEL-139 | chr10-73847589-DEL-124 |
| chr10-73908761-DEL-462 | chr10-73908762-DEL-462 |
| chr10-73934650-DEL-852 | chr10-73934651-DEL-852 |
| chr10-73941951-DEL-55 | chr10-73941952-DEL-55 |
| chr10-73943671-DEL-60 | chr10-73943672-DEL-60 |
| chr10-73956602-DEL-438 | chr10-73956603-DEL-438 |
| chr10-74005203-DEL-74 | chr10-74005204-DEL-74 |
| chr10-74023488-DEL-52 | chr10-74023489-DEL-52 |
| chr10-74067535-DEL-252 | chr10-74067530-DEL-260 |
| chr10-74134132-DEL-64 | chr10-74134133-DEL-64 |
| chr10-74142838-DEL-89 | chr10-74142839-DEL-88 |
| chr10-74144042-DEL-1502 | chr10-74144043-DEL-1502 |
| chr10-74148289-DEL-80 | chr10-74148328-DEL-54 |
| chr10-74155330-DEL-84 | chr10-74155325-DEL-93 |
| chr10-74172900-DEL-401 | chr10-74172901-DEL-401 |
| chr10-74178843-DEL-6262 | chr10-74178844-DEL-6262 |
| chr10-74207500-DEL-80 | chr10-74207459-DEL-78 |
| chr10-74262144-DEL-209 | chr10-74262145-DEL-209 |
| chr10-74292729-DEL-1067 | chr10-74292730-DEL-1067 |
| chr10-74296598-DEL-65 | chr10-74296599-DEL-65 |
| chr10-74357526-DEL-486 | chr10-74357527-DEL-486 |
| chr10-74491197-DEL-6317 | chr10-74491198-DEL-6317 |
| chr10-74517136-DEL-57 | chr10-74517141-DEL-53 |
| chr10-74522415-DEL-3088 | chr10-74522416-DEL-3088 |
| chr10-74526487-DEL-54 | chr10-74526486-DEL-56 |
| chr10-74536289-DEL-1283 | chr10-74536290-DEL-1283 |
| chr10-74550527-DEL-5099 | chr10-74550527-DEL-5099 |
| chr10-74575333-DEL-64 | chr10-74575334-DEL-64 |
| chr10-74582233-DEL-200 | chr10-74582232-DEL-200 |
| chr10-74586102-DEL-388 | chr10-74586105-DEL-386 |
| chr10-74603852-DEL-6626 | chr10-74603852-DEL-6626 |
| chr10-74636527-DEL-1761 | chr10-74636528-DEL-1761 |
| chr10-74660369-DEL-1005 | chr10-74660370-DEL-1005 |
| chr10-74668698-DEL-66 | chr10-74668699-DEL-66 |
| chr10-74693260-DEL-3130 | chr10-74693253-DEL-3128 |
| chr10-74711550-DEL-3679 | chr10-74711548-DEL-3678 |
| chr10-74719023-DEL-3064 | chr10-74719024-DEL-3064 |
| chr10-74722669-DEL-1198 | chr10-74722670-DEL-1198 |
| chr10-74741252-DEL-258 | chr10-74741253-DEL-258 |
| chr10-74762755-DEL-407 | chr10-74762756-DEL-407 |
| chr10-74763919-DEL-53 | chr10-74763920-DEL-53 |
| chr10-74777010-DEL-64 | chr10-74777011-DEL-65 |
| chr10-74844877-DEL-64 | chr10-74844878-DEL-64 |
| chr10-74880833-DEL-148 | chr10-74880828-DEL-143 |
| chr10-74903598-DEL-88 | chr10-74903599-DEL-88 |
| chr10-74909159-DEL-61 | chr10-74909160-DEL-61 |
| chr10-74922940-DEL-582 | chr10-74922941-DEL-582 |
| chr10-74926150-DEL-708 | chr10-74926151-DEL-708 |
| chr10-74937719-DEL-1231 | chr10-74937727-DEL-1224 |
| chr10-74951616-DEL-54 | chr10-74951617-DEL-54 |
| chr10-74977569-DEL-60 | chr10-74977570-DEL-60 |
| chr10-75006584-DEL-71 | chr10-75006588-DEL-69 |
| chr10-75015241-DEL-534 | chr10-75015243-DEL-529 |
| chr10-75033040-DEL-209 | chr10-75033040-DEL-209 |
| chr10-75080717-DEL-53 | chr10-75080718-DEL-53 |
| chr10-75184085-DEL-60 | chr10-75184086-DEL-60 |
| chr10-75215239-DEL-179 | chr10-75215234-DEL-185 |
| chr10-75216335-DEL-875 | chr10-75216336-DEL-875 |
| chr10-75250935-DEL-1637 | chr10-75250936-DEL-1637 |
| chr10-75262096-DEL-383 | chr10-75262096-DEL-383 |
| chr10-75289319-DEL-313 | chr10-75289320-DEL-313 |
| chr10-75309794-DEL-112 | chr10-75309795-DEL-112 |
| chr10-75332438-DEL-159 | chr10-75332439-DEL-159 |
| chr10-75365477-DEL-213 | chr10-75365478-DEL-213 |
| chr10-75410436-DEL-223 | chr10-75410437-DEL-223 |
| chr10-75418612-DEL-86 | chr10-75418613-DEL-86 |
| chr10-75426882-DEL-84 | chr10-75426883-DEL-84 |
| chr10-75454836-DEL-256 | chr10-75454837-DEL-256 |
| chr10-75455763-DEL-60 | chr10-75455764-DEL-60 |

| | |
|---|---|
| chr10-75461063-DEL-275 | chr10-75461064-DEL-275 |
| chr10-75526062-DEL-398 | chr10-75526063-DEL-398 |
| chr10-75542269-DEL-51 | chr10-75542270-DEL-54 |
| chr10-75543647-DEL-145 | chr10-75543696-DEL-145 |
| chr10-75603637-DEL-349 | chr10-75603638-DEL-349 |
| chr10-75604276-DEL-461 | chr10-75604272-DEL-458 |
| chr10-75628811-DEL-54 | chr10-75628811-DEL-54 |
| chr10-75653978-DEL-56 | chr10-75653979-DEL-56 |
| chr10-75711592-DEL-57 | chr10-75711593-DEL-57 |
| chr10-75726567-DEL-203 | chr10-75726567-DEL-202 |
| chr10-75779575-DEL-59 | chr10-75779528-DEL-70 |
| chr10-75791671-DEL-212 | chr10-75791672-DEL-212 |
| chr10-75792490-DEL-272 | chr10-75792491-DEL-272 |
| chr10-75802803-DEL-209 | chr10-75802804-DEL-209 |
| chr10-75807085-DEL-51 | chr10-75807086-DEL-51 |
| chr10-75817546-DEL-1527 | chr10-75817542-DEL-1525 |
| chr10-75830645-DEL-96 | chr10-75830605-DEL-117 |
| chr10-75839905-DEL-54 | chr10-75839906-DEL-54 |
| chr10-75850063-DEL-261 | chr10-75850064-DEL-261 |
| chr10-75854057-DEL-487 | chr10-75854058-DEL-487 |
| chr10-75883964-DEL-918 | chr10-75883965-DEL-918 |
| chr10-75888076-DEL-208 | chr10-75888077-DEL-208 |
| chr10-75894867-DEL-210 | chr10-75894868-DEL-210 |
| chr10-75919710-DEL-210 | chr10-75919711-DEL-210 |
| chr10-75927559-DEL-208 | chr10-75927560-DEL-208 |
| chr10-75960613-DEL-80 | chr10-75960613-DEL-80 |
| chr10-76035164-DEL-88 | chr10-76035165-DEL-88 |
| chr10-76047427-DEL-401 | chr10-76047428-DEL-401 |
| chr10-76050166-DEL-59 | chr10-76050160-DEL-60 |
| chr10-76052142-DEL-206 | chr10-76052143-DEL-206 |
| chr10-76109567-DEL-52 | chr10-76109594-DEL-54 |
| chr10-76160880-DEL-76 | chr10-76160885-DEL-72 |
| chr10-76247226-DEL-121 | chr10-76247227-DEL-122 |
| chr10-76280944-DEL-65 | chr10-76280944-DEL-65 |
| chr10-76297019-DEL-73 | chr10-76297020-DEL-73 |
| chr10-76316550-DEL-220 | chr10-76316551-DEL-220 |
| chr10-76334762-DEL-310 | chr10-76334764-DEL-313 |
| chr10-76341381-DEL-296 | chr10-76341382-DEL-296 |
| chr10-76473887-DEL-150 | chr10-76473888-DEL-150 |
| chr10-76493727-DEL-744 | chr10-76493728-DEL-744 |
| chr10-76530065-DEL-60 | chr10-76530066-DEL-60 |
| chr10-76530622-DEL-100 | chr10-76530623-DEL-100 |
| chr10-76580108-DEL-116 | chr10-76580094-DEL-152 |
| chr10-76583380-DEL-1687 | chr10-76583380-DEL-1687 |
| chr10-76674124-DEL-68 | chr10-76674125-DEL-68 |
| chr10-76753843-DEL-101 | chr10-76753822-DEL-92 |
| chr10-76755844-DEL-3912 | chr10-76755845-DEL-3912 |
| chr10-76764351-DEL-389 | chr10-76764352-DEL-389 |
| chr10-76783242-DEL-216 | chr10-76783243-DEL-216 |
| chr10-76827290-DEL-90 | chr10-76827264-DEL-76 |
| chr10-76841697-DEL-198 | chr10-76841698-DEL-198 |
| chr10-76849017-DEL-771 | chr10-76849016-DEL-771 |
| chr10-76919320-DEL-92 | chr10-76919321-DEL-92 |
| chr10-76937850-DEL-634 | chr10-76937894-DEL-628 |
| chr10-76942644-DEL-179 | chr10-76942663-DEL-161 |
| chr10-76969702-DEL-384 | chr10-76969703-DEL-384 |
| chr10-76970833-DEL-110 | chr10-76970869-DEL-105 |
| chr10-76984470-DEL-214 | chr10-76984471-DEL-214 |
| chr10-76988095-DEL-76 | chr10-76988096-DEL-76 |
| chr10-77002923-DEL-244 | chr10-77002922-DEL-245 |
| chr10-77005124-DEL-210 | chr10-77005120-DEL-215 |
| chr10-77008968-DEL-70 | chr10-77008936-DEL-70 |
| chr10-77040155-DEL-207 | chr10-77040156-DEL-207 |
| chr10-77046091-DEL-59 | chr10-77046092-DEL-58 |
| chr10-77055147-DEL-62 | chr10-77055147-DEL-62 |
| chr10-77067405-DEL-454 | chr10-77067406-DEL-453 |
| chr10-77072966-DEL-434 | chr10-77072967-DEL-434 |
| chr10-77077668-DEL-73 | chr10-77077669-DEL-73 |
| chr10-77117178-DEL-62 | chr10-77117176-DEL-77 |
| chr10-77153559-DEL-765 | chr10-77153576-DEL-765 |
| chr10-77179781-DEL-241 | chr10-77179782-DEL-241 |
| chr10-77185908-DEL-64 | chr10-77185909-DEL-64 |
| chr10-77187841-DEL-215 | chr10-77187842-DEL-215 |
| chr10-77326603-DEL-212 | chr10-77326604-DEL-212 |
| chr10-77327935-DEL-506 | chr10-77327936-DEL-506 |
| chr10-77362101-DEL-387 | chr10-77362102-DEL-387 |
| chr10-77380540-DEL-54 | chr10-77380541-DEL-54 |
| chr10-77438155-DEL-56 | chr10-77438156-DEL-56 |
| chr10-77444842-DEL-158 | chr10-77444843-DEL-197 |
| chr10-77463019-DEL-223 | chr10-77463020-DEL-223 |

| | |
|---|---|
| chr10-77467650-DEL-217 | chr10-77467651-DEL-217 |
| chr10-77503840-DEL-90 | chr10-77503839-DEL-72 |
| chr10-77564548-DEL-74 | chr10-77564549-DEL-74 |
| chr10-77565654-DEL-83 | chr10-77565655-DEL-83 |
| chr10-77568310-DEL-88 | chr10-77568311-DEL-88 |
| chr10-77575683-DEL-271 | chr10-77575684-DEL-271 |
| chr10-77576186-DEL-241 | chr10-77576187-DEL-241 |
| chr10-77582687-DEL-193 | chr10-77582688-DEL-193 |
| chr10-77589331-DEL-209 | chr10-77589332-DEL-209 |
| chr10-77625309-DEL-120 | chr10-77625309-DEL-120 |
| chr10-77643712-DEL-1842 | chr10-77643713-DEL-1842 |
| chr10-77653409-DEL-526 | chr10-77653410-DEL-526 |
| chr10-77655070-DEL-71 | chr10-77655071-DEL-71 |
| chr10-77657245-DEL-55 | chr10-77657246-DEL-55 |
| chr10-77685159-DEL-53 | chr10-77685154-DEL-54 |
| chr10-77691214-DEL-344 | chr10-77691215-DEL-344 |
| chr10-77707796-DEL-592 | chr10-77707797-DEL-592 |
| chr10-77708826-DEL-94 | chr10-77708827-DEL-94 |
| chr10-77711118-DEL-110 | chr10-77711119-DEL-110 |
| chr10-77725497-DEL-54 | chr10-77725501-DEL-51 |
| chr10-77741264-DEL-73 | chr10-77741265-DEL-73 |
| chr10-77768658-DEL-214 | chr10-77768659-DEL-214 |
| chr10-77776229-DEL-64 | chr10-77776230-DEL-64 |
| chr10-77786557-DEL-7160 | chr10-77786558-DEL-7160 |
| chr10-77809463-DEL-192 | chr10-77809464-DEL-192 |
| chr10-77828721-DEL-63 | chr10-77828722-DEL-63 |
| chr10-77856053-DEL-5402 | chr10-77856054-DEL-5402 |
| chr10-77891701-DEL-66 | chr10-77891702-DEL-66 |
| chr10-77901922-DEL-677 | chr10-77901923-DEL-678 |
| chr10-77923534-DEL-95 | chr10-77923535-DEL-95 |
| chr10-77929533-DEL-69 | chr10-77929524-DEL-85 |
| chr10-77942441-DEL-545 | chr10-77942442-DEL-545 |
| chr10-77943331-DEL-90 | chr10-77943343-DEL-83 |
| chr10-77984889-DEL-69 | chr10-77984890-DEL-69 |
| chr10-77986149-DEL-167 | chr10-77986150-DEL-117 |
| chr10-78021611-DEL-67 | chr10-78021603-DEL-62 |
| chr10-78099802-DEL-544 | chr10-78099803-DEL-544 |
| chr10-78109729-DEL-52 | chr10-78109730-DEL-52 |
| chr10-78149779-DEL-68 | chr10-78149780-DEL-68 |
| chr10-78172034-DEL-84 | chr10-78172034-DEL-84 |
| chr10-78177405-DEL-338 | chr10-78177405-DEL-339 |
| chr10-78184479-DEL-112 | chr10-78184480-DEL-112 |
| chr10-78195264-DEL-108 | chr10-78195265-DEL-104 |
| chr10-78229120-DEL-503 | chr10-78229121-DEL-500 |
| chr10-78239668-DEL-78 | chr10-78239669-DEL-78 |
| chr10-78250657-DEL-260 | chr10-78250658-DEL-260 |
| chr10-78251870-DEL-209 | chr10-78251872-DEL-213 |
| chr10-78283059-DEL-395 | chr10-78283060-DEL-395 |
| chr10-78314501-DEL-185 | chr10-78314501-DEL-186 |
| chr10-78335393-DEL-533 | chr10-78335422-DEL-516 |
| chr10-78344206-DEL-90 | chr10-78344207-DEL-90 |
| chr10-78377033-DEL-588 | chr10-78377034-DEL-588 |
| chr10-78408884-DEL-89 | chr10-78408885-DEL-89 |
| chr10-78452743-DEL-80 | chr10-78452772-DEL-63 |
| chr10-78471508-DEL-199 | chr10-78471509-DEL-199 |
| chr10-78479245-DEL-1164 | chr10-78479246-DEL-1164 |
| chr10-78486402-DEL-6167 | chr10-78486403-DEL-6167 |
| chr10-78509131-DEL-1248 | chr10-78509131-DEL-1248 |
| chr10-78511736-DEL-156 | chr10-78511737-DEL-156 |
| chr10-78517125-DEL-997 | chr10-78517124-DEL-991 |
| chr10-78539920-DEL-122 | chr10-78539921-DEL-122 |
| chr10-78559067-DEL-51 | chr10-78559062-DEL-57 |
| chr10-78562347-DEL-56 | chr10-78562348-DEL-56 |
| chr10-78623066-DEL-259 | chr10-78623067-DEL-259 |
| chr10-78687467-DEL-126 | chr10-78687468-DEL-126 |
| chr10-78690074-DEL-844 | chr10-78690075-DEL-844 |
| chr10-78690479-DEL-2139 | chr10-78690479-DEL-2139 |
| chr10-78691053-DEL-1048 | chr10-78691053-DEL-1048 |
| chr10-78695291-DEL-8141 | chr10-78695291-DEL-8141 |
| chr10-78736483-DEL-4260 | chr10-78736483-DEL-4260 |
| chr10-78743317-DEL-498 | chr10-78743318-DEL-498 |
| chr10-78789344-DEL-1144 | chr10-78789344-DEL-1144 |
| chr10-78838265-DEL-2557 | chr10-78838266-DEL-2557 |
| chr10-78876565-DEL-523 | chr10-78876566-DEL-523 |
| chr10-78889663-DEL-317 | chr10-78889664-DEL-317 |
| chr10-78893394-DEL-57 | chr10-78893395-DEL-57 |
| chr10-78895092-DEL-150 | chr10-78895093-DEL-150 |
| chr10-78905784-DEL-1020 | chr10-78905785-DEL-1020 |
| chr10-78916085-DEL-6239 | chr10-78916086-DEL-6239 |
| chr10-78980560-DEL-153 | chr10-78980560-DEL-150 |

| | |
|---|---|
| chr10-79002162-DEL-106 | chr10-79002163-DEL-106 |
| chr10-79051706-DEL-804 | chr10-79051707-DEL-804 |
| chr10-79062141-DEL-150 | chr10-79062141-DEL-150 |
| chr10-79106814-DEL-94 | chr10-79106815-DEL-94 |
| chr10-79200153-DEL-66 | chr10-79200136-DEL-60 |
| chr10-79265042-DEL-209 | chr10-79265043-DEL-209 |
| chr10-79302112-DEL-228 | chr10-79302068-DEL-226 |
| chr10-79407981-DEL-930 | chr10-79407982-DEL-930 |
| chr10-79417745-DEL-52 | chr10-79417746-DEL-52 |
| chr10-79435283-DEL-52 | chr10-79435283-DEL-52 |
| chr10-79513217-DEL-120 | chr10-79513218-DEL-120 |
| chr10-79536166-DEL-215 | chr10-79536167-DEL-215 |
| chr10-79539107-DEL-200 | chr10-79539108-DEL-199 |
| chr10-79607536-DEL-69 | chr10-79607527-DEL-64 |
| chr10-79634593-DEL-71 | chr10-79634593-DEL-71 |
| chr10-79636190-DEL-70 | chr10-79636191-DEL-70 |
| chr10-79640353-DEL-161 | chr10-79640338-DEL-161 |
| chr10-79677756-DEL-54 | chr10-79677757-DEL-54 |
| chr10-79683009-DEL-115 | chr10-79683001-DEL-128 |
| chr10-79706685-DEL-212 | chr10-79706686-DEL-212 |
| chr10-79730627-DEL-100 | chr10-79730630-DEL-100 |
| chr10-79757119-DEL-171 | chr10-79757120-DEL-171 |
| chr10-79817994-DEL-68 | chr10-79817995-DEL-68 |
| chr10-79823536-DEL-51 | chr10-79823535-DEL-52 |
| chr10-79854091-DEL-72 | chr10-79854104-DEL-60 |
| chr10-79881128-DEL-218 | chr10-79881129-DEL-218 |
| chr10-79901498-DEL-174 | chr10-79901499-DEL-173 |
| chr10-80034761-DEL-225 | chr10-80034762-DEL-225 |
| chr10-80047318-DEL-228 | chr10-80047319-DEL-228 |
| chr10-80049888-DEL-207 | chr10-80049891-DEL-205 |
| chr10-80053508-DEL-245 | chr10-80053509-DEL-245 |
| chr10-80080888-DEL-113 | chr10-80080889-DEL-113 |
| chr10-80108682-DEL-165 | chr10-80108646-DEL-201 |
| chr10-80160058-DEL-208 | chr10-80160059-DEL-208 |
| chr10-80272246-DEL-52 | chr10-80272247-DEL-52 |
| chr10-80275380-DEL-86 | chr10-80275381-DEL-86 |
| chr10-80300143-DEL-60 | chr10-80300141-DEL-64 |
| chr10-80315601-DEL-204 | chr10-80315602-DEL-204 |
| chr10-80320204-DEL-92 | chr10-80320205-DEL-90 |
| chr10-80391362-DEL-365 | chr10-80391375-DEL-361 |
| chr10-80471935-DEL-1492 | chr10-80471936-DEL-1492 |
| chr10-80495890-DEL-631 | chr10-80495891-DEL-631 |
| chr10-80497760-DEL-221 | chr10-80497761-DEL-221 |
| chr10-80533050-DEL-204 | chr10-80533051-DEL-204 |
| chr10-80635075-DEL-467 | chr10-80635076-DEL-467 |
| chr10-80649837-DEL-215 | chr10-80649835-DEL-206 |
| chr10-80719034-DEL-120 | chr10-80719035-DEL-120 |
| chr10-80722785-DEL-198 | chr10-80722786-DEL-198 |
| chr10-80742691-DEL-264 | chr10-80742683-DEL-265 |
| chr10-80749147-DEL-217 | chr10-80749148-DEL-217 |
| chr10-80752934-DEL-92 | chr10-80752935-DEL-84 |
| chr10-80760276-DEL-69 | chr10-80760282-DEL-55 |
| chr10-80795572-DEL-65 | chr10-80795573-DEL-65 |
| chr10-80806121-DEL-409 | chr10-80806122-DEL-409 |
| chr10-80811864-DEL-4821 | chr10-80811865-DEL-4821 |
| chr10-80825944-DEL-181 | chr10-80825899-DEL-205 |
| chr10-80872389-DEL-189 | chr10-80872390-DEL-189 |
| chr10-80891869-DEL-64 | chr10-80891870-DEL-64 |
| chr10-80896374-DEL-200 | chr10-80896387-DEL-201 |
| chr10-80902933-DEL-115 | chr10-80902934-DEL-113 |
| chr10-80952674-DEL-228 | chr10-80952675-DEL-228 |
| chr10-80995338-DEL-182 | chr10-80995339-DEL-182 |
| chr10-81005115-DEL-194 | chr10-81005110-DEL-223 |
| chr10-81008154-DEL-210 | chr10-81008155-DEL-210 |
| chr10-81025307-DEL-178 | chr10-81025308-DEL-178 |
| chr10-81059290-DEL-395 | chr10-81059291-DEL-395 |
| chr10-81315680-DEL-78 | chr10-81315681-DEL-78 |
| chr10-81340663-DEL-118 | chr10-81340666-DEL-92 |
| chr10-81344581-DEL-366 | chr10-81344582-DEL-366 |
| chr10-81356989-DEL-104 | chr10-81356943-DEL-106 |
| chr10-81357985-DEL-217 | chr10-81357986-DEL-217 |
| chr10-81459399-DEL-65 | chr10-81459442-DEL-50 |
| chr10-81490326-DEL-220 | chr10-81490327-DEL-220 |
| chr10-81493717-DEL-96 | chr10-81493718-DEL-63 |
| chr10-81500342-DEL-201 | chr10-81500343-DEL-201 |
| chr10-81515285-DEL-184 | chr10-81515269-DEL-185 |
| chr10-81552995-DEL-199 | chr10-81552996-DEL-199 |
| chr10-81587376-DEL-1066 | chr10-81587377-DEL-1066 |
| chr10-81818669-DEL-949 | chr10-81818670-DEL-949 |
| chr10-81867486-DEL-90 | chr10-81867483-DEL-91 |

| | |
|---|---|
| chr10-81873319-DEL-205 | chr10-81873320-DEL-205 |
| chr10-81879412-DEL-209 | chr10-81879413-DEL-209 |
| chr10-81901414-DEL-210 | chr10-81901415-DEL-210 |
| chr10-81976229-DEL-401 | chr10-81976230-DEL-401 |
| chr10-81977551-DEL-400 | chr10-81977545-DEL-400 |
| chr10-81980592-DEL-981 | chr10-81980593-DEL-981 |
| chr10-81981696-DEL-112 | chr10-81981702-DEL-91 |
| chr10-81996722-DEL-1469 | chr10-81996723-DEL-1469 |
| chr10-82003137-DEL-581 | chr10-82003138-DEL-581 |
| chr10-82007613-DEL-56 | chr10-82007614-DEL-56 |
| chr10-82011933-DEL-199 | chr10-82011934-DEL-199 |
| chr10-82041366-DEL-91 | chr10-82041362-DEL-92 |
| chr10-82056641-DEL-66 | chr10-82056642-DEL-66 |
| chr10-82089313-DEL-99 | chr10-82089314-DEL-99 |
| chr10-82166982-DEL-1073 | chr10-82166983-DEL-1073 |
| chr10-82181758-DEL-115 | chr10-82181803-DEL-65 |
| chr10-82200201-DEL-949 | chr10-82200202-DEL-949 |
| chr10-82213344-DEL-83 | chr10-82213337-DEL-84 |
| chr10-82224929-DEL-6470 | chr10-82224930-DEL-6470 |
| chr10-82243648-DEL-2129 | chr10-82243649-DEL-2129 |
| chr10-82259728-DEL-72 | chr10-82259729-DEL-72 |
| chr10-82270254-DEL-178 | chr10-82270222-DEL-212 |
| chr10-82285657-DEL-161 | chr10-82285658-DEL-161 |
| chr10-82299203-DEL-3252 | chr10-82299204-DEL-3252 |
| chr10-82363149-DEL-147 | chr10-82363150-DEL-147 |
| chr10-82364657-DEL-4508 | chr10-82364658-DEL-4508 |
| chr10-82400907-DEL-58 | chr10-82400907-DEL-58 |
| chr10-82431931-DEL-7855 | chr10-82431930-DEL-7856 |
| chr10-82432318-DEL-57 | chr10-82432319-DEL-57 |
| chr10-82435902-DEL-1521 | chr10-82435903-DEL-1521 |
| chr10-82439413-DEL-2567 | chr10-82439414-DEL-2567 |
| chr10-82449529-DEL-61 | chr10-82449530-DEL-59 |
| chr10-82454691-DEL-452 | chr10-82454692-DEL-451 |
| chr10-82473228-DEL-90 | chr10-82473259-DEL-60 |
| chr10-82478435-DEL-120 | chr10-82478462-DEL-96 |
| chr10-82481249-DEL-74 | chr10-82481250-DEL-74 |
| chr10-82492661-DEL-60 | chr10-82492661-DEL-55 |
| chr10-82525126-DEL-64 | chr10-82525127-DEL-50 |
| chr10-82606829-DEL-325 | chr10-82606830-DEL-325 |
| chr10-82619054-DEL-104 | chr10-82619035-DEL-90 |
| chr10-82634461-DEL-122 | chr10-82634414-DEL-126 |
| chr10-82646937-DEL-814 | chr10-82646942-DEL-810 |
| chr10-82661013-DEL-499 | chr10-82660994-DEL-500 |
| chr10-82680866-DEL-78 | chr10-82680867-DEL-107 |
| chr10-82692628-DEL-186 | chr10-82692629-DEL-186 |
| chr10-82698626-DEL-57 | chr10-82698632-DEL-100 |
| chr10-82726714-DEL-124 | chr10-82726715-DEL-124 |
| chr10-82756606-DEL-54 | chr10-82756631-DEL-52 |
| chr10-82764706-DEL-209 | chr10-82764707-DEL-209 |
| chr10-82774922-DEL-2947 | chr10-82774923-DEL-2947 |
| chr10-82775696-DEL-75 | chr10-82775697-DEL-75 |
| chr10-82787157-DEL-61 | chr10-82787158-DEL-61 |
| chr10-82795888-DEL-160 | chr10-82795889-DEL-160 |
| chr10-82800920-DEL-66 | chr10-82800941-DEL-56 |
| chr10-82817857-DEL-54 | chr10-82817858-DEL-54 |
| chr10-82854347-DEL-80 | chr10-82854346-DEL-82 |
| chr10-82894295-DEL-110 | chr10-82894296-DEL-110 |
| chr10-82924917-DEL-327 | chr10-82924918-DEL-327 |
| chr10-82949557-DEL-1314 | chr10-82949544-DEL-1314 |
| chr10-82986386-DEL-57 | chr10-82986389-DEL-55 |
| chr10-83082618-DEL-135 | chr10-83082619-DEL-135 |
| chr10-83085354-DEL-60 | chr10-83085355-DEL-60 |
| chr10-83095515-DEL-60 | chr10-83095513-DEL-61 |
| chr10-83096190-DEL-72 | chr10-83096191-DEL-72 |
| chr10-83098317-DEL-80 | chr10-83098318-DEL-58 |
| chr10-83118317-DEL-193 | chr10-83118318-DEL-193 |
| chr10-83174297-DEL-241 | chr10-83174298-DEL-242 |
| chr10-83187876-DEL-200 | chr10-83187866-DEL-201 |
| chr10-83194012-DEL-311 | chr10-83194013-DEL-311 |
| chr10-83246226-DEL-15747 | chr10-83246226-DEL-15747 |
| chr10-83274172-DEL-2731 | chr10-83274173-DEL-2731 |
| chr10-83317069-DEL-1746 | chr10-83317068-DEL-1746 |
| chr10-83322279-DEL-80 | chr10-83322269-DEL-76 |
| chr10-83334888-DEL-59 | chr10-83334888-DEL-59 |
| chr10-83395441-DEL-76 | chr10-83395474-DEL-91 |
| chr10-83414860-DEL-231 | chr10-83414861-DEL-231 |
| chr10-83431732-DEL-203 | chr10-83431733-DEL-203 |
| chr10-83442374-DEL-205 | chr10-83442375-DEL-205 |
| chr10-83488153-DEL-349 | chr10-83488154-DEL-349 |
| chr10-83506198-DEL-7763 | chr10-83506199-DEL-7763 |

| | |
|---|---|
| chr10-83518330-DEL-91 | chr10-83518331-DEL-91 |
| chr10-83521030-DEL-56 | chr10-83521031-DEL-56 |
| chr10-83604216-DEL-6843 | chr10-83604217-DEL-6843 |
| chr10-83614340-DEL-56 | chr10-83614341-DEL-56 |
| chr10-83622376-DEL-185 | chr10-83622377-DEL-185 |
| chr10-83623333-DEL-5130 | chr10-83623333-DEL-5130 |
| chr10-83633963-DEL-1483 | chr10-83633963-DEL-1483 |
| chr10-83641979-DEL-51 | chr10-83641980-DEL-51 |
| chr10-83645659-DEL-52 | chr10-83645662-DEL-50 |
| chr10-83662744-DEL-168 | chr10-83662745-DEL-168 |
| chr10-83665719-DEL-4433 | chr10-83665720-DEL-4433 |
| chr10-83670631-DEL-1389 | chr10-83670632-DEL-1389 |
| chr10-83695179-DEL-102 | chr10-83695160-DEL-122 |
| chr10-83711069-DEL-65 | chr10-83711073-DEL-61 |
| chr10-83711578-DEL-63 | chr10-83711585-DEL-55 |
| chr10-83730232-DEL-53 | chr10-83730212-DEL-80 |
| chr10-83749054-DEL-710 | chr10-83749053-DEL-708 |
| chr10-83786382-DEL-71 | chr10-83786382-DEL-71 |
| chr10-83789054-DEL-1192 | chr10-83789052-DEL-1192 |
| chr10-83817768-DEL-450 | chr10-83817769-DEL-450 |
| chr10-83819745-DEL-69 | chr10-83819719-DEL-64 |
| chr10-83824507-DEL-453 | chr10-83824508-DEL-453 |
| chr10-83825297-DEL-82 | chr10-83825307-DEL-83 |
| chr10-83841994-DEL-1209 | chr10-83841995-DEL-1209 |
| chr10-83887645-DEL-144 | chr10-83887646-DEL-144 |
| chr10-83904191-DEL-711 | chr10-83904192-DEL-711 |
| chr10-83917475-DEL-51 | chr10-83917476-DEL-51 |
| chr10-83931616-DEL-66 | chr10-83931617-DEL-66 |
| chr10-83984053-DEL-270 | chr10-83984054-DEL-270 |
| chr10-83991069-DEL-6547 | chr10-83991070-DEL-6547 |
| chr10-83999149-DEL-183 | chr10-83999160-DEL-173 |
| chr10-84005905-DEL-324 | chr10-84005906-DEL-324 |
| chr10-84010238-DEL-6442 | chr10-84010239-DEL-6442 |
| chr10-84073150-DEL-1340 | chr10-84073151-DEL-1340 |
| chr10-84087659-DEL-326 | chr10-84087660-DEL-326 |
| chr10-84103406-DEL-56 | chr10-84103407-DEL-56 |
| chr10-84111099-DEL-58 | chr10-84111094-DEL-62 |
| chr10-84143605-DEL-58 | chr10-84143606-DEL-58 |
| chr10-84162749-DEL-583 | chr10-84162749-DEL-583 |
| chr10-84186614-DEL-1861 | chr10-84186615-DEL-1861 |
| chr10-84212540-DEL-66 | chr10-84212540-DEL-86 |
| chr10-84256941-DEL-119 | chr10-84256942-DEL-119 |
| chr10-84264908-DEL-440 | chr10-84264909-DEL-440 |
| chr10-84307608-DEL-1888 | chr10-84307609-DEL-1888 |
| chr10-84311179-DEL-56 | chr10-84311180-DEL-84 |
| chr10-84318422-DEL-449 | chr10-84318423-DEL-449 |
| chr10-84321324-DEL-66 | chr10-84321325-DEL-65 |
| chr10-84331873-DEL-172 | chr10-84331872-DEL-172 |
| chr10-84350595-DEL-58 | chr10-84350586-DEL-53 |
| chr10-84377682-DEL-81 | chr10-84377683-DEL-81 |
| chr10-84407607-DEL-175 | chr10-84407608-DEL-163 |
| chr10-84429966-DEL-189 | chr10-84429967-DEL-189 |
| chr10-84433790-DEL-195 | chr10-84433791-DEL-195 |
| chr10-84434008-DEL-805 | chr10-84434007-DEL-806 |
| chr10-84479297-DEL-173 | chr10-84479298-DEL-173 |
| chr10-84492331-DEL-74 | chr10-84492332-DEL-74 |
| chr10-84500706-DEL-222 | chr10-84500707-DEL-185 |
| chr10-84501167-DEL-208 | chr10-84501168-DEL-208 |
| chr10-84523300-DEL-203 | chr10-84523325-DEL-184 |
| chr10-84524384-DEL-75 | chr10-84524385-DEL-75 |
| chr10-84530870-DEL-216 | chr10-84530871-DEL-216 |
| chr10-84559271-DEL-167 | chr10-84559272-DEL-167 |
| chr10-84577510-DEL-351 | chr10-84577511-DEL-351 |
| chr10-84580447-DEL-5360 | chr10-84580448-DEL-5360 |
| chr10-84602808-DEL-54 | chr10-84602808-DEL-54 |
| chr10-84605932-DEL-205 | chr10-84605933-DEL-205 |
| chr10-84678807-DEL-115 | chr10-84678776-DEL-117 |
| chr10-84679649-DEL-58 | chr10-84679649-DEL-57 |
| chr10-84691305-DEL-442 | chr10-84691308-DEL-440 |
| chr10-84708752-DEL-101 | chr10-84708753-DEL-101 |
| chr10-84728238-DEL-57 | chr10-84728237-DEL-62 |
| chr10-84746146-DEL-149 | chr10-84746147-DEL-149 |
| chr10-84822054-DEL-102 | chr10-84822055-DEL-102 |
| chr10-84879083-DEL-201 | chr10-84879084-DEL-201 |
| chr10-84923358-DEL-232 | chr10-84923359-DEL-232 |
| chr10-84980448-DEL-83 | chr10-84980442-DEL-91 |
| chr10-84985979-DEL-76 | chr10-84985980-DEL-70 |
| chr10-84991085-DEL-187 | chr10-84991086-DEL-187 |
| chr10-84993590-DEL-215 | chr10-84993591-DEL-215 |
| chr10-85007815-DEL-272 | chr10-85007816-DEL-272 |

| | |
|---|---|
| chr10-85025447-DEL-150 | chr10-85025448-DEL-150 |
| chr10-85031993-DEL-3236 | chr10-85031994-DEL-3236 |
| chr10-85054123-DEL-275 | chr10-85054124-DEL-275 |
| chr10-85059315-DEL-170 | chr10-85059316-DEL-167 |
| chr10-85082646-DEL-147 | chr10-85082692-DEL-100 |
| chr10-85105299-DEL-60 | chr10-85105300-DEL-60 |
| chr10-85163641-DEL-1224 | chr10-85163642-DEL-1224 |
| chr10-85167318-DEL-77 | chr10-85167317-DEL-78 |
| chr10-85180057-DEL-60 | chr10-85180058-DEL-60 |
| chr10-85200214-DEL-834 | chr10-85200229-DEL-820 |
| chr10-85201943-DEL-51 | chr10-85201944-DEL-51 |
| chr10-85213479-DEL-112 | chr10-85213480-DEL-120 |
| chr10-85263438-DEL-165 | chr10-85263405-DEL-166 |
| chr10-85365370-DEL-229 | chr10-85365371-DEL-229 |
| chr10-85368237-DEL-723 | chr10-85368242-DEL-719 |
| chr10-85378547-DEL-108 | chr10-85378545-DEL-111 |
| chr10-85417997-DEL-82 | chr10-85417994-DEL-68 |
| chr10-85425089-DEL-63 | chr10-85425082-DEL-78 |
| chr10-85425424-DEL-479 | chr10-85425425-DEL-479 |
| chr10-85427540-DEL-54 | chr10-85427545-DEL-50 |
| chr10-85433657-DEL-90 | chr10-85433692-DEL-50 |
| chr10-85549852-DEL-8040 | chr10-85549852-DEL-8040 |
| chr10-85604283-DEL-4807 | chr10-85604284-DEL-4807 |
| chr10-85623557-DEL-514 | chr10-85623558-DEL-514 |
| chr10-85658030-DEL-15197 | chr10-85658031-DEL-15197 |
| chr10-85675505-DEL-3215 | chr10-85675505-DEL-3215 |
| chr10-85691396-DEL-6431 | chr10-85691397-DEL-6431 |
| chr10-85719474-DEL-82 | chr10-85719475-DEL-82 |
| chr10-85733313-DEL-90 | chr10-85733300-DEL-90 |
| chr10-85761251-DEL-78 | chr10-85761252-DEL-68 |
| chr10-85791873-DEL-180 | chr10-85791874-DEL-180 |
| chr10-85875941-DEL-143 | chr10-85875942-DEL-143 |
| chr10-85906138-DEL-51 | chr10-85906139-DEL-51 |
| chr10-85931002-DEL-608 | chr10-85931003-DEL-606 |
| chr10-85978708-DEL-60 | chr10-85978708-DEL-63 |
| chr10-85982901-DEL-65 | chr10-85982902-DEL-65 |
| chr10-86022834-DEL-69 | chr10-86022834-DEL-69 |
| chr10-86033965-DEL-2132 | chr10-86033966-DEL-2132 |
| chr10-86048747-DEL-51 | chr10-86048748-DEL-51 |
| chr10-86069912-DEL-7099 | chr10-86069913-DEL-7099 |
| chr10-86087923-DEL-62 | chr10-86087924-DEL-62 |
| chr10-86229048-DEL-211 | chr10-86229049-DEL-211 |
| chr10-86237914-DEL-1475 | chr10-86237915-DEL-1475 |
| chr10-86238106-DEL-175 | chr10-86238107-DEL-175 |
| chr10-86247211-DEL-654 | chr10-86247212-DEL-654 |
| chr10-86289541-DEL-252 | chr10-86289542-DEL-252 |
| chr10-86302355-DEL-114 | chr10-86302356-DEL-114 |
| chr10-86338909-DEL-353 | chr10-86338910-DEL-353 |
| chr10-86397457-DEL-4122 | chr10-86397455-DEL-4122 |
| chr10-86419929-DEL-3214 | chr10-86419930-DEL-3212 |
| chr10-86444566-DEL-628 | chr10-86444566-DEL-631 |
| chr10-86547437-DEL-258 | chr10-86547430-DEL-266 |
| chr10-86551410-DEL-1856 | chr10-86551411-DEL-1856 |
| chr10-86560866-DEL-143 | chr10-86560867-DEL-143 |
| chr10-86565073-DEL-60 | chr10-86565062-DEL-72 |
| chr10-86573443-DEL-220 | chr10-86573444-DEL-220 |
| chr10-86577754-DEL-1337 | chr10-86577755-DEL-1337 |
| chr10-86581105-DEL-88 | chr10-86581106-DEL-88 |
| chr10-86632239-DEL-132 | chr10-86632209-DEL-117 |
| chr10-86632688-DEL-6466 | chr10-86632689-DEL-6466 |
| chr10-86650596-DEL-60 | chr10-86650597-DEL-60 |
| chr10-86681265-DEL-294 | chr10-86681266-DEL-294 |
| chr10-86695261-DEL-106 | chr10-86695262-DEL-106 |
| chr10-86699737-DEL-88 | chr10-86699744-DEL-82 |
| chr10-86721938-DEL-987 | chr10-86721939-DEL-987 |
| chr10-86825086-DEL-1928 | chr10-86825087-DEL-1928 |
| chr10-86829485-DEL-604 | chr10-86829486-DEL-604 |
| chr10-86862454-DEL-61 | chr10-86862455-DEL-61 |
| chr10-86900730-DEL-6659 | chr10-86900720-DEL-6660 |
| chr10-86907030-DEL-212 | chr10-86907031-DEL-212 |
| chr10-86944714-DEL-142 | chr10-86944699-DEL-158 |
| chr10-86978730-DEL-78 | chr10-86978731-DEL-78 |
| chr10-86981514-DEL-52 | chr10-86981515-DEL-52 |
| chr10-86986306-DEL-74 | chr10-86986307-DEL-74 |
| chr10-86990575-DEL-55 | chr10-86990575-DEL-50 |
| chr10-86997999-DEL-68 | chr10-86998000-DEL-68 |
| chr10-87010097-DEL-6602 | chr10-87010098-DEL-6602 |
| chr10-87016136-DEL-402 | chr10-87016136-DEL-402 |
| chr10-87029731-DEL-54 | chr10-87029732-DEL-54 |
| chr10-87082003-DEL-884 | chr10-87082004-DEL-884 |

| | |
|---|---|
| chr10-87105327-DEL-75 | chr10-87105328-DEL-75 |
| chr10-87114847-DEL-850 | chr10-87114848-DEL-850 |
| chr10-87138187-DEL-2658 | chr10-87138188-DEL-2658 |
| chr10-87156384-DEL-1624 | chr10-87156385-DEL-1624 |
| chr10-87190100-DEL-74 | chr10-87190101-DEL-74 |
| chr10-87197094-DEL-2913 | chr10-87197095-DEL-2913 |
| chr10-87233392-DEL-76 | chr10-87233393-DEL-76 |
| chr10-87234357-DEL-58 | chr10-87234359-DEL-56 |
| chr10-87234776-DEL-162 | chr10-87234777-DEL-162 |
| chr10-87235962-DEL-9129 | chr10-87235963-DEL-9129 |
| chr10-87241125-DEL-2910 | chr10-87241126-DEL-2910 |
| chr10-87303475-DEL-2079 | chr10-87303476-DEL-2061 |
| chr10-87303731-DEL-927 | chr10-87303732-DEL-927 |
| chr10-87313354-DEL-84 | chr10-87313355-DEL-84 |
| chr10-87344998-DEL-628 | chr10-87344999-DEL-628 |
| chr10-87352810-DEL-203 | chr10-87352811-DEL-203 |
| chr10-87366478-DEL-438 | chr10-87366479-DEL-438 |
| chr10-87370511-DEL-399 | chr10-87370512-DEL-399 |
| chr10-87376629-DEL-131 | chr10-87376621-DEL-132 |
| chr10-87395182-DEL-210 | chr10-87395183-DEL-210 |
| chr10-87467648-DEL-3421 | chr10-87467664-DEL-3419 |
| chr10-87478087-DEL-326 | chr10-87478088-DEL-326 |
| chr10-87523071-DEL-446 | chr10-87523073-DEL-444 |
| chr10-87540413-DEL-147 | chr10-87540415-DEL-132 |
| chr10-87568044-DEL-372 | chr10-87568050-DEL-364 |
| chr10-87635728-DEL-2076 | chr10-87635729-DEL-2076 |
| chr10-87637003-DEL-1589 | chr10-87637004-DEL-1589 |
| chr10-87671785-DEL-118 | chr10-87671786-DEL-118 |
| chr10-87676358-DEL-68 | chr10-87676377-DEL-56 |
| chr10-87690598-DEL-150 | chr10-87690619-DEL-130 |
| chr10-87774437-DEL-323 | chr10-87774438-DEL-323 |
| chr10-87794999-DEL-195 | chr10-87795000-DEL-177 |
| chr10-87808403-DEL-70 | chr10-87808404-DEL-70 |
| chr10-87814597-DEL-410 | chr10-87814598-DEL-410 |
| chr10-87826078-DEL-2400 | chr10-87826079-DEL-2400 |
| chr10-87829132-DEL-3556 | chr10-87829133-DEL-3556 |
| chr10-87834587-DEL-168 | chr10-87834588-DEL-168 |
| chr10-87857029-DEL-117 | chr10-87856991-DEL-156 |
| chr10-87858310-DEL-284 | chr10-87858298-DEL-285 |
| chr10-87866913-DEL-66 | chr10-87866952-DEL-53 |
| chr10-87879200-DEL-100 | chr10-87879201-DEL-100 |
| chr10-87885652-DEL-1890 | chr10-87885653-DEL-1890 |
| chr10-87906862-DEL-573 | chr10-87906863-DEL-573 |
| chr10-87908050-DEL-1891 | chr10-87908064-DEL-1861 |
| chr10-87912384-DEL-203 | chr10-87912385-DEL-203 |
| chr10-87926034-DEL-58 | chr10-87926022-DEL-61 |
| chr10-87931116-DEL-6545 | chr10-87931117-DEL-6545 |
| chr10-87939936-DEL-143 | chr10-87939937-DEL-143 |
| chr10-87944057-DEL-201 | chr10-87944055-DEL-202 |
| chr10-87960058-DEL-209 | chr10-87960059-DEL-207 |
| chr10-87964654-DEL-748 | chr10-87964655-DEL-748 |
| chr10-87990367-DEL-52 | chr10-87990404-DEL-56 |
| chr10-88023457-DEL-4113 | chr10-88023456-DEL-4110 |
| chr10-88035510-DEL-213 | chr10-88035511-DEL-213 |
| chr10-88070579-DEL-126 | chr10-88070580-DEL-126 |
| chr10-88079377-DEL-498 | chr10-88079378-DEL-498 |
| chr10-88183668-DEL-90 | chr10-88183673-DEL-135 |
| chr10-88197630-DEL-201 | chr10-88197631-DEL-201 |
| chr10-88208812-DEL-477 | chr10-88208813-DEL-478 |
| chr10-88259923-DEL-1826 | chr10-88259924-DEL-1826 |
| chr10-88288999-DEL-120 | chr10-88289000-DEL-120 |
| chr10-88291156-DEL-52 | chr10-88291145-DEL-64 |
| chr10-88318456-DEL-284 | chr10-88318457-DEL-284 |
| chr10-88319417-DEL-208 | chr10-88319418-DEL-208 |
| chr10-88333415-DEL-1171 | chr10-88333416-DEL-1171 |
| chr10-88370741-DEL-179 | chr10-88370742-DEL-179 |
| chr10-88375898-DEL-215 | chr10-88375899-DEL-215 |
| chr10-88384254-DEL-85 | chr10-88384255-DEL-66 |
| chr10-88391869-DEL-925 | chr10-88391870-DEL-925 |
| chr10-88398591-DEL-52 | chr10-88398592-DEL-52 |
| chr10-88443717-DEL-64 | chr10-88443698-DEL-65 |
| chr10-88453588-DEL-394 | chr10-88453589-DEL-394 |
| chr10-88454150-DEL-480 | chr10-88454151-DEL-480 |
| chr10-88459800-DEL-53 | chr10-88459800-DEL-53 |
| chr10-88461135-DEL-433 | chr10-88461136-DEL-433 |
| chr10-88461915-DEL-80 | chr10-88461916-DEL-77 |
| chr10-88472597-DEL-193 | chr10-88472598-DEL-193 |
| chr10-88486768-DEL-7166 | chr10-88486769-DEL-7166 |
| chr10-88498770-DEL-762 | chr10-88498771-DEL-762 |
| chr10-88500474-DEL-203 | chr10-88500491-DEL-183 |

| | |
|---|---|
| chr10-88506521-DEL-1251 | chr10-88506522-DEL-1251 |
| chr10-88544095-DEL-81 | chr10-88544096-DEL-81 |
| chr10-88553421-DEL-108 | chr10-88553457-DEL-72 |
| chr10-88554026-DEL-53 | chr10-88554028-DEL-54 |
| chr10-88582096-DEL-210 | chr10-88582097-DEL-210 |
| chr10-88584645-DEL-112 | chr10-88584616-DEL-133 |
| chr10-88599356-DEL-648 | chr10-88599356-DEL-648 |
| chr10-88625877-DEL-181 | chr10-88625884-DEL-165 |
| chr10-88640261-DEL-66 | chr10-88640261-DEL-66 |
| chr10-88649349-DEL-62 | chr10-88649350-DEL-56 |
| chr10-88727314-DEL-181 | chr10-88727315-DEL-181 |
| chr10-88758445-DEL-91 | chr10-88758428-DEL-74 |
| chr10-88764045-DEL-6548 | chr10-88764043-DEL-6548 |
| chr10-88794380-DEL-665 | chr10-88794381-DEL-665 |
| chr10-88799734-DEL-4459 | chr10-88799734-DEL-4459 |
| chr10-88806608-DEL-58 | chr10-88806593-DEL-59 |
| chr10-88817712-DEL-252 | chr10-88817713-DEL-250 |
| chr10-88818335-DEL-530 | chr10-88818336-DEL-530 |
| chr10-88823084-DEL-86 | chr10-88823074-DEL-114 |
| chr10-88826827-DEL-1178 | chr10-88826828-DEL-1177 |
| chr10-88881122-DEL-887 | chr10-88881123-DEL-887 |
| chr10-88894703-DEL-531 | chr10-88894704-DEL-531 |
| chr10-88909001-DEL-165 | chr10-88909002-DEL-165 |
| chr10-88922988-DEL-54 | chr10-88922965-DEL-52 |
| chr10-88926303-DEL-72 | chr10-88926304-DEL-72 |
| chr10-88931787-DEL-84 | chr10-88931788-DEL-84 |
| chr10-88939682-DEL-71 | chr10-88939683-DEL-71 |
| chr10-88951619-DEL-289 | chr10-88951620-DEL-289 |
| chr10-88992968-DEL-52 | chr10-88992980-DEL-57 |
| chr10-89007342-DEL-434 | chr10-89007343-DEL-434 |
| chr10-89018106-DEL-175 | chr10-89018137-DEL-133 |
| chr10-89035231-DEL-3850 | chr10-89035232-DEL-3850 |
| chr10-89045881-DEL-807 | chr10-89045882-DEL-807 |
| chr10-89098352-DEL-94 | chr10-89098353-DEL-94 |
| chr10-89166527-DEL-219 | chr10-89166528-DEL-219 |
| chr10-89220336-DEL-293 | chr10-89220337-DEL-293 |
| chr10-89223426-DEL-968 | chr10-89223427-DEL-968 |
| chr10-89225432-DEL-66 | chr10-89225433-DEL-50 |
| chr10-89232454-DEL-54 | chr10-89232455-DEL-54 |
| chr10-89234745-DEL-1275 | chr10-89234743-DEL-1275 |
| chr10-89245616-DEL-207 | chr10-89245617-DEL-207 |
| chr10-89259097-DEL-69 | chr10-89259098-DEL-69 |
| chr10-89261351-DEL-305 | chr10-89261351-DEL-306 |
| chr10-89268336-DEL-75 | chr10-89268337-DEL-75 |
| chr10-89285273-DEL-799 | chr10-89285274-DEL-799 |
| chr10-89304913-DEL-574 | chr10-89304914-DEL-574 |
| chr10-89332106-DEL-116 | chr10-89332107-DEL-116 |
| chr10-89342980-DEL-60 | chr10-89342981-DEL-62 |
| chr10-89346358-DEL-56 | chr10-89346359-DEL-56 |
| chr10-89349171-DEL-64 | chr10-89349172-DEL-64 |
| chr10-89371358-DEL-1845 | chr10-89371359-DEL-1845 |
| chr10-89373432-DEL-680 | chr10-89373433-DEL-680 |
| chr10-89374547-DEL-64 | chr10-89374548-DEL-64 |
| chr10-89392838-DEL-124 | chr10-89392839-DEL-124 |
| chr10-89396686-DEL-619 | chr10-89396685-DEL-620 |
| chr10-89406403-DEL-85 | chr10-89406402-DEL-63 |
| chr10-89413393-DEL-350 | chr10-89413394-DEL-350 |
| chr10-89414731-DEL-93 | chr10-89414732-DEL-93 |
| chr10-89444887-DEL-284 | chr10-89444888-DEL-284 |
| chr10-89475031-DEL-60 | chr10-89475032-DEL-60 |
| chr10-89506240-DEL-53 | chr10-89506241-DEL-53 |
| chr10-89517597-DEL-84 | chr10-89517597-DEL-84 |
| chr10-89525744-DEL-1083 | chr10-89525745-DEL-1083 |
| chr10-89534416-DEL-2284 | chr10-89534417-DEL-2284 |
| chr10-89543151-DEL-194 | chr10-89543167-DEL-179 |
| chr10-89544165-DEL-51 | chr10-89544166-DEL-51 |
| chr10-89553704-DEL-987 | chr10-89553705-DEL-987 |
| chr10-89588371-DEL-229 | chr10-89588372-DEL-229 |
| chr10-89600359-DEL-94 | chr10-89600360-DEL-94 |
| chr10-89605495-DEL-127 | chr10-89605492-DEL-131 |
| chr10-89637650-DEL-202 | chr10-89637651-DEL-202 |
| chr10-89638322-DEL-60 | chr10-89638323-DEL-60 |
| chr10-89660622-DEL-157 | chr10-89660610-DEL-157 |
| chr10-89671543-DEL-59 | chr10-89671541-DEL-65 |
| chr10-89681113-DEL-539 | chr10-89681114-DEL-539 |
| chr10-89690448-DEL-204 | chr10-89690449-DEL-204 |
| chr10-89712186-DEL-669 | chr10-89712187-DEL-669 |
| chr10-89714097-DEL-363 | chr10-89714098-DEL-363 |
| chr10-89716995-DEL-75 | chr10-89716996-DEL-75 |
| chr10-89742029-DEL-56 | chr10-89742030-DEL-56 |

| | |
|---|---|
| chr10-89759265-DEL-107 | chr10-89759265-DEL-106 |
| chr10-89771266-DEL-6134 | chr10-89771267-DEL-6134 |
| chr10-89776521-DEL-514 | chr10-89776525-DEL-512 |
| chr10-89804196-DEL-85 | chr10-89804197-DEL-85 |
| chr10-89815394-DEL-174 | chr10-89815395-DEL-174 |
| chr10-89815919-DEL-82 | chr10-89815888-DEL-75 |
| chr10-89825039-DEL-3119 | chr10-89825040-DEL-3119 |
| chr10-89836612-DEL-72 | chr10-89836623-DEL-60 |
| chr10-89839212-DEL-112 | chr10-89839213-DEL-112 |
| chr10-89844393-DEL-1430 | chr10-89844394-DEL-1430 |
| chr10-89848262-DEL-206 | chr10-89848252-DEL-206 |
| chr10-89851641-DEL-404 | chr10-89851642-DEL-403 |
| chr10-89854626-DEL-74 | chr10-89854624-DEL-70 |
| chr10-89870041-DEL-51 | chr10-89870032-DEL-52 |
| chr10-89873050-DEL-395 | chr10-89873059-DEL-385 |
| chr10-89890849-DEL-5258 | chr10-89890852-DEL-5258 |
| chr10-89903925-DEL-83 | chr10-89903929-DEL-124 |
| chr10-89937254-DEL-59 | chr10-89937254-DEL-59 |
| chr10-89971139-DEL-99 | chr10-89971140-DEL-99 |
| chr10-89986522-DEL-56 | chr10-89986523-DEL-56 |
| chr10-89996373-DEL-601 | chr10-89996374-DEL-601 |
| chr10-90004270-DEL-63 | chr10-90004271-DEL-62 |
| chr10-90028290-DEL-571 | chr10-90028290-DEL-569 |
| chr10-90034931-DEL-93 | chr10-90034932-DEL-93 |
| chr10-90039723-DEL-73 | chr10-90039736-DEL-61 |
| chr10-90051423-DEL-3430 | chr10-90051424-DEL-3430 |
| chr10-90084804-DEL-6449 | chr10-90084805-DEL-6449 |
| chr10-90084838-DEL-212 | chr10-90084839-DEL-212 |
| chr10-90093034-DEL-50 | chr10-90093035-DEL-50 |
| chr10-90111665-DEL-498 | chr10-90111666-DEL-498 |
| chr10-90123371-DEL-1892 | chr10-90123371-DEL-1893 |
| chr10-90135938-DEL-455 | chr10-90135939-DEL-455 |
| chr10-90160484-DEL-5109 | chr10-90160485-DEL-5109 |
| chr10-90190752-DEL-928 | chr10-90190753-DEL-928 |
| chr10-90199050-DEL-60 | chr10-90199051-DEL-60 |
| chr10-90227389-DEL-59 | chr10-90227390-DEL-59 |
| chr10-90229817-DEL-160 | chr10-90229818-DEL-160 |
| chr10-90232656-DEL-90 | chr10-90232657-DEL-90 |
| chr10-90233823-DEL-1717 | chr10-90233824-DEL-1717 |
| chr10-90257722-DEL-867 | chr10-90257723-DEL-867 |
| chr10-90299203-DEL-1776 | chr10-90299204-DEL-1776 |
| chr10-90307026-DEL-3031 | chr10-90307027-DEL-3031 |
| chr10-90323334-DEL-415 | chr10-90323335-DEL-415 |
| chr10-90327448-DEL-95 | chr10-90327480-DEL-59 |
| chr10-90359880-DEL-90 | chr10-90359881-DEL-90 |
| chr10-90375981-DEL-64 | chr10-90375982-DEL-64 |
| chr10-90382234-DEL-80 | chr10-90382222-DEL-84 |
| chr10-90386819-DEL-1627 | chr10-90386811-DEL-1628 |
| chr10-90407644-DEL-279 | chr10-90407645-DEL-279 |
| chr10-90436654-DEL-1691 | chr10-90436655-DEL-1691 |
| chr10-90440543-DEL-322 | chr10-90440548-DEL-318 |
| chr10-90441319-DEL-72 | chr10-90441316-DEL-72 |
| chr10-90449503-DEL-126 | chr10-90449504-DEL-128 |
| chr10-90462629-DEL-100 | chr10-90462630-DEL-100 |
| chr10-90529646-DEL-488 | chr10-90529648-DEL-487 |
| chr10-90549137-DEL-76 | chr10-90549138-DEL-76 |
| chr10-90603007-DEL-56 | chr10-90603008-DEL-56 |
| chr10-90614015-DEL-54 | chr10-90614016-DEL-54 |
| chr10-90627383-DEL-56 | chr10-90627384-DEL-56 |
| chr10-90724930-DEL-1194 | chr10-90724931-DEL-1194 |
| chr10-90772331-DEL-90 | chr10-90772318-DEL-103 |
| chr10-90842804-DEL-65 | chr10-90842785-DEL-84 |
| chr10-90859771-DEL-3508 | chr10-90859772-DEL-3508 |
| chr10-90865593-DEL-70 | chr10-90865604-DEL-67 |
| chr10-90876207-DEL-74 | chr10-90876208-DEL-74 |
| chr10-90890577-DEL-72 | chr10-90890578-DEL-72 |
| chr10-90905842-DEL-64 | chr10-90905843-DEL-66 |
| chr10-90921976-DEL-164 | chr10-90921977-DEL-164 |
| chr10-90926179-DEL-200 | chr10-90926180-DEL-200 |
| chr10-90949307-DEL-163 | chr10-90949307-DEL-164 |
| chr10-90961751-DEL-250 | chr10-90961730-DEL-258 |
| chr10-90967845-DEL-173 | chr10-90967845-DEL-173 |
| chr10-90973969-DEL-200 | chr10-90973970-DEL-200 |
| chr10-90978227-DEL-200 | chr10-90978231-DEL-203 |
| chr10-91004641-DEL-336 | chr10-91004642-DEL-336 |
| chr10-91010016-DEL-65 | chr10-91010017-DEL-65 |
| chr10-91040650-DEL-190 | chr10-91040651-DEL-190 |
| chr10-91045706-DEL-238 | chr10-91045706-DEL-238 |
| chr10-91056075-DEL-140 | chr10-91056076-DEL-140 |
| chr10-91087386-DEL-71 | chr10-91087391-DEL-67 |

| | |
|---|---|
| chr10-91095229-DEL-328 | chr10-91095225-DEL-327 |
| chr10-91123523-DEL-453 | chr10-91123524-DEL-453 |
| chr10-91163499-DEL-448 | chr10-91163500-DEL-447 |
| chr10-91168022-DEL-6805 | chr10-91168023-DEL-6804 |
| chr10-91168464-DEL-208 | chr10-91168465-DEL-208 |
| chr10-91184915-DEL-100 | chr10-91184916-DEL-100 |
| chr10-91230556-DEL-606 | chr10-91230557-DEL-606 |
| chr10-91255616-DEL-60 | chr10-91255611-DEL-66 |
| chr10-91277346-DEL-90 | chr10-91277347-DEL-90 |
| chr10-91286657-DEL-100 | chr10-91286658-DEL-100 |
| chr10-91291656-DEL-102 | chr10-91291665-DEL-94 |
| chr10-91297917-DEL-80 | chr10-91297918-DEL-80 |
| chr10-91306202-DEL-349 | chr10-91306203-DEL-349 |
| chr10-91332321-DEL-140 | chr10-91332321-DEL-140 |
| chr10-91361350-DEL-145 | chr10-91361351-DEL-145 |
| chr10-91362337-DEL-96 | chr10-91362327-DEL-62 |
| chr10-91389947-DEL-1190 | chr10-91389948-DEL-1190 |
| chr10-91401678-DEL-90 | chr10-91401679-DEL-90 |
| chr10-91421051-DEL-2395 | chr10-91421052-DEL-2395 |
| chr10-91429762-DEL-313 | chr10-91429763-DEL-313 |
| chr10-91435541-DEL-208 | chr10-91435541-DEL-208 |
| chr10-91466125-DEL-188 | chr10-91466128-DEL-186 |
| chr10-91530518-DEL-347 | chr10-91530519-DEL-347 |
| chr10-91548783-DEL-90 | chr10-91548784-DEL-88 |
| chr10-91589865-DEL-55 | chr10-91589871-DEL-50 |
| chr10-91602618-DEL-803 | chr10-91602619-DEL-803 |
| chr10-91617801-DEL-64 | chr10-91617802-DEL-64 |
| chr10-91677059-DEL-60 | chr10-91677060-DEL-60 |
| chr10-91687038-DEL-63 | chr10-91687039-DEL-63 |
| chr10-91709079-DEL-309 | chr10-91709080-DEL-309 |
| chr10-91735378-DEL-71 | chr10-91735379-DEL-71 |
| chr10-91764634-DEL-6456 | chr10-91764635-DEL-6456 |
| chr10-91785210-DEL-77 | chr10-91785228-DEL-52 |
| chr10-91790949-DEL-56 | chr10-91790949-DEL-56 |
| chr10-91827443-DEL-65 | chr10-91827444-DEL-65 |
| chr10-91828781-DEL-53 | chr10-91828782-DEL-53 |
| chr10-91833878-DEL-76 | chr10-91833921-DEL-64 |
| chr10-91843425-DEL-199 | chr10-91843426-DEL-199 |
| chr10-91873465-DEL-1594 | chr10-91873466-DEL-1594 |
| chr10-91894044-DEL-734 | chr10-91894045-DEL-734 |
| chr10-91900349-DEL-52 | chr10-91900350-DEL-52 |
| chr10-91908746-DEL-55 | chr10-91908743-DEL-59 |
| chr10-91920580-DEL-1236 | chr10-91920581-DEL-1236 |
| chr10-91924847-DEL-75 | chr10-91924866-DEL-77 |
| chr10-91930081-DEL-82 | chr10-91930084-DEL-80 |
| chr10-91943945-DEL-74 | chr10-91943946-DEL-74 |
| chr10-91958294-DEL-69 | chr10-91958294-DEL-73 |
| chr10-91973482-DEL-72 | chr10-91973483-DEL-56 |
| chr10-91985137-DEL-2061 | chr10-91985138-DEL-2061 |
| chr10-92020473-DEL-201 | chr10-92020474-DEL-201 |
| chr10-92023166-DEL-92 | chr10-92023165-DEL-91 |
| chr10-92023870-DEL-197 | chr10-92023871-DEL-197 |
| chr10-92114991-DEL-88 | chr10-92114996-DEL-84 |
| chr10-92120076-DEL-52 | chr10-92120077-DEL-52 |
| chr10-92127413-DEL-52 | chr10-92127414-DEL-52 |
| chr10-92219278-DEL-60 | chr10-92219279-DEL-50 |
| chr10-92224029-DEL-118 | chr10-92224030-DEL-118 |
| chr10-92230202-DEL-246 | chr10-92230203-DEL-250 |
| chr10-92231667-DEL-347 | chr10-92231666-DEL-348 |
| chr10-92263467-DEL-67 | chr10-92263478-DEL-57 |
| chr10-92269623-DEL-116 | chr10-92269617-DEL-123 |
| chr10-92272570-DEL-52 | chr10-92272568-DEL-63 |
| chr10-92276579-DEL-52 | chr10-92276568-DEL-56 |
| chr10-92311307-DEL-1168 | chr10-92311308-DEL-1168 |
| chr10-92354657-DEL-2850 | chr10-92354658-DEL-2850 |
| chr10-92381597-DEL-1312 | chr10-92381585-DEL-1313 |
| chr10-92410087-DEL-52 | chr10-92410082-DEL-58 |
| chr10-92444733-DEL-312 | chr10-92444738-DEL-300 |
| chr10-92546115-DEL-59 | chr10-92546116-DEL-59 |
| chr10-92556846-DEL-76 | chr10-92556847-DEL-74 |
| chr10-92579969-DEL-238 | chr10-92579970-DEL-238 |
| chr10-92585805-DEL-288 | chr10-92585806-DEL-288 |
| chr10-92586396-DEL-3617 | chr10-92586397-DEL-3617 |
| chr10-92606539-DEL-336 | chr10-92606540-DEL-336 |
| chr10-92620745-DEL-355 | chr10-92620753-DEL-345 |
| chr10-92639321-DEL-2152 | chr10-92639321-DEL-2152 |
| chr10-92744960-DEL-6388 | chr10-92744961-DEL-6388 |
| chr10-92755723-DEL-105 | chr10-92755727-DEL-106 |
| chr10-92763790-DEL-83 | chr10-92763791-DEL-83 |
| chr10-92827805-DEL-83 | chr10-92827806-DEL-83 |

| | |
|---|---|
| chr10-92841530-DEL-1451 | chr10-92841531-DEL-1451 |
| chr10-92841949-DEL-58 | chr10-92841950-DEL-58 |
| chr10-92867082-DEL-110 | chr10-92867093-DEL-92 |
| chr10-92890022-DEL-6410 | chr10-92890023-DEL-6410 |
| chr10-92899006-DEL-63 | chr10-92899007-DEL-63 |
| chr10-92910430-DEL-92 | chr10-92910431-DEL-87 |
| chr10-92915012-DEL-1065 | chr10-92915013-DEL-1065 |
| chr10-92915206-DEL-189 | chr10-92915207-DEL-189 |
| chr10-92935482-DEL-52 | chr10-92935484-DEL-51 |
| chr10-92949475-DEL-63 | chr10-92949476-DEL-63 |
| chr10-92966790-DEL-881 | chr10-92966791-DEL-881 |
| chr10-92969556-DEL-1085 | chr10-92969557-DEL-1085 |
| chr10-92984058-DEL-63 | chr10-92984064-DEL-58 |
| chr10-92998731-DEL-205 | chr10-92998732-DEL-205 |
| chr10-93008596-DEL-56 | chr10-93008597-DEL-56 |
| chr10-93022653-DEL-385 | chr10-93022654-DEL-385 |
| chr10-93037466-DEL-60 | chr10-93037477-DEL-50 |
| chr10-93065550-DEL-200 | chr10-93065551-DEL-200 |
| chr10-93069026-DEL-204 | chr10-93069027-DEL-240 |
| chr10-93096051-DEL-64 | chr10-93096052-DEL-64 |
| chr10-93096765-DEL-72 | chr10-93096766-DEL-72 |
| chr10-93098677-DEL-135 | chr10-93098673-DEL-149 |
| chr10-93109919-DEL-79 | chr10-93109911-DEL-82 |
| chr10-93115398-DEL-128 | chr10-93115399-DEL-128 |
| chr10-93128011-DEL-124 | chr10-93128009-DEL-115 |
| chr10-93174303-DEL-135 | chr10-93174301-DEL-138 |
| chr10-93181687-DEL-1463 | chr10-93181688-DEL-1463 |
| chr10-93187807-DEL-60 | chr10-93187808-DEL-60 |
| chr10-93190155-DEL-53 | chr10-93190153-DEL-54 |
| chr10-93213158-DEL-2778 | chr10-93213159-DEL-2778 |
| chr10-93221633-DEL-99 | chr10-93221644-DEL-54 |
| chr10-93228421-DEL-128 | chr10-93228403-DEL-124 |
| chr10-93237135-DEL-400 | chr10-93237136-DEL-400 |
| chr10-93238476-DEL-138 | chr10-93238477-DEL-138 |
| chr10-93251545-DEL-197 | chr10-93251545-DEL-197 |
| chr10-93257419-DEL-75 | chr10-93257420-DEL-75 |
| chr10-93262337-DEL-78 | chr10-93262314-DEL-73 |
| chr10-93268743-DEL-210 | chr10-93268744-DEL-210 |
| chr10-93271119-DEL-102 | chr10-93271120-DEL-102 |
| chr10-93280906-DEL-908 | chr10-93280905-DEL-901 |
| chr10-93339758-DEL-69 | chr10-93339759-DEL-69 |
| chr10-93366853-DEL-80 | chr10-93366840-DEL-84 |
| chr10-93419674-DEL-64 | chr10-93419675-DEL-64 |
| chr10-93428212-DEL-142 | chr10-93428213-DEL-142 |
| chr10-93435897-DEL-2069 | chr10-93435925-DEL-2063 |
| chr10-93446670-DEL-6285 | chr10-93446670-DEL-6285 |
| chr10-93448425-DEL-171 | chr10-93448427-DEL-170 |
| chr10-93455544-DEL-71 | chr10-93455545-DEL-75 |
| chr10-93457777-DEL-102 | chr10-93457777-DEL-102 |
| chr10-93479689-DEL-195 | chr10-93479690-DEL-195 |
| chr10-93506468-DEL-60 | chr10-93506465-DEL-52 |
| chr10-93509716-DEL-78 | chr10-93509715-DEL-71 |
| chr10-93519492-DEL-53 | chr10-93519493-DEL-53 |
| chr10-93521262-DEL-66 | chr10-93521263-DEL-66 |
| chr10-93522767-DEL-118 | chr10-93522764-DEL-122 |
| chr10-93524147-DEL-64 | chr10-93524136-DEL-61 |
| chr10-93528782-DEL-1621 | chr10-93528779-DEL-1619 |
| chr10-93540942-DEL-219 | chr10-93540943-DEL-219 |
| chr10-93550762-DEL-56 | chr10-93550764-DEL-55 |
| chr10-93556712-DEL-109 | chr10-93556713-DEL-109 |
| chr10-93567831-DEL-111 | chr10-93567809-DEL-89 |
| chr10-93568279-DEL-177 | chr10-93568273-DEL-182 |
| chr10-93580694-DEL-62 | chr10-93580695-DEL-62 |
| chr10-93598602-DEL-303 | chr10-93598603-DEL-303 |
| chr10-93609772-DEL-356 | chr10-93609771-DEL-357 |
| chr10-93614456-DEL-1195 | chr10-93614457-DEL-1195 |
| chr10-93618142-DEL-53 | chr10-93618135-DEL-51 |
| chr10-93621207-DEL-7425 | chr10-93621208-DEL-7425 |
| chr10-93634226-DEL-56 | chr10-93634231-DEL-52 |
| chr10-93643146-DEL-499 | chr10-93643147-DEL-499 |
| chr10-93659213-DEL-121 | chr10-93659214-DEL-121 |
| chr10-93663011-DEL-115 | chr10-93663012-DEL-115 |
| chr10-93664608-DEL-60 | chr10-93664609-DEL-60 |
| chr10-93688059-DEL-82 | chr10-93688060-DEL-82 |
| chr10-93688817-DEL-207 | chr10-93688818-DEL-207 |
| chr10-93730996-DEL-62 | chr10-93730966-DEL-93 |
| chr10-93734608-DEL-64 | chr10-93734616-DEL-68 |
| chr10-93740617-DEL-56 | chr10-93740618-DEL-56 |
| chr10-93745403-DEL-397 | chr10-93745404-DEL-397 |
| chr10-93760483-DEL-93 | chr10-93760484-DEL-92 |

| | |
|---|---|
| chr10-93778671-DEL-585 | chr10-93778672-DEL-585 |
| chr10-93779571-DEL-289 | chr10-93779572-DEL-289 |
| chr10-93792317-DEL-120 | chr10-93792318-DEL-120 |
| chr10-93793769-DEL-593 | chr10-93793770-DEL-593 |
| chr10-93803098-DEL-60 | chr10-93803102-DEL-60 |
| chr10-93813585-DEL-328 | chr10-93813578-DEL-329 |
| chr10-93818154-DEL-56 | chr10-93818155-DEL-56 |
| chr10-93827415-DEL-134 | chr10-93827416-DEL-134 |
| chr10-93828352-DEL-88 | chr10-93828355-DEL-86 |
| chr10-93852153-DEL-114 | chr10-93852154-DEL-114 |
| chr10-93852734-DEL-101 | chr10-93852742-DEL-96 |
| chr10-93887434-DEL-55 | chr10-93887435-DEL-55 |
| chr10-93890518-DEL-80 | chr10-93890519-DEL-80 |
| chr10-93894164-DEL-175 | chr10-93894165-DEL-175 |
| chr10-93922548-DEL-286 | chr10-93922549-DEL-286 |
| chr10-93923609-DEL-58 | chr10-93923607-DEL-60 |
| chr10-93938554-DEL-56 | chr10-93938554-DEL-56 |
| chr10-93945334-DEL-75 | chr10-93945335-DEL-75 |
| chr10-93946282-DEL-181 | chr10-93946283-DEL-181 |
| chr10-93955577-DEL-207 | chr10-93955578-DEL-207 |
| chr10-93959210-DEL-96 | chr10-93959211-DEL-96 |
| chr10-93964802-DEL-86 | chr10-93964803-DEL-86 |
| chr10-93967845-DEL-52 | chr10-93967846-DEL-52 |
| chr10-93997440-DEL-82 | chr10-93997441-DEL-82 |
| chr10-94025032-DEL-1121 | chr10-94025033-DEL-1121 |
| chr10-94040167-DEL-668 | chr10-94040168-DEL-668 |
| chr10-94054497-DEL-52 | chr10-94054498-DEL-52 |
| chr10-94056353-DEL-59 | chr10-94056354-DEL-59 |
| chr10-94074496-DEL-65 | chr10-94074513-DEL-66 |
| chr10-94109154-DEL-56 | chr10-94109149-DEL-62 |
| chr10-94119345-DEL-333 | chr10-94119346-DEL-333 |
| chr10-94120595-DEL-2079 | chr10-94120597-DEL-2088 |
| chr10-94124493-DEL-51 | chr10-94124493-DEL-51 |
| chr10-94198443-DEL-6669 | chr10-94198444-DEL-6669 |
| chr10-94253954-DEL-238 | chr10-94253955-DEL-238 |
| chr10-94273258-DEL-70 | chr10-94273259-DEL-70 |
| chr10-94348302-DEL-57 | chr10-94348323-DEL-69 |
| chr10-94357057-DEL-300 | chr10-94357058-DEL-300 |
| chr10-94372068-DEL-411 | chr10-94372069-DEL-411 |
| chr10-94400994-DEL-131 | chr10-94400984-DEL-108 |
| chr10-94409709-DEL-258 | chr10-94409714-DEL-246 |
| chr10-94428296-DEL-90 | chr10-94428297-DEL-90 |
| chr10-94435591-DEL-189 | chr10-94435592-DEL-189 |
| chr10-94454836-DEL-424 | chr10-94454837-DEL-424 |
| chr10-94478786-DEL-230 | chr10-94478787-DEL-230 |
| chr10-94490256-DEL-51 | chr10-94490257-DEL-51 |
| chr10-94513741-DEL-192 | chr10-94513742-DEL-192 |
| chr10-94515748-DEL-830 | chr10-94515751-DEL-828 |
| chr10-94519593-DEL-785 | chr10-94519594-DEL-785 |
| chr10-94526898-DEL-185 | chr10-94526899-DEL-185 |
| chr10-94550980-DEL-997 | chr10-94551006-DEL-967 |
| chr10-94686031-DEL-110 | chr10-94686032-DEL-110 |
| chr10-94704609-DEL-150 | chr10-94704612-DEL-153 |
| chr10-94713718-DEL-64 | chr10-94713709-DEL-82 |
| chr10-94732973-DEL-457 | chr10-94732974-DEL-457 |
| chr10-94766462-DEL-179 | chr10-94766463-DEL-179 |
| chr10-94788878-DEL-57 | chr10-94788879-DEL-57 |
| chr10-94790777-DEL-1249 | chr10-94790777-DEL-1249 |
| chr10-94801707-DEL-1365 | chr10-94801710-DEL-1364 |
| chr10-94807546-DEL-91 | chr10-94807547-DEL-91 |
| chr10-94811425-DEL-189 | chr10-94811429-DEL-184 |
| chr10-94819310-DEL-70 | chr10-94819311-DEL-70 |
| chr10-94820174-DEL-1219 | chr10-94820175-DEL-1219 |
| chr10-94824111-DEL-54 | chr10-94824112-DEL-54 |
| chr10-94833266-DEL-1433 | chr10-94833267-DEL-1433 |
| chr10-94854906-DEL-175 | chr10-94854907-DEL-175 |
| chr10-94891796-DEL-54 | chr10-94891795-DEL-54 |
| chr10-94920794-DEL-277 | chr10-94920792-DEL-271 |
| chr10-94927942-DEL-74 | chr10-94927935-DEL-74 |
| chr10-94940879-DEL-828 | chr10-94940868-DEL-817 |
| chr10-94966466-DEL-92 | chr10-94966457-DEL-80 |
| chr10-94972859-DEL-1656 | chr10-94972860-DEL-1656 |
| chr10-95056791-DEL-403 | chr10-95056793-DEL-389 |
| chr10-95076888-DEL-273 | chr10-95076889-DEL-273 |
| chr10-95124018-DEL-94 | chr10-95124019-DEL-94 |
| chr10-95140768-DEL-52 | chr10-95140769-DEL-52 |
| chr10-95144469-DEL-240 | chr10-95144470-DEL-240 |
| chr10-95183504-DEL-202 | chr10-95183505-DEL-202 |
| chr10-95193697-DEL-53 | chr10-95193678-DEL-51 |
| chr10-95209992-DEL-431 | chr10-95209993-DEL-431 |

| | |
|---|---|
| chr10-95213984-DEL-60 | chr10-95213985-DEL-60 |
| chr10-95238905-DEL-52 | chr10-95238906-DEL-52 |
| chr10-95273013-DEL-148 | chr10-95273014-DEL-148 |
| chr10-95289523-DEL-68 | chr10-95289525-DEL-67 |
| chr10-95326808-DEL-2360 | chr10-95326809-DEL-2360 |
| chr10-95333182-DEL-212 | chr10-95333183-DEL-212 |
| chr10-95334233-DEL-253 | chr10-95334234-DEL-253 |
| chr10-95340388-DEL-107 | chr10-95340389-DEL-107 |
| chr10-95340896-DEL-220 | chr10-95340897-DEL-220 |
| chr10-95342232-DEL-118 | chr10-95342233-DEL-118 |
| chr10-95389296-DEL-61 | chr10-95389297-DEL-61 |
| chr10-95396955-DEL-220 | chr10-95396955-DEL-220 |
| chr10-95406229-DEL-62 | chr10-95406230-DEL-66 |
| chr10-95418205-DEL-114 | chr10-95418206-DEL-114 |
| chr10-95432483-DEL-76 | chr10-95432484-DEL-76 |
| chr10-95441792-DEL-75 | chr10-95441763-DEL-95 |
| chr10-95450780-DEL-99 | chr10-95450785-DEL-93 |
| chr10-95462899-DEL-242 | chr10-95462900-DEL-242 |
| chr10-95468483-DEL-164 | chr10-95468484-DEL-164 |
| chr10-95469621-DEL-406 | chr10-95469622-DEL-406 |
| chr10-95496178-DEL-112 | chr10-95496179-DEL-112 |
| chr10-95552781-DEL-112 | chr10-95552768-DEL-103 |
| chr10-95568670-DEL-381 | chr10-95568671-DEL-381 |
| chr10-95573468-DEL-178 | chr10-95573466-DEL-179 |
| chr10-95587371-DEL-295 | chr10-95587363-DEL-296 |
| chr10-95587908-DEL-196 | chr10-95587909-DEL-196 |
| chr10-95652436-DEL-86 | chr10-95652437-DEL-90 |
| chr10-95691099-DEL-200 | chr10-95691100-DEL-200 |
| chr10-95705075-DEL-199 | chr10-95705075-DEL-199 |
| chr10-95715085-DEL-778 | chr10-95715086-DEL-778 |
| chr10-95716034-DEL-56 | chr10-95716035-DEL-56 |
| chr10-95722714-DEL-3201 | chr10-95722715-DEL-3201 |
| chr10-95735931-DEL-166 | chr10-95735953-DEL-176 |
| chr10-95742108-DEL-239 | chr10-95742110-DEL-237 |
| chr10-95742640-DEL-468 | chr10-95742641-DEL-468 |
| chr10-95757003-DEL-285 | chr10-95757004-DEL-283 |
| chr10-95764018-DEL-54 | chr10-95764019-DEL-54 |
| chr10-95794976-DEL-56 | chr10-95794977-DEL-56 |
| chr10-95806553-DEL-63 | chr10-95806554-DEL-59 |
| chr10-95817744-DEL-70 | chr10-95817747-DEL-62 |
| chr10-95913377-DEL-426 | chr10-95913378-DEL-431 |
| chr10-95920384-DEL-130 | chr10-95920384-DEL-135 |
| chr10-95962860-DEL-115 | chr10-95962861-DEL-115 |
| chr10-95968942-DEL-451 | chr10-95968943-DEL-451 |
| chr10-95991724-DEL-128 | chr10-95991725-DEL-128 |
| chr10-95993077-DEL-5229 | chr10-95993078-DEL-5229 |
| chr10-96007767-DEL-2441 | chr10-96007768-DEL-2441 |
| chr10-96015323-DEL-152 | chr10-96015359-DEL-166 |
| chr10-96028391-DEL-7173 | chr10-96028389-DEL-7173 |
| chr10-96036237-DEL-86 | chr10-96036239-DEL-101 |
| chr10-96042239-DEL-1628 | chr10-96042240-DEL-1628 |
| chr10-96053795-DEL-73 | chr10-96053796-DEL-73 |
| chr10-96080047-DEL-81 | chr10-96080050-DEL-79 |
| chr10-96100435-DEL-51 | chr10-96100436-DEL-51 |
| chr10-96112333-DEL-241 | chr10-96112376-DEL-199 |
| chr10-96114759-DEL-156 | chr10-96114760-DEL-156 |
| chr10-96120382-DEL-56 | chr10-96120383-DEL-56 |
| chr10-96124791-DEL-99 | chr10-96124792-DEL-99 |
| chr10-96137293-DEL-211 | chr10-96137294-DEL-211 |
| chr10-96171390-DEL-231 | chr10-96171391-DEL-231 |
| chr10-96189761-DEL-164 | chr10-96189725-DEL-152 |
| chr10-96207685-DEL-231 | chr10-96207686-DEL-231 |
| chr10-96219712-DEL-196 | chr10-96219727-DEL-192 |
| chr10-96250429-DEL-331 | chr10-96250430-DEL-331 |
| chr10-96269763-DEL-117 | chr10-96269764-DEL-117 |
| chr10-96339734-DEL-81 | chr10-96339735-DEL-77 |
| chr10-96341080-DEL-7017 | chr10-96341081-DEL-7017 |
| chr10-96364937-DEL-2187 | chr10-96364938-DEL-2187 |
| chr10-96383188-DEL-1484 | chr10-96383192-DEL-1489 |
| chr10-96389606-DEL-190 | chr10-96389603-DEL-191 |
| chr10-96420789-DEL-476 | chr10-96420792-DEL-474 |
| chr10-96423595-DEL-54 | chr10-96423586-DEL-68 |
| chr10-96436798-DEL-60 | chr10-96436787-DEL-72 |
| chr10-96503552-DEL-6500 | chr10-96503553-DEL-6500 |
| chr10-96549246-DEL-69 | chr10-96549247-DEL-69 |
| chr10-96589211-DEL-100 | chr10-96589220-DEL-66 |
| chr10-96606855-DEL-68 | chr10-96606856-DEL-68 |
| chr10-96607665-DEL-198 | chr10-96607682-DEL-198 |
| chr10-96611978-DEL-54 | chr10-96611979-DEL-54 |
| chr10-96720874-DEL-1671 | chr10-96720875-DEL-1671 |

| | |
|---|---|
| chr10-96723441-DEL-7364 | chr10-96723442-DEL-7364 |
| chr10-96753254-DEL-1586 | chr10-96753254-DEL-1586 |
| chr10-96835682-DEL-678 | chr10-96835683-DEL-678 |
| chr10-96872836-DEL-81 | chr10-96872837-DEL-81 |
| chr10-96891287-DEL-88 | chr10-96891288-DEL-88 |
| chr10-96931804-DEL-417 | chr10-96931800-DEL-418 |
| chr10-96936759-DEL-77 | chr10-96936760-DEL-77 |
| chr10-96983982-DEL-2328 | chr10-96983983-DEL-2328 |
| chr10-96984115-DEL-566 | chr10-96984116-DEL-565 |
| chr10-97007518-DEL-1020 | chr10-97007519-DEL-1020 |
| chr10-97015901-DEL-6223 | chr10-97015902-DEL-6223 |
| chr10-97030539-DEL-138 | chr10-97030535-DEL-139 |
| chr10-97033169-DEL-310 | chr10-97033151-DEL-309 |
| chr10-97034320-DEL-134 | chr10-97034307-DEL-135 |
| chr10-97038421-DEL-358 | chr10-97038422-DEL-358 |
| chr10-97049774-DEL-196 | chr10-97049775-DEL-196 |
| chr10-97053888-DEL-647 | chr10-97053889-DEL-647 |
| chr10-97130654-DEL-280 | chr10-97130655-DEL-280 |
| chr10-97152188-DEL-509 | chr10-97152188-DEL-509 |
| chr10-97156322-DEL-9274 | chr10-97156323-DEL-9274 |
| chr10-97158875-DEL-1890 | chr10-97158876-DEL-1890 |
| chr10-97206062-DEL-52 | chr10-97206063-DEL-52 |
| chr10-97207336-DEL-150 | chr10-97207337-DEL-150 |
| chr10-97239393-DEL-90 | chr10-97239390-DEL-94 |
| chr10-97246871-DEL-591 | chr10-97246871-DEL-591 |
| chr10-97273410-DEL-91 | chr10-97273411-DEL-91 |
| chr10-97340578-DEL-136 | chr10-97340579-DEL-136 |
| chr10-97361457-DEL-8453 | chr10-97361463-DEL-8460 |
| chr10-97375617-DEL-56 | chr10-97375620-DEL-54 |
| chr10-97377651-DEL-343 | chr10-97377652-DEL-343 |
| chr10-97383059-DEL-184 | chr10-97383090-DEL-180 |
| chr10-97395301-DEL-511 | chr10-97395302-DEL-511 |
| chr10-97398865-DEL-538 | chr10-97398866-DEL-538 |
| chr10-97452842-DEL-1568 | chr10-97452843-DEL-1568 |
| chr10-97455454-DEL-1263 | chr10-97455455-DEL-1263 |
| chr10-97460181-DEL-74 | chr10-97460185-DEL-75 |
| chr10-97463633-DEL-197 | chr10-97463586-DEL-247 |
| chr10-97501565-DEL-195 | chr10-97501566-DEL-195 |
| chr10-97514296-DEL-567 | chr10-97514288-DEL-568 |
| chr10-97516375-DEL-417 | chr10-97516376-DEL-417 |
| chr10-97528314-DEL-68 | chr10-97528315-DEL-66 |
| chr10-97538816-DEL-657 | chr10-97538817-DEL-657 |
| chr10-97541099-DEL-82 | chr10-97541121-DEL-83 |
| chr10-97541499-DEL-199 | chr10-97541502-DEL-208 |
| chr10-97572505-DEL-73 | chr10-97572502-DEL-77 |
| chr10-97590857-DEL-6225 | chr10-97590860-DEL-6223 |
| chr10-97652805-DEL-69 | chr10-97652802-DEL-66 |
| chr10-97661350-DEL-159 | chr10-97661351-DEL-158 |
| chr10-97661852-DEL-58 | chr10-97661887-DEL-51 |
| chr10-97665652-DEL-208 | chr10-97665653-DEL-208 |
| chr10-97688238-DEL-123 | chr10-97688245-DEL-117 |
| chr10-97688275-DEL-742 | chr10-97688276-DEL-739 |
| chr10-97697096-DEL-64 | chr10-97697097-DEL-64 |
| chr10-97704211-DEL-58 | chr10-97704212-DEL-58 |
| chr10-97718564-DEL-1755 | chr10-97718565-DEL-1755 |
| chr10-97732635-DEL-967 | chr10-97732643-DEL-968 |
| chr10-97736281-DEL-74 | chr10-97736282-DEL-74 |
| chr10-97749429-DEL-6832 | chr10-97749430-DEL-6832 |
| chr10-97762779-DEL-2136 | chr10-97762776-DEL-2137 |
| chr10-97781188-DEL-60 | chr10-97781190-DEL-63 |
| chr10-97799593-DEL-145 | chr10-97799594-DEL-145 |
| chr10-97809034-DEL-60 | chr10-97809035-DEL-56 |
| chr10-97833941-DEL-6253 | chr10-97833942-DEL-6253 |
| chr10-97847525-DEL-66 | chr10-97847526-DEL-66 |
| chr10-97849713-DEL-160 | chr10-97849713-DEL-160 |
| chr10-97902292-DEL-410 | chr10-97902292-DEL-410 |
| chr10-97910662-DEL-90 | chr10-97910664-DEL-72 |
| chr10-97949840-DEL-418 | chr10-97949841-DEL-418 |
| chr10-97958028-DEL-89 | chr10-97958031-DEL-52 |
| chr10-97961276-DEL-4728 | chr10-97961277-DEL-4728 |
| chr10-97977240-DEL-354 | chr10-97977241-DEL-354 |
| chr10-97987060-DEL-58 | chr10-97987061-DEL-58 |
| chr10-98006921-DEL-60 | chr10-98006917-DEL-60 |
| chr10-98074989-DEL-1405 | chr10-98074990-DEL-1405 |
| chr10-98076088-DEL-184 | chr10-98076089-DEL-184 |
| chr10-98124743-DEL-53 | chr10-98124744-DEL-51 |
| chr10-98173164-DEL-6395 | chr10-98173146-DEL-6396 |
| chr10-98196529-DEL-118 | chr10-98196530-DEL-118 |
| chr10-98204398-DEL-108 | chr10-98204398-DEL-109 |
| chr10-98251402-DEL-850 | chr10-98251403-DEL-850 |

| | |
|---|---|
| chr10-98264049-DEL-439 | chr10-98264050-DEL-439 |
| chr10-98274051-DEL-6119 | chr10-98274052-DEL-6119 |
| chr10-98318104-DEL-66 | chr10-98318107-DEL-64 |
| chr10-98368863-DEL-1261 | chr10-98368864-DEL-1261 |
| chr10-98376772-DEL-96 | chr10-98376732-DEL-114 |
| chr10-98380248-DEL-6698 | chr10-98380248-DEL-6698 |
| chr10-98415582-DEL-78 | chr10-98415588-DEL-78 |
| chr10-98429085-DEL-196 | chr10-98429086-DEL-196 |
| chr10-98441842-DEL-1291 | chr10-98441813-DEL-1295 |
| chr10-98604157-DEL-239 | chr10-98604149-DEL-240 |
| chr10-98655954-DEL-62 | chr10-98655963-DEL-86 |
| chr10-98658240-DEL-53 | chr10-98658240-DEL-53 |
| chr10-98667289-DEL-63 | chr10-98667290-DEL-63 |
| chr10-98668325-DEL-395 | chr10-98668335-DEL-394 |
| chr10-98669589-DEL-331 | chr10-98669590-DEL-331 |
| chr10-98687831-DEL-144 | chr10-98687832-DEL-144 |
| chr10-98692491-DEL-506 | chr10-98692480-DEL-507 |
| chr10-98694020-DEL-60 | chr10-98694021-DEL-60 |
| chr10-98705780-DEL-400 | chr10-98705781-DEL-400 |
| chr10-98760862-DEL-62 | chr10-98760861-DEL-62 |
| chr10-98788270-DEL-1891 | chr10-98788271-DEL-1891 |
| chr10-98822528-DEL-55 | chr10-98822529-DEL-55 |
| chr10-98872703-DEL-73 | chr10-98872704-DEL-74 |
| chr10-98878425-DEL-170 | chr10-98878426-DEL-169 |
| chr10-98880755-DEL-1695 | chr10-98880756-DEL-1695 |
| chr10-98903696-DEL-6440 | chr10-98903697-DEL-6440 |
| chr10-98934048-DEL-5242 | chr10-98934049-DEL-5242 |
| chr10-98993181-DEL-63 | chr10-98993225-DEL-61 |
| chr10-98998770-DEL-71 | chr10-98998770-DEL-77 |
| chr10-98999302-DEL-195 | chr10-98999303-DEL-195 |
| chr10-99052212-DEL-403 | chr10-99052210-DEL-404 |
| chr10-99059754-DEL-103 | chr10-99059755-DEL-103 |
| chr10-99068219-DEL-72 | chr10-99068208-DEL-64 |
| chr10-99108486-DEL-113 | chr10-99108503-DEL-93 |
| chr10-99118864-DEL-84 | chr10-99118864-DEL-54 |
| chr10-99174894-DEL-7215 | chr10-99174895-DEL-7215 |
| chr10-99190292-DEL-184 | chr10-99190293-DEL-184 |
| chr10-99199050-DEL-83 | chr10-99199047-DEL-97 |
| chr10-99205556-DEL-70 | chr10-99205556-DEL-71 |
| chr10-99239758-DEL-51 | chr10-99239781-DEL-57 |
| chr10-99241406-DEL-180 | chr10-99241407-DEL-180 |
| chr10-99242537-DEL-74 | chr10-99242535-DEL-77 |
| chr10-99260900-DEL-72 | chr10-99260899-DEL-74 |
| chr10-99291460-DEL-407 | chr10-99291461-DEL-407 |
| chr10-99341472-DEL-172 | chr10-99341473-DEL-172 |
| chr10-99342969-DEL-76 | chr10-99342961-DEL-52 |
| chr10-99352990-DEL-327 | chr10-99352991-DEL-327 |
| chr10-99375883-DEL-71 | chr10-99375890-DEL-65 |
| chr10-99381707-DEL-5593 | chr10-99381737-DEL-5549 |
| chr10-99401759-DEL-79 | chr10-99401751-DEL-60 |
| chr10-99422616-DEL-273 | chr10-99422617-DEL-273 |
| chr10-99425511-DEL-65 | chr10-99425512-DEL-65 |
| chr10-99432819-DEL-626 | chr10-99432820-DEL-626 |
| chr10-99433571-DEL-67 | chr10-99433572-DEL-65 |
| chr10-99433850-DEL-470 | chr10-99433851-DEL-470 |
| chr10-99435411-DEL-490 | chr10-99435412-DEL-490 |
| chr10-99440261-DEL-1017 | chr10-99440262-DEL-1017 |
| chr10-99444607-DEL-480 | chr10-99444608-DEL-480 |
| chr10-99447668-DEL-89 | chr10-99447661-DEL-97 |
| chr10-99458992-DEL-2431 | chr10-99458993-DEL-2431 |
| chr10-99464147-DEL-1928 | chr10-99464148-DEL-1928 |
| chr10-99466182-DEL-82 | chr10-99466183-DEL-82 |
| chr10-99497832-DEL-6313 | chr10-99497833-DEL-6313 |
| chr10-99505156-DEL-65 | chr10-99505157-DEL-59 |
| chr10-99524321-DEL-62 | chr10-99524322-DEL-62 |
| chr10-99528955-DEL-55 | chr10-99528956-DEL-55 |
| chr10-99548984-DEL-141 | chr10-99548985-DEL-141 |
| chr10-99553447-DEL-64 | chr10-99553447-DEL-62 |
| chr10-99589677-DEL-69 | chr10-99589674-DEL-112 |
| chr10-99609402-DEL-98 | chr10-99609391-DEL-106 |
| chr10-99616009-DEL-345 | chr10-99616010-DEL-345 |
| chr10-99620821-DEL-289 | chr10-99620822-DEL-289 |
| chr10-99674929-DEL-54 | chr10-99674930-DEL-54 |
| chr10-99692029-DEL-220 | chr10-99692030-DEL-220 |
| chr10-99701742-DEL-72 | chr10-99701743-DEL-72 |
| chr10-99717075-DEL-70 | chr10-99717077-DEL-69 |
| chr10-99720710-DEL-6448 | chr10-99720711-DEL-6448 |
| chr10-99760755-DEL-58 | chr10-99760756-DEL-58 |
| chr10-99765594-DEL-52 | chr10-99765594-DEL-52 |
| chr10-99770848-DEL-59 | chr10-99770849-DEL-59 |

| | |
|---|---|
| chr10-99813959-DEL-390 | chr10-99813959-DEL-390 |
| chr10-99830489-DEL-57 | chr10-99830490-DEL-58 |
| chr10-99841247-DEL-574 | chr10-99841221-DEL-594 |
| chr10-99848101-DEL-287 | chr10-99848102-DEL-287 |
| chr10-99862195-DEL-90 | chr10-99862196-DEL-90 |
| chr10-99865964-DEL-55 | chr10-99865978-DEL-56 |
| chr10-99871374-DEL-470 | chr10-99871375-DEL-470 |
| chr10-99878684-DEL-786 | chr10-99878685-DEL-786 |
| chr10-99881843-DEL-206 | chr10-99881808-DEL-222 |
| chr10-99923395-DEL-58 | chr10-99923396-DEL-58 |
| chr10-99949464-DEL-414 | chr10-99949465-DEL-414 |
| chr10-99966747-DEL-105 | chr10-99966748-DEL-105 |
| chr10-99975145-DEL-5265 | chr10-99975146-DEL-5265 |
| chr10-99982684-DEL-78 | chr10-99982701-DEL-108 |
| chr10-99986119-DEL-137 | chr10-99986120-DEL-137 |
| chr10-100002107-DEL-594 | chr10-100002108-DEL-594 |
| chr10-100026588-DEL-55 | chr10-100026589-DEL-50 |
| chr10-100029747-DEL-1148 | chr10-100029748-DEL-1148 |
| chr10-100039630-DEL-255 | chr10-100039631-DEL-255 |
| chr10-100048364-DEL-6642 | chr10-100048365-DEL-6642 |
| chr10-100064745-DEL-3176 | chr10-100064746-DEL-3176 |
| chr10-100071952-DEL-165 | chr10-100071953-DEL-138 |
| chr10-100142068-DEL-63 | chr10-100142069-DEL-63 |
| chr10-100201755-DEL-54 | chr10-100201754-DEL-54 |
| chr10-100208556-DEL-102 | chr10-100208556-DEL-84 |
| chr10-100222496-DEL-101 | chr10-100222497-DEL-101 |
| chr10-100222754-DEL-563 | chr10-100222755-DEL-563 |
| chr10-100254827-DEL-70 | chr10-100254828-DEL-70 |
| chr10-100268761-DEL-836 | chr10-100268762-DEL-836 |
| chr10-100276191-DEL-2458 | chr10-100276192-DEL-2458 |
| chr10-100314871-DEL-504 | chr10-100314872-DEL-504 |
| chr10-100317359-DEL-84 | chr10-100317361-DEL-82 |
| chr10-100317564-DEL-445 | chr10-100317565-DEL-445 |
| chr10-100320517-DEL-62 | chr10-100320509-DEL-57 |
| chr10-100353865-DEL-200 | chr10-100353859-DEL-201 |
| chr10-100385550-DEL-189 | chr10-100385551-DEL-189 |
| chr10-100386529-DEL-88 | chr10-100386530-DEL-88 |
| chr10-100400933-DEL-134 | chr10-100400934-DEL-134 |
| chr10-100414832-DEL-4790 | chr10-100414833-DEL-4791 |
| chr10-100436267-DEL-208 | chr10-100436268-DEL-208 |
| chr10-100452855-DEL-72 | chr10-100452856-DEL-72 |
| chr10-100466352-DEL-52 | chr10-100466353-DEL-52 |
| chr10-100468582-DEL-115 | chr10-100468629-DEL-114 |
| chr10-100482441-DEL-6368 | chr10-100482442-DEL-6368 |
| chr10-100492307-DEL-55 | chr10-100492308-DEL-55 |
| chr10-100508034-DEL-127 | chr10-100508034-DEL-127 |
| chr10-100510792-DEL-946 | chr10-100510792-DEL-946 |
| chr10-100542774-DEL-59 | chr10-100542738-DEL-60 |
| chr10-100555556-DEL-1376 | chr10-100555557-DEL-1376 |
| chr10-100564644-DEL-99 | chr10-100564645-DEL-99 |
| chr10-100576856-DEL-60 | chr10-100576857-DEL-60 |
| chr10-100635711-DEL-56 | chr10-100635712-DEL-56 |
| chr10-100636950-DEL-68 | chr10-100636951-DEL-68 |
| chr10-100640707-DEL-2884 | chr10-100640706-DEL-2884 |
| chr10-100663375-DEL-92 | chr10-100663378-DEL-90 |
| chr10-100664313-DEL-839 | chr10-100664314-DEL-839 |
| chr10-100668758-DEL-58 | chr10-100668759-DEL-54 |
| chr10-100673784-DEL-54 | chr10-100673793-DEL-50 |
| chr10-100675707-DEL-1361 | chr10-100675708-DEL-1361 |
| chr10-100684773-DEL-156 | chr10-100684786-DEL-154 |
| chr10-100685246-DEL-299 | chr10-100685247-DEL-299 |
| chr10-100744652-DEL-75 | chr10-100744653-DEL-75 |
| chr10-100761713-DEL-422 | chr10-100761714-DEL-422 |
| chr10-100769151-DEL-422 | chr10-100769152-DEL-422 |
| chr10-100773715-DEL-122 | chr10-100773703-DEL-100 |
| chr10-100793567-DEL-55 | chr10-100793568-DEL-55 |
| chr10-100806874-DEL-1167 | chr10-100806870-DEL-1168 |
| chr10-100822150-DEL-56 | chr10-100822150-DEL-56 |
| chr10-100822415-DEL-690 | chr10-100822428-DEL-679 |
| chr10-100830346-DEL-228 | chr10-100830347-DEL-228 |
| chr10-100888644-DEL-98 | chr10-100888660-DEL-118 |
| chr10-100906012-DEL-1452 | chr10-100906012-DEL-1452 |
| chr10-100911932-DEL-1853 | chr10-100911933-DEL-1853 |
| chr10-100920165-DEL-56 | chr10-100920161-DEL-56 |
| chr10-100948873-DEL-148 | chr10-100948874-DEL-148 |
| chr10-100958686-DEL-67 | chr10-100958687-DEL-67 |
| chr10-100960632-DEL-872 | chr10-100960633-DEL-871 |
| chr10-100978243-DEL-87 | chr10-100978246-DEL-96 |
| chr10-100985090-DEL-1732 | chr10-100985077-DEL-1733 |
| chr10-100992170-DEL-125 | chr10-100992146-DEL-150 |

| | |
|---|---|
| chr10-101028056-DEL-392 | chr10-101028057-DEL-392 |
| chr10-101035063-DEL-59 | chr10-101035065-DEL-57 |
| chr10-101042423-DEL-51 | chr10-101042417-DEL-53 |
| chr10-101057241-DEL-487 | chr10-101057242-DEL-487 |
| chr10-101065969-DEL-268 | chr10-101065970-DEL-268 |
| chr10-101068880-DEL-3467 | chr10-101068881-DEL-3467 |
| chr10-101075724-DEL-1764 | chr10-101075725-DEL-1764 |
| chr10-101078153-DEL-862 | chr10-101078154-DEL-862 |
| chr10-101085683-DEL-158 | chr10-101085681-DEL-147 |
| chr10-101150245-DEL-1888 | chr10-101150246-DEL-1888 |
| chr10-101177547-DEL-6576 | chr10-101177563-DEL-6561 |
| chr10-101209691-DEL-2808 | chr10-101209692-DEL-2808 |
| chr10-101215747-DEL-177 | chr10-101215748-DEL-177 |
| chr10-101226133-DEL-151 | chr10-101226134-DEL-151 |
| chr10-101233513-DEL-95 | chr10-101233470-DEL-89 |
| chr10-101259717-DEL-382 | chr10-101259718-DEL-382 |
| chr10-101278714-DEL-1263 | chr10-101278715-DEL-1262 |
| chr10-101281539-DEL-6369 | chr10-101281542-DEL-6370 |
| chr10-101287596-DEL-201 | chr10-101287597-DEL-201 |
| chr10-101335324-DEL-247 | chr10-101335323-DEL-248 |
| chr10-101349593-DEL-345 | chr10-101349594-DEL-345 |
| chr10-101359319-DEL-864 | chr10-101359320-DEL-864 |
| chr10-101402630-DEL-6824 | chr10-101402631-DEL-6824 |
| chr10-101403190-DEL-207 | chr10-101403167-DEL-208 |
| chr10-101411243-DEL-66 | chr10-101411244-DEL-66 |
| chr10-101416232-DEL-516 | chr10-101416233-DEL-516 |
| chr10-101433433-DEL-117 | chr10-101433446-DEL-98 |
| chr10-101441418-DEL-1867 | chr10-101441419-DEL-1867 |
| chr10-101460585-DEL-6466 | chr10-101460585-DEL-6466 |
| chr10-101471549-DEL-61 | chr10-101471544-DEL-66 |
| chr10-101482659-DEL-224 | chr10-101482658-DEL-220 |
| chr10-101487817-DEL-68 | chr10-101487816-DEL-70 |
| chr10-101508122-DEL-449 | chr10-101508123-DEL-449 |
| chr10-101519061-DEL-78 | chr10-101519071-DEL-52 |
| chr10-101534431-DEL-52 | chr10-101534430-DEL-80 |
| chr10-101569399-DEL-536 | chr10-101569391-DEL-535 |
| chr10-101571724-DEL-104 | chr10-101571733-DEL-116 |
| chr10-101580892-DEL-191 | chr10-101580895-DEL-189 |
| chr10-101617011-DEL-86 | chr10-101617012-DEL-66 |
| chr10-101621399-DEL-577 | chr10-101621400-DEL-577 |
| chr10-101623027-DEL-64 | chr10-101623039-DEL-65 |
| chr10-101625198-DEL-1991 | chr10-101625227-DEL-1986 |
| chr10-101655181-DEL-904 | chr10-101655182-DEL-904 |
| chr10-101696692-DEL-62 | chr10-101696693-DEL-62 |
| chr10-101698417-DEL-159 | chr10-101698417-DEL-159 |
| chr10-101714988-DEL-2407 | chr10-101714989-DEL-2407 |
| chr10-101722843-DEL-97 | chr10-101722855-DEL-50 |
| chr10-101741824-DEL-75 | chr10-101741825-DEL-75 |
| chr10-101743334-DEL-134 | chr10-101743336-DEL-123 |
| chr10-101746032-DEL-82 | chr10-101746057-DEL-65 |
| chr10-101771126-DEL-269 | chr10-101771127-DEL-269 |
| chr10-101782531-DEL-70 | chr10-101782532-DEL-70 |
| chr10-101786010-DEL-1314 | chr10-101786012-DEL-1313 |
| chr10-101807649-DEL-432 | chr10-101807649-DEL-432 |
| chr10-101814234-DEL-87 | chr10-101814235-DEL-87 |
| chr10-101816335-DEL-53 | chr10-101816331-DEL-59 |
| chr10-101837982-DEL-66 | chr10-101837983-DEL-62 |
| chr10-101842713-DEL-99 | chr10-101842718-DEL-98 |
| chr10-101844594-DEL-130 | chr10-101844595-DEL-130 |
| chr10-101851674-DEL-372 | chr10-101851686-DEL-352 |
| chr10-101867980-DEL-55 | chr10-101867981-DEL-56 |
| chr10-101876410-DEL-1372 | chr10-101876411-DEL-1372 |
| chr10-101905960-DEL-93 | chr10-101905964-DEL-92 |
| chr10-101909459-DEL-104 | chr10-101909460-DEL-104 |
| chr10-101917377-DEL-1421 | chr10-101917378-DEL-1421 |
| chr10-101941175-DEL-309 | chr10-101941176-DEL-309 |
| chr10-101957485-DEL-639 | chr10-101957486-DEL-639 |
| chr10-102015524-DEL-483 | chr10-102015521-DEL-484 |
| chr10-102024959-DEL-1151 | chr10-102024960-DEL-1151 |
| chr10-102028440-DEL-2373 | chr10-102028441-DEL-2373 |
| chr10-102098211-DEL-121 | chr10-102098212-DEL-121 |
| chr10-102113053-DEL-1191 | chr10-102113054-DEL-1191 |
| chr10-102115290-DEL-106 | chr10-102115291-DEL-106 |
| chr10-102151384-DEL-6380 | chr10-102151385-DEL-6380 |
| chr10-102159436-DEL-6435 | chr10-102159437-DEL-6435 |
| chr10-102188979-DEL-99 | chr10-102188978-DEL-99 |
| chr10-102235405-DEL-72 | chr10-102235358-DEL-116 |
| chr10-102245463-DEL-51 | chr10-102245452-DEL-55 |
| chr10-102255271-DEL-195 | chr10-102255272-DEL-195 |
| chr10-102259226-DEL-876 | chr10-102259226-DEL-876 |

| | |
|---|---|
| chr10-102262696-DEL-6341 | chr10-102262702-DEL-6342 |
| chr10-102271123-DEL-332 | chr10-102271124-DEL-332 |
| chr10-102277585-DEL-65 | chr10-102277586-DEL-65 |
| chr10-102284207-DEL-107 | chr10-102284208-DEL-104 |
| chr10-102298110-DEL-4546 | chr10-102298111-DEL-4546 |
| chr10-102347169-DEL-83 | chr10-102347132-DEL-121 |
| chr10-102348847-DEL-97 | chr10-102348848-DEL-97 |
| chr10-102367238-DEL-234 | chr10-102367248-DEL-235 |
| chr10-102450331-DEL-231 | chr10-102450332-DEL-231 |
| chr10-102532296-DEL-940 | chr10-102532297-DEL-940 |
| chr10-102539976-DEL-88 | chr10-102539979-DEL-86 |
| chr10-102568864-DEL-651 | chr10-102568865-DEL-651 |
| chr10-102583214-DEL-353 | chr10-102583213-DEL-353 |
| chr10-102612049-DEL-71 | chr10-102612058-DEL-63 |
| chr10-102612667-DEL-53 | chr10-102612686-DEL-51 |
| chr10-102656965-DEL-80 | chr10-102656977-DEL-68 |
| chr10-102685931-DEL-399 | chr10-102685932-DEL-399 |
| chr10-102695650-DEL-2319 | chr10-102695651-DEL-2319 |
| chr10-102727080-DEL-1072 | chr10-102727081-DEL-1072 |
| chr10-102732831-DEL-71 | chr10-102732834-DEL-72 |
| chr10-102771540-DEL-359 | chr10-102771541-DEL-359 |
| chr10-102792416-DEL-54 | chr10-102792414-DEL-52 |
| chr10-102889414-DEL-55 | chr10-102889415-DEL-55 |
| chr10-102928612-DEL-52 | chr10-102928614-DEL-50 |
| chr10-102940507-DEL-52 | chr10-102940508-DEL-52 |
| chr10-102961751-DEL-92 | chr10-102961754-DEL-70 |
| chr10-102999535-DEL-224 | chr10-102999535-DEL-220 |
| chr10-103024391-DEL-165 | chr10-103024392-DEL-165 |
| chr10-103025504-DEL-78 | chr10-103025506-DEL-73 |
| chr10-103043365-DEL-6649 | chr10-103043366-DEL-6649 |
| chr10-103068773-DEL-179 | chr10-103068777-DEL-184 |
| chr10-103073359-DEL-3327 | chr10-103073360-DEL-3327 |
| chr10-103076777-DEL-93 | chr10-103076778-DEL-93 |
| chr10-103084699-DEL-207 | chr10-103084682-DEL-243 |
| chr10-103086216-DEL-59 | chr10-103086216-DEL-59 |
| chr10-103109849-DEL-213 | chr10-103109850-DEL-213 |
| chr10-103126133-DEL-1292 | chr10-103126134-DEL-1292 |
| chr10-103133788-DEL-101 | chr10-103133789-DEL-101 |
| chr10-103137810-DEL-79 | chr10-103137807-DEL-83 |
| chr10-103149228-DEL-166 | chr10-103149229-DEL-166 |
| chr10-103163054-DEL-148 | chr10-103163055-DEL-148 |
| chr10-103165780-DEL-646 | chr10-103165781-DEL-646 |
| chr10-103185898-DEL-230 | chr10-103185901-DEL-225 |
| chr10-103193420-DEL-1644 | chr10-103193421-DEL-1644 |
| chr10-103221094-DEL-62 | chr10-103221095-DEL-60 |
| chr10-103249808-DEL-206 | chr10-103249809-DEL-206 |
| chr10-103289816-DEL-158 | chr10-103289817-DEL-145 |
| chr10-103306243-DEL-770 | chr10-103306245-DEL-771 |
| chr10-103308506-DEL-923 | chr10-103308500-DEL-924 |
| chr10-103310874-DEL-174 | chr10-103310875-DEL-173 |
| chr10-103322780-DEL-123 | chr10-103322785-DEL-118 |
| chr10-103324640-DEL-224 | chr10-103324641-DEL-223 |
| chr10-103358648-DEL-1875 | chr10-103358640-DEL-1881 |
| chr10-103362831-DEL-430 | chr10-103362832-DEL-430 |
| chr10-103382723-DEL-1681 | chr10-103382724-DEL-1681 |
| chr10-103389065-DEL-109 | chr10-103389066-DEL-109 |
| chr10-103402963-DEL-390 | chr10-103402964-DEL-390 |
| chr10-103405248-DEL-6206 | chr10-103405248-DEL-6206 |
| chr10-103435915-DEL-1965 | chr10-103435916-DEL-1965 |
| chr10-103436176-DEL-195 | chr10-103436144-DEL-196 |
| chr10-103462050-DEL-62 | chr10-103462050-DEL-60 |
| chr10-103468301-DEL-3760 | chr10-103468301-DEL-3760 |
| chr10-103490929-DEL-6708 | chr10-103490930-DEL-6708 |
| chr10-103499246-DEL-168 | chr10-103499247-DEL-168 |
| chr10-103510188-DEL-1546 | chr10-103510189-DEL-1546 |
| chr10-103513808-DEL-52 | chr10-103513809-DEL-52 |
| chr10-103515762-DEL-145 | chr10-103515763-DEL-145 |
| chr10-103520438-DEL-201 | chr10-103520439-DEL-201 |
| chr10-103531584-DEL-118 | chr10-103531585-DEL-118 |
| chr10-103548645-DEL-74 | chr10-103548646-DEL-76 |
| chr10-103555960-DEL-96 | chr10-103555961-DEL-96 |
| chr10-103556819-DEL-249 | chr10-103556816-DEL-250 |
| chr10-103582788-DEL-165 | chr10-103582789-DEL-165 |
| chr10-103583749-DEL-6675 | chr10-103583750-DEL-6675 |
| chr10-103596774-DEL-1565 | chr10-103596775-DEL-1565 |
| chr10-103631727-DEL-202 | chr10-103631728-DEL-203 |
| chr10-103632837-DEL-2937 | chr10-103632838-DEL-2937 |
| chr10-103634902-DEL-304 | chr10-103634903-DEL-304 |
| chr10-103636768-DEL-63 | chr10-103636770-DEL-59 |
| chr10-103693995-DEL-498 | chr10-103693996-DEL-498 |

| | |
|---|---|
| chr10-103703359-DEL-6220 | chr10-103703361-DEL-6219 |
| chr10-103713765-DEL-225 | chr10-103713766-DEL-225 |
| chr10-103736787-DEL-2039 | chr10-103736788-DEL-2039 |
| chr10-103739329-DEL-1430 | chr10-103739330-DEL-1430 |
| chr10-103743182-DEL-467 | chr10-103743180-DEL-468 |
| chr10-103765866-DEL-177 | chr10-103765867-DEL-177 |
| chr10-103770969-DEL-642 | chr10-103770970-DEL-642 |
| chr10-103789587-DEL-100 | chr10-103789587-DEL-124 |
| chr10-103828967-DEL-4591 | chr10-103828953-DEL-4592 |
| chr10-103837953-DEL-60 | chr10-103837956-DEL-58 |
| chr10-103857958-DEL-69 | chr10-103857959-DEL-69 |
| chr10-103862042-DEL-1453 | chr10-103862040-DEL-1454 |
| chr10-103863907-DEL-53 | chr10-103863900-DEL-54 |
| chr10-103871235-DEL-1229 | chr10-103871236-DEL-1229 |
| chr10-103876291-DEL-383 | chr10-103876291-DEL-383 |
| chr10-103878206-DEL-58 | chr10-103878206-DEL-58 |
| chr10-103901831-DEL-102 | chr10-103901831-DEL-102 |
| chr10-103909467-DEL-2565 | chr10-103909468-DEL-2565 |
| chr10-103939057-DEL-68 | chr10-103939054-DEL-72 |
| chr10-104036777-DEL-352 | chr10-104036787-DEL-353 |
| chr10-104090536-DEL-4739 | chr10-104090534-DEL-4740 |
| chr10-104135116-DEL-5398 | chr10-104135116-DEL-5396 |
| chr10-104156238-DEL-70 | chr10-104156239-DEL-70 |
| chr10-104169032-DEL-191 | chr10-104169033-DEL-189 |
| chr10-104180654-DEL-66 | chr10-104180655-DEL-72 |
| chr10-104212283-DEL-199 | chr10-104212284-DEL-199 |
| chr10-104240516-DEL-324 | chr10-104240517-DEL-324 |
| chr10-104253646-DEL-1843 | chr10-104253648-DEL-1844 |
| chr10-104275236-DEL-52 | chr10-104275240-DEL-53 |
| chr10-104287368-DEL-3067 | chr10-104287368-DEL-3067 |
| chr10-104300748-DEL-73 | chr10-104300749-DEL-73 |
| chr10-104329474-DEL-6384 | chr10-104329475-DEL-6384 |
| chr10-104410860-DEL-2956 | chr10-104410861-DEL-2956 |
| chr10-104413037-DEL-81 | chr10-104413038-DEL-81 |
| chr10-104451688-DEL-6551 | chr10-104451689-DEL-6551 |
| chr10-104475704-DEL-62 | chr10-104475705-DEL-62 |
| chr10-104507417-DEL-231 | chr10-104507418-DEL-231 |
| chr10-104532361-DEL-58 | chr10-104532362-DEL-58 |
| chr10-104554531-DEL-997 | chr10-104554532-DEL-997 |
| chr10-104560733-DEL-2707 | chr10-104560733-DEL-2707 |
| chr10-104578192-DEL-187 | chr10-104578193-DEL-187 |
| chr10-104583946-DEL-6783 | chr10-104583947-DEL-6783 |
| chr10-104697095-DEL-235 | chr10-104697096-DEL-235 |
| chr10-104726106-DEL-6302 | chr10-104726107-DEL-6302 |
| chr10-104744995-DEL-122 | chr10-104744950-DEL-123 |
| chr10-104748230-DEL-55 | chr10-104748191-DEL-56 |
| chr10-104756558-DEL-135 | chr10-104756559-DEL-135 |
| chr10-104760671-DEL-116 | chr10-104760677-DEL-111 |
| chr10-104768596-DEL-104 | chr10-104768599-DEL-102 |
| chr10-104777800-DEL-66 | chr10-104777801-DEL-66 |
| chr10-104801956-DEL-136 | chr10-104801957-DEL-136 |
| chr10-104807037-DEL-96 | chr10-104807037-DEL-96 |
| chr10-104810558-DEL-1490 | chr10-104810559-DEL-1490 |
| chr10-104831175-DEL-6485 | chr10-104831176-DEL-6485 |
| chr10-104845814-DEL-1303 | chr10-104845815-DEL-1303 |
| chr10-104857616-DEL-373 | chr10-104857617-DEL-373 |
| chr10-104863319-DEL-88 | chr10-104863318-DEL-54 |
| chr10-104874632-DEL-172 | chr10-104874633-DEL-172 |
| chr10-104882578-DEL-3137 | chr10-104882579-DEL-3137 |
| chr10-104904910-DEL-92 | chr10-104904910-DEL-92 |
| chr10-104908711-DEL-1709 | chr10-104908712-DEL-1709 |
| chr10-104911543-DEL-66 | chr10-104911544-DEL-66 |
| chr10-104924439-DEL-66 | chr10-104924441-DEL-65 |
| chr10-104945344-DEL-56 | chr10-104945345-DEL-56 |
| chr10-104967020-DEL-470 | chr10-104967021-DEL-470 |
| chr10-104969824-DEL-135 | chr10-104969825-DEL-134 |
| chr10-104971419-DEL-53 | chr10-104971420-DEL-53 |
| chr10-104984601-DEL-1165 | chr10-104984594-DEL-1168 |
| chr10-104993288-DEL-101 | chr10-104993256-DEL-102 |
| chr10-105012975-DEL-4680 | chr10-105012976-DEL-4680 |
| chr10-105083403-DEL-115 | chr10-105083404-DEL-115 |
| chr10-105084185-DEL-456 | chr10-105084212-DEL-446 |
| chr10-105092558-DEL-914 | chr10-105092559-DEL-914 |
| chr10-105164912-DEL-110 | chr10-105164905-DEL-100 |
| chr10-105197756-DEL-56 | chr10-105197757-DEL-56 |
| chr10-105227212-DEL-191 | chr10-105227213-DEL-191 |
| chr10-105251599-DEL-56 | chr10-105251606-DEL-50 |
| chr10-105262986-DEL-60 | chr10-105262986-DEL-60 |
| chr10-105295140-DEL-58 | chr10-105295131-DEL-68 |
| chr10-105348530-DEL-92 | chr10-105348531-DEL-92 |

| | |
|---|---|
| chr10-105367906-DEL-82 | chr10-105367907-DEL-82 |
| chr10-105376712-DEL-102 | chr10-105376713-DEL-82 |
| chr10-105394633-DEL-999 | chr10-105394617-DEL-1015 |
| chr10-105415672-DEL-1000 | chr10-105415673-DEL-1000 |
| chr10-105427385-DEL-434 | chr10-105427386-DEL-434 |
| chr10-105440380-DEL-995 | chr10-105440380-DEL-995 |
| chr10-105454267-DEL-2579 | chr10-105454268-DEL-2579 |
| chr10-105464970-DEL-830 | chr10-105464971-DEL-830 |
| chr10-105471650-DEL-61 | chr10-105471651-DEL-61 |
| chr10-105486414-DEL-414 | chr10-105486427-DEL-404 |
| chr10-105497127-DEL-200 | chr10-105497129-DEL-199 |
| chr10-105504472-DEL-64 | chr10-105504473-DEL-64 |
| chr10-105508456-DEL-5338 | chr10-105508457-DEL-5338 |
| chr10-105523971-DEL-1493 | chr10-105523972-DEL-1493 |
| chr10-105539459-DEL-58 | chr10-105539460-DEL-58 |
| chr10-105549273-DEL-215 | chr10-105549274-DEL-215 |
| chr10-105564774-DEL-106 | chr10-105564775-DEL-106 |
| chr10-105565758-DEL-446 | chr10-105565762-DEL-443 |
| chr10-105568725-DEL-259 | chr10-105568738-DEL-247 |
| chr10-105581448-DEL-173 | chr10-105581449-DEL-173 |
| chr10-105635651-DEL-414 | chr10-105635652-DEL-414 |
| chr10-105644891-DEL-6409 | chr10-105644892-DEL-6409 |
| chr10-105656400-DEL-242 | chr10-105656403-DEL-239 |
| chr10-105674414-DEL-1505 | chr10-105674415-DEL-1505 |
| chr10-105681894-DEL-91 | chr10-105681869-DEL-88 |
| chr10-105710231-DEL-185 | chr10-105710232-DEL-185 |
| chr10-105722881-DEL-1512 | chr10-105722882-DEL-1512 |
| chr10-105725755-DEL-851 | chr10-105725753-DEL-852 |
| chr10-105754696-DEL-6818 | chr10-105754692-DEL-6823 |
| chr10-105767065-DEL-829 | chr10-105767066-DEL-829 |
| chr10-105772057-DEL-82 | chr10-105772085-DEL-54 |
| chr10-105782249-DEL-69 | chr10-105782250-DEL-69 |
| chr10-105782656-DEL-554 | chr10-105782657-DEL-549 |
| chr10-105830152-DEL-99 | chr10-105830170-DEL-65 |
| chr10-105848991-DEL-1442 | chr10-105848977-DEL-1445 |
| chr10-105876630-DEL-212 | chr10-105876629-DEL-212 |
| chr10-105879343-DEL-51 | chr10-105879344-DEL-51 |
| chr10-105913197-DEL-143 | chr10-105913198-DEL-143 |
| chr10-105916013-DEL-6533 | chr10-105916014-DEL-6533 |
| chr10-105923689-DEL-6379 | chr10-105923690-DEL-6379 |
| chr10-105946401-DEL-65 | chr10-105946402-DEL-65 |
| chr10-105974538-DEL-258 | chr10-105974525-DEL-258 |
| chr10-105981437-DEL-124 | chr10-105981438-DEL-124 |
| chr10-106018503-DEL-5644 | chr10-106018504-DEL-5644 |
| chr10-106024741-DEL-3213 | chr10-106024742-DEL-3213 |
| chr10-106035358-DEL-175 | chr10-106035359-DEL-175 |
| chr10-106042366-DEL-7074 | chr10-106042367-DEL-7073 |
| chr10-106042382-DEL-424 | chr10-106042382-DEL-424 |
| chr10-106066869-DEL-79 | chr10-106066870-DEL-79 |
| chr10-106068198-DEL-3404 | chr10-106068199-DEL-3404 |
| chr10-106077469-DEL-510 | chr10-106077502-DEL-472 |
| chr10-106104922-DEL-4194 | chr10-106104934-DEL-4199 |
| chr10-106108509-DEL-74 | chr10-106108498-DEL-84 |
| chr10-106109288-DEL-65 | chr10-106109289-DEL-65 |
| chr10-106116838-DEL-90 | chr10-106116815-DEL-126 |
| chr10-106121463-DEL-560 | chr10-106121464-DEL-559 |
| chr10-106129655-DEL-71 | chr10-106129673-DEL-51 |
| chr10-106148853-DEL-6379 | chr10-106148854-DEL-6379 |
| chr10-106159492-DEL-63 | chr10-106159493-DEL-61 |
| chr10-106182221-DEL-57 | chr10-106182222-DEL-57 |
| chr10-106217068-DEL-460 | chr10-106217079-DEL-461 |
| chr10-106226483-DEL-12458 | chr10-106226484-DEL-12458 |
| chr10-106250480-DEL-62 | chr10-106250481-DEL-50 |
| chr10-106257999-DEL-128 | chr10-106258000-DEL-128 |
| chr10-106271929-DEL-6630 | chr10-106271930-DEL-6630 |
| chr10-106302559-DEL-73 | chr10-106302559-DEL-68 |
| chr10-106326779-DEL-53 | chr10-106326780-DEL-52 |
| chr10-106379771-DEL-396 | chr10-106379772-DEL-396 |
| chr10-106444850-DEL-58 | chr10-106444851-DEL-58 |
| chr10-106492295-DEL-301 | chr10-106492295-DEL-301 |
| chr10-106504038-DEL-70 | chr10-106504039-DEL-70 |
| chr10-106519203-DEL-681 | chr10-106519204-DEL-680 |
| chr10-106531413-DEL-84 | chr10-106531414-DEL-84 |
| chr10-106549290-DEL-238 | chr10-106549293-DEL-236 |
| chr10-106552844-DEL-74 | chr10-106552857-DEL-72 |
| chr10-106560504-DEL-186 | chr10-106560505-DEL-184 |
| chr10-106585472-DEL-80 | chr10-106585473-DEL-85 |
| chr10-106612811-DEL-492 | chr10-106612811-DEL-492 |
| chr10-106614560-DEL-506 | chr10-106614560-DEL-506 |
| chr10-106621155-DEL-1744 | chr10-106621156-DEL-1744 |

| | |
|---|---|
| chr10-106621739-DEL-77 | chr10-106621740-DEL-77 |
| chr10-106646145-DEL-155 | chr10-106646138-DEL-155 |
| chr10-106652082-DEL-87 | chr10-106652083-DEL-87 |
| chr10-106657489-DEL-4634 | chr10-106657490-DEL-4634 |
| chr10-106678811-DEL-4821 | chr10-106678812-DEL-4821 |
| chr10-106687784-DEL-385 | chr10-106687781-DEL-385 |
| chr10-106715541-DEL-84 | chr10-106715542-DEL-84 |
| chr10-106779700-DEL-56 | chr10-106779703-DEL-54 |
| chr10-106804496-DEL-1367 | chr10-106804497-DEL-1364 |
| chr10-106807769-DEL-1128 | chr10-106807770-DEL-1128 |
| chr10-106817183-DEL-109 | chr10-106817184-DEL-109 |
| chr10-106838192-DEL-142 | chr10-106838193-DEL-120 |
| chr10-106841510-DEL-104 | chr10-106841505-DEL-98 |
| chr10-106842328-DEL-104 | chr10-106842329-DEL-100 |
| chr10-106852523-DEL-67 | chr10-106852524-DEL-51 |
| chr10-106862578-DEL-123 | chr10-106862579-DEL-123 |
| chr10-106864523-DEL-547 | chr10-106864524-DEL-547 |
| chr10-106865336-DEL-163 | chr10-106865337-DEL-163 |
| chr10-106868070-DEL-57 | chr10-106868071-DEL-57 |
| chr10-106922778-DEL-113 | chr10-106922797-DEL-95 |
| chr10-106936273-DEL-3160 | chr10-106936274-DEL-3160 |
| chr10-107035676-DEL-168 | chr10-107035677-DEL-165 |
| chr10-107042714-DEL-68 | chr10-107042715-DEL-68 |
| chr10-107053126-DEL-72 | chr10-107053127-DEL-73 |
| chr10-107062361-DEL-2215 | chr10-107062362-DEL-2215 |
| chr10-107072432-DEL-289 | chr10-107072433-DEL-289 |
| chr10-107091837-DEL-390 | chr10-107091838-DEL-390 |
| chr10-107095269-DEL-70 | chr10-107095270-DEL-70 |
| chr10-107129643-DEL-64 | chr10-107129644-DEL-64 |
| chr10-107162510-DEL-6545 | chr10-107162511-DEL-6545 |
| chr10-107193275-DEL-376 | chr10-107193277-DEL-375 |
| chr10-107202324-DEL-6549 | chr10-107202325-DEL-6549 |
| chr10-107202340-DEL-204 | chr10-107202341-DEL-204 |
| chr10-107211346-DEL-838 | chr10-107211347-DEL-838 |
| chr10-107220182-DEL-506 | chr10-107220183-DEL-506 |
| chr10-107280543-DEL-2918 | chr10-107280544-DEL-2918 |
| chr10-107282606-DEL-357 | chr10-107282606-DEL-357 |
| chr10-107299225-DEL-2517 | chr10-107299226-DEL-2517 |
| chr10-107345349-DEL-195 | chr10-107345350-DEL-195 |
| chr10-107376078-DEL-76 | chr10-107376079-DEL-74 |
| chr10-107390408-DEL-6241 | chr10-107390408-DEL-6241 |
| chr10-107400779-DEL-76 | chr10-107400780-DEL-76 |
| chr10-107431500-DEL-334 | chr10-107431501-DEL-333 |
| chr10-107439379-DEL-204 | chr10-107439396-DEL-184 |
| chr10-107445570-DEL-178 | chr10-107445569-DEL-180 |
| chr10-107489854-DEL-73 | chr10-107489855-DEL-71 |
| chr10-107493443-DEL-357 | chr10-107493439-DEL-358 |
| chr10-107493970-DEL-160 | chr10-107493971-DEL-160 |
| chr10-107505672-DEL-171 | chr10-107505695-DEL-129 |
| chr10-107511298-DEL-1050 | chr10-107511299-DEL-1049 |
| chr10-107514865-DEL-71 | chr10-107514878-DEL-77 |
| chr10-107599188-DEL-1115 | chr10-107599189-DEL-1115 |
| chr10-107623593-DEL-303 | chr10-107623594-DEL-303 |
| chr10-107631613-DEL-173 | chr10-107631613-DEL-171 |
| chr10-107643556-DEL-91 | chr10-107643551-DEL-85 |
| chr10-107648950-DEL-102 | chr10-107648951-DEL-102 |
| chr10-107659239-DEL-100 | chr10-107659240-DEL-100 |
| chr10-107666071-DEL-239 | chr10-107666072-DEL-239 |
| chr10-107675260-DEL-60 | chr10-107675284-DEL-53 |
| chr10-107681214-DEL-7632 | chr10-107681215-DEL-7632 |
| chr10-107711548-DEL-76 | chr10-107711549-DEL-82 |
| chr10-107771754-DEL-69 | chr10-107771758-DEL-68 |
| chr10-107780521-DEL-68 | chr10-107780522-DEL-68 |
| chr10-107786677-DEL-336 | chr10-107786678-DEL-336 |
| chr10-107787240-DEL-60 | chr10-107787241-DEL-60 |
| chr10-107787786-DEL-243 | chr10-107787787-DEL-243 |
| chr10-107791665-DEL-393 | chr10-107791666-DEL-393 |
| chr10-107802030-DEL-130 | chr10-107802031-DEL-130 |
| chr10-107814771-DEL-2281 | chr10-107814772-DEL-2281 |
| chr10-107829780-DEL-126 | chr10-107829781-DEL-126 |
| chr10-107835337-DEL-7162 | chr10-107835355-DEL-7151 |
| chr10-107880944-DEL-689 | chr10-107880950-DEL-688 |
| chr10-107888666-DEL-221 | chr10-107888667-DEL-221 |
| chr10-107930135-DEL-3236 | chr10-107930136-DEL-3236 |
| chr10-107939658-DEL-60 | chr10-107939659-DEL-62 |
| chr10-107952550-DEL-67 | chr10-107952551-DEL-67 |
| chr10-108024980-DEL-52 | chr10-108024981-DEL-51 |
| chr10-108031054-DEL-206 | chr10-108031055-DEL-206 |
| chr10-108051770-DEL-131 | chr10-108051771-DEL-131 |
| chr10-108065532-DEL-112 | chr10-108065571-DEL-74 |

| | |
|---|---|
| chr10-108100642-DEL-52 | chr10-108100643-DEL-52 |
| chr10-108116899-DEL-218 | chr10-108116900-DEL-218 |
| chr10-108129391-DEL-207 | chr10-108129392-DEL-207 |
| chr10-108130023-DEL-195 | chr10-108130024-DEL-195 |
| chr10-108140225-DEL-507 | chr10-108140226-DEL-507 |
| chr10-108142404-DEL-219 | chr10-108142405-DEL-219 |
| chr10-108153061-DEL-87 | chr10-108153056-DEL-99 |
| chr10-108163407-DEL-635 | chr10-108163408-DEL-635 |
| chr10-108194454-DEL-175 | chr10-108194444-DEL-176 |
| chr10-108229434-DEL-232 | chr10-108229435-DEL-232 |
| chr10-108233211-DEL-206 | chr10-108233212-DEL-206 |
| chr10-108238179-DEL-84 | chr10-108238180-DEL-84 |
| chr10-108248202-DEL-211 | chr10-108248203-DEL-211 |
| chr10-108294853-DEL-67 | chr10-108294841-DEL-56 |
| chr10-108323375-DEL-72 | chr10-108323376-DEL-72 |
| chr10-108324795-DEL-92 | chr10-108324796-DEL-92 |
| chr10-108329570-DEL-69 | chr10-108329577-DEL-57 |
| chr10-108364985-DEL-65 | chr10-108364986-DEL-50 |
| chr10-108402112-DEL-191 | chr10-108402113-DEL-191 |
| chr10-108420598-DEL-83 | chr10-108420599-DEL-83 |
| chr10-108434738-DEL-657 | chr10-108434739-DEL-657 |
| chr10-108452267-DEL-77 | chr10-108452252-DEL-82 |
| chr10-108493947-DEL-182 | chr10-108493948-DEL-182 |
| chr10-108530133-DEL-748 | chr10-108530134-DEL-749 |
| chr10-108611557-DEL-58 | chr10-108611557-DEL-58 |
| chr10-108630418-DEL-85 | chr10-108630419-DEL-85 |
| chr10-108729357-DEL-61 | chr10-108729310-DEL-60 |
| chr10-108733296-DEL-441 | chr10-108733297-DEL-441 |
| chr10-108748630-DEL-52 | chr10-108748631-DEL-52 |
| chr10-108787589-DEL-116 | chr10-108787589-DEL-116 |
| chr10-108809181-DEL-113 | chr10-108809182-DEL-113 |
| chr10-108822706-DEL-215 | chr10-108822707-DEL-215 |
| chr10-108823826-DEL-624 | chr10-108823820-DEL-625 |
| chr10-108849670-DEL-65 | chr10-108849671-DEL-65 |
| chr10-108919485-DEL-6353 | chr10-108919485-DEL-6365 |
| chr10-108968757-DEL-51 | chr10-108968750-DEL-52 |
| chr10-108983494-DEL-737 | chr10-108983495-DEL-737 |
| chr10-109048376-DEL-6159 | chr10-109048377-DEL-6159 |
| chr10-109151085-DEL-7090 | chr10-109151086-DEL-7090 |
| chr10-109200880-DEL-201 | chr10-109200882-DEL-201 |
| chr10-109208992-DEL-127 | chr10-109208978-DEL-118 |
| chr10-109228748-DEL-56 | chr10-109228700-DEL-105 |
| chr10-109243208-DEL-6220 | chr10-109243206-DEL-6221 |
| chr10-109265486-DEL-325 | chr10-109265487-DEL-325 |
| chr10-109287770-DEL-66 | chr10-109287771-DEL-66 |
| chr10-109289459-DEL-70 | chr10-109289437-DEL-58 |
| chr10-109297105-DEL-2324 | chr10-109297104-DEL-2325 |
| chr10-109317141-DEL-118 | chr10-109317142-DEL-118 |
| chr10-109322122-DEL-64 | chr10-109322127-DEL-60 |
| chr10-109358645-DEL-1127 | chr10-109358646-DEL-1127 |
| chr10-109358726-DEL-446 | chr10-109358727-DEL-446 |
| chr10-109364859-DEL-215 | chr10-109364860-DEL-215 |
| chr10-109370129-DEL-284 | chr10-109370130-DEL-284 |
| chr10-109370915-DEL-1888 | chr10-109370916-DEL-1888 |
| chr10-109377523-DEL-400 | chr10-109377524-DEL-400 |
| chr10-109379168-DEL-62 | chr10-109379163-DEL-52 |
| chr10-109407099-DEL-76 | chr10-109407097-DEL-76 |
| chr10-109422838-DEL-208 | chr10-109422839-DEL-208 |
| chr10-109471675-DEL-862 | chr10-109471676-DEL-862 |
| chr10-109504795-DEL-113 | chr10-109504796-DEL-113 |
| chr10-109523657-DEL-238 | chr10-109523658-DEL-238 |
| chr10-109566547-DEL-55 | chr10-109566540-DEL-56 |
| chr10-109568707-DEL-343 | chr10-109568708-DEL-343 |
| chr10-109573792-DEL-92 | chr10-109573791-DEL-91 |
| chr10-109599800-DEL-107 | chr10-109599801-DEL-107 |
| chr10-109609815-DEL-1337 | chr10-109609816-DEL-1337 |
| chr10-109622535-DEL-76 | chr10-109622550-DEL-50 |
| chr10-109702263-DEL-71 | chr10-109702265-DEL-70 |
| chr10-109773005-DEL-616 | chr10-109773005-DEL-616 |
| chr10-109823055-DEL-92 | chr10-109823056-DEL-92 |
| chr10-109844821-DEL-7711 | chr10-109844822-DEL-7711 |
| chr10-109861434-DEL-53 | chr10-109861435-DEL-53 |
| chr10-109863646-DEL-236 | chr10-109863647-DEL-236 |
| chr10-109890035-DEL-1856 | chr10-109890036-DEL-1856 |
| chr10-109898953-DEL-74 | chr10-109898928-DEL-111 |
| chr10-109918283-DEL-400 | chr10-109918284-DEL-400 |
| chr10-109939282-DEL-322 | chr10-109939283-DEL-322 |
| chr10-109939943-DEL-212 | chr10-109939938-DEL-220 |
| chr10-109940696-DEL-6439 | chr10-109940695-DEL-6440 |
| chr10-109956325-DEL-75 | chr10-109956323-DEL-84 |

| | |
|---|---|
| chr10-109959771-DEL-150 | chr10-109959772-DEL-150 |
| chr10-109962481-DEL-64 | chr10-109962465-DEL-82 |
| chr10-109977426-DEL-301 | chr10-109977427-DEL-301 |
| chr10-110013384-DEL-72 | chr10-110013385-DEL-72 |
| chr10-110033822-DEL-1338 | chr10-110033823-DEL-1338 |
| chr10-110043676-DEL-890 | chr10-110043677-DEL-892 |
| chr10-110055274-DEL-359 | chr10-110055275-DEL-359 |
| chr10-110058236-DEL-290 | chr10-110058237-DEL-298 |
| chr10-110060763-DEL-6414 | chr10-110060764-DEL-6414 |
| chr10-110116747-DEL-58 | chr10-110116748-DEL-58 |
| chr10-110164510-DEL-739 | chr10-110164511-DEL-739 |
| chr10-110182717-DEL-54 | chr10-110182721-DEL-51 |
| chr10-110200171-DEL-6390 | chr10-110200167-DEL-6391 |
| chr10-110214018-DEL-450 | chr10-110214019-DEL-445 |
| chr10-110247573-DEL-845 | chr10-110247574-DEL-845 |
| chr10-110259159-DEL-1233 | chr10-110259160-DEL-1232 |
| chr10-110262948-DEL-951 | chr10-110262949-DEL-951 |
| chr10-110273590-DEL-57 | chr10-110273590-DEL-57 |
| chr10-110285381-DEL-55 | chr10-110285381-DEL-55 |
| chr10-110353324-DEL-1351 | chr10-110353309-DEL-1366 |
| chr10-110382604-DEL-505 | chr10-110382603-DEL-506 |
| chr10-110389266-DEL-54 | chr10-110389267-DEL-54 |
| chr10-110391552-DEL-706 | chr10-110391552-DEL-706 |
| chr10-110396912-DEL-54 | chr10-110396916-DEL-50 |
| chr10-110408674-DEL-70 | chr10-110408675-DEL-70 |
| chr10-110427121-DEL-1458 | chr10-110427122-DEL-1458 |
| chr10-110462607-DEL-7453 | chr10-110462608-DEL-7453 |
| chr10-110473855-DEL-71 | chr10-110473856-DEL-71 |
| chr10-110487974-DEL-498 | chr10-110487975-DEL-498 |
| chr10-110519617-DEL-6464 | chr10-110519618-DEL-6464 |
| chr10-110527806-DEL-63 | chr10-110527807-DEL-63 |
| chr10-110532142-DEL-4518 | chr10-110532143-DEL-4518 |
| chr10-110544886-DEL-213 | chr10-110544887-DEL-213 |
| chr10-110545476-DEL-142 | chr10-110545476-DEL-142 |
| chr10-110546898-DEL-219 | chr10-110546903-DEL-211 |
| chr10-110586150-DEL-65 | chr10-110586145-DEL-71 |
| chr10-110606283-DEL-2875 | chr10-110606284-DEL-2875 |
| chr10-110638218-DEL-186 | chr10-110638219-DEL-186 |
| chr10-110655978-DEL-52 | chr10-110655985-DEL-76 |
| chr10-110657008-DEL-225 | chr10-110657026-DEL-218 |
| chr10-110682727-DEL-7016 | chr10-110682728-DEL-7016 |
| chr10-110708912-DEL-86 | chr10-110708947-DEL-52 |
| chr10-110718451-DEL-64 | chr10-110718452-DEL-64 |
| chr10-110734428-DEL-53 | chr10-110734429-DEL-53 |
| chr10-110737277-DEL-54 | chr10-110737278-DEL-54 |
| chr10-110747255-DEL-291 | chr10-110747286-DEL-289 |
| chr10-110769658-DEL-76 | chr10-110769659-DEL-74 |
| chr10-110772953-DEL-127 | chr10-110772942-DEL-128 |
| chr10-110775556-DEL-93 | chr10-110775557-DEL-93 |
| chr10-110795181-DEL-132 | chr10-110795182-DEL-132 |
| chr10-110803026-DEL-102 | chr10-110802982-DEL-103 |
| chr10-110814006-DEL-70 | chr10-110814007-DEL-70 |
| chr10-110835847-DEL-2938 | chr10-110835846-DEL-2939 |
| chr10-110880351-DEL-194 | chr10-110880352-DEL-194 |
| chr10-110891940-DEL-118 | chr10-110891941-DEL-118 |
| chr10-110920950-DEL-61 | chr10-110920951-DEL-61 |
| chr10-110930160-DEL-60 | chr10-110930161-DEL-56 |
| chr10-110944513-DEL-62 | chr10-110944514-DEL-62 |
| chr10-110949719-DEL-601 | chr10-110949720-DEL-601 |
| chr10-110961631-DEL-951 | chr10-110961632-DEL-951 |
| chr10-110987181-DEL-1829 | chr10-110987183-DEL-1828 |
| chr10-110991815-DEL-62 | chr10-110991803-DEL-62 |
| chr10-111014429-DEL-568 | chr10-111014430-DEL-568 |
| chr10-111024420-DEL-71 | chr10-111024421-DEL-70 |
| chr10-111039860-DEL-129 | chr10-111039862-DEL-128 |
| chr10-111065343-DEL-930 | chr10-111065344-DEL-930 |
| chr10-111068341-DEL-204 | chr10-111068340-DEL-224 |
| chr10-111071765-DEL-266 | chr10-111071766-DEL-266 |
| chr10-111097864-DEL-137 | chr10-111097865-DEL-137 |
| chr10-111109735-DEL-223 | chr10-111109736-DEL-223 |
| chr10-111119877-DEL-219 | chr10-111119878-DEL-219 |
| chr10-111152288-DEL-84 | chr10-111152259-DEL-117 |
| chr10-111160938-DEL-396 | chr10-111160938-DEL-396 |
| chr10-111187079-DEL-54 | chr10-111187079-DEL-54 |
| chr10-111189599-DEL-198 | chr10-111189599-DEL-198 |
| chr10-111217552-DEL-725 | chr10-111217553-DEL-725 |
| chr10-111223418-DEL-178 | chr10-111223419-DEL-178 |
| chr10-111261797-DEL-1314 | chr10-111261798-DEL-1314 |
| chr10-111282704-DEL-192 | chr10-111282705-DEL-191 |
| chr10-111314801-DEL-511 | chr10-111314802-DEL-511 |

| | |
|---|---|
| chr10-111348891-DEL-655 | chr10-111348892-DEL-655 |
| chr10-111354777-DEL-65 | chr10-111354809-DEL-56 |
| chr10-111404000-DEL-181 | chr10-111404001-DEL-181 |
| chr10-111420477-DEL-257 | chr10-111420522-DEL-207 |
| chr10-111426503-DEL-58 | chr10-111426523-DEL-50 |
| chr10-111446965-DEL-338 | chr10-111446965-DEL-339 |
| chr10-111457058-DEL-144 | chr10-111457059-DEL-144 |
| chr10-111463952-DEL-127 | chr10-111463953-DEL-127 |
| chr10-111489624-DEL-210 | chr10-111489625-DEL-210 |
| chr10-111504673-DEL-74 | chr10-111504674-DEL-74 |
| chr10-111548844-DEL-74 | chr10-111548845-DEL-74 |
| chr10-111568063-DEL-76 | chr10-111568064-DEL-75 |
| chr10-111576054-DEL-194 | chr10-111576055-DEL-194 |
| chr10-111583990-DEL-83 | chr10-111583990-DEL-83 |
| chr10-111597566-DEL-1153 | chr10-111597567-DEL-1153 |
| chr10-111619093-DEL-218 | chr10-111619094-DEL-218 |
| chr10-111631144-DEL-60 | chr10-111631145-DEL-60 |
| chr10-111633685-DEL-116 | chr10-111633686-DEL-158 |
| chr10-111644115-DEL-132 | chr10-111644117-DEL-127 |
| chr10-111715081-DEL-328 | chr10-111715082-DEL-328 |
| chr10-111715931-DEL-51 | chr10-111715932-DEL-51 |
| chr10-111747884-DEL-504 | chr10-111747881-DEL-505 |
| chr10-111751542-DEL-252 | chr10-111751543-DEL-252 |
| chr10-111756371-DEL-98 | chr10-111756379-DEL-91 |
| chr10-111756870-DEL-177 | chr10-111756869-DEL-179 |
| chr10-111767968-DEL-305 | chr10-111767969-DEL-305 |
| chr10-111777489-DEL-80 | chr10-111777513-DEL-79 |
| chr10-111799048-DEL-227 | chr10-111799049-DEL-227 |
| chr10-111814440-DEL-255 | chr10-111814400-DEL-256 |
| chr10-111833282-DEL-52 | chr10-111833264-DEL-53 |
| chr10-111859901-DEL-81 | chr10-111859903-DEL-55 |
| chr10-111881218-DEL-906 | chr10-111881202-DEL-906 |
| chr10-111889412-DEL-4891 | chr10-111889413-DEL-4891 |
| chr10-111896866-DEL-473 | chr10-111896864-DEL-471 |
| chr10-111923753-DEL-1727 | chr10-111923751-DEL-1729 |
| chr10-111958698-DEL-1123 | chr10-111958697-DEL-1119 |
| chr10-111975385-DEL-53 | chr10-111975386-DEL-53 |
| chr10-111984390-DEL-4613 | chr10-111984391-DEL-4613 |
| chr10-112030101-DEL-609 | chr10-112030101-DEL-609 |
| chr10-112053218-DEL-1545 | chr10-112053218-DEL-1545 |
| chr10-112077884-DEL-730 | chr10-112077873-DEL-738 |
| chr10-112143212-DEL-198 | chr10-112143213-DEL-198 |
| chr10-112149225-DEL-58 | chr10-112149226-DEL-58 |
| chr10-112209188-DEL-11293 | chr10-112209189-DEL-11293 |
| chr10-112219274-DEL-391 | chr10-112219275-DEL-391 |
| chr10-112223614-DEL-147 | chr10-112223615-DEL-147 |
| chr10-112260963-DEL-6913 | chr10-112260964-DEL-6913 |
| chr10-112261334-DEL-212 | chr10-112261335-DEL-212 |
| chr10-112271855-DEL-677 | chr10-112271856-DEL-677 |
| chr10-112277494-DEL-114 | chr10-112277476-DEL-132 |
| chr10-112311642-DEL-80 | chr10-112311609-DEL-78 |
| chr10-112325255-DEL-743 | chr10-112325245-DEL-744 |
| chr10-112341904-DEL-6397 | chr10-112341905-DEL-6397 |
| chr10-112348545-DEL-6173 | chr10-112348545-DEL-6173 |
| chr10-112360127-DEL-395 | chr10-112360123-DEL-393 |
| chr10-112380865-DEL-72 | chr10-112380866-DEL-66 |
| chr10-112397910-DEL-68 | chr10-112397960-DEL-62 |
| chr10-112422902-DEL-6652 | chr10-112422903-DEL-6652 |
| chr10-112458037-DEL-186 | chr10-112458033-DEL-183 |
| chr10-112475809-DEL-99 | chr10-112475810-DEL-99 |
| chr10-112499940-DEL-75 | chr10-112499898-DEL-117 |
| chr10-112504528-DEL-82 | chr10-112504536-DEL-56 |
| chr10-112511916-DEL-517 | chr10-112511917-DEL-512 |
| chr10-112512585-DEL-305 | chr10-112512586-DEL-305 |
| chr10-112534961-DEL-6865 | chr10-112534962-DEL-6865 |
| chr10-112546217-DEL-193 | chr10-112546218-DEL-193 |
| chr10-112547464-DEL-812 | chr10-112547464-DEL-793 |
| chr10-112555166-DEL-2827 | chr10-112555167-DEL-2827 |
| chr10-112572194-DEL-57 | chr10-112572196-DEL-56 |
| chr10-112573232-DEL-982 | chr10-112573233-DEL-979 |
| chr10-112579976-DEL-207 | chr10-112579977-DEL-198 |
| chr10-112621737-DEL-80 | chr10-112621704-DEL-110 |
| chr10-112640094-DEL-69 | chr10-112640095-DEL-69 |
| chr10-112674002-DEL-1494 | chr10-112674003-DEL-1494 |
| chr10-112701869-DEL-6514 | chr10-112701871-DEL-6513 |
| chr10-112731522-DEL-63 | chr10-112731516-DEL-64 |
| chr10-112742786-DEL-65 | chr10-112742792-DEL-60 |
| chr10-112753315-DEL-68 | chr10-112753356-DEL-58 |
| chr10-112774022-DEL-811 | chr10-112774023-DEL-811 |
| chr10-112823060-DEL-533 | chr10-112823061-DEL-533 |

| | |
|---|---|
| chr10-112825956-DEL-198 | chr10-112825957-DEL-198 |
| chr10-112866759-DEL-612 | chr10-112866760-DEL-612 |
| chr10-112867473-DEL-184 | chr10-112867474-DEL-184 |
| chr10-112876132-DEL-94 | chr10-112876129-DEL-98 |
| chr10-112878808-DEL-313 | chr10-112878809-DEL-313 |
| chr10-112894427-DEL-1647 | chr10-112894428-DEL-1647 |
| chr10-113041413-DEL-216 | chr10-113041414-DEL-216 |
| chr10-113108525-DEL-474 | chr10-113108526-DEL-474 |
| chr10-113110560-DEL-100 | chr10-113110561-DEL-100 |
| chr10-113115632-DEL-54 | chr10-113115637-DEL-50 |
| chr10-113116192-DEL-281 | chr10-113116193-DEL-281 |
| chr10-113127409-DEL-72 | chr10-113127410-DEL-74 |
| chr10-113154436-DEL-79 | chr10-113154437-DEL-79 |
| chr10-113218824-DEL-6431 | chr10-113218825-DEL-6431 |
| chr10-113285228-DEL-108 | chr10-113285235-DEL-102 |
| chr10-113317655-DEL-301 | chr10-113317656-DEL-301 |
| chr10-113401936-DEL-81 | chr10-113401937-DEL-81 |
| chr10-113409691-DEL-3355 | chr10-113409692-DEL-3355 |
| chr10-113414949-DEL-1474 | chr10-113414950-DEL-1474 |
| chr10-113426292-DEL-64 | chr10-113426297-DEL-60 |
| chr10-113514198-DEL-2031 | chr10-113514195-DEL-2032 |
| chr10-113530356-DEL-36082 | chr10-113530356-DEL-36082 |
| chr10-113578954-DEL-52 | chr10-113578955-DEL-54 |
| chr10-113604091-DEL-390 | chr10-113604092-DEL-390 |
| chr10-113611397-DEL-120 | chr10-113611398-DEL-120 |
| chr10-113618887-DEL-343 | chr10-113618888-DEL-343 |
| chr10-113629201-DEL-2306 | chr10-113629201-DEL-2306 |
| chr10-113632935-DEL-62 | chr10-113632936-DEL-62 |
| chr10-113654862-DEL-2163 | chr10-113654863-DEL-2163 |
| chr10-113660039-DEL-64 | chr10-113660040-DEL-64 |
| chr10-113691813-DEL-86 | chr10-113691857-DEL-90 |
| chr10-113700815-DEL-951 | chr10-113700816-DEL-951 |
| chr10-113732718-DEL-112 | chr10-113732718-DEL-112 |
| chr10-113743618-DEL-64 | chr10-113743619-DEL-64 |
| chr10-113748120-DEL-64 | chr10-113748117-DEL-76 |
| chr10-113748828-DEL-578 | chr10-113748829-DEL-578 |
| chr10-113756018-DEL-232 | chr10-113756019-DEL-232 |
| chr10-113756857-DEL-2278 | chr10-113756858-DEL-2278 |
| chr10-113760170-DEL-8261 | chr10-113760171-DEL-8261 |
| chr10-113805609-DEL-173 | chr10-113805610-DEL-173 |
| chr10-113807574-DEL-214 | chr10-113807575-DEL-214 |
| chr10-113843850-DEL-615 | chr10-113843851-DEL-615 |
| chr10-113856750-DEL-684 | chr10-113856751-DEL-684 |
| chr10-113891526-DEL-6391 | chr10-113891517-DEL-6392 |
| chr10-113903798-DEL-56 | chr10-113903803-DEL-54 |
| chr10-113927899-DEL-91 | chr10-113927900-DEL-91 |
| chr10-113971010-DEL-278 | chr10-113971011-DEL-278 |
| chr10-113993592-DEL-510 | chr10-113993592-DEL-510 |
| chr10-114002829-DEL-207 | chr10-114002830-DEL-207 |
| chr10-114009840-DEL-3208 | chr10-114009841-DEL-3208 |
| chr10-114028392-DEL-64 | chr10-114028393-DEL-64 |
| chr10-114038107-DEL-490 | chr10-114038108-DEL-490 |
| chr10-114038750-DEL-113 | chr10-114038751-DEL-113 |
| chr10-114068267-DEL-83 | chr10-114068275-DEL-91 |
| chr10-114100302-DEL-58 | chr10-114100303-DEL-58 |
| chr10-114101196-DEL-96 | chr10-114101209-DEL-84 |
| chr10-114111912-DEL-1546 | chr10-114111913-DEL-1546 |
| chr10-114165794-DEL-6481 | chr10-114165795-DEL-6481 |
| chr10-114258535-DEL-581 | chr10-114258536-DEL-581 |
| chr10-114286905-DEL-171 | chr10-114286906-DEL-141 |
| chr10-114306622-DEL-266 | chr10-114306623-DEL-266 |
| chr10-114319955-DEL-70 | chr10-114319964-DEL-66 |
| chr10-114368168-DEL-198 | chr10-114368169-DEL-198 |
| chr10-114371368-DEL-7011 | chr10-114371369-DEL-7011 |
| chr10-114371614-DEL-424 | chr10-114371615-DEL-424 |
| chr10-114398993-DEL-128 | chr10-114398992-DEL-126 |
| chr10-114399487-DEL-349 | chr10-114399493-DEL-348 |
| chr10-114400494-DEL-127 | chr10-114400495-DEL-127 |
| chr10-114441499-DEL-52 | chr10-114441500-DEL-52 |
| chr10-114463102-DEL-58 | chr10-114463103-DEL-58 |
| chr10-114496844-DEL-556 | chr10-114496845-DEL-556 |
| chr10-114573149-DEL-152 | chr10-114573150-DEL-134 |
| chr10-114598323-DEL-134 | chr10-114598315-DEL-135 |
| chr10-114624673-DEL-4324 | chr10-114624674-DEL-4324 |
| chr10-114631899-DEL-53 | chr10-114631901-DEL-52 |
| chr10-114640238-DEL-55 | chr10-114640236-DEL-84 |
| chr10-114653440-DEL-3783 | chr10-114653441-DEL-3783 |
| chr10-114663561-DEL-394 | chr10-114663560-DEL-394 |
| chr10-114672138-DEL-2489 | chr10-114672139-DEL-2489 |
| chr10-114684098-DEL-55 | chr10-114684091-DEL-63 |

| | |
|---|---|
| chr10-114696498-DEL-6380 | chr10-114696499-DEL-6380 |
| chr10-114725184-DEL-84 | chr10-114725189-DEL-80 |
| chr10-114730490-DEL-181 | chr10-114730491-DEL-181 |
| chr10-114735830-DEL-1296 | chr10-114735831-DEL-1296 |
| chr10-114754832-DEL-82 | chr10-114754833-DEL-82 |
| chr10-114756833-DEL-358 | chr10-114756834-DEL-358 |
| chr10-114793133-DEL-148 | chr10-114793142-DEL-140 |
| chr10-114834048-DEL-665 | chr10-114834049-DEL-665 |
| chr10-114835076-DEL-92 | chr10-114835085-DEL-84 |
| chr10-114866440-DEL-68 | chr10-114866441-DEL-68 |
| chr10-114872025-DEL-6662 | chr10-114872026-DEL-6662 |
| chr10-114878971-DEL-119 | chr10-114878972-DEL-121 |
| chr10-114889825-DEL-973 | chr10-114889826-DEL-970 |
| chr10-114897150-DEL-68 | chr10-114897151-DEL-68 |
| chr10-114901721-DEL-70 | chr10-114901721-DEL-71 |
| chr10-114909326-DEL-206 | chr10-114909327-DEL-206 |
| chr10-114943227-DEL-176 | chr10-114943228-DEL-176 |
| chr10-114945096-DEL-107 | chr10-114945097-DEL-107 |
| chr10-114950554-DEL-397 | chr10-114950555-DEL-397 |
| chr10-114972432-DEL-231 | chr10-114972433-DEL-231 |
| chr10-114976009-DEL-87 | chr10-114976010-DEL-87 |
| chr10-114988166-DEL-55 | chr10-114988167-DEL-55 |
| chr10-115006158-DEL-84 | chr10-115006165-DEL-70 |
| chr10-115025508-DEL-479 | chr10-115025508-DEL-476 |
| chr10-115048381-DEL-63 | chr10-115048381-DEL-63 |
| chr10-115058641-DEL-701 | chr10-115058643-DEL-700 |
| chr10-115075581-DEL-327 | chr10-115075582-DEL-327 |
| chr10-115088838-DEL-218 | chr10-115088839-DEL-218 |
| chr10-115105836-DEL-534 | chr10-115105837-DEL-534 |
| chr10-115113278-DEL-1315 | chr10-115113278-DEL-1315 |
| chr10-115117943-DEL-76 | chr10-115117944-DEL-76 |
| chr10-115121857-DEL-70 | chr10-115121843-DEL-60 |
| chr10-115133790-DEL-167 | chr10-115133791-DEL-167 |
| chr10-115166120-DEL-1289 | chr10-115166121-DEL-1289 |
| chr10-115167916-DEL-107 | chr10-115167927-DEL-77 |
| chr10-115178771-DEL-59 | chr10-115178772-DEL-59 |
| chr10-115182768-DEL-144 | chr10-115182769-DEL-144 |
| chr10-115190032-DEL-90 | chr10-115190033-DEL-90 |
| chr10-115211008-DEL-174 | chr10-115211009-DEL-174 |
| chr10-115265176-DEL-210 | chr10-115265177-DEL-210 |
| chr10-115271338-DEL-581 | chr10-115271330-DEL-548 |
| chr10-115333787-DEL-161 | chr10-115333788-DEL-165 |
| chr10-115334722-DEL-67 | chr10-115334723-DEL-67 |
| chr10-115337104-DEL-74 | chr10-115337096-DEL-83 |
| chr10-115359183-DEL-130 | chr10-115359188-DEL-126 |
| chr10-115363645-DEL-216 | chr10-115363646-DEL-215 |
| chr10-115368907-DEL-192 | chr10-115368907-DEL-148 |
| chr10-115369863-DEL-95 | chr10-115369863-DEL-96 |
| chr10-115371710-DEL-6453 | chr10-115371711-DEL-6453 |
| chr10-115383666-DEL-170 | chr10-115383671-DEL-163 |
| chr10-115410379-DEL-60 | chr10-115410380-DEL-60 |
| chr10-115435654-DEL-512 | chr10-115435655-DEL-512 |
| chr10-115490736-DEL-2637 | chr10-115490737-DEL-2637 |
| chr10-115525459-DEL-539 | chr10-115525460-DEL-539 |
| chr10-115535581-DEL-253 | chr10-115535582-DEL-253 |
| chr10-115540041-DEL-939 | chr10-115540042-DEL-939 |
| chr10-115568376-DEL-1026 | chr10-115568376-DEL-1026 |
| chr10-115620509-DEL-458 | chr10-115620510-DEL-458 |
| chr10-115622402-DEL-60 | chr10-115622401-DEL-62 |
| chr10-115635449-DEL-82 | chr10-115635450-DEL-82 |
| chr10-115644216-DEL-145 | chr10-115644247-DEL-115 |
| chr10-115651183-DEL-55 | chr10-115651196-DEL-72 |
| chr10-115656167-DEL-287 | chr10-115656168-DEL-287 |
| chr10-115673446-DEL-271 | chr10-115673447-DEL-271 |
| chr10-115676239-DEL-567 | chr10-115676237-DEL-568 |
| chr10-115692786-DEL-55 | chr10-115692787-DEL-55 |
| chr10-115740557-DEL-1180 | chr10-115740558-DEL-1180 |
| chr10-115752698-DEL-90 | chr10-115752699-DEL-90 |
| chr10-115761357-DEL-200 | chr10-115761358-DEL-200 |
| chr10-115785564-DEL-72 | chr10-115785565-DEL-72 |
| chr10-115785568-DEL-1199 | chr10-115785569-DEL-1199 |
| chr10-115796383-DEL-70 | chr10-115796389-DEL-78 |
| chr10-115847158-DEL-104 | chr10-115847170-DEL-100 |
| chr10-115860197-DEL-6240 | chr10-115860198-DEL-6240 |
| chr10-115887864-DEL-94 | chr10-115887882-DEL-54 |
| chr10-115901508-DEL-684 | chr10-115901501-DEL-691 |
| chr10-115904839-DEL-62 | chr10-115904836-DEL-56 |
| chr10-115910932-DEL-400 | chr10-115910933-DEL-401 |
| chr10-115925517-DEL-56 | chr10-115925518-DEL-56 |
| chr10-115957202-DEL-74 | chr10-115957211-DEL-65 |

| | |
|---|---|
| chr10-115973056-DEL-326 | chr10-115973057-DEL-326 |
| chr10-115997223-DEL-326 | chr10-115997224-DEL-326 |
| chr10-115998984-DEL-254 | chr10-115998985-DEL-254 |
| chr10-116006907-DEL-2152 | chr10-116006908-DEL-2152 |
| chr10-116049736-DEL-2505 | chr10-116049737-DEL-2505 |
| chr10-116054403-DEL-780 | chr10-116054404-DEL-780 |
| chr10-116060376-DEL-81 | chr10-116060361-DEL-97 |
| chr10-116060888-DEL-818 | chr10-116060889-DEL-818 |
| chr10-116066267-DEL-83 | chr10-116066268-DEL-83 |
| chr10-116076979-DEL-60 | chr10-116076980-DEL-88 |
| chr10-116082101-DEL-840 | chr10-116082102-DEL-840 |
| chr10-116084864-DEL-71 | chr10-116084865-DEL-70 |
| chr10-116114006-DEL-70 | chr10-116114005-DEL-50 |
| chr10-116124976-DEL-1115 | chr10-116124993-DEL-1108 |
| chr10-116172162-DEL-1925 | chr10-116172191-DEL-1882 |
| chr10-116178492-DEL-68 | chr10-116178494-DEL-80 |
| chr10-116183035-DEL-58 | chr10-116183036-DEL-58 |
| chr10-116199691-DEL-123 | chr10-116199679-DEL-126 |
| chr10-116207882-DEL-113 | chr10-116207883-DEL-113 |
| chr10-116209139-DEL-863 | chr10-116209140-DEL-863 |
| chr10-116230114-DEL-776 | chr10-116230115-DEL-776 |
| chr10-116266942-DEL-192 | chr10-116266953-DEL-176 |
| chr10-116301396-DEL-257 | chr10-116301397-DEL-257 |
| chr10-116325708-DEL-360 | chr10-116325709-DEL-405 |
| chr10-116360826-DEL-78 | chr10-116360827-DEL-78 |
| chr10-116393640-DEL-366 | chr10-116393638-DEL-364 |
| chr10-116419564-DEL-76 | chr10-116419565-DEL-76 |
| chr10-116425782-DEL-371 | chr10-116425783-DEL-371 |
| chr10-116442158-DEL-7123 | chr10-116442159-DEL-7123 |
| chr10-116470497-DEL-151 | chr10-116470498-DEL-151 |
| chr10-116487548-DEL-106 | chr10-116487549-DEL-66 |
| chr10-116496945-DEL-66 | chr10-116496942-DEL-63 |
| chr10-116501843-DEL-53 | chr10-116501844-DEL-53 |
| chr10-116520910-DEL-110 | chr10-116520902-DEL-90 |
| chr10-116534518-DEL-204 | chr10-116534519-DEL-204 |
| chr10-116587560-DEL-147 | chr10-116587604-DEL-104 |
| chr10-116595488-DEL-167 | chr10-116595489-DEL-167 |
| chr10-116598096-DEL-1887 | chr10-116598097-DEL-1887 |
| chr10-116601254-DEL-718 | chr10-116601255-DEL-718 |
| chr10-116624796-DEL-167 | chr10-116624787-DEL-181 |
| chr10-116636902-DEL-74 | chr10-116636898-DEL-64 |
| chr10-116676270-DEL-53 | chr10-116676271-DEL-53 |
| chr10-116681589-DEL-112 | chr10-116681590-DEL-112 |
| chr10-116692858-DEL-821 | chr10-116692860-DEL-821 |
| chr10-116714118-DEL-1456 | chr10-116714119-DEL-1456 |
| chr10-116734655-DEL-1492 | chr10-116734656-DEL-1492 |
| chr10-116752211-DEL-59 | chr10-116752212-DEL-59 |
| chr10-116773062-DEL-60 | chr10-116773063-DEL-60 |
| chr10-116782912-DEL-51 | chr10-116782909-DEL-52 |
| chr10-116794804-DEL-74 | chr10-116794804-DEL-71 |
| chr10-116806443-DEL-399 | chr10-116806440-DEL-400 |
| chr10-116807569-DEL-225 | chr10-116807570-DEL-225 |
| chr10-116816535-DEL-190 | chr10-116816536-DEL-190 |
| chr10-116817779-DEL-212 | chr10-116817780-DEL-212 |
| chr10-116833916-DEL-58 | chr10-116833917-DEL-60 |
| chr10-116834561-DEL-65 | chr10-116834562-DEL-65 |
| chr10-116851102-DEL-391 | chr10-116851103-DEL-391 |
| chr10-116857989-DEL-389 | chr10-116857990-DEL-389 |
| chr10-116861738-DEL-399 | chr10-116861728-DEL-387 |
| chr10-116867545-DEL-699 | chr10-116867546-DEL-699 |
| chr10-116879814-DEL-297 | chr10-116879781-DEL-332 |
| chr10-116901334-DEL-174 | chr10-116901336-DEL-172 |
| chr10-116911328-DEL-56 | chr10-116911286-DEL-57 |
| chr10-116968857-DEL-207 | chr10-116968858-DEL-207 |
| chr10-116975939-DEL-51 | chr10-116975935-DEL-54 |
| chr10-117006537-DEL-73 | chr10-117006532-DEL-71 |
| chr10-117037914-DEL-316 | chr10-117037915-DEL-316 |
| chr10-117054871-DEL-416 | chr10-117054872-DEL-416 |
| chr10-117081479-DEL-184 | chr10-117081479-DEL-192 |
| chr10-117127619-DEL-185 | chr10-117127620-DEL-185 |
| chr10-117130285-DEL-818 | chr10-117130286-DEL-818 |
| chr10-117135663-DEL-802 | chr10-117135664-DEL-802 |
| chr10-117202353-DEL-60 | chr10-117202354-DEL-60 |
| chr10-117206471-DEL-57 | chr10-117206472-DEL-57 |
| chr10-117224626-DEL-127 | chr10-117224627-DEL-127 |
| chr10-117225776-DEL-84 | chr10-117225777-DEL-84 |
| chr10-117232980-DEL-53 | chr10-117232981-DEL-53 |
| chr10-117233507-DEL-71 | chr10-117233508-DEL-71 |
| chr10-117235739-DEL-59 | chr10-117235740-DEL-60 |
| chr10-117248889-DEL-210 | chr10-117248890-DEL-210 |

| | |
|---|---|
| chr10-117262781-DEL-2481 | chr10-117262782-DEL-2481 |
| chr10-117266776-DEL-209 | chr10-117266777-DEL-209 |
| chr10-117273824-DEL-519 | chr10-117273825-DEL-511 |
| chr10-117275578-DEL-72 | chr10-117275580-DEL-56 |
| chr10-117303397-DEL-925 | chr10-117303398-DEL-923 |
| chr10-117304662-DEL-203 | chr10-117304663-DEL-203 |
| chr10-117316877-DEL-84 | chr10-117316910-DEL-51 |
| chr10-117319563-DEL-7065 | chr10-117319564-DEL-7065 |
| chr10-117331089-DEL-209 | chr10-117331090-DEL-209 |
| chr10-117341853-DEL-56 | chr10-117341854-DEL-56 |
| chr10-117344605-DEL-196 | chr10-117344604-DEL-197 |
| chr10-117370112-DEL-58 | chr10-117370112-DEL-58 |
| chr10-117374942-DEL-359 | chr10-117374943-DEL-359 |
| chr10-117385059-DEL-137 | chr10-117385060-DEL-137 |
| chr10-117418226-DEL-383 | chr10-117418208-DEL-362 |
| chr10-117436403-DEL-208 | chr10-117436398-DEL-209 |
| chr10-117472564-DEL-65 | chr10-117472565-DEL-65 |
| chr10-117548717-DEL-145 | chr10-117548718-DEL-145 |
| chr10-117563984-DEL-497 | chr10-117563985-DEL-497 |
| chr10-117569714-DEL-54 | chr10-117569713-DEL-53 |
| chr10-117592829-DEL-61 | chr10-117592830-DEL-61 |
| chr10-117607933-DEL-202 | chr10-117607933-DEL-201 |
| chr10-117612860-DEL-129 | chr10-117612859-DEL-128 |
| chr10-117637355-DEL-508 | chr10-117637352-DEL-508 |
| chr10-117670334-DEL-101 | chr10-117670335-DEL-101 |
| chr10-117686747-DEL-209 | chr10-117686748-DEL-209 |
| chr10-117693310-DEL-74 | chr10-117693303-DEL-75 |
| chr10-117697749-DEL-65 | chr10-117697766-DEL-50 |
| chr10-117700479-DEL-56 | chr10-117700479-DEL-56 |
| chr10-117721900-DEL-171 | chr10-117721901-DEL-172 |
| chr10-117743094-DEL-78 | chr10-117743095-DEL-76 |
| chr10-117756843-DEL-80 | chr10-117756844-DEL-80 |
| chr10-117769772-DEL-57 | chr10-117769773-DEL-57 |
| chr10-117805290-DEL-1957 | chr10-117805291-DEL-1957 |
| chr10-117815353-DEL-54 | chr10-117815355-DEL-52 |
| chr10-117821962-DEL-269 | chr10-117821963-DEL-269 |
| chr10-117836871-DEL-54 | chr10-117836872-DEL-54 |
| chr10-117851930-DEL-52 | chr10-117851931-DEL-52 |
| chr10-117860721-DEL-3626 | chr10-117860721-DEL-3626 |
| chr10-117864688-DEL-109 | chr10-117864688-DEL-109 |
| chr10-117865369-DEL-126 | chr10-117865370-DEL-126 |
| chr10-117866289-DEL-2850 | chr10-117866290-DEL-2850 |
| chr10-117873830-DEL-55 | chr10-117873831-DEL-55 |
| chr10-117887152-DEL-63 | chr10-117887153-DEL-63 |
| chr10-117894822-DEL-139 | chr10-117894811-DEL-140 |
| chr10-117906538-DEL-5834 | chr10-117906539-DEL-5834 |
| chr10-117919811-DEL-353 | chr10-117919812-DEL-353 |
| chr10-117925878-DEL-479 | chr10-117925881-DEL-477 |
| chr10-117940700-DEL-56 | chr10-117940701-DEL-56 |
| chr10-117941428-DEL-88 | chr10-117941439-DEL-78 |
| chr10-117954571-DEL-204 | chr10-117954572-DEL-204 |
| chr10-117970826-DEL-414 | chr10-117970828-DEL-411 |
| chr10-117974362-DEL-675 | chr10-117974363-DEL-675 |
| chr10-117989047-DEL-238 | chr10-117989048-DEL-238 |
| chr10-117994770-DEL-73 | chr10-117994768-DEL-72 |
| chr10-118008985-DEL-93 | chr10-118008986-DEL-93 |
| chr10-118012546-DEL-1101 | chr10-118012549-DEL-1096 |
| chr10-118026696-DEL-63 | chr10-118026697-DEL-63 |
| chr10-118035653-DEL-146 | chr10-118035625-DEL-126 |
| chr10-118053560-DEL-624 | chr10-118053561-DEL-624 |
| chr10-118141990-DEL-84 | chr10-118141991-DEL-84 |
| chr10-118146031-DEL-75 | chr10-118146032-DEL-75 |
| chr10-118219870-DEL-968 | chr10-118219871-DEL-968 |
| chr10-118226063-DEL-84 | chr10-118226073-DEL-81 |
| chr10-118228826-DEL-994 | chr10-118228827-DEL-994 |
| chr10-118237207-DEL-75 | chr10-118237208-DEL-75 |
| chr10-118297555-DEL-154 | chr10-118297556-DEL-154 |
| chr10-118302412-DEL-794 | chr10-118302422-DEL-781 |
| chr10-118309921-DEL-2104 | chr10-118309922-DEL-2104 |
| chr10-118345220-DEL-62 | chr10-118345221-DEL-62 |
| chr10-118345809-DEL-53 | chr10-118345781-DEL-52 |
| chr10-118349216-DEL-76 | chr10-118349216-DEL-66 |
| chr10-118364676-DEL-68 | chr10-118364691-DEL-54 |
| chr10-118370701-DEL-199 | chr10-118370702-DEL-199 |
| chr10-118375924-DEL-52 | chr10-118375944-DEL-51 |
| chr10-118408347-DEL-112 | chr10-118408332-DEL-128 |
| chr10-118412009-DEL-70 | chr10-118412010-DEL-70 |
| chr10-118427100-DEL-138 | chr10-118427100-DEL-122 |
| chr10-118430604-DEL-207 | chr10-118430605-DEL-207 |
| chr10-118433110-DEL-748 | chr10-118433111-DEL-748 |

| | |
|---|---|
| chr10-118440433-DEL-59 | chr10-118440436-DEL-57 |
| chr10-118442765-DEL-72 | chr10-118442762-DEL-62 |
| chr10-118444064-DEL-54 | chr10-118444065-DEL-54 |
| chr10-118449947-DEL-231 | chr10-118449948-DEL-231 |
| chr10-118452599-DEL-390 | chr10-118452600-DEL-390 |
| chr10-118502994-DEL-52 | chr10-118502999-DEL-56 |
| chr10-118510746-DEL-52 | chr10-118510747-DEL-52 |
| chr10-118520778-DEL-60 | chr10-118520779-DEL-60 |
| chr10-118524229-DEL-70 | chr10-118524218-DEL-62 |
| chr10-118534200-DEL-107 | chr10-118534201-DEL-107 |
| chr10-118572309-DEL-211 | chr10-118572310-DEL-211 |
| chr10-118594312-DEL-56 | chr10-118594313-DEL-56 |
| chr10-118605437-DEL-109 | chr10-118605438-DEL-109 |
| chr10-118610208-DEL-88 | chr10-118610209-DEL-88 |
| chr10-118612280-DEL-204 | chr10-118612281-DEL-204 |
| chr10-118615675-DEL-616 | chr10-118615676-DEL-616 |
| chr10-118623257-DEL-151 | chr10-118623258-DEL-151 |
| chr10-118653178-DEL-94 | chr10-118653179-DEL-94 |
| chr10-118689341-DEL-53 | chr10-118689342-DEL-54 |
| chr10-118710659-DEL-80 | chr10-118710655-DEL-85 |
| chr10-118736921-DEL-130 | chr10-118736922-DEL-130 |
| chr10-118741162-DEL-80 | chr10-118741163-DEL-80 |
| chr10-118746392-DEL-6626 | chr10-118746393-DEL-6626 |
| chr10-118746910-DEL-201 | chr10-118746911-DEL-201 |
| chr10-118773443-DEL-6440 | chr10-118773444-DEL-6440 |
| chr10-118780704-DEL-158 | chr10-118780705-DEL-158 |
| chr10-118787489-DEL-862 | chr10-118787515-DEL-862 |
| chr10-118789752-DEL-194 | chr10-118789753-DEL-194 |
| chr10-118800163-DEL-87 | chr10-118800164-DEL-87 |
| chr10-118800767-DEL-178 | chr10-118800768-DEL-178 |
| chr10-118813707-DEL-200 | chr10-118813708-DEL-200 |
| chr10-118814589-DEL-192 | chr10-118814590-DEL-152 |
| chr10-118825345-DEL-137 | chr10-118825346-DEL-137 |
| chr10-118834483-DEL-116 | chr10-118834484-DEL-116 |
| chr10-118859829-DEL-100 | chr10-118859818-DEL-103 |
| chr10-118886728-DEL-82 | chr10-118886700-DEL-113 |
| chr10-118906085-DEL-75 | chr10-118906092-DEL-55 |
| chr10-118941787-DEL-866 | chr10-118941788-DEL-866 |
| chr10-118944573-DEL-70 | chr10-118944573-DEL-70 |
| chr10-118954512-DEL-105 | chr10-118954502-DEL-111 |
| chr10-118963557-DEL-264 | chr10-118963558-DEL-264 |
| chr10-118965184-DEL-397 | chr10-118965185-DEL-397 |
| chr10-118992306-DEL-89 | chr10-118992307-DEL-89 |
| chr10-119011328-DEL-513 | chr10-119011325-DEL-512 |
| chr10-119013564-DEL-543 | chr10-119013565-DEL-543 |
| chr10-119016625-DEL-203 | chr10-119016626-DEL-203 |
| chr10-119019353-DEL-58 | chr10-119019354-DEL-58 |
| chr10-119084537-DEL-209 | chr10-119084538-DEL-209 |
| chr10-119089086-DEL-240 | chr10-119089087-DEL-240 |
| chr10-119107361-DEL-68 | chr10-119107362-DEL-68 |
| chr10-119111824-DEL-382 | chr10-119111825-DEL-382 |
| chr10-119117037-DEL-6618 | chr10-119117038-DEL-6618 |
| chr10-119123079-DEL-424 | chr10-119123080-DEL-424 |
| chr10-119141918-DEL-454 | chr10-119141919-DEL-490 |
| chr10-119143766-DEL-198 | chr10-119143767-DEL-198 |
| chr10-119146809-DEL-213 | chr10-119146810-DEL-213 |
| chr10-119170254-DEL-113 | chr10-119170255-DEL-113 |
| chr10-119205751-DEL-59 | chr10-119205751-DEL-59 |
| chr10-119268881-DEL-64 | chr10-119268884-DEL-62 |
| chr10-119292987-DEL-56 | chr10-119292972-DEL-60 |
| chr10-119311654-DEL-155 | chr10-119311658-DEL-154 |
| chr10-119320958-DEL-62 | chr10-119320958-DEL-62 |
| chr10-119338832-DEL-242 | chr10-119338818-DEL-242 |
| chr10-119355508-DEL-507 | chr10-119355509-DEL-507 |
| chr10-119370288-DEL-57 | chr10-119370289-DEL-57 |
| chr10-119381885-DEL-94 | chr10-119381898-DEL-56 |
| chr10-119402681-DEL-527 | chr10-119402682-DEL-527 |
| chr10-119412190-DEL-87 | chr10-119412191-DEL-87 |
| chr10-119425263-DEL-194 | chr10-119425290-DEL-168 |
| chr10-119451613-DEL-152 | chr10-119451608-DEL-130 |
| chr10-119464983-DEL-571 | chr10-119464984-DEL-571 |
| chr10-119530671-DEL-82 | chr10-119530672-DEL-123 |
| chr10-119534738-DEL-2796 | chr10-119534739-DEL-2796 |
| chr10-119545980-DEL-386 | chr10-119545977-DEL-387 |
| chr10-119599878-DEL-6274 | chr10-119599879-DEL-6274 |
| chr10-119603346-DEL-336 | chr10-119603348-DEL-334 |
| chr10-119606872-DEL-159 | chr10-119606873-DEL-158 |
| chr10-119648601-DEL-220 | chr10-119648602-DEL-220 |
| chr10-119677997-DEL-1853 | chr10-119677998-DEL-1853 |
| chr10-119681450-DEL-68 | chr10-119681450-DEL-68 |

| | |
|---|---|
| chr10-119742535-DEL-78 | chr10-119742527-DEL-88 |
| chr10-119751948-DEL-266 | chr10-119751942-DEL-267 |
| chr10-119753172-DEL-1614 | chr10-119753173-DEL-1614 |
| chr10-119755342-DEL-247 | chr10-119755347-DEL-244 |
| chr10-119775148-DEL-2185 | chr10-119775149-DEL-2185 |
| chr10-119809792-DEL-474 | chr10-119809793-DEL-506 |
| chr10-119815012-DEL-56 | chr10-119815012-DEL-56 |
| chr10-119857084-DEL-138 | chr10-119857085-DEL-138 |
| chr10-119917812-DEL-51 | chr10-119917813-DEL-51 |
| chr10-119962698-DEL-60 | chr10-119962699-DEL-60 |
| chr10-119963711-DEL-53 | chr10-119963712-DEL-53 |
| chr10-119970381-DEL-204 | chr10-119970382-DEL-204 |
| chr10-119979812-DEL-84 | chr10-119979813-DEL-84 |
| chr10-119980712-DEL-91 | chr10-119980713-DEL-91 |
| chr10-119989700-DEL-293 | chr10-119989698-DEL-294 |
| chr10-119991218-DEL-1205 | chr10-119991231-DEL-1179 |
| chr10-119994718-DEL-308 | chr10-119994719-DEL-308 |
| chr10-120007547-DEL-332 | chr10-120007548-DEL-332 |
| chr10-120023194-DEL-603 | chr10-120023195-DEL-603 |
| chr10-120039843-DEL-210 | chr10-120039844-DEL-210 |
| chr10-120052985-DEL-80 | chr10-120052986-DEL-80 |
| chr10-120089500-DEL-72 | chr10-120089501-DEL-72 |
| chr10-120102114-DEL-201 | chr10-120102115-DEL-201 |
| chr10-120128682-DEL-695 | chr10-120128707-DEL-676 |
| chr10-120132739-DEL-109 | chr10-120132733-DEL-116 |
| chr10-120154563-DEL-330 | chr10-120154564-DEL-330 |
| chr10-120172078-DEL-200 | chr10-120172079-DEL-200 |
| chr10-120207783-DEL-52 | chr10-120207784-DEL-52 |
| chr10-120209657-DEL-165 | chr10-120209658-DEL-165 |
| chr10-120221407-DEL-109 | chr10-120221408-DEL-109 |
| chr10-120260217-DEL-1611 | chr10-120260218-DEL-1611 |
| chr10-120271376-DEL-51 | chr10-120271377-DEL-51 |
| chr10-120272119-DEL-58 | chr10-120272120-DEL-58 |
| chr10-120281751-DEL-197 | chr10-120281752-DEL-197 |
| chr10-120282787-DEL-176 | chr10-120282802-DEL-176 |
| chr10-120330583-DEL-675 | chr10-120330584-DEL-675 |
| chr10-120338096-DEL-207 | chr10-120338097-DEL-207 |
| chr10-120342894-DEL-164 | chr10-120342899-DEL-160 |
| chr10-120348437-DEL-255 | chr10-120348438-DEL-255 |
| chr10-120349400-DEL-52 | chr10-120349411-DEL-55 |
| chr10-120359645-DEL-163 | chr10-120359646-DEL-163 |
| chr10-120360829-DEL-92 | chr10-120360828-DEL-104 |
| chr10-120399216-DEL-222 | chr10-120399195-DEL-244 |
| chr10-120446495-DEL-201 | chr10-120446496-DEL-201 |
| chr10-120447352-DEL-409 | chr10-120447362-DEL-415 |
| chr10-120450696-DEL-76 | chr10-120450697-DEL-76 |
| chr10-120469988-DEL-536 | chr10-120469989-DEL-536 |
| chr10-120473886-DEL-60 | chr10-120473887-DEL-60 |
| chr10-120481015-DEL-62 | chr10-120481024-DEL-54 |
| chr10-120566939-DEL-52 | chr10-120566940-DEL-78 |
| chr10-120573536-DEL-389 | chr10-120573537-DEL-388 |
| chr10-120596604-DEL-84 | chr10-120596605-DEL-84 |
| chr10-120617714-DEL-64 | chr10-120617715-DEL-57 |
| chr10-120636427-DEL-73 | chr10-120636428-DEL-73 |
| chr10-120656245-DEL-65 | chr10-120656227-DEL-66 |
| chr10-120666951-DEL-76 | chr10-120666948-DEL-72 |
| chr10-120672467-DEL-61 | chr10-120672468-DEL-61 |
| chr10-120696811-DEL-51 | chr10-120696809-DEL-52 |
| chr10-120703547-DEL-828 | chr10-120703548-DEL-828 |
| chr10-120721225-DEL-2764 | chr10-120721226-DEL-2764 |
| chr10-120732218-DEL-119 | chr10-120732219-DEL-119 |
| chr10-120738170-DEL-92 | chr10-120738140-DEL-93 |
| chr10-120766430-DEL-107 | chr10-120766431-DEL-107 |
| chr10-120771970-DEL-206 | chr10-120771971-DEL-206 |
| chr10-120802117-DEL-217 | chr10-120802116-DEL-217 |
| chr10-120819625-DEL-83 | chr10-120819619-DEL-102 |
| chr10-120864245-DEL-54 | chr10-120864246-DEL-54 |
| chr10-120914826-DEL-62 | chr10-120914827-DEL-62 |
| chr10-120920978-DEL-238 | chr10-120920989-DEL-228 |
| chr10-120921525-DEL-1739 | chr10-120921526-DEL-1739 |
| chr10-120928760-DEL-329 | chr10-120928763-DEL-310 |
| chr10-120933020-DEL-139 | chr10-120933021-DEL-139 |
| chr10-120967307-DEL-228 | chr10-120967308-DEL-228 |
| chr10-120969079-DEL-57 | chr10-120969076-DEL-58 |
| chr10-120971959-DEL-7137 | chr10-120971959-DEL-7137 |
| chr10-120990368-DEL-54 | chr10-120990369-DEL-54 |
| chr10-121000647-DEL-4516 | chr10-121000648-DEL-4516 |
| chr10-121011591-DEL-72 | chr10-121011612-DEL-92 |
| chr10-121055856-DEL-69 | chr10-121055857-DEL-69 |
| chr10-121095731-DEL-514 | chr10-121095732-DEL-514 |

| | |
|---|---|
| chr10-121104068-DEL-69 | chr10-121104069-DEL-69 |
| chr10-121117099-DEL-2541 | chr10-121117100-DEL-2541 |
| chr10-121120973-DEL-2277 | chr10-121120975-DEL-2276 |
| chr10-121147474-DEL-65 | chr10-121147475-DEL-57 |
| chr10-121157437-DEL-466 | chr10-121157425-DEL-434 |
| chr10-121164846-DEL-158 | chr10-121164847-DEL-158 |
| chr10-121171740-DEL-1099 | chr10-121171741-DEL-1099 |
| chr10-121192864-DEL-195 | chr10-121192865-DEL-195 |
| chr10-121213523-DEL-73 | chr10-121213524-DEL-73 |
| chr10-121237649-DEL-201 | chr10-121237650-DEL-201 |
| chr10-121241945-DEL-1477 | chr10-121241946-DEL-1477 |
| chr10-121254584-DEL-99 | chr10-121254585-DEL-99 |
| chr10-121258570-DEL-52 | chr10-121258570-DEL-52 |
| chr10-121263053-DEL-58 | chr10-121263053-DEL-57 |
| chr10-121273312-DEL-208 | chr10-121273313-DEL-208 |
| chr10-121276835-DEL-617 | chr10-121276835-DEL-617 |
| chr10-121289416-DEL-64 | chr10-121289417-DEL-64 |
| chr10-121314332-DEL-194 | chr10-121314330-DEL-195 |
| chr10-121333450-DEL-51 | chr10-121333451-DEL-51 |
| chr10-121352500-DEL-90 | chr10-121352497-DEL-84 |
| chr10-121359905-DEL-78 | chr10-121359906-DEL-78 |
| chr10-121361435-DEL-62 | chr10-121361436-DEL-62 |
| chr10-121366247-DEL-76 | chr10-121366248-DEL-76 |
| chr10-121372511-DEL-52 | chr10-121372512-DEL-50 |
| chr10-121392556-DEL-62 | chr10-121392557-DEL-62 |
| chr10-121400508-DEL-1115 | chr10-121400509-DEL-1115 |
| chr10-121408439-DEL-1558 | chr10-121408440-DEL-1558 |
| chr10-121412877-DEL-420 | chr10-121412890-DEL-450 |
| chr10-121417904-DEL-261 | chr10-121417905-DEL-261 |
| chr10-121432772-DEL-127 | chr10-121432773-DEL-127 |
| chr10-121448894-DEL-75 | chr10-121448895-DEL-75 |
| chr10-121474556-DEL-109 | chr10-121474557-DEL-109 |
| chr10-121483589-DEL-163 | chr10-121483590-DEL-163 |
| chr10-121487684-DEL-201 | chr10-121487685-DEL-201 |
| chr10-121496788-DEL-139 | chr10-121496778-DEL-136 |
| chr10-121534201-DEL-59 | chr10-121534202-DEL-59 |
| chr10-121542761-DEL-7748 | chr10-121542762-DEL-7748 |
| chr10-121542775-DEL-201 | chr10-121542776-DEL-201 |
| chr10-121591251-DEL-238 | chr10-121591251-DEL-220 |
| chr10-121603466-DEL-1943 | chr10-121603467-DEL-1943 |
| chr10-121608868-DEL-512 | chr10-121608869-DEL-512 |
| chr10-121633690-DEL-656 | chr10-121633691-DEL-656 |
| chr10-121645846-DEL-60 | chr10-121645847-DEL-60 |
| chr10-121662240-DEL-97 | chr10-121662239-DEL-53 |
| chr10-121668808-DEL-192 | chr10-121668809-DEL-192 |
| chr10-121670457-DEL-58 | chr10-121670463-DEL-50 |
| chr10-121677077-DEL-98 | chr10-121677078-DEL-112 |
| chr10-121685077-DEL-85 | chr10-121685078-DEL-85 |
| chr10-121693903-DEL-72 | chr10-121693880-DEL-87 |
| chr10-121715501-DEL-58 | chr10-121715518-DEL-53 |
| chr10-121738380-DEL-191 | chr10-121738381-DEL-189 |
| chr10-121746484-DEL-203 | chr10-121746485-DEL-203 |
| chr10-121752844-DEL-56 | chr10-121752845-DEL-56 |
| chr10-121760317-DEL-52 | chr10-121760317-DEL-52 |
| chr10-121799788-DEL-79 | chr10-121799789-DEL-79 |
| chr10-121829255-DEL-596 | chr10-121829256-DEL-596 |
| chr10-121830382-DEL-175 | chr10-121830371-DEL-187 |
| chr10-121836651-DEL-51 | chr10-121836653-DEL-52 |
| chr10-121853317-DEL-124 | chr10-121853318-DEL-125 |
| chr10-121872426-DEL-66 | chr10-121872427-DEL-66 |
| chr10-121884369-DEL-337 | chr10-121884370-DEL-337 |
| chr10-121903667-DEL-1061 | chr10-121903668-DEL-1061 |
| chr10-121905154-DEL-341 | chr10-121905155-DEL-341 |
| chr10-121943341-DEL-7235 | chr10-121943342-DEL-7235 |
| chr10-121974820-DEL-54 | chr10-121974792-DEL-89 |
| chr10-121976899-DEL-2654 | chr10-121976901-DEL-2653 |
| chr10-121981462-DEL-60 | chr10-121981466-DEL-57 |
| chr10-121988760-DEL-6390 | chr10-121988760-DEL-6390 |
| chr10-122039099-DEL-148 | chr10-122039100-DEL-148 |
| chr10-122049790-DEL-3124 | chr10-122049773-DEL-3139 |
| chr10-122063294-DEL-72 | chr10-122063295-DEL-52 |
| chr10-122069286-DEL-119 | chr10-122069287-DEL-119 |
| chr10-122070821-DEL-6881 | chr10-122070822-DEL-6881 |
| chr10-122088215-DEL-93 | chr10-122088235-DEL-81 |
| chr10-122095191-DEL-1864 | chr10-122095192-DEL-1864 |
| chr10-122098368-DEL-132 | chr10-122098375-DEL-128 |
| chr10-122105538-DEL-58 | chr10-122105537-DEL-56 |
| chr10-122115441-DEL-3129 | chr10-122115442-DEL-3129 |
| chr10-122125486-DEL-89 | chr10-122125473-DEL-88 |
| chr10-122139426-DEL-5083 | chr10-122139427-DEL-5083 |

| | |
|---|---|
| chr10-122150566-DEL-58 | chr10-122150567-DEL-58 |
| chr10-122175357-DEL-104 | chr10-122175358-DEL-104 |
| chr10-122185579-DEL-1771 | chr10-122185570-DEL-1772 |
| chr10-122201960-DEL-4937 | chr10-122201961-DEL-4937 |
| chr10-122227267-DEL-6870 | chr10-122227268-DEL-6870 |
| chr10-122234281-DEL-72 | chr10-122234281-DEL-72 |
| chr10-122235361-DEL-85 | chr10-122235362-DEL-85 |
| chr10-122239907-DEL-285 | chr10-122239908-DEL-285 |
| chr10-122254623-DEL-6468 | chr10-122254624-DEL-6468 |
| chr10-122299645-DEL-170 | chr10-122299646-DEL-170 |
| chr10-122300538-DEL-84 | chr10-122300539-DEL-84 |
| chr10-122316339-DEL-1335 | chr10-122316339-DEL-1337 |
| chr10-122319156-DEL-60 | chr10-122319157-DEL-60 |
| chr10-122341566-DEL-7182 | chr10-122341567-DEL-7182 |
| chr10-122365770-DEL-400 | chr10-122365771-DEL-400 |
| chr10-122376428-DEL-181 | chr10-122376428-DEL-178 |
| chr10-122380468-DEL-79 | chr10-122380469-DEL-79 |
| chr10-122380685-DEL-365 | chr10-122380686-DEL-365 |
| chr10-122384377-DEL-210 | chr10-122384378-DEL-210 |
| chr10-122403937-DEL-102 | chr10-122403889-DEL-106 |
| chr10-122436251-DEL-135 | chr10-122436252-DEL-135 |
| chr10-122447604-DEL-84 | chr10-122447605-DEL-84 |
| chr10-122462302-DEL-142 | chr10-122462295-DEL-142 |
| chr10-122463665-DEL-124 | chr10-122463648-DEL-154 |
| chr10-122503389-DEL-52 | chr10-122503390-DEL-52 |
| chr10-122516989-DEL-60 | chr10-122516958-DEL-61 |
| chr10-122529736-DEL-54 | chr10-122529735-DEL-66 |
| chr10-122537950-DEL-717 | chr10-122537951-DEL-717 |
| chr10-122570604-DEL-845 | chr10-122570605-DEL-845 |
| chr10-122579229-DEL-185 | chr10-122579230-DEL-185 |
| chr10-122593569-DEL-59 | chr10-122593570-DEL-59 |
| chr10-122598369-DEL-95 | chr10-122598375-DEL-106 |
| chr10-122600766-DEL-151 | chr10-122600767-DEL-151 |
| chr10-122612531-DEL-58 | chr10-122612532-DEL-58 |
| chr10-122627651-DEL-96 | chr10-122627656-DEL-59 |
| chr10-122630722-DEL-71 | chr10-122630723-DEL-71 |
| chr10-122632429-DEL-195 | chr10-122632430-DEL-195 |
| chr10-122634946-DEL-122 | chr10-122634947-DEL-122 |
| chr10-122640111-DEL-147 | chr10-122640112-DEL-147 |
| chr10-122666749-DEL-56 | chr10-122666748-DEL-56 |
| chr10-122668763-DEL-7057 | chr10-122668764-DEL-7057 |
| chr10-122676035-DEL-226 | chr10-122676036-DEL-226 |
| chr10-122676887-DEL-80 | chr10-122676888-DEL-78 |
| chr10-122702464-DEL-624 | chr10-122702465-DEL-624 |
| chr10-122722157-DEL-112 | chr10-122722158-DEL-96 |
| chr10-122742876-DEL-56 | chr10-122742877-DEL-56 |
| chr10-122771194-DEL-1973 | chr10-122771193-DEL-1973 |
| chr10-122776060-DEL-85 | chr10-122776061-DEL-85 |
| chr10-122779222-DEL-565 | chr10-122779223-DEL-565 |
| chr10-122785307-DEL-79 | chr10-122785308-DEL-79 |
| chr10-122789869-DEL-217 | chr10-122789870-DEL-217 |
| chr10-122801287-DEL-389 | chr10-122801307-DEL-380 |
| chr10-122825447-DEL-57 | chr10-122825448-DEL-63 |
| chr10-122828255-DEL-135 | chr10-122828302-DEL-107 |
| chr10-122862652-DEL-51 | chr10-122862653-DEL-51 |
| chr10-122868165-DEL-70 | chr10-122868166-DEL-70 |
| chr10-122870564-DEL-199 | chr10-122870564-DEL-197 |
| chr10-122920489-DEL-68 | chr10-122920490-DEL-68 |
| chr10-122924413-DEL-580 | chr10-122924414-DEL-568 |
| chr10-122925179-DEL-207 | chr10-122925179-DEL-207 |
| chr10-122927160-DEL-452 | chr10-122927161-DEL-468 |
| chr10-122930386-DEL-53 | chr10-122930368-DEL-70 |
| chr10-122931613-DEL-203 | chr10-122931614-DEL-203 |
| chr10-122965148-DEL-126 | chr10-122965149-DEL-126 |
| chr10-122971181-DEL-225 | chr10-122971182-DEL-225 |
| chr10-122975019-DEL-1854 | chr10-122975020-DEL-1854 |
| chr10-123008154-DEL-183 | chr10-123008155-DEL-183 |
| chr10-123022114-DEL-101 | chr10-123022102-DEL-87 |
| chr10-123042381-DEL-315 | chr10-123042382-DEL-315 |
| chr10-123067832-DEL-565 | chr10-123067833-DEL-565 |
| chr10-123086690-DEL-140 | chr10-123086707-DEL-137 |
| chr10-123114824-DEL-467 | chr10-123114831-DEL-468 |
| chr10-123124589-DEL-110 | chr10-123124591-DEL-110 |
| chr10-123125162-DEL-80 | chr10-123125169-DEL-78 |
| chr10-123154358-DEL-521 | chr10-123154359-DEL-521 |
| chr10-123157823-DEL-92 | chr10-123157824-DEL-92 |
| chr10-123159762-DEL-837 | chr10-123159763-DEL-837 |
| chr10-123165909-DEL-88 | chr10-123165900-DEL-89 |
| chr10-123168407-DEL-134 | chr10-123168426-DEL-124 |
| chr10-123184030-DEL-100 | chr10-123184027-DEL-102 |

| | |
|---|---|
| chr10-123195816-DEL-168 | chr10-123195817-DEL-168 |
| chr10-123231741-DEL-110 | chr10-123231742-DEL-110 |
| chr10-123255888-DEL-72 | chr10-123255873-DEL-70 |
| chr10-123282530-DEL-55 | chr10-123282530-DEL-56 |
| chr10-123318451-DEL-583 | chr10-123318451-DEL-583 |
| chr10-123326447-DEL-6623 | chr10-123326448-DEL-6623 |
| chr10-123356824-DEL-55 | chr10-123356824-DEL-55 |
| chr10-123393949-DEL-57 | chr10-123393990-DEL-65 |
| chr10-123402741-DEL-222 | chr10-123402742-DEL-222 |
| chr10-123410461-DEL-156 | chr10-123410459-DEL-155 |
| chr10-123447980-DEL-519 | chr10-123447981-DEL-519 |
| chr10-123457485-DEL-80 | chr10-123457510-DEL-79 |
| chr10-123487315-DEL-475 | chr10-123487316-DEL-475 |
| chr10-123516954-DEL-65 | chr10-123516955-DEL-65 |
| chr10-123567032-DEL-98 | chr10-123567029-DEL-99 |
| chr10-123628502-DEL-4555 | chr10-123628518-DEL-4540 |
| chr10-123641610-DEL-99 | chr10-123641623-DEL-72 |
| chr10-123651252-DEL-516 | chr10-123651253-DEL-516 |
| chr10-123684329-DEL-396 | chr10-123684329-DEL-396 |
| chr10-123694564-DEL-8207 | chr10-123694567-DEL-8205 |
| chr10-123700919-DEL-326 | chr10-123700920-DEL-326 |
| chr10-123715405-DEL-52 | chr10-123715406-DEL-52 |
| chr10-123720477-DEL-58 | chr10-123720476-DEL-59 |
| chr10-123798967-DEL-1867 | chr10-123798968-DEL-1867 |
| chr10-123802615-DEL-1569 | chr10-123802616-DEL-1569 |
| chr10-123863560-DEL-384 | chr10-123863561-DEL-384 |
| chr10-123907398-DEL-219 | chr10-123907399-DEL-219 |
| chr10-123933873-DEL-2203 | chr10-123933874-DEL-2203 |
| chr10-123970569-DEL-664 | chr10-123970550-DEL-684 |
| chr10-123985290-DEL-59 | chr10-123985291-DEL-59 |
| chr10-124028500-DEL-133 | chr10-124028466-DEL-168 |
| chr10-124032790-DEL-1035 | chr10-124032791-DEL-1035 |
| chr10-124033744-DEL-68 | chr10-124033747-DEL-66 |
| chr10-124038261-DEL-142 | chr10-124038261-DEL-148 |
| chr10-124043618-DEL-204 | chr10-124043626-DEL-198 |
| chr10-124096828-DEL-106 | chr10-124096842-DEL-72 |
| chr10-124112504-DEL-100 | chr10-124112516-DEL-103 |
| chr10-124131675-DEL-70 | chr10-124131680-DEL-58 |
| chr10-124145656-DEL-2998 | chr10-124145657-DEL-2998 |
| chr10-124168243-DEL-80 | chr10-124168244-DEL-80 |
| chr10-124187940-DEL-54 | chr10-124187939-DEL-56 |
| chr10-124198614-DEL-64 | chr10-124198615-DEL-62 |
| chr10-124208147-DEL-55 | chr10-124208148-DEL-55 |
| chr10-124222947-DEL-90 | chr10-124222944-DEL-94 |
| chr10-124239243-DEL-852 | chr10-124239259-DEL-837 |
| chr10-124248838-DEL-1895 | chr10-124248839-DEL-1895 |
| chr10-124276535-DEL-68 | chr10-124276528-DEL-77 |
| chr10-124278376-DEL-99 | chr10-124278382-DEL-99 |
| chr10-124288751-DEL-944 | chr10-124288752-DEL-944 |
| chr10-124293080-DEL-64 | chr10-124293081-DEL-64 |
| chr10-124302647-DEL-2298 | chr10-124302644-DEL-2297 |
| chr10-124380565-DEL-2176 | chr10-124380566-DEL-2176 |
| chr10-124412219-DEL-7105 | chr10-124412220-DEL-7105 |
| chr10-124441656-DEL-5186 | chr10-124441657-DEL-5186 |
| chr10-124454646-DEL-78 | chr10-124454645-DEL-69 |
| chr10-124462656-DEL-261 | chr10-124462657-DEL-261 |
| chr10-124467211-DEL-892 | chr10-124467220-DEL-882 |
| chr10-124484032-DEL-2087 | chr10-124484034-DEL-2086 |
| chr10-124490929-DEL-57 | chr10-124490930-DEL-57 |
| chr10-124516361-DEL-5590 | chr10-124516362-DEL-5590 |
| chr10-124527060-DEL-52 | chr10-124527061-DEL-52 |
| chr10-124562507-DEL-134 | chr10-124562508-DEL-134 |
| chr10-124602145-DEL-1544 | chr10-124602146-DEL-1544 |
| chr10-124623405-DEL-1316 | chr10-124623405-DEL-1316 |
| chr10-124630736-DEL-408 | chr10-124630737-DEL-408 |
| chr10-124658706-DEL-59 | chr10-124658685-DEL-63 |
| chr10-124665354-DEL-82 | chr10-124665355-DEL-82 |
| chr10-124666640-DEL-663 | chr10-124666641-DEL-663 |
| chr10-124677461-DEL-354 | chr10-124677462-DEL-354 |
| chr10-124691641-DEL-70 | chr10-124691641-DEL-73 |
| chr10-124748699-DEL-77 | chr10-124748700-DEL-77 |
| chr10-124752400-DEL-436 | chr10-124752393-DEL-437 |
| chr10-124756403-DEL-66 | chr10-124756404-DEL-60 |
| chr10-124766225-DEL-1145 | chr10-124766226-DEL-1145 |
| chr10-124769474-DEL-81 | chr10-124769476-DEL-52 |
| chr10-124774853-DEL-155 | chr10-124774854-DEL-155 |
| chr10-124787255-DEL-53 | chr10-124787256-DEL-67 |
| chr10-124794783-DEL-104 | chr10-124794782-DEL-126 |
| chr10-124805899-DEL-52 | chr10-124805900-DEL-52 |
| chr10-124923684-DEL-79 | chr10-124923664-DEL-70 |

| | |
|---|---|
| chr10-124926899-DEL-54 | chr10-124926875-DEL-58 |
| chr10-124955439-DEL-101 | chr10-124955440-DEL-96 |
| chr10-124961883-DEL-428 | chr10-124961887-DEL-380 |
| chr10-124962933-DEL-65 | chr10-124962930-DEL-73 |
| chr10-124967712-DEL-95 | chr10-124967713-DEL-95 |
| chr10-124991365-DEL-2005 | chr10-124991366-DEL-2005 |
| chr10-124993618-DEL-74 | chr10-124993619-DEL-76 |
| chr10-124997512-DEL-85 | chr10-124997506-DEL-86 |
| chr10-125000256-DEL-1051 | chr10-125000257-DEL-1053 |
| chr10-125001394-DEL-189 | chr10-125001395-DEL-189 |
| chr10-125041156-DEL-134 | chr10-125041156-DEL-135 |
| chr10-125044638-DEL-71 | chr10-125044639-DEL-71 |
| chr10-125111605-DEL-2300 | chr10-125111611-DEL-2301 |
| chr10-125131103-DEL-60 | chr10-125131104-DEL-60 |
| chr10-125172755-DEL-58 | chr10-125172760-DEL-56 |
| chr10-125196254-DEL-90 | chr10-125196254-DEL-120 |
| chr10-125221657-DEL-60 | chr10-125221657-DEL-60 |
| chr10-125226573-DEL-63 | chr10-125226574-DEL-63 |
| chr10-125263287-DEL-58 | chr10-125263283-DEL-62 |
| chr10-125288302-DEL-61 | chr10-125288303-DEL-61 |
| chr10-125288650-DEL-2306 | chr10-125288651-DEL-2306 |
| chr10-125330178-DEL-52 | chr10-125330179-DEL-52 |
| chr10-125335488-DEL-56 | chr10-125335487-DEL-56 |
| chr10-125339924-DEL-3799 | chr10-125339925-DEL-3799 |
| chr10-125359768-DEL-174 | chr10-125359775-DEL-162 |
| chr10-125375330-DEL-81 | chr10-125375331-DEL-81 |
| chr10-125383214-DEL-107 | chr10-125383215-DEL-107 |
| chr10-125389812-DEL-66 | chr10-125389785-DEL-66 |
| chr10-125423609-DEL-1495 | chr10-125423610-DEL-1495 |
| chr10-125434672-DEL-164 | chr10-125434673-DEL-164 |
| chr10-125435678-DEL-58 | chr10-125435679-DEL-58 |
| chr10-125439076-DEL-91 | chr10-125439077-DEL-90 |
| chr10-125454661-DEL-440 | chr10-125454662-DEL-440 |
| chr10-125535689-DEL-59 | chr10-125535690-DEL-60 |
| chr10-125545149-DEL-94 | chr10-125545150-DEL-94 |
| chr10-125565049-DEL-518 | chr10-125565050-DEL-518 |
| chr10-125580269-DEL-198 | chr10-125580270-DEL-198 |
| chr10-125581231-DEL-102 | chr10-125581214-DEL-122 |
| chr10-125586823-DEL-62 | chr10-125586824-DEL-62 |
| chr10-125596365-DEL-54 | chr10-125596366-DEL-54 |
| chr10-125609912-DEL-478 | chr10-125609909-DEL-479 |
| chr10-125611418-DEL-55 | chr10-125611380-DEL-50 |
| chr10-125631070-DEL-153 | chr10-125631071-DEL-153 |
| chr10-125632605-DEL-58 | chr10-125632606-DEL-58 |
| chr10-125649254-DEL-351 | chr10-125649255-DEL-351 |
| chr10-125655379-DEL-6419 | chr10-125655380-DEL-6419 |
| chr10-125677386-DEL-1427 | chr10-125677387-DEL-1427 |
| chr10-125701238-DEL-60 | chr10-125701239-DEL-60 |
| chr10-125742439-DEL-52 | chr10-125742440-DEL-52 |
| chr10-125744621-DEL-52 | chr10-125744622-DEL-52 |
| chr10-125753112-DEL-90 | chr10-125753083-DEL-85 |
| chr10-125754993-DEL-436 | chr10-125754994-DEL-436 |
| chr10-125755656-DEL-64 | chr10-125755657-DEL-64 |
| chr10-125820167-DEL-54 | chr10-125820168-DEL-54 |
| chr10-125834299-DEL-51 | chr10-125834291-DEL-60 |
| chr10-125843167-DEL-146 | chr10-125843168-DEL-146 |
| chr10-125866255-DEL-544 | chr10-125866258-DEL-543 |
| chr10-125934811-DEL-54 | chr10-125934812-DEL-54 |
| chr10-125941978-DEL-65 | chr10-125941960-DEL-66 |
| chr10-125968088-DEL-65 | chr10-125968089-DEL-65 |
| chr10-125974201-DEL-166 | chr10-125974202-DEL-166 |
| chr10-125982927-DEL-83 | chr10-125982927-DEL-74 |
| chr10-126000345-DEL-364 | chr10-126000346-DEL-364 |
| chr10-126000853-DEL-914 | chr10-126000851-DEL-913 |
| chr10-126015651-DEL-122 | chr10-126015652-DEL-122 |
| chr10-126025405-DEL-6311 | chr10-126025406-DEL-6311 |
| chr10-126051241-DEL-56 | chr10-126051239-DEL-59 |
| chr10-126052792-DEL-468 | chr10-126052793-DEL-468 |
| chr10-126054295-DEL-380 | chr10-126054296-DEL-380 |
| chr10-126058186-DEL-617 | chr10-126058183-DEL-618 |
| chr10-126061957-DEL-123 | chr10-126061936-DEL-129 |
| chr10-126077935-DEL-631 | chr10-126077936-DEL-631 |
| chr10-126082519-DEL-1149 | chr10-126082520-DEL-1149 |
| chr10-126098251-DEL-4227 | chr10-126098252-DEL-4227 |
| chr10-126121193-DEL-6444 | chr10-126121194-DEL-6444 |
| chr10-126130892-DEL-1259 | chr10-126130893-DEL-1259 |
| chr10-126137301-DEL-833 | chr10-126137302-DEL-833 |
| chr10-126168737-DEL-162 | chr10-126168738-DEL-162 |
| chr10-126181541-DEL-155 | chr10-126181534-DEL-155 |
| chr10-126244026-DEL-92 | chr10-126244009-DEL-112 |

| | |
|---|---|
| chr10-126257906-DEL-4125 | chr10-126257886-DEL-4129 |
| chr10-126270973-DEL-504 | chr10-126270974-DEL-504 |
| chr10-126272356-DEL-578 | chr10-126272357-DEL-578 |
| chr10-126295214-DEL-1885 | chr10-126295215-DEL-1885 |
| chr10-126307114-DEL-251 | chr10-126307115-DEL-251 |
| chr10-126327624-DEL-84 | chr10-126327625-DEL-84 |
| chr10-126347500-DEL-77 | chr10-126347546-DEL-77 |
| chr10-126364834-DEL-64 | chr10-126364835-DEL-64 |
| chr10-126369639-DEL-436 | chr10-126369640-DEL-436 |
| chr10-126389215-DEL-184 | chr10-126389182-DEL-218 |
| chr10-126398054-DEL-1001 | chr10-126398055-DEL-1001 |
| chr10-126425621-DEL-67 | chr10-126425622-DEL-67 |
| chr10-126432982-DEL-76 | chr10-126432983-DEL-76 |
| chr10-126489093-DEL-1020 | chr10-126489094-DEL-1020 |
| chr10-126491738-DEL-74 | chr10-126491739-DEL-74 |
| chr10-126510418-DEL-1878 | chr10-126510422-DEL-1872 |
| chr10-126515251-DEL-65 | chr10-126515252-DEL-65 |
| chr10-126517041-DEL-165 | chr10-126517016-DEL-165 |
| chr10-126517585-DEL-5555 | chr10-126517586-DEL-5555 |
| chr10-126526942-DEL-77 | chr10-126526902-DEL-78 |
| chr10-126536947-DEL-6480 | chr10-126536948-DEL-6480 |
| chr10-126561208-DEL-92 | chr10-126561241-DEL-104 |
| chr10-126568916-DEL-1221 | chr10-126568917-DEL-1221 |
| chr10-126569097-DEL-53 | chr10-126569098-DEL-53 |
| chr10-126580245-DEL-75 | chr10-126580246-DEL-60 |
| chr10-126581263-DEL-1291 | chr10-126581266-DEL-1290 |
| chr10-126589729-DEL-93 | chr10-126589718-DEL-94 |
| chr10-126590929-DEL-107 | chr10-126590933-DEL-105 |
| chr10-126621513-DEL-1492 | chr10-126621514-DEL-1492 |
| chr10-126632690-DEL-145 | chr10-126632691-DEL-145 |
| chr10-126661394-DEL-64 | chr10-126661403-DEL-56 |
| chr10-126665862-DEL-534 | chr10-126665863-DEL-534 |
| chr10-126683173-DEL-294 | chr10-126683174-DEL-294 |
| chr10-126699902-DEL-211 | chr10-126699903-DEL-211 |
| chr10-126702130-DEL-147 | chr10-126702109-DEL-123 |
| chr10-126710389-DEL-52 | chr10-126710390-DEL-62 |
| chr10-126725472-DEL-932 | chr10-126725469-DEL-933 |
| chr10-126731956-DEL-349 | chr10-126731957-DEL-349 |
| chr10-126743619-DEL-83 | chr10-126743614-DEL-76 |
| chr10-126744664-DEL-193 | chr10-126744665-DEL-191 |
| chr10-126777634-DEL-498 | chr10-126777619-DEL-514 |
| chr10-126835252-DEL-140 | chr10-126835212-DEL-141 |
| chr10-126846495-DEL-83 | chr10-126846496-DEL-83 |
| chr10-126861771-DEL-66 | chr10-126861772-DEL-66 |
| chr10-126868013-DEL-196 | chr10-126868014-DEL-196 |
| chr10-126892849-DEL-168 | chr10-126892850-DEL-168 |
| chr10-126921917-DEL-233 | chr10-126921918-DEL-231 |
| chr10-126938739-DEL-51 | chr10-126938740-DEL-51 |
| chr10-126941534-DEL-212 | chr10-126941535-DEL-212 |
| chr10-126946023-DEL-214 | chr10-126946062-DEL-176 |
| chr10-126984713-DEL-204 | chr10-126984714-DEL-204 |
| chr10-126998563-DEL-221 | chr10-126998564-DEL-221 |
| chr10-127011239-DEL-65 | chr10-127011240-DEL-65 |
| chr10-127043131-DEL-155 | chr10-127043132-DEL-155 |
| chr10-127044072-DEL-72 | chr10-127044073-DEL-72 |
| chr10-127070482-DEL-61 | chr10-127070530-DEL-61 |
| chr10-127071267-DEL-121 | chr10-127071285-DEL-99 |
| chr10-127078152-DEL-62 | chr10-127078153-DEL-62 |
| chr10-127096243-DEL-64 | chr10-127096244-DEL-64 |
| chr10-127129414-DEL-96 | chr10-127129415-DEL-96 |
| chr10-127134594-DEL-188 | chr10-127134595-DEL-188 |
| chr10-127151802-DEL-74 | chr10-127151781-DEL-94 |
| chr10-127169346-DEL-54 | chr10-127169347-DEL-54 |
| chr10-127200111-DEL-216 | chr10-127200112-DEL-216 |
| chr10-127203485-DEL-53 | chr10-127203486-DEL-53 |
| chr10-127217544-DEL-168 | chr10-127217545-DEL-168 |
| chr10-127250434-DEL-226 | chr10-127250435-DEL-226 |
| chr10-127262121-DEL-104 | chr10-127262122-DEL-74 |
| chr10-127263817-DEL-101 | chr10-127263818-DEL-101 |
| chr10-127273843-DEL-206 | chr10-127273844-DEL-206 |
| chr10-127286059-DEL-284 | chr10-127286060-DEL-284 |
| chr10-127291855-DEL-105 | chr10-127291860-DEL-101 |
| chr10-127303212-DEL-275 | chr10-127303213-DEL-275 |
| chr10-127315453-DEL-203 | chr10-127315454-DEL-203 |
| chr10-127349570-DEL-224 | chr10-127349571-DEL-224 |
| chr10-127396139-DEL-51 | chr10-127396140-DEL-51 |
| chr10-127401466-DEL-186 | chr10-127401461-DEL-192 |
| chr10-127406590-DEL-214 | chr10-127406591-DEL-214 |
| chr10-127471427-DEL-70 | chr10-127471433-DEL-65 |
| chr10-127474362-DEL-232 | chr10-127474363-DEL-232 |

| | |
|---|---|
| chr10-127542125-DEL-60 | chr10-127542126-DEL-60 |
| chr10-127582640-DEL-125 | chr10-127582641-DEL-125 |
| chr10-127609789-DEL-324 | chr10-127609790-DEL-324 |
| chr10-127630062-DEL-1646 | chr10-127630063-DEL-1646 |
| chr10-127653123-DEL-5829 | chr10-127653123-DEL-5829 |
| chr10-127662143-DEL-152 | chr10-127662144-DEL-169 |
| chr10-127665381-DEL-410 | chr10-127665382-DEL-410 |
| chr10-127667255-DEL-958 | chr10-127667256-DEL-958 |
| chr10-127744675-DEL-127 | chr10-127744676-DEL-127 |
| chr10-127751549-DEL-76 | chr10-127751550-DEL-76 |
| chr10-127753178-DEL-78 | chr10-127753179-DEL-117 |
| chr10-127754787-DEL-134 | chr10-127754788-DEL-134 |
| chr10-127775703-DEL-55 | chr10-127775701-DEL-56 |
| chr10-127780659-DEL-81 | chr10-127780641-DEL-126 |
| chr10-127793842-DEL-58 | chr10-127793847-DEL-54 |
| chr10-127823509-DEL-535 | chr10-127823510-DEL-535 |
| chr10-127828899-DEL-206 | chr10-127828900-DEL-206 |
| chr10-127853468-DEL-198 | chr10-127853469-DEL-198 |
| chr10-127860564-DEL-177 | chr10-127860565-DEL-177 |
| chr10-127860996-DEL-369 | chr10-127860997-DEL-369 |
| chr10-127883121-DEL-256 | chr10-127883122-DEL-256 |
| chr10-127887228-DEL-198 | chr10-127887229-DEL-198 |
| chr10-127888341-DEL-180 | chr10-127888342-DEL-180 |
| chr10-127892167-DEL-181 | chr10-127892167-DEL-181 |
| chr10-127910353-DEL-65 | chr10-127910354-DEL-65 |
| chr10-127922960-DEL-200 | chr10-127922961-DEL-200 |
| chr10-127992354-DEL-224 | chr10-127992355-DEL-224 |
| chr10-127994345-DEL-174 | chr10-127994343-DEL-175 |
| chr10-128001457-DEL-82 | chr10-128001456-DEL-75 |
| chr10-128002764-DEL-210 | chr10-128002765-DEL-210 |
| chr10-128004321-DEL-84 | chr10-128004322-DEL-84 |
| chr10-128006100-DEL-145 | chr10-128006054-DEL-184 |
| chr10-128013732-DEL-210 | chr10-128013733-DEL-210 |
| chr10-128038150-DEL-54 | chr10-128038153-DEL-52 |
| chr10-128038563-DEL-214 | chr10-128038564-DEL-214 |
| chr10-128054027-DEL-56 | chr10-128054027-DEL-56 |
| chr10-128062224-DEL-123 | chr10-128062228-DEL-120 |
| chr10-128098290-DEL-60 | chr10-128098291-DEL-60 |
| chr10-128109820-DEL-107 | chr10-128109821-DEL-107 |
| chr10-128124948-DEL-66 | chr10-128124949-DEL-66 |
| chr10-128131638-DEL-111 | chr10-128131649-DEL-82 |
| chr10-128160706-DEL-361 | chr10-128160707-DEL-361 |
| chr10-128182651-DEL-208 | chr10-128182652-DEL-208 |
| chr10-128217185-DEL-286 | chr10-128217186-DEL-286 |
| chr10-128240820-DEL-179 | chr10-128240813-DEL-182 |
| chr10-128254507-DEL-79 | chr10-128254540-DEL-55 |
| chr10-128255808-DEL-176 | chr10-128255809-DEL-176 |
| chr10-128263828-DEL-210 | chr10-128263829-DEL-210 |
| chr10-128268734-DEL-241 | chr10-128268735-DEL-241 |
| chr10-128277405-DEL-102 | chr10-128277406-DEL-102 |
| chr10-128286000-DEL-102 | chr10-128285999-DEL-106 |
| chr10-128315245-DEL-190 | chr10-128315246-DEL-190 |
| chr10-128319538-DEL-113 | chr10-128319539-DEL-113 |
| chr10-128339111-DEL-76 | chr10-128339112-DEL-76 |
| chr10-128368258-DEL-180 | chr10-128368259-DEL-180 |
| chr10-128374481-DEL-175 | chr10-128374482-DEL-175 |
| chr10-128443787-DEL-599 | chr10-128443788-DEL-599 |
| chr10-128444567-DEL-102 | chr10-128444568-DEL-102 |
| chr10-128445321-DEL-383 | chr10-128445328-DEL-376 |
| chr10-128447682-DEL-456 | chr10-128447682-DEL-457 |
| chr10-128451901-DEL-224 | chr10-128451902-DEL-224 |
| chr10-128456046-DEL-629 | chr10-128456047-DEL-628 |
| chr10-128457972-DEL-63 | chr10-128457972-DEL-63 |
| chr10-128486782-DEL-247 | chr10-128486783-DEL-247 |
| chr10-128490938-DEL-208 | chr10-128490939-DEL-208 |
| chr10-128497533-DEL-198 | chr10-128497534-DEL-198 |
| chr10-128553271-DEL-212 | chr10-128553272-DEL-212 |
| chr10-128566681-DEL-211 | chr10-128566682-DEL-211 |
| chr10-128575133-DEL-198 | chr10-128575121-DEL-211 |
| chr10-128576883-DEL-213 | chr10-128576882-DEL-214 |
| chr10-128596890-DEL-264 | chr10-128596891-DEL-264 |
| chr10-128603982-DEL-159 | chr10-128604022-DEL-146 |
| chr10-128622475-DEL-136 | chr10-128622476-DEL-108 |
| chr10-128646586-DEL-1891 | chr10-128646587-DEL-1891 |
| chr10-128666944-DEL-223 | chr10-128666945-DEL-223 |
| chr10-128679848-DEL-213 | chr10-128679849-DEL-213 |
| chr10-128710988-DEL-156 | chr10-128710989-DEL-156 |
| chr10-128717198-DEL-131 | chr10-128717199-DEL-131 |
| chr10-128734180-DEL-237 | chr10-128734181-DEL-237 |
| chr10-128737218-DEL-108 | chr10-128737219-DEL-108 |

| | |
|---|---|
| chr10-128787110-DEL-53 | chr10-128787102-DEL-54 |
| chr10-128792208-DEL-51 | chr10-128792209-DEL-51 |
| chr10-128804721-DEL-63 | chr10-128804714-DEL-71 |
| chr10-128811119-DEL-115 | chr10-128811108-DEL-108 |
| chr10-128815587-DEL-59 | chr10-128815590-DEL-58 |
| chr10-128816562-DEL-88 | chr10-128816563-DEL-88 |
| chr10-128822585-DEL-1512 | chr10-128822586-DEL-1512 |
| chr10-128829431-DEL-1653 | chr10-128829432-DEL-1653 |
| chr10-128854800-DEL-77 | chr10-128854801-DEL-77 |
| chr10-128856642-DEL-66 | chr10-128856643-DEL-66 |
| chr10-128878516-DEL-77 | chr10-128878516-DEL-77 |
| chr10-128910269-DEL-198 | chr10-128910270-DEL-198 |
| chr10-128945524-DEL-399 | chr10-128945525-DEL-399 |
| chr10-128987842-DEL-1066 | chr10-128987843-DEL-1062 |
| chr10-128990298-DEL-983 | chr10-128990298-DEL-983 |
| chr10-129001656-DEL-2543 | chr10-129001657-DEL-2543 |
| chr10-129004631-DEL-71 | chr10-129004632-DEL-71 |
| chr10-129007881-DEL-66 | chr10-129007882-DEL-66 |
| chr10-129025310-DEL-6606 | chr10-129025311-DEL-6606 |
| chr10-129057960-DEL-378 | chr10-129057966-DEL-374 |
| chr10-129074976-DEL-2602 | chr10-129074977-DEL-2602 |
| chr10-129080651-DEL-54 | chr10-129080652-DEL-54 |
| chr10-129090369-DEL-84 | chr10-129090354-DEL-103 |
| chr10-129108590-DEL-6328 | chr10-129108581-DEL-6336 |
| chr10-129119468-DEL-1868 | chr10-129119468-DEL-1868 |
| chr10-129121575-DEL-54 | chr10-129121575-DEL-54 |
| chr10-129161869-DEL-7027 | chr10-129161870-DEL-7027 |
| chr10-129175029-DEL-5491 | chr10-129175029-DEL-5491 |
| chr10-129177046-DEL-904 | chr10-129177047-DEL-904 |
| chr10-129181580-DEL-593 | chr10-129181581-DEL-593 |
| chr10-129212741-DEL-64 | chr10-129212742-DEL-61 |
| chr10-129218399-DEL-481 | chr10-129218400-DEL-481 |
| chr10-129224679-DEL-7305 | chr10-129224680-DEL-7305 |
| chr10-129237920-DEL-67 | chr10-129237921-DEL-67 |
| chr10-129249104-DEL-6365 | chr10-129249105-DEL-6365 |
| chr10-129264630-DEL-67 | chr10-129264631-DEL-67 |
| chr10-129277861-DEL-12997 | chr10-129277864-DEL-12996 |
| chr10-129311812-DEL-59 | chr10-129311798-DEL-56 |
| chr10-129312768-DEL-1854 | chr10-129312769-DEL-1854 |
| chr10-129315496-DEL-72 | chr10-129315497-DEL-72 |
| chr10-129321389-DEL-130 | chr10-129321420-DEL-135 |
| chr10-129341831-DEL-229 | chr10-129341832-DEL-229 |
| chr10-129346353-DEL-572 | chr10-129346354-DEL-572 |
| chr10-129348817-DEL-63 | chr10-129348818-DEL-63 |
| chr10-129360022-DEL-723 | chr10-129360019-DEL-720 |
| chr10-129380563-DEL-52 | chr10-129380537-DEL-53 |
| chr10-129397216-DEL-683 | chr10-129397217-DEL-683 |
| chr10-129399695-DEL-429 | chr10-129399696-DEL-429 |
| chr10-129409517-DEL-104 | chr10-129409518-DEL-104 |
| chr10-129420397-DEL-9608 | chr10-129420397-DEL-9608 |
| chr10-129454557-DEL-272 | chr10-129454558-DEL-272 |
| chr10-129486612-DEL-186 | chr10-129486613-DEL-186 |
| chr10-129591737-DEL-200 | chr10-129591738-DEL-200 |
| chr10-129684689-DEL-437 | chr10-129684690-DEL-437 |
| chr10-129694720-DEL-54 | chr10-129694721-DEL-54 |
| chr10-129713547-DEL-1989 | chr10-129713548-DEL-1989 |
| chr10-129718619-DEL-983 | chr10-129718620-DEL-983 |
| chr10-129722096-DEL-205 | chr10-129722097-DEL-205 |
| chr10-129728909-DEL-414 | chr10-129728910-DEL-414 |
| chr10-129734145-DEL-678 | chr10-129734146-DEL-678 |
| chr10-129761133-DEL-4386 | chr10-129761134-DEL-4386 |
| chr10-129770830-DEL-120 | chr10-129770831-DEL-120 |
| chr10-129798285-DEL-70 | chr10-129798287-DEL-73 |
| chr10-129798886-DEL-2385 | chr10-129798887-DEL-2385 |
| chr10-129878241-DEL-16097 | chr10-129878241-DEL-16095 |
| chr10-129924071-DEL-790 | chr10-129924072-DEL-790 |
| chr10-129952250-DEL-993 | chr10-129952251-DEL-993 |
| chr10-129964005-DEL-415 | chr10-129963983-DEL-437 |
| chr10-129968684-DEL-338 | chr10-129968685-DEL-338 |
| chr10-129980431-DEL-1248 | chr10-129980432-DEL-1248 |
| chr10-129986075-DEL-69 | chr10-129986067-DEL-70 |
| chr10-129987446-DEL-1348 | chr10-129987447-DEL-1347 |
| chr10-129993344-DEL-383 | chr10-129993345-DEL-383 |
| chr10-130024727-DEL-2646 | chr10-130024728-DEL-2646 |
| chr10-130060021-DEL-201 | chr10-130060022-DEL-201 |
| chr10-130138994-DEL-1180 | chr10-130138995-DEL-1180 |
| chr10-130144899-DEL-1138 | chr10-130144900-DEL-1138 |
| chr10-130163415-DEL-2950 | chr10-130163430-DEL-2933 |
| chr10-130170448-DEL-766 | chr10-130170449-DEL-766 |
| chr10-130247004-DEL-3099 | chr10-130247005-DEL-3099 |

| | |
|---|---|
| chr10-130267836-DEL-193 | chr10-130267841-DEL-192 |
| chr10-130283371-DEL-212 | chr10-130283372-DEL-212 |
| chr10-130310271-DEL-1476 | chr10-130310276-DEL-1471 |
| chr10-130365600-DEL-398 | chr10-130365602-DEL-396 |
| chr10-130396017-DEL-5965 | chr10-130396018-DEL-5965 |
| chr10-130409895-DEL-336 | chr10-130409896-DEL-336 |
| chr11-3148648-DEL-63 | chr11-3148645-DEL-65 |
| chr11-3208471-DEL-134 | chr11-3208472-DEL-134 |
| chr11-3301497-DEL-200 | chr11-3301498-DEL-200 |
| chr11-3306537-DEL-210 | chr11-3306538-DEL-210 |
| chr11-3318018-DEL-96 | chr11-3318025-DEL-93 |
| chr11-3326936-DEL-123 | chr11-3326937-DEL-123 |
| chr11-3363343-DEL-143 | chr11-3363343-DEL-151 |
| chr11-3424401-DEL-221 | chr11-3424402-DEL-221 |
| chr11-3431345-DEL-210 | chr11-3431346-DEL-210 |
| chr11-3514349-DEL-73 | chr11-3514350-DEL-73 |
| chr11-3548294-DEL-1143 | chr11-3548295-DEL-1143 |
| chr11-3553844-DEL-279 | chr11-3553845-DEL-279 |
| chr11-3564939-DEL-88 | chr11-3564940-DEL-88 |
| chr11-3684931-DEL-72 | chr11-3684932-DEL-72 |
| chr11-3693777-DEL-104 | chr11-3693778-DEL-104 |
| chr11-3759536-DEL-380 | chr11-3759537-DEL-380 |
| chr11-3776948-DEL-201 | chr11-3776949-DEL-201 |
| chr11-3818674-DEL-515 | chr11-3818675-DEL-516 |
| chr11-3899114-DEL-2488 | chr11-3899115-DEL-2488 |
| chr11-3944217-DEL-287 | chr11-3944218-DEL-287 |
| chr11-3957837-DEL-180 | chr11-3957838-DEL-133 |
| chr11-3977736-DEL-3218 | chr11-3977737-DEL-3218 |
| chr11-3988325-DEL-52 | chr11-3988326-DEL-52 |
| chr11-3998943-DEL-3089 | chr11-3998944-DEL-3089 |
| chr11-4018919-DEL-94 | chr11-4018920-DEL-79 |
| chr11-4038201-DEL-2683 | chr11-4038202-DEL-2683 |
| chr11-4104289-DEL-215 | chr11-4104290-DEL-215 |
| chr11-4139670-DEL-104 | chr11-4139671-DEL-104 |
| chr11-4172014-DEL-53 | chr11-4172015-DEL-53 |
| chr11-4175616-DEL-125 | chr11-4175617-DEL-125 |
| chr11-4198045-DEL-138 | chr11-4198046-DEL-138 |
| chr11-4203215-DEL-507 | chr11-4203216-DEL-507 |
| chr11-4222838-DEL-76 | chr11-4222839-DEL-76 |
| chr11-4273941-DEL-314 | chr11-4273951-DEL-317 |
| chr11-4281982-DEL-210 | chr11-4281983-DEL-210 |
| chr11-4313229-DEL-2092 | chr11-4313230-DEL-2092 |
| chr11-4328228-DEL-186 | chr11-4328227-DEL-183 |
| chr11-4335505-DEL-167 | chr11-4335504-DEL-168 |
| chr11-4445962-DEL-60 | chr11-4445963-DEL-60 |
| chr11-4456229-DEL-238 | chr11-4456230-DEL-238 |
| chr11-4460614-DEL-281 | chr11-4460615-DEL-281 |
| chr11-4517894-DEL-64 | chr11-4517895-DEL-64 |
| chr11-4527401-DEL-193 | chr11-4527402-DEL-193 |
| chr11-4531264-DEL-117 | chr11-4531265-DEL-117 |
| chr11-4532467-DEL-80 | chr11-4532468-DEL-80 |
| chr11-4537917-DEL-212 | chr11-4537918-DEL-212 |
| chr11-4590845-DEL-72 | chr11-4590846-DEL-72 |
| chr11-4595415-DEL-253 | chr11-4595416-DEL-253 |
| chr11-4609992-DEL-58 | chr11-4609993-DEL-58 |
| chr11-4614351-DEL-2761 | chr11-4614352-DEL-2761 |
| chr11-4645462-DEL-214 | chr11-4645463-DEL-214 |
| chr11-4649195-DEL-200 | chr11-4649196-DEL-200 |
| chr11-4664183-DEL-204 | chr11-4664184-DEL-204 |
| chr11-4709412-DEL-66 | chr11-4709413-DEL-66 |
| chr11-4709839-DEL-205 | chr11-4709840-DEL-205 |
| chr11-4725372-DEL-741 | chr11-4725373-DEL-741 |
| chr11-4747649-DEL-56 | chr11-4747650-DEL-56 |
| chr11-4776531-DEL-54 | chr11-4776532-DEL-54 |
| chr11-4801416-DEL-215 | chr11-4801417-DEL-215 |
| chr11-4804009-DEL-241 | chr11-4804010-DEL-241 |
| chr11-4806578-DEL-154 | chr11-4806562-DEL-174 |
| chr11-4812194-DEL-147 | chr11-4812195-DEL-147 |
| chr11-4814570-DEL-205 | chr11-4814571-DEL-205 |
| chr11-4814837-DEL-545 | chr11-4814838-DEL-545 |
| chr11-4821155-DEL-90 | chr11-4821155-DEL-78 |
| chr11-4826199-DEL-1242 | chr11-4826200-DEL-1242 |
| chr11-4826982-DEL-189 | chr11-4826983-DEL-189 |
| chr11-4828351-DEL-182 | chr11-4828352-DEL-182 |
| chr11-4829721-DEL-95 | chr11-4829722-DEL-95 |
| chr11-4836206-DEL-182 | chr11-4836206-DEL-182 |
| chr11-4838030-DEL-210 | chr11-4838031-DEL-210 |
| chr11-4844733-DEL-182 | chr11-4844734-DEL-182 |
| chr11-4856955-DEL-232 | chr11-4856976-DEL-212 |
| chr11-4869445-DEL-80 | chr11-4869446-DEL-80 |

| | |
|---|---|
| chr11-4890447-DEL-72 | chr11-4890448-DEL-72 |
| chr11-4923192-DEL-169 | chr11-4923193-DEL-169 |
| chr11-4929909-DEL-60 | chr11-4929910-DEL-60 |
| chr11-4954653-DEL-193 | chr11-4954654-DEL-193 |
| chr11-4960796-DEL-490 | chr11-4960797-DEL-490 |
| chr11-4963429-DEL-241 | chr11-4963430-DEL-241 |
| chr11-4994974-DEL-239 | chr11-4994975-DEL-239 |
| chr11-4995595-DEL-203 | chr11-4995596-DEL-202 |
| chr11-4999371-DEL-60 | chr11-4999347-DEL-95 |
| chr11-5004637-DEL-127 | chr11-5004638-DEL-127 |
| chr11-5023860-DEL-138 | chr11-5023861-DEL-138 |
| chr11-5040419-DEL-284 | chr11-5040420-DEL-284 |
| chr11-5077163-DEL-178 | chr11-5077143-DEL-204 |
| chr11-5103188-DEL-108 | chr11-5103228-DEL-58 |
| chr11-5118266-DEL-1777 | chr11-5118267-DEL-1777 |
| chr11-5146416-DEL-418 | chr11-5146417-DEL-419 |
| chr11-5150203-DEL-336 | chr11-5150204-DEL-336 |
| chr11-5161062-DEL-126 | chr11-5161063-DEL-126 |
| chr11-5176931-DEL-245 | chr11-5176932-DEL-245 |
| chr11-5184884-DEL-82 | chr11-5184898-DEL-50 |
| chr11-5250430-DEL-78 | chr11-5250431-DEL-78 |
| chr11-5259543-DEL-107 | chr11-5259554-DEL-97 |
| chr11-5276477-DEL-92 | chr11-5276478-DEL-92 |
| chr11-5311212-DEL-4158 | chr11-5311212-DEL-4158 |
| chr11-5345846-DEL-834 | chr11-5345847-DEL-834 |
| chr11-5377711-DEL-922 | chr11-5377712-DEL-922 |
| chr11-5377738-DEL-72 | chr11-5377739-DEL-72 |
| chr11-5415967-DEL-74 | chr11-5415968-DEL-74 |
| chr11-5421413-DEL-108 | chr11-5421414-DEL-108 |
| chr11-5425928-DEL-214 | chr11-5425928-DEL-215 |
| chr11-5428182-DEL-72 | chr11-5428183-DEL-72 |
| chr11-5443472-DEL-217 | chr11-5443473-DEL-217 |
| chr11-5453441-DEL-169 | chr11-5453439-DEL-157 |
| chr11-5456546-DEL-208 | chr11-5456562-DEL-209 |
| chr11-5457592-DEL-799 | chr11-5457593-DEL-799 |
| chr11-5460720-DEL-64 | chr11-5460721-DEL-64 |
| chr11-5461216-DEL-84 | chr11-5461217-DEL-84 |
| chr11-5485469-DEL-474 | chr11-5485470-DEL-474 |
| chr11-5490130-DEL-174 | chr11-5490131-DEL-174 |
| chr11-5498719-DEL-2457 | chr11-5498720-DEL-2457 |
| chr11-5502736-DEL-5382 | chr11-5502737-DEL-5382 |
| chr11-5538276-DEL-209 | chr11-5538277-DEL-209 |
| chr11-5563353-DEL-72 | chr11-5563333-DEL-72 |
| chr11-5589716-DEL-221 | chr11-5589717-DEL-221 |
| chr11-5606134-DEL-213 | chr11-5606135-DEL-213 |
| chr11-5616655-DEL-226 | chr11-5616656-DEL-226 |
| chr11-5620406-DEL-201 | chr11-5620407-DEL-201 |
| chr11-5621131-DEL-120 | chr11-5621132-DEL-120 |
| chr11-5628009-DEL-201 | chr11-5628010-DEL-201 |
| chr11-5633398-DEL-210 | chr11-5633399-DEL-210 |
| chr11-5636183-DEL-444 | chr11-5636185-DEL-435 |
| chr11-5644767-DEL-5363 | chr11-5644765-DEL-5363 |
| chr11-5653510-DEL-55 | chr11-5653511-DEL-55 |
| chr11-5657879-DEL-223 | chr11-5657880-DEL-223 |
| chr11-5669919-DEL-198 | chr11-5669920-DEL-198 |
| chr11-5674437-DEL-64 | chr11-5674400-DEL-64 |
| chr11-5686648-DEL-79 | chr11-5686649-DEL-79 |
| chr11-5692964-DEL-78 | chr11-5692965-DEL-78 |
| chr11-5700268-DEL-210 | chr11-5700269-DEL-210 |
| chr11-5708238-DEL-201 | chr11-5708239-DEL-201 |
| chr11-5711177-DEL-182 | chr11-5711169-DEL-189 |
| chr11-5712877-DEL-214 | chr11-5712878-DEL-214 |
| chr11-5724487-DEL-60 | chr11-5724488-DEL-60 |
| chr11-5730545-DEL-84 | chr11-5730546-DEL-84 |
| chr11-5748625-DEL-185 | chr11-5748626-DEL-185 |
| chr11-5768384-DEL-352 | chr11-5768385-DEL-352 |
| chr11-5772590-DEL-167 | chr11-5772591-DEL-167 |
| chr11-5792711-DEL-173 | chr11-5792707-DEL-175 |
| chr11-5838542-DEL-195 | chr11-5838543-DEL-195 |
| chr11-5840295-DEL-53 | chr11-5840298-DEL-51 |
| chr11-5847573-DEL-201 | chr11-5847574-DEL-201 |
| chr11-5893272-DEL-52 | chr11-5893261-DEL-53 |
| chr11-5948509-DEL-136 | chr11-5948510-DEL-136 |
| chr11-5955647-DEL-5868 | chr11-5955648-DEL-5868 |
| chr11-5962516-DEL-78 | chr11-5962517-DEL-78 |
| chr11-5973167-DEL-101 | chr11-5973167-DEL-102 |
| chr11-5998016-DEL-82 | chr11-5998017-DEL-82 |
| chr11-6007052-DEL-58 | chr11-6007053-DEL-58 |
| chr11-6016573-DEL-88 | chr11-6016574-DEL-88 |
| chr11-6021844-DEL-54 | chr11-6021844-DEL-54 |

| | |
|---|---|
| chr11-6032737-DEL-62 | chr11-6032738-DEL-62 |
| chr11-6091505-DEL-7125 | chr11-6091506-DEL-7125 |
| chr11-6159791-DEL-57 | chr11-6159792-DEL-57 |
| chr11-6179694-DEL-148 | chr11-6179695-DEL-149 |
| chr11-6188746-DEL-3037 | chr11-6188747-DEL-3037 |
| chr11-6192802-DEL-60 | chr11-6192811-DEL-52 |
| chr11-6253225-DEL-486 | chr11-6253226-DEL-486 |
| chr11-6256635-DEL-74 | chr11-6256636-DEL-74 |
| chr11-6302792-DEL-206 | chr11-6302793-DEL-206 |
| chr11-6307978-DEL-195 | chr11-6307979-DEL-195 |
| chr11-6308483-DEL-213 | chr11-6308482-DEL-214 |
| chr11-6312039-DEL-193 | chr11-6312045-DEL-188 |
| chr11-6330625-DEL-201 | chr11-6330626-DEL-200 |
| chr11-6331829-DEL-180 | chr11-6331830-DEL-180 |
| chr11-6335040-DEL-200 | chr11-6335041-DEL-200 |
| chr11-6374252-DEL-198 | chr11-6374253-DEL-198 |
| chr11-6375783-DEL-228 | chr11-6375782-DEL-228 |
| chr11-6398367-DEL-3231 | chr11-6398368-DEL-3231 |
| chr11-6409666-DEL-1282 | chr11-6409667-DEL-1282 |
| chr11-6439076-DEL-237 | chr11-6439077-DEL-237 |
| chr11-6441398-DEL-205 | chr11-6441385-DEL-200 |
| chr11-6476888-DEL-416 | chr11-6476889-DEL-416 |
| chr11-6480286-DEL-398 | chr11-6480286-DEL-398 |
| chr11-6487729-DEL-202 | chr11-6487730-DEL-202 |
| chr11-6514893-DEL-206 | chr11-6514893-DEL-206 |
| chr11-6522813-DEL-130 | chr11-6522814-DEL-130 |
| chr11-6527716-DEL-183 | chr11-6527716-DEL-183 |
| chr11-6529069-DEL-228 | chr11-6529070-DEL-228 |
| chr11-6536541-DEL-178 | chr11-6536543-DEL-181 |
| chr11-6544608-DEL-224 | chr11-6544609-DEL-224 |
| chr11-6588137-DEL-65 | chr11-6588137-DEL-65 |
| chr11-6591340-DEL-79 | chr11-6591307-DEL-80 |
| chr11-6610936-DEL-271 | chr11-6610983-DEL-270 |
| chr11-6640755-DEL-198 | chr11-6640756-DEL-198 |
| chr11-6643675-DEL-205 | chr11-6643676-DEL-205 |
| chr11-6687129-DEL-120 | chr11-6687130-DEL-120 |
| chr11-6699417-DEL-8974 | chr11-6699418-DEL-8974 |
| chr11-6741013-DEL-9253 | chr11-6741014-DEL-9253 |
| chr11-6778627-DEL-158 | chr11-6778626-DEL-160 |
| chr11-6783844-DEL-118 | chr11-6783845-DEL-118 |
| chr11-6806125-DEL-231 | chr11-6806126-DEL-231 |
| chr11-6815274-DEL-328 | chr11-6815275-DEL-328 |
| chr11-6816860-DEL-73 | chr11-6816861-DEL-73 |
| chr11-6822570-DEL-71 | chr11-6822571-DEL-71 |
| chr11-6900641-DEL-60 | chr11-6900642-DEL-60 |
| chr11-6942909-DEL-63 | chr11-6942910-DEL-61 |
| chr11-6958798-DEL-85 | chr11-6958799-DEL-85 |
| chr11-6976838-DEL-64 | chr11-6976855-DEL-74 |
| chr11-6983330-DEL-237 | chr11-6983331-DEL-237 |
| chr11-6998836-DEL-330 | chr11-6998837-DEL-330 |
| chr11-7048092-DEL-122 | chr11-7048093-DEL-122 |
| chr11-7056055-DEL-52 | chr11-7056056-DEL-52 |
| chr11-7060065-DEL-2219 | chr11-7060066-DEL-2219 |
| chr11-7067061-DEL-90 | chr11-7067044-DEL-91 |
| chr11-7072683-DEL-396 | chr11-7072679-DEL-399 |
| chr11-7139308-DEL-659 | chr11-7139308-DEL-659 |
| chr11-7149620-DEL-74 | chr11-7149621-DEL-74 |
| chr11-7153096-DEL-199 | chr11-7153097-DEL-199 |
| chr11-7187309-DEL-53 | chr11-7187314-DEL-52 |
| chr11-7199258-DEL-54 | chr11-7199259-DEL-54 |
| chr11-7222940-DEL-71 | chr11-7222941-DEL-71 |
| chr11-7237859-DEL-60 | chr11-7237860-DEL-60 |
| chr11-7277735-DEL-1637 | chr11-7277736-DEL-1635 |
| chr11-7290875-DEL-1236 | chr11-7290876-DEL-1236 |
| chr11-7303265-DEL-545 | chr11-7303266-DEL-545 |
| chr11-7307280-DEL-59 | chr11-7307278-DEL-60 |
| chr11-7321854-DEL-90 | chr11-7321855-DEL-90 |
| chr11-7331874-DEL-82 | chr11-7331877-DEL-80 |
| chr11-7353466-DEL-137 | chr11-7353467-DEL-137 |
| chr11-7376655-DEL-554 | chr11-7376656-DEL-554 |
| chr11-7396893-DEL-57 | chr11-7396894-DEL-57 |
| chr11-7437457-DEL-4926 | chr11-7437458-DEL-4926 |
| chr11-7461101-DEL-126 | chr11-7461102-DEL-126 |
| chr11-7494229-DEL-62 | chr11-7494230-DEL-62 |
| chr11-7525688-DEL-7226 | chr11-7525689-DEL-7226 |
| chr11-7534046-DEL-7920 | chr11-7534047-DEL-7920 |
| chr11-7546780-DEL-276 | chr11-7546777-DEL-288 |
| chr11-7562741-DEL-68 | chr11-7562742-DEL-68 |
| chr11-7567752-DEL-1859 | chr11-7567753-DEL-1859 |
| chr11-7609483-DEL-507 | chr11-7609484-DEL-507 |

| | |
|---|---|
| chr11-7613897-DEL-60 | chr11-7613906-DEL-74 |
| chr11-7659703-DEL-342 | chr11-7659704-DEL-342 |
| chr11-7662273-DEL-80 | chr11-7662274-DEL-80 |
| chr11-7663886-DEL-114 | chr11-7663896-DEL-141 |
| chr11-7664780-DEL-6214 | chr11-7664781-DEL-6214 |
| chr11-7676761-DEL-240 | chr11-7676762-DEL-240 |
| chr11-7686663-DEL-60 | chr11-7686665-DEL-62 |
| chr11-7758298-DEL-55 | chr11-7758299-DEL-55 |
| chr11-7761396-DEL-236 | chr11-7761397-DEL-236 |
| chr11-7772814-DEL-62 | chr11-7772815-DEL-62 |
| chr11-7794332-DEL-141 | chr11-7794333-DEL-141 |
| chr11-7801945-DEL-250 | chr11-7801944-DEL-251 |
| chr11-7810342-DEL-1552 | chr11-7810343-DEL-1552 |
| chr11-7860981-DEL-131 | chr11-7860965-DEL-108 |
| chr11-7870727-DEL-6877 | chr11-7870728-DEL-6877 |
| chr11-7877042-DEL-214 | chr11-7877042-DEL-214 |
| chr11-7880335-DEL-82 | chr11-7880338-DEL-104 |
| chr11-7881645-DEL-342 | chr11-7881646-DEL-342 |
| chr11-7905099-DEL-1454 | chr11-7905085-DEL-1442 |
| chr11-7915561-DEL-1171 | chr11-7915561-DEL-1171 |
| chr11-7936784-DEL-441 | chr11-7936785-DEL-441 |
| chr11-7956612-DEL-309 | chr11-7956613-DEL-309 |
| chr11-7962118-DEL-514 | chr11-7962119-DEL-514 |
| chr11-7965013-DEL-425 | chr11-7965014-DEL-425 |
| chr11-7994415-DEL-756 | chr11-7994416-DEL-756 |
| chr11-8026473-DEL-6452 | chr11-8026474-DEL-6452 |
| chr11-8049685-DEL-372 | chr11-8049686-DEL-372 |
| chr11-8062243-DEL-1130 | chr11-8062244-DEL-1130 |
| chr11-8076676-DEL-401 | chr11-8076677-DEL-401 |
| chr11-8082304-DEL-52 | chr11-8082305-DEL-52 |
| chr11-8099298-DEL-6721 | chr11-8099285-DEL-6722 |
| chr11-8138747-DEL-102 | chr11-8138748-DEL-106 |
| chr11-8169901-DEL-61 | chr11-8169902-DEL-61 |
| chr11-8217841-DEL-974 | chr11-8217842-DEL-974 |
| chr11-8220598-DEL-3662 | chr11-8220599-DEL-3662 |
| chr11-8228205-DEL-69 | chr11-8228206-DEL-70 |
| chr11-8237817-DEL-226 | chr11-8237818-DEL-226 |
| chr11-8272053-DEL-1110 | chr11-8272052-DEL-1110 |
| chr11-8288045-DEL-1642 | chr11-8288046-DEL-1642 |
| chr11-8309981-DEL-82 | chr11-8309949-DEL-68 |
| chr11-8310427-DEL-259 | chr11-8310428-DEL-259 |
| chr11-8319765-DEL-992 | chr11-8319766-DEL-992 |
| chr11-8323773-DEL-96 | chr11-8323774-DEL-96 |
| chr11-8378848-DEL-353 | chr11-8378850-DEL-350 |
| chr11-8404843-DEL-98 | chr11-8404827-DEL-113 |
| chr11-8421552-DEL-668 | chr11-8421553-DEL-668 |
| chr11-8424810-DEL-3168 | chr11-8424811-DEL-3166 |
| chr11-8444565-DEL-133 | chr11-8444566-DEL-133 |
| chr11-8489344-DEL-91 | chr11-8489361-DEL-71 |
| chr11-8493031-DEL-195 | chr11-8493032-DEL-195 |
| chr11-8512497-DEL-388 | chr11-8512498-DEL-388 |
| chr11-8580880-DEL-75 | chr11-8580881-DEL-75 |
| chr11-8583368-DEL-50 | chr11-8583383-DEL-73 |
| chr11-8604352-DEL-1163 | chr11-8604353-DEL-1163 |
| chr11-8683261-DEL-208 | chr11-8683262-DEL-208 |
| chr11-8691474-DEL-2354 | chr11-8691474-DEL-2354 |
| chr11-8694527-DEL-62 | chr11-8694528-DEL-62 |
| chr11-8700915-DEL-65 | chr11-8700927-DEL-59 |
| chr11-8716219-DEL-67 | chr11-8716210-DEL-67 |
| chr11-8724855-DEL-62 | chr11-8724856-DEL-62 |
| chr11-8731242-DEL-67 | chr11-8731243-DEL-67 |
| chr11-8734131-DEL-137 | chr11-8734128-DEL-132 |
| chr11-8762485-DEL-64 | chr11-8762498-DEL-52 |
| chr11-8787866-DEL-5172 | chr11-8787866-DEL-5172 |
| chr11-8823388-DEL-122 | chr11-8823389-DEL-122 |
| chr11-8849115-DEL-53 | chr11-8849112-DEL-53 |
| chr11-8870293-DEL-8984 | chr11-8870294-DEL-8984 |
| chr11-8881771-DEL-53 | chr11-8881758-DEL-51 |
| chr11-8887885-DEL-857 | chr11-8887886-DEL-857 |
| chr11-8896775-DEL-199 | chr11-8896776-DEL-199 |
| chr11-8899006-DEL-197 | chr11-8899007-DEL-197 |
| chr11-8903228-DEL-199 | chr11-8903229-DEL-199 |
| chr11-8917947-DEL-158 | chr11-8917948-DEL-158 |
| chr11-8927477-DEL-794 | chr11-8927475-DEL-797 |
| chr11-8962645-DEL-487 | chr11-8962646-DEL-487 |
| chr11-8978441-DEL-86 | chr11-8978442-DEL-86 |
| chr11-8982184-DEL-156 | chr11-8982185-DEL-156 |
| chr11-9042595-DEL-54 | chr11-9042598-DEL-52 |
| chr11-9080508-DEL-490 | chr11-9080509-DEL-490 |
| chr11-9088017-DEL-627 | chr11-9088018-DEL-627 |

| | |
|---|---|
| chr11-9099442-DEL-1234 | chr11-9099443-DEL-1234 |
| chr11-9122201-DEL-1110 | chr11-9122203-DEL-1109 |
| chr11-9148443-DEL-618 | chr11-9148444-DEL-618 |
| chr11-9162249-DEL-60 | chr11-9162250-DEL-60 |
| chr11-9186313-DEL-307 | chr11-9186314-DEL-307 |
| chr11-9195376-DEL-75 | chr11-9195386-DEL-85 |
| chr11-9199302-DEL-66 | chr11-9199303-DEL-64 |
| chr11-9200137-DEL-107 | chr11-9200159-DEL-108 |
| chr11-9203847-DEL-74 | chr11-9203848-DEL-70 |
| chr11-9205783-DEL-197 | chr11-9205784-DEL-197 |
| chr11-9206805-DEL-214 | chr11-9206806-DEL-214 |
| chr11-9256517-DEL-69 | chr11-9256531-DEL-56 |
| chr11-9269750-DEL-52 | chr11-9269751-DEL-52 |
| chr11-9297783-DEL-53 | chr11-9297784-DEL-53 |
| chr11-9308814-DEL-6667 | chr11-9308815-DEL-6667 |
| chr11-9338317-DEL-131 | chr11-9338318-DEL-133 |
| chr11-9339670-DEL-95 | chr11-9339671-DEL-65 |
| chr11-9356447-DEL-52 | chr11-9356448-DEL-52 |
| chr11-9367284-DEL-769 | chr11-9367283-DEL-770 |
| chr11-9392654-DEL-7986 | chr11-9392654-DEL-7986 |
| chr11-9411637-DEL-78 | chr11-9411650-DEL-68 |
| chr11-9480718-DEL-52 | chr11-9480719-DEL-52 |
| chr11-9497378-DEL-1331 | chr11-9497379-DEL-1331 |
| chr11-9570923-DEL-96 | chr11-9570924-DEL-96 |
| chr11-9605746-DEL-235 | chr11-9605744-DEL-235 |
| chr11-9611799-DEL-231 | chr11-9611800-DEL-231 |
| chr11-9643862-DEL-147 | chr11-9643863-DEL-147 |
| chr11-9657979-DEL-7089 | chr11-9657980-DEL-7089 |
| chr11-9669582-DEL-582 | chr11-9669583-DEL-582 |
| chr11-9671141-DEL-158 | chr11-9671142-DEL-157 |
| chr11-9685572-DEL-70 | chr11-9685573-DEL-70 |
| chr11-9686986-DEL-622 | chr11-9686987-DEL-622 |
| chr11-9700140-DEL-201 | chr11-9700141-DEL-201 |
| chr11-9717822-DEL-4043 | chr11-9717823-DEL-4043 |
| chr11-9722778-DEL-147 | chr11-9722779-DEL-147 |
| chr11-9735675-DEL-62 | chr11-9735676-DEL-54 |
| chr11-9748658-DEL-244 | chr11-9748659-DEL-244 |
| chr11-9769387-DEL-584 | chr11-9769388-DEL-584 |
| chr11-9775205-DEL-54 | chr11-9775206-DEL-54 |
| chr11-9786957-DEL-352 | chr11-9786959-DEL-323 |
| chr11-9895506-DEL-1246 | chr11-9895507-DEL-1246 |
| chr11-9920763-DEL-5543 | chr11-9920764-DEL-5543 |
| chr11-9946527-DEL-475 | chr11-9946528-DEL-475 |
| chr11-9973361-DEL-269 | chr11-9973362-DEL-269 |
| chr11-9981115-DEL-866 | chr11-9981115-DEL-869 |
| chr11-9988249-DEL-344 | chr11-9988250-DEL-344 |
| chr11-10004270-DEL-56 | chr11-10004270-DEL-56 |
| chr11-10015127-DEL-82 | chr11-10015128-DEL-82 |
| chr11-10047747-DEL-52 | chr11-10047748-DEL-52 |
| chr11-10078381-DEL-130 | chr11-10078382-DEL-130 |
| chr11-10095217-DEL-1419 | chr11-10095218-DEL-1419 |
| chr11-10127525-DEL-498 | chr11-10127526-DEL-498 |
| chr11-10148082-DEL-2922 | chr11-10148083-DEL-2922 |
| chr11-10160067-DEL-282 | chr11-10160068-DEL-282 |
| chr11-10166660-DEL-94 | chr11-10166660-DEL-94 |
| chr11-10172146-DEL-4183 | chr11-10172145-DEL-4178 |
| chr11-10228971-DEL-116 | chr11-10228963-DEL-117 |
| chr11-10230011-DEL-68 | chr11-10230034-DEL-73 |
| chr11-10264407-DEL-284 | chr11-10264424-DEL-270 |
| chr11-10269188-DEL-714 | chr11-10269189-DEL-714 |
| chr11-10278860-DEL-430 | chr11-10278861-DEL-430 |
| chr11-10307060-DEL-3724 | chr11-10307060-DEL-3724 |
| chr11-10371845-DEL-817 | chr11-10371846-DEL-817 |
| chr11-10413821-DEL-78 | chr11-10413822-DEL-78 |
| chr11-10433146-DEL-57 | chr11-10433172-DEL-59 |
| chr11-10479292-DEL-6560 | chr11-10479294-DEL-6559 |
| chr11-10488330-DEL-51 | chr11-10488328-DEL-54 |
| chr11-10500757-DEL-76 | chr11-10500761-DEL-77 |
| chr11-10569799-DEL-60 | chr11-10569800-DEL-60 |
| chr11-10593212-DEL-3517 | chr11-10593213-DEL-3517 |
| chr11-10624938-DEL-1579 | chr11-10624939-DEL-1579 |
| chr11-10644867-DEL-231 | chr11-10644868-DEL-231 |
| chr11-10653657-DEL-1008 | chr11-10653658-DEL-1008 |
| chr11-10673667-DEL-325 | chr11-10673668-DEL-325 |
| chr11-10732471-DEL-695 | chr11-10732472-DEL-695 |
| chr11-10764289-DEL-103 | chr11-10764285-DEL-75 |
| chr11-10776926-DEL-2347 | chr11-10776927-DEL-2347 |
| chr11-10801447-DEL-6215 | chr11-10801448-DEL-6215 |
| chr11-10829284-DEL-6239 | chr11-10829286-DEL-6238 |
| chr11-10845775-DEL-169 | chr11-10845776-DEL-169 |

| | |
|---|---|
| chr11-10869498-DEL-644 | chr11-10869497-DEL-643 |
| chr11-10917953-DEL-717 | chr11-10917952-DEL-717 |
| chr11-10984136-DEL-60 | chr11-10984140-DEL-75 |
| chr11-10992817-DEL-6407 | chr11-10992818-DEL-6407 |
| chr11-11005938-DEL-1223 | chr11-11005939-DEL-1223 |
| chr11-11022530-DEL-86 | chr11-11022531-DEL-86 |
| chr11-11028909-DEL-53 | chr11-11028910-DEL-53 |
| chr11-11057702-DEL-787 | chr11-11057703-DEL-787 |
| chr11-11070769-DEL-133 | chr11-11070770-DEL-133 |
| chr11-11073885-DEL-3018 | chr11-11073886-DEL-3018 |
| chr11-11091599-DEL-6478 | chr11-11091600-DEL-6478 |
| chr11-11130180-DEL-403 | chr11-11130181-DEL-403 |
| chr11-11139959-DEL-298 | chr11-11139960-DEL-298 |
| chr11-11173422-DEL-2084 | chr11-11173423-DEL-2084 |
| chr11-11207779-DEL-168 | chr11-11207780-DEL-168 |
| chr11-11224798-DEL-59 | chr11-11224801-DEL-55 |
| chr11-11237209-DEL-7313 | chr11-11237210-DEL-7313 |
| chr11-11255858-DEL-2241 | chr11-11255859-DEL-2241 |
| chr11-11409158-DEL-139 | chr11-11409164-DEL-137 |
| chr11-11428014-DEL-1194 | chr11-11428015-DEL-1195 |
| chr11-11488597-DEL-54 | chr11-11488602-DEL-50 |
| chr11-11491007-DEL-187 | chr11-11491008-DEL-187 |
| chr11-11512536-DEL-432 | chr11-11512537-DEL-432 |
| chr11-11632609-DEL-64 | chr11-11632621-DEL-64 |
| chr11-11695621-DEL-78 | chr11-11695622-DEL-52 |
| chr11-11712655-DEL-207 | chr11-11712656-DEL-207 |
| chr11-11728141-DEL-184 | chr11-11728142-DEL-184 |
| chr11-11742360-DEL-362 | chr11-11742361-DEL-362 |
| chr11-11755248-DEL-173 | chr11-11755249-DEL-173 |
| chr11-11819578-DEL-121 | chr11-11819579-DEL-121 |
| chr11-11825784-DEL-64 | chr11-11825784-DEL-64 |
| chr11-11846011-DEL-361 | chr11-11846014-DEL-328 |
| chr11-11847194-DEL-66 | chr11-11847195-DEL-66 |
| chr11-11848111-DEL-66 | chr11-11848112-DEL-66 |
| chr11-11858355-DEL-213 | chr11-11858356-DEL-213 |
| chr11-11862450-DEL-230 | chr11-11862451-DEL-230 |
| chr11-11879997-DEL-117 | chr11-11879998-DEL-117 |
| chr11-11930390-DEL-485 | chr11-11930390-DEL-482 |
| chr11-11957568-DEL-54 | chr11-11957570-DEL-52 |
| chr11-12045359-DEL-1177 | chr11-12045376-DEL-1178 |
| chr11-12050780-DEL-213 | chr11-12050824-DEL-213 |
| chr11-12126592-DEL-3310 | chr11-12126593-DEL-3310 |
| chr11-12131245-DEL-74 | chr11-12131245-DEL-74 |
| chr11-12180325-DEL-510 | chr11-12180326-DEL-510 |
| chr11-12253541-DEL-233 | chr11-12253542-DEL-233 |
| chr11-12283418-DEL-1365 | chr11-12283419-DEL-1365 |
| chr11-12325025-DEL-193 | chr11-12325026-DEL-193 |
| chr11-12373868-DEL-213 | chr11-12373869-DEL-213 |
| chr11-12436000-DEL-1042 | chr11-12436001-DEL-1042 |
| chr11-12455017-DEL-56 | chr11-12455025-DEL-57 |
| chr11-12457301-DEL-5298 | chr11-12457302-DEL-5298 |
| chr11-12511902-DEL-7127 | chr11-12511903-DEL-7127 |
| chr11-12519458-DEL-467 | chr11-12519459-DEL-467 |
| chr11-12526794-DEL-829 | chr11-12526794-DEL-829 |
| chr11-12532460-DEL-70 | chr11-12532461-DEL-70 |
| chr11-12535075-DEL-155 | chr11-12535076-DEL-155 |
| chr11-12538497-DEL-79 | chr11-12538495-DEL-79 |
| chr11-12541648-DEL-2216 | chr11-12541649-DEL-2216 |
| chr11-12587924-DEL-6274 | chr11-12587925-DEL-6274 |
| chr11-12599987-DEL-570 | chr11-12599988-DEL-570 |
| chr11-12612436-DEL-2418 | chr11-12612437-DEL-2418 |
| chr11-12628320-DEL-374 | chr11-12628321-DEL-374 |
| chr11-12631857-DEL-1166 | chr11-12631858-DEL-1162 |
| chr11-12660435-DEL-552 | chr11-12660435-DEL-552 |
| chr11-12745530-DEL-7170 | chr11-12745531-DEL-7170 |
| chr11-12753139-DEL-586 | chr11-12753140-DEL-586 |
| chr11-12848630-DEL-5485 | chr11-12848629-DEL-5485 |
| chr11-12856651-DEL-135 | chr11-12856652-DEL-135 |
| chr11-12876669-DEL-428 | chr11-12876681-DEL-444 |
| chr11-12904016-DEL-5193 | chr11-12904017-DEL-5193 |
| chr11-12913733-DEL-1124 | chr11-12913734-DEL-1124 |
| chr11-12944783-DEL-108 | chr11-12944782-DEL-109 |
| chr11-12951167-DEL-177 | chr11-12951164-DEL-144 |
| chr11-13090803-DEL-506 | chr11-13090804-DEL-506 |
| chr11-13112988-DEL-1482 | chr11-13112989-DEL-1482 |
| chr11-13113102-DEL-351 | chr11-13113103-DEL-351 |
| chr11-13130824-DEL-400 | chr11-13130825-DEL-400 |
| chr11-13166000-DEL-400 | chr11-13166000-DEL-400 |
| chr11-13189685-DEL-236 | chr11-13189686-DEL-236 |
| chr11-13190996-DEL-52 | chr11-13190992-DEL-56 |

| | |
|---|---|
| chr11-13201889-DEL-1669 | chr11-13201890-DEL-1669 |
| chr11-13220779-DEL-150 | chr11-13220780-DEL-146 |
| chr11-13237617-DEL-6157 | chr11-13237617-DEL-6157 |
| chr11-13259023-DEL-7208 | chr11-13259024-DEL-7208 |
| chr11-13265271-DEL-212 | chr11-13265272-DEL-212 |
| chr11-13305145-DEL-54 | chr11-13305146-DEL-54 |
| chr11-13352575-DEL-184 | chr11-13352576-DEL-184 |
| chr11-13405730-DEL-483 | chr11-13405732-DEL-482 |
| chr11-13415538-DEL-80 | chr11-13415539-DEL-80 |
| chr11-13419204-DEL-109 | chr11-13419221-DEL-83 |
| chr11-13434253-DEL-1002 | chr11-13434301-DEL-954 |
| chr11-13437851-DEL-1795 | chr11-13437830-DEL-1796 |
| chr11-13481323-DEL-68 | chr11-13481324-DEL-68 |
| chr11-13493909-DEL-76 | chr11-13493934-DEL-52 |
| chr11-13495408-DEL-71 | chr11-13495409-DEL-71 |
| chr11-13496221-DEL-6831 | chr11-13496222-DEL-6831 |
| chr11-13511746-DEL-288 | chr11-13511747-DEL-288 |
| chr11-13519861-DEL-5541 | chr11-13519861-DEL-5541 |
| chr11-13536255-DEL-1466 | chr11-13536255-DEL-1466 |
| chr11-13542276-DEL-51 | chr11-13542267-DEL-68 |
| chr11-13545570-DEL-54 | chr11-13545569-DEL-60 |
| chr11-13565240-DEL-7499 | chr11-13565235-DEL-7498 |
| chr11-13577499-DEL-71 | chr11-13577508-DEL-50 |
| chr11-13582341-DEL-52 | chr11-13582340-DEL-54 |
| chr11-13609544-DEL-434 | chr11-13609545-DEL-434 |
| chr11-13620609-DEL-61 | chr11-13620611-DEL-60 |
| chr11-13662614-DEL-525 | chr11-13662615-DEL-525 |
| chr11-13671930-DEL-66 | chr11-13671931-DEL-66 |
| chr11-13677725-DEL-1256 | chr11-13677726-DEL-1256 |
| chr11-13691229-DEL-789 | chr11-13691230-DEL-788 |
| chr11-13726455-DEL-52 | chr11-13726464-DEL-56 |
| chr11-13765937-DEL-7009 | chr11-13765938-DEL-7009 |
| chr11-13778291-DEL-52 | chr11-13778292-DEL-53 |
| chr11-13822863-DEL-2974 | chr11-13822865-DEL-2975 |
| chr11-13849258-DEL-444 | chr11-13849259-DEL-444 |
| chr11-13859400-DEL-123 | chr11-13859395-DEL-123 |
| chr11-13865756-DEL-2639 | chr11-13865757-DEL-2639 |
| chr11-13872185-DEL-7467 | chr11-13872186-DEL-7467 |
| chr11-13881416-DEL-2171 | chr11-13881417-DEL-2171 |
| chr11-13896574-DEL-6945 | chr11-13896575-DEL-6945 |
| chr11-13914083-DEL-212 | chr11-13914084-DEL-212 |
| chr11-13916642-DEL-74 | chr11-13916643-DEL-74 |
| chr11-13943410-DEL-708 | chr11-13943411-DEL-708 |
| chr11-13956640-DEL-51 | chr11-13956641-DEL-51 |
| chr11-13962357-DEL-191 | chr11-13962358-DEL-191 |
| chr11-14012697-DEL-163 | chr11-14012698-DEL-163 |
| chr11-14039999-DEL-64 | chr11-14040000-DEL-64 |
| chr11-14059338-DEL-239 | chr11-14059339-DEL-239 |
| chr11-14062003-DEL-118 | chr11-14062004-DEL-118 |
| chr11-14079545-DEL-140 | chr11-14079545-DEL-140 |
| chr11-14086523-DEL-9611 | chr11-14086524-DEL-9611 |
| chr11-14092412-DEL-1659 | chr11-14092409-DEL-1660 |
| chr11-14153999-DEL-66 | chr11-14153998-DEL-62 |
| chr11-14156960-DEL-79 | chr11-14156959-DEL-78 |
| chr11-14181234-DEL-55 | chr11-14181219-DEL-70 |
| chr11-14187467-DEL-57 | chr11-14187468-DEL-57 |
| chr11-14214571-DEL-258 | chr11-14214573-DEL-247 |
| chr11-14217402-DEL-64 | chr11-14217399-DEL-66 |
| chr11-14220208-DEL-61 | chr11-14220211-DEL-58 |
| chr11-14230186-DEL-2303 | chr11-14230187-DEL-2303 |
| chr11-14259997-DEL-2612 | chr11-14259992-DEL-2613 |
| chr11-14274110-DEL-966 | chr11-14274111-DEL-966 |
| chr11-14297985-DEL-122 | chr11-14297986-DEL-122 |
| chr11-14339933-DEL-4632 | chr11-14339934-DEL-4632 |
| chr11-14345774-DEL-398 | chr11-14345775-DEL-398 |
| chr11-14361152-DEL-1387 | chr11-14361153-DEL-1387 |
| chr11-14381964-DEL-1844 | chr11-14381965-DEL-1842 |
| chr11-14403887-DEL-11592 | chr11-14403888-DEL-11592 |
| chr11-14422965-DEL-3139 | chr11-14422965-DEL-3139 |
| chr11-14431112-DEL-135 | chr11-14431113-DEL-132 |
| chr11-14463709-DEL-6008 | chr11-14463710-DEL-6008 |
| chr11-14471154-DEL-485 | chr11-14471155-DEL-485 |
| chr11-14477607-DEL-64 | chr11-14477605-DEL-86 |
| chr11-14498501-DEL-725 | chr11-14498502-DEL-725 |
| chr11-14535377-DEL-899 | chr11-14535378-DEL-899 |
| chr11-14562150-DEL-602 | chr11-14562151-DEL-602 |
| chr11-14582070-DEL-76 | chr11-14582071-DEL-76 |
| chr11-14597104-DEL-1500 | chr11-14597104-DEL-1500 |
| chr11-14663987-DEL-60 | chr11-14663988-DEL-60 |
| chr11-14665625-DEL-354 | chr11-14665626-DEL-354 |

| | |
|---|---|
| chr11-14678065-DEL-92 | chr11-14678066-DEL-92 |
| chr11-14683447-DEL-199 | chr11-14683448-DEL-199 |
| chr11-14699168-DEL-76 | chr11-14699169-DEL-76 |
| chr11-14746098-DEL-71 | chr11-14746104-DEL-72 |
| chr11-14776368-DEL-136 | chr11-14776369-DEL-136 |
| chr11-14790558-DEL-737 | chr11-14790559-DEL-737 |
| chr11-14807207-DEL-1895 | chr11-14807208-DEL-1895 |
| chr11-14846616-DEL-3448 | chr11-14846613-DEL-3449 |
| chr11-14851370-DEL-438 | chr11-14851371-DEL-438 |
| chr11-14852415-DEL-94 | chr11-14852407-DEL-95 |
| chr11-14865859-DEL-6276 | chr11-14865860-DEL-6276 |
| chr11-14872432-DEL-59 | chr11-14872433-DEL-59 |
| chr11-14876902-DEL-363 | chr11-14876903-DEL-363 |
| chr11-14883333-DEL-211 | chr11-14883334-DEL-211 |
| chr11-14909700-DEL-327 | chr11-14909701-DEL-327 |
| chr11-14924624-DEL-453 | chr11-14924625-DEL-453 |
| chr11-14927042-DEL-1960 | chr11-14927043-DEL-1960 |
| chr11-14936502-DEL-856 | chr11-14936492-DEL-857 |
| chr11-14945698-DEL-62 | chr11-14945699-DEL-62 |
| chr11-14956823-DEL-994 | chr11-14956824-DEL-994 |
| chr11-14971416-DEL-284 | chr11-14971416-DEL-284 |
| chr11-14978351-DEL-984 | chr11-14978352-DEL-984 |
| chr11-14979948-DEL-2116 | chr11-14979949-DEL-2116 |
| chr11-14983091-DEL-870 | chr11-14983092-DEL-870 |
| chr11-15006270-DEL-2869 | chr11-15006270-DEL-2869 |
| chr11-15011770-DEL-709 | chr11-15011771-DEL-709 |
| chr11-15016748-DEL-2512 | chr11-15016749-DEL-2512 |
| chr11-15027787-DEL-2440 | chr11-15027788-DEL-2440 |
| chr11-15035437-DEL-60 | chr11-15035438-DEL-60 |
| chr11-15039653-DEL-2094 | chr11-15039654-DEL-2091 |
| chr11-15046500-DEL-177 | chr11-15046500-DEL-177 |
| chr11-15117913-DEL-5519 | chr11-15117914-DEL-5519 |
| chr11-15159021-DEL-75 | chr11-15159022-DEL-75 |
| chr11-15177685-DEL-7180 | chr11-15177683-DEL-7181 |
| chr11-15194254-DEL-384 | chr11-15194255-DEL-384 |
| chr11-15205439-DEL-755 | chr11-15205440-DEL-755 |
| chr11-15248980-DEL-6176 | chr11-15248981-DEL-6176 |
| chr11-15267690-DEL-80 | chr11-15267691-DEL-80 |
| chr11-15284137-DEL-431 | chr11-15284138-DEL-431 |
| chr11-15295436-DEL-1689 | chr11-15295437-DEL-1689 |
| chr11-15313573-DEL-153 | chr11-15313574-DEL-153 |
| chr11-15341551-DEL-975 | chr11-15341552-DEL-975 |
| chr11-15348051-DEL-3473 | chr11-15348052-DEL-3473 |
| chr11-15373754-DEL-2317 | chr11-15373755-DEL-2317 |
| chr11-15375411-DEL-342 | chr11-15375412-DEL-342 |
| chr11-15380184-DEL-55 | chr11-15380186-DEL-56 |
| chr11-15425302-DEL-438 | chr11-15425302-DEL-437 |
| chr11-15430914-DEL-52 | chr11-15430915-DEL-54 |
| chr11-15454363-DEL-331 | chr11-15454364-DEL-331 |
| chr11-15470830-DEL-55 | chr11-15470830-DEL-56 |
| chr11-15513666-DEL-87 | chr11-15513657-DEL-68 |
| chr11-15515668-DEL-56 | chr11-15515669-DEL-56 |
| chr11-15558076-DEL-66 | chr11-15558077-DEL-66 |
| chr11-15596467-DEL-52 | chr11-15596469-DEL-78 |
| chr11-15639377-DEL-78 | chr11-15639378-DEL-78 |
| chr11-15642308-DEL-689 | chr11-15642308-DEL-689 |
| chr11-15644744-DEL-3159 | chr11-15644745-DEL-3159 |
| chr11-15668093-DEL-839 | chr11-15668094-DEL-839 |
| chr11-15679264-DEL-6382 | chr11-15679265-DEL-6383 |
| chr11-15685920-DEL-76 | chr11-15685921-DEL-66 |
| chr11-15688340-DEL-59 | chr11-15688353-DEL-63 |
| chr11-15699373-DEL-682 | chr11-15699373-DEL-682 |
| chr11-15704522-DEL-181 | chr11-15704522-DEL-181 |
| chr11-15728125-DEL-6457 | chr11-15728126-DEL-6457 |
| chr11-15747794-DEL-91 | chr11-15747794-DEL-91 |
| chr11-15748458-DEL-2523 | chr11-15748458-DEL-2523 |
| chr11-15752734-DEL-687 | chr11-15752734-DEL-688 |
| chr11-15755369-DEL-357 | chr11-15755368-DEL-357 |
| chr11-15773221-DEL-1420 | chr11-15773222-DEL-1420 |
| chr11-15777359-DEL-2567 | chr11-15777360-DEL-2567 |
| chr11-15778617-DEL-6606 | chr11-15778618-DEL-6606 |
| chr11-15786964-DEL-3891 | chr11-15786965-DEL-3891 |
| chr11-15805100-DEL-9142 | chr11-15805101-DEL-9142 |
| chr11-15829489-DEL-445 | chr11-15829483-DEL-437 |
| chr11-15833671-DEL-4129 | chr11-15833672-DEL-4129 |
| chr11-15841086-DEL-409 | chr11-15841086-DEL-410 |
| chr11-15846904-DEL-65 | chr11-15846904-DEL-65 |
| chr11-15896279-DEL-99 | chr11-15896280-DEL-99 |
| chr11-15902419-DEL-195 | chr11-15902420-DEL-195 |
| chr11-15951785-DEL-64 | chr11-15951786-DEL-64 |

| | |
|---|---|
| chr11-16045892-DEL-6194 | chr11-16045893-DEL-6194 |
| chr11-16060565-DEL-106 | chr11-16060578-DEL-82 |
| chr11-16062516-DEL-81 | chr11-16062521-DEL-76 |
| chr11-16113761-DEL-2409 | chr11-16113762-DEL-2409 |
| chr11-16120363-DEL-90 | chr11-16120348-DEL-60 |
| chr11-16138520-DEL-3927 | chr11-16138521-DEL-3927 |
| chr11-16146412-DEL-54 | chr11-16146417-DEL-53 |
| chr11-16154255-DEL-2663 | chr11-16154256-DEL-2663 |
| chr11-16158272-DEL-1095 | chr11-16158273-DEL-1095 |
| chr11-16160187-DEL-60 | chr11-16160188-DEL-54 |
| chr11-16160967-DEL-174 | chr11-16160968-DEL-174 |
| chr11-16178386-DEL-4409 | chr11-16178387-DEL-4409 |
| chr11-16183857-DEL-3274 | chr11-16183858-DEL-3274 |
| chr11-16226536-DEL-56 | chr11-16226537-DEL-56 |
| chr11-16230737-DEL-79 | chr11-16230740-DEL-76 |
| chr11-16255656-DEL-86 | chr11-16255657-DEL-86 |
| chr11-16272911-DEL-373 | chr11-16272912-DEL-373 |
| chr11-16273585-DEL-57 | chr11-16273586-DEL-57 |
| chr11-16273822-DEL-1133 | chr11-16273823-DEL-1133 |
| chr11-16303576-DEL-889 | chr11-16303577-DEL-889 |
| chr11-16309098-DEL-324 | chr11-16309099-DEL-324 |
| chr11-16317921-DEL-61 | chr11-16317921-DEL-61 |
| chr11-16332766-DEL-1827 | chr11-16332776-DEL-1827 |
| chr11-16335688-DEL-6663 | chr11-16335689-DEL-6663 |
| chr11-16365337-DEL-238 | chr11-16365338-DEL-238 |
| chr11-16384672-DEL-222 | chr11-16384673-DEL-222 |
| chr11-16423091-DEL-530 | chr11-16423092-DEL-530 |
| chr11-16424054-DEL-187 | chr11-16424057-DEL-181 |
| chr11-16429123-DEL-5622 | chr11-16429111-DEL-5634 |
| chr11-16436602-DEL-745 | chr11-16436603-DEL-745 |
| chr11-16487633-DEL-73 | chr11-16487643-DEL-65 |
| chr11-16493330-DEL-1845 | chr11-16493331-DEL-1845 |
| chr11-16508501-DEL-283 | chr11-16508504-DEL-281 |
| chr11-16517016-DEL-5983 | chr11-16517017-DEL-5983 |
| chr11-16523844-DEL-210 | chr11-16523845-DEL-211 |
| chr11-16528024-DEL-73 | chr11-16528025-DEL-73 |
| chr11-16584068-DEL-2077 | chr11-16584069-DEL-2077 |
| chr11-16586641-DEL-111 | chr11-16586642-DEL-111 |
| chr11-16597019-DEL-306 | chr11-16597020-DEL-306 |
| chr11-16606550-DEL-104 | chr11-16606550-DEL-80 |
| chr11-16612936-DEL-65 | chr11-16612936-DEL-65 |
| chr11-16615605-DEL-861 | chr11-16615608-DEL-859 |
| chr11-16618456-DEL-63 | chr11-16618456-DEL-63 |
| chr11-16638902-DEL-5802 | chr11-16638903-DEL-5802 |
| chr11-16699741-DEL-206 | chr11-16699742-DEL-206 |
| chr11-16730050-DEL-130 | chr11-16730048-DEL-131 |
| chr11-16765121-DEL-87 | chr11-16765122-DEL-86 |
| chr11-16780083-DEL-56 | chr11-16780084-DEL-56 |
| chr11-16797008-DEL-409 | chr11-16797009-DEL-409 |
| chr11-16822598-DEL-73 | chr11-16822574-DEL-74 |
| chr11-16841306-DEL-52 | chr11-16841307-DEL-52 |
| chr11-16874219-DEL-397 | chr11-16874220-DEL-397 |
| chr11-16911112-DEL-87 | chr11-16911113-DEL-87 |
| chr11-16911477-DEL-480 | chr11-16911478-DEL-480 |
| chr11-16956385-DEL-233 | chr11-16956386-DEL-233 |
| chr11-16961241-DEL-75 | chr11-16961242-DEL-75 |
| chr11-16982254-DEL-1553 | chr11-16982255-DEL-1553 |
| chr11-17008481-DEL-3624 | chr11-17008482-DEL-3624 |
| chr11-17014907-DEL-62 | chr11-17014908-DEL-62 |
| chr11-17042647-DEL-404 | chr11-17042648-DEL-404 |
| chr11-17048972-DEL-6232 | chr11-17048973-DEL-6232 |
| chr11-17071697-DEL-157 | chr11-17071697-DEL-157 |
| chr11-17072653-DEL-789 | chr11-17072663-DEL-794 |
| chr11-17079127-DEL-179 | chr11-17079130-DEL-132 |
| chr11-17101627-DEL-2256 | chr11-17101628-DEL-2256 |
| chr11-17104463-DEL-188 | chr11-17104464-DEL-188 |
| chr11-17125066-DEL-79 | chr11-17125080-DEL-68 |
| chr11-17128806-DEL-1456 | chr11-17128807-DEL-1456 |
| chr11-17167032-DEL-243 | chr11-17167058-DEL-199 |
| chr11-17184379-DEL-544 | chr11-17184380-DEL-544 |
| chr11-17190174-DEL-1009 | chr11-17190175-DEL-1009 |
| chr11-17192437-DEL-1450 | chr11-17192438-DEL-1450 |
| chr11-17203875-DEL-1263 | chr11-17203876-DEL-1263 |
| chr11-17204220-DEL-461 | chr11-17204221-DEL-461 |
| chr11-17205835-DEL-183 | chr11-17205830-DEL-184 |
| chr11-17224200-DEL-91 | chr11-17224211-DEL-63 |
| chr11-17227119-DEL-61 | chr11-17227119-DEL-66 |
| chr11-17228526-DEL-202 | chr11-17228511-DEL-198 |
| chr11-17232746-DEL-1259 | chr11-17232758-DEL-1272 |
| chr11-17234079-DEL-54 | chr11-17234080-DEL-54 |

| | |
|---|---|
| chr11-17240022-DEL-394 | chr11-17240023-DEL-394 |
| chr11-17244017-DEL-6418 | chr11-17244019-DEL-6417 |
| chr11-17252836-DEL-66 | chr11-17252837-DEL-66 |
| chr11-17266209-DEL-912 | chr11-17266210-DEL-912 |
| chr11-17267176-DEL-153 | chr11-17267177-DEL-153 |
| chr11-17323642-DEL-219 | chr11-17323644-DEL-206 |
| chr11-17324770-DEL-431 | chr11-17324771-DEL-432 |
| chr11-17330890-DEL-272 | chr11-17330891-DEL-272 |
| chr11-17351320-DEL-6417 | chr11-17351321-DEL-6417 |
| chr11-17358142-DEL-180 | chr11-17358143-DEL-180 |
| chr11-17369170-DEL-278 | chr11-17369171-DEL-278 |
| chr11-17393093-DEL-60 | chr11-17393093-DEL-62 |
| chr11-17407089-DEL-75 | chr11-17407090-DEL-75 |
| chr11-17434893-DEL-64 | chr11-17434890-DEL-71 |
| chr11-17461613-DEL-79 | chr11-17461614-DEL-79 |
| chr11-17504381-DEL-132 | chr11-17504382-DEL-132 |
| chr11-17519707-DEL-58 | chr11-17519726-DEL-52 |
| chr11-17532444-DEL-2438 | chr11-17532444-DEL-2438 |
| chr11-17539097-DEL-70 | chr11-17539098-DEL-70 |
| chr11-17543358-DEL-58 | chr11-17543363-DEL-63 |
| chr11-17560195-DEL-746 | chr11-17560196-DEL-746 |
| chr11-17580129-DEL-62 | chr11-17580132-DEL-60 |
| chr11-17617682-DEL-7258 | chr11-17617683-DEL-7258 |
| chr11-17627840-DEL-66 | chr11-17627841-DEL-66 |
| chr11-17640694-DEL-312 | chr11-17640695-DEL-312 |
| chr11-17645711-DEL-144 | chr11-17645712-DEL-144 |
| chr11-17652472-DEL-331 | chr11-17652473-DEL-331 |
| chr11-17680665-DEL-172 | chr11-17680666-DEL-172 |
| chr11-17709997-DEL-135 | chr11-17709998-DEL-135 |
| chr11-17753449-DEL-1541 | chr11-17753450-DEL-1541 |
| chr11-17767920-DEL-66 | chr11-17767921-DEL-66 |
| chr11-17817030-DEL-92 | chr11-17817031-DEL-92 |
| chr11-17839310-DEL-513 | chr11-17839311-DEL-513 |
| chr11-17844365-DEL-250 | chr11-17844366-DEL-250 |
| chr11-17852593-DEL-55 | chr11-17852593-DEL-56 |
| chr11-17854921-DEL-60 | chr11-17854920-DEL-58 |
| chr11-17889093-DEL-91 | chr11-17889094-DEL-91 |
| chr11-17899549-DEL-223 | chr11-17899550-DEL-223 |
| chr11-17900195-DEL-75 | chr11-17900196-DEL-70 |
| chr11-17907893-DEL-177 | chr11-17907894-DEL-177 |
| chr11-17913785-DEL-200 | chr11-17913760-DEL-201 |
| chr11-17923016-DEL-550 | chr11-17923017-DEL-550 |
| chr11-17925334-DEL-241 | chr11-17925335-DEL-241 |
| chr11-17947919-DEL-62 | chr11-17947920-DEL-62 |
| chr11-17955155-DEL-240 | chr11-17955156-DEL-240 |
| chr11-17956480-DEL-917 | chr11-17956481-DEL-917 |
| chr11-17978458-DEL-51 | chr11-17978459-DEL-54 |
| chr11-17990265-DEL-63 | chr11-17990265-DEL-63 |
| chr11-17995989-DEL-6732 | chr11-17995990-DEL-6732 |
| chr11-18028469-DEL-6145 | chr11-18028467-DEL-6146 |
| chr11-18056589-DEL-2767 | chr11-18056590-DEL-2767 |
| chr11-18079998-DEL-158 | chr11-18079999-DEL-157 |
| chr11-18088464-DEL-417 | chr11-18088465-DEL-417 |
| chr11-18099879-DEL-177 | chr11-18099880-DEL-177 |
| chr11-18109756-DEL-84 | chr11-18109757-DEL-84 |
| chr11-18131992-DEL-1429 | chr11-18131993-DEL-1429 |
| chr11-18135459-DEL-7018 | chr11-18135458-DEL-7018 |
| chr11-18148133-DEL-1162 | chr11-18148134-DEL-1162 |
| chr11-18163856-DEL-51 | chr11-18163857-DEL-51 |
| chr11-18179886-DEL-54 | chr11-18179891-DEL-52 |
| chr11-18205666-DEL-1274 | chr11-18205667-DEL-1273 |
| chr11-18207445-DEL-248 | chr11-18207434-DEL-240 |
| chr11-18208954-DEL-94 | chr11-18208955-DEL-94 |
| chr11-18208957-DEL-1287 | chr11-18208958-DEL-1287 |
| chr11-18231651-DEL-80 | chr11-18231652-DEL-80 |
| chr11-18235672-DEL-1079 | chr11-18235673-DEL-1079 |
| chr11-18259390-DEL-1298 | chr11-18259391-DEL-1298 |
| chr11-18270285-DEL-58 | chr11-18270290-DEL-54 |
| chr11-18281363-DEL-71 | chr11-18281336-DEL-75 |
| chr11-18286952-DEL-6540 | chr11-18286953-DEL-6540 |
| chr11-18328858-DEL-1553 | chr11-18328859-DEL-1553 |
| chr11-18334231-DEL-265 | chr11-18334231-DEL-265 |
| chr11-18341084-DEL-2560 | chr11-18341085-DEL-2560 |
| chr11-18369969-DEL-110 | chr11-18369970-DEL-110 |
| chr11-18396896-DEL-72 | chr11-18396897-DEL-72 |
| chr11-18408233-DEL-970 | chr11-18408234-DEL-970 |
| chr11-18438512-DEL-96 | chr11-18438513-DEL-97 |
| chr11-18450751-DEL-182 | chr11-18450748-DEL-182 |
| chr11-18474043-DEL-100 | chr11-18474044-DEL-100 |
| chr11-18534057-DEL-54 | chr11-18534058-DEL-54 |

| | |
|---|---|
| chr11-18560576-DEL-379 | chr11-18560577-DEL-379 |
| chr11-18611697-DEL-146 | chr11-18611698-DEL-146 |
| chr11-18635437-DEL-3026 | chr11-18635438-DEL-3026 |
| chr11-18638857-DEL-102 | chr11-18638849-DEL-63 |
| chr11-18655583-DEL-2264 | chr11-18655584-DEL-2264 |
| chr11-18661271-DEL-51 | chr11-18661242-DEL-76 |
| chr11-18691424-DEL-413 | chr11-18691425-DEL-413 |
| chr11-18692124-DEL-56 | chr11-18692125-DEL-56 |
| chr11-18723590-DEL-6483 | chr11-18723591-DEL-6483 |
| chr11-18744678-DEL-126 | chr11-18744679-DEL-126 |
| chr11-18751461-DEL-1804 | chr11-18751462-DEL-1804 |
| chr11-18756332-DEL-7430 | chr11-18756333-DEL-7430 |
| chr11-18781638-DEL-280 | chr11-18781639-DEL-280 |
| chr11-18784449-DEL-521 | chr11-18784450-DEL-521 |
| chr11-18796976-DEL-142 | chr11-18796977-DEL-142 |
| chr11-18815798-DEL-66 | chr11-18815799-DEL-66 |
| chr11-18859199-DEL-58 | chr11-18859196-DEL-62 |
| chr11-18886751-DEL-165 | chr11-18886752-DEL-165 |
| chr11-18910282-DEL-58 | chr11-18910283-DEL-58 |
| chr11-18920959-DEL-65 | chr11-18920959-DEL-66 |
| chr11-18981134-DEL-124 | chr11-18981147-DEL-96 |
| chr11-18995775-DEL-69 | chr11-18995782-DEL-62 |
| chr11-19024166-DEL-65 | chr11-19024167-DEL-65 |
| chr11-19032566-DEL-6490 | chr11-19032567-DEL-6490 |
| chr11-19046025-DEL-487 | chr11-19046026-DEL-486 |
| chr11-19050580-DEL-527 | chr11-19050581-DEL-528 |
| chr11-19073205-DEL-69 | chr11-19073205-DEL-66 |
| chr11-19080093-DEL-305 | chr11-19080094-DEL-305 |
| chr11-19101570-DEL-96 | chr11-19101561-DEL-84 |
| chr11-19140578-DEL-139 | chr11-19140579-DEL-139 |
| chr11-19141724-DEL-75 | chr11-19141677-DEL-90 |
| chr11-19172525-DEL-162 | chr11-19172526-DEL-162 |
| chr11-19212152-DEL-133 | chr11-19212153-DEL-133 |
| chr11-19244152-DEL-53 | chr11-19244153-DEL-53 |
| chr11-19278232-DEL-558 | chr11-19278233-DEL-558 |
| chr11-19289158-DEL-150 | chr11-19289159-DEL-150 |
| chr11-19352315-DEL-139 | chr11-19352316-DEL-140 |
| chr11-19359127-DEL-7082 | chr11-19359128-DEL-7082 |
| chr11-19387749-DEL-60 | chr11-19387720-DEL-62 |
| chr11-19421817-DEL-92 | chr11-19421816-DEL-92 |
| chr11-19434450-DEL-538 | chr11-19434451-DEL-538 |
| chr11-19460880-DEL-89 | chr11-19460881-DEL-89 |
| chr11-19465828-DEL-2363 | chr11-19465829-DEL-2363 |
| chr11-19478458-DEL-140 | chr11-19478492-DEL-113 |
| chr11-19508194-DEL-91 | chr11-19508195-DEL-91 |
| chr11-19515314-DEL-497 | chr11-19515315-DEL-497 |
| chr11-19516128-DEL-4449 | chr11-19516129-DEL-4449 |
| chr11-19539544-DEL-104 | chr11-19539520-DEL-112 |
| chr11-19542317-DEL-227 | chr11-19542318-DEL-227 |
| chr11-19571752-DEL-192 | chr11-19571753-DEL-192 |
| chr11-19577503-DEL-78 | chr11-19577504-DEL-78 |
| chr11-19593426-DEL-275 | chr11-19593427-DEL-275 |
| chr11-19619250-DEL-237 | chr11-19619251-DEL-249 |
| chr11-19627157-DEL-371 | chr11-19627156-DEL-372 |
| chr11-19652830-DEL-321 | chr11-19652812-DEL-333 |
| chr11-19654394-DEL-298 | chr11-19654395-DEL-298 |
| chr11-19674279-DEL-184 | chr11-19674280-DEL-184 |
| chr11-19701993-DEL-74 | chr11-19701994-DEL-78 |
| chr11-19705159-DEL-555 | chr11-19705160-DEL-555 |
| chr11-19706720-DEL-77 | chr11-19706717-DEL-78 |
| chr11-19725420-DEL-60 | chr11-19725421-DEL-60 |
| chr11-19731994-DEL-85 | chr11-19732001-DEL-55 |
| chr11-19738626-DEL-5069 | chr11-19738627-DEL-5069 |
| chr11-19752938-DEL-411 | chr11-19752939-DEL-411 |
| chr11-19753651-DEL-79 | chr11-19753663-DEL-79 |
| chr11-19758952-DEL-485 | chr11-19758953-DEL-485 |
| chr11-19773080-DEL-90 | chr11-19773080-DEL-90 |
| chr11-19787347-DEL-486 | chr11-19787348-DEL-486 |
| chr11-19809334-DEL-218 | chr11-19809335-DEL-218 |
| chr11-19812366-DEL-432 | chr11-19812367-DEL-432 |
| chr11-19818377-DEL-3425 | chr11-19818378-DEL-3425 |
| chr11-19840578-DEL-66 | chr11-19840599-DEL-58 |
| chr11-19842840-DEL-72 | chr11-19842841-DEL-72 |
| chr11-19857649-DEL-152 | chr11-19857636-DEL-152 |
| chr11-19859906-DEL-154 | chr11-19859906-DEL-154 |
| chr11-19863792-DEL-100 | chr11-19863800-DEL-62 |
| chr11-19866504-DEL-163 | chr11-19866514-DEL-162 |
| chr11-19884452-DEL-213 | chr11-19884453-DEL-213 |
| chr11-19916672-DEL-229 | chr11-19916673-DEL-229 |
| chr11-19940451-DEL-169 | chr11-19940452-DEL-169 |

| | |
|---|---|
| chr11-19949686-DEL-412 | chr11-19949687-DEL-412 |
| chr11-19979635-DEL-197 | chr11-19979636-DEL-197 |
| chr11-19982026-DEL-230 | chr11-19982006-DEL-262 |
| chr11-19982389-DEL-680 | chr11-19982390-DEL-720 |
| chr11-19999434-DEL-68 | chr11-19999435-DEL-68 |
| chr11-20001652-DEL-212 | chr11-20001653-DEL-209 |
| chr11-20004676-DEL-57 | chr11-20004673-DEL-58 |
| chr11-20011555-DEL-79 | chr11-20011536-DEL-79 |
| chr11-20019039-DEL-66 | chr11-20019040-DEL-66 |
| chr11-20020764-DEL-248 | chr11-20020765-DEL-248 |
| chr11-20025203-DEL-176 | chr11-20025204-DEL-176 |
| chr11-20027642-DEL-212 | chr11-20027643-DEL-212 |
| chr11-20031904-DEL-72 | chr11-20031905-DEL-72 |
| chr11-20036851-DEL-214 | chr11-20036852-DEL-214 |
| chr11-20041688-DEL-200 | chr11-20041689-DEL-200 |
| chr11-20064156-DEL-125 | chr11-20064157-DEL-125 |
| chr11-20089151-DEL-515 | chr11-20089147-DEL-516 |
| chr11-20091145-DEL-1370 | chr11-20091146-DEL-1370 |
| chr11-20104541-DEL-67 | chr11-20104542-DEL-67 |
| chr11-20111523-DEL-154 | chr11-20111527-DEL-148 |
| chr11-20124097-DEL-127 | chr11-20124102-DEL-123 |
| chr11-20130685-DEL-154 | chr11-20130688-DEL-155 |
| chr11-20135232-DEL-234 | chr11-20135233-DEL-234 |
| chr11-20140286-DEL-116 | chr11-20140281-DEL-110 |
| chr11-20143727-DEL-174 | chr11-20143728-DEL-174 |
| chr11-20171896-DEL-225 | chr11-20171897-DEL-225 |
| chr11-20181912-DEL-186 | chr11-20181913-DEL-186 |
| chr11-20201982-DEL-55 | chr11-20201983-DEL-56 |
| chr11-20213312-DEL-54 | chr11-20213320-DEL-62 |
| chr11-20219108-DEL-198 | chr11-20219109-DEL-198 |
| chr11-20226469-DEL-183 | chr11-20226470-DEL-183 |
| chr11-20234422-DEL-69 | chr11-20234423-DEL-69 |
| chr11-20246888-DEL-338 | chr11-20246889-DEL-337 |
| chr11-20265848-DEL-264 | chr11-20265849-DEL-264 |
| chr11-20284808-DEL-1285 | chr11-20284801-DEL-1286 |
| chr11-20288302-DEL-70 | chr11-20288307-DEL-60 |
| chr11-20296890-DEL-80 | chr11-20296860-DEL-78 |
| chr11-20299091-DEL-67 | chr11-20299092-DEL-65 |
| chr11-20300356-DEL-317 | chr11-20300357-DEL-317 |
| chr11-20306025-DEL-722 | chr11-20306024-DEL-723 |
| chr11-20327489-DEL-4203 | chr11-20327490-DEL-4203 |
| chr11-20355345-DEL-56 | chr11-20355346-DEL-56 |
| chr11-20358281-DEL-217 | chr11-20358282-DEL-209 |
| chr11-20369240-DEL-88 | chr11-20369241-DEL-88 |
| chr11-20372030-DEL-187 | chr11-20372031-DEL-187 |
| chr11-20373267-DEL-61 | chr11-20373274-DEL-55 |
| chr11-20374788-DEL-179 | chr11-20374789-DEL-179 |
| chr11-20395363-DEL-59 | chr11-20395364-DEL-59 |
| chr11-20398389-DEL-225 | chr11-20398390-DEL-225 |
| chr11-20401021-DEL-219 | chr11-20401021-DEL-219 |
| chr11-20402388-DEL-197 | chr11-20402389-DEL-197 |
| chr11-20408794-DEL-286 | chr11-20408795-DEL-286 |
| chr11-20430003-DEL-204 | chr11-20429988-DEL-216 |
| chr11-20446326-DEL-165 | chr11-20446327-DEL-165 |
| chr11-20447761-DEL-69 | chr11-20447760-DEL-61 |
| chr11-20454348-DEL-227 | chr11-20454349-DEL-227 |
| chr11-20458738-DEL-182 | chr11-20458738-DEL-182 |
| chr11-20459101-DEL-2780 | chr11-20459100-DEL-2781 |
| chr11-20466389-DEL-4802 | chr11-20466390-DEL-4802 |
| chr11-20476988-DEL-64 | chr11-20476989-DEL-64 |
| chr11-20545829-DEL-2294 | chr11-20545830-DEL-2294 |
| chr11-20560376-DEL-67 | chr11-20560392-DEL-52 |
| chr11-20577679-DEL-60 | chr11-20577682-DEL-58 |
| chr11-20629907-DEL-105 | chr11-20629908-DEL-105 |
| chr11-20649178-DEL-96 | chr11-20649179-DEL-96 |
| chr11-20739721-DEL-212 | chr11-20739718-DEL-213 |
| chr11-20756590-DEL-1617 | chr11-20756591-DEL-1617 |
| chr11-20768372-DEL-604 | chr11-20768376-DEL-602 |
| chr11-20821395-DEL-181 | chr11-20821396-DEL-181 |
| chr11-20835909-DEL-75 | chr11-20835902-DEL-76 |
| chr11-20852563-DEL-283 | chr11-20852564-DEL-277 |
| chr11-20864321-DEL-121 | chr11-20864322-DEL-121 |
| chr11-20875383-DEL-124 | chr11-20875384-DEL-124 |
| chr11-20923508-DEL-208 | chr11-20923509-DEL-208 |
| chr11-20955806-DEL-111 | chr11-20955807-DEL-111 |
| chr11-21003980-DEL-62 | chr11-21003977-DEL-66 |
| chr11-21007265-DEL-53 | chr11-21007272-DEL-91 |
| chr11-21017030-DEL-2700 | chr11-21017031-DEL-2700 |
| chr11-21033370-DEL-130 | chr11-21033371-DEL-94 |
| chr11-21064729-DEL-127 | chr11-21064730-DEL-127 |

| | |
|---|---|
| chr11-21139406-DEL-68 | chr11-21139407-DEL-68 |
| chr11-21147468-DEL-147 | chr11-21147469-DEL-147 |
| chr11-21181224-DEL-96 | chr11-21181230-DEL-90 |
| chr11-21191188-DEL-66 | chr11-21191189-DEL-66 |
| chr11-21207940-DEL-823 | chr11-21207941-DEL-823 |
| chr11-21236379-DEL-87 | chr11-21236379-DEL-87 |
| chr11-21260235-DEL-219 | chr11-21260236-DEL-218 |
| chr11-21267667-DEL-61 | chr11-21267668-DEL-61 |
| chr11-21285141-DEL-475 | chr11-21285142-DEL-462 |
| chr11-21324596-DEL-534 | chr11-21324585-DEL-534 |
| chr11-21403971-DEL-1633 | chr11-21403972-DEL-1633 |
| chr11-21424311-DEL-1181 | chr11-21424312-DEL-1181 |
| chr11-21426175-DEL-207 | chr11-21426176-DEL-207 |
| chr11-21427173-DEL-197 | chr11-21427174-DEL-197 |
| chr11-21429171-DEL-201 | chr11-21429172-DEL-201 |
| chr11-21491319-DEL-370 | chr11-21491322-DEL-369 |
| chr11-21524466-DEL-120 | chr11-21524464-DEL-116 |
| chr11-21534732-DEL-96 | chr11-21534738-DEL-93 |
| chr11-21546943-DEL-429 | chr11-21546944-DEL-429 |
| chr11-21554864-DEL-52 | chr11-21554865-DEL-52 |
| chr11-21560916-DEL-3469 | chr11-21560916-DEL-3469 |
| chr11-21604578-DEL-4418 | chr11-21604579-DEL-4418 |
| chr11-21618072-DEL-204 | chr11-21618073-DEL-204 |
| chr11-21620088-DEL-393 | chr11-21620110-DEL-366 |
| chr11-21660028-DEL-60 | chr11-21660020-DEL-80 |
| chr11-21715431-DEL-1671 | chr11-21715432-DEL-1671 |
| chr11-21729469-DEL-401 | chr11-21729470-DEL-401 |
| chr11-21730820-DEL-373 | chr11-21730821-DEL-374 |
| chr11-21848043-DEL-311 | chr11-21848050-DEL-311 |
| chr11-21856972-DEL-6450 | chr11-21856973-DEL-6450 |
| chr11-21857102-DEL-211 | chr11-21857079-DEL-212 |
| chr11-21871304-DEL-85 | chr11-21871299-DEL-91 |
| chr11-21895059-DEL-69 | chr11-21895057-DEL-84 |
| chr11-21896645-DEL-54 | chr11-21896643-DEL-55 |
| chr11-21912339-DEL-207 | chr11-21912340-DEL-207 |
| chr11-22071801-DEL-72 | chr11-22071802-DEL-72 |
| chr11-22118827-DEL-96 | chr11-22118828-DEL-96 |
| chr11-22132859-DEL-1557 | chr11-22132860-DEL-1557 |
| chr11-22193416-DEL-54 | chr11-22193417-DEL-66 |
| chr11-22227746-DEL-427 | chr11-22227747-DEL-427 |
| chr11-22260476-DEL-1868 | chr11-22260476-DEL-1859 |
| chr11-22271945-DEL-1163 | chr11-22271946-DEL-1163 |
| chr11-22281054-DEL-123 | chr11-22281055-DEL-123 |
| chr11-22285217-DEL-5318 | chr11-22285218-DEL-5318 |
| chr11-22302844-DEL-1756 | chr11-22302845-DEL-1756 |
| chr11-22345416-DEL-6403 | chr11-22345417-DEL-6403 |
| chr11-22374050-DEL-65 | chr11-22374050-DEL-55 |
| chr11-22377114-DEL-3631 | chr11-22377115-DEL-3631 |
| chr11-22389581-DEL-400 | chr11-22389582-DEL-400 |
| chr11-22416293-DEL-445 | chr11-22416294-DEL-445 |
| chr11-22424033-DEL-67 | chr11-22424018-DEL-93 |
| chr11-22426356-DEL-612 | chr11-22426352-DEL-616 |
| chr11-22435122-DEL-414 | chr11-22435123-DEL-414 |
| chr11-22464899-DEL-176 | chr11-22464900-DEL-168 |
| chr11-22466167-DEL-74 | chr11-22466168-DEL-74 |
| chr11-22469174-DEL-916 | chr11-22469175-DEL-916 |
| chr11-22473795-DEL-191 | chr11-22473796-DEL-191 |
| chr11-22481065-DEL-154 | chr11-22481066-DEL-154 |
| chr11-22484191-DEL-225 | chr11-22484190-DEL-227 |
| chr11-22484880-DEL-1893 | chr11-22484881-DEL-1893 |
| chr11-22492222-DEL-84 | chr11-22492228-DEL-72 |
| chr11-22509898-DEL-222 | chr11-22509898-DEL-222 |
| chr11-22517033-DEL-239 | chr11-22517034-DEL-239 |
| chr11-22518257-DEL-1886 | chr11-22518258-DEL-1886 |
| chr11-22527700-DEL-5338 | chr11-22527701-DEL-5333 |
| chr11-22553620-DEL-87 | chr11-22553620-DEL-86 |
| chr11-22583598-DEL-53 | chr11-22583599-DEL-52 |
| chr11-22591608-DEL-66 | chr11-22591609-DEL-66 |
| chr11-22610143-DEL-7486 | chr11-22610144-DEL-7486 |
| chr11-22614441-DEL-193 | chr11-22614437-DEL-199 |
| chr11-22623375-DEL-208 | chr11-22623376-DEL-208 |
| chr11-22636221-DEL-493 | chr11-22636222-DEL-493 |
| chr11-22652053-DEL-257 | chr11-22652054-DEL-257 |
| chr11-22655724-DEL-108 | chr11-22655725-DEL-112 |
| chr11-22657221-DEL-173 | chr11-22657222-DEL-173 |
| chr11-22699359-DEL-1484 | chr11-22699360-DEL-1484 |
| chr11-22704584-DEL-60 | chr11-22704607-DEL-50 |
| chr11-22711891-DEL-72 | chr11-22711899-DEL-84 |
| chr11-22747782-DEL-51 | chr11-22747783-DEL-56 |
| chr11-22748566-DEL-162 | chr11-22748567-DEL-162 |

| | |
|---|---|
| chr11-22775510-DEL-61 | chr11-22775515-DEL-57 |
| chr11-22865022-DEL-58 | chr11-22865031-DEL-50 |
| chr11-22889142-DEL-112 | chr11-22889143-DEL-112 |
| chr11-22889833-DEL-4839 | chr11-22889834-DEL-4839 |
| chr11-22911912-DEL-125 | chr11-22911925-DEL-93 |
| chr11-22968967-DEL-55 | chr11-22968932-DEL-56 |
| chr11-23019204-DEL-361 | chr11-23019205-DEL-361 |
| chr11-23032737-DEL-76 | chr11-23032730-DEL-84 |
| chr11-23035661-DEL-52 | chr11-23035661-DEL-52 |
| chr11-23036849-DEL-236 | chr11-23036849-DEL-236 |
| chr11-23047463-DEL-195 | chr11-23047464-DEL-195 |
| chr11-23070034-DEL-1086 | chr11-23070025-DEL-1096 |
| chr11-23072920-DEL-87 | chr11-23072921-DEL-58 |
| chr11-23093542-DEL-112 | chr11-23093543-DEL-112 |
| chr11-23103718-DEL-472 | chr11-23103719-DEL-472 |
| chr11-23107050-DEL-214 | chr11-23107051-DEL-214 |
| chr11-23116760-DEL-574 | chr11-23116782-DEL-551 |
| chr11-23120342-DEL-53 | chr11-23120343-DEL-53 |
| chr11-23144024-DEL-190 | chr11-23144024-DEL-190 |
| chr11-23161837-DEL-196 | chr11-23161836-DEL-202 |
| chr11-23168576-DEL-7098 | chr11-23168577-DEL-7098 |
| chr11-23192791-DEL-4480 | chr11-23192792-DEL-4480 |
| chr11-23209845-DEL-204 | chr11-23209834-DEL-203 |
| chr11-23244398-DEL-131 | chr11-23244399-DEL-131 |
| chr11-23265051-DEL-80 | chr11-23265068-DEL-52 |
| chr11-23286207-DEL-3135 | chr11-23286208-DEL-3135 |
| chr11-23308560-DEL-192 | chr11-23308609-DEL-144 |
| chr11-23316713-DEL-182 | chr11-23316714-DEL-182 |
| chr11-23350059-DEL-84 | chr11-23350069-DEL-65 |
| chr11-23389433-DEL-222 | chr11-23389434-DEL-222 |
| chr11-23391441-DEL-1610 | chr11-23391442-DEL-1610 |
| chr11-23410873-DEL-119 | chr11-23410897-DEL-87 |
| chr11-23452026-DEL-230 | chr11-23452027-DEL-230 |
| chr11-23452629-DEL-192 | chr11-23452615-DEL-193 |
| chr11-23454332-DEL-857 | chr11-23454333-DEL-857 |
| chr11-23460975-DEL-72 | chr11-23460969-DEL-70 |
| chr11-23468413-DEL-54 | chr11-23468438-DEL-57 |
| chr11-23480468-DEL-200 | chr11-23480455-DEL-201 |
| chr11-23491180-DEL-148 | chr11-23491181-DEL-148 |
| chr11-23496395-DEL-6440 | chr11-23496396-DEL-6440 |
| chr11-23513693-DEL-113 | chr11-23513692-DEL-114 |
| chr11-23532068-DEL-62 | chr11-23532069-DEL-62 |
| chr11-23549601-DEL-242 | chr11-23549602-DEL-242 |
| chr11-23557507-DEL-218 | chr11-23557508-DEL-218 |
| chr11-23567333-DEL-463 | chr11-23567334-DEL-463 |
| chr11-23585574-DEL-188 | chr11-23585606-DEL-156 |
| chr11-23617504-DEL-361 | chr11-23617505-DEL-361 |
| chr11-23673692-DEL-51 | chr11-23673691-DEL-53 |
| chr11-23674607-DEL-209 | chr11-23674601-DEL-168 |
| chr11-23717587-DEL-199 | chr11-23717588-DEL-199 |
| chr11-23733931-DEL-163 | chr11-23733932-DEL-163 |
| chr11-23769492-DEL-166 | chr11-23769491-DEL-166 |
| chr11-23773895-DEL-210 | chr11-23773896-DEL-210 |
| chr11-23784732-DEL-6459 | chr11-23784733-DEL-6459 |
| chr11-23857146-DEL-155 | chr11-23857142-DEL-152 |
| chr11-23908171-DEL-227 | chr11-23908172-DEL-227 |
| chr11-23990830-DEL-164 | chr11-23990831-DEL-164 |
| chr11-24055878-DEL-214 | chr11-24055879-DEL-214 |
| chr11-24107417-DEL-66 | chr11-24107418-DEL-66 |
| chr11-24109661-DEL-98 | chr11-24109662-DEL-98 |
| chr11-24219797-DEL-94 | chr11-24219798-DEL-94 |
| chr11-24228201-DEL-428 | chr11-24228202-DEL-428 |
| chr11-24277144-DEL-60 | chr11-24277145-DEL-60 |
| chr11-24288475-DEL-70 | chr11-24288476-DEL-70 |
| chr11-24342533-DEL-269 | chr11-24342534-DEL-269 |
| chr11-24397054-DEL-55 | chr11-24397055-DEL-55 |
| chr11-24409742-DEL-499 | chr11-24409743-DEL-499 |
| chr11-24439603-DEL-507 | chr11-24439604-DEL-507 |
| chr11-24477489-DEL-1987 | chr11-24477490-DEL-1987 |
| chr11-24483512-DEL-401 | chr11-24483505-DEL-387 |
| chr11-24506316-DEL-52 | chr11-24506312-DEL-53 |
| chr11-24510654-DEL-2305 | chr11-24510655-DEL-2305 |
| chr11-24534141-DEL-60 | chr11-24534142-DEL-60 |
| chr11-24540654-DEL-6345 | chr11-24540654-DEL-6345 |
| chr11-24543629-DEL-1268 | chr11-24543630-DEL-1268 |
| chr11-24546572-DEL-199 | chr11-24546573-DEL-199 |
| chr11-24715586-DEL-63 | chr11-24715587-DEL-63 |
| chr11-24755460-DEL-6446 | chr11-24755461-DEL-6446 |
| chr11-24765297-DEL-325 | chr11-24765300-DEL-310 |
| chr11-24823336-DEL-1560 | chr11-24823337-DEL-1558 |

| | |
|---|---|
| chr11-24825650-DEL-6624 | chr11-24825633-DEL-6622 |
| chr11-24841859-DEL-54 | chr11-24841860-DEL-54 |
| chr11-24845870-DEL-402 | chr11-24845867-DEL-402 |
| chr11-24861869-DEL-56 | chr11-24861869-DEL-56 |
| chr11-24872943-DEL-90 | chr11-24872944-DEL-90 |
| chr11-24881707-DEL-53 | chr11-24881689-DEL-51 |
| chr11-24905983-DEL-56 | chr11-24905984-DEL-56 |
| chr11-24906859-DEL-2094 | chr11-24906861-DEL-2095 |
| chr11-24966739-DEL-1192 | chr11-24966740-DEL-1192 |
| chr11-24977236-DEL-67 | chr11-24977248-DEL-55 |
| chr11-24983212-DEL-57 | chr11-24983213-DEL-75 |
| chr11-25010982-DEL-64 | chr11-25010987-DEL-56 |
| chr11-25014642-DEL-5142 | chr11-25014643-DEL-5142 |
| chr11-25016508-DEL-136 | chr11-25016509-DEL-136 |
| chr11-25088787-DEL-52 | chr11-25088788-DEL-56 |
| chr11-25123795-DEL-66 | chr11-25123796-DEL-66 |
| chr11-25146575-DEL-6057 | chr11-25146576-DEL-6057 |
| chr11-25166218-DEL-1106 | chr11-25166219-DEL-1106 |
| chr11-25175214-DEL-80 | chr11-25175211-DEL-80 |
| chr11-25177021-DEL-130 | chr11-25177022-DEL-130 |
| chr11-25217757-DEL-72 | chr11-25217758-DEL-72 |
| chr11-25248116-DEL-172 | chr11-25248117-DEL-172 |
| chr11-25250029-DEL-60 | chr11-25250030-DEL-60 |
| chr11-25289351-DEL-461 | chr11-25289352-DEL-459 |
| chr11-25398112-DEL-100 | chr11-25398113-DEL-100 |
| chr11-25433598-DEL-438 | chr11-25433599-DEL-438 |
| chr11-25447230-DEL-61 | chr11-25447232-DEL-50 |
| chr11-25507129-DEL-6320 | chr11-25507130-DEL-6320 |
| chr11-25515663-DEL-690 | chr11-25515664-DEL-689 |
| chr11-25554297-DEL-58 | chr11-25554298-DEL-58 |
| chr11-25560231-DEL-2822 | chr11-25560232-DEL-2822 |
| chr11-25664058-DEL-280 | chr11-25664059-DEL-274 |
| chr11-25667383-DEL-139 | chr11-25667384-DEL-138 |
| chr11-25688090-DEL-56 | chr11-25688091-DEL-56 |
| chr11-25745339-DEL-6762 | chr11-25745340-DEL-6762 |
| chr11-25819455-DEL-110 | chr11-25819461-DEL-105 |
| chr11-25825903-DEL-1184 | chr11-25825904-DEL-1184 |
| chr11-25831044-DEL-95 | chr11-25831045-DEL-95 |
| chr11-25843629-DEL-352 | chr11-25843630-DEL-352 |
| chr11-25920263-DEL-6228 | chr11-25920264-DEL-6228 |
| chr11-25947150-DEL-58 | chr11-25947151-DEL-58 |
| chr11-25958316-DEL-65 | chr11-25958315-DEL-66 |
| chr11-25961198-DEL-54 | chr11-25961199-DEL-54 |
| chr11-25977808-DEL-433 | chr11-25977809-DEL-433 |
| chr11-25980271-DEL-3477 | chr11-25980272-DEL-3477 |
| chr11-26009380-DEL-62 | chr11-26009381-DEL-62 |
| chr11-26048659-DEL-57 | chr11-26048656-DEL-61 |
| chr11-26099632-DEL-102 | chr11-26099633-DEL-102 |
| chr11-26121461-DEL-4772 | chr11-26121462-DEL-4772 |
| chr11-26133639-DEL-55 | chr11-26133640-DEL-55 |
| chr11-26152456-DEL-124 | chr11-26152457-DEL-124 |
| chr11-26154928-DEL-951 | chr11-26154928-DEL-944 |
| chr11-26155066-DEL-91 | chr11-26155067-DEL-91 |
| chr11-26164196-DEL-4586 | chr11-26164197-DEL-4586 |
| chr11-26186456-DEL-94 | chr11-26186457-DEL-94 |
| chr11-26213371-DEL-74 | chr11-26213371-DEL-75 |
| chr11-26225901-DEL-231 | chr11-26225902-DEL-231 |
| chr11-26244614-DEL-202 | chr11-26244615-DEL-202 |
| chr11-26249892-DEL-386 | chr11-26249893-DEL-386 |
| chr11-26262696-DEL-153 | chr11-26262697-DEL-153 |
| chr11-26275808-DEL-76 | chr11-26275809-DEL-76 |
| chr11-26276554-DEL-59 | chr11-26276555-DEL-59 |
| chr11-26289831-DEL-233 | chr11-26289819-DEL-246 |
| chr11-26298794-DEL-55 | chr11-26298795-DEL-55 |
| chr11-26363102-DEL-52 | chr11-26363103-DEL-52 |
| chr11-26376516-DEL-6861 | chr11-26376517-DEL-6861 |
| chr11-26405753-DEL-74 | chr11-26405754-DEL-74 |
| chr11-26411447-DEL-6394 | chr11-26411448-DEL-6394 |
| chr11-26447101-DEL-147 | chr11-26447102-DEL-147 |
| chr11-26449274-DEL-185 | chr11-26449289-DEL-155 |
| chr11-26453341-DEL-1341 | chr11-26453342-DEL-1341 |
| chr11-26462461-DEL-66 | chr11-26462413-DEL-90 |
| chr11-26469765-DEL-278 | chr11-26469766-DEL-278 |
| chr11-26479215-DEL-75 | chr11-26479193-DEL-60 |
| chr11-26486726-DEL-76 | chr11-26486717-DEL-86 |
| chr11-26505439-DEL-5345 | chr11-26505440-DEL-5345 |
| chr11-26556722-DEL-172 | chr11-26556723-DEL-172 |
| chr11-26579290-DEL-6249 | chr11-26579291-DEL-6249 |
| chr11-26619395-DEL-214 | chr11-26619396-DEL-214 |
| chr11-26661300-DEL-110 | chr11-26661301-DEL-110 |

| | |
|---|---|
| chr11-26682836-DEL-6443 | chr11-26682837-DEL-6443 |
| chr11-26736550-DEL-1295 | chr11-26736551-DEL-1291 |
| chr11-26792373-DEL-59 | chr11-26792374-DEL-59 |
| chr11-26895311-DEL-270 | chr11-26895312-DEL-270 |
| chr11-26919987-DEL-374 | chr11-26919988-DEL-374 |
| chr11-26921812-DEL-66 | chr11-26921813-DEL-66 |
| chr11-26927842-DEL-247 | chr11-26927843-DEL-247 |
| chr11-26959093-DEL-64 | chr11-26959094-DEL-65 |
| chr11-26961107-DEL-1416 | chr11-26961112-DEL-1415 |
| chr11-26977478-DEL-140 | chr11-26977479-DEL-140 |
| chr11-26994088-DEL-476 | chr11-26994089-DEL-476 |
| chr11-27009227-DEL-58 | chr11-27009228-DEL-58 |
| chr11-27013216-DEL-198 | chr11-27013217-DEL-198 |
| chr11-27014923-DEL-828 | chr11-27014924-DEL-828 |
| chr11-27016062-DEL-1120 | chr11-27016063-DEL-1120 |
| chr11-27026107-DEL-586 | chr11-27026108-DEL-586 |
| chr11-27034107-DEL-733 | chr11-27034107-DEL-733 |
| chr11-27037168-DEL-343 | chr11-27037169-DEL-342 |
| chr11-27044776-DEL-466 | chr11-27044777-DEL-466 |
| chr11-27046982-DEL-85 | chr11-27046978-DEL-86 |
| chr11-27049237-DEL-97 | chr11-27049238-DEL-97 |
| chr11-27049984-DEL-83 | chr11-27049985-DEL-83 |
| chr11-27067428-DEL-52 | chr11-27067429-DEL-52 |
| chr11-27068427-DEL-454 | chr11-27068428-DEL-454 |
| chr11-27068948-DEL-128 | chr11-27068949-DEL-127 |
| chr11-27083703-DEL-4397 | chr11-27083704-DEL-4397 |
| chr11-27098053-DEL-84 | chr11-27098054-DEL-84 |
| chr11-27111801-DEL-84 | chr11-27111802-DEL-80 |
| chr11-27114246-DEL-66 | chr11-27114246-DEL-66 |
| chr11-27124130-DEL-7025 | chr11-27124131-DEL-7025 |
| chr11-27140853-DEL-130 | chr11-27140854-DEL-130 |
| chr11-27144587-DEL-134 | chr11-27144589-DEL-128 |
| chr11-27164331-DEL-923 | chr11-27164332-DEL-923 |
| chr11-27228140-DEL-168 | chr11-27228141-DEL-168 |
| chr11-27250654-DEL-1331 | chr11-27250655-DEL-1331 |
| chr11-27303824-DEL-108 | chr11-27303825-DEL-108 |
| chr11-27306883-DEL-6180 | chr11-27306884-DEL-6180 |
| chr11-27318145-DEL-110 | chr11-27318146-DEL-110 |
| chr11-27322295-DEL-96 | chr11-27322296-DEL-96 |
| chr11-27327668-DEL-476 | chr11-27327669-DEL-476 |
| chr11-27332587-DEL-6916 | chr11-27332588-DEL-6916 |
| chr11-27365062-DEL-2662 | chr11-27365063-DEL-2662 |
| chr11-27376015-DEL-91 | chr11-27376016-DEL-91 |
| chr11-27393571-DEL-71 | chr11-27393572-DEL-71 |
| chr11-27423112-DEL-1663 | chr11-27423113-DEL-1661 |
| chr11-27436137-DEL-918 | chr11-27436138-DEL-918 |
| chr11-27444454-DEL-123 | chr11-27444455-DEL-123 |
| chr11-27453891-DEL-56 | chr11-27453883-DEL-54 |
| chr11-27485059-DEL-6245 | chr11-27485060-DEL-6245 |
| chr11-27526618-DEL-3406 | chr11-27526618-DEL-3410 |
| chr11-27551655-DEL-220 | chr11-27551656-DEL-220 |
| chr11-27558899-DEL-7486 | chr11-27558892-DEL-7487 |
| chr11-27591186-DEL-1886 | chr11-27591186-DEL-1886 |
| chr11-27609308-DEL-460 | chr11-27609309-DEL-460 |
| chr11-27613042-DEL-71 | chr11-27613026-DEL-72 |
| chr11-27624000-DEL-276 | chr11-27624001-DEL-276 |
| chr11-27643423-DEL-1838 | chr11-27643421-DEL-1838 |
| chr11-27652162-DEL-415 | chr11-27652163-DEL-415 |
| chr11-27656952-DEL-83 | chr11-27656955-DEL-81 |
| chr11-27658626-DEL-566 | chr11-27658627-DEL-566 |
| chr11-27730794-DEL-63 | chr11-27730770-DEL-62 |
| chr11-27738290-DEL-482 | chr11-27738291-DEL-482 |
| chr11-27744514-DEL-59 | chr11-27744515-DEL-58 |
| chr11-27772603-DEL-1376 | chr11-27772604-DEL-1376 |
| chr11-27798202-DEL-232 | chr11-27798203-DEL-232 |
| chr11-27803664-DEL-1770 | chr11-27803665-DEL-1770 |
| chr11-27811324-DEL-61 | chr11-27811319-DEL-61 |
| chr11-27886878-DEL-358 | chr11-27886879-DEL-358 |
| chr11-27895318-DEL-1547 | chr11-27895317-DEL-1547 |
| chr11-27901061-DEL-5997 | chr11-27901062-DEL-5997 |
| chr11-27909451-DEL-413 | chr11-27909452-DEL-413 |
| chr11-27923966-DEL-330 | chr11-27923961-DEL-331 |
| chr11-27927582-DEL-547 | chr11-27927583-DEL-547 |
| chr11-27973278-DEL-60 | chr11-27973279-DEL-60 |
| chr11-28002451-DEL-392 | chr11-28002452-DEL-392 |
| chr11-28015786-DEL-452 | chr11-28015787-DEL-452 |
| chr11-28029595-DEL-502 | chr11-28029596-DEL-502 |
| chr11-28031928-DEL-2040 | chr11-28031937-DEL-2055 |
| chr11-28047278-DEL-7125 | chr11-28047279-DEL-7125 |
| chr11-28062033-DEL-60 | chr11-28062034-DEL-60 |

| | |
|---|---|
| chr11-28076362-DEL-532 | chr11-28076362-DEL-532 |
| chr11-28081893-DEL-1212 | chr11-28081894-DEL-1212 |
| chr11-28084769-DEL-2163 | chr11-28084771-DEL-2160 |
| chr11-28090999-DEL-4992 | chr11-28090999-DEL-4992 |
| chr11-28102849-DEL-54 | chr11-28102854-DEL-50 |
| chr11-28130920-DEL-1396 | chr11-28130920-DEL-1396 |
| chr11-28140753-DEL-856 | chr11-28140744-DEL-855 |
| chr11-28180726-DEL-3329 | chr11-28180740-DEL-3321 |
| chr11-28209707-DEL-60 | chr11-28209712-DEL-50 |
| chr11-28238842-DEL-325 | chr11-28238843-DEL-325 |
| chr11-28242594-DEL-6232 | chr11-28242595-DEL-6232 |
| chr11-28252282-DEL-2031 | chr11-28252283-DEL-2031 |
| chr11-28257720-DEL-1386 | chr11-28257721-DEL-1386 |
| chr11-28257720-DEL-57 | chr11-28257722-DEL-54 |
| chr11-28279934-DEL-52 | chr11-28279935-DEL-52 |
| chr11-28281927-DEL-109 | chr11-28281927-DEL-109 |
| chr11-28294533-DEL-54 | chr11-28294519-DEL-57 |
| chr11-28343405-DEL-83 | chr11-28343406-DEL-83 |
| chr11-28355397-DEL-60 | chr11-28355350-DEL-70 |
| chr11-28370038-DEL-148 | chr11-28370039-DEL-148 |
| chr11-28385388-DEL-55 | chr11-28385389-DEL-55 |
| chr11-28407693-DEL-79 | chr11-28407694-DEL-79 |
| chr11-28409578-DEL-147 | chr11-28409579-DEL-147 |
| chr11-28411333-DEL-1673 | chr11-28411334-DEL-1673 |
| chr11-28412591-DEL-1223 | chr11-28412591-DEL-1223 |
| chr11-28413368-DEL-1068 | chr11-28413369-DEL-1068 |
| chr11-28423830-DEL-3009 | chr11-28423831-DEL-3009 |
| chr11-28435066-DEL-154 | chr11-28435067-DEL-154 |
| chr11-28470965-DEL-466 | chr11-28470964-DEL-467 |
| chr11-28475912-DEL-65 | chr11-28475913-DEL-65 |
| chr11-28479278-DEL-52 | chr11-28479279-DEL-52 |
| chr11-28479782-DEL-970 | chr11-28479783-DEL-970 |
| chr11-28505462-DEL-141 | chr11-28505463-DEL-141 |
| chr11-28529895-DEL-1784 | chr11-28529896-DEL-1784 |
| chr11-28548673-DEL-58 | chr11-28548649-DEL-59 |
| chr11-28555632-DEL-69 | chr11-28555633-DEL-69 |
| chr11-28559811-DEL-65 | chr11-28559774-DEL-66 |
| chr11-28627885-DEL-567 | chr11-28627886-DEL-567 |
| chr11-28638540-DEL-472 | chr11-28638541-DEL-472 |
| chr11-28704786-DEL-423 | chr11-28704787-DEL-423 |
| chr11-28709506-DEL-113 | chr11-28709516-DEL-104 |
| chr11-28716903-DEL-88 | chr11-28716904-DEL-88 |
| chr11-28772732-DEL-56 | chr11-28772733-DEL-56 |
| chr11-28795725-DEL-207 | chr11-28795726-DEL-206 |
| chr11-28800696-DEL-592 | chr11-28800697-DEL-592 |
| chr11-28880318-DEL-313 | chr11-28880319-DEL-313 |
| chr11-28905950-DEL-814 | chr11-28905951-DEL-814 |
| chr11-28909563-DEL-761 | chr11-28909564-DEL-761 |
| chr11-28910906-DEL-186 | chr11-28910907-DEL-184 |
| chr11-28953677-DEL-117 | chr11-28953678-DEL-117 |
| chr11-28975090-DEL-5739 | chr11-28975091-DEL-5739 |
| chr11-29074550-DEL-5227 | chr11-29074551-DEL-5227 |
| chr11-29098865-DEL-68 | chr11-29098866-DEL-68 |
| chr11-29107814-DEL-75 | chr11-29107814-DEL-75 |
| chr11-29113878-DEL-2755 | chr11-29113874-DEL-2760 |
| chr11-29128742-DEL-204 | chr11-29128743-DEL-204 |
| chr11-29166975-DEL-120 | chr11-29166976-DEL-120 |
| chr11-29178612-DEL-52 | chr11-29178613-DEL-52 |
| chr11-29185441-DEL-417 | chr11-29185442-DEL-417 |
| chr11-29224243-DEL-53 | chr11-29224244-DEL-53 |
| chr11-29229839-DEL-4024 | chr11-29229840-DEL-4024 |
| chr11-29250196-DEL-52 | chr11-29250197-DEL-52 |
| chr11-29254190-DEL-93 | chr11-29254191-DEL-93 |
| chr11-29260790-DEL-461 | chr11-29260791-DEL-461 |
| chr11-29273274-DEL-284 | chr11-29273272-DEL-297 |
| chr11-29352712-DEL-122 | chr11-29352713-DEL-122 |
| chr11-29388710-DEL-6475 | chr11-29388711-DEL-6475 |
| chr11-29516417-DEL-233 | chr11-29516417-DEL-233 |
| chr11-29523350-DEL-87 | chr11-29523351-DEL-87 |
| chr11-29523890-DEL-77 | chr11-29523899-DEL-52 |
| chr11-29535982-DEL-6402 | chr11-29535983-DEL-6402 |
| chr11-29554277-DEL-61 | chr11-29554278-DEL-61 |
| chr11-29573956-DEL-180 | chr11-29573957-DEL-180 |
| chr11-29579841-DEL-2393 | chr11-29579842-DEL-2393 |
| chr11-29580164-DEL-211 | chr11-29580165-DEL-211 |
| chr11-29607653-DEL-519 | chr11-29607652-DEL-520 |
| chr11-29613143-DEL-200 | chr11-29613148-DEL-196 |
| chr11-29613744-DEL-197 | chr11-29613745-DEL-197 |
| chr11-29616728-DEL-219 | chr11-29616729-DEL-219 |
| chr11-29634197-DEL-54 | chr11-29634198-DEL-54 |

| | |
|---|---|
| chr11-29646365-DEL-79 | chr11-29646366-DEL-79 |
| chr11-29661839-DEL-192 | chr11-29661840-DEL-192 |
| chr11-29666013-DEL-96 | chr11-29666005-DEL-104 |
| chr11-29679545-DEL-835 | chr11-29679546-DEL-835 |
| chr11-29700561-DEL-114 | chr11-29700561-DEL-114 |
| chr11-29710718-DEL-213 | chr11-29710719-DEL-213 |
| chr11-29717280-DEL-91 | chr11-29717281-DEL-91 |
| chr11-29730305-DEL-946 | chr11-29730306-DEL-946 |
| chr11-29764264-DEL-80 | chr11-29764264-DEL-80 |
| chr11-29787889-DEL-223 | chr11-29787890-DEL-223 |
| chr11-29791984-DEL-171 | chr11-29791983-DEL-174 |
| chr11-29801781-DEL-232 | chr11-29801782-DEL-232 |
| chr11-29866442-DEL-68 | chr11-29866443-DEL-68 |
| chr11-29884606-DEL-131 | chr11-29884607-DEL-131 |
| chr11-29898291-DEL-647 | chr11-29898292-DEL-648 |
| chr11-29916859-DEL-166 | chr11-29916860-DEL-166 |
| chr11-29958233-DEL-56 | chr11-29958234-DEL-56 |
| chr11-29965386-DEL-84 | chr11-29965387-DEL-84 |
| chr11-29989454-DEL-65 | chr11-29989455-DEL-65 |
| chr11-29993097-DEL-1669 | chr11-29993097-DEL-1669 |
| chr11-29993550-DEL-112 | chr11-29993551-DEL-111 |
| chr11-30025769-DEL-63 | chr11-30025770-DEL-63 |
| chr11-30073416-DEL-58 | chr11-30073417-DEL-58 |
| chr11-30078759-DEL-213 | chr11-30078760-DEL-213 |
| chr11-30096155-DEL-75 | chr11-30096139-DEL-90 |
| chr11-30119314-DEL-915 | chr11-30119315-DEL-915 |
| chr11-30131518-DEL-80 | chr11-30131519-DEL-80 |
| chr11-30155948-DEL-64 | chr11-30155949-DEL-64 |
| chr11-30178645-DEL-277 | chr11-30178646-DEL-276 |
| chr11-30182590-DEL-68 | chr11-30182591-DEL-66 |
| chr11-30187870-DEL-57 | chr11-30187871-DEL-57 |
| chr11-30210039-DEL-388 | chr11-30210045-DEL-383 |
| chr11-30211321-DEL-76 | chr11-30211322-DEL-76 |
| chr11-30220009-DEL-99 | chr11-30220010-DEL-100 |
| chr11-30232419-DEL-253 | chr11-30232423-DEL-250 |
| chr11-30255479-DEL-54 | chr11-30255480-DEL-50 |
| chr11-30273579-DEL-860 | chr11-30273573-DEL-861 |
| chr11-30296716-DEL-360 | chr11-30296717-DEL-360 |
| chr11-30311583-DEL-838 | chr11-30311584-DEL-838 |
| chr11-30314093-DEL-52 | chr11-30314097-DEL-67 |
| chr11-30332262-DEL-91 | chr11-30332263-DEL-91 |
| chr11-30339349-DEL-65 | chr11-30339350-DEL-65 |
| chr11-30364064-DEL-205 | chr11-30364065-DEL-205 |
| chr11-30383849-DEL-3932 | chr11-30383850-DEL-3932 |
| chr11-30413535-DEL-2403 | chr11-30413536-DEL-2403 |
| chr11-30432674-DEL-149 | chr11-30432675-DEL-149 |
| chr11-30437995-DEL-503 | chr11-30437996-DEL-503 |
| chr11-30479021-DEL-165 | chr11-30479022-DEL-162 |
| chr11-30484160-DEL-1025 | chr11-30484161-DEL-1025 |
| chr11-30509841-DEL-107 | chr11-30509827-DEL-117 |
| chr11-30513322-DEL-1592 | chr11-30513323-DEL-1592 |
| chr11-30516065-DEL-76 | chr11-30516066-DEL-76 |
| chr11-30518373-DEL-221 | chr11-30518374-DEL-221 |
| chr11-30518951-DEL-1584 | chr11-30518958-DEL-1578 |
| chr11-30532082-DEL-426 | chr11-30532083-DEL-426 |
| chr11-30537764-DEL-1807 | chr11-30537757-DEL-1808 |
| chr11-30554855-DEL-53 | chr11-30554847-DEL-54 |
| chr11-30562200-DEL-78 | chr11-30562201-DEL-78 |
| chr11-30574602-DEL-200 | chr11-30574603-DEL-200 |
| chr11-30579292-DEL-55 | chr11-30579292-DEL-56 |
| chr11-30580069-DEL-104 | chr11-30580070-DEL-104 |
| chr11-30586280-DEL-6487 | chr11-30586281-DEL-6487 |
| chr11-30590745-DEL-398 | chr11-30590746-DEL-398 |
| chr11-30598329-DEL-53 | chr11-30598319-DEL-64 |
| chr11-30601899-DEL-196 | chr11-30601899-DEL-196 |
| chr11-30602751-DEL-51 | chr11-30602751-DEL-52 |
| chr11-30606608-DEL-199 | chr11-30606598-DEL-200 |
| chr11-30618841-DEL-51 | chr11-30618842-DEL-51 |
| chr11-30628455-DEL-111 | chr11-30628460-DEL-109 |
| chr11-30629719-DEL-3330 | chr11-30629720-DEL-3330 |
| chr11-30636667-DEL-1235 | chr11-30636667-DEL-1235 |
| chr11-30641929-DEL-52 | chr11-30641930-DEL-52 |
| chr11-30643277-DEL-3053 | chr11-30643277-DEL-3053 |
| chr11-30668947-DEL-539 | chr11-30668948-DEL-539 |
| chr11-30692411-DEL-355 | chr11-30692412-DEL-355 |
| chr11-30696338-DEL-154 | chr11-30696355-DEL-142 |
| chr11-30698323-DEL-205 | chr11-30698324-DEL-205 |
| chr11-30701241-DEL-75 | chr11-30701222-DEL-70 |
| chr11-30747849-DEL-208 | chr11-30747850-DEL-208 |
| chr11-30790157-DEL-549 | chr11-30790158-DEL-549 |

| | |
|---|---|
| chr11-30821576-DEL-52 | chr11-30821577-DEL-52 |
| chr11-30853764-DEL-60 | chr11-30853765-DEL-60 |
| chr11-30904145-DEL-57 | chr11-30904145-DEL-72 |
| chr11-30919140-DEL-100 | chr11-30919135-DEL-97 |
| chr11-30932870-DEL-61 | chr11-30932874-DEL-58 |
| chr11-30939589-DEL-892 | chr11-30939590-DEL-892 |
| chr11-30946378-DEL-143 | chr11-30946379-DEL-143 |
| chr11-30980203-DEL-545 | chr11-30980204-DEL-545 |
| chr11-30981667-DEL-181 | chr11-30981668-DEL-181 |
| chr11-30982563-DEL-1341 | chr11-30982564-DEL-1341 |
| chr11-30985662-DEL-6713 | chr11-30985660-DEL-6726 |
| chr11-31004946-DEL-204 | chr11-31004947-DEL-204 |
| chr11-31011144-DEL-149 | chr11-31011149-DEL-144 |
| chr11-31011747-DEL-892 | chr11-31011748-DEL-892 |
| chr11-31025240-DEL-390 | chr11-31025241-DEL-390 |
| chr11-31078211-DEL-337 | chr11-31078212-DEL-337 |
| chr11-31089410-DEL-121 | chr11-31089411-DEL-121 |
| chr11-31099223-DEL-58 | chr11-31099224-DEL-57 |
| chr11-31099694-DEL-6198 | chr11-31099695-DEL-6198 |
| chr11-31106510-DEL-54 | chr11-31106511-DEL-54 |
| chr11-31108374-DEL-251 | chr11-31108380-DEL-243 |
| chr11-31111493-DEL-507 | chr11-31111494-DEL-505 |
| chr11-31114739-DEL-56 | chr11-31114739-DEL-56 |
| chr11-31123230-DEL-4495 | chr11-31123234-DEL-4494 |
| chr11-31147042-DEL-399 | chr11-31147043-DEL-399 |
| chr11-31151661-DEL-54 | chr11-31151662-DEL-54 |
| chr11-31156879-DEL-69 | chr11-31156880-DEL-65 |
| chr11-31159986-DEL-5890 | chr11-31159987-DEL-5890 |
| chr11-31167700-DEL-155 | chr11-31167701-DEL-155 |
| chr11-31180489-DEL-84 | chr11-31180490-DEL-84 |
| chr11-31195048-DEL-1478 | chr11-31195049-DEL-1478 |
| chr11-31225702-DEL-212 | chr11-31225703-DEL-212 |
| chr11-31231745-DEL-262 | chr11-31231745-DEL-262 |
| chr11-31237918-DEL-65 | chr11-31237923-DEL-60 |
| chr11-31238792-DEL-80 | chr11-31238793-DEL-80 |
| chr11-31241269-DEL-185 | chr11-31241269-DEL-185 |
| chr11-31244339-DEL-392 | chr11-31244340-DEL-392 |
| chr11-31273669-DEL-73 | chr11-31273670-DEL-73 |
| chr11-31280771-DEL-52 | chr11-31280766-DEL-58 |
| chr11-31291355-DEL-83 | chr11-31291353-DEL-76 |
| chr11-31320323-DEL-114 | chr11-31320302-DEL-156 |
| chr11-31354736-DEL-263 | chr11-31354737-DEL-263 |
| chr11-31360446-DEL-197 | chr11-31360446-DEL-197 |
| chr11-31361781-DEL-1988 | chr11-31361782-DEL-1988 |
| chr11-31364707-DEL-2960 | chr11-31364708-DEL-2960 |
| chr11-31375113-DEL-56 | chr11-31375094-DEL-52 |
| chr11-31382685-DEL-199 | chr11-31382685-DEL-199 |
| chr11-31385605-DEL-4266 | chr11-31385605-DEL-4266 |
| chr11-31397433-DEL-7406 | chr11-31397434-DEL-7406 |
| chr11-31405204-DEL-99 | chr11-31405205-DEL-105 |
| chr11-31432737-DEL-60 | chr11-31432738-DEL-60 |
| chr11-31435428-DEL-164 | chr11-31435429-DEL-164 |
| chr11-31436533-DEL-181 | chr11-31436534-DEL-181 |
| chr11-31452014-DEL-92 | chr11-31452014-DEL-92 |
| chr11-31482243-DEL-160 | chr11-31482244-DEL-160 |
| chr11-31486296-DEL-561 | chr11-31486297-DEL-561 |
| chr11-31546421-DEL-3075 | chr11-31546422-DEL-3075 |
| chr11-31555695-DEL-58 | chr11-31555704-DEL-50 |
| chr11-31583103-DEL-896 | chr11-31583104-DEL-896 |
| chr11-31586337-DEL-1350 | chr11-31586338-DEL-1350 |
| chr11-31588585-DEL-63 | chr11-31588586-DEL-63 |
| chr11-31594398-DEL-1480 | chr11-31594399-DEL-1480 |
| chr11-31621217-DEL-100 | chr11-31621238-DEL-80 |
| chr11-31635230-DEL-160 | chr11-31635231-DEL-160 |
| chr11-31640225-DEL-2129 | chr11-31640226-DEL-2129 |
| chr11-31645837-DEL-53 | chr11-31645796-DEL-64 |
| chr11-31658559-DEL-52 | chr11-31658556-DEL-52 |
| chr11-31661123-DEL-255 | chr11-31661139-DEL-240 |
| chr11-31704326-DEL-54 | chr11-31704327-DEL-54 |
| chr11-31724153-DEL-3018 | chr11-31724154-DEL-3018 |
| chr11-31739008-DEL-2077 | chr11-31739009-DEL-2077 |
| chr11-31745791-DEL-62 | chr11-31745792-DEL-62 |
| chr11-31754864-DEL-56 | chr11-31754865-DEL-56 |
| chr11-31756407-DEL-476 | chr11-31756407-DEL-476 |
| chr11-31792163-DEL-60 | chr11-31792164-DEL-60 |
| chr11-31803388-DEL-1307 | chr11-31803389-DEL-1307 |
| chr11-31807362-DEL-79 | chr11-31807363-DEL-79 |
| chr11-31808512-DEL-407 | chr11-31808513-DEL-407 |
| chr11-31860742-DEL-323 | chr11-31860743-DEL-323 |
| chr11-31872589-DEL-200 | chr11-31872590-DEL-200 |

| | |
|---|---|
| chr11-31902022-DEL-1588 | chr11-31902068-DEL-1543 |
| chr11-31958373-DEL-55 | chr11-31958374-DEL-55 |
| chr11-31984187-DEL-52 | chr11-31984212-DEL-57 |
| chr11-31989262-DEL-187 | chr11-31989308-DEL-182 |
| chr11-32021456-DEL-78 | chr11-32021457-DEL-78 |
| chr11-32053546-DEL-330 | chr11-32053547-DEL-330 |
| chr11-32059478-DEL-66 | chr11-32059479-DEL-66 |
| chr11-32071116-DEL-73 | chr11-32071089-DEL-95 |
| chr11-32072813-DEL-111 | chr11-32072814-DEL-111 |
| chr11-32101349-DEL-51 | chr11-32101350-DEL-51 |
| chr11-32112651-DEL-1431 | chr11-32112652-DEL-1431 |
| chr11-32115175-DEL-155 | chr11-32115176-DEL-155 |
| chr11-32118056-DEL-60 | chr11-32118051-DEL-62 |
| chr11-32128751-DEL-3613 | chr11-32128752-DEL-3613 |
| chr11-32130447-DEL-367 | chr11-32130448-DEL-386 |
| chr11-32142189-DEL-65 | chr11-32142186-DEL-61 |
| chr11-32209663-DEL-349 | chr11-32209649-DEL-350 |
| chr11-32223592-DEL-66 | chr11-32223593-DEL-66 |
| chr11-32238085-DEL-64 | chr11-32238086-DEL-64 |
| chr11-32249672-DEL-75 | chr11-32249665-DEL-75 |
| chr11-32254542-DEL-677 | chr11-32254543-DEL-677 |
| chr11-32262305-DEL-102 | chr11-32262317-DEL-86 |
| chr11-32265091-DEL-3578 | chr11-32265092-DEL-3578 |
| chr11-32288335-DEL-132 | chr11-32288361-DEL-89 |
| chr11-32306436-DEL-165 | chr11-32306437-DEL-165 |
| chr11-32326229-DEL-71 | chr11-32326230-DEL-71 |
| chr11-32335112-DEL-108 | chr11-32335116-DEL-116 |
| chr11-32338496-DEL-92 | chr11-32338497-DEL-92 |
| chr11-32367289-DEL-737 | chr11-32367290-DEL-737 |
| chr11-32368189-DEL-91 | chr11-32368153-DEL-92 |
| chr11-32426505-DEL-277 | chr11-32426506-DEL-277 |
| chr11-32445044-DEL-747 | chr11-32445044-DEL-712 |
| chr11-32454376-DEL-138 | chr11-32454377-DEL-138 |
| chr11-32505812-DEL-106 | chr11-32505829-DEL-100 |
| chr11-32522025-DEL-55 | chr11-32522026-DEL-55 |
| chr11-32529557-DEL-315 | chr11-32529558-DEL-315 |
| chr11-32547246-DEL-197 | chr11-32547247-DEL-196 |
| chr11-32565232-DEL-154 | chr11-32565247-DEL-154 |
| chr11-32567567-DEL-228 | chr11-32567557-DEL-229 |
| chr11-32579330-DEL-212 | chr11-32579331-DEL-212 |
| chr11-32622319-DEL-1876 | chr11-32622320-DEL-1876 |
| chr11-32636624-DEL-320 | chr11-32636625-DEL-320 |
| chr11-32639603-DEL-1042 | chr11-32639603-DEL-1043 |
| chr11-32675495-DEL-1377 | chr11-32675496-DEL-1377 |
| chr11-32707396-DEL-76 | chr11-32707397-DEL-74 |
| chr11-32728728-DEL-53 | chr11-32728764-DEL-53 |
| chr11-32750423-DEL-141 | chr11-32750423-DEL-141 |
| chr11-32755681-DEL-275 | chr11-32755682-DEL-275 |
| chr11-32777701-DEL-108 | chr11-32777703-DEL-82 |
| chr11-32804964-DEL-2368 | chr11-32804965-DEL-2368 |
| chr11-32810931-DEL-76 | chr11-32810932-DEL-76 |
| chr11-32822631-DEL-86 | chr11-32822635-DEL-62 |
| chr11-32823084-DEL-265 | chr11-32823085-DEL-265 |
| chr11-32841282-DEL-410 | chr11-32841283-DEL-410 |
| chr11-32859246-DEL-1844 | chr11-32859247-DEL-1844 |
| chr11-32880179-DEL-73 | chr11-32880180-DEL-73 |
| chr11-32886449-DEL-179 | chr11-32886433-DEL-169 |
| chr11-32908915-DEL-61 | chr11-32908916-DEL-61 |
| chr11-32914792-DEL-142 | chr11-32914804-DEL-152 |
| chr11-32918153-DEL-95 | chr11-32918154-DEL-95 |
| chr11-32935439-DEL-114 | chr11-32935440-DEL-114 |
| chr11-32984472-DEL-189 | chr11-32984473-DEL-189 |
| chr11-32986178-DEL-164 | chr11-32986179-DEL-164 |
| chr11-32998729-DEL-498 | chr11-32998730-DEL-498 |
| chr11-33012300-DEL-6439 | chr11-33012301-DEL-6440 |
| chr11-33021099-DEL-104 | chr11-33021069-DEL-110 |
| chr11-33033876-DEL-136 | chr11-33033901-DEL-112 |
| chr11-33046625-DEL-52 | chr11-33046593-DEL-54 |
| chr11-33059368-DEL-51 | chr11-33059369-DEL-51 |
| chr11-33063725-DEL-111 | chr11-33063726-DEL-111 |
| chr11-33116283-DEL-64 | chr11-33116284-DEL-62 |
| chr11-33119405-DEL-216 | chr11-33119406-DEL-216 |
| chr11-33131768-DEL-56 | chr11-33131769-DEL-56 |
| chr11-33133265-DEL-52 | chr11-33133260-DEL-53 |
| chr11-33134493-DEL-58 | chr11-33134502-DEL-52 |
| chr11-33144672-DEL-69 | chr11-33144657-DEL-84 |
| chr11-33201831-DEL-223 | chr11-33201832-DEL-223 |
| chr11-33202788-DEL-61 | chr11-33202789-DEL-61 |
| chr11-33216592-DEL-94 | chr11-33216613-DEL-112 |
| chr11-33235546-DEL-63 | chr11-33235547-DEL-63 |

| | |
|---|---|
| chr11-33315463-DEL-2341 | chr11-33315464-DEL-2341 |
| chr11-33315585-DEL-336 | chr11-33315586-DEL-336 |
| chr11-33408579-DEL-52 | chr11-33408580-DEL-52 |
| chr11-33421370-DEL-60 | chr11-33421371-DEL-60 |
| chr11-33422675-DEL-57 | chr11-33422676-DEL-57 |
| chr11-33455932-DEL-207 | chr11-33455933-DEL-207 |
| chr11-33457769-DEL-656 | chr11-33457770-DEL-656 |
| chr11-33475448-DEL-59 | chr11-33475440-DEL-58 |
| chr11-33489796-DEL-97 | chr11-33489797-DEL-97 |
| chr11-33494818-DEL-204 | chr11-33494819-DEL-204 |
| chr11-33511716-DEL-806 | chr11-33511717-DEL-806 |
| chr11-33512345-DEL-96 | chr11-33512346-DEL-96 |
| chr11-33518820-DEL-211 | chr11-33518821-DEL-211 |
| chr11-33522154-DEL-84 | chr11-33522155-DEL-84 |
| chr11-33530711-DEL-54 | chr11-33530699-DEL-66 |
| chr11-33543296-DEL-51 | chr11-33543297-DEL-51 |
| chr11-33553975-DEL-125 | chr11-33553971-DEL-130 |
| chr11-33555951-DEL-90 | chr11-33555952-DEL-90 |
| chr11-33563144-DEL-94 | chr11-33563124-DEL-111 |
| chr11-33609993-DEL-51 | chr11-33609994-DEL-51 |
| chr11-33616991-DEL-247 | chr11-33616964-DEL-275 |
| chr11-33628485-DEL-72 | chr11-33628505-DEL-58 |
| chr11-33659369-DEL-68 | chr11-33659376-DEL-62 |
| chr11-33662451-DEL-400 | chr11-33662452-DEL-400 |
| chr11-33674721-DEL-263 | chr11-33674725-DEL-260 |
| chr11-33677923-DEL-189 | chr11-33677875-DEL-194 |
| chr11-33764997-DEL-679 | chr11-33764998-DEL-679 |
| chr11-33766006-DEL-54 | chr11-33766007-DEL-54 |
| chr11-33813304-DEL-76 | chr11-33813333-DEL-68 |
| chr11-33834213-DEL-106 | chr11-33834221-DEL-124 |
| chr11-33881985-DEL-156 | chr11-33882021-DEL-118 |
| chr11-33937173-DEL-173 | chr11-33937174-DEL-173 |
| chr11-33959243-DEL-70 | chr11-33959241-DEL-70 |
| chr11-33961200-DEL-248 | chr11-33961201-DEL-248 |
| chr11-33963874-DEL-6299 | chr11-33963875-DEL-6299 |
| chr11-33972258-DEL-92 | chr11-33972259-DEL-92 |
| chr11-33985523-DEL-86 | chr11-33985524-DEL-84 |
| chr11-33986984-DEL-104 | chr11-33986985-DEL-108 |
| chr11-34035443-DEL-80 | chr11-34035444-DEL-80 |
| chr11-34039919-DEL-58 | chr11-34039920-DEL-87 |
| chr11-34041268-DEL-1144 | chr11-34041269-DEL-1144 |
| chr11-34064478-DEL-719 | chr11-34064472-DEL-720 |
| chr11-34067530-DEL-88 | chr11-34067530-DEL-88 |
| chr11-34075701-DEL-204 | chr11-34075702-DEL-204 |
| chr11-34113817-DEL-177 | chr11-34113818-DEL-177 |
| chr11-34114888-DEL-106 | chr11-34114889-DEL-106 |
| chr11-34122040-DEL-58 | chr11-34122041-DEL-58 |
| chr11-34165904-DEL-435 | chr11-34165905-DEL-435 |
| chr11-34180368-DEL-51 | chr11-34180371-DEL-52 |
| chr11-34228056-DEL-109 | chr11-34228057-DEL-109 |
| chr11-34275089-DEL-90 | chr11-34275126-DEL-54 |
| chr11-34298100-DEL-62 | chr11-34298098-DEL-82 |
| chr11-34332718-DEL-84 | chr11-34332719-DEL-84 |
| chr11-34360257-DEL-134 | chr11-34360258-DEL-134 |
| chr11-34365038-DEL-272 | chr11-34365044-DEL-267 |
| chr11-34394346-DEL-54 | chr11-34394351-DEL-50 |
| chr11-34411410-DEL-95 | chr11-34411411-DEL-95 |
| chr11-34571446-DEL-384 | chr11-34571447-DEL-384 |
| chr11-34581203-DEL-62 | chr11-34581204-DEL-62 |
| chr11-34609099-DEL-79 | chr11-34609099-DEL-80 |
| chr11-34618483-DEL-921 | chr11-34618484-DEL-921 |
| chr11-34636461-DEL-129 | chr11-34636462-DEL-129 |
| chr11-34676855-DEL-176 | chr11-34676856-DEL-176 |
| chr11-34678086-DEL-201 | chr11-34678087-DEL-201 |
| chr11-34692982-DEL-322 | chr11-34692983-DEL-322 |
| chr11-34740765-DEL-6235 | chr11-34740766-DEL-6235 |
| chr11-34798025-DEL-325 | chr11-34798028-DEL-323 |
| chr11-34814243-DEL-131 | chr11-34814251-DEL-126 |
| chr11-34849466-DEL-89 | chr11-34849473-DEL-89 |
| chr11-34857577-DEL-417 | chr11-34857578-DEL-417 |
| chr11-34936758-DEL-1171 | chr11-34936759-DEL-1171 |
| chr11-34950797-DEL-154 | chr11-34950798-DEL-154 |
| chr11-34958866-DEL-54 | chr11-34958867-DEL-54 |
| chr11-34962738-DEL-58 | chr11-34962749-DEL-50 |
| chr11-34967360-DEL-153 | chr11-34967361-DEL-153 |
| chr11-35015340-DEL-51 | chr11-35015341-DEL-51 |
| chr11-35020378-DEL-165 | chr11-35020379-DEL-166 |
| chr11-35021355-DEL-94 | chr11-35021355-DEL-94 |
| chr11-35077162-DEL-239 | chr11-35077163-DEL-238 |
| chr11-35105431-DEL-99 | chr11-35105432-DEL-99 |

| | |
|---|---|
| chr11-35109751-DEL-88 | chr11-35109758-DEL-56 |
| chr11-35138261-DEL-554 | chr11-35138262-DEL-554 |
| chr11-35221179-DEL-54 | chr11-35221180-DEL-54 |
| chr11-35245362-DEL-375 | chr11-35245363-DEL-375 |
| chr11-35317283-DEL-1163 | chr11-35317284-DEL-1163 |
| chr11-35320086-DEL-89 | chr11-35320087-DEL-89 |
| chr11-35354782-DEL-93 | chr11-35354783-DEL-91 |
| chr11-35415557-DEL-78 | chr11-35415558-DEL-78 |
| chr11-35446437-DEL-73 | chr11-35446438-DEL-73 |
| chr11-35489118-DEL-220 | chr11-35489119-DEL-220 |
| chr11-35490899-DEL-72 | chr11-35490900-DEL-68 |
| chr11-35491999-DEL-70 | chr11-35492000-DEL-70 |
| chr11-35501109-DEL-78 | chr11-35501110-DEL-78 |
| chr11-35510791-DEL-51 | chr11-35510792-DEL-51 |
| chr11-35530394-DEL-107 | chr11-35530395-DEL-107 |
| chr11-35537146-DEL-208 | chr11-35537147-DEL-208 |
| chr11-35546539-DEL-60 | chr11-35546519-DEL-50 |
| chr11-35554209-DEL-60 | chr11-35554210-DEL-60 |
| chr11-35575735-DEL-364 | chr11-35575736-DEL-364 |
| chr11-35581271-DEL-80 | chr11-35581272-DEL-80 |
| chr11-35590354-DEL-74 | chr11-35590355-DEL-74 |
| chr11-35621859-DEL-276 | chr11-35621860-DEL-276 |
| chr11-35622712-DEL-321 | chr11-35622674-DEL-344 |
| chr11-35624063-DEL-120 | chr11-35624064-DEL-111 |
| chr11-35627413-DEL-185 | chr11-35627414-DEL-185 |
| chr11-35636945-DEL-223 | chr11-35636984-DEL-177 |
| chr11-35646958-DEL-212 | chr11-35646959-DEL-212 |
| chr11-35655425-DEL-256 | chr11-35655424-DEL-256 |
| chr11-35664295-DEL-159 | chr11-35664296-DEL-159 |
| chr11-35733276-DEL-416 | chr11-35733277-DEL-416 |
| chr11-35736395-DEL-999 | chr11-35736396-DEL-999 |
| chr11-35739461-DEL-52 | chr11-35739462-DEL-51 |
| chr11-35750369-DEL-207 | chr11-35750370-DEL-207 |
| chr11-35764028-DEL-161 | chr11-35764028-DEL-203 |
| chr11-35777221-DEL-194 | chr11-35777222-DEL-194 |
| chr11-35835408-DEL-156 | chr11-35835409-DEL-156 |
| chr11-35843377-DEL-56 | chr11-35843394-DEL-54 |
| chr11-35856289-DEL-141 | chr11-35856290-DEL-141 |
| chr11-35865243-DEL-80 | chr11-35865244-DEL-80 |
| chr11-35872077-DEL-64 | chr11-35872078-DEL-64 |
| chr11-35909850-DEL-60 | chr11-35909851-DEL-60 |
| chr11-35910420-DEL-58 | chr11-35910384-DEL-101 |
| chr11-35915948-DEL-76 | chr11-35915961-DEL-52 |
| chr11-35961574-DEL-51 | chr11-35961575-DEL-51 |
| chr11-35966600-DEL-63 | chr11-35966630-DEL-55 |
| chr11-35998242-DEL-54 | chr11-35998201-DEL-52 |
| chr11-36011113-DEL-98 | chr11-36011114-DEL-98 |
| chr11-36046949-DEL-314 | chr11-36046950-DEL-314 |
| chr11-36050113-DEL-196 | chr11-36050099-DEL-197 |
| chr11-36079326-DEL-73 | chr11-36079331-DEL-73 |
| chr11-36084340-DEL-141 | chr11-36084341-DEL-141 |
| chr11-36088772-DEL-742 | chr11-36088757-DEL-758 |
| chr11-36112065-DEL-57 | chr11-36112065-DEL-57 |
| chr11-36128547-DEL-550 | chr11-36128548-DEL-550 |
| chr11-36233262-DEL-6415 | chr11-36233253-DEL-6416 |
| chr11-36311471-DEL-86 | chr11-36311488-DEL-70 |
| chr11-36320234-DEL-51 | chr11-36320235-DEL-51 |
| chr11-36345638-DEL-207 | chr11-36345639-DEL-207 |
| chr11-36367340-DEL-52 | chr11-36367341-DEL-52 |
| chr11-36376679-DEL-71 | chr11-36376676-DEL-59 |
| chr11-36387986-DEL-517 | chr11-36387987-DEL-517 |
| chr11-36389471-DEL-62 | chr11-36389440-DEL-80 |
| chr11-36446634-DEL-181 | chr11-36446635-DEL-181 |
| chr11-36450249-DEL-96 | chr11-36450250-DEL-96 |
| chr11-36460725-DEL-55 | chr11-36460726-DEL-55 |
| chr11-36468098-DEL-1022 | chr11-36468099-DEL-1022 |
| chr11-36510834-DEL-90 | chr11-36510834-DEL-90 |
| chr11-36624142-DEL-1843 | chr11-36624143-DEL-1843 |
| chr11-36644120-DEL-100 | chr11-36644121-DEL-100 |
| chr11-36654494-DEL-293 | chr11-36654495-DEL-293 |
| chr11-36662600-DEL-72 | chr11-36662614-DEL-88 |
| chr11-36711091-DEL-140 | chr11-36711074-DEL-141 |
| chr11-36719462-DEL-74 | chr11-36719449-DEL-54 |
| chr11-36744813-DEL-240 | chr11-36744814-DEL-240 |
| chr11-36758032-DEL-823 | chr11-36758033-DEL-823 |
| chr11-36800278-DEL-56 | chr11-36800280-DEL-54 |
| chr11-36870500-DEL-480 | chr11-36870500-DEL-480 |
| chr11-36914353-DEL-80 | chr11-36914330-DEL-122 |
| chr11-36949439-DEL-60 | chr11-36949485-DEL-64 |
| chr11-36949889-DEL-889 | chr11-36949890-DEL-889 |

| | |
|---|---|
| chr11-36977904-DEL-123 | chr11-36977905-DEL-123 |
| chr11-36992290-DEL-123 | chr11-36992291-DEL-90 |
| chr11-36992674-DEL-517 | chr11-36992675-DEL-517 |
| chr11-37074751-DEL-82 | chr11-37074749-DEL-60 |
| chr11-37079046-DEL-60 | chr11-37079047-DEL-58 |
| chr11-37159667-DEL-182 | chr11-37159668-DEL-182 |
| chr11-37171624-DEL-168 | chr11-37171627-DEL-166 |
| chr11-37202118-DEL-239 | chr11-37202118-DEL-240 |
| chr11-37262987-DEL-149 | chr11-37262988-DEL-149 |
| chr11-37275052-DEL-511 | chr11-37275055-DEL-512 |
| chr11-37308645-DEL-134 | chr11-37308646-DEL-134 |
| chr11-37372612-DEL-3285 | chr11-37372613-DEL-3285 |
| chr11-37397858-DEL-396 | chr11-37397859-DEL-396 |
| chr11-37411367-DEL-56 | chr11-37411368-DEL-56 |
| chr11-37417565-DEL-159 | chr11-37417566-DEL-159 |
| chr11-37524712-DEL-6584 | chr11-37524713-DEL-6584 |
| chr11-37579352-DEL-106 | chr11-37579347-DEL-103 |
| chr11-37583812-DEL-517 | chr11-37583813-DEL-517 |
| chr11-37602940-DEL-997 | chr11-37602940-DEL-997 |
| chr11-37607925-DEL-1854 | chr11-37607926-DEL-1854 |
| chr11-37612956-DEL-359 | chr11-37612951-DEL-357 |
| chr11-37643254-DEL-7104 | chr11-37643255-DEL-7104 |
| chr11-37643270-DEL-425 | chr11-37643271-DEL-425 |
| chr11-37669674-DEL-577 | chr11-37669675-DEL-577 |
| chr11-37680867-DEL-1749 | chr11-37680868-DEL-1749 |
| chr11-37723904-DEL-70 | chr11-37723905-DEL-70 |
| chr11-37725566-DEL-102 | chr11-37725586-DEL-116 |
| chr11-37746147-DEL-88 | chr11-37746157-DEL-60 |
| chr11-37754571-DEL-178 | chr11-37754572-DEL-173 |
| chr11-37761171-DEL-122 | chr11-37761172-DEL-122 |
| chr11-37763540-DEL-204 | chr11-37763551-DEL-197 |
| chr11-37765237-DEL-73 | chr11-37765238-DEL-73 |
| chr11-37811766-DEL-2880 | chr11-37811760-DEL-2881 |
| chr11-37828322-DEL-548 | chr11-37828323-DEL-548 |
| chr11-37904059-DEL-6445 | chr11-37904093-DEL-6413 |
| chr11-37916357-DEL-687 | chr11-37916361-DEL-675 |
| chr11-37987611-DEL-5321 | chr11-37987617-DEL-5318 |
| chr11-37990240-DEL-136 | chr11-37990241-DEL-136 |
| chr11-37990859-DEL-237 | chr11-37990860-DEL-237 |
| chr11-37995896-DEL-132 | chr11-37995897-DEL-132 |
| chr11-38011393-DEL-162 | chr11-38011395-DEL-161 |
| chr11-38013931-DEL-62 | chr11-38013929-DEL-61 |
| chr11-38021047-DEL-462 | chr11-38021048-DEL-462 |
| chr11-38137894-DEL-7090 | chr11-38137895-DEL-7090 |
| chr11-38221814-DEL-66 | chr11-38221815-DEL-66 |
| chr11-38257612-DEL-185 | chr11-38257613-DEL-185 |
| chr11-38372356-DEL-5796 | chr11-38372357-DEL-5796 |
| chr11-38394964-DEL-57 | chr11-38394927-DEL-57 |
| chr11-38397687-DEL-111 | chr11-38397696-DEL-107 |
| chr11-38453567-DEL-73 | chr11-38453568-DEL-73 |
| chr11-38466960-DEL-56 | chr11-38466963-DEL-54 |
| chr11-38484039-DEL-61 | chr11-38484038-DEL-62 |
| chr11-38486198-DEL-76 | chr11-38486199-DEL-64 |
| chr11-38508578-DEL-82 | chr11-38508581-DEL-54 |
| chr11-38510846-DEL-148 | chr11-38510845-DEL-150 |
| chr11-38553302-DEL-956 | chr11-38553303-DEL-956 |
| chr11-38593482-DEL-84 | chr11-38593483-DEL-84 |
| chr11-38628035-DEL-70 | chr11-38628058-DEL-94 |
| chr11-38632529-DEL-2024 | chr11-38632530-DEL-2024 |
| chr11-38668092-DEL-1310 | chr11-38668093-DEL-1310 |
| chr11-38678418-DEL-53 | chr11-38678417-DEL-54 |
| chr11-38687118-DEL-51 | chr11-38687121-DEL-69 |
| chr11-38713089-DEL-80 | chr11-38713115-DEL-91 |
| chr11-38714570-DEL-61 | chr11-38714571-DEL-61 |
| chr11-38721911-DEL-1464 | chr11-38721912-DEL-1464 |
| chr11-38735329-DEL-69 | chr11-38735330-DEL-69 |
| chr11-38767412-DEL-2900 | chr11-38767413-DEL-2900 |
| chr11-38775384-DEL-482 | chr11-38775384-DEL-482 |
| chr11-38793448-DEL-7025 | chr11-38793449-DEL-7025 |
| chr11-38817582-DEL-247 | chr11-38817583-DEL-247 |
| chr11-38870782-DEL-6307 | chr11-38870792-DEL-6300 |
| chr11-38889011-DEL-267 | chr11-38889012-DEL-267 |
| chr11-38889972-DEL-580 | chr11-38889973-DEL-580 |
| chr11-38897461-DEL-91 | chr11-38897462-DEL-91 |
| chr11-38907326-DEL-1899 | chr11-38907327-DEL-1899 |
| chr11-38909195-DEL-293 | chr11-38909196-DEL-293 |
| chr11-38909905-DEL-86 | chr11-38909906-DEL-86 |
| chr11-38963136-DEL-68 | chr11-38963139-DEL-66 |
| chr11-38994506-DEL-1476 | chr11-38994507-DEL-1476 |
| chr11-39023920-DEL-68 | chr11-39023921-DEL-68 |

| | |
|---|---|
| chr11-39026261-DEL-744 | chr11-39026262-DEL-744 |
| chr11-39029350-DEL-65 | chr11-39029351-DEL-65 |
| chr11-39032916-DEL-228 | chr11-39032917-DEL-227 |
| chr11-39083642-DEL-544 | chr11-39083643-DEL-544 |
| chr11-39104087-DEL-1923 | chr11-39104088-DEL-1923 |
| chr11-39112819-DEL-683 | chr11-39112820-DEL-683 |
| chr11-39143020-DEL-823 | chr11-39143021-DEL-823 |
| chr11-39145261-DEL-67 | chr11-39145248-DEL-56 |
| chr11-39151441-DEL-120 | chr11-39151442-DEL-120 |
| chr11-39196156-DEL-326 | chr11-39196157-DEL-326 |
| chr11-39196783-DEL-701 | chr11-39196781-DEL-702 |
| chr11-39207388-DEL-962 | chr11-39207389-DEL-962 |
| chr11-39219541-DEL-252 | chr11-39219542-DEL-252 |
| chr11-39251065-DEL-66 | chr11-39251055-DEL-52 |
| chr11-39274294-DEL-155 | chr11-39274295-DEL-155 |
| chr11-39329238-DEL-80 | chr11-39329195-DEL-127 |
| chr11-39396965-DEL-245 | chr11-39396966-DEL-245 |
| chr11-39420938-DEL-464 | chr11-39420939-DEL-464 |
| chr11-39432684-DEL-198 | chr11-39432685-DEL-198 |
| chr11-39443371-DEL-84 | chr11-39443372-DEL-77 |
| chr11-39463949-DEL-86 | chr11-39463950-DEL-86 |
| chr11-39465750-DEL-140 | chr11-39465751-DEL-140 |
| chr11-39490226-DEL-69 | chr11-39490231-DEL-57 |
| chr11-39505829-DEL-177 | chr11-39505830-DEL-177 |
| chr11-39525181-DEL-142 | chr11-39525180-DEL-143 |
| chr11-39535800-DEL-467 | chr11-39535803-DEL-466 |
| chr11-39591951-DEL-2003 | chr11-39591952-DEL-2003 |
| chr11-39692012-DEL-549 | chr11-39692013-DEL-549 |
| chr11-39703179-DEL-60 | chr11-39703180-DEL-60 |
| chr11-39704031-DEL-2332 | chr11-39704032-DEL-2332 |
| chr11-39708674-DEL-932 | chr11-39708675-DEL-930 |
| chr11-39709747-DEL-398 | chr11-39709744-DEL-414 |
| chr11-39748930-DEL-63 | chr11-39748931-DEL-63 |
| chr11-39750706-DEL-53 | chr11-39750703-DEL-51 |
| chr11-39794067-DEL-73 | chr11-39794068-DEL-73 |
| chr11-39831778-DEL-73 | chr11-39831779-DEL-73 |
| chr11-39870511-DEL-533 | chr11-39870512-DEL-533 |
| chr11-39874537-DEL-75 | chr11-39874538-DEL-75 |
| chr11-39882407-DEL-6155 | chr11-39882408-DEL-6155 |
| chr11-39909899-DEL-67 | chr11-39909897-DEL-65 |
| chr11-39928252-DEL-54 | chr11-39928247-DEL-72 |
| chr11-39933465-DEL-1189 | chr11-39933455-DEL-1183 |
| chr11-39954564-DEL-7086 | chr11-39954565-DEL-7086 |
| chr11-39969322-DEL-113 | chr11-39969344-DEL-89 |
| chr11-39979214-DEL-74 | chr11-39979215-DEL-74 |
| chr11-39983308-DEL-119 | chr11-39983309-DEL-119 |
| chr11-40000449-DEL-50 | chr11-40000471-DEL-52 |
| chr11-40013869-DEL-1069 | chr11-40013869-DEL-1069 |
| chr11-40060516-DEL-887 | chr11-40060517-DEL-887 |
| chr11-40121518-DEL-6579 | chr11-40121519-DEL-6579 |
| chr11-40139800-DEL-214 | chr11-40139801-DEL-214 |
| chr11-40154605-DEL-136 | chr11-40154594-DEL-117 |
| chr11-40173603-DEL-115 | chr11-40173604-DEL-115 |
| chr11-40206640-DEL-53 | chr11-40206641-DEL-53 |
| chr11-40232955-DEL-7208 | chr11-40232956-DEL-7208 |
| chr11-40255515-DEL-171 | chr11-40255516-DEL-171 |
| chr11-40266749-DEL-107 | chr11-40266750-DEL-107 |
| chr11-40268532-DEL-70 | chr11-40268509-DEL-62 |
| chr11-40277351-DEL-389 | chr11-40277352-DEL-389 |
| chr11-40277437-DEL-2776 | chr11-40277438-DEL-2776 |
| chr11-40294570-DEL-121 | chr11-40294571-DEL-121 |
| chr11-40295217-DEL-153 | chr11-40295220-DEL-152 |
| chr11-40304275-DEL-7073 | chr11-40304276-DEL-7073 |
| chr11-40327558-DEL-63 | chr11-40327565-DEL-58 |
| chr11-40336564-DEL-110 | chr11-40336565-DEL-110 |
| chr11-40393170-DEL-56 | chr11-40393171-DEL-56 |
| chr11-40395281-DEL-119 | chr11-40395300-DEL-101 |
| chr11-40406543-DEL-51 | chr11-40406540-DEL-52 |
| chr11-40516343-DEL-6442 | chr11-40516344-DEL-6442 |
| chr11-40524706-DEL-6498 | chr11-40524707-DEL-6498 |
| chr11-40543953-DEL-74 | chr11-40543954-DEL-74 |
| chr11-40558796-DEL-65 | chr11-40558796-DEL-65 |
| chr11-40563348-DEL-1373 | chr11-40563349-DEL-1373 |
| chr11-40579127-DEL-66 | chr11-40579128-DEL-66 |
| chr11-40592460-DEL-344 | chr11-40592461-DEL-344 |
| chr11-40594021-DEL-213 | chr11-40594022-DEL-213 |
| chr11-40603458-DEL-1077 | chr11-40603459-DEL-1077 |
| chr11-40603852-DEL-56 | chr11-40603853-DEL-56 |
| chr11-40614713-DEL-100 | chr11-40614714-DEL-100 |
| chr11-40631320-DEL-82 | chr11-40631326-DEL-77 |

| | |
|---|---|
| chr11-40669721-DEL-193 | chr11-40669723-DEL-192 |
| chr11-40685001-DEL-265 | chr11-40685002-DEL-265 |
| chr11-40685833-DEL-2005 | chr11-40685834-DEL-2005 |
| chr11-40729574-DEL-6813 | chr11-40729575-DEL-6813 |
| chr11-40738262-DEL-2221 | chr11-40738263-DEL-2221 |
| chr11-40800308-DEL-189 | chr11-40800309-DEL-189 |
| chr11-40804766-DEL-6730 | chr11-40804767-DEL-6730 |
| chr11-40847169-DEL-15790 | chr11-40847156-DEL-15776 |
| chr11-40876307-DEL-6339 | chr11-40876314-DEL-6333 |
| chr11-40891308-DEL-148 | chr11-40891309-DEL-148 |
| chr11-40907077-DEL-6594 | chr11-40907078-DEL-6594 |
| chr11-40959842-DEL-63 | chr11-40959843-DEL-63 |
| chr11-40979332-DEL-3222 | chr11-40979333-DEL-3222 |
| chr11-40988673-DEL-397 | chr11-40988674-DEL-397 |
| chr11-40989174-DEL-981 | chr11-40989175-DEL-981 |
| chr11-40996084-DEL-6652 | chr11-40996085-DEL-6652 |
| chr11-41012284-DEL-53 | chr11-41012285-DEL-53 |
| chr11-41030166-DEL-721 | chr11-41030167-DEL-721 |
| chr11-41032469-DEL-2276 | chr11-41032470-DEL-2276 |
| chr11-41041128-DEL-210 | chr11-41041125-DEL-213 |
| chr11-41053744-DEL-53 | chr11-41053775-DEL-50 |
| chr11-41054960-DEL-68 | chr11-41054961-DEL-66 |
| chr11-41089535-DEL-92 | chr11-41089557-DEL-52 |
| chr11-41109519-DEL-5646 | chr11-41109520-DEL-5646 |
| chr11-41146848-DEL-429 | chr11-41146849-DEL-429 |
| chr11-41160318-DEL-828 | chr11-41160319-DEL-828 |
| chr11-41173804-DEL-530 | chr11-41173805-DEL-530 |
| chr11-41177507-DEL-119 | chr11-41177508-DEL-119 |
| chr11-41182847-DEL-718 | chr11-41182849-DEL-717 |
| chr11-41186558-DEL-66 | chr11-41186559-DEL-66 |
| chr11-41222619-DEL-314 | chr11-41222620-DEL-314 |
| chr11-41240037-DEL-217 | chr11-41240038-DEL-217 |
| chr11-41248298-DEL-183 | chr11-41248299-DEL-183 |
| chr11-41271710-DEL-6438 | chr11-41271697-DEL-6439 |
| chr11-41271808-DEL-207 | chr11-41271767-DEL-208 |
| chr11-41287951-DEL-310 | chr11-41287952-DEL-310 |
| chr11-41300164-DEL-758 | chr11-41300163-DEL-758 |
| chr11-41335596-DEL-331 | chr11-41335597-DEL-331 |
| chr11-41340035-DEL-72 | chr11-41340043-DEL-62 |
| chr11-41389468-DEL-2272 | chr11-41389468-DEL-2272 |
| chr11-41428462-DEL-1186 | chr11-41428463-DEL-1186 |
| chr11-41435442-DEL-51 | chr11-41435442-DEL-51 |
| chr11-41438396-DEL-64 | chr11-41438397-DEL-64 |
| chr11-41439793-DEL-1777 | chr11-41439794-DEL-1777 |
| chr11-41445776-DEL-210 | chr11-41445777-DEL-210 |
| chr11-41449473-DEL-6255 | chr11-41449465-DEL-6256 |
| chr11-41457365-DEL-1379 | chr11-41457366-DEL-1379 |
| chr11-41464751-DEL-555 | chr11-41464752-DEL-555 |
| chr11-41476551-DEL-938 | chr11-41476552-DEL-938 |
| chr11-41480794-DEL-520 | chr11-41480795-DEL-520 |
| chr11-41525479-DEL-70 | chr11-41525480-DEL-70 |
| chr11-41563478-DEL-73 | chr11-41563474-DEL-73 |
| chr11-41575011-DEL-63 | chr11-41575012-DEL-63 |
| chr11-41586629-DEL-66 | chr11-41586630-DEL-66 |
| chr11-41594733-DEL-6592 | chr11-41594734-DEL-6592 |
| chr11-41626041-DEL-66 | chr11-41626039-DEL-67 |
| chr11-41636361-DEL-62 | chr11-41636362-DEL-62 |
| chr11-41687344-DEL-127 | chr11-41687345-DEL-127 |
| chr11-41718301-DEL-935 | chr11-41718302-DEL-935 |
| chr11-41731666-DEL-3287 | chr11-41731669-DEL-3288 |
| chr11-41750962-DEL-324 | chr11-41750963-DEL-324 |
| chr11-41764217-DEL-1168 | chr11-41764218-DEL-1168 |
| chr11-41767383-DEL-369 | chr11-41767384-DEL-369 |
| chr11-41768366-DEL-7595 | chr11-41768367-DEL-7595 |
| chr11-41791595-DEL-139 | chr11-41791596-DEL-139 |
| chr11-41796862-DEL-58 | chr11-41796863-DEL-58 |
| chr11-41814410-DEL-211 | chr11-41814411-DEL-211 |
| chr11-41831164-DEL-260 | chr11-41831165-DEL-260 |
| chr11-41833503-DEL-54 | chr11-41833496-DEL-64 |
| chr11-41879464-DEL-61 | chr11-41879465-DEL-61 |
| chr11-41947519-DEL-86 | chr11-41947520-DEL-86 |
| chr11-41966331-DEL-52 | chr11-41966332-DEL-52 |
| chr11-41980892-DEL-83 | chr11-41980893-DEL-83 |
| chr11-41985755-DEL-80 | chr11-41985767-DEL-71 |
| chr11-41991516-DEL-1537 | chr11-41991517-DEL-1537 |
| chr11-42037593-DEL-134 | chr11-42037594-DEL-134 |
| chr11-42071203-DEL-68 | chr11-42071204-DEL-68 |
| chr11-42076476-DEL-97 | chr11-42076477-DEL-97 |
| chr11-42099458-DEL-466 | chr11-42099459-DEL-466 |
| chr11-42119832-DEL-142 | chr11-42119833-DEL-142 |

| | |
|---|---|
| chr11-42120900-DEL-164 | chr11-42120899-DEL-144 |
| chr11-42137572-DEL-219 | chr11-42137573-DEL-219 |
| chr11-42141955-DEL-78 | chr11-42141956-DEL-78 |
| chr11-42151603-DEL-213 | chr11-42151604-DEL-213 |
| chr11-42159414-DEL-62 | chr11-42159425-DEL-60 |
| chr11-42179239-DEL-3459 | chr11-42179240-DEL-3459 |
| chr11-42198705-DEL-65 | chr11-42198708-DEL-62 |
| chr11-42238020-DEL-147 | chr11-42238017-DEL-148 |
| chr11-42292599-DEL-83 | chr11-42292587-DEL-84 |
| chr11-42332933-DEL-253 | chr11-42332937-DEL-221 |
| chr11-42359810-DEL-83 | chr11-42359808-DEL-79 |
| chr11-42474610-DEL-517 | chr11-42474611-DEL-517 |
| chr11-42547017-DEL-78 | chr11-42547018-DEL-78 |
| chr11-42548292-DEL-95 | chr11-42548293-DEL-95 |
| chr11-42561843-DEL-1221 | chr11-42561844-DEL-1221 |
| chr11-42590972-DEL-810 | chr11-42590973-DEL-810 |
| chr11-42622427-DEL-413 | chr11-42622428-DEL-413 |
| chr11-42658024-DEL-1451 | chr11-42658025-DEL-1451 |
| chr11-42670021-DEL-6241 | chr11-42670022-DEL-6241 |
| chr11-42689007-DEL-427 | chr11-42689008-DEL-427 |
| chr11-42721791-DEL-351 | chr11-42721792-DEL-351 |
| chr11-42751798-DEL-422 | chr11-42751799-DEL-422 |
| chr11-42756985-DEL-2416 | chr11-42756986-DEL-2416 |
| chr11-42779476-DEL-1252 | chr11-42779477-DEL-1253 |
| chr11-42783350-DEL-1086 | chr11-42783351-DEL-1086 |
| chr11-42787850-DEL-52 | chr11-42787845-DEL-52 |
| chr11-42813887-DEL-760 | chr11-42813888-DEL-760 |
| chr11-42826572-DEL-85 | chr11-42826573-DEL-85 |
| chr11-42829322-DEL-124 | chr11-42829323-DEL-124 |
| chr11-42860706-DEL-6379 | chr11-42860707-DEL-6379 |
| chr11-42874141-DEL-2708 | chr11-42874142-DEL-2708 |
| chr11-42876952-DEL-56 | chr11-42876958-DEL-56 |
| chr11-42878715-DEL-1109 | chr11-42878716-DEL-1109 |
| chr11-42892863-DEL-62 | chr11-42892868-DEL-54 |
| chr11-42904483-DEL-72 | chr11-42904484-DEL-64 |
| chr11-42920619-DEL-54 | chr11-42920620-DEL-54 |
| chr11-42947960-DEL-66 | chr11-42947961-DEL-66 |
| chr11-42955257-DEL-1064 | chr11-42955258-DEL-1064 |
| chr11-42967383-DEL-302 | chr11-42967384-DEL-302 |
| chr11-42969279-DEL-455 | chr11-42969280-DEL-456 |
| chr11-42982189-DEL-112 | chr11-42982190-DEL-112 |
| chr11-43013251-DEL-75 | chr11-43013252-DEL-75 |
| chr11-43016441-DEL-136 | chr11-43016428-DEL-150 |
| chr11-43018554-DEL-64 | chr11-43018554-DEL-65 |
| chr11-43035603-DEL-53 | chr11-43035601-DEL-83 |
| chr11-43047244-DEL-536 | chr11-43047245-DEL-536 |
| chr11-43064368-DEL-52 | chr11-43064369-DEL-52 |
| chr11-43082344-DEL-309 | chr11-43082342-DEL-312 |
| chr11-43108866-DEL-199 | chr11-43108867-DEL-199 |
| chr11-43128172-DEL-1864 | chr11-43128173-DEL-1864 |
| chr11-43167686-DEL-68 | chr11-43167707-DEL-60 |
| chr11-43176005-DEL-1020 | chr11-43176006-DEL-1020 |
| chr11-43194535-DEL-6630 | chr11-43194536-DEL-6630 |
| chr11-43218751-DEL-1842 | chr11-43218752-DEL-1842 |
| chr11-43236601-DEL-52 | chr11-43236596-DEL-56 |
| chr11-43246100-DEL-93 | chr11-43246101-DEL-92 |
| chr11-43255760-DEL-85 | chr11-43255761-DEL-85 |
| chr11-43297397-DEL-199 | chr11-43297398-DEL-199 |
| chr11-43376471-DEL-59 | chr11-43376472-DEL-59 |
| chr11-43379518-DEL-1607 | chr11-43379519-DEL-1603 |
| chr11-43381125-DEL-76 | chr11-43381126-DEL-86 |
| chr11-43389375-DEL-1690 | chr11-43389376-DEL-1690 |
| chr11-43390513-DEL-197 | chr11-43390494-DEL-198 |
| chr11-43392498-DEL-140 | chr11-43392499-DEL-152 |
| chr11-43394778-DEL-428 | chr11-43394776-DEL-429 |
| chr11-43441958-DEL-181 | chr11-43441959-DEL-181 |
| chr11-43444239-DEL-172 | chr11-43444240-DEL-172 |
| chr11-43498696-DEL-60 | chr11-43498693-DEL-60 |
| chr11-43511214-DEL-181 | chr11-43511176-DEL-177 |
| chr11-43521686-DEL-501 | chr11-43521682-DEL-502 |
| chr11-43523314-DEL-55 | chr11-43523320-DEL-51 |
| chr11-43535735-DEL-1579 | chr11-43535736-DEL-1576 |
| chr11-43539826-DEL-62 | chr11-43539826-DEL-57 |
| chr11-43550544-DEL-4100 | chr11-43550545-DEL-4100 |
| chr11-43560752-DEL-66 | chr11-43560756-DEL-64 |
| chr11-43565707-DEL-194 | chr11-43565708-DEL-194 |
| chr11-43571894-DEL-52 | chr11-43571895-DEL-52 |
| chr11-43579584-DEL-73 | chr11-43579560-DEL-73 |
| chr11-43620147-DEL-97 | chr11-43620148-DEL-97 |
| chr11-43668586-DEL-259 | chr11-43668624-DEL-222 |

| | |
|---|---|
| chr11-43705211-DEL-77 | chr11-43705164-DEL-78 |
| chr11-43713443-DEL-78 | chr11-43713444-DEL-78 |
| chr11-43714129-DEL-52 | chr11-43714128-DEL-52 |
| chr11-43716841-DEL-97 | chr11-43716843-DEL-98 |
| chr11-43750538-DEL-159 | chr11-43750539-DEL-159 |
| chr11-43757386-DEL-77 | chr11-43757387-DEL-77 |
| chr11-43761168-DEL-62 | chr11-43761169-DEL-62 |
| chr11-43778328-DEL-69 | chr11-43778329-DEL-69 |
| chr11-43797328-DEL-134 | chr11-43797332-DEL-131 |
| chr11-43803563-DEL-186 | chr11-43803564-DEL-186 |
| chr11-43828344-DEL-375 | chr11-43828344-DEL-375 |
| chr11-43833020-DEL-78 | chr11-43833017-DEL-56 |
| chr11-43853890-DEL-728 | chr11-43853891-DEL-728 |
| chr11-43910952-DEL-63 | chr11-43910953-DEL-63 |
| chr11-43935155-DEL-192 | chr11-43935156-DEL-144 |
| chr11-43968635-DEL-714 | chr11-43968636-DEL-715 |
| chr11-43976331-DEL-432 | chr11-43976332-DEL-432 |
| chr11-44008590-DEL-1168 | chr11-44008591-DEL-1168 |
| chr11-44013319-DEL-66 | chr11-44013320-DEL-66 |
| chr11-44031185-DEL-58 | chr11-44031191-DEL-98 |
| chr11-44057170-DEL-403 | chr11-44057158-DEL-404 |
| chr11-44078408-DEL-51 | chr11-44078412-DEL-50 |
| chr11-44165990-DEL-2203 | chr11-44165990-DEL-2203 |
| chr11-44169052-DEL-76 | chr11-44169055-DEL-74 |
| chr11-44183504-DEL-693 | chr11-44183505-DEL-693 |
| chr11-44185628-DEL-115 | chr11-44185644-DEL-119 |
| chr11-44196891-DEL-6458 | chr11-44196892-DEL-6458 |
| chr11-44227071-DEL-141 | chr11-44227072-DEL-141 |
| chr11-44258394-DEL-59 | chr11-44258388-DEL-60 |
| chr11-44299130-DEL-74 | chr11-44299135-DEL-70 |
| chr11-44322246-DEL-352 | chr11-44322247-DEL-352 |
| chr11-44327186-DEL-170 | chr11-44327187-DEL-170 |
| chr11-44330140-DEL-1279 | chr11-44330132-DEL-1277 |
| chr11-44330163-DEL-71 | chr11-44330164-DEL-71 |
| chr11-44331011-DEL-586 | chr11-44331012-DEL-586 |
| chr11-44332337-DEL-284 | chr11-44332338-DEL-284 |
| chr11-44339883-DEL-58 | chr11-44339884-DEL-58 |
| chr11-44351352-DEL-108 | chr11-44351353-DEL-108 |
| chr11-44372998-DEL-624 | chr11-44372999-DEL-624 |
| chr11-44388656-DEL-90 | chr11-44388655-DEL-80 |
| chr11-44397391-DEL-879 | chr11-44397391-DEL-877 |
| chr11-44419356-DEL-95 | chr11-44419357-DEL-95 |
| chr11-44433903-DEL-70 | chr11-44433904-DEL-70 |
| chr11-44447425-DEL-208 | chr11-44447429-DEL-210 |
| chr11-44474751-DEL-1211 | chr11-44474752-DEL-1211 |
| chr11-44477384-DEL-88 | chr11-44477385-DEL-88 |
| chr11-44480229-DEL-60 | chr11-44480230-DEL-60 |
| chr11-44488272-DEL-210 | chr11-44488273-DEL-210 |
| chr11-44490340-DEL-202 | chr11-44490341-DEL-202 |
| chr11-44505589-DEL-61 | chr11-44505590-DEL-64 |
| chr11-44549868-DEL-60 | chr11-44549866-DEL-61 |
| chr11-44562171-DEL-174 | chr11-44562182-DEL-139 |
| chr11-44581403-DEL-64 | chr11-44581404-DEL-64 |
| chr11-44615783-DEL-400 | chr11-44615784-DEL-400 |
| chr11-44677460-DEL-54 | chr11-44677460-DEL-54 |
| chr11-44698005-DEL-144 | chr11-44698006-DEL-144 |
| chr11-44701139-DEL-273 | chr11-44701140-DEL-267 |
| chr11-44712340-DEL-102 | chr11-44712341-DEL-102 |
| chr11-44731092-DEL-232 | chr11-44731093-DEL-232 |
| chr11-44747700-DEL-1318 | chr11-44747701-DEL-1318 |
| chr11-44778926-DEL-65 | chr11-44778941-DEL-51 |
| chr11-44805120-DEL-60 | chr11-44805116-DEL-64 |
| chr11-44810139-DEL-168 | chr11-44810139-DEL-171 |
| chr11-44820902-DEL-85 | chr11-44820903-DEL-85 |
| chr11-44828199-DEL-98 | chr11-44828222-DEL-98 |
| chr11-44841067-DEL-56 | chr11-44841068-DEL-56 |
| chr11-44899404-DEL-60 | chr11-44899396-DEL-70 |
| chr11-44904080-DEL-68 | chr11-44904046-DEL-54 |
| chr11-44929420-DEL-64 | chr11-44929443-DEL-69 |
| chr11-44931305-DEL-75 | chr11-44931306-DEL-75 |
| chr11-44936100-DEL-330 | chr11-44936107-DEL-324 |
| chr11-44984287-DEL-51 | chr11-44984288-DEL-51 |
| chr11-44986624-DEL-6225 | chr11-44986622-DEL-6226 |
| chr11-44999305-DEL-58 | chr11-44999306-DEL-56 |
| chr11-45008295-DEL-58 | chr11-45008296-DEL-58 |
| chr11-45096439-DEL-55 | chr11-45096438-DEL-56 |
| chr11-45118510-DEL-54 | chr11-45118511-DEL-54 |
| chr11-45171411-DEL-368 | chr11-45171412-DEL-368 |
| chr11-45183168-DEL-78 | chr11-45183169-DEL-78 |
| chr11-45189728-DEL-52 | chr11-45189728-DEL-52 |

| | |
|---|---|
| chr11-45191239-DEL-75 | chr11-45191253-DEL-53 |
| chr11-45203414-DEL-54 | chr11-45203415-DEL-54 |
| chr11-45210122-DEL-58 | chr11-45210108-DEL-85 |
| chr11-45251738-DEL-1142 | chr11-45251739-DEL-1142 |
| chr11-45258596-DEL-6867 | chr11-45258597-DEL-6867 |
| chr11-45305766-DEL-82 | chr11-45305759-DEL-84 |
| chr11-45315011-DEL-3228 | chr11-45315012-DEL-3228 |
| chr11-45331643-DEL-90 | chr11-45331607-DEL-88 |
| chr11-45342032-DEL-57 | chr11-45342019-DEL-58 |
| chr11-45343359-DEL-223 | chr11-45343355-DEL-223 |
| chr11-45358849-DEL-106 | chr11-45358850-DEL-106 |
| chr11-45378973-DEL-427 | chr11-45378974-DEL-427 |
| chr11-45383670-DEL-65 | chr11-45383656-DEL-79 |
| chr11-45453634-DEL-172 | chr11-45453635-DEL-172 |
| chr11-45471352-DEL-681 | chr11-45471353-DEL-681 |
| chr11-45473748-DEL-6205 | chr11-45473749-DEL-6205 |
| chr11-45516737-DEL-352 | chr11-45516738-DEL-352 |
| chr11-45522097-DEL-144 | chr11-45522104-DEL-110 |
| chr11-45531810-DEL-175 | chr11-45531810-DEL-176 |
| chr11-45541037-DEL-5382 | chr11-45541038-DEL-5382 |
| chr11-45606578-DEL-69 | chr11-45606581-DEL-67 |
| chr11-45608021-DEL-136 | chr11-45608022-DEL-136 |
| chr11-45614426-DEL-62 | chr11-45614427-DEL-62 |
| chr11-45615252-DEL-7912 | chr11-45615253-DEL-7912 |
| chr11-45616749-DEL-220 | chr11-45616749-DEL-220 |
| chr11-45634867-DEL-309 | chr11-45634868-DEL-308 |
| chr11-45640728-DEL-53 | chr11-45640728-DEL-53 |
| chr11-45660357-DEL-7495 | chr11-45660357-DEL-7494 |
| chr11-45677417-DEL-51 | chr11-45677418-DEL-51 |
| chr11-45677942-DEL-56 | chr11-45677939-DEL-66 |
| chr11-45679042-DEL-96 | chr11-45679043-DEL-96 |
| chr11-45682276-DEL-193 | chr11-45682265-DEL-194 |
| chr11-45716969-DEL-59 | chr11-45716970-DEL-59 |
| chr11-45718948-DEL-514 | chr11-45718950-DEL-514 |
| chr11-45719927-DEL-942 | chr11-45719928-DEL-942 |
| chr11-45745571-DEL-77 | chr11-45745572-DEL-77 |
| chr11-45747655-DEL-1062 | chr11-45747654-DEL-1062 |
| chr11-45768708-DEL-213 | chr11-45768709-DEL-213 |
| chr11-45810178-DEL-219 | chr11-45810179-DEL-219 |
| chr11-45830699-DEL-872 | chr11-45830699-DEL-873 |
| chr11-45848339-DEL-98 | chr11-45848342-DEL-112 |
| chr11-45855168-DEL-751 | chr11-45855169-DEL-751 |
| chr11-45927541-DEL-110 | chr11-45927542-DEL-110 |
| chr11-45937626-DEL-230 | chr11-45937607-DEL-211 |
| chr11-45942064-DEL-50 | chr11-45942059-DEL-55 |
| chr11-45942179-DEL-998 | chr11-45942179-DEL-998 |
| chr11-45952201-DEL-60 | chr11-45952198-DEL-64 |
| chr11-45992457-DEL-170 | chr11-45992458-DEL-170 |
| chr11-46043742-DEL-230 | chr11-46043741-DEL-211 |
| chr11-46054399-DEL-97 | chr11-46054400-DEL-97 |
| chr11-46065326-DEL-198 | chr11-46065327-DEL-198 |
| chr11-46075250-DEL-84 | chr11-46075251-DEL-84 |
| chr11-46103986-DEL-434 | chr11-46103987-DEL-434 |
| chr11-46133398-DEL-109 | chr11-46133424-DEL-76 |
| chr11-46186330-DEL-348 | chr11-46186331-DEL-348 |
| chr11-46190548-DEL-228 | chr11-46190549-DEL-228 |
| chr11-46229840-DEL-84 | chr11-46229804-DEL-83 |
| chr11-46240605-DEL-53 | chr11-46240606-DEL-53 |
| chr11-46241692-DEL-55 | chr11-46241670-DEL-52 |
| chr11-46247439-DEL-71 | chr11-46247440-DEL-71 |
| chr11-46255192-DEL-236 | chr11-46255193-DEL-237 |
| chr11-46273664-DEL-197 | chr11-46273665-DEL-193 |
| chr11-46297545-DEL-63 | chr11-46297545-DEL-63 |
| chr11-46300367-DEL-55 | chr11-46300368-DEL-55 |
| chr11-46303043-DEL-2991 | chr11-46303044-DEL-2991 |
| chr11-46350033-DEL-175 | chr11-46350034-DEL-175 |
| chr11-46356640-DEL-302 | chr11-46356641-DEL-302 |
| chr11-46357362-DEL-71 | chr11-46357363-DEL-71 |
| chr11-46358245-DEL-202 | chr11-46358246-DEL-202 |
| chr11-46377932-DEL-84 | chr11-46377932-DEL-88 |
| chr11-46383540-DEL-12240 | chr11-46383541-DEL-12240 |
| chr11-46386683-DEL-5023 | chr11-46386683-DEL-5023 |
| chr11-46394827-DEL-208 | chr11-46394828-DEL-208 |
| chr11-46403608-DEL-2719 | chr11-46403609-DEL-2719 |
| chr11-46406537-DEL-60 | chr11-46406577-DEL-55 |
| chr11-46429079-DEL-635 | chr11-46429080-DEL-635 |
| chr11-46440789-DEL-66 | chr11-46440790-DEL-66 |
| chr11-46458302-DEL-470 | chr11-46458303-DEL-470 |
| chr11-46467249-DEL-106 | chr11-46467251-DEL-105 |
| chr11-46469093-DEL-201 | chr11-46469100-DEL-199 |

| | |
|---|---|
| chr11-46481889-DEL-802 | chr11-46481890-DEL-802 |
| chr11-46482473-DEL-363 | chr11-46482474-DEL-363 |
| chr11-46489343-DEL-6653 | chr11-46489344-DEL-6653 |
| chr11-46495646-DEL-208 | chr11-46495647-DEL-208 |
| chr11-46550333-DEL-473 | chr11-46550334-DEL-473 |
| chr11-46557280-DEL-3041 | chr11-46557281-DEL-3041 |
| chr11-46569186-DEL-607 | chr11-46569187-DEL-607 |
| chr11-46581334-DEL-244 | chr11-46581335-DEL-244 |
| chr11-46582581-DEL-52 | chr11-46582582-DEL-52 |
| chr11-46612244-DEL-7283 | chr11-46612245-DEL-7282 |
| chr11-46649796-DEL-401 | chr11-46649797-DEL-401 |
| chr11-46650357-DEL-7078 | chr11-46650358-DEL-7078 |
| chr11-46668746-DEL-68 | chr11-46668747-DEL-68 |
| chr11-46669382-DEL-132 | chr11-46669383-DEL-132 |
| chr11-46693949-DEL-67 | chr11-46693968-DEL-51 |
| chr11-46694584-DEL-212 | chr11-46694585-DEL-212 |
| chr11-46728899-DEL-53 | chr11-46728900-DEL-53 |
| chr11-46746962-DEL-151 | chr11-46746963-DEL-151 |
| chr11-46771080-DEL-190 | chr11-46771081-DEL-190 |
| chr11-46773699-DEL-126 | chr11-46773700-DEL-126 |
| chr11-46784965-DEL-223 | chr11-46784966-DEL-223 |
| chr11-46790740-DEL-104 | chr11-46790751-DEL-94 |
| chr11-46795691-DEL-6388 | chr11-46795692-DEL-6388 |
| chr11-46845675-DEL-138 | chr11-46845676-DEL-138 |
| chr11-46854157-DEL-1264 | chr11-46854158-DEL-1264 |
| chr11-46891506-DEL-53 | chr11-46891507-DEL-53 |
| chr11-46901255-DEL-1104 | chr11-46901256-DEL-1104 |
| chr11-46906500-DEL-56 | chr11-46906513-DEL-58 |
| chr11-47043401-DEL-85 | chr11-47043402-DEL-85 |
| chr11-47067433-DEL-79 | chr11-47067423-DEL-80 |
| chr11-47100159-DEL-128 | chr11-47100160-DEL-128 |
| chr11-47118289-DEL-126 | chr11-47118284-DEL-127 |
| chr11-47120808-DEL-58 | chr11-47120809-DEL-58 |
| chr11-47206737-DEL-883 | chr11-47206738-DEL-883 |
| chr11-47216341-DEL-54 | chr11-47216342-DEL-54 |
| chr11-47232075-DEL-197 | chr11-47232076-DEL-197 |
| chr11-47251249-DEL-177 | chr11-47251250-DEL-177 |
| chr11-47352770-DEL-6474 | chr11-47352771-DEL-6474 |
| chr11-47423952-DEL-52 | chr11-47423953-DEL-52 |
| chr11-47504780-DEL-135 | chr11-47504781-DEL-135 |
| chr11-47518100-DEL-772 | chr11-47518101-DEL-772 |
| chr11-47531732-DEL-147 | chr11-47531733-DEL-147 |
| chr11-47542044-DEL-145 | chr11-47542045-DEL-145 |
| chr11-47553661-DEL-53 | chr11-47553659-DEL-54 |
| chr11-47616664-DEL-203 | chr11-47616665-DEL-203 |
| chr11-47652642-DEL-870 | chr11-47652643-DEL-870 |
| chr11-47684978-DEL-6376 | chr11-47684979-DEL-6376 |
| chr11-47698035-DEL-60 | chr11-47698036-DEL-60 |
| chr11-47738382-DEL-13368 | chr11-47738383-DEL-13368 |
| chr11-47753966-DEL-482 | chr11-47753967-DEL-482 |
| chr11-47777764-DEL-52 | chr11-47777764-DEL-52 |
| chr11-47778717-DEL-129 | chr11-47778715-DEL-129 |
| chr11-47804629-DEL-1613 | chr11-47804630-DEL-1613 |
| chr11-47811898-DEL-201 | chr11-47811899-DEL-201 |
| chr11-47842322-DEL-99 | chr11-47842323-DEL-99 |
| chr11-47845122-DEL-121 | chr11-47845128-DEL-113 |
| chr11-47847964-DEL-56 | chr11-47847957-DEL-68 |
| chr11-47852873-DEL-698 | chr11-47852874-DEL-698 |
| chr11-47872807-DEL-55 | chr11-47872808-DEL-55 |
| chr11-47911974-DEL-69 | chr11-47911993-DEL-50 |
| chr11-47914170-DEL-162 | chr11-47914171-DEL-162 |
| chr11-47954571-DEL-66 | chr11-47954572-DEL-66 |
| chr11-47979645-DEL-399 | chr11-47979646-DEL-399 |
| chr11-47999772-DEL-703 | chr11-47999773-DEL-703 |
| chr11-48011944-DEL-56 | chr11-48011944-DEL-56 |
| chr11-48049576-DEL-136 | chr11-48049577-DEL-130 |
| chr11-48068660-DEL-284 | chr11-48068661-DEL-284 |
| chr11-48072669-DEL-174 | chr11-48072669-DEL-175 |
| chr11-48082270-DEL-66 | chr11-48082272-DEL-58 |
| chr11-48122635-DEL-2983 | chr11-48122635-DEL-2983 |
| chr11-48144868-DEL-58 | chr11-48144881-DEL-50 |
| chr11-48180301-DEL-843 | chr11-48180302-DEL-843 |
| chr11-48183196-DEL-275 | chr11-48183197-DEL-275 |
| chr11-48188861-DEL-490 | chr11-48188863-DEL-488 |
| chr11-48199021-DEL-78 | chr11-48199034-DEL-66 |
| chr11-48207551-DEL-4106 | chr11-48207552-DEL-4106 |
| chr11-48254564-DEL-56 | chr11-48254561-DEL-60 |
| chr11-48258614-DEL-6391 | chr11-48258615-DEL-6391 |
| chr11-48278933-DEL-504 | chr11-48278934-DEL-504 |
| chr11-48281287-DEL-51 | chr11-48281283-DEL-67 |

| | |
|---|---|
| chr11-48308822-DEL-72 | chr11-48308833-DEL-58 |
| chr11-48312448-DEL-404 | chr11-48312449-DEL-404 |
| chr11-48336692-DEL-400 | chr11-48336693-DEL-400 |
| chr11-48355333-DEL-220 | chr11-48355341-DEL-183 |
| chr11-48356195-DEL-397 | chr11-48356196-DEL-398 |
| chr11-48357884-DEL-9653 | chr11-48357885-DEL-9653 |
| chr11-48380345-DEL-1691 | chr11-48380346-DEL-1691 |
| chr11-48390065-DEL-97 | chr11-48390064-DEL-98 |
| chr11-48406097-DEL-297 | chr11-48406098-DEL-297 |
| chr11-48414111-DEL-144 | chr11-48414112-DEL-144 |
| chr11-48419555-DEL-7423 | chr11-48419556-DEL-7423 |
| chr11-48443985-DEL-141 | chr11-48443986-DEL-141 |
| chr11-48474617-DEL-402 | chr11-48474618-DEL-402 |
| chr11-48480660-DEL-80 | chr11-48480652-DEL-79 |
| chr11-48491820-DEL-216 | chr11-48491821-DEL-216 |
| chr11-48510674-DEL-247 | chr11-48510675-DEL-247 |
| chr11-48515123-DEL-2919 | chr11-48515124-DEL-2919 |
| chr11-48597474-DEL-70 | chr11-48597475-DEL-70 |
| chr11-48598606-DEL-73 | chr11-48598599-DEL-69 |
| chr11-48607064-DEL-6393 | chr11-48607065-DEL-6393 |
| chr11-48621581-DEL-1586 | chr11-48621582-DEL-1586 |
| chr11-48641589-DEL-846 | chr11-48641590-DEL-846 |
| chr11-48681064-DEL-110 | chr11-48681065-DEL-110 |
| chr11-48685238-DEL-486 | chr11-48685239-DEL-486 |
| chr11-48716159-DEL-1411 | chr11-48716160-DEL-1411 |
| chr11-48777075-DEL-62 | chr11-48777076-DEL-61 |
| chr11-48792235-DEL-66 | chr11-48792235-DEL-75 |
| chr11-48794226-DEL-425 | chr11-48794227-DEL-425 |
| chr11-48810891-DEL-94 | chr11-48810892-DEL-94 |
| chr11-48983519-DEL-1568 | chr11-48983520-DEL-1568 |
| chr11-48988333-DEL-92 | chr11-48988334-DEL-92 |
| chr11-49007923-DEL-80 | chr11-49007936-DEL-84 |
| chr11-49013402-DEL-108 | chr11-49013403-DEL-108 |
| chr11-49019099-DEL-94 | chr11-49019099-DEL-74 |
| chr11-49033935-DEL-445 | chr11-49033936-DEL-444 |
| chr11-49049619-DEL-88 | chr11-49049618-DEL-90 |
| chr11-49085291-DEL-481 | chr11-49085292-DEL-481 |
| chr11-49119543-DEL-175 | chr11-49119536-DEL-176 |
| chr11-49134157-DEL-75 | chr11-49134158-DEL-75 |
| chr11-49159678-DEL-731 | chr11-49159677-DEL-731 |
| chr11-49178534-DEL-232 | chr11-49178535-DEL-232 |
| chr11-49187425-DEL-77 | chr11-49187412-DEL-81 |
| chr11-49188961-DEL-1280 | chr11-49188959-DEL-1278 |
| chr11-49233899-DEL-6465 | chr11-49233900-DEL-6465 |
| chr11-49255627-DEL-3923 | chr11-49255628-DEL-3923 |
| chr11-49272535-DEL-576 | chr11-49272536-DEL-576 |
| chr11-49273878-DEL-80 | chr11-49273878-DEL-80 |
| chr11-49293260-DEL-1617 | chr11-49293261-DEL-1617 |
| chr11-49319802-DEL-51 | chr11-49319803-DEL-51 |
| chr11-49336003-DEL-7482 | chr11-49336004-DEL-7482 |
| chr11-49353507-DEL-70 | chr11-49353516-DEL-70 |
| chr11-49479083-DEL-72 | chr11-49479084-DEL-72 |
| chr11-49489559-DEL-683 | chr11-49489560-DEL-683 |
| chr11-49529528-DEL-296 | chr11-49529529-DEL-296 |
| chr11-49569462-DEL-504 | chr11-49569465-DEL-501 |
| chr11-49576373-DEL-213 | chr11-49576374-DEL-213 |
| chr11-49610116-DEL-195 | chr11-49610117-DEL-195 |
| chr11-49639698-DEL-206 | chr11-49639699-DEL-206 |
| chr11-49658579-DEL-209 | chr11-49658580-DEL-209 |
| chr11-49668807-DEL-101 | chr11-49668808-DEL-101 |
| chr11-49672559-DEL-144 | chr11-49672560-DEL-144 |
| chr11-49675262-DEL-54 | chr11-49675263-DEL-54 |
| chr11-49698028-DEL-98 | chr11-49698038-DEL-89 |
| chr11-49710136-DEL-62 | chr11-49710137-DEL-62 |
| chr11-49756554-DEL-944 | chr11-49756555-DEL-944 |
| chr11-49771434-DEL-402 | chr11-49771435-DEL-402 |
| chr11-49777346-DEL-56 | chr11-49777347-DEL-56 |
| chr11-49777504-DEL-854 | chr11-49777500-DEL-855 |
| chr11-49789105-DEL-128 | chr11-49789104-DEL-124 |
| chr11-49791222-DEL-222 | chr11-49791224-DEL-214 |
| chr11-49825005-DEL-86 | chr11-49825008-DEL-66 |
| chr11-49852604-DEL-62 | chr11-49852605-DEL-62 |
| chr11-49858537-DEL-506 | chr11-49858538-DEL-506 |
| chr11-49870017-DEL-80 | chr11-49870018-DEL-80 |
| chr11-49885810-DEL-152 | chr11-49885812-DEL-126 |
| chr11-49889272-DEL-86 | chr11-49889273-DEL-86 |
| chr11-49898469-DEL-166 | chr11-49898470-DEL-166 |
| chr11-49948717-DEL-146 | chr11-49948718-DEL-146 |
| chr11-50018747-DEL-78 | chr11-50018761-DEL-56 |
| chr11-50019331-DEL-60 | chr11-50019332-DEL-60 |

| | |
|---|---|
| chr11-50037622-DEL-203 | chr11-50037623-DEL-202 |
| chr11-50057215-DEL-72 | chr11-50057216-DEL-72 |
| chr11-50083962-DEL-95 | chr11-50083947-DEL-94 |
| chr11-50089530-DEL-184 | chr11-50089531-DEL-184 |
| chr11-50130788-DEL-210 | chr11-50130789-DEL-210 |
| chr11-50133652-DEL-76 | chr11-50133653-DEL-62 |
| chr11-50137745-DEL-79 | chr11-50137746-DEL-79 |
| chr11-50139219-DEL-87 | chr11-50139248-DEL-95 |
| chr11-50162713-DEL-65 | chr11-50162703-DEL-65 |
| chr11-50164764-DEL-52 | chr11-50164765-DEL-52 |
| chr11-50172795-DEL-224 | chr11-50172796-DEL-224 |
| chr11-50194119-DEL-173 | chr11-50194120-DEL-173 |
| chr11-50234641-DEL-79 | chr11-50234640-DEL-79 |
| chr11-50240081-DEL-1894 | chr11-50240096-DEL-1860 |
| chr11-50243947-DEL-198 | chr11-50243948-DEL-198 |
| chr11-50251663-DEL-120 | chr11-50251664-DEL-120 |
| chr11-50263382-DEL-86 | chr11-50263382-DEL-87 |
| chr11-50281572-DEL-194 | chr11-50281573-DEL-194 |
| chr11-50286420-DEL-2022 | chr11-50286421-DEL-2022 |
| chr11-50286931-DEL-212 | chr11-50286932-DEL-212 |
| chr11-50302507-DEL-1916 | chr11-50302513-DEL-1910 |
| chr11-50336005-DEL-3164 | chr11-50336005-DEL-3164 |
| chr11-50354467-DEL-5639 | chr11-50354467-DEL-5636 |
| chr11-50374291-DEL-190 | chr11-50374292-DEL-152 |
| chr11-50376233-DEL-64 | chr11-50376234-DEL-62 |
| chr11-50399472-DEL-57 | chr11-50399485-DEL-54 |
| chr11-50402763-DEL-464 | chr11-50402764-DEL-464 |
| chr11-50404770-DEL-58 | chr11-50404785-DEL-61 |
| chr11-50412638-DEL-110 | chr11-50412639-DEL-110 |
| chr11-50413421-DEL-78 | chr11-50413422-DEL-78 |
| chr11-50429683-DEL-982 | chr11-50429684-DEL-982 |
| chr11-50445887-DEL-67 | chr11-50445867-DEL-69 |
| chr11-50457584-DEL-92 | chr11-50457575-DEL-102 |
| chr11-50488501-DEL-99 | chr11-50488488-DEL-67 |
| chr11-50517287-DEL-96 | chr11-50517288-DEL-96 |
| chr11-50527485-DEL-70 | chr11-50527485-DEL-70 |
| chr11-50536309-DEL-82 | chr11-50536310-DEL-82 |
| chr11-50575821-DEL-82 | chr11-50575822-DEL-82 |
| chr11-50631439-DEL-82 | chr11-50631440-DEL-82 |
| chr11-50650983-DEL-69 | chr11-50650984-DEL-69 |
| chr11-50675950-DEL-98 | chr11-50675951-DEL-98 |
| chr11-50680911-DEL-78 | chr11-50680902-DEL-81 |
| chr11-50691194-DEL-75 | chr11-50691199-DEL-71 |
| chr11-50698579-DEL-53 | chr11-50698600-DEL-53 |
| chr11-50712636-DEL-316 | chr11-50712637-DEL-316 |
| chr11-50718893-DEL-933 | chr11-50718894-DEL-933 |
| chr11-50746220-DEL-126 | chr11-50746207-DEL-140 |
| chr11-50759978-DEL-59 | chr11-50759973-DEL-98 |
| chr11-50809176-DEL-56 | chr11-50809177-DEL-56 |
| chr11-50824291-DEL-89 | chr11-50824292-DEL-89 |
| chr11-50825139-DEL-1071 | chr11-50825139-DEL-1071 |
| chr11-50829961-DEL-62 | chr11-50829962-DEL-62 |
| chr11-50870558-DEL-1419 | chr11-50870559-DEL-1419 |
| chr11-50871952-DEL-84 | chr11-50871996-DEL-60 |
| chr11-50873684-DEL-99 | chr11-50873685-DEL-99 |
| chr11-50879833-DEL-1095 | chr11-50879835-DEL-1093 |
| chr11-50885025-DEL-72 | chr11-50885026-DEL-72 |
| chr11-50915002-DEL-208 | chr11-50915003-DEL-208 |
| chr11-50922331-DEL-343 | chr11-50922331-DEL-343 |
| chr11-50952556-DEL-58 | chr11-50952557-DEL-58 |
| chr11-50975315-DEL-2703 | chr11-50975316-DEL-2703 |
| chr11-50990559-DEL-52 | chr11-50990541-DEL-72 |
| chr11-51025324-DEL-201 | chr11-51025324-DEL-202 |
| chr11-51028703-DEL-88 | chr11-51028708-DEL-84 |
| chr11-51032970-DEL-63 | chr11-51032977-DEL-55 |
| chr11-51044077-DEL-74 | chr11-51044078-DEL-74 |
| chr11-51069558-DEL-66 | chr11-51069559-DEL-66 |
| chr11-51078702-DEL-79 | chr11-51078667-DEL-112 |
| chr11-51137858-DEL-86 | chr11-51137859-DEL-86 |
| chr11-51139015-DEL-812 | chr11-51139016-DEL-813 |
| chr11-51168439-DEL-213 | chr11-51168440-DEL-213 |
| chr11-51170308-DEL-400 | chr11-51170309-DEL-400 |
| chr11-51215967-DEL-74 | chr11-51215936-DEL-78 |
| chr11-51218691-DEL-114 | chr11-51218691-DEL-114 |
| chr11-51253291-DEL-56 | chr11-51253292-DEL-56 |
| chr11-51264336-DEL-303 | chr11-51264335-DEL-304 |
| chr11-51328536-DEL-4345 | chr11-51328537-DEL-4345 |
| chr11-51336277-DEL-555 | chr11-51336278-DEL-555 |
| chr11-51346440-DEL-146 | chr11-51346441-DEL-146 |
| chr11-51346999-DEL-53 | chr11-51346974-DEL-68 |

| | |
|---|---|
| chr11-51380479-DEL-96 | chr11-51380480-DEL-96 |
| chr11-51410499-DEL-451 | chr11-51410498-DEL-451 |
| chr11-51501334-DEL-147 | chr11-51501335-DEL-147 |
| chr11-51508299-DEL-193 | chr11-51508300-DEL-193 |
| chr11-51543908-DEL-54 | chr11-51543891-DEL-55 |
| chr11-51553205-DEL-221 | chr11-51553206-DEL-188 |
| chr11-51555866-DEL-5179 | chr11-51555867-DEL-5179 |
| chr11-51573163-DEL-52 | chr11-51573158-DEL-54 |
| chr11-51574242-DEL-209 | chr11-51574243-DEL-209 |
| chr11-51581358-DEL-207 | chr11-51581359-DEL-207 |
| chr11-51586226-DEL-230 | chr11-51586227-DEL-230 |
| chr11-51606172-DEL-205 | chr11-51606173-DEL-205 |
| chr11-51609325-DEL-78 | chr11-51609326-DEL-78 |
| chr11-51615986-DEL-82 | chr11-51615985-DEL-84 |
| chr11-51618897-DEL-88 | chr11-51618898-DEL-88 |
| chr11-51625140-DEL-129 | chr11-51625141-DEL-129 |
| chr11-51628750-DEL-208 | chr11-51628751-DEL-208 |
| chr11-51631197-DEL-52 | chr11-51631198-DEL-52 |
| chr11-51642447-DEL-163 | chr11-51642448-DEL-163 |
| chr11-51642991-DEL-52 | chr11-51642990-DEL-54 |
| chr11-51648849-DEL-200 | chr11-51648850-DEL-200 |
| chr11-51653170-DEL-72 | chr11-51653168-DEL-72 |
| chr11-51680833-DEL-607 | chr11-51680834-DEL-607 |
| chr11-51688873-DEL-2399 | chr11-51688874-DEL-2399 |
| chr11-51691080-DEL-83 | chr11-51691129-DEL-60 |
| chr11-51697711-DEL-205 | chr11-51697737-DEL-180 |
| chr11-51699516-DEL-235 | chr11-51699514-DEL-242 |
| chr11-51731367-DEL-90 | chr11-51731397-DEL-78 |
| chr11-51760010-DEL-2559 | chr11-51760011-DEL-2559 |
| chr11-51778132-DEL-186 | chr11-51778133-DEL-185 |
| chr11-51782080-DEL-107 | chr11-51782100-DEL-77 |
| chr11-51859376-DEL-198 | chr11-51859377-DEL-198 |
| chr11-51882491-DEL-235 | chr11-51882492-DEL-235 |
| chr11-51906611-DEL-133 | chr11-51906575-DEL-134 |
| chr11-51911044-DEL-392 | chr11-51911045-DEL-392 |
| chr11-51911777-DEL-60 | chr11-51911760-DEL-51 |
| chr11-51921366-DEL-150 | chr11-51921367-DEL-150 |
| chr11-51929931-DEL-89 | chr11-51929932-DEL-89 |
| chr11-51932440-DEL-253 | chr11-51932441-DEL-253 |
| chr11-51945770-DEL-119 | chr11-51945771-DEL-119 |
| chr11-51957252-DEL-811 | chr11-51957253-DEL-811 |
| chr11-51963482-DEL-156 | chr11-51963483-DEL-156 |
| chr11-51981832-DEL-232 | chr11-51981833-DEL-232 |
| chr11-51994857-DEL-116 | chr11-51994858-DEL-116 |
| chr11-52000941-DEL-61 | chr11-52000941-DEL-57 |
| chr11-52042395-DEL-1502 | chr11-52042396-DEL-1502 |
| chr11-52076701-DEL-86 | chr11-52076715-DEL-58 |
| chr11-52105905-DEL-326 | chr11-52105906-DEL-326 |
| chr11-52110242-DEL-139 | chr11-52110216-DEL-141 |
| chr11-52161977-DEL-206 | chr11-52161978-DEL-206 |
| chr11-52181216-DEL-202 | chr11-52181217-DEL-202 |
| chr11-52186197-DEL-173 | chr11-52186197-DEL-173 |
| chr11-52192313-DEL-76 | chr11-52192309-DEL-66 |
| chr11-52199040-DEL-86 | chr11-52199047-DEL-80 |
| chr11-52227797-DEL-66 | chr11-52227798-DEL-66 |
| chr11-52237992-DEL-60 | chr11-52237977-DEL-63 |
| chr11-52247371-DEL-203 | chr11-52247372-DEL-203 |
| chr11-52248859-DEL-52 | chr11-52248856-DEL-56 |
| chr11-52274118-DEL-201 | chr11-52274119-DEL-201 |
| chr11-52281713-DEL-197 | chr11-52281732-DEL-193 |
| chr11-52300309-DEL-1802 | chr11-52300310-DEL-1802 |
| chr11-52314555-DEL-73 | chr11-52314556-DEL-71 |
| chr11-52315798-DEL-94 | chr11-52315825-DEL-86 |
| chr11-52340480-DEL-92 | chr11-52340481-DEL-92 |
| chr11-52342669-DEL-56 | chr11-52342670-DEL-56 |
| chr11-52364897-DEL-147 | chr11-52364898-DEL-147 |
| chr11-52374148-DEL-826 | chr11-52374149-DEL-826 |
| chr11-52388223-DEL-234 | chr11-52388222-DEL-232 |
| chr11-52392598-DEL-201 | chr11-52392555-DEL-201 |
| chr11-52434686-DEL-218 | chr11-52434687-DEL-218 |
| chr11-52462814-DEL-108 | chr11-52462815-DEL-108 |
| chr11-52496915-DEL-497 | chr11-52496916-DEL-497 |
| chr11-52505516-DEL-101 | chr11-52505555-DEL-76 |
| chr11-52525097-DEL-53 | chr11-52525096-DEL-52 |
| chr11-52527262-DEL-132 | chr11-52527264-DEL-130 |
| chr11-52534531-DEL-348 | chr11-52534532-DEL-347 |
| chr11-52596265-DEL-79 | chr11-52596266-DEL-79 |
| chr11-52621581-DEL-78 | chr11-52621582-DEL-78 |
| chr11-52667400-DEL-127 | chr11-52667401-DEL-80 |
| chr11-52702809-DEL-86 | chr11-52702842-DEL-82 |

| | |
|---|---|
| chr11-52747802-DEL-440 | chr11-52747803-DEL-440 |
| chr11-52772559-DEL-106 | chr11-52772560-DEL-106 |
| chr11-52795920-DEL-105 | chr11-52795921-DEL-105 |
| chr11-52798112-DEL-130 | chr11-52798113-DEL-130 |
| chr11-52813520-DEL-170 | chr11-52813521-DEL-170 |
| chr11-52864980-DEL-51 | chr11-52864981-DEL-51 |
| chr11-52872007-DEL-80 | chr11-52872008-DEL-80 |
| chr11-52879441-DEL-174 | chr11-52879441-DEL-170 |
| chr11-52935774-DEL-198 | chr11-52935775-DEL-198 |
| chr11-52938103-DEL-88 | chr11-52938138-DEL-54 |
| chr11-53003877-DEL-54 | chr11-53003878-DEL-54 |
| chr11-53011684-DEL-197 | chr11-53011687-DEL-197 |
| chr11-53014045-DEL-104 | chr11-53014046-DEL-104 |
| chr11-53048326-DEL-82 | chr11-53048327-DEL-82 |
| chr11-53066000-DEL-1008 | chr11-53066001-DEL-1008 |
| chr11-53121829-DEL-105 | chr11-53121834-DEL-101 |
| chr11-53128052-DEL-486 | chr11-53128053-DEL-486 |
| chr11-53133096-DEL-104 | chr11-53133097-DEL-104 |
| chr11-53194504-DEL-199 | chr11-53194505-DEL-199 |
| chr11-53201235-DEL-195 | chr11-53201235-DEL-196 |
| chr11-53203941-DEL-206 | chr11-53203966-DEL-182 |
| chr11-53209155-DEL-188 | chr11-53209156-DEL-188 |
| chr11-53220965-DEL-182 | chr11-53220966-DEL-182 |
| chr11-53238552-DEL-206 | chr11-53238553-DEL-206 |
| chr11-53244292-DEL-227 | chr11-53244293-DEL-227 |
| chr11-53247621-DEL-168 | chr11-53247622-DEL-168 |
| chr11-53269370-DEL-218 | chr11-53269371-DEL-218 |
| chr11-53278385-DEL-180 | chr11-53278386-DEL-180 |
| chr11-53280632-DEL-215 | chr11-53280633-DEL-215 |
| chr11-53286836-DEL-83 | chr11-53286810-DEL-76 |
| chr11-53317473-DEL-176 | chr11-53317474-DEL-176 |
| chr11-53325431-DEL-195 | chr11-53325428-DEL-202 |
| chr11-53341506-DEL-127 | chr11-53341507-DEL-127 |
| chr11-53362048-DEL-84 | chr11-53362049-DEL-84 |
| chr11-53375563-DEL-199 | chr11-53375564-DEL-199 |
| chr11-53378342-DEL-65 | chr11-53378342-DEL-65 |
| chr11-53381975-DEL-86 | chr11-53381976-DEL-86 |
| chr11-53388878-DEL-131 | chr11-53388879-DEL-131 |
| chr11-53429624-DEL-79 | chr11-53429624-DEL-53 |
| chr11-53444575-DEL-218 | chr11-53444575-DEL-218 |
| chr11-53471936-DEL-393 | chr11-53471937-DEL-393 |
| chr11-53486222-DEL-208 | chr11-53486223-DEL-208 |
| chr11-53486948-DEL-146 | chr11-53486971-DEL-142 |
| chr11-53492676-DEL-145 | chr11-53492680-DEL-144 |
| chr11-53501343-DEL-76 | chr11-53501345-DEL-68 |
| chr11-53502311-DEL-440 | chr11-53502310-DEL-450 |
| chr11-53507211-DEL-82 | chr11-53507171-DEL-123 |
| chr11-53516831-DEL-138 | chr11-53516836-DEL-134 |
| chr11-53520231-DEL-56 | chr11-53520263-DEL-50 |
| chr11-53561810-DEL-52 | chr11-53561814-DEL-60 |
| chr11-53569028-DEL-246 | chr11-53569029-DEL-246 |
| chr11-53578594-DEL-168 | chr11-53578595-DEL-168 |
| chr11-53579626-DEL-207 | chr11-53579627-DEL-207 |
| chr11-53580031-DEL-2370 | chr11-53580032-DEL-2370 |
| chr11-53600048-DEL-511 | chr11-53600049-DEL-511 |
| chr11-53613665-DEL-81 | chr11-53613666-DEL-83 |
| chr11-53627176-DEL-187 | chr11-53627177-DEL-187 |
| chr11-53646334-DEL-209 | chr11-53646335-DEL-209 |
| chr11-53650684-DEL-78 | chr11-53650685-DEL-78 |
| chr11-53656084-DEL-99 | chr11-53656085-DEL-99 |
| chr11-53657158-DEL-207 | chr11-53657159-DEL-207 |
| chr11-53689485-DEL-179 | chr11-53689467-DEL-164 |
| chr11-53722680-DEL-2068 | chr11-53722683-DEL-2067 |
| chr11-53728194-DEL-164 | chr11-53728205-DEL-148 |
| chr11-53735934-DEL-180 | chr11-53735935-DEL-180 |
| chr11-53738497-DEL-202 | chr11-53738498-DEL-202 |
| chr11-53752047-DEL-213 | chr11-53752048-DEL-213 |
| chr11-53768757-DEL-195 | chr11-53768758-DEL-195 |
| chr11-53790851-DEL-441 | chr11-53790852-DEL-441 |
| chr11-53793396-DEL-60 | chr11-53793407-DEL-50 |
| chr11-53799989-DEL-1521 | chr11-53799990-DEL-1521 |
| chr11-53878949-DEL-74 | chr11-53878950-DEL-74 |
| chr11-53896036-DEL-82 | chr11-53896037-DEL-82 |
| chr11-53897211-DEL-152 | chr11-53897212-DEL-152 |
| chr11-53901411-DEL-70 | chr11-53901412-DEL-70 |
| chr11-53908282-DEL-141 | chr11-53908270-DEL-166 |
| chr11-53932200-DEL-72 | chr11-53932201-DEL-60 |
| chr11-53970257-DEL-54 | chr11-53970267-DEL-50 |
| chr11-53974905-DEL-57 | chr11-53974906-DEL-56 |
| chr11-53989400-DEL-498 | chr11-53989401-DEL-498 |

| | |
|---|---|
| chr11-53995452-DEL-497 | chr11-53995453-DEL-497 |
| chr11-54026248-DEL-3808 | chr11-54026249-DEL-3808 |
| chr11-54030191-DEL-52 | chr11-54030192-DEL-52 |
| chr11-54037010-DEL-107 | chr11-54037011-DEL-107 |
| chr11-54043806-DEL-69 | chr11-54043807-DEL-69 |
| chr11-54054307-DEL-65 | chr11-54054308-DEL-65 |
| chr11-54059331-DEL-130 | chr11-54059358-DEL-104 |
| chr11-54073510-DEL-54 | chr11-54073511-DEL-54 |
| chr11-54165015-DEL-75 | chr11-54165019-DEL-72 |
| chr11-54177089-DEL-80 | chr11-54177088-DEL-81 |
| chr11-54180692-DEL-52 | chr11-54180693-DEL-52 |
| chr11-54184361-DEL-116 | chr11-54184366-DEL-112 |
| chr11-54217199-DEL-260 | chr11-54217200-DEL-260 |
| chr11-54240406-DEL-109 | chr11-54240407-DEL-109 |
| chr11-54272232-DEL-7140 | chr11-54272233-DEL-7140 |
| chr11-54278645-DEL-636 | chr11-54278646-DEL-636 |
| chr11-54291761-DEL-1624 | chr11-54291762-DEL-1624 |
| chr11-54295604-DEL-104 | chr11-54295605-DEL-104 |
| chr11-54320276-DEL-172 | chr11-54320277-DEL-172 |
| chr11-54321433-DEL-675 | chr11-54321434-DEL-676 |
| chr11-54337937-DEL-126 | chr11-54337938-DEL-126 |
| chr11-54339494-DEL-622 | chr11-54339495-DEL-622 |
| chr11-54341741-DEL-120 | chr11-54341742-DEL-80 |
| chr11-54372777-DEL-205 | chr11-54372778-DEL-205 |
| chr11-54444732-DEL-178 | chr11-54444733-DEL-178 |
| chr11-54464328-DEL-336 | chr11-54464329-DEL-336 |
| chr11-54467399-DEL-7125 | chr11-54467400-DEL-7125 |
| chr11-54564338-DEL-208 | chr11-54564339-DEL-208 |
| chr11-54626683-DEL-115 | chr11-54626682-DEL-114 |
| chr11-54689573-DEL-256 | chr11-54689574-DEL-256 |
| chr11-54697847-DEL-58 | chr11-54697848-DEL-58 |
| chr11-54715169-DEL-172 | chr11-54715170-DEL-172 |
| chr11-54721228-DEL-218 | chr11-54721247-DEL-242 |
| chr11-54734763-DEL-217 | chr11-54734762-DEL-213 |
| chr11-54738897-DEL-444 | chr11-54738898-DEL-444 |
| chr11-54744830-DEL-2907 | chr11-54744831-DEL-2907 |
| chr11-54755214-DEL-63 | chr11-54755215-DEL-63 |
| chr11-54772523-DEL-103 | chr11-54772524-DEL-103 |
| chr11-54776877-DEL-85 | chr11-54776878-DEL-85 |
| chr11-54784275-DEL-163 | chr11-54784226-DEL-195 |
| chr11-54840898-DEL-58 | chr11-54840889-DEL-56 |
| chr11-54892916-DEL-262 | chr11-54892917-DEL-262 |
| chr11-54921342-DEL-60 | chr11-54921342-DEL-54 |
| chr11-54935930-DEL-54 | chr11-54935933-DEL-50 |
| chr11-54939654-DEL-199 | chr11-54939655-DEL-199 |
| chr11-54958970-DEL-220 | chr11-54958978-DEL-212 |
| chr11-55007301-DEL-208 | chr11-55007301-DEL-208 |
| chr11-55013643-DEL-54 | chr11-55013644-DEL-54 |
| chr11-55083454-DEL-481 | chr11-55083455-DEL-481 |
| chr11-55118302-DEL-97 | chr11-55118298-DEL-109 |
| chr11-55121810-DEL-56 | chr11-55121815-DEL-52 |
| chr11-55131745-DEL-7110 | chr11-55131746-DEL-7110 |
| chr11-55156218-DEL-66 | chr11-55156219-DEL-66 |
| chr11-55185919-DEL-70 | chr11-55185906-DEL-71 |
| chr11-55188114-DEL-355 | chr11-55188069-DEL-392 |
| chr11-55212737-DEL-51 | chr11-55212746-DEL-51 |
| chr11-55248744-DEL-60 | chr11-55248745-DEL-58 |
| chr11-55263859-DEL-72 | chr11-55263870-DEL-75 |
| chr11-55270070-DEL-73 | chr11-55270071-DEL-73 |
| chr11-55282629-DEL-70 | chr11-55282630-DEL-70 |
| chr11-55287131-DEL-329 | chr11-55287122-DEL-329 |
| chr11-55291521-DEL-60 | chr11-55291538-DEL-54 |
| chr11-55311623-DEL-86 | chr11-55311625-DEL-85 |
| chr11-55318472-DEL-64 | chr11-55318472-DEL-68 |
| chr11-55322482-DEL-115 | chr11-55322483-DEL-115 |
| chr11-55331509-DEL-164 | chr11-55331510-DEL-162 |
| chr11-55339095-DEL-393 | chr11-55339091-DEL-394 |
| chr11-55356649-DEL-433 | chr11-55356662-DEL-421 |
| chr11-55364773-DEL-153 | chr11-55364774-DEL-153 |
| chr11-55382640-DEL-78 | chr11-55382637-DEL-70 |
| chr11-55402171-DEL-58 | chr11-55402170-DEL-86 |
| chr11-55408562-DEL-52 | chr11-55408562-DEL-52 |
| chr11-55419597-DEL-53 | chr11-55419598-DEL-53 |
| chr11-55515361-DEL-120 | chr11-55515370-DEL-112 |
| chr11-55519945-DEL-182 | chr11-55519946-DEL-182 |
| chr11-55541246-DEL-501 | chr11-55541249-DEL-499 |
| chr11-55554816-DEL-6774 | chr11-55554817-DEL-6774 |
| chr11-55561337-DEL-208 | chr11-55561338-DEL-208 |
| chr11-55578317-DEL-324 | chr11-55578317-DEL-324 |
| chr11-55579410-DEL-344 | chr11-55579411-DEL-344 |

| | |
|---|---|
| chr11-55579961-DEL-103 | chr11-55579951-DEL-101 |
| chr11-55585153-DEL-2267 | chr11-55585154-DEL-2267 |
| chr11-55639759-DEL-761 | chr11-55639762-DEL-757 |
| chr11-55644750-DEL-461 | chr11-55644751-DEL-461 |
| chr11-55713697-DEL-969 | chr11-55713698-DEL-969 |
| chr11-55719396-DEL-75 | chr11-55719397-DEL-75 |
| chr11-55722137-DEL-6213 | chr11-55722137-DEL-6213 |
| chr11-55738283-DEL-803 | chr11-55738284-DEL-803 |
| chr11-55745283-DEL-55 | chr11-55745283-DEL-55 |
| chr11-55749361-DEL-69 | chr11-55749362-DEL-69 |
| chr11-55790607-DEL-368 | chr11-55790604-DEL-364 |
| chr11-55811576-DEL-137 | chr11-55811580-DEL-149 |
| chr11-55814444-DEL-88 | chr11-55814447-DEL-89 |
| chr11-55841204-DEL-57 | chr11-55841204-DEL-58 |
| chr11-55842370-DEL-91 | chr11-55842371-DEL-91 |
| chr11-55843442-DEL-279 | chr11-55843442-DEL-274 |
| chr11-55850344-DEL-52 | chr11-55850345-DEL-52 |
| chr11-55871745-DEL-1718 | chr11-55871744-DEL-1718 |
| chr11-55886296-DEL-292 | chr11-55886297-DEL-292 |
| chr11-55901015-DEL-51 | chr11-55901015-DEL-51 |
| chr11-55909848-DEL-93 | chr11-55909849-DEL-93 |
| chr11-55923608-DEL-90 | chr11-55923628-DEL-78 |
| chr11-55954169-DEL-88 | chr11-55954170-DEL-88 |
| chr11-55970627-DEL-52 | chr11-55970628-DEL-52 |
| chr11-56001647-DEL-273 | chr11-56001648-DEL-273 |
| chr11-56004269-DEL-2020 | chr11-56004270-DEL-2020 |
| chr11-56022781-DEL-2469 | chr11-56022782-DEL-2469 |
| chr11-56023560-DEL-54 | chr11-56023561-DEL-54 |
| chr11-56026792-DEL-136 | chr11-56026793-DEL-132 |
| chr11-56035876-DEL-69 | chr11-56035877-DEL-69 |
| chr11-56063994-DEL-78 | chr11-56064001-DEL-72 |
| chr11-56067932-DEL-6402 | chr11-56067933-DEL-6402 |
| chr11-56094625-DEL-7529 | chr11-56094626-DEL-7529 |
| chr11-56117199-DEL-6648 | chr11-56117200-DEL-6648 |
| chr11-56125186-DEL-60 | chr11-56125193-DEL-52 |
| chr11-56125351-DEL-5007 | chr11-56125352-DEL-5007 |
| chr11-56131485-DEL-1476 | chr11-56131486-DEL-1476 |
| chr11-56150100-DEL-57 | chr11-56150098-DEL-60 |
| chr11-56154055-DEL-107 | chr11-56154056-DEL-107 |
| chr11-56160293-DEL-189 | chr11-56160294-DEL-189 |
| chr11-56161962-DEL-459 | chr11-56161963-DEL-459 |
| chr11-56162572-DEL-58 | chr11-56162573-DEL-58 |
| chr11-56202964-DEL-810 | chr11-56202965-DEL-810 |
| chr11-56213918-DEL-342 | chr11-56213919-DEL-342 |
| chr11-56219084-DEL-53 | chr11-56219097-DEL-62 |
| chr11-56261375-DEL-54 | chr11-56261376-DEL-54 |
| chr11-56277695-DEL-51 | chr11-56277696-DEL-51 |
| chr11-56278309-DEL-73 | chr11-56278310-DEL-73 |
| chr11-56285034-DEL-6245 | chr11-56285035-DEL-6245 |
| chr11-56317739-DEL-162 | chr11-56317740-DEL-162 |
| chr11-56350135-DEL-3281 | chr11-56350150-DEL-3281 |
| chr11-56368157-DEL-429 | chr11-56368158-DEL-429 |
| chr11-56369805-DEL-90 | chr11-56369806-DEL-90 |
| chr11-56370574-DEL-497 | chr11-56370575-DEL-497 |
| chr11-56380706-DEL-6369 | chr11-56380707-DEL-6369 |
| chr11-56394345-DEL-73 | chr11-56394349-DEL-61 |
| chr11-56398442-DEL-147 | chr11-56398491-DEL-106 |
| chr11-56429388-DEL-85 | chr11-56429386-DEL-85 |
| chr11-56455427-DEL-646 | chr11-56455428-DEL-646 |
| chr11-56458841-DEL-58 | chr11-56458842-DEL-58 |
| chr11-56463033-DEL-77 | chr11-56463034-DEL-77 |
| chr11-56465328-DEL-685 | chr11-56465329-DEL-685 |
| chr11-56469666-DEL-986 | chr11-56469643-DEL-987 |
| chr11-56475862-DEL-51 | chr11-56475863-DEL-51 |
| chr11-56478334-DEL-6197 | chr11-56478335-DEL-6197 |
| chr11-56509080-DEL-398 | chr11-56509081-DEL-398 |
| chr11-56510461-DEL-66 | chr11-56510452-DEL-50 |
| chr11-56518171-DEL-72 | chr11-56518153-DEL-73 |
| chr11-56523794-DEL-128 | chr11-56523807-DEL-118 |
| chr11-56568337-DEL-118 | chr11-56568338-DEL-118 |
| chr11-56586298-DEL-518 | chr11-56586299-DEL-518 |
| chr11-56594898-DEL-1626 | chr11-56594898-DEL-1626 |
| chr11-56597362-DEL-480 | chr11-56597363-DEL-480 |
| chr11-56598832-DEL-203 | chr11-56598833-DEL-203 |
| chr11-56623143-DEL-6470 | chr11-56623144-DEL-6470 |
| chr11-56623259-DEL-214 | chr11-56623260-DEL-214 |
| chr11-56664283-DEL-56 | chr11-56664284-DEL-56 |
| chr11-56691265-DEL-6922 | chr11-56691266-DEL-6922 |
| chr11-56699027-DEL-53 | chr11-56699029-DEL-50 |
| chr11-56734890-DEL-4456 | chr11-56734891-DEL-4456 |

| | |
|---|---|
| chr11-56737337-DEL-6089 | chr11-56737338-DEL-6090 |
| chr11-56755238-DEL-2312 | chr11-56755239-DEL-2312 |
| chr11-56776389-DEL-70 | chr11-56776390-DEL-70 |
| chr11-56807309-DEL-123 | chr11-56807309-DEL-123 |
| chr11-56835758-DEL-123 | chr11-56835759-DEL-123 |
| chr11-56841883-DEL-79 | chr11-56841884-DEL-79 |
| chr11-56862862-DEL-90 | chr11-56862863-DEL-90 |
| chr11-56886480-DEL-205 | chr11-56886481-DEL-205 |
| chr11-56887712-DEL-54 | chr11-56887713-DEL-54 |
| chr11-56894152-DEL-82 | chr11-56894124-DEL-70 |
| chr11-56914473-DEL-1570 | chr11-56914473-DEL-1571 |
| chr11-56922847-DEL-269 | chr11-56922823-DEL-270 |
| chr11-56969926-DEL-72 | chr11-56969951-DEL-56 |
| chr11-57010418-DEL-56 | chr11-57010419-DEL-56 |
| chr11-57022539-DEL-144 | chr11-57022538-DEL-141 |
| chr11-57038746-DEL-247 | chr11-57038747-DEL-247 |
| chr11-57066894-DEL-88 | chr11-57066899-DEL-72 |
| chr11-57073152-DEL-533 | chr11-57073153-DEL-533 |
| chr11-57111194-DEL-97 | chr11-57111189-DEL-98 |
| chr11-57223511-DEL-52 | chr11-57223512-DEL-52 |
| chr11-57227740-DEL-91 | chr11-57227751-DEL-81 |
| chr11-57236374-DEL-6475 | chr11-57236375-DEL-6475 |
| chr11-57263684-DEL-126 | chr11-57263685-DEL-126 |
| chr11-57277110-DEL-1894 | chr11-57277111-DEL-1894 |
| chr11-57292047-DEL-76 | chr11-57292048-DEL-76 |
| chr11-57313525-DEL-331 | chr11-57313526-DEL-331 |
| chr11-57314307-DEL-1445 | chr11-57314308-DEL-1445 |
| chr11-57343009-DEL-58 | chr11-57343010-DEL-58 |
| chr11-57344797-DEL-91 | chr11-57344798-DEL-90 |
| chr11-57355470-DEL-242 | chr11-57355471-DEL-242 |
| chr11-57363064-DEL-89 | chr11-57363064-DEL-89 |
| chr11-57391572-DEL-167 | chr11-57391573-DEL-167 |
| chr11-57427974-DEL-1529 | chr11-57427975-DEL-1529 |
| chr11-57433655-DEL-549 | chr11-57433656-DEL-549 |
| chr11-57438287-DEL-121 | chr11-57438288-DEL-121 |
| chr11-57445546-DEL-5751 | chr11-57445547-DEL-5751 |
| chr11-57466445-DEL-212 | chr11-57466446-DEL-212 |
| chr11-57469442-DEL-167 | chr11-57469443-DEL-167 |
| chr11-57474821-DEL-3066 | chr11-57474818-DEL-3067 |
| chr11-57484129-DEL-52 | chr11-57484129-DEL-52 |
| chr11-57485908-DEL-58 | chr11-57485861-DEL-106 |
| chr11-57503144-DEL-171 | chr11-57503145-DEL-171 |
| chr11-57505000-DEL-739 | chr11-57505001-DEL-739 |
| chr11-57525771-DEL-56 | chr11-57525772-DEL-56 |
| chr11-57556511-DEL-62 | chr11-57556505-DEL-63 |
| chr11-57585145-DEL-70 | chr11-57585146-DEL-70 |
| chr11-57593625-DEL-56 | chr11-57593628-DEL-54 |
| chr11-57596593-DEL-52 | chr11-57596594-DEL-52 |
| chr11-57608749-DEL-202 | chr11-57608750-DEL-202 |
| chr11-57625463-DEL-54 | chr11-57625439-DEL-56 |
| chr11-57625686-DEL-353 | chr11-57625687-DEL-353 |
| chr11-57631956-DEL-65 | chr11-57631908-DEL-66 |
| chr11-57641363-DEL-59 | chr11-57641364-DEL-59 |
| chr11-57652675-DEL-1655 | chr11-57652665-DEL-1648 |
| chr11-57667081-DEL-91 | chr11-57667082-DEL-91 |
| chr11-57688261-DEL-76 | chr11-57688266-DEL-68 |
| chr11-57716457-DEL-107 | chr11-57716458-DEL-107 |
| chr11-57743923-DEL-55 | chr11-57743917-DEL-52 |
| chr11-57762534-DEL-82 | chr11-57762535-DEL-82 |
| chr11-57771363-DEL-54 | chr11-57771364-DEL-52 |
| chr11-57793414-DEL-78 | chr11-57793440-DEL-73 |
| chr11-57814902-DEL-58 | chr11-57814903-DEL-58 |
| chr11-57817765-DEL-73 | chr11-57817766-DEL-73 |
| chr11-57830759-DEL-125 | chr11-57830760-DEL-125 |
| chr11-57839051-DEL-63 | chr11-57839052-DEL-63 |
| chr11-57847463-DEL-77 | chr11-57847464-DEL-76 |
| chr11-57857232-DEL-203 | chr11-57857233-DEL-203 |
| chr11-57873217-DEL-171 | chr11-57873218-DEL-171 |
| chr11-57895658-DEL-56 | chr11-57895665-DEL-50 |
| chr11-57913880-DEL-56 | chr11-57913881-DEL-56 |
| chr11-57929283-DEL-451 | chr11-57929281-DEL-450 |
| chr11-57965453-DEL-200 | chr11-57965458-DEL-195 |
| chr11-57974258-DEL-216 | chr11-57974259-DEL-216 |
| chr11-57977835-DEL-226 | chr11-57977836-DEL-226 |
| chr11-57979130-DEL-1301 | chr11-57979131-DEL-1301 |
| chr11-57992810-DEL-549 | chr11-57992811-DEL-549 |
| chr11-58009999-DEL-590 | chr11-58010000-DEL-590 |
| chr11-58032916-DEL-304 | chr11-58032923-DEL-300 |
| chr11-58049131-DEL-240 | chr11-58049131-DEL-237 |
| chr11-58050379-DEL-51 | chr11-58050380-DEL-52 |

| | |
|---|---|
| chr11-58061347-DEL-200 | chr11-58061348-DEL-200 |
| chr11-58071380-DEL-185 | chr11-58071381-DEL-185 |
| chr11-58093132-DEL-118 | chr11-58093133-DEL-118 |
| chr11-58115714-DEL-89 | chr11-58115715-DEL-89 |
| chr11-58136244-DEL-144 | chr11-58136239-DEL-168 |
| chr11-58141044-DEL-218 | chr11-58141045-DEL-218 |
| chr11-58141740-DEL-86 | chr11-58141741-DEL-90 |
| chr11-58143175-DEL-65 | chr11-58143182-DEL-80 |
| chr11-58146453-DEL-215 | chr11-58146454-DEL-215 |
| chr11-58148316-DEL-52 | chr11-58148316-DEL-52 |
| chr11-58162856-DEL-85 | chr11-58162856-DEL-105 |
| chr11-58167560-DEL-356 | chr11-58167561-DEL-356 |
| chr11-58174722-DEL-328 | chr11-58174706-DEL-332 |
| chr11-58175629-DEL-58 | chr11-58175630-DEL-58 |
| chr11-58180359-DEL-73 | chr11-58180360-DEL-73 |
| chr11-58182617-DEL-62 | chr11-58182612-DEL-60 |
| chr11-58190853-DEL-235 | chr11-58190860-DEL-225 |
| chr11-58194378-DEL-53 | chr11-58194379-DEL-53 |
| chr11-58195274-DEL-93 | chr11-58195286-DEL-86 |
| chr11-58231225-DEL-94 | chr11-58231226-DEL-94 |
| chr11-58258872-DEL-71 | chr11-58258892-DEL-72 |
| chr11-58294834-DEL-62 | chr11-58294832-DEL-50 |
| chr11-58321277-DEL-70 | chr11-58321278-DEL-70 |
| chr11-58347425-DEL-51 | chr11-58347422-DEL-52 |
| chr11-58355784-DEL-647 | chr11-58355785-DEL-647 |
| chr11-58359782-DEL-6562 | chr11-58359769-DEL-6561 |
| chr11-58388221-DEL-528 | chr11-58388222-DEL-528 |
| chr11-58409395-DEL-3665 | chr11-58409396-DEL-3665 |
| chr11-58463361-DEL-66 | chr11-58463362-DEL-66 |
| chr11-58473886-DEL-96 | chr11-58473887-DEL-96 |
| chr11-58480261-DEL-187 | chr11-58480262-DEL-187 |
| chr11-58482028-DEL-350 | chr11-58482029-DEL-350 |
| chr11-58486336-DEL-54 | chr11-58486337-DEL-54 |
| chr11-58487267-DEL-248 | chr11-58487268-DEL-248 |
| chr11-58501128-DEL-63 | chr11-58501128-DEL-63 |
| chr11-58519495-DEL-102 | chr11-58519521-DEL-81 |
| chr11-58541522-DEL-94 | chr11-58541523-DEL-94 |
| chr11-58543013-DEL-68 | chr11-58543014-DEL-68 |
| chr11-58552278-DEL-52 | chr11-58552279-DEL-52 |
| chr11-58553524-DEL-102 | chr11-58553525-DEL-102 |
| chr11-58564621-DEL-7009 | chr11-58564622-DEL-7009 |
| chr11-58564656-DEL-416 | chr11-58564656-DEL-416 |
| chr11-58594386-DEL-3678 | chr11-58594387-DEL-3696 |
| chr11-58598021-DEL-62 | chr11-58598031-DEL-62 |
| chr11-58598904-DEL-2966 | chr11-58598905-DEL-2966 |
| chr11-58600197-DEL-422 | chr11-58600190-DEL-424 |
| chr11-58601682-DEL-265 | chr11-58601683-DEL-265 |
| chr11-58649768-DEL-1940 | chr11-58649769-DEL-1940 |
| chr11-58650785-DEL-126 | chr11-58650784-DEL-128 |
| chr11-58658403-DEL-95 | chr11-58658404-DEL-95 |
| chr11-58668163-DEL-100 | chr11-58668164-DEL-100 |
| chr11-58689178-DEL-52 | chr11-58689182-DEL-56 |
| chr11-58706864-DEL-117 | chr11-58706865-DEL-104 |
| chr11-58738882-DEL-186 | chr11-58738883-DEL-186 |
| chr11-58753784-DEL-73 | chr11-58753789-DEL-69 |
| chr11-58761850-DEL-1434 | chr11-58761851-DEL-1434 |
| chr11-58766411-DEL-60 | chr11-58766400-DEL-58 |
| chr11-58797864-DEL-210 | chr11-58797865-DEL-210 |
| chr11-58798792-DEL-211 | chr11-58798793-DEL-210 |
| chr11-58848984-DEL-120 | chr11-58848990-DEL-115 |
| chr11-58874189-DEL-222 | chr11-58874202-DEL-240 |
| chr11-58895048-DEL-83 | chr11-58895035-DEL-84 |
| chr11-58924133-DEL-63 | chr11-58924134-DEL-63 |
| chr11-58949265-DEL-56 | chr11-58949266-DEL-56 |
| chr11-58982869-DEL-970 | chr11-58982866-DEL-968 |
| chr11-58986976-DEL-1136 | chr11-58986977-DEL-1136 |
| chr11-59013984-DEL-88 | chr11-59013985-DEL-78 |
| chr11-59019842-DEL-52 | chr11-59019843-DEL-52 |
| chr11-59037575-DEL-72 | chr11-59037578-DEL-70 |
| chr11-59080306-DEL-130 | chr11-59080307-DEL-130 |
| chr11-59081655-DEL-52 | chr11-59081656-DEL-52 |
| chr11-59090759-DEL-235 | chr11-59090728-DEL-236 |
| chr11-59098380-DEL-65 | chr11-59098381-DEL-65 |
| chr11-59156834-DEL-192 | chr11-59156835-DEL-192 |
| chr11-59176622-DEL-54 | chr11-59176623-DEL-54 |
| chr11-59236670-DEL-62 | chr11-59236659-DEL-63 |
| chr11-59240662-DEL-101 | chr11-59240663-DEL-101 |
| chr11-59249940-DEL-393 | chr11-59249941-DEL-393 |
| chr11-59265718-DEL-62 | chr11-59265719-DEL-62 |
| chr11-59268025-DEL-97 | chr11-59268026-DEL-97 |

| | |
|---|---|
| chr11-59273248-DEL-84 | chr11-59273255-DEL-80 |
| chr11-59286676-DEL-121 | chr11-59286692-DEL-85 |
| chr11-59326056-DEL-209 | chr11-59326057-DEL-209 |
| chr11-59332862-DEL-312 | chr11-59332863-DEL-312 |
| chr11-59336969-DEL-200 | chr11-59336974-DEL-196 |
| chr11-59402942-DEL-200 | chr11-59402943-DEL-200 |
| chr11-59404336-DEL-90 | chr11-59404333-DEL-83 |
| chr11-59409234-DEL-61 | chr11-59409234-DEL-61 |
| chr11-59418740-DEL-54 | chr11-59418765-DEL-76 |
| chr11-59431560-DEL-85 | chr11-59431561-DEL-85 |
| chr11-59438093-DEL-66 | chr11-59438094-DEL-66 |
| chr11-59454298-DEL-514 | chr11-59454299-DEL-514 |
| chr11-59455195-DEL-73 | chr11-59455196-DEL-73 |
| chr11-59456134-DEL-89 | chr11-59456135-DEL-89 |
| chr11-59467101-DEL-63 | chr11-59467105-DEL-58 |
| chr11-59468066-DEL-66 | chr11-59468067-DEL-66 |
| chr11-59486105-DEL-114 | chr11-59486106-DEL-114 |
| chr11-59498067-DEL-337 | chr11-59498068-DEL-337 |
| chr11-59521263-DEL-175 | chr11-59521264-DEL-175 |
| chr11-59531402-DEL-287 | chr11-59531382-DEL-267 |
| chr11-59599542-DEL-87 | chr11-59599543-DEL-93 |
| chr11-59600981-DEL-166 | chr11-59600982-DEL-166 |
| chr11-59663556-DEL-85 | chr11-59663514-DEL-128 |
| chr11-59728543-DEL-194 | chr11-59728545-DEL-193 |
| chr11-59772917-DEL-1232 | chr11-59772918-DEL-1232 |
| chr11-59777717-DEL-64 | chr11-59777718-DEL-64 |
| chr11-59778243-DEL-498 | chr11-59778244-DEL-498 |
| chr11-59792286-DEL-196 | chr11-59792303-DEL-166 |
| chr11-59833072-DEL-260 | chr11-59833073-DEL-260 |
| chr11-59872745-DEL-199 | chr11-59872746-DEL-199 |
| chr11-59881614-DEL-271 | chr11-59881615-DEL-271 |
| chr11-59883055-DEL-224 | chr11-59883056-DEL-224 |
| chr11-59889644-DEL-56 | chr11-59889645-DEL-56 |
| chr11-59904365-DEL-51 | chr11-59904366-DEL-51 |
| chr11-59905275-DEL-132 | chr11-59905276-DEL-132 |
| chr11-59912611-DEL-80 | chr11-59912612-DEL-80 |
| chr11-59946927-DEL-100 | chr11-59946928-DEL-100 |
| chr11-60019726-DEL-60 | chr11-60019727-DEL-60 |
| chr11-60056291-DEL-259 | chr11-60056292-DEL-259 |
| chr11-60093276-DEL-127 | chr11-60093277-DEL-127 |
| chr11-60171596-DEL-55 | chr11-60171591-DEL-56 |
| chr11-60172414-DEL-370 | chr11-60172415-DEL-369 |
| chr11-60184371-DEL-218 | chr11-60184372-DEL-218 |
| chr11-60216693-DEL-216 | chr11-60216694-DEL-216 |
| chr11-60264595-DEL-218 | chr11-60264596-DEL-218 |
| chr11-60360528-DEL-109 | chr11-60360529-DEL-109 |
| chr11-60391485-DEL-179 | chr11-60391486-DEL-179 |
| chr11-60414305-DEL-63 | chr11-60414305-DEL-64 |
| chr11-60418337-DEL-199 | chr11-60418338-DEL-199 |
| chr11-60419770-DEL-1660 | chr11-60419771-DEL-1660 |
| chr11-60424753-DEL-149 | chr11-60424754-DEL-149 |
| chr11-60487308-DEL-81 | chr11-60487300-DEL-93 |
| chr11-60505697-DEL-89 | chr11-60505730-DEL-70 |
| chr11-60529264-DEL-101 | chr11-60529263-DEL-101 |
| chr11-60537456-DEL-86 | chr11-60537437-DEL-96 |
| chr11-60553049-DEL-93 | chr11-60553050-DEL-95 |
| chr11-60557945-DEL-727 | chr11-60557937-DEL-729 |
| chr11-60613614-DEL-81 | chr11-60613615-DEL-81 |
| chr11-60626365-DEL-201 | chr11-60626360-DEL-202 |
| chr11-60648234-DEL-89 | chr11-60648235-DEL-89 |
| chr11-60651483-DEL-150 | chr11-60651484-DEL-150 |
| chr11-60655769-DEL-60 | chr11-60655746-DEL-56 |
| chr11-60660510-DEL-116 | chr11-60660511-DEL-116 |
| chr11-60673787-DEL-222 | chr11-60673788-DEL-222 |
| chr11-60684217-DEL-210 | chr11-60684218-DEL-209 |
| chr11-60704767-DEL-198 | chr11-60704768-DEL-196 |
| chr11-60705135-DEL-1890 | chr11-60705136-DEL-1890 |
| chr11-60709491-DEL-76 | chr11-60709492-DEL-76 |
| chr11-60726093-DEL-1643 | chr11-60726094-DEL-1643 |
| chr11-60760467-DEL-1065 | chr11-60760454-DEL-1069 |
| chr11-60764725-DEL-147 | chr11-60764732-DEL-143 |
| chr11-60773591-DEL-217 | chr11-60773592-DEL-217 |
| chr11-60778058-DEL-211 | chr11-60778059-DEL-211 |
| chr11-60785312-DEL-595 | chr11-60785313-DEL-595 |
| chr11-60801749-DEL-266 | chr11-60801750-DEL-265 |
| chr11-60850500-DEL-63 | chr11-60850501-DEL-84 |
| chr11-60889217-DEL-85 | chr11-60889218-DEL-85 |
| chr11-60938729-DEL-396 | chr11-60938730-DEL-396 |
| chr11-60979167-DEL-627 | chr11-60979168-DEL-627 |
| chr11-60990364-DEL-108 | chr11-60990365-DEL-108 |

| | |
|---|---|
| chr11-61007452-DEL-140 | chr11-61007453-DEL-140 |
| chr11-61019364-DEL-89 | chr11-61019365-DEL-89 |
| chr11-61083191-DEL-506 | chr11-61083192-DEL-506 |
| chr11-61087592-DEL-57 | chr11-61087593-DEL-57 |
| chr11-61137441-DEL-189 | chr11-61137442-DEL-189 |
| chr11-61166679-DEL-97 | chr11-61166680-DEL-97 |
| chr11-61175998-DEL-198 | chr11-61175998-DEL-190 |
| chr11-61182485-DEL-209 | chr11-61182486-DEL-209 |
| chr11-61188744-DEL-52 | chr11-61188745-DEL-52 |
| chr11-61195571-DEL-332 | chr11-61195563-DEL-341 |
| chr11-61200968-DEL-95 | chr11-61200970-DEL-94 |
| chr11-61202752-DEL-166 | chr11-61202753-DEL-166 |
| chr11-61205525-DEL-221 | chr11-61205526-DEL-221 |
| chr11-61275667-DEL-58 | chr11-61275684-DEL-50 |
| chr11-61278956-DEL-53 | chr11-61278957-DEL-52 |
| chr11-61306443-DEL-88 | chr11-61306431-DEL-88 |
| chr11-61311188-DEL-7217 | chr11-61311189-DEL-7217 |
| chr11-61328414-DEL-92 | chr11-61328415-DEL-92 |
| chr11-61345472-DEL-76 | chr11-61345499-DEL-50 |
| chr11-61439561-DEL-745 | chr11-61439562-DEL-745 |
| chr11-61484444-DEL-2813 | chr11-61484445-DEL-2813 |
| chr11-61544840-DEL-445 | chr11-61544840-DEL-445 |
| chr11-61546119-DEL-60 | chr11-61546120-DEL-60 |
| chr11-61609380-DEL-61 | chr11-61609380-DEL-61 |
| chr11-61617247-DEL-1280 | chr11-61617248-DEL-1280 |
| chr11-61717684-DEL-54 | chr11-61717685-DEL-54 |
| chr11-61720424-DEL-98 | chr11-61720425-DEL-98 |
| chr11-61726328-DEL-212 | chr11-61726329-DEL-212 |
| chr11-61738965-DEL-52 | chr11-61738966-DEL-52 |
| chr11-61754702-DEL-84 | chr11-61754703-DEL-84 |
| chr11-61764061-DEL-1057 | chr11-61764062-DEL-1057 |
| chr11-61781530-DEL-163 | chr11-61781531-DEL-163 |
| chr11-61831431-DEL-198 | chr11-61831432-DEL-198 |
| chr11-61849635-DEL-52 | chr11-61849635-DEL-52 |
| chr11-61864457-DEL-60 | chr11-61864469-DEL-70 |
| chr11-61865985-DEL-59 | chr11-61865985-DEL-59 |
| chr11-61869506-DEL-56 | chr11-61869510-DEL-52 |
| chr11-61873456-DEL-223 | chr11-61873457-DEL-223 |
| chr11-61881923-DEL-965 | chr11-61881929-DEL-949 |
| chr11-61887535-DEL-54 | chr11-61887527-DEL-54 |
| chr11-61891413-DEL-306 | chr11-61891415-DEL-307 |
| chr11-61898622-DEL-419 | chr11-61898623-DEL-419 |
| chr11-61903144-DEL-312 | chr11-61903145-DEL-312 |
| chr11-61907527-DEL-91 | chr11-61907508-DEL-92 |
| chr11-61908763-DEL-125 | chr11-61908764-DEL-125 |
| chr11-61911628-DEL-107 | chr11-61911582-DEL-108 |
| chr11-61931417-DEL-1689 | chr11-61931418-DEL-1689 |
| chr11-61946599-DEL-55 | chr11-61946568-DEL-56 |
| chr11-62005920-DEL-114 | chr11-62005921-DEL-113 |
| chr11-62109035-DEL-122 | chr11-62109036-DEL-126 |
| chr11-62109539-DEL-81 | chr11-62109540-DEL-81 |
| chr11-62110475-DEL-79 | chr11-62110476-DEL-79 |
| chr11-62121682-DEL-105 | chr11-62121672-DEL-111 |
| chr11-62211419-DEL-211 | chr11-62211420-DEL-211 |
| chr11-62303073-DEL-195 | chr11-62303074-DEL-195 |
| chr11-62307945-DEL-198 | chr11-62307946-DEL-198 |
| chr11-62320510-DEL-2457 | chr11-62320501-DEL-2470 |
| chr11-62342585-DEL-213 | chr11-62342586-DEL-213 |
| chr11-62345051-DEL-207 | chr11-62345055-DEL-202 |
| chr11-62376950-DEL-176 | chr11-62376951-DEL-176 |
| chr11-62382274-DEL-245 | chr11-62382275-DEL-245 |
| chr11-62392182-DEL-96 | chr11-62392168-DEL-101 |
| chr11-62399025-DEL-92 | chr11-62399024-DEL-93 |
| chr11-62410990-DEL-375 | chr11-62410991-DEL-375 |
| chr11-62411780-DEL-1001 | chr11-62411781-DEL-1001 |
| chr11-62423670-DEL-445 | chr11-62423671-DEL-445 |
| chr11-62432579-DEL-1179 | chr11-62432580-DEL-1179 |
| chr11-62448822-DEL-835 | chr11-62448821-DEL-837 |
| chr11-62458840-DEL-204 | chr11-62458841-DEL-204 |
| chr11-62459867-DEL-75 | chr11-62459868-DEL-60 |
| chr11-62464526-DEL-285 | chr11-62464527-DEL-285 |
| chr11-62470774-DEL-52 | chr11-62470775-DEL-52 |
| chr11-62487179-DEL-195 | chr11-62487180-DEL-195 |
| chr11-62491160-DEL-176 | chr11-62491161-DEL-176 |
| chr11-62507346-DEL-374 | chr11-62507347-DEL-374 |
| chr11-62518237-DEL-1087 | chr11-62518238-DEL-1087 |
| chr11-62547375-DEL-77 | chr11-62547376-DEL-77 |
| chr11-62563435-DEL-70 | chr11-62563436-DEL-70 |
| chr11-62564635-DEL-233 | chr11-62564623-DEL-231 |
| chr11-62583043-DEL-226 | chr11-62583044-DEL-226 |

| | |
|---|---|
| chr11-62584539-DEL-230 | chr11-62584540-DEL-230 |
| chr11-62593110-DEL-210 | chr11-62593111-DEL-210 |
| chr11-62594447-DEL-198 | chr11-62594448-DEL-198 |
| chr11-62607237-DEL-546 | chr11-62607238-DEL-546 |
| chr11-62607903-DEL-430 | chr11-62607904-DEL-430 |
| chr11-62610667-DEL-195 | chr11-62610668-DEL-195 |
| chr11-62653737-DEL-204 | chr11-62653738-DEL-204 |
| chr11-62654817-DEL-261 | chr11-62654818-DEL-261 |
| chr11-62664365-DEL-193 | chr11-62664366-DEL-194 |
| chr11-62666687-DEL-51 | chr11-62666683-DEL-52 |
| chr11-62703799-DEL-236 | chr11-62703800-DEL-236 |
| chr11-62719444-DEL-55 | chr11-62719432-DEL-65 |
| chr11-62728732-DEL-96 | chr11-62728733-DEL-96 |
| chr11-62746953-DEL-66 | chr11-62746963-DEL-74 |
| chr11-62752827-DEL-94 | chr11-62752828-DEL-96 |
| chr11-62770792-DEL-220 | chr11-62770793-DEL-220 |
| chr11-62775250-DEL-327 | chr11-62775251-DEL-327 |
| chr11-62784183-DEL-58 | chr11-62784183-DEL-58 |
| chr11-62787511-DEL-52 | chr11-62787512-DEL-52 |
| chr11-62790724-DEL-1229 | chr11-62790725-DEL-1229 |
| chr11-62792049-DEL-197 | chr11-62792050-DEL-197 |
| chr11-62818258-DEL-64 | chr11-62818259-DEL-62 |
| chr11-62819005-DEL-162 | chr11-62819006-DEL-162 |
| chr11-62822556-DEL-116 | chr11-62822557-DEL-116 |
| chr11-62843251-DEL-54 | chr11-62843252-DEL-54 |
| chr11-62848022-DEL-475 | chr11-62848023-DEL-475 |
| chr11-62856473-DEL-103 | chr11-62856474-DEL-103 |
| chr11-62859159-DEL-6367 | chr11-62859160-DEL-6367 |
| chr11-62901565-DEL-66 | chr11-62901570-DEL-62 |
| chr11-62960656-DEL-56 | chr11-62960656-DEL-56 |
| chr11-62966047-DEL-214 | chr11-62966048-DEL-214 |
| chr11-62974204-DEL-55 | chr11-62974205-DEL-55 |
| chr11-63011280-DEL-129 | chr11-63011281-DEL-129 |
| chr11-63085473-DEL-6498 | chr11-63085474-DEL-6498 |
| chr11-63096536-DEL-786 | chr11-63096537-DEL-786 |
| chr11-63103434-DEL-54 | chr11-63103435-DEL-54 |
| chr11-63130206-DEL-219 | chr11-63130206-DEL-223 |
| chr11-63139185-DEL-64 | chr11-63139186-DEL-64 |
| chr11-63162989-DEL-104 | chr11-63163000-DEL-94 |
| chr11-63164652-DEL-98 | chr11-63164657-DEL-63 |
| chr11-63171836-DEL-71 | chr11-63171835-DEL-71 |
| chr11-63202907-DEL-1303 | chr11-63202908-DEL-1303 |
| chr11-63217368-DEL-263 | chr11-63217369-DEL-263 |
| chr11-63219810-DEL-192 | chr11-63219795-DEL-192 |
| chr11-63224654-DEL-116 | chr11-63224659-DEL-112 |
| chr11-63265595-DEL-70 | chr11-63265587-DEL-72 |
| chr11-63271736-DEL-52 | chr11-63271737-DEL-52 |
| chr11-63307986-DEL-127 | chr11-63307987-DEL-127 |
| chr11-63347096-DEL-82 | chr11-63347099-DEL-52 |
| chr11-63358536-DEL-7181 | chr11-63358537-DEL-7181 |
| chr11-63366504-DEL-367 | chr11-63366505-DEL-367 |
| chr11-63368753-DEL-71 | chr11-63368757-DEL-72 |
| chr11-63370740-DEL-160 | chr11-63370741-DEL-120 |
| chr11-63381659-DEL-1948 | chr11-63381660-DEL-1948 |
| chr11-63410262-DEL-53 | chr11-63410262-DEL-53 |
| chr11-63443207-DEL-1262 | chr11-63443218-DEL-1259 |
| chr11-63484580-DEL-2489 | chr11-63484581-DEL-2481 |
| chr11-63498827-DEL-704 | chr11-63498828-DEL-704 |
| chr11-63502403-DEL-1087 | chr11-63502404-DEL-1087 |
| chr11-63514659-DEL-498 | chr11-63514660-DEL-498 |
| chr11-63525172-DEL-62 | chr11-63525179-DEL-56 |
| chr11-63537724-DEL-6690 | chr11-63537725-DEL-6690 |
| chr11-63566641-DEL-504 | chr11-63566684-DEL-471 |
| chr11-63577810-DEL-122 | chr11-63577826-DEL-113 |
| chr11-63590716-DEL-68 | chr11-63590714-DEL-70 |
| chr11-63631452-DEL-401 | chr11-63631453-DEL-401 |
| chr11-63678730-DEL-272 | chr11-63678731-DEL-272 |
| chr11-63690120-DEL-6362 | chr11-63690121-DEL-6362 |
| chr11-63701733-DEL-98 | chr11-63701769-DEL-84 |
| chr11-63705270-DEL-2152 | chr11-63705265-DEL-2145 |
| chr11-63710182-DEL-80 | chr11-63710183-DEL-80 |
| chr11-63728837-DEL-1440 | chr11-63728838-DEL-1440 |
| chr11-63730758-DEL-123 | chr11-63730759-DEL-123 |
| chr11-63736888-DEL-885 | chr11-63736889-DEL-885 |
| chr11-63753512-DEL-64 | chr11-63753513-DEL-64 |
| chr11-63756611-DEL-146 | chr11-63756612-DEL-146 |
| chr11-63759476-DEL-52 | chr11-63759471-DEL-53 |
| chr11-63780624-DEL-1093 | chr11-63780625-DEL-1094 |
| chr11-63798570-DEL-52 | chr11-63798566-DEL-56 |
| chr11-63799753-DEL-126 | chr11-63799754-DEL-123 |

| | |
|---|---|
| chr11-63804024-DEL-128 | chr11-63804056-DEL-97 |
| chr11-63831170-DEL-557 | chr11-63831171-DEL-557 |
| chr11-63835137-DEL-72 | chr11-63835136-DEL-97 |
| chr11-63845193-DEL-56 | chr11-63845194-DEL-56 |
| chr11-63866688-DEL-152 | chr11-63866691-DEL-153 |
| chr11-63874790-DEL-201 | chr11-63874791-DEL-201 |
| chr11-63896655-DEL-211 | chr11-63896656-DEL-211 |
| chr11-63907991-DEL-54 | chr11-63907977-DEL-72 |
| chr11-63989757-DEL-92 | chr11-63989758-DEL-69 |
| chr11-63998151-DEL-64 | chr11-63998155-DEL-59 |
| chr11-64013877-DEL-131 | chr11-64013878-DEL-131 |
| chr11-64028522-DEL-63 | chr11-64028523-DEL-63 |
| chr11-64029593-DEL-101 | chr11-64029594-DEL-101 |
| chr11-64046640-DEL-725 | chr11-64046641-DEL-725 |
| chr11-64089927-DEL-1386 | chr11-64089928-DEL-1386 |
| chr11-64093844-DEL-532 | chr11-64093845-DEL-532 |
| chr11-64126639-DEL-58 | chr11-64126639-DEL-58 |
| chr11-64137520-DEL-74 | chr11-64137520-DEL-74 |
| chr11-64142050-DEL-54 | chr11-64142051-DEL-54 |
| chr11-64162798-DEL-63 | chr11-64162839-DEL-60 |
| chr11-64178935-DEL-974 | chr11-64178936-DEL-974 |
| chr11-64186603-DEL-53 | chr11-64186604-DEL-53 |
| chr11-64189957-DEL-6861 | chr11-64189958-DEL-6861 |
| chr11-64220527-DEL-72 | chr11-64220536-DEL-57 |
| chr11-64228206-DEL-56 | chr11-64228207-DEL-56 |
| chr11-64233542-DEL-863 | chr11-64233543-DEL-863 |
| chr11-64252798-DEL-432 | chr11-64252799-DEL-432 |
| chr11-64265347-DEL-788 | chr11-64265347-DEL-788 |
| chr11-64271330-DEL-79 | chr11-64271328-DEL-82 |
| chr11-64278589-DEL-85 | chr11-64278590-DEL-85 |
| chr11-64347663-DEL-1864 | chr11-64347664-DEL-1864 |
| chr11-64360268-DEL-108 | chr11-64360233-DEL-144 |
| chr11-64365669-DEL-120 | chr11-64365650-DEL-137 |
| chr11-64420504-DEL-291 | chr11-64420505-DEL-291 |
| chr11-64426855-DEL-990 | chr11-64426858-DEL-988 |
| chr11-64458837-DEL-6209 | chr11-64458838-DEL-6209 |
| chr11-64480787-DEL-72 | chr11-64480793-DEL-52 |
| chr11-64510845-DEL-52 | chr11-64510846-DEL-52 |
| chr11-64522534-DEL-208 | chr11-64522535-DEL-208 |
| chr11-64560716-DEL-588 | chr11-64560717-DEL-588 |
| chr11-64586230-DEL-238 | chr11-64586232-DEL-237 |
| chr11-64604287-DEL-839 | chr11-64604297-DEL-830 |
| chr11-64615278-DEL-255 | chr11-64615279-DEL-255 |
| chr11-64616587-DEL-145 | chr11-64616591-DEL-141 |
| chr11-64666562-DEL-159 | chr11-64666563-DEL-159 |
| chr11-64668497-DEL-108 | chr11-64668497-DEL-109 |
| chr11-64682863-DEL-5415 | chr11-64682864-DEL-5415 |
| chr11-64698681-DEL-1238 | chr11-64698682-DEL-1238 |
| chr11-64704537-DEL-429 | chr11-64704536-DEL-430 |
| chr11-64705305-DEL-51 | chr11-64705306-DEL-51 |
| chr11-64710980-DEL-70 | chr11-64710981-DEL-70 |
| chr11-64716290-DEL-1854 | chr11-64716291-DEL-1854 |
| chr11-64742303-DEL-510 | chr11-64742304-DEL-510 |
| chr11-64759988-DEL-112 | chr11-64759989-DEL-112 |
| chr11-64767253-DEL-165 | chr11-64767254-DEL-165 |
| chr11-64784527-DEL-352 | chr11-64784528-DEL-352 |
| chr11-64802167-DEL-52 | chr11-64802174-DEL-52 |
| chr11-64810797-DEL-112 | chr11-64810826-DEL-84 |
| chr11-64833383-DEL-1137 | chr11-64833384-DEL-1137 |
| chr11-64837931-DEL-1092 | chr11-64837941-DEL-1083 |
| chr11-64886524-DEL-801 | chr11-64886525-DEL-801 |
| chr11-64948015-DEL-104 | chr11-64948016-DEL-104 |
| chr11-64966391-DEL-53 | chr11-64966392-DEL-53 |
| chr11-64972018-DEL-60 | chr11-64972018-DEL-50 |
| chr11-64993529-DEL-307 | chr11-64993530-DEL-307 |
| chr11-65011399-DEL-71 | chr11-65011400-DEL-71 |
| chr11-65017982-DEL-123 | chr11-65017983-DEL-123 |
| chr11-65093794-DEL-166 | chr11-65093795-DEL-166 |
| chr11-65104048-DEL-68 | chr11-65104063-DEL-83 |
| chr11-65159052-DEL-87 | chr11-65159053-DEL-87 |
| chr11-65173055-DEL-52 | chr11-65173055-DEL-52 |
| chr11-65199463-DEL-759 | chr11-65199465-DEL-758 |
| chr11-65236228-DEL-179 | chr11-65236229-DEL-179 |
| chr11-65241096-DEL-1163 | chr11-65241097-DEL-1163 |
| chr11-65244270-DEL-59 | chr11-65244271-DEL-59 |
| chr11-65269133-DEL-159 | chr11-65269134-DEL-159 |
| chr11-65269904-DEL-67 | chr11-65269905-DEL-67 |
| chr11-65319942-DEL-324 | chr11-65319943-DEL-324 |
| chr11-65320031-DEL-810 | chr11-65320032-DEL-810 |
| chr11-65334118-DEL-81 | chr11-65334159-DEL-82 |

| | |
|---|---|
| chr11-65337847-DEL-111 | chr11-65337848-DEL-112 |
| chr11-65342604-DEL-7248 | chr11-65342601-DEL-7249 |
| chr11-65379210-DEL-74 | chr11-65379186-DEL-93 |
| chr11-65387452-DEL-690 | chr11-65387457-DEL-688 |
| chr11-65391745-DEL-57 | chr11-65391746-DEL-57 |
| chr11-65463736-DEL-94 | chr11-65463737-DEL-92 |
| chr11-65514969-DEL-63 | chr11-65514973-DEL-57 |
| chr11-65517203-DEL-376 | chr11-65517204-DEL-376 |
| chr11-65545895-DEL-57 | chr11-65545896-DEL-58 |
| chr11-65564501-DEL-163 | chr11-65564502-DEL-163 |
| chr11-65569369-DEL-285 | chr11-65569370-DEL-284 |
| chr11-65653586-DEL-116 | chr11-65653587-DEL-116 |
| chr11-65673178-DEL-92 | chr11-65673178-DEL-89 |
| chr11-65677361-DEL-75 | chr11-65677377-DEL-73 |
| chr11-65688139-DEL-132 | chr11-65688140-DEL-132 |
| chr11-65706011-DEL-51 | chr11-65706012-DEL-51 |
| chr11-65736842-DEL-102 | chr11-65736843-DEL-102 |
| chr11-65737579-DEL-750 | chr11-65737580-DEL-750 |
| chr11-65775750-DEL-199 | chr11-65775768-DEL-191 |
| chr11-65782252-DEL-199 | chr11-65782259-DEL-199 |
| chr11-65838056-DEL-92 | chr11-65838085-DEL-65 |
| chr11-65840264-DEL-72 | chr11-65840265-DEL-72 |
| chr11-65868528-DEL-56 | chr11-65868553-DEL-58 |
| chr11-65901704-DEL-244 | chr11-65901704-DEL-244 |
| chr11-65904257-DEL-113 | chr11-65904258-DEL-112 |
| chr11-65912455-DEL-846 | chr11-65912456-DEL-846 |
| chr11-65935866-DEL-118 | chr11-65935867-DEL-115 |
| chr11-65941996-DEL-723 | chr11-65941997-DEL-723 |
| chr11-65944956-DEL-60 | chr11-65944959-DEL-58 |
| chr11-65979674-DEL-199 | chr11-65979663-DEL-235 |
| chr11-65990251-DEL-58 | chr11-65990252-DEL-56 |
| chr11-66083818-DEL-6194 | chr11-66083819-DEL-6194 |
| chr11-66111208-DEL-102 | chr11-66111209-DEL-102 |
| chr11-66170189-DEL-58 | chr11-66170194-DEL-52 |
| chr11-66174914-DEL-590 | chr11-66174915-DEL-590 |
| chr11-66206114-DEL-1004 | chr11-66206115-DEL-1004 |
| chr11-66208774-DEL-57 | chr11-66208775-DEL-57 |
| chr11-66223297-DEL-1890 | chr11-66223298-DEL-1890 |
| chr11-66223534-DEL-255 | chr11-66223535-DEL-255 |
| chr11-66228378-DEL-69 | chr11-66228379-DEL-69 |
| chr11-66233741-DEL-53 | chr11-66233738-DEL-61 |
| chr11-66237601-DEL-62 | chr11-66237602-DEL-62 |
| chr11-66241100-DEL-826 | chr11-66241101-DEL-826 |
| chr11-66245713-DEL-69 | chr11-66245710-DEL-69 |
| chr11-66262998-DEL-292 | chr11-66262986-DEL-285 |
| chr11-66298976-DEL-58 | chr11-66298959-DEL-76 |
| chr11-66343874-DEL-582 | chr11-66343875-DEL-582 |
| chr11-66346275-DEL-57 | chr11-66346279-DEL-71 |
| chr11-66358921-DEL-65 | chr11-66358922-DEL-65 |
| chr11-66386239-DEL-389 | chr11-66386240-DEL-389 |
| chr11-66412171-DEL-208 | chr11-66412172-DEL-208 |
| chr11-66433813-DEL-278 | chr11-66433834-DEL-261 |
| chr11-66468754-DEL-67 | chr11-66468755-DEL-67 |
| chr11-66472366-DEL-1442 | chr11-66472367-DEL-1442 |
| chr11-66492737-DEL-343 | chr11-66492738-DEL-343 |
| chr11-66500059-DEL-63 | chr11-66500073-DEL-63 |
| chr11-66509097-DEL-266 | chr11-66509104-DEL-260 |
| chr11-66511099-DEL-354 | chr11-66511094-DEL-330 |
| chr11-66519100-DEL-6188 | chr11-66519091-DEL-6189 |
| chr11-66536715-DEL-155 | chr11-66536733-DEL-131 |
| chr11-66558809-DEL-1477 | chr11-66558810-DEL-1477 |
| chr11-66583750-DEL-107 | chr11-66583751-DEL-107 |
| chr11-66597004-DEL-6228 | chr11-66597005-DEL-6228 |
| chr11-66628442-DEL-393 | chr11-66628443-DEL-393 |
| chr11-66660152-DEL-100 | chr11-66660153-DEL-100 |
| chr11-66666675-DEL-589 | chr11-66666673-DEL-590 |
| chr11-66674933-DEL-2745 | chr11-66674934-DEL-2745 |
| chr11-66684380-DEL-3421 | chr11-66684380-DEL-3421 |
| chr11-66718610-DEL-328 | chr11-66718611-DEL-328 |
| chr11-66747310-DEL-1369 | chr11-66747311-DEL-1369 |
| chr11-66751428-DEL-96 | chr11-66751429-DEL-96 |
| chr11-66755985-DEL-576 | chr11-66755987-DEL-571 |
| chr11-66785188-DEL-206 | chr11-66785203-DEL-196 |
| chr11-66791400-DEL-65 | chr11-66791401-DEL-65 |
| chr11-66807446-DEL-54 | chr11-66807446-DEL-54 |
| chr11-66831144-DEL-55 | chr11-66831145-DEL-55 |
| chr11-66838662-DEL-6519 | chr11-66838663-DEL-6519 |
| chr11-66885651-DEL-83 | chr11-66885639-DEL-81 |
| chr11-66926241-DEL-51 | chr11-66926216-DEL-74 |
| chr11-66941358-DEL-306 | chr11-66941359-DEL-306 |

| | |
|---|---|
| chr11-66960395-DEL-146 | chr11-66960426-DEL-104 |
| chr11-66988300-DEL-198 | chr11-66988301-DEL-198 |
| chr11-67020530-DEL-2669 | chr11-67020531-DEL-2669 |
| chr11-67045891-DEL-177 | chr11-67045892-DEL-177 |
| chr11-67058455-DEL-84 | chr11-67058460-DEL-80 |
| chr11-67077741-DEL-53 | chr11-67077742-DEL-53 |
| chr11-67078258-DEL-82 | chr11-67078259-DEL-82 |
| chr11-67178603-DEL-60 | chr11-67178608-DEL-50 |
| chr11-67184813-DEL-51 | chr11-67184814-DEL-51 |
| chr11-67186820-DEL-469 | chr11-67186821-DEL-469 |
| chr11-67195867-DEL-76 | chr11-67195868-DEL-72 |
| chr11-67230152-DEL-54 | chr11-67230155-DEL-55 |
| chr11-67245978-DEL-78 | chr11-67245979-DEL-78 |
| chr11-67282564-DEL-52 | chr11-67282565-DEL-52 |
| chr11-67293774-DEL-268 | chr11-67293775-DEL-268 |
| chr11-67310091-DEL-108 | chr11-67310109-DEL-78 |
| chr11-67353006-DEL-842 | chr11-67353007-DEL-842 |
| chr11-67353905-DEL-299 | chr11-67353906-DEL-299 |
| chr11-67383167-DEL-455 | chr11-67383168-DEL-455 |
| chr11-67397764-DEL-178 | chr11-67397765-DEL-179 |
| chr11-67412622-DEL-51 | chr11-67412623-DEL-51 |
| chr11-67430134-DEL-102 | chr11-67430135-DEL-102 |
| chr11-67453032-DEL-57 | chr11-67453033-DEL-57 |
| chr11-67526671-DEL-94 | chr11-67526709-DEL-102 |
| chr11-67568460-DEL-266 | chr11-67568467-DEL-258 |
| chr11-67569238-DEL-708 | chr11-67569220-DEL-714 |
| chr11-67577782-DEL-56 | chr11-67577783-DEL-56 |
| chr11-67580174-DEL-117 | chr11-67580175-DEL-117 |
| chr11-67585525-DEL-62 | chr11-67585530-DEL-58 |
| chr11-67602213-DEL-313 | chr11-67602225-DEL-322 |
| chr11-67654236-DEL-567 | chr11-67654237-DEL-567 |
| chr11-67658830-DEL-100 | chr11-67658876-DEL-55 |
| chr11-67671848-DEL-96 | chr11-67671849-DEL-96 |
| chr11-67707265-DEL-98 | chr11-67707251-DEL-105 |
| chr11-67729768-DEL-62 | chr11-67729753-DEL-54 |
| chr11-67734278-DEL-91 | chr11-67734279-DEL-91 |
| chr11-67740696-DEL-100 | chr11-67740690-DEL-103 |
| chr11-67750097-DEL-2722 | chr11-67750098-DEL-2722 |
| chr11-67757167-DEL-6401 | chr11-67757168-DEL-6401 |
| chr11-67771260-DEL-58 | chr11-67771261-DEL-58 |
| chr11-67783751-DEL-58 | chr11-67783748-DEL-60 |
| chr11-67808382-DEL-192 | chr11-67808383-DEL-192 |
| chr11-67819019-DEL-88 | chr11-67819020-DEL-88 |
| chr11-67883344-DEL-99 | chr11-67883345-DEL-99 |
| chr11-67897804-DEL-172 | chr11-67897795-DEL-182 |
| chr11-67921117-DEL-204 | chr11-67921118-DEL-204 |
| chr11-67970130-DEL-72 | chr11-67970131-DEL-72 |
| chr11-67975867-DEL-112 | chr11-67975868-DEL-112 |
| chr11-67985092-DEL-174 | chr11-67985093-DEL-174 |
| chr11-68006258-DEL-210 | chr11-68006259-DEL-210 |
| chr11-68023612-DEL-153 | chr11-68023613-DEL-153 |
| chr11-68042536-DEL-207 | chr11-68042537-DEL-206 |
| chr11-68055634-DEL-223 | chr11-68055634-DEL-223 |
| chr11-68072580-DEL-201 | chr11-68072581-DEL-201 |
| chr11-68088742-DEL-53 | chr11-68088725-DEL-71 |
| chr11-68149421-DEL-93 | chr11-68149422-DEL-94 |
| chr11-68175082-DEL-191 | chr11-68175083-DEL-191 |
| chr11-68188447-DEL-445 | chr11-68188448-DEL-445 |
| chr11-68190920-DEL-96 | chr11-68190921-DEL-96 |
| chr11-68284140-DEL-212 | chr11-68284141-DEL-212 |
| chr11-68308947-DEL-78 | chr11-68308948-DEL-78 |
| chr11-68351736-DEL-59 | chr11-68351717-DEL-91 |
| chr11-68373658-DEL-174 | chr11-68373665-DEL-142 |
| chr11-68376789-DEL-144 | chr11-68376790-DEL-144 |
| chr11-68379679-DEL-52 | chr11-68379680-DEL-52 |
| chr11-68397166-DEL-107 | chr11-68397166-DEL-107 |
| chr11-68398162-DEL-344 | chr11-68398163-DEL-344 |
| chr11-68402864-DEL-190 | chr11-68402863-DEL-189 |
| chr11-68430916-DEL-77 | chr11-68430917-DEL-77 |
| chr11-68449369-DEL-231 | chr11-68449370-DEL-231 |
| chr11-68457331-DEL-53 | chr11-68457325-DEL-54 |
| chr11-68460004-DEL-162 | chr11-68460005-DEL-162 |
| chr11-68464322-DEL-66 | chr11-68464323-DEL-66 |
| chr11-68467034-DEL-406 | chr11-68467035-DEL-406 |
| chr11-68469727-DEL-52 | chr11-68469728-DEL-52 |
| chr11-68472988-DEL-206 | chr11-68472989-DEL-206 |
| chr11-68473968-DEL-58 | chr11-68473965-DEL-62 |
| chr11-68483737-DEL-62 | chr11-68483750-DEL-52 |
| chr11-68486163-DEL-767 | chr11-68486164-DEL-767 |
| chr11-68512174-DEL-207 | chr11-68512175-DEL-207 |

| | |
|---|---|
| chr11-68519122-DEL-89 | chr11-68519123-DEL-89 |
| chr11-68542660-DEL-1134 | chr11-68542661-DEL-1132 |
| chr11-68551240-DEL-64 | chr11-68551249-DEL-62 |
| chr11-68593876-DEL-93 | chr11-68593877-DEL-93 |
| chr11-68600081-DEL-139 | chr11-68600082-DEL-139 |
| chr11-68602095-DEL-137 | chr11-68602095-DEL-138 |
| chr11-68624259-DEL-176 | chr11-68624260-DEL-220 |
| chr11-68650370-DEL-142 | chr11-68650371-DEL-142 |
| chr11-68662291-DEL-93 | chr11-68662292-DEL-93 |
| chr11-68675454-DEL-163 | chr11-68675455-DEL-163 |
| chr11-68677737-DEL-66 | chr11-68677738-DEL-66 |
| chr11-68686296-DEL-473 | chr11-68686297-DEL-473 |
| chr11-68697597-DEL-63 | chr11-68697603-DEL-64 |
| chr11-68709676-DEL-95 | chr11-68709677-DEL-95 |
| chr11-68718689-DEL-227 | chr11-68718690-DEL-227 |
| chr11-68731434-DEL-204 | chr11-68731431-DEL-203 |
| chr11-68756413-DEL-53 | chr11-68756414-DEL-53 |
| chr11-68757194-DEL-191 | chr11-68757195-DEL-191 |
| chr11-68777488-DEL-155 | chr11-68777489-DEL-155 |
| chr11-68778445-DEL-133 | chr11-68778446-DEL-133 |
| chr11-68805728-DEL-110 | chr11-68805729-DEL-110 |
| chr11-68808334-DEL-53 | chr11-68808310-DEL-70 |
| chr11-68808561-DEL-1860 | chr11-68808561-DEL-1861 |
| chr11-68811806-DEL-95 | chr11-68811807-DEL-95 |
| chr11-68851832-DEL-174 | chr11-68851833-DEL-174 |
| chr11-68868559-DEL-221 | chr11-68868560-DEL-221 |
| chr11-68876222-DEL-68 | chr11-68876223-DEL-68 |
| chr11-68900863-DEL-56 | chr11-68900864-DEL-56 |
| chr11-68905514-DEL-66 | chr11-68905515-DEL-66 |
| chr11-68925834-DEL-87 | chr11-68925835-DEL-85 |
| chr11-68966006-DEL-211 | chr11-68966007-DEL-211 |
| chr11-68966531-DEL-214 | chr11-68966532-DEL-214 |
| chr11-68969268-DEL-201 | chr11-68969269-DEL-201 |
| chr11-68974853-DEL-653 | chr11-68974854-DEL-653 |
| chr11-69018983-DEL-171 | chr11-69018984-DEL-171 |
| chr11-69043783-DEL-90 | chr11-69043780-DEL-90 |
| chr11-69050468-DEL-91 | chr11-69050460-DEL-92 |
| chr11-69062937-DEL-111 | chr11-69062938-DEL-111 |
| chr11-69064083-DEL-57 | chr11-69064084-DEL-57 |
| chr11-69073510-DEL-249 | chr11-69073511-DEL-249 |
| chr11-69131076-DEL-64 | chr11-69131076-DEL-64 |
| chr11-69154132-DEL-306 | chr11-69154149-DEL-298 |
| chr11-69165228-DEL-843 | chr11-69165229-DEL-843 |
| chr11-69170074-DEL-166 | chr11-69170059-DEL-141 |
| chr11-69181770-DEL-393 | chr11-69181771-DEL-393 |
| chr11-69209938-DEL-56 | chr11-69209939-DEL-56 |
| chr11-69227259-DEL-55 | chr11-69227260-DEL-55 |
| chr11-69246759-DEL-170 | chr11-69246762-DEL-171 |
| chr11-69284835-DEL-218 | chr11-69284836-DEL-218 |
| chr11-69293323-DEL-615 | chr11-69293324-DEL-615 |
| chr11-69328996-DEL-54 | chr11-69328997-DEL-62 |
| chr11-69376369-DEL-72 | chr11-69376369-DEL-71 |
| chr11-69379164-DEL-198 | chr11-69379166-DEL-197 |
| chr11-69387761-DEL-90 | chr11-69387762-DEL-90 |
| chr11-69413605-DEL-211 | chr11-69413606-DEL-211 |
| chr11-69432473-DEL-220 | chr11-69432474-DEL-220 |
| chr11-69509736-DEL-1129 | chr11-69509736-DEL-1123 |
| chr11-69515369-DEL-197 | chr11-69515370-DEL-197 |
| chr11-69550726-DEL-769 | chr11-69550746-DEL-752 |
| chr11-69580434-DEL-93 | chr11-69580435-DEL-93 |
| chr11-69593059-DEL-196 | chr11-69593060-DEL-196 |
| chr11-69593576-DEL-221 | chr11-69593577-DEL-221 |
| chr11-69594437-DEL-213 | chr11-69594438-DEL-213 |
| chr11-69595672-DEL-83 | chr11-69595678-DEL-78 |
| chr11-69608642-DEL-238 | chr11-69608643-DEL-238 |
| chr11-69610571-DEL-63 | chr11-69610570-DEL-63 |
| chr11-69614850-DEL-193 | chr11-69614851-DEL-193 |
| chr11-69619097-DEL-201 | chr11-69619098-DEL-201 |
| chr11-69644996-DEL-648 | chr11-69644997-DEL-648 |
| chr11-69654214-DEL-147 | chr11-69654216-DEL-145 |
| chr11-69655750-DEL-400 | chr11-69655779-DEL-368 |
| chr11-69706985-DEL-206 | chr11-69706986-DEL-206 |
| chr11-69751611-DEL-150 | chr11-69751612-DEL-102 |
| chr11-69754901-DEL-144 | chr11-69754902-DEL-144 |
| chr11-69779096-DEL-257 | chr11-69779094-DEL-215 |
| chr11-69861869-DEL-252 | chr11-69861870-DEL-252 |
| chr11-69876792-DEL-215 | chr11-69876800-DEL-208 |
| chr11-69913685-DEL-82 | chr11-69913686-DEL-64 |
| chr11-69931904-DEL-54 | chr11-69931922-DEL-86 |
| chr11-69957336-DEL-99 | chr11-69957304-DEL-94 |

| | |
|---|---|
| chr11-69958355-DEL-80 | chr11-69958356-DEL-80 |
| chr11-69959818-DEL-195 | chr11-69959819-DEL-195 |
| chr11-69978262-DEL-61 | chr11-69978263-DEL-61 |
| chr11-70004017-DEL-129 | chr11-70004018-DEL-129 |
| chr11-70015033-DEL-95 | chr11-70015034-DEL-95 |
| chr11-70018631-DEL-63 | chr11-70018632-DEL-63 |
| chr11-70026844-DEL-53 | chr11-70026890-DEL-53 |
| chr11-70039288-DEL-154 | chr11-70039289-DEL-154 |
| chr11-70046656-DEL-6463 | chr11-70046657-DEL-6463 |
| chr11-70078058-DEL-89 | chr11-70078071-DEL-77 |
| chr11-70102350-DEL-193 | chr11-70102351-DEL-193 |
| chr11-70152308-DEL-52 | chr11-70152309-DEL-52 |
| chr11-70190988-DEL-93 | chr11-70190989-DEL-92 |
| chr11-70198601-DEL-549 | chr11-70198601-DEL-547 |
| chr11-70201713-DEL-213 | chr11-70201714-DEL-213 |
| chr11-70217226-DEL-438 | chr11-70217227-DEL-438 |
| chr11-70240179-DEL-64 | chr11-70240180-DEL-64 |
| chr11-70243174-DEL-56 | chr11-70243175-DEL-50 |
| chr11-70254661-DEL-320 | chr11-70254662-DEL-320 |
| chr11-70299841-DEL-79 | chr11-70299842-DEL-79 |
| chr11-70302059-DEL-151 | chr11-70302060-DEL-151 |
| chr11-70364023-DEL-214 | chr11-70364024-DEL-214 |
| chr11-70365439-DEL-547 | chr11-70365440-DEL-547 |
| chr11-70372610-DEL-206 | chr11-70372611-DEL-206 |
| chr11-70407901-DEL-224 | chr11-70407902-DEL-224 |
| chr11-70419870-DEL-965 | chr11-70419871-DEL-965 |
| chr11-70511518-DEL-199 | chr11-70511519-DEL-199 |
| chr11-70524102-DEL-326 | chr11-70524103-DEL-326 |
| chr11-70527927-DEL-123 | chr11-70527928-DEL-123 |
| chr11-70586933-DEL-112 | chr11-70586961-DEL-102 |
| chr11-70615343-DEL-64 | chr11-70615344-DEL-64 |
| chr11-70629267-DEL-212 | chr11-70629268-DEL-212 |
| chr11-70650444-DEL-75 | chr11-70650445-DEL-74 |
| chr11-70661476-DEL-69 | chr11-70661468-DEL-70 |
| chr11-70663920-DEL-118 | chr11-70663908-DEL-146 |
| chr11-70686804-DEL-919 | chr11-70686805-DEL-919 |
| chr11-70708903-DEL-425 | chr11-70708904-DEL-425 |
| chr11-70721151-DEL-54 | chr11-70721151-DEL-54 |
| chr11-70723738-DEL-200 | chr11-70723739-DEL-200 |
| chr11-70801637-DEL-233 | chr11-70801638-DEL-233 |
| chr11-70827611-DEL-74 | chr11-70827612-DEL-74 |
| chr11-70860172-DEL-66 | chr11-70860165-DEL-74 |
| chr11-70896222-DEL-196 | chr11-70896246-DEL-214 |
| chr11-70931085-DEL-4460 | chr11-70931084-DEL-4460 |
| chr11-70943837-DEL-655 | chr11-70943838-DEL-655 |
| chr11-70949490-DEL-717 | chr11-70949489-DEL-712 |
| chr11-70953579-DEL-989 | chr11-70953580-DEL-989 |
| chr11-70970502-DEL-11404 | chr11-70970502-DEL-11404 |
| chr11-71034945-DEL-2721 | chr11-71034946-DEL-2721 |
| chr11-71049842-DEL-59 | chr11-71049843-DEL-67 |
| chr11-71060123-DEL-238 | chr11-71060122-DEL-196 |
| chr11-71064114-DEL-3807 | chr11-71064114-DEL-3807 |
| chr11-71076835-DEL-170 | chr11-71076833-DEL-170 |
| chr11-71086140-DEL-1151 | chr11-71086141-DEL-1151 |
| chr11-71093183-DEL-68 | chr11-71093172-DEL-80 |
| chr11-71142205-DEL-113 | chr11-71142176-DEL-128 |
| chr11-71160960-DEL-2191 | chr11-71160953-DEL-2181 |
| chr11-71231858-DEL-175 | chr11-71231859-DEL-175 |
| chr11-71237999-DEL-742 | chr11-71238000-DEL-742 |
| chr11-71249694-DEL-268 | chr11-71249694-DEL-268 |
| chr11-71267702-DEL-7984 | chr11-71267703-DEL-7984 |
| chr11-71274832-DEL-208 | chr11-71274833-DEL-208 |
| chr11-71339897-DEL-6173 | chr11-71339898-DEL-6173 |
| chr11-71348784-DEL-68 | chr11-71348785-DEL-68 |
| chr11-71357746-DEL-212 | chr11-71357747-DEL-212 |
| chr11-71390391-DEL-133 | chr11-71390418-DEL-90 |
| chr11-71396562-DEL-6411 | chr11-71396563-DEL-6411 |
| chr11-71408784-DEL-51 | chr11-71408785-DEL-51 |
| chr11-71420687-DEL-716 | chr11-71420698-DEL-719 |
| chr11-71425235-DEL-68 | chr11-71425236-DEL-70 |
| chr11-71449023-DEL-78 | chr11-71449024-DEL-78 |
| chr11-71464667-DEL-270 | chr11-71464668-DEL-270 |
| chr11-71524579-DEL-60 | chr11-71524579-DEL-60 |
| chr11-71549714-DEL-328 | chr11-71549715-DEL-328 |
| chr11-71550705-DEL-147 | chr11-71550706-DEL-143 |
| chr11-71600282-DEL-226 | chr11-71600284-DEL-225 |
| chr11-71610792-DEL-375 | chr11-71610793-DEL-375 |
| chr11-71612326-DEL-246 | chr11-71612327-DEL-246 |
| chr11-71623473-DEL-70 | chr11-71623474-DEL-70 |
| chr11-71648438-DEL-54 | chr11-71648439-DEL-50 |

| | |
|---|---|
| chr11-71668689-DEL-118 | chr11-71668690-DEL-118 |
| chr11-71693438-DEL-86 | chr11-71693439-DEL-86 |
| chr11-71712463-DEL-66 | chr11-71712464-DEL-63 |
| chr11-71736516-DEL-170 | chr11-71736510-DEL-168 |
| chr11-71752290-DEL-453 | chr11-71752289-DEL-452 |
| chr11-71766058-DEL-6399 | chr11-71766059-DEL-6399 |
| chr11-71818790-DEL-343 | chr11-71818791-DEL-343 |
| chr11-71820347-DEL-74 | chr11-71820366-DEL-56 |
| chr11-71833069-DEL-51 | chr11-71833045-DEL-62 |
| chr11-71834881-DEL-58 | chr11-71834882-DEL-58 |
| chr11-71845730-DEL-52 | chr11-71845731-DEL-52 |
| chr11-71854909-DEL-54 | chr11-71854906-DEL-57 |
| chr11-71883339-DEL-106 | chr11-71883361-DEL-84 |
| chr11-71963678-DEL-51 | chr11-71963679-DEL-51 |
| chr11-72062146-DEL-206 | chr11-72062147-DEL-206 |
| chr11-72091274-DEL-64 | chr11-72091258-DEL-64 |
| chr11-72167422-DEL-2881 | chr11-72167423-DEL-2881 |
| chr11-72183492-DEL-3773 | chr11-72183482-DEL-3783 |
| chr11-72217358-DEL-6432 | chr11-72217359-DEL-6432 |
| chr11-72226893-DEL-450 | chr11-72226894-DEL-451 |
| chr11-72233492-DEL-330 | chr11-72233493-DEL-330 |
| chr11-72266441-DEL-120 | chr11-72266442-DEL-120 |
| chr11-72348440-DEL-295 | chr11-72348441-DEL-295 |
| chr11-72367223-DEL-126 | chr11-72367224-DEL-126 |
| chr11-72386874-DEL-55 | chr11-72386874-DEL-55 |
| chr11-72446175-DEL-203 | chr11-72446176-DEL-203 |
| chr11-72450764-DEL-60 | chr11-72450765-DEL-68 |
| chr11-72459370-DEL-201 | chr11-72459348-DEL-202 |
| chr11-72461877-DEL-292 | chr11-72461878-DEL-292 |
| chr11-72469676-DEL-211 | chr11-72469676-DEL-212 |
| chr11-72477830-DEL-166 | chr11-72477816-DEL-167 |
| chr11-72487650-DEL-238 | chr11-72487651-DEL-238 |
| chr11-72532989-DEL-252 | chr11-72532990-DEL-252 |
| chr11-72547697-DEL-214 | chr11-72547712-DEL-202 |
| chr11-72590474-DEL-597 | chr11-72590475-DEL-597 |
| chr11-72617693-DEL-218 | chr11-72617694-DEL-218 |
| chr11-72655968-DEL-1753 | chr11-72655983-DEL-1728 |
| chr11-72662700-DEL-201 | chr11-72662701-DEL-201 |
| chr11-72705501-DEL-80 | chr11-72705502-DEL-80 |
| chr11-72711012-DEL-126 | chr11-72711013-DEL-126 |
| chr11-72724878-DEL-58 | chr11-72724879-DEL-58 |
| chr11-72734164-DEL-295 | chr11-72734165-DEL-295 |
| chr11-72843961-DEL-54 | chr11-72843962-DEL-54 |
| chr11-72884188-DEL-628 | chr11-72884189-DEL-628 |
| chr11-72925143-DEL-772 | chr11-72925148-DEL-772 |
| chr11-72952901-DEL-161 | chr11-72952902-DEL-161 |
| chr11-72954283-DEL-409 | chr11-72954284-DEL-409 |
| chr11-72956567-DEL-758 | chr11-72956553-DEL-759 |
| chr11-72965234-DEL-62 | chr11-72965235-DEL-62 |
| chr11-72974199-DEL-227 | chr11-72974200-DEL-227 |
| chr11-72990062-DEL-68 | chr11-72990073-DEL-79 |
| chr11-73009870-DEL-120 | chr11-73009871-DEL-120 |
| chr11-73017229-DEL-197 | chr11-73017230-DEL-197 |
| chr11-73030132-DEL-599 | chr11-73030100-DEL-600 |
| chr11-73033376-DEL-204 | chr11-73033377-DEL-204 |
| chr11-73152170-DEL-76 | chr11-73152171-DEL-76 |
| chr11-73225734-DEL-70 | chr11-73225745-DEL-78 |
| chr11-73228447-DEL-1251 | chr11-73228448-DEL-1251 |
| chr11-73253041-DEL-2239 | chr11-73253041-DEL-2239 |
| chr11-73299878-DEL-166 | chr11-73299879-DEL-166 |
| chr11-73311549-DEL-75 | chr11-73311550-DEL-59 |
| chr11-73327976-DEL-1897 | chr11-73327977-DEL-1897 |
| chr11-73334749-DEL-1033 | chr11-73334750-DEL-1033 |
| chr11-73382708-DEL-208 | chr11-73382709-DEL-208 |
| chr11-73389472-DEL-768 | chr11-73389472-DEL-768 |
| chr11-73400411-DEL-80 | chr11-73400417-DEL-65 |
| chr11-73421544-DEL-3881 | chr11-73421545-DEL-3881 |
| chr11-73451268-DEL-3044 | chr11-73451269-DEL-3044 |
| chr11-73495514-DEL-52 | chr11-73495515-DEL-52 |
| chr11-73501586-DEL-7420 | chr11-73501587-DEL-7420 |
| chr11-73507822-DEL-208 | chr11-73507823-DEL-208 |
| chr11-73586716-DEL-169 | chr11-73586717-DEL-169 |
| chr11-73593088-DEL-124 | chr11-73593089-DEL-124 |
| chr11-73626777-DEL-4858 | chr11-73626778-DEL-4858 |
| chr11-73699006-DEL-60 | chr11-73699007-DEL-52 |
| chr11-73709115-DEL-2764 | chr11-73709116-DEL-2764 |
| chr11-73717319-DEL-6541 | chr11-73717320-DEL-6541 |
| chr11-73723545-DEL-205 | chr11-73723538-DEL-206 |
| chr11-73756683-DEL-351 | chr11-73756684-DEL-351 |
| chr11-73769147-DEL-3001 | chr11-73769148-DEL-3001 |

| | |
|---|---|
| chr11-73837012-DEL-3861 | chr11-73837013-DEL-3861 |
| chr11-73842721-DEL-176 | chr11-73842717-DEL-177 |
| chr11-74011761-DEL-1350 | chr11-74011762-DEL-1350 |
| chr11-74019727-DEL-7135 | chr11-74019728-DEL-7135 |
| chr11-74055812-DEL-1313 | chr11-74055816-DEL-1312 |
| chr11-74110511-DEL-7056 | chr11-74110512-DEL-7056 |
| chr11-74192663-DEL-13588 | chr11-74192664-DEL-13588 |
| chr11-74249481-DEL-179 | chr11-74249478-DEL-179 |
| chr11-74263870-DEL-90 | chr11-74263856-DEL-105 |
| chr11-74273855-DEL-55 | chr11-74273856-DEL-55 |
| chr11-74310115-DEL-219 | chr11-74310116-DEL-219 |
| chr11-74338298-DEL-54 | chr11-74338299-DEL-64 |
| chr11-74353453-DEL-254 | chr11-74353454-DEL-254 |
| chr11-74362129-DEL-890 | chr11-74362130-DEL-890 |
| chr11-74362267-DEL-100 | chr11-74362268-DEL-100 |
| chr11-74399623-DEL-944 | chr11-74399613-DEL-945 |
| chr11-74416948-DEL-114 | chr11-74416949-DEL-118 |
| chr11-74429127-DEL-51 | chr11-74429092-DEL-60 |
| chr11-74431000-DEL-5225 | chr11-74430997-DEL-5226 |
| chr11-74463998-DEL-55 | chr11-74463999-DEL-55 |
| chr11-74466346-DEL-82 | chr11-74466347-DEL-82 |
| chr11-74472454-DEL-218 | chr11-74472434-DEL-219 |
| chr11-74486660-DEL-105 | chr11-74486661-DEL-105 |
| chr11-74491585-DEL-155 | chr11-74491586-DEL-155 |
| chr11-74497710-DEL-179 | chr11-74497711-DEL-179 |
| chr11-74503661-DEL-74 | chr11-74503662-DEL-74 |
| chr11-74509156-DEL-184 | chr11-74509157-DEL-184 |
| chr11-74526163-DEL-198 | chr11-74526164-DEL-198 |
| chr11-74527595-DEL-178 | chr11-74527604-DEL-174 |
| chr11-74578660-DEL-210 | chr11-74578661-DEL-210 |
| chr11-74597148-DEL-181 | chr11-74597149-DEL-180 |
| chr11-74600693-DEL-225 | chr11-74600683-DEL-226 |
| chr11-74607656-DEL-67 | chr11-74607657-DEL-67 |
| chr11-74608856-DEL-249 | chr11-74608857-DEL-249 |
| chr11-74650740-DEL-80 | chr11-74650761-DEL-74 |
| chr11-74685182-DEL-209 | chr11-74685183-DEL-209 |
| chr11-74771536-DEL-189 | chr11-74771537-DEL-189 |
| chr11-74775230-DEL-7099 | chr11-74775231-DEL-7099 |
| chr11-74787575-DEL-66 | chr11-74787576-DEL-64 |
| chr11-74823814-DEL-53 | chr11-74823815-DEL-53 |
| chr11-74846594-DEL-173 | chr11-74846595-DEL-173 |
| chr11-74858648-DEL-718 | chr11-74858651-DEL-716 |
| chr11-74871797-DEL-159 | chr11-74871798-DEL-159 |
| chr11-74877124-DEL-206 | chr11-74877125-DEL-206 |
| chr11-74966749-DEL-166 | chr11-74966748-DEL-166 |
| chr11-74974759-DEL-174 | chr11-74974755-DEL-173 |
| chr11-74999014-DEL-219 | chr11-74999015-DEL-219 |
| chr11-75004844-DEL-227 | chr11-75004845-DEL-227 |
| chr11-75013951-DEL-195 | chr11-75013945-DEL-196 |
| chr11-75104073-DEL-51 | chr11-75104054-DEL-71 |
| chr11-75113093-DEL-206 | chr11-75113094-DEL-206 |
| chr11-75144425-DEL-209 | chr11-75144426-DEL-209 |
| chr11-75146788-DEL-301 | chr11-75146789-DEL-300 |
| chr11-75164885-DEL-208 | chr11-75164885-DEL-208 |
| chr11-75175034-DEL-81 | chr11-75175035-DEL-81 |
| chr11-75194717-DEL-207 | chr11-75194718-DEL-207 |
| chr11-75247658-DEL-8803 | chr11-75247659-DEL-8802 |
| chr11-75248792-DEL-140 | chr11-75248793-DEL-140 |
| chr11-75261159-DEL-99 | chr11-75261160-DEL-99 |
| chr11-75271144-DEL-155 | chr11-75271140-DEL-151 |
| chr11-75274450-DEL-80 | chr11-75274410-DEL-80 |
| chr11-75276911-DEL-78 | chr11-75276897-DEL-59 |
| chr11-75296855-DEL-102 | chr11-75296871-DEL-136 |
| chr11-75317661-DEL-165 | chr11-75317662-DEL-165 |
| chr11-75330701-DEL-78 | chr11-75330708-DEL-72 |
| chr11-75360925-DEL-401 | chr11-75360926-DEL-401 |
| chr11-75369367-DEL-204 | chr11-75369368-DEL-204 |
| chr11-75409265-DEL-130 | chr11-75409264-DEL-134 |
| chr11-75427933-DEL-93 | chr11-75427917-DEL-93 |
| chr11-75430540-DEL-201 | chr11-75430541-DEL-201 |
| chr11-75484898-DEL-181 | chr11-75484899-DEL-181 |
| chr11-75488300-DEL-469 | chr11-75488301-DEL-469 |
| chr11-75490447-DEL-427 | chr11-75490448-DEL-427 |
| chr11-75509175-DEL-225 | chr11-75509176-DEL-225 |
| chr11-75537054-DEL-208 | chr11-75537055-DEL-208 |
| chr11-75582434-DEL-216 | chr11-75582431-DEL-217 |
| chr11-75587183-DEL-105 | chr11-75587184-DEL-105 |
| chr11-75602916-DEL-200 | chr11-75602917-DEL-200 |
| chr11-75603986-DEL-155 | chr11-75603987-DEL-155 |
| chr11-75616029-DEL-193 | chr11-75616030-DEL-193 |

| | |
|---|---|
| chr11-75618518-DEL-1232 | chr11-75618524-DEL-1228 |
| chr11-75636516-DEL-210 | chr11-75636517-DEL-210 |
| chr11-75678564-DEL-219 | chr11-75678565-DEL-219 |
| chr11-75681859-DEL-54 | chr11-75681862-DEL-60 |
| chr11-75707623-DEL-74 | chr11-75707648-DEL-53 |
| chr11-75734956-DEL-56 | chr11-75734953-DEL-50 |
| chr11-75776534-DEL-363 | chr11-75776535-DEL-363 |
| chr11-75824076-DEL-53 | chr11-75824077-DEL-53 |
| chr11-75896996-DEL-208 | chr11-75896997-DEL-208 |
| chr11-75898385-DEL-77 | chr11-75898386-DEL-77 |
| chr11-75913225-DEL-200 | chr11-75913226-DEL-200 |
| chr11-75920818-DEL-56 | chr11-75920811-DEL-64 |
| chr11-75927647-DEL-70 | chr11-75927648-DEL-70 |
| chr11-75946365-DEL-200 | chr11-75946366-DEL-200 |
| chr11-76001719-DEL-209 | chr11-76001722-DEL-206 |
| chr11-76036596-DEL-219 | chr11-76036597-DEL-219 |
| chr11-76040647-DEL-194 | chr11-76040648-DEL-194 |
| chr11-76085281-DEL-180 | chr11-76085282-DEL-180 |
| chr11-76120213-DEL-206 | chr11-76120214-DEL-206 |
| chr11-76172679-DEL-222 | chr11-76172680-DEL-222 |
| chr11-76187324-DEL-351 | chr11-76187349-DEL-323 |
| chr11-76214646-DEL-73 | chr11-76214651-DEL-72 |
| chr11-76236826-DEL-220 | chr11-76236827-DEL-220 |
| chr11-76247242-DEL-178 | chr11-76247243-DEL-178 |
| chr11-76282472-DEL-213 | chr11-76282473-DEL-213 |
| chr11-76298828-DEL-103 | chr11-76298827-DEL-115 |
| chr11-76302238-DEL-221 | chr11-76302239-DEL-221 |
| chr11-76366829-DEL-218 | chr11-76366830-DEL-217 |
| chr11-76402291-DEL-101 | chr11-76402299-DEL-100 |
| chr11-76415378-DEL-90 | chr11-76415379-DEL-90 |
| chr11-76417280-DEL-2068 | chr11-76417275-DEL-2066 |
| chr11-76439631-DEL-9049 | chr11-76439632-DEL-9049 |
| chr11-76450441-DEL-97 | chr11-76450412-DEL-78 |
| chr11-76473278-DEL-94 | chr11-76473279-DEL-94 |
| chr11-76492092-DEL-411 | chr11-76492093-DEL-411 |
| chr11-76511304-DEL-64 | chr11-76511305-DEL-64 |
| chr11-76534307-DEL-82 | chr11-76534312-DEL-80 |
| chr11-76538964-DEL-89 | chr11-76538965-DEL-89 |
| chr11-76543374-DEL-68 | chr11-76543375-DEL-68 |
| chr11-76553174-DEL-144 | chr11-76553175-DEL-144 |
| chr11-76559097-DEL-63 | chr11-76559098-DEL-63 |
| chr11-76570296-DEL-66 | chr11-76570297-DEL-54 |
| chr11-76575390-DEL-224 | chr11-76575391-DEL-224 |
| chr11-76583321-DEL-103 | chr11-76583321-DEL-103 |
| chr11-76590314-DEL-74 | chr11-76590315-DEL-74 |
| chr11-76622445-DEL-76 | chr11-76622446-DEL-76 |
| chr11-76638338-DEL-244 | chr11-76638339-DEL-244 |
| chr11-76662107-DEL-365 | chr11-76662108-DEL-365 |
| chr11-76663115-DEL-60 | chr11-76663116-DEL-60 |
| chr11-76687795-DEL-144 | chr11-76687791-DEL-144 |
| chr11-76718945-DEL-6441 | chr11-76718946-DEL-6441 |
| chr11-76753126-DEL-6390 | chr11-76753126-DEL-6382 |
| chr11-76766354-DEL-417 | chr11-76766355-DEL-417 |
| chr11-76798892-DEL-62 | chr11-76798890-DEL-61 |
| chr11-76822955-DEL-160 | chr11-76822956-DEL-160 |
| chr11-76858860-DEL-56 | chr11-76858861-DEL-56 |
| chr11-76885091-DEL-66 | chr11-76885091-DEL-66 |
| chr11-76898619-DEL-82 | chr11-76898619-DEL-81 |
| chr11-76911586-DEL-92 | chr11-76911587-DEL-92 |
| chr11-76999505-DEL-119 | chr11-76999506-DEL-119 |
| chr11-77050483-DEL-143 | chr11-77050482-DEL-145 |
| chr11-77060830-DEL-55 | chr11-77060832-DEL-50 |
| chr11-77110305-DEL-8605 | chr11-77110306-DEL-8604 |
| chr11-77136810-DEL-202 | chr11-77136811-DEL-202 |
| chr11-77138736-DEL-176 | chr11-77138737-DEL-176 |
| chr11-77145971-DEL-1916 | chr11-77145973-DEL-1897 |
| chr11-77158815-DEL-443 | chr11-77158816-DEL-443 |
| chr11-77175996-DEL-103 | chr11-77175996-DEL-104 |
| chr11-77184251-DEL-2999 | chr11-77184252-DEL-2999 |
| chr11-77207624-DEL-56 | chr11-77207625-DEL-57 |
| chr11-77259350-DEL-104 | chr11-77259351-DEL-104 |
| chr11-77261563-DEL-91 | chr11-77261564-DEL-91 |
| chr11-77324340-DEL-170 | chr11-77324341-DEL-170 |
| chr11-77339232-DEL-110 | chr11-77339233-DEL-110 |
| chr11-77352471-DEL-72 | chr11-77352472-DEL-72 |
| chr11-77365492-DEL-400 | chr11-77365493-DEL-400 |
| chr11-77401866-DEL-216 | chr11-77401867-DEL-216 |
| chr11-77431327-DEL-56 | chr11-77431336-DEL-55 |
| chr11-77458442-DEL-200 | chr11-77458443-DEL-200 |
| chr11-77512692-DEL-190 | chr11-77512693-DEL-190 |

| | |
|---|---|
| chr11-77530057-DEL-194 | chr11-77530058-DEL-194 |
| chr11-77530891-DEL-220 | chr11-77530906-DEL-200 |
| chr11-77532185-DEL-61 | chr11-77532185-DEL-61 |
| chr11-77542024-DEL-180 | chr11-77542025-DEL-180 |
| chr11-77543219-DEL-229 | chr11-77543220-DEL-229 |
| chr11-77545475-DEL-131 | chr11-77545468-DEL-138 |
| chr11-77559215-DEL-319 | chr11-77559216-DEL-319 |
| chr11-77586805-DEL-218 | chr11-77586803-DEL-219 |
| chr11-77587351-DEL-218 | chr11-77587352-DEL-218 |
| chr11-77588263-DEL-982 | chr11-77588261-DEL-983 |
| chr11-77593271-DEL-346 | chr11-77593272-DEL-346 |
| chr11-77599160-DEL-177 | chr11-77599161-DEL-177 |
| chr11-77623057-DEL-404 | chr11-77623058-DEL-404 |
| chr11-77626031-DEL-223 | chr11-77626032-DEL-223 |
| chr11-77634414-DEL-125 | chr11-77634414-DEL-131 |
| chr11-77637922-DEL-174 | chr11-77637923-DEL-174 |
| chr11-77647284-DEL-215 | chr11-77647285-DEL-215 |
| chr11-77648162-DEL-202 | chr11-77648163-DEL-202 |
| chr11-77679010-DEL-150 | chr11-77679011-DEL-150 |
| chr11-77680127-DEL-213 | chr11-77680128-DEL-213 |
| chr11-77703988-DEL-183 | chr11-77703989-DEL-183 |
| chr11-77728620-DEL-77 | chr11-77728612-DEL-62 |
| chr11-77740407-DEL-122 | chr11-77740408-DEL-131 |
| chr11-77765086-DEL-155 | chr11-77765087-DEL-155 |
| chr11-77832295-DEL-100 | chr11-77832296-DEL-75 |
| chr11-77843498-DEL-87 | chr11-77843484-DEL-102 |
| chr11-77857807-DEL-176 | chr11-77857808-DEL-176 |
| chr11-77895662-DEL-214 | chr11-77895663-DEL-214 |
| chr11-77915991-DEL-213 | chr11-77915992-DEL-213 |
| chr11-77926724-DEL-86 | chr11-77926725-DEL-86 |
| chr11-77969452-DEL-199 | chr11-77969453-DEL-198 |
| chr11-77972404-DEL-228 | chr11-77972405-DEL-228 |
| chr11-77984763-DEL-198 | chr11-77984764-DEL-198 |
| chr11-78002962-DEL-58 | chr11-78002963-DEL-58 |
| chr11-78028048-DEL-206 | chr11-78028049-DEL-206 |
| chr11-78033572-DEL-210 | chr11-78033573-DEL-210 |
| chr11-78066474-DEL-184 | chr11-78066475-DEL-184 |
| chr11-78086130-DEL-64 | chr11-78086132-DEL-62 |
| chr11-78129835-DEL-222 | chr11-78129836-DEL-215 |
| chr11-78143667-DEL-221 | chr11-78143668-DEL-221 |
| chr11-78167501-DEL-99 | chr11-78167502-DEL-99 |
| chr11-78179364-DEL-211 | chr11-78179365-DEL-211 |
| chr11-78202931-DEL-208 | chr11-78202931-DEL-208 |
| chr11-78218674-DEL-214 | chr11-78218675-DEL-214 |
| chr11-78220832-DEL-498 | chr11-78220833-DEL-500 |
| chr11-78222887-DEL-630 | chr11-78222888-DEL-630 |
| chr11-78245060-DEL-207 | chr11-78245061-DEL-207 |
| chr11-78270618-DEL-75 | chr11-78270619-DEL-75 |
| chr11-78281891-DEL-191 | chr11-78281892-DEL-191 |
| chr11-78288849-DEL-54 | chr11-78288850-DEL-54 |
| chr11-78338896-DEL-194 | chr11-78338897-DEL-194 |
| chr11-78340035-DEL-73 | chr11-78340036-DEL-71 |
| chr11-78340425-DEL-188 | chr11-78340426-DEL-188 |
| chr11-78350527-DEL-578 | chr11-78350536-DEL-552 |
| chr11-78382951-DEL-199 | chr11-78382952-DEL-199 |
| chr11-78419457-DEL-145 | chr11-78419424-DEL-179 |
| chr11-78422318-DEL-153 | chr11-78422319-DEL-153 |
| chr11-78423444-DEL-1459 | chr11-78423445-DEL-1459 |
| chr11-78456063-DEL-210 | chr11-78456064-DEL-210 |
| chr11-78465068-DEL-245 | chr11-78465069-DEL-245 |
| chr11-78514407-DEL-710 | chr11-78514406-DEL-711 |
| chr11-78562132-DEL-204 | chr11-78562133-DEL-204 |
| chr11-78595535-DEL-114 | chr11-78595537-DEL-113 |
| chr11-78616330-DEL-206 | chr11-78616331-DEL-206 |
| chr11-78634764-DEL-171 | chr11-78634781-DEL-154 |
| chr11-78635641-DEL-207 | chr11-78635642-DEL-207 |
| chr11-78670137-DEL-79 | chr11-78670143-DEL-80 |
| chr11-78688312-DEL-136 | chr11-78688313-DEL-136 |
| chr11-78790270-DEL-54 | chr11-78790275-DEL-50 |
| chr11-78797431-DEL-195 | chr11-78797432-DEL-195 |
| chr11-78804291-DEL-206 | chr11-78804292-DEL-206 |
| chr11-78836990-DEL-79 | chr11-78836991-DEL-79 |
| chr11-78853120-DEL-68 | chr11-78853120-DEL-68 |
| chr11-78869822-DEL-174 | chr11-78869824-DEL-176 |
| chr11-78962757-DEL-100 | chr11-78962758-DEL-100 |
| chr11-78994383-DEL-551 | chr11-78994384-DEL-551 |
| chr11-79015633-DEL-260 | chr11-79015634-DEL-256 |
| chr11-79038887-DEL-208 | chr11-79038888-DEL-208 |
| chr11-79059727-DEL-662 | chr11-79059722-DEL-653 |
| chr11-79064355-DEL-154 | chr11-79064356-DEL-132 |

| | |
|---|---|
| chr11-79077507-DEL-222 | chr11-79077496-DEL-234 |
| chr11-79096678-DEL-70 | chr11-79096679-DEL-68 |
| chr11-79099008-DEL-141 | chr11-79099009-DEL-117 |
| chr11-79123493-DEL-4437 | chr11-79123494-DEL-4437 |
| chr11-79141375-DEL-351 | chr11-79141376-DEL-351 |
| chr11-79151153-DEL-179 | chr11-79151154-DEL-179 |
| chr11-79162950-DEL-206 | chr11-79162950-DEL-207 |
| chr11-79175666-DEL-60 | chr11-79175667-DEL-60 |
| chr11-79187434-DEL-179 | chr11-79187435-DEL-179 |
| chr11-79203954-DEL-199 | chr11-79203955-DEL-199 |
| chr11-79217531-DEL-201 | chr11-79217530-DEL-202 |
| chr11-79243185-DEL-204 | chr11-79243186-DEL-204 |
| chr11-79246941-DEL-173 | chr11-79246942-DEL-173 |
| chr11-79251462-DEL-64 | chr11-79251463-DEL-64 |
| chr11-79253270-DEL-114 | chr11-79253271-DEL-114 |
| chr11-79259578-DEL-59 | chr11-79259579-DEL-59 |
| chr11-79263066-DEL-129 | chr11-79263067-DEL-129 |
| chr11-79266982-DEL-170 | chr11-79266967-DEL-182 |
| chr11-79276454-DEL-207 | chr11-79276455-DEL-207 |
| chr11-79322364-DEL-83 | chr11-79322370-DEL-81 |
| chr11-79346571-DEL-70 | chr11-79346570-DEL-60 |
| chr11-79423521-DEL-632 | chr11-79423520-DEL-628 |
| chr11-79515967-DEL-94 | chr11-79515970-DEL-92 |
| chr11-79521069-DEL-85 | chr11-79521069-DEL-85 |
| chr11-79528125-DEL-60 | chr11-79528078-DEL-66 |
| chr11-79589583-DEL-205 | chr11-79589584-DEL-205 |
| chr11-79631286-DEL-157 | chr11-79631287-DEL-157 |
| chr11-79632311-DEL-4394 | chr11-79632312-DEL-4394 |
| chr11-79659430-DEL-199 | chr11-79659431-DEL-199 |
| chr11-79663267-DEL-388 | chr11-79663268-DEL-388 |
| chr11-79687870-DEL-63 | chr11-79687872-DEL-62 |
| chr11-79700836-DEL-83 | chr11-79700837-DEL-83 |
| chr11-79710916-DEL-593 | chr11-79710917-DEL-593 |
| chr11-79723885-DEL-62 | chr11-79723886-DEL-62 |
| chr11-79736089-DEL-1893 | chr11-79736090-DEL-1893 |
| chr11-79744153-DEL-99 | chr11-79744154-DEL-99 |
| chr11-79763482-DEL-4393 | chr11-79763489-DEL-4397 |
| chr11-79808103-DEL-1888 | chr11-79808104-DEL-1888 |
| chr11-79828818-DEL-195 | chr11-79828819-DEL-195 |
| chr11-79860908-DEL-181 | chr11-79860909-DEL-181 |
| chr11-79878568-DEL-265 | chr11-79878560-DEL-266 |
| chr11-79879857-DEL-204 | chr11-79879858-DEL-204 |
| chr11-79898615-DEL-216 | chr11-79898616-DEL-216 |
| chr11-79902148-DEL-139 | chr11-79902155-DEL-147 |
| chr11-79903310-DEL-407 | chr11-79903311-DEL-407 |
| chr11-79907577-DEL-218 | chr11-79907588-DEL-216 |
| chr11-79909756-DEL-471 | chr11-79909757-DEL-471 |
| chr11-79930254-DEL-178 | chr11-79930254-DEL-179 |
| chr11-79930906-DEL-174 | chr11-79930907-DEL-174 |
| chr11-79960251-DEL-188 | chr11-79960252-DEL-188 |
| chr11-79983658-DEL-211 | chr11-79983659-DEL-211 |
| chr11-79990885-DEL-192 | chr11-79990883-DEL-191 |
| chr11-79994656-DEL-74 | chr11-79994657-DEL-74 |
| chr11-80005526-DEL-181 | chr11-80005519-DEL-182 |
| chr11-80015022-DEL-140 | chr11-80015023-DEL-140 |
| chr11-80017684-DEL-228 | chr11-80017685-DEL-228 |
| chr11-80019020-DEL-232 | chr11-80019018-DEL-233 |
| chr11-80021455-DEL-208 | chr11-80021456-DEL-208 |
| chr11-80029142-DEL-186 | chr11-80029143-DEL-186 |
| chr11-80038974-DEL-222 | chr11-80038975-DEL-222 |
| chr11-80040042-DEL-182 | chr11-80040037-DEL-188 |
| chr11-80041647-DEL-213 | chr11-80041648-DEL-213 |
| chr11-80042973-DEL-51 | chr11-80042958-DEL-52 |
| chr11-80071062-DEL-174 | chr11-80071063-DEL-174 |
| chr11-80078071-DEL-1583 | chr11-80078072-DEL-1583 |
| chr11-80109293-DEL-191 | chr11-80109294-DEL-191 |
| chr11-80121126-DEL-191 | chr11-80121127-DEL-193 |
| chr11-80135341-DEL-54 | chr11-80135342-DEL-54 |
| chr11-80144829-DEL-222 | chr11-80144830-DEL-222 |
| chr11-80154319-DEL-68 | chr11-80154320-DEL-68 |
| chr11-80159749-DEL-155 | chr11-80159763-DEL-135 |
| chr11-80218899-DEL-54 | chr11-80218899-DEL-62 |
| chr11-80269982-DEL-188 | chr11-80269983-DEL-188 |
| chr11-80347763-DEL-80 | chr11-80347764-DEL-80 |
| chr11-80458343-DEL-55 | chr11-80458343-DEL-55 |
| chr11-80464848-DEL-53 | chr11-80464849-DEL-53 |
| chr11-80487986-DEL-120 | chr11-80488026-DEL-92 |
| chr11-80499805-DEL-220 | chr11-80499803-DEL-221 |
| chr11-80503339-DEL-739 | chr11-80503340-DEL-739 |
| chr11-80514952-DEL-224 | chr11-80514953-DEL-224 |

| | |
|---|---|
| chr11-80533833-DEL-64 | chr11-80533834-DEL-64 |
| chr11-80559904-DEL-558 | chr11-80559905-DEL-559 |
| chr11-80627919-DEL-2172 | chr11-80627918-DEL-2173 |
| chr11-80723854-DEL-1517 | chr11-80723855-DEL-1517 |
| chr11-80727306-DEL-823 | chr11-80727307-DEL-823 |
| chr11-80730745-DEL-152 | chr11-80730745-DEL-110 |
| chr11-80737555-DEL-68 | chr11-80737556-DEL-68 |
| chr11-80806709-DEL-245 | chr11-80806710-DEL-245 |
| chr11-80808476-DEL-95 | chr11-80808477-DEL-95 |
| chr11-80811500-DEL-104 | chr11-80811478-DEL-98 |
| chr11-80830267-DEL-89 | chr11-80830268-DEL-89 |
| chr11-80847556-DEL-667 | chr11-80847555-DEL-668 |
| chr11-80863039-DEL-265 | chr11-80863040-DEL-265 |
| chr11-80913615-DEL-2224 | chr11-80913616-DEL-2224 |
| chr11-80966052-DEL-237 | chr11-80966053-DEL-237 |
| chr11-80975553-DEL-172 | chr11-80975554-DEL-172 |
| chr11-81020784-DEL-331 | chr11-81020785-DEL-331 |
| chr11-81061925-DEL-85 | chr11-81061926-DEL-85 |
| chr11-81086391-DEL-187 | chr11-81086390-DEL-182 |
| chr11-81093955-DEL-56 | chr11-81093956-DEL-56 |
| chr11-81129485-DEL-76 | chr11-81129486-DEL-76 |
| chr11-81132165-DEL-318 | chr11-81132166-DEL-318 |
| chr11-81238998-DEL-144 | chr11-81238999-DEL-144 |
| chr11-81279203-DEL-84 | chr11-81279204-DEL-84 |
| chr11-81290548-DEL-102 | chr11-81290549-DEL-102 |
| chr11-81313888-DEL-78 | chr11-81313889-DEL-78 |
| chr11-81321694-DEL-6414 | chr11-81321695-DEL-6414 |
| chr11-81342493-DEL-118 | chr11-81342498-DEL-124 |
| chr11-81356091-DEL-67 | chr11-81356092-DEL-67 |
| chr11-81373123-DEL-87 | chr11-81373127-DEL-84 |
| chr11-81385791-DEL-622 | chr11-81385792-DEL-623 |
| chr11-81419007-DEL-2239 | chr11-81419008-DEL-2239 |
| chr11-81422301-DEL-74 | chr11-81422292-DEL-94 |
| chr11-81423981-DEL-94 | chr11-81423979-DEL-94 |
| chr11-81436299-DEL-7217 | chr11-81436300-DEL-7217 |
| chr11-81450895-DEL-175 | chr11-81450938-DEL-173 |
| chr11-81458083-DEL-119 | chr11-81458084-DEL-119 |
| chr11-81459868-DEL-801 | chr11-81459869-DEL-801 |
| chr11-81520302-DEL-246 | chr11-81520303-DEL-246 |
| chr11-81558774-DEL-365 | chr11-81558777-DEL-362 |
| chr11-81629925-DEL-55 | chr11-81629926-DEL-55 |
| chr11-81668634-DEL-139 | chr11-81668638-DEL-127 |
| chr11-81686696-DEL-105 | chr11-81686702-DEL-97 |
| chr11-81749558-DEL-54 | chr11-81749558-DEL-55 |
| chr11-81785954-DEL-1079 | chr11-81785954-DEL-1079 |
| chr11-81843209-DEL-62 | chr11-81843186-DEL-50 |
| chr11-81853427-DEL-55 | chr11-81853427-DEL-56 |
| chr11-81854616-DEL-92 | chr11-81854617-DEL-92 |
| chr11-81908344-DEL-677 | chr11-81908345-DEL-678 |
| chr11-81972996-DEL-66 | chr11-81973000-DEL-56 |
| chr11-81975058-DEL-66 | chr11-81975089-DEL-61 |
| chr11-82019117-DEL-603 | chr11-82019116-DEL-603 |
| chr11-82070075-DEL-66 | chr11-82070040-DEL-102 |
| chr11-82115899-DEL-57 | chr11-82115907-DEL-51 |
| chr11-82169805-DEL-72 | chr11-82169806-DEL-72 |
| chr11-82193397-DEL-53 | chr11-82193409-DEL-78 |
| chr11-82224837-DEL-4317 | chr11-82224838-DEL-4317 |
| chr11-82304143-DEL-85 | chr11-82304144-DEL-85 |
| chr11-82359466-DEL-400 | chr11-82359467-DEL-400 |
| chr11-82360049-DEL-91 | chr11-82360067-DEL-108 |
| chr11-82368611-DEL-78 | chr11-82368612-DEL-78 |
| chr11-82393025-DEL-1177 | chr11-82393026-DEL-1177 |
| chr11-82421731-DEL-601 | chr11-82421732-DEL-601 |
| chr11-82435784-DEL-88 | chr11-82435766-DEL-89 |
| chr11-82443715-DEL-174 | chr11-82443693-DEL-197 |
| chr11-82472567-DEL-451 | chr11-82472568-DEL-451 |
| chr11-82476049-DEL-145 | chr11-82476050-DEL-145 |
| chr11-82507090-DEL-53 | chr11-82507093-DEL-51 |
| chr11-82559450-DEL-203 | chr11-82559453-DEL-201 |
| chr11-82605051-DEL-195 | chr11-82605052-DEL-195 |
| chr11-82631067-DEL-200 | chr11-82631068-DEL-200 |
| chr11-82639107-DEL-457 | chr11-82639108-DEL-457 |
| chr11-82649829-DEL-79 | chr11-82649834-DEL-68 |
| chr11-82707777-DEL-288 | chr11-82707778-DEL-288 |
| chr11-82771152-DEL-372 | chr11-82771144-DEL-373 |
| chr11-82772202-DEL-84 | chr11-82772203-DEL-84 |
| chr11-82781626-DEL-51 | chr11-82781627-DEL-51 |
| chr11-82787509-DEL-694 | chr11-82787509-DEL-695 |
| chr11-82826235-DEL-53 | chr11-82826230-DEL-59 |
| chr11-82854588-DEL-234 | chr11-82854589-DEL-234 |

| | |
|---|---|
| chr11-82859261-DEL-268 | chr11-82859262-DEL-268 |
| chr11-82865500-DEL-107 | chr11-82865499-DEL-153 |
| chr11-82883056-DEL-57 | chr11-82883057-DEL-57 |
| chr11-82898677-DEL-122 | chr11-82898677-DEL-122 |
| chr11-82900554-DEL-340 | chr11-82900554-DEL-340 |
| chr11-82906284-DEL-974 | chr11-82906285-DEL-974 |
| chr11-82913868-DEL-99 | chr11-82913853-DEL-96 |
| chr11-82918244-DEL-609 | chr11-82918245-DEL-608 |
| chr11-82930610-DEL-77 | chr11-82930611-DEL-77 |
| chr11-82941203-DEL-6401 | chr11-82941204-DEL-6401 |
| chr11-82951072-DEL-76 | chr11-82951073-DEL-76 |
| chr11-82961828-DEL-5940 | chr11-82961827-DEL-5940 |
| chr11-82979239-DEL-506 | chr11-82979240-DEL-507 |
| chr11-82985728-DEL-211 | chr11-82985728-DEL-211 |
| chr11-83085988-DEL-223 | chr11-83085989-DEL-223 |
| chr11-83123045-DEL-5447 | chr11-83123037-DEL-5448 |
| chr11-83151683-DEL-87 | chr11-83151684-DEL-87 |
| chr11-83156779-DEL-213 | chr11-83156773-DEL-214 |
| chr11-83207727-DEL-76 | chr11-83207728-DEL-76 |
| chr11-83209783-DEL-194 | chr11-83209784-DEL-194 |
| chr11-83229240-DEL-67 | chr11-83229262-DEL-63 |
| chr11-83235368-DEL-58 | chr11-83235364-DEL-59 |
| chr11-83239424-DEL-203 | chr11-83239425-DEL-203 |
| chr11-83265836-DEL-207 | chr11-83265837-DEL-207 |
| chr11-83314270-DEL-88 | chr11-83314245-DEL-80 |
| chr11-83360448-DEL-877 | chr11-83360449-DEL-877 |
| chr11-83418899-DEL-95 | chr11-83418911-DEL-95 |
| chr11-83437326-DEL-1389 | chr11-83437324-DEL-1409 |
| chr11-83442944-DEL-382 | chr11-83442945-DEL-382 |
| chr11-83446105-DEL-822 | chr11-83446102-DEL-826 |
| chr11-83450021-DEL-124 | chr11-83450022-DEL-124 |
| chr11-83456701-DEL-794 | chr11-83456702-DEL-794 |
| chr11-83487517-DEL-170 | chr11-83487518-DEL-175 |
| chr11-83519246-DEL-212 | chr11-83519247-DEL-212 |
| chr11-83585462-DEL-217 | chr11-83585463-DEL-217 |
| chr11-83588409-DEL-3211 | chr11-83588409-DEL-3211 |
| chr11-83612020-DEL-67 | chr11-83612021-DEL-67 |
| chr11-83614445-DEL-89 | chr11-83614446-DEL-89 |
| chr11-83615210-DEL-1974 | chr11-83615211-DEL-1974 |
| chr11-83634814-DEL-59 | chr11-83634815-DEL-59 |
| chr11-83655157-DEL-304 | chr11-83655158-DEL-304 |
| chr11-83680787-DEL-103 | chr11-83680788-DEL-103 |
| chr11-83681852-DEL-60 | chr11-83681851-DEL-54 |
| chr11-83713843-DEL-56 | chr11-83713835-DEL-54 |
| chr11-83736257-DEL-178 | chr11-83736296-DEL-186 |
| chr11-83739826-DEL-59 | chr11-83739843-DEL-51 |
| chr11-83811690-DEL-147 | chr11-83811691-DEL-147 |
| chr11-83818180-DEL-765 | chr11-83818181-DEL-765 |
| chr11-83826060-DEL-67 | chr11-83826061-DEL-67 |
| chr11-83840942-DEL-132 | chr11-83840942-DEL-132 |
| chr11-83841522-DEL-205 | chr11-83841523-DEL-205 |
| chr11-83926669-DEL-565 | chr11-83926670-DEL-565 |
| chr11-83955775-DEL-203 | chr11-83955776-DEL-203 |
| chr11-83981199-DEL-72 | chr11-83981225-DEL-80 |
| chr11-84008752-DEL-52 | chr11-84008753-DEL-52 |
| chr11-84033615-DEL-1459 | chr11-84033616-DEL-1459 |
| chr11-84034360-DEL-51 | chr11-84034404-DEL-50 |
| chr11-84040065-DEL-71 | chr11-84040066-DEL-71 |
| chr11-84052572-DEL-193 | chr11-84052565-DEL-196 |
| chr11-84078039-DEL-200 | chr11-84078040-DEL-200 |
| chr11-84101664-DEL-964 | chr11-84101665-DEL-964 |
| chr11-84113186-DEL-52 | chr11-84113187-DEL-52 |
| chr11-84117779-DEL-232 | chr11-84117780-DEL-232 |
| chr11-84123324-DEL-55 | chr11-84123355-DEL-62 |
| chr11-84126362-DEL-400 | chr11-84126363-DEL-400 |
| chr11-84130485-DEL-773 | chr11-84130486-DEL-773 |
| chr11-84147069-DEL-62 | chr11-84147074-DEL-56 |
| chr11-84198399-DEL-2993 | chr11-84198400-DEL-2993 |
| chr11-84202590-DEL-214 | chr11-84202591-DEL-214 |
| chr11-84298322-DEL-221 | chr11-84298323-DEL-221 |
| chr11-84308489-DEL-210 | chr11-84308490-DEL-210 |
| chr11-84309887-DEL-202 | chr11-84309888-DEL-202 |
| chr11-84360291-DEL-108 | chr11-84360292-DEL-108 |
| chr11-84380581-DEL-513 | chr11-84380581-DEL-513 |
| chr11-84399802-DEL-366 | chr11-84399803-DEL-366 |
| chr11-84401653-DEL-235 | chr11-84401654-DEL-235 |
| chr11-84411719-DEL-252 | chr11-84411720-DEL-252 |
| chr11-84433497-DEL-83 | chr11-84433498-DEL-82 |
| chr11-84505633-DEL-54 | chr11-84505636-DEL-58 |
| chr11-84544084-DEL-72 | chr11-84544085-DEL-66 |

| | |
|---|---|
| chr11-84568692-DEL-187 | chr11-84568655-DEL-204 |
| chr11-84576170-DEL-59 | chr11-84576171-DEL-59 |
| chr11-84583307-DEL-3647 | chr11-84583308-DEL-3647 |
| chr11-84590400-DEL-179 | chr11-84590401-DEL-179 |
| chr11-84606005-DEL-202 | chr11-84606006-DEL-202 |
| chr11-84668923-DEL-733 | chr11-84668928-DEL-731 |
| chr11-84683552-DEL-98 | chr11-84683553-DEL-98 |
| chr11-84696282-DEL-178 | chr11-84696278-DEL-165 |
| chr11-84779825-DEL-190 | chr11-84779830-DEL-189 |
| chr11-84795141-DEL-52 | chr11-84795134-DEL-52 |
| chr11-84832952-DEL-197 | chr11-84832952-DEL-196 |
| chr11-84878394-DEL-213 | chr11-84878395-DEL-213 |
| chr11-84913690-DEL-195 | chr11-84913691-DEL-195 |
| chr11-84952267-DEL-6427 | chr11-84952268-DEL-6427 |
| chr11-85000358-DEL-680 | chr11-85000359-DEL-680 |
| chr11-85012208-DEL-1184 | chr11-85012209-DEL-1184 |
| chr11-85015297-DEL-202 | chr11-85015298-DEL-202 |
| chr11-85020279-DEL-199 | chr11-85020280-DEL-199 |
| chr11-85032326-DEL-150 | chr11-85032327-DEL-150 |
| chr11-85046323-DEL-55 | chr11-85046315-DEL-52 |
| chr11-85054098-DEL-150 | chr11-85054099-DEL-150 |
| chr11-85067232-DEL-486 | chr11-85067233-DEL-486 |
| chr11-85074641-DEL-202 | chr11-85074642-DEL-202 |
| chr11-85077755-DEL-218 | chr11-85077756-DEL-218 |
| chr11-85142649-DEL-183 | chr11-85142650-DEL-183 |
| chr11-85157038-DEL-87 | chr11-85157061-DEL-64 |
| chr11-85165303-DEL-121 | chr11-85165304-DEL-122 |
| chr11-85173862-DEL-205 | chr11-85173863-DEL-205 |
| chr11-85176390-DEL-7116 | chr11-85176391-DEL-7116 |
| chr11-85184620-DEL-371 | chr11-85184621-DEL-371 |
| chr11-85191133-DEL-215 | chr11-85191134-DEL-215 |
| chr11-85261420-DEL-201 | chr11-85261421-DEL-201 |
| chr11-85276976-DEL-451 | chr11-85276977-DEL-451 |
| chr11-85285820-DEL-317 | chr11-85285821-DEL-317 |
| chr11-85310635-DEL-1125 | chr11-85310636-DEL-1125 |
| chr11-85312245-DEL-291 | chr11-85312246-DEL-291 |
| chr11-85345404-DEL-363 | chr11-85345405-DEL-363 |
| chr11-85366755-DEL-222 | chr11-85366756-DEL-222 |
| chr11-85403155-DEL-2604 | chr11-85403156-DEL-2604 |
| chr11-85429991-DEL-60 | chr11-85429980-DEL-52 |
| chr11-85455400-DEL-56 | chr11-85455397-DEL-66 |
| chr11-85525845-DEL-93 | chr11-85525846-DEL-93 |
| chr11-85556145-DEL-1155 | chr11-85556146-DEL-1155 |
| chr11-85598810-DEL-349 | chr11-85598811-DEL-349 |
| chr11-85678974-DEL-220 | chr11-85678975-DEL-220 |
| chr11-85743902-DEL-57 | chr11-85743903-DEL-57 |
| chr11-85744554-DEL-57 | chr11-85744543-DEL-69 |
| chr11-85748231-DEL-219 | chr11-85748232-DEL-219 |
| chr11-85800653-DEL-67 | chr11-85800654-DEL-67 |
| chr11-85939155-DEL-161 | chr11-85939156-DEL-161 |
| chr11-85945733-DEL-836 | chr11-85945734-DEL-836 |
| chr11-85967908-DEL-79 | chr11-85967931-DEL-79 |
| chr11-85969688-DEL-203 | chr11-85969686-DEL-202 |
| chr11-85983162-DEL-202 | chr11-85983163-DEL-202 |
| chr11-85996040-DEL-397 | chr11-85996040-DEL-397 |
| chr11-85996752-DEL-166 | chr11-85996752-DEL-166 |
| chr11-86032156-DEL-394 | chr11-86032157-DEL-394 |
| chr11-86065760-DEL-66 | chr11-86065752-DEL-60 |
| chr11-86068272-DEL-7194 | chr11-86068273-DEL-7194 |
| chr11-86072721-DEL-222 | chr11-86072722-DEL-222 |
| chr11-86078973-DEL-60 | chr11-86078974-DEL-60 |
| chr11-86110565-DEL-194 | chr11-86110566-DEL-194 |
| chr11-86112639-DEL-209 | chr11-86112640-DEL-209 |
| chr11-86125287-DEL-104 | chr11-86125287-DEL-104 |
| chr11-86142584-DEL-58 | chr11-86142570-DEL-57 |
| chr11-86145711-DEL-205 | chr11-86145712-DEL-205 |
| chr11-86152685-DEL-2691 | chr11-86152686-DEL-2691 |
| chr11-86155501-DEL-954 | chr11-86155501-DEL-955 |
| chr11-86232422-DEL-68 | chr11-86232423-DEL-68 |
| chr11-86254548-DEL-162 | chr11-86254549-DEL-162 |
| chr11-86268494-DEL-71 | chr11-86268495-DEL-71 |
| chr11-86269173-DEL-209 | chr11-86269174-DEL-209 |
| chr11-86318338-DEL-120 | chr11-86318338-DEL-120 |
| chr11-86324001-DEL-234 | chr11-86324002-DEL-234 |
| chr11-86345217-DEL-207 | chr11-86345218-DEL-207 |
| chr11-86351067-DEL-178 | chr11-86351068-DEL-178 |
| chr11-86354523-DEL-196 | chr11-86354524-DEL-196 |
| chr11-86357338-DEL-55 | chr11-86357339-DEL-55 |
| chr11-86417479-DEL-225 | chr11-86417480-DEL-225 |
| chr11-86428225-DEL-1162 | chr11-86428226-DEL-1162 |

| | |
|---|---|
| chr11-86442308-DEL-205 | chr11-86442309-DEL-205 |
| chr11-86455184-DEL-87 | chr11-86455185-DEL-87 |
| chr11-86472801-DEL-132 | chr11-86472802-DEL-132 |
| chr11-86505274-DEL-220 | chr11-86505275-DEL-220 |
| chr11-86524972-DEL-221 | chr11-86524973-DEL-221 |
| chr11-86551152-DEL-207 | chr11-86551153-DEL-207 |
| chr11-86569040-DEL-214 | chr11-86569048-DEL-190 |
| chr11-86576867-DEL-52 | chr11-86576868-DEL-52 |
| chr11-86606794-DEL-218 | chr11-86606795-DEL-218 |
| chr11-86656980-DEL-402 | chr11-86656981-DEL-402 |
| chr11-86668946-DEL-95 | chr11-86668947-DEL-95 |
| chr11-86669812-DEL-200 | chr11-86669813-DEL-200 |
| chr11-86670993-DEL-169 | chr11-86670992-DEL-125 |
| chr11-86701000-DEL-93 | chr11-86701001-DEL-93 |
| chr11-86702563-DEL-4658 | chr11-86702564-DEL-4658 |
| chr11-86715515-DEL-334 | chr11-86715516-DEL-334 |
| chr11-86726154-DEL-875 | chr11-86726155-DEL-875 |
| chr11-86734243-DEL-207 | chr11-86734244-DEL-207 |
| chr11-86736462-DEL-4521 | chr11-86736463-DEL-4521 |
| chr11-86773137-DEL-9045 | chr11-86773138-DEL-9045 |
| chr11-86851851-DEL-62 | chr11-86851858-DEL-50 |
| chr11-86861435-DEL-158 | chr11-86861436-DEL-119 |
| chr11-86869436-DEL-145 | chr11-86869437-DEL-145 |
| chr11-86889329-DEL-366 | chr11-86889330-DEL-366 |
| chr11-86913986-DEL-223 | chr11-86913987-DEL-223 |
| chr11-86915108-DEL-322 | chr11-86915109-DEL-322 |
| chr11-86918481-DEL-186 | chr11-86918482-DEL-186 |
| chr11-86920110-DEL-53 | chr11-86920110-DEL-53 |
| chr11-86963574-DEL-230 | chr11-86963575-DEL-230 |
| chr11-86990729-DEL-204 | chr11-86990730-DEL-204 |
| chr11-87007195-DEL-863 | chr11-87007196-DEL-863 |
| chr11-87026866-DEL-212 | chr11-87026867-DEL-212 |
| chr11-87067339-DEL-200 | chr11-87067340-DEL-200 |
| chr11-87093883-DEL-97 | chr11-87093884-DEL-97 |
| chr11-87153979-DEL-118 | chr11-87153980-DEL-118 |
| chr11-87159403-DEL-329 | chr11-87159404-DEL-329 |
| chr11-87195350-DEL-58 | chr11-87195351-DEL-58 |
| chr11-87238159-DEL-186 | chr11-87238160-DEL-186 |
| chr11-87287804-DEL-203 | chr11-87287805-DEL-203 |
| chr11-87289993-DEL-419 | chr11-87289992-DEL-420 |
| chr11-87321998-DEL-472 | chr11-87321999-DEL-472 |
| chr11-87360058-DEL-213 | chr11-87360059-DEL-213 |
| chr11-87362478-DEL-582 | chr11-87362478-DEL-582 |
| chr11-87382209-DEL-209 | chr11-87382209-DEL-209 |
| chr11-87387909-DEL-219 | chr11-87387910-DEL-219 |
| chr11-87401486-DEL-97 | chr11-87401487-DEL-97 |
| chr11-87436657-DEL-229 | chr11-87436631-DEL-206 |
| chr11-87518395-DEL-51 | chr11-87518396-DEL-51 |
| chr11-87571032-DEL-422 | chr11-87571033-DEL-422 |
| chr11-87586787-DEL-53 | chr11-87586793-DEL-54 |
| chr11-87594203-DEL-52 | chr11-87594207-DEL-52 |
| chr11-87604420-DEL-615 | chr11-87604421-DEL-615 |
| chr11-87608350-DEL-112 | chr11-87608351-DEL-112 |
| chr11-87718122-DEL-129 | chr11-87718089-DEL-115 |
| chr11-87740594-DEL-106 | chr11-87740595-DEL-106 |
| chr11-87743194-DEL-51 | chr11-87743195-DEL-51 |
| chr11-87761483-DEL-69 | chr11-87761484-DEL-69 |
| chr11-87793437-DEL-198 | chr11-87793438-DEL-198 |
| chr11-87798949-DEL-164 | chr11-87798950-DEL-164 |
| chr11-87806649-DEL-170 | chr11-87806650-DEL-170 |
| chr11-87814688-DEL-74 | chr11-87814689-DEL-70 |
| chr11-87822255-DEL-130 | chr11-87822256-DEL-130 |
| chr11-87831703-DEL-342 | chr11-87831704-DEL-342 |
| chr11-87833684-DEL-140 | chr11-87833662-DEL-135 |
| chr11-87834186-DEL-58 | chr11-87834187-DEL-58 |
| chr11-87841469-DEL-92 | chr11-87841469-DEL-98 |
| chr11-87853647-DEL-206 | chr11-87853648-DEL-206 |
| chr11-87859094-DEL-99 | chr11-87859095-DEL-99 |
| chr11-87862829-DEL-90 | chr11-87862828-DEL-69 |
| chr11-87863502-DEL-90 | chr11-87863503-DEL-95 |
| chr11-87865788-DEL-55 | chr11-87865793-DEL-51 |
| chr11-87911692-DEL-628 | chr11-87911692-DEL-627 |
| chr11-87912377-DEL-486 | chr11-87912370-DEL-486 |
| chr11-87923226-DEL-282 | chr11-87923227-DEL-282 |
| chr11-87929728-DEL-209 | chr11-87929729-DEL-209 |
| chr11-87932013-DEL-204 | chr11-87932014-DEL-204 |
| chr11-87953735-DEL-246 | chr11-87953735-DEL-246 |
| chr11-87985511-DEL-1124 | chr11-87985510-DEL-1125 |
| chr11-87987415-DEL-119 | chr11-87987416-DEL-119 |
| chr11-88015557-DEL-132 | chr11-88015579-DEL-131 |

| | |
|---|---|
| chr11-88022019-DEL-200 | chr11-88022020-DEL-200 |
| chr11-88027848-DEL-101 | chr11-88027849-DEL-95 |
| chr11-88053402-DEL-124 | chr11-88053400-DEL-132 |
| chr11-88065476-DEL-2282 | chr11-88065476-DEL-2282 |
| chr11-88067167-DEL-400 | chr11-88067168-DEL-400 |
| chr11-88069713-DEL-267 | chr11-88069714-DEL-267 |
| chr11-88081122-DEL-106 | chr11-88081115-DEL-102 |
| chr11-88121530-DEL-411 | chr11-88121539-DEL-403 |
| chr11-88126165-DEL-92 | chr11-88126166-DEL-92 |
| chr11-88136855-DEL-218 | chr11-88136856-DEL-218 |
| chr11-88157780-DEL-52 | chr11-88157783-DEL-50 |
| chr11-88177477-DEL-65 | chr11-88177478-DEL-65 |
| chr11-88219217-DEL-140 | chr11-88219220-DEL-131 |
| chr11-88261359-DEL-105 | chr11-88261360-DEL-105 |
| chr11-88301929-DEL-81 | chr11-88301928-DEL-82 |
| chr11-88324055-DEL-182 | chr11-88324056-DEL-182 |
| chr11-88347102-DEL-176 | chr11-88347103-DEL-176 |
| chr11-88369808-DEL-83 | chr11-88369809-DEL-83 |
| chr11-88390176-DEL-194 | chr11-88390174-DEL-196 |
| chr11-88394541-DEL-99 | chr11-88394544-DEL-100 |
| chr11-88421412-DEL-1189 | chr11-88421413-DEL-1189 |
| chr11-88424129-DEL-233 | chr11-88424129-DEL-233 |
| chr11-88447751-DEL-66 | chr11-88447752-DEL-66 |
| chr11-88459992-DEL-52 | chr11-88459993-DEL-52 |
| chr11-88466877-DEL-670 | chr11-88466878-DEL-670 |
| chr11-88485190-DEL-86 | chr11-88485188-DEL-100 |
| chr11-88540938-DEL-113 | chr11-88540939-DEL-113 |
| chr11-88567751-DEL-101 | chr11-88567752-DEL-101 |
| chr11-88583607-DEL-90 | chr11-88583604-DEL-80 |
| chr11-88616394-DEL-63 | chr11-88616404-DEL-50 |
| chr11-88630778-DEL-933 | chr11-88630779-DEL-933 |
| chr11-88641176-DEL-57 | chr11-88641183-DEL-51 |
| chr11-88670668-DEL-243 | chr11-88670677-DEL-250 |
| chr11-88690911-DEL-1840 | chr11-88690912-DEL-1840 |
| chr11-88695129-DEL-120 | chr11-88695130-DEL-120 |
| chr11-88696556-DEL-76 | chr11-88696555-DEL-60 |
| chr11-88702449-DEL-53 | chr11-88702451-DEL-54 |
| chr11-88718773-DEL-65 | chr11-88718774-DEL-65 |
| chr11-88723700-DEL-57 | chr11-88723707-DEL-51 |
| chr11-88726559-DEL-94 | chr11-88726578-DEL-50 |
| chr11-88748868-DEL-185 | chr11-88748869-DEL-185 |
| chr11-88758575-DEL-78 | chr11-88758575-DEL-78 |
| chr11-88781001-DEL-6791 | chr11-88781002-DEL-6791 |
| chr11-88801775-DEL-7296 | chr11-88801779-DEL-7297 |
| chr11-88813277-DEL-341 | chr11-88813252-DEL-327 |
| chr11-88814499-DEL-64 | chr11-88814500-DEL-62 |
| chr11-88820414-DEL-108 | chr11-88820392-DEL-92 |
| chr11-88823272-DEL-168 | chr11-88823273-DEL-168 |
| chr11-88827337-DEL-121 | chr11-88827338-DEL-109 |
| chr11-88836150-DEL-100 | chr11-88836151-DEL-100 |
| chr11-88844196-DEL-52 | chr11-88844197-DEL-52 |
| chr11-88852579-DEL-201 | chr11-88852580-DEL-201 |
| chr11-88854225-DEL-208 | chr11-88854226-DEL-208 |
| chr11-88864228-DEL-341 | chr11-88864229-DEL-341 |
| chr11-88869490-DEL-213 | chr11-88869491-DEL-213 |
| chr11-88871526-DEL-205 | chr11-88871527-DEL-205 |
| chr11-88883705-DEL-209 | chr11-88883706-DEL-209 |
| chr11-88900198-DEL-63 | chr11-88900199-DEL-62 |
| chr11-88922829-DEL-53 | chr11-88922830-DEL-53 |
| chr11-88932474-DEL-51 | chr11-88932475-DEL-52 |
| chr11-88936589-DEL-380 | chr11-88936590-DEL-380 |
| chr11-88947866-DEL-203 | chr11-88947867-DEL-199 |
| chr11-88971237-DEL-118 | chr11-88971247-DEL-79 |
| chr11-88972516-DEL-389 | chr11-88972513-DEL-390 |
| chr11-88979863-DEL-1557 | chr11-88979864-DEL-1556 |
| chr11-88982200-DEL-563 | chr11-88982201-DEL-563 |
| chr11-88990908-DEL-91 | chr11-88990909-DEL-111 |
| chr11-89001921-DEL-99 | chr11-89001889-DEL-132 |
| chr11-89030241-DEL-190 | chr11-89030238-DEL-178 |
| chr11-89050575-DEL-179 | chr11-89050576-DEL-178 |
| chr11-89082615-DEL-58 | chr11-89082616-DEL-58 |
| chr11-89085554-DEL-215 | chr11-89085555-DEL-215 |
| chr11-89087689-DEL-257 | chr11-89087690-DEL-258 |
| chr11-89096747-DEL-82 | chr11-89096754-DEL-92 |
| chr11-89118101-DEL-61 | chr11-89118102-DEL-61 |
| chr11-89121948-DEL-492 | chr11-89121944-DEL-493 |
| chr11-89169185-DEL-76 | chr11-89169186-DEL-76 |
| chr11-89215157-DEL-56 | chr11-89215158-DEL-56 |
| chr11-89238588-DEL-50 | chr11-89238588-DEL-62 |
| chr11-89240918-DEL-82 | chr11-89240919-DEL-82 |

| | |
|---|---|
| chr11-89260076-DEL-124 | chr11-89260077-DEL-124 |
| chr11-89268892-DEL-62 | chr11-89268875-DEL-81 |
| chr11-89275910-DEL-71 | chr11-89275904-DEL-68 |
| chr11-89326414-DEL-58 | chr11-89326415-DEL-58 |
| chr11-89344657-DEL-62 | chr11-89344658-DEL-62 |
| chr11-89353948-DEL-60 | chr11-89353949-DEL-60 |
| chr11-89378627-DEL-551 | chr11-89378628-DEL-551 |
| chr11-89386173-DEL-90 | chr11-89386174-DEL-90 |
| chr11-89391211-DEL-86 | chr11-89391212-DEL-85 |
| chr11-89408310-DEL-252 | chr11-89408311-DEL-257 |
| chr11-89439379-DEL-56 | chr11-89439380-DEL-56 |
| chr11-89444893-DEL-6537 | chr11-89444893-DEL-6537 |
| chr11-89462484-DEL-53 | chr11-89462485-DEL-53 |
| chr11-89496413-DEL-74 | chr11-89496404-DEL-84 |
| chr11-89518505-DEL-76 | chr11-89518492-DEL-77 |
| chr11-89555603-DEL-66 | chr11-89555604-DEL-66 |
| chr11-89572573-DEL-63 | chr11-89572574-DEL-63 |
| chr11-89573653-DEL-7155 | chr11-89573654-DEL-7155 |
| chr11-89580549-DEL-52 | chr11-89580550-DEL-52 |
| chr11-89593354-DEL-75 | chr11-89593356-DEL-73 |
| chr11-89602725-DEL-115 | chr11-89602762-DEL-111 |
| chr11-89603394-DEL-54 | chr11-89603395-DEL-54 |
| chr11-89612806-DEL-60 | chr11-89612807-DEL-60 |
| chr11-89620936-DEL-69 | chr11-89620937-DEL-69 |
| chr11-89621171-DEL-1121 | chr11-89621172-DEL-1121 |
| chr11-89636311-DEL-1917 | chr11-89636304-DEL-1918 |
| chr11-89644019-DEL-206 | chr11-89644020-DEL-206 |
| chr11-89652581-DEL-125 | chr11-89652588-DEL-126 |
| chr11-89670457-DEL-104 | chr11-89670462-DEL-100 |
| chr11-89682085-DEL-212 | chr11-89682073-DEL-213 |
| chr11-89713621-DEL-950 | chr11-89713622-DEL-950 |
| chr11-89735575-DEL-93 | chr11-89735555-DEL-113 |
| chr11-89737150-DEL-72 | chr11-89737151-DEL-72 |
| chr11-89742771-DEL-124 | chr11-89742772-DEL-124 |
| chr11-89754536-DEL-51 | chr11-89754537-DEL-52 |
| chr11-89759161-DEL-108 | chr11-89759162-DEL-108 |
| chr11-89773567-DEL-193 | chr11-89773568-DEL-193 |
| chr11-89778011-DEL-191 | chr11-89778012-DEL-191 |
| chr11-89794882-DEL-6481 | chr11-89794883-DEL-6481 |
| chr11-89812500-DEL-58 | chr11-89812501-DEL-58 |
| chr11-89816212-DEL-2195 | chr11-89816213-DEL-2195 |
| chr11-89846294-DEL-80 | chr11-89846295-DEL-80 |
| chr11-89851046-DEL-8434 | chr11-89851047-DEL-8434 |
| chr11-89868145-DEL-52 | chr11-89868146-DEL-54 |
| chr11-89892315-DEL-56 | chr11-89892316-DEL-56 |
| chr11-89899796-DEL-1258 | chr11-89899797-DEL-1258 |
| chr11-89913224-DEL-1035 | chr11-89913224-DEL-1036 |
| chr11-89945068-DEL-60 | chr11-89945040-DEL-55 |
| chr11-89962945-DEL-63 | chr11-89962946-DEL-63 |
| chr11-89981010-DEL-68 | chr11-89981008-DEL-69 |
| chr11-89988427-DEL-58 | chr11-89988414-DEL-58 |
| chr11-89995965-DEL-51 | chr11-89995954-DEL-52 |
| chr11-90004308-DEL-196 | chr11-90004300-DEL-194 |
| chr11-90039325-DEL-55 | chr11-90039326-DEL-50 |
| chr11-90043062-DEL-741 | chr11-90043066-DEL-742 |
| chr11-90047728-DEL-2693 | chr11-90047728-DEL-2693 |
| chr11-90050625-DEL-1898 | chr11-90050625-DEL-1898 |
| chr11-90063957-DEL-57 | chr11-90063958-DEL-57 |
| chr11-90086393-DEL-60 | chr11-90086443-DEL-53 |
| chr11-90153353-DEL-98 | chr11-90153354-DEL-98 |
| chr11-90196617-DEL-311 | chr11-90196618-DEL-311 |
| chr11-90201203-DEL-2072 | chr11-90201204-DEL-2072 |
| chr11-90237646-DEL-61 | chr11-90237646-DEL-60 |
| chr11-90240116-DEL-70 | chr11-90240117-DEL-70 |
| chr11-90267401-DEL-94 | chr11-90267402-DEL-94 |
| chr11-90270250-DEL-94 | chr11-90270251-DEL-94 |
| chr11-90272370-DEL-59 | chr11-90272371-DEL-59 |
| chr11-90283936-DEL-459 | chr11-90283939-DEL-459 |
| chr11-90318497-DEL-4302 | chr11-90318498-DEL-4302 |
| chr11-90329261-DEL-176 | chr11-90329262-DEL-176 |
| chr11-90335082-DEL-137 | chr11-90335083-DEL-137 |
| chr11-90344092-DEL-54 | chr11-90344093-DEL-54 |
| chr11-90350945-DEL-76 | chr11-90350946-DEL-76 |
| chr11-90393863-DEL-2832 | chr11-90393864-DEL-2832 |
| chr11-90422627-DEL-56 | chr11-90422628-DEL-56 |
| chr11-90469925-DEL-6130 | chr11-90469926-DEL-6130 |
| chr11-90498387-DEL-92 | chr11-90498381-DEL-93 |
| chr11-90502721-DEL-298 | chr11-90502722-DEL-298 |
| chr11-90533659-DEL-51 | chr11-90533660-DEL-51 |
| chr11-90549581-DEL-187 | chr11-90549597-DEL-172 |

| | |
|---|---|
| chr11-90570262-DEL-98 | chr11-90570263-DEL-98 |
| chr11-90574534-DEL-659 | chr11-90574535-DEL-659 |
| chr11-90589781-DEL-51 | chr11-90589782-DEL-51 |
| chr11-90615824-DEL-62 | chr11-90615825-DEL-62 |
| chr11-90629096-DEL-150 | chr11-90629099-DEL-157 |
| chr11-90659980-DEL-6224 | chr11-90659981-DEL-6224 |
| chr11-90674588-DEL-314 | chr11-90674589-DEL-314 |
| chr11-90720327-DEL-5433 | chr11-90720328-DEL-5433 |
| chr11-90755638-DEL-2577 | chr11-90755639-DEL-2577 |
| chr11-90764884-DEL-1197 | chr11-90764885-DEL-1197 |
| chr11-90777593-DEL-52 | chr11-90777610-DEL-56 |
| chr11-90783000-DEL-385 | chr11-90783001-DEL-385 |
| chr11-90792546-DEL-6537 | chr11-90792547-DEL-6537 |
| chr11-90831834-DEL-164 | chr11-90831835-DEL-164 |
| chr11-90834286-DEL-918 | chr11-90834287-DEL-918 |
| chr11-90849318-DEL-15509 | chr11-90849319-DEL-15509 |
| chr11-90851007-DEL-7092 | chr11-90851008-DEL-7092 |
| chr11-90878046-DEL-2117 | chr11-90878046-DEL-2117 |
| chr11-90880530-DEL-136 | chr11-90880531-DEL-136 |
| chr11-90883448-DEL-63 | chr11-90883457-DEL-51 |
| chr11-90888192-DEL-6183 | chr11-90888193-DEL-6183 |
| chr11-90909117-DEL-168 | chr11-90909118-DEL-168 |
| chr11-90911301-DEL-6494 | chr11-90911302-DEL-6494 |
| chr11-90931542-DEL-2790 | chr11-90931542-DEL-2790 |
| chr11-90934685-DEL-1009 | chr11-90934686-DEL-1009 |
| chr11-90938170-DEL-1858 | chr11-90938171-DEL-1857 |
| chr11-90971454-DEL-54 | chr11-90971455-DEL-54 |
| chr11-90976801-DEL-65 | chr11-90976801-DEL-57 |
| chr11-90990743-DEL-60 | chr11-90990744-DEL-60 |
| chr11-91008128-DEL-300 | chr11-91008129-DEL-300 |
| chr11-91022499-DEL-1594 | chr11-91022500-DEL-1594 |
| chr11-91029725-DEL-60 | chr11-91029726-DEL-78 |
| chr11-91034023-DEL-52 | chr11-91034023-DEL-53 |
| chr11-91045658-DEL-2413 | chr11-91045659-DEL-2413 |
| chr11-91059030-DEL-306 | chr11-91059031-DEL-306 |
| chr11-91064751-DEL-58 | chr11-91064740-DEL-59 |
| chr11-91078773-DEL-496 | chr11-91078774-DEL-496 |
| chr11-91079701-DEL-63 | chr11-91079702-DEL-63 |
| chr11-91096616-DEL-98 | chr11-91096618-DEL-104 |
| chr11-91120400-DEL-826 | chr11-91120401-DEL-826 |
| chr11-91137674-DEL-76 | chr11-91137675-DEL-76 |
| chr11-91144937-DEL-54 | chr11-91144934-DEL-54 |
| chr11-91198600-DEL-274 | chr11-91198600-DEL-273 |
| chr11-91215035-DEL-64 | chr11-91214996-DEL-64 |
| chr11-91220370-DEL-2375 | chr11-91220371-DEL-2375 |
| chr11-91252889-DEL-429 | chr11-91252890-DEL-429 |
| chr11-91290623-DEL-2484 | chr11-91290624-DEL-2484 |
| chr11-91328815-DEL-52 | chr11-91328816-DEL-52 |
| chr11-91329648-DEL-1507 | chr11-91329649-DEL-1507 |
| chr11-91332518-DEL-53 | chr11-91332516-DEL-54 |
| chr11-91333555-DEL-4862 | chr11-91333556-DEL-4862 |
| chr11-91363849-DEL-74 | chr11-91363850-DEL-70 |
| chr11-91378719-DEL-52 | chr11-91378719-DEL-53 |
| chr11-91408004-DEL-73 | chr11-91407973-DEL-54 |
| chr11-91424287-DEL-249 | chr11-91424288-DEL-249 |
| chr11-91452834-DEL-225 | chr11-91452845-DEL-217 |
| chr11-91462058-DEL-57 | chr11-91462059-DEL-57 |
| chr11-91474949-DEL-98 | chr11-91474948-DEL-98 |
| chr11-91491631-DEL-67 | chr11-91491631-DEL-61 |
| chr11-91504951-DEL-310 | chr11-91504952-DEL-310 |
| chr11-91509603-DEL-191 | chr11-91509604-DEL-191 |
| chr11-91527725-DEL-6985 | chr11-91527726-DEL-6985 |
| chr11-91535254-DEL-969 | chr11-91535255-DEL-969 |
| chr11-91558605-DEL-442 | chr11-91558606-DEL-442 |
| chr11-91564791-DEL-60 | chr11-91564791-DEL-60 |
| chr11-91572298-DEL-1528 | chr11-91572299-DEL-1528 |
| chr11-91608304-DEL-3099 | chr11-91608304-DEL-3099 |
| chr11-91623772-DEL-56 | chr11-91623773-DEL-56 |
| chr11-91629187-DEL-1453 | chr11-91629188-DEL-1453 |
| chr11-91655417-DEL-2104 | chr11-91655418-DEL-2104 |
| chr11-91672457-DEL-132 | chr11-91672458-DEL-132 |
| chr11-91683614-DEL-85 | chr11-91683615-DEL-85 |
| chr11-91772789-DEL-127 | chr11-91772789-DEL-127 |
| chr11-91800459-DEL-330 | chr11-91800460-DEL-330 |
| chr11-91803129-DEL-1219 | chr11-91803130-DEL-1219 |
| chr11-91811307-DEL-7238 | chr11-91811308-DEL-7238 |
| chr11-91812277-DEL-412 | chr11-91812318-DEL-372 |
| chr11-91821092-DEL-536 | chr11-91821090-DEL-537 |
| chr11-91840962-DEL-263 | chr11-91840963-DEL-263 |
| chr11-91845566-DEL-514 | chr11-91845567-DEL-514 |

| | |
|---|---|
| chr11-91865526-DEL-101 | chr11-91865527-DEL-101 |
| chr11-91903437-DEL-178 | chr11-91903438-DEL-178 |
| chr11-91910806-DEL-303 | chr11-91910807-DEL-303 |
| chr11-91916822-DEL-386 | chr11-91916823-DEL-386 |
| chr11-91928321-DEL-91 | chr11-91928322-DEL-91 |
| chr11-91941489-DEL-140 | chr11-91941515-DEL-114 |
| chr11-91947799-DEL-243 | chr11-91947800-DEL-243 |
| chr11-91950402-DEL-324 | chr11-91950403-DEL-324 |
| chr11-91984939-DEL-880 | chr11-91984939-DEL-880 |
| chr11-91989860-DEL-7183 | chr11-91989869-DEL-7182 |
| chr11-91997378-DEL-1721 | chr11-91997368-DEL-1722 |
| chr11-92002727-DEL-3053 | chr11-92002728-DEL-3053 |
| chr11-92019257-DEL-2819 | chr11-92019255-DEL-2820 |
| chr11-92063727-DEL-1120 | chr11-92063728-DEL-1120 |
| chr11-92067218-DEL-2461 | chr11-92067219-DEL-2461 |
| chr11-92077276-DEL-58 | chr11-92077288-DEL-58 |
| chr11-92078291-DEL-245 | chr11-92078292-DEL-245 |
| chr11-92101032-DEL-1510 | chr11-92101033-DEL-1510 |
| chr11-92101041-DEL-60 | chr11-92101042-DEL-60 |
| chr11-92103398-DEL-155 | chr11-92103399-DEL-155 |
| chr11-92104845-DEL-477 | chr11-92104846-DEL-477 |
| chr11-92119250-DEL-155 | chr11-92119251-DEL-155 |
| chr11-92173569-DEL-717 | chr11-92173570-DEL-717 |
| chr11-92189678-DEL-1178 | chr11-92189679-DEL-1178 |
| chr11-92242404-DEL-210 | chr11-92242400-DEL-198 |
| chr11-92262738-DEL-76 | chr11-92262739-DEL-76 |
| chr11-92272513-DEL-54 | chr11-92272514-DEL-54 |
| chr11-92288625-DEL-80 | chr11-92288627-DEL-81 |
| chr11-92348506-DEL-62 | chr11-92348507-DEL-62 |
| chr11-92364587-DEL-62 | chr11-92364588-DEL-62 |
| chr11-92403512-DEL-93 | chr11-92403513-DEL-93 |
| chr11-92464648-DEL-202 | chr11-92464649-DEL-202 |
| chr11-92467709-DEL-56 | chr11-92467710-DEL-56 |
| chr11-92476345-DEL-6436 | chr11-92476346-DEL-6436 |
| chr11-92491482-DEL-2842 | chr11-92491488-DEL-2843 |
| chr11-92503669-DEL-52 | chr11-92503670-DEL-54 |
| chr11-92577263-DEL-136 | chr11-92577256-DEL-186 |
| chr11-92585041-DEL-7216 | chr11-92585042-DEL-7216 |
| chr11-92630308-DEL-167 | chr11-92630312-DEL-164 |
| chr11-92631364-DEL-552 | chr11-92631367-DEL-550 |
| chr11-92653660-DEL-65 | chr11-92653656-DEL-70 |
| chr11-92661254-DEL-342 | chr11-92661255-DEL-342 |
| chr11-92714862-DEL-1889 | chr11-92714863-DEL-1889 |
| chr11-92734603-DEL-622 | chr11-92734602-DEL-622 |
| chr11-92746426-DEL-1135 | chr11-92746427-DEL-1135 |
| chr11-92764591-DEL-67 | chr11-92764591-DEL-67 |
| chr11-92768238-DEL-1618 | chr11-92768239-DEL-1618 |
| chr11-92807658-DEL-2498 | chr11-92807659-DEL-2498 |
| chr11-92811163-DEL-93 | chr11-92811164-DEL-93 |
| chr11-92819868-DEL-163 | chr11-92819869-DEL-163 |
| chr11-92834282-DEL-86 | chr11-92834283-DEL-86 |
| chr11-92848605-DEL-769 | chr11-92848613-DEL-770 |
| chr11-92852815-DEL-80 | chr11-92852816-DEL-80 |
| chr11-92852972-DEL-636 | chr11-92852973-DEL-636 |
| chr11-92858834-DEL-189 | chr11-92858835-DEL-189 |
| chr11-92859618-DEL-327 | chr11-92859617-DEL-328 |
| chr11-92875416-DEL-58 | chr11-92875385-DEL-60 |
| chr11-92878374-DEL-67 | chr11-92878375-DEL-67 |
| chr11-92882221-DEL-111 | chr11-92882224-DEL-126 |
| chr11-92888286-DEL-179 | chr11-92888288-DEL-180 |
| chr11-92922416-DEL-468 | chr11-92922417-DEL-468 |
| chr11-93011851-DEL-82 | chr11-93011872-DEL-82 |
| chr11-93072540-DEL-746 | chr11-93072541-DEL-746 |
| chr11-93081703-DEL-149 | chr11-93081704-DEL-149 |
| chr11-93106270-DEL-81 | chr11-93106271-DEL-81 |
| chr11-93116971-DEL-118 | chr11-93116973-DEL-118 |
| chr11-93135944-DEL-60 | chr11-93135960-DEL-52 |
| chr11-93157092-DEL-69 | chr11-93157093-DEL-66 |
| chr11-93159887-DEL-54 | chr11-93159888-DEL-54 |
| chr11-93178667-DEL-65 | chr11-93178659-DEL-50 |
| chr11-93206194-DEL-52 | chr11-93206191-DEL-56 |
| chr11-93209416-DEL-175 | chr11-93209417-DEL-175 |
| chr11-93229624-DEL-890 | chr11-93229625-DEL-890 |
| chr11-93240452-DEL-115 | chr11-93240453-DEL-115 |
| chr11-93282393-DEL-66 | chr11-93282394-DEL-66 |
| chr11-93289232-DEL-181 | chr11-93289220-DEL-182 |
| chr11-93309509-DEL-274 | chr11-93309510-DEL-274 |
| chr11-93345655-DEL-62 | chr11-93345656-DEL-62 |
| chr11-93356179-DEL-66 | chr11-93356189-DEL-51 |
| chr11-93360839-DEL-76 | chr11-93360841-DEL-75 |

| | |
|---|---|
| chr11-93366512-DEL-58 | chr11-93366513-DEL-58 |
| chr11-93375859-DEL-92 | chr11-93375860-DEL-91 |
| chr11-93376899-DEL-54 | chr11-93376900-DEL-54 |
| chr11-93431132-DEL-4528 | chr11-93431133-DEL-4528 |
| chr11-93471901-DEL-100 | chr11-93471902-DEL-100 |
| chr11-93501692-DEL-216 | chr11-93501691-DEL-215 |
| chr11-93508431-DEL-75 | chr11-93508432-DEL-75 |
| chr11-93509934-DEL-68 | chr11-93509935-DEL-68 |
| chr11-93527844-DEL-532 | chr11-93527845-DEL-532 |
| chr11-93582481-DEL-198 | chr11-93582482-DEL-198 |
| chr11-93590905-DEL-54 | chr11-93590906-DEL-50 |
| chr11-93601520-DEL-6242 | chr11-93601506-DEL-6235 |
| chr11-93646772-DEL-51 | chr11-93646768-DEL-55 |
| chr11-93677771-DEL-168 | chr11-93677772-DEL-168 |
| chr11-93696021-DEL-462 | chr11-93696022-DEL-462 |
| chr11-93697638-DEL-3581 | chr11-93697639-DEL-3581 |
| chr11-93702551-DEL-225 | chr11-93702552-DEL-225 |
| chr11-93712230-DEL-54 | chr11-93712231-DEL-54 |
| chr11-93724907-DEL-60 | chr11-93724908-DEL-60 |
| chr11-93740538-DEL-400 | chr11-93740539-DEL-400 |
| chr11-93741712-DEL-60 | chr11-93741713-DEL-60 |
| chr11-93794152-DEL-57 | chr11-93794159-DEL-53 |
| chr11-93799296-DEL-60 | chr11-93799297-DEL-60 |
| chr11-93821878-DEL-168 | chr11-93821879-DEL-168 |
| chr11-93826781-DEL-94 | chr11-93826782-DEL-95 |
| chr11-93854159-DEL-118 | chr11-93854160-DEL-118 |
| chr11-93858552-DEL-64 | chr11-93858553-DEL-64 |
| chr11-93861797-DEL-189 | chr11-93861798-DEL-189 |
| chr11-93869233-DEL-1344 | chr11-93869234-DEL-1344 |
| chr11-93870956-DEL-72 | chr11-93870957-DEL-72 |
| chr11-93871597-DEL-1063 | chr11-93871598-DEL-1066 |
| chr11-93872579-DEL-58 | chr11-93872580-DEL-58 |
| chr11-93880443-DEL-178 | chr11-93880444-DEL-178 |
| chr11-93905100-DEL-688 | chr11-93905101-DEL-688 |
| chr11-93905156-DEL-232 | chr11-93905157-DEL-232 |
| chr11-93927948-DEL-556 | chr11-93927949-DEL-556 |
| chr11-93949445-DEL-214 | chr11-93949446-DEL-214 |
| chr11-93951481-DEL-631 | chr11-93951482-DEL-631 |
| chr11-93955317-DEL-274 | chr11-93955318-DEL-274 |
| chr11-93965468-DEL-320 | chr11-93965469-DEL-320 |
| chr11-93967664-DEL-89 | chr11-93967665-DEL-89 |
| chr11-93983797-DEL-103 | chr11-93983798-DEL-103 |
| chr11-94005634-DEL-77 | chr11-94005629-DEL-77 |
| chr11-94018244-DEL-453 | chr11-94018245-DEL-453 |
| chr11-94048842-DEL-85 | chr11-94048843-DEL-85 |
| chr11-94061918-DEL-272 | chr11-94061919-DEL-272 |
| chr11-94076403-DEL-188 | chr11-94076404-DEL-188 |
| chr11-94108565-DEL-219 | chr11-94108572-DEL-197 |
| chr11-94131675-DEL-412 | chr11-94131676-DEL-412 |
| chr11-94158655-DEL-158 | chr11-94158656-DEL-158 |
| chr11-94195430-DEL-178 | chr11-94195434-DEL-179 |
| chr11-94198142-DEL-215 | chr11-94198143-DEL-215 |
| chr11-94247421-DEL-55 | chr11-94247414-DEL-56 |
| chr11-94338366-DEL-75 | chr11-94338370-DEL-72 |
| chr11-94419131-DEL-72 | chr11-94419132-DEL-72 |
| chr11-94419956-DEL-239 | chr11-94419957-DEL-239 |
| chr11-94446942-DEL-90 | chr11-94446943-DEL-90 |
| chr11-94473997-DEL-60 | chr11-94474000-DEL-54 |
| chr11-94479104-DEL-88 | chr11-94479105-DEL-88 |
| chr11-94479853-DEL-214 | chr11-94479854-DEL-213 |
| chr11-94502776-DEL-65 | chr11-94502766-DEL-87 |
| chr11-94510022-DEL-165 | chr11-94510023-DEL-165 |
| chr11-94513740-DEL-6401 | chr11-94513741-DEL-6401 |
| chr11-94542309-DEL-216 | chr11-94542310-DEL-216 |
| chr11-94547025-DEL-56 | chr11-94547032-DEL-56 |
| chr11-94552752-DEL-220 | chr11-94552753-DEL-220 |
| chr11-94561667-DEL-72 | chr11-94561668-DEL-72 |
| chr11-94571126-DEL-56 | chr11-94571127-DEL-56 |
| chr11-94576656-DEL-108 | chr11-94576657-DEL-108 |
| chr11-94582329-DEL-463 | chr11-94582330-DEL-461 |
| chr11-94587655-DEL-60 | chr11-94587656-DEL-60 |
| chr11-94593357-DEL-121 | chr11-94593358-DEL-109 |
| chr11-94595665-DEL-95 | chr11-94595666-DEL-95 |
| chr11-94619537-DEL-101 | chr11-94619538-DEL-101 |
| chr11-94638318-DEL-99 | chr11-94638319-DEL-99 |
| chr11-94714759-DEL-53 | chr11-94714759-DEL-53 |
| chr11-94731820-DEL-229 | chr11-94731821-DEL-229 |
| chr11-94774479-DEL-87 | chr11-94774480-DEL-87 |
| chr11-94788855-DEL-188 | chr11-94788856-DEL-188 |
| chr11-94794207-DEL-217 | chr11-94794208-DEL-217 |

| | |
|---|---|
| chr11-94810674-DEL-1294 | chr11-94810679-DEL-1290 |
| chr11-94810686-DEL-90 | chr11-94810685-DEL-92 |
| chr11-94811532-DEL-84 | chr11-94811537-DEL-86 |
| chr11-94814331-DEL-374 | chr11-94814332-DEL-374 |
| chr11-94860767-DEL-216 | chr11-94860768-DEL-216 |
| chr11-94946071-DEL-196 | chr11-94946079-DEL-178 |
| chr11-94972183-DEL-197 | chr11-94972184-DEL-197 |
| chr11-94975490-DEL-208 | chr11-94975491-DEL-208 |
| chr11-94976041-DEL-118 | chr11-94976042-DEL-118 |
| chr11-94980841-DEL-60 | chr11-94980842-DEL-60 |
| chr11-95064962-DEL-70 | chr11-95064963-DEL-70 |
| chr11-95097981-DEL-328 | chr11-95097982-DEL-328 |
| chr11-95120249-DEL-189 | chr11-95120250-DEL-189 |
| chr11-95141795-DEL-185 | chr11-95141796-DEL-185 |
| chr11-95147061-DEL-339 | chr11-95147062-DEL-339 |
| chr11-95176149-DEL-192 | chr11-95176147-DEL-192 |
| chr11-95186449-DEL-190 | chr11-95186450-DEL-190 |
| chr11-95188720-DEL-136 | chr11-95188721-DEL-136 |
| chr11-95205059-DEL-78 | chr11-95205087-DEL-63 |
| chr11-95207985-DEL-666 | chr11-95207986-DEL-666 |
| chr11-95208783-DEL-69 | chr11-95208786-DEL-70 |
| chr11-95225036-DEL-172 | chr11-95225037-DEL-172 |
| chr11-95262397-DEL-193 | chr11-95262398-DEL-193 |
| chr11-95266200-DEL-124 | chr11-95266203-DEL-120 |
| chr11-95273439-DEL-211 | chr11-95273456-DEL-212 |
| chr11-95291961-DEL-89 | chr11-95291961-DEL-90 |
| chr11-95311633-DEL-222 | chr11-95311634-DEL-222 |
| chr11-95356562-DEL-1802 | chr11-95356563-DEL-1802 |
| chr11-95357532-DEL-69 | chr11-95357533-DEL-66 |
| chr11-95431579-DEL-63 | chr11-95431575-DEL-64 |
| chr11-95448498-DEL-74 | chr11-95448499-DEL-74 |
| chr11-95452003-DEL-141 | chr11-95451966-DEL-145 |
| chr11-95464760-DEL-112 | chr11-95464719-DEL-130 |
| chr11-95498080-DEL-193 | chr11-95498081-DEL-192 |
| chr11-95509239-DEL-211 | chr11-95509240-DEL-211 |
| chr11-95511370-DEL-207 | chr11-95511371-DEL-207 |
| chr11-95563218-DEL-178 | chr11-95563219-DEL-178 |
| chr11-95569752-DEL-730 | chr11-95569753-DEL-730 |
| chr11-95587679-DEL-306 | chr11-95587682-DEL-307 |
| chr11-95624062-DEL-208 | chr11-95624063-DEL-208 |
| chr11-95630117-DEL-215 | chr11-95630118-DEL-215 |
| chr11-95631299-DEL-808 | chr11-95631300-DEL-808 |
| chr11-95646222-DEL-860 | chr11-95646223-DEL-860 |
| chr11-95652059-DEL-53 | chr11-95652060-DEL-53 |
| chr11-95670604-DEL-278 | chr11-95670618-DEL-274 |
| chr11-95670961-DEL-3100 | chr11-95670962-DEL-3100 |
| chr11-95671676-DEL-229 | chr11-95671665-DEL-222 |
| chr11-95688763-DEL-118 | chr11-95688773-DEL-100 |
| chr11-95690310-DEL-134 | chr11-95690299-DEL-116 |
| chr11-95717062-DEL-155 | chr11-95717055-DEL-152 |
| chr11-95753855-DEL-173 | chr11-95753851-DEL-174 |
| chr11-95757887-DEL-172 | chr11-95757879-DEL-170 |
| chr11-95768230-DEL-200 | chr11-95768231-DEL-200 |
| chr11-95771327-DEL-52 | chr11-95771327-DEL-52 |
| chr11-95784347-DEL-205 | chr11-95784348-DEL-205 |
| chr11-95795818-DEL-186 | chr11-95795819-DEL-186 |
| chr11-95813645-DEL-1604 | chr11-95813646-DEL-1604 |
| chr11-95814113-DEL-323 | chr11-95814114-DEL-323 |
| chr11-95816830-DEL-6421 | chr11-95816845-DEL-6406 |
| chr11-95822900-DEL-219 | chr11-95822901-DEL-219 |
| chr11-95826863-DEL-3290 | chr11-95826864-DEL-3294 |
| chr11-95827937-DEL-68 | chr11-95827927-DEL-69 |
| chr11-95831825-DEL-1726 | chr11-95831826-DEL-1726 |
| chr11-95840346-DEL-211 | chr11-95840347-DEL-211 |
| chr11-95843636-DEL-85 | chr11-95843631-DEL-91 |
| chr11-95846229-DEL-90 | chr11-95846226-DEL-88 |
| chr11-95863098-DEL-200 | chr11-95863099-DEL-200 |
| chr11-95892196-DEL-53 | chr11-95892197-DEL-53 |
| chr11-95904285-DEL-216 | chr11-95904286-DEL-216 |
| chr11-95910513-DEL-111 | chr11-95910514-DEL-111 |
| chr11-95912890-DEL-321 | chr11-95912891-DEL-321 |
| chr11-95918870-DEL-52 | chr11-95918871-DEL-52 |
| chr11-95930534-DEL-216 | chr11-95930535-DEL-215 |
| chr11-95932200-DEL-208 | chr11-95932201-DEL-208 |
| chr11-95933361-DEL-100 | chr11-95933362-DEL-100 |
| chr11-96007720-DEL-205 | chr11-96007721-DEL-205 |
| chr11-96008798-DEL-392 | chr11-96008799-DEL-392 |
| chr11-96014288-DEL-110 | chr11-96014289-DEL-74 |
| chr11-96041452-DEL-54 | chr11-96041453-DEL-54 |
| chr11-96042081-DEL-85 | chr11-96042082-DEL-85 |

| | |
|---|---|
| chr11-96070362-DEL-56 | chr11-96070363-DEL-56 |
| chr11-96093012-DEL-164 | chr11-96093013-DEL-164 |
| chr11-96099110-DEL-57 | chr11-96099111-DEL-57 |
| chr11-96111950-DEL-946 | chr11-96111952-DEL-937 |
| chr11-96113028-DEL-206 | chr11-96113029-DEL-206 |
| chr11-96124987-DEL-210 | chr11-96124988-DEL-210 |
| chr11-96131414-DEL-67 | chr11-96131415-DEL-67 |
| chr11-96139408-DEL-663 | chr11-96139409-DEL-663 |
| chr11-96139441-DEL-73 | chr11-96139442-DEL-73 |
| chr11-96179927-DEL-192 | chr11-96179930-DEL-190 |
| chr11-96250884-DEL-203 | chr11-96250885-DEL-203 |
| chr11-96253417-DEL-60 | chr11-96253405-DEL-64 |
| chr11-96268636-DEL-60 | chr11-96268637-DEL-57 |
| chr11-96282895-DEL-650 | chr11-96282896-DEL-650 |
| chr11-96292328-DEL-81 | chr11-96292329-DEL-81 |
| chr11-96308941-DEL-194 | chr11-96308942-DEL-194 |
| chr11-96326634-DEL-61 | chr11-96326635-DEL-61 |
| chr11-96349236-DEL-58 | chr11-96349237-DEL-58 |
| chr11-96363057-DEL-319 | chr11-96363060-DEL-317 |
| chr11-96365500-DEL-56 | chr11-96365501-DEL-52 |
| chr11-96382408-DEL-84 | chr11-96382424-DEL-60 |
| chr11-96396200-DEL-51 | chr11-96396205-DEL-54 |
| chr11-96400108-DEL-62 | chr11-96400109-DEL-62 |
| chr11-96404625-DEL-82 | chr11-96404613-DEL-82 |
| chr11-96410165-DEL-2686 | chr11-96410166-DEL-2686 |
| chr11-96417892-DEL-64 | chr11-96417893-DEL-64 |
| chr11-96453582-DEL-90 | chr11-96453583-DEL-90 |
| chr11-96461475-DEL-65 | chr11-96461476-DEL-65 |
| chr11-96530126-DEL-126 | chr11-96530127-DEL-126 |
| chr11-96542118-DEL-291 | chr11-96542124-DEL-290 |
| chr11-96543444-DEL-317 | chr11-96543445-DEL-317 |
| chr11-96581154-DEL-84 | chr11-96581155-DEL-84 |
| chr11-96605925-DEL-228 | chr11-96605926-DEL-228 |
| chr11-96610553-DEL-64 | chr11-96610553-DEL-64 |
| chr11-96631508-DEL-55 | chr11-96631509-DEL-55 |
| chr11-96654375-DEL-207 | chr11-96654376-DEL-207 |
| chr11-96676228-DEL-205 | chr11-96676229-DEL-205 |
| chr11-96722913-DEL-213 | chr11-96722912-DEL-213 |
| chr11-96726669-DEL-488 | chr11-96726670-DEL-488 |
| chr11-96743016-DEL-141 | chr11-96743015-DEL-142 |
| chr11-96761985-DEL-99 | chr11-96761980-DEL-103 |
| chr11-96793217-DEL-70 | chr11-96793218-DEL-70 |
| chr11-96795254-DEL-62 | chr11-96795248-DEL-104 |
| chr11-96805311-DEL-202 | chr11-96805308-DEL-203 |
| chr11-96812833-DEL-219 | chr11-96812834-DEL-219 |
| chr11-96849117-DEL-52 | chr11-96849118-DEL-52 |
| chr11-96865186-DEL-158 | chr11-96865187-DEL-158 |
| chr11-96881539-DEL-202 | chr11-96881551-DEL-193 |
| chr11-96905703-DEL-59 | chr11-96905704-DEL-59 |
| chr11-96943968-DEL-218 | chr11-96943969-DEL-218 |
| chr11-96947960-DEL-531 | chr11-96947938-DEL-532 |
| chr11-96962444-DEL-60 | chr11-96962445-DEL-60 |
| chr11-96969059-DEL-208 | chr11-96969060-DEL-208 |
| chr11-96976307-DEL-199 | chr11-96976308-DEL-199 |
| chr11-96986325-DEL-208 | chr11-96986326-DEL-208 |
| chr11-96991613-DEL-100 | chr11-96991647-DEL-96 |
| chr11-97024494-DEL-69 | chr11-97024476-DEL-66 |
| chr11-97081433-DEL-220 | chr11-97081434-DEL-220 |
| chr11-97086935-DEL-102 | chr11-97086935-DEL-102 |
| chr11-97093323-DEL-163 | chr11-97093324-DEL-163 |
| chr11-97095347-DEL-1169 | chr11-97095343-DEL-1170 |
| chr11-97115744-DEL-209 | chr11-97115744-DEL-209 |
| chr11-97122877-DEL-181 | chr11-97122878-DEL-181 |
| chr11-97182028-DEL-138 | chr11-97182028-DEL-138 |
| chr11-97183296-DEL-145 | chr11-97183297-DEL-145 |
| chr11-97187243-DEL-180 | chr11-97187244-DEL-180 |
| chr11-97189489-DEL-206 | chr11-97189490-DEL-206 |
| chr11-97199296-DEL-232 | chr11-97199297-DEL-232 |
| chr11-97202730-DEL-209 | chr11-97202731-DEL-209 |
| chr11-97203934-DEL-277 | chr11-97203935-DEL-277 |
| chr11-97209510-DEL-206 | chr11-97209511-DEL-206 |
| chr11-97213561-DEL-208 | chr11-97213562-DEL-208 |
| chr11-97215698-DEL-180 | chr11-97215699-DEL-180 |
| chr11-97232671-DEL-205 | chr11-97232672-DEL-205 |
| chr11-97248383-DEL-225 | chr11-97248384-DEL-225 |
| chr11-97249451-DEL-225 | chr11-97249453-DEL-224 |
| chr11-97271959-DEL-193 | chr11-97271960-DEL-193 |
| chr11-97300742-DEL-211 | chr11-97300743-DEL-211 |
| chr11-97311074-DEL-211 | chr11-97311074-DEL-211 |
| chr11-97317359-DEL-213 | chr11-97317359-DEL-213 |

| | |
|---|---|
| chr11-97381424-DEL-165 | chr11-97381425-DEL-165 |
| chr11-97385613-DEL-183 | chr11-97385614-DEL-184 |
| chr11-97405051-DEL-56 | chr11-97405051-DEL-56 |
| chr11-97511276-DEL-206 | chr11-97511277-DEL-206 |
| chr11-97533957-DEL-58 | chr11-97533958-DEL-58 |
| chr11-97536735-DEL-129 | chr11-97536736-DEL-129 |
| chr11-97538606-DEL-204 | chr11-97538607-DEL-204 |
| chr11-97578786-DEL-54 | chr11-97578787-DEL-54 |
| chr11-97615944-DEL-58 | chr11-97615927-DEL-72 |
| chr11-97620980-DEL-56 | chr11-97620982-DEL-59 |
| chr11-97625561-DEL-90 | chr11-97625575-DEL-78 |
| chr11-97645187-DEL-174 | chr11-97645188-DEL-174 |
| chr11-97675696-DEL-606 | chr11-97675697-DEL-606 |
| chr11-97683044-DEL-55 | chr11-97683045-DEL-55 |
| chr11-97711512-DEL-76 | chr11-97711513-DEL-76 |
| chr11-97722320-DEL-192 | chr11-97722322-DEL-190 |
| chr11-97737583-DEL-246 | chr11-97737583-DEL-263 |
| chr11-97786138-DEL-65 | chr11-97786139-DEL-60 |
| chr11-97789539-DEL-60 | chr11-97789540-DEL-60 |
| chr11-97792244-DEL-92 | chr11-97792243-DEL-60 |
| chr11-97800059-DEL-177 | chr11-97800060-DEL-177 |
| chr11-97805913-DEL-172 | chr11-97805914-DEL-172 |
| chr11-97823955-DEL-161 | chr11-97823956-DEL-161 |
| chr11-97836357-DEL-210 | chr11-97836358-DEL-210 |
| chr11-97843948-DEL-540 | chr11-97843949-DEL-540 |
| chr11-97855506-DEL-208 | chr11-97855537-DEL-188 |
| chr11-97860144-DEL-75 | chr11-97860145-DEL-75 |
| chr11-97864180-DEL-62 | chr11-97864181-DEL-62 |
| chr11-97869780-DEL-58 | chr11-97869781-DEL-58 |
| chr11-97873358-DEL-67 | chr11-97873359-DEL-67 |
| chr11-97884677-DEL-125 | chr11-97884678-DEL-119 |
| chr11-97903998-DEL-64 | chr11-97903999-DEL-64 |
| chr11-97923841-DEL-108 | chr11-97923842-DEL-108 |
| chr11-97925309-DEL-210 | chr11-97925310-DEL-210 |
| chr11-97972951-DEL-116 | chr11-97972952-DEL-116 |
| chr11-97978622-DEL-142 | chr11-97978623-DEL-142 |
| chr11-97989159-DEL-55 | chr11-97989160-DEL-55 |
| chr11-97996639-DEL-70 | chr11-97996640-DEL-70 |
| chr11-98013764-DEL-178 | chr11-98013765-DEL-178 |
| chr11-98053234-DEL-191 | chr11-98053248-DEL-191 |
| chr11-98068810-DEL-210 | chr11-98068811-DEL-210 |
| chr11-98076365-DEL-207 | chr11-98076366-DEL-207 |
| chr11-98077629-DEL-213 | chr11-98077630-DEL-213 |
| chr11-98119163-DEL-210 | chr11-98119164-DEL-210 |
| chr11-98159957-DEL-216 | chr11-98159958-DEL-216 |
| chr11-98259145-DEL-207 | chr11-98259146-DEL-207 |
| chr11-98264609-DEL-77 | chr11-98264596-DEL-69 |
| chr11-98266137-DEL-91 | chr11-98266138-DEL-91 |
| chr11-98272273-DEL-144 | chr11-98272292-DEL-126 |
| chr11-98273256-DEL-50 | chr11-98273242-DEL-64 |
| chr11-98289976-DEL-110 | chr11-98289977-DEL-110 |
| chr11-98293439-DEL-108 | chr11-98293440-DEL-108 |
| chr11-98299017-DEL-252 | chr11-98299018-DEL-252 |
| chr11-98385872-DEL-91 | chr11-98385921-DEL-89 |
| chr11-98402261-DEL-809 | chr11-98402262-DEL-809 |
| chr11-98421853-DEL-54 | chr11-98421826-DEL-56 |
| chr11-98425841-DEL-120 | chr11-98425842-DEL-120 |
| chr11-98435402-DEL-55 | chr11-98435403-DEL-55 |
| chr11-98456151-DEL-1015 | chr11-98456152-DEL-1015 |
| chr11-98462292-DEL-72 | chr11-98462286-DEL-78 |
| chr11-98498400-DEL-107 | chr11-98498395-DEL-112 |
| chr11-98500058-DEL-469 | chr11-98500059-DEL-469 |
| chr11-98502158-DEL-196 | chr11-98502159-DEL-196 |
| chr11-98504097-DEL-7105 | chr11-98504098-DEL-7105 |
| chr11-98543917-DEL-204 | chr11-98543918-DEL-204 |
| chr11-98549052-DEL-2432 | chr11-98549053-DEL-2432 |
| chr11-98579066-DEL-115 | chr11-98579072-DEL-110 |
| chr11-98579666-DEL-406 | chr11-98579667-DEL-406 |
| chr11-98581119-DEL-101 | chr11-98581120-DEL-101 |
| chr11-98602746-DEL-210 | chr11-98602743-DEL-211 |
| chr11-98668723-DEL-55 | chr11-98668724-DEL-55 |
| chr11-98669390-DEL-56 | chr11-98669391-DEL-56 |
| chr11-98703910-DEL-215 | chr11-98703911-DEL-215 |
| chr11-98739764-DEL-125 | chr11-98739765-DEL-125 |
| chr11-98757560-DEL-57 | chr11-98757561-DEL-57 |
| chr11-98766994-DEL-207 | chr11-98766995-DEL-207 |
| chr11-98770463-DEL-403 | chr11-98770465-DEL-402 |
| chr11-98789426-DEL-75 | chr11-98789427-DEL-72 |
| chr11-98849408-DEL-52 | chr11-98849409-DEL-52 |
| chr11-98881202-DEL-1411 | chr11-98881203-DEL-1411 |

| | |
|---|---|
| chr11-98889033-DEL-86 | chr11-98889034-DEL-86 |
| chr11-98917520-DEL-211 | chr11-98917521-DEL-211 |
| chr11-98923193-DEL-135 | chr11-98923224-DEL-132 |
| chr11-98927722-DEL-1053 | chr11-98927723-DEL-1053 |
| chr11-98992114-DEL-59 | chr11-98992095-DEL-60 |
| chr11-98998805-DEL-206 | chr11-98998806-DEL-206 |
| chr11-99019744-DEL-52 | chr11-99019745-DEL-52 |
| chr11-99031447-DEL-58 | chr11-99031447-DEL-58 |
| chr11-99064355-DEL-177 | chr11-99064368-DEL-177 |
| chr11-99064854-DEL-1750 | chr11-99064855-DEL-1750 |
| chr11-99096058-DEL-744 | chr11-99096060-DEL-743 |
| chr11-99101433-DEL-177 | chr11-99101432-DEL-177 |
| chr11-99102099-DEL-208 | chr11-99102100-DEL-208 |
| chr11-99126210-DEL-120 | chr11-99126218-DEL-104 |
| chr11-99131620-DEL-55 | chr11-99131621-DEL-55 |
| chr11-99142623-DEL-59 | chr11-99142612-DEL-66 |
| chr11-99152357-DEL-413 | chr11-99152356-DEL-405 |
| chr11-99157087-DEL-62 | chr11-99157119-DEL-64 |
| chr11-99180266-DEL-421 | chr11-99180267-DEL-421 |
| chr11-99229763-DEL-59 | chr11-99229764-DEL-59 |
| chr11-99245371-DEL-72 | chr11-99245368-DEL-72 |
| chr11-99248663-DEL-158 | chr11-99248664-DEL-158 |
| chr11-99253755-DEL-413 | chr11-99253761-DEL-408 |
| chr11-99280929-DEL-78 | chr11-99280930-DEL-78 |
| chr11-99283981-DEL-120 | chr11-99283982-DEL-120 |
| chr11-99324542-DEL-103 | chr11-99324543-DEL-103 |
| chr11-99347720-DEL-1035 | chr11-99347721-DEL-1035 |
| chr11-99363528-DEL-617 | chr11-99363524-DEL-614 |
| chr11-99367797-DEL-184 | chr11-99367790-DEL-185 |
| chr11-99426342-DEL-95 | chr11-99426341-DEL-96 |
| chr11-99445432-DEL-383 | chr11-99445433-DEL-381 |
| chr11-99475100-DEL-56 | chr11-99475101-DEL-56 |
| chr11-99520497-DEL-224 | chr11-99520500-DEL-223 |
| chr11-99532781-DEL-106 | chr11-99532782-DEL-88 |
| chr11-99545894-DEL-510 | chr11-99545894-DEL-511 |
| chr11-99551590-DEL-56 | chr11-99551552-DEL-62 |
| chr11-99603539-DEL-4501 | chr11-99603540-DEL-4501 |
| chr11-99616620-DEL-64 | chr11-99616617-DEL-86 |
| chr11-99626493-DEL-353 | chr11-99626491-DEL-354 |
| chr11-99641427-DEL-1166 | chr11-99641428-DEL-1166 |
| chr11-99662881-DEL-225 | chr11-99662882-DEL-225 |
| chr11-99667762-DEL-114 | chr11-99667763-DEL-110 |
| chr11-99676257-DEL-132 | chr11-99676258-DEL-132 |
| chr11-99715614-DEL-8352 | chr11-99715614-DEL-8352 |
| chr11-99715632-DEL-848 | chr11-99715631-DEL-848 |
| chr11-99752589-DEL-313 | chr11-99752590-DEL-313 |
| chr11-99773702-DEL-7248 | chr11-99773703-DEL-7248 |
| chr11-99782937-DEL-143 | chr11-99782938-DEL-143 |
| chr11-99812420-DEL-6206 | chr11-99812421-DEL-6206 |
| chr11-99824151-DEL-520 | chr11-99824152-DEL-520 |
| chr11-99862990-DEL-104 | chr11-99863003-DEL-92 |
| chr11-99919665-DEL-124 | chr11-99919666-DEL-124 |
| chr11-99960386-DEL-57 | chr11-99960388-DEL-56 |
| chr11-99967229-DEL-698 | chr11-99967230-DEL-698 |
| chr11-99985192-DEL-110 | chr11-99985203-DEL-110 |
| chr11-100005646-DEL-559 | chr11-100005647-DEL-559 |
| chr11-100029268-DEL-57 | chr11-100029264-DEL-58 |
| chr11-100045775-DEL-152 | chr11-100045776-DEL-152 |
| chr11-100049295-DEL-152 | chr11-100049296-DEL-152 |
| chr11-100051824-DEL-162 | chr11-100051869-DEL-162 |
| chr11-100053820-DEL-370 | chr11-100053821-DEL-370 |
| chr11-100055829-DEL-59 | chr11-100055830-DEL-59 |
| chr11-100062041-DEL-60 | chr11-100062042-DEL-60 |
| chr11-100069083-DEL-64 | chr11-100069084-DEL-64 |
| chr11-100071810-DEL-3637 | chr11-100071805-DEL-3638 |
| chr11-100091523-DEL-120 | chr11-100091524-DEL-118 |
| chr11-100093206-DEL-73 | chr11-100093205-DEL-68 |
| chr11-100103247-DEL-143 | chr11-100103248-DEL-143 |
| chr11-100145732-DEL-53 | chr11-100145733-DEL-53 |
| chr11-100166047-DEL-401 | chr11-100166048-DEL-401 |
| chr11-100174209-DEL-61 | chr11-100174237-DEL-56 |
| chr11-100192902-DEL-344 | chr11-100192903-DEL-344 |
| chr11-100198320-DEL-70 | chr11-100198321-DEL-70 |
| chr11-100206138-DEL-201 | chr11-100206139-DEL-201 |
| chr11-100212721-DEL-77 | chr11-100212722-DEL-77 |
| chr11-100241725-DEL-56 | chr11-100241726-DEL-56 |
| chr11-100251753-DEL-1229 | chr11-100251754-DEL-1229 |
| chr11-100258876-DEL-80 | chr11-100258877-DEL-80 |
| chr11-100270465-DEL-101 | chr11-100270466-DEL-101 |
| chr11-100323343-DEL-52 | chr11-100323344-DEL-52 |

| | |
|---|---|
| chr11-100333847-DEL-176 | chr11-100333842-DEL-182 |
| chr11-100339286-DEL-97 | chr11-100339287-DEL-97 |
| chr11-100339657-DEL-1179 | chr11-100339658-DEL-1179 |
| chr11-100341703-DEL-853 | chr11-100341704-DEL-853 |
| chr11-100385016-DEL-434 | chr11-100385017-DEL-434 |
| chr11-100398087-DEL-224 | chr11-100398044-DEL-224 |
| chr11-100458456-DEL-207 | chr11-100458457-DEL-207 |
| chr11-100539872-DEL-144 | chr11-100539837-DEL-184 |
| chr11-100546315-DEL-178 | chr11-100546316-DEL-178 |
| chr11-100553721-DEL-6629 | chr11-100553715-DEL-6633 |
| chr11-100591323-DEL-182 | chr11-100591324-DEL-182 |
| chr11-100625385-DEL-212 | chr11-100625386-DEL-212 |
| chr11-100674831-DEL-62 | chr11-100674853-DEL-54 |
| chr11-100685240-DEL-52 | chr11-100685241-DEL-52 |
| chr11-100704038-DEL-212 | chr11-100704039-DEL-212 |
| chr11-100728632-DEL-175 | chr11-100728633-DEL-175 |
| chr11-100729997-DEL-184 | chr11-100729999-DEL-183 |
| chr11-100766928-DEL-144 | chr11-100766909-DEL-132 |
| chr11-100768659-DEL-76 | chr11-100768658-DEL-84 |
| chr11-100800207-DEL-212 | chr11-100800208-DEL-212 |
| chr11-100837568-DEL-741 | chr11-100837569-DEL-741 |
| chr11-100843967-DEL-79 | chr11-100843968-DEL-79 |
| chr11-100845696-DEL-52 | chr11-100845696-DEL-52 |
| chr11-100862821-DEL-294 | chr11-100862822-DEL-294 |
| chr11-100869558-DEL-208 | chr11-100869559-DEL-208 |
| chr11-100873652-DEL-64 | chr11-100873653-DEL-64 |
| chr11-100885509-DEL-63 | chr11-100885510-DEL-63 |
| chr11-100887657-DEL-60 | chr11-100887650-DEL-54 |
| chr11-100893278-DEL-63 | chr11-100893278-DEL-52 |
| chr11-100894719-DEL-243 | chr11-100894707-DEL-241 |
| chr11-100898705-DEL-101 | chr11-100898706-DEL-101 |
| chr11-100950064-DEL-207 | chr11-100950059-DEL-201 |
| chr11-100966255-DEL-220 | chr11-100966256-DEL-220 |
| chr11-101016752-DEL-68 | chr11-101016753-DEL-68 |
| chr11-101020733-DEL-93 | chr11-101020742-DEL-85 |
| chr11-101023212-DEL-171 | chr11-101023213-DEL-171 |
| chr11-101024585-DEL-184 | chr11-101024586-DEL-184 |
| chr11-101047833-DEL-224 | chr11-101047834-DEL-224 |
| chr11-101048344-DEL-111 | chr11-101048345-DEL-111 |
| chr11-101129883-DEL-806 | chr11-101129884-DEL-806 |
| chr11-101140560-DEL-60 | chr11-101140561-DEL-61 |
| chr11-101146064-DEL-187 | chr11-101146064-DEL-188 |
| chr11-101158641-DEL-193 | chr11-101158642-DEL-193 |
| chr11-101231249-DEL-212 | chr11-101231250-DEL-212 |
| chr11-101242009-DEL-120 | chr11-101242010-DEL-120 |
| chr11-101289829-DEL-401 | chr11-101289830-DEL-401 |
| chr11-101304003-DEL-67 | chr11-101304003-DEL-67 |
| chr11-101318809-DEL-210 | chr11-101318811-DEL-209 |
| chr11-101329039-DEL-242 | chr11-101329040-DEL-242 |
| chr11-101329949-DEL-195 | chr11-101329950-DEL-195 |
| chr11-101330723-DEL-212 | chr11-101330723-DEL-212 |
| chr11-101331684-DEL-204 | chr11-101331685-DEL-204 |
| chr11-101337489-DEL-199 | chr11-101337490-DEL-199 |
| chr11-101338023-DEL-227 | chr11-101338024-DEL-227 |
| chr11-101346933-DEL-64 | chr11-101346933-DEL-55 |
| chr11-101360474-DEL-223 | chr11-101360475-DEL-223 |
| chr11-101366837-DEL-214 | chr11-101366838-DEL-214 |
| chr11-101381440-DEL-196 | chr11-101381441-DEL-196 |
| chr11-101384471-DEL-205 | chr11-101384472-DEL-205 |
| chr11-101401023-DEL-60 | chr11-101401024-DEL-60 |
| chr11-101429577-DEL-196 | chr11-101429565-DEL-199 |
| chr11-101457599-DEL-206 | chr11-101457600-DEL-206 |
| chr11-101458158-DEL-221 | chr11-101458159-DEL-221 |
| chr11-101470068-DEL-150 | chr11-101470069-DEL-150 |
| chr11-101478483-DEL-51 | chr11-101478484-DEL-51 |
| chr11-101516578-DEL-177 | chr11-101516586-DEL-174 |
| chr11-101534800-DEL-215 | chr11-101534801-DEL-215 |
| chr11-101535740-DEL-232 | chr11-101535756-DEL-195 |
| chr11-101538164-DEL-229 | chr11-101538165-DEL-229 |
| chr11-101560919-DEL-215 | chr11-101560920-DEL-215 |
| chr11-101567070-DEL-192 | chr11-101567071-DEL-192 |
| chr11-101589198-DEL-96 | chr11-101589199-DEL-96 |
| chr11-101601427-DEL-199 | chr11-101601428-DEL-199 |
| chr11-101612434-DEL-53 | chr11-101612433-DEL-54 |
| chr11-101639336-DEL-69 | chr11-101639337-DEL-69 |
| chr11-101655846-DEL-78 | chr11-101655861-DEL-62 |
| chr11-101719396-DEL-364 | chr11-101719396-DEL-364 |
| chr11-101736011-DEL-194 | chr11-101736012-DEL-194 |
| chr11-101759658-DEL-59 | chr11-101759669-DEL-50 |
| chr11-101786494-DEL-89 | chr11-101786495-DEL-89 |

| | |
|---|---|
| chr11-101808088-DEL-60 | chr11-101808089-DEL-60 |
| chr11-101813919-DEL-250 | chr11-101813920-DEL-250 |
| chr11-101840532-DEL-550 | chr11-101840533-DEL-550 |
| chr11-101893514-DEL-195 | chr11-101893526-DEL-191 |
| chr11-101905593-DEL-113 | chr11-101905594-DEL-113 |
| chr11-101953719-DEL-113 | chr11-101953715-DEL-114 |
| chr11-101981217-DEL-68 | chr11-101981217-DEL-69 |
| chr11-102000288-DEL-176 | chr11-102000289-DEL-176 |
| chr11-102014860-DEL-642 | chr11-102014861-DEL-642 |
| chr11-102016462-DEL-184 | chr11-102016463-DEL-184 |
| chr11-102020343-DEL-62 | chr11-102020342-DEL-64 |
| chr11-102022977-DEL-76 | chr11-102022939-DEL-119 |
| chr11-102032737-DEL-216 | chr11-102032738-DEL-216 |
| chr11-102035340-DEL-343 | chr11-102035341-DEL-343 |
| chr11-102048894-DEL-52 | chr11-102048891-DEL-56 |
| chr11-102134721-DEL-165 | chr11-102134727-DEL-159 |
| chr11-102141096-DEL-60 | chr11-102141095-DEL-65 |
| chr11-102156522-DEL-91 | chr11-102156523-DEL-91 |
| chr11-102173543-DEL-621 | chr11-102173545-DEL-618 |
| chr11-102216068-DEL-209 | chr11-102216069-DEL-209 |
| chr11-102230864-DEL-207 | chr11-102230865-DEL-207 |
| chr11-102231718-DEL-92 | chr11-102231719-DEL-92 |
| chr11-102262649-DEL-60 | chr11-102262654-DEL-56 |
| chr11-102268342-DEL-153 | chr11-102268343-DEL-153 |
| chr11-102280462-DEL-219 | chr11-102280463-DEL-219 |
| chr11-102362815-DEL-1594 | chr11-102362816-DEL-1594 |
| chr11-102394917-DEL-1818 | chr11-102394918-DEL-1818 |
| chr11-102414061-DEL-184 | chr11-102414062-DEL-184 |
| chr11-102423798-DEL-55 | chr11-102423799-DEL-55 |
| chr11-102431872-DEL-234 | chr11-102431873-DEL-234 |
| chr11-102460161-DEL-145 | chr11-102460162-DEL-145 |
| chr11-102478186-DEL-251 | chr11-102478187-DEL-251 |
| chr11-102484205-DEL-269 | chr11-102484200-DEL-270 |
| chr11-102489357-DEL-431 | chr11-102489358-DEL-431 |
| chr11-102507104-DEL-3434 | chr11-102507108-DEL-3432 |
| chr11-102568118-DEL-80 | chr11-102568119-DEL-80 |
| chr11-102582755-DEL-69 | chr11-102582763-DEL-61 |
| chr11-102621151-DEL-72 | chr11-102621152-DEL-72 |
| chr11-102635312-DEL-216 | chr11-102635313-DEL-216 |
| chr11-102636293-DEL-67 | chr11-102636293-DEL-79 |
| chr11-102640482-DEL-79 | chr11-102640448-DEL-80 |
| chr11-102675122-DEL-2772 | chr11-102675125-DEL-2770 |
| chr11-102675542-DEL-95 | chr11-102675543-DEL-95 |
| chr11-102677963-DEL-339 | chr11-102677964-DEL-339 |
| chr11-102689036-DEL-201 | chr11-102689037-DEL-201 |
| chr11-102692890-DEL-160 | chr11-102692890-DEL-160 |
| chr11-102702762-DEL-56 | chr11-102702764-DEL-55 |
| chr11-102712529-DEL-54 | chr11-102712552-DEL-64 |
| chr11-102717785-DEL-62 | chr11-102717794-DEL-53 |
| chr11-102723930-DEL-63 | chr11-102723911-DEL-63 |
| chr11-102726531-DEL-75 | chr11-102726512-DEL-58 |
| chr11-102733184-DEL-70 | chr11-102733203-DEL-71 |
| chr11-102748562-DEL-102 | chr11-102748563-DEL-102 |
| chr11-102758064-DEL-75 | chr11-102758065-DEL-75 |
| chr11-102763459-DEL-1144 | chr11-102763490-DEL-1102 |
| chr11-102781655-DEL-104 | chr11-102781655-DEL-104 |
| chr11-102883498-DEL-116 | chr11-102883499-DEL-116 |
| chr11-102894899-DEL-64 | chr11-102894900-DEL-64 |
| chr11-102900319-DEL-204 | chr11-102900319-DEL-204 |
| chr11-102908278-DEL-103 | chr11-102908279-DEL-103 |
| chr11-102910002-DEL-171 | chr11-102910002-DEL-171 |
| chr11-102913272-DEL-202 | chr11-102913227-DEL-240 |
| chr11-102920533-DEL-174 | chr11-102920534-DEL-174 |
| chr11-102938720-DEL-1772 | chr11-102938721-DEL-1772 |
| chr11-102941886-DEL-203 | chr11-102941887-DEL-203 |
| chr11-102974483-DEL-9047 | chr11-102974484-DEL-9047 |
| chr11-102983571-DEL-603 | chr11-102983571-DEL-603 |
| chr11-102999537-DEL-53 | chr11-102999536-DEL-50 |
| chr11-103011712-DEL-887 | chr11-103011713-DEL-887 |
| chr11-103033041-DEL-50 | chr11-103033057-DEL-59 |
| chr11-103040339-DEL-52 | chr11-103040338-DEL-53 |
| chr11-103078284-DEL-194 | chr11-103078287-DEL-192 |
| chr11-103097164-DEL-381 | chr11-103097165-DEL-381 |
| chr11-103120895-DEL-105 | chr11-103120896-DEL-105 |
| chr11-103206693-DEL-486 | chr11-103206694-DEL-486 |
| chr11-103222074-DEL-69 | chr11-103222075-DEL-69 |
| chr11-103237377-DEL-109 | chr11-103237378-DEL-109 |
| chr11-103238277-DEL-86 | chr11-103238276-DEL-88 |
| chr11-103239040-DEL-62 | chr11-103239041-DEL-62 |
| chr11-103246443-DEL-60 | chr11-103246444-DEL-60 |

| | |
|---|---|
| chr11-103284347-DEL-69 | chr11-103284348-DEL-69 |
| chr11-103295593-DEL-72 | chr11-103295594-DEL-72 |
| chr11-103296424-DEL-212 | chr11-103296425-DEL-212 |
| chr11-103311786-DEL-125 | chr11-103311787-DEL-125 |
| chr11-103318905-DEL-75 | chr11-103318858-DEL-76 |
| chr11-103359078-DEL-72 | chr11-103359079-DEL-72 |
| chr11-103363403-DEL-232 | chr11-103363404-DEL-232 |
| chr11-103401624-DEL-54 | chr11-103401625-DEL-54 |
| chr11-103411837-DEL-218 | chr11-103411816-DEL-222 |
| chr11-103430821-DEL-525 | chr11-103430822-DEL-525 |
| chr11-103457360-DEL-218 | chr11-103457361-DEL-218 |
| chr11-103460485-DEL-126 | chr11-103460506-DEL-104 |
| chr11-103474246-DEL-69 | chr11-103474241-DEL-55 |
| chr11-103494911-DEL-71 | chr11-103494912-DEL-71 |
| chr11-103576962-DEL-65 | chr11-103576963-DEL-65 |
| chr11-103608155-DEL-71 | chr11-103608156-DEL-71 |
| chr11-103618068-DEL-60 | chr11-103618069-DEL-60 |
| chr11-103709654-DEL-181 | chr11-103709662-DEL-174 |
| chr11-103717482-DEL-332 | chr11-103717484-DEL-331 |
| chr11-103725102-DEL-217 | chr11-103725103-DEL-217 |
| chr11-103733864-DEL-78 | chr11-103733816-DEL-99 |
| chr11-103769406-DEL-162 | chr11-103769407-DEL-162 |
| chr11-103798403-DEL-59 | chr11-103798427-DEL-58 |
| chr11-103823019-DEL-248 | chr11-103823019-DEL-248 |
| chr11-103830865-DEL-108 | chr11-103830866-DEL-108 |
| chr11-103856676-DEL-197 | chr11-103856677-DEL-197 |
| chr11-103881084-DEL-118 | chr11-103881071-DEL-119 |
| chr11-103907130-DEL-54 | chr11-103907119-DEL-52 |
| chr11-103911287-DEL-113 | chr11-103911288-DEL-112 |
| chr11-103929974-DEL-802 | chr11-103929975-DEL-802 |
| chr11-103937381-DEL-1753 | chr11-103937382-DEL-1753 |
| chr11-103963490-DEL-472 | chr11-103963491-DEL-472 |
| chr11-103964220-DEL-409 | chr11-103964221-DEL-409 |
| chr11-103973310-DEL-61 | chr11-103973311-DEL-61 |
| chr11-104002075-DEL-62 | chr11-104002090-DEL-53 |
| chr11-104012219-DEL-78 | chr11-104012220-DEL-78 |
| chr11-104028625-DEL-52 | chr11-104028626-DEL-52 |
| chr11-104099522-DEL-119 | chr11-104099524-DEL-120 |
| chr11-104105210-DEL-192 | chr11-104105211-DEL-192 |
| chr11-104129835-DEL-3638 | chr11-104129832-DEL-3639 |
| chr11-104151342-DEL-203 | chr11-104151336-DEL-204 |
| chr11-104174152-DEL-192 | chr11-104174153-DEL-192 |
| chr11-104174944-DEL-64 | chr11-104174945-DEL-64 |
| chr11-104193895-DEL-72 | chr11-104193896-DEL-72 |
| chr11-104203539-DEL-199 | chr11-104203528-DEL-200 |
| chr11-104211000-DEL-153 | chr11-104211001-DEL-153 |
| chr11-104214109-DEL-198 | chr11-104214110-DEL-198 |
| chr11-104215311-DEL-206 | chr11-104215312-DEL-206 |
| chr11-104227701-DEL-212 | chr11-104227702-DEL-212 |
| chr11-104324761-DEL-1261 | chr11-104324762-DEL-1261 |
| chr11-104380799-DEL-56 | chr11-104380800-DEL-56 |
| chr11-104401353-DEL-64 | chr11-104401354-DEL-64 |
| chr11-104433444-DEL-203 | chr11-104433445-DEL-203 |
| chr11-104464351-DEL-2152 | chr11-104464352-DEL-2152 |
| chr11-104540040-DEL-6369 | chr11-104540041-DEL-6369 |
| chr11-104563238-DEL-225 | chr11-104563239-DEL-225 |
| chr11-104586699-DEL-70 | chr11-104586700-DEL-70 |
| chr11-104590289-DEL-220 | chr11-104590290-DEL-220 |
| chr11-104596387-DEL-207 | chr11-104596388-DEL-207 |
| chr11-104629294-DEL-97 | chr11-104629343-DEL-91 |
| chr11-104655453-DEL-662 | chr11-104655454-DEL-662 |
| chr11-104668552-DEL-76 | chr11-104668553-DEL-76 |
| chr11-104668865-DEL-704 | chr11-104668866-DEL-704 |
| chr11-104679835-DEL-78 | chr11-104679836-DEL-78 |
| chr11-104680675-DEL-432 | chr11-104680681-DEL-412 |
| chr11-104698892-DEL-6212 | chr11-104698889-DEL-6213 |
| chr11-104704842-DEL-197 | chr11-104704843-DEL-197 |
| chr11-104728582-DEL-1183 | chr11-104728583-DEL-1183 |
| chr11-104729093-DEL-121 | chr11-104729094-DEL-121 |
| chr11-104750434-DEL-2721 | chr11-104750435-DEL-2721 |
| chr11-104794345-DEL-206 | chr11-104794346-DEL-206 |
| chr11-104843763-DEL-73 | chr11-104843766-DEL-69 |
| chr11-104883711-DEL-709 | chr11-104883712-DEL-709 |
| chr11-104893616-DEL-541 | chr11-104893617-DEL-541 |
| chr11-104923338-DEL-226 | chr11-104923339-DEL-226 |
| chr11-104945068-DEL-1508 | chr11-104945069-DEL-1508 |
| chr11-104954639-DEL-74 | chr11-104954637-DEL-77 |
| chr11-104969695-DEL-587 | chr11-104969696-DEL-587 |
| chr11-104980224-DEL-1758 | chr11-104980225-DEL-1758 |
| chr11-104994510-DEL-218 | chr11-104994511-DEL-218 |

| | |
|---|---|
| chr11-104996804-DEL-180 | chr11-104996805-DEL-180 |
| chr11-105003412-DEL-68 | chr11-105003380-DEL-69 |
| chr11-105050468-DEL-95 | chr11-105050470-DEL-94 |
| chr11-105051820-DEL-221 | chr11-105051821-DEL-221 |
| chr11-105066104-DEL-214 | chr11-105066105-DEL-214 |
| chr11-105096650-DEL-244 | chr11-105096642-DEL-243 |
| chr11-105150482-DEL-135 | chr11-105150438-DEL-180 |
| chr11-105167969-DEL-58 | chr11-105167969-DEL-58 |
| chr11-105207647-DEL-78 | chr11-105207648-DEL-78 |
| chr11-105210543-DEL-180 | chr11-105210544-DEL-180 |
| chr11-105215889-DEL-55 | chr11-105215890-DEL-55 |
| chr11-105248663-DEL-66 | chr11-105248701-DEL-76 |
| chr11-105268262-DEL-53 | chr11-105268263-DEL-53 |
| chr11-105295480-DEL-67 | chr11-105295481-DEL-67 |
| chr11-105297303-DEL-274 | chr11-105297304-DEL-274 |
| chr11-105319000-DEL-6440 | chr11-105319001-DEL-6440 |
| chr11-105351463-DEL-96 | chr11-105351464-DEL-96 |
| chr11-105359003-DEL-152 | chr11-105359004-DEL-152 |
| chr11-105430034-DEL-93 | chr11-105430035-DEL-93 |
| chr11-105447320-DEL-91 | chr11-105447321-DEL-91 |
| chr11-105488460-DEL-54 | chr11-105488461-DEL-54 |
| chr11-105548753-DEL-389 | chr11-105548754-DEL-389 |
| chr11-105561845-DEL-122 | chr11-105561846-DEL-122 |
| chr11-105596350-DEL-6315 | chr11-105596361-DEL-6316 |
| chr11-105754067-DEL-67 | chr11-105754068-DEL-67 |
| chr11-105788922-DEL-239 | chr11-105788923-DEL-239 |
| chr11-105791521-DEL-180 | chr11-105791522-DEL-180 |
| chr11-105855921-DEL-359 | chr11-105855921-DEL-324 |
| chr11-105882183-DEL-70 | chr11-105882171-DEL-51 |
| chr11-105919065-DEL-5321 | chr11-105919066-DEL-5321 |
| chr11-105960089-DEL-97 | chr11-105960090-DEL-97 |
| chr11-105969241-DEL-225 | chr11-105969242-DEL-225 |
| chr11-106017733-DEL-236 | chr11-106017730-DEL-237 |
| chr11-106062534-DEL-198 | chr11-106062535-DEL-198 |
| chr11-106110078-DEL-66 | chr11-106110079-DEL-66 |
| chr11-106179603-DEL-346 | chr11-106179604-DEL-346 |
| chr11-106183531-DEL-4878 | chr11-106183532-DEL-4878 |
| chr11-106193686-DEL-95 | chr11-106193687-DEL-96 |
| chr11-106247851-DEL-120 | chr11-106247866-DEL-133 |
| chr11-106250937-DEL-645 | chr11-106250938-DEL-645 |
| chr11-106306211-DEL-324 | chr11-106306212-DEL-324 |
| chr11-106333326-DEL-207 | chr11-106333327-DEL-207 |
| chr11-106340428-DEL-207 | chr11-106340429-DEL-207 |
| chr11-106344769-DEL-205 | chr11-106344761-DEL-177 |
| chr11-106396638-DEL-71 | chr11-106396639-DEL-71 |
| chr11-106452585-DEL-220 | chr11-106452586-DEL-220 |
| chr11-106456654-DEL-235 | chr11-106456655-DEL-235 |
| chr11-106475118-DEL-158 | chr11-106475083-DEL-156 |
| chr11-106524938-DEL-217 | chr11-106524939-DEL-217 |
| chr11-106532519-DEL-555 | chr11-106532520-DEL-555 |
| chr11-106554441-DEL-108 | chr11-106554442-DEL-108 |
| chr11-106584780-DEL-498 | chr11-106584781-DEL-498 |
| chr11-106631808-DEL-172 | chr11-106631809-DEL-172 |
| chr11-106635104-DEL-75 | chr11-106635106-DEL-78 |
| chr11-106636166-DEL-709 | chr11-106636167-DEL-709 |
| chr11-106637141-DEL-358 | chr11-106637142-DEL-358 |
| chr11-106662713-DEL-204 | chr11-106662714-DEL-204 |
| chr11-106667243-DEL-150 | chr11-106667244-DEL-150 |
| chr11-106668084-DEL-72 | chr11-106668082-DEL-64 |
| chr11-106682403-DEL-54 | chr11-106682404-DEL-54 |
| chr11-106692407-DEL-197 | chr11-106692408-DEL-197 |
| chr11-106696788-DEL-224 | chr11-106696789-DEL-224 |
| chr11-106698183-DEL-214 | chr11-106698184-DEL-214 |
| chr11-106782620-DEL-105 | chr11-106782620-DEL-105 |
| chr11-106799789-DEL-59 | chr11-106799772-DEL-70 |
| chr11-106821974-DEL-203 | chr11-106821975-DEL-203 |
| chr11-106827302-DEL-175 | chr11-106827303-DEL-175 |
| chr11-106832947-DEL-912 | chr11-106832924-DEL-904 |
| chr11-106846084-DEL-139 | chr11-106846067-DEL-136 |
| chr11-106853518-DEL-220 | chr11-106853519-DEL-220 |
| chr11-106856057-DEL-351 | chr11-106856058-DEL-351 |
| chr11-106868606-DEL-196 | chr11-106868607-DEL-196 |
| chr11-106872204-DEL-174 | chr11-106872205-DEL-174 |
| chr11-106884558-DEL-243 | chr11-106884559-DEL-243 |
| chr11-106891174-DEL-182 | chr11-106891175-DEL-182 |
| chr11-106895878-DEL-234 | chr11-106895879-DEL-234 |
| chr11-106896835-DEL-104 | chr11-106896836-DEL-104 |
| chr11-106900905-DEL-195 | chr11-106900906-DEL-195 |
| chr11-106906611-DEL-247 | chr11-106906598-DEL-211 |
| chr11-106910761-DEL-203 | chr11-106910762-DEL-203 |

| | |
|---|---|
| chr11-106914426-DEL-215 | chr11-106914427-DEL-215 |
| chr11-106940311-DEL-578 | chr11-106940311-DEL-578 |
| chr11-106955159-DEL-209 | chr11-106955160-DEL-209 |
| chr11-107016346-DEL-205 | chr11-107016347-DEL-205 |
| chr11-107025302-DEL-212 | chr11-107025303-DEL-212 |
| chr11-107035072-DEL-237 | chr11-107035073-DEL-237 |
| chr11-107049701-DEL-89 | chr11-107049740-DEL-59 |
| chr11-107053767-DEL-563 | chr11-107053768-DEL-563 |
| chr11-107070129-DEL-171 | chr11-107070130-DEL-171 |
| chr11-107089246-DEL-240 | chr11-107089279-DEL-240 |
| chr11-107126234-DEL-358 | chr11-107126235-DEL-358 |
| chr11-107160797-DEL-165 | chr11-107160798-DEL-165 |
| chr11-107180052-DEL-198 | chr11-107180058-DEL-192 |
| chr11-107185128-DEL-101 | chr11-107185129-DEL-101 |
| chr11-107192028-DEL-236 | chr11-107192029-DEL-236 |
| chr11-107194029-DEL-170 | chr11-107194030-DEL-170 |
| chr11-107197612-DEL-199 | chr11-107197613-DEL-199 |
| chr11-107234471-DEL-462 | chr11-107234472-DEL-462 |
| chr11-107264183-DEL-514 | chr11-107264183-DEL-515 |
| chr11-107269670-DEL-178 | chr11-107269671-DEL-178 |
| chr11-107271392-DEL-203 | chr11-107271393-DEL-203 |
| chr11-107282421-DEL-206 | chr11-107282422-DEL-206 |
| chr11-107303400-DEL-359 | chr11-107303399-DEL-352 |
| chr11-107314710-DEL-78 | chr11-107314712-DEL-79 |
| chr11-107318030-DEL-117 | chr11-107318031-DEL-117 |
| chr11-107332217-DEL-227 | chr11-107332213-DEL-232 |
| chr11-107347011-DEL-196 | chr11-107347012-DEL-196 |
| chr11-107377924-DEL-217 | chr11-107377925-DEL-217 |
| chr11-107395452-DEL-180 | chr11-107395445-DEL-181 |
| chr11-107411505-DEL-98 | chr11-107411506-DEL-98 |
| chr11-107423386-DEL-132 | chr11-107423387-DEL-132 |
| chr11-107424179-DEL-209 | chr11-107424179-DEL-209 |
| chr11-107431616-DEL-204 | chr11-107431617-DEL-204 |
| chr11-107449652-DEL-94 | chr11-107449653-DEL-94 |
| chr11-107472567-DEL-217 | chr11-107472568-DEL-217 |
| chr11-107486685-DEL-209 | chr11-107486686-DEL-209 |
| chr11-107497634-DEL-4826 | chr11-107497635-DEL-4826 |
| chr11-107500050-DEL-207 | chr11-107500051-DEL-207 |
| chr11-107506814-DEL-214 | chr11-107506815-DEL-214 |
| chr11-107549094-DEL-102 | chr11-107549095-DEL-102 |
| chr11-107632482-DEL-83 | chr11-107632483-DEL-83 |
| chr11-107638655-DEL-97 | chr11-107638670-DEL-75 |
| chr11-107641761-DEL-67 | chr11-107641752-DEL-66 |
| chr11-107643370-DEL-77 | chr11-107643371-DEL-77 |
| chr11-107655188-DEL-98 | chr11-107655200-DEL-62 |
| chr11-107675461-DEL-232 | chr11-107675462-DEL-232 |
| chr11-107682012-DEL-74 | chr11-107682000-DEL-87 |
| chr11-107779024-DEL-463 | chr11-107779025-DEL-463 |
| chr11-107845560-DEL-200 | chr11-107845561-DEL-200 |
| chr11-107860545-DEL-51 | chr11-107860546-DEL-51 |
| chr11-107888630-DEL-90 | chr11-107888631-DEL-90 |
| chr11-107927340-DEL-349 | chr11-107927341-DEL-349 |
| chr11-107954717-DEL-132 | chr11-107954751-DEL-114 |
| chr11-107966328-DEL-56 | chr11-107966329-DEL-56 |
| chr11-107996671-DEL-6667 | chr11-107996672-DEL-6667 |
| chr11-108016064-DEL-62 | chr11-108016037-DEL-65 |
| chr11-108023794-DEL-53 | chr11-108023795-DEL-57 |
| chr11-108025583-DEL-107 | chr11-108025584-DEL-97 |
| chr11-108068352-DEL-98 | chr11-108068353-DEL-98 |
| chr11-108218983-DEL-106 | chr11-108218994-DEL-96 |
| chr11-108228166-DEL-483 | chr11-108228167-DEL-483 |
| chr11-108251233-DEL-163 | chr11-108251231-DEL-163 |
| chr11-108290011-DEL-5260 | chr11-108290016-DEL-5256 |
| chr11-108301352-DEL-500 | chr11-108301353-DEL-500 |
| chr11-108307145-DEL-60 | chr11-108307146-DEL-60 |
| chr11-108333394-DEL-51 | chr11-108333395-DEL-51 |
| chr11-108348392-DEL-54 | chr11-108348393-DEL-54 |
| chr11-108351126-DEL-168 | chr11-108351127-DEL-168 |
| chr11-108365846-DEL-498 | chr11-108365847-DEL-498 |
| chr11-108405958-DEL-260 | chr11-108405959-DEL-260 |
| chr11-108407471-DEL-51 | chr11-108407472-DEL-51 |
| chr11-108411664-DEL-1762 | chr11-108411665-DEL-1762 |
| chr11-108443058-DEL-81 | chr11-108443059-DEL-81 |
| chr11-108506763-DEL-88 | chr11-108506764-DEL-74 |
| chr11-108509977-DEL-643 | chr11-108509978-DEL-644 |
| chr11-108540719-DEL-5345 | chr11-108540720-DEL-5345 |
| chr11-108547131-DEL-54 | chr11-108547132-DEL-54 |
| chr11-108550954-DEL-52 | chr11-108550953-DEL-54 |
| chr11-108587945-DEL-550 | chr11-108587945-DEL-550 |
| chr11-108610257-DEL-293 | chr11-108610258-DEL-293 |

| | |
|---|---|
| chr11-108622872-DEL-59 | chr11-108622873-DEL-59 |
| chr11-108655469-DEL-185 | chr11-108655470-DEL-185 |
| chr11-108689701-DEL-75 | chr11-108689701-DEL-75 |
| chr11-108692960-DEL-298 | chr11-108692961-DEL-298 |
| chr11-108715209-DEL-2023 | chr11-108715209-DEL-2023 |
| chr11-108718825-DEL-175 | chr11-108718826-DEL-175 |
| chr11-108778314-DEL-498 | chr11-108778319-DEL-493 |
| chr11-108783238-DEL-51 | chr11-108783239-DEL-51 |
| chr11-108784875-DEL-75 | chr11-108784876-DEL-75 |
| chr11-108826706-DEL-53 | chr11-108826707-DEL-53 |
| chr11-108859064-DEL-621 | chr11-108859065-DEL-621 |
| chr11-108884765-DEL-349 | chr11-108884766-DEL-349 |
| chr11-108919041-DEL-544 | chr11-108919072-DEL-561 |
| chr11-108941873-DEL-2550 | chr11-108941874-DEL-2550 |
| chr11-108947011-DEL-265 | chr11-108947012-DEL-265 |
| chr11-108990205-DEL-111 | chr11-108990205-DEL-112 |
| chr11-109005182-DEL-120 | chr11-109005183-DEL-120 |
| chr11-109012452-DEL-108 | chr11-109012453-DEL-108 |
| chr11-109017941-DEL-82 | chr11-109017942-DEL-78 |
| chr11-109025600-DEL-69 | chr11-109025607-DEL-68 |
| chr11-109031807-DEL-87 | chr11-109031808-DEL-87 |
| chr11-109038594-DEL-116 | chr11-109038594-DEL-113 |
| chr11-109053735-DEL-52 | chr11-109053735-DEL-52 |
| chr11-109057854-DEL-60 | chr11-109057860-DEL-54 |
| chr11-109073891-DEL-347 | chr11-109073892-DEL-347 |
| chr11-109077022-DEL-110 | chr11-109077023-DEL-110 |
| chr11-109111335-DEL-216 | chr11-109111363-DEL-171 |
| chr11-109116653-DEL-60 | chr11-109116654-DEL-59 |
| chr11-109118638-DEL-79 | chr11-109118639-DEL-79 |
| chr11-109146766-DEL-81 | chr11-109146765-DEL-82 |
| chr11-109148570-DEL-106 | chr11-109148571-DEL-106 |
| chr11-109153092-DEL-96 | chr11-109153078-DEL-100 |
| chr11-109173770-DEL-1602 | chr11-109173770-DEL-1602 |
| chr11-109180444-DEL-64 | chr11-109180445-DEL-64 |
| chr11-109187469-DEL-52 | chr11-109187470-DEL-52 |
| chr11-109197984-DEL-215 | chr11-109197985-DEL-215 |
| chr11-109209470-DEL-122 | chr11-109209471-DEL-122 |
| chr11-109212697-DEL-111 | chr11-109212677-DEL-160 |
| chr11-109233768-DEL-60 | chr11-109233770-DEL-58 |
| chr11-109242187-DEL-65 | chr11-109242188-DEL-65 |
| chr11-109281381-DEL-167 | chr11-109281382-DEL-167 |
| chr11-109287785-DEL-84 | chr11-109287786-DEL-84 |
| chr11-109300153-DEL-54 | chr11-109300154-DEL-54 |
| chr11-109308593-DEL-60 | chr11-109308594-DEL-60 |
| chr11-109313585-DEL-248 | chr11-109313585-DEL-248 |
| chr11-109360822-DEL-51 | chr11-109360822-DEL-52 |
| chr11-109383843-DEL-139 | chr11-109383837-DEL-167 |
| chr11-109404271-DEL-72 | chr11-109404263-DEL-108 |
| chr11-109439068-DEL-226 | chr11-109439069-DEL-226 |
| chr11-109442359-DEL-112 | chr11-109442360-DEL-112 |
| chr11-109488249-DEL-78 | chr11-109488250-DEL-78 |
| chr11-109498917-DEL-112 | chr11-109498924-DEL-106 |
| chr11-109516362-DEL-136 | chr11-109516365-DEL-140 |
| chr11-109534680-DEL-394 | chr11-109534681-DEL-394 |
| chr11-109535465-DEL-601 | chr11-109535466-DEL-601 |
| chr11-109536205-DEL-62 | chr11-109536206-DEL-56 |
| chr11-109538434-DEL-196 | chr11-109538435-DEL-196 |
| chr11-109597155-DEL-65 | chr11-109597161-DEL-61 |
| chr11-109602203-DEL-102 | chr11-109602216-DEL-72 |
| chr11-109608805-DEL-66 | chr11-109608779-DEL-66 |
| chr11-109621001-DEL-395 | chr11-109621002-DEL-395 |
| chr11-109630838-DEL-1836 | chr11-109630839-DEL-1836 |
| chr11-109677596-DEL-202 | chr11-109677597-DEL-202 |
| chr11-109681044-DEL-97 | chr11-109681043-DEL-97 |
| chr11-109715446-DEL-75 | chr11-109715446-DEL-72 |
| chr11-109733639-DEL-60 | chr11-109733631-DEL-60 |
| chr11-109743169-DEL-1374 | chr11-109743170-DEL-1374 |
| chr11-109801578-DEL-105 | chr11-109801565-DEL-75 |
| chr11-109805669-DEL-119 | chr11-109805685-DEL-104 |
| chr11-109823923-DEL-68 | chr11-109823924-DEL-68 |
| chr11-109885753-DEL-60 | chr11-109885754-DEL-60 |
| chr11-109930267-DEL-97 | chr11-109930269-DEL-95 |
| chr11-109955174-DEL-633 | chr11-109955175-DEL-626 |
| chr11-109957603-DEL-61 | chr11-109957604-DEL-61 |
| chr11-109960420-DEL-85 | chr11-109960421-DEL-85 |
| chr11-110004252-DEL-54 | chr11-110004226-DEL-54 |
| chr11-110050253-DEL-115 | chr11-110050254-DEL-115 |
| chr11-110055797-DEL-152 | chr11-110055798-DEL-152 |
| chr11-110061275-DEL-585 | chr11-110061276-DEL-584 |
| chr11-110062525-DEL-1259 | chr11-110062526-DEL-1259 |

| | |
|---|---|
| chr11-110083017-DEL-55 | chr11-110083018-DEL-55 |
| chr11-110083843-DEL-187 | chr11-110083844-DEL-187 |
| chr11-110095067-DEL-209 | chr11-110095068-DEL-209 |
| chr11-110097423-DEL-119 | chr11-110097420-DEL-113 |
| chr11-110151528-DEL-147 | chr11-110151529-DEL-147 |
| chr11-110152773-DEL-66 | chr11-110152774-DEL-66 |
| chr11-110154744-DEL-1806 | chr11-110154745-DEL-1806 |
| chr11-110185819-DEL-61 | chr11-110185818-DEL-61 |
| chr11-110198352-DEL-588 | chr11-110198351-DEL-589 |
| chr11-110201922-DEL-62 | chr11-110201923-DEL-62 |
| chr11-110207314-DEL-709 | chr11-110207315-DEL-709 |
| chr11-110226666-DEL-60 | chr11-110226667-DEL-60 |
| chr11-110231368-DEL-2155 | chr11-110231369-DEL-2155 |
| chr11-110256362-DEL-194 | chr11-110256368-DEL-193 |
| chr11-110266855-DEL-175 | chr11-110266856-DEL-175 |
| chr11-110269644-DEL-206 | chr11-110269645-DEL-206 |
| chr11-110275265-DEL-62 | chr11-110275276-DEL-52 |
| chr11-110294212-DEL-87 | chr11-110294213-DEL-87 |
| chr11-110332601-DEL-55 | chr11-110332602-DEL-55 |
| chr11-110347003-DEL-223 | chr11-110347004-DEL-223 |
| chr11-110356108-DEL-104 | chr11-110356147-DEL-61 |
| chr11-110363894-DEL-71 | chr11-110363895-DEL-71 |
| chr11-110420178-DEL-209 | chr11-110420179-DEL-209 |
| chr11-110424108-DEL-56 | chr11-110424109-DEL-56 |
| chr11-110428638-DEL-79 | chr11-110428639-DEL-79 |
| chr11-110439098-DEL-1113 | chr11-110439099-DEL-1113 |
| chr11-110471714-DEL-584 | chr11-110471730-DEL-563 |
| chr11-110497198-DEL-138 | chr11-110497206-DEL-141 |
| chr11-110516479-DEL-56 | chr11-110516480-DEL-60 |
| chr11-110531852-DEL-510 | chr11-110531852-DEL-510 |
| chr11-110550284-DEL-54 | chr11-110550285-DEL-54 |
| chr11-110551005-DEL-151 | chr11-110551009-DEL-147 |
| chr11-110717011-DEL-54 | chr11-110717011-DEL-54 |
| chr11-110743745-DEL-57 | chr11-110743744-DEL-55 |
| chr11-110748730-DEL-65 | chr11-110748744-DEL-56 |
| chr11-110751682-DEL-101 | chr11-110751687-DEL-97 |
| chr11-110753908-DEL-224 | chr11-110753956-DEL-176 |
| chr11-110775240-DEL-607 | chr11-110775241-DEL-607 |
| chr11-110777565-DEL-273 | chr11-110777566-DEL-273 |
| chr11-110826395-DEL-80 | chr11-110826403-DEL-52 |
| chr11-110861778-DEL-60 | chr11-110861774-DEL-65 |
| chr11-110936673-DEL-68 | chr11-110936674-DEL-68 |
| chr11-110983136-DEL-327 | chr11-110983137-DEL-327 |
| chr11-110988727-DEL-55 | chr11-110988728-DEL-50 |
| chr11-110993994-DEL-259 | chr11-110993995-DEL-259 |
| chr11-111007956-DEL-59 | chr11-111007909-DEL-60 |
| chr11-111063882-DEL-810 | chr11-111063883-DEL-810 |
| chr11-111073330-DEL-7113 | chr11-111073331-DEL-7113 |
| chr11-111082363-DEL-2988 | chr11-111082364-DEL-2987 |
| chr11-111106479-DEL-7137 | chr11-111106480-DEL-7137 |
| chr11-111121524-DEL-54 | chr11-111121525-DEL-54 |
| chr11-111132932-DEL-80 | chr11-111132882-DEL-84 |
| chr11-111159707-DEL-79 | chr11-111159708-DEL-79 |
| chr11-111199659-DEL-1675 | chr11-111199660-DEL-1675 |
| chr11-111201424-DEL-74 | chr11-111201427-DEL-60 |
| chr11-111205856-DEL-104 | chr11-111205905-DEL-60 |
| chr11-111244301-DEL-7107 | chr11-111244296-DEL-7112 |
| chr11-111259562-DEL-769 | chr11-111259563-DEL-769 |
| chr11-111260901-DEL-197 | chr11-111260902-DEL-197 |
| chr11-111310588-DEL-437 | chr11-111310589-DEL-437 |
| chr11-111344156-DEL-6215 | chr11-111344150-DEL-6222 |
| chr11-111361177-DEL-174 | chr11-111361178-DEL-174 |
| chr11-111400106-DEL-52 | chr11-111400107-DEL-50 |
| chr11-111400859-DEL-169 | chr11-111400837-DEL-192 |
| chr11-111438938-DEL-88 | chr11-111438939-DEL-88 |
| chr11-111445462-DEL-54 | chr11-111445463-DEL-54 |
| chr11-111461544-DEL-170 | chr11-111461545-DEL-170 |
| chr11-111495180-DEL-87 | chr11-111495202-DEL-54 |
| chr11-111507427-DEL-100 | chr11-111507428-DEL-100 |
| chr11-111536623-DEL-525 | chr11-111536624-DEL-525 |
| chr11-111561693-DEL-50 | chr11-111561652-DEL-91 |
| chr11-111570453-DEL-68 | chr11-111570454-DEL-72 |
| chr11-111594380-DEL-224 | chr11-111594374-DEL-227 |
| chr11-111603872-DEL-53 | chr11-111603873-DEL-51 |
| chr11-111613706-DEL-57 | chr11-111613691-DEL-59 |
| chr11-111615255-DEL-1193 | chr11-111615255-DEL-1193 |
| chr11-111617236-DEL-718 | chr11-111617237-DEL-718 |
| chr11-111648982-DEL-129 | chr11-111648981-DEL-130 |
| chr11-111652641-DEL-55 | chr11-111652644-DEL-56 |
| chr11-111691288-DEL-1117 | chr11-111691289-DEL-1117 |

| | |
|---|---|
| chr11-111705314-DEL-54 | chr11-111705315-DEL-54 |
| chr11-111746195-DEL-1463 | chr11-111746196-DEL-1463 |
| chr11-111746728-DEL-570 | chr11-111746729-DEL-570 |
| chr11-111825264-DEL-61 | chr11-111825267-DEL-59 |
| chr11-111831368-DEL-106 | chr11-111831369-DEL-90 |
| chr11-111847109-DEL-151 | chr11-111847110-DEL-151 |
| chr11-111856289-DEL-548 | chr11-111856283-DEL-554 |
| chr11-111880837-DEL-285 | chr11-111880838-DEL-285 |
| chr11-111892154-DEL-83 | chr11-111892155-DEL-83 |
| chr11-111901771-DEL-6139 | chr11-111901775-DEL-6136 |
| chr11-111913526-DEL-70 | chr11-111913538-DEL-90 |
| chr11-111940546-DEL-81 | chr11-111940547-DEL-81 |
| chr11-111944928-DEL-177 | chr11-111944916-DEL-178 |
| chr11-111949454-DEL-52 | chr11-111949444-DEL-62 |
| chr11-111951817-DEL-772 | chr11-111951815-DEL-769 |
| chr11-111953229-DEL-81 | chr11-111953230-DEL-81 |
| chr11-111960156-DEL-72 | chr11-111960155-DEL-96 |
| chr11-111965786-DEL-50 | chr11-111965786-DEL-50 |
| chr11-111968666-DEL-117 | chr11-111968667-DEL-117 |
| chr11-111973454-DEL-56 | chr11-111973455-DEL-56 |
| chr11-112013928-DEL-94 | chr11-112013956-DEL-105 |
| chr11-112022396-DEL-63 | chr11-112022388-DEL-82 |
| chr11-112055217-DEL-119 | chr11-112055218-DEL-118 |
| chr11-112058589-DEL-52 | chr11-112058590-DEL-52 |
| chr11-112075432-DEL-768 | chr11-112075433-DEL-768 |
| chr11-112093211-DEL-400 | chr11-112093212-DEL-400 |
| chr11-112114579-DEL-114 | chr11-112114601-DEL-93 |
| chr11-112119005-DEL-123 | chr11-112119007-DEL-124 |
| chr11-112133440-DEL-69 | chr11-112133429-DEL-81 |
| chr11-112148136-DEL-634 | chr11-112148137-DEL-634 |
| chr11-112187156-DEL-169 | chr11-112187157-DEL-169 |
| chr11-112187811-DEL-60 | chr11-112187812-DEL-60 |
| chr11-112192208-DEL-92 | chr11-112192209-DEL-92 |
| chr11-112243258-DEL-56 | chr11-112243259-DEL-56 |
| chr11-112244953-DEL-277 | chr11-112244953-DEL-274 |
| chr11-112250436-DEL-51 | chr11-112250436-DEL-51 |
| chr11-112256417-DEL-289 | chr11-112256418-DEL-289 |
| chr11-112267697-DEL-2998 | chr11-112267698-DEL-2998 |
| chr11-112294004-DEL-82 | chr11-112294052-DEL-58 |
| chr11-112320769-DEL-3256 | chr11-112320770-DEL-3256 |
| chr11-112329627-DEL-179 | chr11-112329628-DEL-179 |
| chr11-112336613-DEL-50 | chr11-112336581-DEL-59 |
| chr11-112350475-DEL-348 | chr11-112350456-DEL-359 |
| chr11-112351167-DEL-51 | chr11-112351168-DEL-51 |
| chr11-112377965-DEL-647 | chr11-112377966-DEL-647 |
| chr11-112383939-DEL-306 | chr11-112383940-DEL-306 |
| chr11-112404472-DEL-1120 | chr11-112404473-DEL-1120 |
| chr11-112412257-DEL-54 | chr11-112412258-DEL-54 |
| chr11-112438522-DEL-1088 | chr11-112438523-DEL-1088 |
| chr11-112483051-DEL-78 | chr11-112483054-DEL-68 |
| chr11-112492357-DEL-63 | chr11-112492358-DEL-63 |
| chr11-112503306-DEL-71 | chr11-112503307-DEL-50 |
| chr11-112519202-DEL-375 | chr11-112519203-DEL-375 |
| chr11-112528442-DEL-179 | chr11-112528440-DEL-173 |
| chr11-112529600-DEL-72 | chr11-112529607-DEL-66 |
| chr11-112564721-DEL-212 | chr11-112564722-DEL-212 |
| chr11-112620299-DEL-207 | chr11-112620300-DEL-207 |
| chr11-112620951-DEL-78 | chr11-112620952-DEL-78 |
| chr11-112636523-DEL-1106 | chr11-112636524-DEL-1106 |
| chr11-112642677-DEL-53 | chr11-112642678-DEL-53 |
| chr11-112653991-DEL-102 | chr11-112653977-DEL-96 |
| chr11-112682530-DEL-81 | chr11-112682559-DEL-56 |
| chr11-112697903-DEL-76 | chr11-112697895-DEL-71 |
| chr11-112749638-DEL-82 | chr11-112749639-DEL-82 |
| chr11-112772840-DEL-129 | chr11-112772839-DEL-129 |
| chr11-112791768-DEL-82 | chr11-112791769-DEL-82 |
| chr11-112800524-DEL-113 | chr11-112800525-DEL-110 |
| chr11-112812410-DEL-350 | chr11-112812411-DEL-350 |
| chr11-112813952-DEL-70 | chr11-112813953-DEL-70 |
| chr11-112824259-DEL-51 | chr11-112824260-DEL-51 |
| chr11-112867438-DEL-58 | chr11-112867441-DEL-56 |
| chr11-112944726-DEL-1595 | chr11-112944727-DEL-1595 |
| chr11-112952796-DEL-109 | chr11-112952820-DEL-108 |
| chr11-112997000-DEL-276 | chr11-112997001-DEL-276 |
| chr11-113015648-DEL-227 | chr11-113015649-DEL-227 |
| chr11-113025001-DEL-94 | chr11-113025015-DEL-93 |
| chr11-113032349-DEL-91 | chr11-113032350-DEL-90 |
| chr11-113035613-DEL-579 | chr11-113035614-DEL-579 |
| chr11-113045658-DEL-74 | chr11-113045659-DEL-74 |
| chr11-113057619-DEL-216 | chr11-113057620-DEL-216 |

| | |
|---|---|
| chr11-113059692-DEL-52 | chr11-113059693-DEL-52 |
| chr11-113062222-DEL-287 | chr11-113062210-DEL-293 |
| chr11-113063836-DEL-479 | chr11-113063837-DEL-479 |
| chr11-113064472-DEL-252 | chr11-113064473-DEL-252 |
| chr11-113093619-DEL-76 | chr11-113093622-DEL-74 |
| chr11-113110584-DEL-60 | chr11-113110578-DEL-51 |
| chr11-113156908-DEL-76 | chr11-113156928-DEL-57 |
| chr11-113174036-DEL-54 | chr11-113174037-DEL-54 |
| chr11-113178180-DEL-70 | chr11-113178181-DEL-70 |
| chr11-113183639-DEL-128 | chr11-113183643-DEL-84 |
| chr11-113197238-DEL-164 | chr11-113197217-DEL-151 |
| chr11-113216503-DEL-72 | chr11-113216504-DEL-72 |
| chr11-113274665-DEL-172 | chr11-113274666-DEL-172 |
| chr11-113375295-DEL-55 | chr11-113375296-DEL-56 |
| chr11-113392020-DEL-56 | chr11-113391987-DEL-57 |
| chr11-113418644-DEL-415 | chr11-113418645-DEL-415 |
| chr11-113431131-DEL-315 | chr11-113431132-DEL-315 |
| chr11-113436102-DEL-104 | chr11-113436127-DEL-64 |
| chr11-113438759-DEL-1493 | chr11-113438760-DEL-1493 |
| chr11-113460516-DEL-60 | chr11-113460517-DEL-60 |
| chr11-113507192-DEL-79 | chr11-113507192-DEL-79 |
| chr11-113517101-DEL-621 | chr11-113517101-DEL-621 |
| chr11-113519907-DEL-144 | chr11-113519938-DEL-102 |
| chr11-113521295-DEL-60 | chr11-113521320-DEL-51 |
| chr11-113529804-DEL-296 | chr11-113529805-DEL-296 |
| chr11-113533225-DEL-54 | chr11-113533226-DEL-54 |
| chr11-113542203-DEL-70 | chr11-113542204-DEL-70 |
| chr11-113550486-DEL-177 | chr11-113550487-DEL-177 |
| chr11-113551313-DEL-196 | chr11-113551314-DEL-196 |
| chr11-113555684-DEL-1105 | chr11-113555685-DEL-1105 |
| chr11-113571430-DEL-79 | chr11-113571431-DEL-79 |
| chr11-113582267-DEL-80 | chr11-113582221-DEL-79 |
| chr11-113589746-DEL-306 | chr11-113589747-DEL-306 |
| chr11-113595321-DEL-186 | chr11-113595320-DEL-187 |
| chr11-113606712-DEL-87 | chr11-113606713-DEL-87 |
| chr11-113635215-DEL-84 | chr11-113635216-DEL-84 |
| chr11-113635746-DEL-214 | chr11-113635747-DEL-214 |
| chr11-113644352-DEL-199 | chr11-113644353-DEL-199 |
| chr11-113648600-DEL-54 | chr11-113648603-DEL-50 |
| chr11-113659410-DEL-491 | chr11-113659411-DEL-491 |
| chr11-113662783-DEL-206 | chr11-113662784-DEL-206 |
| chr11-113663973-DEL-95 | chr11-113663975-DEL-94 |
| chr11-113673822-DEL-887 | chr11-113673823-DEL-886 |
| chr11-113715159-DEL-69 | chr11-113715160-DEL-69 |
| chr11-113728109-DEL-64 | chr11-113728110-DEL-64 |
| chr11-113739170-DEL-381 | chr11-113739172-DEL-379 |
| chr11-113767350-DEL-226 | chr11-113767351-DEL-226 |
| chr11-113803003-DEL-7124 | chr11-113803004-DEL-7124 |
| chr11-113811957-DEL-198 | chr11-113811958-DEL-198 |
| chr11-113845688-DEL-90 | chr11-113845689-DEL-90 |
| chr11-113901206-DEL-130 | chr11-113901227-DEL-114 |
| chr11-113932449-DEL-79 | chr11-113932436-DEL-90 |
| chr11-113969487-DEL-77 | chr11-113969502-DEL-76 |
| chr11-113987213-DEL-53 | chr11-113987172-DEL-98 |
| chr11-113999848-DEL-52 | chr11-113999830-DEL-70 |
| chr11-114004397-DEL-102 | chr11-114004398-DEL-102 |
| chr11-114021864-DEL-196 | chr11-114021853-DEL-178 |
| chr11-114096715-DEL-146 | chr11-114096715-DEL-146 |
| chr11-114104994-DEL-470 | chr11-114104995-DEL-470 |
| chr11-114159707-DEL-163 | chr11-114159692-DEL-192 |
| chr11-114187034-DEL-106 | chr11-114187019-DEL-121 |
| chr11-114223551-DEL-505 | chr11-114223552-DEL-505 |
| chr11-114275578-DEL-503 | chr11-114275579-DEL-503 |
| chr11-114279295-DEL-233 | chr11-114279296-DEL-233 |
| chr11-114283076-DEL-267 | chr11-114283077-DEL-267 |
| chr11-114303425-DEL-97 | chr11-114303426-DEL-97 |
| chr11-114325211-DEL-63 | chr11-114325208-DEL-64 |
| chr11-114343728-DEL-357 | chr11-114343729-DEL-357 |
| chr11-114347127-DEL-6436 | chr11-114347128-DEL-6414 |
| chr11-114356796-DEL-72 | chr11-114356797-DEL-72 |
| chr11-114360397-DEL-61 | chr11-114360398-DEL-61 |
| chr11-114387090-DEL-62 | chr11-114387091-DEL-62 |
| chr11-114421815-DEL-160 | chr11-114421816-DEL-160 |
| chr11-114433820-DEL-78 | chr11-114433801-DEL-97 |
| chr11-114460146-DEL-100 | chr11-114460136-DEL-98 |
| chr11-114462985-DEL-1894 | chr11-114462980-DEL-1904 |
| chr11-114465462-DEL-647 | chr11-114465463-DEL-647 |
| chr11-114472747-DEL-88 | chr11-114472748-DEL-88 |
| chr11-114497269-DEL-213 | chr11-114497270-DEL-213 |
| chr11-114504607-DEL-202 | chr11-114504608-DEL-202 |

| | |
|---|---|
| chr11-114510980-DEL-411 | chr11-114510981-DEL-411 |
| chr11-114514582-DEL-120 | chr11-114514583-DEL-120 |
| chr11-114523689-DEL-166 | chr11-114523670-DEL-186 |
| chr11-114531233-DEL-58 | chr11-114531234-DEL-58 |
| chr11-114541739-DEL-83 | chr11-114541740-DEL-83 |
| chr11-114544128-DEL-210 | chr11-114544129-DEL-210 |
| chr11-114550227-DEL-2820 | chr11-114550228-DEL-2820 |
| chr11-114558326-DEL-197 | chr11-114558327-DEL-197 |
| chr11-114565578-DEL-186 | chr11-114565579-DEL-186 |
| chr11-114578415-DEL-81 | chr11-114578415-DEL-81 |
| chr11-114596798-DEL-194 | chr11-114596800-DEL-188 |
| chr11-114605146-DEL-170 | chr11-114605147-DEL-170 |
| chr11-114624327-DEL-72 | chr11-114624328-DEL-75 |
| chr11-114695297-DEL-634 | chr11-114695298-DEL-634 |
| chr11-114715522-DEL-499 | chr11-114715523-DEL-499 |
| chr11-114717930-DEL-123 | chr11-114717931-DEL-123 |
| chr11-114748153-DEL-446 | chr11-114748154-DEL-446 |
| chr11-114750327-DEL-205 | chr11-114750328-DEL-205 |
| chr11-114752391-DEL-97 | chr11-114752392-DEL-96 |
| chr11-114753265-DEL-108 | chr11-114753266-DEL-108 |
| chr11-114792960-DEL-470 | chr11-114792961-DEL-470 |
| chr11-114800600-DEL-228 | chr11-114800601-DEL-228 |
| chr11-114800992-DEL-377 | chr11-114800993-DEL-377 |
| chr11-114832875-DEL-1018 | chr11-114832866-DEL-1019 |
| chr11-114835040-DEL-569 | chr11-114835041-DEL-569 |
| chr11-114838260-DEL-562 | chr11-114838261-DEL-562 |
| chr11-114847214-DEL-64 | chr11-114847215-DEL-64 |
| chr11-114860007-DEL-383 | chr11-114860008-DEL-383 |
| chr11-114865070-DEL-5109 | chr11-114865070-DEL-5109 |
| chr11-114907885-DEL-77 | chr11-114907886-DEL-77 |
| chr11-114920149-DEL-52 | chr11-114920134-DEL-50 |
| chr11-114931161-DEL-77 | chr11-114931144-DEL-78 |
| chr11-114947643-DEL-191 | chr11-114947657-DEL-187 |
| chr11-114953320-DEL-1310 | chr11-114953321-DEL-1310 |
| chr11-114957523-DEL-99 | chr11-114957524-DEL-99 |
| chr11-114998047-DEL-104 | chr11-114998048-DEL-104 |
| chr11-115043475-DEL-79 | chr11-115043476-DEL-79 |
| chr11-115068357-DEL-56 | chr11-115068358-DEL-56 |
| chr11-115086632-DEL-69 | chr11-115086608-DEL-99 |
| chr11-115121980-DEL-93 | chr11-115121980-DEL-93 |
| chr11-115135637-DEL-122 | chr11-115135638-DEL-122 |
| chr11-115192902-DEL-51 | chr11-115192903-DEL-51 |
| chr11-115194459-DEL-239 | chr11-115194445-DEL-205 |
| chr11-115231930-DEL-82 | chr11-115231955-DEL-60 |
| chr11-115326185-DEL-254 | chr11-115326186-DEL-254 |
| chr11-115349802-DEL-2574 | chr11-115349803-DEL-2574 |
| chr11-115357596-DEL-80 | chr11-115357625-DEL-68 |
| chr11-115376592-DEL-200 | chr11-115376593-DEL-200 |
| chr11-115386468-DEL-106 | chr11-115386469-DEL-106 |
| chr11-115387056-DEL-248 | chr11-115387057-DEL-248 |
| chr11-115394497-DEL-65 | chr11-115394498-DEL-65 |
| chr11-115395297-DEL-210 | chr11-115395298-DEL-210 |
| chr11-115400072-DEL-238 | chr11-115400073-DEL-238 |
| chr11-115419818-DEL-219 | chr11-115419819-DEL-219 |
| chr11-115431013-DEL-216 | chr11-115431014-DEL-216 |
| chr11-115444791-DEL-54 | chr11-115444792-DEL-54 |
| chr11-115451327-DEL-220 | chr11-115451328-DEL-220 |
| chr11-115508820-DEL-56 | chr11-115508821-DEL-56 |
| chr11-115520276-DEL-172 | chr11-115520282-DEL-170 |
| chr11-115526784-DEL-150 | chr11-115526790-DEL-145 |
| chr11-115547976-DEL-408 | chr11-115547977-DEL-408 |
| chr11-115556680-DEL-199 | chr11-115556681-DEL-199 |
| chr11-115559175-DEL-56 | chr11-115559172-DEL-58 |
| chr11-115566639-DEL-1880 | chr11-115566640-DEL-1880 |
| chr11-115585820-DEL-66 | chr11-115585821-DEL-60 |
| chr11-115602369-DEL-53 | chr11-115602371-DEL-54 |
| chr11-115606121-DEL-497 | chr11-115606122-DEL-497 |
| chr11-115615085-DEL-577 | chr11-115615086-DEL-577 |
| chr11-115619553-DEL-391 | chr11-115619534-DEL-421 |
| chr11-115620904-DEL-171 | chr11-115620908-DEL-153 |
| chr11-115621523-DEL-261 | chr11-115621524-DEL-261 |
| chr11-115623064-DEL-224 | chr11-115623064-DEL-224 |
| chr11-115628101-DEL-186 | chr11-115628102-DEL-186 |
| chr11-115633102-DEL-202 | chr11-115633102-DEL-202 |
| chr11-115639727-DEL-219 | chr11-115639743-DEL-208 |
| chr11-115641398-DEL-52 | chr11-115641399-DEL-60 |
| chr11-115642448-DEL-67 | chr11-115642468-DEL-52 |
| chr11-115645718-DEL-7109 | chr11-115645719-DEL-7109 |
| chr11-115661555-DEL-69 | chr11-115661549-DEL-70 |
| chr11-115696693-DEL-79 | chr11-115696694-DEL-79 |

| | |
|---|---|
| chr11-115706836-DEL-87 | chr11-115706837-DEL-87 |
| chr11-115710035-DEL-508 | chr11-115710036-DEL-508 |
| chr11-115754789-DEL-180 | chr11-115754790-DEL-180 |
| chr11-115761411-DEL-78 | chr11-115761412-DEL-78 |
| chr11-115813561-DEL-207 | chr11-115813562-DEL-207 |
| chr11-115814892-DEL-54 | chr11-115814893-DEL-54 |
| chr11-115831887-DEL-436 | chr11-115831888-DEL-436 |
| chr11-115849390-DEL-102 | chr11-115849419-DEL-74 |
| chr11-115871260-DEL-52 | chr11-115871252-DEL-56 |
| chr11-115875623-DEL-150 | chr11-115875624-DEL-150 |
| chr11-115891847-DEL-788 | chr11-115891848-DEL-788 |
| chr11-115893294-DEL-66 | chr11-115893295-DEL-70 |
| chr11-115916647-DEL-213 | chr11-115916648-DEL-213 |
| chr11-115925197-DEL-63 | chr11-115925198-DEL-62 |
| chr11-115926188-DEL-209 | chr11-115926189-DEL-209 |
| chr11-115930246-DEL-199 | chr11-115930247-DEL-184 |
| chr11-115953889-DEL-318 | chr11-115953890-DEL-318 |
| chr11-115957049-DEL-435 | chr11-115957050-DEL-435 |
| chr11-115966202-DEL-233 | chr11-115966203-DEL-233 |
| chr11-115981177-DEL-209 | chr11-115981178-DEL-209 |
| chr11-115982082-DEL-2396 | chr11-115982083-DEL-2396 |
| chr11-115983999-DEL-196 | chr11-115983994-DEL-195 |
| chr11-115986065-DEL-206 | chr11-115986066-DEL-206 |
| chr11-116003378-DEL-208 | chr11-116003379-DEL-208 |
| chr11-116007862-DEL-59 | chr11-116007863-DEL-59 |
| chr11-116031794-DEL-288 | chr11-116031762-DEL-289 |
| chr11-116053561-DEL-168 | chr11-116053562-DEL-168 |
| chr11-116060561-DEL-211 | chr11-116060562-DEL-211 |
| chr11-116067102-DEL-208 | chr11-116067103-DEL-209 |
| chr11-116072082-DEL-152 | chr11-116072083-DEL-152 |
| chr11-116143877-DEL-82 | chr11-116143878-DEL-82 |
| chr11-116167218-DEL-206 | chr11-116167219-DEL-206 |
| chr11-116170124-DEL-208 | chr11-116170125-DEL-208 |
| chr11-116177438-DEL-533 | chr11-116177439-DEL-533 |
| chr11-116256098-DEL-127 | chr11-116256093-DEL-125 |
| chr11-116259566-DEL-193 | chr11-116259567-DEL-193 |
| chr11-116265551-DEL-214 | chr11-116265552-DEL-214 |
| chr11-116269887-DEL-206 | chr11-116269888-DEL-206 |
| chr11-116280223-DEL-5311 | chr11-116280224-DEL-5311 |
| chr11-116289791-DEL-154 | chr11-116289792-DEL-154 |
| chr11-116290363-DEL-203 | chr11-116290364-DEL-203 |
| chr11-116310528-DEL-114 | chr11-116310529-DEL-114 |
| chr11-116360279-DEL-87 | chr11-116360280-DEL-87 |
| chr11-116364603-DEL-1243 | chr11-116364604-DEL-1243 |
| chr11-116390871-DEL-120 | chr11-116390892-DEL-100 |
| chr11-116400880-DEL-763 | chr11-116400880-DEL-763 |
| chr11-116407132-DEL-108 | chr11-116407141-DEL-92 |
| chr11-116416116-DEL-60 | chr11-116416117-DEL-60 |
| chr11-116447580-DEL-1198 | chr11-116447576-DEL-1199 |
| chr11-116460131-DEL-162 | chr11-116460132-DEL-162 |
| chr11-116481518-DEL-114 | chr11-116481517-DEL-115 |
| chr11-116501736-DEL-193 | chr11-116501737-DEL-193 |
| chr11-116524385-DEL-256 | chr11-116524386-DEL-256 |
| chr11-116532399-DEL-80 | chr11-116532366-DEL-108 |
| chr11-116546184-DEL-204 | chr11-116546185-DEL-204 |
| chr11-116557747-DEL-218 | chr11-116557748-DEL-218 |
| chr11-116582637-DEL-245 | chr11-116582638-DEL-245 |
| chr11-116598717-DEL-205 | chr11-116598718-DEL-205 |
| chr11-116599529-DEL-110 | chr11-116599530-DEL-125 |
| chr11-116607125-DEL-198 | chr11-116607115-DEL-201 |
| chr11-116614737-DEL-201 | chr11-116614738-DEL-201 |
| chr11-116616846-DEL-66 | chr11-116616870-DEL-88 |
| chr11-116623779-DEL-229 | chr11-116623775-DEL-229 |
| chr11-116636759-DEL-150 | chr11-116636800-DEL-108 |
| chr11-116640648-DEL-115 | chr11-116640649-DEL-115 |
| chr11-116650116-DEL-5105 | chr11-116650116-DEL-5105 |
| chr11-116662973-DEL-168 | chr11-116662992-DEL-176 |
| chr11-116676177-DEL-7379 | chr11-116676177-DEL-7379 |
| chr11-116688871-DEL-100 | chr11-116688872-DEL-100 |
| chr11-116690786-DEL-211 | chr11-116690787-DEL-211 |
| chr11-116694928-DEL-154 | chr11-116694928-DEL-154 |
| chr11-116698237-DEL-126 | chr11-116698237-DEL-126 |
| chr11-116746226-DEL-188 | chr11-116746227-DEL-188 |
| chr11-116766282-DEL-215 | chr11-116766283-DEL-215 |
| chr11-116769572-DEL-109 | chr11-116769576-DEL-108 |
| chr11-116770984-DEL-200 | chr11-116770984-DEL-198 |
| chr11-116772617-DEL-315 | chr11-116772638-DEL-294 |
| chr11-116773886-DEL-205 | chr11-116773885-DEL-206 |
| chr11-116779544-DEL-90 | chr11-116779545-DEL-90 |
| chr11-116784819-DEL-74 | chr11-116784820-DEL-74 |

| | |
|---|---|
| chr11-116786013-DEL-195 | chr11-116786014-DEL-195 |
| chr11-116790189-DEL-319 | chr11-116790190-DEL-319 |
| chr11-116796500-DEL-327 | chr11-116796501-DEL-327 |
| chr11-116800572-DEL-209 | chr11-116800573-DEL-209 |
| chr11-116831132-DEL-166 | chr11-116831133-DEL-166 |
| chr11-116893656-DEL-116 | chr11-116893652-DEL-110 |
| chr11-116896725-DEL-123 | chr11-116896726-DEL-123 |
| chr11-116929635-DEL-215 | chr11-116929636-DEL-215 |
| chr11-116944135-DEL-80 | chr11-116944136-DEL-80 |
| chr11-116946347-DEL-190 | chr11-116946348-DEL-190 |
| chr11-116956358-DEL-242 | chr11-116956359-DEL-242 |
| chr11-116959767-DEL-208 | chr11-116959768-DEL-208 |
| chr11-116960614-DEL-197 | chr11-116960615-DEL-197 |
| chr11-116965830-DEL-431 | chr11-116965831-DEL-431 |
| chr11-116970284-DEL-264 | chr11-116970286-DEL-262 |
| chr11-116977606-DEL-207 | chr11-116977607-DEL-207 |
| chr11-116991587-DEL-190 | chr11-116991588-DEL-190 |
| chr11-116998890-DEL-50 | chr11-116998900-DEL-67 |
| chr11-117002669-DEL-179 | chr11-117002670-DEL-179 |
| chr11-117016939-DEL-111 | chr11-117016940-DEL-111 |
| chr11-117028550-DEL-211 | chr11-117028551-DEL-211 |
| chr11-117042850-DEL-177 | chr11-117042851-DEL-177 |
| chr11-117054882-DEL-1135 | chr11-117054883-DEL-1135 |
| chr11-117105395-DEL-84 | chr11-117105389-DEL-80 |
| chr11-117112601-DEL-134 | chr11-117112639-DEL-96 |
| chr11-117127850-DEL-189 | chr11-117127851-DEL-189 |
| chr11-117180142-DEL-200 | chr11-117180143-DEL-196 |
| chr11-117193572-DEL-66 | chr11-117193572-DEL-62 |
| chr11-117212690-DEL-52 | chr11-117212691-DEL-52 |
| chr11-117214322-DEL-175 | chr11-117214332-DEL-166 |
| chr11-117219415-DEL-191 | chr11-117219425-DEL-191 |
| chr11-117227643-DEL-52 | chr11-117227643-DEL-52 |
| chr11-117244853-DEL-81 | chr11-117244853-DEL-80 |
| chr11-117248937-DEL-244 | chr11-117248938-DEL-244 |
| chr11-117260565-DEL-91 | chr11-117260566-DEL-91 |
| chr11-117274197-DEL-54 | chr11-117274220-DEL-52 |
| chr11-117307938-DEL-199 | chr11-117307939-DEL-199 |
| chr11-117333280-DEL-268 | chr11-117333281-DEL-262 |
| chr11-117336951-DEL-55 | chr11-117336952-DEL-55 |
| chr11-117340598-DEL-80 | chr11-117340598-DEL-80 |
| chr11-117380892-DEL-88 | chr11-117380875-DEL-83 |
| chr11-117402158-DEL-98 | chr11-117402159-DEL-98 |
| chr11-117418420-DEL-324 | chr11-117418421-DEL-324 |
| chr11-117422051-DEL-186 | chr11-117422052-DEL-186 |
| chr11-117423916-DEL-64 | chr11-117423917-DEL-64 |
| chr11-117424650-DEL-207 | chr11-117424651-DEL-207 |
| chr11-117440775-DEL-88 | chr11-117440776-DEL-88 |
| chr11-117444681-DEL-213 | chr11-117444681-DEL-213 |
| chr11-117456519-DEL-58 | chr11-117456524-DEL-66 |
| chr11-117467303-DEL-109 | chr11-117467304-DEL-109 |
| chr11-117473211-DEL-184 | chr11-117473226-DEL-180 |
| chr11-117473985-DEL-204 | chr11-117473986-DEL-204 |
| chr11-117475623-DEL-202 | chr11-117475624-DEL-202 |
| chr11-117484928-DEL-206 | chr11-117484929-DEL-206 |
| chr11-117514423-DEL-211 | chr11-117514424-DEL-211 |
| chr11-117526559-DEL-64 | chr11-117526539-DEL-82 |
| chr11-117528107-DEL-68 | chr11-117528089-DEL-62 |
| chr11-117536148-DEL-100 | chr11-117536132-DEL-107 |
| chr11-117555434-DEL-70 | chr11-117555437-DEL-84 |
| chr11-117647935-DEL-58 | chr11-117647936-DEL-58 |
| chr11-117649987-DEL-61 | chr11-117649946-DEL-63 |
| chr11-117684608-DEL-203 | chr11-117684609-DEL-203 |
| chr11-117687410-DEL-387 | chr11-117687413-DEL-387 |
| chr11-117699167-DEL-52 | chr11-117699168-DEL-50 |
| chr11-117707066-DEL-107 | chr11-117707067-DEL-103 |
| chr11-117724280-DEL-219 | chr11-117724281-DEL-219 |
| chr11-117728640-DEL-214 | chr11-117728641-DEL-214 |
| chr11-117787385-DEL-78 | chr11-117787386-DEL-78 |
| chr11-117788241-DEL-3333 | chr11-117788242-DEL-3333 |
| chr11-117794649-DEL-57 | chr11-117794650-DEL-57 |
| chr11-117835329-DEL-78 | chr11-117835335-DEL-77 |
| chr11-117839047-DEL-236 | chr11-117839044-DEL-240 |
| chr11-117852194-DEL-412 | chr11-117852184-DEL-410 |
| chr11-117893571-DEL-215 | chr11-117893572-DEL-215 |
| chr11-117944343-DEL-144 | chr11-117944344-DEL-109 |
| chr11-117969904-DEL-215 | chr11-117969905-DEL-215 |
| chr11-117982685-DEL-220 | chr11-117982686-DEL-220 |
| chr11-118049336-DEL-335 | chr11-118049337-DEL-335 |
| chr11-118086717-DEL-372 | chr11-118086718-DEL-372 |
| chr11-118146755-DEL-68 | chr11-118146756-DEL-68 |

| | |
|---|---|
| chr11-118205331-DEL-109 | chr11-118205323-DEL-111 |
| chr11-118227845-DEL-214 | chr11-118227846-DEL-214 |
| chr11-118247445-DEL-252 | chr11-118247454-DEL-241 |
| chr11-118332747-DEL-167 | chr11-118332748-DEL-167 |
| chr11-118370492-DEL-168 | chr11-118370493-DEL-168 |
| chr11-118490358-DEL-119 | chr11-118490322-DEL-115 |
| chr11-118493236-DEL-156 | chr11-118493218-DEL-175 |
| chr11-118577112-DEL-100 | chr11-118577082-DEL-140 |
| chr11-118610015-DEL-90 | chr11-118610016-DEL-90 |
| chr11-118634375-DEL-54 | chr11-118634370-DEL-60 |
| chr11-118645037-DEL-91 | chr11-118645060-DEL-51 |
| chr11-118691242-DEL-100 | chr11-118691243-DEL-100 |
| chr11-118707932-DEL-66 | chr11-118707933-DEL-66 |
| chr11-118737304-DEL-340 | chr11-118737305-DEL-340 |
| chr11-118761125-DEL-117 | chr11-118761109-DEL-154 |
| chr11-118764830-DEL-102 | chr11-118764831-DEL-103 |
| chr11-118778792-DEL-72 | chr11-118778793-DEL-72 |
| chr11-118804499-DEL-68 | chr11-118804500-DEL-68 |
| chr11-118826635-DEL-96 | chr11-118826635-DEL-96 |
| chr11-118829236-DEL-118 | chr11-118829236-DEL-118 |
| chr11-118856580-DEL-69 | chr11-118856587-DEL-69 |
| chr11-118866111-DEL-65 | chr11-118866111-DEL-65 |
| chr11-118871069-DEL-158 | chr11-118871046-DEL-142 |
| chr11-118880101-DEL-86 | chr11-118880100-DEL-87 |
| chr11-118922112-DEL-193 | chr11-118922113-DEL-193 |
| chr11-118997735-DEL-89 | chr11-118997710-DEL-90 |
| chr11-119000922-DEL-123 | chr11-119000950-DEL-96 |
| chr11-119065494-DEL-85 | chr11-119065535-DEL-85 |
| chr11-119085296-DEL-117 | chr11-119085269-DEL-118 |
| chr11-119095368-DEL-83 | chr11-119095369-DEL-83 |
| chr11-119097056-DEL-272 | chr11-119097057-DEL-272 |
| chr11-119105749-DEL-93 | chr11-119105750-DEL-93 |
| chr11-119109104-DEL-143 | chr11-119109104-DEL-143 |
| chr11-119116553-DEL-378 | chr11-119116554-DEL-378 |
| chr11-119154587-DEL-80 | chr11-119154588-DEL-86 |
| chr11-119180922-DEL-193 | chr11-119180888-DEL-212 |
| chr11-119199947-DEL-87 | chr11-119199973-DEL-83 |
| chr11-119202559-DEL-1992 | chr11-119202560-DEL-1992 |
| chr11-119207107-DEL-112 | chr11-119207108-DEL-112 |
| chr11-119208466-DEL-55 | chr11-119208467-DEL-55 |
| chr11-119210462-DEL-1873 | chr11-119210468-DEL-1846 |
| chr11-119257469-DEL-230 | chr11-119257472-DEL-227 |
| chr11-119276544-DEL-198 | chr11-119276545-DEL-198 |
| chr11-119281695-DEL-106 | chr11-119281696-DEL-106 |
| chr11-119291498-DEL-211 | chr11-119291499-DEL-211 |
| chr11-119350178-DEL-150 | chr11-119350179-DEL-150 |
| chr11-119359349-DEL-189 | chr11-119359350-DEL-189 |
| chr11-119384735-DEL-56 | chr11-119384736-DEL-56 |
| chr11-119397031-DEL-166 | chr11-119396993-DEL-205 |
| chr11-119415264-DEL-90 | chr11-119415265-DEL-90 |
| chr11-119420551-DEL-65 | chr11-119420552-DEL-65 |
| chr11-119426324-DEL-115 | chr11-119426348-DEL-69 |
| chr11-119461711-DEL-96 | chr11-119461695-DEL-84 |
| chr11-119466995-DEL-168 | chr11-119467000-DEL-164 |
| chr11-119473841-DEL-392 | chr11-119473842-DEL-392 |
| chr11-119545167-DEL-2113 | chr11-119545171-DEL-2110 |
| chr11-119571919-DEL-71 | chr11-119571920-DEL-71 |
| chr11-119584591-DEL-171 | chr11-119584592-DEL-171 |
| chr11-119589891-DEL-6436 | chr11-119589892-DEL-6436 |
| chr11-119613919-DEL-76 | chr11-119613920-DEL-76 |
| chr11-119616405-DEL-270 | chr11-119616406-DEL-270 |
| chr11-119669563-DEL-465 | chr11-119669564-DEL-465 |
| chr11-119706780-DEL-68 | chr11-119706781-DEL-102 |
| chr11-119752528-DEL-132 | chr11-119752505-DEL-156 |
| chr11-119757039-DEL-129 | chr11-119757040-DEL-129 |
| chr11-119774892-DEL-84 | chr11-119774893-DEL-78 |
| chr11-119818859-DEL-68 | chr11-119818871-DEL-83 |
| chr11-119837044-DEL-68 | chr11-119837045-DEL-68 |
| chr11-119918406-DEL-54 | chr11-119918407-DEL-54 |
| chr11-119943616-DEL-748 | chr11-119943619-DEL-746 |
| chr11-119964213-DEL-68 | chr11-119964214-DEL-68 |
| chr11-119992694-DEL-175 | chr11-119992695-DEL-175 |
| chr11-120005441-DEL-78 | chr11-120005442-DEL-78 |
| chr11-120044553-DEL-219 | chr11-120044552-DEL-218 |
| chr11-120102250-DEL-194 | chr11-120102251-DEL-194 |
| chr11-120140684-DEL-52 | chr11-120140684-DEL-53 |
| chr11-120198718-DEL-55 | chr11-120198728-DEL-58 |
| chr11-120200405-DEL-143 | chr11-120200403-DEL-143 |
| chr11-120208971-DEL-187 | chr11-120208972-DEL-187 |
| chr11-120211895-DEL-7193 | chr11-120211896-DEL-7193 |

| | |
|---|---|
| chr11-120232192-DEL-221 | chr11-120232193-DEL-221 |
| chr11-120281343-DEL-813 | chr11-120281344-DEL-813 |
| chr11-120304680-DEL-210 | chr11-120304681-DEL-210 |
| chr11-120315041-DEL-213 | chr11-120315042-DEL-213 |
| chr11-120341109-DEL-63 | chr11-120341116-DEL-96 |
| chr11-120351236-DEL-178 | chr11-120351236-DEL-180 |
| chr11-120365599-DEL-207 | chr11-120365600-DEL-207 |
| chr11-120366669-DEL-200 | chr11-120366670-DEL-200 |
| chr11-120401917-DEL-66 | chr11-120401916-DEL-68 |
| chr11-120447070-DEL-416 | chr11-120447071-DEL-416 |
| chr11-120447654-DEL-146 | chr11-120447655-DEL-146 |
| chr11-120450667-DEL-61 | chr11-120450668-DEL-61 |
| chr11-120460122-DEL-204 | chr11-120460123-DEL-204 |
| chr11-120475890-DEL-631 | chr11-120475891-DEL-631 |
| chr11-120626776-DEL-211 | chr11-120626777-DEL-211 |
| chr11-120628241-DEL-170 | chr11-120628242-DEL-170 |
| chr11-120646660-DEL-286 | chr11-120646661-DEL-286 |
| chr11-120647644-DEL-268 | chr11-120647645-DEL-268 |
| chr11-120653603-DEL-1641 | chr11-120653604-DEL-1641 |
| chr11-120661725-DEL-1539 | chr11-120661726-DEL-1539 |
| chr11-120672220-DEL-134 | chr11-120672261-DEL-135 |
| chr11-120716615-DEL-426 | chr11-120716616-DEL-426 |
| chr11-120746447-DEL-60 | chr11-120746447-DEL-61 |
| chr11-120754679-DEL-56 | chr11-120754680-DEL-56 |
| chr11-120799205-DEL-5088 | chr11-120799206-DEL-5088 |
| chr11-120817618-DEL-209 | chr11-120817613-DEL-210 |
| chr11-120905311-DEL-65 | chr11-120905312-DEL-65 |
| chr11-120933990-DEL-147 | chr11-120933991-DEL-147 |
| chr11-120951478-DEL-208 | chr11-120951478-DEL-208 |
| chr11-120960732-DEL-5000 | chr11-120960733-DEL-5000 |
| chr11-120982995-DEL-59 | chr11-120983004-DEL-51 |
| chr11-120986535-DEL-383 | chr11-120986527-DEL-384 |
| chr11-120994788-DEL-1829 | chr11-120994789-DEL-1829 |
| chr11-121016258-DEL-164 | chr11-121016259-DEL-164 |
| chr11-121017059-DEL-1226 | chr11-121017060-DEL-1226 |
| chr11-121041856-DEL-953 | chr11-121041857-DEL-953 |
| chr11-121077356-DEL-52 | chr11-121077357-DEL-52 |
| chr11-121096382-DEL-1888 | chr11-121096383-DEL-1888 |
| chr11-121102560-DEL-55 | chr11-121102561-DEL-55 |
| chr11-121105507-DEL-257 | chr11-121105507-DEL-223 |
| chr11-121108581-DEL-190 | chr11-121108582-DEL-190 |
| chr11-121138174-DEL-226 | chr11-121138175-DEL-226 |
| chr11-121176950-DEL-52 | chr11-121176951-DEL-52 |
| chr11-121181947-DEL-64 | chr11-121181948-DEL-64 |
| chr11-121201862-DEL-8975 | chr11-121201863-DEL-8975 |
| chr11-121208424-DEL-208 | chr11-121208425-DEL-208 |
| chr11-121215158-DEL-221 | chr11-121215159-DEL-221 |
| chr11-121235437-DEL-204 | chr11-121235438-DEL-204 |
| chr11-121258688-DEL-211 | chr11-121258689-DEL-211 |
| chr11-121259319-DEL-489 | chr11-121259320-DEL-489 |
| chr11-121287047-DEL-7110 | chr11-121287048-DEL-7110 |
| chr11-121357566-DEL-63 | chr11-121357532-DEL-94 |
| chr11-121387009-DEL-98 | chr11-121387029-DEL-79 |
| chr11-121452481-DEL-1533 | chr11-121452477-DEL-1534 |
| chr11-121476089-DEL-202 | chr11-121476090-DEL-202 |
| chr11-121502771-DEL-1864 | chr11-121502772-DEL-1864 |
| chr11-121540092-DEL-497 | chr11-121540093-DEL-497 |
| chr11-121602716-DEL-445 | chr11-121602717-DEL-445 |
| chr11-121615172-DEL-545 | chr11-121615173-DEL-545 |
| chr11-121654475-DEL-64 | chr11-121654476-DEL-64 |
| chr11-121670210-DEL-324 | chr11-121670211-DEL-324 |
| chr11-121742664-DEL-141 | chr11-121742665-DEL-141 |
| chr11-121766789-DEL-211 | chr11-121766790-DEL-211 |
| chr11-121780735-DEL-1305 | chr11-121780736-DEL-1305 |
| chr11-121781603-DEL-205 | chr11-121781604-DEL-205 |
| chr11-121787375-DEL-3470 | chr11-121787376-DEL-3471 |
| chr11-121788462-DEL-638 | chr11-121788463-DEL-638 |
| chr11-121794823-DEL-1277 | chr11-121794824-DEL-1277 |
| chr11-121799194-DEL-2501 | chr11-121799196-DEL-2500 |
| chr11-121803910-DEL-79 | chr11-121803911-DEL-79 |
| chr11-121820810-DEL-50 | chr11-121820810-DEL-50 |
| chr11-121841032-DEL-79 | chr11-121841033-DEL-79 |
| chr12-3142897-DEL-60 | chr12-3142898-DEL-60 |
| chr12-3214738-DEL-75 | chr12-3214742-DEL-72 |
| chr12-3308551-DEL-2201 | chr12-3308557-DEL-2197 |
| chr12-3371377-DEL-118 | chr12-3371378-DEL-118 |
| chr12-3380815-DEL-7122 | chr12-3380816-DEL-7122 |
| chr12-3390987-DEL-62 | chr12-3390988-DEL-62 |
| chr12-3463100-DEL-206 | chr12-3463101-DEL-206 |
| chr12-3518632-DEL-96 | chr12-3518601-DEL-124 |

| | |
|---|---|
| chr12-3520402-DEL-75 | chr12-3520383-DEL-57 |
| chr12-3542350-DEL-199 | chr12-3542351-DEL-199 |
| chr12-3561884-DEL-237 | chr12-3561885-DEL-237 |
| chr12-3582391-DEL-57 | chr12-3582389-DEL-54 |
| chr12-3585323-DEL-1074 | chr12-3585324-DEL-1074 |
| chr12-3591710-DEL-59 | chr12-3591711-DEL-59 |
| chr12-3594306-DEL-211 | chr12-3594307-DEL-211 |
| chr12-3600755-DEL-149 | chr12-3600756-DEL-149 |
| chr12-3609869-DEL-244 | chr12-3609870-DEL-244 |
| chr12-3615173-DEL-78 | chr12-3615174-DEL-78 |
| chr12-3654405-DEL-233 | chr12-3654407-DEL-232 |
| chr12-3667912-DEL-180 | chr12-3667913-DEL-180 |
| chr12-3685669-DEL-159 | chr12-3685670-DEL-159 |
| chr12-3708128-DEL-51 | chr12-3708129-DEL-51 |
| chr12-3757889-DEL-669 | chr12-3757890-DEL-669 |
| chr12-3761094-DEL-209 | chr12-3761095-DEL-209 |
| chr12-3791895-DEL-119 | chr12-3791896-DEL-119 |
| chr12-3795124-DEL-181 | chr12-3795125-DEL-181 |
| chr12-3839330-DEL-54 | chr12-3839331-DEL-54 |
| chr12-3877238-DEL-214 | chr12-3877239-DEL-214 |
| chr12-3879279-DEL-168 | chr12-3879304-DEL-126 |
| chr12-3926948-DEL-275 | chr12-3926949-DEL-275 |
| chr12-3944780-DEL-81 | chr12-3944781-DEL-81 |
| chr12-3950857-DEL-205 | chr12-3950878-DEL-185 |
| chr12-3969577-DEL-76 | chr12-3969578-DEL-76 |
| chr12-3980318-DEL-140 | chr12-3980319-DEL-140 |
| chr12-4000438-DEL-202 | chr12-4000439-DEL-202 |
| chr12-4011104-DEL-235 | chr12-4011105-DEL-235 |
| chr12-4017347-DEL-60 | chr12-4017347-DEL-60 |
| chr12-4051314-DEL-2819 | chr12-4051316-DEL-2820 |
| chr12-4096501-DEL-84 | chr12-4096504-DEL-82 |
| chr12-4106386-DEL-177 | chr12-4106396-DEL-168 |
| chr12-4107060-DEL-87 | chr12-4107061-DEL-87 |
| chr12-4126487-DEL-59 | chr12-4126488-DEL-59 |
| chr12-4134863-DEL-201 | chr12-4134864-DEL-201 |
| chr12-4136062-DEL-499 | chr12-4136060-DEL-500 |
| chr12-4136601-DEL-60 | chr12-4136602-DEL-61 |
| chr12-4162680-DEL-66 | chr12-4162693-DEL-58 |
| chr12-4169211-DEL-97 | chr12-4169212-DEL-97 |
| chr12-4191477-DEL-92 | chr12-4191478-DEL-92 |
| chr12-4198849-DEL-438 | chr12-4198850-DEL-438 |
| chr12-4224481-DEL-194 | chr12-4224482-DEL-194 |
| chr12-4226937-DEL-216 | chr12-4226938-DEL-216 |
| chr12-4235054-DEL-172 | chr12-4235055-DEL-172 |
| chr12-4305149-DEL-72 | chr12-4305150-DEL-72 |
| chr12-4306571-DEL-53 | chr12-4306572-DEL-53 |
| chr12-4326417-DEL-476 | chr12-4326421-DEL-457 |
| chr12-4335048-DEL-117 | chr12-4335022-DEL-114 |
| chr12-4342687-DEL-76 | chr12-4342688-DEL-76 |
| chr12-4348532-DEL-54 | chr12-4348530-DEL-51 |
| chr12-4351038-DEL-221 | chr12-4351039-DEL-221 |
| chr12-4353580-DEL-217 | chr12-4353581-DEL-217 |
| chr12-4378769-DEL-1916 | chr12-4378770-DEL-1915 |
| chr12-4407408-DEL-421 | chr12-4407409-DEL-421 |
| chr12-4418086-DEL-55 | chr12-4418066-DEL-60 |
| chr12-4422698-DEL-75 | chr12-4422699-DEL-75 |
| chr12-4436981-DEL-236 | chr12-4436982-DEL-236 |
| chr12-4443987-DEL-237 | chr12-4443982-DEL-237 |
| chr12-4455913-DEL-203 | chr12-4455914-DEL-203 |
| chr12-4465082-DEL-609 | chr12-4465084-DEL-608 |
| chr12-4495170-DEL-77 | chr12-4495152-DEL-62 |
| chr12-4508632-DEL-64 | chr12-4508633-DEL-64 |
| chr12-4552837-DEL-86 | chr12-4552851-DEL-73 |
| chr12-4568537-DEL-70 | chr12-4568538-DEL-70 |
| chr12-4586844-DEL-72 | chr12-4586863-DEL-56 |
| chr12-4602450-DEL-284 | chr12-4602451-DEL-284 |
| chr12-4630101-DEL-214 | chr12-4630102-DEL-214 |
| chr12-4713680-DEL-69 | chr12-4713681-DEL-69 |
| chr12-4738929-DEL-204 | chr12-4738930-DEL-204 |
| chr12-4775339-DEL-178 | chr12-4775322-DEL-178 |
| chr12-4778774-DEL-183 | chr12-4778773-DEL-181 |
| chr12-4808531-DEL-1716 | chr12-4808532-DEL-1716 |
| chr12-4811607-DEL-187 | chr12-4811608-DEL-187 |
| chr12-4815617-DEL-66 | chr12-4815618-DEL-76 |
| chr12-4822059-DEL-123 | chr12-4822060-DEL-123 |
| chr12-4837758-DEL-2446 | chr12-4837759-DEL-2446 |
| chr12-4843960-DEL-55 | chr12-4843961-DEL-55 |
| chr12-4850784-DEL-8823 | chr12-4850785-DEL-8823 |
| chr12-4861943-DEL-68 | chr12-4861944-DEL-68 |
| chr12-4875445-DEL-248 | chr12-4875450-DEL-244 |

| | |
|---|---|
| chr12-4878835-DEL-69 | chr12-4878868-DEL-73 |
| chr12-4880475-DEL-86 | chr12-4880472-DEL-87 |
| chr12-4895647-DEL-1258 | chr12-4895648-DEL-1258 |
| chr12-4895887-DEL-158 | chr12-4895888-DEL-158 |
| chr12-4904576-DEL-58 | chr12-4904575-DEL-64 |
| chr12-4928694-DEL-83 | chr12-4928695-DEL-83 |
| chr12-4965932-DEL-66 | chr12-4965914-DEL-85 |
| chr12-4966360-DEL-426 | chr12-4966401-DEL-386 |
| chr12-5000850-DEL-57 | chr12-5000851-DEL-57 |
| chr12-5008946-DEL-158 | chr12-5008947-DEL-158 |
| chr12-5010631-DEL-103 | chr12-5010632-DEL-103 |
| chr12-5059818-DEL-191 | chr12-5059813-DEL-191 |
| chr12-5063420-DEL-78 | chr12-5063421-DEL-78 |
| chr12-5072703-DEL-93 | chr12-5072704-DEL-93 |
| chr12-5077874-DEL-201 | chr12-5077875-DEL-201 |
| chr12-5131796-DEL-90 | chr12-5131793-DEL-94 |
| chr12-5167997-DEL-185 | chr12-5167978-DEL-162 |
| chr12-5228463-DEL-51 | chr12-5228464-DEL-51 |
| chr12-5329367-DEL-102 | chr12-5329384-DEL-116 |
| chr12-5375386-DEL-87 | chr12-5375387-DEL-116 |
| chr12-5394707-DEL-459 | chr12-5394708-DEL-459 |
| chr12-5398825-DEL-68 | chr12-5398821-DEL-69 |
| chr12-5504284-DEL-70 | chr12-5504285-DEL-70 |
| chr12-5505072-DEL-158 | chr12-5505072-DEL-158 |
| chr12-5529102-DEL-6431 | chr12-5529103-DEL-6431 |
| chr12-5540431-DEL-136 | chr12-5540432-DEL-136 |
| chr12-5573619-DEL-56 | chr12-5573620-DEL-55 |
| chr12-5633972-DEL-57 | chr12-5633973-DEL-65 |
| chr12-5648844-DEL-62 | chr12-5648844-DEL-62 |
| chr12-5663770-DEL-4167 | chr12-5663771-DEL-4167 |
| chr12-5725535-DEL-90 | chr12-5725540-DEL-82 |
| chr12-5737938-DEL-230 | chr12-5737937-DEL-231 |
| chr12-5767019-DEL-641 | chr12-5767020-DEL-641 |
| chr12-5794709-DEL-52 | chr12-5794720-DEL-64 |
| chr12-5847689-DEL-325 | chr12-5847690-DEL-325 |
| chr12-5865694-DEL-138 | chr12-5865694-DEL-136 |
| chr12-5878509-DEL-6208 | chr12-5878510-DEL-6208 |
| chr12-5896465-DEL-190 | chr12-5896467-DEL-188 |
| chr12-5941841-DEL-427 | chr12-5941842-DEL-427 |
| chr12-6005022-DEL-92 | chr12-6005012-DEL-102 |
| chr12-6014353-DEL-1472 | chr12-6014352-DEL-1473 |
| chr12-6016282-DEL-101 | chr12-6016286-DEL-89 |
| chr12-6063003-DEL-6569 | chr12-6063004-DEL-6569 |
| chr12-6127459-DEL-6682 | chr12-6127460-DEL-6682 |
| chr12-6213123-DEL-100 | chr12-6213122-DEL-100 |
| chr12-6229713-DEL-62 | chr12-6229714-DEL-60 |
| chr12-6285103-DEL-214 | chr12-6285104-DEL-214 |
| chr12-6288627-DEL-1489 | chr12-6288628-DEL-1489 |
| chr12-6295247-DEL-52 | chr12-6295248-DEL-52 |
| chr12-6324768-DEL-59 | chr12-6324767-DEL-61 |
| chr12-6473738-DEL-4219 | chr12-6473739-DEL-4219 |
| chr12-6479282-DEL-69 | chr12-6479283-DEL-69 |
| chr12-6481570-DEL-6346 | chr12-6481571-DEL-6346 |
| chr12-6487662-DEL-214 | chr12-6487663-DEL-214 |
| chr12-6489100-DEL-68 | chr12-6489101-DEL-68 |
| chr12-6533160-DEL-569 | chr12-6533161-DEL-569 |
| chr12-6566465-DEL-7212 | chr12-6566466-DEL-7212 |
| chr12-6588339-DEL-96 | chr12-6588357-DEL-66 |
| chr12-6653994-DEL-68 | chr12-6653995-DEL-68 |
| chr12-6656471-DEL-6186 | chr12-6656472-DEL-6186 |
| chr12-6716577-DEL-6247 | chr12-6716578-DEL-6247 |
| chr12-6750583-DEL-92 | chr12-6750584-DEL-92 |
| chr12-6791310-DEL-138 | chr12-6791311-DEL-138 |
| chr12-6812673-DEL-287 | chr12-6812674-DEL-287 |
| chr12-6813346-DEL-110 | chr12-6813347-DEL-64 |
| chr12-6820937-DEL-6521 | chr12-6820938-DEL-6521 |
| chr12-6853016-DEL-7109 | chr12-6853017-DEL-7109 |
| chr12-6965669-DEL-88 | chr12-6965679-DEL-50 |
| chr12-6968187-DEL-71 | chr12-6968192-DEL-71 |
| chr12-7044124-DEL-483 | chr12-7044120-DEL-483 |
| chr12-7045011-DEL-303 | chr12-7045009-DEL-296 |
| chr12-7047293-DEL-564 | chr12-7047294-DEL-564 |
| chr12-7055272-DEL-52 | chr12-7055275-DEL-69 |
| chr12-7088911-DEL-419 | chr12-7088910-DEL-421 |
| chr12-7105039-DEL-71 | chr12-7105041-DEL-70 |
| chr12-7161614-DEL-6584 | chr12-7161615-DEL-6584 |
| chr12-7215325-DEL-117 | chr12-7215318-DEL-123 |
| chr12-7218672-DEL-1454 | chr12-7218673-DEL-1454 |
| chr12-7224982-DEL-62 | chr12-7224964-DEL-80 |
| chr12-7305561-DEL-88 | chr12-7305545-DEL-94 |

| | |
|---|---|
| chr12-7322389-DEL-96 | chr12-7322436-DEL-87 |
| chr12-7344565-DEL-458 | chr12-7344566-DEL-458 |
| chr12-7370444-DEL-58 | chr12-7370475-DEL-55 |
| chr12-7395014-DEL-168 | chr12-7395015-DEL-168 |
| chr12-7454463-DEL-2227 | chr12-7454464-DEL-2227 |
| chr12-7484403-DEL-239 | chr12-7484404-DEL-239 |
| chr12-7502858-DEL-518 | chr12-7502859-DEL-518 |
| chr12-7522042-DEL-59 | chr12-7522031-DEL-56 |
| chr12-7560575-DEL-186 | chr12-7560576-DEL-186 |
| chr12-7608041-DEL-1077 | chr12-7608042-DEL-1077 |
| chr12-7639084-DEL-3027 | chr12-7639085-DEL-3027 |
| chr12-7670882-DEL-972 | chr12-7670883-DEL-967 |
| chr12-7676851-DEL-3286 | chr12-7676852-DEL-3286 |
| chr12-7697344-DEL-160 | chr12-7697346-DEL-120 |
| chr12-7761547-DEL-129 | chr12-7761546-DEL-130 |
| chr12-7763222-DEL-375 | chr12-7763223-DEL-375 |
| chr12-7767204-DEL-149 | chr12-7767205-DEL-149 |
| chr12-7808171-DEL-199 | chr12-7808172-DEL-199 |
| chr12-7810803-DEL-1058 | chr12-7810804-DEL-1058 |
| chr12-7849618-DEL-437 | chr12-7849618-DEL-437 |
| chr12-7851843-DEL-6683 | chr12-7851844-DEL-6683 |
| chr12-7898107-DEL-248 | chr12-7898107-DEL-245 |
| chr12-7922945-DEL-70 | chr12-7922974-DEL-52 |
| chr12-7937117-DEL-493 | chr12-7937118-DEL-493 |
| chr12-8015379-DEL-69 | chr12-8015380-DEL-69 |
| chr12-8026524-DEL-204 | chr12-8026525-DEL-204 |
| chr12-8091097-DEL-197 | chr12-8091098-DEL-197 |
| chr12-8105242-DEL-52 | chr12-8105243-DEL-52 |
| chr12-8140682-DEL-97 | chr12-8140656-DEL-97 |
| chr12-8149301-DEL-72 | chr12-8149266-DEL-72 |
| chr12-8166994-DEL-60 | chr12-8166995-DEL-60 |
| chr12-8173875-DEL-6641 | chr12-8173876-DEL-6641 |
| chr12-8173892-DEL-212 | chr12-8173893-DEL-212 |
| chr12-8187078-DEL-76 | chr12-8187065-DEL-80 |
| chr12-8201044-DEL-96 | chr12-8201045-DEL-96 |
| chr12-8232343-DEL-216 | chr12-8232344-DEL-216 |
| chr12-8245319-DEL-4627 | chr12-8245300-DEL-4628 |
| chr12-8254821-DEL-587 | chr12-8254822-DEL-587 |
| chr12-8306436-DEL-64 | chr12-8306424-DEL-92 |
| chr12-8318892-DEL-326 | chr12-8318893-DEL-326 |
| chr12-8320371-DEL-1047 | chr12-8320372-DEL-1047 |
| chr12-8380735-DEL-685 | chr12-8380736-DEL-685 |
| chr12-8382745-DEL-96 | chr12-8382746-DEL-110 |
| chr12-8391298-DEL-58 | chr12-8391294-DEL-59 |
| chr12-8405461-DEL-60 | chr12-8405462-DEL-52 |
| chr12-8453169-DEL-5290 | chr12-8453170-DEL-5290 |
| chr12-8465336-DEL-305 | chr12-8465335-DEL-305 |
| chr12-8520023-DEL-56 | chr12-8520024-DEL-56 |
| chr12-8536219-DEL-208 | chr12-8536220-DEL-207 |
| chr12-8540654-DEL-196 | chr12-8540655-DEL-196 |
| chr12-8545354-DEL-252 | chr12-8545355-DEL-252 |
| chr12-8605000-DEL-240 | chr12-8605001-DEL-240 |
| chr12-8626972-DEL-11726 | chr12-8626973-DEL-11726 |
| chr12-8632841-DEL-4735 | chr12-8632842-DEL-4735 |
| chr12-8664355-DEL-53 | chr12-8664356-DEL-53 |
| chr12-8671008-DEL-7118 | chr12-8671008-DEL-7118 |
| chr12-8684027-DEL-70 | chr12-8684028-DEL-70 |
| chr12-8685564-DEL-64 | chr12-8685567-DEL-62 |
| chr12-8702519-DEL-199 | chr12-8702527-DEL-201 |
| chr12-8716886-DEL-216 | chr12-8716887-DEL-216 |
| chr12-8733205-DEL-145 | chr12-8733206-DEL-145 |
| chr12-8736401-DEL-179 | chr12-8736403-DEL-178 |
| chr12-8753124-DEL-195 | chr12-8753119-DEL-196 |
| chr12-8818337-DEL-199 | chr12-8818338-DEL-199 |
| chr12-8863911-DEL-250 | chr12-8863912-DEL-250 |
| chr12-8872213-DEL-217 | chr12-8872214-DEL-217 |
| chr12-8893692-DEL-233 | chr12-8893693-DEL-225 |
| chr12-8898798-DEL-278 | chr12-8898799-DEL-278 |
| chr12-8953917-DEL-4074 | chr12-8953918-DEL-4074 |
| chr12-8959290-DEL-1894 | chr12-8959291-DEL-1894 |
| chr12-8976697-DEL-543 | chr12-8976698-DEL-543 |
| chr12-8978392-DEL-93 | chr12-8978385-DEL-101 |
| chr12-8984296-DEL-209 | chr12-8984297-DEL-209 |
| chr12-9014204-DEL-258 | chr12-9014204-DEL-258 |
| chr12-9088965-DEL-77 | chr12-9088966-DEL-77 |
| chr12-9121100-DEL-84 | chr12-9121097-DEL-86 |
| chr12-9132561-DEL-201 | chr12-9132562-DEL-201 |
| chr12-9157147-DEL-64 | chr12-9157148-DEL-65 |
| chr12-9189500-DEL-3081 | chr12-9189491-DEL-3091 |
| chr12-9196744-DEL-64 | chr12-9196726-DEL-58 |

| | |
|---|---|
| chr12-9255742-DEL-345 | chr12-9255733-DEL-345 |
| chr12-9315316-DEL-55 | chr12-9315295-DEL-65 |
| chr12-9322537-DEL-56 | chr12-9322538-DEL-56 |
| chr12-9351100-DEL-1677 | chr12-9351101-DEL-1677 |
| chr12-9354933-DEL-68 | chr12-9354934-DEL-68 |
| chr12-9366813-DEL-60 | chr12-9366853-DEL-51 |
| chr12-9370700-DEL-109 | chr12-9370701-DEL-109 |
| chr12-9399376-DEL-66 | chr12-9399377-DEL-66 |
| chr12-9427551-DEL-80 | chr12-9427552-DEL-80 |
| chr12-9483909-DEL-2041 | chr12-9483910-DEL-2041 |
| chr12-9493585-DEL-270 | chr12-9493585-DEL-282 |
| chr12-9501973-DEL-76 | chr12-9501974-DEL-76 |
| chr12-9504607-DEL-1820 | chr12-9504608-DEL-1820 |
| chr12-9506740-DEL-51 | chr12-9506726-DEL-66 |
| chr12-9523599-DEL-64 | chr12-9523600-DEL-64 |
| chr12-9553529-DEL-6413 | chr12-9553530-DEL-6413 |
| chr12-9563990-DEL-1853 | chr12-9563991-DEL-1853 |
| chr12-9603506-DEL-1357 | chr12-9603506-DEL-1357 |
| chr12-9676721-DEL-6509 | chr12-9676721-DEL-6509 |
| chr12-9704407-DEL-959 | chr12-9704408-DEL-959 |
| chr12-9727063-DEL-6641 | chr12-9727064-DEL-6641 |
| chr12-9742782-DEL-64 | chr12-9742783-DEL-64 |
| chr12-9743690-DEL-57 | chr12-9743690-DEL-57 |
| chr12-9769117-DEL-897 | chr12-9769118-DEL-897 |
| chr12-9803213-DEL-68 | chr12-9803213-DEL-68 |
| chr12-9807031-DEL-55 | chr12-9807021-DEL-54 |
| chr12-9822320-DEL-64 | chr12-9822321-DEL-64 |
| chr12-9834677-DEL-204 | chr12-9834679-DEL-206 |
| chr12-9844365-DEL-56 | chr12-9844370-DEL-52 |
| chr12-9869586-DEL-735 | chr12-9869587-DEL-735 |
| chr12-9880820-DEL-429 | chr12-9880820-DEL-429 |
| chr12-9891265-DEL-52 | chr12-9891262-DEL-68 |
| chr12-9986649-DEL-66 | chr12-9986651-DEL-56 |
| chr12-10000381-DEL-90 | chr12-10000382-DEL-90 |
| chr12-10006708-DEL-55 | chr12-10006709-DEL-65 |
| chr12-10089009-DEL-62 | chr12-10089010-DEL-62 |
| chr12-10147164-DEL-52 | chr12-10147164-DEL-52 |
| chr12-10158214-DEL-106 | chr12-10158215-DEL-98 |
| chr12-10160712-DEL-1619 | chr12-10160716-DEL-1618 |
| chr12-10166632-DEL-196 | chr12-10166633-DEL-196 |
| chr12-10171912-DEL-65 | chr12-10171913-DEL-65 |
| chr12-10196732-DEL-69 | chr12-10196734-DEL-68 |
| chr12-10197005-DEL-509 | chr12-10197006-DEL-509 |
| chr12-10200148-DEL-1522 | chr12-10200149-DEL-1522 |
| chr12-10202122-DEL-8107 | chr12-10202123-DEL-8107 |
| chr12-10214129-DEL-64 | chr12-10214120-DEL-106 |
| chr12-10216711-DEL-215 | chr12-10216751-DEL-175 |
| chr12-10245854-DEL-135 | chr12-10245855-DEL-135 |
| chr12-10268050-DEL-243 | chr12-10268051-DEL-243 |
| chr12-10288591-DEL-52 | chr12-10288588-DEL-54 |
| chr12-10364853-DEL-692 | chr12-10364854-DEL-692 |
| chr12-10385258-DEL-257 | chr12-10385259-DEL-257 |
| chr12-10393994-DEL-185 | chr12-10393995-DEL-185 |
| chr12-10398450-DEL-225 | chr12-10398451-DEL-225 |
| chr12-10413801-DEL-120 | chr12-10413846-DEL-75 |
| chr12-10446619-DEL-75 | chr12-10446624-DEL-74 |
| chr12-10483279-DEL-183 | chr12-10483280-DEL-183 |
| chr12-10531460-DEL-62 | chr12-10531461-DEL-62 |
| chr12-10543073-DEL-54 | chr12-10543080-DEL-56 |
| chr12-10545973-DEL-497 | chr12-10545974-DEL-497 |
| chr12-10655121-DEL-1323 | chr12-10655122-DEL-1323 |
| chr12-10658131-DEL-53 | chr12-10658132-DEL-53 |
| chr12-10667028-DEL-97 | chr12-10667031-DEL-95 |
| chr12-10676380-DEL-5962 | chr12-10676381-DEL-5962 |
| chr12-10689581-DEL-499 | chr12-10689582-DEL-499 |
| chr12-10696962-DEL-75 | chr12-10697002-DEL-64 |
| chr12-10724235-DEL-640 | chr12-10724236-DEL-640 |
| chr12-10747909-DEL-6398 | chr12-10747910-DEL-6398 |
| chr12-10754783-DEL-1281 | chr12-10754784-DEL-1281 |
| chr12-10768817-DEL-61 | chr12-10768818-DEL-56 |
| chr12-10780414-DEL-52 | chr12-10780415-DEL-52 |
| chr12-10785942-DEL-55 | chr12-10785957-DEL-52 |
| chr12-10808765-DEL-52 | chr12-10808766-DEL-50 |
| chr12-10847238-DEL-2835 | chr12-10847239-DEL-2835 |
| chr12-10879365-DEL-6125 | chr12-10879366-DEL-6125 |
| chr12-10890631-DEL-56 | chr12-10890595-DEL-59 |
| chr12-10894185-DEL-168 | chr12-10894186-DEL-168 |
| chr12-10928576-DEL-84 | chr12-10928580-DEL-81 |
| chr12-10935304-DEL-6230 | chr12-10935305-DEL-6230 |
| chr12-10948542-DEL-1775 | chr12-10948543-DEL-1775 |

| | |
|---|---|
| chr12-10965450-DEL-8525 | chr12-10965449-DEL-8525 |
| chr12-10996321-DEL-1224 | chr12-10996322-DEL-1224 |
| chr12-10999892-DEL-184 | chr12-10999893-DEL-150 |
| chr12-11045993-DEL-116 | chr12-11045972-DEL-88 |
| chr12-11081415-DEL-6553 | chr12-11081416-DEL-6553 |
| chr12-11106973-DEL-54 | chr12-11106974-DEL-54 |
| chr12-11109193-DEL-161 | chr12-11109197-DEL-159 |
| chr12-11123048-DEL-2675 | chr12-11123049-DEL-2675 |
| chr12-11132223-DEL-541 | chr12-11132224-DEL-541 |
| chr12-11136802-DEL-66 | chr12-11136789-DEL-104 |
| chr12-11150844-DEL-573 | chr12-11150845-DEL-573 |
| chr12-11153653-DEL-7085 | chr12-11153654-DEL-7085 |
| chr12-11165388-DEL-54 | chr12-11165389-DEL-54 |
| chr12-11169068-DEL-108 | chr12-11169069-DEL-108 |
| chr12-11174204-DEL-200 | chr12-11174205-DEL-200 |
| chr12-11183672-DEL-134 | chr12-11183673-DEL-134 |
| chr12-11197933-DEL-206 | chr12-11197934-DEL-206 |
| chr12-11213959-DEL-74 | chr12-11213962-DEL-72 |
| chr12-11226806-DEL-343 | chr12-11226807-DEL-343 |
| chr12-11237521-DEL-6628 | chr12-11237522-DEL-6628 |
| chr12-11250170-DEL-358 | chr12-11250172-DEL-356 |
| chr12-11251080-DEL-1019 | chr12-11251081-DEL-1019 |
| chr12-11273607-DEL-70 | chr12-11273611-DEL-62 |
| chr12-11275399-DEL-218 | chr12-11275400-DEL-218 |
| chr12-11311331-DEL-533 | chr12-11311332-DEL-533 |
| chr12-11374379-DEL-107 | chr12-11374380-DEL-107 |
| chr12-11380797-DEL-132 | chr12-11380798-DEL-132 |
| chr12-11387071-DEL-72 | chr12-11387072-DEL-72 |
| chr12-11392330-DEL-60 | chr12-11392301-DEL-55 |
| chr12-11424972-DEL-725 | chr12-11424973-DEL-724 |
| chr12-11436592-DEL-69 | chr12-11436600-DEL-60 |
| chr12-11447301-DEL-166 | chr12-11447302-DEL-166 |
| chr12-11463640-DEL-75 | chr12-11463641-DEL-75 |
| chr12-11484288-DEL-60 | chr12-11484306-DEL-58 |
| chr12-11537776-DEL-204 | chr12-11537804-DEL-204 |
| chr12-11565807-DEL-56 | chr12-11565807-DEL-56 |
| chr12-11592314-DEL-6439 | chr12-11592315-DEL-6439 |
| chr12-11636166-DEL-125 | chr12-11636167-DEL-126 |
| chr12-11659964-DEL-54 | chr12-11659964-DEL-54 |
| chr12-11680015-DEL-124 | chr12-11680016-DEL-124 |
| chr12-11684806-DEL-182 | chr12-11684800-DEL-178 |
| chr12-11691780-DEL-2520 | chr12-11691766-DEL-2532 |
| chr12-11702301-DEL-5770 | chr12-11702302-DEL-5770 |
| chr12-11707441-DEL-604 | chr12-11707442-DEL-604 |
| chr12-11711476-DEL-1113 | chr12-11711477-DEL-1113 |
| chr12-11772063-DEL-761 | chr12-11772064-DEL-761 |
| chr12-11776354-DEL-51 | chr12-11776343-DEL-99 |
| chr12-11778301-DEL-73 | chr12-11778302-DEL-73 |
| chr12-11781481-DEL-2605 | chr12-11781482-DEL-2605 |
| chr12-11784244-DEL-579 | chr12-11784245-DEL-579 |
| chr12-11789557-DEL-62 | chr12-11789558-DEL-62 |
| chr12-11791588-DEL-92 | chr12-11791562-DEL-119 |
| chr12-11805502-DEL-64 | chr12-11805495-DEL-61 |
| chr12-11818599-DEL-54 | chr12-11818602-DEL-52 |
| chr12-11833371-DEL-207 | chr12-11833372-DEL-207 |
| chr12-11843186-DEL-52 | chr12-11843187-DEL-52 |
| chr12-11843955-DEL-399 | chr12-11843956-DEL-399 |
| chr12-11857500-DEL-1095 | chr12-11857501-DEL-1095 |
| chr12-11874918-DEL-93 | chr12-11874919-DEL-93 |
| chr12-11880730-DEL-56 | chr12-11880730-DEL-56 |
| chr12-11896839-DEL-62 | chr12-11896840-DEL-62 |
| chr12-11924995-DEL-52 | chr12-11924996-DEL-52 |
| chr12-11930567-DEL-198 | chr12-11930568-DEL-198 |
| chr12-11966086-DEL-788 | chr12-11966089-DEL-787 |
| chr12-11966388-DEL-5605 | chr12-11966389-DEL-5605 |
| chr12-12003738-DEL-6418 | chr12-12003739-DEL-6418 |
| chr12-12011040-DEL-52 | chr12-12011041-DEL-53 |
| chr12-12020425-DEL-2444 | chr12-12020443-DEL-2426 |
| chr12-12027258-DEL-741 | chr12-12027259-DEL-741 |
| chr12-12031923-DEL-64 | chr12-12031924-DEL-64 |
| chr12-12053722-DEL-395 | chr12-12053721-DEL-394 |
| chr12-12061918-DEL-319 | chr12-12061919-DEL-319 |
| chr12-12069667-DEL-52 | chr12-12069668-DEL-52 |
| chr12-12072905-DEL-73 | chr12-12072906-DEL-73 |
| chr12-12072994-DEL-741 | chr12-12072982-DEL-742 |
| chr12-12090640-DEL-79 | chr12-12090641-DEL-79 |
| chr12-12091538-DEL-213 | chr12-12091539-DEL-213 |
| chr12-12142500-DEL-6495 | chr12-12142501-DEL-6495 |
| chr12-12153328-DEL-674 | chr12-12153348-DEL-665 |
| chr12-12157929-DEL-603 | chr12-12157930-DEL-603 |

| | |
|---|---|
| chr12-12195739-DEL-274 | chr12-12195740-DEL-274 |
| chr12-12203342-DEL-6980 | chr12-12203343-DEL-6980 |
| chr12-12208114-DEL-922 | chr12-12208115-DEL-922 |
| chr12-12250202-DEL-51 | chr12-12250197-DEL-52 |
| chr12-12263955-DEL-100 | chr12-12263970-DEL-86 |
| chr12-12271463-DEL-81 | chr12-12271416-DEL-81 |
| chr12-12329993-DEL-266 | chr12-12329994-DEL-266 |
| chr12-12330555-DEL-95 | chr12-12330556-DEL-95 |
| chr12-12374511-DEL-60 | chr12-12374494-DEL-106 |
| chr12-12377426-DEL-68 | chr12-12377427-DEL-68 |
| chr12-12395281-DEL-2028 | chr12-12395282-DEL-2028 |
| chr12-12413184-DEL-140 | chr12-12413189-DEL-136 |
| chr12-12420858-DEL-82 | chr12-12420843-DEL-102 |
| chr12-12434712-DEL-315 | chr12-12434713-DEL-315 |
| chr12-12440435-DEL-328 | chr12-12440436-DEL-328 |
| chr12-12442524-DEL-168 | chr12-12442514-DEL-168 |
| chr12-12446083-DEL-64 | chr12-12446084-DEL-64 |
| chr12-12468832-DEL-111 | chr12-12468838-DEL-110 |
| chr12-12500761-DEL-64 | chr12-12500770-DEL-56 |
| chr12-12516293-DEL-209 | chr12-12516323-DEL-193 |
| chr12-12517440-DEL-6180 | chr12-12517441-DEL-6180 |
| chr12-12554284-DEL-222 | chr12-12554285-DEL-222 |
| chr12-12563431-DEL-58 | chr12-12563432-DEL-62 |
| chr12-12603844-DEL-696 | chr12-12603845-DEL-696 |
| chr12-12613821-DEL-327 | chr12-12613822-DEL-327 |
| chr12-12641502-DEL-74 | chr12-12641520-DEL-56 |
| chr12-12650942-DEL-129 | chr12-12650950-DEL-126 |
| chr12-12654654-DEL-140 | chr12-12654655-DEL-140 |
| chr12-12676799-DEL-203 | chr12-12676800-DEL-203 |
| chr12-12679586-DEL-64 | chr12-12679562-DEL-53 |
| chr12-12708724-DEL-4519 | chr12-12708725-DEL-4519 |
| chr12-12716658-DEL-394 | chr12-12716662-DEL-396 |
| chr12-12754264-DEL-60 | chr12-12754265-DEL-57 |
| chr12-12758318-DEL-92 | chr12-12758319-DEL-69 |
| chr12-12767396-DEL-59 | chr12-12767392-DEL-63 |
| chr12-12779097-DEL-60 | chr12-12779087-DEL-62 |
| chr12-12784082-DEL-60 | chr12-12784083-DEL-60 |
| chr12-12800290-DEL-70 | chr12-12800301-DEL-50 |
| chr12-12802967-DEL-537 | chr12-12802968-DEL-537 |
| chr12-12805775-DEL-105 | chr12-12805800-DEL-74 |
| chr12-12823102-DEL-798 | chr12-12823103-DEL-798 |
| chr12-12844658-DEL-72 | chr12-12844659-DEL-72 |
| chr12-12848327-DEL-220 | chr12-12848326-DEL-222 |
| chr12-12850901-DEL-108 | chr12-12850902-DEL-108 |
| chr12-12874946-DEL-51 | chr12-12874947-DEL-51 |
| chr12-12882358-DEL-356 | chr12-12882359-DEL-356 |
| chr12-12909433-DEL-6327 | chr12-12909432-DEL-6325 |
| chr12-12918887-DEL-74 | chr12-12918888-DEL-74 |
| chr12-12928709-DEL-72 | chr12-12928710-DEL-72 |
| chr12-12976893-DEL-1809 | chr12-12976888-DEL-1811 |
| chr12-13022015-DEL-85 | chr12-13022016-DEL-90 |
| chr12-13040804-DEL-68 | chr12-13040813-DEL-60 |
| chr12-13051547-DEL-208 | chr12-13051548-DEL-208 |
| chr12-13054946-DEL-1211 | chr12-13054947-DEL-1211 |
| chr12-13065383-DEL-58 | chr12-13065388-DEL-56 |
| chr12-13083587-DEL-1011 | chr12-13083588-DEL-1011 |
| chr12-13107045-DEL-664 | chr12-13107046-DEL-664 |
| chr12-13107888-DEL-651 | chr12-13107889-DEL-651 |
| chr12-13109882-DEL-69 | chr12-13109876-DEL-50 |
| chr12-13147773-DEL-1360 | chr12-13147774-DEL-1360 |
| chr12-13163470-DEL-167 | chr12-13163463-DEL-168 |
| chr12-13184133-DEL-51 | chr12-13184134-DEL-51 |
| chr12-13211061-DEL-390 | chr12-13211062-DEL-390 |
| chr12-13239711-DEL-180 | chr12-13239712-DEL-174 |
| chr12-13270847-DEL-2019 | chr12-13270848-DEL-2019 |
| chr12-13302442-DEL-59 | chr12-13302443-DEL-59 |
| chr12-13308028-DEL-68 | chr12-13308029-DEL-68 |
| chr12-13308707-DEL-689 | chr12-13308709-DEL-688 |
| chr12-13310726-DEL-424 | chr12-13310727-DEL-424 |
| chr12-13311778-DEL-193 | chr12-13311779-DEL-193 |
| chr12-13332905-DEL-108 | chr12-13332906-DEL-72 |
| chr12-13336551-DEL-163 | chr12-13336543-DEL-164 |
| chr12-13364683-DEL-65 | chr12-13364684-DEL-65 |
| chr12-13409051-DEL-449 | chr12-13409052-DEL-449 |
| chr12-13469452-DEL-6467 | chr12-13469453-DEL-6467 |
| chr12-13479224-DEL-60 | chr12-13479238-DEL-74 |
| chr12-13483570-DEL-422 | chr12-13483571-DEL-422 |
| chr12-13490443-DEL-60 | chr12-13490460-DEL-58 |
| chr12-13535883-DEL-112 | chr12-13535884-DEL-112 |
| chr12-13537844-DEL-74 | chr12-13537845-DEL-70 |

| | |
|---|---|
| chr12-13552323-DEL-59 | chr12-13552324-DEL-59 |
| chr12-13553716-DEL-84 | chr12-13553718-DEL-103 |
| chr12-13556978-DEL-991 | chr12-13556979-DEL-991 |
| chr12-13566626-DEL-91 | chr12-13566627-DEL-91 |
| chr12-13567306-DEL-65 | chr12-13567307-DEL-65 |
| chr12-13575657-DEL-76 | chr12-13575642-DEL-70 |
| chr12-13641948-DEL-1197 | chr12-13641949-DEL-1197 |
| chr12-13653194-DEL-273 | chr12-13653182-DEL-286 |
| chr12-13670601-DEL-555 | chr12-13670560-DEL-589 |
| chr12-13728677-DEL-52 | chr12-13728678-DEL-52 |
| chr12-13751471-DEL-294 | chr12-13751468-DEL-295 |
| chr12-13807629-DEL-6288 | chr12-13807630-DEL-6288 |
| chr12-13835787-DEL-1484 | chr12-13835788-DEL-1484 |
| chr12-13839485-DEL-2057 | chr12-13839486-DEL-2057 |
| chr12-13857049-DEL-699 | chr12-13857050-DEL-699 |
| chr12-13866069-DEL-167 | chr12-13866030-DEL-168 |
| chr12-13904343-DEL-1021 | chr12-13904344-DEL-1020 |
| chr12-13913262-DEL-225 | chr12-13913263-DEL-225 |
| chr12-13953770-DEL-1867 | chr12-13953770-DEL-1867 |
| chr12-13961455-DEL-6245 | chr12-13961454-DEL-6248 |
| chr12-13981511-DEL-104 | chr12-13981510-DEL-105 |
| chr12-14002327-DEL-2773 | chr12-14002328-DEL-2773 |
| chr12-14006959-DEL-486 | chr12-14006960-DEL-486 |
| chr12-14030780-DEL-934 | chr12-14030781-DEL-934 |
| chr12-14041386-DEL-7085 | chr12-14041387-DEL-7085 |
| chr12-14047938-DEL-212 | chr12-14047939-DEL-212 |
| chr12-14066865-DEL-1039 | chr12-14066868-DEL-1037 |
| chr12-14110315-DEL-54 | chr12-14110316-DEL-54 |
| chr12-14139580-DEL-388 | chr12-14139581-DEL-388 |
| chr12-14235424-DEL-284 | chr12-14235425-DEL-284 |
| chr12-14253029-DEL-51 | chr12-14253066-DEL-54 |
| chr12-14253973-DEL-346 | chr12-14253986-DEL-346 |
| chr12-14292622-DEL-581 | chr12-14292616-DEL-582 |
| chr12-14328865-DEL-274 | chr12-14328890-DEL-247 |
| chr12-14394082-DEL-430 | chr12-14394083-DEL-430 |
| chr12-14433585-DEL-69 | chr12-14433584-DEL-70 |
| chr12-14438326-DEL-6550 | chr12-14438327-DEL-6550 |
| chr12-14490332-DEL-3731 | chr12-14490334-DEL-3736 |
| chr12-14499698-DEL-6252 | chr12-14499699-DEL-6252 |
| chr12-14517279-DEL-145 | chr12-14517280-DEL-145 |
| chr12-14543589-DEL-1674 | chr12-14543590-DEL-1673 |
| chr12-14589358-DEL-6414 | chr12-14589359-DEL-6414 |
| chr12-14605884-DEL-215 | chr12-14605885-DEL-215 |
| chr12-14610402-DEL-548 | chr12-14610403-DEL-548 |
| chr12-14643094-DEL-108 | chr12-14643097-DEL-62 |
| chr12-14644194-DEL-7257 | chr12-14644195-DEL-7257 |
| chr12-14700925-DEL-217 | chr12-14700922-DEL-221 |
| chr12-14719724-DEL-6252 | chr12-14719725-DEL-6252 |
| chr12-14733241-DEL-88 | chr12-14733287-DEL-52 |
| chr12-14773346-DEL-433 | chr12-14773347-DEL-433 |
| chr12-14792662-DEL-254 | chr12-14792663-DEL-254 |
| chr12-14801856-DEL-492 | chr12-14801857-DEL-492 |
| chr12-14808491-DEL-1455 | chr12-14808492-DEL-1455 |
| chr12-14831211-DEL-90 | chr12-14831212-DEL-90 |
| chr12-14840861-DEL-6798 | chr12-14840862-DEL-6798 |
| chr12-14871288-DEL-67 | chr12-14871289-DEL-67 |
| chr12-14909899-DEL-51 | chr12-14909900-DEL-51 |
| chr12-14945265-DEL-180 | chr12-14945230-DEL-228 |
| chr12-14947971-DEL-441 | chr12-14947972-DEL-441 |
| chr12-14949989-DEL-94 | chr12-14949990-DEL-94 |
| chr12-14963497-DEL-965 | chr12-14963498-DEL-965 |
| chr12-14964854-DEL-1454 | chr12-14964855-DEL-1454 |
| chr12-14969174-DEL-65 | chr12-14969152-DEL-90 |
| chr12-14969825-DEL-419 | chr12-14969826-DEL-419 |
| chr12-14988293-DEL-200 | chr12-14988274-DEL-201 |
| chr12-15007730-DEL-6266 | chr12-15007732-DEL-6265 |
| chr12-15035832-DEL-101 | chr12-15035833-DEL-102 |
| chr12-15046038-DEL-61 | chr12-15046082-DEL-60 |
| chr12-15058935-DEL-185 | chr12-15058936-DEL-185 |
| chr12-15075946-DEL-6748 | chr12-15075947-DEL-6748 |
| chr12-15076223-DEL-140 | chr12-15076223-DEL-141 |
| chr12-15100259-DEL-319 | chr12-15100260-DEL-319 |
| chr12-15105104-DEL-6267 | chr12-15105105-DEL-6267 |
| chr12-15187918-DEL-313 | chr12-15187919-DEL-313 |
| chr12-15199969-DEL-230 | chr12-15199948-DEL-208 |
| chr12-15206112-DEL-1020 | chr12-15206113-DEL-1020 |
| chr12-15220292-DEL-193 | chr12-15220284-DEL-194 |
| chr12-15223530-DEL-427 | chr12-15223546-DEL-409 |
| chr12-15241466-DEL-7199 | chr12-15241467-DEL-7199 |
| chr12-15300361-DEL-59 | chr12-15300362-DEL-55 |

| | |
|---|---|
| chr12-15300879-DEL-97 | chr12-15300883-DEL-96 |
| chr12-15303505-DEL-6176 | chr12-15303506-DEL-6176 |
| chr12-15349774-DEL-2494 | chr12-15349772-DEL-2490 |
| chr12-15367128-DEL-1527 | chr12-15367129-DEL-1527 |
| chr12-15381009-DEL-1765 | chr12-15381010-DEL-1766 |
| chr12-15394729-DEL-181 | chr12-15394730-DEL-181 |
| chr12-15402002-DEL-82 | chr12-15402003-DEL-82 |
| chr12-15418755-DEL-1496 | chr12-15418756-DEL-1496 |
| chr12-15428501-DEL-6641 | chr12-15428502-DEL-6641 |
| chr12-15473410-DEL-6487 | chr12-15473411-DEL-6487 |
| chr12-15493960-DEL-56 | chr12-15493961-DEL-56 |
| chr12-15509117-DEL-81 | chr12-15509119-DEL-82 |
| chr12-15530229-DEL-93 | chr12-15530230-DEL-93 |
| chr12-15553224-DEL-748 | chr12-15553225-DEL-748 |
| chr12-15554533-DEL-441 | chr12-15554535-DEL-442 |
| chr12-15579040-DEL-200 | chr12-15579041-DEL-200 |
| chr12-15580120-DEL-84 | chr12-15580121-DEL-84 |
| chr12-15594812-DEL-323 | chr12-15594813-DEL-323 |
| chr12-15600361-DEL-58 | chr12-15600362-DEL-58 |
| chr12-15625658-DEL-68 | chr12-15625660-DEL-81 |
| chr12-15645197-DEL-66 | chr12-15645198-DEL-66 |
| chr12-15672170-DEL-6478 | chr12-15672172-DEL-6478 |
| chr12-15679036-DEL-80 | chr12-15679045-DEL-79 |
| chr12-15712039-DEL-10481 | chr12-15712040-DEL-10481 |
| chr12-15815216-DEL-533 | chr12-15815217-DEL-533 |
| chr12-15874013-DEL-187 | chr12-15874014-DEL-187 |
| chr12-15885486-DEL-615 | chr12-15885487-DEL-615 |
| chr12-15899545-DEL-58 | chr12-15899546-DEL-58 |
| chr12-15914595-DEL-65 | chr12-15914596-DEL-65 |
| chr12-15915999-DEL-6394 | chr12-15916000-DEL-6394 |
| chr12-15937209-DEL-133 | chr12-15937210-DEL-133 |
| chr12-15941752-DEL-71 | chr12-15941755-DEL-75 |
| chr12-15985933-DEL-96 | chr12-15985934-DEL-96 |
| chr12-16023863-DEL-54 | chr12-16023864-DEL-54 |
| chr12-16062875-DEL-427 | chr12-16062842-DEL-425 |
| chr12-16065493-DEL-62 | chr12-16065494-DEL-62 |
| chr12-16085934-DEL-97 | chr12-16085937-DEL-95 |
| chr12-16088881-DEL-1186 | chr12-16088882-DEL-1186 |
| chr12-16095284-DEL-76 | chr12-16095285-DEL-76 |
| chr12-16106094-DEL-80 | chr12-16106095-DEL-80 |
| chr12-16121981-DEL-1847 | chr12-16121982-DEL-1847 |
| chr12-16163283-DEL-1136 | chr12-16163284-DEL-1136 |
| chr12-16163731-DEL-202 | chr12-16163732-DEL-202 |
| chr12-16164953-DEL-1052 | chr12-16164954-DEL-1052 |
| chr12-16184807-DEL-123 | chr12-16184818-DEL-107 |
| chr12-16217041-DEL-94 | chr12-16217042-DEL-94 |
| chr12-16227842-DEL-1060 | chr12-16227843-DEL-1060 |
| chr12-16238076-DEL-64 | chr12-16238076-DEL-64 |
| chr12-16262421-DEL-56 | chr12-16262422-DEL-52 |
| chr12-16267017-DEL-56 | chr12-16267018-DEL-56 |
| chr12-16274219-DEL-70 | chr12-16274220-DEL-70 |
| chr12-16277265-DEL-123 | chr12-16277266-DEL-123 |
| chr12-16296012-DEL-58 | chr12-16295996-DEL-58 |
| chr12-16297557-DEL-64 | chr12-16297557-DEL-64 |
| chr12-16309666-DEL-63 | chr12-16309667-DEL-63 |
| chr12-16380605-DEL-740 | chr12-16380606-DEL-740 |
| chr12-16388192-DEL-224 | chr12-16388191-DEL-219 |
| chr12-16420716-DEL-73 | chr12-16420725-DEL-66 |
| chr12-16423159-DEL-986 | chr12-16423160-DEL-986 |
| chr12-16424194-DEL-122 | chr12-16424183-DEL-131 |
| chr12-16430827-DEL-52 | chr12-16430811-DEL-84 |
| chr12-16436801-DEL-740 | chr12-16436802-DEL-740 |
| chr12-16458820-DEL-6932 | chr12-16458821-DEL-6932 |
| chr12-16500065-DEL-161 | chr12-16500066-DEL-161 |
| chr12-16503546-DEL-756 | chr12-16503547-DEL-756 |
| chr12-16504591-DEL-91 | chr12-16504592-DEL-91 |
| chr12-16528888-DEL-87 | chr12-16528891-DEL-83 |
| chr12-16536290-DEL-96 | chr12-16536290-DEL-82 |
| chr12-16550124-DEL-59 | chr12-16550126-DEL-60 |
| chr12-16574838-DEL-381 | chr12-16574839-DEL-381 |
| chr12-16596594-DEL-58 | chr12-16596595-DEL-58 |
| chr12-16616054-DEL-58 | chr12-16616055-DEL-58 |
| chr12-16617326-DEL-52 | chr12-16617323-DEL-52 |
| chr12-16624049-DEL-89 | chr12-16624050-DEL-89 |
| chr12-16626647-DEL-146 | chr12-16626648-DEL-146 |
| chr12-16628556-DEL-80 | chr12-16628557-DEL-80 |
| chr12-16672716-DEL-6424 | chr12-16672717-DEL-6424 |
| chr12-16686892-DEL-216 | chr12-16686885-DEL-224 |
| chr12-16687594-DEL-56 | chr12-16687545-DEL-82 |
| chr12-16688045-DEL-353 | chr12-16688043-DEL-355 |

| | |
|---|---|
| chr12-16707405-DEL-56 | chr12-16707406-DEL-56 |
| chr12-16715176-DEL-72 | chr12-16715177-DEL-72 |
| chr12-16728900-DEL-56 | chr12-16728901-DEL-54 |
| chr12-16733437-DEL-108 | chr12-16733436-DEL-110 |
| chr12-16739408-DEL-741 | chr12-16739409-DEL-741 |
| chr12-16745071-DEL-320 | chr12-16745072-DEL-320 |
| chr12-16747739-DEL-57 | chr12-16747739-DEL-57 |
| chr12-16769105-DEL-71 | chr12-16769111-DEL-66 |
| chr12-16797817-DEL-72 | chr12-16797818-DEL-72 |
| chr12-16808080-DEL-51 | chr12-16808075-DEL-52 |
| chr12-16811915-DEL-65 | chr12-16811916-DEL-65 |
| chr12-16812569-DEL-52 | chr12-16812570-DEL-52 |
| chr12-16834594-DEL-280 | chr12-16834600-DEL-281 |
| chr12-16861792-DEL-65 | chr12-16861793-DEL-65 |
| chr12-16880235-DEL-493 | chr12-16880236-DEL-493 |
| chr12-16910246-DEL-210 | chr12-16910247-DEL-210 |
| chr12-16938263-DEL-60 | chr12-16938264-DEL-60 |
| chr12-16972982-DEL-244 | chr12-16972983-DEL-244 |
| chr12-16997536-DEL-76 | chr12-16997537-DEL-76 |
| chr12-17050531-DEL-66 | chr12-17050534-DEL-64 |
| chr12-17058880-DEL-67 | chr12-17058881-DEL-68 |
| chr12-17066095-DEL-224 | chr12-17066112-DEL-196 |
| chr12-17109580-DEL-549 | chr12-17109581-DEL-549 |
| chr12-17112691-DEL-94 | chr12-17112692-DEL-94 |
| chr12-17133768-DEL-436 | chr12-17133769-DEL-428 |
| chr12-17135533-DEL-166 | chr12-17135534-DEL-166 |
| chr12-17136524-DEL-174 | chr12-17136532-DEL-176 |
| chr12-17137145-DEL-348 | chr12-17137146-DEL-348 |
| chr12-17148929-DEL-248 | chr12-17148930-DEL-248 |
| chr12-17152318-DEL-200 | chr12-17152319-DEL-200 |
| chr12-17161163-DEL-102 | chr12-17161164-DEL-102 |
| chr12-17172150-DEL-7206 | chr12-17172151-DEL-7206 |
| chr12-17219245-DEL-72 | chr12-17219246-DEL-72 |
| chr12-17231287-DEL-340 | chr12-17231282-DEL-341 |
| chr12-17232621-DEL-309 | chr12-17232608-DEL-282 |
| chr12-17237259-DEL-389 | chr12-17237248-DEL-400 |
| chr12-17238577-DEL-137 | chr12-17238578-DEL-128 |
| chr12-17256169-DEL-54 | chr12-17256170-DEL-52 |
| chr12-17257616-DEL-251 | chr12-17257617-DEL-251 |
| chr12-17281701-DEL-108 | chr12-17281702-DEL-112 |
| chr12-17293491-DEL-5029 | chr12-17293492-DEL-5029 |
| chr12-17324730-DEL-54 | chr12-17324731-DEL-54 |
| chr12-17336096-DEL-61 | chr12-17336097-DEL-61 |
| chr12-17352877-DEL-52 | chr12-17352878-DEL-52 |
| chr12-17360690-DEL-128 | chr12-17360697-DEL-114 |
| chr12-17390078-DEL-590 | chr12-17390079-DEL-590 |
| chr12-17394305-DEL-101 | chr12-17394306-DEL-99 |
| chr12-17448251-DEL-68 | chr12-17448252-DEL-68 |
| chr12-17456618-DEL-79 | chr12-17456619-DEL-79 |
| chr12-17461140-DEL-54 | chr12-17461141-DEL-54 |
| chr12-17477647-DEL-51 | chr12-17477648-DEL-51 |
| chr12-17511212-DEL-203 | chr12-17511200-DEL-204 |
| chr12-17521959-DEL-54 | chr12-17521958-DEL-52 |
| chr12-17537904-DEL-134 | chr12-17537905-DEL-134 |
| chr12-17541134-DEL-212 | chr12-17541135-DEL-212 |
| chr12-17544006-DEL-54 | chr12-17544007-DEL-54 |
| chr12-17568787-DEL-52 | chr12-17568787-DEL-52 |
| chr12-17572807-DEL-175 | chr12-17572795-DEL-176 |
| chr12-17591322-DEL-74 | chr12-17591337-DEL-67 |
| chr12-17609148-DEL-53 | chr12-17609150-DEL-52 |
| chr12-17639667-DEL-333 | chr12-17639657-DEL-334 |
| chr12-17642687-DEL-217 | chr12-17642688-DEL-217 |
| chr12-17653392-DEL-83 | chr12-17653347-DEL-94 |
| chr12-17657037-DEL-440 | chr12-17657038-DEL-440 |
| chr12-17665854-DEL-77 | chr12-17665816-DEL-115 |
| chr12-17691984-DEL-54 | chr12-17691985-DEL-54 |
| chr12-17720868-DEL-124 | chr12-17720869-DEL-124 |
| chr12-17724120-DEL-166 | chr12-17724123-DEL-164 |
| chr12-17724795-DEL-55 | chr12-17724796-DEL-55 |
| chr12-17725166-DEL-208 | chr12-17725167-DEL-208 |
| chr12-17736083-DEL-87 | chr12-17736084-DEL-87 |
| chr12-17817837-DEL-211 | chr12-17817872-DEL-213 |
| chr12-17821107-DEL-62 | chr12-17821108-DEL-62 |
| chr12-17833295-DEL-71 | chr12-17833296-DEL-71 |
| chr12-17867866-DEL-7112 | chr12-17867867-DEL-7112 |
| chr12-17911960-DEL-102 | chr12-17911961-DEL-102 |
| chr12-17920623-DEL-4937 | chr12-17920624-DEL-4937 |
| chr12-18282736-DEL-873 | chr12-18282737-DEL-873 |
| chr12-18283815-DEL-1414 | chr12-18283822-DEL-1411 |
| chr12-18291940-DEL-1651 | chr12-18291941-DEL-1651 |

| | |
|---|---|
| chr12-18293768-DEL-59 | chr12-18293769-DEL-59 |
| chr12-18314350-DEL-440 | chr12-18314350-DEL-440 |
| chr12-18315480-DEL-164 | chr12-18315478-DEL-165 |
| chr12-18325284-DEL-6942 | chr12-18325284-DEL-6942 |
| chr12-18389501-DEL-6488 | chr12-18389501-DEL-6488 |
| chr12-18395718-DEL-211 | chr12-18395719-DEL-211 |
| chr12-18397003-DEL-56 | chr12-18397002-DEL-56 |
| chr12-18402947-DEL-1894 | chr12-18402948-DEL-1894 |
| chr12-18408238-DEL-198 | chr12-18408237-DEL-198 |
| chr12-18409312-DEL-55 | chr12-18409313-DEL-55 |
| chr12-18414027-DEL-59 | chr12-18414021-DEL-54 |
| chr12-18424895-DEL-61 | chr12-18424896-DEL-61 |
| chr12-18440677-DEL-573 | chr12-18440678-DEL-573 |
| chr12-18459383-DEL-127 | chr12-18459384-DEL-132 |
| chr12-18622836-DEL-198 | chr12-18622837-DEL-198 |
| chr12-18639695-DEL-132 | chr12-18639696-DEL-132 |
| chr12-18680438-DEL-160 | chr12-18680439-DEL-160 |
| chr12-18703944-DEL-62 | chr12-18703925-DEL-60 |
| chr12-18793767-DEL-133 | chr12-18793768-DEL-133 |
| chr12-18986116-DEL-68 | chr12-18986103-DEL-60 |
| chr12-18989620-DEL-56 | chr12-18989613-DEL-60 |
| chr12-18995321-DEL-288 | chr12-18995320-DEL-288 |
| chr12-19230392-DEL-58 | chr12-19230393-DEL-58 |
| chr12-19239218-DEL-202 | chr12-19239201-DEL-202 |
| chr12-19249565-DEL-125 | chr12-19249566-DEL-125 |
| chr12-19270461-DEL-108 | chr12-19270462-DEL-108 |
| chr12-19356525-DEL-209 | chr12-19356526-DEL-209 |
| chr12-19369635-DEL-62 | chr12-19369635-DEL-62 |
| chr12-19483692-DEL-216 | chr12-19483693-DEL-216 |
| chr12-19487975-DEL-52 | chr12-19487970-DEL-53 |
| chr12-19609462-DEL-3351 | chr12-19609461-DEL-3351 |
| chr12-19813468-DEL-350 | chr12-19813468-DEL-350 |
| chr12-19905425-DEL-498 | chr12-19905425-DEL-498 |
| chr12-19921931-DEL-238 | chr12-19921932-DEL-238 |
| chr12-19935730-DEL-136 | chr12-19935739-DEL-137 |
| chr12-19942713-DEL-60 | chr12-19942713-DEL-54 |
| chr12-19956145-DEL-221 | chr12-19956145-DEL-221 |
| chr12-19981727-DEL-6456 | chr12-19981728-DEL-6456 |
| chr12-19995357-DEL-2950 | chr12-19995360-DEL-2951 |
| chr12-20025806-DEL-252 | chr12-20025805-DEL-252 |
| chr12-20029778-DEL-401 | chr12-20029779-DEL-401 |
| chr12-20149374-DEL-213 | chr12-20149375-DEL-213 |
| chr12-20314066-DEL-168 | chr12-20314106-DEL-118 |
| chr12-20323535-DEL-209 | chr12-20323535-DEL-209 |
| chr12-20329154-DEL-190 | chr12-20329176-DEL-184 |
| chr12-20372168-DEL-168 | chr12-20372167-DEL-168 |
| chr12-20471191-DEL-57 | chr12-20471192-DEL-57 |
| chr12-20495149-DEL-1080 | chr12-20495150-DEL-1080 |
| chr12-20512465-DEL-78 | chr12-20512474-DEL-70 |
| chr12-20521463-DEL-66 | chr12-20521462-DEL-66 |
| chr12-20679231-DEL-1460 | chr12-20679232-DEL-1460 |
| chr12-20688482-DEL-553 | chr12-20688482-DEL-553 |
| chr12-20748888-DEL-5765 | chr12-20748889-DEL-5765 |
| chr12-20769806-DEL-60 | chr12-20769806-DEL-52 |
| chr12-20782941-DEL-3190 | chr12-20782938-DEL-3193 |
| chr12-20795963-DEL-999 | chr12-20795964-DEL-999 |
| chr12-20807960-DEL-498 | chr12-20807961-DEL-498 |
| chr12-20877729-DEL-188 | chr12-20877730-DEL-188 |
| chr12-20891091-DEL-181 | chr12-20891092-DEL-181 |
| chr12-20916659-DEL-3786 | chr12-20916648-DEL-3798 |
| chr12-20920075-DEL-66 | chr12-20920076-DEL-66 |
| chr12-21030856-DEL-192 | chr12-21030857-DEL-192 |
| chr12-21055323-DEL-2878 | chr12-21055324-DEL-2878 |
| chr12-21111647-DEL-129 | chr12-21111643-DEL-128 |
| chr12-21113804-DEL-91 | chr12-21113805-DEL-92 |
| chr12-21116892-DEL-199 | chr12-21116893-DEL-199 |
| chr12-21134934-DEL-57 | chr12-21134936-DEL-54 |
| chr12-21208484-DEL-60 | chr12-21208485-DEL-60 |
| chr12-21219860-DEL-55 | chr12-21219861-DEL-55 |
| chr12-21221416-DEL-54 | chr12-21221417-DEL-52 |
| chr12-21242548-DEL-70 | chr12-21242549-DEL-71 |
| chr12-21244931-DEL-1154 | chr12-21244932-DEL-1154 |
| chr12-21250979-DEL-168 | chr12-21250980-DEL-153 |
| chr12-21263851-DEL-166 | chr12-21263852-DEL-166 |
| chr12-21265176-DEL-105 | chr12-21265177-DEL-105 |
| chr12-21269073-DEL-978 | chr12-21269074-DEL-978 |
| chr12-21274983-DEL-178 | chr12-21274984-DEL-178 |
| chr12-21285018-DEL-67 | chr12-21284987-DEL-50 |
| chr12-21286265-DEL-1955 | chr12-21286265-DEL-1953 |
| chr12-21290272-DEL-813 | chr12-21290268-DEL-812 |

| | |
|---|---|
| chr12-21328406-DEL-66 | chr12-21328407-DEL-65 |
| chr12-21334858-DEL-176 | chr12-21334859-DEL-173 |
| chr12-21366148-DEL-119 | chr12-21366149-DEL-119 |
| chr12-21404835-DEL-174 | chr12-21404846-DEL-164 |
| chr12-21406151-DEL-204 | chr12-21406152-DEL-204 |
| chr12-21408175-DEL-592 | chr12-21408176-DEL-592 |
| chr12-21413641-DEL-64 | chr12-21413642-DEL-64 |
| chr12-21415224-DEL-863 | chr12-21415225-DEL-863 |
| chr12-21421263-DEL-179 | chr12-21421264-DEL-179 |
| chr12-21426299-DEL-66 | chr12-21426300-DEL-65 |
| chr12-21431489-DEL-98 | chr12-21431480-DEL-103 |
| chr12-21435268-DEL-2142 | chr12-21435269-DEL-2142 |
| chr12-21436860-DEL-493 | chr12-21436861-DEL-493 |
| chr12-21458155-DEL-211 | chr12-21458156-DEL-211 |
| chr12-21469482-DEL-210 | chr12-21469483-DEL-210 |
| chr12-21478734-DEL-149 | chr12-21478735-DEL-149 |
| chr12-21479572-DEL-282 | chr12-21479564-DEL-285 |
| chr12-21495295-DEL-199 | chr12-21495295-DEL-200 |
| chr12-21498329-DEL-200 | chr12-21498330-DEL-200 |
| chr12-21503370-DEL-56 | chr12-21503375-DEL-56 |
| chr12-21509280-DEL-197 | chr12-21509281-DEL-197 |
| chr12-21520601-DEL-385 | chr12-21520601-DEL-386 |
| chr12-21559260-DEL-83 | chr12-21559261-DEL-83 |
| chr12-21593892-DEL-188 | chr12-21593893-DEL-188 |
| chr12-21626798-DEL-55 | chr12-21626809-DEL-50 |
| chr12-21631812-DEL-57 | chr12-21631778-DEL-90 |
| chr12-21654172-DEL-2814 | chr12-21654173-DEL-2814 |
| chr12-21662993-DEL-588 | chr12-21662996-DEL-586 |
| chr12-21666656-DEL-61 | chr12-21666657-DEL-61 |
| chr12-21793598-DEL-62 | chr12-21793599-DEL-62 |
| chr12-21823252-DEL-58 | chr12-21823253-DEL-57 |
| chr12-21906305-DEL-59 | chr12-21906277-DEL-80 |
| chr12-22052731-DEL-96 | chr12-22052737-DEL-91 |
| chr12-22339031-DEL-137 | chr12-22339030-DEL-137 |
| chr12-22647879-DEL-400 | chr12-22647880-DEL-400 |
| chr12-22676146-DEL-185 | chr12-22676155-DEL-179 |
| chr12-22691407-DEL-201 | chr12-22691408-DEL-201 |
| chr12-22790868-DEL-58 | chr12-22790846-DEL-58 |
| chr12-22792723-DEL-210 | chr12-22792723-DEL-210 |
| chr12-22854060-DEL-69 | chr12-22854060-DEL-74 |
| chr12-23039957-DEL-168 | chr12-23039958-DEL-168 |
| chr12-23053373-DEL-202 | chr12-23053374-DEL-202 |
| chr12-23061527-DEL-203 | chr12-23061526-DEL-203 |
| chr12-23117901-DEL-240 | chr12-23117902-DEL-240 |
| chr12-23121955-DEL-94 | chr12-23121974-DEL-95 |
| chr12-23127545-DEL-66 | chr12-23127544-DEL-55 |
| chr12-23155106-DEL-142 | chr12-23155113-DEL-136 |
| chr12-23259186-DEL-51 | chr12-23259172-DEL-56 |
| chr12-23485490-DEL-52 | chr12-23485487-DEL-56 |
| chr12-23707035-DEL-75 | chr12-23707034-DEL-75 |
| chr12-23782239-DEL-94 | chr12-23782239-DEL-94 |
| chr12-24180705-DEL-1394 | chr12-24180705-DEL-1394 |
| chr12-24185270-DEL-63 | chr12-24185270-DEL-63 |
| chr12-24203300-DEL-2112 | chr12-24203299-DEL-2112 |
| chr12-24210008-DEL-201 | chr12-24210029-DEL-201 |
| chr12-24297533-DEL-116 | chr12-24297561-DEL-115 |
| chr12-24327463-DEL-84 | chr12-24327464-DEL-84 |
| chr12-24331926-DEL-56 | chr12-24331931-DEL-52 |
| chr12-24336049-DEL-53 | chr12-24336018-DEL-54 |
| chr12-24344042-DEL-70 | chr12-24344043-DEL-70 |
| chr12-24344981-DEL-166 | chr12-24344991-DEL-195 |
| chr12-24419633-DEL-76 | chr12-24419622-DEL-88 |
| chr12-24421073-DEL-66 | chr12-24421073-DEL-67 |
| chr12-24423238-DEL-54 | chr12-24423237-DEL-56 |
| chr12-24439912-DEL-52 | chr12-24439913-DEL-52 |
| chr12-24441670-DEL-64 | chr12-24441671-DEL-61 |
| chr12-24445366-DEL-7238 | chr12-24445365-DEL-7239 |
| chr12-24459096-DEL-58 | chr12-24459097-DEL-58 |
| chr12-24467419-DEL-29701 | chr12-24467419-DEL-29701 |
| chr12-24575269-DEL-80 | chr12-24575270-DEL-80 |
| chr12-24610991-DEL-114 | chr12-24610992-DEL-114 |
| chr12-24681975-DEL-182 | chr12-24681976-DEL-182 |
| chr12-24687952-DEL-1112 | chr12-24687953-DEL-1112 |
| chr12-24696660-DEL-110 | chr12-24696660-DEL-111 |
| chr12-24699202-DEL-55 | chr12-24699203-DEL-55 |
| chr12-24719974-DEL-135 | chr12-24719975-DEL-135 |
| chr12-24727266-DEL-55 | chr12-24727267-DEL-55 |
| chr12-24733719-DEL-69 | chr12-24733720-DEL-69 |
| chr12-24739546-DEL-203 | chr12-24739547-DEL-203 |
| chr12-24741546-DEL-399 | chr12-24741547-DEL-399 |

| | |
|---|---|
| chr12-24742074-DEL-4012 | chr12-24742075-DEL-4012 |
| chr12-24752205-DEL-125 | chr12-24752206-DEL-125 |
| chr12-24765578-DEL-102 | chr12-24765579-DEL-102 |
| chr12-24768116-DEL-53 | chr12-24768113-DEL-53 |
| chr12-24771650-DEL-58 | chr12-24771651-DEL-52 |
| chr12-24774782-DEL-2805 | chr12-24774783-DEL-2805 |
| chr12-24793496-DEL-701 | chr12-24793497-DEL-701 |
| chr12-24799698-DEL-205 | chr12-24799699-DEL-205 |
| chr12-24802971-DEL-373 | chr12-24802974-DEL-372 |
| chr12-24807800-DEL-199 | chr12-24807801-DEL-199 |
| chr12-24813495-DEL-185 | chr12-24813496-DEL-185 |
| chr12-24846649-DEL-77 | chr12-24846650-DEL-77 |
| chr12-24864457-DEL-223 | chr12-24864415-DEL-224 |
| chr12-24884528-DEL-125 | chr12-24884502-DEL-128 |
| chr12-24928750-DEL-96 | chr12-24928751-DEL-96 |
| chr12-24938247-DEL-810 | chr12-24938249-DEL-809 |
| chr12-24950182-DEL-372 | chr12-24950183-DEL-372 |
| chr12-24951910-DEL-198 | chr12-24951911-DEL-198 |
| chr12-24955665-DEL-168 | chr12-24955659-DEL-169 |
| chr12-24987292-DEL-78 | chr12-24987293-DEL-78 |
| chr12-24991481-DEL-813 | chr12-24991486-DEL-811 |
| chr12-25041602-DEL-55 | chr12-25041569-DEL-53 |
| chr12-25057556-DEL-85 | chr12-25057557-DEL-85 |
| chr12-25066573-DEL-84 | chr12-25066574-DEL-84 |
| chr12-25070557-DEL-640 | chr12-25070558-DEL-640 |
| chr12-25071347-DEL-205 | chr12-25071348-DEL-205 |
| chr12-25075818-DEL-124 | chr12-25075814-DEL-121 |
| chr12-25099755-DEL-241 | chr12-25099756-DEL-241 |
| chr12-25116774-DEL-60 | chr12-25116781-DEL-53 |
| chr12-25125684-DEL-75 | chr12-25125685-DEL-95 |
| chr12-25132393-DEL-217 | chr12-25132394-DEL-218 |
| chr12-25152177-DEL-202 | chr12-25152178-DEL-202 |
| chr12-25153347-DEL-62 | chr12-25153333-DEL-68 |
| chr12-25165963-DEL-56 | chr12-25165934-DEL-86 |
| chr12-25178318-DEL-636 | chr12-25178319-DEL-636 |
| chr12-25212255-DEL-731 | chr12-25212256-DEL-731 |
| chr12-25234987-DEL-54 | chr12-25234986-DEL-56 |
| chr12-25247710-DEL-58 | chr12-25247676-DEL-92 |
| chr12-25310083-DEL-2451 | chr12-25310084-DEL-2451 |
| chr12-25335468-DEL-66 | chr12-25335469-DEL-66 |
| chr12-25340415-DEL-81 | chr12-25340416-DEL-81 |
| chr12-25352490-DEL-55 | chr12-25352492-DEL-53 |
| chr12-25361242-DEL-164 | chr12-25361243-DEL-164 |
| chr12-25375711-DEL-355 | chr12-25375712-DEL-355 |
| chr12-25387319-DEL-408 | chr12-25387320-DEL-408 |
| chr12-25395343-DEL-119 | chr12-25395344-DEL-91 |
| chr12-25407316-DEL-86 | chr12-25407327-DEL-68 |
| chr12-25407673-DEL-772 | chr12-25407674-DEL-772 |
| chr12-25422052-DEL-56 | chr12-25422053-DEL-56 |
| chr12-25433713-DEL-100 | chr12-25433710-DEL-96 |
| chr12-25438522-DEL-93 | chr12-25438523-DEL-93 |
| chr12-25467142-DEL-55 | chr12-25467128-DEL-70 |
| chr12-25499482-DEL-250 | chr12-25499483-DEL-248 |
| chr12-25508901-DEL-2197 | chr12-25508900-DEL-2197 |
| chr12-25528940-DEL-114 | chr12-25528924-DEL-115 |
| chr12-25541232-DEL-60 | chr12-25541236-DEL-57 |
| chr12-25588862-DEL-55 | chr12-25588866-DEL-52 |
| chr12-25596155-DEL-1941 | chr12-25596151-DEL-1946 |
| chr12-25638704-DEL-176 | chr12-25638705-DEL-176 |
| chr12-25661725-DEL-310 | chr12-25661726-DEL-310 |
| chr12-25719355-DEL-400 | chr12-25719356-DEL-400 |
| chr12-25739166-DEL-497 | chr12-25739167-DEL-497 |
| chr12-25744655-DEL-165 | chr12-25744656-DEL-171 |
| chr12-25753852-DEL-54 | chr12-25753849-DEL-54 |
| chr12-25792170-DEL-51 | chr12-25792169-DEL-51 |
| chr12-25793964-DEL-148 | chr12-25793965-DEL-148 |
| chr12-25797089-DEL-6370 | chr12-25797090-DEL-6370 |
| chr12-25818121-DEL-62 | chr12-25818093-DEL-103 |
| chr12-25862838-DEL-314 | chr12-25862839-DEL-314 |
| chr12-25907257-DEL-416 | chr12-25907258-DEL-416 |
| chr12-25953286-DEL-53 | chr12-25953286-DEL-56 |
| chr12-25980722-DEL-197 | chr12-25980723-DEL-197 |
| chr12-25985637-DEL-70 | chr12-25985680-DEL-66 |
| chr12-26075465-DEL-91 | chr12-26075466-DEL-91 |
| chr12-26084893-DEL-6568 | chr12-26084906-DEL-6551 |
| chr12-26098244-DEL-7517 | chr12-26098245-DEL-7517 |
| chr12-26111655-DEL-189 | chr12-26111656-DEL-189 |
| chr12-26131396-DEL-66 | chr12-26131409-DEL-54 |
| chr12-26156446-DEL-2746 | chr12-26156447-DEL-2746 |
| chr12-26209445-DEL-223 | chr12-26209446-DEL-223 |

| | |
|---|---|
| chr12-26218426-DEL-273 | chr12-26218427-DEL-273 |
| chr12-26245347-DEL-99 | chr12-26245369-DEL-78 |
| chr12-26283736-DEL-368 | chr12-26283737-DEL-368 |
| chr12-26301720-DEL-88 | chr12-26301721-DEL-84 |
| chr12-26306303-DEL-399 | chr12-26306304-DEL-399 |
| chr12-26346028-DEL-56 | chr12-26346029-DEL-54 |
| chr12-26353936-DEL-70 | chr12-26353937-DEL-62 |
| chr12-26384264-DEL-77 | chr12-26384287-DEL-87 |
| chr12-26397265-DEL-228 | chr12-26397266-DEL-228 |
| chr12-26405587-DEL-64 | chr12-26405588-DEL-64 |
| chr12-26411677-DEL-712 | chr12-26411678-DEL-712 |
| chr12-26450627-DEL-58 | chr12-26450628-DEL-58 |
| chr12-26476700-DEL-99 | chr12-26476701-DEL-99 |
| chr12-26518395-DEL-52 | chr12-26518372-DEL-66 |
| chr12-26536107-DEL-79 | chr12-26536108-DEL-79 |
| chr12-26549209-DEL-373 | chr12-26549210-DEL-373 |
| chr12-26569044-DEL-4837 | chr12-26569045-DEL-4837 |
| chr12-26581332-DEL-600 | chr12-26581333-DEL-600 |
| chr12-26585666-DEL-66 | chr12-26585667-DEL-66 |
| chr12-26590366-DEL-194 | chr12-26590367-DEL-194 |
| chr12-26591997-DEL-60 | chr12-26591991-DEL-66 |
| chr12-26598238-DEL-349 | chr12-26598239-DEL-349 |
| chr12-26642659-DEL-64 | chr12-26642651-DEL-68 |
| chr12-26660807-DEL-58 | chr12-26660808-DEL-58 |
| chr12-26662188-DEL-54 | chr12-26662205-DEL-62 |
| chr12-26664635-DEL-94 | chr12-26664629-DEL-74 |
| chr12-26682799-DEL-194 | chr12-26682800-DEL-194 |
| chr12-26686322-DEL-52 | chr12-26686322-DEL-52 |
| chr12-26688710-DEL-135 | chr12-26688709-DEL-135 |
| chr12-26708654-DEL-829 | chr12-26708658-DEL-827 |
| chr12-26752277-DEL-846 | chr12-26752278-DEL-846 |
| chr12-26762228-DEL-901 | chr12-26762228-DEL-902 |
| chr12-26788782-DEL-55 | chr12-26788782-DEL-56 |
| chr12-26794556-DEL-2784 | chr12-26794557-DEL-2785 |
| chr12-26808746-DEL-3303 | chr12-26808747-DEL-3303 |
| chr12-26810439-DEL-63 | chr12-26810435-DEL-64 |
| chr12-26812195-DEL-53 | chr12-26812191-DEL-54 |
| chr12-26827956-DEL-52 | chr12-26827957-DEL-52 |
| chr12-26836381-DEL-80 | chr12-26836382-DEL-80 |
| chr12-26842327-DEL-52 | chr12-26842327-DEL-52 |
| chr12-26852355-DEL-186 | chr12-26852352-DEL-190 |
| chr12-26883296-DEL-586 | chr12-26883297-DEL-586 |
| chr12-26902244-DEL-63 | chr12-26902247-DEL-97 |
| chr12-26919110-DEL-128 | chr12-26919094-DEL-137 |
| chr12-26960989-DEL-182 | chr12-26961010-DEL-162 |
| chr12-26964465-DEL-3972 | chr12-26964466-DEL-3972 |
| chr12-26986339-DEL-54 | chr12-26986340-DEL-54 |
| chr12-26995027-DEL-1278 | chr12-26995028-DEL-1278 |
| chr12-27004418-DEL-672 | chr12-27004419-DEL-672 |
| chr12-27016606-DEL-66 | chr12-27016607-DEL-66 |
| chr12-27048545-DEL-54 | chr12-27048546-DEL-54 |
| chr12-27060100-DEL-686 | chr12-27060101-DEL-686 |
| chr12-27149676-DEL-208 | chr12-27149681-DEL-222 |
| chr12-27160703-DEL-80 | chr12-27160702-DEL-50 |
| chr12-27171869-DEL-54 | chr12-27171870-DEL-54 |
| chr12-27185497-DEL-52 | chr12-27185497-DEL-52 |
| chr12-27215430-DEL-81 | chr12-27215445-DEL-84 |
| chr12-27216483-DEL-738 | chr12-27216482-DEL-738 |
| chr12-27275955-DEL-62 | chr12-27275956-DEL-62 |
| chr12-27281949-DEL-3068 | chr12-27281950-DEL-3068 |
| chr12-27307994-DEL-79 | chr12-27308007-DEL-67 |
| chr12-27312400-DEL-128 | chr12-27312375-DEL-128 |
| chr12-27323464-DEL-58 | chr12-27323465-DEL-52 |
| chr12-27327606-DEL-92 | chr12-27327607-DEL-92 |
| chr12-27327732-DEL-986 | chr12-27327732-DEL-986 |
| chr12-27331396-DEL-52 | chr12-27331397-DEL-52 |
| chr12-27337015-DEL-204 | chr12-27337014-DEL-204 |
| chr12-27380958-DEL-134 | chr12-27380959-DEL-134 |
| chr12-27430890-DEL-97 | chr12-27430869-DEL-135 |
| chr12-27447059-DEL-62 | chr12-27447060-DEL-62 |
| chr12-27488380-DEL-128 | chr12-27488381-DEL-120 |
| chr12-27500158-DEL-116 | chr12-27500159-DEL-117 |
| chr12-27508356-DEL-64 | chr12-27508361-DEL-60 |
| chr12-27528474-DEL-56 | chr12-27528475-DEL-54 |
| chr12-27532056-DEL-53 | chr12-27532072-DEL-53 |
| chr12-27564172-DEL-52 | chr12-27564173-DEL-52 |
| chr12-27568892-DEL-55 | chr12-27568888-DEL-62 |
| chr12-27570353-DEL-57 | chr12-27570353-DEL-58 |
| chr12-27601704-DEL-124 | chr12-27601709-DEL-128 |
| chr12-27606730-DEL-1063 | chr12-27606731-DEL-1063 |

| | |
|---|---|
| chr12-27609563-DEL-54 | chr12-27609564-DEL-84 |
| chr12-27628556-DEL-354 | chr12-27628557-DEL-354 |
| chr12-27641792-DEL-514 | chr12-27641793-DEL-514 |
| chr12-27668871-DEL-449 | chr12-27668872-DEL-449 |
| chr12-27676869-DEL-114 | chr12-27676871-DEL-115 |
| chr12-27684440-DEL-55 | chr12-27684460-DEL-70 |
| chr12-27685182-DEL-124 | chr12-27685183-DEL-124 |
| chr12-27730267-DEL-309 | chr12-27730268-DEL-309 |
| chr12-27741756-DEL-6270 | chr12-27741755-DEL-6270 |
| chr12-27755443-DEL-54 | chr12-27755444-DEL-54 |
| chr12-27761627-DEL-158 | chr12-27761630-DEL-147 |
| chr12-27770908-DEL-61 | chr12-27770907-DEL-62 |
| chr12-27813582-DEL-75 | chr12-27813580-DEL-76 |
| chr12-27828683-DEL-1889 | chr12-27828684-DEL-1889 |
| chr12-27840846-DEL-78 | chr12-27840849-DEL-75 |
| chr12-27851651-DEL-104 | chr12-27851652-DEL-104 |
| chr12-27854596-DEL-54 | chr12-27854597-DEL-54 |
| chr12-27875331-DEL-62 | chr12-27875332-DEL-62 |
| chr12-27885159-DEL-1884 | chr12-27885160-DEL-1884 |
| chr12-27893567-DEL-94 | chr12-27893569-DEL-93 |
| chr12-27917962-DEL-1205 | chr12-27917963-DEL-1205 |
| chr12-27924733-DEL-150 | chr12-27924732-DEL-146 |
| chr12-27941557-DEL-453 | chr12-27941558-DEL-453 |
| chr12-27946123-DEL-7661 | chr12-27946124-DEL-7661 |
| chr12-27982207-DEL-82 | chr12-27982208-DEL-82 |
| chr12-28018835-DEL-71 | chr12-28018804-DEL-72 |
| chr12-28038567-DEL-56 | chr12-28038549-DEL-74 |
| chr12-28085076-DEL-95 | chr12-28085057-DEL-114 |
| chr12-28089952-DEL-474 | chr12-28089959-DEL-468 |
| chr12-28115545-DEL-299 | chr12-28115546-DEL-299 |
| chr12-28117421-DEL-102 | chr12-28117404-DEL-103 |
| chr12-28123626-DEL-109 | chr12-28123625-DEL-108 |
| chr12-28127702-DEL-52 | chr12-28127703-DEL-52 |
| chr12-28211272-DEL-6841 | chr12-28211273-DEL-6841 |
| chr12-28224407-DEL-55 | chr12-28224407-DEL-56 |
| chr12-28233558-DEL-56 | chr12-28233559-DEL-56 |
| chr12-28241304-DEL-6460 | chr12-28241304-DEL-6459 |
| chr12-28293527-DEL-4042 | chr12-28293528-DEL-4042 |
| chr12-28317055-DEL-93 | chr12-28317056-DEL-93 |
| chr12-28341411-DEL-105 | chr12-28341424-DEL-87 |
| chr12-28344512-DEL-140 | chr12-28344513-DEL-140 |
| chr12-28403861-DEL-185 | chr12-28403862-DEL-185 |
| chr12-28411074-DEL-102 | chr12-28411075-DEL-86 |
| chr12-28418803-DEL-279 | chr12-28418804-DEL-279 |
| chr12-28440711-DEL-54 | chr12-28440706-DEL-60 |
| chr12-28481590-DEL-66 | chr12-28481599-DEL-62 |
| chr12-28487952-DEL-498 | chr12-28487953-DEL-498 |
| chr12-28498882-DEL-56 | chr12-28498882-DEL-56 |
| chr12-28515099-DEL-393 | chr12-28515100-DEL-393 |
| chr12-28516174-DEL-75 | chr12-28516200-DEL-69 |
| chr12-28516702-DEL-128 | chr12-28516703-DEL-128 |
| chr12-28519154-DEL-94 | chr12-28519154-DEL-50 |
| chr12-28520376-DEL-60 | chr12-28520377-DEL-60 |
| chr12-28522994-DEL-100 | chr12-28522981-DEL-112 |
| chr12-28530234-DEL-103 | chr12-28530235-DEL-115 |
| chr12-28539596-DEL-4488 | chr12-28539597-DEL-4488 |
| chr12-28558842-DEL-58 | chr12-28558843-DEL-50 |
| chr12-28567086-DEL-156 | chr12-28567087-DEL-144 |
| chr12-28583974-DEL-464 | chr12-28583990-DEL-455 |
| chr12-28586677-DEL-333 | chr12-28586700-DEL-311 |
| chr12-28587879-DEL-56 | chr12-28587882-DEL-54 |
| chr12-28603003-DEL-73 | chr12-28603003-DEL-75 |
| chr12-28616671-DEL-148 | chr12-28616672-DEL-148 |
| chr12-28621754-DEL-119 | chr12-28621754-DEL-111 |
| chr12-28622907-DEL-64 | chr12-28622921-DEL-63 |
| chr12-28624899-DEL-156 | chr12-28624894-DEL-157 |
| chr12-28637600-DEL-113 | chr12-28637593-DEL-110 |
| chr12-28653377-DEL-53 | chr12-28653379-DEL-50 |
| chr12-28659325-DEL-94 | chr12-28659326-DEL-94 |
| chr12-28666324-DEL-223 | chr12-28666325-DEL-223 |
| chr12-28688194-DEL-1343 | chr12-28688195-DEL-1343 |
| chr12-28695042-DEL-52 | chr12-28695043-DEL-52 |
| chr12-28702396-DEL-67 | chr12-28702397-DEL-67 |
| chr12-28708434-DEL-79 | chr12-28708435-DEL-79 |
| chr12-28719432-DEL-202 | chr12-28719433-DEL-203 |
| chr12-28725814-DEL-289 | chr12-28725815-DEL-289 |
| chr12-28762155-DEL-67 | chr12-28762156-DEL-67 |
| chr12-28807513-DEL-1890 | chr12-28807514-DEL-1890 |
| chr12-28818819-DEL-59 | chr12-28818821-DEL-54 |
| chr12-28823374-DEL-72 | chr12-28823341-DEL-90 |

| | |
|---|---|
| chr12-28829065-DEL-72 | chr12-28829066-DEL-72 |
| chr12-28870559-DEL-274 | chr12-28870569-DEL-252 |
| chr12-28874027-DEL-193 | chr12-28874028-DEL-193 |
| chr12-28880868-DEL-207 | chr12-28880869-DEL-207 |
| chr12-28890068-DEL-460 | chr12-28890069-DEL-460 |
| chr12-28893382-DEL-441 | chr12-28893383-DEL-441 |
| chr12-28903546-DEL-617 | chr12-28903547-DEL-617 |
| chr12-28917530-DEL-285 | chr12-28917531-DEL-285 |
| chr12-28924326-DEL-108 | chr12-28924327-DEL-108 |
| chr12-28961294-DEL-63 | chr12-28961321-DEL-57 |
| chr12-28985029-DEL-76 | chr12-28985030-DEL-72 |
| chr12-29018456-DEL-52 | chr12-29018455-DEL-62 |
| chr12-29076524-DEL-542 | chr12-29076525-DEL-542 |
| chr12-29086329-DEL-89 | chr12-29086320-DEL-85 |
| chr12-29090511-DEL-852 | chr12-29090509-DEL-853 |
| chr12-29101551-DEL-1277 | chr12-29101552-DEL-1277 |
| chr12-29106868-DEL-82 | chr12-29106870-DEL-83 |
| chr12-29107823-DEL-1184 | chr12-29107824-DEL-1184 |
| chr12-29128389-DEL-210 | chr12-29128389-DEL-214 |
| chr12-29155438-DEL-103 | chr12-29155439-DEL-102 |
| chr12-29163923-DEL-3524 | chr12-29163924-DEL-3524 |
| chr12-29190418-DEL-60 | chr12-29190418-DEL-60 |
| chr12-29194487-DEL-54 | chr12-29194475-DEL-62 |
| chr12-29202412-DEL-89 | chr12-29202412-DEL-87 |
| chr12-29203294-DEL-2335 | chr12-29203294-DEL-2335 |
| chr12-29223674-DEL-868 | chr12-29223675-DEL-868 |
| chr12-29279876-DEL-396 | chr12-29279878-DEL-395 |
| chr12-29295430-DEL-277 | chr12-29295431-DEL-277 |
| chr12-29317831-DEL-58 | chr12-29317832-DEL-58 |
| chr12-29345576-DEL-568 | chr12-29345577-DEL-568 |
| chr12-29351891-DEL-202 | chr12-29351892-DEL-202 |
| chr12-29376836-DEL-91 | chr12-29376837-DEL-91 |
| chr12-29400468-DEL-55 | chr12-29400469-DEL-55 |
| chr12-29412451-DEL-97 | chr12-29412452-DEL-97 |
| chr12-29429362-DEL-10404 | chr12-29429363-DEL-10404 |
| chr12-29455612-DEL-54 | chr12-29455613-DEL-54 |
| chr12-29484272-DEL-385 | chr12-29484273-DEL-385 |
| chr12-29485983-DEL-70 | chr12-29485984-DEL-70 |
| chr12-29515467-DEL-193 | chr12-29515497-DEL-193 |
| chr12-29519193-DEL-108 | chr12-29519194-DEL-108 |
| chr12-29528907-DEL-59 | chr12-29528907-DEL-60 |
| chr12-29539023-DEL-8361 | chr12-29539024-DEL-8361 |
| chr12-29608800-DEL-54 | chr12-29608801-DEL-54 |
| chr12-29647264-DEL-67 | chr12-29647265-DEL-67 |
| chr12-29660090-DEL-115 | chr12-29660090-DEL-115 |
| chr12-29712714-DEL-149 | chr12-29712715-DEL-149 |
| chr12-29714031-DEL-58 | chr12-29714032-DEL-58 |
| chr12-29723554-DEL-88 | chr12-29723555-DEL-88 |
| chr12-29742624-DEL-59 | chr12-29742625-DEL-55 |
| chr12-29766400-DEL-52 | chr12-29766401-DEL-52 |
| chr12-29856117-DEL-51 | chr12-29856118-DEL-51 |
| chr12-29897300-DEL-115 | chr12-29897306-DEL-110 |
| chr12-29902286-DEL-364 | chr12-29902287-DEL-364 |
| chr12-29947095-DEL-175 | chr12-29947096-DEL-175 |
| chr12-29958106-DEL-428 | chr12-29958107-DEL-428 |
| chr12-29965732-DEL-55 | chr12-29965733-DEL-55 |
| chr12-29985078-DEL-133 | chr12-29985079-DEL-133 |
| chr12-29993035-DEL-120 | chr12-29993036-DEL-120 |
| chr12-29999948-DEL-202 | chr12-29999949-DEL-202 |
| chr12-30003261-DEL-78 | chr12-30003262-DEL-73 |
| chr12-30005280-DEL-1193 | chr12-30005281-DEL-1193 |
| chr12-30016618-DEL-195 | chr12-30016619-DEL-195 |
| chr12-30020779-DEL-92 | chr12-30020783-DEL-82 |
| chr12-30066276-DEL-53 | chr12-30066277-DEL-53 |
| chr12-30092689-DEL-177 | chr12-30092690-DEL-176 |
| chr12-30106131-DEL-1128 | chr12-30106132-DEL-1128 |
| chr12-30125222-DEL-365 | chr12-30125223-DEL-365 |
| chr12-30134988-DEL-6473 | chr12-30134989-DEL-6473 |
| chr12-30149282-DEL-355 | chr12-30149283-DEL-355 |
| chr12-30155640-DEL-144 | chr12-30155641-DEL-144 |
| chr12-30193940-DEL-263 | chr12-30193941-DEL-263 |
| chr12-30217490-DEL-124 | chr12-30217491-DEL-124 |
| chr12-30225461-DEL-971 | chr12-30225462-DEL-971 |
| chr12-30227134-DEL-66 | chr12-30227135-DEL-66 |
| chr12-30230165-DEL-56 | chr12-30230166-DEL-56 |
| chr12-30237309-DEL-6280 | chr12-30237310-DEL-6280 |
| chr12-30266446-DEL-94 | chr12-30266447-DEL-94 |
| chr12-30267919-DEL-120 | chr12-30267920-DEL-120 |
| chr12-30270834-DEL-128 | chr12-30270844-DEL-119 |
| chr12-30311963-DEL-51 | chr12-30311984-DEL-63 |

| | |
|---|---|
| chr12-30318372-DEL-83 | chr12-30318373-DEL-82 |
| chr12-30330532-DEL-101 | chr12-30330544-DEL-100 |
| chr12-30350651-DEL-643 | chr12-30350652-DEL-643 |
| chr12-30365920-DEL-100 | chr12-30365921-DEL-100 |
| chr12-30377489-DEL-738 | chr12-30377490-DEL-738 |
| chr12-30402894-DEL-50 | chr12-30402890-DEL-54 |
| chr12-30412217-DEL-99 | chr12-30412224-DEL-94 |
| chr12-30435721-DEL-66 | chr12-30435713-DEL-66 |
| chr12-30439592-DEL-104 | chr12-30439593-DEL-104 |
| chr12-30465765-DEL-830 | chr12-30465762-DEL-831 |
| chr12-30564074-DEL-2076 | chr12-30564075-DEL-2076 |
| chr12-30583917-DEL-108 | chr12-30583918-DEL-102 |
| chr12-30585000-DEL-160 | chr12-30585001-DEL-160 |
| chr12-30585625-DEL-1895 | chr12-30585626-DEL-1895 |
| chr12-30640422-DEL-375 | chr12-30640423-DEL-375 |
| chr12-30672026-DEL-320 | chr12-30672027-DEL-320 |
| chr12-30677469-DEL-53 | chr12-30677471-DEL-71 |
| chr12-30688982-DEL-1378 | chr12-30688983-DEL-1378 |
| chr12-30705662-DEL-693 | chr12-30705663-DEL-693 |
| chr12-30706227-DEL-805 | chr12-30706225-DEL-806 |
| chr12-30719846-DEL-100 | chr12-30719847-DEL-100 |
| chr12-30723298-DEL-56 | chr12-30723305-DEL-56 |
| chr12-30730615-DEL-6263 | chr12-30730616-DEL-6263 |
| chr12-30750294-DEL-190 | chr12-30750295-DEL-190 |
| chr12-30750903-DEL-313 | chr12-30750904-DEL-313 |
| chr12-30759492-DEL-569 | chr12-30759493-DEL-569 |
| chr12-30819778-DEL-2529 | chr12-30819779-DEL-2529 |
| chr12-30826977-DEL-72 | chr12-30826978-DEL-72 |
| chr12-30831313-DEL-62 | chr12-30831314-DEL-62 |
| chr12-30838440-DEL-266 | chr12-30838441-DEL-266 |
| chr12-30847172-DEL-60 | chr12-30847173-DEL-60 |
| chr12-30851057-DEL-1894 | chr12-30851058-DEL-1894 |
| chr12-30870524-DEL-6426 | chr12-30870525-DEL-6426 |
| chr12-30913976-DEL-196 | chr12-30913977-DEL-196 |
| chr12-30963913-DEL-70 | chr12-30963914-DEL-70 |
| chr12-30984363-DEL-57 | chr12-30984365-DEL-56 |
| chr12-30984730-DEL-285 | chr12-30984731-DEL-284 |
| chr12-31041409-DEL-6703 | chr12-31041410-DEL-6703 |
| chr12-31041426-DEL-212 | chr12-31041426-DEL-212 |
| chr12-31072369-DEL-90 | chr12-31072370-DEL-90 |
| chr12-31078077-DEL-119 | chr12-31078096-DEL-84 |
| chr12-31092034-DEL-2006 | chr12-31092035-DEL-2003 |
| chr12-31092769-DEL-58 | chr12-31092770-DEL-58 |
| chr12-31095633-DEL-6466 | chr12-31095634-DEL-6466 |
| chr12-31107636-DEL-1304 | chr12-31107637-DEL-1304 |
| chr12-31110209-DEL-66 | chr12-31110212-DEL-64 |
| chr12-31142036-DEL-54 | chr12-31142031-DEL-60 |
| chr12-31146192-DEL-210 | chr12-31146193-DEL-210 |
| chr12-31152868-DEL-1000 | chr12-31152869-DEL-1000 |
| chr12-31154693-DEL-96 | chr12-31154694-DEL-96 |
| chr12-31155559-DEL-6342 | chr12-31155560-DEL-6344 |
| chr12-31205699-DEL-76 | chr12-31205683-DEL-84 |
| chr12-31212691-DEL-1886 | chr12-31212692-DEL-1886 |
| chr12-31221594-DEL-67 | chr12-31221595-DEL-67 |
| chr12-31230389-DEL-218 | chr12-31230390-DEL-218 |
| chr12-31232172-DEL-53 | chr12-31232173-DEL-53 |
| chr12-31248590-DEL-275 | chr12-31248591-DEL-275 |
| chr12-31255641-DEL-52 | chr12-31255642-DEL-52 |
| chr12-31267088-DEL-205 | chr12-31267083-DEL-209 |
| chr12-31269164-DEL-196 | chr12-31269165-DEL-196 |
| chr12-31307736-DEL-144 | chr12-31307737-DEL-144 |
| chr12-31308588-DEL-480 | chr12-31308592-DEL-484 |
| chr12-31387338-DEL-197 | chr12-31387339-DEL-197 |
| chr12-31394039-DEL-195 | chr12-31394040-DEL-194 |
| chr12-31397866-DEL-363 | chr12-31397867-DEL-363 |
| chr12-31408149-DEL-7074 | chr12-31408150-DEL-7074 |
| chr12-31463537-DEL-67 | chr12-31463553-DEL-62 |
| chr12-31540803-DEL-83 | chr12-31540805-DEL-80 |
| chr12-31588518-DEL-62 | chr12-31588519-DEL-60 |
| chr12-31623080-DEL-199 | chr12-31623081-DEL-199 |
| chr12-31626983-DEL-100 | chr12-31626984-DEL-100 |
| chr12-31630374-DEL-71 | chr12-31630375-DEL-89 |
| chr12-31653498-DEL-73 | chr12-31653499-DEL-73 |
| chr12-31800442-DEL-203 | chr12-31800443-DEL-203 |
| chr12-31817625-DEL-199 | chr12-31817626-DEL-199 |
| chr12-31832442-DEL-82 | chr12-31832443-DEL-86 |
| chr12-31850658-DEL-214 | chr12-31850659-DEL-214 |
| chr12-31858393-DEL-280 | chr12-31858394-DEL-280 |
| chr12-31863470-DEL-183 | chr12-31863471-DEL-183 |
| chr12-31876324-DEL-74 | chr12-31876323-DEL-72 |

| | |
|---|---|
| chr12-31877088-DEL-197 | chr12-31877089-DEL-197 |
| chr12-31881359-DEL-80 | chr12-31881360-DEL-80 |
| chr12-31916621-DEL-1113 | chr12-31916622-DEL-1114 |
| chr12-31921829-DEL-210 | chr12-31921830-DEL-240 |
| chr12-31944437-DEL-80 | chr12-31944422-DEL-96 |
| chr12-31950018-DEL-258 | chr12-31950019-DEL-258 |
| chr12-31951648-DEL-54 | chr12-31951649-DEL-54 |
| chr12-31960833-DEL-96 | chr12-31960834-DEL-96 |
| chr12-31965610-DEL-287 | chr12-31965611-DEL-287 |
| chr12-32013193-DEL-654 | chr12-32013194-DEL-654 |
| chr12-32024306-DEL-361 | chr12-32024296-DEL-353 |
| chr12-32037523-DEL-127 | chr12-32037524-DEL-127 |
| chr12-32044756-DEL-64 | chr12-32044757-DEL-64 |
| chr12-32059200-DEL-107 | chr12-32059201-DEL-107 |
| chr12-32059890-DEL-180 | chr12-32059900-DEL-179 |
| chr12-32061999-DEL-2472 | chr12-32062000-DEL-2472 |
| chr12-32078013-DEL-4368 | chr12-32078014-DEL-4368 |
| chr12-32135708-DEL-57 | chr12-32135709-DEL-61 |
| chr12-32145704-DEL-75 | chr12-32145705-DEL-75 |
| chr12-32156702-DEL-595 | chr12-32156703-DEL-595 |
| chr12-32181739-DEL-69 | chr12-32181709-DEL-73 |
| chr12-32187239-DEL-95 | chr12-32187239-DEL-95 |
| chr12-32200146-DEL-53 | chr12-32200150-DEL-50 |
| chr12-32200767-DEL-111 | chr12-32200766-DEL-120 |
| chr12-32241932-DEL-64 | chr12-32241933-DEL-64 |
| chr12-32259731-DEL-91 | chr12-32259732-DEL-91 |
| chr12-32267681-DEL-1882 | chr12-32267682-DEL-1882 |
| chr12-32308138-DEL-467 | chr12-32308139-DEL-467 |
| chr12-32311718-DEL-82 | chr12-32311717-DEL-83 |
| chr12-32360255-DEL-181 | chr12-32360252-DEL-158 |
| chr12-32382384-DEL-200 | chr12-32382356-DEL-198 |
| chr12-32388975-DEL-62 | chr12-32388958-DEL-62 |
| chr12-32420092-DEL-3700 | chr12-32420093-DEL-3700 |
| chr12-32480790-DEL-172 | chr12-32480791-DEL-172 |
| chr12-32549014-DEL-298 | chr12-32549015-DEL-298 |
| chr12-32583366-DEL-325 | chr12-32583367-DEL-325 |
| chr12-32585112-DEL-7383 | chr12-32585113-DEL-7383 |
| chr12-32592660-DEL-1487 | chr12-32592660-DEL-1487 |
| chr12-32610188-DEL-144 | chr12-32610189-DEL-144 |
| chr12-32646479-DEL-4073 | chr12-32646480-DEL-4073 |
| chr12-32652195-DEL-58 | chr12-32652196-DEL-58 |
| chr12-32658356-DEL-87 | chr12-32658357-DEL-87 |
| chr12-32720952-DEL-252 | chr12-32720974-DEL-231 |
| chr12-32782525-DEL-2114 | chr12-32782526-DEL-2114 |
| chr12-32801890-DEL-170 | chr12-32801889-DEL-172 |
| chr12-32802871-DEL-183 | chr12-32802879-DEL-185 |
| chr12-32830329-DEL-1452 | chr12-32830330-DEL-1452 |
| chr12-32845270-DEL-92 | chr12-32845277-DEL-87 |
| chr12-32854250-DEL-328 | chr12-32854249-DEL-329 |
| chr12-32883330-DEL-176 | chr12-32883331-DEL-176 |
| chr12-32898194-DEL-374 | chr12-32898191-DEL-374 |
| chr12-32923171-DEL-107 | chr12-32923172-DEL-107 |
| chr12-32971725-DEL-217 | chr12-32971726-DEL-217 |
| chr12-33029542-DEL-176 | chr12-33029543-DEL-176 |
| chr12-33049726-DEL-86 | chr12-33049727-DEL-86 |
| chr12-33052970-DEL-116 | chr12-33052971-DEL-117 |
| chr12-33061075-DEL-275 | chr12-33061076-DEL-275 |
| chr12-33101348-DEL-68 | chr12-33101349-DEL-68 |
| chr12-33133184-DEL-68 | chr12-33133167-DEL-51 |
| chr12-33147960-DEL-148 | chr12-33147960-DEL-150 |
| chr12-33154988-DEL-403 | chr12-33154989-DEL-403 |
| chr12-33179860-DEL-51 | chr12-33179861-DEL-52 |
| chr12-33190209-DEL-204 | chr12-33190210-DEL-204 |
| chr12-33190897-DEL-61 | chr12-33190898-DEL-62 |
| chr12-33223439-DEL-125 | chr12-33223400-DEL-167 |
| chr12-33257719-DEL-357 | chr12-33257720-DEL-357 |
| chr12-33273780-DEL-53 | chr12-33273812-DEL-50 |
| chr12-33278655-DEL-6403 | chr12-33278656-DEL-6403 |
| chr12-33344709-DEL-165 | chr12-33344675-DEL-164 |
| chr12-33362696-DEL-100 | chr12-33362697-DEL-95 |
| chr12-33383944-DEL-97 | chr12-33383945-DEL-97 |
| chr12-33460443-DEL-711 | chr12-33460444-DEL-711 |
| chr12-33493292-DEL-1475 | chr12-33493293-DEL-1475 |
| chr12-33496186-DEL-50 | chr12-33496192-DEL-57 |
| chr12-33498430-DEL-155 | chr12-33498431-DEL-155 |
| chr12-33499790-DEL-254 | chr12-33499791-DEL-254 |
| chr12-33508127-DEL-313 | chr12-33508128-DEL-313 |
| chr12-33553318-DEL-2798 | chr12-33553319-DEL-2798 |
| chr12-33601084-DEL-327 | chr12-33601108-DEL-328 |
| chr12-33662705-DEL-77 | chr12-33662706-DEL-77 |

| | |
|---|---|
| chr12-33668628-DEL-78 | chr12-33668629-DEL-78 |
| chr12-33707524-DEL-1573 | chr12-33707525-DEL-1573 |
| chr12-33714366-DEL-6438 | chr12-33714367-DEL-6438 |
| chr12-33747248-DEL-107 | chr12-33747249-DEL-107 |
| chr12-33756587-DEL-197 | chr12-33756588-DEL-197 |
| chr12-33797263-DEL-2754 | chr12-33797264-DEL-2754 |
| chr12-33816497-DEL-6418 | chr12-33816498-DEL-6418 |
| chr12-33832492-DEL-120 | chr12-33832493-DEL-120 |
| chr12-33855172-DEL-811 | chr12-33855184-DEL-799 |
| chr12-33857691-DEL-461 | chr12-33857692-DEL-460 |
| chr12-33860283-DEL-322 | chr12-33860284-DEL-322 |
| chr12-33862609-DEL-76 | chr12-33862610-DEL-76 |
| chr12-33914782-DEL-557 | chr12-33914783-DEL-557 |
| chr12-33919170-DEL-1558 | chr12-33919178-DEL-1554 |
| chr12-33937710-DEL-59 | chr12-33937685-DEL-101 |
| chr12-33964975-DEL-53 | chr12-33964976-DEL-53 |
| chr12-33979653-DEL-1001 | chr12-33979654-DEL-1001 |
| chr12-33995749-DEL-56 | chr12-33995750-DEL-56 |
| chr12-34021992-DEL-194 | chr12-34021993-DEL-194 |
| chr12-34043967-DEL-6570 | chr12-34043968-DEL-6570 |
| chr12-34063821-DEL-609 | chr12-34063822-DEL-609 |
| chr12-34074357-DEL-6160 | chr12-34074364-DEL-6144 |
| chr12-34098619-DEL-66 | chr12-34098620-DEL-66 |
| chr12-34106768-DEL-1271 | chr12-34106769-DEL-1271 |
| chr12-34116914-DEL-6899 | chr12-34116915-DEL-6899 |
| chr12-34117096-DEL-213 | chr12-34117097-DEL-213 |
| chr12-34132356-DEL-366 | chr12-34132370-DEL-360 |
| chr12-34138042-DEL-398 | chr12-34138043-DEL-398 |
| chr12-34178657-DEL-3361 | chr12-34178658-DEL-3361 |
| chr12-34188171-DEL-1756 | chr12-34188172-DEL-1756 |
| chr12-34211297-DEL-2587 | chr12-34211298-DEL-2587 |
| chr12-34240202-DEL-461 | chr12-34240203-DEL-461 |
| chr12-34286465-DEL-70 | chr12-34286466-DEL-70 |
| chr12-34304308-DEL-699 | chr12-34304309-DEL-699 |
| chr12-34331777-DEL-55 | chr12-34331778-DEL-55 |
| chr12-34359356-DEL-84 | chr12-34359357-DEL-64 |
| chr12-34369073-DEL-1507 | chr12-34369073-DEL-1507 |
| chr12-34573428-DEL-66 | chr12-34573429-DEL-66 |
| chr12-34587459-DEL-52 | chr12-34587460-DEL-52 |
| chr12-34603316-DEL-289 | chr12-34603322-DEL-284 |
| chr12-34607105-DEL-612 | chr12-34607103-DEL-612 |
| chr12-34672951-DEL-5985 | chr12-34672952-DEL-5985 |
| chr12-34688718-DEL-56 | chr12-34688760-DEL-56 |
| chr12-34691666-DEL-56 | chr12-34691667-DEL-56 |
| chr12-34693627-DEL-78 | chr12-34693608-DEL-52 |
| chr12-34694944-DEL-987 | chr12-34694945-DEL-987 |
| chr12-34703166-DEL-95 | chr12-34703167-DEL-87 |
| chr12-34711360-DEL-97 | chr12-34711361-DEL-97 |
| chr12-34731463-DEL-51 | chr12-34731464-DEL-51 |
| chr12-34743659-DEL-62 | chr12-34743660-DEL-62 |
| chr12-34743813-DEL-771 | chr12-34743814-DEL-771 |
| chr12-34783387-DEL-6517 | chr12-34783388-DEL-6517 |
| chr12-34815704-DEL-171 | chr12-34815705-DEL-171 |
| chr12-34834426-DEL-54 | chr12-34834427-DEL-54 |
| chr12-34853146-DEL-77 | chr12-34853147-DEL-77 |
| chr12-34855917-DEL-68 | chr12-34855918-DEL-68 |
| chr12-34898735-DEL-161 | chr12-34898746-DEL-193 |
| chr12-34918502-DEL-640 | chr12-34918503-DEL-640 |
| chr12-34934731-DEL-118 | chr12-34934732-DEL-118 |
| chr12-34968171-DEL-127 | chr12-34968172-DEL-127 |
| chr12-34972456-DEL-65 | chr12-34972445-DEL-66 |
| chr12-34972732-DEL-375 | chr12-34972766-DEL-342 |
| chr12-34974882-DEL-400 | chr12-34974883-DEL-400 |
| chr12-34975623-DEL-639 | chr12-34975634-DEL-646 |
| chr12-35004557-DEL-64 | chr12-35004545-DEL-60 |
| chr12-35057058-DEL-168 | chr12-35057059-DEL-168 |
| chr12-35079033-DEL-343 | chr12-35079034-DEL-343 |
| chr12-35080892-DEL-1383 | chr12-35080893-DEL-1383 |
| chr12-35084960-DEL-77 | chr12-35084975-DEL-92 |
| chr12-35125014-DEL-1098 | chr12-35125012-DEL-1100 |
| chr12-35130473-DEL-52 | chr12-35130474-DEL-52 |
| chr12-35177867-DEL-751 | chr12-35177868-DEL-751 |
| chr12-35202523-DEL-358 | chr12-35202524-DEL-358 |
| chr12-35261791-DEL-808 | chr12-35261791-DEL-806 |
| chr12-35275426-DEL-53 | chr12-35275410-DEL-70 |
| chr12-35276783-DEL-56 | chr12-35276784-DEL-56 |
| chr12-35286831-DEL-52 | chr12-35286832-DEL-52 |
| chr12-35323436-DEL-6233 | chr12-35323437-DEL-6233 |
| chr12-35353137-DEL-84 | chr12-35353137-DEL-84 |
| chr12-35358946-DEL-162 | chr12-35358923-DEL-153 |

| | |
|---|---|
| chr12-35360627-DEL-201 | chr12-35360628-DEL-201 |
| chr12-35375922-DEL-51 | chr12-35375905-DEL-54 |
| chr12-35385324-DEL-142 | chr12-35385325-DEL-142 |
| chr12-35397174-DEL-6131 | chr12-35397175-DEL-6131 |
| chr12-35405912-DEL-57 | chr12-35405878-DEL-60 |
| chr12-35408965-DEL-97 | chr12-35408966-DEL-97 |
| chr12-35435279-DEL-51 | chr12-35435280-DEL-51 |
| chr12-35465386-DEL-78 | chr12-35465387-DEL-78 |
| chr12-35476740-DEL-77 | chr12-35476741-DEL-77 |
| chr12-35499707-DEL-60 | chr12-35499708-DEL-55 |
| chr12-35515545-DEL-127 | chr12-35515546-DEL-125 |
| chr12-35520109-DEL-1198 | chr12-35520110-DEL-1198 |
| chr12-35534711-DEL-250 | chr12-35534712-DEL-250 |
| chr12-35567311-DEL-1470 | chr12-35567312-DEL-1470 |
| chr12-35595951-DEL-59 | chr12-35595929-DEL-59 |
| chr12-35630497-DEL-80 | chr12-35630497-DEL-78 |
| chr12-35643563-DEL-1242 | chr12-35643564-DEL-1242 |
| chr12-35665327-DEL-2234 | chr12-35665328-DEL-2234 |
| chr12-35671009-DEL-108 | chr12-35671010-DEL-108 |
| chr12-35700489-DEL-80 | chr12-35700490-DEL-82 |
| chr12-35730719-DEL-251 | chr12-35730720-DEL-251 |
| chr12-35745879-DEL-237 | chr12-35745880-DEL-237 |
| chr12-35789583-DEL-971 | chr12-35789584-DEL-971 |
| chr12-35793317-DEL-977 | chr12-35793318-DEL-977 |
| chr12-35802669-DEL-52 | chr12-35802670-DEL-52 |
| chr12-35847222-DEL-192 | chr12-35847223-DEL-192 |
| chr12-35852841-DEL-214 | chr12-35852842-DEL-214 |
| chr12-35904160-DEL-7491 | chr12-35904161-DEL-7491 |
| chr12-35924816-DEL-208 | chr12-35924817-DEL-208 |
| chr12-35937592-DEL-71 | chr12-35937593-DEL-67 |
| chr12-35948207-DEL-1158 | chr12-35948208-DEL-1158 |
| chr12-35948929-DEL-132 | chr12-35948930-DEL-132 |
| chr12-35952793-DEL-6254 | chr12-35952794-DEL-6254 |
| chr12-35968606-DEL-399 | chr12-35968601-DEL-400 |
| chr12-35975857-DEL-150 | chr12-35975858-DEL-150 |
| chr12-35993454-DEL-157 | chr12-35993458-DEL-152 |
| chr12-36006227-DEL-171 | chr12-36006228-DEL-171 |
| chr12-36030824-DEL-102 | chr12-36030868-DEL-102 |
| chr12-36057972-DEL-55 | chr12-36057972-DEL-55 |
| chr12-36064178-DEL-204 | chr12-36064179-DEL-204 |
| chr12-36066027-DEL-86 | chr12-36066028-DEL-86 |
| chr12-36104297-DEL-210 | chr12-36104293-DEL-215 |
| chr12-36119619-DEL-182 | chr12-36119617-DEL-194 |
| chr12-36149364-DEL-2804 | chr12-36149365-DEL-2804 |
| chr12-36174726-DEL-393 | chr12-36174727-DEL-393 |
| chr12-36177132-DEL-416 | chr12-36177133-DEL-416 |
| chr12-36185268-DEL-97 | chr12-36185269-DEL-97 |
| chr12-36201604-DEL-719 | chr12-36201605-DEL-719 |
| chr12-36222344-DEL-56 | chr12-36222295-DEL-57 |
| chr12-36249263-DEL-6414 | chr12-36249262-DEL-6415 |
| chr12-36258741-DEL-90 | chr12-36258742-DEL-90 |
| chr12-36309495-DEL-86 | chr12-36309493-DEL-87 |
| chr12-36328962-DEL-6450 | chr12-36328963-DEL-6450 |
| chr12-36338118-DEL-66 | chr12-36338119-DEL-66 |
| chr12-36344939-DEL-324 | chr12-36344940-DEL-324 |
| chr12-36386140-DEL-6902 | chr12-36386141-DEL-6902 |
| chr12-36410355-DEL-117 | chr12-36410356-DEL-117 |
| chr12-36422188-DEL-2151 | chr12-36422190-DEL-2152 |
| chr12-36423977-DEL-69 | chr12-36423942-DEL-70 |
| chr12-36426049-DEL-2368 | chr12-36426047-DEL-2369 |
| chr12-36463331-DEL-7549 | chr12-36463332-DEL-7549 |
| chr12-36510064-DEL-330 | chr12-36510065-DEL-330 |
| chr12-36525569-DEL-163 | chr12-36525570-DEL-163 |
| chr12-36536146-DEL-6970 | chr12-36536147-DEL-6970 |
| chr12-36543076-DEL-830 | chr12-36543076-DEL-830 |
| chr12-36555583-DEL-74 | chr12-36555584-DEL-74 |
| chr12-36589293-DEL-655 | chr12-36589294-DEL-655 |
| chr12-36605859-DEL-157 | chr12-36605860-DEL-157 |
| chr12-36665963-DEL-2628 | chr12-36665964-DEL-2628 |
| chr12-36679093-DEL-1162 | chr12-36679089-DEL-1166 |
| chr12-36715625-DEL-222 | chr12-36715670-DEL-196 |
| chr12-36762288-DEL-76 | chr12-36762289-DEL-76 |
| chr12-36795543-DEL-432 | chr12-36795544-DEL-432 |
| chr12-36801660-DEL-71 | chr12-36801651-DEL-83 |
| chr12-36802901-DEL-66 | chr12-36802902-DEL-66 |
| chr12-36826743-DEL-77 | chr12-36826764-DEL-56 |
| chr12-36836153-DEL-525 | chr12-36836154-DEL-525 |
| chr12-36838230-DEL-209 | chr12-36838231-DEL-209 |
| chr12-36854429-DEL-420 | chr12-36854430-DEL-420 |
| chr12-36862887-DEL-59 | chr12-36862888-DEL-59 |

| | |
|---|---|
| chr12-36908235-DEL-63 | chr12-36908236-DEL-66 |
| chr12-36919427-DEL-86 | chr12-36919417-DEL-69 |
| chr12-36935511-DEL-74 | chr12-36935512-DEL-74 |
| chr12-36953982-DEL-111 | chr12-36953983-DEL-111 |
| chr12-36954573-DEL-292 | chr12-36954574-DEL-292 |
| chr12-37001690-DEL-6443 | chr12-37001691-DEL-6443 |
| chr12-37001783-DEL-213 | chr12-37001757-DEL-214 |
| chr12-37008995-DEL-2624 | chr12-37008996-DEL-2624 |
| chr12-37020869-DEL-194 | chr12-37020871-DEL-193 |
| chr12-37034305-DEL-618 | chr12-37034306-DEL-618 |
| chr12-37045078-DEL-1690 | chr12-37045079-DEL-1690 |
| chr12-37085622-DEL-399 | chr12-37085623-DEL-399 |
| chr12-37112209-DEL-227 | chr12-37112210-DEL-227 |
| chr12-37114701-DEL-6430 | chr12-37114702-DEL-6430 |
| chr12-37140470-DEL-82 | chr12-37140454-DEL-83 |
| chr12-37197766-DEL-4249 | chr12-37197767-DEL-4249 |
| chr12-37204756-DEL-6808 | chr12-37204757-DEL-6808 |
| chr12-37230284-DEL-504 | chr12-37230285-DEL-504 |
| chr12-37233689-DEL-78 | chr12-37233684-DEL-56 |
| chr12-37239846-DEL-57 | chr12-37239847-DEL-57 |
| chr12-37240668-DEL-7196 | chr12-37240669-DEL-7196 |
| chr12-37263109-DEL-338 | chr12-37263110-DEL-338 |
| chr12-37263918-DEL-1562 | chr12-37263919-DEL-1562 |
| chr12-37266070-DEL-1691 | chr12-37266071-DEL-1691 |
| chr12-37303842-DEL-56 | chr12-37303847-DEL-62 |
| chr12-37311316-DEL-258 | chr12-37311309-DEL-259 |
| chr12-37422178-DEL-884 | chr12-37422179-DEL-884 |
| chr12-37424368-DEL-140 | chr12-37424376-DEL-112 |
| chr12-37468407-DEL-60 | chr12-37468408-DEL-60 |
| chr12-37472629-DEL-52 | chr12-37472630-DEL-52 |
| chr12-37480799-DEL-81 | chr12-37480767-DEL-75 |
| chr12-37546287-DEL-1068 | chr12-37546289-DEL-1067 |
| chr12-37550811-DEL-2597 | chr12-37550812-DEL-2596 |
| chr12-37551035-DEL-485 | chr12-37551036-DEL-485 |
| chr12-37581062-DEL-6123 | chr12-37581063-DEL-6123 |
| chr12-37612535-DEL-1595 | chr12-37612536-DEL-1595 |
| chr12-37623672-DEL-252 | chr12-37623673-DEL-252 |
| chr12-37625140-DEL-743 | chr12-37625141-DEL-743 |
| chr12-37647250-DEL-744 | chr12-37647251-DEL-744 |
| chr12-37648590-DEL-51 | chr12-37648591-DEL-51 |
| chr12-37652915-DEL-5257 | chr12-37652916-DEL-5257 |
| chr12-37718112-DEL-6211 | chr12-37718113-DEL-6211 |
| chr12-37725152-DEL-55 | chr12-37725153-DEL-55 |
| chr12-37737353-DEL-58 | chr12-37737354-DEL-56 |
| chr12-37747018-DEL-59 | chr12-37747019-DEL-63 |
| chr12-37811513-DEL-1950 | chr12-37811514-DEL-1950 |
| chr12-37821550-DEL-205 | chr12-37821551-DEL-205 |
| chr12-37828196-DEL-4681 | chr12-37828197-DEL-4681 |
| chr12-37839038-DEL-1473 | chr12-37839039-DEL-1473 |
| chr12-37893148-DEL-102 | chr12-37893149-DEL-102 |
| chr12-37910831-DEL-443 | chr12-37910832-DEL-443 |
| chr12-37913676-DEL-70 | chr12-37913677-DEL-70 |
| chr12-37923723-DEL-777 | chr12-37923724-DEL-777 |
| chr12-37932742-DEL-100 | chr12-37932739-DEL-104 |
| chr12-37943825-DEL-78 | chr12-37943825-DEL-78 |
| chr12-37986105-DEL-60 | chr12-37986091-DEL-75 |
| chr12-38011463-DEL-61 | chr12-38011465-DEL-58 |
| chr12-38053670-DEL-57 | chr12-38053647-DEL-55 |
| chr12-38097595-DEL-7346 | chr12-38097596-DEL-7346 |
| chr12-38111639-DEL-1284 | chr12-38111640-DEL-1284 |
| chr12-38118848-DEL-92 | chr12-38118825-DEL-97 |
| chr12-38124613-DEL-6482 | chr12-38124614-DEL-6482 |
| chr12-38186066-DEL-59 | chr12-38186058-DEL-68 |
| chr12-38187658-DEL-449 | chr12-38187659-DEL-449 |
| chr12-38192962-DEL-70 | chr12-38192975-DEL-68 |
| chr12-38206702-DEL-54 | chr12-38206710-DEL-52 |
| chr12-38242931-DEL-646 | chr12-38242932-DEL-646 |
| chr12-38245649-DEL-10264 | chr12-38245650-DEL-10264 |
| chr12-38252981-DEL-808 | chr12-38252985-DEL-807 |
| chr12-38293987-DEL-84 | chr12-38293988-DEL-84 |
| chr12-38305143-DEL-111 | chr12-38305144-DEL-111 |
| chr12-38325772-DEL-1817 | chr12-38325773-DEL-1817 |
| chr12-38383215-DEL-502 | chr12-38383216-DEL-502 |
| chr12-38453730-DEL-57 | chr12-38453731-DEL-57 |
| chr12-38513928-DEL-1834 | chr12-38513930-DEL-1836 |
| chr12-38522545-DEL-1053 | chr12-38522546-DEL-1053 |
| chr12-38536890-DEL-1637 | chr12-38536891-DEL-1637 |
| chr12-38594461-DEL-330 | chr12-38594462-DEL-330 |
| chr12-38630657-DEL-293 | chr12-38630658-DEL-293 |
| chr12-38642868-DEL-274 | chr12-38642869-DEL-274 |

| | |
|---|---|
| chr12-38781746-DEL-52 | chr12-38781747-DEL-52 |
| chr12-38789565-DEL-274 | chr12-38789566-DEL-274 |
| chr12-38795902-DEL-132 | chr12-38795893-DEL-147 |
| chr12-38796821-DEL-2994 | chr12-38796822-DEL-2994 |
| chr12-38861270-DEL-200 | chr12-38861271-DEL-200 |
| chr12-38867138-DEL-156 | chr12-38867093-DEL-172 |
| chr12-38918319-DEL-52 | chr12-38918299-DEL-100 |
| chr12-38922377-DEL-70 | chr12-38922385-DEL-68 |
| chr12-38940478-DEL-95 | chr12-38940479-DEL-95 |
| chr12-38944540-DEL-275 | chr12-38944541-DEL-275 |
| chr12-38946618-DEL-52 | chr12-38946619-DEL-52 |
| chr12-38950864-DEL-65 | chr12-38950865-DEL-65 |
| chr12-38960945-DEL-8327 | chr12-38960946-DEL-8327 |
| chr12-39002778-DEL-84 | chr12-39002799-DEL-65 |
| chr12-39027750-DEL-1221 | chr12-39027751-DEL-1221 |
| chr12-39029230-DEL-3111 | chr12-39029231-DEL-3111 |
| chr12-39039072-DEL-3805 | chr12-39039073-DEL-3805 |
| chr12-39047032-DEL-129 | chr12-39047034-DEL-127 |
| chr12-39054605-DEL-101 | chr12-39054604-DEL-102 |
| chr12-39068092-DEL-62 | chr12-39068093-DEL-62 |
| chr12-39089386-DEL-431 | chr12-39089384-DEL-432 |
| chr12-39093843-DEL-439 | chr12-39093844-DEL-437 |
| chr12-39114386-DEL-84 | chr12-39114387-DEL-60 |
| chr12-39138910-DEL-339 | chr12-39138911-DEL-339 |
| chr12-39209695-DEL-58 | chr12-39209695-DEL-52 |
| chr12-39224259-DEL-221 | chr12-39224260-DEL-221 |
| chr12-39236629-DEL-51 | chr12-39236630-DEL-51 |
| chr12-39247368-DEL-1019 | chr12-39247369-DEL-1019 |
| chr12-39286827-DEL-88 | chr12-39286836-DEL-83 |
| chr12-39326710-DEL-75 | chr12-39326711-DEL-75 |
| chr12-39330143-DEL-848 | chr12-39330144-DEL-848 |
| chr12-39350153-DEL-99 | chr12-39350154-DEL-99 |
| chr12-39359343-DEL-66 | chr12-39359344-DEL-66 |
| chr12-39361533-DEL-608 | chr12-39361534-DEL-608 |
| chr12-39405476-DEL-631 | chr12-39405478-DEL-632 |
| chr12-39409684-DEL-6132 | chr12-39409685-DEL-6132 |
| chr12-39481643-DEL-4570 | chr12-39481644-DEL-4570 |
| chr12-39588745-DEL-6465 | chr12-39588746-DEL-6465 |
| chr12-39588901-DEL-208 | chr12-39588902-DEL-208 |
| chr12-39614450-DEL-9201 | chr12-39614453-DEL-9197 |
| chr12-39666920-DEL-291 | chr12-39666921-DEL-291 |
| chr12-39672689-DEL-4126 | chr12-39672690-DEL-4126 |
| chr12-39677471-DEL-433 | chr12-39677471-DEL-433 |
| chr12-39695004-DEL-53 | chr12-39695005-DEL-53 |
| chr12-39750742-DEL-1165 | chr12-39750743-DEL-1165 |
| chr12-39754876-DEL-622 | chr12-39754906-DEL-608 |
| chr12-39767614-DEL-294 | chr12-39767615-DEL-294 |
| chr12-39772091-DEL-64 | chr12-39772098-DEL-58 |
| chr12-39810746-DEL-53 | chr12-39810743-DEL-54 |
| chr12-39811177-DEL-184 | chr12-39811178-DEL-184 |
| chr12-39835668-DEL-1319 | chr12-39835646-DEL-1320 |
| chr12-39882721-DEL-417 | chr12-39882722-DEL-417 |
| chr12-39898866-DEL-114 | chr12-39898909-DEL-88 |
| chr12-39949509-DEL-6565 | chr12-39949510-DEL-6565 |
| chr12-39998452-DEL-666 | chr12-39998419-DEL-684 |
| chr12-40000051-DEL-209 | chr12-40000052-DEL-209 |
| chr12-40035122-DEL-967 | chr12-40035123-DEL-967 |
| chr12-40042528-DEL-99 | chr12-40042529-DEL-99 |
| chr12-40110254-DEL-83 | chr12-40110257-DEL-87 |
| chr12-40117392-DEL-607 | chr12-40117393-DEL-607 |
| chr12-40130724-DEL-67 | chr12-40130724-DEL-58 |
| chr12-40142851-DEL-138 | chr12-40142852-DEL-138 |
| chr12-40160750-DEL-1947 | chr12-40160740-DEL-1944 |
| chr12-40184886-DEL-201 | chr12-40184887-DEL-201 |
| chr12-40192703-DEL-930 | chr12-40192704-DEL-930 |
| chr12-40201261-DEL-171 | chr12-40201262-DEL-171 |
| chr12-40203576-DEL-56 | chr12-40203577-DEL-56 |
| chr12-40206702-DEL-64 | chr12-40206699-DEL-50 |
| chr12-40217128-DEL-66 | chr12-40217129-DEL-66 |
| chr12-40231490-DEL-2831 | chr12-40231490-DEL-2831 |
| chr12-40232054-DEL-78 | chr12-40232055-DEL-78 |
| chr12-40236360-DEL-1891 | chr12-40236355-DEL-1892 |
| chr12-40244387-DEL-1895 | chr12-40244385-DEL-1895 |
| chr12-40334960-DEL-213 | chr12-40334961-DEL-213 |
| chr12-40350019-DEL-872 | chr12-40350020-DEL-872 |
| chr12-40416149-DEL-444 | chr12-40416150-DEL-444 |
| chr12-40420799-DEL-6709 | chr12-40420800-DEL-6709 |
| chr12-40449045-DEL-59 | chr12-40449046-DEL-59 |
| chr12-40452084-DEL-7116 | chr12-40452085-DEL-7116 |
| chr12-40492317-DEL-116 | chr12-40492318-DEL-116 |

| | |
|---|---|
| chr12-40518931-DEL-92 | chr12-40518932-DEL-92 |
| chr12-40547477-DEL-82 | chr12-40547478-DEL-82 |
| chr12-40560961-DEL-124 | chr12-40560962-DEL-124 |
| chr12-40574129-DEL-494 | chr12-40574130-DEL-494 |
| chr12-40579065-DEL-86 | chr12-40579066-DEL-86 |
| chr12-40586694-DEL-208 | chr12-40586695-DEL-208 |
| chr12-40629270-DEL-212 | chr12-40629271-DEL-212 |
| chr12-40697768-DEL-94 | chr12-40697734-DEL-97 |
| chr12-40698589-DEL-137 | chr12-40698577-DEL-136 |
| chr12-40736070-DEL-868 | chr12-40736071-DEL-868 |
| chr12-40770257-DEL-486 | chr12-40770258-DEL-486 |
| chr12-40784410-DEL-243 | chr12-40784411-DEL-243 |
| chr12-40786038-DEL-210 | chr12-40786039-DEL-210 |
| chr12-40822275-DEL-64 | chr12-40822276-DEL-64 |
| chr12-40829456-DEL-54 | chr12-40829457-DEL-54 |
| chr12-40854674-DEL-465 | chr12-40854675-DEL-465 |
| chr12-40919023-DEL-205 | chr12-40919024-DEL-205 |
| chr12-40960325-DEL-198 | chr12-40960347-DEL-177 |
| chr12-40990854-DEL-459 | chr12-40990855-DEL-459 |
| chr12-41005501-DEL-127 | chr12-41005502-DEL-127 |
| chr12-41041157-DEL-104 | chr12-41041117-DEL-103 |
| chr12-41049988-DEL-52 | chr12-41049962-DEL-53 |
| chr12-41058218-DEL-451 | chr12-41058219-DEL-451 |
| chr12-41072104-DEL-53 | chr12-41072089-DEL-54 |
| chr12-41078837-DEL-51 | chr12-41078836-DEL-53 |
| chr12-41080742-DEL-67 | chr12-41080741-DEL-64 |
| chr12-41084976-DEL-184 | chr12-41084968-DEL-227 |
| chr12-41114838-DEL-1017 | chr12-41114839-DEL-1017 |
| chr12-41136830-DEL-273 | chr12-41136831-DEL-273 |
| chr12-41240042-DEL-98 | chr12-41240043-DEL-98 |
| chr12-41271302-DEL-90 | chr12-41271303-DEL-90 |
| chr12-41280531-DEL-1228 | chr12-41280532-DEL-1228 |
| chr12-41305454-DEL-60 | chr12-41305482-DEL-61 |
| chr12-41315401-DEL-8412 | chr12-41315402-DEL-8412 |
| chr12-41335105-DEL-53 | chr12-41335106-DEL-53 |
| chr12-41346063-DEL-400 | chr12-41346068-DEL-399 |
| chr12-41372426-DEL-101 | chr12-41372464-DEL-103 |
| chr12-41413536-DEL-7090 | chr12-41413537-DEL-7090 |
| chr12-41446371-DEL-323 | chr12-41446372-DEL-323 |
| chr12-41447759-DEL-696 | chr12-41447760-DEL-696 |
| chr12-41463634-DEL-98 | chr12-41463677-DEL-68 |
| chr12-41464135-DEL-745 | chr12-41464136-DEL-742 |
| chr12-41472541-DEL-639 | chr12-41472542-DEL-639 |
| chr12-41552534-DEL-5970 | chr12-41552535-DEL-5970 |
| chr12-41565113-DEL-69 | chr12-41565114-DEL-63 |
| chr12-41569783-DEL-80 | chr12-41569792-DEL-68 |
| chr12-41578990-DEL-1671 | chr12-41578991-DEL-1671 |
| chr12-41592141-DEL-1005 | chr12-41592142-DEL-1005 |
| chr12-41618782-DEL-598 | chr12-41618785-DEL-596 |
| chr12-41652486-DEL-1714 | chr12-41652487-DEL-1714 |
| chr12-41738588-DEL-119 | chr12-41738589-DEL-119 |
| chr12-41784975-DEL-2171 | chr12-41784976-DEL-2171 |
| chr12-41791582-DEL-724 | chr12-41791583-DEL-724 |
| chr12-41799021-DEL-146 | chr12-41799014-DEL-118 |
| chr12-41824559-DEL-786 | chr12-41824560-DEL-786 |
| chr12-41836440-DEL-62 | chr12-41836441-DEL-62 |
| chr12-41840178-DEL-66 | chr12-41840179-DEL-66 |
| chr12-41843950-DEL-60 | chr12-41843951-DEL-60 |
| chr12-41855378-DEL-388 | chr12-41855371-DEL-402 |
| chr12-41866858-DEL-58 | chr12-41866809-DEL-84 |
| chr12-41880560-DEL-96 | chr12-41880561-DEL-96 |
| chr12-41881788-DEL-1725 | chr12-41881789-DEL-1725 |
| chr12-41904542-DEL-1234 | chr12-41904543-DEL-1234 |
| chr12-41906694-DEL-2013 | chr12-41906695-DEL-2013 |
| chr12-41915605-DEL-1465 | chr12-41915606-DEL-1465 |
| chr12-41962217-DEL-3311 | chr12-41962218-DEL-3311 |
| chr12-42003372-DEL-301 | chr12-42003373-DEL-301 |
| chr12-42004239-DEL-74 | chr12-42004226-DEL-64 |
| chr12-42023699-DEL-6000 | chr12-42023700-DEL-6000 |
| chr12-42107567-DEL-452 | chr12-42107568-DEL-451 |
| chr12-42119857-DEL-233 | chr12-42119858-DEL-233 |
| chr12-42198337-DEL-388 | chr12-42198338-DEL-388 |
| chr12-42231285-DEL-820 | chr12-42231286-DEL-820 |
| chr12-42258615-DEL-344 | chr12-42258616-DEL-344 |
| chr12-42262466-DEL-61 | chr12-42262467-DEL-61 |
| chr12-42287434-DEL-7206 | chr12-42287435-DEL-7206 |
| chr12-42288006-DEL-624 | chr12-42288007-DEL-624 |
| chr12-42322231-DEL-74 | chr12-42322228-DEL-75 |
| chr12-42331580-DEL-67 | chr12-42331581-DEL-67 |
| chr12-42333700-DEL-178 | chr12-42333701-DEL-175 |

| | |
|---|---|
| chr12-42363806-DEL-1014 | chr12-42363805-DEL-1015 |
| chr12-42373748-DEL-407 | chr12-42373749-DEL-407 |
| chr12-42378923-DEL-6264 | chr12-42378924-DEL-6264 |
| chr12-42399071-DEL-6334 | chr12-42399072-DEL-6335 |
| chr12-42426601-DEL-99 | chr12-42426602-DEL-99 |
| chr12-42488870-DEL-521 | chr12-42488871-DEL-521 |
| chr12-42501190-DEL-52 | chr12-42501188-DEL-88 |
| chr12-42512159-DEL-251 | chr12-42512154-DEL-252 |
| chr12-42525812-DEL-845 | chr12-42525813-DEL-845 |
| chr12-42554259-DEL-946 | chr12-42554260-DEL-946 |
| chr12-42558502-DEL-74 | chr12-42558503-DEL-74 |
| chr12-42563265-DEL-681 | chr12-42563265-DEL-681 |
| chr12-42564591-DEL-365 | chr12-42564592-DEL-365 |
| chr12-42566070-DEL-51 | chr12-42566051-DEL-73 |
| chr12-42589841-DEL-199 | chr12-42589842-DEL-199 |
| chr12-42606792-DEL-6827 | chr12-42606793-DEL-6827 |
| chr12-42620060-DEL-97 | chr12-42620061-DEL-97 |
| chr12-42638389-DEL-7436 | chr12-42638389-DEL-7436 |
| chr12-42670160-DEL-6648 | chr12-42670161-DEL-6648 |
| chr12-42713570-DEL-6492 | chr12-42713553-DEL-6495 |
| chr12-42721268-DEL-70 | chr12-42721269-DEL-70 |
| chr12-42727566-DEL-210 | chr12-42727566-DEL-210 |
| chr12-42782561-DEL-125 | chr12-42782562-DEL-125 |
| chr12-42786089-DEL-6872 | chr12-42786090-DEL-6872 |
| chr12-42792237-DEL-424 | chr12-42792238-DEL-424 |
| chr12-42822547-DEL-6125 | chr12-42822548-DEL-6125 |
| chr12-42828165-DEL-111 | chr12-42828166-DEL-111 |
| chr12-42838471-DEL-55 | chr12-42838451-DEL-62 |
| chr12-42865680-DEL-640 | chr12-42865680-DEL-640 |
| chr12-42869508-DEL-5636 | chr12-42869509-DEL-5636 |
| chr12-42917163-DEL-5302 | chr12-42917156-DEL-5287 |
| chr12-42944685-DEL-57 | chr12-42944672-DEL-58 |
| chr12-42949944-DEL-65 | chr12-42949945-DEL-65 |
| chr12-42968587-DEL-1419 | chr12-42968588-DEL-1418 |
| chr12-42972460-DEL-868 | chr12-42972460-DEL-868 |
| chr12-42978402-DEL-191 | chr12-42978403-DEL-191 |
| chr12-43080946-DEL-187 | chr12-43080947-DEL-187 |
| chr12-43081561-DEL-610 | chr12-43081562-DEL-610 |
| chr12-43083385-DEL-7303 | chr12-43083386-DEL-7303 |
| chr12-43109497-DEL-1363 | chr12-43109498-DEL-1363 |
| chr12-43117957-DEL-59 | chr12-43117958-DEL-59 |
| chr12-43160337-DEL-59 | chr12-43160338-DEL-59 |
| chr12-43168358-DEL-462 | chr12-43168359-DEL-462 |
| chr12-43205009-DEL-52 | chr12-43205010-DEL-52 |
| chr12-43222050-DEL-6452 | chr12-43222051-DEL-6452 |
| chr12-43244868-DEL-1939 | chr12-43244869-DEL-1939 |
| chr12-43262971-DEL-427 | chr12-43262978-DEL-428 |
| chr12-43268528-DEL-1095 | chr12-43268570-DEL-1085 |
| chr12-43272176-DEL-151 | chr12-43272177-DEL-151 |
| chr12-43282251-DEL-63 | chr12-43282252-DEL-63 |
| chr12-43295571-DEL-407 | chr12-43295574-DEL-408 |
| chr12-43304694-DEL-129 | chr12-43304695-DEL-129 |
| chr12-43314802-DEL-66 | chr12-43314766-DEL-68 |
| chr12-43327786-DEL-85 | chr12-43327787-DEL-85 |
| chr12-43370017-DEL-59 | chr12-43370022-DEL-55 |
| chr12-43370784-DEL-548 | chr12-43370785-DEL-548 |
| chr12-43382277-DEL-1617 | chr12-43382278-DEL-1617 |
| chr12-43390083-DEL-6486 | chr12-43390084-DEL-6486 |
| chr12-43465730-DEL-875 | chr12-43465731-DEL-875 |
| chr12-43470388-DEL-64 | chr12-43470389-DEL-64 |
| chr12-43523035-DEL-1246 | chr12-43523037-DEL-1205 |
| chr12-43546561-DEL-2844 | chr12-43546561-DEL-2844 |
| chr12-43591101-DEL-387 | chr12-43591102-DEL-387 |
| chr12-43596581-DEL-995 | chr12-43596582-DEL-994 |
| chr12-43609332-DEL-547 | chr12-43609333-DEL-547 |
| chr12-43614422-DEL-3207 | chr12-43614423-DEL-3207 |
| chr12-43624918-DEL-4771 | chr12-43624919-DEL-4771 |
| chr12-43628594-DEL-479 | chr12-43628595-DEL-479 |
| chr12-43636507-DEL-308 | chr12-43636508-DEL-308 |
| chr12-43637451-DEL-1082 | chr12-43637457-DEL-1077 |
| chr12-43646411-DEL-653 | chr12-43646411-DEL-652 |
| chr12-43660394-DEL-127 | chr12-43660395-DEL-127 |
| chr12-43661503-DEL-900 | chr12-43661504-DEL-899 |
| chr12-43716821-DEL-8768 | chr12-43716818-DEL-8769 |
| chr12-43726413-DEL-86 | chr12-43726414-DEL-86 |
| chr12-43730243-DEL-187 | chr12-43730244-DEL-187 |
| chr12-43748536-DEL-54 | chr12-43748537-DEL-54 |
| chr12-43761021-DEL-301 | chr12-43761022-DEL-301 |
| chr12-43766628-DEL-845 | chr12-43766629-DEL-845 |
| chr12-43771002-DEL-6454 | chr12-43771003-DEL-6454 |

| | |
|---|---|
| chr12-43795536-DEL-6545 | chr12-43795536-DEL-6545 |
| chr12-43802524-DEL-962 | chr12-43802524-DEL-962 |
| chr12-43916097-DEL-175 | chr12-43916098-DEL-175 |
| chr12-43931213-DEL-72 | chr12-43931220-DEL-71 |
| chr12-43933219-DEL-6416 | chr12-43933220-DEL-6416 |
| chr12-43975694-DEL-214 | chr12-43975695-DEL-214 |
| chr12-44024653-DEL-402 | chr12-44024654-DEL-401 |
| chr12-44032482-DEL-311 | chr12-44032511-DEL-283 |
| chr12-44035747-DEL-403 | chr12-44035748-DEL-403 |
| chr12-44036352-DEL-280 | chr12-44036353-DEL-280 |
| chr12-44042769-DEL-1292 | chr12-44042770-DEL-1292 |
| chr12-44073843-DEL-524 | chr12-44073844-DEL-524 |
| chr12-44110754-DEL-52 | chr12-44110755-DEL-52 |
| chr12-44163269-DEL-185 | chr12-44163270-DEL-182 |
| chr12-44166853-DEL-199 | chr12-44166854-DEL-199 |
| chr12-44173426-DEL-1884 | chr12-44173426-DEL-1884 |
| chr12-44198452-DEL-4983 | chr12-44198453-DEL-4983 |
| chr12-44207674-DEL-485 | chr12-44207675-DEL-485 |
| chr12-44278083-DEL-6599 | chr12-44278085-DEL-6599 |
| chr12-44285398-DEL-382 | chr12-44285400-DEL-381 |
| chr12-44290911-DEL-84 | chr12-44290913-DEL-83 |
| chr12-44297678-DEL-63 | chr12-44297678-DEL-61 |
| chr12-44302223-DEL-6506 | chr12-44302224-DEL-6506 |
| chr12-44317772-DEL-809 | chr12-44317773-DEL-809 |
| chr12-44318353-DEL-202 | chr12-44318354-DEL-202 |
| chr12-44339539-DEL-3033 | chr12-44339540-DEL-3033 |
| chr12-44410431-DEL-952 | chr12-44410424-DEL-953 |
| chr12-44432529-DEL-157 | chr12-44432559-DEL-160 |
| chr12-44440765-DEL-75 | chr12-44440763-DEL-76 |
| chr12-44493722-DEL-59 | chr12-44493738-DEL-60 |
| chr12-44524756-DEL-938 | chr12-44524754-DEL-913 |
| chr12-44529852-DEL-9088 | chr12-44529853-DEL-9088 |
| chr12-44541090-DEL-2376 | chr12-44541091-DEL-2376 |
| chr12-44583061-DEL-104 | chr12-44583062-DEL-104 |
| chr12-44588933-DEL-85 | chr12-44588934-DEL-84 |
| chr12-44590517-DEL-914 | chr12-44590518-DEL-914 |
| chr12-44591245-DEL-74 | chr12-44591216-DEL-54 |
| chr12-44607721-DEL-317 | chr12-44607722-DEL-317 |
| chr12-44659629-DEL-1209 | chr12-44659630-DEL-1209 |
| chr12-44661278-DEL-91 | chr12-44661294-DEL-84 |
| chr12-44671724-DEL-106 | chr12-44671725-DEL-106 |
| chr12-44682415-DEL-704 | chr12-44682417-DEL-703 |
| chr12-44684065-DEL-789 | chr12-44684066-DEL-789 |
| chr12-44693293-DEL-6220 | chr12-44693281-DEL-6221 |
| chr12-44712332-DEL-52 | chr12-44712333-DEL-52 |
| chr12-44720624-DEL-832 | chr12-44720625-DEL-832 |
| chr12-44723141-DEL-6410 | chr12-44723142-DEL-6410 |
| chr12-44747244-DEL-487 | chr12-44747245-DEL-487 |
| chr12-44758947-DEL-76 | chr12-44758944-DEL-80 |
| chr12-44784424-DEL-214 | chr12-44784425-DEL-214 |
| chr12-44786440-DEL-938 | chr12-44786441-DEL-938 |
| chr12-44789734-DEL-1869 | chr12-44789735-DEL-1869 |
| chr12-44842380-DEL-214 | chr12-44842380-DEL-214 |
| chr12-44843193-DEL-61 | chr12-44843194-DEL-61 |
| chr12-44880137-DEL-938 | chr12-44880138-DEL-938 |
| chr12-44887781-DEL-132 | chr12-44887782-DEL-132 |
| chr12-44892539-DEL-753 | chr12-44892539-DEL-753 |
| chr12-44893398-DEL-3682 | chr12-44893402-DEL-3670 |
| chr12-44921827-DEL-164 | chr12-44921828-DEL-164 |
| chr12-44937333-DEL-2270 | chr12-44937334-DEL-2270 |
| chr12-44950625-DEL-156 | chr12-44950626-DEL-156 |
| chr12-44965984-DEL-70 | chr12-44965985-DEL-70 |
| chr12-44970380-DEL-292 | chr12-44970381-DEL-292 |
| chr12-44982603-DEL-280 | chr12-44982602-DEL-281 |
| chr12-45014607-DEL-297 | chr12-45014608-DEL-297 |
| chr12-45033579-DEL-51 | chr12-45033580-DEL-52 |
| chr12-45037087-DEL-150 | chr12-45037088-DEL-150 |
| chr12-45064826-DEL-899 | chr12-45064824-DEL-896 |
| chr12-45097806-DEL-429 | chr12-45097837-DEL-399 |
| chr12-45129414-DEL-74 | chr12-45129415-DEL-111 |
| chr12-45139899-DEL-146 | chr12-45139900-DEL-146 |
| chr12-45149936-DEL-54 | chr12-45149910-DEL-61 |
| chr12-45158225-DEL-172 | chr12-45158226-DEL-171 |
| chr12-45164581-DEL-58 | chr12-45164581-DEL-58 |
| chr12-45165739-DEL-1121 | chr12-45165740-DEL-1120 |
| chr12-45173549-DEL-82 | chr12-45173549-DEL-83 |
| chr12-45187895-DEL-60 | chr12-45187896-DEL-60 |
| chr12-45217104-DEL-54 | chr12-45217105-DEL-54 |
| chr12-45217918-DEL-400 | chr12-45217919-DEL-400 |
| chr12-45232627-DEL-88 | chr12-45232628-DEL-100 |

| | |
|---|---|
| chr12-45240779-DEL-3617 | chr12-45240780-DEL-3617 |
| chr12-45277410-DEL-116 | chr12-45277411-DEL-116 |
| chr12-45280277-DEL-93 | chr12-45280278-DEL-88 |
| chr12-45283051-DEL-71 | chr12-45283052-DEL-71 |
| chr12-45330779-DEL-107 | chr12-45330780-DEL-107 |
| chr12-45333475-DEL-82 | chr12-45333462-DEL-69 |
| chr12-45333546-DEL-510 | chr12-45333563-DEL-494 |
| chr12-45334426-DEL-52 | chr12-45334427-DEL-52 |
| chr12-45339584-DEL-779 | chr12-45339584-DEL-779 |
| chr12-45341640-DEL-362 | chr12-45341640-DEL-362 |
| chr12-45354030-DEL-518 | chr12-45354040-DEL-503 |
| chr12-45355469-DEL-1139 | chr12-45355469-DEL-1139 |
| chr12-45360587-DEL-1206 | chr12-45360587-DEL-1206 |
| chr12-45363957-DEL-344 | chr12-45363957-DEL-344 |
| chr12-45380445-DEL-2129 | chr12-45380446-DEL-2129 |
| chr12-45386443-DEL-740 | chr12-45386444-DEL-740 |
| chr12-45387802-DEL-77 | chr12-45387803-DEL-80 |
| chr12-45424639-DEL-233 | chr12-45424640-DEL-230 |
| chr12-45449109-DEL-62 | chr12-45449110-DEL-62 |
| chr12-45454789-DEL-660 | chr12-45454789-DEL-660 |
| chr12-45459445-DEL-56 | chr12-45459445-DEL-56 |
| chr12-45468067-DEL-486 | chr12-45468067-DEL-486 |
| chr12-45519307-DEL-1590 | chr12-45519308-DEL-1590 |
| chr12-45520438-DEL-54 | chr12-45520439-DEL-54 |
| chr12-45526939-DEL-936 | chr12-45526940-DEL-936 |
| chr12-45543458-DEL-54 | chr12-45543459-DEL-54 |
| chr12-45552585-DEL-56 | chr12-45552586-DEL-56 |
| chr12-45645679-DEL-197 | chr12-45645680-DEL-197 |
| chr12-45646856-DEL-53 | chr12-45646857-DEL-53 |
| chr12-45647367-DEL-1890 | chr12-45647368-DEL-1890 |
| chr12-45756415-DEL-53 | chr12-45756416-DEL-53 |
| chr12-45803071-DEL-3570 | chr12-45803072-DEL-3570 |
| chr12-45814483-DEL-7722 | chr12-45814484-DEL-7722 |
| chr12-45819123-DEL-1448 | chr12-45819124-DEL-1448 |
| chr12-45831844-DEL-57 | chr12-45831852-DEL-68 |
| chr12-45834477-DEL-143 | chr12-45834478-DEL-143 |
| chr12-45836188-DEL-235 | chr12-45836168-DEL-209 |
| chr12-45845416-DEL-121 | chr12-45845419-DEL-119 |
| chr12-45853020-DEL-935 | chr12-45853021-DEL-935 |
| chr12-45863373-DEL-1505 | chr12-45863374-DEL-1505 |
| chr12-45903529-DEL-335 | chr12-45903530-DEL-335 |
| chr12-45935062-DEL-1521 | chr12-45935063-DEL-1521 |
| chr12-45936691-DEL-435 | chr12-45936691-DEL-435 |
| chr12-45941437-DEL-333 | chr12-45941437-DEL-333 |
| chr12-45958536-DEL-101 | chr12-45958567-DEL-94 |
| chr12-45968911-DEL-63 | chr12-45968913-DEL-62 |
| chr12-45995957-DEL-13779 | chr12-45995958-DEL-13779 |
| chr12-46030495-DEL-258 | chr12-46030496-DEL-258 |
| chr12-46060830-DEL-926 | chr12-46060831-DEL-926 |
| chr12-46091378-DEL-7005 | chr12-46091371-DEL-7006 |
| chr12-46091460-DEL-402 | chr12-46091461-DEL-402 |
| chr12-46104621-DEL-63 | chr12-46104621-DEL-63 |
| chr12-46110578-DEL-633 | chr12-46110579-DEL-633 |
| chr12-46118450-DEL-2811 | chr12-46118451-DEL-2811 |
| chr12-46180795-DEL-62 | chr12-46180789-DEL-63 |
| chr12-46182896-DEL-78 | chr12-46182903-DEL-72 |
| chr12-46202308-DEL-1181 | chr12-46202307-DEL-1181 |
| chr12-46230927-DEL-1174 | chr12-46230928-DEL-1174 |
| chr12-46236208-DEL-59 | chr12-46236209-DEL-59 |
| chr12-46272161-DEL-51 | chr12-46272161-DEL-51 |
| chr12-46298717-DEL-3109 | chr12-46298718-DEL-3109 |
| chr12-46317264-DEL-2820 | chr12-46317265-DEL-2820 |
| chr12-46357874-DEL-210 | chr12-46357875-DEL-210 |
| chr12-46369577-DEL-55 | chr12-46369578-DEL-55 |
| chr12-46391079-DEL-298 | chr12-46391080-DEL-298 |
| chr12-46409900-DEL-92 | chr12-46409909-DEL-104 |
| chr12-46452980-DEL-2445 | chr12-46452981-DEL-2445 |
| chr12-46456856-DEL-436 | chr12-46456853-DEL-436 |
| chr12-46459604-DEL-1140 | chr12-46459605-DEL-1140 |
| chr12-46464800-DEL-57 | chr12-46464801-DEL-57 |
| chr12-46469986-DEL-62 | chr12-46469987-DEL-64 |
| chr12-46472411-DEL-116 | chr12-46472411-DEL-117 |
| chr12-46481066-DEL-297 | chr12-46481067-DEL-297 |
| chr12-46486480-DEL-2076 | chr12-46486481-DEL-2076 |
| chr12-46497386-DEL-60 | chr12-46497345-DEL-57 |
| chr12-46499173-DEL-1987 | chr12-46499174-DEL-1987 |
| chr12-46577244-DEL-60 | chr12-46577243-DEL-61 |
| chr12-46685104-DEL-532 | chr12-46685105-DEL-532 |
| chr12-46687621-DEL-83 | chr12-46687616-DEL-82 |
| chr12-46754311-DEL-980 | chr12-46754313-DEL-981 |

| | |
|---|---|
| chr12-46778647-DEL-422 | chr12-46778648-DEL-422 |
| chr12-46785367-DEL-62 | chr12-46785374-DEL-58 |
| chr12-46857414-DEL-112 | chr12-46857415-DEL-112 |
| chr12-46867084-DEL-795 | chr12-46867085-DEL-795 |
| chr12-46894590-DEL-223 | chr12-46894591-DEL-223 |
| chr12-46905048-DEL-93 | chr12-46905045-DEL-57 |
| chr12-46915997-DEL-70 | chr12-46915998-DEL-70 |
| chr12-46979152-DEL-68 | chr12-46979153-DEL-68 |
| chr12-46990593-DEL-308 | chr12-46990595-DEL-307 |
| chr12-47016618-DEL-374 | chr12-47016619-DEL-373 |
| chr12-47030597-DEL-72 | chr12-47030602-DEL-68 |
| chr12-47037494-DEL-62 | chr12-47037502-DEL-55 |
| chr12-47117853-DEL-2816 | chr12-47117854-DEL-2816 |
| chr12-47219692-DEL-69 | chr12-47219679-DEL-69 |
| chr12-47226443-DEL-125 | chr12-47226444-DEL-125 |
| chr12-47276874-DEL-5836 | chr12-47276875-DEL-5836 |
| chr12-47290233-DEL-56 | chr12-47290233-DEL-60 |
| chr12-47296967-DEL-400 | chr12-47296968-DEL-400 |
| chr12-47301782-DEL-65 | chr12-47301790-DEL-67 |
| chr12-47304160-DEL-63 | chr12-47304161-DEL-63 |
| chr12-47308638-DEL-88 | chr12-47308639-DEL-88 |
| chr12-47317125-DEL-254 | chr12-47317126-DEL-254 |
| chr12-47326207-DEL-78 | chr12-47326208-DEL-78 |
| chr12-47326800-DEL-6894 | chr12-47326801-DEL-6894 |
| chr12-47341632-DEL-116 | chr12-47341633-DEL-116 |
| chr12-47346121-DEL-1075 | chr12-47346118-DEL-1078 |
| chr12-47348000-DEL-1554 | chr12-47348001-DEL-1554 |
| chr12-47350416-DEL-936 | chr12-47350416-DEL-936 |
| chr12-47370294-DEL-58 | chr12-47370302-DEL-64 |
| chr12-47371975-DEL-295 | chr12-47371976-DEL-295 |
| chr12-47375323-DEL-3413 | chr12-47375324-DEL-3413 |
| chr12-47382387-DEL-377 | chr12-47382388-DEL-377 |
| chr12-47385752-DEL-2772 | chr12-47385749-DEL-2773 |
| chr12-47419745-DEL-52 | chr12-47419746-DEL-52 |
| chr12-47425763-DEL-390 | chr12-47425764-DEL-390 |
| chr12-47430791-DEL-6439 | chr12-47430792-DEL-6439 |
| chr12-47464497-DEL-327 | chr12-47464498-DEL-327 |
| chr12-47479864-DEL-325 | chr12-47479865-DEL-325 |
| chr12-47484718-DEL-430 | chr12-47484719-DEL-430 |
| chr12-47506308-DEL-62 | chr12-47506333-DEL-50 |
| chr12-47541406-DEL-1176 | chr12-47541407-DEL-1176 |
| chr12-47557252-DEL-486 | chr12-47557253-DEL-486 |
| chr12-47569957-DEL-61 | chr12-47569959-DEL-60 |
| chr12-47572810-DEL-6400 | chr12-47572811-DEL-6400 |
| chr12-47601378-DEL-209 | chr12-47601379-DEL-209 |
| chr12-47625689-DEL-6743 | chr12-47625690-DEL-6743 |
| chr12-47747899-DEL-69 | chr12-47747910-DEL-78 |
| chr12-47761809-DEL-4658 | chr12-47761810-DEL-4658 |
| chr12-47802787-DEL-203 | chr12-47802788-DEL-203 |
| chr12-47806286-DEL-102 | chr12-47806287-DEL-102 |
| chr12-47814924-DEL-209 | chr12-47814925-DEL-209 |
| chr12-47816362-DEL-79 | chr12-47816364-DEL-78 |
| chr12-47847988-DEL-95 | chr12-47847987-DEL-99 |
| chr12-47859653-DEL-534 | chr12-47859654-DEL-534 |
| chr12-47930618-DEL-428 | chr12-47930619-DEL-428 |
| chr12-47962089-DEL-51 | chr12-47962092-DEL-50 |
| chr12-47982232-DEL-80 | chr12-47982249-DEL-81 |
| chr12-47996237-DEL-100 | chr12-47996272-DEL-54 |
| chr12-48030485-DEL-409 | chr12-48030486-DEL-409 |
| chr12-48048431-DEL-506 | chr12-48048462-DEL-505 |
| chr12-48089345-DEL-7321 | chr12-48089346-DEL-7321 |
| chr12-48090276-DEL-496 | chr12-48090277-DEL-496 |
| chr12-48111647-DEL-1115 | chr12-48111650-DEL-1114 |
| chr12-48160118-DEL-828 | chr12-48160119-DEL-828 |
| chr12-48162475-DEL-332 | chr12-48162476-DEL-333 |
| chr12-48177195-DEL-658 | chr12-48177191-DEL-659 |
| chr12-48181390-DEL-78 | chr12-48181413-DEL-98 |
| chr12-48210941-DEL-1891 | chr12-48210942-DEL-1891 |
| chr12-48229567-DEL-1507 | chr12-48229568-DEL-1507 |
| chr12-48237985-DEL-55 | chr12-48237981-DEL-70 |
| chr12-48243537-DEL-54 | chr12-48243531-DEL-66 |
| chr12-48264766-DEL-158 | chr12-48264767-DEL-158 |
| chr12-48298722-DEL-65 | chr12-48298723-DEL-65 |
| chr12-48316168-DEL-2115 | chr12-48316169-DEL-2115 |
| chr12-48347601-DEL-71 | chr12-48347602-DEL-82 |
| chr12-48350363-DEL-77 | chr12-48350364-DEL-77 |
| chr12-48352465-DEL-209 | chr12-48352478-DEL-197 |
| chr12-48387124-DEL-6096 | chr12-48387125-DEL-6096 |
| chr12-48397052-DEL-58 | chr12-48397052-DEL-58 |
| chr12-48406542-DEL-73 | chr12-48406543-DEL-73 |

| | |
|---|---|
| chr12-48421532-DEL-1888 | chr12-48421533-DEL-1888 |
| chr12-48427352-DEL-2223 | chr12-48427353-DEL-2223 |
| chr12-48440241-DEL-74 | chr12-48440242-DEL-74 |
| chr12-48453421-DEL-1886 | chr12-48453422-DEL-1886 |
| chr12-48471972-DEL-201 | chr12-48471973-DEL-201 |
| chr12-48475612-DEL-53 | chr12-48475592-DEL-54 |
| chr12-48486188-DEL-149 | chr12-48486189-DEL-149 |
| chr12-48490330-DEL-70 | chr12-48490331-DEL-70 |
| chr12-48511497-DEL-70 | chr12-48511500-DEL-68 |
| chr12-48516386-DEL-67 | chr12-48516387-DEL-67 |
| chr12-48548974-DEL-901 | chr12-48548975-DEL-901 |
| chr12-48575760-DEL-90 | chr12-48575762-DEL-60 |
| chr12-48578959-DEL-7141 | chr12-48578960-DEL-7141 |
| chr12-48600935-DEL-5830 | chr12-48600935-DEL-5830 |
| chr12-48615209-DEL-218 | chr12-48615210-DEL-218 |
| chr12-48717418-DEL-581 | chr12-48717421-DEL-582 |
| chr12-48776958-DEL-114 | chr12-48776959-DEL-114 |
| chr12-48778295-DEL-186 | chr12-48778300-DEL-185 |
| chr12-48781361-DEL-72 | chr12-48781362-DEL-56 |
| chr12-48825841-DEL-7504 | chr12-48825842-DEL-7504 |
| chr12-48846978-DEL-69 | chr12-48846949-DEL-90 |
| chr12-48852786-DEL-146 | chr12-48852780-DEL-110 |
| chr12-48861118-DEL-259 | chr12-48861119-DEL-259 |
| chr12-48879653-DEL-54 | chr12-48879654-DEL-54 |
| chr12-48882276-DEL-315 | chr12-48882277-DEL-315 |
| chr12-48887291-DEL-712 | chr12-48887289-DEL-713 |
| chr12-48888379-DEL-58 | chr12-48888390-DEL-74 |
| chr12-48889719-DEL-6649 | chr12-48889718-DEL-6651 |
| chr12-48904529-DEL-52 | chr12-48904518-DEL-72 |
| chr12-48945578-DEL-54 | chr12-48945579-DEL-54 |
| chr12-48958775-DEL-5618 | chr12-48958775-DEL-5618 |
| chr12-48976826-DEL-112 | chr12-48976827-DEL-112 |
| chr12-48996983-DEL-53 | chr12-48996993-DEL-53 |
| chr12-49029396-DEL-1944 | chr12-49029397-DEL-1944 |
| chr12-49038372-DEL-185 | chr12-49038373-DEL-185 |
| chr12-49041567-DEL-7107 | chr12-49041568-DEL-7107 |
| chr12-49097914-DEL-6840 | chr12-49097915-DEL-6840 |
| chr12-49123101-DEL-214 | chr12-49123102-DEL-214 |
| chr12-49189278-DEL-345 | chr12-49189279-DEL-345 |
| chr12-49195259-DEL-7104 | chr12-49195260-DEL-7104 |
| chr12-49209262-DEL-646 | chr12-49209263-DEL-646 |
| chr12-49213194-DEL-68 | chr12-49213188-DEL-56 |
| chr12-49239497-DEL-6751 | chr12-49239498-DEL-6751 |
| chr12-49246993-DEL-85 | chr12-49246994-DEL-85 |
| chr12-49253916-DEL-63 | chr12-49253921-DEL-57 |
| chr12-49260904-DEL-9224 | chr12-49260905-DEL-9224 |
| chr12-49266532-DEL-3064 | chr12-49266533-DEL-3064 |
| chr12-49277174-DEL-1257 | chr12-49277175-DEL-1257 |
| chr12-49290177-DEL-51 | chr12-49290172-DEL-66 |
| chr12-49314160-DEL-86 | chr12-49314197-DEL-60 |
| chr12-49318750-DEL-1396 | chr12-49318751-DEL-1396 |
| chr12-49325622-DEL-68 | chr12-49325614-DEL-58 |
| chr12-49328767-DEL-202 | chr12-49328767-DEL-202 |
| chr12-49344884-DEL-1573 | chr12-49344885-DEL-1573 |
| chr12-49349375-DEL-1251 | chr12-49349376-DEL-1238 |
| chr12-49351132-DEL-3562 | chr12-49351133-DEL-3562 |
| chr12-49368640-DEL-193 | chr12-49368641-DEL-193 |
| chr12-49392759-DEL-939 | chr12-49392760-DEL-939 |
| chr12-49409610-DEL-4931 | chr12-49409611-DEL-4931 |
| chr12-49452322-DEL-400 | chr12-49452323-DEL-400 |
| chr12-49545433-DEL-59 | chr12-49545434-DEL-59 |
| chr12-49547358-DEL-2142 | chr12-49547359-DEL-2142 |
| chr12-49548399-DEL-6461 | chr12-49548400-DEL-6461 |
| chr12-49573835-DEL-6365 | chr12-49573836-DEL-6365 |
| chr12-49582258-DEL-362 | chr12-49582259-DEL-362 |
| chr12-49586080-DEL-64 | chr12-49586080-DEL-64 |
| chr12-49593787-DEL-675 | chr12-49593792-DEL-672 |
| chr12-49595413-DEL-2831 | chr12-49595414-DEL-2831 |
| chr12-49602034-DEL-287 | chr12-49602034-DEL-287 |
| chr12-49614534-DEL-535 | chr12-49614548-DEL-522 |
| chr12-49619870-DEL-64 | chr12-49619871-DEL-64 |
| chr12-49627875-DEL-272 | chr12-49627876-DEL-272 |
| chr12-49640423-DEL-72 | chr12-49640430-DEL-50 |
| chr12-49646779-DEL-57 | chr12-49646766-DEL-72 |
| chr12-49652893-DEL-6457 | chr12-49652894-DEL-6457 |
| chr12-49664417-DEL-61 | chr12-49664417-DEL-60 |
| chr12-49664651-DEL-839 | chr12-49664652-DEL-839 |
| chr12-49748621-DEL-6660 | chr12-49748619-DEL-6656 |
| chr12-49748651-DEL-212 | chr12-49748652-DEL-212 |
| chr12-49757264-DEL-360 | chr12-49757265-DEL-360 |

| | |
|---|---|
| chr12-49762640-DEL-246 | chr12-49762641-DEL-246 |
| chr12-49789287-DEL-230 | chr12-49789288-DEL-230 |
| chr12-49813058-DEL-291 | chr12-49813059-DEL-291 |
| chr12-49837024-DEL-931 | chr12-49837024-DEL-932 |
| chr12-49872668-DEL-63 | chr12-49872693-DEL-72 |
| chr12-49917496-DEL-79 | chr12-49917499-DEL-75 |
| chr12-49926788-DEL-2790 | chr12-49926789-DEL-2790 |
| chr12-49928164-DEL-222 | chr12-49928165-DEL-222 |
| chr12-49929808-DEL-861 | chr12-49929809-DEL-861 |
| chr12-49951235-DEL-400 | chr12-49951236-DEL-400 |
| chr12-49960602-DEL-91 | chr12-49960598-DEL-110 |
| chr12-49976127-DEL-147 | chr12-49976140-DEL-142 |
| chr12-49985689-DEL-78 | chr12-49985682-DEL-80 |
| chr12-49989020-DEL-461 | chr12-49989020-DEL-461 |
| chr12-50068642-DEL-1798 | chr12-50068643-DEL-1798 |
| chr12-50072433-DEL-430 | chr12-50072434-DEL-430 |
| chr12-50090083-DEL-63 | chr12-50090073-DEL-55 |
| chr12-50114525-DEL-1234 | chr12-50114526-DEL-1233 |
| chr12-50161053-DEL-72 | chr12-50161054-DEL-72 |
| chr12-50180229-DEL-1855 | chr12-50180230-DEL-1855 |
| chr12-50190672-DEL-64 | chr12-50190673-DEL-64 |
| chr12-50282041-DEL-656 | chr12-50282042-DEL-656 |
| chr12-50288971-DEL-6795 | chr12-50288972-DEL-6795 |
| chr12-50300720-DEL-3989 | chr12-50300717-DEL-3985 |
| chr12-50329704-DEL-102 | chr12-50329705-DEL-99 |
| chr12-50363276-DEL-1007 | chr12-50363277-DEL-1007 |
| chr12-50372366-DEL-71 | chr12-50372367-DEL-71 |
| chr12-50386106-DEL-416 | chr12-50386107-DEL-416 |
| chr12-50401143-DEL-1004 | chr12-50401144-DEL-1004 |
| chr12-50422985-DEL-54 | chr12-50422985-DEL-54 |
| chr12-50469734-DEL-96 | chr12-50469734-DEL-92 |
| chr12-50482160-DEL-135 | chr12-50482161-DEL-135 |
| chr12-50487125-DEL-216 | chr12-50487126-DEL-216 |
| chr12-50488209-DEL-171 | chr12-50488203-DEL-151 |
| chr12-50520926-DEL-224 | chr12-50520927-DEL-224 |
| chr12-50545446-DEL-71 | chr12-50545447-DEL-71 |
| chr12-50564477-DEL-82 | chr12-50564478-DEL-82 |
| chr12-50566110-DEL-196 | chr12-50566111-DEL-196 |
| chr12-50607298-DEL-236 | chr12-50607299-DEL-236 |
| chr12-50627807-DEL-6372 | chr12-50627809-DEL-6358 |
| chr12-50634673-DEL-249 | chr12-50634674-DEL-249 |
| chr12-50663134-DEL-130 | chr12-50663157-DEL-84 |
| chr12-50670017-DEL-503 | chr12-50670018-DEL-503 |
| chr12-50732706-DEL-6147 | chr12-50732707-DEL-6147 |
| chr12-50752354-DEL-460 | chr12-50752355-DEL-460 |
| chr12-50769257-DEL-75 | chr12-50769258-DEL-75 |
| chr12-50773065-DEL-57 | chr12-50773065-DEL-57 |
| chr12-50789427-DEL-912 | chr12-50789428-DEL-912 |
| chr12-50813082-DEL-57 | chr12-50813083-DEL-57 |
| chr12-50813151-DEL-782 | chr12-50813152-DEL-782 |
| chr12-50825315-DEL-208 | chr12-50825316-DEL-208 |
| chr12-50831383-DEL-5371 | chr12-50831384-DEL-5371 |
| chr12-50854399-DEL-567 | chr12-50854374-DEL-557 |
| chr12-50910411-DEL-1329 | chr12-50910412-DEL-1329 |
| chr12-50913343-DEL-1636 | chr12-50913344-DEL-1636 |
| chr12-50930917-DEL-330 | chr12-50930916-DEL-334 |
| chr12-50942542-DEL-82 | chr12-50942543-DEL-82 |
| chr12-50962896-DEL-73 | chr12-50962897-DEL-73 |
| chr12-50967417-DEL-352 | chr12-50967418-DEL-352 |
| chr12-50992685-DEL-71 | chr12-50992686-DEL-71 |
| chr12-51022005-DEL-5553 | chr12-51022006-DEL-5553 |
| chr12-51035949-DEL-76 | chr12-51035964-DEL-50 |
| chr12-51039364-DEL-364 | chr12-51039365-DEL-364 |
| chr12-51080003-DEL-3397 | chr12-51080004-DEL-3397 |
| chr12-51080303-DEL-1498 | chr12-51080302-DEL-1498 |
| chr12-51084665-DEL-152 | chr12-51084621-DEL-174 |
| chr12-51095086-DEL-822 | chr12-51095087-DEL-822 |
| chr12-51096276-DEL-1016 | chr12-51096277-DEL-1016 |
| chr12-51116846-DEL-52 | chr12-51116847-DEL-52 |
| chr12-51133322-DEL-127 | chr12-51133323-DEL-127 |
| chr12-51138404-DEL-108 | chr12-51138405-DEL-108 |
| chr12-51147424-DEL-181 | chr12-51147408-DEL-181 |
| chr12-51161151-DEL-504 | chr12-51161152-DEL-504 |
| chr12-51166092-DEL-2024 | chr12-51166092-DEL-2024 |
| chr12-51183527-DEL-395 | chr12-51183528-DEL-395 |
| chr12-51214834-DEL-6375 | chr12-51214835-DEL-6375 |
| chr12-51251115-DEL-3766 | chr12-51251116-DEL-3766 |
| chr12-51264992-DEL-127 | chr12-51264993-DEL-127 |
| chr12-51270667-DEL-1747 | chr12-51270668-DEL-1747 |
| chr12-51272121-DEL-1477 | chr12-51272122-DEL-1477 |

| | |
|---|---|
| chr12-51334806-DEL-114 | chr12-51334796-DEL-93 |
| chr12-51375342-DEL-206 | chr12-51375343-DEL-206 |
| chr12-51384340-DEL-1287 | chr12-51384308-DEL-1320 |
| chr12-51396843-DEL-106 | chr12-51396844-DEL-106 |
| chr12-51426867-DEL-927 | chr12-51426868-DEL-927 |
| chr12-51454030-DEL-2677 | chr12-51454031-DEL-2677 |
| chr12-51471589-DEL-62 | chr12-51471590-DEL-62 |
| chr12-51472002-DEL-461 | chr12-51472003-DEL-461 |
| chr12-51500321-DEL-340 | chr12-51500322-DEL-340 |
| chr12-51513435-DEL-2061 | chr12-51513436-DEL-2061 |
| chr12-51525891-DEL-187 | chr12-51525892-DEL-187 |
| chr12-51533043-DEL-5406 | chr12-51533044-DEL-5406 |
| chr12-51561549-DEL-117 | chr12-51561549-DEL-116 |
| chr12-51564977-DEL-74 | chr12-51564978-DEL-74 |
| chr12-51572973-DEL-934 | chr12-51572974-DEL-934 |
| chr12-51574554-DEL-78 | chr12-51574543-DEL-79 |
| chr12-51578205-DEL-205 | chr12-51578206-DEL-205 |
| chr12-51591483-DEL-320 | chr12-51591484-DEL-320 |
| chr12-51604811-DEL-125 | chr12-51604812-DEL-125 |
| chr12-51623623-DEL-177 | chr12-51623624-DEL-177 |
| chr12-51629206-DEL-627 | chr12-51629207-DEL-627 |
| chr12-51633548-DEL-118 | chr12-51633549-DEL-118 |
| chr12-51643969-DEL-51 | chr12-51643970-DEL-51 |
| chr12-51670733-DEL-2158 | chr12-51670734-DEL-2158 |
| chr12-51704556-DEL-64 | chr12-51704557-DEL-64 |
| chr12-51730009-DEL-62 | chr12-51730010-DEL-62 |
| chr12-51760851-DEL-237 | chr12-51760852-DEL-237 |
| chr12-51774915-DEL-196 | chr12-51774916-DEL-196 |
| chr12-51778485-DEL-423 | chr12-51778486-DEL-423 |
| chr12-51796860-DEL-54 | chr12-51796860-DEL-54 |
| chr12-51811979-DEL-62 | chr12-51811980-DEL-62 |
| chr12-51820059-DEL-68 | chr12-51820060-DEL-68 |
| chr12-51831421-DEL-772 | chr12-51831425-DEL-771 |
| chr12-51842658-DEL-64 | chr12-51842665-DEL-58 |
| chr12-51863045-DEL-126 | chr12-51863046-DEL-126 |
| chr12-51900569-DEL-442 | chr12-51900570-DEL-442 |
| chr12-51906781-DEL-66 | chr12-51906739-DEL-90 |
| chr12-51915303-DEL-198 | chr12-51915304-DEL-198 |
| chr12-51932307-DEL-193 | chr12-51932308-DEL-193 |
| chr12-51945344-DEL-66 | chr12-51945345-DEL-66 |
| chr12-51953482-DEL-194 | chr12-51953483-DEL-194 |
| chr12-51988058-DEL-211 | chr12-51988059-DEL-211 |
| chr12-51996313-DEL-56 | chr12-51996314-DEL-56 |
| chr12-52011892-DEL-55 | chr12-52011842-DEL-103 |
| chr12-52013868-DEL-535 | chr12-52013869-DEL-535 |
| chr12-52020370-DEL-250 | chr12-52020371-DEL-250 |
| chr12-52059296-DEL-64 | chr12-52059297-DEL-64 |
| chr12-52068082-DEL-68 | chr12-52068079-DEL-72 |
| chr12-52086579-DEL-7960 | chr12-52086580-DEL-7960 |
| chr12-52089098-DEL-96 | chr12-52089111-DEL-95 |
| chr12-52105564-DEL-58 | chr12-52105596-DEL-62 |
| chr12-52115536-DEL-84 | chr12-52115567-DEL-54 |
| chr12-52125663-DEL-772 | chr12-52125664-DEL-772 |
| chr12-52127612-DEL-7435 | chr12-52127613-DEL-7435 |
| chr12-52156572-DEL-6660 | chr12-52156573-DEL-6660 |
| chr12-52173406-DEL-75 | chr12-52173407-DEL-75 |
| chr12-52219447-DEL-3985 | chr12-52219448-DEL-3985 |
| chr12-52247872-DEL-643 | chr12-52247872-DEL-643 |
| chr12-52253990-DEL-204 | chr12-52253991-DEL-204 |
| chr12-52278322-DEL-60 | chr12-52278323-DEL-60 |
| chr12-52308133-DEL-643 | chr12-52308134-DEL-643 |
| chr12-52347495-DEL-58 | chr12-52347496-DEL-58 |
| chr12-52357819-DEL-268 | chr12-52357820-DEL-268 |
| chr12-52380498-DEL-258 | chr12-52380499-DEL-226 |
| chr12-52420898-DEL-135 | chr12-52420899-DEL-135 |
| chr12-52463651-DEL-1168 | chr12-52463651-DEL-1167 |
| chr12-52474336-DEL-113 | chr12-52474336-DEL-116 |
| chr12-52506037-DEL-2129 | chr12-52506036-DEL-2129 |
| chr12-52528730-DEL-74 | chr12-52528731-DEL-58 |
| chr12-52533440-DEL-64 | chr12-52533444-DEL-62 |
| chr12-52537727-DEL-114 | chr12-52537728-DEL-114 |
| chr12-52561592-DEL-100 | chr12-52561593-DEL-100 |
| chr12-52598536-DEL-113 | chr12-52598537-DEL-113 |
| chr12-52621440-DEL-535 | chr12-52621441-DEL-535 |
| chr12-52631505-DEL-79 | chr12-52631506-DEL-79 |
| chr12-52648239-DEL-311 | chr12-52648240-DEL-311 |
| chr12-52672004-DEL-56 | chr12-52672001-DEL-63 |
| chr12-52681483-DEL-80 | chr12-52681484-DEL-80 |
| chr12-52698152-DEL-943 | chr12-52698153-DEL-943 |
| chr12-52699212-DEL-1021 | chr12-52699213-DEL-1021 |

| | |
|---|---|
| chr12-52719548-DEL-115 | chr12-52719549-DEL-115 |
| chr12-52734032-DEL-56 | chr12-52734033-DEL-56 |
| chr12-52735493-DEL-53 | chr12-52735494-DEL-53 |
| chr12-52759583-DEL-1177 | chr12-52759584-DEL-1177 |
| chr12-52763875-DEL-140 | chr12-52763876-DEL-140 |
| chr12-52770364-DEL-57 | chr12-52770365-DEL-51 |
| chr12-52771195-DEL-208 | chr12-52771196-DEL-208 |
| chr12-52776881-DEL-5211 | chr12-52776882-DEL-5211 |
| chr12-52794360-DEL-1030 | chr12-52794379-DEL-1012 |
| chr12-52812494-DEL-1727 | chr12-52812495-DEL-1727 |
| chr12-52814749-DEL-93 | chr12-52814741-DEL-72 |
| chr12-52815287-DEL-92 | chr12-52815288-DEL-82 |
| chr12-52822457-DEL-60 | chr12-52822457-DEL-66 |
| chr12-52840500-DEL-57 | chr12-52840501-DEL-57 |
| chr12-52844848-DEL-187 | chr12-52844849-DEL-187 |
| chr12-52849247-DEL-81 | chr12-52849250-DEL-79 |
| chr12-52855038-DEL-1910 | chr12-52855039-DEL-1910 |
| chr12-52872180-DEL-863 | chr12-52872181-DEL-863 |
| chr12-52874646-DEL-69 | chr12-52874649-DEL-68 |
| chr12-52884260-DEL-93 | chr12-52884261-DEL-93 |
| chr12-52889177-DEL-192 | chr12-52889178-DEL-192 |
| chr12-52890602-DEL-2253 | chr12-52890603-DEL-2253 |
| chr12-52891713-DEL-911 | chr12-52891714-DEL-911 |
| chr12-52902381-DEL-79 | chr12-52902392-DEL-76 |
| chr12-52905871-DEL-1031 | chr12-52905901-DEL-999 |
| chr12-52907815-DEL-51 | chr12-52907816-DEL-51 |
| chr12-52953126-DEL-74 | chr12-52953127-DEL-74 |
| chr12-52977587-DEL-85 | chr12-52977588-DEL-85 |
| chr12-52980700-DEL-62 | chr12-52980701-DEL-62 |
| chr12-52987307-DEL-54 | chr12-52987307-DEL-54 |
| chr12-52989179-DEL-93 | chr12-52989180-DEL-93 |
| chr12-52997998-DEL-1545 | chr12-52997982-DEL-1556 |
| chr12-53000395-DEL-761 | chr12-53000395-DEL-762 |
| chr12-53025900-DEL-75 | chr12-53025901-DEL-75 |
| chr12-53151617-DEL-213 | chr12-53151618-DEL-213 |
| chr12-53156822-DEL-609 | chr12-53156823-DEL-609 |
| chr12-53178279-DEL-75 | chr12-53178280-DEL-75 |
| chr12-53198895-DEL-87 | chr12-53198896-DEL-87 |
| chr12-53228447-DEL-2220 | chr12-53228450-DEL-2217 |
| chr12-53237212-DEL-527 | chr12-53237213-DEL-527 |
| chr12-53240578-DEL-84 | chr12-53240579-DEL-84 |
| chr12-53244899-DEL-135 | chr12-53244900-DEL-135 |
| chr12-53330944-DEL-98 | chr12-53330945-DEL-97 |
| chr12-53395738-DEL-148 | chr12-53395739-DEL-148 |
| chr12-53511661-DEL-70 | chr12-53511662-DEL-70 |
| chr12-53531141-DEL-58 | chr12-53531142-DEL-58 |
| chr12-53532571-DEL-52 | chr12-53532569-DEL-50 |
| chr12-53536295-DEL-163 | chr12-53536296-DEL-161 |
| chr12-53539174-DEL-52 | chr12-53539175-DEL-52 |
| chr12-53563412-DEL-208 | chr12-53563419-DEL-170 |
| chr12-53629106-DEL-67 | chr12-53629106-DEL-68 |
| chr12-53675807-DEL-135 | chr12-53675803-DEL-139 |
| chr12-53704215-DEL-488 | chr12-53704216-DEL-488 |
| chr12-53743315-DEL-72 | chr12-53743316-DEL-72 |
| chr12-53761155-DEL-4713 | chr12-53761156-DEL-4713 |
| chr12-53765180-DEL-61 | chr12-53765180-DEL-60 |
| chr12-53768042-DEL-60 | chr12-53768043-DEL-60 |
| chr12-53772578-DEL-333 | chr12-53772579-DEL-333 |
| chr12-53809942-DEL-58 | chr12-53809943-DEL-58 |
| chr12-53811045-DEL-63 | chr12-53811046-DEL-63 |
| chr12-53830713-DEL-55 | chr12-53830714-DEL-55 |
| chr12-53868800-DEL-56 | chr12-53868801-DEL-56 |
| chr12-53882893-DEL-52 | chr12-53882894-DEL-50 |
| chr12-53944977-DEL-997 | chr12-53944978-DEL-997 |
| chr12-53949419-DEL-1721 | chr12-53949420-DEL-1721 |
| chr12-53967674-DEL-447 | chr12-53967675-DEL-447 |
| chr12-53997478-DEL-129 | chr12-53997479-DEL-129 |
| chr12-54044693-DEL-58 | chr12-54044694-DEL-58 |
| chr12-54048541-DEL-124 | chr12-54048546-DEL-123 |
| chr12-54086437-DEL-199 | chr12-54086438-DEL-199 |
| chr12-54088455-DEL-73 | chr12-54088458-DEL-71 |
| chr12-54112888-DEL-99 | chr12-54112850-DEL-111 |
| chr12-54138717-DEL-70 | chr12-54138713-DEL-68 |
| chr12-54153447-DEL-416 | chr12-54153448-DEL-416 |
| chr12-54159886-DEL-540 | chr12-54159887-DEL-540 |
| chr12-54166716-DEL-285 | chr12-54166717-DEL-284 |
| chr12-54204214-DEL-62 | chr12-54204177-DEL-80 |
| chr12-54304904-DEL-154 | chr12-54304903-DEL-108 |
| chr12-54316146-DEL-942 | chr12-54316147-DEL-942 |
| chr12-54336327-DEL-434 | chr12-54336328-DEL-434 |

| | |
|---|---|
| chr12-54344838-DEL-195 | chr12-54344839-DEL-197 |
| chr12-54360792-DEL-56 | chr12-54360792-DEL-56 |
| chr12-54361852-DEL-119 | chr12-54361853-DEL-119 |
| chr12-54368528-DEL-498 | chr12-54368530-DEL-498 |
| chr12-54373504-DEL-60 | chr12-54373505-DEL-60 |
| chr12-54395996-DEL-168 | chr12-54395997-DEL-168 |
| chr12-54405287-DEL-3015 | chr12-54405288-DEL-3015 |
| chr12-54409826-DEL-514 | chr12-54409827-DEL-514 |
| chr12-54411959-DEL-4113 | chr12-54411960-DEL-4113 |
| chr12-54426034-DEL-52 | chr12-54426035-DEL-52 |
| chr12-54458188-DEL-60 | chr12-54458189-DEL-60 |
| chr12-54481678-DEL-450 | chr12-54481687-DEL-451 |
| chr12-54505391-DEL-155 | chr12-54505392-DEL-155 |
| chr12-54523707-DEL-54 | chr12-54523708-DEL-54 |
| chr12-54527156-DEL-215 | chr12-54527157-DEL-215 |
| chr12-54554269-DEL-95 | chr12-54554266-DEL-90 |
| chr12-54572778-DEL-238 | chr12-54572780-DEL-189 |
| chr12-54579770-DEL-299 | chr12-54579771-DEL-299 |
| chr12-54594844-DEL-108 | chr12-54594829-DEL-78 |
| chr12-54625473-DEL-328 | chr12-54625474-DEL-328 |
| chr12-54628610-DEL-1074 | chr12-54628611-DEL-1074 |
| chr12-54635958-DEL-332 | chr12-54635959-DEL-332 |
| chr12-54637225-DEL-54 | chr12-54637226-DEL-54 |
| chr12-54645447-DEL-431 | chr12-54645448-DEL-431 |
| chr12-54652577-DEL-193 | chr12-54652578-DEL-193 |
| chr12-54668533-DEL-9447 | chr12-54668534-DEL-9447 |
| chr12-54686310-DEL-337 | chr12-54686311-DEL-337 |
| chr12-54718633-DEL-216 | chr12-54718634-DEL-216 |
| chr12-54743791-DEL-70 | chr12-54743792-DEL-70 |
| chr12-54752761-DEL-182 | chr12-54752761-DEL-182 |
| chr12-54765705-DEL-60 | chr12-54765706-DEL-60 |
| chr12-54776420-DEL-65 | chr12-54776420-DEL-65 |
| chr12-54789555-DEL-173 | chr12-54789556-DEL-173 |
| chr12-54797167-DEL-215 | chr12-54797161-DEL-216 |
| chr12-54819112-DEL-176 | chr12-54819113-DEL-176 |
| chr12-54821004-DEL-8986 | chr12-54821005-DEL-8986 |
| chr12-54833128-DEL-76 | chr12-54833144-DEL-60 |
| chr12-54835336-DEL-478 | chr12-54835337-DEL-478 |
| chr12-54856804-DEL-63 | chr12-54856805-DEL-63 |
| chr12-54868661-DEL-89 | chr12-54868666-DEL-80 |
| chr12-54876895-DEL-73 | chr12-54876896-DEL-73 |
| chr12-54880649-DEL-206 | chr12-54880650-DEL-206 |
| chr12-54894896-DEL-228 | chr12-54894897-DEL-228 |
| chr12-54900045-DEL-58 | chr12-54900046-DEL-58 |
| chr12-54912103-DEL-60 | chr12-54912112-DEL-52 |
| chr12-54915846-DEL-207 | chr12-54915847-DEL-207 |
| chr12-54917712-DEL-1268 | chr12-54917713-DEL-1268 |
| chr12-54921271-DEL-64 | chr12-54921225-DEL-86 |
| chr12-54928878-DEL-207 | chr12-54928879-DEL-207 |
| chr12-54930343-DEL-164 | chr12-54930344-DEL-164 |
| chr12-54936984-DEL-320 | chr12-54936985-DEL-320 |
| chr12-54938079-DEL-112 | chr12-54938080-DEL-112 |
| chr12-54939315-DEL-382 | chr12-54939319-DEL-378 |
| chr12-54962275-DEL-215 | chr12-54962276-DEL-215 |
| chr12-54981280-DEL-230 | chr12-54981281-DEL-230 |
| chr12-54990163-DEL-74 | chr12-54990164-DEL-74 |
| chr12-55004295-DEL-158 | chr12-55004296-DEL-158 |
| chr12-55013563-DEL-76 | chr12-55013564-DEL-76 |
| chr12-55028763-DEL-320 | chr12-55028764-DEL-320 |
| chr12-55039120-DEL-269 | chr12-55039121-DEL-269 |
| chr12-55062032-DEL-190 | chr12-55062033-DEL-190 |
| chr12-55120813-DEL-215 | chr12-55120814-DEL-215 |
| chr12-55128939-DEL-131 | chr12-55128939-DEL-131 |
| chr12-55359903-DEL-214 | chr12-55359904-DEL-214 |
| chr12-55367489-DEL-171 | chr12-55367522-DEL-148 |
| chr12-55391053-DEL-1022 | chr12-55391054-DEL-1022 |
| chr12-55444716-DEL-70 | chr12-55444717-DEL-70 |
| chr12-55449340-DEL-101 | chr12-55449341-DEL-101 |
| chr12-55457721-DEL-88 | chr12-55457728-DEL-82 |
| chr12-55502385-DEL-164 | chr12-55502387-DEL-163 |
| chr12-55548861-DEL-305 | chr12-55548862-DEL-305 |
| chr12-55596116-DEL-51 | chr12-55596117-DEL-51 |
| chr12-55597398-DEL-152 | chr12-55597399-DEL-152 |
| chr12-55600690-DEL-88 | chr12-55600677-DEL-59 |
| chr12-55602171-DEL-554 | chr12-55602172-DEL-554 |
| chr12-55609767-DEL-210 | chr12-55609768-DEL-210 |
| chr12-55663724-DEL-247 | chr12-55663725-DEL-247 |
| chr12-55679383-DEL-7086 | chr12-55679384-DEL-7086 |
| chr12-55709911-DEL-197 | chr12-55709912-DEL-197 |
| chr12-55800262-DEL-56 | chr12-55800263-DEL-56 |

| | |
|---|---|
| chr12-55826751-DEL-821 | chr12-55826751-DEL-819 |
| chr12-55836506-DEL-56 | chr12-55836506-DEL-60 |
| chr12-55839124-DEL-1106 | chr12-55839125-DEL-1106 |
| chr12-55863818-DEL-140 | chr12-55863801-DEL-158 |
| chr12-55878558-DEL-53 | chr12-55878559-DEL-54 |
| chr12-55912368-DEL-94 | chr12-55912369-DEL-94 |
| chr12-55948865-DEL-312 | chr12-55948866-DEL-312 |
| chr12-55963774-DEL-75 | chr12-55963747-DEL-57 |
| chr12-56006822-DEL-416 | chr12-56006824-DEL-415 |
| chr12-56018315-DEL-144 | chr12-56018316-DEL-116 |
| chr12-56023565-DEL-66 | chr12-56023580-DEL-52 |
| chr12-56043730-DEL-52 | chr12-56043727-DEL-58 |
| chr12-56047815-DEL-138 | chr12-56047816-DEL-138 |
| chr12-56095091-DEL-5013 | chr12-56095092-DEL-5013 |
| chr12-56102635-DEL-70 | chr12-56102643-DEL-102 |
| chr12-56150956-DEL-6440 | chr12-56150957-DEL-6440 |
| chr12-56219183-DEL-91 | chr12-56219184-DEL-91 |
| chr12-56243192-DEL-635 | chr12-56243193-DEL-635 |
| chr12-56257635-DEL-119 | chr12-56257640-DEL-117 |
| chr12-56318114-DEL-161 | chr12-56318115-DEL-161 |
| chr12-56325061-DEL-56 | chr12-56325060-DEL-52 |
| chr12-56345599-DEL-68 | chr12-56345582-DEL-54 |
| chr12-56346960-DEL-237 | chr12-56346961-DEL-237 |
| chr12-56350983-DEL-1041 | chr12-56350984-DEL-1041 |
| chr12-56356251-DEL-182 | chr12-56356261-DEL-170 |
| chr12-56359684-DEL-355 | chr12-56359685-DEL-355 |
| chr12-56373124-DEL-178 | chr12-56373125-DEL-178 |
| chr12-56412346-DEL-217 | chr12-56412355-DEL-219 |
| chr12-56431450-DEL-56 | chr12-56431451-DEL-56 |
| chr12-56476422-DEL-678 | chr12-56476423-DEL-678 |
| chr12-56479656-DEL-340 | chr12-56479657-DEL-340 |
| chr12-56571963-DEL-357 | chr12-56571964-DEL-357 |
| chr12-56597949-DEL-52 | chr12-56597950-DEL-52 |
| chr12-56603414-DEL-108 | chr12-56603450-DEL-90 |
| chr12-56605018-DEL-68 | chr12-56605020-DEL-83 |
| chr12-56624710-DEL-351 | chr12-56624710-DEL-351 |
| chr12-56625101-DEL-6390 | chr12-56625102-DEL-6390 |
| chr12-56672966-DEL-3619 | chr12-56672967-DEL-3619 |
| chr12-56673684-DEL-231 | chr12-56673685-DEL-231 |
| chr12-56675333-DEL-494 | chr12-56675334-DEL-494 |
| chr12-56743183-DEL-52 | chr12-56743184-DEL-52 |
| chr12-56794633-DEL-4327 | chr12-56794634-DEL-4327 |
| chr12-56809457-DEL-581 | chr12-56809458-DEL-581 |
| chr12-56832891-DEL-191 | chr12-56832892-DEL-191 |
| chr12-56835424-DEL-287 | chr12-56835425-DEL-301 |
| chr12-56872637-DEL-54 | chr12-56872638-DEL-74 |
| chr12-56905875-DEL-241 | chr12-56905885-DEL-232 |
| chr12-56934284-DEL-64 | chr12-56934285-DEL-64 |
| chr12-56942399-DEL-58 | chr12-56942401-DEL-57 |
| chr12-56985624-DEL-114 | chr12-56985600-DEL-138 |
| chr12-56993676-DEL-1463 | chr12-56993660-DEL-1464 |
| chr12-57013023-DEL-1180 | chr12-57013024-DEL-1180 |
| chr12-57062001-DEL-62 | chr12-57061997-DEL-67 |
| chr12-57065010-DEL-75 | chr12-57065021-DEL-71 |
| chr12-57088731-DEL-79 | chr12-57088732-DEL-79 |
| chr12-57145502-DEL-245 | chr12-57145503-DEL-245 |
| chr12-57198420-DEL-59 | chr12-57198418-DEL-60 |
| chr12-57211669-DEL-575 | chr12-57211670-DEL-575 |
| chr12-57218258-DEL-109 | chr12-57218259-DEL-108 |
| chr12-57220547-DEL-1352 | chr12-57220536-DEL-1353 |
| chr12-57234103-DEL-1246 | chr12-57234104-DEL-1246 |
| chr12-57245287-DEL-54 | chr12-57245288-DEL-54 |
| chr12-57253743-DEL-61 | chr12-57253739-DEL-66 |
| chr12-57255931-DEL-59 | chr12-57255932-DEL-59 |
| chr12-57269108-DEL-189 | chr12-57269109-DEL-189 |
| chr12-57279488-DEL-99 | chr12-57279489-DEL-99 |
| chr12-57295687-DEL-791 | chr12-57295688-DEL-787 |
| chr12-57393410-DEL-188 | chr12-57393411-DEL-188 |
| chr12-57452587-DEL-231 | chr12-57452588-DEL-231 |
| chr12-57466249-DEL-334 | chr12-57466250-DEL-334 |
| chr12-57473196-DEL-504 | chr12-57473197-DEL-504 |
| chr12-57533222-DEL-689 | chr12-57533223-DEL-689 |
| chr12-57605163-DEL-66 | chr12-57605164-DEL-66 |
| chr12-57631306-DEL-94 | chr12-57631308-DEL-92 |
| chr12-57646840-DEL-6101 | chr12-57646841-DEL-6101 |
| chr12-57662390-DEL-65 | chr12-57662340-DEL-66 |
| chr12-57686226-DEL-342 | chr12-57686227-DEL-342 |
| chr12-57714336-DEL-1627 | chr12-57714337-DEL-1627 |
| chr12-57716643-DEL-54 | chr12-57716644-DEL-70 |
| chr12-57742560-DEL-212 | chr12-57742561-DEL-208 |

| | |
|---|---|
| chr12-57746907-DEL-1896 | chr12-57746908-DEL-1896 |
| chr12-57795560-DEL-86 | chr12-57795561-DEL-86 |
| chr12-57862534-DEL-6421 | chr12-57862535-DEL-6421 |
| chr12-57883021-DEL-68 | chr12-57883022-DEL-68 |
| chr12-57898384-DEL-143 | chr12-57898382-DEL-112 |
| chr12-57940711-DEL-104 | chr12-57940692-DEL-89 |
| chr12-57953044-DEL-239 | chr12-57953044-DEL-239 |
| chr12-57957515-DEL-6264 | chr12-57957516-DEL-6264 |
| chr12-57964362-DEL-90 | chr12-57964407-DEL-74 |
| chr12-57972822-DEL-72 | chr12-57972823-DEL-72 |
| chr12-57982458-DEL-6265 | chr12-57982452-DEL-6266 |
| chr12-57989000-DEL-275 | chr12-57989001-DEL-275 |
| chr12-58004699-DEL-1365 | chr12-58004700-DEL-1365 |
| chr12-58009237-DEL-310 | chr12-58009238-DEL-310 |
| chr12-58011007-DEL-409 | chr12-58011008-DEL-409 |
| chr12-58046151-DEL-406 | chr12-58046154-DEL-404 |
| chr12-58064059-DEL-97 | chr12-58064065-DEL-91 |
| chr12-58067810-DEL-595 | chr12-58067811-DEL-595 |
| chr12-58069661-DEL-341 | chr12-58069662-DEL-341 |
| chr12-58112008-DEL-154 | chr12-58112009-DEL-154 |
| chr12-58119037-DEL-456 | chr12-58119038-DEL-456 |
| chr12-58128516-DEL-661 | chr12-58128517-DEL-661 |
| chr12-58138099-DEL-94 | chr12-58138100-DEL-94 |
| chr12-58153772-DEL-54 | chr12-58153773-DEL-56 |
| chr12-58156188-DEL-2802 | chr12-58156191-DEL-2800 |
| chr12-58166555-DEL-8441 | chr12-58166556-DEL-8441 |
| chr12-58186359-DEL-90 | chr12-58186360-DEL-90 |
| chr12-58196176-DEL-830 | chr12-58196177-DEL-830 |
| chr12-58196607-DEL-1149 | chr12-58196608-DEL-1149 |
| chr12-58202572-DEL-66 | chr12-58202573-DEL-66 |
| chr12-58202991-DEL-6815 | chr12-58202992-DEL-6815 |
| chr12-58203155-DEL-416 | chr12-58203156-DEL-416 |
| chr12-58242364-DEL-454 | chr12-58242365-DEL-454 |
| chr12-58294456-DEL-1294 | chr12-58294456-DEL-1294 |
| chr12-58324760-DEL-341 | chr12-58324761-DEL-341 |
| chr12-58337552-DEL-1891 | chr12-58337553-DEL-1891 |
| chr12-58341612-DEL-1890 | chr12-58341613-DEL-1890 |
| chr12-58360880-DEL-2392 | chr12-58360880-DEL-2392 |
| chr12-58360933-DEL-214 | chr12-58360934-DEL-214 |
| chr12-58364102-DEL-2796 | chr12-58364102-DEL-2796 |
| chr12-58385498-DEL-1137 | chr12-58385505-DEL-1141 |
| chr12-58397915-DEL-1522 | chr12-58397916-DEL-1522 |
| chr12-58428781-DEL-935 | chr12-58428782-DEL-935 |
| chr12-58444131-DEL-218 | chr12-58444132-DEL-218 |
| chr12-58454720-DEL-76 | chr12-58454721-DEL-76 |
| chr12-58498500-DEL-54 | chr12-58498473-DEL-50 |
| chr12-58523118-DEL-250 | chr12-58523118-DEL-243 |
| chr12-58525439-DEL-74 | chr12-58525440-DEL-72 |
| chr12-58533674-DEL-84 | chr12-58533676-DEL-83 |
| chr12-58534070-DEL-12586 | chr12-58534071-DEL-12586 |
| chr12-58555192-DEL-1957 | chr12-58555190-DEL-1957 |
| chr12-58633063-DEL-60 | chr12-58633070-DEL-58 |
| chr12-58641916-DEL-825 | chr12-58641917-DEL-825 |
| chr12-58648505-DEL-6635 | chr12-58648506-DEL-6635 |
| chr12-58659819-DEL-62 | chr12-58659822-DEL-60 |
| chr12-58673555-DEL-89 | chr12-58673556-DEL-89 |
| chr12-58706462-DEL-6408 | chr12-58706463-DEL-6408 |
| chr12-58714975-DEL-96 | chr12-58714976-DEL-96 |
| chr12-58724580-DEL-163 | chr12-58724569-DEL-166 |
| chr12-58726393-DEL-1886 | chr12-58726394-DEL-1886 |
| chr12-58733652-DEL-200 | chr12-58733653-DEL-200 |
| chr12-58736386-DEL-1747 | chr12-58736387-DEL-1747 |
| chr12-58759831-DEL-56 | chr12-58759832-DEL-56 |
| chr12-58767109-DEL-1222 | chr12-58767135-DEL-1197 |
| chr12-58786059-DEL-78 | chr12-58786060-DEL-78 |
| chr12-58812806-DEL-92 | chr12-58812801-DEL-98 |
| chr12-58814523-DEL-51 | chr12-58814499-DEL-52 |
| chr12-58816579-DEL-6466 | chr12-58816580-DEL-6466 |
| chr12-58841683-DEL-74 | chr12-58841684-DEL-74 |
| chr12-58847160-DEL-77 | chr12-58847161-DEL-77 |
| chr12-58855920-DEL-225 | chr12-58855921-DEL-237 |
| chr12-58869268-DEL-195 | chr12-58869269-DEL-195 |
| chr12-58869520-DEL-1506 | chr12-58869521-DEL-1506 |
| chr12-58910585-DEL-214 | chr12-58910585-DEL-214 |
| chr12-58929219-DEL-126 | chr12-58929220-DEL-126 |
| chr12-58940220-DEL-80 | chr12-58940221-DEL-80 |
| chr12-58989190-DEL-56 | chr12-58989195-DEL-52 |
| chr12-59008883-DEL-124 | chr12-59008884-DEL-124 |
| chr12-59022046-DEL-213 | chr12-59022046-DEL-213 |
| chr12-59038487-DEL-198 | chr12-59038487-DEL-198 |

| | |
|---|---|
| chr12-59046824-DEL-171 | chr12-59046825-DEL-171 |
| chr12-59049567-DEL-68 | chr12-59049559-DEL-69 |
| chr12-59054312-DEL-187 | chr12-59054313-DEL-187 |
| chr12-59061092-DEL-197 | chr12-59061093-DEL-197 |
| chr12-59061610-DEL-74 | chr12-59061611-DEL-74 |
| chr12-59070337-DEL-207 | chr12-59070338-DEL-207 |
| chr12-59076905-DEL-928 | chr12-59076914-DEL-919 |
| chr12-59094497-DEL-230 | chr12-59094498-DEL-230 |
| chr12-59099546-DEL-111 | chr12-59099547-DEL-106 |
| chr12-59105006-DEL-202 | chr12-59105007-DEL-202 |
| chr12-59111302-DEL-68 | chr12-59111303-DEL-68 |
| chr12-59112546-DEL-65 | chr12-59112538-DEL-60 |
| chr12-59122404-DEL-186 | chr12-59122405-DEL-186 |
| chr12-59123955-DEL-639 | chr12-59123956-DEL-639 |
| chr12-59127764-DEL-976 | chr12-59127772-DEL-978 |
| chr12-59143396-DEL-54 | chr12-59143397-DEL-54 |
| chr12-59147401-DEL-173 | chr12-59147405-DEL-169 |
| chr12-59156525-DEL-60 | chr12-59156526-DEL-62 |
| chr12-59159583-DEL-1054 | chr12-59159580-DEL-1060 |
| chr12-59168077-DEL-3429 | chr12-59168077-DEL-3429 |
| chr12-59234587-DEL-228 | chr12-59234586-DEL-230 |
| chr12-59237537-DEL-249 | chr12-59237537-DEL-228 |
| chr12-59258861-DEL-358 | chr12-59258862-DEL-358 |
| chr12-59269404-DEL-4521 | chr12-59269405-DEL-4521 |
| chr12-59295821-DEL-404 | chr12-59295822-DEL-404 |
| chr12-59316602-DEL-7056 | chr12-59316603-DEL-7056 |
| chr12-59327109-DEL-51 | chr12-59327110-DEL-51 |
| chr12-59328247-DEL-341 | chr12-59328248-DEL-341 |
| chr12-59340508-DEL-553 | chr12-59340509-DEL-553 |
| chr12-59344625-DEL-6452 | chr12-59344626-DEL-6452 |
| chr12-59377559-DEL-56 | chr12-59377560-DEL-56 |
| chr12-59394125-DEL-1212 | chr12-59394126-DEL-1212 |
| chr12-59405836-DEL-490 | chr12-59405837-DEL-490 |
| chr12-59406240-DEL-1248 | chr12-59406241-DEL-1248 |
| chr12-59408555-DEL-2280 | chr12-59408556-DEL-2280 |
| chr12-59414094-DEL-2215 | chr12-59414095-DEL-2214 |
| chr12-59438614-DEL-324 | chr12-59438615-DEL-324 |
| chr12-59483643-DEL-398 | chr12-59483644-DEL-398 |
| chr12-59487137-DEL-7134 | chr12-59487138-DEL-7134 |
| chr12-59525075-DEL-82 | chr12-59525075-DEL-82 |
| chr12-59545277-DEL-6371 | chr12-59545278-DEL-6371 |
| chr12-59569241-DEL-52 | chr12-59569242-DEL-52 |
| chr12-59587656-DEL-686 | chr12-59587657-DEL-686 |
| chr12-59608865-DEL-540 | chr12-59608866-DEL-540 |
| chr12-59646825-DEL-6196 | chr12-59646826-DEL-6196 |
| chr12-59660808-DEL-54 | chr12-59660809-DEL-54 |
| chr12-59690166-DEL-143 | chr12-59690200-DEL-105 |
| chr12-59698951-DEL-336 | chr12-59698952-DEL-336 |
| chr12-59737492-DEL-429 | chr12-59737493-DEL-429 |
| chr12-59775794-DEL-176 | chr12-59775795-DEL-176 |
| chr12-59846210-DEL-186 | chr12-59846211-DEL-186 |
| chr12-59854122-DEL-1101 | chr12-59854121-DEL-1101 |
| chr12-59926261-DEL-247 | chr12-59926261-DEL-246 |
| chr12-59936076-DEL-420 | chr12-59936077-DEL-420 |
| chr12-59952230-DEL-72 | chr12-59952231-DEL-64 |
| chr12-59961313-DEL-785 | chr12-59961314-DEL-785 |
| chr12-59964301-DEL-1042 | chr12-59964296-DEL-1043 |
| chr12-59969572-DEL-238 | chr12-59969573-DEL-238 |
| chr12-59970482-DEL-162 | chr12-59970483-DEL-162 |
| chr12-59994531-DEL-199 | chr12-59994492-DEL-199 |
| chr12-60008219-DEL-1630 | chr12-60008220-DEL-1630 |
| chr12-60012762-DEL-70 | chr12-60012763-DEL-70 |
| chr12-60022181-DEL-71 | chr12-60022179-DEL-69 |
| chr12-60030982-DEL-63 | chr12-60030983-DEL-63 |
| chr12-60033441-DEL-211 | chr12-60033442-DEL-211 |
| chr12-60074927-DEL-65 | chr12-60074927-DEL-74 |
| chr12-60088842-DEL-475 | chr12-60088848-DEL-457 |
| chr12-60108912-DEL-188 | chr12-60108913-DEL-188 |
| chr12-60116197-DEL-1394 | chr12-60116198-DEL-1394 |
| chr12-60119452-DEL-223 | chr12-60119457-DEL-219 |
| chr12-60134162-DEL-149 | chr12-60134163-DEL-149 |
| chr12-60141137-DEL-111 | chr12-60141166-DEL-94 |
| chr12-60147893-DEL-391 | chr12-60147904-DEL-390 |
| chr12-60181301-DEL-52 | chr12-60181302-DEL-52 |
| chr12-60216143-DEL-2138 | chr12-60216144-DEL-2138 |
| chr12-60216676-DEL-799 | chr12-60216677-DEL-799 |
| chr12-60234231-DEL-54 | chr12-60234238-DEL-54 |
| chr12-60243805-DEL-497 | chr12-60243806-DEL-497 |
| chr12-60247467-DEL-58 | chr12-60247468-DEL-58 |
| chr12-60248474-DEL-265 | chr12-60248475-DEL-265 |

| | |
|---|---|
| chr12-60248991-DEL-6169 | chr12-60248978-DEL-6170 |
| chr12-60257717-DEL-195 | chr12-60257718-DEL-195 |
| chr12-60269417-DEL-54 | chr12-60269406-DEL-65 |
| chr12-60274226-DEL-2904 | chr12-60274221-DEL-2899 |
| chr12-60275037-DEL-974 | chr12-60275038-DEL-974 |
| chr12-60301476-DEL-106 | chr12-60301475-DEL-111 |
| chr12-60304228-DEL-1531 | chr12-60304229-DEL-1531 |
| chr12-60311993-DEL-396 | chr12-60311994-DEL-396 |
| chr12-60319267-DEL-6961 | chr12-60319267-DEL-6961 |
| chr12-60347306-DEL-647 | chr12-60347307-DEL-647 |
| chr12-60363767-DEL-133 | chr12-60363768-DEL-133 |
| chr12-60369849-DEL-160 | chr12-60369849-DEL-161 |
| chr12-60395399-DEL-77 | chr12-60395400-DEL-77 |
| chr12-60407423-DEL-2856 | chr12-60407424-DEL-2856 |
| chr12-60430479-DEL-136 | chr12-60430480-DEL-136 |
| chr12-60443280-DEL-8892 | chr12-60443281-DEL-8892 |
| chr12-60450406-DEL-2302 | chr12-60450420-DEL-2289 |
| chr12-60462307-DEL-121 | chr12-60462315-DEL-85 |
| chr12-60516836-DEL-416 | chr12-60516837-DEL-416 |
| chr12-60547914-DEL-4997 | chr12-60547915-DEL-4997 |
| chr12-60559944-DEL-1522 | chr12-60559944-DEL-1522 |
| chr12-60572010-DEL-3145 | chr12-60572011-DEL-3145 |
| chr12-60582598-DEL-2085 | chr12-60582599-DEL-2085 |
| chr12-60602722-DEL-90 | chr12-60602723-DEL-90 |
| chr12-60610370-DEL-341 | chr12-60610371-DEL-341 |
| chr12-60616910-DEL-102 | chr12-60616911-DEL-102 |
| chr12-60665315-DEL-192 | chr12-60665299-DEL-191 |
| chr12-60666039-DEL-832 | chr12-60666037-DEL-833 |
| chr12-60673920-DEL-480 | chr12-60673912-DEL-489 |
| chr12-60687550-DEL-206 | chr12-60687552-DEL-205 |
| chr12-60688522-DEL-110 | chr12-60688522-DEL-126 |
| chr12-60694902-DEL-2238 | chr12-60694903-DEL-2239 |
| chr12-60753715-DEL-7123 | chr12-60753716-DEL-7123 |
| chr12-60783920-DEL-2458 | chr12-60783921-DEL-2458 |
| chr12-60794457-DEL-665 | chr12-60794459-DEL-666 |
| chr12-60801052-DEL-1837 | chr12-60801053-DEL-1837 |
| chr12-60823485-DEL-1496 | chr12-60823485-DEL-1496 |
| chr12-60835745-DEL-2008 | chr12-60835746-DEL-2008 |
| chr12-60850355-DEL-488 | chr12-60850356-DEL-488 |
| chr12-60880824-DEL-621 | chr12-60880824-DEL-621 |
| chr12-60884804-DEL-2301 | chr12-60884805-DEL-2301 |
| chr12-60915979-DEL-65 | chr12-60915980-DEL-65 |
| chr12-60922633-DEL-287 | chr12-60922634-DEL-287 |
| chr12-60946457-DEL-286 | chr12-60946458-DEL-282 |
| chr12-60953902-DEL-72 | chr12-60953901-DEL-73 |
| chr12-60964855-DEL-433 | chr12-60964855-DEL-433 |
| chr12-60970550-DEL-87 | chr12-60970540-DEL-62 |
| chr12-60973786-DEL-356 | chr12-60973787-DEL-356 |
| chr12-60990566-DEL-722 | chr12-60990567-DEL-722 |
| chr12-60997626-DEL-154 | chr12-60997627-DEL-154 |
| chr12-61025030-DEL-239 | chr12-61025022-DEL-240 |
| chr12-61051594-DEL-88 | chr12-61051587-DEL-72 |
| chr12-61064298-DEL-259 | chr12-61064299-DEL-258 |
| chr12-61078646-DEL-1115 | chr12-61078647-DEL-1115 |
| chr12-61082187-DEL-342 | chr12-61082188-DEL-342 |
| chr12-61088018-DEL-155 | chr12-61088019-DEL-155 |
| chr12-61090944-DEL-1135 | chr12-61090945-DEL-1135 |
| chr12-61095306-DEL-51 | chr12-61095256-DEL-95 |
| chr12-61099720-DEL-195 | chr12-61099757-DEL-202 |
| chr12-61108546-DEL-413 | chr12-61108593-DEL-390 |
| chr12-61140514-DEL-810 | chr12-61140515-DEL-810 |
| chr12-61161423-DEL-2717 | chr12-61161424-DEL-2717 |
| chr12-61170281-DEL-75 | chr12-61170283-DEL-68 |
| chr12-61171886-DEL-62 | chr12-61171887-DEL-74 |
| chr12-61175186-DEL-1037 | chr12-61175187-DEL-1034 |
| chr12-61179274-DEL-89 | chr12-61179275-DEL-89 |
| chr12-61186365-DEL-67 | chr12-61186366-DEL-67 |
| chr12-61201266-DEL-6416 | chr12-61201267-DEL-6416 |
| chr12-61226955-DEL-4031 | chr12-61226962-DEL-3982 |
| chr12-61239703-DEL-92 | chr12-61239704-DEL-92 |
| chr12-61242127-DEL-65 | chr12-61242128-DEL-65 |
| chr12-61256843-DEL-93 | chr12-61256836-DEL-50 |
| chr12-61274080-DEL-95 | chr12-61274081-DEL-95 |
| chr12-61285316-DEL-55 | chr12-61285317-DEL-55 |
| chr12-61286505-DEL-52 | chr12-61286505-DEL-52 |
| chr12-61292921-DEL-86 | chr12-61292921-DEL-84 |
| chr12-61333188-DEL-56 | chr12-61333191-DEL-54 |
| chr12-61335915-DEL-4575 | chr12-61335916-DEL-4575 |
| chr12-61349329-DEL-672 | chr12-61349330-DEL-672 |
| chr12-61410286-DEL-6784 | chr12-61410287-DEL-6784 |

| | |
|---|---|
| chr12-61453845-DEL-136 | chr12-61453846-DEL-136 |
| chr12-61464827-DEL-2000 | chr12-61464828-DEL-2000 |
| chr12-61508217-DEL-115 | chr12-61508218-DEL-115 |
| chr12-61510073-DEL-672 | chr12-61510075-DEL-670 |
| chr12-61522025-DEL-61 | chr12-61522013-DEL-72 |
| chr12-61527345-DEL-373 | chr12-61527346-DEL-373 |
| chr12-61584090-DEL-100 | chr12-61584091-DEL-100 |
| chr12-61631168-DEL-72 | chr12-61631174-DEL-67 |
| chr12-61631903-DEL-296 | chr12-61631904-DEL-296 |
| chr12-61640504-DEL-58 | chr12-61640506-DEL-66 |
| chr12-61650208-DEL-58 | chr12-61650209-DEL-58 |
| chr12-61651832-DEL-55 | chr12-61651828-DEL-59 |
| chr12-61657425-DEL-60 | chr12-61657450-DEL-59 |
| chr12-61678749-DEL-9714 | chr12-61678750-DEL-9714 |
| chr12-61708705-DEL-115 | chr12-61708706-DEL-115 |
| chr12-61715845-DEL-378 | chr12-61715846-DEL-378 |
| chr12-61730254-DEL-109 | chr12-61730255-DEL-109 |
| chr12-61742405-DEL-2346 | chr12-61742406-DEL-2346 |
| chr12-61745715-DEL-6680 | chr12-61745716-DEL-6680 |
| chr12-61763256-DEL-57 | chr12-61763251-DEL-60 |
| chr12-61766590-DEL-2293 | chr12-61766590-DEL-2293 |
| chr12-61769389-DEL-86 | chr12-61769390-DEL-85 |
| chr12-61803181-DEL-6992 | chr12-61803182-DEL-6992 |
| chr12-61837740-DEL-61 | chr12-61837741-DEL-61 |
| chr12-61857714-DEL-1370 | chr12-61857715-DEL-1370 |
| chr12-61861465-DEL-2012 | chr12-61861466-DEL-2012 |
| chr12-61870574-DEL-70 | chr12-61870569-DEL-76 |
| chr12-61894765-DEL-392 | chr12-61894766-DEL-392 |
| chr12-61914917-DEL-105 | chr12-61914918-DEL-105 |
| chr12-61918202-DEL-308 | chr12-61918209-DEL-301 |
| chr12-61919375-DEL-563 | chr12-61919376-DEL-563 |
| chr12-61925222-DEL-3013 | chr12-61925223-DEL-3013 |
| chr12-61934130-DEL-638 | chr12-61934131-DEL-638 |
| chr12-61978846-DEL-131 | chr12-61978847-DEL-131 |
| chr12-62000413-DEL-84 | chr12-62000414-DEL-84 |
| chr12-62002252-DEL-60 | chr12-62002253-DEL-60 |
| chr12-62033285-DEL-58 | chr12-62033286-DEL-58 |
| chr12-62074249-DEL-929 | chr12-62074250-DEL-929 |
| chr12-62115038-DEL-54 | chr12-62115037-DEL-50 |
| chr12-62116472-DEL-6592 | chr12-62116473-DEL-6592 |
| chr12-62124726-DEL-122 | chr12-62124727-DEL-122 |
| chr12-62130093-DEL-64 | chr12-62130094-DEL-64 |
| chr12-62154036-DEL-161 | chr12-62154037-DEL-161 |
| chr12-62167559-DEL-616 | chr12-62167560-DEL-616 |
| chr12-62175607-DEL-2577 | chr12-62175608-DEL-2577 |
| chr12-62181861-DEL-1219 | chr12-62181862-DEL-1219 |
| chr12-62191656-DEL-6417 | chr12-62191657-DEL-6417 |
| chr12-62223485-DEL-7525 | chr12-62223486-DEL-7525 |
| chr12-62249703-DEL-2769 | chr12-62249714-DEL-2764 |
| chr12-62263220-DEL-132 | chr12-62263221-DEL-132 |
| chr12-62273580-DEL-5517 | chr12-62273581-DEL-5517 |
| chr12-62304643-DEL-3403 | chr12-62304644-DEL-3403 |
| chr12-62323483-DEL-1674 | chr12-62323466-DEL-1691 |
| chr12-62354799-DEL-6195 | chr12-62354800-DEL-6195 |
| chr12-62457804-DEL-60 | chr12-62457805-DEL-60 |
| chr12-62494628-DEL-192 | chr12-62494629-DEL-192 |
| chr12-62515650-DEL-88 | chr12-62515683-DEL-88 |
| chr12-62555543-DEL-2420 | chr12-62555544-DEL-2420 |
| chr12-62557412-DEL-278 | chr12-62557412-DEL-279 |
| chr12-62581661-DEL-3971 | chr12-62581662-DEL-3971 |
| chr12-62588518-DEL-8145 | chr12-62588519-DEL-8145 |
| chr12-62597869-DEL-114 | chr12-62597866-DEL-118 |
| chr12-62637162-DEL-175 | chr12-62637163-DEL-175 |
| chr12-62645702-DEL-68 | chr12-62645683-DEL-88 |
| chr12-62677326-DEL-54 | chr12-62677330-DEL-51 |
| chr12-62678198-DEL-359 | chr12-62678199-DEL-359 |
| chr12-62682168-DEL-1046 | chr12-62682169-DEL-1046 |
| chr12-62688119-DEL-93 | chr12-62688141-DEL-50 |
| chr12-62688695-DEL-420 | chr12-62688696-DEL-420 |
| chr12-62694432-DEL-66 | chr12-62694461-DEL-76 |
| chr12-62697109-DEL-478 | chr12-62697111-DEL-477 |
| chr12-62730522-DEL-197 | chr12-62730523-DEL-197 |
| chr12-62742142-DEL-124 | chr12-62742143-DEL-124 |
| chr12-62777303-DEL-588 | chr12-62777303-DEL-588 |
| chr12-62782577-DEL-860 | chr12-62782578-DEL-860 |
| chr12-62812329-DEL-894 | chr12-62812330-DEL-894 |
| chr12-62819827-DEL-193 | chr12-62819828-DEL-193 |
| chr12-62845108-DEL-1179 | chr12-62845109-DEL-1179 |
| chr12-62888263-DEL-164 | chr12-62888263-DEL-163 |
| chr12-62889530-DEL-348 | chr12-62889530-DEL-348 |

| | |
|---|---|
| chr12-62938760-DEL-324 | chr12-62938760-DEL-324 |
| chr12-62952901-DEL-144 | chr12-62952911-DEL-105 |
| chr12-62965838-DEL-2568 | chr12-62965839-DEL-2568 |
| chr12-63006501-DEL-7305 | chr12-63006502-DEL-7305 |
| chr12-63064431-DEL-53 | chr12-63064432-DEL-53 |
| chr12-63084327-DEL-433 | chr12-63084328-DEL-433 |
| chr12-63088481-DEL-6604 | chr12-63088483-DEL-6600 |
| chr12-63097067-DEL-1195 | chr12-63097068-DEL-1195 |
| chr12-63099135-DEL-199 | chr12-63099125-DEL-198 |
| chr12-63111813-DEL-1537 | chr12-63111813-DEL-1537 |
| chr12-63127942-DEL-138 | chr12-63127943-DEL-138 |
| chr12-63137686-DEL-1065 | chr12-63137687-DEL-1065 |
| chr12-63156313-DEL-1100 | chr12-63156314-DEL-1100 |
| chr12-63158292-DEL-56 | chr12-63158293-DEL-56 |
| chr12-63163496-DEL-274 | chr12-63163497-DEL-274 |
| chr12-63166052-DEL-7199 | chr12-63166053-DEL-7199 |
| chr12-63176063-DEL-287 | chr12-63176074-DEL-273 |
| chr12-63184188-DEL-1397 | chr12-63184189-DEL-1397 |
| chr12-63188041-DEL-125 | chr12-63188049-DEL-103 |
| chr12-63213513-DEL-58 | chr12-63213518-DEL-60 |
| chr12-63215119-DEL-6388 | chr12-63215120-DEL-6388 |
| chr12-63238902-DEL-442 | chr12-63238903-DEL-442 |
| chr12-63244585-DEL-94 | chr12-63244586-DEL-94 |
| chr12-63249393-DEL-253 | chr12-63249393-DEL-253 |
| chr12-63251824-DEL-438 | chr12-63251824-DEL-438 |
| chr12-63257406-DEL-52 | chr12-63257391-DEL-59 |
| chr12-63259092-DEL-70 | chr12-63259093-DEL-70 |
| chr12-63277055-DEL-106 | chr12-63277060-DEL-98 |
| chr12-63311228-DEL-659 | chr12-63311230-DEL-658 |
| chr12-63319458-DEL-548 | chr12-63319459-DEL-548 |
| chr12-63323652-DEL-56 | chr12-63323653-DEL-56 |
| chr12-63336348-DEL-2397 | chr12-63336349-DEL-2397 |
| chr12-63344245-DEL-4005 | chr12-63344246-DEL-4005 |
| chr12-63397960-DEL-7398 | chr12-63397961-DEL-7398 |
| chr12-63440575-DEL-6664 | chr12-63440576-DEL-6664 |
| chr12-63460429-DEL-1330 | chr12-63460430-DEL-1330 |
| chr12-63478411-DEL-84 | chr12-63478412-DEL-84 |
| chr12-63500954-DEL-284 | chr12-63500955-DEL-284 |
| chr12-63506277-DEL-1775 | chr12-63506278-DEL-1775 |
| chr12-63522228-DEL-1565 | chr12-63522229-DEL-1565 |
| chr12-63528265-DEL-82 | chr12-63528266-DEL-82 |
| chr12-63547699-DEL-1725 | chr12-63547700-DEL-1725 |
| chr12-63597016-DEL-78 | chr12-63597012-DEL-83 |
| chr12-63644299-DEL-76 | chr12-63644300-DEL-76 |
| chr12-63676345-DEL-3960 | chr12-63676346-DEL-3960 |
| chr12-63696007-DEL-97 | chr12-63696008-DEL-70 |
| chr12-63700761-DEL-170 | chr12-63700762-DEL-170 |
| chr12-63709331-DEL-400 | chr12-63709332-DEL-400 |
| chr12-63738324-DEL-6674 | chr12-63738325-DEL-6674 |
| chr12-63738507-DEL-212 | chr12-63738508-DEL-212 |
| chr12-63791827-DEL-250 | chr12-63791828-DEL-250 |
| chr12-63793701-DEL-968 | chr12-63793702-DEL-968 |
| chr12-63830226-DEL-426 | chr12-63830243-DEL-407 |
| chr12-63880628-DEL-636 | chr12-63880629-DEL-636 |
| chr12-63883254-DEL-3772 | chr12-63883255-DEL-3772 |
| chr12-63897536-DEL-1371 | chr12-63897537-DEL-1371 |
| chr12-63903973-DEL-321 | chr12-63903965-DEL-322 |
| chr12-63906897-DEL-58 | chr12-63906891-DEL-54 |
| chr12-63919925-DEL-340 | chr12-63919926-DEL-340 |
| chr12-63936553-DEL-431 | chr12-63936569-DEL-405 |
| chr12-63948959-DEL-476 | chr12-63948960-DEL-476 |
| chr12-63953010-DEL-438 | chr12-63953011-DEL-438 |
| chr12-63955843-DEL-201 | chr12-63955844-DEL-201 |
| chr12-63967785-DEL-133 | chr12-63967834-DEL-85 |
| chr12-64024018-DEL-52 | chr12-64023991-DEL-66 |
| chr12-64033660-DEL-2313 | chr12-64033661-DEL-2313 |
| chr12-64039781-DEL-51 | chr12-64039755-DEL-100 |
| chr12-64077030-DEL-2774 | chr12-64077031-DEL-2774 |
| chr12-64096105-DEL-71 | chr12-64096124-DEL-55 |
| chr12-64142516-DEL-90 | chr12-64142513-DEL-94 |
| chr12-64159915-DEL-57 | chr12-64159916-DEL-57 |
| chr12-64174565-DEL-179 | chr12-64174566-DEL-177 |
| chr12-64178320-DEL-2954 | chr12-64178321-DEL-2954 |
| chr12-64192843-DEL-10741 | chr12-64192844-DEL-10741 |
| chr12-64207680-DEL-190 | chr12-64207681-DEL-190 |
| chr12-64216756-DEL-140 | chr12-64216757-DEL-140 |
| chr12-64221242-DEL-5512 | chr12-64221242-DEL-5512 |
| chr12-64239386-DEL-72 | chr12-64239387-DEL-72 |
| chr12-64253140-DEL-90 | chr12-64253141-DEL-90 |
| chr12-64254188-DEL-113 | chr12-64254184-DEL-114 |

| | |
|---|---|
| chr12-64269715-DEL-60 | chr12-64269716-DEL-60 |
| chr12-64298429-DEL-90 | chr12-64298464-DEL-56 |
| chr12-64318540-DEL-63 | chr12-64318541-DEL-63 |
| chr12-64320921-DEL-96 | chr12-64320922-DEL-76 |
| chr12-64323887-DEL-3052 | chr12-64323873-DEL-3066 |
| chr12-64345809-DEL-6190 | chr12-64345810-DEL-6190 |
| chr12-64358609-DEL-68 | chr12-64358618-DEL-60 |
| chr12-64367168-DEL-690 | chr12-64367169-DEL-690 |
| chr12-64375454-DEL-82 | chr12-64375452-DEL-82 |
| chr12-64402724-DEL-157 | chr12-64402723-DEL-157 |
| chr12-64417434-DEL-56 | chr12-64417435-DEL-56 |
| chr12-64420058-DEL-1187 | chr12-64420060-DEL-1186 |
| chr12-64440373-DEL-2946 | chr12-64440374-DEL-2946 |
| chr12-64447008-DEL-385 | chr12-64446999-DEL-395 |
| chr12-64464087-DEL-2351 | chr12-64464088-DEL-2351 |
| chr12-64480160-DEL-179 | chr12-64480161-DEL-179 |
| chr12-64481776-DEL-440 | chr12-64481777-DEL-440 |
| chr12-64510755-DEL-1888 | chr12-64510756-DEL-1888 |
| chr12-64553811-DEL-607 | chr12-64553812-DEL-607 |
| chr12-64574243-DEL-123 | chr12-64574244-DEL-123 |
| chr12-64617152-DEL-140 | chr12-64617153-DEL-140 |
| chr12-64649505-DEL-6389 | chr12-64649506-DEL-6389 |
| chr12-64661407-DEL-463 | chr12-64661408-DEL-463 |
| chr12-64665847-DEL-85 | chr12-64665857-DEL-79 |
| chr12-64693320-DEL-60 | chr12-64693321-DEL-60 |
| chr12-64695294-DEL-3485 | chr12-64695295-DEL-3485 |
| chr12-64699649-DEL-591 | chr12-64699642-DEL-596 |
| chr12-64712779-DEL-900 | chr12-64712780-DEL-899 |
| chr12-64737016-DEL-54 | chr12-64737018-DEL-52 |
| chr12-64771716-DEL-371 | chr12-64771717-DEL-371 |
| chr12-64772747-DEL-9901 | chr12-64772748-DEL-9901 |
| chr12-64777053-DEL-2881 | chr12-64777053-DEL-2881 |
| chr12-64784094-DEL-60 | chr12-64784095-DEL-60 |
| chr12-64832950-DEL-1542 | chr12-64832951-DEL-1542 |
| chr12-64837956-DEL-1413 | chr12-64837957-DEL-1413 |
| chr12-64844321-DEL-566 | chr12-64844322-DEL-566 |
| chr12-64891574-DEL-67 | chr12-64891575-DEL-67 |
| chr12-64921957-DEL-478 | chr12-64921958-DEL-478 |
| chr12-64929089-DEL-90 | chr12-64929088-DEL-92 |
| chr12-64942133-DEL-102 | chr12-64942134-DEL-100 |
| chr12-64945030-DEL-2868 | chr12-64945038-DEL-2865 |
| chr12-64955559-DEL-525 | chr12-64955560-DEL-525 |
| chr12-65070431-DEL-157 | chr12-65070431-DEL-152 |
| chr12-65073953-DEL-7128 | chr12-65073954-DEL-7128 |
| chr12-65117860-DEL-61 | chr12-65117861-DEL-61 |
| chr12-65206069-DEL-205 | chr12-65206070-DEL-205 |
| chr12-65283797-DEL-100 | chr12-65283798-DEL-100 |
| chr12-65292515-DEL-128 | chr12-65292516-DEL-128 |
| chr12-65306789-DEL-6987 | chr12-65306790-DEL-6987 |
| chr12-65392961-DEL-508 | chr12-65392962-DEL-508 |
| chr12-65434337-DEL-454 | chr12-65434346-DEL-446 |
| chr12-65458838-DEL-559 | chr12-65458839-DEL-559 |
| chr12-65521467-DEL-119 | chr12-65521468-DEL-119 |
| chr12-65529121-DEL-59 | chr12-65529122-DEL-59 |
| chr12-65550252-DEL-89 | chr12-65550249-DEL-89 |
| chr12-65550860-DEL-73 | chr12-65550865-DEL-72 |
| chr12-65553697-DEL-2429 | chr12-65553698-DEL-2429 |
| chr12-65603517-DEL-54 | chr12-65603533-DEL-59 |
| chr12-65635087-DEL-51 | chr12-65635068-DEL-71 |
| chr12-65640106-DEL-66 | chr12-65640106-DEL-63 |
| chr12-65665154-DEL-680 | chr12-65665156-DEL-678 |
| chr12-65668134-DEL-494 | chr12-65668135-DEL-494 |
| chr12-65679975-DEL-7182 | chr12-65679976-DEL-7182 |
| chr12-65707866-DEL-886 | chr12-65707867-DEL-886 |
| chr12-65730065-DEL-2655 | chr12-65730066-DEL-2655 |
| chr12-65751502-DEL-54 | chr12-65751503-DEL-54 |
| chr12-65780156-DEL-58 | chr12-65780159-DEL-56 |
| chr12-65793788-DEL-104 | chr12-65793789-DEL-104 |
| chr12-65834965-DEL-67 | chr12-65834966-DEL-67 |
| chr12-65838536-DEL-420 | chr12-65838537-DEL-420 |
| chr12-65848017-DEL-7079 | chr12-65848018-DEL-7079 |
| chr12-65875084-DEL-240 | chr12-65875085-DEL-240 |
| chr12-65879588-DEL-1738 | chr12-65879589-DEL-1738 |
| chr12-65888524-DEL-1278 | chr12-65888524-DEL-1278 |
| chr12-65890544-DEL-423 | chr12-65890542-DEL-423 |
| chr12-65892982-DEL-52 | chr12-65892983-DEL-62 |
| chr12-65907821-DEL-1530 | chr12-65907822-DEL-1530 |
| chr12-65945200-DEL-3962 | chr12-65945190-DEL-3963 |
| chr12-65961979-DEL-6865 | chr12-65961980-DEL-6864 |
| chr12-65985911-DEL-148 | chr12-65985949-DEL-111 |

| | |
|---|---|
| chr12-66046628-DEL-83 | chr12-66046630-DEL-80 |
| chr12-66052582-DEL-76 | chr12-66052583-DEL-76 |
| chr12-66089436-DEL-52 | chr12-66089437-DEL-52 |
| chr12-66139814-DEL-148 | chr12-66139780-DEL-149 |
| chr12-66150109-DEL-6217 | chr12-66150110-DEL-6217 |
| chr12-66203017-DEL-209 | chr12-66203018-DEL-209 |
| chr12-66211646-DEL-6375 | chr12-66211647-DEL-6375 |
| chr12-66229049-DEL-404 | chr12-66229050-DEL-404 |
| chr12-66234521-DEL-852 | chr12-66234522-DEL-852 |
| chr12-66237329-DEL-226 | chr12-66237330-DEL-226 |
| chr12-66247008-DEL-3164 | chr12-66247009-DEL-3164 |
| chr12-66260084-DEL-1281 | chr12-66260085-DEL-1281 |
| chr12-66293331-DEL-2542 | chr12-66293332-DEL-2542 |
| chr12-66300913-DEL-6271 | chr12-66300914-DEL-6271 |
| chr12-66316563-DEL-405 | chr12-66316564-DEL-399 |
| chr12-66320126-DEL-57 | chr12-66320127-DEL-53 |
| chr12-66321920-DEL-503 | chr12-66321921-DEL-503 |
| chr12-66327252-DEL-499 | chr12-66327253-DEL-499 |
| chr12-66362778-DEL-1182 | chr12-66362779-DEL-1182 |
| chr12-66422228-DEL-69 | chr12-66422229-DEL-69 |
| chr12-66453816-DEL-7071 | chr12-66453817-DEL-7071 |
| chr12-66477007-DEL-1155 | chr12-66477008-DEL-1155 |
| chr12-66491653-DEL-58 | chr12-66491655-DEL-57 |
| chr12-66546970-DEL-353 | chr12-66546971-DEL-353 |
| chr12-66549591-DEL-1966 | chr12-66549592-DEL-1966 |
| chr12-66581792-DEL-154 | chr12-66581793-DEL-154 |
| chr12-66582796-DEL-1357 | chr12-66582797-DEL-1357 |
| chr12-66628577-DEL-227 | chr12-66628589-DEL-224 |
| chr12-66652775-DEL-322 | chr12-66652776-DEL-322 |
| chr12-66669064-DEL-273 | chr12-66669065-DEL-273 |
| chr12-66691794-DEL-122 | chr12-66691795-DEL-122 |
| chr12-66715307-DEL-72 | chr12-66715308-DEL-72 |
| chr12-66717814-DEL-52 | chr12-66717815-DEL-52 |
| chr12-66725767-DEL-133 | chr12-66725768-DEL-133 |
| chr12-66739427-DEL-744 | chr12-66739428-DEL-744 |
| chr12-66744617-DEL-254 | chr12-66744618-DEL-254 |
| chr12-66746071-DEL-55 | chr12-66746072-DEL-55 |
| chr12-66760043-DEL-214 | chr12-66760045-DEL-213 |
| chr12-66776735-DEL-56 | chr12-66776736-DEL-56 |
| chr12-66779016-DEL-80 | chr12-66779017-DEL-80 |
| chr12-66780599-DEL-61 | chr12-66780600-DEL-61 |
| chr12-66798529-DEL-64 | chr12-66798526-DEL-62 |
| chr12-66799618-DEL-225 | chr12-66799619-DEL-225 |
| chr12-66805399-DEL-70 | chr12-66805400-DEL-70 |
| chr12-66818546-DEL-216 | chr12-66818547-DEL-216 |
| chr12-66828232-DEL-204 | chr12-66828233-DEL-204 |
| chr12-66829233-DEL-2520 | chr12-66829234-DEL-2520 |
| chr12-66842241-DEL-202 | chr12-66842243-DEL-201 |
| chr12-66851198-DEL-853 | chr12-66851199-DEL-853 |
| chr12-66853394-DEL-67 | chr12-66853401-DEL-60 |
| chr12-66858497-DEL-563 | chr12-66858498-DEL-563 |
| chr12-66874214-DEL-73 | chr12-66874215-DEL-73 |
| chr12-66917820-DEL-217 | chr12-66917821-DEL-217 |
| chr12-66957380-DEL-668 | chr12-66957381-DEL-675 |
| chr12-66967521-DEL-62 | chr12-66967516-DEL-68 |
| chr12-66978947-DEL-335 | chr12-66978948-DEL-335 |
| chr12-66991248-DEL-2356 | chr12-66991249-DEL-2356 |
| chr12-66996948-DEL-53 | chr12-66996949-DEL-53 |
| chr12-67004598-DEL-1360 | chr12-67004599-DEL-1360 |
| chr12-67064498-DEL-57 | chr12-67064489-DEL-58 |
| chr12-67160223-DEL-115 | chr12-67160224-DEL-115 |
| chr12-67224616-DEL-217 | chr12-67224617-DEL-217 |
| chr12-67244545-DEL-403 | chr12-67244546-DEL-403 |
| chr12-67246043-DEL-133 | chr12-67246044-DEL-133 |
| chr12-67253995-DEL-175 | chr12-67253996-DEL-175 |
| chr12-67275257-DEL-2424 | chr12-67275258-DEL-2424 |
| chr12-67306930-DEL-6413 | chr12-67306931-DEL-6413 |
| chr12-67338018-DEL-70 | chr12-67338018-DEL-70 |
| chr12-67341801-DEL-284 | chr12-67341802-DEL-284 |
| chr12-67342932-DEL-111 | chr12-67342937-DEL-112 |
| chr12-67399593-DEL-1489 | chr12-67399594-DEL-1489 |
| chr12-67401109-DEL-70 | chr12-67401111-DEL-60 |
| chr12-67421726-DEL-158 | chr12-67421727-DEL-158 |
| chr12-67426471-DEL-796 | chr12-67426472-DEL-796 |
| chr12-67451358-DEL-54 | chr12-67451359-DEL-56 |
| chr12-67454732-DEL-110 | chr12-67454733-DEL-110 |
| chr12-67475109-DEL-357 | chr12-67475110-DEL-357 |
| chr12-67478629-DEL-4224 | chr12-67478630-DEL-4224 |
| chr12-67517221-DEL-2882 | chr12-67517222-DEL-2882 |
| chr12-67557478-DEL-678 | chr12-67557479-DEL-678 |

| | |
|---|---|
| chr12-67566018-DEL-1892 | chr12-67566019-DEL-1892 |
| chr12-67572746-DEL-54 | chr12-67572747-DEL-54 |
| chr12-67582170-DEL-2001 | chr12-67582171-DEL-2001 |
| chr12-67601633-DEL-6800 | chr12-67601634-DEL-6800 |
| chr12-67616880-DEL-1510 | chr12-67616881-DEL-1510 |
| chr12-67652017-DEL-168 | chr12-67652018-DEL-168 |
| chr12-67656669-DEL-132 | chr12-67656670-DEL-140 |
| chr12-67735158-DEL-123 | chr12-67735158-DEL-120 |
| chr12-67767468-DEL-2117 | chr12-67767469-DEL-2117 |
| chr12-67793859-DEL-332 | chr12-67793860-DEL-332 |
| chr12-67795310-DEL-126 | chr12-67795311-DEL-126 |
| chr12-67810028-DEL-101 | chr12-67810032-DEL-99 |
| chr12-67819730-DEL-52 | chr12-67819731-DEL-52 |
| chr12-67860783-DEL-145 | chr12-67860784-DEL-145 |
| chr12-67866873-DEL-131 | chr12-67866878-DEL-126 |
| chr12-67869768-DEL-390 | chr12-67869768-DEL-390 |
| chr12-67902297-DEL-155 | chr12-67902291-DEL-162 |
| chr12-67920827-DEL-554 | chr12-67920828-DEL-554 |
| chr12-67923172-DEL-1107 | chr12-67923173-DEL-1107 |
| chr12-67932336-DEL-84 | chr12-67932337-DEL-84 |
| chr12-67967632-DEL-302 | chr12-67967633-DEL-302 |
| chr12-67996912-DEL-7007 | chr12-67996913-DEL-7007 |
| chr12-67996933-DEL-418 | chr12-67996934-DEL-418 |
| chr12-68025904-DEL-60 | chr12-68025905-DEL-60 |
| chr12-68033150-DEL-3107 | chr12-68033151-DEL-3107 |
| chr12-68048347-DEL-522 | chr12-68048348-DEL-522 |
| chr12-68059339-DEL-166 | chr12-68059316-DEL-150 |
| chr12-68064155-DEL-135 | chr12-68064156-DEL-135 |
| chr12-68065115-DEL-60 | chr12-68065118-DEL-58 |
| chr12-68072850-DEL-162 | chr12-68072851-DEL-162 |
| chr12-68083421-DEL-5743 | chr12-68083422-DEL-5743 |
| chr12-68100908-DEL-84 | chr12-68100881-DEL-96 |
| chr12-68156898-DEL-65 | chr12-68156901-DEL-65 |
| chr12-68180260-DEL-105 | chr12-68180261-DEL-103 |
| chr12-68182739-DEL-169 | chr12-68182740-DEL-169 |
| chr12-68200712-DEL-201 | chr12-68200713-DEL-201 |
| chr12-68212650-DEL-238 | chr12-68212651-DEL-238 |
| chr12-68253584-DEL-171 | chr12-68253585-DEL-171 |
| chr12-68258549-DEL-427 | chr12-68258550-DEL-427 |
| chr12-68308390-DEL-117 | chr12-68308391-DEL-117 |
| chr12-68330194-DEL-56 | chr12-68330202-DEL-74 |
| chr12-68351497-DEL-962 | chr12-68351498-DEL-962 |
| chr12-68414920-DEL-66 | chr12-68414919-DEL-68 |
| chr12-68420374-DEL-87 | chr12-68420375-DEL-87 |
| chr12-68452014-DEL-665 | chr12-68452015-DEL-665 |
| chr12-68453528-DEL-7098 | chr12-68453529-DEL-7098 |
| chr12-68472606-DEL-101 | chr12-68472611-DEL-99 |
| chr12-68481630-DEL-6448 | chr12-68481634-DEL-6449 |
| chr12-68540433-DEL-370 | chr12-68540434-DEL-370 |
| chr12-68565461-DEL-230 | chr12-68565462-DEL-230 |
| chr12-68591204-DEL-92 | chr12-68591205-DEL-92 |
| chr12-68597307-DEL-118 | chr12-68597308-DEL-118 |
| chr12-68598087-DEL-783 | chr12-68598088-DEL-783 |
| chr12-68601502-DEL-427 | chr12-68601503-DEL-427 |
| chr12-68604842-DEL-5543 | chr12-68604843-DEL-5543 |
| chr12-68626892-DEL-65 | chr12-68626893-DEL-65 |
| chr12-68661442-DEL-55 | chr12-68661443-DEL-55 |
| chr12-68663012-DEL-80 | chr12-68663013-DEL-80 |
| chr12-68663850-DEL-484 | chr12-68663851-DEL-484 |
| chr12-68668480-DEL-7243 | chr12-68668481-DEL-7243 |
| chr12-68690079-DEL-237 | chr12-68690073-DEL-233 |
| chr12-68712370-DEL-557 | chr12-68712371-DEL-557 |
| chr12-68752218-DEL-6383 | chr12-68752207-DEL-6384 |
| chr12-68779760-DEL-108 | chr12-68779753-DEL-116 |
| chr12-68791944-DEL-5281 | chr12-68791945-DEL-5281 |
| chr12-68802230-DEL-1377 | chr12-68802231-DEL-1377 |
| chr12-68809453-DEL-66 | chr12-68809453-DEL-66 |
| chr12-68826940-DEL-453 | chr12-68826941-DEL-453 |
| chr12-68831579-DEL-51 | chr12-68831578-DEL-72 |
| chr12-68842982-DEL-58 | chr12-68842983-DEL-58 |
| chr12-68870320-DEL-406 | chr12-68870321-DEL-406 |
| chr12-68877393-DEL-510 | chr12-68877394-DEL-510 |
| chr12-68902458-DEL-52 | chr12-68902459-DEL-52 |
| chr12-68937288-DEL-142 | chr12-68937285-DEL-142 |
| chr12-68992717-DEL-54 | chr12-68992724-DEL-52 |
| chr12-69034690-DEL-72 | chr12-69034691-DEL-56 |
| chr12-69068709-DEL-1548 | chr12-69068710-DEL-1548 |
| chr12-69110005-DEL-199 | chr12-69110006-DEL-199 |
| chr12-69116158-DEL-540 | chr12-69116159-DEL-540 |
| chr12-69120921-DEL-100 | chr12-69120922-DEL-100 |

| | |
|---|---|
| chr12-69123530-DEL-52 | chr12-69123529-DEL-52 |
| chr12-69152858-DEL-72 | chr12-69152859-DEL-72 |
| chr12-69165702-DEL-65 | chr12-69165703-DEL-65 |
| chr12-69168745-DEL-162 | chr12-69168746-DEL-162 |
| chr12-69169727-DEL-1020 | chr12-69169732-DEL-1008 |
| chr12-69180275-DEL-266 | chr12-69180277-DEL-265 |
| chr12-69194549-DEL-228 | chr12-69194552-DEL-226 |
| chr12-69195329-DEL-52 | chr12-69195330-DEL-52 |
| chr12-69213860-DEL-216 | chr12-69213861-DEL-216 |
| chr12-69240468-DEL-1114 | chr12-69240469-DEL-1114 |
| chr12-69251761-DEL-88 | chr12-69251768-DEL-50 |
| chr12-69279934-DEL-344 | chr12-69279935-DEL-344 |
| chr12-69296580-DEL-88 | chr12-69296581-DEL-88 |
| chr12-69308426-DEL-79 | chr12-69308425-DEL-81 |
| chr12-69313622-DEL-498 | chr12-69313623-DEL-498 |
| chr12-69316080-DEL-191 | chr12-69316081-DEL-191 |
| chr12-69349587-DEL-206 | chr12-69349588-DEL-206 |
| chr12-69361899-DEL-51 | chr12-69361900-DEL-51 |
| chr12-69365738-DEL-177 | chr12-69365739-DEL-176 |
| chr12-69370887-DEL-68 | chr12-69370888-DEL-66 |
| chr12-69402446-DEL-80 | chr12-69402447-DEL-80 |
| chr12-69410005-DEL-71 | chr12-69410006-DEL-71 |
| chr12-69423467-DEL-64 | chr12-69423468-DEL-64 |
| chr12-69444504-DEL-1606 | chr12-69444505-DEL-1606 |
| chr12-69454342-DEL-454 | chr12-69454343-DEL-454 |
| chr12-69470768-DEL-60 | chr12-69470769-DEL-60 |
| chr12-69482084-DEL-191 | chr12-69482085-DEL-191 |
| chr12-69488364-DEL-106 | chr12-69488329-DEL-151 |
| chr12-69490163-DEL-107 | chr12-69490168-DEL-71 |
| chr12-69506822-DEL-59 | chr12-69506811-DEL-60 |
| chr12-69513279-DEL-70 | chr12-69513280-DEL-70 |
| chr12-69532186-DEL-60 | chr12-69532195-DEL-51 |
| chr12-69538506-DEL-1004 | chr12-69538507-DEL-1004 |
| chr12-69553431-DEL-410 | chr12-69553432-DEL-410 |
| chr12-69561236-DEL-120 | chr12-69561237-DEL-120 |
| chr12-69570416-DEL-392 | chr12-69570417-DEL-392 |
| chr12-69581580-DEL-111 | chr12-69581581-DEL-111 |
| chr12-69593575-DEL-9045 | chr12-69593576-DEL-9045 |
| chr12-69613071-DEL-134 | chr12-69613072-DEL-134 |
| chr12-69653698-DEL-82 | chr12-69653699-DEL-82 |
| chr12-69657937-DEL-641 | chr12-69657930-DEL-615 |
| chr12-69658011-DEL-60 | chr12-69658012-DEL-60 |
| chr12-69659438-DEL-81 | chr12-69659415-DEL-54 |
| chr12-69669491-DEL-452 | chr12-69669491-DEL-453 |
| chr12-69671016-DEL-858 | chr12-69671017-DEL-858 |
| chr12-69680339-DEL-387 | chr12-69680340-DEL-387 |
| chr12-69694416-DEL-65 | chr12-69694417-DEL-65 |
| chr12-69709123-DEL-150 | chr12-69709123-DEL-150 |
| chr12-69710581-DEL-206 | chr12-69710582-DEL-206 |
| chr12-69754189-DEL-99 | chr12-69754190-DEL-99 |
| chr12-69773002-DEL-235 | chr12-69773003-DEL-235 |
| chr12-69793090-DEL-184 | chr12-69793104-DEL-192 |
| chr12-69849169-DEL-203 | chr12-69849157-DEL-204 |
| chr12-69853319-DEL-460 | chr12-69853320-DEL-460 |
| chr12-69868593-DEL-120 | chr12-69868594-DEL-120 |
| chr12-69897402-DEL-168 | chr12-69897403-DEL-169 |
| chr12-69906469-DEL-1454 | chr12-69906470-DEL-1454 |
| chr12-69963846-DEL-2687 | chr12-69963847-DEL-2687 |
| chr12-69975361-DEL-194 | chr12-69975362-DEL-194 |
| chr12-70027931-DEL-51 | chr12-70027913-DEL-50 |
| chr12-70047772-DEL-4196 | chr12-70047773-DEL-4196 |
| chr12-70099059-DEL-78 | chr12-70099060-DEL-78 |
| chr12-70107627-DEL-164 | chr12-70107628-DEL-164 |
| chr12-70128801-DEL-86 | chr12-70128793-DEL-67 |
| chr12-70146794-DEL-98 | chr12-70146795-DEL-98 |
| chr12-70149020-DEL-164 | chr12-70149021-DEL-164 |
| chr12-70174006-DEL-362 | chr12-70174010-DEL-363 |
| chr12-70181085-DEL-92 | chr12-70181086-DEL-92 |
| chr12-70207568-DEL-57 | chr12-70207580-DEL-58 |
| chr12-70231234-DEL-77 | chr12-70231236-DEL-76 |
| chr12-70240200-DEL-55 | chr12-70240201-DEL-55 |
| chr12-70258027-DEL-180 | chr12-70258027-DEL-180 |
| chr12-70262255-DEL-681 | chr12-70262256-DEL-681 |
| chr12-70263519-DEL-1441 | chr12-70263520-DEL-1441 |
| chr12-70346576-DEL-106 | chr12-70346571-DEL-104 |
| chr12-70353042-DEL-824 | chr12-70353043-DEL-824 |
| chr12-70401446-DEL-84 | chr12-70401447-DEL-84 |
| chr12-70406728-DEL-770 | chr12-70406716-DEL-770 |
| chr12-70418337-DEL-810 | chr12-70418338-DEL-810 |
| chr12-70434735-DEL-188 | chr12-70434738-DEL-187 |

| | |
|---|---|
| chr12-70495868-DEL-160 | chr12-70495869-DEL-160 |
| chr12-70503159-DEL-119 | chr12-70503160-DEL-119 |
| chr12-70509609-DEL-100 | chr12-70509610-DEL-100 |
| chr12-70526664-DEL-5986 | chr12-70526665-DEL-5986 |
| chr12-70535518-DEL-826 | chr12-70535519-DEL-826 |
| chr12-70592555-DEL-91 | chr12-70592556-DEL-91 |
| chr12-70598049-DEL-683 | chr12-70598050-DEL-683 |
| chr12-70618494-DEL-94 | chr12-70618495-DEL-94 |
| chr12-70619662-DEL-62 | chr12-70619669-DEL-56 |
| chr12-70628727-DEL-62 | chr12-70628728-DEL-61 |
| chr12-70636664-DEL-102 | chr12-70636695-DEL-84 |
| chr12-70663621-DEL-106 | chr12-70663621-DEL-82 |
| chr12-70707948-DEL-7109 | chr12-70707949-DEL-7109 |
| chr12-70731655-DEL-51 | chr12-70731656-DEL-51 |
| chr12-70795239-DEL-344 | chr12-70795240-DEL-344 |
| chr12-70834041-DEL-213 | chr12-70834042-DEL-213 |
| chr12-70841078-DEL-52 | chr12-70841090-DEL-71 |
| chr12-70849728-DEL-95 | chr12-70849729-DEL-52 |
| chr12-70878831-DEL-70 | chr12-70878831-DEL-70 |
| chr12-70879341-DEL-64 | chr12-70879342-DEL-56 |
| chr12-70918603-DEL-162 | chr12-70918604-DEL-162 |
| chr12-70929611-DEL-277 | chr12-70929625-DEL-280 |
| chr12-70931747-DEL-83 | chr12-70931743-DEL-84 |
| chr12-70962695-DEL-165 | chr12-70962698-DEL-151 |
| chr12-70964774-DEL-7300 | chr12-70964781-DEL-7300 |
| chr12-70977243-DEL-196 | chr12-70977246-DEL-198 |
| chr12-70979935-DEL-210 | chr12-70979936-DEL-210 |
| chr12-71009574-DEL-223 | chr12-71009570-DEL-222 |
| chr12-71020920-DEL-186 | chr12-71020920-DEL-187 |
| chr12-71023928-DEL-213 | chr12-71023929-DEL-213 |
| chr12-71050452-DEL-221 | chr12-71050453-DEL-221 |
| chr12-71075933-DEL-399 | chr12-71075934-DEL-399 |
| chr12-71077366-DEL-160 | chr12-71077367-DEL-160 |
| chr12-71109258-DEL-203 | chr12-71109258-DEL-204 |
| chr12-71111850-DEL-193 | chr12-71111851-DEL-193 |
| chr12-71116046-DEL-184 | chr12-71116047-DEL-184 |
| chr12-71123917-DEL-6442 | chr12-71123918-DEL-6442 |
| chr12-71131171-DEL-244 | chr12-71131125-DEL-242 |
| chr12-71149695-DEL-645 | chr12-71149696-DEL-645 |
| chr12-71151022-DEL-200 | chr12-71151023-DEL-200 |
| chr12-71151590-DEL-3337 | chr12-71151591-DEL-3337 |
| chr12-71177533-DEL-66 | chr12-71177515-DEL-69 |
| chr12-71210588-DEL-326 | chr12-71210588-DEL-286 |
| chr12-71211717-DEL-141 | chr12-71211730-DEL-94 |
| chr12-71215593-DEL-53 | chr12-71215594-DEL-53 |
| chr12-71230274-DEL-210 | chr12-71230275-DEL-210 |
| chr12-71230787-DEL-100 | chr12-71230788-DEL-100 |
| chr12-71238610-DEL-72 | chr12-71238611-DEL-72 |
| chr12-71243256-DEL-77 | chr12-71243257-DEL-102 |
| chr12-71253622-DEL-6608 | chr12-71253623-DEL-6608 |
| chr12-71265658-DEL-116 | chr12-71265658-DEL-116 |
| chr12-71269306-DEL-217 | chr12-71269307-DEL-217 |
| chr12-71284688-DEL-1887 | chr12-71284694-DEL-1882 |
| chr12-71296955-DEL-146 | chr12-71296954-DEL-139 |
| chr12-71375648-DEL-55 | chr12-71375649-DEL-55 |
| chr12-71376412-DEL-722 | chr12-71376413-DEL-711 |
| chr12-71391192-DEL-80 | chr12-71391193-DEL-80 |
| chr12-71396500-DEL-71 | chr12-71396501-DEL-71 |
| chr12-71398902-DEL-386 | chr12-71398903-DEL-386 |
| chr12-71403432-DEL-137 | chr12-71403433-DEL-137 |
| chr12-71452554-DEL-342 | chr12-71452555-DEL-342 |
| chr12-71491540-DEL-212 | chr12-71491529-DEL-213 |
| chr12-71509876-DEL-57 | chr12-71509877-DEL-57 |
| chr12-71525825-DEL-55 | chr12-71525826-DEL-55 |
| chr12-71535473-DEL-188 | chr12-71535474-DEL-188 |
| chr12-71552013-DEL-54 | chr12-71552014-DEL-54 |
| chr12-71570387-DEL-67 | chr12-71570343-DEL-115 |
| chr12-71591039-DEL-72 | chr12-71591048-DEL-64 |
| chr12-71603493-DEL-82 | chr12-71603494-DEL-82 |
| chr12-71636394-DEL-6651 | chr12-71636395-DEL-6651 |
| chr12-71679440-DEL-82 | chr12-71679452-DEL-52 |
| chr12-71685443-DEL-951 | chr12-71685444-DEL-951 |
| chr12-71703919-DEL-94 | chr12-71703920-DEL-94 |
| chr12-71714370-DEL-1288 | chr12-71714371-DEL-1288 |
| chr12-71733296-DEL-196 | chr12-71733285-DEL-197 |
| chr12-71736259-DEL-120 | chr12-71736259-DEL-120 |
| chr12-71737235-DEL-639 | chr12-71737235-DEL-645 |
| chr12-71762197-DEL-74 | chr12-71762217-DEL-95 |
| chr12-71784361-DEL-414 | chr12-71784362-DEL-414 |
| chr12-71786957-DEL-402 | chr12-71786958-DEL-402 |

| | |
|---|---|
| chr12-71809121-DEL-58 | chr12-71809122-DEL-58 |
| chr12-71823004-DEL-1355 | chr12-71823005-DEL-1355 |
| chr12-71829117-DEL-58 | chr12-71829118-DEL-58 |
| chr12-71873149-DEL-2118 | chr12-71873150-DEL-2118 |
| chr12-71897644-DEL-130 | chr12-71897645-DEL-130 |
| chr12-71924386-DEL-683 | chr12-71924387-DEL-683 |
| chr12-71969858-DEL-836 | chr12-71969859-DEL-836 |
| chr12-72009860-DEL-4685 | chr12-72009861-DEL-4685 |
| chr12-72028382-DEL-78 | chr12-72028383-DEL-72 |
| chr12-72029090-DEL-74 | chr12-72029103-DEL-84 |
| chr12-72054031-DEL-70 | chr12-72054032-DEL-60 |
| chr12-72057829-DEL-64 | chr12-72057830-DEL-64 |
| chr12-72059598-DEL-150 | chr12-72059599-DEL-150 |
| chr12-72075383-DEL-175 | chr12-72075381-DEL-200 |
| chr12-72087524-DEL-70 | chr12-72087520-DEL-74 |
| chr12-72210830-DEL-1069 | chr12-72210818-DEL-1054 |
| chr12-72215510-DEL-1985 | chr12-72215511-DEL-1985 |
| chr12-72239148-DEL-60 | chr12-72239149-DEL-60 |
| chr12-72249751-DEL-6321 | chr12-72249752-DEL-6321 |
| chr12-72266516-DEL-257 | chr12-72266519-DEL-258 |
| chr12-72277610-DEL-95 | chr12-72277611-DEL-95 |
| chr12-72287857-DEL-59 | chr12-72287858-DEL-59 |
| chr12-72309112-DEL-649 | chr12-72309113-DEL-648 |
| chr12-72317304-DEL-1892 | chr12-72317305-DEL-1892 |
| chr12-72321255-DEL-80 | chr12-72321289-DEL-52 |
| chr12-72349425-DEL-200 | chr12-72349411-DEL-201 |
| chr12-72364947-DEL-173 | chr12-72364907-DEL-174 |
| chr12-72373788-DEL-229 | chr12-72373799-DEL-218 |
| chr12-72377774-DEL-467 | chr12-72377775-DEL-467 |
| chr12-72380389-DEL-69 | chr12-72380390-DEL-69 |
| chr12-72394428-DEL-56 | chr12-72394429-DEL-56 |
| chr12-72413816-DEL-51 | chr12-72413806-DEL-52 |
| chr12-72444977-DEL-65 | chr12-72444976-DEL-65 |
| chr12-72452084-DEL-58 | chr12-72452085-DEL-58 |
| chr12-72464956-DEL-357 | chr12-72464957-DEL-357 |
| chr12-72471914-DEL-323 | chr12-72471915-DEL-320 |
| chr12-72477027-DEL-52 | chr12-72477028-DEL-52 |
| chr12-72491168-DEL-2344 | chr12-72491169-DEL-2344 |
| chr12-72493275-DEL-59 | chr12-72493276-DEL-59 |
| chr12-72532539-DEL-189 | chr12-72532531-DEL-190 |
| chr12-72559207-DEL-86 | chr12-72559208-DEL-86 |
| chr12-72589637-DEL-278 | chr12-72589638-DEL-278 |
| chr12-72594976-DEL-229 | chr12-72594977-DEL-229 |
| chr12-72604249-DEL-6438 | chr12-72604250-DEL-6438 |
| chr12-72629690-DEL-54 | chr12-72629691-DEL-54 |
| chr12-72634232-DEL-89 | chr12-72634233-DEL-89 |
| chr12-72672278-DEL-286 | chr12-72672279-DEL-286 |
| chr12-72716217-DEL-75 | chr12-72716218-DEL-75 |
| chr12-72722482-DEL-114 | chr12-72722483-DEL-114 |
| chr12-72726771-DEL-132 | chr12-72726772-DEL-132 |
| chr12-72730694-DEL-207 | chr12-72730695-DEL-207 |
| chr12-72739445-DEL-215 | chr12-72739446-DEL-215 |
| chr12-72751240-DEL-76 | chr12-72751241-DEL-76 |
| chr12-72755940-DEL-57 | chr12-72755936-DEL-53 |
| chr12-72794855-DEL-229 | chr12-72794856-DEL-229 |
| chr12-72798814-DEL-53 | chr12-72798815-DEL-53 |
| chr12-72816462-DEL-61 | chr12-72816463-DEL-61 |
| chr12-72817064-DEL-56 | chr12-72817071-DEL-52 |
| chr12-72847969-DEL-155 | chr12-72847970-DEL-124 |
| chr12-72864051-DEL-195 | chr12-72864052-DEL-188 |
| chr12-72891706-DEL-225 | chr12-72891707-DEL-225 |
| chr12-72904559-DEL-70 | chr12-72904560-DEL-70 |
| chr12-72994524-DEL-52 | chr12-72994533-DEL-51 |
| chr12-73013525-DEL-716 | chr12-73013526-DEL-716 |
| chr12-73040959-DEL-206 | chr12-73040950-DEL-205 |
| chr12-73043881-DEL-77 | chr12-73043890-DEL-75 |
| chr12-73114171-DEL-196 | chr12-73114172-DEL-196 |
| chr12-73120319-DEL-67 | chr12-73120320-DEL-66 |
| chr12-73120996-DEL-6442 | chr12-73120997-DEL-6442 |
| chr12-73177399-DEL-138 | chr12-73177400-DEL-138 |
| chr12-73220347-DEL-219 | chr12-73220348-DEL-219 |
| chr12-73233584-DEL-56 | chr12-73233584-DEL-56 |
| chr12-73257762-DEL-191 | chr12-73257763-DEL-191 |
| chr12-73261359-DEL-51 | chr12-73261360-DEL-51 |
| chr12-73276205-DEL-415 | chr12-73276206-DEL-415 |
| chr12-73308729-DEL-221 | chr12-73308730-DEL-221 |
| chr12-73310537-DEL-732 | chr12-73310538-DEL-732 |
| chr12-73325995-DEL-52 | chr12-73325996-DEL-52 |
| chr12-73336899-DEL-449 | chr12-73336901-DEL-448 |
| chr12-73353425-DEL-196 | chr12-73353426-DEL-196 |

| | |
|---|---|
| chr12-73357760-DEL-4874 | chr12-73357761-DEL-4874 |
| chr12-73443718-DEL-94 | chr12-73443719-DEL-94 |
| chr12-73454850-DEL-94 | chr12-73454851-DEL-94 |
| chr12-73482387-DEL-194 | chr12-73482349-DEL-218 |
| chr12-73483714-DEL-54 | chr12-73483715-DEL-54 |
| chr12-73503993-DEL-231 | chr12-73503994-DEL-231 |
| chr12-73585162-DEL-88 | chr12-73585163-DEL-88 |
| chr12-73599888-DEL-192 | chr12-73599889-DEL-192 |
| chr12-73604846-DEL-76 | chr12-73604847-DEL-76 |
| chr12-73643434-DEL-83 | chr12-73643435-DEL-83 |
| chr12-73656062-DEL-504 | chr12-73656063-DEL-504 |
| chr12-73668716-DEL-58 | chr12-73668717-DEL-58 |
| chr12-73692562-DEL-498 | chr12-73692563-DEL-498 |
| chr12-73715912-DEL-441 | chr12-73715913-DEL-441 |
| chr12-73742115-DEL-71 | chr12-73742116-DEL-69 |
| chr12-73756263-DEL-153 | chr12-73756267-DEL-155 |
| chr12-73763761-DEL-51 | chr12-73763762-DEL-59 |
| chr12-73783194-DEL-255 | chr12-73783195-DEL-255 |
| chr12-73785627-DEL-98 | chr12-73785628-DEL-98 |
| chr12-73839816-DEL-74 | chr12-73839837-DEL-66 |
| chr12-73841860-DEL-64 | chr12-73841861-DEL-64 |
| chr12-73865041-DEL-1538 | chr12-73865041-DEL-1538 |
| chr12-73891932-DEL-58 | chr12-73891933-DEL-58 |
| chr12-73937858-DEL-216 | chr12-73937859-DEL-216 |
| chr12-73938402-DEL-2159 | chr12-73938403-DEL-2159 |
| chr12-73940630-DEL-578 | chr12-73940631-DEL-578 |
| chr12-73987952-DEL-180 | chr12-73987953-DEL-180 |
| chr12-74010813-DEL-91 | chr12-74010810-DEL-91 |
| chr12-74016047-DEL-58 | chr12-74016048-DEL-58 |
| chr12-74018182-DEL-72 | chr12-74018183-DEL-72 |
| chr12-74046282-DEL-1070 | chr12-74046283-DEL-1070 |
| chr12-74093918-DEL-2254 | chr12-74093918-DEL-2254 |
| chr12-74112205-DEL-504 | chr12-74112206-DEL-504 |
| chr12-74185631-DEL-767 | chr12-74185621-DEL-766 |
| chr12-74232595-DEL-98 | chr12-74232596-DEL-98 |
| chr12-74254089-DEL-80 | chr12-74254086-DEL-90 |
| chr12-74274625-DEL-92 | chr12-74274625-DEL-92 |
| chr12-74312228-DEL-134 | chr12-74312229-DEL-134 |
| chr12-74328552-DEL-57 | chr12-74328553-DEL-59 |
| chr12-74338458-DEL-109 | chr12-74338459-DEL-109 |
| chr12-74352703-DEL-4657 | chr12-74352704-DEL-4657 |
| chr12-74360902-DEL-69 | chr12-74360906-DEL-66 |
| chr12-74362748-DEL-55 | chr12-74362762-DEL-91 |
| chr12-74394958-DEL-57 | chr12-74394989-DEL-63 |
| chr12-74397057-DEL-202 | chr12-74397057-DEL-202 |
| chr12-74401273-DEL-571 | chr12-74401274-DEL-571 |
| chr12-74413004-DEL-189 | chr12-74413005-DEL-189 |
| chr12-74414036-DEL-390 | chr12-74414037-DEL-390 |
| chr12-74434310-DEL-1209 | chr12-74434311-DEL-1209 |
| chr12-74478808-DEL-54 | chr12-74478819-DEL-54 |
| chr12-74508792-DEL-133 | chr12-74508793-DEL-133 |
| chr12-74511124-DEL-3355 | chr12-74511124-DEL-3355 |
| chr12-74514958-DEL-301 | chr12-74514959-DEL-301 |
| chr12-74611576-DEL-478 | chr12-74611577-DEL-480 |
| chr12-74624813-DEL-1373 | chr12-74624798-DEL-1374 |
| chr12-74625860-DEL-129 | chr12-74625861-DEL-129 |
| chr12-74638701-DEL-451 | chr12-74638702-DEL-451 |
| chr12-74645781-DEL-1097 | chr12-74645782-DEL-1097 |
| chr12-74651965-DEL-66 | chr12-74651966-DEL-66 |
| chr12-74663939-DEL-331 | chr12-74663940-DEL-331 |
| chr12-74667501-DEL-3417 | chr12-74667502-DEL-3417 |
| chr12-74682608-DEL-6265 | chr12-74682610-DEL-6264 |
| chr12-74699616-DEL-134 | chr12-74699617-DEL-134 |
| chr12-74728836-DEL-76 | chr12-74728837-DEL-76 |
| chr12-74731226-DEL-650 | chr12-74731227-DEL-650 |
| chr12-74738885-DEL-245 | chr12-74738886-DEL-245 |
| chr12-74745100-DEL-2772 | chr12-74745101-DEL-2772 |
| chr12-74754869-DEL-237 | chr12-74754870-DEL-237 |
| chr12-74757075-DEL-551 | chr12-74757071-DEL-543 |
| chr12-74796088-DEL-54 | chr12-74796089-DEL-54 |
| chr12-74813431-DEL-160 | chr12-74813432-DEL-160 |
| chr12-74822509-DEL-2457 | chr12-74822511-DEL-2456 |
| chr12-74840841-DEL-70 | chr12-74840838-DEL-74 |
| chr12-74854337-DEL-440 | chr12-74854338-DEL-440 |
| chr12-74861540-DEL-514 | chr12-74861541-DEL-514 |
| chr12-74874761-DEL-79 | chr12-74874786-DEL-54 |
| chr12-74922960-DEL-189 | chr12-74922961-DEL-189 |
| chr12-74926711-DEL-51 | chr12-74926732-DEL-53 |
| chr12-74939466-DEL-94 | chr12-74939480-DEL-68 |
| chr12-74973291-DEL-7163 | chr12-74973292-DEL-7163 |

| | |
|---|---|
| chr12-74986162-DEL-56 | chr12-74986163-DEL-58 |
| chr12-75017510-DEL-56 | chr12-75017511-DEL-56 |
| chr12-75163535-DEL-108 | chr12-75163536-DEL-108 |
| chr12-75220351-DEL-52 | chr12-75220354-DEL-50 |
| chr12-75390605-DEL-70 | chr12-75390606-DEL-70 |
| chr12-75395659-DEL-53 | chr12-75395660-DEL-54 |
| chr12-75426180-DEL-6906 | chr12-75426181-DEL-6906 |
| chr12-75517561-DEL-62 | chr12-75517562-DEL-70 |
| chr12-75544088-DEL-53 | chr12-75544090-DEL-52 |
| chr12-75556783-DEL-56 | chr12-75556783-DEL-57 |
| chr12-75598552-DEL-102 | chr12-75598553-DEL-102 |
| chr12-75656711-DEL-174 | chr12-75656712-DEL-174 |
| chr12-75663403-DEL-215 | chr12-75663404-DEL-215 |
| chr12-75675393-DEL-752 | chr12-75675394-DEL-752 |
| chr12-75677655-DEL-690 | chr12-75677656-DEL-690 |
| chr12-75689184-DEL-171 | chr12-75689185-DEL-171 |
| chr12-75696766-DEL-298 | chr12-75696766-DEL-298 |
| chr12-75736690-DEL-198 | chr12-75736691-DEL-198 |
| chr12-75742561-DEL-81 | chr12-75742541-DEL-75 |
| chr12-75776566-DEL-193 | chr12-75776567-DEL-193 |
| chr12-75798118-DEL-180 | chr12-75798119-DEL-180 |
| chr12-75807801-DEL-531 | chr12-75807802-DEL-531 |
| chr12-75818819-DEL-54 | chr12-75818819-DEL-54 |
| chr12-75834093-DEL-154 | chr12-75834095-DEL-153 |
| chr12-75845214-DEL-179 | chr12-75845186-DEL-227 |
| chr12-75855751-DEL-232 | chr12-75855752-DEL-232 |
| chr12-75857906-DEL-218 | chr12-75857907-DEL-218 |
| chr12-75906434-DEL-373 | chr12-75906435-DEL-373 |
| chr12-75940033-DEL-52 | chr12-75940041-DEL-54 |
| chr12-75960656-DEL-73 | chr12-75960657-DEL-69 |
| chr12-75975602-DEL-84 | chr12-75975608-DEL-79 |
| chr12-75994204-DEL-196 | chr12-75994205-DEL-194 |
| chr12-75995256-DEL-117 | chr12-75995267-DEL-110 |
| chr12-76054513-DEL-205 | chr12-76054514-DEL-205 |
| chr12-76114757-DEL-401 | chr12-76114758-DEL-401 |
| chr12-76122976-DEL-193 | chr12-76122977-DEL-193 |
| chr12-76136997-DEL-113 | chr12-76136998-DEL-113 |
| chr12-76197049-DEL-241 | chr12-76197052-DEL-240 |
| chr12-76204825-DEL-72 | chr12-76204833-DEL-64 |
| chr12-76205946-DEL-123 | chr12-76205937-DEL-133 |
| chr12-76214865-DEL-56 | chr12-76214866-DEL-60 |
| chr12-76219420-DEL-475 | chr12-76219421-DEL-475 |
| chr12-76254176-DEL-207 | chr12-76254176-DEL-207 |
| chr12-76258624-DEL-247 | chr12-76258625-DEL-247 |
| chr12-76268128-DEL-74 | chr12-76268129-DEL-92 |
| chr12-76269112-DEL-7098 | chr12-76269113-DEL-7098 |
| chr12-76314269-DEL-189 | chr12-76314270-DEL-189 |
| chr12-76333158-DEL-400 | chr12-76333138-DEL-386 |
| chr12-76391797-DEL-211 | chr12-76391798-DEL-211 |
| chr12-76397971-DEL-1134 | chr12-76397972-DEL-1134 |
| chr12-76403493-DEL-182 | chr12-76403494-DEL-182 |
| chr12-76420168-DEL-216 | chr12-76420169-DEL-216 |
| chr12-76431292-DEL-55 | chr12-76431293-DEL-55 |
| chr12-76457962-DEL-1274 | chr12-76457961-DEL-1274 |
| chr12-76499323-DEL-199 | chr12-76499324-DEL-199 |
| chr12-76506612-DEL-506 | chr12-76506613-DEL-506 |
| chr12-76515742-DEL-225 | chr12-76515739-DEL-226 |
| chr12-76521419-DEL-108 | chr12-76521420-DEL-64 |
| chr12-76523457-DEL-158 | chr12-76523462-DEL-165 |
| chr12-76567052-DEL-1039 | chr12-76567047-DEL-1042 |
| chr12-76621276-DEL-195 | chr12-76621277-DEL-195 |
| chr12-76631209-DEL-66 | chr12-76631208-DEL-68 |
| chr12-76638328-DEL-79 | chr12-76638342-DEL-60 |
| chr12-76685065-DEL-208 | chr12-76685066-DEL-208 |
| chr12-76700055-DEL-2462 | chr12-76700053-DEL-2462 |
| chr12-76732529-DEL-119 | chr12-76732530-DEL-119 |
| chr12-76764168-DEL-52 | chr12-76764122-DEL-56 |
| chr12-76799716-DEL-77 | chr12-76799717-DEL-77 |
| chr12-76802359-DEL-6442 | chr12-76802357-DEL-6442 |
| chr12-76814886-DEL-192 | chr12-76814887-DEL-192 |
| chr12-76823285-DEL-373 | chr12-76823286-DEL-373 |
| chr12-76825547-DEL-84 | chr12-76825548-DEL-84 |
| chr12-76828766-DEL-52 | chr12-76828767-DEL-52 |
| chr12-76829723-DEL-197 | chr12-76829726-DEL-197 |
| chr12-76855627-DEL-405 | chr12-76855628-DEL-405 |
| chr12-76886494-DEL-70 | chr12-76886495-DEL-70 |
| chr12-76902037-DEL-207 | chr12-76902038-DEL-207 |
| chr12-76904286-DEL-1005 | chr12-76904287-DEL-1005 |
| chr12-76907498-DEL-222 | chr12-76907499-DEL-222 |
| chr12-76915920-DEL-217 | chr12-76915897-DEL-253 |

| | |
|---|---|
| chr12-76924898-DEL-88 | chr12-76924898-DEL-88 |
| chr12-76939790-DEL-92 | chr12-76939791-DEL-138 |
| chr12-76941770-DEL-328 | chr12-76941771-DEL-328 |
| chr12-76962173-DEL-501 | chr12-76962174-DEL-501 |
| chr12-76978319-DEL-198 | chr12-76978320-DEL-198 |
| chr12-76980155-DEL-143 | chr12-76980156-DEL-143 |
| chr12-77031674-DEL-56 | chr12-77031677-DEL-54 |
| chr12-77038308-DEL-219 | chr12-77038307-DEL-241 |
| chr12-77044773-DEL-59 | chr12-77044774-DEL-59 |
| chr12-77062181-DEL-135 | chr12-77062179-DEL-164 |
| chr12-77073649-DEL-65 | chr12-77073601-DEL-81 |
| chr12-77075497-DEL-67 | chr12-77075498-DEL-67 |
| chr12-77084388-DEL-218 | chr12-77084389-DEL-218 |
| chr12-77090633-DEL-189 | chr12-77090634-DEL-189 |
| chr12-77128938-DEL-761 | chr12-77128939-DEL-752 |
| chr12-77143030-DEL-217 | chr12-77143031-DEL-217 |
| chr12-77149575-DEL-83 | chr12-77149576-DEL-83 |
| chr12-77165328-DEL-640 | chr12-77165319-DEL-640 |
| chr12-77197200-DEL-185 | chr12-77197201-DEL-185 |
| chr12-77210204-DEL-873 | chr12-77210199-DEL-869 |
| chr12-77215434-DEL-547 | chr12-77215435-DEL-547 |
| chr12-77224428-DEL-61 | chr12-77224440-DEL-57 |
| chr12-77233365-DEL-97 | chr12-77233366-DEL-97 |
| chr12-77300456-DEL-60 | chr12-77300442-DEL-61 |
| chr12-77316198-DEL-1079 | chr12-77316199-DEL-1079 |
| chr12-77338546-DEL-206 | chr12-77338547-DEL-206 |
| chr12-77350935-DEL-217 | chr12-77350939-DEL-214 |
| chr12-77430116-DEL-6336 | chr12-77430117-DEL-6336 |
| chr12-77437402-DEL-92 | chr12-77437407-DEL-88 |
| chr12-77438762-DEL-823 | chr12-77438763-DEL-823 |
| chr12-77451621-DEL-6463 | chr12-77451622-DEL-6463 |
| chr12-77460772-DEL-330 | chr12-77460773-DEL-330 |
| chr12-77529023-DEL-59 | chr12-77529024-DEL-59 |
| chr12-77560537-DEL-420 | chr12-77560538-DEL-420 |
| chr12-77565954-DEL-130 | chr12-77565967-DEL-129 |
| chr12-77593877-DEL-6671 | chr12-77593878-DEL-6671 |
| chr12-77709439-DEL-174 | chr12-77709440-DEL-174 |
| chr12-77715213-DEL-105 | chr12-77715214-DEL-105 |
| chr12-77780958-DEL-79 | chr12-77780959-DEL-79 |
| chr12-77815705-DEL-6647 | chr12-77815706-DEL-6647 |
| chr12-77848699-DEL-2228 | chr12-77848700-DEL-2228 |
| chr12-77885794-DEL-148 | chr12-77885807-DEL-140 |
| chr12-77893112-DEL-573 | chr12-77893113-DEL-573 |
| chr12-77920878-DEL-99 | chr12-77920853-DEL-108 |
| chr12-77925656-DEL-8453 | chr12-77925647-DEL-8437 |
| chr12-77946031-DEL-1215 | chr12-77946032-DEL-1211 |
| chr12-77965437-DEL-603 | chr12-77965438-DEL-603 |
| chr12-77968122-DEL-498 | chr12-77968123-DEL-498 |
| chr12-77970865-DEL-3425 | chr12-77970865-DEL-3425 |
| chr12-77984492-DEL-187 | chr12-77984491-DEL-157 |
| chr12-77996038-DEL-55 | chr12-77996038-DEL-55 |
| chr12-78009888-DEL-69 | chr12-78009889-DEL-69 |
| chr12-78054160-DEL-1647 | chr12-78054161-DEL-1647 |
| chr12-78056125-DEL-326 | chr12-78056126-DEL-326 |
| chr12-78057989-DEL-1233 | chr12-78057980-DEL-1232 |
| chr12-78076700-DEL-366 | chr12-78076701-DEL-366 |
| chr12-78078584-DEL-130 | chr12-78078585-DEL-130 |
| chr12-78097061-DEL-3558 | chr12-78097062-DEL-3558 |
| chr12-78111209-DEL-75 | chr12-78111215-DEL-75 |
| chr12-78121712-DEL-297 | chr12-78121730-DEL-282 |
| chr12-78123368-DEL-134 | chr12-78123369-DEL-134 |
| chr12-78138056-DEL-96 | chr12-78138056-DEL-96 |
| chr12-78149006-DEL-58 | chr12-78149009-DEL-57 |
| chr12-78161944-DEL-1293 | chr12-78161945-DEL-1293 |
| chr12-78168979-DEL-2293 | chr12-78168980-DEL-2293 |
| chr12-78272462-DEL-52 | chr12-78272465-DEL-53 |
| chr12-78335614-DEL-1397 | chr12-78335615-DEL-1397 |
| chr12-78357926-DEL-117 | chr12-78357927-DEL-117 |
| chr12-78381764-DEL-3885 | chr12-78381765-DEL-3885 |
| chr12-78391564-DEL-4585 | chr12-78391565-DEL-4585 |
| chr12-78392054-DEL-53 | chr12-78392056-DEL-51 |
| chr12-78410277-DEL-199 | chr12-78410278-DEL-199 |
| chr12-78453678-DEL-52 | chr12-78453679-DEL-52 |
| chr12-78514043-DEL-65 | chr12-78514045-DEL-64 |
| chr12-78542014-DEL-704 | chr12-78542015-DEL-704 |
| chr12-78562221-DEL-658 | chr12-78562222-DEL-658 |
| chr12-78572851-DEL-577 | chr12-78572852-DEL-577 |
| chr12-78599211-DEL-1354 | chr12-78599212-DEL-1354 |
| chr12-78656599-DEL-1326 | chr12-78656600-DEL-1324 |
| chr12-78707582-DEL-121 | chr12-78707577-DEL-96 |

| | |
|---|---|
| chr12-78718560-DEL-2788 | chr12-78718561-DEL-2788 |
| chr12-78751618-DEL-205 | chr12-78751619-DEL-204 |
| chr12-78765582-DEL-58 | chr12-78765583-DEL-58 |
| chr12-78809548-DEL-518 | chr12-78809549-DEL-518 |
| chr12-78840022-DEL-212 | chr12-78840023-DEL-212 |
| chr12-78852390-DEL-245 | chr12-78852391-DEL-245 |
| chr12-78874910-DEL-337 | chr12-78874911-DEL-337 |
| chr12-78881031-DEL-214 | chr12-78881031-DEL-214 |
| chr12-78887750-DEL-58 | chr12-78887751-DEL-58 |
| chr12-78892420-DEL-626 | chr12-78892423-DEL-624 |
| chr12-78894879-DEL-96 | chr12-78894883-DEL-115 |
| chr12-78899273-DEL-101 | chr12-78899274-DEL-101 |
| chr12-78902264-DEL-931 | chr12-78902265-DEL-931 |
| chr12-78920382-DEL-124 | chr12-78920383-DEL-124 |
| chr12-78951306-DEL-77 | chr12-78951307-DEL-77 |
| chr12-78960300-DEL-450 | chr12-78960301-DEL-450 |
| chr12-78971320-DEL-121 | chr12-78971321-DEL-121 |
| chr12-78987190-DEL-83 | chr12-78987191-DEL-83 |
| chr12-78992802-DEL-271 | chr12-78992803-DEL-271 |
| chr12-79006976-DEL-66 | chr12-79006976-DEL-67 |
| chr12-79013943-DEL-1544 | chr12-79013944-DEL-1544 |
| chr12-79038896-DEL-90 | chr12-79038897-DEL-138 |
| chr12-79045946-DEL-198 | chr12-79045947-DEL-198 |
| chr12-79074554-DEL-187 | chr12-79074555-DEL-187 |
| chr12-79089142-DEL-654 | chr12-79089143-DEL-653 |
| chr12-79090012-DEL-73 | chr12-79090013-DEL-73 |
| chr12-79103217-DEL-3020 | chr12-79103218-DEL-3020 |
| chr12-79109813-DEL-68 | chr12-79109814-DEL-68 |
| chr12-79140661-DEL-180 | chr12-79140662-DEL-180 |
| chr12-79142001-DEL-406 | chr12-79142002-DEL-406 |
| chr12-79161072-DEL-2341 | chr12-79161074-DEL-2341 |
| chr12-79169958-DEL-6391 | chr12-79169959-DEL-6391 |
| chr12-79185105-DEL-64 | chr12-79185108-DEL-65 |
| chr12-79232226-DEL-59 | chr12-79232213-DEL-78 |
| chr12-79247190-DEL-816 | chr12-79247191-DEL-816 |
| chr12-79271428-DEL-52 | chr12-79271429-DEL-52 |
| chr12-79272738-DEL-52 | chr12-79272739-DEL-52 |
| chr12-79284222-DEL-80 | chr12-79284223-DEL-80 |
| chr12-79302201-DEL-60 | chr12-79302187-DEL-88 |
| chr12-79309386-DEL-59 | chr12-79309385-DEL-67 |
| chr12-79309743-DEL-270 | chr12-79309750-DEL-267 |
| chr12-79335242-DEL-202 | chr12-79335243-DEL-201 |
| chr12-79356938-DEL-62 | chr12-79356939-DEL-62 |
| chr12-79369815-DEL-84 | chr12-79369816-DEL-84 |
| chr12-79421024-DEL-181 | chr12-79421025-DEL-181 |
| chr12-79426194-DEL-726 | chr12-79426195-DEL-726 |
| chr12-79439630-DEL-52 | chr12-79439608-DEL-99 |
| chr12-79442611-DEL-3769 | chr12-79442612-DEL-3769 |
| chr12-79444730-DEL-66 | chr12-79444688-DEL-58 |
| chr12-79455216-DEL-120 | chr12-79455217-DEL-120 |
| chr12-79473543-DEL-440 | chr12-79473544-DEL-440 |
| chr12-79482839-DEL-51 | chr12-79482818-DEL-52 |
| chr12-79492381-DEL-55 | chr12-79492409-DEL-56 |
| chr12-79503295-DEL-265 | chr12-79503296-DEL-265 |
| chr12-79514325-DEL-432 | chr12-79514326-DEL-432 |
| chr12-79527016-DEL-68 | chr12-79527010-DEL-81 |
| chr12-79563607-DEL-72 | chr12-79563582-DEL-50 |
| chr12-79566061-DEL-4286 | chr12-79566062-DEL-4286 |
| chr12-79686391-DEL-60 | chr12-79686392-DEL-60 |
| chr12-79687485-DEL-203 | chr12-79687486-DEL-203 |
| chr12-79714228-DEL-180 | chr12-79714229-DEL-180 |
| chr12-79716830-DEL-265 | chr12-79716831-DEL-265 |
| chr12-79765599-DEL-234 | chr12-79765599-DEL-234 |
| chr12-79767010-DEL-434 | chr12-79767011-DEL-434 |
| chr12-79770556-DEL-227 | chr12-79770557-DEL-227 |
| chr12-79784846-DEL-64 | chr12-79784849-DEL-62 |
| chr12-79802911-DEL-66 | chr12-79802911-DEL-66 |
| chr12-79850814-DEL-75 | chr12-79850815-DEL-75 |
| chr12-79876375-DEL-53 | chr12-79876373-DEL-54 |
| chr12-79978736-DEL-54 | chr12-79978737-DEL-54 |
| chr12-79987570-DEL-64 | chr12-79987563-DEL-78 |
| chr12-79999316-DEL-405 | chr12-79999317-DEL-405 |
| chr12-80006151-DEL-90 | chr12-80006152-DEL-90 |
| chr12-80011391-DEL-147 | chr12-80011403-DEL-112 |
| chr12-80027172-DEL-54 | chr12-80027173-DEL-54 |
| chr12-80039444-DEL-67 | chr12-80039445-DEL-67 |
| chr12-80073819-DEL-54 | chr12-80073820-DEL-54 |
| chr12-80097714-DEL-409 | chr12-80097715-DEL-409 |
| chr12-80196591-DEL-203 | chr12-80196592-DEL-203 |
| chr12-80207471-DEL-204 | chr12-80207472-DEL-204 |

| | |
|---|---|
| chr12-80272036-DEL-159 | chr12-80272037-DEL-159 |
| chr12-80285045-DEL-90 | chr12-80285048-DEL-88 |
| chr12-80310749-DEL-733 | chr12-80310750-DEL-733 |
| chr12-80311797-DEL-215 | chr12-80311802-DEL-207 |
| chr12-80342233-DEL-144 | chr12-80342231-DEL-145 |
| chr12-80352185-DEL-401 | chr12-80352186-DEL-401 |
| chr12-80364498-DEL-514 | chr12-80364499-DEL-514 |
| chr12-80392653-DEL-196 | chr12-80392654-DEL-196 |
| chr12-80399902-DEL-203 | chr12-80399903-DEL-203 |
| chr12-80408535-DEL-60 | chr12-80408536-DEL-60 |
| chr12-80419625-DEL-204 | chr12-80419626-DEL-204 |
| chr12-80425230-DEL-802 | chr12-80425199-DEL-803 |
| chr12-80451627-DEL-2929 | chr12-80451628-DEL-2929 |
| chr12-80461383-DEL-11442 | chr12-80461384-DEL-11442 |
| chr12-80477737-DEL-99 | chr12-80477755-DEL-99 |
| chr12-80484728-DEL-208 | chr12-80484729-DEL-208 |
| chr12-80494583-DEL-1899 | chr12-80494584-DEL-1899 |
| chr12-80503245-DEL-201 | chr12-80503246-DEL-201 |
| chr12-80509349-DEL-64 | chr12-80509350-DEL-92 |
| chr12-80519360-DEL-439 | chr12-80519361-DEL-439 |
| chr12-80525181-DEL-209 | chr12-80525182-DEL-209 |
| chr12-80539827-DEL-183 | chr12-80539828-DEL-183 |
| chr12-80576034-DEL-73 | chr12-80576035-DEL-73 |
| chr12-80598115-DEL-189 | chr12-80598116-DEL-189 |
| chr12-80607945-DEL-222 | chr12-80607946-DEL-222 |
| chr12-80652631-DEL-193 | chr12-80652632-DEL-193 |
| chr12-80668960-DEL-62 | chr12-80668961-DEL-62 |
| chr12-80672711-DEL-173 | chr12-80672712-DEL-173 |
| chr12-80700100-DEL-224 | chr12-80700101-DEL-224 |
| chr12-80749356-DEL-69 | chr12-80749357-DEL-69 |
| chr12-80779162-DEL-50 | chr12-80779180-DEL-58 |
| chr12-80791897-DEL-115 | chr12-80791877-DEL-133 |
| chr12-80811456-DEL-54 | chr12-80811457-DEL-54 |
| chr12-80824049-DEL-184 | chr12-80824050-DEL-184 |
| chr12-80826020-DEL-198 | chr12-80826021-DEL-198 |
| chr12-80831137-DEL-52 | chr12-80831118-DEL-60 |
| chr12-80846117-DEL-51 | chr12-80846113-DEL-56 |
| chr12-80911209-DEL-52 | chr12-80911192-DEL-64 |
| chr12-80933132-DEL-60 | chr12-80933159-DEL-63 |
| chr12-80944382-DEL-132 | chr12-80944383-DEL-132 |
| chr12-80964193-DEL-200 | chr12-80964194-DEL-200 |
| chr12-80966445-DEL-1726 | chr12-80966446-DEL-1726 |
| chr12-80991730-DEL-122 | chr12-80991734-DEL-84 |
| chr12-81003396-DEL-286 | chr12-81003397-DEL-286 |
| chr12-81007704-DEL-89 | chr12-81007705-DEL-87 |
| chr12-81032043-DEL-84 | chr12-81032064-DEL-52 |
| chr12-81041210-DEL-79 | chr12-81041208-DEL-80 |
| chr12-81041594-DEL-789 | chr12-81041590-DEL-790 |
| chr12-81075855-DEL-74 | chr12-81075837-DEL-60 |
| chr12-81095706-DEL-66 | chr12-81095746-DEL-51 |
| chr12-81141261-DEL-68 | chr12-81141290-DEL-52 |
| chr12-81144983-DEL-63 | chr12-81144983-DEL-85 |
| chr12-81161024-DEL-182 | chr12-81161025-DEL-182 |
| chr12-81163776-DEL-89 | chr12-81163766-DEL-66 |
| chr12-81165964-DEL-52 | chr12-81165965-DEL-52 |
| chr12-81175167-DEL-82 | chr12-81175168-DEL-82 |
| chr12-81209220-DEL-84 | chr12-81209221-DEL-84 |
| chr12-81300456-DEL-1670 | chr12-81300455-DEL-1670 |
| chr12-81306108-DEL-365 | chr12-81306109-DEL-365 |
| chr12-81316771-DEL-84 | chr12-81316770-DEL-58 |
| chr12-81324819-DEL-95 | chr12-81324820-DEL-95 |
| chr12-81325400-DEL-382 | chr12-81325401-DEL-382 |
| chr12-81352914-DEL-473 | chr12-81352915-DEL-473 |
| chr12-81443697-DEL-53 | chr12-81443730-DEL-51 |
| chr12-81483279-DEL-423 | chr12-81483280-DEL-423 |
| chr12-81568984-DEL-1649 | chr12-81568985-DEL-1649 |
| chr12-81629862-DEL-4407 | chr12-81629863-DEL-4407 |
| chr12-81635224-DEL-67 | chr12-81635221-DEL-53 |
| chr12-81653018-DEL-54 | chr12-81653022-DEL-58 |
| chr12-81659351-DEL-60 | chr12-81659358-DEL-54 |
| chr12-81669658-DEL-274 | chr12-81669659-DEL-274 |
| chr12-81684972-DEL-480 | chr12-81684973-DEL-480 |
| chr12-81724408-DEL-112 | chr12-81724409-DEL-112 |
| chr12-81738377-DEL-52 | chr12-81738378-DEL-52 |
| chr12-81739536-DEL-875 | chr12-81739537-DEL-875 |
| chr12-81747443-DEL-160 | chr12-81747443-DEL-160 |
| chr12-81752481-DEL-5449 | chr12-81752482-DEL-5449 |
| chr12-81791942-DEL-65 | chr12-81791975-DEL-57 |
| chr12-81799850-DEL-401 | chr12-81799851-DEL-401 |
| chr12-81810022-DEL-207 | chr12-81810022-DEL-208 |

| | |
|---|---|
| chr12-81818602-DEL-78 | chr12-81818603-DEL-78 |
| chr12-81830951-DEL-51 | chr12-81830951-DEL-51 |
| chr12-81833575-DEL-702 | chr12-81833576-DEL-702 |
| chr12-81860469-DEL-64 | chr12-81860438-DEL-59 |
| chr12-81872875-DEL-1889 | chr12-81872876-DEL-1890 |
| chr12-81886577-DEL-81 | chr12-81886578-DEL-81 |
| chr12-81895391-DEL-61 | chr12-81895389-DEL-72 |
| chr12-81925454-DEL-214 | chr12-81925455-DEL-214 |
| chr12-81940652-DEL-397 | chr12-81940653-DEL-398 |
| chr12-81970467-DEL-370 | chr12-81970468-DEL-370 |
| chr12-81989218-DEL-166 | chr12-81989219-DEL-166 |
| chr12-81997387-DEL-59 | chr12-81997386-DEL-60 |
| chr12-82028476-DEL-57 | chr12-82028477-DEL-58 |
| chr12-82121043-DEL-81 | chr12-82121050-DEL-78 |
| chr12-82121918-DEL-96 | chr12-82121946-DEL-54 |
| chr12-82138442-DEL-278 | chr12-82138447-DEL-270 |
| chr12-82177986-DEL-874 | chr12-82178002-DEL-858 |
| chr12-82189792-DEL-203 | chr12-82189801-DEL-203 |
| chr12-82230712-DEL-399 | chr12-82230713-DEL-399 |
| chr12-82256312-DEL-213 | chr12-82256313-DEL-213 |
| chr12-82284136-DEL-90 | chr12-82284137-DEL-90 |
| chr12-82289216-DEL-825 | chr12-82289233-DEL-823 |
| chr12-82327973-DEL-66 | chr12-82327973-DEL-66 |
| chr12-82334297-DEL-173 | chr12-82334297-DEL-165 |
| chr12-82337588-DEL-202 | chr12-82337587-DEL-203 |
| chr12-82355975-DEL-207 | chr12-82355976-DEL-207 |
| chr12-82399802-DEL-101 | chr12-82399807-DEL-94 |
| chr12-82426488-DEL-114 | chr12-82426492-DEL-112 |
| chr12-82460932-DEL-56 | chr12-82460933-DEL-56 |
| chr12-82476968-DEL-1069 | chr12-82476969-DEL-1069 |
| chr12-82500007-DEL-195 | chr12-82500020-DEL-184 |
| chr12-82507787-DEL-5433 | chr12-82507788-DEL-5433 |
| chr12-82537826-DEL-497 | chr12-82537827-DEL-497 |
| chr12-82606831-DEL-52 | chr12-82606869-DEL-58 |
| chr12-82611362-DEL-87 | chr12-82611362-DEL-87 |
| chr12-82636477-DEL-52 | chr12-82636474-DEL-56 |
| chr12-82676166-DEL-395 | chr12-82676167-DEL-395 |
| chr12-82677137-DEL-262 | chr12-82677138-DEL-262 |
| chr12-82680392-DEL-6873 | chr12-82680393-DEL-6873 |
| chr12-82693870-DEL-450 | chr12-82693867-DEL-451 |
| chr12-82696618-DEL-74 | chr12-82696619-DEL-74 |
| chr12-82720446-DEL-176 | chr12-82720447-DEL-176 |
| chr12-82740238-DEL-1484 | chr12-82740239-DEL-1484 |
| chr12-82774671-DEL-113 | chr12-82774672-DEL-113 |
| chr12-82775980-DEL-669 | chr12-82775981-DEL-669 |
| chr12-82786288-DEL-3084 | chr12-82786306-DEL-3067 |
| chr12-82802005-DEL-213 | chr12-82802006-DEL-213 |
| chr12-82817287-DEL-1118 | chr12-82817288-DEL-1118 |
| chr12-82827238-DEL-458 | chr12-82827239-DEL-455 |
| chr12-82830240-DEL-6195 | chr12-82830241-DEL-6195 |
| chr12-82845821-DEL-116 | chr12-82845842-DEL-116 |
| chr12-82898955-DEL-53 | chr12-82898958-DEL-51 |
| chr12-82918642-DEL-72 | chr12-82918670-DEL-72 |
| chr12-82920402-DEL-530 | chr12-82920400-DEL-530 |
| chr12-82927835-DEL-322 | chr12-82927836-DEL-322 |
| chr12-82958683-DEL-58 | chr12-82958684-DEL-58 |
| chr12-83028962-DEL-185 | chr12-83028963-DEL-185 |
| chr12-83102998-DEL-435 | chr12-83103004-DEL-429 |
| chr12-83127718-DEL-138 | chr12-83127719-DEL-138 |
| chr12-83134898-DEL-604 | chr12-83134907-DEL-600 |
| chr12-83175663-DEL-674 | chr12-83175664-DEL-674 |
| chr12-83192961-DEL-54 | chr12-83192963-DEL-54 |
| chr12-83211624-DEL-100 | chr12-83211674-DEL-80 |
| chr12-83212204-DEL-1259 | chr12-83212205-DEL-1259 |
| chr12-83258708-DEL-63 | chr12-83258714-DEL-92 |
| chr12-83284409-DEL-180 | chr12-83284395-DEL-223 |
| chr12-83292462-DEL-485 | chr12-83292463-DEL-485 |
| chr12-83334886-DEL-1565 | chr12-83334885-DEL-1565 |
| chr12-83339543-DEL-78 | chr12-83339544-DEL-78 |
| chr12-83393832-DEL-68 | chr12-83393834-DEL-66 |
| chr12-83412715-DEL-358 | chr12-83412716-DEL-358 |
| chr12-83413553-DEL-58 | chr12-83413554-DEL-58 |
| chr12-83423191-DEL-60 | chr12-83423192-DEL-96 |
| chr12-83449940-DEL-202 | chr12-83449941-DEL-202 |
| chr12-83485196-DEL-656 | chr12-83485197-DEL-656 |
| chr12-83550271-DEL-76 | chr12-83550272-DEL-76 |
| chr12-83551519-DEL-1011 | chr12-83551520-DEL-1011 |
| chr12-83561975-DEL-216 | chr12-83561965-DEL-217 |
| chr12-83563186-DEL-327 | chr12-83563187-DEL-327 |
| chr12-83568186-DEL-69 | chr12-83568187-DEL-69 |

| | |
|---|---|
| chr12-83596421-DEL-502 | chr12-83596422-DEL-500 |
| chr12-83598574-DEL-175 | chr12-83598584-DEL-173 |
| chr12-83607116-DEL-54 | chr12-83607117-DEL-54 |
| chr12-83621969-DEL-217 | chr12-83621970-DEL-217 |
| chr12-83649666-DEL-249 | chr12-83649667-DEL-249 |
| chr12-83655930-DEL-2474 | chr12-83655931-DEL-2474 |
| chr12-83666396-DEL-51 | chr12-83666397-DEL-51 |
| chr12-83671755-DEL-6452 | chr12-83671756-DEL-6452 |
| chr12-83681689-DEL-218 | chr12-83681690-DEL-218 |
| chr12-83693927-DEL-211 | chr12-83693928-DEL-211 |
| chr12-83724802-DEL-2997 | chr12-83724803-DEL-2997 |
| chr12-83751320-DEL-173 | chr12-83751321-DEL-173 |
| chr12-83780130-DEL-193 | chr12-83780131-DEL-193 |
| chr12-83806897-DEL-164 | chr12-83806898-DEL-164 |
| chr12-83812383-DEL-222 | chr12-83812384-DEL-222 |
| chr12-83855654-DEL-69 | chr12-83855615-DEL-92 |
| chr12-83861369-DEL-64 | chr12-83861370-DEL-64 |
| chr12-83865768-DEL-124 | chr12-83865769-DEL-124 |
| chr12-83901663-DEL-65 | chr12-83901648-DEL-60 |
| chr12-83991562-DEL-208 | chr12-83991600-DEL-171 |
| chr12-84007246-DEL-3291 | chr12-84007247-DEL-3291 |
| chr12-84035863-DEL-768 | chr12-84035864-DEL-768 |
| chr12-84044616-DEL-203 | chr12-84044616-DEL-204 |
| chr12-84049794-DEL-417 | chr12-84049795-DEL-417 |
| chr12-84068525-DEL-359 | chr12-84068526-DEL-359 |
| chr12-84112177-DEL-452 | chr12-84112178-DEL-452 |
| chr12-84123427-DEL-55 | chr12-84123427-DEL-55 |
| chr12-84163341-DEL-536 | chr12-84163341-DEL-543 |
| chr12-84174304-DEL-61 | chr12-84174305-DEL-53 |
| chr12-84177741-DEL-59 | chr12-84177742-DEL-59 |
| chr12-84181379-DEL-71 | chr12-84181380-DEL-71 |
| chr12-84184490-DEL-54 | chr12-84184491-DEL-54 |
| chr12-84270361-DEL-179 | chr12-84270362-DEL-179 |
| chr12-84272021-DEL-201 | chr12-84272022-DEL-201 |
| chr12-84283775-DEL-209 | chr12-84283776-DEL-209 |
| chr12-84305017-DEL-746 | chr12-84305018-DEL-746 |
| chr12-84316612-DEL-75 | chr12-84316613-DEL-75 |
| chr12-84359017-DEL-211 | chr12-84359018-DEL-211 |
| chr12-84376960-DEL-202 | chr12-84376961-DEL-202 |
| chr12-84377638-DEL-227 | chr12-84377639-DEL-226 |
| chr12-84405228-DEL-400 | chr12-84405229-DEL-400 |
| chr12-84448247-DEL-200 | chr12-84448231-DEL-197 |
| chr12-84451168-DEL-232 | chr12-84451169-DEL-232 |
| chr12-84468001-DEL-212 | chr12-84468002-DEL-212 |
| chr12-84472106-DEL-66 | chr12-84472105-DEL-71 |
| chr12-84520212-DEL-174 | chr12-84520219-DEL-169 |
| chr12-84539360-DEL-207 | chr12-84539361-DEL-207 |
| chr12-84545332-DEL-66 | chr12-84545333-DEL-66 |
| chr12-84556714-DEL-224 | chr12-84556715-DEL-224 |
| chr12-84560039-DEL-55 | chr12-84560040-DEL-55 |
| chr12-84586559-DEL-318 | chr12-84586560-DEL-318 |
| chr12-84587760-DEL-247 | chr12-84587761-DEL-247 |
| chr12-84607898-DEL-80 | chr12-84607902-DEL-77 |
| chr12-84670136-DEL-204 | chr12-84670137-DEL-204 |
| chr12-84693050-DEL-129 | chr12-84693050-DEL-129 |
| chr12-84706777-DEL-222 | chr12-84706778-DEL-222 |
| chr12-84708069-DEL-807 | chr12-84708069-DEL-806 |
| chr12-84712002-DEL-178 | chr12-84712002-DEL-178 |
| chr12-84717080-DEL-230 | chr12-84717087-DEL-212 |
| chr12-84758137-DEL-205 | chr12-84758138-DEL-204 |
| chr12-84763630-DEL-212 | chr12-84763641-DEL-202 |
| chr12-84782525-DEL-6905 | chr12-84782526-DEL-6905 |
| chr12-84822228-DEL-55 | chr12-84822224-DEL-53 |
| chr12-84889103-DEL-188 | chr12-84889104-DEL-188 |
| chr12-84895975-DEL-108 | chr12-84895976-DEL-108 |
| chr12-84901996-DEL-51 | chr12-84901997-DEL-51 |
| chr12-84914863-DEL-173 | chr12-84914864-DEL-173 |
| chr12-84927939-DEL-138 | chr12-84927936-DEL-139 |
| chr12-84980341-DEL-56 | chr12-84980342-DEL-56 |
| chr12-84988956-DEL-139 | chr12-84988920-DEL-176 |
| chr12-85012653-DEL-57 | chr12-85012654-DEL-57 |
| chr12-85018601-DEL-216 | chr12-85018602-DEL-216 |
| chr12-85021807-DEL-54 | chr12-85021808-DEL-54 |
| chr12-85033800-DEL-203 | chr12-85033769-DEL-231 |
| chr12-85050285-DEL-484 | chr12-85050286-DEL-484 |
| chr12-85055672-DEL-203 | chr12-85055672-DEL-204 |
| chr12-85164042-DEL-507 | chr12-85164043-DEL-507 |
| chr12-85178827-DEL-236 | chr12-85178817-DEL-243 |
| chr12-85203621-DEL-200 | chr12-85203622-DEL-199 |
| chr12-85206961-DEL-203 | chr12-85206962-DEL-203 |

| | |
|---|---|
| chr12-85209201-DEL-74 | chr12-85209204-DEL-73 |
| chr12-85215047-DEL-60 | chr12-85215060-DEL-52 |
| chr12-85254856-DEL-88 | chr12-85254821-DEL-81 |
| chr12-85257305-DEL-61 | chr12-85257306-DEL-61 |
| chr12-85288155-DEL-72 | chr12-85288126-DEL-56 |
| chr12-85296553-DEL-78 | chr12-85296536-DEL-96 |
| chr12-85328757-DEL-2284 | chr12-85328758-DEL-2284 |
| chr12-85404130-DEL-102 | chr12-85404131-DEL-102 |
| chr12-85406311-DEL-262 | chr12-85406310-DEL-263 |
| chr12-85411898-DEL-77 | chr12-85411922-DEL-50 |
| chr12-85417948-DEL-55 | chr12-85417929-DEL-51 |
| chr12-85430407-DEL-202 | chr12-85430408-DEL-202 |
| chr12-85435568-DEL-134 | chr12-85435570-DEL-133 |
| chr12-85447793-DEL-7158 | chr12-85447814-DEL-7138 |
| chr12-85499433-DEL-73 | chr12-85499450-DEL-57 |
| chr12-85538293-DEL-65 | chr12-85538296-DEL-62 |
| chr12-85545374-DEL-229 | chr12-85545339-DEL-251 |
| chr12-85572524-DEL-266 | chr12-85572525-DEL-266 |
| chr12-85575936-DEL-59 | chr12-85575937-DEL-59 |
| chr12-85584580-DEL-429 | chr12-85584581-DEL-429 |
| chr12-85603802-DEL-60 | chr12-85603799-DEL-54 |
| chr12-85667375-DEL-155 | chr12-85667376-DEL-155 |
| chr12-85691654-DEL-254 | chr12-85691655-DEL-254 |
| chr12-85696701-DEL-428 | chr12-85696702-DEL-428 |
| chr12-85704959-DEL-130 | chr12-85704960-DEL-130 |
| chr12-85708207-DEL-563 | chr12-85708208-DEL-563 |
| chr12-85744944-DEL-414 | chr12-85744945-DEL-414 |
| chr12-85760173-DEL-94 | chr12-85760185-DEL-66 |
| chr12-85764033-DEL-91 | chr12-85764036-DEL-89 |
| chr12-85769316-DEL-186 | chr12-85769318-DEL-186 |
| chr12-85779522-DEL-59 | chr12-85779523-DEL-59 |
| chr12-85779853-DEL-266 | chr12-85779854-DEL-266 |
| chr12-85786830-DEL-194 | chr12-85786831-DEL-194 |
| chr12-85790936-DEL-203 | chr12-85790937-DEL-203 |
| chr12-85804455-DEL-181 | chr12-85804456-DEL-181 |
| chr12-85807939-DEL-175 | chr12-85807935-DEL-176 |
| chr12-85809416-DEL-174 | chr12-85809417-DEL-174 |
| chr12-85849373-DEL-91 | chr12-85849384-DEL-112 |
| chr12-85861056-DEL-327 | chr12-85861057-DEL-327 |
| chr12-85879064-DEL-382 | chr12-85879065-DEL-382 |
| chr12-85880717-DEL-400 | chr12-85880718-DEL-400 |
| chr12-85896871-DEL-160 | chr12-85896872-DEL-158 |
| chr12-85914683-DEL-368 | chr12-85914684-DEL-368 |
| chr12-85927623-DEL-172 | chr12-85927626-DEL-173 |
| chr12-85984807-DEL-52 | chr12-85984808-DEL-52 |
| chr12-86022960-DEL-202 | chr12-86022961-DEL-202 |
| chr12-86030276-DEL-75 | chr12-86030277-DEL-75 |
| chr12-86054965-DEL-53 | chr12-86054965-DEL-53 |
| chr12-86098015-DEL-486 | chr12-86098016-DEL-486 |
| chr12-86166757-DEL-469 | chr12-86166758-DEL-469 |
| chr12-86177563-DEL-337 | chr12-86177564-DEL-337 |
| chr12-86188204-DEL-187 | chr12-86188205-DEL-187 |
| chr12-86192518-DEL-51 | chr12-86192518-DEL-52 |
| chr12-86223270-DEL-198 | chr12-86223271-DEL-198 |
| chr12-86229794-DEL-195 | chr12-86229795-DEL-195 |
| chr12-86232818-DEL-75 | chr12-86232819-DEL-75 |
| chr12-86321981-DEL-208 | chr12-86321982-DEL-208 |
| chr12-86389846-DEL-94 | chr12-86389861-DEL-60 |
| chr12-86408959-DEL-85 | chr12-86408960-DEL-97 |
| chr12-86416398-DEL-127 | chr12-86416399-DEL-127 |
| chr12-86427445-DEL-56 | chr12-86427446-DEL-55 |
| chr12-86430997-DEL-99 | chr12-86430968-DEL-79 |
| chr12-86478761-DEL-206 | chr12-86478762-DEL-206 |
| chr12-86513772-DEL-76 | chr12-86513769-DEL-80 |
| chr12-86532229-DEL-61 | chr12-86532221-DEL-50 |
| chr12-86546253-DEL-838 | chr12-86546255-DEL-837 |
| chr12-86551342-DEL-170 | chr12-86551343-DEL-170 |
| chr12-86570948-DEL-129 | chr12-86570965-DEL-113 |
| chr12-86577172-DEL-52 | chr12-86577173-DEL-52 |
| chr12-86583863-DEL-51 | chr12-86583864-DEL-51 |
| chr12-86589676-DEL-892 | chr12-86589674-DEL-893 |
| chr12-86639212-DEL-75 | chr12-86639213-DEL-75 |
| chr12-86692475-DEL-70 | chr12-86692476-DEL-70 |
| chr12-86699897-DEL-60 | chr12-86699897-DEL-60 |
| chr12-86706951-DEL-88 | chr12-86706952-DEL-86 |
| chr12-86717716-DEL-91 | chr12-86717717-DEL-91 |
| chr12-86721821-DEL-63 | chr12-86721818-DEL-61 |
| chr12-86797701-DEL-629 | chr12-86797702-DEL-629 |
| chr12-86801329-DEL-66 | chr12-86801330-DEL-66 |
| chr12-86805929-DEL-178 | chr12-86805929-DEL-179 |

| | |
|---|---|
| chr12-86807381-DEL-68 | chr12-86807382-DEL-68 |
| chr12-86842730-DEL-76 | chr12-86842731-DEL-76 |
| chr12-86844131-DEL-339 | chr12-86844132-DEL-339 |
| chr12-86845366-DEL-172 | chr12-86845367-DEL-172 |
| chr12-86875738-DEL-187 | chr12-86875732-DEL-191 |
| chr12-86884099-DEL-579 | chr12-86884100-DEL-579 |
| chr12-86887938-DEL-221 | chr12-86887939-DEL-221 |
| chr12-86892491-DEL-791 | chr12-86892492-DEL-791 |
| chr12-86949593-DEL-238 | chr12-86949592-DEL-239 |
| chr12-86951534-DEL-539 | chr12-86951535-DEL-539 |
| chr12-86971992-DEL-464 | chr12-86971993-DEL-464 |
| chr12-86975731-DEL-88 | chr12-86975723-DEL-98 |
| chr12-86996010-DEL-63 | chr12-86996011-DEL-63 |
| chr12-87047834-DEL-94 | chr12-87047835-DEL-94 |
| chr12-87099684-DEL-72 | chr12-87099688-DEL-66 |
| chr12-87185161-DEL-66 | chr12-87185162-DEL-66 |
| chr12-87255723-DEL-216 | chr12-87255724-DEL-216 |
| chr12-87306530-DEL-209 | chr12-87306531-DEL-209 |
| chr12-87338979-DEL-179 | chr12-87338980-DEL-179 |
| chr12-87339385-DEL-547 | chr12-87339386-DEL-547 |
| chr12-87353189-DEL-132 | chr12-87353190-DEL-132 |
| chr12-87362718-DEL-215 | chr12-87362719-DEL-215 |
| chr12-87382153-DEL-226 | chr12-87382154-DEL-226 |
| chr12-87413632-DEL-77 | chr12-87413662-DEL-76 |
| chr12-87430141-DEL-204 | chr12-87430142-DEL-204 |
| chr12-87448428-DEL-165 | chr12-87448429-DEL-165 |
| chr12-87465317-DEL-197 | chr12-87465318-DEL-197 |
| chr12-87477009-DEL-85 | chr12-87477010-DEL-85 |
| chr12-87485733-DEL-183 | chr12-87485734-DEL-183 |
| chr12-87508052-DEL-220 | chr12-87508048-DEL-219 |
| chr12-87533635-DEL-136 | chr12-87533636-DEL-136 |
| chr12-87534587-DEL-2443 | chr12-87534588-DEL-2443 |
| chr12-87536675-DEL-217 | chr12-87536675-DEL-217 |
| chr12-87607708-DEL-204 | chr12-87607709-DEL-204 |
| chr12-87624281-DEL-395 | chr12-87624281-DEL-395 |
| chr12-87638826-DEL-6762 | chr12-87638827-DEL-6762 |
| chr12-87752540-DEL-51 | chr12-87752537-DEL-52 |
| chr12-87994244-DEL-181 | chr12-87994245-DEL-181 |
| chr12-88101952-DEL-116 | chr12-88101952-DEL-117 |
| chr12-88147329-DEL-58 | chr12-88147330-DEL-58 |
| chr12-88162423-DEL-77 | chr12-88162424-DEL-77 |
| chr12-88277353-DEL-210 | chr12-88277354-DEL-210 |
| chr12-88340824-DEL-106 | chr12-88340816-DEL-98 |
| chr12-88342697-DEL-179 | chr12-88342698-DEL-179 |
| chr12-88357383-DEL-59 | chr12-88357384-DEL-59 |
| chr12-88365092-DEL-74 | chr12-88365061-DEL-70 |
| chr12-88386411-DEL-63 | chr12-88386412-DEL-63 |
| chr12-88410316-DEL-371 | chr12-88410316-DEL-371 |
| chr12-88438739-DEL-340 | chr12-88438740-DEL-340 |
| chr12-88460075-DEL-1846 | chr12-88460076-DEL-1846 |
| chr12-88478463-DEL-4932 | chr12-88478464-DEL-4932 |
| chr12-88486883-DEL-499 | chr12-88486884-DEL-497 |
| chr12-88509645-DEL-84 | chr12-88509640-DEL-85 |
| chr12-88570805-DEL-52 | chr12-88570823-DEL-61 |
| chr12-88592157-DEL-58 | chr12-88592158-DEL-58 |
| chr12-88606605-DEL-497 | chr12-88606606-DEL-497 |
| chr12-88624054-DEL-3443 | chr12-88624055-DEL-3443 |
| chr12-88634093-DEL-58 | chr12-88634093-DEL-58 |
| chr12-88643929-DEL-827 | chr12-88643929-DEL-827 |
| chr12-88656663-DEL-1897 | chr12-88656664-DEL-1897 |
| chr12-88719963-DEL-428 | chr12-88719964-DEL-428 |
| chr12-88779692-DEL-368 | chr12-88779698-DEL-365 |
| chr12-88780204-DEL-736 | chr12-88780205-DEL-736 |
| chr12-88803299-DEL-243 | chr12-88803300-DEL-243 |
| chr12-88814796-DEL-63 | chr12-88814795-DEL-64 |
| chr12-88853561-DEL-346 | chr12-88853562-DEL-346 |
| chr12-88878662-DEL-974 | chr12-88878663-DEL-974 |
| chr12-88935061-DEL-92 | chr12-88935062-DEL-92 |
| chr12-88959551-DEL-76 | chr12-88959552-DEL-76 |
| chr12-88983952-DEL-480 | chr12-88983953-DEL-480 |
| chr12-89042611-DEL-51 | chr12-89042654-DEL-51 |
| chr12-89107205-DEL-86 | chr12-89107212-DEL-63 |
| chr12-89113430-DEL-1057 | chr12-89113431-DEL-1057 |
| chr12-89151858-DEL-59 | chr12-89151825-DEL-93 |
| chr12-89178524-DEL-152 | chr12-89178525-DEL-152 |
| chr12-89185792-DEL-356 | chr12-89185791-DEL-357 |
| chr12-89192430-DEL-52 | chr12-89192431-DEL-52 |
| chr12-89213030-DEL-2106 | chr12-89213031-DEL-2105 |
| chr12-89218374-DEL-1032 | chr12-89218375-DEL-1032 |
| chr12-89233292-DEL-2284 | chr12-89233293-DEL-2284 |

| | |
|---|---|
| chr12-89252593-DEL-64 | chr12-89252593-DEL-64 |
| chr12-89306769-DEL-223 | chr12-89306770-DEL-223 |
| chr12-89332015-DEL-259 | chr12-89332016-DEL-259 |
| chr12-89352432-DEL-60 | chr12-89352433-DEL-56 |
| chr12-89389875-DEL-488 | chr12-89389863-DEL-489 |
| chr12-89404576-DEL-57 | chr12-89404575-DEL-60 |
| chr12-89457105-DEL-1664 | chr12-89457104-DEL-1663 |
| chr12-89517408-DEL-102 | chr12-89517450-DEL-102 |
| chr12-89561975-DEL-6415 | chr12-89561976-DEL-6415 |
| chr12-89576927-DEL-52 | chr12-89576942-DEL-69 |
| chr12-89625701-DEL-572 | chr12-89625699-DEL-569 |
| chr12-89632946-DEL-170 | chr12-89632947-DEL-170 |
| chr12-89637894-DEL-326 | chr12-89637895-DEL-326 |
| chr12-89683150-DEL-111 | chr12-89683151-DEL-111 |
| chr12-89690438-DEL-66 | chr12-89690439-DEL-67 |
| chr12-89741543-DEL-80 | chr12-89741544-DEL-80 |
| chr12-89804230-DEL-96 | chr12-89804254-DEL-76 |
| chr12-89815254-DEL-80 | chr12-89815235-DEL-81 |
| chr12-89907107-DEL-81 | chr12-89907108-DEL-81 |
| chr12-89919194-DEL-111 | chr12-89919195-DEL-111 |
| chr12-89927376-DEL-383 | chr12-89927377-DEL-383 |
| chr12-89974203-DEL-64 | chr12-89974204-DEL-64 |
| chr12-90021369-DEL-839 | chr12-90021370-DEL-839 |
| chr12-90042215-DEL-6856 | chr12-90042216-DEL-6856 |
| chr12-90054105-DEL-62 | chr12-90054068-DEL-90 |
| chr12-90061166-DEL-150 | chr12-90061165-DEL-150 |
| chr12-90076153-DEL-690 | chr12-90076154-DEL-690 |
| chr12-90088854-DEL-52 | chr12-90088855-DEL-68 |
| chr12-90172239-DEL-123 | chr12-90172240-DEL-123 |
| chr12-90178099-DEL-58 | chr12-90178106-DEL-52 |
| chr12-90205596-DEL-71 | chr12-90205599-DEL-55 |
| chr12-90211327-DEL-889 | chr12-90211328-DEL-889 |
| chr12-90215089-DEL-71 | chr12-90215082-DEL-77 |
| chr12-90219016-DEL-64 | chr12-90219031-DEL-54 |
| chr12-90227474-DEL-2785 | chr12-90227475-DEL-2785 |
| chr12-90348119-DEL-699 | chr12-90348120-DEL-699 |
| chr12-90395729-DEL-238 | chr12-90395730-DEL-237 |
| chr12-90402961-DEL-71 | chr12-90402961-DEL-71 |
| chr12-90404175-DEL-2134 | chr12-90404175-DEL-2134 |
| chr12-90443529-DEL-6170 | chr12-90443515-DEL-6184 |
| chr12-90481625-DEL-191 | chr12-90481626-DEL-191 |
| chr12-90482759-DEL-101 | chr12-90482760-DEL-101 |
| chr12-90485291-DEL-215 | chr12-90485292-DEL-213 |
| chr12-90528085-DEL-498 | chr12-90528086-DEL-498 |
| chr12-90530457-DEL-65 | chr12-90530461-DEL-62 |
| chr12-90531302-DEL-484 | chr12-90531303-DEL-484 |
| chr12-90561626-DEL-729 | chr12-90561627-DEL-729 |
| chr12-90572469-DEL-806 | chr12-90572470-DEL-806 |
| chr12-90586571-DEL-52 | chr12-90586572-DEL-52 |
| chr12-90591068-DEL-1690 | chr12-90591069-DEL-1690 |
| chr12-90601213-DEL-826 | chr12-90601214-DEL-826 |
| chr12-90602133-DEL-90 | chr12-90602130-DEL-60 |
| chr12-90616184-DEL-73 | chr12-90616195-DEL-63 |
| chr12-90628370-DEL-2141 | chr12-90628371-DEL-2141 |
| chr12-90636707-DEL-898 | chr12-90636708-DEL-898 |
| chr12-90639025-DEL-319 | chr12-90639026-DEL-319 |
| chr12-90654674-DEL-6028 | chr12-90654675-DEL-6028 |
| chr12-90681549-DEL-52 | chr12-90681564-DEL-78 |
| chr12-90724125-DEL-202 | chr12-90724124-DEL-203 |
| chr12-90724470-DEL-2239 | chr12-90724471-DEL-2239 |
| chr12-90733789-DEL-4695 | chr12-90733790-DEL-4695 |
| chr12-90763419-DEL-334 | chr12-90763413-DEL-335 |
| chr12-90783992-DEL-329 | chr12-90783993-DEL-329 |
| chr12-90801599-DEL-6599 | chr12-90801600-DEL-6599 |
| chr12-90825350-DEL-248 | chr12-90825351-DEL-248 |
| chr12-90837696-DEL-54 | chr12-90837697-DEL-54 |
| chr12-90902496-DEL-1087 | chr12-90902497-DEL-1087 |
| chr12-90914967-DEL-1155 | chr12-90914968-DEL-1155 |
| chr12-90918516-DEL-108 | chr12-90918517-DEL-108 |
| chr12-90928860-DEL-6220 | chr12-90928861-DEL-6220 |
| chr12-90946503-DEL-72 | chr12-90946504-DEL-60 |
| chr12-90949167-DEL-886 | chr12-90949168-DEL-885 |
| chr12-90971289-DEL-2006 | chr12-90971290-DEL-2006 |
| chr12-90976133-DEL-194 | chr12-90976134-DEL-194 |
| chr12-91019790-DEL-2462 | chr12-91019791-DEL-2462 |
| chr12-91043172-DEL-3246 | chr12-91043173-DEL-3246 |
| chr12-91080224-DEL-191 | chr12-91080223-DEL-187 |
| chr12-91089430-DEL-2048 | chr12-91089431-DEL-2048 |
| chr12-91090063-DEL-86 | chr12-91090066-DEL-84 |
| chr12-91106512-DEL-3022 | chr12-91106513-DEL-3022 |

| | |
|---|---|
| chr12-91124116-DEL-672 | chr12-91124117-DEL-672 |
| chr12-91128721-DEL-518 | chr12-91128722-DEL-518 |
| chr12-91129849-DEL-80 | chr12-91129823-DEL-60 |
| chr12-91130724-DEL-6799 | chr12-91130725-DEL-6799 |
| chr12-91150416-DEL-68 | chr12-91150417-DEL-68 |
| chr12-91177620-DEL-56 | chr12-91177623-DEL-66 |
| chr12-91183527-DEL-262 | chr12-91183528-DEL-262 |
| chr12-91188132-DEL-3145 | chr12-91188127-DEL-3146 |
| chr12-91194311-DEL-2561 | chr12-91194312-DEL-2561 |
| chr12-91202425-DEL-450 | chr12-91202426-DEL-450 |
| chr12-91205124-DEL-60 | chr12-91205125-DEL-60 |
| chr12-91207227-DEL-6112 | chr12-91207228-DEL-6112 |
| chr12-91214660-DEL-434 | chr12-91214649-DEL-427 |
| chr12-91244667-DEL-6195 | chr12-91244668-DEL-6195 |
| chr12-91283671-DEL-66 | chr12-91283672-DEL-66 |
| chr12-91305619-DEL-96 | chr12-91305620-DEL-96 |
| chr12-91340853-DEL-220 | chr12-91340854-DEL-220 |
| chr12-91397505-DEL-59 | chr12-91397486-DEL-60 |
| chr12-91405863-DEL-1972 | chr12-91405864-DEL-1972 |
| chr12-91467641-DEL-74 | chr12-91467642-DEL-69 |
| chr12-91469195-DEL-368 | chr12-91469198-DEL-332 |
| chr12-91485910-DEL-75 | chr12-91485908-DEL-52 |
| chr12-91508348-DEL-2499 | chr12-91508349-DEL-2499 |
| chr12-91520273-DEL-415 | chr12-91520274-DEL-415 |
| chr12-91545414-DEL-216 | chr12-91545415-DEL-216 |
| chr12-91560280-DEL-103 | chr12-91560288-DEL-74 |
| chr12-91568112-DEL-1771 | chr12-91568113-DEL-1771 |
| chr12-91576421-DEL-714 | chr12-91576422-DEL-715 |
| chr12-91617189-DEL-2896 | chr12-91617184-DEL-2897 |
| chr12-91618867-DEL-214 | chr12-91618868-DEL-210 |
| chr12-91625893-DEL-76 | chr12-91625880-DEL-77 |
| chr12-91644389-DEL-141 | chr12-91644391-DEL-140 |
| chr12-91648389-DEL-54 | chr12-91648347-DEL-80 |
| chr12-91650788-DEL-80 | chr12-91650789-DEL-80 |
| chr12-91685116-DEL-416 | chr12-91685117-DEL-416 |
| chr12-91688426-DEL-399 | chr12-91688421-DEL-400 |
| chr12-91690068-DEL-62 | chr12-91690069-DEL-62 |
| chr12-91696896-DEL-295 | chr12-91696897-DEL-295 |
| chr12-91697557-DEL-119 | chr12-91697560-DEL-93 |
| chr12-91755280-DEL-204 | chr12-91755281-DEL-204 |
| chr12-91757700-DEL-107 | chr12-91757701-DEL-107 |
| chr12-91759067-DEL-1063 | chr12-91759068-DEL-1063 |
| chr12-91762574-DEL-89 | chr12-91762575-DEL-89 |
| chr12-91779133-DEL-121 | chr12-91779134-DEL-121 |
| chr12-91817327-DEL-265 | chr12-91817328-DEL-265 |
| chr12-91835786-DEL-761 | chr12-91835787-DEL-761 |
| chr12-91857668-DEL-469 | chr12-91857669-DEL-469 |
| chr12-91901272-DEL-118 | chr12-91901274-DEL-116 |
| chr12-91902957-DEL-7566 | chr12-91902958-DEL-7566 |
| chr12-91961145-DEL-395 | chr12-91961146-DEL-395 |
| chr12-92047272-DEL-60 | chr12-92047273-DEL-60 |
| chr12-92074127-DEL-75 | chr12-92074128-DEL-75 |
| chr12-92086021-DEL-101 | chr12-92086021-DEL-101 |
| chr12-92089802-DEL-63 | chr12-92089803-DEL-63 |
| chr12-92112427-DEL-242 | chr12-92112426-DEL-243 |
| chr12-92124990-DEL-400 | chr12-92124990-DEL-400 |
| chr12-92132341-DEL-7205 | chr12-92132342-DEL-7205 |
| chr12-92142252-DEL-86 | chr12-92142253-DEL-86 |
| chr12-92171533-DEL-145 | chr12-92171534-DEL-145 |
| chr12-92175856-DEL-1805 | chr12-92175857-DEL-1805 |
| chr12-92204435-DEL-56 | chr12-92204449-DEL-57 |
| chr12-92205402-DEL-421 | chr12-92205403-DEL-421 |
| chr12-92229358-DEL-2200 | chr12-92229359-DEL-2200 |
| chr12-92232258-DEL-203 | chr12-92232259-DEL-203 |
| chr12-92276627-DEL-825 | chr12-92276628-DEL-825 |
| chr12-92279248-DEL-356 | chr12-92279249-DEL-356 |
| chr12-92324816-DEL-7310 | chr12-92324817-DEL-7310 |
| chr12-92360893-DEL-1706 | chr12-92360894-DEL-1706 |
| chr12-92380515-DEL-3887 | chr12-92380516-DEL-3887 |
| chr12-92410152-DEL-1017 | chr12-92410153-DEL-1016 |
| chr12-92428886-DEL-105 | chr12-92428887-DEL-105 |
| chr12-92485677-DEL-458 | chr12-92485678-DEL-458 |
| chr12-92506245-DEL-405 | chr12-92506245-DEL-405 |
| chr12-92506745-DEL-1247 | chr12-92506745-DEL-1247 |
| chr12-92529511-DEL-85 | chr12-92529512-DEL-85 |
| chr12-92544146-DEL-79 | chr12-92544147-DEL-79 |
| chr12-92552008-DEL-2708 | chr12-92552009-DEL-2708 |
| chr12-92558062-DEL-6237 | chr12-92558063-DEL-6237 |
| chr12-92564387-DEL-76 | chr12-92564397-DEL-52 |
| chr12-92565762-DEL-318 | chr12-92565763-DEL-318 |

| | |
|---|---|
| chr12-92572286-DEL-53 | chr12-92572287-DEL-53 |
| chr12-92599463-DEL-74 | chr12-92599466-DEL-50 |
| chr12-92602059-DEL-60 | chr12-92602054-DEL-66 |
| chr12-92610802-DEL-133 | chr12-92610803-DEL-133 |
| chr12-92617345-DEL-204 | chr12-92617329-DEL-204 |
| chr12-92671563-DEL-1312 | chr12-92671564-DEL-1312 |
| chr12-92694028-DEL-142 | chr12-92694029-DEL-142 |
| chr12-92699945-DEL-54 | chr12-92699946-DEL-54 |
| chr12-92708586-DEL-6211 | chr12-92708587-DEL-6211 |
| chr12-92740399-DEL-211 | chr12-92740400-DEL-211 |
| chr12-92754917-DEL-111 | chr12-92754918-DEL-111 |
| chr12-92786196-DEL-54 | chr12-92786168-DEL-72 |
| chr12-92798806-DEL-642 | chr12-92798807-DEL-642 |
| chr12-92800984-DEL-769 | chr12-92800985-DEL-769 |
| chr12-92803226-DEL-63 | chr12-92803229-DEL-64 |
| chr12-92828733-DEL-6382 | chr12-92828734-DEL-6382 |
| chr12-92849307-DEL-9907 | chr12-92849308-DEL-9907 |
| chr12-92917431-DEL-364 | chr12-92917432-DEL-364 |
| chr12-92960950-DEL-2085 | chr12-92960951-DEL-2085 |
| chr12-92977778-DEL-68 | chr12-92977788-DEL-61 |
| chr12-92992469-DEL-1061 | chr12-92992470-DEL-1061 |
| chr12-92995948-DEL-68 | chr12-92995953-DEL-64 |
| chr12-93010247-DEL-64 | chr12-93010248-DEL-64 |
| chr12-93041430-DEL-67 | chr12-93041431-DEL-67 |
| chr12-93043470-DEL-1725 | chr12-93043471-DEL-1725 |
| chr12-93069914-DEL-6019 | chr12-93069915-DEL-6019 |
| chr12-93079667-DEL-116 | chr12-93079668-DEL-116 |
| chr12-93105550-DEL-704 | chr12-93105551-DEL-704 |
| chr12-93149686-DEL-130 | chr12-93149684-DEL-130 |
| chr12-93168691-DEL-61 | chr12-93168692-DEL-61 |
| chr12-93183453-DEL-66 | chr12-93183454-DEL-66 |
| chr12-93190161-DEL-82 | chr12-93190170-DEL-73 |
| chr12-93225596-DEL-6875 | chr12-93225596-DEL-6875 |
| chr12-93263740-DEL-6265 | chr12-93263741-DEL-6265 |
| chr12-93284262-DEL-6342 | chr12-93284263-DEL-6342 |
| chr12-93314937-DEL-59 | chr12-93314933-DEL-60 |
| chr12-93326323-DEL-1373 | chr12-93326313-DEL-1370 |
| chr12-93333733-DEL-754 | chr12-93333734-DEL-754 |
| chr12-93336008-DEL-468 | chr12-93336011-DEL-465 |
| chr12-93355481-DEL-171 | chr12-93355494-DEL-172 |
| chr12-93358861-DEL-959 | chr12-93358862-DEL-958 |
| chr12-93382363-DEL-51 | chr12-93382377-DEL-52 |
| chr12-93387571-DEL-95 | chr12-93387572-DEL-95 |
| chr12-93421242-DEL-107 | chr12-93421243-DEL-107 |
| chr12-93424537-DEL-62 | chr12-93424538-DEL-62 |
| chr12-93430051-DEL-464 | chr12-93430052-DEL-464 |
| chr12-93446350-DEL-1597 | chr12-93446351-DEL-1597 |
| chr12-93455173-DEL-4354 | chr12-93455174-DEL-4354 |
| chr12-93460899-DEL-72 | chr12-93460900-DEL-72 |
| chr12-93476282-DEL-7149 | chr12-93476283-DEL-7149 |
| chr12-93485781-DEL-633 | chr12-93485782-DEL-633 |
| chr12-93486739-DEL-992 | chr12-93486729-DEL-993 |
| chr12-93495557-DEL-268 | chr12-93495550-DEL-272 |
| chr12-93496238-DEL-85 | chr12-93496239-DEL-85 |
| chr12-93496683-DEL-563 | chr12-93496682-DEL-564 |
| chr12-93505347-DEL-4171 | chr12-93505348-DEL-4171 |
| chr12-93523583-DEL-164 | chr12-93523584-DEL-164 |
| chr12-93525900-DEL-6872 | chr12-93525901-DEL-6872 |
| chr12-93550375-DEL-52 | chr12-93550376-DEL-52 |
| chr12-93550937-DEL-52 | chr12-93550937-DEL-52 |
| chr12-93551202-DEL-6469 | chr12-93551185-DEL-6486 |
| chr12-93557215-DEL-215 | chr12-93557216-DEL-215 |
| chr12-93647733-DEL-90 | chr12-93647734-DEL-90 |
| chr12-93657366-DEL-2175 | chr12-93657367-DEL-2175 |
| chr12-93660273-DEL-943 | chr12-93660274-DEL-943 |
| chr12-93673843-DEL-95 | chr12-93673834-DEL-96 |
| chr12-93674521-DEL-1342 | chr12-93674521-DEL-1342 |
| chr12-93688015-DEL-587 | chr12-93688016-DEL-587 |
| chr12-93693945-DEL-6146 | chr12-93693946-DEL-6146 |
| chr12-93712528-DEL-459 | chr12-93712571-DEL-417 |
| chr12-93733254-DEL-3041 | chr12-93733255-DEL-3041 |
| chr12-93780728-DEL-244 | chr12-93780735-DEL-219 |
| chr12-93786980-DEL-2329 | chr12-93786981-DEL-2328 |
| chr12-93855630-DEL-62 | chr12-93855631-DEL-60 |
| chr12-93887090-DEL-9115 | chr12-93887090-DEL-9115 |
| chr12-93923563-DEL-62 | chr12-93923566-DEL-60 |
| chr12-93987889-DEL-133 | chr12-93987890-DEL-133 |
| chr12-93990392-DEL-453 | chr12-93990369-DEL-472 |
| chr12-94003198-DEL-7341 | chr12-94003199-DEL-7342 |
| chr12-94020328-DEL-800 | chr12-94020329-DEL-800 |

| | |
|---|---|
| chr12-94039336-DEL-497 | chr12-94039337-DEL-497 |
| chr12-94040623-DEL-865 | chr12-94040624-DEL-865 |
| chr12-94063025-DEL-203 | chr12-94063026-DEL-203 |
| chr12-94064539-DEL-88 | chr12-94064568-DEL-66 |
| chr12-94066321-DEL-112 | chr12-94066322-DEL-112 |
| chr12-94074044-DEL-96 | chr12-94074057-DEL-84 |
| chr12-94113043-DEL-481 | chr12-94113044-DEL-481 |
| chr12-94136919-DEL-53 | chr12-94136919-DEL-57 |
| chr12-94143886-DEL-183 | chr12-94143887-DEL-183 |
| chr12-94195376-DEL-64 | chr12-94195368-DEL-71 |
| chr12-94196328-DEL-345 | chr12-94196329-DEL-345 |
| chr12-94215445-DEL-400 | chr12-94215446-DEL-400 |
| chr12-94218901-DEL-58 | chr12-94218902-DEL-59 |
| chr12-94219580-DEL-62 | chr12-94219581-DEL-62 |
| chr12-94235309-DEL-6632 | chr12-94235310-DEL-6632 |
| chr12-94256058-DEL-53 | chr12-94256063-DEL-72 |
| chr12-94260563-DEL-121 | chr12-94260564-DEL-121 |
| chr12-94266894-DEL-144 | chr12-94266895-DEL-144 |
| chr12-94300217-DEL-117 | chr12-94300218-DEL-117 |
| chr12-94371039-DEL-80 | chr12-94371027-DEL-81 |
| chr12-94400671-DEL-524 | chr12-94400672-DEL-524 |
| chr12-94401250-DEL-77 | chr12-94401251-DEL-77 |
| chr12-94404164-DEL-6084 | chr12-94404165-DEL-6084 |
| chr12-94412302-DEL-61 | chr12-94412302-DEL-65 |
| chr12-94433919-DEL-98 | chr12-94433920-DEL-98 |
| chr12-94458917-DEL-1566 | chr12-94458918-DEL-1566 |
| chr12-94479623-DEL-1914 | chr12-94479624-DEL-1914 |
| chr12-94484663-DEL-375 | chr12-94484664-DEL-375 |
| chr12-94493548-DEL-489 | chr12-94493549-DEL-489 |
| chr12-94502090-DEL-316 | chr12-94502091-DEL-316 |
| chr12-94542326-DEL-678 | chr12-94542327-DEL-678 |
| chr12-94562286-DEL-56 | chr12-94562293-DEL-68 |
| chr12-94584439-DEL-5286 | chr12-94584439-DEL-5286 |
| chr12-94601711-DEL-65 | chr12-94601712-DEL-65 |
| chr12-94605911-DEL-378 | chr12-94605912-DEL-378 |
| chr12-94616095-DEL-68 | chr12-94616095-DEL-60 |
| chr12-94621608-DEL-60 | chr12-94621609-DEL-60 |
| chr12-94628528-DEL-199 | chr12-94628529-DEL-199 |
| chr12-94656234-DEL-118 | chr12-94656235-DEL-118 |
| chr12-94678525-DEL-266 | chr12-94678525-DEL-266 |
| chr12-94679182-DEL-1379 | chr12-94679182-DEL-1379 |
| chr12-94723784-DEL-125 | chr12-94723785-DEL-125 |
| chr12-94736567-DEL-56 | chr12-94736568-DEL-56 |
| chr12-94790571-DEL-76 | chr12-94790570-DEL-76 |
| chr12-94808531-DEL-738 | chr12-94808523-DEL-727 |
| chr12-94815274-DEL-6655 | chr12-94815274-DEL-6655 |
| chr12-94834165-DEL-222 | chr12-94834169-DEL-195 |
| chr12-94850290-DEL-649 | chr12-94850290-DEL-649 |
| chr12-94858571-DEL-2551 | chr12-94858572-DEL-2551 |
| chr12-94865708-DEL-58 | chr12-94865705-DEL-58 |
| chr12-94899975-DEL-1455 | chr12-94899977-DEL-1448 |
| chr12-94943951-DEL-216 | chr12-94943953-DEL-216 |
| chr12-94945025-DEL-1309 | chr12-94945026-DEL-1309 |
| chr12-94945224-DEL-308 | chr12-94945223-DEL-315 |
| chr12-94955540-DEL-484 | chr12-94955557-DEL-484 |
| chr12-94964454-DEL-910 | chr12-94964455-DEL-910 |
| chr12-94972658-DEL-2207 | chr12-94972659-DEL-2207 |
| chr12-94995943-DEL-51 | chr12-94995945-DEL-66 |
| chr12-95032781-DEL-803 | chr12-95032782-DEL-803 |
| chr12-95047437-DEL-379 | chr12-95047438-DEL-379 |
| chr12-95087678-DEL-2025 | chr12-95087679-DEL-2026 |
| chr12-95102814-DEL-246 | chr12-95102815-DEL-246 |
| chr12-95111713-DEL-548 | chr12-95111714-DEL-548 |
| chr12-95124472-DEL-758 | chr12-95124471-DEL-760 |
| chr12-95157041-DEL-677 | chr12-95157042-DEL-677 |
| chr12-95166044-DEL-85 | chr12-95166044-DEL-86 |
| chr12-95170240-DEL-62 | chr12-95170241-DEL-62 |
| chr12-95177561-DEL-434 | chr12-95177562-DEL-434 |
| chr12-95209728-DEL-72 | chr12-95209711-DEL-79 |
| chr12-95215175-DEL-7135 | chr12-95215176-DEL-7135 |
| chr12-95223380-DEL-79 | chr12-95223375-DEL-65 |
| chr12-95225588-DEL-1373 | chr12-95225588-DEL-1373 |
| chr12-95242562-DEL-2172 | chr12-95242578-DEL-2188 |
| chr12-95254914-DEL-2669 | chr12-95254915-DEL-2669 |
| chr12-95290012-DEL-78 | chr12-95290010-DEL-84 |
| chr12-95307305-DEL-6481 | chr12-95307306-DEL-6481 |
| chr12-95324592-DEL-95 | chr12-95324590-DEL-106 |
| chr12-95336394-DEL-85 | chr12-95336387-DEL-93 |
| chr12-95364808-DEL-62 | chr12-95364809-DEL-60 |
| chr12-95367507-DEL-213 | chr12-95367500-DEL-211 |

| | |
|---|---|
| chr12-95370245-DEL-144 | chr12-95370246-DEL-144 |
| chr12-95371245-DEL-75 | chr12-95371246-DEL-75 |
| chr12-95380306-DEL-4510 | chr12-95380307-DEL-4511 |
| chr12-95381866-DEL-390 | chr12-95381867-DEL-390 |
| chr12-95390275-DEL-66 | chr12-95390258-DEL-84 |
| chr12-95403185-DEL-669 | chr12-95403186-DEL-669 |
| chr12-95410596-DEL-1068 | chr12-95410597-DEL-1068 |
| chr12-95436280-DEL-262 | chr12-95436281-DEL-262 |
| chr12-95437041-DEL-60 | chr12-95437042-DEL-50 |
| chr12-95546154-DEL-6642 | chr12-95546155-DEL-6642 |
| chr12-95578170-DEL-6819 | chr12-95578171-DEL-6821 |
| chr12-95586582-DEL-487 | chr12-95586583-DEL-487 |
| chr12-95588610-DEL-90 | chr12-95588611-DEL-90 |
| chr12-95597024-DEL-373 | chr12-95597025-DEL-373 |
| chr12-95602818-DEL-81 | chr12-95602819-DEL-81 |
| chr12-95621907-DEL-52 | chr12-95621908-DEL-52 |
| chr12-95638935-DEL-1133 | chr12-95638936-DEL-1133 |
| chr12-95639657-DEL-207 | chr12-95639658-DEL-207 |
| chr12-95656517-DEL-58 | chr12-95656517-DEL-58 |
| chr12-95706873-DEL-84 | chr12-95706874-DEL-84 |
| chr12-95725094-DEL-168 | chr12-95725095-DEL-168 |
| chr12-95763845-DEL-6377 | chr12-95763846-DEL-6377 |
| chr12-95766556-DEL-203 | chr12-95766557-DEL-203 |
| chr12-95788803-DEL-7936 | chr12-95788804-DEL-7936 |
| chr12-95789275-DEL-212 | chr12-95789276-DEL-212 |
| chr12-95858810-DEL-960 | chr12-95858811-DEL-960 |
| chr12-95897109-DEL-6526 | chr12-95897109-DEL-6527 |
| chr12-95936361-DEL-115 | chr12-95936370-DEL-107 |
| chr12-95941969-DEL-177 | chr12-95941961-DEL-177 |
| chr12-95942844-DEL-4987 | chr12-95942845-DEL-4987 |
| chr12-95948604-DEL-196 | chr12-95948604-DEL-197 |
| chr12-95967079-DEL-6988 | chr12-95967080-DEL-6988 |
| chr12-95973274-DEL-603 | chr12-95973275-DEL-603 |
| chr12-96044082-DEL-75 | chr12-96044072-DEL-76 |
| chr12-96044457-DEL-6636 | chr12-96044458-DEL-6636 |
| chr12-96050551-DEL-208 | chr12-96050552-DEL-208 |
| chr12-96053563-DEL-70 | chr12-96053563-DEL-50 |
| chr12-96087524-DEL-93 | chr12-96087525-DEL-93 |
| chr12-96089873-DEL-71 | chr12-96089873-DEL-71 |
| chr12-96104904-DEL-3369 | chr12-96104905-DEL-3369 |
| chr12-96112077-DEL-52 | chr12-96112049-DEL-50 |
| chr12-96130921-DEL-285 | chr12-96130922-DEL-285 |
| chr12-96148016-DEL-59 | chr12-96148017-DEL-59 |
| chr12-96148719-DEL-56 | chr12-96148718-DEL-56 |
| chr12-96153248-DEL-70 | chr12-96153247-DEL-54 |
| chr12-96157938-DEL-131 | chr12-96157939-DEL-131 |
| chr12-96160763-DEL-66 | chr12-96160769-DEL-90 |
| chr12-96162114-DEL-2401 | chr12-96162114-DEL-2401 |
| chr12-96175427-DEL-93 | chr12-96175428-DEL-93 |
| chr12-96196926-DEL-4380 | chr12-96196927-DEL-4380 |
| chr12-96212055-DEL-1877 | chr12-96212056-DEL-1877 |
| chr12-96224601-DEL-204 | chr12-96224602-DEL-204 |
| chr12-96255359-DEL-2032 | chr12-96255359-DEL-2032 |
| chr12-96258406-DEL-114 | chr12-96258407-DEL-114 |
| chr12-96264487-DEL-400 | chr12-96264488-DEL-400 |
| chr12-96319273-DEL-205 | chr12-96319274-DEL-205 |
| chr12-96336119-DEL-337 | chr12-96336120-DEL-337 |
| chr12-96349970-DEL-53 | chr12-96349971-DEL-53 |
| chr12-96355059-DEL-1996 | chr12-96355060-DEL-1996 |
| chr12-96381253-DEL-175 | chr12-96381254-DEL-175 |
| chr12-96413903-DEL-186 | chr12-96413904-DEL-186 |
| chr12-96426268-DEL-55 | chr12-96426262-DEL-57 |
| chr12-96431867-DEL-92 | chr12-96431868-DEL-92 |
| chr12-96444726-DEL-54 | chr12-96444731-DEL-98 |
| chr12-96447884-DEL-1222 | chr12-96447885-DEL-1222 |
| chr12-96477508-DEL-7048 | chr12-96477509-DEL-7048 |
| chr12-96499152-DEL-66 | chr12-96499153-DEL-66 |
| chr12-96500653-DEL-58 | chr12-96500654-DEL-58 |
| chr12-96508154-DEL-54 | chr12-96508161-DEL-60 |
| chr12-96531611-DEL-54 | chr12-96531612-DEL-54 |
| chr12-96535841-DEL-823 | chr12-96535841-DEL-823 |
| chr12-96536836-DEL-3597 | chr12-96536837-DEL-3597 |
| chr12-96543639-DEL-67 | chr12-96543640-DEL-67 |
| chr12-96550799-DEL-2607 | chr12-96550800-DEL-2607 |
| chr12-96557196-DEL-64 | chr12-96557197-DEL-64 |
| chr12-96566098-DEL-596 | chr12-96566099-DEL-596 |
| chr12-96567388-DEL-1310 | chr12-96567389-DEL-1310 |
| chr12-96568488-DEL-248 | chr12-96568489-DEL-248 |
| chr12-96582226-DEL-56 | chr12-96582227-DEL-65 |
| chr12-96651515-DEL-228 | chr12-96651516-DEL-228 |

| | |
|---|---|
| chr12-96679358-DEL-246 | chr12-96679354-DEL-247 |
| chr12-96725190-DEL-932 | chr12-96725191-DEL-932 |
| chr12-96726737-DEL-185 | chr12-96726738-DEL-185 |
| chr12-96732660-DEL-146 | chr12-96732661-DEL-146 |
| chr12-96734604-DEL-1146 | chr12-96734605-DEL-1146 |
| chr12-96762836-DEL-62 | chr12-96762837-DEL-62 |
| chr12-96784409-DEL-400 | chr12-96784410-DEL-400 |
| chr12-96791649-DEL-173 | chr12-96791650-DEL-173 |
| chr12-96804864-DEL-77 | chr12-96804858-DEL-78 |
| chr12-96808338-DEL-6960 | chr12-96808339-DEL-6960 |
| chr12-96814663-DEL-214 | chr12-96814664-DEL-214 |
| chr12-96818372-DEL-146 | chr12-96818373-DEL-146 |
| chr12-96855886-DEL-209 | chr12-96855886-DEL-208 |
| chr12-96856814-DEL-54 | chr12-96856815-DEL-54 |
| chr12-96881782-DEL-1343 | chr12-96881783-DEL-1343 |
| chr12-96889030-DEL-832 | chr12-96889031-DEL-832 |
| chr12-96898221-DEL-390 | chr12-96898222-DEL-390 |
| chr12-96928059-DEL-1893 | chr12-96928035-DEL-1898 |
| chr12-96933060-DEL-94 | chr12-96933061-DEL-94 |
| chr12-96936496-DEL-53 | chr12-96936497-DEL-53 |
| chr12-96937497-DEL-243 | chr12-96937498-DEL-243 |
| chr12-96965809-DEL-1768 | chr12-96965804-DEL-1760 |
| chr12-96975823-DEL-6290 | chr12-96975824-DEL-6290 |
| chr12-96988520-DEL-250 | chr12-96988521-DEL-250 |
| chr12-96996607-DEL-742 | chr12-96996604-DEL-743 |
| chr12-97003991-DEL-7133 | chr12-97003992-DEL-7133 |
| chr12-97011817-DEL-60 | chr12-97011818-DEL-54 |
| chr12-97023194-DEL-170 | chr12-97023195-DEL-170 |
| chr12-97025241-DEL-60 | chr12-97025242-DEL-60 |
| chr12-97028196-DEL-365 | chr12-97028195-DEL-366 |
| chr12-97029208-DEL-1335 | chr12-97029209-DEL-1335 |
| chr12-97037143-DEL-1790 | chr12-97037144-DEL-1790 |
| chr12-97046717-DEL-65 | chr12-97046737-DEL-85 |
| chr12-97056192-DEL-218 | chr12-97056193-DEL-218 |
| chr12-97099752-DEL-138 | chr12-97099737-DEL-152 |
| chr12-97120474-DEL-521 | chr12-97120475-DEL-521 |
| chr12-97121145-DEL-64 | chr12-97121146-DEL-64 |
| chr12-97138885-DEL-916 | chr12-97138886-DEL-916 |
| chr12-97153173-DEL-1601 | chr12-97153174-DEL-1601 |
| chr12-97172789-DEL-118 | chr12-97172790-DEL-118 |
| chr12-97195824-DEL-82 | chr12-97195825-DEL-82 |
| chr12-97198791-DEL-199 | chr12-97198792-DEL-199 |
| chr12-97214935-DEL-390 | chr12-97214936-DEL-390 |
| chr12-97225887-DEL-110 | chr12-97225888-DEL-82 |
| chr12-97256544-DEL-2467 | chr12-97256545-DEL-2467 |
| chr12-97261245-DEL-97 | chr12-97261244-DEL-59 |
| chr12-97281664-DEL-97 | chr12-97281665-DEL-94 |
| chr12-97285175-DEL-1144 | chr12-97285176-DEL-1144 |
| chr12-97289462-DEL-97 | chr12-97289463-DEL-91 |
| chr12-97295974-DEL-389 | chr12-97295973-DEL-390 |
| chr12-97344430-DEL-6396 | chr12-97344431-DEL-6396 |
| chr12-97356937-DEL-52 | chr12-97356937-DEL-88 |
| chr12-97358187-DEL-239 | chr12-97358188-DEL-239 |
| chr12-97367014-DEL-325 | chr12-97367015-DEL-325 |
| chr12-97392387-DEL-328 | chr12-97392389-DEL-323 |
| chr12-97429342-DEL-315 | chr12-97429343-DEL-315 |
| chr12-97449019-DEL-1265 | chr12-97449020-DEL-1263 |
| chr12-97468677-DEL-149 | chr12-97468678-DEL-149 |
| chr12-97523377-DEL-5114 | chr12-97523378-DEL-5114 |
| chr12-97531348-DEL-477 | chr12-97531349-DEL-477 |
| chr12-97555734-DEL-1791 | chr12-97555735-DEL-1791 |
| chr12-97576589-DEL-62 | chr12-97576573-DEL-60 |
| chr12-97594919-DEL-346 | chr12-97594939-DEL-310 |
| chr12-97601446-DEL-54 | chr12-97601447-DEL-54 |
| chr12-97631383-DEL-451 | chr12-97631384-DEL-451 |
| chr12-97633077-DEL-715 | chr12-97633078-DEL-715 |
| chr12-97636599-DEL-5994 | chr12-97636600-DEL-5994 |
| chr12-97644438-DEL-7816 | chr12-97644441-DEL-7815 |
| chr12-97653772-DEL-78 | chr12-97653772-DEL-78 |
| chr12-97679684-DEL-149 | chr12-97679685-DEL-148 |
| chr12-97693417-DEL-100 | chr12-97693426-DEL-100 |
| chr12-97700101-DEL-59 | chr12-97700102-DEL-58 |
| chr12-97728633-DEL-57 | chr12-97728634-DEL-57 |
| chr12-97731018-DEL-486 | chr12-97731019-DEL-486 |
| chr12-97732849-DEL-3276 | chr12-97732870-DEL-3255 |
| chr12-97780486-DEL-327 | chr12-97780487-DEL-327 |
| chr12-97786101-DEL-7582 | chr12-97786102-DEL-7582 |
| chr12-97880224-DEL-115 | chr12-97880225-DEL-115 |
| chr12-97888779-DEL-2707 | chr12-97888780-DEL-2707 |
| chr12-97893259-DEL-66 | chr12-97893262-DEL-86 |

| | |
|---|---|
| chr12-97899085-DEL-55 | chr12-97899071-DEL-55 |
| chr12-97929746-DEL-54 | chr12-97929747-DEL-54 |
| chr12-97952013-DEL-96 | chr12-97952014-DEL-96 |
| chr12-97985244-DEL-55 | chr12-97985242-DEL-52 |
| chr12-97989108-DEL-66 | chr12-97989109-DEL-66 |
| chr12-98019864-DEL-200 | chr12-98019865-DEL-200 |
| chr12-98048034-DEL-98 | chr12-98048035-DEL-98 |
| chr12-98051882-DEL-99 | chr12-98051883-DEL-98 |
| chr12-98054738-DEL-1618 | chr12-98054739-DEL-1618 |
| chr12-98081955-DEL-6194 | chr12-98081953-DEL-6195 |
| chr12-98095651-DEL-164 | chr12-98095652-DEL-164 |
| chr12-98139965-DEL-2859 | chr12-98139966-DEL-2859 |
| chr12-98177699-DEL-57 | chr12-98177700-DEL-57 |
| chr12-98187487-DEL-814 | chr12-98187488-DEL-814 |
| chr12-98190136-DEL-1098 | chr12-98190136-DEL-1097 |
| chr12-98271977-DEL-876 | chr12-98271978-DEL-876 |
| chr12-98278961-DEL-1741 | chr12-98278962-DEL-1741 |
| chr12-98283018-DEL-68 | chr12-98283019-DEL-56 |
| chr12-98286803-DEL-589 | chr12-98286804-DEL-589 |
| chr12-98289891-DEL-670 | chr12-98289901-DEL-659 |
| chr12-98292492-DEL-339 | chr12-98292493-DEL-339 |
| chr12-98305641-DEL-79 | chr12-98305654-DEL-85 |
| chr12-98321486-DEL-7121 | chr12-98321487-DEL-7121 |
| chr12-98328993-DEL-56 | chr12-98328974-DEL-58 |
| chr12-98338046-DEL-84 | chr12-98338068-DEL-51 |
| chr12-98376107-DEL-863 | chr12-98376108-DEL-863 |
| chr12-98427680-DEL-6434 | chr12-98427681-DEL-6434 |
| chr12-98449676-DEL-5928 | chr12-98449677-DEL-5928 |
| chr12-98465450-DEL-52 | chr12-98465450-DEL-52 |
| chr12-98489960-DEL-55 | chr12-98489961-DEL-55 |
| chr12-98505087-DEL-1615 | chr12-98505088-DEL-1615 |
| chr12-98560892-DEL-240 | chr12-98560893-DEL-240 |
| chr12-98582306-DEL-64 | chr12-98582307-DEL-64 |
| chr12-98593151-DEL-214 | chr12-98593152-DEL-214 |
| chr12-98600984-DEL-224 | chr12-98600985-DEL-224 |
| chr12-98666887-DEL-198 | chr12-98666888-DEL-198 |
| chr12-98689306-DEL-183 | chr12-98689307-DEL-183 |
| chr12-98695182-DEL-66 | chr12-98695185-DEL-64 |
| chr12-98705638-DEL-72 | chr12-98705639-DEL-72 |
| chr12-98788661-DEL-218 | chr12-98788662-DEL-218 |
| chr12-98881100-DEL-68 | chr12-98881101-DEL-68 |
| chr12-98889536-DEL-6251 | chr12-98889537-DEL-6251 |
| chr12-98896667-DEL-519 | chr12-98896668-DEL-519 |
| chr12-98911359-DEL-92 | chr12-98911360-DEL-92 |
| chr12-98916496-DEL-126 | chr12-98916493-DEL-114 |
| chr12-98917891-DEL-63 | chr12-98917860-DEL-64 |
| chr12-98929477-DEL-602 | chr12-98929478-DEL-602 |
| chr12-98937109-DEL-171 | chr12-98937152-DEL-129 |
| chr12-98953903-DEL-532 | chr12-98953904-DEL-532 |
| chr12-99021663-DEL-354 | chr12-99021664-DEL-354 |
| chr12-99033543-DEL-60 | chr12-99033514-DEL-90 |
| chr12-99099324-DEL-109 | chr12-99099325-DEL-109 |
| chr12-99108188-DEL-6734 | chr12-99108189-DEL-6734 |
| chr12-99143565-DEL-205 | chr12-99143566-DEL-205 |
| chr12-99174308-DEL-72 | chr12-99174309-DEL-71 |
| chr12-99221310-DEL-87 | chr12-99221323-DEL-75 |
| chr12-99222917-DEL-76 | chr12-99222930-DEL-68 |
| chr12-99237157-DEL-1711 | chr12-99237158-DEL-1710 |
| chr12-99238487-DEL-150 | chr12-99238488-DEL-150 |
| chr12-99239195-DEL-89 | chr12-99239196-DEL-89 |
| chr12-99368154-DEL-513 | chr12-99368155-DEL-513 |
| chr12-99400910-DEL-238 | chr12-99400911-DEL-238 |
| chr12-99414833-DEL-191 | chr12-99414834-DEL-191 |
| chr12-99420427-DEL-156 | chr12-99420414-DEL-150 |
| chr12-99441436-DEL-56 | chr12-99441437-DEL-56 |
| chr12-99442730-DEL-58 | chr12-99442731-DEL-58 |
| chr12-99458703-DEL-178 | chr12-99458682-DEL-200 |
| chr12-99472491-DEL-170 | chr12-99472479-DEL-147 |
| chr12-99474987-DEL-598 | chr12-99474988-DEL-598 |
| chr12-99479004-DEL-106 | chr12-99478969-DEL-91 |
| chr12-99521990-DEL-105 | chr12-99521991-DEL-105 |
| chr12-99531568-DEL-220 | chr12-99531564-DEL-225 |
| chr12-99534572-DEL-236 | chr12-99534573-DEL-236 |
| chr12-99551021-DEL-64 | chr12-99551029-DEL-56 |
| chr12-99554807-DEL-52 | chr12-99554808-DEL-52 |
| chr12-99557910-DEL-69 | chr12-99557893-DEL-64 |
| chr12-99564023-DEL-602 | chr12-99564021-DEL-597 |
| chr12-99578257-DEL-58 | chr12-99578258-DEL-58 |
| chr12-99585233-DEL-131 | chr12-99585231-DEL-132 |
| chr12-99589086-DEL-202 | chr12-99589087-DEL-202 |

| | |
|---|---|
| chr12-99736474-DEL-59 | chr12-99736475-DEL-59 |
| chr12-99773243-DEL-200 | chr12-99773244-DEL-200 |
| chr12-99784341-DEL-322 | chr12-99784342-DEL-322 |
| chr12-99816279-DEL-60 | chr12-99816280-DEL-60 |
| chr12-99820292-DEL-56 | chr12-99820293-DEL-56 |
| chr12-99835586-DEL-1846 | chr12-99835587-DEL-1845 |
| chr12-99847957-DEL-72 | chr12-99847958-DEL-72 |
| chr12-99854656-DEL-444 | chr12-99854657-DEL-444 |
| chr12-99916667-DEL-190 | chr12-99916668-DEL-190 |
| chr12-99961551-DEL-283 | chr12-99961552-DEL-283 |
| chr12-99987348-DEL-396 | chr12-99987349-DEL-396 |
| chr12-100024679-DEL-62 | chr12-100024679-DEL-60 |
| chr12-100030162-DEL-59 | chr12-100030163-DEL-59 |
| chr12-100031174-DEL-262 | chr12-100031175-DEL-260 |
| chr12-100034827-DEL-55 | chr12-100034828-DEL-55 |
| chr12-100040501-DEL-96 | chr12-100040550-DEL-72 |
| chr12-100049709-DEL-63 | chr12-100049706-DEL-66 |
| chr12-100060112-DEL-504 | chr12-100060113-DEL-504 |
| chr12-100072799-DEL-222 | chr12-100072800-DEL-222 |
| chr12-100079803-DEL-317 | chr12-100079758-DEL-355 |
| chr12-100202243-DEL-97 | chr12-100202244-DEL-97 |
| chr12-100216994-DEL-280 | chr12-100216995-DEL-278 |
| chr12-100226052-DEL-94 | chr12-100226053-DEL-95 |
| chr12-100230940-DEL-69 | chr12-100230941-DEL-69 |
| chr12-100289012-DEL-55 | chr12-100289013-DEL-55 |
| chr12-100357765-DEL-281 | chr12-100357766-DEL-281 |
| chr12-100390122-DEL-208 | chr12-100390123-DEL-208 |
| chr12-100425096-DEL-6391 | chr12-100425090-DEL-6401 |
| chr12-100469444-DEL-69 | chr12-100469453-DEL-66 |
| chr12-100487532-DEL-77 | chr12-100487533-DEL-77 |
| chr12-100527290-DEL-86 | chr12-100527291-DEL-86 |
| chr12-100568873-DEL-193 | chr12-100568874-DEL-193 |
| chr12-100578771-DEL-1926 | chr12-100578772-DEL-1926 |
| chr12-100608388-DEL-554 | chr12-100608389-DEL-554 |
| chr12-100609003-DEL-929 | chr12-100609004-DEL-930 |
| chr12-100639259-DEL-88 | chr12-100639260-DEL-88 |
| chr12-100641212-DEL-696 | chr12-100641213-DEL-696 |
| chr12-100710724-DEL-520 | chr12-100710725-DEL-520 |
| chr12-100749990-DEL-117 | chr12-100749991-DEL-117 |
| chr12-100821989-DEL-273 | chr12-100821990-DEL-273 |
| chr12-100829032-DEL-66 | chr12-100829033-DEL-66 |
| chr12-100836877-DEL-204 | chr12-100836878-DEL-204 |
| chr12-100851012-DEL-203 | chr12-100851013-DEL-203 |
| chr12-100851591-DEL-89 | chr12-100851592-DEL-89 |
| chr12-100861027-DEL-141 | chr12-100861024-DEL-141 |
| chr12-100882422-DEL-405 | chr12-100882423-DEL-405 |
| chr12-100894108-DEL-71 | chr12-100894109-DEL-71 |
| chr12-100898588-DEL-58 | chr12-100898568-DEL-56 |
| chr12-100901018-DEL-129 | chr12-100901019-DEL-129 |
| chr12-100905978-DEL-169 | chr12-100905979-DEL-169 |
| chr12-100915761-DEL-54 | chr12-100915762-DEL-51 |
| chr12-100930108-DEL-161 | chr12-100930109-DEL-161 |
| chr12-100931963-DEL-115 | chr12-100931964-DEL-115 |
| chr12-100953275-DEL-141 | chr12-100953276-DEL-141 |
| chr12-100967901-DEL-67 | chr12-100967902-DEL-67 |
| chr12-100968590-DEL-79 | chr12-100968591-DEL-79 |
| chr12-100996573-DEL-183 | chr12-100996572-DEL-181 |
| chr12-101003297-DEL-224 | chr12-101003298-DEL-224 |
| chr12-101042191-DEL-74 | chr12-101042192-DEL-74 |
| chr12-101052030-DEL-164 | chr12-101052031-DEL-164 |
| chr12-101058324-DEL-732 | chr12-101058325-DEL-732 |
| chr12-101102546-DEL-7530 | chr12-101102547-DEL-7530 |
| chr12-101132756-DEL-58 | chr12-101132757-DEL-58 |
| chr12-101157259-DEL-185 | chr12-101157258-DEL-186 |
| chr12-101171593-DEL-120 | chr12-101171594-DEL-121 |
| chr12-101212811-DEL-72 | chr12-101212812-DEL-68 |
| chr12-101220821-DEL-3285 | chr12-101220823-DEL-3284 |
| chr12-101296975-DEL-92 | chr12-101296976-DEL-108 |
| chr12-101311554-DEL-1287 | chr12-101311554-DEL-1288 |
| chr12-101325404-DEL-606 | chr12-101325405-DEL-606 |
| chr12-101413136-DEL-6692 | chr12-101413137-DEL-6692 |
| chr12-101422609-DEL-73 | chr12-101422610-DEL-73 |
| chr12-101425279-DEL-1209 | chr12-101425303-DEL-1185 |
| chr12-101511632-DEL-781 | chr12-101511633-DEL-781 |
| chr12-101560616-DEL-232 | chr12-101560617-DEL-232 |
| chr12-101570351-DEL-1828 | chr12-101570352-DEL-1828 |
| chr12-101579604-DEL-6406 | chr12-101579605-DEL-6406 |
| chr12-101599096-DEL-79 | chr12-101599097-DEL-79 |
| chr12-101611027-DEL-178 | chr12-101611028-DEL-178 |
| chr12-101644127-DEL-114 | chr12-101644130-DEL-112 |

| | |
|---|---|
| chr12-101648163-DEL-162 | chr12-101648164-DEL-162 |
| chr12-101651062-DEL-2664 | chr12-101651063-DEL-2664 |
| chr12-101668501-DEL-1180 | chr12-101668502-DEL-1180 |
| chr12-101677878-DEL-203 | chr12-101677878-DEL-203 |
| chr12-101689775-DEL-520 | chr12-101689776-DEL-520 |
| chr12-101717831-DEL-54 | chr12-101717831-DEL-54 |
| chr12-101738046-DEL-596 | chr12-101738047-DEL-596 |
| chr12-101743458-DEL-335 | chr12-101743459-DEL-335 |
| chr12-101754097-DEL-93 | chr12-101754147-DEL-55 |
| chr12-101762218-DEL-274 | chr12-101762219-DEL-274 |
| chr12-101770480-DEL-314 | chr12-101770481-DEL-314 |
| chr12-101791596-DEL-165 | chr12-101791599-DEL-153 |
| chr12-101807345-DEL-485 | chr12-101807346-DEL-485 |
| chr12-101808431-DEL-1684 | chr12-101808432-DEL-1684 |
| chr12-101813890-DEL-1892 | chr12-101813891-DEL-1892 |
| chr12-101823962-DEL-214 | chr12-101823931-DEL-215 |
| chr12-101829704-DEL-217 | chr12-101829705-DEL-217 |
| chr12-101855215-DEL-194 | chr12-101855216-DEL-194 |
| chr12-101876279-DEL-206 | chr12-101876280-DEL-206 |
| chr12-101906295-DEL-212 | chr12-101906291-DEL-216 |
| chr12-101907689-DEL-59 | chr12-101907690-DEL-59 |
| chr12-101928374-DEL-70 | chr12-101928375-DEL-66 |
| chr12-101941135-DEL-52 | chr12-101941136-DEL-52 |
| chr12-101946496-DEL-74 | chr12-101946497-DEL-74 |
| chr12-101959233-DEL-205 | chr12-101959234-DEL-205 |
| chr12-101988545-DEL-593 | chr12-101988545-DEL-593 |
| chr12-101998627-DEL-239 | chr12-101998626-DEL-240 |
| chr12-102002313-DEL-431 | chr12-102002317-DEL-386 |
| chr12-102003316-DEL-384 | chr12-102003317-DEL-384 |
| chr12-102021466-DEL-261 | chr12-102021493-DEL-263 |
| chr12-102046498-DEL-60 | chr12-102046498-DEL-60 |
| chr12-102061192-DEL-152 | chr12-102061181-DEL-116 |
| chr12-102063960-DEL-92 | chr12-102063961-DEL-92 |
| chr12-102086097-DEL-6224 | chr12-102086098-DEL-6224 |
| chr12-102118481-DEL-335 | chr12-102118482-DEL-335 |
| chr12-102124972-DEL-146 | chr12-102124973-DEL-146 |
| chr12-102145981-DEL-382 | chr12-102145982-DEL-382 |
| chr12-102150874-DEL-56 | chr12-102150875-DEL-56 |
| chr12-102210595-DEL-60 | chr12-102210606-DEL-50 |
| chr12-102219712-DEL-242 | chr12-102219713-DEL-242 |
| chr12-102245861-DEL-838 | chr12-102245862-DEL-838 |
| chr12-102267042-DEL-90 | chr12-102267011-DEL-75 |
| chr12-102272522-DEL-234 | chr12-102272523-DEL-234 |
| chr12-102302468-DEL-391 | chr12-102302469-DEL-391 |
| chr12-102304179-DEL-66 | chr12-102304180-DEL-78 |
| chr12-102330330-DEL-369 | chr12-102330331-DEL-369 |
| chr12-102348254-DEL-112 | chr12-102348255-DEL-112 |
| chr12-102350243-DEL-182 | chr12-102350244-DEL-182 |
| chr12-102351882-DEL-112 | chr12-102351883-DEL-112 |
| chr12-102353371-DEL-57 | chr12-102353372-DEL-59 |
| chr12-102357909-DEL-59 | chr12-102357910-DEL-59 |
| chr12-102371771-DEL-65 | chr12-102371757-DEL-80 |
| chr12-102372331-DEL-6703 | chr12-102372332-DEL-6703 |
| chr12-102378506-DEL-424 | chr12-102378507-DEL-424 |
| chr12-102392909-DEL-3073 | chr12-102392910-DEL-3073 |
| chr12-102410834-DEL-194 | chr12-102410835-DEL-194 |
| chr12-102443778-DEL-197 | chr12-102443779-DEL-197 |
| chr12-102465667-DEL-725 | chr12-102465668-DEL-725 |
| chr12-102466547-DEL-175 | chr12-102466539-DEL-183 |
| chr12-102481574-DEL-554 | chr12-102481579-DEL-550 |
| chr12-102485473-DEL-444 | chr12-102485492-DEL-426 |
| chr12-102508207-DEL-181 | chr12-102508204-DEL-182 |
| chr12-102543555-DEL-172 | chr12-102543556-DEL-172 |
| chr12-102556615-DEL-66 | chr12-102556616-DEL-66 |
| chr12-102596007-DEL-204 | chr12-102596008-DEL-204 |
| chr12-102597720-DEL-60 | chr12-102597713-DEL-66 |
| chr12-102601193-DEL-480 | chr12-102601194-DEL-480 |
| chr12-102616226-DEL-77 | chr12-102616225-DEL-68 |
| chr12-102704173-DEL-184 | chr12-102704174-DEL-184 |
| chr12-102717185-DEL-133 | chr12-102717183-DEL-109 |
| chr12-102719708-DEL-202 | chr12-102719709-DEL-202 |
| chr12-102741359-DEL-127 | chr12-102741360-DEL-125 |
| chr12-102763458-DEL-63 | chr12-102763459-DEL-63 |
| chr12-102766253-DEL-209 | chr12-102766254-DEL-209 |
| chr12-102770457-DEL-208 | chr12-102770458-DEL-208 |
| chr12-102775766-DEL-117 | chr12-102775767-DEL-117 |
| chr12-102780189-DEL-412 | chr12-102780190-DEL-412 |
| chr12-102782717-DEL-208 | chr12-102782718-DEL-208 |
| chr12-102788116-DEL-215 | chr12-102788117-DEL-215 |
| chr12-102793100-DEL-96 | chr12-102793101-DEL-96 |

| | |
|---|---|
| chr12-102799374-DEL-863 | chr12-102799373-DEL-862 |
| chr12-102849425-DEL-80 | chr12-102849426-DEL-80 |
| chr12-102858727-DEL-128 | chr12-102858728-DEL-128 |
| chr12-102870896-DEL-182 | chr12-102870897-DEL-182 |
| chr12-102932091-DEL-54 | chr12-102932092-DEL-56 |
| chr12-102938600-DEL-87 | chr12-102938601-DEL-87 |
| chr12-102942810-DEL-114 | chr12-102942811-DEL-114 |
| chr12-102950672-DEL-421 | chr12-102950674-DEL-449 |
| chr12-102979975-DEL-72 | chr12-102979976-DEL-72 |
| chr12-102986426-DEL-96 | chr12-102986427-DEL-96 |
| chr12-103000801-DEL-72 | chr12-103000802-DEL-72 |
| chr12-103048117-DEL-157 | chr12-103048143-DEL-129 |
| chr12-103054156-DEL-107 | chr12-103054157-DEL-107 |
| chr12-103141007-DEL-147 | chr12-103141008-DEL-147 |
| chr12-103174998-DEL-240 | chr12-103175038-DEL-222 |
| chr12-103215101-DEL-638 | chr12-103215102-DEL-638 |
| chr12-103226960-DEL-54 | chr12-103226944-DEL-70 |
| chr12-103230310-DEL-390 | chr12-103230311-DEL-410 |
| chr12-103236242-DEL-98 | chr12-103236243-DEL-98 |
| chr12-103237460-DEL-74 | chr12-103237461-DEL-74 |
| chr12-103241614-DEL-52 | chr12-103241616-DEL-51 |
| chr12-103305582-DEL-89 | chr12-103305583-DEL-89 |
| chr12-103308938-DEL-125 | chr12-103308933-DEL-131 |
| chr12-103323128-DEL-78 | chr12-103323129-DEL-115 |
| chr12-103351921-DEL-52 | chr12-103351916-DEL-58 |
| chr12-103486456-DEL-207 | chr12-103486457-DEL-207 |
| chr12-103487736-DEL-107 | chr12-103487760-DEL-105 |
| chr12-103539891-DEL-147 | chr12-103539891-DEL-148 |
| chr12-103558279-DEL-86 | chr12-103558314-DEL-50 |
| chr12-103568131-DEL-58 | chr12-103568127-DEL-59 |
| chr12-103582617-DEL-52 | chr12-103582618-DEL-52 |
| chr12-103649057-DEL-87 | chr12-103649058-DEL-87 |
| chr12-103653792-DEL-52 | chr12-103653793-DEL-54 |
| chr12-103664427-DEL-1797 | chr12-103664428-DEL-1796 |
| chr12-103680328-DEL-62 | chr12-103680331-DEL-60 |
| chr12-103755137-DEL-52 | chr12-103755137-DEL-52 |
| chr12-103760849-DEL-213 | chr12-103760865-DEL-213 |
| chr12-103762281-DEL-478 | chr12-103762280-DEL-479 |
| chr12-103812711-DEL-1734 | chr12-103812712-DEL-1734 |
| chr12-103895912-DEL-2688 | chr12-103895912-DEL-2688 |
| chr12-103922561-DEL-60 | chr12-103922564-DEL-58 |
| chr12-103943710-DEL-62 | chr12-103943711-DEL-60 |
| chr12-103961819-DEL-57 | chr12-103961818-DEL-66 |
| chr12-103977028-DEL-54 | chr12-103977029-DEL-52 |
| chr12-104032127-DEL-52 | chr12-104032127-DEL-52 |
| chr12-104034472-DEL-60 | chr12-104034437-DEL-64 |
| chr12-104051878-DEL-62 | chr12-104051879-DEL-62 |
| chr12-104062686-DEL-1038 | chr12-104062688-DEL-1034 |
| chr12-104095711-DEL-85 | chr12-104095712-DEL-85 |
| chr12-104136619-DEL-3767 | chr12-104136619-DEL-3767 |
| chr12-104167865-DEL-6716 | chr12-104167866-DEL-6716 |
| chr12-104181328-DEL-395 | chr12-104181329-DEL-395 |
| chr12-104200993-DEL-70 | chr12-104200994-DEL-72 |
| chr12-104210064-DEL-634 | chr12-104210065-DEL-634 |
| chr12-104241227-DEL-53 | chr12-104241228-DEL-53 |
| chr12-104248878-DEL-870 | chr12-104248879-DEL-870 |
| chr12-104257288-DEL-51 | chr12-104257289-DEL-51 |
| chr12-104260975-DEL-891 | chr12-104260976-DEL-891 |
| chr12-104268496-DEL-52 | chr12-104268530-DEL-58 |
| chr12-104297450-DEL-200 | chr12-104297451-DEL-200 |
| chr12-104335566-DEL-120 | chr12-104335581-DEL-106 |
| chr12-104344986-DEL-64 | chr12-104344985-DEL-65 |
| chr12-104348611-DEL-391 | chr12-104348608-DEL-391 |
| chr12-104360565-DEL-167 | chr12-104360569-DEL-164 |
| chr12-104369928-DEL-54 | chr12-104369929-DEL-54 |
| chr12-104371316-DEL-315 | chr12-104371317-DEL-315 |
| chr12-104383130-DEL-78 | chr12-104383131-DEL-54 |
| chr12-104401021-DEL-111 | chr12-104401022-DEL-111 |
| chr12-104424450-DEL-110 | chr12-104424457-DEL-102 |
| chr12-104451924-DEL-59 | chr12-104451924-DEL-60 |
| chr12-104466374-DEL-108 | chr12-104466382-DEL-109 |
| chr12-104473493-DEL-274 | chr12-104473494-DEL-274 |
| chr12-104483241-DEL-64 | chr12-104483242-DEL-64 |
| chr12-104495647-DEL-609 | chr12-104495648-DEL-609 |
| chr12-104503883-DEL-142 | chr12-104503884-DEL-142 |
| chr12-104506883-DEL-65 | chr12-104506884-DEL-65 |
| chr12-104510169-DEL-79 | chr12-104510182-DEL-67 |
| chr12-104514236-DEL-59 | chr12-104514232-DEL-54 |
| chr12-104527079-DEL-96 | chr12-104527098-DEL-64 |
| chr12-104575131-DEL-211 | chr12-104575132-DEL-211 |

| | |
|---|---|
| chr12-104628760-DEL-150 | chr12-104628761-DEL-150 |
| chr12-104637019-DEL-251 | chr12-104637019-DEL-251 |
| chr12-104770370-DEL-208 | chr12-104770371-DEL-208 |
| chr12-104792159-DEL-64 | chr12-104792160-DEL-64 |
| chr12-104803743-DEL-94 | chr12-104803744-DEL-94 |
| chr12-104809604-DEL-56 | chr12-104809605-DEL-56 |
| chr12-104854133-DEL-153 | chr12-104854134-DEL-153 |
| chr12-104880810-DEL-282 | chr12-104880809-DEL-282 |
| chr12-104915437-DEL-1219 | chr12-104915438-DEL-1219 |
| chr12-104950552-DEL-82 | chr12-104950545-DEL-90 |
| chr12-104960664-DEL-62 | chr12-104960665-DEL-62 |
| chr12-104987663-DEL-150 | chr12-104987664-DEL-150 |
| chr12-105001834-DEL-82 | chr12-105001835-DEL-82 |
| chr12-105013760-DEL-462 | chr12-105013761-DEL-462 |
| chr12-105043412-DEL-87 | chr12-105043429-DEL-71 |
| chr12-105045969-DEL-1278 | chr12-105045970-DEL-1278 |
| chr12-105046638-DEL-194 | chr12-105046639-DEL-194 |
| chr12-105070280-DEL-3009 | chr12-105070281-DEL-3009 |
| chr12-105122922-DEL-70 | chr12-105122923-DEL-68 |
| chr12-105129097-DEL-84 | chr12-105129095-DEL-84 |
| chr12-105139300-DEL-262 | chr12-105139301-DEL-262 |
| chr12-105157627-DEL-143 | chr12-105157626-DEL-143 |
| chr12-105178032-DEL-184 | chr12-105178033-DEL-178 |
| chr12-105178878-DEL-160 | chr12-105178879-DEL-160 |
| chr12-105185412-DEL-205 | chr12-105185413-DEL-205 |
| chr12-105198053-DEL-76 | chr12-105198054-DEL-76 |
| chr12-105221222-DEL-933 | chr12-105221223-DEL-933 |
| chr12-105261364-DEL-143 | chr12-105261365-DEL-143 |
| chr12-105265841-DEL-52 | chr12-105265821-DEL-62 |
| chr12-105266339-DEL-7461 | chr12-105266339-DEL-7461 |
| chr12-105295367-DEL-59 | chr12-105295373-DEL-60 |
| chr12-105311766-DEL-317 | chr12-105311768-DEL-316 |
| chr12-105355070-DEL-217 | chr12-105355108-DEL-185 |
| chr12-105392473-DEL-60 | chr12-105392471-DEL-66 |
| chr12-105445708-DEL-492 | chr12-105445709-DEL-492 |
| chr12-105447277-DEL-1066 | chr12-105447278-DEL-1066 |
| chr12-105455537-DEL-52 | chr12-105455538-DEL-52 |
| chr12-105499628-DEL-53 | chr12-105499627-DEL-55 |
| chr12-105519744-DEL-270 | chr12-105519744-DEL-270 |
| chr12-105542507-DEL-139 | chr12-105542508-DEL-138 |
| chr12-105552901-DEL-581 | chr12-105552902-DEL-581 |
| chr12-105555102-DEL-834 | chr12-105555103-DEL-834 |
| chr12-105562784-DEL-1445 | chr12-105562785-DEL-1445 |
| chr12-105585371-DEL-1282 | chr12-105585372-DEL-1282 |
| chr12-105594283-DEL-3341 | chr12-105594284-DEL-3341 |
| chr12-105621817-DEL-196 | chr12-105621818-DEL-196 |
| chr12-105638461-DEL-295 | chr12-105638462-DEL-292 |
| chr12-105645502-DEL-203 | chr12-105645503-DEL-203 |
| chr12-105655132-DEL-218 | chr12-105655116-DEL-189 |
| chr12-105661159-DEL-88 | chr12-105661119-DEL-107 |
| chr12-105668801-DEL-60 | chr12-105668802-DEL-60 |
| chr12-105681137-DEL-186 | chr12-105681138-DEL-184 |
| chr12-105682064-DEL-83 | chr12-105682076-DEL-74 |
| chr12-105697937-DEL-203 | chr12-105697934-DEL-207 |
| chr12-105699381-DEL-185 | chr12-105699382-DEL-185 |
| chr12-105700571-DEL-85 | chr12-105700563-DEL-51 |
| chr12-105702683-DEL-188 | chr12-105702684-DEL-188 |
| chr12-105703535-DEL-216 | chr12-105703536-DEL-216 |
| chr12-105712645-DEL-195 | chr12-105712652-DEL-209 |
| chr12-105715032-DEL-3433 | chr12-105715033-DEL-3433 |
| chr12-105718570-DEL-75 | chr12-105718543-DEL-76 |
| chr12-105731368-DEL-226 | chr12-105731364-DEL-225 |
| chr12-105742087-DEL-76 | chr12-105742073-DEL-88 |
| chr12-105849055-DEL-698 | chr12-105849056-DEL-698 |
| chr12-105854004-DEL-76 | chr12-105854000-DEL-52 |
| chr12-105858858-DEL-205 | chr12-105858859-DEL-205 |
| chr12-105882834-DEL-263 | chr12-105882835-DEL-263 |
| chr12-105917449-DEL-204 | chr12-105917450-DEL-204 |
| chr12-105937927-DEL-160 | chr12-105937913-DEL-161 |
| chr12-105940326-DEL-63 | chr12-105940327-DEL-51 |
| chr12-105946376-DEL-550 | chr12-105946369-DEL-533 |
| chr12-105983830-DEL-103 | chr12-105983825-DEL-61 |
| chr12-106003002-DEL-234 | chr12-106003003-DEL-234 |
| chr12-106010682-DEL-217 | chr12-106010681-DEL-217 |
| chr12-106011892-DEL-602 | chr12-106011893-DEL-602 |
| chr12-106075463-DEL-63 | chr12-106075464-DEL-63 |
| chr12-106097399-DEL-223 | chr12-106097400-DEL-223 |
| chr12-106102513-DEL-55 | chr12-106102519-DEL-50 |
| chr12-106111484-DEL-97 | chr12-106111485-DEL-97 |
| chr12-106134150-DEL-98 | chr12-106134151-DEL-98 |

| | |
|---|---|
| chr12-106168081-DEL-52 | chr12-106168107-DEL-60 |
| chr12-106174311-DEL-219 | chr12-106174312-DEL-219 |
| chr12-106184393-DEL-85 | chr12-106184404-DEL-75 |
| chr12-106186086-DEL-146 | chr12-106186126-DEL-107 |
| chr12-106201957-DEL-83 | chr12-106201967-DEL-84 |
| chr12-106203794-DEL-111 | chr12-106203817-DEL-116 |
| chr12-106206400-DEL-66 | chr12-106206400-DEL-58 |
| chr12-106230888-DEL-127 | chr12-106230889-DEL-118 |
| chr12-106248507-DEL-97 | chr12-106248508-DEL-97 |
| chr12-106253412-DEL-67 | chr12-106253403-DEL-77 |
| chr12-106263816-DEL-94 | chr12-106263825-DEL-86 |
| chr12-106291142-DEL-63 | chr12-106291144-DEL-61 |
| chr12-106311167-DEL-1570 | chr12-106311168-DEL-1570 |
| chr12-106339898-DEL-67 | chr12-106339893-DEL-73 |
| chr12-106350170-DEL-161 | chr12-106350171-DEL-161 |
| chr12-106351952-DEL-58 | chr12-106351952-DEL-66 |
| chr12-106356716-DEL-92 | chr12-106356743-DEL-66 |
| chr12-106400413-DEL-52 | chr12-106400408-DEL-53 |
| chr12-106412017-DEL-59 | chr12-106411999-DEL-106 |
| chr12-106425004-DEL-115 | chr12-106425005-DEL-113 |
| chr12-106513552-DEL-69 | chr12-106513551-DEL-70 |
| chr12-106558285-DEL-153 | chr12-106558286-DEL-153 |
| chr12-106559965-DEL-76 | chr12-106559960-DEL-54 |
| chr12-106562866-DEL-78 | chr12-106562867-DEL-78 |
| chr12-106577788-DEL-51 | chr12-106577789-DEL-51 |
| chr12-106606777-DEL-249 | chr12-106606778-DEL-227 |
| chr12-106609565-DEL-60 | chr12-106609544-DEL-50 |
| chr12-106619694-DEL-172 | chr12-106619695-DEL-172 |
| chr12-106623029-DEL-79 | chr12-106623023-DEL-67 |
| chr12-106656250-DEL-110 | chr12-106656251-DEL-110 |
| chr12-106668826-DEL-199 | chr12-106668823-DEL-202 |
| chr12-106679491-DEL-104 | chr12-106679492-DEL-104 |
| chr12-106686354-DEL-79 | chr12-106686349-DEL-76 |
| chr12-106692229-DEL-54 | chr12-106692228-DEL-60 |
| chr12-106707951-DEL-127 | chr12-106707951-DEL-127 |
| chr12-106726638-DEL-1930 | chr12-106726639-DEL-1930 |
| chr12-106793948-DEL-3883 | chr12-106793949-DEL-3883 |
| chr12-106804405-DEL-96 | chr12-106804398-DEL-95 |
| chr12-106805140-DEL-59 | chr12-106805135-DEL-60 |
| chr12-106858108-DEL-117 | chr12-106858109-DEL-117 |
| chr12-106888894-DEL-76 | chr12-106888895-DEL-76 |
| chr12-106891661-DEL-66 | chr12-106891662-DEL-66 |
| chr12-106903167-DEL-61 | chr12-106903178-DEL-51 |
| chr12-106913993-DEL-85 | chr12-106913994-DEL-85 |
| chr12-106923462-DEL-115 | chr12-106923463-DEL-115 |
| chr12-106951057-DEL-143 | chr12-106951058-DEL-143 |
| chr12-106982109-DEL-123 | chr12-106982110-DEL-115 |
| chr12-107024053-DEL-78 | chr12-107024080-DEL-52 |
| chr12-107079947-DEL-58 | chr12-107079948-DEL-58 |
| chr12-107083464-DEL-62 | chr12-107083465-DEL-66 |
| chr12-107087570-DEL-98 | chr12-107087571-DEL-98 |
| chr12-107132749-DEL-1262 | chr12-107132750-DEL-1262 |
| chr12-107157616-DEL-292 | chr12-107157615-DEL-294 |
| chr12-107183106-DEL-480 | chr12-107183107-DEL-477 |
| chr12-107185120-DEL-173 | chr12-107185119-DEL-143 |
| chr12-107221021-DEL-54 | chr12-107221022-DEL-54 |
| chr12-107226818-DEL-65 | chr12-107226818-DEL-66 |
| chr12-107245893-DEL-203 | chr12-107245894-DEL-203 |
| chr12-107265321-DEL-64 | chr12-107265322-DEL-64 |
| chr12-107276373-DEL-401 | chr12-107276374-DEL-401 |
| chr12-107277257-DEL-58 | chr12-107277258-DEL-58 |
| chr12-107302247-DEL-60 | chr12-107302248-DEL-60 |
| chr12-107303194-DEL-96 | chr12-107303178-DEL-86 |
| chr12-107309115-DEL-273 | chr12-107309110-DEL-271 |
| chr12-107310182-DEL-428 | chr12-107310209-DEL-428 |
| chr12-107350953-DEL-228 | chr12-107350954-DEL-228 |
| chr12-107365068-DEL-80 | chr12-107365069-DEL-80 |
| chr12-107374079-DEL-142 | chr12-107374080-DEL-142 |
| chr12-107385404-DEL-284 | chr12-107385405-DEL-283 |
| chr12-107390887-DEL-97 | chr12-107390888-DEL-97 |
| chr12-107396991-DEL-56 | chr12-107396992-DEL-56 |
| chr12-107405701-DEL-400 | chr12-107405702-DEL-400 |
| chr12-107446503-DEL-167 | chr12-107446547-DEL-124 |
| chr12-107450110-DEL-896 | chr12-107450111-DEL-896 |
| chr12-107460472-DEL-52 | chr12-107460473-DEL-52 |
| chr12-107467587-DEL-961 | chr12-107467588-DEL-961 |
| chr12-107499670-DEL-106 | chr12-107499671-DEL-106 |
| chr12-107501171-DEL-94 | chr12-107501209-DEL-60 |
| chr12-107555049-DEL-69 | chr12-107555050-DEL-69 |
| chr12-107581339-DEL-164 | chr12-107581340-DEL-164 |

| | |
|---|---|
| chr12-107597308-DEL-51 | chr12-107597309-DEL-51 |
| chr12-107610627-DEL-236 | chr12-107610618-DEL-246 |
| chr12-107648833-DEL-58 | chr12-107648834-DEL-58 |
| chr12-107681755-DEL-60 | chr12-107681755-DEL-60 |
| chr12-107683497-DEL-89 | chr12-107683496-DEL-90 |
| chr12-107701798-DEL-96 | chr12-107701799-DEL-96 |
| chr12-107708790-DEL-52 | chr12-107708791-DEL-52 |
| chr12-107712818-DEL-91 | chr12-107712823-DEL-87 |
| chr12-107728131-DEL-71 | chr12-107728123-DEL-72 |
| chr12-107743573-DEL-68 | chr12-107743574-DEL-68 |
| chr12-107779849-DEL-81 | chr12-107779850-DEL-81 |
| chr12-107832585-DEL-520 | chr12-107832586-DEL-520 |
| chr12-107836710-DEL-125 | chr12-107836703-DEL-156 |
| chr12-107839937-DEL-340 | chr12-107839958-DEL-320 |
| chr12-107847432-DEL-176 | chr12-107847433-DEL-176 |
| chr12-107848847-DEL-122 | chr12-107848848-DEL-122 |
| chr12-107905217-DEL-68 | chr12-107905209-DEL-59 |
| chr12-107911188-DEL-65 | chr12-107911189-DEL-65 |
| chr12-107914345-DEL-337 | chr12-107914346-DEL-337 |
| chr12-108026093-DEL-437 | chr12-108026094-DEL-437 |
| chr12-108028231-DEL-138 | chr12-108028232-DEL-138 |
| chr12-108029589-DEL-261 | chr12-108029586-DEL-261 |
| chr12-108038315-DEL-200 | chr12-108038316-DEL-200 |
| chr12-108043440-DEL-59 | chr12-108043441-DEL-59 |
| chr12-108045544-DEL-5136 | chr12-108045545-DEL-5136 |
| chr12-108065694-DEL-207 | chr12-108065695-DEL-207 |
| chr12-108091869-DEL-240 | chr12-108091870-DEL-240 |
| chr12-108121086-DEL-56 | chr12-108121087-DEL-56 |
| chr12-108196976-DEL-215 | chr12-108196977-DEL-215 |
| chr12-108213946-DEL-138 | chr12-108213952-DEL-137 |
| chr12-108217974-DEL-74 | chr12-108217982-DEL-58 |
| chr12-108249892-DEL-232 | chr12-108249893-DEL-232 |
| chr12-108266457-DEL-523 | chr12-108266458-DEL-523 |
| chr12-108289025-DEL-84 | chr12-108289026-DEL-84 |
| chr12-108291147-DEL-114 | chr12-108291148-DEL-114 |
| chr12-108314064-DEL-199 | chr12-108314065-DEL-199 |
| chr12-108321216-DEL-198 | chr12-108321217-DEL-198 |
| chr12-108325925-DEL-78 | chr12-108325926-DEL-78 |
| chr12-108364741-DEL-208 | chr12-108364742-DEL-208 |
| chr12-108381958-DEL-782 | chr12-108381960-DEL-781 |
| chr12-108405256-DEL-53 | chr12-108405257-DEL-53 |
| chr12-108405941-DEL-205 | chr12-108405942-DEL-205 |
| chr12-108436247-DEL-180 | chr12-108436248-DEL-180 |
| chr12-108468097-DEL-51 | chr12-108468090-DEL-52 |
| chr12-108484113-DEL-66 | chr12-108484120-DEL-60 |
| chr12-108512060-DEL-124 | chr12-108512043-DEL-94 |
| chr12-108525521-DEL-495 | chr12-108525522-DEL-495 |
| chr12-108533769-DEL-6452 | chr12-108533770-DEL-6452 |
| chr12-108539744-DEL-214 | chr12-108539745-DEL-214 |
| chr12-108541504-DEL-207 | chr12-108541505-DEL-207 |
| chr12-108638548-DEL-181 | chr12-108638549-DEL-181 |
| chr12-108679805-DEL-213 | chr12-108679806-DEL-213 |
| chr12-108684334-DEL-219 | chr12-108684335-DEL-219 |
| chr12-108701582-DEL-103 | chr12-108701583-DEL-103 |
| chr12-108703399-DEL-484 | chr12-108703400-DEL-484 |
| chr12-108710602-DEL-127 | chr12-108710603-DEL-127 |
| chr12-108711880-DEL-202 | chr12-108711881-DEL-202 |
| chr12-108718974-DEL-227 | chr12-108718975-DEL-227 |
| chr12-108736692-DEL-90 | chr12-108736693-DEL-90 |
| chr12-108739999-DEL-209 | chr12-108740000-DEL-209 |
| chr12-108769831-DEL-66 | chr12-108769832-DEL-66 |
| chr12-108773680-DEL-165 | chr12-108773681-DEL-165 |
| chr12-108788803-DEL-201 | chr12-108788804-DEL-201 |
| chr12-108798628-DEL-136 | chr12-108798629-DEL-136 |
| chr12-108806480-DEL-191 | chr12-108806482-DEL-190 |
| chr12-108812298-DEL-929 | chr12-108812298-DEL-929 |
| chr12-108813351-DEL-850 | chr12-108813351-DEL-850 |
| chr12-108814024-DEL-104 | chr12-108814025-DEL-104 |
| chr12-108828089-DEL-73 | chr12-108828054-DEL-105 |
| chr12-108875472-DEL-240 | chr12-108875473-DEL-240 |
| chr12-108887842-DEL-81 | chr12-108887843-DEL-81 |
| chr12-108932414-DEL-186 | chr12-108932415-DEL-186 |
| chr12-108967159-DEL-88 | chr12-108967156-DEL-92 |
| chr12-108968486-DEL-81 | chr12-108968492-DEL-73 |
| chr12-108991993-DEL-84 | chr12-108991993-DEL-88 |
| chr12-109046730-DEL-80 | chr12-109046732-DEL-79 |
| chr12-109048667-DEL-75 | chr12-109048675-DEL-50 |
| chr12-109050253-DEL-55 | chr12-109050249-DEL-60 |
| chr12-109067580-DEL-57 | chr12-109067581-DEL-57 |
| chr12-109069385-DEL-1085 | chr12-109069386-DEL-1085 |

| | |
|---|---|
| chr12-109087950-DEL-21617 | chr12-109087950-DEL-21617 |
| chr12-109110009-DEL-82 | chr12-109110008-DEL-82 |
| chr12-109133724-DEL-108 | chr12-109133727-DEL-106 |
| chr12-109137022-DEL-196 | chr12-109137023-DEL-196 |
| chr12-109140621-DEL-196 | chr12-109140622-DEL-196 |
| chr12-109142351-DEL-815 | chr12-109142352-DEL-815 |
| chr12-109150906-DEL-63 | chr12-109150904-DEL-63 |
| chr12-109151445-DEL-198 | chr12-109151446-DEL-198 |
| chr12-109153599-DEL-1097 | chr12-109153600-DEL-1097 |
| chr12-109157745-DEL-67 | chr12-109157710-DEL-91 |
| chr12-109180267-DEL-472 | chr12-109180268-DEL-472 |
| chr12-109186766-DEL-714 | chr12-109186767-DEL-714 |
| chr12-109197607-DEL-195 | chr12-109197572-DEL-196 |
| chr12-109250999-DEL-3003 | chr12-109251000-DEL-3003 |
| chr12-109330401-DEL-60 | chr12-109330402-DEL-76 |
| chr12-109331338-DEL-228 | chr12-109331329-DEL-205 |
| chr12-109389542-DEL-161 | chr12-109389543-DEL-161 |
| chr12-109443739-DEL-1114 | chr12-109443739-DEL-1114 |
| chr12-109445150-DEL-1090 | chr12-109445145-DEL-1091 |
| chr12-109598014-DEL-484 | chr12-109598015-DEL-484 |
| chr12-109630414-DEL-90 | chr12-109630415-DEL-90 |
| chr12-109645405-DEL-218 | chr12-109645406-DEL-216 |
| chr12-109673653-DEL-60 | chr12-109673654-DEL-60 |
| chr12-109711161-DEL-208 | chr12-109711162-DEL-208 |
| chr12-109712039-DEL-72 | chr12-109712043-DEL-79 |
| chr12-109730488-DEL-84 | chr12-109730492-DEL-85 |
| chr12-109734685-DEL-2535 | chr12-109734686-DEL-2535 |
| chr12-109769320-DEL-87 | chr12-109769319-DEL-58 |
| chr12-109778308-DEL-168 | chr12-109778309-DEL-168 |
| chr12-109781214-DEL-106 | chr12-109781211-DEL-115 |
| chr12-109786936-DEL-2696 | chr12-109786937-DEL-2696 |
| chr12-109792991-DEL-110 | chr12-109793015-DEL-128 |
| chr12-109821408-DEL-70 | chr12-109821409-DEL-70 |
| chr12-109859049-DEL-97 | chr12-109859044-DEL-103 |
| chr12-109860779-DEL-787 | chr12-109860780-DEL-787 |
| chr12-109868392-DEL-7078 | chr12-109868393-DEL-7078 |
| chr12-109895653-DEL-1863 | chr12-109895654-DEL-1863 |
| chr12-109898506-DEL-3006 | chr12-109898507-DEL-3006 |
| chr12-109923686-DEL-74 | chr12-109923697-DEL-64 |
| chr12-109954026-DEL-312 | chr12-109954045-DEL-294 |
| chr12-109959345-DEL-221 | chr12-109959346-DEL-221 |
| chr12-109977744-DEL-386 | chr12-109977745-DEL-386 |
| chr12-109978805-DEL-273 | chr12-109978835-DEL-286 |
| chr12-109982709-DEL-141 | chr12-109982714-DEL-137 |
| chr12-110041389-DEL-462 | chr12-110041390-DEL-462 |
| chr12-110075879-DEL-215 | chr12-110075880-DEL-215 |
| chr12-110085888-DEL-63 | chr12-110085884-DEL-52 |
| chr12-110122439-DEL-79 | chr12-110122440-DEL-75 |
| chr12-110143471-DEL-62 | chr12-110143472-DEL-62 |
| chr12-110164773-DEL-52 | chr12-110164774-DEL-52 |
| chr12-110172474-DEL-104 | chr12-110172475-DEL-104 |
| chr12-110188902-DEL-724 | chr12-110188903-DEL-724 |
| chr12-110205111-DEL-56 | chr12-110205110-DEL-51 |
| chr12-110258003-DEL-84 | chr12-110258004-DEL-84 |
| chr12-110269570-DEL-1673 | chr12-110269593-DEL-1651 |
| chr12-110271543-DEL-3207 | chr12-110271544-DEL-3207 |
| chr12-110282344-DEL-514 | chr12-110282345-DEL-514 |
| chr12-110307076-DEL-345 | chr12-110307077-DEL-344 |
| chr12-110323535-DEL-367 | chr12-110323536-DEL-367 |
| chr12-110327651-DEL-56 | chr12-110327652-DEL-56 |
| chr12-110358754-DEL-108 | chr12-110358767-DEL-106 |
| chr12-110385408-DEL-170 | chr12-110385409-DEL-170 |
| chr12-110386757-DEL-1473 | chr12-110386758-DEL-1467 |
| chr12-110387875-DEL-55 | chr12-110387876-DEL-53 |
| chr12-110408462-DEL-168 | chr12-110408463-DEL-168 |
| chr12-110428088-DEL-77 | chr12-110428091-DEL-75 |
| chr12-110432075-DEL-52 | chr12-110432113-DEL-55 |
| chr12-110457327-DEL-102 | chr12-110457320-DEL-110 |
| chr12-110458693-DEL-252 | chr12-110458694-DEL-252 |
| chr12-110486257-DEL-130 | chr12-110486258-DEL-130 |
| chr12-110487747-DEL-53 | chr12-110487748-DEL-53 |
| chr12-110504408-DEL-3911 | chr12-110504409-DEL-3911 |
| chr12-110505406-DEL-236 | chr12-110505407-DEL-236 |
| chr12-110515428-DEL-52 | chr12-110515429-DEL-52 |
| chr12-110540368-DEL-642 | chr12-110540369-DEL-642 |
| chr12-110553875-DEL-181 | chr12-110553880-DEL-177 |
| chr12-110561486-DEL-173 | chr12-110561486-DEL-173 |
| chr12-110570813-DEL-200 | chr12-110570814-DEL-200 |
| chr12-110610801-DEL-200 | chr12-110610802-DEL-200 |
| chr12-110611339-DEL-212 | chr12-110611340-DEL-212 |

| | |
|---|---|
| chr12-110614927-DEL-58 | chr12-110614928-DEL-58 |
| chr12-110626948-DEL-132 | chr12-110626949-DEL-132 |
| chr12-110645078-DEL-158 | chr12-110645079-DEL-158 |
| chr12-110650302-DEL-204 | chr12-110650303-DEL-204 |
| chr12-110699934-DEL-285 | chr12-110699935-DEL-285 |
| chr12-110716427-DEL-200 | chr12-110716428-DEL-200 |
| chr12-110725261-DEL-182 | chr12-110725262-DEL-182 |
| chr12-110727463-DEL-841 | chr12-110727464-DEL-841 |
| chr12-110778492-DEL-168 | chr12-110778493-DEL-168 |
| chr12-110784757-DEL-78 | chr12-110784758-DEL-78 |
| chr12-110788704-DEL-1349 | chr12-110788705-DEL-1349 |
| chr12-110791637-DEL-67 | chr12-110791644-DEL-62 |
| chr12-110796942-DEL-209 | chr12-110796943-DEL-209 |
| chr12-110798468-DEL-213 | chr12-110798469-DEL-213 |
| chr12-110801996-DEL-162 | chr12-110801997-DEL-162 |
| chr12-110809349-DEL-74 | chr12-110809353-DEL-72 |
| chr12-110824471-DEL-918 | chr12-110824472-DEL-918 |
| chr12-110829111-DEL-214 | chr12-110829112-DEL-238 |
| chr12-110844532-DEL-583 | chr12-110844531-DEL-582 |
| chr12-110887464-DEL-285 | chr12-110887462-DEL-285 |
| chr12-110989239-DEL-78 | chr12-110989240-DEL-72 |
| chr12-110997361-DEL-96 | chr12-110997362-DEL-96 |
| chr12-111028083-DEL-104 | chr12-111028088-DEL-100 |
| chr12-111029072-DEL-215 | chr12-111029073-DEL-215 |
| chr12-111049444-DEL-57 | chr12-111049445-DEL-57 |
| chr12-111051785-DEL-62 | chr12-111051814-DEL-62 |
| chr12-111059634-DEL-214 | chr12-111059635-DEL-214 |
| chr12-111108745-DEL-427 | chr12-111108746-DEL-427 |
| chr12-111125933-DEL-127 | chr12-111125939-DEL-119 |
| chr12-111130361-DEL-227 | chr12-111130362-DEL-227 |
| chr12-111135316-DEL-84 | chr12-111135307-DEL-80 |
| chr12-111153128-DEL-59 | chr12-111153129-DEL-59 |
| chr12-111161114-DEL-216 | chr12-111161115-DEL-216 |
| chr12-111173326-DEL-331 | chr12-111173326-DEL-331 |
| chr12-111174511-DEL-153 | chr12-111174512-DEL-153 |
| chr12-111187383-DEL-934 | chr12-111187384-DEL-934 |
| chr12-111188411-DEL-90 | chr12-111188407-DEL-90 |
| chr12-111224515-DEL-236 | chr12-111224516-DEL-236 |
| chr12-111225087-DEL-215 | chr12-111225088-DEL-215 |
| chr12-111244368-DEL-418 | chr12-111244369-DEL-418 |
| chr12-111247290-DEL-116 | chr12-111247291-DEL-116 |
| chr12-111257574-DEL-904 | chr12-111257575-DEL-904 |
| chr12-111258925-DEL-61 | chr12-111258926-DEL-61 |
| chr12-111267467-DEL-452 | chr12-111267466-DEL-449 |
| chr12-111274962-DEL-218 | chr12-111274963-DEL-218 |
| chr12-111282544-DEL-69 | chr12-111282557-DEL-57 |
| chr12-111308317-DEL-66 | chr12-111308318-DEL-66 |
| chr12-111316669-DEL-84 | chr12-111316670-DEL-87 |
| chr12-111354632-DEL-5218 | chr12-111354633-DEL-5218 |
| chr12-111367112-DEL-52 | chr12-111367113-DEL-52 |
| chr12-111427568-DEL-62 | chr12-111427575-DEL-56 |
| chr12-111438825-DEL-240 | chr12-111438826-DEL-240 |
| chr12-111476940-DEL-2053 | chr12-111476941-DEL-2053 |
| chr12-111479702-DEL-201 | chr12-111479703-DEL-201 |
| chr12-111486094-DEL-366 | chr12-111486095-DEL-366 |
| chr12-111516185-DEL-209 | chr12-111516186-DEL-209 |
| chr12-111522433-DEL-57 | chr12-111522434-DEL-57 |
| chr12-111534538-DEL-214 | chr12-111534533-DEL-220 |
| chr12-111543626-DEL-188 | chr12-111543627-DEL-188 |
| chr12-111544806-DEL-213 | chr12-111544807-DEL-213 |
| chr12-111554586-DEL-72 | chr12-111554587-DEL-72 |
| chr12-111582257-DEL-88 | chr12-111582258-DEL-88 |
| chr12-111586751-DEL-64 | chr12-111586752-DEL-64 |
| chr12-111590159-DEL-80 | chr12-111590154-DEL-81 |
| chr12-111669273-DEL-236 | chr12-111669266-DEL-244 |
| chr12-111674944-DEL-179 | chr12-111674942-DEL-171 |
| chr12-111683043-DEL-66 | chr12-111683044-DEL-66 |
| chr12-111683714-DEL-315 | chr12-111683715-DEL-315 |
| chr12-111689704-DEL-199 | chr12-111689705-DEL-199 |
| chr12-111707135-DEL-81 | chr12-111707119-DEL-74 |
| chr12-111709241-DEL-417 | chr12-111709252-DEL-421 |
| chr12-111718588-DEL-201 | chr12-111718589-DEL-201 |
| chr12-111720544-DEL-207 | chr12-111720545-DEL-207 |
| chr12-111735388-DEL-207 | chr12-111735389-DEL-207 |
| chr12-111736586-DEL-80 | chr12-111736615-DEL-50 |
| chr12-111756507-DEL-51 | chr12-111756508-DEL-51 |
| chr12-111770484-DEL-68 | chr12-111770465-DEL-66 |
| chr12-111776601-DEL-52 | chr12-111776602-DEL-52 |
| chr12-111793568-DEL-84 | chr12-111793569-DEL-84 |
| chr12-111813833-DEL-68 | chr12-111813846-DEL-56 |

| | |
|---|---|
| chr12-111815351-DEL-430 | chr12-111815352-DEL-430 |
| chr12-111816830-DEL-140 | chr12-111816839-DEL-103 |
| chr12-111829607-DEL-64 | chr12-111829624-DEL-50 |
| chr12-111836969-DEL-130 | chr12-111836944-DEL-156 |
| chr12-111844493-DEL-206 | chr12-111844494-DEL-206 |
| chr12-111880824-DEL-52 | chr12-111880825-DEL-52 |
| chr12-111906018-DEL-117 | chr12-111906019-DEL-117 |
| chr12-111920092-DEL-125 | chr12-111920100-DEL-120 |
| chr12-111928185-DEL-57 | chr12-111928186-DEL-57 |
| chr12-111938620-DEL-72 | chr12-111938621-DEL-72 |
| chr12-111976254-DEL-94 | chr12-111976255-DEL-94 |
| chr12-111976765-DEL-112 | chr12-111976796-DEL-82 |
| chr12-111983370-DEL-4398 | chr12-111983371-DEL-4398 |
| chr12-111990079-DEL-111 | chr12-111990047-DEL-107 |
| chr12-112002119-DEL-226 | chr12-112002115-DEL-225 |
| chr12-112014962-DEL-196 | chr12-112014963-DEL-196 |
| chr12-112021109-DEL-6493 | chr12-112021110-DEL-6493 |
| chr12-112036859-DEL-1339 | chr12-112036859-DEL-1337 |
| chr12-112043555-DEL-51 | chr12-112043544-DEL-51 |
| chr12-112047335-DEL-78 | chr12-112047336-DEL-78 |
| chr12-112166247-DEL-74 | chr12-112166229-DEL-75 |
| chr12-112169659-DEL-51 | chr12-112169633-DEL-64 |
| chr12-112170704-DEL-189 | chr12-112170705-DEL-189 |
| chr12-112192825-DEL-112 | chr12-112192830-DEL-108 |
| chr12-112202073-DEL-64 | chr12-112202095-DEL-76 |
| chr12-112224539-DEL-196 | chr12-112224540-DEL-196 |
| chr12-112236124-DEL-72 | chr12-112236118-DEL-70 |
| chr12-112261318-DEL-100 | chr12-112261318-DEL-100 |
| chr12-112263151-DEL-62 | chr12-112263144-DEL-70 |
| chr12-112276270-DEL-52 | chr12-112276221-DEL-72 |
| chr12-112351589-DEL-89 | chr12-112351599-DEL-79 |
| chr12-112359247-DEL-617 | chr12-112359248-DEL-617 |
| chr12-112371332-DEL-62 | chr12-112371333-DEL-62 |
| chr12-112374156-DEL-200 | chr12-112374167-DEL-239 |
| chr12-112380111-DEL-110 | chr12-112380112-DEL-110 |
| chr12-112385717-DEL-91 | chr12-112385718-DEL-90 |
| chr12-112406802-DEL-56 | chr12-112406803-DEL-56 |
| chr12-112410987-DEL-142 | chr12-112410988-DEL-142 |
| chr12-112423732-DEL-70 | chr12-112423741-DEL-62 |
| chr12-112434845-DEL-59 | chr12-112434846-DEL-59 |
| chr12-112443491-DEL-498 | chr12-112443492-DEL-498 |
| chr12-112446987-DEL-56 | chr12-112446988-DEL-56 |
| chr12-112454108-DEL-176 | chr12-112454109-DEL-176 |
| chr12-112461641-DEL-466 | chr12-112461642-DEL-466 |
| chr12-112480051-DEL-56 | chr12-112480008-DEL-68 |
| chr12-112491044-DEL-1038 | chr12-112491045-DEL-1038 |
| chr12-112510295-DEL-104 | chr12-112510296-DEL-104 |
| chr12-112513768-DEL-1218 | chr12-112513768-DEL-1218 |
| chr12-112515584-DEL-198 | chr12-112515585-DEL-198 |
| chr12-112523753-DEL-171 | chr12-112523754-DEL-171 |
| chr12-112525305-DEL-74 | chr12-112525306-DEL-74 |
| chr12-112539006-DEL-290 | chr12-112538988-DEL-332 |
| chr12-112557123-DEL-80 | chr12-112557124-DEL-80 |
| chr12-112563380-DEL-60 | chr12-112563381-DEL-60 |
| chr12-112567157-DEL-175 | chr12-112567158-DEL-175 |
| chr12-112622836-DEL-61 | chr12-112622837-DEL-61 |
| chr12-112651371-DEL-102 | chr12-112651372-DEL-102 |
| chr12-112670199-DEL-200 | chr12-112670200-DEL-200 |
| chr12-112679754-DEL-124 | chr12-112679755-DEL-124 |
| chr12-112713733-DEL-89 | chr12-112713734-DEL-89 |
| chr12-112719306-DEL-205 | chr12-112719307-DEL-205 |
| chr12-112754277-DEL-52 | chr12-112754262-DEL-68 |
| chr12-112790675-DEL-56 | chr12-112790665-DEL-56 |
| chr12-112800724-DEL-457 | chr12-112800725-DEL-457 |
| chr12-112801806-DEL-227 | chr12-112801774-DEL-228 |
| chr12-112803089-DEL-121 | chr12-112803090-DEL-121 |
| chr12-112805222-DEL-56 | chr12-112805223-DEL-72 |
| chr12-112808639-DEL-54 | chr12-112808639-DEL-55 |
| chr12-112810184-DEL-84 | chr12-112810182-DEL-93 |
| chr12-112819789-DEL-79 | chr12-112819813-DEL-55 |
| chr12-112823460-DEL-59 | chr12-112823459-DEL-60 |
| chr12-112825047-DEL-191 | chr12-112825054-DEL-190 |
| chr12-112826260-DEL-297 | chr12-112826254-DEL-309 |
| chr12-112850901-DEL-2935 | chr12-112850902-DEL-2935 |
| chr12-112854053-DEL-423 | chr12-112854054-DEL-423 |
| chr12-112859775-DEL-266 | chr12-112859748-DEL-248 |
| chr12-112863669-DEL-62 | chr12-112863670-DEL-74 |
| chr12-112888706-DEL-700 | chr12-112888707-DEL-700 |
| chr12-112900230-DEL-64 | chr12-112900231-DEL-64 |
| chr12-112900745-DEL-55 | chr12-112900746-DEL-55 |

| | |
|---|---|
| chr12-112959296-DEL-151 | chr12-112959297-DEL-151 |
| chr12-112980146-DEL-55 | chr12-112980147-DEL-55 |
| chr12-112990119-DEL-75 | chr12-112990118-DEL-76 |
| chr12-112993245-DEL-212 | chr12-112993246-DEL-212 |
| chr12-112996847-DEL-66 | chr12-112996848-DEL-66 |
| chr12-113005888-DEL-53 | chr12-113005889-DEL-53 |
| chr12-113019515-DEL-57 | chr12-113019516-DEL-57 |
| chr12-113024986-DEL-119 | chr12-113024987-DEL-119 |
| chr12-113036094-DEL-98 | chr12-113036095-DEL-98 |
| chr12-113040299-DEL-168 | chr12-113040300-DEL-168 |
| chr12-113045170-DEL-56 | chr12-113045154-DEL-72 |
| chr12-113074267-DEL-1120 | chr12-113074268-DEL-1120 |
| chr12-113079293-DEL-259 | chr12-113079294-DEL-259 |
| chr12-113099488-DEL-132 | chr12-113099489-DEL-132 |
| chr12-113132012-DEL-387 | chr12-113132013-DEL-387 |
| chr12-113147970-DEL-124 | chr12-113147971-DEL-124 |
| chr12-113159133-DEL-5627 | chr12-113159167-DEL-5594 |
| chr12-113167067-DEL-103 | chr12-113167068-DEL-103 |
| chr12-113171875-DEL-223 | chr12-113171876-DEL-223 |
| chr12-113173221-DEL-217 | chr12-113173216-DEL-218 |
| chr12-113188960-DEL-285 | chr12-113188961-DEL-285 |
| chr12-113193159-DEL-157 | chr12-113193194-DEL-153 |
| chr12-113203494-DEL-80 | chr12-113203495-DEL-80 |
| chr12-113215817-DEL-65 | chr12-113215818-DEL-65 |
| chr12-113224910-DEL-55 | chr12-113224911-DEL-55 |
| chr12-113225843-DEL-87 | chr12-113225844-DEL-87 |
| chr12-113231587-DEL-65 | chr12-113231588-DEL-65 |
| chr12-113247663-DEL-72 | chr12-113247666-DEL-70 |
| chr12-113257055-DEL-415 | chr12-113257056-DEL-415 |
| chr12-113258101-DEL-104 | chr12-113258102-DEL-104 |
| chr12-113258697-DEL-52 | chr12-113258698-DEL-52 |
| chr12-113261102-DEL-1889 | chr12-113261103-DEL-1889 |
| chr12-113274150-DEL-538 | chr12-113274151-DEL-538 |
| chr12-113282542-DEL-121 | chr12-113282535-DEL-128 |
| chr12-113285358-DEL-110 | chr12-113285359-DEL-110 |
| chr12-113325249-DEL-51 | chr12-113325244-DEL-51 |
| chr12-113328523-DEL-93 | chr12-113328523-DEL-93 |
| chr12-113334331-DEL-415 | chr12-113334332-DEL-415 |
| chr12-113343016-DEL-2270 | chr12-113343017-DEL-2270 |
| chr12-113352052-DEL-87 | chr12-113352053-DEL-87 |
| chr12-113377096-DEL-80 | chr12-113377097-DEL-80 |
| chr12-113410678-DEL-68 | chr12-113410678-DEL-68 |
| chr12-113411613-DEL-56 | chr12-113411615-DEL-54 |
| chr12-113412914-DEL-158 | chr12-113412914-DEL-158 |
| chr12-113417177-DEL-176 | chr12-113417177-DEL-164 |
| chr12-113419335-DEL-511 | chr12-113419335-DEL-511 |
| chr12-113435320-DEL-327 | chr12-113435321-DEL-327 |
| chr12-113436658-DEL-61 | chr12-113436662-DEL-57 |
| chr12-113441047-DEL-85 | chr12-113441050-DEL-66 |
| chr12-113443626-DEL-601 | chr12-113443628-DEL-602 |
| chr12-113473136-DEL-61 | chr12-113473137-DEL-61 |
| chr12-113497050-DEL-60 | chr12-113497051-DEL-60 |
| chr12-113523605-DEL-784 | chr12-113523606-DEL-784 |
| chr12-113540353-DEL-965 | chr12-113540354-DEL-965 |
| chr12-113591559-DEL-3131 | chr12-113591560-DEL-3131 |
| chr12-113603566-DEL-486 | chr12-113603567-DEL-486 |
| chr12-113621333-DEL-56 | chr12-113621339-DEL-50 |
| chr12-113638850-DEL-244 | chr12-113638851-DEL-244 |
| chr12-113650750-DEL-51 | chr12-113650794-DEL-52 |
| chr12-113658418-DEL-55 | chr12-113658417-DEL-55 |
| chr12-113689077-DEL-114 | chr12-113689077-DEL-114 |
| chr12-113696353-DEL-690 | chr12-113696352-DEL-690 |
| chr12-113727836-DEL-335 | chr12-113727836-DEL-335 |
| chr12-113759666-DEL-497 | chr12-113759666-DEL-497 |
| chr12-113765475-DEL-290 | chr12-113765476-DEL-290 |
| chr12-113767831-DEL-1231 | chr12-113767831-DEL-1231 |
| chr12-113771655-DEL-802 | chr12-113771656-DEL-802 |
| chr12-113772219-DEL-68 | chr12-113772216-DEL-69 |
| chr12-113784585-DEL-7721 | chr12-113784586-DEL-7721 |
| chr12-113793931-DEL-408 | chr12-113793932-DEL-408 |
| chr12-113799961-DEL-6102 | chr12-113799961-DEL-6102 |
| chr12-113810879-DEL-612 | chr12-113810879-DEL-612 |
| chr12-113812502-DEL-66 | chr12-113812502-DEL-66 |
| chr12-113812839-DEL-814 | chr12-113812839-DEL-814 |
| chr12-113845656-DEL-56 | chr12-113845657-DEL-56 |
| chr12-113846435-DEL-216 | chr12-113846436-DEL-216 |
| chr12-113847292-DEL-1391 | chr12-113847292-DEL-1391 |
| chr12-113850162-DEL-431 | chr12-113850162-DEL-431 |
| chr12-113883293-DEL-3963 | chr12-113883294-DEL-3963 |
| chr12-113890630-DEL-965 | chr12-113890631-DEL-965 |

| | |
|---|---|
| chr12-113897948-DEL-215 | chr12-113897948-DEL-215 |
| chr12-113917863-DEL-1065 | chr12-113917864-DEL-1065 |
| chr12-113923438-DEL-857 | chr12-113923438-DEL-857 |
| chr12-113924986-DEL-465 | chr12-113924976-DEL-468 |
| chr12-113933817-DEL-711 | chr12-113933817-DEL-711 |
| chr12-113946433-DEL-74 | chr12-113946433-DEL-74 |
| chr12-113966326-DEL-1813 | chr12-113966326-DEL-1813 |
| chr12-113970449-DEL-684 | chr12-113970449-DEL-684 |
| chr12-113971865-DEL-102 | chr12-113971864-DEL-102 |
| chr12-113985252-DEL-54 | chr12-113985253-DEL-54 |
| chr12-113988355-DEL-66 | chr12-113988331-DEL-66 |
| chr12-113994409-DEL-462 | chr12-113994395-DEL-465 |
| chr12-114014498-DEL-75 | chr12-114014497-DEL-75 |
| chr12-114020574-DEL-63 | chr12-114020574-DEL-63 |
| chr12-114032177-DEL-10713 | chr12-114032178-DEL-10713 |
| chr12-114049858-DEL-52 | chr12-114049859-DEL-52 |
| chr12-114124255-DEL-90 | chr12-114124256-DEL-91 |
| chr12-114134014-DEL-71 | chr12-114134007-DEL-71 |
| chr12-114136485-DEL-598 | chr12-114136486-DEL-598 |
| chr12-114142640-DEL-68 | chr12-114142635-DEL-74 |
| chr12-114145845-DEL-76 | chr12-114145846-DEL-76 |
| chr12-114150893-DEL-752 | chr12-114150895-DEL-751 |
| chr12-114162223-DEL-146 | chr12-114162224-DEL-146 |
| chr12-114165505-DEL-871 | chr12-114165506-DEL-871 |
| chr12-114167823-DEL-1765 | chr12-114167823-DEL-1765 |
| chr12-114170880-DEL-418 | chr12-114170881-DEL-418 |
| chr12-114186244-DEL-83 | chr12-114186245-DEL-83 |
| chr12-114198711-DEL-6795 | chr12-114198712-DEL-6795 |
| chr12-114209010-DEL-64 | chr12-114209025-DEL-54 |
| chr12-114224975-DEL-861 | chr12-114224976-DEL-861 |
| chr12-114232417-DEL-1277 | chr12-114232418-DEL-1277 |
| chr12-114232441-DEL-94 | chr12-114232442-DEL-80 |
| chr12-114232952-DEL-424 | chr12-114232948-DEL-425 |
| chr12-114239205-DEL-308 | chr12-114239206-DEL-308 |
| chr12-114240162-DEL-58 | chr12-114240162-DEL-58 |
| chr12-114242036-DEL-3526 | chr12-114242037-DEL-3526 |
| chr12-114255500-DEL-870 | chr12-114255501-DEL-870 |
| chr12-114283098-DEL-675 | chr12-114283099-DEL-675 |
| chr12-114287170-DEL-58 | chr12-114287171-DEL-58 |
| chr12-114302194-DEL-231 | chr12-114302191-DEL-235 |
| chr12-114336388-DEL-105 | chr12-114336385-DEL-109 |
| chr12-114341053-DEL-2115 | chr12-114341054-DEL-2115 |
| chr12-114344309-DEL-64 | chr12-114344293-DEL-106 |
| chr12-114344967-DEL-4012 | chr12-114344968-DEL-4012 |
| chr12-114350811-DEL-586 | chr12-114350811-DEL-586 |
| chr12-114368962-DEL-870 | chr12-114368961-DEL-868 |
| chr12-114377210-DEL-82 | chr12-114377241-DEL-56 |
| chr12-114377928-DEL-53 | chr12-114377930-DEL-52 |
| chr12-114380842-DEL-172 | chr12-114380842-DEL-172 |
| chr12-114383373-DEL-478 | chr12-114383372-DEL-478 |
| chr12-114387601-DEL-113 | chr12-114387601-DEL-99 |
| chr12-114390644-DEL-70 | chr12-114390643-DEL-70 |
| chr12-114400566-DEL-4799 | chr12-114400565-DEL-4799 |
| chr12-114407111-DEL-72 | chr12-114407105-DEL-78 |
| chr12-114408460-DEL-138 | chr12-114408461-DEL-126 |
| chr12-114414794-DEL-171 | chr12-114414795-DEL-171 |
| chr12-114420384-DEL-2539 | chr12-114420385-DEL-2539 |
| chr12-114461478-DEL-3179 | chr12-114461478-DEL-3179 |
| chr12-114471751-DEL-1562 | chr12-114471752-DEL-1562 |
| chr12-114480188-DEL-841 | chr12-114480187-DEL-839 |
| chr12-114483722-DEL-376 | chr12-114483721-DEL-376 |
| chr12-114488108-DEL-2778 | chr12-114488109-DEL-2778 |
| chr12-114491506-DEL-1963 | chr12-114491507-DEL-1963 |
| chr12-114511588-DEL-939 | chr12-114511589-DEL-939 |
| chr12-114512195-DEL-99 | chr12-114512196-DEL-99 |
| chr12-114520196-DEL-124 | chr12-114520196-DEL-124 |
| chr12-114520989-DEL-57 | chr12-114520989-DEL-57 |
| chr12-114525052-DEL-1367 | chr12-114525052-DEL-1367 |
| chr12-114525158-DEL-427 | chr12-114525157-DEL-427 |
| chr12-114526798-DEL-85 | chr12-114526808-DEL-83 |
| chr12-114530796-DEL-1503 | chr12-114530796-DEL-1503 |
| chr12-114549684-DEL-75 | chr12-114549683-DEL-75 |
| chr12-114551291-DEL-73 | chr12-114551293-DEL-64 |
| chr12-114553734-DEL-128 | chr12-114553735-DEL-128 |
| chr12-114554409-DEL-6391 | chr12-114554410-DEL-6391 |
| chr12-114587338-DEL-211 | chr12-114587385-DEL-204 |
| chr12-114596824-DEL-7457 | chr12-114596825-DEL-7457 |
| chr12-114610988-DEL-1153 | chr12-114610988-DEL-1153 |
| chr12-114618602-DEL-134 | chr12-114618602-DEL-120 |
| chr12-114619963-DEL-77 | chr12-114619963-DEL-77 |

| | |
|---|---|
| chr12-114622650-DEL-58 | chr12-114622661-DEL-57 |
| chr12-114625014-DEL-70 | chr12-114625015-DEL-70 |
| chr12-114628845-DEL-107 | chr12-114628846-DEL-107 |
| chr12-114654242-DEL-2034 | chr12-114654242-DEL-2034 |
| chr12-114665594-DEL-947 | chr12-114665594-DEL-947 |
| chr12-114667230-DEL-1811 | chr12-114667231-DEL-1811 |
| chr12-114681918-DEL-203 | chr12-114681919-DEL-203 |
| chr12-114692021-DEL-1545 | chr12-114692020-DEL-1548 |
| chr12-114749730-DEL-1052 | chr12-114749730-DEL-1052 |
| chr12-114751448-DEL-298 | chr12-114751447-DEL-298 |
| chr12-114816662-DEL-2738 | chr12-114816661-DEL-2738 |
| chr12-114830469-DEL-446 | chr12-114830469-DEL-446 |
| chr12-114863891-DEL-1207 | chr12-114863891-DEL-1207 |
| chr12-114882951-DEL-1379 | chr12-114882952-DEL-1379 |
| chr12-114904600-DEL-2969 | chr12-114904597-DEL-2969 |
| chr12-114909158-DEL-152 | chr12-114909158-DEL-152 |
| chr12-114911726-DEL-288 | chr12-114911725-DEL-288 |
| chr12-114922254-DEL-297 | chr12-114922255-DEL-297 |
| chr12-114927824-DEL-524 | chr12-114927824-DEL-524 |
| chr12-114930385-DEL-90 | chr12-114930385-DEL-90 |
| chr12-114931280-DEL-64 | chr12-114931281-DEL-64 |
| chr12-114935503-DEL-663 | chr12-114935504-DEL-663 |
| chr12-114937493-DEL-69 | chr12-114937492-DEL-69 |
| chr12-114942464-DEL-64 | chr12-114942465-DEL-64 |
| chr12-114944861-DEL-57 | chr12-114944844-DEL-58 |
| chr12-114952312-DEL-66 | chr12-114952312-DEL-67 |
| chr12-114972820-DEL-79 | chr12-114972811-DEL-75 |
| chr12-114976133-DEL-1386 | chr12-114976134-DEL-1386 |
| chr12-114977790-DEL-126 | chr12-114977791-DEL-125 |
| chr12-114986013-DEL-83 | chr12-114986028-DEL-75 |
| chr12-114988494-DEL-85 | chr12-114988495-DEL-85 |
| chr12-115005390-DEL-401 | chr12-115005391-DEL-401 |
| chr12-115036307-DEL-1234 | chr12-115036299-DEL-1215 |
| chr12-115041692-DEL-499 | chr12-115041692-DEL-499 |
| chr12-115045702-DEL-367 | chr12-115045702-DEL-367 |
| chr12-115056873-DEL-3204 | chr12-115056874-DEL-3204 |
| chr12-115084316-DEL-477 | chr12-115084319-DEL-477 |
| chr12-115099609-DEL-129 | chr12-115099605-DEL-133 |
| chr12-115116346-DEL-98 | chr12-115116347-DEL-98 |
| chr12-115137566-DEL-1925 | chr12-115137567-DEL-1925 |
| chr12-115144044-DEL-616 | chr12-115144043-DEL-616 |
| chr12-115158108-DEL-105 | chr12-115158108-DEL-105 |
| chr12-115161667-DEL-2187 | chr12-115161666-DEL-2187 |
| chr12-115164476-DEL-2878 | chr12-115164476-DEL-2879 |
| chr12-115171409-DEL-286 | chr12-115171409-DEL-286 |
| chr12-115207797-DEL-367 | chr12-115207801-DEL-372 |
| chr12-115219334-DEL-77 | chr12-115219325-DEL-81 |
| chr12-115222068-DEL-383 | chr12-115222069-DEL-381 |
| chr12-115250589-DEL-672 | chr12-115250589-DEL-672 |
| chr12-115262396-DEL-2182 | chr12-115262396-DEL-2182 |
| chr12-115277501-DEL-2162 | chr12-115277516-DEL-2163 |
| chr12-115294255-DEL-180 | chr12-115294255-DEL-180 |
| chr12-115313082-DEL-498 | chr12-115313083-DEL-498 |
| chr12-115335305-DEL-285 | chr12-115335305-DEL-285 |
| chr12-115340118-DEL-81 | chr12-115340118-DEL-71 |
| chr12-115377342-DEL-54 | chr12-115377343-DEL-54 |
| chr12-115377650-DEL-266 | chr12-115377651-DEL-266 |
| chr12-115396691-DEL-59 | chr12-115396692-DEL-59 |
| chr12-115405791-DEL-59 | chr12-115405791-DEL-59 |
| chr12-115420738-DEL-84 | chr12-115420726-DEL-82 |
| chr12-115445239-DEL-1440 | chr12-115445239-DEL-1441 |
| chr12-115469149-DEL-62 | chr12-115469150-DEL-60 |
| chr12-115483543-DEL-460 | chr12-115483543-DEL-460 |
| chr12-115496752-DEL-62 | chr12-115496753-DEL-62 |
| chr12-115565098-DEL-100 | chr12-115565091-DEL-108 |
| chr12-115587723-DEL-161 | chr12-115587722-DEL-161 |
| chr12-115599276-DEL-1787 | chr12-115599276-DEL-1787 |
| chr12-115602752-DEL-63 | chr12-115602758-DEL-57 |
| chr12-115620450-DEL-1064 | chr12-115620450-DEL-1064 |
| chr12-115672382-DEL-901 | chr12-115672383-DEL-901 |
| chr12-115686099-DEL-60 | chr12-115686100-DEL-60 |
| chr12-115736989-DEL-1050 | chr12-115736993-DEL-1050 |
| chr12-115755195-DEL-72 | chr12-115755194-DEL-72 |
| chr12-115758947-DEL-512 | chr12-115758948-DEL-511 |
| chr12-115787958-DEL-871 | chr12-115787958-DEL-871 |
| chr12-115795218-DEL-303 | chr12-115795218-DEL-303 |
| chr12-115826734-DEL-119 | chr12-115826734-DEL-119 |
| chr12-115845683-DEL-765 | chr12-115845686-DEL-762 |
| chr12-115856406-DEL-4599 | chr12-115856407-DEL-4599 |
| chr12-115868317-DEL-366 | chr12-115868318-DEL-366 |

| | |
|---|---|
| chr12-115878918-DEL-74 | chr12-115878870-DEL-123 |
| chr12-115897169-DEL-431 | chr12-115897170-DEL-431 |
| chr12-115905563-DEL-70 | chr12-115905563-DEL-70 |
| chr12-115924181-DEL-375 | chr12-115924182-DEL-369 |
| chr12-115933062-DEL-2924 | chr12-115933062-DEL-2924 |
| chr12-115936804-DEL-215 | chr12-115936805-DEL-215 |
| chr12-115946057-DEL-9742 | chr12-115946058-DEL-9742 |
| chr12-115964628-DEL-109 | chr12-115964593-DEL-96 |
| chr12-115979626-DEL-3168 | chr12-115979627-DEL-3168 |
| chr12-115980871-DEL-1463 | chr12-115980872-DEL-1463 |
| chr12-115984290-DEL-735 | chr12-115984291-DEL-733 |
| chr12-115992218-DEL-584 | chr12-115992219-DEL-584 |
| chr12-115997926-DEL-337 | chr12-115997923-DEL-333 |
| chr12-116001908-DEL-6214 | chr12-116001909-DEL-6213 |
| chr12-116009987-DEL-615 | chr12-116009988-DEL-615 |
| chr12-116029652-DEL-53 | chr12-116029653-DEL-54 |
| chr12-116036212-DEL-200 | chr12-116036213-DEL-200 |
| chr12-116057998-DEL-59 | chr12-116057999-DEL-59 |
| chr12-116071307-DEL-72 | chr12-116071309-DEL-71 |
| chr12-116091103-DEL-174 | chr12-116091104-DEL-174 |
| chr12-116110278-DEL-113 | chr12-116110279-DEL-113 |
| chr12-116124602-DEL-58 | chr12-116124603-DEL-58 |
| chr12-116136920-DEL-212 | chr12-116136921-DEL-212 |
| chr12-116151275-DEL-833 | chr12-116151276-DEL-833 |
| chr12-116153197-DEL-343 | chr12-116153198-DEL-343 |
| chr12-116159488-DEL-134 | chr12-116159509-DEL-114 |
| chr12-116170475-DEL-56 | chr12-116170476-DEL-56 |
| chr12-116179951-DEL-75 | chr12-116179952-DEL-75 |
| chr12-116184927-DEL-68 | chr12-116184928-DEL-68 |
| chr12-116185884-DEL-123 | chr12-116185885-DEL-123 |
| chr12-116201463-DEL-64 | chr12-116201464-DEL-64 |
| chr12-116212472-DEL-721 | chr12-116212473-DEL-721 |
| chr12-116239079-DEL-121 | chr12-116239066-DEL-114 |
| chr12-116255760-DEL-52 | chr12-116255731-DEL-67 |
| chr12-116268060-DEL-366 | chr12-116268061-DEL-366 |
| chr12-116286778-DEL-51 | chr12-116286779-DEL-51 |
| chr12-116288483-DEL-199 | chr12-116288484-DEL-199 |
| chr12-116292143-DEL-1245 | chr12-116292144-DEL-1245 |
| chr12-116315377-DEL-80 | chr12-116315370-DEL-88 |
| chr12-116348163-DEL-610 | chr12-116348151-DEL-611 |
| chr12-116362179-DEL-51 | chr12-116362180-DEL-51 |
| chr12-116385662-DEL-54 | chr12-116385662-DEL-54 |
| chr12-116399354-DEL-399 | chr12-116399355-DEL-399 |
| chr12-116406843-DEL-68 | chr12-116406844-DEL-68 |
| chr12-116413050-DEL-180 | chr12-116413051-DEL-181 |
| chr12-116417290-DEL-3800 | chr12-116417291-DEL-3800 |
| chr12-116441059-DEL-143 | chr12-116441060-DEL-143 |
| chr12-116461426-DEL-54 | chr12-116461427-DEL-54 |
| chr12-116464729-DEL-131 | chr12-116464717-DEL-108 |
| chr12-116470401-DEL-1174 | chr12-116470402-DEL-1174 |
| chr12-116471919-DEL-259 | chr12-116471920-DEL-259 |
| chr12-116473484-DEL-243 | chr12-116473484-DEL-243 |
| chr12-116480232-DEL-96 | chr12-116480233-DEL-88 |
| chr12-116537016-DEL-224 | chr12-116537017-DEL-224 |
| chr12-116540492-DEL-1080 | chr12-116540492-DEL-1060 |
| chr12-116549895-DEL-285 | chr12-116549896-DEL-285 |
| chr12-116557110-DEL-326 | chr12-116557111-DEL-326 |
| chr12-116569601-DEL-209 | chr12-116569602-DEL-209 |
| chr12-116573322-DEL-3520 | chr12-116573323-DEL-3520 |
| chr12-116581399-DEL-70 | chr12-116581400-DEL-68 |
| chr12-116629099-DEL-72 | chr12-116629101-DEL-72 |
| chr12-116669609-DEL-331 | chr12-116669610-DEL-331 |
| chr12-116675766-DEL-1565 | chr12-116675767-DEL-1565 |
| chr12-116694290-DEL-444 | chr12-116694285-DEL-445 |
| chr12-116701210-DEL-127 | chr12-116701211-DEL-127 |
| chr12-116726376-DEL-440 | chr12-116726377-DEL-440 |
| chr12-116747864-DEL-178 | chr12-116747865-DEL-178 |
| chr12-116748925-DEL-197 | chr12-116748926-DEL-197 |
| chr12-116759736-DEL-6412 | chr12-116759720-DEL-6420 |
| chr12-116765828-DEL-204 | chr12-116765829-DEL-204 |
| chr12-116774870-DEL-1460 | chr12-116774871-DEL-1460 |
| chr12-116777296-DEL-117 | chr12-116777297-DEL-117 |
| chr12-116781249-DEL-68 | chr12-116781253-DEL-56 |
| chr12-116791690-DEL-343 | chr12-116791691-DEL-343 |
| chr12-116809369-DEL-121 | chr12-116809372-DEL-119 |
| chr12-116811050-DEL-6039 | chr12-116811051-DEL-6039 |
| chr12-116818803-DEL-72 | chr12-116818804-DEL-72 |
| chr12-116833613-DEL-500 | chr12-116833614-DEL-500 |
| chr12-116838939-DEL-93 | chr12-116838940-DEL-93 |
| chr12-116848558-DEL-64 | chr12-116848562-DEL-61 |

| | |
|---|---|
| chr12-116902436-DEL-176 | chr12-116902437-DEL-176 |
| chr12-116915151-DEL-90 | chr12-116915113-DEL-91 |
| chr12-116919757-DEL-209 | chr12-116919761-DEL-210 |
| chr12-116950644-DEL-51 | chr12-116950645-DEL-53 |
| chr12-116979393-DEL-457 | chr12-116979394-DEL-457 |
| chr12-117008906-DEL-398 | chr12-117008907-DEL-398 |
| chr12-117029363-DEL-6398 | chr12-117029363-DEL-6398 |
| chr12-117035362-DEL-199 | chr12-117035363-DEL-199 |
| chr12-117039531-DEL-653 | chr12-117039532-DEL-653 |
| chr12-117044784-DEL-121 | chr12-117044785-DEL-121 |
| chr12-117050787-DEL-69 | chr12-117050788-DEL-69 |
| chr12-117081516-DEL-1637 | chr12-117081517-DEL-1637 |
| chr12-117090050-DEL-108 | chr12-117090052-DEL-104 |
| chr12-117129126-DEL-207 | chr12-117129125-DEL-217 |
| chr12-117174008-DEL-132 | chr12-117174010-DEL-131 |
| chr12-117178758-DEL-556 | chr12-117178759-DEL-556 |
| chr12-117179482-DEL-1036 | chr12-117179483-DEL-1036 |
| chr12-117222395-DEL-53 | chr12-117222396-DEL-53 |
| chr12-117240434-DEL-87 | chr12-117240432-DEL-88 |
| chr12-117295487-DEL-100 | chr12-117295488-DEL-100 |
| chr12-117299383-DEL-203 | chr12-117299384-DEL-203 |
| chr12-117307179-DEL-197 | chr12-117307180-DEL-197 |
| chr12-117316908-DEL-51 | chr12-117316909-DEL-51 |
| chr12-117318862-DEL-117 | chr12-117318863-DEL-117 |
| chr12-117332132-DEL-681 | chr12-117332139-DEL-641 |
| chr12-117346116-DEL-127 | chr12-117346117-DEL-127 |
| chr12-117346286-DEL-2468 | chr12-117346286-DEL-2468 |
| chr12-117357949-DEL-1416 | chr12-117357950-DEL-1416 |
| chr12-117361857-DEL-328 | chr12-117361858-DEL-328 |
| chr12-117362434-DEL-1127 | chr12-117362435-DEL-1127 |
| chr12-117366074-DEL-471 | chr12-117366075-DEL-471 |
| chr12-117366735-DEL-169 | chr12-117366737-DEL-168 |
| chr12-117381489-DEL-74 | chr12-117381480-DEL-82 |
| chr12-117387807-DEL-199 | chr12-117387808-DEL-199 |
| chr12-117400285-DEL-64 | chr12-117400286-DEL-64 |
| chr12-117404204-DEL-62 | chr12-117404206-DEL-60 |
| chr12-117406611-DEL-192 | chr12-117406612-DEL-192 |
| chr12-117426956-DEL-51 | chr12-117426949-DEL-69 |
| chr12-117434097-DEL-107 | chr12-117434096-DEL-108 |
| chr12-117486933-DEL-216 | chr12-117486934-DEL-216 |
| chr12-117521196-DEL-65 | chr12-117521197-DEL-65 |
| chr12-117522483-DEL-714 | chr12-117522484-DEL-714 |
| chr12-117525881-DEL-55 | chr12-117525882-DEL-55 |
| chr12-117575041-DEL-53 | chr12-117575040-DEL-53 |
| chr12-117590164-DEL-52 | chr12-117590157-DEL-60 |
| chr12-117628648-DEL-427 | chr12-117628649-DEL-427 |
| chr12-117639467-DEL-627 | chr12-117639468-DEL-627 |
| chr12-117641762-DEL-73 | chr12-117641763-DEL-73 |
| chr12-117643657-DEL-105 | chr12-117643658-DEL-105 |
| chr12-117666505-DEL-204 | chr12-117666492-DEL-207 |
| chr12-117700052-DEL-552 | chr12-117700053-DEL-552 |
| chr12-117705470-DEL-272 | chr12-117705471-DEL-272 |
| chr12-117727213-DEL-66 | chr12-117727214-DEL-66 |
| chr12-117732690-DEL-52 | chr12-117732691-DEL-52 |
| chr12-117739029-DEL-1591 | chr12-117739030-DEL-1591 |
| chr12-117742579-DEL-79 | chr12-117742549-DEL-80 |
| chr12-117752147-DEL-88 | chr12-117752148-DEL-86 |
| chr12-117761783-DEL-6413 | chr12-117761786-DEL-6413 |
| chr12-117821667-DEL-456 | chr12-117821668-DEL-456 |
| chr12-117862085-DEL-545 | chr12-117862086-DEL-545 |
| chr12-117864308-DEL-531 | chr12-117864293-DEL-532 |
| chr12-117869063-DEL-51 | chr12-117869061-DEL-54 |
| chr12-117870828-DEL-432 | chr12-117870823-DEL-403 |
| chr12-117871777-DEL-2736 | chr12-117871778-DEL-2736 |
| chr12-117893714-DEL-846 | chr12-117893715-DEL-846 |
| chr12-117908038-DEL-212 | chr12-117908039-DEL-212 |
| chr12-117912810-DEL-176 | chr12-117912811-DEL-176 |
| chr12-117917304-DEL-178 | chr12-117917305-DEL-178 |
| chr12-117919246-DEL-513 | chr12-117919247-DEL-514 |
| chr12-117957851-DEL-83 | chr12-117957875-DEL-56 |
| chr12-117995340-DEL-3220 | chr12-117995341-DEL-3220 |
| chr12-118037859-DEL-380 | chr12-118037860-DEL-380 |
| chr12-118041044-DEL-83 | chr12-118041015-DEL-84 |
| chr12-118049955-DEL-100 | chr12-118049956-DEL-100 |
| chr12-118095548-DEL-115 | chr12-118095550-DEL-110 |
| chr12-118095814-DEL-6254 | chr12-118095815-DEL-6254 |
| chr12-118104042-DEL-65 | chr12-118104049-DEL-59 |
| chr12-118107683-DEL-435 | chr12-118107684-DEL-435 |
| chr12-118109334-DEL-1768 | chr12-118109335-DEL-1768 |
| chr12-118126164-DEL-1224 | chr12-118126165-DEL-1224 |

| | |
|---|---|
| chr12-118131342-DEL-2348 | chr12-118131339-DEL-2346 |
| chr12-118134652-DEL-75 | chr12-118134653-DEL-75 |
| chr12-118142281-DEL-252 | chr12-118142282-DEL-252 |
| chr12-118180607-DEL-1896 | chr12-118180608-DEL-1896 |
| chr12-118185576-DEL-208 | chr12-118185577-DEL-208 |
| chr12-118281229-DEL-677 | chr12-118281230-DEL-677 |
| chr12-118310413-DEL-117 | chr12-118310414-DEL-117 |
| chr12-118344091-DEL-688 | chr12-118344084-DEL-692 |
| chr12-118345552-DEL-6591 | chr12-118345553-DEL-6591 |
| chr12-118351593-DEL-208 | chr12-118351594-DEL-208 |
| chr12-118353685-DEL-197 | chr12-118353685-DEL-197 |
| chr12-118358174-DEL-124 | chr12-118358158-DEL-125 |
| chr12-118367222-DEL-134 | chr12-118367223-DEL-144 |
| chr12-118376532-DEL-671 | chr12-118376533-DEL-671 |
| chr12-118409499-DEL-228 | chr12-118409500-DEL-228 |
| chr12-118448590-DEL-340 | chr12-118448591-DEL-340 |
| chr12-118458019-DEL-99 | chr12-118457985-DEL-101 |
| chr12-118518608-DEL-193 | chr12-118518609-DEL-193 |
| chr12-118581325-DEL-305 | chr12-118581309-DEL-306 |
| chr12-118602778-DEL-55 | chr12-118602779-DEL-55 |
| chr12-118647134-DEL-6678 | chr12-118647135-DEL-6678 |
| chr12-118669451-DEL-70 | chr12-118669455-DEL-56 |
| chr12-118692787-DEL-975 | chr12-118692788-DEL-975 |
| chr12-118700632-DEL-54 | chr12-118700633-DEL-56 |
| chr12-118720254-DEL-76 | chr12-118720255-DEL-76 |
| chr12-118740420-DEL-103 | chr12-118740411-DEL-108 |
| chr12-118744818-DEL-308 | chr12-118744800-DEL-315 |
| chr12-118788066-DEL-2065 | chr12-118788066-DEL-2065 |
| chr12-118835872-DEL-60 | chr12-118835873-DEL-60 |
| chr12-118839896-DEL-192 | chr12-118839897-DEL-192 |
| chr12-118882861-DEL-497 | chr12-118882864-DEL-493 |
| chr12-118886070-DEL-2046 | chr12-118886071-DEL-2046 |
| chr12-118908137-DEL-68 | chr12-118908138-DEL-64 |
| chr12-118915284-DEL-152 | chr12-118915285-DEL-151 |
| chr12-118918745-DEL-204 | chr12-118918746-DEL-204 |
| chr12-118975460-DEL-6419 | chr12-118975461-DEL-6419 |
| chr12-118975499-DEL-201 | chr12-118975500-DEL-201 |
| chr12-119028498-DEL-1165 | chr12-119028499-DEL-1165 |
| chr12-119055158-DEL-440 | chr12-119055159-DEL-440 |
| chr12-119083845-DEL-435 | chr12-119083846-DEL-435 |
| chr12-119098517-DEL-1033 | chr12-119098518-DEL-1033 |
| chr12-119155542-DEL-97 | chr12-119155543-DEL-93 |
| chr12-119224651-DEL-3403 | chr12-119224652-DEL-3403 |
| chr12-119249304-DEL-6144 | chr12-119249305-DEL-6144 |
| chr12-119260963-DEL-3265 | chr12-119260964-DEL-3265 |
| chr12-119265716-DEL-238 | chr12-119265706-DEL-227 |
| chr12-119300449-DEL-51 | chr12-119300450-DEL-51 |
| chr12-119309933-DEL-6293 | chr12-119309934-DEL-6293 |
| chr12-119372058-DEL-71 | chr12-119372059-DEL-71 |
| chr12-119394233-DEL-681 | chr12-119394234-DEL-681 |
| chr12-119396224-DEL-3280 | chr12-119396225-DEL-3280 |
| chr12-119433138-DEL-151 | chr12-119433122-DEL-152 |
| chr12-119438953-DEL-71 | chr12-119438954-DEL-71 |
| chr12-119442756-DEL-1065 | chr12-119442757-DEL-1065 |
| chr12-119450893-DEL-119 | chr12-119450895-DEL-118 |
| chr12-119477355-DEL-7126 | chr12-119477356-DEL-7126 |
| chr12-119511565-DEL-2356 | chr12-119511566-DEL-2356 |
| chr12-119516332-DEL-4333 | chr12-119516294-DEL-4333 |
| chr12-119536802-DEL-3095 | chr12-119536803-DEL-3095 |
| chr12-119538201-DEL-1341 | chr12-119538201-DEL-1341 |
| chr12-119554451-DEL-7205 | chr12-119554452-DEL-7205 |
| chr12-119580756-DEL-218 | chr12-119580757-DEL-218 |
| chr12-119600996-DEL-2731 | chr12-119600997-DEL-2729 |
| chr12-119624743-DEL-6597 | chr12-119624744-DEL-6597 |
| chr12-119654648-DEL-207 | chr12-119654649-DEL-207 |
| chr12-119675790-DEL-380 | chr12-119675791-DEL-380 |
| chr12-119713235-DEL-151 | chr12-119713236-DEL-151 |
| chr12-119742679-DEL-5932 | chr12-119742680-DEL-5932 |
| chr12-119770676-DEL-122 | chr12-119770677-DEL-122 |
| chr12-119808296-DEL-78 | chr12-119808297-DEL-78 |
| chr12-119822624-DEL-201 | chr12-119822626-DEL-192 |
| chr12-119895643-DEL-91 | chr12-119895644-DEL-90 |
| chr12-119901535-DEL-340 | chr12-119901536-DEL-340 |
| chr12-119903229-DEL-225 | chr12-119903230-DEL-225 |
| chr12-119956757-DEL-205 | chr12-119956757-DEL-205 |
| chr13-3065581-DEL-2766 | chr13-3065582-DEL-2766 |
| chr13-3071008-DEL-6440 | chr13-3071009-DEL-6440 |
| chr13-3139070-DEL-140 | chr13-3139058-DEL-141 |
| chr13-3159363-DEL-7527 | chr13-3159364-DEL-7527 |
| chr13-3159553-DEL-212 | chr13-3159554-DEL-212 |

| | |
|---|---|
| chr13-3167615-DEL-83 | chr13-3167599-DEL-120 |
| chr13-3168262-DEL-119 | chr13-3168263-DEL-119 |
| chr13-3208919-DEL-204 | chr13-3208920-DEL-204 |
| chr13-3222629-DEL-8845 | chr13-3222630-DEL-8845 |
| chr13-3247218-DEL-5465 | chr13-3247220-DEL-5464 |
| chr13-3266764-DEL-116 | chr13-3266761-DEL-120 |
| chr13-3281281-DEL-5492 | chr13-3281282-DEL-5492 |
| chr13-3296272-DEL-1846 | chr13-3296273-DEL-1846 |
| chr13-3311163-DEL-54 | chr13-3311158-DEL-60 |
| chr13-3329266-DEL-558 | chr13-3329267-DEL-558 |
| chr13-3331757-DEL-51 | chr13-3331758-DEL-54 |
| chr13-3397728-DEL-126 | chr13-3397728-DEL-126 |
| chr13-3415184-DEL-191 | chr13-3415182-DEL-192 |
| chr13-3416964-DEL-169 | chr13-3416965-DEL-169 |
| chr13-3417642-DEL-1464 | chr13-3417642-DEL-1464 |
| chr13-3428230-DEL-83 | chr13-3428231-DEL-80 |
| chr13-3436025-DEL-214 | chr13-3436026-DEL-214 |
| chr13-3483965-DEL-92 | chr13-3483919-DEL-91 |
| chr13-3502859-DEL-141 | chr13-3502860-DEL-141 |
| chr13-3518924-DEL-1317 | chr13-3518925-DEL-1317 |
| chr13-3521911-DEL-79 | chr13-3521906-DEL-68 |
| chr13-3527017-DEL-105 | chr13-3527017-DEL-105 |
| chr13-3528395-DEL-86 | chr13-3528396-DEL-86 |
| chr13-3564300-DEL-221 | chr13-3564301-DEL-221 |
| chr13-3596329-DEL-1179 | chr13-3596330-DEL-1179 |
| chr13-3633461-DEL-561 | chr13-3633462-DEL-561 |
| chr13-3664955-DEL-3954 | chr13-3664955-DEL-3954 |
| chr13-3665685-DEL-75 | chr13-3665686-DEL-75 |
| chr13-3671674-DEL-582 | chr13-3671675-DEL-582 |
| chr13-3713551-DEL-324 | chr13-3713552-DEL-324 |
| chr13-3721813-DEL-60 | chr13-3721814-DEL-60 |
| chr13-3755739-DEL-111 | chr13-3755740-DEL-111 |
| chr13-3880051-DEL-50 | chr13-3880050-DEL-52 |
| chr13-3931782-DEL-136 | chr13-3931762-DEL-144 |
| chr13-3948150-DEL-501 | chr13-3948129-DEL-478 |
| chr13-3951257-DEL-1790 | chr13-3951209-DEL-1791 |
| chr13-3963573-DEL-240 | chr13-3963573-DEL-241 |
| chr13-3988123-DEL-363 | chr13-3988124-DEL-363 |
| chr13-4000041-DEL-92 | chr13-4000052-DEL-102 |
| chr13-4007101-DEL-209 | chr13-4007102-DEL-209 |
| chr13-4018489-DEL-550 | chr13-4018490-DEL-550 |
| chr13-4023122-DEL-6156 | chr13-4023123-DEL-6156 |
| chr13-4038291-DEL-3019 | chr13-4038292-DEL-3019 |
| chr13-4069110-DEL-164 | chr13-4069111-DEL-164 |
| chr13-4115503-DEL-10538 | chr13-4115504-DEL-10538 |
| chr13-4220467-DEL-124 | chr13-4220468-DEL-124 |
| chr13-4226737-DEL-444 | chr13-4226738-DEL-444 |
| chr13-4308506-DEL-3717 | chr13-4308507-DEL-3717 |
| chr13-4340580-DEL-234 | chr13-4340581-DEL-234 |
| chr13-4358192-DEL-190 | chr13-4358192-DEL-190 |
| chr13-4388457-DEL-7106 | chr13-4388458-DEL-7106 |
| chr13-4399293-DEL-603 | chr13-4399294-DEL-602 |
| chr13-4431774-DEL-642 | chr13-4431775-DEL-642 |
| chr13-4436306-DEL-422 | chr13-4436307-DEL-422 |
| chr13-4443212-DEL-1866 | chr13-4443213-DEL-1866 |
| chr13-4445840-DEL-284 | chr13-4445841-DEL-284 |
| chr13-4477533-DEL-517 | chr13-4477534-DEL-517 |
| chr13-4479475-DEL-686 | chr13-4479476-DEL-686 |
| chr13-4492933-DEL-438 | chr13-4492934-DEL-438 |
| chr13-4509700-DEL-4997 | chr13-4509701-DEL-4997 |
| chr13-4522380-DEL-2770 | chr13-4522381-DEL-2770 |
| chr13-4530795-DEL-206 | chr13-4530796-DEL-206 |
| chr13-4560328-DEL-55 | chr13-4560329-DEL-55 |
| chr13-4592903-DEL-92 | chr13-4592921-DEL-86 |
| chr13-4600761-DEL-3299 | chr13-4600762-DEL-3299 |
| chr13-4653352-DEL-170 | chr13-4653353-DEL-170 |
| chr13-4663693-DEL-102 | chr13-4663691-DEL-105 |
| chr13-4713180-DEL-85 | chr13-4713190-DEL-75 |
| chr13-4721392-DEL-164 | chr13-4721393-DEL-164 |
| chr13-4736700-DEL-50 | chr13-4736701-DEL-56 |
| chr13-4749588-DEL-83 | chr13-4749589-DEL-83 |
| chr13-4759937-DEL-54 | chr13-4759938-DEL-54 |
| chr13-4791012-DEL-1378 | chr13-4791013-DEL-1378 |
| chr13-4791959-DEL-99 | chr13-4791965-DEL-102 |
| chr13-4802833-DEL-54 | chr13-4802831-DEL-54 |
| chr13-4815063-DEL-105 | chr13-4815064-DEL-105 |
| chr13-4841924-DEL-1283 | chr13-4841925-DEL-1283 |
| chr13-4855574-DEL-155 | chr13-4855575-DEL-155 |
| chr13-4879447-DEL-75 | chr13-4879448-DEL-75 |
| chr13-4917964-DEL-195 | chr13-4917965-DEL-195 |

| | |
|---|---|
| chr13-4920736-DEL-5140 | chr13-4920737-DEL-5140 |
| chr13-4933460-DEL-650 | chr13-4933461-DEL-650 |
| chr13-4936312-DEL-494 | chr13-4936313-DEL-494 |
| chr13-4992645-DEL-476 | chr13-4992646-DEL-476 |
| chr13-5031919-DEL-7955 | chr13-5031920-DEL-7955 |
| chr13-5113882-DEL-6810 | chr13-5113879-DEL-6808 |
| chr13-5142160-DEL-6675 | chr13-5142161-DEL-6675 |
| chr13-5171064-DEL-70 | chr13-5171065-DEL-70 |
| chr13-5189295-DEL-145 | chr13-5189296-DEL-145 |
| chr13-5191306-DEL-6175 | chr13-5191307-DEL-6175 |
| chr13-5201564-DEL-138 | chr13-5201571-DEL-129 |
| chr13-5221413-DEL-7080 | chr13-5221414-DEL-7080 |
| chr13-5244885-DEL-7454 | chr13-5244886-DEL-7454 |
| chr13-5271697-DEL-668 | chr13-5271698-DEL-668 |
| chr13-5316586-DEL-74 | chr13-5316587-DEL-74 |
| chr13-5320748-DEL-64 | chr13-5320749-DEL-64 |
| chr13-5328068-DEL-5972 | chr13-5328070-DEL-5972 |
| chr13-5350405-DEL-199 | chr13-5350406-DEL-199 |
| chr13-5373165-DEL-62 | chr13-5373164-DEL-64 |
| chr13-5379213-DEL-58 | chr13-5379214-DEL-58 |
| chr13-5382267-DEL-816 | chr13-5382268-DEL-816 |
| chr13-5397169-DEL-144 | chr13-5397182-DEL-132 |
| chr13-5429253-DEL-392 | chr13-5429254-DEL-392 |
| chr13-5495888-DEL-1079 | chr13-5495888-DEL-1079 |
| chr13-5520481-DEL-54 | chr13-5520482-DEL-54 |
| chr13-5537134-DEL-1550 | chr13-5537135-DEL-1550 |
| chr13-5578634-DEL-103 | chr13-5578635-DEL-101 |
| chr13-5627686-DEL-62 | chr13-5627687-DEL-62 |
| chr13-5677333-DEL-74 | chr13-5677334-DEL-69 |
| chr13-5688039-DEL-77 | chr13-5688040-DEL-77 |
| chr13-5705568-DEL-88 | chr13-5705569-DEL-88 |
| chr13-5742875-DEL-230 | chr13-5742876-DEL-230 |
| chr13-5774697-DEL-201 | chr13-5774698-DEL-201 |
| chr13-5801319-DEL-532 | chr13-5801320-DEL-532 |
| chr13-5812634-DEL-222 | chr13-5812635-DEL-222 |
| chr13-5816135-DEL-208 | chr13-5816130-DEL-214 |
| chr13-5836305-DEL-121 | chr13-5836266-DEL-123 |
| chr13-5844216-DEL-54 | chr13-5844217-DEL-54 |
| chr13-5925950-DEL-138 | chr13-5925951-DEL-138 |
| chr13-5932572-DEL-2959 | chr13-5932573-DEL-2959 |
| chr13-5990604-DEL-65 | chr13-5990604-DEL-66 |
| chr13-6008034-DEL-53 | chr13-6008034-DEL-54 |
| chr13-6052949-DEL-61 | chr13-6052945-DEL-65 |
| chr13-6063480-DEL-65 | chr13-6063481-DEL-65 |
| chr13-6088685-DEL-92 | chr13-6088686-DEL-92 |
| chr13-6091274-DEL-63 | chr13-6091281-DEL-56 |
| chr13-6184137-DEL-53 | chr13-6184178-DEL-58 |
| chr13-6185424-DEL-154 | chr13-6185425-DEL-154 |
| chr13-6198530-DEL-1179 | chr13-6198531-DEL-1180 |
| chr13-6213834-DEL-235 | chr13-6213835-DEL-235 |
| chr13-6226872-DEL-108 | chr13-6226919-DEL-62 |
| chr13-6234196-DEL-426 | chr13-6234197-DEL-426 |
| chr13-6236762-DEL-55 | chr13-6236763-DEL-55 |
| chr13-6244206-DEL-56 | chr13-6244181-DEL-74 |
| chr13-6262239-DEL-69 | chr13-6262240-DEL-69 |
| chr13-6267554-DEL-70 | chr13-6267534-DEL-90 |
| chr13-6299896-DEL-70 | chr13-6299897-DEL-70 |
| chr13-6316830-DEL-2738 | chr13-6316831-DEL-2738 |
| chr13-6350264-DEL-929 | chr13-6350259-DEL-930 |
| chr13-6358902-DEL-1565 | chr13-6358903-DEL-1565 |
| chr13-6361100-DEL-179 | chr13-6361101-DEL-179 |
| chr13-6378590-DEL-63 | chr13-6378591-DEL-63 |
| chr13-6379242-DEL-468 | chr13-6379243-DEL-468 |
| chr13-6380267-DEL-151 | chr13-6380268-DEL-151 |
| chr13-6383551-DEL-67 | chr13-6383552-DEL-67 |
| chr13-6395144-DEL-74 | chr13-6395106-DEL-64 |
| chr13-6454674-DEL-104 | chr13-6454675-DEL-104 |
| chr13-6458298-DEL-62 | chr13-6458314-DEL-56 |
| chr13-6485732-DEL-6743 | chr13-6485733-DEL-6743 |
| chr13-6486091-DEL-214 | chr13-6486092-DEL-214 |
| chr13-6508182-DEL-1781 | chr13-6508183-DEL-1781 |
| chr13-6539444-DEL-2956 | chr13-6539445-DEL-2956 |
| chr13-6544634-DEL-61 | chr13-6544635-DEL-61 |
| chr13-6548333-DEL-54 | chr13-6548336-DEL-52 |
| chr13-6559208-DEL-150 | chr13-6559209-DEL-150 |
| chr13-6560749-DEL-3876 | chr13-6560750-DEL-3876 |
| chr13-6589363-DEL-87 | chr13-6589364-DEL-86 |
| chr13-6601932-DEL-52 | chr13-6601933-DEL-78 |
| chr13-6623947-DEL-62 | chr13-6623941-DEL-80 |
| chr13-6659880-DEL-2700 | chr13-6659880-DEL-2700 |

| | |
|---|---|
| chr13-6660997-DEL-60 | chr13-6660998-DEL-60 |
| chr13-6706192-DEL-108 | chr13-6706193-DEL-108 |
| chr13-6712516-DEL-494 | chr13-6712517-DEL-494 |
| chr13-6717799-DEL-6876 | chr13-6717800-DEL-6876 |
| chr13-6732711-DEL-71 | chr13-6732712-DEL-71 |
| chr13-6785853-DEL-214 | chr13-6785854-DEL-214 |
| chr13-6798083-DEL-137 | chr13-6798084-DEL-133 |
| chr13-6810861-DEL-751 | chr13-6810862-DEL-750 |
| chr13-6812107-DEL-2477 | chr13-6812108-DEL-2477 |
| chr13-6826793-DEL-110 | chr13-6826794-DEL-110 |
| chr13-6828423-DEL-6895 | chr13-6828424-DEL-6895 |
| chr13-6838987-DEL-119 | chr13-6838985-DEL-97 |
| chr13-6870016-DEL-6255 | chr13-6870017-DEL-6255 |
| chr13-6881475-DEL-381 | chr13-6881476-DEL-381 |
| chr13-6927814-DEL-7065 | chr13-6927815-DEL-7065 |
| chr13-6934223-DEL-212 | chr13-6934224-DEL-212 |
| chr13-6990339-DEL-6413 | chr13-6990340-DEL-6413 |
| chr13-6999616-DEL-80 | chr13-6999618-DEL-66 |
| chr13-7000575-DEL-433 | chr13-7000576-DEL-433 |
| chr13-7006596-DEL-1531 | chr13-7006597-DEL-1531 |
| chr13-7063126-DEL-4220 | chr13-7063126-DEL-4220 |
| chr13-7089537-DEL-56 | chr13-7089542-DEL-56 |
| chr13-7091183-DEL-2458 | chr13-7091182-DEL-2459 |
| chr13-7093968-DEL-484 | chr13-7093969-DEL-484 |
| chr13-7099369-DEL-6552 | chr13-7099370-DEL-6552 |
| chr13-7106300-DEL-641 | chr13-7106300-DEL-641 |
| chr13-7117782-DEL-439 | chr13-7117771-DEL-439 |
| chr13-7118446-DEL-6386 | chr13-7118447-DEL-6386 |
| chr13-7146895-DEL-62 | chr13-7146890-DEL-74 |
| chr13-7154148-DEL-92 | chr13-7154149-DEL-92 |
| chr13-7164115-DEL-68 | chr13-7164116-DEL-60 |
| chr13-7174311-DEL-6617 | chr13-7174311-DEL-6617 |
| chr13-7191073-DEL-202 | chr13-7191074-DEL-202 |
| chr13-7195925-DEL-127 | chr13-7195908-DEL-134 |
| chr13-7202528-DEL-80 | chr13-7202529-DEL-80 |
| chr13-7224661-DEL-268 | chr13-7224662-DEL-268 |
| chr13-7229694-DEL-91 | chr13-7229695-DEL-91 |
| chr13-7247915-DEL-58 | chr13-7247916-DEL-58 |
| chr13-7295780-DEL-213 | chr13-7295779-DEL-213 |
| chr13-7325208-DEL-4416 | chr13-7325209-DEL-4416 |
| chr13-7442563-DEL-67 | chr13-7442556-DEL-55 |
| chr13-7469599-DEL-63 | chr13-7469600-DEL-63 |
| chr13-7485387-DEL-121 | chr13-7485388-DEL-121 |
| chr13-7488448-DEL-1289 | chr13-7488449-DEL-1289 |
| chr13-7490232-DEL-2339 | chr13-7490233-DEL-2339 |
| chr13-7510070-DEL-1539 | chr13-7510071-DEL-1539 |
| chr13-7511944-DEL-1416 | chr13-7511945-DEL-1416 |
| chr13-7518992-DEL-106 | chr13-7518992-DEL-145 |
| chr13-7548091-DEL-2015 | chr13-7548092-DEL-2015 |
| chr13-7570815-DEL-137 | chr13-7570816-DEL-137 |
| chr13-7606416-DEL-87 | chr13-7606374-DEL-90 |
| chr13-7615096-DEL-66 | chr13-7615097-DEL-66 |
| chr13-7621191-DEL-254 | chr13-7621192-DEL-226 |
| chr13-7674285-DEL-70 | chr13-7674286-DEL-70 |
| chr13-7676174-DEL-149 | chr13-7676175-DEL-149 |
| chr13-7685896-DEL-483 | chr13-7685895-DEL-483 |
| chr13-7686543-DEL-211 | chr13-7686544-DEL-211 |
| chr13-7699310-DEL-68 | chr13-7699276-DEL-52 |
| chr13-7702151-DEL-55 | chr13-7702152-DEL-55 |
| chr13-7721139-DEL-202 | chr13-7721140-DEL-202 |
| chr13-7725318-DEL-52 | chr13-7725317-DEL-54 |
| chr13-7764155-DEL-768 | chr13-7764156-DEL-768 |
| chr13-7810439-DEL-399 | chr13-7810440-DEL-399 |
| chr13-7918301-DEL-216 | chr13-7918301-DEL-209 |
| chr13-7936068-DEL-60 | chr13-7936069-DEL-58 |
| chr13-7999170-DEL-7108 | chr13-7999171-DEL-7108 |
| chr13-8007172-DEL-505 | chr13-8007173-DEL-505 |
| chr13-8012599-DEL-400 | chr13-8012600-DEL-400 |
| chr13-8013771-DEL-2672 | chr13-8013776-DEL-2668 |
| chr13-8054169-DEL-85 | chr13-8054170-DEL-85 |
| chr13-8063237-DEL-64 | chr13-8063245-DEL-53 |
| chr13-8071940-DEL-2683 | chr13-8071941-DEL-2683 |
| chr13-8087411-DEL-6701 | chr13-8087412-DEL-6701 |
| chr13-8096948-DEL-64 | chr13-8096949-DEL-64 |
| chr13-8120699-DEL-55 | chr13-8120687-DEL-68 |
| chr13-8139480-DEL-57 | chr13-8139481-DEL-57 |
| chr13-8142821-DEL-69 | chr13-8142812-DEL-91 |
| chr13-8144074-DEL-64 | chr13-8144061-DEL-54 |
| chr13-8189739-DEL-134 | chr13-8189727-DEL-122 |
| chr13-8245661-DEL-76 | chr13-8245662-DEL-76 |

| | |
|---|---|
| chr13-8276117-DEL-58 | chr13-8276118-DEL-58 |
| chr13-8292181-DEL-503 | chr13-8292182-DEL-503 |
| chr13-8317624-DEL-2396 | chr13-8317625-DEL-2396 |
| chr13-8339058-DEL-1111 | chr13-8339059-DEL-1111 |
| chr13-8351561-DEL-958 | chr13-8351562-DEL-958 |
| chr13-8368192-DEL-145 | chr13-8368193-DEL-145 |
| chr13-8379649-DEL-68 | chr13-8379650-DEL-68 |
| chr13-8414990-DEL-502 | chr13-8414991-DEL-502 |
| chr13-8422694-DEL-3059 | chr13-8422695-DEL-3059 |
| chr13-8453647-DEL-153 | chr13-8453648-DEL-153 |
| chr13-8457380-DEL-910 | chr13-8457381-DEL-910 |
| chr13-8464541-DEL-348 | chr13-8464542-DEL-348 |
| chr13-8478469-DEL-710 | chr13-8478470-DEL-710 |
| chr13-8480577-DEL-209 | chr13-8480578-DEL-207 |
| chr13-8491246-DEL-862 | chr13-8491247-DEL-862 |
| chr13-8501045-DEL-251 | chr13-8501046-DEL-251 |
| chr13-8519663-DEL-5214 | chr13-8519664-DEL-5214 |
| chr13-8543952-DEL-198 | chr13-8543953-DEL-198 |
| chr13-8554533-DEL-84 | chr13-8554534-DEL-84 |
| chr13-8576717-DEL-103 | chr13-8576718-DEL-103 |
| chr13-8601017-DEL-95 | chr13-8601018-DEL-95 |
| chr13-8649965-DEL-197 | chr13-8649966-DEL-197 |
| chr13-8677423-DEL-470 | chr13-8677424-DEL-470 |
| chr13-8684827-DEL-3949 | chr13-8684828-DEL-3949 |
| chr13-8714059-DEL-62 | chr13-8714060-DEL-62 |
| chr13-8715737-DEL-2755 | chr13-8715738-DEL-2755 |
| chr13-8767899-DEL-94 | chr13-8767900-DEL-94 |
| chr13-8770955-DEL-140 | chr13-8770969-DEL-100 |
| chr13-8781377-DEL-89 | chr13-8781376-DEL-90 |
| chr13-8805085-DEL-895 | chr13-8805101-DEL-880 |
| chr13-8805848-DEL-63 | chr13-8805809-DEL-103 |
| chr13-8811873-DEL-5544 | chr13-8811874-DEL-5544 |
| chr13-8836291-DEL-921 | chr13-8836292-DEL-921 |
| chr13-8840111-DEL-400 | chr13-8840112-DEL-400 |
| chr13-8854302-DEL-79 | chr13-8854303-DEL-79 |
| chr13-8858597-DEL-7391 | chr13-8858587-DEL-7390 |
| chr13-8887804-DEL-109 | chr13-8887805-DEL-109 |
| chr13-8907383-DEL-638 | chr13-8907384-DEL-638 |
| chr13-8909748-DEL-55 | chr13-8909787-DEL-52 |
| chr13-8910790-DEL-77 | chr13-8910791-DEL-77 |
| chr13-8913814-DEL-228 | chr13-8913815-DEL-228 |
| chr13-8925180-DEL-322 | chr13-8925181-DEL-322 |
| chr13-8943187-DEL-214 | chr13-8943188-DEL-214 |
| chr13-8945315-DEL-7150 | chr13-8945314-DEL-7150 |
| chr13-8968553-DEL-1897 | chr13-8968553-DEL-1898 |
| chr13-8983039-DEL-60 | chr13-8983040-DEL-60 |
| chr13-8985579-DEL-514 | chr13-8985580-DEL-514 |
| chr13-9033886-DEL-220 | chr13-9033887-DEL-220 |
| chr13-9034781-DEL-195 | chr13-9034778-DEL-196 |
| chr13-9055292-DEL-54 | chr13-9055289-DEL-56 |
| chr13-9102095-DEL-264 | chr13-9102091-DEL-263 |
| chr13-9103515-DEL-202 | chr13-9103534-DEL-198 |
| chr13-9135861-DEL-51 | chr13-9135862-DEL-51 |
| chr13-9141042-DEL-146 | chr13-9141043-DEL-146 |
| chr13-9187397-DEL-97 | chr13-9187398-DEL-97 |
| chr13-9256542-DEL-484 | chr13-9256543-DEL-484 |
| chr13-9260353-DEL-1345 | chr13-9260354-DEL-1345 |
| chr13-9266659-DEL-59 | chr13-9266678-DEL-63 |
| chr13-9275079-DEL-1276 | chr13-9275080-DEL-1276 |
| chr13-9285597-DEL-512 | chr13-9285598-DEL-512 |
| chr13-9303882-DEL-216 | chr13-9303883-DEL-216 |
| chr13-9305436-DEL-126 | chr13-9305437-DEL-76 |
| chr13-9310182-DEL-74 | chr13-9310183-DEL-86 |
| chr13-9335057-DEL-257 | chr13-9335058-DEL-257 |
| chr13-9359416-DEL-201 | chr13-9359417-DEL-201 |
| chr13-9377247-DEL-140 | chr13-9377248-DEL-140 |
| chr13-9378617-DEL-112 | chr13-9378618-DEL-112 |
| chr13-9397353-DEL-3117 | chr13-9397354-DEL-3117 |
| chr13-9403397-DEL-226 | chr13-9403398-DEL-226 |
| chr13-9442842-DEL-780 | chr13-9442843-DEL-780 |
| chr13-9448466-DEL-150 | chr13-9448487-DEL-135 |
| chr13-9454366-DEL-362 | chr13-9454367-DEL-362 |
| chr13-9482688-DEL-7242 | chr13-9482689-DEL-7242 |
| chr13-9525371-DEL-2937 | chr13-9525375-DEL-2933 |
| chr13-9530028-DEL-66 | chr13-9530029-DEL-66 |
| chr13-9576814-DEL-257 | chr13-9576814-DEL-257 |
| chr13-9597523-DEL-372 | chr13-9597524-DEL-372 |
| chr13-9598992-DEL-79 | chr13-9598993-DEL-79 |
| chr13-9620317-DEL-86 | chr13-9620318-DEL-86 |
| chr13-9650976-DEL-86 | chr13-9650977-DEL-86 |

| | |
|---|---|
| chr13-9657507-DEL-56 | chr13-9657504-DEL-60 |
| chr13-9661710-DEL-60 | chr13-9661710-DEL-60 |
| chr13-9694330-DEL-459 | chr13-9694331-DEL-459 |
| chr13-9706215-DEL-165 | chr13-9706216-DEL-165 |
| chr13-9721897-DEL-84 | chr13-9721898-DEL-84 |
| chr13-9728769-DEL-2886 | chr13-9728770-DEL-2886 |
| chr13-9749149-DEL-637 | chr13-9749150-DEL-637 |
| chr13-9756058-DEL-60 | chr13-9756020-DEL-52 |
| chr13-9769088-DEL-76 | chr13-9769089-DEL-76 |
| chr13-9782560-DEL-358 | chr13-9782561-DEL-358 |
| chr13-9825378-DEL-105 | chr13-9825379-DEL-103 |
| chr13-9829466-DEL-662 | chr13-9829466-DEL-663 |
| chr13-9833176-DEL-701 | chr13-9833177-DEL-701 |
| chr13-9848839-DEL-205 | chr13-9848840-DEL-205 |
| chr13-9858567-DEL-78 | chr13-9858551-DEL-50 |
| chr13-9865108-DEL-104 | chr13-9865109-DEL-104 |
| chr13-9868410-DEL-2471 | chr13-9868411-DEL-2471 |
| chr13-9871411-DEL-92 | chr13-9871386-DEL-90 |
| chr13-9872109-DEL-76 | chr13-9872110-DEL-76 |
| chr13-9884295-DEL-304 | chr13-9884295-DEL-304 |
| chr13-9921863-DEL-469 | chr13-9921864-DEL-469 |
| chr13-9934912-DEL-155 | chr13-9934913-DEL-155 |
| chr13-9945080-DEL-68 | chr13-9945081-DEL-68 |
| chr13-9974277-DEL-84 | chr13-9974303-DEL-85 |
| chr13-10005946-DEL-1235 | chr13-10005947-DEL-1235 |
| chr13-10045263-DEL-60 | chr13-10045264-DEL-60 |
| chr13-10075832-DEL-62 | chr13-10075833-DEL-62 |
| chr13-10094913-DEL-139 | chr13-10094887-DEL-140 |
| chr13-10099057-DEL-1653 | chr13-10099058-DEL-1653 |
| chr13-10111763-DEL-58 | chr13-10111754-DEL-62 |
| chr13-10122423-DEL-73 | chr13-10122423-DEL-63 |
| chr13-10132650-DEL-55 | chr13-10132651-DEL-55 |
| chr13-10143627-DEL-527 | chr13-10143628-DEL-528 |
| chr13-10176816-DEL-1906 | chr13-10176817-DEL-1906 |
| chr13-10208958-DEL-465 | chr13-10208959-DEL-465 |
| chr13-10214377-DEL-386 | chr13-10214377-DEL-387 |
| chr13-10224808-DEL-6334 | chr13-10224809-DEL-6334 |
| chr13-10255020-DEL-70 | chr13-10255021-DEL-96 |
| chr13-10255304-DEL-667 | chr13-10255306-DEL-666 |
| chr13-10271500-DEL-120 | chr13-10271501-DEL-120 |
| chr13-10281630-DEL-108 | chr13-10281637-DEL-98 |
| chr13-10303804-DEL-513 | chr13-10303816-DEL-514 |
| chr13-10346784-DEL-66 | chr13-10346789-DEL-54 |
| chr13-10357601-DEL-738 | chr13-10357602-DEL-738 |
| chr13-10380770-DEL-149 | chr13-10380771-DEL-149 |
| chr13-10387928-DEL-61 | chr13-10387951-DEL-86 |
| chr13-10411610-DEL-65 | chr13-10411598-DEL-64 |
| chr13-10413484-DEL-116 | chr13-10413485-DEL-119 |
| chr13-10431701-DEL-105 | chr13-10431676-DEL-106 |
| chr13-10437808-DEL-506 | chr13-10437809-DEL-506 |
| chr13-10463020-DEL-54 | chr13-10463024-DEL-50 |
| chr13-10469834-DEL-166 | chr13-10469835-DEL-166 |
| chr13-10471443-DEL-938 | chr13-10471444-DEL-938 |
| chr13-10474807-DEL-584 | chr13-10474801-DEL-589 |
| chr13-10506334-DEL-235 | chr13-10506335-DEL-235 |
| chr13-10513272-DEL-60 | chr13-10513273-DEL-60 |
| chr13-10529449-DEL-96 | chr13-10529450-DEL-96 |
| chr13-10549606-DEL-132 | chr13-10549607-DEL-132 |
| chr13-10568860-DEL-1471 | chr13-10568861-DEL-1471 |
| chr13-10578434-DEL-1094 | chr13-10578435-DEL-1094 |
| chr13-10608299-DEL-933 | chr13-10608300-DEL-933 |
| chr13-10646523-DEL-82 | chr13-10646524-DEL-82 |
| chr13-10652805-DEL-6413 | chr13-10652806-DEL-6412 |
| chr13-10652838-DEL-64 | chr13-10652839-DEL-64 |
| chr13-10659431-DEL-543 | chr13-10659432-DEL-543 |
| chr13-10660046-DEL-2342 | chr13-10660047-DEL-2341 |
| chr13-10739520-DEL-72 | chr13-10739511-DEL-60 |
| chr13-10826972-DEL-2506 | chr13-10826973-DEL-2506 |
| chr13-10851924-DEL-262 | chr13-10851925-DEL-262 |
| chr13-10862015-DEL-374 | chr13-10862016-DEL-374 |
| chr13-10865678-DEL-183 | chr13-10865678-DEL-189 |
| chr13-10870805-DEL-315 | chr13-10870806-DEL-315 |
| chr13-10871305-DEL-68 | chr13-10871306-DEL-68 |
| chr13-10881117-DEL-198 | chr13-10881118-DEL-198 |
| chr13-10893368-DEL-887 | chr13-10893389-DEL-887 |
| chr13-10913135-DEL-196 | chr13-10913160-DEL-196 |
| chr13-10916338-DEL-387 | chr13-10916339-DEL-387 |
| chr13-10926919-DEL-155 | chr13-10926924-DEL-156 |
| chr13-10928232-DEL-138 | chr13-10928233-DEL-138 |
| chr13-10939234-DEL-7124 | chr13-10939235-DEL-7124 |

| | |
|---|---|
| chr13-10966054-DEL-71 | chr13-10966056-DEL-70 |
| chr13-10972525-DEL-1049 | chr13-10972526-DEL-1049 |
| chr13-10986009-DEL-422 | chr13-10986010-DEL-422 |
| chr13-10989050-DEL-714 | chr13-10989051-DEL-714 |
| chr13-11024147-DEL-5881 | chr13-11024148-DEL-5881 |
| chr13-11039685-DEL-58 | chr13-11039686-DEL-58 |
| chr13-11073714-DEL-108 | chr13-11073715-DEL-106 |
| chr13-11074968-DEL-64 | chr13-11074971-DEL-62 |
| chr13-11093129-DEL-318 | chr13-11093122-DEL-324 |
| chr13-11097636-DEL-60 | chr13-11097637-DEL-60 |
| chr13-11123947-DEL-56 | chr13-11123947-DEL-56 |
| chr13-11132498-DEL-635 | chr13-11132499-DEL-635 |
| chr13-11133837-DEL-404 | chr13-11133844-DEL-396 |
| chr13-11134483-DEL-84 | chr13-11134484-DEL-84 |
| chr13-11135307-DEL-170 | chr13-11135308-DEL-170 |
| chr13-11142545-DEL-75 | chr13-11142546-DEL-75 |
| chr13-11156924-DEL-58 | chr13-11156920-DEL-57 |
| chr13-11203595-DEL-1892 | chr13-11203596-DEL-1892 |
| chr13-11263725-DEL-57 | chr13-11263726-DEL-57 |
| chr13-11289713-DEL-248 | chr13-11289714-DEL-248 |
| chr13-11314219-DEL-926 | chr13-11314220-DEL-926 |
| chr13-11334800-DEL-112 | chr13-11334802-DEL-124 |
| chr13-11350411-DEL-1438 | chr13-11350412-DEL-1438 |
| chr13-11368559-DEL-154 | chr13-11368560-DEL-154 |
| chr13-11369290-DEL-369 | chr13-11369291-DEL-369 |
| chr13-11417531-DEL-52 | chr13-11417532-DEL-52 |
| chr13-11442387-DEL-1323 | chr13-11442388-DEL-1323 |
| chr13-11455372-DEL-832 | chr13-11455373-DEL-832 |
| chr13-11463462-DEL-103 | chr13-11463484-DEL-115 |
| chr13-11477437-DEL-1288 | chr13-11477438-DEL-1286 |
| chr13-11504643-DEL-67 | chr13-11504644-DEL-67 |
| chr13-11515977-DEL-7054 | chr13-11515978-DEL-7054 |
| chr13-11531240-DEL-86 | chr13-11531241-DEL-86 |
| chr13-11534263-DEL-130 | chr13-11534264-DEL-130 |
| chr13-11535547-DEL-4821 | chr13-11535548-DEL-4821 |
| chr13-11542739-DEL-159 | chr13-11542740-DEL-159 |
| chr13-11552755-DEL-468 | chr13-11552763-DEL-468 |
| chr13-11563241-DEL-76 | chr13-11563243-DEL-70 |
| chr13-11598128-DEL-58 | chr13-11598129-DEL-58 |
| chr13-11617696-DEL-83 | chr13-11617696-DEL-84 |
| chr13-11619334-DEL-99 | chr13-11619335-DEL-99 |
| chr13-11629844-DEL-248 | chr13-11629845-DEL-248 |
| chr13-11656181-DEL-56 | chr13-11656182-DEL-56 |
| chr13-11674074-DEL-93 | chr13-11674078-DEL-94 |
| chr13-11696424-DEL-817 | chr13-11696425-DEL-817 |
| chr13-11707725-DEL-3725 | chr13-11707726-DEL-3725 |
| chr13-11787864-DEL-67 | chr13-11787865-DEL-67 |
| chr13-11803798-DEL-110 | chr13-11803816-DEL-108 |
| chr13-11807288-DEL-214 | chr13-11807289-DEL-214 |
| chr13-11833082-DEL-53 | chr13-11833083-DEL-53 |
| chr13-11840489-DEL-63 | chr13-11840490-DEL-65 |
| chr13-11841825-DEL-60 | chr13-11841789-DEL-62 |
| chr13-11849865-DEL-58 | chr13-11849866-DEL-58 |
| chr13-11881480-DEL-72 | chr13-11881481-DEL-84 |
| chr13-11942503-DEL-460 | chr13-11942504-DEL-460 |
| chr13-11950271-DEL-1728 | chr13-11950272-DEL-1728 |
| chr13-11955978-DEL-127 | chr13-11955977-DEL-128 |
| chr13-11974891-DEL-132 | chr13-11974892-DEL-132 |
| chr13-12010841-DEL-235 | chr13-12010842-DEL-235 |
| chr13-12016268-DEL-113 | chr13-12016269-DEL-113 |
| chr13-12049637-DEL-110 | chr13-12049638-DEL-110 |
| chr13-12058961-DEL-667 | chr13-12058962-DEL-667 |
| chr13-12079254-DEL-885 | chr13-12079255-DEL-885 |
| chr13-12088965-DEL-208 | chr13-12088966-DEL-208 |
| chr13-12159782-DEL-342 | chr13-12159783-DEL-342 |
| chr13-12182912-DEL-105 | chr13-12182870-DEL-148 |
| chr13-12203262-DEL-513 | chr13-12203263-DEL-513 |
| chr13-12217205-DEL-227 | chr13-12217192-DEL-228 |
| chr13-12218364-DEL-201 | chr13-12218361-DEL-194 |
| chr13-12221837-DEL-186 | chr13-12221838-DEL-186 |
| chr13-12231384-DEL-53 | chr13-12231387-DEL-54 |
| chr13-12234377-DEL-1873 | chr13-12234378-DEL-1873 |
| chr13-12243771-DEL-138 | chr13-12243772-DEL-139 |
| chr13-12258264-DEL-443 | chr13-12258265-DEL-443 |
| chr13-12274349-DEL-350 | chr13-12274350-DEL-350 |
| chr13-12286914-DEL-158 | chr13-12286914-DEL-158 |
| chr13-12325284-DEL-85 | chr13-12325285-DEL-56 |
| chr13-12328065-DEL-227 | chr13-12328066-DEL-227 |
| chr13-12330420-DEL-70 | chr13-12330424-DEL-50 |
| chr13-12331257-DEL-201 | chr13-12331258-DEL-201 |

| | |
|---|---|
| chr13-12332641-DEL-70 | chr13-12332640-DEL-72 |
| chr13-12333403-DEL-65 | chr13-12333388-DEL-66 |
| chr13-12347485-DEL-275 | chr13-12347486-DEL-275 |
| chr13-12368375-DEL-73 | chr13-12368380-DEL-73 |
| chr13-12387695-DEL-96 | chr13-12387696-DEL-96 |
| chr13-12389007-DEL-202 | chr13-12389008-DEL-202 |
| chr13-12406000-DEL-793 | chr13-12406001-DEL-793 |
| chr13-12415480-DEL-169 | chr13-12415481-DEL-169 |
| chr13-12472611-DEL-68 | chr13-12472604-DEL-76 |
| chr13-12487977-DEL-54 | chr13-12487978-DEL-54 |
| chr13-12489984-DEL-103 | chr13-12489988-DEL-102 |
| chr13-12505535-DEL-196 | chr13-12505536-DEL-196 |
| chr13-12534631-DEL-53 | chr13-12534623-DEL-62 |
| chr13-12541704-DEL-55 | chr13-12541711-DEL-62 |
| chr13-12545140-DEL-1037 | chr13-12545141-DEL-1037 |
| chr13-12550990-DEL-82 | chr13-12550991-DEL-82 |
| chr13-12594449-DEL-171 | chr13-12594449-DEL-171 |
| chr13-13592310-DEL-312 | chr13-13592302-DEL-313 |
| chr13-13602741-DEL-92 | chr13-13602742-DEL-92 |
| chr13-13623952-DEL-173 | chr13-13623944-DEL-171 |
| chr13-13641521-DEL-686 | chr13-13641510-DEL-679 |
| chr13-13644847-DEL-133 | chr13-13644848-DEL-133 |
| chr13-13668979-DEL-281 | chr13-13668980-DEL-281 |
| chr13-13702908-DEL-178 | chr13-13702909-DEL-178 |
| chr13-13719533-DEL-79 | chr13-13719534-DEL-79 |
| chr13-13724796-DEL-512 | chr13-13724797-DEL-512 |
| chr13-13773869-DEL-448 | chr13-13773870-DEL-448 |
| chr13-13775812-DEL-110 | chr13-13775813-DEL-110 |
| chr13-13818944-DEL-84 | chr13-13818945-DEL-84 |
| chr13-13827177-DEL-834 | chr13-13827178-DEL-834 |
| chr13-13849207-DEL-55 | chr13-13849208-DEL-55 |
| chr13-13892402-DEL-2376 | chr13-13892401-DEL-2377 |
| chr13-13943961-DEL-4525 | chr13-13943962-DEL-4525 |
| chr13-13974918-DEL-158 | chr13-13974919-DEL-158 |
| chr13-13979867-DEL-54 | chr13-13979868-DEL-54 |
| chr13-14009272-DEL-210 | chr13-14009273-DEL-210 |
| chr13-14024157-DEL-506 | chr13-14024158-DEL-506 |
| chr13-14031908-DEL-61 | chr13-14031926-DEL-62 |
| chr13-14038078-DEL-307 | chr13-14038079-DEL-307 |
| chr13-14048059-DEL-2237 | chr13-14048055-DEL-2238 |
| chr13-14057500-DEL-6253 | chr13-14057501-DEL-6253 |
| chr13-14126399-DEL-921 | chr13-14126400-DEL-921 |
| chr13-14156161-DEL-1736 | chr13-14156162-DEL-1736 |
| chr13-14174281-DEL-57 | chr13-14174282-DEL-57 |
| chr13-14194792-DEL-4858 | chr13-14194793-DEL-4858 |
| chr13-14231027-DEL-503 | chr13-14231015-DEL-504 |
| chr13-14352165-DEL-56 | chr13-14352166-DEL-52 |
| chr13-14382378-DEL-6858 | chr13-14382379-DEL-6858 |
| chr13-14417235-DEL-354 | chr13-14417236-DEL-354 |
| chr13-14423480-DEL-750 | chr13-14423481-DEL-750 |
| chr13-14425530-DEL-76 | chr13-14425543-DEL-64 |
| chr13-14439052-DEL-51 | chr13-14439052-DEL-51 |
| chr13-14440789-DEL-126 | chr13-14440790-DEL-126 |
| chr13-14497914-DEL-489 | chr13-14497915-DEL-484 |
| chr13-14501276-DEL-80 | chr13-14501258-DEL-85 |
| chr13-14507727-DEL-947 | chr13-14507728-DEL-947 |
| chr13-14560411-DEL-920 | chr13-14560412-DEL-920 |
| chr13-14574347-DEL-2544 | chr13-14574348-DEL-2544 |
| chr13-14602000-DEL-2738 | chr13-14601992-DEL-2739 |
| chr13-14620536-DEL-840 | chr13-14620536-DEL-840 |
| chr13-14622384-DEL-3980 | chr13-14622385-DEL-3980 |
| chr13-14633154-DEL-150 | chr13-14633155-DEL-150 |
| chr13-14656186-DEL-54 | chr13-14656193-DEL-58 |
| chr13-14732261-DEL-160 | chr13-14732249-DEL-158 |
| chr13-14774344-DEL-680 | chr13-14774345-DEL-680 |
| chr13-14777204-DEL-195 | chr13-14777200-DEL-195 |
| chr13-14788625-DEL-785 | chr13-14788626-DEL-785 |
| chr13-14797682-DEL-215 | chr13-14797683-DEL-215 |
| chr13-14806832-DEL-300 | chr13-14806833-DEL-299 |
| chr13-14833658-DEL-125 | chr13-14833669-DEL-118 |
| chr13-14844308-DEL-116 | chr13-14844309-DEL-116 |
| chr13-14851843-DEL-58 | chr13-14851844-DEL-58 |
| chr13-14861854-DEL-465 | chr13-14861855-DEL-464 |
| chr13-14865091-DEL-6413 | chr13-14865092-DEL-6413 |
| chr13-14871204-DEL-141 | chr13-14871204-DEL-141 |
| chr13-14882005-DEL-1864 | chr13-14882006-DEL-1864 |
| chr13-14885360-DEL-263 | chr13-14885361-DEL-263 |
| chr13-14910838-DEL-52 | chr13-14910839-DEL-52 |
| chr13-14977331-DEL-7026 | chr13-14977332-DEL-7026 |
| chr13-15053301-DEL-70 | chr13-15053302-DEL-74 |

| | |
|---|---|
| chr13-15079370-DEL-328 | chr13-15079371-DEL-328 |
| chr13-15081977-DEL-1454 | chr13-15081978-DEL-1454 |
| chr13-15116964-DEL-54 | chr13-15116961-DEL-55 |
| chr13-15129127-DEL-251 | chr13-15129127-DEL-250 |
| chr13-15149706-DEL-410 | chr13-15149707-DEL-410 |
| chr13-15201781-DEL-6436 | chr13-15201782-DEL-6436 |
| chr13-15206442-DEL-137 | chr13-15206443-DEL-136 |
| chr13-15212463-DEL-1056 | chr13-15212463-DEL-1056 |
| chr13-15215372-DEL-226 | chr13-15215373-DEL-226 |
| chr13-15273027-DEL-2739 | chr13-15273028-DEL-2739 |
| chr13-15291378-DEL-399 | chr13-15291379-DEL-399 |
| chr13-15312886-DEL-3088 | chr13-15312887-DEL-3088 |
| chr13-15334851-DEL-1695 | chr13-15334851-DEL-1695 |
| chr13-15347981-DEL-120 | chr13-15347985-DEL-117 |
| chr13-15399999-DEL-61 | chr13-15399997-DEL-61 |
| chr13-15421472-DEL-211 | chr13-15421473-DEL-211 |
| chr13-15426395-DEL-6550 | chr13-15426396-DEL-6550 |
| chr13-15449249-DEL-1893 | chr13-15449249-DEL-1893 |
| chr13-15480511-DEL-255 | chr13-15480512-DEL-255 |
| chr13-15492040-DEL-1403 | chr13-15492041-DEL-1403 |
| chr13-15496306-DEL-1891 | chr13-15496307-DEL-1891 |
| chr13-15516390-DEL-112 | chr13-15516391-DEL-112 |
| chr13-15517985-DEL-132 | chr13-15518032-DEL-84 |
| chr13-15602679-DEL-188 | chr13-15602680-DEL-188 |
| chr13-15606045-DEL-7232 | chr13-15606046-DEL-7232 |
| chr13-15622180-DEL-96 | chr13-15622187-DEL-82 |
| chr13-15626827-DEL-647 | chr13-15626828-DEL-647 |
| chr13-15676134-DEL-59 | chr13-15676135-DEL-59 |
| chr13-15692333-DEL-51 | chr13-15692322-DEL-52 |
| chr13-15696303-DEL-1197 | chr13-15696304-DEL-1197 |
| chr13-15769494-DEL-53 | chr13-15769495-DEL-53 |
| chr13-15769843-DEL-828 | chr13-15769844-DEL-828 |
| chr13-15804662-DEL-57 | chr13-15804663-DEL-57 |
| chr13-15867285-DEL-118 | chr13-15867286-DEL-118 |
| chr13-15883770-DEL-180 | chr13-15883769-DEL-196 |
| chr13-15936615-DEL-100 | chr13-15936620-DEL-96 |
| chr13-15951600-DEL-358 | chr13-15951601-DEL-358 |
| chr13-15969794-DEL-336 | chr13-15969795-DEL-336 |
| chr13-15975591-DEL-709 | chr13-15975602-DEL-687 |
| chr13-15983788-DEL-3263 | chr13-15983789-DEL-3263 |
| chr13-15996860-DEL-293 | chr13-15996861-DEL-293 |
| chr13-16004073-DEL-52 | chr13-16004074-DEL-54 |
| chr13-16018038-DEL-563 | chr13-16018039-DEL-563 |
| chr13-16057823-DEL-329 | chr13-16057824-DEL-329 |
| chr13-16087761-DEL-393 | chr13-16087762-DEL-393 |
| chr13-16101041-DEL-1862 | chr13-16101042-DEL-1862 |
| chr13-16105510-DEL-105 | chr13-16105511-DEL-105 |
| chr13-16107614-DEL-58 | chr13-16107622-DEL-50 |
| chr13-16143366-DEL-201 | chr13-16143366-DEL-201 |
| chr13-16202265-DEL-70 | chr13-16202266-DEL-54 |
| chr13-16213751-DEL-1985 | chr13-16213752-DEL-1985 |
| chr13-16220962-DEL-94 | chr13-16220983-DEL-74 |
| chr13-16224444-DEL-611 | chr13-16224445-DEL-611 |
| chr13-16310569-DEL-5592 | chr13-16310570-DEL-5592 |
| chr13-16352806-DEL-61 | chr13-16352807-DEL-61 |
| chr13-16370632-DEL-63 | chr13-16370627-DEL-63 |
| chr13-16425571-DEL-2594 | chr13-16425571-DEL-2594 |
| chr13-16542518-DEL-5547 | chr13-16542519-DEL-5547 |
| chr13-16559791-DEL-6901 | chr13-16559792-DEL-6901 |
| chr13-16565958-DEL-424 | chr13-16565958-DEL-424 |
| chr13-16578464-DEL-65 | chr13-16578465-DEL-65 |
| chr13-16582914-DEL-6229 | chr13-16582915-DEL-6229 |
| chr13-16604902-DEL-1510 | chr13-16604944-DEL-1467 |
| chr13-16612443-DEL-56 | chr13-16612453-DEL-54 |
| chr13-16618797-DEL-63 | chr13-16618797-DEL-62 |
| chr13-16642504-DEL-61 | chr13-16642505-DEL-62 |
| chr13-16660687-DEL-173 | chr13-16660688-DEL-173 |
| chr13-16674432-DEL-7036 | chr13-16674433-DEL-7036 |
| chr13-16691375-DEL-71 | chr13-16691376-DEL-71 |
| chr13-16695159-DEL-125 | chr13-16695160-DEL-125 |
| chr13-16702981-DEL-232 | chr13-16702978-DEL-236 |
| chr13-16722551-DEL-70 | chr13-16722552-DEL-70 |
| chr13-16725205-DEL-827 | chr13-16725207-DEL-828 |
| chr13-16727700-DEL-374 | chr13-16727700-DEL-374 |
| chr13-16746220-DEL-58 | chr13-16746207-DEL-72 |
| chr13-16752651-DEL-62 | chr13-16752644-DEL-70 |
| chr13-16772930-DEL-9676 | chr13-16772931-DEL-9676 |
| chr13-16775907-DEL-2577 | chr13-16775908-DEL-2577 |
| chr13-16782335-DEL-208 | chr13-16782336-DEL-208 |
| chr13-16792208-DEL-384 | chr13-16792209-DEL-384 |

| | |
|---|---|
| chr13-16902412-DEL-2655 | chr13-16902407-DEL-2656 |
| chr13-16912200-DEL-86 | chr13-16912235-DEL-63 |
| chr13-16976148-DEL-1840 | chr13-16976149-DEL-1840 |
| chr13-16979360-DEL-139 | chr13-16979361-DEL-139 |
| chr13-16991182-DEL-52 | chr13-16991185-DEL-50 |
| chr13-17054108-DEL-8774 | chr13-17054109-DEL-8774 |
| chr13-17114562-DEL-6274 | chr13-17114563-DEL-6274 |
| chr13-17159333-DEL-80 | chr13-17159306-DEL-108 |
| chr13-17177984-DEL-159 | chr13-17177985-DEL-159 |
| chr13-17230155-DEL-1140 | chr13-17230156-DEL-1140 |
| chr13-17237111-DEL-456 | chr13-17237112-DEL-456 |
| chr13-17271394-DEL-249 | chr13-17271395-DEL-249 |
| chr13-17291825-DEL-1504 | chr13-17291826-DEL-1504 |
| chr13-17322106-DEL-6404 | chr13-17322092-DEL-6386 |
| chr13-17344657-DEL-1555 | chr13-17344658-DEL-1555 |
| chr13-17374440-DEL-95 | chr13-17374411-DEL-115 |
| chr13-17375115-DEL-3941 | chr13-17375112-DEL-3938 |
| chr13-17435727-DEL-280 | chr13-17435727-DEL-282 |
| chr13-17453147-DEL-1670 | chr13-17453148-DEL-1670 |
| chr13-17468303-DEL-464 | chr13-17468304-DEL-464 |
| chr13-17473508-DEL-144 | chr13-17473509-DEL-136 |
| chr13-17496602-DEL-1033 | chr13-17496603-DEL-1033 |
| chr13-17499660-DEL-241 | chr13-17499661-DEL-241 |
| chr13-17503359-DEL-323 | chr13-17503360-DEL-323 |
| chr13-17503822-DEL-6214 | chr13-17503823-DEL-6214 |
| chr13-17523147-DEL-1539 | chr13-17523148-DEL-1539 |
| chr13-17530657-DEL-350 | chr13-17530658-DEL-350 |
| chr13-17533179-DEL-6221 | chr13-17533180-DEL-6221 |
| chr13-17541933-DEL-69 | chr13-17541910-DEL-93 |
| chr13-17542538-DEL-499 | chr13-17542539-DEL-499 |
| chr13-17557280-DEL-55 | chr13-17557281-DEL-55 |
| chr13-17558082-DEL-226 | chr13-17558083-DEL-226 |
| chr13-17603103-DEL-72 | chr13-17603110-DEL-72 |
| chr13-17615359-DEL-5299 | chr13-17615360-DEL-5299 |
| chr13-17623295-DEL-231 | chr13-17623278-DEL-249 |
| chr13-17626699-DEL-80 | chr13-17626696-DEL-80 |
| chr13-17632412-DEL-1236 | chr13-17632413-DEL-1236 |
| chr13-17636648-DEL-3913 | chr13-17636649-DEL-3913 |
| chr13-17647879-DEL-6562 | chr13-17647880-DEL-6561 |
| chr13-17648316-DEL-443 | chr13-17648317-DEL-443 |
| chr13-17672202-DEL-79 | chr13-17672202-DEL-77 |
| chr13-17674509-DEL-239 | chr13-17674510-DEL-238 |
| chr13-17688149-DEL-299 | chr13-17688150-DEL-299 |
| chr13-17701612-DEL-2906 | chr13-17701613-DEL-2906 |
| chr13-17726841-DEL-52 | chr13-17726838-DEL-56 |
| chr13-17733637-DEL-257 | chr13-17733638-DEL-257 |
| chr13-17816660-DEL-71 | chr13-17816662-DEL-70 |
| chr13-17828035-DEL-571 | chr13-17828030-DEL-572 |
| chr13-17853299-DEL-677 | chr13-17853300-DEL-677 |
| chr13-17860286-DEL-639 | chr13-17860287-DEL-639 |
| chr13-17906486-DEL-237 | chr13-17906487-DEL-237 |
| chr13-17933525-DEL-659 | chr13-17933526-DEL-660 |
| chr13-17946143-DEL-221 | chr13-17946144-DEL-221 |
| chr13-17962854-DEL-3096 | chr13-17962855-DEL-3096 |
| chr13-17966138-DEL-1689 | chr13-17966139-DEL-1689 |
| chr13-17984165-DEL-117 | chr13-17984166-DEL-117 |
| chr13-17985838-DEL-150 | chr13-17985839-DEL-150 |
| chr13-17994320-DEL-212 | chr13-17994321-DEL-212 |
| chr13-18005153-DEL-182 | chr13-18005154-DEL-182 |
| chr13-18008959-DEL-1229 | chr13-18008943-DEL-1249 |
| chr13-18010876-DEL-53 | chr13-18010872-DEL-73 |
| chr13-18022113-DEL-621 | chr13-18022114-DEL-621 |
| chr13-18025470-DEL-55 | chr13-18025470-DEL-55 |
| chr13-18038919-DEL-147 | chr13-18038919-DEL-147 |
| chr13-18091806-DEL-409 | chr13-18091807-DEL-409 |
| chr13-18106665-DEL-6402 | chr13-18106666-DEL-6402 |
| chr13-18133206-DEL-721 | chr13-18133207-DEL-721 |
| chr13-18138939-DEL-1832 | chr13-18138940-DEL-1832 |
| chr13-18144224-DEL-325 | chr13-18144225-DEL-325 |
| chr13-18168275-DEL-407 | chr13-18168276-DEL-407 |
| chr13-18176840-DEL-6387 | chr13-18176841-DEL-6387 |
| chr13-18204889-DEL-1333 | chr13-18204884-DEL-1332 |
| chr13-18216858-DEL-190 | chr13-18216864-DEL-195 |
| chr13-18257690-DEL-315 | chr13-18257691-DEL-315 |
| chr13-18268226-DEL-195 | chr13-18268227-DEL-195 |
| chr13-18333216-DEL-152 | chr13-18333217-DEL-148 |
| chr13-18389833-DEL-51 | chr13-18389834-DEL-51 |
| chr13-18462936-DEL-2264 | chr13-18462932-DEL-2263 |
| chr13-18474878-DEL-67 | chr13-18474854-DEL-51 |
| chr13-18476106-DEL-57 | chr13-18476107-DEL-57 |

| | |
|---|---|
| chr13-18496878-DEL-77 | chr13-18496879-DEL-77 |
| chr13-18613707-DEL-67 | chr13-18613708-DEL-67 |
| chr13-18632724-DEL-209 | chr13-18632725-DEL-209 |
| chr13-18642015-DEL-401 | chr13-18642016-DEL-401 |
| chr13-18676954-DEL-5817 | chr13-18676955-DEL-5817 |
| chr13-18678569-DEL-151 | chr13-18678570-DEL-151 |
| chr13-18694072-DEL-288 | chr13-18694073-DEL-288 |
| chr13-18711370-DEL-225 | chr13-18711371-DEL-225 |
| chr13-18726425-DEL-223 | chr13-18726426-DEL-223 |
| chr13-18738324-DEL-213 | chr13-18738325-DEL-213 |
| chr13-18741760-DEL-900 | chr13-18741761-DEL-900 |
| chr13-18768592-DEL-66 | chr13-18768614-DEL-51 |
| chr13-18782849-DEL-141 | chr13-18782841-DEL-160 |
| chr13-18823405-DEL-6425 | chr13-18823406-DEL-6425 |
| chr13-18837000-DEL-220 | chr13-18837001-DEL-220 |
| chr13-18840741-DEL-148 | chr13-18840742-DEL-148 |
| chr13-18857851-DEL-104 | chr13-18857899-DEL-98 |
| chr13-18860885-DEL-52 | chr13-18860910-DEL-76 |
| chr13-18861926-DEL-1209 | chr13-18861927-DEL-1209 |
| chr13-18866054-DEL-176 | chr13-18866056-DEL-177 |
| chr13-18959349-DEL-168 | chr13-18959350-DEL-126 |
| chr13-18960352-DEL-583 | chr13-18960353-DEL-583 |
| chr13-19009602-DEL-1522 | chr13-19009603-DEL-1522 |
| chr13-19016877-DEL-1530 | chr13-19016878-DEL-1530 |
| chr13-19018518-DEL-1316 | chr13-19018519-DEL-1316 |
| chr13-19057794-DEL-377 | chr13-19057795-DEL-377 |
| chr13-19061104-DEL-299 | chr13-19061105-DEL-299 |
| chr13-19068883-DEL-1895 | chr13-19068883-DEL-1895 |
| chr13-19078399-DEL-356 | chr13-19078400-DEL-356 |
| chr13-19082786-DEL-181 | chr13-19082787-DEL-181 |
| chr13-19087660-DEL-356 | chr13-19087660-DEL-355 |
| chr13-19115517-DEL-8738 | chr13-19115513-DEL-8737 |
| chr13-19142146-DEL-6270 | chr13-19142147-DEL-6270 |
| chr13-19160386-DEL-320 | chr13-19160387-DEL-320 |
| chr13-19166329-DEL-144 | chr13-19166330-DEL-144 |
| chr13-19173477-DEL-106 | chr13-19173478-DEL-106 |
| chr13-19178035-DEL-7213 | chr13-19178036-DEL-7213 |
| chr13-19198153-DEL-254 | chr13-19198154-DEL-254 |
| chr13-19243431-DEL-6505 | chr13-19243432-DEL-6505 |
| chr13-19254073-DEL-166 | chr13-19254074-DEL-166 |
| chr13-19267095-DEL-290 | chr13-19267096-DEL-290 |
| chr13-19299169-DEL-363 | chr13-19299170-DEL-363 |
| chr13-19308682-DEL-146 | chr13-19308723-DEL-102 |
| chr13-19326311-DEL-177 | chr13-19326312-DEL-177 |
| chr13-19382421-DEL-835 | chr13-19382424-DEL-832 |
| chr13-19412999-DEL-91 | chr13-19413000-DEL-91 |
| chr13-19430730-DEL-1181 | chr13-19430731-DEL-1181 |
| chr13-19479490-DEL-378 | chr13-19479491-DEL-378 |
| chr13-19499185-DEL-7646 | chr13-19499186-DEL-7646 |
| chr13-19514135-DEL-120 | chr13-19514136-DEL-120 |
| chr13-19546115-DEL-89 | chr13-19546131-DEL-83 |
| chr13-19547375-DEL-1059 | chr13-19547372-DEL-1065 |
| chr13-19548054-DEL-195 | chr13-19548055-DEL-195 |
| chr13-19620873-DEL-72 | chr13-19620874-DEL-72 |
| chr13-19621484-DEL-68 | chr13-19621485-DEL-66 |
| chr13-19622879-DEL-7461 | chr13-19622880-DEL-7461 |
| chr13-19623106-DEL-238 | chr13-19623107-DEL-238 |
| chr13-19627858-DEL-525 | chr13-19627859-DEL-525 |
| chr13-19659654-DEL-130 | chr13-19659655-DEL-127 |
| chr13-19670752-DEL-6209 | chr13-19670753-DEL-6209 |
| chr13-19679281-DEL-994 | chr13-19679282-DEL-994 |
| chr13-19692246-DEL-1487 | chr13-19692247-DEL-1487 |
| chr13-19703698-DEL-66 | chr13-19703694-DEL-74 |
| chr13-19727274-DEL-234 | chr13-19727275-DEL-234 |
| chr13-19769727-DEL-344 | chr13-19769728-DEL-344 |
| chr13-19794774-DEL-499 | chr13-19794761-DEL-494 |
| chr13-19820248-DEL-475 | chr13-19820249-DEL-475 |
| chr13-19822048-DEL-64 | chr13-19822030-DEL-53 |
| chr13-19830752-DEL-55 | chr13-19830753-DEL-55 |
| chr13-19831050-DEL-1555 | chr13-19831051-DEL-1555 |
| chr13-19834587-DEL-66 | chr13-19834588-DEL-54 |
| chr13-19848360-DEL-202 | chr13-19848324-DEL-203 |
| chr13-19854201-DEL-59 | chr13-19854202-DEL-53 |
| chr13-19870895-DEL-146 | chr13-19870896-DEL-146 |
| chr13-19882426-DEL-76 | chr13-19882427-DEL-64 |
| chr13-19889766-DEL-1458 | chr13-19889767-DEL-1458 |
| chr13-19931809-DEL-144 | chr13-19931834-DEL-156 |
| chr13-19937034-DEL-1370 | chr13-19937035-DEL-1370 |
| chr13-19949254-DEL-86 | chr13-19949269-DEL-78 |
| chr13-19959011-DEL-54 | chr13-19958994-DEL-64 |

| | |
|---|---|
| chr13-19962276-DEL-87 | chr13-19962277-DEL-66 |
| chr13-19968030-DEL-949 | chr13-19968031-DEL-949 |
| chr13-19976566-DEL-170 | chr13-19976529-DEL-175 |
| chr13-20017671-DEL-66 | chr13-20017670-DEL-68 |
| chr13-20028197-DEL-207 | chr13-20028198-DEL-207 |
| chr13-20040344-DEL-214 | chr13-20040337-DEL-215 |
| chr13-20045039-DEL-1687 | chr13-20045040-DEL-1687 |
| chr13-20051701-DEL-4743 | chr13-20051702-DEL-4743 |
| chr13-20065031-DEL-72 | chr13-20064984-DEL-74 |
| chr13-20102516-DEL-75 | chr13-20102517-DEL-75 |
| chr13-20103426-DEL-716 | chr13-20103427-DEL-716 |
| chr13-20114399-DEL-840 | chr13-20114400-DEL-840 |
| chr13-20122634-DEL-2055 | chr13-20122635-DEL-2055 |
| chr13-20126310-DEL-545 | chr13-20126311-DEL-544 |
| chr13-20133317-DEL-180 | chr13-20133318-DEL-170 |
| chr13-20138696-DEL-6270 | chr13-20138697-DEL-6270 |
| chr13-20162310-DEL-150 | chr13-20162311-DEL-150 |
| chr13-20173433-DEL-53 | chr13-20173435-DEL-73 |
| chr13-20178819-DEL-172 | chr13-20178820-DEL-172 |
| chr13-20217537-DEL-199 | chr13-20217538-DEL-199 |
| chr13-20235417-DEL-734 | chr13-20235427-DEL-705 |
| chr13-20273273-DEL-55 | chr13-20273243-DEL-56 |
| chr13-20292917-DEL-90 | chr13-20292918-DEL-90 |
| chr13-20319229-DEL-430 | chr13-20319230-DEL-430 |
| chr13-20334605-DEL-114 | chr13-20334606-DEL-114 |
| chr13-20345858-DEL-2545 | chr13-20345859-DEL-2545 |
| chr13-20353610-DEL-103 | chr13-20353610-DEL-133 |
| chr13-20360533-DEL-445 | chr13-20360534-DEL-445 |
| chr13-20389340-DEL-57 | chr13-20389341-DEL-57 |
| chr13-20407822-DEL-99 | chr13-20407839-DEL-65 |
| chr13-20418374-DEL-106 | chr13-20418375-DEL-106 |
| chr13-20420813-DEL-134 | chr13-20420803-DEL-130 |
| chr13-20434812-DEL-258 | chr13-20434811-DEL-260 |
| chr13-20459844-DEL-92 | chr13-20459845-DEL-92 |
| chr13-20501800-DEL-91 | chr13-20501801-DEL-91 |
| chr13-20530013-DEL-323 | chr13-20530014-DEL-323 |
| chr13-20531724-DEL-146 | chr13-20531685-DEL-162 |
| chr13-20552978-DEL-128 | chr13-20552979-DEL-128 |
| chr13-20557713-DEL-203 | chr13-20557714-DEL-203 |
| chr13-20572120-DEL-125 | chr13-20572119-DEL-125 |
| chr13-20618156-DEL-60 | chr13-20618157-DEL-60 |
| chr13-20669647-DEL-1508 | chr13-20669648-DEL-1508 |
| chr13-20682144-DEL-96 | chr13-20682145-DEL-96 |
| chr13-20714770-DEL-6390 | chr13-20714771-DEL-6390 |
| chr13-20779550-DEL-362 | chr13-20779551-DEL-362 |
| chr13-20801155-DEL-91 | chr13-20801159-DEL-83 |
| chr13-20812394-DEL-74 | chr13-20812395-DEL-74 |
| chr13-20813870-DEL-86 | chr13-20813871-DEL-86 |
| chr13-20862694-DEL-60 | chr13-20862706-DEL-58 |
| chr13-20877618-DEL-90 | chr13-20877618-DEL-86 |
| chr13-20881780-DEL-203 | chr13-20881781-DEL-203 |
| chr13-20887614-DEL-62 | chr13-20887615-DEL-56 |
| chr13-20896818-DEL-6379 | chr13-20896819-DEL-6379 |
| chr13-20906017-DEL-81 | chr13-20906025-DEL-73 |
| chr13-20914523-DEL-82 | chr13-20914524-DEL-82 |
| chr13-20951390-DEL-96 | chr13-20951395-DEL-92 |
| chr13-20955256-DEL-123 | chr13-20955257-DEL-123 |
| chr13-20987202-DEL-77 | chr13-20987192-DEL-75 |
| chr13-21007010-DEL-374 | chr13-21007017-DEL-364 |
| chr13-21009001-DEL-65 | chr13-21009002-DEL-65 |
| chr13-21010745-DEL-113 | chr13-21010734-DEL-125 |
| chr13-21012210-DEL-503 | chr13-21012211-DEL-503 |
| chr13-21016850-DEL-72 | chr13-21016863-DEL-70 |
| chr13-21027434-DEL-2089 | chr13-21027435-DEL-2089 |
| chr13-21114427-DEL-180 | chr13-21114428-DEL-180 |
| chr13-21144075-DEL-133 | chr13-21144086-DEL-84 |
| chr13-21159086-DEL-122 | chr13-21159097-DEL-108 |
| chr13-21183622-DEL-73 | chr13-21183623-DEL-73 |
| chr13-21185775-DEL-110 | chr13-21185776-DEL-110 |
| chr13-21187362-DEL-66 | chr13-21187363-DEL-66 |
| chr13-21226183-DEL-408 | chr13-21226184-DEL-406 |
| chr13-21245273-DEL-244 | chr13-21245274-DEL-244 |
| chr13-21259982-DEL-699 | chr13-21259983-DEL-699 |
| chr13-21291430-DEL-444 | chr13-21291431-DEL-444 |
| chr13-21301491-DEL-125 | chr13-21301498-DEL-124 |
| chr13-21303101-DEL-237 | chr13-21303102-DEL-237 |
| chr13-21322122-DEL-935 | chr13-21322123-DEL-935 |
| chr13-21361653-DEL-71 | chr13-21361656-DEL-72 |
| chr13-21377391-DEL-134 | chr13-21377392-DEL-134 |
| chr13-21377934-DEL-208 | chr13-21377935-DEL-208 |

| | |
|---|---|
| chr13-21392641-DEL-683 | chr13-21392642-DEL-683 |
| chr13-21412646-DEL-441 | chr13-21412647-DEL-441 |
| chr13-21439384-DEL-433 | chr13-21439385-DEL-433 |
| chr13-21440275-DEL-940 | chr13-21440276-DEL-940 |
| chr13-21443754-DEL-71 | chr13-21443765-DEL-65 |
| chr13-21487187-DEL-197 | chr13-21487188-DEL-197 |
| chr13-21513956-DEL-214 | chr13-21513957-DEL-214 |
| chr13-21520395-DEL-68 | chr13-21520365-DEL-72 |
| chr13-21557659-DEL-1803 | chr13-21557660-DEL-1803 |
| chr13-21575892-DEL-172 | chr13-21575893-DEL-172 |
| chr13-21627332-DEL-59 | chr13-21627333-DEL-59 |
| chr13-21641678-DEL-56 | chr13-21641679-DEL-56 |
| chr13-21658468-DEL-458 | chr13-21658469-DEL-458 |
| chr13-21661755-DEL-152 | chr13-21661756-DEL-152 |
| chr13-21671701-DEL-58 | chr13-21671702-DEL-58 |
| chr13-21679386-DEL-1937 | chr13-21679379-DEL-1938 |
| chr13-21685081-DEL-539 | chr13-21685082-DEL-539 |
| chr13-21698189-DEL-1285 | chr13-21698190-DEL-1285 |
| chr13-21722767-DEL-60 | chr13-21722768-DEL-60 |
| chr13-21747872-DEL-88 | chr13-21747915-DEL-54 |
| chr13-21759308-DEL-64 | chr13-21759309-DEL-64 |
| chr13-21783162-DEL-1929 | chr13-21783162-DEL-1929 |
| chr13-21786618-DEL-392 | chr13-21786618-DEL-393 |
| chr13-21790355-DEL-59 | chr13-21790356-DEL-59 |
| chr13-21790512-DEL-2081 | chr13-21790513-DEL-2081 |
| chr13-21792239-DEL-128 | chr13-21792240-DEL-126 |
| chr13-21793816-DEL-64 | chr13-21793817-DEL-63 |
| chr13-21794863-DEL-69 | chr13-21794864-DEL-69 |
| chr13-21801493-DEL-81 | chr13-21801496-DEL-80 |
| chr13-21813981-DEL-7098 | chr13-21813982-DEL-7098 |
| chr13-21863709-DEL-306 | chr13-21863710-DEL-306 |
| chr13-21870837-DEL-3800 | chr13-21870838-DEL-3801 |
| chr13-21883699-DEL-64 | chr13-21883700-DEL-64 |
| chr13-21893085-DEL-183 | chr13-21893086-DEL-183 |
| chr13-21906873-DEL-2118 | chr13-21906874-DEL-2118 |
| chr13-21912599-DEL-511 | chr13-21912600-DEL-511 |
| chr13-21919799-DEL-1269 | chr13-21919800-DEL-1269 |
| chr13-21946960-DEL-5400 | chr13-21946961-DEL-5400 |
| chr13-21957329-DEL-1714 | chr13-21957330-DEL-1714 |
| chr13-21974436-DEL-215 | chr13-21974437-DEL-215 |
| chr13-21985394-DEL-78 | chr13-21985412-DEL-60 |
| chr13-21987842-DEL-2268 | chr13-21987841-DEL-2268 |
| chr13-22023451-DEL-539 | chr13-22023452-DEL-539 |
| chr13-22033099-DEL-902 | chr13-22033099-DEL-902 |
| chr13-22081860-DEL-59 | chr13-22081856-DEL-60 |
| chr13-22144965-DEL-85 | chr13-22144966-DEL-85 |
| chr13-22195225-DEL-78 | chr13-22195232-DEL-72 |
| chr13-22200068-DEL-195 | chr13-22200069-DEL-195 |
| chr13-22216559-DEL-185 | chr13-22216560-DEL-185 |
| chr13-22246490-DEL-927 | chr13-22246481-DEL-940 |
| chr13-22258176-DEL-553 | chr13-22258177-DEL-553 |
| chr13-22267415-DEL-146 | chr13-22267420-DEL-128 |
| chr13-22271687-DEL-84 | chr13-22271688-DEL-84 |
| chr13-22283263-DEL-542 | chr13-22283264-DEL-542 |
| chr13-22294787-DEL-16901 | chr13-22294788-DEL-16901 |
| chr13-22296757-DEL-130 | chr13-22296757-DEL-130 |
| chr13-22298936-DEL-7428 | chr13-22298933-DEL-7431 |
| chr13-22376323-DEL-365 | chr13-22376324-DEL-365 |
| chr13-22387807-DEL-891 | chr13-22387808-DEL-891 |
| chr13-22389085-DEL-6351 | chr13-22389086-DEL-6351 |
| chr13-22412628-DEL-4337 | chr13-22412629-DEL-4337 |
| chr13-22432091-DEL-415 | chr13-22432092-DEL-415 |
| chr13-22444002-DEL-867 | chr13-22444003-DEL-865 |
| chr13-22516446-DEL-3036 | chr13-22516447-DEL-3036 |
| chr13-22525487-DEL-889 | chr13-22525487-DEL-889 |
| chr13-22527730-DEL-112 | chr13-22527768-DEL-67 |
| chr13-22550428-DEL-307 | chr13-22550429-DEL-307 |
| chr13-22561299-DEL-1082 | chr13-22561300-DEL-1082 |
| chr13-22607128-DEL-6910 | chr13-22607129-DEL-6910 |
| chr13-22641692-DEL-204 | chr13-22641701-DEL-204 |
| chr13-22667305-DEL-70 | chr13-22667307-DEL-79 |
| chr13-22723043-DEL-95 | chr13-22723044-DEL-95 |
| chr13-22727269-DEL-7081 | chr13-22727270-DEL-7081 |
| chr13-22734024-DEL-208 | chr13-22734025-DEL-208 |
| chr13-22757169-DEL-555 | chr13-22757170-DEL-555 |
| chr13-22782459-DEL-6210 | chr13-22782460-DEL-6210 |
| chr13-22808310-DEL-1434 | chr13-22808311-DEL-1434 |
| chr13-22839639-DEL-605 | chr13-22839640-DEL-605 |
| chr13-22846479-DEL-198 | chr13-22846480-DEL-198 |
| chr13-22873156-DEL-347 | chr13-22873157-DEL-347 |

| | |
|---|---|
| chr13-22885304-DEL-149 | chr13-22885304-DEL-149 |
| chr13-22887223-DEL-3625 | chr13-22887221-DEL-3626 |
| chr13-22957838-DEL-62 | chr13-22957837-DEL-64 |
| chr13-23072677-DEL-234 | chr13-23072678-DEL-234 |
| chr13-23147712-DEL-108 | chr13-23147761-DEL-74 |
| chr13-23225775-DEL-58 | chr13-23225776-DEL-58 |
| chr13-23340398-DEL-52 | chr13-23340399-DEL-52 |
| chr13-23341812-DEL-428 | chr13-23341813-DEL-428 |
| chr13-23354324-DEL-7077 | chr13-23354325-DEL-7077 |
| chr13-23374652-DEL-214 | chr13-23374653-DEL-214 |
| chr13-23462230-DEL-337 | chr13-23462231-DEL-337 |
| chr13-23508693-DEL-5493 | chr13-23508694-DEL-5493 |
| chr13-23518977-DEL-486 | chr13-23518977-DEL-484 |
| chr13-23529137-DEL-391 | chr13-23529094-DEL-391 |
| chr13-23542839-DEL-1647 | chr13-23542840-DEL-1648 |
| chr13-23598305-DEL-185 | chr13-23598306-DEL-185 |
| chr13-23673585-DEL-27786 | chr13-23673579-DEL-27792 |
| chr13-23732875-DEL-89 | chr13-23732876-DEL-89 |
| chr13-23795054-DEL-328 | chr13-23795055-DEL-328 |
| chr13-23800013-DEL-215 | chr13-23800017-DEL-214 |
| chr13-23822931-DEL-7182 | chr13-23822934-DEL-7181 |
| chr13-23835932-DEL-160 | chr13-23835933-DEL-160 |
| chr13-23852764-DEL-194 | chr13-23852755-DEL-195 |
| chr13-23897014-DEL-94 | chr13-23897017-DEL-92 |
| chr13-23914737-DEL-58 | chr13-23914738-DEL-58 |
| chr13-23976299-DEL-91 | chr13-23976300-DEL-91 |
| chr13-24011196-DEL-95 | chr13-24011197-DEL-95 |
| chr13-24033176-DEL-115 | chr13-24033177-DEL-115 |
| chr13-24040328-DEL-69 | chr13-24040329-DEL-69 |
| chr13-24063389-DEL-126 | chr13-24063390-DEL-126 |
| chr13-24105970-DEL-178 | chr13-24105971-DEL-178 |
| chr13-24125793-DEL-1249 | chr13-24125794-DEL-1249 |
| chr13-24171072-DEL-187 | chr13-24171073-DEL-187 |
| chr13-24212304-DEL-6610 | chr13-24212305-DEL-6610 |
| chr13-24288868-DEL-211 | chr13-24288869-DEL-211 |
| chr13-24313945-DEL-843 | chr13-24313946-DEL-841 |
| chr13-24327817-DEL-81 | chr13-24327818-DEL-81 |
| chr13-24336092-DEL-66 | chr13-24336069-DEL-68 |
| chr13-24351668-DEL-73 | chr13-24351676-DEL-74 |
| chr13-24384370-DEL-561 | chr13-24384371-DEL-561 |
| chr13-24387533-DEL-81 | chr13-24387532-DEL-82 |
| chr13-24410660-DEL-3658 | chr13-24410661-DEL-3658 |
| chr13-24436751-DEL-63 | chr13-24436752-DEL-64 |
| chr13-24484993-DEL-2842 | chr13-24484994-DEL-2842 |
| chr13-24502212-DEL-199 | chr13-24502213-DEL-199 |
| chr13-24507294-DEL-3001 | chr13-24507336-DEL-2959 |
| chr13-24519765-DEL-94 | chr13-24519765-DEL-94 |
| chr13-24521219-DEL-164 | chr13-24521220-DEL-164 |
| chr13-24527296-DEL-204 | chr13-24527297-DEL-204 |
| chr13-24546531-DEL-70 | chr13-24546532-DEL-70 |
| chr13-24555765-DEL-68 | chr13-24555766-DEL-68 |
| chr13-24571791-DEL-3057 | chr13-24571792-DEL-3057 |
| chr13-24575121-DEL-387 | chr13-24575145-DEL-364 |
| chr13-24575665-DEL-58 | chr13-24575668-DEL-56 |
| chr13-24615943-DEL-63 | chr13-24615949-DEL-87 |
| chr13-24625810-DEL-66 | chr13-24625811-DEL-66 |
| chr13-24645607-DEL-85 | chr13-24645608-DEL-85 |
| chr13-24663996-DEL-57 | chr13-24663997-DEL-57 |
| chr13-24667353-DEL-60 | chr13-24667357-DEL-55 |
| chr13-24687554-DEL-58 | chr13-24687568-DEL-60 |
| chr13-24717878-DEL-68 | chr13-24717878-DEL-68 |
| chr13-24725577-DEL-234 | chr13-24725553-DEL-234 |
| chr13-24737921-DEL-575 | chr13-24737924-DEL-573 |
| chr13-24761724-DEL-3320 | chr13-24761725-DEL-3320 |
| chr13-24809530-DEL-64 | chr13-24809531-DEL-64 |
| chr13-24860340-DEL-163 | chr13-24860341-DEL-163 |
| chr13-24865591-DEL-336 | chr13-24865592-DEL-336 |
| chr13-24875435-DEL-209 | chr13-24875433-DEL-210 |
| chr13-24903559-DEL-199 | chr13-24903551-DEL-208 |
| chr13-24910067-DEL-225 | chr13-24910068-DEL-237 |
| chr13-24927838-DEL-111 | chr13-24927863-DEL-91 |
| chr13-24952596-DEL-66 | chr13-24952597-DEL-66 |
| chr13-24955810-DEL-66 | chr13-24955811-DEL-66 |
| chr13-24968173-DEL-82 | chr13-24968174-DEL-82 |
| chr13-25000070-DEL-202 | chr13-25000071-DEL-202 |
| chr13-25011205-DEL-52 | chr13-25011206-DEL-52 |
| chr13-25013233-DEL-166 | chr13-25013234-DEL-166 |
| chr13-25031317-DEL-176 | chr13-25031318-DEL-176 |
| chr13-25038314-DEL-60 | chr13-25038315-DEL-60 |
| chr13-25042956-DEL-64 | chr13-25042957-DEL-64 |

| | |
|---|---|
| chr13-25043821-DEL-325 | chr13-25043822-DEL-325 |
| chr13-25062595-DEL-214 | chr13-25062596-DEL-214 |
| chr13-25068667-DEL-84 | chr13-25068668-DEL-84 |
| chr13-25080265-DEL-179 | chr13-25080266-DEL-179 |
| chr13-25084846-DEL-57 | chr13-25084847-DEL-57 |
| chr13-25097554-DEL-102 | chr13-25097551-DEL-106 |
| chr13-25153895-DEL-763 | chr13-25153896-DEL-763 |
| chr13-25191372-DEL-8326 | chr13-25191373-DEL-8326 |
| chr13-25213595-DEL-6600 | chr13-25213605-DEL-6598 |
| chr13-25230365-DEL-780 | chr13-25230362-DEL-780 |
| chr13-25234392-DEL-192 | chr13-25234426-DEL-177 |
| chr13-25248138-DEL-358 | chr13-25248139-DEL-358 |
| chr13-25273193-DEL-161 | chr13-25273192-DEL-160 |
| chr13-25285346-DEL-62 | chr13-25285340-DEL-68 |
| chr13-25294686-DEL-61 | chr13-25294687-DEL-61 |
| chr13-25320613-DEL-64 | chr13-25320614-DEL-64 |
| chr13-25349628-DEL-206 | chr13-25349630-DEL-188 |
| chr13-25409769-DEL-199 | chr13-25409770-DEL-199 |
| chr13-25419589-DEL-6592 | chr13-25419583-DEL-6599 |
| chr13-25433075-DEL-61 | chr13-25433076-DEL-61 |
| chr13-25454711-DEL-204 | chr13-25454712-DEL-201 |
| chr13-25486338-DEL-132 | chr13-25486340-DEL-140 |
| chr13-25501291-DEL-85 | chr13-25501292-DEL-85 |
| chr13-25526331-DEL-54 | chr13-25526333-DEL-55 |
| chr13-25534642-DEL-7285 | chr13-25534643-DEL-7285 |
| chr13-25565979-DEL-152 | chr13-25565966-DEL-199 |
| chr13-25663337-DEL-96 | chr13-25663338-DEL-96 |
| chr13-25665852-DEL-460 | chr13-25665853-DEL-460 |
| chr13-25672179-DEL-336 | chr13-25672180-DEL-336 |
| chr13-25684236-DEL-273 | chr13-25684238-DEL-274 |
| chr13-25704656-DEL-57 | chr13-25704655-DEL-58 |
| chr13-25711127-DEL-78 | chr13-25711128-DEL-78 |
| chr13-25718254-DEL-266 | chr13-25718259-DEL-265 |
| chr13-25720508-DEL-6676 | chr13-25720509-DEL-6676 |
| chr13-25761815-DEL-101 | chr13-25761816-DEL-101 |
| chr13-25786333-DEL-214 | chr13-25786334-DEL-214 |
| chr13-25788011-DEL-632 | chr13-25788012-DEL-632 |
| chr13-25822392-DEL-299 | chr13-25822393-DEL-299 |
| chr13-25835960-DEL-1164 | chr13-25835961-DEL-1164 |
| chr13-25839067-DEL-3398 | chr13-25839068-DEL-3398 |
| chr13-25850645-DEL-363 | chr13-25850646-DEL-363 |
| chr13-25891439-DEL-322 | chr13-25891439-DEL-323 |
| chr13-25898835-DEL-157 | chr13-25898836-DEL-157 |
| chr13-25906100-DEL-70 | chr13-25906101-DEL-70 |
| chr13-25914283-DEL-301 | chr13-25914284-DEL-301 |
| chr13-25939947-DEL-724 | chr13-25939982-DEL-691 |
| chr13-25949121-DEL-817 | chr13-25949122-DEL-817 |
| chr13-25954275-DEL-110 | chr13-25954256-DEL-118 |
| chr13-25955909-DEL-1027 | chr13-25955910-DEL-1027 |
| chr13-25970013-DEL-369 | chr13-25970013-DEL-362 |
| chr13-25971146-DEL-60 | chr13-25971139-DEL-68 |
| chr13-25988455-DEL-201 | chr13-25988456-DEL-201 |
| chr13-26007300-DEL-1547 | chr13-26007301-DEL-1547 |
| chr13-26048443-DEL-62 | chr13-26048461-DEL-77 |
| chr13-26054394-DEL-619 | chr13-26054395-DEL-619 |
| chr13-26061084-DEL-78 | chr13-26061085-DEL-78 |
| chr13-26085218-DEL-54 | chr13-26085219-DEL-54 |
| chr13-26093374-DEL-672 | chr13-26093375-DEL-672 |
| chr13-26123851-DEL-67 | chr13-26123852-DEL-66 |
| chr13-26133467-DEL-78 | chr13-26133468-DEL-78 |
| chr13-26159352-DEL-1015 | chr13-26159356-DEL-1009 |
| chr13-26171533-DEL-83 | chr13-26171534-DEL-83 |
| chr13-26175308-DEL-52 | chr13-26175309-DEL-52 |
| chr13-26181408-DEL-550 | chr13-26181409-DEL-550 |
| chr13-26211267-DEL-54 | chr13-26211275-DEL-52 |
| chr13-26232743-DEL-6401 | chr13-26232744-DEL-6401 |
| chr13-26243697-DEL-118 | chr13-26243697-DEL-142 |
| chr13-26251035-DEL-111 | chr13-26251036-DEL-111 |
| chr13-26285155-DEL-349 | chr13-26285156-DEL-349 |
| chr13-26308696-DEL-111 | chr13-26308697-DEL-111 |
| chr13-26319243-DEL-626 | chr13-26319244-DEL-626 |
| chr13-26328349-DEL-164 | chr13-26328349-DEL-164 |
| chr13-26332271-DEL-640 | chr13-26332257-DEL-654 |
| chr13-26359995-DEL-6250 | chr13-26359996-DEL-6250 |
| chr13-26423301-DEL-6684 | chr13-26423302-DEL-6684 |
| chr13-26468506-DEL-57 | chr13-26468506-DEL-56 |
| chr13-26479017-DEL-52 | chr13-26479018-DEL-52 |
| chr13-26514711-DEL-63 | chr13-26514711-DEL-63 |
| chr13-26542914-DEL-3583 | chr13-26542906-DEL-3581 |
| chr13-26551104-DEL-7135 | chr13-26551099-DEL-7135 |

| | |
|---|---|
| chr13-26568625-DEL-70 | chr13-26568650-DEL-63 |
| chr13-26574854-DEL-240 | chr13-26574847-DEL-243 |
| chr13-26594992-DEL-200 | chr13-26594990-DEL-200 |
| chr13-26615614-DEL-116 | chr13-26615612-DEL-115 |
| chr13-26624567-DEL-7663 | chr13-26624568-DEL-7663 |
| chr13-26662447-DEL-210 | chr13-26662443-DEL-210 |
| chr13-26669806-DEL-215 | chr13-26669807-DEL-215 |
| chr13-26675205-DEL-149 | chr13-26675206-DEL-149 |
| chr13-26676494-DEL-720 | chr13-26676490-DEL-721 |
| chr13-26690612-DEL-63 | chr13-26690586-DEL-72 |
| chr13-26712220-DEL-372 | chr13-26712221-DEL-372 |
| chr13-26784851-DEL-258 | chr13-26784852-DEL-258 |
| chr13-26810141-DEL-327 | chr13-26810142-DEL-327 |
| chr13-26824538-DEL-84 | chr13-26824525-DEL-85 |
| chr13-26867885-DEL-3031 | chr13-26867886-DEL-3031 |
| chr13-26889081-DEL-158 | chr13-26889082-DEL-158 |
| chr13-26918874-DEL-85 | chr13-26918903-DEL-103 |
| chr13-26993544-DEL-238 | chr13-26993545-DEL-238 |
| chr13-26996628-DEL-400 | chr13-26996629-DEL-400 |
| chr13-27073466-DEL-55 | chr13-27073469-DEL-53 |
| chr13-27082951-DEL-424 | chr13-27082952-DEL-424 |
| chr13-27093725-DEL-452 | chr13-27093726-DEL-450 |
| chr13-27114666-DEL-152 | chr13-27114667-DEL-150 |
| chr13-27117437-DEL-108 | chr13-27117437-DEL-107 |
| chr13-27126241-DEL-252 | chr13-27126235-DEL-252 |
| chr13-27191797-DEL-1228 | chr13-27191798-DEL-1228 |
| chr13-27295581-DEL-963 | chr13-27295581-DEL-963 |
| chr13-27317783-DEL-65 | chr13-27317784-DEL-64 |
| chr13-27347726-DEL-5325 | chr13-27347727-DEL-5325 |
| chr13-27369454-DEL-2989 | chr13-27369454-DEL-2987 |
| chr13-27390205-DEL-60 | chr13-27390206-DEL-60 |
| chr13-27391484-DEL-603 | chr13-27391485-DEL-603 |
| chr13-27393421-DEL-52 | chr13-27393406-DEL-72 |
| chr13-27410880-DEL-3804 | chr13-27410881-DEL-3804 |
| chr13-27419468-DEL-212 | chr13-27419469-DEL-212 |
| chr13-27455353-DEL-759 | chr13-27455354-DEL-759 |
| chr13-27472050-DEL-2699 | chr13-27472051-DEL-2699 |
| chr13-27605441-DEL-80 | chr13-27605442-DEL-80 |
| chr13-27727697-DEL-2018 | chr13-27727698-DEL-2018 |
| chr13-27745930-DEL-53 | chr13-27745918-DEL-65 |
| chr13-27784708-DEL-89 | chr13-27784705-DEL-90 |
| chr13-27842053-DEL-80 | chr13-27842088-DEL-64 |
| chr13-27889904-DEL-2379 | chr13-27889905-DEL-2379 |
| chr13-27930996-DEL-134 | chr13-27931003-DEL-137 |
| chr13-27970763-DEL-4145 | chr13-27970764-DEL-4145 |
| chr13-28005353-DEL-72 | chr13-28005354-DEL-74 |
| chr13-28008076-DEL-6360 | chr13-28008077-DEL-6360 |
| chr13-28083460-DEL-480 | chr13-28083461-DEL-480 |
| chr13-28086580-DEL-91 | chr13-28086587-DEL-74 |
| chr13-28145109-DEL-258 | chr13-28145110-DEL-258 |
| chr13-28152506-DEL-394 | chr13-28152507-DEL-394 |
| chr13-28167259-DEL-214 | chr13-28167260-DEL-214 |
| chr13-28196249-DEL-153 | chr13-28196250-DEL-153 |
| chr13-28233522-DEL-6042 | chr13-28233523-DEL-6042 |
| chr13-28245352-DEL-54 | chr13-28245353-DEL-54 |
| chr13-28248705-DEL-417 | chr13-28248706-DEL-417 |
| chr13-28254694-DEL-579 | chr13-28254695-DEL-579 |
| chr13-28256828-DEL-67 | chr13-28256809-DEL-67 |
| chr13-28259641-DEL-391 | chr13-28259642-DEL-391 |
| chr13-28263474-DEL-6215 | chr13-28263475-DEL-6215 |
| chr13-28286920-DEL-52 | chr13-28286920-DEL-53 |
| chr13-28308397-DEL-7076 | chr13-28308398-DEL-7076 |
| chr13-28321111-DEL-167 | chr13-28321112-DEL-167 |
| chr13-28326282-DEL-67 | chr13-28326286-DEL-68 |
| chr13-28337735-DEL-348 | chr13-28337736-DEL-348 |
| chr13-28354618-DEL-119 | chr13-28354654-DEL-79 |
| chr13-28360611-DEL-90 | chr13-28360612-DEL-90 |
| chr13-28364370-DEL-436 | chr13-28364371-DEL-436 |
| chr13-28365312-DEL-90 | chr13-28365323-DEL-80 |
| chr13-28372364-DEL-68 | chr13-28372365-DEL-68 |
| chr13-28382430-DEL-2904 | chr13-28382431-DEL-2904 |
| chr13-28397266-DEL-3164 | chr13-28397267-DEL-3164 |
| chr13-28413204-DEL-137 | chr13-28413205-DEL-137 |
| chr13-28414423-DEL-374 | chr13-28414424-DEL-374 |
| chr13-28477029-DEL-191 | chr13-28477030-DEL-214 |
| chr13-28512967-DEL-756 | chr13-28512968-DEL-756 |
| chr13-28537714-DEL-154 | chr13-28537715-DEL-153 |
| chr13-28545285-DEL-51 | chr13-28545286-DEL-51 |
| chr13-28566774-DEL-219 | chr13-28566775-DEL-219 |
| chr13-28590991-DEL-84 | chr13-28590980-DEL-96 |

| | |
|---|---|
| chr13-28622041-DEL-1722 | chr13-28622042-DEL-1722 |
| chr13-28653304-DEL-6797 | chr13-28653305-DEL-6797 |
| chr13-28673309-DEL-1131 | chr13-28673310-DEL-1131 |
| chr13-28679482-DEL-3697 | chr13-28679483-DEL-3697 |
| chr13-28684346-DEL-98 | chr13-28684347-DEL-98 |
| chr13-28765143-DEL-855 | chr13-28765144-DEL-855 |
| chr13-28766766-DEL-211 | chr13-28766767-DEL-211 |
| chr13-28804665-DEL-3498 | chr13-28804666-DEL-3498 |
| chr13-28842136-DEL-60 | chr13-28842150-DEL-92 |
| chr13-28859562-DEL-1369 | chr13-28859563-DEL-1369 |
| chr13-28879313-DEL-163 | chr13-28879314-DEL-163 |
| chr13-28936214-DEL-64 | chr13-28936204-DEL-56 |
| chr13-28947656-DEL-174 | chr13-28947657-DEL-174 |
| chr13-28963371-DEL-87 | chr13-28963384-DEL-73 |
| chr13-28972789-DEL-221 | chr13-28972790-DEL-221 |
| chr13-28987038-DEL-977 | chr13-28987039-DEL-977 |
| chr13-28993805-DEL-119 | chr13-28993806-DEL-119 |
| chr13-29020096-DEL-211 | chr13-29020100-DEL-208 |
| chr13-29021470-DEL-52 | chr13-29021440-DEL-53 |
| chr13-29028246-DEL-192 | chr13-29028247-DEL-191 |
| chr13-29052640-DEL-52 | chr13-29052641-DEL-52 |
| chr13-29108806-DEL-58 | chr13-29108806-DEL-58 |
| chr13-29109547-DEL-60 | chr13-29109558-DEL-50 |
| chr13-29114904-DEL-64 | chr13-29114905-DEL-64 |
| chr13-29125615-DEL-193 | chr13-29125616-DEL-193 |
| chr13-29130001-DEL-231 | chr13-29130002-DEL-231 |
| chr13-29148236-DEL-80 | chr13-29148237-DEL-80 |
| chr13-29170910-DEL-106 | chr13-29170936-DEL-90 |
| chr13-29177136-DEL-117 | chr13-29177130-DEL-126 |
| chr13-29229043-DEL-159 | chr13-29229042-DEL-160 |
| chr13-29248794-DEL-8147 | chr13-29248795-DEL-8147 |
| chr13-29266692-DEL-60 | chr13-29266693-DEL-60 |
| chr13-29268612-DEL-83 | chr13-29268613-DEL-83 |
| chr13-29286592-DEL-68 | chr13-29286593-DEL-74 |
| chr13-29294034-DEL-91 | chr13-29294035-DEL-91 |
| chr13-29315267-DEL-1224 | chr13-29315268-DEL-1224 |
| chr13-29319408-DEL-52 | chr13-29319409-DEL-52 |
| chr13-29331001-DEL-265 | chr13-29331002-DEL-265 |
| chr13-29340176-DEL-62 | chr13-29340177-DEL-62 |
| chr13-29375146-DEL-52 | chr13-29375147-DEL-52 |
| chr13-29397743-DEL-1489 | chr13-29397744-DEL-1489 |
| chr13-29420874-DEL-64 | chr13-29420875-DEL-64 |
| chr13-29433019-DEL-6792 | chr13-29433020-DEL-6792 |
| chr13-29449154-DEL-603 | chr13-29449147-DEL-604 |
| chr13-29451553-DEL-1059 | chr13-29451539-DEL-1060 |
| chr13-29478295-DEL-138 | chr13-29478287-DEL-144 |
| chr13-29505712-DEL-88 | chr13-29505713-DEL-88 |
| chr13-29515132-DEL-262 | chr13-29515133-DEL-262 |
| chr13-29526261-DEL-2636 | chr13-29526262-DEL-2636 |
| chr13-29571690-DEL-215 | chr13-29571691-DEL-215 |
| chr13-29623265-DEL-2405 | chr13-29623266-DEL-2405 |
| chr13-29684789-DEL-54 | chr13-29684790-DEL-54 |
| chr13-29698816-DEL-549 | chr13-29698817-DEL-549 |
| chr13-29710186-DEL-106 | chr13-29710181-DEL-106 |
| chr13-29743862-DEL-100 | chr13-29743861-DEL-102 |
| chr13-29751322-DEL-205 | chr13-29751323-DEL-205 |
| chr13-29760219-DEL-102 | chr13-29760220-DEL-102 |
| chr13-29785803-DEL-598 | chr13-29785804-DEL-598 |
| chr13-29822244-DEL-55 | chr13-29822245-DEL-55 |
| chr13-29869054-DEL-404 | chr13-29869055-DEL-404 |
| chr13-29886780-DEL-148 | chr13-29886780-DEL-148 |
| chr13-29888847-DEL-2123 | chr13-29888848-DEL-2123 |
| chr13-29961584-DEL-110 | chr13-29961604-DEL-84 |
| chr13-29964396-DEL-323 | chr13-29964397-DEL-323 |
| chr13-30024240-DEL-56 | chr13-30024241-DEL-56 |
| chr13-30041834-DEL-156 | chr13-30041837-DEL-144 |
| chr13-30053639-DEL-208 | chr13-30053640-DEL-208 |
| chr13-30064624-DEL-6119 | chr13-30064625-DEL-6119 |
| chr13-30078152-DEL-3311 | chr13-30078153-DEL-3311 |
| chr13-30082261-DEL-210 | chr13-30082262-DEL-210 |
| chr13-30116366-DEL-343 | chr13-30116367-DEL-343 |
| chr13-30118819-DEL-96 | chr13-30118809-DEL-72 |
| chr13-30149592-DEL-70 | chr13-30149593-DEL-70 |
| chr13-30239560-DEL-89 | chr13-30239553-DEL-90 |
| chr13-30244137-DEL-217 | chr13-30244141-DEL-215 |
| chr13-30245522-DEL-168 | chr13-30245525-DEL-162 |
| chr13-30257202-DEL-57 | chr13-30257203-DEL-57 |
| chr13-30272576-DEL-309 | chr13-30272577-DEL-309 |
| chr13-30285426-DEL-133 | chr13-30285426-DEL-133 |
| chr13-30292649-DEL-201 | chr13-30292650-DEL-201 |

| | |
|---|---|
| chr13-30299516-DEL-101 | chr13-30299512-DEL-101 |
| chr13-30300838-DEL-6352 | chr13-30300816-DEL-6377 |
| chr13-30306898-DEL-201 | chr13-30306899-DEL-201 |
| chr13-30345228-DEL-62 | chr13-30345207-DEL-50 |
| chr13-30370968-DEL-3597 | chr13-30370969-DEL-3597 |
| chr13-30408922-DEL-195 | chr13-30408914-DEL-196 |
| chr13-30457481-DEL-2745 | chr13-30457479-DEL-2746 |
| chr13-30468644-DEL-51 | chr13-30468645-DEL-51 |
| chr13-30470500-DEL-1650 | chr13-30470501-DEL-1650 |
| chr13-30494512-DEL-326 | chr13-30494513-DEL-326 |
| chr13-30496179-DEL-63 | chr13-30496198-DEL-85 |
| chr13-30498664-DEL-6403 | chr13-30498665-DEL-6403 |
| chr13-30526674-DEL-188 | chr13-30526675-DEL-188 |
| chr13-30554103-DEL-7113 | chr13-30554104-DEL-7113 |
| chr13-30624797-DEL-84 | chr13-30624798-DEL-84 |
| chr13-30633774-DEL-59 | chr13-30633775-DEL-59 |
| chr13-30657922-DEL-1515 | chr13-30657938-DEL-1533 |
| chr13-30659500-DEL-90 | chr13-30659507-DEL-84 |
| chr13-30670636-DEL-4030 | chr13-30670637-DEL-4030 |
| chr13-30683223-DEL-1890 | chr13-30683224-DEL-1890 |
| chr13-30700053-DEL-358 | chr13-30700054-DEL-358 |
| chr13-30719588-DEL-1878 | chr13-30719589-DEL-1878 |
| chr13-30751855-DEL-450 | chr13-30751856-DEL-450 |
| chr13-30780569-DEL-220 | chr13-30780570-DEL-220 |
| chr13-30782087-DEL-188 | chr13-30782055-DEL-221 |
| chr13-30951164-DEL-62 | chr13-30951158-DEL-67 |
| chr13-30965806-DEL-91 | chr13-30965807-DEL-91 |
| chr13-30981555-DEL-1604 | chr13-30981556-DEL-1604 |
| chr13-31018992-DEL-119 | chr13-31018993-DEL-119 |
| chr13-31019868-DEL-54 | chr13-31019869-DEL-54 |
| chr13-31028019-DEL-878 | chr13-31028016-DEL-875 |
| chr13-31034210-DEL-4509 | chr13-31034211-DEL-4509 |
| chr13-31035728-DEL-121 | chr13-31035712-DEL-136 |
| chr13-31048275-DEL-56 | chr13-31048276-DEL-56 |
| chr13-31049962-DEL-164 | chr13-31049964-DEL-160 |
| chr13-31193479-DEL-4291 | chr13-31193480-DEL-4291 |
| chr13-31242284-DEL-78 | chr13-31242293-DEL-94 |
| chr13-31270908-DEL-1459 | chr13-31270909-DEL-1459 |
| chr13-31295553-DEL-6607 | chr13-31295553-DEL-6607 |
| chr13-31300764-DEL-493 | chr13-31300765-DEL-493 |
| chr13-31306176-DEL-52 | chr13-31306177-DEL-64 |
| chr13-31310195-DEL-74 | chr13-31310206-DEL-64 |
| chr13-31324004-DEL-6205 | chr13-31324005-DEL-6205 |
| chr13-31331129-DEL-55 | chr13-31331130-DEL-55 |
| chr13-31335757-DEL-400 | chr13-31335756-DEL-400 |
| chr13-31370530-DEL-54 | chr13-31370525-DEL-66 |
| chr13-31399124-DEL-193 | chr13-31399123-DEL-193 |
| chr13-31406486-DEL-64 | chr13-31406487-DEL-64 |
| chr13-31417343-DEL-7126 | chr13-31417344-DEL-7126 |
| chr13-31438597-DEL-72 | chr13-31438598-DEL-72 |
| chr13-31453483-DEL-78 | chr13-31453484-DEL-78 |
| chr13-31455459-DEL-58 | chr13-31455459-DEL-58 |
| chr13-31456305-DEL-67 | chr13-31456306-DEL-67 |
| chr13-31463267-DEL-3773 | chr13-31463268-DEL-3773 |
| chr13-31481554-DEL-95 | chr13-31481555-DEL-95 |
| chr13-31507568-DEL-301 | chr13-31507569-DEL-301 |
| chr13-31535826-DEL-80 | chr13-31535827-DEL-78 |
| chr13-31544263-DEL-186 | chr13-31544259-DEL-187 |
| chr13-31567020-DEL-6649 | chr13-31567021-DEL-6649 |
| chr13-31591651-DEL-349 | chr13-31591652-DEL-349 |
| chr13-31614682-DEL-50 | chr13-31614683-DEL-55 |
| chr13-31625979-DEL-344 | chr13-31625980-DEL-343 |
| chr13-31637748-DEL-2168 | chr13-31637749-DEL-2168 |
| chr13-31663793-DEL-71 | chr13-31663792-DEL-72 |
| chr13-31666720-DEL-102 | chr13-31666721-DEL-102 |
| chr13-31668920-DEL-319 | chr13-31668921-DEL-319 |
| chr13-31675373-DEL-68 | chr13-31675392-DEL-50 |
| chr13-31688137-DEL-239 | chr13-31688138-DEL-239 |
| chr13-31699037-DEL-62 | chr13-31699040-DEL-60 |
| chr13-31714399-DEL-61 | chr13-31714407-DEL-54 |
| chr13-31727772-DEL-1660 | chr13-31727773-DEL-1660 |
| chr13-31737033-DEL-102 | chr13-31737034-DEL-102 |
| chr13-31770709-DEL-107 | chr13-31770703-DEL-84 |
| chr13-31777866-DEL-3802 | chr13-31777867-DEL-3802 |
| chr13-31792586-DEL-3244 | chr13-31792587-DEL-3240 |
| chr13-31801783-DEL-64 | chr13-31801785-DEL-63 |
| chr13-31803703-DEL-64 | chr13-31803657-DEL-65 |
| chr13-31864791-DEL-112 | chr13-31864792-DEL-112 |
| chr13-31866437-DEL-52 | chr13-31866438-DEL-52 |
| chr13-31885278-DEL-1808 | chr13-31885279-DEL-1808 |

| | |
|---|---|
| chr13-31910069-DEL-59 | chr13-31910070-DEL-59 |
| chr13-31919736-DEL-61 | chr13-31919730-DEL-62 |
| chr13-31922259-DEL-431 | chr13-31922264-DEL-430 |
| chr13-31978534-DEL-459 | chr13-31978535-DEL-458 |
| chr13-32004381-DEL-74 | chr13-32004382-DEL-70 |
| chr13-32026949-DEL-55 | chr13-32026950-DEL-55 |
| chr13-32032001-DEL-108 | chr13-32032004-DEL-106 |
| chr13-32038906-DEL-789 | chr13-32038910-DEL-786 |
| chr13-32068382-DEL-441 | chr13-32068383-DEL-441 |
| chr13-32167957-DEL-209 | chr13-32167958-DEL-209 |
| chr13-32433273-DEL-126 | chr13-32433274-DEL-126 |
| chr13-32456325-DEL-465 | chr13-32456326-DEL-465 |
| chr13-32520037-DEL-60 | chr13-32520038-DEL-60 |
| chr13-32534420-DEL-7225 | chr13-32534421-DEL-7225 |
| chr13-32550567-DEL-1543 | chr13-32550568-DEL-1543 |
| chr13-32560326-DEL-159 | chr13-32560327-DEL-159 |
| chr13-32574569-DEL-51 | chr13-32574570-DEL-51 |
| chr13-32578714-DEL-70 | chr13-32578749-DEL-50 |
| chr13-32682940-DEL-57 | chr13-32682937-DEL-61 |
| chr13-32752184-DEL-64 | chr13-32752185-DEL-64 |
| chr13-32821496-DEL-2811 | chr13-32821497-DEL-2811 |
| chr13-32846537-DEL-154 | chr13-32846538-DEL-132 |
| chr13-32904142-DEL-64 | chr13-32904143-DEL-64 |
| chr13-32904707-DEL-74 | chr13-32904692-DEL-64 |
| chr13-32921890-DEL-76 | chr13-32921891-DEL-76 |
| chr13-32972471-DEL-98 | chr13-32972472-DEL-98 |
| chr13-32983201-DEL-57 | chr13-32983192-DEL-55 |
| chr13-32994171-DEL-287 | chr13-32994172-DEL-287 |
| chr13-33059273-DEL-70 | chr13-33059274-DEL-70 |
| chr13-33100188-DEL-361 | chr13-33100189-DEL-361 |
| chr13-33113606-DEL-2812 | chr13-33113607-DEL-2812 |
| chr13-33124841-DEL-2461 | chr13-33124842-DEL-2461 |
| chr13-33131543-DEL-7118 | chr13-33131544-DEL-7118 |
| chr13-33199225-DEL-106 | chr13-33199228-DEL-105 |
| chr13-33235560-DEL-140 | chr13-33235561-DEL-130 |
| chr13-33239029-DEL-7712 | chr13-33239030-DEL-7712 |
| chr13-33250975-DEL-65 | chr13-33250981-DEL-63 |
| chr13-33272151-DEL-68 | chr13-33272151-DEL-68 |
| chr13-33275028-DEL-76 | chr13-33275006-DEL-72 |
| chr13-33282413-DEL-53 | chr13-33282414-DEL-53 |
| chr13-33286013-DEL-66 | chr13-33286026-DEL-66 |
| chr13-33337911-DEL-191 | chr13-33337912-DEL-191 |
| chr13-33361152-DEL-368 | chr13-33361152-DEL-368 |
| chr13-33363242-DEL-2867 | chr13-33363243-DEL-2867 |
| chr13-33434997-DEL-228 | chr13-33434998-DEL-228 |
| chr13-33536303-DEL-738 | chr13-33536304-DEL-738 |
| chr13-33536909-DEL-229 | chr13-33536910-DEL-229 |
| chr13-33656723-DEL-6232 | chr13-33656724-DEL-6232 |
| chr13-33685586-DEL-2129 | chr13-33685596-DEL-2128 |
| chr13-33702837-DEL-195 | chr13-33702838-DEL-195 |
| chr13-33717171-DEL-821 | chr13-33717173-DEL-817 |
| chr13-33731095-DEL-3388 | chr13-33731097-DEL-3388 |
| chr13-33758840-DEL-10413 | chr13-33758838-DEL-10415 |
| chr13-33773609-DEL-130 | chr13-33773618-DEL-121 |
| chr13-33776224-DEL-486 | chr13-33776224-DEL-486 |
| chr13-33821474-DEL-662 | chr13-33821475-DEL-662 |
| chr13-33822779-DEL-79 | chr13-33822778-DEL-79 |
| chr13-33861280-DEL-6457 | chr13-33861281-DEL-6457 |
| chr13-33877317-DEL-305 | chr13-33877318-DEL-306 |
| chr13-33884613-DEL-390 | chr13-33884614-DEL-390 |
| chr13-33892598-DEL-101 | chr13-33892614-DEL-82 |
| chr13-33910182-DEL-5896 | chr13-33910182-DEL-5896 |
| chr13-33917575-DEL-4975 | chr13-33917574-DEL-4974 |
| chr13-33935792-DEL-1400 | chr13-33935793-DEL-1400 |
| chr13-33937764-DEL-2125 | chr13-33937765-DEL-2125 |
| chr13-33946091-DEL-299 | chr13-33946082-DEL-300 |
| chr13-33985057-DEL-4828 | chr13-33985058-DEL-4828 |
| chr13-34005229-DEL-521 | chr13-34005230-DEL-521 |
| chr13-34013207-DEL-1680 | chr13-34013208-DEL-1680 |
| chr13-34013583-DEL-172 | chr13-34013584-DEL-172 |
| chr13-34039101-DEL-605 | chr13-34039102-DEL-605 |
| chr13-34041306-DEL-126 | chr13-34041301-DEL-124 |
| chr13-34046015-DEL-7091 | chr13-34046011-DEL-7092 |
| chr13-34074472-DEL-2101 | chr13-34074472-DEL-2101 |
| chr13-34126679-DEL-56 | chr13-34126680-DEL-56 |
| chr13-34147666-DEL-386 | chr13-34147667-DEL-386 |
| chr13-34238513-DEL-377 | chr13-34238514-DEL-377 |
| chr13-34279859-DEL-62 | chr13-34279848-DEL-68 |
| chr13-34286927-DEL-1056 | chr13-34286928-DEL-1056 |
| chr13-34286927-DEL-484 | chr13-34286928-DEL-484 |

| | |
|---|---|
| chr13-34323281-DEL-3046 | chr13-34323282-DEL-3046 |
| chr13-34343610-DEL-78 | chr13-34343611-DEL-78 |
| chr13-34350749-DEL-7237 | chr13-34350750-DEL-7237 |
| chr13-34401688-DEL-260 | chr13-34401675-DEL-261 |
| chr13-34405454-DEL-152 | chr13-34405447-DEL-122 |
| chr13-34466556-DEL-1270 | chr13-34466557-DEL-1270 |
| chr13-34477480-DEL-319 | chr13-34477469-DEL-331 |
| chr13-34540262-DEL-89 | chr13-34540263-DEL-89 |
| chr13-34540890-DEL-192 | chr13-34540891-DEL-192 |
| chr13-34626057-DEL-317 | chr13-34626077-DEL-291 |
| chr13-34628168-DEL-78 | chr13-34628169-DEL-75 |
| chr13-34638250-DEL-204 | chr13-34638251-DEL-204 |
| chr13-34645519-DEL-79 | chr13-34645520-DEL-79 |
| chr13-34654852-DEL-6213 | chr13-34654853-DEL-6213 |
| chr13-34678054-DEL-2057 | chr13-34678055-DEL-2057 |
| chr13-34689974-DEL-78 | chr13-34689974-DEL-78 |
| chr13-34697750-DEL-7182 | chr13-34697751-DEL-7182 |
| chr13-34708301-DEL-462 | chr13-34708307-DEL-457 |
| chr13-34721103-DEL-246 | chr13-34721103-DEL-246 |
| chr13-34731563-DEL-1507 | chr13-34731564-DEL-1507 |
| chr13-34731578-DEL-60 | chr13-34731574-DEL-75 |
| chr13-34738935-DEL-54 | chr13-34738936-DEL-54 |
| chr13-34750099-DEL-446 | chr13-34750100-DEL-446 |
| chr13-34757339-DEL-57 | chr13-34757342-DEL-64 |
| chr13-34764655-DEL-511 | chr13-34764656-DEL-511 |
| chr13-34800450-DEL-1024 | chr13-34800450-DEL-1024 |
| chr13-34849098-DEL-82 | chr13-34849099-DEL-82 |
| chr13-34874774-DEL-60 | chr13-34874770-DEL-65 |
| chr13-34904958-DEL-141 | chr13-34904948-DEL-128 |
| chr13-34933340-DEL-6488 | chr13-34933341-DEL-6488 |
| chr13-34975323-DEL-209 | chr13-34975324-DEL-209 |
| chr13-34988005-DEL-115 | chr13-34988006-DEL-80 |
| chr13-34992744-DEL-398 | chr13-34992745-DEL-398 |
| chr13-35003129-DEL-109 | chr13-35003151-DEL-100 |
| chr13-35005366-DEL-89 | chr13-35005338-DEL-90 |
| chr13-35018994-DEL-710 | chr13-35018991-DEL-713 |
| chr13-35036139-DEL-60 | chr13-35036138-DEL-62 |
| chr13-35040369-DEL-56 | chr13-35040369-DEL-57 |
| chr13-35130046-DEL-215 | chr13-35130047-DEL-215 |
| chr13-35146353-DEL-399 | chr13-35146354-DEL-397 |
| chr13-35185281-DEL-68 | chr13-35185282-DEL-68 |
| chr13-35218470-DEL-112 | chr13-35218516-DEL-112 |
| chr13-35222204-DEL-237 | chr13-35222205-DEL-237 |
| chr13-35230470-DEL-6401 | chr13-35230470-DEL-6401 |
| chr13-35240760-DEL-214 | chr13-35240761-DEL-214 |
| chr13-35243107-DEL-51 | chr13-35243108-DEL-51 |
| chr13-35266091-DEL-1432 | chr13-35266092-DEL-1432 |
| chr13-35315039-DEL-164 | chr13-35315040-DEL-164 |
| chr13-35322708-DEL-199 | chr13-35322709-DEL-199 |
| chr13-35341101-DEL-682 | chr13-35341102-DEL-682 |
| chr13-35345540-DEL-113 | chr13-35345541-DEL-113 |
| chr13-35355341-DEL-59 | chr13-35355342-DEL-59 |
| chr13-35363646-DEL-285 | chr13-35363647-DEL-285 |
| chr13-35450900-DEL-1752 | chr13-35450901-DEL-1752 |
| chr13-35477323-DEL-1122 | chr13-35477323-DEL-1102 |
| chr13-35486420-DEL-3048 | chr13-35486420-DEL-3048 |
| chr13-35508355-DEL-499 | chr13-35508356-DEL-499 |
| chr13-35581762-DEL-74 | chr13-35581765-DEL-72 |
| chr13-35595384-DEL-1326 | chr13-35595385-DEL-1326 |
| chr13-35602599-DEL-192 | chr13-35602600-DEL-192 |
| chr13-35633564-DEL-52 | chr13-35633565-DEL-52 |
| chr13-35656918-DEL-70 | chr13-35656919-DEL-70 |
| chr13-35694331-DEL-401 | chr13-35694332-DEL-401 |
| chr13-35727675-DEL-400 | chr13-35727678-DEL-399 |
| chr13-35747445-DEL-498 | chr13-35747446-DEL-498 |
| chr13-35751124-DEL-6841 | chr13-35751125-DEL-6841 |
| chr13-35758564-DEL-3904 | chr13-35758564-DEL-3904 |
| chr13-35863861-DEL-446 | chr13-35863862-DEL-446 |
| chr13-35871058-DEL-211 | chr13-35871059-DEL-211 |
| chr13-36004463-DEL-52 | chr13-36004464-DEL-52 |
| chr13-36028595-DEL-75 | chr13-36028596-DEL-75 |
| chr13-36069582-DEL-62 | chr13-36069583-DEL-65 |
| chr13-36074377-DEL-73 | chr13-36074374-DEL-74 |
| chr13-36074534-DEL-1921 | chr13-36074535-DEL-1921 |
| chr13-36095024-DEL-74 | chr13-36095027-DEL-69 |
| chr13-36102368-DEL-437 | chr13-36102369-DEL-437 |
| chr13-36130403-DEL-108 | chr13-36130390-DEL-90 |
| chr13-36155214-DEL-773 | chr13-36155215-DEL-773 |
| chr13-36168407-DEL-216 | chr13-36168408-DEL-216 |
| chr13-36174174-DEL-1359 | chr13-36174174-DEL-1359 |

| | |
|---|---|
| chr13-36189703-DEL-4051 | chr13-36189704-DEL-4051 |
| chr13-36202468-DEL-84 | chr13-36202469-DEL-84 |
| chr13-36232109-DEL-874 | chr13-36232109-DEL-874 |
| chr13-36268103-DEL-207 | chr13-36268104-DEL-207 |
| chr13-36417528-DEL-963 | chr13-36417529-DEL-963 |
| chr13-36454583-DEL-68 | chr13-36454586-DEL-66 |
| chr13-36483512-DEL-85 | chr13-36483513-DEL-85 |
| chr13-36538248-DEL-1602 | chr13-36538248-DEL-1602 |
| chr13-36541718-DEL-225 | chr13-36541719-DEL-225 |
| chr13-36552532-DEL-78 | chr13-36552537-DEL-74 |
| chr13-36557145-DEL-3948 | chr13-36557146-DEL-3948 |
| chr13-36590532-DEL-1043 | chr13-36590533-DEL-1043 |
| chr13-36628069-DEL-92 | chr13-36628070-DEL-92 |
| chr13-36643072-DEL-210 | chr13-36643073-DEL-210 |
| chr13-36667910-DEL-115 | chr13-36667911-DEL-115 |
| chr13-36716257-DEL-60 | chr13-36716258-DEL-60 |
| chr13-36795545-DEL-79 | chr13-36795546-DEL-79 |
| chr13-36828902-DEL-58 | chr13-36828902-DEL-58 |
| chr13-36842406-DEL-550 | chr13-36842407-DEL-550 |
| chr13-36889715-DEL-53 | chr13-36889716-DEL-53 |
| chr13-36949223-DEL-114 | chr13-36949224-DEL-114 |
| chr13-36958116-DEL-125 | chr13-36958123-DEL-104 |
| chr13-36974224-DEL-102 | chr13-36974225-DEL-102 |
| chr13-37012334-DEL-354 | chr13-37012335-DEL-354 |
| chr13-37027178-DEL-10468 | chr13-37027179-DEL-10468 |
| chr13-37090483-DEL-1251 | chr13-37090484-DEL-1251 |
| chr13-37116592-DEL-222 | chr13-37116582-DEL-214 |
| chr13-37168297-DEL-59 | chr13-37168298-DEL-59 |
| chr13-37201838-DEL-1263 | chr13-37201839-DEL-1263 |
| chr13-37211805-DEL-78 | chr13-37211810-DEL-60 |
| chr13-37220690-DEL-7132 | chr13-37220691-DEL-7132 |
| chr13-37274452-DEL-1001 | chr13-37274453-DEL-1001 |
| chr13-37306543-DEL-129 | chr13-37306544-DEL-129 |
| chr13-37308024-DEL-309 | chr13-37308025-DEL-309 |
| chr13-37330843-DEL-534 | chr13-37330844-DEL-534 |
| chr13-37350680-DEL-89 | chr13-37350685-DEL-87 |
| chr13-37351020-DEL-7394 | chr13-37351021-DEL-7394 |
| chr13-37405656-DEL-2045 | chr13-37405657-DEL-2045 |
| chr13-37418702-DEL-506 | chr13-37418703-DEL-506 |
| chr13-37446063-DEL-414 | chr13-37446064-DEL-414 |
| chr13-37446959-DEL-6484 | chr13-37446960-DEL-6484 |
| chr13-37465066-DEL-54 | chr13-37465067-DEL-54 |
| chr13-37504087-DEL-84 | chr13-37504088-DEL-84 |
| chr13-37559688-DEL-217 | chr13-37559651-DEL-261 |
| chr13-37565055-DEL-211 | chr13-37565056-DEL-211 |
| chr13-37569815-DEL-219 | chr13-37569816-DEL-219 |
| chr13-37597659-DEL-54 | chr13-37597660-DEL-54 |
| chr13-37605062-DEL-84 | chr13-37605063-DEL-84 |
| chr13-37629674-DEL-775 | chr13-37629675-DEL-775 |
| chr13-37632038-DEL-1066 | chr13-37632039-DEL-1066 |
| chr13-37639193-DEL-72 | chr13-37639193-DEL-72 |
| chr13-37649131-DEL-449 | chr13-37649130-DEL-449 |
| chr13-37651053-DEL-353 | chr13-37651054-DEL-353 |
| chr13-37656157-DEL-136 | chr13-37656135-DEL-137 |
| chr13-37657900-DEL-59 | chr13-37657883-DEL-62 |
| chr13-37660832-DEL-62 | chr13-37660833-DEL-62 |
| chr13-37666460-DEL-73 | chr13-37666419-DEL-98 |
| chr13-37677644-DEL-270 | chr13-37677645-DEL-270 |
| chr13-37703561-DEL-227 | chr13-37703562-DEL-227 |
| chr13-37723350-DEL-58 | chr13-37723351-DEL-58 |
| chr13-37739516-DEL-1436 | chr13-37739517-DEL-1436 |
| chr13-37764603-DEL-61 | chr13-37764584-DEL-62 |
| chr13-37790600-DEL-67 | chr13-37790561-DEL-68 |
| chr13-37803537-DEL-51 | chr13-37803537-DEL-51 |
| chr13-37819106-DEL-423 | chr13-37819107-DEL-389 |
| chr13-37836871-DEL-54 | chr13-37836872-DEL-52 |
| chr13-37843893-DEL-61 | chr13-37843887-DEL-88 |
| chr13-37899652-DEL-67 | chr13-37899653-DEL-67 |
| chr13-37923713-DEL-54 | chr13-37923714-DEL-54 |
| chr13-37966045-DEL-54 | chr13-37966045-DEL-72 |
| chr13-37980609-DEL-55 | chr13-37980605-DEL-56 |
| chr13-37981323-DEL-227 | chr13-37981324-DEL-227 |
| chr13-38002003-DEL-61 | chr13-38002004-DEL-61 |
| chr13-38018839-DEL-201 | chr13-38018840-DEL-201 |
| chr13-38032173-DEL-232 | chr13-38032174-DEL-232 |
| chr13-38070331-DEL-215 | chr13-38070332-DEL-215 |
| chr13-38111272-DEL-333 | chr13-38111273-DEL-309 |
| chr13-38183929-DEL-370 | chr13-38183930-DEL-370 |
| chr13-38215276-DEL-307 | chr13-38215277-DEL-307 |
| chr13-38235988-DEL-54 | chr13-38235989-DEL-54 |

| | |
|---|---|
| chr13-38238233-DEL-63 | chr13-38238234-DEL-63 |
| chr13-38262624-DEL-140 | chr13-38262627-DEL-137 |
| chr13-38266414-DEL-72 | chr13-38266454-DEL-76 |
| chr13-38283939-DEL-71 | chr13-38283940-DEL-71 |
| chr13-38284220-DEL-448 | chr13-38284215-DEL-449 |
| chr13-38285339-DEL-770 | chr13-38285340-DEL-771 |
| chr13-38290888-DEL-117 | chr13-38290889-DEL-117 |
| chr13-38293612-DEL-5385 | chr13-38293613-DEL-5385 |
| chr13-38354535-DEL-54 | chr13-38354536-DEL-54 |
| chr13-38413371-DEL-498 | chr13-38413372-DEL-498 |
| chr13-38428620-DEL-74 | chr13-38428621-DEL-74 |
| chr13-38451146-DEL-60 | chr13-38451146-DEL-60 |
| chr13-38457742-DEL-61 | chr13-38457744-DEL-60 |
| chr13-38470625-DEL-193 | chr13-38470626-DEL-193 |
| chr13-38495882-DEL-72 | chr13-38495883-DEL-72 |
| chr13-38668393-DEL-72 | chr13-38668394-DEL-72 |
| chr13-38685896-DEL-75 | chr13-38685897-DEL-75 |
| chr13-38741639-DEL-1007 | chr13-38741640-DEL-1007 |
| chr13-38769704-DEL-2156 | chr13-38769705-DEL-2156 |
| chr13-38811719-DEL-484 | chr13-38811720-DEL-484 |
| chr13-38826876-DEL-242 | chr13-38826849-DEL-243 |
| chr13-38840607-DEL-252 | chr13-38840589-DEL-253 |
| chr13-38844263-DEL-318 | chr13-38844264-DEL-318 |
| chr13-38884670-DEL-332 | chr13-38884671-DEL-332 |
| chr13-38893688-DEL-498 | chr13-38893689-DEL-498 |
| chr13-38950819-DEL-2372 | chr13-38950820-DEL-2372 |
| chr13-38984825-DEL-413 | chr13-38984826-DEL-413 |
| chr13-39005586-DEL-387 | chr13-39005586-DEL-387 |
| chr13-39007436-DEL-153 | chr13-39007437-DEL-153 |
| chr13-39049305-DEL-100 | chr13-39049306-DEL-100 |
| chr13-39050654-DEL-81 | chr13-39050655-DEL-81 |
| chr13-39105868-DEL-56 | chr13-39105869-DEL-56 |
| chr13-39112032-DEL-56 | chr13-39112033-DEL-56 |
| chr13-39125591-DEL-54 | chr13-39125592-DEL-54 |
| chr13-39131544-DEL-117 | chr13-39131545-DEL-117 |
| chr13-39135054-DEL-249 | chr13-39135055-DEL-249 |
| chr13-39220802-DEL-2866 | chr13-39220803-DEL-2866 |
| chr13-39232894-DEL-56 | chr13-39232895-DEL-76 |
| chr13-39233893-DEL-6780 | chr13-39233894-DEL-6780 |
| chr13-39264545-DEL-56 | chr13-39264546-DEL-56 |
| chr13-39279120-DEL-64 | chr13-39279121-DEL-64 |
| chr13-39296424-DEL-1119 | chr13-39296425-DEL-1119 |
| chr13-39306326-DEL-250 | chr13-39306327-DEL-250 |
| chr13-39335467-DEL-51 | chr13-39335467-DEL-51 |
| chr13-39373502-DEL-86 | chr13-39373503-DEL-86 |
| chr13-39397379-DEL-262 | chr13-39397380-DEL-262 |
| chr13-39467138-DEL-286 | chr13-39467137-DEL-288 |
| chr13-39486491-DEL-139 | chr13-39486509-DEL-153 |
| chr13-39511352-DEL-54 | chr13-39511321-DEL-64 |
| chr13-39549258-DEL-82 | chr13-39549257-DEL-88 |
| chr13-39550238-DEL-212 | chr13-39550239-DEL-212 |
| chr13-39551378-DEL-51 | chr13-39551355-DEL-63 |
| chr13-39557224-DEL-450 | chr13-39557225-DEL-450 |
| chr13-39580218-DEL-531 | chr13-39580233-DEL-555 |
| chr13-39597055-DEL-70 | chr13-39597056-DEL-70 |
| chr13-39622667-DEL-54 | chr13-39622668-DEL-54 |
| chr13-39627268-DEL-56 | chr13-39627230-DEL-72 |
| chr13-39650980-DEL-130 | chr13-39651025-DEL-86 |
| chr13-39690454-DEL-8480 | chr13-39690455-DEL-8480 |
| chr13-39711003-DEL-304 | chr13-39711004-DEL-304 |
| chr13-39713256-DEL-99 | chr13-39713257-DEL-99 |
| chr13-39724098-DEL-94 | chr13-39724099-DEL-94 |
| chr13-39727659-DEL-55 | chr13-39727659-DEL-55 |
| chr13-39728495-DEL-6610 | chr13-39728496-DEL-6610 |
| chr13-39778284-DEL-5102 | chr13-39778285-DEL-5102 |
| chr13-39807448-DEL-439 | chr13-39807449-DEL-439 |
| chr13-39858732-DEL-52 | chr13-39858687-DEL-75 |
| chr13-39862418-DEL-2699 | chr13-39862419-DEL-2699 |
| chr13-39870726-DEL-2044 | chr13-39870727-DEL-2044 |
| chr13-39892844-DEL-79 | chr13-39892845-DEL-52 |
| chr13-39894157-DEL-10780 | chr13-39894158-DEL-10780 |
| chr13-39904371-DEL-208 | chr13-39904372-DEL-208 |
| chr13-39943109-DEL-400 | chr13-39943110-DEL-400 |
| chr13-39973870-DEL-77 | chr13-39973871-DEL-82 |
| chr13-40003278-DEL-380 | chr13-40003274-DEL-404 |
| chr13-40068194-DEL-288 | chr13-40068194-DEL-286 |
| chr13-40070107-DEL-1520 | chr13-40070108-DEL-1520 |
| chr13-40079888-DEL-1819 | chr13-40079889-DEL-1819 |
| chr13-40083647-DEL-106 | chr13-40083662-DEL-92 |
| chr13-40113908-DEL-51 | chr13-40113909-DEL-51 |

| | |
|---|---|
| chr13-40199803-DEL-158 | chr13-40199804-DEL-158 |
| chr13-40202895-DEL-6628 | chr13-40202896-DEL-6628 |
| chr13-40324646-DEL-68 | chr13-40324643-DEL-72 |
| chr13-40349344-DEL-451 | chr13-40349345-DEL-451 |
| chr13-40359349-DEL-52 | chr13-40359350-DEL-52 |
| chr13-40364349-DEL-58 | chr13-40364319-DEL-100 |
| chr13-40381368-DEL-101 | chr13-40381369-DEL-101 |
| chr13-40384242-DEL-74 | chr13-40384243-DEL-74 |
| chr13-40419101-DEL-52 | chr13-40419094-DEL-60 |
| chr13-40427163-DEL-131 | chr13-40427164-DEL-131 |
| chr13-40451328-DEL-68 | chr13-40451325-DEL-80 |
| chr13-40464562-DEL-54 | chr13-40464563-DEL-54 |
| chr13-40466788-DEL-124 | chr13-40466791-DEL-124 |
| chr13-40492364-DEL-467 | chr13-40492365-DEL-467 |
| chr13-40505647-DEL-1279 | chr13-40505646-DEL-1277 |
| chr13-40533120-DEL-66 | chr13-40533129-DEL-58 |
| chr13-40542098-DEL-141 | chr13-40542101-DEL-140 |
| chr13-40564315-DEL-1114 | chr13-40564316-DEL-1114 |
| chr13-40635634-DEL-52 | chr13-40635619-DEL-78 |
| chr13-40653414-DEL-255 | chr13-40653405-DEL-254 |
| chr13-40655964-DEL-246 | chr13-40655965-DEL-247 |
| chr13-40699163-DEL-1349 | chr13-40699164-DEL-1349 |
| chr13-40716571-DEL-108 | chr13-40716600-DEL-60 |
| chr13-40741112-DEL-6465 | chr13-40741113-DEL-6465 |
| chr13-40779536-DEL-6624 | chr13-40779537-DEL-6624 |
| chr13-40807560-DEL-114 | chr13-40807551-DEL-124 |
| chr13-40823971-DEL-56 | chr13-40823978-DEL-50 |
| chr13-40829077-DEL-340 | chr13-40829078-DEL-340 |
| chr13-40906148-DEL-456 | chr13-40906149-DEL-446 |
| chr13-40919496-DEL-110 | chr13-40919492-DEL-110 |
| chr13-40921481-DEL-171 | chr13-40921481-DEL-172 |
| chr13-40925636-DEL-406 | chr13-40925637-DEL-406 |
| chr13-40930364-DEL-497 | chr13-40930365-DEL-497 |
| chr13-40959214-DEL-202 | chr13-40959214-DEL-202 |
| chr13-40962882-DEL-171 | chr13-40962874-DEL-172 |
| chr13-40966941-DEL-211 | chr13-40966942-DEL-211 |
| chr13-40978914-DEL-118 | chr13-40978915-DEL-118 |
| chr13-40980401-DEL-3412 | chr13-40980403-DEL-3410 |
| chr13-41003431-DEL-271 | chr13-41003432-DEL-271 |
| chr13-41015598-DEL-997 | chr13-41015599-DEL-997 |
| chr13-41041924-DEL-203 | chr13-41041924-DEL-200 |
| chr13-41073411-DEL-660 | chr13-41073412-DEL-660 |
| chr13-41081625-DEL-84 | chr13-41081626-DEL-84 |
| chr13-41114662-DEL-1031 | chr13-41114661-DEL-1031 |
| chr13-41120063-DEL-506 | chr13-41120064-DEL-506 |
| chr13-41131984-DEL-83 | chr13-41131985-DEL-83 |
| chr13-41134677-DEL-179 | chr13-41134678-DEL-179 |
| chr13-41138250-DEL-190 | chr13-41138251-DEL-190 |
| chr13-41151424-DEL-107 | chr13-41151425-DEL-70 |
| chr13-41211788-DEL-6293 | chr13-41211789-DEL-6293 |
| chr13-41222941-DEL-206 | chr13-41222942-DEL-206 |
| chr13-41227892-DEL-100 | chr13-41227889-DEL-101 |
| chr13-41228801-DEL-511 | chr13-41228802-DEL-511 |
| chr13-41237651-DEL-52 | chr13-41237648-DEL-52 |
| chr13-41270919-DEL-370 | chr13-41270920-DEL-370 |
| chr13-41297114-DEL-52 | chr13-41297115-DEL-52 |
| chr13-41298866-DEL-64 | chr13-41298867-DEL-64 |
| chr13-41319510-DEL-2065 | chr13-41319521-DEL-2066 |
| chr13-41330785-DEL-88 | chr13-41330786-DEL-88 |
| chr13-41358603-DEL-1126 | chr13-41358601-DEL-1127 |
| chr13-41359095-DEL-122 | chr13-41359096-DEL-122 |
| chr13-41367533-DEL-98 | chr13-41367539-DEL-95 |
| chr13-41382381-DEL-121 | chr13-41382382-DEL-121 |
| chr13-41412102-DEL-3105 | chr13-41412103-DEL-3105 |
| chr13-41419920-DEL-199 | chr13-41419921-DEL-199 |
| chr13-41435391-DEL-157 | chr13-41435392-DEL-157 |
| chr13-41440396-DEL-160 | chr13-41440397-DEL-160 |
| chr13-41456049-DEL-54 | chr13-41456050-DEL-54 |
| chr13-41457459-DEL-69 | chr13-41457460-DEL-69 |
| chr13-41481513-DEL-85 | chr13-41481514-DEL-85 |
| chr13-41520369-DEL-133 | chr13-41520360-DEL-143 |
| chr13-41544930-DEL-203 | chr13-41544931-DEL-203 |
| chr13-41589450-DEL-63 | chr13-41589450-DEL-63 |
| chr13-41596749-DEL-183 | chr13-41596750-DEL-183 |
| chr13-41604991-DEL-90 | chr13-41605039-DEL-60 |
| chr13-41622440-DEL-78 | chr13-41622441-DEL-78 |
| chr13-41649349-DEL-743 | chr13-41649350-DEL-743 |
| chr13-41715069-DEL-458 | chr13-41715070-DEL-458 |
| chr13-41823196-DEL-90 | chr13-41823196-DEL-88 |
| chr13-41825404-DEL-85 | chr13-41825405-DEL-85 |

| | |
|---|---|
| chr13-41840222-DEL-260 | chr13-41840223-DEL-260 |
| chr13-41849348-DEL-53 | chr13-41849349-DEL-53 |
| chr13-41868504-DEL-177 | chr13-41868505-DEL-177 |
| chr13-41877596-DEL-990 | chr13-41877597-DEL-990 |
| chr13-41888586-DEL-437 | chr13-41888571-DEL-454 |
| chr13-41937591-DEL-565 | chr13-41937590-DEL-566 |
| chr13-41940840-DEL-460 | chr13-41940841-DEL-460 |
| chr13-41943251-DEL-933 | chr13-41943252-DEL-933 |
| chr13-41947687-DEL-116 | chr13-41947692-DEL-115 |
| chr13-41961852-DEL-188 | chr13-41961853-DEL-188 |
| chr13-41965867-DEL-58 | chr13-41965909-DEL-58 |
| chr13-41971598-DEL-395 | chr13-41971599-DEL-395 |
| chr13-41979943-DEL-64 | chr13-41979914-DEL-67 |
| chr13-41986239-DEL-839 | chr13-41986238-DEL-840 |
| chr13-41998919-DEL-176 | chr13-41998920-DEL-176 |
| chr13-42014236-DEL-69 | chr13-42014237-DEL-69 |
| chr13-42041757-DEL-360 | chr13-42041758-DEL-360 |
| chr13-42046386-DEL-2087 | chr13-42046387-DEL-2087 |
| chr13-42069244-DEL-470 | chr13-42069245-DEL-470 |
| chr13-42077741-DEL-116 | chr13-42077742-DEL-116 |
| chr13-42084494-DEL-66 | chr13-42084495-DEL-66 |
| chr13-42104292-DEL-1873 | chr13-42104293-DEL-1873 |
| chr13-42112509-DEL-4296 | chr13-42112510-DEL-4297 |
| chr13-42133393-DEL-8851 | chr13-42133394-DEL-8851 |
| chr13-42137043-DEL-210 | chr13-42137044-DEL-210 |
| chr13-42160408-DEL-132 | chr13-42160424-DEL-141 |
| chr13-42177196-DEL-488 | chr13-42177193-DEL-498 |
| chr13-42222507-DEL-77 | chr13-42222508-DEL-77 |
| chr13-42230314-DEL-59 | chr13-42230315-DEL-59 |
| chr13-42239417-DEL-120 | chr13-42239418-DEL-120 |
| chr13-42248164-DEL-773 | chr13-42248165-DEL-773 |
| chr13-42262803-DEL-228 | chr13-42262804-DEL-228 |
| chr13-42274226-DEL-92 | chr13-42274227-DEL-78 |
| chr13-42281561-DEL-2927 | chr13-42281562-DEL-2927 |
| chr13-42295238-DEL-84 | chr13-42295239-DEL-84 |
| chr13-42305905-DEL-54 | chr13-42305905-DEL-54 |
| chr13-42323821-DEL-110 | chr13-42323822-DEL-110 |
| chr13-42325023-DEL-70 | chr13-42325023-DEL-70 |
| chr13-42329200-DEL-398 | chr13-42329201-DEL-398 |
| chr13-42385753-DEL-90 | chr13-42385754-DEL-90 |
| chr13-42401982-DEL-60 | chr13-42401953-DEL-55 |
| chr13-42404026-DEL-176 | chr13-42404027-DEL-176 |
| chr13-42404859-DEL-174 | chr13-42404860-DEL-174 |
| chr13-42425461-DEL-64 | chr13-42425462-DEL-60 |
| chr13-42451424-DEL-62 | chr13-42451427-DEL-60 |
| chr13-42483077-DEL-589 | chr13-42483078-DEL-589 |
| chr13-42496312-DEL-61 | chr13-42496306-DEL-67 |
| chr13-42512015-DEL-2209 | chr13-42512011-DEL-2209 |
| chr13-42551276-DEL-52 | chr13-42551276-DEL-52 |
| chr13-42563645-DEL-1470 | chr13-42563646-DEL-1470 |
| chr13-42629598-DEL-57 | chr13-42629599-DEL-57 |
| chr13-42645152-DEL-62 | chr13-42645153-DEL-62 |
| chr13-42701382-DEL-72 | chr13-42701383-DEL-72 |
| chr13-42705246-DEL-126 | chr13-42705294-DEL-87 |
| chr13-42739574-DEL-58 | chr13-42739575-DEL-58 |
| chr13-42749758-DEL-107 | chr13-42749759-DEL-107 |
| chr13-42757592-DEL-80 | chr13-42757590-DEL-83 |
| chr13-42758338-DEL-674 | chr13-42758339-DEL-674 |
| chr13-42827956-DEL-81 | chr13-42827957-DEL-81 |
| chr13-42849190-DEL-52 | chr13-42849191-DEL-52 |
| chr13-42869030-DEL-356 | chr13-42869031-DEL-356 |
| chr13-42919914-DEL-74 | chr13-42919921-DEL-72 |
| chr13-42928486-DEL-325 | chr13-42928487-DEL-325 |
| chr13-42964404-DEL-137 | chr13-42964411-DEL-169 |
| chr13-42966464-DEL-141 | chr13-42966465-DEL-141 |
| chr13-42972133-DEL-1111 | chr13-42972133-DEL-1111 |
| chr13-42993903-DEL-62 | chr13-42993902-DEL-62 |
| chr13-43010450-DEL-68 | chr13-43010451-DEL-68 |
| chr13-43011807-DEL-213 | chr13-43011808-DEL-213 |
| chr13-43030352-DEL-164 | chr13-43030353-DEL-164 |
| chr13-43067584-DEL-689 | chr13-43067585-DEL-689 |
| chr13-43085830-DEL-254 | chr13-43085831-DEL-254 |
| chr13-43126629-DEL-55 | chr13-43126629-DEL-55 |
| chr13-43155608-DEL-499 | chr13-43155609-DEL-499 |
| chr13-43161002-DEL-185 | chr13-43161003-DEL-185 |
| chr13-43193667-DEL-215 | chr13-43193668-DEL-215 |
| chr13-43201144-DEL-142 | chr13-43201137-DEL-143 |
| chr13-43244654-DEL-60 | chr13-43244655-DEL-58 |
| chr13-43252176-DEL-130 | chr13-43252177-DEL-130 |
| chr13-43261021-DEL-225 | chr13-43261022-DEL-225 |

| | |
|---|---|
| chr13-43264159-DEL-143 | chr13-43264144-DEL-144 |
| chr13-43278458-DEL-109 | chr13-43278462-DEL-106 |
| chr13-43295159-DEL-319 | chr13-43295164-DEL-318 |
| chr13-43324080-DEL-67 | chr13-43324081-DEL-67 |
| chr13-43399228-DEL-620 | chr13-43399230-DEL-619 |
| chr13-43431011-DEL-7120 | chr13-43431012-DEL-7120 |
| chr13-43468621-DEL-1825 | chr13-43468622-DEL-1825 |
| chr13-43529096-DEL-76 | chr13-43529099-DEL-86 |
| chr13-43540262-DEL-7089 | chr13-43540270-DEL-7082 |
| chr13-43587879-DEL-77 | chr13-43587880-DEL-77 |
| chr13-43612503-DEL-190 | chr13-43612504-DEL-190 |
| chr13-43616469-DEL-55 | chr13-43616440-DEL-105 |
| chr13-43625620-DEL-411 | chr13-43625621-DEL-411 |
| chr13-43656275-DEL-94 | chr13-43656278-DEL-88 |
| chr13-43673742-DEL-62 | chr13-43673766-DEL-55 |
| chr13-43695431-DEL-1892 | chr13-43695432-DEL-1892 |
| chr13-43726730-DEL-866 | chr13-43726731-DEL-866 |
| chr13-43739556-DEL-1147 | chr13-43739557-DEL-1147 |
| chr13-43740987-DEL-58 | chr13-43740980-DEL-58 |
| chr13-43761070-DEL-60 | chr13-43761071-DEL-60 |
| chr13-43767153-DEL-279 | chr13-43767141-DEL-277 |
| chr13-43817500-DEL-214 | chr13-43817501-DEL-214 |
| chr13-43836355-DEL-400 | chr13-43836356-DEL-400 |
| chr13-43894010-DEL-57 | chr13-43894011-DEL-57 |
| chr13-43981669-DEL-57 | chr13-43981670-DEL-57 |
| chr13-43990866-DEL-71 | chr13-43990867-DEL-71 |
| chr13-43993633-DEL-80 | chr13-43993634-DEL-80 |
| chr13-44110810-DEL-925 | chr13-44110811-DEL-925 |
| chr13-44160183-DEL-1737 | chr13-44160184-DEL-1737 |
| chr13-44211483-DEL-1372 | chr13-44211484-DEL-1372 |
| chr13-44315275-DEL-1567 | chr13-44315276-DEL-1567 |
| chr13-44336666-DEL-122 | chr13-44336667-DEL-122 |
| chr13-44417846-DEL-7091 | chr13-44417847-DEL-7091 |
| chr13-44440770-DEL-76 | chr13-44440771-DEL-76 |
| chr13-44457464-DEL-195 | chr13-44457465-DEL-195 |
| chr13-44473192-DEL-58 | chr13-44473193-DEL-58 |
| chr13-44504045-DEL-2815 | chr13-44504046-DEL-2815 |
| chr13-44517915-DEL-124 | chr13-44517916-DEL-124 |
| chr13-44523334-DEL-53 | chr13-44523332-DEL-52 |
| chr13-44534063-DEL-53 | chr13-44534064-DEL-53 |
| chr13-44539331-DEL-223 | chr13-44539332-DEL-223 |
| chr13-44546787-DEL-75 | chr13-44546788-DEL-75 |
| chr13-44563708-DEL-96 | chr13-44563709-DEL-96 |
| chr13-44578030-DEL-231 | chr13-44578031-DEL-231 |
| chr13-44580611-DEL-110 | chr13-44580612-DEL-110 |
| chr13-44626284-DEL-86 | chr13-44626285-DEL-86 |
| chr13-44633404-DEL-267 | chr13-44633405-DEL-267 |
| chr13-44634667-DEL-98 | chr13-44634697-DEL-68 |
| chr13-44639338-DEL-393 | chr13-44639341-DEL-391 |
| chr13-44641686-DEL-1864 | chr13-44641687-DEL-1864 |
| chr13-44647090-DEL-227 | chr13-44647091-DEL-227 |
| chr13-44666897-DEL-82 | chr13-44666926-DEL-50 |
| chr13-44681864-DEL-53 | chr13-44681865-DEL-63 |
| chr13-44695547-DEL-186 | chr13-44695547-DEL-187 |
| chr13-44768198-DEL-210 | chr13-44768199-DEL-210 |
| chr13-44774366-DEL-218 | chr13-44774367-DEL-218 |
| chr13-44781152-DEL-394 | chr13-44781160-DEL-397 |
| chr13-44797632-DEL-181 | chr13-44797627-DEL-187 |
| chr13-44809888-DEL-380 | chr13-44809885-DEL-381 |
| chr13-44826265-DEL-61 | chr13-44826258-DEL-85 |
| chr13-44828324-DEL-86 | chr13-44828326-DEL-79 |
| chr13-44862344-DEL-176 | chr13-44862345-DEL-176 |
| chr13-44866099-DEL-1887 | chr13-44866100-DEL-1887 |
| chr13-44877822-DEL-63 | chr13-44877823-DEL-63 |
| chr13-44916366-DEL-52 | chr13-44916364-DEL-54 |
| chr13-44927114-DEL-152 | chr13-44927115-DEL-152 |
| chr13-44956104-DEL-210 | chr13-44956105-DEL-210 |
| chr13-44957435-DEL-209 | chr13-44957436-DEL-209 |
| chr13-44965692-DEL-217 | chr13-44965693-DEL-217 |
| chr13-44966619-DEL-90 | chr13-44966620-DEL-90 |
| chr13-44969938-DEL-479 | chr13-44969939-DEL-479 |
| chr13-45056888-DEL-66 | chr13-45056889-DEL-51 |
| chr13-45134333-DEL-73 | chr13-45134334-DEL-73 |
| chr13-45160175-DEL-67 | chr13-45160197-DEL-71 |
| chr13-45164621-DEL-7092 | chr13-45164623-DEL-7091 |
| chr13-45231795-DEL-866 | chr13-45231796-DEL-866 |
| chr13-45251267-DEL-349 | chr13-45251268-DEL-349 |
| chr13-45261167-DEL-1259 | chr13-45261168-DEL-1259 |
| chr13-45291622-DEL-1649 | chr13-45291623-DEL-1649 |
| chr13-45309100-DEL-93 | chr13-45309098-DEL-93 |

| | |
|---|---|
| chr13-45327606-DEL-509 | chr13-45327607-DEL-509 |
| chr13-45328486-DEL-96 | chr13-45328489-DEL-97 |
| chr13-45394342-DEL-103 | chr13-45394343-DEL-103 |
| chr13-45429354-DEL-53 | chr13-45429354-DEL-53 |
| chr13-45491558-DEL-95 | chr13-45491559-DEL-95 |
| chr13-45509516-DEL-431 | chr13-45509517-DEL-431 |
| chr13-45529152-DEL-56 | chr13-45529153-DEL-56 |
| chr13-45585513-DEL-2162 | chr13-45585514-DEL-2162 |
| chr13-45600451-DEL-243 | chr13-45600452-DEL-243 |
| chr13-45618689-DEL-76 | chr13-45618711-DEL-54 |
| chr13-45623634-DEL-998 | chr13-45623635-DEL-998 |
| chr13-45676067-DEL-434 | chr13-45676068-DEL-434 |
| chr13-45679403-DEL-162 | chr13-45679408-DEL-158 |
| chr13-45760638-DEL-58 | chr13-45760639-DEL-58 |
| chr13-45790477-DEL-52 | chr13-45790467-DEL-62 |
| chr13-45800769-DEL-68 | chr13-45800770-DEL-68 |
| chr13-45821556-DEL-565 | chr13-45821557-DEL-565 |
| chr13-45828782-DEL-54 | chr13-45828743-DEL-94 |
| chr13-45965598-DEL-77 | chr13-45965599-DEL-77 |
| chr13-45967696-DEL-54 | chr13-45967691-DEL-60 |
| chr13-45978257-DEL-235 | chr13-45978256-DEL-235 |
| chr13-46080327-DEL-56 | chr13-46080328-DEL-56 |
| chr13-46105589-DEL-84 | chr13-46105590-DEL-84 |
| chr13-46151730-DEL-3028 | chr13-46151731-DEL-3028 |
| chr13-46165653-DEL-60 | chr13-46165635-DEL-104 |
| chr13-46170253-DEL-72 | chr13-46170254-DEL-72 |
| chr13-46173640-DEL-106 | chr13-46173616-DEL-101 |
| chr13-46198841-DEL-275 | chr13-46198837-DEL-291 |
| chr13-46199545-DEL-347 | chr13-46199546-DEL-347 |
| chr13-46217170-DEL-1050 | chr13-46217171-DEL-1050 |
| chr13-46232720-DEL-503 | chr13-46232721-DEL-503 |
| chr13-46296545-DEL-250 | chr13-46296546-DEL-250 |
| chr13-46303087-DEL-99 | chr13-46303088-DEL-99 |
| chr13-46350251-DEL-235 | chr13-46350282-DEL-205 |
| chr13-46351520-DEL-1080 | chr13-46351521-DEL-1080 |
| chr13-46379007-DEL-177 | chr13-46379008-DEL-177 |
| chr13-46379862-DEL-63 | chr13-46379908-DEL-58 |
| chr13-46387464-DEL-104 | chr13-46387465-DEL-104 |
| chr13-46416494-DEL-1198 | chr13-46416495-DEL-1198 |
| chr13-46474384-DEL-3137 | chr13-46474385-DEL-3137 |
| chr13-46529663-DEL-7073 | chr13-46529664-DEL-7073 |
| chr13-46538274-DEL-101 | chr13-46538241-DEL-116 |
| chr13-46579443-DEL-202 | chr13-46579443-DEL-203 |
| chr13-46593500-DEL-1890 | chr13-46593507-DEL-1897 |
| chr13-46607377-DEL-904 | chr13-46607378-DEL-903 |
| chr13-46639047-DEL-83 | chr13-46639006-DEL-84 |
| chr13-46639761-DEL-81 | chr13-46639773-DEL-61 |
| chr13-46647523-DEL-74 | chr13-46647524-DEL-74 |
| chr13-46651752-DEL-66 | chr13-46651753-DEL-66 |
| chr13-46654530-DEL-67 | chr13-46654531-DEL-67 |
| chr13-46666169-DEL-264 | chr13-46666170-DEL-264 |
| chr13-46698519-DEL-1224 | chr13-46698520-DEL-1224 |
| chr13-46707233-DEL-70 | chr13-46707234-DEL-70 |
| chr13-46719781-DEL-57 | chr13-46719791-DEL-52 |
| chr13-46726109-DEL-72 | chr13-46726110-DEL-72 |
| chr13-46739172-DEL-83 | chr13-46739189-DEL-71 |
| chr13-46753852-DEL-69 | chr13-46753853-DEL-69 |
| chr13-46777136-DEL-69 | chr13-46777134-DEL-107 |
| chr13-46786551-DEL-149 | chr13-46786563-DEL-148 |
| chr13-46830945-DEL-100 | chr13-46830946-DEL-100 |
| chr13-46895197-DEL-177 | chr13-46895198-DEL-177 |
| chr13-46929811-DEL-155 | chr13-46929812-DEL-155 |
| chr13-46953372-DEL-118 | chr13-46953373-DEL-118 |
| chr13-46969525-DEL-65 | chr13-46969512-DEL-66 |
| chr13-46981986-DEL-185 | chr13-46981987-DEL-185 |
| chr13-46990607-DEL-224 | chr13-46990608-DEL-224 |
| chr13-46996328-DEL-60 | chr13-46996329-DEL-60 |
| chr13-47001162-DEL-216 | chr13-47001163-DEL-216 |
| chr13-47010384-DEL-55 | chr13-47010381-DEL-52 |
| chr13-47022094-DEL-56 | chr13-47022097-DEL-54 |
| chr13-47040672-DEL-66 | chr13-47040664-DEL-66 |
| chr13-47091941-DEL-465 | chr13-47091942-DEL-465 |
| chr13-47103102-DEL-2052 | chr13-47103100-DEL-2053 |
| chr13-47104341-DEL-182 | chr13-47104342-DEL-182 |
| chr13-47120790-DEL-60 | chr13-47120795-DEL-74 |
| chr13-47130562-DEL-87 | chr13-47130563-DEL-87 |
| chr13-47149271-DEL-1340 | chr13-47149272-DEL-1340 |
| chr13-47158497-DEL-68 | chr13-47158498-DEL-68 |
| chr13-47175227-DEL-734 | chr13-47175229-DEL-732 |
| chr13-47179674-DEL-248 | chr13-47179675-DEL-248 |

| | |
|---|---|
| chr13-47181188-DEL-228 | chr13-47181189-DEL-228 |
| chr13-47183431-DEL-201 | chr13-47183422-DEL-198 |
| chr13-47217815-DEL-88 | chr13-47217816-DEL-88 |
| chr13-47268364-DEL-217 | chr13-47268365-DEL-217 |
| chr13-47269104-DEL-54 | chr13-47269068-DEL-55 |
| chr13-47312071-DEL-55 | chr13-47312072-DEL-55 |
| chr13-47344183-DEL-176 | chr13-47344201-DEL-129 |
| chr13-47358164-DEL-56 | chr13-47358165-DEL-56 |
| chr13-47367812-DEL-62 | chr13-47367814-DEL-61 |
| chr13-47384960-DEL-229 | chr13-47384961-DEL-229 |
| chr13-47444865-DEL-226 | chr13-47444866-DEL-226 |
| chr13-47449242-DEL-486 | chr13-47449243-DEL-486 |
| chr13-47451294-DEL-55 | chr13-47451291-DEL-61 |
| chr13-47457746-DEL-93 | chr13-47457743-DEL-58 |
| chr13-47467072-DEL-61 | chr13-47467059-DEL-50 |
| chr13-47477210-DEL-59 | chr13-47477211-DEL-59 |
| chr13-47494214-DEL-666 | chr13-47494215-DEL-666 |
| chr13-47503833-DEL-63 | chr13-47503830-DEL-67 |
| chr13-47506744-DEL-73 | chr13-47506747-DEL-70 |
| chr13-47535968-DEL-367 | chr13-47535969-DEL-367 |
| chr13-47556230-DEL-52 | chr13-47556231-DEL-52 |
| chr13-47561119-DEL-203 | chr13-47561120-DEL-203 |
| chr13-47577185-DEL-102 | chr13-47577186-DEL-136 |
| chr13-47579755-DEL-104 | chr13-47579780-DEL-95 |
| chr13-47612086-DEL-54 | chr13-47612087-DEL-52 |
| chr13-47622558-DEL-143 | chr13-47622559-DEL-142 |
| chr13-47636402-DEL-908 | chr13-47636403-DEL-908 |
| chr13-47651527-DEL-404 | chr13-47651528-DEL-404 |
| chr13-47662472-DEL-64 | chr13-47662477-DEL-60 |
| chr13-47669597-DEL-312 | chr13-47669598-DEL-312 |
| chr13-47674000-DEL-7934 | chr13-47674001-DEL-7934 |
| chr13-47680679-DEL-416 | chr13-47680680-DEL-416 |
| chr13-47701337-DEL-114 | chr13-47701336-DEL-84 |
| chr13-47747314-DEL-62 | chr13-47747315-DEL-62 |
| chr13-47768563-DEL-317 | chr13-47768564-DEL-320 |
| chr13-47774504-DEL-551 | chr13-47774505-DEL-551 |
| chr13-47785708-DEL-118 | chr13-47785711-DEL-90 |
| chr13-47792358-DEL-52 | chr13-47792358-DEL-52 |
| chr13-47797453-DEL-102 | chr13-47797442-DEL-100 |
| chr13-47857576-DEL-132 | chr13-47857577-DEL-133 |
| chr13-47893449-DEL-6586 | chr13-47893450-DEL-6586 |
| chr13-47931624-DEL-71 | chr13-47931625-DEL-71 |
| chr13-47955414-DEL-70 | chr13-47955414-DEL-70 |
| chr13-47967924-DEL-67 | chr13-47967919-DEL-57 |
| chr13-47979019-DEL-1355 | chr13-47979020-DEL-1355 |
| chr13-48004091-DEL-102 | chr13-48004092-DEL-102 |
| chr13-48017442-DEL-713 | chr13-48017443-DEL-713 |
| chr13-48022777-DEL-79 | chr13-48022778-DEL-79 |
| chr13-48079868-DEL-852 | chr13-48079869-DEL-852 |
| chr13-48214044-DEL-72 | chr13-48214045-DEL-72 |
| chr13-48378618-DEL-56 | chr13-48378619-DEL-56 |
| chr13-48387339-DEL-420 | chr13-48387340-DEL-420 |
| chr13-48450544-DEL-6839 | chr13-48450544-DEL-6839 |
| chr13-48464314-DEL-79 | chr13-48464315-DEL-79 |
| chr13-48514973-DEL-135 | chr13-48514974-DEL-135 |
| chr13-48517041-DEL-567 | chr13-48517039-DEL-554 |
| chr13-48575479-DEL-102 | chr13-48575480-DEL-102 |
| chr13-48624528-DEL-97 | chr13-48624529-DEL-97 |
| chr13-48625621-DEL-1130 | chr13-48625622-DEL-1130 |
| chr13-48628930-DEL-440 | chr13-48628931-DEL-440 |
| chr13-48651536-DEL-55 | chr13-48651537-DEL-55 |
| chr13-48654551-DEL-399 | chr13-48654552-DEL-399 |
| chr13-48755752-DEL-63 | chr13-48755753-DEL-63 |
| chr13-48839446-DEL-82 | chr13-48839409-DEL-71 |
| chr13-48841261-DEL-52 | chr13-48841269-DEL-64 |
| chr13-48870464-DEL-69 | chr13-48870465-DEL-69 |
| chr13-48881998-DEL-63 | chr13-48881999-DEL-62 |
| chr13-48895171-DEL-658 | chr13-48895172-DEL-658 |
| chr13-48926270-DEL-66 | chr13-48926270-DEL-66 |
| chr13-48963397-DEL-67 | chr13-48963398-DEL-67 |
| chr13-49002447-DEL-76 | chr13-49002452-DEL-56 |
| chr13-49006018-DEL-72 | chr13-49006025-DEL-60 |
| chr13-49060613-DEL-54 | chr13-49060614-DEL-54 |
| chr13-49092282-DEL-56 | chr13-49092283-DEL-56 |
| chr13-49141524-DEL-620 | chr13-49141525-DEL-620 |
| chr13-49156249-DEL-51 | chr13-49156240-DEL-52 |
| chr13-49158766-DEL-211 | chr13-49158767-DEL-211 |
| chr13-49175967-DEL-198 | chr13-49175968-DEL-198 |
| chr13-49177019-DEL-181 | chr13-49177020-DEL-181 |
| chr13-49188492-DEL-176 | chr13-49188493-DEL-176 |

| | |
|---|---|
| chr13-49276757-DEL-71 | chr13-49276758-DEL-71 |
| chr13-49278328-DEL-1367 | chr13-49278329-DEL-1367 |
| chr13-49378953-DEL-432 | chr13-49378954-DEL-432 |
| chr13-49426717-DEL-76 | chr13-49426720-DEL-74 |
| chr13-49493852-DEL-55 | chr13-49493854-DEL-53 |
| chr13-49519719-DEL-86 | chr13-49519716-DEL-90 |
| chr13-49599117-DEL-232 | chr13-49599118-DEL-232 |
| chr13-49636335-DEL-116 | chr13-49636358-DEL-70 |
| chr13-49758741-DEL-107 | chr13-49758742-DEL-107 |
| chr13-49759154-DEL-442 | chr13-49759155-DEL-442 |
| chr13-49786967-DEL-1930 | chr13-49786968-DEL-1930 |
| chr13-49799486-DEL-151 | chr13-49799487-DEL-151 |
| chr13-49847266-DEL-6403 | chr13-49847267-DEL-6403 |
| chr13-49866150-DEL-60 | chr13-49866164-DEL-52 |
| chr13-49895695-DEL-64 | chr13-49895703-DEL-57 |
| chr13-49896914-DEL-132 | chr13-49896914-DEL-99 |
| chr13-49913534-DEL-162 | chr13-49913534-DEL-163 |
| chr13-49922999-DEL-340 | chr13-49923000-DEL-340 |
| chr13-49934334-DEL-728 | chr13-49934335-DEL-728 |
| chr13-49945368-DEL-100 | chr13-49945369-DEL-100 |
| chr13-49953065-DEL-2710 | chr13-49953059-DEL-2716 |
| chr13-49970892-DEL-56 | chr13-49970893-DEL-56 |
| chr13-49976212-DEL-81 | chr13-49976213-DEL-81 |
| chr13-50000441-DEL-56 | chr13-50000441-DEL-57 |
| chr13-50003102-DEL-94 | chr13-50003103-DEL-94 |
| chr13-50071379-DEL-2863 | chr13-50071380-DEL-2863 |
| chr13-50123731-DEL-161 | chr13-50123732-DEL-161 |
| chr13-50129300-DEL-57 | chr13-50129301-DEL-57 |
| chr13-50231376-DEL-381 | chr13-50231376-DEL-381 |
| chr13-50259244-DEL-412 | chr13-50259244-DEL-412 |
| chr13-50267931-DEL-109 | chr13-50267932-DEL-109 |
| chr13-50319970-DEL-1658 | chr13-50319971-DEL-1658 |
| chr13-50323527-DEL-3338 | chr13-50323528-DEL-3338 |
| chr13-50376764-DEL-76 | chr13-50376765-DEL-76 |
| chr13-50393973-DEL-78 | chr13-50393974-DEL-78 |
| chr13-50555208-DEL-1091 | chr13-50555209-DEL-1091 |
| chr13-50655528-DEL-3859 | chr13-50655529-DEL-3859 |
| chr13-50792366-DEL-3292 | chr13-50792367-DEL-3292 |
| chr13-50797561-DEL-1652 | chr13-50797562-DEL-1652 |
| chr13-50884907-DEL-4797 | chr13-50884908-DEL-4797 |
| chr13-50967371-DEL-280 | chr13-50967372-DEL-280 |
| chr13-50974663-DEL-384 | chr13-50974712-DEL-336 |
| chr13-50994330-DEL-1386 | chr13-50994331-DEL-1386 |
| chr13-51047389-DEL-6465 | chr13-51047390-DEL-6465 |
| chr13-51067805-DEL-51 | chr13-51067806-DEL-51 |
| chr13-51117017-DEL-88 | chr13-51117018-DEL-88 |
| chr13-51154718-DEL-337 | chr13-51154719-DEL-337 |
| chr13-51155851-DEL-160 | chr13-51155852-DEL-160 |
| chr13-51167656-DEL-246 | chr13-51167657-DEL-247 |
| chr13-51173186-DEL-72 | chr13-51173187-DEL-72 |
| chr13-51181904-DEL-552 | chr13-51181914-DEL-543 |
| chr13-51186311-DEL-3911 | chr13-51186332-DEL-3878 |
| chr13-51202451-DEL-82 | chr13-51202452-DEL-82 |
| chr13-51238548-DEL-221 | chr13-51238549-DEL-221 |
| chr13-51251109-DEL-340 | chr13-51251110-DEL-340 |
| chr13-51296166-DEL-113 | chr13-51296167-DEL-113 |
| chr13-51341999-DEL-192 | chr13-51341983-DEL-205 |
| chr13-51356312-DEL-785 | chr13-51356313-DEL-785 |
| chr13-51364092-DEL-125 | chr13-51364093-DEL-125 |
| chr13-51373989-DEL-1598 | chr13-51373994-DEL-1596 |
| chr13-51406296-DEL-432 | chr13-51406297-DEL-432 |
| chr13-51411913-DEL-86 | chr13-51411883-DEL-117 |
| chr13-51419744-DEL-5265 | chr13-51419745-DEL-5265 |
| chr13-51427786-DEL-785 | chr13-51427787-DEL-785 |
| chr13-51429708-DEL-71 | chr13-51429709-DEL-71 |
| chr13-51448780-DEL-75 | chr13-51448760-DEL-92 |
| chr13-51459209-DEL-63 | chr13-51459178-DEL-63 |
| chr13-51470066-DEL-71 | chr13-51470080-DEL-58 |
| chr13-51499284-DEL-106 | chr13-51499258-DEL-106 |
| chr13-51512100-DEL-263 | chr13-51512101-DEL-263 |
| chr13-51514942-DEL-981 | chr13-51514943-DEL-981 |
| chr13-51529623-DEL-64 | chr13-51529608-DEL-59 |
| chr13-51534073-DEL-184 | chr13-51534087-DEL-180 |
| chr13-51556189-DEL-55 | chr13-51556189-DEL-55 |
| chr13-51595175-DEL-75 | chr13-51595156-DEL-89 |
| chr13-51641079-DEL-103 | chr13-51641080-DEL-101 |
| chr13-51647077-DEL-401 | chr13-51647078-DEL-401 |
| chr13-51658456-DEL-183 | chr13-51658471-DEL-173 |
| chr13-51676468-DEL-91 | chr13-51676495-DEL-55 |
| chr13-51686617-DEL-170 | chr13-51686618-DEL-170 |

| | |
|---|---|
| chr13-51741057-DEL-55 | chr13-51741058-DEL-55 |
| chr13-51747463-DEL-290 | chr13-51747463-DEL-294 |
| chr13-51760779-DEL-72 | chr13-51760753-DEL-81 |
| chr13-51826119-DEL-813 | chr13-51826120-DEL-813 |
| chr13-51826183-DEL-214 | chr13-51826184-DEL-214 |
| chr13-51845948-DEL-215 | chr13-51845949-DEL-215 |
| chr13-51864520-DEL-57 | chr13-51864520-DEL-57 |
| chr13-51880607-DEL-260 | chr13-51880608-DEL-260 |
| chr13-51881671-DEL-202 | chr13-51881692-DEL-190 |
| chr13-51907870-DEL-62 | chr13-51907871-DEL-62 |
| chr13-51923349-DEL-59 | chr13-51923348-DEL-68 |
| chr13-51949569-DEL-164 | chr13-51949570-DEL-164 |
| chr13-51967256-DEL-51 | chr13-51967257-DEL-51 |
| chr13-51971650-DEL-66 | chr13-51971634-DEL-52 |
| chr13-51992459-DEL-135 | chr13-51992460-DEL-135 |
| chr13-51999057-DEL-81 | chr13-51999058-DEL-83 |
| chr13-52010342-DEL-81 | chr13-52010349-DEL-75 |
| chr13-52024237-DEL-493 | chr13-52024238-DEL-493 |
| chr13-52043820-DEL-59 | chr13-52043821-DEL-59 |
| chr13-52044989-DEL-162 | chr13-52044952-DEL-202 |
| chr13-52047778-DEL-3024 | chr13-52047779-DEL-3024 |
| chr13-52064094-DEL-212 | chr13-52064095-DEL-212 |
| chr13-52064915-DEL-54 | chr13-52064916-DEL-54 |
| chr13-52067969-DEL-216 | chr13-52067974-DEL-211 |
| chr13-52068825-DEL-76 | chr13-52068820-DEL-108 |
| chr13-52111845-DEL-134 | chr13-52111846-DEL-134 |
| chr13-52139511-DEL-5269 | chr13-52139512-DEL-5269 |
| chr13-52204476-DEL-3710 | chr13-52204477-DEL-3710 |
| chr13-52209573-DEL-529 | chr13-52209574-DEL-529 |
| chr13-52228480-DEL-971 | chr13-52228481-DEL-971 |
| chr13-52235903-DEL-494 | chr13-52235904-DEL-494 |
| chr13-52241561-DEL-79 | chr13-52241562-DEL-79 |
| chr13-52332530-DEL-87 | chr13-52332531-DEL-87 |
| chr13-52361206-DEL-309 | chr13-52361207-DEL-309 |
| chr13-52362953-DEL-54 | chr13-52362954-DEL-62 |
| chr13-52370402-DEL-54 | chr13-52370403-DEL-54 |
| chr13-52389287-DEL-68 | chr13-52389292-DEL-64 |
| chr13-52402041-DEL-109 | chr13-52402042-DEL-61 |
| chr13-52416311-DEL-95 | chr13-52416312-DEL-95 |
| chr13-52443548-DEL-5400 | chr13-52443549-DEL-5400 |
| chr13-52474449-DEL-111 | chr13-52474462-DEL-99 |
| chr13-52478499-DEL-72 | chr13-52478493-DEL-73 |
| chr13-52504282-DEL-1274 | chr13-52504283-DEL-1274 |
| chr13-52513336-DEL-142 | chr13-52513337-DEL-142 |
| chr13-52520793-DEL-176 | chr13-52520794-DEL-176 |
| chr13-52523340-DEL-53 | chr13-52523338-DEL-53 |
| chr13-52536123-DEL-7097 | chr13-52536124-DEL-7097 |
| chr13-52556080-DEL-82 | chr13-52556097-DEL-66 |
| chr13-52557917-DEL-105 | chr13-52557918-DEL-105 |
| chr13-52571689-DEL-517 | chr13-52571690-DEL-517 |
| chr13-52591416-DEL-70 | chr13-52591417-DEL-68 |
| chr13-52599955-DEL-55 | chr13-52599956-DEL-55 |
| chr13-52607790-DEL-55 | chr13-52607789-DEL-57 |
| chr13-52619670-DEL-88 | chr13-52619671-DEL-88 |
| chr13-52641631-DEL-63 | chr13-52641632-DEL-63 |
| chr13-52647145-DEL-90 | chr13-52647146-DEL-90 |
| chr13-52656119-DEL-140 | chr13-52656120-DEL-140 |
| chr13-52676067-DEL-429 | chr13-52676068-DEL-429 |
| chr13-52691113-DEL-68 | chr13-52691114-DEL-67 |
| chr13-52694819-DEL-1171 | chr13-52694820-DEL-1172 |
| chr13-52713043-DEL-81 | chr13-52713044-DEL-81 |
| chr13-52718433-DEL-120 | chr13-52718474-DEL-80 |
| chr13-52727946-DEL-80 | chr13-52727947-DEL-80 |
| chr13-52745123-DEL-153 | chr13-52745127-DEL-149 |
| chr13-52793040-DEL-82 | chr13-52793041-DEL-82 |
| chr13-52802451-DEL-102 | chr13-52802452-DEL-102 |
| chr13-52803812-DEL-112 | chr13-52803813-DEL-112 |
| chr13-52807588-DEL-56 | chr13-52807589-DEL-56 |
| chr13-52809488-DEL-95 | chr13-52809489-DEL-95 |
| chr13-52826034-DEL-203 | chr13-52826035-DEL-203 |
| chr13-52837300-DEL-332 | chr13-52837302-DEL-328 |
| chr13-52871083-DEL-394 | chr13-52871084-DEL-394 |
| chr13-52873120-DEL-57 | chr13-52873107-DEL-58 |
| chr13-52888233-DEL-56 | chr13-52888234-DEL-56 |
| chr13-52944514-DEL-65 | chr13-52944514-DEL-66 |
| chr13-52983897-DEL-65 | chr13-52983901-DEL-66 |
| chr13-53000534-DEL-54 | chr13-53000535-DEL-54 |
| chr13-53047285-DEL-131 | chr13-53047286-DEL-131 |
| chr13-53051401-DEL-59 | chr13-53051398-DEL-63 |
| chr13-53054142-DEL-60 | chr13-53054143-DEL-60 |

| | |
|---|---|
| chr13-53105851-DEL-53 | chr13-53105852-DEL-53 |
| chr13-53127287-DEL-238 | chr13-53127288-DEL-238 |
| chr13-53149923-DEL-52 | chr13-53149924-DEL-52 |
| chr13-53153211-DEL-51 | chr13-53153212-DEL-51 |
| chr13-53170879-DEL-1205 | chr13-53170880-DEL-1205 |
| chr13-53179732-DEL-99 | chr13-53179733-DEL-99 |
| chr13-53180191-DEL-348 | chr13-53180192-DEL-346 |
| chr13-53190524-DEL-332 | chr13-53190525-DEL-332 |
| chr13-53195017-DEL-204 | chr13-53195018-DEL-204 |
| chr13-53218415-DEL-56 | chr13-53218416-DEL-56 |
| chr13-53257105-DEL-188 | chr13-53257106-DEL-188 |
| chr13-53267574-DEL-138 | chr13-53267575-DEL-147 |
| chr13-53289422-DEL-1870 | chr13-53289423-DEL-1870 |
| chr13-53303992-DEL-59 | chr13-53303973-DEL-81 |
| chr13-53312608-DEL-150 | chr13-53312609-DEL-150 |
| chr13-53429713-DEL-52 | chr13-53429714-DEL-52 |
| chr13-53457818-DEL-170 | chr13-53457819-DEL-170 |
| chr13-53482984-DEL-309 | chr13-53482985-DEL-309 |
| chr13-53534538-DEL-1325 | chr13-53534539-DEL-1325 |
| chr13-53566377-DEL-126 | chr13-53566378-DEL-126 |
| chr13-53585905-DEL-503 | chr13-53585906-DEL-503 |
| chr13-53597264-DEL-197 | chr13-53597266-DEL-196 |
| chr13-53600328-DEL-54 | chr13-53600317-DEL-74 |
| chr13-53605935-DEL-130 | chr13-53605936-DEL-128 |
| chr13-53607793-DEL-71 | chr13-53607795-DEL-65 |
| chr13-53642396-DEL-52 | chr13-53642397-DEL-52 |
| chr13-53658815-DEL-104 | chr13-53658810-DEL-103 |
| chr13-53681629-DEL-254 | chr13-53681630-DEL-254 |
| chr13-53703819-DEL-214 | chr13-53703820-DEL-214 |
| chr13-53705850-DEL-106 | chr13-53705850-DEL-106 |
| chr13-53712154-DEL-179 | chr13-53712151-DEL-165 |
| chr13-53743201-DEL-119 | chr13-53743202-DEL-119 |
| chr13-53746118-DEL-54 | chr13-53746119-DEL-54 |
| chr13-53753917-DEL-146 | chr13-53753930-DEL-134 |
| chr13-53755538-DEL-92 | chr13-53755554-DEL-77 |
| chr13-53777304-DEL-70 | chr13-53777305-DEL-70 |
| chr13-53792427-DEL-169 | chr13-53792428-DEL-169 |
| chr13-53815790-DEL-128 | chr13-53815796-DEL-124 |
| chr13-53847745-DEL-92 | chr13-53847734-DEL-70 |
| chr13-53888543-DEL-767 | chr13-53888544-DEL-767 |
| chr13-53898917-DEL-2129 | chr13-53898918-DEL-2129 |
| chr13-53925214-DEL-153 | chr13-53925215-DEL-139 |
| chr13-53949071-DEL-54 | chr13-53949076-DEL-50 |
| chr13-53954647-DEL-54 | chr13-53954648-DEL-54 |
| chr13-53990323-DEL-96 | chr13-53990324-DEL-64 |
| chr13-54001522-DEL-1189 | chr13-54001523-DEL-1189 |
| chr13-54001607-DEL-498 | chr13-54001608-DEL-498 |
| chr13-54006555-DEL-134 | chr13-54006544-DEL-146 |
| chr13-54027666-DEL-199 | chr13-54027662-DEL-200 |
| chr13-54079874-DEL-193 | chr13-54079875-DEL-193 |
| chr13-54086973-DEL-401 | chr13-54086974-DEL-401 |
| chr13-54087805-DEL-1620 | chr13-54087806-DEL-1620 |
| chr13-54102770-DEL-3068 | chr13-54102771-DEL-3068 |
| chr13-54111387-DEL-6291 | chr13-54111388-DEL-6291 |
| chr13-54120653-DEL-57 | chr13-54120654-DEL-57 |
| chr13-54122660-DEL-341 | chr13-54122644-DEL-353 |
| chr13-54127276-DEL-992 | chr13-54127277-DEL-992 |
| chr13-54132091-DEL-105 | chr13-54132089-DEL-110 |
| chr13-54138051-DEL-844 | chr13-54138052-DEL-844 |
| chr13-54182342-DEL-440 | chr13-54182343-DEL-440 |
| chr13-54196580-DEL-184 | chr13-54196581-DEL-184 |
| chr13-54200939-DEL-216 | chr13-54200940-DEL-216 |
| chr13-54225349-DEL-248 | chr13-54225350-DEL-248 |
| chr13-54235664-DEL-203 | chr13-54235665-DEL-203 |
| chr13-54286777-DEL-382 | chr13-54286778-DEL-382 |
| chr13-54287828-DEL-131 | chr13-54287829-DEL-131 |
| chr13-54301644-DEL-80 | chr13-54301645-DEL-104 |
| chr13-54310721-DEL-373 | chr13-54310722-DEL-373 |
| chr13-54323742-DEL-100 | chr13-54323743-DEL-100 |
| chr13-54337910-DEL-53 | chr13-54337911-DEL-54 |
| chr13-54365808-DEL-76 | chr13-54365813-DEL-72 |
| chr13-54390747-DEL-153 | chr13-54390748-DEL-153 |
| chr13-54399938-DEL-60 | chr13-54399938-DEL-60 |
| chr13-54479613-DEL-102 | chr13-54479614-DEL-102 |
| chr13-54482836-DEL-55 | chr13-54482798-DEL-98 |
| chr13-54504492-DEL-93 | chr13-54504493-DEL-93 |
| chr13-54514999-DEL-67 | chr13-54515000-DEL-67 |
| chr13-54560473-DEL-59 | chr13-54560474-DEL-59 |
| chr13-54568813-DEL-2592 | chr13-54568814-DEL-2592 |
| chr13-54583941-DEL-160 | chr13-54583942-DEL-160 |

| | |
|---|---|
| chr13-54592349-DEL-70 | chr13-54592350-DEL-70 |
| chr13-54626848-DEL-276 | chr13-54626855-DEL-278 |
| chr13-54647937-DEL-199 | chr13-54647938-DEL-199 |
| chr13-54655801-DEL-61 | chr13-54655802-DEL-61 |
| chr13-54692768-DEL-1486 | chr13-54692769-DEL-1486 |
| chr13-54720211-DEL-205 | chr13-54720212-DEL-205 |
| chr13-54726964-DEL-953 | chr13-54726961-DEL-954 |
| chr13-54755409-DEL-86 | chr13-54755423-DEL-81 |
| chr13-54779771-DEL-210 | chr13-54779772-DEL-210 |
| chr13-54795055-DEL-55 | chr13-54795056-DEL-55 |
| chr13-54818723-DEL-69 | chr13-54818732-DEL-61 |
| chr13-54854625-DEL-55 | chr13-54854626-DEL-55 |
| chr13-54875270-DEL-62 | chr13-54875271-DEL-62 |
| chr13-54877440-DEL-493 | chr13-54877441-DEL-493 |
| chr13-54888829-DEL-201 | chr13-54888830-DEL-201 |
| chr13-54918902-DEL-4785 | chr13-54918903-DEL-4785 |
| chr13-54973226-DEL-1882 | chr13-54973227-DEL-1882 |
| chr13-54999074-DEL-58 | chr13-54999073-DEL-58 |
| chr13-55001264-DEL-290 | chr13-55001265-DEL-290 |
| chr13-55004859-DEL-196 | chr13-55004860-DEL-196 |
| chr13-55049795-DEL-176 | chr13-55049796-DEL-176 |
| chr13-55053495-DEL-76 | chr13-55053512-DEL-86 |
| chr13-55246783-DEL-76 | chr13-55246786-DEL-74 |
| chr13-55249916-DEL-64 | chr13-55249915-DEL-66 |
| chr13-55257823-DEL-182 | chr13-55257824-DEL-182 |
| chr13-55275043-DEL-625 | chr13-55275044-DEL-625 |
| chr13-55292151-DEL-771 | chr13-55292152-DEL-771 |
| chr13-55312161-DEL-173 | chr13-55312162-DEL-173 |
| chr13-55313064-DEL-165 | chr13-55313065-DEL-165 |
| chr13-55314383-DEL-173 | chr13-55314384-DEL-173 |
| chr13-55332255-DEL-4798 | chr13-55332256-DEL-4798 |
| chr13-55343192-DEL-67 | chr13-55343200-DEL-60 |
| chr13-55369686-DEL-221 | chr13-55369687-DEL-221 |
| chr13-55372157-DEL-72 | chr13-55372158-DEL-72 |
| chr13-55376483-DEL-150 | chr13-55376484-DEL-150 |
| chr13-55385069-DEL-121 | chr13-55385070-DEL-121 |
| chr13-55436531-DEL-239 | chr13-55436533-DEL-204 |
| chr13-55476852-DEL-75 | chr13-55476861-DEL-58 |
| chr13-55491015-DEL-51 | chr13-55491016-DEL-51 |
| chr13-55537087-DEL-290 | chr13-55537088-DEL-290 |
| chr13-55538000-DEL-394 | chr13-55538001-DEL-394 |
| chr13-55549200-DEL-1253 | chr13-55549201-DEL-1253 |
| chr13-55566758-DEL-204 | chr13-55566759-DEL-204 |
| chr13-55727875-DEL-100 | chr13-55727835-DEL-101 |
| chr13-55734924-DEL-224 | chr13-55734921-DEL-228 |
| chr13-55741931-DEL-177 | chr13-55741932-DEL-177 |
| chr13-55767506-DEL-149 | chr13-55767507-DEL-149 |
| chr13-55775316-DEL-279 | chr13-55775317-DEL-279 |
| chr13-55778191-DEL-219 | chr13-55778192-DEL-219 |
| chr13-55818805-DEL-209 | chr13-55818806-DEL-209 |
| chr13-55836709-DEL-175 | chr13-55836710-DEL-175 |
| chr13-55844043-DEL-180 | chr13-55844044-DEL-180 |
| chr13-55848022-DEL-175 | chr13-55848023-DEL-175 |
| chr13-55855323-DEL-201 | chr13-55855324-DEL-201 |
| chr13-55899723-DEL-209 | chr13-55899724-DEL-209 |
| chr13-55906613-DEL-217 | chr13-55906614-DEL-217 |
| chr13-55926869-DEL-4690 | chr13-55926870-DEL-4690 |
| chr13-55936902-DEL-205 | chr13-55936903-DEL-205 |
| chr13-55962061-DEL-154 | chr13-55962084-DEL-122 |
| chr13-55992303-DEL-1501 | chr13-55992304-DEL-1501 |
| chr13-56031908-DEL-755 | chr13-56031909-DEL-755 |
| chr13-56070616-DEL-53 | chr13-56070617-DEL-53 |
| chr13-56073547-DEL-231 | chr13-56073548-DEL-231 |
| chr13-56086750-DEL-53 | chr13-56086790-DEL-61 |
| chr13-56118117-DEL-60 | chr13-56118118-DEL-60 |
| chr13-56119123-DEL-58 | chr13-56119117-DEL-59 |
| chr13-56139571-DEL-203 | chr13-56139572-DEL-203 |
| chr13-56175742-DEL-192 | chr13-56175743-DEL-192 |
| chr13-56193785-DEL-980 | chr13-56193797-DEL-981 |
| chr13-56194289-DEL-106 | chr13-56194286-DEL-110 |
| chr13-56246380-DEL-835 | chr13-56246381-DEL-835 |
| chr13-56277210-DEL-188 | chr13-56277211-DEL-188 |
| chr13-56288103-DEL-2193 | chr13-56288104-DEL-2193 |
| chr13-56306591-DEL-113 | chr13-56306592-DEL-113 |
| chr13-56322789-DEL-84 | chr13-56322790-DEL-84 |
| chr13-56339547-DEL-86 | chr13-56339548-DEL-86 |
| chr13-56343872-DEL-434 | chr13-56343873-DEL-434 |
| chr13-56353892-DEL-570 | chr13-56353893-DEL-570 |
| chr13-56365399-DEL-56 | chr13-56365387-DEL-63 |
| chr13-56366636-DEL-52 | chr13-56366628-DEL-68 |

| | |
|---|---|
| chr13-56367469-DEL-202 | chr13-56367470-DEL-202 |
| chr13-56383612-DEL-56 | chr13-56383613-DEL-56 |
| chr13-56385101-DEL-54 | chr13-56385103-DEL-52 |
| chr13-56389319-DEL-54 | chr13-56389320-DEL-54 |
| chr13-56407583-DEL-56 | chr13-56407584-DEL-56 |
| chr13-56409542-DEL-53 | chr13-56409543-DEL-53 |
| chr13-56424109-DEL-69 | chr13-56424110-DEL-69 |
| chr13-56433102-DEL-69 | chr13-56433103-DEL-69 |
| chr13-56451779-DEL-345 | chr13-56451780-DEL-345 |
| chr13-56503327-DEL-120 | chr13-56503328-DEL-120 |
| chr13-56509612-DEL-118 | chr13-56509613-DEL-117 |
| chr13-56512496-DEL-131 | chr13-56512503-DEL-127 |
| chr13-56572299-DEL-53 | chr13-56572300-DEL-53 |
| chr13-56599942-DEL-588 | chr13-56599944-DEL-587 |
| chr13-56609803-DEL-507 | chr13-56609797-DEL-497 |
| chr13-56620234-DEL-129 | chr13-56620247-DEL-111 |
| chr13-56621551-DEL-506 | chr13-56621552-DEL-506 |
| chr13-56640623-DEL-2321 | chr13-56640619-DEL-2360 |
| chr13-56651977-DEL-340 | chr13-56651978-DEL-340 |
| chr13-56656092-DEL-441 | chr13-56656093-DEL-441 |
| chr13-56661857-DEL-59 | chr13-56661858-DEL-59 |
| chr13-56688845-DEL-142 | chr13-56688892-DEL-108 |
| chr13-56698767-DEL-64 | chr13-56698758-DEL-63 |
| chr13-56723287-DEL-115 | chr13-56723333-DEL-118 |
| chr13-56764555-DEL-86 | chr13-56764556-DEL-86 |
| chr13-56813527-DEL-105 | chr13-56813528-DEL-105 |
| chr13-56828015-DEL-986 | chr13-56828016-DEL-986 |
| chr13-56835171-DEL-57 | chr13-56835172-DEL-57 |
| chr13-56860687-DEL-7103 | chr13-56860688-DEL-7103 |
| chr13-56902543-DEL-184 | chr13-56902544-DEL-184 |
| chr13-56903485-DEL-90 | chr13-56903486-DEL-90 |
| chr13-56933192-DEL-72 | chr13-56933193-DEL-72 |
| chr13-56936421-DEL-64 | chr13-56936422-DEL-64 |
| chr13-56977489-DEL-98 | chr13-56977490-DEL-128 |
| chr13-56993315-DEL-249 | chr13-56993316-DEL-249 |
| chr13-57013138-DEL-349 | chr13-57013139-DEL-349 |
| chr13-57027110-DEL-65 | chr13-57027121-DEL-55 |
| chr13-57074865-DEL-138 | chr13-57074866-DEL-138 |
| chr13-57077205-DEL-914 | chr13-57077206-DEL-914 |
| chr13-57083528-DEL-94 | chr13-57083529-DEL-94 |
| chr13-57097067-DEL-981 | chr13-57097068-DEL-981 |
| chr13-57125477-DEL-6200 | chr13-57125477-DEL-6201 |
| chr13-57289657-DEL-54 | chr13-57289629-DEL-55 |
| chr13-57304010-DEL-117 | chr13-57304011-DEL-117 |
| chr13-57324961-DEL-79 | chr13-57324962-DEL-79 |
| chr13-57338149-DEL-105 | chr13-57338150-DEL-105 |
| chr13-57349800-DEL-6244 | chr13-57349801-DEL-6244 |
| chr13-57376148-DEL-312 | chr13-57376149-DEL-303 |
| chr13-57382979-DEL-114 | chr13-57382978-DEL-116 |
| chr13-57393658-DEL-68 | chr13-57393659-DEL-68 |
| chr13-57431137-DEL-54 | chr13-57431138-DEL-74 |
| chr13-57447686-DEL-112 | chr13-57447661-DEL-154 |
| chr13-57520093-DEL-76 | chr13-57520094-DEL-76 |
| chr13-57528606-DEL-199 | chr13-57528607-DEL-198 |
| chr13-57530299-DEL-167 | chr13-57530302-DEL-164 |
| chr13-57543902-DEL-76 | chr13-57543903-DEL-76 |
| chr13-57584346-DEL-273 | chr13-57584342-DEL-275 |
| chr13-57618814-DEL-226 | chr13-57618815-DEL-226 |
| chr13-57630795-DEL-194 | chr13-57630796-DEL-194 |
| chr13-57645836-DEL-90 | chr13-57645837-DEL-90 |
| chr13-57696151-DEL-57 | chr13-57696152-DEL-57 |
| chr13-57705191-DEL-1051 | chr13-57705192-DEL-1051 |
| chr13-57716631-DEL-58 | chr13-57716635-DEL-55 |
| chr13-57825916-DEL-113 | chr13-57825915-DEL-114 |
| chr13-57945515-DEL-95 | chr13-57945516-DEL-95 |
| chr13-57966561-DEL-233 | chr13-57966562-DEL-233 |
| chr13-57983856-DEL-839 | chr13-57983856-DEL-839 |
| chr13-58035644-DEL-52 | chr13-58035645-DEL-52 |
| chr13-58058068-DEL-138 | chr13-58058069-DEL-144 |
| chr13-58078406-DEL-72 | chr13-58078414-DEL-70 |
| chr13-58088700-DEL-105 | chr13-58088667-DEL-102 |
| chr13-58137390-DEL-81 | chr13-58137391-DEL-81 |
| chr13-58137987-DEL-69 | chr13-58137988-DEL-69 |
| chr13-58141590-DEL-321 | chr13-58141617-DEL-291 |
| chr13-58149675-DEL-91 | chr13-58149677-DEL-90 |
| chr13-58154898-DEL-163 | chr13-58154899-DEL-122 |
| chr13-58168159-DEL-52 | chr13-58168160-DEL-54 |
| chr13-58176676-DEL-109 | chr13-58176679-DEL-107 |
| chr13-58258209-DEL-112 | chr13-58258210-DEL-112 |
| chr13-58278441-DEL-52 | chr13-58278438-DEL-52 |

| | |
|---|---|
| chr13-58294677-DEL-67 | chr13-58294672-DEL-63 |
| chr13-58299053-DEL-2593 | chr13-58299054-DEL-2593 |
| chr13-58311022-DEL-142 | chr13-58311023-DEL-142 |
| chr13-58319763-DEL-217 | chr13-58319764-DEL-217 |
| chr13-58321271-DEL-76 | chr13-58321272-DEL-76 |
| chr13-58354338-DEL-163 | chr13-58354339-DEL-163 |
| chr13-58411628-DEL-93 | chr13-58411629-DEL-93 |
| chr13-58438638-DEL-61 | chr13-58438639-DEL-61 |
| chr13-58446347-DEL-216 | chr13-58446336-DEL-239 |
| chr13-58449432-DEL-63 | chr13-58449433-DEL-63 |
| chr13-58477717-DEL-94 | chr13-58477718-DEL-94 |
| chr13-58490951-DEL-61 | chr13-58490950-DEL-62 |
| chr13-58493728-DEL-202 | chr13-58493729-DEL-202 |
| chr13-58494203-DEL-1984 | chr13-58494231-DEL-1984 |
| chr13-58503146-DEL-441 | chr13-58503147-DEL-441 |
| chr13-58512802-DEL-53 | chr13-58512803-DEL-53 |
| chr13-58576386-DEL-6401 | chr13-58576387-DEL-6401 |
| chr13-58594123-DEL-207 | chr13-58594124-DEL-207 |
| chr13-58606555-DEL-208 | chr13-58606556-DEL-208 |
| chr13-58640406-DEL-98 | chr13-58640407-DEL-98 |
| chr13-58653928-DEL-62 | chr13-58653929-DEL-62 |
| chr13-58674677-DEL-225 | chr13-58674678-DEL-225 |
| chr13-58681183-DEL-227 | chr13-58681184-DEL-227 |
| chr13-58701551-DEL-172 | chr13-58701552-DEL-172 |
| chr13-58722358-DEL-90 | chr13-58722359-DEL-90 |
| chr13-58731682-DEL-1890 | chr13-58731683-DEL-1890 |
| chr13-58751587-DEL-51 | chr13-58751588-DEL-51 |
| chr13-58763302-DEL-199 | chr13-58763303-DEL-199 |
| chr13-58780301-DEL-185 | chr13-58780302-DEL-185 |
| chr13-58806705-DEL-149 | chr13-58806705-DEL-112 |
| chr13-58807082-DEL-342 | chr13-58807083-DEL-342 |
| chr13-58837674-DEL-72 | chr13-58837675-DEL-72 |
| chr13-58853777-DEL-197 | chr13-58853778-DEL-197 |
| chr13-58860558-DEL-52 | chr13-58860559-DEL-52 |
| chr13-58861143-DEL-1177 | chr13-58861144-DEL-1177 |
| chr13-58876334-DEL-54 | chr13-58876335-DEL-54 |
| chr13-58884715-DEL-96 | chr13-58884748-DEL-96 |
| chr13-58983267-DEL-80 | chr13-58983272-DEL-76 |
| chr13-58985008-DEL-83 | chr13-58985009-DEL-83 |
| chr13-59001239-DEL-278 | chr13-59001240-DEL-278 |
| chr13-59030699-DEL-113 | chr13-59030700-DEL-113 |
| chr13-59060598-DEL-71 | chr13-59060599-DEL-71 |
| chr13-59064038-DEL-56 | chr13-59064039-DEL-56 |
| chr13-59121288-DEL-65 | chr13-59121289-DEL-65 |
| chr13-59125375-DEL-52 | chr13-59125376-DEL-52 |
| chr13-59144876-DEL-75 | chr13-59144882-DEL-55 |
| chr13-59166434-DEL-64 | chr13-59166431-DEL-76 |
| chr13-59172459-DEL-155 | chr13-59172460-DEL-155 |
| chr13-59192494-DEL-224 | chr13-59192495-DEL-222 |
| chr13-59213450-DEL-94 | chr13-59213451-DEL-94 |
| chr13-59244591-DEL-60 | chr13-59244623-DEL-60 |
| chr13-59248675-DEL-64 | chr13-59248683-DEL-50 |
| chr13-59249291-DEL-6473 | chr13-59249292-DEL-6473 |
| chr13-59269300-DEL-3931 | chr13-59269293-DEL-3932 |
| chr13-59285123-DEL-200 | chr13-59285124-DEL-200 |
| chr13-59356006-DEL-351 | chr13-59356008-DEL-350 |
| chr13-59359221-DEL-111 | chr13-59359222-DEL-111 |
| chr13-59365029-DEL-60 | chr13-59365030-DEL-60 |
| chr13-59387565-DEL-172 | chr13-59387566-DEL-172 |
| chr13-59421642-DEL-236 | chr13-59421661-DEL-218 |
| chr13-59443101-DEL-62 | chr13-59443106-DEL-58 |
| chr13-59453407-DEL-61 | chr13-59453408-DEL-61 |
| chr13-59457026-DEL-162 | chr13-59457027-DEL-162 |
| chr13-59473649-DEL-681 | chr13-59473650-DEL-681 |
| chr13-59481017-DEL-51 | chr13-59481018-DEL-51 |
| chr13-59533894-DEL-1134 | chr13-59533895-DEL-1134 |
| chr13-59550950-DEL-53 | chr13-59550950-DEL-55 |
| chr13-59551751-DEL-400 | chr13-59551752-DEL-400 |
| chr13-59559265-DEL-452 | chr13-59559266-DEL-452 |
| chr13-59567802-DEL-1128 | chr13-59567803-DEL-1128 |
| chr13-59571996-DEL-143 | chr13-59571999-DEL-146 |
| chr13-59579816-DEL-160 | chr13-59579847-DEL-152 |
| chr13-59587156-DEL-163 | chr13-59587155-DEL-162 |
| chr13-59612791-DEL-72 | chr13-59612792-DEL-72 |
| chr13-59642786-DEL-62 | chr13-59642787-DEL-62 |
| chr13-59654246-DEL-60 | chr13-59654247-DEL-60 |
| chr13-59791482-DEL-198 | chr13-59791483-DEL-198 |
| chr13-59800716-DEL-62 | chr13-59800717-DEL-62 |
| chr13-59801570-DEL-5653 | chr13-59801571-DEL-5653 |
| chr13-59806766-DEL-320 | chr13-59806767-DEL-320 |

| | |
|---|---|
| chr13-59825079-DEL-516 | chr13-59825103-DEL-507 |
| chr13-59828536-DEL-58 | chr13-59828533-DEL-63 |
| chr13-59899751-DEL-205 | chr13-59899752-DEL-204 |
| chr13-59927932-DEL-61 | chr13-59927933-DEL-60 |
| chr13-59932177-DEL-72 | chr13-59932178-DEL-72 |
| chr13-59982968-DEL-66 | chr13-59983001-DEL-82 |
| chr13-59998586-DEL-63 | chr13-59998587-DEL-63 |
| chr13-60017971-DEL-80 | chr13-60017972-DEL-80 |
| chr13-60023266-DEL-63 | chr13-60023267-DEL-63 |
| chr13-60037926-DEL-62 | chr13-60037927-DEL-62 |
| chr13-60060326-DEL-103 | chr13-60060327-DEL-103 |
| chr13-60069570-DEL-177 | chr13-60069571-DEL-177 |
| chr13-60078242-DEL-899 | chr13-60078222-DEL-892 |
| chr13-60115350-DEL-70 | chr13-60115334-DEL-86 |
| chr13-60126138-DEL-248 | chr13-60126139-DEL-248 |
| chr13-60153416-DEL-82 | chr13-60153417-DEL-82 |
| chr13-60157952-DEL-62 | chr13-60157953-DEL-62 |
| chr13-60168264-DEL-61 | chr13-60168264-DEL-62 |
| chr13-60169431-DEL-7084 | chr13-60169432-DEL-7084 |
| chr13-60177930-DEL-717 | chr13-60177934-DEL-718 |
| chr13-60186677-DEL-66 | chr13-60186678-DEL-66 |
| chr13-60187376-DEL-67 | chr13-60187377-DEL-67 |
| chr13-60189425-DEL-131 | chr13-60189426-DEL-131 |
| chr13-60202714-DEL-60 | chr13-60202715-DEL-60 |
| chr13-60204346-DEL-53 | chr13-60204347-DEL-53 |
| chr13-60214015-DEL-246 | chr13-60214016-DEL-246 |
| chr13-60218239-DEL-548 | chr13-60218240-DEL-548 |
| chr13-60225393-DEL-57 | chr13-60225394-DEL-57 |
| chr13-60229517-DEL-1600 | chr13-60229512-DEL-1562 |
| chr13-60264814-DEL-80 | chr13-60264815-DEL-82 |
| chr13-60297142-DEL-59 | chr13-60297145-DEL-87 |
| chr13-60334036-DEL-563 | chr13-60334037-DEL-563 |
| chr13-60344496-DEL-7207 | chr13-60344496-DEL-7207 |
| chr13-60368350-DEL-1163 | chr13-60368351-DEL-1163 |
| chr13-60377399-DEL-59 | chr13-60377400-DEL-59 |
| chr13-60387832-DEL-342 | chr13-60387833-DEL-342 |
| chr13-60389273-DEL-70 | chr13-60389258-DEL-82 |
| chr13-60399725-DEL-88 | chr13-60399726-DEL-88 |
| chr13-60410851-DEL-55 | chr13-60410862-DEL-56 |
| chr13-60412546-DEL-703 | chr13-60412547-DEL-703 |
| chr13-60414165-DEL-70 | chr13-60414166-DEL-70 |
| chr13-60435174-DEL-55 | chr13-60435181-DEL-76 |
| chr13-60440222-DEL-1140 | chr13-60440223-DEL-1140 |
| chr13-60473428-DEL-2497 | chr13-60473429-DEL-2497 |
| chr13-60476570-DEL-468 | chr13-60476571-DEL-468 |
| chr13-60481659-DEL-158 | chr13-60481662-DEL-144 |
| chr13-60483089-DEL-63 | chr13-60483090-DEL-63 |
| chr13-60489099-DEL-155 | chr13-60489082-DEL-173 |
| chr13-60531198-DEL-202 | chr13-60531198-DEL-202 |
| chr13-60537441-DEL-214 | chr13-60537480-DEL-214 |
| chr13-60543091-DEL-206 | chr13-60543092-DEL-206 |
| chr13-60591557-DEL-84 | chr13-60591558-DEL-84 |
| chr13-60609626-DEL-1492 | chr13-60609579-DEL-1493 |
| chr13-60642075-DEL-2862 | chr13-60642076-DEL-2862 |
| chr13-60679314-DEL-2036 | chr13-60679314-DEL-2036 |
| chr13-60701400-DEL-75 | chr13-60701401-DEL-60 |
| chr13-60704114-DEL-78 | chr13-60704114-DEL-70 |
| chr13-60720296-DEL-54 | chr13-60720297-DEL-54 |
| chr13-60736074-DEL-161 | chr13-60736075-DEL-162 |
| chr13-60758662-DEL-198 | chr13-60758663-DEL-198 |
| chr13-60769726-DEL-2129 | chr13-60769727-DEL-2129 |
| chr13-60789585-DEL-878 | chr13-60789586-DEL-878 |
| chr13-60791392-DEL-90 | chr13-60791393-DEL-90 |
| chr13-60824407-DEL-60 | chr13-60824408-DEL-60 |
| chr13-60845402-DEL-401 | chr13-60845403-DEL-401 |
| chr13-60847491-DEL-65 | chr13-60847502-DEL-50 |
| chr13-60860875-DEL-71 | chr13-60860873-DEL-78 |
| chr13-60863740-DEL-64 | chr13-60863741-DEL-64 |
| chr13-60875849-DEL-67 | chr13-60875849-DEL-69 |
| chr13-60900178-DEL-81 | chr13-60900179-DEL-81 |
| chr13-60916446-DEL-113 | chr13-60916447-DEL-113 |
| chr13-60916888-DEL-204 | chr13-60916889-DEL-204 |
| chr13-60976331-DEL-3046 | chr13-60976332-DEL-3046 |
| chr13-61003644-DEL-6680 | chr13-61003645-DEL-6680 |
| chr13-61022236-DEL-715 | chr13-61022237-DEL-715 |
| chr13-61027772-DEL-9888 | chr13-61027773-DEL-9888 |
| chr13-61047701-DEL-763 | chr13-61047702-DEL-763 |
| chr13-61050515-DEL-1143 | chr13-61050510-DEL-1141 |
| chr13-61094877-DEL-4693 | chr13-61094878-DEL-4693 |
| chr13-61122215-DEL-6376 | chr13-61122207-DEL-6377 |

| | |
|---|---|
| chr13-61145767-DEL-481 | chr13-61145768-DEL-481 |
| chr13-61167336-DEL-2103 | chr13-61167337-DEL-2103 |
| chr13-61222030-DEL-722 | chr13-61222031-DEL-722 |
| chr13-61242286-DEL-241 | chr13-61242286-DEL-241 |
| chr13-61370431-DEL-872 | chr13-61370432-DEL-872 |
| chr13-61402075-DEL-165 | chr13-61402076-DEL-165 |
| chr13-61425049-DEL-567 | chr13-61425050-DEL-567 |
| chr13-61426925-DEL-54 | chr13-61426926-DEL-54 |
| chr13-61448104-DEL-404 | chr13-61448105-DEL-404 |
| chr13-61460083-DEL-51 | chr13-61460083-DEL-51 |
| chr13-61474197-DEL-52 | chr13-61474198-DEL-52 |
| chr13-61488451-DEL-69 | chr13-61488470-DEL-82 |
| chr13-61495774-DEL-56 | chr13-61495775-DEL-56 |
| chr13-61580004-DEL-106 | chr13-61580005-DEL-106 |
| chr13-61603568-DEL-522 | chr13-61603569-DEL-522 |
| chr13-61629265-DEL-58 | chr13-61629266-DEL-58 |
| chr13-61630734-DEL-4529 | chr13-61630735-DEL-4529 |
| chr13-61672792-DEL-1660 | chr13-61672792-DEL-1660 |
| chr13-61706748-DEL-238 | chr13-61706749-DEL-238 |
| chr13-61711740-DEL-138 | chr13-61711741-DEL-134 |
| chr13-61755464-DEL-970 | chr13-61755465-DEL-969 |
| chr13-61757211-DEL-63 | chr13-61757212-DEL-63 |
| chr13-61810678-DEL-89 | chr13-61810678-DEL-89 |
| chr13-61812354-DEL-76 | chr13-61812353-DEL-76 |
| chr13-61912431-DEL-208 | chr13-61912431-DEL-208 |
| chr13-61917643-DEL-205 | chr13-61917644-DEL-205 |
| chr13-61932097-DEL-731 | chr13-61932097-DEL-731 |
| chr13-61983749-DEL-830 | chr13-61983749-DEL-830 |
| chr13-61990374-DEL-486 | chr13-61990374-DEL-486 |
| chr13-62263656-DEL-200 | chr13-62263655-DEL-200 |
| chr13-62282369-DEL-217 | chr13-62282368-DEL-217 |
| chr13-62284570-DEL-412 | chr13-62284569-DEL-412 |
| chr13-62321985-DEL-557 | chr13-62321975-DEL-559 |
| chr13-62361271-DEL-212 | chr13-62361272-DEL-202 |
| chr13-62371497-DEL-2015 | chr13-62371498-DEL-2015 |
| chr13-62387053-DEL-253 | chr13-62387054-DEL-253 |
| chr13-62390926-DEL-53 | chr13-62390927-DEL-53 |
| chr13-62395030-DEL-6805 | chr13-62395031-DEL-6805 |
| chr13-62439495-DEL-4602 | chr13-62439496-DEL-4602 |
| chr13-62461153-DEL-1893 | chr13-62461154-DEL-1893 |
| chr13-62466404-DEL-446 | chr13-62466405-DEL-446 |
| chr13-62501673-DEL-2754 | chr13-62501674-DEL-2754 |
| chr13-62510223-DEL-498 | chr13-62510224-DEL-498 |
| chr13-62520955-DEL-211 | chr13-62520959-DEL-208 |
| chr13-62524397-DEL-208 | chr13-62524398-DEL-208 |
| chr13-62536110-DEL-157 | chr13-62536111-DEL-157 |
| chr13-62538823-DEL-158 | chr13-62538822-DEL-158 |
| chr13-62547574-DEL-203 | chr13-62547575-DEL-203 |
| chr13-62553208-DEL-438 | chr13-62553209-DEL-438 |
| chr13-62592222-DEL-212 | chr13-62592223-DEL-212 |
| chr13-62599347-DEL-839 | chr13-62599348-DEL-839 |
| chr13-62620797-DEL-7166 | chr13-62620798-DEL-7166 |
| chr13-62640405-DEL-187 | chr13-62640406-DEL-187 |
| chr13-62672052-DEL-119 | chr13-62672052-DEL-119 |
| chr13-62690908-DEL-403 | chr13-62690909-DEL-403 |
| chr13-62693323-DEL-217 | chr13-62693324-DEL-217 |
| chr13-62713149-DEL-155 | chr13-62713149-DEL-155 |
| chr13-62715413-DEL-328 | chr13-62715412-DEL-329 |
| chr13-62716338-DEL-774 | chr13-62716337-DEL-775 |
| chr13-62738464-DEL-189 | chr13-62738464-DEL-189 |
| chr13-62739992-DEL-345 | chr13-62739993-DEL-345 |
| chr13-62760460-DEL-201 | chr13-62760461-DEL-201 |
| chr13-62803478-DEL-754 | chr13-62803479-DEL-754 |
| chr13-62807103-DEL-5154 | chr13-62807104-DEL-5154 |
| chr13-62817808-DEL-301 | chr13-62817815-DEL-296 |
| chr13-62823957-DEL-285 | chr13-62823958-DEL-285 |
| chr13-62829765-DEL-205 | chr13-62829766-DEL-205 |
| chr13-62835610-DEL-196 | chr13-62835620-DEL-195 |
| chr13-62857786-DEL-62 | chr13-62857778-DEL-56 |
| chr13-62874304-DEL-6475 | chr13-62874305-DEL-6475 |
| chr13-62922241-DEL-84 | chr13-62922242-DEL-84 |
| chr13-62944812-DEL-209 | chr13-62944813-DEL-209 |
| chr13-62962767-DEL-388 | chr13-62962768-DEL-388 |
| chr13-62977665-DEL-524 | chr13-62977666-DEL-524 |
| chr13-62983970-DEL-51 | chr13-62983971-DEL-51 |
| chr13-62990916-DEL-113 | chr13-62990917-DEL-113 |
| chr13-62993623-DEL-70 | chr13-62993597-DEL-67 |
| chr13-62996463-DEL-56 | chr13-62996463-DEL-57 |
| chr13-63008222-DEL-204 | chr13-63008225-DEL-205 |
| chr13-63016417-DEL-58 | chr13-63016392-DEL-50 |

| | |
|---|---|
| chr13-63024800-DEL-77 | chr13-63024800-DEL-77 |
| chr13-63031158-DEL-58 | chr13-63031176-DEL-69 |
| chr13-63032869-DEL-199 | chr13-63032870-DEL-199 |
| chr13-63167117-DEL-3203 | chr13-63167118-DEL-3203 |
| chr13-63289823-DEL-3198 | chr13-63289824-DEL-3198 |
| chr13-63304701-DEL-6837 | chr13-63304702-DEL-6837 |
| chr13-63327610-DEL-966 | chr13-63327611-DEL-966 |
| chr13-63365808-DEL-5153 | chr13-63365809-DEL-5153 |
| chr13-63382351-DEL-90 | chr13-63382374-DEL-68 |
| chr13-63432118-DEL-153 | chr13-63432135-DEL-114 |
| chr13-63440168-DEL-281 | chr13-63440163-DEL-287 |
| chr13-63487068-DEL-86 | chr13-63487083-DEL-72 |
| chr13-63496981-DEL-56 | chr13-63496982-DEL-56 |
| chr13-63511991-DEL-174 | chr13-63511992-DEL-140 |
| chr13-63516691-DEL-229 | chr13-63516696-DEL-227 |
| chr13-63540749-DEL-64 | chr13-63540764-DEL-50 |
| chr13-63553639-DEL-123 | chr13-63553672-DEL-124 |
| chr13-63564658-DEL-164 | chr13-63564659-DEL-164 |
| chr13-63585443-DEL-6506 | chr13-63585444-DEL-6506 |
| chr13-63594259-DEL-1164 | chr13-63594260-DEL-1164 |
| chr13-63618932-DEL-1186 | chr13-63618933-DEL-1186 |
| chr13-63627835-DEL-437 | chr13-63627836-DEL-437 |
| chr13-63649643-DEL-197 | chr13-63649640-DEL-196 |
| chr13-63668941-DEL-731 | chr13-63668942-DEL-731 |
| chr13-63679998-DEL-55 | chr13-63679998-DEL-55 |
| chr13-63726719-DEL-270 | chr13-63726720-DEL-270 |
| chr13-63774890-DEL-83 | chr13-63774891-DEL-70 |
| chr13-63785788-DEL-2416 | chr13-63785789-DEL-2416 |
| chr13-63842054-DEL-80 | chr13-63842053-DEL-81 |
| chr13-63862117-DEL-1767 | chr13-63862118-DEL-1767 |
| chr13-63865602-DEL-251 | chr13-63865603-DEL-251 |
| chr13-63897453-DEL-60 | chr13-63897462-DEL-55 |
| chr13-63916495-DEL-62 | chr13-63916508-DEL-50 |
| chr13-63923574-DEL-201 | chr13-63923575-DEL-201 |
| chr13-63957146-DEL-63 | chr13-63957147-DEL-63 |
| chr13-64009054-DEL-550 | chr13-64009055-DEL-550 |
| chr13-64058884-DEL-241 | chr13-64058879-DEL-239 |
| chr13-64067921-DEL-63 | chr13-64067922-DEL-63 |
| chr13-64070085-DEL-53 | chr13-64070086-DEL-53 |
| chr13-64075729-DEL-88 | chr13-64075759-DEL-70 |
| chr13-64101885-DEL-70 | chr13-64101878-DEL-78 |
| chr13-64154163-DEL-172 | chr13-64154156-DEL-190 |
| chr13-64160314-DEL-80 | chr13-64160310-DEL-95 |
| chr13-64175015-DEL-53 | chr13-64175016-DEL-53 |
| chr13-64177494-DEL-58 | chr13-64177495-DEL-58 |
| chr13-64189562-DEL-52 | chr13-64189563-DEL-52 |
| chr13-64194398-DEL-188 | chr13-64194397-DEL-195 |
| chr13-64199707-DEL-1327 | chr13-64199708-DEL-1327 |
| chr13-64237682-DEL-74 | chr13-64237683-DEL-74 |
| chr13-64242510-DEL-76 | chr13-64242516-DEL-75 |
| chr13-64263708-DEL-190 | chr13-64263709-DEL-190 |
| chr13-64267772-DEL-145 | chr13-64267773-DEL-145 |
| chr13-64272833-DEL-75 | chr13-64272838-DEL-74 |
| chr13-64280514-DEL-52 | chr13-64280523-DEL-52 |
| chr13-64294238-DEL-750 | chr13-64294241-DEL-749 |
| chr13-64313553-DEL-159 | chr13-64313554-DEL-159 |
| chr13-64319481-DEL-88 | chr13-64319482-DEL-88 |
| chr13-64322198-DEL-101 | chr13-64322199-DEL-101 |
| chr13-64328458-DEL-64 | chr13-64328473-DEL-53 |
| chr13-64352724-DEL-57 | chr13-64352725-DEL-57 |
| chr13-64358919-DEL-890 | chr13-64358920-DEL-890 |
| chr13-64366048-DEL-231 | chr13-64366049-DEL-231 |
| chr13-64370351-DEL-99 | chr13-64370380-DEL-56 |
| chr13-64376201-DEL-62 | chr13-64376192-DEL-51 |
| chr13-64388598-DEL-54 | chr13-64388597-DEL-55 |
| chr13-64401990-DEL-174 | chr13-64401991-DEL-174 |
| chr13-64402515-DEL-78 | chr13-64402516-DEL-78 |
| chr13-64403699-DEL-527 | chr13-64403698-DEL-525 |
| chr13-64409150-DEL-168 | chr13-64409151-DEL-169 |
| chr13-64417804-DEL-134 | chr13-64417804-DEL-138 |
| chr13-64424171-DEL-77 | chr13-64424172-DEL-59 |
| chr13-64429277-DEL-73 | chr13-64429296-DEL-60 |
| chr13-64431842-DEL-56 | chr13-64431843-DEL-56 |
| chr13-64442212-DEL-380 | chr13-64442221-DEL-370 |
| chr13-64448303-DEL-104 | chr13-64448304-DEL-104 |
| chr13-64451909-DEL-190 | chr13-64451910-DEL-190 |
| chr13-64463907-DEL-96 | chr13-64463908-DEL-96 |
| chr13-64478841-DEL-182 | chr13-64478842-DEL-182 |
| chr13-64497746-DEL-252 | chr13-64497747-DEL-249 |
| chr13-64509626-DEL-231 | chr13-64509628-DEL-227 |

| | |
|---|---|
| chr13-64510882-DEL-86 | chr13-64510883-DEL-86 |
| chr13-64531266-DEL-156 | chr13-64531267-DEL-156 |
| chr13-64541445-DEL-405 | chr13-64541435-DEL-406 |
| chr13-64543351-DEL-59 | chr13-64543351-DEL-87 |
| chr13-64544301-DEL-123 | chr13-64544302-DEL-123 |
| chr13-64546634-DEL-52 | chr13-64546635-DEL-52 |
| chr13-64552727-DEL-51 | chr13-64552721-DEL-52 |
| chr13-64566357-DEL-1824 | chr13-64566358-DEL-1824 |
| chr13-64568160-DEL-60 | chr13-64568159-DEL-67 |
| chr13-64581386-DEL-98 | chr13-64581386-DEL-82 |
| chr13-64590295-DEL-55 | chr13-64590271-DEL-88 |
| chr13-64604443-DEL-347 | chr13-64604444-DEL-347 |
| chr13-64617025-DEL-65 | chr13-64617026-DEL-65 |
| chr13-64621111-DEL-184 | chr13-64621112-DEL-184 |
| chr13-64624804-DEL-216 | chr13-64624805-DEL-168 |
| chr13-64640154-DEL-286 | chr13-64640155-DEL-286 |
| chr13-64646306-DEL-2649 | chr13-64646307-DEL-2649 |
| chr13-64655100-DEL-175 | chr13-64655101-DEL-175 |
| chr13-64664428-DEL-691 | chr13-64664429-DEL-691 |
| chr13-64688471-DEL-4330 | chr13-64688471-DEL-4330 |
| chr13-64700711-DEL-316 | chr13-64700738-DEL-284 |
| chr13-64707258-DEL-456 | chr13-64707259-DEL-456 |
| chr13-64730453-DEL-6820 | chr13-64730454-DEL-6820 |
| chr13-64738357-DEL-93 | chr13-64738366-DEL-85 |
| chr13-64750832-DEL-2969 | chr13-64750833-DEL-2969 |
| chr13-64754065-DEL-302 | chr13-64754065-DEL-302 |
| chr13-64754425-DEL-6252 | chr13-64754426-DEL-6252 |
| chr13-64764064-DEL-88 | chr13-64764065-DEL-64 |
| chr13-64778469-DEL-560 | chr13-64778470-DEL-558 |
| chr13-64790439-DEL-6690 | chr13-64790440-DEL-6690 |
| chr13-64838445-DEL-103 | chr13-64838398-DEL-152 |
| chr13-64854737-DEL-665 | chr13-64854738-DEL-665 |
| chr13-64856013-DEL-6022 | chr13-64856014-DEL-6022 |
| chr13-64873167-DEL-952 | chr13-64873168-DEL-952 |
| chr13-64876488-DEL-64 | chr13-64876489-DEL-80 |
| chr13-64882540-DEL-75 | chr13-64882541-DEL-75 |
| chr13-64912206-DEL-64 | chr13-64912207-DEL-64 |
| chr13-64938386-DEL-198 | chr13-64938380-DEL-199 |
| chr13-64952764-DEL-2999 | chr13-64952767-DEL-3000 |
| chr13-64965808-DEL-107 | chr13-64965807-DEL-111 |
| chr13-64971181-DEL-119 | chr13-64971182-DEL-119 |
| chr13-64977768-DEL-5483 | chr13-64977769-DEL-5483 |
| chr13-64990306-DEL-56 | chr13-64990307-DEL-56 |
| chr13-65025866-DEL-7326 | chr13-65025867-DEL-7326 |
| chr13-65058657-DEL-77 | chr13-65058697-DEL-52 |
| chr13-65075726-DEL-1050 | chr13-65075727-DEL-1050 |
| chr13-65079528-DEL-459 | chr13-65079529-DEL-459 |
| chr13-65087668-DEL-59 | chr13-65087669-DEL-55 |
| chr13-65089655-DEL-200 | chr13-65089656-DEL-200 |
| chr13-65098132-DEL-193 | chr13-65098130-DEL-193 |
| chr13-65118761-DEL-343 | chr13-65118762-DEL-343 |
| chr13-65147035-DEL-587 | chr13-65147036-DEL-587 |
| chr13-65159594-DEL-146 | chr13-65159595-DEL-146 |
| chr13-65159810-DEL-3082 | chr13-65159811-DEL-3082 |
| chr13-65200857-DEL-51 | chr13-65200858-DEL-51 |
| chr13-65203284-DEL-2799 | chr13-65203285-DEL-2799 |
| chr13-65239432-DEL-171 | chr13-65239433-DEL-171 |
| chr13-65274323-DEL-68 | chr13-65274324-DEL-68 |
| chr13-65287059-DEL-197 | chr13-65287060-DEL-197 |
| chr13-65302547-DEL-64 | chr13-65302570-DEL-75 |
| chr13-65310176-DEL-60 | chr13-65310177-DEL-60 |
| chr13-65315963-DEL-760 | chr13-65315965-DEL-759 |
| chr13-65352217-DEL-205 | chr13-65352208-DEL-201 |
| chr13-65388357-DEL-52 | chr13-65388358-DEL-52 |
| chr13-65396897-DEL-498 | chr13-65396881-DEL-499 |
| chr13-65400949-DEL-205 | chr13-65400950-DEL-205 |
| chr13-65405757-DEL-88 | chr13-65405769-DEL-86 |
| chr13-65410531-DEL-2055 | chr13-65410532-DEL-2055 |
| chr13-65418632-DEL-202 | chr13-65418633-DEL-202 |
| chr13-65469626-DEL-3107 | chr13-65469627-DEL-3107 |
| chr13-65519744-DEL-227 | chr13-65519743-DEL-227 |
| chr13-65520640-DEL-70 | chr13-65520657-DEL-52 |
| chr13-65625251-DEL-256 | chr13-65625250-DEL-256 |
| chr13-65660909-DEL-84 | chr13-65660929-DEL-82 |
| chr13-65692095-DEL-81 | chr13-65692098-DEL-69 |
| chr13-65703131-DEL-213 | chr13-65703130-DEL-213 |
| chr13-65730689-DEL-220 | chr13-65730689-DEL-220 |
| chr13-65735343-DEL-57 | chr13-65735344-DEL-57 |
| chr13-65746118-DEL-200 | chr13-65746120-DEL-199 |
| chr13-65761706-DEL-76 | chr13-65761707-DEL-76 |

| | |
|---|---|
| chr13-65907604-DEL-165 | chr13-65907604-DEL-165 |
| chr13-65910054-DEL-168 | chr13-65910055-DEL-168 |
| chr13-66012188-DEL-393 | chr13-66012189-DEL-393 |
| chr13-66019217-DEL-65 | chr13-66019209-DEL-73 |
| chr13-66030644-DEL-898 | chr13-66030644-DEL-898 |
| chr13-66054656-DEL-504 | chr13-66054657-DEL-504 |
| chr13-66108878-DEL-199 | chr13-66108878-DEL-199 |
| chr13-66144936-DEL-4022 | chr13-66144937-DEL-4022 |
| chr13-66308872-DEL-171 | chr13-66308872-DEL-171 |
| chr13-66327515-DEL-107 | chr13-66327515-DEL-109 |
| chr13-66415560-DEL-81 | chr13-66415556-DEL-56 |
| chr13-66429579-DEL-84 | chr13-66429532-DEL-84 |
| chr13-66460623-DEL-52 | chr13-66460624-DEL-55 |
| chr13-66466075-DEL-137 | chr13-66466075-DEL-137 |
| chr13-66576523-DEL-89 | chr13-66576505-DEL-96 |
| chr13-66588331-DEL-450 | chr13-66588330-DEL-450 |
| chr13-66703880-DEL-54 | chr13-66703880-DEL-54 |
| chr13-66743840-DEL-195 | chr13-66743840-DEL-195 |
| chr13-66762500-DEL-837 | chr13-66762500-DEL-837 |
| chr13-66767440-DEL-435 | chr13-66767440-DEL-435 |
| chr13-66770272-DEL-74 | chr13-66770290-DEL-84 |
| chr13-66789238-DEL-56 | chr13-66789237-DEL-56 |
| chr13-66927006-DEL-98 | chr13-66926994-DEL-106 |
| chr13-66933728-DEL-217 | chr13-66933728-DEL-217 |
| chr13-66940564-DEL-84 | chr13-66940563-DEL-84 |
| chr13-66948307-DEL-1149 | chr13-66948308-DEL-1149 |
| chr13-66974008-DEL-1109 | chr13-66973995-DEL-1117 |
| chr13-67009400-DEL-202 | chr13-67009401-DEL-202 |
| chr13-67044212-DEL-439 | chr13-67044213-DEL-439 |
| chr13-67090633-DEL-207 | chr13-67090634-DEL-207 |
| chr13-67096325-DEL-67 | chr13-67096326-DEL-67 |
| chr13-67137573-DEL-1879 | chr13-67137574-DEL-1879 |
| chr13-67184538-DEL-516 | chr13-67184539-DEL-516 |
| chr13-67192284-DEL-202 | chr13-67192285-DEL-202 |
| chr13-67216492-DEL-182 | chr13-67216493-DEL-182 |
| chr13-67240595-DEL-2289 | chr13-67240596-DEL-2289 |
| chr13-67286834-DEL-84 | chr13-67286835-DEL-84 |
| chr13-67303125-DEL-6364 | chr13-67303122-DEL-6365 |
| chr13-67322214-DEL-283 | chr13-67322215-DEL-283 |
| chr13-67333310-DEL-196 | chr13-67333311-DEL-196 |
| chr13-67344729-DEL-178 | chr13-67344730-DEL-178 |
| chr13-67346070-DEL-156 | chr13-67346071-DEL-117 |
| chr13-67361653-DEL-60 | chr13-67361654-DEL-60 |
| chr13-67377031-DEL-203 | chr13-67377032-DEL-203 |
| chr13-67379597-DEL-579 | chr13-67379598-DEL-579 |
| chr13-67381960-DEL-5528 | chr13-67381961-DEL-5528 |
| chr13-67388078-DEL-130 | chr13-67388079-DEL-130 |
| chr13-67389348-DEL-62 | chr13-67389347-DEL-52 |
| chr13-67409020-DEL-69 | chr13-67409021-DEL-69 |
| chr13-67417638-DEL-209 | chr13-67417644-DEL-208 |
| chr13-67419733-DEL-6213 | chr13-67419734-DEL-6213 |
| chr13-67432885-DEL-203 | chr13-67432886-DEL-203 |
| chr13-67439813-DEL-873 | chr13-67439814-DEL-873 |
| chr13-67441673-DEL-84 | chr13-67441674-DEL-84 |
| chr13-67445084-DEL-1756 | chr13-67445085-DEL-1756 |
| chr13-67451985-DEL-71 | chr13-67451986-DEL-71 |
| chr13-67476391-DEL-986 | chr13-67476385-DEL-984 |
| chr13-67494849-DEL-308 | chr13-67494850-DEL-308 |
| chr13-67499763-DEL-236 | chr13-67499764-DEL-236 |
| chr13-67517452-DEL-2282 | chr13-67517453-DEL-2282 |
| chr13-67521291-DEL-78 | chr13-67521253-DEL-117 |
| chr13-67544275-DEL-1948 | chr13-67544276-DEL-1948 |
| chr13-67550678-DEL-209 | chr13-67550679-DEL-209 |
| chr13-67558561-DEL-868 | chr13-67558562-DEL-869 |
| chr13-67579173-DEL-176 | chr13-67579174-DEL-176 |
| chr13-67581353-DEL-177 | chr13-67581354-DEL-177 |
| chr13-67588654-DEL-1125 | chr13-67588655-DEL-1125 |
| chr13-67618375-DEL-227 | chr13-67618376-DEL-227 |
| chr13-67625876-DEL-1478 | chr13-67625877-DEL-1478 |
| chr13-67631101-DEL-180 | chr13-67631102-DEL-180 |
| chr13-67634431-DEL-56 | chr13-67634434-DEL-58 |
| chr13-67647038-DEL-195 | chr13-67647039-DEL-195 |
| chr13-67715378-DEL-60 | chr13-67715379-DEL-52 |
| chr13-67731634-DEL-51 | chr13-67731635-DEL-51 |
| chr13-67748010-DEL-2099 | chr13-67748011-DEL-2099 |
| chr13-67756311-DEL-68 | chr13-67756344-DEL-80 |
| chr13-67757973-DEL-189 | chr13-67757974-DEL-189 |
| chr13-67760008-DEL-284 | chr13-67760007-DEL-284 |
| chr13-67770970-DEL-238 | chr13-67770971-DEL-238 |
| chr13-67775061-DEL-388 | chr13-67775062-DEL-388 |

| | |
|---|---|
| chr13-67789705-DEL-9265 | chr13-67789706-DEL-9265 |
| chr13-67812301-DEL-939 | chr13-67812299-DEL-940 |
| chr13-67814254-DEL-330 | chr13-67814255-DEL-330 |
| chr13-67824919-DEL-171 | chr13-67824920-DEL-171 |
| chr13-67853135-DEL-60 | chr13-67853136-DEL-60 |
| chr13-67907996-DEL-161 | chr13-67907997-DEL-161 |
| chr13-67910939-DEL-211 | chr13-67910940-DEL-211 |
| chr13-67912176-DEL-201 | chr13-67912177-DEL-201 |
| chr13-67917027-DEL-54 | chr13-67917027-DEL-50 |
| chr13-67925424-DEL-213 | chr13-67925425-DEL-213 |
| chr13-67939275-DEL-591 | chr13-67939276-DEL-591 |
| chr13-67940345-DEL-85 | chr13-67940346-DEL-85 |
| chr13-67947732-DEL-69 | chr13-67947732-DEL-105 |
| chr13-67999359-DEL-268 | chr13-67999360-DEL-268 |
| chr13-68004266-DEL-485 | chr13-68004267-DEL-483 |
| chr13-68012781-DEL-56 | chr13-68012782-DEL-56 |
| chr13-68018708-DEL-461 | chr13-68018709-DEL-461 |
| chr13-68024428-DEL-532 | chr13-68024429-DEL-532 |
| chr13-68033002-DEL-8691 | chr13-68033003-DEL-8691 |
| chr13-68062461-DEL-8804 | chr13-68062462-DEL-8804 |
| chr13-68075312-DEL-307 | chr13-68075314-DEL-306 |
| chr13-68138587-DEL-205 | chr13-68138587-DEL-205 |
| chr13-68141642-DEL-178 | chr13-68141641-DEL-178 |
| chr13-68142885-DEL-4009 | chr13-68142885-DEL-4009 |
| chr13-68148547-DEL-117 | chr13-68148546-DEL-117 |
| chr13-68186286-DEL-78 | chr13-68186285-DEL-78 |
| chr13-68234669-DEL-53 | chr13-68234668-DEL-53 |
| chr13-68314432-DEL-57 | chr13-68314430-DEL-56 |
| chr13-68322364-DEL-197 | chr13-68322365-DEL-197 |
| chr13-68350242-DEL-116 | chr13-68350243-DEL-116 |
| chr13-68458633-DEL-115 | chr13-68458651-DEL-83 |
| chr13-68548915-DEL-755 | chr13-68548916-DEL-755 |
| chr13-68612489-DEL-82 | chr13-68612489-DEL-82 |
| chr13-68618265-DEL-69 | chr13-68618313-DEL-69 |
| chr13-68621447-DEL-1790 | chr13-68621447-DEL-1790 |
| chr13-68701596-DEL-57 | chr13-68701593-DEL-56 |
| chr13-68704333-DEL-215 | chr13-68704334-DEL-215 |
| chr13-68759018-DEL-76 | chr13-68759023-DEL-74 |
| chr13-68760566-DEL-103 | chr13-68760567-DEL-103 |
| chr13-68763422-DEL-3438 | chr13-68763423-DEL-3438 |
| chr13-68783247-DEL-68 | chr13-68783248-DEL-68 |
| chr13-68791240-DEL-516 | chr13-68791241-DEL-516 |
| chr13-68793322-DEL-54 | chr13-68793323-DEL-54 |
| chr13-68797406-DEL-54 | chr13-68797407-DEL-54 |
| chr13-68810600-DEL-76 | chr13-68810601-DEL-76 |
| chr13-68829865-DEL-111 | chr13-68829864-DEL-110 |
| chr13-68852007-DEL-1189 | chr13-68852008-DEL-1189 |
| chr13-68861908-DEL-100 | chr13-68861906-DEL-100 |
| chr13-68862906-DEL-156 | chr13-68862891-DEL-160 |
| chr13-68868017-DEL-60 | chr13-68868018-DEL-60 |
| chr13-68874392-DEL-56 | chr13-68874393-DEL-56 |
| chr13-68883283-DEL-127 | chr13-68883277-DEL-149 |
| chr13-68901445-DEL-67 | chr13-68901445-DEL-68 |
| chr13-68918074-DEL-1043 | chr13-68918075-DEL-1043 |
| chr13-68959654-DEL-119 | chr13-68959646-DEL-121 |
| chr13-68967263-DEL-321 | chr13-68967262-DEL-322 |
| chr13-68969951-DEL-133 | chr13-68969952-DEL-131 |
| chr13-68977230-DEL-164 | chr13-68977231-DEL-164 |
| chr13-68987576-DEL-52 | chr13-68987577-DEL-52 |
| chr13-69028570-DEL-75 | chr13-69028571-DEL-75 |
| chr13-69041565-DEL-6460 | chr13-69041566-DEL-6460 |
| chr13-69076309-DEL-138 | chr13-69076308-DEL-136 |
| chr13-69088200-DEL-910 | chr13-69088201-DEL-910 |
| chr13-69118588-DEL-157 | chr13-69118560-DEL-166 |
| chr13-69131156-DEL-54 | chr13-69131157-DEL-54 |
| chr13-69187719-DEL-1453 | chr13-69187719-DEL-1453 |
| chr13-69207674-DEL-57 | chr13-69207673-DEL-58 |
| chr13-69218933-DEL-66 | chr13-69218934-DEL-66 |
| chr13-69227741-DEL-6408 | chr13-69227742-DEL-6408 |
| chr13-69235133-DEL-53 | chr13-69235116-DEL-54 |
| chr13-69243555-DEL-203 | chr13-69243553-DEL-204 |
| chr13-69249908-DEL-639 | chr13-69249909-DEL-639 |
| chr13-69264630-DEL-3038 | chr13-69264631-DEL-3038 |
| chr13-69270004-DEL-110 | chr13-69270005-DEL-109 |
| chr13-69300706-DEL-6436 | chr13-69300707-DEL-6436 |
| chr13-69312501-DEL-56 | chr13-69312502-DEL-56 |
| chr13-69319163-DEL-6856 | chr13-69319164-DEL-6856 |
| chr13-69335844-DEL-124 | chr13-69335845-DEL-128 |
| chr13-69341557-DEL-53 | chr13-69341552-DEL-54 |
| chr13-69342209-DEL-1165 | chr13-69342210-DEL-1165 |

| | |
|---|---|
| chr13-69366568-DEL-169 | chr13-69366569-DEL-169 |
| chr13-69385266-DEL-564 | chr13-69385267-DEL-564 |
| chr13-69389000-DEL-55 | chr13-69388991-DEL-50 |
| chr13-69398951-DEL-240 | chr13-69398952-DEL-240 |
| chr13-69424176-DEL-60 | chr13-69424177-DEL-60 |
| chr13-69427552-DEL-118 | chr13-69427553-DEL-118 |
| chr13-69431529-DEL-6384 | chr13-69431530-DEL-6384 |
| chr13-69440058-DEL-283 | chr13-69440059-DEL-283 |
| chr13-69462706-DEL-3794 | chr13-69462705-DEL-3794 |
| chr13-69463720-DEL-96 | chr13-69463721-DEL-96 |
| chr13-69487585-DEL-52 | chr13-69487586-DEL-52 |
| chr13-69502167-DEL-277 | chr13-69502166-DEL-273 |
| chr13-69522263-DEL-233 | chr13-69522264-DEL-233 |
| chr13-69525480-DEL-283 | chr13-69525483-DEL-284 |
| chr13-69547323-DEL-56 | chr13-69547324-DEL-62 |
| chr13-69549348-DEL-60 | chr13-69549349-DEL-60 |
| chr13-69551620-DEL-58 | chr13-69551621-DEL-58 |
| chr13-69567795-DEL-682 | chr13-69567796-DEL-682 |
| chr13-69592208-DEL-112 | chr13-69592209-DEL-112 |
| chr13-69594251-DEL-316 | chr13-69594252-DEL-316 |
| chr13-69597067-DEL-287 | chr13-69597068-DEL-287 |
| chr13-69597887-DEL-116 | chr13-69597894-DEL-117 |
| chr13-69612883-DEL-209 | chr13-69612884-DEL-209 |
| chr13-69615620-DEL-1966 | chr13-69615621-DEL-1966 |
| chr13-69656250-DEL-61 | chr13-69656251-DEL-61 |
| chr13-69659390-DEL-165 | chr13-69659391-DEL-165 |
| chr13-69664200-DEL-144 | chr13-69664200-DEL-145 |
| chr13-69726371-DEL-2220 | chr13-69726372-DEL-2220 |
| chr13-69742908-DEL-114 | chr13-69742909-DEL-76 |
| chr13-69767983-DEL-78 | chr13-69767986-DEL-76 |
| chr13-69795495-DEL-58 | chr13-69795496-DEL-58 |
| chr13-69803009-DEL-213 | chr13-69803010-DEL-213 |
| chr13-69804102-DEL-1842 | chr13-69804103-DEL-1842 |
| chr13-69805559-DEL-180 | chr13-69805551-DEL-174 |
| chr13-69814282-DEL-89 | chr13-69814283-DEL-89 |
| chr13-69825537-DEL-60 | chr13-69825538-DEL-60 |
| chr13-69827549-DEL-200 | chr13-69827561-DEL-200 |
| chr13-69835608-DEL-56 | chr13-69835609-DEL-56 |
| chr13-69848164-DEL-108 | chr13-69848166-DEL-107 |
| chr13-69870717-DEL-97 | chr13-69870727-DEL-88 |
| chr13-69872894-DEL-68 | chr13-69872895-DEL-68 |
| chr13-69897066-DEL-1664 | chr13-69897066-DEL-1662 |
| chr13-69903184-DEL-576 | chr13-69903185-DEL-576 |
| chr13-69915494-DEL-119 | chr13-69915495-DEL-119 |
| chr13-69954565-DEL-485 | chr13-69954566-DEL-485 |
| chr13-69979027-DEL-585 | chr13-69979028-DEL-585 |
| chr13-69985715-DEL-401 | chr13-69985717-DEL-400 |
| chr13-70024985-DEL-441 | chr13-70024986-DEL-441 |
| chr13-70029948-DEL-8725 | chr13-70029949-DEL-8725 |
| chr13-70057378-DEL-148 | chr13-70057375-DEL-152 |
| chr13-70078636-DEL-184 | chr13-70078637-DEL-184 |
| chr13-70112848-DEL-54 | chr13-70112849-DEL-54 |
| chr13-70133342-DEL-98 | chr13-70133343-DEL-98 |
| chr13-70153781-DEL-266 | chr13-70153782-DEL-266 |
| chr13-70154422-DEL-168 | chr13-70154417-DEL-185 |
| chr13-70159632-DEL-214 | chr13-70159633-DEL-214 |
| chr13-70175811-DEL-1101 | chr13-70175824-DEL-1083 |
| chr13-70177693-DEL-862 | chr13-70177694-DEL-862 |
| chr13-70180490-DEL-104 | chr13-70180491-DEL-104 |
| chr13-70183771-DEL-57 | chr13-70183758-DEL-58 |
| chr13-70199867-DEL-209 | chr13-70199868-DEL-209 |
| chr13-70210520-DEL-544 | chr13-70210521-DEL-544 |
| chr13-70215744-DEL-224 | chr13-70215745-DEL-224 |
| chr13-70257157-DEL-2172 | chr13-70257158-DEL-2172 |
| chr13-70258903-DEL-84 | chr13-70258900-DEL-100 |
| chr13-70272079-DEL-52 | chr13-70272080-DEL-52 |
| chr13-70283789-DEL-546 | chr13-70283788-DEL-546 |
| chr13-70313624-DEL-92 | chr13-70313625-DEL-92 |
| chr13-70320483-DEL-88 | chr13-70320492-DEL-80 |
| chr13-70335599-DEL-831 | chr13-70335600-DEL-831 |
| chr13-70347887-DEL-469 | chr13-70347888-DEL-469 |
| chr13-70378339-DEL-66 | chr13-70378340-DEL-88 |
| chr13-70399935-DEL-100 | chr13-70399936-DEL-100 |
| chr13-70414306-DEL-3255 | chr13-70414307-DEL-3255 |
| chr13-70425231-DEL-64 | chr13-70425232-DEL-64 |
| chr13-70434333-DEL-134 | chr13-70434336-DEL-131 |
| chr13-70436394-DEL-102 | chr13-70436395-DEL-102 |
| chr13-70460062-DEL-72 | chr13-70460063-DEL-72 |
| chr13-70463209-DEL-217 | chr13-70463210-DEL-217 |
| chr13-70465279-DEL-64 | chr13-70465280-DEL-64 |

| | |
|---|---|
| chr13-70465485-DEL-1750 | chr13-70465486-DEL-1750 |
| chr13-70476578-DEL-66 | chr13-70476579-DEL-66 |
| chr13-70534590-DEL-178 | chr13-70534587-DEL-178 |
| chr13-70535451-DEL-1561 | chr13-70535452-DEL-1562 |
| chr13-70538646-DEL-5800 | chr13-70538646-DEL-5800 |
| chr13-70545625-DEL-1003 | chr13-70545626-DEL-1003 |
| chr13-70569284-DEL-67 | chr13-70569285-DEL-67 |
| chr13-70595437-DEL-80 | chr13-70595438-DEL-80 |
| chr13-70607687-DEL-291 | chr13-70607685-DEL-292 |
| chr13-70627067-DEL-150 | chr13-70627073-DEL-135 |
| chr13-70631806-DEL-101 | chr13-70631807-DEL-101 |
| chr13-70633311-DEL-3462 | chr13-70633324-DEL-3464 |
| chr13-70640834-DEL-703 | chr13-70640859-DEL-679 |
| chr13-70652814-DEL-208 | chr13-70652814-DEL-208 |
| chr13-70664474-DEL-257 | chr13-70664475-DEL-257 |
| chr13-70668603-DEL-6131 | chr13-70668604-DEL-6131 |
| chr13-70670004-DEL-767 | chr13-70670005-DEL-767 |
| chr13-70674563-DEL-162 | chr13-70674564-DEL-162 |
| chr13-70689208-DEL-61 | chr13-70689209-DEL-61 |
| chr13-70709270-DEL-103 | chr13-70709271-DEL-103 |
| chr13-70776305-DEL-996 | chr13-70776306-DEL-996 |
| chr13-70787011-DEL-198 | chr13-70787012-DEL-198 |
| chr13-70793845-DEL-73 | chr13-70793844-DEL-75 |
| chr13-70832465-DEL-59 | chr13-70832466-DEL-59 |
| chr13-70842683-DEL-210 | chr13-70842684-DEL-210 |
| chr13-70849926-DEL-167 | chr13-70849916-DEL-168 |
| chr13-70860946-DEL-79 | chr13-70860945-DEL-80 |
| chr13-70933675-DEL-401 | chr13-70933676-DEL-401 |
| chr13-70959286-DEL-58 | chr13-70959288-DEL-57 |
| chr13-70970882-DEL-66 | chr13-70970858-DEL-62 |
| chr13-70980057-DEL-203 | chr13-70980058-DEL-203 |
| chr13-71005284-DEL-505 | chr13-71005285-DEL-505 |
| chr13-71009584-DEL-194 | chr13-71009585-DEL-194 |
| chr13-71037835-DEL-62 | chr13-71037836-DEL-52 |
| chr13-71041476-DEL-168 | chr13-71041501-DEL-192 |
| chr13-71045115-DEL-52 | chr13-71045116-DEL-56 |
| chr13-71046640-DEL-1230 | chr13-71046644-DEL-1220 |
| chr13-71049136-DEL-575 | chr13-71049138-DEL-572 |
| chr13-71055240-DEL-817 | chr13-71055241-DEL-817 |
| chr13-71068025-DEL-180 | chr13-71068026-DEL-180 |
| chr13-71072178-DEL-5600 | chr13-71072179-DEL-5600 |
| chr13-71086301-DEL-138 | chr13-71086301-DEL-138 |
| chr13-71151025-DEL-51 | chr13-71151028-DEL-52 |
| chr13-71157377-DEL-362 | chr13-71157378-DEL-362 |
| chr13-71166397-DEL-2395 | chr13-71166398-DEL-2395 |
| chr13-71186877-DEL-54 | chr13-71186877-DEL-54 |
| chr13-71195671-DEL-75 | chr13-71195672-DEL-75 |
| chr13-71233776-DEL-6452 | chr13-71233777-DEL-6452 |
| chr13-71272872-DEL-85 | chr13-71272881-DEL-74 |
| chr13-71285567-DEL-156 | chr13-71285568-DEL-156 |
| chr13-71360833-DEL-57 | chr13-71360834-DEL-57 |
| chr13-71376659-DEL-1013 | chr13-71376659-DEL-1013 |
| chr13-71393103-DEL-63 | chr13-71393108-DEL-65 |
| chr13-71407985-DEL-120 | chr13-71408010-DEL-84 |
| chr13-71435331-DEL-139 | chr13-71435306-DEL-116 |
| chr13-71451192-DEL-192 | chr13-71451193-DEL-192 |
| chr13-71459803-DEL-177 | chr13-71459804-DEL-177 |
| chr13-71491732-DEL-6853 | chr13-71491733-DEL-6853 |
| chr13-71505838-DEL-100 | chr13-71505837-DEL-100 |
| chr13-71517499-DEL-66 | chr13-71517500-DEL-66 |
| chr13-71522039-DEL-60 | chr13-71522040-DEL-60 |
| chr13-71555700-DEL-176 | chr13-71555723-DEL-166 |
| chr13-71556643-DEL-251 | chr13-71556644-DEL-251 |
| chr13-71572614-DEL-224 | chr13-71572615-DEL-224 |
| chr13-71573187-DEL-1981 | chr13-71573188-DEL-1981 |
| chr13-71576380-DEL-6676 | chr13-71576365-DEL-6677 |
| chr13-71584171-DEL-1234 | chr13-71584172-DEL-1234 |
| chr13-71594245-DEL-56 | chr13-71594245-DEL-56 |
| chr13-71627425-DEL-140 | chr13-71627430-DEL-138 |
| chr13-71639749-DEL-6512 | chr13-71639750-DEL-6512 |
| chr13-71676431-DEL-56 | chr13-71676432-DEL-56 |
| chr13-71683929-DEL-567 | chr13-71683930-DEL-567 |
| chr13-71704352-DEL-1835 | chr13-71704353-DEL-1835 |
| chr13-71708551-DEL-257 | chr13-71708554-DEL-252 |
| chr13-71732567-DEL-51 | chr13-71732568-DEL-51 |
| chr13-71743455-DEL-131 | chr13-71743438-DEL-121 |
| chr13-71745744-DEL-86 | chr13-71745745-DEL-86 |
| chr13-71803267-DEL-66 | chr13-71803269-DEL-67 |
| chr13-71812748-DEL-52 | chr13-71812749-DEL-52 |
| chr13-71826594-DEL-60 | chr13-71826595-DEL-60 |

| | |
|---|---|
| chr13-71828787-DEL-569 | chr13-71828788-DEL-569 |
| chr13-71845743-DEL-57 | chr13-71845744-DEL-57 |
| chr13-71851738-DEL-84 | chr13-71851745-DEL-78 |
| chr13-71855421-DEL-54 | chr13-71855422-DEL-54 |
| chr13-71868740-DEL-93 | chr13-71868741-DEL-93 |
| chr13-71877620-DEL-87 | chr13-71877657-DEL-86 |
| chr13-71904837-DEL-254 | chr13-71904838-DEL-254 |
| chr13-71997905-DEL-95 | chr13-71997858-DEL-93 |
| chr13-72018004-DEL-366 | chr13-72018000-DEL-379 |
| chr13-72027616-DEL-6432 | chr13-72027617-DEL-6432 |
| chr13-72044661-DEL-92 | chr13-72044662-DEL-92 |
| chr13-72060473-DEL-60 | chr13-72060474-DEL-60 |
| chr13-72087023-DEL-129 | chr13-72087024-DEL-129 |
| chr13-72094324-DEL-148 | chr13-72094299-DEL-172 |
| chr13-72137476-DEL-523 | chr13-72137477-DEL-521 |
| chr13-72142427-DEL-54 | chr13-72142428-DEL-54 |
| chr13-72143712-DEL-288 | chr13-72143713-DEL-288 |
| chr13-72145598-DEL-103 | chr13-72145599-DEL-103 |
| chr13-72183888-DEL-6544 | chr13-72183889-DEL-6544 |
| chr13-72184089-DEL-204 | chr13-72184090-DEL-204 |
| chr13-72201523-DEL-771 | chr13-72201524-DEL-771 |
| chr13-72306294-DEL-141 | chr13-72306295-DEL-141 |
| chr13-72308984-DEL-92 | chr13-72308965-DEL-82 |
| chr13-72342857-DEL-68 | chr13-72342858-DEL-68 |
| chr13-72352395-DEL-2856 | chr13-72352396-DEL-2856 |
| chr13-72358545-DEL-60 | chr13-72358546-DEL-60 |
| chr13-72374294-DEL-96 | chr13-72374295-DEL-96 |
| chr13-72402843-DEL-114 | chr13-72402844-DEL-115 |
| chr13-72448370-DEL-80 | chr13-72448371-DEL-88 |
| chr13-72462151-DEL-66 | chr13-72462152-DEL-52 |
| chr13-72484794-DEL-3321 | chr13-72484795-DEL-3321 |
| chr13-72507424-DEL-3057 | chr13-72507425-DEL-3057 |
| chr13-72521990-DEL-62 | chr13-72521991-DEL-60 |
| chr13-72529042-DEL-1723 | chr13-72529043-DEL-1723 |
| chr13-72533872-DEL-7097 | chr13-72533873-DEL-7097 |
| chr13-72546268-DEL-608 | chr13-72546269-DEL-608 |
| chr13-72607757-DEL-244 | chr13-72607758-DEL-244 |
| chr13-72608599-DEL-490 | chr13-72608600-DEL-490 |
| chr13-72627164-DEL-229 | chr13-72627187-DEL-217 |
| chr13-72658553-DEL-51 | chr13-72658554-DEL-52 |
| chr13-72682966-DEL-146 | chr13-72682967-DEL-164 |
| chr13-72690371-DEL-51 | chr13-72690371-DEL-51 |
| chr13-72699637-DEL-486 | chr13-72699638-DEL-486 |
| chr13-72716996-DEL-1183 | chr13-72716996-DEL-1183 |
| chr13-72748660-DEL-53 | chr13-72748661-DEL-53 |
| chr13-72760799-DEL-110 | chr13-72760801-DEL-72 |
| chr13-72768410-DEL-69 | chr13-72768399-DEL-81 |
| chr13-72797664-DEL-1493 | chr13-72797665-DEL-1493 |
| chr13-72825448-DEL-6588 | chr13-72825449-DEL-6588 |
| chr13-72863869-DEL-61 | chr13-72863855-DEL-64 |
| chr13-72870503-DEL-118 | chr13-72870504-DEL-118 |
| chr13-72907804-DEL-66 | chr13-72907811-DEL-60 |
| chr13-72935425-DEL-201 | chr13-72935441-DEL-173 |
| chr13-72946907-DEL-72 | chr13-72946916-DEL-64 |
| chr13-72992877-DEL-1281 | chr13-72992878-DEL-1281 |
| chr13-73034014-DEL-511 | chr13-73034015-DEL-511 |
| chr13-73087862-DEL-128 | chr13-73087861-DEL-130 |
| chr13-73090618-DEL-188 | chr13-73090619-DEL-188 |
| chr13-73132452-DEL-113 | chr13-73132453-DEL-113 |
| chr13-73193967-DEL-64 | chr13-73193968-DEL-63 |
| chr13-73250197-DEL-110 | chr13-73250175-DEL-125 |
| chr13-73267600-DEL-1216 | chr13-73267601-DEL-1216 |
| chr13-73272092-DEL-6247 | chr13-73272093-DEL-6247 |
| chr13-73316791-DEL-107 | chr13-73316793-DEL-102 |
| chr13-73351316-DEL-220 | chr13-73351317-DEL-244 |
| chr13-73357563-DEL-168 | chr13-73357564-DEL-168 |
| chr13-73386678-DEL-209 | chr13-73386679-DEL-209 |
| chr13-73390427-DEL-230 | chr13-73390428-DEL-230 |
| chr13-73465938-DEL-338 | chr13-73465930-DEL-339 |
| chr13-73491631-DEL-68 | chr13-73491632-DEL-68 |
| chr13-73492335-DEL-1196 | chr13-73492336-DEL-1196 |
| chr13-73495964-DEL-177 | chr13-73495948-DEL-178 |
| chr13-73529496-DEL-80 | chr13-73529497-DEL-80 |
| chr13-73530262-DEL-62 | chr13-73530263-DEL-62 |
| chr13-73578758-DEL-68 | chr13-73578714-DEL-77 |
| chr13-73600290-DEL-223 | chr13-73600291-DEL-223 |
| chr13-73634519-DEL-116 | chr13-73634506-DEL-114 |
| chr13-73665089-DEL-1655 | chr13-73665089-DEL-1655 |
| chr13-73700465-DEL-55 | chr13-73700464-DEL-54 |
| chr13-73703378-DEL-273 | chr13-73703379-DEL-273 |

| | |
|---|---|
| chr13-73720717-DEL-56 | chr13-73720718-DEL-56 |
| chr13-73773710-DEL-90 | chr13-73773711-DEL-90 |
| chr13-73781012-DEL-134 | chr13-73781013-DEL-134 |
| chr13-73786230-DEL-55 | chr13-73786231-DEL-54 |
| chr13-73791523-DEL-520 | chr13-73791524-DEL-520 |
| chr13-73853051-DEL-145 | chr13-73853081-DEL-116 |
| chr13-73854880-DEL-237 | chr13-73854881-DEL-237 |
| chr13-73870286-DEL-59 | chr13-73870284-DEL-59 |
| chr13-73872942-DEL-477 | chr13-73872943-DEL-437 |
| chr13-73893997-DEL-92 | chr13-73893997-DEL-92 |
| chr13-73908887-DEL-202 | chr13-73908888-DEL-202 |
| chr13-73924949-DEL-212 | chr13-73924950-DEL-212 |
| chr13-73949546-DEL-177 | chr13-73949547-DEL-177 |
| chr13-73975309-DEL-2371 | chr13-73975310-DEL-2371 |
| chr13-74024559-DEL-3296 | chr13-74024560-DEL-3296 |
| chr13-74053049-DEL-163 | chr13-74053050-DEL-163 |
| chr13-74093722-DEL-254 | chr13-74093723-DEL-254 |
| chr13-74154841-DEL-148 | chr13-74154827-DEL-145 |
| chr13-74219443-DEL-111 | chr13-74219444-DEL-111 |
| chr13-74240422-DEL-352 | chr13-74240423-DEL-352 |
| chr13-74297833-DEL-60 | chr13-74297834-DEL-60 |
| chr13-74301464-DEL-299 | chr13-74301465-DEL-299 |
| chr13-74312012-DEL-70 | chr13-74312013-DEL-70 |
| chr13-74346223-DEL-54 | chr13-74346224-DEL-54 |
| chr13-74396542-DEL-54 | chr13-74396540-DEL-54 |
| chr13-74437112-DEL-497 | chr13-74437113-DEL-497 |
| chr13-74441326-DEL-586 | chr13-74441327-DEL-586 |
| chr13-74448075-DEL-534 | chr13-74448076-DEL-534 |
| chr13-74482053-DEL-95 | chr13-74482054-DEL-95 |
| chr13-74491860-DEL-171 | chr13-74491861-DEL-171 |
| chr13-74503190-DEL-52 | chr13-74503190-DEL-52 |
| chr13-74523478-DEL-170 | chr13-74523479-DEL-170 |
| chr13-74554515-DEL-962 | chr13-74554501-DEL-976 |
| chr13-74561123-DEL-18053 | chr13-74561123-DEL-18053 |
| chr13-74588059-DEL-140 | chr13-74588060-DEL-140 |
| chr13-74664598-DEL-255 | chr13-74664581-DEL-275 |
| chr13-74674882-DEL-207 | chr13-74674879-DEL-208 |
| chr13-74707453-DEL-343 | chr13-74707454-DEL-343 |
| chr13-74708799-DEL-66 | chr13-74708800-DEL-66 |
| chr13-74725521-DEL-429 | chr13-74725522-DEL-429 |
| chr13-74757340-DEL-816 | chr13-74757341-DEL-816 |
| chr13-74784131-DEL-1696 | chr13-74784132-DEL-1696 |
| chr13-74809801-DEL-68 | chr13-74809802-DEL-68 |
| chr13-74818676-DEL-72 | chr13-74818667-DEL-72 |
| chr13-74832093-DEL-6196 | chr13-74832094-DEL-6196 |
| chr13-74853702-DEL-6668 | chr13-74853702-DEL-6668 |
| chr13-74904353-DEL-99 | chr13-74904342-DEL-66 |
| chr13-74923872-DEL-204 | chr13-74923873-DEL-204 |
| chr13-74924614-DEL-400 | chr13-74924615-DEL-400 |
| chr13-74932048-DEL-1108 | chr13-74932049-DEL-1108 |
| chr13-74936233-DEL-351 | chr13-74936230-DEL-355 |
| chr13-74944010-DEL-310 | chr13-74944011-DEL-310 |
| chr13-74958863-DEL-2813 | chr13-74958864-DEL-2813 |
| chr13-74971360-DEL-131 | chr13-74971332-DEL-159 |
| chr13-74993724-DEL-395 | chr13-74993725-DEL-395 |
| chr13-75052325-DEL-208 | chr13-75052319-DEL-209 |
| chr13-75094938-DEL-1404 | chr13-75094939-DEL-1404 |
| chr13-75105002-DEL-66 | chr13-75105003-DEL-64 |
| chr13-75118112-DEL-120 | chr13-75118125-DEL-92 |
| chr13-75140556-DEL-875 | chr13-75140557-DEL-875 |
| chr13-75195202-DEL-4309 | chr13-75195201-DEL-4310 |
| chr13-75214030-DEL-400 | chr13-75214031-DEL-400 |
| chr13-75233913-DEL-58 | chr13-75233914-DEL-58 |
| chr13-75266073-DEL-1914 | chr13-75266074-DEL-1914 |
| chr13-75272112-DEL-1086 | chr13-75272113-DEL-1086 |
| chr13-75286471-DEL-196 | chr13-75286472-DEL-196 |
| chr13-75303348-DEL-464 | chr13-75303349-DEL-464 |
| chr13-75310549-DEL-102 | chr13-75310550-DEL-102 |
| chr13-75319749-DEL-10360 | chr13-75319750-DEL-10360 |
| chr13-75319764-DEL-214 | chr13-75319765-DEL-214 |
| chr13-75324657-DEL-3729 | chr13-75324658-DEL-3729 |
| chr13-75354050-DEL-8894 | chr13-75354051-DEL-8894 |
| chr13-75392973-DEL-54 | chr13-75392974-DEL-54 |
| chr13-75449465-DEL-75 | chr13-75449466-DEL-75 |
| chr13-75490476-DEL-152 | chr13-75490499-DEL-150 |
| chr13-75508656-DEL-220 | chr13-75508657-DEL-220 |
| chr13-75512883-DEL-695 | chr13-75512884-DEL-695 |
| chr13-75556330-DEL-203 | chr13-75556324-DEL-204 |
| chr13-75574143-DEL-60 | chr13-75574160-DEL-56 |
| chr13-75611557-DEL-400 | chr13-75611558-DEL-400 |

| | |
|---|---|
| chr13-75615516-DEL-64 | chr13-75615517-DEL-64 |
| chr13-75621924-DEL-1162 | chr13-75621925-DEL-1162 |
| chr13-75664617-DEL-1434 | chr13-75664618-DEL-1434 |
| chr13-75694141-DEL-85 | chr13-75694158-DEL-66 |
| chr13-75700735-DEL-76 | chr13-75700736-DEL-76 |
| chr13-75706165-DEL-607 | chr13-75706154-DEL-608 |
| chr13-75709745-DEL-67 | chr13-75709746-DEL-67 |
| chr13-75723650-DEL-2089 | chr13-75723651-DEL-2089 |
| chr13-75731654-DEL-60 | chr13-75731655-DEL-60 |
| chr13-75792966-DEL-396 | chr13-75792967-DEL-396 |
| chr13-75809792-DEL-483 | chr13-75809793-DEL-483 |
| chr13-75825130-DEL-5355 | chr13-75825131-DEL-5355 |
| chr13-75845348-DEL-2797 | chr13-75845349-DEL-2797 |
| chr13-75860889-DEL-205 | chr13-75860890-DEL-205 |
| chr13-75911473-DEL-60 | chr13-75911474-DEL-60 |
| chr13-75930290-DEL-313 | chr13-75930291-DEL-313 |
| chr13-75938968-DEL-76 | chr13-75938969-DEL-76 |
| chr13-75970433-DEL-204 | chr13-75970433-DEL-205 |
| chr13-75970851-DEL-549 | chr13-75970852-DEL-549 |
| chr13-75975013-DEL-2585 | chr13-75975014-DEL-2585 |
| chr13-75997120-DEL-84 | chr13-75997121-DEL-84 |
| chr13-75998491-DEL-691 | chr13-75998492-DEL-691 |
| chr13-76009984-DEL-3017 | chr13-76009985-DEL-3017 |
| chr13-76017122-DEL-145 | chr13-76017123-DEL-145 |
| chr13-76029903-DEL-182 | chr13-76029903-DEL-182 |
| chr13-76040861-DEL-53 | chr13-76040862-DEL-53 |
| chr13-76074203-DEL-9993 | chr13-76074201-DEL-9993 |
| chr13-76102210-DEL-977 | chr13-76102211-DEL-977 |
| chr13-76103856-DEL-72 | chr13-76103857-DEL-72 |
| chr13-76141634-DEL-86 | chr13-76141665-DEL-56 |
| chr13-76147941-DEL-286 | chr13-76147942-DEL-286 |
| chr13-76150423-DEL-109 | chr13-76150376-DEL-110 |
| chr13-76179882-DEL-522 | chr13-76179883-DEL-522 |
| chr13-76180756-DEL-4070 | chr13-76180757-DEL-4070 |
| chr13-76199626-DEL-87 | chr13-76199626-DEL-87 |
| chr13-76225324-DEL-6457 | chr13-76225325-DEL-6457 |
| chr13-76233919-DEL-207 | chr13-76233920-DEL-207 |
| chr13-76263849-DEL-85 | chr13-76263850-DEL-85 |
| chr13-76289348-DEL-55 | chr13-76289349-DEL-65 |
| chr13-76315571-DEL-2367 | chr13-76315572-DEL-2367 |
| chr13-76347747-DEL-64 | chr13-76347732-DEL-65 |
| chr13-76351925-DEL-6377 | chr13-76351926-DEL-6375 |
| chr13-76380335-DEL-256 | chr13-76380336-DEL-256 |
| chr13-76389838-DEL-470 | chr13-76389839-DEL-470 |
| chr13-76393105-DEL-720 | chr13-76393106-DEL-720 |
| chr13-76405420-DEL-182 | chr13-76405423-DEL-180 |
| chr13-76436047-DEL-184 | chr13-76436048-DEL-184 |
| chr13-76439014-DEL-55 | chr13-76439015-DEL-55 |
| chr13-76453246-DEL-83 | chr13-76453282-DEL-63 |
| chr13-76469403-DEL-70 | chr13-76469404-DEL-70 |
| chr13-76493786-DEL-6344 | chr13-76493787-DEL-6344 |
| chr13-76503376-DEL-571 | chr13-76503377-DEL-571 |
| chr13-76509353-DEL-1619 | chr13-76509354-DEL-1619 |
| chr13-76523089-DEL-78 | chr13-76523115-DEL-55 |
| chr13-76525274-DEL-123 | chr13-76525303-DEL-95 |
| chr13-76528380-DEL-53 | chr13-76528381-DEL-55 |
| chr13-76536393-DEL-167 | chr13-76536394-DEL-167 |
| chr13-76538481-DEL-148 | chr13-76538482-DEL-148 |
| chr13-76546564-DEL-286 | chr13-76546565-DEL-286 |
| chr13-76553697-DEL-97 | chr13-76553698-DEL-97 |
| chr13-76554139-DEL-309 | chr13-76554139-DEL-310 |
| chr13-76574377-DEL-1339 | chr13-76574377-DEL-1339 |
| chr13-76576146-DEL-80 | chr13-76576147-DEL-72 |
| chr13-76583287-DEL-179 | chr13-76583280-DEL-180 |
| chr13-76585877-DEL-177 | chr13-76585878-DEL-177 |
| chr13-76587455-DEL-327 | chr13-76587446-DEL-324 |
| chr13-76597933-DEL-171 | chr13-76597934-DEL-166 |
| chr13-76632043-DEL-88 | chr13-76632043-DEL-88 |
| chr13-76633136-DEL-52 | chr13-76633137-DEL-52 |
| chr13-76636966-DEL-1270 | chr13-76636967-DEL-1270 |
| chr13-76638009-DEL-83 | chr13-76638010-DEL-71 |
| chr13-76642026-DEL-11303 | chr13-76642027-DEL-11303 |
| chr13-76644032-DEL-5342 | chr13-76644033-DEL-5342 |
| chr13-76659649-DEL-1109 | chr13-76659650-DEL-1109 |
| chr13-76662065-DEL-703 | chr13-76662039-DEL-704 |
| chr13-76685234-DEL-563 | chr13-76685235-DEL-563 |
| chr13-76692584-DEL-262 | chr13-76692585-DEL-262 |
| chr13-76730507-DEL-620 | chr13-76730508-DEL-620 |
| chr13-76773580-DEL-2728 | chr13-76773581-DEL-2727 |
| chr13-76775704-DEL-2994 | chr13-76775705-DEL-2994 |

| | |
|---|---|
| chr13-76783479-DEL-2494 | chr13-76783480-DEL-2494 |
| chr13-76793892-DEL-113 | chr13-76793895-DEL-111 |
| chr13-76798135-DEL-316 | chr13-76798136-DEL-316 |
| chr13-76825083-DEL-1394 | chr13-76825084-DEL-1394 |
| chr13-76846798-DEL-70 | chr13-76846799-DEL-70 |
| chr13-76862907-DEL-6114 | chr13-76862908-DEL-6114 |
| chr13-76902081-DEL-907 | chr13-76902082-DEL-907 |
| chr13-76914165-DEL-63 | chr13-76914166-DEL-63 |
| chr13-76916311-DEL-241 | chr13-76916312-DEL-241 |
| chr13-76924552-DEL-315 | chr13-76924553-DEL-315 |
| chr13-76936766-DEL-547 | chr13-76936767-DEL-547 |
| chr13-76940119-DEL-1534 | chr13-76940120-DEL-1535 |
| chr13-76988569-DEL-100 | chr13-76988570-DEL-92 |
| chr13-77012399-DEL-495 | chr13-77012400-DEL-495 |
| chr13-77051263-DEL-6204 | chr13-77051264-DEL-6204 |
| chr13-77069119-DEL-62 | chr13-77069126-DEL-56 |
| chr13-77108794-DEL-2689 | chr13-77108795-DEL-2689 |
| chr13-77156176-DEL-52 | chr13-77156177-DEL-52 |
| chr13-77157998-DEL-847 | chr13-77157999-DEL-847 |
| chr13-77168602-DEL-96 | chr13-77168603-DEL-88 |
| chr13-77169882-DEL-54 | chr13-77169883-DEL-52 |
| chr13-77190116-DEL-407 | chr13-77190117-DEL-407 |
| chr13-77212449-DEL-56 | chr13-77212454-DEL-52 |
| chr13-77242687-DEL-5738 | chr13-77242687-DEL-5738 |
| chr13-77262291-DEL-211 | chr13-77262292-DEL-211 |
| chr13-77280970-DEL-911 | chr13-77280971-DEL-911 |
| chr13-77293041-DEL-84 | chr13-77293042-DEL-78 |
| chr13-77316627-DEL-52 | chr13-77316611-DEL-57 |
| chr13-77317235-DEL-2676 | chr13-77317236-DEL-2676 |
| chr13-77319152-DEL-54 | chr13-77319155-DEL-52 |
| chr13-77346326-DEL-713 | chr13-77346327-DEL-713 |
| chr13-77376113-DEL-7395 | chr13-77376114-DEL-7395 |
| chr13-77402064-DEL-60 | chr13-77402061-DEL-80 |
| chr13-77414560-DEL-1065 | chr13-77414561-DEL-1065 |
| chr13-77446059-DEL-93 | chr13-77446088-DEL-65 |
| chr13-77452161-DEL-62 | chr13-77452133-DEL-62 |
| chr13-77488032-DEL-70 | chr13-77488033-DEL-70 |
| chr13-77501064-DEL-2245 | chr13-77501065-DEL-2245 |
| chr13-77576986-DEL-6358 | chr13-77576987-DEL-6358 |
| chr13-77577051-DEL-214 | chr13-77577052-DEL-214 |
| chr13-77584628-DEL-214 | chr13-77584630-DEL-212 |
| chr13-77587433-DEL-169 | chr13-77587433-DEL-168 |
| chr13-77605967-DEL-2154 | chr13-77605968-DEL-2154 |
| chr13-77645236-DEL-58 | chr13-77645237-DEL-56 |
| chr13-77685933-DEL-6684 | chr13-77685934-DEL-6684 |
| chr13-77735315-DEL-5667 | chr13-77735316-DEL-5667 |
| chr13-77758793-DEL-438 | chr13-77758794-DEL-438 |
| chr13-77762919-DEL-310 | chr13-77762920-DEL-309 |
| chr13-77769321-DEL-2318 | chr13-77769322-DEL-2318 |
| chr13-77783572-DEL-8742 | chr13-77783573-DEL-8742 |
| chr13-77812499-DEL-390 | chr13-77812500-DEL-390 |
| chr13-77826772-DEL-409 | chr13-77826773-DEL-409 |
| chr13-77828864-DEL-2912 | chr13-77828864-DEL-2912 |
| chr13-77833218-DEL-513 | chr13-77833219-DEL-513 |
| chr13-77835960-DEL-202 | chr13-77835961-DEL-202 |
| chr13-77843920-DEL-6424 | chr13-77843921-DEL-6424 |
| chr13-77891902-DEL-2490 | chr13-77891903-DEL-2490 |
| chr13-77919395-DEL-54 | chr13-77919441-DEL-54 |
| chr13-77920898-DEL-1640 | chr13-77920899-DEL-1640 |
| chr13-77928477-DEL-641 | chr13-77928479-DEL-642 |
| chr13-77930579-DEL-498 | chr13-77930580-DEL-498 |
| chr13-77939916-DEL-61 | chr13-77939917-DEL-61 |
| chr13-77943106-DEL-11331 | chr13-77943107-DEL-11331 |
| chr13-77946279-DEL-17407 | chr13-77946280-DEL-17407 |
| chr13-77967850-DEL-292 | chr13-77967851-DEL-292 |
| chr13-78011933-DEL-64 | chr13-78011923-DEL-92 |
| chr13-78060546-DEL-54 | chr13-78060547-DEL-54 |
| chr13-78068675-DEL-108 | chr13-78068676-DEL-108 |
| chr13-78093383-DEL-2743 | chr13-78093384-DEL-2743 |
| chr13-78099389-DEL-199 | chr13-78099390-DEL-199 |
| chr13-78101425-DEL-383 | chr13-78101425-DEL-384 |
| chr13-78111490-DEL-122 | chr13-78111491-DEL-122 |
| chr13-78148435-DEL-817 | chr13-78148436-DEL-817 |
| chr13-78159391-DEL-56 | chr13-78159392-DEL-56 |
| chr13-78235624-DEL-64 | chr13-78235625-DEL-66 |
| chr13-78278910-DEL-56 | chr13-78278911-DEL-56 |
| chr13-78280696-DEL-6514 | chr13-78280697-DEL-6514 |
| chr13-78289902-DEL-785 | chr13-78289901-DEL-785 |
| chr13-78376848-DEL-2233 | chr13-78376845-DEL-2233 |
| chr13-78384874-DEL-71 | chr13-78384875-DEL-71 |

| | |
|---|---|
| chr13-78389143-DEL-63 | chr13-78389144-DEL-63 |
| chr13-78409303-DEL-6281 | chr13-78409304-DEL-6281 |
| chr13-78419418-DEL-1530 | chr13-78419402-DEL-1546 |
| chr13-78426891-DEL-53 | chr13-78426889-DEL-54 |
| chr13-78459433-DEL-85 | chr13-78459399-DEL-88 |
| chr13-78491563-DEL-1127 | chr13-78491564-DEL-1127 |
| chr13-78521900-DEL-5151 | chr13-78521901-DEL-5151 |
| chr13-78535054-DEL-1994 | chr13-78535055-DEL-1994 |
| chr13-78561747-DEL-595 | chr13-78561748-DEL-595 |
| chr13-78563987-DEL-202 | chr13-78563988-DEL-202 |
| chr13-78580156-DEL-372 | chr13-78580157-DEL-372 |
| chr13-78609849-DEL-3328 | chr13-78609850-DEL-3328 |
| chr13-78620505-DEL-82 | chr13-78620506-DEL-79 |
| chr13-78639008-DEL-2852 | chr13-78639009-DEL-2852 |
| chr13-78641799-DEL-117 | chr13-78641799-DEL-118 |
| chr13-78696582-DEL-66 | chr13-78696583-DEL-66 |
| chr13-78824237-DEL-853 | chr13-78824238-DEL-853 |
| chr13-78856086-DEL-139 | chr13-78856087-DEL-139 |
| chr13-78869781-DEL-689 | chr13-78869786-DEL-686 |
| chr13-78900200-DEL-2375 | chr13-78900201-DEL-2375 |
| chr13-78960873-DEL-175 | chr13-78960874-DEL-175 |
| chr13-78973325-DEL-1021 | chr13-78973326-DEL-1021 |
| chr13-78989633-DEL-202 | chr13-78989633-DEL-202 |
| chr13-78996937-DEL-62 | chr13-78996938-DEL-62 |
| chr13-79002464-DEL-93 | chr13-79002465-DEL-93 |
| chr13-79017495-DEL-63 | chr13-79017489-DEL-64 |
| chr13-79028770-DEL-133 | chr13-79028771-DEL-133 |
| chr13-79093799-DEL-6731 | chr13-79093800-DEL-6731 |
| chr13-79093922-DEL-214 | chr13-79093922-DEL-214 |
| chr13-79113901-DEL-117 | chr13-79113902-DEL-117 |
| chr13-79119941-DEL-5951 | chr13-79119942-DEL-5951 |
| chr13-79120729-DEL-52 | chr13-79120730-DEL-52 |
| chr13-79123153-DEL-58 | chr13-79123154-DEL-58 |
| chr13-79124391-DEL-598 | chr13-79124392-DEL-598 |
| chr13-79159183-DEL-148 | chr13-79159184-DEL-148 |
| chr13-79171943-DEL-157 | chr13-79171968-DEL-108 |
| chr13-79210407-DEL-212 | chr13-79210408-DEL-212 |
| chr13-79212432-DEL-125 | chr13-79212433-DEL-125 |
| chr13-79217987-DEL-54 | chr13-79217988-DEL-54 |
| chr13-79230342-DEL-57 | chr13-79230345-DEL-55 |
| chr13-79230900-DEL-1838 | chr13-79230901-DEL-1838 |
| chr13-79264710-DEL-114 | chr13-79264711-DEL-114 |
| chr13-79272488-DEL-110 | chr13-79272489-DEL-110 |
| chr13-79274074-DEL-4489 | chr13-79274075-DEL-4489 |
| chr13-79278863-DEL-100 | chr13-79278903-DEL-64 |
| chr13-79309240-DEL-66 | chr13-79309227-DEL-64 |
| chr13-79329088-DEL-1261 | chr13-79329088-DEL-1261 |
| chr13-79389530-DEL-155 | chr13-79389531-DEL-155 |
| chr13-79390929-DEL-318 | chr13-79390943-DEL-311 |
| chr13-79407726-DEL-71 | chr13-79407727-DEL-71 |
| chr13-79411905-DEL-3091 | chr13-79411906-DEL-3091 |
| chr13-79462670-DEL-80 | chr13-79462671-DEL-80 |
| chr13-79505061-DEL-322 | chr13-79505062-DEL-322 |
| chr13-79516194-DEL-2683 | chr13-79516195-DEL-2683 |
| chr13-79569940-DEL-6177 | chr13-79569941-DEL-6177 |
| chr13-79600404-DEL-113 | chr13-79600413-DEL-108 |
| chr13-79603072-DEL-1391 | chr13-79603073-DEL-1391 |
| chr13-79636851-DEL-2483 | chr13-79636848-DEL-2481 |
| chr13-79637997-DEL-1076 | chr13-79637998-DEL-1076 |
| chr13-79657187-DEL-831 | chr13-79657188-DEL-831 |
| chr13-79684829-DEL-3182 | chr13-79684830-DEL-3182 |
| chr13-79707156-DEL-446 | chr13-79707157-DEL-445 |
| chr13-79711627-DEL-7111 | chr13-79711628-DEL-7111 |
| chr13-79749990-DEL-1859 | chr13-79749991-DEL-1859 |
| chr13-79801262-DEL-191 | chr13-79801263-DEL-191 |
| chr13-79834108-DEL-216 | chr13-79834109-DEL-216 |
| chr13-79869688-DEL-203 | chr13-79869647-DEL-201 |
| chr13-79898908-DEL-780 | chr13-79898897-DEL-810 |
| chr13-79929090-DEL-53 | chr13-79929091-DEL-53 |
| chr13-79950344-DEL-6186 | chr13-79950339-DEL-6187 |
| chr13-79973466-DEL-285 | chr13-79973467-DEL-285 |
| chr13-79987354-DEL-77 | chr13-79987357-DEL-74 |
| chr13-80003693-DEL-1003 | chr13-80003694-DEL-1003 |
| chr13-80051670-DEL-358 | chr13-80051671-DEL-358 |
| chr13-80095282-DEL-194 | chr13-80095283-DEL-194 |
| chr13-80108574-DEL-197 | chr13-80108575-DEL-197 |
| chr13-80159032-DEL-1684 | chr13-80159033-DEL-1684 |
| chr13-80172129-DEL-6904 | chr13-80172130-DEL-6904 |
| chr13-80213918-DEL-581 | chr13-80213919-DEL-581 |
| chr13-80245098-DEL-261 | chr13-80245099-DEL-261 |

| | |
|---|---|
| chr13-80260269-DEL-95 | chr13-80260270-DEL-92 |
| chr13-80266008-DEL-130 | chr13-80266007-DEL-132 |
| chr13-80286987-DEL-389 | chr13-80286987-DEL-389 |
| chr13-80287741-DEL-115 | chr13-80287741-DEL-115 |
| chr13-80293334-DEL-1261 | chr13-80293345-DEL-1260 |
| chr13-80305741-DEL-180 | chr13-80305742-DEL-180 |
| chr13-80352031-DEL-1069 | chr13-80352032-DEL-1069 |
| chr13-80364845-DEL-1599 | chr13-80364846-DEL-1599 |
| chr13-80401628-DEL-120 | chr13-80401629-DEL-120 |
| chr13-80405491-DEL-92 | chr13-80405470-DEL-114 |
| chr13-80416429-DEL-293 | chr13-80416430-DEL-293 |
| chr13-80422123-DEL-83 | chr13-80422124-DEL-83 |
| chr13-80424303-DEL-3271 | chr13-80424304-DEL-3271 |
| chr13-80523529-DEL-1491 | chr13-80523530-DEL-1491 |
| chr13-80541243-DEL-405 | chr13-80541244-DEL-410 |
| chr13-80577018-DEL-6413 | chr13-80577019-DEL-6413 |
| chr13-80596224-DEL-2227 | chr13-80596225-DEL-2227 |
| chr13-80610563-DEL-2060 | chr13-80610564-DEL-2060 |
| chr13-80637206-DEL-3164 | chr13-80637207-DEL-3164 |
| chr13-80660448-DEL-391 | chr13-80660449-DEL-391 |
| chr13-80711727-DEL-66 | chr13-80711728-DEL-66 |
| chr13-80732858-DEL-340 | chr13-80732851-DEL-341 |
| chr13-80820676-DEL-57 | chr13-80820672-DEL-68 |
| chr13-80831175-DEL-6635 | chr13-80831175-DEL-6635 |
| chr13-80848867-DEL-5407 | chr13-80848868-DEL-5407 |
| chr13-80874679-DEL-55 | chr13-80874680-DEL-55 |
| chr13-80878289-DEL-376 | chr13-80878290-DEL-376 |
| chr13-80883327-DEL-70 | chr13-80883331-DEL-75 |
| chr13-80895442-DEL-54 | chr13-80895442-DEL-60 |
| chr13-80942112-DEL-97 | chr13-80942109-DEL-101 |
| chr13-80951675-DEL-418 | chr13-80951676-DEL-418 |
| chr13-80982604-DEL-6266 | chr13-80982605-DEL-6266 |
| chr13-80985408-DEL-838 | chr13-80985409-DEL-838 |
| chr13-80994885-DEL-498 | chr13-80994887-DEL-497 |
| chr13-80995419-DEL-60 | chr13-80995420-DEL-56 |
| chr13-80996077-DEL-59 | chr13-80996076-DEL-61 |
| chr13-80999084-DEL-203 | chr13-80999085-DEL-203 |
| chr13-81014702-DEL-82 | chr13-81014703-DEL-77 |
| chr13-81076814-DEL-58 | chr13-81076815-DEL-58 |
| chr13-81100427-DEL-76 | chr13-81100428-DEL-76 |
| chr13-81109600-DEL-51 | chr13-81109595-DEL-52 |
| chr13-81126385-DEL-106 | chr13-81126386-DEL-146 |
| chr13-81143915-DEL-1141 | chr13-81143916-DEL-1141 |
| chr13-81165133-DEL-89 | chr13-81165177-DEL-50 |
| chr13-81191708-DEL-605 | chr13-81191709-DEL-605 |
| chr13-81197670-DEL-90 | chr13-81197624-DEL-94 |
| chr13-81212707-DEL-1939 | chr13-81212708-DEL-1939 |
| chr13-81219486-DEL-4235 | chr13-81219487-DEL-4235 |
| chr13-81231730-DEL-3105 | chr13-81231711-DEL-3124 |
| chr13-81289304-DEL-74 | chr13-81289304-DEL-74 |
| chr13-81299335-DEL-590 | chr13-81299336-DEL-590 |
| chr13-81312169-DEL-56 | chr13-81312168-DEL-54 |
| chr13-81343048-DEL-9115 | chr13-81343049-DEL-9115 |
| chr13-81346573-DEL-2868 | chr13-81346572-DEL-2868 |
| chr13-81374169-DEL-733 | chr13-81374170-DEL-733 |
| chr13-81387858-DEL-179 | chr13-81387853-DEL-180 |
| chr13-81389034-DEL-51 | chr13-81389035-DEL-51 |
| chr13-81401037-DEL-64 | chr13-81401038-DEL-62 |
| chr13-81412380-DEL-95 | chr13-81412398-DEL-78 |
| chr13-81421247-DEL-159 | chr13-81421247-DEL-159 |
| chr13-81452512-DEL-1293 | chr13-81452513-DEL-1293 |
| chr13-81458269-DEL-379 | chr13-81458276-DEL-371 |
| chr13-81538267-DEL-121 | chr13-81538268-DEL-121 |
| chr13-81550013-DEL-61 | chr13-81550000-DEL-59 |
| chr13-81551937-DEL-132 | chr13-81551947-DEL-132 |
| chr13-81561904-DEL-52 | chr13-81561905-DEL-52 |
| chr13-81570697-DEL-1606 | chr13-81570698-DEL-1606 |
| chr13-81573555-DEL-51 | chr13-81573553-DEL-58 |
| chr13-81588584-DEL-992 | chr13-81588585-DEL-992 |
| chr13-81590310-DEL-53 | chr13-81590311-DEL-53 |
| chr13-81601015-DEL-6248 | chr13-81601016-DEL-6248 |
| chr13-81664699-DEL-76 | chr13-81664722-DEL-89 |
| chr13-81686822-DEL-145 | chr13-81686823-DEL-142 |
| chr13-81691286-DEL-83 | chr13-81691293-DEL-57 |
| chr13-81730863-DEL-123 | chr13-81730902-DEL-122 |
| chr13-81734022-DEL-6259 | chr13-81734023-DEL-6259 |
| chr13-81786442-DEL-68 | chr13-81786443-DEL-68 |
| chr13-81814864-DEL-1492 | chr13-81814865-DEL-1492 |
| chr13-81832788-DEL-580 | chr13-81832789-DEL-580 |
| chr13-81837417-DEL-192 | chr13-81837418-DEL-192 |

| | |
|---|---|
| chr13-81843275-DEL-223 | chr13-81843268-DEL-226 |
| chr13-81846737-DEL-235 | chr13-81846738-DEL-235 |
| chr13-81865240-DEL-78 | chr13-81865248-DEL-70 |
| chr13-81867491-DEL-6190 | chr13-81867492-DEL-6190 |
| chr13-81914626-DEL-75 | chr13-81914627-DEL-75 |
| chr13-81958960-DEL-333 | chr13-81958961-DEL-333 |
| chr13-81962898-DEL-824 | chr13-81962901-DEL-824 |
| chr13-81992441-DEL-804 | chr13-81992442-DEL-804 |
| chr13-82014715-DEL-80 | chr13-82014716-DEL-72 |
| chr13-82039090-DEL-7088 | chr13-82039091-DEL-7088 |
| chr13-82088224-DEL-201 | chr13-82088225-DEL-201 |
| chr13-82090178-DEL-153 | chr13-82090179-DEL-153 |
| chr13-82104586-DEL-72 | chr13-82104587-DEL-72 |
| chr13-82117274-DEL-132 | chr13-82117275-DEL-132 |
| chr13-82132901-DEL-6195 | chr13-82132902-DEL-6194 |
| chr13-82138872-DEL-199 | chr13-82138858-DEL-199 |
| chr13-82166887-DEL-250 | chr13-82166886-DEL-250 |
| chr13-82170174-DEL-532 | chr13-82170174-DEL-532 |
| chr13-82201019-DEL-474 | chr13-82201018-DEL-476 |
| chr13-82259115-DEL-2843 | chr13-82259116-DEL-2843 |
| chr13-82302598-DEL-401 | chr13-82302597-DEL-401 |
| chr13-82309579-DEL-94 | chr13-82309584-DEL-89 |
| chr13-82353466-DEL-4951 | chr13-82353465-DEL-4952 |
| chr13-82397530-DEL-498 | chr13-82397531-DEL-498 |
| chr13-82432235-DEL-218 | chr13-82432236-DEL-218 |
| chr13-82444403-DEL-104 | chr13-82444404-DEL-104 |
| chr13-82473941-DEL-117 | chr13-82473942-DEL-117 |
| chr13-82480462-DEL-105 | chr13-82480463-DEL-104 |
| chr13-82483767-DEL-6720 | chr13-82483768-DEL-6720 |
| chr13-82516465-DEL-1490 | chr13-82516466-DEL-1490 |
| chr13-82523475-DEL-418 | chr13-82523473-DEL-418 |
| chr13-82566817-DEL-88 | chr13-82566813-DEL-61 |
| chr13-82592589-DEL-886 | chr13-82592590-DEL-886 |
| chr13-82612752-DEL-1147 | chr13-82612753-DEL-1147 |
| chr13-82654402-DEL-51 | chr13-82654404-DEL-64 |
| chr13-82688569-DEL-1114 | chr13-82688570-DEL-1114 |
| chr13-82697678-DEL-1370 | chr13-82697663-DEL-1371 |
| chr13-82699068-DEL-337 | chr13-82699056-DEL-335 |
| chr13-82703325-DEL-2594 | chr13-82703326-DEL-2584 |
| chr13-82708205-DEL-82 | chr13-82708206-DEL-82 |
| chr13-82720898-DEL-78 | chr13-82720899-DEL-78 |
| chr13-82729861-DEL-107 | chr13-82729862-DEL-107 |
| chr13-82733815-DEL-66 | chr13-82733816-DEL-66 |
| chr13-82735423-DEL-56 | chr13-82735424-DEL-54 |
| chr13-82792093-DEL-1365 | chr13-82792097-DEL-1361 |
| chr13-82801739-DEL-5800 | chr13-82801740-DEL-5800 |
| chr13-82836265-DEL-6554 | chr13-82836266-DEL-6554 |
| chr13-82848979-DEL-206 | chr13-82848980-DEL-206 |
| chr13-82872612-DEL-54 | chr13-82872611-DEL-56 |
| chr13-82900543-DEL-84 | chr13-82900544-DEL-84 |
| chr13-82918423-DEL-800 | chr13-82918424-DEL-800 |
| chr13-82952960-DEL-1042 | chr13-82952961-DEL-1042 |
| chr13-82965410-DEL-491 | chr13-82965411-DEL-491 |
| chr13-82968481-DEL-6436 | chr13-82968482-DEL-6436 |
| chr13-82968554-DEL-216 | chr13-82968555-DEL-216 |
| chr13-82986983-DEL-68 | chr13-82986984-DEL-68 |
| chr13-82996594-DEL-148 | chr13-82996595-DEL-148 |
| chr13-83012527-DEL-80 | chr13-83012544-DEL-70 |
| chr13-83016228-DEL-241 | chr13-83016229-DEL-241 |
| chr13-83064758-DEL-242 | chr13-83064759-DEL-242 |
| chr13-83129342-DEL-62 | chr13-83129343-DEL-62 |
| chr13-83137988-DEL-114 | chr13-83137986-DEL-113 |
| chr13-83174107-DEL-60 | chr13-83174108-DEL-60 |
| chr13-83184616-DEL-1227 | chr13-83184617-DEL-1227 |
| chr13-83282639-DEL-3191 | chr13-83282639-DEL-3192 |
| chr13-83299475-DEL-82 | chr13-83299476-DEL-84 |
| chr13-83319534-DEL-72 | chr13-83319538-DEL-68 |
| chr13-83328826-DEL-6623 | chr13-83328827-DEL-6623 |
| chr13-83335026-DEL-208 | chr13-83335026-DEL-208 |
| chr13-83348779-DEL-91 | chr13-83348780-DEL-91 |
| chr13-83404852-DEL-6466 | chr13-83404853-DEL-6466 |
| chr13-83470526-DEL-1838 | chr13-83470527-DEL-1838 |
| chr13-83497714-DEL-596 | chr13-83497715-DEL-596 |
| chr13-83567116-DEL-7134 | chr13-83567117-DEL-7134 |
| chr13-83581228-DEL-211 | chr13-83581229-DEL-211 |
| chr13-83589851-DEL-2657 | chr13-83589852-DEL-2657 |
| chr13-83693816-DEL-1689 | chr13-83693817-DEL-1689 |
| chr13-83710247-DEL-86 | chr13-83710248-DEL-86 |
| chr13-83731455-DEL-90 | chr13-83731466-DEL-84 |
| chr13-83738892-DEL-131 | chr13-83738893-DEL-131 |

| | |
|---|---|
| chr13-83776632-DEL-94 | chr13-83776633-DEL-94 |
| chr13-83785527-DEL-66 | chr13-83785528-DEL-56 |
| chr13-83798422-DEL-216 | chr13-83798423-DEL-216 |
| chr13-83825230-DEL-217 | chr13-83825231-DEL-217 |
| chr13-83835300-DEL-455 | chr13-83835301-DEL-455 |
| chr13-83838904-DEL-914 | chr13-83838903-DEL-913 |
| chr13-83844766-DEL-73 | chr13-83844767-DEL-73 |
| chr13-83906520-DEL-53 | chr13-83906515-DEL-83 |
| chr13-83934933-DEL-141 | chr13-83934945-DEL-179 |
| chr13-83947077-DEL-66 | chr13-83947053-DEL-78 |
| chr13-83960860-DEL-6225 | chr13-83960861-DEL-6225 |
| chr13-84070790-DEL-670 | chr13-84070793-DEL-666 |
| chr13-84175135-DEL-2846 | chr13-84175136-DEL-2846 |
| chr13-84191006-DEL-676 | chr13-84191007-DEL-676 |
| chr13-84253922-DEL-6047 | chr13-84253923-DEL-6047 |
| chr13-84261260-DEL-2479 | chr13-84261261-DEL-2479 |
| chr13-84294282-DEL-60 | chr13-84294280-DEL-63 |
| chr13-84295364-DEL-911 | chr13-84295364-DEL-911 |
| chr13-84298750-DEL-58 | chr13-84298762-DEL-62 |
| chr13-84349081-DEL-1520 | chr13-84349082-DEL-1520 |
| chr13-84364865-DEL-65 | chr13-84364866-DEL-65 |
| chr13-84377822-DEL-57 | chr13-84377823-DEL-57 |
| chr13-84435765-DEL-86 | chr13-84435766-DEL-86 |
| chr13-84436618-DEL-111 | chr13-84436622-DEL-107 |
| chr13-84446571-DEL-71 | chr13-84446572-DEL-71 |
| chr13-84469460-DEL-57 | chr13-84469461-DEL-57 |
| chr13-84473209-DEL-81 | chr13-84473212-DEL-73 |
| chr13-84482014-DEL-67 | chr13-84482000-DEL-68 |
| chr13-84506586-DEL-1573 | chr13-84506587-DEL-1573 |
| chr13-84525126-DEL-67 | chr13-84525127-DEL-67 |
| chr13-84560596-DEL-71 | chr13-84560602-DEL-66 |
| chr13-84568999-DEL-2855 | chr13-84569000-DEL-2855 |
| chr13-84589098-DEL-6612 | chr13-84589099-DEL-6612 |
| chr13-84614173-DEL-1077 | chr13-84614169-DEL-1084 |
| chr13-84631516-DEL-7283 | chr13-84631517-DEL-7283 |
| chr13-84651125-DEL-55 | chr13-84651114-DEL-54 |
| chr13-84654620-DEL-718 | chr13-84654614-DEL-719 |
| chr13-84687589-DEL-1142 | chr13-84687590-DEL-1142 |
| chr13-84724943-DEL-180 | chr13-84724944-DEL-180 |
| chr13-84741535-DEL-178 | chr13-84741530-DEL-182 |
| chr13-84783679-DEL-54 | chr13-84783680-DEL-58 |
| chr13-84799805-DEL-61 | chr13-84799786-DEL-81 |
| chr13-84803858-DEL-75 | chr13-84803902-DEL-80 |
| chr13-84813664-DEL-498 | chr13-84813665-DEL-498 |
| chr13-84816974-DEL-2177 | chr13-84816974-DEL-2177 |
| chr13-84819474-DEL-74 | chr13-84819475-DEL-74 |
| chr13-84866082-DEL-154 | chr13-84866083-DEL-154 |
| chr13-84918534-DEL-58 | chr13-84918522-DEL-68 |
| chr13-84982664-DEL-6632 | chr13-84982664-DEL-6632 |
| chr13-84998020-DEL-156 | chr13-84998021-DEL-156 |
| chr13-84999700-DEL-2660 | chr13-84999701-DEL-2660 |
| chr13-85015838-DEL-1744 | chr13-85015839-DEL-1744 |
| chr13-85018085-DEL-412 | chr13-85018086-DEL-412 |
| chr13-85037347-DEL-2194 | chr13-85037348-DEL-2194 |
| chr13-85060586-DEL-624 | chr13-85060589-DEL-618 |
| chr13-85072120-DEL-368 | chr13-85072131-DEL-369 |
| chr13-85085530-DEL-390 | chr13-85085531-DEL-390 |
| chr13-85087298-DEL-1063 | chr13-85087301-DEL-1061 |
| chr13-85093185-DEL-904 | chr13-85093186-DEL-904 |
| chr13-85094576-DEL-1906 | chr13-85094577-DEL-1906 |
| chr13-85104903-DEL-1332 | chr13-85104904-DEL-1332 |
| chr13-85109568-DEL-3404 | chr13-85109569-DEL-3404 |
| chr13-85172074-DEL-1478 | chr13-85172075-DEL-1478 |
| chr13-85176819-DEL-168 | chr13-85176819-DEL-161 |
| chr13-85186664-DEL-448 | chr13-85186665-DEL-448 |
| chr13-85187653-DEL-6287 | chr13-85187654-DEL-6287 |
| chr13-85211614-DEL-204 | chr13-85211615-DEL-204 |
| chr13-85223076-DEL-387 | chr13-85223077-DEL-387 |
| chr13-85253112-DEL-78 | chr13-85253113-DEL-78 |
| chr13-85274672-DEL-975 | chr13-85274665-DEL-975 |
| chr13-85280783-DEL-3200 | chr13-85280784-DEL-3200 |
| chr13-85287526-DEL-407 | chr13-85287527-DEL-407 |
| chr13-85290719-DEL-211 | chr13-85290720-DEL-211 |
| chr13-85291348-DEL-2683 | chr13-85291348-DEL-2683 |
| chr13-85296156-DEL-189 | chr13-85296157-DEL-189 |
| chr13-85297492-DEL-191 | chr13-85297493-DEL-191 |
| chr13-85326500-DEL-6645 | chr13-85326503-DEL-6646 |
| chr13-85343547-DEL-56 | chr13-85343553-DEL-50 |
| chr13-85353626-DEL-56 | chr13-85353621-DEL-62 |
| chr13-85400430-DEL-70 | chr13-85400431-DEL-70 |

| | |
|---|---|
| chr13-85406304-DEL-67 | chr13-85406294-DEL-68 |
| chr13-85471566-DEL-6619 | chr13-85471567-DEL-6619 |
| chr13-85477742-DEL-208 | chr13-85477726-DEL-208 |
| chr13-85499590-DEL-54 | chr13-85499591-DEL-50 |
| chr13-85511059-DEL-445 | chr13-85511060-DEL-445 |
| chr13-85539960-DEL-58 | chr13-85539960-DEL-70 |
| chr13-85544262-DEL-542 | chr13-85544263-DEL-542 |
| chr13-85545837-DEL-112 | chr13-85545835-DEL-99 |
| chr13-85552223-DEL-3659 | chr13-85552222-DEL-3659 |
| chr13-85562666-DEL-736 | chr13-85562667-DEL-736 |
| chr13-85569096-DEL-4639 | chr13-85569097-DEL-4639 |
| chr13-85580923-DEL-70 | chr13-85580924-DEL-70 |
| chr13-85588014-DEL-709 | chr13-85588015-DEL-709 |
| chr13-85593655-DEL-728 | chr13-85593656-DEL-728 |
| chr13-85611110-DEL-7623 | chr13-85611111-DEL-7623 |
| chr13-85625218-DEL-115 | chr13-85625219-DEL-115 |
| chr13-85638263-DEL-5608 | chr13-85638264-DEL-5608 |
| chr13-85642763-DEL-100 | chr13-85642764-DEL-100 |
| chr13-85647768-DEL-118 | chr13-85647769-DEL-118 |
| chr13-85673761-DEL-51 | chr13-85673758-DEL-51 |
| chr13-85689836-DEL-1150 | chr13-85689837-DEL-1150 |
| chr13-85747021-DEL-6577 | chr13-85747022-DEL-6577 |
| chr13-85786599-DEL-591 | chr13-85786600-DEL-591 |
| chr13-85820353-DEL-6555 | chr13-85820354-DEL-6555 |
| chr13-85847441-DEL-71 | chr13-85847434-DEL-62 |
| chr13-85865339-DEL-1458 | chr13-85865340-DEL-1458 |
| chr13-85880759-DEL-201 | chr13-85880760-DEL-201 |
| chr13-85928749-DEL-96 | chr13-85928748-DEL-72 |
| chr13-85946921-DEL-268 | chr13-85946961-DEL-256 |
| chr13-85986113-DEL-7807 | chr13-85986114-DEL-7807 |
| chr13-86007414-DEL-84 | chr13-86007415-DEL-83 |
| chr13-86013502-DEL-62 | chr13-86013462-DEL-99 |
| chr13-86018503-DEL-360 | chr13-86018494-DEL-368 |
| chr13-86021627-DEL-110 | chr13-86021628-DEL-110 |
| chr13-86063941-DEL-1087 | chr13-86063942-DEL-1087 |
| chr13-86097667-DEL-95 | chr13-86097668-DEL-95 |
| chr13-86120410-DEL-244 | chr13-86120411-DEL-244 |
| chr13-86154619-DEL-70 | chr13-86154620-DEL-69 |
| chr13-86162911-DEL-134 | chr13-86162914-DEL-132 |
| chr13-86174208-DEL-381 | chr13-86174209-DEL-381 |
| chr13-86177439-DEL-202 | chr13-86177440-DEL-202 |
| chr13-86229183-DEL-210 | chr13-86229184-DEL-210 |
| chr13-86238105-DEL-81 | chr13-86238109-DEL-78 |
| chr13-86242943-DEL-233 | chr13-86242943-DEL-233 |
| chr13-86251666-DEL-94 | chr13-86251667-DEL-94 |
| chr13-86252263-DEL-873 | chr13-86252264-DEL-873 |
| chr13-86263259-DEL-146 | chr13-86263257-DEL-146 |
| chr13-86265906-DEL-202 | chr13-86265907-DEL-202 |
| chr13-86297683-DEL-2219 | chr13-86297683-DEL-2219 |
| chr13-86304036-DEL-192 | chr13-86304045-DEL-195 |
| chr13-86332104-DEL-53 | chr13-86332105-DEL-53 |
| chr13-86383204-DEL-1142 | chr13-86383205-DEL-1142 |
| chr13-86423993-DEL-53 | chr13-86423994-DEL-53 |
| chr13-86431947-DEL-357 | chr13-86431948-DEL-357 |
| chr13-86446501-DEL-2250 | chr13-86446501-DEL-2250 |
| chr13-86447832-DEL-2251 | chr13-86447832-DEL-2251 |
| chr13-86461370-DEL-641 | chr13-86461370-DEL-641 |
| chr13-86470640-DEL-1549 | chr13-86470641-DEL-1549 |
| chr13-86472874-DEL-390 | chr13-86472875-DEL-390 |
| chr13-86497468-DEL-6658 | chr13-86497469-DEL-6658 |
| chr13-86508819-DEL-777 | chr13-86508820-DEL-777 |
| chr13-86539616-DEL-85 | chr13-86539588-DEL-93 |
| chr13-86551546-DEL-61 | chr13-86551547-DEL-61 |
| chr13-86558507-DEL-105 | chr13-86558517-DEL-75 |
| chr13-86560649-DEL-2416 | chr13-86560650-DEL-2416 |
| chr13-86574325-DEL-71 | chr13-86574326-DEL-71 |
| chr13-86591134-DEL-74 | chr13-86591135-DEL-74 |
| chr13-86595721-DEL-104 | chr13-86595710-DEL-116 |
| chr13-86597082-DEL-579 | chr13-86597083-DEL-579 |
| chr13-86599169-DEL-433 | chr13-86599150-DEL-434 |
| chr13-86601787-DEL-502 | chr13-86601788-DEL-502 |
| chr13-86610885-DEL-60 | chr13-86610884-DEL-60 |
| chr13-86625747-DEL-493 | chr13-86625750-DEL-495 |
| chr13-86650629-DEL-9249 | chr13-86650630-DEL-9249 |
| chr13-86655111-DEL-2996 | chr13-86655112-DEL-2996 |
| chr13-86705655-DEL-280 | chr13-86705656-DEL-280 |
| chr13-86712265-DEL-1937 | chr13-86712266-DEL-1937 |
| chr13-86719760-DEL-104 | chr13-86719761-DEL-104 |
| chr13-86732632-DEL-6435 | chr13-86732618-DEL-6447 |
| chr13-86744025-DEL-261 | chr13-86744026-DEL-261 |

| | |
|---|---|
| chr13-86751922-DEL-65 | chr13-86751922-DEL-66 |
| chr13-86763193-DEL-431 | chr13-86763194-DEL-429 |
| chr13-86775717-DEL-3013 | chr13-86775696-DEL-3013 |
| chr13-86779395-DEL-6811 | chr13-86779396-DEL-6811 |
| chr13-86790275-DEL-2163 | chr13-86790272-DEL-2164 |
| chr13-86795404-DEL-118 | chr13-86795400-DEL-122 |
| chr13-86819196-DEL-445 | chr13-86819197-DEL-445 |
| chr13-86839483-DEL-238 | chr13-86839484-DEL-238 |
| chr13-86846439-DEL-963 | chr13-86846460-DEL-941 |
| chr13-86860942-DEL-7085 | chr13-86860943-DEL-7085 |
| chr13-86882229-DEL-859 | chr13-86882230-DEL-858 |
| chr13-86884391-DEL-208 | chr13-86884394-DEL-209 |
| chr13-86920768-DEL-5399 | chr13-86920769-DEL-5399 |
| chr13-86934303-DEL-123 | chr13-86934304-DEL-123 |
| chr13-86944307-DEL-732 | chr13-86944308-DEL-732 |
| chr13-86956276-DEL-54 | chr13-86956276-DEL-54 |
| chr13-87014066-DEL-895 | chr13-87014067-DEL-895 |
| chr13-87035449-DEL-2382 | chr13-87035450-DEL-2382 |
| chr13-87041900-DEL-1770 | chr13-87041901-DEL-1770 |
| chr13-87042870-DEL-740 | chr13-87042871-DEL-740 |
| chr13-87067692-DEL-5769 | chr13-87067693-DEL-5769 |
| chr13-87088055-DEL-447 | chr13-87088055-DEL-444 |
| chr13-87104553-DEL-184 | chr13-87104547-DEL-185 |
| chr13-87113090-DEL-937 | chr13-87113091-DEL-937 |
| chr13-87119469-DEL-324 | chr13-87119470-DEL-324 |
| chr13-87124259-DEL-72 | chr13-87124260-DEL-72 |
| chr13-87131400-DEL-56 | chr13-87131401-DEL-56 |
| chr13-87211993-DEL-7186 | chr13-87211994-DEL-7186 |
| chr13-87221372-DEL-91 | chr13-87221372-DEL-92 |
| chr13-87223742-DEL-115 | chr13-87223743-DEL-115 |
| chr13-87230797-DEL-436 | chr13-87230794-DEL-434 |
| chr13-87232864-DEL-382 | chr13-87232865-DEL-382 |
| chr13-87235765-DEL-988 | chr13-87235765-DEL-988 |
| chr13-87264952-DEL-196 | chr13-87264938-DEL-198 |
| chr13-87266361-DEL-52 | chr13-87266361-DEL-53 |
| chr13-87269493-DEL-70 | chr13-87269493-DEL-67 |
| chr13-87300805-DEL-197 | chr13-87300806-DEL-197 |
| chr13-87309637-DEL-150 | chr13-87309638-DEL-150 |
| chr13-87312146-DEL-81 | chr13-87312147-DEL-81 |
| chr13-87333778-DEL-4175 | chr13-87333778-DEL-4175 |
| chr13-87338574-DEL-871 | chr13-87338574-DEL-871 |
| chr13-87339709-DEL-567 | chr13-87339709-DEL-567 |
| chr13-87362973-DEL-172 | chr13-87362974-DEL-172 |
| chr13-87401657-DEL-144 | chr13-87401658-DEL-144 |
| chr13-87403538-DEL-5074 | chr13-87403539-DEL-5074 |
| chr13-87420779-DEL-246 | chr13-87420780-DEL-240 |
| chr13-87431891-DEL-456 | chr13-87431892-DEL-457 |
| chr13-87448457-DEL-336 | chr13-87448458-DEL-336 |
| chr13-87451122-DEL-65 | chr13-87451123-DEL-65 |
| chr13-87458852-DEL-2694 | chr13-87458853-DEL-2694 |
| chr13-87463129-DEL-665 | chr13-87463130-DEL-665 |
| chr13-87497047-DEL-5189 | chr13-87497048-DEL-5189 |
| chr13-87528973-DEL-54 | chr13-87528925-DEL-78 |
| chr13-87532499-DEL-417 | chr13-87532500-DEL-417 |
| chr13-87535833-DEL-467 | chr13-87535834-DEL-467 |
| chr13-87580511-DEL-168 | chr13-87580512-DEL-168 |
| chr13-87589247-DEL-200 | chr13-87589234-DEL-201 |
| chr13-87607932-DEL-1477 | chr13-87607933-DEL-1477 |
| chr13-87610271-DEL-77 | chr13-87610275-DEL-76 |
| chr13-87622982-DEL-497 | chr13-87622983-DEL-497 |
| chr13-87624794-DEL-6433 | chr13-87624795-DEL-6433 |
| chr13-87650387-DEL-12510 | chr13-87650388-DEL-12510 |
| chr13-87717560-DEL-7143 | chr13-87717571-DEL-7140 |
| chr13-87727060-DEL-171 | chr13-87727061-DEL-171 |
| chr13-87729481-DEL-805 | chr13-87729482-DEL-805 |
| chr13-87732554-DEL-707 | chr13-87732555-DEL-707 |
| chr13-87736807-DEL-1309 | chr13-87736808-DEL-1309 |
| chr13-87748002-DEL-629 | chr13-87748003-DEL-629 |
| chr13-87760993-DEL-419 | chr13-87760992-DEL-419 |
| chr13-87804823-DEL-391 | chr13-87804818-DEL-392 |
| chr13-87813234-DEL-1176 | chr13-87813235-DEL-1176 |
| chr13-87834367-DEL-199 | chr13-87834358-DEL-188 |
| chr13-87861017-DEL-1199 | chr13-87861018-DEL-1199 |
| chr13-87868593-DEL-120 | chr13-87868595-DEL-72 |
| chr13-87921949-DEL-55 | chr13-87921973-DEL-60 |
| chr13-87925569-DEL-308 | chr13-87925569-DEL-308 |
| chr13-87946324-DEL-66 | chr13-87946350-DEL-58 |
| chr13-87958638-DEL-180 | chr13-87958639-DEL-180 |
| chr13-87983442-DEL-64 | chr13-87983452-DEL-55 |
| chr13-88006795-DEL-400 | chr13-88006795-DEL-400 |

| | |
|---|---|
| chr13-88011633-DEL-79 | chr13-88011640-DEL-67 |
| chr13-88013483-DEL-296 | chr13-88013483-DEL-296 |
| chr13-88014793-DEL-357 | chr13-88014793-DEL-357 |
| chr13-88038033-DEL-6354 | chr13-88038034-DEL-6355 |
| chr13-88045040-DEL-8874 | chr13-88045041-DEL-8874 |
| chr13-88060223-DEL-103 | chr13-88060224-DEL-70 |
| chr13-88078346-DEL-124 | chr13-88078334-DEL-121 |
| chr13-88103161-DEL-71 | chr13-88103162-DEL-71 |
| chr13-88114952-DEL-424 | chr13-88114953-DEL-424 |
| chr13-88137588-DEL-1891 | chr13-88137589-DEL-1891 |
| chr13-88140995-DEL-1642 | chr13-88140996-DEL-1642 |
| chr13-88158910-DEL-259 | chr13-88158911-DEL-259 |
| chr13-88192219-DEL-179 | chr13-88192223-DEL-175 |
| chr13-88192618-DEL-6155 | chr13-88192619-DEL-6155 |
| chr13-88239412-DEL-130 | chr13-88239413-DEL-129 |
| chr13-88240823-DEL-3073 | chr13-88240822-DEL-3073 |
| chr13-88246213-DEL-116 | chr13-88246235-DEL-89 |
| chr13-88248177-DEL-1114 | chr13-88248178-DEL-1114 |
| chr13-88276098-DEL-70 | chr13-88276099-DEL-70 |
| chr13-88289571-DEL-270 | chr13-88289572-DEL-270 |
| chr13-88295579-DEL-896 | chr13-88295580-DEL-896 |
| chr13-88306521-DEL-81 | chr13-88306522-DEL-81 |
| chr13-88308394-DEL-2059 | chr13-88308395-DEL-2059 |
| chr13-88385698-DEL-909 | chr13-88385699-DEL-909 |
| chr13-88396958-DEL-165 | chr13-88396959-DEL-165 |
| chr13-88413912-DEL-82 | chr13-88413913-DEL-82 |
| chr13-88443243-DEL-135 | chr13-88443244-DEL-135 |
| chr13-88456049-DEL-1276 | chr13-88456049-DEL-1276 |
| chr13-88462280-DEL-466 | chr13-88462280-DEL-466 |
| chr13-88471493-DEL-3423 | chr13-88471494-DEL-3423 |
| chr13-88492445-DEL-1893 | chr13-88492445-DEL-1893 |
| chr13-88505735-DEL-759 | chr13-88505736-DEL-759 |
| chr13-88546876-DEL-525 | chr13-88546877-DEL-525 |
| chr13-88548421-DEL-440 | chr13-88548422-DEL-440 |
| chr13-88555888-DEL-147 | chr13-88555885-DEL-148 |
| chr13-88601438-DEL-127 | chr13-88601437-DEL-100 |
| chr13-88637499-DEL-90 | chr13-88637500-DEL-90 |
| chr13-88639800-DEL-200 | chr13-88639801-DEL-200 |
| chr13-88642355-DEL-6220 | chr13-88642355-DEL-6220 |
| chr13-88654946-DEL-54 | chr13-88654925-DEL-86 |
| chr13-88678034-DEL-204 | chr13-88678035-DEL-204 |
| chr13-88683974-DEL-676 | chr13-88683995-DEL-653 |
| chr13-88714054-DEL-74 | chr13-88714075-DEL-50 |
| chr13-88771457-DEL-202 | chr13-88771458-DEL-202 |
| chr13-88782594-DEL-388 | chr13-88782595-DEL-388 |
| chr13-88828734-DEL-2937 | chr13-88828735-DEL-2937 |
| chr13-88868339-DEL-929 | chr13-88868340-DEL-929 |
| chr13-88870407-DEL-114 | chr13-88870408-DEL-114 |
| chr13-88917239-DEL-160 | chr13-88917284-DEL-121 |
| chr13-88919655-DEL-824 | chr13-88919656-DEL-824 |
| chr13-88922120-DEL-2568 | chr13-88922121-DEL-2568 |
| chr13-88986327-DEL-826 | chr13-88986328-DEL-826 |
| chr13-88991150-DEL-286 | chr13-88991151-DEL-286 |
| chr13-88992685-DEL-228 | chr13-88992686-DEL-228 |
| chr13-88994956-DEL-102 | chr13-88994940-DEL-120 |
| chr13-89005184-DEL-1286 | chr13-89005185-DEL-1286 |
| chr13-89065123-DEL-69 | chr13-89065123-DEL-94 |
| chr13-89108134-DEL-3455 | chr13-89108135-DEL-3455 |
| chr13-89114228-DEL-737 | chr13-89114229-DEL-737 |
| chr13-89116330-DEL-76 | chr13-89116331-DEL-76 |
| chr13-89121294-DEL-170 | chr13-89121295-DEL-170 |
| chr13-89128427-DEL-381 | chr13-89128427-DEL-381 |
| chr13-89135053-DEL-6216 | chr13-89135054-DEL-6216 |
| chr13-89197889-DEL-194 | chr13-89197890-DEL-194 |
| chr13-89258114-DEL-230 | chr13-89258116-DEL-229 |
| chr13-89261661-DEL-1258 | chr13-89261662-DEL-1258 |
| chr13-89297608-DEL-241 | chr13-89297609-DEL-241 |
| chr13-89305388-DEL-6778 | chr13-89305389-DEL-6778 |
| chr13-89325673-DEL-393 | chr13-89325674-DEL-393 |
| chr13-89338906-DEL-951 | chr13-89338907-DEL-951 |
| chr13-89356076-DEL-203 | chr13-89356077-DEL-203 |
| chr13-89357758-DEL-58 | chr13-89357759-DEL-57 |
| chr13-89363794-DEL-6289 | chr13-89363795-DEL-6289 |
| chr13-89434695-DEL-120 | chr13-89434696-DEL-120 |
| chr13-89447602-DEL-248 | chr13-89447601-DEL-249 |
| chr13-89477323-DEL-56 | chr13-89477320-DEL-51 |
| chr13-89482883-DEL-287 | chr13-89482884-DEL-287 |
| chr13-89501922-DEL-459 | chr13-89501923-DEL-459 |
| chr13-89508659-DEL-53 | chr13-89508649-DEL-54 |
| chr13-89511665-DEL-301 | chr13-89511666-DEL-301 |

| | |
|---|---|
| chr13-89531909-DEL-1000 | chr13-89531910-DEL-1000 |
| chr13-89532867-DEL-303 | chr13-89532868-DEL-303 |
| chr13-89562601-DEL-64 | chr13-89562602-DEL-64 |
| chr13-89584403-DEL-458 | chr13-89584404-DEL-458 |
| chr13-89588113-DEL-2394 | chr13-89588114-DEL-2394 |
| chr13-89596637-DEL-2663 | chr13-89596638-DEL-2663 |
| chr13-89617459-DEL-82 | chr13-89617460-DEL-82 |
| chr13-89625880-DEL-423 | chr13-89625881-DEL-423 |
| chr13-89686545-DEL-67 | chr13-89686520-DEL-64 |
| chr13-89695383-DEL-56 | chr13-89695384-DEL-56 |
| chr13-89721814-DEL-322 | chr13-89721804-DEL-319 |
| chr13-89727380-DEL-1948 | chr13-89727381-DEL-1948 |
| chr13-89763954-DEL-1238 | chr13-89763955-DEL-1238 |
| chr13-89766433-DEL-110 | chr13-89766434-DEL-110 |
| chr13-89771887-DEL-390 | chr13-89771888-DEL-390 |
| chr13-89776013-DEL-1399 | chr13-89776014-DEL-1399 |
| chr13-89829136-DEL-51 | chr13-89829137-DEL-51 |
| chr13-89865728-DEL-157 | chr13-89865705-DEL-181 |
| chr13-89877071-DEL-62 | chr13-89877072-DEL-62 |
| chr13-89898591-DEL-178 | chr13-89898591-DEL-170 |
| chr13-89936148-DEL-85 | chr13-89936172-DEL-52 |
| chr13-89973121-DEL-54 | chr13-89973127-DEL-51 |
| chr13-89978040-DEL-972 | chr13-89978041-DEL-972 |
| chr13-89982306-DEL-75 | chr13-89982308-DEL-67 |
| chr13-89985004-DEL-85 | chr13-89985001-DEL-80 |
| chr13-90010092-DEL-6462 | chr13-90010093-DEL-6462 |
| chr13-90061016-DEL-60 | chr13-90061015-DEL-52 |
| chr13-90121484-DEL-1816 | chr13-90121485-DEL-1816 |
| chr13-90169284-DEL-917 | chr13-90169285-DEL-917 |
| chr13-90185348-DEL-177 | chr13-90185349-DEL-177 |
| chr13-90191009-DEL-6659 | chr13-90191010-DEL-6659 |
| chr13-90207755-DEL-1884 | chr13-90207756-DEL-1884 |
| chr13-90211671-DEL-63 | chr13-90211690-DEL-60 |
| chr13-90281082-DEL-88 | chr13-90281080-DEL-88 |
| chr13-90322015-DEL-802 | chr13-90322016-DEL-802 |
| chr13-90349333-DEL-193 | chr13-90349334-DEL-193 |
| chr13-90350713-DEL-62 | chr13-90350713-DEL-71 |
| chr13-90369669-DEL-1363 | chr13-90369670-DEL-1363 |
| chr13-90380198-DEL-190 | chr13-90380199-DEL-190 |
| chr13-90405590-DEL-53 | chr13-90405574-DEL-51 |
| chr13-90433959-DEL-5492 | chr13-90433960-DEL-5492 |
| chr13-90445344-DEL-664 | chr13-90445345-DEL-664 |
| chr13-90468005-DEL-2640 | chr13-90468006-DEL-2640 |
| chr13-90474451-DEL-179 | chr13-90474452-DEL-179 |
| chr13-90476511-DEL-1172 | chr13-90476511-DEL-1172 |
| chr13-90525569-DEL-155 | chr13-90525521-DEL-198 |
| chr13-90560281-DEL-2565 | chr13-90560282-DEL-2565 |
| chr13-90563389-DEL-1888 | chr13-90563390-DEL-1888 |
| chr13-90585820-DEL-2336 | chr13-90585821-DEL-2336 |
| chr13-90601555-DEL-73 | chr13-90601556-DEL-73 |
| chr13-90604189-DEL-216 | chr13-90604190-DEL-216 |
| chr13-90619152-DEL-63 | chr13-90619153-DEL-63 |
| chr13-90631259-DEL-54 | chr13-90631260-DEL-54 |
| chr13-90644926-DEL-53 | chr13-90644927-DEL-53 |
| chr13-90688879-DEL-3851 | chr13-90688878-DEL-3852 |
| chr13-90741567-DEL-6496 | chr13-90741568-DEL-6496 |
| chr13-90749276-DEL-597 | chr13-90749277-DEL-597 |
| chr13-90750340-DEL-3663 | chr13-90750341-DEL-3663 |
| chr13-90754499-DEL-2006 | chr13-90754498-DEL-2007 |
| chr13-90782041-DEL-77 | chr13-90782042-DEL-78 |
| chr13-90783046-DEL-60 | chr13-90783042-DEL-64 |
| chr13-90811581-DEL-6588 | chr13-90811582-DEL-6588 |
| chr13-90811598-DEL-208 | chr13-90811599-DEL-208 |
| chr13-90812123-DEL-198 | chr13-90812124-DEL-198 |
| chr13-90852595-DEL-128 | chr13-90852596-DEL-128 |
| chr13-90863558-DEL-843 | chr13-90863558-DEL-844 |
| chr13-90865523-DEL-1018 | chr13-90865524-DEL-1018 |
| chr13-90889753-DEL-274 | chr13-90889754-DEL-274 |
| chr13-90911152-DEL-995 | chr13-90911153-DEL-995 |
| chr13-90913243-DEL-126 | chr13-90913244-DEL-125 |
| chr13-90925722-DEL-934 | chr13-90925723-DEL-934 |
| chr13-90944514-DEL-60 | chr13-90944515-DEL-60 |
| chr13-90952702-DEL-88 | chr13-90952707-DEL-86 |
| chr13-90953931-DEL-99 | chr13-90953932-DEL-99 |
| chr13-90954714-DEL-197 | chr13-90954715-DEL-197 |
| chr13-90973370-DEL-85 | chr13-90973371-DEL-85 |
| chr13-90984070-DEL-198 | chr13-90984066-DEL-199 |
| chr13-90989849-DEL-100 | chr13-90989850-DEL-100 |
| chr13-91057804-DEL-197 | chr13-91057805-DEL-197 |
| chr13-91066742-DEL-133 | chr13-91066743-DEL-133 |

| | |
|---|---|
| chr13-91067521-DEL-1115 | chr13-91067522-DEL-1115 |
| chr13-91070195-DEL-69 | chr13-91070196-DEL-69 |
| chr13-91078883-DEL-349 | chr13-91078882-DEL-348 |
| chr13-91085572-DEL-99 | chr13-91085572-DEL-99 |
| chr13-91175460-DEL-963 | chr13-91175461-DEL-963 |
| chr13-91182686-DEL-65 | chr13-91182687-DEL-65 |
| chr13-91201739-DEL-95 | chr13-91201740-DEL-92 |
| chr13-91215766-DEL-64 | chr13-91215767-DEL-64 |
| chr13-91222153-DEL-1710 | chr13-91222154-DEL-1710 |
| chr13-91228518-DEL-395 | chr13-91228519-DEL-395 |
| chr13-91232893-DEL-203 | chr13-91232894-DEL-203 |
| chr13-91293882-DEL-215 | chr13-91293883-DEL-215 |
| chr13-91333321-DEL-133 | chr13-91333322-DEL-133 |
| chr13-91377996-DEL-326 | chr13-91377997-DEL-326 |
| chr13-91400321-DEL-204 | chr13-91400322-DEL-204 |
| chr13-91401717-DEL-305 | chr13-91401718-DEL-305 |
| chr13-91418109-DEL-231 | chr13-91418106-DEL-235 |
| chr13-91420452-DEL-1310 | chr13-91420453-DEL-1310 |
| chr13-91444510-DEL-173 | chr13-91444511-DEL-173 |
| chr13-91459980-DEL-650 | chr13-91459981-DEL-650 |
| chr13-91475924-DEL-6188 | chr13-91475925-DEL-6188 |
| chr13-91484131-DEL-209 | chr13-91484131-DEL-207 |
| chr13-91493035-DEL-473 | chr13-91493036-DEL-473 |
| chr13-91600237-DEL-72 | chr13-91600240-DEL-52 |
| chr13-91637902-DEL-60 | chr13-91637903-DEL-60 |
| chr13-91643645-DEL-432 | chr13-91643646-DEL-432 |
| chr13-91651609-DEL-578 | chr13-91651610-DEL-578 |
| chr13-91675713-DEL-118 | chr13-91675714-DEL-118 |
| chr13-91810156-DEL-60 | chr13-91810157-DEL-60 |
| chr13-91816174-DEL-1492 | chr13-91816175-DEL-1492 |
| chr13-91816603-DEL-56 | chr13-91816604-DEL-56 |
| chr13-91820926-DEL-401 | chr13-91820927-DEL-401 |
| chr13-91826709-DEL-1363 | chr13-91826711-DEL-1361 |
| chr13-91828479-DEL-129 | chr13-91828480-DEL-129 |
| chr13-91864627-DEL-66 | chr13-91864621-DEL-68 |
| chr13-91876476-DEL-7054 | chr13-91876477-DEL-7054 |
| chr13-91893410-DEL-65 | chr13-91893377-DEL-72 |
| chr13-91896057-DEL-129 | chr13-91896058-DEL-129 |
| chr13-91921500-DEL-1132 | chr13-91921501-DEL-1132 |
| chr13-91939185-DEL-111 | chr13-91939204-DEL-117 |
| chr13-91950309-DEL-2090 | chr13-91950303-DEL-2100 |
| chr13-91963284-DEL-6441 | chr13-91963285-DEL-6441 |
| chr13-91984835-DEL-205 | chr13-91984836-DEL-205 |
| chr13-91994282-DEL-83 | chr13-91994283-DEL-83 |
| chr13-92027486-DEL-388 | chr13-92027487-DEL-388 |
| chr13-92048546-DEL-212 | chr13-92048547-DEL-212 |
| chr13-92059844-DEL-76 | chr13-92059845-DEL-76 |
| chr13-92069013-DEL-83 | chr13-92069012-DEL-84 |
| chr13-92079159-DEL-1762 | chr13-92079160-DEL-1762 |
| chr13-92094411-DEL-782 | chr13-92094412-DEL-782 |
| chr13-92098006-DEL-2096 | chr13-92098007-DEL-2096 |
| chr13-92121197-DEL-86 | chr13-92121198-DEL-69 |
| chr13-92143989-DEL-72 | chr13-92144017-DEL-56 |
| chr13-92146102-DEL-1893 | chr13-92146091-DEL-1905 |
| chr13-92147100-DEL-180 | chr13-92147101-DEL-180 |
| chr13-92193962-DEL-1200 | chr13-92193956-DEL-1201 |
| chr13-92221950-DEL-56 | chr13-92221930-DEL-52 |
| chr13-92227422-DEL-2204 | chr13-92227423-DEL-2204 |
| chr13-92240261-DEL-53 | chr13-92240262-DEL-53 |
| chr13-92259609-DEL-170 | chr13-92259610-DEL-170 |
| chr13-92277982-DEL-170 | chr13-92277983-DEL-170 |
| chr13-92283112-DEL-1993 | chr13-92283113-DEL-1993 |
| chr13-92289362-DEL-512 | chr13-92289363-DEL-512 |
| chr13-92317492-DEL-300 | chr13-92317493-DEL-300 |
| chr13-92320810-DEL-838 | chr13-92320811-DEL-838 |
| chr13-92331405-DEL-84 | chr13-92331406-DEL-63 |
| chr13-92338225-DEL-58 | chr13-92338226-DEL-58 |
| chr13-92345889-DEL-52 | chr13-92345888-DEL-54 |
| chr13-92349546-DEL-182 | chr13-92349547-DEL-182 |
| chr13-92353449-DEL-259 | chr13-92353435-DEL-261 |
| chr13-92385321-DEL-55 | chr13-92385322-DEL-55 |
| chr13-92391349-DEL-112 | chr13-92391350-DEL-112 |
| chr13-92394450-DEL-932 | chr13-92394451-DEL-932 |
| chr13-92395425-DEL-253 | chr13-92395426-DEL-253 |
| chr13-92396525-DEL-52 | chr13-92396526-DEL-52 |
| chr13-92404963-DEL-4341 | chr13-92404964-DEL-4341 |
| chr13-92437258-DEL-64 | chr13-92437259-DEL-64 |
| chr13-92446824-DEL-213 | chr13-92446825-DEL-213 |
| chr13-92448432-DEL-77 | chr13-92448433-DEL-77 |
| chr13-92452122-DEL-632 | chr13-92452123-DEL-632 |

| | |
|---|---|
| chr13-92453053-DEL-648 | chr13-92453054-DEL-648 |
| chr13-92463788-DEL-199 | chr13-92463789-DEL-199 |
| chr13-92473505-DEL-54 | chr13-92473506-DEL-54 |
| chr13-92476281-DEL-94 | chr13-92476282-DEL-94 |
| chr13-92484544-DEL-387 | chr13-92484545-DEL-387 |
| chr13-92542870-DEL-225 | chr13-92542871-DEL-225 |
| chr13-92543242-DEL-507 | chr13-92543243-DEL-507 |
| chr13-92546253-DEL-238 | chr13-92546254-DEL-238 |
| chr13-92558192-DEL-351 | chr13-92558193-DEL-351 |
| chr13-92566733-DEL-180 | chr13-92566774-DEL-140 |
| chr13-92590970-DEL-1087 | chr13-92590960-DEL-1088 |
| chr13-92615495-DEL-500 | chr13-92615496-DEL-500 |
| chr13-92618014-DEL-223 | chr13-92618015-DEL-223 |
| chr13-92625879-DEL-120 | chr13-92625909-DEL-91 |
| chr13-92635040-DEL-498 | chr13-92635041-DEL-498 |
| chr13-92685585-DEL-836 | chr13-92685586-DEL-836 |
| chr13-92707963-DEL-999 | chr13-92707964-DEL-999 |
| chr13-92709259-DEL-465 | chr13-92709251-DEL-466 |
| chr13-92718957-DEL-498 | chr13-92718958-DEL-498 |
| chr13-92722350-DEL-298 | chr13-92722351-DEL-298 |
| chr13-92735489-DEL-6420 | chr13-92735490-DEL-6420 |
| chr13-92750348-DEL-65 | chr13-92750370-DEL-53 |
| chr13-92753427-DEL-8505 | chr13-92753428-DEL-8505 |
| chr13-92765098-DEL-3125 | chr13-92765099-DEL-3125 |
| chr13-92778760-DEL-888 | chr13-92778761-DEL-888 |
| chr13-92780607-DEL-219 | chr13-92780608-DEL-219 |
| chr13-92807834-DEL-441 | chr13-92807835-DEL-441 |
| chr13-92860855-DEL-1357 | chr13-92860856-DEL-1357 |
| chr13-92862936-DEL-678 | chr13-92862937-DEL-678 |
| chr13-92883984-DEL-1304 | chr13-92883985-DEL-1303 |
| chr13-92886109-DEL-492 | chr13-92886110-DEL-492 |
| chr13-92926522-DEL-339 | chr13-92926523-DEL-339 |
| chr13-92953506-DEL-211 | chr13-92953507-DEL-211 |
| chr13-92970445-DEL-6249 | chr13-92970446-DEL-6249 |
| chr13-93007385-DEL-75 | chr13-93007390-DEL-72 |
| chr13-93025303-DEL-58 | chr13-93025304-DEL-58 |
| chr13-93026923-DEL-6007 | chr13-93026916-DEL-6008 |
| chr13-93038710-DEL-54 | chr13-93038711-DEL-54 |
| chr13-93042571-DEL-73 | chr13-93042572-DEL-73 |
| chr13-93069996-DEL-85 | chr13-93069997-DEL-85 |
| chr13-93078252-DEL-84 | chr13-93078253-DEL-84 |
| chr13-93089108-DEL-68 | chr13-93089109-DEL-68 |
| chr13-93111455-DEL-118 | chr13-93111456-DEL-118 |
| chr13-93114706-DEL-112 | chr13-93114751-DEL-66 |
| chr13-93120022-DEL-52 | chr13-93120023-DEL-52 |
| chr13-93148531-DEL-190 | chr13-93148532-DEL-190 |
| chr13-93151682-DEL-53 | chr13-93151682-DEL-51 |
| chr13-93159589-DEL-76 | chr13-93159590-DEL-76 |
| chr13-93172127-DEL-302 | chr13-93172128-DEL-302 |
| chr13-93181349-DEL-884 | chr13-93181350-DEL-884 |
| chr13-93197306-DEL-160 | chr13-93197307-DEL-160 |
| chr13-93212445-DEL-108 | chr13-93212446-DEL-108 |
| chr13-93245171-DEL-7108 | chr13-93245172-DEL-7108 |
| chr13-93259833-DEL-3133 | chr13-93259834-DEL-3133 |
| chr13-93270838-DEL-735 | chr13-93270839-DEL-735 |
| chr13-93317435-DEL-59 | chr13-93317432-DEL-59 |
| chr13-93325292-DEL-248 | chr13-93325298-DEL-243 |
| chr13-93334429-DEL-86 | chr13-93334430-DEL-86 |
| chr13-93335115-DEL-4141 | chr13-93335116-DEL-4141 |
| chr13-93351103-DEL-430 | chr13-93351098-DEL-433 |
| chr13-93355830-DEL-5441 | chr13-93355815-DEL-5415 |
| chr13-93362047-DEL-76 | chr13-93362048-DEL-76 |
| chr13-93399247-DEL-54 | chr13-93399247-DEL-54 |
| chr13-93402831-DEL-207 | chr13-93402832-DEL-207 |
| chr13-93406348-DEL-109 | chr13-93406339-DEL-112 |
| chr13-93415687-DEL-5955 | chr13-93415688-DEL-5955 |
| chr13-93433624-DEL-161 | chr13-93433625-DEL-161 |
| chr13-93466917-DEL-99 | chr13-93466916-DEL-78 |
| chr13-93491959-DEL-188 | chr13-93491960-DEL-188 |
| chr13-93572349-DEL-266 | chr13-93572350-DEL-266 |
| chr13-93611532-DEL-2420 | chr13-93611533-DEL-2420 |
| chr13-93660583-DEL-223 | chr13-93660617-DEL-221 |
| chr13-93668219-DEL-2359 | chr13-93668220-DEL-2358 |
| chr13-93670832-DEL-3665 | chr13-93670833-DEL-3665 |
| chr13-93679734-DEL-374 | chr13-93679735-DEL-374 |
| chr13-93722978-DEL-513 | chr13-93722979-DEL-513 |
| chr13-93725820-DEL-65 | chr13-93725821-DEL-64 |
| chr13-93730746-DEL-197 | chr13-93730747-DEL-197 |
| chr13-93732668-DEL-201 | chr13-93732669-DEL-201 |
| chr13-93735139-DEL-597 | chr13-93735175-DEL-562 |

| | |
|---|---|
| chr13-93751125-DEL-68 | chr13-93751126-DEL-68 |
| chr13-93753061-DEL-84 | chr13-93753062-DEL-84 |
| chr13-93755345-DEL-203 | chr13-93755346-DEL-203 |
| chr13-93756845-DEL-56 | chr13-93756846-DEL-50 |
| chr13-93791831-DEL-787 | chr13-93791832-DEL-787 |
| chr13-93794164-DEL-202 | chr13-93794165-DEL-202 |
| chr13-93797610-DEL-131 | chr13-93797611-DEL-131 |
| chr13-93818318-DEL-183 | chr13-93818320-DEL-182 |
| chr13-93819729-DEL-64 | chr13-93819730-DEL-64 |
| chr13-93834382-DEL-284 | chr13-93834385-DEL-282 |
| chr13-93865414-DEL-54 | chr13-93865414-DEL-54 |
| chr13-93867959-DEL-168 | chr13-93867960-DEL-165 |
| chr13-93898619-DEL-612 | chr13-93898620-DEL-612 |
| chr13-93904587-DEL-307 | chr13-93904602-DEL-308 |
| chr13-93909762-DEL-69 | chr13-93909751-DEL-70 |
| chr13-93934334-DEL-216 | chr13-93934335-DEL-216 |
| chr13-93936693-DEL-6254 | chr13-93936694-DEL-6254 |
| chr13-93963134-DEL-68 | chr13-93963135-DEL-68 |
| chr13-93971620-DEL-84 | chr13-93971621-DEL-82 |
| chr13-93979950-DEL-86 | chr13-93979933-DEL-77 |
| chr13-93993709-DEL-426 | chr13-93993710-DEL-426 |
| chr13-93998128-DEL-252 | chr13-93998129-DEL-252 |
| chr13-94022739-DEL-837 | chr13-94022740-DEL-837 |
| chr13-94034480-DEL-80 | chr13-94034473-DEL-60 |
| chr13-94079689-DEL-141 | chr13-94079692-DEL-136 |
| chr13-94090527-DEL-1299 | chr13-94090528-DEL-1299 |
| chr13-94113365-DEL-170 | chr13-94113366-DEL-170 |
| chr13-94127856-DEL-605 | chr13-94127842-DEL-606 |
| chr13-94143080-DEL-129 | chr13-94143080-DEL-128 |
| chr13-94151170-DEL-6168 | chr13-94151171-DEL-6168 |
| chr13-94157945-DEL-51 | chr13-94157946-DEL-51 |
| chr13-94161234-DEL-500 | chr13-94161234-DEL-500 |
| chr13-94171511-DEL-89 | chr13-94171512-DEL-77 |
| chr13-94283627-DEL-79 | chr13-94283628-DEL-79 |
| chr13-94341755-DEL-62 | chr13-94341756-DEL-60 |
| chr13-94347345-DEL-76 | chr13-94347346-DEL-76 |
| chr13-94351114-DEL-6403 | chr13-94351115-DEL-6403 |
| chr13-94360447-DEL-82 | chr13-94360448-DEL-82 |
| chr13-94362777-DEL-3109 | chr13-94362778-DEL-3109 |
| chr13-94370795-DEL-59 | chr13-94370801-DEL-54 |
| chr13-94372380-DEL-225 | chr13-94372381-DEL-225 |
| chr13-94380384-DEL-56 | chr13-94380385-DEL-56 |
| chr13-94386951-DEL-358 | chr13-94386976-DEL-314 |
| chr13-94391539-DEL-1308 | chr13-94391540-DEL-1308 |
| chr13-94393292-DEL-1026 | chr13-94393293-DEL-1026 |
| chr13-94431226-DEL-54 | chr13-94431217-DEL-62 |
| chr13-94440447-DEL-209 | chr13-94440448-DEL-209 |
| chr13-94449490-DEL-84 | chr13-94449491-DEL-84 |
| chr13-94467662-DEL-196 | chr13-94467679-DEL-202 |
| chr13-94481825-DEL-104 | chr13-94481826-DEL-104 |
| chr13-94519573-DEL-290 | chr13-94519574-DEL-290 |
| chr13-94520716-DEL-61 | chr13-94520714-DEL-61 |
| chr13-94523996-DEL-60 | chr13-94524008-DEL-60 |
| chr13-94552919-DEL-197 | chr13-94552919-DEL-197 |
| chr13-94555833-DEL-273 | chr13-94555834-DEL-273 |
| chr13-94567997-DEL-128 | chr13-94567994-DEL-154 |
| chr13-94568642-DEL-65 | chr13-94568643-DEL-65 |
| chr13-94602328-DEL-72 | chr13-94602331-DEL-70 |
| chr13-94622033-DEL-195 | chr13-94622034-DEL-195 |
| chr13-94624305-DEL-51 | chr13-94624306-DEL-51 |
| chr13-94628493-DEL-72 | chr13-94628486-DEL-96 |
| chr13-94635645-DEL-56 | chr13-94635646-DEL-56 |
| chr13-94650077-DEL-58 | chr13-94650078-DEL-58 |
| chr13-94688132-DEL-6387 | chr13-94688133-DEL-6387 |
| chr13-94717825-DEL-187 | chr13-94717826-DEL-142 |
| chr13-94722499-DEL-474 | chr13-94722500-DEL-474 |
| chr13-94787393-DEL-60 | chr13-94787394-DEL-60 |
| chr13-94788274-DEL-366 | chr13-94788275-DEL-366 |
| chr13-94800387-DEL-91 | chr13-94800386-DEL-92 |
| chr13-94812624-DEL-95 | chr13-94812625-DEL-95 |
| chr13-94834690-DEL-189 | chr13-94834685-DEL-195 |
| chr13-94847660-DEL-72 | chr13-94847661-DEL-108 |
| chr13-94868930-DEL-329 | chr13-94868930-DEL-330 |
| chr13-94872005-DEL-7135 | chr13-94871998-DEL-7142 |
| chr13-94882071-DEL-220 | chr13-94882078-DEL-219 |
| chr13-94886193-DEL-183 | chr13-94886194-DEL-183 |
| chr13-94923280-DEL-84 | chr13-94923281-DEL-84 |
| chr13-94944386-DEL-86 | chr13-94944391-DEL-70 |
| chr13-94950960-DEL-215 | chr13-94950961-DEL-215 |
| chr13-94980652-DEL-185 | chr13-94980653-DEL-185 |

| | |
|---|---|
| chr13-94993580-DEL-340 | chr13-94993583-DEL-338 |
| chr13-94993939-DEL-652 | chr13-94993940-DEL-647 |
| chr13-95046585-DEL-64 | chr13-95046549-DEL-58 |
| chr13-95053197-DEL-91 | chr13-95053197-DEL-100 |
| chr13-95053901-DEL-108 | chr13-95053929-DEL-78 |
| chr13-95055167-DEL-177 | chr13-95055168-DEL-177 |
| chr13-95086874-DEL-140 | chr13-95086881-DEL-155 |
| chr13-95087161-DEL-719 | chr13-95087143-DEL-713 |
| chr13-95088057-DEL-89 | chr13-95088057-DEL-89 |
| chr13-95102193-DEL-64 | chr13-95102194-DEL-64 |
| chr13-95149003-DEL-650 | chr13-95149011-DEL-648 |
| chr13-95240159-DEL-168 | chr13-95240160-DEL-134 |
| chr13-95260761-DEL-251 | chr13-95260762-DEL-251 |
| chr13-95275110-DEL-70 | chr13-95275111-DEL-70 |
| chr13-95280003-DEL-1512 | chr13-95279958-DEL-1516 |
| chr13-95291479-DEL-80 | chr13-95291480-DEL-80 |
| chr13-95316320-DEL-2693 | chr13-95316321-DEL-2693 |
| chr13-95321542-DEL-72 | chr13-95321553-DEL-50 |
| chr13-95335067-DEL-374 | chr13-95335068-DEL-374 |
| chr13-95344040-DEL-835 | chr13-95344041-DEL-835 |
| chr13-95431330-DEL-268 | chr13-95431331-DEL-268 |
| chr13-95447528-DEL-54 | chr13-95447528-DEL-54 |
| chr13-95457333-DEL-180 | chr13-95457374-DEL-173 |
| chr13-95458951-DEL-248 | chr13-95458946-DEL-249 |
| chr13-95499787-DEL-152 | chr13-95499788-DEL-152 |
| chr13-95520437-DEL-64 | chr13-95520453-DEL-56 |
| chr13-95536091-DEL-648 | chr13-95536079-DEL-649 |
| chr13-95541132-DEL-70 | chr13-95541133-DEL-70 |
| chr13-95543459-DEL-59 | chr13-95543459-DEL-59 |
| chr13-95548541-DEL-52 | chr13-95548541-DEL-52 |
| chr13-95550134-DEL-609 | chr13-95550141-DEL-603 |
| chr13-95562924-DEL-64 | chr13-95562925-DEL-64 |
| chr13-95572336-DEL-98 | chr13-95572322-DEL-78 |
| chr13-95590534-DEL-184 | chr13-95590536-DEL-185 |
| chr13-95591702-DEL-65 | chr13-95591703-DEL-65 |
| chr13-95594949-DEL-1055 | chr13-95594950-DEL-1055 |
| chr13-95597021-DEL-66 | chr13-95597014-DEL-74 |
| chr13-95597673-DEL-351 | chr13-95597674-DEL-350 |
| chr13-95616771-DEL-70 | chr13-95616772-DEL-70 |
| chr13-95622878-DEL-362 | chr13-95622881-DEL-360 |
| chr13-95632821-DEL-340 | chr13-95632822-DEL-340 |
| chr13-95637179-DEL-59 | chr13-95637169-DEL-91 |
| chr13-95640624-DEL-59 | chr13-95640624-DEL-60 |
| chr13-95643335-DEL-367 | chr13-95643291-DEL-410 |
| chr13-95647962-DEL-83 | chr13-95647965-DEL-81 |
| chr13-95655164-DEL-76 | chr13-95655125-DEL-66 |
| chr13-95657947-DEL-281 | chr13-95657948-DEL-281 |
| chr13-95665200-DEL-191 | chr13-95665194-DEL-188 |
| chr13-95670004-DEL-81 | chr13-95670003-DEL-79 |
| chr13-95675418-DEL-124 | chr13-95675407-DEL-136 |
| chr13-95676471-DEL-53 | chr13-95676472-DEL-53 |
| chr13-95708021-DEL-109 | chr13-95708022-DEL-109 |
| chr13-95713655-DEL-65 | chr13-95713656-DEL-69 |
| chr13-95714205-DEL-1201 | chr13-95714206-DEL-1201 |
| chr13-95715379-DEL-127 | chr13-95715380-DEL-127 |
| chr13-95716245-DEL-1186 | chr13-95716246-DEL-1186 |
| chr13-95717214-DEL-237 | chr13-95717215-DEL-237 |
| chr13-95721955-DEL-60 | chr13-95721956-DEL-60 |
| chr13-95723170-DEL-85 | chr13-95723171-DEL-85 |
| chr13-95732884-DEL-1352 | chr13-95732885-DEL-1352 |
| chr13-95782444-DEL-59 | chr13-95782445-DEL-59 |
| chr13-95790673-DEL-229 | chr13-95790674-DEL-223 |
| chr13-95792306-DEL-406 | chr13-95792307-DEL-406 |
| chr13-95812693-DEL-166 | chr13-95812694-DEL-166 |
| chr13-95834272-DEL-227 | chr13-95834269-DEL-228 |
| chr13-95856320-DEL-53 | chr13-95856321-DEL-53 |
| chr13-95869180-DEL-51 | chr13-95869181-DEL-51 |
| chr13-95874399-DEL-192 | chr13-95874400-DEL-192 |
| chr13-95894789-DEL-56 | chr13-95894795-DEL-50 |
| chr13-95904085-DEL-186 | chr13-95904086-DEL-186 |
| chr13-95922172-DEL-58 | chr13-95922173-DEL-58 |
| chr13-95948842-DEL-72 | chr13-95948843-DEL-69 |
| chr13-95950198-DEL-1330 | chr13-95950192-DEL-1331 |
| chr13-95967177-DEL-88 | chr13-95967178-DEL-88 |
| chr13-96015952-DEL-76 | chr13-96015953-DEL-76 |
| chr13-96016786-DEL-62 | chr13-96016787-DEL-62 |
| chr13-96027077-DEL-52 | chr13-96027078-DEL-52 |
| chr13-96058809-DEL-58 | chr13-96058810-DEL-58 |
| chr13-96070581-DEL-515 | chr13-96070582-DEL-515 |
| chr13-96097251-DEL-62 | chr13-96097243-DEL-64 |

| | |
|---|---|
| chr13-96126786-DEL-138 | chr13-96126787-DEL-138 |
| chr13-96138121-DEL-57 | chr13-96138122-DEL-57 |
| chr13-96145160-DEL-418 | chr13-96145161-DEL-418 |
| chr13-96150142-DEL-6793 | chr13-96150143-DEL-6793 |
| chr13-96173989-DEL-90 | chr13-96173989-DEL-54 |
| chr13-96191558-DEL-109 | chr13-96191560-DEL-107 |
| chr13-96193801-DEL-106 | chr13-96193802-DEL-104 |
| chr13-96215727-DEL-2166 | chr13-96215717-DEL-2167 |
| chr13-96229671-DEL-63 | chr13-96229672-DEL-63 |
| chr13-96265191-DEL-54 | chr13-96265190-DEL-55 |
| chr13-96316607-DEL-3413 | chr13-96316608-DEL-3413 |
| chr13-96325314-DEL-52 | chr13-96325315-DEL-52 |
| chr13-96343016-DEL-401 | chr13-96343017-DEL-401 |
| chr13-96357574-DEL-1395 | chr13-96357575-DEL-1394 |
| chr13-96359683-DEL-1635 | chr13-96359682-DEL-1635 |
| chr13-96366223-DEL-204 | chr13-96366224-DEL-204 |
| chr13-96382018-DEL-643 | chr13-96382019-DEL-643 |
| chr13-96385323-DEL-690 | chr13-96385317-DEL-691 |
| chr13-96415647-DEL-313 | chr13-96415648-DEL-313 |
| chr13-96428431-DEL-216 | chr13-96428432-DEL-216 |
| chr13-96444354-DEL-3549 | chr13-96444355-DEL-3549 |
| chr13-96462659-DEL-400 | chr13-96462660-DEL-400 |
| chr13-96477412-DEL-175 | chr13-96477413-DEL-175 |
| chr13-96494082-DEL-6797 | chr13-96494083-DEL-6797 |
| chr13-96496102-DEL-231 | chr13-96496103-DEL-231 |
| chr13-96525745-DEL-196 | chr13-96525746-DEL-196 |
| chr13-96589986-DEL-122 | chr13-96589994-DEL-124 |
| chr13-96609927-DEL-292 | chr13-96609916-DEL-288 |
| chr13-96614258-DEL-327 | chr13-96614259-DEL-327 |
| chr13-96616454-DEL-249 | chr13-96616455-DEL-249 |
| chr13-96655269-DEL-179 | chr13-96655270-DEL-179 |
| chr13-96708588-DEL-761 | chr13-96708571-DEL-760 |
| chr13-96722244-DEL-3483 | chr13-96722245-DEL-3483 |
| chr13-96777268-DEL-58 | chr13-96777269-DEL-58 |
| chr13-96780419-DEL-2297 | chr13-96780420-DEL-2297 |
| chr13-96836837-DEL-197 | chr13-96836836-DEL-197 |
| chr13-96842395-DEL-84 | chr13-96842426-DEL-76 |
| chr13-96843211-DEL-2501 | chr13-96843212-DEL-2501 |
| chr13-96875802-DEL-190 | chr13-96875803-DEL-190 |
| chr13-96880012-DEL-121 | chr13-96880013-DEL-121 |
| chr13-96883089-DEL-1247 | chr13-96883090-DEL-1247 |
| chr13-96891134-DEL-95 | chr13-96891135-DEL-95 |
| chr13-96901195-DEL-239 | chr13-96901196-DEL-239 |
| chr13-96920140-DEL-2333 | chr13-96920141-DEL-2333 |
| chr13-96987180-DEL-77 | chr13-96987181-DEL-77 |
| chr13-96995987-DEL-6540 | chr13-96995988-DEL-6540 |
| chr13-97032641-DEL-755 | chr13-97032642-DEL-755 |
| chr13-97078566-DEL-66 | chr13-97078567-DEL-66 |
| chr13-97090237-DEL-56 | chr13-97090238-DEL-56 |
| chr13-97104744-DEL-337 | chr13-97104745-DEL-337 |
| chr13-97110183-DEL-72 | chr13-97110184-DEL-76 |
| chr13-97111591-DEL-84 | chr13-97111556-DEL-104 |
| chr13-97116926-DEL-946 | chr13-97116927-DEL-946 |
| chr13-97129712-DEL-344 | chr13-97129713-DEL-344 |
| chr13-97132340-DEL-208 | chr13-97132341-DEL-208 |
| chr13-97141202-DEL-185 | chr13-97141200-DEL-215 |
| chr13-97145578-DEL-53 | chr13-97145573-DEL-60 |
| chr13-97167821-DEL-1158 | chr13-97167822-DEL-1158 |
| chr13-97172544-DEL-66 | chr13-97172545-DEL-66 |
| chr13-97191493-DEL-97 | chr13-97191500-DEL-78 |
| chr13-97197228-DEL-63 | chr13-97197221-DEL-64 |
| chr13-97215054-DEL-183 | chr13-97215055-DEL-183 |
| chr13-97222790-DEL-190 | chr13-97222791-DEL-190 |
| chr13-97234706-DEL-1143 | chr13-97234707-DEL-1143 |
| chr13-97237162-DEL-206 | chr13-97237163-DEL-206 |
| chr13-97240229-DEL-119 | chr13-97240230-DEL-119 |
| chr13-97253047-DEL-151 | chr13-97253048-DEL-151 |
| chr13-97255280-DEL-209 | chr13-97255281-DEL-209 |
| chr13-97272946-DEL-148 | chr13-97272955-DEL-177 |
| chr13-97275021-DEL-205 | chr13-97275025-DEL-200 |
| chr13-97281026-DEL-197 | chr13-97281027-DEL-197 |
| chr13-97296237-DEL-220 | chr13-97296238-DEL-220 |
| chr13-97304253-DEL-465 | chr13-97304256-DEL-466 |
| chr13-97305054-DEL-169 | chr13-97305055-DEL-169 |
| chr13-97329807-DEL-112 | chr13-97329808-DEL-112 |
| chr13-97331105-DEL-534 | chr13-97331119-DEL-521 |
| chr13-97346868-DEL-52 | chr13-97346869-DEL-52 |
| chr13-97348344-DEL-88 | chr13-97348345-DEL-88 |
| chr13-97362683-DEL-54 | chr13-97362683-DEL-54 |
| chr13-97366171-DEL-291 | chr13-97366172-DEL-291 |

| | |
|---|---|
| chr13-97368133-DEL-85 | chr13-97368134-DEL-84 |
| chr13-97414549-DEL-85 | chr13-97414550-DEL-85 |
| chr13-97426222-DEL-123 | chr13-97426223-DEL-123 |
| chr13-97448321-DEL-56 | chr13-97448321-DEL-57 |
| chr13-97526541-DEL-240 | chr13-97526542-DEL-240 |
| chr13-97559238-DEL-94 | chr13-97559239-DEL-94 |
| chr13-97600226-DEL-345 | chr13-97600227-DEL-345 |
| chr13-97612930-DEL-193 | chr13-97612931-DEL-193 |
| chr13-97658154-DEL-173 | chr13-97658153-DEL-170 |
| chr13-97661164-DEL-206 | chr13-97661165-DEL-206 |
| chr13-97672568-DEL-124 | chr13-97672569-DEL-124 |
| chr13-97680235-DEL-98 | chr13-97680224-DEL-94 |
| chr13-97681081-DEL-54 | chr13-97681082-DEL-57 |
| chr13-97686550-DEL-207 | chr13-97686551-DEL-207 |
| chr13-97721813-DEL-69 | chr13-97721818-DEL-67 |
| chr13-97727304-DEL-147 | chr13-97727305-DEL-147 |
| chr13-97730103-DEL-51 | chr13-97730104-DEL-51 |
| chr13-97753132-DEL-200 | chr13-97753148-DEL-178 |
| chr13-97821149-DEL-78 | chr13-97821143-DEL-76 |
| chr13-97834326-DEL-142 | chr13-97834332-DEL-128 |
| chr13-97848694-DEL-200 | chr13-97848695-DEL-200 |
| chr13-97888481-DEL-1089 | chr13-97888482-DEL-1089 |
| chr13-97896573-DEL-567 | chr13-97896574-DEL-567 |
| chr13-97903883-DEL-100 | chr13-97903900-DEL-56 |
| chr13-97938183-DEL-140 | chr13-97938185-DEL-139 |
| chr13-97957302-DEL-74 | chr13-97957314-DEL-52 |
| chr13-97977655-DEL-58 | chr13-97977664-DEL-62 |
| chr13-97997103-DEL-1306 | chr13-97997104-DEL-1306 |
| chr13-98014291-DEL-411 | chr13-98014292-DEL-411 |
| chr13-98035371-DEL-61 | chr13-98035372-DEL-61 |
| chr13-98112592-DEL-72 | chr13-98112593-DEL-72 |
| chr13-98145755-DEL-60 | chr13-98145755-DEL-60 |
| chr13-98182694-DEL-1109 | chr13-98182695-DEL-1109 |
| chr13-98194589-DEL-398 | chr13-98194590-DEL-398 |
| chr13-98204654-DEL-236 | chr13-98204655-DEL-236 |
| chr13-98219216-DEL-7487 | chr13-98219217-DEL-7487 |
| chr13-98243011-DEL-70 | chr13-98243012-DEL-63 |
| chr13-98267754-DEL-80 | chr13-98267755-DEL-80 |
| chr13-98271548-DEL-56 | chr13-98271553-DEL-50 |
| chr13-98276951-DEL-401 | chr13-98276952-DEL-401 |
| chr13-98290838-DEL-1028 | chr13-98290839-DEL-1028 |
| chr13-98311526-DEL-78 | chr13-98311488-DEL-84 |
| chr13-98328518-DEL-54 | chr13-98328515-DEL-58 |
| chr13-98330804-DEL-71 | chr13-98330806-DEL-74 |
| chr13-98336761-DEL-758 | chr13-98336762-DEL-758 |
| chr13-98351899-DEL-71 | chr13-98351893-DEL-60 |
| chr13-98352822-DEL-63 | chr13-98352840-DEL-56 |
| chr13-98406720-DEL-54 | chr13-98406721-DEL-54 |
| chr13-98413108-DEL-183 | chr13-98413109-DEL-183 |
| chr13-98420121-DEL-176 | chr13-98420122-DEL-176 |
| chr13-98458838-DEL-275 | chr13-98458839-DEL-275 |
| chr13-98460353-DEL-82 | chr13-98460354-DEL-82 |
| chr13-98462210-DEL-189 | chr13-98462210-DEL-189 |
| chr13-98462530-DEL-4425 | chr13-98462530-DEL-4425 |
| chr13-98478538-DEL-60 | chr13-98478539-DEL-60 |
| chr13-98506916-DEL-86 | chr13-98506917-DEL-86 |
| chr13-98522126-DEL-101 | chr13-98522133-DEL-92 |
| chr13-98524239-DEL-62 | chr13-98524270-DEL-59 |
| chr13-98544234-DEL-198 | chr13-98544274-DEL-149 |
| chr13-98554990-DEL-1891 | chr13-98554991-DEL-1891 |
| chr13-98592520-DEL-62 | chr13-98592517-DEL-53 |
| chr13-98594993-DEL-485 | chr13-98594994-DEL-485 |
| chr13-98611911-DEL-83 | chr13-98611912-DEL-83 |
| chr13-98624368-DEL-169 | chr13-98624369-DEL-169 |
| chr13-98625846-DEL-532 | chr13-98625841-DEL-495 |
| chr13-98631482-DEL-1006 | chr13-98631498-DEL-964 |
| chr13-98654081-DEL-54 | chr13-98654082-DEL-54 |
| chr13-98662235-DEL-434 | chr13-98662236-DEL-434 |
| chr13-98743022-DEL-77 | chr13-98743007-DEL-87 |
| chr13-98747293-DEL-68 | chr13-98747294-DEL-56 |
| chr13-98750875-DEL-333 | chr13-98750878-DEL-334 |
| chr13-98755869-DEL-83 | chr13-98755869-DEL-83 |
| chr13-98777941-DEL-392 | chr13-98777942-DEL-392 |
| chr13-98785219-DEL-175 | chr13-98785220-DEL-175 |
| chr13-98787542-DEL-1230 | chr13-98787543-DEL-1230 |
| chr13-98807382-DEL-983 | chr13-98807383-DEL-983 |
| chr13-98820603-DEL-78 | chr13-98820604-DEL-78 |
| chr13-98874541-DEL-55 | chr13-98874537-DEL-63 |
| chr13-98887460-DEL-202 | chr13-98887461-DEL-202 |
| chr13-98893793-DEL-104 | chr13-98893794-DEL-104 |

| | |
|---|---|
| chr13-98900405-DEL-72 | chr13-98900404-DEL-72 |
| chr13-98909418-DEL-441 | chr13-98909419-DEL-441 |
| chr13-98965856-DEL-250 | chr13-98965857-DEL-256 |
| chr13-99010132-DEL-58 | chr13-99010133-DEL-58 |
| chr13-99030266-DEL-52 | chr13-99030267-DEL-52 |
| chr13-99034766-DEL-208 | chr13-99034767-DEL-208 |
| chr13-99040688-DEL-551 | chr13-99040689-DEL-551 |
| chr13-99043274-DEL-211 | chr13-99043275-DEL-211 |
| chr13-99048774-DEL-630 | chr13-99048775-DEL-630 |
| chr13-99051870-DEL-1241 | chr13-99051871-DEL-1241 |
| chr13-99056645-DEL-1704 | chr13-99056646-DEL-1704 |
| chr13-99089128-DEL-1125 | chr13-99089129-DEL-1125 |
| chr13-99122778-DEL-8985 | chr13-99122779-DEL-8985 |
| chr13-99138718-DEL-331 | chr13-99138712-DEL-344 |
| chr13-99149904-DEL-204 | chr13-99149875-DEL-226 |
| chr13-99154310-DEL-68 | chr13-99154311-DEL-72 |
| chr13-99166039-DEL-86 | chr13-99166007-DEL-115 |
| chr13-99225580-DEL-919 | chr13-99225581-DEL-919 |
| chr13-99253362-DEL-432 | chr13-99253360-DEL-433 |
| chr13-99257310-DEL-59 | chr13-99257312-DEL-59 |
| chr13-99259301-DEL-400 | chr13-99259302-DEL-400 |
| chr13-99263709-DEL-458 | chr13-99263710-DEL-458 |
| chr13-99274574-DEL-51 | chr13-99274574-DEL-74 |
| chr13-99293265-DEL-60 | chr13-99293266-DEL-60 |
| chr13-99385315-DEL-132 | chr13-99385316-DEL-132 |
| chr13-99390464-DEL-52 | chr13-99390465-DEL-52 |
| chr13-99411735-DEL-56 | chr13-99411736-DEL-56 |
| chr13-99442306-DEL-56 | chr13-99442309-DEL-54 |
| chr13-99443549-DEL-54 | chr13-99443550-DEL-54 |
| chr13-99453050-DEL-66 | chr13-99453026-DEL-66 |
| chr13-99462785-DEL-100 | chr13-99462786-DEL-100 |
| chr13-99492217-DEL-84 | chr13-99492218-DEL-84 |
| chr13-99507248-DEL-128 | chr13-99507249-DEL-128 |
| chr13-99509154-DEL-1573 | chr13-99509155-DEL-1573 |
| chr13-99525389-DEL-492 | chr13-99525390-DEL-492 |
| chr13-99527754-DEL-53 | chr13-99527715-DEL-87 |
| chr13-99534492-DEL-90 | chr13-99534493-DEL-90 |
| chr13-99537714-DEL-723 | chr13-99537715-DEL-723 |
| chr13-99588318-DEL-70 | chr13-99588310-DEL-78 |
| chr13-99595021-DEL-213 | chr13-99595022-DEL-213 |
| chr13-99633613-DEL-102 | chr13-99633614-DEL-102 |
| chr13-99663868-DEL-96 | chr13-99663859-DEL-106 |
| chr13-99696748-DEL-214 | chr13-99696749-DEL-214 |
| chr13-99711432-DEL-55 | chr13-99711433-DEL-54 |
| chr13-99728478-DEL-60 | chr13-99728479-DEL-60 |
| chr13-99736254-DEL-338 | chr13-99736256-DEL-337 |
| chr13-99737033-DEL-450 | chr13-99737034-DEL-450 |
| chr13-99743002-DEL-54 | chr13-99743003-DEL-54 |
| chr13-99751125-DEL-1896 | chr13-99751126-DEL-1896 |
| chr13-99772029-DEL-405 | chr13-99772030-DEL-405 |
| chr13-99773419-DEL-399 | chr13-99773420-DEL-399 |
| chr13-99800021-DEL-151 | chr13-99800022-DEL-151 |
| chr13-99817008-DEL-1097 | chr13-99817009-DEL-1097 |
| chr13-99830367-DEL-360 | chr13-99830368-DEL-360 |
| chr13-99838989-DEL-2068 | chr13-99838988-DEL-2068 |
| chr13-99845875-DEL-6444 | chr13-99845876-DEL-6444 |
| chr13-99898885-DEL-202 | chr13-99898886-DEL-202 |
| chr13-99912428-DEL-120 | chr13-99912429-DEL-120 |
| chr13-99916265-DEL-88 | chr13-99916266-DEL-88 |
| chr13-99921796-DEL-116 | chr13-99921834-DEL-69 |
| chr13-99924992-DEL-700 | chr13-99924993-DEL-700 |
| chr13-99927426-DEL-233 | chr13-99927428-DEL-233 |
| chr13-99954735-DEL-4725 | chr13-99954736-DEL-4725 |
| chr13-99967165-DEL-248 | chr13-99967166-DEL-248 |
| chr13-99970676-DEL-3022 | chr13-99970676-DEL-3023 |
| chr13-99976607-DEL-1974 | chr13-99976608-DEL-1974 |
| chr13-99984298-DEL-343 | chr13-99984299-DEL-343 |
| chr13-100007177-DEL-60 | chr13-100007186-DEL-58 |
| chr13-100009088-DEL-62 | chr13-100009079-DEL-72 |
| chr13-100038720-DEL-296 | chr13-100038721-DEL-296 |
| chr13-100046724-DEL-291 | chr13-100046725-DEL-291 |
| chr13-100070561-DEL-52 | chr13-100070562-DEL-52 |
| chr13-100074746-DEL-202 | chr13-100074747-DEL-202 |
| chr13-100122971-DEL-511 | chr13-100122972-DEL-511 |
| chr13-100140255-DEL-216 | chr13-100140256-DEL-216 |
| chr13-100165519-DEL-58 | chr13-100165520-DEL-58 |
| chr13-100207252-DEL-815 | chr13-100207253-DEL-815 |
| chr13-100211507-DEL-195 | chr13-100211508-DEL-195 |
| chr13-100235218-DEL-186 | chr13-100235219-DEL-186 |
| chr13-100254234-DEL-6002 | chr13-100254227-DEL-6006 |

| | |
|---|---|
| chr13-100265873-DEL-1040 | chr13-100265876-DEL-1038 |
| chr13-100267749-DEL-266 | chr13-100267750-DEL-266 |
| chr13-100331477-DEL-86 | chr13-100331478-DEL-50 |
| chr13-100334090-DEL-201 | chr13-100334091-DEL-201 |
| chr13-100360466-DEL-3603 | chr13-100360467-DEL-3603 |
| chr13-100397155-DEL-226 | chr13-100397156-DEL-226 |
| chr13-100425414-DEL-3793 | chr13-100425415-DEL-3793 |
| chr13-100471129-DEL-4552 | chr13-100471128-DEL-4552 |
| chr13-100502622-DEL-1583 | chr13-100502623-DEL-1583 |
| chr13-100516545-DEL-6384 | chr13-100516546-DEL-6384 |
| chr13-100532986-DEL-90 | chr13-100532987-DEL-90 |
| chr13-100548056-DEL-1577 | chr13-100548057-DEL-1577 |
| chr13-100556455-DEL-61 | chr13-100556456-DEL-61 |
| chr13-100583991-DEL-324 | chr13-100583992-DEL-324 |
| chr13-100585541-DEL-433 | chr13-100585542-DEL-433 |
| chr13-100662307-DEL-98 | chr13-100662313-DEL-93 |
| chr13-100664701-DEL-146 | chr13-100664702-DEL-146 |
| chr13-100680516-DEL-93 | chr13-100680520-DEL-120 |
| chr13-100681433-DEL-216 | chr13-100681424-DEL-217 |
| chr13-100692700-DEL-114 | chr13-100692714-DEL-101 |
| chr13-100701241-DEL-202 | chr13-100701242-DEL-202 |
| chr13-100704901-DEL-826 | chr13-100704902-DEL-826 |
| chr13-100714816-DEL-808 | chr13-100714813-DEL-809 |
| chr13-100717571-DEL-209 | chr13-100717572-DEL-209 |
| chr13-100725453-DEL-5280 | chr13-100725454-DEL-5280 |
| chr13-100744748-DEL-427 | chr13-100744756-DEL-419 |
| chr13-100745745-DEL-238 | chr13-100745746-DEL-238 |
| chr13-100747389-DEL-202 | chr13-100747390-DEL-202 |
| chr13-100763269-DEL-167 | chr13-100763258-DEL-168 |
| chr13-100785449-DEL-209 | chr13-100785450-DEL-209 |
| chr13-100798840-DEL-815 | chr13-100798841-DEL-815 |
| chr13-100819815-DEL-349 | chr13-100819816-DEL-349 |
| chr13-100823477-DEL-56 | chr13-100823478-DEL-56 |
| chr13-100832301-DEL-65 | chr13-100832302-DEL-65 |
| chr13-100841255-DEL-326 | chr13-100841256-DEL-326 |
| chr13-100845629-DEL-56 | chr13-100845630-DEL-56 |
| chr13-100863008-DEL-54 | chr13-100863009-DEL-54 |
| chr13-100863835-DEL-1198 | chr13-100863836-DEL-1198 |
| chr13-100868409-DEL-218 | chr13-100868410-DEL-218 |
| chr13-100873583-DEL-621 | chr13-100873584-DEL-584 |
| chr13-100886197-DEL-70 | chr13-100886196-DEL-71 |
| chr13-100909880-DEL-155 | chr13-100909881-DEL-155 |
| chr13-100932737-DEL-202 | chr13-100932738-DEL-202 |
| chr13-100938210-DEL-199 | chr13-100938212-DEL-198 |
| chr13-100975763-DEL-237 | chr13-100975758-DEL-238 |
| chr13-100978000-DEL-68 | chr13-100978001-DEL-68 |
| chr13-100991571-DEL-1769 | chr13-100991572-DEL-1769 |
| chr13-101001171-DEL-200 | chr13-101001171-DEL-200 |
| chr13-101032790-DEL-147 | chr13-101032794-DEL-144 |
| chr13-101045532-DEL-892 | chr13-101045533-DEL-892 |
| chr13-101057821-DEL-6590 | chr13-101057822-DEL-6590 |
| chr13-101071138-DEL-1036 | chr13-101071139-DEL-1036 |
| chr13-101080340-DEL-247 | chr13-101080341-DEL-247 |
| chr13-101082377-DEL-2309 | chr13-101082378-DEL-2309 |
| chr13-101116670-DEL-166 | chr13-101116671-DEL-166 |
| chr13-101117874-DEL-5987 | chr13-101117875-DEL-5987 |
| chr13-101128407-DEL-199 | chr13-101128408-DEL-199 |
| chr13-101145864-DEL-474 | chr13-101145865-DEL-474 |
| chr13-101149493-DEL-361 | chr13-101149494-DEL-361 |
| chr13-101174582-DEL-1282 | chr13-101174580-DEL-1283 |
| chr13-101179206-DEL-6690 | chr13-101179207-DEL-6690 |
| chr13-101207977-DEL-72 | chr13-101207977-DEL-92 |
| chr13-101229965-DEL-145 | chr13-101229966-DEL-145 |
| chr13-101254492-DEL-76 | chr13-101254493-DEL-76 |
| chr13-101287703-DEL-649 | chr13-101287704-DEL-644 |
| chr13-101293702-DEL-201 | chr13-101293703-DEL-201 |
| chr13-101298732-DEL-94 | chr13-101298733-DEL-94 |
| chr13-101331992-DEL-101 | chr13-101331993-DEL-101 |
| chr13-101344135-DEL-701 | chr13-101344136-DEL-701 |
| chr13-101346818-DEL-72 | chr13-101346798-DEL-72 |
| chr13-101399345-DEL-244 | chr13-101399346-DEL-244 |
| chr13-101411795-DEL-52 | chr13-101411796-DEL-51 |
| chr13-101415692-DEL-222 | chr13-101415693-DEL-222 |
| chr13-101417810-DEL-203 | chr13-101417811-DEL-203 |
| chr13-101429023-DEL-92 | chr13-101429024-DEL-92 |
| chr13-101440544-DEL-51 | chr13-101440545-DEL-51 |
| chr13-101453303-DEL-62 | chr13-101453304-DEL-62 |
| chr13-101463702-DEL-186 | chr13-101463745-DEL-168 |
| chr13-101472903-DEL-1329 | chr13-101472904-DEL-1329 |
| chr13-101513757-DEL-68 | chr13-101513758-DEL-68 |

| | |
|---|---|
| chr13-101530481-DEL-56 | chr13-101530482-DEL-56 |
| chr13-101543184-DEL-217 | chr13-101543185-DEL-217 |
| chr13-101670317-DEL-123 | chr13-101670318-DEL-119 |
| chr13-101682108-DEL-58 | chr13-101682109-DEL-58 |
| chr13-101767105-DEL-56 | chr13-101767106-DEL-56 |
| chr13-101811835-DEL-148 | chr13-101811836-DEL-148 |
| chr13-101826376-DEL-76 | chr13-101826377-DEL-76 |
| chr13-101846991-DEL-73 | chr13-101846981-DEL-76 |
| chr13-101861079-DEL-122 | chr13-101861097-DEL-78 |
| chr13-101919983-DEL-1236 | chr13-101919984-DEL-1235 |
| chr13-101957320-DEL-280 | chr13-101957321-DEL-280 |
| chr13-102039684-DEL-98 | chr13-102039685-DEL-98 |
| chr13-102097615-DEL-3996 | chr13-102097616-DEL-3996 |
| chr13-102141689-DEL-323 | chr13-102141690-DEL-323 |
| chr13-102161094-DEL-61 | chr13-102161095-DEL-61 |
| chr13-102189193-DEL-902 | chr13-102189194-DEL-902 |
| chr13-102217297-DEL-70 | chr13-102217296-DEL-56 |
| chr13-102240129-DEL-725 | chr13-102240130-DEL-725 |
| chr13-102292571-DEL-1027 | chr13-102292572-DEL-1027 |
| chr13-102322333-DEL-54 | chr13-102322335-DEL-53 |
| chr13-102360683-DEL-2391 | chr13-102360679-DEL-2392 |
| chr13-102391249-DEL-96 | chr13-102391222-DEL-100 |
| chr13-102400183-DEL-6011 | chr13-102400184-DEL-6011 |
| chr13-102425519-DEL-161 | chr13-102425520-DEL-161 |
| chr13-102427435-DEL-3103 | chr13-102427436-DEL-3103 |
| chr13-102430903-DEL-52 | chr13-102430891-DEL-51 |
| chr13-102487865-DEL-178 | chr13-102487866-DEL-178 |
| chr13-102513479-DEL-211 | chr13-102513480-DEL-211 |
| chr13-102514265-DEL-241 | chr13-102514269-DEL-238 |
| chr13-102515756-DEL-76 | chr13-102515756-DEL-98 |
| chr13-102518371-DEL-6415 | chr13-102518363-DEL-6416 |
| chr13-102536035-DEL-90 | chr13-102536036-DEL-90 |
| chr13-102550209-DEL-108 | chr13-102550210-DEL-98 |
| chr13-102551758-DEL-1502 | chr13-102551759-DEL-1502 |
| chr13-102554529-DEL-259 | chr13-102554530-DEL-259 |
| chr13-102557837-DEL-212 | chr13-102557838-DEL-212 |
| chr13-102571840-DEL-2807 | chr13-102571841-DEL-2807 |
| chr13-102592766-DEL-64 | chr13-102592767-DEL-64 |
| chr13-102607124-DEL-244 | chr13-102607125-DEL-244 |
| chr13-102609709-DEL-1589 | chr13-102609710-DEL-1589 |
| chr13-102628927-DEL-6526 | chr13-102628928-DEL-6526 |
| chr13-102659452-DEL-123 | chr13-102659460-DEL-115 |
| chr13-102671682-DEL-857 | chr13-102671684-DEL-858 |
| chr13-102721633-DEL-220 | chr13-102721635-DEL-219 |
| chr13-102735922-DEL-88 | chr13-102735906-DEL-117 |
| chr13-102743476-DEL-6472 | chr13-102743477-DEL-6472 |
| chr13-102750807-DEL-6440 | chr13-102750808-DEL-6440 |
| chr13-102758974-DEL-84 | chr13-102758975-DEL-84 |
| chr13-102765279-DEL-224 | chr13-102765280-DEL-224 |
| chr13-102769781-DEL-63 | chr13-102769782-DEL-63 |
| chr13-102777628-DEL-74 | chr13-102777629-DEL-74 |
| chr13-102779707-DEL-56 | chr13-102779695-DEL-54 |
| chr13-102785927-DEL-69 | chr13-102785920-DEL-77 |
| chr13-102793077-DEL-180 | chr13-102793078-DEL-180 |
| chr13-102834247-DEL-353 | chr13-102834248-DEL-353 |
| chr13-102866392-DEL-60 | chr13-102866393-DEL-60 |
| chr13-102916737-DEL-317 | chr13-102916738-DEL-317 |
| chr13-102934848-DEL-164 | chr13-102934849-DEL-164 |
| chr13-102961957-DEL-182 | chr13-102961980-DEL-160 |
| chr13-102963606-DEL-7160 | chr13-102963607-DEL-7160 |
| chr13-102980681-DEL-113 | chr13-102980673-DEL-86 |
| chr13-103036599-DEL-72 | chr13-103036600-DEL-72 |
| chr13-103054421-DEL-87 | chr13-103054422-DEL-87 |
| chr13-103060341-DEL-484 | chr13-103060342-DEL-484 |
| chr13-103065642-DEL-133 | chr13-103065643-DEL-133 |
| chr13-103087165-DEL-255 | chr13-103087166-DEL-253 |
| chr13-103101198-DEL-55 | chr13-103101199-DEL-55 |
| chr13-103101579-DEL-398 | chr13-103101580-DEL-398 |
| chr13-103102575-DEL-587 | chr13-103102572-DEL-585 |
| chr13-103104373-DEL-400 | chr13-103104374-DEL-400 |
| chr13-103146750-DEL-421 | chr13-103146751-DEL-419 |
| chr13-103152349-DEL-109 | chr13-103152353-DEL-111 |
| chr13-103152540-DEL-1599 | chr13-103152541-DEL-1599 |
| chr13-103167023-DEL-426 | chr13-103167022-DEL-424 |
| chr13-103169328-DEL-59 | chr13-103169333-DEL-53 |
| chr13-103172698-DEL-191 | chr13-103172696-DEL-184 |
| chr13-103197278-DEL-232 | chr13-103197279-DEL-232 |
| chr13-103205278-DEL-55 | chr13-103205279-DEL-55 |
| chr13-103219344-DEL-110 | chr13-103219309-DEL-102 |
| chr13-103227040-DEL-75 | chr13-103227041-DEL-75 |

| | |
|---|---|
| chr13-103236055-DEL-58 | chr13-103236056-DEL-58 |
| chr13-103244269-DEL-1021 | chr13-103244270-DEL-1021 |
| chr13-103283170-DEL-73 | chr13-103283164-DEL-79 |
| chr13-103292979-DEL-89 | chr13-103292980-DEL-65 |
| chr13-103303778-DEL-78 | chr13-103303779-DEL-78 |
| chr13-103326586-DEL-383 | chr13-103326587-DEL-383 |
| chr13-103338618-DEL-74 | chr13-103338619-DEL-66 |
| chr13-103356709-DEL-52 | chr13-103356710-DEL-52 |
| chr13-103367297-DEL-3118 | chr13-103367298-DEL-3118 |
| chr13-103368865-DEL-1082 | chr13-103368866-DEL-1082 |
| chr13-103412524-DEL-142 | chr13-103412525-DEL-135 |
| chr13-103429192-DEL-329 | chr13-103429193-DEL-329 |
| chr13-103435484-DEL-62 | chr13-103435485-DEL-62 |
| chr13-103442982-DEL-190 | chr13-103442983-DEL-190 |
| chr13-103472785-DEL-97 | chr13-103472786-DEL-97 |
| chr13-103480200-DEL-53 | chr13-103480200-DEL-53 |
| chr13-103482565-DEL-117 | chr13-103482566-DEL-117 |
| chr13-103492675-DEL-3061 | chr13-103492676-DEL-3061 |
| chr13-103495800-DEL-230 | chr13-103495801-DEL-230 |
| chr13-103507483-DEL-136 | chr13-103507484-DEL-136 |
| chr13-103508875-DEL-147 | chr13-103508873-DEL-148 |
| chr13-103514875-DEL-128 | chr13-103514876-DEL-128 |
| chr13-103518761-DEL-991 | chr13-103518762-DEL-991 |
| chr13-103533468-DEL-52 | chr13-103533469-DEL-52 |
| chr13-103537232-DEL-6620 | chr13-103537233-DEL-6620 |
| chr13-103573166-DEL-60 | chr13-103573166-DEL-60 |
| chr13-103592494-DEL-80 | chr13-103592495-DEL-81 |
| chr13-103621252-DEL-101 | chr13-103621236-DEL-102 |
| chr13-103669264-DEL-1812 | chr13-103669265-DEL-1812 |
| chr13-103677215-DEL-190 | chr13-103677214-DEL-170 |
| chr13-103680693-DEL-68 | chr13-103680694-DEL-68 |
| chr13-103702302-DEL-52 | chr13-103702303-DEL-52 |
| chr13-103702777-DEL-6693 | chr13-103702778-DEL-6693 |
| chr13-103729241-DEL-181 | chr13-103729212-DEL-181 |
| chr13-103757548-DEL-81 | chr13-103757549-DEL-81 |
| chr13-103773905-DEL-818 | chr13-103773906-DEL-818 |
| chr13-103779357-DEL-66 | chr13-103779361-DEL-62 |
| chr13-103786332-DEL-808 | chr13-103786333-DEL-808 |
| chr13-103790125-DEL-78 | chr13-103790137-DEL-64 |
| chr13-103790983-DEL-115 | chr13-103790982-DEL-117 |
| chr13-103795130-DEL-145 | chr13-103795131-DEL-145 |
| chr13-103810044-DEL-1383 | chr13-103810045-DEL-1383 |
| chr13-103824099-DEL-99 | chr13-103824107-DEL-84 |
| chr13-103830966-DEL-122 | chr13-103830968-DEL-118 |
| chr13-103845152-DEL-541 | chr13-103845153-DEL-542 |
| chr13-103848921-DEL-8432 | chr13-103848956-DEL-8404 |
| chr13-103866996-DEL-720 | chr13-103867033-DEL-703 |
| chr13-103877385-DEL-141 | chr13-103877384-DEL-135 |
| chr13-103884612-DEL-390 | chr13-103884604-DEL-399 |
| chr13-103892679-DEL-276 | chr13-103892724-DEL-268 |
| chr13-103892699-DEL-864 | chr13-103892668-DEL-872 |
| chr13-103914512-DEL-126 | chr13-103914513-DEL-126 |
| chr13-103921264-DEL-198 | chr13-103921257-DEL-199 |
| chr13-103925002-DEL-742 | chr13-103925003-DEL-741 |
| chr13-103928471-DEL-75 | chr13-103928468-DEL-75 |
| chr13-103945558-DEL-457 | chr13-103945559-DEL-457 |
| chr13-103983028-DEL-58 | chr13-103983029-DEL-58 |
| chr13-103995492-DEL-105 | chr13-103995493-DEL-105 |
| chr13-104012554-DEL-99 | chr13-104012556-DEL-98 |
| chr13-104024923-DEL-74 | chr13-104024924-DEL-74 |
| chr13-104027202-DEL-91 | chr13-104027203-DEL-91 |
| chr13-104044984-DEL-162 | chr13-104044983-DEL-163 |
| chr13-104073889-DEL-210 | chr13-104073890-DEL-210 |
| chr13-104079519-DEL-75 | chr13-104079528-DEL-58 |
| chr13-104083498-DEL-2845 | chr13-104083499-DEL-2845 |
| chr13-104092675-DEL-73 | chr13-104092675-DEL-73 |
| chr13-104105533-DEL-62 | chr13-104105518-DEL-74 |
| chr13-104105880-DEL-359 | chr13-104105881-DEL-359 |
| chr13-104124791-DEL-65 | chr13-104124792-DEL-65 |
| chr13-104132526-DEL-6590 | chr13-104132527-DEL-6591 |
| chr13-104146472-DEL-187 | chr13-104146473-DEL-187 |
| chr13-104154673-DEL-83 | chr13-104154682-DEL-65 |
| chr13-104174368-DEL-60 | chr13-104174369-DEL-60 |
| chr13-104193395-DEL-620 | chr13-104193396-DEL-620 |
| chr13-104194530-DEL-168 | chr13-104194531-DEL-168 |
| chr13-104212047-DEL-630 | chr13-104212048-DEL-626 |
| chr13-104217664-DEL-52 | chr13-104217657-DEL-52 |
| chr13-104234996-DEL-102 | chr13-104234988-DEL-102 |
| chr13-104244550-DEL-202 | chr13-104244551-DEL-202 |
| chr13-104257336-DEL-533 | chr13-104257337-DEL-533 |

| | |
|---|---|
| chr13-104282521-DEL-250 | chr13-104282522-DEL-250 |
| chr13-104291814-DEL-62 | chr13-104291815-DEL-62 |
| chr13-104333954-DEL-232 | chr13-104333955-DEL-232 |
| chr13-104351687-DEL-64 | chr13-104351689-DEL-56 |
| chr13-104355685-DEL-52 | chr13-104355686-DEL-52 |
| chr13-104416732-DEL-210 | chr13-104416733-DEL-210 |
| chr13-104471885-DEL-99 | chr13-104471886-DEL-99 |
| chr13-104483960-DEL-1112 | chr13-104483961-DEL-1112 |
| chr13-104485453-DEL-210 | chr13-104485454-DEL-210 |
| chr13-104570500-DEL-232 | chr13-104570501-DEL-232 |
| chr13-104576437-DEL-54 | chr13-104576439-DEL-54 |
| chr13-104621053-DEL-4922 | chr13-104621054-DEL-4922 |
| chr13-104638296-DEL-263 | chr13-104638297-DEL-263 |
| chr13-104663409-DEL-137 | chr13-104663406-DEL-138 |
| chr13-104678137-DEL-76 | chr13-104678138-DEL-76 |
| chr13-104692300-DEL-67 | chr13-104692300-DEL-60 |
| chr13-104698675-DEL-288 | chr13-104698652-DEL-312 |
| chr13-104728469-DEL-161 | chr13-104728470-DEL-159 |
| chr13-104735361-DEL-103 | chr13-104735362-DEL-103 |
| chr13-104745021-DEL-73 | chr13-104745033-DEL-51 |
| chr13-104776229-DEL-172 | chr13-104776230-DEL-172 |
| chr13-104779432-DEL-2456 | chr13-104779433-DEL-2456 |
| chr13-104780634-DEL-82 | chr13-104780635-DEL-82 |
| chr13-104840127-DEL-80 | chr13-104840128-DEL-80 |
| chr13-104859597-DEL-2406 | chr13-104859598-DEL-2406 |
| chr13-104871735-DEL-58 | chr13-104871736-DEL-58 |
| chr13-104879535-DEL-59 | chr13-104879524-DEL-59 |
| chr13-104895029-DEL-82 | chr13-104895028-DEL-79 |
| chr13-104904105-DEL-6274 | chr13-104904099-DEL-6282 |
| chr13-104904369-DEL-56 | chr13-104904370-DEL-56 |
| chr13-104975830-DEL-561 | chr13-104975822-DEL-566 |
| chr13-105001142-DEL-99 | chr13-105001143-DEL-99 |
| chr13-105012384-DEL-84 | chr13-105012359-DEL-84 |
| chr13-105036264-DEL-7412 | chr13-105036265-DEL-7412 |
| chr13-105090593-DEL-366 | chr13-105090594-DEL-366 |
| chr13-105094252-DEL-90 | chr13-105094253-DEL-90 |
| chr13-105127720-DEL-5619 | chr13-105127721-DEL-5619 |
| chr13-105142862-DEL-96 | chr13-105142863-DEL-96 |
| chr13-105151736-DEL-56 | chr13-105151737-DEL-56 |
| chr13-105231492-DEL-964 | chr13-105231493-DEL-964 |
| chr13-105259995-DEL-5184 | chr13-105259996-DEL-5184 |
| chr13-105260003-DEL-2377 | chr13-105260002-DEL-2377 |
| chr13-105268150-DEL-54 | chr13-105268127-DEL-78 |
| chr13-105292219-DEL-1343 | chr13-105292220-DEL-1341 |
| chr13-105301860-DEL-66 | chr13-105301867-DEL-72 |
| chr13-105315843-DEL-80 | chr13-105315844-DEL-80 |
| chr13-105336553-DEL-62 | chr13-105336554-DEL-60 |
| chr13-105368485-DEL-185 | chr13-105368486-DEL-185 |
| chr13-105481539-DEL-3847 | chr13-105481540-DEL-3847 |
| chr13-105485987-DEL-76 | chr13-105485982-DEL-52 |
| chr13-105526692-DEL-64 | chr13-105526676-DEL-113 |
| chr13-105529216-DEL-56 | chr13-105529217-DEL-56 |
| chr13-105563959-DEL-97 | chr13-105563960-DEL-97 |
| chr13-105565045-DEL-54 | chr13-105565023-DEL-50 |
| chr13-105567739-DEL-388 | chr13-105567740-DEL-388 |
| chr13-105593063-DEL-361 | chr13-105593064-DEL-360 |
| chr13-105599021-DEL-216 | chr13-105599022-DEL-216 |
| chr13-105612029-DEL-409 | chr13-105612030-DEL-419 |
| chr13-105627587-DEL-1893 | chr13-105627588-DEL-1893 |
| chr13-105663196-DEL-539 | chr13-105663197-DEL-539 |
| chr13-105691328-DEL-177 | chr13-105691328-DEL-177 |
| chr13-105691826-DEL-971 | chr13-105691827-DEL-971 |
| chr13-105745235-DEL-2410 | chr13-105745236-DEL-2410 |
| chr13-105752832-DEL-1920 | chr13-105752833-DEL-1871 |
| chr13-105779521-DEL-188 | chr13-105779522-DEL-188 |
| chr13-105780430-DEL-1772 | chr13-105780431-DEL-1772 |
| chr13-105823158-DEL-55 | chr13-105823159-DEL-55 |
| chr13-105826199-DEL-6261 | chr13-105826200-DEL-6261 |
| chr13-105845815-DEL-6438 | chr13-105845816-DEL-6438 |
| chr13-105874649-DEL-108 | chr13-105874650-DEL-108 |
| chr13-105876173-DEL-3215 | chr13-105876173-DEL-3215 |
| chr13-105884621-DEL-95 | chr13-105884623-DEL-98 |
| chr13-105888892-DEL-91 | chr13-105888893-DEL-90 |
| chr13-105895682-DEL-68 | chr13-105895689-DEL-70 |
| chr13-105901432-DEL-52 | chr13-105901433-DEL-56 |
| chr13-105915917-DEL-364 | chr13-105915917-DEL-364 |
| chr13-105926998-DEL-114 | chr13-105926999-DEL-114 |
| chr13-105929624-DEL-6464 | chr13-105929625-DEL-6463 |
| chr13-105941075-DEL-439 | chr13-105941076-DEL-439 |
| chr13-105963444-DEL-64 | chr13-105963444-DEL-65 |

| | |
|---|---|
| chr13-105975823-DEL-83 | chr13-105975824-DEL-83 |
| chr13-105992960-DEL-336 | chr13-105992961-DEL-336 |
| chr13-106005651-DEL-3101 | chr13-106005652-DEL-3101 |
| chr13-106025375-DEL-316 | chr13-106025376-DEL-316 |
| chr13-106027184-DEL-1221 | chr13-106027185-DEL-1221 |
| chr13-106030395-DEL-1603 | chr13-106030395-DEL-1603 |
| chr13-106032976-DEL-1005 | chr13-106032975-DEL-1005 |
| chr13-106035811-DEL-126 | chr13-106035809-DEL-127 |
| chr13-106038238-DEL-68 | chr13-106038202-DEL-69 |
| chr13-106039669-DEL-967 | chr13-106039670-DEL-967 |
| chr13-106048898-DEL-1594 | chr13-106048899-DEL-1594 |
| chr13-106064001-DEL-225 | chr13-106064001-DEL-225 |
| chr13-106066860-DEL-564 | chr13-106066861-DEL-564 |
| chr13-106080343-DEL-1212 | chr13-106080344-DEL-1212 |
| chr13-106107417-DEL-8066 | chr13-106107418-DEL-8066 |
| chr13-106110919-DEL-2842 | chr13-106110920-DEL-2842 |
| chr13-106141419-DEL-67 | chr13-106141418-DEL-67 |
| chr13-106142781-DEL-57 | chr13-106142781-DEL-57 |
| chr13-106157465-DEL-887 | chr13-106157465-DEL-887 |
| chr13-106158461-DEL-58 | chr13-106158461-DEL-57 |
| chr13-106169471-DEL-5175 | chr13-106169471-DEL-5176 |
| chr13-106176140-DEL-57 | chr13-106176142-DEL-56 |
| chr13-106203664-DEL-77 | chr13-106203653-DEL-89 |
| chr13-106271695-DEL-56 | chr13-106271693-DEL-72 |
| chr13-106283226-DEL-424 | chr13-106283227-DEL-424 |
| chr13-106287337-DEL-882 | chr13-106287338-DEL-882 |
| chr13-106330764-DEL-598 | chr13-106330765-DEL-598 |
| chr13-106341876-DEL-1188 | chr13-106341877-DEL-1188 |
| chr13-106361274-DEL-6057 | chr13-106361275-DEL-6057 |
| chr13-106421000-DEL-130 | chr13-106421001-DEL-130 |
| chr13-106427553-DEL-9348 | chr13-106427554-DEL-9348 |
| chr13-106453853-DEL-1496 | chr13-106453854-DEL-1496 |
| chr13-106456018-DEL-1250 | chr13-106456019-DEL-1250 |
| chr13-106468892-DEL-56 | chr13-106468894-DEL-54 |
| chr13-106492708-DEL-2502 | chr13-106492719-DEL-2503 |
| chr13-106498942-DEL-62 | chr13-106498943-DEL-62 |
| chr13-106509196-DEL-7429 | chr13-106509197-DEL-7429 |
| chr13-106517980-DEL-6377 | chr13-106517981-DEL-6377 |
| chr13-106560012-DEL-1256 | chr13-106560013-DEL-1256 |
| chr13-106563529-DEL-1642 | chr13-106563530-DEL-1642 |
| chr13-106583460-DEL-62 | chr13-106583461-DEL-62 |
| chr13-106586595-DEL-936 | chr13-106586596-DEL-936 |
| chr13-106592892-DEL-208 | chr13-106592890-DEL-209 |
| chr13-106649225-DEL-369 | chr13-106649226-DEL-369 |
| chr13-106652691-DEL-1479 | chr13-106652692-DEL-1479 |
| chr13-106656813-DEL-70 | chr13-106656814-DEL-70 |
| chr13-106662831-DEL-7038 | chr13-106662832-DEL-7038 |
| chr13-106703838-DEL-133 | chr13-106703839-DEL-133 |
| chr13-106730460-DEL-463 | chr13-106730461-DEL-463 |
| chr13-106732598-DEL-278 | chr13-106732599-DEL-278 |
| chr13-106747615-DEL-211 | chr13-106747616-DEL-211 |
| chr13-106751067-DEL-714 | chr13-106751067-DEL-714 |
| chr13-106776465-DEL-468 | chr13-106776466-DEL-468 |
| chr13-106777441-DEL-91 | chr13-106777441-DEL-92 |
| chr13-106786733-DEL-6590 | chr13-106786734-DEL-6590 |
| chr13-106800119-DEL-147 | chr13-106800120-DEL-147 |
| chr13-106816277-DEL-216 | chr13-106816281-DEL-214 |
| chr13-106831297-DEL-2492 | chr13-106831298-DEL-2492 |
| chr13-106834811-DEL-399 | chr13-106834812-DEL-399 |
| chr13-106861634-DEL-200 | chr13-106861635-DEL-200 |
| chr13-106902406-DEL-98 | chr13-106902375-DEL-94 |
| chr13-106913038-DEL-141 | chr13-106913039-DEL-141 |
| chr13-106921844-DEL-2948 | chr13-106921843-DEL-2955 |
| chr13-106934333-DEL-104 | chr13-106934334-DEL-104 |
| chr13-106978306-DEL-75 | chr13-106978307-DEL-75 |
| chr13-106991749-DEL-225 | chr13-106991750-DEL-225 |
| chr13-106998479-DEL-62 | chr13-106998473-DEL-73 |
| chr13-107021204-DEL-216 | chr13-107021205-DEL-216 |
| chr13-107023408-DEL-84 | chr13-107023409-DEL-72 |
| chr13-107067339-DEL-212 | chr13-107067340-DEL-212 |
| chr13-107080164-DEL-108 | chr13-107080165-DEL-108 |
| chr13-107087470-DEL-202 | chr13-107087471-DEL-202 |
| chr13-107112173-DEL-407 | chr13-107112174-DEL-407 |
| chr13-107143379-DEL-1531 | chr13-107143380-DEL-1531 |
| chr13-107160724-DEL-508 | chr13-107160725-DEL-508 |
| chr13-107167856-DEL-56 | chr13-107167854-DEL-55 |
| chr13-107185590-DEL-1118 | chr13-107185591-DEL-1118 |
| chr13-107206822-DEL-161 | chr13-107206823-DEL-161 |
| chr13-107213000-DEL-97 | chr13-107213001-DEL-97 |
| chr13-107218951-DEL-72 | chr13-107218958-DEL-64 |

| | |
|---|---|
| chr13-107220315-DEL-193 | chr13-107220316-DEL-193 |
| chr13-107245437-DEL-56 | chr13-107245438-DEL-56 |
| chr13-107248599-DEL-379 | chr13-107248600-DEL-363 |
| chr13-107269686-DEL-214 | chr13-107269687-DEL-214 |
| chr13-107274798-DEL-70 | chr13-107274778-DEL-70 |
| chr13-107280311-DEL-397 | chr13-107280312-DEL-397 |
| chr13-107290577-DEL-133 | chr13-107290578-DEL-133 |
| chr13-107309662-DEL-150 | chr13-107309664-DEL-141 |
| chr13-107313005-DEL-60 | chr13-107313005-DEL-55 |
| chr13-107351466-DEL-55 | chr13-107351460-DEL-54 |
| chr13-107376106-DEL-333 | chr13-107376107-DEL-333 |
| chr13-107425688-DEL-66 | chr13-107425689-DEL-66 |
| chr13-107484541-DEL-55 | chr13-107484529-DEL-56 |
| chr13-107490543-DEL-6204 | chr13-107490544-DEL-6204 |
| chr13-107504470-DEL-58 | chr13-107504471-DEL-58 |
| chr13-107550328-DEL-940 | chr13-107550329-DEL-940 |
| chr13-107568043-DEL-140 | chr13-107568038-DEL-141 |
| chr13-107568735-DEL-5475 | chr13-107568736-DEL-5475 |
| chr13-107580429-DEL-354 | chr13-107580430-DEL-354 |
| chr13-107586922-DEL-52 | chr13-107586923-DEL-52 |
| chr13-107609946-DEL-55 | chr13-107609947-DEL-55 |
| chr13-107647787-DEL-392 | chr13-107647788-DEL-392 |
| chr13-107656692-DEL-1274 | chr13-107656693-DEL-1274 |
| chr13-107666137-DEL-172 | chr13-107666157-DEL-153 |
| chr13-107668483-DEL-153 | chr13-107668467-DEL-154 |
| chr13-107686615-DEL-57 | chr13-107686616-DEL-57 |
| chr13-107693849-DEL-372 | chr13-107693830-DEL-401 |
| chr13-107718978-DEL-59 | chr13-107718951-DEL-66 |
| chr13-107746582-DEL-64 | chr13-107746583-DEL-64 |
| chr13-107796581-DEL-943 | chr13-107796582-DEL-943 |
| chr13-107811272-DEL-267 | chr13-107811273-DEL-267 |
| chr13-107878715-DEL-54 | chr13-107878716-DEL-54 |
| chr13-107887211-DEL-69 | chr13-107887245-DEL-59 |
| chr13-107910948-DEL-496 | chr13-107910949-DEL-496 |
| chr13-107955125-DEL-129 | chr13-107955126-DEL-129 |
| chr13-107965855-DEL-106 | chr13-107965852-DEL-101 |
| chr13-107975861-DEL-250 | chr13-107975862-DEL-250 |
| chr13-107977112-DEL-60 | chr13-107977113-DEL-60 |
| chr13-107986624-DEL-57 | chr13-107986625-DEL-57 |
| chr13-107994789-DEL-1716 | chr13-107994771-DEL-1755 |
| chr13-107994789-DEL-312 | chr13-107994790-DEL-312 |
| chr13-108042905-DEL-112 | chr13-108042904-DEL-113 |
| chr13-108049662-DEL-216 | chr13-108049663-DEL-213 |
| chr13-108083114-DEL-85 | chr13-108083115-DEL-85 |
| chr13-108093689-DEL-126 | chr13-108093704-DEL-117 |
| chr13-108101713-DEL-208 | chr13-108101714-DEL-208 |
| chr13-108129175-DEL-87 | chr13-108129190-DEL-96 |
| chr13-108134222-DEL-5378 | chr13-108134223-DEL-5378 |
| chr13-108144384-DEL-59 | chr13-108144365-DEL-61 |
| chr13-108163185-DEL-180 | chr13-108163194-DEL-186 |
| chr13-108166035-DEL-98 | chr13-108166036-DEL-98 |
| chr13-108189181-DEL-53 | chr13-108189182-DEL-53 |
| chr13-108209406-DEL-250 | chr13-108209407-DEL-250 |
| chr13-108232696-DEL-52 | chr13-108232697-DEL-52 |
| chr13-108249357-DEL-341 | chr13-108249354-DEL-342 |
| chr13-108254379-DEL-73 | chr13-108254380-DEL-73 |
| chr13-108271113-DEL-307 | chr13-108271114-DEL-307 |
| chr13-108282825-DEL-403 | chr13-108282828-DEL-395 |
| chr13-108292137-DEL-69 | chr13-108292143-DEL-64 |
| chr13-108329540-DEL-161 | chr13-108329537-DEL-162 |
| chr13-108336851-DEL-92 | chr13-108336852-DEL-92 |
| chr13-108340344-DEL-409 | chr13-108340345-DEL-409 |
| chr13-108347188-DEL-92 | chr13-108347193-DEL-88 |
| chr13-108372682-DEL-86 | chr13-108372683-DEL-88 |
| chr13-108379608-DEL-56 | chr13-108379597-DEL-68 |
| chr13-108425479-DEL-70 | chr13-108425480-DEL-70 |
| chr13-108441765-DEL-65 | chr13-108441751-DEL-79 |
| chr13-108495469-DEL-766 | chr13-108495470-DEL-766 |
| chr13-108499870-DEL-80 | chr13-108499892-DEL-70 |
| chr13-108505224-DEL-64 | chr13-108505225-DEL-64 |
| chr13-108516441-DEL-131 | chr13-108516442-DEL-131 |
| chr13-108526704-DEL-70 | chr13-108526705-DEL-70 |
| chr13-108531044-DEL-6380 | chr13-108531045-DEL-6380 |
| chr13-108546628-DEL-125 | chr13-108546619-DEL-116 |
| chr13-108547680-DEL-53 | chr13-108547681-DEL-53 |
| chr13-108559591-DEL-83 | chr13-108559592-DEL-83 |
| chr13-108567984-DEL-91 | chr13-108567968-DEL-106 |
| chr13-108583404-DEL-105 | chr13-108583405-DEL-105 |
| chr13-108590416-DEL-111 | chr13-108590414-DEL-114 |
| chr13-108592682-DEL-2784 | chr13-108592683-DEL-2784 |

| | |
|---|---|
| chr13-108621071-DEL-113 | chr13-108621098-DEL-102 |
| chr13-108639735-DEL-6677 | chr13-108639736-DEL-6677 |
| chr13-108649715-DEL-5652 | chr13-108649716-DEL-5652 |
| chr13-108658434-DEL-72 | chr13-108658435-DEL-72 |
| chr13-108659032-DEL-60 | chr13-108659013-DEL-95 |
| chr13-108701856-DEL-58 | chr13-108701823-DEL-62 |
| chr13-108704079-DEL-1515 | chr13-108704080-DEL-1515 |
| chr13-108719054-DEL-2330 | chr13-108719055-DEL-2330 |
| chr13-108724847-DEL-443 | chr13-108724848-DEL-443 |
| chr13-108732546-DEL-96 | chr13-108732547-DEL-96 |
| chr13-108742387-DEL-134 | chr13-108742387-DEL-134 |
| chr13-108742568-DEL-6171 | chr13-108742569-DEL-6171 |
| chr13-108751441-DEL-52 | chr13-108751441-DEL-52 |
| chr13-108755787-DEL-126 | chr13-108755788-DEL-126 |
| chr13-108759167-DEL-75 | chr13-108759168-DEL-75 |
| chr13-108768999-DEL-458 | chr13-108769000-DEL-458 |
| chr13-108782694-DEL-192 | chr13-108782677-DEL-190 |
| chr13-108789527-DEL-51 | chr13-108789528-DEL-51 |
| chr13-108806834-DEL-55 | chr13-108806837-DEL-54 |
| chr13-108807096-DEL-886 | chr13-108807097-DEL-886 |
| chr13-108812555-DEL-136 | chr13-108812556-DEL-136 |
| chr13-108815878-DEL-867 | chr13-108815879-DEL-867 |
| chr13-108817648-DEL-68 | chr13-108817649-DEL-68 |
| chr13-108827197-DEL-488 | chr13-108827196-DEL-483 |
| chr13-108850379-DEL-190 | chr13-108850380-DEL-190 |
| chr13-108856060-DEL-60 | chr13-108856061-DEL-60 |
| chr13-108868337-DEL-119 | chr13-108868338-DEL-119 |
| chr13-108878773-DEL-210 | chr13-108878774-DEL-210 |
| chr13-108926634-DEL-6438 | chr13-108926633-DEL-6439 |
| chr13-108948687-DEL-345 | chr13-108948688-DEL-345 |
| chr13-108962732-DEL-1265 | chr13-108962733-DEL-1265 |
| chr13-108978670-DEL-84 | chr13-108978671-DEL-84 |
| chr13-108979223-DEL-2562 | chr13-108979224-DEL-2562 |
| chr13-108989246-DEL-90 | chr13-108989247-DEL-90 |
| chr13-109008295-DEL-210 | chr13-109008296-DEL-210 |
| chr13-109018941-DEL-113 | chr13-109018948-DEL-107 |
| chr13-109039493-DEL-59 | chr13-109039494-DEL-59 |
| chr13-109053869-DEL-395 | chr13-109053870-DEL-395 |
| chr13-109055933-DEL-61 | chr13-109055934-DEL-61 |
| chr13-109132106-DEL-66 | chr13-109132095-DEL-82 |
| chr13-109133420-DEL-83 | chr13-109133421-DEL-83 |
| chr13-109173715-DEL-389 | chr13-109173716-DEL-389 |
| chr13-109178205-DEL-411 | chr13-109178211-DEL-409 |
| chr13-109198078-DEL-473 | chr13-109198079-DEL-473 |
| chr13-109212577-DEL-112 | chr13-109212585-DEL-87 |
| chr13-109213681-DEL-96 | chr13-109213682-DEL-96 |
| chr13-109241865-DEL-1428 | chr13-109241866-DEL-1428 |
| chr13-109247604-DEL-66 | chr13-109247606-DEL-50 |
| chr13-109256504-DEL-65 | chr13-109256505-DEL-63 |
| chr13-109257899-DEL-97 | chr13-109257900-DEL-97 |
| chr13-109281448-DEL-55 | chr13-109281449-DEL-55 |
| chr13-109315205-DEL-60 | chr13-109315206-DEL-60 |
| chr13-109325399-DEL-207 | chr13-109325400-DEL-207 |
| chr13-109337172-DEL-72 | chr13-109337173-DEL-72 |
| chr13-109343698-DEL-281 | chr13-109343699-DEL-281 |
| chr13-109391203-DEL-70 | chr13-109391226-DEL-70 |
| chr13-109396490-DEL-68 | chr13-109396491-DEL-68 |
| chr13-109416261-DEL-159 | chr13-109416262-DEL-159 |
| chr13-109423251-DEL-110 | chr13-109423252-DEL-110 |
| chr13-109433229-DEL-77 | chr13-109433232-DEL-75 |
| chr13-109459121-DEL-403 | chr13-109459122-DEL-403 |
| chr13-109491335-DEL-480 | chr13-109491330-DEL-481 |
| chr13-109496869-DEL-1920 | chr13-109496870-DEL-1920 |
| chr13-109502727-DEL-6701 | chr13-109502728-DEL-6701 |
| chr13-109508739-DEL-221 | chr13-109508740-DEL-221 |
| chr13-109546490-DEL-6424 | chr13-109546491-DEL-6424 |
| chr13-109621517-DEL-136 | chr13-109621493-DEL-170 |
| chr13-109625214-DEL-1050 | chr13-109625215-DEL-1050 |
| chr13-109648891-DEL-118 | chr13-109648892-DEL-118 |
| chr13-109660789-DEL-254 | chr13-109660790-DEL-254 |
| chr13-109666023-DEL-58 | chr13-109666024-DEL-58 |
| chr13-109688769-DEL-118 | chr13-109688770-DEL-118 |
| chr13-109690168-DEL-160 | chr13-109690167-DEL-161 |
| chr13-109724662-DEL-61 | chr13-109724657-DEL-57 |
| chr13-109739802-DEL-215 | chr13-109739803-DEL-215 |
| chr13-109762718-DEL-207 | chr13-109762719-DEL-207 |
| chr13-109794689-DEL-201 | chr13-109794690-DEL-201 |
| chr13-109809212-DEL-2929 | chr13-109809213-DEL-2929 |
| chr13-109821360-DEL-129 | chr13-109821361-DEL-129 |
| chr13-109834035-DEL-186 | chr13-109834037-DEL-185 |

| | |
|---|---|
| chr13-109835649-DEL-115 | chr13-109835650-DEL-115 |
| chr13-109838082-DEL-75 | chr13-109838079-DEL-79 |
| chr13-109848428-DEL-1707 | chr13-109848429-DEL-1707 |
| chr13-109855347-DEL-105 | chr13-109855348-DEL-105 |
| chr13-109898432-DEL-6693 | chr13-109898433-DEL-6693 |
| chr13-109920614-DEL-55 | chr13-109920615-DEL-55 |
| chr13-109929266-DEL-197 | chr13-109929267-DEL-197 |
| chr13-109959801-DEL-64 | chr13-109959802-DEL-64 |
| chr13-109978043-DEL-481 | chr13-109978043-DEL-481 |
| chr13-109986209-DEL-174 | chr13-109986210-DEL-174 |
| chr13-109988219-DEL-7167 | chr13-109988220-DEL-7167 |
| chr13-110000757-DEL-59 | chr13-110000759-DEL-51 |
| chr13-110002549-DEL-72 | chr13-110002550-DEL-72 |
| chr13-110010838-DEL-1445 | chr13-110010839-DEL-1445 |
| chr13-110028522-DEL-291 | chr13-110028536-DEL-269 |
| chr13-110034017-DEL-913 | chr13-110034018-DEL-913 |
| chr13-110037021-DEL-98 | chr13-110037022-DEL-98 |
| chr13-110069693-DEL-135 | chr13-110069738-DEL-90 |
| chr13-110072463-DEL-58 | chr13-110072468-DEL-54 |
| chr13-110124878-DEL-145 | chr13-110124879-DEL-145 |
| chr13-110125575-DEL-976 | chr13-110125576-DEL-976 |
| chr13-110137540-DEL-54 | chr13-110137539-DEL-56 |
| chr13-110164723-DEL-300 | chr13-110164724-DEL-300 |
| chr13-110165693-DEL-380 | chr13-110165694-DEL-380 |
| chr13-110172637-DEL-821 | chr13-110172638-DEL-821 |
| chr13-110178578-DEL-498 | chr13-110178579-DEL-498 |
| chr13-110181445-DEL-52 | chr13-110181476-DEL-52 |
| chr13-110183446-DEL-53 | chr13-110183447-DEL-53 |
| chr13-110184426-DEL-108 | chr13-110184470-DEL-88 |
| chr13-110200921-DEL-2397 | chr13-110200922-DEL-2397 |
| chr13-110229478-DEL-54 | chr13-110229479-DEL-54 |
| chr13-110263263-DEL-86 | chr13-110263264-DEL-86 |
| chr13-110266937-DEL-185 | chr13-110266942-DEL-181 |
| chr13-110282717-DEL-2608 | chr13-110282718-DEL-2608 |
| chr13-110287262-DEL-190 | chr13-110287222-DEL-193 |
| chr13-110289671-DEL-498 | chr13-110289672-DEL-498 |
| chr13-110294836-DEL-199 | chr13-110294837-DEL-199 |
| chr13-110323098-DEL-85 | chr13-110323099-DEL-85 |
| chr13-110352989-DEL-106 | chr13-110352988-DEL-109 |
| chr13-110406200-DEL-54 | chr13-110406159-DEL-68 |
| chr13-110420105-DEL-62 | chr13-110420098-DEL-70 |
| chr13-110440286-DEL-1848 | chr13-110440287-DEL-1848 |
| chr13-110446019-DEL-6734 | chr13-110446020-DEL-6734 |
| chr13-110474839-DEL-2436 | chr13-110474804-DEL-2466 |
| chr13-110485746-DEL-92 | chr13-110485745-DEL-94 |
| chr13-110505468-DEL-572 | chr13-110505469-DEL-572 |
| chr13-110527719-DEL-132 | chr13-110527720-DEL-132 |
| chr13-110538653-DEL-4335 | chr13-110538654-DEL-4335 |
| chr13-110542879-DEL-60 | chr13-110542880-DEL-60 |
| chr13-110557731-DEL-368 | chr13-110557732-DEL-368 |
| chr13-110560490-DEL-1092 | chr13-110560491-DEL-1092 |
| chr13-110574922-DEL-1412 | chr13-110574923-DEL-1412 |
| chr13-110585657-DEL-53 | chr13-110585639-DEL-54 |
| chr13-110653367-DEL-139 | chr13-110653368-DEL-137 |
| chr13-110657381-DEL-400 | chr13-110657382-DEL-400 |
| chr13-110666445-DEL-201 | chr13-110666445-DEL-202 |
| chr13-110727492-DEL-55 | chr13-110727500-DEL-54 |
| chr13-110737617-DEL-60 | chr13-110737614-DEL-52 |
| chr13-110740889-DEL-51 | chr13-110740890-DEL-51 |
| chr13-110745846-DEL-112 | chr13-110745853-DEL-100 |
| chr13-110746616-DEL-6213 | chr13-110746649-DEL-6180 |
| chr13-110756375-DEL-160 | chr13-110756376-DEL-160 |
| chr13-110776127-DEL-131 | chr13-110776128-DEL-131 |
| chr13-110782047-DEL-118 | chr13-110782051-DEL-134 |
| chr13-110787059-DEL-66 | chr13-110787064-DEL-64 |
| chr13-110795332-DEL-130 | chr13-110795333-DEL-130 |
| chr13-110811311-DEL-295 | chr13-110811340-DEL-284 |
| chr13-110839326-DEL-57 | chr13-110839327-DEL-57 |
| chr13-110848484-DEL-79 | chr13-110848502-DEL-57 |
| chr13-110858352-DEL-78 | chr13-110858353-DEL-78 |
| chr13-110866067-DEL-2795 | chr13-110866068-DEL-2795 |
| chr13-110885255-DEL-219 | chr13-110885256-DEL-237 |
| chr13-110932390-DEL-91 | chr13-110932389-DEL-92 |
| chr13-110936156-DEL-74 | chr13-110936155-DEL-74 |
| chr13-110944532-DEL-54 | chr13-110944533-DEL-54 |
| chr13-111003187-DEL-71 | chr13-111003179-DEL-72 |
| chr13-111010310-DEL-74 | chr13-111010311-DEL-74 |
| chr13-111056525-DEL-615 | chr13-111056526-DEL-615 |
| chr13-111091009-DEL-116 | chr13-111091019-DEL-120 |
| chr13-111115615-DEL-104 | chr13-111115612-DEL-74 |

| | |
|---|---|
| chr13-111119445-DEL-69 | chr13-111119445-DEL-69 |
| chr13-111130885-DEL-72 | chr13-111130886-DEL-72 |
| chr13-111134886-DEL-154 | chr13-111134884-DEL-154 |
| chr13-111184415-DEL-218 | chr13-111184414-DEL-219 |
| chr13-111187981-DEL-192 | chr13-111187982-DEL-192 |
| chr13-111198511-DEL-12573 | chr13-111198511-DEL-12573 |
| chr13-111204264-DEL-5450 | chr13-111204264-DEL-5450 |
| chr13-111213611-DEL-539 | chr13-111213612-DEL-539 |
| chr13-111225771-DEL-4118 | chr13-111225770-DEL-4119 |
| chr13-111231263-DEL-514 | chr13-111231264-DEL-514 |
| chr13-111280692-DEL-638 | chr13-111280694-DEL-633 |
| chr13-111311401-DEL-273 | chr13-111311402-DEL-273 |
| chr13-111320151-DEL-78 | chr13-111320152-DEL-78 |
| chr13-111324143-DEL-2055 | chr13-111324144-DEL-2055 |
| chr13-111331518-DEL-54 | chr13-111331519-DEL-54 |
| chr13-111337470-DEL-342 | chr13-111337471-DEL-342 |
| chr13-111344382-DEL-990 | chr13-111344351-DEL-991 |
| chr13-111356518-DEL-4435 | chr13-111356519-DEL-4435 |
| chr13-111367805-DEL-81 | chr13-111367806-DEL-81 |
| chr13-111374375-DEL-62 | chr13-111374375-DEL-63 |
| chr13-111383012-DEL-59 | chr13-111383013-DEL-58 |
| chr13-111415388-DEL-58 | chr13-111415386-DEL-58 |
| chr13-111424893-DEL-1960 | chr13-111424894-DEL-1960 |
| chr13-111446027-DEL-72 | chr13-111445999-DEL-57 |
| chr13-111456330-DEL-60 | chr13-111456331-DEL-60 |
| chr13-111458400-DEL-328 | chr13-111458401-DEL-328 |
| chr13-111467902-DEL-331 | chr13-111467901-DEL-332 |
| chr13-111475530-DEL-313 | chr13-111475531-DEL-313 |
| chr13-111485951-DEL-393 | chr13-111485952-DEL-393 |
| chr13-111487270-DEL-1289 | chr13-111487271-DEL-1289 |
| chr13-111491912-DEL-579 | chr13-111491913-DEL-579 |
| chr13-111510939-DEL-206 | chr13-111510954-DEL-192 |
| chr13-111522367-DEL-117 | chr13-111522366-DEL-107 |
| chr13-111523962-DEL-51 | chr13-111523947-DEL-52 |
| chr13-111524968-DEL-120 | chr13-111524969-DEL-120 |
| chr13-111536895-DEL-2352 | chr13-111536893-DEL-2353 |
| chr13-111544793-DEL-82 | chr13-111544798-DEL-78 |
| chr13-111558061-DEL-53 | chr13-111558062-DEL-53 |
| chr13-111577709-DEL-100 | chr13-111577703-DEL-101 |
| chr13-111594048-DEL-314 | chr13-111594050-DEL-312 |
| chr13-111636181-DEL-6986 | chr13-111636182-DEL-6986 |
| chr13-111650472-DEL-1896 | chr13-111650433-DEL-1936 |
| chr13-111675073-DEL-127 | chr13-111675074-DEL-127 |
| chr13-111679475-DEL-86 | chr13-111679436-DEL-86 |
| chr13-111687919-DEL-102 | chr13-111687926-DEL-96 |
| chr13-111700152-DEL-159 | chr13-111700152-DEL-159 |
| chr13-111706515-DEL-353 | chr13-111706516-DEL-353 |
| chr13-111711681-DEL-64 | chr13-111711682-DEL-64 |
| chr13-111715845-DEL-168 | chr13-111715846-DEL-168 |
| chr13-111729371-DEL-79 | chr13-111729375-DEL-74 |
| chr13-111729780-DEL-406 | chr13-111729781-DEL-406 |
| chr13-111731943-DEL-206 | chr13-111731944-DEL-206 |
| chr13-111733167-DEL-85 | chr13-111733168-DEL-85 |
| chr13-111776108-DEL-200 | chr13-111776122-DEL-207 |
| chr13-111792855-DEL-59 | chr13-111792857-DEL-58 |
| chr13-111818661-DEL-174 | chr13-111818662-DEL-174 |
| chr13-111825320-DEL-142 | chr13-111825322-DEL-141 |
| chr13-111830762-DEL-6594 | chr13-111830763-DEL-6594 |
| chr13-111856596-DEL-82 | chr13-111856597-DEL-82 |
| chr13-111857863-DEL-490 | chr13-111857853-DEL-491 |
| chr13-111865100-DEL-8814 | chr13-111865100-DEL-8815 |
| chr13-111874745-DEL-187 | chr13-111874746-DEL-187 |
| chr13-111879941-DEL-52 | chr13-111879942-DEL-52 |
| chr13-111930322-DEL-228 | chr13-111930323-DEL-228 |
| chr13-111962324-DEL-80 | chr13-111962325-DEL-110 |
| chr13-111976587-DEL-51 | chr13-111976596-DEL-56 |
| chr13-111980115-DEL-170 | chr13-111980106-DEL-171 |
| chr13-112003636-DEL-57 | chr13-112003637-DEL-57 |
| chr13-112018753-DEL-2207 | chr13-112018749-DEL-2205 |
| chr13-112023379-DEL-1305 | chr13-112023380-DEL-1305 |
| chr13-112033814-DEL-129 | chr13-112033859-DEL-108 |
| chr13-112046560-DEL-181 | chr13-112046561-DEL-181 |
| chr13-112059950-DEL-94 | chr13-112059951-DEL-94 |
| chr13-112078571-DEL-643 | chr13-112078572-DEL-643 |
| chr13-112094057-DEL-92 | chr13-112094066-DEL-84 |
| chr13-112099105-DEL-91 | chr13-112099106-DEL-91 |
| chr13-112111573-DEL-51 | chr13-112111574-DEL-51 |
| chr13-112121278-DEL-655 | chr13-112121327-DEL-607 |
| chr13-112128935-DEL-56 | chr13-112128937-DEL-54 |
| chr13-112144050-DEL-91 | chr13-112144008-DEL-126 |

| | |
|---|---|
| chr13-112157762-DEL-86 | chr13-112157763-DEL-86 |
| chr13-112167638-DEL-167 | chr13-112167638-DEL-165 |
| chr13-112211633-DEL-70 | chr13-112211639-DEL-70 |
| chr13-112234979-DEL-84 | chr13-112234980-DEL-84 |
| chr13-112272706-DEL-338 | chr13-112272707-DEL-338 |
| chr13-112274446-DEL-85 | chr13-112274480-DEL-86 |
| chr13-112275331-DEL-1232 | chr13-112275332-DEL-1229 |
| chr13-112278537-DEL-91 | chr13-112278538-DEL-106 |
| chr13-112283888-DEL-2346 | chr13-112283889-DEL-2346 |
| chr13-112287106-DEL-668 | chr13-112287107-DEL-668 |
| chr13-112292823-DEL-163 | chr13-112292824-DEL-163 |
| chr13-112301339-DEL-54 | chr13-112301339-DEL-54 |
| chr13-112307176-DEL-282 | chr13-112307176-DEL-298 |
| chr13-112311812-DEL-375 | chr13-112311813-DEL-375 |
| chr13-112328416-DEL-66 | chr13-112328417-DEL-66 |
| chr13-112342450-DEL-60 | chr13-112342451-DEL-60 |
| chr13-112351074-DEL-604 | chr13-112351075-DEL-604 |
| chr13-112359092-DEL-161 | chr13-112359093-DEL-161 |
| chr13-112361107-DEL-2875 | chr13-112361108-DEL-2875 |
| chr13-112392049-DEL-116 | chr13-112392050-DEL-112 |
| chr13-112415660-DEL-835 | chr13-112415661-DEL-835 |
| chr13-112418213-DEL-440 | chr13-112418214-DEL-440 |
| chr13-112438418-DEL-62 | chr13-112438416-DEL-58 |
| chr13-112440345-DEL-527 | chr13-112440346-DEL-527 |
| chr13-112481627-DEL-54 | chr13-112481628-DEL-54 |
| chr13-112507783-DEL-269 | chr13-112507784-DEL-269 |
| chr13-112515971-DEL-985 | chr13-112515972-DEL-985 |
| chr13-112517097-DEL-72 | chr13-112517098-DEL-72 |
| chr13-112529694-DEL-54 | chr13-112529694-DEL-55 |
| chr13-112537529-DEL-62 | chr13-112537541-DEL-52 |
| chr13-112538137-DEL-565 | chr13-112538138-DEL-565 |
| chr13-112558826-DEL-51 | chr13-112558826-DEL-51 |
| chr13-112564027-DEL-82 | chr13-112564048-DEL-92 |
| chr13-112565450-DEL-64 | chr13-112565468-DEL-78 |
| chr13-112567138-DEL-62 | chr13-112567137-DEL-62 |
| chr13-112568036-DEL-124 | chr13-112568037-DEL-124 |
| chr13-112586973-DEL-76 | chr13-112586972-DEL-76 |
| chr13-112591904-DEL-202 | chr13-112591905-DEL-202 |
| chr13-112594365-DEL-198 | chr13-112594340-DEL-198 |
| chr13-112597228-DEL-68 | chr13-112597229-DEL-68 |
| chr13-112606786-DEL-1154 | chr13-112606787-DEL-1154 |
| chr13-112609407-DEL-150 | chr13-112609424-DEL-100 |
| chr13-112649965-DEL-63 | chr13-112649974-DEL-55 |
| chr13-112690478-DEL-54 | chr13-112690479-DEL-54 |
| chr13-112698944-DEL-54 | chr13-112698948-DEL-52 |
| chr13-112701669-DEL-55 | chr13-112701670-DEL-55 |
| chr13-112730097-DEL-1623 | chr13-112730098-DEL-1623 |
| chr13-112783331-DEL-105 | chr13-112783381-DEL-104 |
| chr13-112810330-DEL-54 | chr13-112810331-DEL-54 |
| chr13-112828832-DEL-200 | chr13-112828821-DEL-212 |
| chr13-112832236-DEL-105 | chr13-112832237-DEL-105 |
| chr13-112848944-DEL-78 | chr13-112848945-DEL-78 |
| chr13-112852765-DEL-211 | chr13-112852766-DEL-211 |
| chr13-112854293-DEL-105 | chr13-112854257-DEL-106 |
| chr13-112929635-DEL-132 | chr13-112929636-DEL-132 |
| chr13-112939104-DEL-71 | chr13-112939107-DEL-69 |
| chr13-112956619-DEL-64 | chr13-112956620-DEL-64 |
| chr13-112960780-DEL-92 | chr13-112960794-DEL-52 |
| chr13-112982799-DEL-364 | chr13-112982800-DEL-364 |
| chr13-113015791-DEL-182 | chr13-113015778-DEL-168 |
| chr13-113018101-DEL-206 | chr13-113018103-DEL-205 |
| chr13-113019464-DEL-52 | chr13-113019464-DEL-68 |
| chr13-113023578-DEL-121 | chr13-113023579-DEL-120 |
| chr13-113029417-DEL-59 | chr13-113029418-DEL-59 |
| chr13-113031088-DEL-64 | chr13-113031089-DEL-64 |
| chr13-113038166-DEL-369 | chr13-113038167-DEL-369 |
| chr13-113058759-DEL-3166 | chr13-113058761-DEL-3165 |
| chr13-113071006-DEL-255 | chr13-113071007-DEL-255 |
| chr13-113072530-DEL-217 | chr13-113072531-DEL-217 |
| chr13-113085934-DEL-59 | chr13-113085935-DEL-59 |
| chr13-113097993-DEL-109 | chr13-113097994-DEL-109 |
| chr13-113121631-DEL-194 | chr13-113121624-DEL-192 |
| chr13-113128599-DEL-243 | chr13-113128621-DEL-213 |
| chr13-113129521-DEL-164 | chr13-113129522-DEL-164 |
| chr13-113131992-DEL-198 | chr13-113131993-DEL-198 |
| chr13-113147797-DEL-259 | chr13-113147795-DEL-258 |
| chr13-113150199-DEL-326 | chr13-113150200-DEL-326 |
| chr13-113156586-DEL-982 | chr13-113156587-DEL-982 |
| chr13-113158442-DEL-326 | chr13-113158443-DEL-326 |
| chr13-113159347-DEL-1825 | chr13-113159364-DEL-1816 |

| | |
|---|---|
| chr13-113162824-DEL-590 | chr13-113162825-DEL-590 |
| chr13-113164873-DEL-57 | chr13-113164876-DEL-57 |
| chr13-113184742-DEL-81 | chr13-113184743-DEL-81 |
| chr13-113191847-DEL-71 | chr13-113191848-DEL-71 |
| chr13-113193304-DEL-209 | chr13-113193284-DEL-210 |
| chr13-113195141-DEL-64 | chr13-113195142-DEL-62 |
| chr13-113204502-DEL-212 | chr13-113204506-DEL-208 |
| chr13-113225461-DEL-76 | chr13-113225460-DEL-74 |
| chr13-113244003-DEL-54 | chr13-113244004-DEL-54 |
| chr13-113260677-DEL-55 | chr13-113260678-DEL-55 |
| chr13-113263747-DEL-56 | chr13-113263742-DEL-58 |
| chr13-113270808-DEL-409 | chr13-113270809-DEL-409 |
| chr13-113274051-DEL-206 | chr13-113274052-DEL-206 |
| chr13-113280969-DEL-184 | chr13-113280970-DEL-184 |
| chr13-113285155-DEL-1000 | chr13-113285156-DEL-1000 |
| chr13-113287695-DEL-211 | chr13-113287696-DEL-211 |
| chr13-113295895-DEL-83 | chr13-113295896-DEL-83 |
| chr13-113298102-DEL-481 | chr13-113298103-DEL-481 |
| chr13-113300448-DEL-72 | chr13-113300449-DEL-72 |
| chr13-113313799-DEL-430 | chr13-113313799-DEL-429 |
| chr13-113373515-DEL-1525 | chr13-113373515-DEL-1525 |
| chr13-113375289-DEL-5998 | chr13-113375290-DEL-5998 |
| chr13-113382195-DEL-2006 | chr13-113382196-DEL-2004 |
| chr13-113402321-DEL-51 | chr13-113402322-DEL-54 |
| chr13-113409079-DEL-13579 | chr13-113409080-DEL-13579 |
| chr13-113433747-DEL-54 | chr13-113433748-DEL-54 |
| chr13-113444456-DEL-3090 | chr13-113444457-DEL-3090 |
| chr13-113450260-DEL-598 | chr13-113450261-DEL-598 |
| chr13-113451840-DEL-64 | chr13-113451841-DEL-64 |
| chr13-113461803-DEL-53 | chr13-113461786-DEL-57 |
| chr13-113471108-DEL-80 | chr13-113471108-DEL-79 |
| chr13-113494728-DEL-400 | chr13-113494729-DEL-400 |
| chr13-113512790-DEL-1158 | chr13-113512791-DEL-1158 |
| chr13-113519993-DEL-552 | chr13-113519994-DEL-552 |
| chr13-113532486-DEL-202 | chr13-113532479-DEL-210 |
| chr13-113542955-DEL-70 | chr13-113542956-DEL-70 |
| chr13-113550722-DEL-68 | chr13-113550722-DEL-68 |
| chr13-113573216-DEL-66 | chr13-113573187-DEL-50 |
| chr13-113593921-DEL-1596 | chr13-113593922-DEL-1596 |
| chr13-113598737-DEL-52 | chr13-113598748-DEL-74 |
| chr13-113606907-DEL-80 | chr13-113606890-DEL-68 |
| chr13-113620527-DEL-70 | chr13-113620528-DEL-70 |
| chr13-113643033-DEL-116 | chr13-113643034-DEL-116 |
| chr13-113648166-DEL-1893 | chr13-113648167-DEL-1893 |
| chr13-113668133-DEL-217 | chr13-113668134-DEL-217 |
| chr13-113694774-DEL-201 | chr13-113694775-DEL-201 |
| chr13-113698316-DEL-4551 | chr13-113698317-DEL-4551 |
| chr13-113759787-DEL-191 | chr13-113759788-DEL-191 |
| chr13-113783002-DEL-104 | chr13-113783003-DEL-104 |
| chr13-113785892-DEL-2852 | chr13-113785893-DEL-2852 |
| chr13-113789339-DEL-168 | chr13-113789344-DEL-167 |
| chr13-113800883-DEL-2623 | chr13-113800883-DEL-2623 |
| chr13-113809156-DEL-216 | chr13-113809155-DEL-216 |
| chr13-113815088-DEL-81 | chr13-113815094-DEL-76 |
| chr13-113818252-DEL-701 | chr13-113818253-DEL-701 |
| chr13-113824390-DEL-80 | chr13-113824391-DEL-80 |
| chr13-113851134-DEL-174 | chr13-113851135-DEL-174 |
| chr13-113854942-DEL-5549 | chr13-113854951-DEL-5545 |
| chr13-113860981-DEL-52 | chr13-113860982-DEL-52 |
| chr13-113871560-DEL-211 | chr13-113871561-DEL-211 |
| chr13-113892666-DEL-72 | chr13-113892667-DEL-96 |
| chr13-113900651-DEL-158 | chr13-113900650-DEL-159 |
| chr13-113903077-DEL-1162 | chr13-113903078-DEL-1162 |
| chr13-113926498-DEL-200 | chr13-113926499-DEL-200 |
| chr13-113927950-DEL-64 | chr13-113927951-DEL-64 |
| chr13-113939673-DEL-4003 | chr13-113939674-DEL-4003 |
| chr13-113994021-DEL-393 | chr13-113994021-DEL-393 |
| chr13-114025806-DEL-83 | chr13-114025807-DEL-83 |
| chr13-114026875-DEL-60 | chr13-114026893-DEL-52 |
| chr13-114041091-DEL-73 | chr13-114041092-DEL-73 |
| chr13-114077239-DEL-90 | chr13-114077240-DEL-54 |
| chr13-114096906-DEL-131 | chr13-114096907-DEL-131 |
| chr13-114097597-DEL-784 | chr13-114097598-DEL-784 |
| chr13-114135886-DEL-214 | chr13-114135887-DEL-214 |
| chr13-114176517-DEL-6439 | chr13-114176518-DEL-6439 |
| chr13-114207267-DEL-217 | chr13-114207268-DEL-217 |
| chr13-114208959-DEL-201 | chr13-114208954-DEL-199 |
| chr13-114259378-DEL-396 | chr13-114259379-DEL-396 |
| chr13-114262257-DEL-372 | chr13-114262258-DEL-372 |
| chr13-114264681-DEL-184 | chr13-114264682-DEL-184 |

| | |
|---|---|
| chr13-114277160-DEL-55 | chr13-114277161-DEL-55 |
| chr13-114291976-DEL-64 | chr13-114291977-DEL-64 |
| chr13-114305770-DEL-63 | chr13-114305769-DEL-64 |
| chr13-114368084-DEL-136 | chr13-114368085-DEL-136 |
| chr13-114380941-DEL-209 | chr13-114380942-DEL-209 |
| chr13-114382797-DEL-71 | chr13-114382782-DEL-80 |
| chr13-114419267-DEL-555 | chr13-114419268-DEL-555 |
| chr13-114494977-DEL-69 | chr13-114494978-DEL-69 |
| chr13-114497681-DEL-188 | chr13-114497682-DEL-188 |
| chr13-114545533-DEL-62 | chr13-114545522-DEL-72 |
| chr13-114553768-DEL-86 | chr13-114553772-DEL-85 |
| chr13-114564281-DEL-277 | chr13-114564234-DEL-313 |
| chr13-114579983-DEL-384 | chr13-114579984-DEL-384 |
| chr13-114601132-DEL-209 | chr13-114601123-DEL-210 |
| chr13-114634750-DEL-81 | chr13-114634751-DEL-81 |
| chr13-114635328-DEL-220 | chr13-114635329-DEL-220 |
| chr13-114675512-DEL-4703 | chr13-114675549-DEL-4660 |
| chr13-114709602-DEL-76 | chr13-114709603-DEL-76 |
| chr13-114726402-DEL-2943 | chr13-114726403-DEL-2943 |
| chr13-114758733-DEL-5287 | chr13-114758734-DEL-5287 |
| chr13-114802532-DEL-2056 | chr13-114802533-DEL-2056 |
| chr13-114807072-DEL-3119 | chr13-114807073-DEL-3119 |
| chr13-114811955-DEL-344 | chr13-114812005-DEL-312 |
| chr13-114813413-DEL-120 | chr13-114813414-DEL-120 |
| chr13-114860131-DEL-990 | chr13-114860131-DEL-987 |
| chr13-114895053-DEL-122 | chr13-114895091-DEL-128 |
| chr13-114914799-DEL-623 | chr13-114914800-DEL-623 |
| chr13-114922815-DEL-80 | chr13-114922816-DEL-84 |
| chr13-114930190-DEL-319 | chr13-114930188-DEL-318 |
| chr13-114935974-DEL-1606 | chr13-114935975-DEL-1606 |
| chr13-114946023-DEL-71 | chr13-114946024-DEL-71 |
| chr13-114946824-DEL-2503 | chr13-114946825-DEL-2503 |
| chr13-114949938-DEL-420 | chr13-114949939-DEL-420 |
| chr13-114956178-DEL-174 | chr13-114956182-DEL-176 |
| chr13-114978321-DEL-731 | chr13-114978322-DEL-731 |
| chr13-114990014-DEL-151 | chr13-114990015-DEL-151 |
| chr13-115047243-DEL-72 | chr13-115047238-DEL-98 |
| chr13-115063046-DEL-337 | chr13-115063046-DEL-335 |
| chr13-115098036-DEL-86 | chr13-115098037-DEL-86 |
| chr13-115176530-DEL-412 | chr13-115176531-DEL-412 |
| chr13-115181429-DEL-218 | chr13-115181430-DEL-218 |
| chr13-115183027-DEL-204 | chr13-115183028-DEL-204 |
| chr13-115199057-DEL-114 | chr13-115199053-DEL-109 |
| chr13-115201334-DEL-53 | chr13-115201335-DEL-53 |
| chr13-115207205-DEL-271 | chr13-115207206-DEL-271 |
| chr13-115229039-DEL-79 | chr13-115229039-DEL-79 |
| chr13-115231857-DEL-350 | chr13-115231858-DEL-350 |
| chr13-115239807-DEL-85 | chr13-115239805-DEL-100 |
| chr13-115304753-DEL-117 | chr13-115304767-DEL-117 |
| chr13-115331237-DEL-65 | chr13-115331238-DEL-65 |
| chr13-115340146-DEL-1909 | chr13-115340145-DEL-1908 |
| chr13-115351631-DEL-52 | chr13-115351623-DEL-76 |
| chr13-115358419-DEL-53 | chr13-115358420-DEL-53 |
| chr13-115361279-DEL-79 | chr13-115361280-DEL-77 |
| chr13-115384161-DEL-69 | chr13-115384201-DEL-76 |
| chr13-115403405-DEL-205 | chr13-115403414-DEL-193 |
| chr13-115411503-DEL-110 | chr13-115411504-DEL-110 |
| chr13-115427553-DEL-246 | chr13-115427554-DEL-246 |
| chr13-115428303-DEL-1870 | chr13-115428304-DEL-1870 |
| chr13-115461083-DEL-62 | chr13-115461085-DEL-63 |
| chr13-115462498-DEL-66 | chr13-115462499-DEL-66 |
| chr13-115471648-DEL-82 | chr13-115471649-DEL-83 |
| chr13-115474314-DEL-432 | chr13-115474315-DEL-432 |
| chr13-115495913-DEL-102 | chr13-115495912-DEL-94 |
| chr13-115502222-DEL-68 | chr13-115502223-DEL-68 |
| chr13-115529691-DEL-72 | chr13-115529692-DEL-72 |
| chr13-115535370-DEL-339 | chr13-115535371-DEL-339 |
| chr13-115643665-DEL-78 | chr13-115643677-DEL-78 |
| chr13-115652908-DEL-100 | chr13-115652907-DEL-94 |
| chr13-115675363-DEL-119 | chr13-115675364-DEL-119 |
| chr13-115690338-DEL-810 | chr13-115690339-DEL-810 |
| chr13-115694258-DEL-204 | chr13-115694259-DEL-204 |
| chr13-115694879-DEL-1791 | chr13-115694880-DEL-1791 |
| chr13-115700537-DEL-62 | chr13-115700537-DEL-62 |
| chr13-115756651-DEL-507 | chr13-115756652-DEL-507 |
| chr13-115758847-DEL-64 | chr13-115758836-DEL-108 |
| chr13-115771163-DEL-65 | chr13-115771164-DEL-65 |
| chr13-115776874-DEL-120 | chr13-115776853-DEL-131 |
| chr13-115798116-DEL-52 | chr13-115798116-DEL-52 |
| chr13-115804276-DEL-70 | chr13-115804263-DEL-75 |

| | |
|---|---|
| chr13-115824643-DEL-96 | chr13-115824644-DEL-96 |
| chr13-115857333-DEL-963 | chr13-115857334-DEL-963 |
| chr13-115898986-DEL-158 | chr13-115898987-DEL-158 |
| chr13-115912230-DEL-54 | chr13-115912207-DEL-94 |
| chr13-115930744-DEL-288 | chr13-115930721-DEL-294 |
| chr13-115966184-DEL-106 | chr13-115966185-DEL-106 |
| chr13-115984241-DEL-400 | chr13-115984242-DEL-400 |
| chr13-115985014-DEL-124 | chr13-115985007-DEL-119 |
| chr13-116013500-DEL-531 | chr13-116013492-DEL-533 |
| chr13-116019024-DEL-255 | chr13-116019023-DEL-256 |
| chr13-116025593-DEL-7464 | chr13-116025592-DEL-7449 |
| chr13-116041816-DEL-86 | chr13-116041817-DEL-86 |
| chr13-116060739-DEL-68 | chr13-116060767-DEL-50 |
| chr13-116062436-DEL-443 | chr13-116062437-DEL-443 |
| chr13-116067288-DEL-52 | chr13-116067281-DEL-75 |
| chr13-116075309-DEL-52 | chr13-116075310-DEL-52 |
| chr13-116078454-DEL-2799 | chr13-116078452-DEL-2800 |
| chr13-116101725-DEL-7102 | chr13-116101722-DEL-7103 |
| chr13-116115640-DEL-939 | chr13-116115641-DEL-939 |
| chr13-116137286-DEL-52 | chr13-116137287-DEL-52 |
| chr13-116176522-DEL-196 | chr13-116176522-DEL-190 |
| chr13-116202767-DEL-1138 | chr13-116202768-DEL-1138 |
| chr13-116208220-DEL-805 | chr13-116208221-DEL-805 |
| chr13-116221128-DEL-328 | chr13-116221149-DEL-285 |
| chr13-116227203-DEL-6466 | chr13-116227204-DEL-6466 |
| chr13-116251956-DEL-590 | chr13-116251957-DEL-590 |
| chr13-116257639-DEL-503 | chr13-116257640-DEL-503 |
| chr13-116262593-DEL-5260 | chr13-116262594-DEL-5260 |
| chr13-116280557-DEL-2364 | chr13-116280558-DEL-2364 |
| chr13-116285710-DEL-647 | chr13-116285711-DEL-647 |
| chr13-116290257-DEL-480 | chr13-116290258-DEL-480 |
| chr13-116319382-DEL-396 | chr13-116319383-DEL-396 |
| chr13-116326329-DEL-51 | chr13-116326330-DEL-51 |
| chr13-116329432-DEL-66 | chr13-116329433-DEL-66 |
| chr13-116355356-DEL-278 | chr13-116355367-DEL-268 |
| chr13-116372510-DEL-89 | chr13-116372512-DEL-86 |
| chr13-116372900-DEL-3664 | chr13-116372901-DEL-3664 |
| chr13-116392083-DEL-56 | chr13-116392090-DEL-57 |
| chr13-116436684-DEL-265 | chr13-116436684-DEL-263 |
| chr13-116480420-DEL-54 | chr13-116480421-DEL-54 |
| chr13-116510948-DEL-78 | chr13-116510949-DEL-78 |
| chr13-116517643-DEL-75 | chr13-116517644-DEL-75 |
| chr13-116518924-DEL-482 | chr13-116518922-DEL-483 |
| chr13-116521507-DEL-192 | chr13-116521508-DEL-192 |
| chr13-116522931-DEL-84 | chr13-116522932-DEL-84 |
| chr13-116542774-DEL-58 | chr13-116542750-DEL-74 |
| chr13-116567857-DEL-1010 | chr13-116567858-DEL-1010 |
| chr13-116569238-DEL-223 | chr13-116569239-DEL-223 |
| chr13-116599393-DEL-251 | chr13-116599394-DEL-251 |
| chr13-116601901-DEL-3517 | chr13-116601902-DEL-3517 |
| chr13-116621743-DEL-92 | chr13-116621742-DEL-90 |
| chr13-116637382-DEL-1299 | chr13-116637383-DEL-1299 |
| chr13-116689897-DEL-66 | chr13-116689898-DEL-66 |
| chr13-116703655-DEL-255 | chr13-116703654-DEL-256 |
| chr13-116714579-DEL-7147 | chr13-116714580-DEL-7147 |
| chr13-116740878-DEL-55 | chr13-116740879-DEL-55 |
| chr13-116745740-DEL-248 | chr13-116745741-DEL-247 |
| chr13-116754041-DEL-232 | chr13-116754039-DEL-233 |
| chr13-116760164-DEL-124 | chr13-116760165-DEL-124 |
| chr13-116768613-DEL-64 | chr13-116768613-DEL-64 |
| chr13-116798799-DEL-115 | chr13-116798836-DEL-119 |
| chr13-116801099-DEL-440 | chr13-116801100-DEL-440 |
| chr13-116802370-DEL-78 | chr13-116802371-DEL-54 |
| chr13-116804624-DEL-58 | chr13-116804625-DEL-58 |
| chr13-116820133-DEL-369 | chr13-116820134-DEL-369 |
| chr13-116832480-DEL-109 | chr13-116832482-DEL-85 |
| chr13-116898083-DEL-210 | chr13-116898084-DEL-210 |
| chr13-116904425-DEL-6080 | chr13-116904426-DEL-6080 |
| chr13-116906700-DEL-104 | chr13-116906717-DEL-97 |
| chr13-116907587-DEL-2289 | chr13-116907588-DEL-2289 |
| chr13-116919507-DEL-60 | chr13-116919498-DEL-65 |
| chr13-116949442-DEL-268 | chr13-116949441-DEL-256 |
| chr13-116973756-DEL-6186 | chr13-116973741-DEL-6200 |
| chr13-117009769-DEL-54 | chr13-117009770-DEL-54 |
| chr13-117009821-DEL-2036 | chr13-117009820-DEL-2037 |
| chr13-117020199-DEL-85 | chr13-117020200-DEL-85 |
| chr13-117027061-DEL-309 | chr13-117027062-DEL-309 |
| chr13-117055527-DEL-758 | chr13-117055528-DEL-758 |
| chr13-117068441-DEL-345 | chr13-117068446-DEL-340 |
| chr13-117073801-DEL-968 | chr13-117073802-DEL-966 |

| | |
|---|---|
| chr13-117103752-DEL-54 | chr13-117103753-DEL-68 |
| chr13-117126447-DEL-206 | chr13-117126448-DEL-201 |
| chr13-117141112-DEL-55 | chr13-117141113-DEL-55 |
| chr13-117169896-DEL-140 | chr13-117169897-DEL-140 |
| chr13-117174026-DEL-2666 | chr13-117174026-DEL-2666 |
| chr13-117186458-DEL-803 | chr13-117186459-DEL-803 |
| chr13-117190034-DEL-51 | chr13-117190028-DEL-63 |
| chr13-117224050-DEL-80 | chr13-117224051-DEL-80 |
| chr13-117228888-DEL-95 | chr13-117228886-DEL-95 |
| chr13-117250788-DEL-211 | chr13-117250789-DEL-210 |
| chr13-117266831-DEL-3299 | chr13-117266832-DEL-3298 |
| chr13-117288899-DEL-77 | chr13-117288900-DEL-56 |
| chr13-117301849-DEL-8452 | chr13-117301850-DEL-8452 |
| chr13-117311906-DEL-63 | chr13-117311907-DEL-63 |
| chr13-117323262-DEL-70 | chr13-117323263-DEL-70 |
| chr13-117351251-DEL-213 | chr13-117351252-DEL-213 |
| chr13-117373106-DEL-52 | chr13-117373107-DEL-52 |
| chr13-117378214-DEL-206 | chr13-117378215-DEL-206 |
| chr13-117387234-DEL-4383 | chr13-117387235-DEL-4383 |
| chr13-117387725-DEL-1780 | chr13-117387726-DEL-1779 |
| chr13-117406209-DEL-123 | chr13-117406210-DEL-123 |
| chr13-117414072-DEL-116 | chr13-117414073-DEL-116 |
| chr13-117422041-DEL-54 | chr13-117422042-DEL-55 |
| chr13-117432318-DEL-7205 | chr13-117432319-DEL-7205 |
| chr13-117451418-DEL-52 | chr13-117451419-DEL-52 |
| chr13-117463466-DEL-55 | chr13-117463475-DEL-51 |
| chr13-117467477-DEL-589 | chr13-117467478-DEL-589 |
| chr13-117487445-DEL-263 | chr13-117487431-DEL-278 |
| chr13-117491053-DEL-247 | chr13-117491052-DEL-244 |
| chr13-117494412-DEL-843 | chr13-117494413-DEL-843 |
| chr13-117507552-DEL-1229 | chr13-117507553-DEL-1229 |
| chr13-117518101-DEL-200 | chr13-117518101-DEL-200 |
| chr13-117527655-DEL-207 | chr13-117527692-DEL-207 |
| chr13-117536814-DEL-58 | chr13-117536816-DEL-56 |
| chr13-117539220-DEL-55 | chr13-117539221-DEL-55 |
| chr13-117552522-DEL-119 | chr13-117552515-DEL-118 |
| chr13-117621976-DEL-194 | chr13-117621977-DEL-194 |
| chr13-117633276-DEL-58 | chr13-117633273-DEL-62 |
| chr13-117641312-DEL-445 | chr13-117641313-DEL-445 |
| chr13-117651124-DEL-64 | chr13-117651125-DEL-54 |
| chr13-117658482-DEL-108 | chr13-117658483-DEL-108 |
| chr13-117683898-DEL-197 | chr13-117683897-DEL-198 |
| chr13-117715623-DEL-344 | chr13-117715619-DEL-341 |
| chr13-117716134-DEL-101 | chr13-117716135-DEL-101 |
| chr13-117727789-DEL-86 | chr13-117727790-DEL-86 |
| chr13-117758806-DEL-6186 | chr13-117758806-DEL-6186 |
| chr13-117780691-DEL-83 | chr13-117780695-DEL-80 |
| chr13-117801160-DEL-51 | chr13-117801161-DEL-55 |
| chr13-117805587-DEL-77 | chr13-117805585-DEL-78 |
| chr13-117830981-DEL-80 | chr13-117830980-DEL-82 |
| chr13-117837601-DEL-160 | chr13-117837602-DEL-160 |
| chr13-117846610-DEL-446 | chr13-117846611-DEL-446 |
| chr13-117860467-DEL-114 | chr13-117860468-DEL-114 |
| chr13-117863716-DEL-97 | chr13-117863716-DEL-98 |
| chr13-117879851-DEL-71 | chr13-117879856-DEL-67 |
| chr13-117881601-DEL-175 | chr13-117881595-DEL-176 |
| chr13-117890632-DEL-140 | chr13-117890632-DEL-140 |
| chr13-117913748-DEL-415 | chr13-117913735-DEL-414 |
| chr13-117941532-DEL-66 | chr13-117941533-DEL-66 |
| chr13-117953627-DEL-1474 | chr13-117953634-DEL-1479 |
| chr13-117985906-DEL-194 | chr13-117985907-DEL-194 |
| chr13-118003189-DEL-94 | chr13-118003192-DEL-92 |
| chr13-118012218-DEL-61 | chr13-118012219-DEL-61 |
| chr13-118022903-DEL-454 | chr13-118022916-DEL-453 |
| chr13-118045394-DEL-185 | chr13-118045395-DEL-185 |
| chr13-118048359-DEL-284 | chr13-118048360-DEL-284 |
| chr13-118055706-DEL-206 | chr13-118055707-DEL-206 |
| chr13-118069293-DEL-1190 | chr13-118069285-DEL-1191 |
| chr13-118076984-DEL-126 | chr13-118076985-DEL-126 |
| chr13-118082607-DEL-59 | chr13-118082609-DEL-58 |
| chr13-118102522-DEL-334 | chr13-118102523-DEL-334 |
| chr13-118144020-DEL-59 | chr13-118144019-DEL-60 |
| chr13-118145234-DEL-275 | chr13-118145235-DEL-275 |
| chr13-118152448-DEL-122 | chr13-118152461-DEL-119 |
| chr13-118155616-DEL-112 | chr13-118155617-DEL-112 |
| chr13-118167036-DEL-108 | chr13-118167075-DEL-70 |
| chr13-118179411-DEL-391 | chr13-118179412-DEL-390 |
| chr13-118180533-DEL-180 | chr13-118180534-DEL-180 |
| chr13-118182087-DEL-115 | chr13-118182088-DEL-115 |
| chr13-118195198-DEL-90 | chr13-118195205-DEL-84 |

| | |
|---|---|
| chr13-118207046-DEL-3845 | chr13-118207047-DEL-3845 |
| chr13-118213232-DEL-290 | chr13-118213233-DEL-290 |
| chr13-118217998-DEL-125 | chr13-118217999-DEL-125 |
| chr13-118220433-DEL-134 | chr13-118220436-DEL-129 |
| chr13-118235612-DEL-71 | chr13-118235612-DEL-71 |
| chr13-118261361-DEL-506 | chr13-118261362-DEL-506 |
| chr13-118271601-DEL-199 | chr13-118271602-DEL-199 |
| chr13-118313273-DEL-167 | chr13-118313274-DEL-167 |
| chr13-118328695-DEL-131 | chr13-118328714-DEL-123 |
| chr13-118352858-DEL-2050 | chr13-118352859-DEL-2049 |
| chr13-118358782-DEL-522 | chr13-118358783-DEL-522 |
| chr13-118387577-DEL-793 | chr13-118387576-DEL-791 |
| chr13-118396817-DEL-2473 | chr13-118396818-DEL-2473 |
| chr13-118405539-DEL-478 | chr13-118405540-DEL-478 |
| chr13-118409914-DEL-1575 | chr13-118409915-DEL-1575 |
| chr13-118429527-DEL-1126 | chr13-118429528-DEL-1126 |
| chr13-118435272-DEL-451 | chr13-118435273-DEL-451 |
| chr13-118453925-DEL-6428 | chr13-118453926-DEL-6428 |
| chr13-118459997-DEL-217 | chr13-118459998-DEL-217 |
| chr13-118469368-DEL-52 | chr13-118469367-DEL-54 |
| chr13-118481777-DEL-1078 | chr13-118481778-DEL-1078 |
| chr13-118489529-DEL-508 | chr13-118489530-DEL-508 |
| chr13-118526568-DEL-138 | chr13-118526569-DEL-138 |
| chr13-118545382-DEL-556 | chr13-118545380-DEL-555 |
| chr13-118586533-DEL-945 | chr13-118586534-DEL-945 |
| chr13-118623242-DEL-51 | chr13-118623243-DEL-51 |
| chr13-118625982-DEL-73 | chr13-118626000-DEL-58 |
| chr13-118636511-DEL-158 | chr13-118636532-DEL-118 |
| chr13-118658878-DEL-6432 | chr13-118658865-DEL-6433 |
| chr13-118669451-DEL-52 | chr13-118669452-DEL-52 |
| chr13-118679875-DEL-1379 | chr13-118679876-DEL-1379 |
| chr13-118697558-DEL-2835 | chr13-118697559-DEL-2835 |
| chr13-118747997-DEL-1726 | chr13-118747998-DEL-1726 |
| chr13-118749833-DEL-309 | chr13-118749832-DEL-310 |
| chr13-118765565-DEL-220 | chr13-118765566-DEL-220 |
| chr13-118788397-DEL-400 | chr13-118788398-DEL-400 |
| chr13-118794356-DEL-192 | chr13-118794356-DEL-195 |
| chr13-118802993-DEL-68 | chr13-118802994-DEL-68 |
| chr13-118816766-DEL-60 | chr13-118816767-DEL-60 |
| chr13-118825254-DEL-336 | chr13-118825255-DEL-331 |
| chr13-118865218-DEL-57 | chr13-118865221-DEL-55 |
| chr13-118868713-DEL-58 | chr13-118868714-DEL-58 |
| chr13-118891627-DEL-56 | chr13-118891627-DEL-56 |
| chr13-118914276-DEL-143 | chr13-118914268-DEL-144 |
| chr13-118915217-DEL-55 | chr13-118915218-DEL-55 |
| chr13-118934373-DEL-63 | chr13-118934374-DEL-63 |
| chr13-118935944-DEL-211 | chr13-118935945-DEL-211 |
| chr13-118957133-DEL-135 | chr13-118957134-DEL-143 |
| chr13-118995567-DEL-616 | chr13-118995573-DEL-612 |
| chr13-119012546-DEL-6086 | chr13-119012547-DEL-6086 |
| chr13-119038972-DEL-457 | chr13-119038975-DEL-456 |
| chr13-119039659-DEL-6167 | chr13-119039660-DEL-6167 |
| chr13-119050761-DEL-147 | chr13-119050762-DEL-147 |
| chr13-119052194-DEL-1173 | chr13-119052195-DEL-1173 |
| chr13-119057760-DEL-826 | chr13-119057761-DEL-826 |
| chr13-119070437-DEL-74 | chr13-119070438-DEL-74 |
| chr13-119072128-DEL-1066 | chr13-119072129-DEL-1066 |
| chr13-119079883-DEL-70 | chr13-119079884-DEL-70 |
| chr13-119080373-DEL-90 | chr13-119080376-DEL-88 |
| chr13-119081629-DEL-2068 | chr13-119081630-DEL-2068 |
| chr13-119095135-DEL-603 | chr13-119095134-DEL-609 |
| chr13-119138479-DEL-95 | chr13-119138480-DEL-95 |
| chr13-119141637-DEL-145 | chr13-119141620-DEL-146 |
| chr13-119150918-DEL-68 | chr13-119150919-DEL-68 |
| chr13-119158638-DEL-95 | chr13-119158627-DEL-93 |
| chr13-119160092-DEL-99 | chr13-119160081-DEL-94 |
| chr13-119182371-DEL-385 | chr13-119182362-DEL-386 |
| chr13-119209821-DEL-58 | chr13-119209822-DEL-58 |
| chr13-119234631-DEL-6343 | chr13-119234632-DEL-6343 |
| chr13-119259029-DEL-287 | chr13-119259030-DEL-287 |
| chr13-119261490-DEL-435 | chr13-119261491-DEL-435 |
| chr13-119264853-DEL-7086 | chr13-119264854-DEL-7086 |
| chr13-119293915-DEL-59 | chr13-119293916-DEL-59 |
| chr13-119309276-DEL-96 | chr13-119309277-DEL-80 |
| chr13-119313524-DEL-60 | chr13-119313516-DEL-56 |
| chr13-119324708-DEL-208 | chr13-119324709-DEL-208 |
| chr13-119366170-DEL-888 | chr13-119366171-DEL-888 |
| chr13-119395177-DEL-72 | chr13-119395178-DEL-72 |
| chr13-119403319-DEL-88 | chr13-119403316-DEL-92 |
| chr13-119425066-DEL-1119 | chr13-119425066-DEL-1119 |

| | |
|---|---|
| chr13-119427486-DEL-907 | chr13-119427485-DEL-908 |
| chr13-119438331-DEL-118 | chr13-119438332-DEL-118 |
| chr13-119512963-DEL-87 | chr13-119512964-DEL-89 |
| chr13-119516585-DEL-212 | chr13-119516586-DEL-212 |
| chr13-119546421-DEL-184 | chr13-119546421-DEL-184 |
| chr13-119575922-DEL-100 | chr13-119575923-DEL-100 |
| chr13-119589249-DEL-87 | chr13-119589251-DEL-86 |
| chr13-119603168-DEL-215 | chr13-119603169-DEL-215 |
| chr13-120093253-DEL-70 | chr13-120093254-DEL-70 |
| chr13-120104765-DEL-58 | chr13-120104766-DEL-58 |
| chr13-120135285-DEL-66 | chr13-120135286-DEL-66 |
| chr13-120147772-DEL-175 | chr13-120147773-DEL-175 |
| chr13-120166390-DEL-157 | chr13-120166390-DEL-157 |
| chr13-120182788-DEL-290 | chr13-120182789-DEL-290 |
| chr13-120212293-DEL-238 | chr13-120212294-DEL-238 |
| chr13-120218664-DEL-247 | chr13-120218665-DEL-247 |
| chr13-120219728-DEL-260 | chr13-120219729-DEL-260 |
| chr13-120223785-DEL-1345 | chr13-120223786-DEL-1345 |
| chr13-120246939-DEL-51 | chr13-120246940-DEL-51 |
| chr13-120249513-DEL-208 | chr13-120249514-DEL-208 |
| chr13-120257413-DEL-183 | chr13-120257413-DEL-183 |
| chr13-120261247-DEL-192 | chr13-120261248-DEL-192 |
| chr13-120264508-DEL-54 | chr13-120264535-DEL-50 |
| chr13-120270577-DEL-170 | chr13-120270581-DEL-167 |
| chr13-120298294-DEL-620 | chr13-120298294-DEL-620 |
| chr13-120343416-DEL-332 | chr13-120343412-DEL-336 |
| chr13-120384936-DEL-101 | chr13-120384937-DEL-101 |
| chr13-120397196-DEL-322 | chr13-120397196-DEL-322 |
| chr13-120480522-DEL-213 | chr13-120480523-DEL-213 |
| chr13-120481134-DEL-213 | chr13-120481135-DEL-213 |
| chr13-120497310-DEL-84 | chr13-120497296-DEL-80 |
| chr13-120501092-DEL-211 | chr13-120501092-DEL-211 |
| chr13-120515276-DEL-236 | chr13-120515277-DEL-236 |
| chr13-120520354-DEL-448 | chr13-120520355-DEL-448 |
| chr13-120644600-DEL-490 | chr13-120644600-DEL-490 |
| chr13-120714561-DEL-101 | chr13-120714562-DEL-101 |
| chr14-3055982-DEL-82 | chr14-3055995-DEL-82 |
| chr14-3093848-DEL-1125 | chr14-3093847-DEL-1123 |
| chr14-3127924-DEL-13549 | chr14-3127925-DEL-13549 |
| chr14-3165379-DEL-214 | chr14-3165380-DEL-214 |
| chr14-3213903-DEL-3985 | chr14-3213904-DEL-3985 |
| chr14-3222141-DEL-53 | chr14-3222142-DEL-53 |
| chr14-3233437-DEL-198 | chr14-3233438-DEL-198 |
| chr14-3238886-DEL-55 | chr14-3238887-DEL-52 |
| chr14-3239780-DEL-84 | chr14-3239753-DEL-85 |
| chr14-3251042-DEL-490 | chr14-3251043-DEL-490 |
| chr14-3292938-DEL-1322 | chr14-3292939-DEL-1322 |
| chr14-3309024-DEL-328 | chr14-3309026-DEL-327 |
| chr14-3320368-DEL-619 | chr14-3320372-DEL-618 |
| chr14-3327469-DEL-1628 | chr14-3327470-DEL-1628 |
| chr14-3378633-DEL-341 | chr14-3378634-DEL-341 |
| chr14-3414546-DEL-54 | chr14-3414547-DEL-54 |
| chr14-3456675-DEL-7152 | chr14-3456676-DEL-7152 |
| chr14-3537344-DEL-6050 | chr14-3537345-DEL-6050 |
| chr14-3603084-DEL-7086 | chr14-3603085-DEL-7086 |
| chr14-3625544-DEL-752 | chr14-3625545-DEL-752 |
| chr14-3655843-DEL-614 | chr14-3655844-DEL-614 |
| chr14-3661504-DEL-835 | chr14-3661505-DEL-835 |
| chr14-3684290-DEL-3790 | chr14-3684291-DEL-3790 |
| chr14-3728328-DEL-1220 | chr14-3728329-DEL-1220 |
| chr14-3746805-DEL-324 | chr14-3746806-DEL-324 |
| chr14-3756048-DEL-74 | chr14-3756067-DEL-50 |
| chr14-3761317-DEL-695 | chr14-3761317-DEL-695 |
| chr14-3775329-DEL-1892 | chr14-3775330-DEL-1892 |
| chr14-3788975-DEL-802 | chr14-3788975-DEL-801 |
| chr14-3806381-DEL-845 | chr14-3806382-DEL-845 |
| chr14-3837559-DEL-56 | chr14-3837560-DEL-56 |
| chr14-3881513-DEL-207 | chr14-3881514-DEL-207 |
| chr14-3894083-DEL-713 | chr14-3894080-DEL-714 |
| chr14-3921353-DEL-3828 | chr14-3921355-DEL-3827 |
| chr14-3922592-DEL-529 | chr14-3922593-DEL-529 |
| chr14-3932322-DEL-198 | chr14-3932323-DEL-198 |
| chr14-3934475-DEL-434 | chr14-3934473-DEL-435 |
| chr14-3948106-DEL-201 | chr14-3948080-DEL-201 |
| chr14-3968025-DEL-56 | chr14-3968025-DEL-56 |
| chr14-4020080-DEL-384 | chr14-4020081-DEL-384 |
| chr14-4029349-DEL-2715 | chr14-4029350-DEL-2715 |
| chr14-4033030-DEL-6191 | chr14-4033031-DEL-6191 |
| chr14-4063619-DEL-56 | chr14-4063612-DEL-62 |
| chr14-4088009-DEL-206 | chr14-4088000-DEL-211 |

| | |
|---|---|
| chr14-4092503-DEL-67 | chr14-4092504-DEL-67 |
| chr14-4124567-DEL-136 | chr14-4124556-DEL-148 |
| chr14-4140078-DEL-72 | chr14-4140078-DEL-72 |
| chr14-4170788-DEL-3508 | chr14-4170789-DEL-3508 |
| chr14-4173229-DEL-1481 | chr14-4173230-DEL-1481 |
| chr14-4194217-DEL-206 | chr14-4194218-DEL-206 |
| chr14-4196502-DEL-470 | chr14-4196503-DEL-470 |
| chr14-4211677-DEL-3971 | chr14-4211678-DEL-3971 |
| chr14-4217253-DEL-1153 | chr14-4217254-DEL-1153 |
| chr14-4222422-DEL-60 | chr14-4222423-DEL-60 |
| chr14-4228255-DEL-60 | chr14-4228256-DEL-60 |
| chr14-4272807-DEL-4309 | chr14-4272808-DEL-4309 |
| chr14-4272917-DEL-200 | chr14-4272918-DEL-200 |
| chr14-4284630-DEL-5693 | chr14-4284631-DEL-5693 |
| chr14-4322866-DEL-7109 | chr14-4322867-DEL-7109 |
| chr14-4352937-DEL-2331 | chr14-4352938-DEL-2331 |
| chr14-4397385-DEL-1682 | chr14-4397384-DEL-1682 |
| chr14-4399090-DEL-1222 | chr14-4399089-DEL-1222 |
| chr14-4411894-DEL-209 | chr14-4411895-DEL-209 |
| chr14-4424222-DEL-538 | chr14-4424223-DEL-536 |
| chr14-4470989-DEL-4268 | chr14-4470990-DEL-4268 |
| chr14-4504965-DEL-56 | chr14-4504966-DEL-56 |
| chr14-4546111-DEL-79 | chr14-4546112-DEL-79 |
| chr14-4547831-DEL-117 | chr14-4547832-DEL-117 |
| chr14-4558882-DEL-360 | chr14-4558883-DEL-360 |
| chr14-4603215-DEL-54 | chr14-4603218-DEL-52 |
| chr14-4678360-DEL-75 | chr14-4678360-DEL-75 |
| chr14-4687912-DEL-272 | chr14-4687882-DEL-273 |
| chr14-4709547-DEL-102 | chr14-4709548-DEL-102 |
| chr14-4730551-DEL-1514 | chr14-4730552-DEL-1514 |
| chr14-4747493-DEL-705 | chr14-4747494-DEL-705 |
| chr14-4789563-DEL-402 | chr14-4789558-DEL-403 |
| chr14-4823027-DEL-367 | chr14-4823028-DEL-367 |
| chr14-4844019-DEL-55 | chr14-4844016-DEL-58 |
| chr14-4852047-DEL-54 | chr14-4852048-DEL-54 |
| chr14-4854129-DEL-111 | chr14-4854098-DEL-104 |
| chr14-4871219-DEL-4272 | chr14-4871220-DEL-4272 |
| chr14-4912695-DEL-99 | chr14-4912648-DEL-99 |
| chr14-4918836-DEL-757 | chr14-4918837-DEL-757 |
| chr14-4922616-DEL-1815 | chr14-4922617-DEL-1815 |
| chr14-4931973-DEL-717 | chr14-4931974-DEL-717 |
| chr14-4976162-DEL-63 | chr14-4976163-DEL-63 |
| chr14-5038999-DEL-420 | chr14-5039000-DEL-420 |
| chr14-5055476-DEL-68 | chr14-5055460-DEL-85 |
| chr14-5080988-DEL-116 | chr14-5080989-DEL-116 |
| chr14-5087662-DEL-3845 | chr14-5087663-DEL-3845 |
| chr14-5099303-DEL-1937 | chr14-5099304-DEL-1937 |
| chr14-5101895-DEL-1896 | chr14-5101896-DEL-1896 |
| chr14-5117131-DEL-100 | chr14-5117132-DEL-100 |
| chr14-5135231-DEL-55 | chr14-5135228-DEL-59 |
| chr14-5143466-DEL-308 | chr14-5143470-DEL-305 |
| chr14-5148553-DEL-6219 | chr14-5148554-DEL-6219 |
| chr14-5164428-DEL-243 | chr14-5164429-DEL-242 |
| chr14-5225526-DEL-6201 | chr14-5225527-DEL-6201 |
| chr14-5246874-DEL-355 | chr14-5246882-DEL-354 |
| chr14-5263666-DEL-89 | chr14-5263667-DEL-89 |
| chr14-5269179-DEL-184 | chr14-5269180-DEL-184 |
| chr14-5286829-DEL-3060 | chr14-5286801-DEL-3059 |
| chr14-5316758-DEL-1039 | chr14-5316759-DEL-1039 |
| chr14-5319868-DEL-436 | chr14-5319869-DEL-436 |
| chr14-5326976-DEL-297 | chr14-5326977-DEL-297 |
| chr14-5328722-DEL-7817 | chr14-5328708-DEL-7817 |
| chr14-5354554-DEL-94 | chr14-5354555-DEL-94 |
| chr14-5360644-DEL-99 | chr14-5360645-DEL-99 |
| chr14-5363249-DEL-185 | chr14-5363244-DEL-181 |
| chr14-5367409-DEL-2079 | chr14-5367410-DEL-2079 |
| chr14-5398515-DEL-205 | chr14-5398516-DEL-205 |
| chr14-5426201-DEL-61 | chr14-5426202-DEL-62 |
| chr14-5437528-DEL-6653 | chr14-5437529-DEL-6653 |
| chr14-5457541-DEL-6219 | chr14-5457542-DEL-6219 |
| chr14-5488899-DEL-77 | chr14-5488900-DEL-77 |
| chr14-5489148-DEL-6466 | chr14-5489149-DEL-6466 |
| chr14-5526991-DEL-374 | chr14-5526992-DEL-374 |
| chr14-5528500-DEL-3147 | chr14-5528501-DEL-3147 |
| chr14-5535431-DEL-1942 | chr14-5535432-DEL-1942 |
| chr14-5549631-DEL-110 | chr14-5549650-DEL-92 |
| chr14-5555678-DEL-6290 | chr14-5555679-DEL-6290 |
| chr14-5560682-DEL-77 | chr14-5560685-DEL-71 |
| chr14-5566309-DEL-56 | chr14-5566310-DEL-72 |
| chr14-5629043-DEL-1678 | chr14-5629044-DEL-1678 |

| | |
|---|---|
| chr14-5643626-DEL-66 | chr14-5643627-DEL-66 |
| chr14-5648214-DEL-687 | chr14-5648215-DEL-687 |
| chr14-5657402-DEL-5298 | chr14-5657403-DEL-5298 |
| chr14-5665688-DEL-6351 | chr14-5665689-DEL-6351 |
| chr14-5689801-DEL-1636 | chr14-5689802-DEL-1636 |
| chr14-5693213-DEL-55 | chr14-5693210-DEL-64 |
| chr14-5703489-DEL-1503 | chr14-5703490-DEL-1503 |
| chr14-5705580-DEL-70 | chr14-5705580-DEL-57 |
| chr14-5785593-DEL-66 | chr14-5785591-DEL-67 |
| chr14-5790325-DEL-96 | chr14-5790326-DEL-96 |
| chr14-5799700-DEL-103 | chr14-5799701-DEL-103 |
| chr14-5810507-DEL-6378 | chr14-5810508-DEL-6378 |
| chr14-5839182-DEL-1879 | chr14-5839182-DEL-1879 |
| chr14-5842875-DEL-53 | chr14-5842879-DEL-54 |
| chr14-5844605-DEL-1036 | chr14-5844606-DEL-1036 |
| chr14-5911690-DEL-1018 | chr14-5911691-DEL-1018 |
| chr14-5929520-DEL-1280 | chr14-5929521-DEL-1280 |
| chr14-5933326-DEL-64 | chr14-5933327-DEL-64 |
| chr14-5954893-DEL-63 | chr14-5954887-DEL-63 |
| chr14-5969877-DEL-446 | chr14-5969878-DEL-446 |
| chr14-5973564-DEL-431 | chr14-5973565-DEL-431 |
| chr14-5978945-DEL-120 | chr14-5978970-DEL-96 |
| chr14-5981182-DEL-1944 | chr14-5981182-DEL-1944 |
| chr14-5982628-DEL-387 | chr14-5982628-DEL-386 |
| chr14-5993755-DEL-1150 | chr14-5993756-DEL-1150 |
| chr14-5997290-DEL-1885 | chr14-5997290-DEL-1886 |
| chr14-6016317-DEL-72 | chr14-6016318-DEL-72 |
| chr14-6108260-DEL-408 | chr14-6108261-DEL-408 |
| chr14-6122227-DEL-192 | chr14-6122228-DEL-192 |
| chr14-6124071-DEL-292 | chr14-6124055-DEL-293 |
| chr14-6124968-DEL-444 | chr14-6124969-DEL-444 |
| chr14-6131624-DEL-204 | chr14-6131625-DEL-204 |
| chr14-6137316-DEL-75 | chr14-6137317-DEL-50 |
| chr14-6181556-DEL-693 | chr14-6181557-DEL-693 |
| chr14-6202610-DEL-185 | chr14-6202611-DEL-185 |
| chr14-6210340-DEL-561 | chr14-6210340-DEL-561 |
| chr14-6239710-DEL-76 | chr14-6239710-DEL-76 |
| chr14-6271055-DEL-1565 | chr14-6271058-DEL-1559 |
| chr14-6283507-DEL-68 | chr14-6283508-DEL-68 |
| chr14-6364623-DEL-162 | chr14-6364609-DEL-163 |
| chr14-6381986-DEL-2298 | chr14-6381987-DEL-2298 |
| chr14-6384345-DEL-440 | chr14-6384346-DEL-440 |
| chr14-6388492-DEL-1888 | chr14-6388493-DEL-1888 |
| chr14-6441072-DEL-85 | chr14-6441073-DEL-85 |
| chr14-6445427-DEL-501 | chr14-6445428-DEL-501 |
| chr14-6446520-DEL-6227 | chr14-6446521-DEL-6227 |
| chr14-6454513-DEL-2734 | chr14-6454514-DEL-2734 |
| chr14-6462411-DEL-70 | chr14-6462418-DEL-95 |
| chr14-6489922-DEL-56 | chr14-6489923-DEL-56 |
| chr14-6504904-DEL-4228 | chr14-6504905-DEL-4228 |
| chr14-6511054-DEL-68 | chr14-6511067-DEL-65 |
| chr14-6512368-DEL-194 | chr14-6512369-DEL-194 |
| chr14-6514170-DEL-407 | chr14-6514171-DEL-407 |
| chr14-6517462-DEL-207 | chr14-6517463-DEL-207 |
| chr14-6596321-DEL-443 | chr14-6596322-DEL-443 |
| chr14-6603649-DEL-100 | chr14-6603650-DEL-100 |
| chr14-6608228-DEL-201 | chr14-6608229-DEL-201 |
| chr14-6668479-DEL-53 | chr14-6668480-DEL-53 |
| chr14-6676252-DEL-261 | chr14-6676253-DEL-261 |
| chr14-6685549-DEL-826 | chr14-6685550-DEL-826 |
| chr14-6711238-DEL-60 | chr14-6711239-DEL-60 |
| chr14-6733231-DEL-95 | chr14-6733233-DEL-96 |
| chr14-6741334-DEL-1860 | chr14-6741335-DEL-1860 |
| chr14-6757251-DEL-590 | chr14-6757252-DEL-588 |
| chr14-6776745-DEL-90 | chr14-6776745-DEL-72 |
| chr14-6782616-DEL-76 | chr14-6782617-DEL-76 |
| chr14-6792173-DEL-66 | chr14-6792174-DEL-66 |
| chr14-6801005-DEL-2580 | chr14-6801006-DEL-2580 |
| chr14-6816900-DEL-59 | chr14-6816901-DEL-59 |
| chr14-6824258-DEL-306 | chr14-6824259-DEL-306 |
| chr14-6832180-DEL-157 | chr14-6832181-DEL-157 |
| chr14-6849670-DEL-54 | chr14-6849669-DEL-55 |
| chr14-6854078-DEL-56 | chr14-6854077-DEL-57 |
| chr14-6925942-DEL-6255 | chr14-6925943-DEL-6255 |
| chr14-7001579-DEL-8223 | chr14-7001580-DEL-8223 |
| chr14-7036071-DEL-180 | chr14-7036088-DEL-170 |
| chr14-7081385-DEL-930 | chr14-7081386-DEL-930 |
| chr14-7083663-DEL-52 | chr14-7083663-DEL-52 |
| chr14-7090127-DEL-106 | chr14-7090128-DEL-106 |
| chr14-7137521-DEL-179 | chr14-7137522-DEL-179 |

| | |
|---|---|
| chr14-7140140-DEL-70 | chr14-7140141-DEL-70 |
| chr14-7162640-DEL-142 | chr14-7162641-DEL-142 |
| chr14-7179908-DEL-59 | chr14-7179909-DEL-60 |
| chr14-7207833-DEL-1227 | chr14-7207834-DEL-1227 |
| chr14-7211888-DEL-318 | chr14-7211889-DEL-318 |
| chr14-7218092-DEL-180 | chr14-7218097-DEL-179 |
| chr14-7223159-DEL-56 | chr14-7223160-DEL-56 |
| chr14-7252053-DEL-198 | chr14-7252054-DEL-198 |
| chr14-7272947-DEL-246 | chr14-7272952-DEL-232 |
| chr14-7342572-DEL-6212 | chr14-7342573-DEL-6212 |
| chr14-7355688-DEL-431 | chr14-7355689-DEL-431 |
| chr14-7357292-DEL-1458 | chr14-7357290-DEL-1459 |
| chr14-7359377-DEL-56 | chr14-7359352-DEL-56 |
| chr14-7373545-DEL-65 | chr14-7373545-DEL-65 |
| chr14-7375590-DEL-6048 | chr14-7375591-DEL-6048 |
| chr14-7392676-DEL-57 | chr14-7392677-DEL-57 |
| chr14-7397125-DEL-204 | chr14-7397126-DEL-204 |
| chr14-7414472-DEL-143 | chr14-7414473-DEL-147 |
| chr14-7422857-DEL-106 | chr14-7422866-DEL-98 |
| chr14-7449878-DEL-69 | chr14-7449847-DEL-68 |
| chr14-7485545-DEL-147 | chr14-7485546-DEL-147 |
| chr14-7496787-DEL-201 | chr14-7496788-DEL-201 |
| chr14-7520854-DEL-223 | chr14-7520855-DEL-223 |
| chr14-7524512-DEL-102 | chr14-7524513-DEL-102 |
| chr14-7539345-DEL-103 | chr14-7539346-DEL-103 |
| chr14-7583352-DEL-57 | chr14-7583353-DEL-55 |
| chr14-7610087-DEL-99 | chr14-7610088-DEL-99 |
| chr14-7614949-DEL-224 | chr14-7614950-DEL-224 |
| chr14-7654279-DEL-1990 | chr14-7654280-DEL-1990 |
| chr14-7666135-DEL-321 | chr14-7666136-DEL-321 |
| chr14-7726147-DEL-87 | chr14-7726139-DEL-88 |
| chr14-7762686-DEL-59 | chr14-7762669-DEL-64 |
| chr14-7781110-DEL-74 | chr14-7781101-DEL-78 |
| chr14-7789626-DEL-1731 | chr14-7789627-DEL-1731 |
| chr14-7793482-DEL-252 | chr14-7793483-DEL-252 |
| chr14-7799865-DEL-116 | chr14-7799866-DEL-116 |
| chr14-7820303-DEL-666 | chr14-7820304-DEL-666 |
| chr14-7841868-DEL-6648 | chr14-7841869-DEL-6648 |
| chr14-7904532-DEL-1435 | chr14-7904533-DEL-1435 |
| chr14-7905764-DEL-1444 | chr14-7905764-DEL-1444 |
| chr14-7909985-DEL-72 | chr14-7909985-DEL-72 |
| chr14-7947358-DEL-54 | chr14-7947358-DEL-54 |
| chr14-7964523-DEL-87 | chr14-7964512-DEL-99 |
| chr14-8034944-DEL-6593 | chr14-8034945-DEL-6593 |
| chr14-8129505-DEL-7150 | chr14-8129506-DEL-7150 |
| chr14-8142274-DEL-118 | chr14-8142275-DEL-118 |
| chr14-8148451-DEL-205 | chr14-8148448-DEL-206 |
| chr14-8177533-DEL-486 | chr14-8177534-DEL-486 |
| chr14-8209790-DEL-5714 | chr14-8209791-DEL-5714 |
| chr14-8255268-DEL-209 | chr14-8255269-DEL-209 |
| chr14-8315083-DEL-103 | chr14-8315084-DEL-103 |
| chr14-8316678-DEL-74 | chr14-8316663-DEL-75 |
| chr14-8319754-DEL-6419 | chr14-8319755-DEL-6419 |
| chr14-8349775-DEL-78 | chr14-8349776-DEL-78 |
| chr14-8370484-DEL-62 | chr14-8370485-DEL-62 |
| chr14-8393065-DEL-1590 | chr14-8393066-DEL-1590 |
| chr14-8411174-DEL-54 | chr14-8411175-DEL-54 |
| chr14-8421136-DEL-1683 | chr14-8421137-DEL-1683 |
| chr14-8437486-DEL-629 | chr14-8437490-DEL-627 |
| chr14-8445540-DEL-61 | chr14-8445532-DEL-62 |
| chr14-8447214-DEL-52 | chr14-8447215-DEL-52 |
| chr14-8464837-DEL-1834 | chr14-8464838-DEL-1834 |
| chr14-8478143-DEL-424 | chr14-8478144-DEL-424 |
| chr14-8478981-DEL-120 | chr14-8478982-DEL-70 |
| chr14-8513636-DEL-132 | chr14-8513637-DEL-132 |
| chr14-8541385-DEL-54 | chr14-8541386-DEL-54 |
| chr14-8546068-DEL-516 | chr14-8546068-DEL-516 |
| chr14-8550512-DEL-211 | chr14-8550513-DEL-212 |
| chr14-8556205-DEL-643 | chr14-8556206-DEL-643 |
| chr14-8558190-DEL-57 | chr14-8558191-DEL-57 |
| chr14-8561419-DEL-204 | chr14-8561420-DEL-204 |
| chr14-8580480-DEL-1304 | chr14-8580480-DEL-1299 |
| chr14-8622887-DEL-216 | chr14-8622888-DEL-216 |
| chr14-8677130-DEL-105 | chr14-8677131-DEL-105 |
| chr14-8690222-DEL-852 | chr14-8690223-DEL-852 |
| chr14-8708414-DEL-5653 | chr14-8708415-DEL-5650 |
| chr14-8715376-DEL-106 | chr14-8715377-DEL-106 |
| chr14-8715796-DEL-4750 | chr14-8715796-DEL-4750 |
| chr14-8738618-DEL-434 | chr14-8738619-DEL-434 |
| chr14-8740930-DEL-1372 | chr14-8740931-DEL-1372 |

| | |
|---|---|
| chr14-8751016-DEL-458 | chr14-8751017-DEL-457 |
| chr14-8769015-DEL-7031 | chr14-8769016-DEL-7031 |
| chr14-8775306-DEL-212 | chr14-8775307-DEL-212 |
| chr14-8780350-DEL-434 | chr14-8780351-DEL-434 |
| chr14-8814751-DEL-401 | chr14-8814752-DEL-401 |
| chr14-8822021-DEL-7491 | chr14-8822022-DEL-7491 |
| chr14-8828153-DEL-411 | chr14-8828154-DEL-411 |
| chr14-8849317-DEL-165 | chr14-8849318-DEL-165 |
| chr14-8871446-DEL-569 | chr14-8871447-DEL-569 |
| chr14-8909932-DEL-201 | chr14-8909929-DEL-202 |
| chr14-8948392-DEL-63 | chr14-8948392-DEL-63 |
| chr14-8951085-DEL-2757 | chr14-8951086-DEL-2757 |
| chr14-8956993-DEL-505 | chr14-8957017-DEL-482 |
| chr14-8958559-DEL-126 | chr14-8958560-DEL-126 |
| chr14-8959643-DEL-53 | chr14-8959644-DEL-53 |
| chr14-8985003-DEL-181 | chr14-8985004-DEL-181 |
| chr14-9033707-DEL-5643 | chr14-9033701-DEL-5644 |
| chr14-9080085-DEL-60 | chr14-9080106-DEL-50 |
| chr14-9139340-DEL-50 | chr14-9139341-DEL-59 |
| chr14-9188656-DEL-316 | chr14-9188657-DEL-316 |
| chr14-9207388-DEL-2394 | chr14-9207389-DEL-2394 |
| chr14-9207965-DEL-141 | chr14-9207966-DEL-141 |
| chr14-9210532-DEL-110 | chr14-9210533-DEL-110 |
| chr14-9244144-DEL-235 | chr14-9244145-DEL-235 |
| chr14-9248624-DEL-203 | chr14-9248625-DEL-203 |
| chr14-9278081-DEL-427 | chr14-9278082-DEL-427 |
| chr14-9325214-DEL-334 | chr14-9325214-DEL-334 |
| chr14-9329268-DEL-75 | chr14-9329269-DEL-54 |
| chr14-9329832-DEL-751 | chr14-9329833-DEL-751 |
| chr14-9330723-DEL-148 | chr14-9330724-DEL-148 |
| chr14-9335281-DEL-2556 | chr14-9335282-DEL-2556 |
| chr14-9346573-DEL-6478 | chr14-9346574-DEL-6478 |
| chr14-9352711-DEL-214 | chr14-9352712-DEL-214 |
| chr14-9358767-DEL-206 | chr14-9358750-DEL-200 |
| chr14-9363761-DEL-504 | chr14-9363761-DEL-504 |
| chr14-9384914-DEL-391 | chr14-9384915-DEL-391 |
| chr14-9392487-DEL-448 | chr14-9392488-DEL-448 |
| chr14-9394745-DEL-219 | chr14-9394746-DEL-219 |
| chr14-9411866-DEL-2358 | chr14-9411867-DEL-2358 |
| chr14-9437896-DEL-55 | chr14-9437899-DEL-55 |
| chr14-9495542-DEL-124 | chr14-9495519-DEL-132 |
| chr14-9497702-DEL-60 | chr14-9497699-DEL-63 |
| chr14-9527157-DEL-211 | chr14-9527164-DEL-195 |
| chr14-9557924-DEL-381 | chr14-9557925-DEL-381 |
| chr14-9560653-DEL-66 | chr14-9560653-DEL-66 |
| chr14-9582559-DEL-198 | chr14-9582560-DEL-198 |
| chr14-9615613-DEL-62 | chr14-9615614-DEL-62 |
| chr14-9640085-DEL-2725 | chr14-9640086-DEL-2725 |
| chr14-9677346-DEL-56 | chr14-9677347-DEL-56 |
| chr14-9719254-DEL-198 | chr14-9719248-DEL-199 |
| chr14-9721940-DEL-324 | chr14-9721937-DEL-312 |
| chr14-9722750-DEL-240 | chr14-9722751-DEL-240 |
| chr14-9740205-DEL-864 | chr14-9740206-DEL-868 |
| chr14-9741464-DEL-88 | chr14-9741465-DEL-88 |
| chr14-9745974-DEL-116 | chr14-9745954-DEL-117 |
| chr14-9807701-DEL-82 | chr14-9807666-DEL-70 |
| chr14-9827747-DEL-72 | chr14-9827749-DEL-71 |
| chr14-9846655-DEL-2026 | chr14-9846656-DEL-2026 |
| chr14-9849839-DEL-2777 | chr14-9849840-DEL-2777 |
| chr14-9853145-DEL-60 | chr14-9853141-DEL-55 |
| chr14-9881638-DEL-335 | chr14-9881639-DEL-335 |
| chr14-9907745-DEL-168 | chr14-9907746-DEL-168 |
| chr14-9925889-DEL-87 | chr14-9925890-DEL-88 |
| chr14-9937437-DEL-204 | chr14-9937438-DEL-204 |
| chr14-9972402-DEL-62 | chr14-9972413-DEL-52 |
| chr14-9974372-DEL-55 | chr14-9974373-DEL-50 |
| chr14-10015844-DEL-494 | chr14-10015845-DEL-493 |
| chr14-10024928-DEL-58 | chr14-10024925-DEL-50 |
| chr14-10027353-DEL-60 | chr14-10027354-DEL-60 |
| chr14-10037138-DEL-77 | chr14-10037140-DEL-75 |
| chr14-10143498-DEL-54 | chr14-10143499-DEL-54 |
| chr14-10179650-DEL-177 | chr14-10179651-DEL-177 |
| chr14-10209619-DEL-13033 | chr14-10209606-DEL-13046 |
| chr14-10229142-DEL-221 | chr14-10229143-DEL-221 |
| chr14-10233851-DEL-223 | chr14-10233852-DEL-223 |
| chr14-10251401-DEL-52 | chr14-10251402-DEL-50 |
| chr14-10257725-DEL-297 | chr14-10257726-DEL-297 |
| chr14-10304696-DEL-446 | chr14-10304697-DEL-446 |
| chr14-10309618-DEL-207 | chr14-10309618-DEL-207 |
| chr14-10318453-DEL-124 | chr14-10318438-DEL-122 |

| | |
|---|---|
| chr14-10351162-DEL-181 | chr14-10351163-DEL-181 |
| chr14-10372641-DEL-504 | chr14-10372642-DEL-504 |
| chr14-10375142-DEL-60 | chr14-10375142-DEL-59 |
| chr14-10381967-DEL-399 | chr14-10381968-DEL-399 |
| chr14-10395740-DEL-87 | chr14-10395741-DEL-87 |
| chr14-10402603-DEL-51 | chr14-10402604-DEL-51 |
| chr14-10407375-DEL-854 | chr14-10407376-DEL-854 |
| chr14-10475334-DEL-202 | chr14-10475335-DEL-202 |
| chr14-10483163-DEL-75 | chr14-10483164-DEL-75 |
| chr14-10500984-DEL-100 | chr14-10500968-DEL-78 |
| chr14-10505318-DEL-202 | chr14-10505319-DEL-202 |
| chr14-10525718-DEL-207 | chr14-10525719-DEL-207 |
| chr14-10530928-DEL-69 | chr14-10530935-DEL-63 |
| chr14-10542792-DEL-1389 | chr14-10542793-DEL-1389 |
| chr14-10564990-DEL-1227 | chr14-10564991-DEL-1227 |
| chr14-10571885-DEL-385 | chr14-10571886-DEL-385 |
| chr14-10598661-DEL-58 | chr14-10598662-DEL-58 |
| chr14-10615009-DEL-441 | chr14-10615011-DEL-439 |
| chr14-10640230-DEL-53 | chr14-10640231-DEL-53 |
| chr14-10642655-DEL-1141 | chr14-10642656-DEL-1141 |
| chr14-10664704-DEL-52 | chr14-10664688-DEL-63 |
| chr14-10681712-DEL-116 | chr14-10681713-DEL-116 |
| chr14-10688694-DEL-80 | chr14-10688695-DEL-80 |
| chr14-10692457-DEL-78 | chr14-10692458-DEL-78 |
| chr14-10738575-DEL-94 | chr14-10738576-DEL-94 |
| chr14-10757664-DEL-214 | chr14-10757680-DEL-187 |
| chr14-10795136-DEL-432 | chr14-10795137-DEL-432 |
| chr14-10800759-DEL-232 | chr14-10800760-DEL-232 |
| chr14-10808672-DEL-169 | chr14-10808673-DEL-169 |
| chr14-10836087-DEL-60 | chr14-10836087-DEL-60 |
| chr14-10852652-DEL-135 | chr14-10852653-DEL-135 |
| chr14-10885974-DEL-162 | chr14-10885975-DEL-162 |
| chr14-10899427-DEL-116 | chr14-10899430-DEL-114 |
| chr14-10910099-DEL-54 | chr14-10910100-DEL-54 |
| chr14-10939606-DEL-204 | chr14-10939607-DEL-204 |
| chr14-10941884-DEL-197 | chr14-10941885-DEL-197 |
| chr14-10945316-DEL-64 | chr14-10945317-DEL-64 |
| chr14-10970397-DEL-90 | chr14-10970398-DEL-90 |
| chr14-10985021-DEL-117 | chr14-10985013-DEL-126 |
| chr14-11029700-DEL-192 | chr14-11029701-DEL-192 |
| chr14-11047760-DEL-90 | chr14-11047761-DEL-90 |
| chr14-11051601-DEL-964 | chr14-11051602-DEL-964 |
| chr14-11075640-DEL-1208 | chr14-11075641-DEL-1208 |
| chr14-11077280-DEL-205 | chr14-11077283-DEL-206 |
| chr14-11079970-DEL-662 | chr14-11079971-DEL-662 |
| chr14-11082391-DEL-123 | chr14-11082391-DEL-123 |
| chr14-11108308-DEL-54 | chr14-11108309-DEL-54 |
| chr14-11114953-DEL-57 | chr14-11114954-DEL-57 |
| chr14-11141516-DEL-448 | chr14-11141519-DEL-449 |
| chr14-11143400-DEL-2834 | chr14-11143400-DEL-2834 |
| chr14-11153341-DEL-352 | chr14-11153342-DEL-352 |
| chr14-11176815-DEL-182 | chr14-11176816-DEL-182 |
| chr14-11187752-DEL-1294 | chr14-11187752-DEL-1294 |
| chr14-11193739-DEL-60 | chr14-11193738-DEL-60 |
| chr14-11197837-DEL-525 | chr14-11197838-DEL-525 |
| chr14-11201210-DEL-398 | chr14-11201211-DEL-398 |
| chr14-11203799-DEL-70 | chr14-11203799-DEL-50 |
| chr14-11209928-DEL-55 | chr14-11209928-DEL-56 |
| chr14-11219633-DEL-65 | chr14-11219628-DEL-71 |
| chr14-11222505-DEL-53 | chr14-11222506-DEL-53 |
| chr14-11235416-DEL-1893 | chr14-11235417-DEL-1893 |
| chr14-11277856-DEL-358 | chr14-11277857-DEL-358 |
| chr14-11283623-DEL-51 | chr14-11283624-DEL-51 |
| chr14-11290004-DEL-249 | chr14-11290007-DEL-250 |
| chr14-11332356-DEL-210 | chr14-11332347-DEL-220 |
| chr14-11364886-DEL-126 | chr14-11364919-DEL-124 |
| chr14-11382347-DEL-318 | chr14-11382348-DEL-318 |
| chr14-11386273-DEL-80 | chr14-11386273-DEL-78 |
| chr14-11390048-DEL-88 | chr14-11390048-DEL-88 |
| chr14-11395641-DEL-2380 | chr14-11395642-DEL-2380 |
| chr14-11426466-DEL-1347 | chr14-11426467-DEL-1347 |
| chr14-11432074-DEL-52 | chr14-11432075-DEL-52 |
| chr14-11476533-DEL-56 | chr14-11476534-DEL-56 |
| chr14-11490208-DEL-59 | chr14-11490209-DEL-59 |
| chr14-11496086-DEL-54 | chr14-11496087-DEL-54 |
| chr14-11538683-DEL-1031 | chr14-11538684-DEL-1031 |
| chr14-11567900-DEL-1344 | chr14-11567901-DEL-1344 |
| chr14-11574248-DEL-60 | chr14-11574247-DEL-60 |
| chr14-11584777-DEL-264 | chr14-11584820-DEL-222 |
| chr14-11587450-DEL-686 | chr14-11587451-DEL-686 |

| | |
|---|---|
| chr14-11600946-DEL-53 | chr14-11600939-DEL-54 |
| chr14-11617940-DEL-733 | chr14-11617941-DEL-733 |
| chr14-11626022-DEL-80 | chr14-11626023-DEL-80 |
| chr14-11631867-DEL-800 | chr14-11631868-DEL-800 |
| chr14-11648440-DEL-284 | chr14-11648441-DEL-284 |
| chr14-11656965-DEL-70 | chr14-11656966-DEL-70 |
| chr14-11658453-DEL-253 | chr14-11658454-DEL-253 |
| chr14-11672584-DEL-264 | chr14-11672585-DEL-288 |
| chr14-11694075-DEL-1768 | chr14-11694075-DEL-1768 |
| chr14-11710623-DEL-57 | chr14-11710628-DEL-53 |
| chr14-11714550-DEL-81 | chr14-11714545-DEL-88 |
| chr14-11724196-DEL-2781 | chr14-11724197-DEL-2781 |
| chr14-11739408-DEL-1225 | chr14-11739408-DEL-1225 |
| chr14-11747521-DEL-66 | chr14-11747522-DEL-66 |
| chr14-11759579-DEL-593 | chr14-11759580-DEL-593 |
| chr14-11769627-DEL-55 | chr14-11769628-DEL-54 |
| chr14-11784952-DEL-354 | chr14-11784953-DEL-354 |
| chr14-11821116-DEL-52 | chr14-11821117-DEL-52 |
| chr14-11831903-DEL-165 | chr14-11831904-DEL-165 |
| chr14-11849319-DEL-57 | chr14-11849320-DEL-57 |
| chr14-11854867-DEL-6440 | chr14-11854868-DEL-6440 |
| chr14-11911140-DEL-879 | chr14-11911140-DEL-879 |
| chr14-11927793-DEL-646 | chr14-11927794-DEL-646 |
| chr14-11947458-DEL-4980 | chr14-11947459-DEL-4980 |
| chr14-11952128-DEL-64 | chr14-11952129-DEL-64 |
| chr14-11953474-DEL-324 | chr14-11953474-DEL-325 |
| chr14-11956251-DEL-93 | chr14-11956272-DEL-73 |
| chr14-11960631-DEL-64 | chr14-11960632-DEL-64 |
| chr14-11966578-DEL-446 | chr14-11966579-DEL-446 |
| chr14-11968796-DEL-726 | chr14-11968797-DEL-726 |
| chr14-11972486-DEL-7597 | chr14-11972487-DEL-7597 |
| chr14-11972642-DEL-428 | chr14-11972642-DEL-428 |
| chr14-11989850-DEL-203 | chr14-11989802-DEL-204 |
| chr14-12034435-DEL-72 | chr14-12034466-DEL-77 |
| chr14-12043884-DEL-445 | chr14-12043885-DEL-445 |
| chr14-12084346-DEL-774 | chr14-12084347-DEL-774 |
| chr14-12097511-DEL-87 | chr14-12097512-DEL-87 |
| chr14-12117081-DEL-84 | chr14-12117082-DEL-84 |
| chr14-12121495-DEL-57 | chr14-12121496-DEL-57 |
| chr14-12155392-DEL-109 | chr14-12155400-DEL-106 |
| chr14-12178994-DEL-52 | chr14-12178995-DEL-52 |
| chr14-12194483-DEL-392 | chr14-12194484-DEL-392 |
| chr14-12238542-DEL-1085 | chr14-12238543-DEL-1085 |
| chr14-12241275-DEL-110 | chr14-12241276-DEL-110 |
| chr14-12270179-DEL-1890 | chr14-12270180-DEL-1890 |
| chr14-12283914-DEL-7101 | chr14-12283906-DEL-7102 |
| chr14-12300130-DEL-58 | chr14-12300131-DEL-58 |
| chr14-12330096-DEL-814 | chr14-12330097-DEL-814 |
| chr14-12388843-DEL-101 | chr14-12388844-DEL-101 |
| chr14-12418393-DEL-211 | chr14-12418394-DEL-211 |
| chr14-12420561-DEL-96 | chr14-12420546-DEL-116 |
| chr14-12450438-DEL-1057 | chr14-12450439-DEL-1057 |
| chr14-12463640-DEL-424 | chr14-12463641-DEL-424 |
| chr14-12471408-DEL-572 | chr14-12471409-DEL-571 |
| chr14-12477310-DEL-57 | chr14-12477311-DEL-57 |
| chr14-12487062-DEL-2667 | chr14-12487063-DEL-2667 |
| chr14-12500787-DEL-567 | chr14-12500788-DEL-567 |
| chr14-12519228-DEL-2184 | chr14-12519229-DEL-2184 |
| chr14-12570798-DEL-708 | chr14-12570799-DEL-708 |
| chr14-12609291-DEL-6458 | chr14-12609291-DEL-6458 |
| chr14-12664427-DEL-116 | chr14-12664422-DEL-116 |
| chr14-12670917-DEL-424 | chr14-12670918-DEL-424 |
| chr14-12677038-DEL-54 | chr14-12677039-DEL-53 |
| chr14-12684677-DEL-13412 | chr14-12684680-DEL-13410 |
| chr14-12698006-DEL-687 | chr14-12698007-DEL-687 |
| chr14-12715668-DEL-71 | chr14-12715703-DEL-74 |
| chr14-12729051-DEL-354 | chr14-12729041-DEL-346 |
| chr14-12733903-DEL-7090 | chr14-12733882-DEL-7112 |
| chr14-12760242-DEL-60 | chr14-12760243-DEL-60 |
| chr14-12783976-DEL-82 | chr14-12783977-DEL-82 |
| chr14-12807155-DEL-85 | chr14-12807157-DEL-84 |
| chr14-12828453-DEL-131 | chr14-12828447-DEL-132 |
| chr14-12856921-DEL-1894 | chr14-12856922-DEL-1894 |
| chr14-12864365-DEL-182 | chr14-12864366-DEL-182 |
| chr14-12903237-DEL-242 | chr14-12903238-DEL-242 |
| chr14-12941124-DEL-103 | chr14-12941111-DEL-104 |
| chr14-12968035-DEL-234 | chr14-12968036-DEL-234 |
| chr14-12982569-DEL-66 | chr14-12982576-DEL-52 |
| chr14-12998284-DEL-340 | chr14-12998276-DEL-304 |
| chr14-13020423-DEL-257 | chr14-13020424-DEL-257 |

| | |
|---|---|
| chr14-13028068-DEL-54 | chr14-13028069-DEL-54 |
| chr14-13036766-DEL-55 | chr14-13036767-DEL-55 |
| chr14-13038613-DEL-59 | chr14-13038614-DEL-69 |
| chr14-13045898-DEL-8696 | chr14-13045899-DEL-8696 |
| chr14-13082358-DEL-78 | chr14-13082359-DEL-78 |
| chr14-13085971-DEL-52 | chr14-13085972-DEL-52 |
| chr14-13091539-DEL-6524 | chr14-13091525-DEL-6525 |
| chr14-13103393-DEL-217 | chr14-13103394-DEL-217 |
| chr14-13118259-DEL-52 | chr14-13118262-DEL-56 |
| chr14-13135752-DEL-156 | chr14-13135753-DEL-156 |
| chr14-13167274-DEL-56 | chr14-13167275-DEL-56 |
| chr14-13183939-DEL-238 | chr14-13183940-DEL-238 |
| chr14-13203321-DEL-557 | chr14-13203322-DEL-557 |
| chr14-13208785-DEL-2488 | chr14-13208786-DEL-2488 |
| chr14-13226729-DEL-75 | chr14-13226730-DEL-75 |
| chr14-13240317-DEL-92 | chr14-13240317-DEL-95 |
| chr14-13249214-DEL-78 | chr14-13249215-DEL-78 |
| chr14-13254804-DEL-571 | chr14-13254805-DEL-571 |
| chr14-13261931-DEL-202 | chr14-13261932-DEL-202 |
| chr14-13264127-DEL-330 | chr14-13264128-DEL-330 |
| chr14-13267689-DEL-417 | chr14-13267690-DEL-417 |
| chr14-13292236-DEL-1295 | chr14-13292237-DEL-1295 |
| chr14-13294447-DEL-1493 | chr14-13294447-DEL-1493 |
| chr14-13298276-DEL-4318 | chr14-13298277-DEL-4319 |
| chr14-13337168-DEL-868 | chr14-13337169-DEL-868 |
| chr14-13338238-DEL-4349 | chr14-13338239-DEL-4349 |
| chr14-13356637-DEL-2487 | chr14-13356638-DEL-2487 |
| chr14-13366879-DEL-6250 | chr14-13366880-DEL-6250 |
| chr14-13386487-DEL-120 | chr14-13386494-DEL-76 |
| chr14-13417712-DEL-7114 | chr14-13417713-DEL-7114 |
| chr14-13443087-DEL-101 | chr14-13443088-DEL-101 |
| chr14-13489660-DEL-60 | chr14-13489661-DEL-60 |
| chr14-13501404-DEL-328 | chr14-13501405-DEL-328 |
| chr14-13502447-DEL-51 | chr14-13502448-DEL-51 |
| chr14-13525463-DEL-503 | chr14-13525464-DEL-503 |
| chr14-13530866-DEL-58 | chr14-13530866-DEL-58 |
| chr14-13540660-DEL-130 | chr14-13540661-DEL-130 |
| chr14-13578779-DEL-3144 | chr14-13578780-DEL-3144 |
| chr14-13585833-DEL-1300 | chr14-13585834-DEL-1300 |
| chr14-13610169-DEL-1321 | chr14-13610170-DEL-1321 |
| chr14-13623506-DEL-100 | chr14-13623507-DEL-100 |
| chr14-13669239-DEL-1237 | chr14-13669240-DEL-1237 |
| chr14-13685323-DEL-52 | chr14-13685324-DEL-50 |
| chr14-13688380-DEL-50 | chr14-13688365-DEL-66 |
| chr14-13713702-DEL-14096 | chr14-13713703-DEL-14096 |
| chr14-13739503-DEL-514 | chr14-13739504-DEL-514 |
| chr14-13781801-DEL-70 | chr14-13781802-DEL-70 |
| chr14-13789083-DEL-56 | chr14-13789084-DEL-56 |
| chr14-13795615-DEL-1085 | chr14-13795616-DEL-1085 |
| chr14-13804434-DEL-201 | chr14-13804435-DEL-198 |
| chr14-13811007-DEL-7098 | chr14-13811008-DEL-7098 |
| chr14-13871991-DEL-948 | chr14-13871992-DEL-948 |
| chr14-13914352-DEL-1127 | chr14-13914353-DEL-1127 |
| chr14-13922374-DEL-2105 | chr14-13922375-DEL-2105 |
| chr14-13930181-DEL-559 | chr14-13930182-DEL-559 |
| chr14-13947627-DEL-258 | chr14-13947628-DEL-259 |
| chr14-13967791-DEL-56 | chr14-13967792-DEL-56 |
| chr14-13973401-DEL-114 | chr14-13973400-DEL-115 |
| chr14-13982048-DEL-312 | chr14-13982049-DEL-312 |
| chr14-13993244-DEL-75 | chr14-13993244-DEL-76 |
| chr14-14035620-DEL-52 | chr14-14035637-DEL-50 |
| chr14-14040749-DEL-60 | chr14-14040730-DEL-84 |
| chr14-14057036-DEL-56 | chr14-14057037-DEL-56 |
| chr14-14116931-DEL-76 | chr14-14116932-DEL-76 |
| chr14-14147097-DEL-654 | chr14-14147098-DEL-654 |
| chr14-14167438-DEL-72 | chr14-14167439-DEL-72 |
| chr14-14176025-DEL-694 | chr14-14176026-DEL-694 |
| chr14-14189772-DEL-589 | chr14-14189773-DEL-589 |
| chr14-14191260-DEL-54 | chr14-14191261-DEL-54 |
| chr14-14244479-DEL-88 | chr14-14244480-DEL-88 |
| chr14-14289962-DEL-2828 | chr14-14289962-DEL-2828 |
| chr14-14293893-DEL-52 | chr14-14293894-DEL-52 |
| chr14-14295682-DEL-212 | chr14-14295683-DEL-212 |
| chr14-14313716-DEL-174 | chr14-14313717-DEL-174 |
| chr14-14339250-DEL-79 | chr14-14339251-DEL-79 |
| chr14-14339694-DEL-721 | chr14-14339695-DEL-720 |
| chr14-14340984-DEL-138 | chr14-14340985-DEL-138 |
| chr14-14381225-DEL-323 | chr14-14381226-DEL-323 |
| chr14-14387148-DEL-82 | chr14-14387149-DEL-82 |
| chr14-14405946-DEL-113 | chr14-14405939-DEL-113 |

| | |
|---|---|
| chr14-14423441-DEL-181 | chr14-14423442-DEL-181 |
| chr14-14431060-DEL-51 | chr14-14431061-DEL-51 |
| chr14-14439985-DEL-56 | chr14-14439986-DEL-56 |
| chr14-14443527-DEL-53 | chr14-14443528-DEL-53 |
| chr14-14444009-DEL-766 | chr14-14444010-DEL-765 |
| chr14-14468082-DEL-52 | chr14-14468083-DEL-52 |
| chr14-14479740-DEL-89 | chr14-14479741-DEL-89 |
| chr14-14515898-DEL-78 | chr14-14515899-DEL-78 |
| chr14-14548361-DEL-135 | chr14-14548392-DEL-87 |
| chr14-14553417-DEL-209 | chr14-14553418-DEL-209 |
| chr14-14573703-DEL-543 | chr14-14573704-DEL-543 |
| chr14-14575668-DEL-74 | chr14-14575668-DEL-72 |
| chr14-14590320-DEL-57 | chr14-14590367-DEL-57 |
| chr14-14595256-DEL-214 | chr14-14595257-DEL-214 |
| chr14-14598045-DEL-1188 | chr14-14598046-DEL-1188 |
| chr14-14610240-DEL-672 | chr14-14610241-DEL-672 |
| chr14-14632115-DEL-63 | chr14-14632116-DEL-63 |
| chr14-14636963-DEL-51 | chr14-14636964-DEL-51 |
| chr14-14645886-DEL-201 | chr14-14645887-DEL-201 |
| chr14-14662160-DEL-65 | chr14-14662161-DEL-56 |
| chr14-14662630-DEL-142 | chr14-14662631-DEL-142 |
| chr14-14670950-DEL-82 | chr14-14670959-DEL-74 |
| chr14-14672905-DEL-94 | chr14-14672900-DEL-95 |
| chr14-14683857-DEL-570 | chr14-14683868-DEL-568 |
| chr14-14685462-DEL-55 | chr14-14685462-DEL-55 |
| chr14-14699940-DEL-89 | chr14-14699931-DEL-58 |
| chr14-14703201-DEL-62 | chr14-14703202-DEL-62 |
| chr14-14758458-DEL-869 | chr14-14758459-DEL-869 |
| chr14-14767549-DEL-7092 | chr14-14767550-DEL-7092 |
| chr14-14783302-DEL-1131 | chr14-14783303-DEL-1131 |
| chr14-14785232-DEL-230 | chr14-14785233-DEL-230 |
| chr14-15504598-DEL-1063 | chr14-15504599-DEL-1063 |
| chr14-15562341-DEL-369 | chr14-15562341-DEL-369 |
| chr14-15912715-DEL-208 | chr14-15912716-DEL-208 |
| chr14-18331163-DEL-369 | chr14-18331163-DEL-369 |
| chr14-18915851-DEL-68 | chr14-18915826-DEL-96 |
| chr14-18920995-DEL-284 | chr14-18920996-DEL-284 |
| chr14-19744759-DEL-204 | chr14-19744760-DEL-204 |
| chr14-19746303-DEL-64 | chr14-19746304-DEL-64 |
| chr14-19751805-DEL-93 | chr14-19751806-DEL-93 |
| chr14-19786278-DEL-68 | chr14-19786279-DEL-68 |
| chr14-19787554-DEL-70 | chr14-19787555-DEL-70 |
| chr14-19810373-DEL-147 | chr14-19810374-DEL-147 |
| chr14-19811066-DEL-166 | chr14-19811067-DEL-166 |
| chr14-19831472-DEL-1005 | chr14-19831475-DEL-1003 |
| chr14-19831780-DEL-103 | chr14-19831737-DEL-104 |
| chr14-19836930-DEL-161 | chr14-19836931-DEL-161 |
| chr14-19837481-DEL-1705 | chr14-19837482-DEL-1705 |
| chr14-19842235-DEL-222 | chr14-19842236-DEL-222 |
| chr14-19847430-DEL-58 | chr14-19847431-DEL-58 |
| chr14-19858488-DEL-435 | chr14-19858489-DEL-435 |
| chr14-19859406-DEL-213 | chr14-19859407-DEL-213 |
| chr14-19880880-DEL-681 | chr14-19880881-DEL-681 |
| chr14-19894755-DEL-1886 | chr14-19894756-DEL-1886 |
| chr14-19896834-DEL-61 | chr14-19896835-DEL-61 |
| chr14-19897461-DEL-76 | chr14-19897462-DEL-76 |
| chr14-19910663-DEL-549 | chr14-19910664-DEL-549 |
| chr14-19913608-DEL-650 | chr14-19913609-DEL-650 |
| chr14-19915028-DEL-335 | chr14-19915029-DEL-335 |
| chr14-19915526-DEL-358 | chr14-19915528-DEL-357 |
| chr14-19916281-DEL-187 | chr14-19916282-DEL-187 |
| chr14-19935706-DEL-135 | chr14-19935707-DEL-135 |
| chr14-20056732-DEL-60 | chr14-20056730-DEL-63 |
| chr14-20073735-DEL-328 | chr14-20073736-DEL-328 |
| chr14-20087756-DEL-3183 | chr14-20087757-DEL-3183 |
| chr14-20111847-DEL-62 | chr14-20111840-DEL-50 |
| chr14-20115248-DEL-113 | chr14-20115249-DEL-113 |
| chr14-20151657-DEL-258 | chr14-20151634-DEL-282 |
| chr14-20158186-DEL-96 | chr14-20158187-DEL-96 |
| chr14-20207923-DEL-98 | chr14-20207924-DEL-98 |
| chr14-20238608-DEL-104 | chr14-20238607-DEL-101 |
| chr14-20264787-DEL-52 | chr14-20264787-DEL-52 |
| chr14-20291710-DEL-74 | chr14-20291698-DEL-75 |
| chr14-20303899-DEL-87 | chr14-20303934-DEL-95 |
| chr14-20315931-DEL-148 | chr14-20315962-DEL-143 |
| chr14-20329971-DEL-7175 | chr14-20329972-DEL-7175 |
| chr14-20344109-DEL-69 | chr14-20344110-DEL-69 |
| chr14-20352203-DEL-205 | chr14-20352196-DEL-205 |
| chr14-20362941-DEL-66 | chr14-20362942-DEL-66 |
| chr14-20372521-DEL-77 | chr14-20372522-DEL-77 |

| | |
|---|---|
| chr14-20393948-DEL-420 | chr14-20393949-DEL-420 |
| chr14-20421210-DEL-187 | chr14-20421211-DEL-187 |
| chr14-20476693-DEL-947 | chr14-20476694-DEL-947 |
| chr14-20483378-DEL-1687 | chr14-20483379-DEL-1687 |
| chr14-20505109-DEL-200 | chr14-20505110-DEL-199 |
| chr14-20601643-DEL-75 | chr14-20601644-DEL-75 |
| chr14-20641258-DEL-60 | chr14-20641259-DEL-60 |
| chr14-20653068-DEL-78 | chr14-20653068-DEL-78 |
| chr14-20673687-DEL-212 | chr14-20673688-DEL-212 |
| chr14-20677349-DEL-204 | chr14-20677350-DEL-204 |
| chr14-20740552-DEL-169 | chr14-20740553-DEL-159 |
| chr14-20753524-DEL-177 | chr14-20753525-DEL-177 |
| chr14-20867374-DEL-84 | chr14-20867379-DEL-77 |
| chr14-20870693-DEL-98 | chr14-20870694-DEL-103 |
| chr14-20907475-DEL-66 | chr14-20907482-DEL-54 |
| chr14-20917642-DEL-56 | chr14-20917642-DEL-56 |
| chr14-20919704-DEL-51 | chr14-20919679-DEL-52 |
| chr14-20937293-DEL-80 | chr14-20937293-DEL-78 |
| chr14-20941344-DEL-196 | chr14-20941345-DEL-197 |
| chr14-20945691-DEL-3115 | chr14-20945692-DEL-3115 |
| chr14-20947096-DEL-200 | chr14-20947097-DEL-200 |
| chr14-20957221-DEL-64 | chr14-20957207-DEL-73 |
| chr14-20969188-DEL-203 | chr14-20969189-DEL-204 |
| chr14-21043954-DEL-69 | chr14-21043951-DEL-74 |
| chr14-21057650-DEL-63 | chr14-21057649-DEL-69 |
| chr14-21060550-DEL-863 | chr14-21060551-DEL-863 |
| chr14-21073984-DEL-57 | chr14-21073975-DEL-58 |
| chr14-21131242-DEL-418 | chr14-21131243-DEL-418 |
| chr14-21183674-DEL-53 | chr14-21183675-DEL-53 |
| chr14-21245219-DEL-156 | chr14-21245220-DEL-156 |
| chr14-21248480-DEL-114 | chr14-21248481-DEL-114 |
| chr14-21250144-DEL-197 | chr14-21250168-DEL-148 |
| chr14-21261370-DEL-161 | chr14-21261371-DEL-161 |
| chr14-21262140-DEL-5333 | chr14-21262141-DEL-5333 |
| chr14-21343818-DEL-833 | chr14-21343819-DEL-833 |
| chr14-21359147-DEL-199 | chr14-21359148-DEL-199 |
| chr14-21371615-DEL-585 | chr14-21371616-DEL-585 |
| chr14-21410439-DEL-5547 | chr14-21410440-DEL-5547 |
| chr14-21444711-DEL-71 | chr14-21444712-DEL-71 |
| chr14-21453975-DEL-750 | chr14-21453971-DEL-751 |
| chr14-21456361-DEL-213 | chr14-21456362-DEL-213 |
| chr14-21465106-DEL-81 | chr14-21465115-DEL-69 |
| chr14-21470174-DEL-66 | chr14-21470175-DEL-66 |
| chr14-21487698-DEL-198 | chr14-21487699-DEL-198 |
| chr14-21527421-DEL-92 | chr14-21527444-DEL-70 |
| chr14-21538148-DEL-567 | chr14-21538149-DEL-567 |
| chr14-21575291-DEL-207 | chr14-21575292-DEL-207 |
| chr14-21576642-DEL-153 | chr14-21576643-DEL-153 |
| chr14-21579597-DEL-445 | chr14-21579598-DEL-444 |
| chr14-21608070-DEL-51 | chr14-21608071-DEL-51 |
| chr14-21615317-DEL-52 | chr14-21615320-DEL-50 |
| chr14-21644040-DEL-2936 | chr14-21644041-DEL-2936 |
| chr14-21688233-DEL-75 | chr14-21688234-DEL-75 |
| chr14-21688999-DEL-460 | chr14-21689000-DEL-460 |
| chr14-21693279-DEL-219 | chr14-21693280-DEL-219 |
| chr14-21715813-DEL-62 | chr14-21715814-DEL-62 |
| chr14-21748676-DEL-112 | chr14-21748677-DEL-112 |
| chr14-21754973-DEL-673 | chr14-21754974-DEL-673 |
| chr14-21765222-DEL-68 | chr14-21765223-DEL-70 |
| chr14-21788345-DEL-77 | chr14-21788361-DEL-62 |
| chr14-21827180-DEL-52 | chr14-21827181-DEL-52 |
| chr14-21863184-DEL-69 | chr14-21863185-DEL-65 |
| chr14-21864540-DEL-184 | chr14-21864541-DEL-184 |
| chr14-21865567-DEL-207 | chr14-21865568-DEL-207 |
| chr14-21886519-DEL-202 | chr14-21886520-DEL-202 |
| chr14-21914129-DEL-132 | chr14-21914130-DEL-132 |
| chr14-21927539-DEL-84 | chr14-21927540-DEL-84 |
| chr14-21928615-DEL-6201 | chr14-21928616-DEL-6201 |
| chr14-21929539-DEL-373 | chr14-21929540-DEL-373 |
| chr14-21949411-DEL-196 | chr14-21949408-DEL-198 |
| chr14-22021212-DEL-70 | chr14-22021192-DEL-90 |
| chr14-22023352-DEL-870 | chr14-22023353-DEL-870 |
| chr14-22053671-DEL-56 | chr14-22053636-DEL-54 |
| chr14-22080839-DEL-379 | chr14-22080840-DEL-379 |
| chr14-22101426-DEL-62 | chr14-22101427-DEL-62 |
| chr14-22127155-DEL-444 | chr14-22127155-DEL-444 |
| chr14-22130643-DEL-225 | chr14-22130681-DEL-177 |
| chr14-22185060-DEL-199 | chr14-22185061-DEL-211 |
| chr14-22215440-DEL-111 | chr14-22215441-DEL-111 |
| chr14-22231469-DEL-75 | chr14-22231440-DEL-76 |

| | |
|---|---|
| chr14-22247434-DEL-189 | chr14-22247435-DEL-189 |
| chr14-22253273-DEL-170 | chr14-22253274-DEL-170 |
| chr14-22263531-DEL-53 | chr14-22263530-DEL-53 |
| chr14-22283017-DEL-182 | chr14-22283018-DEL-182 |
| chr14-22288593-DEL-52 | chr14-22288620-DEL-62 |
| chr14-22325839-DEL-61 | chr14-22325840-DEL-61 |
| chr14-22350462-DEL-52 | chr14-22350463-DEL-52 |
| chr14-22352702-DEL-54 | chr14-22352703-DEL-54 |
| chr14-22361085-DEL-6238 | chr14-22361086-DEL-6238 |
| chr14-22381273-DEL-52 | chr14-22381274-DEL-52 |
| chr14-22399151-DEL-110 | chr14-22399200-DEL-62 |
| chr14-22446802-DEL-64 | chr14-22446803-DEL-64 |
| chr14-22449301-DEL-301 | chr14-22449302-DEL-297 |
| chr14-22518971-DEL-1301 | chr14-22518972-DEL-1301 |
| chr14-22608038-DEL-55 | chr14-22608020-DEL-56 |
| chr14-22609038-DEL-58 | chr14-22609039-DEL-58 |
| chr14-22651922-DEL-55 | chr14-22651923-DEL-55 |
| chr14-22653125-DEL-93 | chr14-22653122-DEL-97 |
| chr14-22657116-DEL-269 | chr14-22657117-DEL-269 |
| chr14-22666611-DEL-106 | chr14-22666612-DEL-98 |
| chr14-22670004-DEL-120 | chr14-22670054-DEL-80 |
| chr14-22673629-DEL-126 | chr14-22673630-DEL-126 |
| chr14-22674539-DEL-93 | chr14-22674550-DEL-83 |
| chr14-22709412-DEL-225 | chr14-22709412-DEL-269 |
| chr14-22728823-DEL-111 | chr14-22728850-DEL-76 |
| chr14-22745325-DEL-158 | chr14-22745326-DEL-154 |
| chr14-22804862-DEL-95 | chr14-22804863-DEL-96 |
| chr14-22818938-DEL-86 | chr14-22818965-DEL-56 |
| chr14-22839025-DEL-1210 | chr14-22839018-DEL-1211 |
| chr14-22853460-DEL-290 | chr14-22853456-DEL-291 |
| chr14-22879406-DEL-7135 | chr14-22879407-DEL-7135 |
| chr14-22892101-DEL-541 | chr14-22892102-DEL-541 |
| chr14-22898368-DEL-59 | chr14-22898369-DEL-59 |
| chr14-22923087-DEL-53 | chr14-22923088-DEL-53 |
| chr14-22929372-DEL-51 | chr14-22929348-DEL-60 |
| chr14-22950815-DEL-626 | chr14-22950803-DEL-627 |
| chr14-22983571-DEL-2341 | chr14-22983572-DEL-2341 |
| chr14-23007769-DEL-462 | chr14-23007769-DEL-462 |
| chr14-23021797-DEL-4518 | chr14-23021798-DEL-4518 |
| chr14-23039675-DEL-1891 | chr14-23039676-DEL-1891 |
| chr14-23066934-DEL-6326 | chr14-23066935-DEL-6326 |
| chr14-23115896-DEL-140 | chr14-23115900-DEL-139 |
| chr14-23147053-DEL-84 | chr14-23147053-DEL-84 |
| chr14-23171813-DEL-6863 | chr14-23171799-DEL-6864 |
| chr14-23226559-DEL-399 | chr14-23226560-DEL-399 |
| chr14-23328627-DEL-52 | chr14-23328612-DEL-56 |
| chr14-23435548-DEL-132 | chr14-23435569-DEL-104 |
| chr14-23459879-DEL-6940 | chr14-23459880-DEL-6940 |
| chr14-23459932-DEL-214 | chr14-23459933-DEL-214 |
| chr14-23513943-DEL-52 | chr14-23513895-DEL-52 |
| chr14-23619399-DEL-115 | chr14-23619400-DEL-116 |
| chr14-23630490-DEL-52 | chr14-23630492-DEL-50 |
| chr14-23666011-DEL-508 | chr14-23666012-DEL-508 |
| chr14-23672455-DEL-292 | chr14-23672505-DEL-249 |
| chr14-23704481-DEL-54 | chr14-23704500-DEL-54 |
| chr14-23802476-DEL-92 | chr14-23802473-DEL-89 |
| chr14-23842557-DEL-68 | chr14-23842592-DEL-60 |
| chr14-23924796-DEL-173 | chr14-23924797-DEL-173 |
| chr14-23943465-DEL-95 | chr14-23943435-DEL-88 |
| chr14-24004551-DEL-51 | chr14-24004552-DEL-51 |
| chr14-24007624-DEL-57 | chr14-24007625-DEL-57 |
| chr14-24023110-DEL-79 | chr14-24023111-DEL-79 |
| chr14-24037331-DEL-62 | chr14-24037324-DEL-70 |
| chr14-24104683-DEL-182 | chr14-24104684-DEL-182 |
| chr14-24150469-DEL-438 | chr14-24150454-DEL-454 |
| chr14-24176412-DEL-65 | chr14-24176413-DEL-65 |
| chr14-24214272-DEL-59 | chr14-24214273-DEL-59 |
| chr14-24233532-DEL-422 | chr14-24233534-DEL-422 |
| chr14-24247581-DEL-64 | chr14-24247582-DEL-64 |
| chr14-24261578-DEL-85 | chr14-24261573-DEL-61 |
| chr14-24266717-DEL-52 | chr14-24266718-DEL-52 |
| chr14-24275121-DEL-2982 | chr14-24275134-DEL-2963 |
| chr14-24327534-DEL-331 | chr14-24327535-DEL-331 |
| chr14-24334295-DEL-182 | chr14-24334296-DEL-182 |
| chr14-24348482-DEL-62 | chr14-24348483-DEL-63 |
| chr14-24358114-DEL-84 | chr14-24358115-DEL-84 |
| chr14-24367350-DEL-7004 | chr14-24367351-DEL-7004 |
| chr14-24390090-DEL-244 | chr14-24390083-DEL-248 |
| chr14-24423151-DEL-107 | chr14-24423153-DEL-107 |
| chr14-24426620-DEL-52 | chr14-24426621-DEL-52 |

| | |
|---|---|
| chr14-24428839-DEL-249 | chr14-24428840-DEL-249 |
| chr14-24429608-DEL-4523 | chr14-24429609-DEL-4523 |
| chr14-24440744-DEL-3085 | chr14-24440745-DEL-3085 |
| chr14-24458064-DEL-706 | chr14-24458068-DEL-700 |
| chr14-24462823-DEL-4437 | chr14-24462824-DEL-4437 |
| chr14-24468258-DEL-199 | chr14-24468259-DEL-199 |
| chr14-24510129-DEL-90 | chr14-24510152-DEL-90 |
| chr14-24539327-DEL-983 | chr14-24539328-DEL-983 |
| chr14-24555507-DEL-56 | chr14-24555508-DEL-56 |
| chr14-24581546-DEL-218 | chr14-24581546-DEL-218 |
| chr14-24613703-DEL-66 | chr14-24613704-DEL-66 |
| chr14-24632330-DEL-156 | chr14-24632321-DEL-166 |
| chr14-24744417-DEL-65 | chr14-24744415-DEL-66 |
| chr14-24746786-DEL-56 | chr14-24746779-DEL-62 |
| chr14-24769548-DEL-52 | chr14-24769544-DEL-72 |
| chr14-24797704-DEL-106 | chr14-24797705-DEL-106 |
| chr14-24853869-DEL-54 | chr14-24853870-DEL-54 |
| chr14-24881981-DEL-143 | chr14-24881974-DEL-137 |
| chr14-24928344-DEL-56 | chr14-24928345-DEL-58 |
| chr14-24929872-DEL-243 | chr14-24929873-DEL-243 |
| chr14-24950230-DEL-137 | chr14-24950231-DEL-137 |
| chr14-24964061-DEL-53 | chr14-24964062-DEL-51 |
| chr14-24989713-DEL-92 | chr14-24989743-DEL-84 |
| chr14-25011630-DEL-62 | chr14-25011631-DEL-56 |
| chr14-25017503-DEL-52 | chr14-25017504-DEL-52 |
| chr14-25037804-DEL-69 | chr14-25037805-DEL-69 |
| chr14-25060248-DEL-68 | chr14-25060249-DEL-68 |
| chr14-25068653-DEL-86 | chr14-25068672-DEL-56 |
| chr14-25126002-DEL-52 | chr14-25126003-DEL-52 |
| chr14-25190057-DEL-90 | chr14-25190058-DEL-90 |
| chr14-25217828-DEL-102 | chr14-25217812-DEL-78 |
| chr14-25232393-DEL-88 | chr14-25232394-DEL-88 |
| chr14-25272628-DEL-62 | chr14-25272640-DEL-50 |
| chr14-25296677-DEL-1887 | chr14-25296678-DEL-1887 |
| chr14-25315098-DEL-62 | chr14-25315090-DEL-64 |
| chr14-25324510-DEL-102 | chr14-25324511-DEL-102 |
| chr14-25341152-DEL-74 | chr14-25341153-DEL-74 |
| chr14-25364129-DEL-172 | chr14-25364138-DEL-164 |
| chr14-25410767-DEL-126 | chr14-25410768-DEL-126 |
| chr14-25428894-DEL-1482 | chr14-25428894-DEL-1482 |
| chr14-25433860-DEL-546 | chr14-25433861-DEL-546 |
| chr14-25507619-DEL-63 | chr14-25507620-DEL-63 |
| chr14-25527704-DEL-64 | chr14-25527705-DEL-64 |
| chr14-25530396-DEL-131 | chr14-25530402-DEL-107 |
| chr14-25566406-DEL-178 | chr14-25566407-DEL-178 |
| chr14-25609772-DEL-55 | chr14-25609775-DEL-51 |
| chr14-25691408-DEL-370 | chr14-25691409-DEL-370 |
| chr14-25704380-DEL-68 | chr14-25704381-DEL-66 |
| chr14-25706928-DEL-566 | chr14-25706929-DEL-566 |
| chr14-25728245-DEL-6404 | chr14-25728246-DEL-6404 |
| chr14-25758761-DEL-118 | chr14-25758762-DEL-118 |
| chr14-25787883-DEL-1686 | chr14-25787884-DEL-1686 |
| chr14-25791924-DEL-180 | chr14-25791925-DEL-180 |
| chr14-25792758-DEL-210 | chr14-25792759-DEL-210 |
| chr14-25808374-DEL-165 | chr14-25808375-DEL-165 |
| chr14-25820222-DEL-78 | chr14-25820223-DEL-78 |
| chr14-25839318-DEL-603 | chr14-25839319-DEL-603 |
| chr14-25841033-DEL-66 | chr14-25841034-DEL-66 |
| chr14-25849065-DEL-219 | chr14-25849066-DEL-219 |
| chr14-25882735-DEL-206 | chr14-25882747-DEL-200 |
| chr14-25907014-DEL-1371 | chr14-25907014-DEL-1369 |
| chr14-25921072-DEL-78 | chr14-25921072-DEL-78 |
| chr14-25959302-DEL-237 | chr14-25959291-DEL-238 |
| chr14-25972526-DEL-517 | chr14-25972528-DEL-518 |
| chr14-25981236-DEL-51 | chr14-25981231-DEL-57 |
| chr14-25982979-DEL-72 | chr14-25982979-DEL-72 |
| chr14-25990963-DEL-173 | chr14-25990964-DEL-173 |
| chr14-25995304-DEL-571 | chr14-25995335-DEL-585 |
| chr14-26013334-DEL-454 | chr14-26013335-DEL-454 |
| chr14-26017113-DEL-54 | chr14-26017114-DEL-54 |
| chr14-26028629-DEL-342 | chr14-26028630-DEL-342 |
| chr14-26049426-DEL-211 | chr14-26049427-DEL-211 |
| chr14-26050810-DEL-170 | chr14-26050810-DEL-170 |
| chr14-26058944-DEL-592 | chr14-26058939-DEL-591 |
| chr14-26076736-DEL-692 | chr14-26076737-DEL-692 |
| chr14-26092108-DEL-139 | chr14-26092109-DEL-139 |
| chr14-26095363-DEL-158 | chr14-26095358-DEL-159 |
| chr14-26096809-DEL-113 | chr14-26096809-DEL-110 |
| chr14-26100559-DEL-117 | chr14-26100560-DEL-117 |
| chr14-26125324-DEL-58 | chr14-26125315-DEL-68 |

| | |
|---|---|
| chr14-26132899-DEL-86 | chr14-26132853-DEL-133 |
| chr14-26154872-DEL-85 | chr14-26154873-DEL-85 |
| chr14-26159999-DEL-56 | chr14-26160000-DEL-56 |
| chr14-26179492-DEL-194 | chr14-26179493-DEL-194 |
| chr14-26193915-DEL-96 | chr14-26193879-DEL-90 |
| chr14-26200223-DEL-1173 | chr14-26200224-DEL-1173 |
| chr14-26213235-DEL-58 | chr14-26213241-DEL-64 |
| chr14-26250547-DEL-58 | chr14-26250548-DEL-58 |
| chr14-26313830-DEL-135 | chr14-26313809-DEL-130 |
| chr14-26323827-DEL-209 | chr14-26323837-DEL-210 |
| chr14-26365065-DEL-198 | chr14-26365059-DEL-199 |
| chr14-26388681-DEL-68 | chr14-26388689-DEL-60 |
| chr14-26408551-DEL-738 | chr14-26408552-DEL-744 |
| chr14-26419407-DEL-6859 | chr14-26419408-DEL-6859 |
| chr14-26433988-DEL-970 | chr14-26433989-DEL-971 |
| chr14-26439674-DEL-212 | chr14-26439675-DEL-212 |
| chr14-26449678-DEL-1379 | chr14-26449679-DEL-1379 |
| chr14-26457386-DEL-803 | chr14-26457387-DEL-803 |
| chr14-26461225-DEL-54 | chr14-26461226-DEL-54 |
| chr14-26464694-DEL-54 | chr14-26464693-DEL-54 |
| chr14-26465834-DEL-880 | chr14-26465834-DEL-880 |
| chr14-26470459-DEL-641 | chr14-26470459-DEL-641 |
| chr14-26471835-DEL-347 | chr14-26471836-DEL-347 |
| chr14-26473118-DEL-81 | chr14-26473119-DEL-81 |
| chr14-26479276-DEL-81 | chr14-26479277-DEL-81 |
| chr14-26479786-DEL-135 | chr14-26479787-DEL-135 |
| chr14-26510648-DEL-2317 | chr14-26510649-DEL-2317 |
| chr14-26573984-DEL-197 | chr14-26573985-DEL-197 |
| chr14-26575139-DEL-64 | chr14-26575140-DEL-64 |
| chr14-26620718-DEL-200 | chr14-26620727-DEL-192 |
| chr14-26627183-DEL-200 | chr14-26627179-DEL-201 |
| chr14-26627540-DEL-879 | chr14-26627541-DEL-879 |
| chr14-26629326-DEL-69 | chr14-26629327-DEL-69 |
| chr14-26634613-DEL-56 | chr14-26634615-DEL-54 |
| chr14-26646416-DEL-155 | chr14-26646417-DEL-155 |
| chr14-26651249-DEL-138 | chr14-26651250-DEL-138 |
| chr14-26655027-DEL-207 | chr14-26655028-DEL-207 |
| chr14-26675704-DEL-207 | chr14-26675705-DEL-207 |
| chr14-26698056-DEL-260 | chr14-26698057-DEL-260 |
| chr14-26707339-DEL-233 | chr14-26707340-DEL-233 |
| chr14-26709893-DEL-70 | chr14-26709894-DEL-70 |
| chr14-26737462-DEL-88 | chr14-26737463-DEL-88 |
| chr14-26738452-DEL-765 | chr14-26738453-DEL-765 |
| chr14-26739560-DEL-176 | chr14-26739561-DEL-176 |
| chr14-26749349-DEL-507 | chr14-26749350-DEL-507 |
| chr14-26758553-DEL-164 | chr14-26758554-DEL-164 |
| chr14-26843533-DEL-84 | chr14-26843534-DEL-84 |
| chr14-26858939-DEL-95 | chr14-26858945-DEL-90 |
| chr14-26865897-DEL-51 | chr14-26865898-DEL-51 |
| chr14-26875538-DEL-68 | chr14-26875539-DEL-68 |
| chr14-26896565-DEL-117 | chr14-26896566-DEL-117 |
| chr14-26929236-DEL-202 | chr14-26929237-DEL-202 |
| chr14-26969201-DEL-181 | chr14-26969202-DEL-181 |
| chr14-26988103-DEL-152 | chr14-26988104-DEL-152 |
| chr14-27013194-DEL-208 | chr14-27013195-DEL-208 |
| chr14-27014793-DEL-211 | chr14-27014794-DEL-211 |
| chr14-27017122-DEL-353 | chr14-27017123-DEL-353 |
| chr14-27024578-DEL-60 | chr14-27024579-DEL-60 |
| chr14-27028424-DEL-196 | chr14-27028425-DEL-196 |
| chr14-27036251-DEL-189 | chr14-27036252-DEL-189 |
| chr14-27048290-DEL-209 | chr14-27048291-DEL-209 |
| chr14-27071421-DEL-74 | chr14-27071394-DEL-62 |
| chr14-27076778-DEL-209 | chr14-27076779-DEL-209 |
| chr14-27080611-DEL-89 | chr14-27080612-DEL-89 |
| chr14-27081209-DEL-205 | chr14-27081210-DEL-205 |
| chr14-27085866-DEL-160 | chr14-27085867-DEL-160 |
| chr14-27106678-DEL-647 | chr14-27106682-DEL-645 |
| chr14-27147222-DEL-52 | chr14-27147223-DEL-52 |
| chr14-27161966-DEL-89 | chr14-27161967-DEL-89 |
| chr14-27242009-DEL-99 | chr14-27242010-DEL-99 |
| chr14-27247393-DEL-199 | chr14-27247394-DEL-199 |
| chr14-27252350-DEL-62 | chr14-27252351-DEL-62 |
| chr14-27270633-DEL-268 | chr14-27270634-DEL-268 |
| chr14-27284110-DEL-139 | chr14-27284111-DEL-139 |
| chr14-27292068-DEL-6026 | chr14-27292069-DEL-6026 |
| chr14-27330359-DEL-71 | chr14-27330360-DEL-71 |
| chr14-27363754-DEL-66 | chr14-27363755-DEL-66 |
| chr14-27382358-DEL-204 | chr14-27382359-DEL-204 |
| chr14-27405337-DEL-7366 | chr14-27405338-DEL-7366 |
| chr14-27424286-DEL-338 | chr14-27424287-DEL-338 |

| | |
|---|---|
| chr14-27453290-DEL-272 | chr14-27453291-DEL-272 |
| chr14-27493670-DEL-127 | chr14-27493671-DEL-127 |
| chr14-27526036-DEL-141 | chr14-27526037-DEL-141 |
| chr14-27554942-DEL-51 | chr14-27554940-DEL-51 |
| chr14-27564175-DEL-1892 | chr14-27564176-DEL-1892 |
| chr14-27577546-DEL-72 | chr14-27577547-DEL-72 |
| chr14-27589749-DEL-237 | chr14-27589750-DEL-237 |
| chr14-27596129-DEL-95 | chr14-27596130-DEL-95 |
| chr14-27607090-DEL-90 | chr14-27607091-DEL-90 |
| chr14-27636654-DEL-304 | chr14-27636655-DEL-304 |
| chr14-27652092-DEL-112 | chr14-27652093-DEL-112 |
| chr14-27680626-DEL-327 | chr14-27680620-DEL-328 |
| chr14-27690355-DEL-1892 | chr14-27690356-DEL-1892 |
| chr14-27695256-DEL-88 | chr14-27695257-DEL-88 |
| chr14-27739334-DEL-249 | chr14-27739335-DEL-249 |
| chr14-27759032-DEL-655 | chr14-27759033-DEL-655 |
| chr14-27811199-DEL-89 | chr14-27811200-DEL-89 |
| chr14-27855871-DEL-132 | chr14-27855872-DEL-132 |
| chr14-27891756-DEL-80 | chr14-27891756-DEL-80 |
| chr14-27904440-DEL-6915 | chr14-27904435-DEL-6915 |
| chr14-27966636-DEL-60 | chr14-27966636-DEL-60 |
| chr14-27967673-DEL-364 | chr14-27967676-DEL-362 |
| chr14-27971610-DEL-463 | chr14-27971611-DEL-463 |
| chr14-28014450-DEL-79 | chr14-28014470-DEL-56 |
| chr14-28095637-DEL-95 | chr14-28095638-DEL-95 |
| chr14-28154625-DEL-59 | chr14-28154626-DEL-59 |
| chr14-28170231-DEL-74 | chr14-28170231-DEL-80 |
| chr14-28217513-DEL-675 | chr14-28217512-DEL-676 |
| chr14-28220053-DEL-154 | chr14-28220054-DEL-154 |
| chr14-28290243-DEL-59 | chr14-28290221-DEL-57 |
| chr14-28299820-DEL-480 | chr14-28299821-DEL-480 |
| chr14-28300631-DEL-76 | chr14-28300646-DEL-62 |
| chr14-28324614-DEL-144 | chr14-28324615-DEL-144 |
| chr14-28353287-DEL-52 | chr14-28353287-DEL-51 |
| chr14-28359673-DEL-105 | chr14-28359661-DEL-106 |
| chr14-28362623-DEL-485 | chr14-28362624-DEL-485 |
| chr14-28366269-DEL-205 | chr14-28366270-DEL-205 |
| chr14-28407238-DEL-65 | chr14-28407239-DEL-65 |
| chr14-28445326-DEL-234 | chr14-28445327-DEL-234 |
| chr14-28491637-DEL-66 | chr14-28491644-DEL-59 |
| chr14-28504077-DEL-216 | chr14-28504077-DEL-216 |
| chr14-28551443-DEL-82 | chr14-28551424-DEL-58 |
| chr14-28562666-DEL-53 | chr14-28562659-DEL-54 |
| chr14-28612316-DEL-104 | chr14-28612307-DEL-105 |
| chr14-28633701-DEL-174 | chr14-28633702-DEL-174 |
| chr14-28654187-DEL-66 | chr14-28654188-DEL-66 |
| chr14-28658956-DEL-467 | chr14-28658957-DEL-467 |
| chr14-28698237-DEL-127 | chr14-28698238-DEL-127 |
| chr14-28722530-DEL-66 | chr14-28722531-DEL-66 |
| chr14-28723223-DEL-1864 | chr14-28723224-DEL-1864 |
| chr14-28792206-DEL-65 | chr14-28792211-DEL-61 |
| chr14-28804867-DEL-304 | chr14-28804868-DEL-304 |
| chr14-28805762-DEL-58 | chr14-28805763-DEL-58 |
| chr14-28870313-DEL-3960 | chr14-28870314-DEL-3960 |
| chr14-28897133-DEL-2481 | chr14-28897134-DEL-2481 |
| chr14-28970837-DEL-6930 | chr14-28970838-DEL-6930 |
| chr14-29021888-DEL-507 | chr14-29021862-DEL-508 |
| chr14-29031066-DEL-947 | chr14-29031067-DEL-947 |
| chr14-29065869-DEL-1829 | chr14-29065870-DEL-1829 |
| chr14-29095899-DEL-170 | chr14-29095900-DEL-170 |
| chr14-29105327-DEL-171 | chr14-29105328-DEL-171 |
| chr14-29107339-DEL-54 | chr14-29107340-DEL-54 |
| chr14-29132321-DEL-56 | chr14-29132307-DEL-85 |
| chr14-29168132-DEL-67 | chr14-29168142-DEL-90 |
| chr14-29170522-DEL-54 | chr14-29170525-DEL-52 |
| chr14-29180516-DEL-256 | chr14-29180514-DEL-216 |
| chr14-29184573-DEL-577 | chr14-29184574-DEL-578 |
| chr14-29208565-DEL-168 | chr14-29208566-DEL-168 |
| chr14-29226384-DEL-66 | chr14-29226385-DEL-66 |
| chr14-29237629-DEL-53 | chr14-29237630-DEL-53 |
| chr14-29264329-DEL-2626 | chr14-29264330-DEL-2626 |
| chr14-29293732-DEL-1184 | chr14-29293733-DEL-1184 |
| chr14-29340054-DEL-64 | chr14-29340055-DEL-64 |
| chr14-29340859-DEL-66 | chr14-29340860-DEL-77 |
| chr14-29343798-DEL-68 | chr14-29343790-DEL-88 |
| chr14-29435182-DEL-52 | chr14-29435183-DEL-52 |
| chr14-29474372-DEL-125 | chr14-29474373-DEL-125 |
| chr14-29483287-DEL-5033 | chr14-29483288-DEL-5033 |
| chr14-29516599-DEL-51 | chr14-29516600-DEL-51 |
| chr14-29542412-DEL-51 | chr14-29542413-DEL-51 |

| | |
|---|---|
| chr14-29609315-DEL-68 | chr14-29609303-DEL-104 |
| chr14-29641443-DEL-6579 | chr14-29641423-DEL-6579 |
| chr14-29665022-DEL-92 | chr14-29665040-DEL-56 |
| chr14-29677126-DEL-124 | chr14-29677127-DEL-124 |
| chr14-29694532-DEL-127 | chr14-29694533-DEL-127 |
| chr14-29783745-DEL-146 | chr14-29783770-DEL-126 |
| chr14-29784214-DEL-208 | chr14-29784215-DEL-208 |
| chr14-29806613-DEL-150 | chr14-29806614-DEL-150 |
| chr14-29825045-DEL-63 | chr14-29825048-DEL-61 |
| chr14-29828189-DEL-199 | chr14-29828184-DEL-200 |
| chr14-29843430-DEL-52 | chr14-29843431-DEL-52 |
| chr14-29881734-DEL-68 | chr14-29881739-DEL-64 |
| chr14-29890515-DEL-56 | chr14-29890516-DEL-56 |
| chr14-29984846-DEL-64 | chr14-29984847-DEL-64 |
| chr14-30006887-DEL-61 | chr14-30006887-DEL-70 |
| chr14-30124243-DEL-78 | chr14-30124225-DEL-75 |
| chr14-30125040-DEL-102 | chr14-30125013-DEL-102 |
| chr14-30134856-DEL-150 | chr14-30134857-DEL-156 |
| chr14-30184112-DEL-221 | chr14-30184116-DEL-221 |
| chr14-30186646-DEL-66 | chr14-30186647-DEL-66 |
| chr14-30203594-DEL-114 | chr14-30203594-DEL-115 |
| chr14-30257132-DEL-106 | chr14-30257133-DEL-106 |
| chr14-30259560-DEL-217 | chr14-30259561-DEL-217 |
| chr14-30269283-DEL-76 | chr14-30269302-DEL-60 |
| chr14-30284729-DEL-378 | chr14-30284730-DEL-378 |
| chr14-30298833-DEL-876 | chr14-30298834-DEL-876 |
| chr14-30300443-DEL-198 | chr14-30300455-DEL-180 |
| chr14-30304644-DEL-139 | chr14-30304648-DEL-135 |
| chr14-30310133-DEL-128 | chr14-30310134-DEL-128 |
| chr14-30501475-DEL-3516 | chr14-30501476-DEL-3516 |
| chr14-30533675-DEL-306 | chr14-30533676-DEL-306 |
| chr14-30554117-DEL-62 | chr14-30554134-DEL-57 |
| chr14-30563585-DEL-56 | chr14-30563583-DEL-59 |
| chr14-30620993-DEL-187 | chr14-30620994-DEL-187 |
| chr14-30655641-DEL-64 | chr14-30655642-DEL-64 |
| chr14-30676228-DEL-431 | chr14-30676228-DEL-430 |
| chr14-30812563-DEL-253 | chr14-30812564-DEL-253 |
| chr14-30849824-DEL-80 | chr14-30849825-DEL-76 |
| chr14-30942135-DEL-103 | chr14-30942135-DEL-100 |
| chr14-30998915-DEL-466 | chr14-30998916-DEL-466 |
| chr14-31034055-DEL-75 | chr14-31034056-DEL-75 |
| chr14-31060838-DEL-219 | chr14-31060839-DEL-219 |
| chr14-31178394-DEL-74 | chr14-31178395-DEL-74 |
| chr14-31193975-DEL-90 | chr14-31194006-DEL-90 |
| chr14-31212327-DEL-205 | chr14-31212328-DEL-205 |
| chr14-31236158-DEL-57 | chr14-31236159-DEL-57 |
| chr14-31264549-DEL-101 | chr14-31264550-DEL-101 |
| chr14-31321372-DEL-192 | chr14-31321377-DEL-190 |
| chr14-31331350-DEL-112 | chr14-31331350-DEL-112 |
| chr14-31333370-DEL-1486 | chr14-31333371-DEL-1486 |
| chr14-31348727-DEL-180 | chr14-31348728-DEL-180 |
| chr14-31358540-DEL-198 | chr14-31358541-DEL-198 |
| chr14-31379208-DEL-60 | chr14-31379209-DEL-60 |
| chr14-31394492-DEL-1253 | chr14-31394493-DEL-1253 |
| chr14-31399594-DEL-78 | chr14-31399595-DEL-78 |
| chr14-31412203-DEL-425 | chr14-31412204-DEL-425 |
| chr14-31573703-DEL-94 | chr14-31573706-DEL-92 |
| chr14-31595121-DEL-173 | chr14-31595122-DEL-173 |
| chr14-31604204-DEL-1880 | chr14-31604205-DEL-1880 |
| chr14-31610093-DEL-1985 | chr14-31610094-DEL-1985 |
| chr14-31612214-DEL-1086 | chr14-31612214-DEL-1086 |
| chr14-31641263-DEL-4713 | chr14-31641264-DEL-4713 |
| chr14-31661868-DEL-52 | chr14-31661869-DEL-52 |
| chr14-31703485-DEL-53 | chr14-31703485-DEL-57 |
| chr14-31708309-DEL-464 | chr14-31708314-DEL-460 |
| chr14-31721896-DEL-81 | chr14-31721907-DEL-83 |
| chr14-31723486-DEL-254 | chr14-31723482-DEL-253 |
| chr14-31728934-DEL-599 | chr14-31728935-DEL-599 |
| chr14-31732784-DEL-54 | chr14-31732791-DEL-50 |
| chr14-31739790-DEL-194 | chr14-31739791-DEL-194 |
| chr14-31755121-DEL-1105 | chr14-31755122-DEL-1105 |
| chr14-31770744-DEL-140 | chr14-31770739-DEL-105 |
| chr14-31784269-DEL-54 | chr14-31784271-DEL-52 |
| chr14-31800715-DEL-197 | chr14-31800716-DEL-197 |
| chr14-31809729-DEL-240 | chr14-31809730-DEL-240 |
| chr14-31823416-DEL-201 | chr14-31823417-DEL-200 |
| chr14-31832227-DEL-184 | chr14-31832267-DEL-144 |
| chr14-31836022-DEL-203 | chr14-31836023-DEL-203 |
| chr14-31837256-DEL-5246 | chr14-31837257-DEL-5246 |
| chr14-32017604-DEL-56 | chr14-32017605-DEL-56 |

| | |
|---|---|
| chr14-32019317-DEL-201 | chr14-32019318-DEL-201 |
| chr14-32050235-DEL-218 | chr14-32050236-DEL-218 |
| chr14-32051757-DEL-301 | chr14-32051778-DEL-281 |
| chr14-32053189-DEL-193 | chr14-32053190-DEL-193 |
| chr14-32073418-DEL-119 | chr14-32073419-DEL-114 |
| chr14-32074753-DEL-57 | chr14-32074753-DEL-57 |
| chr14-32080112-DEL-149 | chr14-32080110-DEL-150 |
| chr14-32082936-DEL-209 | chr14-32082937-DEL-209 |
| chr14-32090460-DEL-81 | chr14-32090461-DEL-54 |
| chr14-32091884-DEL-280 | chr14-32091885-DEL-280 |
| chr14-32115221-DEL-7093 | chr14-32115222-DEL-7093 |
| chr14-32127325-DEL-154 | chr14-32127326-DEL-154 |
| chr14-32138598-DEL-400 | chr14-32138599-DEL-400 |
| chr14-32171586-DEL-68 | chr14-32171587-DEL-68 |
| chr14-32179679-DEL-89 | chr14-32179680-DEL-89 |
| chr14-32232416-DEL-140 | chr14-32232417-DEL-140 |
| chr14-32273716-DEL-52 | chr14-32273717-DEL-52 |
| chr14-32348193-DEL-73 | chr14-32348194-DEL-73 |
| chr14-32359700-DEL-456 | chr14-32359701-DEL-456 |
| chr14-32400645-DEL-1008 | chr14-32400646-DEL-1008 |
| chr14-32403696-DEL-96 | chr14-32403697-DEL-96 |
| chr14-32443957-DEL-77 | chr14-32443957-DEL-77 |
| chr14-32449937-DEL-380 | chr14-32449938-DEL-380 |
| chr14-32464336-DEL-95 | chr14-32464337-DEL-95 |
| chr14-32464817-DEL-214 | chr14-32464818-DEL-214 |
| chr14-32472547-DEL-260 | chr14-32472546-DEL-261 |
| chr14-32479520-DEL-1509 | chr14-32479521-DEL-1509 |
| chr14-32531057-DEL-1066 | chr14-32531058-DEL-1066 |
| chr14-32568693-DEL-6253 | chr14-32568694-DEL-6253 |
| chr14-32599187-DEL-67 | chr14-32599177-DEL-62 |
| chr14-32612656-DEL-1493 | chr14-32612657-DEL-1493 |
| chr14-32620833-DEL-2002 | chr14-32620834-DEL-2002 |
| chr14-32625786-DEL-95 | chr14-32625787-DEL-95 |
| chr14-32692239-DEL-120 | chr14-32692249-DEL-128 |
| chr14-32694931-DEL-464 | chr14-32694932-DEL-464 |
| chr14-32720753-DEL-66 | chr14-32720754-DEL-66 |
| chr14-32751296-DEL-57 | chr14-32751295-DEL-59 |
| chr14-32806433-DEL-60 | chr14-32806422-DEL-72 |
| chr14-32827087-DEL-376 | chr14-32827088-DEL-376 |
| chr14-32879496-DEL-105 | chr14-32879527-DEL-70 |
| chr14-32900342-DEL-80 | chr14-32900343-DEL-80 |
| chr14-32902859-DEL-418 | chr14-32902860-DEL-418 |
| chr14-32903377-DEL-309 | chr14-32903378-DEL-309 |
| chr14-32919688-DEL-194 | chr14-32919688-DEL-195 |
| chr14-32920304-DEL-944 | chr14-32920306-DEL-941 |
| chr14-32927614-DEL-83 | chr14-32927615-DEL-73 |
| chr14-32942693-DEL-4236 | chr14-32942694-DEL-4236 |
| chr14-32974084-DEL-119 | chr14-32974089-DEL-115 |
| chr14-32989097-DEL-1603 | chr14-32989098-DEL-1603 |
| chr14-32991557-DEL-441 | chr14-32991558-DEL-441 |
| chr14-33000245-DEL-1002 | chr14-33000246-DEL-1002 |
| chr14-33014995-DEL-114 | chr14-33014996-DEL-72 |
| chr14-33021774-DEL-56 | chr14-33021793-DEL-56 |
| chr14-33045912-DEL-161 | chr14-33045913-DEL-161 |
| chr14-33049449-DEL-74 | chr14-33049450-DEL-74 |
| chr14-33056580-DEL-171 | chr14-33056581-DEL-171 |
| chr14-33069680-DEL-54 | chr14-33069681-DEL-54 |
| chr14-33089770-DEL-656 | chr14-33089771-DEL-656 |
| chr14-33096791-DEL-73 | chr14-33096782-DEL-83 |
| chr14-33123090-DEL-89 | chr14-33123095-DEL-83 |
| chr14-33179586-DEL-1058 | chr14-33179587-DEL-1058 |
| chr14-33209103-DEL-86 | chr14-33209104-DEL-86 |
| chr14-33289602-DEL-132 | chr14-33289603-DEL-132 |
| chr14-33298985-DEL-516 | chr14-33298986-DEL-516 |
| chr14-33305701-DEL-931 | chr14-33305702-DEL-931 |
| chr14-33315949-DEL-99 | chr14-33315950-DEL-99 |
| chr14-33369789-DEL-2957 | chr14-33369790-DEL-2957 |
| chr14-33376117-DEL-219 | chr14-33376118-DEL-219 |
| chr14-33389259-DEL-2710 | chr14-33389260-DEL-2710 |
| chr14-33408183-DEL-492 | chr14-33408184-DEL-492 |
| chr14-33411599-DEL-70 | chr14-33411600-DEL-70 |
| chr14-33480422-DEL-68 | chr14-33480411-DEL-69 |
| chr14-33484620-DEL-74 | chr14-33484607-DEL-70 |
| chr14-33503611-DEL-119 | chr14-33503612-DEL-119 |
| chr14-33526095-DEL-167 | chr14-33526097-DEL-165 |
| chr14-33613853-DEL-59 | chr14-33613854-DEL-59 |
| chr14-33632203-DEL-198 | chr14-33632204-DEL-198 |
| chr14-33639909-DEL-124 | chr14-33639889-DEL-156 |
| chr14-33664711-DEL-68 | chr14-33664712-DEL-68 |
| chr14-33673537-DEL-392 | chr14-33673538-DEL-392 |

| | |
|---|---|
| chr14-33702247-DEL-7095 | chr14-33702248-DEL-7095 |
| chr14-33744335-DEL-302 | chr14-33744336-DEL-302 |
| chr14-33803219-DEL-1283 | chr14-33803220-DEL-1283 |
| chr14-33816994-DEL-119 | chr14-33816988-DEL-120 |
| chr14-33822816-DEL-320 | chr14-33822817-DEL-320 |
| chr14-33836680-DEL-52 | chr14-33836686-DEL-52 |
| chr14-33861517-DEL-439 | chr14-33861518-DEL-439 |
| chr14-33879920-DEL-1384 | chr14-33879921-DEL-1384 |
| chr14-33888947-DEL-233 | chr14-33888939-DEL-242 |
| chr14-33892584-DEL-561 | chr14-33892585-DEL-561 |
| chr14-33939343-DEL-123 | chr14-33939348-DEL-122 |
| chr14-33947157-DEL-82 | chr14-33947158-DEL-82 |
| chr14-33973696-DEL-131 | chr14-33973697-DEL-131 |
| chr14-33975470-DEL-52 | chr14-33975471-DEL-52 |
| chr14-33981872-DEL-214 | chr14-33981873-DEL-214 |
| chr14-33984192-DEL-61 | chr14-33984196-DEL-60 |
| chr14-33991001-DEL-84 | chr14-33991002-DEL-84 |
| chr14-33993722-DEL-1358 | chr14-33993723-DEL-1358 |
| chr14-34027324-DEL-228 | chr14-34027325-DEL-228 |
| chr14-34029967-DEL-88 | chr14-34029968-DEL-88 |
| chr14-34037675-DEL-196 | chr14-34037676-DEL-196 |
| chr14-34043971-DEL-942 | chr14-34043972-DEL-942 |
| chr14-34062370-DEL-222 | chr14-34062371-DEL-221 |
| chr14-34067223-DEL-417 | chr14-34067224-DEL-417 |
| chr14-34068858-DEL-101 | chr14-34068850-DEL-89 |
| chr14-34084947-DEL-90 | chr14-34084948-DEL-90 |
| chr14-34105805-DEL-5295 | chr14-34105806-DEL-5295 |
| chr14-34115158-DEL-145 | chr14-34115159-DEL-145 |
| chr14-34122424-DEL-206 | chr14-34122454-DEL-166 |
| chr14-34123956-DEL-123 | chr14-34123937-DEL-124 |
| chr14-34156855-DEL-202 | chr14-34156856-DEL-202 |
| chr14-34157830-DEL-239 | chr14-34157831-DEL-239 |
| chr14-34160153-DEL-157 | chr14-34160157-DEL-156 |
| chr14-34164008-DEL-186 | chr14-34164009-DEL-186 |
| chr14-34217020-DEL-205 | chr14-34217021-DEL-205 |
| chr14-34235191-DEL-96 | chr14-34235192-DEL-96 |
| chr14-34263383-DEL-209 | chr14-34263384-DEL-209 |
| chr14-34287228-DEL-61 | chr14-34287229-DEL-61 |
| chr14-34301743-DEL-370 | chr14-34301744-DEL-370 |
| chr14-34328893-DEL-57 | chr14-34328884-DEL-65 |
| chr14-34333964-DEL-51 | chr14-34333965-DEL-51 |
| chr14-34341743-DEL-223 | chr14-34341748-DEL-225 |
| chr14-34360725-DEL-61 | chr14-34360726-DEL-53 |
| chr14-34361672-DEL-1886 | chr14-34361673-DEL-1886 |
| chr14-34368413-DEL-392 | chr14-34368414-DEL-382 |
| chr14-34374172-DEL-707 | chr14-34374173-DEL-707 |
| chr14-34379807-DEL-547 | chr14-34379808-DEL-547 |
| chr14-34413328-DEL-210 | chr14-34413329-DEL-210 |
| chr14-34420685-DEL-230 | chr14-34420686-DEL-230 |
| chr14-34445069-DEL-83 | chr14-34445067-DEL-76 |
| chr14-34449157-DEL-64 | chr14-34449150-DEL-72 |
| chr14-34478855-DEL-401 | chr14-34478856-DEL-401 |
| chr14-34493797-DEL-328 | chr14-34493798-DEL-328 |
| chr14-34505475-DEL-63 | chr14-34505443-DEL-55 |
| chr14-34517302-DEL-11966 | chr14-34517303-DEL-11966 |
| chr14-34536356-DEL-625 | chr14-34536357-DEL-625 |
| chr14-34570036-DEL-572 | chr14-34570037-DEL-572 |
| chr14-34600595-DEL-112 | chr14-34600596-DEL-112 |
| chr14-34620458-DEL-2192 | chr14-34620461-DEL-2193 |
| chr14-34662509-DEL-124 | chr14-34662510-DEL-124 |
| chr14-34697921-DEL-771 | chr14-34697922-DEL-771 |
| chr14-34700115-DEL-221 | chr14-34700116-DEL-221 |
| chr14-34711534-DEL-1658 | chr14-34711535-DEL-1658 |
| chr14-34714975-DEL-225 | chr14-34714976-DEL-225 |
| chr14-34754290-DEL-92 | chr14-34754293-DEL-90 |
| chr14-34797278-DEL-54 | chr14-34797279-DEL-58 |
| chr14-34807505-DEL-99 | chr14-34807500-DEL-100 |
| chr14-34815055-DEL-430 | chr14-34815056-DEL-430 |
| chr14-34871190-DEL-503 | chr14-34871191-DEL-503 |
| chr14-34887551-DEL-86 | chr14-34887560-DEL-55 |
| chr14-34892665-DEL-221 | chr14-34892666-DEL-221 |
| chr14-34921553-DEL-6912 | chr14-34921537-DEL-6928 |
| chr14-34928162-DEL-205 | chr14-34928163-DEL-205 |
| chr14-34930052-DEL-646 | chr14-34930053-DEL-646 |
| chr14-34934569-DEL-4024 | chr14-34934565-DEL-4025 |
| chr14-34945960-DEL-138 | chr14-34945961-DEL-138 |
| chr14-34955920-DEL-264 | chr14-34955921-DEL-264 |
| chr14-34997186-DEL-6276 | chr14-34997187-DEL-6276 |
| chr14-35015983-DEL-86 | chr14-35015964-DEL-74 |
| chr14-35050041-DEL-61 | chr14-35050030-DEL-64 |

| | |
|---|---|
| chr14-35069834-DEL-174 | chr14-35069834-DEL-175 |
| chr14-35092476-DEL-2143 | chr14-35092477-DEL-2143 |
| chr14-35110673-DEL-59 | chr14-35110674-DEL-59 |
| chr14-35119480-DEL-6434 | chr14-35119494-DEL-6432 |
| chr14-35154344-DEL-7318 | chr14-35154345-DEL-7318 |
| chr14-35176039-DEL-57 | chr14-35176040-DEL-57 |
| chr14-35197556-DEL-1382 | chr14-35197557-DEL-1382 |
| chr14-35208833-DEL-76 | chr14-35208834-DEL-76 |
| chr14-35236154-DEL-403 | chr14-35236155-DEL-403 |
| chr14-35249151-DEL-6663 | chr14-35249152-DEL-6663 |
| chr14-35323860-DEL-940 | chr14-35323862-DEL-939 |
| chr14-35410647-DEL-52 | chr14-35410648-DEL-52 |
| chr14-35413394-DEL-210 | chr14-35413395-DEL-210 |
| chr14-35414882-DEL-79 | chr14-35414859-DEL-84 |
| chr14-35431107-DEL-6394 | chr14-35431108-DEL-6394 |
| chr14-35438692-DEL-5443 | chr14-35438693-DEL-5443 |
| chr14-35448459-DEL-72 | chr14-35448451-DEL-80 |
| chr14-35472358-DEL-116 | chr14-35472359-DEL-116 |
| chr14-35518762-DEL-3745 | chr14-35518763-DEL-3745 |
| chr14-35554780-DEL-2419 | chr14-35554781-DEL-2418 |
| chr14-35613841-DEL-6068 | chr14-35613842-DEL-6068 |
| chr14-35624444-DEL-64 | chr14-35624463-DEL-53 |
| chr14-35652777-DEL-66 | chr14-35652778-DEL-64 |
| chr14-35699883-DEL-67 | chr14-35699883-DEL-68 |
| chr14-35703633-DEL-91 | chr14-35703634-DEL-91 |
| chr14-35722391-DEL-84 | chr14-35722396-DEL-80 |
| chr14-35733515-DEL-493 | chr14-35733516-DEL-493 |
| chr14-35734861-DEL-51 | chr14-35734861-DEL-51 |
| chr14-35744894-DEL-62 | chr14-35744895-DEL-62 |
| chr14-35786681-DEL-204 | chr14-35786682-DEL-204 |
| chr14-35825161-DEL-245 | chr14-35825162-DEL-245 |
| chr14-35828324-DEL-56 | chr14-35828325-DEL-52 |
| chr14-35907601-DEL-1099 | chr14-35907602-DEL-1099 |
| chr14-35914986-DEL-65 | chr14-35914987-DEL-65 |
| chr14-35920705-DEL-4338 | chr14-35920706-DEL-4338 |
| chr14-35955834-DEL-83 | chr14-35955834-DEL-83 |
| chr14-35997227-DEL-86 | chr14-35997234-DEL-80 |
| chr14-36007681-DEL-707 | chr14-36007682-DEL-707 |
| chr14-36011701-DEL-1906 | chr14-36011702-DEL-1906 |
| chr14-36021804-DEL-52 | chr14-36021797-DEL-79 |
| chr14-36024850-DEL-4749 | chr14-36024851-DEL-4749 |
| chr14-36064812-DEL-94 | chr14-36064823-DEL-80 |
| chr14-36070603-DEL-554 | chr14-36070604-DEL-554 |
| chr14-36075498-DEL-232 | chr14-36075499-DEL-232 |
| chr14-36090049-DEL-874 | chr14-36090050-DEL-874 |
| chr14-36149667-DEL-6329 | chr14-36149668-DEL-6329 |
| chr14-36159094-DEL-3233 | chr14-36159073-DEL-3235 |
| chr14-36244453-DEL-122 | chr14-36244450-DEL-126 |
| chr14-36261380-DEL-6411 | chr14-36261381-DEL-6411 |
| chr14-36291402-DEL-382 | chr14-36291403-DEL-379 |
| chr14-36318299-DEL-6664 | chr14-36318293-DEL-6665 |
| chr14-36354761-DEL-585 | chr14-36354762-DEL-585 |
| chr14-36393466-DEL-93 | chr14-36393465-DEL-95 |
| chr14-36565690-DEL-891 | chr14-36565691-DEL-891 |
| chr14-36569161-DEL-1345 | chr14-36569162-DEL-1345 |
| chr14-36603321-DEL-51 | chr14-36603354-DEL-51 |
| chr14-36623492-DEL-56 | chr14-36623493-DEL-52 |
| chr14-36632706-DEL-218 | chr14-36632707-DEL-218 |
| chr14-36682016-DEL-57 | chr14-36682015-DEL-58 |
| chr14-36703504-DEL-82 | chr14-36703525-DEL-62 |
| chr14-36719644-DEL-60 | chr14-36719643-DEL-58 |
| chr14-36742270-DEL-3697 | chr14-36742271-DEL-3697 |
| chr14-36766936-DEL-87 | chr14-36766935-DEL-95 |
| chr14-36828859-DEL-54 | chr14-36828848-DEL-66 |
| chr14-36830512-DEL-95 | chr14-36830506-DEL-100 |
| chr14-36837706-DEL-507 | chr14-36837698-DEL-510 |
| chr14-36841690-DEL-86 | chr14-36841691-DEL-86 |
| chr14-36859832-DEL-427 | chr14-36859828-DEL-422 |
| chr14-36864228-DEL-78 | chr14-36864229-DEL-78 |
| chr14-36865149-DEL-401 | chr14-36865150-DEL-401 |
| chr14-36879331-DEL-1823 | chr14-36879332-DEL-1823 |
| chr14-36895446-DEL-52 | chr14-36895446-DEL-50 |
| chr14-36899854-DEL-194 | chr14-36899879-DEL-170 |
| chr14-36908993-DEL-251 | chr14-36908996-DEL-249 |
| chr14-36913645-DEL-6375 | chr14-36913656-DEL-6365 |
| chr14-36919456-DEL-212 | chr14-36919456-DEL-212 |
| chr14-36927667-DEL-143 | chr14-36927668-DEL-143 |
| chr14-36951148-DEL-7062 | chr14-36951149-DEL-7062 |
| chr14-36964349-DEL-419 | chr14-36964350-DEL-419 |
| chr14-36976721-DEL-6397 | chr14-36976722-DEL-6397 |

| | |
|---|---|
| chr14-36990271-DEL-86 | chr14-36990272-DEL-86 |
| chr14-37015046-DEL-66 | chr14-37015054-DEL-58 |
| chr14-37021773-DEL-158 | chr14-37021774-DEL-158 |
| chr14-37025665-DEL-123 | chr14-37025666-DEL-123 |
| chr14-37054435-DEL-98 | chr14-37054436-DEL-98 |
| chr14-37070434-DEL-302 | chr14-37070435-DEL-302 |
| chr14-37085807-DEL-116 | chr14-37085795-DEL-117 |
| chr14-37115465-DEL-6808 | chr14-37115466-DEL-6808 |
| chr14-37132164-DEL-196 | chr14-37132157-DEL-204 |
| chr14-37175709-DEL-58 | chr14-37175710-DEL-58 |
| chr14-37180360-DEL-1873 | chr14-37180361-DEL-1873 |
| chr14-37183021-DEL-5727 | chr14-37183021-DEL-5729 |
| chr14-37192984-DEL-332 | chr14-37192985-DEL-332 |
| chr14-37196001-DEL-73 | chr14-37196002-DEL-73 |
| chr14-37222650-DEL-4376 | chr14-37222651-DEL-4376 |
| chr14-37228611-DEL-117 | chr14-37228612-DEL-117 |
| chr14-37229788-DEL-449 | chr14-37229789-DEL-449 |
| chr14-37258523-DEL-292 | chr14-37258523-DEL-292 |
| chr14-37275132-DEL-68 | chr14-37275133-DEL-68 |
| chr14-37276442-DEL-145 | chr14-37276444-DEL-165 |
| chr14-37295993-DEL-1017 | chr14-37295993-DEL-1018 |
| chr14-37322279-DEL-92 | chr14-37322280-DEL-92 |
| chr14-37386378-DEL-1090 | chr14-37386379-DEL-1090 |
| chr14-37390472-DEL-1595 | chr14-37390466-DEL-1590 |
| chr14-37391292-DEL-1393 | chr14-37391293-DEL-1393 |
| chr14-37393798-DEL-178 | chr14-37393799-DEL-178 |
| chr14-37399596-DEL-460 | chr14-37399597-DEL-460 |
| chr14-37402574-DEL-206 | chr14-37402575-DEL-206 |
| chr14-37403239-DEL-376 | chr14-37403240-DEL-376 |
| chr14-37406826-DEL-52 | chr14-37406827-DEL-52 |
| chr14-37410562-DEL-73 | chr14-37410563-DEL-73 |
| chr14-37420880-DEL-234 | chr14-37420881-DEL-234 |
| chr14-37427494-DEL-1278 | chr14-37427495-DEL-1278 |
| chr14-37429805-DEL-1542 | chr14-37429805-DEL-1542 |
| chr14-37447444-DEL-61 | chr14-37447443-DEL-62 |
| chr14-37450155-DEL-207 | chr14-37450156-DEL-207 |
| chr14-37464502-DEL-184 | chr14-37464503-DEL-183 |
| chr14-37468046-DEL-414 | chr14-37468047-DEL-409 |
| chr14-37483742-DEL-68 | chr14-37483743-DEL-68 |
| chr14-37500484-DEL-6174 | chr14-37500485-DEL-6174 |
| chr14-37511076-DEL-338 | chr14-37511077-DEL-338 |
| chr14-37536966-DEL-124 | chr14-37536967-DEL-124 |
| chr14-37537883-DEL-52 | chr14-37537885-DEL-58 |
| chr14-37538671-DEL-500 | chr14-37538672-DEL-500 |
| chr14-37543490-DEL-58 | chr14-37543491-DEL-58 |
| chr14-37559553-DEL-6379 | chr14-37559554-DEL-6379 |
| chr14-37584412-DEL-87 | chr14-37584412-DEL-81 |
| chr14-37598194-DEL-386 | chr14-37598195-DEL-386 |
| chr14-37614883-DEL-395 | chr14-37614884-DEL-395 |
| chr14-37628817-DEL-1656 | chr14-37628818-DEL-1656 |
| chr14-37641373-DEL-62 | chr14-37641374-DEL-62 |
| chr14-37647345-DEL-424 | chr14-37647346-DEL-424 |
| chr14-37658924-DEL-6411 | chr14-37658925-DEL-6411 |
| chr14-37689093-DEL-53 | chr14-37689094-DEL-53 |
| chr14-37701254-DEL-143 | chr14-37701255-DEL-162 |
| chr14-37714140-DEL-68 | chr14-37714141-DEL-68 |
| chr14-37734431-DEL-284 | chr14-37734432-DEL-284 |
| chr14-37747248-DEL-2694 | chr14-37747249-DEL-2694 |
| chr14-37757889-DEL-3259 | chr14-37757890-DEL-3259 |
| chr14-37805371-DEL-634 | chr14-37805372-DEL-634 |
| chr14-37808706-DEL-1255 | chr14-37808707-DEL-1255 |
| chr14-37820144-DEL-54 | chr14-37820145-DEL-54 |
| chr14-37830579-DEL-9258 | chr14-37830582-DEL-9255 |
| chr14-37845577-DEL-322 | chr14-37845578-DEL-322 |
| chr14-37849498-DEL-894 | chr14-37849499-DEL-894 |
| chr14-37860097-DEL-1681 | chr14-37860098-DEL-1681 |
| chr14-37870614-DEL-3292 | chr14-37870613-DEL-3293 |
| chr14-37891577-DEL-6463 | chr14-37891569-DEL-6464 |
| chr14-37893012-DEL-462 | chr14-37893013-DEL-462 |
| chr14-37922512-DEL-183 | chr14-37922513-DEL-183 |
| chr14-37927318-DEL-1068 | chr14-37927319-DEL-1068 |
| chr14-37993158-DEL-58 | chr14-37993159-DEL-58 |
| chr14-38044563-DEL-523 | chr14-38044564-DEL-523 |
| chr14-38048978-DEL-253 | chr14-38048998-DEL-224 |
| chr14-38068182-DEL-1624 | chr14-38068183-DEL-1623 |
| chr14-38072861-DEL-923 | chr14-38072862-DEL-922 |
| chr14-38077802-DEL-195 | chr14-38077803-DEL-191 |
| chr14-38079593-DEL-53 | chr14-38079590-DEL-57 |
| chr14-38102396-DEL-417 | chr14-38102397-DEL-417 |
| chr14-38146656-DEL-64 | chr14-38146663-DEL-58 |

| | |
|---|---|
| chr14-38149514-DEL-324 | chr14-38149513-DEL-324 |
| chr14-38177052-DEL-7079 | chr14-38177053-DEL-7079 |
| chr14-38189497-DEL-56 | chr14-38189478-DEL-78 |
| chr14-38208979-DEL-78 | chr14-38208948-DEL-127 |
| chr14-38216762-DEL-90 | chr14-38216757-DEL-96 |
| chr14-38219606-DEL-621 | chr14-38219607-DEL-621 |
| chr14-38235265-DEL-72 | chr14-38235268-DEL-72 |
| chr14-38249712-DEL-201 | chr14-38249713-DEL-200 |
| chr14-38251122-DEL-894 | chr14-38251123-DEL-894 |
| chr14-38257957-DEL-7319 | chr14-38257958-DEL-7319 |
| chr14-38303826-DEL-67 | chr14-38303827-DEL-54 |
| chr14-38307668-DEL-766 | chr14-38307668-DEL-766 |
| chr14-38316806-DEL-167 | chr14-38316806-DEL-167 |
| chr14-38327294-DEL-6158 | chr14-38327295-DEL-6158 |
| chr14-38361295-DEL-64 | chr14-38361296-DEL-64 |
| chr14-38361903-DEL-62 | chr14-38361904-DEL-60 |
| chr14-38365150-DEL-447 | chr14-38365151-DEL-447 |
| chr14-38394592-DEL-464 | chr14-38394593-DEL-465 |
| chr14-38396177-DEL-2702 | chr14-38396194-DEL-2698 |
| chr14-38399019-DEL-61 | chr14-38399018-DEL-67 |
| chr14-38415077-DEL-1459 | chr14-38415077-DEL-1459 |
| chr14-38419743-DEL-74 | chr14-38419744-DEL-74 |
| chr14-38468387-DEL-587 | chr14-38468375-DEL-599 |
| chr14-38475947-DEL-2090 | chr14-38475948-DEL-2090 |
| chr14-38489997-DEL-441 | chr14-38489997-DEL-441 |
| chr14-38490740-DEL-132 | chr14-38490741-DEL-132 |
| chr14-38503257-DEL-420 | chr14-38503258-DEL-420 |
| chr14-38508103-DEL-2033 | chr14-38508104-DEL-2033 |
| chr14-38525192-DEL-60 | chr14-38525191-DEL-54 |
| chr14-38529507-DEL-81 | chr14-38529494-DEL-83 |
| chr14-38562180-DEL-497 | chr14-38562182-DEL-496 |
| chr14-38566600-DEL-569 | chr14-38566601-DEL-569 |
| chr14-38584902-DEL-419 | chr14-38584903-DEL-419 |
| chr14-38635512-DEL-60 | chr14-38635493-DEL-60 |
| chr14-38647714-DEL-76 | chr14-38647715-DEL-76 |
| chr14-38683908-DEL-61 | chr14-38683906-DEL-61 |
| chr14-38702190-DEL-1832 | chr14-38702191-DEL-1832 |
| chr14-38724646-DEL-68 | chr14-38724647-DEL-68 |
| chr14-38729636-DEL-186 | chr14-38729633-DEL-186 |
| chr14-38750565-DEL-609 | chr14-38750566-DEL-609 |
| chr14-38786253-DEL-63 | chr14-38786254-DEL-63 |
| chr14-38820066-DEL-56 | chr14-38820069-DEL-54 |
| chr14-38821239-DEL-102 | chr14-38821243-DEL-54 |
| chr14-38878251-DEL-64 | chr14-38878252-DEL-64 |
| chr14-38908610-DEL-141 | chr14-38908611-DEL-141 |
| chr14-38915875-DEL-963 | chr14-38915873-DEL-964 |
| chr14-38931520-DEL-1421 | chr14-38931521-DEL-1421 |
| chr14-38964324-DEL-1759 | chr14-38964325-DEL-1759 |
| chr14-39005081-DEL-6387 | chr14-39005082-DEL-6387 |
| chr14-39202460-DEL-255 | chr14-39202457-DEL-253 |
| chr14-39215315-DEL-3781 | chr14-39215316-DEL-3781 |
| chr14-39226444-DEL-58 | chr14-39226445-DEL-58 |
| chr14-39266161-DEL-482 | chr14-39266162-DEL-482 |
| chr14-39268792-DEL-6791 | chr14-39268793-DEL-6791 |
| chr14-39274840-DEL-416 | chr14-39274841-DEL-416 |
| chr14-39306609-DEL-6469 | chr14-39306610-DEL-6469 |
| chr14-39314379-DEL-389 | chr14-39314380-DEL-389 |
| chr14-39314874-DEL-608 | chr14-39314875-DEL-608 |
| chr14-39315879-DEL-5376 | chr14-39315880-DEL-5376 |
| chr14-39326666-DEL-6195 | chr14-39326667-DEL-6195 |
| chr14-39361090-DEL-4366 | chr14-39361091-DEL-4365 |
| chr14-39369742-DEL-445 | chr14-39369743-DEL-445 |
| chr14-39378063-DEL-128 | chr14-39378026-DEL-120 |
| chr14-39387326-DEL-5685 | chr14-39387327-DEL-5685 |
| chr14-39398766-DEL-149 | chr14-39398767-DEL-149 |
| chr14-39433850-DEL-57 | chr14-39433851-DEL-57 |
| chr14-39442869-DEL-59 | chr14-39442868-DEL-61 |
| chr14-39453386-DEL-5811 | chr14-39453387-DEL-5811 |
| chr14-39460390-DEL-58 | chr14-39460391-DEL-58 |
| chr14-39466322-DEL-54 | chr14-39466322-DEL-54 |
| chr14-39467307-DEL-282 | chr14-39467308-DEL-282 |
| chr14-39468019-DEL-60 | chr14-39468020-DEL-60 |
| chr14-39479461-DEL-7216 | chr14-39479462-DEL-7216 |
| chr14-39498299-DEL-274 | chr14-39498318-DEL-271 |
| chr14-39505979-DEL-95 | chr14-39505980-DEL-94 |
| chr14-39506202-DEL-495 | chr14-39506203-DEL-495 |
| chr14-39516309-DEL-2789 | chr14-39516310-DEL-2789 |
| chr14-39524656-DEL-3495 | chr14-39524657-DEL-3495 |
| chr14-39529034-DEL-71 | chr14-39529014-DEL-78 |
| chr14-39559627-DEL-263 | chr14-39559628-DEL-263 |

| | |
|---|---|
| chr14-39574567-DEL-54 | chr14-39574567-DEL-54 |
| chr14-39627001-DEL-6274 | chr14-39627002-DEL-6274 |
| chr14-39665611-DEL-196 | chr14-39665624-DEL-188 |
| chr14-39701635-DEL-437 | chr14-39701636-DEL-437 |
| chr14-39705689-DEL-277 | chr14-39705690-DEL-277 |
| chr14-39757188-DEL-874 | chr14-39757189-DEL-874 |
| chr14-39765250-DEL-338 | chr14-39765251-DEL-338 |
| chr14-39771563-DEL-83 | chr14-39771562-DEL-84 |
| chr14-39779799-DEL-2795 | chr14-39779800-DEL-2795 |
| chr14-39788002-DEL-6325 | chr14-39788003-DEL-6325 |
| chr14-39818966-DEL-7144 | chr14-39818967-DEL-7144 |
| chr14-39834193-DEL-861 | chr14-39834194-DEL-861 |
| chr14-39835714-DEL-52 | chr14-39835716-DEL-51 |
| chr14-39842926-DEL-354 | chr14-39842927-DEL-354 |
| chr14-39843614-DEL-14815 | chr14-39843615-DEL-14815 |
| chr14-39863302-DEL-67 | chr14-39863303-DEL-67 |
| chr14-39882238-DEL-750 | chr14-39882239-DEL-750 |
| chr14-39898186-DEL-350 | chr14-39898187-DEL-350 |
| chr14-39902751-DEL-211 | chr14-39902752-DEL-211 |
| chr14-39905810-DEL-6439 | chr14-39905811-DEL-6439 |
| chr14-39952914-DEL-1814 | chr14-39952915-DEL-1814 |
| chr14-39995217-DEL-93 | chr14-39995218-DEL-93 |
| chr14-40084695-DEL-89 | chr14-40084696-DEL-89 |
| chr14-40192848-DEL-62 | chr14-40192865-DEL-62 |
| chr14-40217671-DEL-123 | chr14-40217661-DEL-120 |
| chr14-40241806-DEL-764 | chr14-40241808-DEL-758 |
| chr14-40244361-DEL-60 | chr14-40244390-DEL-58 |
| chr14-40265771-DEL-183 | chr14-40265773-DEL-180 |
| chr14-40268549-DEL-7595 | chr14-40268550-DEL-7595 |
| chr14-40286551-DEL-72 | chr14-40286547-DEL-76 |
| chr14-40296627-DEL-75 | chr14-40296627-DEL-70 |
| chr14-40304599-DEL-76 | chr14-40304600-DEL-76 |
| chr14-40343942-DEL-450 | chr14-40343943-DEL-450 |
| chr14-40373013-DEL-114 | chr14-40373014-DEL-114 |
| chr14-40392697-DEL-57 | chr14-40392709-DEL-50 |
| chr14-40410022-DEL-122 | chr14-40410023-DEL-122 |
| chr14-40440189-DEL-68 | chr14-40440190-DEL-68 |
| chr14-40477339-DEL-747 | chr14-40477326-DEL-748 |
| chr14-40483511-DEL-3693 | chr14-40483512-DEL-3693 |
| chr14-40503603-DEL-6206 | chr14-40503604-DEL-6206 |
| chr14-40534091-DEL-206 | chr14-40534092-DEL-206 |
| chr14-40535380-DEL-1203 | chr14-40535381-DEL-1203 |
| chr14-40541410-DEL-54 | chr14-40541411-DEL-54 |
| chr14-40566280-DEL-6820 | chr14-40566276-DEL-6821 |
| chr14-40574015-DEL-54 | chr14-40574016-DEL-54 |
| chr14-40598240-DEL-94 | chr14-40598273-DEL-79 |
| chr14-40620805-DEL-249 | chr14-40620806-DEL-249 |
| chr14-40682788-DEL-459 | chr14-40682789-DEL-459 |
| chr14-40703829-DEL-166 | chr14-40703830-DEL-166 |
| chr14-40738594-DEL-110 | chr14-40738595-DEL-111 |
| chr14-40759074-DEL-116 | chr14-40759075-DEL-116 |
| chr14-40765761-DEL-548 | chr14-40765762-DEL-548 |
| chr14-40775345-DEL-8270 | chr14-40775346-DEL-8270 |
| chr14-40803595-DEL-62 | chr14-40803584-DEL-59 |
| chr14-40818852-DEL-80 | chr14-40818853-DEL-80 |
| chr14-40850732-DEL-426 | chr14-40850733-DEL-426 |
| chr14-40859870-DEL-98 | chr14-40859871-DEL-98 |
| chr14-40863358-DEL-441 | chr14-40863359-DEL-441 |
| chr14-40866730-DEL-246 | chr14-40866723-DEL-247 |
| chr14-40868654-DEL-83 | chr14-40868655-DEL-84 |
| chr14-40870268-DEL-106 | chr14-40870269-DEL-106 |
| chr14-40912253-DEL-402 | chr14-40912254-DEL-402 |
| chr14-40924035-DEL-53 | chr14-40924036-DEL-53 |
| chr14-40954374-DEL-478 | chr14-40954375-DEL-478 |
| chr14-41077492-DEL-167 | chr14-41077479-DEL-169 |
| chr14-41220399-DEL-204 | chr14-41220400-DEL-204 |
| chr14-41233103-DEL-851 | chr14-41233104-DEL-851 |
| chr14-41342746-DEL-68 | chr14-41342724-DEL-50 |
| chr14-41446020-DEL-345 | chr14-41446019-DEL-345 |
| chr14-41460169-DEL-1478 | chr14-41460170-DEL-1478 |
| chr14-41816423-DEL-66 | chr14-41816424-DEL-66 |
| chr14-42031895-DEL-1031 | chr14-42031895-DEL-1031 |
| chr14-42394270-DEL-77 | chr14-42394271-DEL-77 |
| chr14-42542593-DEL-785 | chr14-42542593-DEL-785 |
| chr14-42982136-DEL-145 | chr14-42982140-DEL-98 |
| chr14-43001945-DEL-681 | chr14-43001946-DEL-681 |
| chr14-43776431-DEL-214 | chr14-43776432-DEL-214 |
| chr14-44016199-DEL-529 | chr14-44016200-DEL-529 |
| chr14-44016853-DEL-87 | chr14-44016855-DEL-88 |
| chr14-44117089-DEL-147 | chr14-44117089-DEL-147 |

| | |
|---|---|
| chr14-44223159-DEL-201 | chr14-44223159-DEL-201 |
| chr14-44341079-DEL-482 | chr14-44341078-DEL-478 |
| chr14-44354523-DEL-275 | chr14-44354524-DEL-275 |
| chr14-44946261-DEL-518 | chr14-44946261-DEL-518 |
| chr14-45000936-DEL-473 | chr14-45000937-DEL-473 |
| chr14-45027916-DEL-71 | chr14-45027930-DEL-58 |
| chr14-45037868-DEL-165 | chr14-45037869-DEL-165 |
| chr14-45050249-DEL-82 | chr14-45050250-DEL-82 |
| chr14-45056709-DEL-1656 | chr14-45056710-DEL-1655 |
| chr14-45074228-DEL-228 | chr14-45074214-DEL-227 |
| chr14-45076893-DEL-432 | chr14-45076889-DEL-436 |
| chr14-45114627-DEL-1543 | chr14-45114628-DEL-1543 |
| chr14-45124407-DEL-489 | chr14-45124408-DEL-489 |
| chr14-45127477-DEL-136 | chr14-45127495-DEL-118 |
| chr14-45160706-DEL-66 | chr14-45160707-DEL-66 |
| chr14-45170504-DEL-82 | chr14-45170503-DEL-82 |
| chr14-45171226-DEL-725 | chr14-45171224-DEL-726 |
| chr14-45212543-DEL-687 | chr14-45212555-DEL-675 |
| chr14-45246636-DEL-64 | chr14-45246637-DEL-66 |
| chr14-45251084-DEL-6664 | chr14-45251085-DEL-6664 |
| chr14-45289431-DEL-104 | chr14-45289432-DEL-104 |
| chr14-45300266-DEL-379 | chr14-45300267-DEL-379 |
| chr14-45324854-DEL-1665 | chr14-45324855-DEL-1665 |
| chr14-45330414-DEL-56 | chr14-45330415-DEL-56 |
| chr14-45339223-DEL-56 | chr14-45339224-DEL-56 |
| chr14-45347326-DEL-1687 | chr14-45347327-DEL-1687 |
| chr14-45370862-DEL-216 | chr14-45370863-DEL-216 |
| chr14-45383054-DEL-235 | chr14-45383055-DEL-235 |
| chr14-45405246-DEL-1013 | chr14-45405247-DEL-1013 |
| chr14-45422889-DEL-390 | chr14-45422890-DEL-390 |
| chr14-45480002-DEL-95 | chr14-45480008-DEL-80 |
| chr14-45490103-DEL-206 | chr14-45490104-DEL-206 |
| chr14-45497344-DEL-158 | chr14-45497345-DEL-158 |
| chr14-45501835-DEL-96 | chr14-45501836-DEL-96 |
| chr14-45570660-DEL-133 | chr14-45570661-DEL-133 |
| chr14-45584947-DEL-95 | chr14-45584948-DEL-95 |
| chr14-45587196-DEL-74 | chr14-45587195-DEL-75 |
| chr14-45627368-DEL-191 | chr14-45627369-DEL-191 |
| chr14-45630823-DEL-175 | chr14-45630824-DEL-175 |
| chr14-45652626-DEL-207 | chr14-45652627-DEL-207 |
| chr14-45653443-DEL-180 | chr14-45653444-DEL-180 |
| chr14-45656062-DEL-53 | chr14-45656063-DEL-53 |
| chr14-45658373-DEL-196 | chr14-45658374-DEL-196 |
| chr14-45670510-DEL-79 | chr14-45670511-DEL-79 |
| chr14-45676638-DEL-171 | chr14-45676639-DEL-170 |
| chr14-45679285-DEL-58 | chr14-45679286-DEL-58 |
| chr14-45681792-DEL-174 | chr14-45681793-DEL-174 |
| chr14-45694250-DEL-85 | chr14-45694251-DEL-80 |
| chr14-45803949-DEL-88 | chr14-45803973-DEL-96 |
| chr14-45818895-DEL-169 | chr14-45818896-DEL-169 |
| chr14-45863548-DEL-64 | chr14-45863550-DEL-68 |
| chr14-45873616-DEL-124 | chr14-45873617-DEL-124 |
| chr14-45936941-DEL-205 | chr14-45936955-DEL-195 |
| chr14-45939877-DEL-151 | chr14-45939878-DEL-151 |
| chr14-45950575-DEL-707 | chr14-45950576-DEL-707 |
| chr14-45959313-DEL-180 | chr14-45959314-DEL-180 |
| chr14-45964476-DEL-73 | chr14-45964474-DEL-74 |
| chr14-45970283-DEL-56 | chr14-45970284-DEL-56 |
| chr14-45974449-DEL-313 | chr14-45974410-DEL-317 |
| chr14-45985566-DEL-75 | chr14-45985567-DEL-75 |
| chr14-45985884-DEL-6437 | chr14-45985881-DEL-6438 |
| chr14-45999226-DEL-52 | chr14-45999210-DEL-63 |
| chr14-46007711-DEL-142 | chr14-46007712-DEL-142 |
| chr14-46019971-DEL-60 | chr14-46019982-DEL-50 |
| chr14-46027599-DEL-210 | chr14-46027600-DEL-210 |
| chr14-46060669-DEL-220 | chr14-46060670-DEL-220 |
| chr14-46065863-DEL-79 | chr14-46065864-DEL-79 |
| chr14-46095427-DEL-112 | chr14-46095428-DEL-112 |
| chr14-46097319-DEL-76 | chr14-46097320-DEL-76 |
| chr14-46099902-DEL-2723 | chr14-46099894-DEL-2724 |
| chr14-46112732-DEL-1387 | chr14-46112733-DEL-1387 |
| chr14-46117970-DEL-2188 | chr14-46117971-DEL-2188 |
| chr14-46121706-DEL-114 | chr14-46121707-DEL-114 |
| chr14-46126568-DEL-6522 | chr14-46126569-DEL-6522 |
| chr14-46141735-DEL-57 | chr14-46141733-DEL-58 |
| chr14-46176243-DEL-72 | chr14-46176244-DEL-72 |
| chr14-46203508-DEL-1433 | chr14-46203509-DEL-1433 |
| chr14-46211731-DEL-125 | chr14-46211732-DEL-125 |
| chr14-46217699-DEL-312 | chr14-46217700-DEL-307 |
| chr14-46302511-DEL-466 | chr14-46302512-DEL-466 |

| | |
|---|---|
| chr14-46321474-DEL-965 | chr14-46321475-DEL-965 |
| chr14-46338722-DEL-91 | chr14-46338721-DEL-89 |
| chr14-46346568-DEL-268 | chr14-46346570-DEL-267 |
| chr14-46353360-DEL-61 | chr14-46353357-DEL-57 |
| chr14-46365061-DEL-825 | chr14-46365062-DEL-825 |
| chr14-46367989-DEL-117 | chr14-46367995-DEL-114 |
| chr14-46372121-DEL-85 | chr14-46372122-DEL-85 |
| chr14-46373005-DEL-68 | chr14-46373006-DEL-66 |
| chr14-46379553-DEL-56 | chr14-46379554-DEL-56 |
| chr14-46421508-DEL-984 | chr14-46421509-DEL-984 |
| chr14-46422686-DEL-1343 | chr14-46422687-DEL-1343 |
| chr14-46432230-DEL-74 | chr14-46432231-DEL-74 |
| chr14-46448951-DEL-6807 | chr14-46448952-DEL-6807 |
| chr14-46474702-DEL-54 | chr14-46474703-DEL-54 |
| chr14-46482770-DEL-60 | chr14-46482771-DEL-60 |
| chr14-46505748-DEL-203 | chr14-46505749-DEL-203 |
| chr14-46524163-DEL-125 | chr14-46524156-DEL-129 |
| chr14-46528493-DEL-58 | chr14-46528494-DEL-50 |
| chr14-46573127-DEL-53 | chr14-46573122-DEL-57 |
| chr14-46589731-DEL-2538 | chr14-46589732-DEL-2538 |
| chr14-46602401-DEL-105 | chr14-46602402-DEL-105 |
| chr14-46604800-DEL-152 | chr14-46604801-DEL-148 |
| chr14-46632898-DEL-319 | chr14-46632899-DEL-289 |
| chr14-46649895-DEL-56 | chr14-46649896-DEL-56 |
| chr14-46675190-DEL-54 | chr14-46675189-DEL-52 |
| chr14-46677782-DEL-280 | chr14-46677783-DEL-280 |
| chr14-46738117-DEL-55 | chr14-46738119-DEL-56 |
| chr14-46748099-DEL-202 | chr14-46748100-DEL-202 |
| chr14-46791024-DEL-64 | chr14-46791025-DEL-64 |
| chr14-46795249-DEL-547 | chr14-46795250-DEL-547 |
| chr14-46818036-DEL-234 | chr14-46818037-DEL-234 |
| chr14-46828608-DEL-180 | chr14-46828609-DEL-180 |
| chr14-46851090-DEL-104 | chr14-46851091-DEL-104 |
| chr14-46856410-DEL-545 | chr14-46856411-DEL-545 |
| chr14-46862046-DEL-68 | chr14-46862058-DEL-57 |
| chr14-46874779-DEL-340 | chr14-46874780-DEL-340 |
| chr14-46891065-DEL-252 | chr14-46891066-DEL-252 |
| chr14-46921362-DEL-298 | chr14-46921363-DEL-298 |
| chr14-46968259-DEL-60 | chr14-46968260-DEL-60 |
| chr14-46983235-DEL-6427 | chr14-46983236-DEL-6427 |
| chr14-46996011-DEL-860 | chr14-46996012-DEL-860 |
| chr14-47001310-DEL-72 | chr14-47001311-DEL-72 |
| chr14-47005525-DEL-286 | chr14-47005526-DEL-286 |
| chr14-47037640-DEL-439 | chr14-47037641-DEL-439 |
| chr14-47039909-DEL-50 | chr14-47039907-DEL-50 |
| chr14-47057660-DEL-207 | chr14-47057661-DEL-207 |
| chr14-47074381-DEL-52 | chr14-47074363-DEL-51 |
| chr14-47110407-DEL-98 | chr14-47110428-DEL-86 |
| chr14-47110557-DEL-490 | chr14-47110544-DEL-486 |
| chr14-47127400-DEL-94 | chr14-47127401-DEL-94 |
| chr14-47135130-DEL-57 | chr14-47135137-DEL-51 |
| chr14-47142342-DEL-224 | chr14-47142343-DEL-224 |
| chr14-47185259-DEL-62 | chr14-47185260-DEL-62 |
| chr14-47194738-DEL-84 | chr14-47194739-DEL-84 |
| chr14-47198521-DEL-358 | chr14-47198520-DEL-358 |
| chr14-47226269-DEL-924 | chr14-47226270-DEL-924 |
| chr14-47230561-DEL-52 | chr14-47230562-DEL-52 |
| chr14-47234681-DEL-68 | chr14-47234674-DEL-69 |
| chr14-47240602-DEL-77 | chr14-47240603-DEL-77 |
| chr14-47247931-DEL-80 | chr14-47247932-DEL-80 |
| chr14-47251737-DEL-123 | chr14-47251738-DEL-123 |
| chr14-47264126-DEL-99 | chr14-47264127-DEL-99 |
| chr14-47268139-DEL-1900 | chr14-47268140-DEL-1899 |
| chr14-47305663-DEL-64 | chr14-47305664-DEL-64 |
| chr14-47308033-DEL-52 | chr14-47308034-DEL-52 |
| chr14-47313104-DEL-59 | chr14-47313105-DEL-59 |
| chr14-47347518-DEL-523 | chr14-47347519-DEL-523 |
| chr14-47352004-DEL-53 | chr14-47352004-DEL-53 |
| chr14-47356482-DEL-61 | chr14-47356483-DEL-61 |
| chr14-47376784-DEL-96 | chr14-47376785-DEL-96 |
| chr14-47381334-DEL-95 | chr14-47381335-DEL-133 |
| chr14-47390488-DEL-136 | chr14-47390489-DEL-136 |
| chr14-47404859-DEL-65 | chr14-47404860-DEL-58 |
| chr14-47411470-DEL-679 | chr14-47411470-DEL-678 |
| chr14-47413824-DEL-147 | chr14-47413825-DEL-147 |
| chr14-47473505-DEL-86 | chr14-47473506-DEL-86 |
| chr14-47477728-DEL-212 | chr14-47477729-DEL-212 |
| chr14-47494518-DEL-132 | chr14-47494519-DEL-132 |
| chr14-47500053-DEL-70 | chr14-47500054-DEL-70 |
| chr14-47508049-DEL-203 | chr14-47508050-DEL-203 |

| | |
|---|---|
| chr14-47519291-DEL-63 | chr14-47519292-DEL-63 |
| chr14-47549003-DEL-201 | chr14-47549004-DEL-201 |
| chr14-47616332-DEL-384 | chr14-47616333-DEL-384 |
| chr14-47642753-DEL-86 | chr14-47642754-DEL-86 |
| chr14-47646260-DEL-83 | chr14-47646261-DEL-83 |
| chr14-47687738-DEL-53 | chr14-47687739-DEL-53 |
| chr14-47688425-DEL-174 | chr14-47688426-DEL-174 |
| chr14-47691337-DEL-196 | chr14-47691338-DEL-196 |
| chr14-47691926-DEL-59 | chr14-47691916-DEL-60 |
| chr14-47698051-DEL-1055 | chr14-47698052-DEL-1055 |
| chr14-47708521-DEL-52 | chr14-47708513-DEL-50 |
| chr14-47750002-DEL-296 | chr14-47750003-DEL-296 |
| chr14-47761356-DEL-109 | chr14-47761357-DEL-109 |
| chr14-47785109-DEL-57 | chr14-47785110-DEL-57 |
| chr14-47812143-DEL-52 | chr14-47812144-DEL-52 |
| chr14-47820943-DEL-201 | chr14-47820944-DEL-201 |
| chr14-47840880-DEL-232 | chr14-47840880-DEL-232 |
| chr14-47844877-DEL-3124 | chr14-47844878-DEL-3124 |
| chr14-47879561-DEL-423 | chr14-47879562-DEL-422 |
| chr14-47890201-DEL-66 | chr14-47890202-DEL-66 |
| chr14-47918347-DEL-200 | chr14-47918338-DEL-201 |
| chr14-47935599-DEL-222 | chr14-47935600-DEL-222 |
| chr14-47940780-DEL-52 | chr14-47940771-DEL-60 |
| chr14-47986908-DEL-57 | chr14-47986909-DEL-57 |
| chr14-48000740-DEL-200 | chr14-48000732-DEL-199 |
| chr14-48001919-DEL-63 | chr14-48001920-DEL-63 |
| chr14-48027212-DEL-64 | chr14-48027209-DEL-64 |
| chr14-48039338-DEL-189 | chr14-48039290-DEL-230 |
| chr14-48059935-DEL-451 | chr14-48059936-DEL-451 |
| chr14-48102568-DEL-552 | chr14-48102569-DEL-552 |
| chr14-48127575-DEL-84 | chr14-48127589-DEL-62 |
| chr14-48141096-DEL-228 | chr14-48141097-DEL-228 |
| chr14-48183157-DEL-506 | chr14-48183158-DEL-506 |
| chr14-48185912-DEL-1444 | chr14-48185913-DEL-1444 |
| chr14-48189368-DEL-2900 | chr14-48189369-DEL-2900 |
| chr14-48208808-DEL-2448 | chr14-48208809-DEL-2448 |
| chr14-48238312-DEL-224 | chr14-48238313-DEL-224 |
| chr14-48239902-DEL-948 | chr14-48239903-DEL-948 |
| chr14-48253082-DEL-231 | chr14-48253102-DEL-239 |
| chr14-48256099-DEL-80 | chr14-48256101-DEL-78 |
| chr14-48289804-DEL-72 | chr14-48289805-DEL-74 |
| chr14-48299201-DEL-286 | chr14-48299201-DEL-287 |
| chr14-48301212-DEL-170 | chr14-48301213-DEL-170 |
| chr14-48316553-DEL-716 | chr14-48316554-DEL-716 |
| chr14-48320419-DEL-207 | chr14-48320426-DEL-214 |
| chr14-48323262-DEL-68 | chr14-48323263-DEL-68 |
| chr14-48324135-DEL-66 | chr14-48324136-DEL-66 |
| chr14-48332732-DEL-56 | chr14-48332733-DEL-56 |
| chr14-48341459-DEL-56 | chr14-48341465-DEL-51 |
| chr14-48354956-DEL-75 | chr14-48354957-DEL-75 |
| chr14-48386137-DEL-2116 | chr14-48386141-DEL-2115 |
| chr14-48404104-DEL-167 | chr14-48404105-DEL-167 |
| chr14-48421347-DEL-109 | chr14-48421348-DEL-109 |
| chr14-48443018-DEL-60 | chr14-48443019-DEL-60 |
| chr14-48510866-DEL-163 | chr14-48510823-DEL-190 |
| chr14-48519469-DEL-52 | chr14-48519436-DEL-90 |
| chr14-48523129-DEL-1035 | chr14-48523130-DEL-1035 |
| chr14-48557812-DEL-69 | chr14-48557812-DEL-70 |
| chr14-48562626-DEL-208 | chr14-48562627-DEL-208 |
| chr14-48591548-DEL-86 | chr14-48591549-DEL-86 |
| chr14-48602390-DEL-1113 | chr14-48602391-DEL-1112 |
| chr14-48612745-DEL-54 | chr14-48612746-DEL-54 |
| chr14-48615448-DEL-837 | chr14-48615449-DEL-837 |
| chr14-48664629-DEL-160 | chr14-48664630-DEL-160 |
| chr14-48670628-DEL-152 | chr14-48670629-DEL-152 |
| chr14-48700123-DEL-64 | chr14-48700123-DEL-64 |
| chr14-48715525-DEL-137 | chr14-48715526-DEL-137 |
| chr14-48721147-DEL-56 | chr14-48721148-DEL-54 |
| chr14-48722608-DEL-60 | chr14-48722609-DEL-60 |
| chr14-48767036-DEL-122 | chr14-48767006-DEL-123 |
| chr14-48786810-DEL-631 | chr14-48786811-DEL-631 |
| chr14-48787811-DEL-92 | chr14-48787812-DEL-92 |
| chr14-48798856-DEL-846 | chr14-48798857-DEL-846 |
| chr14-48826879-DEL-82 | chr14-48826880-DEL-82 |
| chr14-48837725-DEL-178 | chr14-48837726-DEL-178 |
| chr14-48848975-DEL-228 | chr14-48848976-DEL-228 |
| chr14-48855875-DEL-131 | chr14-48855883-DEL-123 |
| chr14-48858886-DEL-56 | chr14-48858886-DEL-56 |
| chr14-48859844-DEL-160 | chr14-48859855-DEL-112 |
| chr14-48870373-DEL-148 | chr14-48870374-DEL-148 |

| | |
|---|---|
| chr14-48883207-DEL-67 | chr14-48883207-DEL-67 |
| chr14-48886801-DEL-334 | chr14-48886780-DEL-376 |
| chr14-48939555-DEL-52 | chr14-48939556-DEL-52 |
| chr14-48940193-DEL-93 | chr14-48940194-DEL-93 |
| chr14-48958661-DEL-202 | chr14-48958662-DEL-202 |
| chr14-48968573-DEL-82 | chr14-48968542-DEL-114 |
| chr14-48980292-DEL-64 | chr14-48980293-DEL-70 |
| chr14-48981168-DEL-187 | chr14-48981169-DEL-187 |
| chr14-48996776-DEL-62 | chr14-48996777-DEL-62 |
| chr14-49013476-DEL-103 | chr14-49013477-DEL-107 |
| chr14-49021022-DEL-101 | chr14-49021023-DEL-101 |
| chr14-49053210-DEL-56 | chr14-49053211-DEL-56 |
| chr14-49065848-DEL-157 | chr14-49065849-DEL-157 |
| chr14-49190367-DEL-219 | chr14-49190368-DEL-219 |
| chr14-49194271-DEL-108 | chr14-49194272-DEL-108 |
| chr14-49212658-DEL-54 | chr14-49212659-DEL-50 |
| chr14-49226709-DEL-207 | chr14-49226710-DEL-207 |
| chr14-49266854-DEL-932 | chr14-49266855-DEL-932 |
| chr14-49300699-DEL-56 | chr14-49300700-DEL-56 |
| chr14-49307674-DEL-66 | chr14-49307679-DEL-60 |
| chr14-49309128-DEL-72 | chr14-49309129-DEL-70 |
| chr14-49331826-DEL-78 | chr14-49331827-DEL-78 |
| chr14-49342464-DEL-51 | chr14-49342465-DEL-51 |
| chr14-49375147-DEL-424 | chr14-49375148-DEL-424 |
| chr14-49376162-DEL-85 | chr14-49376120-DEL-86 |
| chr14-49381388-DEL-76 | chr14-49381395-DEL-64 |
| chr14-49394029-DEL-101 | chr14-49394030-DEL-101 |
| chr14-49419598-DEL-146 | chr14-49419600-DEL-145 |
| chr14-49431129-DEL-211 | chr14-49431130-DEL-211 |
| chr14-49455631-DEL-167 | chr14-49455632-DEL-167 |
| chr14-49498240-DEL-84 | chr14-49498235-DEL-87 |
| chr14-49504525-DEL-66 | chr14-49504528-DEL-64 |
| chr14-49515907-DEL-474 | chr14-49515908-DEL-474 |
| chr14-49587132-DEL-64 | chr14-49587133-DEL-64 |
| chr14-49613551-DEL-380 | chr14-49613552-DEL-380 |
| chr14-49633994-DEL-84 | chr14-49633995-DEL-90 |
| chr14-49652511-DEL-1128 | chr14-49652512-DEL-1128 |
| chr14-49669762-DEL-144 | chr14-49669763-DEL-144 |
| chr14-49753786-DEL-68 | chr14-49753763-DEL-54 |
| chr14-49763803-DEL-129 | chr14-49763804-DEL-126 |
| chr14-49801859-DEL-187 | chr14-49801858-DEL-188 |
| chr14-49808088-DEL-54 | chr14-49808089-DEL-50 |
| chr14-49839618-DEL-7079 | chr14-49839619-DEL-7079 |
| chr14-49853234-DEL-1969 | chr14-49853227-DEL-1963 |
| chr14-49865959-DEL-6551 | chr14-49865960-DEL-6551 |
| chr14-49923473-DEL-529 | chr14-49923474-DEL-529 |
| chr14-49932767-DEL-817 | chr14-49932768-DEL-817 |
| chr14-49950462-DEL-99 | chr14-49950452-DEL-102 |
| chr14-49966225-DEL-60 | chr14-49966225-DEL-58 |
| chr14-50013322-DEL-411 | chr14-50013322-DEL-410 |
| chr14-50017090-DEL-67 | chr14-50017090-DEL-67 |
| chr14-50025528-DEL-156 | chr14-50025529-DEL-156 |
| chr14-50034658-DEL-1760 | chr14-50034659-DEL-1760 |
| chr14-50115846-DEL-6736 | chr14-50115847-DEL-6736 |
| chr14-50133426-DEL-1117 | chr14-50133427-DEL-1117 |
| chr14-50140260-DEL-62 | chr14-50140222-DEL-63 |
| chr14-50154310-DEL-6423 | chr14-50154311-DEL-6423 |
| chr14-50177653-DEL-147 | chr14-50177654-DEL-147 |
| chr14-50208760-DEL-253 | chr14-50208762-DEL-252 |
| chr14-50212893-DEL-104 | chr14-50212903-DEL-95 |
| chr14-50215857-DEL-95 | chr14-50215858-DEL-95 |
| chr14-50221847-DEL-221 | chr14-50221848-DEL-221 |
| chr14-50239605-DEL-6704 | chr14-50239604-DEL-6705 |
| chr14-50269479-DEL-149 | chr14-50269480-DEL-149 |
| chr14-50378368-DEL-146 | chr14-50378370-DEL-147 |
| chr14-50392082-DEL-119 | chr14-50392085-DEL-124 |
| chr14-50451126-DEL-233 | chr14-50451115-DEL-245 |
| chr14-50457930-DEL-820 | chr14-50457931-DEL-820 |
| chr14-50486807-DEL-6378 | chr14-50486807-DEL-6378 |
| chr14-50493626-DEL-404 | chr14-50493627-DEL-404 |
| chr14-50544345-DEL-226 | chr14-50544346-DEL-226 |
| chr14-50576480-DEL-102 | chr14-50576481-DEL-102 |
| chr14-50592948-DEL-93 | chr14-50592949-DEL-93 |
| chr14-50609519-DEL-332 | chr14-50609522-DEL-331 |
| chr14-50614778-DEL-68 | chr14-50614779-DEL-68 |
| chr14-50620811-DEL-1288 | chr14-50620812-DEL-1288 |
| chr14-50622550-DEL-71 | chr14-50622552-DEL-69 |
| chr14-50629457-DEL-66 | chr14-50629458-DEL-54 |
| chr14-50634011-DEL-72 | chr14-50634009-DEL-76 |
| chr14-50664454-DEL-791 | chr14-50664454-DEL-791 |

| | |
|---|---|
| chr14-50703338-DEL-2566 | chr14-50703339-DEL-2566 |
| chr14-50726064-DEL-96 | chr14-50726061-DEL-100 |
| chr14-50726560-DEL-3437 | chr14-50726581-DEL-3454 |
| chr14-50735794-DEL-908 | chr14-50735794-DEL-908 |
| chr14-50739799-DEL-70 | chr14-50739800-DEL-70 |
| chr14-50772857-DEL-160 | chr14-50772858-DEL-160 |
| chr14-50817451-DEL-337 | chr14-50817449-DEL-338 |
| chr14-50820528-DEL-6264 | chr14-50820529-DEL-6264 |
| chr14-50830011-DEL-54 | chr14-50830014-DEL-52 |
| chr14-50833220-DEL-1204 | chr14-50833241-DEL-1185 |
| chr14-50866342-DEL-9035 | chr14-50866343-DEL-9035 |
| chr14-50895970-DEL-66 | chr14-50895970-DEL-66 |
| chr14-50896266-DEL-1159 | chr14-50896267-DEL-1159 |
| chr14-50914832-DEL-75 | chr14-50914833-DEL-75 |
| chr14-50925978-DEL-381 | chr14-50925979-DEL-381 |
| chr14-50969260-DEL-205 | chr14-50969261-DEL-205 |
| chr14-50988771-DEL-93 | chr14-50988760-DEL-105 |
| chr14-50992981-DEL-310 | chr14-50992982-DEL-310 |
| chr14-50997365-DEL-687 | chr14-50997366-DEL-687 |
| chr14-50998690-DEL-1626 | chr14-50998691-DEL-1626 |
| chr14-51005827-DEL-108 | chr14-51005828-DEL-108 |
| chr14-51028244-DEL-60 | chr14-51028233-DEL-72 |
| chr14-51046365-DEL-194 | chr14-51046366-DEL-194 |
| chr14-51048861-DEL-193 | chr14-51048862-DEL-193 |
| chr14-51053130-DEL-196 | chr14-51053131-DEL-196 |
| chr14-51063443-DEL-91 | chr14-51063444-DEL-91 |
| chr14-51076223-DEL-121 | chr14-51076179-DEL-122 |
| chr14-51092090-DEL-134 | chr14-51092092-DEL-135 |
| chr14-51094896-DEL-205 | chr14-51094897-DEL-205 |
| chr14-51098922-DEL-72 | chr14-51098923-DEL-68 |
| chr14-51117661-DEL-4657 | chr14-51117662-DEL-4657 |
| chr14-51138827-DEL-707 | chr14-51138828-DEL-707 |
| chr14-51156001-DEL-52 | chr14-51156002-DEL-52 |
| chr14-51198560-DEL-436 | chr14-51198578-DEL-431 |
| chr14-51215617-DEL-799 | chr14-51215618-DEL-799 |
| chr14-51218240-DEL-197 | chr14-51218241-DEL-197 |
| chr14-51226185-DEL-214 | chr14-51226186-DEL-214 |
| chr14-51235736-DEL-114 | chr14-51235737-DEL-114 |
| chr14-51262060-DEL-205 | chr14-51262061-DEL-205 |
| chr14-51312170-DEL-72 | chr14-51312171-DEL-72 |
| chr14-51330611-DEL-52 | chr14-51330612-DEL-52 |
| chr14-51341644-DEL-80 | chr14-51341645-DEL-80 |
| chr14-51359610-DEL-170 | chr14-51359613-DEL-164 |
| chr14-51363163-DEL-59 | chr14-51363164-DEL-59 |
| chr14-51370243-DEL-58 | chr14-51370244-DEL-50 |
| chr14-51412428-DEL-80 | chr14-51412431-DEL-78 |
| chr14-51413116-DEL-106 | chr14-51413099-DEL-142 |
| chr14-51427863-DEL-143 | chr14-51427864-DEL-143 |
| chr14-51447413-DEL-1588 | chr14-51447414-DEL-1588 |
| chr14-51451749-DEL-4342 | chr14-51451748-DEL-4340 |
| chr14-51461341-DEL-63 | chr14-51461342-DEL-63 |
| chr14-51465934-DEL-222 | chr14-51465935-DEL-222 |
| chr14-51475971-DEL-297 | chr14-51475972-DEL-297 |
| chr14-51476523-DEL-97 | chr14-51476523-DEL-97 |
| chr14-51494359-DEL-2750 | chr14-51494360-DEL-2750 |
| chr14-51507447-DEL-1271 | chr14-51507448-DEL-1271 |
| chr14-51525719-DEL-225 | chr14-51525720-DEL-225 |
| chr14-51535790-DEL-5907 | chr14-51535791-DEL-5907 |
| chr14-51547124-DEL-81 | chr14-51547125-DEL-81 |
| chr14-51557437-DEL-2606 | chr14-51557430-DEL-2607 |
| chr14-51577181-DEL-391 | chr14-51577182-DEL-391 |
| chr14-51593351-DEL-957 | chr14-51593352-DEL-957 |
| chr14-51599291-DEL-1070 | chr14-51599292-DEL-1070 |
| chr14-51619535-DEL-3931 | chr14-51619536-DEL-3931 |
| chr14-51623638-DEL-2405 | chr14-51623639-DEL-2405 |
| chr14-51644938-DEL-2298 | chr14-51644939-DEL-2298 |
| chr14-51687793-DEL-1244 | chr14-51687794-DEL-1244 |
| chr14-51692834-DEL-54 | chr14-51692835-DEL-54 |
| chr14-51722599-DEL-116 | chr14-51722599-DEL-116 |
| chr14-51784803-DEL-54 | chr14-51784803-DEL-54 |
| chr14-51791038-DEL-942 | chr14-51791039-DEL-942 |
| chr14-51792946-DEL-551 | chr14-51792947-DEL-551 |
| chr14-51795651-DEL-56 | chr14-51795652-DEL-56 |
| chr14-51812503-DEL-77 | chr14-51812504-DEL-77 |
| chr14-51821738-DEL-156 | chr14-51821739-DEL-156 |
| chr14-51840904-DEL-156 | chr14-51840905-DEL-156 |
| chr14-51860080-DEL-362 | chr14-51860081-DEL-362 |
| chr14-51866444-DEL-232 | chr14-51866444-DEL-232 |
| chr14-51869508-DEL-74 | chr14-51869508-DEL-74 |
| chr14-51900923-DEL-578 | chr14-51900926-DEL-579 |

| | |
|---|---|
| chr14-51940824-DEL-6157 | chr14-51940825-DEL-6157 |
| chr14-51971343-DEL-400 | chr14-51971343-DEL-400 |
| chr14-52003178-DEL-78 | chr14-52003179-DEL-78 |
| chr14-52008125-DEL-118 | chr14-52008126-DEL-116 |
| chr14-52012071-DEL-708 | chr14-52012072-DEL-708 |
| chr14-52025405-DEL-549 | chr14-52025404-DEL-549 |
| chr14-52046053-DEL-221 | chr14-52046054-DEL-221 |
| chr14-52053698-DEL-739 | chr14-52053699-DEL-739 |
| chr14-52054871-DEL-959 | chr14-52054872-DEL-959 |
| chr14-52074566-DEL-201 | chr14-52074567-DEL-201 |
| chr14-52096107-DEL-186 | chr14-52096108-DEL-186 |
| chr14-52111560-DEL-193 | chr14-52111557-DEL-193 |
| chr14-52113662-DEL-7199 | chr14-52113663-DEL-7211 |
| chr14-52120779-DEL-70 | chr14-52120792-DEL-58 |
| chr14-52171829-DEL-60 | chr14-52171830-DEL-60 |
| chr14-52177109-DEL-107 | chr14-52177106-DEL-111 |
| chr14-52187921-DEL-117 | chr14-52187922-DEL-117 |
| chr14-52192981-DEL-3087 | chr14-52192982-DEL-3087 |
| chr14-52195007-DEL-68 | chr14-52195004-DEL-58 |
| chr14-52198202-DEL-52 | chr14-52198188-DEL-52 |
| chr14-52202434-DEL-55 | chr14-52202435-DEL-55 |
| chr14-52231144-DEL-217 | chr14-52231145-DEL-217 |
| chr14-52239083-DEL-202 | chr14-52239084-DEL-202 |
| chr14-52245267-DEL-264 | chr14-52245268-DEL-232 |
| chr14-52280048-DEL-93 | chr14-52280028-DEL-96 |
| chr14-52286842-DEL-143 | chr14-52286804-DEL-138 |
| chr14-52289586-DEL-404 | chr14-52289587-DEL-404 |
| chr14-52294987-DEL-217 | chr14-52294988-DEL-217 |
| chr14-52295339-DEL-64 | chr14-52295331-DEL-65 |
| chr14-52298711-DEL-661 | chr14-52298712-DEL-661 |
| chr14-52302067-DEL-1068 | chr14-52302068-DEL-1068 |
| chr14-52302727-DEL-211 | chr14-52302728-DEL-211 |
| chr14-52316580-DEL-215 | chr14-52316581-DEL-215 |
| chr14-52329364-DEL-207 | chr14-52329365-DEL-207 |
| chr14-52331618-DEL-262 | chr14-52331619-DEL-262 |
| chr14-52340035-DEL-185 | chr14-52340035-DEL-185 |
| chr14-52343183-DEL-56 | chr14-52343180-DEL-60 |
| chr14-52345977-DEL-160 | chr14-52345971-DEL-162 |
| chr14-52346676-DEL-80 | chr14-52346677-DEL-80 |
| chr14-52350421-DEL-197 | chr14-52350422-DEL-197 |
| chr14-52352795-DEL-1507 | chr14-52352796-DEL-1507 |
| chr14-52362941-DEL-235 | chr14-52362942-DEL-235 |
| chr14-52363949-DEL-181 | chr14-52363950-DEL-181 |
| chr14-52365197-DEL-994 | chr14-52365198-DEL-993 |
| chr14-52385435-DEL-77 | chr14-52385436-DEL-77 |
| chr14-52386439-DEL-58 | chr14-52386440-DEL-58 |
| chr14-52391441-DEL-207 | chr14-52391442-DEL-207 |
| chr14-52405308-DEL-89 | chr14-52405343-DEL-55 |
| chr14-52406911-DEL-223 | chr14-52406880-DEL-219 |
| chr14-52423215-DEL-209 | chr14-52423216-DEL-209 |
| chr14-52428102-DEL-156 | chr14-52428130-DEL-129 |
| chr14-52431248-DEL-237 | chr14-52431249-DEL-237 |
| chr14-52462917-DEL-56 | chr14-52462918-DEL-56 |
| chr14-52487866-DEL-967 | chr14-52487867-DEL-967 |
| chr14-52504170-DEL-886 | chr14-52504171-DEL-886 |
| chr14-52507710-DEL-104 | chr14-52507711-DEL-104 |
| chr14-52522415-DEL-58 | chr14-52522424-DEL-54 |
| chr14-52527332-DEL-837 | chr14-52527333-DEL-837 |
| chr14-52542956-DEL-68 | chr14-52542930-DEL-67 |
| chr14-52567526-DEL-178 | chr14-52567527-DEL-157 |
| chr14-52573267-DEL-365 | chr14-52573268-DEL-365 |
| chr14-52574891-DEL-797 | chr14-52574892-DEL-797 |
| chr14-52579614-DEL-55 | chr14-52579602-DEL-67 |
| chr14-52582406-DEL-250 | chr14-52582402-DEL-251 |
| chr14-52593340-DEL-195 | chr14-52593341-DEL-195 |
| chr14-52617351-DEL-234 | chr14-52617350-DEL-197 |
| chr14-52617601-DEL-4370 | chr14-52617602-DEL-4370 |
| chr14-52638287-DEL-425 | chr14-52638288-DEL-422 |
| chr14-52640464-DEL-506 | chr14-52640465-DEL-506 |
| chr14-52649696-DEL-60 | chr14-52649693-DEL-60 |
| chr14-52659628-DEL-183 | chr14-52659629-DEL-183 |
| chr14-52671064-DEL-2131 | chr14-52671065-DEL-2131 |
| chr14-52675976-DEL-1098 | chr14-52675977-DEL-1098 |
| chr14-52685805-DEL-152 | chr14-52685806-DEL-152 |
| chr14-52713717-DEL-3039 | chr14-52713717-DEL-3039 |
| chr14-52720195-DEL-317 | chr14-52720196-DEL-317 |
| chr14-52742898-DEL-98 | chr14-52742899-DEL-98 |
| chr14-52743509-DEL-805 | chr14-52743510-DEL-805 |
| chr14-52765707-DEL-257 | chr14-52765708-DEL-257 |
| chr14-52768966-DEL-2652 | chr14-52768967-DEL-2652 |

| | |
|---|---|
| chr14-52778679-DEL-102 | chr14-52778678-DEL-104 |
| chr14-52807952-DEL-1393 | chr14-52807953-DEL-1393 |
| chr14-52823982-DEL-6376 | chr14-52823983-DEL-6376 |
| chr14-52829984-DEL-201 | chr14-52829985-DEL-201 |
| chr14-52848110-DEL-81 | chr14-52848111-DEL-81 |
| chr14-52856713-DEL-65 | chr14-52856714-DEL-65 |
| chr14-52865023-DEL-6173 | chr14-52865024-DEL-6173 |
| chr14-52879611-DEL-981 | chr14-52879612-DEL-981 |
| chr14-52894110-DEL-162 | chr14-52894111-DEL-162 |
| chr14-52894879-DEL-4314 | chr14-52894880-DEL-4314 |
| chr14-52901884-DEL-117 | chr14-52901885-DEL-117 |
| chr14-52929564-DEL-3951 | chr14-52929565-DEL-3951 |
| chr14-52933516-DEL-2310 | chr14-52933515-DEL-2310 |
| chr14-52940435-DEL-6079 | chr14-52940431-DEL-6080 |
| chr14-52956235-DEL-66 | chr14-52956234-DEL-66 |
| chr14-52962822-DEL-1891 | chr14-52962823-DEL-1891 |
| chr14-52975845-DEL-81 | chr14-52975846-DEL-81 |
| chr14-52995362-DEL-7095 | chr14-52995363-DEL-7096 |
| chr14-53037285-DEL-1681 | chr14-53037276-DEL-1690 |
| chr14-53043676-DEL-375 | chr14-53043676-DEL-375 |
| chr14-53051239-DEL-518 | chr14-53051240-DEL-518 |
| chr14-53061198-DEL-55 | chr14-53061199-DEL-55 |
| chr14-53071099-DEL-103 | chr14-53071074-DEL-102 |
| chr14-53147943-DEL-60 | chr14-53147944-DEL-58 |
| chr14-53277311-DEL-896 | chr14-53277311-DEL-896 |
| chr14-53338338-DEL-78 | chr14-53338345-DEL-90 |
| chr14-53560057-DEL-80 | chr14-53560058-DEL-74 |
| chr14-53566086-DEL-375 | chr14-53566085-DEL-375 |
| chr14-53572798-DEL-101 | chr14-53572799-DEL-101 |
| chr14-53575411-DEL-66 | chr14-53575411-DEL-70 |
| chr14-53587679-DEL-62 | chr14-53587678-DEL-62 |
| chr14-53588047-DEL-230 | chr14-53588046-DEL-230 |
| chr14-53602836-DEL-221 | chr14-53602828-DEL-224 |
| chr14-53609631-DEL-866 | chr14-53609632-DEL-866 |
| chr14-53655716-DEL-74 | chr14-53655721-DEL-70 |
| chr14-53665005-DEL-109 | chr14-53665006-DEL-109 |
| chr14-53676397-DEL-584 | chr14-53676398-DEL-584 |
| chr14-53678574-DEL-501 | chr14-53678574-DEL-501 |
| chr14-53679810-DEL-363 | chr14-53679810-DEL-363 |
| chr14-53712621-DEL-4508 | chr14-53712622-DEL-4508 |
| chr14-53740695-DEL-856 | chr14-53740696-DEL-856 |
| chr14-53741762-DEL-52 | chr14-53741763-DEL-52 |
| chr14-53744065-DEL-125 | chr14-53744029-DEL-126 |
| chr14-53747286-DEL-2352 | chr14-53747287-DEL-2352 |
| chr14-53838850-DEL-110 | chr14-53838879-DEL-93 |
| chr14-53877807-DEL-55 | chr14-53877793-DEL-52 |
| chr14-53880679-DEL-487 | chr14-53880680-DEL-487 |
| chr14-53885723-DEL-72 | chr14-53885740-DEL-54 |
| chr14-53980120-DEL-239 | chr14-53980121-DEL-239 |
| chr14-54011345-DEL-115 | chr14-54011347-DEL-111 |
| chr14-54029413-DEL-93 | chr14-54029414-DEL-93 |
| chr14-54029616-DEL-399 | chr14-54029617-DEL-399 |
| chr14-54031998-DEL-56 | chr14-54031999-DEL-58 |
| chr14-54091999-DEL-385 | chr14-54092000-DEL-385 |
| chr14-54093987-DEL-111 | chr14-54093988-DEL-111 |
| chr14-54097493-DEL-86 | chr14-54097492-DEL-88 |
| chr14-54105666-DEL-52 | chr14-54105667-DEL-52 |
| chr14-54133455-DEL-526 | chr14-54133456-DEL-526 |
| chr14-54136432-DEL-1571 | chr14-54136433-DEL-1571 |
| chr14-54204320-DEL-63 | chr14-54204305-DEL-66 |
| chr14-54213572-DEL-397 | chr14-54213572-DEL-397 |
| chr14-54214089-DEL-221 | chr14-54214090-DEL-216 |
| chr14-54223417-DEL-79 | chr14-54223418-DEL-79 |
| chr14-54241862-DEL-106 | chr14-54241863-DEL-106 |
| chr14-54247991-DEL-108 | chr14-54247992-DEL-108 |
| chr14-54257452-DEL-339 | chr14-54257453-DEL-339 |
| chr14-54274331-DEL-52 | chr14-54274332-DEL-52 |
| chr14-54289557-DEL-108 | chr14-54289562-DEL-68 |
| chr14-54293395-DEL-82 | chr14-54293389-DEL-83 |
| chr14-54338771-DEL-55 | chr14-54338779-DEL-56 |
| chr14-54360861-DEL-62 | chr14-54360862-DEL-62 |
| chr14-54367074-DEL-6741 | chr14-54367075-DEL-6741 |
| chr14-54382348-DEL-55 | chr14-54382349-DEL-55 |
| chr14-54400594-DEL-86 | chr14-54400609-DEL-50 |
| chr14-54424085-DEL-51 | chr14-54424083-DEL-54 |
| chr14-54437543-DEL-220 | chr14-54437544-DEL-220 |
| chr14-54507445-DEL-236 | chr14-54507446-DEL-227 |
| chr14-54528261-DEL-68 | chr14-54528262-DEL-68 |
| chr14-54529181-DEL-205 | chr14-54529182-DEL-205 |
| chr14-54529824-DEL-5548 | chr14-54529825-DEL-5548 |

| | |
|---|---|
| chr14-54542620-DEL-2467 | chr14-54542621-DEL-2467 |
| chr14-54552069-DEL-4173 | chr14-54552070-DEL-4173 |
| chr14-54573335-DEL-52 | chr14-54573362-DEL-58 |
| chr14-54574943-DEL-52 | chr14-54574944-DEL-52 |
| chr14-54589965-DEL-2104 | chr14-54589966-DEL-2104 |
| chr14-54602304-DEL-224 | chr14-54602305-DEL-224 |
| chr14-54608419-DEL-505 | chr14-54608420-DEL-505 |
| chr14-54609758-DEL-64 | chr14-54609758-DEL-60 |
| chr14-54645032-DEL-82 | chr14-54645033-DEL-82 |
| chr14-54653563-DEL-122 | chr14-54653564-DEL-124 |
| chr14-54659386-DEL-62 | chr14-54659387-DEL-68 |
| chr14-54660039-DEL-207 | chr14-54660037-DEL-201 |
| chr14-54675383-DEL-162 | chr14-54675384-DEL-161 |
| chr14-54685295-DEL-147 | chr14-54685296-DEL-147 |
| chr14-54687388-DEL-133 | chr14-54687389-DEL-133 |
| chr14-54688529-DEL-66 | chr14-54688530-DEL-66 |
| chr14-54693526-DEL-324 | chr14-54693526-DEL-321 |
| chr14-54694412-DEL-69 | chr14-54694419-DEL-78 |
| chr14-54699531-DEL-60 | chr14-54699531-DEL-60 |
| chr14-54728016-DEL-803 | chr14-54728017-DEL-803 |
| chr14-54729890-DEL-66 | chr14-54729891-DEL-66 |
| chr14-54738265-DEL-200 | chr14-54738266-DEL-200 |
| chr14-54739209-DEL-108 | chr14-54739210-DEL-109 |
| chr14-54757710-DEL-80 | chr14-54757689-DEL-93 |
| chr14-54758430-DEL-56 | chr14-54758436-DEL-55 |
| chr14-54767817-DEL-7827 | chr14-54767818-DEL-7827 |
| chr14-54798762-DEL-52 | chr14-54798765-DEL-50 |
| chr14-54818259-DEL-200 | chr14-54818260-DEL-200 |
| chr14-54831453-DEL-67 | chr14-54831453-DEL-68 |
| chr14-54837516-DEL-216 | chr14-54837517-DEL-216 |
| chr14-54851185-DEL-356 | chr14-54851186-DEL-356 |
| chr14-54859673-DEL-55 | chr14-54859680-DEL-64 |
| chr14-54865107-DEL-68 | chr14-54865097-DEL-60 |
| chr14-54877912-DEL-62 | chr14-54877947-DEL-53 |
| chr14-54891631-DEL-63 | chr14-54891632-DEL-63 |
| chr14-54917637-DEL-100 | chr14-54917638-DEL-100 |
| chr14-54941910-DEL-579 | chr14-54941912-DEL-562 |
| chr14-54977847-DEL-268 | chr14-54977848-DEL-268 |
| chr14-54985171-DEL-54 | chr14-54985168-DEL-75 |
| chr14-55000349-DEL-1887 | chr14-55000350-DEL-1887 |
| chr14-55014018-DEL-51 | chr14-55014051-DEL-57 |
| chr14-55042820-DEL-231 | chr14-55042821-DEL-231 |
| chr14-55046052-DEL-63 | chr14-55046040-DEL-75 |
| chr14-55047448-DEL-187 | chr14-55047449-DEL-187 |
| chr14-55069042-DEL-210 | chr14-55069043-DEL-210 |
| chr14-55078120-DEL-190 | chr14-55078118-DEL-191 |
| chr14-55098662-DEL-198 | chr14-55098663-DEL-198 |
| chr14-55109064-DEL-218 | chr14-55109065-DEL-218 |
| chr14-55141990-DEL-197 | chr14-55141991-DEL-197 |
| chr14-55160522-DEL-299 | chr14-55160522-DEL-300 |
| chr14-55174567-DEL-206 | chr14-55174568-DEL-206 |
| chr14-55192086-DEL-786 | chr14-55192087-DEL-786 |
| chr14-55203540-DEL-67 | chr14-55203524-DEL-52 |
| chr14-55242647-DEL-58 | chr14-55242648-DEL-58 |
| chr14-55261190-DEL-189 | chr14-55261191-DEL-143 |
| chr14-55280195-DEL-61 | chr14-55280196-DEL-61 |
| chr14-55281409-DEL-181 | chr14-55281410-DEL-181 |
| chr14-55287469-DEL-63 | chr14-55287470-DEL-63 |
| chr14-55290499-DEL-57 | chr14-55290500-DEL-57 |
| chr14-55359145-DEL-211 | chr14-55359146-DEL-211 |
| chr14-55391975-DEL-240 | chr14-55391992-DEL-224 |
| chr14-55461920-DEL-1254 | chr14-55461921-DEL-1254 |
| chr14-55463819-DEL-3927 | chr14-55463820-DEL-3927 |
| chr14-55470972-DEL-394 | chr14-55470973-DEL-394 |
| chr14-55496158-DEL-400 | chr14-55496159-DEL-400 |
| chr14-55496944-DEL-1565 | chr14-55496945-DEL-1565 |
| chr14-55516431-DEL-236 | chr14-55516432-DEL-236 |
| chr14-55527584-DEL-377 | chr14-55527585-DEL-377 |
| chr14-55548672-DEL-4005 | chr14-55548673-DEL-4005 |
| chr14-55560011-DEL-98 | chr14-55560042-DEL-76 |
| chr14-55570323-DEL-137 | chr14-55570324-DEL-137 |
| chr14-55592352-DEL-1590 | chr14-55592353-DEL-1590 |
| chr14-55635075-DEL-315 | chr14-55635076-DEL-315 |
| chr14-55644360-DEL-993 | chr14-55644361-DEL-993 |
| chr14-55683181-DEL-2947 | chr14-55683182-DEL-2947 |
| chr14-55719907-DEL-1334 | chr14-55719908-DEL-1334 |
| chr14-55765704-DEL-573 | chr14-55765705-DEL-573 |
| chr14-55780669-DEL-51 | chr14-55780697-DEL-59 |
| chr14-55803282-DEL-64 | chr14-55803251-DEL-64 |
| chr14-55927177-DEL-105 | chr14-55927178-DEL-105 |

| | |
|---|---|
| chr14-55933725-DEL-208 | chr14-55933721-DEL-209 |
| chr14-55996913-DEL-176 | chr14-55996914-DEL-176 |
| chr14-56027897-DEL-63 | chr14-56027897-DEL-63 |
| chr14-56032464-DEL-465 | chr14-56032464-DEL-465 |
| chr14-56034490-DEL-7445 | chr14-56034484-DEL-7446 |
| chr14-56071848-DEL-52 | chr14-56071848-DEL-52 |
| chr14-56115808-DEL-221 | chr14-56115809-DEL-221 |
| chr14-56139215-DEL-211 | chr14-56139216-DEL-211 |
| chr14-56150772-DEL-62 | chr14-56150773-DEL-62 |
| chr14-56175288-DEL-420 | chr14-56175301-DEL-416 |
| chr14-56186013-DEL-167 | chr14-56186014-DEL-167 |
| chr14-56198248-DEL-119 | chr14-56198282-DEL-70 |
| chr14-56201027-DEL-468 | chr14-56201028-DEL-468 |
| chr14-56216185-DEL-8412 | chr14-56216186-DEL-8412 |
| chr14-56234563-DEL-926 | chr14-56234564-DEL-925 |
| chr14-56238860-DEL-475 | chr14-56238861-DEL-475 |
| chr14-56265412-DEL-123 | chr14-56265413-DEL-123 |
| chr14-56291127-DEL-1070 | chr14-56291128-DEL-1070 |
| chr14-56336346-DEL-153 | chr14-56336347-DEL-153 |
| chr14-56363534-DEL-110 | chr14-56363535-DEL-110 |
| chr14-56371835-DEL-213 | chr14-56371836-DEL-213 |
| chr14-56391036-DEL-369 | chr14-56391037-DEL-369 |
| chr14-56411021-DEL-58 | chr14-56411022-DEL-58 |
| chr14-56425723-DEL-12345 | chr14-56425723-DEL-12345 |
| chr14-56460323-DEL-1890 | chr14-56460324-DEL-1890 |
| chr14-56510945-DEL-224 | chr14-56510946-DEL-224 |
| chr14-56545297-DEL-60 | chr14-56545298-DEL-60 |
| chr14-56567612-DEL-60 | chr14-56567584-DEL-88 |
| chr14-56578405-DEL-421 | chr14-56578406-DEL-421 |
| chr14-56638741-DEL-223 | chr14-56638742-DEL-223 |
| chr14-56653000-DEL-2593 | chr14-56653001-DEL-2593 |
| chr14-56671193-DEL-148 | chr14-56671194-DEL-148 |
| chr14-56798042-DEL-1833 | chr14-56798067-DEL-1809 |
| chr14-56800258-DEL-53 | chr14-56800259-DEL-57 |
| chr14-56801319-DEL-86 | chr14-56801319-DEL-86 |
| chr14-56804519-DEL-298 | chr14-56804520-DEL-298 |
| chr14-56833291-DEL-60 | chr14-56833300-DEL-52 |
| chr14-56885179-DEL-288 | chr14-56885204-DEL-282 |
| chr14-56908204-DEL-191 | chr14-56908213-DEL-190 |
| chr14-56931735-DEL-51 | chr14-56931736-DEL-51 |
| chr14-56973412-DEL-232 | chr14-56973413-DEL-232 |
| chr14-57007451-DEL-138 | chr14-57007452-DEL-138 |
| chr14-57009887-DEL-212 | chr14-57009901-DEL-206 |
| chr14-57044315-DEL-198 | chr14-57044316-DEL-198 |
| chr14-57059063-DEL-613 | chr14-57059063-DEL-613 |
| chr14-57061891-DEL-59 | chr14-57061866-DEL-63 |
| chr14-57097256-DEL-58 | chr14-57097259-DEL-52 |
| chr14-57114744-DEL-507 | chr14-57114745-DEL-507 |
| chr14-57229569-DEL-6669 | chr14-57229570-DEL-6669 |
| chr14-57263346-DEL-198 | chr14-57263346-DEL-198 |
| chr14-57319784-DEL-57 | chr14-57319784-DEL-58 |
| chr14-57327885-DEL-309 | chr14-57327886-DEL-309 |
| chr14-57338935-DEL-100 | chr14-57338936-DEL-92 |
| chr14-57347794-DEL-218 | chr14-57347795-DEL-216 |
| chr14-57382262-DEL-421 | chr14-57382263-DEL-421 |
| chr14-57393376-DEL-72 | chr14-57393377-DEL-72 |
| chr14-57419121-DEL-775 | chr14-57419122-DEL-775 |
| chr14-57430203-DEL-90 | chr14-57430204-DEL-90 |
| chr14-57454490-DEL-86 | chr14-57454493-DEL-84 |
| chr14-57477257-DEL-84 | chr14-57477258-DEL-84 |
| chr14-57478896-DEL-234 | chr14-57478897-DEL-234 |
| chr14-57516178-DEL-1883 | chr14-57516179-DEL-1883 |
| chr14-57518893-DEL-117 | chr14-57518894-DEL-117 |
| chr14-57528048-DEL-195 | chr14-57528049-DEL-195 |
| chr14-57542097-DEL-73 | chr14-57542098-DEL-73 |
| chr14-57545560-DEL-92 | chr14-57545567-DEL-80 |
| chr14-57559190-DEL-1246 | chr14-57559156-DEL-1282 |
| chr14-57560463-DEL-1009 | chr14-57560463-DEL-1009 |
| chr14-57590972-DEL-69 | chr14-57590992-DEL-93 |
| chr14-57597717-DEL-60 | chr14-57597718-DEL-60 |
| chr14-57603034-DEL-205 | chr14-57603035-DEL-205 |
| chr14-57616580-DEL-166 | chr14-57616540-DEL-187 |
| chr14-57628427-DEL-217 | chr14-57628428-DEL-217 |
| chr14-57684375-DEL-201 | chr14-57684376-DEL-201 |
| chr14-57686778-DEL-243 | chr14-57686749-DEL-247 |
| chr14-57688866-DEL-204 | chr14-57688866-DEL-205 |
| chr14-57697301-DEL-57 | chr14-57697302-DEL-57 |
| chr14-57705979-DEL-134 | chr14-57705980-DEL-134 |
| chr14-57707858-DEL-2524 | chr14-57707859-DEL-2522 |
| chr14-57711767-DEL-238 | chr14-57711768-DEL-238 |

| | |
|---|---|
| chr14-57714378-DEL-253 | chr14-57714379-DEL-253 |
| chr14-57723929-DEL-198 | chr14-57723930-DEL-198 |
| chr14-57728547-DEL-215 | chr14-57728548-DEL-215 |
| chr14-57730826-DEL-327 | chr14-57730827-DEL-327 |
| chr14-57747984-DEL-54 | chr14-57747982-DEL-55 |
| chr14-57755775-DEL-97 | chr14-57755776-DEL-97 |
| chr14-57757146-DEL-358 | chr14-57757147-DEL-358 |
| chr14-57791106-DEL-587 | chr14-57791107-DEL-587 |
| chr14-57792182-DEL-150 | chr14-57792183-DEL-125 |
| chr14-57795826-DEL-59 | chr14-57795827-DEL-59 |
| chr14-57801070-DEL-211 | chr14-57801071-DEL-211 |
| chr14-57802583-DEL-191 | chr14-57802584-DEL-191 |
| chr14-57804469-DEL-225 | chr14-57804470-DEL-225 |
| chr14-57809997-DEL-188 | chr14-57809998-DEL-188 |
| chr14-57832367-DEL-557 | chr14-57832368-DEL-557 |
| chr14-57833199-DEL-64 | chr14-57833200-DEL-64 |
| chr14-57857461-DEL-203 | chr14-57857462-DEL-203 |
| chr14-57858094-DEL-113 | chr14-57858095-DEL-113 |
| chr14-57858693-DEL-94 | chr14-57858694-DEL-94 |
| chr14-57892279-DEL-221 | chr14-57892280-DEL-221 |
| chr14-57912111-DEL-180 | chr14-57912112-DEL-180 |
| chr14-57914636-DEL-202 | chr14-57914637-DEL-202 |
| chr14-57916528-DEL-227 | chr14-57916529-DEL-227 |
| chr14-57929877-DEL-195 | chr14-57929878-DEL-195 |
| chr14-57930795-DEL-206 | chr14-57930796-DEL-206 |
| chr14-57941939-DEL-169 | chr14-57941940-DEL-169 |
| chr14-57962902-DEL-78 | chr14-57962903-DEL-78 |
| chr14-58014622-DEL-193 | chr14-58014623-DEL-193 |
| chr14-58040728-DEL-65 | chr14-58040729-DEL-65 |
| chr14-58061571-DEL-72 | chr14-58061573-DEL-73 |
| chr14-58077260-DEL-160 | chr14-58077261-DEL-160 |
| chr14-58086217-DEL-474 | chr14-58086218-DEL-474 |
| chr14-58117725-DEL-216 | chr14-58117725-DEL-213 |
| chr14-58132269-DEL-203 | chr14-58132270-DEL-203 |
| chr14-58142445-DEL-60 | chr14-58142444-DEL-61 |
| chr14-58152864-DEL-75 | chr14-58152865-DEL-75 |
| chr14-58157612-DEL-227 | chr14-58157613-DEL-227 |
| chr14-58163110-DEL-202 | chr14-58163111-DEL-202 |
| chr14-58164418-DEL-212 | chr14-58164419-DEL-212 |
| chr14-58168188-DEL-120 | chr14-58168189-DEL-120 |
| chr14-58203668-DEL-744 | chr14-58203669-DEL-744 |
| chr14-58290040-DEL-7086 | chr14-58290041-DEL-7086 |
| chr14-58350475-DEL-93 | chr14-58350476-DEL-93 |
| chr14-58364565-DEL-70 | chr14-58364565-DEL-69 |
| chr14-58434951-DEL-88 | chr14-58434952-DEL-80 |
| chr14-58443306-DEL-98 | chr14-58443312-DEL-93 |
| chr14-58483882-DEL-6619 | chr14-58483883-DEL-6619 |
| chr14-58490043-DEL-208 | chr14-58490044-DEL-208 |
| chr14-58507234-DEL-419 | chr14-58507235-DEL-419 |
| chr14-58521659-DEL-262 | chr14-58521660-DEL-261 |
| chr14-58550005-DEL-1850 | chr14-58550006-DEL-1850 |
| chr14-58563104-DEL-6008 | chr14-58563105-DEL-6008 |
| chr14-58641975-DEL-6652 | chr14-58641975-DEL-6652 |
| chr14-58650019-DEL-783 | chr14-58650020-DEL-783 |
| chr14-58674367-DEL-153 | chr14-58674368-DEL-153 |
| chr14-58700161-DEL-1862 | chr14-58700162-DEL-1862 |
| chr14-58720591-DEL-762 | chr14-58720592-DEL-762 |
| chr14-58721522-DEL-379 | chr14-58721523-DEL-379 |
| chr14-58766431-DEL-98 | chr14-58766433-DEL-95 |
| chr14-58824800-DEL-6501 | chr14-58824801-DEL-6501 |
| chr14-58849129-DEL-1390 | chr14-58849130-DEL-1390 |
| chr14-58945660-DEL-188 | chr14-58945678-DEL-192 |
| chr14-58961933-DEL-64 | chr14-58961934-DEL-64 |
| chr14-59077376-DEL-1176 | chr14-59077377-DEL-1176 |
| chr14-59197297-DEL-329 | chr14-59197308-DEL-315 |
| chr14-59270982-DEL-59 | chr14-59270984-DEL-52 |
| chr14-59508473-DEL-233 | chr14-59508474-DEL-233 |
| chr14-59648975-DEL-65 | chr14-59648976-DEL-64 |
| chr14-59697090-DEL-1097 | chr14-59697091-DEL-1097 |
| chr14-59756437-DEL-55 | chr14-59756434-DEL-54 |
| chr14-59773985-DEL-333 | chr14-59773987-DEL-332 |
| chr14-59869474-DEL-82 | chr14-59869475-DEL-82 |
| chr14-59956368-DEL-61 | chr14-59956369-DEL-61 |
| chr14-59968334-DEL-120 | chr14-59968304-DEL-138 |
| chr14-60011763-DEL-3401 | chr14-60011764-DEL-3401 |
| chr14-60075186-DEL-547 | chr14-60075205-DEL-532 |
| chr14-60146121-DEL-942 | chr14-60146123-DEL-940 |
| chr14-60208407-DEL-94 | chr14-60208408-DEL-94 |
| chr14-60218162-DEL-6318 | chr14-60218161-DEL-6319 |
| chr14-60230580-DEL-264 | chr14-60230555-DEL-288 |

| | |
|---|---|
| chr14-60230881-DEL-1224 | chr14-60230882-DEL-1224 |
| chr14-60236456-DEL-407 | chr14-60236457-DEL-406 |
| chr14-60246025-DEL-161 | chr14-60246026-DEL-161 |
| chr14-60250890-DEL-111 | chr14-60250888-DEL-112 |
| chr14-60275229-DEL-1008 | chr14-60275230-DEL-1008 |
| chr14-60277014-DEL-2270 | chr14-60277015-DEL-2270 |
| chr14-60284840-DEL-245 | chr14-60284845-DEL-245 |
| chr14-60348066-DEL-4399 | chr14-60348067-DEL-4399 |
| chr14-60392827-DEL-6606 | chr14-60392828-DEL-6606 |
| chr14-60392882-DEL-201 | chr14-60392883-DEL-201 |
| chr14-60439352-DEL-128 | chr14-60439353-DEL-128 |
| chr14-60495937-DEL-1008 | chr14-60495938-DEL-1008 |
| chr14-60520658-DEL-72 | chr14-60520659-DEL-72 |
| chr14-60544255-DEL-70 | chr14-60544256-DEL-70 |
| chr14-60547710-DEL-76 | chr14-60547711-DEL-76 |
| chr14-60565119-DEL-6445 | chr14-60565120-DEL-6445 |
| chr14-60579874-DEL-250 | chr14-60579875-DEL-250 |
| chr14-60588320-DEL-6475 | chr14-60588321-DEL-6475 |
| chr14-60632187-DEL-1390 | chr14-60632187-DEL-1390 |
| chr14-60706229-DEL-56 | chr14-60706230-DEL-56 |
| chr14-60742401-DEL-72 | chr14-60742366-DEL-75 |
| chr14-60744884-DEL-3599 | chr14-60744885-DEL-3599 |
| chr14-60776283-DEL-310 | chr14-60776284-DEL-310 |
| chr14-60801542-DEL-5759 | chr14-60801543-DEL-5759 |
| chr14-60841605-DEL-118 | chr14-60841616-DEL-108 |
| chr14-60843036-DEL-65 | chr14-60843037-DEL-65 |
| chr14-60849283-DEL-164 | chr14-60849284-DEL-164 |
| chr14-60915212-DEL-747 | chr14-60915206-DEL-744 |
| chr14-60922545-DEL-122 | chr14-60922543-DEL-122 |
| chr14-60952233-DEL-133 | chr14-60952234-DEL-133 |
| chr14-60954144-DEL-54 | chr14-60954145-DEL-54 |
| chr14-60962663-DEL-203 | chr14-60962664-DEL-203 |
| chr14-60973460-DEL-52 | chr14-60973461-DEL-52 |
| chr14-61011530-DEL-59 | chr14-61011525-DEL-75 |
| chr14-61015904-DEL-57 | chr14-61015903-DEL-58 |
| chr14-61053393-DEL-137 | chr14-61053394-DEL-137 |
| chr14-61053946-DEL-400 | chr14-61053947-DEL-400 |
| chr14-61057719-DEL-6440 | chr14-61057720-DEL-6440 |
| chr14-61091687-DEL-354 | chr14-61091688-DEL-354 |
| chr14-61109023-DEL-58 | chr14-61109024-DEL-58 |
| chr14-61112904-DEL-6233 | chr14-61112905-DEL-6233 |
| chr14-61138035-DEL-51 | chr14-61138021-DEL-56 |
| chr14-61155606-DEL-93 | chr14-61155607-DEL-93 |
| chr14-61171467-DEL-328 | chr14-61171468-DEL-328 |
| chr14-61197518-DEL-1864 | chr14-61197519-DEL-1864 |
| chr14-61215278-DEL-74 | chr14-61215279-DEL-74 |
| chr14-61232325-DEL-58 | chr14-61232325-DEL-58 |
| chr14-61240702-DEL-378 | chr14-61240703-DEL-378 |
| chr14-61252378-DEL-154 | chr14-61252379-DEL-154 |
| chr14-61276877-DEL-226 | chr14-61276878-DEL-226 |
| chr14-61336346-DEL-207 | chr14-61336347-DEL-207 |
| chr14-61345697-DEL-1214 | chr14-61345698-DEL-1214 |
| chr14-61361139-DEL-247 | chr14-61361140-DEL-247 |
| chr14-61388122-DEL-217 | chr14-61388123-DEL-217 |
| chr14-61414962-DEL-94 | chr14-61414963-DEL-94 |
| chr14-61427693-DEL-52 | chr14-61427694-DEL-52 |
| chr14-61481328-DEL-395 | chr14-61481316-DEL-396 |
| chr14-61485134-DEL-207 | chr14-61485135-DEL-207 |
| chr14-61487600-DEL-328 | chr14-61487601-DEL-328 |
| chr14-61514226-DEL-124 | chr14-61514227-DEL-124 |
| chr14-61519408-DEL-773 | chr14-61519409-DEL-773 |
| chr14-61520743-DEL-325 | chr14-61520744-DEL-325 |
| chr14-61525534-DEL-1246 | chr14-61525535-DEL-1246 |
| chr14-61532572-DEL-67 | chr14-61532573-DEL-67 |
| chr14-61582925-DEL-140 | chr14-61582926-DEL-105 |
| chr14-61592495-DEL-769 | chr14-61592496-DEL-769 |
| chr14-61599337-DEL-254 | chr14-61599348-DEL-258 |
| chr14-61663559-DEL-98 | chr14-61663560-DEL-98 |
| chr14-61664748-DEL-328 | chr14-61664749-DEL-328 |
| chr14-61669777-DEL-56 | chr14-61669770-DEL-52 |
| chr14-61696037-DEL-52 | chr14-61696038-DEL-52 |
| chr14-61705623-DEL-96 | chr14-61705624-DEL-96 |
| chr14-61732519-DEL-485 | chr14-61732520-DEL-485 |
| chr14-61735006-DEL-175 | chr14-61735007-DEL-175 |
| chr14-61755121-DEL-83 | chr14-61755122-DEL-83 |
| chr14-61760009-DEL-934 | chr14-61760010-DEL-934 |
| chr14-61762579-DEL-251 | chr14-61762580-DEL-248 |
| chr14-61774932-DEL-484 | chr14-61774933-DEL-484 |
| chr14-61802426-DEL-55 | chr14-61802428-DEL-54 |
| chr14-61807204-DEL-217 | chr14-61807205-DEL-217 |

| | |
|---|---|
| chr14-61851291-DEL-182 | chr14-61851291-DEL-182 |
| chr14-61863556-DEL-198 | chr14-61863557-DEL-198 |
| chr14-61867670-DEL-209 | chr14-61867671-DEL-209 |
| chr14-61879119-DEL-222 | chr14-61879119-DEL-223 |
| chr14-61943823-DEL-441 | chr14-61943824-DEL-441 |
| chr14-62018875-DEL-64 | chr14-62018876-DEL-52 |
| chr14-62041652-DEL-158 | chr14-62041653-DEL-158 |
| chr14-62059201-DEL-68 | chr14-62059202-DEL-68 |
| chr14-62078081-DEL-217 | chr14-62078082-DEL-217 |
| chr14-62098041-DEL-68 | chr14-62098058-DEL-52 |
| chr14-62104348-DEL-55 | chr14-62104349-DEL-55 |
| chr14-62104953-DEL-53 | chr14-62104954-DEL-53 |
| chr14-62147006-DEL-192 | chr14-62147007-DEL-192 |
| chr14-62198553-DEL-157 | chr14-62198554-DEL-157 |
| chr14-62203566-DEL-380 | chr14-62203567-DEL-380 |
| chr14-62242894-DEL-57 | chr14-62242895-DEL-57 |
| chr14-62302550-DEL-138 | chr14-62302551-DEL-138 |
| chr14-62309504-DEL-75 | chr14-62309521-DEL-59 |
| chr14-62358667-DEL-67 | chr14-62358668-DEL-66 |
| chr14-62362214-DEL-1474 | chr14-62362215-DEL-1474 |
| chr14-62364033-DEL-55 | chr14-62364034-DEL-70 |
| chr14-62368659-DEL-61 | chr14-62368626-DEL-69 |
| chr14-62398500-DEL-464 | chr14-62398501-DEL-464 |
| chr14-62421231-DEL-60 | chr14-62421232-DEL-58 |
| chr14-62463365-DEL-56 | chr14-62463366-DEL-56 |
| chr14-62483370-DEL-100 | chr14-62483371-DEL-100 |
| chr14-62485048-DEL-189 | chr14-62485049-DEL-189 |
| chr14-62492632-DEL-345 | chr14-62492633-DEL-345 |
| chr14-62499020-DEL-6554 | chr14-62499021-DEL-6554 |
| chr14-62524308-DEL-64 | chr14-62524293-DEL-80 |
| chr14-62573689-DEL-280 | chr14-62573690-DEL-280 |
| chr14-62579001-DEL-383 | chr14-62579002-DEL-383 |
| chr14-62588528-DEL-54 | chr14-62588529-DEL-54 |
| chr14-62609297-DEL-159 | chr14-62609298-DEL-159 |
| chr14-62616581-DEL-51 | chr14-62616579-DEL-52 |
| chr14-62620914-DEL-77 | chr14-62620922-DEL-70 |
| chr14-62642423-DEL-93 | chr14-62642424-DEL-93 |
| chr14-62676434-DEL-122 | chr14-62676435-DEL-122 |
| chr14-62742224-DEL-703 | chr14-62742225-DEL-703 |
| chr14-62755032-DEL-80 | chr14-62755035-DEL-78 |
| chr14-62761722-DEL-118 | chr14-62761724-DEL-119 |
| chr14-62762351-DEL-1340 | chr14-62762352-DEL-1340 |
| chr14-62763446-DEL-172 | chr14-62763447-DEL-172 |
| chr14-62804295-DEL-153 | chr14-62804296-DEL-153 |
| chr14-62813163-DEL-138 | chr14-62813164-DEL-138 |
| chr14-62817827-DEL-136 | chr14-62817790-DEL-136 |
| chr14-62877275-DEL-59 | chr14-62877276-DEL-59 |
| chr14-62880054-DEL-4550 | chr14-62880055-DEL-4550 |
| chr14-62949419-DEL-714 | chr14-62949419-DEL-715 |
| chr14-62959151-DEL-524 | chr14-62959152-DEL-524 |
| chr14-62972646-DEL-1577 | chr14-62972647-DEL-1577 |
| chr14-62980917-DEL-75 | chr14-62980918-DEL-75 |
| chr14-63013200-DEL-53 | chr14-63013201-DEL-53 |
| chr14-63048531-DEL-93 | chr14-63048532-DEL-93 |
| chr14-63115735-DEL-51 | chr14-63115736-DEL-51 |
| chr14-63124227-DEL-63 | chr14-63124228-DEL-55 |
| chr14-63182392-DEL-189 | chr14-63182393-DEL-189 |
| chr14-63188781-DEL-60 | chr14-63188784-DEL-58 |
| chr14-63200675-DEL-2373 | chr14-63200676-DEL-2373 |
| chr14-63216141-DEL-84 | chr14-63216142-DEL-84 |
| chr14-63231695-DEL-51 | chr14-63231682-DEL-51 |
| chr14-63251838-DEL-208 | chr14-63251839-DEL-208 |
| chr14-63253756-DEL-400 | chr14-63253757-DEL-400 |
| chr14-63257360-DEL-86 | chr14-63257351-DEL-87 |
| chr14-63288955-DEL-55 | chr14-63288956-DEL-55 |
| chr14-63294582-DEL-185 | chr14-63294583-DEL-185 |
| chr14-63299219-DEL-178 | chr14-63299220-DEL-178 |
| chr14-63301024-DEL-84 | chr14-63301023-DEL-79 |
| chr14-63337088-DEL-80 | chr14-63337089-DEL-80 |
| chr14-63350144-DEL-1669 | chr14-63350149-DEL-1660 |
| chr14-63355425-DEL-96 | chr14-63355426-DEL-78 |
| chr14-63441575-DEL-303 | chr14-63441570-DEL-309 |
| chr14-63455069-DEL-309 | chr14-63455070-DEL-309 |
| chr14-63517978-DEL-186 | chr14-63517978-DEL-186 |
| chr14-63525973-DEL-228 | chr14-63525974-DEL-228 |
| chr14-63528173-DEL-503 | chr14-63528174-DEL-503 |
| chr14-63552052-DEL-398 | chr14-63552053-DEL-398 |
| chr14-63553792-DEL-58 | chr14-63553795-DEL-56 |
| chr14-63574435-DEL-200 | chr14-63574436-DEL-200 |
| chr14-63579051-DEL-150 | chr14-63579074-DEL-141 |

| | |
|---|---|
| chr14-63586361-DEL-101 | chr14-63586361-DEL-101 |
| chr14-63593088-DEL-88 | chr14-63593089-DEL-88 |
| chr14-63633458-DEL-166 | chr14-63633456-DEL-168 |
| chr14-63658856-DEL-154 | chr14-63658861-DEL-150 |
| chr14-63709437-DEL-1488 | chr14-63709438-DEL-1488 |
| chr14-63721401-DEL-428 | chr14-63721402-DEL-428 |
| chr14-63727272-DEL-1344 | chr14-63727273-DEL-1344 |
| chr14-63748239-DEL-79 | chr14-63748240-DEL-79 |
| chr14-63759070-DEL-986 | chr14-63759071-DEL-986 |
| chr14-63765119-DEL-57 | chr14-63765114-DEL-61 |
| chr14-63796654-DEL-208 | chr14-63796655-DEL-208 |
| chr14-63831399-DEL-61 | chr14-63831396-DEL-61 |
| chr14-63847042-DEL-157 | chr14-63847043-DEL-157 |
| chr14-63857092-DEL-133 | chr14-63857093-DEL-133 |
| chr14-63879794-DEL-7223 | chr14-63879795-DEL-7223 |
| chr14-63894944-DEL-207 | chr14-63894945-DEL-207 |
| chr14-63923074-DEL-272 | chr14-63923075-DEL-272 |
| chr14-63927550-DEL-312 | chr14-63927551-DEL-312 |
| chr14-63943337-DEL-5665 | chr14-63943338-DEL-5665 |
| chr14-63962849-DEL-52 | chr14-63962850-DEL-52 |
| chr14-63972592-DEL-58 | chr14-63972593-DEL-58 |
| chr14-64000138-DEL-187 | chr14-64000142-DEL-187 |
| chr14-64014799-DEL-64 | chr14-64014800-DEL-56 |
| chr14-64028467-DEL-64 | chr14-64028467-DEL-56 |
| chr14-64037251-DEL-108 | chr14-64037252-DEL-108 |
| chr14-64043724-DEL-56 | chr14-64043725-DEL-52 |
| chr14-64047629-DEL-223 | chr14-64047646-DEL-207 |
| chr14-64132501-DEL-341 | chr14-64132502-DEL-341 |
| chr14-64195011-DEL-127 | chr14-64195012-DEL-127 |
| chr14-64231397-DEL-57 | chr14-64231394-DEL-58 |
| chr14-64243434-DEL-228 | chr14-64243431-DEL-232 |
| chr14-64261078-DEL-729 | chr14-64261079-DEL-729 |
| chr14-64286156-DEL-124 | chr14-64286139-DEL-130 |
| chr14-64291465-DEL-67 | chr14-64291466-DEL-67 |
| chr14-64324232-DEL-108 | chr14-64324233-DEL-90 |
| chr14-64331911-DEL-59 | chr14-64331903-DEL-56 |
| chr14-64344375-DEL-78 | chr14-64344376-DEL-78 |
| chr14-64364235-DEL-137 | chr14-64364230-DEL-142 |
| chr14-64379783-DEL-405 | chr14-64379784-DEL-405 |
| chr14-64384895-DEL-60 | chr14-64384860-DEL-66 |
| chr14-64397456-DEL-58 | chr14-64397457-DEL-58 |
| chr14-64400016-DEL-75 | chr14-64399988-DEL-65 |
| chr14-64462802-DEL-2785 | chr14-64462803-DEL-2785 |
| chr14-64496155-DEL-752 | chr14-64496156-DEL-752 |
| chr14-64534173-DEL-56 | chr14-64534174-DEL-56 |
| chr14-64543450-DEL-522 | chr14-64543451-DEL-522 |
| chr14-64617586-DEL-180 | chr14-64617587-DEL-180 |
| chr14-64664738-DEL-135 | chr14-64664739-DEL-135 |
| chr14-64685213-DEL-92 | chr14-64685214-DEL-102 |
| chr14-64700822-DEL-5342 | chr14-64700823-DEL-5342 |
| chr14-64708970-DEL-596 | chr14-64708974-DEL-592 |
| chr14-64736631-DEL-188 | chr14-64736632-DEL-188 |
| chr14-64742230-DEL-6413 | chr14-64742231-DEL-6413 |
| chr14-64845406-DEL-115 | chr14-64845407-DEL-115 |
| chr14-64847707-DEL-114 | chr14-64847718-DEL-104 |
| chr14-64915997-DEL-96 | chr14-64916004-DEL-82 |
| chr14-64938632-DEL-105 | chr14-64938668-DEL-70 |
| chr14-64953789-DEL-163 | chr14-64953782-DEL-164 |
| chr14-65078987-DEL-56 | chr14-65078988-DEL-56 |
| chr14-65093870-DEL-203 | chr14-65093871-DEL-203 |
| chr14-65174143-DEL-3671 | chr14-65174136-DEL-3672 |
| chr14-65177808-DEL-196 | chr14-65177809-DEL-196 |
| chr14-65207963-DEL-2623 | chr14-65207964-DEL-2623 |
| chr14-65211398-DEL-655 | chr14-65211400-DEL-652 |
| chr14-65212991-DEL-63 | chr14-65212992-DEL-63 |
| chr14-65224680-DEL-111 | chr14-65224680-DEL-111 |
| chr14-65225355-DEL-111 | chr14-65225318-DEL-111 |
| chr14-65248156-DEL-111 | chr14-65248157-DEL-111 |
| chr14-65255144-DEL-81 | chr14-65255145-DEL-79 |
| chr14-65275633-DEL-941 | chr14-65275633-DEL-942 |
| chr14-65340660-DEL-225 | chr14-65340661-DEL-225 |
| chr14-65354517-DEL-180 | chr14-65354518-DEL-180 |
| chr14-65366286-DEL-6403 | chr14-65366287-DEL-6403 |
| chr14-65407275-DEL-255 | chr14-65407273-DEL-252 |
| chr14-65415946-DEL-5131 | chr14-65415947-DEL-5131 |
| chr14-65451815-DEL-640 | chr14-65451816-DEL-640 |
| chr14-65516173-DEL-6452 | chr14-65516174-DEL-6452 |
| chr14-65554620-DEL-223 | chr14-65554617-DEL-230 |
| chr14-65610116-DEL-903 | chr14-65610116-DEL-904 |
| chr14-65610804-DEL-56 | chr14-65610754-DEL-60 |

| | |
|---|---|
| chr14-65648192-DEL-59 | chr14-65648193-DEL-59 |
| chr14-65689147-DEL-66 | chr14-65689148-DEL-70 |
| chr14-65793413-DEL-70 | chr14-65793414-DEL-70 |
| chr14-65833890-DEL-403 | chr14-65833891-DEL-403 |
| chr14-65865520-DEL-97 | chr14-65865521-DEL-97 |
| chr14-65879968-DEL-58 | chr14-65879967-DEL-60 |
| chr14-65886612-DEL-81 | chr14-65886637-DEL-52 |
| chr14-65947884-DEL-192 | chr14-65947884-DEL-194 |
| chr14-65959004-DEL-56 | chr14-65959005-DEL-56 |
| chr14-65970986-DEL-1068 | chr14-65970987-DEL-1068 |
| chr14-65993766-DEL-399 | chr14-65993767-DEL-399 |
| chr14-66003151-DEL-51 | chr14-66003150-DEL-51 |
| chr14-66026478-DEL-67 | chr14-66026482-DEL-66 |
| chr14-66027946-DEL-328 | chr14-66027947-DEL-328 |
| chr14-66067431-DEL-207 | chr14-66067432-DEL-207 |
| chr14-66072722-DEL-174 | chr14-66072723-DEL-174 |
| chr14-66112962-DEL-117 | chr14-66112961-DEL-118 |
| chr14-66115272-DEL-62 | chr14-66115273-DEL-62 |
| chr14-66123999-DEL-72 | chr14-66124000-DEL-72 |
| chr14-66129526-DEL-202 | chr14-66129527-DEL-202 |
| chr14-66174503-DEL-105 | chr14-66174504-DEL-105 |
| chr14-66195024-DEL-61 | chr14-66195033-DEL-51 |
| chr14-66197394-DEL-1710 | chr14-66197395-DEL-1710 |
| chr14-66199834-DEL-148 | chr14-66199835-DEL-144 |
| chr14-66232188-DEL-52 | chr14-66232189-DEL-52 |
| chr14-66233676-DEL-102 | chr14-66233677-DEL-102 |
| chr14-66247502-DEL-1059 | chr14-66247503-DEL-1059 |
| chr14-66248245-DEL-219 | chr14-66248246-DEL-219 |
| chr14-66268433-DEL-399 | chr14-66268434-DEL-399 |
| chr14-66282374-DEL-1163 | chr14-66282375-DEL-1163 |
| chr14-66364065-DEL-2244 | chr14-66364065-DEL-2244 |
| chr14-66410161-DEL-109 | chr14-66410162-DEL-109 |
| chr14-66440662-DEL-3321 | chr14-66440663-DEL-3321 |
| chr14-66465181-DEL-53 | chr14-66465182-DEL-53 |
| chr14-66481344-DEL-56 | chr14-66481343-DEL-52 |
| chr14-66482843-DEL-180 | chr14-66482844-DEL-180 |
| chr14-66488931-DEL-65 | chr14-66488919-DEL-65 |
| chr14-66505318-DEL-61 | chr14-66505319-DEL-61 |
| chr14-66521705-DEL-223 | chr14-66521706-DEL-223 |
| chr14-66604459-DEL-1966 | chr14-66604459-DEL-1938 |
| chr14-66607591-DEL-168 | chr14-66607592-DEL-168 |
| chr14-66617218-DEL-65 | chr14-66617189-DEL-68 |
| chr14-66642269-DEL-345 | chr14-66642270-DEL-345 |
| chr14-66657111-DEL-74 | chr14-66657106-DEL-80 |
| chr14-66672314-DEL-98 | chr14-66672315-DEL-98 |
| chr14-66690184-DEL-51 | chr14-66690185-DEL-51 |
| chr14-66700131-DEL-7157 | chr14-66700132-DEL-7157 |
| chr14-66745890-DEL-102 | chr14-66745891-DEL-102 |
| chr14-66746601-DEL-65 | chr14-66746602-DEL-65 |
| chr14-66758545-DEL-102 | chr14-66758546-DEL-102 |
| chr14-66772975-DEL-54 | chr14-66772974-DEL-56 |
| chr14-66820684-DEL-58 | chr14-66820637-DEL-103 |
| chr14-66850919-DEL-3580 | chr14-66850920-DEL-3580 |
| chr14-66864935-DEL-519 | chr14-66864936-DEL-519 |
| chr14-66906329-DEL-139 | chr14-66906330-DEL-139 |
| chr14-66913122-DEL-62 | chr14-66913123-DEL-62 |
| chr14-66938471-DEL-3117 | chr14-66938467-DEL-3114 |
| chr14-66944596-DEL-58 | chr14-66944599-DEL-56 |
| chr14-66949253-DEL-291 | chr14-66949254-DEL-291 |
| chr14-66984899-DEL-62 | chr14-66984904-DEL-54 |
| chr14-66991707-DEL-82 | chr14-66991741-DEL-76 |
| chr14-67000775-DEL-60 | chr14-67000775-DEL-60 |
| chr14-67002376-DEL-77 | chr14-67002377-DEL-77 |
| chr14-67022257-DEL-141 | chr14-67022258-DEL-141 |
| chr14-67084495-DEL-72 | chr14-67084496-DEL-72 |
| chr14-67116541-DEL-809 | chr14-67116542-DEL-809 |
| chr14-67121531-DEL-2236 | chr14-67121531-DEL-2236 |
| chr14-67132284-DEL-83 | chr14-67132284-DEL-83 |
| chr14-67205858-DEL-329 | chr14-67205861-DEL-330 |
| chr14-67207439-DEL-1364 | chr14-67207440-DEL-1364 |
| chr14-67262383-DEL-58 | chr14-67262384-DEL-56 |
| chr14-67263832-DEL-184 | chr14-67263833-DEL-184 |
| chr14-67342093-DEL-76 | chr14-67342096-DEL-74 |
| chr14-67382902-DEL-70 | chr14-67382898-DEL-71 |
| chr14-67402953-DEL-88 | chr14-67402961-DEL-80 |
| chr14-67448340-DEL-776 | chr14-67448341-DEL-776 |
| chr14-67459632-DEL-193 | chr14-67459633-DEL-193 |
| chr14-67495497-DEL-109 | chr14-67495515-DEL-92 |
| chr14-67527084-DEL-160 | chr14-67527085-DEL-160 |
| chr14-67572288-DEL-70 | chr14-67572289-DEL-70 |

| | |
|---|---|
| chr14-67577045-DEL-66 | chr14-67577046-DEL-56 |
| chr14-67587797-DEL-104 | chr14-67587814-DEL-75 |
| chr14-67656107-DEL-76 | chr14-67656108-DEL-76 |
| chr14-67659318-DEL-53 | chr14-67659319-DEL-53 |
| chr14-67670653-DEL-127 | chr14-67670654-DEL-127 |
| chr14-67699985-DEL-132 | chr14-67700002-DEL-116 |
| chr14-67715254-DEL-90 | chr14-67715249-DEL-91 |
| chr14-67726300-DEL-443 | chr14-67726327-DEL-462 |
| chr14-67728770-DEL-70 | chr14-67728771-DEL-70 |
| chr14-67748104-DEL-367 | chr14-67748105-DEL-367 |
| chr14-67751193-DEL-197 | chr14-67751192-DEL-195 |
| chr14-67752572-DEL-206 | chr14-67752573-DEL-206 |
| chr14-67758674-DEL-134 | chr14-67758675-DEL-134 |
| chr14-67763245-DEL-60 | chr14-67763246-DEL-60 |
| chr14-67765893-DEL-120 | chr14-67765894-DEL-120 |
| chr14-67777227-DEL-288 | chr14-67777208-DEL-281 |
| chr14-67795357-DEL-211 | chr14-67795358-DEL-222 |
| chr14-67885419-DEL-63 | chr14-67885420-DEL-63 |
| chr14-67887964-DEL-1696 | chr14-67887961-DEL-1682 |
| chr14-67904824-DEL-199 | chr14-67904825-DEL-199 |
| chr14-67961526-DEL-324 | chr14-67961532-DEL-321 |
| chr14-67992671-DEL-56 | chr14-67992672-DEL-56 |
| chr14-68020589-DEL-98 | chr14-68020582-DEL-62 |
| chr14-68108223-DEL-86 | chr14-68108224-DEL-86 |
| chr14-68111118-DEL-93 | chr14-68111077-DEL-136 |
| chr14-68128949-DEL-60 | chr14-68128953-DEL-54 |
| chr14-68133386-DEL-104 | chr14-68133387-DEL-104 |
| chr14-68181303-DEL-118 | chr14-68181304-DEL-118 |
| chr14-68190237-DEL-1092 | chr14-68190238-DEL-1092 |
| chr14-68199779-DEL-60 | chr14-68199780-DEL-60 |
| chr14-68216234-DEL-620 | chr14-68216218-DEL-636 |
| chr14-68224719-DEL-203 | chr14-68224720-DEL-203 |
| chr14-68239128-DEL-2203 | chr14-68239129-DEL-2203 |
| chr14-68244284-DEL-7130 | chr14-68244276-DEL-7132 |
| chr14-68252513-DEL-197 | chr14-68252514-DEL-197 |
| chr14-68254098-DEL-149 | chr14-68254097-DEL-150 |
| chr14-68267129-DEL-574 | chr14-68267130-DEL-574 |
| chr14-68275330-DEL-88 | chr14-68275331-DEL-88 |
| chr14-68302339-DEL-59 | chr14-68302340-DEL-59 |
| chr14-68312968-DEL-73 | chr14-68312948-DEL-80 |
| chr14-68319228-DEL-72 | chr14-68319221-DEL-68 |
| chr14-68340729-DEL-95 | chr14-68340730-DEL-95 |
| chr14-68343095-DEL-58 | chr14-68343096-DEL-58 |
| chr14-68354094-DEL-549 | chr14-68354095-DEL-549 |
| chr14-68359128-DEL-56 | chr14-68359111-DEL-57 |
| chr14-68380625-DEL-56 | chr14-68380627-DEL-52 |
| chr14-68437488-DEL-98 | chr14-68437489-DEL-98 |
| chr14-68439035-DEL-907 | chr14-68439036-DEL-907 |
| chr14-68444117-DEL-59 | chr14-68444118-DEL-59 |
| chr14-68447731-DEL-59 | chr14-68447690-DEL-72 |
| chr14-68514868-DEL-411 | chr14-68514869-DEL-411 |
| chr14-68518374-DEL-52 | chr14-68518375-DEL-52 |
| chr14-68530179-DEL-65 | chr14-68530180-DEL-68 |
| chr14-68540127-DEL-72 | chr14-68540128-DEL-72 |
| chr14-68559733-DEL-60 | chr14-68559734-DEL-60 |
| chr14-68561499-DEL-68 | chr14-68561505-DEL-50 |
| chr14-68567654-DEL-247 | chr14-68567655-DEL-247 |
| chr14-68572315-DEL-141 | chr14-68572338-DEL-127 |
| chr14-68606309-DEL-52 | chr14-68606310-DEL-52 |
| chr14-68617170-DEL-356 | chr14-68617171-DEL-356 |
| chr14-68661094-DEL-365 | chr14-68661089-DEL-366 |
| chr14-68673409-DEL-401 | chr14-68673410-DEL-401 |
| chr14-68707127-DEL-8401 | chr14-68707128-DEL-8401 |
| chr14-68733718-DEL-311 | chr14-68733719-DEL-311 |
| chr14-68752725-DEL-739 | chr14-68752726-DEL-739 |
| chr14-68786528-DEL-6653 | chr14-68786529-DEL-6653 |
| chr14-68799003-DEL-196 | chr14-68799004-DEL-196 |
| chr14-68811859-DEL-73 | chr14-68811860-DEL-74 |
| chr14-68819782-DEL-66 | chr14-68819783-DEL-66 |
| chr14-68832445-DEL-123 | chr14-68832448-DEL-120 |
| chr14-68881444-DEL-78 | chr14-68881445-DEL-78 |
| chr14-68882978-DEL-401 | chr14-68882979-DEL-401 |
| chr14-68913402-DEL-7087 | chr14-68913403-DEL-7087 |
| chr14-68924526-DEL-70 | chr14-68924527-DEL-70 |
| chr14-68945720-DEL-5845 | chr14-68945723-DEL-5842 |
| chr14-68962704-DEL-820 | chr14-68962690-DEL-809 |
| chr14-68972896-DEL-437 | chr14-68972898-DEL-436 |
| chr14-68974183-DEL-321 | chr14-68974184-DEL-321 |
| chr14-68986159-DEL-6272 | chr14-68986160-DEL-6272 |
| chr14-69011481-DEL-1033 | chr14-69011482-DEL-1033 |

| | |
|---|---|
| chr14-69047722-DEL-2739 | chr14-69047723-DEL-2739 |
| chr14-69055155-DEL-90 | chr14-69055158-DEL-88 |
| chr14-69062534-DEL-80 | chr14-69062530-DEL-104 |
| chr14-69088265-DEL-6186 | chr14-69088266-DEL-6186 |
| chr14-69116709-DEL-61 | chr14-69116710-DEL-61 |
| chr14-69138006-DEL-62 | chr14-69138010-DEL-61 |
| chr14-69146039-DEL-404 | chr14-69146034-DEL-391 |
| chr14-69150512-DEL-1807 | chr14-69150513-DEL-1807 |
| chr14-69163589-DEL-225 | chr14-69163590-DEL-225 |
| chr14-69196009-DEL-95 | chr14-69196010-DEL-89 |
| chr14-69197665-DEL-411 | chr14-69197666-DEL-411 |
| chr14-69199769-DEL-308 | chr14-69199777-DEL-307 |
| chr14-69202344-DEL-64 | chr14-69202344-DEL-64 |
| chr14-69230968-DEL-108 | chr14-69230965-DEL-112 |
| chr14-69245517-DEL-1310 | chr14-69245518-DEL-1310 |
| chr14-69347026-DEL-7068 | chr14-69347026-DEL-7069 |
| chr14-69418561-DEL-133 | chr14-69418562-DEL-133 |
| chr14-69459554-DEL-466 | chr14-69459555-DEL-466 |
| chr14-69495194-DEL-68 | chr14-69495195-DEL-68 |
| chr14-69495463-DEL-309 | chr14-69495464-DEL-309 |
| chr14-69925563-DEL-57 | chr14-69925564-DEL-57 |
| chr14-69932943-DEL-205 | chr14-69932944-DEL-205 |
| chr14-69941687-DEL-135 | chr14-69941660-DEL-136 |
| chr14-69964858-DEL-113 | chr14-69964858-DEL-113 |
| chr14-69973970-DEL-511 | chr14-69973970-DEL-511 |
| chr14-69993274-DEL-1305 | chr14-69993275-DEL-1305 |
| chr14-69995712-DEL-369 | chr14-69995713-DEL-369 |
| chr14-70018677-DEL-414 | chr14-70018678-DEL-397 |
| chr14-70044794-DEL-165 | chr14-70044795-DEL-165 |
| chr14-70059373-DEL-111 | chr14-70059404-DEL-120 |
| chr14-70064737-DEL-62 | chr14-70064738-DEL-62 |
| chr14-70084783-DEL-112 | chr14-70084775-DEL-80 |
| chr14-70091695-DEL-59 | chr14-70091696-DEL-59 |
| chr14-70101811-DEL-230 | chr14-70101812-DEL-230 |
| chr14-70130928-DEL-175 | chr14-70130929-DEL-175 |
| chr14-70156353-DEL-202 | chr14-70156354-DEL-201 |
| chr14-70217360-DEL-99 | chr14-70217361-DEL-99 |
| chr14-70341713-DEL-1882 | chr14-70341714-DEL-1882 |
| chr14-70348457-DEL-62 | chr14-70348436-DEL-63 |
| chr14-70355528-DEL-217 | chr14-70355529-DEL-217 |
| chr14-70500032-DEL-514 | chr14-70500033-DEL-514 |
| chr14-70558485-DEL-56 | chr14-70558486-DEL-52 |
| chr14-70578273-DEL-117 | chr14-70578274-DEL-117 |
| chr14-70597441-DEL-1502 | chr14-70597441-DEL-1503 |
| chr14-70618985-DEL-58 | chr14-70618966-DEL-58 |
| chr14-70628437-DEL-201 | chr14-70628438-DEL-201 |
| chr14-70633667-DEL-460 | chr14-70633672-DEL-457 |
| chr14-70663445-DEL-280 | chr14-70663446-DEL-280 |
| chr14-70689851-DEL-63 | chr14-70689852-DEL-63 |
| chr14-70697340-DEL-60 | chr14-70697341-DEL-60 |
| chr14-70700109-DEL-58 | chr14-70700110-DEL-58 |
| chr14-70806220-DEL-146 | chr14-70806226-DEL-116 |
| chr14-70821113-DEL-128 | chr14-70821114-DEL-128 |
| chr14-70833717-DEL-52 | chr14-70833718-DEL-52 |
| chr14-70857398-DEL-248 | chr14-70857399-DEL-248 |
| chr14-70914823-DEL-211 | chr14-70914824-DEL-211 |
| chr14-70941604-DEL-327 | chr14-70941605-DEL-327 |
| chr14-70965268-DEL-6766 | chr14-70965269-DEL-6766 |
| chr14-70987078-DEL-196 | chr14-70987079-DEL-196 |
| chr14-71001752-DEL-181 | chr14-71001753-DEL-181 |
| chr14-71008325-DEL-201 | chr14-71008325-DEL-201 |
| chr14-71033040-DEL-52 | chr14-71033045-DEL-52 |
| chr14-71038497-DEL-206 | chr14-71038498-DEL-206 |
| chr14-71062167-DEL-198 | chr14-71062168-DEL-198 |
| chr14-71071418-DEL-101 | chr14-71071418-DEL-101 |
| chr14-71075910-DEL-500 | chr14-71075910-DEL-500 |
| chr14-71078380-DEL-124 | chr14-71078380-DEL-124 |
| chr14-71100785-DEL-54 | chr14-71100786-DEL-54 |
| chr14-71154370-DEL-1390 | chr14-71154371-DEL-1390 |
| chr14-71155264-DEL-60 | chr14-71155265-DEL-60 |
| chr14-71186603-DEL-63 | chr14-71186604-DEL-63 |
| chr14-71194730-DEL-110 | chr14-71194730-DEL-111 |
| chr14-71211859-DEL-1445 | chr14-71211860-DEL-1445 |
| chr14-71225780-DEL-50 | chr14-71225788-DEL-59 |
| chr14-71263247-DEL-78 | chr14-71263248-DEL-78 |
| chr14-71281343-DEL-256 | chr14-71281344-DEL-256 |
| chr14-71295327-DEL-1927 | chr14-71295327-DEL-1927 |
| chr14-71337061-DEL-341 | chr14-71337062-DEL-341 |
| chr14-71358632-DEL-5502 | chr14-71358633-DEL-5502 |
| chr14-71375018-DEL-64 | chr14-71374983-DEL-65 |

| | |
|---|---|
| chr14-71430647-DEL-56 | chr14-71430648-DEL-56 |
| chr14-71456141-DEL-3926 | chr14-71456142-DEL-3926 |
| chr14-71470116-DEL-77 | chr14-71470136-DEL-52 |
| chr14-71482139-DEL-343 | chr14-71482140-DEL-343 |
| chr14-71523200-DEL-384 | chr14-71523201-DEL-384 |
| chr14-71568444-DEL-231 | chr14-71568445-DEL-231 |
| chr14-71581687-DEL-771 | chr14-71581688-DEL-771 |
| chr14-71620758-DEL-903 | chr14-71620759-DEL-903 |
| chr14-71628849-DEL-6665 | chr14-71628850-DEL-6665 |
| chr14-71635604-DEL-6427 | chr14-71635604-DEL-6427 |
| chr14-71648513-DEL-362 | chr14-71648553-DEL-339 |
| chr14-71674191-DEL-793 | chr14-71674192-DEL-793 |
| chr14-71679816-DEL-957 | chr14-71679811-DEL-958 |
| chr14-71687062-DEL-6006 | chr14-71687063-DEL-6006 |
| chr14-71696398-DEL-454 | chr14-71696399-DEL-454 |
| chr14-71791326-DEL-594 | chr14-71791327-DEL-594 |
| chr14-71798725-DEL-53 | chr14-71798726-DEL-53 |
| chr14-71812124-DEL-54 | chr14-71812125-DEL-54 |
| chr14-71821191-DEL-156 | chr14-71821221-DEL-140 |
| chr14-71858593-DEL-6381 | chr14-71858594-DEL-6381 |
| chr14-71866085-DEL-4219 | chr14-71866086-DEL-4219 |
| chr14-71932256-DEL-421 | chr14-71932255-DEL-421 |
| chr14-71957130-DEL-296 | chr14-71957131-DEL-296 |
| chr14-71967734-DEL-218 | chr14-71967735-DEL-227 |
| chr14-71974911-DEL-76 | chr14-71974911-DEL-76 |
| chr14-71979843-DEL-308 | chr14-71979844-DEL-308 |
| chr14-72007317-DEL-56 | chr14-72007318-DEL-56 |
| chr14-72038507-DEL-2940 | chr14-72038508-DEL-2940 |
| chr14-72049211-DEL-86 | chr14-72049212-DEL-86 |
| chr14-72083907-DEL-897 | chr14-72083908-DEL-897 |
| chr14-72092679-DEL-53 | chr14-72092680-DEL-53 |
| chr14-72094584-DEL-88 | chr14-72094567-DEL-82 |
| chr14-72110646-DEL-54 | chr14-72110647-DEL-50 |
| chr14-72113714-DEL-59 | chr14-72113719-DEL-56 |
| chr14-72166410-DEL-670 | chr14-72166402-DEL-671 |
| chr14-72198418-DEL-873 | chr14-72198419-DEL-873 |
| chr14-72204433-DEL-97 | chr14-72204434-DEL-97 |
| chr14-72206862-DEL-60 | chr14-72206863-DEL-61 |
| chr14-72220644-DEL-216 | chr14-72220645-DEL-216 |
| chr14-72270512-DEL-285 | chr14-72270513-DEL-285 |
| chr14-72288502-DEL-6615 | chr14-72288488-DEL-6615 |
| chr14-72294731-DEL-214 | chr14-72294732-DEL-214 |
| chr14-72334881-DEL-1895 | chr14-72334882-DEL-1895 |
| chr14-72337629-DEL-166 | chr14-72337630-DEL-166 |
| chr14-72361511-DEL-64 | chr14-72361514-DEL-62 |
| chr14-72365020-DEL-3649 | chr14-72365022-DEL-3646 |
| chr14-72372403-DEL-442 | chr14-72372404-DEL-442 |
| chr14-72397660-DEL-202 | chr14-72397660-DEL-203 |
| chr14-72422224-DEL-60 | chr14-72422215-DEL-70 |
| chr14-72456062-DEL-122 | chr14-72456063-DEL-122 |
| chr14-72525779-DEL-771 | chr14-72525779-DEL-771 |
| chr14-72544690-DEL-400 | chr14-72544691-DEL-400 |
| chr14-72545783-DEL-391 | chr14-72545783-DEL-391 |
| chr14-72548489-DEL-6595 | chr14-72548490-DEL-6595 |
| chr14-72556475-DEL-61 | chr14-72556497-DEL-54 |
| chr14-72570617-DEL-797 | chr14-72570618-DEL-797 |
| chr14-72589324-DEL-997 | chr14-72589324-DEL-997 |
| chr14-72596657-DEL-56 | chr14-72596658-DEL-56 |
| chr14-72647520-DEL-6704 | chr14-72647521-DEL-6704 |
| chr14-72661009-DEL-932 | chr14-72661010-DEL-932 |
| chr14-72668755-DEL-64 | chr14-72668756-DEL-64 |
| chr14-72681872-DEL-208 | chr14-72681873-DEL-208 |
| chr14-72685019-DEL-119 | chr14-72685019-DEL-119 |
| chr14-72688369-DEL-126 | chr14-72688370-DEL-126 |
| chr14-72710592-DEL-58 | chr14-72710593-DEL-58 |
| chr14-72725200-DEL-397 | chr14-72725201-DEL-397 |
| chr14-72731961-DEL-500 | chr14-72731962-DEL-500 |
| chr14-72735124-DEL-1078 | chr14-72735125-DEL-1054 |
| chr14-72764168-DEL-227 | chr14-72764169-DEL-227 |
| chr14-72774708-DEL-51 | chr14-72774709-DEL-51 |
| chr14-72778662-DEL-88 | chr14-72778663-DEL-88 |
| chr14-72819785-DEL-125 | chr14-72819786-DEL-125 |
| chr14-72829827-DEL-351 | chr14-72829828-DEL-351 |
| chr14-72852655-DEL-5621 | chr14-72852656-DEL-5621 |
| chr14-72912007-DEL-762 | chr14-72912007-DEL-762 |
| chr14-72919584-DEL-1097 | chr14-72919568-DEL-1065 |
| chr14-73004096-DEL-1805 | chr14-73004097-DEL-1805 |
| chr14-73007227-DEL-53 | chr14-73007228-DEL-53 |
| chr14-73021770-DEL-54 | chr14-73021773-DEL-52 |
| chr14-73022644-DEL-60 | chr14-73022645-DEL-60 |

| | |
|---|---|
| chr14-73030306-DEL-6243 | chr14-73030307-DEL-6243 |
| chr14-73039267-DEL-72 | chr14-73039268-DEL-72 |
| chr14-73056847-DEL-793 | chr14-73056848-DEL-793 |
| chr14-73073224-DEL-1411 | chr14-73073225-DEL-1411 |
| chr14-73098985-DEL-435 | chr14-73098986-DEL-435 |
| chr14-73103841-DEL-54 | chr14-73103842-DEL-54 |
| chr14-73117835-DEL-3841 | chr14-73117836-DEL-3841 |
| chr14-73137506-DEL-1115 | chr14-73137507-DEL-1115 |
| chr14-73146766-DEL-7610 | chr14-73146774-DEL-7635 |
| chr14-73157323-DEL-159 | chr14-73157324-DEL-159 |
| chr14-73170342-DEL-678 | chr14-73170343-DEL-678 |
| chr14-73181045-DEL-83 | chr14-73181045-DEL-61 |
| chr14-73184335-DEL-158 | chr14-73184336-DEL-158 |
| chr14-73185442-DEL-12239 | chr14-73185443-DEL-12239 |
| chr14-73212819-DEL-85 | chr14-73212820-DEL-85 |
| chr14-73219166-DEL-416 | chr14-73219167-DEL-416 |
| chr14-73224412-DEL-60 | chr14-73224413-DEL-60 |
| chr14-73248579-DEL-68 | chr14-73248580-DEL-70 |
| chr14-73263921-DEL-1585 | chr14-73263922-DEL-1585 |
| chr14-73293160-DEL-208 | chr14-73293161-DEL-208 |
| chr14-73339355-DEL-93 | chr14-73339356-DEL-93 |
| chr14-73352371-DEL-194 | chr14-73352372-DEL-194 |
| chr14-73353545-DEL-60 | chr14-73353554-DEL-58 |
| chr14-73354680-DEL-85 | chr14-73354681-DEL-85 |
| chr14-73364827-DEL-322 | chr14-73364828-DEL-322 |
| chr14-73399098-DEL-66 | chr14-73399099-DEL-66 |
| chr14-73413828-DEL-64 | chr14-73413848-DEL-61 |
| chr14-73424695-DEL-7484 | chr14-73424695-DEL-7485 |
| chr14-73456862-DEL-57 | chr14-73456862-DEL-54 |
| chr14-73466914-DEL-180 | chr14-73466915-DEL-180 |
| chr14-73470981-DEL-401 | chr14-73470982-DEL-401 |
| chr14-73502722-DEL-91 | chr14-73502723-DEL-91 |
| chr14-73537171-DEL-107 | chr14-73537172-DEL-107 |
| chr14-73538291-DEL-160 | chr14-73538245-DEL-206 |
| chr14-73575748-DEL-94 | chr14-73575749-DEL-90 |
| chr14-73604368-DEL-77 | chr14-73604369-DEL-77 |
| chr14-73652597-DEL-75 | chr14-73652598-DEL-57 |
| chr14-73683931-DEL-69 | chr14-73683938-DEL-63 |
| chr14-73688335-DEL-118 | chr14-73688336-DEL-118 |
| chr14-73691562-DEL-145 | chr14-73691559-DEL-166 |
| chr14-73701843-DEL-58 | chr14-73701844-DEL-58 |
| chr14-73711459-DEL-105 | chr14-73711460-DEL-105 |
| chr14-73721725-DEL-866 | chr14-73721733-DEL-853 |
| chr14-73765364-DEL-508 | chr14-73765376-DEL-497 |
| chr14-73766428-DEL-120 | chr14-73766429-DEL-120 |
| chr14-73767737-DEL-64 | chr14-73767738-DEL-64 |
| chr14-73782615-DEL-81 | chr14-73782615-DEL-81 |
| chr14-73798419-DEL-170 | chr14-73798420-DEL-170 |
| chr14-73827894-DEL-57 | chr14-73827901-DEL-56 |
| chr14-73877775-DEL-196 | chr14-73877776-DEL-210 |
| chr14-73885457-DEL-1929 | chr14-73885457-DEL-1929 |
| chr14-73889501-DEL-66 | chr14-73889514-DEL-54 |
| chr14-73896938-DEL-199 | chr14-73896942-DEL-196 |
| chr14-73911210-DEL-6218 | chr14-73911211-DEL-6218 |
| chr14-73916534-DEL-412 | chr14-73916534-DEL-412 |
| chr14-73923048-DEL-196 | chr14-73923049-DEL-196 |
| chr14-73928755-DEL-708 | chr14-73928756-DEL-708 |
| chr14-73954179-DEL-1847 | chr14-73954180-DEL-1847 |
| chr14-74033921-DEL-851 | chr14-74033922-DEL-851 |
| chr14-74121587-DEL-53 | chr14-74121588-DEL-53 |
| chr14-74143229-DEL-1015 | chr14-74143232-DEL-1013 |
| chr14-74146008-DEL-62 | chr14-74145964-DEL-64 |
| chr14-74178570-DEL-56 | chr14-74178564-DEL-50 |
| chr14-74181130-DEL-74 | chr14-74181131-DEL-74 |
| chr14-74191322-DEL-1743 | chr14-74191323-DEL-1743 |
| chr14-74194649-DEL-3032 | chr14-74194650-DEL-3032 |
| chr14-74202691-DEL-1044 | chr14-74202692-DEL-1044 |
| chr14-74211912-DEL-54 | chr14-74211910-DEL-55 |
| chr14-74226885-DEL-72 | chr14-74226890-DEL-60 |
| chr14-74238578-DEL-110 | chr14-74238571-DEL-120 |
| chr14-74240234-DEL-196 | chr14-74240235-DEL-196 |
| chr14-74261011-DEL-890 | chr14-74261012-DEL-890 |
| chr14-74280325-DEL-146 | chr14-74280326-DEL-140 |
| chr14-74292527-DEL-6776 | chr14-74292528-DEL-6777 |
| chr14-74303088-DEL-305 | chr14-74303089-DEL-287 |
| chr14-74360075-DEL-153 | chr14-74360084-DEL-130 |
| chr14-74361146-DEL-484 | chr14-74361147-DEL-484 |
| chr14-74361777-DEL-6565 | chr14-74361778-DEL-6565 |
| chr14-74369138-DEL-6868 | chr14-74369139-DEL-6868 |
| chr14-74375271-DEL-425 | chr14-74375224-DEL-426 |

| | |
|---|---|
| chr14-74499231-DEL-1009 | chr14-74499236-DEL-1005 |
| chr14-74507400-DEL-5565 | chr14-74507401-DEL-5565 |
| chr14-74530179-DEL-97 | chr14-74530170-DEL-105 |
| chr14-74535765-DEL-53 | chr14-74535766-DEL-53 |
| chr14-74545533-DEL-60 | chr14-74545534-DEL-60 |
| chr14-74603096-DEL-110 | chr14-74603097-DEL-108 |
| chr14-74642595-DEL-203 | chr14-74642596-DEL-203 |
| chr14-74647916-DEL-94 | chr14-74647909-DEL-98 |
| chr14-74654135-DEL-62 | chr14-74654136-DEL-62 |
| chr14-74671440-DEL-208 | chr14-74671441-DEL-208 |
| chr14-74671842-DEL-14606 | chr14-74671842-DEL-14607 |
| chr14-74678429-DEL-6809 | chr14-74678429-DEL-6809 |
| chr14-74678493-DEL-208 | chr14-74678493-DEL-208 |
| chr14-74722967-DEL-52 | chr14-74722968-DEL-52 |
| chr14-74761488-DEL-589 | chr14-74761489-DEL-575 |
| chr14-74770056-DEL-325 | chr14-74770057-DEL-325 |
| chr14-74784369-DEL-116 | chr14-74784395-DEL-100 |
| chr14-74808016-DEL-326 | chr14-74808017-DEL-326 |
| chr14-74808621-DEL-79 | chr14-74808622-DEL-79 |
| chr14-74814374-DEL-655 | chr14-74814375-DEL-655 |
| chr14-74922046-DEL-157 | chr14-74922047-DEL-157 |
| chr14-74927775-DEL-77 | chr14-74927776-DEL-77 |
| chr14-74949745-DEL-57 | chr14-74949746-DEL-57 |
| chr14-74952630-DEL-827 | chr14-74952631-DEL-827 |
| chr14-74986175-DEL-56 | chr14-74986176-DEL-54 |
| chr14-74988744-DEL-55 | chr14-74988745-DEL-55 |
| chr14-75003968-DEL-60 | chr14-75004007-DEL-62 |
| chr14-75014438-DEL-6285 | chr14-75014439-DEL-6285 |
| chr14-75027033-DEL-227 | chr14-75027034-DEL-227 |
| chr14-75033059-DEL-132 | chr14-75033060-DEL-132 |
| chr14-75045768-DEL-89 | chr14-75045769-DEL-89 |
| chr14-75046677-DEL-81 | chr14-75046674-DEL-83 |
| chr14-75088776-DEL-54 | chr14-75088784-DEL-55 |
| chr14-75095843-DEL-58 | chr14-75095844-DEL-58 |
| chr14-75097975-DEL-308 | chr14-75097976-DEL-308 |
| chr14-75105695-DEL-57 | chr14-75105696-DEL-53 |
| chr14-75130847-DEL-120 | chr14-75130833-DEL-100 |
| chr14-75145605-DEL-205 | chr14-75145606-DEL-205 |
| chr14-75168439-DEL-137 | chr14-75168433-DEL-115 |
| chr14-75177312-DEL-115 | chr14-75177313-DEL-115 |
| chr14-75198379-DEL-157 | chr14-75198378-DEL-153 |
| chr14-75249885-DEL-1941 | chr14-75249886-DEL-1941 |
| chr14-75257499-DEL-352 | chr14-75257500-DEL-352 |
| chr14-75275394-DEL-85 | chr14-75275395-DEL-85 |
| chr14-75284280-DEL-82 | chr14-75284281-DEL-82 |
| chr14-75302118-DEL-72 | chr14-75302119-DEL-72 |
| chr14-75304467-DEL-405 | chr14-75304468-DEL-405 |
| chr14-75331688-DEL-456 | chr14-75331689-DEL-456 |
| chr14-75335366-DEL-52 | chr14-75335351-DEL-54 |
| chr14-75339671-DEL-60 | chr14-75339672-DEL-60 |
| chr14-75342638-DEL-82 | chr14-75342639-DEL-82 |
| chr14-75381642-DEL-217 | chr14-75381643-DEL-217 |
| chr14-75396351-DEL-102 | chr14-75396352-DEL-102 |
| chr14-75404660-DEL-82 | chr14-75404659-DEL-76 |
| chr14-75444620-DEL-62 | chr14-75444611-DEL-73 |
| chr14-75454739-DEL-125 | chr14-75454740-DEL-125 |
| chr14-75457582-DEL-169 | chr14-75457583-DEL-169 |
| chr14-75478720-DEL-689 | chr14-75478721-DEL-689 |
| chr14-75495820-DEL-678 | chr14-75495821-DEL-678 |
| chr14-75514790-DEL-105 | chr14-75514791-DEL-105 |
| chr14-75519843-DEL-93 | chr14-75519829-DEL-108 |
| chr14-75580829-DEL-131 | chr14-75580830-DEL-131 |
| chr14-75591786-DEL-3720 | chr14-75591787-DEL-3720 |
| chr14-75596410-DEL-132 | chr14-75596411-DEL-132 |
| chr14-75596933-DEL-88 | chr14-75596925-DEL-87 |
| chr14-75601407-DEL-53 | chr14-75601408-DEL-53 |
| chr14-75627363-DEL-122 | chr14-75627364-DEL-122 |
| chr14-75639820-DEL-60 | chr14-75639825-DEL-56 |
| chr14-75648684-DEL-79 | chr14-75648705-DEL-85 |
| chr14-75651793-DEL-727 | chr14-75651790-DEL-728 |
| chr14-75655120-DEL-86 | chr14-75655141-DEL-65 |
| chr14-75662756-DEL-5977 | chr14-75662757-DEL-5977 |
| chr14-75678598-DEL-204 | chr14-75678599-DEL-204 |
| chr14-75715427-DEL-348 | chr14-75715428-DEL-348 |
| chr14-75746910-DEL-486 | chr14-75746911-DEL-486 |
| chr14-75793724-DEL-549 | chr14-75793725-DEL-549 |
| chr14-75794755-DEL-128 | chr14-75794756-DEL-128 |
| chr14-75824338-DEL-60 | chr14-75824338-DEL-60 |
| chr14-75836896-DEL-55 | chr14-75836899-DEL-90 |
| chr14-75838421-DEL-84 | chr14-75838422-DEL-84 |

| | |
|---|---|
| chr14-75868496-DEL-319 | chr14-75868497-DEL-319 |
| chr14-75883877-DEL-79 | chr14-75883878-DEL-79 |
| chr14-75906852-DEL-56 | chr14-75906853-DEL-56 |
| chr14-75955245-DEL-70 | chr14-75955246-DEL-70 |
| chr14-75974987-DEL-70 | chr14-75974970-DEL-96 |
| chr14-76003193-DEL-699 | chr14-76003194-DEL-699 |
| chr14-76004185-DEL-92 | chr14-76004182-DEL-100 |
| chr14-76013814-DEL-62 | chr14-76013850-DEL-63 |
| chr14-76022205-DEL-232 | chr14-76022206-DEL-232 |
| chr14-76035888-DEL-1864 | chr14-76035889-DEL-1864 |
| chr14-76038560-DEL-192 | chr14-76038561-DEL-201 |
| chr14-76060522-DEL-56 | chr14-76060523-DEL-56 |
| chr14-76077897-DEL-1440 | chr14-76077898-DEL-1440 |
| chr14-76115532-DEL-190 | chr14-76115533-DEL-190 |
| chr14-76117384-DEL-220 | chr14-76117384-DEL-220 |
| chr14-76121016-DEL-54 | chr14-76121017-DEL-54 |
| chr14-76122999-DEL-179 | chr14-76123000-DEL-179 |
| chr14-76124264-DEL-331 | chr14-76124304-DEL-317 |
| chr14-76146988-DEL-60 | chr14-76146989-DEL-60 |
| chr14-76167159-DEL-229 | chr14-76167160-DEL-229 |
| chr14-76188599-DEL-391 | chr14-76188600-DEL-391 |
| chr14-76193661-DEL-349 | chr14-76193660-DEL-351 |
| chr14-76194676-DEL-161 | chr14-76194677-DEL-161 |
| chr14-76209127-DEL-357 | chr14-76209128-DEL-357 |
| chr14-76257040-DEL-51 | chr14-76257041-DEL-51 |
| chr14-76375673-DEL-198 | chr14-76375674-DEL-198 |
| chr14-76398019-DEL-170 | chr14-76398020-DEL-170 |
| chr14-76421562-DEL-190 | chr14-76421563-DEL-190 |
| chr14-76438002-DEL-409 | chr14-76438003-DEL-409 |
| chr14-76448537-DEL-62 | chr14-76448538-DEL-62 |
| chr14-76449577-DEL-494 | chr14-76449578-DEL-494 |
| chr14-76461352-DEL-121 | chr14-76461353-DEL-121 |
| chr14-76461957-DEL-59 | chr14-76461958-DEL-59 |
| chr14-76466901-DEL-4308 | chr14-76466902-DEL-4308 |
| chr14-76474039-DEL-972 | chr14-76474040-DEL-972 |
| chr14-76477015-DEL-75 | chr14-76477015-DEL-75 |
| chr14-76515034-DEL-52 | chr14-76515034-DEL-52 |
| chr14-76525327-DEL-1230 | chr14-76525328-DEL-1230 |
| chr14-76525586-DEL-132 | chr14-76525587-DEL-131 |
| chr14-76529631-DEL-2385 | chr14-76529632-DEL-2385 |
| chr14-76542604-DEL-378 | chr14-76542605-DEL-378 |
| chr14-76558079-DEL-83 | chr14-76558080-DEL-97 |
| chr14-76568593-DEL-57 | chr14-76568589-DEL-62 |
| chr14-76575130-DEL-51 | chr14-76575151-DEL-53 |
| chr14-76586552-DEL-944 | chr14-76586553-DEL-944 |
| chr14-76588465-DEL-226 | chr14-76588465-DEL-230 |
| chr14-76591515-DEL-614 | chr14-76591516-DEL-614 |
| chr14-76594155-DEL-564 | chr14-76594156-DEL-564 |
| chr14-76643747-DEL-71 | chr14-76643748-DEL-71 |
| chr14-76715892-DEL-108 | chr14-76715893-DEL-108 |
| chr14-76721706-DEL-330 | chr14-76721711-DEL-326 |
| chr14-76729077-DEL-55 | chr14-76729074-DEL-58 |
| chr14-76736331-DEL-187 | chr14-76736327-DEL-188 |
| chr14-76757664-DEL-128 | chr14-76757669-DEL-124 |
| chr14-76760091-DEL-400 | chr14-76760092-DEL-400 |
| chr14-76762572-DEL-88 | chr14-76762573-DEL-88 |
| chr14-76836797-DEL-123 | chr14-76836802-DEL-119 |
| chr14-76844475-DEL-154 | chr14-76844476-DEL-154 |
| chr14-76857020-DEL-177 | chr14-76857021-DEL-177 |
| chr14-76861279-DEL-414 | chr14-76861280-DEL-414 |
| chr14-76881840-DEL-121 | chr14-76881841-DEL-121 |
| chr14-76896298-DEL-170 | chr14-76896299-DEL-170 |
| chr14-76905292-DEL-1115 | chr14-76905291-DEL-1115 |
| chr14-77031178-DEL-279 | chr14-77031179-DEL-279 |
| chr14-77040646-DEL-90 | chr14-77040669-DEL-59 |
| chr14-77043713-DEL-3109 | chr14-77043714-DEL-3109 |
| chr14-77047188-DEL-186 | chr14-77047189-DEL-186 |
| chr14-77060827-DEL-459 | chr14-77060828-DEL-459 |
| chr14-77072113-DEL-119 | chr14-77072114-DEL-119 |
| chr14-77079317-DEL-376 | chr14-77079318-DEL-376 |
| chr14-77157349-DEL-1001 | chr14-77157350-DEL-1001 |
| chr14-77194369-DEL-55 | chr14-77194369-DEL-55 |
| chr14-77202895-DEL-1133 | chr14-77202896-DEL-1133 |
| chr14-77277626-DEL-60 | chr14-77277633-DEL-54 |
| chr14-77292586-DEL-148 | chr14-77292610-DEL-139 |
| chr14-77302236-DEL-184 | chr14-77302237-DEL-184 |
| chr14-77308079-DEL-7006 | chr14-77308080-DEL-7006 |
| chr14-77364082-DEL-208 | chr14-77364083-DEL-208 |
| chr14-77449136-DEL-281 | chr14-77449137-DEL-281 |
| chr14-77471411-DEL-3183 | chr14-77471412-DEL-3183 |

| | |
|---|---|
| chr14-77489226-DEL-545 | chr14-77489238-DEL-525 |
| chr14-77506967-DEL-101 | chr14-77506968-DEL-101 |
| chr14-77536784-DEL-6465 | chr14-77536810-DEL-6440 |
| chr14-77546265-DEL-75 | chr14-77546284-DEL-71 |
| chr14-77552088-DEL-67 | chr14-77552089-DEL-76 |
| chr14-77604898-DEL-69 | chr14-77604899-DEL-69 |
| chr14-77649010-DEL-395 | chr14-77649011-DEL-395 |
| chr14-77683049-DEL-95 | chr14-77683050-DEL-95 |
| chr14-77725874-DEL-211 | chr14-77725875-DEL-211 |
| chr14-77740191-DEL-179 | chr14-77740192-DEL-179 |
| chr14-77749691-DEL-551 | chr14-77749692-DEL-551 |
| chr14-77777140-DEL-1355 | chr14-77777133-DEL-1356 |
| chr14-77786134-DEL-201 | chr14-77786135-DEL-201 |
| chr14-77790378-DEL-3531 | chr14-77790379-DEL-3531 |
| chr14-77802744-DEL-621 | chr14-77802745-DEL-621 |
| chr14-77810965-DEL-190 | chr14-77810966-DEL-190 |
| chr14-77834031-DEL-894 | chr14-77834032-DEL-894 |
| chr14-77887484-DEL-125 | chr14-77887482-DEL-123 |
| chr14-77919687-DEL-478 | chr14-77919688-DEL-478 |
| chr14-77945693-DEL-52 | chr14-77945694-DEL-52 |
| chr14-77985177-DEL-432 | chr14-77985178-DEL-432 |
| chr14-78018134-DEL-437 | chr14-78018135-DEL-437 |
| chr14-78034028-DEL-271 | chr14-78034029-DEL-271 |
| chr14-78041551-DEL-98 | chr14-78041552-DEL-98 |
| chr14-78073277-DEL-394 | chr14-78073279-DEL-392 |
| chr14-78100204-DEL-494 | chr14-78100204-DEL-494 |
| chr14-78109470-DEL-188 | chr14-78109471-DEL-188 |
| chr14-78115282-DEL-178 | chr14-78115283-DEL-178 |
| chr14-78150210-DEL-281 | chr14-78150211-DEL-281 |
| chr14-78154619-DEL-81 | chr14-78154620-DEL-81 |
| chr14-78167518-DEL-188 | chr14-78167507-DEL-189 |
| chr14-78182780-DEL-53 | chr14-78182780-DEL-53 |
| chr14-78189959-DEL-84 | chr14-78189960-DEL-123 |
| chr14-78203975-DEL-85 | chr14-78203980-DEL-81 |
| chr14-78209372-DEL-60 | chr14-78209401-DEL-60 |
| chr14-78212552-DEL-81 | chr14-78212553-DEL-81 |
| chr14-78227307-DEL-1168 | chr14-78227308-DEL-1168 |
| chr14-78246379-DEL-58 | chr14-78246380-DEL-58 |
| chr14-78249033-DEL-62 | chr14-78249034-DEL-62 |
| chr14-78251751-DEL-120 | chr14-78251752-DEL-120 |
| chr14-78260039-DEL-134 | chr14-78260032-DEL-136 |
| chr14-78327857-DEL-111 | chr14-78327858-DEL-111 |
| chr14-78328075-DEL-971 | chr14-78328076-DEL-971 |
| chr14-78368081-DEL-69 | chr14-78368078-DEL-76 |
| chr14-78370045-DEL-58 | chr14-78370046-DEL-58 |
| chr14-78399520-DEL-6207 | chr14-78399521-DEL-6207 |
| chr14-78437265-DEL-384 | chr14-78437266-DEL-384 |
| chr14-78439660-DEL-87 | chr14-78439673-DEL-75 |
| chr14-78445701-DEL-6498 | chr14-78445694-DEL-6492 |
| chr14-78468755-DEL-197 | chr14-78468756-DEL-197 |
| chr14-78532397-DEL-81 | chr14-78532394-DEL-82 |
| chr14-78546523-DEL-54 | chr14-78546505-DEL-85 |
| chr14-78572191-DEL-5362 | chr14-78572192-DEL-5362 |
| chr14-78589902-DEL-104 | chr14-78589903-DEL-104 |
| chr14-78638416-DEL-70 | chr14-78638417-DEL-70 |
| chr14-78649999-DEL-54 | chr14-78650000-DEL-50 |
| chr14-78659951-DEL-69 | chr14-78659955-DEL-68 |
| chr14-78660999-DEL-829 | chr14-78661000-DEL-829 |
| chr14-78666571-DEL-739 | chr14-78666572-DEL-739 |
| chr14-78699036-DEL-52 | chr14-78699025-DEL-53 |
| chr14-78700371-DEL-73 | chr14-78700369-DEL-108 |
| chr14-78709284-DEL-110 | chr14-78709285-DEL-110 |
| chr14-78710284-DEL-123 | chr14-78710276-DEL-138 |
| chr14-78718442-DEL-122 | chr14-78718443-DEL-122 |
| chr14-78722919-DEL-1282 | chr14-78722920-DEL-1282 |
| chr14-78788943-DEL-68 | chr14-78788946-DEL-66 |
| chr14-78792515-DEL-379 | chr14-78792516-DEL-379 |
| chr14-78808141-DEL-68 | chr14-78808140-DEL-64 |
| chr14-78831518-DEL-240 | chr14-78831519-DEL-240 |
| chr14-78835401-DEL-143 | chr14-78835402-DEL-143 |
| chr14-78851416-DEL-56 | chr14-78851417-DEL-56 |
| chr14-78871499-DEL-117 | chr14-78871500-DEL-117 |
| chr14-78882012-DEL-2370 | chr14-78882013-DEL-2370 |
| chr14-78901396-DEL-2741 | chr14-78901397-DEL-2741 |
| chr14-78904689-DEL-377 | chr14-78904690-DEL-377 |
| chr14-78914767-DEL-144 | chr14-78914763-DEL-143 |
| chr14-78943457-DEL-63 | chr14-78943458-DEL-63 |
| chr14-78953322-DEL-114 | chr14-78953295-DEL-114 |
| chr14-78956642-DEL-123 | chr14-78956643-DEL-123 |
| chr14-78969918-DEL-57 | chr14-78969919-DEL-57 |

| | |
|---|---|
| chr14-78995402-DEL-80 | chr14-78995403-DEL-100 |
| chr14-78999008-DEL-194 | chr14-78999044-DEL-158 |
| chr14-79005637-DEL-224 | chr14-79005638-DEL-224 |
| chr14-79022072-DEL-5240 | chr14-79022073-DEL-5240 |
| chr14-79030775-DEL-57 | chr14-79030776-DEL-57 |
| chr14-79067864-DEL-207 | chr14-79067865-DEL-207 |
| chr14-79074518-DEL-521 | chr14-79074519-DEL-521 |
| chr14-79090332-DEL-73 | chr14-79090333-DEL-73 |
| chr14-79112036-DEL-253 | chr14-79112037-DEL-249 |
| chr14-79125049-DEL-5922 | chr14-79125050-DEL-5922 |
| chr14-79132562-DEL-175 | chr14-79132535-DEL-175 |
| chr14-79135409-DEL-69 | chr14-79135410-DEL-69 |
| chr14-79164227-DEL-3513 | chr14-79164212-DEL-3514 |
| chr14-79183168-DEL-86 | chr14-79183169-DEL-86 |
| chr14-79232802-DEL-341 | chr14-79232803-DEL-341 |
| chr14-79234409-DEL-6368 | chr14-79234409-DEL-6368 |
| chr14-79242271-DEL-865 | chr14-79242272-DEL-865 |
| chr14-79244249-DEL-211 | chr14-79244250-DEL-211 |
| chr14-79304401-DEL-99 | chr14-79304402-DEL-99 |
| chr14-79329337-DEL-71 | chr14-79329338-DEL-71 |
| chr14-79330257-DEL-199 | chr14-79330258-DEL-199 |
| chr14-79343148-DEL-692 | chr14-79343151-DEL-693 |
| chr14-79350024-DEL-364 | chr14-79350025-DEL-364 |
| chr14-79370663-DEL-62 | chr14-79370664-DEL-62 |
| chr14-79380149-DEL-434 | chr14-79380150-DEL-434 |
| chr14-79382682-DEL-6454 | chr14-79382683-DEL-6454 |
| chr14-79427388-DEL-6483 | chr14-79427389-DEL-6483 |
| chr14-79427478-DEL-204 | chr14-79427479-DEL-204 |
| chr14-79457742-DEL-1115 | chr14-79457743-DEL-1115 |
| chr14-79493565-DEL-144 | chr14-79493566-DEL-144 |
| chr14-79508188-DEL-145 | chr14-79508189-DEL-145 |
| chr14-79547427-DEL-432 | chr14-79547428-DEL-432 |
| chr14-79554543-DEL-197 | chr14-79554544-DEL-197 |
| chr14-79556199-DEL-470 | chr14-79556200-DEL-470 |
| chr14-79563600-DEL-75 | chr14-79563601-DEL-80 |
| chr14-79575936-DEL-227 | chr14-79575937-DEL-226 |
| chr14-79597717-DEL-103 | chr14-79597718-DEL-103 |
| chr14-79602404-DEL-198 | chr14-79602405-DEL-199 |
| chr14-79613171-DEL-197 | chr14-79613172-DEL-197 |
| chr14-79628525-DEL-162 | chr14-79628526-DEL-162 |
| chr14-79628822-DEL-400 | chr14-79628824-DEL-399 |
| chr14-79630484-DEL-171 | chr14-79630485-DEL-171 |
| chr14-79673919-DEL-636 | chr14-79673912-DEL-643 |
| chr14-79675252-DEL-54 | chr14-79675253-DEL-54 |
| chr14-79714599-DEL-202 | chr14-79714600-DEL-202 |
| chr14-79734598-DEL-90 | chr14-79734592-DEL-122 |
| chr14-79766955-DEL-635 | chr14-79766956-DEL-635 |
| chr14-79776881-DEL-176 | chr14-79776882-DEL-176 |
| chr14-79777653-DEL-401 | chr14-79777654-DEL-401 |
| chr14-79799588-DEL-187 | chr14-79799589-DEL-187 |
| chr14-79808640-DEL-54 | chr14-79808641-DEL-54 |
| chr14-79811105-DEL-96 | chr14-79811106-DEL-96 |
| chr14-79820052-DEL-52 | chr14-79820054-DEL-52 |
| chr14-79840361-DEL-148 | chr14-79840362-DEL-148 |
| chr14-79843699-DEL-222 | chr14-79843702-DEL-226 |
| chr14-79847497-DEL-70 | chr14-79847498-DEL-70 |
| chr14-79853011-DEL-215 | chr14-79853012-DEL-215 |
| chr14-79859992-DEL-564 | chr14-79859993-DEL-564 |
| chr14-79861897-DEL-200 | chr14-79861898-DEL-200 |
| chr14-79868337-DEL-75 | chr14-79868338-DEL-76 |
| chr14-79895124-DEL-60 | chr14-79895125-DEL-60 |
| chr14-79926209-DEL-367 | chr14-79926210-DEL-367 |
| chr14-79948256-DEL-132 | chr14-79948257-DEL-130 |
| chr14-80032922-DEL-4356 | chr14-80032923-DEL-4356 |
| chr14-80052899-DEL-1421 | chr14-80052900-DEL-1421 |
| chr14-80066211-DEL-7538 | chr14-80066212-DEL-7538 |
| chr14-80072287-DEL-636 | chr14-80072286-DEL-636 |
| chr14-80097901-DEL-2656 | chr14-80097902-DEL-2656 |
| chr14-80114962-DEL-55 | chr14-80114928-DEL-54 |
| chr14-80115348-DEL-280 | chr14-80115349-DEL-279 |
| chr14-80153168-DEL-130 | chr14-80153132-DEL-128 |
| chr14-80155875-DEL-53 | chr14-80155876-DEL-53 |
| chr14-80156894-DEL-6377 | chr14-80156895-DEL-6377 |
| chr14-80173762-DEL-87 | chr14-80173742-DEL-107 |
| chr14-80197872-DEL-79 | chr14-80197899-DEL-79 |
| chr14-80202520-DEL-753 | chr14-80202521-DEL-753 |
| chr14-80206279-DEL-1067 | chr14-80206280-DEL-1066 |
| chr14-80217536-DEL-499 | chr14-80217537-DEL-499 |
| chr14-80237125-DEL-181 | chr14-80237126-DEL-178 |
| chr14-80239128-DEL-80 | chr14-80239129-DEL-80 |

| | |
|---|---|
| chr14-80254380-DEL-56 | chr14-80254376-DEL-55 |
| chr14-80275543-DEL-177 | chr14-80275545-DEL-178 |
| chr14-80294718-DEL-358 | chr14-80294719-DEL-358 |
| chr14-80321976-DEL-505 | chr14-80321977-DEL-505 |
| chr14-80326563-DEL-7607 | chr14-80326564-DEL-7607 |
| chr14-80348077-DEL-607 | chr14-80348078-DEL-607 |
| chr14-80385147-DEL-3154 | chr14-80385148-DEL-3154 |
| chr14-80416473-DEL-6510 | chr14-80416473-DEL-6510 |
| chr14-80423020-DEL-1457 | chr14-80423020-DEL-1457 |
| chr14-80471651-DEL-725 | chr14-80471654-DEL-723 |
| chr14-80481156-DEL-74 | chr14-80481157-DEL-74 |
| chr14-80499249-DEL-60 | chr14-80499250-DEL-54 |
| chr14-80512312-DEL-1227 | chr14-80512313-DEL-1227 |
| chr14-80586502-DEL-7069 | chr14-80586493-DEL-7070 |
| chr14-80659166-DEL-199 | chr14-80659167-DEL-199 |
| chr14-80661382-DEL-199 | chr14-80661383-DEL-199 |
| chr14-80673697-DEL-3001 | chr14-80673698-DEL-3001 |
| chr14-80686226-DEL-1096 | chr14-80686227-DEL-1096 |
| chr14-80719788-DEL-55 | chr14-80719789-DEL-55 |
| chr14-80731686-DEL-444 | chr14-80731687-DEL-444 |
| chr14-80804403-DEL-6403 | chr14-80804404-DEL-6403 |
| chr14-80810546-DEL-64 | chr14-80810547-DEL-64 |
| chr14-80819139-DEL-2802 | chr14-80819140-DEL-2802 |
| chr14-80847649-DEL-2785 | chr14-80847650-DEL-2785 |
| chr14-80857267-DEL-1968 | chr14-80857268-DEL-1966 |
| chr14-80860033-DEL-5143 | chr14-80860034-DEL-5143 |
| chr14-80941208-DEL-867 | chr14-80941209-DEL-867 |
| chr14-80953998-DEL-160 | chr14-80953999-DEL-160 |
| chr14-80973682-DEL-5086 | chr14-80973683-DEL-5086 |
| chr14-80985124-DEL-131 | chr14-80985125-DEL-131 |
| chr14-81014078-DEL-64 | chr14-81014042-DEL-54 |
| chr14-81041849-DEL-71 | chr14-81041848-DEL-76 |
| chr14-81043868-DEL-3381 | chr14-81043869-DEL-3381 |
| chr14-81126692-DEL-1722 | chr14-81126694-DEL-1720 |
| chr14-81159643-DEL-482 | chr14-81159644-DEL-482 |
| chr14-81166590-DEL-3781 | chr14-81166591-DEL-3781 |
| chr14-81192767-DEL-2763 | chr14-81192767-DEL-2763 |
| chr14-81276862-DEL-6816 | chr14-81276863-DEL-6816 |
| chr14-81283175-DEL-402 | chr14-81283176-DEL-402 |
| chr14-81323517-DEL-9334 | chr14-81323518-DEL-9334 |
| chr14-81351143-DEL-50 | chr14-81351103-DEL-65 |
| chr14-81356246-DEL-872 | chr14-81356247-DEL-872 |
| chr14-81366526-DEL-423 | chr14-81366527-DEL-423 |
| chr14-81388810-DEL-3954 | chr14-81388811-DEL-3954 |
| chr14-81406095-DEL-54 | chr14-81406096-DEL-54 |
| chr14-81408781-DEL-1070 | chr14-81408773-DEL-1070 |
| chr14-81412083-DEL-695 | chr14-81412078-DEL-700 |
| chr14-81413228-DEL-633 | chr14-81413228-DEL-633 |
| chr14-81425035-DEL-93 | chr14-81425036-DEL-93 |
| chr14-81462656-DEL-1909 | chr14-81462657-DEL-1909 |
| chr14-81480056-DEL-1599 | chr14-81480057-DEL-1599 |
| chr14-81482416-DEL-939 | chr14-81482417-DEL-937 |
| chr14-81499505-DEL-65 | chr14-81499506-DEL-65 |
| chr14-81564092-DEL-54 | chr14-81564093-DEL-54 |
| chr14-81569156-DEL-5989 | chr14-81569157-DEL-5989 |
| chr14-81572182-DEL-1696 | chr14-81572183-DEL-1696 |
| chr14-81590647-DEL-96 | chr14-81590646-DEL-97 |
| chr14-81610346-DEL-170 | chr14-81610347-DEL-170 |
| chr14-81639370-DEL-72 | chr14-81639390-DEL-53 |
| chr14-81645077-DEL-195 | chr14-81645078-DEL-195 |
| chr14-81645604-DEL-182 | chr14-81645605-DEL-182 |
| chr14-81652965-DEL-1190 | chr14-81652979-DEL-1166 |
| chr14-81662933-DEL-55 | chr14-81662929-DEL-60 |
| chr14-81677447-DEL-64 | chr14-81677448-DEL-64 |
| chr14-81681465-DEL-74 | chr14-81681466-DEL-74 |
| chr14-81711400-DEL-201 | chr14-81711401-DEL-201 |
| chr14-81723935-DEL-60 | chr14-81723905-DEL-95 |
| chr14-81732744-DEL-6496 | chr14-81732745-DEL-6496 |
| chr14-81758393-DEL-207 | chr14-81758394-DEL-207 |
| chr14-81792135-DEL-6440 | chr14-81792125-DEL-6441 |
| chr14-81823674-DEL-52 | chr14-81823675-DEL-52 |
| chr14-81825149-DEL-64 | chr14-81825123-DEL-50 |
| chr14-81828551-DEL-1964 | chr14-81828552-DEL-1964 |
| chr14-81866624-DEL-2992 | chr14-81866624-DEL-2992 |
| chr14-81873764-DEL-1527 | chr14-81873765-DEL-1527 |
| chr14-81887745-DEL-993 | chr14-81887746-DEL-993 |
| chr14-81894016-DEL-210 | chr14-81894017-DEL-210 |
| chr14-81908880-DEL-54 | chr14-81908894-DEL-70 |
| chr14-81927677-DEL-143 | chr14-81927678-DEL-143 |
| chr14-81945904-DEL-54 | chr14-81945907-DEL-52 |

| | |
|---|---|
| chr14-81964379-DEL-116 | chr14-81964376-DEL-120 |
| chr14-81968848-DEL-140 | chr14-81968849-DEL-140 |
| chr14-81988308-DEL-148 | chr14-81988309-DEL-148 |
| chr14-81991520-DEL-215 | chr14-81991519-DEL-193 |
| chr14-82034832-DEL-206 | chr14-82034833-DEL-206 |
| chr14-82060633-DEL-161 | chr14-82060634-DEL-161 |
| chr14-82066222-DEL-199 | chr14-82066232-DEL-190 |
| chr14-82103246-DEL-475 | chr14-82103247-DEL-475 |
| chr14-82104994-DEL-100 | chr14-82105000-DEL-101 |
| chr14-82122087-DEL-84 | chr14-82122088-DEL-84 |
| chr14-82138251-DEL-5436 | chr14-82138233-DEL-5420 |
| chr14-82155775-DEL-1025 | chr14-82155776-DEL-1025 |
| chr14-82163018-DEL-90 | chr14-82163019-DEL-84 |
| chr14-82172337-DEL-2009 | chr14-82172338-DEL-2009 |
| chr14-82205292-DEL-883 | chr14-82205293-DEL-883 |
| chr14-82206934-DEL-128 | chr14-82206935-DEL-128 |
| chr14-82209951-DEL-536 | chr14-82209952-DEL-536 |
| chr14-82211444-DEL-567 | chr14-82211445-DEL-567 |
| chr14-82228662-DEL-63 | chr14-82228663-DEL-66 |
| chr14-82259800-DEL-132 | chr14-82259807-DEL-120 |
| chr14-82274804-DEL-6507 | chr14-82274796-DEL-6508 |
| chr14-82314982-DEL-7301 | chr14-82314983-DEL-7301 |
| chr14-82328163-DEL-74 | chr14-82328164-DEL-74 |
| chr14-82333660-DEL-64 | chr14-82333661-DEL-64 |
| chr14-82336225-DEL-5862 | chr14-82336226-DEL-5862 |
| chr14-82342170-DEL-60 | chr14-82342148-DEL-58 |
| chr14-82347195-DEL-54 | chr14-82347183-DEL-70 |
| chr14-82357700-DEL-493 | chr14-82357701-DEL-493 |
| chr14-82362310-DEL-122 | chr14-82362311-DEL-122 |
| chr14-82366241-DEL-1635 | chr14-82366242-DEL-1635 |
| chr14-82394835-DEL-67 | chr14-82394835-DEL-67 |
| chr14-82397264-DEL-73 | chr14-82397257-DEL-80 |
| chr14-82415640-DEL-3827 | chr14-82415641-DEL-3827 |
| chr14-82441894-DEL-1164 | chr14-82441895-DEL-1164 |
| chr14-82459015-DEL-1099 | chr14-82459016-DEL-1099 |
| chr14-82467486-DEL-1491 | chr14-82467487-DEL-1491 |
| chr14-82500561-DEL-6406 | chr14-82500562-DEL-6406 |
| chr14-82509008-DEL-56 | chr14-82509009-DEL-56 |
| chr14-82531831-DEL-52 | chr14-82531832-DEL-52 |
| chr14-82595844-DEL-7002 | chr14-82595845-DEL-7002 |
| chr14-82601973-DEL-416 | chr14-82601974-DEL-416 |
| chr14-82606383-DEL-1605 | chr14-82606383-DEL-1605 |
| chr14-82623280-DEL-103 | chr14-82623281-DEL-103 |
| chr14-82625617-DEL-2264 | chr14-82625618-DEL-2264 |
| chr14-82629376-DEL-62 | chr14-82629373-DEL-62 |
| chr14-82646676-DEL-6252 | chr14-82646677-DEL-6252 |
| chr14-82658687-DEL-90 | chr14-82658688-DEL-90 |
| chr14-82775577-DEL-322 | chr14-82775574-DEL-309 |
| chr14-82792690-DEL-1633 | chr14-82792690-DEL-1637 |
| chr14-82797427-DEL-312 | chr14-82797428-DEL-312 |
| chr14-82866590-DEL-276 | chr14-82866591-DEL-276 |
| chr14-82905406-DEL-52 | chr14-82905407-DEL-52 |
| chr14-82917952-DEL-545 | chr14-82917952-DEL-545 |
| chr14-82938301-DEL-342 | chr14-82938302-DEL-342 |
| chr14-82938306-DEL-2480 | chr14-82938306-DEL-2480 |
| chr14-82990724-DEL-332 | chr14-82990725-DEL-332 |
| chr14-83039145-DEL-314 | chr14-83039146-DEL-314 |
| chr14-83042362-DEL-60 | chr14-83042362-DEL-62 |
| chr14-83044123-DEL-51 | chr14-83044114-DEL-76 |
| chr14-83049301-DEL-198 | chr14-83049302-DEL-198 |
| chr14-83050110-DEL-72 | chr14-83050111-DEL-72 |
| chr14-83057596-DEL-54 | chr14-83057597-DEL-54 |
| chr14-83072170-DEL-288 | chr14-83072171-DEL-288 |
| chr14-83079515-DEL-105 | chr14-83079517-DEL-100 |
| chr14-83084296-DEL-6202 | chr14-83084297-DEL-6202 |
| chr14-83138221-DEL-2563 | chr14-83138222-DEL-2563 |
| chr14-83147516-DEL-5680 | chr14-83147517-DEL-5680 |
| chr14-83195117-DEL-6440 | chr14-83195118-DEL-6440 |
| chr14-83253732-DEL-78 | chr14-83253733-DEL-78 |
| chr14-83260895-DEL-134 | chr14-83260896-DEL-134 |
| chr14-83261320-DEL-5543 | chr14-83261321-DEL-5543 |
| chr14-83285467-DEL-930 | chr14-83285467-DEL-930 |
| chr14-83290602-DEL-68 | chr14-83290601-DEL-69 |
| chr14-83307834-DEL-220 | chr14-83307835-DEL-220 |
| chr14-83316604-DEL-123 | chr14-83316606-DEL-122 |
| chr14-83333697-DEL-163 | chr14-83333698-DEL-163 |
| chr14-83336859-DEL-119 | chr14-83336906-DEL-72 |
| chr14-83379166-DEL-84 | chr14-83379148-DEL-69 |
| chr14-83402531-DEL-1799 | chr14-83402533-DEL-1799 |
| chr14-83405214-DEL-880 | chr14-83405214-DEL-880 |

| | |
|---|---|
| chr14-83434901-DEL-6143 | chr14-83434902-DEL-6143 |
| chr14-83444288-DEL-492 | chr14-83444289-DEL-492 |
| chr14-83455610-DEL-205 | chr14-83455609-DEL-205 |
| chr14-83489521-DEL-56 | chr14-83489522-DEL-56 |
| chr14-83501528-DEL-919 | chr14-83501524-DEL-920 |
| chr14-83509948-DEL-166 | chr14-83509949-DEL-161 |
| chr14-83618973-DEL-207 | chr14-83618974-DEL-207 |
| chr14-83620045-DEL-61 | chr14-83620046-DEL-61 |
| chr14-83632286-DEL-326 | chr14-83632287-DEL-326 |
| chr14-83643587-DEL-501 | chr14-83643588-DEL-501 |
| chr14-83659166-DEL-547 | chr14-83659167-DEL-547 |
| chr14-83660502-DEL-2178 | chr14-83660503-DEL-2178 |
| chr14-83673869-DEL-63 | chr14-83673870-DEL-63 |
| chr14-83731925-DEL-84 | chr14-83731895-DEL-69 |
| chr14-83738892-DEL-52 | chr14-83738893-DEL-52 |
| chr14-83748316-DEL-54 | chr14-83748317-DEL-53 |
| chr14-83752588-DEL-203 | chr14-83752589-DEL-203 |
| chr14-83792198-DEL-2469 | chr14-83792199-DEL-2469 |
| chr14-83854253-DEL-513 | chr14-83854254-DEL-513 |
| chr14-83880097-DEL-1904 | chr14-83880098-DEL-1902 |
| chr14-83934346-DEL-4785 | chr14-83934347-DEL-4785 |
| chr14-83962885-DEL-53 | chr14-83962880-DEL-58 |
| chr14-83965391-DEL-6270 | chr14-83965392-DEL-6270 |
| chr14-83980892-DEL-77 | chr14-83980893-DEL-77 |
| chr14-83999156-DEL-1599 | chr14-83999157-DEL-1599 |
| chr14-84003084-DEL-75 | chr14-84003085-DEL-75 |
| chr14-84020778-DEL-176 | chr14-84020778-DEL-176 |
| chr14-84040192-DEL-57 | chr14-84040192-DEL-58 |
| chr14-84060839-DEL-194 | chr14-84060840-DEL-194 |
| chr14-84062026-DEL-927 | chr14-84062027-DEL-927 |
| chr14-84067429-DEL-66 | chr14-84067430-DEL-66 |
| chr14-84077745-DEL-5880 | chr14-84077746-DEL-5880 |
| chr14-84078408-DEL-2639 | chr14-84078411-DEL-2639 |
| chr14-84088626-DEL-114 | chr14-84088664-DEL-112 |
| chr14-84103504-DEL-7956 | chr14-84103505-DEL-7956 |
| chr14-84124507-DEL-62 | chr14-84124511-DEL-58 |
| chr14-84173381-DEL-147 | chr14-84173382-DEL-147 |
| chr14-84175990-DEL-96 | chr14-84175991-DEL-96 |
| chr14-84221577-DEL-185 | chr14-84221578-DEL-185 |
| chr14-84229374-DEL-196 | chr14-84229358-DEL-217 |
| chr14-84269178-DEL-60 | chr14-84269181-DEL-58 |
| chr14-84317655-DEL-74 | chr14-84317656-DEL-74 |
| chr14-84331156-DEL-58 | chr14-84331157-DEL-58 |
| chr14-84343388-DEL-582 | chr14-84343389-DEL-582 |
| chr14-84362388-DEL-298 | chr14-84362389-DEL-298 |
| chr14-84377588-DEL-71 | chr14-84377589-DEL-71 |
| chr14-84386179-DEL-787 | chr14-84386180-DEL-787 |
| chr14-84395594-DEL-2586 | chr14-84395595-DEL-2586 |
| chr14-84401887-DEL-1038 | chr14-84401891-DEL-1037 |
| chr14-84409080-DEL-110 | chr14-84409081-DEL-110 |
| chr14-84411671-DEL-80 | chr14-84411672-DEL-80 |
| chr14-84414787-DEL-52 | chr14-84414788-DEL-52 |
| chr14-84424141-DEL-4298 | chr14-84424142-DEL-4298 |
| chr14-84430984-DEL-206 | chr14-84430976-DEL-207 |
| chr14-84431859-DEL-100 | chr14-84431858-DEL-66 |
| chr14-84456853-DEL-95 | chr14-84456857-DEL-76 |
| chr14-84502234-DEL-53 | chr14-84502235-DEL-53 |
| chr14-84514430-DEL-90 | chr14-84514431-DEL-90 |
| chr14-84525856-DEL-57 | chr14-84525857-DEL-58 |
| chr14-84535552-DEL-53 | chr14-84535553-DEL-53 |
| chr14-84547461-DEL-6209 | chr14-84547453-DEL-6210 |
| chr14-84573802-DEL-1031 | chr14-84573802-DEL-1031 |
| chr14-84584736-DEL-59 | chr14-84584737-DEL-59 |
| chr14-84590191-DEL-413 | chr14-84590192-DEL-413 |
| chr14-84590797-DEL-2631 | chr14-84590798-DEL-2631 |
| chr14-84666030-DEL-6339 | chr14-84666031-DEL-6339 |
| chr14-84739522-DEL-54 | chr14-84739523-DEL-54 |
| chr14-84788392-DEL-189 | chr14-84788393-DEL-187 |
| chr14-84792648-DEL-959 | chr14-84792647-DEL-960 |
| chr14-84819123-DEL-204 | chr14-84819124-DEL-204 |
| chr14-84838848-DEL-6362 | chr14-84838849-DEL-6362 |
| chr14-84851869-DEL-238 | chr14-84851870-DEL-238 |
| chr14-84867018-DEL-458 | chr14-84867019-DEL-458 |
| chr14-84931965-DEL-1828 | chr14-84931966-DEL-1828 |
| chr14-84948442-DEL-223 | chr14-84948443-DEL-223 |
| chr14-84966795-DEL-5431 | chr14-84966796-DEL-5431 |
| chr14-84985483-DEL-954 | chr14-84985484-DEL-952 |
| chr14-84988151-DEL-369 | chr14-84988151-DEL-369 |
| chr14-84989449-DEL-697 | chr14-84989450-DEL-697 |
| chr14-84994559-DEL-175 | chr14-84994593-DEL-173 |

| | |
|---|---|
| chr14-84995494-DEL-63 | chr14-84995490-DEL-64 |
| chr14-85000111-DEL-1756 | chr14-85000112-DEL-1756 |
| chr14-85017447-DEL-809 | chr14-85017448-DEL-809 |
| chr14-85020265-DEL-112 | chr14-85020266-DEL-112 |
| chr14-85072510-DEL-4459 | chr14-85072511-DEL-4459 |
| chr14-85080272-DEL-138 | chr14-85080273-DEL-138 |
| chr14-85092582-DEL-7117 | chr14-85092583-DEL-7117 |
| chr14-85120957-DEL-2442 | chr14-85120958-DEL-2442 |
| chr14-85154164-DEL-1114 | chr14-85154165-DEL-1114 |
| chr14-85175145-DEL-6380 | chr14-85175146-DEL-6380 |
| chr14-85184995-DEL-128 | chr14-85184994-DEL-130 |
| chr14-85203004-DEL-64 | chr14-85203005-DEL-64 |
| chr14-85240029-DEL-2704 | chr14-85240030-DEL-2704 |
| chr14-85269220-DEL-52 | chr14-85269220-DEL-52 |
| chr14-85294999-DEL-101 | chr14-85295000-DEL-101 |
| chr14-85306377-DEL-1043 | chr14-85306378-DEL-1043 |
| chr14-85310684-DEL-200 | chr14-85310685-DEL-200 |
| chr14-85343011-DEL-2116 | chr14-85343012-DEL-2116 |
| chr14-85415156-DEL-4535 | chr14-85415157-DEL-4535 |
| chr14-85494322-DEL-97 | chr14-85494327-DEL-98 |
| chr14-85499547-DEL-6187 | chr14-85499548-DEL-6187 |
| chr14-85522044-DEL-53 | chr14-85522045-DEL-53 |
| chr14-85534190-DEL-248 | chr14-85534191-DEL-248 |
| chr14-85561303-DEL-1814 | chr14-85561304-DEL-1814 |
| chr14-85567334-DEL-69 | chr14-85567334-DEL-68 |
| chr14-85582756-DEL-2140 | chr14-85582754-DEL-2140 |
| chr14-85619677-DEL-122 | chr14-85619677-DEL-123 |
| chr14-85647933-DEL-191 | chr14-85647932-DEL-190 |
| chr14-85648115-DEL-6171 | chr14-85648116-DEL-6171 |
| chr14-85658295-DEL-857 | chr14-85658296-DEL-857 |
| chr14-85681126-DEL-52 | chr14-85681140-DEL-58 |
| chr14-85745258-DEL-54 | chr14-85745257-DEL-54 |
| chr14-85758607-DEL-4353 | chr14-85758608-DEL-4353 |
| chr14-85765453-DEL-7123 | chr14-85765454-DEL-7123 |
| chr14-85765588-DEL-66 | chr14-85765589-DEL-66 |
| chr14-85779657-DEL-1912 | chr14-85779658-DEL-1912 |
| chr14-85826298-DEL-177 | chr14-85826299-DEL-177 |
| chr14-85857864-DEL-62 | chr14-85857865-DEL-62 |
| chr14-85892216-DEL-72 | chr14-85892217-DEL-70 |
| chr14-85896894-DEL-2823 | chr14-85896895-DEL-2823 |
| chr14-85908980-DEL-65 | chr14-85908985-DEL-66 |
| chr14-85972348-DEL-603 | chr14-85972349-DEL-603 |
| chr14-85985860-DEL-74 | chr14-85985865-DEL-72 |
| chr14-85994043-DEL-59 | chr14-85994029-DEL-60 |
| chr14-85997147-DEL-71 | chr14-85997155-DEL-106 |
| chr14-86003080-DEL-797 | chr14-86003081-DEL-797 |
| chr14-86106830-DEL-498 | chr14-86106831-DEL-498 |
| chr14-86108636-DEL-64 | chr14-86108637-DEL-64 |
| chr14-86126529-DEL-7118 | chr14-86126530-DEL-7118 |
| chr14-86180893-DEL-56 | chr14-86180894-DEL-56 |
| chr14-86188945-DEL-278 | chr14-86188946-DEL-278 |
| chr14-86189845-DEL-62 | chr14-86189846-DEL-62 |
| chr14-86190718-DEL-6300 | chr14-86190718-DEL-6300 |
| chr14-86209882-DEL-90 | chr14-86209883-DEL-90 |
| chr14-86287131-DEL-61 | chr14-86287132-DEL-61 |
| chr14-86298324-DEL-60 | chr14-86298325-DEL-60 |
| chr14-86311244-DEL-2216 | chr14-86311245-DEL-2216 |
| chr14-86324341-DEL-65 | chr14-86324342-DEL-53 |
| chr14-86368971-DEL-4507 | chr14-86368972-DEL-4507 |
| chr14-86376781-DEL-172 | chr14-86376782-DEL-172 |
| chr14-86436436-DEL-147 | chr14-86436435-DEL-147 |
| chr14-86437136-DEL-60 | chr14-86437132-DEL-51 |
| chr14-86459120-DEL-108 | chr14-86459129-DEL-100 |
| chr14-86462077-DEL-121 | chr14-86462078-DEL-121 |
| chr14-86503561-DEL-109 | chr14-86503562-DEL-109 |
| chr14-86557598-DEL-685 | chr14-86557593-DEL-681 |
| chr14-86578962-DEL-54 | chr14-86578963-DEL-54 |
| chr14-86582005-DEL-1321 | chr14-86582006-DEL-1321 |
| chr14-86604185-DEL-54 | chr14-86604186-DEL-54 |
| chr14-86609673-DEL-102 | chr14-86609674-DEL-102 |
| chr14-86616695-DEL-199 | chr14-86616696-DEL-199 |
| chr14-86620676-DEL-820 | chr14-86620677-DEL-820 |
| chr14-86625987-DEL-55 | chr14-86625988-DEL-50 |
| chr14-86635545-DEL-484 | chr14-86635548-DEL-485 |
| chr14-86696359-DEL-2827 | chr14-86696360-DEL-2827 |
| chr14-86709722-DEL-70 | chr14-86709717-DEL-78 |
| chr14-86716487-DEL-58 | chr14-86716492-DEL-59 |
| chr14-86720486-DEL-58 | chr14-86720487-DEL-58 |
| chr14-86762215-DEL-82 | chr14-86762204-DEL-78 |
| chr14-86776130-DEL-60 | chr14-86776131-DEL-60 |

| | |
|---|---|
| chr14-86821182-DEL-261 | chr14-86821183-DEL-261 |
| chr14-86823665-DEL-90 | chr14-86823664-DEL-92 |
| chr14-86827950-DEL-7085 | chr14-86827951-DEL-7085 |
| chr14-86847208-DEL-1052 | chr14-86847209-DEL-1051 |
| chr14-86860710-DEL-298 | chr14-86860711-DEL-298 |
| chr14-86883441-DEL-71 | chr14-86883442-DEL-71 |
| chr14-86885753-DEL-109 | chr14-86885753-DEL-110 |
| chr14-86919094-DEL-137 | chr14-86919079-DEL-184 |
| chr14-86924247-DEL-322 | chr14-86924248-DEL-322 |
| chr14-87002411-DEL-53 | chr14-87002394-DEL-79 |
| chr14-87011797-DEL-227 | chr14-87011798-DEL-227 |
| chr14-87020436-DEL-1176 | chr14-87020433-DEL-1172 |
| chr14-87095322-DEL-1711 | chr14-87095323-DEL-1711 |
| chr14-87109661-DEL-4350 | chr14-87109662-DEL-4351 |
| chr14-87135569-DEL-200 | chr14-87135562-DEL-207 |
| chr14-87142089-DEL-95 | chr14-87142071-DEL-96 |
| chr14-87151300-DEL-3426 | chr14-87151301-DEL-3426 |
| chr14-87163517-DEL-2563 | chr14-87163514-DEL-2558 |
| chr14-87168769-DEL-52 | chr14-87168769-DEL-52 |
| chr14-87169942-DEL-738 | chr14-87169943-DEL-738 |
| chr14-87177864-DEL-6504 | chr14-87177849-DEL-6505 |
| chr14-87190625-DEL-73 | chr14-87190626-DEL-73 |
| chr14-87209400-DEL-78 | chr14-87209393-DEL-59 |
| chr14-87217087-DEL-149 | chr14-87217094-DEL-129 |
| chr14-87217835-DEL-379 | chr14-87217831-DEL-376 |
| chr14-87250589-DEL-62 | chr14-87250584-DEL-67 |
| chr14-87259683-DEL-797 | chr14-87259684-DEL-797 |
| chr14-87288530-DEL-1785 | chr14-87288531-DEL-1785 |
| chr14-87299696-DEL-131 | chr14-87299697-DEL-131 |
| chr14-87305790-DEL-325 | chr14-87305791-DEL-325 |
| chr14-87340085-DEL-74 | chr14-87340086-DEL-74 |
| chr14-87346932-DEL-65 | chr14-87346962-DEL-81 |
| chr14-87376249-DEL-156 | chr14-87376250-DEL-156 |
| chr14-87377084-DEL-120 | chr14-87377127-DEL-75 |
| chr14-87391032-DEL-52 | chr14-87391032-DEL-52 |
| chr14-87401824-DEL-69 | chr14-87401825-DEL-69 |
| chr14-87402918-DEL-670 | chr14-87402919-DEL-670 |
| chr14-87404480-DEL-70 | chr14-87404475-DEL-68 |
| chr14-87416278-DEL-104 | chr14-87416256-DEL-114 |
| chr14-87425538-DEL-355 | chr14-87425539-DEL-355 |
| chr14-87433676-DEL-1059 | chr14-87433677-DEL-1059 |
| chr14-87436784-DEL-56 | chr14-87436781-DEL-52 |
| chr14-87467733-DEL-1574 | chr14-87467734-DEL-1574 |
| chr14-87470291-DEL-58 | chr14-87470294-DEL-51 |
| chr14-87479723-DEL-2507 | chr14-87479724-DEL-2481 |
| chr14-87487137-DEL-582 | chr14-87487146-DEL-579 |
| chr14-87498552-DEL-86 | chr14-87498553-DEL-86 |
| chr14-87510602-DEL-90 | chr14-87510603-DEL-90 |
| chr14-87520488-DEL-60 | chr14-87520488-DEL-60 |
| chr14-87536660-DEL-60 | chr14-87536660-DEL-60 |
| chr14-87562795-DEL-75 | chr14-87562796-DEL-70 |
| chr14-87566402-DEL-1405 | chr14-87566403-DEL-1405 |
| chr14-87572337-DEL-105 | chr14-87572338-DEL-123 |
| chr14-87583506-DEL-56 | chr14-87583503-DEL-50 |
| chr14-87586620-DEL-231 | chr14-87586621-DEL-231 |
| chr14-87590491-DEL-962 | chr14-87590492-DEL-962 |
| chr14-87591957-DEL-6378 | chr14-87591958-DEL-6378 |
| chr14-87608845-DEL-1579 | chr14-87608846-DEL-1579 |
| chr14-87615063-DEL-95 | chr14-87615064-DEL-95 |
| chr14-87615300-DEL-840 | chr14-87615304-DEL-845 |
| chr14-87624063-DEL-5497 | chr14-87624054-DEL-5504 |
| chr14-87652421-DEL-197 | chr14-87652422-DEL-197 |
| chr14-87672830-DEL-278 | chr14-87672831-DEL-278 |
| chr14-87681563-DEL-241 | chr14-87681564-DEL-242 |
| chr14-87684937-DEL-200 | chr14-87684952-DEL-197 |
| chr14-87694039-DEL-60 | chr14-87694036-DEL-61 |
| chr14-87694681-DEL-123 | chr14-87694682-DEL-123 |
| chr14-87716705-DEL-333 | chr14-87716706-DEL-333 |
| chr14-87718812-DEL-65 | chr14-87718813-DEL-65 |
| chr14-87721604-DEL-55 | chr14-87721637-DEL-55 |
| chr14-87731048-DEL-58 | chr14-87731049-DEL-66 |
| chr14-87739803-DEL-112 | chr14-87739804-DEL-112 |
| chr14-87764071-DEL-286 | chr14-87764072-DEL-286 |
| chr14-87781081-DEL-880 | chr14-87781082-DEL-880 |
| chr14-87798010-DEL-1863 | chr14-87798011-DEL-1863 |
| chr14-87841372-DEL-876 | chr14-87841373-DEL-876 |
| chr14-87842850-DEL-3252 | chr14-87842851-DEL-3252 |
| chr14-87859877-DEL-264 | chr14-87859878-DEL-264 |
| chr14-87908748-DEL-6109 | chr14-87908749-DEL-6109 |
| chr14-87917016-DEL-671 | chr14-87917017-DEL-671 |

| | |
|---|---|
| chr14-87962162-DEL-360 | chr14-87962163-DEL-360 |
| chr14-88023208-DEL-786 | chr14-88023209-DEL-786 |
| chr14-88046790-DEL-2706 | chr14-88046791-DEL-2706 |
| chr14-88050997-DEL-5830 | chr14-88050998-DEL-5830 |
| chr14-88087014-DEL-125 | chr14-88087015-DEL-119 |
| chr14-88087960-DEL-6354 | chr14-88087961-DEL-6354 |
| chr14-88104605-DEL-675 | chr14-88104606-DEL-675 |
| chr14-88129982-DEL-246 | chr14-88130017-DEL-244 |
| chr14-88165528-DEL-160 | chr14-88165528-DEL-160 |
| chr14-88167107-DEL-910 | chr14-88167108-DEL-910 |
| chr14-88199526-DEL-75 | chr14-88199521-DEL-69 |
| chr14-88216435-DEL-285 | chr14-88216436-DEL-285 |
| chr14-88224919-DEL-162 | chr14-88224920-DEL-162 |
| chr14-88271164-DEL-7060 | chr14-88271165-DEL-7060 |
| chr14-88278786-DEL-550 | chr14-88278787-DEL-550 |
| chr14-88284887-DEL-66 | chr14-88284888-DEL-66 |
| chr14-88325787-DEL-144 | chr14-88325787-DEL-130 |
| chr14-88337766-DEL-1273 | chr14-88337767-DEL-1273 |
| chr14-88347463-DEL-64 | chr14-88347464-DEL-64 |
| chr14-88351812-DEL-923 | chr14-88351813-DEL-923 |
| chr14-88360379-DEL-890 | chr14-88360380-DEL-890 |
| chr14-88362794-DEL-52 | chr14-88362795-DEL-52 |
| chr14-88365775-DEL-92 | chr14-88365769-DEL-77 |
| chr14-88372630-DEL-1554 | chr14-88372631-DEL-1554 |
| chr14-88375546-DEL-68 | chr14-88375547-DEL-72 |
| chr14-88399666-DEL-275 | chr14-88399667-DEL-275 |
| chr14-88413824-DEL-151 | chr14-88413823-DEL-151 |
| chr14-88451239-DEL-3326 | chr14-88451240-DEL-3326 |
| chr14-88451247-DEL-1268 | chr14-88451248-DEL-1268 |
| chr14-88453379-DEL-2830 | chr14-88453380-DEL-2830 |
| chr14-88480871-DEL-52 | chr14-88480872-DEL-52 |
| chr14-88503446-DEL-613 | chr14-88503447-DEL-613 |
| chr14-88505234-DEL-1737 | chr14-88505235-DEL-1737 |
| chr14-88544312-DEL-704 | chr14-88544313-DEL-704 |
| chr14-88557021-DEL-1033 | chr14-88557022-DEL-1033 |
| chr14-88571846-DEL-70 | chr14-88571848-DEL-69 |
| chr14-88582549-DEL-60 | chr14-88582562-DEL-52 |
| chr14-88605522-DEL-6594 | chr14-88605523-DEL-6594 |
| chr14-88674735-DEL-136 | chr14-88674736-DEL-136 |
| chr14-88675227-DEL-1255 | chr14-88675228-DEL-1255 |
| chr14-88683630-DEL-53 | chr14-88683631-DEL-53 |
| chr14-88687565-DEL-3540 | chr14-88687566-DEL-3540 |
| chr14-88696088-DEL-417 | chr14-88696089-DEL-417 |
| chr14-88715032-DEL-358 | chr14-88715033-DEL-358 |
| chr14-88722264-DEL-436 | chr14-88722265-DEL-436 |
| chr14-88750636-DEL-399 | chr14-88750629-DEL-400 |
| chr14-88774155-DEL-132 | chr14-88774156-DEL-132 |
| chr14-88777020-DEL-610 | chr14-88777020-DEL-610 |
| chr14-88778212-DEL-56 | chr14-88778213-DEL-56 |
| chr14-88847257-DEL-60 | chr14-88847258-DEL-54 |
| chr14-88850782-DEL-6607 | chr14-88850783-DEL-6607 |
| chr14-88864094-DEL-645 | chr14-88864095-DEL-645 |
| chr14-88891897-DEL-95 | chr14-88891891-DEL-79 |
| chr14-88901806-DEL-1804 | chr14-88901806-DEL-1804 |
| chr14-88907357-DEL-829 | chr14-88907358-DEL-829 |
| chr14-88941086-DEL-59 | chr14-88941087-DEL-58 |
| chr14-88942176-DEL-77 | chr14-88942174-DEL-61 |
| chr14-88975078-DEL-101 | chr14-88975079-DEL-101 |
| chr14-89008574-DEL-99 | chr14-89008617-DEL-52 |
| chr14-89024462-DEL-113 | chr14-89024463-DEL-113 |
| chr14-89029960-DEL-2974 | chr14-89029961-DEL-2974 |
| chr14-89031402-DEL-62 | chr14-89031403-DEL-62 |
| chr14-89052234-DEL-300 | chr14-89052235-DEL-300 |
| chr14-89052941-DEL-1395 | chr14-89052942-DEL-1395 |
| chr14-89055505-DEL-70 | chr14-89055506-DEL-70 |
| chr14-89088422-DEL-86 | chr14-89088423-DEL-86 |
| chr14-89099547-DEL-75 | chr14-89099550-DEL-73 |
| chr14-89103374-DEL-2911 | chr14-89103375-DEL-2911 |
| chr14-89111715-DEL-115 | chr14-89111686-DEL-100 |
| chr14-89120681-DEL-375 | chr14-89120682-DEL-375 |
| chr14-89122099-DEL-945 | chr14-89122100-DEL-945 |
| chr14-89131244-DEL-377 | chr14-89131244-DEL-378 |
| chr14-89137581-DEL-70 | chr14-89137582-DEL-55 |
| chr14-89173925-DEL-1657 | chr14-89173926-DEL-1657 |
| chr14-89175779-DEL-2679 | chr14-89175780-DEL-2679 |
| chr14-89192726-DEL-213 | chr14-89192728-DEL-212 |
| chr14-89206927-DEL-2839 | chr14-89206912-DEL-2854 |
| chr14-89213782-DEL-54 | chr14-89213785-DEL-52 |
| chr14-89216113-DEL-251 | chr14-89216114-DEL-251 |
| chr14-89227798-DEL-399 | chr14-89227799-DEL-399 |

| | |
|---|---|
| chr14-89242607-DEL-478 | chr14-89242608-DEL-478 |
| chr14-89260337-DEL-52 | chr14-89260332-DEL-53 |
| chr14-89262806-DEL-550 | chr14-89262807-DEL-550 |
| chr14-89264514-DEL-143 | chr14-89264515-DEL-143 |
| chr14-89275096-DEL-2605 | chr14-89275097-DEL-2605 |
| chr14-89305397-DEL-88 | chr14-89305404-DEL-82 |
| chr14-89344746-DEL-675 | chr14-89344747-DEL-675 |
| chr14-89351306-DEL-171 | chr14-89351300-DEL-162 |
| chr14-89357014-DEL-70 | chr14-89357015-DEL-70 |
| chr14-89398218-DEL-160 | chr14-89398217-DEL-161 |
| chr14-89423672-DEL-102 | chr14-89423673-DEL-102 |
| chr14-89464569-DEL-124 | chr14-89464569-DEL-124 |
| chr14-89468237-DEL-62 | chr14-89468238-DEL-62 |
| chr14-89480447-DEL-55 | chr14-89480448-DEL-55 |
| chr14-89491508-DEL-5044 | chr14-89491509-DEL-5044 |
| chr14-89494926-DEL-342 | chr14-89494927-DEL-342 |
| chr14-89502472-DEL-110 | chr14-89502473-DEL-110 |
| chr14-89511110-DEL-300 | chr14-89511111-DEL-300 |
| chr14-89511669-DEL-77 | chr14-89511686-DEL-79 |
| chr14-89551786-DEL-226 | chr14-89551787-DEL-222 |
| chr14-89579209-DEL-241 | chr14-89579210-DEL-239 |
| chr14-89608453-DEL-2394 | chr14-89608454-DEL-2394 |
| chr14-89622131-DEL-277 | chr14-89622117-DEL-270 |
| chr14-89657844-DEL-1191 | chr14-89657845-DEL-1191 |
| chr14-89659737-DEL-601 | chr14-89659738-DEL-601 |
| chr14-89661616-DEL-597 | chr14-89661609-DEL-598 |
| chr14-89688439-DEL-3556 | chr14-89688440-DEL-3556 |
| chr14-89723780-DEL-6195 | chr14-89723781-DEL-6195 |
| chr14-89731504-DEL-511 | chr14-89731506-DEL-509 |
| chr14-89735802-DEL-115 | chr14-89735803-DEL-115 |
| chr14-89743422-DEL-175 | chr14-89743423-DEL-175 |
| chr14-89743845-DEL-713 | chr14-89743846-DEL-713 |
| chr14-89755486-DEL-217 | chr14-89755487-DEL-217 |
| chr14-89766571-DEL-95 | chr14-89766568-DEL-90 |
| chr14-89769189-DEL-2207 | chr14-89769190-DEL-2207 |
| chr14-89792121-DEL-258 | chr14-89792122-DEL-258 |
| chr14-89821238-DEL-75 | chr14-89821238-DEL-75 |
| chr14-89841947-DEL-380 | chr14-89841948-DEL-380 |
| chr14-89843157-DEL-318 | chr14-89843158-DEL-318 |
| chr14-89847048-DEL-582 | chr14-89847049-DEL-582 |
| chr14-89922156-DEL-88 | chr14-89922161-DEL-84 |
| chr14-89930154-DEL-1513 | chr14-89930154-DEL-1513 |
| chr14-89951429-DEL-646 | chr14-89951445-DEL-627 |
| chr14-89979381-DEL-658 | chr14-89979382-DEL-658 |
| chr14-90005620-DEL-455 | chr14-90005621-DEL-455 |
| chr14-90018987-DEL-460 | chr14-90018988-DEL-460 |
| chr14-90022794-DEL-577 | chr14-90022795-DEL-577 |
| chr14-90032062-DEL-101 | chr14-90032063-DEL-101 |
| chr14-90070933-DEL-58 | chr14-90070934-DEL-58 |
| chr14-90079107-DEL-71 | chr14-90079070-DEL-96 |
| chr14-90084497-DEL-426 | chr14-90084498-DEL-426 |
| chr14-90107982-DEL-402 | chr14-90107983-DEL-402 |
| chr14-90127774-DEL-608 | chr14-90127775-DEL-608 |
| chr14-90133639-DEL-3096 | chr14-90133639-DEL-3096 |
| chr14-90181709-DEL-382 | chr14-90181710-DEL-382 |
| chr14-90204991-DEL-305 | chr14-90204992-DEL-304 |
| chr14-90206876-DEL-938 | chr14-90206877-DEL-938 |
| chr14-90221750-DEL-80 | chr14-90221751-DEL-80 |
| chr14-90225068-DEL-58 | chr14-90225068-DEL-58 |
| chr14-90248749-DEL-932 | chr14-90248748-DEL-926 |
| chr14-90250484-DEL-143 | chr14-90250485-DEL-143 |
| chr14-90266082-DEL-52 | chr14-90266083-DEL-52 |
| chr14-90275544-DEL-119 | chr14-90275544-DEL-117 |
| chr14-90279228-DEL-226 | chr14-90279229-DEL-226 |
| chr14-90286014-DEL-246 | chr14-90286015-DEL-246 |
| chr14-90307301-DEL-759 | chr14-90307302-DEL-759 |
| chr14-90315780-DEL-7245 | chr14-90315805-DEL-7232 |
| chr14-90342335-DEL-82 | chr14-90342335-DEL-70 |
| chr14-90370617-DEL-177 | chr14-90370618-DEL-177 |
| chr14-90380154-DEL-65 | chr14-90380155-DEL-65 |
| chr14-90421093-DEL-58 | chr14-90421094-DEL-58 |
| chr14-90427553-DEL-51 | chr14-90427554-DEL-51 |
| chr14-90432679-DEL-1417 | chr14-90432680-DEL-1417 |
| chr14-90443643-DEL-60 | chr14-90443638-DEL-90 |
| chr14-90451109-DEL-2964 | chr14-90451110-DEL-2954 |
| chr14-90471298-DEL-307 | chr14-90471298-DEL-307 |
| chr14-90473783-DEL-144 | chr14-90473784-DEL-144 |
| chr14-90486223-DEL-170 | chr14-90486226-DEL-168 |
| chr14-90532078-DEL-7221 | chr14-90532079-DEL-7221 |
| chr14-90550973-DEL-56 | chr14-90550972-DEL-58 |

| | |
|---|---|
| chr14-90574569-DEL-64 | chr14-90574570-DEL-64 |
| chr14-90611816-DEL-84 | chr14-90611830-DEL-73 |
| chr14-90624673-DEL-641 | chr14-90624674-DEL-641 |
| chr14-90651069-DEL-115 | chr14-90651070-DEL-91 |
| chr14-90654517-DEL-1408 | chr14-90654518-DEL-1408 |
| chr14-90664984-DEL-188 | chr14-90664984-DEL-188 |
| chr14-90682433-DEL-52 | chr14-90682425-DEL-64 |
| chr14-90705021-DEL-53 | chr14-90705020-DEL-54 |
| chr14-90712301-DEL-63 | chr14-90712301-DEL-64 |
| chr14-90719437-DEL-723 | chr14-90719438-DEL-723 |
| chr14-90756788-DEL-1657 | chr14-90756789-DEL-1657 |
| chr14-90758573-DEL-52 | chr14-90758574-DEL-52 |
| chr14-90786988-DEL-2283 | chr14-90786984-DEL-2284 |
| chr14-90788532-DEL-64 | chr14-90788541-DEL-56 |
| chr14-90800353-DEL-977 | chr14-90800352-DEL-977 |
| chr14-90835878-DEL-55 | chr14-90835885-DEL-53 |
| chr14-90850795-DEL-134 | chr14-90850796-DEL-134 |
| chr14-90873313-DEL-58 | chr14-90873314-DEL-58 |
| chr14-90875464-DEL-1096 | chr14-90875464-DEL-1094 |
| chr14-90876804-DEL-77 | chr14-90876806-DEL-78 |
| chr14-90884241-DEL-500 | chr14-90884242-DEL-500 |
| chr14-90894790-DEL-81 | chr14-90894791-DEL-81 |
| chr14-90920550-DEL-7262 | chr14-90920551-DEL-7262 |
| chr14-90974921-DEL-261 | chr14-90974922-DEL-261 |
| chr14-90975758-DEL-1466 | chr14-90975759-DEL-1466 |
| chr14-90976215-DEL-604 | chr14-90976216-DEL-603 |
| chr14-91008005-DEL-1059 | chr14-91008006-DEL-1059 |
| chr14-91011317-DEL-4070 | chr14-91011315-DEL-4066 |
| chr14-91032537-DEL-402 | chr14-91032530-DEL-400 |
| chr14-91045656-DEL-94 | chr14-91045657-DEL-94 |
| chr14-91055663-DEL-255 | chr14-91055664-DEL-255 |
| chr14-91067495-DEL-76 | chr14-91067496-DEL-76 |
| chr14-91106979-DEL-124 | chr14-91106988-DEL-116 |
| chr14-91121533-DEL-1307 | chr14-91121534-DEL-1307 |
| chr14-91139689-DEL-57 | chr14-91139689-DEL-58 |
| chr14-91141708-DEL-1313 | chr14-91141709-DEL-1313 |
| chr14-91161999-DEL-489 | chr14-91162000-DEL-489 |
| chr14-91171232-DEL-6877 | chr14-91171233-DEL-6877 |
| chr14-91191236-DEL-881 | chr14-91191237-DEL-881 |
| chr14-91210198-DEL-8366 | chr14-91210195-DEL-8367 |
| chr14-91219485-DEL-1574 | chr14-91219486-DEL-1574 |
| chr14-91265590-DEL-2178 | chr14-91265591-DEL-2178 |
| chr14-91311328-DEL-57 | chr14-91311327-DEL-59 |
| chr14-91322810-DEL-1755 | chr14-91322811-DEL-1755 |
| chr14-91340014-DEL-158 | chr14-91340015-DEL-158 |
| chr14-91363238-DEL-6399 | chr14-91363239-DEL-6399 |
| chr14-91370611-DEL-55 | chr14-91370616-DEL-54 |
| chr14-91374454-DEL-60 | chr14-91374455-DEL-60 |
| chr14-91375143-DEL-872 | chr14-91375144-DEL-872 |
| chr14-91376224-DEL-321 | chr14-91376225-DEL-321 |
| chr14-91380897-DEL-63 | chr14-91380897-DEL-63 |
| chr14-91390700-DEL-277 | chr14-91390701-DEL-277 |
| chr14-91406093-DEL-6440 | chr14-91406094-DEL-6440 |
| chr14-91413667-DEL-2219 | chr14-91413668-DEL-2219 |
| chr14-91416426-DEL-82 | chr14-91416427-DEL-82 |
| chr14-91448932-DEL-394 | chr14-91448933-DEL-393 |
| chr14-91466393-DEL-63 | chr14-91466391-DEL-66 |
| chr14-91476319-DEL-1321 | chr14-91476321-DEL-1320 |
| chr14-91479280-DEL-199 | chr14-91479263-DEL-200 |
| chr14-91503794-DEL-366 | chr14-91503795-DEL-366 |
| chr14-91505656-DEL-377 | chr14-91505660-DEL-370 |
| chr14-91516968-DEL-6148 | chr14-91516969-DEL-6148 |
| chr14-91577707-DEL-1183 | chr14-91577708-DEL-1182 |
| chr14-91587185-DEL-1913 | chr14-91587186-DEL-1913 |
| chr14-91593313-DEL-901 | chr14-91593325-DEL-892 |
| chr14-91594360-DEL-108 | chr14-91594361-DEL-108 |
| chr14-91620757-DEL-1754 | chr14-91620761-DEL-1751 |
| chr14-91624573-DEL-53 | chr14-91624575-DEL-51 |
| chr14-91639227-DEL-235 | chr14-91639228-DEL-235 |
| chr14-91642378-DEL-60 | chr14-91642373-DEL-60 |
| chr14-91651838-DEL-203 | chr14-91651839-DEL-203 |
| chr14-91652778-DEL-529 | chr14-91652779-DEL-529 |
| chr14-91664430-DEL-64 | chr14-91664433-DEL-62 |
| chr14-91702986-DEL-52 | chr14-91702987-DEL-52 |
| chr14-91707106-DEL-459 | chr14-91707107-DEL-459 |
| chr14-91738160-DEL-311 | chr14-91738161-DEL-311 |
| chr14-91748184-DEL-4328 | chr14-91748185-DEL-4328 |
| chr14-91756440-DEL-144 | chr14-91756441-DEL-144 |
| chr14-91802626-DEL-365 | chr14-91802627-DEL-365 |
| chr14-91826998-DEL-1174 | chr14-91826999-DEL-1174 |

| | |
|---|---|
| chr14-91857250-DEL-1532 | chr14-91857251-DEL-1531 |
| chr14-91864517-DEL-82 | chr14-91864518-DEL-82 |
| chr14-91878416-DEL-51 | chr14-91878417-DEL-51 |
| chr14-91884750-DEL-573 | chr14-91884751-DEL-573 |
| chr14-91899305-DEL-215 | chr14-91899306-DEL-240 |
| chr14-91908512-DEL-6465 | chr14-91908513-DEL-6465 |
| chr14-91925134-DEL-157 | chr14-91925135-DEL-157 |
| chr14-91928595-DEL-217 | chr14-91928595-DEL-217 |
| chr14-91954105-DEL-59 | chr14-91954119-DEL-51 |
| chr14-91989506-DEL-1495 | chr14-91989507-DEL-1495 |
| chr14-92006397-DEL-194 | chr14-92006398-DEL-194 |
| chr14-92033858-DEL-378 | chr14-92033859-DEL-378 |
| chr14-92060823-DEL-1740 | chr14-92060824-DEL-1740 |
| chr14-92069017-DEL-830 | chr14-92069018-DEL-830 |
| chr14-92084127-DEL-76 | chr14-92084128-DEL-76 |
| chr14-92090121-DEL-101 | chr14-92090150-DEL-84 |
| chr14-92094616-DEL-141 | chr14-92094639-DEL-158 |
| chr14-92106244-DEL-58 | chr14-92106245-DEL-58 |
| chr14-92113086-DEL-150 | chr14-92113087-DEL-144 |
| chr14-92180094-DEL-537 | chr14-92180095-DEL-537 |
| chr14-92190634-DEL-66 | chr14-92190635-DEL-66 |
| chr14-92191492-DEL-201 | chr14-92191470-DEL-243 |
| chr14-92198471-DEL-3180 | chr14-92198472-DEL-3180 |
| chr14-92207581-DEL-6510 | chr14-92207582-DEL-6510 |
| chr14-92221004-DEL-1762 | chr14-92221005-DEL-1762 |
| chr14-92229540-DEL-3457 | chr14-92229541-DEL-3457 |
| chr14-92244838-DEL-193 | chr14-92244838-DEL-193 |
| chr14-92269269-DEL-9214 | chr14-92269270-DEL-9214 |
| chr14-92283243-DEL-379 | chr14-92283244-DEL-379 |
| chr14-92286593-DEL-2997 | chr14-92286594-DEL-2997 |
| chr14-92312468-DEL-313 | chr14-92312511-DEL-271 |
| chr14-92367071-DEL-399 | chr14-92367072-DEL-399 |
| chr14-92367788-DEL-66 | chr14-92367789-DEL-66 |
| chr14-92391115-DEL-70 | chr14-92391112-DEL-71 |
| chr14-92400362-DEL-68 | chr14-92400363-DEL-68 |
| chr14-92422364-DEL-555 | chr14-92422365-DEL-555 |
| chr14-92443547-DEL-390 | chr14-92443548-DEL-390 |
| chr14-92454447-DEL-124 | chr14-92454448-DEL-124 |
| chr14-92465666-DEL-6262 | chr14-92465667-DEL-6262 |
| chr14-92481665-DEL-118 | chr14-92481701-DEL-96 |
| chr14-92487595-DEL-69 | chr14-92487596-DEL-69 |
| chr14-92520420-DEL-1846 | chr14-92520421-DEL-1854 |
| chr14-92530210-DEL-3096 | chr14-92530211-DEL-3097 |
| chr14-92557289-DEL-60 | chr14-92557285-DEL-60 |
| chr14-92587608-DEL-183 | chr14-92587609-DEL-183 |
| chr14-92600215-DEL-105 | chr14-92600231-DEL-103 |
| chr14-92601683-DEL-554 | chr14-92601684-DEL-554 |
| chr14-92603922-DEL-361 | chr14-92603923-DEL-361 |
| chr14-92620956-DEL-6375 | chr14-92620957-DEL-6375 |
| chr14-92657293-DEL-95 | chr14-92657294-DEL-95 |
| chr14-92660721-DEL-56 | chr14-92660722-DEL-56 |
| chr14-92679192-DEL-306 | chr14-92679193-DEL-306 |
| chr14-92680877-DEL-66 | chr14-92680886-DEL-58 |
| chr14-92696041-DEL-58 | chr14-92696040-DEL-59 |
| chr14-92702675-DEL-60 | chr14-92702676-DEL-60 |
| chr14-92708592-DEL-67 | chr14-92708593-DEL-65 |
| chr14-92714452-DEL-92 | chr14-92714453-DEL-88 |
| chr14-92732761-DEL-95 | chr14-92732762-DEL-97 |
| chr14-92749270-DEL-956 | chr14-92749271-DEL-956 |
| chr14-92753215-DEL-643 | chr14-92753216-DEL-642 |
| chr14-92773930-DEL-843 | chr14-92773931-DEL-843 |
| chr14-92793746-DEL-198 | chr14-92793747-DEL-198 |
| chr14-92809213-DEL-289 | chr14-92809214-DEL-289 |
| chr14-92817988-DEL-68 | chr14-92817995-DEL-69 |
| chr14-92833374-DEL-6365 | chr14-92833375-DEL-6365 |
| chr14-92840938-DEL-1154 | chr14-92840939-DEL-1154 |
| chr14-92842641-DEL-147 | chr14-92842642-DEL-147 |
| chr14-92895968-DEL-1842 | chr14-92895969-DEL-1842 |
| chr14-92899191-DEL-52 | chr14-92899192-DEL-52 |
| chr14-92901839-DEL-627 | chr14-92901842-DEL-625 |
| chr14-92933491-DEL-501 | chr14-92933492-DEL-501 |
| chr14-92937429-DEL-92 | chr14-92937430-DEL-92 |
| chr14-92942518-DEL-112 | chr14-92942519-DEL-112 |
| chr14-92950058-DEL-59 | chr14-92950053-DEL-65 |
| chr14-93009295-DEL-91 | chr14-93009285-DEL-78 |
| chr14-93020850-DEL-1639 | chr14-93020851-DEL-1639 |
| chr14-93026495-DEL-1048 | chr14-93026496-DEL-1048 |
| chr14-93029937-DEL-817 | chr14-93029937-DEL-818 |
| chr14-93032280-DEL-3781 | chr14-93032253-DEL-3781 |
| chr14-93040166-DEL-239 | chr14-93040130-DEL-232 |

| | |
|---|---|
| chr14-93041909-DEL-247 | chr14-93041912-DEL-248 |
| chr14-93048786-DEL-65 | chr14-93048787-DEL-65 |
| chr14-93053675-DEL-155 | chr14-93053676-DEL-154 |
| chr14-93055804-DEL-62 | chr14-93055803-DEL-66 |
| chr14-93070138-DEL-97 | chr14-93070139-DEL-97 |
| chr14-93081782-DEL-6286 | chr14-93081783-DEL-6286 |
| chr14-93093793-DEL-93 | chr14-93093794-DEL-91 |
| chr14-93094025-DEL-753 | chr14-93094026-DEL-753 |
| chr14-93100990-DEL-107 | chr14-93100991-DEL-107 |
| chr14-93123830-DEL-80 | chr14-93123830-DEL-80 |
| chr14-93140149-DEL-184 | chr14-93140166-DEL-168 |
| chr14-93182644-DEL-56 | chr14-93182636-DEL-57 |
| chr14-93190845-DEL-1221 | chr14-93190846-DEL-1221 |
| chr14-93211517-DEL-55 | chr14-93211518-DEL-50 |
| chr14-93213414-DEL-52 | chr14-93213437-DEL-73 |
| chr14-93216911-DEL-75 | chr14-93216912-DEL-75 |
| chr14-93230201-DEL-52 | chr14-93230202-DEL-52 |
| chr14-93254915-DEL-5262 | chr14-93254916-DEL-5262 |
| chr14-93279011-DEL-56 | chr14-93279012-DEL-56 |
| chr14-93283371-DEL-4757 | chr14-93283372-DEL-4757 |
| chr14-93302192-DEL-71 | chr14-93302195-DEL-53 |
| chr14-93354930-DEL-76 | chr14-93354887-DEL-126 |
| chr14-93387649-DEL-359 | chr14-93387675-DEL-364 |
| chr14-93394643-DEL-54 | chr14-93394644-DEL-52 |
| chr14-93402784-DEL-74 | chr14-93402785-DEL-74 |
| chr14-93439342-DEL-72 | chr14-93439343-DEL-68 |
| chr14-93444912-DEL-458 | chr14-93444913-DEL-458 |
| chr14-93446454-DEL-7462 | chr14-93446455-DEL-7462 |
| chr14-93488625-DEL-575 | chr14-93488626-DEL-575 |
| chr14-93494747-DEL-89 | chr14-93494748-DEL-89 |
| chr14-93497265-DEL-60 | chr14-93497233-DEL-85 |
| chr14-93521301-DEL-54 | chr14-93521303-DEL-52 |
| chr14-93540032-DEL-371 | chr14-93540033-DEL-371 |
| chr14-93546591-DEL-52 | chr14-93546550-DEL-50 |
| chr14-93609210-DEL-53 | chr14-93609198-DEL-66 |
| chr14-93618510-DEL-76 | chr14-93618511-DEL-76 |
| chr14-93627237-DEL-56 | chr14-93627238-DEL-56 |
| chr14-93646396-DEL-87 | chr14-93646432-DEL-87 |
| chr14-93660616-DEL-93 | chr14-93660615-DEL-93 |
| chr14-93661657-DEL-845 | chr14-93661658-DEL-846 |
| chr14-93691450-DEL-160 | chr14-93691429-DEL-182 |
| chr14-93709367-DEL-130 | chr14-93709368-DEL-130 |
| chr14-93728783-DEL-62 | chr14-93728785-DEL-60 |
| chr14-93803587-DEL-334 | chr14-93803588-DEL-333 |
| chr14-93852900-DEL-86 | chr14-93852914-DEL-78 |
| chr14-93869957-DEL-1024 | chr14-93869954-DEL-1025 |
| chr14-93875353-DEL-51 | chr14-93875348-DEL-57 |
| chr14-93879455-DEL-117 | chr14-93879457-DEL-117 |
| chr14-93897965-DEL-758 | chr14-93897966-DEL-758 |
| chr14-93941785-DEL-201 | chr14-93941786-DEL-201 |
| chr14-93959036-DEL-58 | chr14-93959034-DEL-58 |
| chr14-93969312-DEL-207 | chr14-93969313-DEL-207 |
| chr14-93998365-DEL-51 | chr14-93998368-DEL-52 |
| chr14-93998454-DEL-1282 | chr14-93998452-DEL-1275 |
| chr14-94003821-DEL-56 | chr14-94003804-DEL-57 |
| chr14-94018517-DEL-52 | chr14-94018518-DEL-52 |
| chr14-94053471-DEL-228 | chr14-94053472-DEL-228 |
| chr14-94133290-DEL-75 | chr14-94133261-DEL-64 |
| chr14-94142988-DEL-87 | chr14-94142989-DEL-87 |
| chr14-94148466-DEL-63 | chr14-94148466-DEL-63 |
| chr14-94161233-DEL-720 | chr14-94161233-DEL-721 |
| chr14-94168361-DEL-6613 | chr14-94168362-DEL-6613 |
| chr14-94168711-DEL-2104 | chr14-94168712-DEL-2104 |
| chr14-94186939-DEL-51 | chr14-94186940-DEL-51 |
| chr14-94196328-DEL-116 | chr14-94196330-DEL-115 |
| chr14-94206797-DEL-147 | chr14-94206798-DEL-147 |
| chr14-94267592-DEL-270 | chr14-94267593-DEL-270 |
| chr14-94287562-DEL-246 | chr14-94287563-DEL-246 |
| chr14-94288940-DEL-6073 | chr14-94288941-DEL-6073 |
| chr14-94299021-DEL-548 | chr14-94299028-DEL-542 |
| chr14-94320401-DEL-128 | chr14-94320402-DEL-128 |
| chr14-94323708-DEL-1096 | chr14-94323708-DEL-1096 |
| chr14-94334907-DEL-712 | chr14-94334908-DEL-712 |
| chr14-94355857-DEL-201 | chr14-94355858-DEL-201 |
| chr14-94374659-DEL-57 | chr14-94374660-DEL-52 |
| chr14-94386902-DEL-71 | chr14-94386898-DEL-55 |
| chr14-94398023-DEL-60 | chr14-94398024-DEL-60 |
| chr14-94415178-DEL-549 | chr14-94415179-DEL-549 |
| chr14-94419295-DEL-53 | chr14-94419289-DEL-59 |
| chr14-94420216-DEL-236 | chr14-94420217-DEL-236 |

| | |
|---|---|
| chr14-94423592-DEL-3987 | chr14-94423595-DEL-3985 |
| chr14-94449306-DEL-52 | chr14-94449307-DEL-52 |
| chr14-94452651-DEL-1643 | chr14-94452652-DEL-1643 |
| chr14-94483291-DEL-430 | chr14-94483287-DEL-410 |
| chr14-94487973-DEL-82 | chr14-94487974-DEL-76 |
| chr14-94492129-DEL-73 | chr14-94492130-DEL-73 |
| chr14-94493890-DEL-85 | chr14-94493870-DEL-96 |
| chr14-94508570-DEL-80 | chr14-94508571-DEL-80 |
| chr14-94527340-DEL-206 | chr14-94527341-DEL-206 |
| chr14-94538294-DEL-52 | chr14-94538295-DEL-52 |
| chr14-94542422-DEL-76 | chr14-94542423-DEL-80 |
| chr14-94548923-DEL-1539 | chr14-94548924-DEL-1539 |
| chr14-94554490-DEL-382 | chr14-94554491-DEL-382 |
| chr14-94571639-DEL-3827 | chr14-94571640-DEL-3827 |
| chr14-94604688-DEL-63 | chr14-94604690-DEL-61 |
| chr14-94607261-DEL-98 | chr14-94607262-DEL-98 |
| chr14-94615419-DEL-2056 | chr14-94615420-DEL-2056 |
| chr14-94635385-DEL-80 | chr14-94635386-DEL-80 |
| chr14-94655101-DEL-54 | chr14-94655102-DEL-54 |
| chr14-94660632-DEL-445 | chr14-94660633-DEL-451 |
| chr14-94682682-DEL-53 | chr14-94682683-DEL-53 |
| chr14-94685430-DEL-2272 | chr14-94685431-DEL-2272 |
| chr14-94691988-DEL-2703 | chr14-94691988-DEL-2703 |
| chr14-94710557-DEL-60 | chr14-94710558-DEL-60 |
| chr14-94721879-DEL-81 | chr14-94721877-DEL-84 |
| chr14-94744610-DEL-653 | chr14-94744611-DEL-653 |
| chr14-94756252-DEL-183 | chr14-94756299-DEL-164 |
| chr14-94802773-DEL-713 | chr14-94802774-DEL-713 |
| chr14-94827913-DEL-1352 | chr14-94827908-DEL-1352 |
| chr14-94872679-DEL-2686 | chr14-94872680-DEL-2686 |
| chr14-94876208-DEL-437 | chr14-94876209-DEL-437 |
| chr14-94880231-DEL-3263 | chr14-94880232-DEL-3263 |
| chr14-94892429-DEL-53 | chr14-94892424-DEL-57 |
| chr14-94910050-DEL-56 | chr14-94910051-DEL-56 |
| chr14-94933332-DEL-167 | chr14-94933311-DEL-172 |
| chr14-94934585-DEL-56 | chr14-94934586-DEL-68 |
| chr14-94944021-DEL-2203 | chr14-94944022-DEL-2203 |
| chr14-94946367-DEL-614 | chr14-94946368-DEL-614 |
| chr14-94964298-DEL-64 | chr14-94964299-DEL-56 |
| chr14-94973488-DEL-396 | chr14-94973489-DEL-396 |
| chr14-94987226-DEL-90 | chr14-94987227-DEL-90 |
| chr14-94990916-DEL-100 | chr14-94990921-DEL-96 |
| chr14-94993841-DEL-60 | chr14-94993842-DEL-60 |
| chr14-94995575-DEL-52 | chr14-94995576-DEL-54 |
| chr14-95016613-DEL-187 | chr14-95016614-DEL-187 |
| chr14-95047316-DEL-4127 | chr14-95047317-DEL-4127 |
| chr14-95061262-DEL-86 | chr14-95061265-DEL-84 |
| chr14-95079722-DEL-6273 | chr14-95079723-DEL-6273 |
| chr14-95098916-DEL-62 | chr14-95098917-DEL-62 |
| chr14-95127329-DEL-52 | chr14-95127330-DEL-52 |
| chr14-95138727-DEL-1576 | chr14-95138728-DEL-1576 |
| chr14-95143901-DEL-283 | chr14-95143895-DEL-289 |
| chr14-95152345-DEL-2661 | chr14-95152345-DEL-2661 |
| chr14-95169827-DEL-188 | chr14-95169828-DEL-188 |
| chr14-95190298-DEL-265 | chr14-95190299-DEL-265 |
| chr14-95202806-DEL-111 | chr14-95202807-DEL-111 |
| chr14-95207668-DEL-196 | chr14-95207669-DEL-196 |
| chr14-95218861-DEL-6371 | chr14-95218862-DEL-6371 |
| chr14-95243407-DEL-81 | chr14-95243408-DEL-81 |
| chr14-95266048-DEL-192 | chr14-95266049-DEL-192 |
| chr14-95271188-DEL-58 | chr14-95271189-DEL-58 |
| chr14-95288574-DEL-1731 | chr14-95288561-DEL-1729 |
| chr14-95292685-DEL-6176 | chr14-95292686-DEL-6176 |
| chr14-95309585-DEL-78 | chr14-95309586-DEL-74 |
| chr14-95322845-DEL-230 | chr14-95322846-DEL-230 |
| chr14-95329155-DEL-224 | chr14-95329162-DEL-218 |
| chr14-95345964-DEL-560 | chr14-95345963-DEL-561 |
| chr14-95384016-DEL-1339 | chr14-95384017-DEL-1339 |
| chr14-95387827-DEL-1441 | chr14-95387828-DEL-1441 |
| chr14-95389424-DEL-434 | chr14-95389425-DEL-434 |
| chr14-95401449-DEL-124 | chr14-95401450-DEL-124 |
| chr14-95415196-DEL-199 | chr14-95415196-DEL-200 |
| chr14-95415854-DEL-98 | chr14-95415855-DEL-108 |
| chr14-95446230-DEL-72 | chr14-95446231-DEL-72 |
| chr14-95452283-DEL-6368 | chr14-95452284-DEL-6368 |
| chr14-95461662-DEL-60 | chr14-95461663-DEL-56 |
| chr14-95476391-DEL-66 | chr14-95476400-DEL-54 |
| chr14-95488952-DEL-62 | chr14-95488947-DEL-68 |
| chr14-95491422-DEL-555 | chr14-95491423-DEL-555 |
| chr14-95507859-DEL-2410 | chr14-95507860-DEL-2410 |

| | |
|---|---|
| chr14-95512511-DEL-66 | chr14-95512512-DEL-66 |
| chr14-95515439-DEL-1376 | chr14-95515440-DEL-1376 |
| chr14-95516868-DEL-383 | chr14-95516869-DEL-383 |
| chr14-95521964-DEL-1161 | chr14-95521965-DEL-1161 |
| chr14-95523541-DEL-72 | chr14-95523562-DEL-52 |
| chr14-95552416-DEL-3924 | chr14-95552416-DEL-3924 |
| chr14-95562422-DEL-60 | chr14-95562423-DEL-50 |
| chr14-95569359-DEL-450 | chr14-95569360-DEL-450 |
| chr14-95576383-DEL-80 | chr14-95576384-DEL-80 |
| chr14-95622477-DEL-845 | chr14-95622478-DEL-845 |
| chr14-95647748-DEL-53 | chr14-95647768-DEL-54 |
| chr14-95650363-DEL-513 | chr14-95650364-DEL-513 |
| chr14-95652571-DEL-62 | chr14-95652576-DEL-58 |
| chr14-95655806-DEL-108 | chr14-95655807-DEL-108 |
| chr14-95667054-DEL-529 | chr14-95667055-DEL-529 |
| chr14-95685846-DEL-3014 | chr14-95685847-DEL-3014 |
| chr14-95721556-DEL-307 | chr14-95721529-DEL-316 |
| chr14-95737343-DEL-197 | chr14-95737344-DEL-197 |
| chr14-95778014-DEL-63 | chr14-95778015-DEL-63 |
| chr14-95797349-DEL-54 | chr14-95797350-DEL-54 |
| chr14-95798948-DEL-7082 | chr14-95798949-DEL-7082 |
| chr14-95809504-DEL-124 | chr14-95809509-DEL-120 |
| chr14-95818304-DEL-353 | chr14-95818306-DEL-354 |
| chr14-95822498-DEL-154 | chr14-95822501-DEL-152 |
| chr14-95851949-DEL-53 | chr14-95851968-DEL-60 |
| chr14-95857093-DEL-1129 | chr14-95857094-DEL-1129 |
| chr14-95876111-DEL-63 | chr14-95876113-DEL-53 |
| chr14-95936425-DEL-60 | chr14-95936426-DEL-60 |
| chr14-95937523-DEL-639 | chr14-95937524-DEL-639 |
| chr14-95947291-DEL-829 | chr14-95947292-DEL-829 |
| chr14-95955922-DEL-2832 | chr14-95955923-DEL-2832 |
| chr14-95960767-DEL-76 | chr14-95960768-DEL-76 |
| chr14-95970073-DEL-363 | chr14-95970072-DEL-369 |
| chr14-95971870-DEL-56 | chr14-95971864-DEL-55 |
| chr14-95977573-DEL-1494 | chr14-95977574-DEL-1494 |
| chr14-95997027-DEL-185 | chr14-95997028-DEL-185 |
| chr14-96054293-DEL-77 | chr14-96054298-DEL-78 |
| chr14-96060847-DEL-373 | chr14-96060848-DEL-373 |
| chr14-96065705-DEL-1489 | chr14-96065706-DEL-1489 |
| chr14-96120866-DEL-270 | chr14-96120867-DEL-270 |
| chr14-96131474-DEL-1415 | chr14-96131475-DEL-1414 |
| chr14-96152973-DEL-446 | chr14-96152974-DEL-446 |
| chr14-96198432-DEL-164 | chr14-96198413-DEL-164 |
| chr14-96218226-DEL-160 | chr14-96218226-DEL-160 |
| chr14-96242036-DEL-116 | chr14-96242037-DEL-116 |
| chr14-96244363-DEL-260 | chr14-96244364-DEL-257 |
| chr14-96285351-DEL-6399 | chr14-96285352-DEL-6399 |
| chr14-96330978-DEL-2632 | chr14-96330978-DEL-2632 |
| chr14-96337826-DEL-2788 | chr14-96337827-DEL-2788 |
| chr14-96341929-DEL-57 | chr14-96341930-DEL-57 |
| chr14-96347433-DEL-6385 | chr14-96347434-DEL-6385 |
| chr14-96368271-DEL-56 | chr14-96368272-DEL-56 |
| chr14-96370809-DEL-475 | chr14-96370810-DEL-475 |
| chr14-96375258-DEL-326 | chr14-96375259-DEL-326 |
| chr14-96388573-DEL-56 | chr14-96388576-DEL-58 |
| chr14-96394979-DEL-629 | chr14-96394979-DEL-627 |
| chr14-96409458-DEL-162 | chr14-96409459-DEL-162 |
| chr14-96412266-DEL-2043 | chr14-96412267-DEL-2043 |
| chr14-96418449-DEL-14919 | chr14-96418479-DEL-14889 |
| chr14-96426053-DEL-6753 | chr14-96426052-DEL-6754 |
| chr14-96439332-DEL-82 | chr14-96439330-DEL-79 |
| chr14-96457409-DEL-68 | chr14-96457412-DEL-66 |
| chr14-96464339-DEL-352 | chr14-96464340-DEL-352 |
| chr14-96484603-DEL-60 | chr14-96484604-DEL-60 |
| chr14-96485315-DEL-64 | chr14-96485316-DEL-64 |
| chr14-96486437-DEL-2237 | chr14-96486438-DEL-2237 |
| chr14-96493081-DEL-109 | chr14-96493082-DEL-109 |
| chr14-96514697-DEL-100 | chr14-96514698-DEL-100 |
| chr14-96521354-DEL-7078 | chr14-96521355-DEL-7078 |
| chr14-96545677-DEL-400 | chr14-96545678-DEL-400 |
| chr14-96558248-DEL-59 | chr14-96558249-DEL-59 |
| chr14-96565705-DEL-56 | chr14-96565705-DEL-56 |
| chr14-96566822-DEL-129 | chr14-96566823-DEL-129 |
| chr14-96579449-DEL-60 | chr14-96579450-DEL-60 |
| chr14-96583604-DEL-230 | chr14-96583604-DEL-229 |
| chr14-96600745-DEL-1643 | chr14-96600745-DEL-1644 |
| chr14-96605647-DEL-67 | chr14-96605649-DEL-68 |
| chr14-96622970-DEL-51 | chr14-96622971-DEL-51 |
| chr14-96627065-DEL-292 | chr14-96627064-DEL-292 |
| chr14-96633145-DEL-2162 | chr14-96633146-DEL-2162 |

| | |
|---|---|
| chr14-96641657-DEL-83 | chr14-96641673-DEL-68 |
| chr14-96642428-DEL-6317 | chr14-96642428-DEL-6317 |
| chr14-96655979-DEL-678 | chr14-96655980-DEL-678 |
| chr14-96676861-DEL-52 | chr14-96676860-DEL-53 |
| chr14-96679133-DEL-108 | chr14-96679134-DEL-108 |
| chr14-96680153-DEL-2011 | chr14-96680154-DEL-2011 |
| chr14-96688356-DEL-399 | chr14-96688357-DEL-399 |
| chr14-96694259-DEL-81 | chr14-96694260-DEL-81 |
| chr14-96696249-DEL-95 | chr14-96696250-DEL-95 |
| chr14-96705226-DEL-432 | chr14-96705227-DEL-432 |
| chr14-96715319-DEL-1794 | chr14-96715320-DEL-1794 |
| chr14-96720011-DEL-54 | chr14-96720012-DEL-54 |
| chr14-96742430-DEL-110 | chr14-96742435-DEL-106 |
| chr14-96747090-DEL-307 | chr14-96747091-DEL-307 |
| chr14-96749036-DEL-95 | chr14-96749046-DEL-86 |
| chr14-96759051-DEL-59 | chr14-96759052-DEL-59 |
| chr14-96759527-DEL-116 | chr14-96759534-DEL-110 |
| chr14-96774971-DEL-109 | chr14-96774972-DEL-109 |
| chr14-96787719-DEL-54 | chr14-96787722-DEL-56 |
| chr14-96790078-DEL-742 | chr14-96790090-DEL-743 |
| chr14-96804451-DEL-52 | chr14-96804450-DEL-54 |
| chr14-96821836-DEL-55 | chr14-96821837-DEL-55 |
| chr14-96838595-DEL-312 | chr14-96838617-DEL-304 |
| chr14-96839302-DEL-1077 | chr14-96839303-DEL-1076 |
| chr14-96847620-DEL-376 | chr14-96847621-DEL-376 |
| chr14-96850987-DEL-955 | chr14-96850988-DEL-954 |
| chr14-96871794-DEL-65 | chr14-96871795-DEL-65 |
| chr14-96894083-DEL-280 | chr14-96894084-DEL-280 |
| chr14-96909244-DEL-87 | chr14-96909245-DEL-87 |
| chr14-96910946-DEL-64 | chr14-96910943-DEL-68 |
| chr14-96918087-DEL-54 | chr14-96918088-DEL-54 |
| chr14-96923064-DEL-102 | chr14-96923065-DEL-102 |
| chr14-96937939-DEL-2647 | chr14-96937941-DEL-2645 |
| chr14-96943152-DEL-861 | chr14-96943153-DEL-861 |
| chr14-96950528-DEL-1090 | chr14-96950529-DEL-1090 |
| chr14-96956937-DEL-84 | chr14-96956945-DEL-76 |
| chr14-96967786-DEL-55 | chr14-96967790-DEL-54 |
| chr14-96976229-DEL-68 | chr14-96976230-DEL-68 |
| chr14-97039059-DEL-1623 | chr14-97039060-DEL-1623 |
| chr14-97040537-DEL-664 | chr14-97040538-DEL-664 |
| chr14-97070030-DEL-274 | chr14-97070031-DEL-274 |
| chr14-97087677-DEL-6362 | chr14-97087678-DEL-6362 |
| chr14-97096276-DEL-827 | chr14-97096277-DEL-827 |
| chr14-97133321-DEL-6942 | chr14-97133322-DEL-6942 |
| chr14-97134940-DEL-477 | chr14-97134941-DEL-477 |
| chr14-97135618-DEL-57 | chr14-97135619-DEL-57 |
| chr14-97162644-DEL-73 | chr14-97162653-DEL-71 |
| chr14-97166708-DEL-2851 | chr14-97166709-DEL-2851 |
| chr14-97223493-DEL-92 | chr14-97223520-DEL-102 |
| chr14-97236011-DEL-350 | chr14-97236012-DEL-350 |
| chr14-97242052-DEL-1213 | chr14-97242053-DEL-1213 |
| chr14-97268506-DEL-6932 | chr14-97268500-DEL-6925 |
| chr14-97277757-DEL-54 | chr14-97277758-DEL-54 |
| chr14-97285334-DEL-409 | chr14-97285335-DEL-409 |
| chr14-97297974-DEL-99 | chr14-97297976-DEL-98 |
| chr14-97305364-DEL-195 | chr14-97305365-DEL-195 |
| chr14-97312853-DEL-72 | chr14-97312854-DEL-72 |
| chr14-97317839-DEL-59 | chr14-97317840-DEL-59 |
| chr14-97327424-DEL-60 | chr14-97327425-DEL-60 |
| chr14-97354026-DEL-6279 | chr14-97354027-DEL-6279 |
| chr14-97377029-DEL-6421 | chr14-97377029-DEL-6421 |
| chr14-97410903-DEL-61 | chr14-97410904-DEL-61 |
| chr14-97425101-DEL-2189 | chr14-97425102-DEL-2189 |
| chr14-97506243-DEL-416 | chr14-97506244-DEL-416 |
| chr14-97537832-DEL-670 | chr14-97537827-DEL-670 |
| chr14-97552583-DEL-99 | chr14-97552604-DEL-78 |
| chr14-97560663-DEL-2915 | chr14-97560664-DEL-2915 |
| chr14-97597118-DEL-496 | chr14-97597119-DEL-496 |
| chr14-97599378-DEL-56 | chr14-97599379-DEL-56 |
| chr14-97602093-DEL-1420 | chr14-97602094-DEL-1420 |
| chr14-97605038-DEL-434 | chr14-97605039-DEL-426 |
| chr14-97636219-DEL-71 | chr14-97636220-DEL-71 |
| chr14-97664031-DEL-133 | chr14-97664048-DEL-94 |
| chr14-97699420-DEL-73 | chr14-97699421-DEL-73 |
| chr14-97701177-DEL-88 | chr14-97701178-DEL-88 |
| chr14-97713086-DEL-116 | chr14-97713087-DEL-116 |
| chr14-97731530-DEL-2249 | chr14-97731531-DEL-2249 |
| chr14-97750604-DEL-733 | chr14-97750605-DEL-733 |
| chr14-97752162-DEL-358 | chr14-97752163-DEL-358 |
| chr14-97766046-DEL-105 | chr14-97766047-DEL-105 |

| | |
|---|---|
| chr14-97767732-DEL-773 | chr14-97767733-DEL-773 |
| chr14-97772481-DEL-56 | chr14-97772482-DEL-56 |
| chr14-97775966-DEL-824 | chr14-97775967-DEL-824 |
| chr14-97796916-DEL-50 | chr14-97796930-DEL-54 |
| chr14-97810503-DEL-162 | chr14-97810504-DEL-162 |
| chr14-97826517-DEL-98 | chr14-97826511-DEL-96 |
| chr14-97891496-DEL-66 | chr14-97891494-DEL-67 |
| chr14-97910617-DEL-959 | chr14-97910618-DEL-959 |
| chr14-97913488-DEL-60 | chr14-97913481-DEL-56 |
| chr14-97928344-DEL-1595 | chr14-97928345-DEL-1595 |
| chr14-97946433-DEL-78 | chr14-97946434-DEL-78 |
| chr14-97952308-DEL-6128 | chr14-97952309-DEL-6128 |
| chr14-97964362-DEL-380 | chr14-97964363-DEL-380 |
| chr14-97973488-DEL-1762 | chr14-97973489-DEL-1762 |
| chr14-97981280-DEL-1059 | chr14-97981281-DEL-1059 |
| chr14-98015799-DEL-203 | chr14-98015800-DEL-203 |
| chr14-98038345-DEL-206 | chr14-98038346-DEL-206 |
| chr14-98039746-DEL-58 | chr14-98039747-DEL-58 |
| chr14-98046457-DEL-518 | chr14-98046458-DEL-518 |
| chr14-98056964-DEL-76 | chr14-98056962-DEL-92 |
| chr14-98058845-DEL-197 | chr14-98058872-DEL-214 |
| chr14-98062670-DEL-52 | chr14-98062671-DEL-52 |
| chr14-98082535-DEL-497 | chr14-98082536-DEL-497 |
| chr14-98096327-DEL-240 | chr14-98096338-DEL-234 |
| chr14-98118379-DEL-1021 | chr14-98118380-DEL-1021 |
| chr14-98142446-DEL-60 | chr14-98142448-DEL-58 |
| chr14-98143173-DEL-85 | chr14-98143174-DEL-85 |
| chr14-98196233-DEL-88 | chr14-98196215-DEL-76 |
| chr14-98227009-DEL-54 | chr14-98227012-DEL-52 |
| chr14-98257379-DEL-61 | chr14-98257380-DEL-61 |
| chr14-98275404-DEL-54 | chr14-98275405-DEL-54 |
| chr14-98281650-DEL-54 | chr14-98281651-DEL-54 |
| chr14-98305927-DEL-62 | chr14-98305928-DEL-62 |
| chr14-98370238-DEL-56 | chr14-98370259-DEL-58 |
| chr14-98377889-DEL-67 | chr14-98377888-DEL-68 |
| chr14-98387748-DEL-54 | chr14-98387749-DEL-54 |
| chr14-98435524-DEL-62 | chr14-98435525-DEL-62 |
| chr14-98462474-DEL-17155 | chr14-98462475-DEL-17155 |
| chr14-98545159-DEL-327 | chr14-98545160-DEL-327 |
| chr14-98564925-DEL-52 | chr14-98564925-DEL-52 |
| chr14-98583220-DEL-258 | chr14-98583221-DEL-258 |
| chr14-98588817-DEL-1939 | chr14-98588818-DEL-1939 |
| chr14-98594450-DEL-378 | chr14-98594451-DEL-378 |
| chr14-98632379-DEL-370 | chr14-98632380-DEL-370 |
| chr14-98687977-DEL-210 | chr14-98687978-DEL-210 |
| chr14-98704939-DEL-111 | chr14-98704934-DEL-77 |
| chr14-98715221-DEL-54 | chr14-98715222-DEL-54 |
| chr14-98717836-DEL-69 | chr14-98717841-DEL-50 |
| chr14-98781741-DEL-78 | chr14-98781752-DEL-56 |
| chr14-98785839-DEL-76 | chr14-98785840-DEL-50 |
| chr14-98788233-DEL-4681 | chr14-98788234-DEL-4681 |
| chr14-98788839-DEL-67 | chr14-98788840-DEL-67 |
| chr14-98819117-DEL-338 | chr14-98819118-DEL-338 |
| chr14-98820022-DEL-93 | chr14-98820022-DEL-94 |
| chr14-98820834-DEL-98 | chr14-98820830-DEL-62 |
| chr14-98835436-DEL-204 | chr14-98835437-DEL-204 |
| chr14-98875176-DEL-130 | chr14-98875177-DEL-130 |
| chr14-98891260-DEL-133 | chr14-98891279-DEL-96 |
| chr14-98899665-DEL-92 | chr14-98899666-DEL-92 |
| chr14-98903896-DEL-1905 | chr14-98903878-DEL-1917 |
| chr14-98910561-DEL-88 | chr14-98910562-DEL-100 |
| chr14-98914866-DEL-111 | chr14-98914867-DEL-111 |
| chr14-98933437-DEL-90 | chr14-98933438-DEL-86 |
| chr14-98934464-DEL-669 | chr14-98934465-DEL-669 |
| chr14-98969412-DEL-6611 | chr14-98969413-DEL-6611 |
| chr14-98992351-DEL-2704 | chr14-98992352-DEL-2704 |
| chr14-98996573-DEL-127 | chr14-98996573-DEL-128 |
| chr14-99008170-DEL-92 | chr14-99008180-DEL-64 |
| chr14-99072853-DEL-156 | chr14-99072854-DEL-156 |
| chr14-99105302-DEL-76 | chr14-99105302-DEL-88 |
| chr14-99140712-DEL-69 | chr14-99140713-DEL-69 |
| chr14-99198094-DEL-89 | chr14-99198095-DEL-89 |
| chr14-99205334-DEL-1447 | chr14-99205335-DEL-1447 |
| chr14-99238477-DEL-55 | chr14-99238478-DEL-55 |
| chr14-99258209-DEL-95 | chr14-99258210-DEL-95 |
| chr14-99271178-DEL-159 | chr14-99271179-DEL-159 |
| chr14-99271696-DEL-91 | chr14-99271746-DEL-54 |
| chr14-99285586-DEL-100 | chr14-99285560-DEL-127 |
| chr14-99293215-DEL-106 | chr14-99293216-DEL-100 |
| chr14-99299998-DEL-194 | chr14-99299980-DEL-215 |

| | |
|---|---|
| chr14-99308985-DEL-205 | chr14-99308986-DEL-205 |
| chr14-99330219-DEL-1539 | chr14-99330220-DEL-1539 |
| chr14-99331875-DEL-454 | chr14-99331876-DEL-454 |
| chr14-99339813-DEL-72 | chr14-99339806-DEL-120 |
| chr14-99341403-DEL-290 | chr14-99341404-DEL-290 |
| chr14-99360563-DEL-2473 | chr14-99360576-DEL-2461 |
| chr14-99408839-DEL-212 | chr14-99408840-DEL-212 |
| chr14-99413830-DEL-165 | chr14-99413831-DEL-132 |
| chr14-99429030-DEL-66 | chr14-99429031-DEL-66 |
| chr14-99444477-DEL-1180 | chr14-99444478-DEL-1180 |
| chr14-99449527-DEL-123 | chr14-99449528-DEL-123 |
| chr14-99481700-DEL-193 | chr14-99481701-DEL-193 |
| chr14-99487121-DEL-217 | chr14-99487122-DEL-217 |
| chr14-99504557-DEL-54 | chr14-99504560-DEL-52 |
| chr14-99508948-DEL-54 | chr14-99508949-DEL-54 |
| chr14-99510110-DEL-63 | chr14-99510111-DEL-63 |
| chr14-99510648-DEL-199 | chr14-99510649-DEL-199 |
| chr14-99520599-DEL-1129 | chr14-99520603-DEL-1117 |
| chr14-99531294-DEL-77 | chr14-99531316-DEL-56 |
| chr14-99553446-DEL-221 | chr14-99553447-DEL-221 |
| chr14-99561567-DEL-55 | chr14-99561562-DEL-56 |
| chr14-99594468-DEL-226 | chr14-99594469-DEL-226 |
| chr14-99595510-DEL-180 | chr14-99595511-DEL-180 |
| chr14-99598798-DEL-206 | chr14-99598799-DEL-206 |
| chr14-99607501-DEL-198 | chr14-99607502-DEL-198 |
| chr14-99609379-DEL-60 | chr14-99609368-DEL-80 |
| chr14-99615062-DEL-60 | chr14-99615063-DEL-60 |
| chr14-99621173-DEL-166 | chr14-99621174-DEL-172 |
| chr14-99627859-DEL-57 | chr14-99627864-DEL-85 |
| chr14-99634637-DEL-211 | chr14-99634638-DEL-211 |
| chr14-99643149-DEL-62 | chr14-99643150-DEL-62 |
| chr14-99646465-DEL-109 | chr14-99646466-DEL-109 |
| chr14-99648082-DEL-73 | chr14-99648083-DEL-73 |
| chr14-99664911-DEL-80 | chr14-99664912-DEL-79 |
| chr14-99665719-DEL-201 | chr14-99665719-DEL-202 |
| chr14-99679567-DEL-124 | chr14-99679568-DEL-124 |
| chr14-99683173-DEL-1394 | chr14-99683174-DEL-1394 |
| chr14-99688852-DEL-91 | chr14-99688853-DEL-91 |
| chr14-99697042-DEL-90 | chr14-99697042-DEL-81 |
| chr14-99699071-DEL-160 | chr14-99699072-DEL-160 |
| chr14-99717426-DEL-87 | chr14-99717444-DEL-102 |
| chr14-99758661-DEL-60 | chr14-99758662-DEL-60 |
| chr14-99776990-DEL-52 | chr14-99776991-DEL-52 |
| chr14-99792507-DEL-200 | chr14-99792508-DEL-200 |
| chr14-99808982-DEL-117 | chr14-99808985-DEL-115 |
| chr14-99865348-DEL-1950 | chr14-99865349-DEL-1950 |
| chr14-99878649-DEL-52 | chr14-99878650-DEL-52 |
| chr14-99884845-DEL-62 | chr14-99884846-DEL-62 |
| chr14-99888374-DEL-168 | chr14-99888373-DEL-169 |
| chr14-99896404-DEL-169 | chr14-99896405-DEL-169 |
| chr14-99903851-DEL-622 | chr14-99903852-DEL-622 |
| chr14-99909677-DEL-180 | chr14-99909678-DEL-180 |
| chr14-99935554-DEL-171 | chr14-99935555-DEL-177 |
| chr14-99976909-DEL-56 | chr14-99976919-DEL-70 |
| chr14-99999407-DEL-71 | chr14-99999408-DEL-71 |
| chr14-99999946-DEL-91 | chr14-99999947-DEL-91 |
| chr14-100002134-DEL-215 | chr14-100002135-DEL-215 |
| chr14-100004451-DEL-388 | chr14-100004452-DEL-388 |
| chr14-100008212-DEL-315 | chr14-100008213-DEL-315 |
| chr14-100045096-DEL-473 | chr14-100045097-DEL-474 |
| chr14-100052920-DEL-4849 | chr14-100052921-DEL-4849 |
| chr14-100053398-DEL-2172 | chr14-100053397-DEL-2173 |
| chr14-100085396-DEL-138 | chr14-100085397-DEL-138 |
| chr14-100100721-DEL-198 | chr14-100100711-DEL-192 |
| chr14-100116836-DEL-296 | chr14-100116837-DEL-296 |
| chr14-100168623-DEL-62 | chr14-100168624-DEL-62 |
| chr14-100252044-DEL-138 | chr14-100252045-DEL-138 |
| chr14-100265695-DEL-315 | chr14-100265696-DEL-315 |
| chr14-100284582-DEL-66 | chr14-100284583-DEL-66 |
| chr14-100285815-DEL-52 | chr14-100285805-DEL-53 |
| chr14-100292701-DEL-6699 | chr14-100292689-DEL-6700 |
| chr14-100360465-DEL-95 | chr14-100360465-DEL-97 |
| chr14-100368850-DEL-84 | chr14-100368851-DEL-84 |
| chr14-100398624-DEL-109 | chr14-100398625-DEL-108 |
| chr14-100402553-DEL-289 | chr14-100402554-DEL-289 |
| chr14-100456806-DEL-403 | chr14-100456825-DEL-385 |
| chr14-100482347-DEL-66 | chr14-100482348-DEL-66 |
| chr14-100486030-DEL-7113 | chr14-100486031-DEL-7113 |
| chr14-100494231-DEL-1457 | chr14-100494232-DEL-1457 |
| chr14-100496242-DEL-1680 | chr14-100496243-DEL-1680 |

| | |
|---|---|
| chr14-100509667-DEL-69 | chr14-100509670-DEL-70 |
| chr14-100538152-DEL-1190 | chr14-100538153-DEL-1190 |
| chr14-100565900-DEL-56 | chr14-100565893-DEL-62 |
| chr14-100584643-DEL-367 | chr14-100584644-DEL-367 |
| chr14-100592972-DEL-89 | chr14-100592973-DEL-89 |
| chr14-100616516-DEL-10721 | chr14-100616517-DEL-10721 |
| chr14-100642682-DEL-75 | chr14-100642682-DEL-75 |
| chr14-100686939-DEL-200 | chr14-100686940-DEL-200 |
| chr14-100692664-DEL-70 | chr14-100692665-DEL-70 |
| chr14-100707075-DEL-176 | chr14-100707076-DEL-176 |
| chr14-100736985-DEL-1848 | chr14-100736986-DEL-1849 |
| chr14-100755685-DEL-510 | chr14-100755686-DEL-510 |
| chr14-100825889-DEL-158 | chr14-100825880-DEL-124 |
| chr14-100832881-DEL-120 | chr14-100832850-DEL-152 |
| chr14-100848355-DEL-92 | chr14-100848360-DEL-88 |
| chr14-100863018-DEL-268 | chr14-100863019-DEL-268 |
| chr14-100886541-DEL-6803 | chr14-100886542-DEL-6803 |
| chr14-100907585-DEL-590 | chr14-100907586-DEL-590 |
| chr14-100913669-DEL-9974 | chr14-100913670-DEL-9974 |
| chr14-100922370-DEL-132 | chr14-100922371-DEL-128 |
| chr14-100930472-DEL-2992 | chr14-100930473-DEL-2992 |
| chr14-100933784-DEL-965 | chr14-100933785-DEL-965 |
| chr14-100942614-DEL-239 | chr14-100942615-DEL-239 |
| chr14-100960363-DEL-96 | chr14-100960364-DEL-96 |
| chr14-100974752-DEL-6600 | chr14-100974751-DEL-6601 |
| chr14-101013587-DEL-2220 | chr14-101013588-DEL-2220 |
| chr14-101032569-DEL-7065 | chr14-101032570-DEL-7065 |
| chr14-101038704-DEL-212 | chr14-101038704-DEL-212 |
| chr14-101052515-DEL-345 | chr14-101052510-DEL-343 |
| chr14-101067098-DEL-53 | chr14-101067099-DEL-51 |
| chr14-101069116-DEL-4319 | chr14-101069117-DEL-4319 |
| chr14-101093357-DEL-3195 | chr14-101093358-DEL-3195 |
| chr14-101148284-DEL-150 | chr14-101148288-DEL-105 |
| chr14-101199614-DEL-420 | chr14-101199615-DEL-420 |
| chr14-101223311-DEL-52 | chr14-101223312-DEL-52 |
| chr14-101230618-DEL-1822 | chr14-101230619-DEL-1822 |
| chr14-101242956-DEL-97 | chr14-101242957-DEL-97 |
| chr14-101250655-DEL-54 | chr14-101250656-DEL-60 |
| chr14-101251934-DEL-69 | chr14-101251966-DEL-56 |
| chr14-101262075-DEL-356 | chr14-101262074-DEL-350 |
| chr14-101313820-DEL-6452 | chr14-101313821-DEL-6452 |
| chr14-101320405-DEL-72 | chr14-101320403-DEL-54 |
| chr14-101345872-DEL-1611 | chr14-101345876-DEL-1599 |
| chr14-101372998-DEL-109 | chr14-101372999-DEL-110 |
| chr14-101387333-DEL-81 | chr14-101387332-DEL-83 |
| chr14-101387569-DEL-994 | chr14-101387569-DEL-994 |
| chr14-101397301-DEL-97 | chr14-101397302-DEL-97 |
| chr14-101399186-DEL-4862 | chr14-101399187-DEL-4862 |
| chr14-101425277-DEL-298 | chr14-101425278-DEL-298 |
| chr14-101452131-DEL-99 | chr14-101452132-DEL-99 |
| chr14-101482241-DEL-138 | chr14-101482236-DEL-144 |
| chr14-101493128-DEL-71 | chr14-101493128-DEL-72 |
| chr14-101494215-DEL-78 | chr14-101494180-DEL-82 |
| chr14-101495920-DEL-366 | chr14-101495921-DEL-366 |
| chr14-101502763-DEL-887 | chr14-101502764-DEL-887 |
| chr14-101511522-DEL-391 | chr14-101511523-DEL-391 |
| chr14-101552748-DEL-72 | chr14-101552749-DEL-72 |
| chr14-101593534-DEL-88 | chr14-101593530-DEL-78 |
| chr14-101594914-DEL-60 | chr14-101594923-DEL-52 |
| chr14-101597137-DEL-53 | chr14-101597138-DEL-53 |
| chr14-101601048-DEL-3012 | chr14-101601049-DEL-3012 |
| chr14-101602723-DEL-952 | chr14-101602724-DEL-952 |
| chr14-101671233-DEL-399 | chr14-101671228-DEL-400 |
| chr14-101674722-DEL-75 | chr14-101674723-DEL-75 |
| chr14-101676390-DEL-324 | chr14-101676391-DEL-324 |
| chr14-101701127-DEL-464 | chr14-101701132-DEL-496 |
| chr14-101735140-DEL-543 | chr14-101735141-DEL-543 |
| chr14-101818590-DEL-3053 | chr14-101818591-DEL-3053 |
| chr14-101843178-DEL-180 | chr14-101843179-DEL-180 |
| chr14-101851611-DEL-54 | chr14-101851594-DEL-51 |
| chr14-101880288-DEL-242 | chr14-101880289-DEL-242 |
| chr14-101893768-DEL-136 | chr14-101893769-DEL-136 |
| chr14-101897282-DEL-52 | chr14-101897283-DEL-52 |
| chr14-101898117-DEL-211 | chr14-101898118-DEL-211 |
| chr14-101932719-DEL-244 | chr14-101932721-DEL-243 |
| chr14-101959558-DEL-279 | chr14-101959567-DEL-274 |
| chr14-101972353-DEL-54 | chr14-101972356-DEL-51 |
| chr14-101985246-DEL-54 | chr14-101985247-DEL-54 |
| chr14-101991671-DEL-105 | chr14-101991635-DEL-140 |
| chr14-101992488-DEL-68 | chr14-101992494-DEL-82 |

| | |
|---|---|
| chr14-101992899-DEL-364 | chr14-101992900-DEL-364 |
| chr14-101995282-DEL-196 | chr14-101995283-DEL-196 |
| chr14-102007251-DEL-105 | chr14-102007230-DEL-151 |
| chr14-102008014-DEL-64 | chr14-102007998-DEL-65 |
| chr14-102025862-DEL-88 | chr14-102025877-DEL-90 |
| chr14-102026871-DEL-93 | chr14-102026872-DEL-93 |
| chr14-102035086-DEL-60 | chr14-102035095-DEL-60 |
| chr14-102046984-DEL-6434 | chr14-102047010-DEL-6409 |
| chr14-102066747-DEL-278 | chr14-102066745-DEL-279 |
| chr14-102082495-DEL-335 | chr14-102082505-DEL-327 |
| chr14-102096234-DEL-206 | chr14-102096235-DEL-206 |
| chr14-102100819-DEL-2291 | chr14-102100820-DEL-2290 |
| chr14-102103278-DEL-116 | chr14-102103279-DEL-116 |
| chr14-102195123-DEL-6245 | chr14-102195124-DEL-6245 |
| chr14-102208002-DEL-100 | chr14-102208047-DEL-56 |
| chr14-102208758-DEL-493 | chr14-102208759-DEL-493 |
| chr14-102219361-DEL-905 | chr14-102219355-DEL-906 |
| chr14-102227345-DEL-52 | chr14-102227346-DEL-52 |
| chr14-102259868-DEL-215 | chr14-102259869-DEL-215 |
| chr14-102279594-DEL-227 | chr14-102279598-DEL-224 |
| chr14-102284309-DEL-75 | chr14-102284319-DEL-66 |
| chr14-102289656-DEL-1839 | chr14-102289657-DEL-1839 |
| chr14-102300052-DEL-133 | chr14-102300053-DEL-133 |
| chr14-102387843-DEL-52 | chr14-102387844-DEL-52 |
| chr14-102402001-DEL-167 | chr14-102402002-DEL-167 |
| chr14-102424535-DEL-688 | chr14-102424536-DEL-688 |
| chr14-102513230-DEL-90 | chr14-102513231-DEL-90 |
| chr14-102542798-DEL-765 | chr14-102542799-DEL-765 |
| chr14-102559346-DEL-175 | chr14-102559347-DEL-175 |
| chr14-102572329-DEL-107 | chr14-102572328-DEL-99 |
| chr14-102578903-DEL-456 | chr14-102578904-DEL-456 |
| chr14-102639365-DEL-78 | chr14-102639366-DEL-78 |
| chr14-102668629-DEL-197 | chr14-102668630-DEL-197 |
| chr14-102706125-DEL-229 | chr14-102706126-DEL-229 |
| chr14-102711433-DEL-1006 | chr14-102711434-DEL-1003 |
| chr14-102715014-DEL-571 | chr14-102715015-DEL-571 |
| chr14-102724110-DEL-68 | chr14-102724111-DEL-68 |
| chr14-102735622-DEL-237 | chr14-102735622-DEL-237 |
| chr14-102767616-DEL-300 | chr14-102767617-DEL-300 |
| chr14-102808093-DEL-612 | chr14-102808094-DEL-612 |
| chr14-102814193-DEL-80 | chr14-102814194-DEL-80 |
| chr14-102836625-DEL-68 | chr14-102836602-DEL-75 |
| chr14-102844038-DEL-959 | chr14-102844039-DEL-959 |
| chr14-102845085-DEL-160 | chr14-102845082-DEL-160 |
| chr14-102847357-DEL-253 | chr14-102847358-DEL-253 |
| chr14-102899080-DEL-51 | chr14-102899062-DEL-52 |
| chr14-102903167-DEL-1040 | chr14-102903168-DEL-1040 |
| chr14-102915143-DEL-299 | chr14-102915144-DEL-299 |
| chr14-102920452-DEL-1750 | chr14-102920453-DEL-1750 |
| chr14-102939325-DEL-206 | chr14-102939326-DEL-206 |
| chr14-102946741-DEL-687 | chr14-102946742-DEL-687 |
| chr14-102950936-DEL-60 | chr14-102950937-DEL-60 |
| chr14-102967214-DEL-215 | chr14-102967215-DEL-215 |
| chr14-102987981-DEL-320 | chr14-102987982-DEL-320 |
| chr14-102999200-DEL-792 | chr14-102999200-DEL-792 |
| chr14-103024557-DEL-106 | chr14-103024558-DEL-106 |
| chr14-103050173-DEL-516 | chr14-103050174-DEL-516 |
| chr14-103050989-DEL-2875 | chr14-103050991-DEL-2863 |
| chr14-103082328-DEL-6382 | chr14-103082329-DEL-6382 |
| chr14-103098770-DEL-104 | chr14-103098787-DEL-72 |
| chr14-103123263-DEL-383 | chr14-103123264-DEL-383 |
| chr14-103195177-DEL-183 | chr14-103195178-DEL-183 |
| chr14-103233243-DEL-6723 | chr14-103233244-DEL-6723 |
| chr14-103300225-DEL-71 | chr14-103300219-DEL-78 |
| chr14-103300753-DEL-1595 | chr14-103300753-DEL-1595 |
| chr14-103321821-DEL-104 | chr14-103321822-DEL-104 |
| chr14-103385788-DEL-54 | chr14-103385789-DEL-54 |
| chr14-103400232-DEL-74 | chr14-103400233-DEL-74 |
| chr14-103451730-DEL-376 | chr14-103451731-DEL-376 |
| chr14-103456044-DEL-246 | chr14-103456045-DEL-246 |
| chr14-103456870-DEL-197 | chr14-103456871-DEL-197 |
| chr14-103466056-DEL-60 | chr14-103466057-DEL-60 |
| chr14-103476333-DEL-303 | chr14-103476334-DEL-303 |
| chr14-103480615-DEL-99 | chr14-103480616-DEL-99 |
| chr14-103503248-DEL-70 | chr14-103503248-DEL-70 |
| chr14-103543211-DEL-287 | chr14-103543213-DEL-286 |
| chr14-103570221-DEL-62 | chr14-103570222-DEL-62 |
| chr14-103607681-DEL-1273 | chr14-103607682-DEL-1273 |
| chr14-103632028-DEL-70 | chr14-103632029-DEL-64 |
| chr14-103635729-DEL-86 | chr14-103635720-DEL-94 |

| | |
|---|---|
| chr14-103643601-DEL-613 | chr14-103643602-DEL-612 |
| chr14-103644965-DEL-6500 | chr14-103644966-DEL-6500 |
| chr14-103694084-DEL-72 | chr14-103694084-DEL-50 |
| chr14-103710801-DEL-74 | chr14-103710798-DEL-72 |
| chr14-103713913-DEL-743 | chr14-103713914-DEL-743 |
| chr14-103719535-DEL-603 | chr14-103719536-DEL-602 |
| chr14-103722441-DEL-275 | chr14-103722441-DEL-273 |
| chr14-103763481-DEL-76 | chr14-103763481-DEL-72 |
| chr14-103822203-DEL-136 | chr14-103822204-DEL-136 |
| chr14-103857948-DEL-1036 | chr14-103857949-DEL-1036 |
| chr14-103872722-DEL-7073 | chr14-103872723-DEL-7073 |
| chr14-103913966-DEL-229 | chr14-103913967-DEL-229 |
| chr14-103929335-DEL-93 | chr14-103929336-DEL-93 |
| chr14-103933616-DEL-88 | chr14-103933617-DEL-88 |
| chr14-103934220-DEL-1854 | chr14-103934221-DEL-1854 |
| chr14-103947817-DEL-211 | chr14-103947818-DEL-210 |
| chr14-103950085-DEL-52 | chr14-103950090-DEL-79 |
| chr14-103972218-DEL-6440 | chr14-103972218-DEL-6440 |
| chr14-104018063-DEL-1591 | chr14-104018064-DEL-1589 |
| chr14-104031275-DEL-54 | chr14-104031272-DEL-58 |
| chr14-104161906-DEL-453 | chr14-104161925-DEL-435 |
| chr14-104165180-DEL-418 | chr14-104165180-DEL-418 |
| chr14-104168755-DEL-200 | chr14-104168755-DEL-200 |
| chr14-104193034-DEL-120 | chr14-104193053-DEL-102 |
| chr14-104197030-DEL-1348 | chr14-104197031-DEL-1348 |
| chr14-104200400-DEL-103 | chr14-104200397-DEL-127 |
| chr14-104235164-DEL-2382 | chr14-104235165-DEL-2382 |
| chr14-104236823-DEL-1198 | chr14-104236824-DEL-1186 |
| chr14-104250214-DEL-109 | chr14-104250215-DEL-109 |
| chr14-104256710-DEL-6219 | chr14-104256711-DEL-6219 |
| chr14-104292898-DEL-1420 | chr14-104292899-DEL-1420 |
| chr14-104302643-DEL-6885 | chr14-104302644-DEL-6885 |
| chr14-104315568-DEL-62 | chr14-104315569-DEL-62 |
| chr14-104359135-DEL-6560 | chr14-104359136-DEL-6560 |
| chr14-104370292-DEL-159 | chr14-104370281-DEL-165 |
| chr14-104373768-DEL-70 | chr14-104373766-DEL-54 |
| chr14-104377386-DEL-74 | chr14-104377390-DEL-71 |
| chr14-104391249-DEL-92 | chr14-104391251-DEL-80 |
| chr14-104416264-DEL-813 | chr14-104416266-DEL-812 |
| chr14-104457845-DEL-562 | chr14-104457846-DEL-562 |
| chr14-104468117-DEL-283 | chr14-104468118-DEL-283 |
| chr14-104519941-DEL-1105 | chr14-104519942-DEL-1105 |
| chr14-104523275-DEL-56 | chr14-104523272-DEL-60 |
| chr14-104554874-DEL-63 | chr14-104554875-DEL-63 |
| chr14-104555417-DEL-80 | chr14-104555420-DEL-78 |
| chr14-104560540-DEL-68 | chr14-104560541-DEL-68 |
| chr14-104574427-DEL-90 | chr14-104574415-DEL-106 |
| chr14-104584655-DEL-183 | chr14-104584636-DEL-178 |
| chr14-104634364-DEL-81 | chr14-104634358-DEL-82 |
| chr14-104649590-DEL-340 | chr14-104649593-DEL-338 |
| chr14-104664019-DEL-217 | chr14-104664020-DEL-217 |
| chr14-104723068-DEL-75 | chr14-104723069-DEL-75 |
| chr14-104733579-DEL-99 | chr14-104733581-DEL-98 |
| chr14-104746325-DEL-73 | chr14-104746326-DEL-73 |
| chr14-104778687-DEL-52 | chr14-104778640-DEL-52 |
| chr14-104791787-DEL-5446 | chr14-104791788-DEL-5446 |
| chr14-104820046-DEL-7110 | chr14-104820047-DEL-7110 |
| chr14-104900993-DEL-54 | chr14-104900994-DEL-54 |
| chr14-104920363-DEL-90 | chr14-104920364-DEL-90 |
| chr14-104938011-DEL-197 | chr14-104938012-DEL-181 |
| chr14-104942323-DEL-7547 | chr14-104942324-DEL-7547 |
| chr14-104948613-DEL-425 | chr14-104948605-DEL-426 |
| chr14-104953188-DEL-331 | chr14-104953189-DEL-331 |
| chr14-104962178-DEL-1036 | chr14-104962174-DEL-1041 |
| chr14-104973022-DEL-583 | chr14-104973023-DEL-583 |
| chr14-104975518-DEL-352 | chr14-104975518-DEL-351 |
| chr14-105009884-DEL-74 | chr14-105009896-DEL-51 |
| chr14-105017089-DEL-400 | chr14-105017089-DEL-400 |
| chr14-105032890-DEL-56 | chr14-105032890-DEL-56 |
| chr14-105038629-DEL-92 | chr14-105038634-DEL-108 |
| chr14-105041829-DEL-56 | chr14-105041832-DEL-54 |
| chr14-105043290-DEL-56 | chr14-105043291-DEL-56 |
| chr14-105046904-DEL-52 | chr14-105046905-DEL-52 |
| chr14-105083269-DEL-576 | chr14-105083270-DEL-576 |
| chr14-105089828-DEL-223 | chr14-105089829-DEL-223 |
| chr14-105106590-DEL-64 | chr14-105106580-DEL-107 |
| chr14-105115578-DEL-405 | chr14-105115579-DEL-405 |
| chr14-105127656-DEL-352 | chr14-105127640-DEL-336 |
| chr14-105185047-DEL-106 | chr14-105185049-DEL-108 |
| chr14-105214806-DEL-94 | chr14-105214809-DEL-92 |

| | |
|---|---|
| chr14-105224343-DEL-6239 | chr14-105224354-DEL-6239 |
| chr14-105282589-DEL-6282 | chr14-105282590-DEL-6282 |
| chr14-105311332-DEL-103 | chr14-105311333-DEL-103 |
| chr14-105329259-DEL-109 | chr14-105329260-DEL-109 |
| chr14-105337604-DEL-135 | chr14-105337625-DEL-115 |
| chr14-105420711-DEL-187 | chr14-105420712-DEL-186 |
| chr14-105422838-DEL-2121 | chr14-105422839-DEL-2121 |
| chr14-105431001-DEL-714 | chr14-105431002-DEL-714 |
| chr14-105445067-DEL-6634 | chr14-105445068-DEL-6634 |
| chr14-105465342-DEL-2717 | chr14-105465343-DEL-2717 |
| chr14-105486115-DEL-90 | chr14-105486116-DEL-90 |
| chr14-105515536-DEL-171 | chr14-105515537-DEL-171 |
| chr14-105534394-DEL-66 | chr14-105534395-DEL-66 |
| chr14-105536074-DEL-196 | chr14-105536075-DEL-196 |
| chr14-105541812-DEL-121 | chr14-105541813-DEL-121 |
| chr14-105574706-DEL-574 | chr14-105574697-DEL-575 |
| chr14-105582026-DEL-633 | chr14-105582027-DEL-633 |
| chr14-105584800-DEL-716 | chr14-105584801-DEL-716 |
| chr14-105593263-DEL-276 | chr14-105593244-DEL-296 |
| chr14-105610216-DEL-1499 | chr14-105610217-DEL-1499 |
| chr14-105612928-DEL-86 | chr14-105612909-DEL-60 |
| chr14-105620187-DEL-444 | chr14-105620188-DEL-444 |
| chr14-105639191-DEL-358 | chr14-105639192-DEL-358 |
| chr14-105707756-DEL-152 | chr14-105707755-DEL-153 |
| chr14-105732295-DEL-184 | chr14-105732313-DEL-190 |
| chr14-105772598-DEL-78 | chr14-105772599-DEL-60 |
| chr14-105781748-DEL-161 | chr14-105781749-DEL-161 |
| chr14-105786623-DEL-2396 | chr14-105786624-DEL-2396 |
| chr14-105848877-DEL-91 | chr14-105848878-DEL-91 |
| chr14-105857065-DEL-77 | chr14-105857048-DEL-78 |
| chr14-105919245-DEL-415 | chr14-105919246-DEL-415 |
| chr14-105986904-DEL-61 | chr14-105986905-DEL-61 |
| chr14-105994826-DEL-7046 | chr14-105994826-DEL-7046 |
| chr14-106000193-DEL-369 | chr14-106000194-DEL-369 |
| chr14-106022706-DEL-106 | chr14-106022707-DEL-106 |
| chr14-106043709-DEL-2504 | chr14-106043710-DEL-2504 |
| chr14-106054541-DEL-191 | chr14-106054542-DEL-191 |
| chr14-106060439-DEL-59 | chr14-106060440-DEL-59 |
| chr14-106067488-DEL-80 | chr14-106067485-DEL-84 |
| chr14-106072366-DEL-177 | chr14-106072367-DEL-177 |
| chr14-106072760-DEL-466 | chr14-106072761-DEL-466 |
| chr14-106111215-DEL-600 | chr14-106111216-DEL-600 |
| chr14-106112500-DEL-418 | chr14-106112501-DEL-408 |
| chr14-106118611-DEL-527 | chr14-106118612-DEL-527 |
| chr14-106142008-DEL-116 | chr14-106142009-DEL-116 |
| chr14-106143871-DEL-57 | chr14-106143872-DEL-57 |
| chr14-106171102-DEL-128 | chr14-106171103-DEL-128 |
| chr14-106216917-DEL-72 | chr14-106216918-DEL-72 |
| chr14-106240120-DEL-3025 | chr14-106240121-DEL-3025 |
| chr14-106256269-DEL-1608 | chr14-106256270-DEL-1608 |
| chr14-106332123-DEL-987 | chr14-106332123-DEL-986 |
| chr14-106335510-DEL-331 | chr14-106335511-DEL-331 |
| chr14-106336473-DEL-200 | chr14-106336474-DEL-200 |
| chr14-106341499-DEL-61 | chr14-106341500-DEL-61 |
| chr14-106355245-DEL-358 | chr14-106355246-DEL-358 |
| chr14-106369974-DEL-69 | chr14-106369979-DEL-62 |
| chr14-106417818-DEL-191 | chr14-106417835-DEL-173 |
| chr14-106424575-DEL-73 | chr14-106424554-DEL-74 |
| chr14-106478150-DEL-54 | chr14-106478151-DEL-54 |
| chr14-106501140-DEL-104 | chr14-106501126-DEL-114 |
| chr14-106514316-DEL-999 | chr14-106514317-DEL-999 |
| chr14-106516458-DEL-52 | chr14-106516418-DEL-52 |
| chr14-106538287-DEL-6557 | chr14-106538288-DEL-6557 |
| chr14-106561819-DEL-62 | chr14-106561820-DEL-62 |
| chr14-106567091-DEL-105 | chr14-106567067-DEL-150 |
| chr14-106568894-DEL-7804 | chr14-106568895-DEL-7804 |
| chr14-106621408-DEL-475 | chr14-106621409-DEL-475 |
| chr14-106639364-DEL-54 | chr14-106639365-DEL-54 |
| chr14-106644147-DEL-64 | chr14-106644148-DEL-64 |
| chr14-106645485-DEL-295 | chr14-106645486-DEL-295 |
| chr14-106733239-DEL-60 | chr14-106733244-DEL-60 |
| chr14-106735987-DEL-696 | chr14-106735988-DEL-696 |
| chr14-106737257-DEL-93 | chr14-106737258-DEL-93 |
| chr14-106749687-DEL-76 | chr14-106749688-DEL-76 |
| chr14-106770698-DEL-69 | chr14-106770699-DEL-69 |
| chr14-106776962-DEL-63 | chr14-106776957-DEL-64 |
| chr14-106783713-DEL-893 | chr14-106783714-DEL-893 |
| chr14-106812798-DEL-196 | chr14-106812799-DEL-196 |
| chr14-106851416-DEL-208 | chr14-106851415-DEL-205 |
| chr14-106856975-DEL-848 | chr14-106856973-DEL-851 |

| | |
|---|---|
| chr14-106878660-DEL-2616 | chr14-106878660-DEL-2616 |
| chr14-106878677-DEL-56 | chr14-106878677-DEL-57 |
| chr14-106888746-DEL-308 | chr14-106888747-DEL-308 |
| chr14-106897900-DEL-357 | chr14-106897901-DEL-357 |
| chr14-106914625-DEL-104 | chr14-106914626-DEL-104 |
| chr14-106958341-DEL-7836 | chr14-106958342-DEL-7836 |
| chr14-106979863-DEL-135 | chr14-106979850-DEL-136 |
| chr14-106991657-DEL-70 | chr14-106991658-DEL-70 |
| chr14-107019360-DEL-3633 | chr14-107019361-DEL-3633 |
| chr14-107020875-DEL-450 | chr14-107020876-DEL-450 |
| chr14-107029554-DEL-59 | chr14-107029555-DEL-65 |
| chr14-107031322-DEL-75 | chr14-107031323-DEL-75 |
| chr14-107054440-DEL-647 | chr14-107054441-DEL-647 |
| chr14-107063134-DEL-2651 | chr14-107063135-DEL-2651 |
| chr14-107093254-DEL-3126 | chr14-107093255-DEL-3126 |
| chr14-107101722-DEL-102 | chr14-107101772-DEL-60 |
| chr14-107114548-DEL-64 | chr14-107114549-DEL-62 |
| chr14-107116439-DEL-108 | chr14-107116440-DEL-108 |
| chr14-107123409-DEL-130 | chr14-107123410-DEL-130 |
| chr14-107132718-DEL-1542 | chr14-107132719-DEL-1542 |
| chr14-107163132-DEL-6800 | chr14-107163133-DEL-6800 |
| chr14-107163339-DEL-416 | chr14-107163340-DEL-416 |
| chr14-107173447-DEL-976 | chr14-107173446-DEL-977 |
| chr14-107179847-DEL-437 | chr14-107179848-DEL-437 |
| chr14-107211020-DEL-205 | chr14-107211021-DEL-205 |
| chr14-107270232-DEL-1373 | chr14-107270233-DEL-1373 |
| chr14-107360830-DEL-10496 | chr14-107360831-DEL-10496 |
| chr14-107364313-DEL-919 | chr14-107364314-DEL-919 |
| chr14-107377279-DEL-804 | chr14-107377280-DEL-804 |
| chr14-107389715-DEL-6403 | chr14-107389716-DEL-6403 |
| chr14-107396888-DEL-7221 | chr14-107396883-DEL-7222 |
| chr14-107431461-DEL-247 | chr14-107431462-DEL-247 |
| chr14-107486783-DEL-73 | chr14-107486792-DEL-61 |
| chr14-107494124-DEL-72 | chr14-107494125-DEL-72 |
| chr14-107499342-DEL-1181 | chr14-107499343-DEL-1181 |
| chr14-107512226-DEL-53 | chr14-107512223-DEL-54 |
| chr14-107543716-DEL-401 | chr14-107543728-DEL-395 |
| chr14-107616102-DEL-187 | chr14-107616103-DEL-187 |
| chr14-107630033-DEL-72 | chr14-107630018-DEL-80 |
| chr14-107651313-DEL-452 | chr14-107651314-DEL-452 |
| chr14-107656614-DEL-532 | chr14-107656615-DEL-532 |
| chr14-107666955-DEL-1052 | chr14-107666956-DEL-1053 |
| chr14-107673312-DEL-62 | chr14-107673323-DEL-57 |
| chr14-107675946-DEL-143 | chr14-107675947-DEL-143 |
| chr14-107685185-DEL-94 | chr14-107685186-DEL-94 |
| chr14-107717123-DEL-134 | chr14-107717124-DEL-134 |
| chr14-107721900-DEL-2812 | chr14-107721901-DEL-2812 |
| chr14-107746452-DEL-66 | chr14-107746453-DEL-66 |
| chr14-107759101-DEL-68 | chr14-107759102-DEL-68 |
| chr14-107788893-DEL-128 | chr14-107788894-DEL-128 |
| chr14-107796489-DEL-776 | chr14-107796489-DEL-776 |
| chr14-107816630-DEL-66 | chr14-107816633-DEL-52 |
| chr14-107862394-DEL-1251 | chr14-107862395-DEL-1251 |
| chr14-107953013-DEL-206 | chr14-107953019-DEL-205 |
| chr14-107976568-DEL-108 | chr14-107976569-DEL-108 |
| chr14-107983488-DEL-98 | chr14-107983489-DEL-98 |
| chr14-107994062-DEL-6674 | chr14-107994063-DEL-6674 |
| chr14-107994527-DEL-211 | chr14-107994528-DEL-211 |
| chr14-108085465-DEL-164 | chr14-108085463-DEL-161 |
| chr14-108091286-DEL-842 | chr14-108091288-DEL-841 |
| chr14-108100750-DEL-218 | chr14-108100751-DEL-218 |
| chr14-108115118-DEL-574 | chr14-108115119-DEL-574 |
| chr14-108160707-DEL-1128 | chr14-108160708-DEL-1128 |
| chr14-108185330-DEL-160 | chr14-108185331-DEL-160 |
| chr14-108245746-DEL-475 | chr14-108245749-DEL-473 |
| chr14-108247335-DEL-302 | chr14-108247336-DEL-302 |
| chr14-108250746-DEL-108 | chr14-108250747-DEL-108 |
| chr14-108270307-DEL-488 | chr14-108270308-DEL-488 |
| chr14-108277914-DEL-909 | chr14-108277915-DEL-909 |
| chr14-108300169-DEL-652 | chr14-108300170-DEL-652 |
| chr14-108324792-DEL-57 | chr14-108324808-DEL-52 |
| chr14-108359174-DEL-68 | chr14-108359175-DEL-68 |
| chr14-108401768-DEL-1596 | chr14-108401768-DEL-1596 |
| chr14-108403853-DEL-2575 | chr14-108403850-DEL-2576 |
| chr14-108406837-DEL-5475 | chr14-108406838-DEL-5475 |
| chr14-108447582-DEL-397 | chr14-108447583-DEL-397 |
| chr14-108457246-DEL-144 | chr14-108457247-DEL-136 |
| chr14-108503630-DEL-368 | chr14-108503630-DEL-368 |
| chr14-108506985-DEL-526 | chr14-108506986-DEL-526 |
| chr14-108554466-DEL-391 | chr14-108554467-DEL-391 |

| | |
|---|---|
| chr14-108564305-DEL-527 | chr14-108564306-DEL-527 |
| chr14-108568619-DEL-312 | chr14-108568619-DEL-313 |
| chr14-108575439-DEL-2772 | chr14-108575464-DEL-2748 |
| chr14-108589269-DEL-124 | chr14-108589241-DEL-124 |
| chr14-108621008-DEL-84 | chr14-108621008-DEL-64 |
| chr14-108637796-DEL-79 | chr14-108637788-DEL-64 |
| chr14-108648309-DEL-104 | chr14-108648310-DEL-104 |
| chr14-108648446-DEL-912 | chr14-108648494-DEL-878 |
| chr14-108657164-DEL-1281 | chr14-108657161-DEL-1282 |
| chr14-108658547-DEL-66 | chr14-108658532-DEL-66 |
| chr14-108669034-DEL-149 | chr14-108669035-DEL-149 |
| chr14-108691274-DEL-93 | chr14-108691275-DEL-93 |
| chr14-108693384-DEL-64 | chr14-108693385-DEL-64 |
| chr14-108697612-DEL-58 | chr14-108697613-DEL-58 |
| chr14-108736979-DEL-256 | chr14-108736980-DEL-256 |
| chr14-108757627-DEL-411 | chr14-108757628-DEL-411 |
| chr14-108769445-DEL-182 | chr14-108769446-DEL-181 |
| chr14-108776239-DEL-64 | chr14-108776243-DEL-61 |
| chr14-108780707-DEL-122 | chr14-108780724-DEL-106 |
| chr14-108786089-DEL-1577 | chr14-108786090-DEL-1577 |
| chr14-108797729-DEL-604 | chr14-108797730-DEL-604 |
| chr14-108808425-DEL-72 | chr14-108808425-DEL-72 |
| chr14-108809907-DEL-853 | chr14-108809907-DEL-853 |
| chr14-108814454-DEL-710 | chr14-108814455-DEL-710 |
| chr14-108816248-DEL-58 | chr14-108816247-DEL-59 |
| chr14-108817214-DEL-58 | chr14-108817204-DEL-74 |
| chr14-108832875-DEL-922 | chr14-108832876-DEL-922 |
| chr14-108841992-DEL-199 | chr14-108841993-DEL-199 |
| chr14-108843670-DEL-102 | chr14-108843680-DEL-96 |
| chr14-108858868-DEL-61 | chr14-108858869-DEL-61 |
| chr14-108877544-DEL-760 | chr14-108877544-DEL-770 |
| chr14-108884266-DEL-114 | chr14-108884267-DEL-114 |
| chr14-109001974-DEL-52 | chr14-109001975-DEL-50 |
| chr14-109008309-DEL-348 | chr14-109008310-DEL-348 |
| chr14-109063192-DEL-70 | chr14-109063192-DEL-74 |
| chr14-109066893-DEL-192 | chr14-109066894-DEL-192 |
| chr14-109069013-DEL-218 | chr14-109069014-DEL-218 |
| chr14-109073522-DEL-128 | chr14-109073510-DEL-126 |
| chr14-109079019-DEL-458 | chr14-109079019-DEL-458 |
| chr14-109108933-DEL-90 | chr14-109108940-DEL-56 |
| chr14-109139305-DEL-60 | chr14-109139273-DEL-89 |
| chr14-109143073-DEL-101 | chr14-109143074-DEL-101 |
| chr14-109179852-DEL-268 | chr14-109179856-DEL-268 |
| chr14-109187900-DEL-6204 | chr14-109187901-DEL-6204 |
| chr14-109195719-DEL-79 | chr14-109195734-DEL-59 |
| chr14-109202065-DEL-171 | chr14-109202066-DEL-169 |
| chr14-109204452-DEL-56 | chr14-109204453-DEL-54 |
| chr14-109213063-DEL-880 | chr14-109213064-DEL-880 |
| chr14-109232541-DEL-110 | chr14-109232537-DEL-78 |
| chr14-109258810-DEL-499 | chr14-109258811-DEL-499 |
| chr14-109275054-DEL-491 | chr14-109275055-DEL-491 |
| chr14-109285854-DEL-1395 | chr14-109285855-DEL-1395 |
| chr14-109301649-DEL-186 | chr14-109301650-DEL-186 |
| chr14-109316773-DEL-60 | chr14-109316774-DEL-58 |
| chr14-109320322-DEL-122 | chr14-109320323-DEL-122 |
| chr14-109332121-DEL-65 | chr14-109332115-DEL-66 |
| chr14-109363201-DEL-324 | chr14-109363201-DEL-324 |
| chr14-109381045-DEL-66 | chr14-109381049-DEL-54 |
| chr14-109388005-DEL-54 | chr14-109388005-DEL-54 |
| chr14-109405638-DEL-982 | chr14-109405639-DEL-982 |
| chr14-109414286-DEL-211 | chr14-109414287-DEL-211 |
| chr14-109416787-DEL-616 | chr14-109416788-DEL-616 |
| chr14-109428709-DEL-2065 | chr14-109428710-DEL-2065 |
| chr14-109431635-DEL-1489 | chr14-109431636-DEL-1489 |
| chr14-109487477-DEL-60 | chr14-109487478-DEL-60 |
| chr14-109514241-DEL-500 | chr14-109514242-DEL-500 |
| chr14-109539840-DEL-1039 | chr14-109539841-DEL-1039 |
| chr14-109540834-DEL-185 | chr14-109540835-DEL-185 |
| chr14-109543279-DEL-189 | chr14-109543280-DEL-189 |
| chr14-109554768-DEL-1236 | chr14-109554769-DEL-1232 |
| chr14-109612174-DEL-204 | chr14-109612171-DEL-208 |
| chr14-109624608-DEL-1124 | chr14-109624611-DEL-1122 |
| chr14-109637015-DEL-487 | chr14-109637016-DEL-487 |
| chr14-109646150-DEL-1465 | chr14-109646151-DEL-1465 |
| chr14-109654691-DEL-63 | chr14-109654692-DEL-63 |
| chr14-109675984-DEL-188 | chr14-109675986-DEL-187 |
| chr14-109681135-DEL-68 | chr14-109681136-DEL-68 |
| chr14-109683971-DEL-823 | chr14-109683972-DEL-823 |
| chr14-109690475-DEL-80 | chr14-109690476-DEL-80 |
| chr14-109713534-DEL-76 | chr14-109713535-DEL-76 |

| | |
|---|---|
| chr14-109730246-DEL-536 | chr14-109730247-DEL-536 |
| chr14-109768724-DEL-445 | chr14-109768725-DEL-445 |
| chr14-109791140-DEL-269 | chr14-109791141-DEL-269 |
| chr14-109796803-DEL-311 | chr14-109796804-DEL-311 |
| chr14-109805141-DEL-55 | chr14-109805142-DEL-55 |
| chr14-109811340-DEL-140 | chr14-109811341-DEL-140 |
| chr14-109818504-DEL-2055 | chr14-109818505-DEL-2055 |
| chr14-109841282-DEL-3654 | chr14-109841283-DEL-3654 |
| chr14-109849424-DEL-276 | chr14-109849419-DEL-288 |
| chr14-109851312-DEL-180 | chr14-109851313-DEL-180 |
| chr14-109865205-DEL-395 | chr14-109865206-DEL-395 |
| chr14-109868049-DEL-3409 | chr14-109868050-DEL-3409 |
| chr14-109881281-DEL-1887 | chr14-109881282-DEL-1883 |
| chr14-109887178-DEL-56 | chr14-109887179-DEL-54 |
| chr14-109899547-DEL-120 | chr14-109899547-DEL-120 |
| chr14-109906461-DEL-810 | chr14-109906462-DEL-810 |
| chr14-109909926-DEL-738 | chr14-109909927-DEL-738 |
| chr14-109913660-DEL-66 | chr14-109913659-DEL-66 |
| chr14-109914230-DEL-472 | chr14-109914231-DEL-472 |
| chr14-109917438-DEL-1218 | chr14-109917439-DEL-1218 |
| chr14-109945933-DEL-90 | chr14-109945937-DEL-87 |
| chr14-109950352-DEL-1664 | chr14-109950353-DEL-1663 |
| chr14-109970351-DEL-170 | chr14-109970352-DEL-170 |
| chr14-109978304-DEL-6204 | chr14-109978305-DEL-6204 |
| chr14-110008579-DEL-108 | chr14-110008584-DEL-104 |
| chr14-110064781-DEL-7116 | chr14-110064782-DEL-7116 |
| chr14-110086890-DEL-306 | chr14-110086891-DEL-306 |
| chr14-110109733-DEL-632 | chr14-110109734-DEL-632 |
| chr14-110144461-DEL-140 | chr14-110144462-DEL-140 |
| chr14-110148990-DEL-6381 | chr14-110148991-DEL-6381 |
| chr14-110188827-DEL-124 | chr14-110188824-DEL-125 |
| chr14-110199577-DEL-2759 | chr14-110199578-DEL-2759 |
| chr14-110205366-DEL-2307 | chr14-110205367-DEL-2307 |
| chr14-110224088-DEL-63 | chr14-110224089-DEL-63 |
| chr14-110247534-DEL-693 | chr14-110247535-DEL-693 |
| chr14-110267995-DEL-59 | chr14-110267992-DEL-63 |
| chr14-110268201-DEL-686 | chr14-110268202-DEL-686 |
| chr14-110278452-DEL-511 | chr14-110278453-DEL-511 |
| chr14-110281242-DEL-195 | chr14-110281241-DEL-196 |
| chr14-110291060-DEL-148 | chr14-110291061-DEL-148 |
| chr14-110292466-DEL-7160 | chr14-110292467-DEL-7160 |
| chr14-110312875-DEL-289 | chr14-110312876-DEL-289 |
| chr14-110314171-DEL-725 | chr14-110314171-DEL-725 |
| chr14-110317124-DEL-7075 | chr14-110317125-DEL-7075 |
| chr14-110333587-DEL-62 | chr14-110333585-DEL-71 |
| chr14-110335779-DEL-87 | chr14-110335780-DEL-87 |
| chr14-110339238-DEL-6400 | chr14-110339239-DEL-6400 |
| chr14-110348747-DEL-100 | chr14-110348753-DEL-95 |
| chr14-110360489-DEL-79 | chr14-110360490-DEL-79 |
| chr14-110373270-DEL-6044 | chr14-110373270-DEL-6043 |
| chr14-110381519-DEL-112 | chr14-110381520-DEL-112 |
| chr14-110384170-DEL-639 | chr14-110384171-DEL-639 |
| chr14-110384904-DEL-770 | chr14-110384905-DEL-770 |
| chr14-110441853-DEL-324 | chr14-110441854-DEL-324 |
| chr14-110445248-DEL-85 | chr14-110445249-DEL-85 |
| chr14-110480833-DEL-778 | chr14-110480834-DEL-778 |
| chr14-110498454-DEL-1004 | chr14-110498455-DEL-1004 |
| chr14-110502301-DEL-59 | chr14-110502302-DEL-59 |
| chr14-110520947-DEL-54 | chr14-110520954-DEL-50 |
| chr14-110527713-DEL-65 | chr14-110527714-DEL-65 |
| chr14-110531332-DEL-510 | chr14-110531333-DEL-510 |
| chr14-110533902-DEL-200 | chr14-110533903-DEL-200 |
| chr14-110537097-DEL-90 | chr14-110537094-DEL-94 |
| chr14-110537834-DEL-1893 | chr14-110537835-DEL-1893 |
| chr14-110541600-DEL-464 | chr14-110541600-DEL-464 |
| chr14-110542367-DEL-1501 | chr14-110542367-DEL-1500 |
| chr14-110545747-DEL-2746 | chr14-110545748-DEL-2746 |
| chr14-110555336-DEL-287 | chr14-110555337-DEL-287 |
| chr14-110557690-DEL-196 | chr14-110557691-DEL-196 |
| chr14-110590772-DEL-287 | chr14-110590773-DEL-287 |
| chr14-110593243-DEL-9713 | chr14-110593243-DEL-9713 |
| chr14-110608539-DEL-456 | chr14-110608539-DEL-453 |
| chr14-110639915-DEL-3588 | chr14-110639916-DEL-3588 |
| chr14-110645447-DEL-1626 | chr14-110645448-DEL-1626 |
| chr14-110682506-DEL-130 | chr14-110682509-DEL-128 |
| chr14-110688405-DEL-64 | chr14-110688406-DEL-64 |
| chr14-110693402-DEL-1789 | chr14-110693403-DEL-1789 |
| chr14-110704072-DEL-85 | chr14-110704076-DEL-82 |
| chr14-110729544-DEL-4227 | chr14-110729545-DEL-4227 |
| chr14-110738700-DEL-67 | chr14-110738701-DEL-67 |

| | |
|---|---|
| chr14-110758058-DEL-1490 | chr14-110758059-DEL-1490 |
| chr14-110784686-DEL-6121 | chr14-110784679-DEL-6120 |
| chr14-110791927-DEL-80 | chr14-110791928-DEL-80 |
| chr14-110799858-DEL-154 | chr14-110799859-DEL-154 |
| chr14-110834981-DEL-173 | chr14-110834982-DEL-173 |
| chr14-110867594-DEL-2694 | chr14-110867595-DEL-2694 |
| chr14-110868880-DEL-1316 | chr14-110868880-DEL-1316 |
| chr14-110884410-DEL-1296 | chr14-110884412-DEL-1293 |
| chr14-110891943-DEL-53 | chr14-110891934-DEL-53 |
| chr14-110956200-DEL-128 | chr14-110956204-DEL-125 |
| chr14-110976411-DEL-168 | chr14-110976412-DEL-168 |
| chr14-110978550-DEL-85 | chr14-110978560-DEL-72 |
| chr14-110991898-DEL-60 | chr14-110991899-DEL-60 |
| chr14-111005288-DEL-54 | chr14-111005291-DEL-50 |
| chr14-111006056-DEL-53 | chr14-111006070-DEL-50 |
| chr14-111011793-DEL-53 | chr14-111011794-DEL-53 |
| chr14-111025543-DEL-74 | chr14-111025544-DEL-74 |
| chr14-111031264-DEL-2992 | chr14-111031265-DEL-2992 |
| chr14-111046991-DEL-471 | chr14-111046992-DEL-471 |
| chr14-111128004-DEL-614 | chr14-111128005-DEL-614 |
| chr14-111141777-DEL-6272 | chr14-111141778-DEL-6272 |
| chr14-111162823-DEL-417 | chr14-111162817-DEL-418 |
| chr14-111163534-DEL-1336 | chr14-111163535-DEL-1336 |
| chr14-111190499-DEL-758 | chr14-111190500-DEL-758 |
| chr14-111220130-DEL-848 | chr14-111220127-DEL-844 |
| chr14-111222405-DEL-1481 | chr14-111222406-DEL-1481 |
| chr14-111232963-DEL-190 | chr14-111232964-DEL-190 |
| chr14-111264513-DEL-74 | chr14-111264532-DEL-65 |
| chr14-111284013-DEL-72 | chr14-111284014-DEL-72 |
| chr14-111290186-DEL-59 | chr14-111290187-DEL-59 |
| chr14-111297400-DEL-230 | chr14-111297396-DEL-227 |
| chr14-111297945-DEL-5980 | chr14-111297970-DEL-5978 |
| chr14-111308081-DEL-74 | chr14-111308082-DEL-70 |
| chr14-111317917-DEL-2664 | chr14-111317918-DEL-2664 |
| chr14-111334655-DEL-790 | chr14-111334644-DEL-791 |
| chr14-111341463-DEL-74 | chr14-111341464-DEL-74 |
| chr14-111345169-DEL-1545 | chr14-111345169-DEL-1545 |
| chr14-111347851-DEL-180 | chr14-111347855-DEL-176 |
| chr14-111394100-DEL-2058 | chr14-111394101-DEL-2058 |
| chr14-111397254-DEL-141 | chr14-111397255-DEL-141 |
| chr14-111414721-DEL-77 | chr14-111414722-DEL-77 |
| chr14-111422097-DEL-165 | chr14-111422098-DEL-165 |
| chr14-111435192-DEL-76 | chr14-111435193-DEL-76 |
| chr14-111437812-DEL-206 | chr14-111437813-DEL-206 |
| chr14-111461532-DEL-2155 | chr14-111461533-DEL-2155 |
| chr14-111497684-DEL-68 | chr14-111497687-DEL-50 |
| chr14-111502976-DEL-2034 | chr14-111502977-DEL-2034 |
| chr14-111503123-DEL-773 | chr14-111503124-DEL-773 |
| chr14-111504415-DEL-353 | chr14-111504420-DEL-342 |
| chr14-111508263-DEL-2627 | chr14-111508264-DEL-2627 |
| chr14-111524794-DEL-80 | chr14-111524795-DEL-80 |
| chr14-111526120-DEL-210 | chr14-111526121-DEL-210 |
| chr14-111541455-DEL-211 | chr14-111541456-DEL-211 |
| chr14-111551287-DEL-60 | chr14-111551288-DEL-60 |
| chr14-111605090-DEL-63 | chr14-111605091-DEL-63 |
| chr14-111612905-DEL-390 | chr14-111612905-DEL-390 |
| chr14-111617291-DEL-4981 | chr14-111617292-DEL-4981 |
| chr14-111644618-DEL-775 | chr14-111644621-DEL-804 |
| chr14-111646657-DEL-1002 | chr14-111646693-DEL-967 |
| chr14-111649396-DEL-377 | chr14-111649412-DEL-372 |
| chr14-111652281-DEL-797 | chr14-111652280-DEL-798 |
| chr14-111661974-DEL-2174 | chr14-111661975-DEL-2170 |
| chr14-111666901-DEL-114 | chr14-111666902-DEL-114 |
| chr14-111693604-DEL-343 | chr14-111693614-DEL-339 |
| chr14-111699264-DEL-137 | chr14-111699265-DEL-137 |
| chr14-111729498-DEL-560 | chr14-111729499-DEL-560 |
| chr14-111766268-DEL-60 | chr14-111766269-DEL-60 |
| chr14-111784281-DEL-89 | chr14-111784284-DEL-87 |
| chr14-111827747-DEL-327 | chr14-111827747-DEL-327 |
| chr14-111841177-DEL-101 | chr14-111841178-DEL-101 |
| chr14-111846472-DEL-57 | chr14-111846473-DEL-57 |
| chr14-111859320-DEL-1447 | chr14-111859320-DEL-1447 |
| chr14-111897169-DEL-54 | chr14-111897170-DEL-54 |
| chr14-111902374-DEL-5085 | chr14-111902377-DEL-5082 |
| chr14-112034259-DEL-370 | chr14-112034249-DEL-361 |
| chr14-112043703-DEL-68 | chr14-112043683-DEL-68 |
| chr14-112070147-DEL-5929 | chr14-112070148-DEL-5929 |
| chr14-112077910-DEL-100 | chr14-112077935-DEL-68 |
| chr14-112080033-DEL-2450 | chr14-112080034-DEL-2450 |
| chr14-112082661-DEL-848 | chr14-112082662-DEL-849 |

| | |
|---|---|
| chr14-112111232-DEL-2946 | chr14-112111233-DEL-2946 |
| chr14-112127382-DEL-105 | chr14-112127383-DEL-105 |
| chr14-112140615-DEL-6764 | chr14-112140616-DEL-6764 |
| chr14-112151975-DEL-580 | chr14-112151976-DEL-580 |
| chr14-112172888-DEL-3063 | chr14-112172889-DEL-3062 |
| chr14-112182613-DEL-216 | chr14-112182614-DEL-216 |
| chr14-112250446-DEL-294 | chr14-112250441-DEL-278 |
| chr14-112272753-DEL-649 | chr14-112272754-DEL-649 |
| chr14-112293175-DEL-607 | chr14-112293176-DEL-607 |
| chr14-112298499-DEL-767 | chr14-112298495-DEL-765 |
| chr14-112312106-DEL-152 | chr14-112312131-DEL-108 |
| chr14-112338334-DEL-351 | chr14-112338334-DEL-351 |
| chr14-112362438-DEL-203 | chr14-112362432-DEL-204 |
| chr14-112365138-DEL-6444 | chr14-112365139-DEL-6444 |
| chr14-112380645-DEL-567 | chr14-112380694-DEL-520 |
| chr14-112386663-DEL-111 | chr14-112386664-DEL-111 |
| chr14-112388585-DEL-56 | chr14-112388586-DEL-54 |
| chr14-112409857-DEL-54 | chr14-112409858-DEL-57 |
| chr14-112415238-DEL-1828 | chr14-112415246-DEL-1822 |
| chr14-112447627-DEL-6279 | chr14-112447628-DEL-6279 |
| chr14-112486915-DEL-104 | chr14-112486919-DEL-101 |
| chr14-112492274-DEL-817 | chr14-112492275-DEL-817 |
| chr14-112500117-DEL-2329 | chr14-112500118-DEL-2329 |
| chr14-112510422-DEL-243 | chr14-112510423-DEL-241 |
| chr14-112518135-DEL-920 | chr14-112518136-DEL-920 |
| chr14-112521448-DEL-8366 | chr14-112521443-DEL-8367 |
| chr14-112524910-DEL-1892 | chr14-112524911-DEL-1892 |
| chr14-112570245-DEL-7137 | chr14-112570246-DEL-7137 |
| chr14-112603563-DEL-5542 | chr14-112603562-DEL-5543 |
| chr14-112628319-DEL-5842 | chr14-112628320-DEL-5842 |
| chr14-112642767-DEL-110 | chr14-112642765-DEL-112 |
| chr14-112663774-DEL-6756 | chr14-112663775-DEL-6756 |
| chr14-112676220-DEL-901 | chr14-112676221-DEL-901 |
| chr14-112680379-DEL-52 | chr14-112680382-DEL-50 |
| chr14-112687600-DEL-3439 | chr14-112687601-DEL-3439 |
| chr14-112699292-DEL-536 | chr14-112699293-DEL-536 |
| chr14-112718221-DEL-5866 | chr14-112718198-DEL-5892 |
| chr14-112722301-DEL-423 | chr14-112722302-DEL-423 |
| chr14-112727689-DEL-1020 | chr14-112727689-DEL-1017 |
| chr14-112732586-DEL-129 | chr14-112732587-DEL-129 |
| chr14-112735309-DEL-444 | chr14-112735310-DEL-444 |
| chr14-112742320-DEL-2063 | chr14-112742321-DEL-2063 |
| chr14-112745006-DEL-56 | chr14-112744991-DEL-72 |
| chr14-112756326-DEL-69 | chr14-112756327-DEL-69 |
| chr14-112769436-DEL-180 | chr14-112769437-DEL-180 |
| chr14-112775500-DEL-1278 | chr14-112775501-DEL-1278 |
| chr14-112776896-DEL-63 | chr14-112776897-DEL-63 |
| chr14-112782487-DEL-393 | chr14-112782481-DEL-394 |
| chr14-112786524-DEL-336 | chr14-112786537-DEL-337 |
| chr14-112820765-DEL-51 | chr14-112820766-DEL-51 |
| chr14-112833622-DEL-7644 | chr14-112833623-DEL-7644 |
| chr14-112834892-DEL-435 | chr14-112834892-DEL-435 |
| chr14-112847892-DEL-60 | chr14-112847893-DEL-60 |
| chr14-112852933-DEL-56 | chr14-112852934-DEL-52 |
| chr14-112856462-DEL-71 | chr14-112856463-DEL-71 |
| chr14-112865182-DEL-7089 | chr14-112865183-DEL-7089 |
| chr14-112896258-DEL-7801 | chr14-112896259-DEL-7801 |
| chr14-112904601-DEL-224 | chr14-112904601-DEL-217 |
| chr14-112960751-DEL-1447 | chr14-112960754-DEL-1448 |
| chr14-113012472-DEL-1343 | chr14-113012472-DEL-1343 |
| chr14-113034907-DEL-57 | chr14-113034908-DEL-57 |
| chr14-113043293-DEL-97 | chr14-113043294-DEL-97 |
| chr14-113051449-DEL-3284 | chr14-113051451-DEL-3283 |
| chr14-113055286-DEL-336 | chr14-113055287-DEL-336 |
| chr14-113126855-DEL-106 | chr14-113126858-DEL-72 |
| chr14-113136739-DEL-650 | chr14-113136740-DEL-650 |
| chr14-113141078-DEL-497 | chr14-113141071-DEL-498 |
| chr14-113199702-DEL-4009 | chr14-113199703-DEL-4009 |
| chr14-113210890-DEL-58 | chr14-113210893-DEL-56 |
| chr14-113255892-DEL-7205 | chr14-113255893-DEL-7205 |
| chr14-113271740-DEL-4544 | chr14-113271741-DEL-4544 |
| chr14-113280938-DEL-52 | chr14-113280935-DEL-56 |
| chr14-113294127-DEL-1144 | chr14-113294125-DEL-1147 |
| chr14-113299656-DEL-68 | chr14-113299629-DEL-95 |
| chr14-113365517-DEL-84 | chr14-113365521-DEL-75 |
| chr14-113411352-DEL-144 | chr14-113411353-DEL-144 |
| chr14-113416265-DEL-56 | chr14-113416266-DEL-56 |
| chr14-113419961-DEL-1783 | chr14-113419962-DEL-1783 |
| chr14-113425881-DEL-185 | chr14-113425882-DEL-185 |
| chr14-113444798-DEL-847 | chr14-113444799-DEL-847 |

| | |
|---|---|
| chr14-113449424-DEL-72 | chr14-113449425-DEL-72 |
| chr14-113481546-DEL-519 | chr14-113481547-DEL-519 |
| chr14-113487571-DEL-9229 | chr14-113487572-DEL-9229 |
| chr14-113493826-DEL-208 | chr14-113493827-DEL-208 |
| chr14-113525520-DEL-5291 | chr14-113525521-DEL-5291 |
| chr14-113552041-DEL-2130 | chr14-113552025-DEL-2144 |
| chr14-113573721-DEL-535 | chr14-113573722-DEL-563 |
| chr14-113574879-DEL-136 | chr14-113574881-DEL-135 |
| chr14-113608049-DEL-398 | chr14-113608050-DEL-398 |
| chr14-113640894-DEL-88 | chr14-113640894-DEL-88 |
| chr14-113642472-DEL-348 | chr14-113642467-DEL-340 |
| chr14-113740989-DEL-56 | chr14-113740990-DEL-56 |
| chr14-113751559-DEL-1681 | chr14-113751560-DEL-1681 |
| chr14-113761889-DEL-430 | chr14-113761890-DEL-430 |
| chr14-113775806-DEL-2911 | chr14-113775807-DEL-2911 |
| chr14-113803188-DEL-91 | chr14-113803156-DEL-128 |
| chr14-113822696-DEL-66 | chr14-113822697-DEL-66 |
| chr14-113830171-DEL-3507 | chr14-113830172-DEL-3507 |
| chr14-113831029-DEL-1691 | chr14-113831029-DEL-1691 |
| chr14-113838220-DEL-440 | chr14-113838221-DEL-440 |
| chr14-113940125-DEL-81 | chr14-113940106-DEL-89 |
| chr14-113946099-DEL-100 | chr14-113946100-DEL-100 |
| chr14-113991468-DEL-742 | chr14-113991469-DEL-742 |
| chr14-114017174-DEL-909 | chr14-114017175-DEL-909 |
| chr14-114022001-DEL-3111 | chr14-114021986-DEL-3115 |
| chr14-114022307-DEL-99 | chr14-114022339-DEL-66 |
| chr14-114058786-DEL-518 | chr14-114058787-DEL-518 |
| chr14-114059712-DEL-368 | chr14-114059739-DEL-390 |
| chr14-114076282-DEL-3883 | chr14-114076283-DEL-3883 |
| chr14-114076652-DEL-304 | chr14-114076653-DEL-304 |
| chr14-114102853-DEL-336 | chr14-114102854-DEL-336 |
| chr14-114109261-DEL-6212 | chr14-114109263-DEL-6211 |
| chr14-114119400-DEL-1396 | chr14-114119401-DEL-1396 |
| chr14-114135510-DEL-353 | chr14-114135508-DEL-356 |
| chr14-114150304-DEL-54 | chr14-114150310-DEL-54 |
| chr14-114212297-DEL-7822 | chr14-114212298-DEL-7822 |
| chr14-114238469-DEL-110 | chr14-114238471-DEL-62 |
| chr14-114253831-DEL-2688 | chr14-114253832-DEL-2688 |
| chr14-114332345-DEL-7987 | chr14-114332345-DEL-7987 |
| chr14-114362318-DEL-63 | chr14-114362317-DEL-64 |
| chr14-114400155-DEL-65 | chr14-114400156-DEL-65 |
| chr14-114441052-DEL-7166 | chr14-114441053-DEL-7166 |
| chr14-114452989-DEL-76 | chr14-114452990-DEL-76 |
| chr14-114474654-DEL-335 | chr14-114474655-DEL-335 |
| chr14-114475119-DEL-2757 | chr14-114475114-DEL-2756 |
| chr14-114535953-DEL-928 | chr14-114535954-DEL-928 |
| chr14-114538487-DEL-55 | chr14-114538488-DEL-55 |
| chr14-114546477-DEL-69 | chr14-114546478-DEL-69 |
| chr14-114556068-DEL-214 | chr14-114556068-DEL-214 |
| chr14-114563362-DEL-313 | chr14-114563363-DEL-313 |
| chr14-114575026-DEL-825 | chr14-114575027-DEL-825 |
| chr14-114585332-DEL-61 | chr14-114585333-DEL-61 |
| chr14-114587203-DEL-498 | chr14-114587204-DEL-498 |
| chr14-114614161-DEL-83 | chr14-114614176-DEL-95 |
| chr14-114614349-DEL-418 | chr14-114614350-DEL-419 |
| chr14-114639313-DEL-74 | chr14-114639315-DEL-75 |
| chr14-114657739-DEL-4117 | chr14-114657740-DEL-4117 |
| chr14-114696097-DEL-1777 | chr14-114696098-DEL-1777 |
| chr14-114729683-DEL-291 | chr14-114729684-DEL-289 |
| chr14-114739782-DEL-175 | chr14-114739783-DEL-175 |
| chr14-114743244-DEL-7152 | chr14-114743245-DEL-7152 |
| chr14-114753209-DEL-55 | chr14-114753196-DEL-56 |
| chr14-114754958-DEL-172 | chr14-114754962-DEL-173 |
| chr14-114762426-DEL-457 | chr14-114762427-DEL-457 |
| chr14-114807029-DEL-74 | chr14-114807022-DEL-80 |
| chr14-114816104-DEL-1870 | chr14-114816109-DEL-1867 |
| chr14-114836218-DEL-235 | chr14-114836215-DEL-236 |
| chr14-114847742-DEL-564 | chr14-114847743-DEL-564 |
| chr14-114853905-DEL-53 | chr14-114853906-DEL-53 |
| chr14-114898161-DEL-5342 | chr14-114898162-DEL-5342 |
| chr14-114904997-DEL-7047 | chr14-114904998-DEL-7047 |
| chr14-114922835-DEL-4111 | chr14-114922836-DEL-4111 |
| chr14-114950898-DEL-345 | chr14-114950899-DEL-345 |
| chr14-114981712-DEL-2017 | chr14-114981713-DEL-2017 |
| chr14-115020410-DEL-110 | chr14-115020411-DEL-110 |
| chr14-115029931-DEL-68 | chr14-115029932-DEL-96 |
| chr14-115046970-DEL-378 | chr14-115046971-DEL-378 |
| chr14-115059484-DEL-1862 | chr14-115059485-DEL-1862 |
| chr14-115069212-DEL-54 | chr14-115069213-DEL-52 |
| chr14-115080882-DEL-124 | chr14-115080876-DEL-125 |

| | |
|---|---|
| chr14-115105061-DEL-334 | chr14-115105062-DEL-334 |
| chr14-115134483-DEL-6883 | chr14-115134483-DEL-6883 |
| chr14-115171268-DEL-701 | chr14-115171269-DEL-701 |
| chr14-115172257-DEL-420 | chr14-115172258-DEL-420 |
| chr14-115175257-DEL-1894 | chr14-115175258-DEL-1894 |
| chr14-115180815-DEL-87 | chr14-115180814-DEL-88 |
| chr14-115211777-DEL-372 | chr14-115211778-DEL-372 |
| chr14-115233665-DEL-838 | chr14-115233666-DEL-838 |
| chr14-115237117-DEL-101 | chr14-115237118-DEL-101 |
| chr14-115347053-DEL-7841 | chr14-115347042-DEL-7842 |
| chr14-115368533-DEL-54 | chr14-115368534-DEL-54 |
| chr14-115377090-DEL-62 | chr14-115377063-DEL-80 |
| chr14-115419816-DEL-76 | chr14-115419829-DEL-64 |
| chr14-115430155-DEL-1472 | chr14-115430156-DEL-1472 |
| chr14-115445205-DEL-697 | chr14-115445178-DEL-710 |
| chr14-115461624-DEL-1957 | chr14-115461625-DEL-1957 |
| chr14-115467837-DEL-98 | chr14-115467862-DEL-72 |
| chr14-115502722-DEL-66 | chr14-115502725-DEL-67 |
| chr14-115509279-DEL-9438 | chr14-115509280-DEL-9438 |
| chr14-115517280-DEL-214 | chr14-115517281-DEL-214 |
| chr14-115520063-DEL-124 | chr14-115520064-DEL-124 |
| chr14-115526479-DEL-96 | chr14-115526486-DEL-66 |
| chr14-115550429-DEL-88 | chr14-115550430-DEL-88 |
| chr14-115560677-DEL-90 | chr14-115560677-DEL-90 |
| chr14-115666573-DEL-468 | chr14-115666563-DEL-469 |
| chr14-115687969-DEL-77 | chr14-115687954-DEL-81 |
| chr14-115691321-DEL-82 | chr14-115691322-DEL-88 |
| chr14-115736838-DEL-60 | chr14-115736843-DEL-56 |
| chr14-115771548-DEL-134 | chr14-115771549-DEL-134 |
| chr14-115809191-DEL-239 | chr14-115809192-DEL-239 |
| chr14-115822342-DEL-91 | chr14-115822343-DEL-91 |
| chr14-115822860-DEL-223 | chr14-115822861-DEL-224 |
| chr14-115863260-DEL-512 | chr14-115863261-DEL-512 |
| chr14-115887437-DEL-6354 | chr14-115887438-DEL-6354 |
| chr14-115896796-DEL-757 | chr14-115896797-DEL-757 |
| chr14-115931902-DEL-221 | chr14-115931904-DEL-238 |
| chr14-115981056-DEL-139 | chr14-115981057-DEL-140 |
| chr14-115994680-DEL-578 | chr14-115994684-DEL-577 |
| chr14-116001940-DEL-212 | chr14-116001941-DEL-212 |
| chr14-116002762-DEL-83 | chr14-116002763-DEL-83 |
| chr14-116045582-DEL-1873 | chr14-116045583-DEL-1873 |
| chr14-116099722-DEL-93 | chr14-116099755-DEL-65 |
| chr14-116109544-DEL-89 | chr14-116109557-DEL-87 |
| chr14-116112309-DEL-381 | chr14-116112310-DEL-381 |
| chr14-116114548-DEL-135 | chr14-116114549-DEL-135 |
| chr14-116121967-DEL-6064 | chr14-116121968-DEL-6064 |
| chr14-116130769-DEL-798 | chr14-116130770-DEL-798 |
| chr14-116156615-DEL-300 | chr14-116156637-DEL-296 |
| chr14-116172243-DEL-55 | chr14-116172236-DEL-52 |
| chr14-116195418-DEL-7068 | chr14-116195419-DEL-7068 |
| chr14-116229770-DEL-93 | chr14-116229771-DEL-93 |
| chr14-116235695-DEL-315 | chr14-116235731-DEL-296 |
| chr14-116243001-DEL-377 | chr14-116243002-DEL-377 |
| chr14-116267504-DEL-278 | chr14-116267505-DEL-278 |
| chr14-116300803-DEL-736 | chr14-116300804-DEL-736 |
| chr14-116311681-DEL-3073 | chr14-116311682-DEL-3073 |
| chr14-116336242-DEL-1313 | chr14-116336243-DEL-1313 |
| chr14-116343838-DEL-78 | chr14-116343835-DEL-60 |
| chr14-116361278-DEL-1539 | chr14-116361279-DEL-1539 |
| chr14-116405133-DEL-141 | chr14-116405134-DEL-141 |
| chr14-116413112-DEL-78 | chr14-116413113-DEL-78 |
| chr14-116464324-DEL-7099 | chr14-116464325-DEL-7099 |
| chr14-116504769-DEL-497 | chr14-116504770-DEL-497 |
| chr14-116524575-DEL-64 | chr14-116524576-DEL-64 |
| chr14-116525082-DEL-110 | chr14-116525076-DEL-142 |
| chr14-116562862-DEL-6313 | chr14-116562863-DEL-6313 |
| chr14-116582736-DEL-214 | chr14-116582737-DEL-214 |
| chr14-116604395-DEL-1767 | chr14-116604396-DEL-1767 |
| chr14-116611404-DEL-77 | chr14-116611405-DEL-78 |
| chr14-116612101-DEL-52 | chr14-116612094-DEL-62 |
| chr14-116640050-DEL-4991 | chr14-116640051-DEL-4991 |
| chr14-116658647-DEL-118 | chr14-116658648-DEL-118 |
| chr14-116659539-DEL-285 | chr14-116659540-DEL-285 |
| chr14-116672927-DEL-6382 | chr14-116672928-DEL-6382 |
| chr14-116682016-DEL-189 | chr14-116682017-DEL-189 |
| chr14-116694570-DEL-69 | chr14-116694547-DEL-93 |
| chr14-116717137-DEL-76 | chr14-116717132-DEL-81 |
| chr14-116725007-DEL-198 | chr14-116725014-DEL-230 |
| chr14-116736725-DEL-80 | chr14-116736720-DEL-104 |
| chr14-116744943-DEL-6450 | chr14-116744944-DEL-6450 |

| | |
|---|---|
| chr14-116755307-DEL-89 | chr14-116755314-DEL-62 |
| chr14-116799952-DEL-66 | chr14-116799953-DEL-66 |
| chr14-116805765-DEL-73 | chr14-116805758-DEL-62 |
| chr14-116808245-DEL-6209 | chr14-116808246-DEL-6209 |
| chr14-116817099-DEL-487 | chr14-116817100-DEL-487 |
| chr14-116840617-DEL-100 | chr14-116840624-DEL-58 |
| chr14-116861508-DEL-389 | chr14-116861509-DEL-389 |
| chr14-116872545-DEL-6437 | chr14-116872545-DEL-6437 |
| chr14-116886958-DEL-240 | chr14-116886959-DEL-240 |
| chr14-116889252-DEL-61 | chr14-116889253-DEL-59 |
| chr14-116915833-DEL-1877 | chr14-116915834-DEL-1877 |
| chr14-116934599-DEL-223 | chr14-116934600-DEL-223 |
| chr14-116949346-DEL-867 | chr14-116949347-DEL-867 |
| chr14-116967614-DEL-648 | chr14-116967608-DEL-643 |
| chr14-117025832-DEL-6683 | chr14-117025833-DEL-6683 |
| chr14-117036017-DEL-54 | chr14-117036022-DEL-50 |
| chr14-117117278-DEL-349 | chr14-117117278-DEL-353 |
| chr14-117119739-DEL-186 | chr14-117119740-DEL-186 |
| chr14-117121711-DEL-65 | chr14-117121712-DEL-65 |
| chr14-117139258-DEL-3316 | chr14-117139259-DEL-3316 |
| chr14-117143807-DEL-61 | chr14-117143808-DEL-61 |
| chr14-117172234-DEL-60 | chr14-117172239-DEL-55 |
| chr14-117235466-DEL-59 | chr14-117235467-DEL-59 |
| chr14-117241706-DEL-83 | chr14-117241703-DEL-83 |
| chr14-117260448-DEL-1470 | chr14-117260449-DEL-1470 |
| chr14-117268143-DEL-51 | chr14-117268144-DEL-51 |
| chr14-117276129-DEL-1704 | chr14-117276126-DEL-1704 |
| chr14-117316380-DEL-4561 | chr14-117316381-DEL-4561 |
| chr14-117324685-DEL-104 | chr14-117324702-DEL-64 |
| chr14-117365675-DEL-97 | chr14-117365676-DEL-97 |
| chr14-117393256-DEL-67 | chr14-117393254-DEL-67 |
| chr14-117416297-DEL-6687 | chr14-117416338-DEL-6660 |
| chr14-117446742-DEL-378 | chr14-117446745-DEL-376 |
| chr14-117460638-DEL-188 | chr14-117460639-DEL-188 |
| chr14-117495618-DEL-6450 | chr14-117495619-DEL-6450 |
| chr14-117495768-DEL-212 | chr14-117495769-DEL-212 |
| chr14-117512558-DEL-987 | chr14-117512562-DEL-984 |
| chr14-117522273-DEL-680 | chr14-117522289-DEL-671 |
| chr14-117523159-DEL-7123 | chr14-117523160-DEL-7123 |
| chr14-117551265-DEL-54 | chr14-117551266-DEL-52 |
| chr14-117559634-DEL-241 | chr14-117559635-DEL-241 |
| chr14-117585619-DEL-122 | chr14-117585620-DEL-122 |
| chr14-117683955-DEL-1239 | chr14-117683956-DEL-1239 |
| chr14-117689440-DEL-69 | chr14-117689434-DEL-74 |
| chr14-117730592-DEL-1137 | chr14-117730593-DEL-1137 |
| chr14-117763439-DEL-63 | chr14-117763425-DEL-53 |
| chr14-117777638-DEL-7868 | chr14-117777639-DEL-7868 |
| chr14-117783933-DEL-1013 | chr14-117783932-DEL-1013 |
| chr14-117790330-DEL-877 | chr14-117790331-DEL-877 |
| chr14-117812766-DEL-225 | chr14-117812767-DEL-225 |
| chr14-117815718-DEL-80 | chr14-117815734-DEL-60 |
| chr14-117822538-DEL-79 | chr14-117822539-DEL-79 |
| chr14-117825267-DEL-1558 | chr14-117825268-DEL-1558 |
| chr14-117842822-DEL-136 | chr14-117842827-DEL-132 |
| chr14-117850897-DEL-3888 | chr14-117850899-DEL-3888 |
| chr14-117853158-DEL-200 | chr14-117853159-DEL-200 |
| chr14-117855241-DEL-52 | chr14-117855242-DEL-52 |
| chr14-117855366-DEL-658 | chr14-117855367-DEL-658 |
| chr14-117900805-DEL-78 | chr14-117900816-DEL-68 |
| chr14-117933520-DEL-256 | chr14-117933521-DEL-256 |
| chr14-117952587-DEL-56 | chr14-117952587-DEL-57 |
| chr14-117958917-DEL-122 | chr14-117958918-DEL-122 |
| chr14-117959413-DEL-305 | chr14-117959414-DEL-305 |
| chr14-117970308-DEL-60 | chr14-117970309-DEL-60 |
| chr14-117983198-DEL-91 | chr14-117983206-DEL-84 |
| chr14-117987196-DEL-66 | chr14-117987235-DEL-64 |
| chr14-118002928-DEL-98 | chr14-118002929-DEL-98 |
| chr14-118007307-DEL-312 | chr14-118007307-DEL-310 |
| chr14-118049430-DEL-180 | chr14-118049431-DEL-180 |
| chr14-118056147-DEL-1338 | chr14-118056147-DEL-1331 |
| chr14-118084118-DEL-950 | chr14-118084119-DEL-950 |
| chr14-118093582-DEL-6988 | chr14-118093583-DEL-6988 |
| chr14-118148798-DEL-482 | chr14-118148797-DEL-483 |
| chr14-118166345-DEL-522 | chr14-118166344-DEL-523 |
| chr14-118170405-DEL-197 | chr14-118170406-DEL-197 |
| chr14-118187103-DEL-102 | chr14-118187104-DEL-102 |
| chr14-118193483-DEL-54 | chr14-118193484-DEL-54 |
| chr14-118214289-DEL-157 | chr14-118214306-DEL-153 |
| chr14-118221991-DEL-84 | chr14-118221991-DEL-84 |
| chr14-118226512-DEL-366 | chr14-118226523-DEL-359 |

| | |
|---|---|
| chr14-118233986-DEL-388 | chr14-118233983-DEL-392 |
| chr14-118236730-DEL-821 | chr14-118236731-DEL-821 |
| chr14-118248820-DEL-481 | chr14-118248821-DEL-481 |
| chr14-118249956-DEL-193 | chr14-118249955-DEL-192 |
| chr14-118251341-DEL-538 | chr14-118251342-DEL-538 |
| chr14-118264400-DEL-207 | chr14-118264397-DEL-207 |
| chr14-118270992-DEL-77 | chr14-118270993-DEL-77 |
| chr14-118336885-DEL-513 | chr14-118336880-DEL-470 |
| chr14-118356974-DEL-55 | chr14-118356974-DEL-55 |
| chr14-118362995-DEL-217 | chr14-118363000-DEL-213 |
| chr14-118380251-DEL-57 | chr14-118380252-DEL-57 |
| chr14-118382555-DEL-1167 | chr14-118382545-DEL-1168 |
| chr14-118394062-DEL-62 | chr14-118394063-DEL-62 |
| chr14-118396733-DEL-82 | chr14-118396769-DEL-82 |
| chr14-118401826-DEL-161 | chr14-118401828-DEL-160 |
| chr14-118402371-DEL-3619 | chr14-118402372-DEL-3619 |
| chr14-118412752-DEL-72 | chr14-118412751-DEL-64 |
| chr14-118420925-DEL-580 | chr14-118420926-DEL-580 |
| chr14-118437279-DEL-142 | chr14-118437280-DEL-142 |
| chr14-118443011-DEL-144 | chr14-118443000-DEL-145 |
| chr14-118450483-DEL-396 | chr14-118450484-DEL-396 |
| chr14-118488025-DEL-53 | chr14-118488026-DEL-53 |
| chr14-118494283-DEL-361 | chr14-118494286-DEL-362 |
| chr14-118504614-DEL-580 | chr14-118504615-DEL-580 |
| chr14-118523712-DEL-183 | chr14-118523713-DEL-183 |
| chr14-118534713-DEL-56 | chr14-118534713-DEL-56 |
| chr14-118579141-DEL-242 | chr14-118579134-DEL-248 |
| chr14-118580958-DEL-58 | chr14-118580933-DEL-50 |
| chr14-118584939-DEL-67 | chr14-118584942-DEL-65 |
| chr14-118614928-DEL-489 | chr14-118614929-DEL-489 |
| chr14-118620762-DEL-198 | chr14-118620773-DEL-188 |
| chr14-118624138-DEL-206 | chr14-118624138-DEL-192 |
| chr14-118647199-DEL-439 | chr14-118647200-DEL-439 |
| chr14-118656498-DEL-67 | chr14-118656499-DEL-67 |
| chr14-118687854-DEL-176 | chr14-118687854-DEL-176 |
| chr14-118696959-DEL-327 | chr14-118696960-DEL-327 |
| chr14-118707434-DEL-112 | chr14-118707431-DEL-112 |
| chr14-118749181-DEL-125 | chr14-118749182-DEL-125 |
| chr14-118750318-DEL-60 | chr14-118750318-DEL-60 |
| chr14-118758243-DEL-106 | chr14-118758247-DEL-107 |
| chr14-118760906-DEL-92 | chr14-118760907-DEL-92 |
| chr14-118767476-DEL-420 | chr14-118767477-DEL-420 |
| chr14-118781020-DEL-50 | chr14-118780978-DEL-72 |
| chr14-118785206-DEL-69 | chr14-118785189-DEL-105 |
| chr14-118791689-DEL-58 | chr14-118791698-DEL-50 |
| chr14-118800549-DEL-129 | chr14-118800550-DEL-129 |
| chr14-118809165-DEL-456 | chr14-118809166-DEL-456 |
| chr14-118831394-DEL-55 | chr14-118831395-DEL-55 |
| chr14-118839053-DEL-115 | chr14-118839054-DEL-115 |
| chr14-118890212-DEL-54 | chr14-118890213-DEL-54 |
| chr14-118892720-DEL-416 | chr14-118892721-DEL-415 |
| chr14-118903978-DEL-847 | chr14-118903979-DEL-847 |
| chr14-118914189-DEL-152 | chr14-118914190-DEL-152 |
| chr14-118989342-DEL-64 | chr14-118989343-DEL-64 |
| chr14-119044445-DEL-56 | chr14-119044446-DEL-72 |
| chr14-119049060-DEL-1316 | chr14-119049061-DEL-1316 |
| chr14-119055193-DEL-112 | chr14-119055194-DEL-112 |
| chr14-119060636-DEL-66 | chr14-119060637-DEL-52 |
| chr14-119132609-DEL-72 | chr14-119132610-DEL-72 |
| chr14-119149638-DEL-118 | chr14-119149639-DEL-118 |
| chr14-119191082-DEL-414 | chr14-119191077-DEL-413 |
| chr14-119194454-DEL-80 | chr14-119194469-DEL-79 |
| chr14-119200306-DEL-1620 | chr14-119200307-DEL-1620 |
| chr14-119201765-DEL-72 | chr14-119201766-DEL-72 |
| chr14-119219638-DEL-393 | chr14-119219639-DEL-393 |
| chr14-119227466-DEL-710 | chr14-119227458-DEL-711 |
| chr14-119228764-DEL-170 | chr14-119228765-DEL-170 |
| chr14-119234439-DEL-51 | chr14-119234406-DEL-62 |
| chr14-119243084-DEL-204 | chr14-119243085-DEL-204 |
| chr14-119244047-DEL-60 | chr14-119244049-DEL-59 |
| chr14-119248785-DEL-123 | chr14-119248786-DEL-123 |
| chr14-119261694-DEL-102 | chr14-119261695-DEL-102 |
| chr14-119281060-DEL-56 | chr14-119281063-DEL-54 |
| chr14-119282553-DEL-147 | chr14-119282554-DEL-147 |
| chr14-119283172-DEL-188 | chr14-119283173-DEL-188 |
| chr14-119285098-DEL-795 | chr14-119285099-DEL-795 |
| chr14-119312180-DEL-58 | chr14-119312195-DEL-50 |
| chr14-119318178-DEL-489 | chr14-119318174-DEL-490 |
| chr14-119340162-DEL-549 | chr14-119340163-DEL-549 |
| chr14-119360743-DEL-76 | chr14-119360744-DEL-76 |

| | |
|---|---|
| chr14-119364903-DEL-146 | chr14-119364904-DEL-146 |
| chr14-119367262-DEL-64 | chr14-119367263-DEL-64 |
| chr14-119370740-DEL-52 | chr14-119370741-DEL-56 |
| chr14-119394032-DEL-146 | chr14-119394033-DEL-146 |
| chr14-119430733-DEL-58 | chr14-119430734-DEL-58 |
| chr14-119432904-DEL-71 | chr14-119432904-DEL-72 |
| chr14-119464902-DEL-59 | chr14-119464903-DEL-59 |
| chr14-119469695-DEL-78 | chr14-119469696-DEL-78 |
| chr14-119477145-DEL-1264 | chr14-119477146-DEL-1263 |
| chr14-119489461-DEL-84 | chr14-119489479-DEL-57 |
| chr14-119506288-DEL-66 | chr14-119506295-DEL-60 |
| chr14-119515683-DEL-2637 | chr14-119515684-DEL-2637 |
| chr14-119551338-DEL-56 | chr14-119551343-DEL-52 |
| chr14-119559253-DEL-143 | chr14-119559254-DEL-143 |
| chr14-119569595-DEL-219 | chr14-119569631-DEL-209 |
| chr14-119589832-DEL-2233 | chr14-119589833-DEL-2233 |
| chr14-119598267-DEL-69 | chr14-119598268-DEL-69 |
| chr14-119612945-DEL-2626 | chr14-119612946-DEL-2626 |
| chr14-119636093-DEL-961 | chr14-119636094-DEL-961 |
| chr14-119664153-DEL-142 | chr14-119664154-DEL-142 |
| chr14-119678888-DEL-3781 | chr14-119678889-DEL-3781 |
| chr14-119695807-DEL-65 | chr14-119695808-DEL-65 |
| chr14-119716031-DEL-4922 | chr14-119716032-DEL-4922 |
| chr14-119737279-DEL-487 | chr14-119737280-DEL-487 |
| chr14-119738178-DEL-462 | chr14-119738179-DEL-462 |
| chr14-119791901-DEL-406 | chr14-119791902-DEL-406 |
| chr14-119801166-DEL-1730 | chr14-119801167-DEL-1730 |
| chr14-119835173-DEL-1814 | chr14-119835174-DEL-1814 |
| chr14-119854794-DEL-384 | chr14-119854795-DEL-384 |
| chr14-119877252-DEL-80 | chr14-119877253-DEL-76 |
| chr14-119882649-DEL-3900 | chr14-119882650-DEL-3900 |
| chr14-119907511-DEL-134 | chr14-119907512-DEL-134 |
| chr14-119925373-DEL-95 | chr14-119925389-DEL-63 |
| chr14-119929204-DEL-6170 | chr14-119929205-DEL-6170 |
| chr14-119938799-DEL-59 | chr14-119938800-DEL-59 |
| chr14-119956363-DEL-52 | chr14-119956364-DEL-52 |
| chr14-119998663-DEL-813 | chr14-119998676-DEL-809 |
| chr14-120022510-DEL-70 | chr14-120022511-DEL-84 |
| chr14-120023214-DEL-1626 | chr14-120023215-DEL-1626 |
| chr14-120031096-DEL-96 | chr14-120031097-DEL-96 |
| chr14-120062008-DEL-107 | chr14-120061991-DEL-108 |
| chr14-120064885-DEL-677 | chr14-120064885-DEL-677 |
| chr14-120136058-DEL-293 | chr14-120136093-DEL-250 |
| chr14-120171123-DEL-6485 | chr14-120171124-DEL-6485 |
| chr14-120174657-DEL-650 | chr14-120174656-DEL-650 |
| chr14-120193483-DEL-7337 | chr14-120193484-DEL-7337 |
| chr14-120236645-DEL-516 | chr14-120236646-DEL-516 |
| chr14-120246007-DEL-56 | chr14-120246000-DEL-52 |
| chr14-120254564-DEL-534 | chr14-120254565-DEL-534 |
| chr14-120261677-DEL-109 | chr14-120261678-DEL-103 |
| chr14-120271765-DEL-243 | chr14-120271766-DEL-243 |
| chr14-120294508-DEL-349 | chr14-120294509-DEL-349 |
| chr14-120299321-DEL-6285 | chr14-120299318-DEL-6269 |
| chr14-120319834-DEL-339 | chr14-120319835-DEL-339 |
| chr14-120325791-DEL-120 | chr14-120325794-DEL-118 |
| chr14-120342134-DEL-917 | chr14-120342135-DEL-916 |
| chr14-120346695-DEL-81 | chr14-120346696-DEL-80 |
| chr14-120351312-DEL-80 | chr14-120351325-DEL-99 |
| chr14-120394110-DEL-63 | chr14-120394107-DEL-66 |
| chr14-120404274-DEL-286 | chr14-120404275-DEL-285 |
| chr14-120416194-DEL-7033 | chr14-120416195-DEL-7033 |
| chr14-120416369-DEL-409 | chr14-120416370-DEL-409 |
| chr14-120434775-DEL-53 | chr14-120434776-DEL-53 |
| chr14-120447058-DEL-681 | chr14-120447059-DEL-681 |
| chr14-120472236-DEL-55 | chr14-120472236-DEL-55 |
| chr14-120555110-DEL-144 | chr14-120555111-DEL-144 |
| chr14-120570843-DEL-52 | chr14-120570844-DEL-52 |
| chr14-120588602-DEL-491 | chr14-120588603-DEL-491 |
| chr14-120589630-DEL-57 | chr14-120589631-DEL-57 |
| chr14-120595408-DEL-280 | chr14-120595409-DEL-280 |
| chr14-120651090-DEL-151 | chr14-120651060-DEL-152 |
| chr14-120673996-DEL-343 | chr14-120673997-DEL-343 |
| chr14-120693959-DEL-1287 | chr14-120693960-DEL-1287 |
| chr14-120699905-DEL-348 | chr14-120699896-DEL-349 |
| chr14-120728580-DEL-3336 | chr14-120728581-DEL-3336 |
| chr14-120760132-DEL-303 | chr14-120760133-DEL-303 |
| chr14-120767990-DEL-60 | chr14-120768011-DEL-64 |
| chr14-120787026-DEL-54 | chr14-120787027-DEL-54 |
| chr14-120787991-DEL-6136 | chr14-120787992-DEL-6136 |
| chr14-120802838-DEL-222 | chr14-120802839-DEL-222 |

| | |
|---|---|
| chr14-120808264-DEL-557 | chr14-120808275-DEL-559 |
| chr14-120837993-DEL-2197 | chr14-120837994-DEL-2197 |
| chr14-120859806-DEL-1353 | chr14-120859807-DEL-1353 |
| chr14-120868807-DEL-164 | chr14-120868789-DEL-165 |
| chr14-120889156-DEL-104 | chr14-120889156-DEL-104 |
| chr14-120911402-DEL-316 | chr14-120911403-DEL-316 |
| chr14-120913948-DEL-151 | chr14-120913949-DEL-151 |
| chr14-120920615-DEL-62 | chr14-120920630-DEL-66 |
| chr14-120921051-DEL-1156 | chr14-120921052-DEL-1156 |
| chr14-120953220-DEL-1019 | chr14-120953221-DEL-1019 |
| chr14-120963028-DEL-61 | chr14-120962998-DEL-66 |
| chr14-120966891-DEL-95 | chr14-120966897-DEL-51 |
| chr14-120971613-DEL-66 | chr14-120971614-DEL-66 |
| chr14-120977170-DEL-57 | chr14-120977171-DEL-57 |
| chr14-120979965-DEL-7329 | chr14-120979967-DEL-7328 |
| chr14-121010370-DEL-204 | chr14-121010371-DEL-204 |
| chr14-121014826-DEL-6617 | chr14-121014826-DEL-6616 |
| chr14-121022624-DEL-1957 | chr14-121022628-DEL-1956 |
| chr14-121030915-DEL-2939 | chr14-121030916-DEL-2939 |
| chr14-121044224-DEL-488 | chr14-121044232-DEL-481 |
| chr14-121045978-DEL-619 | chr14-121045976-DEL-618 |
| chr14-121048172-DEL-369 | chr14-121048173-DEL-369 |
| chr14-121051102-DEL-58 | chr14-121051087-DEL-74 |
| chr14-121053846-DEL-67 | chr14-121053846-DEL-73 |
| chr14-121062694-DEL-51 | chr14-121062695-DEL-51 |
| chr14-121066981-DEL-110 | chr14-121066943-DEL-148 |
| chr14-121074622-DEL-1273 | chr14-121074623-DEL-1273 |
| chr14-121075871-DEL-214 | chr14-121075872-DEL-214 |
| chr14-121085488-DEL-98 | chr14-121085489-DEL-98 |
| chr14-121087149-DEL-376 | chr14-121087150-DEL-336 |
| chr14-121097493-DEL-66 | chr14-121097518-DEL-52 |
| chr14-121125035-DEL-186 | chr14-121125052-DEL-165 |
| chr14-121127695-DEL-552 | chr14-121127728-DEL-530 |
| chr14-121138870-DEL-115 | chr14-121138871-DEL-115 |
| chr14-121194046-DEL-358 | chr14-121194047-DEL-358 |
| chr14-121197478-DEL-57 | chr14-121197476-DEL-58 |
| chr14-121200354-DEL-812 | chr14-121200351-DEL-813 |
| chr14-121203456-DEL-70 | chr14-121203457-DEL-70 |
| chr14-121219649-DEL-201 | chr14-121219650-DEL-201 |
| chr14-121225432-DEL-98 | chr14-121225428-DEL-99 |
| chr14-121232819-DEL-148 | chr14-121232820-DEL-148 |
| chr14-121237040-DEL-131 | chr14-121237045-DEL-127 |
| chr14-121238792-DEL-569 | chr14-121238793-DEL-569 |
| chr14-121241085-DEL-62 | chr14-121241082-DEL-80 |
| chr14-121244500-DEL-4330 | chr14-121244501-DEL-4330 |
| chr14-121254393-DEL-427 | chr14-121254394-DEL-427 |
| chr14-121257261-DEL-1405 | chr14-121257278-DEL-1391 |
| chr14-121261924-DEL-230 | chr14-121261927-DEL-228 |
| chr14-121271065-DEL-163 | chr14-121271066-DEL-163 |
| chr14-121289376-DEL-51 | chr14-121289373-DEL-54 |
| chr14-121303758-DEL-3217 | chr14-121303766-DEL-3216 |
| chr14-121337854-DEL-165 | chr14-121337842-DEL-144 |
| chr14-121359884-DEL-154 | chr14-121359885-DEL-154 |
| chr14-121367700-DEL-59 | chr14-121367700-DEL-66 |
| chr14-121376519-DEL-156 | chr14-121376474-DEL-170 |
| chr14-121383476-DEL-75 | chr14-121383481-DEL-71 |
| chr14-121399188-DEL-2185 | chr14-121399189-DEL-2185 |
| chr14-121403749-DEL-108 | chr14-121403750-DEL-108 |
| chr14-121416909-DEL-170 | chr14-121416909-DEL-171 |
| chr14-121422454-DEL-194 | chr14-121422455-DEL-194 |
| chr14-121439329-DEL-64 | chr14-121439330-DEL-64 |
| chr14-121441703-DEL-86 | chr14-121441704-DEL-86 |
| chr14-121448679-DEL-411 | chr14-121448680-DEL-411 |
| chr14-121460390-DEL-496 | chr14-121460385-DEL-497 |
| chr14-121467664-DEL-182 | chr14-121467665-DEL-181 |
| chr14-121471409-DEL-109 | chr14-121471449-DEL-70 |
| chr14-121533807-DEL-371 | chr14-121533808-DEL-371 |
| chr14-121565217-DEL-1164 | chr14-121565201-DEL-1180 |
| chr14-121594532-DEL-3098 | chr14-121594533-DEL-3098 |
| chr14-121623666-DEL-65 | chr14-121623667-DEL-65 |
| chr14-121626269-DEL-519 | chr14-121626270-DEL-519 |
| chr14-121627781-DEL-176 | chr14-121627782-DEL-176 |
| chr14-121650841-DEL-420 | chr14-121650842-DEL-420 |
| chr14-121656130-DEL-53 | chr14-121656131-DEL-53 |
| chr14-121674236-DEL-72 | chr14-121674237-DEL-72 |
| chr14-121675510-DEL-60 | chr14-121675511-DEL-60 |
| chr14-121690832-DEL-1351 | chr14-121690833-DEL-1351 |
| chr14-121700690-DEL-54 | chr14-121700691-DEL-54 |
| chr14-121702866-DEL-60 | chr14-121702867-DEL-60 |
| chr14-121707673-DEL-66 | chr14-121707683-DEL-54 |

| | |
|---|---|
| chr14-121713655-DEL-495 | chr14-121713656-DEL-495 |
| chr14-121718857-DEL-64 | chr14-121718858-DEL-64 |
| chr14-121754946-DEL-179 | chr14-121754970-DEL-182 |
| chr14-121756354-DEL-424 | chr14-121756355-DEL-424 |
| chr14-121763316-DEL-206 | chr14-121763317-DEL-206 |
| chr14-121798775-DEL-663 | chr14-121798776-DEL-663 |
| chr14-121813456-DEL-128 | chr14-121813457-DEL-128 |
| chr14-121818550-DEL-76 | chr14-121818547-DEL-76 |
| chr14-121838988-DEL-225 | chr14-121838989-DEL-225 |
| chr14-121852802-DEL-113 | chr14-121852803-DEL-113 |
| chr14-121857817-DEL-462 | chr14-121857818-DEL-462 |
| chr14-121873774-DEL-70 | chr14-121873775-DEL-70 |
| chr14-121919063-DEL-336 | chr14-121919060-DEL-337 |
| chr14-121922053-DEL-387 | chr14-121922054-DEL-387 |
| chr14-121925410-DEL-120 | chr14-121925406-DEL-132 |
| chr14-121930458-DEL-62 | chr14-121930459-DEL-62 |
| chr14-121936562-DEL-572 | chr14-121936563-DEL-572 |
| chr14-121992345-DEL-244 | chr14-121992346-DEL-244 |
| chr14-122009434-DEL-714 | chr14-122009435-DEL-714 |
| chr14-122015913-DEL-51 | chr14-122015914-DEL-51 |
| chr14-122026514-DEL-125 | chr14-122026513-DEL-123 |
| chr14-122049339-DEL-240 | chr14-122049340-DEL-240 |
| chr14-122058254-DEL-202 | chr14-122058255-DEL-203 |
| chr14-122078102-DEL-111 | chr14-122078103-DEL-111 |
| chr14-122081080-DEL-55 | chr14-122081062-DEL-66 |
| chr14-122098165-DEL-189 | chr14-122098166-DEL-189 |
| chr14-122099122-DEL-66 | chr14-122099126-DEL-63 |
| chr14-122107251-DEL-54 | chr14-122107252-DEL-54 |
| chr14-122126294-DEL-68 | chr14-122126291-DEL-68 |
| chr14-122151331-DEL-214 | chr14-122151332-DEL-214 |
| chr14-122159342-DEL-100 | chr14-122159352-DEL-110 |
| chr14-122183974-DEL-94 | chr14-122183975-DEL-82 |
| chr14-122214989-DEL-1426 | chr14-122214990-DEL-1426 |
| chr14-122245610-DEL-81 | chr14-122245611-DEL-81 |
| chr14-122268389-DEL-1317 | chr14-122268390-DEL-1317 |
| chr14-122284550-DEL-184 | chr14-122284551-DEL-185 |
| chr14-122302210-DEL-99 | chr14-122302211-DEL-99 |
| chr14-122320648-DEL-104 | chr14-122320649-DEL-104 |
| chr14-122323168-DEL-81 | chr14-122323169-DEL-81 |
| chr14-122328447-DEL-474 | chr14-122328448-DEL-475 |
| chr14-122334072-DEL-661 | chr14-122334073-DEL-661 |
| chr14-122337312-DEL-180 | chr14-122337311-DEL-180 |
| chr14-122339038-DEL-414 | chr14-122339037-DEL-415 |
| chr14-122355282-DEL-1533 | chr14-122355283-DEL-1533 |
| chr14-122415075-DEL-313 | chr14-122415076-DEL-313 |
| chr14-122419686-DEL-429 | chr14-122419687-DEL-429 |
| chr14-122429649-DEL-210 | chr14-122429650-DEL-210 |
| chr14-122432633-DEL-55 | chr14-122432634-DEL-55 |
| chr14-122443682-DEL-70 | chr14-122443664-DEL-69 |
| chr14-122445833-DEL-70 | chr14-122445830-DEL-62 |
| chr14-122453440-DEL-186 | chr14-122453441-DEL-186 |
| chr14-122455453-DEL-130 | chr14-122455454-DEL-130 |
| chr14-122456542-DEL-84 | chr14-122456543-DEL-84 |
| chr14-122491872-DEL-196 | chr14-122491873-DEL-196 |
| chr14-122513112-DEL-72 | chr14-122513111-DEL-72 |
| chr14-122514979-DEL-71 | chr14-122514980-DEL-71 |
| chr14-122528761-DEL-60 | chr14-122528760-DEL-62 |
| chr14-122546282-DEL-58 | chr14-122546284-DEL-60 |
| chr14-122587872-DEL-285 | chr14-122587869-DEL-286 |
| chr14-122655656-DEL-762 | chr14-122655654-DEL-763 |
| chr14-122658590-DEL-70 | chr14-122658591-DEL-70 |
| chr14-122672835-DEL-61 | chr14-122672835-DEL-61 |
| chr14-122755549-DEL-7865 | chr14-122755550-DEL-7865 |
| chr14-122786882-DEL-955 | chr14-122786883-DEL-955 |
| chr14-122787929-DEL-206 | chr14-122787930-DEL-206 |
| chr14-122794079-DEL-52 | chr14-122794080-DEL-52 |
| chr14-122798408-DEL-132 | chr14-122798409-DEL-132 |
| chr14-122806386-DEL-66 | chr14-122806386-DEL-65 |
| chr14-122821425-DEL-68 | chr14-122821430-DEL-52 |
| chr14-122821770-DEL-418 | chr14-122821772-DEL-417 |
| chr14-122822651-DEL-387 | chr14-122822652-DEL-387 |
| chr14-122831845-DEL-156 | chr14-122831894-DEL-129 |
| chr14-122859512-DEL-6465 | chr14-122859513-DEL-6465 |
| chr14-122869446-DEL-168 | chr14-122869447-DEL-168 |
| chr14-122902703-DEL-399 | chr14-122902704-DEL-399 |
| chr14-122915221-DEL-209 | chr14-122915222-DEL-209 |
| chr14-122959810-DEL-74 | chr14-122959811-DEL-74 |
| chr14-122969510-DEL-2918 | chr14-122969511-DEL-2918 |
| chr14-122978683-DEL-1174 | chr14-122978684-DEL-1174 |
| chr14-123001194-DEL-1592 | chr14-123001195-DEL-1592 |

| | |
|---|---|
| chr14-123008128-DEL-6855 | chr14-123008129-DEL-6855 |
| chr14-123129681-DEL-304 | chr14-123129682-DEL-304 |
| chr14-123135979-DEL-57 | chr14-123135980-DEL-57 |
| chr14-123243797-DEL-1856 | chr14-123243798-DEL-1856 |
| chr14-123286568-DEL-120 | chr14-123286557-DEL-124 |
| chr14-123304435-DEL-201 | chr14-123304436-DEL-201 |
| chr14-123329080-DEL-115 | chr14-123329098-DEL-92 |
| chr14-123338893-DEL-6788 | chr14-123338894-DEL-6788 |
| chr14-123339453-DEL-208 | chr14-123339454-DEL-208 |
| chr14-123363162-DEL-4255 | chr14-123363163-DEL-4255 |
| chr14-123368288-DEL-864 | chr14-123368289-DEL-864 |
| chr14-123374895-DEL-328 | chr14-123374896-DEL-328 |
| chr14-123386562-DEL-66 | chr14-123386563-DEL-66 |
| chr14-123459190-DEL-89 | chr14-123459190-DEL-90 |
| chr14-123468215-DEL-1344 | chr14-123468216-DEL-1344 |
| chr14-123487921-DEL-75 | chr14-123487922-DEL-75 |
| chr14-123495245-DEL-150 | chr14-123495246-DEL-150 |
| chr14-123536673-DEL-81 | chr14-123536674-DEL-81 |
| chr14-123586855-DEL-1893 | chr14-123586856-DEL-1893 |
| chr14-123597806-DEL-556 | chr14-123597842-DEL-558 |
| chr14-123619507-DEL-246 | chr14-123619508-DEL-246 |
| chr14-123621378-DEL-364 | chr14-123621379-DEL-364 |
| chr14-123638622-DEL-136 | chr14-123638623-DEL-136 |
| chr14-123643796-DEL-86 | chr14-123643797-DEL-86 |
| chr14-123648621-DEL-145 | chr14-123648616-DEL-144 |
| chr14-123649354-DEL-8190 | chr14-123649351-DEL-8191 |
| chr14-123670394-DEL-424 | chr14-123670395-DEL-424 |
| chr14-123670819-DEL-636 | chr14-123670819-DEL-636 |
| chr14-123675247-DEL-64 | chr14-123675248-DEL-64 |
| chr14-123704167-DEL-51 | chr14-123704168-DEL-51 |
| chr14-123717842-DEL-2789 | chr14-123717843-DEL-2789 |
| chr14-123727142-DEL-68 | chr14-123727143-DEL-68 |
| chr14-123781920-DEL-6979 | chr14-123781921-DEL-6979 |
| chr14-123782054-DEL-409 | chr14-123782055-DEL-409 |
| chr14-123790090-DEL-58 | chr14-123790091-DEL-58 |
| chr14-123806975-DEL-186 | chr14-123806976-DEL-186 |
| chr14-123822376-DEL-515 | chr14-123822377-DEL-515 |
| chr14-123846731-DEL-6329 | chr14-123846748-DEL-6330 |
| chr14-123869283-DEL-615 | chr14-123869284-DEL-615 |
| chr14-123892512-DEL-2653 | chr14-123892513-DEL-2653 |
| chr14-123908911-DEL-171 | chr14-123908912-DEL-171 |
| chr14-123926794-DEL-6814 | chr14-123926795-DEL-6814 |
| chr14-123933113-DEL-409 | chr14-123933114-DEL-409 |
| chr14-123954755-DEL-1611 | chr14-123954756-DEL-1611 |
| chr14-123960413-DEL-12521 | chr14-123960412-DEL-12522 |
| chr14-123977824-DEL-208 | chr14-123977825-DEL-208 |
| chr14-123988169-DEL-113 | chr14-123988170-DEL-113 |
| chr14-124012430-DEL-4742 | chr14-124012431-DEL-4742 |
| chr14-124024060-DEL-6818 | chr14-124024051-DEL-6819 |
| chr14-124042252-DEL-6399 | chr14-124042253-DEL-6399 |
| chr14-124074618-DEL-143 | chr14-124074619-DEL-143 |
| chr14-124087160-DEL-112 | chr14-124087161-DEL-112 |
| chr14-124111227-DEL-7002 | chr14-124111228-DEL-7002 |
| chr14-124117443-DEL-416 | chr14-124117444-DEL-416 |
| chr14-124186374-DEL-3571 | chr14-124186375-DEL-3571 |
| chr14-124195057-DEL-982 | chr14-124195058-DEL-982 |
| chr14-124234224-DEL-76 | chr14-124234225-DEL-76 |
| chr14-124234734-DEL-3291 | chr14-124234735-DEL-3291 |
| chr14-124277560-DEL-255 | chr14-124277561-DEL-255 |
| chr14-124307764-DEL-404 | chr14-124307763-DEL-404 |
| chr14-124439542-DEL-146 | chr14-124439543-DEL-146 |
| chr14-124471535-DEL-520 | chr14-124471536-DEL-520 |
| chr14-124474997-DEL-196 | chr14-124474998-DEL-196 |
| chr14-124504728-DEL-6780 | chr14-124504729-DEL-6780 |
| chr14-124530722-DEL-52 | chr14-124530712-DEL-53 |
| chr14-124547900-DEL-207 | chr14-124547901-DEL-207 |
| chr14-124549383-DEL-2357 | chr14-124549384-DEL-2357 |
| chr14-124571244-DEL-201 | chr14-124571245-DEL-201 |
| chr14-124595462-DEL-334 | chr14-124595463-DEL-334 |
| chr14-124604837-DEL-50 | chr14-124604837-DEL-50 |
| chr14-124621164-DEL-172 | chr14-124621165-DEL-172 |
| chr14-124641092-DEL-732 | chr14-124641093-DEL-732 |
| chr14-124652566-DEL-659 | chr14-124652567-DEL-659 |
| chr14-124662665-DEL-6380 | chr14-124662666-DEL-6380 |
| chr14-124662681-DEL-208 | chr14-124662681-DEL-208 |
| chr14-124669584-DEL-65 | chr14-124669585-DEL-65 |
| chr14-124688718-DEL-71 | chr14-124688719-DEL-70 |
| chr14-124728745-DEL-6968 | chr14-124728746-DEL-6968 |
| chr14-124791792-DEL-360 | chr14-124791793-DEL-360 |
| chr14-124848679-DEL-716 | chr14-124848680-DEL-716 |

| | |
|---|---|
| chr14-124890838-DEL-1000 | chr14-124890839-DEL-1000 |
| chr14-124891628-DEL-7074 | chr14-124891629-DEL-7074 |
| chr14-124891819-DEL-636 | chr14-124891820-DEL-636 |
| chr14-124892396-DEL-241 | chr14-124892397-DEL-241 |
| chr14-124905740-DEL-198 | chr14-124905741-DEL-198 |
| chr14-124921839-DEL-7386 | chr14-124921840-DEL-7386 |
| chr14-124955278-DEL-195 | chr14-124955279-DEL-195 |
| chr14-124988776-DEL-201 | chr14-124988777-DEL-201 |
| chr14-124999129-DEL-208 | chr14-124999130-DEL-208 |
| chr14-125002683-DEL-54 | chr14-125002684-DEL-54 |
| chr15-3138156-DEL-81 | chr15-3138191-DEL-81 |
| chr15-3222698-DEL-422 | chr15-3222699-DEL-422 |
| chr15-3245491-DEL-105 | chr15-3245492-DEL-100 |
| chr15-3275785-DEL-65 | chr15-3275786-DEL-65 |
| chr15-3294567-DEL-1178 | chr15-3294568-DEL-1178 |
| chr15-3315260-DEL-53 | chr15-3315256-DEL-58 |
| chr15-3332153-DEL-68 | chr15-3332162-DEL-70 |
| chr15-3343663-DEL-123 | chr15-3343634-DEL-124 |
| chr15-3390122-DEL-3119 | chr15-3390123-DEL-3119 |
| chr15-3416864-DEL-72 | chr15-3416865-DEL-72 |
| chr15-3507406-DEL-3929 | chr15-3507407-DEL-3929 |
| chr15-3531580-DEL-76 | chr15-3531581-DEL-76 |
| chr15-3544970-DEL-56 | chr15-3544971-DEL-56 |
| chr15-3571285-DEL-176 | chr15-3571285-DEL-176 |
| chr15-3577404-DEL-3170 | chr15-3577405-DEL-3170 |
| chr15-3591421-DEL-2969 | chr15-3591422-DEL-2969 |
| chr15-3646813-DEL-60 | chr15-3646808-DEL-54 |
| chr15-3697704-DEL-74 | chr15-3697713-DEL-50 |
| chr15-3724202-DEL-880 | chr15-3724203-DEL-880 |
| chr15-3767930-DEL-64 | chr15-3767931-DEL-64 |
| chr15-3777416-DEL-52 | chr15-3777417-DEL-52 |
| chr15-3797783-DEL-72 | chr15-3797784-DEL-72 |
| chr15-3830589-DEL-54 | chr15-3830586-DEL-58 |
| chr15-3858566-DEL-408 | chr15-3858567-DEL-408 |
| chr15-3871909-DEL-56 | chr15-3871903-DEL-58 |
| chr15-3890943-DEL-75 | chr15-3890946-DEL-73 |
| chr15-3911225-DEL-6402 | chr15-3911226-DEL-6402 |
| chr15-3923990-DEL-410 | chr15-3923991-DEL-407 |
| chr15-3947784-DEL-308 | chr15-3947785-DEL-308 |
| chr15-3949782-DEL-6869 | chr15-3949783-DEL-6869 |
| chr15-3950149-DEL-212 | chr15-3950150-DEL-212 |
| chr15-3959453-DEL-672 | chr15-3959454-DEL-672 |
| chr15-3962572-DEL-57 | chr15-3962573-DEL-57 |
| chr15-3964787-DEL-822 | chr15-3964788-DEL-822 |
| chr15-3985868-DEL-344 | chr15-3985869-DEL-344 |
| chr15-3995349-DEL-789 | chr15-3995350-DEL-789 |
| chr15-4002275-DEL-420 | chr15-4002276-DEL-421 |
| chr15-4003438-DEL-54 | chr15-4003439-DEL-54 |
| chr15-4052093-DEL-51 | chr15-4052068-DEL-51 |
| chr15-4066864-DEL-100 | chr15-4066865-DEL-72 |
| chr15-4099867-DEL-420 | chr15-4099868-DEL-420 |
| chr15-4100934-DEL-2010 | chr15-4100940-DEL-2005 |
| chr15-4113066-DEL-70 | chr15-4113067-DEL-70 |
| chr15-4143868-DEL-3656 | chr15-4143869-DEL-3656 |
| chr15-4156098-DEL-458 | chr15-4156099-DEL-458 |
| chr15-4163018-DEL-642 | chr15-4163019-DEL-641 |
| chr15-4164046-DEL-78 | chr15-4164047-DEL-78 |
| chr15-4173846-DEL-173 | chr15-4173847-DEL-173 |
| chr15-4193185-DEL-3325 | chr15-4193186-DEL-3325 |
| chr15-4207793-DEL-100 | chr15-4207781-DEL-88 |
| chr15-4208297-DEL-212 | chr15-4208298-DEL-212 |
| chr15-4221869-DEL-92 | chr15-4221876-DEL-86 |
| chr15-4230598-DEL-56 | chr15-4230605-DEL-57 |
| chr15-4244359-DEL-3627 | chr15-4244360-DEL-3627 |
| chr15-4252870-DEL-76 | chr15-4252866-DEL-77 |
| chr15-4278316-DEL-62 | chr15-4278317-DEL-58 |
| chr15-4309684-DEL-6320 | chr15-4309685-DEL-6320 |
| chr15-4343523-DEL-70 | chr15-4343524-DEL-70 |
| chr15-4397862-DEL-61 | chr15-4397863-DEL-61 |
| chr15-4413319-DEL-4048 | chr15-4413319-DEL-4048 |
| chr15-4443347-DEL-6920 | chr15-4443348-DEL-6920 |
| chr15-4464451-DEL-178 | chr15-4464452-DEL-178 |
| chr15-4475774-DEL-56 | chr15-4475773-DEL-56 |
| chr15-4488503-DEL-1079 | chr15-4488504-DEL-1079 |
| chr15-4505427-DEL-110 | chr15-4505428-DEL-110 |
| chr15-4517948-DEL-56 | chr15-4517939-DEL-66 |
| chr15-4524159-DEL-348 | chr15-4524160-DEL-348 |
| chr15-4527014-DEL-322 | chr15-4527015-DEL-322 |
| chr15-4581128-DEL-73 | chr15-4581107-DEL-56 |
| chr15-4589018-DEL-1705 | chr15-4589019-DEL-1705 |

| | |
|---|---|
| chr15-4613305-DEL-6396 | chr15-4613306-DEL-6396 |
| chr15-4643526-DEL-8030 | chr15-4643526-DEL-8031 |
| chr15-4647516-DEL-99 | chr15-4647516-DEL-99 |
| chr15-4695564-DEL-6922 | chr15-4695562-DEL-6923 |
| chr15-4701830-DEL-212 | chr15-4701834-DEL-211 |
| chr15-4710010-DEL-55 | chr15-4710011-DEL-55 |
| chr15-4798559-DEL-6888 | chr15-4798560-DEL-6888 |
| chr15-4830829-DEL-434 | chr15-4830830-DEL-434 |
| chr15-4864261-DEL-713 | chr15-4864259-DEL-714 |
| chr15-4888859-DEL-11969 | chr15-4888860-DEL-11969 |
| chr15-4902423-DEL-79 | chr15-4902423-DEL-91 |
| chr15-4914516-DEL-1280 | chr15-4914517-DEL-1280 |
| chr15-4922320-DEL-1930 | chr15-4922321-DEL-1930 |
| chr15-4924252-DEL-74 | chr15-4924255-DEL-72 |
| chr15-4957208-DEL-164 | chr15-4957208-DEL-162 |
| chr15-4958016-DEL-66 | chr15-4958017-DEL-66 |
| chr15-4960988-DEL-74 | chr15-4960989-DEL-74 |
| chr15-4969538-DEL-86 | chr15-4969532-DEL-62 |
| chr15-4993879-DEL-2117 | chr15-4993880-DEL-2117 |
| chr15-5009863-DEL-7066 | chr15-5009864-DEL-7066 |
| chr15-5026689-DEL-275 | chr15-5026690-DEL-275 |
| chr15-5036751-DEL-78 | chr15-5036752-DEL-51 |
| chr15-5072026-DEL-195 | chr15-5072027-DEL-195 |
| chr15-5075869-DEL-114 | chr15-5075870-DEL-114 |
| chr15-5121956-DEL-762 | chr15-5121957-DEL-762 |
| chr15-5132679-DEL-178 | chr15-5132666-DEL-206 |
| chr15-5150829-DEL-1036 | chr15-5150830-DEL-1036 |
| chr15-5153160-DEL-193 | chr15-5153161-DEL-193 |
| chr15-5165595-DEL-512 | chr15-5165596-DEL-512 |
| chr15-5180819-DEL-338 | chr15-5180820-DEL-338 |
| chr15-5197022-DEL-108 | chr15-5197013-DEL-106 |
| chr15-5255035-DEL-1551 | chr15-5255036-DEL-1551 |
| chr15-5296326-DEL-6862 | chr15-5296327-DEL-6862 |
| chr15-5296842-DEL-212 | chr15-5296843-DEL-212 |
| chr15-5309091-DEL-82 | chr15-5309131-DEL-62 |
| chr15-5313832-DEL-1829 | chr15-5313833-DEL-1829 |
| chr15-5335349-DEL-1839 | chr15-5335350-DEL-1839 |
| chr15-5360115-DEL-100 | chr15-5360116-DEL-100 |
| chr15-5386985-DEL-729 | chr15-5386959-DEL-755 |
| chr15-5441035-DEL-2168 | chr15-5441036-DEL-2168 |
| chr15-5444266-DEL-283 | chr15-5444267-DEL-283 |
| chr15-5463646-DEL-584 | chr15-5463647-DEL-584 |
| chr15-5468712-DEL-1407 | chr15-5468712-DEL-1407 |
| chr15-5521212-DEL-125 | chr15-5521262-DEL-76 |
| chr15-5571108-DEL-1416 | chr15-5571109-DEL-1416 |
| chr15-5572944-DEL-3262 | chr15-5572945-DEL-3262 |
| chr15-5633589-DEL-66 | chr15-5633584-DEL-58 |
| chr15-5678874-DEL-67 | chr15-5678886-DEL-62 |
| chr15-5692034-DEL-179 | chr15-5692035-DEL-179 |
| chr15-5711224-DEL-6423 | chr15-5711225-DEL-6423 |
| chr15-5796291-DEL-73 | chr15-5796290-DEL-71 |
| chr15-5799790-DEL-1845 | chr15-5799791-DEL-1845 |
| chr15-5816801-DEL-60 | chr15-5816802-DEL-60 |
| chr15-5818255-DEL-7600 | chr15-5818256-DEL-7600 |
| chr15-5871322-DEL-340 | chr15-5871318-DEL-341 |
| chr15-5881758-DEL-1339 | chr15-5881759-DEL-1339 |
| chr15-5895080-DEL-162 | chr15-5895081-DEL-162 |
| chr15-5899332-DEL-100 | chr15-5899333-DEL-100 |
| chr15-5911725-DEL-5054 | chr15-5911726-DEL-5054 |
| chr15-5930205-DEL-776 | chr15-5930206-DEL-776 |
| chr15-5936148-DEL-92 | chr15-5936193-DEL-56 |
| chr15-5941297-DEL-70 | chr15-5941300-DEL-68 |
| chr15-5954900-DEL-400 | chr15-5954901-DEL-400 |
| chr15-5995059-DEL-198 | chr15-5995060-DEL-198 |
| chr15-5995869-DEL-445 | chr15-5995870-DEL-445 |
| chr15-6015983-DEL-1439 | chr15-6015982-DEL-1440 |
| chr15-6037349-DEL-614 | chr15-6037350-DEL-614 |
| chr15-6044786-DEL-872 | chr15-6044787-DEL-872 |
| chr15-6049781-DEL-111 | chr15-6049776-DEL-111 |
| chr15-6052245-DEL-346 | chr15-6052246-DEL-346 |
| chr15-6060155-DEL-512 | chr15-6060156-DEL-512 |
| chr15-6099195-DEL-178 | chr15-6099196-DEL-178 |
| chr15-6114650-DEL-1892 | chr15-6114651-DEL-1892 |
| chr15-6139737-DEL-2128 | chr15-6139738-DEL-2128 |
| chr15-6162452-DEL-56 | chr15-6162449-DEL-54 |
| chr15-6173163-DEL-2537 | chr15-6173164-DEL-2537 |
| chr15-6176219-DEL-443 | chr15-6176220-DEL-443 |
| chr15-6178624-DEL-360 | chr15-6178625-DEL-360 |
| chr15-6211892-DEL-102 | chr15-6211893-DEL-102 |
| chr15-6216373-DEL-130 | chr15-6216356-DEL-116 |

| | |
|---|---|
| chr15-6222471-DEL-6990 | chr15-6222467-DEL-6983 |
| chr15-6251047-DEL-54 | chr15-6251048-DEL-54 |
| chr15-6255601-DEL-3091 | chr15-6255602-DEL-3091 |
| chr15-6280557-DEL-1072 | chr15-6280558-DEL-1072 |
| chr15-6299367-DEL-6380 | chr15-6299368-DEL-6380 |
| chr15-6305294-DEL-140 | chr15-6305295-DEL-140 |
| chr15-6321529-DEL-106 | chr15-6321519-DEL-80 |
| chr15-6330633-DEL-161 | chr15-6330633-DEL-162 |
| chr15-6347321-DEL-52 | chr15-6347322-DEL-52 |
| chr15-6349427-DEL-55 | chr15-6349428-DEL-55 |
| chr15-6382043-DEL-51 | chr15-6382043-DEL-51 |
| chr15-6388586-DEL-54 | chr15-6388587-DEL-54 |
| chr15-6403416-DEL-74 | chr15-6403417-DEL-50 |
| chr15-6446706-DEL-52 | chr15-6446707-DEL-52 |
| chr15-6462761-DEL-72 | chr15-6462767-DEL-54 |
| chr15-6465562-DEL-79 | chr15-6465585-DEL-83 |
| chr15-6481164-DEL-708 | chr15-6481165-DEL-708 |
| chr15-6482211-DEL-66 | chr15-6482193-DEL-55 |
| chr15-6485475-DEL-74 | chr15-6485482-DEL-80 |
| chr15-6491083-DEL-63 | chr15-6491084-DEL-63 |
| chr15-6497739-DEL-68 | chr15-6497769-DEL-60 |
| chr15-6517197-DEL-66 | chr15-6517198-DEL-66 |
| chr15-6522995-DEL-72 | chr15-6522996-DEL-72 |
| chr15-6551476-DEL-117 | chr15-6551477-DEL-117 |
| chr15-6563272-DEL-67 | chr15-6563273-DEL-67 |
| chr15-6611551-DEL-332 | chr15-6611552-DEL-332 |
| chr15-6612694-DEL-57 | chr15-6612695-DEL-57 |
| chr15-6637382-DEL-154 | chr15-6637383-DEL-154 |
| chr15-6642080-DEL-876 | chr15-6642081-DEL-876 |
| chr15-6645104-DEL-417 | chr15-6645105-DEL-417 |
| chr15-6648269-DEL-245 | chr15-6648270-DEL-245 |
| chr15-6677705-DEL-62 | chr15-6677698-DEL-70 |
| chr15-6683322-DEL-1044 | chr15-6683322-DEL-1044 |
| chr15-6753135-DEL-312 | chr15-6753136-DEL-309 |
| chr15-6834852-DEL-2169 | chr15-6834853-DEL-2170 |
| chr15-6848159-DEL-73 | chr15-6848130-DEL-87 |
| chr15-6848735-DEL-99 | chr15-6848740-DEL-96 |
| chr15-6922093-DEL-6567 | chr15-6922094-DEL-6567 |
| chr15-6954670-DEL-74 | chr15-6954693-DEL-50 |
| chr15-6961537-DEL-312 | chr15-6961573-DEL-295 |
| chr15-6965473-DEL-2471 | chr15-6965473-DEL-2472 |
| chr15-6998252-DEL-198 | chr15-6998253-DEL-198 |
| chr15-7007167-DEL-202 | chr15-7007176-DEL-194 |
| chr15-7046397-DEL-102 | chr15-7046393-DEL-95 |
| chr15-7059845-DEL-288 | chr15-7059846-DEL-288 |
| chr15-7110998-DEL-80 | chr15-7110999-DEL-80 |
| chr15-7131966-DEL-55 | chr15-7131952-DEL-57 |
| chr15-7140350-DEL-106 | chr15-7140363-DEL-94 |
| chr15-7144555-DEL-81 | chr15-7144556-DEL-81 |
| chr15-7222162-DEL-66 | chr15-7222191-DEL-66 |
| chr15-7223231-DEL-64 | chr15-7223228-DEL-64 |
| chr15-7245013-DEL-108 | chr15-7245014-DEL-108 |
| chr15-7260937-DEL-107 | chr15-7260938-DEL-107 |
| chr15-7273037-DEL-61 | chr15-7273038-DEL-61 |
| chr15-7302850-DEL-6332 | chr15-7302834-DEL-6347 |
| chr15-7336305-DEL-76 | chr15-7336324-DEL-58 |
| chr15-7343045-DEL-56 | chr15-7343046-DEL-56 |
| chr15-7353904-DEL-818 | chr15-7353905-DEL-818 |
| chr15-7361980-DEL-253 | chr15-7361981-DEL-253 |
| chr15-7372748-DEL-258 | chr15-7372749-DEL-258 |
| chr15-7381425-DEL-64 | chr15-7381436-DEL-78 |
| chr15-7383986-DEL-90 | chr15-7383978-DEL-87 |
| chr15-7389858-DEL-6896 | chr15-7389859-DEL-6896 |
| chr15-7398638-DEL-57 | chr15-7398639-DEL-57 |
| chr15-7406129-DEL-65 | chr15-7406120-DEL-72 |
| chr15-7418803-DEL-56 | chr15-7418804-DEL-56 |
| chr15-7429503-DEL-198 | chr15-7429504-DEL-193 |
| chr15-7430423-DEL-1169 | chr15-7430424-DEL-1169 |
| chr15-7485924-DEL-1674 | chr15-7485925-DEL-1674 |
| chr15-7499291-DEL-106 | chr15-7499292-DEL-106 |
| chr15-7512633-DEL-89 | chr15-7512641-DEL-77 |
| chr15-7525565-DEL-115 | chr15-7525551-DEL-165 |
| chr15-7562091-DEL-82 | chr15-7562092-DEL-74 |
| chr15-7568519-DEL-67 | chr15-7568522-DEL-71 |
| chr15-7569451-DEL-8455 | chr15-7569452-DEL-8455 |
| chr15-7620028-DEL-53 | chr15-7620031-DEL-51 |
| chr15-7622348-DEL-2281 | chr15-7622349-DEL-2281 |
| chr15-7626676-DEL-68 | chr15-7626677-DEL-68 |
| chr15-7637574-DEL-517 | chr15-7637575-DEL-517 |
| chr15-7657902-DEL-210 | chr15-7657903-DEL-209 |

| | |
|---|---|
| chr15-7677665-DEL-65 | chr15-7677673-DEL-64 |
| chr15-7692535-DEL-517 | chr15-7692528-DEL-521 |
| chr15-7695414-DEL-722 | chr15-7695415-DEL-722 |
| chr15-7771144-DEL-163 | chr15-7771137-DEL-163 |
| chr15-7772687-DEL-238 | chr15-7772688-DEL-238 |
| chr15-7830569-DEL-1666 | chr15-7830569-DEL-1666 |
| chr15-7869067-DEL-245 | chr15-7869043-DEL-256 |
| chr15-7878833-DEL-7338 | chr15-7878834-DEL-7338 |
| chr15-7912426-DEL-446 | chr15-7912427-DEL-446 |
| chr15-7935339-DEL-268 | chr15-7935340-DEL-268 |
| chr15-7940351-DEL-7131 | chr15-7940352-DEL-7131 |
| chr15-7955699-DEL-3298 | chr15-7955700-DEL-3298 |
| chr15-8030208-DEL-410 | chr15-8030209-DEL-410 |
| chr15-8035412-DEL-94 | chr15-8035413-DEL-141 |
| chr15-8036200-DEL-211 | chr15-8036201-DEL-211 |
| chr15-8063279-DEL-216 | chr15-8063280-DEL-216 |
| chr15-8098069-DEL-167 | chr15-8098070-DEL-167 |
| chr15-8106563-DEL-212 | chr15-8106564-DEL-212 |
| chr15-8115328-DEL-230 | chr15-8115340-DEL-231 |
| chr15-8131345-DEL-545 | chr15-8131346-DEL-545 |
| chr15-8135844-DEL-195 | chr15-8135845-DEL-195 |
| chr15-8155576-DEL-205 | chr15-8155577-DEL-205 |
| chr15-8158122-DEL-60 | chr15-8158123-DEL-60 |
| chr15-8174455-DEL-194 | chr15-8174456-DEL-194 |
| chr15-8192766-DEL-90 | chr15-8192767-DEL-90 |
| chr15-8214193-DEL-211 | chr15-8214194-DEL-211 |
| chr15-8224437-DEL-2413 | chr15-8224438-DEL-2413 |
| chr15-8227295-DEL-245 | chr15-8227296-DEL-245 |
| chr15-8254522-DEL-55 | chr15-8254482-DEL-59 |
| chr15-8264344-DEL-216 | chr15-8264335-DEL-217 |
| chr15-8272450-DEL-515 | chr15-8272451-DEL-515 |
| chr15-8286409-DEL-72 | chr15-8286408-DEL-71 |
| chr15-8289517-DEL-52 | chr15-8289518-DEL-52 |
| chr15-8290485-DEL-208 | chr15-8290486-DEL-208 |
| chr15-8296215-DEL-2630 | chr15-8296216-DEL-2630 |
| chr15-8302967-DEL-4673 | chr15-8302968-DEL-4673 |
| chr15-8304794-DEL-146 | chr15-8304795-DEL-146 |
| chr15-8408354-DEL-539 | chr15-8408355-DEL-539 |
| chr15-8488114-DEL-52 | chr15-8488115-DEL-52 |
| chr15-8510663-DEL-169 | chr15-8510662-DEL-171 |
| chr15-8540462-DEL-203 | chr15-8540463-DEL-203 |
| chr15-8561624-DEL-186 | chr15-8561625-DEL-186 |
| chr15-8569462-DEL-69 | chr15-8569463-DEL-69 |
| chr15-8577515-DEL-4986 | chr15-8577516-DEL-4986 |
| chr15-8591749-DEL-118 | chr15-8591750-DEL-118 |
| chr15-8638746-DEL-1773 | chr15-8638747-DEL-1773 |
| chr15-8674752-DEL-69 | chr15-8674784-DEL-51 |
| chr15-8678771-DEL-55 | chr15-8678772-DEL-55 |
| chr15-8719840-DEL-178 | chr15-8719841-DEL-178 |
| chr15-8726232-DEL-137 | chr15-8726233-DEL-135 |
| chr15-8729261-DEL-105 | chr15-8729285-DEL-78 |
| chr15-8753923-DEL-3100 | chr15-8753924-DEL-3100 |
| chr15-8769422-DEL-201 | chr15-8769441-DEL-183 |
| chr15-8779977-DEL-244 | chr15-8779978-DEL-244 |
| chr15-8808575-DEL-107 | chr15-8808562-DEL-108 |
| chr15-8812378-DEL-142 | chr15-8812414-DEL-99 |
| chr15-8843472-DEL-215 | chr15-8843473-DEL-215 |
| chr15-8850731-DEL-1712 | chr15-8850725-DEL-1718 |
| chr15-8907335-DEL-60 | chr15-8907326-DEL-70 |
| chr15-8911038-DEL-125 | chr15-8911039-DEL-125 |
| chr15-8935120-DEL-629 | chr15-8935120-DEL-629 |
| chr15-8940257-DEL-207 | chr15-8940257-DEL-207 |
| chr15-8951316-DEL-4708 | chr15-8951317-DEL-4708 |
| chr15-9008393-DEL-387 | chr15-9008394-DEL-387 |
| chr15-9027503-DEL-178 | chr15-9027504-DEL-178 |
| chr15-9039752-DEL-68 | chr15-9039753-DEL-68 |
| chr15-9045518-DEL-6348 | chr15-9045519-DEL-6348 |
| chr15-9119166-DEL-55 | chr15-9119167-DEL-55 |
| chr15-9170213-DEL-206 | chr15-9170214-DEL-206 |
| chr15-9176551-DEL-184 | chr15-9176552-DEL-184 |
| chr15-9191353-DEL-223 | chr15-9191354-DEL-223 |
| chr15-9219119-DEL-7493 | chr15-9219120-DEL-7493 |
| chr15-9234269-DEL-104 | chr15-9234270-DEL-104 |
| chr15-9244792-DEL-467 | chr15-9244793-DEL-467 |
| chr15-9394608-DEL-6237 | chr15-9394608-DEL-6237 |
| chr15-9437176-DEL-6473 | chr15-9437177-DEL-6473 |
| chr15-9453235-DEL-361 | chr15-9453236-DEL-361 |
| chr15-9456572-DEL-106 | chr15-9456570-DEL-100 |
| chr15-9499243-DEL-3798 | chr15-9499244-DEL-3798 |
| chr15-9526959-DEL-220 | chr15-9526960-DEL-220 |

| | |
|---|---|
| chr15-9530829-DEL-515 | chr15-9530830-DEL-515 |
| chr15-9536489-DEL-6418 | chr15-9536490-DEL-6418 |
| chr15-9550475-DEL-3753 | chr15-9550476-DEL-3753 |
| chr15-9557843-DEL-90 | chr15-9557816-DEL-73 |
| chr15-9590602-DEL-118 | chr15-9590619-DEL-90 |
| chr15-9671230-DEL-58 | chr15-9671231-DEL-58 |
| chr15-9695103-DEL-389 | chr15-9695104-DEL-389 |
| chr15-9721260-DEL-91 | chr15-9721260-DEL-91 |
| chr15-9738313-DEL-129 | chr15-9738338-DEL-105 |
| chr15-9749110-DEL-306 | chr15-9749097-DEL-307 |
| chr15-9765948-DEL-361 | chr15-9765949-DEL-361 |
| chr15-9811161-DEL-1823 | chr15-9811162-DEL-1823 |
| chr15-9812501-DEL-79 | chr15-9812502-DEL-79 |
| chr15-9813391-DEL-426 | chr15-9813392-DEL-426 |
| chr15-9813396-DEL-1336 | chr15-9813397-DEL-1336 |
| chr15-9824103-DEL-120 | chr15-9824055-DEL-134 |
| chr15-9825335-DEL-3407 | chr15-9825336-DEL-3407 |
| chr15-9831722-DEL-64 | chr15-9831723-DEL-64 |
| chr15-9844287-DEL-261 | chr15-9844286-DEL-262 |
| chr15-9859271-DEL-110 | chr15-9859283-DEL-107 |
| chr15-9899870-DEL-208 | chr15-9899871-DEL-208 |
| chr15-9903210-DEL-608 | chr15-9903211-DEL-608 |
| chr15-9964239-DEL-6309 | chr15-9964240-DEL-6309 |
| chr15-9991797-DEL-6612 | chr15-9991798-DEL-6612 |
| chr15-10008295-DEL-437 | chr15-10008293-DEL-438 |
| chr15-10019396-DEL-2973 | chr15-10019397-DEL-2973 |
| chr15-10079399-DEL-204 | chr15-10079400-DEL-204 |
| chr15-10095683-DEL-409 | chr15-10095685-DEL-408 |
| chr15-10119413-DEL-90 | chr15-10119390-DEL-86 |
| chr15-10156355-DEL-400 | chr15-10156356-DEL-400 |
| chr15-10157773-DEL-90 | chr15-10157774-DEL-120 |
| chr15-10168087-DEL-181 | chr15-10168088-DEL-181 |
| chr15-10169660-DEL-401 | chr15-10169661-DEL-401 |
| chr15-10270391-DEL-462 | chr15-10270392-DEL-462 |
| chr15-10351719-DEL-6743 | chr15-10351725-DEL-6742 |
| chr15-10365439-DEL-687 | chr15-10365436-DEL-688 |
| chr15-10368324-DEL-426 | chr15-10368325-DEL-426 |
| chr15-10413114-DEL-60 | chr15-10413115-DEL-60 |
| chr15-10430566-DEL-156 | chr15-10430564-DEL-153 |
| chr15-10432417-DEL-89 | chr15-10432407-DEL-89 |
| chr15-10434705-DEL-305 | chr15-10434706-DEL-305 |
| chr15-10435715-DEL-109 | chr15-10435719-DEL-110 |
| chr15-10472865-DEL-190 | chr15-10472866-DEL-190 |
| chr15-10473766-DEL-298 | chr15-10473767-DEL-298 |
| chr15-10479848-DEL-200 | chr15-10479849-DEL-200 |
| chr15-10493479-DEL-81 | chr15-10493479-DEL-81 |
| chr15-10494954-DEL-51 | chr15-10494952-DEL-53 |
| chr15-10498389-DEL-358 | chr15-10498390-DEL-358 |
| chr15-10515704-DEL-55 | chr15-10515705-DEL-55 |
| chr15-10537980-DEL-247 | chr15-10537980-DEL-248 |
| chr15-10544945-DEL-56 | chr15-10544946-DEL-56 |
| chr15-10559794-DEL-608 | chr15-10559795-DEL-605 |
| chr15-10560510-DEL-54 | chr15-10560511-DEL-54 |
| chr15-10579660-DEL-2099 | chr15-10579649-DEL-2100 |
| chr15-10627457-DEL-60 | chr15-10627458-DEL-60 |
| chr15-10648900-DEL-184 | chr15-10648903-DEL-182 |
| chr15-10649470-DEL-135 | chr15-10649471-DEL-135 |
| chr15-10655551-DEL-132 | chr15-10655552-DEL-99 |
| chr15-10728404-DEL-157 | chr15-10728452-DEL-120 |
| chr15-10730473-DEL-3125 | chr15-10730474-DEL-3125 |
| chr15-10754643-DEL-192 | chr15-10754644-DEL-191 |
| chr15-10763675-DEL-744 | chr15-10763676-DEL-744 |
| chr15-10772022-DEL-6965 | chr15-10772023-DEL-6965 |
| chr15-10781283-DEL-200 | chr15-10781284-DEL-200 |
| chr15-10792704-DEL-82 | chr15-10792694-DEL-92 |
| chr15-10806585-DEL-6239 | chr15-10806586-DEL-6239 |
| chr15-10835291-DEL-1893 | chr15-10835292-DEL-1893 |
| chr15-10890293-DEL-1433 | chr15-10890294-DEL-1433 |
| chr15-10904166-DEL-54 | chr15-10904167-DEL-52 |
| chr15-10913810-DEL-1300 | chr15-10913811-DEL-1300 |
| chr15-10917921-DEL-112 | chr15-10917922-DEL-96 |
| chr15-10933930-DEL-52 | chr15-10933905-DEL-55 |
| chr15-10949297-DEL-60 | chr15-10949284-DEL-64 |
| chr15-10999684-DEL-80 | chr15-10999685-DEL-80 |
| chr15-11010375-DEL-600 | chr15-11010376-DEL-600 |
| chr15-11013406-DEL-1245 | chr15-11013407-DEL-1240 |
| chr15-11015685-DEL-54 | chr15-11015700-DEL-74 |
| chr15-11045883-DEL-6221 | chr15-11045884-DEL-6221 |
| chr15-11079591-DEL-400 | chr15-11079592-DEL-400 |
| chr15-11094240-DEL-54 | chr15-11094241-DEL-54 |

| | |
|---|---|
| chr15-11157748-DEL-102 | chr15-11157748-DEL-101 |
| chr15-11159856-DEL-64 | chr15-11159857-DEL-62 |
| chr15-11167710-DEL-1088 | chr15-11167711-DEL-1088 |
| chr15-11174520-DEL-532 | chr15-11174521-DEL-532 |
| chr15-11194169-DEL-114 | chr15-11194170-DEL-114 |
| chr15-11212301-DEL-200 | chr15-11212302-DEL-200 |
| chr15-11224408-DEL-6758 | chr15-11224409-DEL-6758 |
| chr15-11259600-DEL-697 | chr15-11259601-DEL-697 |
| chr15-11273738-DEL-72 | chr15-11273739-DEL-72 |
| chr15-11302811-DEL-1129 | chr15-11302812-DEL-1129 |
| chr15-11302880-DEL-62 | chr15-11302881-DEL-68 |
| chr15-11320529-DEL-60 | chr15-11320520-DEL-61 |
| chr15-11352489-DEL-304 | chr15-11352490-DEL-304 |
| chr15-11354556-DEL-1080 | chr15-11354556-DEL-1077 |
| chr15-11363059-DEL-1630 | chr15-11363059-DEL-1630 |
| chr15-11384632-DEL-53 | chr15-11384633-DEL-53 |
| chr15-11410455-DEL-64 | chr15-11410437-DEL-55 |
| chr15-11416626-DEL-74 | chr15-11416627-DEL-74 |
| chr15-11428887-DEL-594 | chr15-11428903-DEL-551 |
| chr15-11436070-DEL-60 | chr15-11436098-DEL-64 |
| chr15-11437528-DEL-54 | chr15-11437529-DEL-54 |
| chr15-11442949-DEL-1350 | chr15-11442950-DEL-1351 |
| chr15-11462357-DEL-345 | chr15-11462358-DEL-343 |
| chr15-11483917-DEL-86 | chr15-11483921-DEL-70 |
| chr15-11555894-DEL-77 | chr15-11555890-DEL-78 |
| chr15-11559163-DEL-100 | chr15-11559164-DEL-100 |
| chr15-11565398-DEL-71 | chr15-11565399-DEL-71 |
| chr15-11580905-DEL-2137 | chr15-11580878-DEL-2137 |
| chr15-11583932-DEL-751 | chr15-11583930-DEL-752 |
| chr15-11632260-DEL-400 | chr15-11632261-DEL-400 |
| chr15-11634852-DEL-1962 | chr15-11634853-DEL-1962 |
| chr15-11646604-DEL-363 | chr15-11646605-DEL-364 |
| chr15-11654092-DEL-450 | chr15-11654093-DEL-450 |
| chr15-11673994-DEL-1143 | chr15-11673995-DEL-1143 |
| chr15-11680298-DEL-518 | chr15-11680294-DEL-519 |
| chr15-11681338-DEL-58 | chr15-11681346-DEL-50 |
| chr15-11705843-DEL-400 | chr15-11705844-DEL-400 |
| chr15-11710743-DEL-826 | chr15-11710744-DEL-826 |
| chr15-11715346-DEL-1165 | chr15-11715347-DEL-1165 |
| chr15-11757589-DEL-88 | chr15-11757590-DEL-88 |
| chr15-11787466-DEL-468 | chr15-11787510-DEL-425 |
| chr15-11814925-DEL-54 | chr15-11814926-DEL-54 |
| chr15-11852400-DEL-70 | chr15-11852401-DEL-52 |
| chr15-11886272-DEL-231 | chr15-11886273-DEL-231 |
| chr15-11929543-DEL-85 | chr15-11929544-DEL-84 |
| chr15-12009179-DEL-110 | chr15-12009179-DEL-109 |
| chr15-12056009-DEL-7169 | chr15-12056010-DEL-7169 |
| chr15-12063493-DEL-425 | chr15-12063494-DEL-422 |
| chr15-12065075-DEL-83 | chr15-12065074-DEL-76 |
| chr15-12081922-DEL-99 | chr15-12081922-DEL-99 |
| chr15-12088713-DEL-88 | chr15-12088710-DEL-86 |
| chr15-12107018-DEL-84 | chr15-12107018-DEL-84 |
| chr15-12188949-DEL-1088 | chr15-12188950-DEL-1088 |
| chr15-12245374-DEL-70 | chr15-12245389-DEL-56 |
| chr15-12254814-DEL-378 | chr15-12254815-DEL-378 |
| chr15-12265158-DEL-54 | chr15-12265159-DEL-54 |
| chr15-12314582-DEL-60 | chr15-12314583-DEL-60 |
| chr15-12365018-DEL-135 | chr15-12365004-DEL-126 |
| chr15-12368929-DEL-63 | chr15-12368936-DEL-63 |
| chr15-12398826-DEL-65 | chr15-12398827-DEL-65 |
| chr15-12430730-DEL-281 | chr15-12430729-DEL-282 |
| chr15-12441341-DEL-61 | chr15-12441342-DEL-61 |
| chr15-12449812-DEL-75 | chr15-12449813-DEL-75 |
| chr15-12507245-DEL-53 | chr15-12507234-DEL-58 |
| chr15-12526853-DEL-5052 | chr15-12526854-DEL-5052 |
| chr15-12543450-DEL-156 | chr15-12543451-DEL-156 |
| chr15-12562841-DEL-98 | chr15-12562841-DEL-98 |
| chr15-12566303-DEL-432 | chr15-12566304-DEL-432 |
| chr15-12572284-DEL-559 | chr15-12572285-DEL-559 |
| chr15-12574887-DEL-1289 | chr15-12574888-DEL-1290 |
| chr15-12578582-DEL-1560 | chr15-12578583-DEL-1560 |
| chr15-12595839-DEL-78 | chr15-12595858-DEL-60 |
| chr15-12602100-DEL-650 | chr15-12602101-DEL-650 |
| chr15-12607998-DEL-96 | chr15-12607999-DEL-96 |
| chr15-12609184-DEL-805 | chr15-12609185-DEL-805 |
| chr15-12612040-DEL-54 | chr15-12612041-DEL-54 |
| chr15-12621009-DEL-56 | chr15-12620962-DEL-56 |
| chr15-12632145-DEL-355 | chr15-12632132-DEL-369 |
| chr15-12648119-DEL-158 | chr15-12648120-DEL-158 |
| chr15-12656905-DEL-499 | chr15-12656921-DEL-494 |

| | |
|---|---|
| chr15-12685526-DEL-63 | chr15-12685527-DEL-63 |
| chr15-12715958-DEL-970 | chr15-12715959-DEL-970 |
| chr15-12752665-DEL-75 | chr15-12752678-DEL-75 |
| chr15-12762127-DEL-238 | chr15-12762128-DEL-238 |
| chr15-12782317-DEL-80 | chr15-12782311-DEL-55 |
| chr15-12783907-DEL-100 | chr15-12783908-DEL-55 |
| chr15-12796612-DEL-66 | chr15-12796613-DEL-66 |
| chr15-12830498-DEL-2136 | chr15-12830499-DEL-2136 |
| chr15-12891412-DEL-66 | chr15-12891412-DEL-64 |
| chr15-12892523-DEL-587 | chr15-12892524-DEL-587 |
| chr15-12906014-DEL-69 | chr15-12906016-DEL-67 |
| chr15-12932361-DEL-1890 | chr15-12932362-DEL-1890 |
| chr15-12951679-DEL-551 | chr15-12951680-DEL-551 |
| chr15-12956342-DEL-54 | chr15-12956338-DEL-58 |
| chr15-12961514-DEL-632 | chr15-12961515-DEL-632 |
| chr15-12965736-DEL-62 | chr15-12965738-DEL-62 |
| chr15-12970519-DEL-188 | chr15-12970528-DEL-180 |
| chr15-12978947-DEL-83 | chr15-12978956-DEL-75 |
| chr15-13002776-DEL-459 | chr15-13002776-DEL-459 |
| chr15-13005434-DEL-1058 | chr15-13005435-DEL-1058 |
| chr15-13072234-DEL-541 | chr15-13072235-DEL-541 |
| chr15-13073201-DEL-109 | chr15-13073202-DEL-109 |
| chr15-13079934-DEL-170 | chr15-13079932-DEL-171 |
| chr15-13146368-DEL-183 | chr15-13146369-DEL-183 |
| chr15-13160687-DEL-132 | chr15-13160688-DEL-132 |
| chr15-13167555-DEL-60 | chr15-13167560-DEL-66 |
| chr15-13170025-DEL-103 | chr15-13170064-DEL-56 |
| chr15-13206540-DEL-285 | chr15-13206541-DEL-285 |
| chr15-13220495-DEL-68 | chr15-13220496-DEL-68 |
| chr15-13242233-DEL-64 | chr15-13242234-DEL-64 |
| chr15-13255519-DEL-54 | chr15-13255520-DEL-54 |
| chr15-13260181-DEL-61 | chr15-13260182-DEL-61 |
| chr15-13291902-DEL-86 | chr15-13291902-DEL-87 |
| chr15-13313093-DEL-70 | chr15-13313094-DEL-70 |
| chr15-13326369-DEL-56 | chr15-13326370-DEL-56 |
| chr15-13347007-DEL-103 | chr15-13347008-DEL-103 |
| chr15-13355972-DEL-60 | chr15-13355973-DEL-60 |
| chr15-13384033-DEL-103 | chr15-13384040-DEL-97 |
| chr15-13390374-DEL-80 | chr15-13390375-DEL-80 |
| chr15-13401895-DEL-886 | chr15-13401896-DEL-886 |
| chr15-13448377-DEL-140 | chr15-13448406-DEL-112 |
| chr15-13484185-DEL-280 | chr15-13484186-DEL-280 |
| chr15-13485094-DEL-164 | chr15-13485089-DEL-180 |
| chr15-13494476-DEL-78 | chr15-13494488-DEL-79 |
| chr15-13507460-DEL-867 | chr15-13507477-DEL-862 |
| chr15-13510183-DEL-57 | chr15-13510184-DEL-57 |
| chr15-13510803-DEL-328 | chr15-13510804-DEL-326 |
| chr15-13592431-DEL-85 | chr15-13592419-DEL-86 |
| chr15-13596517-DEL-77 | chr15-13596477-DEL-106 |
| chr15-13597242-DEL-117 | chr15-13597243-DEL-118 |
| chr15-13614085-DEL-1301 | chr15-13614087-DEL-1300 |
| chr15-13637319-DEL-88 | chr15-13637320-DEL-104 |
| chr15-13662060-DEL-55 | chr15-13662060-DEL-50 |
| chr15-13676776-DEL-249 | chr15-13676777-DEL-249 |
| chr15-13687791-DEL-4150 | chr15-13687792-DEL-4150 |
| chr15-13728922-DEL-97 | chr15-13728923-DEL-97 |
| chr15-13734151-DEL-59 | chr15-13734158-DEL-53 |
| chr15-13750334-DEL-50 | chr15-13750333-DEL-62 |
| chr15-13797668-DEL-239 | chr15-13797668-DEL-240 |
| chr15-13806077-DEL-52 | chr15-13806079-DEL-54 |
| chr15-13815562-DEL-428 | chr15-13815563-DEL-428 |
| chr15-13880760-DEL-6356 | chr15-13880761-DEL-6356 |
| chr15-13934405-DEL-95 | chr15-13934410-DEL-92 |
| chr15-13939618-DEL-267 | chr15-13939619-DEL-267 |
| chr15-13946417-DEL-4544 | chr15-13946418-DEL-4544 |
| chr15-13958390-DEL-68 | chr15-13958391-DEL-68 |
| chr15-13968905-DEL-124 | chr15-13968906-DEL-124 |
| chr15-14047330-DEL-55 | chr15-14047331-DEL-55 |
| chr15-14047639-DEL-824 | chr15-14047640-DEL-824 |
| chr15-14087192-DEL-1915 | chr15-14087193-DEL-1915 |
| chr15-14128599-DEL-146 | chr15-14128600-DEL-146 |
| chr15-14136569-DEL-55 | chr15-14136570-DEL-71 |
| chr15-14141074-DEL-1949 | chr15-14141075-DEL-1949 |
| chr15-14169199-DEL-3652 | chr15-14169186-DEL-3653 |
| chr15-14169947-DEL-212 | chr15-14169948-DEL-212 |
| chr15-14176647-DEL-108 | chr15-14176675-DEL-69 |
| chr15-14182446-DEL-82 | chr15-14182442-DEL-86 |
| chr15-14212969-DEL-399 | chr15-14212970-DEL-399 |
| chr15-14223950-DEL-137 | chr15-14223951-DEL-137 |
| chr15-14235439-DEL-84 | chr15-14235440-DEL-84 |

| | |
|---|---|
| chr15-14261213-DEL-94 | chr15-14261214-DEL-94 |
| chr15-14264424-DEL-72 | chr15-14264441-DEL-52 |
| chr15-14286479-DEL-5419 | chr15-14286480-DEL-5419 |
| chr15-14326095-DEL-241 | chr15-14326123-DEL-214 |
| chr15-14361712-DEL-90 | chr15-14361714-DEL-91 |
| chr15-14368177-DEL-337 | chr15-14368178-DEL-337 |
| chr15-14394325-DEL-227 | chr15-14394326-DEL-227 |
| chr15-14395591-DEL-401 | chr15-14395592-DEL-401 |
| chr15-14403023-DEL-52 | chr15-14403024-DEL-52 |
| chr15-14414660-DEL-1888 | chr15-14414661-DEL-1888 |
| chr15-14432790-DEL-6336 | chr15-14432791-DEL-6336 |
| chr15-14444712-DEL-60 | chr15-14444713-DEL-60 |
| chr15-14470700-DEL-677 | chr15-14470701-DEL-675 |
| chr15-14474464-DEL-66 | chr15-14474463-DEL-67 |
| chr15-14499238-DEL-58 | chr15-14499238-DEL-58 |
| chr15-14505816-DEL-1134 | chr15-14505816-DEL-1134 |
| chr15-14514066-DEL-333 | chr15-14514067-DEL-333 |
| chr15-14519467-DEL-52 | chr15-14519466-DEL-52 |
| chr15-14536301-DEL-497 | chr15-14536301-DEL-497 |
| chr15-14581345-DEL-472 | chr15-14581346-DEL-472 |
| chr15-14635520-DEL-102 | chr15-14635521-DEL-102 |
| chr15-14656653-DEL-175 | chr15-14656654-DEL-175 |
| chr15-14661913-DEL-64 | chr15-14661914-DEL-63 |
| chr15-14673749-DEL-415 | chr15-14673750-DEL-415 |
| chr15-14684040-DEL-220 | chr15-14684041-DEL-228 |
| chr15-14706691-DEL-352 | chr15-14706692-DEL-352 |
| chr15-14718854-DEL-57 | chr15-14718854-DEL-51 |
| chr15-14722167-DEL-11399 | chr15-14722168-DEL-11399 |
| chr15-14738526-DEL-2949 | chr15-14738527-DEL-2949 |
| chr15-14760889-DEL-440 | chr15-14760890-DEL-440 |
| chr15-14776346-DEL-110 | chr15-14776347-DEL-110 |
| chr15-14799720-DEL-1078 | chr15-14799719-DEL-1079 |
| chr15-14805628-DEL-621 | chr15-14805629-DEL-621 |
| chr15-14811507-DEL-1307 | chr15-14811508-DEL-1307 |
| chr15-14813046-DEL-372 | chr15-14813047-DEL-372 |
| chr15-14842269-DEL-2267 | chr15-14842269-DEL-2267 |
| chr15-14845014-DEL-8177 | chr15-14845014-DEL-8177 |
| chr15-14857267-DEL-1020 | chr15-14857268-DEL-1020 |
| chr15-14873040-DEL-5800 | chr15-14873040-DEL-5800 |
| chr15-14903725-DEL-87 | chr15-14903724-DEL-86 |
| chr15-14913055-DEL-314 | chr15-14913056-DEL-314 |
| chr15-14942455-DEL-2888 | chr15-14942440-DEL-2903 |
| chr15-14956014-DEL-348 | chr15-14956013-DEL-348 |
| chr15-14967478-DEL-56 | chr15-14967479-DEL-56 |
| chr15-14977193-DEL-2021 | chr15-14977194-DEL-2020 |
| chr15-14985238-DEL-78 | chr15-14985239-DEL-54 |
| chr15-15004036-DEL-1908 | chr15-15004022-DEL-1909 |
| chr15-15024510-DEL-206 | chr15-15024511-DEL-206 |
| chr15-15027599-DEL-86 | chr15-15027600-DEL-86 |
| chr15-15030289-DEL-105 | chr15-15030290-DEL-105 |
| chr15-15130535-DEL-132 | chr15-15130536-DEL-132 |
| chr15-15158897-DEL-894 | chr15-15158902-DEL-891 |
| chr15-15215251-DEL-58 | chr15-15215252-DEL-58 |
| chr15-15288004-DEL-169 | chr15-15288005-DEL-170 |
| chr15-15314819-DEL-163 | chr15-15314820-DEL-163 |
| chr15-15320383-DEL-454 | chr15-15320384-DEL-454 |
| chr15-15320894-DEL-736 | chr15-15320895-DEL-736 |
| chr15-15370013-DEL-529 | chr15-15370014-DEL-529 |
| chr15-15387562-DEL-644 | chr15-15387554-DEL-652 |
| chr15-15433107-DEL-54 | chr15-15433112-DEL-68 |
| chr15-15445960-DEL-560 | chr15-15445961-DEL-560 |
| chr15-15461043-DEL-504 | chr15-15461044-DEL-504 |
| chr15-15465552-DEL-1765 | chr15-15465553-DEL-1764 |
| chr15-15476704-DEL-814 | chr15-15476705-DEL-814 |
| chr15-15502554-DEL-308 | chr15-15502555-DEL-308 |
| chr15-15503147-DEL-475 | chr15-15503132-DEL-493 |
| chr15-15512451-DEL-78 | chr15-15512472-DEL-50 |
| chr15-15530288-DEL-199 | chr15-15530275-DEL-198 |
| chr15-15547974-DEL-4261 | chr15-15547975-DEL-4261 |
| chr15-15569928-DEL-104 | chr15-15569925-DEL-108 |
| chr15-15580920-DEL-66 | chr15-15580921-DEL-54 |
| chr15-15585167-DEL-701 | chr15-15585168-DEL-701 |
| chr15-15640534-DEL-6462 | chr15-15640535-DEL-6462 |
| chr15-15646477-DEL-201 | chr15-15646478-DEL-201 |
| chr15-15708262-DEL-56 | chr15-15708251-DEL-56 |
| chr15-15720551-DEL-415 | chr15-15720552-DEL-415 |
| chr15-15775405-DEL-57 | chr15-15775405-DEL-57 |
| chr15-15779803-DEL-390 | chr15-15779804-DEL-390 |
| chr15-15784854-DEL-4503 | chr15-15784855-DEL-4503 |
| chr15-15802802-DEL-685 | chr15-15802803-DEL-685 |

| | |
|---|---|
| chr15-15803741-DEL-85 | chr15-15803742-DEL-85 |
| chr15-15831504-DEL-60 | chr15-15831469-DEL-60 |
| chr15-15902320-DEL-6455 | chr15-15902321-DEL-6455 |
| chr15-15950623-DEL-109 | chr15-15950624-DEL-109 |
| chr15-15954140-DEL-60 | chr15-15954142-DEL-76 |
| chr15-15965093-DEL-55 | chr15-15965104-DEL-65 |
| chr15-15971931-DEL-419 | chr15-15971932-DEL-419 |
| chr15-15994443-DEL-1085 | chr15-15994444-DEL-1085 |
| chr15-16015782-DEL-68 | chr15-16015788-DEL-62 |
| chr15-16036189-DEL-52 | chr15-16036190-DEL-52 |
| chr15-16075489-DEL-172 | chr15-16075489-DEL-172 |
| chr15-16086205-DEL-51 | chr15-16086206-DEL-51 |
| chr15-16088976-DEL-63 | chr15-16088958-DEL-75 |
| chr15-16101160-DEL-1385 | chr15-16101161-DEL-1385 |
| chr15-16115154-DEL-6672 | chr15-16115155-DEL-6672 |
| chr15-16117533-DEL-426 | chr15-16117534-DEL-426 |
| chr15-16137323-DEL-401 | chr15-16137324-DEL-401 |
| chr15-16139917-DEL-617 | chr15-16139918-DEL-617 |
| chr15-16162628-DEL-355 | chr15-16162629-DEL-355 |
| chr15-16181575-DEL-4938 | chr15-16181577-DEL-4939 |
| chr15-16198705-DEL-235 | chr15-16198706-DEL-235 |
| chr15-16232081-DEL-276 | chr15-16232082-DEL-274 |
| chr15-16233623-DEL-482 | chr15-16233624-DEL-482 |
| chr15-16248993-DEL-2130 | chr15-16248994-DEL-2130 |
| chr15-16256613-DEL-115 | chr15-16256614-DEL-115 |
| chr15-16280934-DEL-1589 | chr15-16280934-DEL-1589 |
| chr15-16285151-DEL-100 | chr15-16285152-DEL-90 |
| chr15-16291509-DEL-2293 | chr15-16291508-DEL-2290 |
| chr15-16295124-DEL-410 | chr15-16295125-DEL-410 |
| chr15-16299936-DEL-2346 | chr15-16299937-DEL-2345 |
| chr15-16300546-DEL-246 | chr15-16300547-DEL-246 |
| chr15-16303448-DEL-6210 | chr15-16303449-DEL-6210 |
| chr15-16365335-DEL-58 | chr15-16365336-DEL-58 |
| chr15-16387485-DEL-6282 | chr15-16387486-DEL-6282 |
| chr15-16450235-DEL-214 | chr15-16450236-DEL-214 |
| chr15-16454949-DEL-2479 | chr15-16454950-DEL-2479 |
| chr15-16502511-DEL-184 | chr15-16502512-DEL-184 |
| chr15-16526484-DEL-364 | chr15-16526485-DEL-364 |
| chr15-16528268-DEL-381 | chr15-16528269-DEL-381 |
| chr15-16608350-DEL-217 | chr15-16608348-DEL-208 |
| chr15-16612054-DEL-2850 | chr15-16612055-DEL-2850 |
| chr15-16618135-DEL-660 | chr15-16618136-DEL-660 |
| chr15-16619071-DEL-939 | chr15-16619072-DEL-939 |
| chr15-16624466-DEL-55 | chr15-16624467-DEL-55 |
| chr15-16644129-DEL-65 | chr15-16644145-DEL-98 |
| chr15-16663622-DEL-1457 | chr15-16663617-DEL-1458 |
| chr15-16675220-DEL-86 | chr15-16675220-DEL-86 |
| chr15-16692846-DEL-56 | chr15-16692848-DEL-55 |
| chr15-16725068-DEL-94 | chr15-16725069-DEL-94 |
| chr15-16739708-DEL-1658 | chr15-16739709-DEL-1658 |
| chr15-16805470-DEL-108 | chr15-16805471-DEL-108 |
| chr15-16815135-DEL-168 | chr15-16815141-DEL-180 |
| chr15-16822344-DEL-465 | chr15-16822345-DEL-464 |
| chr15-16901413-DEL-6171 | chr15-16901420-DEL-6173 |
| chr15-16934730-DEL-106 | chr15-16934731-DEL-106 |
| chr15-16948865-DEL-6401 | chr15-16948866-DEL-6401 |
| chr15-16970852-DEL-63 | chr15-16970830-DEL-64 |
| chr15-16972903-DEL-81 | chr15-16972916-DEL-69 |
| chr15-17028910-DEL-1954 | chr15-17028911-DEL-1954 |
| chr15-17068267-DEL-76 | chr15-17068268-DEL-76 |
| chr15-17087265-DEL-66 | chr15-17087266-DEL-66 |
| chr15-17096855-DEL-57 | chr15-17096855-DEL-57 |
| chr15-17119320-DEL-68 | chr15-17119321-DEL-68 |
| chr15-17138444-DEL-2573 | chr15-17138445-DEL-2573 |
| chr15-17141148-DEL-178 | chr15-17141148-DEL-179 |
| chr15-17155803-DEL-13885 | chr15-17155804-DEL-13885 |
| chr15-17182430-DEL-977 | chr15-17182431-DEL-977 |
| chr15-17203727-DEL-113 | chr15-17203728-DEL-112 |
| chr15-17225005-DEL-202 | chr15-17225006-DEL-202 |
| chr15-17239945-DEL-52 | chr15-17239946-DEL-52 |
| chr15-17248605-DEL-4629 | chr15-17248606-DEL-4629 |
| chr15-17266481-DEL-7268 | chr15-17266482-DEL-7268 |
| chr15-17325756-DEL-64 | chr15-17325759-DEL-62 |
| chr15-17336983-DEL-607 | chr15-17336984-DEL-607 |
| chr15-17339232-DEL-95 | chr15-17339233-DEL-95 |
| chr15-17343188-DEL-450 | chr15-17343189-DEL-450 |
| chr15-17397526-DEL-64 | chr15-17397530-DEL-53 |
| chr15-17402568-DEL-100 | chr15-17402569-DEL-100 |
| chr15-17404127-DEL-732 | chr15-17404128-DEL-732 |
| chr15-17424455-DEL-112 | chr15-17424467-DEL-78 |

| | |
|---|---|
| chr15-17428984-DEL-451 | chr15-17428985-DEL-451 |
| chr15-17444551-DEL-224 | chr15-17444552-DEL-180 |
| chr15-17464711-DEL-111 | chr15-17464720-DEL-103 |
| chr15-17483711-DEL-67 | chr15-17483706-DEL-51 |
| chr15-17541275-DEL-143 | chr15-17541276-DEL-143 |
| chr15-17551341-DEL-2112 | chr15-17551342-DEL-2112 |
| chr15-17576343-DEL-1851 | chr15-17576341-DEL-1849 |
| chr15-17590449-DEL-68 | chr15-17590450-DEL-68 |
| chr15-17591668-DEL-54 | chr15-17591668-DEL-54 |
| chr15-17612868-DEL-91 | chr15-17612869-DEL-91 |
| chr15-17661500-DEL-144 | chr15-17661501-DEL-144 |
| chr15-17685097-DEL-63 | chr15-17685098-DEL-63 |
| chr15-17685603-DEL-249 | chr15-17685604-DEL-249 |
| chr15-17691690-DEL-959 | chr15-17691691-DEL-959 |
| chr15-17707205-DEL-474 | chr15-17707206-DEL-472 |
| chr15-17726609-DEL-2036 | chr15-17726610-DEL-2036 |
| chr15-17738121-DEL-65 | chr15-17738118-DEL-60 |
| chr15-17752780-DEL-1265 | chr15-17752782-DEL-1262 |
| chr15-17763859-DEL-313 | chr15-17763863-DEL-310 |
| chr15-17772232-DEL-60 | chr15-17772233-DEL-60 |
| chr15-17772989-DEL-1503 | chr15-17772990-DEL-1503 |
| chr15-17777653-DEL-4835 | chr15-17777654-DEL-4835 |
| chr15-17822901-DEL-392 | chr15-17822902-DEL-392 |
| chr15-17825685-DEL-247 | chr15-17825686-DEL-248 |
| chr15-17863431-DEL-1202 | chr15-17863432-DEL-1202 |
| chr15-17864148-DEL-132 | chr15-17864148-DEL-182 |
| chr15-17876350-DEL-486 | chr15-17876351-DEL-486 |
| chr15-17880796-DEL-98 | chr15-17880797-DEL-98 |
| chr15-17882105-DEL-800 | chr15-17882106-DEL-800 |
| chr15-17900531-DEL-150 | chr15-17900532-DEL-150 |
| chr15-17905509-DEL-1527 | chr15-17905510-DEL-1527 |
| chr15-17919023-DEL-214 | chr15-17919024-DEL-214 |
| chr15-17920660-DEL-119 | chr15-17920661-DEL-119 |
| chr15-17928172-DEL-660 | chr15-17928173-DEL-661 |
| chr15-17937790-DEL-1083 | chr15-17937791-DEL-1083 |
| chr15-17938200-DEL-395 | chr15-17938201-DEL-395 |
| chr15-17941981-DEL-1089 | chr15-17941981-DEL-1089 |
| chr15-17966866-DEL-54 | chr15-17966863-DEL-55 |
| chr15-17980820-DEL-1192 | chr15-17980821-DEL-1192 |
| chr15-17983315-DEL-80 | chr15-17983334-DEL-51 |
| chr15-18001756-DEL-221 | chr15-18001757-DEL-221 |
| chr15-18053324-DEL-103 | chr15-18053325-DEL-103 |
| chr15-18056923-DEL-506 | chr15-18056946-DEL-498 |
| chr15-18097074-DEL-53 | chr15-18097074-DEL-53 |
| chr15-18107460-DEL-543 | chr15-18107460-DEL-543 |
| chr15-18125104-DEL-2842 | chr15-18125105-DEL-2842 |
| chr15-18175036-DEL-52 | chr15-18175037-DEL-50 |
| chr15-18175606-DEL-1370 | chr15-18175606-DEL-1370 |
| chr15-18195915-DEL-568 | chr15-18195916-DEL-568 |
| chr15-18199122-DEL-630 | chr15-18199123-DEL-625 |
| chr15-18212624-DEL-52 | chr15-18212624-DEL-52 |
| chr15-18313672-DEL-98 | chr15-18313673-DEL-98 |
| chr15-18314594-DEL-6518 | chr15-18314595-DEL-6518 |
| chr15-18321057-DEL-233 | chr15-18321058-DEL-233 |
| chr15-18328952-DEL-1651 | chr15-18328953-DEL-1651 |
| chr15-18338919-DEL-52 | chr15-18338920-DEL-52 |
| chr15-18343430-DEL-3041 | chr15-18343431-DEL-3041 |
| chr15-18405005-DEL-6078 | chr15-18405006-DEL-6078 |
| chr15-18430550-DEL-756 | chr15-18430592-DEL-757 |
| chr15-18497507-DEL-93 | chr15-18497508-DEL-93 |
| chr15-18503103-DEL-115 | chr15-18503104-DEL-115 |
| chr15-18519818-DEL-7456 | chr15-18519819-DEL-7456 |
| chr15-18554562-DEL-272 | chr15-18554563-DEL-272 |
| chr15-18554565-DEL-1174 | chr15-18554566-DEL-1174 |
| chr15-18566919-DEL-544 | chr15-18566920-DEL-544 |
| chr15-18664462-DEL-2280 | chr15-18664462-DEL-2280 |
| chr15-18731342-DEL-193 | chr15-18731343-DEL-193 |
| chr15-18750845-DEL-54 | chr15-18750846-DEL-54 |
| chr15-18781329-DEL-57 | chr15-18781332-DEL-55 |
| chr15-18835910-DEL-2094 | chr15-18835911-DEL-2094 |
| chr15-18892321-DEL-84 | chr15-18892322-DEL-84 |
| chr15-19045554-DEL-52 | chr15-19045555-DEL-52 |
| chr15-19045956-DEL-344 | chr15-19045957-DEL-344 |
| chr15-19049239-DEL-4542 | chr15-19049240-DEL-4542 |
| chr15-19057405-DEL-6452 | chr15-19057406-DEL-6452 |
| chr15-19067821-DEL-503 | chr15-19067816-DEL-505 |
| chr15-19081036-DEL-897 | chr15-19081034-DEL-900 |
| chr15-19083237-DEL-282 | chr15-19083238-DEL-282 |
| chr15-19085565-DEL-111 | chr15-19085566-DEL-111 |
| chr15-19089181-DEL-845 | chr15-19089182-DEL-845 |

| | |
|---|---|
| chr15-19103481-DEL-1272 | chr15-19103474-DEL-1273 |
| chr15-19118483-DEL-60 | chr15-19118484-DEL-60 |
| chr15-19139259-DEL-52 | chr15-19139260-DEL-52 |
| chr15-19174990-DEL-80 | chr15-19174991-DEL-80 |
| chr15-19200853-DEL-76 | chr15-19200854-DEL-76 |
| chr15-19226809-DEL-78 | chr15-19226793-DEL-70 |
| chr15-19232364-DEL-1171 | chr15-19232365-DEL-1171 |
| chr15-19232605-DEL-69 | chr15-19232602-DEL-70 |
| chr15-19258196-DEL-56 | chr15-19258192-DEL-53 |
| chr15-19264224-DEL-485 | chr15-19264225-DEL-485 |
| chr15-19274187-DEL-678 | chr15-19274233-DEL-632 |
| chr15-19279137-DEL-203 | chr15-19279138-DEL-203 |
| chr15-19312883-DEL-669 | chr15-19312884-DEL-669 |
| chr15-19355594-DEL-85 | chr15-19355601-DEL-84 |
| chr15-19361845-DEL-53 | chr15-19361846-DEL-53 |
| chr15-19363799-DEL-65 | chr15-19363795-DEL-77 |
| chr15-19365488-DEL-1005 | chr15-19365489-DEL-1005 |
| chr15-19369798-DEL-196 | chr15-19369799-DEL-196 |
| chr15-19370252-DEL-519 | chr15-19370253-DEL-519 |
| chr15-19376320-DEL-238 | chr15-19376313-DEL-239 |
| chr15-19376875-DEL-213 | chr15-19376887-DEL-211 |
| chr15-19378442-DEL-324 | chr15-19378443-DEL-324 |
| chr15-19382020-DEL-97 | chr15-19382021-DEL-97 |
| chr15-19402672-DEL-2937 | chr15-19402673-DEL-2937 |
| chr15-19411272-DEL-60 | chr15-19411273-DEL-60 |
| chr15-19418044-DEL-1473 | chr15-19418045-DEL-1473 |
| chr15-19421700-DEL-399 | chr15-19421701-DEL-399 |
| chr15-19440454-DEL-306 | chr15-19440455-DEL-306 |
| chr15-19503259-DEL-105 | chr15-19503241-DEL-95 |
| chr15-19517177-DEL-225 | chr15-19517167-DEL-226 |
| chr15-19518991-DEL-1573 | chr15-19518992-DEL-1573 |
| chr15-19520649-DEL-204 | chr15-19520632-DEL-204 |
| chr15-19524327-DEL-9163 | chr15-19524328-DEL-9163 |
| chr15-19551590-DEL-203 | chr15-19551600-DEL-191 |
| chr15-19574068-DEL-214 | chr15-19574069-DEL-214 |
| chr15-19577533-DEL-112 | chr15-19577534-DEL-112 |
| chr15-19595178-DEL-52 | chr15-19595179-DEL-50 |
| chr15-19626004-DEL-6348 | chr15-19626005-DEL-6348 |
| chr15-19669261-DEL-55 | chr15-19669260-DEL-54 |
| chr15-19678094-DEL-7837 | chr15-19678078-DEL-7838 |
| chr15-19686407-DEL-99 | chr15-19686408-DEL-99 |
| chr15-19698126-DEL-162 | chr15-19698127-DEL-162 |
| chr15-19708941-DEL-104 | chr15-19708942-DEL-104 |
| chr15-19714076-DEL-138 | chr15-19714095-DEL-138 |
| chr15-19718564-DEL-100 | chr15-19718599-DEL-55 |
| chr15-19719120-DEL-117 | chr15-19719121-DEL-117 |
| chr15-19721387-DEL-17123 | chr15-19721390-DEL-17091 |
| chr15-19738480-DEL-6238 | chr15-19738478-DEL-6236 |
| chr15-19774456-DEL-72 | chr15-19774463-DEL-66 |
| chr15-19776916-DEL-74 | chr15-19776917-DEL-74 |
| chr15-19796794-DEL-249 | chr15-19796795-DEL-249 |
| chr15-19800962-DEL-220 | chr15-19800965-DEL-218 |
| chr15-19828574-DEL-51 | chr15-19828567-DEL-50 |
| chr15-19857262-DEL-493 | chr15-19857263-DEL-493 |
| chr15-19877922-DEL-619 | chr15-19877923-DEL-618 |
| chr15-19879889-DEL-238 | chr15-19879890-DEL-238 |
| chr15-19882687-DEL-72 | chr15-19882710-DEL-92 |
| chr15-19883713-DEL-1467 | chr15-19883714-DEL-1467 |
| chr15-19905325-DEL-991 | chr15-19905326-DEL-991 |
| chr15-19932743-DEL-146 | chr15-19932744-DEL-146 |
| chr15-19933942-DEL-120 | chr15-19933943-DEL-119 |
| chr15-19935490-DEL-60 | chr15-19935491-DEL-60 |
| chr15-19956208-DEL-86 | chr15-19956208-DEL-86 |
| chr15-19966729-DEL-946 | chr15-19966730-DEL-946 |
| chr15-20036570-DEL-54 | chr15-20036573-DEL-52 |
| chr15-20040984-DEL-337 | chr15-20040985-DEL-337 |
| chr15-20042070-DEL-210 | chr15-20042071-DEL-210 |
| chr15-20043108-DEL-76 | chr15-20043121-DEL-64 |
| chr15-20080914-DEL-82 | chr15-20080915-DEL-82 |
| chr15-20097945-DEL-1809 | chr15-20097946-DEL-1809 |
| chr15-20128119-DEL-57 | chr15-20128131-DEL-56 |
| chr15-20137685-DEL-90 | chr15-20137686-DEL-88 |
| chr15-20143741-DEL-1551 | chr15-20143742-DEL-1551 |
| chr15-20194333-DEL-64 | chr15-20194341-DEL-56 |
| chr15-20205177-DEL-219 | chr15-20205178-DEL-219 |
| chr15-20316630-DEL-6872 | chr15-20316631-DEL-6872 |
| chr15-20323816-DEL-57 | chr15-20323817-DEL-57 |
| chr15-20336635-DEL-190 | chr15-20336635-DEL-190 |
| chr15-20338724-DEL-673 | chr15-20338722-DEL-655 |
| chr15-20349222-DEL-6578 | chr15-20349223-DEL-6578 |

| | |
|---|---|
| chr15-20384194-DEL-604 | chr15-20384195-DEL-604 |
| chr15-20398752-DEL-1425 | chr15-20398753-DEL-1425 |
| chr15-20408304-DEL-53 | chr15-20408305-DEL-53 |
| chr15-20430823-DEL-86 | chr15-20430824-DEL-90 |
| chr15-20442157-DEL-57 | chr15-20442158-DEL-57 |
| chr15-20474870-DEL-6611 | chr15-20474871-DEL-6611 |
| chr15-20497137-DEL-66 | chr15-20497111-DEL-50 |
| chr15-20499139-DEL-6219 | chr15-20499140-DEL-6219 |
| chr15-20507396-DEL-2799 | chr15-20507397-DEL-2799 |
| chr15-20526097-DEL-3145 | chr15-20526102-DEL-3141 |
| chr15-20531427-DEL-6422 | chr15-20531428-DEL-6422 |
| chr15-20542612-DEL-511 | chr15-20542613-DEL-511 |
| chr15-20545131-DEL-207 | chr15-20545133-DEL-205 |
| chr15-20579383-DEL-298 | chr15-20579384-DEL-298 |
| chr15-20581383-DEL-57 | chr15-20581384-DEL-57 |
| chr15-20597301-DEL-1160 | chr15-20597301-DEL-1160 |
| chr15-20614535-DEL-60 | chr15-20614524-DEL-66 |
| chr15-20620066-DEL-3888 | chr15-20620067-DEL-3888 |
| chr15-20645167-DEL-397 | chr15-20645185-DEL-351 |
| chr15-20647297-DEL-3207 | chr15-20647298-DEL-3207 |
| chr15-20668068-DEL-58 | chr15-20668069-DEL-58 |
| chr15-20728142-DEL-75 | chr15-20728144-DEL-77 |
| chr15-20790059-DEL-477 | chr15-20790060-DEL-477 |
| chr15-20799747-DEL-157 | chr15-20799748-DEL-157 |
| chr15-20808778-DEL-145 | chr15-20808778-DEL-143 |
| chr15-20821518-DEL-6369 | chr15-20821516-DEL-6370 |
| chr15-20849755-DEL-181 | chr15-20849756-DEL-181 |
| chr15-20870955-DEL-3035 | chr15-20870956-DEL-3035 |
| chr15-20892117-DEL-299 | chr15-20892120-DEL-297 |
| chr15-20909852-DEL-51 | chr15-20909853-DEL-51 |
| chr15-20910480-DEL-377 | chr15-20910472-DEL-377 |
| chr15-20960090-DEL-69 | chr15-20960080-DEL-70 |
| chr15-20971645-DEL-887 | chr15-20971643-DEL-888 |
| chr15-20972419-DEL-97 | chr15-20972444-DEL-73 |
| chr15-20972941-DEL-5317 | chr15-20972935-DEL-5318 |
| chr15-21013612-DEL-498 | chr15-21013613-DEL-497 |
| chr15-21057324-DEL-93 | chr15-21057325-DEL-93 |
| chr15-21101836-DEL-561 | chr15-21101837-DEL-561 |
| chr15-21193617-DEL-875 | chr15-21193618-DEL-875 |
| chr15-21262108-DEL-6478 | chr15-21262109-DEL-6478 |
| chr15-21286707-DEL-1027 | chr15-21286709-DEL-1028 |
| chr15-21322773-DEL-51 | chr15-21322774-DEL-51 |
| chr15-21335495-DEL-7166 | chr15-21335496-DEL-7166 |
| chr15-21367032-DEL-89 | chr15-21367033-DEL-89 |
| chr15-21369587-DEL-412 | chr15-21369588-DEL-412 |
| chr15-21373997-DEL-64 | chr15-21373998-DEL-64 |
| chr15-21444220-DEL-199 | chr15-21444221-DEL-199 |
| chr15-21450138-DEL-4281 | chr15-21450137-DEL-4281 |
| chr15-21454212-DEL-205 | chr15-21454213-DEL-205 |
| chr15-21617237-DEL-51 | chr15-21617238-DEL-51 |
| chr15-21682299-DEL-1417 | chr15-21682300-DEL-1415 |
| chr15-21699332-DEL-56 | chr15-21699319-DEL-60 |
| chr15-21719565-DEL-2781 | chr15-21719566-DEL-2781 |
| chr15-21725369-DEL-57 | chr15-21725364-DEL-57 |
| chr15-21726540-DEL-1527 | chr15-21726541-DEL-1527 |
| chr15-21814100-DEL-60 | chr15-21814103-DEL-58 |
| chr15-21829635-DEL-438 | chr15-21829631-DEL-439 |
| chr15-21839424-DEL-2222 | chr15-21839425-DEL-2222 |
| chr15-21887803-DEL-53 | chr15-21887804-DEL-53 |
| chr15-21889402-DEL-2998 | chr15-21889403-DEL-2998 |
| chr15-21901156-DEL-83 | chr15-21901157-DEL-83 |
| chr15-21905877-DEL-1293 | chr15-21905878-DEL-1293 |
| chr15-21971048-DEL-60 | chr15-21971048-DEL-72 |
| chr15-22023083-DEL-87 | chr15-22023083-DEL-87 |
| chr15-22055106-DEL-762 | chr15-22055107-DEL-762 |
| chr15-22068136-DEL-69 | chr15-22068136-DEL-69 |
| chr15-22093404-DEL-56 | chr15-22093405-DEL-50 |
| chr15-22172325-DEL-1884 | chr15-22172326-DEL-1884 |
| chr15-22197333-DEL-777 | chr15-22197334-DEL-777 |
| chr15-22213401-DEL-86 | chr15-22213402-DEL-86 |
| chr15-22235409-DEL-62 | chr15-22235410-DEL-62 |
| chr15-22257603-DEL-153 | chr15-22257604-DEL-153 |
| chr15-22265086-DEL-426 | chr15-22265087-DEL-426 |
| chr15-22268034-DEL-6384 | chr15-22268035-DEL-6384 |
| chr15-22275569-DEL-484 | chr15-22275570-DEL-484 |
| chr15-22315198-DEL-51 | chr15-22315197-DEL-52 |
| chr15-22326627-DEL-59 | chr15-22326594-DEL-56 |
| chr15-22360747-DEL-433 | chr15-22360748-DEL-433 |
| chr15-22362540-DEL-4648 | chr15-22362547-DEL-4648 |
| chr15-22394974-DEL-445 | chr15-22394975-DEL-445 |

| | |
|---|---|
| chr15-22396727-DEL-519 | chr15-22396728-DEL-519 |
| chr15-22479011-DEL-52 | chr15-22479012-DEL-50 |
| chr15-22485361-DEL-62 | chr15-22485357-DEL-57 |
| chr15-22488114-DEL-214 | chr15-22488115-DEL-214 |
| chr15-22536914-DEL-83 | chr15-22536915-DEL-83 |
| chr15-22549814-DEL-142 | chr15-22549815-DEL-142 |
| chr15-22586117-DEL-53 | chr15-22586120-DEL-52 |
| chr15-22621380-DEL-63 | chr15-22621392-DEL-58 |
| chr15-22623294-DEL-358 | chr15-22623295-DEL-358 |
| chr15-22628268-DEL-217 | chr15-22628269-DEL-217 |
| chr15-22645214-DEL-156 | chr15-22645215-DEL-156 |
| chr15-22648809-DEL-284 | chr15-22648810-DEL-284 |
| chr15-22649675-DEL-173 | chr15-22649676-DEL-173 |
| chr15-22672745-DEL-94 | chr15-22672746-DEL-94 |
| chr15-22717419-DEL-66 | chr15-22717426-DEL-90 |
| chr15-22743749-DEL-163 | chr15-22743750-DEL-163 |
| chr15-22751542-DEL-5242 | chr15-22751549-DEL-5234 |
| chr15-22769844-DEL-58 | chr15-22769876-DEL-57 |
| chr15-22773486-DEL-411 | chr15-22773487-DEL-411 |
| chr15-22776942-DEL-56 | chr15-22776943-DEL-56 |
| chr15-22802370-DEL-3432 | chr15-22802370-DEL-3440 |
| chr15-22818525-DEL-1267 | chr15-22818526-DEL-1267 |
| chr15-22833687-DEL-51 | chr15-22833695-DEL-76 |
| chr15-22834812-DEL-942 | chr15-22834813-DEL-942 |
| chr15-22835337-DEL-79 | chr15-22835338-DEL-79 |
| chr15-22849000-DEL-80 | chr15-22849001-DEL-80 |
| chr15-22862249-DEL-64 | chr15-22862250-DEL-64 |
| chr15-22893110-DEL-282 | chr15-22893110-DEL-282 |
| chr15-22962429-DEL-1253 | chr15-22962430-DEL-1253 |
| chr15-22974269-DEL-60 | chr15-22974270-DEL-60 |
| chr15-22976112-DEL-171 | chr15-22976113-DEL-171 |
| chr15-23064587-DEL-125 | chr15-23064589-DEL-123 |
| chr15-23070333-DEL-188 | chr15-23070334-DEL-188 |
| chr15-23114824-DEL-506 | chr15-23114825-DEL-506 |
| chr15-23115569-DEL-1211 | chr15-23115570-DEL-1211 |
| chr15-23117562-DEL-721 | chr15-23117563-DEL-721 |
| chr15-23130283-DEL-335 | chr15-23130284-DEL-335 |
| chr15-23154278-DEL-279 | chr15-23154279-DEL-279 |
| chr15-23184306-DEL-5397 | chr15-23184307-DEL-5397 |
| chr15-23201022-DEL-150 | chr15-23201024-DEL-154 |
| chr15-23214703-DEL-1555 | chr15-23214704-DEL-1555 |
| chr15-23232150-DEL-92 | chr15-23232151-DEL-92 |
| chr15-23234900-DEL-71 | chr15-23234901-DEL-71 |
| chr15-23240461-DEL-7088 | chr15-23240462-DEL-7088 |
| chr15-23250771-DEL-4942 | chr15-23250772-DEL-4942 |
| chr15-23264527-DEL-777 | chr15-23264528-DEL-777 |
| chr15-23291337-DEL-6682 | chr15-23291338-DEL-6682 |
| chr15-23305960-DEL-75 | chr15-23305921-DEL-66 |
| chr15-23315591-DEL-152 | chr15-23315600-DEL-144 |
| chr15-23349380-DEL-82 | chr15-23349350-DEL-82 |
| chr15-23358876-DEL-7561 | chr15-23358877-DEL-7561 |
| chr15-23429446-DEL-649 | chr15-23429447-DEL-649 |
| chr15-23460920-DEL-146 | chr15-23460921-DEL-146 |
| chr15-23493451-DEL-157 | chr15-23493452-DEL-157 |
| chr15-23494148-DEL-175 | chr15-23494149-DEL-175 |
| chr15-23495466-DEL-56 | chr15-23495467-DEL-56 |
| chr15-23528512-DEL-54 | chr15-23528513-DEL-54 |
| chr15-23557962-DEL-183 | chr15-23557963-DEL-183 |
| chr15-23593311-DEL-215 | chr15-23593312-DEL-215 |
| chr15-23614827-DEL-58 | chr15-23614828-DEL-58 |
| chr15-23729298-DEL-6427 | chr15-23729299-DEL-6427 |
| chr15-23729539-DEL-208 | chr15-23729540-DEL-208 |
| chr15-23765593-DEL-1211 | chr15-23765594-DEL-1211 |
| chr15-23784646-DEL-53 | chr15-23784647-DEL-53 |
| chr15-23854587-DEL-876 | chr15-23854588-DEL-876 |
| chr15-23870453-DEL-359 | chr15-23870454-DEL-359 |
| chr15-23886074-DEL-1888 | chr15-23886075-DEL-1888 |
| chr15-23894072-DEL-1463 | chr15-23894073-DEL-1463 |
| chr15-23911628-DEL-102 | chr15-23911629-DEL-102 |
| chr15-23917575-DEL-832 | chr15-23917576-DEL-832 |
| chr15-23937025-DEL-95 | chr15-23937026-DEL-75 |
| chr15-23938345-DEL-1370 | chr15-23938346-DEL-1370 |
| chr15-23958952-DEL-78 | chr15-23958953-DEL-78 |
| chr15-23984637-DEL-76 | chr15-23984638-DEL-76 |
| chr15-23997211-DEL-398 | chr15-23997212-DEL-398 |
| chr15-24038490-DEL-1383 | chr15-24038491-DEL-1383 |
| chr15-24060648-DEL-158 | chr15-24060675-DEL-167 |
| chr15-24062972-DEL-200 | chr15-24062973-DEL-200 |
| chr15-24064376-DEL-199 | chr15-24064375-DEL-199 |
| chr15-24095519-DEL-128 | chr15-24095520-DEL-134 |

| | |
|---|---|
| chr15-24097905-DEL-68 | chr15-24097890-DEL-80 |
| chr15-24136435-DEL-56 | chr15-24136436-DEL-56 |
| chr15-24137272-DEL-1373 | chr15-24137273-DEL-1373 |
| chr15-24175069-DEL-2201 | chr15-24175070-DEL-2201 |
| chr15-24176981-DEL-115 | chr15-24176982-DEL-115 |
| chr15-24246243-DEL-229 | chr15-24246244-DEL-229 |
| chr15-24248480-DEL-99 | chr15-24248480-DEL-91 |
| chr15-24298440-DEL-58 | chr15-24298441-DEL-58 |
| chr15-24310356-DEL-6436 | chr15-24310357-DEL-6436 |
| chr15-24319079-DEL-349 | chr15-24319080-DEL-349 |
| chr15-24321252-DEL-4633 | chr15-24321253-DEL-4633 |
| chr15-24334138-DEL-120 | chr15-24334140-DEL-120 |
| chr15-24411669-DEL-6395 | chr15-24411670-DEL-6395 |
| chr15-24424070-DEL-210 | chr15-24424071-DEL-210 |
| chr15-24439503-DEL-56 | chr15-24439504-DEL-56 |
| chr15-24462901-DEL-1257 | chr15-24462902-DEL-1257 |
| chr15-24470753-DEL-992 | chr15-24470754-DEL-992 |
| chr15-24477996-DEL-80 | chr15-24477997-DEL-52 |
| chr15-24506601-DEL-4731 | chr15-24506602-DEL-4731 |
| chr15-24511461-DEL-1573 | chr15-24511462-DEL-1573 |
| chr15-24539364-DEL-2191 | chr15-24539365-DEL-2191 |
| chr15-24542421-DEL-961 | chr15-24542422-DEL-957 |
| chr15-24618152-DEL-989 | chr15-24618153-DEL-989 |
| chr15-24635440-DEL-81 | chr15-24635441-DEL-81 |
| chr15-24662943-DEL-6423 | chr15-24662944-DEL-6423 |
| chr15-24670263-DEL-6164 | chr15-24670264-DEL-6164 |
| chr15-24694632-DEL-447 | chr15-24694633-DEL-447 |
| chr15-24695520-DEL-442 | chr15-24695521-DEL-442 |
| chr15-24698606-DEL-53 | chr15-24698607-DEL-53 |
| chr15-24717613-DEL-291 | chr15-24717612-DEL-292 |
| chr15-24760665-DEL-6187 | chr15-24760666-DEL-6187 |
| chr15-24787836-DEL-1285 | chr15-24787837-DEL-1285 |
| chr15-24806262-DEL-1569 | chr15-24806263-DEL-1569 |
| chr15-24818144-DEL-198 | chr15-24818145-DEL-198 |
| chr15-24831607-DEL-54 | chr15-24831607-DEL-90 |
| chr15-24834940-DEL-2205 | chr15-24834941-DEL-2205 |
| chr15-24853592-DEL-137 | chr15-24853593-DEL-137 |
| chr15-24868258-DEL-1131 | chr15-24868259-DEL-1129 |
| chr15-24880802-DEL-1972 | chr15-24880803-DEL-1972 |
| chr15-24894471-DEL-698 | chr15-24894457-DEL-688 |
| chr15-24896655-DEL-1862 | chr15-24896656-DEL-1862 |
| chr15-24925312-DEL-372 | chr15-24925309-DEL-372 |
| chr15-24945618-DEL-90 | chr15-24945617-DEL-91 |
| chr15-24995808-DEL-1258 | chr15-24995809-DEL-1258 |
| chr15-25004913-DEL-58 | chr15-25004914-DEL-58 |
| chr15-25006808-DEL-64 | chr15-25006809-DEL-64 |
| chr15-25010244-DEL-507 | chr15-25010245-DEL-507 |
| chr15-25014803-DEL-131 | chr15-25014802-DEL-122 |
| chr15-25031346-DEL-365 | chr15-25031347-DEL-365 |
| chr15-25048730-DEL-166 | chr15-25048730-DEL-166 |
| chr15-25051567-DEL-4291 | chr15-25051568-DEL-4291 |
| chr15-25062466-DEL-2735 | chr15-25062467-DEL-2735 |
| chr15-25087144-DEL-177 | chr15-25087145-DEL-177 |
| chr15-25106105-DEL-56 | chr15-25106106-DEL-56 |
| chr15-25110981-DEL-6503 | chr15-25110982-DEL-6503 |
| chr15-25115586-DEL-1783 | chr15-25115586-DEL-1783 |
| chr15-25124701-DEL-545 | chr15-25124702-DEL-545 |
| chr15-25127438-DEL-81 | chr15-25127433-DEL-65 |
| chr15-25131784-DEL-375 | chr15-25131785-DEL-375 |
| chr15-25140666-DEL-206 | chr15-25140664-DEL-206 |
| chr15-25160135-DEL-6022 | chr15-25160136-DEL-6022 |
| chr15-25177328-DEL-264 | chr15-25177329-DEL-264 |
| chr15-25177734-DEL-2964 | chr15-25177735-DEL-2964 |
| chr15-25194201-DEL-135 | chr15-25194202-DEL-126 |
| chr15-25200379-DEL-201 | chr15-25200380-DEL-201 |
| chr15-25202112-DEL-90 | chr15-25202110-DEL-108 |
| chr15-25233232-DEL-338 | chr15-25233227-DEL-339 |
| chr15-25245761-DEL-55 | chr15-25245762-DEL-55 |
| chr15-25246035-DEL-6168 | chr15-25246035-DEL-6168 |
| chr15-25255295-DEL-709 | chr15-25255296-DEL-709 |
| chr15-25261281-DEL-178 | chr15-25261278-DEL-178 |
| chr15-25269406-DEL-404 | chr15-25269407-DEL-404 |
| chr15-25324692-DEL-59 | chr15-25324696-DEL-92 |
| chr15-25350424-DEL-1369 | chr15-25350425-DEL-1363 |
| chr15-25359876-DEL-69 | chr15-25359892-DEL-54 |
| chr15-25380774-DEL-329 | chr15-25380777-DEL-328 |
| chr15-25398474-DEL-70 | chr15-25398474-DEL-68 |
| chr15-25419197-DEL-170 | chr15-25419216-DEL-147 |
| chr15-25436362-DEL-884 | chr15-25436363-DEL-884 |
| chr15-25492776-DEL-54 | chr15-25492814-DEL-54 |

| | |
|---|---|
| chr15-25506589-DEL-102 | chr15-25506590-DEL-102 |
| chr15-25508149-DEL-103 | chr15-25508150-DEL-102 |
| chr15-25516738-DEL-53 | chr15-25516707-DEL-51 |
| chr15-25526884-DEL-365 | chr15-25526885-DEL-365 |
| chr15-25529208-DEL-1200 | chr15-25529208-DEL-1200 |
| chr15-25545389-DEL-190 | chr15-25545390-DEL-190 |
| chr15-25547026-DEL-1268 | chr15-25547027-DEL-1268 |
| chr15-25551555-DEL-104 | chr15-25551556-DEL-100 |
| chr15-25553388-DEL-403 | chr15-25553389-DEL-403 |
| chr15-25560165-DEL-71 | chr15-25560166-DEL-67 |
| chr15-25571584-DEL-188 | chr15-25571572-DEL-189 |
| chr15-25587460-DEL-59 | chr15-25587425-DEL-51 |
| chr15-25596969-DEL-96 | chr15-25596970-DEL-96 |
| chr15-25661710-DEL-53 | chr15-25661710-DEL-53 |
| chr15-25698800-DEL-66 | chr15-25698801-DEL-66 |
| chr15-25704275-DEL-197 | chr15-25704276-DEL-197 |
| chr15-25718101-DEL-222 | chr15-25718107-DEL-219 |
| chr15-25741333-DEL-202 | chr15-25741334-DEL-202 |
| chr15-25748911-DEL-838 | chr15-25748912-DEL-838 |
| chr15-25750151-DEL-229 | chr15-25750150-DEL-221 |
| chr15-25757097-DEL-373 | chr15-25757098-DEL-373 |
| chr15-25776663-DEL-61 | chr15-25776654-DEL-64 |
| chr15-25791464-DEL-171 | chr15-25791465-DEL-171 |
| chr15-25808420-DEL-188 | chr15-25808421-DEL-188 |
| chr15-25825883-DEL-129 | chr15-25825887-DEL-127 |
| chr15-25849434-DEL-86 | chr15-25849459-DEL-61 |
| chr15-25855746-DEL-54 | chr15-25855747-DEL-54 |
| chr15-25898810-DEL-174 | chr15-25898811-DEL-174 |
| chr15-25927678-DEL-409 | chr15-25927679-DEL-409 |
| chr15-25929385-DEL-172 | chr15-25929386-DEL-172 |
| chr15-25937079-DEL-945 | chr15-25937080-DEL-945 |
| chr15-25959233-DEL-322 | chr15-25959234-DEL-322 |
| chr15-26067069-DEL-303 | chr15-26067069-DEL-303 |
| chr15-26082829-DEL-6225 | chr15-26082830-DEL-6225 |
| chr15-26154451-DEL-88 | chr15-26154490-DEL-50 |
| chr15-26156023-DEL-949 | chr15-26156024-DEL-949 |
| chr15-26169854-DEL-8310 | chr15-26169856-DEL-8323 |
| chr15-26180180-DEL-340 | chr15-26180181-DEL-340 |
| chr15-26211909-DEL-65 | chr15-26211910-DEL-65 |
| chr15-26216375-DEL-103 | chr15-26216377-DEL-115 |
| chr15-26242055-DEL-6497 | chr15-26242056-DEL-6497 |
| chr15-26248805-DEL-1075 | chr15-26248806-DEL-1075 |
| chr15-26266969-DEL-54 | chr15-26266974-DEL-50 |
| chr15-26280311-DEL-6678 | chr15-26280312-DEL-6678 |
| chr15-26370908-DEL-51 | chr15-26370909-DEL-51 |
| chr15-26376147-DEL-100 | chr15-26376147-DEL-98 |
| chr15-26385843-DEL-118 | chr15-26385844-DEL-118 |
| chr15-26394454-DEL-217 | chr15-26394440-DEL-218 |
| chr15-26399206-DEL-53 | chr15-26399203-DEL-54 |
| chr15-26418310-DEL-574 | chr15-26418311-DEL-574 |
| chr15-26418990-DEL-54 | chr15-26418991-DEL-54 |
| chr15-26444338-DEL-210 | chr15-26444339-DEL-210 |
| chr15-26449259-DEL-57 | chr15-26449260-DEL-57 |
| chr15-26500045-DEL-62 | chr15-26500046-DEL-61 |
| chr15-26549273-DEL-408 | chr15-26549264-DEL-409 |
| chr15-26593295-DEL-60 | chr15-26593300-DEL-56 |
| chr15-26630679-DEL-72 | chr15-26630680-DEL-72 |
| chr15-26651137-DEL-1039 | chr15-26651138-DEL-1039 |
| chr15-26674909-DEL-7085 | chr15-26674910-DEL-7085 |
| chr15-26687499-DEL-88 | chr15-26687515-DEL-103 |
| chr15-26723952-DEL-469 | chr15-26723953-DEL-469 |
| chr15-26728344-DEL-279 | chr15-26728348-DEL-276 |
| chr15-26737367-DEL-773 | chr15-26737372-DEL-776 |
| chr15-26739774-DEL-394 | chr15-26739775-DEL-394 |
| chr15-26771723-DEL-53 | chr15-26771724-DEL-53 |
| chr15-26782426-DEL-550 | chr15-26782417-DEL-551 |
| chr15-26790808-DEL-598 | chr15-26790809-DEL-598 |
| chr15-26803251-DEL-80 | chr15-26803252-DEL-80 |
| chr15-26880044-DEL-102 | chr15-26880045-DEL-102 |
| chr15-26882960-DEL-68 | chr15-26882967-DEL-50 |
| chr15-26905128-DEL-4844 | chr15-26905129-DEL-4844 |
| chr15-26912427-DEL-399 | chr15-26912428-DEL-399 |
| chr15-26919920-DEL-542 | chr15-26919921-DEL-542 |
| chr15-26929002-DEL-255 | chr15-26929003-DEL-255 |
| chr15-26938219-DEL-509 | chr15-26938222-DEL-508 |
| chr15-26939460-DEL-697 | chr15-26939461-DEL-697 |
| chr15-26984826-DEL-94 | chr15-26984827-DEL-94 |
| chr15-26988823-DEL-56 | chr15-26988824-DEL-56 |
| chr15-27007372-DEL-154 | chr15-27007373-DEL-154 |
| chr15-27033794-DEL-75 | chr15-27033795-DEL-75 |

| | |
|---|---|
| chr15-27036010-DEL-6005 | chr15-27036011-DEL-6005 |
| chr15-27044812-DEL-681 | chr15-27044803-DEL-682 |
| chr15-27053110-DEL-2622 | chr15-27053111-DEL-2622 |
| chr15-27058198-DEL-566 | chr15-27058199-DEL-566 |
| chr15-27154456-DEL-363 | chr15-27154457-DEL-363 |
| chr15-27156719-DEL-59 | chr15-27156719-DEL-59 |
| chr15-27184078-DEL-210 | chr15-27184079-DEL-210 |
| chr15-27185927-DEL-171 | chr15-27185928-DEL-171 |
| chr15-27230259-DEL-101 | chr15-27230262-DEL-114 |
| chr15-27243826-DEL-142 | chr15-27243822-DEL-107 |
| chr15-27272847-DEL-205 | chr15-27272848-DEL-205 |
| chr15-27339965-DEL-103 | chr15-27339962-DEL-101 |
| chr15-27363514-DEL-203 | chr15-27363508-DEL-201 |
| chr15-27368318-DEL-547 | chr15-27368319-DEL-544 |
| chr15-27372581-DEL-1399 | chr15-27372582-DEL-1399 |
| chr15-27373085-DEL-400 | chr15-27373086-DEL-400 |
| chr15-27380279-DEL-64 | chr15-27380280-DEL-64 |
| chr15-27384229-DEL-6102 | chr15-27384215-DEL-6067 |
| chr15-27391575-DEL-1014 | chr15-27391576-DEL-1014 |
| chr15-27477090-DEL-51 | chr15-27477091-DEL-51 |
| chr15-27484328-DEL-60 | chr15-27484331-DEL-58 |
| chr15-27513017-DEL-231 | chr15-27513018-DEL-232 |
| chr15-27547004-DEL-60 | chr15-27547005-DEL-60 |
| chr15-27556702-DEL-58 | chr15-27556703-DEL-58 |
| chr15-27610062-DEL-54 | chr15-27610063-DEL-54 |
| chr15-27612556-DEL-81 | chr15-27612549-DEL-97 |
| chr15-27637127-DEL-66 | chr15-27637129-DEL-65 |
| chr15-27638879-DEL-862 | chr15-27638880-DEL-862 |
| chr15-27658500-DEL-70 | chr15-27658522-DEL-84 |
| chr15-27686775-DEL-894 | chr15-27686776-DEL-894 |
| chr15-27695958-DEL-66 | chr15-27695958-DEL-66 |
| chr15-27721232-DEL-2619 | chr15-27721233-DEL-2619 |
| chr15-27738657-DEL-57 | chr15-27738658-DEL-57 |
| chr15-27741853-DEL-68 | chr15-27741854-DEL-68 |
| chr15-27761085-DEL-70 | chr15-27761074-DEL-76 |
| chr15-27775565-DEL-108 | chr15-27775566-DEL-108 |
| chr15-27777136-DEL-62 | chr15-27777137-DEL-62 |
| chr15-27806597-DEL-52 | chr15-27806610-DEL-53 |
| chr15-27825621-DEL-74 | chr15-27825621-DEL-74 |
| chr15-27828071-DEL-136 | chr15-27828072-DEL-136 |
| chr15-27848114-DEL-93 | chr15-27848119-DEL-90 |
| chr15-27857615-DEL-628 | chr15-27857616-DEL-628 |
| chr15-27896824-DEL-84 | chr15-27896825-DEL-83 |
| chr15-27899292-DEL-592 | chr15-27899293-DEL-592 |
| chr15-27915139-DEL-126 | chr15-27915140-DEL-126 |
| chr15-28006000-DEL-167 | chr15-28005997-DEL-168 |
| chr15-28035170-DEL-217 | chr15-28035171-DEL-217 |
| chr15-28044360-DEL-139 | chr15-28044362-DEL-136 |
| chr15-28046386-DEL-743 | chr15-28046387-DEL-743 |
| chr15-28101338-DEL-206 | chr15-28101338-DEL-206 |
| chr15-28112594-DEL-400 | chr15-28112595-DEL-400 |
| chr15-28115190-DEL-135 | chr15-28115191-DEL-105 |
| chr15-28139640-DEL-278 | chr15-28139641-DEL-277 |
| chr15-28143445-DEL-404 | chr15-28143446-DEL-404 |
| chr15-28158093-DEL-179 | chr15-28158094-DEL-179 |
| chr15-28160170-DEL-121 | chr15-28160171-DEL-121 |
| chr15-28168092-DEL-677 | chr15-28168093-DEL-677 |
| chr15-28175924-DEL-489 | chr15-28175916-DEL-489 |
| chr15-28192865-DEL-111 | chr15-28192866-DEL-111 |
| chr15-28226035-DEL-58 | chr15-28226036-DEL-58 |
| chr15-28252896-DEL-6650 | chr15-28252897-DEL-6650 |
| chr15-28280619-DEL-6281 | chr15-28280620-DEL-6281 |
| chr15-28317140-DEL-130 | chr15-28317128-DEL-109 |
| chr15-28332807-DEL-79 | chr15-28332816-DEL-50 |
| chr15-28359400-DEL-169 | chr15-28359401-DEL-169 |
| chr15-28374247-DEL-264 | chr15-28374248-DEL-264 |
| chr15-28376722-DEL-80 | chr15-28376723-DEL-80 |
| chr15-28384835-DEL-135 | chr15-28384836-DEL-135 |
| chr15-28423896-DEL-332 | chr15-28423902-DEL-321 |
| chr15-28425148-DEL-9025 | chr15-28425152-DEL-9022 |
| chr15-28429912-DEL-812 | chr15-28429913-DEL-812 |
| chr15-28481785-DEL-2857 | chr15-28481787-DEL-2856 |
| chr15-28508193-DEL-251 | chr15-28508194-DEL-251 |
| chr15-28510333-DEL-156 | chr15-28510334-DEL-156 |
| chr15-28532368-DEL-336 | chr15-28532369-DEL-336 |
| chr15-28538952-DEL-1037 | chr15-28538953-DEL-1037 |
| chr15-28541358-DEL-133 | chr15-28541359-DEL-133 |
| chr15-28579640-DEL-838 | chr15-28579641-DEL-838 |
| chr15-28595796-DEL-68 | chr15-28595829-DEL-60 |
| chr15-28610582-DEL-931 | chr15-28610585-DEL-926 |

| | |
|---|---|
| chr15-28612283-DEL-54 | chr15-28612284-DEL-52 |
| chr15-28615144-DEL-172 | chr15-28615145-DEL-172 |
| chr15-28639380-DEL-267 | chr15-28639355-DEL-293 |
| chr15-28648686-DEL-653 | chr15-28648687-DEL-653 |
| chr15-28676012-DEL-528 | chr15-28676049-DEL-499 |
| chr15-28684818-DEL-172 | chr15-28684819-DEL-172 |
| chr15-28727164-DEL-74 | chr15-28727165-DEL-50 |
| chr15-28727467-DEL-2039 | chr15-28727468-DEL-2039 |
| chr15-28730695-DEL-196 | chr15-28730696-DEL-196 |
| chr15-28754539-DEL-160 | chr15-28754540-DEL-160 |
| chr15-28755609-DEL-56 | chr15-28755644-DEL-60 |
| chr15-28777368-DEL-3580 | chr15-28777371-DEL-3578 |
| chr15-28803726-DEL-1207 | chr15-28803727-DEL-1207 |
| chr15-28852913-DEL-370 | chr15-28852914-DEL-370 |
| chr15-28917273-DEL-80 | chr15-28917274-DEL-80 |
| chr15-28935113-DEL-136 | chr15-28935129-DEL-122 |
| chr15-28973536-DEL-480 | chr15-28973535-DEL-481 |
| chr15-28977617-DEL-594 | chr15-28977618-DEL-594 |
| chr15-28980116-DEL-540 | chr15-28980117-DEL-540 |
| chr15-28982270-DEL-4510 | chr15-28982284-DEL-4497 |
| chr15-28994159-DEL-67 | chr15-28994160-DEL-67 |
| chr15-28997130-DEL-52 | chr15-28997130-DEL-53 |
| chr15-29015256-DEL-462 | chr15-29015257-DEL-462 |
| chr15-29042692-DEL-313 | chr15-29042693-DEL-313 |
| chr15-29075298-DEL-65 | chr15-29075299-DEL-65 |
| chr15-29079072-DEL-97 | chr15-29079073-DEL-98 |
| chr15-29114531-DEL-59 | chr15-29114520-DEL-71 |
| chr15-29125046-DEL-6314 | chr15-29125047-DEL-6314 |
| chr15-29126696-DEL-1954 | chr15-29126699-DEL-1955 |
| chr15-29130966-DEL-3100 | chr15-29130967-DEL-3100 |
| chr15-29133154-DEL-75 | chr15-29133155-DEL-75 |
| chr15-29167247-DEL-2896 | chr15-29167256-DEL-2895 |
| chr15-29204049-DEL-65 | chr15-29204061-DEL-62 |
| chr15-29212701-DEL-95 | chr15-29212702-DEL-95 |
| chr15-29233042-DEL-256 | chr15-29233036-DEL-263 |
| chr15-29236073-DEL-520 | chr15-29236074-DEL-520 |
| chr15-29238078-DEL-94 | chr15-29238079-DEL-94 |
| chr15-29238562-DEL-555 | chr15-29238560-DEL-555 |
| chr15-29276746-DEL-74 | chr15-29276751-DEL-70 |
| chr15-29283488-DEL-205 | chr15-29283489-DEL-205 |
| chr15-29297091-DEL-136 | chr15-29297092-DEL-136 |
| chr15-29303441-DEL-286 | chr15-29303442-DEL-286 |
| chr15-29347156-DEL-1527 | chr15-29347156-DEL-1527 |
| chr15-29387095-DEL-84 | chr15-29387111-DEL-102 |
| chr15-29389357-DEL-6414 | chr15-29389358-DEL-6414 |
| chr15-29400329-DEL-245 | chr15-29400310-DEL-250 |
| chr15-29405660-DEL-54 | chr15-29405661-DEL-82 |
| chr15-29421308-DEL-343 | chr15-29421309-DEL-343 |
| chr15-29447482-DEL-72 | chr15-29447522-DEL-71 |
| chr15-29463936-DEL-6531 | chr15-29463937-DEL-6531 |
| chr15-29520037-DEL-272 | chr15-29519997-DEL-288 |
| chr15-29571881-DEL-310 | chr15-29571878-DEL-311 |
| chr15-29591950-DEL-6368 | chr15-29591951-DEL-6368 |
| chr15-29610049-DEL-82 | chr15-29610049-DEL-82 |
| chr15-29622257-DEL-2999 | chr15-29622258-DEL-2999 |
| chr15-29627849-DEL-60 | chr15-29627843-DEL-61 |
| chr15-29660437-DEL-781 | chr15-29660438-DEL-779 |
| chr15-29663917-DEL-66 | chr15-29663912-DEL-68 |
| chr15-29672772-DEL-52 | chr15-29672773-DEL-52 |
| chr15-29693219-DEL-67 | chr15-29693219-DEL-67 |
| chr15-29694433-DEL-1919 | chr15-29694434-DEL-1919 |
| chr15-29698972-DEL-1965 | chr15-29698973-DEL-1965 |
| chr15-29700461-DEL-66 | chr15-29700462-DEL-66 |
| chr15-29723478-DEL-72 | chr15-29723499-DEL-52 |
| chr15-29763745-DEL-211 | chr15-29763746-DEL-211 |
| chr15-29778084-DEL-51 | chr15-29778076-DEL-57 |
| chr15-29796709-DEL-314 | chr15-29796709-DEL-314 |
| chr15-29819440-DEL-1594 | chr15-29819441-DEL-1593 |
| chr15-29842007-DEL-80 | chr15-29842008-DEL-80 |
| chr15-29866553-DEL-1963 | chr15-29866593-DEL-1961 |
| chr15-29870849-DEL-79 | chr15-29870850-DEL-79 |
| chr15-29889197-DEL-1839 | chr15-29889198-DEL-1839 |
| chr15-29934502-DEL-419 | chr15-29934503-DEL-419 |
| chr15-29958314-DEL-80 | chr15-29958317-DEL-78 |
| chr15-29971624-DEL-1800 | chr15-29971625-DEL-1800 |
| chr15-29983696-DEL-7083 | chr15-29983697-DEL-7083 |
| chr15-29992499-DEL-194 | chr15-29992500-DEL-194 |
| chr15-30022073-DEL-439 | chr15-30022074-DEL-439 |
| chr15-30042573-DEL-7230 | chr15-30042585-DEL-7233 |
| chr15-30077144-DEL-60 | chr15-30077145-DEL-54 |

| | |
|---|---|
| chr15-30087478-DEL-8103 | chr15-30087479-DEL-8103 |
| chr15-30093993-DEL-206 | chr15-30093994-DEL-206 |
| chr15-30095805-DEL-220 | chr15-30095806-DEL-220 |
| chr15-30098819-DEL-180 | chr15-30098820-DEL-180 |
| chr15-30104025-DEL-439 | chr15-30104026-DEL-439 |
| chr15-30114744-DEL-150 | chr15-30114745-DEL-150 |
| chr15-30122935-DEL-3590 | chr15-30122936-DEL-3590 |
| chr15-30149611-DEL-52 | chr15-30149629-DEL-56 |
| chr15-30162137-DEL-90 | chr15-30162129-DEL-96 |
| chr15-30182079-DEL-70 | chr15-30182080-DEL-70 |
| chr15-30193743-DEL-79 | chr15-30193743-DEL-76 |
| chr15-30216899-DEL-6267 | chr15-30216920-DEL-6246 |
| chr15-30224321-DEL-1286 | chr15-30224322-DEL-1286 |
| chr15-30235863-DEL-79 | chr15-30235864-DEL-80 |
| chr15-30242630-DEL-3753 | chr15-30242631-DEL-3753 |
| chr15-30244812-DEL-70 | chr15-30244813-DEL-70 |
| chr15-30251608-DEL-6442 | chr15-30251609-DEL-6442 |
| chr15-30258936-DEL-396 | chr15-30258937-DEL-396 |
| chr15-30299504-DEL-100 | chr15-30299501-DEL-99 |
| chr15-30310527-DEL-108 | chr15-30310528-DEL-108 |
| chr15-30348963-DEL-3332 | chr15-30348964-DEL-3332 |
| chr15-30370895-DEL-72 | chr15-30370896-DEL-72 |
| chr15-30397385-DEL-52 | chr15-30397388-DEL-50 |
| chr15-30397971-DEL-225 | chr15-30397972-DEL-223 |
| chr15-30398579-DEL-647 | chr15-30398580-DEL-647 |
| chr15-30402337-DEL-101 | chr15-30402338-DEL-101 |
| chr15-30413857-DEL-95 | chr15-30413858-DEL-95 |
| chr15-30418323-DEL-58 | chr15-30418323-DEL-58 |
| chr15-30431098-DEL-592 | chr15-30431099-DEL-592 |
| chr15-30432713-DEL-265 | chr15-30432714-DEL-265 |
| chr15-30441562-DEL-6647 | chr15-30441563-DEL-6647 |
| chr15-30482098-DEL-484 | chr15-30482099-DEL-484 |
| chr15-30483488-DEL-1277 | chr15-30483486-DEL-1278 |
| chr15-30488528-DEL-107 | chr15-30488556-DEL-78 |
| chr15-30496701-DEL-249 | chr15-30496702-DEL-249 |
| chr15-30513921-DEL-114 | chr15-30513922-DEL-114 |
| chr15-30516696-DEL-2643 | chr15-30516697-DEL-2643 |
| chr15-30530798-DEL-165 | chr15-30530799-DEL-165 |
| chr15-30531462-DEL-982 | chr15-30531463-DEL-983 |
| chr15-30544207-DEL-260 | chr15-30544208-DEL-260 |
| chr15-30557254-DEL-72 | chr15-30557253-DEL-89 |
| chr15-30577302-DEL-118 | chr15-30577308-DEL-113 |
| chr15-30579282-DEL-6343 | chr15-30579271-DEL-6344 |
| chr15-30579557-DEL-64 | chr15-30579558-DEL-64 |
| chr15-30597623-DEL-133 | chr15-30597625-DEL-131 |
| chr15-30620877-DEL-51 | chr15-30620847-DEL-56 |
| chr15-30632763-DEL-371 | chr15-30632764-DEL-371 |
| chr15-30670467-DEL-801 | chr15-30670468-DEL-801 |
| chr15-30690476-DEL-204 | chr15-30690477-DEL-204 |
| chr15-30729754-DEL-95 | chr15-30729742-DEL-96 |
| chr15-30730813-DEL-499 | chr15-30730814-DEL-499 |
| chr15-30736366-DEL-410 | chr15-30736367-DEL-410 |
| chr15-30747885-DEL-7159 | chr15-30747886-DEL-7159 |
| chr15-30778873-DEL-56 | chr15-30778874-DEL-56 |
| chr15-30780226-DEL-74 | chr15-30780227-DEL-74 |
| chr15-30800229-DEL-411 | chr15-30800230-DEL-411 |
| chr15-30803817-DEL-86 | chr15-30803818-DEL-86 |
| chr15-30827995-DEL-245 | chr15-30827998-DEL-243 |
| chr15-30849355-DEL-51 | chr15-30849356-DEL-51 |
| chr15-30850241-DEL-105 | chr15-30850242-DEL-105 |
| chr15-30860262-DEL-1556 | chr15-30860263-DEL-1556 |
| chr15-30872450-DEL-197 | chr15-30872493-DEL-163 |
| chr15-30887567-DEL-66 | chr15-30887565-DEL-66 |
| chr15-30892987-DEL-1186 | chr15-30892988-DEL-1186 |
| chr15-30898599-DEL-95 | chr15-30898600-DEL-95 |
| chr15-30909786-DEL-97 | chr15-30909787-DEL-96 |
| chr15-30910313-DEL-497 | chr15-30910307-DEL-498 |
| chr15-30931436-DEL-643 | chr15-30931437-DEL-643 |
| chr15-30933042-DEL-118 | chr15-30933043-DEL-118 |
| chr15-30943504-DEL-940 | chr15-30943505-DEL-940 |
| chr15-30950827-DEL-686 | chr15-30950821-DEL-681 |
| chr15-30955739-DEL-493 | chr15-30955740-DEL-493 |
| chr15-30973327-DEL-70 | chr15-30973328-DEL-70 |
| chr15-31002963-DEL-214 | chr15-31002963-DEL-214 |
| chr15-31022016-DEL-80 | chr15-31022017-DEL-80 |
| chr15-31066855-DEL-1326 | chr15-31066856-DEL-1326 |
| chr15-31072347-DEL-11527 | chr15-31072348-DEL-11516 |
| chr15-31073015-DEL-5522 | chr15-31073013-DEL-5522 |
| chr15-31095020-DEL-74 | chr15-31095021-DEL-74 |
| chr15-31097110-DEL-51 | chr15-31097111-DEL-51 |

| | |
|---|---|
| chr15-31111057-DEL-194 | chr15-31111058-DEL-194 |
| chr15-31140977-DEL-55 | chr15-31140978-DEL-55 |
| chr15-31142388-DEL-109 | chr15-31142388-DEL-109 |
| chr15-31144917-DEL-133 | chr15-31144918-DEL-133 |
| chr15-31175902-DEL-290 | chr15-31175903-DEL-265 |
| chr15-31181830-DEL-2453 | chr15-31181831-DEL-2453 |
| chr15-31189590-DEL-638 | chr15-31189591-DEL-638 |
| chr15-31192524-DEL-126 | chr15-31192541-DEL-104 |
| chr15-31200833-DEL-6606 | chr15-31200826-DEL-6607 |
| chr15-31213314-DEL-104 | chr15-31213315-DEL-104 |
| chr15-31217900-DEL-351 | chr15-31217901-DEL-351 |
| chr15-31239519-DEL-100 | chr15-31239498-DEL-98 |
| chr15-31249670-DEL-93 | chr15-31249671-DEL-78 |
| chr15-31270698-DEL-1173 | chr15-31270699-DEL-1172 |
| chr15-31289540-DEL-194 | chr15-31289541-DEL-194 |
| chr15-31308191-DEL-52 | chr15-31308172-DEL-57 |
| chr15-31312931-DEL-72 | chr15-31312932-DEL-72 |
| chr15-31334879-DEL-200 | chr15-31334881-DEL-199 |
| chr15-31338944-DEL-77 | chr15-31338945-DEL-77 |
| chr15-31351546-DEL-60 | chr15-31351547-DEL-60 |
| chr15-31354747-DEL-627 | chr15-31354747-DEL-628 |
| chr15-31356155-DEL-179 | chr15-31356156-DEL-179 |
| chr15-31357492-DEL-68 | chr15-31357493-DEL-68 |
| chr15-31400913-DEL-106 | chr15-31400914-DEL-106 |
| chr15-31423499-DEL-223 | chr15-31423500-DEL-223 |
| chr15-31427542-DEL-53 | chr15-31427542-DEL-53 |
| chr15-31431826-DEL-141 | chr15-31431827-DEL-141 |
| chr15-31445769-DEL-1208 | chr15-31445770-DEL-1208 |
| chr15-31472571-DEL-443 | chr15-31472572-DEL-442 |
| chr15-31486870-DEL-189 | chr15-31486871-DEL-189 |
| chr15-31488659-DEL-207 | chr15-31488660-DEL-207 |
| chr15-31515313-DEL-198 | chr15-31515313-DEL-199 |
| chr15-31522142-DEL-141 | chr15-31522143-DEL-141 |
| chr15-31551710-DEL-204 | chr15-31551711-DEL-204 |
| chr15-31558301-DEL-501 | chr15-31558302-DEL-501 |
| chr15-31562913-DEL-167 | chr15-31562909-DEL-168 |
| chr15-31566960-DEL-85 | chr15-31566961-DEL-79 |
| chr15-31573337-DEL-100 | chr15-31573338-DEL-100 |
| chr15-31588458-DEL-198 | chr15-31588459-DEL-198 |
| chr15-31606365-DEL-90 | chr15-31606366-DEL-90 |
| chr15-31619095-DEL-222 | chr15-31619096-DEL-222 |
| chr15-31627130-DEL-91 | chr15-31627162-DEL-105 |
| chr15-31654828-DEL-904 | chr15-31654829-DEL-904 |
| chr15-31683676-DEL-83 | chr15-31683672-DEL-84 |
| chr15-31704222-DEL-678 | chr15-31704224-DEL-678 |
| chr15-31721944-DEL-53 | chr15-31721933-DEL-73 |
| chr15-31726697-DEL-856 | chr15-31726700-DEL-852 |
| chr15-31727910-DEL-56 | chr15-31727911-DEL-56 |
| chr15-31733165-DEL-273 | chr15-31733166-DEL-273 |
| chr15-31736503-DEL-280 | chr15-31736504-DEL-280 |
| chr15-31750257-DEL-117 | chr15-31750256-DEL-113 |
| chr15-31755999-DEL-2857 | chr15-31756000-DEL-2857 |
| chr15-31774531-DEL-56 | chr15-31774532-DEL-56 |
| chr15-31775055-DEL-87 | chr15-31775067-DEL-82 |
| chr15-31785639-DEL-66 | chr15-31785652-DEL-54 |
| chr15-31813886-DEL-83 | chr15-31813887-DEL-83 |
| chr15-31814706-DEL-74 | chr15-31814714-DEL-52 |
| chr15-31828741-DEL-330 | chr15-31828742-DEL-330 |
| chr15-31847217-DEL-536 | chr15-31847215-DEL-535 |
| chr15-31848740-DEL-748 | chr15-31848741-DEL-748 |
| chr15-31868741-DEL-65 | chr15-31868742-DEL-65 |
| chr15-31872784-DEL-53 | chr15-31872785-DEL-53 |
| chr15-31876964-DEL-772 | chr15-31876965-DEL-772 |
| chr15-31888066-DEL-94 | chr15-31888067-DEL-94 |
| chr15-31915205-DEL-72 | chr15-31915206-DEL-54 |
| chr15-31940727-DEL-6485 | chr15-31940728-DEL-6485 |
| chr15-31946989-DEL-213 | chr15-31946959-DEL-214 |
| chr15-31954774-DEL-84 | chr15-31954775-DEL-84 |
| chr15-31994343-DEL-342 | chr15-31994344-DEL-342 |
| chr15-32013883-DEL-217 | chr15-32013878-DEL-218 |
| chr15-32019009-DEL-379 | chr15-32019002-DEL-381 |
| chr15-32021474-DEL-2118 | chr15-32021475-DEL-2118 |
| chr15-32084959-DEL-54 | chr15-32084958-DEL-56 |
| chr15-32127492-DEL-58 | chr15-32127494-DEL-64 |
| chr15-32140248-DEL-107 | chr15-32140241-DEL-111 |
| chr15-32142949-DEL-70 | chr15-32142950-DEL-50 |
| chr15-32148366-DEL-213 | chr15-32148384-DEL-203 |
| chr15-32157378-DEL-976 | chr15-32157379-DEL-976 |
| chr15-32171219-DEL-66 | chr15-32171220-DEL-66 |
| chr15-32208894-DEL-6099 | chr15-32208895-DEL-6099 |

| | |
|---|---|
| chr15-32285753-DEL-1595 | chr15-32285754-DEL-1595 |
| chr15-32325452-DEL-135 | chr15-32325453-DEL-135 |
| chr15-32336646-DEL-273 | chr15-32336647-DEL-273 |
| chr15-32345072-DEL-297 | chr15-32345073-DEL-297 |
| chr15-32359406-DEL-297 | chr15-32359407-DEL-297 |
| chr15-32363622-DEL-62 | chr15-32363623-DEL-62 |
| chr15-32364836-DEL-874 | chr15-32364837-DEL-874 |
| chr15-32375222-DEL-831 | chr15-32375223-DEL-831 |
| chr15-32389548-DEL-6658 | chr15-32389549-DEL-6658 |
| chr15-32395902-DEL-212 | chr15-32395903-DEL-212 |
| chr15-32416122-DEL-172 | chr15-32416132-DEL-165 |
| chr15-32435745-DEL-51 | chr15-32435744-DEL-53 |
| chr15-32470057-DEL-212 | chr15-32470058-DEL-212 |
| chr15-32507617-DEL-61 | chr15-32507623-DEL-55 |
| chr15-32531523-DEL-53 | chr15-32531507-DEL-54 |
| chr15-32564319-DEL-78 | chr15-32564320-DEL-78 |
| chr15-32566438-DEL-672 | chr15-32566439-DEL-672 |
| chr15-32603990-DEL-80 | chr15-32603990-DEL-80 |
| chr15-32648688-DEL-54 | chr15-32648689-DEL-54 |
| chr15-32705967-DEL-509 | chr15-32705968-DEL-509 |
| chr15-32716546-DEL-137 | chr15-32716547-DEL-137 |
| chr15-32735346-DEL-160 | chr15-32735375-DEL-132 |
| chr15-32741336-DEL-318 | chr15-32741341-DEL-317 |
| chr15-32763348-DEL-141 | chr15-32763349-DEL-141 |
| chr15-32774048-DEL-86 | chr15-32774049-DEL-86 |
| chr15-32784118-DEL-2690 | chr15-32784118-DEL-2690 |
| chr15-32792142-DEL-1432 | chr15-32792143-DEL-1432 |
| chr15-32797838-DEL-343 | chr15-32797837-DEL-343 |
| chr15-32801023-DEL-6184 | chr15-32801024-DEL-6184 |
| chr15-32817125-DEL-64 | chr15-32817126-DEL-64 |
| chr15-32822206-DEL-2380 | chr15-32822207-DEL-2380 |
| chr15-32824265-DEL-3359 | chr15-32824265-DEL-3359 |
| chr15-32826537-DEL-52 | chr15-32826538-DEL-52 |
| chr15-32830502-DEL-62 | chr15-32830503-DEL-62 |
| chr15-32882996-DEL-126 | chr15-32882997-DEL-120 |
| chr15-32946390-DEL-63 | chr15-32946391-DEL-63 |
| chr15-32969385-DEL-1888 | chr15-32969386-DEL-1888 |
| chr15-33001397-DEL-4259 | chr15-33001398-DEL-4259 |
| chr15-33029728-DEL-1504 | chr15-33029729-DEL-1504 |
| chr15-33059517-DEL-1323 | chr15-33059518-DEL-1323 |
| chr15-33076494-DEL-2292 | chr15-33076495-DEL-2292 |
| chr15-33120254-DEL-307 | chr15-33120255-DEL-307 |
| chr15-33122133-DEL-51 | chr15-33122134-DEL-51 |
| chr15-33138777-DEL-2881 | chr15-33138778-DEL-2881 |
| chr15-33153429-DEL-358 | chr15-33153430-DEL-358 |
| chr15-33155760-DEL-608 | chr15-33155761-DEL-608 |
| chr15-33158251-DEL-54 | chr15-33158252-DEL-54 |
| chr15-33168734-DEL-6275 | chr15-33168735-DEL-6275 |
| chr15-33175786-DEL-1198 | chr15-33175786-DEL-1198 |
| chr15-33181766-DEL-52 | chr15-33181767-DEL-52 |
| chr15-33198331-DEL-98 | chr15-33198332-DEL-98 |
| chr15-33200321-DEL-966 | chr15-33200319-DEL-967 |
| chr15-33217198-DEL-1332 | chr15-33217199-DEL-1332 |
| chr15-33221527-DEL-2285 | chr15-33221528-DEL-2285 |
| chr15-33246894-DEL-1734 | chr15-33246895-DEL-1734 |
| chr15-33256359-DEL-533 | chr15-33256358-DEL-533 |
| chr15-33264073-DEL-79 | chr15-33264118-DEL-80 |
| chr15-33267430-DEL-1518 | chr15-33267431-DEL-1518 |
| chr15-33269984-DEL-6382 | chr15-33269985-DEL-6382 |
| chr15-33308929-DEL-1892 | chr15-33308930-DEL-1892 |
| chr15-33332547-DEL-583 | chr15-33332548-DEL-583 |
| chr15-33333684-DEL-908 | chr15-33333685-DEL-908 |
| chr15-33356432-DEL-54 | chr15-33356433-DEL-54 |
| chr15-33377753-DEL-411 | chr15-33377754-DEL-411 |
| chr15-33450689-DEL-4082 | chr15-33450690-DEL-4082 |
| chr15-33453049-DEL-1230 | chr15-33453050-DEL-1230 |
| chr15-33458271-DEL-72 | chr15-33458272-DEL-72 |
| chr15-33486990-DEL-6495 | chr15-33486991-DEL-6495 |
| chr15-33501253-DEL-727 | chr15-33501253-DEL-726 |
| chr15-33510380-DEL-400 | chr15-33510381-DEL-400 |
| chr15-33519882-DEL-57 | chr15-33519883-DEL-57 |
| chr15-33527624-DEL-1458 | chr15-33527625-DEL-1458 |
| chr15-33529648-DEL-634 | chr15-33529649-DEL-633 |
| chr15-33537310-DEL-1036 | chr15-33537311-DEL-1036 |
| chr15-33541735-DEL-111 | chr15-33541732-DEL-112 |
| chr15-33544693-DEL-66 | chr15-33544687-DEL-68 |
| chr15-33544882-DEL-942 | chr15-33544900-DEL-943 |
| chr15-33560355-DEL-6155 | chr15-33560356-DEL-6155 |
| chr15-33568778-DEL-3195 | chr15-33568779-DEL-3195 |
| chr15-33578096-DEL-1601 | chr15-33578097-DEL-1601 |

| | |
|---|---|
| chr15-33597160-DEL-454 | chr15-33597161-DEL-454 |
| chr15-33604601-DEL-145 | chr15-33604602-DEL-155 |
| chr15-33606810-DEL-6486 | chr15-33606810-DEL-6486 |
| chr15-33612968-DEL-214 | chr15-33612969-DEL-214 |
| chr15-33621261-DEL-188 | chr15-33621261-DEL-188 |
| chr15-33635676-DEL-377 | chr15-33635677-DEL-377 |
| chr15-33638938-DEL-1660 | chr15-33638939-DEL-1660 |
| chr15-33659151-DEL-2652 | chr15-33659152-DEL-2652 |
| chr15-33663160-DEL-1167 | chr15-33663161-DEL-1167 |
| chr15-33676302-DEL-94 | chr15-33676305-DEL-92 |
| chr15-33676923-DEL-62 | chr15-33676924-DEL-62 |
| chr15-33694936-DEL-182 | chr15-33694937-DEL-182 |
| chr15-33703942-DEL-459 | chr15-33703943-DEL-459 |
| chr15-33720218-DEL-2352 | chr15-33720219-DEL-2352 |
| chr15-33730890-DEL-15108 | chr15-33730891-DEL-15108 |
| chr15-33735611-DEL-97 | chr15-33735612-DEL-97 |
| chr15-33738958-DEL-6942 | chr15-33738959-DEL-6942 |
| chr15-33761184-DEL-100 | chr15-33761185-DEL-100 |
| chr15-33762972-DEL-1488 | chr15-33762956-DEL-1489 |
| chr15-33765220-DEL-270 | chr15-33765221-DEL-270 |
| chr15-33798017-DEL-158 | chr15-33798018-DEL-158 |
| chr15-33810770-DEL-1766 | chr15-33810771-DEL-1766 |
| chr15-33831579-DEL-7054 | chr15-33831580-DEL-7054 |
| chr15-33839106-DEL-1232 | chr15-33839107-DEL-1232 |
| chr15-33888111-DEL-215 | chr15-33888112-DEL-215 |
| chr15-33931150-DEL-276 | chr15-33931151-DEL-276 |
| chr15-33934650-DEL-127 | chr15-33934651-DEL-127 |
| chr15-33942216-DEL-1891 | chr15-33942217-DEL-1891 |
| chr15-33952524-DEL-112 | chr15-33952525-DEL-112 |
| chr15-33978235-DEL-56 | chr15-33978235-DEL-56 |
| chr15-33978482-DEL-3049 | chr15-33978483-DEL-3049 |
| chr15-33985041-DEL-2897 | chr15-33985042-DEL-2897 |
| chr15-33990361-DEL-84 | chr15-33990384-DEL-98 |
| chr15-33992395-DEL-211 | chr15-33992396-DEL-211 |
| chr15-33993233-DEL-853 | chr15-33993234-DEL-853 |
| chr15-34020515-DEL-82 | chr15-34020516-DEL-82 |
| chr15-34035907-DEL-236 | chr15-34035925-DEL-228 |
| chr15-34130267-DEL-401 | chr15-34130268-DEL-401 |
| chr15-34157768-DEL-160 | chr15-34157769-DEL-160 |
| chr15-34162994-DEL-161 | chr15-34162995-DEL-161 |
| chr15-34170564-DEL-157 | chr15-34170573-DEL-149 |
| chr15-34183029-DEL-76 | chr15-34183030-DEL-76 |
| chr15-34207904-DEL-102 | chr15-34207903-DEL-103 |
| chr15-34212804-DEL-61 | chr15-34212805-DEL-60 |
| chr15-34226748-DEL-3332 | chr15-34226749-DEL-3332 |
| chr15-34244733-DEL-200 | chr15-34244734-DEL-200 |
| chr15-34275027-DEL-284 | chr15-34275028-DEL-284 |
| chr15-34279448-DEL-216 | chr15-34279449-DEL-216 |
| chr15-34286358-DEL-191 | chr15-34286359-DEL-191 |
| chr15-34295308-DEL-52 | chr15-34295309-DEL-52 |
| chr15-34315758-DEL-418 | chr15-34315759-DEL-418 |
| chr15-34327615-DEL-191 | chr15-34327616-DEL-191 |
| chr15-34337342-DEL-133 | chr15-34337343-DEL-86 |
| chr15-34355057-DEL-52 | chr15-34355034-DEL-52 |
| chr15-34360750-DEL-56 | chr15-34360751-DEL-56 |
| chr15-34374374-DEL-90 | chr15-34374378-DEL-78 |
| chr15-34392373-DEL-82 | chr15-34392349-DEL-80 |
| chr15-34401319-DEL-208 | chr15-34401316-DEL-204 |
| chr15-34445266-DEL-175 | chr15-34445267-DEL-175 |
| chr15-34450759-DEL-194 | chr15-34450760-DEL-194 |
| chr15-34451516-DEL-189 | chr15-34451517-DEL-189 |
| chr15-34474537-DEL-64 | chr15-34474538-DEL-64 |
| chr15-34481718-DEL-59 | chr15-34481719-DEL-55 |
| chr15-34500954-DEL-1036 | chr15-34500954-DEL-1036 |
| chr15-34552506-DEL-332 | chr15-34552507-DEL-332 |
| chr15-34555250-DEL-2682 | chr15-34555251-DEL-2682 |
| chr15-34570668-DEL-142 | chr15-34570687-DEL-128 |
| chr15-34600689-DEL-57 | chr15-34600689-DEL-53 |
| chr15-34601207-DEL-526 | chr15-34601208-DEL-526 |
| chr15-34616107-DEL-192 | chr15-34616108-DEL-192 |
| chr15-34630007-DEL-120 | chr15-34630008-DEL-120 |
| chr15-34689679-DEL-614 | chr15-34689680-DEL-614 |
| chr15-34719293-DEL-210 | chr15-34719294-DEL-210 |
| chr15-34724047-DEL-59 | chr15-34724048-DEL-62 |
| chr15-34727417-DEL-52 | chr15-34727418-DEL-52 |
| chr15-34751636-DEL-86 | chr15-34751637-DEL-86 |
| chr15-34760505-DEL-58 | chr15-34760506-DEL-55 |
| chr15-34763233-DEL-6408 | chr15-34763234-DEL-6408 |
| chr15-34793659-DEL-141 | chr15-34793660-DEL-141 |
| chr15-34798184-DEL-199 | chr15-34798185-DEL-199 |

| | |
|---|---|
| chr15-34807552-DEL-380 | chr15-34807553-DEL-380 |
| chr15-34832107-DEL-90 | chr15-34832108-DEL-90 |
| chr15-34917450-DEL-56 | chr15-34917451-DEL-54 |
| chr15-34922929-DEL-62 | chr15-34922930-DEL-62 |
| chr15-34923620-DEL-240 | chr15-34923621-DEL-239 |
| chr15-34932219-DEL-159 | chr15-34932220-DEL-159 |
| chr15-34935852-DEL-650 | chr15-34935853-DEL-650 |
| chr15-34938117-DEL-6693 | chr15-34938118-DEL-6693 |
| chr15-34987578-DEL-6342 | chr15-34987579-DEL-6342 |
| chr15-35015103-DEL-92 | chr15-35015103-DEL-142 |
| chr15-35071805-DEL-132 | chr15-35071804-DEL-134 |
| chr15-35091032-DEL-204 | chr15-35091033-DEL-204 |
| chr15-35102275-DEL-249 | chr15-35102276-DEL-249 |
| chr15-35116659-DEL-7834 | chr15-35116660-DEL-7834 |
| chr15-35138602-DEL-105 | chr15-35138603-DEL-105 |
| chr15-35144036-DEL-66 | chr15-35144037-DEL-66 |
| chr15-35155280-DEL-54 | chr15-35155281-DEL-54 |
| chr15-35164542-DEL-53 | chr15-35164543-DEL-53 |
| chr15-35178885-DEL-401 | chr15-35178886-DEL-401 |
| chr15-35220691-DEL-53 | chr15-35220692-DEL-53 |
| chr15-35242604-DEL-127 | chr15-35242605-DEL-127 |
| chr15-35288265-DEL-72 | chr15-35288266-DEL-72 |
| chr15-35325089-DEL-171 | chr15-35325090-DEL-171 |
| chr15-35363297-DEL-213 | chr15-35363298-DEL-213 |
| chr15-35405365-DEL-401 | chr15-35405366-DEL-401 |
| chr15-35419927-DEL-400 | chr15-35419928-DEL-400 |
| chr15-35441951-DEL-70 | chr15-35441952-DEL-70 |
| chr15-35468489-DEL-61 | chr15-35468490-DEL-61 |
| chr15-35506950-DEL-200 | chr15-35506951-DEL-200 |
| chr15-35531978-DEL-52 | chr15-35531978-DEL-52 |
| chr15-35563614-DEL-4797 | chr15-35563615-DEL-4795 |
| chr15-35583940-DEL-6446 | chr15-35583933-DEL-6447 |
| chr15-35590066-DEL-211 | chr15-35590055-DEL-212 |
| chr15-35655845-DEL-127 | chr15-35655846-DEL-122 |
| chr15-35685232-DEL-142 | chr15-35685233-DEL-142 |
| chr15-35698413-DEL-193 | chr15-35698414-DEL-193 |
| chr15-35707178-DEL-68 | chr15-35707179-DEL-68 |
| chr15-35720941-DEL-101 | chr15-35720942-DEL-101 |
| chr15-35783917-DEL-3232 | chr15-35783918-DEL-3232 |
| chr15-35817984-DEL-1014 | chr15-35817957-DEL-1015 |
| chr15-35819857-DEL-1221 | chr15-35819866-DEL-1211 |
| chr15-35836593-DEL-59 | chr15-35836593-DEL-59 |
| chr15-35855144-DEL-492 | chr15-35855135-DEL-493 |
| chr15-35866725-DEL-55 | chr15-35866726-DEL-55 |
| chr15-35909280-DEL-52 | chr15-35909281-DEL-52 |
| chr15-35944465-DEL-1259 | chr15-35944466-DEL-1259 |
| chr15-35954526-DEL-60 | chr15-35954527-DEL-60 |
| chr15-35991526-DEL-6418 | chr15-35991527-DEL-6418 |
| chr15-36050668-DEL-1505 | chr15-36050670-DEL-1506 |
| chr15-36055735-DEL-52 | chr15-36055736-DEL-50 |
| chr15-36062611-DEL-417 | chr15-36062602-DEL-415 |
| chr15-36103364-DEL-280 | chr15-36103365-DEL-280 |
| chr15-36126988-DEL-185 | chr15-36126989-DEL-185 |
| chr15-36152806-DEL-64 | chr15-36152807-DEL-64 |
| chr15-36167651-DEL-189 | chr15-36167646-DEL-193 |
| chr15-36169317-DEL-63 | chr15-36169300-DEL-61 |
| chr15-36171306-DEL-1342 | chr15-36171307-DEL-1340 |
| chr15-36185008-DEL-57 | chr15-36185009-DEL-57 |
| chr15-36189969-DEL-123 | chr15-36189970-DEL-123 |
| chr15-36219313-DEL-842 | chr15-36219314-DEL-842 |
| chr15-36226424-DEL-51 | chr15-36226392-DEL-52 |
| chr15-36253021-DEL-76 | chr15-36253022-DEL-76 |
| chr15-36273199-DEL-54 | chr15-36273204-DEL-50 |
| chr15-36288037-DEL-762 | chr15-36288038-DEL-762 |
| chr15-36290238-DEL-919 | chr15-36290239-DEL-919 |
| chr15-36294198-DEL-51 | chr15-36294199-DEL-51 |
| chr15-36316595-DEL-153 | chr15-36316596-DEL-153 |
| chr15-36319930-DEL-212 | chr15-36319931-DEL-212 |
| chr15-36328047-DEL-115 | chr15-36328048-DEL-115 |
| chr15-36337034-DEL-597 | chr15-36337035-DEL-597 |
| chr15-36343676-DEL-63 | chr15-36343677-DEL-63 |
| chr15-36347034-DEL-133 | chr15-36347035-DEL-133 |
| chr15-36368646-DEL-70 | chr15-36368646-DEL-71 |
| chr15-36370422-DEL-64 | chr15-36370423-DEL-64 |
| chr15-36381438-DEL-230 | chr15-36381439-DEL-230 |
| chr15-36398135-DEL-104 | chr15-36398116-DEL-97 |
| chr15-36398457-DEL-295 | chr15-36398460-DEL-293 |
| chr15-36413927-DEL-94 | chr15-36413928-DEL-94 |
| chr15-36439991-DEL-65 | chr15-36439992-DEL-65 |
| chr15-36444248-DEL-104 | chr15-36444251-DEL-103 |

| | |
|---|---|
| chr15-36445795-DEL-54 | chr15-36445796-DEL-54 |
| chr15-36458997-DEL-3033 | chr15-36458998-DEL-3033 |
| chr15-36468907-DEL-178 | chr15-36468908-DEL-179 |
| chr15-36481361-DEL-2586 | chr15-36481362-DEL-2586 |
| chr15-36484607-DEL-52 | chr15-36484608-DEL-50 |
| chr15-36485302-DEL-109 | chr15-36485322-DEL-103 |
| chr15-36506191-DEL-133 | chr15-36506192-DEL-133 |
| chr15-36508085-DEL-58 | chr15-36508086-DEL-57 |
| chr15-36511739-DEL-184 | chr15-36511740-DEL-184 |
| chr15-36514480-DEL-57 | chr15-36514481-DEL-57 |
| chr15-36531304-DEL-477 | chr15-36531317-DEL-465 |
| chr15-36533164-DEL-152 | chr15-36533127-DEL-190 |
| chr15-36534936-DEL-60 | chr15-36534898-DEL-68 |
| chr15-36539857-DEL-1055 | chr15-36539858-DEL-1055 |
| chr15-36549090-DEL-218 | chr15-36549091-DEL-218 |
| chr15-36554494-DEL-213 | chr15-36554495-DEL-213 |
| chr15-36592875-DEL-474 | chr15-36592875-DEL-474 |
| chr15-36624443-DEL-72 | chr15-36624443-DEL-72 |
| chr15-36643630-DEL-51 | chr15-36643665-DEL-60 |
| chr15-36646908-DEL-407 | chr15-36646909-DEL-407 |
| chr15-36648644-DEL-68 | chr15-36648645-DEL-68 |
| chr15-36666551-DEL-59 | chr15-36666532-DEL-60 |
| chr15-36673614-DEL-6211 | chr15-36673614-DEL-6210 |
| chr15-36684521-DEL-54 | chr15-36684522-DEL-54 |
| chr15-36703165-DEL-364 | chr15-36703166-DEL-364 |
| chr15-36725297-DEL-499 | chr15-36725292-DEL-500 |
| chr15-36730536-DEL-126 | chr15-36730548-DEL-113 |
| chr15-36733090-DEL-84 | chr15-36733091-DEL-84 |
| chr15-36735414-DEL-1290 | chr15-36735414-DEL-1311 |
| chr15-36746154-DEL-78 | chr15-36746175-DEL-71 |
| chr15-36752212-DEL-238 | chr15-36752213-DEL-238 |
| chr15-36812527-DEL-92 | chr15-36812552-DEL-68 |
| chr15-36831103-DEL-78 | chr15-36831122-DEL-69 |
| chr15-36854263-DEL-205 | chr15-36854264-DEL-205 |
| chr15-36859750-DEL-1502 | chr15-36859751-DEL-1502 |
| chr15-36863359-DEL-64 | chr15-36863356-DEL-74 |
| chr15-36866896-DEL-112 | chr15-36866897-DEL-122 |
| chr15-36888346-DEL-228 | chr15-36888347-DEL-228 |
| chr15-36897504-DEL-136 | chr15-36897505-DEL-136 |
| chr15-36916993-DEL-52 | chr15-36916994-DEL-52 |
| chr15-36929947-DEL-216 | chr15-36929962-DEL-234 |
| chr15-36951206-DEL-506 | chr15-36951207-DEL-506 |
| chr15-36954588-DEL-124 | chr15-36954587-DEL-123 |
| chr15-36968328-DEL-769 | chr15-36968329-DEL-769 |
| chr15-36976461-DEL-866 | chr15-36976462-DEL-866 |
| chr15-36977176-DEL-249 | chr15-36977136-DEL-250 |
| chr15-36990818-DEL-92 | chr15-36990819-DEL-92 |
| chr15-36995802-DEL-1799 | chr15-36995805-DEL-1800 |
| chr15-37023381-DEL-499 | chr15-37023383-DEL-499 |
| chr15-37023956-DEL-183 | chr15-37023957-DEL-183 |
| chr15-37032257-DEL-62 | chr15-37032257-DEL-62 |
| chr15-37034697-DEL-170 | chr15-37034698-DEL-170 |
| chr15-37038412-DEL-453 | chr15-37038413-DEL-453 |
| chr15-37055638-DEL-207 | chr15-37055639-DEL-207 |
| chr15-37072486-DEL-104 | chr15-37072487-DEL-104 |
| chr15-37073106-DEL-7406 | chr15-37073107-DEL-7406 |
| chr15-37092278-DEL-5389 | chr15-37092279-DEL-5389 |
| chr15-37105940-DEL-212 | chr15-37105931-DEL-234 |
| chr15-37116631-DEL-3794 | chr15-37116632-DEL-3794 |
| chr15-37121659-DEL-149 | chr15-37121647-DEL-162 |
| chr15-37125934-DEL-476 | chr15-37125933-DEL-477 |
| chr15-37127643-DEL-137 | chr15-37127644-DEL-137 |
| chr15-37143857-DEL-242 | chr15-37143858-DEL-242 |
| chr15-37149108-DEL-1891 | chr15-37149109-DEL-1891 |
| chr15-37172845-DEL-215 | chr15-37172846-DEL-198 |
| chr15-37202910-DEL-54 | chr15-37202911-DEL-54 |
| chr15-37203551-DEL-80 | chr15-37203552-DEL-80 |
| chr15-37212906-DEL-64 | chr15-37212896-DEL-66 |
| chr15-37226480-DEL-151 | chr15-37226469-DEL-162 |
| chr15-37228643-DEL-174 | chr15-37228644-DEL-174 |
| chr15-37282637-DEL-220 | chr15-37282638-DEL-220 |
| chr15-37286120-DEL-243 | chr15-37286121-DEL-243 |
| chr15-37290367-DEL-66 | chr15-37290368-DEL-64 |
| chr15-37321093-DEL-95 | chr15-37321088-DEL-103 |
| chr15-37335806-DEL-86 | chr15-37335807-DEL-86 |
| chr15-37340266-DEL-1559 | chr15-37340267-DEL-1559 |
| chr15-37342161-DEL-1044 | chr15-37342162-DEL-1044 |
| chr15-37349796-DEL-65 | chr15-37349797-DEL-65 |
| chr15-37394745-DEL-98 | chr15-37394724-DEL-93 |
| chr15-37398011-DEL-75 | chr15-37398008-DEL-76 |

| | |
|---|---|
| chr15-37399779-DEL-79 | chr15-37399730-DEL-120 |
| chr15-37431274-DEL-60 | chr15-37431275-DEL-60 |
| chr15-37465289-DEL-6520 | chr15-37465290-DEL-6520 |
| chr15-37484207-DEL-646 | chr15-37484208-DEL-646 |
| chr15-37491982-DEL-86 | chr15-37491981-DEL-86 |
| chr15-37505296-DEL-400 | chr15-37505297-DEL-400 |
| chr15-37514927-DEL-72 | chr15-37514928-DEL-72 |
| chr15-37523860-DEL-66 | chr15-37523861-DEL-66 |
| chr15-37529970-DEL-380 | chr15-37529971-DEL-380 |
| chr15-37538162-DEL-200 | chr15-37538163-DEL-200 |
| chr15-37566789-DEL-1516 | chr15-37566790-DEL-1516 |
| chr15-37570883-DEL-64 | chr15-37570880-DEL-62 |
| chr15-37605797-DEL-696 | chr15-37605798-DEL-696 |
| chr15-37635191-DEL-1968 | chr15-37635192-DEL-1968 |
| chr15-37648264-DEL-740 | chr15-37648266-DEL-738 |
| chr15-37684413-DEL-1057 | chr15-37684414-DEL-1057 |
| chr15-37696148-DEL-281 | chr15-37696149-DEL-281 |
| chr15-37702069-DEL-53 | chr15-37702059-DEL-71 |
| chr15-37702871-DEL-71 | chr15-37702871-DEL-75 |
| chr15-37738669-DEL-793 | chr15-37738670-DEL-793 |
| chr15-37744551-DEL-492 | chr15-37744552-DEL-492 |
| chr15-37759720-DEL-494 | chr15-37759721-DEL-494 |
| chr15-37762783-DEL-56 | chr15-37762784-DEL-56 |
| chr15-37773743-DEL-68 | chr15-37773751-DEL-60 |
| chr15-37776039-DEL-125 | chr15-37776040-DEL-125 |
| chr15-37785939-DEL-140 | chr15-37785940-DEL-140 |
| chr15-37816857-DEL-2181 | chr15-37816858-DEL-2181 |
| chr15-37827023-DEL-52 | chr15-37827030-DEL-50 |
| chr15-37872884-DEL-54 | chr15-37872885-DEL-54 |
| chr15-37877918-DEL-3831 | chr15-37877919-DEL-3831 |
| chr15-37911841-DEL-390 | chr15-37911842-DEL-390 |
| chr15-37918373-DEL-132 | chr15-37918374-DEL-133 |
| chr15-37932239-DEL-158 | chr15-37932240-DEL-158 |
| chr15-37943481-DEL-559 | chr15-37943482-DEL-559 |
| chr15-37976998-DEL-112 | chr15-37976999-DEL-112 |
| chr15-38073283-DEL-205 | chr15-38073284-DEL-205 |
| chr15-38093692-DEL-87 | chr15-38093693-DEL-87 |
| chr15-38118656-DEL-77 | chr15-38118696-DEL-80 |
| chr15-38119938-DEL-476 | chr15-38119939-DEL-476 |
| chr15-38151319-DEL-169 | chr15-38151307-DEL-170 |
| chr15-38165585-DEL-67 | chr15-38165566-DEL-87 |
| chr15-38177802-DEL-88 | chr15-38177807-DEL-65 |
| chr15-38186097-DEL-203 | chr15-38186098-DEL-203 |
| chr15-38190863-DEL-560 | chr15-38190864-DEL-560 |
| chr15-38244078-DEL-72 | chr15-38244077-DEL-60 |
| chr15-38247612-DEL-216 | chr15-38247613-DEL-216 |
| chr15-38253289-DEL-89 | chr15-38253288-DEL-75 |
| chr15-38254371-DEL-78 | chr15-38254372-DEL-78 |
| chr15-38262157-DEL-56 | chr15-38262162-DEL-52 |
| chr15-38280875-DEL-67 | chr15-38280876-DEL-67 |
| chr15-38285051-DEL-68 | chr15-38285052-DEL-68 |
| chr15-38287892-DEL-81 | chr15-38287893-DEL-81 |
| chr15-38288603-DEL-72 | chr15-38288598-DEL-78 |
| chr15-38305905-DEL-224 | chr15-38305912-DEL-218 |
| chr15-38308967-DEL-402 | chr15-38308934-DEL-396 |
| chr15-38310833-DEL-56 | chr15-38310833-DEL-52 |
| chr15-38316383-DEL-361 | chr15-38316384-DEL-361 |
| chr15-38422683-DEL-70 | chr15-38422684-DEL-70 |
| chr15-38446037-DEL-200 | chr15-38446038-DEL-200 |
| chr15-38451821-DEL-149 | chr15-38451815-DEL-147 |
| chr15-38459838-DEL-79 | chr15-38459822-DEL-77 |
| chr15-38462663-DEL-2587 | chr15-38462664-DEL-2587 |
| chr15-38479030-DEL-4388 | chr15-38479031-DEL-4388 |
| chr15-38509164-DEL-110 | chr15-38509165-DEL-110 |
| chr15-38533823-DEL-318 | chr15-38533804-DEL-314 |
| chr15-38535363-DEL-52 | chr15-38535364-DEL-52 |
| chr15-38541831-DEL-111 | chr15-38541855-DEL-112 |
| chr15-38559139-DEL-980 | chr15-38559140-DEL-980 |
| chr15-38565785-DEL-87 | chr15-38565786-DEL-87 |
| chr15-38576578-DEL-501 | chr15-38576578-DEL-500 |
| chr15-38585209-DEL-176 | chr15-38585210-DEL-176 |
| chr15-38601620-DEL-52 | chr15-38601621-DEL-52 |
| chr15-38622614-DEL-471 | chr15-38622615-DEL-471 |
| chr15-38643254-DEL-196 | chr15-38643255-DEL-196 |
| chr15-38649062-DEL-111 | chr15-38649063-DEL-111 |
| chr15-38654447-DEL-290 | chr15-38654448-DEL-290 |
| chr15-38683852-DEL-608 | chr15-38683853-DEL-608 |
| chr15-38701504-DEL-80 | chr15-38701505-DEL-80 |
| chr15-38705673-DEL-225 | chr15-38705674-DEL-225 |
| chr15-38721309-DEL-85 | chr15-38721310-DEL-85 |

| | |
|---|---|
| chr15-38727590-DEL-91 | chr15-38727595-DEL-89 |
| chr15-38730484-DEL-505 | chr15-38730485-DEL-505 |
| chr15-38748030-DEL-182 | chr15-38748067-DEL-146 |
| chr15-38749620-DEL-56 | chr15-38749621-DEL-52 |
| chr15-38780303-DEL-172 | chr15-38780304-DEL-172 |
| chr15-38809788-DEL-152 | chr15-38809781-DEL-160 |
| chr15-38819587-DEL-62 | chr15-38819588-DEL-62 |
| chr15-38838589-DEL-150 | chr15-38838559-DEL-179 |
| chr15-38844872-DEL-71 | chr15-38844873-DEL-71 |
| chr15-38853005-DEL-341 | chr15-38853006-DEL-341 |
| chr15-38858305-DEL-211 | chr15-38858322-DEL-197 |
| chr15-38879706-DEL-1761 | chr15-38879707-DEL-1761 |
| chr15-38885422-DEL-57 | chr15-38885423-DEL-51 |
| chr15-38885956-DEL-90 | chr15-38885957-DEL-90 |
| chr15-38896061-DEL-114 | chr15-38896031-DEL-111 |
| chr15-38920915-DEL-53 | chr15-38920916-DEL-53 |
| chr15-38930136-DEL-202 | chr15-38930137-DEL-202 |
| chr15-38951626-DEL-1024 | chr15-38951627-DEL-1024 |
| chr15-39029433-DEL-347 | chr15-39029434-DEL-347 |
| chr15-39036430-DEL-5222 | chr15-39036431-DEL-5222 |
| chr15-39045839-DEL-204 | chr15-39045840-DEL-204 |
| chr15-39074715-DEL-142 | chr15-39074752-DEL-106 |
| chr15-39085793-DEL-214 | chr15-39085793-DEL-214 |
| chr15-39087113-DEL-201 | chr15-39087114-DEL-201 |
| chr15-39090892-DEL-69 | chr15-39090882-DEL-70 |
| chr15-39094599-DEL-84 | chr15-39094600-DEL-96 |
| chr15-39097430-DEL-55 | chr15-39097431-DEL-55 |
| chr15-39112982-DEL-480 | chr15-39112983-DEL-480 |
| chr15-39162574-DEL-203 | chr15-39162575-DEL-203 |
| chr15-39167947-DEL-889 | chr15-39167948-DEL-889 |
| chr15-39176595-DEL-52 | chr15-39176595-DEL-53 |
| chr15-39176743-DEL-1209 | chr15-39176744-DEL-1209 |
| chr15-39185907-DEL-386 | chr15-39185908-DEL-386 |
| chr15-39199105-DEL-4105 | chr15-39199095-DEL-4149 |
| chr15-39228671-DEL-259 | chr15-39228672-DEL-259 |
| chr15-39241139-DEL-644 | chr15-39241136-DEL-645 |
| chr15-39273395-DEL-176 | chr15-39273396-DEL-176 |
| chr15-39276010-DEL-1936 | chr15-39276011-DEL-1936 |
| chr15-39280127-DEL-140 | chr15-39280128-DEL-140 |
| chr15-39290401-DEL-1694 | chr15-39290402-DEL-1694 |
| chr15-39343273-DEL-57 | chr15-39343274-DEL-57 |
| chr15-39359414-DEL-3371 | chr15-39359415-DEL-3371 |
| chr15-39385003-DEL-1102 | chr15-39385004-DEL-1102 |
| chr15-39409039-DEL-54 | chr15-39409040-DEL-54 |
| chr15-39442918-DEL-206 | chr15-39442919-DEL-206 |
| chr15-39468283-DEL-804 | chr15-39468284-DEL-804 |
| chr15-39483901-DEL-929 | chr15-39483902-DEL-929 |
| chr15-39495248-DEL-159 | chr15-39495249-DEL-159 |
| chr15-39505973-DEL-183 | chr15-39505974-DEL-183 |
| chr15-39510204-DEL-60 | chr15-39510205-DEL-60 |
| chr15-39511161-DEL-73 | chr15-39511162-DEL-73 |
| chr15-39558692-DEL-8675 | chr15-39558674-DEL-8697 |
| chr15-39577357-DEL-4552 | chr15-39577358-DEL-4552 |
| chr15-39644334-DEL-52 | chr15-39644335-DEL-51 |
| chr15-39655836-DEL-302 | chr15-39655837-DEL-302 |
| chr15-39665076-DEL-72 | chr15-39665077-DEL-72 |
| chr15-39678968-DEL-58 | chr15-39678969-DEL-58 |
| chr15-39722277-DEL-63 | chr15-39722255-DEL-81 |
| chr15-39724549-DEL-62 | chr15-39724550-DEL-62 |
| chr15-39730394-DEL-84 | chr15-39730395-DEL-98 |
| chr15-39791277-DEL-184 | chr15-39791279-DEL-153 |
| chr15-39827015-DEL-74 | chr15-39827016-DEL-74 |
| chr15-39846360-DEL-6492 | chr15-39846362-DEL-6493 |
| chr15-39855095-DEL-617 | chr15-39855099-DEL-618 |
| chr15-39870011-DEL-209 | chr15-39870023-DEL-210 |
| chr15-39871307-DEL-58 | chr15-39871308-DEL-58 |
| chr15-39871815-DEL-129 | chr15-39871816-DEL-129 |
| chr15-39901810-DEL-135 | chr15-39901794-DEL-136 |
| chr15-39913009-DEL-6421 | chr15-39913010-DEL-6421 |
| chr15-39925621-DEL-101 | chr15-39925638-DEL-77 |
| chr15-39950046-DEL-6464 | chr15-39950047-DEL-6464 |
| chr15-39958291-DEL-6564 | chr15-39958292-DEL-6564 |
| chr15-39991981-DEL-1032 | chr15-39991982-DEL-1032 |
| chr15-40014021-DEL-65 | chr15-40014015-DEL-98 |
| chr15-40015091-DEL-126 | chr15-40015092-DEL-126 |
| chr15-40024090-DEL-445 | chr15-40024091-DEL-445 |
| chr15-40059755-DEL-62 | chr15-40059735-DEL-66 |
| chr15-40067114-DEL-276 | chr15-40067115-DEL-276 |
| chr15-40068274-DEL-6408 | chr15-40068280-DEL-6406 |
| chr15-40079295-DEL-161 | chr15-40079297-DEL-157 |

| | |
|---|---|
| chr15-40081344-DEL-103 | chr15-40081345-DEL-75 |
| chr15-40087832-DEL-219 | chr15-40087833-DEL-219 |
| chr15-40159887-DEL-55 | chr15-40159888-DEL-55 |
| chr15-40165940-DEL-57 | chr15-40165941-DEL-57 |
| chr15-40181339-DEL-2953 | chr15-40181340-DEL-2953 |
| chr15-40184767-DEL-3772 | chr15-40184768-DEL-3772 |
| chr15-40189111-DEL-92 | chr15-40189113-DEL-90 |
| chr15-40190549-DEL-152 | chr15-40190550-DEL-152 |
| chr15-40228684-DEL-317 | chr15-40228685-DEL-317 |
| chr15-40263489-DEL-2754 | chr15-40263490-DEL-2754 |
| chr15-40272051-DEL-54 | chr15-40272031-DEL-53 |
| chr15-40275776-DEL-76 | chr15-40275803-DEL-50 |
| chr15-40279618-DEL-92 | chr15-40279633-DEL-78 |
| chr15-40296612-DEL-1721 | chr15-40296613-DEL-1721 |
| chr15-40304923-DEL-6632 | chr15-40304922-DEL-6633 |
| chr15-40306399-DEL-227 | chr15-40306400-DEL-227 |
| chr15-40438992-DEL-89 | chr15-40438988-DEL-85 |
| chr15-40462401-DEL-104 | chr15-40462402-DEL-104 |
| chr15-40466411-DEL-62 | chr15-40466424-DEL-50 |
| chr15-40491909-DEL-267 | chr15-40491910-DEL-267 |
| chr15-40499777-DEL-188 | chr15-40499778-DEL-188 |
| chr15-40500469-DEL-55 | chr15-40500470-DEL-55 |
| chr15-40522437-DEL-52 | chr15-40522438-DEL-52 |
| chr15-40533708-DEL-857 | chr15-40533709-DEL-857 |
| chr15-40579056-DEL-54 | chr15-40579057-DEL-54 |
| chr15-40582079-DEL-58 | chr15-40582080-DEL-58 |
| chr15-40584914-DEL-92 | chr15-40584913-DEL-72 |
| chr15-40615885-DEL-60 | chr15-40615876-DEL-70 |
| chr15-40634027-DEL-2034 | chr15-40634028-DEL-2034 |
| chr15-40728814-DEL-585 | chr15-40728815-DEL-585 |
| chr15-40742773-DEL-6811 | chr15-40742774-DEL-6811 |
| chr15-40751625-DEL-2113 | chr15-40751626-DEL-2113 |
| chr15-40799979-DEL-52 | chr15-40799980-DEL-59 |
| chr15-40800379-DEL-536 | chr15-40800380-DEL-536 |
| chr15-40828303-DEL-55 | chr15-40828301-DEL-50 |
| chr15-40867149-DEL-107 | chr15-40867150-DEL-107 |
| chr15-40941708-DEL-477 | chr15-40941709-DEL-477 |
| chr15-40943777-DEL-119 | chr15-40943778-DEL-119 |
| chr15-40970338-DEL-75 | chr15-40970332-DEL-76 |
| chr15-41000686-DEL-51 | chr15-41000687-DEL-51 |
| chr15-41014257-DEL-81 | chr15-41014256-DEL-89 |
| chr15-41014645-DEL-260 | chr15-41014656-DEL-250 |
| chr15-41016311-DEL-128 | chr15-41016310-DEL-129 |
| chr15-41024075-DEL-1295 | chr15-41024076-DEL-1295 |
| chr15-41066002-DEL-81 | chr15-41066003-DEL-81 |
| chr15-41081039-DEL-112 | chr15-41081072-DEL-117 |
| chr15-41099140-DEL-547 | chr15-41099141-DEL-547 |
| chr15-41105928-DEL-56 | chr15-41105935-DEL-50 |
| chr15-41115537-DEL-215 | chr15-41115539-DEL-214 |
| chr15-41137386-DEL-101 | chr15-41137387-DEL-101 |
| chr15-41148734-DEL-201 | chr15-41148735-DEL-201 |
| chr15-41179287-DEL-602 | chr15-41179288-DEL-602 |
| chr15-41211914-DEL-1857 | chr15-41211905-DEL-1861 |
| chr15-41257823-DEL-397 | chr15-41257824-DEL-397 |
| chr15-41258894-DEL-310 | chr15-41258895-DEL-310 |
| chr15-41263987-DEL-54 | chr15-41263988-DEL-54 |
| chr15-41282800-DEL-823 | chr15-41282801-DEL-823 |
| chr15-41289255-DEL-370 | chr15-41289256-DEL-370 |
| chr15-41290605-DEL-54 | chr15-41290606-DEL-54 |
| chr15-41350385-DEL-158 | chr15-41350386-DEL-158 |
| chr15-41360076-DEL-628 | chr15-41360077-DEL-627 |
| chr15-41360866-DEL-114 | chr15-41360867-DEL-114 |
| chr15-41390504-DEL-2130 | chr15-41390505-DEL-2130 |
| chr15-41419237-DEL-1622 | chr15-41419239-DEL-1621 |
| chr15-41421620-DEL-393 | chr15-41421621-DEL-393 |
| chr15-41424171-DEL-74 | chr15-41424188-DEL-58 |
| chr15-41437231-DEL-494 | chr15-41437232-DEL-487 |
| chr15-41437924-DEL-478 | chr15-41437921-DEL-477 |
| chr15-41442437-DEL-144 | chr15-41442442-DEL-140 |
| chr15-41455286-DEL-202 | chr15-41455281-DEL-203 |
| chr15-41499297-DEL-52 | chr15-41499298-DEL-52 |
| chr15-41524017-DEL-206 | chr15-41524018-DEL-206 |
| chr15-41557721-DEL-206 | chr15-41557722-DEL-206 |
| chr15-41579206-DEL-640 | chr15-41579207-DEL-640 |
| chr15-41602293-DEL-56 | chr15-41602294-DEL-56 |
| chr15-41606291-DEL-82 | chr15-41606292-DEL-70 |
| chr15-41645394-DEL-208 | chr15-41645395-DEL-208 |
| chr15-41658920-DEL-51 | chr15-41658921-DEL-51 |
| chr15-41664546-DEL-54 | chr15-41664534-DEL-54 |
| chr15-41739285-DEL-101 | chr15-41739286-DEL-101 |

| | |
|---|---|
| chr15-41750645-DEL-2083 | chr15-41750646-DEL-2083 |
| chr15-41753767-DEL-6454 | chr15-41753768-DEL-6454 |
| chr15-41764275-DEL-51 | chr15-41764271-DEL-56 |
| chr15-41775718-DEL-292 | chr15-41775735-DEL-276 |
| chr15-41785420-DEL-1520 | chr15-41785421-DEL-1520 |
| chr15-41789871-DEL-6695 | chr15-41789872-DEL-6695 |
| chr15-41797824-DEL-1193 | chr15-41797815-DEL-1211 |
| chr15-41810462-DEL-234 | chr15-41810463-DEL-234 |
| chr15-41845192-DEL-96 | chr15-41845159-DEL-98 |
| chr15-41865633-DEL-1397 | chr15-41865634-DEL-1397 |
| chr15-41896487-DEL-1312 | chr15-41896488-DEL-1312 |
| chr15-41900135-DEL-921 | chr15-41900136-DEL-921 |
| chr15-41934364-DEL-52 | chr15-41934365-DEL-52 |
| chr15-41935532-DEL-390 | chr15-41935533-DEL-388 |
| chr15-41952687-DEL-318 | chr15-41952688-DEL-317 |
| chr15-41953601-DEL-1105 | chr15-41953602-DEL-1105 |
| chr15-42002914-DEL-441 | chr15-42002915-DEL-441 |
| chr15-42011688-DEL-51 | chr15-42011687-DEL-52 |
| chr15-42016311-DEL-66 | chr15-42016300-DEL-62 |
| chr15-42135662-DEL-80 | chr15-42135663-DEL-80 |
| chr15-42139365-DEL-92 | chr15-42139365-DEL-94 |
| chr15-42151231-DEL-675 | chr15-42151232-DEL-675 |
| chr15-42176279-DEL-370 | chr15-42176280-DEL-369 |
| chr15-42181009-DEL-2731 | chr15-42181010-DEL-2731 |
| chr15-42237234-DEL-72 | chr15-42237235-DEL-72 |
| chr15-42256755-DEL-340 | chr15-42256756-DEL-340 |
| chr15-42273755-DEL-72 | chr15-42273756-DEL-72 |
| chr15-42308113-DEL-695 | chr15-42308114-DEL-695 |
| chr15-42324419-DEL-120 | chr15-42324387-DEL-155 |
| chr15-42374065-DEL-144 | chr15-42374066-DEL-144 |
| chr15-42395301-DEL-169 | chr15-42395306-DEL-170 |
| chr15-42395885-DEL-52 | chr15-42395886-DEL-52 |
| chr15-42434331-DEL-56 | chr15-42434331-DEL-56 |
| chr15-42439047-DEL-1047 | chr15-42439048-DEL-1047 |
| chr15-42451239-DEL-63 | chr15-42451240-DEL-54 |
| chr15-42462504-DEL-2878 | chr15-42462505-DEL-2878 |
| chr15-42507739-DEL-93 | chr15-42507740-DEL-90 |
| chr15-42507819-DEL-6668 | chr15-42507820-DEL-6668 |
| chr15-42557157-DEL-155 | chr15-42557158-DEL-155 |
| chr15-42579428-DEL-388 | chr15-42579429-DEL-388 |
| chr15-42580123-DEL-733 | chr15-42580124-DEL-733 |
| chr15-42585114-DEL-58 | chr15-42585115-DEL-58 |
| chr15-42592902-DEL-66 | chr15-42592903-DEL-66 |
| chr15-42610203-DEL-76 | chr15-42610204-DEL-76 |
| chr15-42612557-DEL-2708 | chr15-42612558-DEL-2708 |
| chr15-42667536-DEL-97 | chr15-42667537-DEL-97 |
| chr15-42674960-DEL-359 | chr15-42674961-DEL-359 |
| chr15-42683455-DEL-1540 | chr15-42683456-DEL-1540 |
| chr15-42687148-DEL-1631 | chr15-42687149-DEL-1631 |
| chr15-42689826-DEL-89 | chr15-42689829-DEL-77 |
| chr15-42706780-DEL-68 | chr15-42706781-DEL-68 |
| chr15-42740238-DEL-6991 | chr15-42740239-DEL-6991 |
| chr15-42779326-DEL-86 | chr15-42779323-DEL-54 |
| chr15-42782331-DEL-58 | chr15-42782336-DEL-54 |
| chr15-42787805-DEL-66 | chr15-42787806-DEL-66 |
| chr15-42791581-DEL-14535 | chr15-42791582-DEL-14535 |
| chr15-42795334-DEL-6585 | chr15-42795334-DEL-6585 |
| chr15-42825028-DEL-62 | chr15-42825029-DEL-62 |
| chr15-42904230-DEL-76 | chr15-42904262-DEL-77 |
| chr15-42913586-DEL-76 | chr15-42913600-DEL-67 |
| chr15-42929071-DEL-468 | chr15-42929072-DEL-466 |
| chr15-42931877-DEL-283 | chr15-42931878-DEL-283 |
| chr15-43004776-DEL-6727 | chr15-43004774-DEL-6728 |
| chr15-43004805-DEL-212 | chr15-43004806-DEL-212 |
| chr15-43141226-DEL-59 | chr15-43141227-DEL-59 |
| chr15-43194008-DEL-5287 | chr15-43194009-DEL-5287 |
| chr15-43242123-DEL-54 | chr15-43242124-DEL-54 |
| chr15-43250980-DEL-170 | chr15-43250981-DEL-178 |
| chr15-43299158-DEL-1671 | chr15-43299159-DEL-1671 |
| chr15-43305551-DEL-340 | chr15-43305552-DEL-340 |
| chr15-43327546-DEL-3228 | chr15-43327547-DEL-3228 |
| chr15-43380347-DEL-6806 | chr15-43380348-DEL-6806 |
| chr15-43436132-DEL-156 | chr15-43436134-DEL-153 |
| chr15-43446073-DEL-205 | chr15-43446074-DEL-205 |
| chr15-43455843-DEL-291 | chr15-43455844-DEL-291 |
| chr15-43456476-DEL-6249 | chr15-43456477-DEL-6249 |
| chr15-43483095-DEL-101 | chr15-43483096-DEL-101 |
| chr15-43503808-DEL-105 | chr15-43503809-DEL-105 |
| chr15-43508538-DEL-2632 | chr15-43508539-DEL-2632 |
| chr15-43516813-DEL-1424 | chr15-43516813-DEL-1424 |

| | |
|---|---|
| chr15-43572059-DEL-133 | chr15-43572058-DEL-133 |
| chr15-43586193-DEL-826 | chr15-43586194-DEL-826 |
| chr15-43592053-DEL-71 | chr15-43592054-DEL-71 |
| chr15-43622177-DEL-168 | chr15-43622170-DEL-176 |
| chr15-43632687-DEL-278 | chr15-43632688-DEL-279 |
| chr15-43645247-DEL-62 | chr15-43645248-DEL-62 |
| chr15-43649414-DEL-91 | chr15-43649415-DEL-91 |
| chr15-43655400-DEL-1617 | chr15-43655401-DEL-1617 |
| chr15-43676538-DEL-68 | chr15-43676539-DEL-68 |
| chr15-43679061-DEL-205 | chr15-43679062-DEL-205 |
| chr15-43681841-DEL-5961 | chr15-43681842-DEL-5961 |
| chr15-43728514-DEL-86 | chr15-43728549-DEL-54 |
| chr15-43735298-DEL-1003 | chr15-43735299-DEL-1003 |
| chr15-43742719-DEL-5623 | chr15-43742720-DEL-5623 |
| chr15-43761643-DEL-92 | chr15-43761644-DEL-92 |
| chr15-43835261-DEL-578 | chr15-43835262-DEL-578 |
| chr15-43873197-DEL-7324 | chr15-43873198-DEL-7324 |
| chr15-43879199-DEL-424 | chr15-43879200-DEL-424 |
| chr15-43880804-DEL-92 | chr15-43880791-DEL-105 |
| chr15-43902520-DEL-61 | chr15-43902520-DEL-74 |
| chr15-43935936-DEL-74 | chr15-43935937-DEL-74 |
| chr15-43950150-DEL-6247 | chr15-43950151-DEL-6247 |
| chr15-43957790-DEL-282 | chr15-43957791-DEL-282 |
| chr15-43960385-DEL-206 | chr15-43960386-DEL-206 |
| chr15-43963039-DEL-76 | chr15-43963040-DEL-76 |
| chr15-44000215-DEL-3218 | chr15-44000216-DEL-3218 |
| chr15-44020798-DEL-3200 | chr15-44020799-DEL-3200 |
| chr15-44046810-DEL-304 | chr15-44046811-DEL-304 |
| chr15-44074280-DEL-151 | chr15-44074281-DEL-151 |
| chr15-44076729-DEL-103 | chr15-44076730-DEL-103 |
| chr15-44087666-DEL-143 | chr15-44087667-DEL-143 |
| chr15-44179937-DEL-99 | chr15-44179938-DEL-99 |
| chr15-44193918-DEL-319 | chr15-44193919-DEL-319 |
| chr15-44222624-DEL-497 | chr15-44222625-DEL-497 |
| chr15-44225553-DEL-85 | chr15-44225554-DEL-97 |
| chr15-44229446-DEL-52 | chr15-44229446-DEL-52 |
| chr15-44236851-DEL-788 | chr15-44236852-DEL-788 |
| chr15-44247677-DEL-51 | chr15-44247678-DEL-51 |
| chr15-44252797-DEL-626 | chr15-44252798-DEL-626 |
| chr15-44258018-DEL-237 | chr15-44258019-DEL-237 |
| chr15-44261904-DEL-56 | chr15-44261905-DEL-56 |
| chr15-44276847-DEL-241 | chr15-44276848-DEL-241 |
| chr15-44287988-DEL-467 | chr15-44287988-DEL-468 |
| chr15-44303807-DEL-143 | chr15-44303807-DEL-106 |
| chr15-44329243-DEL-371 | chr15-44329244-DEL-371 |
| chr15-44330405-DEL-63 | chr15-44330406-DEL-63 |
| chr15-44332503-DEL-7235 | chr15-44332504-DEL-7235 |
| chr15-44339029-DEL-208 | chr15-44338984-DEL-208 |
| chr15-44382760-DEL-56 | chr15-44382761-DEL-56 |
| chr15-44385015-DEL-120 | chr15-44385016-DEL-120 |
| chr15-44385812-DEL-75 | chr15-44385811-DEL-76 |
| chr15-44442349-DEL-125 | chr15-44442348-DEL-125 |
| chr15-44517674-DEL-4760 | chr15-44517675-DEL-4760 |
| chr15-44541566-DEL-197 | chr15-44541567-DEL-197 |
| chr15-44577538-DEL-1250 | chr15-44577539-DEL-1250 |
| chr15-44586511-DEL-4881 | chr15-44586512-DEL-4881 |
| chr15-44624232-DEL-73 | chr15-44624227-DEL-57 |
| chr15-44653070-DEL-6396 | chr15-44653071-DEL-6396 |
| chr15-44663451-DEL-92 | chr15-44663452-DEL-89 |
| chr15-44674803-DEL-108 | chr15-44674804-DEL-108 |
| chr15-44691627-DEL-83 | chr15-44691612-DEL-84 |
| chr15-44693362-DEL-61 | chr15-44693363-DEL-61 |
| chr15-44707680-DEL-54 | chr15-44707681-DEL-54 |
| chr15-44719441-DEL-162 | chr15-44719442-DEL-162 |
| chr15-44724391-DEL-6847 | chr15-44724392-DEL-6847 |
| chr15-44730580-DEL-416 | chr15-44730581-DEL-416 |
| chr15-44734553-DEL-193 | chr15-44734552-DEL-194 |
| chr15-44773299-DEL-538 | chr15-44773296-DEL-538 |
| chr15-44832654-DEL-63 | chr15-44832655-DEL-60 |
| chr15-44839940-DEL-6197 | chr15-44839941-DEL-6197 |
| chr15-44936105-DEL-1810 | chr15-44936106-DEL-1810 |
| chr15-44988331-DEL-67 | chr15-44988313-DEL-60 |
| chr15-45045147-DEL-52 | chr15-45045147-DEL-52 |
| chr15-45048237-DEL-177 | chr15-45048238-DEL-177 |
| chr15-45052021-DEL-950 | chr15-45052022-DEL-950 |
| chr15-45057870-DEL-2690 | chr15-45057880-DEL-2662 |
| chr15-45073055-DEL-107 | chr15-45073056-DEL-107 |
| chr15-45079639-DEL-262 | chr15-45079640-DEL-262 |
| chr15-45087628-DEL-77 | chr15-45087629-DEL-77 |
| chr15-45097854-DEL-1485 | chr15-45097855-DEL-1485 |

| | |
|---|---|
| chr15-45105132-DEL-126 | chr15-45105133-DEL-126 |
| chr15-45187930-DEL-63 | chr15-45187934-DEL-59 |
| chr15-45224569-DEL-59 | chr15-45224552-DEL-60 |
| chr15-45263926-DEL-636 | chr15-45263927-DEL-636 |
| chr15-45268589-DEL-70 | chr15-45268590-DEL-70 |
| chr15-45281464-DEL-84 | chr15-45281465-DEL-84 |
| chr15-45295358-DEL-52 | chr15-45295358-DEL-52 |
| chr15-45301248-DEL-66 | chr15-45301248-DEL-66 |
| chr15-45323322-DEL-63 | chr15-45323353-DEL-51 |
| chr15-45358788-DEL-1223 | chr15-45358789-DEL-1222 |
| chr15-45381532-DEL-247 | chr15-45381528-DEL-248 |
| chr15-45387385-DEL-1135 | chr15-45387386-DEL-1135 |
| chr15-45400643-DEL-494 | chr15-45400644-DEL-494 |
| chr15-45439679-DEL-3617 | chr15-45439674-DEL-3622 |
| chr15-45460733-DEL-282 | chr15-45460734-DEL-282 |
| chr15-45468742-DEL-1129 | chr15-45468743-DEL-1129 |
| chr15-45470680-DEL-3080 | chr15-45470681-DEL-3079 |
| chr15-45500606-DEL-3478 | chr15-45500607-DEL-3478 |
| chr15-45500869-DEL-595 | chr15-45500870-DEL-595 |
| chr15-45530456-DEL-390 | chr15-45530457-DEL-390 |
| chr15-45532766-DEL-1009 | chr15-45532767-DEL-1009 |
| chr15-45558434-DEL-6517 | chr15-45558429-DEL-6518 |
| chr15-45564547-DEL-214 | chr15-45564546-DEL-214 |
| chr15-45565695-DEL-231 | chr15-45565696-DEL-231 |
| chr15-45572019-DEL-167 | chr15-45572020-DEL-167 |
| chr15-45575490-DEL-905 | chr15-45575491-DEL-905 |
| chr15-45636387-DEL-157 | chr15-45636389-DEL-158 |
| chr15-45636912-DEL-6215 | chr15-45636913-DEL-6215 |
| chr15-45636993-DEL-199 | chr15-45636993-DEL-200 |
| chr15-45652433-DEL-81 | chr15-45652415-DEL-100 |
| chr15-45658725-DEL-1449 | chr15-45658726-DEL-1449 |
| chr15-45662274-DEL-210 | chr15-45662262-DEL-212 |
| chr15-45670869-DEL-440 | chr15-45670870-DEL-440 |
| chr15-45680443-DEL-65 | chr15-45680443-DEL-65 |
| chr15-45728911-DEL-80 | chr15-45728935-DEL-80 |
| chr15-45757671-DEL-1674 | chr15-45757672-DEL-1675 |
| chr15-45770798-DEL-2852 | chr15-45770799-DEL-2853 |
| chr15-45782914-DEL-491 | chr15-45782915-DEL-491 |
| chr15-45792741-DEL-3184 | chr15-45792741-DEL-3184 |
| chr15-45800945-DEL-402 | chr15-45800946-DEL-402 |
| chr15-45809151-DEL-91 | chr15-45809153-DEL-90 |
| chr15-45840198-DEL-62 | chr15-45840201-DEL-60 |
| chr15-45890410-DEL-111 | chr15-45890411-DEL-111 |
| chr15-45894419-DEL-89 | chr15-45894420-DEL-89 |
| chr15-45912400-DEL-278 | chr15-45912401-DEL-278 |
| chr15-45940794-DEL-51 | chr15-45940782-DEL-54 |
| chr15-45945816-DEL-3292 | chr15-45945817-DEL-3292 |
| chr15-45947486-DEL-366 | chr15-45947487-DEL-366 |
| chr15-45957640-DEL-1617 | chr15-45957641-DEL-1617 |
| chr15-45979160-DEL-198 | chr15-45979161-DEL-198 |
| chr15-45983276-DEL-62 | chr15-45983280-DEL-52 |
| chr15-45989846-DEL-398 | chr15-45989847-DEL-398 |
| chr15-46001328-DEL-3804 | chr15-46001329-DEL-3804 |
| chr15-46015070-DEL-64 | chr15-46015071-DEL-64 |
| chr15-46018113-DEL-6219 | chr15-46018114-DEL-6219 |
| chr15-46042515-DEL-81 | chr15-46042516-DEL-81 |
| chr15-46052487-DEL-6175 | chr15-46052488-DEL-6175 |
| chr15-46079169-DEL-208 | chr15-46079170-DEL-208 |
| chr15-46084017-DEL-142 | chr15-46084017-DEL-142 |
| chr15-46095986-DEL-491 | chr15-46095986-DEL-491 |
| chr15-46119308-DEL-1878 | chr15-46119309-DEL-1878 |
| chr15-46146567-DEL-617 | chr15-46146568-DEL-616 |
| chr15-46156801-DEL-700 | chr15-46156802-DEL-700 |
| chr15-46166853-DEL-64 | chr15-46166850-DEL-64 |
| chr15-46167301-DEL-263 | chr15-46167301-DEL-263 |
| chr15-46198363-DEL-59 | chr15-46198365-DEL-83 |
| chr15-46206290-DEL-170 | chr15-46206290-DEL-170 |
| chr15-46219096-DEL-1891 | chr15-46219097-DEL-1891 |
| chr15-46226877-DEL-6249 | chr15-46226878-DEL-6249 |
| chr15-46235429-DEL-598 | chr15-46235429-DEL-598 |
| chr15-46237729-DEL-380 | chr15-46237729-DEL-380 |
| chr15-46242028-DEL-328 | chr15-46242028-DEL-328 |
| chr15-46243616-DEL-398 | chr15-46243616-DEL-398 |
| chr15-46247648-DEL-52 | chr15-46247642-DEL-80 |
| chr15-46265591-DEL-68 | chr15-46265592-DEL-68 |
| chr15-46299536-DEL-743 | chr15-46299537-DEL-743 |
| chr15-46307541-DEL-786 | chr15-46307541-DEL-786 |
| chr15-46321318-DEL-866 | chr15-46321302-DEL-882 |
| chr15-46337637-DEL-1332 | chr15-46337639-DEL-1330 |
| chr15-46340661-DEL-1052 | chr15-46340662-DEL-1052 |

| | |
|---|---|
| chr15-46364871-DEL-354 | chr15-46364878-DEL-358 |
| chr15-46365686-DEL-4382 | chr15-46365687-DEL-4382 |
| chr15-46389250-DEL-84 | chr15-46389250-DEL-84 |
| chr15-46419841-DEL-206 | chr15-46419841-DEL-206 |
| chr15-46426040-DEL-4216 | chr15-46426041-DEL-4216 |
| chr15-46435282-DEL-220 | chr15-46435283-DEL-220 |
| chr15-46438449-DEL-2749 | chr15-46438450-DEL-2749 |
| chr15-46440042-DEL-82 | chr15-46440043-DEL-82 |
| chr15-46455428-DEL-668 | chr15-46455431-DEL-667 |
| chr15-46469911-DEL-1023 | chr15-46469912-DEL-1023 |
| chr15-46483044-DEL-1914 | chr15-46483045-DEL-1914 |
| chr15-46502723-DEL-1666 | chr15-46502724-DEL-1666 |
| chr15-46514160-DEL-93 | chr15-46514161-DEL-93 |
| chr15-46527524-DEL-1580 | chr15-46527525-DEL-1580 |
| chr15-46541768-DEL-332 | chr15-46541769-DEL-332 |
| chr15-46547422-DEL-330 | chr15-46547408-DEL-345 |
| chr15-46579814-DEL-185 | chr15-46579815-DEL-185 |
| chr15-46584180-DEL-299 | chr15-46584181-DEL-299 |
| chr15-46592294-DEL-1240 | chr15-46592295-DEL-1240 |
| chr15-46652071-DEL-2152 | chr15-46652072-DEL-2152 |
| chr15-46670868-DEL-117 | chr15-46670868-DEL-117 |
| chr15-46715986-DEL-90 | chr15-46715987-DEL-90 |
| chr15-46731942-DEL-6811 | chr15-46731943-DEL-6811 |
| chr15-46751798-DEL-6573 | chr15-46751799-DEL-6573 |
| chr15-46766794-DEL-4226 | chr15-46766801-DEL-4218 |
| chr15-46775664-DEL-55 | chr15-46775666-DEL-53 |
| chr15-46782304-DEL-845 | chr15-46782305-DEL-845 |
| chr15-46790736-DEL-68 | chr15-46790737-DEL-68 |
| chr15-46823042-DEL-68 | chr15-46823043-DEL-68 |
| chr15-46847950-DEL-214 | chr15-46847951-DEL-214 |
| chr15-46886075-DEL-445 | chr15-46886076-DEL-445 |
| chr15-46892602-DEL-1971 | chr15-46892603-DEL-1971 |
| chr15-46916631-DEL-212 | chr15-46916601-DEL-224 |
| chr15-46927338-DEL-61 | chr15-46927339-DEL-61 |
| chr15-46964760-DEL-108 | chr15-46964761-DEL-108 |
| chr15-46977001-DEL-76 | chr15-46977002-DEL-76 |
| chr15-47043981-DEL-6649 | chr15-47043982-DEL-6649 |
| chr15-47113110-DEL-6346 | chr15-47113111-DEL-6346 |
| chr15-47133345-DEL-828 | chr15-47133346-DEL-828 |
| chr15-47144093-DEL-382 | chr15-47144094-DEL-382 |
| chr15-47156525-DEL-196 | chr15-47156525-DEL-195 |
| chr15-47160961-DEL-80 | chr15-47160962-DEL-80 |
| chr15-47205317-DEL-102 | chr15-47205366-DEL-91 |
| chr15-47219900-DEL-7203 | chr15-47219901-DEL-7203 |
| chr15-47243699-DEL-1435 | chr15-47243696-DEL-1436 |
| chr15-47265994-DEL-2790 | chr15-47265995-DEL-2790 |
| chr15-47286259-DEL-103 | chr15-47286260-DEL-103 |
| chr15-47305868-DEL-141 | chr15-47305869-DEL-141 |
| chr15-47310670-DEL-79 | chr15-47310658-DEL-60 |
| chr15-47329870-DEL-669 | chr15-47329868-DEL-670 |
| chr15-47411840-DEL-6896 | chr15-47411841-DEL-6896 |
| chr15-47433107-DEL-53 | chr15-47433111-DEL-52 |
| chr15-47472867-DEL-1030 | chr15-47472866-DEL-1031 |
| chr15-47555517-DEL-203 | chr15-47555518-DEL-203 |
| chr15-47599224-DEL-51 | chr15-47599225-DEL-51 |
| chr15-47621285-DEL-933 | chr15-47621297-DEL-934 |
| chr15-47657054-DEL-1029 | chr15-47657055-DEL-1029 |
| chr15-47677182-DEL-140 | chr15-47677183-DEL-140 |
| chr15-47692053-DEL-176 | chr15-47692054-DEL-176 |
| chr15-47712317-DEL-280 | chr15-47712336-DEL-237 |
| chr15-47776053-DEL-115 | chr15-47776053-DEL-115 |
| chr15-47842148-DEL-916 | chr15-47842149-DEL-916 |
| chr15-47856061-DEL-68 | chr15-47856062-DEL-64 |
| chr15-47862952-DEL-497 | chr15-47862953-DEL-497 |
| chr15-47917274-DEL-939 | chr15-47917275-DEL-939 |
| chr15-47945603-DEL-2345 | chr15-47945604-DEL-2345 |
| chr15-47951209-DEL-60 | chr15-47951196-DEL-80 |
| chr15-47986472-DEL-4284 | chr15-47986473-DEL-4284 |
| chr15-48067422-DEL-115 | chr15-48067423-DEL-115 |
| chr15-48125332-DEL-4261 | chr15-48125333-DEL-4261 |
| chr15-48187186-DEL-470 | chr15-48187187-DEL-470 |
| chr15-48261153-DEL-6643 | chr15-48261154-DEL-6643 |
| chr15-48261353-DEL-211 | chr15-48261338-DEL-212 |
| chr15-48288682-DEL-319 | chr15-48288683-DEL-318 |
| chr15-48300183-DEL-125 | chr15-48300227-DEL-99 |
| chr15-48326173-DEL-84 | chr15-48326174-DEL-84 |
| chr15-48349929-DEL-122 | chr15-48349930-DEL-122 |
| chr15-48370604-DEL-141 | chr15-48370605-DEL-141 |
| chr15-48376963-DEL-6126 | chr15-48376964-DEL-6126 |
| chr15-48418029-DEL-1957 | chr15-48418030-DEL-1957 |

| | |
|---|---|
| chr15-48490751-DEL-1513 | chr15-48490752-DEL-1513 |
| chr15-48503944-DEL-51 | chr15-48503931-DEL-61 |
| chr15-48507237-DEL-170 | chr15-48507235-DEL-171 |
| chr15-48552718-DEL-400 | chr15-48552718-DEL-398 |
| chr15-48628426-DEL-472 | chr15-48628427-DEL-472 |
| chr15-48679046-DEL-636 | chr15-48679047-DEL-636 |
| chr15-48708535-DEL-359 | chr15-48708536-DEL-359 |
| chr15-48708958-DEL-6228 | chr15-48708959-DEL-6228 |
| chr15-48713436-DEL-65 | chr15-48713439-DEL-60 |
| chr15-48795325-DEL-626 | chr15-48795326-DEL-626 |
| chr15-48807790-DEL-64 | chr15-48807771-DEL-60 |
| chr15-48822242-DEL-66 | chr15-48822243-DEL-66 |
| chr15-48939659-DEL-259 | chr15-48939658-DEL-260 |
| chr15-48988096-DEL-864 | chr15-48988104-DEL-862 |
| chr15-49001521-DEL-403 | chr15-49001522-DEL-403 |
| chr15-49025913-DEL-217 | chr15-49025914-DEL-217 |
| chr15-49030314-DEL-1008 | chr15-49030315-DEL-1008 |
| chr15-49032252-DEL-8778 | chr15-49032253-DEL-8778 |
| chr15-49037977-DEL-217 | chr15-49037978-DEL-217 |
| chr15-49097533-DEL-51 | chr15-49097534-DEL-51 |
| chr15-49112197-DEL-7044 | chr15-49112198-DEL-7044 |
| chr15-49182087-DEL-83 | chr15-49182088-DEL-83 |
| chr15-49186023-DEL-4596 | chr15-49186024-DEL-4596 |
| chr15-49195041-DEL-56 | chr15-49195037-DEL-56 |
| chr15-49197344-DEL-54 | chr15-49197345-DEL-72 |
| chr15-49235666-DEL-7416 | chr15-49235667-DEL-7416 |
| chr15-49241910-DEL-208 | chr15-49241911-DEL-208 |
| chr15-49284492-DEL-58 | chr15-49284493-DEL-58 |
| chr15-49286191-DEL-56 | chr15-49286171-DEL-56 |
| chr15-49292562-DEL-1647 | chr15-49292560-DEL-1646 |
| chr15-49332280-DEL-205 | chr15-49332281-DEL-205 |
| chr15-49354156-DEL-63 | chr15-49354157-DEL-63 |
| chr15-49376006-DEL-89 | chr15-49376007-DEL-89 |
| chr15-49381124-DEL-51 | chr15-49381125-DEL-51 |
| chr15-49382342-DEL-2466 | chr15-49382343-DEL-2466 |
| chr15-49390275-DEL-3511 | chr15-49390276-DEL-3511 |
| chr15-49394633-DEL-63 | chr15-49394634-DEL-63 |
| chr15-49399193-DEL-124 | chr15-49399193-DEL-122 |
| chr15-49403234-DEL-799 | chr15-49403235-DEL-799 |
| chr15-49428585-DEL-119 | chr15-49428577-DEL-121 |
| chr15-49439884-DEL-1428 | chr15-49439885-DEL-1428 |
| chr15-49485158-DEL-5158 | chr15-49485167-DEL-5149 |
| chr15-49485159-DEL-117 | chr15-49485157-DEL-105 |
| chr15-49509982-DEL-128 | chr15-49509981-DEL-126 |
| chr15-49519587-DEL-64 | chr15-49519588-DEL-54 |
| chr15-49532143-DEL-493 | chr15-49532144-DEL-493 |
| chr15-49536261-DEL-60 | chr15-49536264-DEL-50 |
| chr15-49556479-DEL-71 | chr15-49556480-DEL-71 |
| chr15-49570383-DEL-58 | chr15-49570384-DEL-58 |
| chr15-49577213-DEL-128 | chr15-49577209-DEL-136 |
| chr15-49597070-DEL-4158 | chr15-49597071-DEL-4158 |
| chr15-49604426-DEL-110 | chr15-49604427-DEL-110 |
| chr15-49611684-DEL-191 | chr15-49611706-DEL-188 |
| chr15-49614310-DEL-516 | chr15-49614311-DEL-516 |
| chr15-49624404-DEL-469 | chr15-49624405-DEL-469 |
| chr15-49663683-DEL-2529 | chr15-49663684-DEL-2529 |
| chr15-49713074-DEL-140 | chr15-49713072-DEL-168 |
| chr15-49731737-DEL-62 | chr15-49731741-DEL-58 |
| chr15-49744798-DEL-56 | chr15-49744799-DEL-56 |
| chr15-49771144-DEL-8252 | chr15-49771145-DEL-8252 |
| chr15-49788884-DEL-175 | chr15-49788885-DEL-175 |
| chr15-49790289-DEL-5769 | chr15-49790290-DEL-5769 |
| chr15-49811527-DEL-1253 | chr15-49811528-DEL-1253 |
| chr15-49813484-DEL-361 | chr15-49813485-DEL-361 |
| chr15-49842815-DEL-2232 | chr15-49842816-DEL-2232 |
| chr15-49893702-DEL-52 | chr15-49893702-DEL-56 |
| chr15-49919713-DEL-60 | chr15-49919714-DEL-60 |
| chr15-49948506-DEL-86 | chr15-49948509-DEL-66 |
| chr15-49950259-DEL-58 | chr15-49950260-DEL-58 |
| chr15-49967158-DEL-72 | chr15-49967159-DEL-73 |
| chr15-49992609-DEL-6609 | chr15-49992610-DEL-6609 |
| chr15-49992820-DEL-212 | chr15-49992821-DEL-212 |
| chr15-50015623-DEL-449 | chr15-50015624-DEL-449 |
| chr15-50090472-DEL-547 | chr15-50090468-DEL-564 |
| chr15-50093126-DEL-1112 | chr15-50093126-DEL-1112 |
| chr15-50105768-DEL-93 | chr15-50105769-DEL-93 |
| chr15-50108388-DEL-429 | chr15-50108389-DEL-429 |
| chr15-50116959-DEL-398 | chr15-50116960-DEL-398 |
| chr15-50122294-DEL-474 | chr15-50122295-DEL-473 |
| chr15-50154930-DEL-91 | chr15-50154931-DEL-91 |

| | |
|---|---|
| chr15-50184565-DEL-56 | chr15-50184566-DEL-56 |
| chr15-50187145-DEL-258 | chr15-50187146-DEL-258 |
| chr15-50197141-DEL-142 | chr15-50197142-DEL-142 |
| chr15-50217152-DEL-112 | chr15-50217153-DEL-112 |
| chr15-50227844-DEL-6166 | chr15-50227845-DEL-6166 |
| chr15-50285080-DEL-420 | chr15-50285081-DEL-420 |
| chr15-50301557-DEL-6314 | chr15-50301558-DEL-6314 |
| chr15-50346315-DEL-8026 | chr15-50346316-DEL-8026 |
| chr15-50346421-DEL-208 | chr15-50346422-DEL-208 |
| chr15-50355740-DEL-325 | chr15-50355741-DEL-325 |
| chr15-50361495-DEL-59 | chr15-50361492-DEL-60 |
| chr15-50384246-DEL-568 | chr15-50384247-DEL-568 |
| chr15-50387696-DEL-176 | chr15-50387689-DEL-194 |
| chr15-50413659-DEL-208 | chr15-50413656-DEL-209 |
| chr15-50436045-DEL-6269 | chr15-50436046-DEL-6268 |
| chr15-50442883-DEL-81 | chr15-50442884-DEL-81 |
| chr15-50458016-DEL-9649 | chr15-50458017-DEL-9649 |
| chr15-50469826-DEL-58 | chr15-50469827-DEL-58 |
| chr15-50488563-DEL-7216 | chr15-50488564-DEL-7216 |
| chr15-50502753-DEL-5704 | chr15-50502754-DEL-5704 |
| chr15-50592236-DEL-54 | chr15-50592236-DEL-54 |
| chr15-50603809-DEL-95 | chr15-50603810-DEL-95 |
| chr15-50660357-DEL-1029 | chr15-50660358-DEL-1029 |
| chr15-50672689-DEL-54 | chr15-50672690-DEL-54 |
| chr15-50701109-DEL-52 | chr15-50701110-DEL-52 |
| chr15-50701893-DEL-80 | chr15-50701894-DEL-80 |
| chr15-50766872-DEL-59 | chr15-50766873-DEL-59 |
| chr15-50794063-DEL-201 | chr15-50794064-DEL-201 |
| chr15-50868656-DEL-618 | chr15-50868657-DEL-618 |
| chr15-50890601-DEL-1160 | chr15-50890602-DEL-1160 |
| chr15-50892639-DEL-90 | chr15-50892640-DEL-69 |
| chr15-50923819-DEL-82 | chr15-50923820-DEL-82 |
| chr15-50946260-DEL-77 | chr15-50946261-DEL-77 |
| chr15-50975032-DEL-6171 | chr15-50975033-DEL-6171 |
| chr15-51030227-DEL-75 | chr15-51030203-DEL-91 |
| chr15-51057612-DEL-340 | chr15-51057613-DEL-340 |
| chr15-51129729-DEL-54 | chr15-51129736-DEL-56 |
| chr15-51143827-DEL-724 | chr15-51143828-DEL-724 |
| chr15-51165201-DEL-62 | chr15-51165202-DEL-62 |
| chr15-51185616-DEL-56 | chr15-51185607-DEL-54 |
| chr15-51188501-DEL-88 | chr15-51188502-DEL-70 |
| chr15-51204779-DEL-60 | chr15-51204772-DEL-74 |
| chr15-51239642-DEL-52 | chr15-51239643-DEL-52 |
| chr15-51243392-DEL-94 | chr15-51243392-DEL-91 |
| chr15-51258354-DEL-164 | chr15-51258351-DEL-164 |
| chr15-51267603-DEL-63 | chr15-51267574-DEL-50 |
| chr15-51319184-DEL-99 | chr15-51319185-DEL-99 |
| chr15-51353861-DEL-936 | chr15-51353862-DEL-936 |
| chr15-51364274-DEL-897 | chr15-51364275-DEL-897 |
| chr15-51378880-DEL-69 | chr15-51378868-DEL-66 |
| chr15-51395297-DEL-1893 | chr15-51395300-DEL-1893 |
| chr15-51400618-DEL-58 | chr15-51400618-DEL-59 |
| chr15-51407004-DEL-199 | chr15-51407005-DEL-195 |
| chr15-51417823-DEL-439 | chr15-51417825-DEL-440 |
| chr15-51448261-DEL-54 | chr15-51448256-DEL-50 |
| chr15-51453341-DEL-508 | chr15-51453342-DEL-508 |
| chr15-51469866-DEL-68 | chr15-51469874-DEL-60 |
| chr15-51470765-DEL-5661 | chr15-51470748-DEL-5678 |
| chr15-51503423-DEL-78 | chr15-51503424-DEL-78 |
| chr15-51561889-DEL-1211 | chr15-51561878-DEL-1206 |
| chr15-51566992-DEL-117 | chr15-51566993-DEL-117 |
| chr15-51571791-DEL-135 | chr15-51571792-DEL-135 |
| chr15-51652423-DEL-83 | chr15-51652424-DEL-83 |
| chr15-51692360-DEL-197 | chr15-51692327-DEL-199 |
| chr15-51736390-DEL-197 | chr15-51736391-DEL-197 |
| chr15-51769900-DEL-193 | chr15-51769895-DEL-194 |
| chr15-51774071-DEL-97 | chr15-51774113-DEL-56 |
| chr15-51778778-DEL-78 | chr15-51778749-DEL-75 |
| chr15-51791268-DEL-58 | chr15-51791269-DEL-58 |
| chr15-51791817-DEL-181 | chr15-51791818-DEL-181 |
| chr15-51797103-DEL-694 | chr15-51797104-DEL-694 |
| chr15-51798174-DEL-56 | chr15-51798174-DEL-56 |
| chr15-51799153-DEL-435 | chr15-51799151-DEL-435 |
| chr15-51816345-DEL-463 | chr15-51816342-DEL-467 |
| chr15-51840006-DEL-214 | chr15-51840007-DEL-214 |
| chr15-51852820-DEL-97 | chr15-51852821-DEL-97 |
| chr15-51885936-DEL-5591 | chr15-51885937-DEL-5591 |
| chr15-51896287-DEL-210 | chr15-51896288-DEL-210 |
| chr15-51919861-DEL-1670 | chr15-51919860-DEL-1671 |
| chr15-51945374-DEL-2830 | chr15-51945377-DEL-2828 |

| | |
|---|---|
| chr15-51949558-DEL-1331 | chr15-51949559-DEL-1331 |
| chr15-51952646-DEL-791 | chr15-51952647-DEL-792 |
| chr15-51964019-DEL-2366 | chr15-51964020-DEL-2366 |
| chr15-51988170-DEL-216 | chr15-51988170-DEL-216 |
| chr15-51988577-DEL-6643 | chr15-51988578-DEL-6643 |
| chr15-52035577-DEL-104 | chr15-52035600-DEL-89 |
| chr15-52050566-DEL-156 | chr15-52050567-DEL-156 |
| chr15-52055911-DEL-69 | chr15-52055906-DEL-106 |
| chr15-52081246-DEL-53 | chr15-52081247-DEL-51 |
| chr15-52085689-DEL-753 | chr15-52085690-DEL-753 |
| chr15-52095348-DEL-339 | chr15-52095349-DEL-339 |
| chr15-52096231-DEL-709 | chr15-52096232-DEL-709 |
| chr15-52107078-DEL-139 | chr15-52107054-DEL-126 |
| chr15-52130990-DEL-201 | chr15-52130990-DEL-201 |
| chr15-52152479-DEL-249 | chr15-52152475-DEL-254 |
| chr15-52171407-DEL-2317 | chr15-52171408-DEL-2317 |
| chr15-52192901-DEL-114 | chr15-52192902-DEL-114 |
| chr15-52193211-DEL-1495 | chr15-52193212-DEL-1495 |
| chr15-52203753-DEL-55 | chr15-52203753-DEL-55 |
| chr15-52214419-DEL-3213 | chr15-52214420-DEL-3213 |
| chr15-52232577-DEL-830 | chr15-52232578-DEL-830 |
| chr15-52235782-DEL-64 | chr15-52235795-DEL-74 |
| chr15-52286329-DEL-55 | chr15-52286330-DEL-56 |
| chr15-52320560-DEL-84 | chr15-52320537-DEL-60 |
| chr15-52340941-DEL-826 | chr15-52340942-DEL-826 |
| chr15-52411107-DEL-570 | chr15-52411108-DEL-570 |
| chr15-52491440-DEL-6381 | chr15-52491441-DEL-6381 |
| chr15-52538425-DEL-53 | chr15-52538421-DEL-58 |
| chr15-52585035-DEL-85 | chr15-52585036-DEL-85 |
| chr15-52586926-DEL-181 | chr15-52586927-DEL-181 |
| chr15-52587501-DEL-126 | chr15-52587502-DEL-113 |
| chr15-52594204-DEL-6443 | chr15-52594206-DEL-6442 |
| chr15-52606870-DEL-480 | chr15-52606871-DEL-480 |
| chr15-52644455-DEL-418 | chr15-52644455-DEL-418 |
| chr15-52658765-DEL-997 | chr15-52658766-DEL-997 |
| chr15-52678589-DEL-658 | chr15-52678590-DEL-658 |
| chr15-52700958-DEL-7758 | chr15-52700959-DEL-7758 |
| chr15-52727480-DEL-636 | chr15-52727481-DEL-636 |
| chr15-52793197-DEL-83 | chr15-52793198-DEL-83 |
| chr15-52834891-DEL-654 | chr15-52834891-DEL-654 |
| chr15-52881854-DEL-136 | chr15-52881855-DEL-136 |
| chr15-52902727-DEL-4801 | chr15-52902728-DEL-4801 |
| chr15-52917936-DEL-62 | chr15-52917937-DEL-62 |
| chr15-52938932-DEL-213 | chr15-52938933-DEL-213 |
| chr15-52949188-DEL-86 | chr15-52949189-DEL-86 |
| chr15-52971199-DEL-223 | chr15-52971200-DEL-223 |
| chr15-52985782-DEL-223 | chr15-52985783-DEL-223 |
| chr15-53001450-DEL-202 | chr15-53001451-DEL-202 |
| chr15-53043855-DEL-77 | chr15-53043849-DEL-88 |
| chr15-53106352-DEL-216 | chr15-53106353-DEL-216 |
| chr15-53201554-DEL-136 | chr15-53201540-DEL-111 |
| chr15-53213270-DEL-832 | chr15-53213271-DEL-832 |
| chr15-53230074-DEL-228 | chr15-53230061-DEL-200 |
| chr15-53239253-DEL-3938 | chr15-53239252-DEL-3939 |
| chr15-53261694-DEL-54 | chr15-53261694-DEL-54 |
| chr15-53264671-DEL-191 | chr15-53264672-DEL-191 |
| chr15-53285561-DEL-84 | chr15-53285562-DEL-88 |
| chr15-53302312-DEL-75 | chr15-53302313-DEL-75 |
| chr15-53317763-DEL-60 | chr15-53317763-DEL-60 |
| chr15-53329880-DEL-207 | chr15-53329876-DEL-208 |
| chr15-53351337-DEL-52 | chr15-53351338-DEL-52 |
| chr15-53358592-DEL-1139 | chr15-53358593-DEL-1139 |
| chr15-53374589-DEL-56 | chr15-53374590-DEL-56 |
| chr15-53437545-DEL-6436 | chr15-53437546-DEL-6436 |
| chr15-53465949-DEL-59 | chr15-53465948-DEL-60 |
| chr15-53474321-DEL-84 | chr15-53474353-DEL-66 |
| chr15-53501947-DEL-96 | chr15-53501948-DEL-96 |
| chr15-53512650-DEL-1435 | chr15-53512651-DEL-1435 |
| chr15-53587389-DEL-84 | chr15-53587387-DEL-85 |
| chr15-53635864-DEL-302 | chr15-53635865-DEL-302 |
| chr15-53658096-DEL-62 | chr15-53658077-DEL-76 |
| chr15-53712895-DEL-177 | chr15-53712896-DEL-177 |
| chr15-53729701-DEL-703 | chr15-53729702-DEL-703 |
| chr15-53731428-DEL-73 | chr15-53731428-DEL-62 |
| chr15-53781497-DEL-2201 | chr15-53781498-DEL-2201 |
| chr15-53810543-DEL-494 | chr15-53810544-DEL-493 |
| chr15-53895510-DEL-92 | chr15-53895472-DEL-122 |
| chr15-53900944-DEL-6436 | chr15-53900929-DEL-6436 |
| chr15-53908156-DEL-116 | chr15-53908157-DEL-116 |
| chr15-53959223-DEL-59 | chr15-53959224-DEL-59 |

| | |
|---|---|
| chr15-53971475-DEL-934 | chr15-53971476-DEL-934 |
| chr15-53980007-DEL-725 | chr15-53980008-DEL-725 |
| chr15-54044221-DEL-7133 | chr15-54044222-DEL-7133 |
| chr15-54054195-DEL-968 | chr15-54054196-DEL-968 |
| chr15-54068070-DEL-6988 | chr15-54068071-DEL-6988 |
| chr15-54074171-DEL-295 | chr15-54074172-DEL-295 |
| chr15-54146107-DEL-72 | chr15-54146083-DEL-72 |
| chr15-54188571-DEL-67 | chr15-54188557-DEL-109 |
| chr15-54199191-DEL-5698 | chr15-54199192-DEL-5698 |
| chr15-54205680-DEL-171 | chr15-54205677-DEL-138 |
| chr15-54208640-DEL-768 | chr15-54208640-DEL-768 |
| chr15-54282693-DEL-721 | chr15-54282692-DEL-722 |
| chr15-54288108-DEL-251 | chr15-54288109-DEL-251 |
| chr15-54307693-DEL-6476 | chr15-54307694-DEL-6476 |
| chr15-54316324-DEL-54 | chr15-54316313-DEL-61 |
| chr15-54368160-DEL-87 | chr15-54368161-DEL-87 |
| chr15-54375015-DEL-76 | chr15-54375016-DEL-76 |
| chr15-54389535-DEL-2773 | chr15-54389535-DEL-2773 |
| chr15-54410836-DEL-5665 | chr15-54410837-DEL-5665 |
| chr15-54417958-DEL-171 | chr15-54417959-DEL-171 |
| chr15-54449412-DEL-245 | chr15-54449413-DEL-245 |
| chr15-54451136-DEL-95 | chr15-54451119-DEL-123 |
| chr15-54452007-DEL-51 | chr15-54452008-DEL-51 |
| chr15-54453559-DEL-128 | chr15-54453560-DEL-128 |
| chr15-54469706-DEL-1016 | chr15-54469707-DEL-1016 |
| chr15-54495809-DEL-63 | chr15-54495810-DEL-63 |
| chr15-54498853-DEL-6599 | chr15-54498854-DEL-6599 |
| chr15-54527361-DEL-222 | chr15-54527362-DEL-222 |
| chr15-54534661-DEL-6261 | chr15-54534662-DEL-6261 |
| chr15-54546434-DEL-1028 | chr15-54546435-DEL-1028 |
| chr15-54567167-DEL-210 | chr15-54567168-DEL-210 |
| chr15-54577478-DEL-108 | chr15-54577478-DEL-108 |
| chr15-54594915-DEL-61 | chr15-54594916-DEL-61 |
| chr15-54600738-DEL-51 | chr15-54600733-DEL-57 |
| chr15-54616277-DEL-68 | chr15-54616278-DEL-68 |
| chr15-54642497-DEL-82 | chr15-54642498-DEL-82 |
| chr15-54658509-DEL-76 | chr15-54658510-DEL-75 |
| chr15-54685620-DEL-208 | chr15-54685621-DEL-208 |
| chr15-54755160-DEL-670 | chr15-54755161-DEL-670 |
| chr15-54769602-DEL-68 | chr15-54769602-DEL-68 |
| chr15-54778110-DEL-144 | chr15-54778111-DEL-152 |
| chr15-54788052-DEL-213 | chr15-54788053-DEL-213 |
| chr15-54792445-DEL-81 | chr15-54792446-DEL-81 |
| chr15-54802931-DEL-98 | chr15-54802932-DEL-98 |
| chr15-54859285-DEL-75 | chr15-54859286-DEL-75 |
| chr15-54868395-DEL-418 | chr15-54868389-DEL-419 |
| chr15-54898142-DEL-100 | chr15-54898143-DEL-100 |
| chr15-54921199-DEL-58 | chr15-54921200-DEL-54 |
| chr15-54936464-DEL-567 | chr15-54936465-DEL-567 |
| chr15-54942745-DEL-113 | chr15-54942743-DEL-114 |
| chr15-54958252-DEL-204 | chr15-54958253-DEL-204 |
| chr15-54958823-DEL-109 | chr15-54958824-DEL-109 |
| chr15-55010154-DEL-225 | chr15-55010155-DEL-225 |
| chr15-55024829-DEL-391 | chr15-55024790-DEL-391 |
| chr15-55029758-DEL-115 | chr15-55029759-DEL-115 |
| chr15-55050233-DEL-81 | chr15-55050234-DEL-81 |
| chr15-55055530-DEL-181 | chr15-55055531-DEL-181 |
| chr15-55064303-DEL-97 | chr15-55064313-DEL-126 |
| chr15-55072663-DEL-200 | chr15-55072664-DEL-200 |
| chr15-55074560-DEL-94 | chr15-55074561-DEL-94 |
| chr15-55080089-DEL-995 | chr15-55080090-DEL-995 |
| chr15-55089076-DEL-112 | chr15-55089077-DEL-116 |
| chr15-55112166-DEL-210 | chr15-55112167-DEL-210 |
| chr15-55123142-DEL-616 | chr15-55123143-DEL-616 |
| chr15-55131187-DEL-142 | chr15-55131188-DEL-142 |
| chr15-55132663-DEL-139 | chr15-55132664-DEL-139 |
| chr15-55140780-DEL-176 | chr15-55140781-DEL-176 |
| chr15-55141940-DEL-60 | chr15-55141984-DEL-60 |
| chr15-55156082-DEL-1291 | chr15-55156059-DEL-1292 |
| chr15-55167036-DEL-378 | chr15-55167037-DEL-378 |
| chr15-55173783-DEL-57 | chr15-55173784-DEL-57 |
| chr15-55217774-DEL-74 | chr15-55217795-DEL-56 |
| chr15-55228594-DEL-364 | chr15-55228595-DEL-364 |
| chr15-55232022-DEL-468 | chr15-55232023-DEL-468 |
| chr15-55240510-DEL-73 | chr15-55240511-DEL-73 |
| chr15-55259495-DEL-7032 | chr15-55259496-DEL-7032 |
| chr15-55282583-DEL-84 | chr15-55282558-DEL-77 |
| chr15-55292594-DEL-547 | chr15-55292595-DEL-540 |
| chr15-55333920-DEL-547 | chr15-55333921-DEL-547 |
| chr15-55357664-DEL-173 | chr15-55357665-DEL-173 |

| | |
|---|---|
| chr15-55371088-DEL-204 | chr15-55371090-DEL-202 |
| chr15-55372433-DEL-458 | chr15-55372434-DEL-458 |
| chr15-55415227-DEL-206 | chr15-55415228-DEL-205 |
| chr15-55433344-DEL-1193 | chr15-55433345-DEL-1193 |
| chr15-55451102-DEL-398 | chr15-55451103-DEL-398 |
| chr15-55457187-DEL-2066 | chr15-55457190-DEL-2073 |
| chr15-55533448-DEL-6115 | chr15-55533449-DEL-6115 |
| chr15-55540977-DEL-7289 | chr15-55540978-DEL-7289 |
| chr15-55611523-DEL-53 | chr15-55611524-DEL-53 |
| chr15-55674538-DEL-1281 | chr15-55674539-DEL-1281 |
| chr15-55734858-DEL-5543 | chr15-55734859-DEL-5543 |
| chr15-55745818-DEL-382 | chr15-55745819-DEL-382 |
| chr15-55762337-DEL-114 | chr15-55762353-DEL-90 |
| chr15-55774157-DEL-102 | chr15-55774159-DEL-100 |
| chr15-55777602-DEL-60 | chr15-55777602-DEL-66 |
| chr15-55797371-DEL-191 | chr15-55797378-DEL-193 |
| chr15-55807553-DEL-54 | chr15-55807522-DEL-92 |
| chr15-55812755-DEL-264 | chr15-55812755-DEL-264 |
| chr15-55822702-DEL-70 | chr15-55822695-DEL-102 |
| chr15-55825134-DEL-138 | chr15-55825135-DEL-138 |
| chr15-55835213-DEL-52 | chr15-55835208-DEL-57 |
| chr15-55835793-DEL-525 | chr15-55835794-DEL-525 |
| chr15-55886116-DEL-199 | chr15-55886117-DEL-199 |
| chr15-55886617-DEL-589 | chr15-55886618-DEL-589 |
| chr15-55895952-DEL-5492 | chr15-55895953-DEL-5492 |
| chr15-55917357-DEL-66 | chr15-55917358-DEL-66 |
| chr15-55947807-DEL-1242 | chr15-55947808-DEL-1242 |
| chr15-55968251-DEL-85 | chr15-55968276-DEL-62 |
| chr15-55983668-DEL-6230 | chr15-55983669-DEL-6230 |
| chr15-56039192-DEL-7232 | chr15-56039193-DEL-7232 |
| chr15-56084411-DEL-68 | chr15-56084412-DEL-78 |
| chr15-56092480-DEL-183 | chr15-56092481-DEL-182 |
| chr15-56127587-DEL-186 | chr15-56127588-DEL-186 |
| chr15-56146201-DEL-652 | chr15-56146202-DEL-652 |
| chr15-56152495-DEL-1987 | chr15-56152496-DEL-1987 |
| chr15-56168430-DEL-2790 | chr15-56168425-DEL-2791 |
| chr15-56173857-DEL-95 | chr15-56173848-DEL-111 |
| chr15-56217018-DEL-127 | chr15-56217019-DEL-127 |
| chr15-56219352-DEL-84 | chr15-56219362-DEL-64 |
| chr15-56231394-DEL-90 | chr15-56231386-DEL-97 |
| chr15-56244664-DEL-100 | chr15-56244665-DEL-100 |
| chr15-56247359-DEL-557 | chr15-56247360-DEL-557 |
| chr15-56263875-DEL-1336 | chr15-56263876-DEL-1336 |
| chr15-56293267-DEL-1821 | chr15-56293268-DEL-1821 |
| chr15-56322291-DEL-2828 | chr15-56322292-DEL-2828 |
| chr15-56359450-DEL-61 | chr15-56359452-DEL-57 |
| chr15-56376763-DEL-441 | chr15-56376766-DEL-439 |
| chr15-56380725-DEL-155 | chr15-56380726-DEL-155 |
| chr15-56390638-DEL-155 | chr15-56390639-DEL-155 |
| chr15-56459340-DEL-353 | chr15-56459341-DEL-353 |
| chr15-56460564-DEL-358 | chr15-56460565-DEL-358 |
| chr15-56499661-DEL-364 | chr15-56499662-DEL-364 |
| chr15-56517404-DEL-88 | chr15-56517451-DEL-64 |
| chr15-56536574-DEL-199 | chr15-56536575-DEL-199 |
| chr15-56550676-DEL-396 | chr15-56550672-DEL-413 |
| chr15-56573469-DEL-90 | chr15-56573500-DEL-69 |
| chr15-56575560-DEL-1082 | chr15-56575561-DEL-1082 |
| chr15-56577524-DEL-228 | chr15-56577525-DEL-228 |
| chr15-56589784-DEL-185 | chr15-56589785-DEL-185 |
| chr15-56596698-DEL-62 | chr15-56596700-DEL-56 |
| chr15-56612243-DEL-103 | chr15-56612277-DEL-70 |
| chr15-56666319-DEL-2715 | chr15-56666320-DEL-2715 |
| chr15-56671366-DEL-1436 | chr15-56671367-DEL-1436 |
| chr15-56706776-DEL-163 | chr15-56706780-DEL-165 |
| chr15-56737205-DEL-3167 | chr15-56737206-DEL-3158 |
| chr15-56790131-DEL-124 | chr15-56790132-DEL-124 |
| chr15-56804870-DEL-7475 | chr15-56804871-DEL-7475 |
| chr15-56821478-DEL-98 | chr15-56821479-DEL-98 |
| chr15-56851337-DEL-208 | chr15-56851337-DEL-208 |
| chr15-56911872-DEL-157 | chr15-56911882-DEL-148 |
| chr15-56923860-DEL-340 | chr15-56923861-DEL-340 |
| chr15-56987282-DEL-1883 | chr15-56987283-DEL-1883 |
| chr15-56993443-DEL-6083 | chr15-56993443-DEL-6083 |
| chr15-57001558-DEL-132 | chr15-57001572-DEL-150 |
| chr15-57028458-DEL-685 | chr15-57028459-DEL-679 |
| chr15-57041377-DEL-214 | chr15-57041376-DEL-214 |
| chr15-57041849-DEL-154 | chr15-57041850-DEL-111 |
| chr15-57045530-DEL-2423 | chr15-57045531-DEL-2423 |
| chr15-57051189-DEL-60 | chr15-57051190-DEL-50 |
| chr15-57069801-DEL-493 | chr15-57069802-DEL-493 |

| | |
|---|---|
| chr15-57070885-DEL-2035 | chr15-57070886-DEL-2035 |
| chr15-57079391-DEL-72 | chr15-57079364-DEL-67 |
| chr15-57119350-DEL-64 | chr15-57119335-DEL-50 |
| chr15-57154166-DEL-58 | chr15-57154167-DEL-58 |
| chr15-57154677-DEL-202 | chr15-57154678-DEL-202 |
| chr15-57165772-DEL-60 | chr15-57165772-DEL-60 |
| chr15-57177121-DEL-961 | chr15-57177122-DEL-961 |
| chr15-57209285-DEL-102 | chr15-57209286-DEL-102 |
| chr15-57214587-DEL-670 | chr15-57214590-DEL-670 |
| chr15-57275392-DEL-15124 | chr15-57275393-DEL-15124 |
| chr15-57280594-DEL-7168 | chr15-57280595-DEL-7168 |
| chr15-57290936-DEL-214 | chr15-57290937-DEL-214 |
| chr15-57292591-DEL-54 | chr15-57292592-DEL-54 |
| chr15-57296586-DEL-62 | chr15-57296577-DEL-63 |
| chr15-57303425-DEL-676 | chr15-57303426-DEL-676 |
| chr15-57304726-DEL-1927 | chr15-57304727-DEL-1927 |
| chr15-57338634-DEL-57 | chr15-57338635-DEL-57 |
| chr15-57374523-DEL-414 | chr15-57374524-DEL-413 |
| chr15-57493280-DEL-112 | chr15-57493281-DEL-112 |
| chr15-57514281-DEL-112 | chr15-57514292-DEL-104 |
| chr15-57526391-DEL-84 | chr15-57526393-DEL-83 |
| chr15-57533278-DEL-58 | chr15-57533279-DEL-58 |
| chr15-57540431-DEL-325 | chr15-57540432-DEL-325 |
| chr15-57569283-DEL-99 | chr15-57569284-DEL-99 |
| chr15-57653048-DEL-65 | chr15-57653049-DEL-65 |
| chr15-57700038-DEL-401 | chr15-57700039-DEL-401 |
| chr15-57707655-DEL-64 | chr15-57707656-DEL-66 |
| chr15-57763406-DEL-2639 | chr15-57763411-DEL-2635 |
| chr15-57772535-DEL-211 | chr15-57772536-DEL-211 |
| chr15-57793776-DEL-86 | chr15-57793775-DEL-66 |
| chr15-57796081-DEL-173 | chr15-57796082-DEL-173 |
| chr15-57808154-DEL-76 | chr15-57808121-DEL-110 |
| chr15-57812773-DEL-80 | chr15-57812820-DEL-68 |
| chr15-57831948-DEL-54 | chr15-57831949-DEL-54 |
| chr15-57835462-DEL-63 | chr15-57835464-DEL-52 |
| chr15-57851876-DEL-59 | chr15-57851860-DEL-66 |
| chr15-57889504-DEL-58 | chr15-57889505-DEL-58 |
| chr15-57899641-DEL-3631 | chr15-57899640-DEL-3631 |
| chr15-57904737-DEL-62 | chr15-57904731-DEL-63 |
| chr15-57954911-DEL-218 | chr15-57954912-DEL-218 |
| chr15-57957048-DEL-64 | chr15-57957049-DEL-64 |
| chr15-58006170-DEL-1469 | chr15-58006171-DEL-1469 |
| chr15-58012176-DEL-192 | chr15-58012177-DEL-192 |
| chr15-58024960-DEL-63 | chr15-58024975-DEL-54 |
| chr15-58035141-DEL-198 | chr15-58035142-DEL-198 |
| chr15-58047786-DEL-90 | chr15-58047787-DEL-126 |
| chr15-58122053-DEL-122 | chr15-58122054-DEL-122 |
| chr15-58147703-DEL-2264 | chr15-58147704-DEL-2264 |
| chr15-58177465-DEL-265 | chr15-58177466-DEL-265 |
| chr15-58178745-DEL-56 | chr15-58178714-DEL-56 |
| chr15-58181286-DEL-61 | chr15-58181288-DEL-62 |
| chr15-58218785-DEL-88 | chr15-58218785-DEL-88 |
| chr15-58233390-DEL-89 | chr15-58233391-DEL-89 |
| chr15-58244901-DEL-208 | chr15-58244902-DEL-208 |
| chr15-58259615-DEL-60 | chr15-58259601-DEL-64 |
| chr15-58280134-DEL-52 | chr15-58280135-DEL-52 |
| chr15-58306667-DEL-64 | chr15-58306668-DEL-64 |
| chr15-58331427-DEL-65 | chr15-58331418-DEL-60 |
| chr15-58342631-DEL-122 | chr15-58342632-DEL-122 |
| chr15-58346897-DEL-188 | chr15-58346898-DEL-188 |
| chr15-58353605-DEL-73 | chr15-58353602-DEL-53 |
| chr15-58359310-DEL-585 | chr15-58359311-DEL-585 |
| chr15-58363211-DEL-242 | chr15-58363212-DEL-242 |
| chr15-58376835-DEL-90 | chr15-58376836-DEL-90 |
| chr15-58394276-DEL-92 | chr15-58394284-DEL-81 |
| chr15-58404623-DEL-61 | chr15-58404624-DEL-61 |
| chr15-58434458-DEL-108 | chr15-58434459-DEL-98 |
| chr15-58439754-DEL-1186 | chr15-58439755-DEL-1186 |
| chr15-58442900-DEL-206 | chr15-58442900-DEL-207 |
| chr15-58445969-DEL-175 | chr15-58445970-DEL-175 |
| chr15-58479173-DEL-6680 | chr15-58479174-DEL-6680 |
| chr15-58485343-DEL-212 | chr15-58485344-DEL-212 |
| chr15-58537925-DEL-382 | chr15-58537926-DEL-382 |
| chr15-58539177-DEL-788 | chr15-58539178-DEL-788 |
| chr15-58563980-DEL-401 | chr15-58563981-DEL-401 |
| chr15-58564678-DEL-192 | chr15-58564676-DEL-195 |
| chr15-58565345-DEL-4706 | chr15-58565346-DEL-4706 |
| chr15-58575786-DEL-243 | chr15-58575784-DEL-244 |
| chr15-58595920-DEL-144 | chr15-58595921-DEL-144 |
| chr15-58630991-DEL-202 | chr15-58630992-DEL-202 |

| | |
|---|---|
| chr15-58638304-DEL-196 | chr15-58638305-DEL-196 |
| chr15-58645410-DEL-115 | chr15-58645436-DEL-90 |
| chr15-58693102-DEL-131 | chr15-58693103-DEL-131 |
| chr15-58723770-DEL-100 | chr15-58723771-DEL-100 |
| chr15-58727022-DEL-64 | chr15-58727020-DEL-67 |
| chr15-58729375-DEL-492 | chr15-58729375-DEL-488 |
| chr15-58742509-DEL-60 | chr15-58742510-DEL-60 |
| chr15-58783309-DEL-1894 | chr15-58783310-DEL-1894 |
| chr15-58844447-DEL-93 | chr15-58844482-DEL-93 |
| chr15-58937460-DEL-58 | chr15-58937461-DEL-58 |
| chr15-58971870-DEL-526 | chr15-58971876-DEL-504 |
| chr15-58982286-DEL-103 | chr15-58982291-DEL-99 |
| chr15-58991111-DEL-201 | chr15-58991112-DEL-201 |
| chr15-58995606-DEL-1084 | chr15-58995607-DEL-1084 |
| chr15-59011462-DEL-58 | chr15-59011463-DEL-58 |
| chr15-59012028-DEL-92 | chr15-59012029-DEL-92 |
| chr15-59015582-DEL-368 | chr15-59015583-DEL-368 |
| chr15-59033550-DEL-5743 | chr15-59033538-DEL-5761 |
| chr15-59052609-DEL-748 | chr15-59052628-DEL-730 |
| chr15-59098350-DEL-60 | chr15-59098351-DEL-54 |
| chr15-59174992-DEL-182 | chr15-59174993-DEL-182 |
| chr15-59179022-DEL-206 | chr15-59179023-DEL-206 |
| chr15-59184215-DEL-196 | chr15-59184216-DEL-196 |
| chr15-59203227-DEL-588 | chr15-59203228-DEL-588 |
| chr15-59234522-DEL-57 | chr15-59234502-DEL-57 |
| chr15-59262274-DEL-167 | chr15-59262275-DEL-166 |
| chr15-59293213-DEL-2123 | chr15-59293204-DEL-2124 |
| chr15-59317893-DEL-163 | chr15-59317894-DEL-163 |
| chr15-59355081-DEL-68 | chr15-59355082-DEL-68 |
| chr15-59358502-DEL-85 | chr15-59358503-DEL-75 |
| chr15-59369112-DEL-52 | chr15-59369112-DEL-52 |
| chr15-59378508-DEL-90 | chr15-59378509-DEL-90 |
| chr15-59390847-DEL-392 | chr15-59390848-DEL-392 |
| chr15-59416661-DEL-216 | chr15-59416662-DEL-216 |
| chr15-59433740-DEL-52 | chr15-59433741-DEL-52 |
| chr15-59443343-DEL-172 | chr15-59443345-DEL-173 |
| chr15-59450897-DEL-200 | chr15-59450898-DEL-200 |
| chr15-59456956-DEL-56 | chr15-59456945-DEL-68 |
| chr15-59512954-DEL-72 | chr15-59512960-DEL-62 |
| chr15-59542098-DEL-231 | chr15-59542099-DEL-231 |
| chr15-59544652-DEL-60 | chr15-59544653-DEL-60 |
| chr15-59569880-DEL-204 | chr15-59569881-DEL-204 |
| chr15-59571981-DEL-204 | chr15-59571982-DEL-204 |
| chr15-59599743-DEL-52 | chr15-59599744-DEL-52 |
| chr15-59616828-DEL-274 | chr15-59616829-DEL-274 |
| chr15-59631122-DEL-51 | chr15-59631121-DEL-52 |
| chr15-59636112-DEL-541 | chr15-59636113-DEL-542 |
| chr15-59661701-DEL-84 | chr15-59661685-DEL-72 |
| chr15-59672329-DEL-1891 | chr15-59672330-DEL-1891 |
| chr15-59689481-DEL-110 | chr15-59689482-DEL-110 |
| chr15-59712153-DEL-82 | chr15-59712154-DEL-82 |
| chr15-59742538-DEL-4441 | chr15-59742542-DEL-4438 |
| chr15-59771035-DEL-1571 | chr15-59771036-DEL-1571 |
| chr15-59784875-DEL-117 | chr15-59784860-DEL-133 |
| chr15-59785937-DEL-2655 | chr15-59785938-DEL-2655 |
| chr15-59793976-DEL-87 | chr15-59794001-DEL-82 |
| chr15-59800216-DEL-1434 | chr15-59800217-DEL-1434 |
| chr15-59807134-DEL-80 | chr15-59807135-DEL-80 |
| chr15-59807740-DEL-88 | chr15-59807737-DEL-120 |
| chr15-59824299-DEL-88 | chr15-59824294-DEL-70 |
| chr15-59894708-DEL-462 | chr15-59894712-DEL-426 |
| chr15-59916408-DEL-2696 | chr15-59916409-DEL-2696 |
| chr15-59923107-DEL-53 | chr15-59923108-DEL-53 |
| chr15-59928338-DEL-63 | chr15-59928339-DEL-63 |
| chr15-59930742-DEL-6571 | chr15-59930746-DEL-6570 |
| chr15-59954984-DEL-73 | chr15-59954985-DEL-71 |
| chr15-59992442-DEL-778 | chr15-59992443-DEL-778 |
| chr15-59998311-DEL-446 | chr15-59998312-DEL-446 |
| chr15-59999117-DEL-939 | chr15-59999118-DEL-939 |
| chr15-60020180-DEL-54 | chr15-60020184-DEL-50 |
| chr15-60028130-DEL-1990 | chr15-60028131-DEL-1990 |
| chr15-60052559-DEL-4058 | chr15-60052560-DEL-4058 |
| chr15-60072131-DEL-3696 | chr15-60072132-DEL-3696 |
| chr15-60092324-DEL-636 | chr15-60092325-DEL-636 |
| chr15-60104487-DEL-6291 | chr15-60104488-DEL-6291 |
| chr15-60130739-DEL-547 | chr15-60130742-DEL-543 |
| chr15-60165179-DEL-55 | chr15-60165180-DEL-55 |
| chr15-60192276-DEL-187 | chr15-60192277-DEL-187 |
| chr15-60200653-DEL-2026 | chr15-60200654-DEL-2026 |
| chr15-60219524-DEL-6676 | chr15-60219525-DEL-6676 |

| | |
|---|---|
| chr15-60230171-DEL-162 | chr15-60230172-DEL-162 |
| chr15-60243044-DEL-68 | chr15-60243045-DEL-66 |
| chr15-60249046-DEL-210 | chr15-60249047-DEL-210 |
| chr15-60261621-DEL-52 | chr15-60261622-DEL-52 |
| chr15-60269931-DEL-6467 | chr15-60269932-DEL-6467 |
| chr15-60310360-DEL-5285 | chr15-60310361-DEL-5285 |
| chr15-60360787-DEL-371 | chr15-60360788-DEL-371 |
| chr15-60365481-DEL-200 | chr15-60365482-DEL-200 |
| chr15-60372575-DEL-504 | chr15-60372576-DEL-504 |
| chr15-60380610-DEL-150 | chr15-60380611-DEL-149 |
| chr15-60400175-DEL-1504 | chr15-60400176-DEL-1504 |
| chr15-60403090-DEL-235 | chr15-60403091-DEL-235 |
| chr15-60406108-DEL-57 | chr15-60406108-DEL-54 |
| chr15-60443963-DEL-581 | chr15-60443966-DEL-582 |
| chr15-60463861-DEL-2986 | chr15-60463862-DEL-2986 |
| chr15-60472622-DEL-79 | chr15-60472623-DEL-79 |
| chr15-60481158-DEL-58 | chr15-60481159-DEL-58 |
| chr15-60482722-DEL-192 | chr15-60482754-DEL-153 |
| chr15-60493854-DEL-1513 | chr15-60493855-DEL-1513 |
| chr15-60495864-DEL-54 | chr15-60495833-DEL-58 |
| chr15-60533775-DEL-325 | chr15-60533776-DEL-325 |
| chr15-60538819-DEL-994 | chr15-60538836-DEL-995 |
| chr15-60576375-DEL-57 | chr15-60576375-DEL-50 |
| chr15-60591793-DEL-104 | chr15-60591794-DEL-100 |
| chr15-60608522-DEL-100 | chr15-60608473-DEL-150 |
| chr15-60629493-DEL-143 | chr15-60629494-DEL-143 |
| chr15-60644997-DEL-1699 | chr15-60644998-DEL-1699 |
| chr15-60647975-DEL-1193 | chr15-60647976-DEL-1193 |
| chr15-60664053-DEL-873 | chr15-60664054-DEL-873 |
| chr15-60672787-DEL-7418 | chr15-60672788-DEL-7418 |
| chr15-60713377-DEL-159 | chr15-60713378-DEL-159 |
| chr15-60715219-DEL-463 | chr15-60715220-DEL-463 |
| chr15-60729279-DEL-111 | chr15-60729280-DEL-111 |
| chr15-60753655-DEL-196 | chr15-60753656-DEL-196 |
| chr15-60779698-DEL-79 | chr15-60779697-DEL-76 |
| chr15-60812745-DEL-52 | chr15-60812745-DEL-52 |
| chr15-60856655-DEL-1892 | chr15-60856656-DEL-1892 |
| chr15-60868909-DEL-220 | chr15-60868910-DEL-220 |
| chr15-60876252-DEL-61 | chr15-60876253-DEL-62 |
| chr15-60898210-DEL-6784 | chr15-60898211-DEL-6784 |
| chr15-60898342-DEL-208 | chr15-60898342-DEL-208 |
| chr15-60919307-DEL-55 | chr15-60919303-DEL-56 |
| chr15-60927781-DEL-56 | chr15-60927782-DEL-62 |
| chr15-60941242-DEL-62 | chr15-60941242-DEL-62 |
| chr15-60949864-DEL-6157 | chr15-60949865-DEL-6157 |
| chr15-60969886-DEL-87 | chr15-60969892-DEL-93 |
| chr15-61013407-DEL-107 | chr15-61013408-DEL-107 |
| chr15-61020427-DEL-2056 | chr15-61020428-DEL-2056 |
| chr15-61043366-DEL-86 | chr15-61043367-DEL-86 |
| chr15-61049454-DEL-54 | chr15-61049443-DEL-68 |
| chr15-61052326-DEL-55 | chr15-61052327-DEL-55 |
| chr15-61053885-DEL-73 | chr15-61053869-DEL-70 |
| chr15-61064792-DEL-141 | chr15-61064793-DEL-141 |
| chr15-61115472-DEL-74 | chr15-61115473-DEL-68 |
| chr15-61152110-DEL-358 | chr15-61152111-DEL-358 |
| chr15-61160288-DEL-89 | chr15-61160289-DEL-89 |
| chr15-61164936-DEL-55 | chr15-61164935-DEL-56 |
| chr15-61194911-DEL-185 | chr15-61194912-DEL-185 |
| chr15-61206708-DEL-56 | chr15-61206709-DEL-56 |
| chr15-61245969-DEL-78 | chr15-61245970-DEL-78 |
| chr15-61246466-DEL-2186 | chr15-61246467-DEL-2186 |
| chr15-61251310-DEL-234 | chr15-61251308-DEL-235 |
| chr15-61262930-DEL-345 | chr15-61262921-DEL-334 |
| chr15-61273745-DEL-1321 | chr15-61273746-DEL-1321 |
| chr15-61274500-DEL-446 | chr15-61274501-DEL-446 |
| chr15-61342802-DEL-69 | chr15-61342803-DEL-69 |
| chr15-61377767-DEL-351 | chr15-61377768-DEL-351 |
| chr15-61383682-DEL-342 | chr15-61383683-DEL-342 |
| chr15-61417300-DEL-85 | chr15-61417301-DEL-86 |
| chr15-61418294-DEL-191 | chr15-61418295-DEL-190 |
| chr15-61489362-DEL-171 | chr15-61489363-DEL-171 |
| chr15-61515113-DEL-201 | chr15-61515114-DEL-201 |
| chr15-61534274-DEL-2591 | chr15-61534275-DEL-2591 |
| chr15-61538330-DEL-178 | chr15-61538331-DEL-178 |
| chr15-61568826-DEL-75 | chr15-61568827-DEL-75 |
| chr15-61615567-DEL-71 | chr15-61615568-DEL-81 |
| chr15-61628981-DEL-2481 | chr15-61628982-DEL-2481 |
| chr15-61634636-DEL-777 | chr15-61634637-DEL-777 |
| chr15-61635868-DEL-131 | chr15-61635864-DEL-132 |
| chr15-61703302-DEL-7049 | chr15-61703303-DEL-7049 |

| | |
|---|---|
| chr15-61709566-DEL-624 | chr15-61709567-DEL-624 |
| chr15-61775309-DEL-794 | chr15-61775310-DEL-794 |
| chr15-61784869-DEL-313 | chr15-61784870-DEL-313 |
| chr15-61786300-DEL-1042 | chr15-61786301-DEL-1042 |
| chr15-61788261-DEL-506 | chr15-61788262-DEL-506 |
| chr15-61817484-DEL-11565 | chr15-61817485-DEL-11565 |
| chr15-61819109-DEL-198 | chr15-61819110-DEL-198 |
| chr15-61823278-DEL-91 | chr15-61823304-DEL-66 |
| chr15-61826782-DEL-70 | chr15-61826783-DEL-70 |
| chr15-61832447-DEL-52 | chr15-61832444-DEL-53 |
| chr15-61836359-DEL-405 | chr15-61836360-DEL-405 |
| chr15-61887721-DEL-202 | chr15-61887722-DEL-202 |
| chr15-61892607-DEL-216 | chr15-61892608-DEL-216 |
| chr15-61895054-DEL-1807 | chr15-61895055-DEL-1807 |
| chr15-61900971-DEL-108 | chr15-61900928-DEL-128 |
| chr15-61906168-DEL-104 | chr15-61906160-DEL-106 |
| chr15-61907565-DEL-79 | chr15-61907566-DEL-79 |
| chr15-61918460-DEL-196 | chr15-61918449-DEL-232 |
| chr15-61924219-DEL-331 | chr15-61924220-DEL-331 |
| chr15-61942645-DEL-354 | chr15-61942646-DEL-354 |
| chr15-61949398-DEL-207 | chr15-61949399-DEL-207 |
| chr15-61983102-DEL-51 | chr15-61983103-DEL-51 |
| chr15-62015556-DEL-68 | chr15-62015557-DEL-68 |
| chr15-62028696-DEL-61 | chr15-62028701-DEL-60 |
| chr15-62046840-DEL-70 | chr15-62046863-DEL-80 |
| chr15-62057654-DEL-67 | chr15-62057655-DEL-67 |
| chr15-62089574-DEL-170 | chr15-62089575-DEL-170 |
| chr15-62148747-DEL-230 | chr15-62148772-DEL-206 |
| chr15-62189162-DEL-1137 | chr15-62189163-DEL-1137 |
| chr15-62229660-DEL-488 | chr15-62229661-DEL-488 |
| chr15-62246016-DEL-214 | chr15-62246015-DEL-216 |
| chr15-62281551-DEL-92 | chr15-62281552-DEL-92 |
| chr15-62282961-DEL-70 | chr15-62282970-DEL-66 |
| chr15-62343777-DEL-677 | chr15-62343778-DEL-677 |
| chr15-62358796-DEL-71 | chr15-62358789-DEL-69 |
| chr15-62387454-DEL-6730 | chr15-62387454-DEL-6730 |
| chr15-62402167-DEL-53 | chr15-62402150-DEL-70 |
| chr15-62408427-DEL-157 | chr15-62408440-DEL-145 |
| chr15-62412167-DEL-304 | chr15-62412170-DEL-305 |
| chr15-62457772-DEL-63 | chr15-62457787-DEL-52 |
| chr15-62459643-DEL-67 | chr15-62459644-DEL-67 |
| chr15-62475843-DEL-208 | chr15-62475844-DEL-208 |
| chr15-62490865-DEL-3430 | chr15-62490866-DEL-3430 |
| chr15-62504057-DEL-3009 | chr15-62504058-DEL-3009 |
| chr15-62522387-DEL-821 | chr15-62522388-DEL-821 |
| chr15-62531987-DEL-429 | chr15-62531988-DEL-429 |
| chr15-62534506-DEL-107 | chr15-62534507-DEL-107 |
| chr15-62558386-DEL-7238 | chr15-62558387-DEL-7238 |
| chr15-62575176-DEL-81 | chr15-62575177-DEL-81 |
| chr15-62589388-DEL-3061 | chr15-62589389-DEL-3061 |
| chr15-62595730-DEL-66 | chr15-62595731-DEL-66 |
| chr15-62609572-DEL-200 | chr15-62609573-DEL-200 |
| chr15-62631109-DEL-111 | chr15-62631110-DEL-111 |
| chr15-62635913-DEL-6813 | chr15-62635913-DEL-6811 |
| chr15-62655646-DEL-529 | chr15-62655646-DEL-529 |
| chr15-62677306-DEL-263 | chr15-62677302-DEL-265 |
| chr15-62682530-DEL-235 | chr15-62682531-DEL-235 |
| chr15-62698326-DEL-88 | chr15-62698336-DEL-81 |
| chr15-62699775-DEL-377 | chr15-62699776-DEL-377 |
| chr15-62729711-DEL-1901 | chr15-62729712-DEL-1901 |
| chr15-62753752-DEL-178 | chr15-62753753-DEL-178 |
| chr15-62764675-DEL-118 | chr15-62764678-DEL-119 |
| chr15-62781213-DEL-92 | chr15-62781220-DEL-81 |
| chr15-62811656-DEL-87 | chr15-62811652-DEL-91 |
| chr15-62815052-DEL-60 | chr15-62815045-DEL-52 |
| chr15-62850245-DEL-258 | chr15-62850246-DEL-258 |
| chr15-62877970-DEL-66 | chr15-62877956-DEL-68 |
| chr15-62894050-DEL-3499 | chr15-62894048-DEL-3500 |
| chr15-62897764-DEL-132 | chr15-62897754-DEL-142 |
| chr15-62903461-DEL-1924 | chr15-62903462-DEL-1924 |
| chr15-62916292-DEL-56 | chr15-62916293-DEL-55 |
| chr15-62927853-DEL-88 | chr15-62927854-DEL-86 |
| chr15-62963482-DEL-89 | chr15-62963483-DEL-89 |
| chr15-62999294-DEL-7021 | chr15-62999295-DEL-7021 |
| chr15-63017179-DEL-528 | chr15-63017180-DEL-528 |
| chr15-63077375-DEL-239 | chr15-63077376-DEL-239 |
| chr15-63102560-DEL-64 | chr15-63102571-DEL-54 |
| chr15-63115515-DEL-1875 | chr15-63115516-DEL-1875 |
| chr15-63131015-DEL-56 | chr15-63131016-DEL-56 |
| chr15-63162496-DEL-120 | chr15-63162495-DEL-106 |

| | |
|---|---|
| chr15-63189397-DEL-62 | chr15-63189395-DEL-64 |
| chr15-63200069-DEL-460 | chr15-63200070-DEL-460 |
| chr15-63206540-DEL-214 | chr15-63206541-DEL-214 |
| chr15-63251986-DEL-335 | chr15-63251987-DEL-335 |
| chr15-63253101-DEL-1313 | chr15-63253103-DEL-1312 |
| chr15-63264982-DEL-306 | chr15-63264983-DEL-304 |
| chr15-63292075-DEL-1598 | chr15-63292076-DEL-1598 |
| chr15-63297371-DEL-6351 | chr15-63297372-DEL-6351 |
| chr15-63307374-DEL-65 | chr15-63307375-DEL-65 |
| chr15-63330013-DEL-625 | chr15-63330014-DEL-625 |
| chr15-63342125-DEL-152 | chr15-63342126-DEL-152 |
| chr15-63352790-DEL-219 | chr15-63352789-DEL-217 |
| chr15-63374806-DEL-218 | chr15-63374807-DEL-218 |
| chr15-63381174-DEL-158 | chr15-63381175-DEL-154 |
| chr15-63432371-DEL-146 | chr15-63432372-DEL-146 |
| chr15-63432729-DEL-6350 | chr15-63432730-DEL-6350 |
| chr15-63443083-DEL-75 | chr15-63443084-DEL-75 |
| chr15-63452928-DEL-424 | chr15-63452929-DEL-424 |
| chr15-63458757-DEL-62 | chr15-63458766-DEL-54 |
| chr15-63465579-DEL-56 | chr15-63465580-DEL-50 |
| chr15-63467111-DEL-65 | chr15-63467112-DEL-65 |
| chr15-63470689-DEL-155 | chr15-63470690-DEL-155 |
| chr15-63489912-DEL-96 | chr15-63489955-DEL-54 |
| chr15-63490784-DEL-85 | chr15-63490785-DEL-85 |
| chr15-63506678-DEL-118 | chr15-63506679-DEL-158 |
| chr15-63525028-DEL-337 | chr15-63525029-DEL-337 |
| chr15-63562827-DEL-575 | chr15-63562828-DEL-575 |
| chr15-63603599-DEL-54 | chr15-63603612-DEL-78 |
| chr15-63608137-DEL-75 | chr15-63608138-DEL-75 |
| chr15-63611752-DEL-110 | chr15-63611753-DEL-94 |
| chr15-63624986-DEL-59 | chr15-63624970-DEL-60 |
| chr15-63663132-DEL-52 | chr15-63663133-DEL-52 |
| chr15-63683601-DEL-159 | chr15-63683602-DEL-159 |
| chr15-63847285-DEL-173 | chr15-63847286-DEL-176 |
| chr15-63881249-DEL-218 | chr15-63881250-DEL-218 |
| chr15-63904527-DEL-494 | chr15-63904528-DEL-494 |
| chr15-63935611-DEL-320 | chr15-63935595-DEL-292 |
| chr15-63964177-DEL-57 | chr15-63964180-DEL-55 |
| chr15-64002421-DEL-89 | chr15-64002420-DEL-88 |
| chr15-64048110-DEL-195 | chr15-64048111-DEL-195 |
| chr15-64060321-DEL-55 | chr15-64060322-DEL-55 |
| chr15-64072896-DEL-76 | chr15-64072897-DEL-76 |
| chr15-64112848-DEL-57 | chr15-64112849-DEL-57 |
| chr15-64136042-DEL-339 | chr15-64136041-DEL-340 |
| chr15-64147576-DEL-60 | chr15-64147577-DEL-60 |
| chr15-64148965-DEL-593 | chr15-64148966-DEL-593 |
| chr15-64168751-DEL-177 | chr15-64168748-DEL-169 |
| chr15-64182198-DEL-87 | chr15-64182170-DEL-96 |
| chr15-64188370-DEL-54 | chr15-64188383-DEL-54 |
| chr15-64289105-DEL-62 | chr15-64289106-DEL-62 |
| chr15-64290394-DEL-324 | chr15-64290395-DEL-324 |
| chr15-64310714-DEL-354 | chr15-64310715-DEL-354 |
| chr15-64321421-DEL-6199 | chr15-64321422-DEL-6199 |
| chr15-64406309-DEL-722 | chr15-64406310-DEL-722 |
| chr15-64410869-DEL-1508 | chr15-64410869-DEL-1508 |
| chr15-64420451-DEL-131 | chr15-64420448-DEL-132 |
| chr15-64435564-DEL-108 | chr15-64435569-DEL-104 |
| chr15-64440636-DEL-60 | chr15-64440640-DEL-61 |
| chr15-64463588-DEL-6472 | chr15-64463598-DEL-6463 |
| chr15-64506121-DEL-400 | chr15-64506122-DEL-400 |
| chr15-64508409-DEL-86 | chr15-64508410-DEL-84 |
| chr15-64534483-DEL-52 | chr15-64534455-DEL-68 |
| chr15-64542537-DEL-52 | chr15-64542538-DEL-52 |
| chr15-64544354-DEL-10733 | chr15-64544355-DEL-10733 |
| chr15-64582866-DEL-471 | chr15-64582867-DEL-471 |
| chr15-64630821-DEL-201 | chr15-64630808-DEL-204 |
| chr15-64662388-DEL-368 | chr15-64662389-DEL-368 |
| chr15-64671441-DEL-494 | chr15-64671442-DEL-494 |
| chr15-64674270-DEL-844 | chr15-64674271-DEL-844 |
| chr15-64684851-DEL-1644 | chr15-64684852-DEL-1644 |
| chr15-64689434-DEL-4399 | chr15-64689435-DEL-4399 |
| chr15-64708157-DEL-1561 | chr15-64708158-DEL-1561 |
| chr15-64714726-DEL-1003 | chr15-64714727-DEL-1003 |
| chr15-64718022-DEL-349 | chr15-64718022-DEL-346 |
| chr15-64721599-DEL-392 | chr15-64721600-DEL-392 |
| chr15-64721738-DEL-2651 | chr15-64721737-DEL-2652 |
| chr15-64766250-DEL-80 | chr15-64766251-DEL-80 |
| chr15-64777676-DEL-74 | chr15-64777679-DEL-69 |
| chr15-64786802-DEL-68 | chr15-64786803-DEL-72 |
| chr15-64799274-DEL-76 | chr15-64799273-DEL-78 |

| | |
|---|---|
| chr15-64810971-DEL-333 | chr15-64810972-DEL-333 |
| chr15-64831474-DEL-649 | chr15-64831475-DEL-649 |
| chr15-64849983-DEL-66 | chr15-64849983-DEL-66 |
| chr15-64859801-DEL-90 | chr15-64859819-DEL-69 |
| chr15-64860028-DEL-2210 | chr15-64860029-DEL-2210 |
| chr15-64867549-DEL-562 | chr15-64867550-DEL-562 |
| chr15-64874224-DEL-1353 | chr15-64874225-DEL-1353 |
| chr15-64902816-DEL-69 | chr15-64902817-DEL-69 |
| chr15-64910943-DEL-56 | chr15-64910944-DEL-56 |
| chr15-64949490-DEL-1131 | chr15-64949491-DEL-1131 |
| chr15-64964697-DEL-912 | chr15-64964698-DEL-912 |
| chr15-64987466-DEL-6413 | chr15-64987467-DEL-6413 |
| chr15-65004360-DEL-203 | chr15-65004361-DEL-203 |
| chr15-65009997-DEL-78 | chr15-65009998-DEL-78 |
| chr15-65049942-DEL-3118 | chr15-65049943-DEL-3118 |
| chr15-65083787-DEL-65 | chr15-65083812-DEL-55 |
| chr15-65093494-DEL-200 | chr15-65093495-DEL-200 |
| chr15-65095540-DEL-1553 | chr15-65095540-DEL-1553 |
| chr15-65101459-DEL-2030 | chr15-65101460-DEL-2030 |
| chr15-65144546-DEL-6895 | chr15-65144547-DEL-6895 |
| chr15-65158166-DEL-671 | chr15-65158167-DEL-671 |
| chr15-65171394-DEL-69 | chr15-65171395-DEL-69 |
| chr15-65201698-DEL-143 | chr15-65201699-DEL-143 |
| chr15-65203712-DEL-538 | chr15-65203713-DEL-538 |
| chr15-65235983-DEL-6857 | chr15-65235983-DEL-6857 |
| chr15-65242112-DEL-411 | chr15-65242113-DEL-411 |
| chr15-65265868-DEL-83 | chr15-65265869-DEL-83 |
| chr15-65305859-DEL-321 | chr15-65305860-DEL-321 |
| chr15-65377356-DEL-59 | chr15-65377358-DEL-67 |
| chr15-65382632-DEL-6193 | chr15-65382633-DEL-6193 |
| chr15-65422244-DEL-84 | chr15-65422245-DEL-76 |
| chr15-65443122-DEL-6718 | chr15-65443123-DEL-6718 |
| chr15-65458203-DEL-82 | chr15-65458190-DEL-96 |
| chr15-65470312-DEL-6194 | chr15-65470315-DEL-6192 |
| chr15-65478069-DEL-1200 | chr15-65478069-DEL-1200 |
| chr15-65494995-DEL-242 | chr15-65494996-DEL-238 |
| chr15-65498262-DEL-60 | chr15-65498263-DEL-60 |
| chr15-65538515-DEL-7081 | chr15-65538515-DEL-7081 |
| chr15-65578907-DEL-95 | chr15-65578908-DEL-95 |
| chr15-65587826-DEL-6414 | chr15-65587827-DEL-6414 |
| chr15-65600909-DEL-1478 | chr15-65600910-DEL-1478 |
| chr15-65609402-DEL-6853 | chr15-65609403-DEL-6853 |
| chr15-65621165-DEL-62 | chr15-65621166-DEL-77 |
| chr15-65640363-DEL-92 | chr15-65640364-DEL-92 |
| chr15-65646756-DEL-1669 | chr15-65646757-DEL-1669 |
| chr15-65704055-DEL-1146 | chr15-65704056-DEL-1146 |
| chr15-65725743-DEL-284 | chr15-65725744-DEL-284 |
| chr15-65757060-DEL-2007 | chr15-65757061-DEL-2007 |
| chr15-65770432-DEL-63 | chr15-65770454-DEL-67 |
| chr15-65808649-DEL-1921 | chr15-65808650-DEL-1921 |
| chr15-65837835-DEL-86 | chr15-65837864-DEL-56 |
| chr15-65839366-DEL-58 | chr15-65839367-DEL-59 |
| chr15-65850541-DEL-130 | chr15-65850542-DEL-130 |
| chr15-65917052-DEL-4926 | chr15-65917053-DEL-4926 |
| chr15-65917060-DEL-2204 | chr15-65917059-DEL-2204 |
| chr15-65919832-DEL-81 | chr15-65919833-DEL-81 |
| chr15-65928943-DEL-7228 | chr15-65928943-DEL-7228 |
| chr15-65949505-DEL-85 | chr15-65949505-DEL-85 |
| chr15-65959201-DEL-175 | chr15-65959202-DEL-175 |
| chr15-65967257-DEL-635 | chr15-65967258-DEL-635 |
| chr15-66002267-DEL-56 | chr15-66002268-DEL-56 |
| chr15-66025408-DEL-66 | chr15-66025409-DEL-72 |
| chr15-66025989-DEL-290 | chr15-66025978-DEL-291 |
| chr15-66027010-DEL-79 | chr15-66027029-DEL-61 |
| chr15-66074089-DEL-257 | chr15-66074090-DEL-257 |
| chr15-66080147-DEL-1890 | chr15-66080148-DEL-1890 |
| chr15-66089162-DEL-2347 | chr15-66089163-DEL-2347 |
| chr15-66154826-DEL-51 | chr15-66154825-DEL-53 |
| chr15-66214434-DEL-98 | chr15-66214435-DEL-97 |
| chr15-66221970-DEL-468 | chr15-66221971-DEL-468 |
| chr15-66229499-DEL-595 | chr15-66229500-DEL-593 |
| chr15-66242797-DEL-72 | chr15-66242810-DEL-60 |
| chr15-66260810-DEL-89 | chr15-66260823-DEL-50 |
| chr15-66340891-DEL-74 | chr15-66340936-DEL-54 |
| chr15-66354759-DEL-1892 | chr15-66354760-DEL-1892 |
| chr15-66377143-DEL-374 | chr15-66377141-DEL-373 |
| chr15-66391505-DEL-1412 | chr15-66391506-DEL-1412 |
| chr15-66393950-DEL-210 | chr15-66393951-DEL-210 |
| chr15-66446978-DEL-66 | chr15-66446979-DEL-66 |
| chr15-66499798-DEL-594 | chr15-66499799-DEL-594 |

| | |
|---|---|
| chr15-66534916-DEL-58 | chr15-66534917-DEL-58 |
| chr15-66536177-DEL-110 | chr15-66536178-DEL-110 |
| chr15-66546400-DEL-68 | chr15-66546405-DEL-64 |
| chr15-66568966-DEL-53 | chr15-66568967-DEL-51 |
| chr15-66593886-DEL-108 | chr15-66593887-DEL-108 |
| chr15-66625695-DEL-72 | chr15-66625689-DEL-73 |
| chr15-66632862-DEL-887 | chr15-66632863-DEL-887 |
| chr15-66680081-DEL-92 | chr15-66680070-DEL-68 |
| chr15-66722929-DEL-64 | chr15-66722964-DEL-70 |
| chr15-66784306-DEL-91 | chr15-66784307-DEL-91 |
| chr15-66806425-DEL-67 | chr15-66806426-DEL-65 |
| chr15-66820529-DEL-54 | chr15-66820561-DEL-60 |
| chr15-66854862-DEL-140 | chr15-66854860-DEL-130 |
| chr15-66860922-DEL-70 | chr15-66860923-DEL-70 |
| chr15-66861427-DEL-712 | chr15-66861428-DEL-712 |
| chr15-66871604-DEL-6413 | chr15-66871605-DEL-6413 |
| chr15-66877802-DEL-200 | chr15-66877759-DEL-201 |
| chr15-66899152-DEL-232 | chr15-66899138-DEL-234 |
| chr15-66921993-DEL-4637 | chr15-66921994-DEL-4637 |
| chr15-66932904-DEL-206 | chr15-66932905-DEL-206 |
| chr15-66939297-DEL-98 | chr15-66939298-DEL-95 |
| chr15-66941143-DEL-3514 | chr15-66941144-DEL-3514 |
| chr15-67039766-DEL-368 | chr15-67039767-DEL-368 |
| chr15-67060688-DEL-86 | chr15-67060684-DEL-71 |
| chr15-67084602-DEL-822 | chr15-67084603-DEL-822 |
| chr15-67113232-DEL-1573 | chr15-67113233-DEL-1573 |
| chr15-67129568-DEL-2942 | chr15-67129569-DEL-2942 |
| chr15-67137687-DEL-6491 | chr15-67137688-DEL-6491 |
| chr15-67147601-DEL-1310 | chr15-67147601-DEL-1310 |
| chr15-67168683-DEL-197 | chr15-67168684-DEL-197 |
| chr15-67174268-DEL-415 | chr15-67174282-DEL-401 |
| chr15-67178004-DEL-5375 | chr15-67178005-DEL-5375 |
| chr15-67195229-DEL-533 | chr15-67195228-DEL-533 |
| chr15-67212737-DEL-206 | chr15-67212738-DEL-206 |
| chr15-67244597-DEL-92 | chr15-67244598-DEL-90 |
| chr15-67250578-DEL-64 | chr15-67250579-DEL-64 |
| chr15-67255419-DEL-2161 | chr15-67255420-DEL-2161 |
| chr15-67278457-DEL-645 | chr15-67278458-DEL-645 |
| chr15-67344678-DEL-54 | chr15-67344677-DEL-55 |
| chr15-67383071-DEL-1314 | chr15-67383072-DEL-1314 |
| chr15-67438491-DEL-1446 | chr15-67438492-DEL-1446 |
| chr15-67459861-DEL-997 | chr15-67459862-DEL-997 |
| chr15-67462201-DEL-138 | chr15-67462202-DEL-138 |
| chr15-67471585-DEL-511 | chr15-67471586-DEL-511 |
| chr15-67514696-DEL-103 | chr15-67514697-DEL-87 |
| chr15-67516439-DEL-4620 | chr15-67516440-DEL-4620 |
| chr15-67523254-DEL-92 | chr15-67523255-DEL-92 |
| chr15-67534918-DEL-229 | chr15-67534919-DEL-229 |
| chr15-67536390-DEL-6175 | chr15-67536391-DEL-6175 |
| chr15-67537706-DEL-2947 | chr15-67537707-DEL-2947 |
| chr15-67556503-DEL-90 | chr15-67556513-DEL-93 |
| chr15-67560367-DEL-1746 | chr15-67560368-DEL-1746 |
| chr15-67566824-DEL-1534 | chr15-67566825-DEL-1534 |
| chr15-67590047-DEL-52 | chr15-67590038-DEL-56 |
| chr15-67605665-DEL-3000 | chr15-67605666-DEL-3000 |
| chr15-67617986-DEL-355 | chr15-67617986-DEL-355 |
| chr15-67635728-DEL-1016 | chr15-67635721-DEL-1017 |
| chr15-67640887-DEL-364 | chr15-67640888-DEL-364 |
| chr15-67669198-DEL-997 | chr15-67669199-DEL-997 |
| chr15-67699673-DEL-7640 | chr15-67699675-DEL-7639 |
| chr15-67701898-DEL-1251 | chr15-67701899-DEL-1251 |
| chr15-67706904-DEL-198 | chr15-67706905-DEL-199 |
| chr15-67763049-DEL-60 | chr15-67763050-DEL-60 |
| chr15-67782117-DEL-8853 | chr15-67782118-DEL-8853 |
| chr15-67788280-DEL-1345 | chr15-67788281-DEL-1345 |
| chr15-67851882-DEL-167 | chr15-67851883-DEL-167 |
| chr15-67852780-DEL-68 | chr15-67852781-DEL-68 |
| chr15-67860447-DEL-72 | chr15-67860424-DEL-100 |
| chr15-67869785-DEL-66 | chr15-67869786-DEL-66 |
| chr15-67870395-DEL-2644 | chr15-67870395-DEL-2644 |
| chr15-67875490-DEL-563 | chr15-67875491-DEL-563 |
| chr15-67904060-DEL-6387 | chr15-67904061-DEL-6387 |
| chr15-67934887-DEL-78 | chr15-67934887-DEL-78 |
| chr15-68026908-DEL-351 | chr15-68026909-DEL-351 |
| chr15-68038869-DEL-60 | chr15-68038877-DEL-62 |
| chr15-68048323-DEL-90 | chr15-68048324-DEL-135 |
| chr15-68059696-DEL-236 | chr15-68059697-DEL-236 |
| chr15-68064976-DEL-76 | chr15-68064977-DEL-76 |
| chr15-68078438-DEL-271 | chr15-68078439-DEL-271 |
| chr15-68080302-DEL-53 | chr15-68080301-DEL-53 |

| | |
|---|---|
| chr15-68102988-DEL-785 | chr15-68102990-DEL-783 |
| chr15-68108909-DEL-223 | chr15-68108910-DEL-223 |
| chr15-68114290-DEL-83 | chr15-68114289-DEL-84 |
| chr15-68116499-DEL-328 | chr15-68116496-DEL-358 |
| chr15-68169505-DEL-913 | chr15-68169506-DEL-913 |
| chr15-68181620-DEL-576 | chr15-68181620-DEL-576 |
| chr15-68203665-DEL-1137 | chr15-68203666-DEL-1137 |
| chr15-68206151-DEL-58 | chr15-68206152-DEL-58 |
| chr15-68239995-DEL-6506 | chr15-68239996-DEL-6506 |
| chr15-68252921-DEL-74 | chr15-68252916-DEL-84 |
| chr15-68293244-DEL-57 | chr15-68293239-DEL-63 |
| chr15-68331298-DEL-111 | chr15-68331299-DEL-111 |
| chr15-68398563-DEL-2931 | chr15-68398556-DEL-2932 |
| chr15-68445686-DEL-510 | chr15-68445687-DEL-510 |
| chr15-68463711-DEL-55 | chr15-68463712-DEL-55 |
| chr15-68499277-DEL-6527 | chr15-68499263-DEL-6534 |
| chr15-68510570-DEL-55 | chr15-68510573-DEL-59 |
| chr15-68527589-DEL-69 | chr15-68527590-DEL-69 |
| chr15-68527746-DEL-735 | chr15-68527746-DEL-735 |
| chr15-68543251-DEL-5234 | chr15-68543252-DEL-5235 |
| chr15-68554659-DEL-344 | chr15-68554660-DEL-344 |
| chr15-68556715-DEL-1504 | chr15-68556715-DEL-1504 |
| chr15-68575700-DEL-1751 | chr15-68575701-DEL-1751 |
| chr15-68583665-DEL-144 | chr15-68583666-DEL-144 |
| chr15-68613622-DEL-197 | chr15-68613630-DEL-191 |
| chr15-68620498-DEL-65 | chr15-68620496-DEL-66 |
| chr15-68647225-DEL-779 | chr15-68647226-DEL-779 |
| chr15-68658838-DEL-693 | chr15-68658839-DEL-693 |
| chr15-68659844-DEL-54 | chr15-68659845-DEL-54 |
| chr15-68672907-DEL-371 | chr15-68672908-DEL-371 |
| chr15-68713437-DEL-60 | chr15-68713438-DEL-60 |
| chr15-68715475-DEL-107 | chr15-68715473-DEL-104 |
| chr15-68722514-DEL-715 | chr15-68722515-DEL-715 |
| chr15-68725990-DEL-94 | chr15-68725991-DEL-94 |
| chr15-68731525-DEL-57 | chr15-68731526-DEL-56 |
| chr15-68757718-DEL-6434 | chr15-68757719-DEL-6434 |
| chr15-68782813-DEL-90 | chr15-68782818-DEL-78 |
| chr15-68785660-DEL-88 | chr15-68785661-DEL-88 |
| chr15-68822523-DEL-93 | chr15-68822510-DEL-93 |
| chr15-68851748-DEL-6658 | chr15-68851749-DEL-6658 |
| chr15-68862353-DEL-208 | chr15-68862354-DEL-208 |
| chr15-68874344-DEL-411 | chr15-68874345-DEL-408 |
| chr15-68993767-DEL-7264 | chr15-68993768-DEL-7264 |
| chr15-69015711-DEL-6187 | chr15-69015712-DEL-6187 |
| chr15-69067207-DEL-249 | chr15-69067208-DEL-249 |
| chr15-69078386-DEL-61 | chr15-69078391-DEL-57 |
| chr15-69103301-DEL-136 | chr15-69103287-DEL-156 |
| chr15-69137400-DEL-553 | chr15-69137401-DEL-553 |
| chr15-69144966-DEL-65 | chr15-69144944-DEL-74 |
| chr15-69164958-DEL-57 | chr15-69164959-DEL-57 |
| chr15-69201289-DEL-84 | chr15-69201305-DEL-63 |
| chr15-69206440-DEL-84 | chr15-69206449-DEL-76 |
| chr15-69234232-DEL-361 | chr15-69234233-DEL-361 |
| chr15-69247305-DEL-533 | chr15-69247306-DEL-533 |
| chr15-69262859-DEL-796 | chr15-69262860-DEL-796 |
| chr15-69268171-DEL-200 | chr15-69268172-DEL-200 |
| chr15-69275607-DEL-498 | chr15-69275608-DEL-498 |
| chr15-69288125-DEL-368 | chr15-69288126-DEL-368 |
| chr15-69299651-DEL-397 | chr15-69299648-DEL-398 |
| chr15-69343768-DEL-56 | chr15-69343769-DEL-56 |
| chr15-69347484-DEL-751 | chr15-69347485-DEL-751 |
| chr15-69358954-DEL-701 | chr15-69358955-DEL-702 |
| chr15-69428490-DEL-1490 | chr15-69428491-DEL-1490 |
| chr15-69468749-DEL-571 | chr15-69468749-DEL-571 |
| chr15-69513895-DEL-6214 | chr15-69513896-DEL-6214 |
| chr15-69517802-DEL-135 | chr15-69517803-DEL-135 |
| chr15-69535391-DEL-90 | chr15-69535392-DEL-90 |
| chr15-69565554-DEL-90 | chr15-69565564-DEL-86 |
| chr15-69579490-DEL-83 | chr15-69579491-DEL-83 |
| chr15-69590945-DEL-1901 | chr15-69590946-DEL-1901 |
| chr15-69595347-DEL-68 | chr15-69595347-DEL-68 |
| chr15-69613414-DEL-3922 | chr15-69613415-DEL-3922 |
| chr15-69646104-DEL-56 | chr15-69646105-DEL-56 |
| chr15-69708445-DEL-280 | chr15-69708446-DEL-280 |
| chr15-69767664-DEL-96 | chr15-69767667-DEL-94 |
| chr15-69776089-DEL-203 | chr15-69776090-DEL-203 |
| chr15-69777704-DEL-52 | chr15-69777704-DEL-52 |
| chr15-69786586-DEL-1619 | chr15-69786587-DEL-1619 |
| chr15-69799218-DEL-101 | chr15-69799232-DEL-112 |
| chr15-69830118-DEL-620 | chr15-69830119-DEL-620 |

| | |
|---|---|
| chr15-69890160-DEL-2750 | chr15-69890161-DEL-2749 |
| chr15-69899856-DEL-209 | chr15-69899857-DEL-210 |
| chr15-69946103-DEL-56 | chr15-69946104-DEL-56 |
| chr15-69954579-DEL-6583 | chr15-69954579-DEL-6583 |
| chr15-69972494-DEL-52 | chr15-69972495-DEL-52 |
| chr15-69975572-DEL-1445 | chr15-69975573-DEL-1445 |
| chr15-69986777-DEL-157 | chr15-69986778-DEL-157 |
| chr15-69990438-DEL-110 | chr15-69990469-DEL-70 |
| chr15-69997293-DEL-60 | chr15-69997294-DEL-56 |
| chr15-70004007-DEL-204 | chr15-70004007-DEL-204 |
| chr15-70026451-DEL-218 | chr15-70026452-DEL-218 |
| chr15-70026464-DEL-1023 | chr15-70026465-DEL-1023 |
| chr15-70030571-DEL-627 | chr15-70030572-DEL-624 |
| chr15-70034496-DEL-72 | chr15-70034502-DEL-54 |
| chr15-70039841-DEL-948 | chr15-70039842-DEL-948 |
| chr15-70041880-DEL-117 | chr15-70041881-DEL-117 |
| chr15-70077226-DEL-4505 | chr15-70077227-DEL-4505 |
| chr15-70099896-DEL-62 | chr15-70099897-DEL-62 |
| chr15-70107351-DEL-144 | chr15-70107352-DEL-144 |
| chr15-70117632-DEL-2323 | chr15-70117633-DEL-2323 |
| chr15-70126751-DEL-83 | chr15-70126752-DEL-83 |
| chr15-70155902-DEL-70 | chr15-70155903-DEL-70 |
| chr15-70164004-DEL-58 | chr15-70164005-DEL-58 |
| chr15-70171033-DEL-6214 | chr15-70171034-DEL-6214 |
| chr15-70201357-DEL-65 | chr15-70201358-DEL-65 |
| chr15-70202526-DEL-195 | chr15-70202527-DEL-195 |
| chr15-70203893-DEL-6453 | chr15-70203894-DEL-6453 |
| chr15-70221131-DEL-426 | chr15-70221132-DEL-427 |
| chr15-70233400-DEL-60 | chr15-70233370-DEL-104 |
| chr15-70235914-DEL-1372 | chr15-70235914-DEL-1372 |
| chr15-70243608-DEL-392 | chr15-70243609-DEL-392 |
| chr15-70244563-DEL-70 | chr15-70244563-DEL-50 |
| chr15-70337575-DEL-268 | chr15-70337576-DEL-268 |
| chr15-70355830-DEL-60 | chr15-70355831-DEL-60 |
| chr15-70357374-DEL-3494 | chr15-70357375-DEL-3494 |
| chr15-70376845-DEL-1301 | chr15-70376846-DEL-1301 |
| chr15-70389817-DEL-2210 | chr15-70389817-DEL-2210 |
| chr15-70500747-DEL-68 | chr15-70500748-DEL-68 |
| chr15-70519535-DEL-69 | chr15-70519537-DEL-58 |
| chr15-70525471-DEL-6207 | chr15-70525472-DEL-6207 |
| chr15-70566719-DEL-2855 | chr15-70566720-DEL-2855 |
| chr15-70634984-DEL-52 | chr15-70634985-DEL-50 |
| chr15-70640232-DEL-206 | chr15-70640233-DEL-206 |
| chr15-70652921-DEL-109 | chr15-70652922-DEL-109 |
| chr15-70656986-DEL-203 | chr15-70656987-DEL-203 |
| chr15-70663552-DEL-573 | chr15-70663553-DEL-573 |
| chr15-70666707-DEL-458 | chr15-70666708-DEL-458 |
| chr15-70672559-DEL-126 | chr15-70672560-DEL-126 |
| chr15-70720585-DEL-57 | chr15-70720584-DEL-58 |
| chr15-70722856-DEL-6424 | chr15-70722857-DEL-6424 |
| chr15-70735273-DEL-181 | chr15-70735274-DEL-181 |
| chr15-70746426-DEL-2276 | chr15-70746427-DEL-2276 |
| chr15-70758162-DEL-817 | chr15-70758163-DEL-817 |
| chr15-70761141-DEL-837 | chr15-70761142-DEL-837 |
| chr15-70765699-DEL-72 | chr15-70765713-DEL-70 |
| chr15-70766863-DEL-6263 | chr15-70766864-DEL-6261 |
| chr15-70782758-DEL-3547 | chr15-70782759-DEL-3547 |
| chr15-70799188-DEL-1142 | chr15-70799189-DEL-1142 |
| chr15-70900878-DEL-340 | chr15-70900879-DEL-340 |
| chr15-70943099-DEL-703 | chr15-70943100-DEL-703 |
| chr15-70976394-DEL-6158 | chr15-70976395-DEL-6158 |
| chr15-70996751-DEL-504 | chr15-70996752-DEL-504 |
| chr15-71023456-DEL-54 | chr15-71023456-DEL-54 |
| chr15-71062522-DEL-668 | chr15-71062524-DEL-637 |
| chr15-71073232-DEL-60 | chr15-71073233-DEL-60 |
| chr15-71076253-DEL-345 | chr15-71076254-DEL-345 |
| chr15-71077671-DEL-8194 | chr15-71077672-DEL-8194 |
| chr15-71102809-DEL-6623 | chr15-71102810-DEL-6623 |
| chr15-71145277-DEL-55 | chr15-71145292-DEL-50 |
| chr15-71157934-DEL-663 | chr15-71157935-DEL-663 |
| chr15-71176300-DEL-509 | chr15-71176315-DEL-503 |
| chr15-71196457-DEL-653 | chr15-71196458-DEL-653 |
| chr15-71201485-DEL-725 | chr15-71201486-DEL-725 |
| chr15-71204662-DEL-726 | chr15-71204663-DEL-726 |
| chr15-71218508-DEL-1964 | chr15-71218509-DEL-1964 |
| chr15-71233441-DEL-955 | chr15-71233438-DEL-959 |
| chr15-71258734-DEL-52 | chr15-71258692-DEL-95 |
| chr15-71276076-DEL-3147 | chr15-71276077-DEL-3147 |
| chr15-71292538-DEL-67 | chr15-71292539-DEL-67 |
| chr15-71303388-DEL-87 | chr15-71303389-DEL-87 |

| | |
|---|---|
| chr15-71305990-DEL-135 | chr15-71306006-DEL-141 |
| chr15-71330606-DEL-270 | chr15-71330607-DEL-270 |
| chr15-71366407-DEL-95 | chr15-71366408-DEL-95 |
| chr15-71444676-DEL-120 | chr15-71444678-DEL-119 |
| chr15-71447831-DEL-1842 | chr15-71447832-DEL-1842 |
| chr15-71495289-DEL-131 | chr15-71495283-DEL-122 |
| chr15-71516284-DEL-223 | chr15-71516285-DEL-223 |
| chr15-71521233-DEL-13363 | chr15-71521234-DEL-13363 |
| chr15-71618119-DEL-690 | chr15-71618120-DEL-690 |
| chr15-71660380-DEL-788 | chr15-71660381-DEL-788 |
| chr15-71664804-DEL-914 | chr15-71664805-DEL-914 |
| chr15-71666022-DEL-1587 | chr15-71666023-DEL-1587 |
| chr15-71689837-DEL-277 | chr15-71689838-DEL-281 |
| chr15-71694554-DEL-342 | chr15-71694555-DEL-342 |
| chr15-71750598-DEL-1199 | chr15-71750599-DEL-1199 |
| chr15-71750646-DEL-74 | chr15-71750645-DEL-76 |
| chr15-71755637-DEL-54 | chr15-71755621-DEL-52 |
| chr15-71791414-DEL-134 | chr15-71791415-DEL-134 |
| chr15-71796246-DEL-80 | chr15-71796247-DEL-80 |
| chr15-71819120-DEL-72 | chr15-71819138-DEL-51 |
| chr15-71832434-DEL-439 | chr15-71832435-DEL-439 |
| chr15-71837999-DEL-136 | chr15-71838019-DEL-116 |
| chr15-71841459-DEL-548 | chr15-71841460-DEL-548 |
| chr15-71867667-DEL-622 | chr15-71867668-DEL-622 |
| chr15-71881499-DEL-143 | chr15-71881456-DEL-144 |
| chr15-71893800-DEL-89 | chr15-71893814-DEL-96 |
| chr15-71919987-DEL-1222 | chr15-71919978-DEL-1226 |
| chr15-71932298-DEL-125 | chr15-71932299-DEL-125 |
| chr15-71937919-DEL-644 | chr15-71937914-DEL-660 |
| chr15-71960123-DEL-7071 | chr15-71960124-DEL-7071 |
| chr15-71966230-DEL-636 | chr15-71966231-DEL-636 |
| chr15-71971563-DEL-733 | chr15-71971564-DEL-733 |
| chr15-71980174-DEL-60 | chr15-71980174-DEL-50 |
| chr15-72001146-DEL-1131 | chr15-72001147-DEL-1131 |
| chr15-72018311-DEL-58 | chr15-72018312-DEL-58 |
| chr15-72084969-DEL-148 | chr15-72084970-DEL-148 |
| chr15-72094651-DEL-68 | chr15-72094652-DEL-68 |
| chr15-72167095-DEL-372 | chr15-72167096-DEL-372 |
| chr15-72169632-DEL-54 | chr15-72169633-DEL-54 |
| chr15-72175414-DEL-60 | chr15-72175415-DEL-68 |
| chr15-72190846-DEL-58 | chr15-72190847-DEL-58 |
| chr15-72211849-DEL-426 | chr15-72211873-DEL-403 |
| chr15-72252346-DEL-590 | chr15-72252347-DEL-590 |
| chr15-72275112-DEL-85 | chr15-72275097-DEL-101 |
| chr15-72291611-DEL-56 | chr15-72291614-DEL-54 |
| chr15-72300983-DEL-6436 | chr15-72300984-DEL-6436 |
| chr15-72312638-DEL-54 | chr15-72312639-DEL-54 |
| chr15-72345375-DEL-123 | chr15-72345418-DEL-126 |
| chr15-72350286-DEL-56 | chr15-72350287-DEL-56 |
| chr15-72387339-DEL-226 | chr15-72387340-DEL-226 |
| chr15-72394529-DEL-916 | chr15-72394529-DEL-916 |
| chr15-72395973-DEL-101 | chr15-72395968-DEL-102 |
| chr15-72427449-DEL-80 | chr15-72427450-DEL-88 |
| chr15-72439417-DEL-63 | chr15-72439400-DEL-81 |
| chr15-72502805-DEL-70 | chr15-72502804-DEL-72 |
| chr15-72642412-DEL-131 | chr15-72642392-DEL-115 |
| chr15-72668498-DEL-595 | chr15-72668499-DEL-595 |
| chr15-72725061-DEL-230 | chr15-72725062-DEL-230 |
| chr15-72729502-DEL-108 | chr15-72729503-DEL-108 |
| chr15-72753224-DEL-76 | chr15-72753225-DEL-50 |
| chr15-72825957-DEL-67 | chr15-72825952-DEL-67 |
| chr15-72835582-DEL-17150 | chr15-72835583-DEL-17150 |
| chr15-72862966-DEL-76 | chr15-72862967-DEL-76 |
| chr15-72868643-DEL-864 | chr15-72868642-DEL-864 |
| chr15-72906318-DEL-154 | chr15-72906318-DEL-153 |
| chr15-72936472-DEL-60 | chr15-72936473-DEL-60 |
| chr15-72948395-DEL-1288 | chr15-72948396-DEL-1288 |
| chr15-72949422-DEL-196 | chr15-72949429-DEL-192 |
| chr15-72953253-DEL-6547 | chr15-72953254-DEL-6547 |
| chr15-72984645-DEL-74 | chr15-72984646-DEL-74 |
| chr15-72987338-DEL-54 | chr15-72987323-DEL-70 |
| chr15-73000923-DEL-201 | chr15-73000924-DEL-201 |
| chr15-73001657-DEL-66 | chr15-73001658-DEL-56 |
| chr15-73034210-DEL-60 | chr15-73034211-DEL-60 |
| chr15-73052486-DEL-184 | chr15-73052487-DEL-184 |
| chr15-73086248-DEL-191 | chr15-73086235-DEL-203 |
| chr15-73089165-DEL-1409 | chr15-73089166-DEL-1409 |
| chr15-73095491-DEL-209 | chr15-73095492-DEL-209 |
| chr15-73099512-DEL-58 | chr15-73099513-DEL-58 |
| chr15-73122467-DEL-849 | chr15-73122468-DEL-849 |

| | |
|---|---|
| chr15-73171533-DEL-145 | chr15-73171534-DEL-140 |
| chr15-73179162-DEL-1668 | chr15-73179163-DEL-1668 |
| chr15-73204334-DEL-52 | chr15-73204334-DEL-50 |
| chr15-73245074-DEL-4955 | chr15-73245075-DEL-4956 |
| chr15-73259937-DEL-1206 | chr15-73259938-DEL-1206 |
| chr15-73269425-DEL-946 | chr15-73269426-DEL-946 |
| chr15-73299875-DEL-1010 | chr15-73299876-DEL-1010 |
| chr15-73322659-DEL-207 | chr15-73322660-DEL-207 |
| chr15-73328744-DEL-56 | chr15-73328745-DEL-60 |
| chr15-73340307-DEL-422 | chr15-73340308-DEL-422 |
| chr15-73356432-DEL-222 | chr15-73356433-DEL-222 |
| chr15-73372812-DEL-231 | chr15-73372823-DEL-216 |
| chr15-73388373-DEL-69 | chr15-73388374-DEL-69 |
| chr15-73388381-DEL-1355 | chr15-73388382-DEL-1355 |
| chr15-73392035-DEL-248 | chr15-73392036-DEL-248 |
| chr15-73399207-DEL-81 | chr15-73399207-DEL-81 |
| chr15-73402535-DEL-61 | chr15-73402522-DEL-53 |
| chr15-73403841-DEL-52 | chr15-73403842-DEL-52 |
| chr15-73413232-DEL-437 | chr15-73413244-DEL-437 |
| chr15-73434356-DEL-216 | chr15-73434357-DEL-216 |
| chr15-73441900-DEL-98 | chr15-73441901-DEL-98 |
| chr15-73470196-DEL-169 | chr15-73470197-DEL-169 |
| chr15-73474981-DEL-214 | chr15-73474982-DEL-214 |
| chr15-73505118-DEL-174 | chr15-73505119-DEL-174 |
| chr15-73520007-DEL-234 | chr15-73520008-DEL-234 |
| chr15-73523246-DEL-69 | chr15-73523247-DEL-69 |
| chr15-73541898-DEL-54 | chr15-73541899-DEL-54 |
| chr15-73545096-DEL-108 | chr15-73545097-DEL-108 |
| chr15-73559860-DEL-54 | chr15-73559861-DEL-54 |
| chr15-73560325-DEL-193 | chr15-73560326-DEL-193 |
| chr15-73570542-DEL-1344 | chr15-73570543-DEL-1344 |
| chr15-73573849-DEL-359 | chr15-73573850-DEL-359 |
| chr15-73582542-DEL-105 | chr15-73582529-DEL-106 |
| chr15-73589580-DEL-64 | chr15-73589580-DEL-64 |
| chr15-73653439-DEL-52 | chr15-73653440-DEL-52 |
| chr15-73669819-DEL-155 | chr15-73669820-DEL-155 |
| chr15-73681278-DEL-504 | chr15-73681279-DEL-504 |
| chr15-73704593-DEL-217 | chr15-73704585-DEL-218 |
| chr15-73734355-DEL-65 | chr15-73734356-DEL-65 |
| chr15-73745001-DEL-7169 | chr15-73745002-DEL-7169 |
| chr15-73812044-DEL-53 | chr15-73812045-DEL-53 |
| chr15-73819057-DEL-53 | chr15-73819057-DEL-53 |
| chr15-73858013-DEL-235 | chr15-73858014-DEL-235 |
| chr15-73874020-DEL-344 | chr15-73874021-DEL-344 |
| chr15-73940438-DEL-56 | chr15-73940439-DEL-52 |
| chr15-73951810-DEL-1595 | chr15-73951811-DEL-1595 |
| chr15-73964993-DEL-1320 | chr15-73964992-DEL-1322 |
| chr15-74059015-DEL-150 | chr15-74059016-DEL-150 |
| chr15-74084728-DEL-405 | chr15-74084729-DEL-405 |
| chr15-74104993-DEL-116 | chr15-74104994-DEL-114 |
| chr15-74112911-DEL-429 | chr15-74112912-DEL-429 |
| chr15-74138221-DEL-1014 | chr15-74138222-DEL-1012 |
| chr15-74176905-DEL-72 | chr15-74176905-DEL-72 |
| chr15-74186350-DEL-154 | chr15-74186351-DEL-154 |
| chr15-74206625-DEL-86 | chr15-74206626-DEL-86 |
| chr15-74212444-DEL-264 | chr15-74212427-DEL-256 |
| chr15-74312718-DEL-292 | chr15-74312719-DEL-292 |
| chr15-74335739-DEL-401 | chr15-74335740-DEL-401 |
| chr15-74382276-DEL-59 | chr15-74382277-DEL-59 |
| chr15-74412037-DEL-62 | chr15-74412038-DEL-62 |
| chr15-74421590-DEL-60 | chr15-74421593-DEL-58 |
| chr15-74427971-DEL-101 | chr15-74427972-DEL-101 |
| chr15-74474742-DEL-580 | chr15-74474743-DEL-580 |
| chr15-74540361-DEL-66 | chr15-74540361-DEL-66 |
| chr15-74607100-DEL-56 | chr15-74607109-DEL-50 |
| chr15-74608402-DEL-484 | chr15-74608403-DEL-484 |
| chr15-74613941-DEL-120 | chr15-74613942-DEL-120 |
| chr15-74657531-DEL-60 | chr15-74657529-DEL-54 |
| chr15-74771825-DEL-394 | chr15-74771826-DEL-394 |
| chr15-74782282-DEL-1223 | chr15-74782283-DEL-1223 |
| chr15-74786441-DEL-188 | chr15-74786442-DEL-188 |
| chr15-74788897-DEL-199 | chr15-74788894-DEL-200 |
| chr15-74805561-DEL-535 | chr15-74805562-DEL-535 |
| chr15-74811464-DEL-7975 | chr15-74811465-DEL-7975 |
| chr15-74903097-DEL-506 | chr15-74903098-DEL-506 |
| chr15-74904572-DEL-2470 | chr15-74904573-DEL-2470 |
| chr15-74933304-DEL-80 | chr15-74933305-DEL-80 |
| chr15-75004815-DEL-56 | chr15-75004812-DEL-52 |
| chr15-75048968-DEL-3027 | chr15-75048968-DEL-3027 |
| chr15-75062059-DEL-80 | chr15-75062060-DEL-79 |

| | |
|---|---|
| chr15-75063375-DEL-85 | chr15-75063376-DEL-85 |
| chr15-75077503-DEL-7089 | chr15-75077504-DEL-7089 |
| chr15-75105653-DEL-414 | chr15-75105654-DEL-414 |
| chr15-75114709-DEL-67 | chr15-75114710-DEL-67 |
| chr15-75130122-DEL-2139 | chr15-75130116-DEL-2152 |
| chr15-75136422-DEL-347 | chr15-75136423-DEL-347 |
| chr15-75199694-DEL-62 | chr15-75199659-DEL-58 |
| chr15-75204738-DEL-6781 | chr15-75204746-DEL-6780 |
| chr15-75231043-DEL-117 | chr15-75231044-DEL-117 |
| chr15-75247666-DEL-1966 | chr15-75247667-DEL-1966 |
| chr15-75251368-DEL-68 | chr15-75251349-DEL-69 |
| chr15-75258306-DEL-140 | chr15-75258307-DEL-140 |
| chr15-75279550-DEL-967 | chr15-75279551-DEL-967 |
| chr15-75302477-DEL-69 | chr15-75302478-DEL-69 |
| chr15-75320967-DEL-102 | chr15-75320964-DEL-102 |
| chr15-75337158-DEL-65 | chr15-75337158-DEL-65 |
| chr15-75376232-DEL-199 | chr15-75376225-DEL-194 |
| chr15-75657870-DEL-55 | chr15-75657874-DEL-56 |
| chr15-75664891-DEL-1338 | chr15-75664892-DEL-1338 |
| chr15-75676410-DEL-221 | chr15-75676411-DEL-221 |
| chr15-75718836-DEL-76 | chr15-75718837-DEL-76 |
| chr15-75722428-DEL-208 | chr15-75722429-DEL-208 |
| chr15-75742015-DEL-80 | chr15-75742016-DEL-85 |
| chr15-75745695-DEL-215 | chr15-75745718-DEL-193 |
| chr15-75749449-DEL-77 | chr15-75749416-DEL-82 |
| chr15-75802588-DEL-538 | chr15-75802589-DEL-538 |
| chr15-75803710-DEL-91 | chr15-75803711-DEL-91 |
| chr15-75807174-DEL-326 | chr15-75807175-DEL-326 |
| chr15-75813794-DEL-74 | chr15-75813795-DEL-74 |
| chr15-75829595-DEL-429 | chr15-75829603-DEL-383 |
| chr15-75835280-DEL-484 | chr15-75835282-DEL-464 |
| chr15-75849068-DEL-133 | chr15-75849069-DEL-133 |
| chr15-76028474-DEL-332 | chr15-76028475-DEL-332 |
| chr15-76034467-DEL-618 | chr15-76034468-DEL-618 |
| chr15-76049742-DEL-78 | chr15-76049743-DEL-102 |
| chr15-76105493-DEL-55 | chr15-76105494-DEL-55 |
| chr15-76106337-DEL-75 | chr15-76106338-DEL-76 |
| chr15-76107474-DEL-190 | chr15-76107475-DEL-190 |
| chr15-76160352-DEL-55 | chr15-76160353-DEL-55 |
| chr15-76171979-DEL-477 | chr15-76171980-DEL-477 |
| chr15-76184153-DEL-115 | chr15-76184154-DEL-115 |
| chr15-76193161-DEL-54 | chr15-76193144-DEL-71 |
| chr15-76299776-DEL-82 | chr15-76299777-DEL-82 |
| chr15-76435371-DEL-85 | chr15-76435372-DEL-85 |
| chr15-76444012-DEL-7190 | chr15-76444013-DEL-7190 |
| chr15-76503571-DEL-65 | chr15-76503572-DEL-60 |
| chr15-76549851-DEL-219 | chr15-76549852-DEL-219 |
| chr15-76560766-DEL-188 | chr15-76560765-DEL-189 |
| chr15-76584864-DEL-99 | chr15-76584865-DEL-99 |
| chr15-76592063-DEL-231 | chr15-76592064-DEL-231 |
| chr15-76628757-DEL-201 | chr15-76628758-DEL-201 |
| chr15-76629802-DEL-78 | chr15-76629806-DEL-89 |
| chr15-76636390-DEL-205 | chr15-76636391-DEL-205 |
| chr15-76655707-DEL-51 | chr15-76655707-DEL-51 |
| chr15-76660341-DEL-1893 | chr15-76660342-DEL-1893 |
| chr15-76697528-DEL-98 | chr15-76697489-DEL-98 |
| chr15-76698500-DEL-98 | chr15-76698501-DEL-98 |
| chr15-76708129-DEL-72 | chr15-76708170-DEL-63 |
| chr15-76711827-DEL-140 | chr15-76711828-DEL-140 |
| chr15-76719811-DEL-1930 | chr15-76719812-DEL-1930 |
| chr15-76742255-DEL-1613 | chr15-76742265-DEL-1589 |
| chr15-76748576-DEL-278 | chr15-76748577-DEL-278 |
| chr15-76751487-DEL-54 | chr15-76751488-DEL-54 |
| chr15-76792577-DEL-203 | chr15-76792578-DEL-203 |
| chr15-76793414-DEL-196 | chr15-76793415-DEL-196 |
| chr15-76797959-DEL-183 | chr15-76797960-DEL-183 |
| chr15-76804318-DEL-204 | chr15-76804319-DEL-204 |
| chr15-76816950-DEL-168 | chr15-76816951-DEL-168 |
| chr15-76819000-DEL-207 | chr15-76819001-DEL-206 |
| chr15-76821643-DEL-189 | chr15-76821644-DEL-189 |
| chr15-76823505-DEL-231 | chr15-76823506-DEL-231 |
| chr15-76826643-DEL-204 | chr15-76826644-DEL-204 |
| chr15-76830758-DEL-1164 | chr15-76830759-DEL-1164 |
| chr15-76837869-DEL-321 | chr15-76837874-DEL-321 |
| chr15-76848735-DEL-401 | chr15-76848735-DEL-401 |
| chr15-76849173-DEL-6401 | chr15-76849174-DEL-6401 |
| chr15-76875443-DEL-64 | chr15-76875442-DEL-68 |
| chr15-76885697-DEL-81 | chr15-76885698-DEL-81 |
| chr15-76886563-DEL-432 | chr15-76886564-DEL-432 |
| chr15-76891070-DEL-727 | chr15-76891071-DEL-727 |

| | |
|---|---|
| chr15-76908188-DEL-60 | chr15-76908188-DEL-60 |
| chr15-76917045-DEL-72 | chr15-76917046-DEL-72 |
| chr15-76949246-DEL-150 | chr15-76949247-DEL-150 |
| chr15-76951977-DEL-339 | chr15-76951978-DEL-339 |
| chr15-76957613-DEL-133 | chr15-76957604-DEL-146 |
| chr15-77000623-DEL-75 | chr15-77000624-DEL-75 |
| chr15-77079277-DEL-2505 | chr15-77079280-DEL-2506 |
| chr15-77084468-DEL-119 | chr15-77084468-DEL-120 |
| chr15-77219316-DEL-224 | chr15-77219312-DEL-225 |
| chr15-77220013-DEL-70 | chr15-77220014-DEL-70 |
| chr15-77230943-DEL-233 | chr15-77230944-DEL-233 |
| chr15-77234744-DEL-1972 | chr15-77234745-DEL-1972 |
| chr15-77244690-DEL-216 | chr15-77244691-DEL-216 |
| chr15-77256127-DEL-84 | chr15-77256128-DEL-84 |
| chr15-77260276-DEL-367 | chr15-77260277-DEL-367 |
| chr15-77296077-DEL-204 | chr15-77296076-DEL-204 |
| chr15-77297135-DEL-1389 | chr15-77297136-DEL-1389 |
| chr15-77323841-DEL-71 | chr15-77323841-DEL-71 |
| chr15-77335824-DEL-230 | chr15-77335825-DEL-230 |
| chr15-77347582-DEL-435 | chr15-77347583-DEL-435 |
| chr15-77366217-DEL-202 | chr15-77366218-DEL-202 |
| chr15-77369477-DEL-60 | chr15-77369478-DEL-60 |
| chr15-77373858-DEL-530 | chr15-77373859-DEL-530 |
| chr15-77428241-DEL-108 | chr15-77428241-DEL-108 |
| chr15-77430380-DEL-244 | chr15-77430381-DEL-244 |
| chr15-77448214-DEL-107 | chr15-77448204-DEL-127 |
| chr15-77464039-DEL-22526 | chr15-77464041-DEL-22546 |
| chr15-77487543-DEL-132 | chr15-77487544-DEL-132 |
| chr15-77488372-DEL-1504 | chr15-77488371-DEL-1504 |
| chr15-77501412-DEL-590 | chr15-77501406-DEL-590 |
| chr15-77509049-DEL-63 | chr15-77509050-DEL-63 |
| chr15-77515941-DEL-68 | chr15-77515942-DEL-68 |
| chr15-77599882-DEL-1573 | chr15-77599883-DEL-1573 |
| chr15-77612284-DEL-202 | chr15-77612284-DEL-202 |
| chr15-77635987-DEL-124 | chr15-77635984-DEL-128 |
| chr15-77649801-DEL-76 | chr15-77649802-DEL-76 |
| chr15-77658903-DEL-70 | chr15-77658904-DEL-70 |
| chr15-77663908-DEL-163 | chr15-77663909-DEL-163 |
| chr15-77717107-DEL-217 | chr15-77717104-DEL-216 |
| chr15-77728054-DEL-207 | chr15-77728055-DEL-207 |
| chr15-77738089-DEL-238 | chr15-77738090-DEL-238 |
| chr15-77749171-DEL-407 | chr15-77749172-DEL-407 |
| chr15-77755060-DEL-205 | chr15-77755061-DEL-205 |
| chr15-77765184-DEL-534 | chr15-77765185-DEL-534 |
| chr15-77768678-DEL-111 | chr15-77768679-DEL-109 |
| chr15-77769954-DEL-230 | chr15-77769955-DEL-230 |
| chr15-77779400-DEL-7157 | chr15-77779401-DEL-7157 |
| chr15-77805682-DEL-201 | chr15-77805683-DEL-201 |
| chr15-77809895-DEL-230 | chr15-77809896-DEL-230 |
| chr15-77869399-DEL-63 | chr15-77869403-DEL-72 |
| chr15-77882977-DEL-53 | chr15-77882968-DEL-68 |
| chr15-77885147-DEL-68 | chr15-77885160-DEL-56 |
| chr15-77887840-DEL-280 | chr15-77887841-DEL-280 |
| chr15-77903618-DEL-350 | chr15-77903619-DEL-350 |
| chr15-77907681-DEL-62 | chr15-77907682-DEL-54 |
| chr15-77918579-DEL-175 | chr15-77918580-DEL-175 |
| chr15-77924677-DEL-55 | chr15-77924678-DEL-55 |
| chr15-77937609-DEL-76 | chr15-77937610-DEL-76 |
| chr15-77944540-DEL-76 | chr15-77944549-DEL-54 |
| chr15-77962297-DEL-67 | chr15-77962295-DEL-68 |
| chr15-77972203-DEL-145 | chr15-77972209-DEL-147 |
| chr15-78013182-DEL-61 | chr15-78013183-DEL-61 |
| chr15-78035562-DEL-110 | chr15-78035561-DEL-112 |
| chr15-78115368-DEL-66 | chr15-78115352-DEL-52 |
| chr15-78118092-DEL-63 | chr15-78118095-DEL-60 |
| chr15-78149696-DEL-75 | chr15-78149697-DEL-75 |
| chr15-78188927-DEL-90 | chr15-78188904-DEL-70 |
| chr15-78193787-DEL-104 | chr15-78193794-DEL-98 |
| chr15-78194717-DEL-1079 | chr15-78194718-DEL-1079 |
| chr15-78208865-DEL-640 | chr15-78208866-DEL-640 |
| chr15-78260522-DEL-200 | chr15-78260523-DEL-200 |
| chr15-78267254-DEL-422 | chr15-78267277-DEL-396 |
| chr15-78269796-DEL-76 | chr15-78269797-DEL-73 |
| chr15-78341079-DEL-136 | chr15-78341080-DEL-136 |
| chr15-78346841-DEL-60 | chr15-78346850-DEL-52 |
| chr15-78358750-DEL-75 | chr15-78358754-DEL-72 |
| chr15-78403049-DEL-168 | chr15-78403050-DEL-168 |
| chr15-78430819-DEL-80 | chr15-78430819-DEL-80 |
| chr15-78468699-DEL-207 | chr15-78468700-DEL-207 |
| chr15-78495386-DEL-51 | chr15-78495387-DEL-51 |

| | |
|---|---|
| chr15-78528583-DEL-55 | chr15-78528584-DEL-55 |
| chr15-78535918-DEL-648 | chr15-78535919-DEL-648 |
| chr15-78559952-DEL-52 | chr15-78559953-DEL-52 |
| chr15-78610066-DEL-3506 | chr15-78610067-DEL-3506 |
| chr15-78623124-DEL-365 | chr15-78623125-DEL-365 |
| chr15-78636211-DEL-208 | chr15-78636212-DEL-208 |
| chr15-78660119-DEL-66 | chr15-78660120-DEL-66 |
| chr15-78660164-DEL-550 | chr15-78660143-DEL-572 |
| chr15-78662618-DEL-52 | chr15-78662619-DEL-58 |
| chr15-78696523-DEL-181 | chr15-78696524-DEL-180 |
| chr15-78708880-DEL-224 | chr15-78708881-DEL-224 |
| chr15-78721481-DEL-82 | chr15-78721482-DEL-70 |
| chr15-78722531-DEL-203 | chr15-78722532-DEL-202 |
| chr15-78782356-DEL-77 | chr15-78782357-DEL-77 |
| chr15-78783791-DEL-72 | chr15-78783794-DEL-70 |
| chr15-78793519-DEL-214 | chr15-78793520-DEL-214 |
| chr15-78828318-DEL-924 | chr15-78828319-DEL-924 |
| chr15-78828901-DEL-63 | chr15-78828902-DEL-75 |
| chr15-78836625-DEL-1445 | chr15-78836626-DEL-1445 |
| chr15-78839164-DEL-52 | chr15-78839165-DEL-52 |
| chr15-78848425-DEL-201 | chr15-78848426-DEL-201 |
| chr15-78851186-DEL-72 | chr15-78851187-DEL-72 |
| chr15-78896709-DEL-74 | chr15-78896720-DEL-78 |
| chr15-78933179-DEL-191 | chr15-78933180-DEL-191 |
| chr15-78957693-DEL-225 | chr15-78957694-DEL-225 |
| chr15-78968972-DEL-235 | chr15-78968973-DEL-235 |
| chr15-78971265-DEL-103 | chr15-78971242-DEL-141 |
| chr15-78971880-DEL-213 | chr15-78971881-DEL-213 |
| chr15-78974684-DEL-177 | chr15-78974685-DEL-177 |
| chr15-78977371-DEL-55 | chr15-78977372-DEL-55 |
| chr15-78979107-DEL-166 | chr15-78979095-DEL-164 |
| chr15-78981327-DEL-380 | chr15-78981328-DEL-380 |
| chr15-79009675-DEL-54 | chr15-79009676-DEL-54 |
| chr15-79027128-DEL-435 | chr15-79027133-DEL-434 |
| chr15-79034584-DEL-219 | chr15-79034585-DEL-219 |
| chr15-79068337-DEL-179 | chr15-79068338-DEL-179 |
| chr15-79103368-DEL-236 | chr15-79103357-DEL-214 |
| chr15-79111704-DEL-69 | chr15-79111705-DEL-69 |
| chr15-79121243-DEL-67 | chr15-79121244-DEL-67 |
| chr15-79128207-DEL-90 | chr15-79128208-DEL-65 |
| chr15-79159121-DEL-225 | chr15-79159122-DEL-225 |
| chr15-79172240-DEL-78 | chr15-79172241-DEL-78 |
| chr15-79174380-DEL-112 | chr15-79174375-DEL-88 |
| chr15-79177589-DEL-80 | chr15-79177590-DEL-80 |
| chr15-79186437-DEL-325 | chr15-79186405-DEL-326 |
| chr15-79204920-DEL-57 | chr15-79204921-DEL-57 |
| chr15-79214053-DEL-217 | chr15-79214054-DEL-217 |
| chr15-79219498-DEL-58 | chr15-79219499-DEL-58 |
| chr15-79221727-DEL-236 | chr15-79221728-DEL-236 |
| chr15-79230106-DEL-151 | chr15-79230150-DEL-106 |
| chr15-79264920-DEL-66 | chr15-79264921-DEL-66 |
| chr15-79296584-DEL-66 | chr15-79296585-DEL-66 |
| chr15-79300142-DEL-191 | chr15-79300143-DEL-191 |
| chr15-79350325-DEL-213 | chr15-79350326-DEL-213 |
| chr15-79354122-DEL-87 | chr15-79354122-DEL-87 |
| chr15-79425443-DEL-52 | chr15-79425442-DEL-68 |
| chr15-79433241-DEL-240 | chr15-79433242-DEL-240 |
| chr15-79438551-DEL-74 | chr15-79438552-DEL-74 |
| chr15-79509459-DEL-941 | chr15-79509460-DEL-941 |
| chr15-79512593-DEL-238 | chr15-79512594-DEL-238 |
| chr15-79541885-DEL-205 | chr15-79541886-DEL-205 |
| chr15-79548126-DEL-64 | chr15-79548126-DEL-64 |
| chr15-79554552-DEL-126 | chr15-79554553-DEL-126 |
| chr15-79555636-DEL-208 | chr15-79555637-DEL-208 |
| chr15-79566303-DEL-113 | chr15-79566304-DEL-113 |
| chr15-79584016-DEL-6228 | chr15-79584017-DEL-6228 |
| chr15-79620541-DEL-96 | chr15-79620540-DEL-67 |
| chr15-79642667-DEL-160 | chr15-79642667-DEL-158 |
| chr15-79644685-DEL-66 | chr15-79644686-DEL-66 |
| chr15-79687089-DEL-57 | chr15-79687053-DEL-60 |
| chr15-79696950-DEL-72 | chr15-79696914-DEL-66 |
| chr15-79703906-DEL-210 | chr15-79703907-DEL-210 |
| chr15-79705350-DEL-87 | chr15-79705349-DEL-88 |
| chr15-79725100-DEL-336 | chr15-79725101-DEL-336 |
| chr15-79736592-DEL-185 | chr15-79736593-DEL-185 |
| chr15-79742858-DEL-204 | chr15-79742859-DEL-204 |
| chr15-79750969-DEL-203 | chr15-79750970-DEL-203 |
| chr15-79752354-DEL-597 | chr15-79752396-DEL-582 |
| chr15-79753972-DEL-3134 | chr15-79753973-DEL-3134 |
| chr15-79771378-DEL-197 | chr15-79771379-DEL-197 |

| | |
|---|---|
| chr15-79778403-DEL-197 | chr15-79778400-DEL-198 |
| chr15-79779759-DEL-535 | chr15-79779760-DEL-535 |
| chr15-79781450-DEL-73 | chr15-79781453-DEL-75 |
| chr15-79787972-DEL-75 | chr15-79787973-DEL-75 |
| chr15-79806141-DEL-1451 | chr15-79806142-DEL-1451 |
| chr15-79807108-DEL-198 | chr15-79807109-DEL-198 |
| chr15-79815971-DEL-52 | chr15-79815972-DEL-52 |
| chr15-79820442-DEL-226 | chr15-79820443-DEL-226 |
| chr15-79823937-DEL-202 | chr15-79823938-DEL-202 |
| chr15-79826234-DEL-307 | chr15-79826235-DEL-307 |
| chr15-79850093-DEL-182 | chr15-79850094-DEL-182 |
| chr15-79850589-DEL-188 | chr15-79850590-DEL-188 |
| chr15-79858818-DEL-3726 | chr15-79858819-DEL-3726 |
| chr15-79863294-DEL-57 | chr15-79863295-DEL-57 |
| chr15-79866161-DEL-93 | chr15-79866162-DEL-93 |
| chr15-79871500-DEL-103 | chr15-79871501-DEL-103 |
| chr15-79872627-DEL-80 | chr15-79872628-DEL-80 |
| chr15-79873946-DEL-205 | chr15-79873947-DEL-205 |
| chr15-79894387-DEL-58 | chr15-79894388-DEL-58 |
| chr15-79914045-DEL-203 | chr15-79914046-DEL-203 |
| chr15-79916147-DEL-226 | chr15-79916148-DEL-226 |
| chr15-79918251-DEL-51 | chr15-79918244-DEL-51 |
| chr15-79930562-DEL-181 | chr15-79930521-DEL-184 |
| chr15-79931460-DEL-131 | chr15-79931461-DEL-131 |
| chr15-79943145-DEL-156 | chr15-79943146-DEL-156 |
| chr15-79953148-DEL-209 | chr15-79953149-DEL-209 |
| chr15-80005066-DEL-212 | chr15-80005055-DEL-213 |
| chr15-80006412-DEL-202 | chr15-80006432-DEL-202 |
| chr15-80010875-DEL-228 | chr15-80010876-DEL-228 |
| chr15-80016533-DEL-159 | chr15-80016534-DEL-159 |
| chr15-80081199-DEL-197 | chr15-80081200-DEL-197 |
| chr15-80094503-DEL-85 | chr15-80094502-DEL-81 |
| chr15-80104497-DEL-267 | chr15-80104498-DEL-267 |
| chr15-80105906-DEL-172 | chr15-80105907-DEL-170 |
| chr15-80116808-DEL-398 | chr15-80116809-DEL-398 |
| chr15-80121894-DEL-164 | chr15-80121895-DEL-164 |
| chr15-80126825-DEL-183 | chr15-80126826-DEL-183 |
| chr15-80184498-DEL-75 | chr15-80184493-DEL-81 |
| chr15-80194550-DEL-132 | chr15-80194551-DEL-126 |
| chr15-80222226-DEL-1234 | chr15-80222227-DEL-1234 |
| chr15-80237835-DEL-53 | chr15-80237830-DEL-60 |
| chr15-80300592-DEL-72 | chr15-80300591-DEL-74 |
| chr15-80317660-DEL-591 | chr15-80317661-DEL-591 |
| chr15-80318591-DEL-53 | chr15-80318590-DEL-54 |
| chr15-80332119-DEL-78 | chr15-80332120-DEL-78 |
| chr15-80345998-DEL-3483 | chr15-80345999-DEL-3483 |
| chr15-80399581-DEL-79 | chr15-80399581-DEL-80 |
| chr15-80408685-DEL-172 | chr15-80408686-DEL-172 |
| chr15-80414193-DEL-82 | chr15-80414194-DEL-82 |
| chr15-80464670-DEL-54 | chr15-80464671-DEL-54 |
| chr15-80512191-DEL-51 | chr15-80512192-DEL-51 |
| chr15-80538158-DEL-632 | chr15-80538159-DEL-632 |
| chr15-80540580-DEL-274 | chr15-80540581-DEL-274 |
| chr15-80588189-DEL-208 | chr15-80588190-DEL-208 |
| chr15-80620135-DEL-210 | chr15-80620136-DEL-210 |
| chr15-80633016-DEL-89 | chr15-80633017-DEL-89 |
| chr15-80642742-DEL-195 | chr15-80642743-DEL-195 |
| chr15-80685804-DEL-68 | chr15-80685808-DEL-61 |
| chr15-80693112-DEL-64 | chr15-80693111-DEL-82 |
| chr15-80715907-DEL-208 | chr15-80715908-DEL-208 |
| chr15-80803593-DEL-1636 | chr15-80803594-DEL-1632 |
| chr15-80805050-DEL-179 | chr15-80805051-DEL-179 |
| chr15-80836083-DEL-235 | chr15-80836084-DEL-235 |
| chr15-80866515-DEL-207 | chr15-80866516-DEL-207 |
| chr15-80878752-DEL-212 | chr15-80878753-DEL-212 |
| chr15-80885208-DEL-207 | chr15-80885203-DEL-213 |
| chr15-80886550-DEL-144 | chr15-80886573-DEL-121 |
| chr15-80909323-DEL-217 | chr15-80909324-DEL-217 |
| chr15-80951673-DEL-218 | chr15-80951662-DEL-218 |
| chr15-80958624-DEL-58 | chr15-80958625-DEL-58 |
| chr15-80994048-DEL-198 | chr15-80994049-DEL-198 |
| chr15-81003087-DEL-52 | chr15-81003088-DEL-52 |
| chr15-81012147-DEL-57 | chr15-81012130-DEL-74 |
| chr15-81019445-DEL-203 | chr15-81019446-DEL-203 |
| chr15-81021049-DEL-51 | chr15-81021050-DEL-51 |
| chr15-81029663-DEL-6403 | chr15-81029664-DEL-6403 |
| chr15-81068744-DEL-240 | chr15-81068745-DEL-240 |
| chr15-81078503-DEL-222 | chr15-81078504-DEL-222 |
| chr15-81092794-DEL-54 | chr15-81092798-DEL-75 |
| chr15-81096034-DEL-98 | chr15-81096033-DEL-99 |

| | |
|---|---|
| chr15-81096571-DEL-275 | chr15-81096572-DEL-275 |
| chr15-81102793-DEL-108 | chr15-81102779-DEL-127 |
| chr15-81149489-DEL-80 | chr15-81149488-DEL-81 |
| chr15-81150246-DEL-307 | chr15-81150247-DEL-279 |
| chr15-81161612-DEL-394 | chr15-81161613-DEL-394 |
| chr15-81166007-DEL-331 | chr15-81166008-DEL-331 |
| chr15-81190273-DEL-2329 | chr15-81190274-DEL-2329 |
| chr15-81193451-DEL-271 | chr15-81193441-DEL-272 |
| chr15-81197993-DEL-223 | chr15-81197979-DEL-224 |
| chr15-81262223-DEL-129 | chr15-81262224-DEL-129 |
| chr15-81268443-DEL-196 | chr15-81268444-DEL-196 |
| chr15-81343082-DEL-168 | chr15-81343079-DEL-178 |
| chr15-81352734-DEL-51 | chr15-81352735-DEL-51 |
| chr15-81354681-DEL-234 | chr15-81354682-DEL-234 |
| chr15-81367454-DEL-208 | chr15-81367444-DEL-213 |
| chr15-81376600-DEL-4777 | chr15-81376601-DEL-4777 |
| chr15-81395382-DEL-66 | chr15-81395383-DEL-66 |
| chr15-81462753-DEL-71 | chr15-81462795-DEL-60 |
| chr15-81518981-DEL-255 | chr15-81518981-DEL-254 |
| chr15-81543910-DEL-195 | chr15-81543911-DEL-195 |
| chr15-81550889-DEL-216 | chr15-81550890-DEL-216 |
| chr15-81563976-DEL-138 | chr15-81563974-DEL-138 |
| chr15-81585676-DEL-210 | chr15-81585629-DEL-210 |
| chr15-81597187-DEL-210 | chr15-81597188-DEL-210 |
| chr15-81598863-DEL-98 | chr15-81598815-DEL-98 |
| chr15-81611639-DEL-117 | chr15-81611640-DEL-117 |
| chr15-81626817-DEL-211 | chr15-81626818-DEL-211 |
| chr15-81645337-DEL-256 | chr15-81645387-DEL-217 |
| chr15-81655322-DEL-316 | chr15-81655323-DEL-316 |
| chr15-81703996-DEL-166 | chr15-81703997-DEL-166 |
| chr15-81738586-DEL-174 | chr15-81738587-DEL-174 |
| chr15-81765835-DEL-157 | chr15-81765838-DEL-190 |
| chr15-81840001-DEL-130 | chr15-81840002-DEL-130 |
| chr15-81848018-DEL-205 | chr15-81848019-DEL-205 |
| chr15-81890491-DEL-74 | chr15-81890471-DEL-71 |
| chr15-81895904-DEL-201 | chr15-81895905-DEL-201 |
| chr15-81901134-DEL-137 | chr15-81901135-DEL-137 |
| chr15-81904566-DEL-60 | chr15-81904566-DEL-60 |
| chr15-81929264-DEL-207 | chr15-81929266-DEL-207 |
| chr15-81963827-DEL-110 | chr15-81963828-DEL-110 |
| chr15-81977300-DEL-72 | chr15-81977301-DEL-70 |
| chr15-81992267-DEL-111 | chr15-81992268-DEL-111 |
| chr15-82002331-DEL-205 | chr15-82002332-DEL-205 |
| chr15-82027565-DEL-213 | chr15-82027566-DEL-213 |
| chr15-82057907-DEL-212 | chr15-82057896-DEL-212 |
| chr15-82133364-DEL-540 | chr15-82133413-DEL-540 |
| chr15-82137071-DEL-97 | chr15-82137033-DEL-98 |
| chr15-82137495-DEL-344 | chr15-82137496-DEL-344 |
| chr15-82143359-DEL-177 | chr15-82143360-DEL-177 |
| chr15-82171744-DEL-168 | chr15-82171745-DEL-168 |
| chr15-82187028-DEL-296 | chr15-82187005-DEL-314 |
| chr15-82191659-DEL-53 | chr15-82191660-DEL-53 |
| chr15-82200428-DEL-398 | chr15-82200429-DEL-398 |
| chr15-82246731-DEL-5472 | chr15-82246732-DEL-5472 |
| chr15-82255052-DEL-52 | chr15-82255053-DEL-52 |
| chr15-82280936-DEL-399 | chr15-82280936-DEL-399 |
| chr15-82281531-DEL-403 | chr15-82281531-DEL-404 |
| chr15-82299814-DEL-194 | chr15-82299815-DEL-194 |
| chr15-82354842-DEL-200 | chr15-82354832-DEL-201 |
| chr15-82358288-DEL-3925 | chr15-82358289-DEL-3925 |
| chr15-82390986-DEL-2147 | chr15-82390986-DEL-2147 |
| chr15-82448254-DEL-824 | chr15-82448255-DEL-824 |
| chr15-82453024-DEL-1200 | chr15-82453025-DEL-1200 |
| chr15-82457473-DEL-211 | chr15-82457474-DEL-211 |
| chr15-82461300-DEL-827 | chr15-82461301-DEL-827 |
| chr15-82474416-DEL-205 | chr15-82474417-DEL-205 |
| chr15-82474991-DEL-7172 | chr15-82474992-DEL-7172 |
| chr15-82489307-DEL-3873 | chr15-82489303-DEL-3874 |
| chr15-82543411-DEL-180 | chr15-82543400-DEL-181 |
| chr15-82548374-DEL-228 | chr15-82548375-DEL-228 |
| chr15-82559986-DEL-1524 | chr15-82559987-DEL-1524 |
| chr15-82609674-DEL-6297 | chr15-82609675-DEL-6297 |
| chr15-82621579-DEL-175 | chr15-82621580-DEL-175 |
| chr15-82635265-DEL-228 | chr15-82635266-DEL-228 |
| chr15-82655743-DEL-1694 | chr15-82655744-DEL-1694 |
| chr15-82707763-DEL-75 | chr15-82707782-DEL-57 |
| chr15-82717453-DEL-98 | chr15-82717476-DEL-72 |
| chr15-82719218-DEL-203 | chr15-82719219-DEL-203 |
| chr15-82755811-DEL-228 | chr15-82755812-DEL-228 |
| chr15-82757528-DEL-220 | chr15-82757529-DEL-220 |

| | |
|---|---|
| chr15-82762760-DEL-340 | chr15-82762747-DEL-357 |
| chr15-82798474-DEL-55 | chr15-82798475-DEL-65 |
| chr15-82854267-DEL-727 | chr15-82854268-DEL-727 |
| chr15-82857814-DEL-52 | chr15-82857814-DEL-52 |
| chr15-82889071-DEL-434 | chr15-82889072-DEL-434 |
| chr15-82892359-DEL-56 | chr15-82892360-DEL-56 |
| chr15-82899900-DEL-405 | chr15-82899901-DEL-405 |
| chr15-82949213-DEL-3627 | chr15-82949214-DEL-3627 |
| chr15-82954580-DEL-66 | chr15-82954565-DEL-82 |
| chr15-82957178-DEL-204 | chr15-82957179-DEL-204 |
| chr15-82962192-DEL-664 | chr15-82962195-DEL-648 |
| chr15-82980758-DEL-86 | chr15-82980769-DEL-79 |
| chr15-83011488-DEL-107 | chr15-83011472-DEL-108 |
| chr15-83070692-DEL-1141 | chr15-83070693-DEL-1141 |
| chr15-83073020-DEL-203 | chr15-83073014-DEL-201 |
| chr15-83075153-DEL-184 | chr15-83075149-DEL-185 |
| chr15-83077024-DEL-354 | chr15-83077047-DEL-359 |
| chr15-83087873-DEL-345 | chr15-83087874-DEL-345 |
| chr15-83099351-DEL-55 | chr15-83099314-DEL-99 |
| chr15-83126784-DEL-1699 | chr15-83126785-DEL-1699 |
| chr15-83127821-DEL-196 | chr15-83127802-DEL-206 |
| chr15-83148543-DEL-72 | chr15-83148544-DEL-68 |
| chr15-83158515-DEL-125 | chr15-83158516-DEL-125 |
| chr15-83224178-DEL-52 | chr15-83224175-DEL-56 |
| chr15-83289229-DEL-1186 | chr15-83289230-DEL-1186 |
| chr15-83300167-DEL-198 | chr15-83300168-DEL-198 |
| chr15-83309992-DEL-94 | chr15-83310012-DEL-56 |
| chr15-83376046-DEL-87 | chr15-83376047-DEL-87 |
| chr15-83388360-DEL-303 | chr15-83388335-DEL-300 |
| chr15-83391000-DEL-345 | chr15-83391044-DEL-346 |
| chr15-83444788-DEL-193 | chr15-83444789-DEL-193 |
| chr15-83457358-DEL-123 | chr15-83457392-DEL-111 |
| chr15-83459656-DEL-141 | chr15-83459657-DEL-141 |
| chr15-83472102-DEL-207 | chr15-83472103-DEL-207 |
| chr15-83481818-DEL-86 | chr15-83481819-DEL-86 |
| chr15-83482489-DEL-192 | chr15-83482490-DEL-192 |
| chr15-83506834-DEL-364 | chr15-83506835-DEL-364 |
| chr15-83540699-DEL-59 | chr15-83540700-DEL-59 |
| chr15-83557367-DEL-84 | chr15-83557368-DEL-84 |
| chr15-83559534-DEL-61 | chr15-83559535-DEL-61 |
| chr15-83613491-DEL-201 | chr15-83613492-DEL-201 |
| chr15-83622293-DEL-126 | chr15-83622294-DEL-126 |
| chr15-83660749-DEL-63 | chr15-83660750-DEL-63 |
| chr15-83689314-DEL-1498 | chr15-83689315-DEL-1498 |
| chr15-83694852-DEL-56 | chr15-83694851-DEL-58 |
| chr15-83699729-DEL-95 | chr15-83699730-DEL-95 |
| chr15-83709732-DEL-337 | chr15-83709716-DEL-340 |
| chr15-83714753-DEL-208 | chr15-83714715-DEL-226 |
| chr15-83722412-DEL-118 | chr15-83722413-DEL-116 |
| chr15-83736641-DEL-59 | chr15-83736642-DEL-59 |
| chr15-83762770-DEL-168 | chr15-83762741-DEL-169 |
| chr15-83768917-DEL-107 | chr15-83768956-DEL-69 |
| chr15-83800240-DEL-511 | chr15-83800241-DEL-511 |
| chr15-83814312-DEL-204 | chr15-83814313-DEL-204 |
| chr15-83875345-DEL-166 | chr15-83875358-DEL-157 |
| chr15-83878662-DEL-83 | chr15-83878662-DEL-83 |
| chr15-83897334-DEL-202 | chr15-83897335-DEL-202 |
| chr15-83907232-DEL-71 | chr15-83907230-DEL-68 |
| chr15-83908657-DEL-153 | chr15-83908658-DEL-153 |
| chr15-83926455-DEL-340 | chr15-83926456-DEL-340 |
| chr15-83934620-DEL-510 | chr15-83934611-DEL-508 |
| chr15-83937007-DEL-121 | chr15-83937008-DEL-121 |
| chr15-83951826-DEL-180 | chr15-83951827-DEL-180 |
| chr15-83957060-DEL-61 | chr15-83957062-DEL-63 |
| chr15-83983995-DEL-204 | chr15-83983996-DEL-204 |
| chr15-83984783-DEL-1978 | chr15-83984784-DEL-1978 |
| chr15-84034844-DEL-198 | chr15-84034845-DEL-198 |
| chr15-84050743-DEL-216 | chr15-84050736-DEL-217 |
| chr15-84067388-DEL-76 | chr15-84067358-DEL-77 |
| chr15-84127784-DEL-202 | chr15-84127785-DEL-202 |
| chr15-84157902-DEL-228 | chr15-84157903-DEL-228 |
| chr15-84160840-DEL-203 | chr15-84160841-DEL-203 |
| chr15-84182670-DEL-83 | chr15-84182671-DEL-83 |
| chr15-84190465-DEL-220 | chr15-84190466-DEL-220 |
| chr15-84192948-DEL-54 | chr15-84192949-DEL-54 |
| chr15-84226039-DEL-229 | chr15-84226040-DEL-229 |
| chr15-84253098-DEL-201 | chr15-84253098-DEL-202 |
| chr15-84253806-DEL-845 | chr15-84253807-DEL-845 |
| chr15-84254804-DEL-444 | chr15-84254805-DEL-444 |
| chr15-84285433-DEL-129 | chr15-84285434-DEL-129 |

| | |
|---|---|
| chr15-84351425-DEL-1262 | chr15-84351426-DEL-1262 |
| chr15-84355134-DEL-53 | chr15-84355132-DEL-54 |
| chr15-84360089-DEL-126 | chr15-84360090-DEL-126 |
| chr15-84378354-DEL-95 | chr15-84378353-DEL-96 |
| chr15-84386052-DEL-98 | chr15-84386053-DEL-97 |
| chr15-84408923-DEL-329 | chr15-84408898-DEL-330 |
| chr15-84491341-DEL-645 | chr15-84491342-DEL-645 |
| chr15-84505353-DEL-140 | chr15-84505354-DEL-140 |
| chr15-84508239-DEL-51 | chr15-84508240-DEL-51 |
| chr15-84514857-DEL-227 | chr15-84514858-DEL-222 |
| chr15-84518060-DEL-98 | chr15-84518073-DEL-83 |
| chr15-84528832-DEL-129 | chr15-84528832-DEL-123 |
| chr15-84534335-DEL-312 | chr15-84534328-DEL-328 |
| chr15-84545380-DEL-192 | chr15-84545380-DEL-193 |
| chr15-84557462-DEL-68 | chr15-84557465-DEL-66 |
| chr15-84594271-DEL-124 | chr15-84594272-DEL-124 |
| chr15-84599889-DEL-134 | chr15-84599885-DEL-139 |
| chr15-84601870-DEL-317 | chr15-84601871-DEL-317 |
| chr15-84628737-DEL-985 | chr15-84628738-DEL-985 |
| chr15-84630767-DEL-6401 | chr15-84630768-DEL-6401 |
| chr15-84652341-DEL-487 | chr15-84652342-DEL-487 |
| chr15-84694220-DEL-1111 | chr15-84694221-DEL-1111 |
| chr15-84705903-DEL-1238 | chr15-84705902-DEL-1228 |
| chr15-84723085-DEL-80 | chr15-84723094-DEL-72 |
| chr15-84749703-DEL-198 | chr15-84749704-DEL-198 |
| chr15-84779140-DEL-186 | chr15-84779141-DEL-186 |
| chr15-84779963-DEL-57 | chr15-84779964-DEL-57 |
| chr15-84801732-DEL-210 | chr15-84801733-DEL-210 |
| chr15-84841309-DEL-232 | chr15-84841310-DEL-232 |
| chr15-84841867-DEL-60 | chr15-84841868-DEL-61 |
| chr15-84856870-DEL-52 | chr15-84856871-DEL-52 |
| chr15-84861858-DEL-53 | chr15-84861859-DEL-53 |
| chr15-84866649-DEL-212 | chr15-84866655-DEL-211 |
| chr15-84906229-DEL-177 | chr15-84906230-DEL-177 |
| chr15-84938634-DEL-222 | chr15-84938635-DEL-222 |
| chr15-84962568-DEL-53 | chr15-84962555-DEL-54 |
| chr15-84963236-DEL-6457 | chr15-84963235-DEL-6458 |
| chr15-84978185-DEL-739 | chr15-84978198-DEL-725 |
| chr15-84998037-DEL-191 | chr15-84998038-DEL-191 |
| chr15-85013453-DEL-72 | chr15-85013456-DEL-70 |
| chr15-85060606-DEL-57 | chr15-85060607-DEL-57 |
| chr15-85074394-DEL-6401 | chr15-85074395-DEL-6401 |
| chr15-85089973-DEL-125 | chr15-85089974-DEL-125 |
| chr15-85112303-DEL-64 | chr15-85112304-DEL-64 |
| chr15-85119806-DEL-54 | chr15-85119807-DEL-54 |
| chr15-85201994-DEL-145 | chr15-85201984-DEL-156 |
| chr15-85212491-DEL-60 | chr15-85212492-DEL-72 |
| chr15-85235512-DEL-181 | chr15-85235513-DEL-181 |
| chr15-85250226-DEL-525 | chr15-85250227-DEL-525 |
| chr15-85252857-DEL-453 | chr15-85252858-DEL-453 |
| chr15-85262486-DEL-1479 | chr15-85262487-DEL-1479 |
| chr15-85278541-DEL-76 | chr15-85278518-DEL-94 |
| chr15-85318046-DEL-220 | chr15-85318047-DEL-220 |
| chr15-85353662-DEL-100 | chr15-85353663-DEL-100 |
| chr15-85368775-DEL-564 | chr15-85368769-DEL-558 |
| chr15-85386080-DEL-199 | chr15-85386081-DEL-199 |
| chr15-85402400-DEL-55 | chr15-85402398-DEL-60 |
| chr15-85434751-DEL-52 | chr15-85434706-DEL-96 |
| chr15-85491255-DEL-208 | chr15-85491256-DEL-208 |
| chr15-85513746-DEL-87 | chr15-85513747-DEL-87 |
| chr15-85527651-DEL-54 | chr15-85527664-DEL-62 |
| chr15-85548206-DEL-171 | chr15-85548207-DEL-171 |
| chr15-85566954-DEL-66 | chr15-85566955-DEL-66 |
| chr15-85580767-DEL-208 | chr15-85580768-DEL-208 |
| chr15-85582192-DEL-69 | chr15-85582192-DEL-69 |
| chr15-85598774-DEL-52 | chr15-85598775-DEL-52 |
| chr15-85614859-DEL-60 | chr15-85614860-DEL-60 |
| chr15-85615590-DEL-302 | chr15-85615591-DEL-303 |
| chr15-85622442-DEL-68 | chr15-85622460-DEL-73 |
| chr15-85637568-DEL-307 | chr15-85637569-DEL-307 |
| chr15-85646260-DEL-51 | chr15-85646262-DEL-59 |
| chr15-85671790-DEL-67 | chr15-85671791-DEL-67 |
| chr15-85771201-DEL-848 | chr15-85771194-DEL-849 |
| chr15-85830882-DEL-62 | chr15-85830889-DEL-56 |
| chr15-85843591-DEL-417 | chr15-85843592-DEL-417 |
| chr15-85871755-DEL-123 | chr15-85871756-DEL-123 |
| chr15-85882070-DEL-70 | chr15-85882071-DEL-70 |
| chr15-85887686-DEL-106 | chr15-85887687-DEL-106 |
| chr15-85909895-DEL-75 | chr15-85909896-DEL-75 |
| chr15-85921200-DEL-221 | chr15-85921202-DEL-220 |

| | |
|---|---|
| chr15-85968189-DEL-126 | chr15-85968190-DEL-126 |
| chr15-85981032-DEL-203 | chr15-85981033-DEL-203 |
| chr15-86022521-DEL-108 | chr15-86022514-DEL-92 |
| chr15-86045560-DEL-204 | chr15-86045561-DEL-204 |
| chr15-86056695-DEL-54 | chr15-86056700-DEL-50 |
| chr15-86061761-DEL-237 | chr15-86061762-DEL-237 |
| chr15-86072025-DEL-172 | chr15-86072026-DEL-172 |
| chr15-86076444-DEL-195 | chr15-86076458-DEL-196 |
| chr15-86096853-DEL-209 | chr15-86096858-DEL-207 |
| chr15-86136538-DEL-55 | chr15-86136537-DEL-57 |
| chr15-86196458-DEL-5336 | chr15-86196459-DEL-5336 |
| chr15-86211476-DEL-211 | chr15-86211477-DEL-211 |
| chr15-86240947-DEL-52 | chr15-86240947-DEL-54 |
| chr15-86293926-DEL-538 | chr15-86293927-DEL-538 |
| chr15-86296381-DEL-60 | chr15-86296382-DEL-60 |
| chr15-86308913-DEL-69 | chr15-86308913-DEL-70 |
| chr15-86372110-DEL-102 | chr15-86372070-DEL-108 |
| chr15-86431190-DEL-56 | chr15-86431191-DEL-56 |
| chr15-86438687-DEL-64 | chr15-86438688-DEL-64 |
| chr15-86443966-DEL-446 | chr15-86443967-DEL-446 |
| chr15-86491701-DEL-62 | chr15-86491702-DEL-62 |
| chr15-86510108-DEL-472 | chr15-86510109-DEL-472 |
| chr15-86512372-DEL-80 | chr15-86512373-DEL-80 |
| chr15-86547714-DEL-51 | chr15-86547715-DEL-51 |
| chr15-86551674-DEL-52 | chr15-86551675-DEL-60 |
| chr15-86589601-DEL-98 | chr15-86589614-DEL-86 |
| chr15-86604080-DEL-218 | chr15-86604081-DEL-218 |
| chr15-86621170-DEL-64 | chr15-86621171-DEL-64 |
| chr15-86676211-DEL-182 | chr15-86676212-DEL-182 |
| chr15-86688526-DEL-105 | chr15-86688524-DEL-108 |
| chr15-86689041-DEL-88 | chr15-86689042-DEL-88 |
| chr15-86702815-DEL-294 | chr15-86702837-DEL-273 |
| chr15-86715486-DEL-120 | chr15-86715515-DEL-139 |
| chr15-86730778-DEL-62 | chr15-86730771-DEL-70 |
| chr15-86768898-DEL-100 | chr15-86768876-DEL-119 |
| chr15-86770279-DEL-52 | chr15-86770280-DEL-91 |
| chr15-86798460-DEL-72 | chr15-86798471-DEL-76 |
| chr15-86802137-DEL-59 | chr15-86802134-DEL-59 |
| chr15-86819192-DEL-1200 | chr15-86819189-DEL-1200 |
| chr15-86847356-DEL-70 | chr15-86847357-DEL-70 |
| chr15-86866868-DEL-52 | chr15-86866868-DEL-54 |
| chr15-86873416-DEL-141 | chr15-86873421-DEL-137 |
| chr15-86882017-DEL-59 | chr15-86882018-DEL-59 |
| chr15-86893606-DEL-741 | chr15-86893607-DEL-741 |
| chr15-86898324-DEL-58 | chr15-86898325-DEL-58 |
| chr15-86925401-DEL-74 | chr15-86925402-DEL-74 |
| chr15-86959697-DEL-182 | chr15-86959698-DEL-182 |
| chr15-86966444-DEL-614 | chr15-86966445-DEL-614 |
| chr15-86967737-DEL-55 | chr15-86967749-DEL-56 |
| chr15-87012440-DEL-522 | chr15-87012430-DEL-523 |
| chr15-87025021-DEL-74 | chr15-87025022-DEL-74 |
| chr15-87027380-DEL-6206 | chr15-87027381-DEL-6206 |
| chr15-87042431-DEL-136 | chr15-87042448-DEL-100 |
| chr15-87066922-DEL-241 | chr15-87066923-DEL-245 |
| chr15-87090278-DEL-76 | chr15-87090279-DEL-60 |
| chr15-87105941-DEL-85 | chr15-87105942-DEL-85 |
| chr15-87127083-DEL-54 | chr15-87127084-DEL-54 |
| chr15-87149727-DEL-60 | chr15-87149751-DEL-62 |
| chr15-87178173-DEL-7091 | chr15-87178158-DEL-7107 |
| chr15-87200223-DEL-81 | chr15-87200224-DEL-81 |
| chr15-87250549-DEL-67 | chr15-87250550-DEL-67 |
| chr15-87279798-DEL-557 | chr15-87279799-DEL-555 |
| chr15-87342242-DEL-55 | chr15-87342243-DEL-55 |
| chr15-87347536-DEL-54 | chr15-87347537-DEL-54 |
| chr15-87351224-DEL-200 | chr15-87351225-DEL-200 |
| chr15-87352894-DEL-56 | chr15-87352937-DEL-60 |
| chr15-87377074-DEL-65 | chr15-87377075-DEL-65 |
| chr15-87379093-DEL-57 | chr15-87379094-DEL-57 |
| chr15-87398759-DEL-2603 | chr15-87398760-DEL-2603 |
| chr15-87409192-DEL-58 | chr15-87409194-DEL-56 |
| chr15-87420128-DEL-58 | chr15-87420129-DEL-58 |
| chr15-87425643-DEL-89 | chr15-87425693-DEL-51 |
| chr15-87444599-DEL-56 | chr15-87444598-DEL-63 |
| chr15-87452674-DEL-53 | chr15-87452662-DEL-54 |
| chr15-87464931-DEL-126 | chr15-87464932-DEL-126 |
| chr15-87465915-DEL-239 | chr15-87465908-DEL-239 |
| chr15-87498831-DEL-451 | chr15-87498832-DEL-451 |
| chr15-87513400-DEL-162 | chr15-87513401-DEL-158 |
| chr15-87535989-DEL-72 | chr15-87535995-DEL-68 |
| chr15-87542646-DEL-3656 | chr15-87542647-DEL-3656 |

| | |
|---|---|
| chr15-87591641-DEL-72 | chr15-87591642-DEL-72 |
| chr15-87595962-DEL-60 | chr15-87595963-DEL-60 |
| chr15-87609473-DEL-97 | chr15-87609470-DEL-98 |
| chr15-87639018-DEL-196 | chr15-87639019-DEL-196 |
| chr15-87639102-DEL-1049 | chr15-87639103-DEL-1049 |
| chr15-87688945-DEL-90 | chr15-87688943-DEL-64 |
| chr15-87695814-DEL-66 | chr15-87695815-DEL-72 |
| chr15-87723371-DEL-2676 | chr15-87723372-DEL-2676 |
| chr15-87733746-DEL-60 | chr15-87733747-DEL-64 |
| chr15-87748304-DEL-1866 | chr15-87748305-DEL-1866 |
| chr15-87750637-DEL-96 | chr15-87750604-DEL-118 |
| chr15-87800318-DEL-149 | chr15-87800335-DEL-117 |
| chr15-87830395-DEL-61 | chr15-87830391-DEL-88 |
| chr15-87833707-DEL-445 | chr15-87833708-DEL-445 |
| chr15-87846134-DEL-1004 | chr15-87846129-DEL-1008 |
| chr15-87850187-DEL-63 | chr15-87850189-DEL-62 |
| chr15-87947733-DEL-2673 | chr15-87947734-DEL-2673 |
| chr15-87985383-DEL-135 | chr15-87985386-DEL-133 |
| chr15-88096124-DEL-75 | chr15-88096143-DEL-54 |
| chr15-88104310-DEL-137 | chr15-88104321-DEL-105 |
| chr15-88117051-DEL-64 | chr15-88117052-DEL-64 |
| chr15-88130620-DEL-79 | chr15-88130621-DEL-79 |
| chr15-88142425-DEL-137 | chr15-88142404-DEL-146 |
| chr15-88151390-DEL-352 | chr15-88151391-DEL-352 |
| chr15-88193814-DEL-106 | chr15-88193815-DEL-106 |
| chr15-88210586-DEL-220 | chr15-88210589-DEL-172 |
| chr15-88265557-DEL-85 | chr15-88265558-DEL-93 |
| chr15-88272027-DEL-2446 | chr15-88272029-DEL-2445 |
| chr15-88277436-DEL-60 | chr15-88277437-DEL-60 |
| chr15-88278073-DEL-116 | chr15-88278074-DEL-116 |
| chr15-88292074-DEL-79 | chr15-88292069-DEL-85 |
| chr15-88333547-DEL-56 | chr15-88333548-DEL-56 |
| chr15-88342052-DEL-106 | chr15-88342053-DEL-106 |
| chr15-88350947-DEL-498 | chr15-88350948-DEL-498 |
| chr15-88408105-DEL-119 | chr15-88408106-DEL-119 |
| chr15-88409646-DEL-394 | chr15-88409660-DEL-378 |
| chr15-88444725-DEL-327 | chr15-88444726-DEL-327 |
| chr15-88451812-DEL-450 | chr15-88451813-DEL-450 |
| chr15-88459356-DEL-1617 | chr15-88459343-DEL-1618 |
| chr15-88461760-DEL-77 | chr15-88461747-DEL-78 |
| chr15-88473285-DEL-4151 | chr15-88473286-DEL-4151 |
| chr15-88504797-DEL-359 | chr15-88504798-DEL-359 |
| chr15-88515669-DEL-68 | chr15-88515670-DEL-68 |
| chr15-88540967-DEL-111 | chr15-88540969-DEL-110 |
| chr15-88548131-DEL-1199 | chr15-88548132-DEL-1199 |
| chr15-88553835-DEL-89 | chr15-88553833-DEL-90 |
| chr15-88557618-DEL-761 | chr15-88557619-DEL-761 |
| chr15-88575271-DEL-51 | chr15-88575271-DEL-51 |
| chr15-88584721-DEL-165 | chr15-88584719-DEL-167 |
| chr15-88604597-DEL-203 | chr15-88604591-DEL-202 |
| chr15-88610128-DEL-421 | chr15-88610111-DEL-460 |
| chr15-88611282-DEL-190 | chr15-88611282-DEL-190 |
| chr15-88623031-DEL-182 | chr15-88623032-DEL-182 |
| chr15-88639974-DEL-388 | chr15-88639975-DEL-388 |
| chr15-88650856-DEL-211 | chr15-88650857-DEL-211 |
| chr15-88674179-DEL-131 | chr15-88674167-DEL-155 |
| chr15-88678875-DEL-215 | chr15-88678876-DEL-215 |
| chr15-88686026-DEL-495 | chr15-88686027-DEL-495 |
| chr15-88693557-DEL-734 | chr15-88693558-DEL-728 |
| chr15-88719487-DEL-267 | chr15-88719488-DEL-267 |
| chr15-88795346-DEL-395 | chr15-88795348-DEL-409 |
| chr15-88804171-DEL-68 | chr15-88804172-DEL-68 |
| chr15-88859238-DEL-923 | chr15-88859210-DEL-924 |
| chr15-88896185-DEL-99 | chr15-88896206-DEL-66 |
| chr15-88897918-DEL-84 | chr15-88897919-DEL-53 |
| chr15-88920563-DEL-3898 | chr15-88920564-DEL-3898 |
| chr15-88931715-DEL-80 | chr15-88931719-DEL-81 |
| chr15-89017800-DEL-155 | chr15-89017801-DEL-155 |
| chr15-89036977-DEL-60 | chr15-89036978-DEL-56 |
| chr15-89097427-DEL-192 | chr15-89097428-DEL-192 |
| chr15-89105232-DEL-231 | chr15-89105233-DEL-231 |
| chr15-89113664-DEL-420 | chr15-89113665-DEL-420 |
| chr15-89212428-DEL-73 | chr15-89212429-DEL-73 |
| chr15-89222349-DEL-435 | chr15-89222350-DEL-435 |
| chr15-89233098-DEL-193 | chr15-89233131-DEL-167 |
| chr15-89259918-DEL-179 | chr15-89259917-DEL-181 |
| chr15-89278981-DEL-154 | chr15-89278982-DEL-153 |
| chr15-89287917-DEL-61 | chr15-89287919-DEL-62 |
| chr15-89300160-DEL-216 | chr15-89300161-DEL-216 |
| chr15-89305178-DEL-98 | chr15-89305179-DEL-98 |

| | |
|---|---|
| chr15-89328155-DEL-62 | chr15-89328156-DEL-62 |
| chr15-89332060-DEL-513 | chr15-89332061-DEL-513 |
| chr15-89410527-DEL-253 | chr15-89410537-DEL-244 |
| chr15-89413154-DEL-82 | chr15-89413155-DEL-82 |
| chr15-89453238-DEL-126 | chr15-89453239-DEL-126 |
| chr15-89463825-DEL-205 | chr15-89463826-DEL-205 |
| chr15-89499302-DEL-213 | chr15-89499303-DEL-213 |
| chr15-89499797-DEL-6538 | chr15-89499798-DEL-6538 |
| chr15-89516842-DEL-194 | chr15-89516843-DEL-194 |
| chr15-89520602-DEL-211 | chr15-89520603-DEL-211 |
| chr15-89521673-DEL-343 | chr15-89521675-DEL-346 |
| chr15-89543836-DEL-6599 | chr15-89543831-DEL-6600 |
| chr15-89594464-DEL-352 | chr15-89594465-DEL-352 |
| chr15-89594995-DEL-3942 | chr15-89594996-DEL-3942 |
| chr15-89616346-DEL-148 | chr15-89616347-DEL-147 |
| chr15-89632432-DEL-1882 | chr15-89632425-DEL-1883 |
| chr15-89648299-DEL-89 | chr15-89648300-DEL-89 |
| chr15-89649515-DEL-3963 | chr15-89649516-DEL-3963 |
| chr15-89652520-DEL-398 | chr15-89652521-DEL-398 |
| chr15-89656866-DEL-93 | chr15-89656867-DEL-93 |
| chr15-89669771-DEL-1168 | chr15-89669772-DEL-1168 |
| chr15-89672201-DEL-76 | chr15-89672201-DEL-76 |
| chr15-89692421-DEL-293 | chr15-89692422-DEL-293 |
| chr15-89700553-DEL-3947 | chr15-89700554-DEL-3947 |
| chr15-89703339-DEL-71 | chr15-89703339-DEL-72 |
| chr15-89720713-DEL-60 | chr15-89720720-DEL-54 |
| chr15-89741465-DEL-1154 | chr15-89741466-DEL-1154 |
| chr15-89742907-DEL-87 | chr15-89742896-DEL-128 |
| chr15-89746758-DEL-706 | chr15-89746757-DEL-707 |
| chr15-89760642-DEL-671 | chr15-89760643-DEL-671 |
| chr15-89768732-DEL-2983 | chr15-89768733-DEL-2983 |
| chr15-89769217-DEL-167 | chr15-89769218-DEL-167 |
| chr15-89799969-DEL-158 | chr15-89799971-DEL-157 |
| chr15-89827198-DEL-8456 | chr15-89827199-DEL-8456 |
| chr15-89853930-DEL-171 | chr15-89853933-DEL-169 |
| chr15-89857297-DEL-93 | chr15-89857293-DEL-91 |
| chr15-89863895-DEL-162 | chr15-89863896-DEL-162 |
| chr15-89876911-DEL-64 | chr15-89876912-DEL-64 |
| chr15-89882726-DEL-88 | chr15-89882733-DEL-82 |
| chr15-89895355-DEL-210 | chr15-89895356-DEL-210 |
| chr15-89896974-DEL-64 | chr15-89896962-DEL-107 |
| chr15-89958622-DEL-2190 | chr15-89958623-DEL-2190 |
| chr15-89964238-DEL-202 | chr15-89964239-DEL-202 |
| chr15-90005014-DEL-174 | chr15-90005013-DEL-175 |
| chr15-90026335-DEL-403 | chr15-90026323-DEL-404 |
| chr15-90029867-DEL-2041 | chr15-90029868-DEL-2041 |
| chr15-90035708-DEL-696 | chr15-90035709-DEL-696 |
| chr15-90068238-DEL-138 | chr15-90068244-DEL-110 |
| chr15-90074073-DEL-2233 | chr15-90074074-DEL-2233 |
| chr15-90074282-DEL-107 | chr15-90074283-DEL-107 |
| chr15-90076864-DEL-6683 | chr15-90076865-DEL-6683 |
| chr15-90076992-DEL-212 | chr15-90076992-DEL-212 |
| chr15-90168070-DEL-88 | chr15-90168051-DEL-122 |
| chr15-90170331-DEL-339 | chr15-90170332-DEL-339 |
| chr15-90184905-DEL-1405 | chr15-90184905-DEL-1405 |
| chr15-90198801-DEL-272 | chr15-90198802-DEL-272 |
| chr15-90208065-DEL-132 | chr15-90208069-DEL-134 |
| chr15-90222389-DEL-58 | chr15-90222390-DEL-58 |
| chr15-90226062-DEL-68 | chr15-90226062-DEL-68 |
| chr15-90261979-DEL-62 | chr15-90261988-DEL-66 |
| chr15-90262548-DEL-6359 | chr15-90262550-DEL-6369 |
| chr15-90278398-DEL-1333 | chr15-90278399-DEL-1333 |
| chr15-90294325-DEL-387 | chr15-90294340-DEL-390 |
| chr15-90296008-DEL-837 | chr15-90296009-DEL-837 |
| chr15-90298636-DEL-2352 | chr15-90298637-DEL-2352 |
| chr15-90303564-DEL-140 | chr15-90303562-DEL-142 |
| chr15-90310845-DEL-54 | chr15-90310865-DEL-58 |
| chr15-90322076-DEL-326 | chr15-90322039-DEL-332 |
| chr15-90339236-DEL-597 | chr15-90339235-DEL-598 |
| chr15-90341644-DEL-424 | chr15-90341645-DEL-424 |
| chr15-90357936-DEL-2046 | chr15-90357937-DEL-2046 |
| chr15-90375766-DEL-82 | chr15-90375767-DEL-82 |
| chr15-90405062-DEL-119 | chr15-90405063-DEL-119 |
| chr15-90409386-DEL-180 | chr15-90409387-DEL-177 |
| chr15-90416829-DEL-196 | chr15-90416830-DEL-196 |
| chr15-90435767-DEL-198 | chr15-90435768-DEL-198 |
| chr15-90450387-DEL-192 | chr15-90450388-DEL-192 |
| chr15-90514940-DEL-410 | chr15-90514937-DEL-411 |
| chr15-90520664-DEL-7160 | chr15-90520665-DEL-7160 |
| chr15-90536854-DEL-95 | chr15-90536855-DEL-98 |

| | |
|---|---|
| chr15-90539550-DEL-66 | chr15-90539550-DEL-66 |
| chr15-90566298-DEL-56 | chr15-90566299-DEL-56 |
| chr15-90576574-DEL-257 | chr15-90576575-DEL-257 |
| chr15-90597749-DEL-55 | chr15-90597750-DEL-55 |
| chr15-90619647-DEL-220 | chr15-90619647-DEL-220 |
| chr15-90620874-DEL-500 | chr15-90620875-DEL-500 |
| chr15-90648410-DEL-538 | chr15-90648411-DEL-538 |
| chr15-90660619-DEL-276 | chr15-90660620-DEL-276 |
| chr15-90667966-DEL-95 | chr15-90667967-DEL-95 |
| chr15-90685326-DEL-76 | chr15-90685352-DEL-50 |
| chr15-90687010-DEL-72 | chr15-90687011-DEL-72 |
| chr15-90693632-DEL-255 | chr15-90693633-DEL-255 |
| chr15-90705234-DEL-179 | chr15-90705256-DEL-177 |
| chr15-90710093-DEL-947 | chr15-90710094-DEL-947 |
| chr15-90725958-DEL-385 | chr15-90725959-DEL-385 |
| chr15-90728025-DEL-60 | chr15-90728015-DEL-64 |
| chr15-90733531-DEL-78 | chr15-90733550-DEL-78 |
| chr15-90737926-DEL-903 | chr15-90737927-DEL-903 |
| chr15-90796555-DEL-137 | chr15-90796556-DEL-137 |
| chr15-90815025-DEL-155 | chr15-90814992-DEL-192 |
| chr15-90829520-DEL-1120 | chr15-90829506-DEL-1121 |
| chr15-90831882-DEL-159 | chr15-90831883-DEL-159 |
| chr15-90886788-DEL-84 | chr15-90886789-DEL-84 |
| chr15-90927266-DEL-115 | chr15-90927267-DEL-115 |
| chr15-90927469-DEL-822 | chr15-90927470-DEL-822 |
| chr15-90928980-DEL-91 | chr15-90928981-DEL-91 |
| chr15-90945376-DEL-81 | chr15-90945377-DEL-81 |
| chr15-90950197-DEL-118 | chr15-90950198-DEL-118 |
| chr15-90954755-DEL-89 | chr15-90954762-DEL-83 |
| chr15-90958570-DEL-51 | chr15-90958570-DEL-51 |
| chr15-90964125-DEL-7481 | chr15-90964125-DEL-7481 |
| chr15-90976101-DEL-270 | chr15-90976102-DEL-270 |
| chr15-90986781-DEL-189 | chr15-90986782-DEL-189 |
| chr15-90990304-DEL-54 | chr15-90990305-DEL-54 |
| chr15-90994579-DEL-216 | chr15-90994577-DEL-217 |
| chr15-90999054-DEL-7109 | chr15-90999055-DEL-7109 |
| chr15-91012459-DEL-7994 | chr15-91012460-DEL-7994 |
| chr15-91025130-DEL-385 | chr15-91025131-DEL-385 |
| chr15-91025742-DEL-2414 | chr15-91025743-DEL-2414 |
| chr15-91059354-DEL-2623 | chr15-91059355-DEL-2623 |
| chr15-91099052-DEL-719 | chr15-91099048-DEL-720 |
| chr15-91119767-DEL-52 | chr15-91119753-DEL-71 |
| chr15-91120388-DEL-294 | chr15-91120389-DEL-294 |
| chr15-91159270-DEL-268 | chr15-91159271-DEL-268 |
| chr15-91170953-DEL-551 | chr15-91170957-DEL-550 |
| chr15-91175439-DEL-187 | chr15-91175439-DEL-187 |
| chr15-91176505-DEL-184 | chr15-91176506-DEL-184 |
| chr15-91179439-DEL-232 | chr15-91179440-DEL-232 |
| chr15-91194500-DEL-188 | chr15-91194501-DEL-188 |
| chr15-91222232-DEL-52 | chr15-91222217-DEL-60 |
| chr15-91279044-DEL-182 | chr15-91279045-DEL-182 |
| chr15-91303680-DEL-2725 | chr15-91303681-DEL-2725 |
| chr15-91318068-DEL-51 | chr15-91318060-DEL-94 |
| chr15-91327303-DEL-116 | chr15-91327304-DEL-116 |
| chr15-91344533-DEL-80 | chr15-91344534-DEL-80 |
| chr15-91368969-DEL-111 | chr15-91368964-DEL-80 |
| chr15-91372785-DEL-985 | chr15-91372786-DEL-985 |
| chr15-91394519-DEL-51 | chr15-91394495-DEL-65 |
| chr15-91396185-DEL-85 | chr15-91396186-DEL-85 |
| chr15-91396735-DEL-210 | chr15-91396736-DEL-204 |
| chr15-91440773-DEL-61 | chr15-91440774-DEL-61 |
| chr15-91460017-DEL-59 | chr15-91460015-DEL-50 |
| chr15-91474967-DEL-58 | chr15-91474974-DEL-52 |
| chr15-91522868-DEL-1374 | chr15-91522869-DEL-1374 |
| chr15-91546978-DEL-443 | chr15-91546979-DEL-443 |
| chr15-91642788-DEL-57 | chr15-91642789-DEL-57 |
| chr15-91658186-DEL-120 | chr15-91658187-DEL-120 |
| chr15-91667332-DEL-58 | chr15-91667333-DEL-58 |
| chr15-91713618-DEL-122 | chr15-91713619-DEL-122 |
| chr15-91718936-DEL-64 | chr15-91718937-DEL-64 |
| chr15-91730390-DEL-72 | chr15-91730409-DEL-54 |
| chr15-91737441-DEL-54 | chr15-91737442-DEL-54 |
| chr15-91741129-DEL-114 | chr15-91741130-DEL-114 |
| chr15-91753222-DEL-58 | chr15-91753223-DEL-58 |
| chr15-91757843-DEL-323 | chr15-91757844-DEL-323 |
| chr15-91833953-DEL-3939 | chr15-91833954-DEL-3939 |
| chr15-91841329-DEL-95 | chr15-91841330-DEL-95 |
| chr15-91868610-DEL-52 | chr15-91868609-DEL-54 |
| chr15-91876722-DEL-2646 | chr15-91876723-DEL-2646 |
| chr15-91876945-DEL-68 | chr15-91876946-DEL-68 |

| | |
|---|---|
| chr15-91882696-DEL-9276 | chr15-91882697-DEL-9276 |
| chr15-91923859-DEL-2144 | chr15-91923860-DEL-2144 |
| chr15-91932174-DEL-57 | chr15-91932175-DEL-57 |
| chr15-91937389-DEL-100 | chr15-91937384-DEL-86 |
| chr15-91937814-DEL-1153 | chr15-91937815-DEL-1153 |
| chr15-91939742-DEL-84 | chr15-91939746-DEL-83 |
| chr15-91977864-DEL-201 | chr15-91977865-DEL-201 |
| chr15-91999812-DEL-77 | chr15-91999821-DEL-69 |
| chr15-92007239-DEL-168 | chr15-92007240-DEL-168 |
| chr15-92009282-DEL-61 | chr15-92009281-DEL-60 |
| chr15-92019742-DEL-63 | chr15-92019747-DEL-64 |
| chr15-92035624-DEL-100 | chr15-92035623-DEL-102 |
| chr15-92068211-DEL-206 | chr15-92068212-DEL-206 |
| chr15-92101871-DEL-67 | chr15-92101872-DEL-67 |
| chr15-92115097-DEL-74 | chr15-92115131-DEL-85 |
| chr15-92126115-DEL-244 | chr15-92126118-DEL-242 |
| chr15-92131981-DEL-58 | chr15-92131982-DEL-54 |
| chr15-92132744-DEL-82 | chr15-92132745-DEL-82 |
| chr15-92141459-DEL-58 | chr15-92141460-DEL-56 |
| chr15-92148066-DEL-62 | chr15-92148067-DEL-62 |
| chr15-92148981-DEL-814 | chr15-92148982-DEL-814 |
| chr15-92213343-DEL-87 | chr15-92213345-DEL-86 |
| chr15-92223238-DEL-53 | chr15-92223239-DEL-53 |
| chr15-92242983-DEL-168 | chr15-92242984-DEL-168 |
| chr15-92263052-DEL-82 | chr15-92263061-DEL-80 |
| chr15-92284627-DEL-88 | chr15-92284628-DEL-88 |
| chr15-92290596-DEL-66 | chr15-92290546-DEL-78 |
| chr15-92294147-DEL-54 | chr15-92294150-DEL-53 |
| chr15-92319440-DEL-93 | chr15-92319435-DEL-95 |
| chr15-92325837-DEL-58 | chr15-92325844-DEL-52 |
| chr15-92337249-DEL-53 | chr15-92337249-DEL-53 |
| chr15-92337714-DEL-288 | chr15-92337715-DEL-276 |
| chr15-92355555-DEL-144 | chr15-92355556-DEL-144 |
| chr15-92358422-DEL-203 | chr15-92358420-DEL-204 |
| chr15-92360436-DEL-52 | chr15-92360437-DEL-52 |
| chr15-92368085-DEL-4853 | chr15-92368086-DEL-4853 |
| chr15-92410647-DEL-82 | chr15-92410648-DEL-79 |
| chr15-92421744-DEL-71 | chr15-92421745-DEL-71 |
| chr15-92442744-DEL-1684 | chr15-92442745-DEL-1684 |
| chr15-92458047-DEL-51 | chr15-92458048-DEL-51 |
| chr15-92574395-DEL-55 | chr15-92574395-DEL-52 |
| chr15-92604090-DEL-470 | chr15-92604091-DEL-470 |
| chr15-92620575-DEL-7470 | chr15-92620576-DEL-7470 |
| chr15-92637163-DEL-179 | chr15-92637169-DEL-172 |
| chr15-92638516-DEL-247 | chr15-92638517-DEL-247 |
| chr15-92660414-DEL-1621 | chr15-92660413-DEL-1622 |
| chr15-92678628-DEL-51 | chr15-92678629-DEL-51 |
| chr15-92694027-DEL-2739 | chr15-92694028-DEL-2739 |
| chr15-92711429-DEL-6652 | chr15-92711419-DEL-6653 |
| chr15-92722011-DEL-94 | chr15-92722012-DEL-94 |
| chr15-92737938-DEL-65 | chr15-92737939-DEL-65 |
| chr15-92791979-DEL-162 | chr15-92791980-DEL-161 |
| chr15-92820432-DEL-194 | chr15-92820433-DEL-194 |
| chr15-92836977-DEL-207 | chr15-92836977-DEL-207 |
| chr15-92846586-DEL-76 | chr15-92846587-DEL-76 |
| chr15-92855528-DEL-152 | chr15-92855529-DEL-152 |
| chr15-92856437-DEL-410 | chr15-92856438-DEL-410 |
| chr15-92885452-DEL-157 | chr15-92885416-DEL-163 |
| chr15-92911976-DEL-498 | chr15-92911977-DEL-498 |
| chr15-92921283-DEL-83 | chr15-92921279-DEL-84 |
| chr15-92930929-DEL-78 | chr15-92930945-DEL-54 |
| chr15-92941936-DEL-6595 | chr15-92941937-DEL-6595 |
| chr15-93008680-DEL-199 | chr15-93008672-DEL-200 |
| chr15-93012515-DEL-78 | chr15-93012515-DEL-79 |
| chr15-93035446-DEL-357 | chr15-93035447-DEL-357 |
| chr15-93039673-DEL-347 | chr15-93039665-DEL-342 |
| chr15-93064809-DEL-294 | chr15-93064810-DEL-294 |
| chr15-93067608-DEL-6477 | chr15-93067608-DEL-6477 |
| chr15-93087160-DEL-81 | chr15-93087154-DEL-82 |
| chr15-93094387-DEL-1302 | chr15-93094388-DEL-1302 |
| chr15-93132226-DEL-255 | chr15-93132227-DEL-255 |
| chr15-93144388-DEL-203 | chr15-93144389-DEL-203 |
| chr15-93170395-DEL-51 | chr15-93170395-DEL-51 |
| chr15-93232203-DEL-74 | chr15-93232172-DEL-61 |
| chr15-93241167-DEL-592 | chr15-93241168-DEL-592 |
| chr15-93265930-DEL-293 | chr15-93265931-DEL-293 |
| chr15-93270455-DEL-874 | chr15-93270456-DEL-874 |
| chr15-93271037-DEL-210 | chr15-93271037-DEL-210 |
| chr15-93277167-DEL-52 | chr15-93277168-DEL-54 |
| chr15-93281151-DEL-52 | chr15-93281152-DEL-52 |

| | |
|---|---|
| chr15-93281747-DEL-1759 | chr15-93281748-DEL-1759 |
| chr15-93283085-DEL-54 | chr15-93283086-DEL-54 |
| chr15-93287162-DEL-145 | chr15-93287157-DEL-146 |
| chr15-93289914-DEL-2388 | chr15-93289915-DEL-2388 |
| chr15-93320965-DEL-236 | chr15-93320966-DEL-236 |
| chr15-93342610-DEL-446 | chr15-93342611-DEL-446 |
| chr15-93351718-DEL-52 | chr15-93351726-DEL-58 |
| chr15-93357121-DEL-214 | chr15-93357122-DEL-214 |
| chr15-93404166-DEL-218 | chr15-93404169-DEL-218 |
| chr15-93423928-DEL-231 | chr15-93423929-DEL-231 |
| chr15-93433307-DEL-207 | chr15-93433308-DEL-207 |
| chr15-93444658-DEL-68 | chr15-93444659-DEL-68 |
| chr15-93462264-DEL-76 | chr15-93462265-DEL-76 |
| chr15-93476054-DEL-59 | chr15-93476049-DEL-60 |
| chr15-93483301-DEL-71 | chr15-93483289-DEL-69 |
| chr15-93484633-DEL-66 | chr15-93484635-DEL-65 |
| chr15-93504631-DEL-204 | chr15-93504620-DEL-199 |
| chr15-93522373-DEL-72 | chr15-93522374-DEL-72 |
| chr15-93537317-DEL-68 | chr15-93537318-DEL-68 |
| chr15-93576472-DEL-1440 | chr15-93576472-DEL-1439 |
| chr15-93589478-DEL-2900 | chr15-93589479-DEL-2900 |
| chr15-93595403-DEL-56 | chr15-93595404-DEL-56 |
| chr15-93631052-DEL-54 | chr15-93631053-DEL-54 |
| chr15-93648042-DEL-84 | chr15-93648040-DEL-84 |
| chr15-93700579-DEL-540 | chr15-93700580-DEL-540 |
| chr15-93710241-DEL-148 | chr15-93710242-DEL-148 |
| chr15-93720506-DEL-63 | chr15-93720507-DEL-63 |
| chr15-93751899-DEL-398 | chr15-93751900-DEL-398 |
| chr15-93764925-DEL-6496 | chr15-93764926-DEL-6496 |
| chr15-93776626-DEL-52 | chr15-93776627-DEL-51 |
| chr15-93779096-DEL-54 | chr15-93779099-DEL-52 |
| chr15-93781564-DEL-98 | chr15-93781575-DEL-85 |
| chr15-93784739-DEL-400 | chr15-93784740-DEL-400 |
| chr15-93807719-DEL-614 | chr15-93807719-DEL-614 |
| chr15-93809394-DEL-1006 | chr15-93809395-DEL-1006 |
| chr15-93816518-DEL-109 | chr15-93816555-DEL-73 |
| chr15-93821271-DEL-53 | chr15-93821263-DEL-54 |
| chr15-93821992-DEL-54 | chr15-93821987-DEL-67 |
| chr15-93824446-DEL-243 | chr15-93824403-DEL-278 |
| chr15-93832885-DEL-100 | chr15-93832886-DEL-100 |
| chr15-93859042-DEL-104 | chr15-93859043-DEL-104 |
| chr15-93864769-DEL-94 | chr15-93864791-DEL-107 |
| chr15-93866116-DEL-901 | chr15-93866117-DEL-901 |
| chr15-93897918-DEL-56 | chr15-93897919-DEL-56 |
| chr15-93910802-DEL-64 | chr15-93910803-DEL-64 |
| chr15-93911587-DEL-446 | chr15-93911588-DEL-446 |
| chr15-93948832-DEL-3567 | chr15-93948833-DEL-3567 |
| chr15-93955319-DEL-5008 | chr15-93955320-DEL-5008 |
| chr15-93984537-DEL-398 | chr15-93984538-DEL-398 |
| chr15-93996103-DEL-2569 | chr15-93996104-DEL-2569 |
| chr15-94007339-DEL-62 | chr15-94007338-DEL-63 |
| chr15-94027673-DEL-86 | chr15-94027694-DEL-84 |
| chr15-94044630-DEL-2556 | chr15-94044631-DEL-2556 |
| chr15-94048326-DEL-6430 | chr15-94048327-DEL-6430 |
| chr15-94068540-DEL-69 | chr15-94068495-DEL-115 |
| chr15-94100657-DEL-118 | chr15-94100668-DEL-108 |
| chr15-94170287-DEL-201 | chr15-94170294-DEL-200 |
| chr15-94193311-DEL-6625 | chr15-94193312-DEL-6625 |
| chr15-94203135-DEL-197 | chr15-94203136-DEL-197 |
| chr15-94204156-DEL-676 | chr15-94204157-DEL-676 |
| chr15-94207890-DEL-125 | chr15-94207891-DEL-125 |
| chr15-94223005-DEL-206 | chr15-94223012-DEL-158 |
| chr15-94229620-DEL-106 | chr15-94229626-DEL-111 |
| chr15-94242589-DEL-67 | chr15-94242594-DEL-65 |
| chr15-94246064-DEL-73 | chr15-94246067-DEL-77 |
| chr15-94254367-DEL-240 | chr15-94254371-DEL-240 |
| chr15-94257138-DEL-852 | chr15-94257139-DEL-852 |
| chr15-94260754-DEL-53 | chr15-94260755-DEL-53 |
| chr15-94270864-DEL-6562 | chr15-94270865-DEL-6562 |
| chr15-94287897-DEL-1445 | chr15-94287898-DEL-1444 |
| chr15-94297563-DEL-191 | chr15-94297601-DEL-154 |
| chr15-94319705-DEL-55 | chr15-94319711-DEL-56 |
| chr15-94321999-DEL-1417 | chr15-94322001-DEL-1414 |
| chr15-94335219-DEL-57 | chr15-94335220-DEL-57 |
| chr15-94336996-DEL-3407 | chr15-94337005-DEL-3406 |
| chr15-94337950-DEL-627 | chr15-94337950-DEL-627 |
| chr15-94371902-DEL-856 | chr15-94371903-DEL-856 |
| chr15-94395814-DEL-509 | chr15-94395815-DEL-509 |
| chr15-94397464-DEL-358 | chr15-94397450-DEL-354 |
| chr15-94400090-DEL-135 | chr15-94400090-DEL-135 |

| | |
|---|---|
| chr15-94414504-DEL-54 | chr15-94414505-DEL-54 |
| chr15-94437512-DEL-62 | chr15-94437513-DEL-62 |
| chr15-94469724-DEL-164 | chr15-94469725-DEL-164 |
| chr15-94481850-DEL-56 | chr15-94481851-DEL-56 |
| chr15-94502788-DEL-60 | chr15-94502814-DEL-60 |
| chr15-94506509-DEL-7118 | chr15-94506510-DEL-7118 |
| chr15-94515205-DEL-552 | chr15-94515205-DEL-551 |
| chr15-94517263-DEL-88 | chr15-94517264-DEL-88 |
| chr15-94530225-DEL-139 | chr15-94530212-DEL-147 |
| chr15-94532914-DEL-470 | chr15-94532918-DEL-469 |
| chr15-94536989-DEL-52 | chr15-94536967-DEL-67 |
| chr15-94543070-DEL-268 | chr15-94543071-DEL-268 |
| chr15-94544242-DEL-82 | chr15-94544223-DEL-82 |
| chr15-94563489-DEL-56 | chr15-94563490-DEL-56 |
| chr15-94612400-DEL-126 | chr15-94612401-DEL-126 |
| chr15-94617208-DEL-131 | chr15-94617206-DEL-132 |
| chr15-94617637-DEL-309 | chr15-94617639-DEL-309 |
| chr15-94633399-DEL-118 | chr15-94633409-DEL-138 |
| chr15-94644413-DEL-86 | chr15-94644415-DEL-72 |
| chr15-94661925-DEL-711 | chr15-94661926-DEL-711 |
| chr15-94676100-DEL-156 | chr15-94676071-DEL-194 |
| chr15-94694886-DEL-162 | chr15-94694887-DEL-162 |
| chr15-94713314-DEL-2300 | chr15-94713315-DEL-2300 |
| chr15-94720864-DEL-68 | chr15-94720865-DEL-68 |
| chr15-94733183-DEL-72 | chr15-94733195-DEL-96 |
| chr15-94741603-DEL-90 | chr15-94741604-DEL-90 |
| chr15-94752974-DEL-191 | chr15-94752974-DEL-192 |
| chr15-94803716-DEL-59 | chr15-94803715-DEL-67 |
| chr15-94804231-DEL-60 | chr15-94804232-DEL-60 |
| chr15-94816958-DEL-88 | chr15-94816967-DEL-80 |
| chr15-94852589-DEL-103 | chr15-94852590-DEL-103 |
| chr15-94861507-DEL-155 | chr15-94861508-DEL-155 |
| chr15-94891185-DEL-89 | chr15-94891171-DEL-89 |
| chr15-94891978-DEL-146 | chr15-94892001-DEL-114 |
| chr15-94906156-DEL-1117 | chr15-94906157-DEL-1117 |
| chr15-94935003-DEL-52 | chr15-94935004-DEL-52 |
| chr15-94975199-DEL-61 | chr15-94975200-DEL-52 |
| chr15-94985400-DEL-6147 | chr15-94985401-DEL-6147 |
| chr15-95011513-DEL-95 | chr15-95011509-DEL-68 |
| chr15-95049808-DEL-2140 | chr15-95049809-DEL-2140 |
| chr15-95058058-DEL-386 | chr15-95058059-DEL-386 |
| chr15-95065067-DEL-1128 | chr15-95065068-DEL-1128 |
| chr15-95070363-DEL-91 | chr15-95070369-DEL-90 |
| chr15-95075626-DEL-179 | chr15-95075627-DEL-179 |
| chr15-95105829-DEL-69 | chr15-95105829-DEL-69 |
| chr15-95106938-DEL-62 | chr15-95106939-DEL-62 |
| chr15-95159751-DEL-170 | chr15-95159752-DEL-170 |
| chr15-95177650-DEL-56 | chr15-95177651-DEL-56 |
| chr15-95181430-DEL-219 | chr15-95181431-DEL-219 |
| chr15-95203852-DEL-6369 | chr15-95203853-DEL-6369 |
| chr15-95227027-DEL-65 | chr15-95227026-DEL-66 |
| chr15-95229675-DEL-75 | chr15-95229676-DEL-75 |
| chr15-95273088-DEL-278 | chr15-95273089-DEL-272 |
| chr15-95309357-DEL-94 | chr15-95309358-DEL-94 |
| chr15-95366883-DEL-74 | chr15-95366874-DEL-83 |
| chr15-95431210-DEL-52 | chr15-95431211-DEL-52 |
| chr15-95444315-DEL-89 | chr15-95444316-DEL-89 |
| chr15-95455986-DEL-72 | chr15-95455987-DEL-72 |
| chr15-95475195-DEL-2073 | chr15-95475196-DEL-2073 |
| chr15-95476407-DEL-2285 | chr15-95476405-DEL-2286 |
| chr15-95492933-DEL-54 | chr15-95492934-DEL-54 |
| chr15-95510324-DEL-172 | chr15-95510347-DEL-140 |
| chr15-95567602-DEL-2639 | chr15-95567603-DEL-2639 |
| chr15-95570410-DEL-86 | chr15-95570411-DEL-86 |
| chr15-95582815-DEL-1282 | chr15-95582816-DEL-1282 |
| chr15-95592197-DEL-81 | chr15-95592198-DEL-81 |
| chr15-95592852-DEL-500 | chr15-95592853-DEL-501 |
| chr15-95610787-DEL-90 | chr15-95610806-DEL-98 |
| chr15-95616261-DEL-73 | chr15-95616262-DEL-73 |
| chr15-95626353-DEL-6995 | chr15-95626354-DEL-6995 |
| chr15-95651189-DEL-419 | chr15-95651190-DEL-419 |
| chr15-95652160-DEL-2187 | chr15-95652161-DEL-2187 |
| chr15-95661111-DEL-165 | chr15-95661106-DEL-171 |
| chr15-95665436-DEL-56 | chr15-95665437-DEL-56 |
| chr15-95669306-DEL-64 | chr15-95669307-DEL-64 |
| chr15-95687311-DEL-326 | chr15-95687312-DEL-326 |
| chr15-95726360-DEL-56 | chr15-95726360-DEL-58 |
| chr15-95731195-DEL-371 | chr15-95731196-DEL-371 |
| chr15-95750044-DEL-199 | chr15-95750044-DEL-200 |
| chr15-95750765-DEL-1078 | chr15-95750758-DEL-1078 |

| | |
|---|---|
| chr15-95771496-DEL-105 | chr15-95771497-DEL-105 |
| chr15-95789746-DEL-204 | chr15-95789747-DEL-204 |
| chr15-95815572-DEL-782 | chr15-95815573-DEL-782 |
| chr15-95864709-DEL-223 | chr15-95864710-DEL-223 |
| chr15-95876660-DEL-1383 | chr15-95876652-DEL-1381 |
| chr15-95901954-DEL-2247 | chr15-95901957-DEL-2213 |
| chr15-95913745-DEL-431 | chr15-95913744-DEL-432 |
| chr15-95915594-DEL-58 | chr15-95915595-DEL-58 |
| chr15-95925516-DEL-52 | chr15-95925517-DEL-52 |
| chr15-95969160-DEL-148 | chr15-95969196-DEL-98 |
| chr15-96028274-DEL-75 | chr15-96028275-DEL-75 |
| chr15-96038743-DEL-184 | chr15-96038745-DEL-182 |
| chr15-96047213-DEL-138 | chr15-96047214-DEL-138 |
| chr15-96048273-DEL-861 | chr15-96048274-DEL-861 |
| chr15-96055701-DEL-82 | chr15-96055701-DEL-50 |
| chr15-96084424-DEL-152 | chr15-96084425-DEL-152 |
| chr15-96086458-DEL-685 | chr15-96086459-DEL-685 |
| chr15-96096628-DEL-66 | chr15-96096629-DEL-66 |
| chr15-96103032-DEL-69 | chr15-96103033-DEL-69 |
| chr15-96105159-DEL-60 | chr15-96105160-DEL-60 |
| chr15-96116906-DEL-79 | chr15-96116907-DEL-79 |
| chr15-96128875-DEL-231 | chr15-96128868-DEL-232 |
| chr15-96134943-DEL-112 | chr15-96134944-DEL-112 |
| chr15-96145113-DEL-97 | chr15-96145114-DEL-97 |
| chr15-96163435-DEL-55 | chr15-96163436-DEL-55 |
| chr15-96192239-DEL-51 | chr15-96192240-DEL-51 |
| chr15-96218981-DEL-235 | chr15-96218982-DEL-235 |
| chr15-96229364-DEL-94 | chr15-96229365-DEL-94 |
| chr15-96274924-DEL-65 | chr15-96274925-DEL-65 |
| chr15-96299337-DEL-55 | chr15-96299339-DEL-54 |
| chr15-96306006-DEL-185 | chr15-96306007-DEL-185 |
| chr15-96321628-DEL-221 | chr15-96321629-DEL-221 |
| chr15-96347875-DEL-232 | chr15-96347876-DEL-232 |
| chr15-96350361-DEL-231 | chr15-96350362-DEL-231 |
| chr15-96372959-DEL-197 | chr15-96372960-DEL-197 |
| chr15-96375388-DEL-77 | chr15-96375373-DEL-93 |
| chr15-96382490-DEL-180 | chr15-96382486-DEL-181 |
| chr15-96386765-DEL-57 | chr15-96386766-DEL-57 |
| chr15-96421726-DEL-60 | chr15-96421726-DEL-60 |
| chr15-96438905-DEL-399 | chr15-96438906-DEL-399 |
| chr15-96450900-DEL-6442 | chr15-96450901-DEL-6442 |
| chr15-96459412-DEL-516 | chr15-96459413-DEL-516 |
| chr15-96478473-DEL-2775 | chr15-96478474-DEL-2775 |
| chr15-96486148-DEL-89 | chr15-96486149-DEL-89 |
| chr15-96503255-DEL-59 | chr15-96503256-DEL-55 |
| chr15-96510564-DEL-73 | chr15-96510565-DEL-73 |
| chr15-96522761-DEL-101 | chr15-96522762-DEL-101 |
| chr15-96555939-DEL-55 | chr15-96555940-DEL-55 |
| chr15-96561107-DEL-392 | chr15-96561108-DEL-392 |
| chr15-96611858-DEL-201 | chr15-96611859-DEL-201 |
| chr15-96632779-DEL-122 | chr15-96632776-DEL-123 |
| chr15-96638149-DEL-165 | chr15-96638150-DEL-165 |
| chr15-96650415-DEL-60 | chr15-96650415-DEL-60 |
| chr15-96658689-DEL-105 | chr15-96658722-DEL-69 |
| chr15-96704644-DEL-645 | chr15-96704645-DEL-645 |
| chr15-96717480-DEL-82 | chr15-96717481-DEL-84 |
| chr15-96739522-DEL-75 | chr15-96739491-DEL-110 |
| chr15-96782433-DEL-58 | chr15-96782434-DEL-58 |
| chr15-96786391-DEL-219 | chr15-96786392-DEL-219 |
| chr15-96786842-DEL-7179 | chr15-96786843-DEL-7179 |
| chr15-96802642-DEL-186 | chr15-96802643-DEL-170 |
| chr15-96807536-DEL-76 | chr15-96807537-DEL-76 |
| chr15-96808734-DEL-74 | chr15-96808751-DEL-58 |
| chr15-96821327-DEL-216 | chr15-96821328-DEL-216 |
| chr15-96849278-DEL-70 | chr15-96849283-DEL-65 |
| chr15-96875948-DEL-5374 | chr15-96875949-DEL-5374 |
| chr15-96890503-DEL-199 | chr15-96890504-DEL-199 |
| chr15-96892488-DEL-115 | chr15-96892489-DEL-115 |
| chr15-96961867-DEL-88 | chr15-96961832-DEL-90 |
| chr15-96978480-DEL-199 | chr15-96978481-DEL-199 |
| chr15-96979439-DEL-187 | chr15-96979440-DEL-187 |
| chr15-97022226-DEL-134 | chr15-97022273-DEL-88 |
| chr15-97040505-DEL-57 | chr15-97040506-DEL-57 |
| chr15-97047315-DEL-180 | chr15-97047316-DEL-180 |
| chr15-97054548-DEL-525 | chr15-97054549-DEL-525 |
| chr15-97056210-DEL-66 | chr15-97056241-DEL-62 |
| chr15-97069442-DEL-51 | chr15-97069439-DEL-52 |
| chr15-97070079-DEL-54 | chr15-97070082-DEL-55 |
| chr15-97090759-DEL-101 | chr15-97090775-DEL-67 |
| chr15-97097187-DEL-75 | chr15-97097188-DEL-75 |

| | |
|---|---|
| chr15-97106436-DEL-198 | chr15-97106437-DEL-198 |
| chr15-97107276-DEL-485 | chr15-97107277-DEL-485 |
| chr15-97113803-DEL-178 | chr15-97113804-DEL-178 |
| chr15-97161675-DEL-58 | chr15-97161676-DEL-50 |
| chr15-97165997-DEL-56 | chr15-97165997-DEL-56 |
| chr15-97179036-DEL-64 | chr15-97179037-DEL-64 |
| chr15-97223041-DEL-75 | chr15-97223042-DEL-75 |
| chr15-97225823-DEL-174 | chr15-97225824-DEL-174 |
| chr15-97234947-DEL-76 | chr15-97234941-DEL-82 |
| chr15-97239204-DEL-206 | chr15-97239193-DEL-205 |
| chr15-97252013-DEL-68 | chr15-97252014-DEL-68 |
| chr15-97252785-DEL-218 | chr15-97252788-DEL-238 |
| chr15-97256841-DEL-55 | chr15-97256842-DEL-55 |
| chr15-97261338-DEL-866 | chr15-97261339-DEL-866 |
| chr15-97261980-DEL-187 | chr15-97261981-DEL-187 |
| chr15-97263461-DEL-208 | chr15-97263462-DEL-208 |
| chr15-97290451-DEL-72 | chr15-97290452-DEL-72 |
| chr15-97291180-DEL-679 | chr15-97291181-DEL-679 |
| chr15-97304487-DEL-261 | chr15-97304488-DEL-261 |
| chr15-97312339-DEL-249 | chr15-97312340-DEL-248 |
| chr15-97325566-DEL-206 | chr15-97325567-DEL-206 |
| chr15-97349180-DEL-90 | chr15-97349181-DEL-90 |
| chr15-97349703-DEL-116 | chr15-97349704-DEL-116 |
| chr15-97349869-DEL-1147 | chr15-97349870-DEL-1147 |
| chr15-97354996-DEL-184 | chr15-97354998-DEL-183 |
| chr15-97370788-DEL-5447 | chr15-97370789-DEL-5447 |
| chr15-97377906-DEL-7250 | chr15-97377907-DEL-7250 |
| chr15-97387385-DEL-171 | chr15-97387386-DEL-171 |
| chr15-97390064-DEL-549 | chr15-97390065-DEL-550 |
| chr15-97417590-DEL-226 | chr15-97417591-DEL-226 |
| chr15-97424690-DEL-203 | chr15-97424691-DEL-203 |
| chr15-97435528-DEL-95 | chr15-97435528-DEL-81 |
| chr15-97448003-DEL-7421 | chr15-97448004-DEL-7421 |
| chr15-97499765-DEL-50 | chr15-97499744-DEL-72 |
| chr15-97508123-DEL-167 | chr15-97508124-DEL-167 |
| chr15-97515066-DEL-107 | chr15-97515094-DEL-80 |
| chr15-97515777-DEL-63 | chr15-97515778-DEL-63 |
| chr15-97518964-DEL-92 | chr15-97519014-DEL-50 |
| chr15-97520073-DEL-76 | chr15-97520074-DEL-76 |
| chr15-97521731-DEL-89 | chr15-97521732-DEL-89 |
| chr15-97544965-DEL-104 | chr15-97544965-DEL-104 |
| chr15-97546992-DEL-390 | chr15-97546993-DEL-390 |
| chr15-97569113-DEL-729 | chr15-97569114-DEL-729 |
| chr15-97577476-DEL-56 | chr15-97577476-DEL-56 |
| chr15-97582554-DEL-218 | chr15-97582555-DEL-218 |
| chr15-97590097-DEL-58 | chr15-97590100-DEL-56 |
| chr15-97631395-DEL-82 | chr15-97631396-DEL-82 |
| chr15-97631907-DEL-2684 | chr15-97631908-DEL-2684 |
| chr15-97642568-DEL-54 | chr15-97642567-DEL-52 |
| chr15-97661436-DEL-126 | chr15-97661437-DEL-125 |
| chr15-97663927-DEL-66 | chr15-97663924-DEL-70 |
| chr15-97666927-DEL-611 | chr15-97666928-DEL-611 |
| chr15-97672549-DEL-108 | chr15-97672555-DEL-96 |
| chr15-97701184-DEL-174 | chr15-97701179-DEL-172 |
| chr15-97701873-DEL-168 | chr15-97701874-DEL-168 |
| chr15-97763967-DEL-254 | chr15-97763968-DEL-254 |
| chr15-97784916-DEL-194 | chr15-97784917-DEL-194 |
| chr15-97810173-DEL-77 | chr15-97810173-DEL-77 |
| chr15-97822222-DEL-88 | chr15-97822223-DEL-90 |
| chr15-97840927-DEL-99 | chr15-97840928-DEL-99 |
| chr15-97844666-DEL-1026 | chr15-97844667-DEL-1026 |
| chr15-97888520-DEL-56 | chr15-97888535-DEL-63 |
| chr15-97929907-DEL-60 | chr15-97929908-DEL-60 |
| chr15-97935469-DEL-151 | chr15-97935423-DEL-152 |
| chr15-97949016-DEL-232 | chr15-97949017-DEL-232 |
| chr15-97980805-DEL-86 | chr15-97980806-DEL-86 |
| chr15-97998169-DEL-1880 | chr15-97998170-DEL-1880 |
| chr15-98028506-DEL-53 | chr15-98028507-DEL-53 |
| chr15-98039366-DEL-68 | chr15-98039367-DEL-68 |
| chr15-98041467-DEL-210 | chr15-98041468-DEL-210 |
| chr15-98044302-DEL-547 | chr15-98044303-DEL-547 |
| chr15-98051865-DEL-65 | chr15-98051866-DEL-63 |
| chr15-98077855-DEL-79 | chr15-98077835-DEL-95 |
| chr15-98095569-DEL-552 | chr15-98095570-DEL-538 |
| chr15-98097457-DEL-132 | chr15-98097458-DEL-132 |
| chr15-98121338-DEL-55 | chr15-98121339-DEL-55 |
| chr15-98137531-DEL-1147 | chr15-98137532-DEL-1147 |
| chr15-98143713-DEL-60 | chr15-98143671-DEL-60 |
| chr15-98158827-DEL-72 | chr15-98158831-DEL-69 |
| chr15-98163175-DEL-2698 | chr15-98163176-DEL-2698 |

| | |
|---|---|
| chr15-98164381-DEL-59 | chr15-98164369-DEL-54 |
| chr15-98165244-DEL-72 | chr15-98165245-DEL-72 |
| chr15-98205871-DEL-915 | chr15-98205871-DEL-915 |
| chr15-98226976-DEL-450 | chr15-98226977-DEL-450 |
| chr15-98252380-DEL-166 | chr15-98252344-DEL-160 |
| chr15-98253331-DEL-2688 | chr15-98253332-DEL-2688 |
| chr15-98265842-DEL-233 | chr15-98265841-DEL-235 |
| chr15-98269928-DEL-63 | chr15-98269932-DEL-70 |
| chr15-98270571-DEL-1066 | chr15-98270571-DEL-1064 |
| chr15-98283865-DEL-128 | chr15-98283873-DEL-121 |
| chr15-98291522-DEL-384 | chr15-98291523-DEL-384 |
| chr15-98293689-DEL-401 | chr15-98293690-DEL-401 |
| chr15-98300643-DEL-2140 | chr15-98300644-DEL-2140 |
| chr15-98305410-DEL-416 | chr15-98305411-DEL-404 |
| chr15-98309527-DEL-83 | chr15-98309528-DEL-83 |
| chr15-98315426-DEL-524 | chr15-98315427-DEL-524 |
| chr15-98316589-DEL-3975 | chr15-98316588-DEL-3975 |
| chr15-98322114-DEL-2052 | chr15-98322115-DEL-2052 |
| chr15-98327997-DEL-89 | chr15-98327997-DEL-90 |
| chr15-98332587-DEL-108 | chr15-98332582-DEL-94 |
| chr15-98339270-DEL-146 | chr15-98339282-DEL-133 |
| chr15-98339876-DEL-100 | chr15-98339879-DEL-98 |
| chr15-98384880-DEL-468 | chr15-98384881-DEL-468 |
| chr15-98393264-DEL-202 | chr15-98393265-DEL-202 |
| chr15-98394961-DEL-110 | chr15-98394962-DEL-110 |
| chr15-98395993-DEL-318 | chr15-98395994-DEL-318 |
| chr15-98408326-DEL-76 | chr15-98408340-DEL-57 |
| chr15-98412154-DEL-58 | chr15-98412155-DEL-58 |
| chr15-98421656-DEL-54 | chr15-98421657-DEL-54 |
| chr15-98481047-DEL-85 | chr15-98481048-DEL-85 |
| chr15-98483727-DEL-53 | chr15-98483734-DEL-53 |
| chr15-98484379-DEL-64 | chr15-98484381-DEL-63 |
| chr15-98536039-DEL-215 | chr15-98536040-DEL-215 |
| chr15-98558042-DEL-204 | chr15-98558043-DEL-204 |
| chr15-98561103-DEL-189 | chr15-98561109-DEL-195 |
| chr15-98566637-DEL-251 | chr15-98566638-DEL-251 |
| chr15-98569944-DEL-57 | chr15-98569945-DEL-57 |
| chr15-98591953-DEL-114 | chr15-98591963-DEL-110 |
| chr15-98610102-DEL-400 | chr15-98610103-DEL-400 |
| chr15-98615969-DEL-210 | chr15-98615970-DEL-210 |
| chr15-98681611-DEL-203 | chr15-98681612-DEL-203 |
| chr15-98712748-DEL-75 | chr15-98712749-DEL-75 |
| chr15-98748886-DEL-60 | chr15-98748887-DEL-60 |
| chr15-98772278-DEL-182 | chr15-98772279-DEL-182 |
| chr15-98785828-DEL-208 | chr15-98785829-DEL-208 |
| chr15-98788185-DEL-218 | chr15-98788186-DEL-218 |
| chr15-98797585-DEL-196 | chr15-98797586-DEL-196 |
| chr15-98823598-DEL-441 | chr15-98823599-DEL-441 |
| chr15-98826305-DEL-83 | chr15-98826309-DEL-80 |
| chr15-98834517-DEL-78 | chr15-98834525-DEL-56 |
| chr15-98843404-DEL-342 | chr15-98843405-DEL-342 |
| chr15-98859987-DEL-177 | chr15-98859988-DEL-177 |
| chr15-98862436-DEL-480 | chr15-98862437-DEL-480 |
| chr15-98865853-DEL-202 | chr15-98865854-DEL-202 |
| chr15-98892669-DEL-256 | chr15-98892670-DEL-256 |
| chr15-98907915-DEL-68 | chr15-98907916-DEL-68 |
| chr15-98911947-DEL-373 | chr15-98911948-DEL-373 |
| chr15-98918523-DEL-395 | chr15-98918524-DEL-395 |
| chr15-98920541-DEL-192 | chr15-98920553-DEL-196 |
| chr15-98923952-DEL-99 | chr15-98923953-DEL-99 |
| chr15-98944447-DEL-205 | chr15-98944448-DEL-205 |
| chr15-98959261-DEL-236 | chr15-98959262-DEL-236 |
| chr15-98968496-DEL-209 | chr15-98968490-DEL-207 |
| chr15-98974076-DEL-197 | chr15-98974075-DEL-198 |
| chr15-98992106-DEL-373 | chr15-98992124-DEL-374 |
| chr15-99001377-DEL-196 | chr15-99001378-DEL-196 |
| chr15-99003692-DEL-173 | chr15-99003700-DEL-166 |
| chr15-99004391-DEL-491 | chr15-99004392-DEL-491 |
| chr15-99013261-DEL-171 | chr15-99013262-DEL-171 |
| chr15-99017019-DEL-564 | chr15-99017020-DEL-564 |
| chr15-99018032-DEL-72 | chr15-99018033-DEL-69 |
| chr15-99045993-DEL-76 | chr15-99045978-DEL-92 |
| chr15-99056310-DEL-660 | chr15-99056313-DEL-658 |
| chr15-99057269-DEL-205 | chr15-99057270-DEL-205 |
| chr15-99060546-DEL-51 | chr15-99060547-DEL-51 |
| chr15-99066533-DEL-230 | chr15-99066534-DEL-230 |
| chr15-99083621-DEL-77 | chr15-99083614-DEL-65 |
| chr15-99087886-DEL-69 | chr15-99087887-DEL-69 |
| chr15-99091646-DEL-60 | chr15-99091647-DEL-60 |
| chr15-99114985-DEL-199 | chr15-99114986-DEL-199 |

| | |
|---|---|
| chr15-99116953-DEL-106 | chr15-99116954-DEL-106 |
| chr15-99128988-DEL-81 | chr15-99128997-DEL-73 |
| chr15-99150952-DEL-207 | chr15-99150953-DEL-207 |
| chr15-99157704-DEL-215 | chr15-99157740-DEL-190 |
| chr15-99161669-DEL-426 | chr15-99161670-DEL-426 |
| chr15-99172577-DEL-129 | chr15-99172578-DEL-129 |
| chr15-99181198-DEL-211 | chr15-99181199-DEL-211 |
| chr15-99268774-DEL-181 | chr15-99268775-DEL-181 |
| chr15-99271916-DEL-161 | chr15-99271901-DEL-177 |
| chr15-99283688-DEL-204 | chr15-99283689-DEL-204 |
| chr15-99292001-DEL-144 | chr15-99292002-DEL-143 |
| chr15-99305435-DEL-52 | chr15-99305444-DEL-59 |
| chr15-99326440-DEL-68 | chr15-99326441-DEL-68 |
| chr15-99344325-DEL-214 | chr15-99344326-DEL-214 |
| chr15-99348907-DEL-319 | chr15-99348908-DEL-319 |
| chr15-99357060-DEL-51 | chr15-99357061-DEL-51 |
| chr15-99358679-DEL-62 | chr15-99358680-DEL-62 |
| chr15-99359901-DEL-341 | chr15-99359901-DEL-342 |
| chr15-99374480-DEL-141 | chr15-99374481-DEL-141 |
| chr15-99384938-DEL-332 | chr15-99384939-DEL-332 |
| chr15-99431747-DEL-168 | chr15-99431748-DEL-168 |
| chr15-99446564-DEL-79 | chr15-99446565-DEL-79 |
| chr15-99450341-DEL-475 | chr15-99450360-DEL-472 |
| chr15-99474572-DEL-51 | chr15-99474573-DEL-66 |
| chr15-99507459-DEL-162 | chr15-99507460-DEL-162 |
| chr15-99514346-DEL-60 | chr15-99514347-DEL-60 |
| chr15-99525144-DEL-1279 | chr15-99525145-DEL-1279 |
| chr15-99546834-DEL-53 | chr15-99546835-DEL-53 |
| chr15-99561481-DEL-200 | chr15-99561482-DEL-200 |
| chr15-99577320-DEL-76 | chr15-99577337-DEL-80 |
| chr15-99593855-DEL-95 | chr15-99593869-DEL-82 |
| chr15-99603186-DEL-230 | chr15-99603187-DEL-230 |
| chr15-99604337-DEL-391 | chr15-99604338-DEL-391 |
| chr15-99606014-DEL-64 | chr15-99606015-DEL-64 |
| chr15-99620912-DEL-141 | chr15-99620913-DEL-141 |
| chr15-99625170-DEL-218 | chr15-99625171-DEL-218 |
| chr15-99627583-DEL-156 | chr15-99627583-DEL-156 |
| chr15-99634379-DEL-60 | chr15-99634380-DEL-60 |
| chr15-99639138-DEL-126 | chr15-99639139-DEL-126 |
| chr15-99648558-DEL-276 | chr15-99648559-DEL-276 |
| chr15-99692093-DEL-114 | chr15-99692094-DEL-114 |
| chr15-99692754-DEL-251 | chr15-99692755-DEL-251 |
| chr15-99706870-DEL-3209 | chr15-99706871-DEL-3209 |
| chr15-99711790-DEL-52 | chr15-99711796-DEL-51 |
| chr15-99712404-DEL-79 | chr15-99712401-DEL-80 |
| chr15-99719842-DEL-677 | chr15-99719843-DEL-677 |
| chr15-99743087-DEL-73 | chr15-99743088-DEL-73 |
| chr15-99753647-DEL-214 | chr15-99753648-DEL-214 |
| chr15-99756944-DEL-210 | chr15-99756945-DEL-210 |
| chr15-99759249-DEL-2073 | chr15-99759250-DEL-2073 |
| chr15-99772055-DEL-72 | chr15-99772054-DEL-68 |
| chr15-99775263-DEL-208 | chr15-99775264-DEL-208 |
| chr15-99776148-DEL-194 | chr15-99776150-DEL-193 |
| chr15-99804633-DEL-206 | chr15-99804591-DEL-212 |
| chr15-99810053-DEL-99 | chr15-99810058-DEL-94 |
| chr15-99818804-DEL-209 | chr15-99818805-DEL-209 |
| chr15-99822523-DEL-128 | chr15-99822524-DEL-128 |
| chr15-99842910-DEL-58 | chr15-99842874-DEL-57 |
| chr15-99915811-DEL-191 | chr15-99915803-DEL-192 |
| chr15-99924513-DEL-74 | chr15-99924522-DEL-66 |
| chr15-99947955-DEL-236 | chr15-99947956-DEL-236 |
| chr15-99952409-DEL-202 | chr15-99952410-DEL-202 |
| chr15-99989661-DEL-63 | chr15-99989662-DEL-63 |
| chr15-100002892-DEL-66 | chr15-100002893-DEL-66 |
| chr15-100008651-DEL-57 | chr15-100008625-DEL-88 |
| chr15-100014107-DEL-106 | chr15-100014103-DEL-106 |
| chr15-100076887-DEL-347 | chr15-100076906-DEL-334 |
| chr15-100170249-DEL-332 | chr15-100170250-DEL-332 |
| chr15-100219598-DEL-58 | chr15-100219610-DEL-59 |
| chr15-100225210-DEL-103 | chr15-100225211-DEL-103 |
| chr15-100228003-DEL-139 | chr15-100228001-DEL-139 |
| chr15-100256126-DEL-54 | chr15-100256127-DEL-54 |
| chr15-100279229-DEL-177 | chr15-100279268-DEL-176 |
| chr15-100279914-DEL-74 | chr15-100279885-DEL-68 |
| chr15-100286365-DEL-121 | chr15-100286362-DEL-123 |
| chr15-100295857-DEL-61 | chr15-100295858-DEL-61 |
| chr15-100311577-DEL-52 | chr15-100311571-DEL-58 |
| chr15-100331684-DEL-315 | chr15-100331685-DEL-315 |
| chr15-100341427-DEL-149 | chr15-100341428-DEL-149 |
| chr15-100364963-DEL-245 | chr15-100364964-DEL-245 |

| | |
|---|---|
| chr15-100385628-DEL-134 | chr15-100385629-DEL-134 |
| chr15-100456835-DEL-6457 | chr15-100456836-DEL-6457 |
| chr15-100469706-DEL-401 | chr15-100469682-DEL-401 |
| chr15-100476912-DEL-98 | chr15-100476913-DEL-98 |
| chr15-100495700-DEL-73 | chr15-100495701-DEL-73 |
| chr15-100507134-DEL-57 | chr15-100507115-DEL-74 |
| chr15-100513324-DEL-53 | chr15-100513326-DEL-54 |
| chr15-100522067-DEL-212 | chr15-100522068-DEL-212 |
| chr15-100526571-DEL-57 | chr15-100526585-DEL-63 |
| chr15-100530919-DEL-90 | chr15-100530922-DEL-77 |
| chr15-100537406-DEL-237 | chr15-100537407-DEL-237 |
| chr15-100547220-DEL-69 | chr15-100547221-DEL-69 |
| chr15-100565458-DEL-55 | chr15-100565459-DEL-55 |
| chr15-100604852-DEL-82 | chr15-100604873-DEL-62 |
| chr15-100607732-DEL-205 | chr15-100607733-DEL-205 |
| chr15-100608555-DEL-112 | chr15-100608556-DEL-112 |
| chr15-100611756-DEL-127 | chr15-100611757-DEL-127 |
| chr15-100645158-DEL-84 | chr15-100645180-DEL-63 |
| chr15-100646624-DEL-73 | chr15-100646625-DEL-73 |
| chr15-100666466-DEL-213 | chr15-100666467-DEL-213 |
| chr15-100731965-DEL-67 | chr15-100731966-DEL-67 |
| chr15-100755384-DEL-126 | chr15-100755385-DEL-126 |
| chr15-100755946-DEL-2607 | chr15-100755945-DEL-2608 |
| chr15-100756944-DEL-237 | chr15-100756965-DEL-217 |
| chr15-100793763-DEL-126 | chr15-100793785-DEL-105 |
| chr15-100827040-DEL-84 | chr15-100827062-DEL-63 |
| chr15-100886770-DEL-378 | chr15-100886771-DEL-378 |
| chr15-100981151-DEL-54 | chr15-100981152-DEL-54 |
| chr15-100987298-DEL-99 | chr15-100987303-DEL-94 |
| chr15-101002420-DEL-94 | chr15-101002408-DEL-112 |
| chr15-101083110-DEL-52 | chr15-101083088-DEL-52 |
| chr15-101105562-DEL-108 | chr15-101105555-DEL-108 |
| chr15-101166249-DEL-74 | chr15-101166249-DEL-74 |
| chr15-101178796-DEL-229 | chr15-101178797-DEL-229 |
| chr15-101362417-DEL-112 | chr15-101362458-DEL-72 |
| chr15-101367481-DEL-122 | chr15-101367458-DEL-126 |
| chr15-101403806-DEL-143 | chr15-101403807-DEL-143 |
| chr15-101417070-DEL-69 | chr15-101417071-DEL-69 |
| chr15-101445667-DEL-63 | chr15-101445685-DEL-52 |
| chr15-101452063-DEL-61 | chr15-101452064-DEL-61 |
| chr15-101529543-DEL-287 | chr15-101529544-DEL-287 |
| chr15-101535571-DEL-19127 | chr15-101535529-DEL-19128 |
| chr15-101539364-DEL-88 | chr15-101539366-DEL-89 |
| chr15-101542870-DEL-74 | chr15-101542873-DEL-52 |
| chr15-101558158-DEL-12705 | chr15-101558159-DEL-12706 |
| chr15-101576604-DEL-668 | chr15-101576605-DEL-668 |
| chr15-101583245-DEL-136 | chr15-101583228-DEL-137 |
| chr15-101643965-DEL-103 | chr15-101643966-DEL-103 |
| chr15-101655475-DEL-111 | chr15-101655476-DEL-111 |
| chr15-101659415-DEL-571 | chr15-101659416-DEL-571 |
| chr15-101663788-DEL-1039 | chr15-101663789-DEL-1039 |
| chr15-101667878-DEL-52 | chr15-101667879-DEL-52 |
| chr15-101682337-DEL-59 | chr15-101682338-DEL-59 |
| chr15-101690400-DEL-63 | chr15-101690401-DEL-63 |
| chr15-101696361-DEL-215 | chr15-101696362-DEL-215 |
| chr15-101706094-DEL-160 | chr15-101706118-DEL-120 |
| chr15-101715424-DEL-159 | chr15-101715416-DEL-160 |
| chr15-101716321-DEL-1515 | chr15-101716322-DEL-1515 |
| chr15-101720892-DEL-128 | chr15-101720893-DEL-128 |
| chr15-101753186-DEL-112 | chr15-101753187-DEL-98 |
| chr15-101783023-DEL-74 | chr15-101783024-DEL-74 |
| chr15-101809552-DEL-502 | chr15-101809551-DEL-504 |
| chr15-101814554-DEL-81 | chr15-101814555-DEL-81 |
| chr15-101852326-DEL-63 | chr15-101852327-DEL-63 |
| chr15-101865120-DEL-80 | chr15-101865133-DEL-68 |
| chr15-101878026-DEL-54 | chr15-101878023-DEL-66 |
| chr15-101890315-DEL-110 | chr15-101890316-DEL-110 |
| chr15-101897297-DEL-157 | chr15-101897298-DEL-157 |
| chr15-101908286-DEL-207 | chr15-101908287-DEL-207 |
| chr15-101910115-DEL-115 | chr15-101910112-DEL-104 |
| chr15-101925726-DEL-71 | chr15-101925732-DEL-69 |
| chr15-101962834-DEL-1833 | chr15-101962835-DEL-1833 |
| chr15-101966971-DEL-142 | chr15-101966974-DEL-140 |
| chr15-101998723-DEL-52 | chr15-101998724-DEL-52 |
| chr15-102030872-DEL-218 | chr15-102030873-DEL-218 |
| chr15-102031922-DEL-903 | chr15-102031923-DEL-903 |
| chr15-102034061-DEL-363 | chr15-102034062-DEL-363 |
| chr15-102054949-DEL-391 | chr15-102054950-DEL-391 |
| chr15-102082925-DEL-69 | chr15-102082926-DEL-69 |
| chr15-102090893-DEL-213 | chr15-102090894-DEL-213 |

| | |
|---|---|
| chr15-102103863-DEL-901 | chr15-102103864-DEL-901 |
| chr15-102108431-DEL-160 | chr15-102108433-DEL-188 |
| chr15-102128395-DEL-197 | chr15-102128396-DEL-197 |
| chr15-102176431-DEL-150 | chr15-102176432-DEL-150 |
| chr15-102200826-DEL-67 | chr15-102200827-DEL-67 |
| chr15-102219438-DEL-222 | chr15-102219439-DEL-222 |
| chr15-102233926-DEL-490 | chr15-102233927-DEL-490 |
| chr15-102257892-DEL-207 | chr15-102257893-DEL-207 |
| chr15-102261714-DEL-54 | chr15-102261715-DEL-54 |
| chr15-102270595-DEL-56 | chr15-102270596-DEL-56 |
| chr15-102288140-DEL-191 | chr15-102288141-DEL-191 |
| chr15-102299722-DEL-315 | chr15-102299723-DEL-315 |
| chr15-102325357-DEL-1089 | chr15-102325327-DEL-1075 |
| chr15-102328073-DEL-52 | chr15-102328074-DEL-52 |
| chr15-102328450-DEL-244 | chr15-102328435-DEL-235 |
| chr15-102363323-DEL-176 | chr15-102363336-DEL-164 |
| chr15-102388393-DEL-218 | chr15-102388394-DEL-218 |
| chr15-102441493-DEL-269 | chr15-102441494-DEL-269 |
| chr15-102448200-DEL-199 | chr15-102448201-DEL-199 |
| chr15-102470136-DEL-62 | chr15-102470133-DEL-65 |
| chr15-102478950-DEL-211 | chr15-102478951-DEL-211 |
| chr15-102481071-DEL-196 | chr15-102481072-DEL-196 |
| chr15-102483888-DEL-131 | chr15-102483889-DEL-131 |
| chr15-102487989-DEL-175 | chr15-102487990-DEL-175 |
| chr15-102505218-DEL-6403 | chr15-102505219-DEL-6403 |
| chr15-102513073-DEL-211 | chr15-102513074-DEL-211 |
| chr15-102543400-DEL-387 | chr15-102543401-DEL-387 |
| chr15-102548986-DEL-85 | chr15-102548988-DEL-125 |
| chr15-102558213-DEL-171 | chr15-102558220-DEL-165 |
| chr15-102559134-DEL-141 | chr15-102559143-DEL-133 |
| chr15-102560614-DEL-213 | chr15-102560603-DEL-210 |
| chr15-102584048-DEL-69 | chr15-102584070-DEL-62 |
| chr15-102599473-DEL-56 | chr15-102599442-DEL-67 |
| chr15-102609037-DEL-121 | chr15-102609057-DEL-120 |
| chr15-102612399-DEL-118 | chr15-102612403-DEL-125 |
| chr15-102634828-DEL-195 | chr15-102634829-DEL-195 |
| chr15-102635636-DEL-189 | chr15-102635637-DEL-189 |
| chr15-102637339-DEL-229 | chr15-102637340-DEL-229 |
| chr15-102662613-DEL-61 | chr15-102662614-DEL-61 |
| chr15-102723562-DEL-62 | chr15-102723565-DEL-57 |
| chr15-102737610-DEL-705 | chr15-102737611-DEL-705 |
| chr15-102809561-DEL-257 | chr15-102809562-DEL-257 |
| chr15-102813034-DEL-65 | chr15-102813033-DEL-67 |
| chr15-102973290-DEL-55 | chr15-102973296-DEL-50 |
| chr15-103136570-DEL-210 | chr15-103136571-DEL-210 |
| chr15-103142566-DEL-167 | chr15-103142567-DEL-167 |
| chr15-103168041-DEL-156 | chr15-103168034-DEL-160 |
| chr15-103177198-DEL-67 | chr15-103177201-DEL-68 |
| chr15-103184651-DEL-213 | chr15-103184652-DEL-213 |
| chr15-103202721-DEL-218 | chr15-103202722-DEL-217 |
| chr15-103203562-DEL-52 | chr15-103203563-DEL-52 |
| chr15-103228152-DEL-169 | chr15-103228153-DEL-167 |
| chr15-103239372-DEL-232 | chr15-103239391-DEL-214 |
| chr15-103270649-DEL-221 | chr15-103270650-DEL-221 |
| chr15-103313976-DEL-206 | chr15-103313977-DEL-206 |
| chr15-103319037-DEL-7940 | chr15-103319038-DEL-7940 |
| chr15-103374273-DEL-277 | chr15-103374274-DEL-277 |
| chr15-103444706-DEL-205 | chr15-103444707-DEL-205 |
| chr15-103515112-DEL-4388 | chr15-103515113-DEL-4388 |
| chr15-103545354-DEL-58 | chr15-103545355-DEL-58 |
| chr15-103563900-DEL-6907 | chr15-103563901-DEL-6907 |
| chr15-103590829-DEL-201 | chr15-103590830-DEL-201 |
| chr15-103632465-DEL-360 | chr15-103632466-DEL-360 |
| chr15-103638381-DEL-199 | chr15-103638396-DEL-201 |
| chr15-103741687-DEL-1158 | chr15-103741688-DEL-1158 |
| chr15-103750281-DEL-84 | chr15-103750281-DEL-84 |
| chr15-103843705-DEL-105 | chr15-103843705-DEL-105 |
| chr15-103849420-DEL-3224 | chr15-103849419-DEL-3226 |
| chr15-103859005-DEL-3239 | chr15-103859005-DEL-3239 |
| chr15-103878405-DEL-4193 | chr15-103878406-DEL-4193 |
| chr15-103883165-DEL-338 | chr15-103883166-DEL-338 |
| chr15-103895432-DEL-510 | chr15-103895433-DEL-510 |
| chr15-103906121-DEL-54 | chr15-103906122-DEL-54 |
| chr15-103907449-DEL-85 | chr15-103907450-DEL-85 |
| chr15-103963718-DEL-62 | chr15-103963718-DEL-62 |
| chr16-3064289-DEL-54 | chr16-3064313-DEL-53 |
| chr16-3153168-DEL-324 | chr16-3153167-DEL-324 |
| chr16-3163340-DEL-58 | chr16-3163339-DEL-58 |
| chr16-3185004-DEL-51 | chr16-3185004-DEL-51 |
| chr16-3202128-DEL-7475 | chr16-3202129-DEL-7475 |

| | |
|---|---|
| chr16-3274307-DEL-2933 | chr16-3274307-DEL-2933 |
| chr16-3281059-DEL-13178 | chr16-3281056-DEL-13190 |
| chr16-3335624-DEL-1894 | chr16-3335625-DEL-1894 |
| chr16-3387937-DEL-203 | chr16-3387938-DEL-203 |
| chr16-3466923-DEL-183 | chr16-3466924-DEL-183 |
| chr16-3478582-DEL-1826 | chr16-3478582-DEL-1826 |
| chr16-3480711-DEL-109 | chr16-3480712-DEL-109 |
| chr16-3484299-DEL-725 | chr16-3484300-DEL-725 |
| chr16-3507835-DEL-117 | chr16-3507836-DEL-117 |
| chr16-3520783-DEL-807 | chr16-3520784-DEL-807 |
| chr16-3534194-DEL-200 | chr16-3534195-DEL-200 |
| chr16-3551350-DEL-58 | chr16-3551351-DEL-58 |
| chr16-3582104-DEL-6401 | chr16-3582105-DEL-6401 |
| chr16-3615155-DEL-895 | chr16-3615156-DEL-895 |
| chr16-3638778-DEL-2943 | chr16-3638779-DEL-2943 |
| chr16-3642482-DEL-219 | chr16-3642483-DEL-219 |
| chr16-3646862-DEL-118 | chr16-3646863-DEL-118 |
| chr16-3670571-DEL-58 | chr16-3670571-DEL-58 |
| chr16-3704406-DEL-195 | chr16-3704407-DEL-195 |
| chr16-3759021-DEL-207 | chr16-3759022-DEL-207 |
| chr16-3764270-DEL-288 | chr16-3764271-DEL-288 |
| chr16-3779152-DEL-58 | chr16-3779153-DEL-58 |
| chr16-3792462-DEL-99 | chr16-3792463-DEL-99 |
| chr16-3841611-DEL-172 | chr16-3841612-DEL-172 |
| chr16-3849622-DEL-75 | chr16-3849622-DEL-76 |
| chr16-3850686-DEL-208 | chr16-3850687-DEL-208 |
| chr16-3875219-DEL-196 | chr16-3875220-DEL-196 |
| chr16-3904783-DEL-208 | chr16-3904784-DEL-208 |
| chr16-3931279-DEL-103 | chr16-3931280-DEL-103 |
| chr16-4008783-DEL-51 | chr16-4008784-DEL-51 |
| chr16-4057725-DEL-154 | chr16-4057726-DEL-112 |
| chr16-4094892-DEL-171 | chr16-4094893-DEL-171 |
| chr16-4119855-DEL-52 | chr16-4119856-DEL-52 |
| chr16-4131259-DEL-135 | chr16-4131260-DEL-135 |
| chr16-4161065-DEL-8455 | chr16-4161066-DEL-8455 |
| chr16-4205889-DEL-69 | chr16-4205902-DEL-57 |
| chr16-4259333-DEL-112 | chr16-4259334-DEL-112 |
| chr16-4292091-DEL-187 | chr16-4292091-DEL-187 |
| chr16-4334403-DEL-71 | chr16-4334403-DEL-71 |
| chr16-4337525-DEL-191 | chr16-4337526-DEL-191 |
| chr16-4364320-DEL-222 | chr16-4364321-DEL-222 |
| chr16-4368048-DEL-211 | chr16-4368048-DEL-196 |
| chr16-4386495-DEL-194 | chr16-4386496-DEL-194 |
| chr16-4387826-DEL-180 | chr16-4387827-DEL-180 |
| chr16-4561268-DEL-7080 | chr16-4561269-DEL-7080 |
| chr16-4573787-DEL-1177 | chr16-4573788-DEL-1175 |
| chr16-4588837-DEL-197 | chr16-4588838-DEL-197 |
| chr16-4619965-DEL-2262 | chr16-4619966-DEL-2262 |
| chr16-4623723-DEL-196 | chr16-4623724-DEL-196 |
| chr16-4638179-DEL-186 | chr16-4638180-DEL-159 |
| chr16-4675677-DEL-2832 | chr16-4675678-DEL-2832 |
| chr16-4766505-DEL-240 | chr16-4766506-DEL-240 |
| chr16-4767047-DEL-231 | chr16-4767048-DEL-231 |
| chr16-4802325-DEL-211 | chr16-4802326-DEL-211 |
| chr16-4856481-DEL-72 | chr16-4856464-DEL-52 |
| chr16-4860864-DEL-215 | chr16-4860865-DEL-215 |
| chr16-4863469-DEL-78 | chr16-4863470-DEL-78 |
| chr16-4951698-DEL-62 | chr16-4951726-DEL-51 |
| chr16-4975686-DEL-205 | chr16-4975687-DEL-205 |
| chr16-4983913-DEL-490 | chr16-4983920-DEL-512 |
| chr16-5060970-DEL-62 | chr16-5060971-DEL-62 |
| chr16-5064166-DEL-430 | chr16-5064167-DEL-430 |
| chr16-5074442-DEL-569 | chr16-5074443-DEL-569 |
| chr16-5077727-DEL-198 | chr16-5077728-DEL-198 |
| chr16-5088143-DEL-54 | chr16-5088154-DEL-57 |
| chr16-5096664-DEL-57 | chr16-5096665-DEL-57 |
| chr16-5104647-DEL-340 | chr16-5104648-DEL-340 |
| chr16-5118003-DEL-71 | chr16-5118004-DEL-71 |
| chr16-5131558-DEL-194 | chr16-5131551-DEL-195 |
| chr16-5132946-DEL-360 | chr16-5132947-DEL-360 |
| chr16-5150147-DEL-57 | chr16-5150154-DEL-51 |
| chr16-5161585-DEL-64 | chr16-5161588-DEL-62 |
| chr16-5189232-DEL-82 | chr16-5189233-DEL-82 |
| chr16-5192254-DEL-171 | chr16-5192254-DEL-144 |
| chr16-5221341-DEL-75 | chr16-5221338-DEL-56 |
| chr16-5261479-DEL-538 | chr16-5261480-DEL-536 |
| chr16-5262211-DEL-60 | chr16-5262221-DEL-50 |
| chr16-5265749-DEL-1557 | chr16-5265750-DEL-1557 |
| chr16-5274701-DEL-57 | chr16-5274702-DEL-57 |
| chr16-5302310-DEL-60 | chr16-5302311-DEL-60 |

| | |
|---|---|
| chr16-5343702-DEL-146 | chr16-5343703-DEL-146 |
| chr16-5357196-DEL-129 | chr16-5357184-DEL-130 |
| chr16-5397819-DEL-389 | chr16-5397820-DEL-389 |
| chr16-5407057-DEL-216 | chr16-5407058-DEL-216 |
| chr16-5452321-DEL-2951 | chr16-5452322-DEL-2951 |
| chr16-5460298-DEL-3008 | chr16-5460299-DEL-3008 |
| chr16-5465964-DEL-67 | chr16-5465964-DEL-67 |
| chr16-5488053-DEL-84 | chr16-5488054-DEL-84 |
| chr16-5496339-DEL-330 | chr16-5496349-DEL-321 |
| chr16-5502065-DEL-113 | chr16-5502072-DEL-107 |
| chr16-5530195-DEL-385 | chr16-5530196-DEL-385 |
| chr16-5544112-DEL-186 | chr16-5544111-DEL-184 |
| chr16-5564057-DEL-53 | chr16-5564035-DEL-54 |
| chr16-5567066-DEL-1085 | chr16-5567067-DEL-1085 |
| chr16-5610528-DEL-96 | chr16-5610528-DEL-93 |
| chr16-5612368-DEL-87 | chr16-5612386-DEL-70 |
| chr16-5634534-DEL-112 | chr16-5634535-DEL-112 |
| chr16-5645657-DEL-72 | chr16-5645658-DEL-70 |
| chr16-5664247-DEL-54 | chr16-5664248-DEL-54 |
| chr16-5678844-DEL-841 | chr16-5678845-DEL-841 |
| chr16-5713973-DEL-480 | chr16-5713974-DEL-480 |
| chr16-5729590-DEL-100 | chr16-5729591-DEL-100 |
| chr16-5737412-DEL-51 | chr16-5737388-DEL-52 |
| chr16-5792449-DEL-94 | chr16-5792450-DEL-94 |
| chr16-5814453-DEL-52 | chr16-5814454-DEL-52 |
| chr16-5842385-DEL-76 | chr16-5842386-DEL-76 |
| chr16-5847818-DEL-96 | chr16-5847816-DEL-100 |
| chr16-5873274-DEL-110 | chr16-5873275-DEL-110 |
| chr16-5898066-DEL-100 | chr16-5898020-DEL-101 |
| chr16-5922333-DEL-145 | chr16-5922334-DEL-145 |
| chr16-5929739-DEL-58 | chr16-5929740-DEL-58 |
| chr16-5941002-DEL-54 | chr16-5941003-DEL-54 |
| chr16-6025001-DEL-53 | chr16-6025002-DEL-53 |
| chr16-6030451-DEL-54 | chr16-6030454-DEL-52 |
| chr16-6043985-DEL-605 | chr16-6043982-DEL-606 |
| chr16-6079863-DEL-54 | chr16-6079863-DEL-54 |
| chr16-6083205-DEL-72 | chr16-6083211-DEL-60 |
| chr16-6134836-DEL-61 | chr16-6134837-DEL-61 |
| chr16-6154103-DEL-61 | chr16-6154104-DEL-61 |
| chr16-6160219-DEL-539 | chr16-6160220-DEL-539 |
| chr16-6165504-DEL-70 | chr16-6165505-DEL-54 |
| chr16-6181630-DEL-68 | chr16-6181631-DEL-68 |
| chr16-6190876-DEL-55 | chr16-6190894-DEL-50 |
| chr16-6210974-DEL-465 | chr16-6210975-DEL-465 |
| chr16-6216622-DEL-204 | chr16-6216608-DEL-199 |
| chr16-6246545-DEL-1084 | chr16-6246546-DEL-1084 |
| chr16-6250441-DEL-948 | chr16-6250442-DEL-946 |
| chr16-6254932-DEL-68 | chr16-6254933-DEL-68 |
| chr16-6267735-DEL-167 | chr16-6267738-DEL-168 |
| chr16-6294631-DEL-228 | chr16-6294632-DEL-228 |
| chr16-6331129-DEL-53 | chr16-6331114-DEL-54 |
| chr16-6342668-DEL-51 | chr16-6342669-DEL-51 |
| chr16-6345886-DEL-468 | chr16-6345887-DEL-468 |
| chr16-6381034-DEL-50 | chr16-6381032-DEL-52 |
| chr16-6404505-DEL-105 | chr16-6404506-DEL-105 |
| chr16-6418591-DEL-155 | chr16-6418592-DEL-155 |
| chr16-6432540-DEL-1722 | chr16-6432541-DEL-1722 |
| chr16-6469204-DEL-74 | chr16-6469205-DEL-74 |
| chr16-6501448-DEL-370 | chr16-6501427-DEL-372 |
| chr16-6509081-DEL-59 | chr16-6509082-DEL-59 |
| chr16-6509992-DEL-104 | chr16-6509993-DEL-104 |
| chr16-6537880-DEL-80 | chr16-6537908-DEL-76 |
| chr16-6540946-DEL-184 | chr16-6540938-DEL-181 |
| chr16-6543802-DEL-179 | chr16-6543803-DEL-179 |
| chr16-6545682-DEL-62 | chr16-6545684-DEL-60 |
| chr16-6591186-DEL-131 | chr16-6591196-DEL-132 |
| chr16-6605454-DEL-616 | chr16-6605455-DEL-616 |
| chr16-6622373-DEL-174 | chr16-6622374-DEL-174 |
| chr16-6753402-DEL-108 | chr16-6753403-DEL-116 |
| chr16-6768424-DEL-56 | chr16-6768424-DEL-56 |
| chr16-6778356-DEL-94 | chr16-6778358-DEL-92 |
| chr16-6814612-DEL-96 | chr16-6814612-DEL-96 |
| chr16-6844641-DEL-243 | chr16-6844641-DEL-243 |
| chr16-6853178-DEL-126 | chr16-6853179-DEL-126 |
| chr16-6873174-DEL-67 | chr16-6873163-DEL-79 |
| chr16-6877862-DEL-104 | chr16-6877863-DEL-104 |
| chr16-6884572-DEL-112 | chr16-6884579-DEL-110 |
| chr16-6895189-DEL-54 | chr16-6895190-DEL-54 |
| chr16-6919036-DEL-161 | chr16-6919037-DEL-161 |
| chr16-6973196-DEL-51 | chr16-6973197-DEL-51 |

| | |
|---|---|
| chr16-6988151-DEL-1601 | chr16-6988152-DEL-1601 |
| chr16-6989859-DEL-60 | chr16-6989860-DEL-60 |
| chr16-7035832-DEL-58 | chr16-7035833-DEL-54 |
| chr16-7114084-DEL-1921 | chr16-7114085-DEL-1921 |
| chr16-7120861-DEL-622 | chr16-7120862-DEL-627 |
| chr16-7144571-DEL-260 | chr16-7144566-DEL-263 |
| chr16-7163976-DEL-71 | chr16-7163977-DEL-71 |
| chr16-7194144-DEL-58 | chr16-7194144-DEL-58 |
| chr16-7199157-DEL-56 | chr16-7199158-DEL-56 |
| chr16-7242059-DEL-70 | chr16-7242060-DEL-70 |
| chr16-7278595-DEL-2846 | chr16-7278596-DEL-2846 |
| chr16-7292068-DEL-84 | chr16-7292069-DEL-84 |
| chr16-7308137-DEL-213 | chr16-7308138-DEL-213 |
| chr16-7328228-DEL-52 | chr16-7328222-DEL-70 |
| chr16-7356463-DEL-102 | chr16-7356489-DEL-105 |
| chr16-7421741-DEL-2535 | chr16-7421742-DEL-2535 |
| chr16-7473034-DEL-59 | chr16-7473035-DEL-59 |
| chr16-7491736-DEL-160 | chr16-7491689-DEL-161 |
| chr16-7500275-DEL-66 | chr16-7500276-DEL-66 |
| chr16-7540746-DEL-2322 | chr16-7540747-DEL-2322 |
| chr16-7555773-DEL-3245 | chr16-7555773-DEL-3245 |
| chr16-7571835-DEL-6463 | chr16-7571836-DEL-6463 |
| chr16-7593136-DEL-63 | chr16-7593137-DEL-63 |
| chr16-7604098-DEL-60 | chr16-7604101-DEL-58 |
| chr16-7652324-DEL-65 | chr16-7652325-DEL-65 |
| chr16-7657495-DEL-69 | chr16-7657470-DEL-68 |
| chr16-7676812-DEL-56 | chr16-7676813-DEL-56 |
| chr16-7677450-DEL-3583 | chr16-7677449-DEL-3584 |
| chr16-7687349-DEL-572 | chr16-7687350-DEL-572 |
| chr16-7689183-DEL-853 | chr16-7689184-DEL-853 |
| chr16-7716818-DEL-56 | chr16-7716819-DEL-56 |
| chr16-7729308-DEL-117 | chr16-7729348-DEL-102 |
| chr16-7730745-DEL-1280 | chr16-7730745-DEL-1280 |
| chr16-7732093-DEL-1549 | chr16-7732093-DEL-1549 |
| chr16-7756066-DEL-573 | chr16-7756067-DEL-573 |
| chr16-7759512-DEL-186 | chr16-7759513-DEL-186 |
| chr16-7770136-DEL-6228 | chr16-7770137-DEL-6228 |
| chr16-7776849-DEL-112 | chr16-7776850-DEL-110 |
| chr16-7799823-DEL-57 | chr16-7799815-DEL-54 |
| chr16-7817801-DEL-65 | chr16-7817802-DEL-65 |
| chr16-7851876-DEL-7219 | chr16-7851877-DEL-7219 |
| chr16-7933666-DEL-184 | chr16-7933667-DEL-184 |
| chr16-7936962-DEL-76 | chr16-7936963-DEL-76 |
| chr16-7941583-DEL-96 | chr16-7941584-DEL-96 |
| chr16-7967844-DEL-7705 | chr16-7967845-DEL-7705 |
| chr16-7978389-DEL-62 | chr16-7978371-DEL-60 |
| chr16-7989979-DEL-1991 | chr16-7989980-DEL-1991 |
| chr16-8023563-DEL-55 | chr16-8023564-DEL-55 |
| chr16-8067783-DEL-68 | chr16-8067784-DEL-68 |
| chr16-8071943-DEL-6231 | chr16-8071944-DEL-6231 |
| chr16-8090413-DEL-1025 | chr16-8090414-DEL-1025 |
| chr16-8099264-DEL-6223 | chr16-8099265-DEL-6223 |
| chr16-8111746-DEL-54 | chr16-8111751-DEL-50 |
| chr16-8115141-DEL-54 | chr16-8115142-DEL-54 |
| chr16-8121868-DEL-54 | chr16-8121869-DEL-54 |
| chr16-8138309-DEL-70 | chr16-8138310-DEL-70 |
| chr16-8149115-DEL-158 | chr16-8149148-DEL-153 |
| chr16-8196824-DEL-60 | chr16-8196788-DEL-54 |
| chr16-8209806-DEL-136 | chr16-8209845-DEL-98 |
| chr16-8225596-DEL-68 | chr16-8225597-DEL-68 |
| chr16-8232153-DEL-386 | chr16-8232154-DEL-386 |
| chr16-8235789-DEL-57 | chr16-8235794-DEL-56 |
| chr16-8277888-DEL-173 | chr16-8277889-DEL-173 |
| chr16-8278645-DEL-6835 | chr16-8278646-DEL-6835 |
| chr16-8292425-DEL-1044 | chr16-8292426-DEL-1044 |
| chr16-8333056-DEL-55 | chr16-8333056-DEL-55 |
| chr16-8398928-DEL-269 | chr16-8398942-DEL-222 |
| chr16-8420992-DEL-203 | chr16-8420992-DEL-204 |
| chr16-8441548-DEL-1599 | chr16-8441549-DEL-1599 |
| chr16-8442625-DEL-195 | chr16-8442626-DEL-196 |
| chr16-8542271-DEL-77 | chr16-8542272-DEL-77 |
| chr16-8550246-DEL-213 | chr16-8550250-DEL-212 |
| chr16-8563348-DEL-351 | chr16-8563348-DEL-351 |
| chr16-8566571-DEL-234 | chr16-8566572-DEL-234 |
| chr16-8590554-DEL-80 | chr16-8590555-DEL-80 |
| chr16-8595485-DEL-163 | chr16-8595486-DEL-163 |
| chr16-8597513-DEL-67 | chr16-8597514-DEL-67 |
| chr16-8617303-DEL-176 | chr16-8617304-DEL-174 |
| chr16-8622770-DEL-210 | chr16-8622771-DEL-210 |
| chr16-8623905-DEL-394 | chr16-8623904-DEL-395 |

| | |
|---|---|
| chr16-8627268-DEL-80 | chr16-8627268-DEL-110 |
| chr16-8633115-DEL-199 | chr16-8633115-DEL-199 |
| chr16-8637524-DEL-79 | chr16-8637522-DEL-81 |
| chr16-8638380-DEL-101 | chr16-8638372-DEL-100 |
| chr16-8642448-DEL-80 | chr16-8642449-DEL-84 |
| chr16-8644173-DEL-63 | chr16-8644174-DEL-62 |
| chr16-8654207-DEL-1217 | chr16-8654208-DEL-1217 |
| chr16-8663089-DEL-74 | chr16-8663090-DEL-74 |
| chr16-8678685-DEL-249 | chr16-8678686-DEL-249 |
| chr16-8702591-DEL-52 | chr16-8702575-DEL-53 |
| chr16-8722943-DEL-70 | chr16-8722944-DEL-70 |
| chr16-8729396-DEL-2652 | chr16-8729397-DEL-2652 |
| chr16-8735514-DEL-54 | chr16-8735515-DEL-54 |
| chr16-8739861-DEL-447 | chr16-8739861-DEL-447 |
| chr16-8741943-DEL-242 | chr16-8741944-DEL-242 |
| chr16-8753559-DEL-261 | chr16-8753560-DEL-261 |
| chr16-8767859-DEL-1415 | chr16-8767861-DEL-1413 |
| chr16-8787173-DEL-508 | chr16-8787174-DEL-508 |
| chr16-8809347-DEL-98 | chr16-8809340-DEL-122 |
| chr16-8814406-DEL-220 | chr16-8814407-DEL-219 |
| chr16-8829100-DEL-101 | chr16-8829114-DEL-132 |
| chr16-8834174-DEL-72 | chr16-8834175-DEL-78 |
| chr16-8858760-DEL-90 | chr16-8858791-DEL-60 |
| chr16-8878730-DEL-526 | chr16-8878731-DEL-526 |
| chr16-8881577-DEL-6618 | chr16-8881578-DEL-6618 |
| chr16-8924833-DEL-132 | chr16-8924834-DEL-132 |
| chr16-8944168-DEL-289 | chr16-8944169-DEL-289 |
| chr16-8982025-DEL-215 | chr16-8982012-DEL-216 |
| chr16-8999077-DEL-200 | chr16-8999078-DEL-200 |
| chr16-9024045-DEL-130 | chr16-9024046-DEL-130 |
| chr16-9034500-DEL-154 | chr16-9034489-DEL-134 |
| chr16-9037093-DEL-68 | chr16-9037111-DEL-68 |
| chr16-9037732-DEL-1958 | chr16-9037733-DEL-1956 |
| chr16-9039187-DEL-52 | chr16-9039188-DEL-50 |
| chr16-9044035-DEL-1887 | chr16-9044036-DEL-1887 |
| chr16-9059168-DEL-827 | chr16-9059169-DEL-827 |
| chr16-9067701-DEL-102 | chr16-9067734-DEL-68 |
| chr16-9100835-DEL-829 | chr16-9100836-DEL-829 |
| chr16-9110785-DEL-3041 | chr16-9110786-DEL-3041 |
| chr16-9162683-DEL-56 | chr16-9162680-DEL-60 |
| chr16-9191258-DEL-74 | chr16-9191259-DEL-74 |
| chr16-9210567-DEL-62 | chr16-9210565-DEL-65 |
| chr16-9228969-DEL-1217 | chr16-9228970-DEL-1216 |
| chr16-9248673-DEL-1104 | chr16-9248677-DEL-1105 |
| chr16-9253780-DEL-186 | chr16-9253781-DEL-186 |
| chr16-9256718-DEL-92 | chr16-9256719-DEL-81 |
| chr16-9278123-DEL-71 | chr16-9278110-DEL-64 |
| chr16-9280178-DEL-85 | chr16-9280189-DEL-75 |
| chr16-9377533-DEL-87 | chr16-9377534-DEL-87 |
| chr16-9382443-DEL-125 | chr16-9382444-DEL-126 |
| chr16-9466617-DEL-175 | chr16-9466618-DEL-175 |
| chr16-9486284-DEL-789 | chr16-9486285-DEL-789 |
| chr16-9495816-DEL-67 | chr16-9495817-DEL-67 |
| chr16-9537833-DEL-102 | chr16-9537831-DEL-104 |
| chr16-9542745-DEL-523 | chr16-9542732-DEL-529 |
| chr16-9555867-DEL-60 | chr16-9555868-DEL-60 |
| chr16-9563082-DEL-50 | chr16-9563084-DEL-61 |
| chr16-9599057-DEL-92 | chr16-9599042-DEL-93 |
| chr16-9616860-DEL-78 | chr16-9616861-DEL-66 |
| chr16-9685708-DEL-4881 | chr16-9685709-DEL-4881 |
| chr16-9752373-DEL-465 | chr16-9752374-DEL-465 |
| chr16-9774073-DEL-56 | chr16-9774074-DEL-56 |
| chr16-9784232-DEL-349 | chr16-9784233-DEL-348 |
| chr16-9785171-DEL-68 | chr16-9785156-DEL-92 |
| chr16-9796584-DEL-400 | chr16-9796585-DEL-400 |
| chr16-9802418-DEL-56 | chr16-9802403-DEL-76 |
| chr16-9858417-DEL-412 | chr16-9858418-DEL-412 |
| chr16-9882099-DEL-51 | chr16-9882100-DEL-51 |
| chr16-9902242-DEL-52 | chr16-9902243-DEL-52 |
| chr16-9917624-DEL-341 | chr16-9917625-DEL-341 |
| chr16-9935919-DEL-66 | chr16-9935920-DEL-66 |
| chr16-9959342-DEL-52 | chr16-9959343-DEL-52 |
| chr16-9965926-DEL-161 | chr16-9965927-DEL-161 |
| chr16-9973091-DEL-294 | chr16-9973092-DEL-294 |
| chr16-10035602-DEL-51 | chr16-10035603-DEL-51 |
| chr16-10038961-DEL-201 | chr16-10038962-DEL-201 |
| chr16-10048623-DEL-434 | chr16-10048624-DEL-434 |
| chr16-10083940-DEL-1072 | chr16-10083943-DEL-1072 |
| chr16-10089718-DEL-207 | chr16-10089714-DEL-212 |
| chr16-10096544-DEL-110 | chr16-10096565-DEL-85 |

| | |
|---|---|
| chr16-10157378-DEL-60 | chr16-10157379-DEL-60 |
| chr16-10164307-DEL-52 | chr16-10164308-DEL-52 |
| chr16-10191775-DEL-58 | chr16-10191776-DEL-58 |
| chr16-10206720-DEL-57 | chr16-10206721-DEL-57 |
| chr16-10209355-DEL-73 | chr16-10209356-DEL-73 |
| chr16-10217994-DEL-6317 | chr16-10217995-DEL-6317 |
| chr16-10287521-DEL-102 | chr16-10287522-DEL-102 |
| chr16-10303127-DEL-1344 | chr16-10303128-DEL-1344 |
| chr16-10309099-DEL-78 | chr16-10309100-DEL-117 |
| chr16-10310713-DEL-120 | chr16-10310714-DEL-120 |
| chr16-10324900-DEL-93 | chr16-10324901-DEL-93 |
| chr16-10426315-DEL-144 | chr16-10426277-DEL-194 |
| chr16-10430943-DEL-402 | chr16-10430944-DEL-402 |
| chr16-10440513-DEL-110 | chr16-10440557-DEL-109 |
| chr16-10449104-DEL-1875 | chr16-10449105-DEL-1875 |
| chr16-10459674-DEL-213 | chr16-10459675-DEL-213 |
| chr16-10468983-DEL-182 | chr16-10468984-DEL-182 |
| chr16-10535125-DEL-69 | chr16-10535126-DEL-69 |
| chr16-10536047-DEL-60 | chr16-10536048-DEL-60 |
| chr16-10547830-DEL-148 | chr16-10547831-DEL-142 |
| chr16-10564412-DEL-82 | chr16-10564415-DEL-80 |
| chr16-10571012-DEL-159 | chr16-10571013-DEL-156 |
| chr16-10571335-DEL-863 | chr16-10571333-DEL-864 |
| chr16-10579608-DEL-148 | chr16-10579609-DEL-148 |
| chr16-10623021-DEL-94 | chr16-10623022-DEL-94 |
| chr16-10638906-DEL-1355 | chr16-10638907-DEL-1355 |
| chr16-10704243-DEL-308 | chr16-10704244-DEL-308 |
| chr16-10707983-DEL-71 | chr16-10707985-DEL-72 |
| chr16-10776244-DEL-115 | chr16-10776245-DEL-116 |
| chr16-10792398-DEL-491 | chr16-10792399-DEL-491 |
| chr16-10820971-DEL-244 | chr16-10820967-DEL-242 |
| chr16-10831280-DEL-374 | chr16-10831281-DEL-374 |
| chr16-10834902-DEL-353 | chr16-10834903-DEL-353 |
| chr16-10838758-DEL-276 | chr16-10838759-DEL-276 |
| chr16-10840517-DEL-766 | chr16-10840518-DEL-766 |
| chr16-10844263-DEL-76 | chr16-10844264-DEL-76 |
| chr16-10854160-DEL-51 | chr16-10854160-DEL-51 |
| chr16-10858025-DEL-84 | chr16-10858026-DEL-84 |
| chr16-10863947-DEL-346 | chr16-10863948-DEL-346 |
| chr16-10870961-DEL-644 | chr16-10870962-DEL-644 |
| chr16-10929449-DEL-784 | chr16-10929450-DEL-784 |
| chr16-10941750-DEL-97 | chr16-10941720-DEL-131 |
| chr16-10943918-DEL-1107 | chr16-10943932-DEL-1094 |
| chr16-10948802-DEL-448 | chr16-10948803-DEL-448 |
| chr16-10961774-DEL-410 | chr16-10961774-DEL-410 |
| chr16-10962296-DEL-100 | chr16-10962293-DEL-110 |
| chr16-10975803-DEL-229 | chr16-10975804-DEL-229 |
| chr16-10997028-DEL-53 | chr16-10997028-DEL-53 |
| chr16-10999258-DEL-114 | chr16-10999284-DEL-86 |
| chr16-11002429-DEL-1760 | chr16-11002440-DEL-1759 |
| chr16-11009318-DEL-329 | chr16-11009325-DEL-329 |
| chr16-11013280-DEL-67 | chr16-11013281-DEL-67 |
| chr16-11023741-DEL-110 | chr16-11023742-DEL-110 |
| chr16-11066495-DEL-894 | chr16-11066494-DEL-895 |
| chr16-11104721-DEL-171 | chr16-11104722-DEL-171 |
| chr16-11107008-DEL-196 | chr16-11107000-DEL-197 |
| chr16-11115558-DEL-56 | chr16-11115558-DEL-56 |
| chr16-11133602-DEL-64 | chr16-11133603-DEL-64 |
| chr16-11144351-DEL-173 | chr16-11144352-DEL-173 |
| chr16-11151206-DEL-6413 | chr16-11151207-DEL-6413 |
| chr16-11188856-DEL-136 | chr16-11188857-DEL-136 |
| chr16-11191897-DEL-101 | chr16-11191895-DEL-77 |
| chr16-11202179-DEL-66 | chr16-11202171-DEL-74 |
| chr16-11209161-DEL-450 | chr16-11209162-DEL-450 |
| chr16-11211571-DEL-655 | chr16-11211572-DEL-655 |
| chr16-11215388-DEL-82 | chr16-11215389-DEL-82 |
| chr16-11250274-DEL-115 | chr16-11250274-DEL-109 |
| chr16-11294663-DEL-157 | chr16-11294663-DEL-158 |
| chr16-11300705-DEL-6797 | chr16-11300706-DEL-6797 |
| chr16-11320207-DEL-937 | chr16-11320208-DEL-937 |
| chr16-11331561-DEL-193 | chr16-11331561-DEL-193 |
| chr16-11332978-DEL-54 | chr16-11332979-DEL-54 |
| chr16-11334602-DEL-77 | chr16-11334603-DEL-77 |
| chr16-11337064-DEL-64 | chr16-11337063-DEL-64 |
| chr16-11340218-DEL-71 | chr16-11340219-DEL-71 |
| chr16-11358651-DEL-86 | chr16-11358652-DEL-86 |
| chr16-11378331-DEL-876 | chr16-11378334-DEL-877 |
| chr16-11383959-DEL-248 | chr16-11383960-DEL-248 |
| chr16-11387155-DEL-99 | chr16-11387156-DEL-99 |
| chr16-11408556-DEL-126 | chr16-11408557-DEL-126 |

| | |
|---|---|
| chr16-11414667-DEL-110 | chr16-11414670-DEL-111 |
| chr16-11416282-DEL-229 | chr16-11416282-DEL-230 |
| chr16-11436627-DEL-53 | chr16-11436628-DEL-53 |
| chr16-11443403-DEL-67 | chr16-11443412-DEL-59 |
| chr16-11465666-DEL-121 | chr16-11465667-DEL-97 |
| chr16-11479217-DEL-505 | chr16-11479218-DEL-505 |
| chr16-11497871-DEL-5301 | chr16-11497872-DEL-5301 |
| chr16-11504323-DEL-360 | chr16-11504322-DEL-360 |
| chr16-11519974-DEL-111 | chr16-11519975-DEL-111 |
| chr16-11549477-DEL-294 | chr16-11549478-DEL-294 |
| chr16-11587632-DEL-505 | chr16-11587633-DEL-505 |
| chr16-11614473-DEL-134 | chr16-11614474-DEL-140 |
| chr16-11627955-DEL-1031 | chr16-11627946-DEL-1032 |
| chr16-11677488-DEL-6856 | chr16-11677489-DEL-6856 |
| chr16-11690096-DEL-1124 | chr16-11690097-DEL-1124 |
| chr16-11713159-DEL-360 | chr16-11713160-DEL-360 |
| chr16-11744826-DEL-935 | chr16-11744827-DEL-935 |
| chr16-11755227-DEL-60 | chr16-11755228-DEL-60 |
| chr16-11759571-DEL-54 | chr16-11759572-DEL-54 |
| chr16-11788514-DEL-1568 | chr16-11788514-DEL-1568 |
| chr16-11795943-DEL-88 | chr16-11795914-DEL-88 |
| chr16-11865434-DEL-195 | chr16-11865435-DEL-195 |
| chr16-11889979-DEL-177 | chr16-11889980-DEL-177 |
| chr16-11915427-DEL-707 | chr16-11915428-DEL-707 |
| chr16-11918963-DEL-63 | chr16-11918992-DEL-63 |
| chr16-11944329-DEL-81 | chr16-11944330-DEL-81 |
| chr16-11958552-DEL-274 | chr16-11958553-DEL-274 |
| chr16-12036674-DEL-648 | chr16-12036672-DEL-649 |
| chr16-12054206-DEL-147 | chr16-12054207-DEL-147 |
| chr16-12133655-DEL-51 | chr16-12133655-DEL-51 |
| chr16-12139572-DEL-73 | chr16-12139572-DEL-71 |
| chr16-12142586-DEL-22321 | chr16-12142586-DEL-22321 |
| chr16-12164633-DEL-64 | chr16-12164634-DEL-64 |
| chr16-12193635-DEL-552 | chr16-12193635-DEL-551 |
| chr16-12195539-DEL-701 | chr16-12195538-DEL-700 |
| chr16-12212909-DEL-59 | chr16-12212900-DEL-55 |
| chr16-12216817-DEL-504 | chr16-12216818-DEL-503 |
| chr16-12235106-DEL-6342 | chr16-12235107-DEL-6342 |
| chr16-12252023-DEL-259 | chr16-12252024-DEL-259 |
| chr16-12280245-DEL-7062 | chr16-12280246-DEL-7062 |
| chr16-12280263-DEL-212 | chr16-12280263-DEL-212 |
| chr16-12305880-DEL-97 | chr16-12305881-DEL-97 |
| chr16-12354743-DEL-310 | chr16-12354744-DEL-310 |
| chr16-12407966-DEL-6599 | chr16-12407967-DEL-6599 |
| chr16-12414085-DEL-208 | chr16-12414086-DEL-208 |
| chr16-12418827-DEL-506 | chr16-12418828-DEL-507 |
| chr16-12476427-DEL-325 | chr16-12476395-DEL-307 |
| chr16-12477376-DEL-233 | chr16-12477377-DEL-234 |
| chr16-12501078-DEL-190 | chr16-12501079-DEL-190 |
| chr16-12505013-DEL-58 | chr16-12505008-DEL-64 |
| chr16-12520555-DEL-69 | chr16-12520605-DEL-69 |
| chr16-12646669-DEL-2122 | chr16-12646658-DEL-2123 |
| chr16-12647848-DEL-842 | chr16-12647849-DEL-842 |
| chr16-12657233-DEL-332 | chr16-12657234-DEL-332 |
| chr16-12664112-DEL-107 | chr16-12664113-DEL-107 |
| chr16-12671239-DEL-178 | chr16-12671240-DEL-178 |
| chr16-12674540-DEL-79 | chr16-12674551-DEL-75 |
| chr16-12684167-DEL-2263 | chr16-12684167-DEL-2264 |
| chr16-12695652-DEL-73 | chr16-12695623-DEL-72 |
| chr16-12718285-DEL-68 | chr16-12718286-DEL-68 |
| chr16-12729190-DEL-71 | chr16-12729191-DEL-55 |
| chr16-12744711-DEL-173 | chr16-12744712-DEL-173 |
| chr16-12753832-DEL-64 | chr16-12753833-DEL-64 |
| chr16-12760655-DEL-188 | chr16-12760656-DEL-188 |
| chr16-12765133-DEL-93 | chr16-12765132-DEL-89 |
| chr16-12775089-DEL-1774 | chr16-12775090-DEL-1774 |
| chr16-12794185-DEL-5290 | chr16-12794186-DEL-5290 |
| chr16-12804631-DEL-356 | chr16-12804632-DEL-356 |
| chr16-12860235-DEL-111 | chr16-12860236-DEL-111 |
| chr16-12865956-DEL-102 | chr16-12865957-DEL-102 |
| chr16-12874410-DEL-215 | chr16-12874411-DEL-215 |
| chr16-12951502-DEL-6401 | chr16-12951503-DEL-6401 |
| chr16-12990765-DEL-156 | chr16-12990761-DEL-149 |
| chr16-12994279-DEL-6197 | chr16-12994280-DEL-6197 |
| chr16-13019310-DEL-71 | chr16-13019311-DEL-71 |
| chr16-13031328-DEL-115 | chr16-13031329-DEL-115 |
| chr16-13037981-DEL-54 | chr16-13037982-DEL-54 |
| chr16-13064660-DEL-1059 | chr16-13064661-DEL-1059 |
| chr16-13065597-DEL-229 | chr16-13065598-DEL-229 |
| chr16-13088280-DEL-452 | chr16-13088281-DEL-452 |

| | |
|---|---|
| chr16-13130541-DEL-124 | chr16-13130492-DEL-123 |
| chr16-13140331-DEL-786 | chr16-13140332-DEL-786 |
| chr16-13161487-DEL-71 | chr16-13161488-DEL-71 |
| chr16-13223296-DEL-60 | chr16-13223297-DEL-60 |
| chr16-13313523-DEL-110 | chr16-13313480-DEL-154 |
| chr16-13327257-DEL-174 | chr16-13327256-DEL-176 |
| chr16-13340730-DEL-175 | chr16-13340731-DEL-175 |
| chr16-13359151-DEL-61 | chr16-13359169-DEL-60 |
| chr16-13361024-DEL-156 | chr16-13361025-DEL-156 |
| chr16-13362076-DEL-138 | chr16-13362077-DEL-138 |
| chr16-13382108-DEL-194 | chr16-13382109-DEL-194 |
| chr16-13394111-DEL-215 | chr16-13394112-DEL-215 |
| chr16-13404471-DEL-349 | chr16-13404475-DEL-350 |
| chr16-13409271-DEL-180 | chr16-13409284-DEL-168 |
| chr16-13410521-DEL-84 | chr16-13410522-DEL-84 |
| chr16-13416549-DEL-1431 | chr16-13416550-DEL-1431 |
| chr16-13419568-DEL-226 | chr16-13419569-DEL-226 |
| chr16-13432624-DEL-208 | chr16-13432625-DEL-208 |
| chr16-13452612-DEL-1773 | chr16-13452613-DEL-1773 |
| chr16-13459797-DEL-74 | chr16-13459798-DEL-74 |
| chr16-13502769-DEL-68 | chr16-13502770-DEL-68 |
| chr16-13513374-DEL-180 | chr16-13513332-DEL-180 |
| chr16-13523815-DEL-374 | chr16-13523816-DEL-374 |
| chr16-13538046-DEL-279 | chr16-13538047-DEL-279 |
| chr16-13539773-DEL-151 | chr16-13539803-DEL-155 |
| chr16-13553234-DEL-184 | chr16-13553235-DEL-184 |
| chr16-13577309-DEL-1162 | chr16-13577310-DEL-1141 |
| chr16-13577737-DEL-315 | chr16-13577738-DEL-315 |
| chr16-13580878-DEL-204 | chr16-13580879-DEL-204 |
| chr16-13583311-DEL-208 | chr16-13583312-DEL-208 |
| chr16-13585058-DEL-197 | chr16-13585058-DEL-198 |
| chr16-13592538-DEL-1366 | chr16-13592539-DEL-1366 |
| chr16-13594597-DEL-208 | chr16-13594584-DEL-209 |
| chr16-13603869-DEL-1904 | chr16-13603870-DEL-1904 |
| chr16-13619884-DEL-52 | chr16-13619885-DEL-52 |
| chr16-13688053-DEL-206 | chr16-13688044-DEL-194 |
| chr16-13691465-DEL-213 | chr16-13691466-DEL-213 |
| chr16-13726226-DEL-5926 | chr16-13726224-DEL-5926 |
| chr16-13741903-DEL-1893 | chr16-13741890-DEL-1901 |
| chr16-13747855-DEL-483 | chr16-13747856-DEL-483 |
| chr16-13767013-DEL-227 | chr16-13767014-DEL-227 |
| chr16-13785517-DEL-1045 | chr16-13785518-DEL-1045 |
| chr16-13795835-DEL-215 | chr16-13795836-DEL-215 |
| chr16-13813418-DEL-54 | chr16-13813419-DEL-54 |
| chr16-13815863-DEL-233 | chr16-13815864-DEL-233 |
| chr16-13819668-DEL-116 | chr16-13819657-DEL-117 |
| chr16-13828144-DEL-183 | chr16-13828145-DEL-183 |
| chr16-13831764-DEL-64 | chr16-13831769-DEL-64 |
| chr16-13855767-DEL-94 | chr16-13855725-DEL-126 |
| chr16-13859480-DEL-2855 | chr16-13859481-DEL-2856 |
| chr16-13865364-DEL-1064 | chr16-13865365-DEL-1064 |
| chr16-13866672-DEL-62 | chr16-13866673-DEL-62 |
| chr16-13890825-DEL-180 | chr16-13890812-DEL-194 |
| chr16-13931446-DEL-408 | chr16-13931447-DEL-408 |
| chr16-13933757-DEL-55 | chr16-13933770-DEL-54 |
| chr16-13996645-DEL-205 | chr16-13996646-DEL-205 |
| chr16-14007569-DEL-339 | chr16-14007570-DEL-339 |
| chr16-14024001-DEL-610 | chr16-14024002-DEL-610 |
| chr16-14036349-DEL-198 | chr16-14036350-DEL-198 |
| chr16-14045475-DEL-67 | chr16-14045476-DEL-67 |
| chr16-14062947-DEL-116 | chr16-14062948-DEL-116 |
| chr16-14083921-DEL-68 | chr16-14083895-DEL-70 |
| chr16-14091246-DEL-191 | chr16-14091237-DEL-189 |
| chr16-14103546-DEL-967 | chr16-14103547-DEL-971 |
| chr16-14122996-DEL-287 | chr16-14122995-DEL-287 |
| chr16-14151757-DEL-7121 | chr16-14151758-DEL-7121 |
| chr16-14167786-DEL-90 | chr16-14167736-DEL-105 |
| chr16-14193454-DEL-71 | chr16-14193446-DEL-80 |
| chr16-14199982-DEL-756 | chr16-14199981-DEL-755 |
| chr16-14259522-DEL-140 | chr16-14259523-DEL-150 |
| chr16-14270838-DEL-83 | chr16-14270839-DEL-83 |
| chr16-14276420-DEL-60 | chr16-14276422-DEL-59 |
| chr16-14286395-DEL-63 | chr16-14286396-DEL-63 |
| chr16-14294755-DEL-278 | chr16-14294756-DEL-278 |
| chr16-14327400-DEL-172 | chr16-14327401-DEL-172 |
| chr16-14350075-DEL-123 | chr16-14350052-DEL-147 |
| chr16-14367646-DEL-833 | chr16-14367647-DEL-833 |
| chr16-14371878-DEL-965 | chr16-14371879-DEL-965 |
| chr16-14372968-DEL-78 | chr16-14372969-DEL-78 |
| chr16-14387632-DEL-6403 | chr16-14387633-DEL-6403 |

| | |
|---|---|
| chr16-14433829-DEL-138 | chr16-14433843-DEL-108 |
| chr16-14462975-DEL-2682 | chr16-14462976-DEL-2682 |
| chr16-14493637-DEL-14564 | chr16-14493638-DEL-14564 |
| chr16-14559224-DEL-64 | chr16-14559239-DEL-50 |
| chr16-14566573-DEL-6442 | chr16-14566574-DEL-6442 |
| chr16-14572699-DEL-53 | chr16-14572700-DEL-53 |
| chr16-14623792-DEL-459 | chr16-14623793-DEL-459 |
| chr16-14624737-DEL-396 | chr16-14624738-DEL-396 |
| chr16-14627009-DEL-6438 | chr16-14627010-DEL-6438 |
| chr16-14633617-DEL-68 | chr16-14633618-DEL-68 |
| chr16-14639737-DEL-134 | chr16-14639738-DEL-134 |
| chr16-14674116-DEL-165 | chr16-14674117-DEL-165 |
| chr16-14676863-DEL-111 | chr16-14676864-DEL-111 |
| chr16-14683217-DEL-166 | chr16-14683218-DEL-166 |
| chr16-14690581-DEL-6198 | chr16-14690582-DEL-6198 |
| chr16-14835479-DEL-56 | chr16-14835493-DEL-50 |
| chr16-14838645-DEL-1279 | chr16-14838646-DEL-1279 |
| chr16-14863807-DEL-1875 | chr16-14863816-DEL-1860 |
| chr16-14899124-DEL-661 | chr16-14899125-DEL-661 |
| chr16-14988665-DEL-829 | chr16-14988666-DEL-829 |
| chr16-15014011-DEL-388 | chr16-15014012-DEL-388 |
| chr16-15019080-DEL-505 | chr16-15019071-DEL-506 |
| chr16-15072804-DEL-1934 | chr16-15072805-DEL-1934 |
| chr16-15107092-DEL-66 | chr16-15107093-DEL-66 |
| chr16-15133241-DEL-6654 | chr16-15133242-DEL-6654 |
| chr16-15160737-DEL-343 | chr16-15160738-DEL-343 |
| chr16-15222141-DEL-6583 | chr16-15222142-DEL-6583 |
| chr16-15242185-DEL-219 | chr16-15242186-DEL-219 |
| chr16-15251959-DEL-204 | chr16-15251998-DEL-204 |
| chr16-15257297-DEL-1224 | chr16-15257298-DEL-1224 |
| chr16-15289835-DEL-91 | chr16-15289836-DEL-92 |
| chr16-15306252-DEL-99 | chr16-15306253-DEL-99 |
| chr16-15320668-DEL-7079 | chr16-15320669-DEL-7079 |
| chr16-15336379-DEL-198 | chr16-15336389-DEL-186 |
| chr16-15346810-DEL-963 | chr16-15346811-DEL-963 |
| chr16-15381481-DEL-510 | chr16-15381482-DEL-510 |
| chr16-15401543-DEL-212 | chr16-15401544-DEL-212 |
| chr16-15413950-DEL-194 | chr16-15413951-DEL-194 |
| chr16-15417263-DEL-113 | chr16-15417277-DEL-73 |
| chr16-15435963-DEL-177 | chr16-15435964-DEL-177 |
| chr16-15440366-DEL-923 | chr16-15440367-DEL-923 |
| chr16-15445783-DEL-216 | chr16-15445784-DEL-216 |
| chr16-15456459-DEL-122 | chr16-15456460-DEL-121 |
| chr16-15465167-DEL-61 | chr16-15465168-DEL-61 |
| chr16-15468119-DEL-188 | chr16-15468128-DEL-187 |
| chr16-15471284-DEL-131 | chr16-15471285-DEL-131 |
| chr16-15476985-DEL-188 | chr16-15476983-DEL-191 |
| chr16-15483488-DEL-230 | chr16-15483442-DEL-231 |
| chr16-15489080-DEL-570 | chr16-15489075-DEL-576 |
| chr16-15505919-DEL-982 | chr16-15505920-DEL-982 |
| chr16-15512718-DEL-99 | chr16-15512719-DEL-99 |
| chr16-15549206-DEL-85 | chr16-15549207-DEL-85 |
| chr16-15553591-DEL-860 | chr16-15553592-DEL-860 |
| chr16-15559162-DEL-10581 | chr16-15559163-DEL-10581 |
| chr16-15589053-DEL-225 | chr16-15589054-DEL-225 |
| chr16-15617962-DEL-887 | chr16-15617963-DEL-887 |
| chr16-15641142-DEL-2956 | chr16-15641143-DEL-2956 |
| chr16-15661825-DEL-424 | chr16-15661826-DEL-424 |
| chr16-15662356-DEL-1709 | chr16-15662357-DEL-1709 |
| chr16-15669792-DEL-1306 | chr16-15669793-DEL-1306 |
| chr16-15683393-DEL-351 | chr16-15683394-DEL-351 |
| chr16-15697134-DEL-60 | chr16-15697135-DEL-60 |
| chr16-15769693-DEL-76 | chr16-15769694-DEL-76 |
| chr16-15777659-DEL-185 | chr16-15777660-DEL-185 |
| chr16-15792163-DEL-211 | chr16-15792164-DEL-211 |
| chr16-15816298-DEL-255 | chr16-15816298-DEL-258 |
| chr16-15843119-DEL-2067 | chr16-15843120-DEL-2067 |
| chr16-15860201-DEL-5396 | chr16-15860200-DEL-5397 |
| chr16-15877906-DEL-58 | chr16-15877907-DEL-59 |
| chr16-15880230-DEL-64 | chr16-15880220-DEL-65 |
| chr16-15895989-DEL-105 | chr16-15895990-DEL-105 |
| chr16-15898642-DEL-6566 | chr16-15898643-DEL-6566 |
| chr16-15908733-DEL-614 | chr16-15908743-DEL-605 |
| chr16-15911093-DEL-732 | chr16-15911094-DEL-730 |
| chr16-15927966-DEL-70 | chr16-15927966-DEL-70 |
| chr16-15943854-DEL-2202 | chr16-15943855-DEL-2202 |
| chr16-15955313-DEL-807 | chr16-15955314-DEL-807 |
| chr16-15963414-DEL-98 | chr16-15963422-DEL-97 |
| chr16-15970562-DEL-167 | chr16-15970563-DEL-167 |
| chr16-16001523-DEL-198 | chr16-16001510-DEL-199 |

| | |
|---|---|
| chr16-16055874-DEL-1223 | chr16-16055875-DEL-1223 |
| chr16-16109810-DEL-381 | chr16-16109802-DEL-394 |
| chr16-16119629-DEL-441 | chr16-16119673-DEL-404 |
| chr16-16123446-DEL-302 | chr16-16123447-DEL-302 |
| chr16-16126720-DEL-57 | chr16-16126721-DEL-57 |
| chr16-16182726-DEL-169 | chr16-16182727-DEL-169 |
| chr16-16195283-DEL-308 | chr16-16195284-DEL-304 |
| chr16-16210467-DEL-6255 | chr16-16210495-DEL-6249 |
| chr16-16260219-DEL-66 | chr16-16260220-DEL-66 |
| chr16-16263754-DEL-1706 | chr16-16263755-DEL-1706 |
| chr16-16273316-DEL-350 | chr16-16273330-DEL-324 |
| chr16-16292381-DEL-211 | chr16-16292382-DEL-211 |
| chr16-16324911-DEL-7314 | chr16-16324912-DEL-7314 |
| chr16-16325264-DEL-212 | chr16-16325264-DEL-212 |
| chr16-16362743-DEL-204 | chr16-16362744-DEL-204 |
| chr16-16403630-DEL-510 | chr16-16403631-DEL-510 |
| chr16-16418984-DEL-221 | chr16-16418985-DEL-221 |
| chr16-16440233-DEL-1856 | chr16-16440234-DEL-1856 |
| chr16-16470647-DEL-104 | chr16-16470611-DEL-112 |
| chr16-16478542-DEL-86 | chr16-16478543-DEL-86 |
| chr16-16486058-DEL-1384 | chr16-16486059-DEL-1384 |
| chr16-16513674-DEL-3344 | chr16-16513674-DEL-3344 |
| chr16-16541337-DEL-489 | chr16-16541338-DEL-489 |
| chr16-16542551-DEL-993 | chr16-16542557-DEL-994 |
| chr16-16553629-DEL-55 | chr16-16553615-DEL-56 |
| chr16-16581773-DEL-56 | chr16-16581751-DEL-52 |
| chr16-16602532-DEL-81 | chr16-16602532-DEL-78 |
| chr16-16637562-DEL-492 | chr16-16637553-DEL-493 |
| chr16-16641564-DEL-1405 | chr16-16641564-DEL-1405 |
| chr16-16642941-DEL-78 | chr16-16642942-DEL-52 |
| chr16-16667548-DEL-201 | chr16-16667549-DEL-201 |
| chr16-16737837-DEL-89 | chr16-16737838-DEL-89 |
| chr16-16751711-DEL-6394 | chr16-16751712-DEL-6394 |
| chr16-16760215-DEL-149 | chr16-16760216-DEL-149 |
| chr16-16765134-DEL-220 | chr16-16765135-DEL-220 |
| chr16-16766492-DEL-646 | chr16-16766493-DEL-646 |
| chr16-16793850-DEL-341 | chr16-16793851-DEL-341 |
| chr16-16798517-DEL-198 | chr16-16798518-DEL-198 |
| chr16-16805673-DEL-214 | chr16-16805674-DEL-214 |
| chr16-16813343-DEL-1030 | chr16-16813344-DEL-1030 |
| chr16-16834341-DEL-197 | chr16-16834342-DEL-197 |
| chr16-16897796-DEL-58 | chr16-16897797-DEL-58 |
| chr16-16911330-DEL-651 | chr16-16911348-DEL-633 |
| chr16-16916191-DEL-111 | chr16-16916191-DEL-111 |
| chr16-16926412-DEL-206 | chr16-16926413-DEL-206 |
| chr16-16994994-DEL-66 | chr16-16994994-DEL-66 |
| chr16-16998226-DEL-217 | chr16-16998226-DEL-217 |
| chr16-17002039-DEL-190 | chr16-17002043-DEL-191 |
| chr16-17004184-DEL-217 | chr16-17004185-DEL-217 |
| chr16-17009417-DEL-283 | chr16-17009418-DEL-283 |
| chr16-17015156-DEL-212 | chr16-17015157-DEL-212 |
| chr16-17019917-DEL-2904 | chr16-17019918-DEL-2904 |
| chr16-17039856-DEL-204 | chr16-17039839-DEL-201 |
| chr16-17040634-DEL-125 | chr16-17040635-DEL-125 |
| chr16-17049623-DEL-213 | chr16-17049624-DEL-213 |
| chr16-17087327-DEL-307 | chr16-17087286-DEL-308 |
| chr16-17144696-DEL-2813 | chr16-17144697-DEL-2813 |
| chr16-17156092-DEL-274 | chr16-17156093-DEL-274 |
| chr16-17172616-DEL-57 | chr16-17172617-DEL-56 |
| chr16-17178626-DEL-470 | chr16-17178627-DEL-423 |
| chr16-17186772-DEL-232 | chr16-17186773-DEL-232 |
| chr16-17201963-DEL-217 | chr16-17201964-DEL-217 |
| chr16-17205577-DEL-234 | chr16-17205578-DEL-234 |
| chr16-17209960-DEL-399 | chr16-17209961-DEL-399 |
| chr16-17231678-DEL-60 | chr16-17231679-DEL-60 |
| chr16-17239521-DEL-220 | chr16-17239522-DEL-220 |
| chr16-17248808-DEL-450 | chr16-17248809-DEL-450 |
| chr16-17293467-DEL-193 | chr16-17293468-DEL-193 |
| chr16-17342685-DEL-213 | chr16-17342686-DEL-213 |
| chr16-17365043-DEL-100 | chr16-17365044-DEL-100 |
| chr16-17406696-DEL-201 | chr16-17406697-DEL-201 |
| chr16-17429716-DEL-2731 | chr16-17429717-DEL-2731 |
| chr16-17508361-DEL-208 | chr16-17508362-DEL-208 |
| chr16-17513256-DEL-286 | chr16-17513257-DEL-282 |
| chr16-17528256-DEL-910 | chr16-17528257-DEL-910 |
| chr16-17542202-DEL-130 | chr16-17542203-DEL-130 |
| chr16-17559572-DEL-64 | chr16-17559566-DEL-77 |
| chr16-17572111-DEL-167 | chr16-17572111-DEL-167 |
| chr16-17581362-DEL-54 | chr16-17581377-DEL-56 |
| chr16-17673740-DEL-206 | chr16-17673741-DEL-206 |

| | |
|---|---|
| chr16-17684617-DEL-207 | chr16-17684618-DEL-207 |
| chr16-17691655-DEL-835 | chr16-17691648-DEL-847 |
| chr16-17693212-DEL-79 | chr16-17693213-DEL-79 |
| chr16-17769314-DEL-72 | chr16-17769319-DEL-68 |
| chr16-17770396-DEL-3875 | chr16-17770397-DEL-3875 |
| chr16-17775967-DEL-71 | chr16-17775957-DEL-87 |
| chr16-17783346-DEL-176 | chr16-17783347-DEL-176 |
| chr16-17816122-DEL-1437 | chr16-17816124-DEL-1438 |
| chr16-17829758-DEL-5163 | chr16-17829757-DEL-5163 |
| chr16-17852092-DEL-122 | chr16-17852093-DEL-122 |
| chr16-17885484-DEL-226 | chr16-17885485-DEL-226 |
| chr16-17907894-DEL-62 | chr16-17907890-DEL-78 |
| chr16-17931369-DEL-51 | chr16-17931370-DEL-51 |
| chr16-18034799-DEL-206 | chr16-18034800-DEL-206 |
| chr16-18076730-DEL-52 | chr16-18076731-DEL-52 |
| chr16-18081158-DEL-498 | chr16-18081159-DEL-498 |
| chr16-18082772-DEL-214 | chr16-18082773-DEL-214 |
| chr16-18133304-DEL-116 | chr16-18133305-DEL-116 |
| chr16-18137980-DEL-207 | chr16-18137975-DEL-206 |
| chr16-18140382-DEL-136 | chr16-18140422-DEL-137 |
| chr16-18152948-DEL-346 | chr16-18152949-DEL-344 |
| chr16-18154438-DEL-1830 | chr16-18154439-DEL-1830 |
| chr16-18162842-DEL-203 | chr16-18162843-DEL-202 |
| chr16-18176493-DEL-508 | chr16-18176494-DEL-508 |
| chr16-18183257-DEL-312 | chr16-18183258-DEL-312 |
| chr16-18226048-DEL-230 | chr16-18226058-DEL-237 |
| chr16-18255078-DEL-58 | chr16-18255079-DEL-58 |
| chr16-18257766-DEL-244 | chr16-18257766-DEL-245 |
| chr16-18263748-DEL-199 | chr16-18263752-DEL-196 |
| chr16-18285141-DEL-139 | chr16-18285142-DEL-139 |
| chr16-18315651-DEL-173 | chr16-18315652-DEL-173 |
| chr16-18337823-DEL-53 | chr16-18337824-DEL-53 |
| chr16-18339664-DEL-194 | chr16-18339641-DEL-214 |
| chr16-18349016-DEL-201 | chr16-18349017-DEL-201 |
| chr16-18350924-DEL-1181 | chr16-18350925-DEL-1181 |
| chr16-18397668-DEL-333 | chr16-18397669-DEL-333 |
| chr16-18452893-DEL-436 | chr16-18452895-DEL-427 |
| chr16-18468887-DEL-62 | chr16-18468888-DEL-62 |
| chr16-18469372-DEL-149 | chr16-18469373-DEL-149 |
| chr16-18517908-DEL-223 | chr16-18517909-DEL-223 |
| chr16-18574945-DEL-96 | chr16-18574946-DEL-96 |
| chr16-18591791-DEL-55 | chr16-18591795-DEL-58 |
| chr16-18606256-DEL-253 | chr16-18606257-DEL-253 |
| chr16-18621485-DEL-853 | chr16-18621486-DEL-853 |
| chr16-18648531-DEL-137 | chr16-18648530-DEL-137 |
| chr16-18656656-DEL-172 | chr16-18656657-DEL-172 |
| chr16-18660722-DEL-1658 | chr16-18660723-DEL-1658 |
| chr16-18675518-DEL-97 | chr16-18675519-DEL-97 |
| chr16-18723604-DEL-61 | chr16-18723605-DEL-61 |
| chr16-18776822-DEL-1275 | chr16-18776823-DEL-1275 |
| chr16-18782147-DEL-114 | chr16-18782148-DEL-114 |
| chr16-18795922-DEL-1647 | chr16-18795923-DEL-1647 |
| chr16-18829369-DEL-91 | chr16-18829370-DEL-91 |
| chr16-18858061-DEL-185 | chr16-18858062-DEL-185 |
| chr16-18865908-DEL-3429 | chr16-18865909-DEL-3429 |
| chr16-18870085-DEL-276 | chr16-18870085-DEL-276 |
| chr16-18872400-DEL-2913 | chr16-18872401-DEL-2913 |
| chr16-18958426-DEL-6238 | chr16-18958426-DEL-6238 |
| chr16-18979518-DEL-326 | chr16-18979519-DEL-326 |
| chr16-18988469-DEL-2300 | chr16-18988470-DEL-2300 |
| chr16-18993094-DEL-2059 | chr16-18993096-DEL-2058 |
| chr16-18995980-DEL-261 | chr16-18995980-DEL-262 |
| chr16-19066307-DEL-53 | chr16-19066308-DEL-53 |
| chr16-19087903-DEL-58 | chr16-19087903-DEL-58 |
| chr16-19197770-DEL-58 | chr16-19197778-DEL-64 |
| chr16-19207145-DEL-1003 | chr16-19207146-DEL-1003 |
| chr16-19231804-DEL-6638 | chr16-19231805-DEL-6638 |
| chr16-19232192-DEL-212 | chr16-19232192-DEL-212 |
| chr16-19249154-DEL-52 | chr16-19249155-DEL-52 |
| chr16-19255166-DEL-160 | chr16-19255158-DEL-160 |
| chr16-19292832-DEL-964 | chr16-19292833-DEL-964 |
| chr16-19304538-DEL-87 | chr16-19304539-DEL-87 |
| chr16-19310170-DEL-178 | chr16-19310171-DEL-178 |
| chr16-19313616-DEL-196 | chr16-19313615-DEL-197 |
| chr16-19317095-DEL-362 | chr16-19317096-DEL-362 |
| chr16-19317777-DEL-401 | chr16-19317778-DEL-401 |
| chr16-19318327-DEL-114 | chr16-19318328-DEL-113 |
| chr16-19345348-DEL-1090 | chr16-19345349-DEL-1090 |
| chr16-19357925-DEL-330 | chr16-19357926-DEL-330 |
| chr16-19396391-DEL-76 | chr16-19396392-DEL-76 |

| | |
|---|---|
| chr16-19406584-DEL-5876 | chr16-19406585-DEL-5876 |
| chr16-19413055-DEL-8494 | chr16-19413056-DEL-8494 |
| chr16-19440680-DEL-326 | chr16-19440681-DEL-326 |
| chr16-19460365-DEL-319 | chr16-19460382-DEL-324 |
| chr16-19463354-DEL-116 | chr16-19463350-DEL-121 |
| chr16-19472681-DEL-85 | chr16-19472681-DEL-85 |
| chr16-19491790-DEL-56 | chr16-19491777-DEL-70 |
| chr16-19503717-DEL-821 | chr16-19503722-DEL-817 |
| chr16-19511500-DEL-54 | chr16-19511498-DEL-57 |
| chr16-19530139-DEL-6399 | chr16-19530140-DEL-6399 |
| chr16-19550497-DEL-382 | chr16-19550498-DEL-375 |
| chr16-19638390-DEL-923 | chr16-19638390-DEL-923 |
| chr16-19647607-DEL-53 | chr16-19647608-DEL-53 |
| chr16-19676865-DEL-265 | chr16-19676866-DEL-264 |
| chr16-19719920-DEL-249 | chr16-19719921-DEL-249 |
| chr16-19733879-DEL-110 | chr16-19733874-DEL-152 |
| chr16-19787293-DEL-98 | chr16-19787283-DEL-130 |
| chr16-19813349-DEL-170 | chr16-19813347-DEL-171 |
| chr16-19855339-DEL-7145 | chr16-19855340-DEL-7145 |
| chr16-19888098-DEL-212 | chr16-19888099-DEL-212 |
| chr16-19896250-DEL-1469 | chr16-19896250-DEL-1467 |
| chr16-19931320-DEL-160 | chr16-19931321-DEL-160 |
| chr16-19946737-DEL-197 | chr16-19946738-DEL-197 |
| chr16-19994846-DEL-1052 | chr16-19994847-DEL-1052 |
| chr16-20006595-DEL-215 | chr16-20006593-DEL-216 |
| chr16-20009715-DEL-210 | chr16-20009716-DEL-209 |
| chr16-20035515-DEL-77 | chr16-20035517-DEL-76 |
| chr16-20065942-DEL-1017 | chr16-20065943-DEL-1017 |
| chr16-20068519-DEL-199 | chr16-20068520-DEL-199 |
| chr16-20078869-DEL-2739 | chr16-20078870-DEL-2739 |
| chr16-20089897-DEL-199 | chr16-20089898-DEL-199 |
| chr16-20095199-DEL-119 | chr16-20095200-DEL-119 |
| chr16-20097076-DEL-180 | chr16-20097077-DEL-180 |
| chr16-20098192-DEL-51 | chr16-20098193-DEL-51 |
| chr16-20110087-DEL-171 | chr16-20110088-DEL-171 |
| chr16-20126121-DEL-234 | chr16-20126122-DEL-234 |
| chr16-20142567-DEL-57 | chr16-20142556-DEL-69 |
| chr16-20145890-DEL-1134 | chr16-20145891-DEL-1134 |
| chr16-20158931-DEL-218 | chr16-20158932-DEL-214 |
| chr16-20170928-DEL-209 | chr16-20170927-DEL-210 |
| chr16-20171317-DEL-498 | chr16-20171318-DEL-498 |
| chr16-20208476-DEL-170 | chr16-20208516-DEL-151 |
| chr16-20228872-DEL-591 | chr16-20228873-DEL-591 |
| chr16-20253108-DEL-214 | chr16-20253109-DEL-214 |
| chr16-20256118-DEL-909 | chr16-20256122-DEL-907 |
| chr16-20258042-DEL-156 | chr16-20258036-DEL-143 |
| chr16-20273412-DEL-183 | chr16-20273413-DEL-183 |
| chr16-20283094-DEL-194 | chr16-20283095-DEL-194 |
| chr16-20284486-DEL-138 | chr16-20284487-DEL-138 |
| chr16-20301600-DEL-207 | chr16-20301601-DEL-207 |
| chr16-20302733-DEL-180 | chr16-20302748-DEL-176 |
| chr16-20308829-DEL-199 | chr16-20308830-DEL-199 |
| chr16-20341421-DEL-197 | chr16-20341422-DEL-197 |
| chr16-20351291-DEL-102 | chr16-20351292-DEL-90 |
| chr16-20389302-DEL-175 | chr16-20389295-DEL-183 |
| chr16-20391616-DEL-7224 | chr16-20391617-DEL-7224 |
| chr16-20444979-DEL-52 | chr16-20444980-DEL-52 |
| chr16-20541890-DEL-1102 | chr16-20541893-DEL-1100 |
| chr16-20579693-DEL-199 | chr16-20579694-DEL-199 |
| chr16-20580232-DEL-50 | chr16-20580232-DEL-50 |
| chr16-20582480-DEL-207 | chr16-20582481-DEL-207 |
| chr16-20588890-DEL-214 | chr16-20588891-DEL-214 |
| chr16-20590713-DEL-204 | chr16-20590714-DEL-204 |
| chr16-20600481-DEL-90 | chr16-20600482-DEL-90 |
| chr16-20624237-DEL-207 | chr16-20624238-DEL-207 |
| chr16-20647461-DEL-407 | chr16-20647462-DEL-407 |
| chr16-20650815-DEL-204 | chr16-20650816-DEL-203 |
| chr16-20680639-DEL-529 | chr16-20680640-DEL-529 |
| chr16-20709526-DEL-85 | chr16-20709527-DEL-85 |
| chr16-20733610-DEL-7101 | chr16-20733611-DEL-7101 |
| chr16-20843278-DEL-139 | chr16-20843279-DEL-139 |
| chr16-20864343-DEL-694 | chr16-20864343-DEL-694 |
| chr16-20865586-DEL-68 | chr16-20865586-DEL-68 |
| chr16-20907442-DEL-1287 | chr16-20907443-DEL-1287 |
| chr16-20932265-DEL-201 | chr16-20932265-DEL-201 |
| chr16-20948193-DEL-211 | chr16-20948193-DEL-211 |
| chr16-20950454-DEL-159 | chr16-20950454-DEL-159 |
| chr16-20997623-DEL-186 | chr16-20997624-DEL-186 |
| chr16-21007670-DEL-131 | chr16-21007671-DEL-131 |
| chr16-21012525-DEL-198 | chr16-21012526-DEL-198 |

| | |
|---|---|
| chr16-21050279-DEL-207 | chr16-21050280-DEL-207 |
| chr16-21060132-DEL-198 | chr16-21060133-DEL-198 |
| chr16-21172821-DEL-186 | chr16-21172816-DEL-192 |
| chr16-21257502-DEL-80 | chr16-21257503-DEL-80 |
| chr16-21280315-DEL-143 | chr16-21280316-DEL-143 |
| chr16-21285591-DEL-1856 | chr16-21285592-DEL-1856 |
| chr16-21307644-DEL-1833 | chr16-21307645-DEL-1833 |
| chr16-21313336-DEL-202 | chr16-21313337-DEL-202 |
| chr16-21323348-DEL-223 | chr16-21323349-DEL-223 |
| chr16-21398729-DEL-416 | chr16-21398730-DEL-416 |
| chr16-21404224-DEL-245 | chr16-21404225-DEL-245 |
| chr16-21411065-DEL-264 | chr16-21411060-DEL-265 |
| chr16-21416329-DEL-6384 | chr16-21416330-DEL-6384 |
| chr16-21431765-DEL-266 | chr16-21431766-DEL-266 |
| chr16-21436648-DEL-488 | chr16-21436648-DEL-487 |
| chr16-21442212-DEL-122 | chr16-21442195-DEL-98 |
| chr16-21502387-DEL-552 | chr16-21502388-DEL-552 |
| chr16-21529656-DEL-91 | chr16-21529683-DEL-73 |
| chr16-21532654-DEL-201 | chr16-21532656-DEL-200 |
| chr16-21554631-DEL-87 | chr16-21554615-DEL-53 |
| chr16-21604312-DEL-201 | chr16-21604313-DEL-201 |
| chr16-21608928-DEL-196 | chr16-21608929-DEL-196 |
| chr16-21618828-DEL-58 | chr16-21618829-DEL-58 |
| chr16-21624444-DEL-797 | chr16-21624445-DEL-795 |
| chr16-21629475-DEL-370 | chr16-21629476-DEL-370 |
| chr16-21647042-DEL-168 | chr16-21647043-DEL-168 |
| chr16-21664461-DEL-69 | chr16-21664462-DEL-67 |
| chr16-21681898-DEL-189 | chr16-21681899-DEL-189 |
| chr16-21682987-DEL-124 | chr16-21682988-DEL-124 |
| chr16-21707158-DEL-204 | chr16-21707159-DEL-204 |
| chr16-21715170-DEL-762 | chr16-21715171-DEL-762 |
| chr16-21727518-DEL-234 | chr16-21727519-DEL-195 |
| chr16-21757581-DEL-6410 | chr16-21757582-DEL-6410 |
| chr16-21776197-DEL-766 | chr16-21776198-DEL-766 |
| chr16-21777752-DEL-53 | chr16-21777753-DEL-55 |
| chr16-21782552-DEL-101 | chr16-21782553-DEL-101 |
| chr16-21785516-DEL-1131 | chr16-21785517-DEL-1131 |
| chr16-21790004-DEL-52 | chr16-21790006-DEL-51 |
| chr16-21803233-DEL-175 | chr16-21803230-DEL-176 |
| chr16-21804693-DEL-204 | chr16-21804693-DEL-179 |
| chr16-21841044-DEL-113 | chr16-21841045-DEL-113 |
| chr16-21852124-DEL-2573 | chr16-21852125-DEL-2573 |
| chr16-21855499-DEL-201 | chr16-21855500-DEL-201 |
| chr16-21868416-DEL-328 | chr16-21868419-DEL-326 |
| chr16-21939285-DEL-80 | chr16-21939292-DEL-57 |
| chr16-21947182-DEL-210 | chr16-21947183-DEL-210 |
| chr16-21948348-DEL-255 | chr16-21948344-DEL-257 |
| chr16-21965867-DEL-65 | chr16-21965836-DEL-63 |
| chr16-21976345-DEL-418 | chr16-21976351-DEL-413 |
| chr16-21992073-DEL-227 | chr16-21992086-DEL-214 |
| chr16-21992814-DEL-171 | chr16-21992797-DEL-187 |
| chr16-22015808-DEL-60 | chr16-22015809-DEL-60 |
| chr16-22029507-DEL-201 | chr16-22029512-DEL-199 |
| chr16-22033963-DEL-198 | chr16-22033962-DEL-198 |
| chr16-22035335-DEL-211 | chr16-22035334-DEL-207 |
| chr16-22035720-DEL-412 | chr16-22035721-DEL-412 |
| chr16-22057399-DEL-191 | chr16-22057400-DEL-191 |
| chr16-22073570-DEL-56 | chr16-22073571-DEL-56 |
| chr16-22096111-DEL-338 | chr16-22096112-DEL-338 |
| chr16-22097954-DEL-69 | chr16-22097955-DEL-69 |
| chr16-22098704-DEL-473 | chr16-22098705-DEL-473 |
| chr16-22113706-DEL-519 | chr16-22113707-DEL-519 |
| chr16-22126855-DEL-64 | chr16-22126862-DEL-58 |
| chr16-22128318-DEL-516 | chr16-22128319-DEL-516 |
| chr16-22188683-DEL-171 | chr16-22188685-DEL-167 |
| chr16-22223113-DEL-200 | chr16-22223107-DEL-198 |
| chr16-22267813-DEL-51 | chr16-22267814-DEL-51 |
| chr16-22296957-DEL-287 | chr16-22296956-DEL-287 |
| chr16-22387451-DEL-200 | chr16-22387452-DEL-200 |
| chr16-22400499-DEL-52 | chr16-22400501-DEL-58 |
| chr16-22408949-DEL-67 | chr16-22408914-DEL-64 |
| chr16-22532887-DEL-91 | chr16-22532888-DEL-91 |
| chr16-22547522-DEL-75 | chr16-22547540-DEL-51 |
| chr16-22577039-DEL-264 | chr16-22577040-DEL-264 |
| chr16-22591812-DEL-304 | chr16-22591813-DEL-304 |
| chr16-22646873-DEL-4838 | chr16-22646874-DEL-4838 |
| chr16-22647794-DEL-80 | chr16-22647795-DEL-80 |
| chr16-22651196-DEL-504 | chr16-22651197-DEL-504 |
| chr16-22661296-DEL-61 | chr16-22661273-DEL-69 |
| chr16-22720222-DEL-52 | chr16-22720223-DEL-52 |

| | |
|---|---|
| chr16-22734528-DEL-1302 | chr16-22734529-DEL-1302 |
| chr16-22741547-DEL-6251 | chr16-22741548-DEL-6251 |
| chr16-22753548-DEL-232 | chr16-22753549-DEL-232 |
| chr16-22762570-DEL-433 | chr16-22762569-DEL-433 |
| chr16-22784723-DEL-152 | chr16-22784724-DEL-148 |
| chr16-22787460-DEL-980 | chr16-22787461-DEL-980 |
| chr16-22798789-DEL-3051 | chr16-22798790-DEL-3051 |
| chr16-22824422-DEL-1516 | chr16-22824423-DEL-1516 |
| chr16-22835326-DEL-204 | chr16-22835327-DEL-204 |
| chr16-22854318-DEL-593 | chr16-22854319-DEL-593 |
| chr16-22861829-DEL-637 | chr16-22861824-DEL-638 |
| chr16-22862625-DEL-120 | chr16-22862626-DEL-120 |
| chr16-22879996-DEL-212 | chr16-22879997-DEL-212 |
| chr16-22881378-DEL-119 | chr16-22881379-DEL-94 |
| chr16-22899005-DEL-160 | chr16-22898994-DEL-172 |
| chr16-22923670-DEL-88 | chr16-22923671-DEL-89 |
| chr16-22941686-DEL-77 | chr16-22941681-DEL-52 |
| chr16-22950998-DEL-875 | chr16-22950999-DEL-875 |
| chr16-22962250-DEL-149 | chr16-22962258-DEL-135 |
| chr16-22971980-DEL-225 | chr16-22971987-DEL-219 |
| chr16-22978071-DEL-277 | chr16-22978072-DEL-277 |
| chr16-22993208-DEL-210 | chr16-22993209-DEL-210 |
| chr16-23019438-DEL-86 | chr16-23019439-DEL-86 |
| chr16-23058038-DEL-58 | chr16-23058022-DEL-64 |
| chr16-23070665-DEL-59 | chr16-23070664-DEL-56 |
| chr16-23076663-DEL-63 | chr16-23076655-DEL-72 |
| chr16-23093950-DEL-212 | chr16-23093951-DEL-212 |
| chr16-23100756-DEL-698 | chr16-23100757-DEL-698 |
| chr16-23101505-DEL-352 | chr16-23101506-DEL-352 |
| chr16-23102055-DEL-102 | chr16-23102056-DEL-102 |
| chr16-23124931-DEL-51 | chr16-23124908-DEL-78 |
| chr16-23129624-DEL-716 | chr16-23129625-DEL-716 |
| chr16-23141606-DEL-123 | chr16-23141602-DEL-124 |
| chr16-23145476-DEL-1264 | chr16-23145477-DEL-1264 |
| chr16-23153718-DEL-104 | chr16-23153719-DEL-104 |
| chr16-23160538-DEL-437 | chr16-23160539-DEL-437 |
| chr16-23162371-DEL-252 | chr16-23162372-DEL-252 |
| chr16-23199894-DEL-338 | chr16-23199899-DEL-334 |
| chr16-23201098-DEL-56 | chr16-23201099-DEL-52 |
| chr16-23215606-DEL-56 | chr16-23215606-DEL-52 |
| chr16-23227519-DEL-1210 | chr16-23227520-DEL-1209 |
| chr16-23229019-DEL-75 | chr16-23229031-DEL-64 |
| chr16-23253547-DEL-51 | chr16-23253548-DEL-51 |
| chr16-23261065-DEL-446 | chr16-23261066-DEL-446 |
| chr16-23274325-DEL-57 | chr16-23274326-DEL-73 |
| chr16-23298177-DEL-90 | chr16-23298178-DEL-90 |
| chr16-23367503-DEL-86 | chr16-23367516-DEL-62 |
| chr16-23394995-DEL-52 | chr16-23394988-DEL-84 |
| chr16-23412028-DEL-2401 | chr16-23412029-DEL-2401 |
| chr16-23424151-DEL-548 | chr16-23424152-DEL-548 |
| chr16-23496011-DEL-179 | chr16-23496012-DEL-179 |
| chr16-23504996-DEL-199 | chr16-23504997-DEL-199 |
| chr16-23533787-DEL-1241 | chr16-23533788-DEL-1241 |
| chr16-23557128-DEL-1128 | chr16-23557129-DEL-1128 |
| chr16-23567916-DEL-6231 | chr16-23567917-DEL-6231 |
| chr16-23568093-DEL-64 | chr16-23568094-DEL-64 |
| chr16-23582073-DEL-180 | chr16-23582029-DEL-181 |
| chr16-23595736-DEL-53 | chr16-23595736-DEL-52 |
| chr16-23612136-DEL-64 | chr16-23612137-DEL-64 |
| chr16-23650962-DEL-71 | chr16-23650963-DEL-71 |
| chr16-23653382-DEL-143 | chr16-23653429-DEL-115 |
| chr16-23680306-DEL-232 | chr16-23680343-DEL-196 |
| chr16-23747021-DEL-182 | chr16-23747022-DEL-182 |
| chr16-23763003-DEL-90 | chr16-23762959-DEL-112 |
| chr16-23771363-DEL-66 | chr16-23771371-DEL-56 |
| chr16-23920082-DEL-1268 | chr16-23920083-DEL-1268 |
| chr16-23967095-DEL-55 | chr16-23967096-DEL-60 |
| chr16-23975180-DEL-84 | chr16-23975202-DEL-52 |
| chr16-23989258-DEL-296 | chr16-23989259-DEL-296 |
| chr16-24054481-DEL-290 | chr16-24054482-DEL-290 |
| chr16-24058517-DEL-70 | chr16-24058518-DEL-70 |
| chr16-24085863-DEL-89 | chr16-24085864-DEL-89 |
| chr16-24092297-DEL-1165 | chr16-24092320-DEL-1139 |
| chr16-24103289-DEL-54 | chr16-24103264-DEL-55 |
| chr16-24151271-DEL-626 | chr16-24151272-DEL-626 |
| chr16-24159470-DEL-52 | chr16-24159471-DEL-52 |
| chr16-24171469-DEL-64 | chr16-24171470-DEL-64 |
| chr16-24198252-DEL-1028 | chr16-24198252-DEL-1031 |
| chr16-24202096-DEL-228 | chr16-24202097-DEL-228 |
| chr16-24204014-DEL-2432 | chr16-24204015-DEL-2432 |

| | |
|---|---|
| chr16-24220881-DEL-70 | chr16-24220894-DEL-68 |
| chr16-24230381-DEL-101 | chr16-24230375-DEL-102 |
| chr16-24265148-DEL-84 | chr16-24265141-DEL-84 |
| chr16-24348051-DEL-1729 | chr16-24348052-DEL-1729 |
| chr16-24355499-DEL-299 | chr16-24355500-DEL-299 |
| chr16-24370088-DEL-1237 | chr16-24370089-DEL-1237 |
| chr16-24375256-DEL-1891 | chr16-24375257-DEL-1891 |
| chr16-24406320-DEL-311 | chr16-24406321-DEL-311 |
| chr16-24481017-DEL-157 | chr16-24481018-DEL-157 |
| chr16-24521490-DEL-143 | chr16-24521491-DEL-143 |
| chr16-24552874-DEL-58 | chr16-24552875-DEL-58 |
| chr16-24603006-DEL-72 | chr16-24603028-DEL-51 |
| chr16-24670904-DEL-93 | chr16-24670905-DEL-93 |
| chr16-24772268-DEL-205 | chr16-24772269-DEL-205 |
| chr16-24816176-DEL-56 | chr16-24816179-DEL-54 |
| chr16-24837326-DEL-185 | chr16-24837327-DEL-185 |
| chr16-24850236-DEL-103 | chr16-24850240-DEL-99 |
| chr16-24856446-DEL-847 | chr16-24856447-DEL-847 |
| chr16-24894850-DEL-239 | chr16-24894851-DEL-239 |
| chr16-24932839-DEL-52 | chr16-24932840-DEL-52 |
| chr16-24984649-DEL-182 | chr16-24984653-DEL-177 |
| chr16-24990326-DEL-679 | chr16-24990327-DEL-679 |
| chr16-24993456-DEL-235 | chr16-24993457-DEL-235 |
| chr16-24995265-DEL-326 | chr16-24995266-DEL-326 |
| chr16-25009615-DEL-6256 | chr16-25009616-DEL-6256 |
| chr16-25109574-DEL-591 | chr16-25109575-DEL-591 |
| chr16-25130955-DEL-75 | chr16-25130956-DEL-75 |
| chr16-25145357-DEL-928 | chr16-25145358-DEL-928 |
| chr16-25152019-DEL-104 | chr16-25152040-DEL-88 |
| chr16-25159610-DEL-59 | chr16-25159590-DEL-85 |
| chr16-25160871-DEL-937 | chr16-25160872-DEL-937 |
| chr16-25169589-DEL-56 | chr16-25169590-DEL-56 |
| chr16-25183269-DEL-105 | chr16-25183234-DEL-96 |
| chr16-25245364-DEL-125 | chr16-25245365-DEL-125 |
| chr16-25250454-DEL-330 | chr16-25250455-DEL-330 |
| chr16-25267113-DEL-65 | chr16-25267099-DEL-80 |
| chr16-25274311-DEL-62 | chr16-25274324-DEL-50 |
| chr16-25274943-DEL-91 | chr16-25274950-DEL-83 |
| chr16-25278980-DEL-229 | chr16-25278981-DEL-229 |
| chr16-25285521-DEL-98 | chr16-25285520-DEL-114 |
| chr16-25303515-DEL-238 | chr16-25303516-DEL-238 |
| chr16-25316522-DEL-5310 | chr16-25316523-DEL-5310 |
| chr16-25325538-DEL-748 | chr16-25325539-DEL-748 |
| chr16-25336483-DEL-5533 | chr16-25336483-DEL-5534 |
| chr16-25358558-DEL-429 | chr16-25358559-DEL-429 |
| chr16-25384424-DEL-6658 | chr16-25384414-DEL-6659 |
| chr16-25397063-DEL-336 | chr16-25397064-DEL-335 |
| chr16-25407438-DEL-52 | chr16-25407429-DEL-60 |
| chr16-25433372-DEL-72 | chr16-25433375-DEL-82 |
| chr16-25471265-DEL-801 | chr16-25471254-DEL-802 |
| chr16-25475925-DEL-66 | chr16-25475926-DEL-66 |
| chr16-25490828-DEL-116 | chr16-25490835-DEL-110 |
| chr16-25502259-DEL-52 | chr16-25502260-DEL-52 |
| chr16-25521136-DEL-277 | chr16-25521137-DEL-277 |
| chr16-25543033-DEL-75 | chr16-25543034-DEL-75 |
| chr16-25631833-DEL-100 | chr16-25631834-DEL-100 |
| chr16-25642929-DEL-129 | chr16-25642930-DEL-129 |
| chr16-25655913-DEL-99 | chr16-25655914-DEL-99 |
| chr16-25688850-DEL-189 | chr16-25688851-DEL-189 |
| chr16-25733079-DEL-74 | chr16-25733070-DEL-84 |
| chr16-25826072-DEL-1120 | chr16-25826073-DEL-1120 |
| chr16-25840512-DEL-52 | chr16-25840513-DEL-52 |
| chr16-25856407-DEL-400 | chr16-25856408-DEL-400 |
| chr16-25874108-DEL-58 | chr16-25874113-DEL-54 |
| chr16-25940812-DEL-6396 | chr16-25940813-DEL-6396 |
| chr16-25973217-DEL-1845 | chr16-25973218-DEL-1845 |
| chr16-26002949-DEL-56 | chr16-26002950-DEL-56 |
| chr16-26013613-DEL-100 | chr16-26013613-DEL-102 |
| chr16-26037820-DEL-9121 | chr16-26037821-DEL-9121 |
| chr16-26039544-DEL-2898 | chr16-26039545-DEL-2898 |
| chr16-26059149-DEL-629 | chr16-26059150-DEL-629 |
| chr16-26075269-DEL-6380 | chr16-26075270-DEL-6380 |
| chr16-26136098-DEL-171 | chr16-26136086-DEL-165 |
| chr16-26180767-DEL-59 | chr16-26180768-DEL-77 |
| chr16-26193346-DEL-118 | chr16-26193347-DEL-118 |
| chr16-26195774-DEL-121 | chr16-26195777-DEL-119 |
| chr16-26219077-DEL-65 | chr16-26219080-DEL-51 |
| chr16-26221264-DEL-79 | chr16-26221245-DEL-95 |
| chr16-26264450-DEL-72 | chr16-26264489-DEL-58 |
| chr16-26273533-DEL-613 | chr16-26273534-DEL-613 |

| | |
|---|---|
| chr16-26307338-DEL-188 | chr16-26307339-DEL-188 |
| chr16-26315325-DEL-3334 | chr16-26315326-DEL-3334 |
| chr16-26330108-DEL-70 | chr16-26330109-DEL-70 |
| chr16-26360633-DEL-89 | chr16-26360634-DEL-86 |
| chr16-26380104-DEL-72 | chr16-26380105-DEL-114 |
| chr16-26425194-DEL-89 | chr16-26425195-DEL-89 |
| chr16-26452893-DEL-172 | chr16-26452903-DEL-173 |
| chr16-26474453-DEL-6185 | chr16-26474454-DEL-6185 |
| chr16-26494566-DEL-206 | chr16-26494567-DEL-206 |
| chr16-26544254-DEL-73 | chr16-26544253-DEL-74 |
| chr16-26561530-DEL-1263 | chr16-26561523-DEL-1264 |
| chr16-26577313-DEL-70 | chr16-26577314-DEL-70 |
| chr16-26588423-DEL-124 | chr16-26588387-DEL-134 |
| chr16-26593629-DEL-200 | chr16-26593630-DEL-200 |
| chr16-26600262-DEL-64 | chr16-26600263-DEL-64 |
| chr16-26619762-DEL-176 | chr16-26619763-DEL-176 |
| chr16-26637054-DEL-90 | chr16-26637055-DEL-90 |
| chr16-26670909-DEL-226 | chr16-26670910-DEL-226 |
| chr16-26685182-DEL-7163 | chr16-26685183-DEL-7163 |
| chr16-26698924-DEL-636 | chr16-26698925-DEL-636 |
| chr16-26709373-DEL-57 | chr16-26709374-DEL-57 |
| chr16-26716308-DEL-92 | chr16-26716293-DEL-98 |
| chr16-26717283-DEL-123 | chr16-26717294-DEL-139 |
| chr16-26732491-DEL-60 | chr16-26732492-DEL-72 |
| chr16-26735580-DEL-2147 | chr16-26735581-DEL-2147 |
| chr16-26748322-DEL-229 | chr16-26748323-DEL-229 |
| chr16-26754008-DEL-296 | chr16-26754009-DEL-296 |
| chr16-26755991-DEL-6075 | chr16-26755992-DEL-6075 |
| chr16-26795412-DEL-68 | chr16-26795413-DEL-68 |
| chr16-26797852-DEL-502 | chr16-26797853-DEL-502 |
| chr16-26812156-DEL-99 | chr16-26812163-DEL-63 |
| chr16-26819608-DEL-182 | chr16-26819609-DEL-182 |
| chr16-26840447-DEL-67 | chr16-26840448-DEL-67 |
| chr16-26844623-DEL-112 | chr16-26844668-DEL-76 |
| chr16-26877222-DEL-97 | chr16-26877223-DEL-97 |
| chr16-26884073-DEL-72 | chr16-26884074-DEL-71 |
| chr16-26890041-DEL-16727 | chr16-26890026-DEL-16740 |
| chr16-26933497-DEL-1706 | chr16-26933497-DEL-1706 |
| chr16-26949056-DEL-534 | chr16-26949057-DEL-534 |
| chr16-26950445-DEL-2698 | chr16-26950446-DEL-2698 |
| chr16-26954960-DEL-1879 | chr16-26954961-DEL-1879 |
| chr16-26982053-DEL-336 | chr16-26982054-DEL-336 |
| chr16-26986058-DEL-552 | chr16-26986057-DEL-530 |
| chr16-27002239-DEL-486 | chr16-27002240-DEL-486 |
| chr16-27019888-DEL-1694 | chr16-27019889-DEL-1694 |
| chr16-27022191-DEL-55 | chr16-27022190-DEL-54 |
| chr16-27063586-DEL-179 | chr16-27063587-DEL-179 |
| chr16-27074036-DEL-157 | chr16-27074037-DEL-157 |
| chr16-27092614-DEL-654 | chr16-27092615-DEL-654 |
| chr16-27114879-DEL-2063 | chr16-27114880-DEL-2063 |
| chr16-27122845-DEL-580 | chr16-27122846-DEL-580 |
| chr16-27126141-DEL-72 | chr16-27126142-DEL-74 |
| chr16-27131322-DEL-191 | chr16-27131323-DEL-191 |
| chr16-27146674-DEL-253 | chr16-27146675-DEL-253 |
| chr16-27154912-DEL-596 | chr16-27154912-DEL-596 |
| chr16-27158002-DEL-84 | chr16-27157995-DEL-117 |
| chr16-27159393-DEL-1444 | chr16-27159393-DEL-1444 |
| chr16-27178043-DEL-197 | chr16-27178044-DEL-197 |
| chr16-27183625-DEL-93 | chr16-27183628-DEL-92 |
| chr16-27242063-DEL-7846 | chr16-27242064-DEL-7846 |
| chr16-27270343-DEL-58 | chr16-27270344-DEL-58 |
| chr16-27292887-DEL-869 | chr16-27292888-DEL-869 |
| chr16-27326371-DEL-97 | chr16-27326364-DEL-89 |
| chr16-27391961-DEL-1239 | chr16-27391962-DEL-1239 |
| chr16-27420927-DEL-6621 | chr16-27420928-DEL-6621 |
| chr16-27429134-DEL-790 | chr16-27429135-DEL-790 |
| chr16-27449662-DEL-198 | chr16-27449663-DEL-198 |
| chr16-27457096-DEL-77 | chr16-27457097-DEL-77 |
| chr16-27525094-DEL-848 | chr16-27525095-DEL-848 |
| chr16-27538121-DEL-1793 | chr16-27538122-DEL-1793 |
| chr16-27554462-DEL-6548 | chr16-27554463-DEL-6548 |
| chr16-27592837-DEL-330 | chr16-27592838-DEL-330 |
| chr16-27667037-DEL-327 | chr16-27667038-DEL-327 |
| chr16-27670627-DEL-181 | chr16-27670628-DEL-221 |
| chr16-27679347-DEL-533 | chr16-27679348-DEL-533 |
| chr16-27685401-DEL-1890 | chr16-27685402-DEL-1890 |
| chr16-27724682-DEL-114 | chr16-27724683-DEL-114 |
| chr16-27737046-DEL-2228 | chr16-27737047-DEL-2228 |
| chr16-27762181-DEL-159 | chr16-27762184-DEL-157 |
| chr16-27795726-DEL-57 | chr16-27795730-DEL-50 |

| | |
|---|---|
| chr16-27804277-DEL-283 | chr16-27804278-DEL-283 |
| chr16-27809489-DEL-4341 | chr16-27809490-DEL-4341 |
| chr16-27825378-DEL-148 | chr16-27825379-DEL-148 |
| chr16-27881204-DEL-86 | chr16-27881220-DEL-71 |
| chr16-27925416-DEL-105 | chr16-27925456-DEL-66 |
| chr16-27964959-DEL-1276 | chr16-27964960-DEL-1276 |
| chr16-27988530-DEL-77 | chr16-27988554-DEL-75 |
| chr16-27989269-DEL-6479 | chr16-27989270-DEL-6479 |
| chr16-27998575-DEL-783 | chr16-27998576-DEL-783 |
| chr16-28009896-DEL-6436 | chr16-28009897-DEL-6436 |
| chr16-28033639-DEL-77 | chr16-28033640-DEL-77 |
| chr16-28034310-DEL-171 | chr16-28034311-DEL-171 |
| chr16-28087186-DEL-212 | chr16-28087187-DEL-212 |
| chr16-28093967-DEL-63 | chr16-28093968-DEL-63 |
| chr16-28109360-DEL-196 | chr16-28109361-DEL-196 |
| chr16-28111658-DEL-8453 | chr16-28111659-DEL-8452 |
| chr16-28112723-DEL-416 | chr16-28112723-DEL-416 |
| chr16-28113268-DEL-208 | chr16-28113269-DEL-208 |
| chr16-28121841-DEL-53 | chr16-28121842-DEL-53 |
| chr16-28153240-DEL-352 | chr16-28153241-DEL-352 |
| chr16-28181781-DEL-90 | chr16-28181782-DEL-90 |
| chr16-28185912-DEL-53 | chr16-28185913-DEL-53 |
| chr16-28186906-DEL-903 | chr16-28186907-DEL-903 |
| chr16-28188371-DEL-201 | chr16-28188372-DEL-201 |
| chr16-28189070-DEL-54 | chr16-28189071-DEL-54 |
| chr16-28199077-DEL-78 | chr16-28199078-DEL-78 |
| chr16-28207149-DEL-88 | chr16-28207150-DEL-88 |
| chr16-28290136-DEL-1109 | chr16-28290137-DEL-1109 |
| chr16-28303697-DEL-84 | chr16-28303698-DEL-84 |
| chr16-28311701-DEL-51 | chr16-28311701-DEL-51 |
| chr16-28319007-DEL-7004 | chr16-28319008-DEL-7004 |
| chr16-28346090-DEL-775 | chr16-28346091-DEL-775 |
| chr16-28362535-DEL-97 | chr16-28362546-DEL-94 |
| chr16-28370305-DEL-80 | chr16-28370322-DEL-64 |
| chr16-28404438-DEL-60 | chr16-28404432-DEL-66 |
| chr16-28409120-DEL-54 | chr16-28409121-DEL-90 |
| chr16-28413646-DEL-162 | chr16-28413647-DEL-162 |
| chr16-28452956-DEL-59 | chr16-28452949-DEL-54 |
| chr16-28471860-DEL-1777 | chr16-28471861-DEL-1777 |
| chr16-28510276-DEL-196 | chr16-28510277-DEL-196 |
| chr16-28513492-DEL-281 | chr16-28513491-DEL-282 |
| chr16-28514327-DEL-68 | chr16-28514328-DEL-68 |
| chr16-28596690-DEL-201 | chr16-28596691-DEL-201 |
| chr16-28604614-DEL-60 | chr16-28604615-DEL-55 |
| chr16-28606467-DEL-76 | chr16-28606478-DEL-60 |
| chr16-28609075-DEL-64 | chr16-28609075-DEL-64 |
| chr16-28628403-DEL-527 | chr16-28628404-DEL-527 |
| chr16-28641080-DEL-1239 | chr16-28641081-DEL-1239 |
| chr16-28660518-DEL-95 | chr16-28660499-DEL-83 |
| chr16-28672887-DEL-3846 | chr16-28672888-DEL-3846 |
| chr16-28728619-DEL-70 | chr16-28728620-DEL-70 |
| chr16-28767237-DEL-60 | chr16-28767238-DEL-58 |
| chr16-28770551-DEL-60 | chr16-28770552-DEL-60 |
| chr16-28778882-DEL-58 | chr16-28778882-DEL-54 |
| chr16-28797298-DEL-635 | chr16-28797309-DEL-630 |
| chr16-28802119-DEL-65 | chr16-28802120-DEL-67 |
| chr16-28806704-DEL-56 | chr16-28806705-DEL-56 |
| chr16-28828236-DEL-140 | chr16-28828237-DEL-152 |
| chr16-28848031-DEL-55 | chr16-28848039-DEL-52 |
| chr16-28868460-DEL-56 | chr16-28868461-DEL-56 |
| chr16-28894638-DEL-6326 | chr16-28894629-DEL-6312 |
| chr16-28939953-DEL-537 | chr16-28939954-DEL-537 |
| chr16-28952575-DEL-72 | chr16-28952576-DEL-72 |
| chr16-28969801-DEL-538 | chr16-28969802-DEL-538 |
| chr16-28979273-DEL-302 | chr16-28979274-DEL-302 |
| chr16-28984196-DEL-998 | chr16-28984197-DEL-998 |
| chr16-29010618-DEL-90 | chr16-29010619-DEL-90 |
| chr16-29031480-DEL-75 | chr16-29031490-DEL-60 |
| chr16-29040140-DEL-79 | chr16-29040141-DEL-79 |
| chr16-29073826-DEL-399 | chr16-29073827-DEL-399 |
| chr16-29086241-DEL-6229 | chr16-29086242-DEL-6229 |
| chr16-29097548-DEL-65 | chr16-29097552-DEL-66 |
| chr16-29103859-DEL-3819 | chr16-29103860-DEL-3819 |
| chr16-29111369-DEL-218 | chr16-29111364-DEL-219 |
| chr16-29214442-DEL-58 | chr16-29214443-DEL-58 |
| chr16-29242693-DEL-6364 | chr16-29242694-DEL-6364 |
| chr16-29257518-DEL-86 | chr16-29257519-DEL-86 |
| chr16-29261792-DEL-6561 | chr16-29261793-DEL-6560 |
| chr16-29276587-DEL-106 | chr16-29276584-DEL-133 |
| chr16-29334965-DEL-7212 | chr16-29334966-DEL-7212 |

| | |
|---|---|
| chr16-29342175-DEL-117 | chr16-29342176-DEL-117 |
| chr16-29351846-DEL-110 | chr16-29351847-DEL-111 |
| chr16-29356856-DEL-56 | chr16-29356857-DEL-56 |
| chr16-29374031-DEL-280 | chr16-29374032-DEL-280 |
| chr16-29381898-DEL-56 | chr16-29381908-DEL-60 |
| chr16-29388287-DEL-1176 | chr16-29388288-DEL-1176 |
| chr16-29391166-DEL-498 | chr16-29391167-DEL-498 |
| chr16-29417434-DEL-56 | chr16-29417435-DEL-56 |
| chr16-29425839-DEL-70 | chr16-29425846-DEL-58 |
| chr16-29441092-DEL-150 | chr16-29441063-DEL-180 |
| chr16-29442376-DEL-222 | chr16-29442377-DEL-222 |
| chr16-29449506-DEL-104 | chr16-29449520-DEL-98 |
| chr16-29484453-DEL-224 | chr16-29484454-DEL-224 |
| chr16-29497226-DEL-164 | chr16-29497227-DEL-164 |
| chr16-29510236-DEL-157 | chr16-29510237-DEL-157 |
| chr16-29511419-DEL-335 | chr16-29511393-DEL-341 |
| chr16-29543359-DEL-207 | chr16-29543360-DEL-207 |
| chr16-29575453-DEL-132 | chr16-29575465-DEL-132 |
| chr16-29583775-DEL-248 | chr16-29583776-DEL-248 |
| chr16-29607738-DEL-55 | chr16-29607739-DEL-55 |
| chr16-29650561-DEL-386 | chr16-29650562-DEL-386 |
| chr16-29676292-DEL-273 | chr16-29676293-DEL-273 |
| chr16-29686507-DEL-52 | chr16-29686508-DEL-52 |
| chr16-29706469-DEL-56 | chr16-29706486-DEL-54 |
| chr16-29712158-DEL-161 | chr16-29712157-DEL-159 |
| chr16-29720266-DEL-209 | chr16-29720267-DEL-209 |
| chr16-29786404-DEL-64 | chr16-29786410-DEL-58 |
| chr16-29832171-DEL-62 | chr16-29832172-DEL-62 |
| chr16-29894162-DEL-214 | chr16-29894163-DEL-214 |
| chr16-29934397-DEL-186 | chr16-29934398-DEL-186 |
| chr16-29954562-DEL-66 | chr16-29954563-DEL-66 |
| chr16-29963211-DEL-80 | chr16-29963212-DEL-80 |
| chr16-29964003-DEL-92 | chr16-29964004-DEL-92 |
| chr16-29965193-DEL-68 | chr16-29965194-DEL-68 |
| chr16-29973637-DEL-62 | chr16-29973671-DEL-59 |
| chr16-29978890-DEL-204 | chr16-29978892-DEL-202 |
| chr16-30039230-DEL-677 | chr16-30039231-DEL-677 |
| chr16-30053378-DEL-56 | chr16-30053374-DEL-57 |
| chr16-30066233-DEL-122 | chr16-30066234-DEL-122 |
| chr16-30069492-DEL-206 | chr16-30069493-DEL-206 |
| chr16-30083517-DEL-56 | chr16-30083518-DEL-56 |
| chr16-30109275-DEL-59 | chr16-30109237-DEL-60 |
| chr16-30110498-DEL-217 | chr16-30110499-DEL-217 |
| chr16-30200906-DEL-228 | chr16-30200907-DEL-228 |
| chr16-30227705-DEL-255 | chr16-30227706-DEL-255 |
| chr16-30231107-DEL-65 | chr16-30231087-DEL-85 |
| chr16-30233779-DEL-60 | chr16-30233780-DEL-64 |
| chr16-30248574-DEL-462 | chr16-30248575-DEL-462 |
| chr16-30271125-DEL-142 | chr16-30271126-DEL-142 |
| chr16-30289331-DEL-215 | chr16-30289332-DEL-215 |
| chr16-30300137-DEL-102 | chr16-30300138-DEL-102 |
| chr16-30306435-DEL-663 | chr16-30306436-DEL-663 |
| chr16-30309541-DEL-98 | chr16-30309542-DEL-98 |
| chr16-30325166-DEL-228 | chr16-30325167-DEL-228 |
| chr16-30346665-DEL-1191 | chr16-30346665-DEL-1191 |
| chr16-30352163-DEL-92 | chr16-30352162-DEL-92 |
| chr16-30363050-DEL-474 | chr16-30363051-DEL-474 |
| chr16-30370960-DEL-52 | chr16-30370961-DEL-52 |
| chr16-30384994-DEL-54 | chr16-30384995-DEL-54 |
| chr16-30388813-DEL-217 | chr16-30388804-DEL-217 |
| chr16-30429632-DEL-72 | chr16-30429633-DEL-72 |
| chr16-30458714-DEL-169 | chr16-30458715-DEL-169 |
| chr16-30475118-DEL-90 | chr16-30475151-DEL-58 |
| chr16-30483074-DEL-77 | chr16-30483075-DEL-77 |
| chr16-30513989-DEL-418 | chr16-30513990-DEL-418 |
| chr16-30537559-DEL-281 | chr16-30537551-DEL-282 |
| chr16-30552918-DEL-1115 | chr16-30552919-DEL-1115 |
| chr16-30600515-DEL-114 | chr16-30600516-DEL-114 |
| chr16-30632964-DEL-265 | chr16-30632965-DEL-265 |
| chr16-30636666-DEL-51 | chr16-30636654-DEL-78 |
| chr16-30648389-DEL-72 | chr16-30648408-DEL-54 |
| chr16-30672533-DEL-514 | chr16-30672536-DEL-512 |
| chr16-30684542-DEL-328 | chr16-30684543-DEL-328 |
| chr16-30685071-DEL-254 | chr16-30685072-DEL-254 |
| chr16-30687834-DEL-207 | chr16-30687835-DEL-207 |
| chr16-30690365-DEL-237 | chr16-30690366-DEL-237 |
| chr16-30707116-DEL-217 | chr16-30707138-DEL-202 |
| chr16-30707861-DEL-3180 | chr16-30707862-DEL-3180 |
| chr16-30724735-DEL-7482 | chr16-30724736-DEL-7482 |
| chr16-30759238-DEL-189 | chr16-30759239-DEL-192 |

| | |
|---|---|
| chr16-30782011-DEL-495 | chr16-30782012-DEL-495 |
| chr16-30784861-DEL-393 | chr16-30784862-DEL-393 |
| chr16-30790874-DEL-777 | chr16-30790873-DEL-777 |
| chr16-30805661-DEL-80 | chr16-30805657-DEL-78 |
| chr16-30811969-DEL-234 | chr16-30811970-DEL-234 |
| chr16-30826727-DEL-91 | chr16-30826728-DEL-91 |
| chr16-30844441-DEL-89 | chr16-30844442-DEL-89 |
| chr16-30894396-DEL-57 | chr16-30894397-DEL-58 |
| chr16-30904206-DEL-4771 | chr16-30904207-DEL-4771 |
| chr16-30909882-DEL-1058 | chr16-30909883-DEL-1057 |
| chr16-30924176-DEL-63 | chr16-30924177-DEL-63 |
| chr16-30981310-DEL-594 | chr16-30981336-DEL-595 |
| chr16-31000708-DEL-82 | chr16-31000709-DEL-82 |
| chr16-31008536-DEL-469 | chr16-31008537-DEL-469 |
| chr16-31033842-DEL-169 | chr16-31033849-DEL-163 |
| chr16-31037240-DEL-8776 | chr16-31037241-DEL-8776 |
| chr16-31038213-DEL-204 | chr16-31038214-DEL-204 |
| chr16-31114800-DEL-255 | chr16-31114801-DEL-255 |
| chr16-31121639-DEL-202 | chr16-31121640-DEL-202 |
| chr16-31232327-DEL-212 | chr16-31232328-DEL-212 |
| chr16-31259651-DEL-413 | chr16-31259652-DEL-413 |
| chr16-31270650-DEL-2028 | chr16-31270651-DEL-2028 |
| chr16-31281255-DEL-99 | chr16-31281256-DEL-73 |
| chr16-31332196-DEL-217 | chr16-31332197-DEL-217 |
| chr16-31337579-DEL-54 | chr16-31337580-DEL-54 |
| chr16-31367365-DEL-352 | chr16-31367367-DEL-353 |
| chr16-31379873-DEL-207 | chr16-31379874-DEL-207 |
| chr16-31388053-DEL-82 | chr16-31388054-DEL-82 |
| chr16-31390921-DEL-201 | chr16-31390922-DEL-201 |
| chr16-31392755-DEL-60 | chr16-31392736-DEL-54 |
| chr16-31396746-DEL-1199 | chr16-31396701-DEL-1207 |
| chr16-31407042-DEL-177 | chr16-31407043-DEL-181 |
| chr16-31413260-DEL-2798 | chr16-31413261-DEL-2798 |
| chr16-31441638-DEL-151 | chr16-31441639-DEL-151 |
| chr16-31442898-DEL-104 | chr16-31442899-DEL-104 |
| chr16-31452577-DEL-342 | chr16-31452578-DEL-342 |
| chr16-31459873-DEL-1117 | chr16-31459874-DEL-1117 |
| chr16-31510819-DEL-65 | chr16-31510778-DEL-85 |
| chr16-31556089-DEL-3406 | chr16-31556090-DEL-3406 |
| chr16-31636420-DEL-1156 | chr16-31636420-DEL-1156 |
| chr16-31637570-DEL-6410 | chr16-31637571-DEL-6410 |
| chr16-31670656-DEL-214 | chr16-31670657-DEL-214 |
| chr16-31731474-DEL-94 | chr16-31731475-DEL-100 |
| chr16-31742949-DEL-71 | chr16-31742950-DEL-73 |
| chr16-31771927-DEL-252 | chr16-31771928-DEL-252 |
| chr16-31790893-DEL-2009 | chr16-31790892-DEL-2010 |
| chr16-31795034-DEL-181 | chr16-31795035-DEL-181 |
| chr16-31803766-DEL-56 | chr16-31803767-DEL-56 |
| chr16-31811538-DEL-210 | chr16-31811539-DEL-210 |
| chr16-31828254-DEL-199 | chr16-31828251-DEL-200 |
| chr16-31845648-DEL-198 | chr16-31845649-DEL-198 |
| chr16-31848333-DEL-195 | chr16-31848334-DEL-195 |
| chr16-31861533-DEL-386 | chr16-31861534-DEL-386 |
| chr16-31880383-DEL-332 | chr16-31880391-DEL-331 |
| chr16-31888682-DEL-397 | chr16-31888683-DEL-397 |
| chr16-31907890-DEL-70 | chr16-31907891-DEL-70 |
| chr16-31923099-DEL-202 | chr16-31923100-DEL-202 |
| chr16-31941714-DEL-176 | chr16-31941713-DEL-177 |
| chr16-31950381-DEL-194 | chr16-31950382-DEL-194 |
| chr16-31966897-DEL-5252 | chr16-31966889-DEL-5269 |
| chr16-32000065-DEL-181 | chr16-32000066-DEL-181 |
| chr16-32014348-DEL-4809 | chr16-32014349-DEL-4809 |
| chr16-32021943-DEL-865 | chr16-32021944-DEL-865 |
| chr16-32025860-DEL-225 | chr16-32025853-DEL-224 |
| chr16-32099729-DEL-196 | chr16-32099737-DEL-193 |
| chr16-32123869-DEL-201 | chr16-32123870-DEL-201 |
| chr16-32127221-DEL-191 | chr16-32127223-DEL-190 |
| chr16-32140279-DEL-192 | chr16-32140280-DEL-193 |
| chr16-32147162-DEL-79 | chr16-32147133-DEL-73 |
| chr16-32212675-DEL-200 | chr16-32212676-DEL-200 |
| chr16-32229007-DEL-182 | chr16-32229008-DEL-182 |
| chr16-32230333-DEL-54 | chr16-32230358-DEL-60 |
| chr16-32262430-DEL-90 | chr16-32262431-DEL-98 |
| chr16-32305601-DEL-111 | chr16-32305600-DEL-112 |
| chr16-32360728-DEL-662 | chr16-32360729-DEL-662 |
| chr16-32398043-DEL-7162 | chr16-32398044-DEL-7162 |
| chr16-32414779-DEL-1888 | chr16-32414780-DEL-1888 |
| chr16-32417281-DEL-264 | chr16-32417282-DEL-282 |
| chr16-32418868-DEL-472 | chr16-32418869-DEL-472 |
| chr16-32459453-DEL-54 | chr16-32459454-DEL-72 |

| | |
|---|---|
| chr16-32471032-DEL-315 | chr16-32471035-DEL-316 |
| chr16-32473177-DEL-70 | chr16-32473175-DEL-74 |
| chr16-32482849-DEL-792 | chr16-32482849-DEL-792 |
| chr16-32496441-DEL-188 | chr16-32496444-DEL-186 |
| chr16-32506452-DEL-156 | chr16-32506452-DEL-157 |
| chr16-32509217-DEL-1086 | chr16-32509218-DEL-1086 |
| chr16-32513085-DEL-61 | chr16-32513088-DEL-60 |
| chr16-32515189-DEL-120 | chr16-32515200-DEL-118 |
| chr16-32517332-DEL-56 | chr16-32517333-DEL-56 |
| chr16-32520497-DEL-120 | chr16-32520495-DEL-135 |
| chr16-32541263-DEL-888 | chr16-32541264-DEL-888 |
| chr16-32549032-DEL-65 | chr16-32549035-DEL-61 |
| chr16-32600581-DEL-63 | chr16-32600582-DEL-63 |
| chr16-32639015-DEL-62 | chr16-32639016-DEL-62 |
| chr16-32654484-DEL-51 | chr16-32654485-DEL-51 |
| chr16-32686404-DEL-139 | chr16-32686428-DEL-148 |
| chr16-32730265-DEL-54 | chr16-32730266-DEL-54 |
| chr16-32759968-DEL-112 | chr16-32759969-DEL-112 |
| chr16-32768550-DEL-91 | chr16-32768551-DEL-91 |
| chr16-32788669-DEL-486 | chr16-32788670-DEL-486 |
| chr16-32823007-DEL-372 | chr16-32823008-DEL-366 |
| chr16-32830703-DEL-52 | chr16-32830704-DEL-52 |
| chr16-32838888-DEL-76 | chr16-32838889-DEL-76 |
| chr16-32843478-DEL-173 | chr16-32843479-DEL-173 |
| chr16-32848759-DEL-197 | chr16-32848746-DEL-198 |
| chr16-32850562-DEL-187 | chr16-32850563-DEL-187 |
| chr16-32864481-DEL-268 | chr16-32864482-DEL-268 |
| chr16-32881909-DEL-52 | chr16-32881895-DEL-51 |
| chr16-32885120-DEL-71 | chr16-32885121-DEL-71 |
| chr16-32891380-DEL-66 | chr16-32891383-DEL-64 |
| chr16-32898830-DEL-938 | chr16-32898827-DEL-937 |
| chr16-32904998-DEL-158 | chr16-32904998-DEL-124 |
| chr16-32923392-DEL-230 | chr16-32923393-DEL-230 |
| chr16-32924666-DEL-68 | chr16-32924617-DEL-69 |
| chr16-32929084-DEL-86 | chr16-32929085-DEL-86 |
| chr16-32959376-DEL-208 | chr16-32959371-DEL-209 |
| chr16-32967208-DEL-56 | chr16-32967209-DEL-56 |
| chr16-32969818-DEL-916 | chr16-32969819-DEL-916 |
| chr16-32971446-DEL-208 | chr16-32971447-DEL-208 |
| chr16-32973997-DEL-182 | chr16-32974002-DEL-180 |
| chr16-32988325-DEL-84 | chr16-32988288-DEL-72 |
| chr16-32991642-DEL-97 | chr16-32991647-DEL-93 |
| chr16-33006880-DEL-173 | chr16-33006881-DEL-173 |
| chr16-33018856-DEL-183 | chr16-33018857-DEL-183 |
| chr16-33028004-DEL-484 | chr16-33027991-DEL-502 |
| chr16-33038115-DEL-238 | chr16-33038116-DEL-230 |
| chr16-33040330-DEL-196 | chr16-33040329-DEL-195 |
| chr16-33041378-DEL-852 | chr16-33041379-DEL-852 |
| chr16-33079339-DEL-213 | chr16-33079331-DEL-214 |
| chr16-33092519-DEL-216 | chr16-33092516-DEL-216 |
| chr16-33122942-DEL-404 | chr16-33122942-DEL-404 |
| chr16-33132317-DEL-187 | chr16-33132314-DEL-185 |
| chr16-33135131-DEL-66 | chr16-33135124-DEL-74 |
| chr16-33168974-DEL-181 | chr16-33168975-DEL-181 |
| chr16-33183268-DEL-270 | chr16-33183269-DEL-270 |
| chr16-33193253-DEL-184 | chr16-33193255-DEL-176 |
| chr16-33206224-DEL-212 | chr16-33206225-DEL-212 |
| chr16-33226853-DEL-113 | chr16-33226854-DEL-109 |
| chr16-33230338-DEL-73 | chr16-33230339-DEL-73 |
| chr16-33251050-DEL-99 | chr16-33251035-DEL-98 |
| chr16-33259858-DEL-72 | chr16-33259859-DEL-72 |
| chr16-33357148-DEL-81 | chr16-33357149-DEL-81 |
| chr16-33375916-DEL-1552 | chr16-33375915-DEL-1549 |
| chr16-33387429-DEL-69 | chr16-33387438-DEL-51 |
| chr16-33398786-DEL-61 | chr16-33398740-DEL-62 |
| chr16-33403573-DEL-397 | chr16-33403574-DEL-397 |
| chr16-33408904-DEL-55 | chr16-33408903-DEL-66 |
| chr16-33438621-DEL-51 | chr16-33438605-DEL-78 |
| chr16-33442243-DEL-64 | chr16-33442242-DEL-65 |
| chr16-33446307-DEL-500 | chr16-33446305-DEL-500 |
| chr16-33465520-DEL-51 | chr16-33465521-DEL-51 |
| chr16-33483090-DEL-120 | chr16-33483091-DEL-120 |
| chr16-33549972-DEL-93 | chr16-33549973-DEL-93 |
| chr16-33579601-DEL-140 | chr16-33579602-DEL-140 |
| chr16-33584206-DEL-181 | chr16-33584207-DEL-181 |
| chr16-33611474-DEL-52 | chr16-33611475-DEL-52 |
| chr16-33618142-DEL-206 | chr16-33618143-DEL-206 |
| chr16-33628828-DEL-100 | chr16-33628829-DEL-100 |
| chr16-33632929-DEL-206 | chr16-33632920-DEL-207 |
| chr16-33638207-DEL-199 | chr16-33638208-DEL-200 |

| | |
|---|---|
| chr16-33645307-DEL-173 | chr16-33645322-DEL-162 |
| chr16-33649202-DEL-62 | chr16-33649203-DEL-61 |
| chr16-33660224-DEL-107 | chr16-33660223-DEL-108 |
| chr16-33667608-DEL-5350 | chr16-33667609-DEL-5350 |
| chr16-33674086-DEL-176 | chr16-33674099-DEL-188 |
| chr16-33688571-DEL-207 | chr16-33688572-DEL-207 |
| chr16-33714873-DEL-86 | chr16-33714884-DEL-78 |
| chr16-33717071-DEL-178 | chr16-33717072-DEL-178 |
| chr16-33721250-DEL-212 | chr16-33721250-DEL-212 |
| chr16-33839103-DEL-7171 | chr16-33839104-DEL-7171 |
| chr16-33872445-DEL-64 | chr16-33872440-DEL-62 |
| chr16-33884259-DEL-174 | chr16-33884260-DEL-174 |
| chr16-33917682-DEL-189 | chr16-33917666-DEL-175 |
| chr16-33938578-DEL-84 | chr16-33938579-DEL-84 |
| chr16-33960967-DEL-821 | chr16-33960968-DEL-821 |
| chr16-34019200-DEL-83 | chr16-34019214-DEL-50 |
| chr16-34054326-DEL-84 | chr16-34054327-DEL-84 |
| chr16-34101302-DEL-65 | chr16-34101306-DEL-72 |
| chr16-34136007-DEL-619 | chr16-34136008-DEL-619 |
| chr16-34169128-DEL-85 | chr16-34169126-DEL-74 |
| chr16-34216650-DEL-94 | chr16-34216651-DEL-94 |
| chr16-34245068-DEL-236 | chr16-34245069-DEL-236 |
| chr16-34254560-DEL-1810 | chr16-34254561-DEL-1810 |
| chr16-34257320-DEL-84 | chr16-34257321-DEL-84 |
| chr16-34276354-DEL-214 | chr16-34276355-DEL-214 |
| chr16-34308987-DEL-134 | chr16-34308988-DEL-134 |
| chr16-34312596-DEL-472 | chr16-34312597-DEL-472 |
| chr16-34355103-DEL-66 | chr16-34355103-DEL-67 |
| chr16-34402670-DEL-7257 | chr16-34402671-DEL-7257 |
| chr16-34420025-DEL-72 | chr16-34420041-DEL-57 |
| chr16-34423832-DEL-132 | chr16-34423833-DEL-132 |
| chr16-34428828-DEL-80 | chr16-34428829-DEL-80 |
| chr16-34429056-DEL-398 | chr16-34429057-DEL-398 |
| chr16-34446717-DEL-402 | chr16-34446718-DEL-402 |
| chr16-34480982-DEL-188 | chr16-34480983-DEL-188 |
| chr16-34520944-DEL-122 | chr16-34520933-DEL-133 |
| chr16-34546840-DEL-186 | chr16-34546841-DEL-186 |
| chr16-34562277-DEL-146 | chr16-34562278-DEL-146 |
| chr16-34562845-DEL-353 | chr16-34562846-DEL-353 |
| chr16-34566876-DEL-137 | chr16-34566877-DEL-137 |
| chr16-34580787-DEL-54 | chr16-34580788-DEL-54 |
| chr16-34606050-DEL-210 | chr16-34606051-DEL-210 |
| chr16-34633505-DEL-52 | chr16-34633517-DEL-87 |
| chr16-34650610-DEL-248 | chr16-34650611-DEL-248 |
| chr16-34655666-DEL-218 | chr16-34655667-DEL-218 |
| chr16-34656786-DEL-2196 | chr16-34656787-DEL-2196 |
| chr16-34707953-DEL-6421 | chr16-34707954-DEL-6421 |
| chr16-34785214-DEL-631 | chr16-34785210-DEL-629 |
| chr16-34793414-DEL-56 | chr16-34793416-DEL-102 |
| chr16-34833232-DEL-61 | chr16-34833233-DEL-61 |
| chr16-34859012-DEL-60 | chr16-34859004-DEL-85 |
| chr16-34894662-DEL-61 | chr16-34894652-DEL-62 |
| chr16-35101077-DEL-68 | chr16-35101078-DEL-68 |
| chr16-35158642-DEL-76 | chr16-35158645-DEL-70 |
| chr16-35164049-DEL-1032 | chr16-35164049-DEL-1029 |
| chr16-35195458-DEL-70 | chr16-35195459-DEL-70 |
| chr16-35250554-DEL-75 | chr16-35250555-DEL-75 |
| chr16-35262716-DEL-6393 | chr16-35262717-DEL-6393 |
| chr16-35279187-DEL-573 | chr16-35279181-DEL-564 |
| chr16-35323842-DEL-118 | chr16-35323843-DEL-118 |
| chr16-35348324-DEL-197 | chr16-35348325-DEL-197 |
| chr16-35385972-DEL-159 | chr16-35385973-DEL-159 |
| chr16-35389585-DEL-52 | chr16-35389586-DEL-52 |
| chr16-35393631-DEL-825 | chr16-35393632-DEL-825 |
| chr16-35398862-DEL-89 | chr16-35398833-DEL-89 |
| chr16-35408692-DEL-73 | chr16-35408693-DEL-73 |
| chr16-35411804-DEL-52 | chr16-35411786-DEL-70 |
| chr16-35425297-DEL-116 | chr16-35425285-DEL-83 |
| chr16-35488279-DEL-1038 | chr16-35488278-DEL-1037 |
| chr16-35493339-DEL-181 | chr16-35493340-DEL-181 |
| chr16-35509406-DEL-209 | chr16-35509417-DEL-199 |
| chr16-35540430-DEL-401 | chr16-35540431-DEL-401 |
| chr16-35547961-DEL-397 | chr16-35547970-DEL-389 |
| chr16-35550087-DEL-57 | chr16-35550073-DEL-58 |
| chr16-35573559-DEL-70 | chr16-35573562-DEL-58 |
| chr16-35582340-DEL-52 | chr16-35582339-DEL-50 |
| chr16-35585660-DEL-60 | chr16-35585657-DEL-64 |
| chr16-35605720-DEL-143 | chr16-35605723-DEL-141 |
| chr16-35618848-DEL-199 | chr16-35618849-DEL-199 |
| chr16-35676561-DEL-226 | chr16-35676562-DEL-226 |

| | |
|---|---|
| chr16-35686185-DEL-58 | chr16-35686147-DEL-74 |
| chr16-35693911-DEL-197 | chr16-35693912-DEL-197 |
| chr16-35704284-DEL-57 | chr16-35704284-DEL-57 |
| chr16-35769128-DEL-150 | chr16-35769129-DEL-150 |
| chr16-35777755-DEL-657 | chr16-35777756-DEL-657 |
| chr16-35781174-DEL-178 | chr16-35781164-DEL-179 |
| chr16-35782303-DEL-368 | chr16-35782304-DEL-368 |
| chr16-35810363-DEL-173 | chr16-35810364-DEL-173 |
| chr16-35811231-DEL-209 | chr16-35811232-DEL-209 |
| chr16-35819207-DEL-186 | chr16-35819208-DEL-186 |
| chr16-35829242-DEL-55 | chr16-35829246-DEL-55 |
| chr16-35848730-DEL-395 | chr16-35848731-DEL-395 |
| chr16-35876131-DEL-389 | chr16-35876132-DEL-389 |
| chr16-35886723-DEL-1181 | chr16-35886723-DEL-1181 |
| chr16-35901895-DEL-174 | chr16-35901896-DEL-174 |
| chr16-35971746-DEL-2670 | chr16-35971773-DEL-2673 |
| chr16-36004936-DEL-214 | chr16-36004937-DEL-214 |
| chr16-36009247-DEL-190 | chr16-36009260-DEL-178 |
| chr16-36039446-DEL-202 | chr16-36039447-DEL-202 |
| chr16-36063196-DEL-409 | chr16-36063196-DEL-409 |
| chr16-36116160-DEL-4320 | chr16-36116161-DEL-4320 |
| chr16-36168441-DEL-1853 | chr16-36168442-DEL-1853 |
| chr16-36180286-DEL-693 | chr16-36180287-DEL-693 |
| chr16-36184967-DEL-88 | chr16-36184969-DEL-122 |
| chr16-36282024-DEL-1214 | chr16-36282025-DEL-1214 |
| chr16-36300983-DEL-518 | chr16-36300984-DEL-518 |
| chr16-36308537-DEL-503 | chr16-36308538-DEL-503 |
| chr16-36342074-DEL-55 | chr16-36342075-DEL-55 |
| chr16-36350070-DEL-515 | chr16-36350071-DEL-515 |
| chr16-36393337-DEL-80 | chr16-36393338-DEL-80 |
| chr16-36408975-DEL-53 | chr16-36408976-DEL-62 |
| chr16-36473810-DEL-191 | chr16-36473811-DEL-191 |
| chr16-36585410-DEL-5304 | chr16-36585411-DEL-5304 |
| chr16-36639047-DEL-94 | chr16-36639048-DEL-94 |
| chr16-36764387-DEL-245 | chr16-36764388-DEL-245 |
| chr16-36791504-DEL-6146 | chr16-36791505-DEL-6147 |
| chr16-36816501-DEL-78 | chr16-36816502-DEL-78 |
| chr16-36883430-DEL-78 | chr16-36883473-DEL-50 |
| chr16-36924115-DEL-199 | chr16-36924116-DEL-199 |
| chr16-36938641-DEL-3396 | chr16-36938642-DEL-3396 |
| chr16-36948741-DEL-331 | chr16-36948742-DEL-331 |
| chr16-36982035-DEL-3437 | chr16-36982036-DEL-3437 |
| chr16-36986527-DEL-528 | chr16-36986528-DEL-528 |
| chr16-36988791-DEL-175 | chr16-36988795-DEL-183 |
| chr16-37026122-DEL-1120 | chr16-37026123-DEL-1120 |
| chr16-37089223-DEL-477 | chr16-37089233-DEL-478 |
| chr16-37156019-DEL-65 | chr16-37156027-DEL-58 |
| chr16-37169968-DEL-267 | chr16-37169969-DEL-267 |
| chr16-37230232-DEL-6141 | chr16-37230233-DEL-6141 |
| chr16-37238462-DEL-185 | chr16-37238463-DEL-185 |
| chr16-37248843-DEL-63 | chr16-37248839-DEL-60 |
| chr16-37280839-DEL-59 | chr16-37280875-DEL-50 |
| chr16-37337512-DEL-86 | chr16-37337517-DEL-82 |
| chr16-37344139-DEL-154 | chr16-37344140-DEL-154 |
| chr16-37374015-DEL-602 | chr16-37374016-DEL-602 |
| chr16-37406753-DEL-490 | chr16-37406753-DEL-491 |
| chr16-37477840-DEL-2052 | chr16-37477841-DEL-2052 |
| chr16-37516344-DEL-81 | chr16-37516345-DEL-81 |
| chr16-37562368-DEL-146 | chr16-37562327-DEL-168 |
| chr16-37584876-DEL-117 | chr16-37584877-DEL-117 |
| chr16-37616942-DEL-3147 | chr16-37616943-DEL-3147 |
| chr16-37780342-DEL-392 | chr16-37780340-DEL-392 |
| chr16-37783062-DEL-188 | chr16-37783062-DEL-188 |
| chr16-37790080-DEL-60 | chr16-37790081-DEL-60 |
| chr16-37793766-DEL-92 | chr16-37793763-DEL-95 |
| chr16-37819486-DEL-221 | chr16-37819487-DEL-221 |
| chr16-37883647-DEL-198 | chr16-37883648-DEL-198 |
| chr16-37901775-DEL-81 | chr16-37901776-DEL-81 |
| chr16-37924360-DEL-277 | chr16-37924361-DEL-277 |
| chr16-37955439-DEL-119 | chr16-37955440-DEL-119 |
| chr16-37993335-DEL-141 | chr16-37993336-DEL-141 |
| chr16-38054614-DEL-105 | chr16-38054615-DEL-105 |
| chr16-38082294-DEL-522 | chr16-38082294-DEL-520 |
| chr16-38093482-DEL-3986 | chr16-38093483-DEL-3987 |
| chr16-38099304-DEL-111 | chr16-38099305-DEL-102 |
| chr16-38133526-DEL-221 | chr16-38133527-DEL-221 |
| chr16-38157816-DEL-52 | chr16-38157817-DEL-52 |
| chr16-38159716-DEL-90 | chr16-38159717-DEL-90 |
| chr16-38163953-DEL-160 | chr16-38163954-DEL-160 |
| chr16-38164974-DEL-56 | chr16-38164975-DEL-56 |

| | |
|---|---|
| chr16-38198174-DEL-178 | chr16-38198175-DEL-178 |
| chr16-38208560-DEL-66 | chr16-38208560-DEL-66 |
| chr16-38211226-DEL-990 | chr16-38211227-DEL-990 |
| chr16-38256962-DEL-199 | chr16-38256963-DEL-199 |
| chr16-38317868-DEL-176 | chr16-38317869-DEL-174 |
| chr16-38324781-DEL-90 | chr16-38324812-DEL-60 |
| chr16-38344603-DEL-530 | chr16-38344604-DEL-530 |
| chr16-38350425-DEL-1253 | chr16-38350426-DEL-1253 |
| chr16-38424503-DEL-178 | chr16-38424504-DEL-178 |
| chr16-38434781-DEL-222 | chr16-38434782-DEL-222 |
| chr16-38451457-DEL-86 | chr16-38451479-DEL-90 |
| chr16-38455663-DEL-295 | chr16-38455664-DEL-295 |
| chr16-38476664-DEL-1222 | chr16-38476665-DEL-1222 |
| chr16-38534727-DEL-7110 | chr16-38534728-DEL-7110 |
| chr16-38554127-DEL-102 | chr16-38554128-DEL-102 |
| chr16-38563804-DEL-68 | chr16-38563815-DEL-58 |
| chr16-38616889-DEL-76 | chr16-38616894-DEL-72 |
| chr16-38658663-DEL-6544 | chr16-38658668-DEL-6544 |
| chr16-38667313-DEL-99 | chr16-38667314-DEL-99 |
| chr16-38675007-DEL-86 | chr16-38675003-DEL-134 |
| chr16-38686411-DEL-2249 | chr16-38686414-DEL-2242 |
| chr16-38690030-DEL-1756 | chr16-38690031-DEL-1756 |
| chr16-38692892-DEL-76 | chr16-38692893-DEL-76 |
| chr16-38717136-DEL-60 | chr16-38717137-DEL-57 |
| chr16-38744067-DEL-134 | chr16-38744022-DEL-143 |
| chr16-38748329-DEL-118 | chr16-38748330-DEL-118 |
| chr16-38753016-DEL-201 | chr16-38753017-DEL-201 |
| chr16-38756842-DEL-74 | chr16-38756843-DEL-74 |
| chr16-38794127-DEL-53 | chr16-38794125-DEL-59 |
| chr16-38804690-DEL-54 | chr16-38804691-DEL-54 |
| chr16-38856232-DEL-60 | chr16-38856233-DEL-60 |
| chr16-38863314-DEL-52 | chr16-38863315-DEL-50 |
| chr16-38868257-DEL-118 | chr16-38868264-DEL-126 |
| chr16-38873037-DEL-88 | chr16-38873058-DEL-56 |
| chr16-38881727-DEL-327 | chr16-38881728-DEL-327 |
| chr16-38890873-DEL-498 | chr16-38890874-DEL-498 |
| chr16-38901358-DEL-327 | chr16-38901377-DEL-309 |
| chr16-38904170-DEL-331 | chr16-38904170-DEL-331 |
| chr16-38905235-DEL-6369 | chr16-38905236-DEL-6369 |
| chr16-38923490-DEL-72 | chr16-38923502-DEL-74 |
| chr16-38928119-DEL-68 | chr16-38928076-DEL-88 |
| chr16-38929451-DEL-65 | chr16-38929446-DEL-70 |
| chr16-38937433-DEL-171 | chr16-38937434-DEL-171 |
| chr16-38991404-DEL-97 | chr16-38991399-DEL-109 |
| chr16-38999885-DEL-62 | chr16-38999886-DEL-62 |
| chr16-39003131-DEL-1294 | chr16-39003132-DEL-1294 |
| chr16-39005277-DEL-62 | chr16-39005238-DEL-71 |
| chr16-39074695-DEL-929 | chr16-39074697-DEL-928 |
| chr16-39081519-DEL-118 | chr16-39081515-DEL-119 |
| chr16-39086632-DEL-63 | chr16-39086633-DEL-63 |
| chr16-39095225-DEL-63 | chr16-39095226-DEL-59 |
| chr16-39106012-DEL-143 | chr16-39106013-DEL-143 |
| chr16-39109176-DEL-4109 | chr16-39109177-DEL-4109 |
| chr16-39138776-DEL-70 | chr16-39138736-DEL-66 |
| chr16-39144679-DEL-642 | chr16-39144680-DEL-642 |
| chr16-39196188-DEL-68 | chr16-39196189-DEL-74 |
| chr16-39229333-DEL-558 | chr16-39229334-DEL-558 |
| chr16-39237698-DEL-768 | chr16-39237699-DEL-768 |
| chr16-39255636-DEL-6254 | chr16-39255637-DEL-6254 |
| chr16-39327971-DEL-57 | chr16-39327972-DEL-51 |
| chr16-39338850-DEL-596 | chr16-39338851-DEL-596 |
| chr16-39342325-DEL-298 | chr16-39342327-DEL-293 |
| chr16-39349931-DEL-296 | chr16-39349932-DEL-296 |
| chr16-39356056-DEL-3203 | chr16-39356057-DEL-3203 |
| chr16-39365543-DEL-54 | chr16-39365544-DEL-54 |
| chr16-39372661-DEL-96 | chr16-39372662-DEL-96 |
| chr16-39376381-DEL-372 | chr16-39376382-DEL-372 |
| chr16-39377013-DEL-103 | chr16-39377014-DEL-103 |
| chr16-39377796-DEL-188 | chr16-39377797-DEL-188 |
| chr16-39382455-DEL-6194 | chr16-39382456-DEL-6194 |
| chr16-39403247-DEL-111 | chr16-39403250-DEL-107 |
| chr16-39403428-DEL-507 | chr16-39403381-DEL-528 |
| chr16-39474933-DEL-322 | chr16-39474934-DEL-322 |
| chr16-39590199-DEL-58 | chr16-39590200-DEL-58 |
| chr16-39600024-DEL-3058 | chr16-39600025-DEL-3058 |
| chr16-39632936-DEL-6801 | chr16-39632932-DEL-6802 |
| chr16-39638889-DEL-624 | chr16-39638890-DEL-624 |
| chr16-39649227-DEL-54 | chr16-39649228-DEL-54 |
| chr16-39698993-DEL-744 | chr16-39698994-DEL-744 |
| chr16-39709678-DEL-6358 | chr16-39709679-DEL-6358 |

| | |
|---|---|
| chr16-39744860-DEL-82 | chr16-39744861-DEL-82 |
| chr16-39779852-DEL-672 | chr16-39779853-DEL-672 |
| chr16-39818734-DEL-928 | chr16-39818735-DEL-928 |
| chr16-39845835-DEL-274 | chr16-39845785-DEL-278 |
| chr16-39847389-DEL-67 | chr16-39847385-DEL-68 |
| chr16-39849138-DEL-821 | chr16-39849139-DEL-821 |
| chr16-39854094-DEL-53 | chr16-39854095-DEL-54 |
| chr16-39860850-DEL-56 | chr16-39860851-DEL-56 |
| chr16-39894009-DEL-175 | chr16-39894017-DEL-170 |
| chr16-39925619-DEL-1187 | chr16-39925619-DEL-1184 |
| chr16-39961818-DEL-345 | chr16-39961819-DEL-345 |
| chr16-39971124-DEL-61 | chr16-39971125-DEL-61 |
| chr16-40014300-DEL-282 | chr16-40014301-DEL-282 |
| chr16-40042762-DEL-387 | chr16-40042766-DEL-382 |
| chr16-40061968-DEL-953 | chr16-40061969-DEL-953 |
| chr16-40066990-DEL-200 | chr16-40066991-DEL-200 |
| chr16-40070132-DEL-305 | chr16-40070133-DEL-302 |
| chr16-40076275-DEL-56 | chr16-40076276-DEL-56 |
| chr16-40087548-DEL-52 | chr16-40087549-DEL-52 |
| chr16-40106989-DEL-1355 | chr16-40106990-DEL-1355 |
| chr16-40131336-DEL-731 | chr16-40131337-DEL-731 |
| chr16-40148374-DEL-144 | chr16-40148375-DEL-144 |
| chr16-40188841-DEL-79 | chr16-40188842-DEL-79 |
| chr16-40263609-DEL-120 | chr16-40263653-DEL-97 |
| chr16-40290363-DEL-53 | chr16-40290364-DEL-53 |
| chr16-40306376-DEL-384 | chr16-40306377-DEL-384 |
| chr16-40348941-DEL-3427 | chr16-40348942-DEL-3427 |
| chr16-40378801-DEL-690 | chr16-40378802-DEL-690 |
| chr16-40396096-DEL-134 | chr16-40396097-DEL-134 |
| chr16-40430720-DEL-4354 | chr16-40430721-DEL-4354 |
| chr16-40480299-DEL-6803 | chr16-40480300-DEL-6803 |
| chr16-40634584-DEL-6278 | chr16-40634584-DEL-6278 |
| chr16-40693576-DEL-3458 | chr16-40693577-DEL-3458 |
| chr16-40760424-DEL-876 | chr16-40760425-DEL-876 |
| chr16-40821235-DEL-93 | chr16-40821238-DEL-85 |
| chr16-40874614-DEL-6635 | chr16-40874620-DEL-6633 |
| chr16-40926582-DEL-50 | chr16-40926583-DEL-52 |
| chr16-41077365-DEL-400 | chr16-41077366-DEL-400 |
| chr16-41128649-DEL-1911 | chr16-41128650-DEL-1911 |
| chr16-41142919-DEL-403 | chr16-41142920-DEL-403 |
| chr16-41172536-DEL-4825 | chr16-41172537-DEL-4825 |
| chr16-41292671-DEL-66 | chr16-41292672-DEL-66 |
| chr16-41302637-DEL-52 | chr16-41302638-DEL-51 |
| chr16-41311256-DEL-4330 | chr16-41311257-DEL-4330 |
| chr16-41322270-DEL-77 | chr16-41322271-DEL-77 |
| chr16-41336409-DEL-104 | chr16-41336410-DEL-104 |
| chr16-41374770-DEL-88 | chr16-41374771-DEL-88 |
| chr16-41395403-DEL-7521 | chr16-41395404-DEL-7521 |
| chr16-41434711-DEL-271 | chr16-41434710-DEL-272 |
| chr16-41436914-DEL-6394 | chr16-41436915-DEL-6394 |
| chr16-41451271-DEL-60 | chr16-41451272-DEL-60 |
| chr16-41482362-DEL-53 | chr16-41482363-DEL-53 |
| chr16-41525326-DEL-1179 | chr16-41525327-DEL-1179 |
| chr16-41604958-DEL-539 | chr16-41604959-DEL-539 |
| chr16-41641754-DEL-61 | chr16-41641755-DEL-61 |
| chr16-41675158-DEL-6388 | chr16-41675159-DEL-6389 |
| chr16-41718484-DEL-390 | chr16-41718485-DEL-390 |
| chr16-41823629-DEL-452 | chr16-41823630-DEL-452 |
| chr16-41832260-DEL-479 | chr16-41832261-DEL-478 |
| chr16-41843483-DEL-53 | chr16-41843484-DEL-53 |
| chr16-41887902-DEL-111 | chr16-41887903-DEL-111 |
| chr16-41946259-DEL-206 | chr16-41946259-DEL-207 |
| chr16-42137594-DEL-1399 | chr16-42137595-DEL-1399 |
| chr16-42236443-DEL-362 | chr16-42236444-DEL-362 |
| chr16-42254127-DEL-1582 | chr16-42254128-DEL-1582 |
| chr16-42267719-DEL-89 | chr16-42267746-DEL-62 |
| chr16-42293361-DEL-60 | chr16-42293362-DEL-61 |
| chr16-42313263-DEL-68 | chr16-42313283-DEL-50 |
| chr16-42390131-DEL-412 | chr16-42390132-DEL-412 |
| chr16-42445375-DEL-1152 | chr16-42445376-DEL-1152 |
| chr16-42448534-DEL-401 | chr16-42448535-DEL-401 |
| chr16-42451050-DEL-388 | chr16-42451050-DEL-388 |
| chr16-42484275-DEL-333 | chr16-42484276-DEL-333 |
| chr16-42489184-DEL-58 | chr16-42489185-DEL-58 |
| chr16-42499183-DEL-240 | chr16-42499184-DEL-240 |
| chr16-42502003-DEL-106 | chr16-42502015-DEL-123 |
| chr16-42547278-DEL-70 | chr16-42547279-DEL-70 |
| chr16-42551736-DEL-196 | chr16-42551737-DEL-196 |
| chr16-42567306-DEL-76 | chr16-42567311-DEL-72 |
| chr16-42593510-DEL-6578 | chr16-42593511-DEL-6578 |

| | |
|---|---|
| chr16-42607447-DEL-185 | chr16-42607448-DEL-185 |
| chr16-42633721-DEL-6284 | chr16-42633740-DEL-6293 |
| chr16-42648264-DEL-95 | chr16-42648265-DEL-95 |
| chr16-42662646-DEL-12848 | chr16-42662647-DEL-12845 |
| chr16-42768748-DEL-62 | chr16-42768749-DEL-62 |
| chr16-42775907-DEL-528 | chr16-42775908-DEL-528 |
| chr16-42799175-DEL-7012 | chr16-42799176-DEL-7012 |
| chr16-42862810-DEL-629 | chr16-42862811-DEL-629 |
| chr16-42906636-DEL-2429 | chr16-42906637-DEL-2429 |
| chr16-42932472-DEL-500 | chr16-42932473-DEL-500 |
| chr16-42942534-DEL-66 | chr16-42942539-DEL-58 |
| chr16-42957066-DEL-6554 | chr16-42957067-DEL-6554 |
| chr16-43254297-DEL-76 | chr16-43254297-DEL-84 |
| chr16-43260149-DEL-1375 | chr16-43260150-DEL-1375 |
| chr16-43283998-DEL-121 | chr16-43283999-DEL-121 |
| chr16-43304094-DEL-60 | chr16-43304095-DEL-60 |
| chr16-43356947-DEL-74 | chr16-43356948-DEL-74 |
| chr16-43378264-DEL-705 | chr16-43378265-DEL-705 |
| chr16-43394293-DEL-96 | chr16-43394306-DEL-84 |
| chr16-43405376-DEL-355 | chr16-43405377-DEL-355 |
| chr16-43427035-DEL-53 | chr16-43427036-DEL-53 |
| chr16-43469229-DEL-60 | chr16-43469230-DEL-60 |
| chr16-43470180-DEL-648 | chr16-43470181-DEL-648 |
| chr16-43477364-DEL-58 | chr16-43477365-DEL-58 |
| chr16-43483634-DEL-54 | chr16-43483626-DEL-55 |
| chr16-43548152-DEL-54 | chr16-43548153-DEL-50 |
| chr16-43549276-DEL-462 | chr16-43549277-DEL-462 |
| chr16-43560289-DEL-804 | chr16-43560290-DEL-804 |
| chr16-43584446-DEL-7433 | chr16-43584447-DEL-7433 |
| chr16-43615850-DEL-64 | chr16-43615851-DEL-64 |
| chr16-43618334-DEL-498 | chr16-43618335-DEL-498 |
| chr16-43628874-DEL-65 | chr16-43628875-DEL-65 |
| chr16-43654559-DEL-66 | chr16-43654558-DEL-64 |
| chr16-43684998-DEL-2477 | chr16-43684999-DEL-2477 |
| chr16-43693186-DEL-2470 | chr16-43693187-DEL-2470 |
| chr16-43805483-DEL-421 | chr16-43805484-DEL-421 |
| chr16-43809871-DEL-60 | chr16-43809872-DEL-60 |
| chr16-43824751-DEL-192 | chr16-43824743-DEL-193 |
| chr16-43856817-DEL-658 | chr16-43856818-DEL-658 |
| chr16-43860896-DEL-5703 | chr16-43860897-DEL-5703 |
| chr16-43884023-DEL-197 | chr16-43884024-DEL-197 |
| chr16-43899951-DEL-64 | chr16-43899952-DEL-64 |
| chr16-43901807-DEL-73 | chr16-43901808-DEL-73 |
| chr16-43917400-DEL-52 | chr16-43917401-DEL-52 |
| chr16-43926005-DEL-574 | chr16-43926006-DEL-574 |
| chr16-43930292-DEL-356 | chr16-43930293-DEL-356 |
| chr16-43950482-DEL-194 | chr16-43950483-DEL-194 |
| chr16-43955540-DEL-223 | chr16-43955541-DEL-223 |
| chr16-44014389-DEL-53 | chr16-44014389-DEL-53 |
| chr16-44050436-DEL-615 | chr16-44050436-DEL-615 |
| chr16-44069860-DEL-460 | chr16-44069861-DEL-452 |
| chr16-44085897-DEL-104 | chr16-44085892-DEL-108 |
| chr16-44088537-DEL-221 | chr16-44088538-DEL-221 |
| chr16-44094593-DEL-246 | chr16-44094594-DEL-246 |
| chr16-44099517-DEL-177 | chr16-44099518-DEL-177 |
| chr16-44100328-DEL-68 | chr16-44100329-DEL-68 |
| chr16-44120021-DEL-56 | chr16-44120022-DEL-56 |
| chr16-44123051-DEL-87 | chr16-44123052-DEL-87 |
| chr16-44125419-DEL-69 | chr16-44125420-DEL-69 |
| chr16-44130402-DEL-197 | chr16-44130403-DEL-197 |
| chr16-44160170-DEL-447 | chr16-44160171-DEL-447 |
| chr16-44170502-DEL-1268 | chr16-44170503-DEL-1268 |
| chr16-44187199-DEL-75 | chr16-44187200-DEL-75 |
| chr16-44210620-DEL-204 | chr16-44210621-DEL-204 |
| chr16-44220332-DEL-209 | chr16-44220333-DEL-209 |
| chr16-44270640-DEL-100 | chr16-44270641-DEL-100 |
| chr16-44277006-DEL-331 | chr16-44277008-DEL-327 |
| chr16-44280094-DEL-112 | chr16-44280095-DEL-112 |
| chr16-44283564-DEL-57 | chr16-44283556-DEL-52 |
| chr16-44284408-DEL-244 | chr16-44284409-DEL-244 |
| chr16-44285341-DEL-116 | chr16-44285326-DEL-126 |
| chr16-44294339-DEL-134 | chr16-44294338-DEL-135 |
| chr16-44332869-DEL-72 | chr16-44332869-DEL-72 |
| chr16-44390391-DEL-196 | chr16-44390392-DEL-196 |
| chr16-44393473-DEL-201 | chr16-44393474-DEL-201 |
| chr16-44403807-DEL-346 | chr16-44403808-DEL-346 |
| chr16-44423819-DEL-55 | chr16-44423826-DEL-55 |
| chr16-44457997-DEL-424 | chr16-44457998-DEL-424 |
| chr16-44476783-DEL-65 | chr16-44476784-DEL-65 |
| chr16-44523088-DEL-342 | chr16-44523089-DEL-342 |

| | |
|---|---|
| chr16-44530093-DEL-6440 | chr16-44530094-DEL-6440 |
| chr16-44602912-DEL-554 | chr16-44602913-DEL-554 |
| chr16-44610667-DEL-491 | chr16-44610669-DEL-489 |
| chr16-44615423-DEL-1543 | chr16-44615423-DEL-1543 |
| chr16-44656543-DEL-102 | chr16-44656526-DEL-102 |
| chr16-44714364-DEL-6256 | chr16-44714365-DEL-6256 |
| chr16-44729293-DEL-2505 | chr16-44729291-DEL-2506 |
| chr16-44742997-DEL-1768 | chr16-44742998-DEL-1771 |
| chr16-44753816-DEL-1553 | chr16-44753817-DEL-1553 |
| chr16-44765062-DEL-52 | chr16-44765062-DEL-52 |
| chr16-44767690-DEL-3388 | chr16-44767691-DEL-3388 |
| chr16-44797995-DEL-134 | chr16-44797993-DEL-135 |
| chr16-44809799-DEL-452 | chr16-44809800-DEL-452 |
| chr16-44822662-DEL-887 | chr16-44822648-DEL-886 |
| chr16-44859962-DEL-56 | chr16-44859941-DEL-51 |
| chr16-44870559-DEL-6181 | chr16-44870560-DEL-6181 |
| chr16-44882383-DEL-60 | chr16-44882384-DEL-60 |
| chr16-44907163-DEL-124 | chr16-44907164-DEL-124 |
| chr16-44911360-DEL-946 | chr16-44911361-DEL-947 |
| chr16-44920785-DEL-97 | chr16-44920762-DEL-125 |
| chr16-44986439-DEL-117 | chr16-44986440-DEL-117 |
| chr16-45014650-DEL-58 | chr16-45014651-DEL-58 |
| chr16-45028328-DEL-52 | chr16-45028329-DEL-52 |
| chr16-45030247-DEL-2206 | chr16-45030236-DEL-2207 |
| chr16-45058201-DEL-1005 | chr16-45058211-DEL-1007 |
| chr16-45069568-DEL-94 | chr16-45069569-DEL-94 |
| chr16-45078473-DEL-55 | chr16-45078474-DEL-55 |
| chr16-45083165-DEL-116 | chr16-45083158-DEL-117 |
| chr16-45090077-DEL-57 | chr16-45090078-DEL-57 |
| chr16-45122505-DEL-60 | chr16-45122473-DEL-108 |
| chr16-45129218-DEL-401 | chr16-45129201-DEL-419 |
| chr16-45174500-DEL-1090 | chr16-45174501-DEL-1090 |
| chr16-45188049-DEL-1680 | chr16-45188050-DEL-1680 |
| chr16-45198399-DEL-94 | chr16-45198400-DEL-94 |
| chr16-45230340-DEL-80 | chr16-45230342-DEL-79 |
| chr16-45243103-DEL-1266 | chr16-45243104-DEL-1266 |
| chr16-45258502-DEL-200 | chr16-45258503-DEL-200 |
| chr16-45266336-DEL-2661 | chr16-45266336-DEL-2661 |
| chr16-45284657-DEL-4705 | chr16-45284658-DEL-4705 |
| chr16-45320398-DEL-66 | chr16-45320397-DEL-67 |
| chr16-45332941-DEL-6392 | chr16-45332942-DEL-6392 |
| chr16-45339728-DEL-154 | chr16-45339695-DEL-145 |
| chr16-45342644-DEL-128 | chr16-45342645-DEL-128 |
| chr16-45349167-DEL-2207 | chr16-45349168-DEL-2207 |
| chr16-45360944-DEL-81 | chr16-45360952-DEL-100 |
| chr16-45372462-DEL-63 | chr16-45372462-DEL-63 |
| chr16-45385220-DEL-6383 | chr16-45385221-DEL-6383 |
| chr16-45409654-DEL-57 | chr16-45409655-DEL-57 |
| chr16-45414915-DEL-56 | chr16-45414916-DEL-56 |
| chr16-45444264-DEL-180 | chr16-45444265-DEL-180 |
| chr16-45471801-DEL-112 | chr16-45471798-DEL-115 |
| chr16-45489273-DEL-75 | chr16-45489273-DEL-69 |
| chr16-45499080-DEL-60 | chr16-45499105-DEL-54 |
| chr16-45518403-DEL-109 | chr16-45518404-DEL-109 |
| chr16-45526995-DEL-199 | chr16-45526996-DEL-199 |
| chr16-45547223-DEL-58 | chr16-45547224-DEL-58 |
| chr16-45556919-DEL-191 | chr16-45556920-DEL-191 |
| chr16-45578762-DEL-52 | chr16-45578759-DEL-53 |
| chr16-45602768-DEL-132 | chr16-45602769-DEL-132 |
| chr16-45641440-DEL-348 | chr16-45641441-DEL-348 |
| chr16-45692574-DEL-56 | chr16-45692575-DEL-56 |
| chr16-45694803-DEL-951 | chr16-45694804-DEL-951 |
| chr16-45717665-DEL-62 | chr16-45717670-DEL-58 |
| chr16-45739051-DEL-624 | chr16-45739052-DEL-624 |
| chr16-45749791-DEL-104 | chr16-45749792-DEL-104 |
| chr16-45752154-DEL-200 | chr16-45752155-DEL-199 |
| chr16-45756206-DEL-68 | chr16-45756212-DEL-68 |
| chr16-45764842-DEL-151 | chr16-45764843-DEL-151 |
| chr16-45797584-DEL-68 | chr16-45797587-DEL-60 |
| chr16-45798664-DEL-1348 | chr16-45798662-DEL-1345 |
| chr16-45802006-DEL-214 | chr16-45802001-DEL-216 |
| chr16-45805875-DEL-54 | chr16-45805876-DEL-78 |
| chr16-45808753-DEL-2045 | chr16-45808754-DEL-2045 |
| chr16-45826260-DEL-92 | chr16-45826261-DEL-92 |
| chr16-45860589-DEL-470 | chr16-45860590-DEL-470 |
| chr16-45897585-DEL-6135 | chr16-45897586-DEL-6135 |
| chr16-45903733-DEL-2862 | chr16-45903715-DEL-2883 |
| chr16-45955215-DEL-58 | chr16-45955218-DEL-56 |
| chr16-45973315-DEL-2125 | chr16-45973316-DEL-2125 |
| chr16-46002270-DEL-72 | chr16-46002280-DEL-51 |

| | |
|---|---|
| chr16-46012564-DEL-82 | chr16-46012565-DEL-82 |
| chr16-46013543-DEL-96 | chr16-46013544-DEL-96 |
| chr16-46027286-DEL-74 | chr16-46027289-DEL-73 |
| chr16-46036904-DEL-59 | chr16-46036905-DEL-59 |
| chr16-46052701-DEL-207 | chr16-46052708-DEL-201 |
| chr16-46088322-DEL-707 | chr16-46088323-DEL-707 |
| chr16-46095697-DEL-5302 | chr16-46095698-DEL-5302 |
| chr16-46109533-DEL-242 | chr16-46109534-DEL-242 |
| chr16-46115011-DEL-60 | chr16-46115012-DEL-60 |
| chr16-46138296-DEL-70 | chr16-46138297-DEL-70 |
| chr16-46147201-DEL-2420 | chr16-46147201-DEL-2416 |
| chr16-46181666-DEL-120 | chr16-46181667-DEL-120 |
| chr16-46183535-DEL-6149 | chr16-46183549-DEL-6146 |
| chr16-46223236-DEL-1212 | chr16-46223236-DEL-1212 |
| chr16-46225928-DEL-55 | chr16-46225929-DEL-55 |
| chr16-46228878-DEL-76 | chr16-46228877-DEL-82 |
| chr16-46283738-DEL-63 | chr16-46283739-DEL-63 |
| chr16-46289949-DEL-797 | chr16-46289950-DEL-797 |
| chr16-46328841-DEL-72 | chr16-46328842-DEL-72 |
| chr16-46415458-DEL-2847 | chr16-46415456-DEL-2848 |
| chr16-46439327-DEL-364 | chr16-46439328-DEL-364 |
| chr16-46462922-DEL-7791 | chr16-46462921-DEL-7791 |
| chr16-46470625-DEL-57 | chr16-46470626-DEL-57 |
| chr16-46474633-DEL-432 | chr16-46474634-DEL-432 |
| chr16-46488868-DEL-52 | chr16-46488863-DEL-69 |
| chr16-46534743-DEL-193 | chr16-46534744-DEL-193 |
| chr16-46538111-DEL-162 | chr16-46538112-DEL-162 |
| chr16-46543614-DEL-424 | chr16-46543615-DEL-424 |
| chr16-46552731-DEL-75 | chr16-46552732-DEL-91 |
| chr16-46577581-DEL-6467 | chr16-46577582-DEL-6467 |
| chr16-46587128-DEL-96 | chr16-46587129-DEL-97 |
| chr16-46601644-DEL-243 | chr16-46601645-DEL-243 |
| chr16-46654851-DEL-1873 | chr16-46654852-DEL-1873 |
| chr16-46685979-DEL-730 | chr16-46685980-DEL-730 |
| chr16-46687498-DEL-61 | chr16-46687498-DEL-51 |
| chr16-46698113-DEL-63 | chr16-46698110-DEL-63 |
| chr16-46761160-DEL-506 | chr16-46761161-DEL-506 |
| chr16-46795961-DEL-6438 | chr16-46795962-DEL-6438 |
| chr16-46875130-DEL-60 | chr16-46875130-DEL-60 |
| chr16-46882321-DEL-6188 | chr16-46882322-DEL-6188 |
| chr16-46904383-DEL-72 | chr16-46904384-DEL-72 |
| chr16-46905308-DEL-132 | chr16-46905309-DEL-132 |
| chr16-46907093-DEL-3140 | chr16-46907094-DEL-3140 |
| chr16-46907470-DEL-1359 | chr16-46907471-DEL-1359 |
| chr16-46918005-DEL-184 | chr16-46917997-DEL-185 |
| chr16-46933508-DEL-1808 | chr16-46933509-DEL-1808 |
| chr16-46940300-DEL-4633 | chr16-46940301-DEL-4633 |
| chr16-46947574-DEL-10799 | chr16-46947575-DEL-10799 |
| chr16-46968948-DEL-197 | chr16-46968948-DEL-197 |
| chr16-46972142-DEL-51 | chr16-46972141-DEL-53 |
| chr16-47003702-DEL-360 | chr16-47003703-DEL-360 |
| chr16-47008279-DEL-57 | chr16-47008280-DEL-57 |
| chr16-47019464-DEL-327 | chr16-47019465-DEL-327 |
| chr16-47022985-DEL-672 | chr16-47022985-DEL-670 |
| chr16-47030851-DEL-260 | chr16-47030852-DEL-260 |
| chr16-47037602-DEL-51 | chr16-47037603-DEL-51 |
| chr16-47083212-DEL-58 | chr16-47083213-DEL-58 |
| chr16-47149766-DEL-631 | chr16-47149766-DEL-631 |
| chr16-47196082-DEL-6879 | chr16-47196082-DEL-6879 |
| chr16-47210739-DEL-118 | chr16-47210740-DEL-118 |
| chr16-47212874-DEL-67 | chr16-47212845-DEL-50 |
| chr16-47240992-DEL-1892 | chr16-47240993-DEL-1892 |
| chr16-47292938-DEL-76 | chr16-47292945-DEL-80 |
| chr16-47295285-DEL-55 | chr16-47295286-DEL-55 |
| chr16-47298409-DEL-740 | chr16-47298413-DEL-741 |
| chr16-47308074-DEL-235 | chr16-47308075-DEL-234 |
| chr16-47311339-DEL-159 | chr16-47311340-DEL-159 |
| chr16-47315503-DEL-455 | chr16-47315504-DEL-452 |
| chr16-47317833-DEL-744 | chr16-47317834-DEL-744 |
| chr16-47320573-DEL-72 | chr16-47320573-DEL-72 |
| chr16-47322970-DEL-106 | chr16-47322971-DEL-106 |
| chr16-47375230-DEL-776 | chr16-47375231-DEL-776 |
| chr16-47377898-DEL-6633 | chr16-47377899-DEL-6633 |
| chr16-47386022-DEL-254 | chr16-47386023-DEL-254 |
| chr16-47400744-DEL-52 | chr16-47400745-DEL-52 |
| chr16-47411992-DEL-147 | chr16-47411993-DEL-147 |
| chr16-47440147-DEL-88 | chr16-47440148-DEL-88 |
| chr16-47489847-DEL-80 | chr16-47489848-DEL-80 |
| chr16-47494710-DEL-204 | chr16-47494711-DEL-204 |
| chr16-47496796-DEL-97 | chr16-47496797-DEL-97 |

| | |
|---|---|
| chr16-47508643-DEL-993 | chr16-47508645-DEL-992 |
| chr16-47511346-DEL-58 | chr16-47511348-DEL-57 |
| chr16-47532937-DEL-66 | chr16-47532937-DEL-66 |
| chr16-47534505-DEL-938 | chr16-47534511-DEL-937 |
| chr16-47537804-DEL-3897 | chr16-47537806-DEL-3896 |
| chr16-47567906-DEL-6473 | chr16-47567907-DEL-6473 |
| chr16-47610296-DEL-86 | chr16-47610297-DEL-86 |
| chr16-47619217-DEL-4373 | chr16-47619218-DEL-4374 |
| chr16-47674563-DEL-55 | chr16-47674564-DEL-54 |
| chr16-47686685-DEL-535 | chr16-47686686-DEL-533 |
| chr16-47702796-DEL-82 | chr16-47702833-DEL-86 |
| chr16-47727156-DEL-6785 | chr16-47727157-DEL-6785 |
| chr16-47737682-DEL-86 | chr16-47737689-DEL-80 |
| chr16-47755302-DEL-64 | chr16-47755299-DEL-68 |
| chr16-47755946-DEL-67 | chr16-47755956-DEL-66 |
| chr16-47771202-DEL-101 | chr16-47771203-DEL-101 |
| chr16-47775770-DEL-185 | chr16-47775771-DEL-185 |
| chr16-47801643-DEL-6650 | chr16-47801644-DEL-6650 |
| chr16-47811834-DEL-57 | chr16-47811835-DEL-57 |
| chr16-47833043-DEL-856 | chr16-47833043-DEL-856 |
| chr16-47834054-DEL-3623 | chr16-47834054-DEL-3623 |
| chr16-47838842-DEL-56 | chr16-47838835-DEL-64 |
| chr16-47841955-DEL-6672 | chr16-47841960-DEL-6668 |
| chr16-47876228-DEL-138 | chr16-47876196-DEL-144 |
| chr16-47877897-DEL-4111 | chr16-47877898-DEL-4113 |
| chr16-47897354-DEL-170 | chr16-47897359-DEL-165 |
| chr16-47898289-DEL-325 | chr16-47898290-DEL-325 |
| chr16-47933421-DEL-4913 | chr16-47933422-DEL-4913 |
| chr16-47953204-DEL-220 | chr16-47953205-DEL-220 |
| chr16-47956495-DEL-3416 | chr16-47956496-DEL-3416 |
| chr16-47959513-DEL-112 | chr16-47959514-DEL-112 |
| chr16-47963605-DEL-5272 | chr16-47963606-DEL-5272 |
| chr16-47975760-DEL-76 | chr16-47975761-DEL-76 |
| chr16-47988682-DEL-193 | chr16-47988677-DEL-194 |
| chr16-48005148-DEL-4054 | chr16-48005149-DEL-4054 |
| chr16-48017297-DEL-218 | chr16-48017298-DEL-218 |
| chr16-48026661-DEL-263 | chr16-48026624-DEL-311 |
| chr16-48037324-DEL-363 | chr16-48037325-DEL-363 |
| chr16-48040672-DEL-249 | chr16-48040666-DEL-250 |
| chr16-48044166-DEL-611 | chr16-48044164-DEL-611 |
| chr16-48045958-DEL-6200 | chr16-48045959-DEL-6200 |
| chr16-48057324-DEL-207 | chr16-48057325-DEL-207 |
| chr16-48062175-DEL-85 | chr16-48062173-DEL-85 |
| chr16-48069652-DEL-130 | chr16-48069653-DEL-130 |
| chr16-48070535-DEL-90 | chr16-48070533-DEL-75 |
| chr16-48077674-DEL-507 | chr16-48077675-DEL-507 |
| chr16-48082513-DEL-134 | chr16-48082512-DEL-87 |
| chr16-48087820-DEL-70 | chr16-48087821-DEL-68 |
| chr16-48093730-DEL-819 | chr16-48093731-DEL-819 |
| chr16-48102832-DEL-225 | chr16-48102833-DEL-225 |
| chr16-48120612-DEL-59 | chr16-48120613-DEL-60 |
| chr16-48128023-DEL-208 | chr16-48128024-DEL-208 |
| chr16-48139282-DEL-70 | chr16-48139283-DEL-70 |
| chr16-48149133-DEL-663 | chr16-48149134-DEL-663 |
| chr16-48150137-DEL-79 | chr16-48150138-DEL-79 |
| chr16-48154125-DEL-339 | chr16-48154126-DEL-339 |
| chr16-48185345-DEL-65 | chr16-48185346-DEL-65 |
| chr16-48201516-DEL-232 | chr16-48201517-DEL-232 |
| chr16-48202572-DEL-195 | chr16-48202559-DEL-179 |
| chr16-48218824-DEL-72 | chr16-48218826-DEL-56 |
| chr16-48235406-DEL-78 | chr16-48235407-DEL-78 |
| chr16-48236632-DEL-209 | chr16-48236633-DEL-209 |
| chr16-48245847-DEL-390 | chr16-48245848-DEL-390 |
| chr16-48246849-DEL-499 | chr16-48246850-DEL-499 |
| chr16-48248283-DEL-156 | chr16-48248303-DEL-150 |
| chr16-48310995-DEL-7380 | chr16-48310996-DEL-7380 |
| chr16-48333579-DEL-108 | chr16-48333556-DEL-124 |
| chr16-48343109-DEL-2148 | chr16-48343110-DEL-2148 |
| chr16-48346139-DEL-156 | chr16-48346140-DEL-156 |
| chr16-48350906-DEL-488 | chr16-48350907-DEL-488 |
| chr16-48386787-DEL-325 | chr16-48386788-DEL-325 |
| chr16-48398090-DEL-180 | chr16-48398091-DEL-180 |
| chr16-48408155-DEL-419 | chr16-48408156-DEL-419 |
| chr16-48424554-DEL-218 | chr16-48424555-DEL-218 |
| chr16-48425271-DEL-610 | chr16-48425265-DEL-611 |
| chr16-48451564-DEL-191 | chr16-48451565-DEL-191 |
| chr16-48455836-DEL-499 | chr16-48455837-DEL-499 |
| chr16-48476537-DEL-6271 | chr16-48476538-DEL-6270 |
| chr16-48496367-DEL-192 | chr16-48496368-DEL-192 |
| chr16-48520540-DEL-60 | chr16-48520541-DEL-60 |

| | |
|---|---|
| chr16-48556693-DEL-440 | chr16-48556694-DEL-440 |
| chr16-48572847-DEL-1154 | chr16-48572848-DEL-1154 |
| chr16-48587338-DEL-532 | chr16-48587338-DEL-531 |
| chr16-48596160-DEL-56 | chr16-48596161-DEL-56 |
| chr16-48627965-DEL-1112 | chr16-48627960-DEL-1100 |
| chr16-48636263-DEL-803 | chr16-48636264-DEL-803 |
| chr16-48652694-DEL-853 | chr16-48652695-DEL-853 |
| chr16-48676060-DEL-6732 | chr16-48676061-DEL-6732 |
| chr16-48698849-DEL-8568 | chr16-48698850-DEL-8568 |
| chr16-48725318-DEL-331 | chr16-48725319-DEL-331 |
| chr16-48783187-DEL-8475 | chr16-48783188-DEL-8475 |
| chr16-48783343-DEL-424 | chr16-48783344-DEL-424 |
| chr16-48797525-DEL-627 | chr16-48797526-DEL-627 |
| chr16-48814978-DEL-63 | chr16-48814985-DEL-57 |
| chr16-48845378-DEL-6327 | chr16-48845379-DEL-6327 |
| chr16-48885653-DEL-479 | chr16-48885654-DEL-477 |
| chr16-48887649-DEL-533 | chr16-48887648-DEL-534 |
| chr16-48894040-DEL-199 | chr16-48894041-DEL-199 |
| chr16-48900048-DEL-52 | chr16-48900051-DEL-50 |
| chr16-48905245-DEL-63 | chr16-48905246-DEL-63 |
| chr16-48918498-DEL-62 | chr16-48918501-DEL-60 |
| chr16-48939456-DEL-91 | chr16-48939452-DEL-128 |
| chr16-48998703-DEL-2812 | chr16-48998704-DEL-2812 |
| chr16-49007558-DEL-210 | chr16-49007557-DEL-210 |
| chr16-49015663-DEL-75 | chr16-49015664-DEL-73 |
| chr16-49020674-DEL-6228 | chr16-49020668-DEL-6229 |
| chr16-49028942-DEL-100 | chr16-49028943-DEL-100 |
| chr16-49055039-DEL-150 | chr16-49055040-DEL-150 |
| chr16-49077221-DEL-3779 | chr16-49077222-DEL-3779 |
| chr16-49095465-DEL-598 | chr16-49095466-DEL-598 |
| chr16-49103084-DEL-54 | chr16-49103049-DEL-54 |
| chr16-49119374-DEL-76 | chr16-49119375-DEL-76 |
| chr16-49130195-DEL-414 | chr16-49130196-DEL-414 |
| chr16-49132322-DEL-54 | chr16-49132292-DEL-52 |
| chr16-49137176-DEL-70 | chr16-49137177-DEL-65 |
| chr16-49137929-DEL-833 | chr16-49137930-DEL-833 |
| chr16-49163405-DEL-134 | chr16-49163429-DEL-130 |
| chr16-49180182-DEL-3028 | chr16-49180183-DEL-3028 |
| chr16-49234766-DEL-1111 | chr16-49234765-DEL-1100 |
| chr16-49267980-DEL-6444 | chr16-49267993-DEL-6433 |
| chr16-49278412-DEL-1231 | chr16-49278413-DEL-1231 |
| chr16-49286752-DEL-6568 | chr16-49286753-DEL-6568 |
| chr16-49305107-DEL-4650 | chr16-49305107-DEL-4650 |
| chr16-49331169-DEL-68 | chr16-49331170-DEL-58 |
| chr16-49372122-DEL-5967 | chr16-49372122-DEL-5968 |
| chr16-49382347-DEL-358 | chr16-49382348-DEL-358 |
| chr16-49413055-DEL-144 | chr16-49413055-DEL-145 |
| chr16-49509249-DEL-837 | chr16-49509250-DEL-837 |
| chr16-49516872-DEL-96 | chr16-49516861-DEL-102 |
| chr16-49552622-DEL-1772 | chr16-49552623-DEL-1772 |
| chr16-49565110-DEL-51 | chr16-49565111-DEL-51 |
| chr16-49576541-DEL-161 | chr16-49576542-DEL-161 |
| chr16-49592302-DEL-92 | chr16-49592303-DEL-92 |
| chr16-49593329-DEL-72 | chr16-49593330-DEL-72 |
| chr16-49608403-DEL-499 | chr16-49608404-DEL-499 |
| chr16-49618192-DEL-54 | chr16-49618193-DEL-54 |
| chr16-49621681-DEL-432 | chr16-49621682-DEL-432 |
| chr16-49752199-DEL-509 | chr16-49752200-DEL-509 |
| chr16-49806503-DEL-53 | chr16-49806504-DEL-53 |
| chr16-49824452-DEL-2329 | chr16-49824453-DEL-2329 |
| chr16-49833958-DEL-57 | chr16-49833959-DEL-57 |
| chr16-49846188-DEL-60 | chr16-49846189-DEL-60 |
| chr16-49850849-DEL-303 | chr16-49850836-DEL-279 |
| chr16-49900653-DEL-1610 | chr16-49900654-DEL-1610 |
| chr16-49905346-DEL-4396 | chr16-49905347-DEL-4396 |
| chr16-49962829-DEL-54 | chr16-49962830-DEL-54 |
| chr16-49963547-DEL-255 | chr16-49963548-DEL-255 |
| chr16-49974078-DEL-340 | chr16-49974079-DEL-340 |
| chr16-49981249-DEL-86 | chr16-49981250-DEL-86 |
| chr16-49985373-DEL-66 | chr16-49985374-DEL-66 |
| chr16-49987166-DEL-163 | chr16-49987167-DEL-163 |
| chr16-50001474-DEL-86 | chr16-50001475-DEL-86 |
| chr16-50036308-DEL-52 | chr16-50036335-DEL-52 |
| chr16-50056003-DEL-106 | chr16-50056004-DEL-106 |
| chr16-50059765-DEL-64 | chr16-50059766-DEL-64 |
| chr16-50092331-DEL-58 | chr16-50092334-DEL-56 |
| chr16-50117620-DEL-280 | chr16-50117621-DEL-280 |
| chr16-50123849-DEL-378 | chr16-50123850-DEL-378 |
| chr16-50154528-DEL-3023 | chr16-50154529-DEL-3023 |
| chr16-50163498-DEL-118 | chr16-50163498-DEL-118 |

| | |
|---|---|
| chr16-50213732-DEL-99 | chr16-50213733-DEL-99 |
| chr16-50233143-DEL-227 | chr16-50233144-DEL-227 |
| chr16-50278899-DEL-66 | chr16-50278900-DEL-66 |
| chr16-50280248-DEL-60 | chr16-50280239-DEL-60 |
| chr16-50291866-DEL-124 | chr16-50291882-DEL-126 |
| chr16-50308742-DEL-195 | chr16-50308751-DEL-189 |
| chr16-50341942-DEL-1196 | chr16-50341943-DEL-1196 |
| chr16-50358684-DEL-194 | chr16-50358685-DEL-194 |
| chr16-50377902-DEL-328 | chr16-50377903-DEL-328 |
| chr16-50383119-DEL-52 | chr16-50383141-DEL-54 |
| chr16-50400052-DEL-511 | chr16-50400053-DEL-511 |
| chr16-50413390-DEL-279 | chr16-50413391-DEL-279 |
| chr16-50466385-DEL-115 | chr16-50466381-DEL-120 |
| chr16-50470412-DEL-114 | chr16-50470413-DEL-110 |
| chr16-50474817-DEL-72 | chr16-50474818-DEL-72 |
| chr16-50476418-DEL-52 | chr16-50476419-DEL-52 |
| chr16-50500715-DEL-71 | chr16-50500716-DEL-71 |
| chr16-50515035-DEL-54 | chr16-50515055-DEL-57 |
| chr16-50529216-DEL-433 | chr16-50529217-DEL-433 |
| chr16-50629611-DEL-1889 | chr16-50629612-DEL-1889 |
| chr16-50647687-DEL-2832 | chr16-50647688-DEL-2832 |
| chr16-50649137-DEL-1078 | chr16-50649138-DEL-1078 |
| chr16-50657747-DEL-58 | chr16-50657748-DEL-58 |
| chr16-50658733-DEL-798 | chr16-50658733-DEL-798 |
| chr16-50718280-DEL-76 | chr16-50718281-DEL-76 |
| chr16-50730092-DEL-5398 | chr16-50730093-DEL-5398 |
| chr16-50751550-DEL-309 | chr16-50751551-DEL-309 |
| chr16-50784723-DEL-80 | chr16-50784754-DEL-58 |
| chr16-50788289-DEL-57 | chr16-50788290-DEL-57 |
| chr16-50790336-DEL-186 | chr16-50790335-DEL-187 |
| chr16-50798721-DEL-172 | chr16-50798721-DEL-173 |
| chr16-50801635-DEL-7097 | chr16-50801636-DEL-7098 |
| chr16-50801671-DEL-52 | chr16-50801671-DEL-52 |
| chr16-50810298-DEL-1890 | chr16-50810299-DEL-1890 |
| chr16-50818623-DEL-96 | chr16-50818591-DEL-127 |
| chr16-50843234-DEL-220 | chr16-50843233-DEL-221 |
| chr16-50849375-DEL-448 | chr16-50849376-DEL-448 |
| chr16-50859581-DEL-242 | chr16-50859582-DEL-242 |
| chr16-50920144-DEL-441 | chr16-50920145-DEL-441 |
| chr16-50933860-DEL-52 | chr16-50933861-DEL-52 |
| chr16-50935744-DEL-319 | chr16-50935745-DEL-319 |
| chr16-50949689-DEL-6960 | chr16-50949690-DEL-6960 |
| chr16-50998355-DEL-811 | chr16-50998356-DEL-811 |
| chr16-50999456-DEL-747 | chr16-50999457-DEL-747 |
| chr16-51013967-DEL-204 | chr16-51013968-DEL-204 |
| chr16-51027586-DEL-659 | chr16-51027587-DEL-659 |
| chr16-51029011-DEL-395 | chr16-51029010-DEL-396 |
| chr16-51032472-DEL-1726 | chr16-51032473-DEL-1726 |
| chr16-51034437-DEL-654 | chr16-51034438-DEL-654 |
| chr16-51047525-DEL-100 | chr16-51047525-DEL-100 |
| chr16-51055467-DEL-2243 | chr16-51055467-DEL-2243 |
| chr16-51142103-DEL-4931 | chr16-51142104-DEL-4931 |
| chr16-51152030-DEL-76 | chr16-51152031-DEL-76 |
| chr16-51195065-DEL-66 | chr16-51195065-DEL-66 |
| chr16-51204284-DEL-204 | chr16-51204285-DEL-204 |
| chr16-51251866-DEL-398 | chr16-51251867-DEL-398 |
| chr16-51263730-DEL-52 | chr16-51263731-DEL-52 |
| chr16-51323650-DEL-6222 | chr16-51323651-DEL-6222 |
| chr16-51333536-DEL-329 | chr16-51333518-DEL-341 |
| chr16-51333978-DEL-644 | chr16-51333979-DEL-644 |
| chr16-51348035-DEL-230 | chr16-51348036-DEL-230 |
| chr16-51358221-DEL-189 | chr16-51358222-DEL-189 |
| chr16-51401586-DEL-417 | chr16-51401587-DEL-417 |
| chr16-51414478-DEL-242 | chr16-51414479-DEL-242 |
| chr16-51434489-DEL-54 | chr16-51434490-DEL-56 |
| chr16-51436464-DEL-141 | chr16-51436500-DEL-115 |
| chr16-51445865-DEL-4299 | chr16-51445866-DEL-4299 |
| chr16-51462868-DEL-549 | chr16-51462869-DEL-549 |
| chr16-51465096-DEL-69 | chr16-51465095-DEL-78 |
| chr16-51478368-DEL-1497 | chr16-51478368-DEL-1497 |
| chr16-51496067-DEL-63 | chr16-51496065-DEL-63 |
| chr16-51549901-DEL-6743 | chr16-51549901-DEL-6743 |
| chr16-51550078-DEL-214 | chr16-51550078-DEL-214 |
| chr16-51598804-DEL-102 | chr16-51598799-DEL-103 |
| chr16-51600194-DEL-59 | chr16-51600176-DEL-60 |
| chr16-51629819-DEL-1066 | chr16-51629820-DEL-1066 |
| chr16-51653781-DEL-1532 | chr16-51653781-DEL-1532 |
| chr16-51688379-DEL-1443 | chr16-51688379-DEL-1445 |
| chr16-51757306-DEL-349 | chr16-51757304-DEL-350 |
| chr16-51766283-DEL-7092 | chr16-51766284-DEL-7092 |

| | |
|---|---|
| chr16-51777477-DEL-79 | chr16-51777479-DEL-78 |
| chr16-51784881-DEL-995 | chr16-51784882-DEL-995 |
| chr16-51788482-DEL-6432 | chr16-51788483-DEL-6432 |
| chr16-51796228-DEL-4335 | chr16-51796229-DEL-4335 |
| chr16-51811575-DEL-324 | chr16-51811576-DEL-324 |
| chr16-51912969-DEL-80 | chr16-51912970-DEL-84 |
| chr16-51918413-DEL-1397 | chr16-51918414-DEL-1397 |
| chr16-51937856-DEL-69 | chr16-51937857-DEL-69 |
| chr16-51970624-DEL-110 | chr16-51970625-DEL-110 |
| chr16-52023519-DEL-1124 | chr16-52023520-DEL-1124 |
| chr16-52050276-DEL-189 | chr16-52050277-DEL-189 |
| chr16-52063617-DEL-55 | chr16-52063617-DEL-55 |
| chr16-52101352-DEL-6576 | chr16-52101321-DEL-6614 |
| chr16-52113682-DEL-599 | chr16-52113683-DEL-599 |
| chr16-52120284-DEL-86 | chr16-52120281-DEL-90 |
| chr16-52133326-DEL-73 | chr16-52133334-DEL-66 |
| chr16-52137660-DEL-54 | chr16-52137661-DEL-57 |
| chr16-52189232-DEL-1710 | chr16-52189240-DEL-1708 |
| chr16-52193037-DEL-1221 | chr16-52193038-DEL-1221 |
| chr16-52244570-DEL-127 | chr16-52244591-DEL-88 |
| chr16-52246777-DEL-213 | chr16-52246810-DEL-169 |
| chr16-52282794-DEL-258 | chr16-52282795-DEL-258 |
| chr16-52326529-DEL-1976 | chr16-52326530-DEL-1976 |
| chr16-52335098-DEL-6205 | chr16-52335099-DEL-6205 |
| chr16-52346199-DEL-183 | chr16-52346200-DEL-183 |
| chr16-52355744-DEL-740 | chr16-52355745-DEL-740 |
| chr16-52358251-DEL-1426 | chr16-52358259-DEL-1425 |
| chr16-52374177-DEL-5561 | chr16-52374177-DEL-5561 |
| chr16-52381583-DEL-6198 | chr16-52381584-DEL-6198 |
| chr16-52398111-DEL-522 | chr16-52398112-DEL-522 |
| chr16-52424235-DEL-195 | chr16-52424236-DEL-195 |
| chr16-52436010-DEL-913 | chr16-52436011-DEL-913 |
| chr16-52442190-DEL-544 | chr16-52442191-DEL-544 |
| chr16-52454020-DEL-68 | chr16-52454021-DEL-68 |
| chr16-52484332-DEL-3518 | chr16-52484333-DEL-3518 |
| chr16-52501295-DEL-869 | chr16-52501294-DEL-870 |
| chr16-52554864-DEL-1998 | chr16-52554865-DEL-1991 |
| chr16-52557154-DEL-69 | chr16-52557155-DEL-69 |
| chr16-52572720-DEL-192 | chr16-52572721-DEL-192 |
| chr16-52577453-DEL-388 | chr16-52577454-DEL-388 |
| chr16-52580663-DEL-125 | chr16-52580664-DEL-125 |
| chr16-52581748-DEL-52 | chr16-52581749-DEL-52 |
| chr16-52597177-DEL-55 | chr16-52597174-DEL-59 |
| chr16-52674067-DEL-82 | chr16-52674071-DEL-79 |
| chr16-52691939-DEL-102 | chr16-52691940-DEL-102 |
| chr16-52705426-DEL-439 | chr16-52705427-DEL-439 |
| chr16-52714296-DEL-82 | chr16-52714297-DEL-62 |
| chr16-52732461-DEL-2192 | chr16-52732462-DEL-2192 |
| chr16-52737218-DEL-84 | chr16-52737219-DEL-84 |
| chr16-52769337-DEL-400 | chr16-52769333-DEL-399 |
| chr16-52777921-DEL-6379 | chr16-52777923-DEL-6378 |
| chr16-52808541-DEL-56 | chr16-52808540-DEL-50 |
| chr16-52833870-DEL-4443 | chr16-52833871-DEL-4443 |
| chr16-52867262-DEL-160 | chr16-52867281-DEL-112 |
| chr16-52909544-DEL-770 | chr16-52909545-DEL-770 |
| chr16-52920567-DEL-79 | chr16-52920568-DEL-79 |
| chr16-52924875-DEL-1180 | chr16-52924876-DEL-1180 |
| chr16-52955044-DEL-66 | chr16-52955070-DEL-66 |
| chr16-52969003-DEL-2733 | chr16-52969004-DEL-2733 |
| chr16-52997401-DEL-436 | chr16-52997402-DEL-436 |
| chr16-52998445-DEL-588 | chr16-52998446-DEL-588 |
| chr16-53002942-DEL-165 | chr16-53002943-DEL-165 |
| chr16-53018616-DEL-359 | chr16-53018617-DEL-359 |
| chr16-53047153-DEL-61 | chr16-53047154-DEL-59 |
| chr16-53091784-DEL-200 | chr16-53091785-DEL-198 |
| chr16-53111050-DEL-96 | chr16-53111083-DEL-52 |
| chr16-53125124-DEL-60 | chr16-53125125-DEL-60 |
| chr16-53140434-DEL-755 | chr16-53140434-DEL-755 |
| chr16-53168164-DEL-6552 | chr16-53168165-DEL-6552 |
| chr16-53188011-DEL-52 | chr16-53188012-DEL-52 |
| chr16-53201264-DEL-652 | chr16-53201269-DEL-652 |
| chr16-53205252-DEL-224 | chr16-53205253-DEL-224 |
| chr16-53242963-DEL-6337 | chr16-53242963-DEL-6337 |
| chr16-53246023-DEL-77 | chr16-53246024-DEL-77 |
| chr16-53285563-DEL-393 | chr16-53285564-DEL-393 |
| chr16-53320992-DEL-847 | chr16-53320993-DEL-847 |
| chr16-53341002-DEL-210 | chr16-53341003-DEL-210 |
| chr16-53346656-DEL-67 | chr16-53346659-DEL-65 |
| chr16-53350512-DEL-6634 | chr16-53350506-DEL-6635 |
| chr16-53418917-DEL-158 | chr16-53418911-DEL-164 |

| | |
|---|---|
| chr16-53421599-DEL-63 | chr16-53421597-DEL-67 |
| chr16-53443758-DEL-264 | chr16-53443759-DEL-264 |
| chr16-53445605-DEL-703 | chr16-53445606-DEL-703 |
| chr16-53457828-DEL-451 | chr16-53457829-DEL-451 |
| chr16-53467931-DEL-6997 | chr16-53467932-DEL-6997 |
| chr16-53475468-DEL-96 | chr16-53475469-DEL-96 |
| chr16-53491838-DEL-52 | chr16-53491839-DEL-50 |
| chr16-53496392-DEL-2447 | chr16-53496393-DEL-2447 |
| chr16-53501937-DEL-2200 | chr16-53501938-DEL-2200 |
| chr16-53518779-DEL-83 | chr16-53518780-DEL-83 |
| chr16-53546378-DEL-93 | chr16-53546379-DEL-93 |
| chr16-53564880-DEL-60 | chr16-53564880-DEL-54 |
| chr16-53631439-DEL-612 | chr16-53631440-DEL-612 |
| chr16-53638301-DEL-94 | chr16-53638302-DEL-94 |
| chr16-53644822-DEL-64 | chr16-53644823-DEL-64 |
| chr16-53645290-DEL-8170 | chr16-53645286-DEL-8171 |
| chr16-53657461-DEL-231 | chr16-53657462-DEL-231 |
| chr16-53659136-DEL-174 | chr16-53659137-DEL-174 |
| chr16-53685123-DEL-108 | chr16-53685083-DEL-104 |
| chr16-53697373-DEL-51 | chr16-53697355-DEL-64 |
| chr16-53719259-DEL-98 | chr16-53719272-DEL-86 |
| chr16-53741361-DEL-1083 | chr16-53741362-DEL-1083 |
| chr16-53760522-DEL-572 | chr16-53760523-DEL-571 |
| chr16-53768178-DEL-3253 | chr16-53768179-DEL-3253 |
| chr16-53787001-DEL-72 | chr16-53787002-DEL-72 |
| chr16-53792570-DEL-101 | chr16-53792557-DEL-107 |
| chr16-53804181-DEL-2396 | chr16-53804175-DEL-2402 |
| chr16-53811739-DEL-904 | chr16-53811740-DEL-904 |
| chr16-53823661-DEL-6219 | chr16-53823662-DEL-6219 |
| chr16-53853686-DEL-348 | chr16-53853687-DEL-348 |
| chr16-53855719-DEL-111 | chr16-53855703-DEL-120 |
| chr16-53904842-DEL-52 | chr16-53904810-DEL-90 |
| chr16-53928168-DEL-442 | chr16-53928169-DEL-439 |
| chr16-53928952-DEL-544 | chr16-53928953-DEL-544 |
| chr16-53973839-DEL-122 | chr16-53973840-DEL-122 |
| chr16-53980475-DEL-68 | chr16-53980476-DEL-68 |
| chr16-54012582-DEL-202 | chr16-54012583-DEL-202 |
| chr16-54021582-DEL-2260 | chr16-54021583-DEL-2260 |
| chr16-54024635-DEL-561 | chr16-54024636-DEL-561 |
| chr16-54027326-DEL-76 | chr16-54027327-DEL-76 |
| chr16-54039090-DEL-196 | chr16-54039091-DEL-196 |
| chr16-54042057-DEL-466 | chr16-54042050-DEL-461 |
| chr16-54050837-DEL-124 | chr16-54050838-DEL-124 |
| chr16-54060912-DEL-254 | chr16-54060913-DEL-254 |
| chr16-54074460-DEL-644 | chr16-54074461-DEL-645 |
| chr16-54118731-DEL-6447 | chr16-54118732-DEL-6447 |
| chr16-54179305-DEL-366 | chr16-54179306-DEL-362 |
| chr16-54233051-DEL-196 | chr16-54233052-DEL-196 |
| chr16-54236433-DEL-1396 | chr16-54236434-DEL-1396 |
| chr16-54238543-DEL-84 | chr16-54238544-DEL-84 |
| chr16-54243481-DEL-15386 | chr16-54243478-DEL-15388 |
| chr16-54255766-DEL-386 | chr16-54255767-DEL-387 |
| chr16-54276637-DEL-96 | chr16-54276637-DEL-96 |
| chr16-54285243-DEL-208 | chr16-54285244-DEL-208 |
| chr16-54287232-DEL-2486 | chr16-54287233-DEL-2486 |
| chr16-54293507-DEL-477 | chr16-54293508-DEL-477 |
| chr16-54297380-DEL-112 | chr16-54297381-DEL-112 |
| chr16-54332086-DEL-554 | chr16-54332087-DEL-554 |
| chr16-54338229-DEL-6802 | chr16-54338244-DEL-6801 |
| chr16-54358259-DEL-4727 | chr16-54358260-DEL-4727 |
| chr16-54383448-DEL-3500 | chr16-54383449-DEL-3500 |
| chr16-54391556-DEL-588 | chr16-54391555-DEL-588 |
| chr16-54392959-DEL-61 | chr16-54392960-DEL-61 |
| chr16-54404246-DEL-528 | chr16-54404247-DEL-528 |
| chr16-54409727-DEL-629 | chr16-54409728-DEL-629 |
| chr16-54413834-DEL-197 | chr16-54413835-DEL-197 |
| chr16-54417165-DEL-62 | chr16-54417166-DEL-62 |
| chr16-54422818-DEL-851 | chr16-54422819-DEL-851 |
| chr16-54423755-DEL-68 | chr16-54423760-DEL-68 |
| chr16-54432059-DEL-84 | chr16-54432060-DEL-84 |
| chr16-54432722-DEL-939 | chr16-54432721-DEL-940 |
| chr16-54439636-DEL-713 | chr16-54439637-DEL-713 |
| chr16-54443144-DEL-65 | chr16-54443145-DEL-65 |
| chr16-54470804-DEL-1576 | chr16-54470805-DEL-1576 |
| chr16-54508733-DEL-239 | chr16-54508734-DEL-239 |
| chr16-54537309-DEL-3427 | chr16-54537310-DEL-3427 |
| chr16-54540067-DEL-120 | chr16-54540029-DEL-141 |
| chr16-54566663-DEL-688 | chr16-54566664-DEL-688 |
| chr16-54579109-DEL-1321 | chr16-54579110-DEL-1321 |
| chr16-54590318-DEL-645 | chr16-54590319-DEL-643 |

| | |
|---|---|
| chr16-54643747-DEL-2150 | chr16-54643748-DEL-2150 |
| chr16-54648387-DEL-483 | chr16-54648383-DEL-485 |
| chr16-54649455-DEL-152 | chr16-54649456-DEL-152 |
| chr16-54657688-DEL-224 | chr16-54657689-DEL-224 |
| chr16-54662588-DEL-1058 | chr16-54662589-DEL-1058 |
| chr16-54674546-DEL-81 | chr16-54674547-DEL-81 |
| chr16-54687849-DEL-635 | chr16-54687850-DEL-635 |
| chr16-54716671-DEL-2603 | chr16-54716672-DEL-2603 |
| chr16-54743096-DEL-196 | chr16-54743077-DEL-216 |
| chr16-54758666-DEL-188 | chr16-54758667-DEL-188 |
| chr16-54779426-DEL-6402 | chr16-54779427-DEL-6402 |
| chr16-54806513-DEL-54 | chr16-54806514-DEL-54 |
| chr16-54841847-DEL-78 | chr16-54841847-DEL-80 |
| chr16-54850924-DEL-326 | chr16-54850925-DEL-326 |
| chr16-54872895-DEL-4372 | chr16-54872896-DEL-4372 |
| chr16-54904614-DEL-92 | chr16-54904615-DEL-92 |
| chr16-54935212-DEL-6413 | chr16-54935213-DEL-6413 |
| chr16-54941219-DEL-201 | chr16-54941219-DEL-201 |
| chr16-54966525-DEL-475 | chr16-54966526-DEL-475 |
| chr16-54970056-DEL-205 | chr16-54970057-DEL-204 |
| chr16-54973891-DEL-1121 | chr16-54973892-DEL-1121 |
| chr16-54975357-DEL-6248 | chr16-54975358-DEL-6248 |
| chr16-54975379-DEL-199 | chr16-54975380-DEL-199 |
| chr16-54983359-DEL-75 | chr16-54983360-DEL-75 |
| chr16-54990073-DEL-1536 | chr16-54990074-DEL-1536 |
| chr16-55002631-DEL-114 | chr16-55002632-DEL-114 |
| chr16-55004767-DEL-283 | chr16-55004768-DEL-283 |
| chr16-55055515-DEL-56 | chr16-55055516-DEL-58 |
| chr16-55078093-DEL-1091 | chr16-55078098-DEL-1086 |
| chr16-55109255-DEL-661 | chr16-55109256-DEL-661 |
| chr16-55127683-DEL-3674 | chr16-55127684-DEL-3674 |
| chr16-55143094-DEL-3145 | chr16-55143095-DEL-3145 |
| chr16-55147185-DEL-56 | chr16-55147186-DEL-56 |
| chr16-55158950-DEL-2498 | chr16-55158951-DEL-2498 |
| chr16-55239755-DEL-294 | chr16-55239754-DEL-295 |
| chr16-55240309-DEL-105 | chr16-55240310-DEL-91 |
| chr16-55249458-DEL-1556 | chr16-55249459-DEL-1556 |
| chr16-55274271-DEL-520 | chr16-55274272-DEL-520 |
| chr16-55282067-DEL-60 | chr16-55282068-DEL-60 |
| chr16-55292911-DEL-106 | chr16-55292907-DEL-107 |
| chr16-55297468-DEL-134 | chr16-55297469-DEL-134 |
| chr16-55310347-DEL-503 | chr16-55310348-DEL-503 |
| chr16-55320383-DEL-1268 | chr16-55320384-DEL-1268 |
| chr16-55336006-DEL-182 | chr16-55336007-DEL-182 |
| chr16-55340682-DEL-251 | chr16-55340683-DEL-251 |
| chr16-55351616-DEL-53 | chr16-55351616-DEL-53 |
| chr16-55396029-DEL-264 | chr16-55396030-DEL-264 |
| chr16-55405595-DEL-181 | chr16-55405596-DEL-181 |
| chr16-55441145-DEL-104 | chr16-55441146-DEL-104 |
| chr16-55453906-DEL-274 | chr16-55453906-DEL-274 |
| chr16-55455928-DEL-304 | chr16-55455929-DEL-304 |
| chr16-55468055-DEL-6390 | chr16-55468056-DEL-6390 |
| chr16-55493976-DEL-8094 | chr16-55493977-DEL-8094 |
| chr16-55529648-DEL-74 | chr16-55529649-DEL-74 |
| chr16-55572339-DEL-776 | chr16-55572340-DEL-776 |
| chr16-55628565-DEL-53 | chr16-55628546-DEL-54 |
| chr16-55667105-DEL-194 | chr16-55667106-DEL-194 |
| chr16-55672444-DEL-128 | chr16-55672445-DEL-128 |
| chr16-55680988-DEL-52 | chr16-55680994-DEL-67 |
| chr16-55685572-DEL-338 | chr16-55685573-DEL-338 |
| chr16-55728126-DEL-2699 | chr16-55728127-DEL-2699 |
| chr16-55736782-DEL-142 | chr16-55736783-DEL-140 |
| chr16-55747376-DEL-162 | chr16-55747413-DEL-157 |
| chr16-55762018-DEL-336 | chr16-55762019-DEL-336 |
| chr16-55774115-DEL-100 | chr16-55774124-DEL-100 |
| chr16-55775193-DEL-8403 | chr16-55775194-DEL-8403 |
| chr16-55799235-DEL-984 | chr16-55799240-DEL-978 |
| chr16-55804167-DEL-856 | chr16-55804168-DEL-856 |
| chr16-55842267-DEL-97 | chr16-55842268-DEL-97 |
| chr16-55853912-DEL-94 | chr16-55853913-DEL-94 |
| chr16-55858786-DEL-14714 | chr16-55858787-DEL-14714 |
| chr16-55862446-DEL-6452 | chr16-55862447-DEL-6452 |
| chr16-55874527-DEL-209 | chr16-55874520-DEL-210 |
| chr16-55879011-DEL-2048 | chr16-55879012-DEL-2048 |
| chr16-55883027-DEL-1464 | chr16-55883027-DEL-1464 |
| chr16-55885886-DEL-187 | chr16-55885886-DEL-161 |
| chr16-55886762-DEL-113 | chr16-55886763-DEL-113 |
| chr16-55890618-DEL-54 | chr16-55890619-DEL-54 |
| chr16-55912477-DEL-339 | chr16-55912478-DEL-339 |
| chr16-55919883-DEL-133 | chr16-55919884-DEL-133 |

| | |
|---|---|
| chr16-55987468-DEL-353 | chr16-55987469-DEL-353 |
| chr16-55997491-DEL-1248 | chr16-55997492-DEL-1248 |
| chr16-56002149-DEL-52 | chr16-56002147-DEL-53 |
| chr16-56016381-DEL-790 | chr16-56016382-DEL-790 |
| chr16-56021797-DEL-193 | chr16-56021798-DEL-193 |
| chr16-56022665-DEL-121 | chr16-56022666-DEL-121 |
| chr16-56022935-DEL-1307 | chr16-56022936-DEL-1307 |
| chr16-56026686-DEL-203 | chr16-56026687-DEL-203 |
| chr16-56037091-DEL-1201 | chr16-56037091-DEL-1201 |
| chr16-56050902-DEL-157 | chr16-56050891-DEL-197 |
| chr16-56062278-DEL-1021 | chr16-56062275-DEL-1022 |
| chr16-56063620-DEL-84 | chr16-56063621-DEL-74 |
| chr16-56067090-DEL-1055 | chr16-56067091-DEL-1055 |
| chr16-56107859-DEL-6401 | chr16-56107860-DEL-6401 |
| chr16-56146611-DEL-202 | chr16-56146612-DEL-202 |
| chr16-56170741-DEL-500 | chr16-56170742-DEL-500 |
| chr16-56180498-DEL-426 | chr16-56180486-DEL-427 |
| chr16-56204552-DEL-61 | chr16-56204553-DEL-61 |
| chr16-56208491-DEL-146 | chr16-56208492-DEL-146 |
| chr16-56210876-DEL-1462 | chr16-56210877-DEL-1462 |
| chr16-56214239-DEL-199 | chr16-56214240-DEL-199 |
| chr16-56238907-DEL-616 | chr16-56238906-DEL-617 |
| chr16-56241769-DEL-273 | chr16-56241770-DEL-273 |
| chr16-56246671-DEL-143 | chr16-56246672-DEL-143 |
| chr16-56247816-DEL-6207 | chr16-56247817-DEL-6207 |
| chr16-56289867-DEL-1299 | chr16-56289868-DEL-1299 |
| chr16-56291910-DEL-383 | chr16-56291911-DEL-383 |
| chr16-56358343-DEL-331 | chr16-56358344-DEL-331 |
| chr16-56364740-DEL-3018 | chr16-56364741-DEL-3018 |
| chr16-56365004-DEL-841 | chr16-56365004-DEL-841 |
| chr16-56381677-DEL-194 | chr16-56381678-DEL-194 |
| chr16-56387539-DEL-77 | chr16-56387540-DEL-77 |
| chr16-56412263-DEL-384 | chr16-56412264-DEL-384 |
| chr16-56446509-DEL-220 | chr16-56446510-DEL-220 |
| chr16-56461834-DEL-52 | chr16-56461835-DEL-52 |
| chr16-56469824-DEL-305 | chr16-56469825-DEL-305 |
| chr16-56487624-DEL-72 | chr16-56487596-DEL-81 |
| chr16-56490646-DEL-63 | chr16-56490647-DEL-63 |
| chr16-56593086-DEL-72 | chr16-56593087-DEL-72 |
| chr16-56628473-DEL-210 | chr16-56628435-DEL-258 |
| chr16-56644700-DEL-493 | chr16-56644701-DEL-493 |
| chr16-56659303-DEL-54 | chr16-56659304-DEL-54 |
| chr16-56668751-DEL-367 | chr16-56668752-DEL-367 |
| chr16-56673990-DEL-100 | chr16-56673996-DEL-95 |
| chr16-56710744-DEL-59 | chr16-56710745-DEL-62 |
| chr16-56723710-DEL-190 | chr16-56723715-DEL-189 |
| chr16-56746113-DEL-60 | chr16-56746114-DEL-60 |
| chr16-56794551-DEL-1907 | chr16-56794552-DEL-1907 |
| chr16-56806870-DEL-235 | chr16-56806871-DEL-236 |
| chr16-56823033-DEL-6040 | chr16-56823034-DEL-6040 |
| chr16-56844826-DEL-887 | chr16-56844827-DEL-887 |
| chr16-56878570-DEL-3836 | chr16-56878571-DEL-3836 |
| chr16-56884070-DEL-559 | chr16-56884066-DEL-557 |
| chr16-56888218-DEL-102 | chr16-56888219-DEL-102 |
| chr16-56898633-DEL-416 | chr16-56898633-DEL-417 |
| chr16-56930609-DEL-84 | chr16-56930610-DEL-84 |
| chr16-56936530-DEL-73 | chr16-56936547-DEL-51 |
| chr16-56946202-DEL-282 | chr16-56946203-DEL-279 |
| chr16-56955554-DEL-136 | chr16-56955555-DEL-136 |
| chr16-57016052-DEL-543 | chr16-57016053-DEL-543 |
| chr16-57021337-DEL-6442 | chr16-57021338-DEL-6442 |
| chr16-57038349-DEL-208 | chr16-57038350-DEL-208 |
| chr16-57042148-DEL-58 | chr16-57042149-DEL-58 |
| chr16-57043448-DEL-470 | chr16-57043449-DEL-470 |
| chr16-57052497-DEL-212 | chr16-57052498-DEL-212 |
| chr16-57060326-DEL-212 | chr16-57060327-DEL-211 |
| chr16-57076858-DEL-102 | chr16-57076856-DEL-103 |
| chr16-57099807-DEL-204 | chr16-57099808-DEL-204 |
| chr16-57116660-DEL-1798 | chr16-57116661-DEL-1798 |
| chr16-57188861-DEL-74 | chr16-57188862-DEL-74 |
| chr16-57211484-DEL-557 | chr16-57211484-DEL-557 |
| chr16-57284544-DEL-226 | chr16-57284545-DEL-226 |
| chr16-57285175-DEL-383 | chr16-57285176-DEL-383 |
| chr16-57299964-DEL-58 | chr16-57299964-DEL-58 |
| chr16-57399499-DEL-357 | chr16-57399500-DEL-357 |
| chr16-57406427-DEL-7102 | chr16-57406428-DEL-7102 |
| chr16-57412521-DEL-846 | chr16-57412521-DEL-846 |
| chr16-57430012-DEL-151 | chr16-57430013-DEL-151 |
| chr16-57499184-DEL-515 | chr16-57499185-DEL-515 |
| chr16-57516516-DEL-417 | chr16-57516517-DEL-417 |

| | |
|---|---|
| chr16-57530281-DEL-54 | chr16-57530282-DEL-66 |
| chr16-57544033-DEL-100 | chr16-57544070-DEL-64 |
| chr16-57546699-DEL-52 | chr16-57546700-DEL-52 |
| chr16-57547750-DEL-54 | chr16-57547751-DEL-54 |
| chr16-57625581-DEL-6475 | chr16-57625582-DEL-6475 |
| chr16-57675422-DEL-1058 | chr16-57675422-DEL-1059 |
| chr16-57698846-DEL-440 | chr16-57698847-DEL-440 |
| chr16-57722407-DEL-246 | chr16-57722405-DEL-239 |
| chr16-57754342-DEL-84 | chr16-57754327-DEL-76 |
| chr16-57782111-DEL-6369 | chr16-57782112-DEL-6369 |
| chr16-57789424-DEL-72 | chr16-57789424-DEL-56 |
| chr16-57795124-DEL-1811 | chr16-57795125-DEL-1811 |
| chr16-57808274-DEL-275 | chr16-57808275-DEL-276 |
| chr16-57812788-DEL-170 | chr16-57812789-DEL-170 |
| chr16-57814872-DEL-342 | chr16-57814871-DEL-342 |
| chr16-57819195-DEL-6452 | chr16-57819195-DEL-6452 |
| chr16-57834869-DEL-73 | chr16-57834872-DEL-69 |
| chr16-57836180-DEL-346 | chr16-57836181-DEL-346 |
| chr16-57879845-DEL-130 | chr16-57879846-DEL-130 |
| chr16-57912920-DEL-473 | chr16-57912923-DEL-474 |
| chr16-57925461-DEL-62 | chr16-57925448-DEL-80 |
| chr16-57931247-DEL-143 | chr16-57931208-DEL-141 |
| chr16-57935682-DEL-58 | chr16-57935683-DEL-58 |
| chr16-57936094-DEL-4871 | chr16-57936095-DEL-4870 |
| chr16-57964677-DEL-52 | chr16-57964678-DEL-52 |
| chr16-58015830-DEL-193 | chr16-58015831-DEL-193 |
| chr16-58027862-DEL-128 | chr16-58027863-DEL-128 |
| chr16-58034832-DEL-604 | chr16-58034833-DEL-604 |
| chr16-58069777-DEL-6329 | chr16-58069778-DEL-6329 |
| chr16-58097483-DEL-231 | chr16-58097484-DEL-232 |
| chr16-58100160-DEL-66 | chr16-58100155-DEL-80 |
| chr16-58156143-DEL-7253 | chr16-58156144-DEL-7253 |
| chr16-58179336-DEL-56 | chr16-58179334-DEL-56 |
| chr16-58184348-DEL-4102 | chr16-58184349-DEL-4102 |
| chr16-58191420-DEL-536 | chr16-58191421-DEL-536 |
| chr16-58192683-DEL-3597 | chr16-58192684-DEL-3597 |
| chr16-58205735-DEL-59 | chr16-58205707-DEL-60 |
| chr16-58235939-DEL-62 | chr16-58235940-DEL-62 |
| chr16-58238034-DEL-1932 | chr16-58238035-DEL-1932 |
| chr16-58242844-DEL-124 | chr16-58242845-DEL-124 |
| chr16-58310682-DEL-55 | chr16-58310683-DEL-55 |
| chr16-58316067-DEL-419 | chr16-58316068-DEL-419 |
| chr16-58336500-DEL-154 | chr16-58336501-DEL-154 |
| chr16-58400427-DEL-53 | chr16-58400426-DEL-53 |
| chr16-58470674-DEL-5167 | chr16-58470675-DEL-5167 |
| chr16-58516073-DEL-237 | chr16-58516074-DEL-237 |
| chr16-58518166-DEL-120 | chr16-58518166-DEL-79 |
| chr16-58577174-DEL-56 | chr16-58577175-DEL-56 |
| chr16-58589682-DEL-1541 | chr16-58589683-DEL-1541 |
| chr16-58595412-DEL-4910 | chr16-58595413-DEL-4910 |
| chr16-58611054-DEL-69 | chr16-58611058-DEL-66 |
| chr16-58630545-DEL-472 | chr16-58630546-DEL-472 |
| chr16-58635116-DEL-378 | chr16-58635117-DEL-378 |
| chr16-58667498-DEL-195 | chr16-58667497-DEL-196 |
| chr16-58682811-DEL-6331 | chr16-58682812-DEL-6330 |
| chr16-58734006-DEL-63 | chr16-58734000-DEL-64 |
| chr16-58792051-DEL-1393 | chr16-58792051-DEL-1393 |
| chr16-58832548-DEL-356 | chr16-58832562-DEL-348 |
| chr16-58868002-DEL-646 | chr16-58868003-DEL-646 |
| chr16-58875696-DEL-52 | chr16-58875692-DEL-60 |
| chr16-58921603-DEL-56 | chr16-58921604-DEL-56 |
| chr16-58942276-DEL-7121 | chr16-58942277-DEL-7121 |
| chr16-58981144-DEL-60 | chr16-58981145-DEL-60 |
| chr16-58991910-DEL-7096 | chr16-58991911-DEL-7096 |
| chr16-59024394-DEL-62 | chr16-59024395-DEL-56 |
| chr16-59110847-DEL-436 | chr16-59110848-DEL-436 |
| chr16-59114815-DEL-55 | chr16-59114816-DEL-56 |
| chr16-59155068-DEL-7376 | chr16-59155069-DEL-7376 |
| chr16-59172420-DEL-3033 | chr16-59172421-DEL-3033 |
| chr16-59192924-DEL-186 | chr16-59192925-DEL-186 |
| chr16-59198934-DEL-951 | chr16-59198935-DEL-951 |
| chr16-59236898-DEL-155 | chr16-59236899-DEL-154 |
| chr16-59273202-DEL-1509 | chr16-59273203-DEL-1507 |
| chr16-59277937-DEL-3232 | chr16-59277938-DEL-3232 |
| chr16-59306246-DEL-87 | chr16-59306247-DEL-87 |
| chr16-59327708-DEL-78 | chr16-59327719-DEL-72 |
| chr16-59341097-DEL-1108 | chr16-59341098-DEL-1108 |
| chr16-59355282-DEL-76 | chr16-59355279-DEL-80 |
| chr16-59400398-DEL-565 | chr16-59400397-DEL-564 |
| chr16-59402112-DEL-55 | chr16-59402113-DEL-55 |

| | |
|---|---|
| chr16-59404920-DEL-71 | chr16-59404923-DEL-93 |
| chr16-59414212-DEL-140 | chr16-59414213-DEL-140 |
| chr16-59418949-DEL-54 | chr16-59418950-DEL-54 |
| chr16-59424136-DEL-62 | chr16-59424133-DEL-66 |
| chr16-59464599-DEL-163 | chr16-59464600-DEL-163 |
| chr16-59467310-DEL-68 | chr16-59467311-DEL-68 |
| chr16-59477188-DEL-84 | chr16-59477189-DEL-84 |
| chr16-59484924-DEL-168 | chr16-59484925-DEL-168 |
| chr16-59487727-DEL-55 | chr16-59487728-DEL-55 |
| chr16-59495408-DEL-62 | chr16-59495410-DEL-69 |
| chr16-59518577-DEL-6581 | chr16-59518566-DEL-6579 |
| chr16-59546824-DEL-62 | chr16-59546825-DEL-62 |
| chr16-59598135-DEL-6649 | chr16-59598136-DEL-6649 |
| chr16-59607322-DEL-150 | chr16-59607323-DEL-150 |
| chr16-59614713-DEL-3485 | chr16-59614714-DEL-3485 |
| chr16-59627777-DEL-80 | chr16-59627778-DEL-80 |
| chr16-59664294-DEL-495 | chr16-59664294-DEL-495 |
| chr16-59714321-DEL-6771 | chr16-59714322-DEL-6771 |
| chr16-59730384-DEL-102 | chr16-59730385-DEL-102 |
| chr16-59756837-DEL-1024 | chr16-59756831-DEL-1027 |
| chr16-59765933-DEL-51 | chr16-59765920-DEL-75 |
| chr16-59791709-DEL-184 | chr16-59791710-DEL-184 |
| chr16-59802716-DEL-76 | chr16-59802717-DEL-76 |
| chr16-59817168-DEL-1972 | chr16-59817169-DEL-1972 |
| chr16-59833291-DEL-74 | chr16-59833303-DEL-60 |
| chr16-59836627-DEL-6365 | chr16-59836628-DEL-6365 |
| chr16-59854447-DEL-1145 | chr16-59854448-DEL-1145 |
| chr16-59920355-DEL-64 | chr16-59920355-DEL-64 |
| chr16-59953923-DEL-2184 | chr16-59953923-DEL-2184 |
| chr16-59957595-DEL-67 | chr16-59957596-DEL-57 |
| chr16-59971158-DEL-658 | chr16-59971159-DEL-658 |
| chr16-59979153-DEL-1992 | chr16-59979154-DEL-1992 |
| chr16-59984196-DEL-945 | chr16-59984197-DEL-945 |
| chr16-59993432-DEL-295 | chr16-59993433-DEL-295 |
| chr16-59998760-DEL-314 | chr16-59998761-DEL-314 |
| chr16-60012896-DEL-80 | chr16-60012897-DEL-80 |
| chr16-60013809-DEL-54 | chr16-60013810-DEL-57 |
| chr16-60014806-DEL-66 | chr16-60014807-DEL-66 |
| chr16-60058248-DEL-62 | chr16-60058206-DEL-62 |
| chr16-60069939-DEL-57 | chr16-60069940-DEL-57 |
| chr16-60075181-DEL-2210 | chr16-60075182-DEL-2210 |
| chr16-60095541-DEL-157 | chr16-60095532-DEL-158 |
| chr16-60132345-DEL-54 | chr16-60132357-DEL-54 |
| chr16-60141228-DEL-390 | chr16-60141229-DEL-390 |
| chr16-60170599-DEL-1055 | chr16-60170590-DEL-1056 |
| chr16-60184552-DEL-1649 | chr16-60184553-DEL-1649 |
| chr16-60189216-DEL-523 | chr16-60189217-DEL-523 |
| chr16-60202310-DEL-92 | chr16-60202308-DEL-95 |
| chr16-60228248-DEL-55 | chr16-60228244-DEL-56 |
| chr16-60238143-DEL-72 | chr16-60238144-DEL-64 |
| chr16-60250635-DEL-603 | chr16-60250636-DEL-603 |
| chr16-60251310-DEL-1258 | chr16-60251311-DEL-1258 |
| chr16-60256886-DEL-10511 | chr16-60256887-DEL-10511 |
| chr16-60260326-DEL-692 | chr16-60260327-DEL-692 |
| chr16-60261899-DEL-4222 | chr16-60261898-DEL-4222 |
| chr16-60273905-DEL-315 | chr16-60273906-DEL-315 |
| chr16-60276690-DEL-1558 | chr16-60276691-DEL-1558 |
| chr16-60282834-DEL-116 | chr16-60282835-DEL-116 |
| chr16-60299809-DEL-207 | chr16-60299810-DEL-207 |
| chr16-60302472-DEL-134 | chr16-60302473-DEL-132 |
| chr16-60314565-DEL-2715 | chr16-60314566-DEL-2715 |
| chr16-60344691-DEL-70 | chr16-60344700-DEL-56 |
| chr16-60351613-DEL-57 | chr16-60351614-DEL-57 |
| chr16-60368368-DEL-299 | chr16-60368369-DEL-299 |
| chr16-60370429-DEL-83 | chr16-60370440-DEL-71 |
| chr16-60372948-DEL-6215 | chr16-60372949-DEL-6215 |
| chr16-60383803-DEL-3223 | chr16-60383804-DEL-3223 |
| chr16-60395806-DEL-499 | chr16-60395807-DEL-500 |
| chr16-60412885-DEL-56 | chr16-60412886-DEL-50 |
| chr16-60429897-DEL-476 | chr16-60429898-DEL-476 |
| chr16-60457779-DEL-312 | chr16-60457774-DEL-313 |
| chr16-60459063-DEL-823 | chr16-60459063-DEL-823 |
| chr16-60469991-DEL-74 | chr16-60469992-DEL-81 |
| chr16-60561373-DEL-7353 | chr16-60561374-DEL-7353 |
| chr16-60587526-DEL-207 | chr16-60587527-DEL-207 |
| chr16-60590149-DEL-228 | chr16-60590150-DEL-228 |
| chr16-60593386-DEL-6603 | chr16-60593387-DEL-6603 |
| chr16-60610629-DEL-125 | chr16-60610628-DEL-89 |
| chr16-60617371-DEL-762 | chr16-60617357-DEL-776 |
| chr16-60646909-DEL-6360 | chr16-60646910-DEL-6360 |

| | |
|---|---|
| chr16-60656933-DEL-574 | chr16-60656934-DEL-574 |
| chr16-60673711-DEL-1948 | chr16-60673712-DEL-1948 |
| chr16-60681359-DEL-739 | chr16-60681360-DEL-739 |
| chr16-60707941-DEL-63 | chr16-60707942-DEL-63 |
| chr16-60718343-DEL-953 | chr16-60718344-DEL-953 |
| chr16-60725070-DEL-64 | chr16-60725071-DEL-64 |
| chr16-60746125-DEL-6739 | chr16-60746125-DEL-6739 |
| chr16-60754876-DEL-254 | chr16-60754877-DEL-254 |
| chr16-60779839-DEL-1364 | chr16-60779840-DEL-1362 |
| chr16-60813476-DEL-187 | chr16-60813478-DEL-186 |
| chr16-60818041-DEL-199 | chr16-60818042-DEL-199 |
| chr16-60848315-DEL-260 | chr16-60848316-DEL-260 |
| chr16-60880316-DEL-369 | chr16-60880314-DEL-370 |
| chr16-60881547-DEL-69 | chr16-60881548-DEL-69 |
| chr16-60913758-DEL-75 | chr16-60913749-DEL-99 |
| chr16-60916140-DEL-869 | chr16-60916142-DEL-868 |
| chr16-60918195-DEL-1212 | chr16-60918195-DEL-1212 |
| chr16-60964504-DEL-3140 | chr16-60964505-DEL-3142 |
| chr16-60969909-DEL-1261 | chr16-60969910-DEL-1261 |
| chr16-61003025-DEL-3319 | chr16-61003026-DEL-3319 |
| chr16-61013152-DEL-567 | chr16-61013153-DEL-567 |
| chr16-61071168-DEL-8727 | chr16-61071169-DEL-8727 |
| chr16-61082311-DEL-70 | chr16-61082312-DEL-68 |
| chr16-61104096-DEL-62 | chr16-61104095-DEL-64 |
| chr16-61132942-DEL-4165 | chr16-61132943-DEL-4165 |
| chr16-61144459-DEL-3936 | chr16-61144460-DEL-3936 |
| chr16-61150512-DEL-2628 | chr16-61150513-DEL-2628 |
| chr16-61154254-DEL-6230 | chr16-61154255-DEL-6230 |
| chr16-61162530-DEL-459 | chr16-61162535-DEL-457 |
| chr16-61164407-DEL-96 | chr16-61164410-DEL-81 |
| chr16-61166636-DEL-1081 | chr16-61166638-DEL-1080 |
| chr16-61184000-DEL-6536 | chr16-61183999-DEL-6537 |
| chr16-61190022-DEL-64 | chr16-61190023-DEL-64 |
| chr16-61197417-DEL-2683 | chr16-61197418-DEL-2683 |
| chr16-61239289-DEL-62 | chr16-61239290-DEL-63 |
| chr16-61252279-DEL-53 | chr16-61252280-DEL-53 |
| chr16-61262065-DEL-152 | chr16-61262034-DEL-168 |
| chr16-61307553-DEL-170 | chr16-61307554-DEL-170 |
| chr16-61334724-DEL-52 | chr16-61334724-DEL-56 |
| chr16-61350840-DEL-1982 | chr16-61350841-DEL-1982 |
| chr16-61360081-DEL-1068 | chr16-61360071-DEL-1069 |
| chr16-61364485-DEL-580 | chr16-61364486-DEL-580 |
| chr16-61369866-DEL-62 | chr16-61369867-DEL-62 |
| chr16-61386159-DEL-123 | chr16-61386160-DEL-123 |
| chr16-61388969-DEL-52 | chr16-61388970-DEL-52 |
| chr16-61398823-DEL-1993 | chr16-61398824-DEL-1993 |
| chr16-61402092-DEL-1318 | chr16-61402093-DEL-1318 |
| chr16-61465867-DEL-285 | chr16-61465868-DEL-285 |
| chr16-61475681-DEL-460 | chr16-61475682-DEL-460 |
| chr16-61493980-DEL-197 | chr16-61493981-DEL-197 |
| chr16-61530220-DEL-6784 | chr16-61530221-DEL-6784 |
| chr16-61536297-DEL-402 | chr16-61536298-DEL-402 |
| chr16-61552755-DEL-142 | chr16-61552756-DEL-142 |
| chr16-61554283-DEL-1350 | chr16-61554280-DEL-1353 |
| chr16-61567980-DEL-1012 | chr16-61567981-DEL-1012 |
| chr16-61594952-DEL-376 | chr16-61594953-DEL-376 |
| chr16-61608235-DEL-1070 | chr16-61608238-DEL-1072 |
| chr16-61630904-DEL-102 | chr16-61630905-DEL-102 |
| chr16-61673596-DEL-6648 | chr16-61673597-DEL-6648 |
| chr16-61690255-DEL-79 | chr16-61690265-DEL-70 |
| chr16-61712363-DEL-119 | chr16-61712374-DEL-103 |
| chr16-61736326-DEL-162 | chr16-61736327-DEL-163 |
| chr16-61779106-DEL-51 | chr16-61779111-DEL-64 |
| chr16-61788010-DEL-187 | chr16-61788011-DEL-187 |
| chr16-61789104-DEL-584 | chr16-61789105-DEL-584 |
| chr16-61791251-DEL-223 | chr16-61791252-DEL-223 |
| chr16-61793201-DEL-170 | chr16-61793202-DEL-170 |
| chr16-61794478-DEL-56 | chr16-61794479-DEL-54 |
| chr16-61794758-DEL-566 | chr16-61794759-DEL-566 |
| chr16-61800747-DEL-1134 | chr16-61800751-DEL-1128 |
| chr16-61893392-DEL-63 | chr16-61893393-DEL-62 |
| chr16-61944554-DEL-160 | chr16-61944558-DEL-157 |
| chr16-61985205-DEL-5285 | chr16-61985201-DEL-5283 |
| chr16-61996898-DEL-371 | chr16-61996899-DEL-371 |
| chr16-62018747-DEL-6392 | chr16-62018747-DEL-6392 |
| chr16-62035497-DEL-695 | chr16-62035495-DEL-696 |
| chr16-62053497-DEL-639 | chr16-62053497-DEL-640 |
| chr16-62080549-DEL-2380 | chr16-62080546-DEL-2378 |
| chr16-62103516-DEL-972 | chr16-62103517-DEL-972 |
| chr16-62112629-DEL-511 | chr16-62112629-DEL-511 |

| | |
|---|---|
| chr16-62119153-DEL-203 | chr16-62119154-DEL-203 |
| chr16-62131641-DEL-6277 | chr16-62131652-DEL-6276 |
| chr16-62132986-DEL-1908 | chr16-62132987-DEL-1908 |
| chr16-62158608-DEL-84 | chr16-62158609-DEL-84 |
| chr16-62165590-DEL-6473 | chr16-62165590-DEL-6473 |
| chr16-62170425-DEL-314 | chr16-62170426-DEL-313 |
| chr16-62171799-DEL-485 | chr16-62171800-DEL-485 |
| chr16-62185283-DEL-2403 | chr16-62185284-DEL-2403 |
| chr16-62187683-DEL-162 | chr16-62187683-DEL-163 |
| chr16-62198854-DEL-2703 | chr16-62198855-DEL-2703 |
| chr16-62200936-DEL-78 | chr16-62200962-DEL-78 |
| chr16-62215123-DEL-203 | chr16-62215124-DEL-203 |
| chr16-62251665-DEL-87 | chr16-62251657-DEL-88 |
| chr16-62292241-DEL-402 | chr16-62292242-DEL-381 |
| chr16-62295520-DEL-1284 | chr16-62295521-DEL-1284 |
| chr16-62295535-DEL-56 | chr16-62295533-DEL-57 |
| chr16-62378379-DEL-6106 | chr16-62378380-DEL-6105 |
| chr16-62384712-DEL-6476 | chr16-62384696-DEL-6490 |
| chr16-62423194-DEL-65 | chr16-62423195-DEL-69 |
| chr16-62426838-DEL-296 | chr16-62426839-DEL-284 |
| chr16-62484705-DEL-536 | chr16-62484706-DEL-535 |
| chr16-62493287-DEL-95 | chr16-62493287-DEL-95 |
| chr16-62494186-DEL-51 | chr16-62494187-DEL-51 |
| chr16-62524402-DEL-7315 | chr16-62524403-DEL-7315 |
| chr16-62537302-DEL-1893 | chr16-62537303-DEL-1893 |
| chr16-62557619-DEL-2075 | chr16-62557620-DEL-2075 |
| chr16-62583626-DEL-62 | chr16-62583627-DEL-62 |
| chr16-62611296-DEL-386 | chr16-62611297-DEL-386 |
| chr16-62641358-DEL-53 | chr16-62641359-DEL-53 |
| chr16-62648792-DEL-148 | chr16-62648793-DEL-148 |
| chr16-62688892-DEL-1589 | chr16-62688893-DEL-1589 |
| chr16-62692165-DEL-119 | chr16-62692167-DEL-109 |
| chr16-62696272-DEL-4329 | chr16-62696273-DEL-4329 |
| chr16-62700694-DEL-605 | chr16-62700694-DEL-605 |
| chr16-62710852-DEL-60 | chr16-62710852-DEL-63 |
| chr16-62717128-DEL-54 | chr16-62717129-DEL-54 |
| chr16-62732162-DEL-431 | chr16-62732163-DEL-431 |
| chr16-62735092-DEL-198 | chr16-62735093-DEL-198 |
| chr16-62753285-DEL-502 | chr16-62753286-DEL-502 |
| chr16-62754765-DEL-198 | chr16-62754771-DEL-183 |
| chr16-62795625-DEL-64 | chr16-62795625-DEL-72 |
| chr16-62840108-DEL-265 | chr16-62840106-DEL-228 |
| chr16-62844729-DEL-1977 | chr16-62844730-DEL-1977 |
| chr16-62848191-DEL-70 | chr16-62848192-DEL-70 |
| chr16-62905813-DEL-718 | chr16-62905813-DEL-718 |
| chr16-62914358-DEL-112 | chr16-62914351-DEL-120 |
| chr16-62921723-DEL-60 | chr16-62921729-DEL-51 |
| chr16-62927165-DEL-129 | chr16-62927192-DEL-95 |
| chr16-62928241-DEL-1229 | chr16-62928233-DEL-1228 |
| chr16-62929767-DEL-281 | chr16-62929768-DEL-281 |
| chr16-62930856-DEL-53 | chr16-62930857-DEL-52 |
| chr16-62934602-DEL-2137 | chr16-62934603-DEL-2137 |
| chr16-62944606-DEL-64 | chr16-62944604-DEL-65 |
| chr16-62948416-DEL-6382 | chr16-62948417-DEL-6382 |
| chr16-62968030-DEL-100 | chr16-62968031-DEL-100 |
| chr16-62982424-DEL-52 | chr16-62982412-DEL-76 |
| chr16-62984028-DEL-125 | chr16-62984029-DEL-125 |
| chr16-63005812-DEL-59 | chr16-63005812-DEL-60 |
| chr16-63016675-DEL-6442 | chr16-63016676-DEL-6442 |
| chr16-63054665-DEL-650 | chr16-63054666-DEL-650 |
| chr16-63072621-DEL-538 | chr16-63072625-DEL-537 |
| chr16-63075292-DEL-179 | chr16-63075293-DEL-179 |
| chr16-63078046-DEL-5636 | chr16-63078047-DEL-5636 |
| chr16-63094539-DEL-295 | chr16-63094540-DEL-295 |
| chr16-63186536-DEL-1011 | chr16-63186525-DEL-1012 |
| chr16-63202067-DEL-1782 | chr16-63202068-DEL-1782 |
| chr16-63207685-DEL-57 | chr16-63207679-DEL-54 |
| chr16-63259529-DEL-6488 | chr16-63259530-DEL-6488 |
| chr16-63269615-DEL-72 | chr16-63269621-DEL-58 |
| chr16-63273225-DEL-3781 | chr16-63273226-DEL-3781 |
| chr16-63279098-DEL-388 | chr16-63279099-DEL-388 |
| chr16-63281516-DEL-116 | chr16-63281544-DEL-87 |
| chr16-63336651-DEL-54 | chr16-63336652-DEL-54 |
| chr16-63369622-DEL-62 | chr16-63369637-DEL-67 |
| chr16-63375593-DEL-82 | chr16-63375631-DEL-62 |
| chr16-63386078-DEL-92 | chr16-63386079-DEL-92 |
| chr16-63413017-DEL-118 | chr16-63413010-DEL-130 |
| chr16-63414709-DEL-214 | chr16-63414710-DEL-214 |
| chr16-63451020-DEL-1538 | chr16-63451021-DEL-1538 |
| chr16-63470436-DEL-519 | chr16-63470433-DEL-520 |

| | |
|---|---|
| chr16-63517925-DEL-60 | chr16-63517926-DEL-60 |
| chr16-63542344-DEL-6352 | chr16-63542345-DEL-6352 |
| chr16-63604401-DEL-897 | chr16-63604402-DEL-897 |
| chr16-63644007-DEL-2175 | chr16-63643993-DEL-2161 |
| chr16-63700510-DEL-348 | chr16-63700511-DEL-348 |
| chr16-63705502-DEL-60 | chr16-63705503-DEL-58 |
| chr16-63718894-DEL-132 | chr16-63718895-DEL-132 |
| chr16-63728707-DEL-6218 | chr16-63728708-DEL-6218 |
| chr16-63783086-DEL-57 | chr16-63783087-DEL-57 |
| chr16-63795419-DEL-230 | chr16-63795420-DEL-230 |
| chr16-63797202-DEL-208 | chr16-63797203-DEL-207 |
| chr16-63814488-DEL-70 | chr16-63814489-DEL-70 |
| chr16-63822966-DEL-6585 | chr16-63822967-DEL-6585 |
| chr16-63823158-DEL-194 | chr16-63823159-DEL-194 |
| chr16-63834721-DEL-96 | chr16-63834714-DEL-98 |
| chr16-63841959-DEL-155 | chr16-63841960-DEL-155 |
| chr16-63862452-DEL-63 | chr16-63862453-DEL-63 |
| chr16-63921608-DEL-108 | chr16-63921615-DEL-102 |
| chr16-63961170-DEL-71 | chr16-63961168-DEL-52 |
| chr16-63982557-DEL-518 | chr16-63982593-DEL-475 |
| chr16-64080967-DEL-94 | chr16-64080976-DEL-74 |
| chr16-64082716-DEL-86 | chr16-64082715-DEL-86 |
| chr16-64090366-DEL-90 | chr16-64090367-DEL-90 |
| chr16-64104991-DEL-90 | chr16-64105008-DEL-74 |
| chr16-64117116-DEL-231 | chr16-64117141-DEL-198 |
| chr16-64122866-DEL-201 | chr16-64122867-DEL-201 |
| chr16-64157809-DEL-185 | chr16-64157810-DEL-185 |
| chr16-64162082-DEL-60 | chr16-64162083-DEL-60 |
| chr16-64181204-DEL-412 | chr16-64181205-DEL-412 |
| chr16-64200280-DEL-73 | chr16-64200281-DEL-73 |
| chr16-64204936-DEL-1953 | chr16-64204937-DEL-1953 |
| chr16-64217156-DEL-6803 | chr16-64217157-DEL-6803 |
| chr16-64240848-DEL-324 | chr16-64240849-DEL-324 |
| chr16-64268287-DEL-133 | chr16-64268269-DEL-126 |
| chr16-64277040-DEL-297 | chr16-64277041-DEL-297 |
| chr16-64292919-DEL-70 | chr16-64292904-DEL-117 |
| chr16-64377496-DEL-92 | chr16-64377497-DEL-92 |
| chr16-64391062-DEL-630 | chr16-64391063-DEL-630 |
| chr16-64392617-DEL-6631 | chr16-64392618-DEL-6631 |
| chr16-64400600-DEL-296 | chr16-64400601-DEL-296 |
| chr16-64408370-DEL-439 | chr16-64408371-DEL-439 |
| chr16-64417927-DEL-220 | chr16-64417928-DEL-220 |
| chr16-64463656-DEL-1527 | chr16-64463657-DEL-1527 |
| chr16-64466519-DEL-53 | chr16-64466520-DEL-53 |
| chr16-64473014-DEL-1890 | chr16-64473017-DEL-1889 |
| chr16-64516566-DEL-829 | chr16-64516565-DEL-830 |
| chr16-64558462-DEL-400 | chr16-64558463-DEL-400 |
| chr16-64613382-DEL-163 | chr16-64613383-DEL-163 |
| chr16-64626600-DEL-6800 | chr16-64626601-DEL-6800 |
| chr16-64634229-DEL-205 | chr16-64634230-DEL-205 |
| chr16-64636210-DEL-123 | chr16-64636211-DEL-123 |
| chr16-64638150-DEL-208 | chr16-64638151-DEL-208 |
| chr16-64645751-DEL-134 | chr16-64645752-DEL-134 |
| chr16-64682456-DEL-77 | chr16-64682457-DEL-77 |
| chr16-64683557-DEL-2526 | chr16-64683558-DEL-2526 |
| chr16-64709699-DEL-1329 | chr16-64709700-DEL-1329 |
| chr16-64710869-DEL-4366 | chr16-64710870-DEL-4366 |
| chr16-64712842-DEL-211 | chr16-64712843-DEL-211 |
| chr16-64726888-DEL-219 | chr16-64726889-DEL-219 |
| chr16-64785442-DEL-64 | chr16-64785441-DEL-64 |
| chr16-64786288-DEL-4579 | chr16-64786289-DEL-4579 |
| chr16-64812766-DEL-421 | chr16-64812767-DEL-421 |
| chr16-64814299-DEL-6265 | chr16-64814299-DEL-6265 |
| chr16-64861185-DEL-6887 | chr16-64861186-DEL-6887 |
| chr16-64907111-DEL-462 | chr16-64907112-DEL-462 |
| chr16-64908450-DEL-890 | chr16-64908440-DEL-891 |
| chr16-64910964-DEL-161 | chr16-64910965-DEL-162 |
| chr16-64929375-DEL-91 | chr16-64929376-DEL-91 |
| chr16-64967812-DEL-824 | chr16-64967813-DEL-824 |
| chr16-64974600-DEL-566 | chr16-64974601-DEL-566 |
| chr16-64997358-DEL-4313 | chr16-64997359-DEL-4313 |
| chr16-65009046-DEL-54 | chr16-65009047-DEL-54 |
| chr16-65011843-DEL-420 | chr16-65011844-DEL-420 |
| chr16-65045764-DEL-1370 | chr16-65045765-DEL-1370 |
| chr16-65093140-DEL-65 | chr16-65093140-DEL-65 |
| chr16-65137290-DEL-481 | chr16-65137291-DEL-481 |
| chr16-65137915-DEL-188 | chr16-65137916-DEL-188 |
| chr16-65209660-DEL-121 | chr16-65209662-DEL-122 |
| chr16-65222043-DEL-1392 | chr16-65222044-DEL-1392 |
| chr16-65247052-DEL-609 | chr16-65247053-DEL-609 |

| | |
|---|---|
| chr16-65252174-DEL-82 | chr16-65252175-DEL-82 |
| chr16-65253912-DEL-1055 | chr16-65253913-DEL-1055 |
| chr16-65265629-DEL-133 | chr16-65265629-DEL-133 |
| chr16-65270979-DEL-638 | chr16-65270977-DEL-596 |
| chr16-65288955-DEL-1782 | chr16-65288952-DEL-1783 |
| chr16-65299226-DEL-1208 | chr16-65299227-DEL-1208 |
| chr16-65305095-DEL-178 | chr16-65305096-DEL-178 |
| chr16-65321181-DEL-144 | chr16-65321182-DEL-144 |
| chr16-65331115-DEL-344 | chr16-65331112-DEL-345 |
| chr16-65350818-DEL-301 | chr16-65350819-DEL-301 |
| chr16-65399410-DEL-1974 | chr16-65399411-DEL-1974 |
| chr16-65403726-DEL-416 | chr16-65403727-DEL-416 |
| chr16-65432258-DEL-82 | chr16-65432253-DEL-85 |
| chr16-65505196-DEL-751 | chr16-65505198-DEL-749 |
| chr16-65506042-DEL-273 | chr16-65506052-DEL-286 |
| chr16-65514681-DEL-84 | chr16-65514682-DEL-83 |
| chr16-65520371-DEL-72 | chr16-65520414-DEL-50 |
| chr16-65534247-DEL-3167 | chr16-65534248-DEL-3167 |
| chr16-65541908-DEL-6573 | chr16-65541909-DEL-6573 |
| chr16-65553517-DEL-1920 | chr16-65553518-DEL-1920 |
| chr16-65573293-DEL-165 | chr16-65573294-DEL-165 |
| chr16-65574139-DEL-106 | chr16-65574153-DEL-96 |
| chr16-65575162-DEL-72 | chr16-65575163-DEL-72 |
| chr16-65579947-DEL-886 | chr16-65579948-DEL-886 |
| chr16-65583769-DEL-4890 | chr16-65583770-DEL-4890 |
| chr16-65646126-DEL-78 | chr16-65646077-DEL-78 |
| chr16-65674525-DEL-1609 | chr16-65674526-DEL-1609 |
| chr16-65710544-DEL-1213 | chr16-65710545-DEL-1213 |
| chr16-65714001-DEL-339 | chr16-65714002-DEL-339 |
| chr16-65725938-DEL-6384 | chr16-65725955-DEL-6371 |
| chr16-65730863-DEL-172 | chr16-65730864-DEL-172 |
| chr16-65733253-DEL-2065 | chr16-65733253-DEL-2062 |
| chr16-65753649-DEL-105 | chr16-65753650-DEL-105 |
| chr16-65790600-DEL-80 | chr16-65790601-DEL-56 |
| chr16-65805482-DEL-680 | chr16-65805483-DEL-680 |
| chr16-65840835-DEL-919 | chr16-65840836-DEL-919 |
| chr16-65848002-DEL-203 | chr16-65848005-DEL-203 |
| chr16-65869143-DEL-51 | chr16-65869144-DEL-51 |
| chr16-65877159-DEL-93 | chr16-65877160-DEL-93 |
| chr16-65877360-DEL-1569 | chr16-65877350-DEL-1571 |
| chr16-65885535-DEL-84 | chr16-65885517-DEL-59 |
| chr16-65896789-DEL-68 | chr16-65896787-DEL-76 |
| chr16-65900208-DEL-132 | chr16-65900198-DEL-135 |
| chr16-65911305-DEL-213 | chr16-65911306-DEL-213 |
| chr16-65936508-DEL-2977 | chr16-65936509-DEL-2977 |
| chr16-65950298-DEL-204 | chr16-65950299-DEL-204 |
| chr16-65958821-DEL-60 | chr16-65958822-DEL-60 |
| chr16-65984459-DEL-697 | chr16-65984460-DEL-696 |
| chr16-65989038-DEL-1002 | chr16-65989039-DEL-1002 |
| chr16-66067266-DEL-1613 | chr16-66067267-DEL-1613 |
| chr16-66076973-DEL-105 | chr16-66076974-DEL-105 |
| chr16-66082391-DEL-651 | chr16-66082392-DEL-651 |
| chr16-66108682-DEL-6470 | chr16-66108667-DEL-6447 |
| chr16-66150948-DEL-56 | chr16-66150947-DEL-53 |
| chr16-66177707-DEL-490 | chr16-66177708-DEL-489 |
| chr16-66182564-DEL-73 | chr16-66182565-DEL-73 |
| chr16-66186417-DEL-87 | chr16-66186370-DEL-86 |
| chr16-66189457-DEL-167 | chr16-66189464-DEL-173 |
| chr16-66213417-DEL-58 | chr16-66213418-DEL-58 |
| chr16-66229360-DEL-64 | chr16-66229361-DEL-64 |
| chr16-66268003-DEL-375 | chr16-66268004-DEL-375 |
| chr16-66295180-DEL-2128 | chr16-66295181-DEL-2128 |
| chr16-66303640-DEL-365 | chr16-66303641-DEL-365 |
| chr16-66314944-DEL-6688 | chr16-66314945-DEL-6688 |
| chr16-66332544-DEL-105 | chr16-66332545-DEL-105 |
| chr16-66341886-DEL-70 | chr16-66341905-DEL-52 |
| chr16-66343793-DEL-52 | chr16-66343800-DEL-52 |
| chr16-66356384-DEL-390 | chr16-66356385-DEL-390 |
| chr16-66384649-DEL-854 | chr16-66384650-DEL-854 |
| chr16-66390392-DEL-145 | chr16-66390393-DEL-145 |
| chr16-66407805-DEL-53 | chr16-66407790-DEL-89 |
| chr16-66410236-DEL-299 | chr16-66410225-DEL-302 |
| chr16-66410867-DEL-699 | chr16-66410868-DEL-699 |
| chr16-66412838-DEL-256 | chr16-66412839-DEL-256 |
| chr16-66464954-DEL-527 | chr16-66464953-DEL-528 |
| chr16-66469878-DEL-95 | chr16-66469876-DEL-88 |
| chr16-66497092-DEL-2688 | chr16-66497093-DEL-2688 |
| chr16-66532512-DEL-1090 | chr16-66532513-DEL-1090 |
| chr16-66540946-DEL-70 | chr16-66540947-DEL-70 |
| chr16-66547100-DEL-124 | chr16-66547101-DEL-124 |

| | |
|---|---|
| chr16-66551153-DEL-6648 | chr16-66551154-DEL-6648 |
| chr16-66592517-DEL-696 | chr16-66592518-DEL-696 |
| chr16-66642927-DEL-64 | chr16-66642928-DEL-64 |
| chr16-66672037-DEL-92 | chr16-66672029-DEL-60 |
| chr16-66686126-DEL-60 | chr16-66686143-DEL-54 |
| chr16-66725009-DEL-688 | chr16-66725010-DEL-688 |
| chr16-66731963-DEL-82 | chr16-66731963-DEL-83 |
| chr16-66737649-DEL-2565 | chr16-66737650-DEL-2565 |
| chr16-66813172-DEL-60 | chr16-66813177-DEL-56 |
| chr16-66814747-DEL-598 | chr16-66814748-DEL-598 |
| chr16-66846981-DEL-78 | chr16-66846974-DEL-80 |
| chr16-66861489-DEL-588 | chr16-66861490-DEL-588 |
| chr16-66862478-DEL-418 | chr16-66862479-DEL-418 |
| chr16-66886491-DEL-64 | chr16-66886490-DEL-62 |
| chr16-66908989-DEL-70 | chr16-66908986-DEL-66 |
| chr16-66911444-DEL-795 | chr16-66911445-DEL-795 |
| chr16-66923444-DEL-1001 | chr16-66923445-DEL-1001 |
| chr16-66960770-DEL-456 | chr16-66960771-DEL-456 |
| chr16-66969666-DEL-94 | chr16-66969667-DEL-99 |
| chr16-67040224-DEL-230 | chr16-67040225-DEL-230 |
| chr16-67072344-DEL-339 | chr16-67072345-DEL-339 |
| chr16-67125101-DEL-6413 | chr16-67125102-DEL-6413 |
| chr16-67171419-DEL-54 | chr16-67171420-DEL-54 |
| chr16-67174723-DEL-601 | chr16-67174724-DEL-601 |
| chr16-67182015-DEL-6289 | chr16-67182016-DEL-6289 |
| chr16-67197084-DEL-68 | chr16-67197085-DEL-68 |
| chr16-67198521-DEL-75 | chr16-67198522-DEL-75 |
| chr16-67248184-DEL-87 | chr16-67248185-DEL-87 |
| chr16-67262869-DEL-88 | chr16-67262870-DEL-88 |
| chr16-67275522-DEL-162 | chr16-67275523-DEL-161 |
| chr16-67276717-DEL-771 | chr16-67276718-DEL-771 |
| chr16-67359560-DEL-6194 | chr16-67359561-DEL-6194 |
| chr16-67391206-DEL-70 | chr16-67391207-DEL-70 |
| chr16-67392898-DEL-71 | chr16-67392901-DEL-69 |
| chr16-67426802-DEL-7075 | chr16-67426802-DEL-7077 |
| chr16-67434167-DEL-2185 | chr16-67434168-DEL-2185 |
| chr16-67447265-DEL-61 | chr16-67447272-DEL-55 |
| chr16-67524711-DEL-89 | chr16-67524704-DEL-86 |
| chr16-67542950-DEL-410 | chr16-67542952-DEL-406 |
| chr16-67555411-DEL-312 | chr16-67555412-DEL-312 |
| chr16-67560197-DEL-1802 | chr16-67560200-DEL-1797 |
| chr16-67591552-DEL-195 | chr16-67591553-DEL-195 |
| chr16-67595738-DEL-76 | chr16-67595743-DEL-70 |
| chr16-67607373-DEL-1407 | chr16-67607373-DEL-1407 |
| chr16-67618896-DEL-194 | chr16-67618897-DEL-194 |
| chr16-67619840-DEL-395 | chr16-67619841-DEL-395 |
| chr16-67654009-DEL-1068 | chr16-67654010-DEL-1068 |
| chr16-67694406-DEL-146 | chr16-67694427-DEL-118 |
| chr16-67729701-DEL-11439 | chr16-67729702-DEL-11439 |
| chr16-67755388-DEL-223 | chr16-67755387-DEL-223 |
| chr16-67769085-DEL-60 | chr16-67769086-DEL-52 |
| chr16-67789223-DEL-60 | chr16-67789224-DEL-60 |
| chr16-67799075-DEL-768 | chr16-67799076-DEL-768 |
| chr16-67801495-DEL-116 | chr16-67801496-DEL-72 |
| chr16-67812059-DEL-129 | chr16-67812086-DEL-93 |
| chr16-67870409-DEL-555 | chr16-67870409-DEL-555 |
| chr16-67891211-DEL-6423 | chr16-67891212-DEL-6423 |
| chr16-67928624-DEL-1322 | chr16-67928625-DEL-1322 |
| chr16-67949041-DEL-880 | chr16-67949045-DEL-879 |
| chr16-68028825-DEL-64 | chr16-68028826-DEL-64 |
| chr16-68047313-DEL-2318 | chr16-68047313-DEL-2318 |
| chr16-68050956-DEL-97 | chr16-68050956-DEL-97 |
| chr16-68079360-DEL-97 | chr16-68079361-DEL-97 |
| chr16-68084285-DEL-978 | chr16-68084286-DEL-978 |
| chr16-68090627-DEL-234 | chr16-68090633-DEL-227 |
| chr16-68097666-DEL-62 | chr16-68097618-DEL-63 |
| chr16-68115516-DEL-84 | chr16-68115519-DEL-82 |
| chr16-68247808-DEL-75 | chr16-68247809-DEL-75 |
| chr16-68272737-DEL-61 | chr16-68272738-DEL-61 |
| chr16-68282804-DEL-102 | chr16-68282805-DEL-100 |
| chr16-68356876-DEL-54 | chr16-68356879-DEL-52 |
| chr16-68382487-DEL-1804 | chr16-68382488-DEL-1799 |
| chr16-68398832-DEL-96 | chr16-68398833-DEL-96 |
| chr16-68419255-DEL-4555 | chr16-68419256-DEL-4555 |
| chr16-68453214-DEL-94 | chr16-68453207-DEL-96 |
| chr16-68464411-DEL-613 | chr16-68464410-DEL-615 |
| chr16-68524795-DEL-89 | chr16-68524796-DEL-89 |
| chr16-68534870-DEL-7662 | chr16-68534871-DEL-7662 |
| chr16-68537613-DEL-1303 | chr16-68537614-DEL-1303 |
| chr16-68588124-DEL-225 | chr16-68588125-DEL-225 |

| | |
|---|---|
| chr16-68591996-DEL-59 | chr16-68592012-DEL-54 |
| chr16-68626463-DEL-1857 | chr16-68626464-DEL-1857 |
| chr16-68643156-DEL-564 | chr16-68643157-DEL-563 |
| chr16-68663114-DEL-670 | chr16-68663115-DEL-670 |
| chr16-68690202-DEL-164 | chr16-68690203-DEL-164 |
| chr16-68695426-DEL-312 | chr16-68695427-DEL-312 |
| chr16-68753021-DEL-60 | chr16-68753024-DEL-56 |
| chr16-68776179-DEL-78 | chr16-68776173-DEL-74 |
| chr16-68779829-DEL-781 | chr16-68779830-DEL-781 |
| chr16-68785125-DEL-459 | chr16-68785126-DEL-459 |
| chr16-68791544-DEL-395 | chr16-68791545-DEL-395 |
| chr16-68797943-DEL-2660 | chr16-68797944-DEL-2660 |
| chr16-68838084-DEL-3480 | chr16-68838085-DEL-3480 |
| chr16-68851798-DEL-326 | chr16-68851799-DEL-326 |
| chr16-68854645-DEL-226 | chr16-68854646-DEL-226 |
| chr16-68891681-DEL-4609 | chr16-68891682-DEL-4609 |
| chr16-68900702-DEL-78 | chr16-68900703-DEL-78 |
| chr16-68919366-DEL-892 | chr16-68919367-DEL-892 |
| chr16-68925967-DEL-2315 | chr16-68925962-DEL-2316 |
| chr16-68939088-DEL-390 | chr16-68939089-DEL-390 |
| chr16-68941234-DEL-116 | chr16-68941235-DEL-116 |
| chr16-68945387-DEL-622 | chr16-68945388-DEL-622 |
| chr16-68959552-DEL-2183 | chr16-68959553-DEL-2183 |
| chr16-68975881-DEL-867 | chr16-68975901-DEL-859 |
| chr16-68980706-DEL-620 | chr16-68980707-DEL-620 |
| chr16-69001093-DEL-68 | chr16-69001103-DEL-67 |
| chr16-69005551-DEL-6894 | chr16-69005552-DEL-6894 |
| chr16-69005989-DEL-424 | chr16-69005990-DEL-424 |
| chr16-69012555-DEL-5642 | chr16-69012556-DEL-5642 |
| chr16-69025884-DEL-73 | chr16-69025889-DEL-69 |
| chr16-69088662-DEL-136 | chr16-69088665-DEL-132 |
| chr16-69100095-DEL-61 | chr16-69100096-DEL-61 |
| chr16-69102605-DEL-54 | chr16-69102598-DEL-55 |
| chr16-69112287-DEL-83 | chr16-69112288-DEL-83 |
| chr16-69121040-DEL-4623 | chr16-69121041-DEL-4623 |
| chr16-69150236-DEL-1216 | chr16-69150237-DEL-1216 |
| chr16-69157738-DEL-151 | chr16-69157738-DEL-157 |
| chr16-69172700-DEL-109 | chr16-69172702-DEL-107 |
| chr16-69205308-DEL-6462 | chr16-69205309-DEL-6462 |
| chr16-69222994-DEL-90 | chr16-69222994-DEL-90 |
| chr16-69237822-DEL-324 | chr16-69237823-DEL-324 |
| chr16-69302434-DEL-267 | chr16-69302435-DEL-267 |
| chr16-69334587-DEL-2162 | chr16-69334588-DEL-2162 |
| chr16-69398811-DEL-59 | chr16-69398812-DEL-63 |
| chr16-69402419-DEL-1548 | chr16-69402434-DEL-1518 |
| chr16-69405549-DEL-161 | chr16-69405550-DEL-161 |
| chr16-69491754-DEL-269 | chr16-69491755-DEL-269 |
| chr16-69530861-DEL-2257 | chr16-69530862-DEL-2257 |
| chr16-69540502-DEL-833 | chr16-69540503-DEL-833 |
| chr16-69562023-DEL-241 | chr16-69562028-DEL-237 |
| chr16-69568990-DEL-831 | chr16-69568987-DEL-833 |
| chr16-69579311-DEL-118 | chr16-69579312-DEL-118 |
| chr16-69585587-DEL-99 | chr16-69585588-DEL-99 |
| chr16-69587954-DEL-450 | chr16-69587955-DEL-450 |
| chr16-69590599-DEL-118 | chr16-69590602-DEL-116 |
| chr16-69591207-DEL-199 | chr16-69591208-DEL-199 |
| chr16-69593757-DEL-106 | chr16-69593758-DEL-106 |
| chr16-69661800-DEL-4189 | chr16-69661801-DEL-4189 |
| chr16-69689796-DEL-7091 | chr16-69689797-DEL-7091 |
| chr16-69690062-DEL-212 | chr16-69690063-DEL-212 |
| chr16-69697766-DEL-365 | chr16-69697734-DEL-375 |
| chr16-69706576-DEL-374 | chr16-69706577-DEL-374 |
| chr16-69710278-DEL-55 | chr16-69710279-DEL-50 |
| chr16-69723956-DEL-195 | chr16-69723957-DEL-195 |
| chr16-69759051-DEL-3923 | chr16-69759052-DEL-3923 |
| chr16-69783946-DEL-109 | chr16-69783947-DEL-109 |
| chr16-69791498-DEL-84 | chr16-69791499-DEL-84 |
| chr16-69865327-DEL-86 | chr16-69865344-DEL-54 |
| chr16-69902605-DEL-60 | chr16-69902602-DEL-64 |
| chr16-69918822-DEL-639 | chr16-69918823-DEL-639 |
| chr16-69920185-DEL-75 | chr16-69920186-DEL-75 |
| chr16-69941720-DEL-231 | chr16-69941719-DEL-231 |
| chr16-69944214-DEL-80 | chr16-69944215-DEL-80 |
| chr16-69963288-DEL-83 | chr16-69963295-DEL-102 |
| chr16-69966577-DEL-3559 | chr16-69966578-DEL-3559 |
| chr16-69979572-DEL-125 | chr16-69979570-DEL-126 |
| chr16-69984087-DEL-68 | chr16-69984080-DEL-76 |
| chr16-69999228-DEL-75 | chr16-69999229-DEL-90 |
| chr16-70035230-DEL-67 | chr16-70035231-DEL-67 |
| chr16-70052490-DEL-60 | chr16-70052489-DEL-62 |

| | |
|---|---|
| chr16-70065972-DEL-9407 | chr16-70065973-DEL-9407 |
| chr16-70111900-DEL-78 | chr16-70111901-DEL-78 |
| chr16-70128933-DEL-72 | chr16-70128928-DEL-74 |
| chr16-70141996-DEL-124 | chr16-70141999-DEL-122 |
| chr16-70147676-DEL-78 | chr16-70147675-DEL-66 |
| chr16-70148208-DEL-52 | chr16-70148209-DEL-52 |
| chr16-70161665-DEL-74 | chr16-70161666-DEL-74 |
| chr16-70163548-DEL-6916 | chr16-70163530-DEL-6935 |
| chr16-70170899-DEL-3183 | chr16-70170899-DEL-3183 |
| chr16-70178752-DEL-2771 | chr16-70178753-DEL-2771 |
| chr16-70206043-DEL-52 | chr16-70206044-DEL-52 |
| chr16-70215017-DEL-150 | chr16-70215017-DEL-120 |
| chr16-70228432-DEL-125 | chr16-70228427-DEL-110 |
| chr16-70230960-DEL-56 | chr16-70230956-DEL-52 |
| chr16-70235724-DEL-68 | chr16-70235759-DEL-58 |
| chr16-70255051-DEL-399 | chr16-70255052-DEL-399 |
| chr16-70264929-DEL-141 | chr16-70264930-DEL-141 |
| chr16-70294457-DEL-188 | chr16-70294459-DEL-185 |
| chr16-70300565-DEL-174 | chr16-70300566-DEL-171 |
| chr16-70308973-DEL-214 | chr16-70308974-DEL-214 |
| chr16-70311463-DEL-467 | chr16-70311464-DEL-467 |
| chr16-70336307-DEL-832 | chr16-70336308-DEL-832 |
| chr16-70339325-DEL-5355 | chr16-70339326-DEL-5355 |
| chr16-70361012-DEL-63 | chr16-70361012-DEL-64 |
| chr16-70367284-DEL-843 | chr16-70367285-DEL-843 |
| chr16-70385187-DEL-88 | chr16-70385188-DEL-88 |
| chr16-70430870-DEL-5677 | chr16-70430864-DEL-5688 |
| chr16-70464277-DEL-6385 | chr16-70464278-DEL-6385 |
| chr16-70490394-DEL-87 | chr16-70490395-DEL-86 |
| chr16-70506492-DEL-6216 | chr16-70506493-DEL-6216 |
| chr16-70558562-DEL-56 | chr16-70558563-DEL-56 |
| chr16-70574276-DEL-642 | chr16-70574308-DEL-650 |
| chr16-70578861-DEL-3123 | chr16-70578862-DEL-3123 |
| chr16-70596316-DEL-556 | chr16-70596317-DEL-556 |
| chr16-70600410-DEL-180 | chr16-70600411-DEL-180 |
| chr16-70606460-DEL-888 | chr16-70606461-DEL-888 |
| chr16-70614433-DEL-6813 | chr16-70614434-DEL-6812 |
| chr16-70623223-DEL-430 | chr16-70623229-DEL-420 |
| chr16-70668412-DEL-68 | chr16-70668413-DEL-68 |
| chr16-70669380-DEL-384 | chr16-70669366-DEL-385 |
| chr16-70673025-DEL-105 | chr16-70673026-DEL-63 |
| chr16-70684993-DEL-77 | chr16-70684992-DEL-79 |
| chr16-70689246-DEL-6243 | chr16-70689233-DEL-6244 |
| chr16-70737721-DEL-7371 | chr16-70737722-DEL-7371 |
| chr16-70756981-DEL-842 | chr16-70756991-DEL-837 |
| chr16-70766647-DEL-133 | chr16-70766650-DEL-131 |
| chr16-70771029-DEL-74 | chr16-70771030-DEL-70 |
| chr16-70773782-DEL-6477 | chr16-70773783-DEL-6477 |
| chr16-70820830-DEL-63 | chr16-70820832-DEL-62 |
| chr16-70902302-DEL-748 | chr16-70902303-DEL-748 |
| chr16-70928958-DEL-62 | chr16-70928939-DEL-58 |
| chr16-70933567-DEL-1892 | chr16-70933568-DEL-1892 |
| chr16-70944536-DEL-6624 | chr16-70944532-DEL-6625 |
| chr16-70958718-DEL-721 | chr16-70958719-DEL-721 |
| chr16-70960135-DEL-74 | chr16-70960150-DEL-60 |
| chr16-70967553-DEL-4262 | chr16-70967554-DEL-4262 |
| chr16-70970136-DEL-8745 | chr16-70970134-DEL-8745 |
| chr16-70976353-DEL-5962 | chr16-70976354-DEL-5962 |
| chr16-71000421-DEL-53 | chr16-71000423-DEL-52 |
| chr16-71014599-DEL-108 | chr16-71014600-DEL-108 |
| chr16-71019282-DEL-276 | chr16-71019281-DEL-277 |
| chr16-71043959-DEL-564 | chr16-71043960-DEL-564 |
| chr16-71073310-DEL-6750 | chr16-71073310-DEL-6750 |
| chr16-71084702-DEL-703 | chr16-71084703-DEL-703 |
| chr16-71109818-DEL-63 | chr16-71109819-DEL-63 |
| chr16-71111845-DEL-335 | chr16-71111846-DEL-335 |
| chr16-71117159-DEL-377 | chr16-71117188-DEL-356 |
| chr16-71168784-DEL-2430 | chr16-71168785-DEL-2430 |
| chr16-71235380-DEL-338 | chr16-71235379-DEL-335 |
| chr16-71242833-DEL-124 | chr16-71242834-DEL-124 |
| chr16-71249138-DEL-65 | chr16-71249151-DEL-54 |
| chr16-71266933-DEL-6413 | chr16-71266934-DEL-6414 |
| chr16-71275971-DEL-57 | chr16-71275944-DEL-57 |
| chr16-71299273-DEL-2076 | chr16-71299260-DEL-2072 |
| chr16-71309311-DEL-1946 | chr16-71309312-DEL-1946 |
| chr16-71341766-DEL-84 | chr16-71341767-DEL-84 |
| chr16-71346890-DEL-6385 | chr16-71346891-DEL-6385 |
| chr16-71366531-DEL-306 | chr16-71366528-DEL-302 |
| chr16-71380995-DEL-1668 | chr16-71380996-DEL-1665 |
| chr16-71394313-DEL-54 | chr16-71394316-DEL-57 |

| | |
|---|---|
| chr16-71407201-DEL-89 | chr16-71407202-DEL-89 |
| chr16-71430092-DEL-62 | chr16-71430093-DEL-62 |
| chr16-71448372-DEL-333 | chr16-71448370-DEL-332 |
| chr16-71468255-DEL-67 | chr16-71468255-DEL-67 |
| chr16-71483903-DEL-4357 | chr16-71483904-DEL-4357 |
| chr16-71489461-DEL-82 | chr16-71489462-DEL-82 |
| chr16-71551241-DEL-6594 | chr16-71551242-DEL-6594 |
| chr16-71575216-DEL-65 | chr16-71575217-DEL-65 |
| chr16-71578570-DEL-214 | chr16-71578571-DEL-214 |
| chr16-71579294-DEL-965 | chr16-71579295-DEL-965 |
| chr16-71582444-DEL-7110 | chr16-71582445-DEL-7110 |
| chr16-71612547-DEL-85 | chr16-71612548-DEL-85 |
| chr16-71651556-DEL-59 | chr16-71651556-DEL-59 |
| chr16-71668895-DEL-54 | chr16-71668894-DEL-56 |
| chr16-71674431-DEL-527 | chr16-71674432-DEL-527 |
| chr16-71684345-DEL-418 | chr16-71684343-DEL-419 |
| chr16-71726859-DEL-113 | chr16-71726859-DEL-113 |
| chr16-71753226-DEL-92 | chr16-71753235-DEL-68 |
| chr16-71754129-DEL-109 | chr16-71754128-DEL-111 |
| chr16-71778845-DEL-429 | chr16-71778846-DEL-429 |
| chr16-71793173-DEL-1036 | chr16-71793174-DEL-1042 |
| chr16-71816310-DEL-4390 | chr16-71816311-DEL-4390 |
| chr16-71837811-DEL-833 | chr16-71837812-DEL-833 |
| chr16-71849474-DEL-61 | chr16-71849475-DEL-62 |
| chr16-71849998-DEL-80 | chr16-71850001-DEL-70 |
| chr16-71853025-DEL-6161 | chr16-71853026-DEL-6161 |
| chr16-71875192-DEL-156 | chr16-71875193-DEL-156 |
| chr16-71876371-DEL-347 | chr16-71876372-DEL-347 |
| chr16-71901593-DEL-177 | chr16-71901594-DEL-176 |
| chr16-71921983-DEL-334 | chr16-71921984-DEL-335 |
| chr16-71925906-DEL-1074 | chr16-71925907-DEL-1074 |
| chr16-71928392-DEL-490 | chr16-71928393-DEL-490 |
| chr16-71963584-DEL-72 | chr16-71963586-DEL-71 |
| chr16-72033252-DEL-99 | chr16-72033253-DEL-99 |
| chr16-72042083-DEL-64 | chr16-72042072-DEL-59 |
| chr16-72046943-DEL-198 | chr16-72046944-DEL-198 |
| chr16-72053660-DEL-138 | chr16-72053661-DEL-138 |
| chr16-72055060-DEL-105 | chr16-72055061-DEL-105 |
| chr16-72078128-DEL-51 | chr16-72078129-DEL-51 |
| chr16-72084452-DEL-81 | chr16-72084453-DEL-81 |
| chr16-72097766-DEL-1823 | chr16-72097767-DEL-1823 |
| chr16-72099702-DEL-1356 | chr16-72099703-DEL-1356 |
| chr16-72109733-DEL-51 | chr16-72109729-DEL-52 |
| chr16-72149723-DEL-62 | chr16-72149724-DEL-62 |
| chr16-72165916-DEL-1220 | chr16-72165917-DEL-1220 |
| chr16-72169095-DEL-174 | chr16-72169096-DEL-174 |
| chr16-72169331-DEL-929 | chr16-72169333-DEL-928 |
| chr16-72192860-DEL-690 | chr16-72192861-DEL-690 |
| chr16-72230394-DEL-52 | chr16-72230395-DEL-52 |
| chr16-72234558-DEL-80 | chr16-72234571-DEL-68 |
| chr16-72248530-DEL-482 | chr16-72248531-DEL-482 |
| chr16-72251673-DEL-125 | chr16-72251671-DEL-126 |
| chr16-72261805-DEL-73 | chr16-72261805-DEL-73 |
| chr16-72299399-DEL-75 | chr16-72299400-DEL-123 |
| chr16-72304677-DEL-120 | chr16-72304678-DEL-120 |
| chr16-72310521-DEL-148 | chr16-72310534-DEL-136 |
| chr16-72313604-DEL-211 | chr16-72313606-DEL-210 |
| chr16-72333690-DEL-830 | chr16-72333691-DEL-830 |
| chr16-72343093-DEL-87 | chr16-72343094-DEL-87 |
| chr16-72393381-DEL-6274 | chr16-72393382-DEL-6274 |
| chr16-72402899-DEL-499 | chr16-72402900-DEL-499 |
| chr16-72413605-DEL-118 | chr16-72413606-DEL-118 |
| chr16-72420535-DEL-6873 | chr16-72420536-DEL-6873 |
| chr16-72430473-DEL-134 | chr16-72430474-DEL-134 |
| chr16-72464584-DEL-203 | chr16-72464585-DEL-203 |
| chr16-72470102-DEL-59 | chr16-72470103-DEL-59 |
| chr16-72508734-DEL-675 | chr16-72508780-DEL-674 |
| chr16-72521446-DEL-6440 | chr16-72521447-DEL-6440 |
| chr16-72559046-DEL-64 | chr16-72559047-DEL-66 |
| chr16-72560288-DEL-133 | chr16-72560289-DEL-133 |
| chr16-72565777-DEL-78 | chr16-72565776-DEL-80 |
| chr16-72610891-DEL-358 | chr16-72610892-DEL-358 |
| chr16-72640104-DEL-58 | chr16-72640109-DEL-52 |
| chr16-72650826-DEL-52 | chr16-72650827-DEL-52 |
| chr16-72651574-DEL-78 | chr16-72651575-DEL-78 |
| chr16-72656205-DEL-412 | chr16-72656206-DEL-412 |
| chr16-72662844-DEL-57 | chr16-72662845-DEL-57 |
| chr16-72713372-DEL-1896 | chr16-72713373-DEL-1896 |
| chr16-72719707-DEL-110 | chr16-72719708-DEL-110 |
| chr16-72783312-DEL-88 | chr16-72783293-DEL-122 |

| | |
|---|---|
| chr16-72789913-DEL-86 | chr16-72789913-DEL-70 |
| chr16-72821544-DEL-58 | chr16-72821545-DEL-58 |
| chr16-72865639-DEL-68 | chr16-72865640-DEL-68 |
| chr16-72912407-DEL-682 | chr16-72912408-DEL-682 |
| chr16-72923309-DEL-664 | chr16-72923291-DEL-682 |
| chr16-72935589-DEL-261 | chr16-72935590-DEL-261 |
| chr16-72936692-DEL-910 | chr16-72936692-DEL-908 |
| chr16-72938054-DEL-697 | chr16-72938055-DEL-697 |
| chr16-72945527-DEL-296 | chr16-72945500-DEL-297 |
| chr16-72967316-DEL-116 | chr16-72967317-DEL-116 |
| chr16-72991964-DEL-56 | chr16-72991965-DEL-64 |
| chr16-73001310-DEL-94 | chr16-73001280-DEL-98 |
| chr16-73001861-DEL-58 | chr16-73001901-DEL-52 |
| chr16-73003349-DEL-507 | chr16-73003333-DEL-508 |
| chr16-73016923-DEL-404 | chr16-73016924-DEL-404 |
| chr16-73051741-DEL-186 | chr16-73051742-DEL-186 |
| chr16-73089153-DEL-1402 | chr16-73089154-DEL-1402 |
| chr16-73096638-DEL-82 | chr16-73096638-DEL-82 |
| chr16-73121477-DEL-163 | chr16-73121502-DEL-139 |
| chr16-73136554-DEL-164 | chr16-73136549-DEL-183 |
| chr16-73167313-DEL-467 | chr16-73167314-DEL-467 |
| chr16-73178391-DEL-197 | chr16-73178392-DEL-197 |
| chr16-73195737-DEL-445 | chr16-73195738-DEL-445 |
| chr16-73198009-DEL-61 | chr16-73198010-DEL-61 |
| chr16-73213327-DEL-302 | chr16-73213328-DEL-302 |
| chr16-73285914-DEL-68 | chr16-73285927-DEL-74 |
| chr16-73318062-DEL-58 | chr16-73318063-DEL-58 |
| chr16-73319808-DEL-108 | chr16-73319797-DEL-108 |
| chr16-73344045-DEL-6972 | chr16-73344046-DEL-6972 |
| chr16-73390657-DEL-1104 | chr16-73390658-DEL-1104 |
| chr16-73453609-DEL-6536 | chr16-73453610-DEL-6536 |
| chr16-73453687-DEL-213 | chr16-73453679-DEL-214 |
| chr16-73461587-DEL-1463 | chr16-73461587-DEL-1425 |
| chr16-73478001-DEL-7676 | chr16-73478002-DEL-7676 |
| chr16-73478234-DEL-424 | chr16-73478235-DEL-424 |
| chr16-73527073-DEL-6891 | chr16-73527074-DEL-6891 |
| chr16-73570968-DEL-87 | chr16-73570969-DEL-87 |
| chr16-73594641-DEL-90 | chr16-73594642-DEL-96 |
| chr16-73607014-DEL-988 | chr16-73607038-DEL-964 |
| chr16-73616952-DEL-52 | chr16-73616948-DEL-54 |
| chr16-73642080-DEL-170 | chr16-73642081-DEL-170 |
| chr16-73644736-DEL-747 | chr16-73644737-DEL-747 |
| chr16-73668004-DEL-169 | chr16-73668005-DEL-169 |
| chr16-73698516-DEL-87 | chr16-73698517-DEL-87 |
| chr16-73705069-DEL-1610 | chr16-73705070-DEL-1610 |
| chr16-73723338-DEL-54 | chr16-73723343-DEL-50 |
| chr16-73765353-DEL-264 | chr16-73765354-DEL-264 |
| chr16-73777932-DEL-107 | chr16-73777933-DEL-107 |
| chr16-73788744-DEL-6684 | chr16-73788745-DEL-6684 |
| chr16-73872254-DEL-76 | chr16-73872258-DEL-88 |
| chr16-73950613-DEL-3256 | chr16-73950614-DEL-3256 |
| chr16-73958049-DEL-70 | chr16-73958038-DEL-73 |
| chr16-73994564-DEL-51 | chr16-73994563-DEL-54 |
| chr16-73996982-DEL-54 | chr16-73996983-DEL-54 |
| chr16-74002704-DEL-6455 | chr16-74002705-DEL-6455 |
| chr16-74012833-DEL-344 | chr16-74012835-DEL-342 |
| chr16-74063892-DEL-76 | chr16-74063897-DEL-72 |
| chr16-74065923-DEL-64 | chr16-74065936-DEL-52 |
| chr16-74091980-DEL-88 | chr16-74091981-DEL-86 |
| chr16-74096966-DEL-76 | chr16-74096947-DEL-68 |
| chr16-74118292-DEL-100 | chr16-74118312-DEL-89 |
| chr16-74128554-DEL-410 | chr16-74128555-DEL-410 |
| chr16-74146393-DEL-397 | chr16-74146394-DEL-397 |
| chr16-74154271-DEL-6219 | chr16-74154272-DEL-6219 |
| chr16-74165714-DEL-110 | chr16-74165713-DEL-107 |
| chr16-74225746-DEL-323 | chr16-74225747-DEL-323 |
| chr16-74243040-DEL-99 | chr16-74243033-DEL-85 |
| chr16-74262062-DEL-57 | chr16-74262062-DEL-69 |
| chr16-74282862-DEL-78 | chr16-74282856-DEL-82 |
| chr16-74321118-DEL-1608 | chr16-74321117-DEL-1609 |
| chr16-74346510-DEL-6218 | chr16-74346511-DEL-6218 |
| chr16-74371129-DEL-411 | chr16-74371130-DEL-411 |
| chr16-74374699-DEL-2588 | chr16-74374700-DEL-2588 |
| chr16-74374947-DEL-491 | chr16-74374946-DEL-492 |
| chr16-74385356-DEL-764 | chr16-74385338-DEL-773 |
| chr16-74416330-DEL-250 | chr16-74416331-DEL-250 |
| chr16-74449973-DEL-343 | chr16-74449974-DEL-343 |
| chr16-74481027-DEL-789 | chr16-74481028-DEL-789 |
| chr16-74483796-DEL-3531 | chr16-74483797-DEL-3531 |
| chr16-74507593-DEL-172 | chr16-74507594-DEL-172 |

| | |
|---|---|
| chr16-74528492-DEL-384 | chr16-74528493-DEL-384 |
| chr16-74565078-DEL-126 | chr16-74565079-DEL-126 |
| chr16-74568760-DEL-7127 | chr16-74568761-DEL-7127 |
| chr16-74576671-DEL-82 | chr16-74576675-DEL-80 |
| chr16-74577424-DEL-102 | chr16-74577425-DEL-102 |
| chr16-74580221-DEL-97 | chr16-74580222-DEL-96 |
| chr16-74583542-DEL-66 | chr16-74583542-DEL-67 |
| chr16-74602446-DEL-94 | chr16-74602457-DEL-86 |
| chr16-74651908-DEL-67 | chr16-74651909-DEL-67 |
| chr16-74684873-DEL-113 | chr16-74684874-DEL-113 |
| chr16-74709492-DEL-2117 | chr16-74709493-DEL-2117 |
| chr16-74740555-DEL-64 | chr16-74740569-DEL-65 |
| chr16-74753285-DEL-2065 | chr16-74753286-DEL-2065 |
| chr16-74768422-DEL-6246 | chr16-74768423-DEL-6246 |
| chr16-74809622-DEL-365 | chr16-74809623-DEL-365 |
| chr16-74823344-DEL-1047 | chr16-74823345-DEL-1047 |
| chr16-74825734-DEL-80 | chr16-74825719-DEL-74 |
| chr16-74827424-DEL-52 | chr16-74827416-DEL-52 |
| chr16-74906055-DEL-285 | chr16-74906054-DEL-286 |
| chr16-74965444-DEL-397 | chr16-74965445-DEL-397 |
| chr16-74966594-DEL-86 | chr16-74966595-DEL-86 |
| chr16-74972906-DEL-51 | chr16-74972905-DEL-50 |
| chr16-74991839-DEL-89 | chr16-74991840-DEL-89 |
| chr16-75025439-DEL-1298 | chr16-75025440-DEL-1298 |
| chr16-75039190-DEL-848 | chr16-75039154-DEL-884 |
| chr16-75067610-DEL-1103 | chr16-75067611-DEL-1103 |
| chr16-75074553-DEL-59 | chr16-75074554-DEL-59 |
| chr16-75080286-DEL-64 | chr16-75080287-DEL-66 |
| chr16-75091105-DEL-839 | chr16-75091106-DEL-839 |
| chr16-75099338-DEL-714 | chr16-75099339-DEL-714 |
| chr16-75103649-DEL-95 | chr16-75103649-DEL-95 |
| chr16-75123014-DEL-94 | chr16-75123047-DEL-62 |
| chr16-75125703-DEL-92 | chr16-75125716-DEL-53 |
| chr16-75127329-DEL-64 | chr16-75127330-DEL-52 |
| chr16-75128375-DEL-588 | chr16-75128376-DEL-588 |
| chr16-75130948-DEL-52 | chr16-75130949-DEL-80 |
| chr16-75153473-DEL-430 | chr16-75153474-DEL-429 |
| chr16-75160228-DEL-102 | chr16-75160229-DEL-102 |
| chr16-75168404-DEL-716 | chr16-75168405-DEL-716 |
| chr16-75174967-DEL-56 | chr16-75174968-DEL-56 |
| chr16-75179378-DEL-98 | chr16-75179376-DEL-95 |
| chr16-75190363-DEL-2992 | chr16-75190364-DEL-2992 |
| chr16-75210406-DEL-934 | chr16-75210407-DEL-934 |
| chr16-75218606-DEL-169 | chr16-75218604-DEL-169 |
| chr16-75219815-DEL-150 | chr16-75219816-DEL-150 |
| chr16-75226599-DEL-498 | chr16-75226600-DEL-498 |
| chr16-75227311-DEL-1065 | chr16-75227312-DEL-1065 |
| chr16-75229461-DEL-60 | chr16-75229458-DEL-71 |
| chr16-75255067-DEL-90 | chr16-75255068-DEL-86 |
| chr16-75259723-DEL-56 | chr16-75259724-DEL-56 |
| chr16-75274655-DEL-71 | chr16-75274656-DEL-71 |
| chr16-75281506-DEL-219 | chr16-75281507-DEL-219 |
| chr16-75349007-DEL-780 | chr16-75349008-DEL-780 |
| chr16-75375168-DEL-98 | chr16-75375168-DEL-98 |
| chr16-75387939-DEL-281 | chr16-75387940-DEL-281 |
| chr16-75420174-DEL-727 | chr16-75420175-DEL-727 |
| chr16-75431543-DEL-7152 | chr16-75431544-DEL-7152 |
| chr16-75443456-DEL-107 | chr16-75443456-DEL-107 |
| chr16-75451377-DEL-240 | chr16-75451377-DEL-240 |
| chr16-75457406-DEL-400 | chr16-75457405-DEL-400 |
| chr16-75480204-DEL-392 | chr16-75480205-DEL-392 |
| chr16-75482890-DEL-5923 | chr16-75482890-DEL-5922 |
| chr16-75519733-DEL-138 | chr16-75519733-DEL-139 |
| chr16-75565551-DEL-681 | chr16-75565550-DEL-681 |
| chr16-75595767-DEL-213 | chr16-75595768-DEL-213 |
| chr16-75610651-DEL-132 | chr16-75610652-DEL-132 |
| chr16-75624906-DEL-586 | chr16-75624907-DEL-586 |
| chr16-75670202-DEL-54 | chr16-75670203-DEL-54 |
| chr16-75681405-DEL-492 | chr16-75681406-DEL-492 |
| chr16-75687677-DEL-70 | chr16-75687678-DEL-70 |
| chr16-75709891-DEL-98 | chr16-75709887-DEL-60 |
| chr16-75720966-DEL-63 | chr16-75720966-DEL-52 |
| chr16-75739042-DEL-651 | chr16-75739043-DEL-651 |
| chr16-75749257-DEL-6191 | chr16-75749263-DEL-6192 |
| chr16-75769306-DEL-51 | chr16-75769308-DEL-54 |
| chr16-75788113-DEL-6246 | chr16-75788114-DEL-6246 |
| chr16-75810101-DEL-3409 | chr16-75810102-DEL-3409 |
| chr16-75832896-DEL-1281 | chr16-75832881-DEL-1282 |
| chr16-75853897-DEL-7857 | chr16-75853898-DEL-7857 |
| chr16-75889064-DEL-291 | chr16-75889063-DEL-291 |

| | |
|---|---|
| chr16-75898908-DEL-624 | chr16-75898908-DEL-624 |
| chr16-75913278-DEL-365 | chr16-75913279-DEL-365 |
| chr16-75915358-DEL-1897 | chr16-75915359-DEL-1897 |
| chr16-75931926-DEL-8986 | chr16-75931927-DEL-8986 |
| chr16-75942791-DEL-339 | chr16-75942792-DEL-339 |
| chr16-75966486-DEL-7069 | chr16-75966487-DEL-7069 |
| chr16-75975400-DEL-144 | chr16-75975401-DEL-144 |
| chr16-75992683-DEL-56 | chr16-75992667-DEL-70 |
| chr16-76005464-DEL-58 | chr16-76005483-DEL-59 |
| chr16-76068031-DEL-1891 | chr16-76068032-DEL-1891 |
| chr16-76072990-DEL-82 | chr16-76073013-DEL-57 |
| chr16-76074830-DEL-986 | chr16-76074831-DEL-986 |
| chr16-76082294-DEL-88 | chr16-76082299-DEL-84 |
| chr16-76111146-DEL-85 | chr16-76111147-DEL-85 |
| chr16-76112265-DEL-95 | chr16-76112266-DEL-95 |
| chr16-76146832-DEL-54 | chr16-76146800-DEL-51 |
| chr16-76164946-DEL-592 | chr16-76164947-DEL-592 |
| chr16-76174407-DEL-130 | chr16-76174408-DEL-130 |
| chr16-76200069-DEL-531 | chr16-76200070-DEL-531 |
| chr16-76208594-DEL-853 | chr16-76208589-DEL-853 |
| chr16-76217304-DEL-86 | chr16-76217305-DEL-86 |
| chr16-76241895-DEL-177 | chr16-76241896-DEL-177 |
| chr16-76263555-DEL-56 | chr16-76263545-DEL-63 |
| chr16-76279772-DEL-96 | chr16-76279767-DEL-67 |
| chr16-76287554-DEL-6248 | chr16-76287555-DEL-6248 |
| chr16-76310040-DEL-6487 | chr16-76310040-DEL-6487 |
| chr16-76316194-DEL-212 | chr16-76316195-DEL-212 |
| chr16-76326015-DEL-63 | chr16-76326016-DEL-63 |
| chr16-76332931-DEL-118 | chr16-76332932-DEL-118 |
| chr16-76353186-DEL-67 | chr16-76353174-DEL-78 |
| chr16-76432430-DEL-6776 | chr16-76432431-DEL-6776 |
| chr16-76439493-DEL-1117 | chr16-76439458-DEL-1153 |
| chr16-76460229-DEL-705 | chr16-76460230-DEL-705 |
| chr16-76498474-DEL-219 | chr16-76498490-DEL-197 |
| chr16-76557672-DEL-56 | chr16-76557671-DEL-58 |
| chr16-76585918-DEL-6327 | chr16-76585919-DEL-6327 |
| chr16-76594509-DEL-139 | chr16-76594510-DEL-139 |
| chr16-76604652-DEL-123 | chr16-76604613-DEL-116 |
| chr16-76642678-DEL-1030 | chr16-76642680-DEL-1029 |
| chr16-76662969-DEL-106 | chr16-76662999-DEL-56 |
| chr16-76666358-DEL-72 | chr16-76666359-DEL-72 |
| chr16-76669169-DEL-115 | chr16-76669170-DEL-115 |
| chr16-76672862-DEL-2474 | chr16-76672863-DEL-2474 |
| chr16-76736497-DEL-68 | chr16-76736488-DEL-54 |
| chr16-76737258-DEL-5945 | chr16-76737259-DEL-5945 |
| chr16-76777809-DEL-115 | chr16-76777812-DEL-114 |
| chr16-76779973-DEL-56 | chr16-76779980-DEL-54 |
| chr16-76781308-DEL-83 | chr16-76781308-DEL-67 |
| chr16-76782884-DEL-258 | chr16-76782885-DEL-258 |
| chr16-76789788-DEL-110 | chr16-76789789-DEL-112 |
| chr16-76799823-DEL-51 | chr16-76799823-DEL-55 |
| chr16-76804317-DEL-82 | chr16-76804318-DEL-82 |
| chr16-76843350-DEL-68 | chr16-76843351-DEL-68 |
| chr16-76851202-DEL-79 | chr16-76851204-DEL-78 |
| chr16-76883565-DEL-85 | chr16-76883566-DEL-85 |
| chr16-76889658-DEL-96 | chr16-76889658-DEL-94 |
| chr16-76908434-DEL-66 | chr16-76908435-DEL-60 |
| chr16-76913845-DEL-4887 | chr16-76913846-DEL-4881 |
| chr16-76938584-DEL-52 | chr16-76938553-DEL-64 |
| chr16-76939991-DEL-306 | chr16-76939991-DEL-306 |
| chr16-76952134-DEL-6469 | chr16-76952135-DEL-6469 |
| chr16-76983704-DEL-6646 | chr16-76983695-DEL-6647 |
| chr16-77008977-DEL-203 | chr16-77008980-DEL-201 |
| chr16-77009626-DEL-57 | chr16-77009623-DEL-60 |
| chr16-77010673-DEL-386 | chr16-77010674-DEL-386 |
| chr16-77082012-DEL-165 | chr16-77082013-DEL-166 |
| chr16-77098136-DEL-2401 | chr16-77098138-DEL-2397 |
| chr16-77110484-DEL-1215 | chr16-77110485-DEL-1215 |
| chr16-77134611-DEL-52 | chr16-77134612-DEL-52 |
| chr16-77147059-DEL-434 | chr16-77147060-DEL-434 |
| chr16-77184752-DEL-83 | chr16-77184769-DEL-67 |
| chr16-77248619-DEL-82 | chr16-77248620-DEL-82 |
| chr16-77254135-DEL-441 | chr16-77254136-DEL-441 |
| chr16-77356931-DEL-319 | chr16-77356932-DEL-319 |
| chr16-77415728-DEL-52 | chr16-77415729-DEL-50 |
| chr16-77427166-DEL-122 | chr16-77427170-DEL-121 |
| chr16-77488672-DEL-78 | chr16-77488673-DEL-78 |
| chr16-77521444-DEL-553 | chr16-77521445-DEL-553 |
| chr16-77541352-DEL-324 | chr16-77541353-DEL-324 |
| chr16-77578248-DEL-208 | chr16-77578249-DEL-208 |

| | |
|---|---|
| chr16-77601504-DEL-81 | chr16-77601497-DEL-89 |
| chr16-77607062-DEL-313 | chr16-77607063-DEL-313 |
| chr16-77607905-DEL-4322 | chr16-77607906-DEL-4322 |
| chr16-77609849-DEL-1716 | chr16-77609850-DEL-1715 |
| chr16-77653429-DEL-60 | chr16-77653430-DEL-54 |
| chr16-77659106-DEL-101 | chr16-77659105-DEL-73 |
| chr16-77670036-DEL-95 | chr16-77670037-DEL-95 |
| chr16-77674904-DEL-76 | chr16-77674861-DEL-120 |
| chr16-77681195-DEL-101 | chr16-77681191-DEL-96 |
| chr16-77696945-DEL-1891 | chr16-77696946-DEL-1891 |
| chr16-77704248-DEL-358 | chr16-77704249-DEL-358 |
| chr16-77715042-DEL-6420 | chr16-77715043-DEL-6420 |
| chr16-77743782-DEL-335 | chr16-77743783-DEL-335 |
| chr16-77759636-DEL-72 | chr16-77759637-DEL-72 |
| chr16-77762012-DEL-87 | chr16-77761998-DEL-99 |
| chr16-77781399-DEL-61 | chr16-77781400-DEL-61 |
| chr16-77784899-DEL-423 | chr16-77784900-DEL-423 |
| chr16-77788728-DEL-258 | chr16-77788756-DEL-227 |
| chr16-77797159-DEL-84 | chr16-77797141-DEL-102 |
| chr16-77823477-DEL-3653 | chr16-77823478-DEL-3654 |
| chr16-77942905-DEL-215 | chr16-77942878-DEL-243 |
| chr16-77960053-DEL-727 | chr16-77960054-DEL-727 |
| chr16-77972258-DEL-405 | chr16-77972259-DEL-405 |
| chr16-77982991-DEL-578 | chr16-77982985-DEL-578 |
| chr16-78004481-DEL-87 | chr16-78004481-DEL-89 |
| chr16-78013530-DEL-52 | chr16-78013527-DEL-56 |
| chr16-78015548-DEL-489 | chr16-78015534-DEL-505 |
| chr16-78022235-DEL-157 | chr16-78022235-DEL-157 |
| chr16-78025149-DEL-340 | chr16-78025150-DEL-340 |
| chr16-78029279-DEL-912 | chr16-78029280-DEL-912 |
| chr16-78036022-DEL-185 | chr16-78036023-DEL-185 |
| chr16-78040395-DEL-1468 | chr16-78040396-DEL-1468 |
| chr16-78042003-DEL-52 | chr16-78042004-DEL-52 |
| chr16-78066176-DEL-173 | chr16-78066177-DEL-173 |
| chr16-78069563-DEL-184 | chr16-78069564-DEL-184 |
| chr16-78075061-DEL-52 | chr16-78075012-DEL-65 |
| chr16-78086168-DEL-1930 | chr16-78086165-DEL-1930 |
| chr16-78112820-DEL-359 | chr16-78112821-DEL-359 |
| chr16-78147070-DEL-7109 | chr16-78147071-DEL-7109 |
| chr16-78166273-DEL-61 | chr16-78166274-DEL-61 |
| chr16-78201922-DEL-377 | chr16-78201922-DEL-377 |
| chr16-78219627-DEL-129 | chr16-78219628-DEL-127 |
| chr16-78235244-DEL-131 | chr16-78235244-DEL-131 |
| chr16-78241600-DEL-4573 | chr16-78241599-DEL-4579 |
| chr16-78250216-DEL-149 | chr16-78250181-DEL-185 |
| chr16-78287869-DEL-1046 | chr16-78287869-DEL-1044 |
| chr16-78289822-DEL-6614 | chr16-78289823-DEL-6614 |
| chr16-78289930-DEL-201 | chr16-78289931-DEL-201 |
| chr16-78299144-DEL-104 | chr16-78299145-DEL-104 |
| chr16-78313368-DEL-122 | chr16-78313369-DEL-122 |
| chr16-78332335-DEL-98 | chr16-78332336-DEL-92 |
| chr16-78362600-DEL-6442 | chr16-78362601-DEL-6442 |
| chr16-78390634-DEL-138 | chr16-78390635-DEL-162 |
| chr16-78397173-DEL-54 | chr16-78397159-DEL-55 |
| chr16-78397701-DEL-746 | chr16-78397702-DEL-746 |
| chr16-78398782-DEL-202 | chr16-78398783-DEL-202 |
| chr16-78399906-DEL-155 | chr16-78399914-DEL-110 |
| chr16-78404098-DEL-7839 | chr16-78404099-DEL-7839 |
| chr16-78430993-DEL-52 | chr16-78430994-DEL-52 |
| chr16-78433128-DEL-196 | chr16-78433129-DEL-196 |
| chr16-78457671-DEL-262 | chr16-78457672-DEL-262 |
| chr16-78462737-DEL-6273 | chr16-78462738-DEL-6273 |
| chr16-78471138-DEL-773 | chr16-78471139-DEL-773 |
| chr16-78475100-DEL-1915 | chr16-78475101-DEL-1915 |
| chr16-78496719-DEL-349 | chr16-78496732-DEL-345 |
| chr16-78516418-DEL-73 | chr16-78516417-DEL-84 |
| chr16-78522328-DEL-51 | chr16-78522329-DEL-51 |
| chr16-78547393-DEL-3930 | chr16-78547394-DEL-3930 |
| chr16-78567552-DEL-1444 | chr16-78567553-DEL-1444 |
| chr16-78570317-DEL-2501 | chr16-78570328-DEL-2499 |
| chr16-78576575-DEL-1794 | chr16-78576576-DEL-1794 |
| chr16-78594273-DEL-1707 | chr16-78594275-DEL-1706 |
| chr16-78601480-DEL-5022 | chr16-78601488-DEL-5025 |
| chr16-78611264-DEL-5271 | chr16-78611265-DEL-5271 |
| chr16-78615359-DEL-52 | chr16-78615360-DEL-52 |
| chr16-78643711-DEL-3042 | chr16-78643712-DEL-3042 |
| chr16-78647089-DEL-64 | chr16-78647090-DEL-64 |
| chr16-78660508-DEL-60 | chr16-78660517-DEL-52 |
| chr16-78666890-DEL-54 | chr16-78666891-DEL-54 |
| chr16-78681709-DEL-9095 | chr16-78681697-DEL-9089 |

| | |
|---|---|
| chr16-78704424-DEL-58 | chr16-78704424-DEL-58 |
| chr16-78710059-DEL-389 | chr16-78710060-DEL-389 |
| chr16-78726592-DEL-64 | chr16-78726593-DEL-64 |
| chr16-78773911-DEL-1779 | chr16-78773912-DEL-1779 |
| chr16-78777590-DEL-405 | chr16-78777591-DEL-405 |
| chr16-78784995-DEL-6701 | chr16-78784996-DEL-6701 |
| chr16-78804044-DEL-6450 | chr16-78804045-DEL-6450 |
| chr16-78835316-DEL-6420 | chr16-78835317-DEL-6420 |
| chr16-78844756-DEL-59 | chr16-78844757-DEL-59 |
| chr16-78846890-DEL-2023 | chr16-78846880-DEL-2024 |
| chr16-78858857-DEL-66 | chr16-78858865-DEL-59 |
| chr16-78870256-DEL-312 | chr16-78870256-DEL-312 |
| chr16-78899254-DEL-66 | chr16-78899255-DEL-66 |
| chr16-78923015-DEL-92 | chr16-78923015-DEL-93 |
| chr16-78950401-DEL-1047 | chr16-78950403-DEL-1045 |
| chr16-78956560-DEL-77 | chr16-78956561-DEL-77 |
| chr16-78991719-DEL-3517 | chr16-78991717-DEL-3518 |
| chr16-78997473-DEL-750 | chr16-78997502-DEL-737 |
| chr16-79007577-DEL-70 | chr16-79007605-DEL-88 |
| chr16-79010611-DEL-637 | chr16-79010612-DEL-637 |
| chr16-79033290-DEL-58 | chr16-79033290-DEL-59 |
| chr16-79033506-DEL-507 | chr16-79033531-DEL-476 |
| chr16-79076450-DEL-2836 | chr16-79076451-DEL-2836 |
| chr16-79080958-DEL-198 | chr16-79080959-DEL-198 |
| chr16-79081737-DEL-579 | chr16-79081738-DEL-579 |
| chr16-79091234-DEL-132 | chr16-79091233-DEL-133 |
| chr16-79095212-DEL-2028 | chr16-79095213-DEL-2028 |
| chr16-79106386-DEL-65 | chr16-79106347-DEL-67 |
| chr16-79116491-DEL-1258 | chr16-79116492-DEL-1258 |
| chr16-79122893-DEL-66 | chr16-79122898-DEL-56 |
| chr16-79126963-DEL-52 | chr16-79126964-DEL-52 |
| chr16-79137095-DEL-553 | chr16-79137097-DEL-525 |
| chr16-79143624-DEL-583 | chr16-79143620-DEL-583 |
| chr16-79165578-DEL-7280 | chr16-79165579-DEL-7280 |
| chr16-79186017-DEL-256 | chr16-79186018-DEL-256 |
| chr16-79190545-DEL-53 | chr16-79190539-DEL-54 |
| chr16-79234858-DEL-69 | chr16-79234859-DEL-69 |
| chr16-79248860-DEL-6053 | chr16-79248861-DEL-6053 |
| chr16-79254372-DEL-61 | chr16-79254373-DEL-61 |
| chr16-79255369-DEL-67 | chr16-79255370-DEL-62 |
| chr16-79262604-DEL-50 | chr16-79262604-DEL-50 |
| chr16-79266470-DEL-83 | chr16-79266476-DEL-57 |
| chr16-79268793-DEL-150 | chr16-79268794-DEL-150 |
| chr16-79269787-DEL-133 | chr16-79269777-DEL-138 |
| chr16-79274302-DEL-410 | chr16-79274303-DEL-410 |
| chr16-79290259-DEL-343 | chr16-79290260-DEL-343 |
| chr16-79293986-DEL-135 | chr16-79293977-DEL-145 |
| chr16-79294644-DEL-1557 | chr16-79294645-DEL-1557 |
| chr16-79296494-DEL-1854 | chr16-79296495-DEL-1854 |
| chr16-79302069-DEL-1893 | chr16-79302071-DEL-1892 |
| chr16-79310417-DEL-117 | chr16-79310417-DEL-117 |
| chr16-79319735-DEL-51 | chr16-79319720-DEL-54 |
| chr16-79339313-DEL-4494 | chr16-79339314-DEL-4494 |
| chr16-79359603-DEL-86 | chr16-79359604-DEL-86 |
| chr16-79375355-DEL-1158 | chr16-79375355-DEL-1158 |
| chr16-79378035-DEL-2772 | chr16-79378036-DEL-2772 |
| chr16-79408162-DEL-643 | chr16-79408163-DEL-643 |
| chr16-79430189-DEL-155 | chr16-79430190-DEL-155 |
| chr16-79445188-DEL-537 | chr16-79445217-DEL-530 |
| chr16-79458244-DEL-9503 | chr16-79458245-DEL-9503 |
| chr16-79463896-DEL-3011 | chr16-79463897-DEL-3011 |
| chr16-79480720-DEL-921 | chr16-79480721-DEL-921 |
| chr16-79485409-DEL-247 | chr16-79485409-DEL-248 |
| chr16-79497170-DEL-1545 | chr16-79497171-DEL-1545 |
| chr16-79500951-DEL-62 | chr16-79500952-DEL-62 |
| chr16-79517022-DEL-52 | chr16-79517023-DEL-52 |
| chr16-79525467-DEL-6501 | chr16-79525468-DEL-6501 |
| chr16-79531644-DEL-212 | chr16-79531645-DEL-212 |
| chr16-79540566-DEL-394 | chr16-79540568-DEL-385 |
| chr16-79578572-DEL-6293 | chr16-79578573-DEL-6293 |
| chr16-79602267-DEL-64 | chr16-79602265-DEL-65 |
| chr16-79609586-DEL-936 | chr16-79609587-DEL-936 |
| chr16-79614685-DEL-68 | chr16-79614689-DEL-72 |
| chr16-79615288-DEL-198 | chr16-79615289-DEL-198 |
| chr16-79625090-DEL-5258 | chr16-79625091-DEL-5258 |
| chr16-79630789-DEL-178 | chr16-79630789-DEL-178 |
| chr16-79633715-DEL-1511 | chr16-79633749-DEL-1472 |
| chr16-79649698-DEL-2855 | chr16-79649699-DEL-2855 |
| chr16-79673716-DEL-70 | chr16-79673734-DEL-66 |
| chr16-79711793-DEL-3362 | chr16-79711794-DEL-3362 |

| | |
|---|---|
| chr16-79738471-DEL-72 | chr16-79738472-DEL-72 |
| chr16-79739956-DEL-5422 | chr16-79739957-DEL-5422 |
| chr16-79763792-DEL-69 | chr16-79763775-DEL-64 |
| chr16-79765947-DEL-328 | chr16-79765948-DEL-328 |
| chr16-79790298-DEL-66 | chr16-79790335-DEL-74 |
| chr16-79794551-DEL-120 | chr16-79794552-DEL-120 |
| chr16-79800779-DEL-441 | chr16-79800780-DEL-441 |
| chr16-79869907-DEL-94 | chr16-79869950-DEL-52 |
| chr16-79879989-DEL-1301 | chr16-79879990-DEL-1301 |
| chr16-79886034-DEL-157 | chr16-79886035-DEL-157 |
| chr16-79898061-DEL-6383 | chr16-79898062-DEL-6383 |
| chr16-79917169-DEL-999 | chr16-79917169-DEL-983 |
| chr16-79925197-DEL-82 | chr16-79925198-DEL-82 |
| chr16-79983971-DEL-84 | chr16-79983972-DEL-84 |
| chr16-79990414-DEL-144 | chr16-79990417-DEL-142 |
| chr16-80002415-DEL-1703 | chr16-80002419-DEL-1702 |
| chr16-80020298-DEL-134 | chr16-80020299-DEL-134 |
| chr16-80047291-DEL-62 | chr16-80047292-DEL-62 |
| chr16-80117511-DEL-211 | chr16-80117497-DEL-226 |
| chr16-80121750-DEL-321 | chr16-80121751-DEL-321 |
| chr16-80123551-DEL-87 | chr16-80123552-DEL-87 |
| chr16-80141655-DEL-88 | chr16-80141688-DEL-50 |
| chr16-80160643-DEL-206 | chr16-80160644-DEL-206 |
| chr16-80217019-DEL-360 | chr16-80217020-DEL-360 |
| chr16-80251669-DEL-148 | chr16-80251670-DEL-148 |
| chr16-80259907-DEL-52 | chr16-80259908-DEL-52 |
| chr16-80272289-DEL-1786 | chr16-80272286-DEL-1787 |
| chr16-80284772-DEL-764 | chr16-80284776-DEL-761 |
| chr16-80292814-DEL-56 | chr16-80292815-DEL-56 |
| chr16-80297441-DEL-517 | chr16-80297442-DEL-517 |
| chr16-80325696-DEL-79 | chr16-80325699-DEL-77 |
| chr16-80335509-DEL-64 | chr16-80335512-DEL-62 |
| chr16-80355685-DEL-58 | chr16-80355674-DEL-60 |
| chr16-80387513-DEL-456 | chr16-80387514-DEL-456 |
| chr16-80392184-DEL-59 | chr16-80392185-DEL-59 |
| chr16-80404401-DEL-68 | chr16-80404441-DEL-50 |
| chr16-80503801-DEL-53 | chr16-80503780-DEL-62 |
| chr16-80509528-DEL-5943 | chr16-80509528-DEL-5962 |
| chr16-80515296-DEL-901 | chr16-80515297-DEL-901 |
| chr16-80518407-DEL-58 | chr16-80518406-DEL-60 |
| chr16-80526756-DEL-324 | chr16-80526757-DEL-324 |
| chr16-80542041-DEL-1379 | chr16-80542042-DEL-1379 |
| chr16-80544202-DEL-4286 | chr16-80544202-DEL-4286 |
| chr16-80554734-DEL-200 | chr16-80554734-DEL-200 |
| chr16-80556780-DEL-7076 | chr16-80556781-DEL-7076 |
| chr16-80565330-DEL-158 | chr16-80565347-DEL-108 |
| chr16-80586095-DEL-428 | chr16-80586096-DEL-428 |
| chr16-80610650-DEL-119 | chr16-80610639-DEL-131 |
| chr16-80630433-DEL-201 | chr16-80630434-DEL-201 |
| chr16-80649492-DEL-670 | chr16-80649493-DEL-670 |
| chr16-80692489-DEL-2863 | chr16-80692490-DEL-2863 |
| chr16-80736238-DEL-6444 | chr16-80736257-DEL-6443 |
| chr16-80784284-DEL-54 | chr16-80784285-DEL-54 |
| chr16-80786022-DEL-110 | chr16-80786039-DEL-76 |
| chr16-80792409-DEL-6376 | chr16-80792410-DEL-6376 |
| chr16-80808859-DEL-114 | chr16-80808858-DEL-115 |
| chr16-80812498-DEL-4163 | chr16-80812465-DEL-4196 |
| chr16-80822285-DEL-11455 | chr16-80822286-DEL-11455 |
| chr16-80835731-DEL-102 | chr16-80835732-DEL-102 |
| chr16-80844902-DEL-380 | chr16-80844895-DEL-381 |
| chr16-80868554-DEL-7440 | chr16-80868555-DEL-7440 |
| chr16-80901597-DEL-92 | chr16-80901598-DEL-92 |
| chr16-80930589-DEL-145 | chr16-80930591-DEL-144 |
| chr16-80937443-DEL-1138 | chr16-80937445-DEL-1136 |
| chr16-80961566-DEL-107 | chr16-80961561-DEL-103 |
| chr16-80974815-DEL-1654 | chr16-80974822-DEL-1644 |
| chr16-80982543-DEL-1209 | chr16-80982544-DEL-1209 |
| chr16-81040761-DEL-365 | chr16-81040762-DEL-365 |
| chr16-81059407-DEL-444 | chr16-81059408-DEL-444 |
| chr16-81066719-DEL-80 | chr16-81066719-DEL-80 |
| chr16-81089880-DEL-78 | chr16-81089881-DEL-105 |
| chr16-81103259-DEL-52 | chr16-81103246-DEL-64 |
| chr16-81145104-DEL-265 | chr16-81145105-DEL-265 |
| chr16-81182196-DEL-54 | chr16-81182197-DEL-62 |
| chr16-81218773-DEL-85 | chr16-81218736-DEL-102 |
| chr16-81230760-DEL-75 | chr16-81230761-DEL-75 |
| chr16-81254207-DEL-71 | chr16-81254208-DEL-50 |
| chr16-81308768-DEL-624 | chr16-81308769-DEL-624 |
| chr16-81336494-DEL-186 | chr16-81336494-DEL-186 |
| chr16-81343953-DEL-325 | chr16-81343953-DEL-325 |

| | |
|---|---|
| chr16-81347161-DEL-66 | chr16-81347162-DEL-66 |
| chr16-81369224-DEL-224 | chr16-81369225-DEL-225 |
| chr16-81380554-DEL-2540 | chr16-81380555-DEL-2540 |
| chr16-81394611-DEL-230 | chr16-81394611-DEL-230 |
| chr16-81461441-DEL-225 | chr16-81461442-DEL-225 |
| chr16-81490450-DEL-91 | chr16-81490451-DEL-84 |
| chr16-81544441-DEL-5521 | chr16-81544442-DEL-5521 |
| chr16-81572646-DEL-70 | chr16-81572647-DEL-80 |
| chr16-81576842-DEL-1978 | chr16-81576843-DEL-1978 |
| chr16-81605570-DEL-89 | chr16-81605574-DEL-62 |
| chr16-81641290-DEL-187 | chr16-81641291-DEL-187 |
| chr16-81656077-DEL-6470 | chr16-81656078-DEL-6470 |
| chr16-81662809-DEL-779 | chr16-81662806-DEL-780 |
| chr16-81681247-DEL-719 | chr16-81681248-DEL-719 |
| chr16-81733766-DEL-176 | chr16-81733766-DEL-177 |
| chr16-81758788-DEL-389 | chr16-81758793-DEL-389 |
| chr16-81784709-DEL-904 | chr16-81784710-DEL-904 |
| chr16-81876140-DEL-17546 | chr16-81876131-DEL-17553 |
| chr16-81895248-DEL-1468 | chr16-81895241-DEL-1473 |
| chr16-81904096-DEL-650 | chr16-81904097-DEL-650 |
| chr16-81907500-DEL-754 | chr16-81907501-DEL-752 |
| chr16-81980127-DEL-61 | chr16-81980128-DEL-60 |
| chr16-81984623-DEL-61 | chr16-81984624-DEL-61 |
| chr16-81985078-DEL-3328 | chr16-81985079-DEL-3328 |
| chr16-82004518-DEL-4555 | chr16-82004519-DEL-4555 |
| chr16-82009534-DEL-60 | chr16-82009527-DEL-78 |
| chr16-82011401-DEL-2528 | chr16-82011402-DEL-2528 |
| chr16-82022992-DEL-60 | chr16-82022993-DEL-60 |
| chr16-82034801-DEL-5162 | chr16-82034802-DEL-5162 |
| chr16-82045164-DEL-154 | chr16-82045165-DEL-154 |
| chr16-82059860-DEL-59 | chr16-82059850-DEL-60 |
| chr16-82077641-DEL-74 | chr16-82077652-DEL-64 |
| chr16-82116252-DEL-51 | chr16-82116251-DEL-51 |
| chr16-82117880-DEL-146 | chr16-82117881-DEL-146 |
| chr16-82140785-DEL-51 | chr16-82140797-DEL-51 |
| chr16-82147831-DEL-126 | chr16-82147832-DEL-126 |
| chr16-82162251-DEL-110 | chr16-82162252-DEL-110 |
| chr16-82162742-DEL-561 | chr16-82162743-DEL-561 |
| chr16-82183477-DEL-54 | chr16-82183478-DEL-54 |
| chr16-82208631-DEL-51 | chr16-82208630-DEL-52 |
| chr16-82224407-DEL-341 | chr16-82224408-DEL-341 |
| chr16-82232805-DEL-373 | chr16-82232806-DEL-373 |
| chr16-82238278-DEL-51 | chr16-82238276-DEL-52 |
| chr16-82250535-DEL-959 | chr16-82250531-DEL-961 |
| chr16-82280140-DEL-414 | chr16-82280141-DEL-414 |
| chr16-82304513-DEL-6532 | chr16-82304514-DEL-6532 |
| chr16-82340103-DEL-2895 | chr16-82340103-DEL-2895 |
| chr16-82358443-DEL-56 | chr16-82358444-DEL-56 |
| chr16-82380352-DEL-7135 | chr16-82380353-DEL-7135 |
| chr16-82382106-DEL-114 | chr16-82382107-DEL-114 |
| chr16-82402064-DEL-538 | chr16-82402065-DEL-538 |
| chr16-82416713-DEL-486 | chr16-82416714-DEL-486 |
| chr16-82430731-DEL-6855 | chr16-82430730-DEL-6855 |
| chr16-82441570-DEL-130 | chr16-82441571-DEL-130 |
| chr16-82447837-DEL-75 | chr16-82447836-DEL-77 |
| chr16-82473287-DEL-213 | chr16-82473288-DEL-213 |
| chr16-82474468-DEL-760 | chr16-82474469-DEL-760 |
| chr16-82476373-DEL-233 | chr16-82476374-DEL-233 |
| chr16-82502685-DEL-350 | chr16-82502683-DEL-351 |
| chr16-82551594-DEL-64 | chr16-82551595-DEL-64 |
| chr16-82574154-DEL-143 | chr16-82574161-DEL-139 |
| chr16-82606521-DEL-188 | chr16-82606522-DEL-188 |
| chr16-82623721-DEL-6158 | chr16-82623722-DEL-6158 |
| chr16-82663813-DEL-153 | chr16-82663814-DEL-153 |
| chr16-82668910-DEL-526 | chr16-82668911-DEL-526 |
| chr16-82674114-DEL-3733 | chr16-82674115-DEL-3733 |
| chr16-82682754-DEL-3609 | chr16-82682754-DEL-3609 |
| chr16-82686184-DEL-2986 | chr16-82686185-DEL-2986 |
| chr16-82692849-DEL-192 | chr16-82692850-DEL-192 |
| chr16-82698752-DEL-304 | chr16-82698753-DEL-299 |
| chr16-82706477-DEL-535 | chr16-82706478-DEL-536 |
| chr16-82709041-DEL-6750 | chr16-82709042-DEL-6750 |
| chr16-82739248-DEL-222 | chr16-82739249-DEL-222 |
| chr16-82746352-DEL-6385 | chr16-82746353-DEL-6385 |
| chr16-82807696-DEL-56 | chr16-82807697-DEL-56 |
| chr16-82823208-DEL-389 | chr16-82823209-DEL-389 |
| chr16-82834431-DEL-1421 | chr16-82834432-DEL-1421 |
| chr16-82837532-DEL-6410 | chr16-82837533-DEL-6410 |
| chr16-82859796-DEL-335 | chr16-82859797-DEL-335 |
| chr16-82874044-DEL-6302 | chr16-82874032-DEL-6304 |

| | |
|---|---|
| chr16-82884880-DEL-128 | chr16-82884881-DEL-126 |
| chr16-82910118-DEL-497 | chr16-82910119-DEL-497 |
| chr16-82963868-DEL-477 | chr16-82963865-DEL-478 |
| chr16-82968669-DEL-359 | chr16-82968670-DEL-358 |
| chr16-82974214-DEL-319 | chr16-82974216-DEL-276 |
| chr16-82984892-DEL-106 | chr16-82984884-DEL-114 |
| chr16-82993901-DEL-170 | chr16-82993902-DEL-170 |
| chr16-83031429-DEL-5056 | chr16-83031429-DEL-5056 |
| chr16-83037114-DEL-136 | chr16-83037077-DEL-174 |
| chr16-83047739-DEL-59 | chr16-83047745-DEL-60 |
| chr16-83051581-DEL-79 | chr16-83051582-DEL-79 |
| chr16-83052856-DEL-329 | chr16-83052857-DEL-329 |
| chr16-83058855-DEL-6562 | chr16-83058856-DEL-6562 |
| chr16-83079780-DEL-1309 | chr16-83079781-DEL-1309 |
| chr16-83083285-DEL-63 | chr16-83083290-DEL-64 |
| chr16-83085116-DEL-268 | chr16-83085122-DEL-267 |
| chr16-83100129-DEL-355 | chr16-83100129-DEL-355 |
| chr16-83114549-DEL-6231 | chr16-83114541-DEL-6229 |
| chr16-83132264-DEL-62 | chr16-83132264-DEL-62 |
| chr16-83134698-DEL-2928 | chr16-83134699-DEL-2928 |
| chr16-83136834-DEL-81 | chr16-83136835-DEL-81 |
| chr16-83157597-DEL-7194 | chr16-83157598-DEL-7194 |
| chr16-83177506-DEL-57 | chr16-83177490-DEL-63 |
| chr16-83195899-DEL-4923 | chr16-83195900-DEL-4923 |
| chr16-83202934-DEL-77 | chr16-83202936-DEL-76 |
| chr16-83242664-DEL-187 | chr16-83242664-DEL-187 |
| chr16-83246235-DEL-7045 | chr16-83246236-DEL-7045 |
| chr16-83246717-DEL-201 | chr16-83246718-DEL-201 |
| chr16-83259882-DEL-1185 | chr16-83259884-DEL-1184 |
| chr16-83265287-DEL-2165 | chr16-83265288-DEL-2165 |
| chr16-83280931-DEL-1998 | chr16-83280932-DEL-1998 |
| chr16-83289786-DEL-400 | chr16-83289786-DEL-400 |
| chr16-83290629-DEL-1316 | chr16-83290624-DEL-1317 |
| chr16-83294983-DEL-1963 | chr16-83294984-DEL-1963 |
| chr16-83301238-DEL-139 | chr16-83301233-DEL-140 |
| chr16-83308115-DEL-616 | chr16-83308111-DEL-616 |
| chr16-83310644-DEL-74 | chr16-83310637-DEL-81 |
| chr16-83316469-DEL-2062 | chr16-83316470-DEL-2062 |
| chr16-83364287-DEL-566 | chr16-83364288-DEL-566 |
| chr16-83376220-DEL-1226 | chr16-83376221-DEL-1226 |
| chr16-83393384-DEL-61 | chr16-83393377-DEL-69 |
| chr16-83400272-DEL-64 | chr16-83400273-DEL-64 |
| chr16-83433111-DEL-1856 | chr16-83433112-DEL-1856 |
| chr16-83442758-DEL-406 | chr16-83442759-DEL-406 |
| chr16-83452212-DEL-301 | chr16-83452205-DEL-302 |
| chr16-83468843-DEL-58 | chr16-83468844-DEL-62 |
| chr16-83488676-DEL-50 | chr16-83488667-DEL-60 |
| chr16-83500409-DEL-52 | chr16-83500406-DEL-54 |
| chr16-83503937-DEL-67 | chr16-83503938-DEL-67 |
| chr16-83534270-DEL-70 | chr16-83534264-DEL-71 |
| chr16-83536075-DEL-377 | chr16-83536076-DEL-377 |
| chr16-83545154-DEL-1069 | chr16-83545155-DEL-1069 |
| chr16-83546789-DEL-1927 | chr16-83546790-DEL-1927 |
| chr16-83553022-DEL-202 | chr16-83553038-DEL-164 |
| chr16-83597512-DEL-60 | chr16-83597513-DEL-60 |
| chr16-83603203-DEL-5350 | chr16-83603204-DEL-5350 |
| chr16-83626183-DEL-335 | chr16-83626184-DEL-335 |
| chr16-83659686-DEL-217 | chr16-83659687-DEL-216 |
| chr16-83662874-DEL-247 | chr16-83662875-DEL-247 |
| chr16-83671323-DEL-3191 | chr16-83671306-DEL-3208 |
| chr16-83671977-DEL-895 | chr16-83671964-DEL-896 |
| chr16-83673820-DEL-156 | chr16-83673821-DEL-156 |
| chr16-83676379-DEL-72 | chr16-83676380-DEL-72 |
| chr16-83697506-DEL-96 | chr16-83697517-DEL-106 |
| chr16-83708324-DEL-162 | chr16-83708318-DEL-136 |
| chr16-83717116-DEL-75 | chr16-83717112-DEL-78 |
| chr16-83724424-DEL-52 | chr16-83724432-DEL-61 |
| chr16-83728223-DEL-1277 | chr16-83728224-DEL-1277 |
| chr16-83730929-DEL-1243 | chr16-83730930-DEL-1243 |
| chr16-83743028-DEL-11163 | chr16-83743029-DEL-11163 |
| chr16-83779104-DEL-431 | chr16-83779105-DEL-431 |
| chr16-83856984-DEL-916 | chr16-83856985-DEL-916 |
| chr16-83918649-DEL-75 | chr16-83918658-DEL-61 |
| chr16-83922438-DEL-418 | chr16-83922439-DEL-418 |
| chr16-83924440-DEL-2311 | chr16-83924441-DEL-2311 |
| chr16-83954805-DEL-354 | chr16-83954806-DEL-354 |
| chr16-83978475-DEL-63 | chr16-83978478-DEL-61 |
| chr16-83978628-DEL-1126 | chr16-83978626-DEL-1155 |
| chr16-84054987-DEL-4335 | chr16-84054988-DEL-4335 |
| chr16-84057314-DEL-1903 | chr16-84057315-DEL-1903 |

| | |
|---|---|
| chr16-84073255-DEL-7041 | chr16-84073256-DEL-7041 |
| chr16-84073270-DEL-214 | chr16-84073271-DEL-214 |
| chr16-84105323-DEL-227 | chr16-84105324-DEL-227 |
| chr16-84123676-DEL-2781 | chr16-84123677-DEL-2781 |
| chr16-84124565-DEL-68 | chr16-84124566-DEL-68 |
| chr16-84127197-DEL-1107 | chr16-84127198-DEL-1107 |
| chr16-84128824-DEL-1274 | chr16-84128827-DEL-1273 |
| chr16-84147249-DEL-112 | chr16-84147250-DEL-112 |
| chr16-84147846-DEL-85 | chr16-84147846-DEL-85 |
| chr16-84177423-DEL-6582 | chr16-84177424-DEL-6582 |
| chr16-84200933-DEL-87 | chr16-84200934-DEL-87 |
| chr16-84204945-DEL-77 | chr16-84204903-DEL-78 |
| chr16-84211501-DEL-77 | chr16-84211502-DEL-77 |
| chr16-84214953-DEL-146 | chr16-84214954-DEL-146 |
| chr16-84234235-DEL-54 | chr16-84234228-DEL-62 |
| chr16-84235693-DEL-85 | chr16-84235692-DEL-87 |
| chr16-84236597-DEL-6317 | chr16-84236598-DEL-6317 |
| chr16-84274210-DEL-65 | chr16-84274207-DEL-66 |
| chr16-84302659-DEL-121 | chr16-84302660-DEL-121 |
| chr16-84341044-DEL-888 | chr16-84341045-DEL-888 |
| chr16-84346496-DEL-87 | chr16-84346521-DEL-63 |
| chr16-84355369-DEL-52 | chr16-84355369-DEL-52 |
| chr16-84369717-DEL-393 | chr16-84369766-DEL-393 |
| chr16-84369767-DEL-896 | chr16-84369768-DEL-896 |
| chr16-84371738-DEL-2906 | chr16-84371739-DEL-2906 |
| chr16-84375031-DEL-470 | chr16-84375032-DEL-470 |
| chr16-84378025-DEL-1090 | chr16-84378026-DEL-1090 |
| chr16-84380696-DEL-70 | chr16-84380685-DEL-82 |
| chr16-84398807-DEL-187 | chr16-84398808-DEL-187 |
| chr16-84432920-DEL-1890 | chr16-84432922-DEL-1889 |
| chr16-84451543-DEL-1829 | chr16-84451544-DEL-1829 |
| chr16-84471837-DEL-216 | chr16-84471837-DEL-216 |
| chr16-84507685-DEL-56 | chr16-84507674-DEL-68 |
| chr16-84512269-DEL-237 | chr16-84512270-DEL-237 |
| chr16-84515980-DEL-651 | chr16-84515981-DEL-650 |
| chr16-84526581-DEL-238 | chr16-84526582-DEL-238 |
| chr16-84537233-DEL-82 | chr16-84537226-DEL-117 |
| chr16-84537892-DEL-1836 | chr16-84537893-DEL-1836 |
| chr16-84556086-DEL-111 | chr16-84556114-DEL-112 |
| chr16-84556711-DEL-199 | chr16-84556710-DEL-199 |
| chr16-84557521-DEL-796 | chr16-84557522-DEL-796 |
| chr16-84585787-DEL-183 | chr16-84585788-DEL-183 |
| chr16-84597234-DEL-172 | chr16-84597235-DEL-172 |
| chr16-84613638-DEL-53 | chr16-84613639-DEL-53 |
| chr16-84619246-DEL-51 | chr16-84619247-DEL-51 |
| chr16-84639180-DEL-64 | chr16-84639167-DEL-78 |
| chr16-84659880-DEL-51 | chr16-84659881-DEL-51 |
| chr16-84661614-DEL-1015 | chr16-84661615-DEL-1015 |
| chr16-84663236-DEL-206 | chr16-84663237-DEL-206 |
| chr16-84668152-DEL-132 | chr16-84668138-DEL-147 |
| chr16-84679049-DEL-226 | chr16-84679061-DEL-225 |
| chr16-84690915-DEL-88 | chr16-84690916-DEL-88 |
| chr16-84704498-DEL-920 | chr16-84704499-DEL-920 |
| chr16-84726195-DEL-1369 | chr16-84726196-DEL-1369 |
| chr16-84732663-DEL-7286 | chr16-84732664-DEL-7286 |
| chr16-84775387-DEL-60 | chr16-84775388-DEL-60 |
| chr16-84829846-DEL-192 | chr16-84829847-DEL-192 |
| chr16-84842842-DEL-202 | chr16-84842832-DEL-203 |
| chr16-84894995-DEL-100 | chr16-84894996-DEL-92 |
| chr16-84919710-DEL-1029 | chr16-84919711-DEL-1029 |
| chr16-84933505-DEL-346 | chr16-84933508-DEL-342 |
| chr16-84943380-DEL-245 | chr16-84943381-DEL-245 |
| chr16-84946839-DEL-52 | chr16-84946840-DEL-52 |
| chr16-84961584-DEL-93 | chr16-84961584-DEL-93 |
| chr16-84972743-DEL-86 | chr16-84972744-DEL-86 |
| chr16-84991530-DEL-100 | chr16-84991531-DEL-94 |
| chr16-84999749-DEL-421 | chr16-84999750-DEL-421 |
| chr16-85023920-DEL-2109 | chr16-85023921-DEL-2109 |
| chr16-85034539-DEL-7460 | chr16-85034540-DEL-7460 |
| chr16-85076519-DEL-102 | chr16-85076502-DEL-112 |
| chr16-85106038-DEL-127 | chr16-85106068-DEL-93 |
| chr16-85119402-DEL-500 | chr16-85119402-DEL-500 |
| chr16-85128217-DEL-93 | chr16-85128218-DEL-89 |
| chr16-85129279-DEL-114 | chr16-85129280-DEL-114 |
| chr16-85156680-DEL-814 | chr16-85156678-DEL-815 |
| chr16-85232884-DEL-70 | chr16-85232900-DEL-55 |
| chr16-85236405-DEL-257 | chr16-85236406-DEL-257 |
| chr16-85244581-DEL-64 | chr16-85244581-DEL-64 |
| chr16-85257763-DEL-399 | chr16-85257764-DEL-399 |
| chr16-85273864-DEL-75 | chr16-85273865-DEL-75 |

| | |
|---|---|
| chr16-85294945-DEL-78 | chr16-85294969-DEL-52 |
| chr16-85295507-DEL-56 | chr16-85295508-DEL-56 |
| chr16-85297377-DEL-60 | chr16-85297380-DEL-58 |
| chr16-85306902-DEL-440 | chr16-85306903-DEL-440 |
| chr16-85315511-DEL-63 | chr16-85315487-DEL-59 |
| chr16-85326215-DEL-6255 | chr16-85326216-DEL-6255 |
| chr16-85348825-DEL-2002 | chr16-85348826-DEL-2002 |
| chr16-85400243-DEL-60 | chr16-85400245-DEL-57 |
| chr16-85401511-DEL-65 | chr16-85401512-DEL-65 |
| chr16-85413513-DEL-1149 | chr16-85413514-DEL-1149 |
| chr16-85437426-DEL-161 | chr16-85437439-DEL-136 |
| chr16-85439087-DEL-78 | chr16-85439125-DEL-78 |
| chr16-85441762-DEL-54 | chr16-85441763-DEL-54 |
| chr16-85444908-DEL-154 | chr16-85444909-DEL-154 |
| chr16-85447703-DEL-637 | chr16-85447707-DEL-630 |
| chr16-85451362-DEL-74 | chr16-85451363-DEL-74 |
| chr16-85458354-DEL-652 | chr16-85458355-DEL-652 |
| chr16-85480032-DEL-616 | chr16-85480035-DEL-614 |
| chr16-85510307-DEL-6678 | chr16-85510308-DEL-6678 |
| chr16-85520094-DEL-635 | chr16-85520095-DEL-635 |
| chr16-85532032-DEL-556 | chr16-85532032-DEL-557 |
| chr16-85571148-DEL-396 | chr16-85571149-DEL-396 |
| chr16-85578748-DEL-184 | chr16-85578748-DEL-184 |
| chr16-85579703-DEL-2081 | chr16-85579704-DEL-2081 |
| chr16-85581199-DEL-585 | chr16-85581200-DEL-585 |
| chr16-85582372-DEL-208 | chr16-85582373-DEL-208 |
| chr16-85621894-DEL-363 | chr16-85621894-DEL-363 |
| chr16-85636490-DEL-1516 | chr16-85636491-DEL-1526 |
| chr16-85646165-DEL-1512 | chr16-85646166-DEL-1512 |
| chr16-85660702-DEL-475 | chr16-85660703-DEL-475 |
| chr16-85707014-DEL-78 | chr16-85707045-DEL-96 |
| chr16-85711212-DEL-539 | chr16-85711213-DEL-539 |
| chr16-85712261-DEL-2039 | chr16-85712262-DEL-2039 |
| chr16-85739353-DEL-69 | chr16-85739354-DEL-69 |
| chr16-85755355-DEL-239 | chr16-85755356-DEL-239 |
| chr16-85768330-DEL-497 | chr16-85768331-DEL-497 |
| chr16-85786584-DEL-62 | chr16-85786585-DEL-62 |
| chr16-85797435-DEL-60 | chr16-85797436-DEL-66 |
| chr16-85798217-DEL-2146 | chr16-85798218-DEL-2146 |
| chr16-85830993-DEL-53 | chr16-85830994-DEL-54 |
| chr16-85836530-DEL-367 | chr16-85836531-DEL-378 |
| chr16-85837647-DEL-75 | chr16-85837648-DEL-75 |
| chr16-85846313-DEL-1932 | chr16-85846314-DEL-1932 |
| chr16-85849804-DEL-152 | chr16-85849785-DEL-152 |
| chr16-85853099-DEL-1039 | chr16-85853100-DEL-1039 |
| chr16-85871592-DEL-7063 | chr16-85871593-DEL-7063 |
| chr16-85879495-DEL-261 | chr16-85879497-DEL-260 |
| chr16-85879872-DEL-1317 | chr16-85879877-DEL-1315 |
| chr16-85901528-DEL-65 | chr16-85901527-DEL-80 |
| chr16-85902178-DEL-1912 | chr16-85902179-DEL-1911 |
| chr16-85936654-DEL-156 | chr16-85936655-DEL-155 |
| chr16-85947231-DEL-555 | chr16-85947232-DEL-555 |
| chr16-85960786-DEL-546 | chr16-85960787-DEL-547 |
| chr16-85966906-DEL-51 | chr16-85966894-DEL-64 |
| chr16-85984865-DEL-237 | chr16-85984866-DEL-237 |
| chr16-85992898-DEL-3314 | chr16-85992899-DEL-3314 |
| chr16-86006000-DEL-103 | chr16-86006001-DEL-103 |
| chr16-86013991-DEL-343 | chr16-86013991-DEL-343 |
| chr16-86037185-DEL-71 | chr16-86037186-DEL-71 |
| chr16-86075521-DEL-107 | chr16-86075522-DEL-107 |
| chr16-86094425-DEL-585 | chr16-86094426-DEL-585 |
| chr16-86101781-DEL-348 | chr16-86101781-DEL-348 |
| chr16-86103602-DEL-2300 | chr16-86103603-DEL-2298 |
| chr16-86128597-DEL-6198 | chr16-86128598-DEL-6198 |
| chr16-86171886-DEL-55 | chr16-86171891-DEL-54 |
| chr16-86172411-DEL-376 | chr16-86172412-DEL-376 |
| chr16-86202753-DEL-437 | chr16-86202754-DEL-437 |
| chr16-86213062-DEL-487 | chr16-86213063-DEL-487 |
| chr16-86213954-DEL-2284 | chr16-86213955-DEL-2284 |
| chr16-86220614-DEL-60 | chr16-86220615-DEL-60 |
| chr16-86246936-DEL-864 | chr16-86246921-DEL-865 |
| chr16-86249854-DEL-281 | chr16-86249855-DEL-281 |
| chr16-86252872-DEL-61 | chr16-86252873-DEL-57 |
| chr16-86254028-DEL-895 | chr16-86254029-DEL-895 |
| chr16-86279113-DEL-982 | chr16-86279114-DEL-982 |
| chr16-86286064-DEL-76 | chr16-86286035-DEL-74 |
| chr16-86308119-DEL-1533 | chr16-86308121-DEL-1527 |
| chr16-86321767-DEL-1008 | chr16-86321806-DEL-970 |
| chr16-86331617-DEL-918 | chr16-86331618-DEL-918 |
| chr16-86336351-DEL-2880 | chr16-86336352-DEL-2880 |

| | |
|---|---|
| chr16-86347425-DEL-1629 | chr16-86347422-DEL-1625 |
| chr16-86357399-DEL-102 | chr16-86357400-DEL-102 |
| chr16-86364910-DEL-2289 | chr16-86364911-DEL-2289 |
| chr16-86420845-DEL-470 | chr16-86420846-DEL-470 |
| chr16-86457604-DEL-3481 | chr16-86457605-DEL-3481 |
| chr16-86492467-DEL-310 | chr16-86492468-DEL-310 |
| chr16-86522150-DEL-6396 | chr16-86522151-DEL-6396 |
| chr16-86598423-DEL-1312 | chr16-86598424-DEL-1312 |
| chr16-86602273-DEL-71 | chr16-86602264-DEL-72 |
| chr16-86617669-DEL-428 | chr16-86617670-DEL-428 |
| chr16-86622744-DEL-66 | chr16-86622748-DEL-74 |
| chr16-86626914-DEL-62 | chr16-86626915-DEL-62 |
| chr16-86718871-DEL-198 | chr16-86718872-DEL-198 |
| chr16-86747701-DEL-7291 | chr16-86747702-DEL-7291 |
| chr16-86760092-DEL-2683 | chr16-86760091-DEL-2684 |
| chr16-86772295-DEL-6346 | chr16-86772296-DEL-6346 |
| chr16-86781462-DEL-3962 | chr16-86781463-DEL-3962 |
| chr16-86783965-DEL-184 | chr16-86783965-DEL-185 |
| chr16-86786792-DEL-113 | chr16-86786793-DEL-113 |
| chr16-86793400-DEL-2001 | chr16-86793401-DEL-2001 |
| chr16-86806605-DEL-63 | chr16-86806576-DEL-64 |
| chr16-86825093-DEL-85 | chr16-86825094-DEL-85 |
| chr16-86848790-DEL-346 | chr16-86848791-DEL-346 |
| chr16-86852180-DEL-4698 | chr16-86852180-DEL-4698 |
| chr16-86878738-DEL-82 | chr16-86878739-DEL-82 |
| chr16-86883930-DEL-314 | chr16-86883931-DEL-314 |
| chr16-86888340-DEL-171 | chr16-86888339-DEL-172 |
| chr16-86908992-DEL-6507 | chr16-86908993-DEL-6507 |
| chr16-86921235-DEL-80 | chr16-86921236-DEL-80 |
| chr16-86927708-DEL-59 | chr16-86927673-DEL-60 |
| chr16-86932428-DEL-109 | chr16-86932430-DEL-108 |
| chr16-86946134-DEL-6807 | chr16-86946121-DEL-6806 |
| chr16-86956860-DEL-72 | chr16-86956861-DEL-72 |
| chr16-86959091-DEL-176 | chr16-86959092-DEL-175 |
| chr16-86961731-DEL-160 | chr16-86961732-DEL-160 |
| chr16-86969069-DEL-1125 | chr16-86969070-DEL-1125 |
| chr16-86977513-DEL-4226 | chr16-86977513-DEL-4226 |
| chr16-86984887-DEL-60 | chr16-86984892-DEL-56 |
| chr16-86990094-DEL-6486 | chr16-86990095-DEL-6486 |
| chr16-86998563-DEL-476 | chr16-86998564-DEL-476 |
| chr16-87007366-DEL-538 | chr16-87007365-DEL-538 |
| chr16-87010513-DEL-1487 | chr16-87010523-DEL-1484 |
| chr16-87015441-DEL-116 | chr16-87015441-DEL-116 |
| chr16-87051388-DEL-60 | chr16-87051389-DEL-60 |
| chr16-87073076-DEL-949 | chr16-87073077-DEL-949 |
| chr16-87073086-DEL-192 | chr16-87073087-DEL-192 |
| chr16-87079757-DEL-116 | chr16-87079758-DEL-116 |
| chr16-87084303-DEL-3930 | chr16-87084304-DEL-3930 |
| chr16-87093224-DEL-6407 | chr16-87093225-DEL-6407 |
| chr16-87109477-DEL-53 | chr16-87109478-DEL-53 |
| chr16-87113198-DEL-138 | chr16-87113197-DEL-132 |
| chr16-87149042-DEL-66 | chr16-87148998-DEL-63 |
| chr16-87162115-DEL-214 | chr16-87162097-DEL-206 |
| chr16-87169326-DEL-3774 | chr16-87169348-DEL-3752 |
| chr16-87204701-DEL-201 | chr16-87204702-DEL-201 |
| chr16-87215220-DEL-66 | chr16-87215221-DEL-66 |
| chr16-87220678-DEL-128 | chr16-87220679-DEL-128 |
| chr16-87244917-DEL-582 | chr16-87244918-DEL-582 |
| chr16-87281852-DEL-397 | chr16-87281851-DEL-398 |
| chr16-87295134-DEL-80 | chr16-87295151-DEL-64 |
| chr16-87303688-DEL-74 | chr16-87303690-DEL-72 |
| chr16-87305052-DEL-369 | chr16-87305048-DEL-370 |
| chr16-87312920-DEL-3374 | chr16-87312916-DEL-3375 |
| chr16-87360199-DEL-59 | chr16-87360159-DEL-60 |
| chr16-87367487-DEL-62 | chr16-87367472-DEL-63 |
| chr16-87371596-DEL-74 | chr16-87371597-DEL-74 |
| chr16-87374426-DEL-168 | chr16-87374435-DEL-155 |
| chr16-87386267-DEL-98 | chr16-87386268-DEL-98 |
| chr16-87399015-DEL-64 | chr16-87399016-DEL-64 |
| chr16-87417447-DEL-913 | chr16-87417448-DEL-913 |
| chr16-87433798-DEL-223 | chr16-87433799-DEL-223 |
| chr16-87441778-DEL-54 | chr16-87441771-DEL-62 |
| chr16-87475696-DEL-343 | chr16-87475697-DEL-343 |
| chr16-87499354-DEL-93 | chr16-87499355-DEL-93 |
| chr16-87602350-DEL-1353 | chr16-87602351-DEL-1353 |
| chr16-87624266-DEL-208 | chr16-87624274-DEL-178 |
| chr16-87659445-DEL-1544 | chr16-87659446-DEL-1544 |
| chr16-87669840-DEL-80 | chr16-87669841-DEL-80 |
| chr16-87679995-DEL-551 | chr16-87679996-DEL-551 |
| chr16-87684149-DEL-609 | chr16-87684150-DEL-609 |

| | |
|---|---|
| chr16-87685115-DEL-387 | chr16-87685116-DEL-387 |
| chr16-87696480-DEL-83 | chr16-87696452-DEL-80 |
| chr16-87763684-DEL-54 | chr16-87763683-DEL-54 |
| chr16-87787253-DEL-60 | chr16-87787254-DEL-60 |
| chr16-87789111-DEL-146 | chr16-87789116-DEL-142 |
| chr16-87795185-DEL-683 | chr16-87795186-DEL-683 |
| chr16-87807419-DEL-73 | chr16-87807419-DEL-73 |
| chr16-87819825-DEL-6894 | chr16-87819826-DEL-6894 |
| chr16-87850397-DEL-68 | chr16-87850369-DEL-67 |
| chr16-87857091-DEL-81 | chr16-87857104-DEL-69 |
| chr16-87859197-DEL-6194 | chr16-87859197-DEL-6194 |
| chr16-87859292-DEL-199 | chr16-87859293-DEL-199 |
| chr16-87889353-DEL-98 | chr16-87889363-DEL-117 |
| chr16-87890357-DEL-212 | chr16-87890358-DEL-212 |
| chr16-87900873-DEL-6651 | chr16-87900874-DEL-6651 |
| chr16-87921004-DEL-57 | chr16-87921005-DEL-57 |
| chr16-87924505-DEL-68 | chr16-87924506-DEL-66 |
| chr16-87933497-DEL-74 | chr16-87933498-DEL-74 |
| chr16-87958633-DEL-114 | chr16-87958634-DEL-114 |
| chr16-87974837-DEL-90 | chr16-87974831-DEL-100 |
| chr16-88045295-DEL-153 | chr16-88045297-DEL-151 |
| chr16-88059107-DEL-56 | chr16-88059094-DEL-70 |
| chr16-88079990-DEL-51 | chr16-88079986-DEL-52 |
| chr16-88103539-DEL-944 | chr16-88103540-DEL-944 |
| chr16-88138015-DEL-732 | chr16-88138016-DEL-732 |
| chr16-88140644-DEL-509 | chr16-88140645-DEL-509 |
| chr16-88149566-DEL-824 | chr16-88149567-DEL-824 |
| chr16-88155528-DEL-200 | chr16-88155529-DEL-200 |
| chr16-88192439-DEL-209 | chr16-88192440-DEL-209 |
| chr16-88207184-DEL-424 | chr16-88207192-DEL-431 |
| chr16-88242442-DEL-271 | chr16-88242443-DEL-271 |
| chr16-88250257-DEL-677 | chr16-88250258-DEL-677 |
| chr16-88293443-DEL-203 | chr16-88293421-DEL-250 |
| chr16-88299427-DEL-1052 | chr16-88299428-DEL-1052 |
| chr16-88299926-DEL-66 | chr16-88299907-DEL-70 |
| chr16-88324296-DEL-6652 | chr16-88324297-DEL-6652 |
| chr16-88324517-DEL-212 | chr16-88324518-DEL-212 |
| chr16-88374536-DEL-401 | chr16-88374537-DEL-401 |
| chr16-88381393-DEL-56 | chr16-88381402-DEL-54 |
| chr16-88384559-DEL-52 | chr16-88384560-DEL-52 |
| chr16-88387115-DEL-429 | chr16-88387116-DEL-429 |
| chr16-88410138-DEL-72 | chr16-88410119-DEL-72 |
| chr16-88411838-DEL-82 | chr16-88411853-DEL-68 |
| chr16-88459002-DEL-2280 | chr16-88459003-DEL-2280 |
| chr16-88476362-DEL-58 | chr16-88476363-DEL-58 |
| chr16-88486432-DEL-60 | chr16-88486433-DEL-60 |
| chr16-88492563-DEL-59 | chr16-88492563-DEL-59 |
| chr16-88501618-DEL-680 | chr16-88501618-DEL-680 |
| chr16-88513639-DEL-56 | chr16-88513640-DEL-56 |
| chr16-88526883-DEL-56 | chr16-88526884-DEL-56 |
| chr16-88563203-DEL-282 | chr16-88563204-DEL-282 |
| chr16-88565755-DEL-55 | chr16-88565730-DEL-56 |
| chr16-88570803-DEL-60 | chr16-88570804-DEL-60 |
| chr16-88583250-DEL-77 | chr16-88583245-DEL-71 |
| chr16-88587519-DEL-7201 | chr16-88587520-DEL-7201 |
| chr16-88609930-DEL-530 | chr16-88609936-DEL-528 |
| chr16-88630720-DEL-888 | chr16-88630721-DEL-885 |
| chr16-88670779-DEL-153 | chr16-88670780-DEL-153 |
| chr16-88678818-DEL-263 | chr16-88678819-DEL-263 |
| chr16-88680507-DEL-52 | chr16-88680490-DEL-52 |
| chr16-88684397-DEL-463 | chr16-88684398-DEL-463 |
| chr16-88718902-DEL-142 | chr16-88718903-DEL-142 |
| chr16-88724476-DEL-233 | chr16-88724485-DEL-230 |
| chr16-88733336-DEL-200 | chr16-88733337-DEL-200 |
| chr16-88738325-DEL-947 | chr16-88738329-DEL-947 |
| chr16-88751087-DEL-6183 | chr16-88751087-DEL-6183 |
| chr16-88762096-DEL-3894 | chr16-88762094-DEL-3895 |
| chr16-88762213-DEL-62 | chr16-88762214-DEL-62 |
| chr16-88782228-DEL-73 | chr16-88782229-DEL-73 |
| chr16-88815402-DEL-80 | chr16-88815403-DEL-80 |
| chr16-88820120-DEL-3142 | chr16-88820121-DEL-3142 |
| chr16-88857201-DEL-70 | chr16-88857213-DEL-94 |
| chr16-88922485-DEL-778 | chr16-88922486-DEL-778 |
| chr16-88924593-DEL-60 | chr16-88924592-DEL-62 |
| chr16-88928894-DEL-74 | chr16-88928895-DEL-64 |
| chr16-88934840-DEL-1343 | chr16-88934841-DEL-1343 |
| chr16-88942123-DEL-4537 | chr16-88942124-DEL-4537 |
| chr16-88950521-DEL-118 | chr16-88950553-DEL-87 |
| chr16-88958847-DEL-7116 | chr16-88958848-DEL-7116 |
| chr16-88986567-DEL-7810 | chr16-88986551-DEL-7826 |

| | |
|---|---|
| chr16-88997682-DEL-1321 | chr16-88997666-DEL-1337 |
| chr16-89013856-DEL-910 | chr16-89013857-DEL-910 |
| chr16-89085345-DEL-72 | chr16-89085362-DEL-81 |
| chr16-89098619-DEL-78 | chr16-89098613-DEL-63 |
| chr16-89117213-DEL-4625 | chr16-89117214-DEL-4625 |
| chr16-89126010-DEL-7507 | chr16-89126011-DEL-7507 |
| chr16-89135380-DEL-400 | chr16-89135381-DEL-400 |
| chr16-89143520-DEL-241 | chr16-89143521-DEL-241 |
| chr16-89148960-DEL-116 | chr16-89148981-DEL-96 |
| chr16-89150570-DEL-1015 | chr16-89150571-DEL-1015 |
| chr16-89178547-DEL-154 | chr16-89178552-DEL-155 |
| chr16-89205985-DEL-65 | chr16-89205989-DEL-64 |
| chr16-89208349-DEL-528 | chr16-89208350-DEL-528 |
| chr16-89209535-DEL-631 | chr16-89209536-DEL-631 |
| chr16-89216723-DEL-120 | chr16-89216724-DEL-120 |
| chr16-89223308-DEL-730 | chr16-89223309-DEL-729 |
| chr16-89228644-DEL-3030 | chr16-89228645-DEL-3030 |
| chr16-89239789-DEL-5364 | chr16-89239790-DEL-5364 |
| chr16-89250707-DEL-1491 | chr16-89250671-DEL-1491 |
| chr16-89253717-DEL-3572 | chr16-89253718-DEL-3572 |
| chr16-89254851-DEL-598 | chr16-89254865-DEL-582 |
| chr16-89311837-DEL-481 | chr16-89311838-DEL-481 |
| chr16-89315464-DEL-59 | chr16-89315465-DEL-59 |
| chr16-89320135-DEL-60 | chr16-89320136-DEL-60 |
| chr16-89330329-DEL-87 | chr16-89330332-DEL-88 |
| chr16-89331348-DEL-53 | chr16-89331347-DEL-54 |
| chr16-89336448-DEL-131 | chr16-89336449-DEL-131 |
| chr16-89355001-DEL-84 | chr16-89355002-DEL-84 |
| chr16-89355580-DEL-135 | chr16-89355581-DEL-135 |
| chr16-89379192-DEL-647 | chr16-89379212-DEL-622 |
| chr16-89384831-DEL-60 | chr16-89384832-DEL-60 |
| chr16-89415739-DEL-66 | chr16-89415742-DEL-63 |
| chr16-89430601-DEL-98 | chr16-89430596-DEL-71 |
| chr16-89432992-DEL-214 | chr16-89432993-DEL-214 |
| chr16-89445019-DEL-195 | chr16-89445020-DEL-195 |
| chr16-89482685-DEL-82 | chr16-89482686-DEL-82 |
| chr16-89505874-DEL-71 | chr16-89505875-DEL-71 |
| chr16-89522268-DEL-80 | chr16-89522269-DEL-80 |
| chr16-89529934-DEL-60 | chr16-89529935-DEL-60 |
| chr16-89564077-DEL-212 | chr16-89564078-DEL-212 |
| chr16-89591837-DEL-126 | chr16-89591838-DEL-126 |
| chr16-89597939-DEL-212 | chr16-89597940-DEL-212 |
| chr16-89599635-DEL-55 | chr16-89599636-DEL-55 |
| chr16-89600619-DEL-450 | chr16-89600620-DEL-450 |
| chr16-89639533-DEL-96 | chr16-89639533-DEL-132 |
| chr16-89651011-DEL-80 | chr16-89651012-DEL-80 |
| chr16-89665338-DEL-83 | chr16-89665339-DEL-83 |
| chr16-89701637-DEL-314 | chr16-89701625-DEL-316 |
| chr16-89726232-DEL-3777 | chr16-89726233-DEL-3777 |
| chr16-89760140-DEL-1300 | chr16-89760141-DEL-1300 |
| chr16-89767453-DEL-1063 | chr16-89767424-DEL-1064 |
| chr16-89793432-DEL-2735 | chr16-89793433-DEL-2735 |
| chr16-89796512-DEL-870 | chr16-89796513-DEL-870 |
| chr16-89825737-DEL-222 | chr16-89825738-DEL-222 |
| chr16-89853174-DEL-7260 | chr16-89853175-DEL-7260 |
| chr16-89862423-DEL-3616 | chr16-89862424-DEL-3616 |
| chr16-89903048-DEL-195 | chr16-89903049-DEL-195 |
| chr16-89924946-DEL-423 | chr16-89924946-DEL-423 |
| chr16-89927668-DEL-236 | chr16-89927658-DEL-242 |
| chr16-89934104-DEL-64 | chr16-89934088-DEL-60 |
| chr16-89943350-DEL-2069 | chr16-89943351-DEL-2069 |
| chr16-89949392-DEL-216 | chr16-89949378-DEL-208 |
| chr16-89953178-DEL-144 | chr16-89953201-DEL-122 |
| chr16-89971824-DEL-218 | chr16-89971863-DEL-206 |
| chr16-89974889-DEL-92 | chr16-89974890-DEL-69 |
| chr16-89979161-DEL-75 | chr16-89979168-DEL-69 |
| chr16-89987277-DEL-121 | chr16-89987294-DEL-109 |
| chr16-90013796-DEL-220 | chr16-90013797-DEL-220 |
| chr16-90038627-DEL-81 | chr16-90038614-DEL-70 |
| chr16-90056252-DEL-71 | chr16-90056253-DEL-71 |
| chr16-90070125-DEL-206 | chr16-90070126-DEL-206 |
| chr16-90089787-DEL-955 | chr16-90089788-DEL-955 |
| chr16-90091336-DEL-264 | chr16-90091337-DEL-265 |
| chr16-90091562-DEL-2479 | chr16-90091562-DEL-2479 |
| chr16-90093832-DEL-208 | chr16-90093833-DEL-208 |
| chr16-90097081-DEL-189 | chr16-90097082-DEL-189 |
| chr16-90098202-DEL-437 | chr16-90098203-DEL-437 |
| chr16-90105780-DEL-394 | chr16-90105781-DEL-394 |
| chr16-90109197-DEL-212 | chr16-90109198-DEL-211 |
| chr16-90119387-DEL-206 | chr16-90119388-DEL-206 |

| | |
|---|---|
| chr16-90138425-DEL-8335 | chr16-90138426-DEL-8335 |
| chr16-90152689-DEL-113 | chr16-90152721-DEL-82 |
| chr16-90166063-DEL-112 | chr16-90166062-DEL-113 |
| chr16-90172478-DEL-52 | chr16-90172475-DEL-54 |
| chr16-90195692-DEL-90 | chr16-90195693-DEL-90 |
| chr16-90200719-DEL-144 | chr16-90200720-DEL-144 |
| chr16-90235435-DEL-2023 | chr16-90235436-DEL-2023 |
| chr16-90245888-DEL-71 | chr16-90245888-DEL-71 |
| chr16-90287387-DEL-93 | chr16-90287368-DEL-102 |
| chr16-90287960-DEL-127 | chr16-90287968-DEL-128 |
| chr16-90292448-DEL-54 | chr16-90292449-DEL-54 |
| chr16-90313225-DEL-124 | chr16-90313238-DEL-126 |
| chr16-90361654-DEL-234 | chr16-90361655-DEL-234 |
| chr16-90385761-DEL-137 | chr16-90385762-DEL-137 |
| chr16-90386034-DEL-547 | chr16-90386036-DEL-548 |
| chr16-90426469-DEL-65 | chr16-90426469-DEL-65 |
| chr16-90430004-DEL-228 | chr16-90430002-DEL-216 |
| chr16-90439931-DEL-186 | chr16-90439926-DEL-192 |
| chr16-90466196-DEL-74 | chr16-90466197-DEL-74 |
| chr16-90472073-DEL-401 | chr16-90472071-DEL-402 |
| chr16-90479128-DEL-99 | chr16-90479129-DEL-99 |
| chr16-90487771-DEL-203 | chr16-90487772-DEL-203 |
| chr16-90546871-DEL-52 | chr16-90546872-DEL-52 |
| chr16-90547576-DEL-75 | chr16-90547577-DEL-75 |
| chr16-90576819-DEL-131 | chr16-90576818-DEL-131 |
| chr16-90590300-DEL-75 | chr16-90590301-DEL-75 |
| chr16-90636319-DEL-1013 | chr16-90636320-DEL-1013 |
| chr16-90643442-DEL-74 | chr16-90643443-DEL-74 |
| chr16-90667342-DEL-210 | chr16-90667343-DEL-210 |
| chr16-90670995-DEL-83 | chr16-90670995-DEL-83 |
| chr16-90686414-DEL-67 | chr16-90686416-DEL-50 |
| chr16-90691851-DEL-56 | chr16-90691848-DEL-60 |
| chr16-90692549-DEL-289 | chr16-90692550-DEL-289 |
| chr16-90693485-DEL-58 | chr16-90693464-DEL-90 |
| chr16-90704371-DEL-65 | chr16-90704372-DEL-65 |
| chr16-90708009-DEL-115 | chr16-90708010-DEL-115 |
| chr16-90729001-DEL-53 | chr16-90729002-DEL-53 |
| chr16-90737598-DEL-280 | chr16-90737599-DEL-280 |
| chr16-90750881-DEL-55 | chr16-90750882-DEL-55 |
| chr16-90781676-DEL-216 | chr16-90781677-DEL-216 |
| chr16-90793076-DEL-225 | chr16-90793077-DEL-225 |
| chr16-90834608-DEL-138 | chr16-90834634-DEL-98 |
| chr16-90870401-DEL-56 | chr16-90870381-DEL-70 |
| chr16-90874418-DEL-609 | chr16-90874419-DEL-609 |
| chr16-90899887-DEL-68 | chr16-90899888-DEL-68 |
| chr16-90908617-DEL-174 | chr16-90908633-DEL-148 |
| chr16-90926488-DEL-103 | chr16-90926515-DEL-72 |
| chr16-90946289-DEL-63 | chr16-90946289-DEL-59 |
| chr16-90961835-DEL-116 | chr16-90961836-DEL-116 |
| chr16-90963470-DEL-408 | chr16-90963471-DEL-408 |
| chr16-90982401-DEL-144 | chr16-90982402-DEL-144 |
| chr16-90992367-DEL-108 | chr16-90992402-DEL-68 |
| chr16-90994728-DEL-191 | chr16-90994729-DEL-191 |
| chr16-90999574-DEL-86 | chr16-90999575-DEL-86 |
| chr16-91011780-DEL-1021 | chr16-91011781-DEL-1021 |
| chr16-91028747-DEL-134 | chr16-91028750-DEL-132 |
| chr16-91044510-DEL-130 | chr16-91044511-DEL-130 |
| chr16-91045223-DEL-7483 | chr16-91045224-DEL-7483 |
| chr16-91101632-DEL-75 | chr16-91101636-DEL-78 |
| chr16-91122467-DEL-56 | chr16-91122468-DEL-56 |
| chr16-91130303-DEL-89 | chr16-91130300-DEL-87 |
| chr16-91143932-DEL-52 | chr16-91143923-DEL-62 |
| chr16-91147649-DEL-149 | chr16-91147650-DEL-151 |
| chr16-91168937-DEL-138 | chr16-91168938-DEL-138 |
| chr16-91199887-DEL-51 | chr16-91199888-DEL-51 |
| chr16-91217871-DEL-272 | chr16-91217890-DEL-270 |
| chr16-91232526-DEL-60 | chr16-91232529-DEL-74 |
| chr16-91234874-DEL-206 | chr16-91234875-DEL-206 |
| chr16-91235434-DEL-168 | chr16-91235455-DEL-186 |
| chr16-91253859-DEL-495 | chr16-91253860-DEL-495 |
| chr16-91260297-DEL-53 | chr16-91260297-DEL-59 |
| chr16-91276529-DEL-462 | chr16-91276530-DEL-462 |
| chr16-91305562-DEL-1065 | chr16-91305563-DEL-1065 |
| chr16-91312177-DEL-180 | chr16-91312177-DEL-184 |
| chr16-91314609-DEL-113 | chr16-91314610-DEL-113 |
| chr16-91315103-DEL-204 | chr16-91315104-DEL-204 |
| chr16-91347612-DEL-216 | chr16-91347613-DEL-216 |
| chr16-91380413-DEL-418 | chr16-91380414-DEL-418 |
| chr16-91442215-DEL-211 | chr16-91442216-DEL-211 |
| chr16-91477315-DEL-204 | chr16-91477317-DEL-195 |

| | |
|---|---|
| chr16-91490345-DEL-120 | chr16-91490346-DEL-120 |
| chr16-91508051-DEL-203 | chr16-91508052-DEL-203 |
| chr16-91508627-DEL-193 | chr16-91508616-DEL-194 |
| chr16-91512943-DEL-215 | chr16-91512944-DEL-215 |
| chr16-91535109-DEL-1253 | chr16-91535107-DEL-1254 |
| chr16-91543992-DEL-76 | chr16-91544023-DEL-70 |
| chr16-91604483-DEL-70 | chr16-91604442-DEL-68 |
| chr16-91614873-DEL-63 | chr16-91614874-DEL-63 |
| chr16-91620387-DEL-853 | chr16-91620388-DEL-853 |
| chr16-91621303-DEL-402 | chr16-91621304-DEL-377 |
| chr16-91627394-DEL-81 | chr16-91627395-DEL-81 |
| chr16-91675028-DEL-62 | chr16-91675025-DEL-63 |
| chr16-91687804-DEL-56 | chr16-91687805-DEL-56 |
| chr16-91699330-DEL-1126 | chr16-91699331-DEL-1126 |
| chr16-91718147-DEL-72 | chr16-91718148-DEL-72 |
| chr16-91728928-DEL-152 | chr16-91728929-DEL-152 |
| chr16-91730318-DEL-143 | chr16-91730319-DEL-143 |
| chr16-91769976-DEL-194 | chr16-91769977-DEL-194 |
| chr16-91770813-DEL-212 | chr16-91770814-DEL-212 |
| chr16-91783701-DEL-115 | chr16-91783703-DEL-90 |
| chr16-91799779-DEL-65 | chr16-91799816-DEL-65 |
| chr16-91807135-DEL-67 | chr16-91807136-DEL-67 |
| chr16-91845510-DEL-85 | chr16-91845511-DEL-85 |
| chr16-91861764-DEL-60 | chr16-91861790-DEL-57 |
| chr16-91900019-DEL-246 | chr16-91900020-DEL-246 |
| chr16-91905975-DEL-74 | chr16-91905959-DEL-87 |
| chr16-91915310-DEL-201 | chr16-91915311-DEL-201 |
| chr16-91927657-DEL-87 | chr16-91927658-DEL-87 |
| chr16-91934426-DEL-63 | chr16-91934427-DEL-63 |
| chr16-91941147-DEL-1888 | chr16-91941148-DEL-1888 |
| chr16-91969430-DEL-70 | chr16-91969431-DEL-70 |
| chr16-91984754-DEL-143 | chr16-91984755-DEL-143 |
| chr16-92004406-DEL-63 | chr16-92004407-DEL-63 |
| chr16-92007191-DEL-94 | chr16-92007194-DEL-92 |
| chr16-92029095-DEL-205 | chr16-92029096-DEL-205 |
| chr16-92054319-DEL-1890 | chr16-92054320-DEL-1890 |
| chr16-92078230-DEL-1328 | chr16-92078231-DEL-1328 |
| chr16-92134624-DEL-88 | chr16-92134625-DEL-88 |
| chr16-92147163-DEL-1375 | chr16-92147164-DEL-1373 |
| chr16-92163291-DEL-497 | chr16-92163292-DEL-497 |
| chr16-92169863-DEL-557 | chr16-92169864-DEL-557 |
| chr16-92186625-DEL-124 | chr16-92186642-DEL-112 |
| chr16-92210797-DEL-1175 | chr16-92210798-DEL-1175 |
| chr16-92272836-DEL-176 | chr16-92272837-DEL-176 |
| chr16-92275012-DEL-380 | chr16-92275013-DEL-380 |
| chr16-92309615-DEL-632 | chr16-92309611-DEL-632 |
| chr16-92311859-DEL-132 | chr16-92311860-DEL-132 |
| chr16-92316424-DEL-51 | chr16-92316425-DEL-51 |
| chr16-92318143-DEL-81 | chr16-92318144-DEL-81 |
| chr16-92371238-DEL-216 | chr16-92371239-DEL-216 |
| chr16-92403097-DEL-110 | chr16-92403098-DEL-110 |
| chr16-92457720-DEL-58 | chr16-92457721-DEL-58 |
| chr16-92499385-DEL-51 | chr16-92499364-DEL-72 |
| chr16-92506227-DEL-53 | chr16-92506228-DEL-53 |
| chr16-92519209-DEL-64 | chr16-92519210-DEL-64 |
| chr16-92520250-DEL-345 | chr16-92520251-DEL-345 |
| chr16-92562318-DEL-54 | chr16-92562319-DEL-54 |
| chr16-92587190-DEL-344 | chr16-92587191-DEL-344 |
| chr16-92599572-DEL-84 | chr16-92599573-DEL-52 |
| chr16-92605540-DEL-160 | chr16-92605532-DEL-158 |
| chr16-92609103-DEL-159 | chr16-92609104-DEL-159 |
| chr16-92619919-DEL-58 | chr16-92619920-DEL-56 |
| chr16-92663292-DEL-53 | chr16-92663293-DEL-53 |
| chr16-92685956-DEL-141 | chr16-92685957-DEL-141 |
| chr16-92714216-DEL-1440 | chr16-92714217-DEL-1440 |
| chr16-92795167-DEL-76 | chr16-92795153-DEL-86 |
| chr16-92796018-DEL-194 | chr16-92796019-DEL-194 |
| chr16-92827711-DEL-200 | chr16-92827712-DEL-200 |
| chr16-92855862-DEL-512 | chr16-92855866-DEL-511 |
| chr16-92864112-DEL-337 | chr16-92864113-DEL-337 |
| chr16-92866104-DEL-125 | chr16-92866102-DEL-128 |
| chr16-92890426-DEL-102 | chr16-92890427-DEL-102 |
| chr16-92922695-DEL-62 | chr16-92922741-DEL-54 |
| chr16-92924421-DEL-60 | chr16-92924397-DEL-66 |
| chr16-92931939-DEL-558 | chr16-92931940-DEL-558 |
| chr16-92933754-DEL-65 | chr16-92933755-DEL-65 |
| chr16-92943203-DEL-386 | chr16-92943240-DEL-372 |
| chr16-92967526-DEL-129 | chr16-92967506-DEL-126 |
| chr16-93042077-DEL-120 | chr16-93042074-DEL-124 |
| chr16-93088384-DEL-131 | chr16-93088364-DEL-161 |

| | |
|---|---|
| chr16-93105859-DEL-83 | chr16-93105858-DEL-84 |
| chr16-93110704-DEL-2055 | chr16-93110705-DEL-2055 |
| chr16-93185838-DEL-72 | chr16-93185853-DEL-70 |
| chr16-93190851-DEL-51 | chr16-93190852-DEL-51 |
| chr16-93195784-DEL-58 | chr16-93195784-DEL-53 |
| chr16-93202303-DEL-59 | chr16-93202304-DEL-58 |
| chr16-93217245-DEL-64 | chr16-93217246-DEL-64 |
| chr16-93234016-DEL-180 | chr16-93234065-DEL-132 |
| chr16-93276813-DEL-99 | chr16-93276842-DEL-100 |
| chr16-93281183-DEL-53 | chr16-93281184-DEL-53 |
| chr16-93286812-DEL-187 | chr16-93286813-DEL-186 |
| chr16-93297713-DEL-105 | chr16-93297714-DEL-105 |
| chr16-93310069-DEL-95 | chr16-93310070-DEL-95 |
| chr16-93378442-DEL-1892 | chr16-93378443-DEL-1892 |
| chr16-93413824-DEL-6030 | chr16-93413826-DEL-6029 |
| chr16-93422959-DEL-104 | chr16-93422959-DEL-104 |
| chr16-93432728-DEL-4036 | chr16-93432728-DEL-4036 |
| chr16-93446138-DEL-1122 | chr16-93446139-DEL-1122 |
| chr16-93448514-DEL-125 | chr16-93448515-DEL-125 |
| chr16-93459438-DEL-2945 | chr16-93459439-DEL-2945 |
| chr16-93465773-DEL-182 | chr16-93465777-DEL-181 |
| chr16-93473911-DEL-153 | chr16-93473895-DEL-196 |
| chr16-93477389-DEL-72 | chr16-93477390-DEL-72 |
| chr16-93486069-DEL-62 | chr16-93486070-DEL-62 |
| chr16-93494344-DEL-8284 | chr16-93494344-DEL-8284 |
| chr16-93571435-DEL-865 | chr16-93571436-DEL-865 |
| chr16-93581691-DEL-69 | chr16-93581695-DEL-67 |
| chr16-93586135-DEL-141 | chr16-93586118-DEL-159 |
| chr16-93587233-DEL-64 | chr16-93587234-DEL-64 |
| chr16-93602793-DEL-66 | chr16-93602794-DEL-66 |
| chr16-93609293-DEL-106 | chr16-93609273-DEL-127 |
| chr16-93621046-DEL-667 | chr16-93621047-DEL-667 |
| chr16-93623055-DEL-178 | chr16-93623056-DEL-178 |
| chr16-93643815-DEL-100 | chr16-93643816-DEL-100 |
| chr16-93655110-DEL-206 | chr16-93655102-DEL-210 |
| chr16-93673213-DEL-219 | chr16-93673214-DEL-219 |
| chr16-93693112-DEL-58 | chr16-93693115-DEL-59 |
| chr16-93703984-DEL-616 | chr16-93703985-DEL-616 |
| chr16-93704966-DEL-183 | chr16-93704967-DEL-183 |
| chr16-93708570-DEL-153 | chr16-93708571-DEL-152 |
| chr16-93711107-DEL-83 | chr16-93711108-DEL-83 |
| chr16-93721881-DEL-95 | chr16-93721882-DEL-95 |
| chr16-93729104-DEL-53 | chr16-93729097-DEL-61 |
| chr16-93740847-DEL-62 | chr16-93740848-DEL-62 |
| chr16-93769281-DEL-82 | chr16-93769282-DEL-82 |
| chr16-93784871-DEL-468 | chr16-93784872-DEL-468 |
| chr16-93814315-DEL-100 | chr16-93814316-DEL-50 |
| chr16-93823603-DEL-52 | chr16-93823604-DEL-52 |
| chr16-93825718-DEL-203 | chr16-93825719-DEL-203 |
| chr16-93843424-DEL-676 | chr16-93843425-DEL-676 |
| chr16-93873154-DEL-87 | chr16-93873187-DEL-86 |
| chr16-93910678-DEL-52 | chr16-93910679-DEL-52 |
| chr16-93966522-DEL-160 | chr16-93966520-DEL-159 |
| chr16-94001155-DEL-66 | chr16-94001156-DEL-66 |
| chr16-94031076-DEL-205 | chr16-94031077-DEL-203 |
| chr16-94040957-DEL-81 | chr16-94040958-DEL-78 |
| chr16-94042405-DEL-1847 | chr16-94042406-DEL-1847 |
| chr16-94045760-DEL-74 | chr16-94045760-DEL-71 |
| chr16-94054356-DEL-204 | chr16-94054357-DEL-204 |
| chr16-94075770-DEL-7047 | chr16-94075767-DEL-7050 |
| chr16-94094374-DEL-161 | chr16-94094375-DEL-161 |
| chr16-94140555-DEL-338 | chr16-94140556-DEL-338 |
| chr16-94158049-DEL-2112 | chr16-94158047-DEL-2114 |
| chr16-94159099-DEL-74 | chr16-94159134-DEL-76 |
| chr16-94162805-DEL-54 | chr16-94162806-DEL-54 |
| chr16-94192455-DEL-212 | chr16-94192456-DEL-212 |
| chr16-94233110-DEL-86 | chr16-94233111-DEL-86 |
| chr16-94251055-DEL-79 | chr16-94251056-DEL-79 |
| chr16-94261570-DEL-189 | chr16-94261556-DEL-203 |
| chr16-94274803-DEL-168 | chr16-94274804-DEL-164 |
| chr16-94281864-DEL-1871 | chr16-94281865-DEL-1871 |
| chr16-94291011-DEL-222 | chr16-94291012-DEL-222 |
| chr16-94309979-DEL-68 | chr16-94309980-DEL-68 |
| chr16-94325422-DEL-191 | chr16-94325423-DEL-191 |
| chr16-94328767-DEL-2010 | chr16-94328768-DEL-2010 |
| chr16-94332469-DEL-549 | chr16-94332470-DEL-549 |
| chr16-94337000-DEL-91 | chr16-94337001-DEL-91 |
| chr16-94344107-DEL-264 | chr16-94344115-DEL-259 |
| chr16-94355937-DEL-197 | chr16-94355934-DEL-198 |
| chr16-94409058-DEL-55 | chr16-94409057-DEL-56 |

| | |
|---|---|
| chr16-94426878-DEL-258 | chr16-94426879-DEL-258 |
| chr16-94429651-DEL-82 | chr16-94429652-DEL-82 |
| chr16-94475920-DEL-207 | chr16-94475921-DEL-207 |
| chr16-94477402-DEL-240 | chr16-94477403-DEL-240 |
| chr16-94501285-DEL-2340 | chr16-94501286-DEL-2340 |
| chr16-94520726-DEL-88 | chr16-94520749-DEL-66 |
| chr16-94528101-DEL-129 | chr16-94528117-DEL-128 |
| chr16-94539041-DEL-568 | chr16-94539042-DEL-568 |
| chr16-94628698-DEL-263 | chr16-94628699-DEL-263 |
| chr16-94670339-DEL-54 | chr16-94670320-DEL-55 |
| chr16-94672227-DEL-58 | chr16-94672228-DEL-50 |
| chr16-94680990-DEL-58 | chr16-94680991-DEL-58 |
| chr16-94693094-DEL-147 | chr16-94693095-DEL-147 |
| chr16-94713137-DEL-88 | chr16-94713136-DEL-107 |
| chr16-94731116-DEL-1121 | chr16-94731117-DEL-1121 |
| chr16-94742691-DEL-54 | chr16-94742683-DEL-63 |
| chr16-94744836-DEL-576 | chr16-94744837-DEL-576 |
| chr16-94750982-DEL-466 | chr16-94750974-DEL-467 |
| chr16-94760391-DEL-120 | chr16-94760392-DEL-120 |
| chr16-94811883-DEL-6267 | chr16-94811884-DEL-6267 |
| chr16-94833534-DEL-78 | chr16-94833535-DEL-78 |
| chr16-94854724-DEL-62 | chr16-94854716-DEL-51 |
| chr16-94860188-DEL-78 | chr16-94860235-DEL-66 |
| chr16-94873613-DEL-537 | chr16-94873614-DEL-537 |
| chr16-94883116-DEL-6497 | chr16-94883117-DEL-6497 |
| chr16-94891552-DEL-417 | chr16-94891553-DEL-417 |
| chr16-94898232-DEL-271 | chr16-94898233-DEL-271 |
| chr16-94902986-DEL-312 | chr16-94902970-DEL-341 |
| chr16-94910371-DEL-80 | chr16-94910372-DEL-80 |
| chr16-94928189-DEL-85 | chr16-94928190-DEL-85 |
| chr16-94931499-DEL-131 | chr16-94931500-DEL-131 |
| chr16-94945984-DEL-92 | chr16-94945964-DEL-116 |
| chr16-94972285-DEL-670 | chr16-94972286-DEL-670 |
| chr16-94973130-DEL-58 | chr16-94973131-DEL-58 |
| chr16-94988665-DEL-2159 | chr16-94988666-DEL-2159 |
| chr16-94990074-DEL-61 | chr16-94990075-DEL-61 |
| chr16-94995466-DEL-70 | chr16-94995467-DEL-70 |
| chr16-95039237-DEL-66 | chr16-95039238-DEL-66 |
| chr16-95054963-DEL-53 | chr16-95054964-DEL-53 |
| chr16-95058696-DEL-105 | chr16-95058697-DEL-105 |
| chr16-95089956-DEL-62 | chr16-95089957-DEL-62 |
| chr16-95126800-DEL-211 | chr16-95126801-DEL-211 |
| chr16-95134425-DEL-57 | chr16-95134426-DEL-57 |
| chr16-95145299-DEL-72 | chr16-95145299-DEL-72 |
| chr16-95193212-DEL-350 | chr16-95193209-DEL-350 |
| chr16-95194781-DEL-1909 | chr16-95194782-DEL-1909 |
| chr16-95202082-DEL-959 | chr16-95202082-DEL-960 |
| chr16-95209600-DEL-53 | chr16-95209601-DEL-53 |
| chr16-95245990-DEL-508 | chr16-95245991-DEL-508 |
| chr16-95271772-DEL-2653 | chr16-95271773-DEL-2653 |
| chr16-95304880-DEL-80 | chr16-95304884-DEL-66 |
| chr16-95309791-DEL-8082 | chr16-95309792-DEL-8082 |
| chr16-95348554-DEL-74 | chr16-95348555-DEL-74 |
| chr16-95354097-DEL-156 | chr16-95354064-DEL-195 |
| chr16-95362510-DEL-3596 | chr16-95362511-DEL-3596 |
| chr16-95369295-DEL-102 | chr16-95369296-DEL-102 |
| chr16-95401143-DEL-60 | chr16-95401103-DEL-100 |
| chr16-95513932-DEL-802 | chr16-95513933-DEL-802 |
| chr16-95539413-DEL-110 | chr16-95539414-DEL-110 |
| chr16-95557449-DEL-586 | chr16-95557450-DEL-586 |
| chr16-95578094-DEL-208 | chr16-95578095-DEL-208 |
| chr16-95583680-DEL-6454 | chr16-95583681-DEL-6454 |
| chr16-95611533-DEL-75 | chr16-95611533-DEL-76 |
| chr16-95620194-DEL-200 | chr16-95620193-DEL-218 |
| chr16-95628263-DEL-77 | chr16-95628252-DEL-89 |
| chr16-95666526-DEL-108 | chr16-95666527-DEL-108 |
| chr16-95667602-DEL-56 | chr16-95667603-DEL-56 |
| chr16-95675799-DEL-136 | chr16-95675800-DEL-136 |
| chr16-95707191-DEL-6937 | chr16-95707191-DEL-6937 |
| chr16-95713427-DEL-609 | chr16-95713428-DEL-609 |
| chr16-95721570-DEL-75 | chr16-95721571-DEL-75 |
| chr16-95725626-DEL-156 | chr16-95725622-DEL-161 |
| chr16-95744113-DEL-190 | chr16-95744103-DEL-201 |
| chr16-95745491-DEL-126 | chr16-95745492-DEL-126 |
| chr16-95752505-DEL-181 | chr16-95752491-DEL-182 |
| chr16-95760700-DEL-64 | chr16-95760707-DEL-65 |
| chr16-95761716-DEL-211 | chr16-95761716-DEL-211 |
| chr16-95765534-DEL-66 | chr16-95765519-DEL-68 |
| chr16-95812139-DEL-396 | chr16-95812140-DEL-396 |
| chr16-95814273-DEL-54 | chr16-95814269-DEL-52 |

| | |
|---|---|
| chr16-95823456-DEL-211 | chr16-95823467-DEL-210 |
| chr16-95861889-DEL-74 | chr16-95861890-DEL-74 |
| chr16-95872449-DEL-52 | chr16-95872444-DEL-58 |
| chr16-95875320-DEL-198 | chr16-95875321-DEL-198 |
| chr16-95904036-DEL-157 | chr16-95904033-DEL-163 |
| chr16-95912691-DEL-204 | chr16-95912692-DEL-204 |
| chr16-95933393-DEL-211 | chr16-95933394-DEL-169 |
| chr16-95933902-DEL-96 | chr16-95933924-DEL-56 |
| chr16-95938324-DEL-54 | chr16-95938323-DEL-54 |
| chr16-95946032-DEL-213 | chr16-95946033-DEL-213 |
| chr16-95968322-DEL-205 | chr16-95968323-DEL-205 |
| chr16-95990742-DEL-338 | chr16-95990743-DEL-338 |
| chr16-96005245-DEL-60 | chr16-96005245-DEL-60 |
| chr16-96025904-DEL-171 | chr16-96025905-DEL-171 |
| chr16-96027170-DEL-177 | chr16-96027171-DEL-177 |
| chr16-96058094-DEL-158 | chr16-96058097-DEL-151 |
| chr16-96095185-DEL-98 | chr16-96095189-DEL-99 |
| chr16-96099774-DEL-1080 | chr16-96099775-DEL-1080 |
| chr16-96108397-DEL-56 | chr16-96108398-DEL-56 |
| chr16-96140588-DEL-136 | chr16-96140615-DEL-110 |
| chr16-96150858-DEL-66 | chr16-96150859-DEL-66 |
| chr16-96160786-DEL-180 | chr16-96160787-DEL-180 |
| chr16-96167773-DEL-274 | chr16-96167774-DEL-274 |
| chr16-96168418-DEL-2132 | chr16-96168419-DEL-2132 |
| chr16-96222779-DEL-90 | chr16-96222804-DEL-66 |
| chr16-96244426-DEL-52 | chr16-96244427-DEL-52 |
| chr16-96252171-DEL-336 | chr16-96252172-DEL-336 |
| chr16-96265111-DEL-62 | chr16-96265098-DEL-79 |
| chr16-96286075-DEL-90 | chr16-96286076-DEL-90 |
| chr16-96307965-DEL-101 | chr16-96307938-DEL-102 |
| chr16-96321427-DEL-243 | chr16-96321457-DEL-216 |
| chr16-96470179-DEL-94 | chr16-96470180-DEL-94 |
| chr16-96499747-DEL-526 | chr16-96499748-DEL-526 |
| chr16-96512170-DEL-80 | chr16-96512155-DEL-95 |
| chr16-96522938-DEL-837 | chr16-96522939-DEL-837 |
| chr16-96538548-DEL-2022 | chr16-96538549-DEL-2022 |
| chr16-96542286-DEL-1388 | chr16-96542287-DEL-1388 |
| chr16-96545417-DEL-55 | chr16-96545418-DEL-55 |
| chr16-96576825-DEL-71 | chr16-96576836-DEL-69 |
| chr16-96640031-DEL-977 | chr16-96640032-DEL-977 |
| chr16-96641173-DEL-1378 | chr16-96641174-DEL-1378 |
| chr16-96675452-DEL-1482 | chr16-96675453-DEL-1482 |
| chr16-96682375-DEL-1265 | chr16-96682376-DEL-1265 |
| chr16-96686019-DEL-853 | chr16-96686020-DEL-853 |
| chr16-96704542-DEL-132 | chr16-96704571-DEL-96 |
| chr16-96734578-DEL-53 | chr16-96734579-DEL-53 |
| chr16-96743217-DEL-354 | chr16-96743218-DEL-354 |
| chr16-96767750-DEL-1858 | chr16-96767744-DEL-1859 |
| chr16-96781334-DEL-6431 | chr16-96781334-DEL-6423 |
| chr16-96781464-DEL-201 | chr16-96781465-DEL-201 |
| chr16-96851205-DEL-515 | chr16-96851206-DEL-515 |
| chr16-96855244-DEL-197 | chr16-96855245-DEL-197 |
| chr16-96865377-DEL-398 | chr16-96865370-DEL-393 |
| chr16-96892242-DEL-1961 | chr16-96892243-DEL-1961 |
| chr16-96947200-DEL-162 | chr16-96947201-DEL-162 |
| chr16-96957153-DEL-66 | chr16-96957150-DEL-65 |
| chr16-97001352-DEL-326 | chr16-97001353-DEL-326 |
| chr16-97003488-DEL-1271 | chr16-97003489-DEL-1271 |
| chr16-97015135-DEL-60 | chr16-97015114-DEL-60 |
| chr16-97027101-DEL-54 | chr16-97027102-DEL-54 |
| chr16-97061586-DEL-730 | chr16-97061587-DEL-730 |
| chr16-97131637-DEL-86 | chr16-97131636-DEL-55 |
| chr16-97147561-DEL-6219 | chr16-97147562-DEL-6219 |
| chr16-97165036-DEL-1175 | chr16-97165037-DEL-1175 |
| chr16-97194192-DEL-258 | chr16-97194193-DEL-258 |
| chr16-97272858-DEL-55 | chr16-97272858-DEL-50 |
| chr16-97285506-DEL-56 | chr16-97285507-DEL-56 |
| chr16-97286484-DEL-164 | chr16-97286485-DEL-164 |
| chr16-97291179-DEL-1884 | chr16-97291180-DEL-1884 |
| chr16-97296616-DEL-56 | chr16-97296611-DEL-100 |
| chr16-97303565-DEL-420 | chr16-97303566-DEL-420 |
| chr16-97311718-DEL-288 | chr16-97311718-DEL-286 |
| chr16-97330978-DEL-63 | chr16-97330978-DEL-78 |
| chr16-97350066-DEL-58 | chr16-97350067-DEL-62 |
| chr16-97404001-DEL-113 | chr16-97404007-DEL-110 |
| chr16-97418067-DEL-188 | chr16-97418068-DEL-188 |
| chr16-97426619-DEL-107 | chr16-97426620-DEL-107 |
| chr16-97455953-DEL-68 | chr16-97455949-DEL-107 |
| chr16-97497815-DEL-69 | chr16-97497814-DEL-70 |
| chr16-97522619-DEL-2378 | chr16-97522620-DEL-2378 |

| | |
|---|---|
| chr16-97571717-DEL-7238 | chr16-97571718-DEL-7238 |
| chr16-97581902-DEL-58 | chr16-97581903-DEL-58 |
| chr16-97585417-DEL-763 | chr16-97585418-DEL-761 |
| chr16-97624325-DEL-296 | chr16-97624326-DEL-296 |
| chr16-97653748-DEL-58 | chr16-97653749-DEL-58 |
| chr16-97709694-DEL-836 | chr16-97709695-DEL-836 |
| chr16-97813537-DEL-3370 | chr16-97813537-DEL-3370 |
| chr16-97899448-DEL-221 | chr16-97899449-DEL-221 |
| chr17-3078466-DEL-63 | chr17-3078467-DEL-63 |
| chr17-3079173-DEL-4370 | chr17-3079173-DEL-4370 |
| chr17-3103854-DEL-1817 | chr17-3103857-DEL-1818 |
| chr17-3110717-DEL-81 | chr17-3110718-DEL-81 |
| chr17-3159109-DEL-206 | chr17-3159110-DEL-206 |
| chr17-3164722-DEL-57 | chr17-3164723-DEL-57 |
| chr17-3190970-DEL-52 | chr17-3190971-DEL-52 |
| chr17-3207637-DEL-623 | chr17-3207638-DEL-623 |
| chr17-3242438-DEL-172 | chr17-3242439-DEL-172 |
| chr17-3267740-DEL-271 | chr17-3267741-DEL-271 |
| chr17-3286342-DEL-79 | chr17-3286341-DEL-80 |
| chr17-3311786-DEL-206 | chr17-3311786-DEL-206 |
| chr17-3350761-DEL-192 | chr17-3350762-DEL-192 |
| chr17-3355321-DEL-454 | chr17-3355322-DEL-454 |
| chr17-3379977-DEL-226 | chr17-3379978-DEL-226 |
| chr17-3399258-DEL-162 | chr17-3399259-DEL-158 |
| chr17-3433363-DEL-101 | chr17-3433361-DEL-102 |
| chr17-3436388-DEL-205 | chr17-3436389-DEL-205 |
| chr17-3450368-DEL-62 | chr17-3450369-DEL-62 |
| chr17-3494781-DEL-214 | chr17-3494782-DEL-214 |
| chr17-3501501-DEL-70 | chr17-3501502-DEL-70 |
| chr17-3507348-DEL-146 | chr17-3507349-DEL-146 |
| chr17-3514118-DEL-87 | chr17-3514119-DEL-81 |
| chr17-3530225-DEL-70 | chr17-3530226-DEL-70 |
| chr17-3572103-DEL-204 | chr17-3572104-DEL-204 |
| chr17-3586839-DEL-976 | chr17-3586840-DEL-976 |
| chr17-3602628-DEL-58 | chr17-3602629-DEL-58 |
| chr17-3623894-DEL-116 | chr17-3623887-DEL-128 |
| chr17-3658557-DEL-110 | chr17-3658557-DEL-77 |
| chr17-3684110-DEL-71 | chr17-3684118-DEL-68 |
| chr17-3712461-DEL-56 | chr17-3712462-DEL-56 |
| chr17-3738175-DEL-132 | chr17-3738176-DEL-132 |
| chr17-3753010-DEL-86 | chr17-3753011-DEL-84 |
| chr17-3788421-DEL-54 | chr17-3788422-DEL-56 |
| chr17-3805138-DEL-54 | chr17-3805139-DEL-54 |
| chr17-3819462-DEL-183 | chr17-3819508-DEL-168 |
| chr17-3834588-DEL-619 | chr17-3834607-DEL-594 |
| chr17-3839198-DEL-536 | chr17-3839199-DEL-536 |
| chr17-3860440-DEL-748 | chr17-3860441-DEL-748 |
| chr17-3867498-DEL-328 | chr17-3867499-DEL-328 |
| chr17-3873059-DEL-6171 | chr17-3873060-DEL-6171 |
| chr17-3909492-DEL-54 | chr17-3909493-DEL-54 |
| chr17-3935637-DEL-114 | chr17-3935638-DEL-114 |
| chr17-3954953-DEL-88 | chr17-3954954-DEL-124 |
| chr17-3972662-DEL-940 | chr17-3972663-DEL-940 |
| chr17-3975818-DEL-600 | chr17-3975819-DEL-600 |
| chr17-3977814-DEL-267 | chr17-3977815-DEL-267 |
| chr17-4013372-DEL-67 | chr17-4013373-DEL-67 |
| chr17-4057631-DEL-65 | chr17-4057632-DEL-65 |
| chr17-4078487-DEL-58 | chr17-4078488-DEL-58 |
| chr17-4084753-DEL-333 | chr17-4084754-DEL-333 |
| chr17-4098512-DEL-1074 | chr17-4098513-DEL-1074 |
| chr17-4113278-DEL-367 | chr17-4113280-DEL-366 |
| chr17-4132355-DEL-6219 | chr17-4132355-DEL-6219 |
| chr17-4154009-DEL-141 | chr17-4154014-DEL-137 |
| chr17-4161382-DEL-55 | chr17-4161372-DEL-69 |
| chr17-4195003-DEL-51 | chr17-4194983-DEL-83 |
| chr17-4208511-DEL-56 | chr17-4208512-DEL-53 |
| chr17-4211453-DEL-306 | chr17-4211461-DEL-297 |
| chr17-4215884-DEL-2972 | chr17-4215883-DEL-2967 |
| chr17-4218214-DEL-477 | chr17-4218215-DEL-477 |
| chr17-4254619-DEL-1698 | chr17-4254620-DEL-1698 |
| chr17-4321496-DEL-64 | chr17-4321497-DEL-64 |
| chr17-4338134-DEL-202 | chr17-4338135-DEL-202 |
| chr17-4344288-DEL-161 | chr17-4344289-DEL-161 |
| chr17-4345069-DEL-88 | chr17-4345070-DEL-88 |
| chr17-4364352-DEL-62 | chr17-4364353-DEL-62 |
| chr17-4380439-DEL-652 | chr17-4380440-DEL-652 |
| chr17-4385974-DEL-192 | chr17-4385975-DEL-192 |
| chr17-4390651-DEL-80 | chr17-4390646-DEL-91 |
| chr17-4392451-DEL-6398 | chr17-4392452-DEL-6398 |
| chr17-4400404-DEL-234 | chr17-4400405-DEL-234 |

| | |
|---|---|
| chr17-4404937-DEL-621 | chr17-4404938-DEL-621 |
| chr17-4436049-DEL-82 | chr17-4436050-DEL-82 |
| chr17-4447759-DEL-2393 | chr17-4447760-DEL-2393 |
| chr17-4449505-DEL-4620 | chr17-4449506-DEL-4620 |
| chr17-4472533-DEL-196 | chr17-4472534-DEL-196 |
| chr17-4493756-DEL-132 | chr17-4493732-DEL-133 |
| chr17-4510738-DEL-64 | chr17-4510739-DEL-64 |
| chr17-4535987-DEL-58 | chr17-4535988-DEL-58 |
| chr17-4549658-DEL-80 | chr17-4549659-DEL-80 |
| chr17-4587349-DEL-63 | chr17-4587350-DEL-52 |
| chr17-4601949-DEL-272 | chr17-4601950-DEL-272 |
| chr17-4624171-DEL-218 | chr17-4624167-DEL-217 |
| chr17-4632730-DEL-620 | chr17-4632731-DEL-620 |
| chr17-4637419-DEL-923 | chr17-4637419-DEL-923 |
| chr17-4638980-DEL-788 | chr17-4638981-DEL-788 |
| chr17-4650993-DEL-134 | chr17-4650994-DEL-134 |
| chr17-4658148-DEL-63 | chr17-4658151-DEL-64 |
| chr17-4676461-DEL-385 | chr17-4676462-DEL-385 |
| chr17-4710944-DEL-78 | chr17-4710930-DEL-76 |
| chr17-4864841-DEL-121 | chr17-4864838-DEL-125 |
| chr17-4872850-DEL-58 | chr17-4872832-DEL-56 |
| chr17-4878553-DEL-994 | chr17-4878554-DEL-994 |
| chr17-4895410-DEL-163 | chr17-4895401-DEL-164 |
| chr17-4897837-DEL-72 | chr17-4897838-DEL-72 |
| chr17-4902783-DEL-2157 | chr17-4902784-DEL-2157 |
| chr17-4903169-DEL-417 | chr17-4903170-DEL-417 |
| chr17-4966770-DEL-64 | chr17-4966771-DEL-64 |
| chr17-4981827-DEL-58 | chr17-4981828-DEL-58 |
| chr17-5002543-DEL-1922 | chr17-5002544-DEL-1922 |
| chr17-5003436-DEL-108 | chr17-5003437-DEL-108 |
| chr17-5007988-DEL-86 | chr17-5007997-DEL-56 |
| chr17-5033574-DEL-193 | chr17-5033575-DEL-193 |
| chr17-5055880-DEL-60 | chr17-5055881-DEL-60 |
| chr17-5058984-DEL-59 | chr17-5058985-DEL-59 |
| chr17-5084686-DEL-415 | chr17-5084687-DEL-415 |
| chr17-5088530-DEL-114 | chr17-5088569-DEL-76 |
| chr17-5133512-DEL-79 | chr17-5133513-DEL-79 |
| chr17-5187717-DEL-54 | chr17-5187718-DEL-54 |
| chr17-5190600-DEL-142 | chr17-5190601-DEL-142 |
| chr17-5243509-DEL-75 | chr17-5243475-DEL-76 |
| chr17-5300578-DEL-187 | chr17-5300579-DEL-187 |
| chr17-5303606-DEL-52 | chr17-5303583-DEL-56 |
| chr17-5306835-DEL-75 | chr17-5306836-DEL-75 |
| chr17-5309468-DEL-57 | chr17-5309469-DEL-57 |
| chr17-5316963-DEL-76 | chr17-5316964-DEL-76 |
| chr17-5326740-DEL-201 | chr17-5326741-DEL-201 |
| chr17-5329214-DEL-94 | chr17-5329215-DEL-94 |
| chr17-5343630-DEL-106 | chr17-5343631-DEL-102 |
| chr17-5358904-DEL-82 | chr17-5358905-DEL-82 |
| chr17-5363477-DEL-185 | chr17-5363478-DEL-185 |
| chr17-5375859-DEL-56 | chr17-5375860-DEL-56 |
| chr17-5390397-DEL-70 | chr17-5390398-DEL-70 |
| chr17-5398711-DEL-51 | chr17-5398683-DEL-52 |
| chr17-5406499-DEL-250 | chr17-5406531-DEL-224 |
| chr17-5417207-DEL-166 | chr17-5417208-DEL-166 |
| chr17-5428509-DEL-121 | chr17-5428510-DEL-121 |
| chr17-5471665-DEL-407 | chr17-5471666-DEL-407 |
| chr17-5478468-DEL-375 | chr17-5478469-DEL-375 |
| chr17-5497813-DEL-90 | chr17-5497814-DEL-90 |
| chr17-5508753-DEL-685 | chr17-5508745-DEL-692 |
| chr17-5513190-DEL-873 | chr17-5513188-DEL-876 |
| chr17-5553011-DEL-60 | chr17-5553012-DEL-60 |
| chr17-5574943-DEL-636 | chr17-5574944-DEL-636 |
| chr17-5587026-DEL-61 | chr17-5587027-DEL-61 |
| chr17-5596437-DEL-525 | chr17-5596438-DEL-525 |
| chr17-5672077-DEL-234 | chr17-5672078-DEL-195 |
| chr17-5705080-DEL-184 | chr17-5705074-DEL-184 |
| chr17-5706462-DEL-341 | chr17-5706463-DEL-341 |
| chr17-5712036-DEL-1588 | chr17-5712037-DEL-1588 |
| chr17-5750498-DEL-176 | chr17-5750532-DEL-131 |
| chr17-5810580-DEL-339 | chr17-5810581-DEL-339 |
| chr17-5811667-DEL-192 | chr17-5811698-DEL-176 |
| chr17-5814134-DEL-129 | chr17-5814141-DEL-86 |
| chr17-5831310-DEL-353 | chr17-5831311-DEL-353 |
| chr17-5842557-DEL-96 | chr17-5842558-DEL-96 |
| chr17-5864528-DEL-94 | chr17-5864482-DEL-100 |
| chr17-5866304-DEL-680 | chr17-5866296-DEL-681 |
| chr17-5874019-DEL-196 | chr17-5874020-DEL-196 |
| chr17-5898652-DEL-66 | chr17-5898626-DEL-63 |
| chr17-5910575-DEL-76 | chr17-5910580-DEL-76 |

| | |
|---|---|
| chr17-5936506-DEL-84 | chr17-5936521-DEL-63 |
| chr17-5953167-DEL-178 | chr17-5953138-DEL-208 |
| chr17-5961040-DEL-210 | chr17-5961041-DEL-210 |
| chr17-5967191-DEL-59 | chr17-5967233-DEL-57 |
| chr17-5993978-DEL-519 | chr17-5993979-DEL-519 |
| chr17-6014495-DEL-94 | chr17-6014505-DEL-60 |
| chr17-6018330-DEL-65 | chr17-6018374-DEL-65 |
| chr17-6040699-DEL-808 | chr17-6040700-DEL-808 |
| chr17-6071860-DEL-93 | chr17-6071861-DEL-93 |
| chr17-6125899-DEL-126 | chr17-6125900-DEL-126 |
| chr17-6144156-DEL-2486 | chr17-6144157-DEL-2486 |
| chr17-6153404-DEL-53 | chr17-6153406-DEL-53 |
| chr17-6181920-DEL-968 | chr17-6181921-DEL-968 |
| chr17-6209488-DEL-126 | chr17-6209494-DEL-129 |
| chr17-6220728-DEL-534 | chr17-6220729-DEL-534 |
| chr17-6237496-DEL-68 | chr17-6237497-DEL-68 |
| chr17-6270165-DEL-592 | chr17-6270158-DEL-583 |
| chr17-6298372-DEL-55 | chr17-6298373-DEL-55 |
| chr17-6300698-DEL-126 | chr17-6300698-DEL-106 |
| chr17-6319250-DEL-60 | chr17-6319230-DEL-100 |
| chr17-6351792-DEL-79 | chr17-6351793-DEL-79 |
| chr17-6666216-DEL-1498 | chr17-6666216-DEL-1498 |
| chr17-6716915-DEL-937 | chr17-6716916-DEL-937 |
| chr17-6759977-DEL-661 | chr17-6759977-DEL-661 |
| chr17-6764806-DEL-144 | chr17-6764794-DEL-182 |
| chr17-6776353-DEL-73 | chr17-6776363-DEL-72 |
| chr17-6818911-DEL-223 | chr17-6818910-DEL-223 |
| chr17-6886839-DEL-1503 | chr17-6886840-DEL-1503 |
| chr17-6888676-DEL-937 | chr17-6888677-DEL-937 |
| chr17-6926975-DEL-70 | chr17-6926976-DEL-70 |
| chr17-6950686-DEL-90 | chr17-6950685-DEL-90 |
| chr17-6957074-DEL-149 | chr17-6957032-DEL-152 |
| chr17-6962054-DEL-214 | chr17-6962055-DEL-214 |
| chr17-6973004-DEL-224 | chr17-6973005-DEL-224 |
| chr17-6997689-DEL-93 | chr17-6997690-DEL-93 |
| chr17-7012119-DEL-79 | chr17-7012120-DEL-79 |
| chr17-7047355-DEL-205 | chr17-7047356-DEL-205 |
| chr17-7058042-DEL-82 | chr17-7058057-DEL-56 |
| chr17-7147642-DEL-203 | chr17-7147643-DEL-203 |
| chr17-7162413-DEL-461 | chr17-7162414-DEL-461 |
| chr17-7164450-DEL-96 | chr17-7164451-DEL-96 |
| chr17-7167455-DEL-70 | chr17-7167473-DEL-53 |
| chr17-7172654-DEL-128 | chr17-7172655-DEL-127 |
| chr17-7220990-DEL-60 | chr17-7220991-DEL-60 |
| chr17-7235607-DEL-189 | chr17-7235608-DEL-189 |
| chr17-7381951-DEL-205 | chr17-7381952-DEL-205 |
| chr17-7426960-DEL-60 | chr17-7426961-DEL-60 |
| chr17-7442918-DEL-178 | chr17-7442920-DEL-176 |
| chr17-7447231-DEL-1309 | chr17-7447232-DEL-1309 |
| chr17-7453156-DEL-62 | chr17-7453156-DEL-63 |
| chr17-7458397-DEL-240 | chr17-7458398-DEL-240 |
| chr17-7459410-DEL-244 | chr17-7459411-DEL-244 |
| chr17-7460450-DEL-95 | chr17-7460418-DEL-96 |
| chr17-7480602-DEL-54 | chr17-7480603-DEL-54 |
| chr17-7519276-DEL-113 | chr17-7519277-DEL-113 |
| chr17-7571478-DEL-424 | chr17-7571479-DEL-424 |
| chr17-7574726-DEL-940 | chr17-7574727-DEL-940 |
| chr17-7586520-DEL-69 | chr17-7586521-DEL-69 |
| chr17-7680443-DEL-72 | chr17-7680444-DEL-72 |
| chr17-7688662-DEL-1530 | chr17-7688629-DEL-1564 |
| chr17-7699042-DEL-74 | chr17-7699045-DEL-72 |
| chr17-7707979-DEL-66 | chr17-7707980-DEL-66 |
| chr17-7722202-DEL-62 | chr17-7722203-DEL-62 |
| chr17-7730978-DEL-77 | chr17-7730979-DEL-77 |
| chr17-7735852-DEL-162 | chr17-7735853-DEL-162 |
| chr17-7737761-DEL-156 | chr17-7737762-DEL-156 |
| chr17-7739388-DEL-735 | chr17-7739389-DEL-735 |
| chr17-7747505-DEL-200 | chr17-7747506-DEL-200 |
| chr17-7778606-DEL-563 | chr17-7778607-DEL-563 |
| chr17-7779608-DEL-2579 | chr17-7779609-DEL-2579 |
| chr17-7787290-DEL-127 | chr17-7787291-DEL-126 |
| chr17-7807866-DEL-440 | chr17-7807867-DEL-440 |
| chr17-7808795-DEL-211 | chr17-7808796-DEL-211 |
| chr17-7856524-DEL-440 | chr17-7856525-DEL-440 |
| chr17-7892907-DEL-57 | chr17-7892910-DEL-55 |
| chr17-7911562-DEL-62 | chr17-7911557-DEL-68 |
| chr17-7919799-DEL-69 | chr17-7919816-DEL-69 |
| chr17-7941533-DEL-2673 | chr17-7941534-DEL-2673 |
| chr17-7946260-DEL-2998 | chr17-7946274-DEL-2996 |
| chr17-7962537-DEL-84 | chr17-7962538-DEL-84 |

| | |
|---|---|
| chr17-7970223-DEL-106 | chr17-7970224-DEL-106 |
| chr17-7979398-DEL-90 | chr17-7979399-DEL-90 |
| chr17-7985838-DEL-112 | chr17-7985839-DEL-112 |
| chr17-7995231-DEL-164 | chr17-7995280-DEL-157 |
| chr17-8003726-DEL-905 | chr17-8003768-DEL-861 |
| chr17-8033704-DEL-80 | chr17-8033705-DEL-50 |
| chr17-8056971-DEL-56 | chr17-8056955-DEL-74 |
| chr17-8071993-DEL-474 | chr17-8071994-DEL-474 |
| chr17-8104795-DEL-61 | chr17-8104773-DEL-62 |
| chr17-8105357-DEL-56 | chr17-8105356-DEL-56 |
| chr17-8129161-DEL-594 | chr17-8129159-DEL-597 |
| chr17-8136482-DEL-1582 | chr17-8136483-DEL-1582 |
| chr17-8149253-DEL-324 | chr17-8149254-DEL-324 |
| chr17-8165526-DEL-67 | chr17-8165527-DEL-67 |
| chr17-8209017-DEL-3431 | chr17-8209018-DEL-3431 |
| chr17-8223626-DEL-67 | chr17-8223626-DEL-67 |
| chr17-8278562-DEL-373 | chr17-8278567-DEL-369 |
| chr17-8354968-DEL-72 | chr17-8354976-DEL-73 |
| chr17-8396476-DEL-87 | chr17-8396477-DEL-87 |
| chr17-8418741-DEL-66 | chr17-8418742-DEL-66 |
| chr17-8422430-DEL-287 | chr17-8422431-DEL-287 |
| chr17-8424215-DEL-121 | chr17-8424216-DEL-121 |
| chr17-8434676-DEL-7433 | chr17-8434677-DEL-7433 |
| chr17-8447471-DEL-166 | chr17-8447472-DEL-166 |
| chr17-8455946-DEL-54 | chr17-8455947-DEL-54 |
| chr17-8470385-DEL-96 | chr17-8470386-DEL-96 |
| chr17-8507033-DEL-210 | chr17-8507034-DEL-210 |
| chr17-8509404-DEL-187 | chr17-8509405-DEL-187 |
| chr17-8550071-DEL-6400 | chr17-8550072-DEL-6400 |
| chr17-8583275-DEL-54 | chr17-8583300-DEL-57 |
| chr17-8598046-DEL-67 | chr17-8598047-DEL-67 |
| chr17-8598511-DEL-215 | chr17-8598511-DEL-215 |
| chr17-8656823-DEL-91 | chr17-8656776-DEL-91 |
| chr17-8665091-DEL-77 | chr17-8665092-DEL-77 |
| chr17-8719668-DEL-72 | chr17-8719641-DEL-100 |
| chr17-8788539-DEL-330 | chr17-8788540-DEL-330 |
| chr17-8790008-DEL-126 | chr17-8790034-DEL-83 |
| chr17-8797231-DEL-338 | chr17-8797232-DEL-338 |
| chr17-8806109-DEL-108 | chr17-8806124-DEL-130 |
| chr17-8809263-DEL-66 | chr17-8809264-DEL-66 |
| chr17-8911158-DEL-4707 | chr17-8911159-DEL-4707 |
| chr17-8944197-DEL-55 | chr17-8944197-DEL-55 |
| chr17-8970485-DEL-82 | chr17-8970486-DEL-83 |
| chr17-8977530-DEL-51 | chr17-8977531-DEL-54 |
| chr17-9008091-DEL-74 | chr17-9008096-DEL-70 |
| chr17-9038348-DEL-230 | chr17-9038349-DEL-230 |
| chr17-9052500-DEL-125 | chr17-9052501-DEL-125 |
| chr17-9067889-DEL-123 | chr17-9067890-DEL-123 |
| chr17-9087579-DEL-72 | chr17-9087580-DEL-72 |
| chr17-9124935-DEL-1001 | chr17-9124936-DEL-1001 |
| chr17-9131276-DEL-2541 | chr17-9131277-DEL-2541 |
| chr17-9135787-DEL-1489 | chr17-9135788-DEL-1489 |
| chr17-9156063-DEL-64 | chr17-9156070-DEL-66 |
| chr17-9157583-DEL-79 | chr17-9157580-DEL-68 |
| chr17-9214985-DEL-95 | chr17-9214964-DEL-116 |
| chr17-9238030-DEL-128 | chr17-9237983-DEL-176 |
| chr17-9256627-DEL-333 | chr17-9256628-DEL-333 |
| chr17-9257640-DEL-393 | chr17-9257639-DEL-394 |
| chr17-9273081-DEL-248 | chr17-9273080-DEL-248 |
| chr17-9289925-DEL-53 | chr17-9289924-DEL-55 |
| chr17-9355644-DEL-68 | chr17-9355671-DEL-70 |
| chr17-9375592-DEL-731 | chr17-9375593-DEL-731 |
| chr17-9402119-DEL-76 | chr17-9402120-DEL-76 |
| chr17-9411212-DEL-2680 | chr17-9411213-DEL-2680 |
| chr17-9505102-DEL-58 | chr17-9505105-DEL-59 |
| chr17-9517231-DEL-64 | chr17-9517232-DEL-64 |
| chr17-9526829-DEL-111 | chr17-9526830-DEL-111 |
| chr17-9537576-DEL-104 | chr17-9537577-DEL-104 |
| chr17-9564259-DEL-6253 | chr17-9564260-DEL-6253 |
| chr17-9638157-DEL-58 | chr17-9638158-DEL-50 |
| chr17-9689737-DEL-1437 | chr17-9689734-DEL-1438 |
| chr17-9701412-DEL-78 | chr17-9701373-DEL-76 |
| chr17-9708877-DEL-6358 | chr17-9708878-DEL-6358 |
| chr17-9742499-DEL-129 | chr17-9742511-DEL-129 |
| chr17-9750514-DEL-67 | chr17-9750515-DEL-67 |
| chr17-9790053-DEL-5286 | chr17-9790054-DEL-5286 |
| chr17-9814020-DEL-53 | chr17-9814021-DEL-53 |
| chr17-9820975-DEL-74 | chr17-9820964-DEL-86 |
| chr17-9829084-DEL-122 | chr17-9829085-DEL-118 |
| chr17-9833033-DEL-463 | chr17-9833034-DEL-463 |

| | |
|---|---|
| chr17-9906430-DEL-524 | chr17-9906431-DEL-524 |
| chr17-9914291-DEL-52 | chr17-9914292-DEL-52 |
| chr17-9915851-DEL-74 | chr17-9915856-DEL-70 |
| chr17-9936528-DEL-2404 | chr17-9936529-DEL-2404 |
| chr17-9949181-DEL-4348 | chr17-9949182-DEL-4348 |
| chr17-9967570-DEL-330 | chr17-9967571-DEL-330 |
| chr17-9982523-DEL-61 | chr17-9982516-DEL-59 |
| chr17-9999501-DEL-7115 | chr17-9999502-DEL-7115 |
| chr17-10024211-DEL-6726 | chr17-10024212-DEL-6726 |
| chr17-10057000-DEL-84 | chr17-10056975-DEL-83 |
| chr17-10077408-DEL-1242 | chr17-10077409-DEL-1242 |
| chr17-10120415-DEL-186 | chr17-10120419-DEL-178 |
| chr17-10128240-DEL-125 | chr17-10128238-DEL-114 |
| chr17-10166319-DEL-52 | chr17-10166320-DEL-52 |
| chr17-10177386-DEL-78 | chr17-10177387-DEL-78 |
| chr17-10218829-DEL-82 | chr17-10218830-DEL-82 |
| chr17-10226466-DEL-54 | chr17-10226467-DEL-54 |
| chr17-10239135-DEL-59 | chr17-10239136-DEL-59 |
| chr17-10246917-DEL-54 | chr17-10246869-DEL-55 |
| chr17-10266695-DEL-54 | chr17-10266696-DEL-54 |
| chr17-10364019-DEL-52 | chr17-10364020-DEL-52 |
| chr17-10369363-DEL-60 | chr17-10369364-DEL-60 |
| chr17-10403934-DEL-56 | chr17-10403911-DEL-55 |
| chr17-10439809-DEL-58 | chr17-10439830-DEL-50 |
| chr17-10563428-DEL-90 | chr17-10563401-DEL-114 |
| chr17-10594120-DEL-2166 | chr17-10594121-DEL-2166 |
| chr17-10603559-DEL-54 | chr17-10603560-DEL-54 |
| chr17-10718530-DEL-216 | chr17-10718542-DEL-215 |
| chr17-10745064-DEL-69 | chr17-10745065-DEL-69 |
| chr17-10837279-DEL-537 | chr17-10837280-DEL-537 |
| chr17-10866347-DEL-468 | chr17-10866348-DEL-468 |
| chr17-10880614-DEL-59 | chr17-10880619-DEL-55 |
| chr17-10889811-DEL-1420 | chr17-10889812-DEL-1420 |
| chr17-11022985-DEL-314 | chr17-11022986-DEL-314 |
| chr17-11041049-DEL-70 | chr17-11041050-DEL-70 |
| chr17-11056713-DEL-72 | chr17-11056710-DEL-50 |
| chr17-11170214-DEL-52 | chr17-11170215-DEL-52 |
| chr17-11185884-DEL-7717 | chr17-11185885-DEL-7717 |
| chr17-11206815-DEL-5134 | chr17-11206816-DEL-5134 |
| chr17-11212118-DEL-66 | chr17-11212119-DEL-66 |
| chr17-11212772-DEL-101 | chr17-11212773-DEL-101 |
| chr17-11283177-DEL-80 | chr17-11283178-DEL-80 |
| chr17-11316725-DEL-7177 | chr17-11316728-DEL-7167 |
| chr17-11350113-DEL-54 | chr17-11350114-DEL-52 |
| chr17-11360409-DEL-54 | chr17-11360410-DEL-54 |
| chr17-11364567-DEL-1782 | chr17-11364568-DEL-1782 |
| chr17-11385495-DEL-809 | chr17-11385500-DEL-806 |
| chr17-11387153-DEL-60 | chr17-11387154-DEL-60 |
| chr17-11443099-DEL-258 | chr17-11443100-DEL-258 |
| chr17-11507093-DEL-440 | chr17-11507094-DEL-440 |
| chr17-11517546-DEL-201 | chr17-11517547-DEL-201 |
| chr17-11518437-DEL-378 | chr17-11518438-DEL-378 |
| chr17-11547948-DEL-99 | chr17-11547943-DEL-100 |
| chr17-11553939-DEL-66 | chr17-11553954-DEL-61 |
| chr17-11581899-DEL-52 | chr17-11581900-DEL-52 |
| chr17-11674448-DEL-191 | chr17-11674449-DEL-191 |
| chr17-11712408-DEL-6879 | chr17-11712409-DEL-6879 |
| chr17-11775930-DEL-213 | chr17-11775941-DEL-216 |
| chr17-11834275-DEL-839 | chr17-11834276-DEL-839 |
| chr17-11872442-DEL-76 | chr17-11872467-DEL-52 |
| chr17-11878181-DEL-247 | chr17-11878182-DEL-238 |
| chr17-11904855-DEL-63 | chr17-11904855-DEL-63 |
| chr17-11918096-DEL-538 | chr17-11918097-DEL-538 |
| chr17-11941505-DEL-1896 | chr17-11941506-DEL-1896 |
| chr17-11952404-DEL-81 | chr17-11952405-DEL-81 |
| chr17-11973381-DEL-6092 | chr17-11973382-DEL-6092 |
| chr17-12011962-DEL-166 | chr17-12011963-DEL-118 |
| chr17-12031792-DEL-117 | chr17-12031783-DEL-114 |
| chr17-12047706-DEL-158 | chr17-12047707-DEL-158 |
| chr17-12069557-DEL-70 | chr17-12069558-DEL-70 |
| chr17-12091123-DEL-2062 | chr17-12091124-DEL-2062 |
| chr17-12094278-DEL-62 | chr17-12094276-DEL-60 |
| chr17-12124596-DEL-71 | chr17-12124593-DEL-71 |
| chr17-12134509-DEL-82 | chr17-12134510-DEL-82 |
| chr17-12136180-DEL-87 | chr17-12136181-DEL-87 |
| chr17-12188887-DEL-80 | chr17-12188868-DEL-64 |
| chr17-12190611-DEL-193 | chr17-12190612-DEL-193 |
| chr17-12204922-DEL-217 | chr17-12204923-DEL-217 |
| chr17-12205971-DEL-74 | chr17-12206005-DEL-74 |
| chr17-12215405-DEL-61 | chr17-12215406-DEL-61 |

| | |
|---|---|
| chr17-12216035-DEL-83 | chr17-12216036-DEL-83 |
| chr17-12229162-DEL-9062 | chr17-12229163-DEL-9062 |
| chr17-12285478-DEL-75 | chr17-12285478-DEL-75 |
| chr17-12306401-DEL-3108 | chr17-12306402-DEL-3108 |
| chr17-12329877-DEL-1967 | chr17-12329878-DEL-1967 |
| chr17-12336028-DEL-71 | chr17-12336029-DEL-71 |
| chr17-12346113-DEL-103 | chr17-12346114-DEL-103 |
| chr17-12347809-DEL-169 | chr17-12347810-DEL-169 |
| chr17-12352064-DEL-911 | chr17-12352065-DEL-911 |
| chr17-12356911-DEL-376 | chr17-12356912-DEL-376 |
| chr17-12377330-DEL-60 | chr17-12377331-DEL-60 |
| chr17-12391707-DEL-112 | chr17-12391708-DEL-112 |
| chr17-12402758-DEL-59 | chr17-12402759-DEL-59 |
| chr17-12409072-DEL-4519 | chr17-12409073-DEL-4519 |
| chr17-12412741-DEL-165 | chr17-12412742-DEL-164 |
| chr17-12437641-DEL-179 | chr17-12437642-DEL-179 |
| chr17-12442148-DEL-172 | chr17-12442149-DEL-172 |
| chr17-12442705-DEL-132 | chr17-12442706-DEL-132 |
| chr17-12443429-DEL-187 | chr17-12443430-DEL-187 |
| chr17-12452113-DEL-53 | chr17-12452114-DEL-53 |
| chr17-12453403-DEL-242 | chr17-12453416-DEL-238 |
| chr17-12464531-DEL-301 | chr17-12464569-DEL-302 |
| chr17-12465778-DEL-75 | chr17-12465779-DEL-75 |
| chr17-12500326-DEL-59 | chr17-12500327-DEL-59 |
| chr17-12507236-DEL-291 | chr17-12507237-DEL-291 |
| chr17-12527743-DEL-1884 | chr17-12527744-DEL-1884 |
| chr17-12539795-DEL-120 | chr17-12539796-DEL-120 |
| chr17-12544790-DEL-96 | chr17-12544791-DEL-96 |
| chr17-12563921-DEL-536 | chr17-12563922-DEL-536 |
| chr17-12584434-DEL-198 | chr17-12584435-DEL-198 |
| chr17-12601646-DEL-276 | chr17-12601647-DEL-276 |
| chr17-12690062-DEL-185 | chr17-12690063-DEL-185 |
| chr17-12722905-DEL-51 | chr17-12722906-DEL-51 |
| chr17-12743437-DEL-384 | chr17-12743478-DEL-336 |
| chr17-12743895-DEL-760 | chr17-12743896-DEL-760 |
| chr17-12755810-DEL-222 | chr17-12755796-DEL-236 |
| chr17-12865598-DEL-52 | chr17-12865599-DEL-52 |
| chr17-12913728-DEL-232 | chr17-12913729-DEL-233 |
| chr17-12915863-DEL-53 | chr17-12915863-DEL-53 |
| chr17-12948618-DEL-799 | chr17-12948619-DEL-799 |
| chr17-13067260-DEL-224 | chr17-13067261-DEL-224 |
| chr17-13183202-DEL-211 | chr17-13183203-DEL-211 |
| chr17-13195833-DEL-2245 | chr17-13195834-DEL-2244 |
| chr17-13402730-DEL-90 | chr17-13402705-DEL-89 |
| chr17-13585413-DEL-105 | chr17-13585441-DEL-78 |
| chr17-13701210-DEL-4560 | chr17-13701211-DEL-4560 |
| chr17-13805271-DEL-78 | chr17-13805281-DEL-62 |
| chr17-13901113-DEL-157 | chr17-13901115-DEL-156 |
| chr17-13949959-DEL-65 | chr17-13949959-DEL-65 |
| chr17-13951805-DEL-168 | chr17-13951806-DEL-168 |
| chr17-14013074-DEL-94 | chr17-14013075-DEL-94 |
| chr17-14064486-DEL-636 | chr17-14064487-DEL-636 |
| chr17-14088453-DEL-78 | chr17-14088454-DEL-78 |
| chr17-14155104-DEL-60 | chr17-14155109-DEL-56 |
| chr17-14213882-DEL-160 | chr17-14213876-DEL-163 |
| chr17-14253535-DEL-1958 | chr17-14253536-DEL-1958 |
| chr17-14293159-DEL-5228 | chr17-14293160-DEL-5228 |
| chr17-14328096-DEL-737 | chr17-14328097-DEL-737 |
| chr17-14381028-DEL-683 | chr17-14381028-DEL-684 |
| chr17-14400018-DEL-1888 | chr17-14400019-DEL-1888 |
| chr17-14403023-DEL-138 | chr17-14403024-DEL-138 |
| chr17-14441820-DEL-58 | chr17-14441815-DEL-59 |
| chr17-14452174-DEL-67 | chr17-14452187-DEL-50 |
| chr17-14458606-DEL-1909 | chr17-14458607-DEL-1908 |
| chr17-14479434-DEL-1510 | chr17-14479435-DEL-1510 |
| chr17-14494485-DEL-201 | chr17-14494486-DEL-201 |
| chr17-14530629-DEL-92 | chr17-14530629-DEL-92 |
| chr17-14535526-DEL-79 | chr17-14535527-DEL-79 |
| chr17-14569147-DEL-545 | chr17-14569148-DEL-545 |
| chr17-14572015-DEL-204 | chr17-14572016-DEL-204 |
| chr17-14578813-DEL-55 | chr17-14578781-DEL-76 |
| chr17-14653828-DEL-74 | chr17-14653829-DEL-74 |
| chr17-14657077-DEL-52 | chr17-14657077-DEL-52 |
| chr17-14741306-DEL-68 | chr17-14741323-DEL-52 |
| chr17-14757052-DEL-848 | chr17-14757053-DEL-848 |
| chr17-14778982-DEL-678 | chr17-14778983-DEL-678 |
| chr17-14800857-DEL-6429 | chr17-14800853-DEL-6430 |
| chr17-14833823-DEL-821 | chr17-14833824-DEL-821 |
| chr17-14840250-DEL-108 | chr17-14840251-DEL-108 |
| chr17-14860089-DEL-76 | chr17-14860085-DEL-51 |

| | |
|---|---|
| chr17-14921395-DEL-52 | chr17-14921396-DEL-52 |
| chr17-14936835-DEL-1605 | chr17-14936836-DEL-1605 |
| chr17-14941986-DEL-55 | chr17-14941987-DEL-56 |
| chr17-14978010-DEL-6168 | chr17-14978011-DEL-6168 |
| chr17-15020803-DEL-6288 | chr17-15020804-DEL-6288 |
| chr17-15037907-DEL-3112 | chr17-15037908-DEL-3112 |
| chr17-15067864-DEL-198 | chr17-15067865-DEL-198 |
| chr17-15085055-DEL-6742 | chr17-15085056-DEL-6742 |
| chr17-15093761-DEL-656 | chr17-15093761-DEL-656 |
| chr17-15111069-DEL-420 | chr17-15111070-DEL-420 |
| chr17-15160338-DEL-240 | chr17-15160339-DEL-240 |
| chr17-15304857-DEL-313 | chr17-15304858-DEL-313 |
| chr17-15328063-DEL-181 | chr17-15328064-DEL-181 |
| chr17-15330966-DEL-64 | chr17-15330967-DEL-64 |
| chr17-15334756-DEL-138 | chr17-15334757-DEL-138 |
| chr17-15336510-DEL-64 | chr17-15336511-DEL-64 |
| chr17-15350945-DEL-51 | chr17-15350946-DEL-51 |
| chr17-15390993-DEL-134 | chr17-15390992-DEL-135 |
| chr17-15484367-DEL-293 | chr17-15484368-DEL-293 |
| chr17-15523008-DEL-88 | chr17-15523009-DEL-88 |
| chr17-15523735-DEL-3438 | chr17-15523736-DEL-3438 |
| chr17-15629963-DEL-162 | chr17-15629964-DEL-162 |
| chr17-15642363-DEL-117 | chr17-15642362-DEL-118 |
| chr17-15684591-DEL-194 | chr17-15684592-DEL-194 |
| chr17-15736453-DEL-201 | chr17-15736454-DEL-201 |
| chr17-15800633-DEL-216 | chr17-15800634-DEL-216 |
| chr17-15812906-DEL-632 | chr17-15812907-DEL-632 |
| chr17-15814763-DEL-54 | chr17-15814763-DEL-54 |
| chr17-15823104-DEL-7155 | chr17-15823105-DEL-7155 |
| chr17-15867638-DEL-55 | chr17-15867652-DEL-57 |
| chr17-15876364-DEL-859 | chr17-15876365-DEL-859 |
| chr17-15881836-DEL-218 | chr17-15881837-DEL-218 |
| chr17-15886143-DEL-200 | chr17-15886144-DEL-200 |
| chr17-15889824-DEL-530 | chr17-15889825-DEL-530 |
| chr17-15891392-DEL-335 | chr17-15891395-DEL-327 |
| chr17-15903105-DEL-209 | chr17-15903106-DEL-209 |
| chr17-15940709-DEL-154 | chr17-15940710-DEL-154 |
| chr17-15944364-DEL-212 | chr17-15944365-DEL-212 |
| chr17-15997453-DEL-441 | chr17-15997454-DEL-441 |
| chr17-16003128-DEL-73 | chr17-16003128-DEL-71 |
| chr17-16004926-DEL-613 | chr17-16004927-DEL-613 |
| chr17-16013288-DEL-84 | chr17-16013306-DEL-66 |
| chr17-16017774-DEL-63 | chr17-16017774-DEL-64 |
| chr17-16064244-DEL-946 | chr17-16064226-DEL-946 |
| chr17-16067605-DEL-61 | chr17-16067606-DEL-61 |
| chr17-16071864-DEL-70 | chr17-16071900-DEL-58 |
| chr17-16083990-DEL-527 | chr17-16083991-DEL-527 |
| chr17-16101623-DEL-178 | chr17-16101624-DEL-178 |
| chr17-16117740-DEL-909 | chr17-16117741-DEL-909 |
| chr17-16129613-DEL-1223 | chr17-16129614-DEL-1223 |
| chr17-16133103-DEL-1381 | chr17-16133104-DEL-1381 |
| chr17-16148124-DEL-2618 | chr17-16148125-DEL-2618 |
| chr17-16184554-DEL-73 | chr17-16184554-DEL-71 |
| chr17-16238260-DEL-138 | chr17-16238279-DEL-104 |
| chr17-16242989-DEL-1045 | chr17-16242990-DEL-1045 |
| chr17-16253252-DEL-56 | chr17-16253219-DEL-58 |
| chr17-16265226-DEL-587 | chr17-16265227-DEL-587 |
| chr17-16281269-DEL-509 | chr17-16281270-DEL-509 |
| chr17-16288291-DEL-611 | chr17-16288292-DEL-611 |
| chr17-16328371-DEL-750 | chr17-16328369-DEL-745 |
| chr17-16337589-DEL-72 | chr17-16337590-DEL-66 |
| chr17-16377078-DEL-59 | chr17-16377079-DEL-59 |
| chr17-16420003-DEL-56 | chr17-16420004-DEL-56 |
| chr17-16424722-DEL-108 | chr17-16424723-DEL-108 |
| chr17-16438589-DEL-52 | chr17-16438602-DEL-52 |
| chr17-16464547-DEL-63 | chr17-16464557-DEL-50 |
| chr17-16480515-DEL-6204 | chr17-16480512-DEL-6205 |
| chr17-16489146-DEL-545 | chr17-16489147-DEL-541 |
| chr17-16497472-DEL-419 | chr17-16497473-DEL-420 |
| chr17-16509427-DEL-3573 | chr17-16509428-DEL-3573 |
| chr17-16510520-DEL-1493 | chr17-16510521-DEL-1493 |
| chr17-16527135-DEL-634 | chr17-16527136-DEL-634 |
| chr17-16584898-DEL-3170 | chr17-16584899-DEL-3170 |
| chr17-16721262-DEL-279 | chr17-16721263-DEL-279 |
| chr17-16732463-DEL-251 | chr17-16732464-DEL-251 |
| chr17-16743787-DEL-62 | chr17-16743783-DEL-103 |
| chr17-16753279-DEL-173 | chr17-16753284-DEL-171 |
| chr17-16772244-DEL-70 | chr17-16772245-DEL-70 |
| chr17-16777474-DEL-881 | chr17-16777475-DEL-881 |
| chr17-16816277-DEL-101 | chr17-16816278-DEL-101 |

| | |
|---|---|
| chr17-16820714-DEL-58 | chr17-16820715-DEL-50 |
| chr17-16888998-DEL-8036 | chr17-16888999-DEL-8036 |
| chr17-16920683-DEL-673 | chr17-16920672-DEL-685 |
| chr17-16947833-DEL-494 | chr17-16947834-DEL-494 |
| chr17-16971228-DEL-168 | chr17-16971229-DEL-168 |
| chr17-16979678-DEL-2753 | chr17-16979679-DEL-2753 |
| chr17-16990879-DEL-482 | chr17-16990880-DEL-482 |
| chr17-17011873-DEL-5862 | chr17-17011874-DEL-5862 |
| chr17-17028989-DEL-68 | chr17-17028990-DEL-68 |
| chr17-17037872-DEL-62 | chr17-17037872-DEL-59 |
| chr17-17039249-DEL-53 | chr17-17039246-DEL-57 |
| chr17-17072213-DEL-6263 | chr17-17072214-DEL-6263 |
| chr17-17116693-DEL-471 | chr17-17116693-DEL-471 |
| chr17-17117622-DEL-199 | chr17-17117623-DEL-199 |
| chr17-17141923-DEL-114 | chr17-17141923-DEL-114 |
| chr17-17149625-DEL-54 | chr17-17149626-DEL-54 |
| chr17-17156047-DEL-539 | chr17-17156048-DEL-539 |
| chr17-17164956-DEL-569 | chr17-17164957-DEL-569 |
| chr17-17170489-DEL-486 | chr17-17170490-DEL-486 |
| chr17-17230494-DEL-6418 | chr17-17230481-DEL-6419 |
| chr17-17382836-DEL-143 | chr17-17382835-DEL-143 |
| chr17-17387917-DEL-761 | chr17-17387918-DEL-761 |
| chr17-17420147-DEL-150 | chr17-17420146-DEL-150 |
| chr17-17435175-DEL-201 | chr17-17435176-DEL-201 |
| chr17-17461561-DEL-475 | chr17-17461562-DEL-475 |
| chr17-17472025-DEL-1361 | chr17-17472026-DEL-1361 |
| chr17-17496357-DEL-6403 | chr17-17496358-DEL-6403 |
| chr17-17531512-DEL-78 | chr17-17531507-DEL-84 |
| chr17-17561138-DEL-52 | chr17-17561139-DEL-52 |
| chr17-17563888-DEL-343 | chr17-17563889-DEL-343 |
| chr17-17565380-DEL-1103 | chr17-17565381-DEL-1103 |
| chr17-17604635-DEL-54 | chr17-17604647-DEL-51 |
| chr17-17612719-DEL-149 | chr17-17612720-DEL-149 |
| chr17-17644834-DEL-201 | chr17-17644835-DEL-201 |
| chr17-17691366-DEL-1434 | chr17-17691364-DEL-1435 |
| chr17-17693050-DEL-6023 | chr17-17693047-DEL-6021 |
| chr17-17699068-DEL-963 | chr17-17699068-DEL-963 |
| chr17-17715343-DEL-225 | chr17-17715344-DEL-225 |
| chr17-17721260-DEL-1831 | chr17-17721261-DEL-1831 |
| chr17-17723546-DEL-167 | chr17-17723542-DEL-170 |
| chr17-17774356-DEL-7213 | chr17-17774357-DEL-7213 |
| chr17-17809130-DEL-142 | chr17-17809083-DEL-177 |
| chr17-17817424-DEL-58 | chr17-17817425-DEL-58 |
| chr17-17818923-DEL-179 | chr17-17818924-DEL-179 |
| chr17-17831231-DEL-189 | chr17-17831231-DEL-187 |
| chr17-17850687-DEL-1695 | chr17-17850688-DEL-1695 |
| chr17-17858309-DEL-148 | chr17-17858310-DEL-148 |
| chr17-17894117-DEL-114 | chr17-17894118-DEL-114 |
| chr17-17900904-DEL-6392 | chr17-17900905-DEL-6392 |
| chr17-17906895-DEL-201 | chr17-17906896-DEL-201 |
| chr17-17918873-DEL-1881 | chr17-17918874-DEL-1881 |
| chr17-17920831-DEL-451 | chr17-17920828-DEL-441 |
| chr17-17950279-DEL-77 | chr17-17950280-DEL-77 |
| chr17-17984323-DEL-354 | chr17-17984324-DEL-354 |
| chr17-17989378-DEL-178 | chr17-17989381-DEL-176 |
| chr17-17989510-DEL-973 | chr17-17989511-DEL-973 |
| chr17-17992697-DEL-3554 | chr17-17992698-DEL-3554 |
| chr17-18076437-DEL-66 | chr17-18076438-DEL-66 |
| chr17-18087799-DEL-56 | chr17-18087799-DEL-56 |
| chr17-18117311-DEL-273 | chr17-18117314-DEL-271 |
| chr17-18118174-DEL-88 | chr17-18118174-DEL-89 |
| chr17-18120539-DEL-185 | chr17-18120540-DEL-185 |
| chr17-18311017-DEL-380 | chr17-18311018-DEL-380 |
| chr17-18312411-DEL-281 | chr17-18312412-DEL-281 |
| chr17-18314749-DEL-416 | chr17-18314750-DEL-416 |
| chr17-18320068-DEL-4278 | chr17-18320069-DEL-4278 |
| chr17-18340066-DEL-2438 | chr17-18340065-DEL-2438 |
| chr17-18350732-DEL-214 | chr17-18350733-DEL-214 |
| chr17-18481102-DEL-428 | chr17-18481103-DEL-428 |
| chr17-18485905-DEL-504 | chr17-18485906-DEL-504 |
| chr17-18493151-DEL-60 | chr17-18493160-DEL-52 |
| chr17-18500307-DEL-393 | chr17-18500308-DEL-393 |
| chr17-18506651-DEL-88 | chr17-18506636-DEL-67 |
| chr17-18537724-DEL-386 | chr17-18537725-DEL-386 |
| chr17-18586578-DEL-833 | chr17-18586564-DEL-834 |
| chr17-18601231-DEL-8660 | chr17-18601232-DEL-8660 |
| chr17-18618525-DEL-197 | chr17-18618526-DEL-197 |
| chr17-18631675-DEL-62 | chr17-18631676-DEL-53 |
| chr17-18639592-DEL-335 | chr17-18639593-DEL-335 |
| chr17-18688983-DEL-268 | chr17-18688984-DEL-268 |

| | |
|---|---|
| chr17-18717704-DEL-2811 | chr17-18717705-DEL-2811 |
| chr17-18760118-DEL-444 | chr17-18760119-DEL-444 |
| chr17-18768331-DEL-2673 | chr17-18768332-DEL-2673 |
| chr17-18796357-DEL-1181 | chr17-18796358-DEL-1181 |
| chr17-18799877-DEL-98 | chr17-18799875-DEL-99 |
| chr17-18821787-DEL-6553 | chr17-18821788-DEL-6553 |
| chr17-18907965-DEL-269 | chr17-18907966-DEL-269 |
| chr17-18922597-DEL-64 | chr17-18922598-DEL-64 |
| chr17-18957891-DEL-6459 | chr17-18957892-DEL-6459 |
| chr17-18964870-DEL-441 | chr17-18964871-DEL-441 |
| chr17-19020484-DEL-6427 | chr17-19020485-DEL-6427 |
| chr17-19059323-DEL-600 | chr17-19059324-DEL-600 |
| chr17-19061936-DEL-11691 | chr17-19061924-DEL-11698 |
| chr17-19065024-DEL-5103 | chr17-19065025-DEL-5103 |
| chr17-19117815-DEL-5002 | chr17-19117815-DEL-5002 |
| chr17-19151847-DEL-7089 | chr17-19151847-DEL-7089 |
| chr17-19164754-DEL-64 | chr17-19164749-DEL-65 |
| chr17-19214046-DEL-4686 | chr17-19214047-DEL-4686 |
| chr17-19226986-DEL-59 | chr17-19226987-DEL-59 |
| chr17-19236041-DEL-246 | chr17-19236042-DEL-247 |
| chr17-19253669-DEL-2750 | chr17-19253670-DEL-2750 |
| chr17-19324038-DEL-6888 | chr17-19324039-DEL-6888 |
| chr17-19331424-DEL-1164 | chr17-19331425-DEL-1164 |
| chr17-19337993-DEL-233 | chr17-19337994-DEL-233 |
| chr17-19358445-DEL-218 | chr17-19358446-DEL-218 |
| chr17-19378897-DEL-6197 | chr17-19378898-DEL-6197 |
| chr17-19385208-DEL-55 | chr17-19385209-DEL-55 |
| chr17-19604609-DEL-805 | chr17-19604610-DEL-805 |
| chr17-19611374-DEL-1250 | chr17-19611375-DEL-1250 |
| chr17-19629971-DEL-86 | chr17-19629972-DEL-86 |
| chr17-19678072-DEL-54 | chr17-19678073-DEL-54 |
| chr17-19682710-DEL-59 | chr17-19682711-DEL-59 |
| chr17-19684997-DEL-62 | chr17-19684998-DEL-62 |
| chr17-19710502-DEL-940 | chr17-19710503-DEL-940 |
| chr17-19719969-DEL-2503 | chr17-19719970-DEL-2503 |
| chr17-19907355-DEL-54 | chr17-19907360-DEL-50 |
| chr17-19927993-DEL-54 | chr17-19927994-DEL-50 |
| chr17-19959958-DEL-70 | chr17-19959955-DEL-114 |
| chr17-20010853-DEL-569 | chr17-20010854-DEL-569 |
| chr17-20016990-DEL-2108 | chr17-20016991-DEL-2108 |
| chr17-20049250-DEL-1892 | chr17-20049251-DEL-1892 |
| chr17-20074226-DEL-108 | chr17-20074227-DEL-108 |
| chr17-20091738-DEL-305 | chr17-20091739-DEL-305 |
| chr17-20135735-DEL-3595 | chr17-20135736-DEL-3595 |
| chr17-20142895-DEL-350 | chr17-20142895-DEL-350 |
| chr17-20196032-DEL-90 | chr17-20196007-DEL-92 |
| chr17-20222114-DEL-953 | chr17-20222115-DEL-953 |
| chr17-20235349-DEL-101 | chr17-20235350-DEL-101 |
| chr17-20241982-DEL-64 | chr17-20241983-DEL-64 |
| chr17-20242559-DEL-6172 | chr17-20242560-DEL-6172 |
| chr17-20269924-DEL-62 | chr17-20269925-DEL-62 |
| chr17-20270296-DEL-598 | chr17-20270296-DEL-598 |
| chr17-20271389-DEL-476 | chr17-20271389-DEL-476 |
| chr17-20277866-DEL-59 | chr17-20277863-DEL-55 |
| chr17-20279999-DEL-1087 | chr17-20280000-DEL-1087 |
| chr17-20319912-DEL-246 | chr17-20319913-DEL-246 |
| chr17-20335700-DEL-163 | chr17-20335701-DEL-163 |
| chr17-20345362-DEL-1051 | chr17-20345363-DEL-1051 |
| chr17-20359496-DEL-396 | chr17-20359497-DEL-394 |
| chr17-20392353-DEL-350 | chr17-20392354-DEL-350 |
| chr17-20414374-DEL-61 | chr17-20414373-DEL-62 |
| chr17-20532076-DEL-127 | chr17-20532077-DEL-127 |
| chr17-20542737-DEL-460 | chr17-20542738-DEL-460 |
| chr17-20582337-DEL-174 | chr17-20582337-DEL-174 |
| chr17-20589968-DEL-236 | chr17-20589968-DEL-236 |
| chr17-20590895-DEL-71 | chr17-20590895-DEL-71 |
| chr17-20609032-DEL-886 | chr17-20609033-DEL-886 |
| chr17-20651986-DEL-110 | chr17-20651987-DEL-110 |
| chr17-20658570-DEL-64 | chr17-20658569-DEL-108 |
| chr17-20669256-DEL-136 | chr17-20669257-DEL-136 |
| chr17-20746538-DEL-60 | chr17-20746533-DEL-66 |
| chr17-20768723-DEL-328 | chr17-20768724-DEL-328 |
| chr17-20771541-DEL-414 | chr17-20771540-DEL-416 |
| chr17-20833670-DEL-1869 | chr17-20833671-DEL-1869 |
| chr17-20859264-DEL-6680 | chr17-20859265-DEL-6680 |
| chr17-20876582-DEL-65 | chr17-20876576-DEL-53 |
| chr17-20886567-DEL-144 | chr17-20886588-DEL-117 |
| chr17-20888906-DEL-54 | chr17-20888907-DEL-54 |
| chr17-20902148-DEL-635 | chr17-20902145-DEL-636 |
| chr17-20929224-DEL-496 | chr17-20929225-DEL-496 |

| | |
|---|---|
| chr17-20958874-DEL-357 | chr17-20958875-DEL-357 |
| chr17-20962739-DEL-733 | chr17-20962743-DEL-731 |
| chr17-21022742-DEL-1907 | chr17-21022742-DEL-1907 |
| chr17-21026644-DEL-2707 | chr17-21026645-DEL-2707 |
| chr17-21029438-DEL-610 | chr17-21029439-DEL-608 |
| chr17-21030393-DEL-132 | chr17-21030394-DEL-132 |
| chr17-21048868-DEL-247 | chr17-21048869-DEL-247 |
| chr17-21079501-DEL-6647 | chr17-21079498-DEL-6647 |
| chr17-21145064-DEL-489 | chr17-21145065-DEL-489 |
| chr17-21192205-DEL-825 | chr17-21192206-DEL-825 |
| chr17-21194179-DEL-102 | chr17-21194172-DEL-110 |
| chr17-21198729-DEL-5520 | chr17-21198730-DEL-5520 |
| chr17-21204582-DEL-5487 | chr17-21204583-DEL-5487 |
| chr17-21221303-DEL-183 | chr17-21221301-DEL-184 |
| chr17-21228647-DEL-99 | chr17-21228648-DEL-99 |
| chr17-21253256-DEL-165 | chr17-21253257-DEL-165 |
| chr17-21255619-DEL-64 | chr17-21255619-DEL-64 |
| chr17-21262501-DEL-51 | chr17-21262502-DEL-51 |
| chr17-21324283-DEL-680 | chr17-21324284-DEL-680 |
| chr17-21335490-DEL-195 | chr17-21335491-DEL-195 |
| chr17-21341907-DEL-91 | chr17-21341950-DEL-63 |
| chr17-21388456-DEL-73 | chr17-21388436-DEL-74 |
| chr17-21439270-DEL-75 | chr17-21439271-DEL-75 |
| chr17-21447167-DEL-544 | chr17-21447168-DEL-544 |
| chr17-21457336-DEL-15817 | chr17-21457336-DEL-15813 |
| chr17-21472991-DEL-52 | chr17-21472975-DEL-53 |
| chr17-21490304-DEL-76 | chr17-21490305-DEL-69 |
| chr17-21512655-DEL-1705 | chr17-21512656-DEL-1705 |
| chr17-21532695-DEL-2314 | chr17-21532696-DEL-2314 |
| chr17-21559074-DEL-7273 | chr17-21559075-DEL-7273 |
| chr17-21590725-DEL-387 | chr17-21590722-DEL-387 |
| chr17-21609898-DEL-1443 | chr17-21609901-DEL-1442 |
| chr17-21613626-DEL-148 | chr17-21613627-DEL-148 |
| chr17-21618788-DEL-168 | chr17-21618789-DEL-168 |
| chr17-21634333-DEL-4319 | chr17-21634334-DEL-4319 |
| chr17-21640164-DEL-328 | chr17-21640165-DEL-328 |
| chr17-21650950-DEL-212 | chr17-21650951-DEL-212 |
| chr17-21664585-DEL-187 | chr17-21664585-DEL-185 |
| chr17-21695934-DEL-98 | chr17-21695935-DEL-98 |
| chr17-21749972-DEL-177 | chr17-21749973-DEL-177 |
| chr17-21763679-DEL-8298 | chr17-21763680-DEL-8298 |
| chr17-21807272-DEL-99 | chr17-21807273-DEL-99 |
| chr17-21809805-DEL-55 | chr17-21809806-DEL-55 |
| chr17-21818408-DEL-6193 | chr17-21818389-DEL-6210 |
| chr17-21907121-DEL-196 | chr17-21907122-DEL-196 |
| chr17-21971951-DEL-102 | chr17-21971952-DEL-85 |
| chr17-21983905-DEL-1027 | chr17-21983913-DEL-1021 |
| chr17-21987993-DEL-218 | chr17-21987993-DEL-218 |
| chr17-21993597-DEL-450 | chr17-21993598-DEL-450 |
| chr17-22028589-DEL-6509 | chr17-22028590-DEL-6509 |
| chr17-22079026-DEL-56 | chr17-22079027-DEL-56 |
| chr17-22089145-DEL-1417 | chr17-22089146-DEL-1417 |
| chr17-22174473-DEL-65 | chr17-22174441-DEL-54 |
| chr17-22185396-DEL-72 | chr17-22185397-DEL-72 |
| chr17-22264004-DEL-1439 | chr17-22264003-DEL-1439 |
| chr17-22274705-DEL-3087 | chr17-22274706-DEL-3087 |
| chr17-22306446-DEL-405 | chr17-22306447-DEL-405 |
| chr17-22339378-DEL-1170 | chr17-22339379-DEL-1170 |
| chr17-22400025-DEL-333 | chr17-22400024-DEL-334 |
| chr17-22405713-DEL-342 | chr17-22405714-DEL-342 |
| chr17-22409480-DEL-75 | chr17-22409481-DEL-74 |
| chr17-22433896-DEL-7507 | chr17-22433897-DEL-7507 |
| chr17-22450473-DEL-2377 | chr17-22450474-DEL-2377 |
| chr17-22459016-DEL-6377 | chr17-22459017-DEL-6377 |
| chr17-22472369-DEL-3243 | chr17-22472370-DEL-3241 |
| chr17-22489593-DEL-565 | chr17-22489594-DEL-565 |
| chr17-22512839-DEL-117 | chr17-22512840-DEL-117 |
| chr17-22613345-DEL-6372 | chr17-22613346-DEL-6372 |
| chr17-22671316-DEL-181 | chr17-22671317-DEL-181 |
| chr17-22721104-DEL-1888 | chr17-22721105-DEL-1888 |
| chr17-22749644-DEL-267 | chr17-22749645-DEL-267 |
| chr17-22766130-DEL-108 | chr17-22766131-DEL-108 |
| chr17-22803680-DEL-70 | chr17-22803703-DEL-68 |
| chr17-22850676-DEL-4425 | chr17-22850677-DEL-4425 |
| chr17-22902181-DEL-115 | chr17-22902182-DEL-115 |
| chr17-22918433-DEL-686 | chr17-22918434-DEL-686 |
| chr17-22921032-DEL-144 | chr17-22921035-DEL-141 |
| chr17-22921934-DEL-7009 | chr17-22921935-DEL-7009 |
| chr17-22977282-DEL-1741 | chr17-22977283-DEL-1741 |
| chr17-23002078-DEL-118 | chr17-23002079-DEL-118 |

| | |
|---|---|
| chr17-23004290-DEL-70 | chr17-23004291-DEL-70 |
| chr17-23094459-DEL-291 | chr17-23094460-DEL-291 |
| chr17-23095859-DEL-434 | chr17-23095860-DEL-434 |
| chr17-23116161-DEL-164 | chr17-23116162-DEL-164 |
| chr17-23157661-DEL-1128 | chr17-23157660-DEL-1129 |
| chr17-23225347-DEL-236 | chr17-23225348-DEL-236 |
| chr17-23262976-DEL-348 | chr17-23262977-DEL-346 |
| chr17-23370045-DEL-2054 | chr17-23370046-DEL-2054 |
| chr17-23382372-DEL-86 | chr17-23382410-DEL-54 |
| chr17-23412392-DEL-118 | chr17-23412393-DEL-118 |
| chr17-23418804-DEL-349 | chr17-23418805-DEL-349 |
| chr17-23419849-DEL-345 | chr17-23419850-DEL-345 |
| chr17-23421289-DEL-1920 | chr17-23421278-DEL-1906 |
| chr17-23464119-DEL-133 | chr17-23464119-DEL-133 |
| chr17-23480883-DEL-174 | chr17-23480890-DEL-145 |
| chr17-23491174-DEL-454 | chr17-23491175-DEL-454 |
| chr17-23514962-DEL-655 | chr17-23514949-DEL-655 |
| chr17-23535708-DEL-4863 | chr17-23535709-DEL-4863 |
| chr17-23543238-DEL-2159 | chr17-23543239-DEL-2159 |
| chr17-23552447-DEL-385 | chr17-23552448-DEL-385 |
| chr17-23593001-DEL-203 | chr17-23593002-DEL-203 |
| chr17-23594673-DEL-1032 | chr17-23594674-DEL-1032 |
| chr17-23643227-DEL-640 | chr17-23643228-DEL-640 |
| chr17-23645705-DEL-1220 | chr17-23645706-DEL-1220 |
| chr17-23699861-DEL-401 | chr17-23699862-DEL-401 |
| chr17-23703362-DEL-638 | chr17-23703378-DEL-611 |
| chr17-23714277-DEL-7095 | chr17-23714278-DEL-7095 |
| chr17-23752402-DEL-51 | chr17-23752403-DEL-51 |
| chr17-23754860-DEL-51 | chr17-23754861-DEL-51 |
| chr17-23797930-DEL-71 | chr17-23797927-DEL-65 |
| chr17-23812316-DEL-149 | chr17-23812317-DEL-149 |
| chr17-23814205-DEL-52 | chr17-23814206-DEL-52 |
| chr17-23833852-DEL-212 | chr17-23833853-DEL-212 |
| chr17-23847161-DEL-288 | chr17-23847162-DEL-288 |
| chr17-23856791-DEL-82 | chr17-23856792-DEL-82 |
| chr17-23857688-DEL-107 | chr17-23857688-DEL-107 |
| chr17-23869598-DEL-59 | chr17-23869599-DEL-59 |
| chr17-23883140-DEL-204 | chr17-23883141-DEL-204 |
| chr17-23904988-DEL-6336 | chr17-23904989-DEL-6336 |
| chr17-23933980-DEL-52 | chr17-23933981-DEL-50 |
| chr17-23977159-DEL-65 | chr17-23977159-DEL-65 |
| chr17-23985837-DEL-65 | chr17-23985838-DEL-65 |
| chr17-23986617-DEL-195 | chr17-23986618-DEL-195 |
| chr17-23993474-DEL-62 | chr17-23993475-DEL-62 |
| chr17-24000470-DEL-78 | chr17-24000470-DEL-78 |
| chr17-24003071-DEL-62 | chr17-24003072-DEL-62 |
| chr17-24005323-DEL-99 | chr17-24005324-DEL-98 |
| chr17-24016778-DEL-266 | chr17-24016779-DEL-266 |
| chr17-24071532-DEL-191 | chr17-24071533-DEL-191 |
| chr17-24100100-DEL-60 | chr17-24100121-DEL-54 |
| chr17-24107066-DEL-62 | chr17-24107066-DEL-62 |
| chr17-24115314-DEL-94 | chr17-24115315-DEL-79 |
| chr17-24117984-DEL-61 | chr17-24117988-DEL-73 |
| chr17-24121747-DEL-102 | chr17-24121750-DEL-63 |
| chr17-24153132-DEL-301 | chr17-24153133-DEL-299 |
| chr17-24165062-DEL-52 | chr17-24165063-DEL-52 |
| chr17-24171521-DEL-52 | chr17-24171522-DEL-52 |
| chr17-24175267-DEL-59 | chr17-24175267-DEL-59 |
| chr17-24205760-DEL-87 | chr17-24205761-DEL-87 |
| chr17-24206383-DEL-64 | chr17-24206384-DEL-56 |
| chr17-24208362-DEL-203 | chr17-24208373-DEL-202 |
| chr17-24245158-DEL-519 | chr17-24245153-DEL-520 |
| chr17-24250352-DEL-260 | chr17-24250353-DEL-260 |
| chr17-24252076-DEL-210 | chr17-24252077-DEL-210 |
| chr17-24270880-DEL-194 | chr17-24270881-DEL-194 |
| chr17-24278338-DEL-63 | chr17-24278339-DEL-63 |
| chr17-24317853-DEL-220 | chr17-24317854-DEL-220 |
| chr17-24354410-DEL-466 | chr17-24354411-DEL-466 |
| chr17-24406403-DEL-1352 | chr17-24406404-DEL-1352 |
| chr17-24451677-DEL-145 | chr17-24451678-DEL-145 |
| chr17-24464636-DEL-543 | chr17-24464666-DEL-516 |
| chr17-24478675-DEL-2813 | chr17-24478676-DEL-2813 |
| chr17-24482182-DEL-207 | chr17-24482183-DEL-207 |
| chr17-24483536-DEL-208 | chr17-24483538-DEL-207 |
| chr17-24486147-DEL-54 | chr17-24486147-DEL-54 |
| chr17-24487402-DEL-81 | chr17-24487400-DEL-84 |
| chr17-24488857-DEL-51 | chr17-24488857-DEL-51 |
| chr17-24503958-DEL-55 | chr17-24503953-DEL-62 |
| chr17-24515958-DEL-83 | chr17-24515959-DEL-83 |
| chr17-24524725-DEL-203 | chr17-24524726-DEL-203 |

| | |
|---|---|
| chr17-24529135-DEL-172 | chr17-24529136-DEL-172 |
| chr17-24537696-DEL-451 | chr17-24537698-DEL-449 |
| chr17-24559090-DEL-70 | chr17-24559091-DEL-70 |
| chr17-24588804-DEL-562 | chr17-24588805-DEL-562 |
| chr17-24591921-DEL-57 | chr17-24591922-DEL-57 |
| chr17-24616764-DEL-207 | chr17-24616765-DEL-207 |
| chr17-24641899-DEL-171 | chr17-24641900-DEL-171 |
| chr17-24651263-DEL-182 | chr17-24651264-DEL-182 |
| chr17-24681292-DEL-117 | chr17-24681293-DEL-117 |
| chr17-24686413-DEL-450 | chr17-24686414-DEL-450 |
| chr17-24756034-DEL-357 | chr17-24756026-DEL-357 |
| chr17-24761622-DEL-320 | chr17-24761629-DEL-318 |
| chr17-24774279-DEL-112 | chr17-24774280-DEL-112 |
| chr17-24828316-DEL-63 | chr17-24828333-DEL-50 |
| chr17-24830304-DEL-1845 | chr17-24830305-DEL-1845 |
| chr17-24836103-DEL-94 | chr17-24836112-DEL-115 |
| chr17-24860442-DEL-144 | chr17-24860443-DEL-141 |
| chr17-24874592-DEL-56 | chr17-24874625-DEL-65 |
| chr17-24875471-DEL-80 | chr17-24875472-DEL-80 |
| chr17-24905839-DEL-98 | chr17-24905840-DEL-110 |
| chr17-24912595-DEL-556 | chr17-24912596-DEL-555 |
| chr17-24940686-DEL-88 | chr17-24940687-DEL-88 |
| chr17-24944421-DEL-64 | chr17-24944425-DEL-52 |
| chr17-24948607-DEL-82 | chr17-24948605-DEL-53 |
| chr17-24962619-DEL-108 | chr17-24962620-DEL-108 |
| chr17-24978962-DEL-430 | chr17-24978959-DEL-431 |
| chr17-24994948-DEL-59 | chr17-24994949-DEL-59 |
| chr17-25019213-DEL-100 | chr17-25019239-DEL-74 |
| chr17-25028190-DEL-203 | chr17-25028191-DEL-203 |
| chr17-25034769-DEL-184 | chr17-25034769-DEL-185 |
| chr17-25035923-DEL-96 | chr17-25035924-DEL-96 |
| chr17-25041101-DEL-141 | chr17-25041102-DEL-141 |
| chr17-25045390-DEL-730 | chr17-25045391-DEL-730 |
| chr17-25065475-DEL-56 | chr17-25065523-DEL-52 |
| chr17-25078333-DEL-82 | chr17-25078330-DEL-86 |
| chr17-25092339-DEL-56 | chr17-25092320-DEL-68 |
| chr17-25149735-DEL-204 | chr17-25149736-DEL-204 |
| chr17-25186801-DEL-208 | chr17-25186802-DEL-208 |
| chr17-25190693-DEL-316 | chr17-25190694-DEL-316 |
| chr17-25221652-DEL-377 | chr17-25221653-DEL-377 |
| chr17-25257384-DEL-186 | chr17-25257385-DEL-186 |
| chr17-25261496-DEL-695 | chr17-25261497-DEL-696 |
| chr17-25261554-DEL-58 | chr17-25261535-DEL-58 |
| chr17-25262631-DEL-182 | chr17-25262631-DEL-183 |
| chr17-25266017-DEL-240 | chr17-25266018-DEL-240 |
| chr17-25282270-DEL-278 | chr17-25282271-DEL-278 |
| chr17-25336815-DEL-215 | chr17-25336816-DEL-215 |
| chr17-25345146-DEL-108 | chr17-25345147-DEL-108 |
| chr17-25357029-DEL-301 | chr17-25357030-DEL-303 |
| chr17-25455518-DEL-927 | chr17-25455518-DEL-927 |
| chr17-25457513-DEL-106 | chr17-25457514-DEL-106 |
| chr17-25496132-DEL-315 | chr17-25496133-DEL-315 |
| chr17-25500471-DEL-384 | chr17-25500470-DEL-386 |
| chr17-25502905-DEL-4456 | chr17-25502906-DEL-4456 |
| chr17-25523820-DEL-92 | chr17-25523821-DEL-84 |
| chr17-25525313-DEL-981 | chr17-25525318-DEL-982 |
| chr17-25558424-DEL-66 | chr17-25558425-DEL-66 |
| chr17-25574199-DEL-64 | chr17-25574208-DEL-57 |
| chr17-25576798-DEL-66 | chr17-25576799-DEL-66 |
| chr17-25587223-DEL-232 | chr17-25587224-DEL-232 |
| chr17-25597911-DEL-139 | chr17-25597935-DEL-106 |
| chr17-25674004-DEL-63 | chr17-25674005-DEL-63 |
| chr17-25705590-DEL-54 | chr17-25705591-DEL-89 |
| chr17-25732386-DEL-89 | chr17-25732387-DEL-89 |
| chr17-25744518-DEL-54 | chr17-25744519-DEL-54 |
| chr17-25761247-DEL-435 | chr17-25761252-DEL-451 |
| chr17-25769351-DEL-108 | chr17-25769352-DEL-108 |
| chr17-25784023-DEL-158 | chr17-25784053-DEL-110 |
| chr17-25795778-DEL-65 | chr17-25795779-DEL-65 |
| chr17-25858619-DEL-52 | chr17-25858616-DEL-56 |
| chr17-25895146-DEL-197 | chr17-25895147-DEL-197 |
| chr17-25922406-DEL-213 | chr17-25922407-DEL-213 |
| chr17-25927522-DEL-1170 | chr17-25927523-DEL-1170 |
| chr17-25951273-DEL-96 | chr17-25951274-DEL-96 |
| chr17-25981661-DEL-168 | chr17-25981661-DEL-169 |
| chr17-26007246-DEL-175 | chr17-26007247-DEL-175 |
| chr17-26114347-DEL-90 | chr17-26114344-DEL-94 |
| chr17-26124867-DEL-51 | chr17-26124865-DEL-53 |
| chr17-26131559-DEL-571 | chr17-26131560-DEL-571 |
| chr17-26174149-DEL-56 | chr17-26174135-DEL-57 |

| | |
|---|---|
| chr17-26197770-DEL-61 | chr17-26197771-DEL-61 |
| chr17-26210831-DEL-66 | chr17-26210832-DEL-88 |
| chr17-26218667-DEL-896 | chr17-26218667-DEL-896 |
| chr17-26232312-DEL-206 | chr17-26232313-DEL-206 |
| chr17-26316761-DEL-72 | chr17-26316805-DEL-72 |
| chr17-26317812-DEL-419 | chr17-26317813-DEL-419 |
| chr17-26320576-DEL-68 | chr17-26320597-DEL-68 |
| chr17-26325695-DEL-150 | chr17-26325696-DEL-139 |
| chr17-26333330-DEL-214 | chr17-26333331-DEL-214 |
| chr17-26355410-DEL-166 | chr17-26355411-DEL-166 |
| chr17-26374710-DEL-207 | chr17-26374711-DEL-207 |
| chr17-26427843-DEL-162 | chr17-26427844-DEL-162 |
| chr17-26448015-DEL-94 | chr17-26448007-DEL-91 |
| chr17-26468762-DEL-164 | chr17-26468763-DEL-164 |
| chr17-26484011-DEL-194 | chr17-26484012-DEL-194 |
| chr17-26490700-DEL-78 | chr17-26490701-DEL-78 |
| chr17-26491396-DEL-212 | chr17-26491397-DEL-212 |
| chr17-26493099-DEL-194 | chr17-26493100-DEL-194 |
| chr17-26497652-DEL-180 | chr17-26497653-DEL-180 |
| chr17-26518727-DEL-253 | chr17-26518728-DEL-253 |
| chr17-26559815-DEL-538 | chr17-26559816-DEL-538 |
| chr17-26562375-DEL-83 | chr17-26562380-DEL-79 |
| chr17-26598642-DEL-53 | chr17-26598669-DEL-63 |
| chr17-26603529-DEL-374 | chr17-26603530-DEL-374 |
| chr17-26611039-DEL-344 | chr17-26611040-DEL-344 |
| chr17-26626586-DEL-53 | chr17-26626587-DEL-53 |
| chr17-26629218-DEL-99 | chr17-26629230-DEL-92 |
| chr17-26663355-DEL-63 | chr17-26663356-DEL-63 |
| chr17-26686299-DEL-292 | chr17-26686300-DEL-292 |
| chr17-26739937-DEL-115 | chr17-26739938-DEL-115 |
| chr17-26759686-DEL-199 | chr17-26759687-DEL-199 |
| chr17-26766069-DEL-95 | chr17-26766047-DEL-96 |
| chr17-26804601-DEL-219 | chr17-26804602-DEL-219 |
| chr17-26805409-DEL-392 | chr17-26805411-DEL-391 |
| chr17-26809021-DEL-71 | chr17-26809022-DEL-71 |
| chr17-26835047-DEL-56 | chr17-26835048-DEL-56 |
| chr17-26836147-DEL-80 | chr17-26836148-DEL-80 |
| chr17-26848143-DEL-68 | chr17-26848144-DEL-68 |
| chr17-26855525-DEL-210 | chr17-26855526-DEL-224 |
| chr17-26888287-DEL-1307 | chr17-26888285-DEL-1308 |
| chr17-26970573-DEL-619 | chr17-26970574-DEL-619 |
| chr17-26996134-DEL-186 | chr17-26996135-DEL-186 |
| chr17-27019608-DEL-1489 | chr17-27019608-DEL-1489 |
| chr17-27029004-DEL-330 | chr17-27029015-DEL-331 |
| chr17-27044362-DEL-348 | chr17-27044365-DEL-345 |
| chr17-27071886-DEL-98 | chr17-27071908-DEL-114 |
| chr17-27075829-DEL-68 | chr17-27075830-DEL-68 |
| chr17-27087271-DEL-84 | chr17-27087294-DEL-62 |
| chr17-27088734-DEL-59 | chr17-27088765-DEL-56 |
| chr17-27133525-DEL-212 | chr17-27133526-DEL-212 |
| chr17-27210302-DEL-211 | chr17-27210302-DEL-212 |
| chr17-27212481-DEL-225 | chr17-27212482-DEL-226 |
| chr17-27263669-DEL-201 | chr17-27263670-DEL-201 |
| chr17-27286862-DEL-131 | chr17-27286831-DEL-163 |
| chr17-27296136-DEL-150 | chr17-27296137-DEL-150 |
| chr17-27318896-DEL-207 | chr17-27318897-DEL-207 |
| chr17-27398479-DEL-147 | chr17-27398481-DEL-146 |
| chr17-27417507-DEL-206 | chr17-27417508-DEL-206 |
| chr17-27433466-DEL-148 | chr17-27433467-DEL-148 |
| chr17-27439203-DEL-89 | chr17-27439204-DEL-89 |
| chr17-27440816-DEL-66 | chr17-27440817-DEL-66 |
| chr17-27477789-DEL-419 | chr17-27477790-DEL-419 |
| chr17-27483856-DEL-5059 | chr17-27483857-DEL-5059 |
| chr17-27500552-DEL-6474 | chr17-27500553-DEL-6474 |
| chr17-27640004-DEL-99 | chr17-27640004-DEL-98 |
| chr17-27641231-DEL-52 | chr17-27641234-DEL-50 |
| chr17-27695802-DEL-527 | chr17-27695800-DEL-528 |
| chr17-27712719-DEL-59 | chr17-27712720-DEL-59 |
| chr17-27744031-DEL-212 | chr17-27744032-DEL-212 |
| chr17-27751520-DEL-205 | chr17-27751521-DEL-205 |
| chr17-27752245-DEL-170 | chr17-27752247-DEL-169 |
| chr17-27762839-DEL-70 | chr17-27762840-DEL-70 |
| chr17-27791585-DEL-71 | chr17-27791586-DEL-71 |
| chr17-27818294-DEL-165 | chr17-27818337-DEL-122 |
| chr17-27823857-DEL-554 | chr17-27823858-DEL-554 |
| chr17-27847976-DEL-333 | chr17-27847977-DEL-333 |
| chr17-27859168-DEL-75 | chr17-27859169-DEL-75 |
| chr17-27860611-DEL-52 | chr17-27860612-DEL-52 |
| chr17-27861247-DEL-69 | chr17-27861268-DEL-53 |
| chr17-27879996-DEL-60 | chr17-27879996-DEL-58 |

| | |
|---|---|
| chr17-27881836-DEL-169 | chr17-27881837-DEL-169 |
| chr17-27881997-DEL-2131 | chr17-27881992-DEL-2128 |
| chr17-27882394-DEL-213 | chr17-27882395-DEL-213 |
| chr17-27914557-DEL-1882 | chr17-27914558-DEL-1882 |
| chr17-28015793-DEL-53 | chr17-28015753-DEL-53 |
| chr17-28043651-DEL-55 | chr17-28043673-DEL-51 |
| chr17-28052748-DEL-75 | chr17-28052749-DEL-74 |
| chr17-28054662-DEL-63 | chr17-28054663-DEL-63 |
| chr17-28056818-DEL-493 | chr17-28056819-DEL-493 |
| chr17-28072035-DEL-153 | chr17-28072036-DEL-153 |
| chr17-28107768-DEL-126 | chr17-28107759-DEL-136 |
| chr17-28119439-DEL-172 | chr17-28119440-DEL-172 |
| chr17-28147362-DEL-215 | chr17-28147363-DEL-215 |
| chr17-28218880-DEL-147 | chr17-28218930-DEL-98 |
| chr17-28305620-DEL-120 | chr17-28305621-DEL-120 |
| chr17-28312690-DEL-224 | chr17-28312683-DEL-232 |
| chr17-28314623-DEL-216 | chr17-28314622-DEL-215 |
| chr17-28321259-DEL-942 | chr17-28321264-DEL-944 |
| chr17-28372898-DEL-203 | chr17-28372899-DEL-203 |
| chr17-28467228-DEL-55 | chr17-28467228-DEL-56 |
| chr17-28474920-DEL-116 | chr17-28474921-DEL-116 |
| chr17-28487124-DEL-587 | chr17-28487083-DEL-588 |
| chr17-28528909-DEL-784 | chr17-28528910-DEL-784 |
| chr17-28532035-DEL-166 | chr17-28532036-DEL-166 |
| chr17-28533684-DEL-120 | chr17-28533684-DEL-119 |
| chr17-28542536-DEL-202 | chr17-28542536-DEL-203 |
| chr17-28566845-DEL-77 | chr17-28566884-DEL-53 |
| chr17-28589422-DEL-1385 | chr17-28589423-DEL-1385 |
| chr17-28589721-DEL-72 | chr17-28589721-DEL-72 |
| chr17-28607653-DEL-96 | chr17-28607654-DEL-96 |
| chr17-28608912-DEL-105 | chr17-28608912-DEL-97 |
| chr17-28609798-DEL-218 | chr17-28609799-DEL-222 |
| chr17-28692289-DEL-267 | chr17-28692292-DEL-265 |
| chr17-28695293-DEL-249 | chr17-28695294-DEL-249 |
| chr17-28711551-DEL-119 | chr17-28711552-DEL-119 |
| chr17-28717890-DEL-58 | chr17-28717891-DEL-58 |
| chr17-28770903-DEL-111 | chr17-28770904-DEL-106 |
| chr17-28773259-DEL-108 | chr17-28773260-DEL-108 |
| chr17-28778148-DEL-220 | chr17-28778138-DEL-225 |
| chr17-28786061-DEL-207 | chr17-28786068-DEL-200 |
| chr17-28803132-DEL-193 | chr17-28803131-DEL-195 |
| chr17-28808322-DEL-77 | chr17-28808332-DEL-63 |
| chr17-28822620-DEL-241 | chr17-28822621-DEL-241 |
| chr17-28829543-DEL-101 | chr17-28829576-DEL-57 |
| chr17-28830089-DEL-222 | chr17-28830090-DEL-222 |
| chr17-28834018-DEL-387 | chr17-28834019-DEL-387 |
| chr17-28850683-DEL-140 | chr17-28850684-DEL-140 |
| chr17-28853496-DEL-174 | chr17-28853497-DEL-174 |
| chr17-28870015-DEL-56 | chr17-28870016-DEL-56 |
| chr17-28876749-DEL-54 | chr17-28876744-DEL-52 |
| chr17-28888500-DEL-66 | chr17-28888501-DEL-66 |
| chr17-28923393-DEL-1348 | chr17-28923394-DEL-1348 |
| chr17-28936728-DEL-58 | chr17-28936729-DEL-58 |
| chr17-28946445-DEL-326 | chr17-28946446-DEL-326 |
| chr17-28952525-DEL-231 | chr17-28952526-DEL-231 |
| chr17-28970161-DEL-54 | chr17-28970162-DEL-54 |
| chr17-28974798-DEL-248 | chr17-28974784-DEL-293 |
| chr17-28986536-DEL-151 | chr17-28986537-DEL-151 |
| chr17-28996871-DEL-64 | chr17-28996846-DEL-64 |
| chr17-28999486-DEL-3397 | chr17-28999487-DEL-3397 |
| chr17-29000159-DEL-74 | chr17-29000120-DEL-75 |
| chr17-29003111-DEL-206 | chr17-29003100-DEL-207 |
| chr17-29031087-DEL-391 | chr17-29031088-DEL-391 |
| chr17-29064704-DEL-1250 | chr17-29064705-DEL-1250 |
| chr17-29069951-DEL-104 | chr17-29069988-DEL-59 |
| chr17-29091939-DEL-91 | chr17-29091925-DEL-81 |
| chr17-29094946-DEL-207 | chr17-29094947-DEL-207 |
| chr17-29108790-DEL-582 | chr17-29108791-DEL-581 |
| chr17-29110035-DEL-438 | chr17-29110036-DEL-438 |
| chr17-29112074-DEL-323 | chr17-29112075-DEL-334 |
| chr17-29126499-DEL-223 | chr17-29126500-DEL-223 |
| chr17-29131739-DEL-68 | chr17-29131740-DEL-68 |
| chr17-29151090-DEL-57 | chr17-29151091-DEL-57 |
| chr17-29156375-DEL-2781 | chr17-29156376-DEL-2781 |
| chr17-29168569-DEL-204 | chr17-29168570-DEL-204 |
| chr17-29212554-DEL-207 | chr17-29212555-DEL-207 |
| chr17-29221363-DEL-163 | chr17-29221356-DEL-166 |
| chr17-29225351-DEL-109 | chr17-29225352-DEL-109 |
| chr17-29230499-DEL-84 | chr17-29230500-DEL-84 |
| chr17-29236762-DEL-205 | chr17-29236763-DEL-205 |

| | |
|---|---|
| chr17-29239820-DEL-60 | chr17-29239821-DEL-60 |
| chr17-29242767-DEL-261 | chr17-29242768-DEL-261 |
| chr17-29246860-DEL-197 | chr17-29246861-DEL-197 |
| chr17-29254425-DEL-76 | chr17-29254426-DEL-76 |
| chr17-29299094-DEL-180 | chr17-29299095-DEL-179 |
| chr17-29307388-DEL-197 | chr17-29307389-DEL-197 |
| chr17-29316326-DEL-72 | chr17-29316345-DEL-72 |
| chr17-29343936-DEL-223 | chr17-29343935-DEL-224 |
| chr17-29355880-DEL-471 | chr17-29355869-DEL-472 |
| chr17-29411558-DEL-151 | chr17-29411559-DEL-151 |
| chr17-29413730-DEL-205 | chr17-29413731-DEL-205 |
| chr17-29429830-DEL-431 | chr17-29429834-DEL-415 |
| chr17-29475366-DEL-59 | chr17-29475367-DEL-59 |
| chr17-29484169-DEL-217 | chr17-29484170-DEL-217 |
| chr17-29492511-DEL-212 | chr17-29492512-DEL-212 |
| chr17-29504889-DEL-130 | chr17-29504890-DEL-130 |
| chr17-29512426-DEL-56 | chr17-29512427-DEL-56 |
| chr17-29527055-DEL-583 | chr17-29527056-DEL-583 |
| chr17-29563789-DEL-206 | chr17-29563790-DEL-206 |
| chr17-29570509-DEL-3429 | chr17-29570514-DEL-3425 |
| chr17-29630133-DEL-149 | chr17-29630161-DEL-130 |
| chr17-29653988-DEL-123 | chr17-29653989-DEL-123 |
| chr17-29658523-DEL-80 | chr17-29658524-DEL-80 |
| chr17-29662352-DEL-185 | chr17-29662353-DEL-184 |
| chr17-29665882-DEL-204 | chr17-29665883-DEL-204 |
| chr17-29671767-DEL-51 | chr17-29671789-DEL-64 |
| chr17-29673847-DEL-206 | chr17-29673848-DEL-206 |
| chr17-29684963-DEL-234 | chr17-29684964-DEL-234 |
| chr17-29684972-DEL-1352 | chr17-29684973-DEL-1352 |
| chr17-29690182-DEL-937 | chr17-29690183-DEL-937 |
| chr17-29707046-DEL-232 | chr17-29707047-DEL-232 |
| chr17-29725217-DEL-697 | chr17-29725218-DEL-697 |
| chr17-29748793-DEL-184 | chr17-29748794-DEL-184 |
| chr17-29750538-DEL-63 | chr17-29750539-DEL-63 |
| chr17-29758628-DEL-204 | chr17-29758630-DEL-203 |
| chr17-29759259-DEL-191 | chr17-29759260-DEL-191 |
| chr17-29761689-DEL-67 | chr17-29761690-DEL-67 |
| chr17-29780037-DEL-198 | chr17-29780038-DEL-198 |
| chr17-29785948-DEL-105 | chr17-29785949-DEL-105 |
| chr17-29811510-DEL-198 | chr17-29811511-DEL-198 |
| chr17-29838636-DEL-769 | chr17-29838637-DEL-769 |
| chr17-29863484-DEL-614 | chr17-29863509-DEL-615 |
| chr17-29923671-DEL-114 | chr17-29923672-DEL-114 |
| chr17-29941778-DEL-212 | chr17-29941779-DEL-212 |
| chr17-29979448-DEL-54 | chr17-29979449-DEL-54 |
| chr17-30000962-DEL-99 | chr17-30000963-DEL-91 |
| chr17-30012147-DEL-107 | chr17-30012148-DEL-107 |
| chr17-30077760-DEL-52 | chr17-30077761-DEL-52 |
| chr17-30091513-DEL-117 | chr17-30091497-DEL-96 |
| chr17-30099820-DEL-124 | chr17-30099821-DEL-124 |
| chr17-30100681-DEL-57 | chr17-30100682-DEL-57 |
| chr17-30160440-DEL-193 | chr17-30160465-DEL-172 |
| chr17-30179483-DEL-197 | chr17-30179484-DEL-197 |
| chr17-30196217-DEL-238 | chr17-30196218-DEL-238 |
| chr17-30206525-DEL-185 | chr17-30206521-DEL-186 |
| chr17-30210100-DEL-63 | chr17-30210101-DEL-63 |
| chr17-30247627-DEL-978 | chr17-30247628-DEL-978 |
| chr17-30253473-DEL-153 | chr17-30253474-DEL-153 |
| chr17-30258013-DEL-128 | chr17-30258014-DEL-128 |
| chr17-30258148-DEL-2588 | chr17-30258149-DEL-2588 |
| chr17-30285431-DEL-302 | chr17-30285432-DEL-302 |
| chr17-30312370-DEL-133 | chr17-30312371-DEL-133 |
| chr17-30361226-DEL-392 | chr17-30361227-DEL-392 |
| chr17-30410863-DEL-613 | chr17-30410861-DEL-615 |
| chr17-30430638-DEL-237 | chr17-30430639-DEL-237 |
| chr17-30500659-DEL-389 | chr17-30500628-DEL-390 |
| chr17-30574935-DEL-56 | chr17-30574936-DEL-56 |
| chr17-30592093-DEL-94 | chr17-30592094-DEL-94 |
| chr17-30665967-DEL-64 | chr17-30665968-DEL-64 |
| chr17-30674105-DEL-225 | chr17-30674106-DEL-225 |
| chr17-30686034-DEL-80 | chr17-30686035-DEL-79 |
| chr17-30690499-DEL-56 | chr17-30690508-DEL-76 |
| chr17-30724268-DEL-980 | chr17-30724269-DEL-980 |
| chr17-30737889-DEL-346 | chr17-30737890-DEL-346 |
| chr17-30752823-DEL-221 | chr17-30752824-DEL-221 |
| chr17-30773429-DEL-443 | chr17-30773430-DEL-443 |
| chr17-30849807-DEL-3748 | chr17-30849808-DEL-3748 |
| chr17-30863676-DEL-236 | chr17-30863677-DEL-236 |
| chr17-30886051-DEL-116 | chr17-30886043-DEL-111 |
| chr17-30979841-DEL-53 | chr17-30979842-DEL-53 |

| | |
|---|---|
| chr17-30982058-DEL-140 | chr17-30982059-DEL-140 |
| chr17-31016942-DEL-288 | chr17-31016959-DEL-252 |
| chr17-31025949-DEL-72 | chr17-31025950-DEL-72 |
| chr17-31059768-DEL-71 | chr17-31059779-DEL-56 |
| chr17-31076421-DEL-133 | chr17-31076420-DEL-133 |
| chr17-31080473-DEL-76 | chr17-31080474-DEL-96 |
| chr17-31095906-DEL-100 | chr17-31095942-DEL-104 |
| chr17-31119174-DEL-53 | chr17-31119177-DEL-54 |
| chr17-31145366-DEL-71 | chr17-31145367-DEL-71 |
| chr17-31187691-DEL-51 | chr17-31187692-DEL-51 |
| chr17-31191409-DEL-123 | chr17-31191410-DEL-123 |
| chr17-31193770-DEL-59 | chr17-31193771-DEL-59 |
| chr17-31195552-DEL-75 | chr17-31195552-DEL-74 |
| chr17-31204302-DEL-65 | chr17-31204297-DEL-62 |
| chr17-31210335-DEL-126 | chr17-31210336-DEL-126 |
| chr17-31227338-DEL-64 | chr17-31227315-DEL-51 |
| chr17-31233296-DEL-238 | chr17-31233283-DEL-241 |
| chr17-31240740-DEL-244 | chr17-31240741-DEL-244 |
| chr17-31244799-DEL-54 | chr17-31244800-DEL-54 |
| chr17-31265472-DEL-499 | chr17-31265473-DEL-498 |
| chr17-31281793-DEL-225 | chr17-31281794-DEL-225 |
| chr17-31302684-DEL-201 | chr17-31302685-DEL-201 |
| chr17-31314097-DEL-66 | chr17-31314098-DEL-66 |
| chr17-31318805-DEL-76 | chr17-31318806-DEL-76 |
| chr17-31326216-DEL-378 | chr17-31326217-DEL-378 |
| chr17-31356093-DEL-59 | chr17-31356088-DEL-65 |
| chr17-31420664-DEL-2131 | chr17-31420665-DEL-2131 |
| chr17-31430065-DEL-120 | chr17-31430066-DEL-120 |
| chr17-31437595-DEL-163 | chr17-31437596-DEL-163 |
| chr17-31441140-DEL-72 | chr17-31441142-DEL-71 |
| chr17-31448764-DEL-780 | chr17-31448765-DEL-769 |
| chr17-31449934-DEL-301 | chr17-31449935-DEL-301 |
| chr17-31453423-DEL-77 | chr17-31453427-DEL-69 |
| chr17-31463760-DEL-198 | chr17-31463761-DEL-198 |
| chr17-31526431-DEL-211 | chr17-31526432-DEL-211 |
| chr17-31553859-DEL-76 | chr17-31553860-DEL-76 |
| chr17-31581799-DEL-176 | chr17-31581800-DEL-176 |
| chr17-31585709-DEL-77 | chr17-31585710-DEL-77 |
| chr17-31613124-DEL-89 | chr17-31613125-DEL-89 |
| chr17-31656235-DEL-207 | chr17-31656256-DEL-204 |
| chr17-31658321-DEL-210 | chr17-31658322-DEL-210 |
| chr17-31664890-DEL-62 | chr17-31664883-DEL-50 |
| chr17-31669099-DEL-70 | chr17-31669100-DEL-71 |
| chr17-31691389-DEL-100 | chr17-31691390-DEL-100 |
| chr17-31694201-DEL-192 | chr17-31694202-DEL-192 |
| chr17-31695611-DEL-51 | chr17-31695578-DEL-60 |
| chr17-31702735-DEL-1823 | chr17-31702736-DEL-1823 |
| chr17-31742991-DEL-1338 | chr17-31742994-DEL-1336 |
| chr17-31745315-DEL-176 | chr17-31745316-DEL-176 |
| chr17-31751553-DEL-209 | chr17-31751557-DEL-208 |
| chr17-31771387-DEL-1663 | chr17-31771388-DEL-1663 |
| chr17-31780157-DEL-762 | chr17-31780158-DEL-762 |
| chr17-31813259-DEL-68 | chr17-31813245-DEL-86 |
| chr17-31853529-DEL-68 | chr17-31853529-DEL-68 |
| chr17-31861683-DEL-204 | chr17-31861683-DEL-204 |
| chr17-31874194-DEL-205 | chr17-31874190-DEL-206 |
| chr17-31885262-DEL-59 | chr17-31885263-DEL-59 |
| chr17-31893910-DEL-184 | chr17-31893911-DEL-184 |
| chr17-31902487-DEL-138 | chr17-31902488-DEL-138 |
| chr17-31914262-DEL-204 | chr17-31914263-DEL-204 |
| chr17-31916159-DEL-238 | chr17-31916160-DEL-238 |
| chr17-31939766-DEL-58 | chr17-31939753-DEL-58 |
| chr17-31947703-DEL-191 | chr17-31947704-DEL-191 |
| chr17-31955368-DEL-1540 | chr17-31955369-DEL-1540 |
| chr17-31959123-DEL-119 | chr17-31959124-DEL-119 |
| chr17-31978388-DEL-1172 | chr17-31978388-DEL-1173 |
| chr17-31979735-DEL-68 | chr17-31979735-DEL-68 |
| chr17-32005151-DEL-1806 | chr17-32005152-DEL-1806 |
| chr17-32017866-DEL-2879 | chr17-32017867-DEL-2879 |
| chr17-32026857-DEL-78 | chr17-32026864-DEL-66 |
| chr17-32098067-DEL-54 | chr17-32098068-DEL-54 |
| chr17-32123672-DEL-68 | chr17-32123691-DEL-64 |
| chr17-32136725-DEL-82 | chr17-32136726-DEL-82 |
| chr17-32159125-DEL-325 | chr17-32159126-DEL-325 |
| chr17-32175620-DEL-4785 | chr17-32175621-DEL-4785 |
| chr17-32191366-DEL-466 | chr17-32191385-DEL-416 |
| chr17-32206970-DEL-79 | chr17-32206971-DEL-79 |
| chr17-32207789-DEL-346 | chr17-32207790-DEL-346 |
| chr17-32221482-DEL-120 | chr17-32221483-DEL-120 |
| chr17-32236697-DEL-252 | chr17-32236710-DEL-245 |

| | |
|---|---|
| chr17-32254109-DEL-487 | chr17-32254104-DEL-484 |
| chr17-32295049-DEL-68 | chr17-32295066-DEL-52 |
| chr17-32304652-DEL-176 | chr17-32304653-DEL-176 |
| chr17-32306521-DEL-104 | chr17-32306521-DEL-104 |
| chr17-32319339-DEL-576 | chr17-32319340-DEL-573 |
| chr17-32323326-DEL-70 | chr17-32323336-DEL-85 |
| chr17-32331628-DEL-205 | chr17-32331629-DEL-205 |
| chr17-32351862-DEL-194 | chr17-32351863-DEL-194 |
| chr17-32355984-DEL-403 | chr17-32355985-DEL-403 |
| chr17-32381799-DEL-51 | chr17-32381800-DEL-51 |
| chr17-32389001-DEL-66 | chr17-32389002-DEL-66 |
| chr17-32396815-DEL-200 | chr17-32396816-DEL-200 |
| chr17-32406850-DEL-64 | chr17-32406850-DEL-64 |
| chr17-32410647-DEL-90 | chr17-32410648-DEL-90 |
| chr17-32410835-DEL-1315 | chr17-32410838-DEL-1313 |
| chr17-32419490-DEL-71 | chr17-32419491-DEL-71 |
| chr17-32430443-DEL-125 | chr17-32430443-DEL-125 |
| chr17-32431575-DEL-117 | chr17-32431576-DEL-117 |
| chr17-32439302-DEL-60 | chr17-32439303-DEL-54 |
| chr17-32462729-DEL-52 | chr17-32462692-DEL-50 |
| chr17-32466759-DEL-158 | chr17-32466760-DEL-114 |
| chr17-32480348-DEL-208 | chr17-32480349-DEL-208 |
| chr17-32481632-DEL-103 | chr17-32481635-DEL-99 |
| chr17-32482403-DEL-229 | chr17-32482404-DEL-229 |
| chr17-32521421-DEL-168 | chr17-32521423-DEL-167 |
| chr17-32584765-DEL-180 | chr17-32584766-DEL-180 |
| chr17-32588389-DEL-7101 | chr17-32588390-DEL-7101 |
| chr17-32626487-DEL-133 | chr17-32626488-DEL-133 |
| chr17-32642092-DEL-400 | chr17-32642093-DEL-400 |
| chr17-32729787-DEL-1222 | chr17-32729788-DEL-1222 |
| chr17-32733164-DEL-108 | chr17-32733165-DEL-108 |
| chr17-32758643-DEL-76 | chr17-32758627-DEL-100 |
| chr17-32774090-DEL-6007 | chr17-32774091-DEL-6007 |
| chr17-32807286-DEL-954 | chr17-32807287-DEL-954 |
| chr17-32814314-DEL-2574 | chr17-32814315-DEL-2574 |
| chr17-32826286-DEL-72 | chr17-32826299-DEL-52 |
| chr17-32842202-DEL-133 | chr17-32842202-DEL-134 |
| chr17-32851533-DEL-70 | chr17-32851534-DEL-70 |
| chr17-32861455-DEL-168 | chr17-32861456-DEL-167 |
| chr17-32862239-DEL-435 | chr17-32862240-DEL-435 |
| chr17-32871767-DEL-177 | chr17-32871768-DEL-177 |
| chr17-32876339-DEL-560 | chr17-32876340-DEL-560 |
| chr17-32879753-DEL-56 | chr17-32879754-DEL-56 |
| chr17-32891452-DEL-61 | chr17-32891448-DEL-71 |
| chr17-32919522-DEL-84 | chr17-32919524-DEL-82 |
| chr17-32924103-DEL-3422 | chr17-32924104-DEL-3422 |
| chr17-32946225-DEL-207 | chr17-32946226-DEL-207 |
| chr17-32960296-DEL-727 | chr17-32960297-DEL-727 |
| chr17-32983438-DEL-425 | chr17-32983439-DEL-425 |
| chr17-33024466-DEL-58 | chr17-33024459-DEL-59 |
| chr17-33056086-DEL-205 | chr17-33056086-DEL-205 |
| chr17-33066796-DEL-86 | chr17-33066782-DEL-87 |
| chr17-33119420-DEL-719 | chr17-33119421-DEL-719 |
| chr17-33133238-DEL-79 | chr17-33133237-DEL-75 |
| chr17-33170857-DEL-773 | chr17-33170858-DEL-771 |
| chr17-33179926-DEL-83 | chr17-33179930-DEL-74 |
| chr17-33191688-DEL-1845 | chr17-33191689-DEL-1845 |
| chr17-33198118-DEL-1744 | chr17-33198118-DEL-1744 |
| chr17-33214665-DEL-367 | chr17-33214666-DEL-367 |
| chr17-33216378-DEL-190 | chr17-33216379-DEL-190 |
| chr17-33253707-DEL-67 | chr17-33253708-DEL-67 |
| chr17-33257186-DEL-214 | chr17-33257187-DEL-214 |
| chr17-33262913-DEL-8915 | chr17-33262914-DEL-8915 |
| chr17-33273659-DEL-3377 | chr17-33273660-DEL-3377 |
| chr17-33277107-DEL-435 | chr17-33277108-DEL-435 |
| chr17-33336667-DEL-704 | chr17-33336668-DEL-704 |
| chr17-33359918-DEL-927 | chr17-33359919-DEL-927 |
| chr17-33366378-DEL-84 | chr17-33366379-DEL-84 |
| chr17-33425216-DEL-86 | chr17-33425233-DEL-72 |
| chr17-33458332-DEL-72 | chr17-33458332-DEL-68 |
| chr17-33535057-DEL-922 | chr17-33535058-DEL-922 |
| chr17-33541038-DEL-86 | chr17-33541039-DEL-86 |
| chr17-33574270-DEL-134 | chr17-33574271-DEL-134 |
| chr17-33586465-DEL-1049 | chr17-33586466-DEL-1049 |
| chr17-33588749-DEL-80 | chr17-33588782-DEL-60 |
| chr17-33591965-DEL-98 | chr17-33591965-DEL-99 |
| chr17-33636482-DEL-828 | chr17-33636483-DEL-828 |
| chr17-33637128-DEL-2604 | chr17-33637128-DEL-2604 |
| chr17-33639969-DEL-2266 | chr17-33639970-DEL-2266 |
| chr17-33654983-DEL-120 | chr17-33654978-DEL-78 |

| | |
|---|---|
| chr17-33673609-DEL-250 | chr17-33673610-DEL-250 |
| chr17-33674221-DEL-1140 | chr17-33674222-DEL-1140 |
| chr17-33677854-DEL-836 | chr17-33677855-DEL-836 |
| chr17-33724176-DEL-185 | chr17-33724177-DEL-185 |
| chr17-33727154-DEL-66 | chr17-33727155-DEL-66 |
| chr17-33729103-DEL-929 | chr17-33729104-DEL-929 |
| chr17-33773869-DEL-64 | chr17-33773872-DEL-62 |
| chr17-33779176-DEL-1106 | chr17-33779177-DEL-1106 |
| chr17-33787112-DEL-1893 | chr17-33787113-DEL-1893 |
| chr17-33818451-DEL-80 | chr17-33818452-DEL-80 |
| chr17-33837244-DEL-7111 | chr17-33837245-DEL-7111 |
| chr17-33920069-DEL-94 | chr17-33920060-DEL-56 |
| chr17-33946868-DEL-375 | chr17-33946869-DEL-375 |
| chr17-33947600-DEL-100 | chr17-33947601-DEL-100 |
| chr17-33954859-DEL-242 | chr17-33954860-DEL-242 |
| chr17-33957001-DEL-2835 | chr17-33957002-DEL-2835 |
| chr17-33979744-DEL-251 | chr17-33979745-DEL-258 |
| chr17-34023525-DEL-87 | chr17-34023526-DEL-87 |
| chr17-34028865-DEL-213 | chr17-34028866-DEL-213 |
| chr17-34039732-DEL-103 | chr17-34039733-DEL-103 |
| chr17-34050147-DEL-77 | chr17-34050148-DEL-69 |
| chr17-34051232-DEL-120 | chr17-34051233-DEL-120 |
| chr17-34051710-DEL-331 | chr17-34051713-DEL-328 |
| chr17-34052454-DEL-93 | chr17-34052455-DEL-93 |
| chr17-34057398-DEL-86 | chr17-34057399-DEL-86 |
| chr17-34071672-DEL-51 | chr17-34071660-DEL-68 |
| chr17-34074649-DEL-202 | chr17-34074650-DEL-202 |
| chr17-34084849-DEL-56 | chr17-34084849-DEL-56 |
| chr17-34086110-DEL-6475 | chr17-34086111-DEL-6475 |
| chr17-34094221-DEL-220 | chr17-34094222-DEL-220 |
| chr17-34095155-DEL-116 | chr17-34095156-DEL-116 |
| chr17-34096480-DEL-59 | chr17-34096481-DEL-59 |
| chr17-34099531-DEL-117 | chr17-34099532-DEL-117 |
| chr17-34100552-DEL-173 | chr17-34100564-DEL-172 |
| chr17-34113699-DEL-560 | chr17-34113699-DEL-560 |
| chr17-34114348-DEL-3411 | chr17-34114349-DEL-3411 |
| chr17-34114357-DEL-1153 | chr17-34114358-DEL-1153 |
| chr17-34124300-DEL-53 | chr17-34124296-DEL-58 |
| chr17-34153004-DEL-55 | chr17-34153007-DEL-53 |
| chr17-34156962-DEL-238 | chr17-34156963-DEL-238 |
| chr17-34162925-DEL-204 | chr17-34162926-DEL-204 |
| chr17-34186229-DEL-165 | chr17-34186230-DEL-165 |
| chr17-34191225-DEL-95 | chr17-34191226-DEL-95 |
| chr17-34200236-DEL-438 | chr17-34200237-DEL-438 |
| chr17-34213473-DEL-216 | chr17-34213474-DEL-216 |
| chr17-34223077-DEL-77 | chr17-34223077-DEL-77 |
| chr17-34231093-DEL-63 | chr17-34231094-DEL-63 |
| chr17-34268094-DEL-121 | chr17-34268095-DEL-121 |
| chr17-34286969-DEL-141 | chr17-34286998-DEL-103 |
| chr17-34296052-DEL-142 | chr17-34296052-DEL-142 |
| chr17-34327717-DEL-220 | chr17-34327718-DEL-220 |
| chr17-34343338-DEL-183 | chr17-34343339-DEL-183 |
| chr17-34382141-DEL-251 | chr17-34382143-DEL-252 |
| chr17-34382743-DEL-196 | chr17-34382744-DEL-196 |
| chr17-34386436-DEL-198 | chr17-34386437-DEL-198 |
| chr17-34394233-DEL-54 | chr17-34394234-DEL-54 |
| chr17-34422011-DEL-183 | chr17-34422012-DEL-183 |
| chr17-34429373-DEL-78 | chr17-34429374-DEL-78 |
| chr17-34438328-DEL-4797 | chr17-34438329-DEL-4797 |
| chr17-34444727-DEL-7189 | chr17-34444728-DEL-7189 |
| chr17-34462632-DEL-52 | chr17-34462633-DEL-52 |
| chr17-34465492-DEL-60 | chr17-34465489-DEL-60 |
| chr17-34473075-DEL-52 | chr17-34473076-DEL-52 |
| chr17-34484893-DEL-118 | chr17-34484894-DEL-106 |
| chr17-34487041-DEL-55 | chr17-34487042-DEL-55 |
| chr17-34494865-DEL-800 | chr17-34494865-DEL-800 |
| chr17-34518939-DEL-56 | chr17-34518940-DEL-54 |
| chr17-34523154-DEL-62 | chr17-34523155-DEL-62 |
| chr17-34527462-DEL-250 | chr17-34527463-DEL-250 |
| chr17-34543648-DEL-67 | chr17-34543670-DEL-54 |
| chr17-34569947-DEL-96 | chr17-34569948-DEL-96 |
| chr17-34602879-DEL-96 | chr17-34602873-DEL-97 |
| chr17-34605092-DEL-2074 | chr17-34605093-DEL-2042 |
| chr17-34607059-DEL-64 | chr17-34607060-DEL-64 |
| chr17-34609710-DEL-120 | chr17-34609711-DEL-120 |
| chr17-34612276-DEL-1477 | chr17-34612277-DEL-1477 |
| chr17-34625159-DEL-231 | chr17-34625160-DEL-231 |
| chr17-34677402-DEL-313 | chr17-34677403-DEL-313 |
| chr17-34719660-DEL-100 | chr17-34719657-DEL-100 |
| chr17-34728855-DEL-60 | chr17-34728851-DEL-64 |

| | |
|---|---|
| chr17-34748050-DEL-240 | chr17-34748051-DEL-240 |
| chr17-34779216-DEL-243 | chr17-34779217-DEL-243 |
| chr17-34836954-DEL-127 | chr17-34836955-DEL-127 |
| chr17-34905767-DEL-1009 | chr17-34905768-DEL-1009 |
| chr17-34944783-DEL-52 | chr17-34944784-DEL-52 |
| chr17-34987542-DEL-528 | chr17-34987543-DEL-528 |
| chr17-35010230-DEL-208 | chr17-35010231-DEL-204 |
| chr17-35089820-DEL-54 | chr17-35089821-DEL-54 |
| chr17-35109895-DEL-225 | chr17-35109896-DEL-225 |
| chr17-35163779-DEL-51 | chr17-35163780-DEL-51 |
| chr17-35168132-DEL-200 | chr17-35168120-DEL-207 |
| chr17-35168662-DEL-64 | chr17-35168663-DEL-64 |
| chr17-35181449-DEL-54 | chr17-35181450-DEL-52 |
| chr17-35219184-DEL-534 | chr17-35219185-DEL-534 |
| chr17-35225145-DEL-213 | chr17-35225146-DEL-213 |
| chr17-35254375-DEL-231 | chr17-35254376-DEL-231 |
| chr17-35261789-DEL-208 | chr17-35261790-DEL-208 |
| chr17-35328819-DEL-65 | chr17-35328817-DEL-78 |
| chr17-35352404-DEL-151 | chr17-35352405-DEL-151 |
| chr17-35369922-DEL-223 | chr17-35369923-DEL-223 |
| chr17-35411107-DEL-222 | chr17-35411108-DEL-222 |
| chr17-35441791-DEL-53 | chr17-35441791-DEL-53 |
| chr17-35443008-DEL-790 | chr17-35443009-DEL-790 |
| chr17-35449211-DEL-56 | chr17-35449212-DEL-56 |
| chr17-35470612-DEL-214 | chr17-35470613-DEL-214 |
| chr17-35472934-DEL-52 | chr17-35472928-DEL-50 |
| chr17-35499311-DEL-408 | chr17-35499312-DEL-408 |
| chr17-35500960-DEL-646 | chr17-35500961-DEL-646 |
| chr17-35504215-DEL-55 | chr17-35504216-DEL-55 |
| chr17-35506975-DEL-183 | chr17-35506974-DEL-183 |
| chr17-35554824-DEL-1229 | chr17-35554817-DEL-1230 |
| chr17-35561995-DEL-73 | chr17-35561995-DEL-73 |
| chr17-35600248-DEL-59 | chr17-35600249-DEL-59 |
| chr17-35601551-DEL-211 | chr17-35601552-DEL-211 |
| chr17-35605046-DEL-75 | chr17-35605047-DEL-78 |
| chr17-35628228-DEL-1919 | chr17-35628229-DEL-1919 |
| chr17-35633969-DEL-80 | chr17-35633969-DEL-80 |
| chr17-35635649-DEL-208 | chr17-35635650-DEL-208 |
| chr17-35653612-DEL-132 | chr17-35653611-DEL-132 |
| chr17-35742728-DEL-326 | chr17-35742729-DEL-326 |
| chr17-35747540-DEL-132 | chr17-35747541-DEL-132 |
| chr17-35772782-DEL-205 | chr17-35772782-DEL-205 |
| chr17-35779559-DEL-170 | chr17-35779552-DEL-170 |
| chr17-35789101-DEL-4958 | chr17-35789102-DEL-4958 |
| chr17-35797370-DEL-196 | chr17-35797370-DEL-196 |
| chr17-35800074-DEL-61 | chr17-35800094-DEL-60 |
| chr17-35800528-DEL-1951 | chr17-35800529-DEL-1951 |
| chr17-35819436-DEL-201 | chr17-35819437-DEL-199 |
| chr17-35834626-DEL-60 | chr17-35834626-DEL-61 |
| chr17-35852964-DEL-195 | chr17-35852965-DEL-195 |
| chr17-35884821-DEL-80 | chr17-35884822-DEL-60 |
| chr17-35894154-DEL-104 | chr17-35894155-DEL-104 |
| chr17-35894599-DEL-297 | chr17-35894600-DEL-297 |
| chr17-35900617-DEL-173 | chr17-35900619-DEL-172 |
| chr17-35902444-DEL-153 | chr17-35902445-DEL-153 |
| chr17-35904685-DEL-70 | chr17-35904657-DEL-67 |
| chr17-35912503-DEL-96 | chr17-35912506-DEL-88 |
| chr17-35928177-DEL-57 | chr17-35928178-DEL-57 |
| chr17-35930587-DEL-90 | chr17-35930588-DEL-90 |
| chr17-35930601-DEL-1035 | chr17-35930601-DEL-1035 |
| chr17-35932449-DEL-225 | chr17-35932450-DEL-225 |
| chr17-35933004-DEL-270 | chr17-35933005-DEL-270 |
| chr17-35934438-DEL-336 | chr17-35934439-DEL-336 |
| chr17-35938709-DEL-74 | chr17-35938700-DEL-75 |
| chr17-35973206-DEL-532 | chr17-35973207-DEL-532 |
| chr17-35986676-DEL-499 | chr17-35986677-DEL-499 |
| chr17-35987323-DEL-441 | chr17-35987324-DEL-441 |
| chr17-36020439-DEL-586 | chr17-36020440-DEL-586 |
| chr17-36022635-DEL-112 | chr17-36022636-DEL-112 |
| chr17-36024866-DEL-199 | chr17-36024867-DEL-199 |
| chr17-36031332-DEL-619 | chr17-36031333-DEL-619 |
| chr17-36043641-DEL-4085 | chr17-36043642-DEL-4085 |
| chr17-36057086-DEL-116 | chr17-36057085-DEL-117 |
| chr17-36105214-DEL-208 | chr17-36105215-DEL-208 |
| chr17-36108459-DEL-207 | chr17-36108460-DEL-207 |
| chr17-36110818-DEL-84 | chr17-36110804-DEL-93 |
| chr17-36115923-DEL-205 | chr17-36115931-DEL-207 |
| chr17-36129353-DEL-213 | chr17-36129354-DEL-176 |
| chr17-36162225-DEL-213 | chr17-36162226-DEL-213 |
| chr17-36182671-DEL-207 | chr17-36182672-DEL-207 |

| | |
|---|---|
| chr17-36186515-DEL-253 | chr17-36186516-DEL-253 |
| chr17-36198597-DEL-58 | chr17-36198597-DEL-58 |
| chr17-36247577-DEL-132 | chr17-36247578-DEL-132 |
| chr17-36251895-DEL-5575 | chr17-36251895-DEL-5575 |
| chr17-36252635-DEL-77 | chr17-36252636-DEL-77 |
| chr17-36268175-DEL-60 | chr17-36268221-DEL-60 |
| chr17-36274965-DEL-90 | chr17-36274966-DEL-90 |
| chr17-36295543-DEL-108 | chr17-36295542-DEL-109 |
| chr17-36301221-DEL-320 | chr17-36301219-DEL-321 |
| chr17-36313199-DEL-2832 | chr17-36313200-DEL-2832 |
| chr17-36318029-DEL-7105 | chr17-36318030-DEL-7105 |
| chr17-36328782-DEL-702 | chr17-36328783-DEL-702 |
| chr17-36330143-DEL-63 | chr17-36330145-DEL-62 |
| chr17-36357281-DEL-179 | chr17-36357299-DEL-164 |
| chr17-36359905-DEL-7113 | chr17-36359906-DEL-7113 |
| chr17-36444437-DEL-197 | chr17-36444436-DEL-197 |
| chr17-36452629-DEL-193 | chr17-36452629-DEL-193 |
| chr17-36453673-DEL-197 | chr17-36453673-DEL-197 |
| chr17-36458767-DEL-197 | chr17-36458767-DEL-197 |
| chr17-36477645-DEL-62 | chr17-36477637-DEL-67 |
| chr17-36480489-DEL-514 | chr17-36480491-DEL-518 |
| chr17-36485496-DEL-121 | chr17-36485496-DEL-122 |
| chr17-36509945-DEL-263 | chr17-36509946-DEL-263 |
| chr17-36517549-DEL-218 | chr17-36517535-DEL-219 |
| chr17-36524676-DEL-92 | chr17-36524678-DEL-90 |
| chr17-36536860-DEL-501 | chr17-36536860-DEL-501 |
| chr17-36540453-DEL-94 | chr17-36540454-DEL-93 |
| chr17-36604988-DEL-3535 | chr17-36604989-DEL-3535 |
| chr17-36654485-DEL-6093 | chr17-36654486-DEL-6093 |
| chr17-36660407-DEL-64 | chr17-36660427-DEL-64 |
| chr17-36698096-DEL-115 | chr17-36698096-DEL-126 |
| chr17-36716541-DEL-1877 | chr17-36716542-DEL-1877 |
| chr17-36739086-DEL-1096 | chr17-36739081-DEL-1097 |
| chr17-36744951-DEL-139 | chr17-36744952-DEL-139 |
| chr17-36752533-DEL-651 | chr17-36752534-DEL-651 |
| chr17-36764497-DEL-100 | chr17-36764498-DEL-100 |
| chr17-36784987-DEL-6442 | chr17-36784988-DEL-6442 |
| chr17-36795517-DEL-1025 | chr17-36795518-DEL-1025 |
| chr17-36818148-DEL-73 | chr17-36818149-DEL-73 |
| chr17-36831210-DEL-6413 | chr17-36831211-DEL-6413 |
| chr17-36895773-DEL-771 | chr17-36895782-DEL-767 |
| chr17-36902382-DEL-1810 | chr17-36902383-DEL-1810 |
| chr17-36913662-DEL-1819 | chr17-36913663-DEL-1819 |
| chr17-36920796-DEL-6570 | chr17-36920797-DEL-6570 |
| chr17-36937274-DEL-2837 | chr17-36937275-DEL-2837 |
| chr17-37018201-DEL-9814 | chr17-37018202-DEL-9814 |
| chr17-37018217-DEL-212 | chr17-37018217-DEL-212 |
| chr17-37094379-DEL-896 | chr17-37094380-DEL-896 |
| chr17-37110625-DEL-547 | chr17-37110626-DEL-548 |
| chr17-37111882-DEL-644 | chr17-37111895-DEL-643 |
| chr17-37118963-DEL-65 | chr17-37118964-DEL-65 |
| chr17-37129169-DEL-51 | chr17-37129170-DEL-51 |
| chr17-37222410-DEL-190 | chr17-37222411-DEL-190 |
| chr17-37236403-DEL-100 | chr17-37236408-DEL-96 |
| chr17-37244374-DEL-198 | chr17-37244375-DEL-198 |
| chr17-37345034-DEL-164 | chr17-37345030-DEL-160 |
| chr17-37394626-DEL-625 | chr17-37394643-DEL-609 |
| chr17-37400223-DEL-62 | chr17-37400255-DEL-72 |
| chr17-37436074-DEL-938 | chr17-37436075-DEL-938 |
| chr17-37438686-DEL-198 | chr17-37438697-DEL-199 |
| chr17-37439379-DEL-66 | chr17-37439392-DEL-54 |
| chr17-37461036-DEL-71 | chr17-37461037-DEL-71 |
| chr17-37502055-DEL-1250 | chr17-37502056-DEL-1250 |
| chr17-37510840-DEL-191 | chr17-37510841-DEL-191 |
| chr17-37598576-DEL-328 | chr17-37598577-DEL-328 |
| chr17-37640079-DEL-1906 | chr17-37640080-DEL-1906 |
| chr17-37642623-DEL-138 | chr17-37642621-DEL-134 |
| chr17-37690905-DEL-335 | chr17-37690906-DEL-335 |
| chr17-37704498-DEL-235 | chr17-37704499-DEL-235 |
| chr17-37727947-DEL-588 | chr17-37727948-DEL-588 |
| chr17-37749760-DEL-6128 | chr17-37749761-DEL-6128 |
| chr17-37764380-DEL-57 | chr17-37764350-DEL-54 |
| chr17-37767166-DEL-173 | chr17-37767166-DEL-174 |
| chr17-37820822-DEL-981 | chr17-37820818-DEL-984 |
| chr17-37827854-DEL-65 | chr17-37827854-DEL-64 |
| chr17-37860387-DEL-400 | chr17-37860388-DEL-400 |
| chr17-37863123-DEL-6178 | chr17-37863124-DEL-6178 |
| chr17-37880988-DEL-67 | chr17-37880989-DEL-67 |
| chr17-37892489-DEL-60 | chr17-37892495-DEL-55 |
| chr17-37946089-DEL-3957 | chr17-37946090-DEL-3957 |

| | |
|---|---|
| chr17-37946564-DEL-78 | chr17-37946565-DEL-78 |
| chr17-37952425-DEL-990 | chr17-37952427-DEL-991 |
| chr17-37971356-DEL-516 | chr17-37971357-DEL-514 |
| chr17-37981360-DEL-391 | chr17-37981361-DEL-391 |
| chr17-38005677-DEL-56 | chr17-38005678-DEL-56 |
| chr17-38007158-DEL-112 | chr17-38007167-DEL-104 |
| chr17-38023034-DEL-1792 | chr17-38023021-DEL-1812 |
| chr17-38059741-DEL-86 | chr17-38059742-DEL-86 |
| chr17-38121367-DEL-76 | chr17-38121368-DEL-76 |
| chr17-38145095-DEL-72 | chr17-38145088-DEL-68 |
| chr17-38153766-DEL-215 | chr17-38153769-DEL-214 |
| chr17-38164398-DEL-6582 | chr17-38164398-DEL-6582 |
| chr17-38187538-DEL-6297 | chr17-38187539-DEL-6297 |
| chr17-38256613-DEL-327 | chr17-38256614-DEL-327 |
| chr17-38264891-DEL-7074 | chr17-38264892-DEL-7074 |
| chr17-38545007-DEL-788 | chr17-38545008-DEL-788 |
| chr17-38575706-DEL-50 | chr17-38575732-DEL-62 |
| chr17-38578678-DEL-4817 | chr17-38578679-DEL-4817 |
| chr17-38609149-DEL-52 | chr17-38609150-DEL-52 |
| chr17-38709255-DEL-3169 | chr17-38709256-DEL-3169 |
| chr17-38842991-DEL-465 | chr17-38842997-DEL-468 |
| chr17-38847227-DEL-6597 | chr17-38847228-DEL-6597 |
| chr17-38861080-DEL-90 | chr17-38861109-DEL-80 |
| chr17-38881089-DEL-59 | chr17-38881088-DEL-61 |
| chr17-38946564-DEL-159 | chr17-38946607-DEL-130 |
| chr17-38987406-DEL-10940 | chr17-38987406-DEL-10940 |
| chr17-38994026-DEL-208 | chr17-38994027-DEL-208 |
| chr17-38998600-DEL-793 | chr17-38998601-DEL-793 |
| chr17-39133685-DEL-2521 | chr17-39133686-DEL-2521 |
| chr17-39181487-DEL-203 | chr17-39181488-DEL-203 |
| chr17-39201597-DEL-13193 | chr17-39201597-DEL-13193 |
| chr17-39202014-DEL-212 | chr17-39202015-DEL-212 |
| chr17-39278402-DEL-76 | chr17-39278404-DEL-54 |
| chr17-39300842-DEL-75 | chr17-39300843-DEL-75 |
| chr17-39334433-DEL-54 | chr17-39334434-DEL-54 |
| chr17-39394679-DEL-64 | chr17-39394681-DEL-90 |
| chr17-39581949-DEL-4339 | chr17-39581950-DEL-4339 |
| chr17-39607389-DEL-117 | chr17-39607390-DEL-117 |
| chr17-39657525-DEL-5192 | chr17-39657526-DEL-5192 |
| chr17-39728422-DEL-5828 | chr17-39728422-DEL-5828 |
| chr17-39776124-DEL-78 | chr17-39776125-DEL-78 |
| chr17-39875140-DEL-56 | chr17-39875141-DEL-56 |
| chr17-39884812-DEL-57 | chr17-39884812-DEL-57 |
| chr17-39939165-DEL-1532 | chr17-39939166-DEL-1532 |
| chr17-40073339-DEL-6823 | chr17-40073340-DEL-6823 |
| chr17-40096985-DEL-331 | chr17-40096986-DEL-331 |
| chr17-40153875-DEL-6116 | chr17-40153874-DEL-6116 |
| chr17-40206512-DEL-678 | chr17-40206512-DEL-678 |
| chr17-40216607-DEL-1972 | chr17-40216608-DEL-1972 |
| chr17-40263389-DEL-62 | chr17-40263388-DEL-56 |
| chr17-40303260-DEL-6663 | chr17-40303261-DEL-6663 |
| chr17-40431427-DEL-6578 | chr17-40431428-DEL-6578 |
| chr17-40481656-DEL-1299 | chr17-40481657-DEL-1299 |
| chr17-40487145-DEL-373 | chr17-40487145-DEL-373 |
| chr17-40569387-DEL-250 | chr17-40569381-DEL-256 |
| chr17-40590604-DEL-775 | chr17-40590605-DEL-775 |
| chr17-40624655-DEL-3139 | chr17-40624656-DEL-3139 |
| chr17-40692780-DEL-58 | chr17-40692781-DEL-57 |
| chr17-40694327-DEL-611 | chr17-40694330-DEL-609 |
| chr17-40710504-DEL-139 | chr17-40710506-DEL-154 |
| chr17-40720604-DEL-67 | chr17-40720605-DEL-67 |
| chr17-40723208-DEL-59 | chr17-40723182-DEL-59 |
| chr17-40792407-DEL-841 | chr17-40792408-DEL-841 |
| chr17-40804736-DEL-91 | chr17-40804737-DEL-91 |
| chr17-40840722-DEL-61 | chr17-40840723-DEL-61 |
| chr17-41001668-DEL-6975 | chr17-41001669-DEL-6975 |
| chr17-41024784-DEL-275 | chr17-41024781-DEL-276 |
| chr17-41030400-DEL-333 | chr17-41030401-DEL-333 |
| chr17-41032747-DEL-220 | chr17-41032747-DEL-220 |
| chr17-41038118-DEL-184 | chr17-41038119-DEL-183 |
| chr17-41070421-DEL-1476 | chr17-41070422-DEL-1476 |
| chr17-41071902-DEL-671 | chr17-41071902-DEL-667 |
| chr17-41087677-DEL-64 | chr17-41087678-DEL-64 |
| chr17-41106802-DEL-6387 | chr17-41106803-DEL-6387 |
| chr17-41126193-DEL-1430 | chr17-41126194-DEL-1430 |
| chr17-41148913-DEL-161 | chr17-41148897-DEL-139 |
| chr17-41169418-DEL-165 | chr17-41169419-DEL-165 |
| chr17-41197226-DEL-510 | chr17-41197227-DEL-510 |
| chr17-41199765-DEL-213 | chr17-41199766-DEL-213 |
| chr17-41204726-DEL-160 | chr17-41204723-DEL-161 |

| | |
|---|---|
| chr17-41205485-DEL-74 | chr17-41205485-DEL-70 |
| chr17-41212703-DEL-251 | chr17-41212703-DEL-251 |
| chr17-41216106-DEL-74 | chr17-41216107-DEL-74 |
| chr17-41218581-DEL-72 | chr17-41218582-DEL-72 |
| chr17-41219690-DEL-54 | chr17-41219691-DEL-54 |
| chr17-41255309-DEL-101 | chr17-41255310-DEL-101 |
| chr17-41263880-DEL-821 | chr17-41263881-DEL-821 |
| chr17-41268547-DEL-91 | chr17-41268548-DEL-91 |
| chr17-41270273-DEL-78 | chr17-41270274-DEL-78 |
| chr17-41301611-DEL-695 | chr17-41301612-DEL-695 |
| chr17-41308463-DEL-1194 | chr17-41308464-DEL-1195 |
| chr17-41320623-DEL-196 | chr17-41320624-DEL-196 |
| chr17-41333079-DEL-666 | chr17-41333080-DEL-666 |
| chr17-41340033-DEL-1846 | chr17-41340033-DEL-1846 |
| chr17-41351141-DEL-196 | chr17-41351142-DEL-196 |
| chr17-41356490-DEL-8020 | chr17-41356486-DEL-8001 |
| chr17-41357279-DEL-1131 | chr17-41357279-DEL-1131 |
| chr17-41379594-DEL-91 | chr17-41379595-DEL-91 |
| chr17-41394397-DEL-441 | chr17-41394398-DEL-441 |
| chr17-41397237-DEL-623 | chr17-41397238-DEL-623 |
| chr17-41399912-DEL-90 | chr17-41399913-DEL-92 |
| chr17-41401871-DEL-199 | chr17-41401872-DEL-199 |
| chr17-41417795-DEL-146 | chr17-41417796-DEL-146 |
| chr17-41438974-DEL-1336 | chr17-41438958-DEL-1352 |
| chr17-41442136-DEL-266 | chr17-41442136-DEL-266 |
| chr17-41445432-DEL-105 | chr17-41445434-DEL-105 |
| chr17-41488615-DEL-128 | chr17-41488616-DEL-128 |
| chr17-41489386-DEL-466 | chr17-41489387-DEL-466 |
| chr17-41490742-DEL-441 | chr17-41490743-DEL-441 |
| chr17-41495845-DEL-222 | chr17-41495846-DEL-222 |
| chr17-41541023-DEL-255 | chr17-41541024-DEL-255 |
| chr17-41542734-DEL-464 | chr17-41542735-DEL-464 |
| chr17-41550964-DEL-581 | chr17-41550965-DEL-581 |
| chr17-41551522-DEL-557 | chr17-41551522-DEL-558 |
| chr17-41573029-DEL-228 | chr17-41573031-DEL-227 |
| chr17-41581197-DEL-396 | chr17-41581198-DEL-396 |
| chr17-41591319-DEL-1107 | chr17-41591320-DEL-1107 |
| chr17-41600596-DEL-751 | chr17-41600597-DEL-751 |
| chr17-41616427-DEL-6401 | chr17-41616428-DEL-6401 |
| chr17-41632056-DEL-2185 | chr17-41632057-DEL-2185 |
| chr17-41673312-DEL-122 | chr17-41673313-DEL-122 |
| chr17-41768784-DEL-1945 | chr17-41768785-DEL-1945 |
| chr17-41809165-DEL-69 | chr17-41809144-DEL-50 |
| chr17-41849796-DEL-78 | chr17-41849797-DEL-78 |
| chr17-41879857-DEL-739 | chr17-41879858-DEL-739 |
| chr17-41885747-DEL-3644 | chr17-41885748-DEL-3644 |
| chr17-41934789-DEL-409 | chr17-41934790-DEL-409 |
| chr17-41948717-DEL-85 | chr17-41948718-DEL-85 |
| chr17-41952410-DEL-917 | chr17-41952411-DEL-917 |
| chr17-41968680-DEL-6440 | chr17-41968681-DEL-6440 |
| chr17-41980725-DEL-1789 | chr17-41980726-DEL-1789 |
| chr17-41996868-DEL-88 | chr17-41996872-DEL-85 |
| chr17-42003349-DEL-66 | chr17-42003350-DEL-66 |
| chr17-42020813-DEL-517 | chr17-42020814-DEL-515 |
| chr17-42021948-DEL-8249 | chr17-42021949-DEL-8249 |
| chr17-42059779-DEL-58 | chr17-42059781-DEL-58 |
| chr17-42061964-DEL-461 | chr17-42061965-DEL-461 |
| chr17-42067192-DEL-51 | chr17-42067192-DEL-51 |
| chr17-42076201-DEL-174 | chr17-42076202-DEL-174 |
| chr17-42086024-DEL-690 | chr17-42086024-DEL-691 |
| chr17-42091441-DEL-90 | chr17-42091460-DEL-96 |
| chr17-42094781-DEL-5884 | chr17-42094782-DEL-5884 |
| chr17-42103946-DEL-54 | chr17-42103947-DEL-50 |
| chr17-42155013-DEL-70 | chr17-42155014-DEL-70 |
| chr17-42185011-DEL-6273 | chr17-42185011-DEL-6273 |
| chr17-42189089-DEL-1810 | chr17-42189091-DEL-1809 |
| chr17-42242824-DEL-948 | chr17-42242824-DEL-948 |
| chr17-42248994-DEL-1305 | chr17-42248995-DEL-1305 |
| chr17-42251165-DEL-1695 | chr17-42251166-DEL-1695 |
| chr17-42251254-DEL-299 | chr17-42251255-DEL-299 |
| chr17-42254316-DEL-70 | chr17-42254317-DEL-70 |
| chr17-42259135-DEL-58 | chr17-42259136-DEL-58 |
| chr17-42260101-DEL-3216 | chr17-42260102-DEL-3214 |
| chr17-42262339-DEL-121 | chr17-42262347-DEL-125 |
| chr17-42269451-DEL-54 | chr17-42269440-DEL-66 |
| chr17-42278713-DEL-950 | chr17-42278712-DEL-952 |
| chr17-42295506-DEL-976 | chr17-42295507-DEL-976 |
| chr17-42311389-DEL-7223 | chr17-42311390-DEL-7223 |
| chr17-42334360-DEL-66 | chr17-42334360-DEL-78 |
| chr17-42354222-DEL-56 | chr17-42354250-DEL-66 |

| | |
|---|---|
| chr17-42360031-DEL-1172 | chr17-42360032-DEL-1172 |
| chr17-42371841-DEL-60 | chr17-42371842-DEL-60 |
| chr17-42383467-DEL-208 | chr17-42383467-DEL-208 |
| chr17-42447888-DEL-2476 | chr17-42447889-DEL-2476 |
| chr17-42486092-DEL-4620 | chr17-42486093-DEL-4620 |
| chr17-42502084-DEL-91 | chr17-42502077-DEL-96 |
| chr17-42521752-DEL-1615 | chr17-42521752-DEL-1615 |
| chr17-42523702-DEL-3096 | chr17-42523697-DEL-3097 |
| chr17-42524579-DEL-58 | chr17-42524580-DEL-58 |
| chr17-42540355-DEL-466 | chr17-42540356-DEL-466 |
| chr17-42549259-DEL-942 | chr17-42549260-DEL-941 |
| chr17-42554663-DEL-739 | chr17-42554664-DEL-739 |
| chr17-42560231-DEL-164 | chr17-42560232-DEL-164 |
| chr17-42562782-DEL-240 | chr17-42562783-DEL-240 |
| chr17-42599093-DEL-329 | chr17-42599094-DEL-329 |
| chr17-42621699-DEL-64 | chr17-42621704-DEL-55 |
| chr17-42623255-DEL-60 | chr17-42623256-DEL-59 |
| chr17-42652173-DEL-638 | chr17-42652174-DEL-638 |
| chr17-42655571-DEL-173 | chr17-42655572-DEL-173 |
| chr17-42766273-DEL-102 | chr17-42766279-DEL-110 |
| chr17-42769531-DEL-54 | chr17-42769529-DEL-56 |
| chr17-42791897-DEL-54 | chr17-42791889-DEL-62 |
| chr17-42795303-DEL-387 | chr17-42795304-DEL-387 |
| chr17-42803254-DEL-194 | chr17-42803255-DEL-194 |
| chr17-42889636-DEL-125 | chr17-42889637-DEL-125 |
| chr17-42909563-DEL-92 | chr17-42909564-DEL-92 |
| chr17-42911122-DEL-2700 | chr17-42911123-DEL-2700 |
| chr17-42946355-DEL-351 | chr17-42946356-DEL-351 |
| chr17-42960713-DEL-92 | chr17-42960695-DEL-91 |
| chr17-42966926-DEL-107 | chr17-42966924-DEL-100 |
| chr17-42984470-DEL-55 | chr17-42984455-DEL-50 |
| chr17-42993832-DEL-634 | chr17-42993833-DEL-634 |
| chr17-43010179-DEL-142 | chr17-43010185-DEL-135 |
| chr17-43031581-DEL-933 | chr17-43031582-DEL-933 |
| chr17-43054084-DEL-698 | chr17-43054072-DEL-722 |
| chr17-43116544-DEL-70 | chr17-43116537-DEL-94 |
| chr17-43161932-DEL-217 | chr17-43161933-DEL-217 |
| chr17-43163823-DEL-206 | chr17-43163824-DEL-206 |
| chr17-43167071-DEL-71 | chr17-43167072-DEL-71 |
| chr17-43196503-DEL-79 | chr17-43196504-DEL-79 |
| chr17-43200789-DEL-5368 | chr17-43200790-DEL-5368 |
| chr17-43207820-DEL-287 | chr17-43207821-DEL-287 |
| chr17-43208596-DEL-185 | chr17-43208617-DEL-179 |
| chr17-43217293-DEL-87 | chr17-43217294-DEL-87 |
| chr17-43218532-DEL-185 | chr17-43218533-DEL-185 |
| chr17-43249194-DEL-194 | chr17-43249195-DEL-194 |
| chr17-43250042-DEL-60 | chr17-43250045-DEL-58 |
| chr17-43251851-DEL-66 | chr17-43251852-DEL-66 |
| chr17-43258527-DEL-376 | chr17-43258522-DEL-376 |
| chr17-43269274-DEL-605 | chr17-43269275-DEL-605 |
| chr17-43270791-DEL-125 | chr17-43270792-DEL-125 |
| chr17-43272558-DEL-201 | chr17-43272559-DEL-201 |
| chr17-43273629-DEL-54 | chr17-43273630-DEL-54 |
| chr17-43280054-DEL-1231 | chr17-43280055-DEL-1231 |
| chr17-43283028-DEL-1646 | chr17-43283029-DEL-1646 |
| chr17-43290472-DEL-315 | chr17-43290473-DEL-315 |
| chr17-43319057-DEL-339 | chr17-43319058-DEL-339 |
| chr17-43321307-DEL-54 | chr17-43321268-DEL-52 |
| chr17-43326695-DEL-217 | chr17-43326696-DEL-217 |
| chr17-43328161-DEL-51 | chr17-43328162-DEL-51 |
| chr17-43347805-DEL-246 | chr17-43347806-DEL-246 |
| chr17-43362026-DEL-1406 | chr17-43362027-DEL-1406 |
| chr17-43372647-DEL-146 | chr17-43372644-DEL-150 |
| chr17-43377918-DEL-445 | chr17-43377919-DEL-445 |
| chr17-43391434-DEL-142 | chr17-43391435-DEL-142 |
| chr17-43429238-DEL-64 | chr17-43429235-DEL-64 |
| chr17-43435510-DEL-293 | chr17-43435511-DEL-293 |
| chr17-43459961-DEL-54 | chr17-43459919-DEL-60 |
| chr17-43477697-DEL-712 | chr17-43477698-DEL-709 |
| chr17-43481232-DEL-490 | chr17-43481233-DEL-490 |
| chr17-43492180-DEL-6594 | chr17-43492181-DEL-6594 |
| chr17-43498244-DEL-416 | chr17-43498245-DEL-416 |
| chr17-43515298-DEL-2104 | chr17-43515299-DEL-2104 |
| chr17-43536667-DEL-269 | chr17-43536668-DEL-269 |
| chr17-43544444-DEL-54 | chr17-43544447-DEL-54 |
| chr17-43553387-DEL-194 | chr17-43553388-DEL-194 |
| chr17-43554114-DEL-1580 | chr17-43554115-DEL-1580 |
| chr17-43557094-DEL-51 | chr17-43557095-DEL-51 |
| chr17-43577001-DEL-220 | chr17-43577002-DEL-220 |
| chr17-43582794-DEL-70 | chr17-43582799-DEL-66 |

| | |
|---|---|
| chr17-43586268-DEL-56 | chr17-43586269-DEL-56 |
| chr17-43594425-DEL-74 | chr17-43594448-DEL-52 |
| chr17-43608177-DEL-92 | chr17-43608178-DEL-92 |
| chr17-43699520-DEL-60 | chr17-43699521-DEL-60 |
| chr17-43723120-DEL-312 | chr17-43723121-DEL-312 |
| chr17-43729667-DEL-1647 | chr17-43729668-DEL-1647 |
| chr17-43773278-DEL-3098 | chr17-43773279-DEL-3098 |
| chr17-43784249-DEL-72 | chr17-43784238-DEL-96 |
| chr17-43794668-DEL-1258 | chr17-43794669-DEL-1258 |
| chr17-43804001-DEL-500 | chr17-43804002-DEL-500 |
| chr17-43833481-DEL-51 | chr17-43833478-DEL-52 |
| chr17-43836497-DEL-575 | chr17-43836498-DEL-575 |
| chr17-43838909-DEL-411 | chr17-43838910-DEL-411 |
| chr17-43847895-DEL-118 | chr17-43847896-DEL-118 |
| chr17-43863346-DEL-92 | chr17-43863347-DEL-88 |
| chr17-43874147-DEL-197 | chr17-43874148-DEL-197 |
| chr17-43880455-DEL-55 | chr17-43880435-DEL-82 |
| chr17-43894799-DEL-200 | chr17-43894800-DEL-200 |
| chr17-43912323-DEL-54 | chr17-43912324-DEL-54 |
| chr17-43914549-DEL-92 | chr17-43914550-DEL-92 |
| chr17-43915664-DEL-53 | chr17-43915665-DEL-53 |
| chr17-43923486-DEL-56 | chr17-43923486-DEL-53 |
| chr17-43954806-DEL-62 | chr17-43954812-DEL-55 |
| chr17-44004169-DEL-171 | chr17-44004173-DEL-168 |
| chr17-44006631-DEL-2941 | chr17-44006632-DEL-2941 |
| chr17-44011298-DEL-67 | chr17-44011299-DEL-67 |
| chr17-44013169-DEL-72 | chr17-44013170-DEL-72 |
| chr17-44020229-DEL-1557 | chr17-44020230-DEL-1557 |
| chr17-44031195-DEL-1255 | chr17-44031189-DEL-1261 |
| chr17-44039265-DEL-470 | chr17-44039266-DEL-470 |
| chr17-44070940-DEL-3666 | chr17-44070941-DEL-3666 |
| chr17-44074904-DEL-62 | chr17-44074909-DEL-58 |
| chr17-44077500-DEL-54 | chr17-44077501-DEL-54 |
| chr17-44127387-DEL-93 | chr17-44127387-DEL-93 |
| chr17-44134544-DEL-217 | chr17-44134545-DEL-217 |
| chr17-44192233-DEL-443 | chr17-44192241-DEL-440 |
| chr17-44194360-DEL-193 | chr17-44194379-DEL-175 |
| chr17-44196814-DEL-139 | chr17-44196815-DEL-140 |
| chr17-44237152-DEL-2378 | chr17-44237153-DEL-2378 |
| chr17-44239871-DEL-131 | chr17-44239872-DEL-131 |
| chr17-44249260-DEL-6575 | chr17-44249261-DEL-6575 |
| chr17-44278393-DEL-435 | chr17-44278394-DEL-435 |
| chr17-44294956-DEL-958 | chr17-44294956-DEL-958 |
| chr17-44317625-DEL-413 | chr17-44317621-DEL-413 |
| chr17-44325795-DEL-121 | chr17-44325805-DEL-118 |
| chr17-44339363-DEL-68 | chr17-44339364-DEL-68 |
| chr17-44426040-DEL-206 | chr17-44426041-DEL-206 |
| chr17-44446110-DEL-84 | chr17-44446111-DEL-84 |
| chr17-44476402-DEL-62 | chr17-44476407-DEL-88 |
| chr17-44489586-DEL-53 | chr17-44489566-DEL-68 |
| chr17-44501494-DEL-5515 | chr17-44501495-DEL-5515 |
| chr17-44519719-DEL-166 | chr17-44519720-DEL-163 |
| chr17-44527105-DEL-153 | chr17-44527102-DEL-157 |
| chr17-44538815-DEL-377 | chr17-44538816-DEL-377 |
| chr17-44539759-DEL-104 | chr17-44539756-DEL-108 |
| chr17-44564721-DEL-126 | chr17-44564688-DEL-136 |
| chr17-44610372-DEL-60 | chr17-44610373-DEL-60 |
| chr17-44612695-DEL-52 | chr17-44612681-DEL-57 |
| chr17-44614717-DEL-62 | chr17-44614730-DEL-50 |
| chr17-44631579-DEL-62 | chr17-44631578-DEL-56 |
| chr17-44651586-DEL-63 | chr17-44651598-DEL-58 |
| chr17-44661680-DEL-92 | chr17-44661681-DEL-91 |
| chr17-44666683-DEL-73 | chr17-44666684-DEL-73 |
| chr17-44668529-DEL-165 | chr17-44668530-DEL-165 |
| chr17-44688261-DEL-913 | chr17-44688262-DEL-913 |
| chr17-44692001-DEL-410 | chr17-44692001-DEL-410 |
| chr17-44778769-DEL-102 | chr17-44778743-DEL-129 |
| chr17-44780869-DEL-197 | chr17-44780880-DEL-207 |
| chr17-44781658-DEL-425 | chr17-44781659-DEL-425 |
| chr17-44814705-DEL-369 | chr17-44814682-DEL-393 |
| chr17-44833032-DEL-94 | chr17-44833033-DEL-94 |
| chr17-44905960-DEL-447 | chr17-44905961-DEL-447 |
| chr17-44989302-DEL-127 | chr17-44989321-DEL-109 |
| chr17-44995028-DEL-130 | chr17-44995029-DEL-130 |
| chr17-45059883-DEL-98 | chr17-45059879-DEL-118 |
| chr17-45065370-DEL-209 | chr17-45065371-DEL-209 |
| chr17-45138854-DEL-51 | chr17-45138855-DEL-51 |
| chr17-45197504-DEL-56 | chr17-45197505-DEL-56 |
| chr17-45212490-DEL-73 | chr17-45212490-DEL-72 |
| chr17-45227525-DEL-769 | chr17-45227525-DEL-769 |

| | |
|---|---|
| chr17-45232268-DEL-436 | chr17-45232269-DEL-436 |
| chr17-45248893-DEL-57 | chr17-45248893-DEL-57 |
| chr17-45325215-DEL-1012 | chr17-45325216-DEL-1012 |
| chr17-45414224-DEL-7586 | chr17-45414225-DEL-7586 |
| chr17-45434942-DEL-1837 | chr17-45434943-DEL-1837 |
| chr17-45445511-DEL-1893 | chr17-45445512-DEL-1893 |
| chr17-45449806-DEL-93 | chr17-45449807-DEL-92 |
| chr17-45473605-DEL-183 | chr17-45473606-DEL-183 |
| chr17-45480802-DEL-498 | chr17-45480803-DEL-498 |
| chr17-45494851-DEL-211 | chr17-45494852-DEL-211 |
| chr17-45503118-DEL-102 | chr17-45503098-DEL-99 |
| chr17-45507603-DEL-6166 | chr17-45507604-DEL-6166 |
| chr17-45516294-DEL-54 | chr17-45516255-DEL-82 |
| chr17-45522827-DEL-148 | chr17-45522828-DEL-148 |
| chr17-45545361-DEL-56 | chr17-45545362-DEL-56 |
| chr17-45632547-DEL-191 | chr17-45632565-DEL-184 |
| chr17-45661277-DEL-74 | chr17-45661278-DEL-74 |
| chr17-45666894-DEL-82 | chr17-45666909-DEL-68 |
| chr17-45700085-DEL-202 | chr17-45700086-DEL-202 |
| chr17-45711104-DEL-163 | chr17-45711108-DEL-162 |
| chr17-45714941-DEL-431 | chr17-45714941-DEL-431 |
| chr17-45737984-DEL-72 | chr17-45737985-DEL-72 |
| chr17-45781103-DEL-60 | chr17-45781104-DEL-60 |
| chr17-45796080-DEL-194 | chr17-45796082-DEL-198 |
| chr17-45798872-DEL-132 | chr17-45798871-DEL-116 |
| chr17-45802851-DEL-310 | chr17-45802849-DEL-310 |
| chr17-45805983-DEL-58 | chr17-45805984-DEL-58 |
| chr17-45812309-DEL-1201 | chr17-45812310-DEL-1201 |
| chr17-45823641-DEL-289 | chr17-45823642-DEL-289 |
| chr17-45891721-DEL-54 | chr17-45891722-DEL-54 |
| chr17-45918879-DEL-293 | chr17-45918880-DEL-293 |
| chr17-45926263-DEL-191 | chr17-45926264-DEL-191 |
| chr17-45928560-DEL-51 | chr17-45928561-DEL-51 |
| chr17-45948193-DEL-215 | chr17-45948194-DEL-215 |
| chr17-45955566-DEL-5280 | chr17-45955567-DEL-5280 |
| chr17-45966942-DEL-85 | chr17-45966943-DEL-85 |
| chr17-45986689-DEL-119 | chr17-45986690-DEL-119 |
| chr17-45988387-DEL-208 | chr17-45988388-DEL-208 |
| chr17-46000635-DEL-64 | chr17-46000629-DEL-72 |
| chr17-46006929-DEL-215 | chr17-46006934-DEL-211 |
| chr17-46014626-DEL-192 | chr17-46014627-DEL-192 |
| chr17-46020554-DEL-72 | chr17-46020555-DEL-72 |
| chr17-46028788-DEL-135 | chr17-46028789-DEL-136 |
| chr17-46088936-DEL-52 | chr17-46088925-DEL-64 |
| chr17-46090598-DEL-51 | chr17-46090599-DEL-51 |
| chr17-46091531-DEL-204 | chr17-46091532-DEL-204 |
| chr17-46092960-DEL-182 | chr17-46092980-DEL-196 |
| chr17-46129096-DEL-123 | chr17-46129097-DEL-123 |
| chr17-46160389-DEL-76 | chr17-46160392-DEL-74 |
| chr17-46165736-DEL-72 | chr17-46165733-DEL-73 |
| chr17-46191749-DEL-82 | chr17-46191750-DEL-82 |
| chr17-46213345-DEL-97 | chr17-46213365-DEL-54 |
| chr17-46226147-DEL-67 | chr17-46226148-DEL-67 |
| chr17-46291929-DEL-55 | chr17-46291930-DEL-55 |
| chr17-46300761-DEL-90 | chr17-46300792-DEL-60 |
| chr17-46378679-DEL-58 | chr17-46378680-DEL-60 |
| chr17-46411198-DEL-92 | chr17-46411199-DEL-92 |
| chr17-46411256-DEL-568 | chr17-46411257-DEL-568 |
| chr17-46416574-DEL-209 | chr17-46416575-DEL-209 |
| chr17-46451305-DEL-82 | chr17-46451303-DEL-86 |
| chr17-46462897-DEL-333 | chr17-46462898-DEL-333 |
| chr17-46485853-DEL-212 | chr17-46485854-DEL-212 |
| chr17-46491467-DEL-432 | chr17-46491468-DEL-432 |
| chr17-46500459-DEL-110 | chr17-46500466-DEL-104 |
| chr17-46516647-DEL-590 | chr17-46516648-DEL-590 |
| chr17-46522306-DEL-981 | chr17-46522307-DEL-981 |
| chr17-46563357-DEL-105 | chr17-46563358-DEL-105 |
| chr17-46604683-DEL-352 | chr17-46604660-DEL-380 |
| chr17-46611537-DEL-52 | chr17-46611539-DEL-50 |
| chr17-46622176-DEL-282 | chr17-46622177-DEL-282 |
| chr17-46625309-DEL-52 | chr17-46625309-DEL-52 |
| chr17-46628003-DEL-3071 | chr17-46628004-DEL-3071 |
| chr17-46641331-DEL-295 | chr17-46641332-DEL-295 |
| chr17-46685033-DEL-196 | chr17-46685031-DEL-197 |
| chr17-46772839-DEL-229 | chr17-46772840-DEL-229 |
| chr17-46834332-DEL-71 | chr17-46834333-DEL-71 |
| chr17-46847251-DEL-207 | chr17-46847252-DEL-207 |
| chr17-46860108-DEL-119 | chr17-46860109-DEL-119 |
| chr17-46880950-DEL-862 | chr17-46880951-DEL-862 |
| chr17-46883812-DEL-99 | chr17-46883813-DEL-99 |

| | |
|---|---|
| chr17-46894481-DEL-225 | chr17-46894482-DEL-225 |
| chr17-46904163-DEL-371 | chr17-46904164-DEL-371 |
| chr17-46906919-DEL-129 | chr17-46906920-DEL-129 |
| chr17-46913513-DEL-94 | chr17-46913514-DEL-94 |
| chr17-46915043-DEL-343 | chr17-46915043-DEL-343 |
| chr17-46978167-DEL-199 | chr17-46978170-DEL-198 |
| chr17-46993416-DEL-182 | chr17-46993417-DEL-182 |
| chr17-47002685-DEL-128 | chr17-47002686-DEL-128 |
| chr17-47012810-DEL-53 | chr17-47012811-DEL-53 |
| chr17-47042991-DEL-63 | chr17-47042992-DEL-63 |
| chr17-47049087-DEL-201 | chr17-47049088-DEL-201 |
| chr17-47052676-DEL-99 | chr17-47052677-DEL-99 |
| chr17-47077483-DEL-180 | chr17-47077484-DEL-180 |
| chr17-47084812-DEL-145 | chr17-47084813-DEL-145 |
| chr17-47087518-DEL-157 | chr17-47087519-DEL-157 |
| chr17-47094830-DEL-1923 | chr17-47094829-DEL-1924 |
| chr17-47096106-DEL-227 | chr17-47096104-DEL-221 |
| chr17-47105725-DEL-206 | chr17-47105726-DEL-206 |
| chr17-47127853-DEL-413 | chr17-47127853-DEL-414 |
| chr17-47139363-DEL-55 | chr17-47139364-DEL-55 |
| chr17-47146611-DEL-128 | chr17-47146610-DEL-131 |
| chr17-47147980-DEL-53 | chr17-47147981-DEL-53 |
| chr17-47157563-DEL-347 | chr17-47157564-DEL-347 |
| chr17-47159372-DEL-94 | chr17-47159373-DEL-94 |
| chr17-47179842-DEL-60 | chr17-47179843-DEL-60 |
| chr17-47186114-DEL-956 | chr17-47186115-DEL-956 |
| chr17-47211554-DEL-53 | chr17-47211555-DEL-55 |
| chr17-47213442-DEL-172 | chr17-47213440-DEL-173 |
| chr17-47216376-DEL-100 | chr17-47216389-DEL-64 |
| chr17-47217012-DEL-260 | chr17-47217012-DEL-260 |
| chr17-47218380-DEL-199 | chr17-47218382-DEL-198 |
| chr17-47224571-DEL-67 | chr17-47224572-DEL-67 |
| chr17-47233559-DEL-368 | chr17-47233560-DEL-368 |
| chr17-47241553-DEL-122 | chr17-47241552-DEL-124 |
| chr17-47247784-DEL-100 | chr17-47247785-DEL-100 |
| chr17-47300569-DEL-199 | chr17-47300570-DEL-199 |
| chr17-47309512-DEL-1513 | chr17-47309513-DEL-1513 |
| chr17-47328470-DEL-197 | chr17-47328471-DEL-197 |
| chr17-47345704-DEL-120 | chr17-47345705-DEL-120 |
| chr17-47363820-DEL-50 | chr17-47363788-DEL-82 |
| chr17-47377032-DEL-110 | chr17-47377035-DEL-108 |
| chr17-47386811-DEL-129 | chr17-47386812-DEL-129 |
| chr17-47389000-DEL-7073 | chr17-47389007-DEL-7067 |
| chr17-47418795-DEL-58 | chr17-47418796-DEL-58 |
| chr17-47427414-DEL-59 | chr17-47427391-DEL-51 |
| chr17-47430726-DEL-164 | chr17-47430727-DEL-164 |
| chr17-47438955-DEL-196 | chr17-47438961-DEL-194 |
| chr17-47444743-DEL-104 | chr17-47444714-DEL-105 |
| chr17-47517828-DEL-82 | chr17-47517829-DEL-58 |
| chr17-47534610-DEL-77 | chr17-47534598-DEL-78 |
| chr17-47540907-DEL-56 | chr17-47540908-DEL-56 |
| chr17-47552077-DEL-6467 | chr17-47552078-DEL-6467 |
| chr17-47572211-DEL-56 | chr17-47572215-DEL-54 |
| chr17-47594344-DEL-104 | chr17-47594345-DEL-104 |
| chr17-47624460-DEL-157 | chr17-47624461-DEL-157 |
| chr17-47637171-DEL-207 | chr17-47637172-DEL-207 |
| chr17-47645002-DEL-185 | chr17-47645003-DEL-185 |
| chr17-47666801-DEL-2502 | chr17-47666800-DEL-2502 |
| chr17-47668804-DEL-214 | chr17-47668805-DEL-214 |
| chr17-47674667-DEL-82 | chr17-47674625-DEL-81 |
| chr17-47683765-DEL-748 | chr17-47683766-DEL-748 |
| chr17-47689842-DEL-77 | chr17-47689838-DEL-78 |
| chr17-47691113-DEL-339 | chr17-47691114-DEL-339 |
| chr17-47709985-DEL-60 | chr17-47709986-DEL-60 |
| chr17-47713885-DEL-54 | chr17-47713886-DEL-54 |
| chr17-47714899-DEL-102 | chr17-47714900-DEL-102 |
| chr17-47729374-DEL-160 | chr17-47729375-DEL-160 |
| chr17-47733615-DEL-139 | chr17-47733616-DEL-139 |
| chr17-47737458-DEL-53 | chr17-47737459-DEL-53 |
| chr17-47764348-DEL-756 | chr17-47764349-DEL-756 |
| chr17-47785748-DEL-199 | chr17-47785749-DEL-199 |
| chr17-47796428-DEL-99 | chr17-47796433-DEL-60 |
| chr17-47802281-DEL-70 | chr17-47802282-DEL-70 |
| chr17-47810786-DEL-212 | chr17-47810787-DEL-212 |
| chr17-47827597-DEL-56 | chr17-47827600-DEL-54 |
| chr17-47834744-DEL-1168 | chr17-47834745-DEL-1168 |
| chr17-47836655-DEL-555 | chr17-47836656-DEL-555 |
| chr17-47847956-DEL-224 | chr17-47847957-DEL-224 |
| chr17-47851459-DEL-102 | chr17-47851504-DEL-62 |
| chr17-47857773-DEL-161 | chr17-47857771-DEL-162 |

| | |
|---|---|
| chr17-47860161-DEL-83 | chr17-47860162-DEL-104 |
| chr17-47874791-DEL-66 | chr17-47874792-DEL-62 |
| chr17-47893384-DEL-104 | chr17-47893385-DEL-104 |
| chr17-47900279-DEL-64 | chr17-47900280-DEL-64 |
| chr17-47901511-DEL-152 | chr17-47901512-DEL-152 |
| chr17-47914293-DEL-65 | chr17-47914249-DEL-62 |
| chr17-47925796-DEL-106 | chr17-47925796-DEL-106 |
| chr17-47946994-DEL-213 | chr17-47946995-DEL-213 |
| chr17-47963811-DEL-2567 | chr17-47963809-DEL-2570 |
| chr17-47969143-DEL-53 | chr17-47969144-DEL-54 |
| chr17-47973872-DEL-229 | chr17-47973873-DEL-228 |
| chr17-48029709-DEL-821 | chr17-48029710-DEL-821 |
| chr17-48042470-DEL-66 | chr17-48042471-DEL-54 |
| chr17-48075641-DEL-57 | chr17-48075642-DEL-57 |
| chr17-48089145-DEL-106 | chr17-48089146-DEL-106 |
| chr17-48100099-DEL-77 | chr17-48100089-DEL-78 |
| chr17-48149797-DEL-71 | chr17-48149754-DEL-72 |
| chr17-48164128-DEL-99 | chr17-48164129-DEL-99 |
| chr17-48167572-DEL-68 | chr17-48167573-DEL-68 |
| chr17-48170058-DEL-109 | chr17-48170059-DEL-109 |
| chr17-48179857-DEL-51 | chr17-48179858-DEL-51 |
| chr17-48222887-DEL-203 | chr17-48222888-DEL-203 |
| chr17-48223501-DEL-64 | chr17-48223494-DEL-63 |
| chr17-48239787-DEL-222 | chr17-48239788-DEL-222 |
| chr17-48256454-DEL-184 | chr17-48256455-DEL-184 |
| chr17-48277736-DEL-85 | chr17-48277737-DEL-85 |
| chr17-48281315-DEL-84 | chr17-48281316-DEL-84 |
| chr17-48293218-DEL-265 | chr17-48293249-DEL-235 |
| chr17-48296601-DEL-204 | chr17-48296602-DEL-204 |
| chr17-48302045-DEL-74 | chr17-48302034-DEL-96 |
| chr17-48323347-DEL-56 | chr17-48323350-DEL-64 |
| chr17-48335480-DEL-62 | chr17-48335480-DEL-63 |
| chr17-48339075-DEL-64 | chr17-48339076-DEL-64 |
| chr17-48344863-DEL-156 | chr17-48344864-DEL-154 |
| chr17-48381940-DEL-111 | chr17-48381941-DEL-111 |
| chr17-48383636-DEL-6409 | chr17-48383637-DEL-6409 |
| chr17-48394089-DEL-125 | chr17-48394094-DEL-121 |
| chr17-48412368-DEL-173 | chr17-48412369-DEL-173 |
| chr17-48414763-DEL-2768 | chr17-48414764-DEL-2768 |
| chr17-48462508-DEL-85 | chr17-48462509-DEL-50 |
| chr17-48476344-DEL-526 | chr17-48476345-DEL-526 |
| chr17-48500164-DEL-190 | chr17-48500165-DEL-190 |
| chr17-48526920-DEL-72 | chr17-48526922-DEL-70 |
| chr17-48528807-DEL-593 | chr17-48528808-DEL-594 |
| chr17-48559630-DEL-53 | chr17-48559630-DEL-54 |
| chr17-48561509-DEL-74 | chr17-48561512-DEL-72 |
| chr17-48583364-DEL-51 | chr17-48583365-DEL-51 |
| chr17-48585049-DEL-347 | chr17-48585069-DEL-329 |
| chr17-48588233-DEL-481 | chr17-48588228-DEL-487 |
| chr17-48589099-DEL-4477 | chr17-48589100-DEL-4477 |
| chr17-48606693-DEL-68 | chr17-48606711-DEL-64 |
| chr17-48626548-DEL-73 | chr17-48626549-DEL-73 |
| chr17-48634414-DEL-73 | chr17-48634415-DEL-73 |
| chr17-48649946-DEL-55 | chr17-48649943-DEL-56 |
| chr17-48656129-DEL-405 | chr17-48656130-DEL-405 |
| chr17-48663596-DEL-66 | chr17-48663597-DEL-66 |
| chr17-48677906-DEL-207 | chr17-48677907-DEL-207 |
| chr17-48681723-DEL-439 | chr17-48681720-DEL-437 |
| chr17-48685604-DEL-668 | chr17-48685605-DEL-668 |
| chr17-48686484-DEL-421 | chr17-48686485-DEL-421 |
| chr17-48704388-DEL-61 | chr17-48704389-DEL-60 |
| chr17-48710181-DEL-583 | chr17-48710182-DEL-583 |
| chr17-48722626-DEL-207 | chr17-48722627-DEL-207 |
| chr17-48752184-DEL-177 | chr17-48752185-DEL-177 |
| chr17-48760011-DEL-106 | chr17-48759994-DEL-107 |
| chr17-48786615-DEL-92 | chr17-48786612-DEL-112 |
| chr17-48796392-DEL-77 | chr17-48796370-DEL-72 |
| chr17-48800526-DEL-54 | chr17-48800518-DEL-57 |
| chr17-48803317-DEL-54 | chr17-48803318-DEL-54 |
| chr17-48817119-DEL-1116 | chr17-48817120-DEL-1116 |
| chr17-48817227-DEL-62 | chr17-48817228-DEL-62 |
| chr17-48853935-DEL-87 | chr17-48853967-DEL-56 |
| chr17-48857182-DEL-789 | chr17-48857183-DEL-789 |
| chr17-48857685-DEL-66 | chr17-48857696-DEL-56 |
| chr17-48875014-DEL-63 | chr17-48875013-DEL-64 |
| chr17-48876982-DEL-2466 | chr17-48876983-DEL-2466 |
| chr17-48897794-DEL-399 | chr17-48897795-DEL-399 |
| chr17-48918455-DEL-54 | chr17-48918456-DEL-54 |
| chr17-48946151-DEL-64 | chr17-48946152-DEL-64 |
| chr17-48957611-DEL-57 | chr17-48957612-DEL-58 |

| | |
|---|---|
| chr17-48983709-DEL-837 | chr17-48983720-DEL-838 |
| chr17-48988720-DEL-304 | chr17-48988721-DEL-304 |
| chr17-49011902-DEL-80 | chr17-49011903-DEL-80 |
| chr17-49025442-DEL-400 | chr17-49025443-DEL-400 |
| chr17-49032339-DEL-1370 | chr17-49032340-DEL-1367 |
| chr17-49038236-DEL-62 | chr17-49038237-DEL-62 |
| chr17-49064845-DEL-61 | chr17-49064846-DEL-61 |
| chr17-49073103-DEL-209 | chr17-49073104-DEL-209 |
| chr17-49077731-DEL-113 | chr17-49077756-DEL-105 |
| chr17-49135545-DEL-354 | chr17-49135547-DEL-358 |
| chr17-49180456-DEL-58 | chr17-49180457-DEL-58 |
| chr17-49208095-DEL-88 | chr17-49208096-DEL-86 |
| chr17-49216223-DEL-90 | chr17-49216193-DEL-128 |
| chr17-49229187-DEL-54 | chr17-49229194-DEL-52 |
| chr17-49264414-DEL-62 | chr17-49264417-DEL-58 |
| chr17-49279149-DEL-101 | chr17-49279121-DEL-100 |
| chr17-49293081-DEL-76 | chr17-49293082-DEL-75 |
| chr17-49294734-DEL-129 | chr17-49294735-DEL-129 |
| chr17-49298144-DEL-112 | chr17-49298165-DEL-92 |
| chr17-49313541-DEL-281 | chr17-49313542-DEL-281 |
| chr17-49339353-DEL-145 | chr17-49339354-DEL-145 |
| chr17-49357292-DEL-213 | chr17-49357274-DEL-212 |
| chr17-49390487-DEL-90 | chr17-49390488-DEL-86 |
| chr17-49414065-DEL-395 | chr17-49414066-DEL-395 |
| chr17-49424209-DEL-204 | chr17-49424210-DEL-204 |
| chr17-49428302-DEL-56 | chr17-49428328-DEL-60 |
| chr17-49455674-DEL-444 | chr17-49455675-DEL-444 |
| chr17-49464062-DEL-69 | chr17-49464063-DEL-69 |
| chr17-49470143-DEL-62 | chr17-49470144-DEL-62 |
| chr17-49493275-DEL-54 | chr17-49493276-DEL-52 |
| chr17-49504885-DEL-60 | chr17-49504886-DEL-60 |
| chr17-49514172-DEL-60 | chr17-49514136-DEL-96 |
| chr17-49529557-DEL-394 | chr17-49529591-DEL-361 |
| chr17-49567563-DEL-58 | chr17-49567564-DEL-58 |
| chr17-49583196-DEL-52 | chr17-49583197-DEL-52 |
| chr17-49595016-DEL-76 | chr17-49595017-DEL-74 |
| chr17-49614514-DEL-1251 | chr17-49614515-DEL-1251 |
| chr17-49679250-DEL-66 | chr17-49679252-DEL-98 |
| chr17-49731448-DEL-52 | chr17-49731464-DEL-52 |
| chr17-49748216-DEL-7518 | chr17-49748217-DEL-7518 |
| chr17-49758766-DEL-260 | chr17-49758767-DEL-260 |
| chr17-49760794-DEL-106 | chr17-49760795-DEL-106 |
| chr17-49791877-DEL-117 | chr17-49791878-DEL-117 |
| chr17-49815160-DEL-68 | chr17-49815162-DEL-66 |
| chr17-49856133-DEL-57 | chr17-49856137-DEL-54 |
| chr17-49867277-DEL-91 | chr17-49867307-DEL-85 |
| chr17-49909965-DEL-72 | chr17-49909966-DEL-72 |
| chr17-49937721-DEL-1145 | chr17-49937722-DEL-1145 |
| chr17-49971739-DEL-106 | chr17-49971740-DEL-106 |
| chr17-50030740-DEL-90 | chr17-50030741-DEL-88 |
| chr17-50041216-DEL-1568 | chr17-50041217-DEL-1568 |
| chr17-50082877-DEL-108 | chr17-50082916-DEL-69 |
| chr17-50090655-DEL-54 | chr17-50090656-DEL-52 |
| chr17-50112392-DEL-1551 | chr17-50112394-DEL-1552 |
| chr17-50128715-DEL-739 | chr17-50128716-DEL-739 |
| chr17-50160136-DEL-304 | chr17-50160137-DEL-304 |
| chr17-50164980-DEL-52 | chr17-50164981-DEL-52 |
| chr17-50169720-DEL-103 | chr17-50169736-DEL-100 |
| chr17-50178085-DEL-58 | chr17-50178086-DEL-58 |
| chr17-50213115-DEL-90 | chr17-50213116-DEL-90 |
| chr17-50244307-DEL-1206 | chr17-50244308-DEL-1206 |
| chr17-50248364-DEL-90 | chr17-50248382-DEL-72 |
| chr17-50249553-DEL-71 | chr17-50249554-DEL-71 |
| chr17-50262791-DEL-126 | chr17-50262792-DEL-126 |
| chr17-50269209-DEL-390 | chr17-50269210-DEL-390 |
| chr17-50366146-DEL-183 | chr17-50366147-DEL-183 |
| chr17-50378447-DEL-93 | chr17-50378436-DEL-105 |
| chr17-50379022-DEL-180 | chr17-50379023-DEL-180 |
| chr17-50381494-DEL-4353 | chr17-50381495-DEL-4353 |
| chr17-50402409-DEL-60 | chr17-50402410-DEL-60 |
| chr17-50407861-DEL-162 | chr17-50407864-DEL-182 |
| chr17-50408530-DEL-54 | chr17-50408531-DEL-54 |
| chr17-50415190-DEL-182 | chr17-50415191-DEL-182 |
| chr17-50439398-DEL-64 | chr17-50439399-DEL-66 |
| chr17-50470666-DEL-416 | chr17-50470675-DEL-406 |
| chr17-50555814-DEL-4463 | chr17-50555812-DEL-4464 |
| chr17-50570746-DEL-996 | chr17-50570746-DEL-974 |
| chr17-50679170-DEL-101 | chr17-50679172-DEL-100 |
| chr17-50684219-DEL-238 | chr17-50684220-DEL-238 |
| chr17-50687092-DEL-141 | chr17-50687093-DEL-141 |

| | |
|---|---|
| chr17-50701802-DEL-245 | chr17-50701803-DEL-245 |
| chr17-50827527-DEL-371 | chr17-50827528-DEL-371 |
| chr17-50858937-DEL-56 | chr17-50858938-DEL-56 |
| chr17-50877161-DEL-350 | chr17-50877162-DEL-350 |
| chr17-50895428-DEL-537 | chr17-50895429-DEL-537 |
| chr17-50900253-DEL-258 | chr17-50900265-DEL-256 |
| chr17-50910841-DEL-92 | chr17-50910842-DEL-115 |
| chr17-50934852-DEL-90 | chr17-50934871-DEL-72 |
| chr17-50950164-DEL-118 | chr17-50950180-DEL-114 |
| chr17-50953347-DEL-52 | chr17-50953348-DEL-52 |
| chr17-50967618-DEL-5052 | chr17-50967619-DEL-5052 |
| chr17-51005537-DEL-1388 | chr17-51005538-DEL-1388 |
| chr17-51079482-DEL-454 | chr17-51079483-DEL-454 |
| chr17-51096214-DEL-112 | chr17-51096209-DEL-114 |
| chr17-51217162-DEL-261 | chr17-51217163-DEL-261 |
| chr17-51220128-DEL-253 | chr17-51220129-DEL-252 |
| chr17-51233844-DEL-6386 | chr17-51233844-DEL-6386 |
| chr17-51255126-DEL-162 | chr17-51255127-DEL-162 |
| chr17-51264993-DEL-88 | chr17-51264994-DEL-88 |
| chr17-51385820-DEL-200 | chr17-51385821-DEL-200 |
| chr17-51386314-DEL-550 | chr17-51386317-DEL-551 |
| chr17-51410241-DEL-66 | chr17-51410290-DEL-56 |
| chr17-51443476-DEL-76 | chr17-51443477-DEL-60 |
| chr17-51448667-DEL-99 | chr17-51448667-DEL-99 |
| chr17-51501354-DEL-124 | chr17-51501355-DEL-124 |
| chr17-51508122-DEL-98 | chr17-51508123-DEL-98 |
| chr17-51558453-DEL-82 | chr17-51558454-DEL-82 |
| chr17-51584679-DEL-573 | chr17-51584680-DEL-573 |
| chr17-51596940-DEL-386 | chr17-51596941-DEL-386 |
| chr17-51617808-DEL-69 | chr17-51617812-DEL-68 |
| chr17-51631796-DEL-52 | chr17-51631797-DEL-52 |
| chr17-51638162-DEL-1602 | chr17-51638163-DEL-1602 |
| chr17-51648171-DEL-58 | chr17-51648172-DEL-58 |
| chr17-51670938-DEL-1526 | chr17-51670939-DEL-1526 |
| chr17-51696458-DEL-68 | chr17-51696459-DEL-68 |
| chr17-51700464-DEL-56 | chr17-51700465-DEL-56 |
| chr17-51758104-DEL-136 | chr17-51758104-DEL-137 |
| chr17-51761618-DEL-233 | chr17-51761619-DEL-233 |
| chr17-51889223-DEL-64 | chr17-51889224-DEL-64 |
| chr17-51893787-DEL-81 | chr17-51893790-DEL-81 |
| chr17-51897061-DEL-899 | chr17-51897062-DEL-899 |
| chr17-51900167-DEL-86 | chr17-51900162-DEL-60 |
| chr17-51937938-DEL-62 | chr17-51937939-DEL-62 |
| chr17-51946068-DEL-1894 | chr17-51946066-DEL-1895 |
| chr17-51966158-DEL-110 | chr17-51966159-DEL-74 |
| chr17-51978472-DEL-58 | chr17-51978471-DEL-54 |
| chr17-51981965-DEL-82 | chr17-51981963-DEL-83 |
| chr17-51998957-DEL-915 | chr17-51998958-DEL-915 |
| chr17-52007475-DEL-105 | chr17-52007471-DEL-108 |
| chr17-52011089-DEL-120 | chr17-52011090-DEL-120 |
| chr17-52022403-DEL-200 | chr17-52022404-DEL-200 |
| chr17-52057497-DEL-125 | chr17-52057498-DEL-125 |
| chr17-52064839-DEL-128 | chr17-52064840-DEL-128 |
| chr17-52102848-DEL-196 | chr17-52102849-DEL-188 |
| chr17-52164666-DEL-119 | chr17-52164667-DEL-119 |
| chr17-52171459-DEL-6830 | chr17-52171460-DEL-6830 |
| chr17-52190910-DEL-202 | chr17-52190911-DEL-202 |
| chr17-52195426-DEL-178 | chr17-52195427-DEL-178 |
| chr17-52202574-DEL-200 | chr17-52202576-DEL-201 |
| chr17-52248142-DEL-116 | chr17-52248131-DEL-124 |
| chr17-52248850-DEL-68 | chr17-52248851-DEL-69 |
| chr17-52254135-DEL-72 | chr17-52254136-DEL-72 |
| chr17-52255394-DEL-2814 | chr17-52255394-DEL-2814 |
| chr17-52275273-DEL-176 | chr17-52275270-DEL-180 |
| chr17-52275584-DEL-1271 | chr17-52275585-DEL-1271 |
| chr17-52285299-DEL-315 | chr17-52285300-DEL-315 |
| chr17-52303149-DEL-297 | chr17-52303150-DEL-297 |
| chr17-52306268-DEL-76 | chr17-52306265-DEL-80 |
| chr17-52346852-DEL-2140 | chr17-52346853-DEL-2140 |
| chr17-52349252-DEL-1825 | chr17-52349253-DEL-1825 |
| chr17-52384258-DEL-671 | chr17-52384259-DEL-671 |
| chr17-52396399-DEL-7041 | chr17-52396407-DEL-7042 |
| chr17-52402653-DEL-623 | chr17-52402657-DEL-624 |
| chr17-52408361-DEL-95 | chr17-52408357-DEL-96 |
| chr17-52445401-DEL-95 | chr17-52445399-DEL-89 |
| chr17-52479450-DEL-236 | chr17-52479451-DEL-236 |
| chr17-52515209-DEL-525 | chr17-52515210-DEL-525 |
| chr17-52522756-DEL-336 | chr17-52522756-DEL-336 |
| chr17-52550168-DEL-122 | chr17-52550152-DEL-132 |
| chr17-52559323-DEL-872 | chr17-52559324-DEL-872 |

| | |
|---|---|
| chr17-52576977-DEL-51 | chr17-52576978-DEL-51 |
| chr17-52663812-DEL-1805 | chr17-52663813-DEL-1805 |
| chr17-52682032-DEL-6803 | chr17-52682033-DEL-6803 |
| chr17-52715691-DEL-319 | chr17-52715692-DEL-319 |
| chr17-52720952-DEL-85 | chr17-52720947-DEL-53 |
| chr17-52731838-DEL-1888 | chr17-52731839-DEL-1888 |
| chr17-52756275-DEL-60 | chr17-52756275-DEL-60 |
| chr17-52779220-DEL-6675 | chr17-52779221-DEL-6675 |
| chr17-52793632-DEL-3838 | chr17-52793633-DEL-3838 |
| chr17-52889756-DEL-645 | chr17-52889757-DEL-645 |
| chr17-52931644-DEL-68 | chr17-52931644-DEL-72 |
| chr17-52939337-DEL-69 | chr17-52939338-DEL-69 |
| chr17-52948974-DEL-115 | chr17-52948975-DEL-115 |
| chr17-52966061-DEL-57 | chr17-52966062-DEL-57 |
| chr17-53023028-DEL-1654 | chr17-53023029-DEL-1654 |
| chr17-53038931-DEL-66 | chr17-53038932-DEL-66 |
| chr17-53057131-DEL-241 | chr17-53057132-DEL-241 |
| chr17-53092210-DEL-52 | chr17-53092211-DEL-52 |
| chr17-53123411-DEL-139 | chr17-53123411-DEL-140 |
| chr17-53149964-DEL-52 | chr17-53149965-DEL-52 |
| chr17-53199008-DEL-271 | chr17-53199017-DEL-264 |
| chr17-53218217-DEL-64 | chr17-53218218-DEL-64 |
| chr17-53219409-DEL-61 | chr17-53219410-DEL-61 |
| chr17-53220076-DEL-62 | chr17-53220077-DEL-62 |
| chr17-53231043-DEL-4606 | chr17-53231044-DEL-4606 |
| chr17-53252193-DEL-4649 | chr17-53252194-DEL-4649 |
| chr17-53264387-DEL-60 | chr17-53264394-DEL-54 |
| chr17-53273270-DEL-172 | chr17-53273271-DEL-172 |
| chr17-53279675-DEL-161 | chr17-53279676-DEL-161 |
| chr17-53289610-DEL-6438 | chr17-53289611-DEL-6438 |
| chr17-53313692-DEL-4658 | chr17-53313693-DEL-4658 |
| chr17-53323535-DEL-518 | chr17-53323532-DEL-512 |
| chr17-53324417-DEL-483 | chr17-53324417-DEL-483 |
| chr17-53328973-DEL-2432 | chr17-53328974-DEL-2432 |
| chr17-53340180-DEL-199 | chr17-53340181-DEL-199 |
| chr17-53344312-DEL-2506 | chr17-53344313-DEL-2506 |
| chr17-53356723-DEL-592 | chr17-53356724-DEL-592 |
| chr17-53359976-DEL-82 | chr17-53359977-DEL-82 |
| chr17-53361718-DEL-1431 | chr17-53361703-DEL-1446 |
| chr17-53389058-DEL-4286 | chr17-53389059-DEL-4286 |
| chr17-53404265-DEL-443 | chr17-53404263-DEL-445 |
| chr17-53407472-DEL-311 | chr17-53407473-DEL-311 |
| chr17-53453101-DEL-937 | chr17-53453102-DEL-937 |
| chr17-53459552-DEL-62 | chr17-53459541-DEL-74 |
| chr17-53479179-DEL-858 | chr17-53479177-DEL-858 |
| chr17-53482699-DEL-1055 | chr17-53482698-DEL-1057 |
| chr17-53490090-DEL-6369 | chr17-53490091-DEL-6369 |
| chr17-53490125-DEL-214 | chr17-53490126-DEL-214 |
| chr17-53504168-DEL-568 | chr17-53504154-DEL-535 |
| chr17-53522427-DEL-2964 | chr17-53522429-DEL-2963 |
| chr17-53531367-DEL-99 | chr17-53531385-DEL-103 |
| chr17-53545559-DEL-553 | chr17-53545560-DEL-553 |
| chr17-53566858-DEL-2091 | chr17-53566860-DEL-2078 |
| chr17-53581021-DEL-6230 | chr17-53581022-DEL-6230 |
| chr17-53588442-DEL-67 | chr17-53588441-DEL-66 |
| chr17-53618389-DEL-72 | chr17-53618384-DEL-78 |
| chr17-53620464-DEL-6203 | chr17-53620464-DEL-6203 |
| chr17-53632257-DEL-98 | chr17-53632267-DEL-79 |
| chr17-53636036-DEL-64 | chr17-53636037-DEL-64 |
| chr17-53649854-DEL-293 | chr17-53649855-DEL-293 |
| chr17-53661496-DEL-771 | chr17-53661481-DEL-767 |
| chr17-53671333-DEL-77 | chr17-53671334-DEL-77 |
| chr17-53671995-DEL-9920 | chr17-53671995-DEL-9920 |
| chr17-53703217-DEL-56 | chr17-53703210-DEL-52 |
| chr17-53709482-DEL-146 | chr17-53709483-DEL-146 |
| chr17-53713212-DEL-207 | chr17-53713213-DEL-207 |
| chr17-53723255-DEL-201 | chr17-53723256-DEL-201 |
| chr17-53746576-DEL-154 | chr17-53746577-DEL-154 |
| chr17-53798208-DEL-214 | chr17-53798209-DEL-214 |
| chr17-53810402-DEL-207 | chr17-53810403-DEL-207 |
| chr17-53824490-DEL-89 | chr17-53824491-DEL-89 |
| chr17-53837549-DEL-150 | chr17-53837550-DEL-150 |
| chr17-53855477-DEL-441 | chr17-53855478-DEL-441 |
| chr17-53858059-DEL-256 | chr17-53858055-DEL-265 |
| chr17-53860602-DEL-194 | chr17-53860603-DEL-194 |
| chr17-53863942-DEL-342 | chr17-53863943-DEL-342 |
| chr17-53872146-DEL-462 | chr17-53872144-DEL-463 |
| chr17-53922114-DEL-178 | chr17-53922115-DEL-178 |
| chr17-53926373-DEL-95 | chr17-53926356-DEL-77 |
| chr17-53926844-DEL-202 | chr17-53926845-DEL-202 |

| | |
|---|---|
| chr17-53936935-DEL-1346 | chr17-53936951-DEL-1334 |
| chr17-53940708-DEL-61 | chr17-53940708-DEL-76 |
| chr17-53942561-DEL-108 | chr17-53942562-DEL-112 |
| chr17-53948701-DEL-197 | chr17-53948702-DEL-197 |
| chr17-53955356-DEL-76 | chr17-53955383-DEL-50 |
| chr17-54000919-DEL-61 | chr17-54000915-DEL-62 |
| chr17-54017429-DEL-76 | chr17-54017432-DEL-75 |
| chr17-54054703-DEL-6473 | chr17-54054704-DEL-6473 |
| chr17-54097336-DEL-188 | chr17-54097337-DEL-188 |
| chr17-54122681-DEL-7210 | chr17-54122682-DEL-7210 |
| chr17-54155979-DEL-3413 | chr17-54155980-DEL-3413 |
| chr17-54173417-DEL-651 | chr17-54173418-DEL-651 |
| chr17-54192708-DEL-1943 | chr17-54192709-DEL-1943 |
| chr17-54198389-DEL-385 | chr17-54198390-DEL-385 |
| chr17-54209329-DEL-5469 | chr17-54209338-DEL-5468 |
| chr17-54236897-DEL-6850 | chr17-54236898-DEL-6850 |
| chr17-54256753-DEL-132 | chr17-54256754-DEL-127 |
| chr17-54274977-DEL-1526 | chr17-54274978-DEL-1526 |
| chr17-54282264-DEL-4243 | chr17-54282265-DEL-4243 |
| chr17-54285291-DEL-62 | chr17-54285321-DEL-58 |
| chr17-54289418-DEL-2998 | chr17-54289419-DEL-2998 |
| chr17-54320652-DEL-699 | chr17-54320653-DEL-699 |
| chr17-54323928-DEL-654 | chr17-54323926-DEL-652 |
| chr17-54326578-DEL-890 | chr17-54326580-DEL-889 |
| chr17-54329133-DEL-93 | chr17-54329182-DEL-92 |
| chr17-54335619-DEL-144 | chr17-54335620-DEL-144 |
| chr17-54357015-DEL-214 | chr17-54357016-DEL-214 |
| chr17-54371015-DEL-926 | chr17-54371016-DEL-926 |
| chr17-54384949-DEL-60 | chr17-54384938-DEL-78 |
| chr17-54394132-DEL-246 | chr17-54394133-DEL-246 |
| chr17-54408992-DEL-184 | chr17-54408993-DEL-184 |
| chr17-54410082-DEL-61 | chr17-54410086-DEL-60 |
| chr17-54412718-DEL-31738 | chr17-54412718-DEL-31738 |
| chr17-54414419-DEL-199 | chr17-54414420-DEL-199 |
| chr17-54416676-DEL-14632 | chr17-54416674-DEL-14634 |
| chr17-54454394-DEL-123 | chr17-54454378-DEL-139 |
| chr17-54496512-DEL-142 | chr17-54496526-DEL-136 |
| chr17-54500059-DEL-1016 | chr17-54500060-DEL-1016 |
| chr17-54537993-DEL-121 | chr17-54537994-DEL-121 |
| chr17-54543733-DEL-100 | chr17-54543743-DEL-94 |
| chr17-54548573-DEL-60 | chr17-54548574-DEL-60 |
| chr17-54555984-DEL-65 | chr17-54555985-DEL-65 |
| chr17-54564986-DEL-2711 | chr17-54564985-DEL-2712 |
| chr17-54564986-DEL-6030 | chr17-54564986-DEL-6031 |
| chr17-54568405-DEL-3155 | chr17-54568405-DEL-3155 |
| chr17-54590406-DEL-189 | chr17-54590407-DEL-189 |
| chr17-54615067-DEL-92 | chr17-54615074-DEL-66 |
| chr17-54620614-DEL-2408 | chr17-54620615-DEL-2408 |
| chr17-54628026-DEL-60 | chr17-54628027-DEL-60 |
| chr17-54652973-DEL-2128 | chr17-54652974-DEL-2128 |
| chr17-54690083-DEL-52 | chr17-54690084-DEL-52 |
| chr17-54696989-DEL-505 | chr17-54696990-DEL-505 |
| chr17-54700489-DEL-2768 | chr17-54700490-DEL-2768 |
| chr17-54706547-DEL-59 | chr17-54706548-DEL-59 |
| chr17-54716802-DEL-951 | chr17-54716803-DEL-951 |
| chr17-54718928-DEL-186 | chr17-54718929-DEL-186 |
| chr17-54726606-DEL-1676 | chr17-54726607-DEL-1676 |
| chr17-54731877-DEL-2722 | chr17-54731878-DEL-2722 |
| chr17-54736337-DEL-1281 | chr17-54736338-DEL-1281 |
| chr17-54758878-DEL-477 | chr17-54758876-DEL-478 |
| chr17-54769536-DEL-98 | chr17-54769537-DEL-98 |
| chr17-54811086-DEL-2906 | chr17-54811086-DEL-2906 |
| chr17-54816632-DEL-55 | chr17-54816633-DEL-54 |
| chr17-54820371-DEL-53 | chr17-54820349-DEL-65 |
| chr17-54820688-DEL-298 | chr17-54820689-DEL-298 |
| chr17-54833671-DEL-195 | chr17-54833672-DEL-195 |
| chr17-54834359-DEL-3290 | chr17-54834360-DEL-3290 |
| chr17-54842796-DEL-2212 | chr17-54842797-DEL-2212 |
| chr17-54844062-DEL-77 | chr17-54844062-DEL-77 |
| chr17-54862548-DEL-592 | chr17-54862549-DEL-592 |
| chr17-54872182-DEL-2316 | chr17-54872183-DEL-2317 |
| chr17-54879794-DEL-6445 | chr17-54879795-DEL-6445 |
| chr17-54901818-DEL-61 | chr17-54901827-DEL-53 |
| chr17-54914919-DEL-831 | chr17-54914920-DEL-831 |
| chr17-54928594-DEL-901 | chr17-54928594-DEL-901 |
| chr17-54934660-DEL-117 | chr17-54934659-DEL-119 |
| chr17-54956619-DEL-101 | chr17-54956620-DEL-101 |
| chr17-54967054-DEL-928 | chr17-54967069-DEL-914 |
| chr17-54982068-DEL-1934 | chr17-54982069-DEL-1934 |
| chr17-54994850-DEL-1066 | chr17-54994851-DEL-1066 |

| | |
|---|---|
| chr17-55012052-DEL-951 | chr17-55012053-DEL-952 |
| chr17-55013204-DEL-112 | chr17-55013198-DEL-154 |
| chr17-55017872-DEL-387 | chr17-55017869-DEL-385 |
| chr17-55033833-DEL-56 | chr17-55033836-DEL-57 |
| chr17-55034273-DEL-156 | chr17-55034274-DEL-156 |
| chr17-55072140-DEL-53 | chr17-55072141-DEL-53 |
| chr17-55080652-DEL-100 | chr17-55080652-DEL-100 |
| chr17-55082559-DEL-401 | chr17-55082559-DEL-401 |
| chr17-55085889-DEL-187 | chr17-55085889-DEL-187 |
| chr17-55089980-DEL-2193 | chr17-55089981-DEL-2193 |
| chr17-55093813-DEL-121 | chr17-55093813-DEL-121 |
| chr17-55113258-DEL-582 | chr17-55113259-DEL-582 |
| chr17-55118386-DEL-399 | chr17-55118387-DEL-399 |
| chr17-55125489-DEL-854 | chr17-55125490-DEL-854 |
| chr17-55154858-DEL-346 | chr17-55154859-DEL-346 |
| chr17-55192642-DEL-1217 | chr17-55192643-DEL-1217 |
| chr17-55196441-DEL-173 | chr17-55196442-DEL-173 |
| chr17-55205842-DEL-66 | chr17-55205843-DEL-66 |
| chr17-55207182-DEL-383 | chr17-55207183-DEL-383 |
| chr17-55230953-DEL-824 | chr17-55230954-DEL-824 |
| chr17-55234892-DEL-160 | chr17-55234893-DEL-160 |
| chr17-55256539-DEL-6243 | chr17-55256540-DEL-6243 |
| chr17-55256544-DEL-225 | chr17-55256579-DEL-225 |
| chr17-55287044-DEL-57 | chr17-55287045-DEL-57 |
| chr17-55288296-DEL-1968 | chr17-55288296-DEL-1968 |
| chr17-55298473-DEL-13578 | chr17-55298474-DEL-13578 |
| chr17-55339447-DEL-84 | chr17-55339448-DEL-82 |
| chr17-55348620-DEL-70 | chr17-55348621-DEL-70 |
| chr17-55362693-DEL-256 | chr17-55362694-DEL-256 |
| chr17-55370413-DEL-507 | chr17-55370414-DEL-507 |
| chr17-55371876-DEL-59 | chr17-55371883-DEL-63 |
| chr17-55388438-DEL-171 | chr17-55388438-DEL-168 |
| chr17-55411758-DEL-112 | chr17-55411751-DEL-120 |
| chr17-55415617-DEL-6542 | chr17-55415617-DEL-6544 |
| chr17-55436887-DEL-377 | chr17-55436888-DEL-377 |
| chr17-55450696-DEL-69 | chr17-55450697-DEL-69 |
| chr17-55454627-DEL-1291 | chr17-55454628-DEL-1291 |
| chr17-55470823-DEL-944 | chr17-55470824-DEL-944 |
| chr17-55480583-DEL-6775 | chr17-55480584-DEL-6775 |
| chr17-55499806-DEL-913 | chr17-55499807-DEL-914 |
| chr17-55513845-DEL-2784 | chr17-55513846-DEL-2784 |
| chr17-55515746-DEL-320 | chr17-55515746-DEL-320 |
| chr17-55524335-DEL-78 | chr17-55524336-DEL-78 |
| chr17-55528422-DEL-122 | chr17-55528429-DEL-116 |
| chr17-55566702-DEL-57 | chr17-55566677-DEL-58 |
| chr17-55624769-DEL-365 | chr17-55624770-DEL-359 |
| chr17-55647909-DEL-597 | chr17-55647910-DEL-597 |
| chr17-55651454-DEL-700 | chr17-55651455-DEL-700 |
| chr17-55655735-DEL-115 | chr17-55655736-DEL-115 |
| chr17-55656870-DEL-237 | chr17-55656871-DEL-237 |
| chr17-55671102-DEL-503 | chr17-55671103-DEL-503 |
| chr17-55671215-DEL-1069 | chr17-55671216-DEL-1069 |
| chr17-55697797-DEL-6218 | chr17-55697798-DEL-6218 |
| chr17-55707253-DEL-1092 | chr17-55707254-DEL-1092 |
| chr17-55711374-DEL-170 | chr17-55711376-DEL-168 |
| chr17-55712333-DEL-112 | chr17-55712334-DEL-112 |
| chr17-55733298-DEL-978 | chr17-55733299-DEL-978 |
| chr17-55736287-DEL-56 | chr17-55736288-DEL-54 |
| chr17-55778150-DEL-120 | chr17-55778139-DEL-132 |
| chr17-55783659-DEL-56 | chr17-55783660-DEL-52 |
| chr17-55796793-DEL-206 | chr17-55796794-DEL-206 |
| chr17-55798415-DEL-1480 | chr17-55798416-DEL-1480 |
| chr17-55831289-DEL-2661 | chr17-55831290-DEL-2661 |
| chr17-55906831-DEL-2363 | chr17-55906831-DEL-2363 |
| chr17-55909377-DEL-212 | chr17-55909378-DEL-212 |
| chr17-55927127-DEL-1470 | chr17-55927128-DEL-1470 |
| chr17-55937398-DEL-6587 | chr17-55937399-DEL-6587 |
| chr17-55949630-DEL-227 | chr17-55949631-DEL-227 |
| chr17-55952326-DEL-354 | chr17-55952327-DEL-354 |
| chr17-55962047-DEL-64 | chr17-55962043-DEL-64 |
| chr17-55964016-DEL-1627 | chr17-55964031-DEL-1629 |
| chr17-56002389-DEL-195 | chr17-56002390-DEL-195 |
| chr17-56012162-DEL-470 | chr17-56012163-DEL-470 |
| chr17-56019298-DEL-97 | chr17-56019302-DEL-90 |
| chr17-56032776-DEL-93 | chr17-56032765-DEL-105 |
| chr17-56078754-DEL-78 | chr17-56078755-DEL-78 |
| chr17-56141177-DEL-6596 | chr17-56141178-DEL-6596 |
| chr17-56165322-DEL-492 | chr17-56165323-DEL-492 |
| chr17-56195705-DEL-128 | chr17-56195708-DEL-127 |
| chr17-56201677-DEL-256 | chr17-56201678-DEL-256 |

| | |
|---|---|
| chr17-56207097-DEL-175 | chr17-56207098-DEL-175 |
| chr17-56224173-DEL-216 | chr17-56224174-DEL-216 |
| chr17-56230308-DEL-4430 | chr17-56230308-DEL-4430 |
| chr17-56235780-DEL-321 | chr17-56235792-DEL-296 |
| chr17-56236670-DEL-343 | chr17-56236671-DEL-343 |
| chr17-56260306-DEL-137 | chr17-56260307-DEL-137 |
| chr17-56265327-DEL-524 | chr17-56265327-DEL-523 |
| chr17-56284913-DEL-211 | chr17-56284914-DEL-211 |
| chr17-56288650-DEL-2083 | chr17-56288651-DEL-2083 |
| chr17-56298486-DEL-205 | chr17-56298487-DEL-205 |
| chr17-56312838-DEL-64 | chr17-56312839-DEL-64 |
| chr17-56321588-DEL-206 | chr17-56321589-DEL-206 |
| chr17-56390594-DEL-213 | chr17-56390595-DEL-213 |
| chr17-56411430-DEL-99 | chr17-56411430-DEL-99 |
| chr17-56436675-DEL-178 | chr17-56436676-DEL-178 |
| chr17-56450450-DEL-198 | chr17-56450451-DEL-198 |
| chr17-56483126-DEL-56 | chr17-56483127-DEL-56 |
| chr17-56485598-DEL-259 | chr17-56485599-DEL-259 |
| chr17-56491420-DEL-63 | chr17-56491429-DEL-50 |
| chr17-56521090-DEL-210 | chr17-56521091-DEL-210 |
| chr17-56537615-DEL-501 | chr17-56537616-DEL-501 |
| chr17-56539764-DEL-216 | chr17-56539764-DEL-216 |
| chr17-56550530-DEL-231 | chr17-56550529-DEL-233 |
| chr17-56571503-DEL-204 | chr17-56571504-DEL-204 |
| chr17-56574598-DEL-81 | chr17-56574614-DEL-74 |
| chr17-56575654-DEL-205 | chr17-56575655-DEL-205 |
| chr17-56581835-DEL-207 | chr17-56581836-DEL-207 |
| chr17-56590698-DEL-198 | chr17-56590707-DEL-194 |
| chr17-56608545-DEL-208 | chr17-56608551-DEL-209 |
| chr17-56619420-DEL-383 | chr17-56619421-DEL-383 |
| chr17-56635934-DEL-217 | chr17-56635935-DEL-217 |
| chr17-56717408-DEL-232 | chr17-56717437-DEL-192 |
| chr17-56737292-DEL-225 | chr17-56737293-DEL-225 |
| chr17-56763260-DEL-79 | chr17-56763261-DEL-79 |
| chr17-56766677-DEL-280 | chr17-56766678-DEL-280 |
| chr17-56774334-DEL-59 | chr17-56774335-DEL-59 |
| chr17-56790384-DEL-440 | chr17-56790389-DEL-428 |
| chr17-56798613-DEL-62 | chr17-56798614-DEL-62 |
| chr17-56856620-DEL-230 | chr17-56856621-DEL-230 |
| chr17-56866711-DEL-217 | chr17-56866712-DEL-217 |
| chr17-56935277-DEL-211 | chr17-56935278-DEL-211 |
| chr17-56937014-DEL-404 | chr17-56937015-DEL-404 |
| chr17-56953795-DEL-178 | chr17-56953793-DEL-181 |
| chr17-56977453-DEL-196 | chr17-56977454-DEL-196 |
| chr17-57026689-DEL-60 | chr17-57026690-DEL-60 |
| chr17-57051017-DEL-104 | chr17-57051044-DEL-70 |
| chr17-57140701-DEL-199 | chr17-57140702-DEL-199 |
| chr17-57158544-DEL-203 | chr17-57158545-DEL-203 |
| chr17-57170031-DEL-65 | chr17-57170031-DEL-65 |
| chr17-57188476-DEL-81 | chr17-57188477-DEL-81 |
| chr17-57280170-DEL-220 | chr17-57280171-DEL-220 |
| chr17-57291758-DEL-70 | chr17-57291750-DEL-71 |
| chr17-57293761-DEL-204 | chr17-57293762-DEL-204 |
| chr17-57300660-DEL-183 | chr17-57300661-DEL-183 |
| chr17-57335505-DEL-62 | chr17-57335506-DEL-54 |
| chr17-57349188-DEL-281 | chr17-57349189-DEL-281 |
| chr17-57388953-DEL-72 | chr17-57388954-DEL-72 |
| chr17-57419598-DEL-52 | chr17-57419595-DEL-56 |
| chr17-57434894-DEL-338 | chr17-57434895-DEL-338 |
| chr17-57474534-DEL-127 | chr17-57474535-DEL-127 |
| chr17-57506914-DEL-1059 | chr17-57506881-DEL-1060 |
| chr17-57509959-DEL-446 | chr17-57509960-DEL-446 |
| chr17-57536382-DEL-68 | chr17-57536386-DEL-56 |
| chr17-57584800-DEL-216 | chr17-57584801-DEL-216 |
| chr17-57605937-DEL-62 | chr17-57605938-DEL-62 |
| chr17-57607186-DEL-238 | chr17-57607187-DEL-238 |
| chr17-57675374-DEL-6362 | chr17-57675375-DEL-6362 |
| chr17-57681406-DEL-212 | chr17-57681407-DEL-212 |
| chr17-57704465-DEL-174 | chr17-57704466-DEL-174 |
| chr17-57708270-DEL-54 | chr17-57708271-DEL-54 |
| chr17-57710408-DEL-65 | chr17-57710409-DEL-65 |
| chr17-57729931-DEL-198 | chr17-57729931-DEL-198 |
| chr17-57731237-DEL-200 | chr17-57731236-DEL-227 |
| chr17-57784462-DEL-112 | chr17-57784439-DEL-94 |
| chr17-57801448-DEL-133 | chr17-57801449-DEL-133 |
| chr17-57870017-DEL-36851 | chr17-57870015-DEL-36855 |
| chr17-57874539-DEL-150 | chr17-57874541-DEL-147 |
| chr17-57912549-DEL-93 | chr17-57912536-DEL-99 |
| chr17-57922206-DEL-1291 | chr17-57922206-DEL-1291 |
| chr17-57929297-DEL-2681 | chr17-57929298-DEL-2681 |

| | |
|---|---|
| chr17-57934560-DEL-1132 | chr17-57934561-DEL-1132 |
| chr17-57957200-DEL-9100 | chr17-57957184-DEL-9116 |
| chr17-57968044-DEL-51 | chr17-57968060-DEL-51 |
| chr17-57985089-DEL-960 | chr17-57985090-DEL-960 |
| chr17-58006034-DEL-6686 | chr17-58006035-DEL-6686 |
| chr17-58023244-DEL-961 | chr17-58023246-DEL-960 |
| chr17-58027353-DEL-874 | chr17-58027354-DEL-874 |
| chr17-58028402-DEL-876 | chr17-58028403-DEL-877 |
| chr17-58031065-DEL-78 | chr17-58031066-DEL-78 |
| chr17-58031899-DEL-275 | chr17-58031901-DEL-272 |
| chr17-58034706-DEL-3359 | chr17-58034707-DEL-3359 |
| chr17-58039275-DEL-7106 | chr17-58039276-DEL-7106 |
| chr17-58058916-DEL-1799 | chr17-58058916-DEL-1799 |
| chr17-58080970-DEL-776 | chr17-58080971-DEL-776 |
| chr17-58100636-DEL-65 | chr17-58100637-DEL-65 |
| chr17-58105150-DEL-376 | chr17-58105155-DEL-376 |
| chr17-58111687-DEL-80 | chr17-58111687-DEL-80 |
| chr17-58118355-DEL-1268 | chr17-58118356-DEL-1266 |
| chr17-58121387-DEL-3293 | chr17-58121388-DEL-3293 |
| chr17-58145357-DEL-881 | chr17-58145358-DEL-881 |
| chr17-58162919-DEL-685 | chr17-58162920-DEL-685 |
| chr17-58189240-DEL-2374 | chr17-58189241-DEL-2374 |
| chr17-58199482-DEL-1073 | chr17-58199483-DEL-1073 |
| chr17-58216760-DEL-206 | chr17-58216761-DEL-206 |
| chr17-58236792-DEL-58 | chr17-58236801-DEL-50 |
| chr17-58245001-DEL-864 | chr17-58245002-DEL-864 |
| chr17-58265810-DEL-1288 | chr17-58265810-DEL-1288 |
| chr17-58274693-DEL-6374 | chr17-58274694-DEL-6374 |
| chr17-58282773-DEL-65 | chr17-58282774-DEL-65 |
| chr17-58294153-DEL-1023 | chr17-58294154-DEL-1023 |
| chr17-58311944-DEL-524 | chr17-58311945-DEL-524 |
| chr17-58335855-DEL-245 | chr17-58335856-DEL-245 |
| chr17-58355130-DEL-62 | chr17-58355131-DEL-62 |
| chr17-58357985-DEL-801 | chr17-58357986-DEL-801 |
| chr17-58390712-DEL-66 | chr17-58390704-DEL-104 |
| chr17-58419416-DEL-362 | chr17-58419410-DEL-359 |
| chr17-58435446-DEL-5708 | chr17-58435447-DEL-5708 |
| chr17-58437001-DEL-390 | chr17-58437001-DEL-390 |
| chr17-58440737-DEL-70 | chr17-58440739-DEL-69 |
| chr17-58443670-DEL-963 | chr17-58443671-DEL-963 |
| chr17-58459707-DEL-55 | chr17-58459708-DEL-55 |
| chr17-58470341-DEL-91 | chr17-58470370-DEL-81 |
| chr17-58498407-DEL-2546 | chr17-58498408-DEL-2546 |
| chr17-58567784-DEL-104 | chr17-58567785-DEL-104 |
| chr17-58569375-DEL-2704 | chr17-58569376-DEL-2705 |
| chr17-58569999-DEL-573 | chr17-58570000-DEL-573 |
| chr17-58577284-DEL-112 | chr17-58577285-DEL-112 |
| chr17-58582507-DEL-1613 | chr17-58582506-DEL-1613 |
| chr17-58585891-DEL-117 | chr17-58585895-DEL-96 |
| chr17-58587719-DEL-865 | chr17-58587720-DEL-865 |
| chr17-58596043-DEL-1288 | chr17-58596044-DEL-1288 |
| chr17-58607626-DEL-1094 | chr17-58607627-DEL-1094 |
| chr17-58617092-DEL-54 | chr17-58617087-DEL-54 |
| chr17-58625658-DEL-6562 | chr17-58625658-DEL-6562 |
| chr17-58656102-DEL-81 | chr17-58656100-DEL-80 |
| chr17-58656331-DEL-3137 | chr17-58656332-DEL-3137 |
| chr17-58682596-DEL-54 | chr17-58682597-DEL-60 |
| chr17-58690320-DEL-145 | chr17-58690321-DEL-145 |
| chr17-58716595-DEL-241 | chr17-58716596-DEL-241 |
| chr17-58717355-DEL-92 | chr17-58717356-DEL-96 |
| chr17-58722366-DEL-224 | chr17-58722367-DEL-224 |
| chr17-58731995-DEL-68 | chr17-58732011-DEL-62 |
| chr17-58751086-DEL-102 | chr17-58751087-DEL-102 |
| chr17-58772338-DEL-857 | chr17-58772337-DEL-860 |
| chr17-58776522-DEL-2985 | chr17-58776522-DEL-2985 |
| chr17-58787917-DEL-270 | chr17-58787918-DEL-270 |
| chr17-58792023-DEL-126 | chr17-58792048-DEL-102 |
| chr17-58807097-DEL-437 | chr17-58807097-DEL-437 |
| chr17-58808609-DEL-289 | chr17-58808609-DEL-289 |
| chr17-58816873-DEL-118 | chr17-58816880-DEL-111 |
| chr17-58893237-DEL-77 | chr17-58893248-DEL-79 |
| chr17-58900608-DEL-2102 | chr17-58900593-DEL-2103 |
| chr17-58907738-DEL-54 | chr17-58907739-DEL-54 |
| chr17-58914346-DEL-128 | chr17-58914335-DEL-98 |
| chr17-58933019-DEL-551 | chr17-58933020-DEL-551 |
| chr17-58939050-DEL-234 | chr17-58939050-DEL-234 |
| chr17-58951506-DEL-54 | chr17-58951507-DEL-54 |
| chr17-58970180-DEL-1563 | chr17-58970181-DEL-1563 |
| chr17-58974627-DEL-101 | chr17-58974628-DEL-101 |
| chr17-58982164-DEL-972 | chr17-58982165-DEL-972 |

| | |
|---|---|
| chr17-58982885-DEL-51 | chr17-58982885-DEL-51 |
| chr17-58986145-DEL-691 | chr17-58986146-DEL-691 |
| chr17-58988111-DEL-1346 | chr17-58988111-DEL-1346 |
| chr17-58992610-DEL-54 | chr17-58992584-DEL-67 |
| chr17-59002590-DEL-56 | chr17-59002591-DEL-56 |
| chr17-59019064-DEL-256 | chr17-59019065-DEL-256 |
| chr17-59020186-DEL-392 | chr17-59020187-DEL-392 |
| chr17-59020769-DEL-308 | chr17-59020770-DEL-308 |
| chr17-59065093-DEL-6194 | chr17-59065093-DEL-6194 |
| chr17-59065257-DEL-2667 | chr17-59065259-DEL-2666 |
| chr17-59091904-DEL-307 | chr17-59091905-DEL-307 |
| chr17-59096187-DEL-295 | chr17-59096188-DEL-295 |
| chr17-59109939-DEL-79 | chr17-59109942-DEL-77 |
| chr17-59175932-DEL-178 | chr17-59175937-DEL-174 |
| chr17-59185563-DEL-154 | chr17-59185564-DEL-149 |
| chr17-59188179-DEL-54 | chr17-59188180-DEL-54 |
| chr17-59205802-DEL-1715 | chr17-59205803-DEL-1715 |
| chr17-59206364-DEL-87 | chr17-59206364-DEL-87 |
| chr17-59224143-DEL-184 | chr17-59224143-DEL-184 |
| chr17-59242490-DEL-61 | chr17-59242491-DEL-66 |
| chr17-59319968-DEL-74 | chr17-59319969-DEL-74 |
| chr17-59345864-DEL-253 | chr17-59345865-DEL-253 |
| chr17-59351849-DEL-7855 | chr17-59351849-DEL-7855 |
| chr17-59369488-DEL-510 | chr17-59369489-DEL-510 |
| chr17-59383428-DEL-537 | chr17-59383429-DEL-537 |
| chr17-59400996-DEL-84 | chr17-59400997-DEL-84 |
| chr17-59421567-DEL-217 | chr17-59421568-DEL-217 |
| chr17-59467532-DEL-1746 | chr17-59467533-DEL-1745 |
| chr17-59518471-DEL-399 | chr17-59518472-DEL-399 |
| chr17-59522399-DEL-3212 | chr17-59522413-DEL-3211 |
| chr17-59561683-DEL-596 | chr17-59561683-DEL-591 |
| chr17-59563140-DEL-281 | chr17-59563141-DEL-281 |
| chr17-59625288-DEL-1139 | chr17-59625289-DEL-1139 |
| chr17-59701258-DEL-826 | chr17-59701259-DEL-826 |
| chr17-59716249-DEL-90 | chr17-59716249-DEL-56 |
| chr17-59741520-DEL-96 | chr17-59741525-DEL-92 |
| chr17-59756761-DEL-1060 | chr17-59756762-DEL-1060 |
| chr17-59767780-DEL-69 | chr17-59767781-DEL-63 |
| chr17-59774220-DEL-246 | chr17-59774208-DEL-243 |
| chr17-59804314-DEL-6196 | chr17-59804315-DEL-6196 |
| chr17-59866591-DEL-104 | chr17-59866636-DEL-60 |
| chr17-59899734-DEL-60 | chr17-59899724-DEL-70 |
| chr17-59925028-DEL-57 | chr17-59925036-DEL-55 |
| chr17-59942020-DEL-1214 | chr17-59942021-DEL-1214 |
| chr17-59945212-DEL-1547 | chr17-59945213-DEL-1547 |
| chr17-59951603-DEL-1790 | chr17-59951604-DEL-1790 |
| chr17-59979410-DEL-1071 | chr17-59979411-DEL-1071 |
| chr17-60030261-DEL-62 | chr17-60030262-DEL-52 |
| chr17-60036653-DEL-519 | chr17-60036654-DEL-519 |
| chr17-60041407-DEL-162 | chr17-60041406-DEL-159 |
| chr17-60073777-DEL-143 | chr17-60073776-DEL-143 |
| chr17-60083136-DEL-424 | chr17-60083145-DEL-393 |
| chr17-60094108-DEL-97 | chr17-60094139-DEL-52 |
| chr17-60112796-DEL-92 | chr17-60112814-DEL-76 |
| chr17-60115326-DEL-76 | chr17-60115329-DEL-78 |
| chr17-60120046-DEL-157 | chr17-60120047-DEL-157 |
| chr17-60145522-DEL-339 | chr17-60145523-DEL-339 |
| chr17-60165501-DEL-6452 | chr17-60165502-DEL-6452 |
| chr17-60200601-DEL-154 | chr17-60200597-DEL-154 |
| chr17-60215190-DEL-758 | chr17-60215191-DEL-759 |
| chr17-60219105-DEL-1178 | chr17-60219106-DEL-1178 |
| chr17-60221446-DEL-102 | chr17-60221446-DEL-62 |
| chr17-60232067-DEL-95 | chr17-60232063-DEL-92 |
| chr17-60241400-DEL-56 | chr17-60241407-DEL-50 |
| chr17-60253202-DEL-154 | chr17-60253194-DEL-161 |
| chr17-60320631-DEL-248 | chr17-60320632-DEL-247 |
| chr17-60336174-DEL-4759 | chr17-60336175-DEL-4759 |
| chr17-60369541-DEL-87 | chr17-60369542-DEL-87 |
| chr17-60382016-DEL-2757 | chr17-60382016-DEL-2757 |
| chr17-60382341-DEL-704 | chr17-60382341-DEL-704 |
| chr17-60385322-DEL-1766 | chr17-60385322-DEL-1766 |
| chr17-60390350-DEL-232 | chr17-60390350-DEL-232 |
| chr17-60399682-DEL-911 | chr17-60399650-DEL-933 |
| chr17-60419567-DEL-321 | chr17-60419568-DEL-321 |
| chr17-60444601-DEL-717 | chr17-60444601-DEL-717 |
| chr17-60465363-DEL-58 | chr17-60465384-DEL-50 |
| chr17-60465882-DEL-53 | chr17-60465863-DEL-73 |
| chr17-60468859-DEL-57 | chr17-60468860-DEL-57 |
| chr17-60471060-DEL-52 | chr17-60471061-DEL-54 |
| chr17-60472081-DEL-385 | chr17-60472081-DEL-385 |

| | |
|---|---|
| chr17-60473987-DEL-353 | chr17-60473987-DEL-353 |
| chr17-60502888-DEL-493 | chr17-60502889-DEL-493 |
| chr17-60527278-DEL-459 | chr17-60527279-DEL-459 |
| chr17-60529520-DEL-51 | chr17-60529521-DEL-51 |
| chr17-60543442-DEL-428 | chr17-60543492-DEL-412 |
| chr17-60583544-DEL-54 | chr17-60583545-DEL-54 |
| chr17-60600553-DEL-1109 | chr17-60600554-DEL-1109 |
| chr17-60611711-DEL-6932 | chr17-60611711-DEL-6932 |
| chr17-60639728-DEL-72 | chr17-60639732-DEL-73 |
| chr17-60641465-DEL-104 | chr17-60641466-DEL-104 |
| chr17-60663770-DEL-83 | chr17-60663755-DEL-82 |
| chr17-60674019-DEL-2178 | chr17-60674020-DEL-2178 |
| chr17-60683964-DEL-285 | chr17-60683965-DEL-285 |
| chr17-60691164-DEL-211 | chr17-60691164-DEL-211 |
| chr17-60724605-DEL-2678 | chr17-60724606-DEL-2678 |
| chr17-60744327-DEL-6697 | chr17-60744328-DEL-6697 |
| chr17-60760759-DEL-68 | chr17-60760790-DEL-74 |
| chr17-60761935-DEL-214 | chr17-60761936-DEL-214 |
| chr17-60768795-DEL-199 | chr17-60768787-DEL-191 |
| chr17-60819162-DEL-1467 | chr17-60819163-DEL-1467 |
| chr17-60824815-DEL-53 | chr17-60824816-DEL-53 |
| chr17-60829191-DEL-757 | chr17-60829192-DEL-757 |
| chr17-60830661-DEL-2737 | chr17-60830660-DEL-2738 |
| chr17-60834739-DEL-1103 | chr17-60834739-DEL-1103 |
| chr17-60838080-DEL-68 | chr17-60838081-DEL-68 |
| chr17-60854178-DEL-2881 | chr17-60854181-DEL-2881 |
| chr17-60867066-DEL-6326 | chr17-60867050-DEL-6326 |
| chr17-60904288-DEL-51 | chr17-60904289-DEL-51 |
| chr17-60916471-DEL-295 | chr17-60916475-DEL-291 |
| chr17-60922539-DEL-568 | chr17-60922540-DEL-568 |
| chr17-60952807-DEL-52 | chr17-60952808-DEL-52 |
| chr17-60983530-DEL-353 | chr17-60983531-DEL-353 |
| chr17-61054162-DEL-102 | chr17-61054154-DEL-96 |
| chr17-61058271-DEL-566 | chr17-61058272-DEL-566 |
| chr17-61062649-DEL-51 | chr17-61062646-DEL-55 |
| chr17-61071740-DEL-235 | chr17-61071741-DEL-235 |
| chr17-61076849-DEL-196 | chr17-61076850-DEL-196 |
| chr17-61090273-DEL-7494 | chr17-61090274-DEL-7494 |
| chr17-61090722-DEL-212 | chr17-61090723-DEL-212 |
| chr17-61135048-DEL-115 | chr17-61135049-DEL-115 |
| chr17-61141866-DEL-4077 | chr17-61141867-DEL-4077 |
| chr17-61149983-DEL-443 | chr17-61149984-DEL-443 |
| chr17-61156499-DEL-86 | chr17-61156500-DEL-86 |
| chr17-61174963-DEL-1211 | chr17-61174964-DEL-1211 |
| chr17-61193134-DEL-1976 | chr17-61193135-DEL-1976 |
| chr17-61213749-DEL-106 | chr17-61213782-DEL-62 |
| chr17-61243447-DEL-403 | chr17-61243448-DEL-403 |
| chr17-61263062-DEL-145 | chr17-61263063-DEL-145 |
| chr17-61264652-DEL-423 | chr17-61264653-DEL-424 |
| chr17-61268553-DEL-486 | chr17-61268554-DEL-486 |
| chr17-61281993-DEL-57 | chr17-61281992-DEL-58 |
| chr17-61318930-DEL-3002 | chr17-61318931-DEL-3002 |
| chr17-61327379-DEL-7135 | chr17-61327380-DEL-7135 |
| chr17-61334831-DEL-95 | chr17-61334812-DEL-105 |
| chr17-61353445-DEL-54 | chr17-61353418-DEL-68 |
| chr17-61379427-DEL-353 | chr17-61379428-DEL-353 |
| chr17-61388498-DEL-501 | chr17-61388499-DEL-501 |
| chr17-61391269-DEL-6649 | chr17-61391270-DEL-6649 |
| chr17-61427846-DEL-577 | chr17-61427846-DEL-577 |
| chr17-61437188-DEL-131 | chr17-61437188-DEL-131 |
| chr17-61439674-DEL-715 | chr17-61439674-DEL-715 |
| chr17-61471348-DEL-2280 | chr17-61471339-DEL-2282 |
| chr17-61477413-DEL-165 | chr17-61477414-DEL-164 |
| chr17-61512441-DEL-6453 | chr17-61512442-DEL-6453 |
| chr17-61527124-DEL-4668 | chr17-61527125-DEL-4668 |
| chr17-61533818-DEL-396 | chr17-61533819-DEL-394 |
| chr17-61538209-DEL-599 | chr17-61538210-DEL-599 |
| chr17-61540595-DEL-304 | chr17-61540596-DEL-305 |
| chr17-61568936-DEL-950 | chr17-61568937-DEL-950 |
| chr17-61621630-DEL-96 | chr17-61621631-DEL-96 |
| chr17-61641559-DEL-570 | chr17-61641560-DEL-570 |
| chr17-61658509-DEL-80 | chr17-61658510-DEL-80 |
| chr17-61665241-DEL-58 | chr17-61665242-DEL-58 |
| chr17-61676860-DEL-76 | chr17-61676861-DEL-76 |
| chr17-61684607-DEL-6258 | chr17-61684606-DEL-6259 |
| chr17-61719104-DEL-324 | chr17-61719105-DEL-324 |
| chr17-61737338-DEL-2778 | chr17-61737302-DEL-2778 |
| chr17-61757048-DEL-487 | chr17-61757064-DEL-457 |
| chr17-61814449-DEL-69 | chr17-61814450-DEL-69 |
| chr17-61823498-DEL-54 | chr17-61823523-DEL-60 |

| | |
|---|---|
| chr17-61830691-DEL-56 | chr17-61830692-DEL-56 |
| chr17-61845472-DEL-207 | chr17-61845470-DEL-210 |
| chr17-61883695-DEL-769 | chr17-61883696-DEL-769 |
| chr17-61905887-DEL-61 | chr17-61905884-DEL-60 |
| chr17-61915096-DEL-70 | chr17-61915085-DEL-50 |
| chr17-61970252-DEL-906 | chr17-61970253-DEL-906 |
| chr17-62008259-DEL-112 | chr17-62008280-DEL-76 |
| chr17-62010267-DEL-272 | chr17-62010268-DEL-272 |
| chr17-62013833-DEL-3029 | chr17-62013834-DEL-3029 |
| chr17-62030243-DEL-1358 | chr17-62030244-DEL-1358 |
| chr17-62034407-DEL-2288 | chr17-62034408-DEL-2288 |
| chr17-62051240-DEL-134 | chr17-62051241-DEL-134 |
| chr17-62067897-DEL-143 | chr17-62067894-DEL-141 |
| chr17-62100933-DEL-595 | chr17-62100934-DEL-593 |
| chr17-62138795-DEL-455 | chr17-62138796-DEL-455 |
| chr17-62215189-DEL-310 | chr17-62215190-DEL-310 |
| chr17-62254964-DEL-6452 | chr17-62254965-DEL-6452 |
| chr17-62322745-DEL-157 | chr17-62322746-DEL-157 |
| chr17-62386863-DEL-157 | chr17-62386880-DEL-145 |
| chr17-62397114-DEL-56 | chr17-62397115-DEL-56 |
| chr17-62454588-DEL-103 | chr17-62454583-DEL-67 |
| chr17-62512504-DEL-1174 | chr17-62512505-DEL-1174 |
| chr17-62514470-DEL-67 | chr17-62514457-DEL-79 |
| chr17-62543031-DEL-266 | chr17-62543032-DEL-266 |
| chr17-62545943-DEL-62 | chr17-62545944-DEL-62 |
| chr17-62548937-DEL-71 | chr17-62548960-DEL-76 |
| chr17-62583585-DEL-508 | chr17-62583601-DEL-494 |
| chr17-62614629-DEL-56 | chr17-62614630-DEL-56 |
| chr17-62635271-DEL-56 | chr17-62635246-DEL-74 |
| chr17-62692168-DEL-7779 | chr17-62692169-DEL-7779 |
| chr17-62708644-DEL-914 | chr17-62708645-DEL-914 |
| chr17-62803464-DEL-497 | chr17-62803465-DEL-497 |
| chr17-62807399-DEL-195 | chr17-62807400-DEL-195 |
| chr17-62857700-DEL-60 | chr17-62857701-DEL-60 |
| chr17-62974498-DEL-62 | chr17-62974500-DEL-62 |
| chr17-63016575-DEL-172 | chr17-63016576-DEL-172 |
| chr17-63024066-DEL-78 | chr17-63024067-DEL-76 |
| chr17-63044550-DEL-54 | chr17-63044548-DEL-82 |
| chr17-63088595-DEL-84 | chr17-63088596-DEL-78 |
| chr17-63095631-DEL-95 | chr17-63095632-DEL-95 |
| chr17-63221396-DEL-285 | chr17-63221397-DEL-283 |
| chr17-63245560-DEL-198 | chr17-63245561-DEL-198 |
| chr17-63338962-DEL-377 | chr17-63338963-DEL-377 |
| chr17-63353758-DEL-111 | chr17-63353759-DEL-111 |
| chr17-63357690-DEL-69 | chr17-63357648-DEL-115 |
| chr17-63379229-DEL-56 | chr17-63379229-DEL-91 |
| chr17-63406377-DEL-986 | chr17-63406378-DEL-986 |
| chr17-63407988-DEL-67 | chr17-63407954-DEL-68 |
| chr17-63422909-DEL-2089 | chr17-63422910-DEL-2089 |
| chr17-63428672-DEL-234 | chr17-63428673-DEL-234 |
| chr17-63432974-DEL-7112 | chr17-63432975-DEL-7112 |
| chr17-63499793-DEL-5832 | chr17-63499794-DEL-5832 |
| chr17-63554518-DEL-6685 | chr17-63554519-DEL-6685 |
| chr17-63554531-DEL-408 | chr17-63554532-DEL-408 |
| chr17-63577077-DEL-7517 | chr17-63577078-DEL-7517 |
| chr17-63703169-DEL-360 | chr17-63703150-DEL-336 |
| chr17-63730199-DEL-64 | chr17-63730200-DEL-64 |
| chr17-63732385-DEL-2754 | chr17-63732393-DEL-2745 |
| chr17-63745136-DEL-232 | chr17-63745138-DEL-231 |
| chr17-63811544-DEL-2156 | chr17-63811545-DEL-2156 |
| chr17-63824701-DEL-344 | chr17-63824702-DEL-344 |
| chr17-63829368-DEL-99 | chr17-63829369-DEL-99 |
| chr17-63831964-DEL-639 | chr17-63831965-DEL-639 |
| chr17-63851238-DEL-203 | chr17-63851239-DEL-203 |
| chr17-63854260-DEL-202 | chr17-63854261-DEL-202 |
| chr17-63860013-DEL-6315 | chr17-63860014-DEL-6315 |
| chr17-63867855-DEL-170 | chr17-63867854-DEL-170 |
| chr17-63874487-DEL-101 | chr17-63874459-DEL-88 |
| chr17-63885996-DEL-2452 | chr17-63885997-DEL-2452 |
| chr17-63888586-DEL-241 | chr17-63888587-DEL-241 |
| chr17-63894829-DEL-53 | chr17-63894830-DEL-53 |
| chr17-63896441-DEL-498 | chr17-63896442-DEL-498 |
| chr17-63900556-DEL-599 | chr17-63900557-DEL-599 |
| chr17-63910115-DEL-914 | chr17-63910116-DEL-914 |
| chr17-63918745-DEL-6447 | chr17-63918746-DEL-6447 |
| chr17-63947652-DEL-1904 | chr17-63947652-DEL-1904 |
| chr17-63957231-DEL-86 | chr17-63957232-DEL-86 |
| chr17-63972510-DEL-6794 | chr17-63972511-DEL-6794 |
| chr17-63978651-DEL-424 | chr17-63978652-DEL-424 |
| chr17-63989596-DEL-361 | chr17-63989597-DEL-361 |

| | |
|---|---|
| chr17-64005709-DEL-506 | chr17-64005710-DEL-506 |
| chr17-64033757-DEL-874 | chr17-64033758-DEL-874 |
| chr17-64036246-DEL-2614 | chr17-64036247-DEL-2614 |
| chr17-64038993-DEL-688 | chr17-64038994-DEL-688 |
| chr17-64043636-DEL-1105 | chr17-64043637-DEL-1105 |
| chr17-64044796-DEL-289 | chr17-64044795-DEL-289 |
| chr17-64049260-DEL-7139 | chr17-64049261-DEL-7139 |
| chr17-64057858-DEL-83 | chr17-64057859-DEL-83 |
| chr17-64068637-DEL-4916 | chr17-64068646-DEL-4922 |
| chr17-64083770-DEL-166 | chr17-64083771-DEL-166 |
| chr17-64099990-DEL-584 | chr17-64099991-DEL-584 |
| chr17-64124734-DEL-63 | chr17-64124747-DEL-51 |
| chr17-64235510-DEL-697 | chr17-64235511-DEL-697 |
| chr17-64309914-DEL-593 | chr17-64309915-DEL-593 |
| chr17-64366660-DEL-93 | chr17-64366661-DEL-93 |
| chr17-64395573-DEL-394 | chr17-64395572-DEL-408 |
| chr17-64395597-DEL-936 | chr17-64395597-DEL-935 |
| chr17-64417575-DEL-113 | chr17-64417578-DEL-111 |
| chr17-64438400-DEL-389 | chr17-64438401-DEL-389 |
| chr17-64445371-DEL-55 | chr17-64445372-DEL-55 |
| chr17-64459953-DEL-126 | chr17-64459954-DEL-126 |
| chr17-64469214-DEL-2908 | chr17-64469215-DEL-2908 |
| chr17-64478097-DEL-500 | chr17-64478098-DEL-500 |
| chr17-64489068-DEL-136 | chr17-64489069-DEL-136 |
| chr17-64495383-DEL-1492 | chr17-64495384-DEL-1492 |
| chr17-64499758-DEL-536 | chr17-64499758-DEL-534 |
| chr17-64503981-DEL-65 | chr17-64503982-DEL-65 |
| chr17-64517120-DEL-205 | chr17-64517121-DEL-205 |
| chr17-64517852-DEL-64 | chr17-64517853-DEL-64 |
| chr17-64524049-DEL-6314 | chr17-64524050-DEL-6312 |
| chr17-64546406-DEL-799 | chr17-64546407-DEL-799 |
| chr17-64549692-DEL-2005 | chr17-64549693-DEL-2005 |
| chr17-64551334-DEL-66 | chr17-64551334-DEL-66 |
| chr17-64560489-DEL-801 | chr17-64560488-DEL-802 |
| chr17-64579589-DEL-1088 | chr17-64579590-DEL-1088 |
| chr17-64609799-DEL-541 | chr17-64609800-DEL-541 |
| chr17-64612187-DEL-127 | chr17-64612188-DEL-127 |
| chr17-64632489-DEL-62 | chr17-64632490-DEL-62 |
| chr17-64661881-DEL-506 | chr17-64661882-DEL-506 |
| chr17-64680754-DEL-180 | chr17-64680755-DEL-180 |
| chr17-64702488-DEL-68 | chr17-64702489-DEL-68 |
| chr17-64725065-DEL-6227 | chr17-64725055-DEL-6228 |
| chr17-64748437-DEL-96 | chr17-64748438-DEL-96 |
| chr17-64754439-DEL-59 | chr17-64754440-DEL-59 |
| chr17-64777784-DEL-1148 | chr17-64777802-DEL-1118 |
| chr17-64851790-DEL-68 | chr17-64851791-DEL-68 |
| chr17-64884091-DEL-107 | chr17-64884092-DEL-107 |
| chr17-64894698-DEL-372 | chr17-64894723-DEL-347 |
| chr17-64945437-DEL-215 | chr17-64945438-DEL-215 |
| chr17-64977060-DEL-54 | chr17-64977061-DEL-54 |
| chr17-64983063-DEL-125 | chr17-64983064-DEL-125 |
| chr17-64991290-DEL-66 | chr17-64991291-DEL-66 |
| chr17-65003701-DEL-52 | chr17-65003702-DEL-52 |
| chr17-65010334-DEL-62 | chr17-65010335-DEL-62 |
| chr17-65011704-DEL-1147 | chr17-65011705-DEL-1147 |
| chr17-65022277-DEL-950 | chr17-65022277-DEL-950 |
| chr17-65024781-DEL-86 | chr17-65024782-DEL-86 |
| chr17-65063503-DEL-1709 | chr17-65063543-DEL-1674 |
| chr17-65069885-DEL-575 | chr17-65069886-DEL-575 |
| chr17-65088396-DEL-67 | chr17-65088397-DEL-67 |
| chr17-65165946-DEL-57 | chr17-65165934-DEL-58 |
| chr17-65176913-DEL-343 | chr17-65176914-DEL-343 |
| chr17-65205506-DEL-931 | chr17-65205507-DEL-931 |
| chr17-65206563-DEL-172 | chr17-65206564-DEL-172 |
| chr17-65214840-DEL-332 | chr17-65214841-DEL-332 |
| chr17-65233733-DEL-1238 | chr17-65233724-DEL-1239 |
| chr17-65249650-DEL-216 | chr17-65249654-DEL-209 |
| chr17-65250704-DEL-417 | chr17-65250705-DEL-417 |
| chr17-65260683-DEL-133 | chr17-65260670-DEL-147 |
| chr17-65272586-DEL-56 | chr17-65272560-DEL-54 |
| chr17-65293751-DEL-1587 | chr17-65293750-DEL-1588 |
| chr17-65368362-DEL-57 | chr17-65368363-DEL-57 |
| chr17-65384557-DEL-73 | chr17-65384576-DEL-75 |
| chr17-65390000-DEL-9249 | chr17-65390001-DEL-9249 |
| chr17-65398805-DEL-212 | chr17-65398806-DEL-212 |
| chr17-65417338-DEL-310 | chr17-65417339-DEL-310 |
| chr17-65461709-DEL-7079 | chr17-65461710-DEL-7079 |
| chr17-65540461-DEL-126 | chr17-65540460-DEL-108 |
| chr17-65578161-DEL-55 | chr17-65578161-DEL-64 |
| chr17-65586677-DEL-895 | chr17-65586678-DEL-895 |

| | |
|---|---|
| chr17-65590786-DEL-168 | chr17-65590787-DEL-168 |
| chr17-65605296-DEL-65 | chr17-65605296-DEL-65 |
| chr17-65616936-DEL-53 | chr17-65616935-DEL-55 |
| chr17-65638088-DEL-6676 | chr17-65638086-DEL-6679 |
| chr17-65652195-DEL-391 | chr17-65652196-DEL-391 |
| chr17-65682125-DEL-59 | chr17-65682126-DEL-59 |
| chr17-65683276-DEL-88 | chr17-65683277-DEL-88 |
| chr17-65687705-DEL-132 | chr17-65687706-DEL-132 |
| chr17-65697353-DEL-60 | chr17-65697353-DEL-60 |
| chr17-65701810-DEL-208 | chr17-65701811-DEL-208 |
| chr17-65704445-DEL-213 | chr17-65704445-DEL-210 |
| chr17-65706425-DEL-201 | chr17-65706426-DEL-201 |
| chr17-65710683-DEL-108 | chr17-65710679-DEL-113 |
| chr17-65722394-DEL-432 | chr17-65722391-DEL-436 |
| chr17-65744675-DEL-1019 | chr17-65744676-DEL-1019 |
| chr17-65745997-DEL-64 | chr17-65745998-DEL-64 |
| chr17-65751185-DEL-56 | chr17-65751184-DEL-58 |
| chr17-65764016-DEL-18205 | chr17-65764015-DEL-18206 |
| chr17-65768752-DEL-7095 | chr17-65768753-DEL-7095 |
| chr17-65798439-DEL-524 | chr17-65798440-DEL-524 |
| chr17-65823245-DEL-560 | chr17-65823246-DEL-560 |
| chr17-65841084-DEL-78 | chr17-65841073-DEL-87 |
| chr17-65850401-DEL-122 | chr17-65850402-DEL-120 |
| chr17-65857147-DEL-94 | chr17-65857148-DEL-94 |
| chr17-65869086-DEL-54 | chr17-65869086-DEL-54 |
| chr17-65871250-DEL-125 | chr17-65871251-DEL-125 |
| chr17-65888212-DEL-79 | chr17-65888194-DEL-80 |
| chr17-65967081-DEL-1770 | chr17-65967080-DEL-1771 |
| chr17-66004426-DEL-95 | chr17-66004427-DEL-95 |
| chr17-66021632-DEL-129 | chr17-66021633-DEL-129 |
| chr17-66032534-DEL-905 | chr17-66032535-DEL-906 |
| chr17-66056529-DEL-201 | chr17-66056530-DEL-201 |
| chr17-66058285-DEL-147 | chr17-66058286-DEL-147 |
| chr17-66066844-DEL-53 | chr17-66066845-DEL-53 |
| chr17-66069058-DEL-354 | chr17-66069059-DEL-346 |
| chr17-66070504-DEL-213 | chr17-66070505-DEL-213 |
| chr17-66111793-DEL-205 | chr17-66111794-DEL-205 |
| chr17-66114303-DEL-112 | chr17-66114304-DEL-112 |
| chr17-66144415-DEL-204 | chr17-66144416-DEL-204 |
| chr17-66162669-DEL-105 | chr17-66162670-DEL-105 |
| chr17-66201799-DEL-194 | chr17-66201800-DEL-194 |
| chr17-66205172-DEL-187 | chr17-66205173-DEL-187 |
| chr17-66233491-DEL-407 | chr17-66233491-DEL-407 |
| chr17-66235975-DEL-190 | chr17-66235976-DEL-190 |
| chr17-66262264-DEL-206 | chr17-66262265-DEL-207 |
| chr17-66322102-DEL-5374 | chr17-66322103-DEL-5374 |
| chr17-66328404-DEL-59 | chr17-66328405-DEL-59 |
| chr17-66334902-DEL-185 | chr17-66334928-DEL-160 |
| chr17-66339317-DEL-62 | chr17-66339310-DEL-78 |
| chr17-66346289-DEL-187 | chr17-66346290-DEL-187 |
| chr17-66350241-DEL-161 | chr17-66350234-DEL-124 |
| chr17-66365790-DEL-125 | chr17-66365791-DEL-125 |
| chr17-66389126-DEL-483 | chr17-66389127-DEL-483 |
| chr17-66459584-DEL-58 | chr17-66459585-DEL-58 |
| chr17-66493141-DEL-207 | chr17-66493131-DEL-209 |
| chr17-66509384-DEL-81 | chr17-66509385-DEL-81 |
| chr17-66511527-DEL-267 | chr17-66511528-DEL-267 |
| chr17-66526852-DEL-64 | chr17-66526853-DEL-64 |
| chr17-66537516-DEL-301 | chr17-66537517-DEL-266 |
| chr17-66538827-DEL-224 | chr17-66538828-DEL-224 |
| chr17-66552113-DEL-52 | chr17-66552116-DEL-50 |
| chr17-66564800-DEL-84 | chr17-66564806-DEL-83 |
| chr17-66567662-DEL-57 | chr17-66567663-DEL-57 |
| chr17-66581738-DEL-79 | chr17-66581739-DEL-77 |
| chr17-66609816-DEL-8318 | chr17-66609802-DEL-8314 |
| chr17-66626114-DEL-221 | chr17-66626115-DEL-221 |
| chr17-66626550-DEL-654 | chr17-66626551-DEL-654 |
| chr17-66635770-DEL-208 | chr17-66635771-DEL-208 |
| chr17-66671419-DEL-140 | chr17-66671420-DEL-126 |
| chr17-66675010-DEL-112 | chr17-66675019-DEL-107 |
| chr17-66687728-DEL-210 | chr17-66687729-DEL-210 |
| chr17-66706347-DEL-171 | chr17-66706348-DEL-171 |
| chr17-66709689-DEL-68 | chr17-66709690-DEL-68 |
| chr17-66738899-DEL-203 | chr17-66738900-DEL-203 |
| chr17-66769740-DEL-2511 | chr17-66769741-DEL-2511 |
| chr17-66774204-DEL-226 | chr17-66774204-DEL-226 |
| chr17-66774800-DEL-690 | chr17-66774800-DEL-687 |
| chr17-66788608-DEL-383 | chr17-66788609-DEL-384 |
| chr17-66805817-DEL-78 | chr17-66805818-DEL-78 |
| chr17-66809654-DEL-210 | chr17-66809655-DEL-210 |

| | |
|---|---|
| chr17-66811716-DEL-56 | chr17-66811717-DEL-53 |
| chr17-66813131-DEL-233 | chr17-66813159-DEL-231 |
| chr17-66821839-DEL-107 | chr17-66821840-DEL-107 |
| chr17-66834081-DEL-4531 | chr17-66834082-DEL-4531 |
| chr17-66848990-DEL-56 | chr17-66848993-DEL-54 |
| chr17-66898647-DEL-85 | chr17-66898648-DEL-85 |
| chr17-66899487-DEL-207 | chr17-66899488-DEL-207 |
| chr17-66915245-DEL-201 | chr17-66915246-DEL-201 |
| chr17-66919065-DEL-511 | chr17-66919048-DEL-532 |
| chr17-66938137-DEL-900 | chr17-66938145-DEL-892 |
| chr17-66941098-DEL-1093 | chr17-66941099-DEL-1093 |
| chr17-66942531-DEL-435 | chr17-66942538-DEL-435 |
| chr17-66959176-DEL-6218 | chr17-66959202-DEL-6193 |
| chr17-66968614-DEL-217 | chr17-66968615-DEL-217 |
| chr17-66998044-DEL-1314 | chr17-66998045-DEL-1314 |
| chr17-67016823-DEL-66 | chr17-67016824-DEL-66 |
| chr17-67021220-DEL-268 | chr17-67021221-DEL-268 |
| chr17-67039866-DEL-58 | chr17-67039869-DEL-56 |
| chr17-67043059-DEL-79 | chr17-67043060-DEL-79 |
| chr17-67065409-DEL-136 | chr17-67065359-DEL-170 |
| chr17-67110716-DEL-538 | chr17-67110717-DEL-538 |
| chr17-67149242-DEL-66 | chr17-67149234-DEL-74 |
| chr17-67190959-DEL-1418 | chr17-67190960-DEL-1418 |
| chr17-67210923-DEL-60 | chr17-67210924-DEL-56 |
| chr17-67308119-DEL-4989 | chr17-67308122-DEL-4988 |
| chr17-67316528-DEL-452 | chr17-67316529-DEL-452 |
| chr17-67331845-DEL-4587 | chr17-67331846-DEL-4587 |
| chr17-67368107-DEL-145 | chr17-67368084-DEL-149 |
| chr17-67401141-DEL-69 | chr17-67401134-DEL-68 |
| chr17-67421810-DEL-60 | chr17-67421811-DEL-60 |
| chr17-67431810-DEL-124 | chr17-67431810-DEL-124 |
| chr17-67433653-DEL-97 | chr17-67433607-DEL-96 |
| chr17-67443091-DEL-137 | chr17-67443092-DEL-127 |
| chr17-67444500-DEL-302 | chr17-67444511-DEL-311 |
| chr17-67445151-DEL-121 | chr17-67445167-DEL-121 |
| chr17-67478719-DEL-116 | chr17-67478722-DEL-115 |
| chr17-67492193-DEL-52 | chr17-67492187-DEL-50 |
| chr17-67498130-DEL-88 | chr17-67498131-DEL-88 |
| chr17-67541330-DEL-401 | chr17-67541331-DEL-401 |
| chr17-67576817-DEL-138 | chr17-67576818-DEL-138 |
| chr17-67594196-DEL-199 | chr17-67594198-DEL-196 |
| chr17-67608797-DEL-96 | chr17-67608820-DEL-58 |
| chr17-67629149-DEL-88 | chr17-67629150-DEL-88 |
| chr17-67676543-DEL-59 | chr17-67676544-DEL-59 |
| chr17-67692514-DEL-1084 | chr17-67692515-DEL-1084 |
| chr17-67731140-DEL-355 | chr17-67731141-DEL-355 |
| chr17-67733146-DEL-3908 | chr17-67733147-DEL-3908 |
| chr17-67752544-DEL-185 | chr17-67752549-DEL-186 |
| chr17-67772524-DEL-820 | chr17-67772520-DEL-818 |
| chr17-67791242-DEL-115 | chr17-67791268-DEL-95 |
| chr17-67799233-DEL-128 | chr17-67799235-DEL-160 |
| chr17-67824867-DEL-887 | chr17-67824868-DEL-887 |
| chr17-67831803-DEL-154 | chr17-67831804-DEL-154 |
| chr17-67842010-DEL-6442 | chr17-67842011-DEL-6442 |
| chr17-67885665-DEL-2927 | chr17-67885657-DEL-2935 |
| chr17-67898181-DEL-2017 | chr17-67898182-DEL-2017 |
| chr17-67900353-DEL-1621 | chr17-67900354-DEL-1621 |
| chr17-67901817-DEL-198 | chr17-67901817-DEL-198 |
| chr17-67902297-DEL-5343 | chr17-67902298-DEL-5343 |
| chr17-67925981-DEL-183 | chr17-67925982-DEL-183 |
| chr17-67941130-DEL-1528 | chr17-67941138-DEL-1485 |
| chr17-67942529-DEL-137 | chr17-67942530-DEL-137 |
| chr17-67950286-DEL-197 | chr17-67950289-DEL-194 |
| chr17-68027783-DEL-87 | chr17-68027772-DEL-89 |
| chr17-68065858-DEL-185 | chr17-68065859-DEL-185 |
| chr17-68069703-DEL-61 | chr17-68069704-DEL-61 |
| chr17-68081205-DEL-61 | chr17-68081206-DEL-60 |
| chr17-68095400-DEL-100 | chr17-68095383-DEL-126 |
| chr17-68124691-DEL-185 | chr17-68124692-DEL-184 |
| chr17-68131239-DEL-144 | chr17-68131237-DEL-129 |
| chr17-68135522-DEL-6616 | chr17-68135522-DEL-6616 |
| chr17-68152907-DEL-200 | chr17-68152918-DEL-175 |
| chr17-68157790-DEL-52 | chr17-68157791-DEL-52 |
| chr17-68163594-DEL-174 | chr17-68163595-DEL-174 |
| chr17-68167151-DEL-67 | chr17-68167152-DEL-67 |
| chr17-68185896-DEL-80 | chr17-68185899-DEL-93 |
| chr17-68196022-DEL-86 | chr17-68196031-DEL-74 |
| chr17-68197870-DEL-1548 | chr17-68197871-DEL-1548 |
| chr17-68207453-DEL-339 | chr17-68207454-DEL-339 |
| chr17-68220117-DEL-152 | chr17-68220118-DEL-152 |

| | |
|---|---|
| chr17-68221260-DEL-652 | chr17-68221260-DEL-652 |
| chr17-68222049-DEL-88 | chr17-68222053-DEL-84 |
| chr17-68230152-DEL-80 | chr17-68230140-DEL-102 |
| chr17-68233161-DEL-200 | chr17-68233162-DEL-200 |
| chr17-68233420-DEL-464 | chr17-68233421-DEL-464 |
| chr17-68239341-DEL-96 | chr17-68239348-DEL-102 |
| chr17-68246472-DEL-97 | chr17-68246473-DEL-97 |
| chr17-68274447-DEL-53 | chr17-68274448-DEL-53 |
| chr17-68276689-DEL-162 | chr17-68276686-DEL-166 |
| chr17-68290866-DEL-128 | chr17-68290824-DEL-149 |
| chr17-68293287-DEL-159 | chr17-68293288-DEL-159 |
| chr17-68299586-DEL-88 | chr17-68299573-DEL-54 |
| chr17-68318520-DEL-68 | chr17-68318521-DEL-56 |
| chr17-68332467-DEL-211 | chr17-68332468-DEL-211 |
| chr17-68337280-DEL-1656 | chr17-68337281-DEL-1656 |
| chr17-68374669-DEL-52 | chr17-68374655-DEL-67 |
| chr17-68387111-DEL-53 | chr17-68387112-DEL-53 |
| chr17-68412382-DEL-260 | chr17-68412386-DEL-239 |
| chr17-68434127-DEL-1160 | chr17-68434128-DEL-1160 |
| chr17-68435698-DEL-691 | chr17-68435690-DEL-708 |
| chr17-68458762-DEL-538 | chr17-68458763-DEL-538 |
| chr17-68473086-DEL-81 | chr17-68473087-DEL-81 |
| chr17-68494622-DEL-78 | chr17-68494623-DEL-78 |
| chr17-68506936-DEL-1860 | chr17-68506937-DEL-1860 |
| chr17-68528725-DEL-57 | chr17-68528726-DEL-57 |
| chr17-68540914-DEL-1436 | chr17-68540915-DEL-1436 |
| chr17-68571036-DEL-349 | chr17-68571031-DEL-348 |
| chr17-68603656-DEL-7274 | chr17-68603657-DEL-7274 |
| chr17-68614420-DEL-56 | chr17-68614421-DEL-56 |
| chr17-68623034-DEL-315 | chr17-68623035-DEL-315 |
| chr17-68628110-DEL-246 | chr17-68628111-DEL-246 |
| chr17-68668320-DEL-2960 | chr17-68668321-DEL-2960 |
| chr17-68684641-DEL-133 | chr17-68684640-DEL-130 |
| chr17-68708700-DEL-76 | chr17-68708701-DEL-76 |
| chr17-68760684-DEL-491 | chr17-68760685-DEL-491 |
| chr17-68775784-DEL-74 | chr17-68775773-DEL-86 |
| chr17-68781780-DEL-686 | chr17-68781781-DEL-686 |
| chr17-68809801-DEL-7023 | chr17-68809802-DEL-7023 |
| chr17-68835849-DEL-6388 | chr17-68835850-DEL-6386 |
| chr17-68861763-DEL-889 | chr17-68861764-DEL-889 |
| chr17-68875362-DEL-83 | chr17-68875363-DEL-83 |
| chr17-68879239-DEL-114 | chr17-68879253-DEL-72 |
| chr17-68897529-DEL-105 | chr17-68897538-DEL-103 |
| chr17-68914321-DEL-51 | chr17-68914322-DEL-51 |
| chr17-68952900-DEL-1552 | chr17-68952901-DEL-1552 |
| chr17-68957645-DEL-53 | chr17-68957644-DEL-53 |
| chr17-68977434-DEL-58 | chr17-68977435-DEL-52 |
| chr17-68980856-DEL-146 | chr17-68980858-DEL-147 |
| chr17-68997346-DEL-1528 | chr17-68997346-DEL-1528 |
| chr17-69002009-DEL-208 | chr17-69002010-DEL-208 |
| chr17-69012475-DEL-59 | chr17-69012476-DEL-59 |
| chr17-69020560-DEL-54 | chr17-69020561-DEL-54 |
| chr17-69031895-DEL-541 | chr17-69031896-DEL-541 |
| chr17-69036597-DEL-552 | chr17-69036608-DEL-544 |
| chr17-69054496-DEL-2205 | chr17-69054497-DEL-2205 |
| chr17-69073164-DEL-389 | chr17-69073165-DEL-389 |
| chr17-69081721-DEL-94 | chr17-69081722-DEL-94 |
| chr17-69102033-DEL-1005 | chr17-69102034-DEL-1005 |
| chr17-69117996-DEL-333 | chr17-69117985-DEL-345 |
| chr17-69125343-DEL-52 | chr17-69125344-DEL-50 |
| chr17-69160652-DEL-1250 | chr17-69160653-DEL-1250 |
| chr17-69173922-DEL-118 | chr17-69173923-DEL-118 |
| chr17-69194406-DEL-3360 | chr17-69194407-DEL-3360 |
| chr17-69210190-DEL-227 | chr17-69210179-DEL-239 |
| chr17-69214030-DEL-109 | chr17-69214031-DEL-109 |
| chr17-69233274-DEL-107 | chr17-69233303-DEL-96 |
| chr17-69280968-DEL-1451 | chr17-69280970-DEL-1448 |
| chr17-69332197-DEL-69 | chr17-69332198-DEL-69 |
| chr17-69334534-DEL-1071 | chr17-69334531-DEL-1072 |
| chr17-69337768-DEL-2458 | chr17-69337769-DEL-2458 |
| chr17-69358001-DEL-596 | chr17-69358001-DEL-596 |
| chr17-69390902-DEL-104 | chr17-69390903-DEL-104 |
| chr17-69423698-DEL-623 | chr17-69423702-DEL-618 |
| chr17-69429232-DEL-394 | chr17-69429231-DEL-404 |
| chr17-69432802-DEL-58 | chr17-69432803-DEL-58 |
| chr17-69449610-DEL-142 | chr17-69449611-DEL-142 |
| chr17-69477770-DEL-56 | chr17-69477770-DEL-56 |
| chr17-69519077-DEL-381 | chr17-69519076-DEL-381 |
| chr17-69547512-DEL-506 | chr17-69547514-DEL-507 |
| chr17-69553823-DEL-133 | chr17-69553832-DEL-125 |

| | |
|---|---|
| chr17-69558157-DEL-164 | chr17-69558184-DEL-128 |
| chr17-69582022-DEL-1264 | chr17-69582023-DEL-1264 |
| chr17-69603621-DEL-326 | chr17-69603622-DEL-326 |
| chr17-69630152-DEL-15216 | chr17-69630153-DEL-15216 |
| chr17-69630352-DEL-383 | chr17-69630353-DEL-383 |
| chr17-69665918-DEL-581 | chr17-69665919-DEL-581 |
| chr17-69671907-DEL-200 | chr17-69671908-DEL-200 |
| chr17-69741170-DEL-96 | chr17-69741170-DEL-96 |
| chr17-69744317-DEL-66 | chr17-69744344-DEL-56 |
| chr17-69753522-DEL-110 | chr17-69753523-DEL-110 |
| chr17-69757731-DEL-1851 | chr17-69757732-DEL-1851 |
| chr17-69763341-DEL-198 | chr17-69763342-DEL-198 |
| chr17-69800737-DEL-761 | chr17-69800738-DEL-761 |
| chr17-69821502-DEL-430 | chr17-69821502-DEL-454 |
| chr17-69844006-DEL-5271 | chr17-69844007-DEL-5271 |
| chr17-69884787-DEL-2124 | chr17-69884803-DEL-2122 |
| chr17-69894428-DEL-924 | chr17-69894429-DEL-924 |
| chr17-69899851-DEL-935 | chr17-69899852-DEL-935 |
| chr17-69910477-DEL-60 | chr17-69910478-DEL-60 |
| chr17-69934198-DEL-1647 | chr17-69934190-DEL-1647 |
| chr17-69964969-DEL-155 | chr17-69964969-DEL-155 |
| chr17-69973565-DEL-126 | chr17-69973566-DEL-126 |
| chr17-70015166-DEL-135 | chr17-70015167-DEL-135 |
| chr17-70037757-DEL-52 | chr17-70037758-DEL-50 |
| chr17-70045955-DEL-2576 | chr17-70045955-DEL-2576 |
| chr17-70053638-DEL-555 | chr17-70053638-DEL-555 |
| chr17-70089148-DEL-63 | chr17-70089149-DEL-63 |
| chr17-70089993-DEL-158 | chr17-70089994-DEL-158 |
| chr17-70090630-DEL-1097 | chr17-70090630-DEL-1097 |
| chr17-70101868-DEL-338 | chr17-70101868-DEL-338 |
| chr17-70115826-DEL-375 | chr17-70115826-DEL-375 |
| chr17-70135895-DEL-208 | chr17-70135896-DEL-208 |
| chr17-70139214-DEL-73 | chr17-70139215-DEL-73 |
| chr17-70160504-DEL-2006 | chr17-70160505-DEL-2006 |
| chr17-70161975-DEL-112 | chr17-70161976-DEL-112 |
| chr17-70164882-DEL-68 | chr17-70164883-DEL-68 |
| chr17-70166278-DEL-63 | chr17-70166279-DEL-63 |
| chr17-70181180-DEL-333 | chr17-70181181-DEL-333 |
| chr17-70204988-DEL-4666 | chr17-70204989-DEL-4666 |
| chr17-70218382-DEL-139 | chr17-70218383-DEL-139 |
| chr17-70224732-DEL-87 | chr17-70224733-DEL-87 |
| chr17-70236865-DEL-499 | chr17-70236866-DEL-499 |
| chr17-70239883-DEL-227 | chr17-70239884-DEL-227 |
| chr17-70254437-DEL-1746 | chr17-70254438-DEL-1746 |
| chr17-70256690-DEL-91 | chr17-70256690-DEL-89 |
| chr17-70260959-DEL-671 | chr17-70260958-DEL-671 |
| chr17-70267434-DEL-376 | chr17-70267435-DEL-376 |
| chr17-70274707-DEL-1616 | chr17-70274708-DEL-1616 |
| chr17-70316631-DEL-89 | chr17-70316630-DEL-88 |
| chr17-70357561-DEL-6399 | chr17-70357562-DEL-6399 |
| chr17-70368348-DEL-998 | chr17-70368349-DEL-998 |
| chr17-70387713-DEL-4067 | chr17-70387714-DEL-4067 |
| chr17-70417525-DEL-759 | chr17-70417526-DEL-759 |
| chr17-70418450-DEL-6798 | chr17-70418451-DEL-6798 |
| chr17-70440156-DEL-64 | chr17-70440112-DEL-67 |
| chr17-70443891-DEL-295 | chr17-70443892-DEL-295 |
| chr17-70449829-DEL-109 | chr17-70449824-DEL-115 |
| chr17-70450289-DEL-199 | chr17-70450337-DEL-180 |
| chr17-70456949-DEL-7369 | chr17-70456950-DEL-7369 |
| chr17-70536302-DEL-7365 | chr17-70536303-DEL-7365 |
| chr17-70556415-DEL-87 | chr17-70556402-DEL-94 |
| chr17-70566431-DEL-606 | chr17-70566432-DEL-606 |
| chr17-70575776-DEL-481 | chr17-70575779-DEL-481 |
| chr17-70577032-DEL-727 | chr17-70577033-DEL-727 |
| chr17-70606037-DEL-272 | chr17-70606031-DEL-276 |
| chr17-70622039-DEL-117 | chr17-70622040-DEL-74 |
| chr17-70630635-DEL-6437 | chr17-70630636-DEL-6437 |
| chr17-70652686-DEL-94 | chr17-70652688-DEL-60 |
| chr17-70658857-DEL-164 | chr17-70658858-DEL-164 |
| chr17-70666046-DEL-231 | chr17-70666047-DEL-231 |
| chr17-70671728-DEL-78 | chr17-70671728-DEL-78 |
| chr17-70681579-DEL-230 | chr17-70681580-DEL-230 |
| chr17-70700219-DEL-664 | chr17-70700219-DEL-664 |
| chr17-70736503-DEL-56 | chr17-70736494-DEL-62 |
| chr17-70737437-DEL-54 | chr17-70737439-DEL-51 |
| chr17-70741171-DEL-68 | chr17-70741176-DEL-67 |
| chr17-70779564-DEL-855 | chr17-70779565-DEL-856 |
| chr17-70783599-DEL-1505 | chr17-70783605-DEL-1508 |
| chr17-70784524-DEL-1567 | chr17-70784524-DEL-1567 |
| chr17-70797023-DEL-317 | chr17-70797024-DEL-317 |

| | |
|---|---|
| chr17-70802671-DEL-1744 | chr17-70802672-DEL-1744 |
| chr17-70806552-DEL-631 | chr17-70806553-DEL-631 |
| chr17-70821474-DEL-54 | chr17-70821475-DEL-54 |
| chr17-70826949-DEL-231 | chr17-70826950-DEL-231 |
| chr17-70846182-DEL-98 | chr17-70846183-DEL-98 |
| chr17-70851497-DEL-158 | chr17-70851508-DEL-153 |
| chr17-70866301-DEL-130 | chr17-70866306-DEL-126 |
| chr17-70897334-DEL-281 | chr17-70897335-DEL-281 |
| chr17-70916378-DEL-1153 | chr17-70916379-DEL-1153 |
| chr17-70926449-DEL-183 | chr17-70926450-DEL-183 |
| chr17-70959227-DEL-717 | chr17-70959229-DEL-762 |
| chr17-70965449-DEL-1791 | chr17-70965450-DEL-1791 |
| chr17-70966727-DEL-263 | chr17-70966728-DEL-263 |
| chr17-70974782-DEL-55 | chr17-70974783-DEL-55 |
| chr17-70979778-DEL-73 | chr17-70979787-DEL-65 |
| chr17-70984791-DEL-574 | chr17-70984796-DEL-570 |
| chr17-70987409-DEL-68 | chr17-70987419-DEL-59 |
| chr17-70988798-DEL-120 | chr17-70988801-DEL-121 |
| chr17-70995619-DEL-64 | chr17-70995621-DEL-65 |
| chr17-71002117-DEL-140 | chr17-71002124-DEL-134 |
| chr17-71003189-DEL-261 | chr17-71003190-DEL-261 |
| chr17-71004080-DEL-221 | chr17-71004081-DEL-221 |
| chr17-71006495-DEL-72 | chr17-71006495-DEL-72 |
| chr17-71008942-DEL-51 | chr17-71008938-DEL-52 |
| chr17-71019277-DEL-98 | chr17-71019275-DEL-102 |
| chr17-71030320-DEL-162 | chr17-71030321-DEL-162 |
| chr17-71046073-DEL-101 | chr17-71046074-DEL-101 |
| chr17-71064370-DEL-219 | chr17-71064355-DEL-219 |
| chr17-71091577-DEL-177 | chr17-71091580-DEL-175 |
| chr17-71093416-DEL-202 | chr17-71093417-DEL-202 |
| chr17-71095254-DEL-58 | chr17-71095273-DEL-58 |
| chr17-71099271-DEL-206 | chr17-71099272-DEL-206 |
| chr17-71100173-DEL-200 | chr17-71100174-DEL-200 |
| chr17-71102218-DEL-81 | chr17-71102244-DEL-71 |
| chr17-71112417-DEL-75 | chr17-71112418-DEL-75 |
| chr17-71112421-DEL-1060 | chr17-71112422-DEL-1060 |
| chr17-71113942-DEL-52 | chr17-71113943-DEL-52 |
| chr17-71137944-DEL-258 | chr17-71137945-DEL-258 |
| chr17-71164752-DEL-162 | chr17-71164753-DEL-161 |
| chr17-71165343-DEL-66 | chr17-71165344-DEL-66 |
| chr17-71174014-DEL-217 | chr17-71174015-DEL-217 |
| chr17-71187741-DEL-997 | chr17-71187742-DEL-997 |
| chr17-71190409-DEL-297 | chr17-71190423-DEL-284 |
| chr17-71204633-DEL-605 | chr17-71204634-DEL-605 |
| chr17-71222063-DEL-1040 | chr17-71222064-DEL-1041 |
| chr17-71235020-DEL-143 | chr17-71235021-DEL-143 |
| chr17-71238630-DEL-59 | chr17-71238631-DEL-59 |
| chr17-71239357-DEL-284 | chr17-71239358-DEL-284 |
| chr17-71265049-DEL-70 | chr17-71265041-DEL-57 |
| chr17-71268912-DEL-1986 | chr17-71268913-DEL-1986 |
| chr17-71271452-DEL-353 | chr17-71271446-DEL-354 |
| chr17-71292156-DEL-210 | chr17-71292157-DEL-210 |
| chr17-71304691-DEL-77 | chr17-71304691-DEL-77 |
| chr17-71318552-DEL-205 | chr17-71318553-DEL-205 |
| chr17-71324720-DEL-834 | chr17-71324721-DEL-833 |
| chr17-71336714-DEL-53 | chr17-71336715-DEL-53 |
| chr17-71337414-DEL-131 | chr17-71337415-DEL-127 |
| chr17-71385783-DEL-375 | chr17-71385751-DEL-401 |
| chr17-71391859-DEL-51 | chr17-71391860-DEL-51 |
| chr17-71400479-DEL-88 | chr17-71400480-DEL-88 |
| chr17-71404967-DEL-455 | chr17-71404968-DEL-454 |
| chr17-71411415-DEL-183 | chr17-71411416-DEL-183 |
| chr17-71417569-DEL-60 | chr17-71417570-DEL-60 |
| chr17-71471324-DEL-103 | chr17-71471342-DEL-86 |
| chr17-71487299-DEL-157 | chr17-71487298-DEL-157 |
| chr17-71504437-DEL-64 | chr17-71504427-DEL-66 |
| chr17-71525460-DEL-987 | chr17-71525461-DEL-987 |
| chr17-71532810-DEL-379 | chr17-71532811-DEL-379 |
| chr17-71535136-DEL-306 | chr17-71535137-DEL-306 |
| chr17-71536834-DEL-315 | chr17-71536835-DEL-315 |
| chr17-71543368-DEL-165 | chr17-71543369-DEL-165 |
| chr17-71583259-DEL-461 | chr17-71583258-DEL-462 |
| chr17-71586513-DEL-60 | chr17-71586513-DEL-57 |
| chr17-71589010-DEL-150 | chr17-71589054-DEL-108 |
| chr17-71594502-DEL-61 | chr17-71594502-DEL-63 |
| chr17-71601177-DEL-53 | chr17-71601166-DEL-52 |
| chr17-71603907-DEL-84 | chr17-71603884-DEL-80 |
| chr17-71639995-DEL-57 | chr17-71639993-DEL-58 |
| chr17-71687489-DEL-56 | chr17-71687490-DEL-56 |
| chr17-71695960-DEL-211 | chr17-71695961-DEL-211 |

| | |
|---|---|
| chr17-71726823-DEL-1079 | chr17-71726824-DEL-1079 |
| chr17-71731181-DEL-887 | chr17-71731182-DEL-887 |
| chr17-71732404-DEL-731 | chr17-71732405-DEL-731 |
| chr17-71744617-DEL-107 | chr17-71744618-DEL-107 |
| chr17-71749585-DEL-70 | chr17-71749586-DEL-70 |
| chr17-71770035-DEL-58 | chr17-71770047-DEL-86 |
| chr17-71810055-DEL-162 | chr17-71810056-DEL-162 |
| chr17-71823249-DEL-586 | chr17-71823250-DEL-586 |
| chr17-71836528-DEL-396 | chr17-71836529-DEL-396 |
| chr17-71842802-DEL-210 | chr17-71842803-DEL-210 |
| chr17-71847508-DEL-195 | chr17-71847509-DEL-195 |
| chr17-71866577-DEL-54 | chr17-71866578-DEL-54 |
| chr17-71879634-DEL-118 | chr17-71879635-DEL-118 |
| chr17-71930187-DEL-275 | chr17-71930188-DEL-273 |
| chr17-71935803-DEL-196 | chr17-71935814-DEL-197 |
| chr17-72009318-DEL-79 | chr17-72009318-DEL-66 |
| chr17-72011353-DEL-52 | chr17-72011354-DEL-52 |
| chr17-72041325-DEL-339 | chr17-72041326-DEL-339 |
| chr17-72054126-DEL-64 | chr17-72054141-DEL-62 |
| chr17-72093622-DEL-96 | chr17-72093662-DEL-64 |
| chr17-72106739-DEL-175 | chr17-72106740-DEL-175 |
| chr17-72120778-DEL-229 | chr17-72120779-DEL-229 |
| chr17-72122960-DEL-65 | chr17-72122961-DEL-65 |
| chr17-72127615-DEL-85 | chr17-72127616-DEL-89 |
| chr17-72158411-DEL-121 | chr17-72158412-DEL-121 |
| chr17-72159928-DEL-238 | chr17-72159929-DEL-238 |
| chr17-72177193-DEL-55 | chr17-72177194-DEL-55 |
| chr17-72223273-DEL-93 | chr17-72223274-DEL-93 |
| chr17-72253206-DEL-137 | chr17-72253207-DEL-138 |
| chr17-72264843-DEL-574 | chr17-72264844-DEL-574 |
| chr17-72281187-DEL-60 | chr17-72281188-DEL-61 |
| chr17-72304285-DEL-435 | chr17-72304286-DEL-435 |
| chr17-72309162-DEL-1879 | chr17-72309163-DEL-1879 |
| chr17-72312238-DEL-74 | chr17-72312239-DEL-74 |
| chr17-72341352-DEL-66 | chr17-72341353-DEL-66 |
| chr17-72379611-DEL-126 | chr17-72379657-DEL-85 |
| chr17-72384976-DEL-200 | chr17-72384974-DEL-201 |
| chr17-72387760-DEL-78 | chr17-72387761-DEL-78 |
| chr17-72443302-DEL-414 | chr17-72443304-DEL-414 |
| chr17-72482155-DEL-52 | chr17-72482156-DEL-50 |
| chr17-72562816-DEL-92 | chr17-72562814-DEL-71 |
| chr17-72587836-DEL-399 | chr17-72587837-DEL-399 |
| chr17-72651335-DEL-60 | chr17-72651336-DEL-60 |
| chr17-72661636-DEL-515 | chr17-72661636-DEL-515 |
| chr17-72697295-DEL-80 | chr17-72697285-DEL-92 |
| chr17-72709693-DEL-120 | chr17-72709694-DEL-120 |
| chr17-72771971-DEL-288 | chr17-72771984-DEL-294 |
| chr17-72777690-DEL-132 | chr17-72777691-DEL-132 |
| chr17-72783367-DEL-54 | chr17-72783368-DEL-54 |
| chr17-72825998-DEL-106 | chr17-72825999-DEL-106 |
| chr17-72844080-DEL-831 | chr17-72844081-DEL-831 |
| chr17-72857227-DEL-64 | chr17-72857228-DEL-68 |
| chr17-72882015-DEL-4407 | chr17-72882016-DEL-4407 |
| chr17-72904005-DEL-262 | chr17-72904006-DEL-262 |
| chr17-72922258-DEL-4485 | chr17-72922252-DEL-4473 |
| chr17-72928290-DEL-63 | chr17-72928288-DEL-66 |
| chr17-72932028-DEL-51 | chr17-72932025-DEL-54 |
| chr17-72935685-DEL-6964 | chr17-72935686-DEL-6964 |
| chr17-72948253-DEL-1798 | chr17-72948254-DEL-1798 |
| chr17-72959702-DEL-56 | chr17-72959716-DEL-62 |
| chr17-72986427-DEL-256 | chr17-72986433-DEL-269 |
| chr17-73000146-DEL-129 | chr17-73000145-DEL-129 |
| chr17-73002295-DEL-57 | chr17-73002302-DEL-51 |
| chr17-73010085-DEL-811 | chr17-73010086-DEL-811 |
| chr17-73048547-DEL-1283 | chr17-73048548-DEL-1283 |
| chr17-73103449-DEL-60 | chr17-73103450-DEL-60 |
| chr17-73108251-DEL-68 | chr17-73108252-DEL-68 |
| chr17-73117576-DEL-249 | chr17-73117577-DEL-249 |
| chr17-73122558-DEL-781 | chr17-73122559-DEL-781 |
| chr17-73123640-DEL-181 | chr17-73123641-DEL-181 |
| chr17-73131085-DEL-1362 | chr17-73131086-DEL-1362 |
| chr17-73187652-DEL-189 | chr17-73187658-DEL-184 |
| chr17-73194873-DEL-79 | chr17-73194874-DEL-79 |
| chr17-73198211-DEL-171 | chr17-73198212-DEL-171 |
| chr17-73202641-DEL-51 | chr17-73202642-DEL-51 |
| chr17-73206246-DEL-82 | chr17-73206271-DEL-56 |
| chr17-73260580-DEL-57 | chr17-73260587-DEL-51 |
| chr17-73303831-DEL-391 | chr17-73303832-DEL-391 |
| chr17-73311445-DEL-56 | chr17-73311446-DEL-56 |
| chr17-73323578-DEL-492 | chr17-73323579-DEL-492 |

| | |
|---|---|
| chr17-73338508-DEL-58 | chr17-73338512-DEL-61 |
| chr17-73354417-DEL-89 | chr17-73354418-DEL-89 |
| chr17-73378574-DEL-63 | chr17-73378571-DEL-67 |
| chr17-73431098-DEL-111 | chr17-73431099-DEL-111 |
| chr17-73438095-DEL-57 | chr17-73438093-DEL-50 |
| chr17-73460605-DEL-85 | chr17-73460607-DEL-84 |
| chr17-73472021-DEL-67 | chr17-73472022-DEL-67 |
| chr17-73473431-DEL-271 | chr17-73473432-DEL-271 |
| chr17-73486295-DEL-5199 | chr17-73486295-DEL-5199 |
| chr17-73498326-DEL-179 | chr17-73498327-DEL-179 |
| chr17-73521598-DEL-2404 | chr17-73521599-DEL-2404 |
| chr17-73554394-DEL-198 | chr17-73554395-DEL-198 |
| chr17-73561495-DEL-99 | chr17-73561496-DEL-68 |
| chr17-73562233-DEL-199 | chr17-73562234-DEL-199 |
| chr17-73566190-DEL-236 | chr17-73566191-DEL-236 |
| chr17-73589515-DEL-334 | chr17-73589516-DEL-334 |
| chr17-73609457-DEL-878 | chr17-73609458-DEL-878 |
| chr17-73625651-DEL-1861 | chr17-73625647-DEL-1862 |
| chr17-73663931-DEL-1369 | chr17-73663932-DEL-1369 |
| chr17-73686138-DEL-168 | chr17-73686139-DEL-168 |
| chr17-73733172-DEL-105 | chr17-73733173-DEL-105 |
| chr17-73768773-DEL-1037 | chr17-73768771-DEL-1038 |
| chr17-73772367-DEL-54 | chr17-73772368-DEL-54 |
| chr17-73775135-DEL-592 | chr17-73775136-DEL-592 |
| chr17-73830573-DEL-73 | chr17-73830540-DEL-107 |
| chr17-73836656-DEL-52 | chr17-73836657-DEL-52 |
| chr17-73857363-DEL-51 | chr17-73857364-DEL-51 |
| chr17-73866285-DEL-63 | chr17-73866286-DEL-63 |
| chr17-73871112-DEL-62 | chr17-73871113-DEL-62 |
| chr17-73893789-DEL-88 | chr17-73893788-DEL-94 |
| chr17-73913830-DEL-322 | chr17-73913831-DEL-322 |
| chr17-73933055-DEL-62 | chr17-73933054-DEL-64 |
| chr17-73940553-DEL-88 | chr17-73940554-DEL-88 |
| chr17-73960070-DEL-92 | chr17-73960071-DEL-92 |
| chr17-73960686-DEL-56 | chr17-73960690-DEL-57 |
| chr17-74001330-DEL-123 | chr17-74001331-DEL-123 |
| chr17-74054640-DEL-105 | chr17-74054641-DEL-105 |
| chr17-74076983-DEL-165 | chr17-74076984-DEL-165 |
| chr17-74078856-DEL-59 | chr17-74078816-DEL-64 |
| chr17-74089762-DEL-71 | chr17-74089763-DEL-71 |
| chr17-74092315-DEL-103 | chr17-74092314-DEL-104 |
| chr17-74136239-DEL-311 | chr17-74136240-DEL-311 |
| chr17-74147170-DEL-70 | chr17-74147169-DEL-58 |
| chr17-74163128-DEL-7184 | chr17-74163129-DEL-7184 |
| chr17-74174160-DEL-269 | chr17-74174178-DEL-261 |
| chr17-74177833-DEL-219 | chr17-74177847-DEL-212 |
| chr17-74193649-DEL-3121 | chr17-74193650-DEL-3121 |
| chr17-74222884-DEL-348 | chr17-74222876-DEL-342 |
| chr17-74272435-DEL-753 | chr17-74272436-DEL-754 |
| chr17-74317245-DEL-71 | chr17-74317246-DEL-71 |
| chr17-74324142-DEL-60 | chr17-74324144-DEL-58 |
| chr17-74340098-DEL-357 | chr17-74340099-DEL-357 |
| chr17-74342149-DEL-358 | chr17-74342150-DEL-358 |
| chr17-74389275-DEL-55 | chr17-74389276-DEL-55 |
| chr17-74441289-DEL-56 | chr17-74441293-DEL-52 |
| chr17-74443652-DEL-107 | chr17-74443655-DEL-105 |
| chr17-74452427-DEL-78 | chr17-74452429-DEL-63 |
| chr17-74471403-DEL-157 | chr17-74471404-DEL-157 |
| chr17-74497275-DEL-221 | chr17-74497276-DEL-221 |
| chr17-74516772-DEL-159 | chr17-74516773-DEL-159 |
| chr17-74536461-DEL-146 | chr17-74536462-DEL-146 |
| chr17-74556052-DEL-182 | chr17-74556053-DEL-182 |
| chr17-74566808-DEL-256 | chr17-74566809-DEL-256 |
| chr17-74582849-DEL-423 | chr17-74582850-DEL-423 |
| chr17-74587488-DEL-205 | chr17-74587488-DEL-203 |
| chr17-74594025-DEL-213 | chr17-74594026-DEL-213 |
| chr17-74608841-DEL-171 | chr17-74608842-DEL-171 |
| chr17-74609979-DEL-132 | chr17-74609980-DEL-132 |
| chr17-74611886-DEL-160 | chr17-74611887-DEL-160 |
| chr17-74618914-DEL-354 | chr17-74618915-DEL-354 |
| chr17-74619471-DEL-1585 | chr17-74619472-DEL-1585 |
| chr17-74625505-DEL-221 | chr17-74625506-DEL-203 |
| chr17-74652018-DEL-197 | chr17-74652019-DEL-197 |
| chr17-74652972-DEL-307 | chr17-74652972-DEL-304 |
| chr17-74655168-DEL-200 | chr17-74655169-DEL-200 |
| chr17-74666481-DEL-206 | chr17-74666482-DEL-206 |
| chr17-74679047-DEL-68 | chr17-74679048-DEL-68 |
| chr17-74685100-DEL-152 | chr17-74685101-DEL-152 |
| chr17-74708050-DEL-206 | chr17-74708051-DEL-206 |
| chr17-74722350-DEL-132 | chr17-74722351-DEL-134 |

| | |
|---|---|
| chr17-74732729-DEL-183 | chr17-74732730-DEL-183 |
| chr17-74735830-DEL-112 | chr17-74735831-DEL-112 |
| chr17-74740136-DEL-70 | chr17-74740137-DEL-70 |
| chr17-74773541-DEL-175 | chr17-74773542-DEL-175 |
| chr17-74774748-DEL-218 | chr17-74774749-DEL-218 |
| chr17-74782841-DEL-174 | chr17-74782842-DEL-174 |
| chr17-74789227-DEL-180 | chr17-74789228-DEL-180 |
| chr17-74796097-DEL-102 | chr17-74796098-DEL-102 |
| chr17-74817343-DEL-157 | chr17-74817343-DEL-155 |
| chr17-74818039-DEL-204 | chr17-74818040-DEL-204 |
| chr17-74821622-DEL-94 | chr17-74821591-DEL-88 |
| chr17-74828244-DEL-109 | chr17-74828278-DEL-80 |
| chr17-74837670-DEL-68 | chr17-74837671-DEL-68 |
| chr17-74839356-DEL-214 | chr17-74839366-DEL-210 |
| chr17-74842277-DEL-214 | chr17-74842278-DEL-214 |
| chr17-74846052-DEL-295 | chr17-74846066-DEL-286 |
| chr17-74847330-DEL-210 | chr17-74847331-DEL-210 |
| chr17-74856750-DEL-195 | chr17-74856763-DEL-165 |
| chr17-74858714-DEL-213 | chr17-74858715-DEL-213 |
| chr17-74877960-DEL-453 | chr17-74877973-DEL-441 |
| chr17-74880792-DEL-190 | chr17-74880793-DEL-190 |
| chr17-74893880-DEL-249 | chr17-74893881-DEL-249 |
| chr17-74898638-DEL-156 | chr17-74898655-DEL-156 |
| chr17-74972320-DEL-52 | chr17-74972323-DEL-50 |
| chr17-74988414-DEL-580 | chr17-74988415-DEL-580 |
| chr17-74989352-DEL-484 | chr17-74989353-DEL-484 |
| chr17-75000998-DEL-1457 | chr17-75000999-DEL-1457 |
| chr17-75011349-DEL-208 | chr17-75011350-DEL-208 |
| chr17-75011895-DEL-266 | chr17-75011896-DEL-266 |
| chr17-75015295-DEL-219 | chr17-75015296-DEL-219 |
| chr17-75015938-DEL-99 | chr17-75015939-DEL-99 |
| chr17-75016678-DEL-117 | chr17-75016679-DEL-117 |
| chr17-75021786-DEL-196 | chr17-75021787-DEL-196 |
| chr17-75031843-DEL-3126 | chr17-75031844-DEL-3126 |
| chr17-75066059-DEL-7214 | chr17-75066060-DEL-7214 |
| chr17-75114703-DEL-231 | chr17-75114704-DEL-231 |
| chr17-75148513-DEL-63 | chr17-75148514-DEL-63 |
| chr17-75152545-DEL-54 | chr17-75152548-DEL-52 |
| chr17-75168112-DEL-199 | chr17-75168113-DEL-199 |
| chr17-75268930-DEL-304 | chr17-75268931-DEL-304 |
| chr17-75281191-DEL-226 | chr17-75281192-DEL-226 |
| chr17-75284179-DEL-120 | chr17-75284180-DEL-120 |
| chr17-75295449-DEL-305 | chr17-75295450-DEL-305 |
| chr17-75366040-DEL-81 | chr17-75366041-DEL-81 |
| chr17-75370948-DEL-57 | chr17-75370951-DEL-55 |
| chr17-75371871-DEL-138 | chr17-75371874-DEL-136 |
| chr17-75373676-DEL-300 | chr17-75373677-DEL-300 |
| chr17-75376942-DEL-54 | chr17-75376943-DEL-54 |
| chr17-75386129-DEL-257 | chr17-75386134-DEL-258 |
| chr17-75407247-DEL-248 | chr17-75407251-DEL-251 |
| chr17-75430789-DEL-223 | chr17-75430790-DEL-223 |
| chr17-75437772-DEL-120 | chr17-75437773-DEL-120 |
| chr17-75449541-DEL-720 | chr17-75449542-DEL-720 |
| chr17-75452065-DEL-1186 | chr17-75452066-DEL-1186 |
| chr17-75483657-DEL-68 | chr17-75483658-DEL-72 |
| chr17-75491735-DEL-177 | chr17-75491722-DEL-175 |
| chr17-75500221-DEL-64 | chr17-75500222-DEL-64 |
| chr17-75520723-DEL-87 | chr17-75520713-DEL-88 |
| chr17-75523732-DEL-582 | chr17-75523733-DEL-582 |
| chr17-75525672-DEL-77 | chr17-75525677-DEL-73 |
| chr17-75552550-DEL-227 | chr17-75552551-DEL-227 |
| chr17-75574758-DEL-65 | chr17-75574759-DEL-65 |
| chr17-75583834-DEL-84 | chr17-75583837-DEL-82 |
| chr17-75588155-DEL-60 | chr17-75588155-DEL-64 |
| chr17-75604836-DEL-441 | chr17-75604798-DEL-441 |
| chr17-75609836-DEL-58 | chr17-75609837-DEL-58 |
| chr17-75621023-DEL-57 | chr17-75621024-DEL-57 |
| chr17-75637866-DEL-53 | chr17-75637840-DEL-51 |
| chr17-75650785-DEL-389 | chr17-75650786-DEL-389 |
| chr17-75699785-DEL-257 | chr17-75699786-DEL-257 |
| chr17-75706236-DEL-6382 | chr17-75706237-DEL-6382 |
| chr17-75721442-DEL-51 | chr17-75721443-DEL-51 |
| chr17-75731117-DEL-6715 | chr17-75731118-DEL-6715 |
| chr17-75738812-DEL-69 | chr17-75738815-DEL-67 |
| chr17-75749544-DEL-84 | chr17-75749545-DEL-84 |
| chr17-75752198-DEL-84 | chr17-75752199-DEL-84 |
| chr17-75755218-DEL-365 | chr17-75755233-DEL-362 |
| chr17-75760627-DEL-63 | chr17-75760613-DEL-69 |
| chr17-75763253-DEL-86 | chr17-75763254-DEL-116 |
| chr17-75770366-DEL-60 | chr17-75770367-DEL-60 |

| | |
|---|---|
| chr17-75774458-DEL-1275 | chr17-75774459-DEL-1275 |
| chr17-75781997-DEL-52 | chr17-75781998-DEL-52 |
| chr17-75785261-DEL-399 | chr17-75785262-DEL-399 |
| chr17-75840128-DEL-255 | chr17-75840129-DEL-255 |
| chr17-75841196-DEL-126 | chr17-75841197-DEL-126 |
| chr17-75857237-DEL-184 | chr17-75857238-DEL-184 |
| chr17-75866916-DEL-339 | chr17-75866917-DEL-339 |
| chr17-75873443-DEL-397 | chr17-75873448-DEL-389 |
| chr17-75878608-DEL-60 | chr17-75878593-DEL-95 |
| chr17-75889208-DEL-65 | chr17-75889209-DEL-65 |
| chr17-75890806-DEL-419 | chr17-75890807-DEL-419 |
| chr17-75901143-DEL-54 | chr17-75901135-DEL-63 |
| chr17-75911672-DEL-91 | chr17-75911671-DEL-98 |
| chr17-75930441-DEL-249 | chr17-75930442-DEL-249 |
| chr17-75941511-DEL-105 | chr17-75941512-DEL-105 |
| chr17-75944706-DEL-57 | chr17-75944703-DEL-57 |
| chr17-75949663-DEL-215 | chr17-75949664-DEL-215 |
| chr17-75960720-DEL-328 | chr17-75960721-DEL-328 |
| chr17-75998651-DEL-108 | chr17-75998652-DEL-108 |
| chr17-76030817-DEL-4854 | chr17-76030818-DEL-4854 |
| chr17-76073720-DEL-3827 | chr17-76073721-DEL-3827 |
| chr17-76094060-DEL-241 | chr17-76094056-DEL-238 |
| chr17-76103655-DEL-166 | chr17-76103656-DEL-166 |
| chr17-76120408-DEL-183 | chr17-76120409-DEL-183 |
| chr17-76129059-DEL-65 | chr17-76129080-DEL-69 |
| chr17-76140052-DEL-715 | chr17-76140053-DEL-715 |
| chr17-76167079-DEL-72 | chr17-76167092-DEL-56 |
| chr17-76201032-DEL-68 | chr17-76201023-DEL-50 |
| chr17-76203355-DEL-59 | chr17-76203342-DEL-60 |
| chr17-76204385-DEL-5332 | chr17-76204386-DEL-5332 |
| chr17-76221194-DEL-212 | chr17-76221195-DEL-212 |
| chr17-76230078-DEL-2184 | chr17-76230079-DEL-2184 |
| chr17-76239145-DEL-451 | chr17-76239146-DEL-451 |
| chr17-76244770-DEL-6209 | chr17-76244771-DEL-6207 |
| chr17-76259671-DEL-1324 | chr17-76259672-DEL-1324 |
| chr17-76267800-DEL-319 | chr17-76267804-DEL-303 |
| chr17-76273651-DEL-556 | chr17-76273652-DEL-556 |
| chr17-76276341-DEL-455 | chr17-76276342-DEL-455 |
| chr17-76288476-DEL-4516 | chr17-76288477-DEL-4516 |
| chr17-76316287-DEL-935 | chr17-76316291-DEL-936 |
| chr17-76320961-DEL-649 | chr17-76320962-DEL-649 |
| chr17-76329459-DEL-192 | chr17-76329459-DEL-192 |
| chr17-76332328-DEL-183 | chr17-76332291-DEL-183 |
| chr17-76335355-DEL-52 | chr17-76335355-DEL-52 |
| chr17-76342102-DEL-52 | chr17-76342103-DEL-52 |
| chr17-76344897-DEL-7957 | chr17-76344898-DEL-7957 |
| chr17-76347594-DEL-499 | chr17-76347595-DEL-499 |
| chr17-76375316-DEL-6160 | chr17-76375316-DEL-6160 |
| chr17-76382736-DEL-90 | chr17-76382739-DEL-88 |
| chr17-76387097-DEL-60 | chr17-76387103-DEL-60 |
| chr17-76393235-DEL-57 | chr17-76393236-DEL-57 |
| chr17-76396645-DEL-1357 | chr17-76396646-DEL-1357 |
| chr17-76434362-DEL-426 | chr17-76434363-DEL-426 |
| chr17-76461345-DEL-1969 | chr17-76461346-DEL-1969 |
| chr17-76480958-DEL-52 | chr17-76480959-DEL-52 |
| chr17-76517016-DEL-8861 | chr17-76517017-DEL-8861 |
| chr17-76553109-DEL-60 | chr17-76553110-DEL-60 |
| chr17-76554150-DEL-115 | chr17-76554151-DEL-115 |
| chr17-76569343-DEL-345 | chr17-76569344-DEL-345 |
| chr17-76639418-DEL-937 | chr17-76639419-DEL-937 |
| chr17-76645806-DEL-60 | chr17-76645807-DEL-60 |
| chr17-76669579-DEL-334 | chr17-76669580-DEL-334 |
| chr17-76675567-DEL-72 | chr17-76675568-DEL-72 |
| chr17-76695802-DEL-54 | chr17-76695803-DEL-54 |
| chr17-76698516-DEL-190 | chr17-76698517-DEL-190 |
| chr17-76720348-DEL-153 | chr17-76720349-DEL-147 |
| chr17-76741941-DEL-121 | chr17-76741942-DEL-121 |
| chr17-76774555-DEL-57 | chr17-76774556-DEL-57 |
| chr17-76784113-DEL-59 | chr17-76784109-DEL-60 |
| chr17-76806468-DEL-181 | chr17-76806471-DEL-179 |
| chr17-76807224-DEL-117 | chr17-76807225-DEL-117 |
| chr17-76819649-DEL-7138 | chr17-76819650-DEL-7138 |
| chr17-76852883-DEL-68 | chr17-76852884-DEL-68 |
| chr17-76889029-DEL-85 | chr17-76889030-DEL-89 |
| chr17-76889964-DEL-2218 | chr17-76889965-DEL-2218 |
| chr17-76964926-DEL-103 | chr17-76964970-DEL-83 |
| chr17-76971191-DEL-2521 | chr17-76971194-DEL-2522 |
| chr17-76980634-DEL-2788 | chr17-76980635-DEL-2788 |
| chr17-76987599-DEL-61 | chr17-76987595-DEL-54 |
| chr17-77006051-DEL-110 | chr17-77006052-DEL-86 |

| | |
|---|---|
| chr17-77014298-DEL-1863 | chr17-77014299-DEL-1863 |
| chr17-77031821-DEL-92 | chr17-77031814-DEL-96 |
| chr17-77063600-DEL-2411 | chr17-77063601-DEL-2411 |
| chr17-77078439-DEL-116 | chr17-77078450-DEL-73 |
| chr17-77084118-DEL-70 | chr17-77084119-DEL-70 |
| chr17-77096645-DEL-1979 | chr17-77096646-DEL-1979 |
| chr17-77112886-DEL-57 | chr17-77112887-DEL-57 |
| chr17-77138437-DEL-4044 | chr17-77138438-DEL-4044 |
| chr17-77166953-DEL-106 | chr17-77166954-DEL-106 |
| chr17-77169644-DEL-2767 | chr17-77169645-DEL-2767 |
| chr17-77175420-DEL-439 | chr17-77175421-DEL-439 |
| chr17-77185067-DEL-64 | chr17-77185068-DEL-56 |
| chr17-77188537-DEL-85 | chr17-77188538-DEL-85 |
| chr17-77195135-DEL-54 | chr17-77195136-DEL-54 |
| chr17-77195787-DEL-96 | chr17-77195788-DEL-96 |
| chr17-77197700-DEL-112 | chr17-77197701-DEL-112 |
| chr17-77200832-DEL-6364 | chr17-77200833-DEL-6364 |
| chr17-77220802-DEL-389 | chr17-77220803-DEL-389 |
| chr17-77223680-DEL-53 | chr17-77223703-DEL-50 |
| chr17-77286410-DEL-424 | chr17-77286411-DEL-424 |
| chr17-77299504-DEL-2064 | chr17-77299505-DEL-2064 |
| chr17-77301863-DEL-312 | chr17-77301860-DEL-313 |
| chr17-77304430-DEL-2316 | chr17-77304431-DEL-2316 |
| chr17-77308321-DEL-895 | chr17-77308322-DEL-895 |
| chr17-77310167-DEL-60 | chr17-77310164-DEL-64 |
| chr17-77326288-DEL-73 | chr17-77326287-DEL-72 |
| chr17-77334876-DEL-70 | chr17-77334878-DEL-90 |
| chr17-77336703-DEL-57 | chr17-77336704-DEL-57 |
| chr17-77340737-DEL-259 | chr17-77340738-DEL-259 |
| chr17-77344011-DEL-71 | chr17-77344012-DEL-71 |
| chr17-77440283-DEL-113 | chr17-77440308-DEL-89 |
| chr17-77505696-DEL-847 | chr17-77505697-DEL-847 |
| chr17-77510116-DEL-863 | chr17-77510117-DEL-864 |
| chr17-77531769-DEL-58 | chr17-77531770-DEL-57 |
| chr17-77539406-DEL-6677 | chr17-77539407-DEL-6677 |
| chr17-77548671-DEL-84 | chr17-77548672-DEL-84 |
| chr17-77589309-DEL-471 | chr17-77589310-DEL-471 |
| chr17-77607847-DEL-6459 | chr17-77607848-DEL-6459 |
| chr17-77626154-DEL-65 | chr17-77626155-DEL-65 |
| chr17-77651668-DEL-586 | chr17-77651668-DEL-586 |
| chr17-77712037-DEL-52 | chr17-77712038-DEL-52 |
| chr17-77742501-DEL-102 | chr17-77742484-DEL-120 |
| chr17-77760937-DEL-73 | chr17-77760938-DEL-73 |
| chr17-77767449-DEL-433 | chr17-77767450-DEL-433 |
| chr17-77771376-DEL-918 | chr17-77771377-DEL-918 |
| chr17-77778691-DEL-109 | chr17-77778692-DEL-110 |
| chr17-77797759-DEL-1936 | chr17-77797760-DEL-1936 |
| chr17-77805194-DEL-1477 | chr17-77805195-DEL-1477 |
| chr17-77828558-DEL-120 | chr17-77828559-DEL-130 |
| chr17-77853045-DEL-1174 | chr17-77853046-DEL-1174 |
| chr17-77856045-DEL-282 | chr17-77856046-DEL-282 |
| chr17-77860431-DEL-98 | chr17-77860428-DEL-102 |
| chr17-77864521-DEL-5177 | chr17-77864522-DEL-5177 |
| chr17-77895784-DEL-54 | chr17-77895779-DEL-55 |
| chr17-77913393-DEL-178 | chr17-77913394-DEL-178 |
| chr17-77913959-DEL-565 | chr17-77913960-DEL-565 |
| chr17-77915340-DEL-1563 | chr17-77915341-DEL-1563 |
| chr17-77918150-DEL-99 | chr17-77918151-DEL-97 |
| chr17-77959644-DEL-206 | chr17-77959644-DEL-196 |
| chr17-77971956-DEL-3546 | chr17-77971956-DEL-3544 |
| chr17-77976113-DEL-136 | chr17-77976093-DEL-130 |
| chr17-77981001-DEL-692 | chr17-77981001-DEL-693 |
| chr17-77990684-DEL-118 | chr17-77990685-DEL-108 |
| chr17-78005260-DEL-6183 | chr17-78005261-DEL-6183 |
| chr17-78017935-DEL-74 | chr17-78017936-DEL-72 |
| chr17-78032180-DEL-175 | chr17-78032181-DEL-175 |
| chr17-78033247-DEL-51 | chr17-78033250-DEL-61 |
| chr17-78034332-DEL-53 | chr17-78034336-DEL-54 |
| chr17-78049985-DEL-831 | chr17-78049984-DEL-832 |
| chr17-78053921-DEL-251 | chr17-78053922-DEL-251 |
| chr17-78059830-DEL-217 | chr17-78059831-DEL-217 |
| chr17-78077560-DEL-1787 | chr17-78077561-DEL-1787 |
| chr17-78082948-DEL-5946 | chr17-78082949-DEL-5946 |
| chr17-78104585-DEL-3036 | chr17-78104586-DEL-3036 |
| chr17-78112949-DEL-68 | chr17-78112946-DEL-72 |
| chr17-78136463-DEL-6562 | chr17-78136464-DEL-6562 |
| chr17-78146955-DEL-3587 | chr17-78146956-DEL-3587 |
| chr17-78163540-DEL-256 | chr17-78163541-DEL-248 |
| chr17-78174006-DEL-86 | chr17-78174038-DEL-50 |
| chr17-78218106-DEL-381 | chr17-78218103-DEL-382 |

| | |
|---|---|
| chr17-78227902-DEL-4987 | chr17-78227903-DEL-4987 |
| chr17-78236476-DEL-125 | chr17-78236478-DEL-78 |
| chr17-78261187-DEL-1046 | chr17-78261188-DEL-1046 |
| chr17-78262423-DEL-182 | chr17-78262424-DEL-182 |
| chr17-78295347-DEL-79 | chr17-78295348-DEL-79 |
| chr17-78341782-DEL-857 | chr17-78341783-DEL-857 |
| chr17-78368049-DEL-1661 | chr17-78368050-DEL-1661 |
| chr17-78376535-DEL-6704 | chr17-78376536-DEL-6704 |
| chr17-78386884-DEL-78 | chr17-78386885-DEL-78 |
| chr17-78396036-DEL-639 | chr17-78396037-DEL-639 |
| chr17-78477862-DEL-200 | chr17-78477863-DEL-200 |
| chr17-78482326-DEL-108 | chr17-78482327-DEL-108 |
| chr17-78483999-DEL-60 | chr17-78484000-DEL-60 |
| chr17-78513394-DEL-169 | chr17-78513395-DEL-169 |
| chr17-78555503-DEL-204 | chr17-78555482-DEL-226 |
| chr17-78569499-DEL-135 | chr17-78569497-DEL-112 |
| chr17-78582278-DEL-62 | chr17-78582283-DEL-58 |
| chr17-78688295-DEL-372 | chr17-78688303-DEL-365 |
| chr17-78713297-DEL-175 | chr17-78713298-DEL-175 |
| chr17-78737190-DEL-280 | chr17-78737191-DEL-277 |
| chr17-78748376-DEL-72 | chr17-78748377-DEL-72 |
| chr17-78755961-DEL-52 | chr17-78755962-DEL-52 |
| chr17-78773994-DEL-419 | chr17-78773997-DEL-420 |
| chr17-78778120-DEL-5817 | chr17-78778121-DEL-5817 |
| chr17-78798950-DEL-2033 | chr17-78798951-DEL-2033 |
| chr17-78806849-DEL-73 | chr17-78806849-DEL-73 |
| chr17-78843667-DEL-990 | chr17-78843668-DEL-990 |
| chr17-78848161-DEL-66 | chr17-78848136-DEL-74 |
| chr17-78855124-DEL-55 | chr17-78855125-DEL-55 |
| chr17-78869900-DEL-188 | chr17-78869901-DEL-188 |
| chr17-78871826-DEL-56 | chr17-78871814-DEL-57 |
| chr17-78875811-DEL-137 | chr17-78875817-DEL-166 |
| chr17-78877651-DEL-112 | chr17-78877652-DEL-112 |
| chr17-78879781-DEL-56 | chr17-78879731-DEL-92 |
| chr17-78885687-DEL-105 | chr17-78885688-DEL-100 |
| chr17-78891713-DEL-941 | chr17-78891714-DEL-941 |
| chr17-78908008-DEL-116 | chr17-78908005-DEL-120 |
| chr17-78963896-DEL-116 | chr17-78963897-DEL-116 |
| chr17-78975092-DEL-214 | chr17-78975093-DEL-214 |
| chr17-79002350-DEL-52 | chr17-79002351-DEL-52 |
| chr17-79029571-DEL-92 | chr17-79029572-DEL-92 |
| chr17-79046515-DEL-93 | chr17-79046497-DEL-90 |
| chr17-79068190-DEL-120 | chr17-79068193-DEL-114 |
| chr17-79084127-DEL-198 | chr17-79084139-DEL-197 |
| chr17-79091905-DEL-3698 | chr17-79091906-DEL-3698 |
| chr17-79098115-DEL-165 | chr17-79098128-DEL-169 |
| chr17-79111658-DEL-141 | chr17-79111659-DEL-141 |
| chr17-79123959-DEL-147 | chr17-79123930-DEL-137 |
| chr17-79142100-DEL-73 | chr17-79142096-DEL-75 |
| chr17-79166564-DEL-3508 | chr17-79166543-DEL-3527 |
| chr17-79222986-DEL-68 | chr17-79222987-DEL-68 |
| chr17-79232976-DEL-62 | chr17-79232977-DEL-62 |
| chr17-79236106-DEL-82 | chr17-79236106-DEL-83 |
| chr17-79248098-DEL-150 | chr17-79248099-DEL-150 |
| chr17-79269335-DEL-70 | chr17-79269336-DEL-70 |
| chr17-79274577-DEL-80 | chr17-79274570-DEL-72 |
| chr17-79313365-DEL-106 | chr17-79313366-DEL-106 |
| chr17-79329607-DEL-100 | chr17-79329612-DEL-96 |
| chr17-79331600-DEL-66 | chr17-79331603-DEL-64 |
| chr17-79340843-DEL-55 | chr17-79340844-DEL-60 |
| chr17-79351523-DEL-6139 | chr17-79351524-DEL-6139 |
| chr17-79370187-DEL-162 | chr17-79370182-DEL-168 |
| chr17-79389465-DEL-326 | chr17-79389466-DEL-326 |
| chr17-79394759-DEL-1505 | chr17-79394774-DEL-1501 |
| chr17-79421041-DEL-104 | chr17-79421042-DEL-104 |
| chr17-79422158-DEL-62 | chr17-79422158-DEL-60 |
| chr17-79424231-DEL-84 | chr17-79424232-DEL-84 |
| chr17-79428157-DEL-855 | chr17-79428158-DEL-855 |
| chr17-79441268-DEL-60 | chr17-79441269-DEL-60 |
| chr17-79447617-DEL-560 | chr17-79447615-DEL-560 |
| chr17-79466330-DEL-54 | chr17-79466331-DEL-54 |
| chr17-79467464-DEL-58 | chr17-79467465-DEL-58 |
| chr17-79511731-DEL-102 | chr17-79511734-DEL-100 |
| chr17-79515994-DEL-445 | chr17-79516004-DEL-441 |
| chr17-79527749-DEL-1825 | chr17-79527750-DEL-1825 |
| chr17-79530640-DEL-2156 | chr17-79530625-DEL-2171 |
| chr17-79534284-DEL-80 | chr17-79534285-DEL-80 |
| chr17-79542006-DEL-278 | chr17-79542007-DEL-266 |
| chr17-79545690-DEL-62 | chr17-79545691-DEL-62 |
| chr17-79570019-DEL-66 | chr17-79570020-DEL-66 |

| | |
|---|---|
| chr17-79589491-DEL-74 | chr17-79589492-DEL-74 |
| chr17-79593867-DEL-269 | chr17-79593867-DEL-269 |
| chr17-79596122-DEL-52 | chr17-79596122-DEL-66 |
| chr17-79621575-DEL-57 | chr17-79621575-DEL-60 |
| chr17-79622778-DEL-92 | chr17-79622738-DEL-134 |
| chr17-79632703-DEL-67 | chr17-79632706-DEL-65 |
| chr17-79696535-DEL-52 | chr17-79696542-DEL-51 |
| chr17-79697855-DEL-323 | chr17-79697855-DEL-323 |
| chr17-79708096-DEL-10083 | chr17-79708097-DEL-10083 |
| chr17-79723952-DEL-76 | chr17-79723953-DEL-76 |
| chr17-79745267-DEL-58 | chr17-79745268-DEL-56 |
| chr17-79757623-DEL-116 | chr17-79757624-DEL-116 |
| chr17-79780159-DEL-119 | chr17-79780160-DEL-119 |
| chr17-79781772-DEL-856 | chr17-79781773-DEL-856 |
| chr17-79797083-DEL-1978 | chr17-79797084-DEL-1978 |
| chr17-79824596-DEL-2858 | chr17-79824597-DEL-2858 |
| chr17-79835464-DEL-180 | chr17-79835465-DEL-180 |
| chr17-79869862-DEL-60 | chr17-79869863-DEL-60 |
| chr17-79884957-DEL-56 | chr17-79884958-DEL-52 |
| chr17-79894784-DEL-131 | chr17-79894785-DEL-127 |
| chr17-79948851-DEL-6605 | chr17-79948852-DEL-6605 |
| chr17-79975105-DEL-93 | chr17-79975106-DEL-93 |
| chr17-79996388-DEL-827 | chr17-79996389-DEL-827 |
| chr17-80084320-DEL-236 | chr17-80084323-DEL-234 |
| chr17-80086476-DEL-119 | chr17-80086477-DEL-119 |
| chr17-80090344-DEL-113 | chr17-80090343-DEL-114 |
| chr17-80105296-DEL-269 | chr17-80105297-DEL-269 |
| chr17-80119489-DEL-62 | chr17-80119492-DEL-60 |
| chr17-80137681-DEL-84 | chr17-80137643-DEL-76 |
| chr17-80149996-DEL-178 | chr17-80149997-DEL-178 |
| chr17-80189754-DEL-201 | chr17-80189755-DEL-200 |
| chr17-80193130-DEL-198 | chr17-80193131-DEL-198 |
| chr17-80223408-DEL-68 | chr17-80223408-DEL-68 |
| chr17-80233127-DEL-107 | chr17-80233089-DEL-108 |
| chr17-80264782-DEL-207 | chr17-80264783-DEL-207 |
| chr17-80271090-DEL-80 | chr17-80271090-DEL-80 |
| chr17-80282923-DEL-344 | chr17-80282925-DEL-343 |
| chr17-80283998-DEL-116 | chr17-80283999-DEL-116 |
| chr17-80305247-DEL-81 | chr17-80305281-DEL-62 |
| chr17-80311111-DEL-210 | chr17-80311112-DEL-210 |
| chr17-80317391-DEL-87 | chr17-80317392-DEL-87 |
| chr17-80323781-DEL-206 | chr17-80323782-DEL-206 |
| chr17-80324859-DEL-71 | chr17-80324856-DEL-72 |
| chr17-80331729-DEL-1447 | chr17-80331730-DEL-1447 |
| chr17-80358678-DEL-114 | chr17-80358679-DEL-114 |
| chr17-80362859-DEL-208 | chr17-80362860-DEL-208 |
| chr17-80396663-DEL-199 | chr17-80396669-DEL-194 |
| chr17-80413003-DEL-60 | chr17-80413003-DEL-61 |
| chr17-80421773-DEL-605 | chr17-80421778-DEL-601 |
| chr17-80430410-DEL-94 | chr17-80430411-DEL-84 |
| chr17-80431476-DEL-196 | chr17-80431477-DEL-196 |
| chr17-80432143-DEL-203 | chr17-80432154-DEL-200 |
| chr17-80434596-DEL-146 | chr17-80434597-DEL-146 |
| chr17-80448345-DEL-628 | chr17-80448346-DEL-628 |
| chr17-80459709-DEL-80 | chr17-80459710-DEL-80 |
| chr17-80461145-DEL-89 | chr17-80461146-DEL-89 |
| chr17-80461893-DEL-50 | chr17-80461894-DEL-50 |
| chr17-80467368-DEL-99 | chr17-80467369-DEL-99 |
| chr17-80468527-DEL-131 | chr17-80468528-DEL-131 |
| chr17-80470171-DEL-604 | chr17-80470172-DEL-600 |
| chr17-80506217-DEL-116 | chr17-80506218-DEL-116 |
| chr17-80519850-DEL-56 | chr17-80519880-DEL-68 |
| chr17-80546710-DEL-1362 | chr17-80546711-DEL-1362 |
| chr17-80555180-DEL-206 | chr17-80555181-DEL-206 |
| chr17-80574100-DEL-197 | chr17-80574101-DEL-197 |
| chr17-80584081-DEL-216 | chr17-80584082-DEL-216 |
| chr17-80638494-DEL-263 | chr17-80638495-DEL-263 |
| chr17-80650557-DEL-502 | chr17-80650558-DEL-502 |
| chr17-80651752-DEL-77 | chr17-80651753-DEL-77 |
| chr17-80677353-DEL-53 | chr17-80677354-DEL-53 |
| chr17-80686844-DEL-198 | chr17-80686845-DEL-198 |
| chr17-80711102-DEL-376 | chr17-80711129-DEL-340 |
| chr17-80715055-DEL-73 | chr17-80715046-DEL-68 |
| chr17-80728730-DEL-1064 | chr17-80728731-DEL-1064 |
| chr17-80799592-DEL-212 | chr17-80799593-DEL-212 |
| chr17-80824405-DEL-60 | chr17-80824406-DEL-60 |
| chr17-80833160-DEL-233 | chr17-80833161-DEL-233 |
| chr17-80883861-DEL-91 | chr17-80883862-DEL-91 |
| chr17-80893063-DEL-75 | chr17-80893064-DEL-75 |
| chr17-80920252-DEL-195 | chr17-80920253-DEL-195 |

| | |
|---|---|
| chr17-80941254-DEL-480 | chr17-80941255-DEL-480 |
| chr17-80974084-DEL-85 | chr17-80974085-DEL-85 |
| chr17-80976788-DEL-159 | chr17-80976803-DEL-146 |
| chr17-80977248-DEL-424 | chr17-80977249-DEL-424 |
| chr17-81004411-DEL-274 | chr17-81004373-DEL-275 |
| chr17-81006843-DEL-65 | chr17-81006844-DEL-65 |
| chr17-81010164-DEL-53 | chr17-81010164-DEL-53 |
| chr17-81045672-DEL-206 | chr17-81045673-DEL-206 |
| chr17-81063836-DEL-412 | chr17-81063835-DEL-413 |
| chr17-81082104-DEL-51 | chr17-81082105-DEL-51 |
| chr17-81127441-DEL-60 | chr17-81127442-DEL-60 |
| chr17-81130768-DEL-80 | chr17-81130768-DEL-80 |
| chr17-81139140-DEL-485 | chr17-81139141-DEL-485 |
| chr17-81162520-DEL-54 | chr17-81162521-DEL-54 |
| chr17-81186485-DEL-128 | chr17-81186486-DEL-127 |
| chr17-81199955-DEL-804 | chr17-81199956-DEL-804 |
| chr17-81215256-DEL-4624 | chr17-81215256-DEL-4624 |
| chr17-81222773-DEL-317 | chr17-81222774-DEL-317 |
| chr17-81245324-DEL-894 | chr17-81245325-DEL-894 |
| chr17-81261883-DEL-220 | chr17-81261884-DEL-244 |
| chr17-81294983-DEL-52 | chr17-81294984-DEL-52 |
| chr17-81335619-DEL-100 | chr17-81335620-DEL-100 |
| chr17-81351132-DEL-123 | chr17-81351133-DEL-123 |
| chr17-81356297-DEL-220 | chr17-81356298-DEL-220 |
| chr17-81377229-DEL-51 | chr17-81377226-DEL-51 |
| chr17-81401048-DEL-65 | chr17-81401049-DEL-65 |
| chr17-81409529-DEL-66 | chr17-81409522-DEL-52 |
| chr17-81413155-DEL-1199 | chr17-81413156-DEL-1199 |
| chr17-81463837-DEL-7111 | chr17-81463838-DEL-7111 |
| chr17-81496756-DEL-61 | chr17-81496759-DEL-59 |
| chr17-81502539-DEL-53 | chr17-81502544-DEL-54 |
| chr17-81516185-DEL-134 | chr17-81516186-DEL-134 |
| chr17-81539133-DEL-5019 | chr17-81539134-DEL-5019 |
| chr17-81545548-DEL-7786 | chr17-81545549-DEL-7786 |
| chr17-81591843-DEL-108 | chr17-81591871-DEL-112 |
| chr17-81631964-DEL-561 | chr17-81631965-DEL-561 |
| chr17-81641737-DEL-477 | chr17-81641738-DEL-477 |
| chr17-81649987-DEL-51 | chr17-81649982-DEL-60 |
| chr17-81654786-DEL-6887 | chr17-81654787-DEL-6887 |
| chr17-81702079-DEL-874 | chr17-81702081-DEL-869 |
| chr17-81704459-DEL-1818 | chr17-81704460-DEL-1818 |
| chr17-81721846-DEL-93 | chr17-81721847-DEL-93 |
| chr17-81761152-DEL-60 | chr17-81761153-DEL-60 |
| chr17-81774649-DEL-1689 | chr17-81774650-DEL-1689 |
| chr17-81777480-DEL-68 | chr17-81777478-DEL-61 |
| chr17-81780628-DEL-144 | chr17-81780635-DEL-114 |
| chr17-81812755-DEL-2716 | chr17-81812756-DEL-2716 |
| chr17-81817244-DEL-466 | chr17-81817245-DEL-466 |
| chr17-81824078-DEL-801 | chr17-81824079-DEL-802 |
| chr17-81879286-DEL-276 | chr17-81879283-DEL-280 |
| chr17-81890012-DEL-151 | chr17-81890013-DEL-151 |
| chr17-81905480-DEL-84 | chr17-81905482-DEL-62 |
| chr17-81918212-DEL-82 | chr17-81918213-DEL-82 |
| chr17-81966460-DEL-1207 | chr17-81966461-DEL-1207 |
| chr17-81969867-DEL-126 | chr17-81969860-DEL-119 |
| chr17-81981903-DEL-76 | chr17-81981896-DEL-76 |
| chr17-81982359-DEL-506 | chr17-81982360-DEL-506 |
| chr17-81991260-DEL-6656 | chr17-81991261-DEL-6656 |
| chr17-81991674-DEL-212 | chr17-81991675-DEL-212 |
| chr17-81999432-DEL-1350 | chr17-81999433-DEL-1350 |
| chr17-82008709-DEL-144 | chr17-82008722-DEL-132 |
| chr17-82012633-DEL-66 | chr17-82012634-DEL-66 |
| chr17-82023635-DEL-6635 | chr17-82023636-DEL-6635 |
| chr17-82064031-DEL-564 | chr17-82064032-DEL-561 |
| chr17-82081615-DEL-69 | chr17-82081620-DEL-70 |
| chr17-82088402-DEL-289 | chr17-82088393-DEL-285 |
| chr17-82090285-DEL-82 | chr17-82090284-DEL-88 |
| chr17-82116669-DEL-3094 | chr17-82116670-DEL-3094 |
| chr17-82134163-DEL-198 | chr17-82134164-DEL-198 |
| chr17-82138865-DEL-2394 | chr17-82138866-DEL-2394 |
| chr17-82151088-DEL-74 | chr17-82151091-DEL-66 |
| chr17-82165174-DEL-97 | chr17-82165175-DEL-100 |
| chr17-82168541-DEL-130 | chr17-82168551-DEL-129 |
| chr17-82191876-DEL-120 | chr17-82191860-DEL-109 |
| chr17-82196337-DEL-175 | chr17-82196338-DEL-175 |
| chr17-82252410-DEL-68 | chr17-82252423-DEL-56 |
| chr17-82260118-DEL-57 | chr17-82260118-DEL-57 |
| chr17-82262212-DEL-661 | chr17-82262213-DEL-661 |
| chr17-82274323-DEL-7453 | chr17-82274324-DEL-7453 |
| chr17-82284439-DEL-51 | chr17-82284440-DEL-51 |

| | |
|---|---|
| chr17-82325454-DEL-52 | chr17-82325454-DEL-52 |
| chr17-82347993-DEL-78 | chr17-82347994-DEL-78 |
| chr17-82376720-DEL-102 | chr17-82376721-DEL-102 |
| chr17-82389083-DEL-899 | chr17-82389084-DEL-899 |
| chr17-82397351-DEL-630 | chr17-82397352-DEL-630 |
| chr17-82421889-DEL-329 | chr17-82421890-DEL-329 |
| chr17-82433783-DEL-1958 | chr17-82433784-DEL-1958 |
| chr17-82448567-DEL-274 | chr17-82448568-DEL-274 |
| chr17-82455600-DEL-1255 | chr17-82455611-DEL-1258 |
| chr17-82465939-DEL-90 | chr17-82465940-DEL-95 |
| chr17-82467517-DEL-352 | chr17-82467515-DEL-353 |
| chr17-82470757-DEL-6498 | chr17-82470758-DEL-6498 |
| chr17-82477809-DEL-56 | chr17-82477808-DEL-57 |
| chr17-82519351-DEL-61 | chr17-82519352-DEL-61 |
| chr17-82525995-DEL-1016 | chr17-82525996-DEL-1016 |
| chr17-82530032-DEL-1460 | chr17-82530033-DEL-1460 |
| chr17-82545484-DEL-101 | chr17-82545485-DEL-101 |
| chr17-82587163-DEL-1417 | chr17-82587164-DEL-1417 |
| chr17-82603461-DEL-523 | chr17-82603462-DEL-523 |
| chr17-82617464-DEL-549 | chr17-82617465-DEL-549 |
| chr17-82619692-DEL-66 | chr17-82619693-DEL-66 |
| chr17-82623527-DEL-80 | chr17-82623526-DEL-81 |
| chr17-82656490-DEL-64 | chr17-82656491-DEL-64 |
| chr17-82658649-DEL-114 | chr17-82658650-DEL-114 |
| chr17-82675625-DEL-109 | chr17-82675642-DEL-93 |
| chr17-82685174-DEL-94 | chr17-82685175-DEL-94 |
| chr17-82697470-DEL-6161 | chr17-82697471-DEL-6161 |
| chr17-82717265-DEL-1062 | chr17-82717266-DEL-1062 |
| chr17-82722376-DEL-6148 | chr17-82722377-DEL-6148 |
| chr17-82771523-DEL-6702 | chr17-82771524-DEL-6702 |
| chr17-82783555-DEL-384 | chr17-82783554-DEL-384 |
| chr17-82789075-DEL-2349 | chr17-82789076-DEL-2349 |
| chr17-82801375-DEL-1975 | chr17-82801376-DEL-1975 |
| chr17-82819080-DEL-719 | chr17-82819081-DEL-719 |
| chr17-82824212-DEL-208 | chr17-82824213-DEL-208 |
| chr17-82837257-DEL-201 | chr17-82837258-DEL-201 |
| chr17-82840018-DEL-58 | chr17-82840019-DEL-58 |
| chr17-82846573-DEL-1204 | chr17-82846574-DEL-1204 |
| chr17-82851143-DEL-202 | chr17-82851144-DEL-202 |
| chr17-82861159-DEL-110 | chr17-82861156-DEL-84 |
| chr17-82883440-DEL-6130 | chr17-82883441-DEL-6130 |
| chr17-82892400-DEL-72 | chr17-82892401-DEL-72 |
| chr17-82908173-DEL-71 | chr17-82908174-DEL-71 |
| chr17-82948311-DEL-77 | chr17-82948290-DEL-77 |
| chr17-82957531-DEL-51 | chr17-82957532-DEL-51 |
| chr17-82965943-DEL-92 | chr17-82965944-DEL-92 |
| chr17-82983001-DEL-6301 | chr17-82983002-DEL-6301 |
| chr17-82990241-DEL-199 | chr17-82990242-DEL-199 |
| chr17-83016647-DEL-90 | chr17-83016648-DEL-84 |
| chr17-83043472-DEL-116 | chr17-83043473-DEL-116 |
| chr17-83052744-DEL-183 | chr17-83052745-DEL-183 |
| chr17-83057328-DEL-3037 | chr17-83057329-DEL-3037 |
| chr17-83069797-DEL-120 | chr17-83069806-DEL-112 |
| chr17-83086498-DEL-54 | chr17-83086499-DEL-54 |
| chr17-83105386-DEL-6179 | chr17-83105387-DEL-6179 |
| chr17-83126297-DEL-431 | chr17-83126298-DEL-431 |
| chr17-83136427-DEL-228 | chr17-83136428-DEL-228 |
| chr17-83142815-DEL-70 | chr17-83142815-DEL-70 |
| chr17-83157400-DEL-3111 | chr17-83157401-DEL-3111 |
| chr17-83193603-DEL-700 | chr17-83193603-DEL-700 |
| chr17-83216412-DEL-53 | chr17-83216413-DEL-54 |
| chr17-83224748-DEL-1060 | chr17-83224749-DEL-1060 |
| chr17-83250844-DEL-102 | chr17-83250861-DEL-78 |
| chr17-83259804-DEL-1969 | chr17-83259805-DEL-1969 |
| chr17-83263695-DEL-83 | chr17-83263696-DEL-83 |
| chr17-83264696-DEL-66 | chr17-83264697-DEL-66 |
| chr17-83283284-DEL-6690 | chr17-83283285-DEL-6690 |
| chr17-83296287-DEL-2746 | chr17-83296288-DEL-2746 |
| chr17-83300345-DEL-126 | chr17-83300346-DEL-126 |
| chr17-83311218-DEL-73 | chr17-83311208-DEL-50 |
| chr17-83313573-DEL-258 | chr17-83313610-DEL-253 |
| chr17-83314409-DEL-52 | chr17-83314410-DEL-52 |
| chr17-83319492-DEL-490 | chr17-83319493-DEL-490 |
| chr17-83322816-DEL-227 | chr17-83322817-DEL-227 |
| chr17-83322905-DEL-1448 | chr17-83322906-DEL-1448 |
| chr17-83346599-DEL-353 | chr17-83346600-DEL-353 |
| chr17-83355782-DEL-764 | chr17-83355783-DEL-764 |
| chr17-83370647-DEL-85 | chr17-83370648-DEL-85 |
| chr17-83371755-DEL-324 | chr17-83371756-DEL-324 |
| chr17-83410962-DEL-93 | chr17-83411008-DEL-93 |

| | |
|---|---|
| chr17-83411660-DEL-583 | chr17-83411661-DEL-583 |
| chr17-83415264-DEL-94 | chr17-83415267-DEL-67 |
| chr17-83427899-DEL-76 | chr17-83427900-DEL-76 |
| chr17-83447667-DEL-6442 | chr17-83447668-DEL-6442 |
| chr17-83457946-DEL-408 | chr17-83457947-DEL-408 |
| chr17-83460559-DEL-1774 | chr17-83460560-DEL-1774 |
| chr17-83476479-DEL-1002 | chr17-83476479-DEL-1002 |
| chr17-83482700-DEL-110 | chr17-83482701-DEL-110 |
| chr17-83535954-DEL-492 | chr17-83535954-DEL-444 |
| chr17-83563491-DEL-4335 | chr17-83563492-DEL-4335 |
| chr17-83579798-DEL-60 | chr17-83579760-DEL-75 |
| chr17-83613974-DEL-54 | chr17-83613975-DEL-54 |
| chr17-83617352-DEL-92 | chr17-83617393-DEL-80 |
| chr17-83617933-DEL-132 | chr17-83617934-DEL-132 |
| chr17-83640523-DEL-88 | chr17-83640520-DEL-92 |
| chr17-83644690-DEL-54 | chr17-83644691-DEL-54 |
| chr17-83653406-DEL-662 | chr17-83653407-DEL-662 |
| chr17-83671600-DEL-104 | chr17-83671601-DEL-104 |
| chr17-83696478-DEL-465 | chr17-83696479-DEL-434 |
| chr17-83702179-DEL-476 | chr17-83702180-DEL-476 |
| chr17-83708444-DEL-55 | chr17-83708445-DEL-55 |
| chr17-83788592-DEL-231 | chr17-83788593-DEL-231 |
| chr17-83808533-DEL-152 | chr17-83808531-DEL-155 |
| chr17-83810227-DEL-232 | chr17-83810228-DEL-232 |
| chr17-83821141-DEL-139 | chr17-83821142-DEL-139 |
| chr17-83848661-DEL-67 | chr17-83848662-DEL-67 |
| chr17-83851509-DEL-193 | chr17-83851510-DEL-192 |
| chr17-83862219-DEL-88 | chr17-83862220-DEL-88 |
| chr17-83887080-DEL-164 | chr17-83887081-DEL-164 |
| chr17-83901703-DEL-218 | chr17-83901704-DEL-218 |
| chr17-83906797-DEL-253 | chr17-83906798-DEL-253 |
| chr17-83919694-DEL-225 | chr17-83919695-DEL-202 |
| chr17-83929549-DEL-182 | chr17-83929550-DEL-182 |
| chr17-83945822-DEL-105 | chr17-83945824-DEL-104 |
| chr17-83953674-DEL-130 | chr17-83953673-DEL-132 |
| chr17-83954564-DEL-82 | chr17-83954565-DEL-82 |
| chr17-83955641-DEL-256 | chr17-83955642-DEL-256 |
| chr17-83971975-DEL-162 | chr17-83971984-DEL-153 |
| chr17-83980808-DEL-56 | chr17-83980808-DEL-53 |
| chr17-83988378-DEL-926 | chr17-83988379-DEL-926 |
| chr17-83988720-DEL-82 | chr17-83988721-DEL-82 |
| chr17-83994496-DEL-102 | chr17-83994497-DEL-102 |
| chr17-83999486-DEL-201 | chr17-83999487-DEL-201 |
| chr17-84005669-DEL-54 | chr17-84005637-DEL-52 |
| chr17-84024538-DEL-889 | chr17-84024539-DEL-890 |
| chr17-84034065-DEL-82 | chr17-84034066-DEL-82 |
| chr17-84036417-DEL-64 | chr17-84036418-DEL-96 |
| chr17-84044132-DEL-165 | chr17-84044120-DEL-166 |
| chr17-84064103-DEL-76 | chr17-84064104-DEL-76 |
| chr17-84071790-DEL-201 | chr17-84071791-DEL-201 |
| chr17-84087162-DEL-197 | chr17-84087163-DEL-197 |
| chr17-84088179-DEL-211 | chr17-84088180-DEL-211 |
| chr17-84124131-DEL-90 | chr17-84124132-DEL-90 |
| chr17-84142032-DEL-76 | chr17-84142033-DEL-56 |
| chr17-84147090-DEL-84 | chr17-84147097-DEL-76 |
| chr17-84201758-DEL-95 | chr17-84201725-DEL-116 |
| chr17-84239595-DEL-187 | chr17-84239602-DEL-179 |
| chr17-84260404-DEL-60 | chr17-84260404-DEL-60 |
| chr17-84275588-DEL-149 | chr17-84275604-DEL-134 |
| chr17-84280612-DEL-51 | chr17-84280613-DEL-51 |
| chr17-84299758-DEL-348 | chr17-84299759-DEL-347 |
| chr17-84332258-DEL-80 | chr17-84332299-DEL-80 |
| chr17-84337715-DEL-79 | chr17-84337759-DEL-75 |
| chr17-84342303-DEL-60 | chr17-84342304-DEL-60 |
| chr17-84342826-DEL-52 | chr17-84342827-DEL-70 |
| chr17-84377660-DEL-84 | chr17-84377660-DEL-87 |
| chr17-84385488-DEL-77 | chr17-84385489-DEL-77 |
| chr17-84418999-DEL-134 | chr17-84419000-DEL-134 |
| chr17-84452774-DEL-128 | chr17-84452775-DEL-128 |
| chr17-84478241-DEL-261 | chr17-84478240-DEL-262 |
| chr17-84524975-DEL-306 | chr17-84524984-DEL-305 |
| chr17-84525801-DEL-207 | chr17-84525802-DEL-207 |
| chr17-84555625-DEL-63 | chr17-84555626-DEL-63 |
| chr17-84556552-DEL-52 | chr17-84556532-DEL-72 |
| chr17-84569594-DEL-696 | chr17-84569595-DEL-696 |
| chr17-84576482-DEL-168 | chr17-84576441-DEL-196 |
| chr17-84598792-DEL-64 | chr17-84598789-DEL-65 |
| chr17-84607970-DEL-855 | chr17-84607981-DEL-855 |
| chr17-84621136-DEL-52 | chr17-84621137-DEL-52 |
| chr17-84622184-DEL-264 | chr17-84622185-DEL-264 |

| | |
|---|---|
| chr17-84623883-DEL-206 | chr17-84623884-DEL-206 |
| chr17-84659822-DEL-104 | chr17-84659796-DEL-109 |
| chr17-84668824-DEL-90 | chr17-84668840-DEL-74 |
| chr17-84698844-DEL-178 | chr17-84698845-DEL-178 |
| chr17-84749536-DEL-125 | chr17-84749537-DEL-125 |
| chr17-84750765-DEL-211 | chr17-84750766-DEL-211 |
| chr17-84759569-DEL-219 | chr17-84759570-DEL-219 |
| chr17-84775943-DEL-380 | chr17-84775944-DEL-380 |
| chr17-84783503-DEL-113 | chr17-84783504-DEL-113 |
| chr17-84807169-DEL-122 | chr17-84807170-DEL-122 |
| chr17-84814093-DEL-278 | chr17-84814094-DEL-278 |
| chr17-84823675-DEL-401 | chr17-84823676-DEL-401 |
| chr17-84843203-DEL-115 | chr17-84843179-DEL-112 |
| chr17-84858681-DEL-57 | chr17-84858682-DEL-57 |
| chr17-84876807-DEL-119 | chr17-84876809-DEL-116 |
| chr17-84879603-DEL-108 | chr17-84879604-DEL-108 |
| chr17-84885909-DEL-67 | chr17-84885919-DEL-55 |
| chr17-84889991-DEL-92 | chr17-84889972-DEL-118 |
| chr17-84913816-DEL-94 | chr17-84913817-DEL-94 |
| chr17-84920853-DEL-290 | chr17-84920892-DEL-250 |
| chr17-84925759-DEL-191 | chr17-84925760-DEL-191 |
| chr17-84937794-DEL-72 | chr17-84937794-DEL-72 |
| chr17-84974919-DEL-65 | chr17-84974961-DEL-68 |
| chr17-84986478-DEL-76 | chr17-84986467-DEL-88 |
| chr17-85025408-DEL-58 | chr17-85025409-DEL-58 |
| chr17-85043990-DEL-52 | chr17-85043993-DEL-50 |
| chr17-85077235-DEL-51 | chr17-85077236-DEL-51 |
| chr17-85083227-DEL-197 | chr17-85083228-DEL-197 |
| chr17-85084175-DEL-77 | chr17-85084176-DEL-77 |
| chr17-85084716-DEL-74 | chr17-85084738-DEL-82 |
| chr17-85090680-DEL-78 | chr17-85090681-DEL-78 |
| chr17-85106223-DEL-441 | chr17-85106224-DEL-441 |
| chr17-85113652-DEL-59 | chr17-85113610-DEL-54 |
| chr17-85114402-DEL-52 | chr17-85114397-DEL-84 |
| chr17-85115097-DEL-1025 | chr17-85115098-DEL-1025 |
| chr17-85121751-DEL-129 | chr17-85121752-DEL-129 |
| chr17-85132084-DEL-327 | chr17-85132085-DEL-327 |
| chr17-85136465-DEL-201 | chr17-85136466-DEL-201 |
| chr17-85162442-DEL-65 | chr17-85162442-DEL-59 |
| chr17-85165487-DEL-266 | chr17-85165460-DEL-247 |
| chr17-85169904-DEL-82 | chr17-85169914-DEL-98 |
| chr17-85174859-DEL-172 | chr17-85174860-DEL-172 |
| chr17-85176324-DEL-119 | chr17-85176325-DEL-128 |
| chr17-85177285-DEL-237 | chr17-85177292-DEL-230 |
| chr17-85190349-DEL-4903 | chr17-85190350-DEL-4903 |
| chr17-85202021-DEL-57 | chr17-85202022-DEL-57 |
| chr17-85206977-DEL-128 | chr17-85206980-DEL-126 |
| chr17-85208938-DEL-112 | chr17-85208939-DEL-112 |
| chr17-85219998-DEL-70 | chr17-85219999-DEL-70 |
| chr17-85225181-DEL-378 | chr17-85225182-DEL-378 |
| chr17-85227775-DEL-1086 | chr17-85227776-DEL-1086 |
| chr17-85232700-DEL-58 | chr17-85232699-DEL-64 |
| chr17-85243196-DEL-54 | chr17-85243197-DEL-54 |
| chr17-85254494-DEL-245 | chr17-85254495-DEL-245 |
| chr17-85297991-DEL-60 | chr17-85297992-DEL-60 |
| chr17-85344982-DEL-55 | chr17-85344982-DEL-55 |
| chr17-85352385-DEL-52 | chr17-85352386-DEL-52 |
| chr17-85360635-DEL-91 | chr17-85360630-DEL-75 |
| chr17-85367772-DEL-60 | chr17-85367773-DEL-60 |
| chr17-85378403-DEL-112 | chr17-85378404-DEL-112 |
| chr17-85411849-DEL-211 | chr17-85411850-DEL-211 |
| chr17-85439925-DEL-87 | chr17-85439926-DEL-87 |
| chr17-85447344-DEL-448 | chr17-85447345-DEL-448 |
| chr17-85448345-DEL-85 | chr17-85448328-DEL-86 |
| chr17-85453546-DEL-825 | chr17-85453547-DEL-825 |
| chr17-85463000-DEL-132 | chr17-85463001-DEL-132 |
| chr17-85465272-DEL-231 | chr17-85465273-DEL-231 |
| chr17-85468987-DEL-195 | chr17-85468988-DEL-195 |
| chr17-85469558-DEL-271 | chr17-85469559-DEL-271 |
| chr17-85536416-DEL-6430 | chr17-85536417-DEL-6430 |
| chr17-85544706-DEL-184 | chr17-85544712-DEL-203 |
| chr17-85571474-DEL-1706 | chr17-85571475-DEL-1706 |
| chr17-85628359-DEL-80 | chr17-85628360-DEL-80 |
| chr17-85650552-DEL-72 | chr17-85650569-DEL-52 |
| chr17-85658668-DEL-58 | chr17-85658669-DEL-66 |
| chr17-85663490-DEL-99 | chr17-85663492-DEL-77 |
| chr17-85667662-DEL-162 | chr17-85667663-DEL-162 |
| chr17-85705622-DEL-68 | chr17-85705631-DEL-56 |
| chr17-85729491-DEL-259 | chr17-85729492-DEL-258 |
| chr17-85746477-DEL-207 | chr17-85746478-DEL-207 |

| | |
|---|---|
| chr17-85776049-DEL-302 | chr17-85776041-DEL-283 |
| chr17-85785226-DEL-67 | chr17-85785232-DEL-62 |
| chr17-85791948-DEL-56 | chr17-85791949-DEL-56 |
| chr17-85830900-DEL-1463 | chr17-85830905-DEL-1456 |
| chr17-85873094-DEL-175 | chr17-85873095-DEL-175 |
| chr17-85890731-DEL-319 | chr17-85890732-DEL-319 |
| chr17-85943418-DEL-200 | chr17-85943419-DEL-200 |
| chr17-85962845-DEL-69 | chr17-85962846-DEL-69 |
| chr17-85971577-DEL-150 | chr17-85971578-DEL-150 |
| chr17-85973298-DEL-53 | chr17-85973305-DEL-55 |
| chr17-85976819-DEL-109 | chr17-85976820-DEL-109 |
| chr17-85980863-DEL-70 | chr17-85980853-DEL-73 |
| chr17-85988434-DEL-58 | chr17-85988435-DEL-58 |
| chr17-86005473-DEL-114 | chr17-86005492-DEL-85 |
| chr17-86035096-DEL-72 | chr17-86035113-DEL-56 |
| chr17-86036475-DEL-128 | chr17-86036476-DEL-128 |
| chr17-86045377-DEL-1920 | chr17-86045378-DEL-1920 |
| chr17-86051926-DEL-60 | chr17-86051927-DEL-60 |
| chr17-86068876-DEL-137 | chr17-86068877-DEL-137 |
| chr17-86072844-DEL-191 | chr17-86072825-DEL-201 |
| chr17-86082965-DEL-127 | chr17-86082966-DEL-127 |
| chr17-86140470-DEL-445 | chr17-86140471-DEL-445 |
| chr17-86168298-DEL-54 | chr17-86168299-DEL-54 |
| chr17-86245648-DEL-246 | chr17-86245649-DEL-246 |
| chr17-86270500-DEL-351 | chr17-86270501-DEL-351 |
| chr17-86289899-DEL-56 | chr17-86289900-DEL-56 |
| chr17-86303512-DEL-3955 | chr17-86303511-DEL-3957 |
| chr17-86324517-DEL-2247 | chr17-86324518-DEL-2247 |
| chr17-86457609-DEL-156 | chr17-86457610-DEL-156 |
| chr17-86458463-DEL-104 | chr17-86458464-DEL-104 |
| chr17-86463520-DEL-82 | chr17-86463520-DEL-82 |
| chr17-86509865-DEL-460 | chr17-86509860-DEL-461 |
| chr17-86526229-DEL-521 | chr17-86526230-DEL-521 |
| chr17-86552188-DEL-595 | chr17-86552189-DEL-595 |
| chr17-86556698-DEL-192 | chr17-86556699-DEL-192 |
| chr17-86577675-DEL-3938 | chr17-86577676-DEL-3938 |
| chr17-86604352-DEL-54 | chr17-86604352-DEL-54 |
| chr17-86615762-DEL-52 | chr17-86615762-DEL-52 |
| chr17-86683578-DEL-67 | chr17-86683579-DEL-67 |
| chr17-86690146-DEL-77 | chr17-86690149-DEL-75 |
| chr17-86713600-DEL-67 | chr17-86713601-DEL-67 |
| chr17-86732557-DEL-126 | chr17-86732558-DEL-108 |
| chr17-86736311-DEL-225 | chr17-86736312-DEL-225 |
| chr17-86742005-DEL-106 | chr17-86742006-DEL-106 |
| chr17-86763505-DEL-112 | chr17-86763506-DEL-112 |
| chr17-86774148-DEL-58 | chr17-86774149-DEL-58 |
| chr17-86775408-DEL-62 | chr17-86775409-DEL-62 |
| chr17-86791234-DEL-56 | chr17-86791234-DEL-54 |
| chr17-86820172-DEL-53 | chr17-86820167-DEL-59 |
| chr17-86854596-DEL-839 | chr17-86854597-DEL-839 |
| chr17-86876284-DEL-125 | chr17-86876302-DEL-107 |
| chr17-86879995-DEL-57 | chr17-86879996-DEL-57 |
| chr17-86884389-DEL-80 | chr17-86884402-DEL-82 |
| chr17-86885371-DEL-92 | chr17-86885372-DEL-92 |
| chr17-86888073-DEL-290 | chr17-86888074-DEL-290 |
| chr17-86901447-DEL-259 | chr17-86901448-DEL-259 |
| chr17-86904360-DEL-113 | chr17-86904361-DEL-113 |
| chr17-86929612-DEL-124 | chr17-86929613-DEL-155 |
| chr17-86930771-DEL-1131 | chr17-86930772-DEL-1131 |
| chr17-86953188-DEL-79 | chr17-86953191-DEL-77 |
| chr17-86972111-DEL-68 | chr17-86972126-DEL-50 |
| chr17-86996785-DEL-108 | chr17-86996827-DEL-70 |
| chr17-86999278-DEL-118 | chr17-86999279-DEL-118 |
| chr17-87011769-DEL-92 | chr17-87011776-DEL-56 |
| chr17-87023137-DEL-96 | chr17-87023138-DEL-96 |
| chr17-87027665-DEL-190 | chr17-87027668-DEL-156 |
| chr17-87036725-DEL-84 | chr17-87036722-DEL-85 |
| chr17-87047649-DEL-493 | chr17-87047650-DEL-493 |
| chr17-87103891-DEL-344 | chr17-87103892-DEL-344 |
| chr17-87153172-DEL-2273 | chr17-87153173-DEL-2273 |
| chr17-87171201-DEL-115 | chr17-87171202-DEL-115 |
| chr17-87208921-DEL-67 | chr17-87208897-DEL-53 |
| chr17-87253564-DEL-93 | chr17-87253565-DEL-81 |
| chr17-87262396-DEL-555 | chr17-87262397-DEL-555 |
| chr17-87264016-DEL-87 | chr17-87264017-DEL-87 |
| chr17-87264306-DEL-450 | chr17-87264323-DEL-434 |
| chr17-87283800-DEL-349 | chr17-87283801-DEL-349 |
| chr17-87288045-DEL-53 | chr17-87288046-DEL-53 |
| chr17-87299019-DEL-499 | chr17-87299020-DEL-499 |
| chr17-87318293-DEL-91 | chr17-87318265-DEL-92 |

| | |
|---|---|
| chr17-87322908-DEL-697 | chr17-87322911-DEL-699 |
| chr17-87341217-DEL-387 | chr17-87341205-DEL-400 |
| chr17-87385669-DEL-1999 | chr17-87385670-DEL-1999 |
| chr17-87393795-DEL-52 | chr17-87393796-DEL-52 |
| chr17-87410982-DEL-78 | chr17-87410983-DEL-78 |
| chr17-87421976-DEL-54 | chr17-87421977-DEL-54 |
| chr17-87487464-DEL-175 | chr17-87487465-DEL-175 |
| chr17-87489884-DEL-56 | chr17-87489898-DEL-54 |
| chr17-87512913-DEL-63 | chr17-87512912-DEL-64 |
| chr17-87518661-DEL-54 | chr17-87518661-DEL-54 |
| chr17-87519677-DEL-328 | chr17-87519678-DEL-328 |
| chr17-87535766-DEL-318 | chr17-87535771-DEL-314 |
| chr17-87538491-DEL-70 | chr17-87538492-DEL-70 |
| chr17-87541381-DEL-64 | chr17-87541379-DEL-76 |
| chr17-87548310-DEL-101 | chr17-87548311-DEL-101 |
| chr17-87559437-DEL-70 | chr17-87559438-DEL-70 |
| chr17-87565997-DEL-60 | chr17-87565998-DEL-60 |
| chr17-87579406-DEL-72 | chr17-87579407-DEL-72 |
| chr17-87580111-DEL-58 | chr17-87580093-DEL-59 |
| chr17-87584213-DEL-169 | chr17-87584214-DEL-169 |
| chr17-87607043-DEL-211 | chr17-87607044-DEL-211 |
| chr17-87646837-DEL-52 | chr17-87646837-DEL-52 |
| chr17-87656191-DEL-58 | chr17-87656192-DEL-58 |
| chr17-87667243-DEL-166 | chr17-87667244-DEL-166 |
| chr17-87667787-DEL-65 | chr17-87667794-DEL-103 |
| chr17-87775392-DEL-229 | chr17-87775393-DEL-229 |
| chr17-87778958-DEL-60 | chr17-87778959-DEL-60 |
| chr17-87784596-DEL-100 | chr17-87784596-DEL-100 |
| chr17-87794760-DEL-74 | chr17-87794761-DEL-74 |
| chr17-87814433-DEL-71 | chr17-87814434-DEL-71 |
| chr17-87820114-DEL-95 | chr17-87820115-DEL-95 |
| chr17-87833303-DEL-107 | chr17-87833304-DEL-107 |
| chr17-87841118-DEL-1523 | chr17-87841118-DEL-1523 |
| chr17-87846194-DEL-93 | chr17-87846205-DEL-77 |
| chr17-87875442-DEL-199 | chr17-87875425-DEL-242 |
| chr17-87887226-DEL-365 | chr17-87887227-DEL-365 |
| chr17-87890707-DEL-210 | chr17-87890708-DEL-210 |
| chr17-87897536-DEL-519 | chr17-87897537-DEL-519 |
| chr17-87901101-DEL-328 | chr17-87901116-DEL-314 |
| chr17-87901636-DEL-82 | chr17-87901589-DEL-82 |
| chr17-87906658-DEL-397 | chr17-87906659-DEL-397 |
| chr17-87912257-DEL-500 | chr17-87912258-DEL-500 |
| chr17-87929316-DEL-64 | chr17-87929317-DEL-64 |
| chr17-87934953-DEL-202 | chr17-87934954-DEL-202 |
| chr17-87937665-DEL-178 | chr17-87937666-DEL-178 |
| chr17-87959874-DEL-172 | chr17-87959875-DEL-172 |
| chr17-87966217-DEL-173 | chr17-87966218-DEL-173 |
| chr17-87973837-DEL-541 | chr17-87973838-DEL-541 |
| chr17-87976286-DEL-174 | chr17-87976287-DEL-174 |
| chr17-87984518-DEL-142 | chr17-87984503-DEL-143 |
| chr17-88001733-DEL-342 | chr17-88001768-DEL-319 |
| chr17-88023670-DEL-191 | chr17-88023671-DEL-191 |
| chr17-88028841-DEL-70 | chr17-88028829-DEL-50 |
| chr17-88036130-DEL-84 | chr17-88036131-DEL-60 |
| chr17-88039121-DEL-213 | chr17-88039122-DEL-213 |
| chr17-88076238-DEL-218 | chr17-88076239-DEL-218 |
| chr17-88095451-DEL-212 | chr17-88095452-DEL-212 |
| chr17-88096470-DEL-199 | chr17-88096471-DEL-199 |
| chr17-88151396-DEL-65 | chr17-88151397-DEL-65 |
| chr17-88153321-DEL-53 | chr17-88153322-DEL-53 |
| chr17-88160369-DEL-165 | chr17-88160368-DEL-165 |
| chr17-88164152-DEL-209 | chr17-88164153-DEL-209 |
| chr17-88171706-DEL-1284 | chr17-88171707-DEL-1284 |
| chr17-88232091-DEL-433 | chr17-88232092-DEL-433 |
| chr17-88267131-DEL-223 | chr17-88267132-DEL-223 |
| chr17-88271182-DEL-733 | chr17-88271152-DEL-722 |
| chr17-88272772-DEL-57 | chr17-88272773-DEL-57 |
| chr17-88277940-DEL-91 | chr17-88277941-DEL-91 |
| chr17-88323403-DEL-140 | chr17-88323391-DEL-123 |
| chr17-88376881-DEL-62 | chr17-88376882-DEL-62 |
| chr17-88381220-DEL-673 | chr17-88381221-DEL-673 |
| chr17-88399280-DEL-197 | chr17-88399272-DEL-198 |
| chr17-88443398-DEL-52 | chr17-88443398-DEL-64 |
| chr17-88446808-DEL-357 | chr17-88446837-DEL-368 |
| chr17-88456699-DEL-517 | chr17-88456700-DEL-517 |
| chr17-88460235-DEL-195 | chr17-88460238-DEL-194 |
| chr17-88504324-DEL-76 | chr17-88504309-DEL-120 |
| chr17-88518360-DEL-81 | chr17-88518361-DEL-81 |
| chr17-88527991-DEL-52 | chr17-88527992-DEL-56 |
| chr17-88552121-DEL-131 | chr17-88552114-DEL-139 |

| | |
|---|---|
| chr17-88554456-DEL-278 | chr17-88554457-DEL-278 |
| chr17-88555057-DEL-190 | chr17-88555058-DEL-190 |
| chr17-88556866-DEL-272 | chr17-88556867-DEL-272 |
| chr17-88560354-DEL-115 | chr17-88560369-DEL-141 |
| chr17-88574760-DEL-266 | chr17-88574758-DEL-267 |
| chr17-88577500-DEL-327 | chr17-88577501-DEL-326 |
| chr17-88585171-DEL-153 | chr17-88585172-DEL-153 |
| chr17-88606019-DEL-208 | chr17-88606020-DEL-202 |
| chr17-88608916-DEL-206 | chr17-88608917-DEL-206 |
| chr17-88609991-DEL-104 | chr17-88609994-DEL-102 |
| chr17-88639122-DEL-71 | chr17-88639122-DEL-74 |
| chr17-88650058-DEL-329 | chr17-88650059-DEL-329 |
| chr17-88664188-DEL-91 | chr17-88664164-DEL-69 |
| chr17-88669595-DEL-390 | chr17-88669596-DEL-390 |
| chr17-88682717-DEL-57 | chr17-88682728-DEL-51 |
| chr17-88693002-DEL-79 | chr17-88693003-DEL-79 |
| chr17-88699582-DEL-3212 | chr17-88699583-DEL-3208 |
| chr17-88719209-DEL-95 | chr17-88719210-DEL-95 |
| chr17-88730809-DEL-472 | chr17-88730809-DEL-471 |
| chr17-88745352-DEL-79 | chr17-88745353-DEL-79 |
| chr17-88769620-DEL-205 | chr17-88769621-DEL-205 |
| chr17-88771440-DEL-52 | chr17-88771433-DEL-60 |
| chr17-88782586-DEL-54 | chr17-88782566-DEL-55 |
| chr17-88785722-DEL-221 | chr17-88785723-DEL-221 |
| chr17-88793123-DEL-80 | chr17-88793124-DEL-80 |
| chr17-88807069-DEL-250 | chr17-88807076-DEL-240 |
| chr17-88808220-DEL-3346 | chr17-88808225-DEL-3373 |
| chr17-88813202-DEL-167 | chr17-88813203-DEL-167 |
| chr17-88817102-DEL-209 | chr17-88817112-DEL-207 |
| chr17-88823915-DEL-67 | chr17-88823888-DEL-68 |
| chr17-88851599-DEL-385 | chr17-88851600-DEL-385 |
| chr17-88873369-DEL-75 | chr17-88873368-DEL-75 |
| chr17-88881052-DEL-67 | chr17-88881053-DEL-67 |
| chr17-88883202-DEL-69 | chr17-88883203-DEL-69 |
| chr17-88896057-DEL-213 | chr17-88896058-DEL-213 |
| chr17-88898308-DEL-364 | chr17-88898318-DEL-365 |
| chr17-88907629-DEL-67 | chr17-88907630-DEL-67 |
| chr17-88909496-DEL-75 | chr17-88909497-DEL-75 |
| chr17-88909820-DEL-863 | chr17-88909821-DEL-863 |
| chr17-88919852-DEL-3972 | chr17-88919853-DEL-3972 |
| chr17-88922143-DEL-54 | chr17-88922144-DEL-54 |
| chr17-88924865-DEL-3274 | chr17-88924866-DEL-3274 |
| chr17-88953176-DEL-53 | chr17-88953165-DEL-52 |
| chr17-88983876-DEL-572 | chr17-88983877-DEL-572 |
| chr17-88991151-DEL-74 | chr17-88991152-DEL-74 |
| chr17-89000538-DEL-92 | chr17-89000539-DEL-115 |
| chr17-89014045-DEL-212 | chr17-89014046-DEL-212 |
| chr17-89020632-DEL-542 | chr17-89020633-DEL-542 |
| chr17-89052445-DEL-431 | chr17-89052444-DEL-429 |
| chr17-89056703-DEL-358 | chr17-89056712-DEL-345 |
| chr17-89124757-DEL-73 | chr17-89124758-DEL-74 |
| chr17-89181008-DEL-205 | chr17-89181009-DEL-205 |
| chr17-89196550-DEL-89 | chr17-89196521-DEL-95 |
| chr17-89210641-DEL-71 | chr17-89210642-DEL-71 |
| chr17-89215133-DEL-400 | chr17-89215134-DEL-400 |
| chr17-89228377-DEL-203 | chr17-89228378-DEL-203 |
| chr17-89230415-DEL-59 | chr17-89230409-DEL-57 |
| chr17-89233126-DEL-2930 | chr17-89233128-DEL-2929 |
| chr17-89234761-DEL-235 | chr17-89234762-DEL-235 |
| chr17-89236768-DEL-302 | chr17-89236769-DEL-302 |
| chr17-89237551-DEL-87 | chr17-89237566-DEL-78 |
| chr17-89261177-DEL-84 | chr17-89261178-DEL-85 |
| chr17-89270745-DEL-70 | chr17-89270746-DEL-70 |
| chr17-89271569-DEL-462 | chr17-89271570-DEL-462 |
| chr17-89273184-DEL-78 | chr17-89273185-DEL-78 |
| chr17-89274706-DEL-106 | chr17-89274711-DEL-102 |
| chr17-89300698-DEL-293 | chr17-89300699-DEL-293 |
| chr17-89310928-DEL-1738 | chr17-89310929-DEL-1738 |
| chr17-89328427-DEL-79 | chr17-89328428-DEL-79 |
| chr17-89378269-DEL-683 | chr17-89378270-DEL-683 |
| chr17-89441142-DEL-5384 | chr17-89441143-DEL-5384 |
| chr17-89447131-DEL-198 | chr17-89447132-DEL-198 |
| chr17-89455060-DEL-6783 | chr17-89455053-DEL-6781 |
| chr17-89530574-DEL-1506 | chr17-89530567-DEL-1506 |
| chr17-89546517-DEL-389 | chr17-89546518-DEL-389 |
| chr17-89555189-DEL-88 | chr17-89555194-DEL-87 |
| chr17-89564820-DEL-8457 | chr17-89564821-DEL-8457 |
| chr17-89588047-DEL-108 | chr17-89588048-DEL-108 |
| chr17-89668310-DEL-508 | chr17-89668311-DEL-508 |
| chr17-89674867-DEL-70 | chr17-89674871-DEL-66 |

| | |
|---|---|
| chr17-89693480-DEL-466 | chr17-89693483-DEL-465 |
| chr17-89698351-DEL-498 | chr17-89698344-DEL-499 |
| chr17-89711318-DEL-84 | chr17-89711319-DEL-84 |
| chr17-89729352-DEL-55 | chr17-89729353-DEL-55 |
| chr17-89733181-DEL-211 | chr17-89733182-DEL-211 |
| chr17-89748157-DEL-212 | chr17-89748158-DEL-212 |
| chr17-89757958-DEL-55 | chr17-89757959-DEL-55 |
| chr17-89758682-DEL-78 | chr17-89758683-DEL-78 |
| chr17-89761639-DEL-99 | chr17-89761636-DEL-86 |
| chr17-89787297-DEL-7495 | chr17-89787298-DEL-7495 |
| chr17-89799683-DEL-820 | chr17-89799684-DEL-817 |
| chr17-89836713-DEL-1617 | chr17-89836714-DEL-1617 |
| chr17-89865589-DEL-98 | chr17-89865590-DEL-98 |
| chr17-89874736-DEL-123 | chr17-89874737-DEL-123 |
| chr17-89876595-DEL-107 | chr17-89876578-DEL-137 |
| chr17-89878802-DEL-63 | chr17-89878799-DEL-64 |
| chr17-89880243-DEL-54 | chr17-89880244-DEL-54 |
| chr17-89901052-DEL-2053 | chr17-89901053-DEL-2053 |
| chr17-89906156-DEL-134 | chr17-89906176-DEL-120 |
| chr17-89919720-DEL-2641 | chr17-89919714-DEL-2642 |
| chr17-89994894-DEL-67 | chr17-89994895-DEL-67 |
| chr17-90020276-DEL-65 | chr17-90020281-DEL-106 |
| chr17-90024155-DEL-77 | chr17-90024136-DEL-92 |
| chr17-90033095-DEL-555 | chr17-90033095-DEL-555 |
| chr17-90033729-DEL-1000 | chr17-90033729-DEL-1000 |
| chr17-90052040-DEL-83 | chr17-90052054-DEL-54 |
| chr17-90059222-DEL-198 | chr17-90059199-DEL-196 |
| chr17-90089092-DEL-1915 | chr17-90089093-DEL-1915 |
| chr17-90098925-DEL-178 | chr17-90098926-DEL-178 |
| chr17-90100811-DEL-3779 | chr17-90100812-DEL-3779 |
| chr17-90102992-DEL-343 | chr17-90102993-DEL-343 |
| chr17-90123578-DEL-6921 | chr17-90123579-DEL-6921 |
| chr17-90141271-DEL-6459 | chr17-90141272-DEL-6459 |
| chr17-90151905-DEL-584 | chr17-90151906-DEL-584 |
| chr17-90169586-DEL-823 | chr17-90169586-DEL-823 |
| chr17-90179925-DEL-80 | chr17-90179922-DEL-78 |
| chr17-90197476-DEL-544 | chr17-90197477-DEL-544 |
| chr17-90201394-DEL-313 | chr17-90201395-DEL-313 |
| chr17-90201657-DEL-1567 | chr17-90201658-DEL-1567 |
| chr17-90215868-DEL-2002 | chr17-90215869-DEL-2002 |
| chr17-90238961-DEL-102 | chr17-90238962-DEL-102 |
| chr17-90242974-DEL-7144 | chr17-90242975-DEL-7144 |
| chr17-90256286-DEL-54 | chr17-90256287-DEL-54 |
| chr17-90276271-DEL-2649 | chr17-90276272-DEL-2649 |
| chr17-90293548-DEL-93 | chr17-90293537-DEL-68 |
| chr17-90301376-DEL-184 | chr17-90301374-DEL-185 |
| chr17-90302396-DEL-1250 | chr17-90302397-DEL-1250 |
| chr17-90323860-DEL-108 | chr17-90323861-DEL-102 |
| chr17-90330203-DEL-212 | chr17-90330201-DEL-213 |
| chr17-90331381-DEL-5593 | chr17-90331370-DEL-5594 |
| chr17-90338350-DEL-115 | chr17-90338355-DEL-114 |
| chr17-90391138-DEL-115 | chr17-90391139-DEL-115 |
| chr17-90409853-DEL-7119 | chr17-90409854-DEL-7119 |
| chr17-90493519-DEL-1928 | chr17-90493520-DEL-1928 |
| chr17-90518693-DEL-1184 | chr17-90518694-DEL-1184 |
| chr17-90543307-DEL-83 | chr17-90543300-DEL-91 |
| chr17-90546088-DEL-57 | chr17-90546089-DEL-57 |
| chr17-90680352-DEL-60 | chr17-90680349-DEL-64 |
| chr17-90706626-DEL-52 | chr17-90706627-DEL-52 |
| chr17-90717693-DEL-62 | chr17-90717643-DEL-64 |
| chr17-90725368-DEL-368 | chr17-90725370-DEL-367 |
| chr17-90778358-DEL-1476 | chr17-90778359-DEL-1476 |
| chr17-90781373-DEL-206 | chr17-90781374-DEL-206 |
| chr17-90807544-DEL-56 | chr17-90807545-DEL-50 |
| chr17-90853693-DEL-106 | chr17-90853694-DEL-106 |
| chr17-90866718-DEL-233 | chr17-90866718-DEL-233 |
| chr17-90879882-DEL-300 | chr17-90879883-DEL-300 |
| chr17-90880746-DEL-900 | chr17-90880739-DEL-901 |
| chr17-90884932-DEL-146 | chr17-90884933-DEL-146 |
| chr17-90895372-DEL-163 | chr17-90895379-DEL-151 |
| chr17-90900814-DEL-87 | chr17-90900771-DEL-88 |
| chr17-90910420-DEL-551 | chr17-90910421-DEL-551 |
| chr17-90994157-DEL-77 | chr17-90994153-DEL-80 |
| chr17-91021916-DEL-426 | chr17-91021917-DEL-426 |
| chr17-91031626-DEL-268 | chr17-91031627-DEL-268 |
| chr17-91039341-DEL-846 | chr17-91039342-DEL-846 |
| chr17-91084691-DEL-1775 | chr17-91084692-DEL-1775 |
| chr17-91089778-DEL-582 | chr17-91089779-DEL-582 |
| chr17-91091971-DEL-87 | chr17-91091974-DEL-85 |
| chr17-91137689-DEL-473 | chr17-91137684-DEL-474 |

| | |
|---|---|
| chr17-91147411-DEL-54 | chr17-91147412-DEL-54 |
| chr17-91161925-DEL-71 | chr17-91161926-DEL-65 |
| chr17-91176872-DEL-283 | chr17-91176873-DEL-283 |
| chr17-91184939-DEL-63 | chr17-91184940-DEL-63 |
| chr17-91192763-DEL-332 | chr17-91192764-DEL-332 |
| chr17-91223567-DEL-55 | chr17-91223567-DEL-55 |
| chr17-91258736-DEL-98 | chr17-91258737-DEL-98 |
| chr17-91260056-DEL-414 | chr17-91260057-DEL-414 |
| chr17-91339663-DEL-97 | chr17-91339664-DEL-97 |
| chr17-91406838-DEL-63 | chr17-91406837-DEL-60 |
| chr17-91412342-DEL-196 | chr17-91412343-DEL-196 |
| chr17-91425147-DEL-3539 | chr17-91425148-DEL-3539 |
| chr17-91449402-DEL-100 | chr17-91449407-DEL-96 |
| chr17-91454389-DEL-63 | chr17-91454392-DEL-61 |
| chr17-91487534-DEL-60 | chr17-91487535-DEL-60 |
| chr17-91497858-DEL-714 | chr17-91497859-DEL-714 |
| chr17-91500127-DEL-70 | chr17-91500128-DEL-70 |
| chr17-91500689-DEL-893 | chr17-91500689-DEL-893 |
| chr17-91505634-DEL-1484 | chr17-91505635-DEL-1484 |
| chr17-91530229-DEL-484 | chr17-91530230-DEL-484 |
| chr17-91531131-DEL-373 | chr17-91531132-DEL-373 |
| chr17-91544186-DEL-1378 | chr17-91544187-DEL-1378 |
| chr17-91581175-DEL-431 | chr17-91581176-DEL-431 |
| chr17-91591414-DEL-110 | chr17-91591415-DEL-110 |
| chr17-91598558-DEL-76 | chr17-91598559-DEL-76 |
| chr17-91614717-DEL-2108 | chr17-91614718-DEL-2108 |
| chr17-91635081-DEL-111 | chr17-91635064-DEL-106 |
| chr17-91636171-DEL-56 | chr17-91636172-DEL-56 |
| chr17-91652015-DEL-152 | chr17-91652016-DEL-151 |
| chr17-91714812-DEL-1055 | chr17-91714813-DEL-1055 |
| chr17-91717039-DEL-73 | chr17-91717040-DEL-73 |
| chr17-91784345-DEL-300 | chr17-91784345-DEL-300 |
| chr17-91789059-DEL-476 | chr17-91789053-DEL-482 |
| chr17-91804900-DEL-52 | chr17-91804901-DEL-52 |
| chr17-91830761-DEL-62 | chr17-91830762-DEL-62 |
| chr17-91832078-DEL-6225 | chr17-91832079-DEL-6225 |
| chr17-91854877-DEL-5227 | chr17-91854878-DEL-5227 |
| chr17-91879746-DEL-483 | chr17-91879747-DEL-483 |
| chr17-91899688-DEL-122 | chr17-91899689-DEL-122 |
| chr17-92002801-DEL-1969 | chr17-92002802-DEL-1969 |
| chr17-92029888-DEL-151 | chr17-92029889-DEL-151 |
| chr17-92032414-DEL-6197 | chr17-92032415-DEL-6197 |
| chr17-92055489-DEL-222 | chr17-92055490-DEL-222 |
| chr17-92062017-DEL-86 | chr17-92062033-DEL-87 |
| chr17-92079259-DEL-257 | chr17-92079260-DEL-257 |
| chr17-92133363-DEL-479 | chr17-92133365-DEL-478 |
| chr17-92137424-DEL-7082 | chr17-92137425-DEL-7082 |
| chr17-92204629-DEL-51 | chr17-92204630-DEL-51 |
| chr17-92223176-DEL-75 | chr17-92223169-DEL-74 |
| chr17-92263895-DEL-2841 | chr17-92263896-DEL-2841 |
| chr17-92271144-DEL-208 | chr17-92271143-DEL-204 |
| chr17-92315845-DEL-327 | chr17-92315846-DEL-327 |
| chr17-92333661-DEL-776 | chr17-92333662-DEL-776 |
| chr17-92356193-DEL-1754 | chr17-92356194-DEL-1754 |
| chr17-92374469-DEL-435 | chr17-92374470-DEL-435 |
| chr17-92376320-DEL-1064 | chr17-92376324-DEL-1060 |
| chr17-92387882-DEL-3889 | chr17-92387883-DEL-3889 |
| chr17-92391996-DEL-2782 | chr17-92391997-DEL-2782 |
| chr17-92392844-DEL-1121 | chr17-92392845-DEL-1121 |
| chr17-92403884-DEL-51 | chr17-92403885-DEL-57 |
| chr17-92411098-DEL-410 | chr17-92411127-DEL-385 |
| chr17-92414887-DEL-53 | chr17-92414887-DEL-50 |
| chr17-92423107-DEL-5994 | chr17-92423108-DEL-5994 |
| chr17-92434628-DEL-6350 | chr17-92434629-DEL-6350 |
| chr17-92444199-DEL-1133 | chr17-92444198-DEL-1134 |
| chr17-92506221-DEL-755 | chr17-92506222-DEL-755 |
| chr17-92519373-DEL-358 | chr17-92519373-DEL-358 |
| chr17-92547950-DEL-1913 | chr17-92547950-DEL-1913 |
| chr17-92552484-DEL-835 | chr17-92552488-DEL-832 |
| chr17-92565337-DEL-1055 | chr17-92565337-DEL-1055 |
| chr17-92594694-DEL-416 | chr17-92594695-DEL-416 |
| chr17-92601910-DEL-7199 | chr17-92601911-DEL-7199 |
| chr17-92602367-DEL-79 | chr17-92602367-DEL-79 |
| chr17-92619491-DEL-212 | chr17-92619499-DEL-215 |
| chr17-92638999-DEL-55 | chr17-92639000-DEL-55 |
| chr17-92650548-DEL-64 | chr17-92650549-DEL-64 |
| chr17-92657103-DEL-81 | chr17-92657104-DEL-81 |
| chr17-92678225-DEL-196 | chr17-92678226-DEL-196 |
| chr17-92682831-DEL-198 | chr17-92682832-DEL-198 |
| chr17-92682908-DEL-1746 | chr17-92682909-DEL-1746 |

| | |
|---|---|
| chr17-92683956-DEL-386 | chr17-92683957-DEL-385 |
| chr17-92710451-DEL-1497 | chr17-92710446-DEL-1494 |
| chr17-92728493-DEL-14991 | chr17-92728494-DEL-14991 |
| chr17-92729648-DEL-5194 | chr17-92729649-DEL-5194 |
| chr17-92757644-DEL-1801 | chr17-92757645-DEL-1801 |
| chr17-92770240-DEL-78 | chr17-92770241-DEL-76 |
| chr17-92775252-DEL-6188 | chr17-92775253-DEL-6188 |
| chr17-92795407-DEL-1415 | chr17-92795408-DEL-1415 |
| chr17-92814807-DEL-605 | chr17-92814807-DEL-570 |
| chr17-92824428-DEL-6369 | chr17-92824438-DEL-6357 |
| chr17-92888163-DEL-52 | chr17-92888164-DEL-52 |
| chr17-92897745-DEL-58 | chr17-92897702-DEL-61 |
| chr17-92917549-DEL-434 | chr17-92917550-DEL-434 |
| chr17-92964420-DEL-548 | chr17-92964441-DEL-529 |
| chr17-92973565-DEL-116 | chr17-92973610-DEL-72 |
| chr17-92975999-DEL-794 | chr17-92976014-DEL-802 |
| chr17-92985232-DEL-492 | chr17-92985232-DEL-492 |
| chr17-92988634-DEL-335 | chr17-92988634-DEL-335 |
| chr17-93003258-DEL-6455 | chr17-93003259-DEL-6455 |
| chr17-93021570-DEL-404 | chr17-93021571-DEL-404 |
| chr17-93023604-DEL-76 | chr17-93023605-DEL-76 |
| chr17-93027339-DEL-104 | chr17-93027344-DEL-100 |
| chr17-93028746-DEL-2153 | chr17-93028747-DEL-2153 |
| chr17-93061131-DEL-58 | chr17-93061132-DEL-58 |
| chr17-93069143-DEL-52 | chr17-93069094-DEL-76 |
| chr17-93082083-DEL-2280 | chr17-93082084-DEL-2280 |
| chr17-93093655-DEL-66 | chr17-93093654-DEL-66 |
| chr17-93094946-DEL-61 | chr17-93094977-DEL-53 |
| chr17-93122451-DEL-1406 | chr17-93122439-DEL-1407 |
| chr17-93149890-DEL-75 | chr17-93149891-DEL-75 |
| chr17-93162913-DEL-60 | chr17-93162914-DEL-60 |
| chr17-93184447-DEL-64 | chr17-93184448-DEL-64 |
| chr17-93220986-DEL-208 | chr17-93220987-DEL-208 |
| chr17-93269580-DEL-872 | chr17-93269581-DEL-872 |
| chr17-93274274-DEL-334 | chr17-93274275-DEL-334 |
| chr17-93305405-DEL-381 | chr17-93305406-DEL-381 |
| chr17-93323180-DEL-837 | chr17-93323181-DEL-837 |
| chr17-93324397-DEL-6452 | chr17-93324398-DEL-6452 |
| chr17-93349860-DEL-3237 | chr17-93349861-DEL-3237 |
| chr17-93428559-DEL-2555 | chr17-93428560-DEL-2555 |
| chr17-93456128-DEL-3057 | chr17-93456129-DEL-3057 |
| chr17-93522729-DEL-832 | chr17-93522730-DEL-832 |
| chr17-93526171-DEL-253 | chr17-93526172-DEL-253 |
| chr17-93530653-DEL-206 | chr17-93530659-DEL-206 |
| chr17-93552770-DEL-661 | chr17-93552769-DEL-662 |
| chr17-93590212-DEL-6937 | chr17-93590213-DEL-6937 |
| chr17-93596498-DEL-424 | chr17-93596499-DEL-424 |
| chr17-93614188-DEL-698 | chr17-93614189-DEL-698 |
| chr17-93630513-DEL-62 | chr17-93630514-DEL-62 |
| chr17-93644479-DEL-467 | chr17-93644480-DEL-467 |
| chr17-93680255-DEL-104 | chr17-93680246-DEL-98 |
| chr17-93699993-DEL-1212 | chr17-93699994-DEL-1212 |
| chr17-93774654-DEL-398 | chr17-93774655-DEL-398 |
| chr17-93823163-DEL-87 | chr17-93823161-DEL-137 |
| chr17-93840780-DEL-444 | chr17-93840781-DEL-444 |
| chr17-93860664-DEL-853 | chr17-93860665-DEL-853 |
| chr17-93871541-DEL-1746 | chr17-93871542-DEL-1746 |
| chr17-93899116-DEL-70 | chr17-93899117-DEL-70 |
| chr17-93902056-DEL-1541 | chr17-93902057-DEL-1541 |
| chr17-93940591-DEL-537 | chr17-93940595-DEL-537 |
| chr17-93987012-DEL-607 | chr17-93987013-DEL-607 |
| chr17-93988461-DEL-54 | chr17-93988462-DEL-54 |
| chr17-93993552-DEL-717 | chr17-93993544-DEL-718 |
| chr17-93999411-DEL-777 | chr17-93999412-DEL-777 |
| chr17-94018177-DEL-460 | chr17-94018178-DEL-460 |
| chr17-94039611-DEL-97 | chr17-94039610-DEL-98 |
| chr17-94044045-DEL-87 | chr17-94044046-DEL-87 |
| chr17-94050741-DEL-112 | chr17-94050742-DEL-112 |
| chr17-94054807-DEL-4838 | chr17-94054808-DEL-4838 |
| chr17-94071333-DEL-60 | chr17-94071334-DEL-60 |
| chr17-94088123-DEL-78 | chr17-94088124-DEL-78 |
| chr17-94095351-DEL-70 | chr17-94095372-DEL-51 |
| chr17-94098905-DEL-4577 | chr17-94098906-DEL-4577 |
| chr17-94114242-DEL-59 | chr17-94114243-DEL-59 |
| chr17-94121954-DEL-324 | chr17-94121955-DEL-324 |
| chr17-94129721-DEL-256 | chr17-94129721-DEL-256 |
| chr17-94134372-DEL-505 | chr17-94134373-DEL-505 |
| chr17-94190949-DEL-81 | chr17-94190950-DEL-66 |
| chr17-94206333-DEL-53 | chr17-94206334-DEL-53 |
| chr17-94235993-DEL-14720 | chr17-94235994-DEL-14720 |

| | |
|---|---|
| chr17-94241246-DEL-2314 | chr17-94241247-DEL-2314 |
| chr17-94257491-DEL-3511 | chr17-94257492-DEL-3511 |
| chr17-94331632-DEL-2127 | chr17-94331633-DEL-2127 |
| chr17-94334786-DEL-456 | chr17-94334787-DEL-456 |
| chr17-94368533-DEL-74 | chr17-94368532-DEL-64 |
| chr17-94387590-DEL-196 | chr17-94387591-DEL-196 |
| chr17-94392968-DEL-223 | chr17-94392969-DEL-223 |
| chr17-94396588-DEL-80 | chr17-94396573-DEL-78 |
| chr17-94408071-DEL-2201 | chr17-94408072-DEL-2201 |
| chr17-94442091-DEL-6208 | chr17-94442092-DEL-6207 |
| chr17-94497708-DEL-147 | chr17-94497709-DEL-147 |
| chr17-94603782-DEL-7089 | chr17-94603783-DEL-7089 |
| chr17-94625357-DEL-170 | chr17-94625358-DEL-170 |
| chr17-94695672-DEL-738 | chr17-94695673-DEL-738 |
| chr17-94706647-DEL-1436 | chr17-94706648-DEL-1436 |
| chr17-94755964-DEL-115 | chr17-94755965-DEL-115 |
| chr17-94790572-DEL-1963 | chr17-94790572-DEL-1963 |
| chr17-94828417-DEL-311 | chr17-94828418-DEL-311 |
| chr17-94837040-DEL-410 | chr17-94837041-DEL-410 |
| chr17-94838750-DEL-381 | chr17-94838751-DEL-381 |
| chr17-94841810-DEL-8612 | chr17-94841811-DEL-8612 |
| chr17-94865170-DEL-69 | chr17-94865171-DEL-63 |
| chr17-94875565-DEL-65 | chr17-94875566-DEL-65 |
| chr17-94881955-DEL-7534 | chr17-94881956-DEL-7534 |
| chr17-94887218-DEL-1134 | chr17-94887219-DEL-1134 |
| chr17-95004052-DEL-2731 | chr17-95004053-DEL-2731 |
| chr17-95061194-DEL-242 | chr17-95061196-DEL-241 |
| chr17-95065165-DEL-77 | chr17-95065181-DEL-62 |
| chr17-95065631-DEL-394 | chr17-95065632-DEL-394 |
| chr17-95111847-DEL-528 | chr17-95111848-DEL-527 |
| chr17-95112651-DEL-67 | chr17-95112652-DEL-67 |
| chr17-95129395-DEL-193 | chr17-95129396-DEL-193 |
| chr17-95172582-DEL-200 | chr17-95172583-DEL-200 |
| chr18-3144083-DEL-645 | chr18-3144084-DEL-645 |
| chr18-3215588-DEL-194 | chr18-3215589-DEL-194 |
| chr18-3216706-DEL-3000 | chr18-3216707-DEL-3000 |
| chr18-3347560-DEL-224 | chr18-3347561-DEL-224 |
| chr18-3375313-DEL-184 | chr18-3375314-DEL-184 |
| chr18-3384609-DEL-215 | chr18-3384610-DEL-215 |
| chr18-3427342-DEL-76 | chr18-3427392-DEL-73 |
| chr18-3456428-DEL-4922 | chr18-3456429-DEL-4922 |
| chr18-3465885-DEL-638 | chr18-3465909-DEL-635 |
| chr18-3471314-DEL-7375 | chr18-3471315-DEL-7375 |
| chr18-3484410-DEL-516 | chr18-3484411-DEL-516 |
| chr18-3494053-DEL-210 | chr18-3494054-DEL-210 |
| chr18-3515903-DEL-83 | chr18-3515904-DEL-83 |
| chr18-3523602-DEL-84 | chr18-3523620-DEL-83 |
| chr18-3535141-DEL-2022 | chr18-3535142-DEL-2022 |
| chr18-3568204-DEL-1892 | chr18-3568205-DEL-1892 |
| chr18-3580972-DEL-1872 | chr18-3580973-DEL-1872 |
| chr18-3598578-DEL-77 | chr18-3598565-DEL-54 |
| chr18-3608291-DEL-1665 | chr18-3608292-DEL-1665 |
| chr18-3619851-DEL-6944 | chr18-3619852-DEL-6944 |
| chr18-3699045-DEL-2832 | chr18-3699046-DEL-2832 |
| chr18-3721011-DEL-2985 | chr18-3721011-DEL-2985 |
| chr18-3779597-DEL-6191 | chr18-3779598-DEL-6191 |
| chr18-3790308-DEL-205 | chr18-3790309-DEL-205 |
| chr18-3814857-DEL-3110 | chr18-3814858-DEL-3110 |
| chr18-3840243-DEL-203 | chr18-3840244-DEL-203 |
| chr18-3872462-DEL-3301 | chr18-3872463-DEL-3301 |
| chr18-3928815-DEL-734 | chr18-3928780-DEL-769 |
| chr18-3955856-DEL-537 | chr18-3955849-DEL-538 |
| chr18-3984630-DEL-74 | chr18-3984619-DEL-86 |
| chr18-4016068-DEL-72 | chr18-4016069-DEL-72 |
| chr18-4021868-DEL-7521 | chr18-4021869-DEL-7521 |
| chr18-4035298-DEL-1262 | chr18-4035299-DEL-1262 |
| chr18-4058588-DEL-226 | chr18-4058589-DEL-226 |
| chr18-4068608-DEL-179 | chr18-4068609-DEL-179 |
| chr18-4070054-DEL-440 | chr18-4070055-DEL-440 |
| chr18-4071230-DEL-278 | chr18-4071231-DEL-278 |
| chr18-4111306-DEL-479 | chr18-4111307-DEL-479 |
| chr18-4134249-DEL-69 | chr18-4134250-DEL-69 |
| chr18-4171429-DEL-68 | chr18-4171473-DEL-52 |
| chr18-4190027-DEL-59 | chr18-4190028-DEL-59 |
| chr18-4195166-DEL-4170 | chr18-4195167-DEL-4170 |
| chr18-4200540-DEL-94 | chr18-4200509-DEL-120 |
| chr18-4230058-DEL-2027 | chr18-4230059-DEL-2027 |
| chr18-4250203-DEL-51 | chr18-4250204-DEL-51 |
| chr18-4268966-DEL-546 | chr18-4268967-DEL-546 |
| chr18-4281599-DEL-549 | chr18-4281600-DEL-549 |

| | |
|---|---|
| chr18-4319409-DEL-206 | chr18-4319410-DEL-206 |
| chr18-4377631-DEL-78 | chr18-4377632-DEL-78 |
| chr18-4380910-DEL-172 | chr18-4380911-DEL-172 |
| chr18-4409910-DEL-69 | chr18-4409911-DEL-69 |
| chr18-4422786-DEL-1043 | chr18-4422787-DEL-1043 |
| chr18-4447405-DEL-1506 | chr18-4447406-DEL-1506 |
| chr18-4468010-DEL-6348 | chr18-4468011-DEL-6348 |
| chr18-4475750-DEL-201 | chr18-4475751-DEL-201 |
| chr18-4479604-DEL-4263 | chr18-4479605-DEL-4263 |
| chr18-4494048-DEL-8514 | chr18-4494049-DEL-8514 |
| chr18-4532442-DEL-7455 | chr18-4532443-DEL-7455 |
| chr18-4562402-DEL-558 | chr18-4562403-DEL-558 |
| chr18-4658048-DEL-6400 | chr18-4658042-DEL-6401 |
| chr18-4695227-DEL-6581 | chr18-4695231-DEL-6582 |
| chr18-4705144-DEL-79 | chr18-4705120-DEL-89 |
| chr18-4750203-DEL-1530 | chr18-4750204-DEL-1530 |
| chr18-4757165-DEL-94 | chr18-4757166-DEL-94 |
| chr18-4770755-DEL-2365 | chr18-4770770-DEL-2351 |
| chr18-4801276-DEL-66 | chr18-4801277-DEL-66 |
| chr18-4812255-DEL-317 | chr18-4812256-DEL-317 |
| chr18-4841803-DEL-315 | chr18-4841804-DEL-315 |
| chr18-4870706-DEL-192 | chr18-4870707-DEL-192 |
| chr18-4954318-DEL-83 | chr18-4954340-DEL-83 |
| chr18-5003419-DEL-75 | chr18-5003420-DEL-75 |
| chr18-5004192-DEL-100 | chr18-5004193-DEL-100 |
| chr18-5027052-DEL-55 | chr18-5027053-DEL-50 |
| chr18-5036528-DEL-98 | chr18-5036537-DEL-82 |
| chr18-5037813-DEL-52 | chr18-5037814-DEL-52 |
| chr18-5038097-DEL-391 | chr18-5038098-DEL-391 |
| chr18-5048031-DEL-92 | chr18-5048032-DEL-138 |
| chr18-5070311-DEL-216 | chr18-5070312-DEL-216 |
| chr18-5090366-DEL-100 | chr18-5090367-DEL-100 |
| chr18-5092397-DEL-104 | chr18-5092398-DEL-104 |
| chr18-5102529-DEL-82 | chr18-5102530-DEL-82 |
| chr18-5184289-DEL-145 | chr18-5184290-DEL-145 |
| chr18-5203408-DEL-155 | chr18-5203409-DEL-155 |
| chr18-5206520-DEL-177 | chr18-5206521-DEL-177 |
| chr18-5225327-DEL-119 | chr18-5225328-DEL-118 |
| chr18-5231333-DEL-1860 | chr18-5231334-DEL-1860 |
| chr18-5350757-DEL-199 | chr18-5350758-DEL-199 |
| chr18-5352000-DEL-87 | chr18-5352001-DEL-87 |
| chr18-5377737-DEL-65 | chr18-5377737-DEL-65 |
| chr18-5398598-DEL-216 | chr18-5398598-DEL-216 |
| chr18-5435920-DEL-3114 | chr18-5435921-DEL-3114 |
| chr18-5478777-DEL-823 | chr18-5478778-DEL-823 |
| chr18-5499350-DEL-2651 | chr18-5499351-DEL-2651 |
| chr18-5540481-DEL-642 | chr18-5540482-DEL-642 |
| chr18-5636379-DEL-1271 | chr18-5636380-DEL-1271 |
| chr18-5681214-DEL-276 | chr18-5681215-DEL-276 |
| chr18-5686326-DEL-74 | chr18-5686327-DEL-74 |
| chr18-5720617-DEL-1178 | chr18-5720618-DEL-1178 |
| chr18-5751043-DEL-7348 | chr18-5751044-DEL-7348 |
| chr18-5789204-DEL-71 | chr18-5789175-DEL-86 |
| chr18-5808517-DEL-93 | chr18-5808519-DEL-90 |
| chr18-5841223-DEL-98 | chr18-5841224-DEL-98 |
| chr18-5841969-DEL-92 | chr18-5841970-DEL-92 |
| chr18-5868292-DEL-7462 | chr18-5868293-DEL-7462 |
| chr18-5951470-DEL-225 | chr18-5951471-DEL-225 |
| chr18-5955726-DEL-154 | chr18-5955726-DEL-154 |
| chr18-5956658-DEL-78 | chr18-5956656-DEL-70 |
| chr18-5962223-DEL-162 | chr18-5962241-DEL-149 |
| chr18-5990976-DEL-208 | chr18-5990977-DEL-208 |
| chr18-6043277-DEL-2923 | chr18-6043278-DEL-2923 |
| chr18-6046898-DEL-79 | chr18-6046899-DEL-79 |
| chr18-6083045-DEL-3267 | chr18-6083045-DEL-3267 |
| chr18-6084067-DEL-117 | chr18-6084058-DEL-130 |
| chr18-6105326-DEL-132 | chr18-6105327-DEL-132 |
| chr18-6114921-DEL-806 | chr18-6114922-DEL-806 |
| chr18-6147041-DEL-376 | chr18-6147042-DEL-376 |
| chr18-6152261-DEL-696 | chr18-6152262-DEL-696 |
| chr18-6171221-DEL-209 | chr18-6171222-DEL-209 |
| chr18-6190820-DEL-128 | chr18-6190815-DEL-127 |
| chr18-6196140-DEL-3582 | chr18-6196141-DEL-3582 |
| chr18-6202201-DEL-169 | chr18-6202202-DEL-169 |
| chr18-6207823-DEL-215 | chr18-6207824-DEL-215 |
| chr18-6218340-DEL-66 | chr18-6218350-DEL-54 |
| chr18-6243195-DEL-202 | chr18-6243196-DEL-202 |
| chr18-6254473-DEL-193 | chr18-6254474-DEL-193 |
| chr18-6256010-DEL-7150 | chr18-6256011-DEL-7150 |
| chr18-6279382-DEL-78 | chr18-6279379-DEL-81 |

| | |
|---|---|
| chr18-6287556-DEL-52 | chr18-6287551-DEL-57 |
| chr18-6295251-DEL-187 | chr18-6295252-DEL-187 |
| chr18-6296293-DEL-203 | chr18-6296294-DEL-203 |
| chr18-6302276-DEL-52 | chr18-6302267-DEL-69 |
| chr18-6331489-DEL-2497 | chr18-6331490-DEL-2497 |
| chr18-6346356-DEL-9002 | chr18-6346357-DEL-9002 |
| chr18-6356184-DEL-866 | chr18-6356181-DEL-867 |
| chr18-6364886-DEL-382 | chr18-6364887-DEL-382 |
| chr18-6372009-DEL-4326 | chr18-6372010-DEL-4326 |
| chr18-6380880-DEL-213 | chr18-6380878-DEL-214 |
| chr18-6382754-DEL-179 | chr18-6382754-DEL-179 |
| chr18-6396083-DEL-1600 | chr18-6396084-DEL-1600 |
| chr18-6400259-DEL-328 | chr18-6400260-DEL-328 |
| chr18-6410613-DEL-57 | chr18-6410614-DEL-57 |
| chr18-6421722-DEL-924 | chr18-6421723-DEL-923 |
| chr18-6444108-DEL-135 | chr18-6444092-DEL-136 |
| chr18-6482266-DEL-118 | chr18-6482267-DEL-118 |
| chr18-6508760-DEL-1956 | chr18-6508764-DEL-1963 |
| chr18-6525063-DEL-123 | chr18-6525064-DEL-123 |
| chr18-6553170-DEL-66 | chr18-6553171-DEL-66 |
| chr18-6561459-DEL-570 | chr18-6561460-DEL-570 |
| chr18-6582502-DEL-213 | chr18-6582503-DEL-213 |
| chr18-6599098-DEL-53 | chr18-6599099-DEL-53 |
| chr18-6602935-DEL-58 | chr18-6602936-DEL-58 |
| chr18-6621443-DEL-111 | chr18-6621444-DEL-111 |
| chr18-6621704-DEL-822 | chr18-6621751-DEL-785 |
| chr18-6630884-DEL-937 | chr18-6630885-DEL-937 |
| chr18-6632529-DEL-638 | chr18-6632530-DEL-636 |
| chr18-6663823-DEL-5314 | chr18-6663824-DEL-5314 |
| chr18-6683138-DEL-84 | chr18-6683185-DEL-72 |
| chr18-6697116-DEL-52 | chr18-6697121-DEL-72 |
| chr18-6708353-DEL-55 | chr18-6708354-DEL-55 |
| chr18-6714416-DEL-201 | chr18-6714417-DEL-201 |
| chr18-6732889-DEL-63 | chr18-6732890-DEL-63 |
| chr18-6741575-DEL-208 | chr18-6741576-DEL-208 |
| chr18-6763899-DEL-54 | chr18-6763900-DEL-54 |
| chr18-6772953-DEL-1960 | chr18-6772954-DEL-1960 |
| chr18-6802838-DEL-78 | chr18-6802839-DEL-78 |
| chr18-6807173-DEL-208 | chr18-6807174-DEL-208 |
| chr18-6840000-DEL-97 | chr18-6840025-DEL-73 |
| chr18-6861251-DEL-7123 | chr18-6861252-DEL-7123 |
| chr18-6869677-DEL-83 | chr18-6869633-DEL-84 |
| chr18-6871224-DEL-1743 | chr18-6871224-DEL-1743 |
| chr18-6911178-DEL-147 | chr18-6911179-DEL-147 |
| chr18-6912366-DEL-392 | chr18-6912367-DEL-392 |
| chr18-6913699-DEL-525 | chr18-6913700-DEL-525 |
| chr18-6965931-DEL-80 | chr18-6965941-DEL-68 |
| chr18-6983614-DEL-61 | chr18-6983621-DEL-55 |
| chr18-6997871-DEL-131 | chr18-6997872-DEL-131 |
| chr18-7009821-DEL-192 | chr18-7009822-DEL-192 |
| chr18-7099544-DEL-64 | chr18-7099542-DEL-64 |
| chr18-7102271-DEL-2585 | chr18-7102275-DEL-2582 |
| chr18-7104585-DEL-1319 | chr18-7104586-DEL-1319 |
| chr18-7114667-DEL-55 | chr18-7114668-DEL-55 |
| chr18-7123216-DEL-73 | chr18-7123217-DEL-73 |
| chr18-7196258-DEL-72 | chr18-7196257-DEL-70 |
| chr18-7215224-DEL-65 | chr18-7215225-DEL-65 |
| chr18-7275987-DEL-1322 | chr18-7275998-DEL-1320 |
| chr18-7318604-DEL-8334 | chr18-7318605-DEL-8334 |
| chr18-7357754-DEL-137 | chr18-7357755-DEL-137 |
| chr18-7419215-DEL-166 | chr18-7419216-DEL-166 |
| chr18-7442037-DEL-51 | chr18-7442027-DEL-52 |
| chr18-7447306-DEL-960 | chr18-7447307-DEL-960 |
| chr18-7494737-DEL-3127 | chr18-7494738-DEL-3127 |
| chr18-7527579-DEL-7083 | chr18-7527580-DEL-7083 |
| chr18-7540544-DEL-2553 | chr18-7540545-DEL-2553 |
| chr18-7576791-DEL-54 | chr18-7576792-DEL-54 |
| chr18-7584953-DEL-102 | chr18-7584953-DEL-99 |
| chr18-7592405-DEL-88 | chr18-7592406-DEL-88 |
| chr18-7610681-DEL-79 | chr18-7610684-DEL-74 |
| chr18-7613761-DEL-73 | chr18-7613762-DEL-73 |
| chr18-7623410-DEL-63 | chr18-7623410-DEL-63 |
| chr18-7647581-DEL-401 | chr18-7647582-DEL-401 |
| chr18-7661841-DEL-794 | chr18-7661842-DEL-794 |
| chr18-7771354-DEL-3117 | chr18-7771355-DEL-3117 |
| chr18-7779248-DEL-1542 | chr18-7779248-DEL-1541 |
| chr18-7781589-DEL-6253 | chr18-7781590-DEL-6253 |
| chr18-7796360-DEL-2184 | chr18-7796361-DEL-2184 |
| chr18-7816102-DEL-486 | chr18-7816103-DEL-486 |
| chr18-7829115-DEL-3118 | chr18-7829116-DEL-3118 |

| | |
|---|---|
| chr18-7862976-DEL-674 | chr18-7862977-DEL-674 |
| chr18-7866015-DEL-204 | chr18-7866016-DEL-204 |
| chr18-7907838-DEL-203 | chr18-7907837-DEL-203 |
| chr18-7929353-DEL-138 | chr18-7929354-DEL-138 |
| chr18-7959798-DEL-1006 | chr18-7959803-DEL-1001 |
| chr18-7972982-DEL-206 | chr18-7972981-DEL-206 |
| chr18-8011555-DEL-7144 | chr18-8011556-DEL-7144 |
| chr18-8104899-DEL-147 | chr18-8104899-DEL-147 |
| chr18-8117218-DEL-781 | chr18-8117220-DEL-780 |
| chr18-8140459-DEL-6935 | chr18-8140460-DEL-6935 |
| chr18-8150500-DEL-1727 | chr18-8150501-DEL-1727 |
| chr18-8164586-DEL-6142 | chr18-8164579-DEL-6143 |
| chr18-8179056-DEL-155 | chr18-8179057-DEL-155 |
| chr18-8180314-DEL-3182 | chr18-8180315-DEL-3182 |
| chr18-8199747-DEL-1113 | chr18-8199748-DEL-1113 |
| chr18-8208420-DEL-75 | chr18-8208421-DEL-75 |
| chr18-8209772-DEL-6278 | chr18-8209773-DEL-6278 |
| chr18-8215750-DEL-199 | chr18-8215751-DEL-199 |
| chr18-8223125-DEL-640 | chr18-8223126-DEL-640 |
| chr18-8236447-DEL-457 | chr18-8236448-DEL-457 |
| chr18-8237866-DEL-50 | chr18-8237862-DEL-54 |
| chr18-8240283-DEL-12839 | chr18-8240284-DEL-12839 |
| chr18-8245947-DEL-3215 | chr18-8245948-DEL-3215 |
| chr18-8268614-DEL-3191 | chr18-8268615-DEL-3191 |
| chr18-8292071-DEL-1890 | chr18-8292076-DEL-1892 |
| chr18-8298092-DEL-172 | chr18-8298092-DEL-173 |
| chr18-8304380-DEL-60 | chr18-8304376-DEL-62 |
| chr18-8320975-DEL-101 | chr18-8320976-DEL-101 |
| chr18-8354950-DEL-1091 | chr18-8354951-DEL-1091 |
| chr18-8359729-DEL-146 | chr18-8359776-DEL-129 |
| chr18-8371713-DEL-732 | chr18-8371703-DEL-733 |
| chr18-8414995-DEL-2636 | chr18-8414996-DEL-2636 |
| chr18-8431883-DEL-59 | chr18-8431884-DEL-59 |
| chr18-8432759-DEL-1056 | chr18-8432760-DEL-1056 |
| chr18-8457790-DEL-276 | chr18-8457791-DEL-276 |
| chr18-8460649-DEL-63 | chr18-8460650-DEL-57 |
| chr18-8551276-DEL-84 | chr18-8551275-DEL-56 |
| chr18-8578025-DEL-81 | chr18-8578028-DEL-80 |
| chr18-8600201-DEL-603 | chr18-8600198-DEL-607 |
| chr18-8605214-DEL-52 | chr18-8605207-DEL-72 |
| chr18-8626141-DEL-856 | chr18-8626142-DEL-856 |
| chr18-8636843-DEL-903 | chr18-8636844-DEL-903 |
| chr18-8672938-DEL-201 | chr18-8672939-DEL-201 |
| chr18-8674820-DEL-74 | chr18-8674821-DEL-73 |
| chr18-8678230-DEL-60 | chr18-8678195-DEL-58 |
| chr18-8693733-DEL-3419 | chr18-8693734-DEL-3419 |
| chr18-8721860-DEL-876 | chr18-8721861-DEL-876 |
| chr18-8771528-DEL-4925 | chr18-8771529-DEL-4925 |
| chr18-8782081-DEL-6086 | chr18-8782082-DEL-6086 |
| chr18-8874412-DEL-56 | chr18-8874401-DEL-68 |
| chr18-8889741-DEL-80 | chr18-8889746-DEL-72 |
| chr18-8895379-DEL-439 | chr18-8895380-DEL-439 |
| chr18-8907578-DEL-392 | chr18-8907579-DEL-392 |
| chr18-8946587-DEL-492 | chr18-8946588-DEL-492 |
| chr18-8978695-DEL-165 | chr18-8978683-DEL-159 |
| chr18-8978839-DEL-66 | chr18-8978830-DEL-75 |
| chr18-8983197-DEL-400 | chr18-8983198-DEL-400 |
| chr18-8989640-DEL-62 | chr18-8989641-DEL-62 |
| chr18-9010078-DEL-65 | chr18-9010105-DEL-59 |
| chr18-9022990-DEL-56 | chr18-9022991-DEL-56 |
| chr18-9052091-DEL-52 | chr18-9052124-DEL-62 |
| chr18-9091636-DEL-1749 | chr18-9091637-DEL-1749 |
| chr18-9095153-DEL-2164 | chr18-9095154-DEL-2164 |
| chr18-9111546-DEL-6385 | chr18-9111547-DEL-6385 |
| chr18-9123226-DEL-205 | chr18-9123227-DEL-205 |
| chr18-9125508-DEL-1877 | chr18-9125509-DEL-1877 |
| chr18-9151208-DEL-1844 | chr18-9151209-DEL-1844 |
| chr18-9173034-DEL-5244 | chr18-9173035-DEL-5244 |
| chr18-9178918-DEL-55 | chr18-9178919-DEL-55 |
| chr18-9202437-DEL-6277 | chr18-9202438-DEL-6277 |
| chr18-9225877-DEL-1802 | chr18-9225878-DEL-1802 |
| chr18-9228686-DEL-2087 | chr18-9228687-DEL-2085 |
| chr18-9247673-DEL-2939 | chr18-9247674-DEL-2939 |
| chr18-9263098-DEL-6515 | chr18-9263099-DEL-6515 |
| chr18-9309070-DEL-732 | chr18-9309071-DEL-732 |
| chr18-9320090-DEL-86 | chr18-9320090-DEL-86 |
| chr18-9340632-DEL-760 | chr18-9340633-DEL-760 |
| chr18-9360883-DEL-3142 | chr18-9360884-DEL-3142 |
| chr18-9402602-DEL-3099 | chr18-9402603-DEL-3099 |
| chr18-9420045-DEL-3804 | chr18-9420046-DEL-3804 |

| | |
|---|---|
| chr18-9446564-DEL-53 | chr18-9446565-DEL-53 |
| chr18-9466208-DEL-5331 | chr18-9466209-DEL-5331 |
| chr18-9471974-DEL-264 | chr18-9471975-DEL-264 |
| chr18-9473359-DEL-206 | chr18-9473360-DEL-206 |
| chr18-9504742-DEL-94 | chr18-9504743-DEL-94 |
| chr18-9508464-DEL-194 | chr18-9508448-DEL-180 |
| chr18-9509815-DEL-179 | chr18-9509816-DEL-179 |
| chr18-9525862-DEL-55 | chr18-9525863-DEL-55 |
| chr18-9542779-DEL-2391 | chr18-9542780-DEL-2391 |
| chr18-9545366-DEL-1700 | chr18-9545367-DEL-1700 |
| chr18-9549813-DEL-234 | chr18-9549814-DEL-234 |
| chr18-9573110-DEL-84 | chr18-9573125-DEL-54 |
| chr18-9574334-DEL-78 | chr18-9574336-DEL-77 |
| chr18-9576306-DEL-460 | chr18-9576307-DEL-460 |
| chr18-9586371-DEL-83 | chr18-9586372-DEL-83 |
| chr18-9592382-DEL-5653 | chr18-9592383-DEL-5653 |
| chr18-9677595-DEL-364 | chr18-9677596-DEL-364 |
| chr18-9679083-DEL-1382 | chr18-9679084-DEL-1382 |
| chr18-9699782-DEL-293 | chr18-9699783-DEL-290 |
| chr18-9703518-DEL-1891 | chr18-9703519-DEL-1891 |
| chr18-9725370-DEL-274 | chr18-9725371-DEL-274 |
| chr18-9734442-DEL-68 | chr18-9734442-DEL-68 |
| chr18-9755309-DEL-57 | chr18-9755310-DEL-57 |
| chr18-9758936-DEL-819 | chr18-9758937-DEL-819 |
| chr18-9761783-DEL-58 | chr18-9761784-DEL-58 |
| chr18-9762001-DEL-1861 | chr18-9762002-DEL-1861 |
| chr18-9774630-DEL-81 | chr18-9774630-DEL-81 |
| chr18-9836613-DEL-279 | chr18-9836613-DEL-280 |
| chr18-9865663-DEL-60 | chr18-9865666-DEL-61 |
| chr18-9924020-DEL-1392 | chr18-9924020-DEL-1393 |
| chr18-9942017-DEL-205 | chr18-9942013-DEL-179 |
| chr18-9952517-DEL-212 | chr18-9952518-DEL-212 |
| chr18-9964834-DEL-208 | chr18-9964833-DEL-208 |
| chr18-9972376-DEL-78 | chr18-9972373-DEL-64 |
| chr18-9979484-DEL-213 | chr18-9979485-DEL-213 |
| chr18-10005860-DEL-431 | chr18-10005861-DEL-431 |
| chr18-10015438-DEL-122 | chr18-10015439-DEL-122 |
| chr18-10019347-DEL-499 | chr18-10019348-DEL-499 |
| chr18-10037632-DEL-67 | chr18-10037634-DEL-68 |
| chr18-10061267-DEL-522 | chr18-10061268-DEL-522 |
| chr18-10111005-DEL-101 | chr18-10111006-DEL-101 |
| chr18-10123459-DEL-165 | chr18-10123460-DEL-165 |
| chr18-10230046-DEL-6182 | chr18-10230047-DEL-6182 |
| chr18-10242929-DEL-51 | chr18-10242930-DEL-51 |
| chr18-10258344-DEL-146 | chr18-10258345-DEL-143 |
| chr18-10264995-DEL-82 | chr18-10265000-DEL-89 |
| chr18-10265505-DEL-97 | chr18-10265487-DEL-118 |
| chr18-10302810-DEL-710 | chr18-10302811-DEL-710 |
| chr18-10311950-DEL-209 | chr18-10311951-DEL-209 |
| chr18-10320873-DEL-223 | chr18-10320874-DEL-223 |
| chr18-10333043-DEL-416 | chr18-10333044-DEL-416 |
| chr18-10352721-DEL-76 | chr18-10352709-DEL-119 |
| chr18-10353590-DEL-51 | chr18-10353540-DEL-51 |
| chr18-10356698-DEL-1556 | chr18-10356699-DEL-1556 |
| chr18-10365819-DEL-185 | chr18-10365820-DEL-171 |
| chr18-10374236-DEL-58 | chr18-10374225-DEL-56 |
| chr18-10394186-DEL-52 | chr18-10394187-DEL-52 |
| chr18-10416149-DEL-112 | chr18-10416174-DEL-84 |
| chr18-10431671-DEL-118 | chr18-10431680-DEL-110 |
| chr18-10468191-DEL-52 | chr18-10468192-DEL-52 |
| chr18-10481084-DEL-2774 | chr18-10481085-DEL-2774 |
| chr18-10482636-DEL-908 | chr18-10482637-DEL-908 |
| chr18-10488469-DEL-176 | chr18-10488470-DEL-176 |
| chr18-10496813-DEL-155 | chr18-10496802-DEL-154 |
| chr18-10516299-DEL-74 | chr18-10516299-DEL-90 |
| chr18-10526735-DEL-664 | chr18-10526737-DEL-662 |
| chr18-10526735-DEL-97 | chr18-10526737-DEL-95 |
| chr18-10528967-DEL-122 | chr18-10528965-DEL-122 |
| chr18-10548253-DEL-451 | chr18-10548254-DEL-410 |
| chr18-10548815-DEL-202 | chr18-10548816-DEL-202 |
| chr18-10569719-DEL-126 | chr18-10569719-DEL-84 |
| chr18-10612894-DEL-80 | chr18-10612895-DEL-80 |
| chr18-10625020-DEL-541 | chr18-10625022-DEL-540 |
| chr18-10630210-DEL-174 | chr18-10630211-DEL-174 |
| chr18-10722091-DEL-2293 | chr18-10722092-DEL-2293 |
| chr18-10725779-DEL-57 | chr18-10725780-DEL-57 |
| chr18-10733240-DEL-112 | chr18-10733241-DEL-112 |
| chr18-10743619-DEL-52 | chr18-10743620-DEL-52 |
| chr18-10753209-DEL-56 | chr18-10753210-DEL-54 |
| chr18-10828264-DEL-131 | chr18-10828257-DEL-123 |

| | |
|---|---|
| chr18-10853349-DEL-95 | chr18-10853350-DEL-95 |
| chr18-10875457-DEL-7015 | chr18-10875458-DEL-7015 |
| chr18-10881802-DEL-212 | chr18-10881803-DEL-212 |
| chr18-10887918-DEL-257 | chr18-10887919-DEL-257 |
| chr18-10896028-DEL-74 | chr18-10896029-DEL-74 |
| chr18-10918444-DEL-140 | chr18-10918445-DEL-140 |
| chr18-10929116-DEL-218 | chr18-10929117-DEL-216 |
| chr18-10940470-DEL-94 | chr18-10940471-DEL-94 |
| chr18-10990461-DEL-78 | chr18-10990445-DEL-87 |
| chr18-10994214-DEL-57 | chr18-10994214-DEL-54 |
| chr18-11002766-DEL-72 | chr18-11002767-DEL-72 |
| chr18-11013109-DEL-64 | chr18-11013110-DEL-64 |
| chr18-11022397-DEL-56 | chr18-11022398-DEL-56 |
| chr18-11026342-DEL-84 | chr18-11026391-DEL-68 |
| chr18-11031175-DEL-78 | chr18-11031176-DEL-78 |
| chr18-11033313-DEL-265 | chr18-11033316-DEL-262 |
| chr18-11039768-DEL-160 | chr18-11039769-DEL-160 |
| chr18-11077550-DEL-71 | chr18-11077545-DEL-77 |
| chr18-11091611-DEL-84 | chr18-11091612-DEL-83 |
| chr18-11110050-DEL-52 | chr18-11110051-DEL-52 |
| chr18-11110981-DEL-62 | chr18-11110960-DEL-84 |
| chr18-11118634-DEL-89 | chr18-11118642-DEL-58 |
| chr18-11136215-DEL-52 | chr18-11136216-DEL-52 |
| chr18-11138284-DEL-1245 | chr18-11138285-DEL-1245 |
| chr18-11160489-DEL-212 | chr18-11160490-DEL-212 |
| chr18-11168829-DEL-1198 | chr18-11168830-DEL-1198 |
| chr18-11172376-DEL-268 | chr18-11172377-DEL-268 |
| chr18-11185858-DEL-6296 | chr18-11185859-DEL-6296 |
| chr18-11255373-DEL-124 | chr18-11255398-DEL-100 |
| chr18-11287736-DEL-84 | chr18-11287736-DEL-84 |
| chr18-11291148-DEL-3317 | chr18-11291149-DEL-3317 |
| chr18-11303460-DEL-135 | chr18-11303481-DEL-115 |
| chr18-11318879-DEL-96 | chr18-11318880-DEL-96 |
| chr18-11361622-DEL-185 | chr18-11361617-DEL-186 |
| chr18-11376855-DEL-1407 | chr18-11376856-DEL-1407 |
| chr18-11388434-DEL-6961 | chr18-11388434-DEL-6961 |
| chr18-11399026-DEL-375 | chr18-11399027-DEL-375 |
| chr18-11410122-DEL-57 | chr18-11410123-DEL-57 |
| chr18-11412163-DEL-131 | chr18-11412164-DEL-131 |
| chr18-11423374-DEL-533 | chr18-11423375-DEL-533 |
| chr18-11437829-DEL-1965 | chr18-11437830-DEL-1965 |
| chr18-11504678-DEL-468 | chr18-11504679-DEL-468 |
| chr18-11517457-DEL-175 | chr18-11517449-DEL-176 |
| chr18-11522931-DEL-441 | chr18-11522932-DEL-441 |
| chr18-11545351-DEL-234 | chr18-11545352-DEL-234 |
| chr18-11579190-DEL-296 | chr18-11579190-DEL-294 |
| chr18-11582453-DEL-518 | chr18-11582451-DEL-516 |
| chr18-11595091-DEL-243 | chr18-11595092-DEL-243 |
| chr18-11611952-DEL-141 | chr18-11611930-DEL-142 |
| chr18-11625735-DEL-136 | chr18-11625736-DEL-136 |
| chr18-11629606-DEL-11825 | chr18-11629607-DEL-11825 |
| chr18-11645229-DEL-446 | chr18-11645230-DEL-446 |
| chr18-11655595-DEL-271 | chr18-11655596-DEL-271 |
| chr18-11662164-DEL-53 | chr18-11662165-DEL-53 |
| chr18-11673891-DEL-859 | chr18-11673892-DEL-859 |
| chr18-11675354-DEL-121 | chr18-11675355-DEL-118 |
| chr18-11679436-DEL-148 | chr18-11679437-DEL-148 |
| chr18-11688501-DEL-98 | chr18-11688502-DEL-98 |
| chr18-11727829-DEL-58 | chr18-11727830-DEL-58 |
| chr18-11728212-DEL-206 | chr18-11728213-DEL-206 |
| chr18-11744680-DEL-303 | chr18-11744681-DEL-303 |
| chr18-11759111-DEL-5257 | chr18-11759112-DEL-5257 |
| chr18-11775595-DEL-5414 | chr18-11775595-DEL-5414 |
| chr18-11786491-DEL-140 | chr18-11786492-DEL-140 |
| chr18-11804816-DEL-203 | chr18-11804817-DEL-203 |
| chr18-11815086-DEL-166 | chr18-11815087-DEL-166 |
| chr18-11822197-DEL-3580 | chr18-11822199-DEL-3581 |
| chr18-11828605-DEL-969 | chr18-11828617-DEL-965 |
| chr18-11832227-DEL-211 | chr18-11832228-DEL-211 |
| chr18-11887277-DEL-124 | chr18-11887278-DEL-124 |
| chr18-11889551-DEL-54 | chr18-11889529-DEL-59 |
| chr18-11901103-DEL-60 | chr18-11901104-DEL-60 |
| chr18-11901806-DEL-144 | chr18-11901785-DEL-142 |
| chr18-11910635-DEL-316 | chr18-11910636-DEL-316 |
| chr18-11912289-DEL-202 | chr18-11912290-DEL-202 |
| chr18-11942829-DEL-405 | chr18-11942830-DEL-405 |
| chr18-11967325-DEL-59 | chr18-11967322-DEL-63 |
| chr18-11978575-DEL-67 | chr18-11978576-DEL-67 |
| chr18-12003272-DEL-62 | chr18-12003275-DEL-60 |
| chr18-12009318-DEL-138 | chr18-12009319-DEL-138 |

| | |
|---|---|
| chr18-12025632-DEL-177 | chr18-12025644-DEL-163 |
| chr18-12031665-DEL-72 | chr18-12031666-DEL-72 |
| chr18-12033328-DEL-60 | chr18-12033329-DEL-60 |
| chr18-12062850-DEL-206 | chr18-12062851-DEL-206 |
| chr18-12075878-DEL-99 | chr18-12075876-DEL-99 |
| chr18-12089828-DEL-173 | chr18-12089829-DEL-173 |
| chr18-12092559-DEL-181 | chr18-12092560-DEL-181 |
| chr18-12124595-DEL-221 | chr18-12124596-DEL-221 |
| chr18-12133615-DEL-202 | chr18-12133616-DEL-202 |
| chr18-12138890-DEL-84 | chr18-12138891-DEL-84 |
| chr18-12163043-DEL-69 | chr18-12163044-DEL-69 |
| chr18-12163455-DEL-265 | chr18-12163456-DEL-265 |
| chr18-12171694-DEL-98 | chr18-12171695-DEL-98 |
| chr18-12176810-DEL-100 | chr18-12176814-DEL-50 |
| chr18-12198941-DEL-112 | chr18-12198953-DEL-111 |
| chr18-12213590-DEL-93 | chr18-12213593-DEL-91 |
| chr18-12224858-DEL-1267 | chr18-12224857-DEL-1267 |
| chr18-12232414-DEL-110 | chr18-12232415-DEL-110 |
| chr18-12235707-DEL-64 | chr18-12235704-DEL-62 |
| chr18-12245389-DEL-70 | chr18-12245394-DEL-67 |
| chr18-12245854-DEL-126 | chr18-12245855-DEL-126 |
| chr18-12249944-DEL-210 | chr18-12249945-DEL-210 |
| chr18-12250573-DEL-122 | chr18-12250574-DEL-122 |
| chr18-12271318-DEL-199 | chr18-12271319-DEL-199 |
| chr18-12283441-DEL-715 | chr18-12283442-DEL-715 |
| chr18-12292051-DEL-538 | chr18-12292052-DEL-538 |
| chr18-12293234-DEL-718 | chr18-12293235-DEL-718 |
| chr18-12296604-DEL-211 | chr18-12296605-DEL-211 |
| chr18-12298017-DEL-1049 | chr18-12298018-DEL-1049 |
| chr18-12298726-DEL-204 | chr18-12298727-DEL-204 |
| chr18-12307054-DEL-350 | chr18-12307055-DEL-350 |
| chr18-12310437-DEL-75 | chr18-12310434-DEL-76 |
| chr18-12370802-DEL-484 | chr18-12370803-DEL-484 |
| chr18-12370983-DEL-2942 | chr18-12370984-DEL-2942 |
| chr18-12431656-DEL-56 | chr18-12431655-DEL-52 |
| chr18-12444836-DEL-53 | chr18-12444837-DEL-53 |
| chr18-12458364-DEL-1955 | chr18-12458364-DEL-1955 |
| chr18-12484666-DEL-6218 | chr18-12484667-DEL-6218 |
| chr18-12498636-DEL-1463 | chr18-12498637-DEL-1463 |
| chr18-12514503-DEL-102 | chr18-12514492-DEL-76 |
| chr18-12519190-DEL-615 | chr18-12519191-DEL-615 |
| chr18-12533776-DEL-59 | chr18-12533796-DEL-72 |
| chr18-12548236-DEL-58 | chr18-12548237-DEL-58 |
| chr18-12564813-DEL-96 | chr18-12564832-DEL-78 |
| chr18-12570452-DEL-92 | chr18-12570453-DEL-92 |
| chr18-12603582-DEL-232 | chr18-12603578-DEL-232 |
| chr18-12604734-DEL-64 | chr18-12604735-DEL-64 |
| chr18-12616499-DEL-136 | chr18-12616500-DEL-136 |
| chr18-12627276-DEL-835 | chr18-12627277-DEL-835 |
| chr18-12628143-DEL-80 | chr18-12628144-DEL-80 |
| chr18-12683167-DEL-74 | chr18-12683168-DEL-74 |
| chr18-12688614-DEL-117 | chr18-12688615-DEL-117 |
| chr18-12730655-DEL-57 | chr18-12730656-DEL-57 |
| chr18-12742245-DEL-74 | chr18-12742246-DEL-74 |
| chr18-12748699-DEL-58 | chr18-12748696-DEL-74 |
| chr18-12755247-DEL-1802 | chr18-12755248-DEL-1802 |
| chr18-12759437-DEL-199 | chr18-12759438-DEL-199 |
| chr18-12796849-DEL-62 | chr18-12796844-DEL-62 |
| chr18-12845128-DEL-1884 | chr18-12845119-DEL-1887 |
| chr18-12853186-DEL-80 | chr18-12853187-DEL-80 |
| chr18-12855754-DEL-185 | chr18-12855768-DEL-167 |
| chr18-12858146-DEL-139 | chr18-12858151-DEL-93 |
| chr18-12862483-DEL-78 | chr18-12862483-DEL-79 |
| chr18-12864944-DEL-271 | chr18-12864924-DEL-288 |
| chr18-12865428-DEL-304 | chr18-12865429-DEL-304 |
| chr18-12878320-DEL-202 | chr18-12878321-DEL-202 |
| chr18-12910933-DEL-56 | chr18-12910934-DEL-56 |
| chr18-12912298-DEL-80 | chr18-12912299-DEL-80 |
| chr18-12914835-DEL-54 | chr18-12914836-DEL-54 |
| chr18-12922519-DEL-54 | chr18-12922520-DEL-54 |
| chr18-12963169-DEL-1938 | chr18-12963170-DEL-1938 |
| chr18-13009571-DEL-245 | chr18-13009572-DEL-245 |
| chr18-13011659-DEL-111 | chr18-13011660-DEL-111 |
| chr18-13016393-DEL-53 | chr18-13016394-DEL-53 |
| chr18-13031438-DEL-731 | chr18-13031439-DEL-731 |
| chr18-13036486-DEL-54 | chr18-13036487-DEL-54 |
| chr18-13037491-DEL-97 | chr18-13037496-DEL-93 |
| chr18-13051973-DEL-54 | chr18-13052008-DEL-55 |
| chr18-13057178-DEL-93 | chr18-13057169-DEL-85 |
| chr18-13058969-DEL-135 | chr18-13058961-DEL-135 |

| | |
|---|---|
| chr18-13080396-DEL-189 | chr18-13080386-DEL-190 |
| chr18-13107241-DEL-60 | chr18-13107242-DEL-60 |
| chr18-13127471-DEL-166 | chr18-13127468-DEL-172 |
| chr18-13161483-DEL-60 | chr18-13161484-DEL-60 |
| chr18-13171014-DEL-642 | chr18-13171015-DEL-641 |
| chr18-13175903-DEL-204 | chr18-13175904-DEL-204 |
| chr18-13183250-DEL-56 | chr18-13183251-DEL-56 |
| chr18-13187430-DEL-61 | chr18-13187431-DEL-61 |
| chr18-13239138-DEL-486 | chr18-13239139-DEL-486 |
| chr18-13278968-DEL-87 | chr18-13278969-DEL-87 |
| chr18-13305983-DEL-6260 | chr18-13305984-DEL-6260 |
| chr18-13347247-DEL-1086 | chr18-13347248-DEL-1086 |
| chr18-13352322-DEL-84 | chr18-13352323-DEL-84 |
| chr18-13381591-DEL-208 | chr18-13381592-DEL-208 |
| chr18-13381973-DEL-2129 | chr18-13381945-DEL-2130 |
| chr18-13382590-DEL-355 | chr18-13382591-DEL-355 |
| chr18-13394325-DEL-229 | chr18-13394326-DEL-229 |
| chr18-13449575-DEL-816 | chr18-13449576-DEL-816 |
| chr18-13456710-DEL-106 | chr18-13456711-DEL-106 |
| chr18-13512684-DEL-854 | chr18-13512685-DEL-854 |
| chr18-13542710-DEL-71 | chr18-13542714-DEL-54 |
| chr18-13551555-DEL-57 | chr18-13551556-DEL-57 |
| chr18-13558632-DEL-70 | chr18-13558633-DEL-70 |
| chr18-13578162-DEL-62 | chr18-13578163-DEL-62 |
| chr18-13646239-DEL-4029 | chr18-13646240-DEL-4029 |
| chr18-13664935-DEL-681 | chr18-13664936-DEL-681 |
| chr18-13670740-DEL-156 | chr18-13670741-DEL-156 |
| chr18-13697775-DEL-92 | chr18-13697742-DEL-85 |
| chr18-13701701-DEL-75 | chr18-13701666-DEL-90 |
| chr18-13709153-DEL-6386 | chr18-13709154-DEL-6382 |
| chr18-13756826-DEL-52 | chr18-13756826-DEL-52 |
| chr18-13762577-DEL-151 | chr18-13762578-DEL-115 |
| chr18-13768227-DEL-552 | chr18-13768232-DEL-552 |
| chr18-13832849-DEL-58 | chr18-13832850-DEL-58 |
| chr18-13833690-DEL-85 | chr18-13833691-DEL-85 |
| chr18-13843917-DEL-60 | chr18-13843918-DEL-60 |
| chr18-13864748-DEL-927 | chr18-13864748-DEL-926 |
| chr18-13874048-DEL-170 | chr18-13874049-DEL-170 |
| chr18-13910841-DEL-53 | chr18-13910819-DEL-53 |
| chr18-14069101-DEL-213 | chr18-14069099-DEL-211 |
| chr18-14082730-DEL-210 | chr18-14082731-DEL-210 |
| chr18-14090061-DEL-325 | chr18-14090062-DEL-325 |
| chr18-14097013-DEL-54 | chr18-14097028-DEL-52 |
| chr18-14111512-DEL-54 | chr18-14111513-DEL-54 |
| chr18-14136585-DEL-241 | chr18-14136596-DEL-231 |
| chr18-14166866-DEL-60 | chr18-14166867-DEL-60 |
| chr18-14200032-DEL-51 | chr18-14200033-DEL-51 |
| chr18-14208325-DEL-67 | chr18-14208298-DEL-68 |
| chr18-14225209-DEL-1007 | chr18-14225210-DEL-1007 |
| chr18-14236252-DEL-212 | chr18-14236253-DEL-212 |
| chr18-14253906-DEL-419 | chr18-14253907-DEL-419 |
| chr18-14344068-DEL-58 | chr18-14344069-DEL-57 |
| chr18-14359451-DEL-773 | chr18-14359454-DEL-771 |
| chr18-14416586-DEL-109 | chr18-14416587-DEL-101 |
| chr18-14421807-DEL-96 | chr18-14421812-DEL-92 |
| chr18-14440359-DEL-280 | chr18-14440360-DEL-280 |
| chr18-14448766-DEL-68 | chr18-14448766-DEL-69 |
| chr18-14497706-DEL-175 | chr18-14497709-DEL-173 |
| chr18-14501970-DEL-524 | chr18-14501971-DEL-524 |
| chr18-14577791-DEL-1494 | chr18-14577792-DEL-1494 |
| chr18-14603599-DEL-72 | chr18-14603600-DEL-72 |
| chr18-14619971-DEL-67 | chr18-14619971-DEL-67 |
| chr18-14654045-DEL-52 | chr18-14654046-DEL-52 |
| chr18-14679068-DEL-84 | chr18-14679064-DEL-84 |
| chr18-14682926-DEL-1509 | chr18-14682927-DEL-1509 |
| chr18-14687472-DEL-55 | chr18-14687457-DEL-66 |
| chr18-14690176-DEL-400 | chr18-14690177-DEL-400 |
| chr18-14693586-DEL-86 | chr18-14693587-DEL-86 |
| chr18-14698597-DEL-53 | chr18-14698596-DEL-55 |
| chr18-14703152-DEL-1625 | chr18-14703144-DEL-1626 |
| chr18-14715907-DEL-1831 | chr18-14715908-DEL-1828 |
| chr18-14720628-DEL-88 | chr18-14720629-DEL-86 |
| chr18-14744426-DEL-795 | chr18-14744427-DEL-795 |
| chr18-14752560-DEL-176 | chr18-14752561-DEL-176 |
| chr18-14762937-DEL-60 | chr18-14762937-DEL-60 |
| chr18-14809062-DEL-216 | chr18-14809063-DEL-216 |
| chr18-14821688-DEL-52 | chr18-14821689-DEL-52 |
| chr18-14835629-DEL-238 | chr18-14835629-DEL-238 |
| chr18-14865861-DEL-6275 | chr18-14865862-DEL-6275 |
| chr18-14904361-DEL-134 | chr18-14904362-DEL-134 |

| | |
|---|---|
| chr18-15041828-DEL-57 | chr18-15041824-DEL-64 |
| chr18-15085289-DEL-400 | chr18-15085290-DEL-400 |
| chr18-15132253-DEL-68 | chr18-15132249-DEL-72 |
| chr18-15133222-DEL-75 | chr18-15133246-DEL-74 |
| chr18-15172209-DEL-57 | chr18-15172212-DEL-56 |
| chr18-15177126-DEL-111 | chr18-15177101-DEL-99 |
| chr18-15187369-DEL-543 | chr18-15187370-DEL-531 |
| chr18-15188887-DEL-78 | chr18-15188887-DEL-78 |
| chr18-15213459-DEL-88 | chr18-15213460-DEL-88 |
| chr18-15221746-DEL-64 | chr18-15221735-DEL-50 |
| chr18-15241813-DEL-85 | chr18-15241812-DEL-86 |
| chr18-15256385-DEL-67 | chr18-15256387-DEL-66 |
| chr18-15273191-DEL-57 | chr18-15273190-DEL-57 |
| chr18-15276282-DEL-58 | chr18-15276283-DEL-58 |
| chr18-15319412-DEL-6394 | chr18-15319413-DEL-6394 |
| chr18-15325543-DEL-201 | chr18-15325543-DEL-201 |
| chr18-15326767-DEL-108 | chr18-15326816-DEL-91 |
| chr18-15358585-DEL-7090 | chr18-15358586-DEL-7090 |
| chr18-15411875-DEL-78 | chr18-15411900-DEL-54 |
| chr18-15426820-DEL-132 | chr18-15426821-DEL-132 |
| chr18-15434477-DEL-705 | chr18-15434478-DEL-705 |
| chr18-15439793-DEL-80 | chr18-15439794-DEL-80 |
| chr18-15469175-DEL-145 | chr18-15469176-DEL-145 |
| chr18-15487332-DEL-72 | chr18-15487345-DEL-72 |
| chr18-15499530-DEL-6197 | chr18-15499531-DEL-6197 |
| chr18-15514305-DEL-204 | chr18-15514306-DEL-204 |
| chr18-15593009-DEL-52 | chr18-15593010-DEL-52 |
| chr18-15616536-DEL-6438 | chr18-15616537-DEL-6438 |
| chr18-15635777-DEL-364 | chr18-15635778-DEL-364 |
| chr18-15643143-DEL-109 | chr18-15643180-DEL-92 |
| chr18-15656471-DEL-628 | chr18-15656472-DEL-628 |
| chr18-15689678-DEL-117 | chr18-15689664-DEL-102 |
| chr18-15702850-DEL-340 | chr18-15702851-DEL-334 |
| chr18-15720943-DEL-184 | chr18-15720962-DEL-164 |
| chr18-15730814-DEL-4526 | chr18-15730815-DEL-4526 |
| chr18-15828495-DEL-208 | chr18-15828496-DEL-208 |
| chr18-15863043-DEL-7632 | chr18-15863044-DEL-7632 |
| chr18-15887012-DEL-232 | chr18-15887013-DEL-232 |
| chr18-15911693-DEL-1366 | chr18-15911694-DEL-1366 |
| chr18-15930312-DEL-62 | chr18-15930311-DEL-74 |
| chr18-15946064-DEL-91 | chr18-15946078-DEL-80 |
| chr18-15965813-DEL-619 | chr18-15965814-DEL-619 |
| chr18-15968102-DEL-2421 | chr18-15968103-DEL-2421 |
| chr18-16010885-DEL-98 | chr18-16010886-DEL-98 |
| chr18-16028423-DEL-1676 | chr18-16028424-DEL-1676 |
| chr18-16073560-DEL-3068 | chr18-16073561-DEL-3068 |
| chr18-16077286-DEL-204 | chr18-16077287-DEL-204 |
| chr18-16101638-DEL-264 | chr18-16101639-DEL-264 |
| chr18-16114064-DEL-7277 | chr18-16114065-DEL-7277 |
| chr18-16122516-DEL-208 | chr18-16122517-DEL-208 |
| chr18-16124481-DEL-209 | chr18-16124482-DEL-209 |
| chr18-16125359-DEL-358 | chr18-16125360-DEL-358 |
| chr18-16139119-DEL-56 | chr18-16139145-DEL-65 |
| chr18-16174795-DEL-64 | chr18-16174794-DEL-52 |
| chr18-16175981-DEL-90 | chr18-16175978-DEL-91 |
| chr18-16180859-DEL-63 | chr18-16180860-DEL-63 |
| chr18-16193229-DEL-129 | chr18-16193230-DEL-129 |
| chr18-16193942-DEL-57 | chr18-16193943-DEL-57 |
| chr18-16205708-DEL-74 | chr18-16205709-DEL-74 |
| chr18-16207464-DEL-192 | chr18-16207502-DEL-192 |
| chr18-16211541-DEL-236 | chr18-16211542-DEL-236 |
| chr18-16336172-DEL-92 | chr18-16336174-DEL-90 |
| chr18-16337300-DEL-481 | chr18-16337301-DEL-480 |
| chr18-16368057-DEL-401 | chr18-16368058-DEL-401 |
| chr18-16407009-DEL-189 | chr18-16407010-DEL-189 |
| chr18-16407543-DEL-531 | chr18-16407544-DEL-531 |
| chr18-16416026-DEL-6226 | chr18-16416027-DEL-6226 |
| chr18-16431795-DEL-174 | chr18-16431796-DEL-173 |
| chr18-16467316-DEL-1965 | chr18-16467317-DEL-1965 |
| chr18-16471634-DEL-979 | chr18-16471633-DEL-980 |
| chr18-16511179-DEL-1240 | chr18-16511180-DEL-1240 |
| chr18-16531361-DEL-463 | chr18-16531362-DEL-463 |
| chr18-16534529-DEL-80 | chr18-16534530-DEL-80 |
| chr18-16565036-DEL-92 | chr18-16565037-DEL-92 |
| chr18-16626358-DEL-58 | chr18-16626359-DEL-58 |
| chr18-16628306-DEL-135 | chr18-16628305-DEL-137 |
| chr18-16640021-DEL-572 | chr18-16640022-DEL-572 |
| chr18-16644744-DEL-442 | chr18-16644745-DEL-442 |
| chr18-16648089-DEL-232 | chr18-16648059-DEL-232 |
| chr18-16658380-DEL-241 | chr18-16658381-DEL-241 |

| | |
|---|---|
| chr18-16665000-DEL-64 | chr18-16665001-DEL-64 |
| chr18-16712097-DEL-619 | chr18-16712098-DEL-619 |
| chr18-16720455-DEL-53 | chr18-16720456-DEL-53 |
| chr18-16776246-DEL-71 | chr18-16776245-DEL-72 |
| chr18-16785550-DEL-62 | chr18-16785559-DEL-54 |
| chr18-16852867-DEL-72 | chr18-16852868-DEL-72 |
| chr18-16854026-DEL-1833 | chr18-16854027-DEL-1833 |
| chr18-16883064-DEL-95 | chr18-16883045-DEL-100 |
| chr18-16888367-DEL-80 | chr18-16888372-DEL-76 |
| chr18-16904278-DEL-102 | chr18-16904279-DEL-102 |
| chr18-16904813-DEL-255 | chr18-16904814-DEL-255 |
| chr18-16983427-DEL-4556 | chr18-16983428-DEL-4556 |
| chr18-16995604-DEL-106 | chr18-16995653-DEL-82 |
| chr18-17010271-DEL-78 | chr18-17010286-DEL-53 |
| chr18-17028937-DEL-1014 | chr18-17028938-DEL-1014 |
| chr18-17043988-DEL-142 | chr18-17043969-DEL-143 |
| chr18-17094950-DEL-6457 | chr18-17094951-DEL-6457 |
| chr18-17121650-DEL-6623 | chr18-17121651-DEL-6623 |
| chr18-17142826-DEL-5235 | chr18-17142827-DEL-5235 |
| chr18-17170548-DEL-198 | chr18-17170549-DEL-198 |
| chr18-17186632-DEL-59 | chr18-17186623-DEL-56 |
| chr18-17197407-DEL-66 | chr18-17197406-DEL-68 |
| chr18-17203887-DEL-1611 | chr18-17203888-DEL-1611 |
| chr18-17249096-DEL-3620 | chr18-17249097-DEL-3620 |
| chr18-17320630-DEL-192 | chr18-17320631-DEL-192 |
| chr18-17351166-DEL-8347 | chr18-17351167-DEL-8347 |
| chr18-17378182-DEL-1709 | chr18-17378183-DEL-1709 |
| chr18-17405874-DEL-237 | chr18-17405875-DEL-237 |
| chr18-17435484-DEL-52 | chr18-17435482-DEL-54 |
| chr18-17445255-DEL-1316 | chr18-17445256-DEL-1316 |
| chr18-17479003-DEL-6726 | chr18-17479004-DEL-6726 |
| chr18-17487571-DEL-985 | chr18-17487572-DEL-985 |
| chr18-17583438-DEL-435 | chr18-17583439-DEL-435 |
| chr18-17590717-DEL-955 | chr18-17590718-DEL-955 |
| chr18-17597639-DEL-6757 | chr18-17597640-DEL-6757 |
| chr18-17625751-DEL-117 | chr18-17625752-DEL-117 |
| chr18-17631014-DEL-921 | chr18-17631015-DEL-921 |
| chr18-17644882-DEL-6368 | chr18-17644883-DEL-6368 |
| chr18-17674255-DEL-6384 | chr18-17674256-DEL-6384 |
| chr18-17686727-DEL-106 | chr18-17686742-DEL-95 |
| chr18-17691001-DEL-1257 | chr18-17691002-DEL-1257 |
| chr18-17708095-DEL-342 | chr18-17708096-DEL-342 |
| chr18-17726341-DEL-6230 | chr18-17726342-DEL-6230 |
| chr18-17738804-DEL-66 | chr18-17738803-DEL-66 |
| chr18-17763762-DEL-4257 | chr18-17763763-DEL-4257 |
| chr18-17788508-DEL-4633 | chr18-17788509-DEL-4633 |
| chr18-17794209-DEL-6366 | chr18-17794210-DEL-6366 |
| chr18-17833146-DEL-62 | chr18-17833148-DEL-82 |
| chr18-17853121-DEL-66 | chr18-17853122-DEL-66 |
| chr18-17909811-DEL-78 | chr18-17909812-DEL-78 |
| chr18-17927510-DEL-239 | chr18-17927511-DEL-238 |
| chr18-17927745-DEL-634 | chr18-17927745-DEL-635 |
| chr18-17928486-DEL-3176 | chr18-17928487-DEL-3176 |
| chr18-17948040-DEL-398 | chr18-17948041-DEL-398 |
| chr18-17953900-DEL-6736 | chr18-17953901-DEL-6736 |
| chr18-17961878-DEL-460 | chr18-17961865-DEL-451 |
| chr18-17976910-DEL-65 | chr18-17976910-DEL-65 |
| chr18-17991312-DEL-4287 | chr18-17991313-DEL-4287 |
| chr18-18007134-DEL-69 | chr18-18007135-DEL-69 |
| chr18-18016020-DEL-69 | chr18-18016018-DEL-70 |
| chr18-18017045-DEL-953 | chr18-18017046-DEL-953 |
| chr18-18019363-DEL-820 | chr18-18019364-DEL-820 |
| chr18-18043837-DEL-157 | chr18-18043838-DEL-157 |
| chr18-18062112-DEL-480 | chr18-18062113-DEL-479 |
| chr18-18071329-DEL-7279 | chr18-18071330-DEL-7279 |
| chr18-18081397-DEL-233 | chr18-18081398-DEL-233 |
| chr18-18097736-DEL-743 | chr18-18097737-DEL-743 |
| chr18-18130819-DEL-185 | chr18-18130820-DEL-185 |
| chr18-18176751-DEL-1547 | chr18-18176752-DEL-1547 |
| chr18-18194326-DEL-302 | chr18-18194327-DEL-304 |
| chr18-18216853-DEL-130 | chr18-18216854-DEL-130 |
| chr18-18224312-DEL-329 | chr18-18224312-DEL-330 |
| chr18-18241370-DEL-1152 | chr18-18241371-DEL-1152 |
| chr18-18274004-DEL-2003 | chr18-18274004-DEL-2001 |
| chr18-18276816-DEL-54 | chr18-18276830-DEL-52 |
| chr18-18293041-DEL-57 | chr18-18293042-DEL-57 |
| chr18-18304575-DEL-399 | chr18-18304576-DEL-399 |
| chr18-18305865-DEL-6185 | chr18-18305866-DEL-6185 |
| chr18-18324715-DEL-54 | chr18-18324716-DEL-54 |
| chr18-18326221-DEL-58 | chr18-18326222-DEL-58 |

| | |
|---|---|
| chr18-18337204-DEL-60 | chr18-18337205-DEL-60 |
| chr18-18403818-DEL-110 | chr18-18403823-DEL-106 |
| chr18-18433349-DEL-3485 | chr18-18433350-DEL-3485 |
| chr18-18475349-DEL-414 | chr18-18475350-DEL-414 |
| chr18-18480799-DEL-750 | chr18-18480800-DEL-750 |
| chr18-18486271-DEL-5170 | chr18-18486267-DEL-5168 |
| chr18-18521685-DEL-846 | chr18-18521686-DEL-846 |
| chr18-18528318-DEL-131 | chr18-18528319-DEL-129 |
| chr18-18555792-DEL-6418 | chr18-18555793-DEL-6418 |
| chr18-18566444-DEL-198 | chr18-18566445-DEL-198 |
| chr18-18596583-DEL-980 | chr18-18596584-DEL-980 |
| chr18-18747333-DEL-52 | chr18-18747334-DEL-54 |
| chr18-18764361-DEL-52 | chr18-18764350-DEL-64 |
| chr18-18768546-DEL-3569 | chr18-18768549-DEL-3570 |
| chr18-18797530-DEL-6470 | chr18-18797531-DEL-6470 |
| chr18-18809774-DEL-3303 | chr18-18809775-DEL-3303 |
| chr18-18818053-DEL-57 | chr18-18818054-DEL-57 |
| chr18-18832387-DEL-737 | chr18-18832388-DEL-737 |
| chr18-18834142-DEL-81 | chr18-18834143-DEL-81 |
| chr18-18836542-DEL-438 | chr18-18836543-DEL-438 |
| chr18-18863139-DEL-472 | chr18-18863140-DEL-472 |
| chr18-18871050-DEL-19549 | chr18-18871051-DEL-19549 |
| chr18-18934921-DEL-51 | chr18-18934921-DEL-51 |
| chr18-18941079-DEL-280 | chr18-18941080-DEL-280 |
| chr18-18950876-DEL-686 | chr18-18950877-DEL-686 |
| chr18-18970809-DEL-332 | chr18-18970810-DEL-332 |
| chr18-18987665-DEL-66 | chr18-18987666-DEL-66 |
| chr18-19094995-DEL-108 | chr18-19095031-DEL-72 |
| chr18-19095264-DEL-410 | chr18-19095265-DEL-410 |
| chr18-19097060-DEL-59 | chr18-19097061-DEL-59 |
| chr18-19099298-DEL-171 | chr18-19099298-DEL-171 |
| chr18-19171559-DEL-982 | chr18-19171568-DEL-995 |
| chr18-19177485-DEL-62 | chr18-19177486-DEL-62 |
| chr18-19180649-DEL-126 | chr18-19180650-DEL-126 |
| chr18-19217129-DEL-105 | chr18-19217130-DEL-105 |
| chr18-19224883-DEL-93 | chr18-19224931-DEL-62 |
| chr18-19233194-DEL-382 | chr18-19233194-DEL-382 |
| chr18-19249324-DEL-60 | chr18-19249325-DEL-56 |
| chr18-19366895-DEL-282 | chr18-19366895-DEL-283 |
| chr18-19373690-DEL-6230 | chr18-19373677-DEL-6217 |
| chr18-19415624-DEL-6595 | chr18-19415625-DEL-6595 |
| chr18-19432709-DEL-138 | chr18-19432712-DEL-136 |
| chr18-19435148-DEL-1048 | chr18-19435149-DEL-1048 |
| chr18-19451574-DEL-613 | chr18-19451575-DEL-613 |
| chr18-19478825-DEL-6490 | chr18-19478826-DEL-6490 |
| chr18-19493289-DEL-1118 | chr18-19493294-DEL-1116 |
| chr18-19517487-DEL-3874 | chr18-19517488-DEL-3874 |
| chr18-19544502-DEL-734 | chr18-19544502-DEL-734 |
| chr18-19546289-DEL-2396 | chr18-19546289-DEL-2396 |
| chr18-19560539-DEL-479 | chr18-19560540-DEL-479 |
| chr18-19563552-DEL-4349 | chr18-19563553-DEL-4349 |
| chr18-19575069-DEL-6118 | chr18-19575070-DEL-6118 |
| chr18-19582140-DEL-852 | chr18-19582142-DEL-849 |
| chr18-19586674-DEL-82 | chr18-19586676-DEL-83 |
| chr18-19587530-DEL-63 | chr18-19587531-DEL-63 |
| chr18-19625032-DEL-289 | chr18-19625033-DEL-289 |
| chr18-19638063-DEL-2953 | chr18-19638064-DEL-2953 |
| chr18-19650363-DEL-57 | chr18-19650358-DEL-63 |
| chr18-19651490-DEL-321 | chr18-19651490-DEL-321 |
| chr18-19652763-DEL-522 | chr18-19652763-DEL-522 |
| chr18-19663951-DEL-84 | chr18-19663952-DEL-84 |
| chr18-19687041-DEL-6259 | chr18-19687042-DEL-6259 |
| chr18-19745346-DEL-6600 | chr18-19745347-DEL-6600 |
| chr18-19757244-DEL-207 | chr18-19757245-DEL-207 |
| chr18-19759242-DEL-573 | chr18-19759242-DEL-573 |
| chr18-19761357-DEL-888 | chr18-19761359-DEL-881 |
| chr18-19764943-DEL-1893 | chr18-19764943-DEL-1881 |
| chr18-19766698-DEL-1511 | chr18-19766699-DEL-1511 |
| chr18-19769446-DEL-100 | chr18-19769447-DEL-100 |
| chr18-19787689-DEL-381 | chr18-19787690-DEL-381 |
| chr18-19812691-DEL-1890 | chr18-19812692-DEL-1890 |
| chr18-19854222-DEL-69 | chr18-19854189-DEL-102 |
| chr18-19855852-DEL-1469 | chr18-19855853-DEL-1469 |
| chr18-19859015-DEL-1526 | chr18-19859016-DEL-1526 |
| chr18-19923735-DEL-278 | chr18-19923735-DEL-278 |
| chr18-19926658-DEL-402 | chr18-19926659-DEL-402 |
| chr18-19939329-DEL-442 | chr18-19939330-DEL-442 |
| chr18-19942923-DEL-60 | chr18-19942924-DEL-58 |
| chr18-19945029-DEL-6195 | chr18-19945030-DEL-6195 |
| chr18-19968626-DEL-954 | chr18-19968596-DEL-963 |

| | |
|---|---|
| chr18-19970382-DEL-311 | chr18-19970383-DEL-311 |
| chr18-19995112-DEL-6677 | chr18-19995113-DEL-6677 |
| chr18-19995279-DEL-425 | chr18-19995280-DEL-425 |
| chr18-20049791-DEL-683 | chr18-20049792-DEL-683 |
| chr18-20075721-DEL-148 | chr18-20075722-DEL-164 |
| chr18-20101140-DEL-85 | chr18-20101141-DEL-85 |
| chr18-20110912-DEL-91 | chr18-20110913-DEL-91 |
| chr18-20156338-DEL-939 | chr18-20156339-DEL-939 |
| chr18-20262988-DEL-70 | chr18-20262989-DEL-70 |
| chr18-20330203-DEL-141 | chr18-20330204-DEL-153 |
| chr18-20451234-DEL-85 | chr18-20451191-DEL-86 |
| chr18-20477034-DEL-78 | chr18-20477035-DEL-78 |
| chr18-20479275-DEL-158 | chr18-20479272-DEL-162 |
| chr18-20496085-DEL-6802 | chr18-20496086-DEL-6802 |
| chr18-20514519-DEL-705 | chr18-20514520-DEL-705 |
| chr18-20546051-DEL-2123 | chr18-20546052-DEL-2123 |
| chr18-20552040-DEL-1032 | chr18-20552041-DEL-1032 |
| chr18-20607572-DEL-83 | chr18-20607573-DEL-83 |
| chr18-20649342-DEL-539 | chr18-20649343-DEL-539 |
| chr18-20661895-DEL-66 | chr18-20661900-DEL-62 |
| chr18-20668781-DEL-220 | chr18-20668782-DEL-220 |
| chr18-20708585-DEL-1296 | chr18-20708586-DEL-1296 |
| chr18-20736591-DEL-55 | chr18-20736592-DEL-55 |
| chr18-20748136-DEL-3228 | chr18-20748137-DEL-3228 |
| chr18-20754401-DEL-241 | chr18-20754402-DEL-241 |
| chr18-20755323-DEL-220 | chr18-20755324-DEL-220 |
| chr18-20755783-DEL-2197 | chr18-20755782-DEL-2197 |
| chr18-20756856-DEL-172 | chr18-20756848-DEL-173 |
| chr18-20763650-DEL-51 | chr18-20763650-DEL-51 |
| chr18-20765725-DEL-66 | chr18-20765726-DEL-61 |
| chr18-20767526-DEL-1625 | chr18-20767527-DEL-1623 |
| chr18-20774547-DEL-92 | chr18-20774548-DEL-92 |
| chr18-20779124-DEL-108 | chr18-20779125-DEL-100 |
| chr18-20782836-DEL-201 | chr18-20782837-DEL-201 |
| chr18-20815588-DEL-472 | chr18-20815589-DEL-472 |
| chr18-20826257-DEL-171 | chr18-20826258-DEL-171 |
| chr18-20832871-DEL-238 | chr18-20832872-DEL-238 |
| chr18-20861714-DEL-567 | chr18-20861715-DEL-567 |
| chr18-20892917-DEL-853 | chr18-20892918-DEL-853 |
| chr18-20897287-DEL-189 | chr18-20897288-DEL-189 |
| chr18-20904122-DEL-669 | chr18-20904128-DEL-661 |
| chr18-20909933-DEL-507 | chr18-20909934-DEL-507 |
| chr18-20912399-DEL-68 | chr18-20912400-DEL-68 |
| chr18-20918955-DEL-59 | chr18-20918935-DEL-59 |
| chr18-21018961-DEL-516 | chr18-21018972-DEL-506 |
| chr18-21035850-DEL-7107 | chr18-21035851-DEL-7107 |
| chr18-21056739-DEL-85 | chr18-21056740-DEL-85 |
| chr18-21060228-DEL-140 | chr18-21060229-DEL-140 |
| chr18-21061487-DEL-52 | chr18-21061488-DEL-52 |
| chr18-21068484-DEL-653 | chr18-21068485-DEL-653 |
| chr18-21075312-DEL-194 | chr18-21075313-DEL-194 |
| chr18-21080522-DEL-84 | chr18-21080523-DEL-84 |
| chr18-21095642-DEL-76 | chr18-21095624-DEL-53 |
| chr18-21099283-DEL-489 | chr18-21099284-DEL-489 |
| chr18-21129047-DEL-69 | chr18-21129046-DEL-78 |
| chr18-21130648-DEL-56 | chr18-21130636-DEL-70 |
| chr18-21144150-DEL-199 | chr18-21144151-DEL-199 |
| chr18-21168708-DEL-204 | chr18-21168709-DEL-204 |
| chr18-21182232-DEL-185 | chr18-21182231-DEL-181 |
| chr18-21189205-DEL-81 | chr18-21189206-DEL-81 |
| chr18-21190069-DEL-819 | chr18-21190070-DEL-819 |
| chr18-21212750-DEL-242 | chr18-21212751-DEL-242 |
| chr18-21236978-DEL-65 | chr18-21236979-DEL-65 |
| chr18-21296545-DEL-1356 | chr18-21296546-DEL-1356 |
| chr18-21308117-DEL-1034 | chr18-21308118-DEL-1034 |
| chr18-21329991-DEL-205 | chr18-21329992-DEL-205 |
| chr18-21340958-DEL-938 | chr18-21340959-DEL-938 |
| chr18-21394821-DEL-7121 | chr18-21394822-DEL-7121 |
| chr18-21435540-DEL-445 | chr18-21435541-DEL-445 |
| chr18-21447952-DEL-2562 | chr18-21447953-DEL-2562 |
| chr18-21474753-DEL-2168 | chr18-21474754-DEL-2168 |
| chr18-21513220-DEL-508 | chr18-21513221-DEL-508 |
| chr18-21520164-DEL-352 | chr18-21520165-DEL-352 |
| chr18-21527476-DEL-1086 | chr18-21527464-DEL-1101 |
| chr18-21532908-DEL-3453 | chr18-21532909-DEL-3453 |
| chr18-21540120-DEL-118 | chr18-21540121-DEL-118 |
| chr18-21559334-DEL-486 | chr18-21559335-DEL-486 |
| chr18-21596770-DEL-452 | chr18-21596771-DEL-452 |
| chr18-21600973-DEL-3457 | chr18-21600974-DEL-3457 |
| chr18-21628325-DEL-84 | chr18-21628326-DEL-84 |

| | |
|---|---|
| chr18-21648880-DEL-324 | chr18-21648881-DEL-322 |
| chr18-21669391-DEL-6294 | chr18-21669392-DEL-6294 |
| chr18-21687532-DEL-79 | chr18-21687551-DEL-57 |
| chr18-21729328-DEL-78 | chr18-21729328-DEL-79 |
| chr18-21739722-DEL-189 | chr18-21739723-DEL-189 |
| chr18-21747946-DEL-144 | chr18-21747947-DEL-144 |
| chr18-21837300-DEL-207 | chr18-21837285-DEL-207 |
| chr18-21837836-DEL-95 | chr18-21837837-DEL-95 |
| chr18-21896627-DEL-116 | chr18-21896588-DEL-117 |
| chr18-21900642-DEL-534 | chr18-21900643-DEL-534 |
| chr18-21912300-DEL-62 | chr18-21912307-DEL-70 |
| chr18-21913158-DEL-100 | chr18-21913159-DEL-100 |
| chr18-21914682-DEL-441 | chr18-21914683-DEL-441 |
| chr18-21922152-DEL-3674 | chr18-21922158-DEL-3670 |
| chr18-21927394-DEL-832 | chr18-21927395-DEL-832 |
| chr18-21978785-DEL-56 | chr18-21978785-DEL-63 |
| chr18-21999975-DEL-173 | chr18-21999976-DEL-171 |
| chr18-22034089-DEL-60 | chr18-22034089-DEL-60 |
| chr18-22037655-DEL-135 | chr18-22037656-DEL-135 |
| chr18-22043060-DEL-181 | chr18-22043061-DEL-181 |
| chr18-22060836-DEL-149 | chr18-22060837-DEL-149 |
| chr18-22076306-DEL-1875 | chr18-22076307-DEL-1875 |
| chr18-22086988-DEL-64 | chr18-22086989-DEL-60 |
| chr18-22283587-DEL-202 | chr18-22283588-DEL-202 |
| chr18-22369348-DEL-356 | chr18-22369346-DEL-359 |
| chr18-22369865-DEL-446 | chr18-22369866-DEL-446 |
| chr18-22381286-DEL-210 | chr18-22381287-DEL-210 |
| chr18-22391784-DEL-100 | chr18-22391785-DEL-100 |
| chr18-22395859-DEL-122 | chr18-22395860-DEL-122 |
| chr18-22413869-DEL-502 | chr18-22413870-DEL-502 |
| chr18-22418521-DEL-83 | chr18-22418534-DEL-83 |
| chr18-22430424-DEL-6729 | chr18-22430413-DEL-6730 |
| chr18-22452579-DEL-74 | chr18-22452580-DEL-74 |
| chr18-22457817-DEL-733 | chr18-22457818-DEL-733 |
| chr18-22487117-DEL-60 | chr18-22487118-DEL-60 |
| chr18-22545479-DEL-481 | chr18-22545480-DEL-481 |
| chr18-22572394-DEL-7011 | chr18-22572395-DEL-7011 |
| chr18-22580242-DEL-52 | chr18-22580243-DEL-52 |
| chr18-22605843-DEL-104 | chr18-22605858-DEL-86 |
| chr18-22628245-DEL-52 | chr18-22628246-DEL-52 |
| chr18-22698556-DEL-6275 | chr18-22698557-DEL-6275 |
| chr18-22722280-DEL-321 | chr18-22722284-DEL-319 |
| chr18-22724083-DEL-203 | chr18-22724084-DEL-203 |
| chr18-22727479-DEL-210 | chr18-22727480-DEL-210 |
| chr18-22734168-DEL-304 | chr18-22734167-DEL-304 |
| chr18-22736034-DEL-665 | chr18-22736035-DEL-665 |
| chr18-22762463-DEL-54 | chr18-22762472-DEL-64 |
| chr18-22767263-DEL-6404 | chr18-22767263-DEL-6404 |
| chr18-22788425-DEL-484 | chr18-22788426-DEL-484 |
| chr18-22798084-DEL-63 | chr18-22798085-DEL-63 |
| chr18-22802910-DEL-202 | chr18-22802911-DEL-202 |
| chr18-22828561-DEL-201 | chr18-22828562-DEL-201 |
| chr18-22850795-DEL-133 | chr18-22850806-DEL-114 |
| chr18-22905850-DEL-714 | chr18-22905851-DEL-714 |
| chr18-22914387-DEL-515 | chr18-22914388-DEL-515 |
| chr18-22924780-DEL-3503 | chr18-22924781-DEL-3503 |
| chr18-22926690-DEL-239 | chr18-22926691-DEL-239 |
| chr18-22934730-DEL-1524 | chr18-22934731-DEL-1524 |
| chr18-22936982-DEL-2984 | chr18-22936983-DEL-2984 |
| chr18-22940809-DEL-1646 | chr18-22940810-DEL-1646 |
| chr18-22943120-DEL-55 | chr18-22943117-DEL-54 |
| chr18-23010664-DEL-6686 | chr18-23010665-DEL-6686 |
| chr18-23034249-DEL-584 | chr18-23034257-DEL-585 |
| chr18-23089137-DEL-60 | chr18-23089118-DEL-80 |
| chr18-23096129-DEL-6399 | chr18-23096130-DEL-6399 |
| chr18-23122036-DEL-7505 | chr18-23122037-DEL-7505 |
| chr18-23122433-DEL-212 | chr18-23122434-DEL-212 |
| chr18-23146110-DEL-193 | chr18-23146111-DEL-193 |
| chr18-23150660-DEL-614 | chr18-23150661-DEL-613 |
| chr18-23166560-DEL-1290 | chr18-23166561-DEL-1290 |
| chr18-23186934-DEL-6268 | chr18-23186935-DEL-6268 |
| chr18-23198371-DEL-58 | chr18-23198369-DEL-60 |
| chr18-23217206-DEL-92 | chr18-23217249-DEL-72 |
| chr18-23247009-DEL-2672 | chr18-23247005-DEL-2673 |
| chr18-23265809-DEL-101 | chr18-23265859-DEL-70 |
| chr18-23303838-DEL-174 | chr18-23303839-DEL-174 |
| chr18-23314519-DEL-210 | chr18-23314566-DEL-211 |
| chr18-23339479-DEL-1266 | chr18-23339480-DEL-1266 |
| chr18-23381949-DEL-130 | chr18-23381949-DEL-130 |
| chr18-23421315-DEL-3274 | chr18-23421313-DEL-3289 |

| | |
|---|---|
| chr18-23479433-DEL-3445 | chr18-23479436-DEL-3436 |
| chr18-23575005-DEL-103 | chr18-23575005-DEL-103 |
| chr18-23585294-DEL-101 | chr18-23585328-DEL-68 |
| chr18-23595216-DEL-58 | chr18-23595197-DEL-54 |
| chr18-23640981-DEL-107 | chr18-23640982-DEL-107 |
| chr18-23654595-DEL-118 | chr18-23654601-DEL-114 |
| chr18-23669956-DEL-3436 | chr18-23669957-DEL-3436 |
| chr18-23694735-DEL-549 | chr18-23694736-DEL-549 |
| chr18-23727078-DEL-120 | chr18-23727079-DEL-120 |
| chr18-23753981-DEL-64 | chr18-23753984-DEL-62 |
| chr18-23775327-DEL-58 | chr18-23775328-DEL-58 |
| chr18-23827021-DEL-2066 | chr18-23827022-DEL-2066 |
| chr18-23830224-DEL-618 | chr18-23830224-DEL-619 |
| chr18-23835091-DEL-997 | chr18-23835104-DEL-964 |
| chr18-23836572-DEL-97 | chr18-23836573-DEL-97 |
| chr18-23840213-DEL-54 | chr18-23840214-DEL-54 |
| chr18-23841461-DEL-338 | chr18-23841462-DEL-338 |
| chr18-23857897-DEL-931 | chr18-23857898-DEL-931 |
| chr18-23918374-DEL-806 | chr18-23918375-DEL-806 |
| chr18-23921390-DEL-7076 | chr18-23921391-DEL-7076 |
| chr18-24006440-DEL-94 | chr18-24006441-DEL-94 |
| chr18-24048894-DEL-202 | chr18-24048895-DEL-202 |
| chr18-24060218-DEL-65 | chr18-24060226-DEL-58 |
| chr18-24073837-DEL-52 | chr18-24073838-DEL-52 |
| chr18-24077970-DEL-740 | chr18-24077971-DEL-740 |
| chr18-24081375-DEL-3373 | chr18-24081375-DEL-3373 |
| chr18-24111456-DEL-75 | chr18-24111441-DEL-71 |
| chr18-24114470-DEL-3868 | chr18-24114471-DEL-3868 |
| chr18-24166150-DEL-3841 | chr18-24166151-DEL-3841 |
| chr18-24182333-DEL-60 | chr18-24182336-DEL-58 |
| chr18-24183514-DEL-167 | chr18-24183515-DEL-179 |
| chr18-24186375-DEL-52 | chr18-24186376-DEL-52 |
| chr18-24201345-DEL-196 | chr18-24201346-DEL-196 |
| chr18-24204138-DEL-576 | chr18-24204139-DEL-576 |
| chr18-24205772-DEL-89 | chr18-24205773-DEL-89 |
| chr18-24246274-DEL-184 | chr18-24246275-DEL-184 |
| chr18-24256825-DEL-137 | chr18-24256824-DEL-137 |
| chr18-24271674-DEL-76 | chr18-24271662-DEL-75 |
| chr18-24280684-DEL-60 | chr18-24280685-DEL-90 |
| chr18-24289015-DEL-140 | chr18-24289013-DEL-122 |
| chr18-24289792-DEL-55 | chr18-24289793-DEL-55 |
| chr18-24304606-DEL-192 | chr18-24304607-DEL-192 |
| chr18-24324254-DEL-54 | chr18-24324255-DEL-54 |
| chr18-24334960-DEL-727 | chr18-24334961-DEL-727 |
| chr18-24345005-DEL-97 | chr18-24345026-DEL-77 |
| chr18-24351056-DEL-1910 | chr18-24351057-DEL-1910 |
| chr18-24353692-DEL-170 | chr18-24353693-DEL-170 |
| chr18-24364921-DEL-223 | chr18-24364922-DEL-223 |
| chr18-24381709-DEL-109 | chr18-24381710-DEL-109 |
| chr18-24399535-DEL-67 | chr18-24399536-DEL-67 |
| chr18-24428536-DEL-135 | chr18-24428532-DEL-138 |
| chr18-24445365-DEL-1196 | chr18-24445366-DEL-1196 |
| chr18-24451802-DEL-95 | chr18-24451803-DEL-95 |
| chr18-24452461-DEL-825 | chr18-24452462-DEL-825 |
| chr18-24458672-DEL-52 | chr18-24458671-DEL-54 |
| chr18-24489028-DEL-821 | chr18-24489029-DEL-821 |
| chr18-24497295-DEL-72 | chr18-24497296-DEL-60 |
| chr18-24578839-DEL-217 | chr18-24578845-DEL-221 |
| chr18-24594255-DEL-329 | chr18-24594256-DEL-329 |
| chr18-24621759-DEL-571 | chr18-24621760-DEL-571 |
| chr18-24628253-DEL-332 | chr18-24628240-DEL-340 |
| chr18-24654111-DEL-399 | chr18-24654112-DEL-399 |
| chr18-24702268-DEL-213 | chr18-24702269-DEL-213 |
| chr18-24702925-DEL-6954 | chr18-24702926-DEL-6954 |
| chr18-24714452-DEL-184 | chr18-24714453-DEL-184 |
| chr18-24763385-DEL-104 | chr18-24763384-DEL-108 |
| chr18-24778830-DEL-97 | chr18-24778833-DEL-95 |
| chr18-24788719-DEL-987 | chr18-24788720-DEL-987 |
| chr18-24811684-DEL-52 | chr18-24811703-DEL-51 |
| chr18-24830707-DEL-66 | chr18-24830708-DEL-66 |
| chr18-24833352-DEL-201 | chr18-24833353-DEL-201 |
| chr18-24851644-DEL-202 | chr18-24851645-DEL-202 |
| chr18-24883001-DEL-586 | chr18-24883002-DEL-586 |
| chr18-24909300-DEL-76 | chr18-24909301-DEL-76 |
| chr18-24929006-DEL-809 | chr18-24929007-DEL-809 |
| chr18-24932361-DEL-1128 | chr18-24932362-DEL-1128 |
| chr18-24944708-DEL-332 | chr18-24944709-DEL-332 |
| chr18-24948059-DEL-197 | chr18-24948061-DEL-217 |
| chr18-24955637-DEL-84 | chr18-24955638-DEL-84 |
| chr18-24958957-DEL-545 | chr18-24958958-DEL-545 |

| | |
|---|---|
| chr18-24961692-DEL-3322 | chr18-24961693-DEL-3322 |
| chr18-24964395-DEL-1584 | chr18-24964396-DEL-1584 |
| chr18-24964748-DEL-94 | chr18-24964763-DEL-87 |
| chr18-24968264-DEL-56 | chr18-24968270-DEL-51 |
| chr18-25005193-DEL-60 | chr18-25005194-DEL-59 |
| chr18-25019762-DEL-125 | chr18-25019763-DEL-125 |
| chr18-25086272-DEL-62 | chr18-25086272-DEL-64 |
| chr18-25087298-DEL-100 | chr18-25087299-DEL-100 |
| chr18-25091952-DEL-72 | chr18-25091944-DEL-108 |
| chr18-25106220-DEL-54 | chr18-25106221-DEL-60 |
| chr18-25121505-DEL-70 | chr18-25121472-DEL-91 |
| chr18-25150283-DEL-74 | chr18-25150284-DEL-74 |
| chr18-25167562-DEL-172 | chr18-25167563-DEL-172 |
| chr18-25178252-DEL-2153 | chr18-25178253-DEL-2153 |
| chr18-25236211-DEL-347 | chr18-25236212-DEL-347 |
| chr18-25250016-DEL-188 | chr18-25250015-DEL-188 |
| chr18-25251070-DEL-113 | chr18-25251071-DEL-113 |
| chr18-25264598-DEL-70 | chr18-25264586-DEL-83 |
| chr18-25269295-DEL-59 | chr18-25269296-DEL-59 |
| chr18-25272662-DEL-286 | chr18-25272663-DEL-286 |
| chr18-25291745-DEL-220 | chr18-25291746-DEL-220 |
| chr18-25317273-DEL-490 | chr18-25317274-DEL-490 |
| chr18-25318691-DEL-177 | chr18-25318692-DEL-177 |
| chr18-25331634-DEL-91 | chr18-25331635-DEL-91 |
| chr18-25346872-DEL-426 | chr18-25346872-DEL-426 |
| chr18-25347506-DEL-5562 | chr18-25347507-DEL-5562 |
| chr18-25358003-DEL-2466 | chr18-25358004-DEL-2466 |
| chr18-25378923-DEL-52 | chr18-25378924-DEL-52 |
| chr18-25391984-DEL-64 | chr18-25391985-DEL-64 |
| chr18-25410002-DEL-80 | chr18-25410003-DEL-80 |
| chr18-25425771-DEL-126 | chr18-25425761-DEL-151 |
| chr18-25446781-DEL-92 | chr18-25446782-DEL-100 |
| chr18-25451548-DEL-1133 | chr18-25451549-DEL-1133 |
| chr18-25493841-DEL-6194 | chr18-25493842-DEL-6194 |
| chr18-25546315-DEL-6258 | chr18-25546316-DEL-6258 |
| chr18-25564881-DEL-2435 | chr18-25564882-DEL-2435 |
| chr18-25571346-DEL-56 | chr18-25571348-DEL-54 |
| chr18-25574812-DEL-435 | chr18-25574813-DEL-435 |
| chr18-25585314-DEL-56 | chr18-25585320-DEL-51 |
| chr18-25636867-DEL-119 | chr18-25636868-DEL-116 |
| chr18-25683559-DEL-498 | chr18-25683560-DEL-498 |
| chr18-25695913-DEL-201 | chr18-25695905-DEL-215 |
| chr18-25702301-DEL-75 | chr18-25702298-DEL-61 |
| chr18-25708203-DEL-52 | chr18-25708204-DEL-52 |
| chr18-25739606-DEL-144 | chr18-25739607-DEL-108 |
| chr18-25744465-DEL-182 | chr18-25744424-DEL-184 |
| chr18-25778158-DEL-79 | chr18-25778159-DEL-79 |
| chr18-25782728-DEL-100 | chr18-25782729-DEL-102 |
| chr18-25788981-DEL-68 | chr18-25788977-DEL-69 |
| chr18-25890111-DEL-58 | chr18-25890112-DEL-58 |
| chr18-25913928-DEL-6510 | chr18-25913929-DEL-6510 |
| chr18-25926784-DEL-991 | chr18-25926785-DEL-991 |
| chr18-25928761-DEL-120 | chr18-25928762-DEL-120 |
| chr18-25935920-DEL-1115 | chr18-25935921-DEL-1115 |
| chr18-25946567-DEL-1889 | chr18-25946568-DEL-1889 |
| chr18-26012880-DEL-76 | chr18-26012881-DEL-76 |
| chr18-26024765-DEL-56 | chr18-26024766-DEL-56 |
| chr18-26087032-DEL-55 | chr18-26087028-DEL-54 |
| chr18-26114127-DEL-188 | chr18-26114128-DEL-188 |
| chr18-26114790-DEL-189 | chr18-26114790-DEL-187 |
| chr18-26136077-DEL-5134 | chr18-26136078-DEL-5134 |
| chr18-26153785-DEL-89 | chr18-26153812-DEL-106 |
| chr18-26174219-DEL-1197 | chr18-26174220-DEL-1197 |
| chr18-26183075-DEL-97 | chr18-26183056-DEL-118 |
| chr18-26205816-DEL-89 | chr18-26205817-DEL-89 |
| chr18-26206752-DEL-644 | chr18-26206742-DEL-652 |
| chr18-26215925-DEL-464 | chr18-26215926-DEL-464 |
| chr18-26267086-DEL-80 | chr18-26267087-DEL-80 |
| chr18-26272590-DEL-166 | chr18-26272591-DEL-166 |
| chr18-26284146-DEL-6399 | chr18-26284147-DEL-6399 |
| chr18-26329231-DEL-53 | chr18-26329232-DEL-54 |
| chr18-26356034-DEL-52 | chr18-26356035-DEL-52 |
| chr18-26357028-DEL-1331 | chr18-26357029-DEL-1331 |
| chr18-26364392-DEL-328 | chr18-26364413-DEL-309 |
| chr18-26385677-DEL-66 | chr18-26385678-DEL-66 |
| chr18-26404723-DEL-1079 | chr18-26404724-DEL-1079 |
| chr18-26480829-DEL-196 | chr18-26480812-DEL-192 |
| chr18-26513138-DEL-2416 | chr18-26513139-DEL-2416 |
| chr18-26533601-DEL-54 | chr18-26533603-DEL-53 |
| chr18-26567558-DEL-156 | chr18-26567590-DEL-130 |

| | |
|---|---|
| chr18-26584122-DEL-112 | chr18-26584123-DEL-112 |
| chr18-26597147-DEL-222 | chr18-26597160-DEL-228 |
| chr18-26618765-DEL-170 | chr18-26618766-DEL-170 |
| chr18-26646329-DEL-81 | chr18-26646329-DEL-77 |
| chr18-26677885-DEL-85 | chr18-26677913-DEL-54 |
| chr18-26693456-DEL-2145 | chr18-26693457-DEL-2145 |
| chr18-26715677-DEL-52 | chr18-26715702-DEL-73 |
| chr18-26785405-DEL-358 | chr18-26785406-DEL-358 |
| chr18-26799935-DEL-1888 | chr18-26799936-DEL-1888 |
| chr18-26829499-DEL-6488 | chr18-26829499-DEL-6488 |
| chr18-26855308-DEL-433 | chr18-26855309-DEL-433 |
| chr18-26863945-DEL-856 | chr18-26863946-DEL-856 |
| chr18-26881106-DEL-74 | chr18-26881107-DEL-74 |
| chr18-26884944-DEL-61 | chr18-26884980-DEL-62 |
| chr18-26900044-DEL-6066 | chr18-26900045-DEL-6066 |
| chr18-26919896-DEL-379 | chr18-26919897-DEL-379 |
| chr18-26962760-DEL-7243 | chr18-26962761-DEL-7243 |
| chr18-26979693-DEL-98 | chr18-26979694-DEL-98 |
| chr18-26988749-DEL-131 | chr18-26988750-DEL-129 |
| chr18-27022268-DEL-270 | chr18-27022269-DEL-270 |
| chr18-27029329-DEL-266 | chr18-27029329-DEL-267 |
| chr18-27051754-DEL-73 | chr18-27051755-DEL-73 |
| chr18-27064176-DEL-97 | chr18-27064177-DEL-97 |
| chr18-27072546-DEL-70 | chr18-27072554-DEL-50 |
| chr18-27080801-DEL-68 | chr18-27080801-DEL-67 |
| chr18-27131379-DEL-7130 | chr18-27131380-DEL-7130 |
| chr18-27156962-DEL-75 | chr18-27156963-DEL-75 |
| chr18-27195029-DEL-6454 | chr18-27195024-DEL-6455 |
| chr18-27201963-DEL-3701 | chr18-27201963-DEL-3701 |
| chr18-27215268-DEL-1405 | chr18-27215269-DEL-1405 |
| chr18-27226863-DEL-109 | chr18-27226864-DEL-109 |
| chr18-27267412-DEL-1024 | chr18-27267413-DEL-1024 |
| chr18-27268886-DEL-1261 | chr18-27268887-DEL-1261 |
| chr18-27319950-DEL-426 | chr18-27319951-DEL-426 |
| chr18-27366755-DEL-7149 | chr18-27366756-DEL-7149 |
| chr18-27422560-DEL-119 | chr18-27422561-DEL-119 |
| chr18-27434549-DEL-2785 | chr18-27434550-DEL-2785 |
| chr18-27441776-DEL-60 | chr18-27441801-DEL-64 |
| chr18-27458405-DEL-267 | chr18-27458406-DEL-267 |
| chr18-27458912-DEL-426 | chr18-27458913-DEL-426 |
| chr18-27460589-DEL-72 | chr18-27460589-DEL-72 |
| chr18-27499183-DEL-150 | chr18-27499184-DEL-150 |
| chr18-27518128-DEL-107 | chr18-27518129-DEL-107 |
| chr18-27556835-DEL-301 | chr18-27556836-DEL-301 |
| chr18-27589320-DEL-6874 | chr18-27589321-DEL-6874 |
| chr18-27595694-DEL-212 | chr18-27595695-DEL-212 |
| chr18-27641455-DEL-301 | chr18-27641463-DEL-292 |
| chr18-27668530-DEL-200 | chr18-27668529-DEL-201 |
| chr18-27713680-DEL-64 | chr18-27713681-DEL-64 |
| chr18-27714561-DEL-54 | chr18-27714562-DEL-54 |
| chr18-27731626-DEL-1906 | chr18-27731627-DEL-1906 |
| chr18-27732446-DEL-153 | chr18-27732446-DEL-153 |
| chr18-27820972-DEL-68 | chr18-27820941-DEL-56 |
| chr18-27841904-DEL-258 | chr18-27841905-DEL-258 |
| chr18-27864468-DEL-901 | chr18-27864469-DEL-901 |
| chr18-27897435-DEL-88 | chr18-27897440-DEL-84 |
| chr18-27903069-DEL-71 | chr18-27903070-DEL-71 |
| chr18-28018866-DEL-974 | chr18-28018867-DEL-974 |
| chr18-28028542-DEL-77 | chr18-28028539-DEL-75 |
| chr18-28093453-DEL-60 | chr18-28093454-DEL-60 |
| chr18-28124088-DEL-73 | chr18-28124089-DEL-73 |
| chr18-28142644-DEL-6436 | chr18-28142645-DEL-6436 |
| chr18-28213609-DEL-51 | chr18-28213610-DEL-52 |
| chr18-28234177-DEL-62 | chr18-28234186-DEL-58 |
| chr18-28236250-DEL-547 | chr18-28236251-DEL-547 |
| chr18-28242658-DEL-81 | chr18-28242659-DEL-81 |
| chr18-28245895-DEL-56 | chr18-28245896-DEL-56 |
| chr18-28284229-DEL-1688 | chr18-28284230-DEL-1688 |
| chr18-28291487-DEL-132 | chr18-28291488-DEL-132 |
| chr18-28333241-DEL-105 | chr18-28333242-DEL-105 |
| chr18-28338327-DEL-595 | chr18-28338331-DEL-592 |
| chr18-28339393-DEL-221 | chr18-28339394-DEL-221 |
| chr18-28367173-DEL-1539 | chr18-28367174-DEL-1539 |
| chr18-28368732-DEL-180 | chr18-28368733-DEL-180 |
| chr18-28384781-DEL-82 | chr18-28384782-DEL-88 |
| chr18-28390776-DEL-126 | chr18-28390753-DEL-150 |
| chr18-28436317-DEL-76 | chr18-28436314-DEL-64 |
| chr18-28445616-DEL-3085 | chr18-28445616-DEL-3085 |
| chr18-28447182-DEL-383 | chr18-28447183-DEL-383 |
| chr18-28472921-DEL-391 | chr18-28472922-DEL-391 |

| | |
|---|---|
| chr18-28474406-DEL-845 | chr18-28474406-DEL-845 |
| chr18-28481096-DEL-6802 | chr18-28481097-DEL-6802 |
| chr18-28496658-DEL-57 | chr18-28496660-DEL-66 |
| chr18-28503319-DEL-96 | chr18-28503302-DEL-141 |
| chr18-28518170-DEL-110 | chr18-28518171-DEL-108 |
| chr18-28533167-DEL-89 | chr18-28533140-DEL-90 |
| chr18-28541143-DEL-4802 | chr18-28541144-DEL-4802 |
| chr18-28557108-DEL-83 | chr18-28557109-DEL-83 |
| chr18-28574765-DEL-4796 | chr18-28574772-DEL-4790 |
| chr18-28591958-DEL-323 | chr18-28591959-DEL-323 |
| chr18-28615940-DEL-2641 | chr18-28615941-DEL-2641 |
| chr18-28631114-DEL-62 | chr18-28631115-DEL-62 |
| chr18-28647434-DEL-549 | chr18-28647435-DEL-549 |
| chr18-28661572-DEL-5341 | chr18-28661573-DEL-5341 |
| chr18-28681118-DEL-93 | chr18-28681082-DEL-90 |
| chr18-28704540-DEL-200 | chr18-28704541-DEL-200 |
| chr18-28720295-DEL-6620 | chr18-28720296-DEL-6620 |
| chr18-28732823-DEL-2794 | chr18-28732824-DEL-2794 |
| chr18-28732901-DEL-485 | chr18-28732902-DEL-485 |
| chr18-28765891-DEL-93 | chr18-28765892-DEL-91 |
| chr18-28808652-DEL-347 | chr18-28808657-DEL-343 |
| chr18-28878613-DEL-6456 | chr18-28878614-DEL-6456 |
| chr18-28891052-DEL-52 | chr18-28891048-DEL-53 |
| chr18-28916306-DEL-270 | chr18-28916307-DEL-270 |
| chr18-28929403-DEL-2643 | chr18-28929404-DEL-2643 |
| chr18-28929527-DEL-153 | chr18-28929536-DEL-145 |
| chr18-28940426-DEL-62 | chr18-28940407-DEL-64 |
| chr18-28946592-DEL-6163 | chr18-28946593-DEL-6163 |
| chr18-29042991-DEL-385 | chr18-29042992-DEL-385 |
| chr18-29068258-DEL-57 | chr18-29068258-DEL-57 |
| chr18-29093285-DEL-5454 | chr18-29093286-DEL-5454 |
| chr18-29108908-DEL-103 | chr18-29108909-DEL-103 |
| chr18-29109654-DEL-126 | chr18-29109631-DEL-126 |
| chr18-29138749-DEL-57 | chr18-29138754-DEL-53 |
| chr18-29146593-DEL-8503 | chr18-29146594-DEL-8503 |
| chr18-29209489-DEL-188 | chr18-29209489-DEL-188 |
| chr18-29213257-DEL-98 | chr18-29213239-DEL-114 |
| chr18-29215249-DEL-811 | chr18-29215250-DEL-811 |
| chr18-29221986-DEL-62 | chr18-29221953-DEL-65 |
| chr18-29227436-DEL-508 | chr18-29227437-DEL-508 |
| chr18-29228339-DEL-100 | chr18-29228320-DEL-120 |
| chr18-29230500-DEL-432 | chr18-29230501-DEL-432 |
| chr18-29244146-DEL-162 | chr18-29244147-DEL-161 |
| chr18-29246521-DEL-399 | chr18-29246522-DEL-399 |
| chr18-29263129-DEL-57 | chr18-29263130-DEL-56 |
| chr18-29283554-DEL-195 | chr18-29283554-DEL-195 |
| chr18-29285704-DEL-80 | chr18-29285705-DEL-80 |
| chr18-29297158-DEL-445 | chr18-29297159-DEL-445 |
| chr18-29304283-DEL-777 | chr18-29304284-DEL-777 |
| chr18-29305023-DEL-92 | chr18-29305024-DEL-92 |
| chr18-29325971-DEL-11550 | chr18-29325972-DEL-11550 |
| chr18-29358725-DEL-60 | chr18-29358714-DEL-54 |
| chr18-29370818-DEL-60 | chr18-29370819-DEL-60 |
| chr18-29376908-DEL-57 | chr18-29376909-DEL-57 |
| chr18-29378903-DEL-56 | chr18-29378904-DEL-56 |
| chr18-29389967-DEL-106 | chr18-29389968-DEL-106 |
| chr18-29394828-DEL-82 | chr18-29394831-DEL-80 |
| chr18-29408811-DEL-936 | chr18-29408812-DEL-936 |
| chr18-29420135-DEL-199 | chr18-29420136-DEL-199 |
| chr18-29456945-DEL-347 | chr18-29456946-DEL-347 |
| chr18-29459468-DEL-143 | chr18-29459434-DEL-178 |
| chr18-29534798-DEL-988 | chr18-29534799-DEL-987 |
| chr18-29582025-DEL-160 | chr18-29582020-DEL-192 |
| chr18-29601245-DEL-567 | chr18-29601245-DEL-567 |
| chr18-29608871-DEL-97 | chr18-29608878-DEL-91 |
| chr18-29615509-DEL-2201 | chr18-29615510-DEL-2201 |
| chr18-29622486-DEL-85 | chr18-29622476-DEL-102 |
| chr18-29626734-DEL-58 | chr18-29626735-DEL-58 |
| chr18-29629601-DEL-267 | chr18-29629597-DEL-268 |
| chr18-29648462-DEL-697 | chr18-29648464-DEL-696 |
| chr18-29686577-DEL-204 | chr18-29686577-DEL-204 |
| chr18-29690189-DEL-62 | chr18-29690190-DEL-62 |
| chr18-29792732-DEL-56 | chr18-29792751-DEL-62 |
| chr18-29823738-DEL-728 | chr18-29823739-DEL-728 |
| chr18-29846130-DEL-6262 | chr18-29846131-DEL-6262 |
| chr18-29872222-DEL-187 | chr18-29872223-DEL-186 |
| chr18-29929493-DEL-56 | chr18-29929494-DEL-56 |
| chr18-29954685-DEL-746 | chr18-29954686-DEL-746 |
| chr18-29976357-DEL-74 | chr18-29976347-DEL-90 |
| chr18-29989015-DEL-522 | chr18-29989013-DEL-521 |

| | |
|---|---|
| chr18-29989948-DEL-62 | chr18-29989951-DEL-60 |
| chr18-29991635-DEL-739 | chr18-29991636-DEL-739 |
| chr18-30003385-DEL-101 | chr18-30003422-DEL-65 |
| chr18-30004224-DEL-54 | chr18-30004224-DEL-54 |
| chr18-30008492-DEL-52 | chr18-30008493-DEL-52 |
| chr18-30015381-DEL-100 | chr18-30015357-DEL-125 |
| chr18-30044181-DEL-259 | chr18-30044182-DEL-259 |
| chr18-30103897-DEL-6692 | chr18-30103898-DEL-6692 |
| chr18-30135170-DEL-66 | chr18-30135128-DEL-60 |
| chr18-30139903-DEL-61 | chr18-30139902-DEL-110 |
| chr18-30144539-DEL-4554 | chr18-30144537-DEL-4554 |
| chr18-30153380-DEL-54 | chr18-30153381-DEL-54 |
| chr18-30154494-DEL-58 | chr18-30154495-DEL-58 |
| chr18-30182626-DEL-55 | chr18-30182626-DEL-55 |
| chr18-30239548-DEL-257 | chr18-30239543-DEL-255 |
| chr18-30263528-DEL-1171 | chr18-30263529-DEL-1171 |
| chr18-30311440-DEL-221 | chr18-30311441-DEL-221 |
| chr18-30329660-DEL-54 | chr18-30329655-DEL-70 |
| chr18-30333754-DEL-279 | chr18-30333755-DEL-279 |
| chr18-30344687-DEL-54 | chr18-30344688-DEL-54 |
| chr18-30373616-DEL-96 | chr18-30373617-DEL-96 |
| chr18-30399560-DEL-5693 | chr18-30399561-DEL-5693 |
| chr18-30424646-DEL-115 | chr18-30424660-DEL-87 |
| chr18-30445837-DEL-1938 | chr18-30445838-DEL-1938 |
| chr18-30481659-DEL-70 | chr18-30481660-DEL-71 |
| chr18-30485361-DEL-106 | chr18-30485362-DEL-106 |
| chr18-30500897-DEL-211 | chr18-30500898-DEL-211 |
| chr18-30509829-DEL-433 | chr18-30509830-DEL-433 |
| chr18-30535426-DEL-78 | chr18-30535427-DEL-72 |
| chr18-30545588-DEL-276 | chr18-30545589-DEL-276 |
| chr18-30550578-DEL-318 | chr18-30550579-DEL-318 |
| chr18-30552657-DEL-506 | chr18-30552658-DEL-506 |
| chr18-30554396-DEL-937 | chr18-30554397-DEL-937 |
| chr18-30566612-DEL-503 | chr18-30566613-DEL-504 |
| chr18-30576830-DEL-201 | chr18-30576785-DEL-224 |
| chr18-30599584-DEL-2067 | chr18-30599585-DEL-2067 |
| chr18-30617097-DEL-119 | chr18-30617098-DEL-119 |
| chr18-30635629-DEL-119 | chr18-30635630-DEL-119 |
| chr18-30638717-DEL-342 | chr18-30638718-DEL-342 |
| chr18-30691769-DEL-1707 | chr18-30691781-DEL-1694 |
| chr18-30697167-DEL-60 | chr18-30697168-DEL-60 |
| chr18-30723740-DEL-206 | chr18-30723741-DEL-206 |
| chr18-30724442-DEL-1128 | chr18-30724439-DEL-1122 |
| chr18-30758411-DEL-81 | chr18-30758403-DEL-59 |
| chr18-30798787-DEL-324 | chr18-30798788-DEL-324 |
| chr18-30811848-DEL-543 | chr18-30811846-DEL-543 |
| chr18-30815694-DEL-204 | chr18-30815697-DEL-203 |
| chr18-30818304-DEL-475 | chr18-30818305-DEL-475 |
| chr18-30826689-DEL-75 | chr18-30826693-DEL-71 |
| chr18-30840608-DEL-54 | chr18-30840608-DEL-54 |
| chr18-30849516-DEL-468 | chr18-30849517-DEL-468 |
| chr18-30871869-DEL-76 | chr18-30871907-DEL-60 |
| chr18-30958505-DEL-5831 | chr18-30958506-DEL-5831 |
| chr18-30971814-DEL-129 | chr18-30971813-DEL-130 |
| chr18-30995012-DEL-170 | chr18-30995013-DEL-170 |
| chr18-31027914-DEL-392 | chr18-31027915-DEL-392 |
| chr18-31041990-DEL-524 | chr18-31041981-DEL-534 |
| chr18-31058319-DEL-71 | chr18-31058320-DEL-71 |
| chr18-31087157-DEL-359 | chr18-31087158-DEL-359 |
| chr18-31091053-DEL-53 | chr18-31091053-DEL-53 |
| chr18-31118572-DEL-2873 | chr18-31118573-DEL-2873 |
| chr18-31130626-DEL-89 | chr18-31130649-DEL-65 |
| chr18-31165326-DEL-11059 | chr18-31165327-DEL-11059 |
| chr18-31165884-DEL-810 | chr18-31165887-DEL-811 |
| chr18-31167099-DEL-197 | chr18-31167100-DEL-197 |
| chr18-31213802-DEL-123 | chr18-31213802-DEL-124 |
| chr18-31218821-DEL-80 | chr18-31218825-DEL-92 |
| chr18-31238495-DEL-595 | chr18-31238496-DEL-595 |
| chr18-31245286-DEL-1104 | chr18-31245334-DEL-1056 |
| chr18-31259164-DEL-1005 | chr18-31259165-DEL-1005 |
| chr18-31326269-DEL-868 | chr18-31326270-DEL-868 |
| chr18-31329486-DEL-191 | chr18-31329491-DEL-187 |
| chr18-31337838-DEL-106 | chr18-31337839-DEL-106 |
| chr18-31344808-DEL-362 | chr18-31344809-DEL-362 |
| chr18-31348108-DEL-58 | chr18-31348109-DEL-58 |
| chr18-31377225-DEL-1550 | chr18-31377225-DEL-1551 |
| chr18-31487538-DEL-6577 | chr18-31487539-DEL-6577 |
| chr18-31501004-DEL-4075 | chr18-31501005-DEL-4075 |
| chr18-31509875-DEL-201 | chr18-31509876-DEL-200 |
| chr18-31512529-DEL-1009 | chr18-31512530-DEL-1009 |

| | |
|---|---|
| chr18-31517830-DEL-203 | chr18-31517830-DEL-203 |
| chr18-31528553-DEL-144 | chr18-31528554-DEL-143 |
| chr18-31537666-DEL-55 | chr18-31537667-DEL-55 |
| chr18-31605912-DEL-358 | chr18-31605913-DEL-358 |
| chr18-31625276-DEL-97 | chr18-31625323-DEL-63 |
| chr18-31634751-DEL-198 | chr18-31634752-DEL-198 |
| chr18-31640736-DEL-638 | chr18-31640737-DEL-638 |
| chr18-31652532-DEL-76 | chr18-31652526-DEL-79 |
| chr18-31666623-DEL-1600 | chr18-31666607-DEL-1600 |
| chr18-31668918-DEL-2131 | chr18-31668919-DEL-2131 |
| chr18-31725675-DEL-192 | chr18-31725676-DEL-192 |
| chr18-31747088-DEL-51 | chr18-31747089-DEL-51 |
| chr18-31749173-DEL-2576 | chr18-31749174-DEL-2576 |
| chr18-31778073-DEL-52 | chr18-31778070-DEL-56 |
| chr18-31790872-DEL-592 | chr18-31790875-DEL-590 |
| chr18-31826438-DEL-475 | chr18-31826438-DEL-475 |
| chr18-31828619-DEL-74 | chr18-31828620-DEL-74 |
| chr18-31839172-DEL-1698 | chr18-31839173-DEL-1698 |
| chr18-31848920-DEL-3693 | chr18-31848921-DEL-3693 |
| chr18-31856463-DEL-61 | chr18-31856448-DEL-50 |
| chr18-31869852-DEL-176 | chr18-31869852-DEL-176 |
| chr18-31873967-DEL-198 | chr18-31873968-DEL-193 |
| chr18-31875999-DEL-116 | chr18-31875999-DEL-116 |
| chr18-31885310-DEL-199 | chr18-31885331-DEL-198 |
| chr18-31887366-DEL-380 | chr18-31887367-DEL-380 |
| chr18-31908467-DEL-51 | chr18-31908448-DEL-90 |
| chr18-31954243-DEL-298 | chr18-31954244-DEL-298 |
| chr18-31973133-DEL-61 | chr18-31973134-DEL-61 |
| chr18-31988432-DEL-64 | chr18-31988436-DEL-59 |
| chr18-31994075-DEL-142 | chr18-31994042-DEL-143 |
| chr18-32029077-DEL-204 | chr18-32029078-DEL-204 |
| chr18-32157281-DEL-58 | chr18-32157237-DEL-87 |
| chr18-32158922-DEL-498 | chr18-32158923-DEL-498 |
| chr18-32165569-DEL-99 | chr18-32165570-DEL-99 |
| chr18-32173428-DEL-743 | chr18-32173416-DEL-739 |
| chr18-32175358-DEL-9172 | chr18-32175359-DEL-9172 |
| chr18-32205792-DEL-436 | chr18-32205793-DEL-436 |
| chr18-32267227-DEL-99 | chr18-32267221-DEL-100 |
| chr18-32277618-DEL-56 | chr18-32277586-DEL-72 |
| chr18-32290369-DEL-55 | chr18-32290370-DEL-55 |
| chr18-32314405-DEL-336 | chr18-32314406-DEL-336 |
| chr18-32352261-DEL-70 | chr18-32352254-DEL-72 |
| chr18-32396743-DEL-268 | chr18-32396744-DEL-268 |
| chr18-32404513-DEL-229 | chr18-32404514-DEL-229 |
| chr18-32417879-DEL-223 | chr18-32417880-DEL-223 |
| chr18-32418580-DEL-86 | chr18-32418581-DEL-86 |
| chr18-32422280-DEL-53 | chr18-32422281-DEL-53 |
| chr18-32423803-DEL-75 | chr18-32423804-DEL-75 |
| chr18-32460285-DEL-201 | chr18-32460276-DEL-202 |
| chr18-32490736-DEL-51 | chr18-32490736-DEL-52 |
| chr18-32536618-DEL-219 | chr18-32536617-DEL-220 |
| chr18-32558691-DEL-129 | chr18-32558692-DEL-129 |
| chr18-32572280-DEL-133 | chr18-32572281-DEL-133 |
| chr18-32595870-DEL-245 | chr18-32595871-DEL-245 |
| chr18-32601278-DEL-4634 | chr18-32601278-DEL-4632 |
| chr18-32621296-DEL-266 | chr18-32621297-DEL-266 |
| chr18-32628755-DEL-1246 | chr18-32628756-DEL-1246 |
| chr18-32635480-DEL-245 | chr18-32635481-DEL-245 |
| chr18-32645609-DEL-90 | chr18-32645610-DEL-90 |
| chr18-32651106-DEL-59 | chr18-32651106-DEL-70 |
| chr18-32652518-DEL-436 | chr18-32652519-DEL-436 |
| chr18-32679838-DEL-67 | chr18-32679839-DEL-67 |
| chr18-32688107-DEL-59 | chr18-32688108-DEL-59 |
| chr18-32707535-DEL-239 | chr18-32707536-DEL-239 |
| chr18-32711129-DEL-6432 | chr18-32711130-DEL-6432 |
| chr18-32730271-DEL-213 | chr18-32730272-DEL-213 |
| chr18-32731750-DEL-118 | chr18-32731767-DEL-102 |
| chr18-32781803-DEL-54 | chr18-32781804-DEL-57 |
| chr18-32790614-DEL-596 | chr18-32790614-DEL-597 |
| chr18-32793077-DEL-1254 | chr18-32793077-DEL-1254 |
| chr18-32836132-DEL-54 | chr18-32836098-DEL-88 |
| chr18-32901265-DEL-93 | chr18-32901266-DEL-93 |
| chr18-32924190-DEL-928 | chr18-32924191-DEL-928 |
| chr18-32928547-DEL-65 | chr18-32928544-DEL-68 |
| chr18-32943195-DEL-1122 | chr18-32943196-DEL-1122 |
| chr18-32959826-DEL-340 | chr18-32959828-DEL-341 |
| chr18-32966809-DEL-240 | chr18-32966810-DEL-240 |
| chr18-32992748-DEL-62 | chr18-32992749-DEL-62 |
| chr18-32997548-DEL-106 | chr18-32997549-DEL-106 |
| chr18-33010225-DEL-340 | chr18-33010226-DEL-340 |

| | |
|---|---|
| chr18-33036769-DEL-563 | chr18-33036766-DEL-558 |
| chr18-33109720-DEL-1938 | chr18-33109721-DEL-1938 |
| chr18-33112208-DEL-2570 | chr18-33112209-DEL-2570 |
| chr18-33115353-DEL-498 | chr18-33115354-DEL-498 |
| chr18-33211849-DEL-181 | chr18-33211850-DEL-181 |
| chr18-33219831-DEL-711 | chr18-33219832-DEL-711 |
| chr18-33224085-DEL-412 | chr18-33224086-DEL-412 |
| chr18-33276992-DEL-143 | chr18-33276993-DEL-143 |
| chr18-33352629-DEL-1831 | chr18-33352630-DEL-1831 |
| chr18-33396921-DEL-65 | chr18-33396922-DEL-65 |
| chr18-33419804-DEL-1151 | chr18-33419789-DEL-1152 |
| chr18-33434997-DEL-112 | chr18-33434998-DEL-112 |
| chr18-33439809-DEL-6175 | chr18-33439810-DEL-6175 |
| chr18-33462969-DEL-214 | chr18-33462961-DEL-214 |
| chr18-33500672-DEL-53 | chr18-33500673-DEL-53 |
| chr18-33505219-DEL-55 | chr18-33505220-DEL-55 |
| chr18-33508640-DEL-343 | chr18-33508641-DEL-343 |
| chr18-33523767-DEL-182 | chr18-33523768-DEL-182 |
| chr18-33527935-DEL-6707 | chr18-33527936-DEL-6707 |
| chr18-33556774-DEL-109 | chr18-33556775-DEL-109 |
| chr18-33601661-DEL-1066 | chr18-33601662-DEL-1066 |
| chr18-33605214-DEL-110 | chr18-33605255-DEL-67 |
| chr18-33615286-DEL-68 | chr18-33615287-DEL-66 |
| chr18-33619940-DEL-290 | chr18-33619941-DEL-290 |
| chr18-33621324-DEL-55 | chr18-33621325-DEL-55 |
| chr18-33637373-DEL-55 | chr18-33637372-DEL-55 |
| chr18-33731580-DEL-51 | chr18-33731581-DEL-51 |
| chr18-33736287-DEL-504 | chr18-33736288-DEL-504 |
| chr18-33743495-DEL-191 | chr18-33743496-DEL-191 |
| chr18-33761222-DEL-467 | chr18-33761223-DEL-467 |
| chr18-33762704-DEL-55 | chr18-33762705-DEL-55 |
| chr18-33763327-DEL-385 | chr18-33763328-DEL-383 |
| chr18-33768165-DEL-6431 | chr18-33768166-DEL-6431 |
| chr18-33793313-DEL-58 | chr18-33793314-DEL-58 |
| chr18-33830465-DEL-6454 | chr18-33830466-DEL-6454 |
| chr18-33854661-DEL-1022 | chr18-33854662-DEL-1022 |
| chr18-33857862-DEL-102 | chr18-33857863-DEL-102 |
| chr18-33859420-DEL-88 | chr18-33859421-DEL-88 |
| chr18-33863331-DEL-139 | chr18-33863352-DEL-118 |
| chr18-33888347-DEL-88 | chr18-33888348-DEL-88 |
| chr18-33902470-DEL-134 | chr18-33902443-DEL-135 |
| chr18-33921347-DEL-55 | chr18-33921352-DEL-53 |
| chr18-33926690-DEL-187 | chr18-33926694-DEL-184 |
| chr18-33932501-DEL-78 | chr18-33932510-DEL-60 |
| chr18-33975224-DEL-82 | chr18-33975225-DEL-82 |
| chr18-33985742-DEL-54 | chr18-33985743-DEL-54 |
| chr18-33992928-DEL-128 | chr18-33992929-DEL-128 |
| chr18-33994407-DEL-89 | chr18-33994408-DEL-89 |
| chr18-34003256-DEL-323 | chr18-34003206-DEL-331 |
| chr18-34015313-DEL-96 | chr18-34015314-DEL-96 |
| chr18-34016065-DEL-102 | chr18-34016066-DEL-102 |
| chr18-34017494-DEL-407 | chr18-34017494-DEL-394 |
| chr18-34038232-DEL-88 | chr18-34038225-DEL-89 |
| chr18-34043733-DEL-56 | chr18-34043734-DEL-56 |
| chr18-34046096-DEL-112 | chr18-34046089-DEL-120 |
| chr18-34108667-DEL-124 | chr18-34108667-DEL-124 |
| chr18-34116110-DEL-199 | chr18-34116111-DEL-199 |
| chr18-34127921-DEL-450 | chr18-34127924-DEL-444 |
| chr18-34154962-DEL-576 | chr18-34154963-DEL-576 |
| chr18-34159144-DEL-268 | chr18-34159141-DEL-272 |
| chr18-34162401-DEL-435 | chr18-34162402-DEL-435 |
| chr18-34171917-DEL-266 | chr18-34171918-DEL-266 |
| chr18-34174086-DEL-112 | chr18-34174107-DEL-109 |
| chr18-34194457-DEL-55 | chr18-34194458-DEL-55 |
| chr18-34195034-DEL-228 | chr18-34195035-DEL-228 |
| chr18-34203989-DEL-220 | chr18-34203990-DEL-220 |
| chr18-34205212-DEL-207 | chr18-34205213-DEL-208 |
| chr18-34205446-DEL-961 | chr18-34205447-DEL-961 |
| chr18-34216729-DEL-4128 | chr18-34216730-DEL-4128 |
| chr18-34277540-DEL-197 | chr18-34277541-DEL-197 |
| chr18-34316413-DEL-309 | chr18-34316414-DEL-305 |
| chr18-34349018-DEL-529 | chr18-34349019-DEL-529 |
| chr18-34359729-DEL-1778 | chr18-34359730-DEL-1777 |
| chr18-34406940-DEL-455 | chr18-34406941-DEL-455 |
| chr18-34407727-DEL-512 | chr18-34407728-DEL-512 |
| chr18-34408325-DEL-495 | chr18-34408326-DEL-495 |
| chr18-34412532-DEL-88 | chr18-34412533-DEL-88 |
| chr18-34417121-DEL-219 | chr18-34417122-DEL-219 |
| chr18-34420470-DEL-69 | chr18-34420471-DEL-69 |
| chr18-34440109-DEL-66 | chr18-34440136-DEL-50 |

| | |
|---|---|
| chr18-34488893-DEL-255 | chr18-34488940-DEL-208 |
| chr18-34499665-DEL-120 | chr18-34499666-DEL-120 |
| chr18-34521643-DEL-69 | chr18-34521644-DEL-69 |
| chr18-34524324-DEL-1056 | chr18-34524325-DEL-1056 |
| chr18-34525635-DEL-202 | chr18-34525636-DEL-202 |
| chr18-34530633-DEL-829 | chr18-34530635-DEL-828 |
| chr18-34551155-DEL-62 | chr18-34551113-DEL-62 |
| chr18-34603815-DEL-422 | chr18-34603816-DEL-422 |
| chr18-34611504-DEL-153 | chr18-34611505-DEL-153 |
| chr18-34647731-DEL-205 | chr18-34647732-DEL-205 |
| chr18-34662033-DEL-2335 | chr18-34662034-DEL-2335 |
| chr18-34667740-DEL-207 | chr18-34667741-DEL-207 |
| chr18-34685397-DEL-83 | chr18-34685398-DEL-83 |
| chr18-34686484-DEL-212 | chr18-34686485-DEL-212 |
| chr18-34689799-DEL-184 | chr18-34689800-DEL-184 |
| chr18-34690103-DEL-594 | chr18-34690103-DEL-594 |
| chr18-34701595-DEL-56 | chr18-34701587-DEL-71 |
| chr18-34705289-DEL-230 | chr18-34705290-DEL-230 |
| chr18-34708727-DEL-205 | chr18-34708728-DEL-205 |
| chr18-34732618-DEL-215 | chr18-34732619-DEL-215 |
| chr18-34746503-DEL-351 | chr18-34746504-DEL-351 |
| chr18-34751961-DEL-58 | chr18-34751962-DEL-58 |
| chr18-34752920-DEL-513 | chr18-34752921-DEL-513 |
| chr18-34763881-DEL-84 | chr18-34763882-DEL-84 |
| chr18-34768859-DEL-222 | chr18-34768860-DEL-222 |
| chr18-34802111-DEL-52 | chr18-34802132-DEL-52 |
| chr18-34807037-DEL-205 | chr18-34807038-DEL-205 |
| chr18-34834214-DEL-736 | chr18-34834215-DEL-736 |
| chr18-34852287-DEL-56 | chr18-34852288-DEL-56 |
| chr18-34862633-DEL-177 | chr18-34862634-DEL-177 |
| chr18-34870379-DEL-711 | chr18-34870406-DEL-699 |
| chr18-34891061-DEL-180 | chr18-34891062-DEL-180 |
| chr18-34918212-DEL-185 | chr18-34918213-DEL-185 |
| chr18-34931477-DEL-208 | chr18-34931478-DEL-208 |
| chr18-34953251-DEL-718 | chr18-34953252-DEL-718 |
| chr18-34959654-DEL-339 | chr18-34959655-DEL-339 |
| chr18-35009087-DEL-653 | chr18-35009088-DEL-653 |
| chr18-35014253-DEL-214 | chr18-35014254-DEL-214 |
| chr18-35024989-DEL-91 | chr18-35024990-DEL-91 |
| chr18-35027865-DEL-68 | chr18-35027866-DEL-80 |
| chr18-35067729-DEL-202 | chr18-35067730-DEL-202 |
| chr18-35096789-DEL-124 | chr18-35096790-DEL-124 |
| chr18-35122683-DEL-132 | chr18-35122687-DEL-103 |
| chr18-35133647-DEL-75 | chr18-35133621-DEL-74 |
| chr18-35145867-DEL-231 | chr18-35145868-DEL-231 |
| chr18-35173600-DEL-440 | chr18-35173601-DEL-440 |
| chr18-35182798-DEL-244 | chr18-35182799-DEL-244 |
| chr18-35191542-DEL-942 | chr18-35191543-DEL-942 |
| chr18-35213274-DEL-252 | chr18-35213275-DEL-252 |
| chr18-35229775-DEL-62 | chr18-35229792-DEL-62 |
| chr18-35260893-DEL-169 | chr18-35260894-DEL-169 |
| chr18-35270466-DEL-54 | chr18-35270467-DEL-54 |
| chr18-35299896-DEL-174 | chr18-35299891-DEL-180 |
| chr18-35304712-DEL-187 | chr18-35304713-DEL-187 |
| chr18-35320471-DEL-497 | chr18-35320475-DEL-496 |
| chr18-35322443-DEL-217 | chr18-35322444-DEL-217 |
| chr18-35378355-DEL-994 | chr18-35378356-DEL-994 |
| chr18-35412358-DEL-64 | chr18-35412359-DEL-64 |
| chr18-35440323-DEL-174 | chr18-35440324-DEL-174 |
| chr18-35451497-DEL-1890 | chr18-35451498-DEL-1890 |
| chr18-35464539-DEL-152 | chr18-35464540-DEL-152 |
| chr18-35499132-DEL-52 | chr18-35499133-DEL-52 |
| chr18-35562605-DEL-205 | chr18-35562606-DEL-205 |
| chr18-35570728-DEL-331 | chr18-35570755-DEL-305 |
| chr18-35583793-DEL-58 | chr18-35583794-DEL-58 |
| chr18-35599322-DEL-221 | chr18-35599323-DEL-221 |
| chr18-35605243-DEL-153 | chr18-35605244-DEL-153 |
| chr18-35624618-DEL-941 | chr18-35624619-DEL-941 |
| chr18-35726067-DEL-225 | chr18-35726068-DEL-225 |
| chr18-35761482-DEL-904 | chr18-35761483-DEL-904 |
| chr18-35772709-DEL-211 | chr18-35772709-DEL-200 |
| chr18-35777622-DEL-207 | chr18-35777648-DEL-168 |
| chr18-35792247-DEL-120 | chr18-35792248-DEL-120 |
| chr18-35796468-DEL-51 | chr18-35796468-DEL-51 |
| chr18-35800703-DEL-298 | chr18-35800704-DEL-298 |
| chr18-35821601-DEL-201 | chr18-35821602-DEL-201 |
| chr18-35824270-DEL-61 | chr18-35824271-DEL-61 |
| chr18-35831418-DEL-66 | chr18-35831391-DEL-70 |
| chr18-35890309-DEL-212 | chr18-35890310-DEL-212 |
| chr18-35896142-DEL-52 | chr18-35896143-DEL-52 |

| | |
|---|---|
| chr18-35897294-DEL-210 | chr18-35897279-DEL-214 |
| chr18-35927898-DEL-215 | chr18-35927899-DEL-215 |
| chr18-36040034-DEL-119 | chr18-36040035-DEL-119 |
| chr18-36047340-DEL-51 | chr18-36047341-DEL-51 |
| chr18-36057900-DEL-202 | chr18-36057901-DEL-202 |
| chr18-36061132-DEL-90 | chr18-36061134-DEL-89 |
| chr18-36068632-DEL-218 | chr18-36068633-DEL-218 |
| chr18-36073657-DEL-87 | chr18-36073658-DEL-78 |
| chr18-36079750-DEL-60 | chr18-36079751-DEL-60 |
| chr18-36115507-DEL-55 | chr18-36115511-DEL-58 |
| chr18-36137075-DEL-63 | chr18-36137074-DEL-65 |
| chr18-36170376-DEL-188 | chr18-36170377-DEL-188 |
| chr18-36223453-DEL-214 | chr18-36223454-DEL-214 |
| chr18-36265306-DEL-92 | chr18-36265307-DEL-92 |
| chr18-36349133-DEL-51 | chr18-36349124-DEL-93 |
| chr18-36359442-DEL-85 | chr18-36359440-DEL-87 |
| chr18-36367343-DEL-132 | chr18-36367347-DEL-129 |
| chr18-36389652-DEL-73 | chr18-36389653-DEL-57 |
| chr18-36462697-DEL-63 | chr18-36462692-DEL-91 |
| chr18-36471584-DEL-208 | chr18-36471585-DEL-208 |
| chr18-36526804-DEL-185 | chr18-36526805-DEL-185 |
| chr18-36558795-DEL-277 | chr18-36558796-DEL-277 |
| chr18-36572353-DEL-797 | chr18-36572354-DEL-797 |
| chr18-36607027-DEL-7124 | chr18-36607028-DEL-7124 |
| chr18-36629229-DEL-61 | chr18-36629230-DEL-61 |
| chr18-36637451-DEL-55 | chr18-36637452-DEL-55 |
| chr18-36653787-DEL-90 | chr18-36653769-DEL-138 |
| chr18-36670556-DEL-453 | chr18-36670597-DEL-433 |
| chr18-36681181-DEL-53 | chr18-36681188-DEL-67 |
| chr18-36686867-DEL-64 | chr18-36686868-DEL-64 |
| chr18-36695926-DEL-274 | chr18-36695927-DEL-274 |
| chr18-36778852-DEL-66 | chr18-36778853-DEL-66 |
| chr18-36806630-DEL-68 | chr18-36806631-DEL-68 |
| chr18-36845488-DEL-269 | chr18-36845489-DEL-269 |
| chr18-36850402-DEL-202 | chr18-36850403-DEL-202 |
| chr18-36852167-DEL-54 | chr18-36852168-DEL-56 |
| chr18-36861828-DEL-55 | chr18-36861829-DEL-55 |
| chr18-36873103-DEL-235 | chr18-36873104-DEL-235 |
| chr18-36878889-DEL-69 | chr18-36878890-DEL-69 |
| chr18-36918166-DEL-202 | chr18-36918167-DEL-202 |
| chr18-36940236-DEL-334 | chr18-36940237-DEL-334 |
| chr18-36941788-DEL-104 | chr18-36941803-DEL-86 |
| chr18-36947559-DEL-208 | chr18-36947560-DEL-208 |
| chr18-36965993-DEL-2008 | chr18-36965992-DEL-2009 |
| chr18-36969004-DEL-213 | chr18-36969005-DEL-213 |
| chr18-36972222-DEL-268 | chr18-36972223-DEL-268 |
| chr18-36990304-DEL-4332 | chr18-36990305-DEL-4332 |
| chr18-36991598-DEL-222 | chr18-36991599-DEL-222 |
| chr18-37021896-DEL-347 | chr18-37021897-DEL-347 |
| chr18-37023003-DEL-365 | chr18-37023004-DEL-365 |
| chr18-37024636-DEL-76 | chr18-37024637-DEL-76 |
| chr18-37032177-DEL-456 | chr18-37032178-DEL-456 |
| chr18-37038936-DEL-793 | chr18-37038940-DEL-790 |
| chr18-37039839-DEL-111 | chr18-37039840-DEL-111 |
| chr18-37049187-DEL-68 | chr18-37049188-DEL-62 |
| chr18-37088355-DEL-69 | chr18-37088356-DEL-69 |
| chr18-37098637-DEL-89 | chr18-37098633-DEL-90 |
| chr18-37114576-DEL-7128 | chr18-37114577-DEL-7128 |
| chr18-37155937-DEL-1112 | chr18-37155937-DEL-1113 |
| chr18-37164855-DEL-6476 | chr18-37164856-DEL-6476 |
| chr18-37179112-DEL-202 | chr18-37179113-DEL-202 |
| chr18-37243135-DEL-6452 | chr18-37243136-DEL-6452 |
| chr18-37406004-DEL-200 | chr18-37406005-DEL-200 |
| chr18-37409141-DEL-2539 | chr18-37409141-DEL-2539 |
| chr18-37413938-DEL-407 | chr18-37413938-DEL-407 |
| chr18-37418361-DEL-995 | chr18-37418362-DEL-995 |
| chr18-37418566-DEL-118 | chr18-37418604-DEL-74 |
| chr18-37425892-DEL-77 | chr18-37425888-DEL-82 |
| chr18-37437164-DEL-106 | chr18-37437165-DEL-106 |
| chr18-37463003-DEL-510 | chr18-37463004-DEL-508 |
| chr18-37497141-DEL-7225 | chr18-37497142-DEL-7225 |
| chr18-37511753-DEL-290 | chr18-37511754-DEL-290 |
| chr18-37519478-DEL-204 | chr18-37519479-DEL-204 |
| chr18-37532946-DEL-53 | chr18-37532953-DEL-75 |
| chr18-37540213-DEL-1466 | chr18-37540214-DEL-1466 |
| chr18-37565605-DEL-625 | chr18-37565608-DEL-623 |
| chr18-37636473-DEL-53 | chr18-37636474-DEL-53 |
| chr18-37654868-DEL-100 | chr18-37654917-DEL-70 |
| chr18-37753258-DEL-584 | chr18-37753245-DEL-584 |
| chr18-37753275-DEL-54 | chr18-37753275-DEL-54 |

| | |
|---|---|
| chr18-37763250-DEL-6964 | chr18-37763251-DEL-6964 |
| chr18-37763366-DEL-212 | chr18-37763367-DEL-212 |
| chr18-37785421-DEL-623 | chr18-37785441-DEL-603 |
| chr18-37798339-DEL-915 | chr18-37798340-DEL-915 |
| chr18-37828370-DEL-145 | chr18-37828371-DEL-145 |
| chr18-37852907-DEL-60 | chr18-37852896-DEL-72 |
| chr18-37879372-DEL-53 | chr18-37879373-DEL-53 |
| chr18-37892767-DEL-96 | chr18-37892774-DEL-87 |
| chr18-37893849-DEL-347 | chr18-37893850-DEL-347 |
| chr18-37911327-DEL-201 | chr18-37911314-DEL-202 |
| chr18-37919955-DEL-89 | chr18-37919956-DEL-89 |
| chr18-37961274-DEL-53 | chr18-37961274-DEL-53 |
| chr18-38032868-DEL-239 | chr18-38032889-DEL-216 |
| chr18-38085140-DEL-210 | chr18-38085141-DEL-210 |
| chr18-38109530-DEL-55 | chr18-38109503-DEL-83 |
| chr18-38247245-DEL-54 | chr18-38247246-DEL-54 |
| chr18-38258887-DEL-54 | chr18-38258888-DEL-58 |
| chr18-38263440-DEL-427 | chr18-38263430-DEL-425 |
| chr18-38271098-DEL-1890 | chr18-38271097-DEL-1891 |
| chr18-38288709-DEL-148 | chr18-38288710-DEL-148 |
| chr18-38369444-DEL-144 | chr18-38369410-DEL-144 |
| chr18-38370568-DEL-209 | chr18-38370569-DEL-209 |
| chr18-38394359-DEL-114 | chr18-38394360-DEL-114 |
| chr18-38403725-DEL-458 | chr18-38403726-DEL-458 |
| chr18-38404922-DEL-84 | chr18-38404913-DEL-66 |
| chr18-38423487-DEL-93 | chr18-38423487-DEL-93 |
| chr18-38445131-DEL-86 | chr18-38445133-DEL-85 |
| chr18-38457693-DEL-90 | chr18-38457703-DEL-75 |
| chr18-38476018-DEL-207 | chr18-38476019-DEL-207 |
| chr18-38479193-DEL-353 | chr18-38479194-DEL-353 |
| chr18-38507113-DEL-892 | chr18-38507112-DEL-893 |
| chr18-38520680-DEL-60 | chr18-38520681-DEL-66 |
| chr18-38561451-DEL-493 | chr18-38561452-DEL-493 |
| chr18-38602635-DEL-194 | chr18-38602636-DEL-194 |
| chr18-38609659-DEL-228 | chr18-38609660-DEL-228 |
| chr18-38666406-DEL-53 | chr18-38666407-DEL-53 |
| chr18-38668053-DEL-214 | chr18-38668054-DEL-214 |
| chr18-38670806-DEL-209 | chr18-38670807-DEL-209 |
| chr18-38673321-DEL-52 | chr18-38673304-DEL-54 |
| chr18-38693859-DEL-99 | chr18-38693868-DEL-74 |
| chr18-38707607-DEL-85 | chr18-38707608-DEL-85 |
| chr18-38708719-DEL-169 | chr18-38708720-DEL-169 |
| chr18-38737368-DEL-153 | chr18-38737365-DEL-157 |
| chr18-38763444-DEL-156 | chr18-38763445-DEL-156 |
| chr18-38786293-DEL-135 | chr18-38786294-DEL-135 |
| chr18-38792292-DEL-101 | chr18-38792271-DEL-123 |
| chr18-38794672-DEL-54 | chr18-38794656-DEL-54 |
| chr18-38796512-DEL-1723 | chr18-38796511-DEL-1724 |
| chr18-38810529-DEL-52 | chr18-38810532-DEL-50 |
| chr18-38816528-DEL-212 | chr18-38816529-DEL-212 |
| chr18-38838654-DEL-87 | chr18-38838611-DEL-88 |
| chr18-38839474-DEL-192 | chr18-38839475-DEL-192 |
| chr18-38877137-DEL-233 | chr18-38877138-DEL-233 |
| chr18-38954321-DEL-66 | chr18-38954332-DEL-56 |
| chr18-38976304-DEL-498 | chr18-38976305-DEL-498 |
| chr18-38993511-DEL-576 | chr18-38993512-DEL-576 |
| chr18-39037951-DEL-194 | chr18-39037952-DEL-194 |
| chr18-39040258-DEL-87 | chr18-39040259-DEL-87 |
| chr18-39057189-DEL-85 | chr18-39057190-DEL-85 |
| chr18-39060381-DEL-55 | chr18-39060382-DEL-55 |
| chr18-39153062-DEL-768 | chr18-39153063-DEL-768 |
| chr18-39228750-DEL-183 | chr18-39228751-DEL-183 |
| chr18-39256875-DEL-72 | chr18-39256875-DEL-76 |
| chr18-39273788-DEL-2613 | chr18-39273789-DEL-2613 |
| chr18-39284479-DEL-72 | chr18-39284480-DEL-72 |
| chr18-39312094-DEL-82 | chr18-39312094-DEL-82 |
| chr18-39353298-DEL-829 | chr18-39353300-DEL-825 |
| chr18-39360913-DEL-213 | chr18-39360914-DEL-213 |
| chr18-39369510-DEL-64 | chr18-39369511-DEL-64 |
| chr18-39411158-DEL-70 | chr18-39411159-DEL-70 |
| chr18-39465723-DEL-95 | chr18-39465723-DEL-52 |
| chr18-39536443-DEL-301 | chr18-39536444-DEL-301 |
| chr18-39611515-DEL-114 | chr18-39611516-DEL-114 |
| chr18-39633142-DEL-51 | chr18-39633112-DEL-84 |
| chr18-39645629-DEL-64 | chr18-39645676-DEL-60 |
| chr18-39683018-DEL-79 | chr18-39683045-DEL-64 |
| chr18-39731757-DEL-6400 | chr18-39731758-DEL-6400 |
| chr18-39739813-DEL-56 | chr18-39739816-DEL-54 |
| chr18-39770312-DEL-383 | chr18-39770313-DEL-383 |
| chr18-39801993-DEL-96 | chr18-39802004-DEL-96 |

| | |
|---|---|
| chr18-39817188-DEL-455 | chr18-39817189-DEL-455 |
| chr18-39872842-DEL-62 | chr18-39872843-DEL-62 |
| chr18-39913694-DEL-75 | chr18-39913695-DEL-75 |
| chr18-40063616-DEL-84 | chr18-40063622-DEL-72 |
| chr18-40068863-DEL-70 | chr18-40068828-DEL-119 |
| chr18-40075056-DEL-142 | chr18-40075073-DEL-102 |
| chr18-40144530-DEL-1245 | chr18-40144530-DEL-1245 |
| chr18-40161755-DEL-421 | chr18-40161754-DEL-422 |
| chr18-40229808-DEL-1379 | chr18-40229809-DEL-1379 |
| chr18-40237406-DEL-447 | chr18-40237407-DEL-447 |
| chr18-40238217-DEL-279 | chr18-40238218-DEL-279 |
| chr18-40288272-DEL-117 | chr18-40288273-DEL-117 |
| chr18-40297283-DEL-6608 | chr18-40297284-DEL-6605 |
| chr18-40319921-DEL-192 | chr18-40319922-DEL-192 |
| chr18-40331071-DEL-202 | chr18-40331072-DEL-202 |
| chr18-40377355-DEL-424 | chr18-40377352-DEL-425 |
| chr18-40401000-DEL-320 | chr18-40401006-DEL-315 |
| chr18-40475817-DEL-572 | chr18-40475818-DEL-572 |
| chr18-40487452-DEL-806 | chr18-40487453-DEL-806 |
| chr18-40515212-DEL-648 | chr18-40515213-DEL-648 |
| chr18-40524299-DEL-114 | chr18-40524296-DEL-117 |
| chr18-40526779-DEL-638 | chr18-40526780-DEL-638 |
| chr18-40535676-DEL-631 | chr18-40535677-DEL-631 |
| chr18-40550985-DEL-781 | chr18-40550986-DEL-781 |
| chr18-40592842-DEL-2172 | chr18-40592843-DEL-2172 |
| chr18-40638655-DEL-210 | chr18-40638656-DEL-210 |
| chr18-40734821-DEL-6178 | chr18-40734822-DEL-6178 |
| chr18-40749040-DEL-213 | chr18-40749037-DEL-214 |
| chr18-40783468-DEL-583 | chr18-40783468-DEL-583 |
| chr18-40783890-DEL-2560 | chr18-40783891-DEL-2560 |
| chr18-40793952-DEL-2624 | chr18-40793953-DEL-2624 |
| chr18-40816955-DEL-540 | chr18-40816956-DEL-540 |
| chr18-40841670-DEL-4990 | chr18-40841671-DEL-4990 |
| chr18-40857916-DEL-66 | chr18-40857909-DEL-74 |
| chr18-40885430-DEL-1410 | chr18-40885428-DEL-1363 |
| chr18-40895588-DEL-3685 | chr18-40895587-DEL-3685 |
| chr18-40923931-DEL-69 | chr18-40923932-DEL-71 |
| chr18-40925107-DEL-728 | chr18-40925108-DEL-728 |
| chr18-40942571-DEL-94 | chr18-40942542-DEL-124 |
| chr18-40993484-DEL-4944 | chr18-40993484-DEL-4944 |
| chr18-40999988-DEL-193 | chr18-40999989-DEL-193 |
| chr18-41002615-DEL-92 | chr18-41002620-DEL-88 |
| chr18-41025023-DEL-483 | chr18-41025020-DEL-483 |
| chr18-41035506-DEL-1285 | chr18-41035503-DEL-1286 |
| chr18-41050685-DEL-190 | chr18-41050686-DEL-190 |
| chr18-41086695-DEL-218 | chr18-41086696-DEL-218 |
| chr18-41112850-DEL-502 | chr18-41112851-DEL-502 |
| chr18-41140749-DEL-90 | chr18-41140751-DEL-75 |
| chr18-41171420-DEL-72 | chr18-41171429-DEL-69 |
| chr18-41196792-DEL-325 | chr18-41196811-DEL-308 |
| chr18-41253092-DEL-56 | chr18-41253093-DEL-56 |
| chr18-41312033-DEL-62 | chr18-41312015-DEL-60 |
| chr18-41319734-DEL-60 | chr18-41319752-DEL-53 |
| chr18-41327284-DEL-310 | chr18-41327285-DEL-310 |
| chr18-41334119-DEL-88 | chr18-41334124-DEL-79 |
| chr18-41342768-DEL-3993 | chr18-41342769-DEL-3993 |
| chr18-41347287-DEL-639 | chr18-41347288-DEL-639 |
| chr18-41391995-DEL-663 | chr18-41391996-DEL-663 |
| chr18-41462768-DEL-3238 | chr18-41462769-DEL-3237 |
| chr18-41468599-DEL-2823 | chr18-41468589-DEL-2834 |
| chr18-41476649-DEL-83 | chr18-41476654-DEL-79 |
| chr18-41477524-DEL-163 | chr18-41477525-DEL-163 |
| chr18-41514577-DEL-6223 | chr18-41514578-DEL-6223 |
| chr18-41577449-DEL-258 | chr18-41577450-DEL-258 |
| chr18-41589903-DEL-71 | chr18-41589904-DEL-71 |
| chr18-41597478-DEL-7180 | chr18-41597479-DEL-7168 |
| chr18-41642882-DEL-6609 | chr18-41642883-DEL-6609 |
| chr18-41677480-DEL-59 | chr18-41677484-DEL-53 |
| chr18-41684208-DEL-6405 | chr18-41684209-DEL-6405 |
| chr18-41723341-DEL-1811 | chr18-41723340-DEL-1812 |
| chr18-41736479-DEL-1717 | chr18-41736480-DEL-1717 |
| chr18-41744825-DEL-475 | chr18-41744826-DEL-475 |
| chr18-41786091-DEL-57 | chr18-41786092-DEL-57 |
| chr18-41789749-DEL-1345 | chr18-41789750-DEL-1345 |
| chr18-41819514-DEL-7063 | chr18-41819515-DEL-7063 |
| chr18-41833670-DEL-193 | chr18-41833671-DEL-193 |
| chr18-41846051-DEL-76 | chr18-41846064-DEL-64 |
| chr18-41858493-DEL-449 | chr18-41858494-DEL-449 |
| chr18-41878265-DEL-52 | chr18-41878266-DEL-52 |
| chr18-41879666-DEL-70 | chr18-41879667-DEL-70 |

| | |
|---|---|
| chr18-41892733-DEL-120 | chr18-41892736-DEL-102 |
| chr18-41897543-DEL-95 | chr18-41897538-DEL-101 |
| chr18-41911108-DEL-385 | chr18-41911109-DEL-385 |
| chr18-41937602-DEL-218 | chr18-41937603-DEL-218 |
| chr18-41953154-DEL-6272 | chr18-41953155-DEL-6272 |
| chr18-41965394-DEL-64 | chr18-41965397-DEL-50 |
| chr18-42001373-DEL-6651 | chr18-42001374-DEL-6651 |
| chr18-42007443-DEL-211 | chr18-42007430-DEL-212 |
| chr18-42016211-DEL-92 | chr18-42016212-DEL-92 |
| chr18-42019327-DEL-199 | chr18-42019328-DEL-199 |
| chr18-42052814-DEL-1923 | chr18-42052815-DEL-1923 |
| chr18-42092719-DEL-6442 | chr18-42092720-DEL-6442 |
| chr18-42116635-DEL-655 | chr18-42116638-DEL-656 |
| chr18-42128228-DEL-56 | chr18-42128228-DEL-56 |
| chr18-42138384-DEL-63 | chr18-42138341-DEL-56 |
| chr18-42148583-DEL-78 | chr18-42148586-DEL-76 |
| chr18-42149374-DEL-74 | chr18-42149354-DEL-65 |
| chr18-42153857-DEL-754 | chr18-42153858-DEL-754 |
| chr18-42164466-DEL-1097 | chr18-42164467-DEL-1097 |
| chr18-42183568-DEL-142 | chr18-42183564-DEL-148 |
| chr18-42197594-DEL-227 | chr18-42197597-DEL-227 |
| chr18-42207134-DEL-400 | chr18-42207135-DEL-400 |
| chr18-42210336-DEL-10972 | chr18-42210337-DEL-10972 |
| chr18-42216878-DEL-2034 | chr18-42216879-DEL-2035 |
| chr18-42264083-DEL-841 | chr18-42264084-DEL-841 |
| chr18-42270585-DEL-162 | chr18-42270585-DEL-162 |
| chr18-42281466-DEL-97 | chr18-42281467-DEL-97 |
| chr18-42281505-DEL-1050 | chr18-42281506-DEL-1050 |
| chr18-42316170-DEL-3053 | chr18-42316171-DEL-3053 |
| chr18-42320573-DEL-2090 | chr18-42320574-DEL-2090 |
| chr18-42328229-DEL-787 | chr18-42328230-DEL-787 |
| chr18-42333202-DEL-206 | chr18-42333203-DEL-206 |
| chr18-42341565-DEL-176 | chr18-42341566-DEL-172 |
| chr18-42356481-DEL-198 | chr18-42356482-DEL-198 |
| chr18-42376967-DEL-486 | chr18-42376968-DEL-487 |
| chr18-42388219-DEL-70 | chr18-42388220-DEL-80 |
| chr18-42392010-DEL-191 | chr18-42392016-DEL-186 |
| chr18-42392507-DEL-226 | chr18-42392508-DEL-226 |
| chr18-42404811-DEL-92 | chr18-42404812-DEL-92 |
| chr18-42421827-DEL-144 | chr18-42421828-DEL-144 |
| chr18-42478735-DEL-242 | chr18-42478736-DEL-242 |
| chr18-42500192-DEL-165 | chr18-42500193-DEL-165 |
| chr18-42521085-DEL-434 | chr18-42521086-DEL-434 |
| chr18-42539838-DEL-152 | chr18-42539839-DEL-152 |
| chr18-42555101-DEL-78 | chr18-42555086-DEL-83 |
| chr18-42555753-DEL-732 | chr18-42555754-DEL-732 |
| chr18-42556988-DEL-198 | chr18-42556989-DEL-198 |
| chr18-42572792-DEL-214 | chr18-42572793-DEL-214 |
| chr18-42573525-DEL-198 | chr18-42573526-DEL-198 |
| chr18-42589629-DEL-63 | chr18-42589630-DEL-63 |
| chr18-42605840-DEL-1292 | chr18-42605841-DEL-1292 |
| chr18-42608757-DEL-8778 | chr18-42608758-DEL-8778 |
| chr18-42642323-DEL-72 | chr18-42642286-DEL-85 |
| chr18-42642671-DEL-387 | chr18-42642672-DEL-387 |
| chr18-42650081-DEL-212 | chr18-42650082-DEL-212 |
| chr18-42651937-DEL-194 | chr18-42651938-DEL-192 |
| chr18-42679189-DEL-415 | chr18-42679190-DEL-415 |
| chr18-42690972-DEL-971 | chr18-42690970-DEL-950 |
| chr18-42741673-DEL-51 | chr18-42741674-DEL-51 |
| chr18-42747183-DEL-136 | chr18-42747208-DEL-112 |
| chr18-42754278-DEL-202 | chr18-42754279-DEL-202 |
| chr18-42757150-DEL-207 | chr18-42757151-DEL-207 |
| chr18-42763341-DEL-103 | chr18-42763349-DEL-95 |
| chr18-42776693-DEL-371 | chr18-42776686-DEL-379 |
| chr18-42786398-DEL-65 | chr18-42786399-DEL-65 |
| chr18-42805684-DEL-80 | chr18-42805685-DEL-80 |
| chr18-42835320-DEL-56 | chr18-42835321-DEL-56 |
| chr18-42871639-DEL-62 | chr18-42871640-DEL-62 |
| chr18-42901438-DEL-162 | chr18-42901457-DEL-168 |
| chr18-42943613-DEL-1498 | chr18-42943614-DEL-1498 |
| chr18-42962907-DEL-52 | chr18-42962907-DEL-52 |
| chr18-42970431-DEL-92 | chr18-42970432-DEL-92 |
| chr18-42980630-DEL-248 | chr18-42980631-DEL-248 |
| chr18-42981625-DEL-1240 | chr18-42981626-DEL-1240 |
| chr18-42988564-DEL-123 | chr18-42988565-DEL-123 |
| chr18-43004398-DEL-52 | chr18-43004398-DEL-52 |
| chr18-43062539-DEL-227 | chr18-43062540-DEL-227 |
| chr18-43072466-DEL-65 | chr18-43072463-DEL-61 |
| chr18-43074332-DEL-225 | chr18-43074333-DEL-225 |
| chr18-43084933-DEL-203 | chr18-43084934-DEL-203 |

| | |
|---|---|
| chr18-43108826-DEL-281 | chr18-43108810-DEL-282 |
| chr18-43118918-DEL-56 | chr18-43118919-DEL-56 |
| chr18-43119254-DEL-319 | chr18-43119255-DEL-319 |
| chr18-43129762-DEL-7334 | chr18-43129763-DEL-7334 |
| chr18-43186495-DEL-342 | chr18-43186495-DEL-342 |
| chr18-43193791-DEL-355 | chr18-43193795-DEL-353 |
| chr18-43216920-DEL-447 | chr18-43216921-DEL-447 |
| chr18-43233814-DEL-78 | chr18-43233807-DEL-79 |
| chr18-43241220-DEL-211 | chr18-43241221-DEL-211 |
| chr18-43274071-DEL-56 | chr18-43274041-DEL-59 |
| chr18-43288153-DEL-262 | chr18-43288155-DEL-248 |
| chr18-43388014-DEL-111 | chr18-43387976-DEL-153 |
| chr18-43433137-DEL-605 | chr18-43433138-DEL-605 |
| chr18-43444298-DEL-100 | chr18-43444299-DEL-100 |
| chr18-43447318-DEL-87 | chr18-43447319-DEL-87 |
| chr18-43449958-DEL-54 | chr18-43449959-DEL-54 |
| chr18-43460244-DEL-95 | chr18-43460245-DEL-95 |
| chr18-43463941-DEL-1007 | chr18-43463942-DEL-1007 |
| chr18-43483818-DEL-131 | chr18-43483819-DEL-131 |
| chr18-43484365-DEL-126 | chr18-43484366-DEL-126 |
| chr18-43532030-DEL-56 | chr18-43532031-DEL-56 |
| chr18-43574200-DEL-52 | chr18-43574201-DEL-52 |
| chr18-43593560-DEL-2497 | chr18-43593561-DEL-2497 |
| chr18-43597528-DEL-732 | chr18-43597529-DEL-732 |
| chr18-43598214-DEL-56 | chr18-43598215-DEL-56 |
| chr18-43605702-DEL-5221 | chr18-43605703-DEL-5221 |
| chr18-43641424-DEL-360 | chr18-43641425-DEL-360 |
| chr18-43661107-DEL-62 | chr18-43661108-DEL-60 |
| chr18-43666325-DEL-196 | chr18-43666326-DEL-196 |
| chr18-43680447-DEL-198 | chr18-43680448-DEL-194 |
| chr18-43706867-DEL-422 | chr18-43706868-DEL-421 |
| chr18-43708694-DEL-647 | chr18-43708695-DEL-647 |
| chr18-43778303-DEL-601 | chr18-43778304-DEL-601 |
| chr18-43823472-DEL-60 | chr18-43823473-DEL-60 |
| chr18-43832976-DEL-863 | chr18-43832977-DEL-863 |
| chr18-43834708-DEL-934 | chr18-43834709-DEL-934 |
| chr18-43843303-DEL-6536 | chr18-43843299-DEL-6537 |
| chr18-43853901-DEL-502 | chr18-43853902-DEL-502 |
| chr18-43949091-DEL-85 | chr18-43949091-DEL-77 |
| chr18-43953994-DEL-52 | chr18-43953994-DEL-52 |
| chr18-43962879-DEL-6524 | chr18-43962877-DEL-6525 |
| chr18-43971343-DEL-734 | chr18-43971347-DEL-731 |
| chr18-43974226-DEL-501 | chr18-43974225-DEL-503 |
| chr18-44022256-DEL-12655 | chr18-44022257-DEL-12655 |
| chr18-44065463-DEL-115 | chr18-44065464-DEL-115 |
| chr18-44074718-DEL-2407 | chr18-44074719-DEL-2407 |
| chr18-44165911-DEL-2797 | chr18-44165912-DEL-2797 |
| chr18-44188770-DEL-54 | chr18-44188771-DEL-54 |
| chr18-44228545-DEL-56 | chr18-44228546-DEL-56 |
| chr18-44242118-DEL-3827 | chr18-44242119-DEL-3827 |
| chr18-44250772-DEL-67 | chr18-44250773-DEL-67 |
| chr18-44260808-DEL-4719 | chr18-44260809-DEL-4719 |
| chr18-44337154-DEL-90 | chr18-44337155-DEL-90 |
| chr18-44347892-DEL-774 | chr18-44347893-DEL-774 |
| chr18-44423005-DEL-214 | chr18-44423005-DEL-214 |
| chr18-44429479-DEL-110 | chr18-44429480-DEL-110 |
| chr18-44446693-DEL-209 | chr18-44446694-DEL-209 |
| chr18-44462483-DEL-58 | chr18-44462484-DEL-58 |
| chr18-44494572-DEL-1403 | chr18-44494573-DEL-1403 |
| chr18-44498966-DEL-520 | chr18-44498967-DEL-520 |
| chr18-44499772-DEL-55 | chr18-44499773-DEL-55 |
| chr18-44501684-DEL-6598 | chr18-44501685-DEL-6598 |
| chr18-44507803-DEL-81 | chr18-44507804-DEL-81 |
| chr18-44539129-DEL-293 | chr18-44539130-DEL-293 |
| chr18-44544534-DEL-308 | chr18-44544535-DEL-308 |
| chr18-44559165-DEL-60 | chr18-44559144-DEL-54 |
| chr18-44583181-DEL-287 | chr18-44583182-DEL-287 |
| chr18-44591180-DEL-54 | chr18-44591179-DEL-52 |
| chr18-44614113-DEL-119 | chr18-44614117-DEL-97 |
| chr18-44615659-DEL-678 | chr18-44615660-DEL-678 |
| chr18-44635779-DEL-2954 | chr18-44635793-DEL-2951 |
| chr18-44688559-DEL-94 | chr18-44688560-DEL-94 |
| chr18-44713835-DEL-140 | chr18-44713808-DEL-120 |
| chr18-44715957-DEL-71 | chr18-44715958-DEL-71 |
| chr18-44732870-DEL-214 | chr18-44732871-DEL-214 |
| chr18-44788590-DEL-71 | chr18-44788586-DEL-72 |
| chr18-44807068-DEL-76 | chr18-44807071-DEL-77 |
| chr18-44813355-DEL-143 | chr18-44813339-DEL-136 |
| chr18-44822904-DEL-174 | chr18-44822920-DEL-159 |
| chr18-44829399-DEL-90 | chr18-44829394-DEL-78 |

| | |
|---|---|
| chr18-44864803-DEL-65 | chr18-44864806-DEL-63 |
| chr18-44868338-DEL-5289 | chr18-44868339-DEL-5289 |
| chr18-44887656-DEL-73 | chr18-44887678-DEL-52 |
| chr18-44888711-DEL-551 | chr18-44888712-DEL-551 |
| chr18-44909605-DEL-2138 | chr18-44909606-DEL-2137 |
| chr18-44912832-DEL-731 | chr18-44912853-DEL-704 |
| chr18-44949646-DEL-368 | chr18-44949647-DEL-368 |
| chr18-44971765-DEL-202 | chr18-44971766-DEL-202 |
| chr18-45006246-DEL-1493 | chr18-45006247-DEL-1493 |
| chr18-45033841-DEL-185 | chr18-45033842-DEL-185 |
| chr18-45034162-DEL-411 | chr18-45034163-DEL-411 |
| chr18-45039561-DEL-2816 | chr18-45039562-DEL-2816 |
| chr18-45060147-DEL-60 | chr18-45060148-DEL-60 |
| chr18-45086187-DEL-60 | chr18-45086188-DEL-78 |
| chr18-45093659-DEL-979 | chr18-45093660-DEL-979 |
| chr18-45113990-DEL-5992 | chr18-45113979-DEL-6020 |
| chr18-45130882-DEL-180 | chr18-45130883-DEL-180 |
| chr18-45138941-DEL-176 | chr18-45138942-DEL-176 |
| chr18-45149895-DEL-7962 | chr18-45149896-DEL-7962 |
| chr18-45159612-DEL-6372 | chr18-45159613-DEL-6372 |
| chr18-45180858-DEL-6178 | chr18-45180859-DEL-6177 |
| chr18-45191637-DEL-127 | chr18-45191638-DEL-127 |
| chr18-45192425-DEL-56 | chr18-45192398-DEL-84 |
| chr18-45205613-DEL-77 | chr18-45205614-DEL-77 |
| chr18-45206125-DEL-1828 | chr18-45206125-DEL-1828 |
| chr18-45212458-DEL-101 | chr18-45212459-DEL-101 |
| chr18-45251788-DEL-455 | chr18-45251795-DEL-449 |
| chr18-45252705-DEL-75 | chr18-45252706-DEL-75 |
| chr18-45254343-DEL-704 | chr18-45254344-DEL-704 |
| chr18-45264401-DEL-2496 | chr18-45264397-DEL-2497 |
| chr18-45296505-DEL-2631 | chr18-45296522-DEL-2630 |
| chr18-45304948-DEL-6185 | chr18-45304949-DEL-6185 |
| chr18-45338070-DEL-777 | chr18-45338070-DEL-777 |
| chr18-45339240-DEL-8390 | chr18-45339241-DEL-8390 |
| chr18-45401088-DEL-62 | chr18-45401089-DEL-60 |
| chr18-45418172-DEL-64 | chr18-45418176-DEL-62 |
| chr18-45424855-DEL-4519 | chr18-45424856-DEL-4519 |
| chr18-45434302-DEL-1048 | chr18-45434308-DEL-1038 |
| chr18-45437213-DEL-914 | chr18-45437214-DEL-914 |
| chr18-45440398-DEL-2616 | chr18-45440404-DEL-2610 |
| chr18-45443251-DEL-4122 | chr18-45443252-DEL-4122 |
| chr18-45456229-DEL-60 | chr18-45456232-DEL-50 |
| chr18-45458839-DEL-94 | chr18-45458849-DEL-52 |
| chr18-45460202-DEL-244 | chr18-45460201-DEL-241 |
| chr18-45470727-DEL-54 | chr18-45470728-DEL-54 |
| chr18-45484594-DEL-63 | chr18-45484595-DEL-63 |
| chr18-45524521-DEL-6272 | chr18-45524534-DEL-6265 |
| chr18-45555245-DEL-135 | chr18-45555246-DEL-135 |
| chr18-45555951-DEL-1804 | chr18-45555951-DEL-1804 |
| chr18-45572649-DEL-462 | chr18-45572648-DEL-460 |
| chr18-45582450-DEL-133 | chr18-45582451-DEL-133 |
| chr18-45584393-DEL-343 | chr18-45584394-DEL-343 |
| chr18-45589906-DEL-322 | chr18-45589907-DEL-322 |
| chr18-45596520-DEL-395 | chr18-45596521-DEL-395 |
| chr18-45616652-DEL-304 | chr18-45616653-DEL-304 |
| chr18-45624612-DEL-236 | chr18-45624590-DEL-255 |
| chr18-45626889-DEL-71 | chr18-45626890-DEL-71 |
| chr18-45640015-DEL-6443 | chr18-45640016-DEL-6443 |
| chr18-45654187-DEL-357 | chr18-45654188-DEL-357 |
| chr18-45670642-DEL-6002 | chr18-45670643-DEL-6002 |
| chr18-45719512-DEL-75 | chr18-45719528-DEL-72 |
| chr18-45736020-DEL-70 | chr18-45736016-DEL-75 |
| chr18-45767343-DEL-437 | chr18-45767331-DEL-437 |
| chr18-45797312-DEL-110 | chr18-45797313-DEL-110 |
| chr18-45856204-DEL-191 | chr18-45856205-DEL-191 |
| chr18-45881384-DEL-67 | chr18-45881371-DEL-70 |
| chr18-45899871-DEL-498 | chr18-45899872-DEL-498 |
| chr18-45932048-DEL-54 | chr18-45932053-DEL-53 |
| chr18-45944255-DEL-86 | chr18-45944256-DEL-86 |
| chr18-45983842-DEL-1278 | chr18-45983843-DEL-1278 |
| chr18-45987900-DEL-196 | chr18-45987901-DEL-196 |
| chr18-45993048-DEL-58 | chr18-45993010-DEL-102 |
| chr18-45996075-DEL-7061 | chr18-45996076-DEL-7061 |
| chr18-45996129-DEL-428 | chr18-45996130-DEL-428 |
| chr18-46022847-DEL-400 | chr18-46022848-DEL-400 |
| chr18-46047522-DEL-54 | chr18-46047523-DEL-54 |
| chr18-46050071-DEL-150 | chr18-46050074-DEL-148 |
| chr18-46061133-DEL-403 | chr18-46061134-DEL-400 |
| chr18-46082010-DEL-87 | chr18-46082003-DEL-94 |
| chr18-46089817-DEL-106 | chr18-46089816-DEL-102 |

| | |
|---|---|
| chr18-46091046-DEL-1041 | chr18-46091047-DEL-1041 |
| chr18-46097556-DEL-1526 | chr18-46097550-DEL-1529 |
| chr18-46098905-DEL-182 | chr18-46098906-DEL-180 |
| chr18-46132795-DEL-120 | chr18-46132796-DEL-120 |
| chr18-46137309-DEL-6166 | chr18-46137310-DEL-6166 |
| chr18-46153738-DEL-77 | chr18-46153739-DEL-77 |
| chr18-46162975-DEL-1891 | chr18-46162981-DEL-1878 |
| chr18-46190231-DEL-405 | chr18-46190232-DEL-405 |
| chr18-46223578-DEL-746 | chr18-46223579-DEL-746 |
| chr18-46230013-DEL-4492 | chr18-46230014-DEL-4492 |
| chr18-46239678-DEL-419 | chr18-46239679-DEL-419 |
| chr18-46240508-DEL-345 | chr18-46240509-DEL-345 |
| chr18-46244434-DEL-80 | chr18-46244416-DEL-117 |
| chr18-46271008-DEL-398 | chr18-46271009-DEL-398 |
| chr18-46273805-DEL-399 | chr18-46273806-DEL-399 |
| chr18-46288410-DEL-62 | chr18-46288411-DEL-62 |
| chr18-46304791-DEL-70 | chr18-46304810-DEL-50 |
| chr18-46384495-DEL-63 | chr18-46384496-DEL-65 |
| chr18-46389000-DEL-62 | chr18-46389001-DEL-62 |
| chr18-46418443-DEL-104 | chr18-46418444-DEL-104 |
| chr18-46428204-DEL-251 | chr18-46428205-DEL-251 |
| chr18-46464517-DEL-66 | chr18-46464518-DEL-52 |
| chr18-46467688-DEL-54 | chr18-46467689-DEL-54 |
| chr18-46493718-DEL-67 | chr18-46493719-DEL-67 |
| chr18-46495039-DEL-88 | chr18-46495040-DEL-86 |
| chr18-46528307-DEL-87 | chr18-46528305-DEL-88 |
| chr18-46553935-DEL-59 | chr18-46553939-DEL-60 |
| chr18-46572155-DEL-4894 | chr18-46572156-DEL-4894 |
| chr18-46586749-DEL-54 | chr18-46586746-DEL-60 |
| chr18-46613116-DEL-168 | chr18-46613117-DEL-168 |
| chr18-46618533-DEL-756 | chr18-46618531-DEL-759 |
| chr18-46625157-DEL-227 | chr18-46625158-DEL-227 |
| chr18-46644402-DEL-115 | chr18-46644403-DEL-115 |
| chr18-46650557-DEL-212 | chr18-46650558-DEL-212 |
| chr18-46663300-DEL-406 | chr18-46663301-DEL-406 |
| chr18-46667100-DEL-206 | chr18-46667100-DEL-206 |
| chr18-46672194-DEL-58 | chr18-46672195-DEL-58 |
| chr18-46684772-DEL-766 | chr18-46684773-DEL-760 |
| chr18-46686983-DEL-3274 | chr18-46686984-DEL-3274 |
| chr18-46696390-DEL-321 | chr18-46696391-DEL-321 |
| chr18-46723097-DEL-405 | chr18-46723098-DEL-405 |
| chr18-46733505-DEL-206 | chr18-46733506-DEL-206 |
| chr18-46752224-DEL-249 | chr18-46752218-DEL-250 |
| chr18-46753493-DEL-186 | chr18-46753494-DEL-184 |
| chr18-46758213-DEL-1905 | chr18-46758214-DEL-1905 |
| chr18-46776065-DEL-123 | chr18-46776066-DEL-123 |
| chr18-46799980-DEL-91 | chr18-46799952-DEL-119 |
| chr18-46801465-DEL-79 | chr18-46801481-DEL-75 |
| chr18-46803770-DEL-54 | chr18-46803772-DEL-56 |
| chr18-46819518-DEL-6479 | chr18-46819519-DEL-6479 |
| chr18-46827930-DEL-247 | chr18-46827931-DEL-247 |
| chr18-46879579-DEL-159 | chr18-46879580-DEL-159 |
| chr18-46882704-DEL-210 | chr18-46882705-DEL-210 |
| chr18-46905726-DEL-974 | chr18-46905719-DEL-982 |
| chr18-46906887-DEL-371 | chr18-46906888-DEL-371 |
| chr18-46931311-DEL-72 | chr18-46931302-DEL-79 |
| chr18-46936572-DEL-188 | chr18-46936573-DEL-188 |
| chr18-46966458-DEL-64 | chr18-46966459-DEL-64 |
| chr18-46970936-DEL-142 | chr18-46970927-DEL-152 |
| chr18-46975799-DEL-327 | chr18-46975800-DEL-327 |
| chr18-46979230-DEL-81 | chr18-46979239-DEL-69 |
| chr18-46996082-DEL-86 | chr18-46996079-DEL-93 |
| chr18-47004760-DEL-217 | chr18-47004761-DEL-216 |
| chr18-47007913-DEL-238 | chr18-47007914-DEL-238 |
| chr18-47020507-DEL-324 | chr18-47020508-DEL-324 |
| chr18-47062042-DEL-210 | chr18-47062035-DEL-211 |
| chr18-47071628-DEL-98 | chr18-47071629-DEL-98 |
| chr18-47074642-DEL-614 | chr18-47074643-DEL-614 |
| chr18-47096288-DEL-106 | chr18-47096289-DEL-103 |
| chr18-47102134-DEL-66 | chr18-47102134-DEL-66 |
| chr18-47115441-DEL-2228 | chr18-47115442-DEL-2228 |
| chr18-47116201-DEL-211 | chr18-47116187-DEL-212 |
| chr18-47117462-DEL-66 | chr18-47117463-DEL-66 |
| chr18-47129609-DEL-211 | chr18-47129610-DEL-211 |
| chr18-47245192-DEL-553 | chr18-47245193-DEL-553 |
| chr18-47254920-DEL-144 | chr18-47254921-DEL-144 |
| chr18-47307867-DEL-70 | chr18-47307852-DEL-70 |
| chr18-47315458-DEL-6941 | chr18-47315459-DEL-6941 |
| chr18-47353574-DEL-62 | chr18-47353575-DEL-63 |
| chr18-47356247-DEL-1170 | chr18-47356248-DEL-1170 |

| | |
|---|---|
| chr18-47367495-DEL-1072 | chr18-47367495-DEL-1072 |
| chr18-47408602-DEL-96 | chr18-47408603-DEL-100 |
| chr18-47417445-DEL-1577 | chr18-47417446-DEL-1577 |
| chr18-47512194-DEL-60 | chr18-47512195-DEL-60 |
| chr18-47514729-DEL-57 | chr18-47514731-DEL-57 |
| chr18-47519795-DEL-8496 | chr18-47519796-DEL-8496 |
| chr18-47524201-DEL-1013 | chr18-47524202-DEL-1013 |
| chr18-47526566-DEL-1515 | chr18-47526566-DEL-1515 |
| chr18-47530782-DEL-80 | chr18-47530781-DEL-81 |
| chr18-47546295-DEL-243 | chr18-47546290-DEL-244 |
| chr18-47551417-DEL-124 | chr18-47551418-DEL-124 |
| chr18-47562456-DEL-162 | chr18-47562458-DEL-159 |
| chr18-47563392-DEL-60 | chr18-47563387-DEL-57 |
| chr18-47599243-DEL-57 | chr18-47599244-DEL-57 |
| chr18-47612662-DEL-179 | chr18-47612663-DEL-179 |
| chr18-47630236-DEL-604 | chr18-47630232-DEL-605 |
| chr18-47649291-DEL-191 | chr18-47649292-DEL-191 |
| chr18-47673181-DEL-389 | chr18-47673182-DEL-389 |
| chr18-47688757-DEL-77 | chr18-47688746-DEL-65 |
| chr18-47695794-DEL-78 | chr18-47695795-DEL-78 |
| chr18-47699272-DEL-205 | chr18-47699273-DEL-205 |
| chr18-47728460-DEL-211 | chr18-47728461-DEL-211 |
| chr18-47735016-DEL-397 | chr18-47735017-DEL-397 |
| chr18-47760563-DEL-83 | chr18-47760564-DEL-83 |
| chr18-47781813-DEL-5207 | chr18-47781814-DEL-5207 |
| chr18-47799995-DEL-53 | chr18-47799995-DEL-53 |
| chr18-47815773-DEL-53 | chr18-47815774-DEL-53 |
| chr18-47816752-DEL-162 | chr18-47816753-DEL-162 |
| chr18-47836194-DEL-194 | chr18-47836171-DEL-218 |
| chr18-47850155-DEL-4050 | chr18-47850156-DEL-4050 |
| chr18-47856350-DEL-1102 | chr18-47856351-DEL-1102 |
| chr18-47864891-DEL-311 | chr18-47864892-DEL-311 |
| chr18-47870272-DEL-7179 | chr18-47870273-DEL-7179 |
| chr18-47903074-DEL-7123 | chr18-47903075-DEL-7123 |
| chr18-47932498-DEL-66 | chr18-47932483-DEL-64 |
| chr18-47960532-DEL-1381 | chr18-47960533-DEL-1381 |
| chr18-47969328-DEL-55 | chr18-47969322-DEL-56 |
| chr18-48020034-DEL-128 | chr18-48020027-DEL-100 |
| chr18-48026790-DEL-756 | chr18-48026791-DEL-756 |
| chr18-48048433-DEL-56 | chr18-48048434-DEL-56 |
| chr18-48062183-DEL-76 | chr18-48062195-DEL-77 |
| chr18-48119392-DEL-64 | chr18-48119393-DEL-64 |
| chr18-48125903-DEL-99 | chr18-48125924-DEL-58 |
| chr18-48143746-DEL-54 | chr18-48143749-DEL-52 |
| chr18-48165222-DEL-1680 | chr18-48165223-DEL-1680 |
| chr18-48179699-DEL-1779 | chr18-48179700-DEL-1779 |
| chr18-48217605-DEL-73 | chr18-48217606-DEL-73 |
| chr18-48223779-DEL-106 | chr18-48223780-DEL-106 |
| chr18-48226508-DEL-3984 | chr18-48226509-DEL-3984 |
| chr18-48274576-DEL-399 | chr18-48274577-DEL-399 |
| chr18-48279196-DEL-203 | chr18-48279197-DEL-203 |
| chr18-48296219-DEL-345 | chr18-48296220-DEL-345 |
| chr18-48315710-DEL-65 | chr18-48315711-DEL-65 |
| chr18-48323452-DEL-4122 | chr18-48323453-DEL-4122 |
| chr18-48343670-DEL-7309 | chr18-48343671-DEL-7309 |
| chr18-48352034-DEL-1318 | chr18-48352035-DEL-1318 |
| chr18-48363249-DEL-496 | chr18-48363250-DEL-496 |
| chr18-48390068-DEL-412 | chr18-48390064-DEL-416 |
| chr18-48437174-DEL-110 | chr18-48437174-DEL-108 |
| chr18-48447309-DEL-3328 | chr18-48447307-DEL-3329 |
| chr18-48449747-DEL-52 | chr18-48449750-DEL-54 |
| chr18-48520022-DEL-479 | chr18-48520023-DEL-479 |
| chr18-48525007-DEL-348 | chr18-48525008-DEL-348 |
| chr18-48563311-DEL-480 | chr18-48563312-DEL-480 |
| chr18-48576629-DEL-140 | chr18-48576630-DEL-140 |
| chr18-48577941-DEL-56 | chr18-48577940-DEL-58 |
| chr18-48596731-DEL-81 | chr18-48596734-DEL-83 |
| chr18-48600509-DEL-70 | chr18-48600510-DEL-70 |
| chr18-48607695-DEL-73 | chr18-48607694-DEL-75 |
| chr18-48634514-DEL-593 | chr18-48634516-DEL-594 |
| chr18-48635506-DEL-6188 | chr18-48635506-DEL-6188 |
| chr18-48661876-DEL-3818 | chr18-48661877-DEL-3818 |
| chr18-48668661-DEL-6371 | chr18-48668662-DEL-6371 |
| chr18-48687664-DEL-56 | chr18-48687665-DEL-55 |
| chr18-48691793-DEL-585 | chr18-48691794-DEL-585 |
| chr18-48709374-DEL-977 | chr18-48709375-DEL-977 |
| chr18-48755071-DEL-1067 | chr18-48755072-DEL-1067 |
| chr18-48760557-DEL-1241 | chr18-48760558-DEL-1241 |
| chr18-48777103-DEL-57 | chr18-48777084-DEL-58 |
| chr18-48799667-DEL-84 | chr18-48799667-DEL-83 |

| | |
|---|---|
| chr18-48806139-DEL-6275 | chr18-48806140-DEL-6275 |
| chr18-48813893-DEL-130 | chr18-48813894-DEL-130 |
| chr18-48815781-DEL-63 | chr18-48815742-DEL-63 |
| chr18-48818539-DEL-55 | chr18-48818540-DEL-55 |
| chr18-48838966-DEL-7091 | chr18-48838967-DEL-7091 |
| chr18-48871083-DEL-1464 | chr18-48871084-DEL-1464 |
| chr18-48883541-DEL-386 | chr18-48883542-DEL-386 |
| chr18-48891056-DEL-82 | chr18-48891056-DEL-74 |
| chr18-48906442-DEL-791 | chr18-48906442-DEL-792 |
| chr18-49010259-DEL-72 | chr18-49010295-DEL-68 |
| chr18-49041332-DEL-67 | chr18-49041357-DEL-50 |
| chr18-49101411-DEL-236 | chr18-49101412-DEL-236 |
| chr18-49113079-DEL-1893 | chr18-49113080-DEL-1893 |
| chr18-49117306-DEL-443 | chr18-49117307-DEL-443 |
| chr18-49124792-DEL-76 | chr18-49124794-DEL-75 |
| chr18-49124942-DEL-498 | chr18-49124943-DEL-498 |
| chr18-49130442-DEL-456 | chr18-49130443-DEL-456 |
| chr18-49134914-DEL-1060 | chr18-49134915-DEL-1060 |
| chr18-49167365-DEL-54 | chr18-49167366-DEL-54 |
| chr18-49175873-DEL-4382 | chr18-49175874-DEL-4382 |
| chr18-49177992-DEL-54 | chr18-49177984-DEL-57 |
| chr18-49205647-DEL-1110 | chr18-49205648-DEL-1110 |
| chr18-49220669-DEL-786 | chr18-49220690-DEL-784 |
| chr18-49226889-DEL-52 | chr18-49226890-DEL-52 |
| chr18-49252315-DEL-1098 | chr18-49252316-DEL-1098 |
| chr18-49268743-DEL-60 | chr18-49268743-DEL-60 |
| chr18-49280385-DEL-5559 | chr18-49280386-DEL-5559 |
| chr18-49290127-DEL-251 | chr18-49290163-DEL-264 |
| chr18-49299632-DEL-75 | chr18-49299628-DEL-80 |
| chr18-49307222-DEL-769 | chr18-49307223-DEL-769 |
| chr18-49309301-DEL-2941 | chr18-49309301-DEL-2942 |
| chr18-49320802-DEL-224 | chr18-49320795-DEL-224 |
| chr18-49321907-DEL-63 | chr18-49321911-DEL-60 |
| chr18-49323954-DEL-397 | chr18-49323955-DEL-398 |
| chr18-49328889-DEL-73 | chr18-49328890-DEL-81 |
| chr18-49335745-DEL-505 | chr18-49335746-DEL-505 |
| chr18-49350060-DEL-285 | chr18-49350064-DEL-284 |
| chr18-49418320-DEL-777 | chr18-49418321-DEL-777 |
| chr18-49420538-DEL-311 | chr18-49420539-DEL-311 |
| chr18-49467899-DEL-791 | chr18-49467900-DEL-791 |
| chr18-49468982-DEL-76 | chr18-49468983-DEL-76 |
| chr18-49476100-DEL-54 | chr18-49476101-DEL-54 |
| chr18-49524326-DEL-6579 | chr18-49524327-DEL-6579 |
| chr18-49559452-DEL-386 | chr18-49559453-DEL-386 |
| chr18-49582910-DEL-540 | chr18-49582902-DEL-532 |
| chr18-49674521-DEL-269 | chr18-49674522-DEL-269 |
| chr18-49687154-DEL-206 | chr18-49687155-DEL-206 |
| chr18-49694039-DEL-194 | chr18-49694040-DEL-194 |
| chr18-49728859-DEL-353 | chr18-49728858-DEL-344 |
| chr18-49731362-DEL-180 | chr18-49731363-DEL-180 |
| chr18-49760382-DEL-57 | chr18-49760375-DEL-54 |
| chr18-49769799-DEL-57 | chr18-49769784-DEL-60 |
| chr18-49779213-DEL-90 | chr18-49779214-DEL-87 |
| chr18-49867549-DEL-347 | chr18-49867550-DEL-347 |
| chr18-49871895-DEL-333 | chr18-49871896-DEL-333 |
| chr18-49922005-DEL-438 | chr18-49922006-DEL-438 |
| chr18-49926866-DEL-530 | chr18-49926867-DEL-530 |
| chr18-49931374-DEL-5202 | chr18-49931375-DEL-5202 |
| chr18-49933968-DEL-215 | chr18-49933969-DEL-215 |
| chr18-49953535-DEL-173 | chr18-49953552-DEL-174 |
| chr18-49954517-DEL-190 | chr18-49954518-DEL-190 |
| chr18-49961411-DEL-352 | chr18-49961412-DEL-352 |
| chr18-49994546-DEL-220 | chr18-49994547-DEL-220 |
| chr18-50066665-DEL-306 | chr18-50066666-DEL-306 |
| chr18-50067854-DEL-69 | chr18-50067842-DEL-70 |
| chr18-50088322-DEL-358 | chr18-50088323-DEL-358 |
| chr18-50100099-DEL-54 | chr18-50100100-DEL-54 |
| chr18-50107100-DEL-52 | chr18-50107101-DEL-52 |
| chr18-50110032-DEL-1560 | chr18-50110033-DEL-1560 |
| chr18-50119531-DEL-212 | chr18-50119532-DEL-212 |
| chr18-50147301-DEL-79 | chr18-50147302-DEL-79 |
| chr18-50179900-DEL-58 | chr18-50179913-DEL-56 |
| chr18-50209117-DEL-541 | chr18-50209143-DEL-526 |
| chr18-50215901-DEL-139 | chr18-50215896-DEL-167 |
| chr18-50272823-DEL-60 | chr18-50272824-DEL-60 |
| chr18-50297151-DEL-76 | chr18-50297192-DEL-80 |
| chr18-50307789-DEL-181 | chr18-50307791-DEL-180 |
| chr18-50313341-DEL-60 | chr18-50313342-DEL-60 |
| chr18-50318886-DEL-163 | chr18-50318886-DEL-161 |
| chr18-50321019-DEL-130 | chr18-50321020-DEL-130 |

| | |
|---|---|
| chr18-50336180-DEL-1021 | chr18-50336181-DEL-1021 |
| chr18-50336361-DEL-203 | chr18-50336362-DEL-203 |
| chr18-50346290-DEL-6201 | chr18-50346291-DEL-6201 |
| chr18-50347416-DEL-1664 | chr18-50347417-DEL-1664 |
| chr18-50395022-DEL-6434 | chr18-50395023-DEL-6434 |
| chr18-50425094-DEL-3117 | chr18-50425095-DEL-3117 |
| chr18-50503713-DEL-252 | chr18-50503714-DEL-252 |
| chr18-50518249-DEL-1834 | chr18-50518250-DEL-1834 |
| chr18-50555176-DEL-431 | chr18-50555177-DEL-431 |
| chr18-50566386-DEL-214 | chr18-50566377-DEL-224 |
| chr18-50570455-DEL-7763 | chr18-50570456-DEL-7763 |
| chr18-50621917-DEL-65 | chr18-50621905-DEL-66 |
| chr18-50639205-DEL-82 | chr18-50639206-DEL-82 |
| chr18-50641246-DEL-400 | chr18-50641247-DEL-400 |
| chr18-50665005-DEL-7206 | chr18-50665006-DEL-7206 |
| chr18-50696002-DEL-52 | chr18-50696003-DEL-52 |
| chr18-50703517-DEL-322 | chr18-50703518-DEL-322 |
| chr18-50729529-DEL-344 | chr18-50729530-DEL-344 |
| chr18-50732244-DEL-56 | chr18-50732245-DEL-56 |
| chr18-50741233-DEL-177 | chr18-50741234-DEL-177 |
| chr18-50765276-DEL-4320 | chr18-50765277-DEL-4320 |
| chr18-50773525-DEL-459 | chr18-50773523-DEL-459 |
| chr18-50783351-DEL-1666 | chr18-50783349-DEL-1667 |
| chr18-50791740-DEL-7046 | chr18-50791741-DEL-7046 |
| chr18-50828375-DEL-272 | chr18-50828364-DEL-282 |
| chr18-50842447-DEL-114 | chr18-50842448-DEL-114 |
| chr18-50846334-DEL-153 | chr18-50846333-DEL-154 |
| chr18-50858568-DEL-75 | chr18-50858569-DEL-75 |
| chr18-50879385-DEL-2879 | chr18-50879386-DEL-2879 |
| chr18-50885596-DEL-55 | chr18-50885599-DEL-53 |
| chr18-50915937-DEL-6679 | chr18-50915938-DEL-6679 |
| chr18-50916280-DEL-212 | chr18-50916281-DEL-212 |
| chr18-50928381-DEL-588 | chr18-50928382-DEL-588 |
| chr18-50935928-DEL-52 | chr18-50935929-DEL-52 |
| chr18-50953867-DEL-141 | chr18-50953868-DEL-141 |
| chr18-50969141-DEL-77 | chr18-50969142-DEL-77 |
| chr18-50974268-DEL-172 | chr18-50974272-DEL-169 |
| chr18-50992448-DEL-498 | chr18-50992449-DEL-498 |
| chr18-51000042-DEL-662 | chr18-51000043-DEL-662 |
| chr18-51011825-DEL-216 | chr18-51011829-DEL-214 |
| chr18-51014727-DEL-79 | chr18-51014735-DEL-72 |
| chr18-51016429-DEL-954 | chr18-51016427-DEL-955 |
| chr18-51042088-DEL-351 | chr18-51042089-DEL-351 |
| chr18-51054899-DEL-1032 | chr18-51054900-DEL-1032 |
| chr18-51060450-DEL-2456 | chr18-51060451-DEL-2456 |
| chr18-51060512-DEL-498 | chr18-51060513-DEL-498 |
| chr18-51134478-DEL-201 | chr18-51134479-DEL-201 |
| chr18-51195398-DEL-1743 | chr18-51195399-DEL-1743 |
| chr18-51205156-DEL-5357 | chr18-51205157-DEL-5357 |
| chr18-51221057-DEL-62 | chr18-51221062-DEL-58 |
| chr18-51268546-DEL-225 | chr18-51268547-DEL-225 |
| chr18-51284293-DEL-56 | chr18-51284292-DEL-56 |
| chr18-51290321-DEL-115 | chr18-51290322-DEL-110 |
| chr18-51322384-DEL-55 | chr18-51322384-DEL-55 |
| chr18-51326686-DEL-181 | chr18-51326687-DEL-181 |
| chr18-51342425-DEL-1022 | chr18-51342425-DEL-1022 |
| chr18-51348855-DEL-58 | chr18-51348856-DEL-58 |
| chr18-51363687-DEL-1503 | chr18-51363688-DEL-1496 |
| chr18-51386711-DEL-59 | chr18-51386712-DEL-59 |
| chr18-51388919-DEL-595 | chr18-51388920-DEL-595 |
| chr18-51395187-DEL-538 | chr18-51395188-DEL-538 |
| chr18-51413721-DEL-697 | chr18-51413722-DEL-697 |
| chr18-51416901-DEL-654 | chr18-51416902-DEL-654 |
| chr18-51455670-DEL-158 | chr18-51455671-DEL-158 |
| chr18-51467544-DEL-199 | chr18-51467545-DEL-199 |
| chr18-51490996-DEL-1633 | chr18-51490997-DEL-1633 |
| chr18-51499107-DEL-51 | chr18-51499108-DEL-51 |
| chr18-51527293-DEL-6219 | chr18-51527294-DEL-6219 |
| chr18-51534702-DEL-3876 | chr18-51534703-DEL-3876 |
| chr18-51565632-DEL-182 | chr18-51565633-DEL-182 |
| chr18-51588016-DEL-84 | chr18-51588016-DEL-120 |
| chr18-51626887-DEL-58 | chr18-51626888-DEL-58 |
| chr18-51638553-DEL-1684 | chr18-51638554-DEL-1684 |
| chr18-51642168-DEL-680 | chr18-51642169-DEL-680 |
| chr18-51740740-DEL-269 | chr18-51740741-DEL-269 |
| chr18-51741414-DEL-241 | chr18-51741412-DEL-244 |
| chr18-51745443-DEL-64 | chr18-51745444-DEL-64 |
| chr18-51746903-DEL-233 | chr18-51746903-DEL-233 |
| chr18-51802309-DEL-56 | chr18-51802300-DEL-62 |
| chr18-51806394-DEL-1790 | chr18-51806395-DEL-1790 |

| | |
|---|---|
| chr18-51808427-DEL-270 | chr18-51808428-DEL-270 |
| chr18-51810707-DEL-66 | chr18-51810714-DEL-63 |
| chr18-51826071-DEL-6180 | chr18-51826063-DEL-6181 |
| chr18-51827186-DEL-2841 | chr18-51827186-DEL-2841 |
| chr18-51837891-DEL-2110 | chr18-51837892-DEL-2110 |
| chr18-51856728-DEL-194 | chr18-51856729-DEL-194 |
| chr18-51916623-DEL-90 | chr18-51916624-DEL-90 |
| chr18-51950933-DEL-90 | chr18-51950932-DEL-50 |
| chr18-51975020-DEL-154 | chr18-51975021-DEL-154 |
| chr18-51992922-DEL-434 | chr18-51992923-DEL-434 |
| chr18-51993866-DEL-330 | chr18-51993857-DEL-320 |
| chr18-51996982-DEL-61 | chr18-51996975-DEL-55 |
| chr18-51999349-DEL-274 | chr18-51999350-DEL-274 |
| chr18-52005142-DEL-90 | chr18-52005135-DEL-52 |
| chr18-52022829-DEL-91 | chr18-52022830-DEL-91 |
| chr18-52061425-DEL-106 | chr18-52061426-DEL-106 |
| chr18-52082255-DEL-3465 | chr18-52082251-DEL-3469 |
| chr18-52097114-DEL-497 | chr18-52097115-DEL-497 |
| chr18-52105752-DEL-199 | chr18-52105753-DEL-199 |
| chr18-52125338-DEL-223 | chr18-52125361-DEL-241 |
| chr18-52152166-DEL-7662 | chr18-52152167-DEL-7662 |
| chr18-52209808-DEL-51 | chr18-52209809-DEL-51 |
| chr18-52278802-DEL-90 | chr18-52278803-DEL-54 |
| chr18-52288630-DEL-114 | chr18-52288630-DEL-117 |
| chr18-52315330-DEL-76 | chr18-52315363-DEL-56 |
| chr18-52315862-DEL-58 | chr18-52315861-DEL-59 |
| chr18-52332793-DEL-58 | chr18-52332794-DEL-58 |
| chr18-52333849-DEL-4577 | chr18-52333850-DEL-4577 |
| chr18-52340069-DEL-1926 | chr18-52340070-DEL-1926 |
| chr18-52388671-DEL-6653 | chr18-52388672-DEL-6653 |
| chr18-52405631-DEL-52 | chr18-52405624-DEL-60 |
| chr18-52454093-DEL-1704 | chr18-52454094-DEL-1704 |
| chr18-52474135-DEL-146 | chr18-52474136-DEL-146 |
| chr18-52490317-DEL-90 | chr18-52490318-DEL-90 |
| chr18-52508759-DEL-238 | chr18-52508754-DEL-229 |
| chr18-52511841-DEL-1604 | chr18-52511842-DEL-1604 |
| chr18-52520974-DEL-7129 | chr18-52520975-DEL-7129 |
| chr18-52532879-DEL-1145 | chr18-52532880-DEL-1145 |
| chr18-52539175-DEL-382 | chr18-52539176-DEL-382 |
| chr18-52545025-DEL-3311 | chr18-52545026-DEL-3311 |
| chr18-52548652-DEL-3416 | chr18-52548653-DEL-3416 |
| chr18-52548658-DEL-1612 | chr18-52548657-DEL-1614 |
| chr18-52564398-DEL-498 | chr18-52564399-DEL-498 |
| chr18-52570056-DEL-1348 | chr18-52570057-DEL-1348 |
| chr18-52579746-DEL-8102 | chr18-52579730-DEL-8117 |
| chr18-52600510-DEL-305 | chr18-52600511-DEL-305 |
| chr18-52612303-DEL-625 | chr18-52612304-DEL-625 |
| chr18-52624795-DEL-61 | chr18-52624796-DEL-61 |
| chr18-52626059-DEL-70 | chr18-52626060-DEL-68 |
| chr18-52645145-DEL-60 | chr18-52645145-DEL-60 |
| chr18-52648227-DEL-51 | chr18-52648228-DEL-51 |
| chr18-52655363-DEL-456 | chr18-52655364-DEL-456 |
| chr18-52674592-DEL-402 | chr18-52674593-DEL-402 |
| chr18-52675165-DEL-67 | chr18-52675166-DEL-67 |
| chr18-52676321-DEL-6421 | chr18-52676322-DEL-6421 |
| chr18-52707959-DEL-1684 | chr18-52707960-DEL-1684 |
| chr18-52708578-DEL-98 | chr18-52708579-DEL-97 |
| chr18-52740982-DEL-761 | chr18-52740983-DEL-761 |
| chr18-52752215-DEL-205 | chr18-52752216-DEL-205 |
| chr18-52760928-DEL-166 | chr18-52760929-DEL-167 |
| chr18-52767182-DEL-55 | chr18-52767187-DEL-50 |
| chr18-52781565-DEL-113 | chr18-52781566-DEL-113 |
| chr18-52784939-DEL-84 | chr18-52784985-DEL-78 |
| chr18-52855946-DEL-73 | chr18-52855952-DEL-68 |
| chr18-52871348-DEL-252 | chr18-52871388-DEL-204 |
| chr18-52872516-DEL-330 | chr18-52872517-DEL-330 |
| chr18-52888620-DEL-51 | chr18-52888621-DEL-51 |
| chr18-52910134-DEL-82 | chr18-52910105-DEL-122 |
| chr18-52957169-DEL-62 | chr18-52957170-DEL-62 |
| chr18-52961834-DEL-65 | chr18-52961835-DEL-65 |
| chr18-52999993-DEL-277 | chr18-52999994-DEL-277 |
| chr18-53009342-DEL-6796 | chr18-53009343-DEL-6796 |
| chr18-53071259-DEL-281 | chr18-53071260-DEL-281 |
| chr18-53096598-DEL-107 | chr18-53096598-DEL-107 |
| chr18-53145541-DEL-1929 | chr18-53145542-DEL-1929 |
| chr18-53145637-DEL-76 | chr18-53145638-DEL-76 |
| chr18-53153758-DEL-6535 | chr18-53153759-DEL-6535 |
| chr18-53219167-DEL-89 | chr18-53219168-DEL-89 |
| chr18-53286349-DEL-322 | chr18-53286350-DEL-322 |
| chr18-53311335-DEL-220 | chr18-53311336-DEL-220 |

| | |
|---|---|
| chr18-53333879-DEL-1762 | chr18-53333881-DEL-1772 |
| chr18-53337486-DEL-1236 | chr18-53337487-DEL-1236 |
| chr18-53394969-DEL-56 | chr18-53394970-DEL-56 |
| chr18-53402099-DEL-80 | chr18-53402100-DEL-80 |
| chr18-53410434-DEL-2266 | chr18-53410435-DEL-2266 |
| chr18-53415016-DEL-86 | chr18-53415017-DEL-86 |
| chr18-53431341-DEL-105 | chr18-53431342-DEL-105 |
| chr18-53456213-DEL-71 | chr18-53456217-DEL-64 |
| chr18-53469582-DEL-476 | chr18-53469583-DEL-476 |
| chr18-53476011-DEL-398 | chr18-53476012-DEL-398 |
| chr18-53506019-DEL-58 | chr18-53506020-DEL-58 |
| chr18-53508305-DEL-74 | chr18-53508306-DEL-68 |
| chr18-53567361-DEL-86 | chr18-53567357-DEL-78 |
| chr18-53568126-DEL-51 | chr18-53568127-DEL-51 |
| chr18-53568587-DEL-1085 | chr18-53568588-DEL-1085 |
| chr18-53578164-DEL-349 | chr18-53578164-DEL-350 |
| chr18-53618519-DEL-86 | chr18-53618502-DEL-108 |
| chr18-53621537-DEL-250 | chr18-53621538-DEL-250 |
| chr18-53626227-DEL-143 | chr18-53626228-DEL-144 |
| chr18-53654068-DEL-64 | chr18-53654040-DEL-74 |
| chr18-53658743-DEL-62 | chr18-53658755-DEL-56 |
| chr18-53662367-DEL-116 | chr18-53662368-DEL-116 |
| chr18-53663734-DEL-91 | chr18-53663735-DEL-91 |
| chr18-53693557-DEL-6258 | chr18-53693558-DEL-6258 |
| chr18-53747022-DEL-53 | chr18-53747022-DEL-53 |
| chr18-53749115-DEL-65 | chr18-53749086-DEL-86 |
| chr18-53763796-DEL-67 | chr18-53763797-DEL-67 |
| chr18-53789020-DEL-1007 | chr18-53789021-DEL-1007 |
| chr18-53791595-DEL-728 | chr18-53791596-DEL-728 |
| chr18-53799271-DEL-82 | chr18-53799272-DEL-82 |
| chr18-53805166-DEL-77 | chr18-53805167-DEL-77 |
| chr18-53820635-DEL-166 | chr18-53820636-DEL-170 |
| chr18-53834927-DEL-144 | chr18-53834928-DEL-144 |
| chr18-53849494-DEL-114 | chr18-53849495-DEL-114 |
| chr18-53858593-DEL-66 | chr18-53858608-DEL-52 |
| chr18-53864876-DEL-51 | chr18-53864869-DEL-52 |
| chr18-53867034-DEL-159 | chr18-53867035-DEL-159 |
| chr18-53867729-DEL-400 | chr18-53867730-DEL-400 |
| chr18-53897352-DEL-206 | chr18-53897341-DEL-207 |
| chr18-53910634-DEL-59 | chr18-53910635-DEL-59 |
| chr18-53923577-DEL-183 | chr18-53923578-DEL-183 |
| chr18-53927609-DEL-220 | chr18-53927610-DEL-220 |
| chr18-53938327-DEL-173 | chr18-53938328-DEL-170 |
| chr18-53966479-DEL-6452 | chr18-53966480-DEL-6452 |
| chr18-53984675-DEL-221 | chr18-53984676-DEL-221 |
| chr18-54002496-DEL-7157 | chr18-54002497-DEL-7157 |
| chr18-54022867-DEL-215 | chr18-54022868-DEL-215 |
| chr18-54093046-DEL-920 | chr18-54093044-DEL-921 |
| chr18-54111054-DEL-912 | chr18-54111043-DEL-913 |
| chr18-54130551-DEL-65 | chr18-54130552-DEL-65 |
| chr18-54132147-DEL-263 | chr18-54132148-DEL-263 |
| chr18-54142472-DEL-176 | chr18-54142473-DEL-176 |
| chr18-54156066-DEL-423 | chr18-54156064-DEL-403 |
| chr18-54156827-DEL-6348 | chr18-54156828-DEL-6348 |
| chr18-54196900-DEL-1717 | chr18-54196901-DEL-1717 |
| chr18-54235380-DEL-60 | chr18-54235381-DEL-60 |
| chr18-54256077-DEL-78 | chr18-54256078-DEL-78 |
| chr18-54282318-DEL-60 | chr18-54282319-DEL-60 |
| chr18-54282806-DEL-567 | chr18-54282807-DEL-567 |
| chr18-54291287-DEL-1851 | chr18-54291288-DEL-1851 |
| chr18-54302255-DEL-1589 | chr18-54302255-DEL-1588 |
| chr18-54337078-DEL-326 | chr18-54337079-DEL-326 |
| chr18-54359499-DEL-6579 | chr18-54359500-DEL-6579 |
| chr18-54370793-DEL-395 | chr18-54370794-DEL-395 |
| chr18-54377441-DEL-54 | chr18-54377442-DEL-54 |
| chr18-54421087-DEL-3067 | chr18-54421083-DEL-3068 |
| chr18-54452894-DEL-62 | chr18-54452895-DEL-62 |
| chr18-54459576-DEL-7473 | chr18-54459577-DEL-7473 |
| chr18-54502766-DEL-75 | chr18-54502802-DEL-60 |
| chr18-54538087-DEL-172 | chr18-54538088-DEL-172 |
| chr18-54541932-DEL-69 | chr18-54541933-DEL-69 |
| chr18-54550152-DEL-1537 | chr18-54550153-DEL-1537 |
| chr18-54568131-DEL-102 | chr18-54568091-DEL-104 |
| chr18-54599414-DEL-201 | chr18-54599415-DEL-201 |
| chr18-54611134-DEL-109 | chr18-54611135-DEL-109 |
| chr18-54622594-DEL-191 | chr18-54622591-DEL-193 |
| chr18-54636262-DEL-66 | chr18-54636265-DEL-64 |
| chr18-54636837-DEL-465 | chr18-54636838-DEL-465 |
| chr18-54710141-DEL-61 | chr18-54710140-DEL-63 |
| chr18-54717154-DEL-430 | chr18-54717155-DEL-430 |

| | |
|---|---|
| chr18-54779896-DEL-76 | chr18-54779897-DEL-76 |
| chr18-54816718-DEL-147 | chr18-54816730-DEL-131 |
| chr18-54820045-DEL-61 | chr18-54820046-DEL-61 |
| chr18-54831940-DEL-68 | chr18-54831945-DEL-57 |
| chr18-54900219-DEL-166 | chr18-54900225-DEL-177 |
| chr18-54908508-DEL-2289 | chr18-54908509-DEL-2289 |
| chr18-54921585-DEL-728 | chr18-54921586-DEL-728 |
| chr18-54941803-DEL-209 | chr18-54941804-DEL-208 |
| chr18-54948437-DEL-56 | chr18-54948438-DEL-56 |
| chr18-55025438-DEL-90 | chr18-55025439-DEL-84 |
| chr18-55054132-DEL-65 | chr18-55054110-DEL-54 |
| chr18-55072965-DEL-54 | chr18-55072975-DEL-52 |
| chr18-55097030-DEL-72 | chr18-55097031-DEL-72 |
| chr18-55169945-DEL-156 | chr18-55169942-DEL-154 |
| chr18-55174889-DEL-124 | chr18-55174890-DEL-124 |
| chr18-55271928-DEL-210 | chr18-55271929-DEL-210 |
| chr18-55275739-DEL-115 | chr18-55275730-DEL-116 |
| chr18-55293612-DEL-563 | chr18-55293613-DEL-563 |
| chr18-55322417-DEL-55 | chr18-55322425-DEL-50 |
| chr18-55335215-DEL-328 | chr18-55335216-DEL-328 |
| chr18-55344539-DEL-83 | chr18-55344520-DEL-96 |
| chr18-55351454-DEL-601 | chr18-55351455-DEL-601 |
| chr18-55355526-DEL-1890 | chr18-55355527-DEL-1890 |
| chr18-55362543-DEL-82 | chr18-55362520-DEL-106 |
| chr18-55373909-DEL-75 | chr18-55373911-DEL-74 |
| chr18-55383971-DEL-60 | chr18-55383966-DEL-60 |
| chr18-55408497-DEL-52 | chr18-55408478-DEL-64 |
| chr18-55412953-DEL-60 | chr18-55412926-DEL-55 |
| chr18-55423601-DEL-61 | chr18-55423602-DEL-61 |
| chr18-55444181-DEL-10002 | chr18-55444182-DEL-10002 |
| chr18-55495265-DEL-1931 | chr18-55495266-DEL-1931 |
| chr18-55547882-DEL-564 | chr18-55547889-DEL-558 |
| chr18-55552120-DEL-1891 | chr18-55552121-DEL-1891 |
| chr18-55562742-DEL-6454 | chr18-55562743-DEL-6454 |
| chr18-55569882-DEL-143 | chr18-55569891-DEL-105 |
| chr18-55614690-DEL-125 | chr18-55614710-DEL-105 |
| chr18-55619593-DEL-259 | chr18-55619594-DEL-259 |
| chr18-55651104-DEL-133 | chr18-55651105-DEL-133 |
| chr18-55684983-DEL-88 | chr18-55684984-DEL-88 |
| chr18-55705054-DEL-5956 | chr18-55705055-DEL-5956 |
| chr18-55728997-DEL-66 | chr18-55728998-DEL-66 |
| chr18-55747646-DEL-183 | chr18-55747647-DEL-183 |
| chr18-55765920-DEL-54 | chr18-55765921-DEL-50 |
| chr18-55806615-DEL-64 | chr18-55806616-DEL-64 |
| chr18-55832556-DEL-435 | chr18-55832557-DEL-414 |
| chr18-55837270-DEL-615 | chr18-55837271-DEL-615 |
| chr18-55843194-DEL-972 | chr18-55843195-DEL-972 |
| chr18-55874113-DEL-76 | chr18-55874114-DEL-76 |
| chr18-55888751-DEL-200 | chr18-55888752-DEL-200 |
| chr18-55906205-DEL-693 | chr18-55906205-DEL-693 |
| chr18-55913753-DEL-507 | chr18-55913754-DEL-507 |
| chr18-55914321-DEL-398 | chr18-55914322-DEL-398 |
| chr18-55917457-DEL-84 | chr18-55917458-DEL-84 |
| chr18-56021100-DEL-54 | chr18-56021101-DEL-54 |
| chr18-56030103-DEL-6051 | chr18-56030104-DEL-6051 |
| chr18-56053048-DEL-60 | chr18-56053037-DEL-56 |
| chr18-56061653-DEL-84 | chr18-56061665-DEL-62 |
| chr18-56063341-DEL-510 | chr18-56063342-DEL-510 |
| chr18-56065502-DEL-52 | chr18-56065473-DEL-82 |
| chr18-56093475-DEL-76 | chr18-56093478-DEL-74 |
| chr18-56159570-DEL-322 | chr18-56159571-DEL-322 |
| chr18-56207657-DEL-1652 | chr18-56207658-DEL-1652 |
| chr18-56213056-DEL-4472 | chr18-56213057-DEL-4472 |
| chr18-56219097-DEL-1091 | chr18-56219098-DEL-1091 |
| chr18-56232891-DEL-805 | chr18-56232892-DEL-805 |
| chr18-56249864-DEL-68 | chr18-56249872-DEL-82 |
| chr18-56266796-DEL-72 | chr18-56266797-DEL-72 |
| chr18-56275237-DEL-3888 | chr18-56275238-DEL-3888 |
| chr18-56281185-DEL-84 | chr18-56281186-DEL-84 |
| chr18-56292763-DEL-1011 | chr18-56292763-DEL-1011 |
| chr18-56293326-DEL-182 | chr18-56293324-DEL-184 |
| chr18-56299351-DEL-60 | chr18-56299331-DEL-61 |
| chr18-56302263-DEL-92 | chr18-56302274-DEL-82 |
| chr18-56317431-DEL-1471 | chr18-56317427-DEL-1468 |
| chr18-56349724-DEL-2762 | chr18-56349738-DEL-2756 |
| chr18-56353414-DEL-58 | chr18-56353415-DEL-56 |
| chr18-56355858-DEL-7474 | chr18-56355859-DEL-7474 |
| chr18-56433172-DEL-108 | chr18-56433173-DEL-108 |
| chr18-56436402-DEL-372 | chr18-56436403-DEL-372 |
| chr18-56458878-DEL-202 | chr18-56458879-DEL-202 |

| | |
|---|---|
| chr18-56473920-DEL-5026 | chr18-56473921-DEL-5026 |
| chr18-56483632-DEL-125 | chr18-56483624-DEL-118 |
| chr18-56494320-DEL-890 | chr18-56494321-DEL-890 |
| chr18-56503392-DEL-68 | chr18-56503393-DEL-68 |
| chr18-56535891-DEL-76 | chr18-56535864-DEL-76 |
| chr18-56542680-DEL-72 | chr18-56542669-DEL-84 |
| chr18-56584631-DEL-221 | chr18-56584632-DEL-221 |
| chr18-56600843-DEL-68 | chr18-56600844-DEL-68 |
| chr18-56603868-DEL-60 | chr18-56603869-DEL-60 |
| chr18-56616810-DEL-72 | chr18-56616807-DEL-76 |
| chr18-56648210-DEL-60 | chr18-56648210-DEL-60 |
| chr18-56650365-DEL-141 | chr18-56650363-DEL-132 |
| chr18-56654929-DEL-69 | chr18-56654930-DEL-69 |
| chr18-56673945-DEL-60 | chr18-56673946-DEL-60 |
| chr18-56675030-DEL-55 | chr18-56675027-DEL-55 |
| chr18-56698739-DEL-205 | chr18-56698740-DEL-205 |
| chr18-56715366-DEL-208 | chr18-56715367-DEL-208 |
| chr18-56720629-DEL-120 | chr18-56720630-DEL-120 |
| chr18-56737304-DEL-202 | chr18-56737305-DEL-190 |
| chr18-56739312-DEL-63 | chr18-56739307-DEL-63 |
| chr18-56744313-DEL-179 | chr18-56744314-DEL-179 |
| chr18-56748830-DEL-181 | chr18-56748831-DEL-181 |
| chr18-56749716-DEL-165 | chr18-56749717-DEL-165 |
| chr18-56751390-DEL-771 | chr18-56751391-DEL-771 |
| chr18-56754884-DEL-405 | chr18-56754876-DEL-389 |
| chr18-56761570-DEL-90 | chr18-56761571-DEL-90 |
| chr18-56764797-DEL-52 | chr18-56764798-DEL-52 |
| chr18-56771192-DEL-187 | chr18-56771193-DEL-187 |
| chr18-56775714-DEL-155 | chr18-56775715-DEL-155 |
| chr18-56777302-DEL-217 | chr18-56777303-DEL-217 |
| chr18-56779608-DEL-159 | chr18-56779609-DEL-151 |
| chr18-56784655-DEL-203 | chr18-56784656-DEL-203 |
| chr18-56787959-DEL-400 | chr18-56787960-DEL-400 |
| chr18-56795123-DEL-212 | chr18-56795122-DEL-212 |
| chr18-56796064-DEL-168 | chr18-56796065-DEL-168 |
| chr18-56807507-DEL-224 | chr18-56807508-DEL-224 |
| chr18-56813084-DEL-588 | chr18-56813084-DEL-588 |
| chr18-56814647-DEL-93 | chr18-56814648-DEL-72 |
| chr18-56830128-DEL-228 | chr18-56830130-DEL-226 |
| chr18-56832766-DEL-191 | chr18-56832767-DEL-191 |
| chr18-56839462-DEL-172 | chr18-56839463-DEL-172 |
| chr18-56843528-DEL-223 | chr18-56843529-DEL-223 |
| chr18-56856284-DEL-210 | chr18-56856285-DEL-210 |
| chr18-56870264-DEL-120 | chr18-56870265-DEL-120 |
| chr18-56871405-DEL-224 | chr18-56871406-DEL-224 |
| chr18-56889791-DEL-594 | chr18-56889792-DEL-594 |
| chr18-56960009-DEL-3236 | chr18-56960010-DEL-3236 |
| chr18-56967229-DEL-168 | chr18-56967221-DEL-169 |
| chr18-56985474-DEL-1594 | chr18-56985474-DEL-1583 |
| chr18-57013881-DEL-75 | chr18-57013849-DEL-72 |
| chr18-57017250-DEL-166 | chr18-57017251-DEL-166 |
| chr18-57024923-DEL-54 | chr18-57024924-DEL-54 |
| chr18-57034190-DEL-195 | chr18-57034191-DEL-195 |
| chr18-57050752-DEL-60 | chr18-57050753-DEL-60 |
| chr18-57062122-DEL-191 | chr18-57062123-DEL-191 |
| chr18-57063556-DEL-193 | chr18-57063557-DEL-193 |
| chr18-57093497-DEL-56 | chr18-57093498-DEL-56 |
| chr18-57094394-DEL-68 | chr18-57094395-DEL-68 |
| chr18-57137555-DEL-62 | chr18-57137556-DEL-62 |
| chr18-57140937-DEL-6412 | chr18-57140938-DEL-6412 |
| chr18-57161496-DEL-104 | chr18-57161490-DEL-84 |
| chr18-57171307-DEL-440 | chr18-57171308-DEL-440 |
| chr18-57203204-DEL-52 | chr18-57203204-DEL-52 |
| chr18-57204748-DEL-79 | chr18-57204749-DEL-79 |
| chr18-57241868-DEL-75 | chr18-57241877-DEL-63 |
| chr18-57290003-DEL-69 | chr18-57290004-DEL-69 |
| chr18-57338694-DEL-78 | chr18-57338693-DEL-80 |
| chr18-57347447-DEL-6125 | chr18-57347432-DEL-6126 |
| chr18-57355718-DEL-132 | chr18-57355728-DEL-123 |
| chr18-57364639-DEL-61 | chr18-57364640-DEL-61 |
| chr18-57372880-DEL-609 | chr18-57372881-DEL-609 |
| chr18-57374260-DEL-125 | chr18-57374261-DEL-125 |
| chr18-57411277-DEL-105 | chr18-57411274-DEL-103 |
| chr18-57475066-DEL-171 | chr18-57475067-DEL-171 |
| chr18-57483952-DEL-416 | chr18-57483953-DEL-416 |
| chr18-57499223-DEL-443 | chr18-57499241-DEL-426 |
| chr18-57581309-DEL-71 | chr18-57581310-DEL-71 |
| chr18-57599550-DEL-239 | chr18-57599551-DEL-239 |
| chr18-57623087-DEL-441 | chr18-57623088-DEL-441 |
| chr18-57627959-DEL-127 | chr18-57627960-DEL-127 |

| | |
|---|---|
| chr18-57638005-DEL-337 | chr18-57638006-DEL-328 |
| chr18-57638846-DEL-62 | chr18-57638847-DEL-62 |
| chr18-57641239-DEL-1207 | chr18-57641240-DEL-1207 |
| chr18-57652600-DEL-352 | chr18-57652600-DEL-353 |
| chr18-57685037-DEL-54 | chr18-57685041-DEL-56 |
| chr18-57693574-DEL-1866 | chr18-57693575-DEL-1866 |
| chr18-57785336-DEL-196 | chr18-57785337-DEL-195 |
| chr18-57830183-DEL-126 | chr18-57830184-DEL-126 |
| chr18-57842160-DEL-99 | chr18-57842161-DEL-99 |
| chr18-57867846-DEL-70 | chr18-57867847-DEL-70 |
| chr18-57883513-DEL-59 | chr18-57883514-DEL-59 |
| chr18-57892490-DEL-105 | chr18-57892491-DEL-105 |
| chr18-57898733-DEL-322 | chr18-57898734-DEL-322 |
| chr18-57904459-DEL-92 | chr18-57904430-DEL-90 |
| chr18-57915501-DEL-61 | chr18-57915510-DEL-52 |
| chr18-57925584-DEL-6368 | chr18-57925585-DEL-6368 |
| chr18-57934489-DEL-1525 | chr18-57934490-DEL-1525 |
| chr18-57945152-DEL-1488 | chr18-57945152-DEL-1488 |
| chr18-57947097-DEL-62 | chr18-57947096-DEL-64 |
| chr18-57951163-DEL-128 | chr18-57951164-DEL-128 |
| chr18-57962673-DEL-58 | chr18-57962674-DEL-58 |
| chr18-57980212-DEL-1041 | chr18-57980213-DEL-1041 |
| chr18-58035058-DEL-171 | chr18-58035059-DEL-171 |
| chr18-58041157-DEL-233 | chr18-58041158-DEL-233 |
| chr18-58074495-DEL-126 | chr18-58074473-DEL-117 |
| chr18-58081105-DEL-166 | chr18-58081106-DEL-166 |
| chr18-58097419-DEL-64 | chr18-58097421-DEL-63 |
| chr18-58109668-DEL-73 | chr18-58109680-DEL-75 |
| chr18-58115860-DEL-2633 | chr18-58115861-DEL-2633 |
| chr18-58143969-DEL-154 | chr18-58143970-DEL-154 |
| chr18-58155809-DEL-65 | chr18-58155798-DEL-77 |
| chr18-58164521-DEL-54 | chr18-58164522-DEL-54 |
| chr18-58254768-DEL-91 | chr18-58254769-DEL-91 |
| chr18-58276643-DEL-95 | chr18-58276644-DEL-95 |
| chr18-58289779-DEL-58 | chr18-58289779-DEL-58 |
| chr18-58302573-DEL-87 | chr18-58302574-DEL-87 |
| chr18-58303532-DEL-82 | chr18-58303530-DEL-83 |
| chr18-58320454-DEL-95 | chr18-58320455-DEL-95 |
| chr18-58372501-DEL-785 | chr18-58372502-DEL-785 |
| chr18-58375558-DEL-301 | chr18-58375559-DEL-301 |
| chr18-58384382-DEL-614 | chr18-58384383-DEL-614 |
| chr18-58391279-DEL-320 | chr18-58391280-DEL-320 |
| chr18-58397407-DEL-73 | chr18-58397408-DEL-73 |
| chr18-58417587-DEL-4080 | chr18-58417588-DEL-4080 |
| chr18-58429013-DEL-1262 | chr18-58429013-DEL-1262 |
| chr18-58429022-DEL-238 | chr18-58429023-DEL-238 |
| chr18-58450849-DEL-4989 | chr18-58450850-DEL-4989 |
| chr18-58498470-DEL-6224 | chr18-58498491-DEL-6204 |
| chr18-58562811-DEL-400 | chr18-58562812-DEL-400 |
| chr18-58583609-DEL-404 | chr18-58583610-DEL-404 |
| chr18-58611842-DEL-60 | chr18-58611843-DEL-60 |
| chr18-58634827-DEL-398 | chr18-58634824-DEL-372 |
| chr18-58637709-DEL-80 | chr18-58637710-DEL-80 |
| chr18-58640317-DEL-405 | chr18-58640318-DEL-405 |
| chr18-58662415-DEL-91 | chr18-58662385-DEL-137 |
| chr18-58671431-DEL-335 | chr18-58671432-DEL-335 |
| chr18-58695552-DEL-90 | chr18-58695533-DEL-111 |
| chr18-58705357-DEL-88 | chr18-58705361-DEL-83 |
| chr18-58729246-DEL-190 | chr18-58729247-DEL-190 |
| chr18-58779704-DEL-62 | chr18-58779666-DEL-88 |
| chr18-58786437-DEL-62 | chr18-58786438-DEL-62 |
| chr18-58808416-DEL-62 | chr18-58808417-DEL-62 |
| chr18-58815660-DEL-59 | chr18-58815657-DEL-58 |
| chr18-58821762-DEL-199 | chr18-58821763-DEL-199 |
| chr18-58853235-DEL-84 | chr18-58853240-DEL-82 |
| chr18-58896693-DEL-74 | chr18-58896696-DEL-72 |
| chr18-58927535-DEL-56 | chr18-58927536-DEL-56 |
| chr18-58937374-DEL-6259 | chr18-58937375-DEL-6259 |
| chr18-58973416-DEL-56 | chr18-58973398-DEL-72 |
| chr18-58986708-DEL-6192 | chr18-58986708-DEL-6192 |
| chr18-58994552-DEL-63 | chr18-58994553-DEL-63 |
| chr18-59000202-DEL-131 | chr18-59000203-DEL-119 |
| chr18-59009080-DEL-147 | chr18-59009081-DEL-147 |
| chr18-59041731-DEL-294 | chr18-59041732-DEL-293 |
| chr18-59046338-DEL-571 | chr18-59046339-DEL-571 |
| chr18-59049353-DEL-6788 | chr18-59049354-DEL-6788 |
| chr18-59088925-DEL-169 | chr18-59088926-DEL-169 |
| chr18-59121072-DEL-103 | chr18-59121079-DEL-97 |
| chr18-59125903-DEL-6442 | chr18-59125904-DEL-6442 |
| chr18-59134357-DEL-70 | chr18-59134364-DEL-89 |

| | |
|---|---|
| chr18-59146751-DEL-80 | chr18-59146772-DEL-60 |
| chr18-59217052-DEL-205 | chr18-59217050-DEL-206 |
| chr18-59219482-DEL-299 | chr18-59219483-DEL-299 |
| chr18-59221692-DEL-283 | chr18-59221692-DEL-283 |
| chr18-59248418-DEL-6605 | chr18-59248419-DEL-6605 |
| chr18-59248503-DEL-212 | chr18-59248504-DEL-212 |
| chr18-59262137-DEL-74 | chr18-59262139-DEL-54 |
| chr18-59295922-DEL-1836 | chr18-59295923-DEL-1836 |
| chr18-59299773-DEL-69 | chr18-59299774-DEL-67 |
| chr18-59398193-DEL-3037 | chr18-59398194-DEL-3037 |
| chr18-59418349-DEL-320 | chr18-59418350-DEL-320 |
| chr18-59429678-DEL-2733 | chr18-59429679-DEL-2733 |
| chr18-59450781-DEL-90 | chr18-59450776-DEL-106 |
| chr18-59498682-DEL-94 | chr18-59498659-DEL-118 |
| chr18-59525121-DEL-79 | chr18-59525126-DEL-66 |
| chr18-59534546-DEL-1098 | chr18-59534547-DEL-1098 |
| chr18-59535685-DEL-414 | chr18-59535680-DEL-409 |
| chr18-59560150-DEL-469 | chr18-59560151-DEL-469 |
| chr18-59580191-DEL-1812 | chr18-59580192-DEL-1812 |
| chr18-59596279-DEL-584 | chr18-59596280-DEL-584 |
| chr18-59674903-DEL-60 | chr18-59674904-DEL-56 |
| chr18-59675789-DEL-170 | chr18-59675790-DEL-170 |
| chr18-59680472-DEL-629 | chr18-59680473-DEL-629 |
| chr18-59715506-DEL-72 | chr18-59715507-DEL-72 |
| chr18-59719966-DEL-68 | chr18-59719967-DEL-68 |
| chr18-59750366-DEL-1482 | chr18-59750341-DEL-1483 |
| chr18-59755173-DEL-123 | chr18-59755185-DEL-112 |
| chr18-59766491-DEL-193 | chr18-59766492-DEL-193 |
| chr18-59772898-DEL-68 | chr18-59772892-DEL-77 |
| chr18-59793027-DEL-658 | chr18-59793029-DEL-659 |
| chr18-59832756-DEL-427 | chr18-59832755-DEL-428 |
| chr18-59856247-DEL-86 | chr18-59856260-DEL-57 |
| chr18-59862510-DEL-506 | chr18-59862510-DEL-506 |
| chr18-59863044-DEL-7731 | chr18-59863045-DEL-7731 |
| chr18-59876887-DEL-69 | chr18-59876889-DEL-71 |
| chr18-59887563-DEL-776 | chr18-59887564-DEL-776 |
| chr18-59910131-DEL-443 | chr18-59910132-DEL-442 |
| chr18-59924130-DEL-489 | chr18-59924131-DEL-489 |
| chr18-59932094-DEL-60 | chr18-59932086-DEL-54 |
| chr18-59932581-DEL-1617 | chr18-59932582-DEL-1617 |
| chr18-59935412-DEL-944 | chr18-59935413-DEL-944 |
| chr18-59943756-DEL-151 | chr18-59943757-DEL-151 |
| chr18-59960239-DEL-70 | chr18-59960241-DEL-70 |
| chr18-59961174-DEL-171 | chr18-59961175-DEL-161 |
| chr18-59964089-DEL-81 | chr18-59964090-DEL-81 |
| chr18-59974477-DEL-52 | chr18-59974478-DEL-52 |
| chr18-60004461-DEL-181 | chr18-60004464-DEL-179 |
| chr18-60007245-DEL-1193 | chr18-60007246-DEL-1193 |
| chr18-60046872-DEL-64 | chr18-60046873-DEL-64 |
| chr18-60071124-DEL-717 | chr18-60071125-DEL-717 |
| chr18-60080855-DEL-7303 | chr18-60080856-DEL-7303 |
| chr18-60086946-DEL-848 | chr18-60086945-DEL-848 |
| chr18-60119972-DEL-6253 | chr18-60119973-DEL-6253 |
| chr18-60136747-DEL-1089 | chr18-60136750-DEL-1090 |
| chr18-60149717-DEL-92 | chr18-60149724-DEL-80 |
| chr18-60163129-DEL-51 | chr18-60163130-DEL-51 |
| chr18-60175547-DEL-1886 | chr18-60175548-DEL-1886 |
| chr18-60204054-DEL-1704 | chr18-60204055-DEL-1704 |
| chr18-60205336-DEL-1002 | chr18-60205337-DEL-1002 |
| chr18-60209346-DEL-1319 | chr18-60209347-DEL-1319 |
| chr18-60300105-DEL-352 | chr18-60300106-DEL-352 |
| chr18-60306201-DEL-878 | chr18-60306202-DEL-878 |
| chr18-60309830-DEL-138 | chr18-60309831-DEL-138 |
| chr18-60321784-DEL-489 | chr18-60321785-DEL-489 |
| chr18-60324890-DEL-90 | chr18-60324890-DEL-90 |
| chr18-60368850-DEL-2792 | chr18-60368851-DEL-2792 |
| chr18-60403398-DEL-797 | chr18-60403399-DEL-797 |
| chr18-60535170-DEL-4247 | chr18-60535170-DEL-4247 |
| chr18-60538857-DEL-4285 | chr18-60538856-DEL-4285 |
| chr18-60545484-DEL-258 | chr18-60545485-DEL-258 |
| chr18-60559223-DEL-1117 | chr18-60559224-DEL-1117 |
| chr18-60595189-DEL-100 | chr18-60595190-DEL-100 |
| chr18-60612430-DEL-234 | chr18-60612431-DEL-234 |
| chr18-60619134-DEL-256 | chr18-60619135-DEL-256 |
| chr18-60623614-DEL-60 | chr18-60623619-DEL-69 |
| chr18-60651477-DEL-63 | chr18-60651478-DEL-63 |
| chr18-60661432-DEL-5361 | chr18-60661433-DEL-5361 |
| chr18-60709194-DEL-58 | chr18-60709194-DEL-58 |
| chr18-60715140-DEL-1382 | chr18-60715141-DEL-1382 |
| chr18-60716624-DEL-317 | chr18-60716626-DEL-315 |

| | |
|---|---|
| chr18-60732654-DEL-125 | chr18-60732655-DEL-125 |
| chr18-60792289-DEL-173 | chr18-60792290-DEL-173 |
| chr18-60795534-DEL-208 | chr18-60795535-DEL-208 |
| chr18-60796484-DEL-83 | chr18-60796462-DEL-84 |
| chr18-60866918-DEL-56 | chr18-60866919-DEL-56 |
| chr18-60898570-DEL-115 | chr18-60898617-DEL-89 |
| chr18-60904094-DEL-166 | chr18-60904095-DEL-166 |
| chr18-60923134-DEL-89 | chr18-60923135-DEL-89 |
| chr18-60927156-DEL-209 | chr18-60927157-DEL-209 |
| chr18-60929015-DEL-172 | chr18-60929019-DEL-182 |
| chr18-60937352-DEL-492 | chr18-60937353-DEL-492 |
| chr18-60947464-DEL-60 | chr18-60947465-DEL-60 |
| chr18-60962750-DEL-176 | chr18-60962733-DEL-176 |
| chr18-60985120-DEL-198 | chr18-60985121-DEL-198 |
| chr18-60994512-DEL-60 | chr18-60994513-DEL-60 |
| chr18-61040566-DEL-209 | chr18-61040567-DEL-209 |
| chr18-61047768-DEL-66 | chr18-61047779-DEL-56 |
| chr18-61072664-DEL-80 | chr18-61072665-DEL-80 |
| chr18-61106548-DEL-151 | chr18-61106549-DEL-151 |
| chr18-61171134-DEL-302 | chr18-61171133-DEL-303 |
| chr18-61226741-DEL-52 | chr18-61226742-DEL-52 |
| chr18-61227635-DEL-216 | chr18-61227636-DEL-216 |
| chr18-61230157-DEL-55 | chr18-61230158-DEL-55 |
| chr18-61262190-DEL-71 | chr18-61262191-DEL-72 |
| chr18-61272481-DEL-112 | chr18-61272482-DEL-112 |
| chr18-61315484-DEL-165 | chr18-61315451-DEL-168 |
| chr18-61316294-DEL-720 | chr18-61316295-DEL-720 |
| chr18-61352946-DEL-68 | chr18-61352931-DEL-84 |
| chr18-61375964-DEL-62 | chr18-61375980-DEL-56 |
| chr18-61392072-DEL-987 | chr18-61392073-DEL-987 |
| chr18-61405376-DEL-488 | chr18-61405377-DEL-488 |
| chr18-61406659-DEL-559 | chr18-61406661-DEL-559 |
| chr18-61421431-DEL-55 | chr18-61421432-DEL-55 |
| chr18-61476677-DEL-77 | chr18-61476678-DEL-77 |
| chr18-61497614-DEL-64 | chr18-61497615-DEL-64 |
| chr18-61555859-DEL-55 | chr18-61555860-DEL-55 |
| chr18-61574701-DEL-70 | chr18-61574694-DEL-78 |
| chr18-61634754-DEL-73 | chr18-61634755-DEL-73 |
| chr18-61644118-DEL-152 | chr18-61644140-DEL-150 |
| chr18-61653291-DEL-53 | chr18-61653291-DEL-50 |
| chr18-61655927-DEL-317 | chr18-61655928-DEL-317 |
| chr18-61667114-DEL-125 | chr18-61667139-DEL-101 |
| chr18-61670541-DEL-208 | chr18-61670542-DEL-208 |
| chr18-61732286-DEL-178 | chr18-61732284-DEL-179 |
| chr18-61734768-DEL-207 | chr18-61734769-DEL-207 |
| chr18-61817237-DEL-369 | chr18-61817238-DEL-368 |
| chr18-61826269-DEL-89 | chr18-61826260-DEL-68 |
| chr18-61894709-DEL-1399 | chr18-61894710-DEL-1399 |
| chr18-61929322-DEL-401 | chr18-61929323-DEL-401 |
| chr18-61950856-DEL-60 | chr18-61950874-DEL-58 |
| chr18-61981715-DEL-54 | chr18-61981716-DEL-54 |
| chr18-62029515-DEL-212 | chr18-62029516-DEL-209 |
| chr18-62037316-DEL-53 | chr18-62037317-DEL-53 |
| chr18-62057594-DEL-213 | chr18-62057595-DEL-218 |
| chr18-62066020-DEL-252 | chr18-62066021-DEL-252 |
| chr18-62078433-DEL-211 | chr18-62078434-DEL-211 |
| chr18-62105174-DEL-536 | chr18-62105172-DEL-533 |
| chr18-62105918-DEL-381 | chr18-62105919-DEL-381 |
| chr18-62168893-DEL-108 | chr18-62168894-DEL-126 |
| chr18-62169943-DEL-52 | chr18-62169943-DEL-52 |
| chr18-62178409-DEL-65 | chr18-62178414-DEL-57 |
| chr18-62191891-DEL-124 | chr18-62191894-DEL-120 |
| chr18-62193431-DEL-288 | chr18-62193431-DEL-256 |
| chr18-62203536-DEL-69 | chr18-62203537-DEL-69 |
| chr18-62206954-DEL-541 | chr18-62206955-DEL-541 |
| chr18-62290038-DEL-58 | chr18-62290028-DEL-86 |
| chr18-62310065-DEL-52 | chr18-62310107-DEL-51 |
| chr18-62379663-DEL-1465 | chr18-62379664-DEL-1465 |
| chr18-62393006-DEL-76 | chr18-62393007-DEL-76 |
| chr18-62393969-DEL-54 | chr18-62393970-DEL-54 |
| chr18-62399772-DEL-55 | chr18-62399773-DEL-50 |
| chr18-62406828-DEL-3657 | chr18-62406828-DEL-3657 |
| chr18-62419459-DEL-6856 | chr18-62419460-DEL-6856 |
| chr18-62442459-DEL-61 | chr18-62442458-DEL-50 |
| chr18-62460453-DEL-84 | chr18-62460438-DEL-100 |
| chr18-62488340-DEL-70 | chr18-62488335-DEL-64 |
| chr18-62495030-DEL-265 | chr18-62495031-DEL-265 |
| chr18-62515458-DEL-383 | chr18-62515451-DEL-382 |
| chr18-62525020-DEL-130 | chr18-62525053-DEL-98 |
| chr18-62534239-DEL-60 | chr18-62534240-DEL-60 |

| | |
|---|---|
| chr18-62583384-DEL-1138 | chr18-62583385-DEL-1138 |
| chr18-62617922-DEL-71 | chr18-62617921-DEL-60 |
| chr18-62631103-DEL-78 | chr18-62631103-DEL-78 |
| chr18-62651156-DEL-55 | chr18-62651157-DEL-55 |
| chr18-62668661-DEL-217 | chr18-62668662-DEL-217 |
| chr18-62701824-DEL-505 | chr18-62701825-DEL-505 |
| chr18-62705662-DEL-787 | chr18-62705663-DEL-787 |
| chr18-62714148-DEL-316 | chr18-62714149-DEL-316 |
| chr18-62733397-DEL-461 | chr18-62733398-DEL-461 |
| chr18-62734574-DEL-396 | chr18-62734575-DEL-396 |
| chr18-62742707-DEL-276 | chr18-62742708-DEL-276 |
| chr18-62751107-DEL-454 | chr18-62751108-DEL-454 |
| chr18-62752729-DEL-6364 | chr18-62752730-DEL-6364 |
| chr18-62762924-DEL-548 | chr18-62762925-DEL-548 |
| chr18-62766009-DEL-126 | chr18-62766008-DEL-126 |
| chr18-62800803-DEL-299 | chr18-62800804-DEL-299 |
| chr18-62810360-DEL-62 | chr18-62810363-DEL-60 |
| chr18-62820922-DEL-76 | chr18-62820923-DEL-80 |
| chr18-62830938-DEL-841 | chr18-62830939-DEL-841 |
| chr18-62840309-DEL-150 | chr18-62840318-DEL-140 |
| chr18-62876773-DEL-166 | chr18-62876740-DEL-188 |
| chr18-62886603-DEL-768 | chr18-62886614-DEL-726 |
| chr18-62906599-DEL-210 | chr18-62906606-DEL-198 |
| chr18-62917260-DEL-468 | chr18-62917261-DEL-468 |
| chr18-62937186-DEL-104 | chr18-62937187-DEL-104 |
| chr18-62948578-DEL-182 | chr18-62948579-DEL-182 |
| chr18-62957904-DEL-150 | chr18-62957936-DEL-156 |
| chr18-62960098-DEL-1888 | chr18-62960099-DEL-1888 |
| chr18-62968222-DEL-58 | chr18-62968223-DEL-58 |
| chr18-63009020-DEL-1718 | chr18-63009022-DEL-1716 |
| chr18-63010707-DEL-85 | chr18-63010708-DEL-85 |
| chr18-63018387-DEL-80 | chr18-63018387-DEL-80 |
| chr18-63024021-DEL-108 | chr18-63024012-DEL-76 |
| chr18-63074703-DEL-933 | chr18-63074704-DEL-933 |
| chr18-63092780-DEL-82 | chr18-63092793-DEL-50 |
| chr18-63099543-DEL-58 | chr18-63099538-DEL-57 |
| chr18-63133229-DEL-164 | chr18-63133232-DEL-162 |
| chr18-63138372-DEL-79 | chr18-63138373-DEL-79 |
| chr18-63140520-DEL-64 | chr18-63140545-DEL-50 |
| chr18-63158893-DEL-1486 | chr18-63158894-DEL-1486 |
| chr18-63166712-DEL-75 | chr18-63166733-DEL-62 |
| chr18-63167334-DEL-6224 | chr18-63167335-DEL-6224 |
| chr18-63191578-DEL-102 | chr18-63191597-DEL-60 |
| chr18-63210329-DEL-54 | chr18-63210330-DEL-54 |
| chr18-63223214-DEL-204 | chr18-63223215-DEL-204 |
| chr18-63262555-DEL-168 | chr18-63262556-DEL-168 |
| chr18-63271843-DEL-82 | chr18-63271844-DEL-79 |
| chr18-63274474-DEL-62 | chr18-63274475-DEL-62 |
| chr18-63306625-DEL-52 | chr18-63306608-DEL-68 |
| chr18-63325211-DEL-3804 | chr18-63325212-DEL-3804 |
| chr18-63367450-DEL-76 | chr18-63367453-DEL-52 |
| chr18-63370807-DEL-112 | chr18-63370822-DEL-98 |
| chr18-63423958-DEL-338 | chr18-63423959-DEL-338 |
| chr18-63444644-DEL-70 | chr18-63444645-DEL-65 |
| chr18-63456655-DEL-538 | chr18-63456656-DEL-538 |
| chr18-63468181-DEL-66 | chr18-63468180-DEL-65 |
| chr18-63469733-DEL-350 | chr18-63469736-DEL-348 |
| chr18-63492052-DEL-116 | chr18-63492041-DEL-152 |
| chr18-63512350-DEL-110 | chr18-63512321-DEL-104 |
| chr18-63524398-DEL-54 | chr18-63524399-DEL-54 |
| chr18-63526379-DEL-4869 | chr18-63526380-DEL-4869 |
| chr18-63531721-DEL-508 | chr18-63531722-DEL-508 |
| chr18-63535683-DEL-456 | chr18-63535684-DEL-456 |
| chr18-63540305-DEL-239 | chr18-63540305-DEL-238 |
| chr18-63556888-DEL-1993 | chr18-63556889-DEL-1993 |
| chr18-63559072-DEL-991 | chr18-63559073-DEL-991 |
| chr18-63576221-DEL-6646 | chr18-63576222-DEL-6646 |
| chr18-63599403-DEL-72 | chr18-63599404-DEL-72 |
| chr18-63605668-DEL-351 | chr18-63605668-DEL-348 |
| chr18-63607921-DEL-6410 | chr18-63607922-DEL-6410 |
| chr18-63614565-DEL-82 | chr18-63614564-DEL-84 |
| chr18-63625861-DEL-51 | chr18-63625862-DEL-51 |
| chr18-63638038-DEL-53 | chr18-63638034-DEL-52 |
| chr18-63639213-DEL-64 | chr18-63639217-DEL-54 |
| chr18-63667709-DEL-6982 | chr18-63667710-DEL-6982 |
| chr18-63683862-DEL-100 | chr18-63683862-DEL-100 |
| chr18-63692887-DEL-758 | chr18-63692888-DEL-758 |
| chr18-63704257-DEL-278 | chr18-63704258-DEL-278 |
| chr18-63716927-DEL-180 | chr18-63716966-DEL-159 |
| chr18-63728048-DEL-52 | chr18-63728049-DEL-52 |

| | |
|---|---|
| chr18-63759999-DEL-1046 | chr18-63760000-DEL-1045 |
| chr18-63777435-DEL-99 | chr18-63777463-DEL-66 |
| chr18-63791605-DEL-403 | chr18-63791606-DEL-403 |
| chr18-63815399-DEL-162 | chr18-63815400-DEL-161 |
| chr18-63826468-DEL-68 | chr18-63826473-DEL-50 |
| chr18-63845362-DEL-423 | chr18-63845344-DEL-420 |
| chr18-63857410-DEL-113 | chr18-63857420-DEL-129 |
| chr18-63867707-DEL-56 | chr18-63867704-DEL-52 |
| chr18-63905412-DEL-226 | chr18-63905417-DEL-222 |
| chr18-63932580-DEL-211 | chr18-63932581-DEL-212 |
| chr18-63965064-DEL-975 | chr18-63965065-DEL-959 |
| chr18-64011177-DEL-118 | chr18-64011178-DEL-118 |
| chr18-64013719-DEL-328 | chr18-64013720-DEL-328 |
| chr18-64044283-DEL-6329 | chr18-64044284-DEL-6329 |
| chr18-64053507-DEL-1611 | chr18-64053508-DEL-1611 |
| chr18-64078896-DEL-766 | chr18-64078897-DEL-766 |
| chr18-64081676-DEL-638 | chr18-64081677-DEL-638 |
| chr18-64116944-DEL-211 | chr18-64116945-DEL-211 |
| chr18-64130569-DEL-72 | chr18-64130577-DEL-65 |
| chr18-64167039-DEL-3475 | chr18-64167040-DEL-3475 |
| chr18-64181744-DEL-7270 | chr18-64181745-DEL-7270 |
| chr18-64182254-DEL-205 | chr18-64182252-DEL-203 |
| chr18-64215614-DEL-59 | chr18-64215581-DEL-55 |
| chr18-64225565-DEL-50 | chr18-64225573-DEL-58 |
| chr18-64260377-DEL-966 | chr18-64260377-DEL-966 |
| chr18-64262205-DEL-59 | chr18-64262206-DEL-59 |
| chr18-64262950-DEL-506 | chr18-64262951-DEL-506 |
| chr18-64269454-DEL-394 | chr18-64269455-DEL-394 |
| chr18-64275659-DEL-209 | chr18-64275690-DEL-179 |
| chr18-64345523-DEL-80 | chr18-64345546-DEL-80 |
| chr18-64359683-DEL-169 | chr18-64359684-DEL-169 |
| chr18-64376214-DEL-52 | chr18-64376215-DEL-52 |
| chr18-64440123-DEL-966 | chr18-64440110-DEL-966 |
| chr18-64461395-DEL-206 | chr18-64461396-DEL-206 |
| chr18-64511687-DEL-64 | chr18-64511688-DEL-64 |
| chr18-64581163-DEL-1923 | chr18-64581164-DEL-1923 |
| chr18-64610449-DEL-70 | chr18-64610464-DEL-74 |
| chr18-64660084-DEL-190 | chr18-64660085-DEL-190 |
| chr18-64682231-DEL-1017 | chr18-64682232-DEL-1017 |
| chr18-64713624-DEL-80 | chr18-64713625-DEL-80 |
| chr18-64728677-DEL-244 | chr18-64728678-DEL-244 |
| chr18-64747122-DEL-56 | chr18-64747125-DEL-60 |
| chr18-64769457-DEL-142 | chr18-64769458-DEL-142 |
| chr18-64775804-DEL-203 | chr18-64775805-DEL-203 |
| chr18-64785709-DEL-1526 | chr18-64785710-DEL-1526 |
| chr18-64832420-DEL-201 | chr18-64832421-DEL-201 |
| chr18-64833663-DEL-282 | chr18-64833665-DEL-285 |
| chr18-64835958-DEL-483 | chr18-64835959-DEL-483 |
| chr18-64843341-DEL-71 | chr18-64843342-DEL-71 |
| chr18-64847394-DEL-848 | chr18-64847395-DEL-848 |
| chr18-64857992-DEL-249 | chr18-64857990-DEL-249 |
| chr18-64859073-DEL-62 | chr18-64859074-DEL-64 |
| chr18-64876039-DEL-61 | chr18-64876040-DEL-61 |
| chr18-64889577-DEL-68 | chr18-64889580-DEL-66 |
| chr18-64907786-DEL-60 | chr18-64907773-DEL-64 |
| chr18-64928756-DEL-60 | chr18-64928756-DEL-60 |
| chr18-64956994-DEL-248 | chr18-64956995-DEL-248 |
| chr18-64973071-DEL-87 | chr18-64973057-DEL-88 |
| chr18-65000412-DEL-190 | chr18-65000413-DEL-190 |
| chr18-65005271-DEL-54 | chr18-65005272-DEL-54 |
| chr18-65026986-DEL-221 | chr18-65026987-DEL-221 |
| chr18-65042204-DEL-88 | chr18-65042218-DEL-57 |
| chr18-65053109-DEL-207 | chr18-65053110-DEL-207 |
| chr18-65057985-DEL-74 | chr18-65057988-DEL-72 |
| chr18-65062199-DEL-197 | chr18-65062184-DEL-179 |
| chr18-65066043-DEL-100 | chr18-65066035-DEL-129 |
| chr18-65071221-DEL-232 | chr18-65071222-DEL-232 |
| chr18-65079936-DEL-81 | chr18-65079937-DEL-80 |
| chr18-65080664-DEL-210 | chr18-65080665-DEL-210 |
| chr18-65086719-DEL-90 | chr18-65086720-DEL-90 |
| chr18-65091903-DEL-141 | chr18-65091903-DEL-142 |
| chr18-65100444-DEL-237 | chr18-65100437-DEL-234 |
| chr18-65109186-DEL-62 | chr18-65109187-DEL-62 |
| chr18-65116764-DEL-56 | chr18-65116764-DEL-77 |
| chr18-65147397-DEL-154 | chr18-65147398-DEL-154 |
| chr18-65162551-DEL-6291 | chr18-65162552-DEL-6291 |
| chr18-65201146-DEL-81 | chr18-65201147-DEL-81 |
| chr18-65239971-DEL-59 | chr18-65239972-DEL-59 |
| chr18-65260138-DEL-61 | chr18-65260139-DEL-61 |
| chr18-65262146-DEL-54 | chr18-65262147-DEL-58 |

| | |
|---|---|
| chr18-65360294-DEL-535 | chr18-65360294-DEL-505 |
| chr18-65362686-DEL-97 | chr18-65362687-DEL-97 |
| chr18-65374288-DEL-106 | chr18-65374289-DEL-106 |
| chr18-65397415-DEL-59 | chr18-65397416-DEL-59 |
| chr18-65398084-DEL-54 | chr18-65398087-DEL-52 |
| chr18-65412242-DEL-60 | chr18-65412265-DEL-56 |
| chr18-65414412-DEL-84 | chr18-65414413-DEL-84 |
| chr18-65415966-DEL-716 | chr18-65415967-DEL-716 |
| chr18-65418913-DEL-2721 | chr18-65418914-DEL-2721 |
| chr18-65436199-DEL-171 | chr18-65436199-DEL-171 |
| chr18-65444197-DEL-54 | chr18-65444198-DEL-52 |
| chr18-65476822-DEL-245 | chr18-65476823-DEL-245 |
| chr18-65477185-DEL-458 | chr18-65477186-DEL-458 |
| chr18-65477885-DEL-202 | chr18-65477886-DEL-202 |
| chr18-65494581-DEL-161 | chr18-65494582-DEL-161 |
| chr18-65496173-DEL-204 | chr18-65496174-DEL-204 |
| chr18-65499551-DEL-102 | chr18-65499552-DEL-102 |
| chr18-65507087-DEL-73 | chr18-65507088-DEL-73 |
| chr18-65522731-DEL-247 | chr18-65522732-DEL-247 |
| chr18-65523736-DEL-95 | chr18-65523737-DEL-95 |
| chr18-65532005-DEL-78 | chr18-65532007-DEL-78 |
| chr18-65533238-DEL-4889 | chr18-65533239-DEL-4889 |
| chr18-65540288-DEL-206 | chr18-65540297-DEL-199 |
| chr18-65616746-DEL-205 | chr18-65616747-DEL-205 |
| chr18-65621248-DEL-203 | chr18-65621249-DEL-204 |
| chr18-65625381-DEL-2375 | chr18-65625397-DEL-2368 |
| chr18-65628592-DEL-56 | chr18-65628587-DEL-50 |
| chr18-65630456-DEL-66 | chr18-65630457-DEL-66 |
| chr18-65643165-DEL-154 | chr18-65643166-DEL-154 |
| chr18-65666771-DEL-66 | chr18-65666771-DEL-65 |
| chr18-65676534-DEL-375 | chr18-65676535-DEL-375 |
| chr18-65677301-DEL-57 | chr18-65677304-DEL-55 |
| chr18-65693711-DEL-54 | chr18-65693712-DEL-54 |
| chr18-65695235-DEL-1373 | chr18-65695236-DEL-1373 |
| chr18-65712008-DEL-62 | chr18-65712009-DEL-62 |
| chr18-65732268-DEL-203 | chr18-65732269-DEL-203 |
| chr18-65760197-DEL-91 | chr18-65760198-DEL-91 |
| chr18-65762996-DEL-399 | chr18-65762997-DEL-399 |
| chr18-65770673-DEL-203 | chr18-65770674-DEL-203 |
| chr18-65799253-DEL-65 | chr18-65799253-DEL-65 |
| chr18-65804051-DEL-53 | chr18-65804052-DEL-53 |
| chr18-65822996-DEL-216 | chr18-65822997-DEL-216 |
| chr18-65830342-DEL-68 | chr18-65830353-DEL-67 |
| chr18-65849352-DEL-3118 | chr18-65849353-DEL-3118 |
| chr18-65893370-DEL-92 | chr18-65893371-DEL-92 |
| chr18-65896268-DEL-883 | chr18-65896269-DEL-883 |
| chr18-65920602-DEL-387 | chr18-65920604-DEL-386 |
| chr18-65926153-DEL-991 | chr18-65926154-DEL-991 |
| chr18-65970318-DEL-231 | chr18-65970319-DEL-231 |
| chr18-66021512-DEL-604 | chr18-66021496-DEL-622 |
| chr18-66032326-DEL-108 | chr18-66032327-DEL-108 |
| chr18-66104308-DEL-339 | chr18-66104309-DEL-339 |
| chr18-66144114-DEL-66 | chr18-66144115-DEL-66 |
| chr18-66161732-DEL-384 | chr18-66161735-DEL-388 |
| chr18-66168983-DEL-180 | chr18-66168984-DEL-180 |
| chr18-66176836-DEL-362 | chr18-66176837-DEL-362 |
| chr18-66198315-DEL-173 | chr18-66198316-DEL-173 |
| chr18-66281200-DEL-2326 | chr18-66281201-DEL-2326 |
| chr18-66284104-DEL-1666 | chr18-66284105-DEL-1666 |
| chr18-66286270-DEL-1336 | chr18-66286271-DEL-1336 |
| chr18-66296343-DEL-435 | chr18-66296344-DEL-435 |
| chr18-66311020-DEL-54 | chr18-66310979-DEL-74 |
| chr18-66405850-DEL-116 | chr18-66405851-DEL-106 |
| chr18-66412366-DEL-883 | chr18-66412367-DEL-891 |
| chr18-66440641-DEL-97 | chr18-66440646-DEL-74 |
| chr18-66463580-DEL-396 | chr18-66463581-DEL-396 |
| chr18-66464180-DEL-419 | chr18-66464181-DEL-419 |
| chr18-66467405-DEL-425 | chr18-66467394-DEL-426 |
| chr18-66476271-DEL-60 | chr18-66476272-DEL-60 |
| chr18-66486876-DEL-168 | chr18-66486878-DEL-169 |
| chr18-66487911-DEL-68 | chr18-66487917-DEL-64 |
| chr18-66488656-DEL-424 | chr18-66488656-DEL-424 |
| chr18-66491702-DEL-1508 | chr18-66491698-DEL-1509 |
| chr18-66494447-DEL-182 | chr18-66494448-DEL-182 |
| chr18-66504010-DEL-54 | chr18-66504011-DEL-54 |
| chr18-66516001-DEL-171 | chr18-66515984-DEL-189 |
| chr18-66523787-DEL-102 | chr18-66523788-DEL-102 |
| chr18-66557310-DEL-1621 | chr18-66557311-DEL-1621 |
| chr18-66564546-DEL-209 | chr18-66564547-DEL-209 |
| chr18-66575375-DEL-72 | chr18-66575388-DEL-79 |

| | |
|---|---|
| chr18-66585892-DEL-214 | chr18-66585893-DEL-190 |
| chr18-66645241-DEL-384 | chr18-66645242-DEL-384 |
| chr18-66647854-DEL-51 | chr18-66647855-DEL-51 |
| chr18-66655690-DEL-89 | chr18-66655692-DEL-89 |
| chr18-66659792-DEL-97 | chr18-66659765-DEL-125 |
| chr18-66677940-DEL-130 | chr18-66677941-DEL-130 |
| chr18-66704255-DEL-707 | chr18-66704253-DEL-707 |
| chr18-66705077-DEL-148 | chr18-66705078-DEL-148 |
| chr18-66712583-DEL-220 | chr18-66712581-DEL-229 |
| chr18-66714329-DEL-197 | chr18-66714332-DEL-196 |
| chr18-66720178-DEL-2682 | chr18-66720179-DEL-2682 |
| chr18-66726544-DEL-83 | chr18-66726545-DEL-83 |
| chr18-66753869-DEL-82 | chr18-66753870-DEL-82 |
| chr18-66768014-DEL-523 | chr18-66768015-DEL-523 |
| chr18-66768997-DEL-58 | chr18-66769003-DEL-53 |
| chr18-66776808-DEL-61 | chr18-66776809-DEL-61 |
| chr18-66792745-DEL-62 | chr18-66792730-DEL-58 |
| chr18-66813428-DEL-209 | chr18-66813429-DEL-209 |
| chr18-66826456-DEL-5996 | chr18-66826457-DEL-5996 |
| chr18-66847882-DEL-540 | chr18-66847883-DEL-540 |
| chr18-66854173-DEL-402 | chr18-66854174-DEL-402 |
| chr18-66860520-DEL-76 | chr18-66860521-DEL-76 |
| chr18-66860523-DEL-8089 | chr18-66860524-DEL-8089 |
| chr18-66883164-DEL-5321 | chr18-66883165-DEL-5321 |
| chr18-66898809-DEL-473 | chr18-66898810-DEL-473 |
| chr18-66907749-DEL-198 | chr18-66907797-DEL-176 |
| chr18-66961001-DEL-2050 | chr18-66961002-DEL-2050 |
| chr18-66962084-DEL-563 | chr18-66962088-DEL-558 |
| chr18-66985289-DEL-69 | chr18-66985290-DEL-69 |
| chr18-67014040-DEL-199 | chr18-67014040-DEL-199 |
| chr18-67034589-DEL-268 | chr18-67034594-DEL-264 |
| chr18-67072923-DEL-6798 | chr18-67072936-DEL-6797 |
| chr18-67113493-DEL-70 | chr18-67113494-DEL-70 |
| chr18-67117751-DEL-185 | chr18-67117752-DEL-185 |
| chr18-67144280-DEL-1011 | chr18-67144282-DEL-1012 |
| chr18-67167180-DEL-55 | chr18-67167180-DEL-55 |
| chr18-67170213-DEL-79 | chr18-67170214-DEL-79 |
| chr18-67247209-DEL-51 | chr18-67247200-DEL-61 |
| chr18-67255563-DEL-89 | chr18-67255584-DEL-69 |
| chr18-67280550-DEL-337 | chr18-67280553-DEL-353 |
| chr18-67284169-DEL-202 | chr18-67284165-DEL-194 |
| chr18-67367566-DEL-79 | chr18-67367567-DEL-79 |
| chr18-67368431-DEL-207 | chr18-67368432-DEL-207 |
| chr18-67375550-DEL-92 | chr18-67375551-DEL-92 |
| chr18-67377394-DEL-127 | chr18-67377395-DEL-127 |
| chr18-67386408-DEL-88 | chr18-67386408-DEL-64 |
| chr18-67391863-DEL-64 | chr18-67391876-DEL-65 |
| chr18-67394565-DEL-75 | chr18-67394599-DEL-50 |
| chr18-67400989-DEL-2110 | chr18-67400990-DEL-2110 |
| chr18-67421572-DEL-268 | chr18-67421573-DEL-268 |
| chr18-67424638-DEL-60 | chr18-67424633-DEL-52 |
| chr18-67438599-DEL-116 | chr18-67438600-DEL-116 |
| chr18-67492267-DEL-177 | chr18-67492271-DEL-174 |
| chr18-67495656-DEL-254 | chr18-67495657-DEL-254 |
| chr18-67498612-DEL-262 | chr18-67498613-DEL-262 |
| chr18-67501352-DEL-56 | chr18-67501353-DEL-56 |
| chr18-67502600-DEL-118 | chr18-67502601-DEL-116 |
| chr18-67504548-DEL-264 | chr18-67504549-DEL-264 |
| chr18-67514101-DEL-215 | chr18-67514102-DEL-214 |
| chr18-67531823-DEL-196 | chr18-67531824-DEL-196 |
| chr18-67545435-DEL-244 | chr18-67545436-DEL-244 |
| chr18-67555742-DEL-91 | chr18-67555740-DEL-92 |
| chr18-67585962-DEL-180 | chr18-67585963-DEL-180 |
| chr18-67593210-DEL-90 | chr18-67593199-DEL-102 |
| chr18-67600574-DEL-202 | chr18-67600575-DEL-202 |
| chr18-67618924-DEL-1990 | chr18-67618925-DEL-1990 |
| chr18-67643242-DEL-201 | chr18-67643243-DEL-201 |
| chr18-67644158-DEL-194 | chr18-67644159-DEL-194 |
| chr18-67663805-DEL-302 | chr18-67663806-DEL-302 |
| chr18-67666741-DEL-186 | chr18-67666742-DEL-186 |
| chr18-67674655-DEL-103 | chr18-67674656-DEL-103 |
| chr18-67684501-DEL-67 | chr18-67684502-DEL-67 |
| chr18-67689292-DEL-200 | chr18-67689293-DEL-200 |
| chr18-67695381-DEL-668 | chr18-67695382-DEL-668 |
| chr18-67724298-DEL-89 | chr18-67724315-DEL-86 |
| chr18-67735932-DEL-222 | chr18-67735933-DEL-222 |
| chr18-67738695-DEL-204 | chr18-67738696-DEL-204 |
| chr18-67758239-DEL-209 | chr18-67758227-DEL-210 |
| chr18-67771044-DEL-162 | chr18-67771038-DEL-168 |
| chr18-67791151-DEL-189 | chr18-67791152-DEL-189 |

| | |
|---|---|
| chr18-67794845-DEL-199 | chr18-67794853-DEL-199 |
| chr18-67822813-DEL-184 | chr18-67822808-DEL-179 |
| chr18-67835027-DEL-215 | chr18-67835028-DEL-215 |
| chr18-67859814-DEL-1744 | chr18-67859815-DEL-1744 |
| chr18-67862733-DEL-58 | chr18-67862734-DEL-58 |
| chr18-67875552-DEL-381 | chr18-67875553-DEL-381 |
| chr18-67886449-DEL-57 | chr18-67886438-DEL-58 |
| chr18-67887104-DEL-178 | chr18-67887105-DEL-178 |
| chr18-67890480-DEL-2051 | chr18-67890481-DEL-2051 |
| chr18-67935251-DEL-206 | chr18-67935252-DEL-206 |
| chr18-67958125-DEL-215 | chr18-67958126-DEL-215 |
| chr18-67973582-DEL-220 | chr18-67973583-DEL-220 |
| chr18-67984931-DEL-64 | chr18-67984932-DEL-64 |
| chr18-68008710-DEL-154 | chr18-68008711-DEL-154 |
| chr18-68046782-DEL-1894 | chr18-68046783-DEL-1894 |
| chr18-68058714-DEL-86 | chr18-68058736-DEL-64 |
| chr18-68076933-DEL-194 | chr18-68076934-DEL-194 |
| chr18-68079229-DEL-58 | chr18-68079230-DEL-58 |
| chr18-68084938-DEL-55 | chr18-68084939-DEL-55 |
| chr18-68090844-DEL-250 | chr18-68090860-DEL-263 |
| chr18-68154465-DEL-56 | chr18-68154466-DEL-56 |
| chr18-68159150-DEL-1282 | chr18-68159151-DEL-1282 |
| chr18-68170077-DEL-68 | chr18-68170090-DEL-56 |
| chr18-68174038-DEL-910 | chr18-68174039-DEL-910 |
| chr18-68192832-DEL-176 | chr18-68192833-DEL-176 |
| chr18-68196053-DEL-194 | chr18-68196053-DEL-194 |
| chr18-68205926-DEL-144 | chr18-68205927-DEL-144 |
| chr18-68226245-DEL-216 | chr18-68226241-DEL-217 |
| chr18-68254606-DEL-62 | chr18-68254611-DEL-62 |
| chr18-68260394-DEL-63 | chr18-68260395-DEL-63 |
| chr18-68270935-DEL-52 | chr18-68270938-DEL-50 |
| chr18-68309480-DEL-145 | chr18-68309485-DEL-141 |
| chr18-68337432-DEL-466 | chr18-68337433-DEL-466 |
| chr18-68338340-DEL-466 | chr18-68338341-DEL-466 |
| chr18-68341970-DEL-194 | chr18-68341971-DEL-194 |
| chr18-68344119-DEL-572 | chr18-68344120-DEL-572 |
| chr18-68349229-DEL-52 | chr18-68349189-DEL-89 |
| chr18-68403816-DEL-477 | chr18-68403817-DEL-477 |
| chr18-68422596-DEL-93 | chr18-68422597-DEL-93 |
| chr18-68435974-DEL-76 | chr18-68435975-DEL-76 |
| chr18-68539047-DEL-179 | chr18-68539033-DEL-170 |
| chr18-68543384-DEL-77 | chr18-68543385-DEL-77 |
| chr18-68545308-DEL-7159 | chr18-68545309-DEL-7159 |
| chr18-68572796-DEL-69 | chr18-68572797-DEL-69 |
| chr18-68574473-DEL-298 | chr18-68574472-DEL-299 |
| chr18-68576877-DEL-96 | chr18-68576878-DEL-96 |
| chr18-68608318-DEL-6638 | chr18-68608318-DEL-6638 |
| chr18-68622661-DEL-830 | chr18-68622662-DEL-830 |
| chr18-68730815-DEL-64 | chr18-68730816-DEL-64 |
| chr18-68737028-DEL-56 | chr18-68737028-DEL-59 |
| chr18-68752780-DEL-119 | chr18-68752790-DEL-96 |
| chr18-68758369-DEL-186 | chr18-68758370-DEL-186 |
| chr18-68783066-DEL-1189 | chr18-68783067-DEL-1189 |
| chr18-68803029-DEL-62 | chr18-68803030-DEL-62 |
| chr18-68809856-DEL-733 | chr18-68809857-DEL-733 |
| chr18-68856169-DEL-55 | chr18-68856170-DEL-55 |
| chr18-68860414-DEL-85 | chr18-68860415-DEL-85 |
| chr18-68899583-DEL-109 | chr18-68899584-DEL-109 |
| chr18-68915115-DEL-66 | chr18-68915116-DEL-66 |
| chr18-68940240-DEL-83 | chr18-68940241-DEL-83 |
| chr18-68971644-DEL-70 | chr18-68971645-DEL-70 |
| chr18-69024445-DEL-1490 | chr18-69024446-DEL-1489 |
| chr18-69029005-DEL-928 | chr18-69029006-DEL-928 |
| chr18-69029603-DEL-62 | chr18-69029605-DEL-60 |
| chr18-69035524-DEL-56 | chr18-69035524-DEL-68 |
| chr18-69061867-DEL-218 | chr18-69061868-DEL-218 |
| chr18-69075034-DEL-60 | chr18-69075035-DEL-60 |
| chr18-69093688-DEL-817 | chr18-69093689-DEL-817 |
| chr18-69106354-DEL-55 | chr18-69106355-DEL-55 |
| chr18-69118021-DEL-2601 | chr18-69118022-DEL-2601 |
| chr18-69138995-DEL-52 | chr18-69138996-DEL-52 |
| chr18-69143408-DEL-62 | chr18-69143415-DEL-98 |
| chr18-69170159-DEL-4130 | chr18-69170160-DEL-4130 |
| chr18-69177066-DEL-56 | chr18-69177068-DEL-72 |
| chr18-69191307-DEL-56 | chr18-69191308-DEL-58 |
| chr18-69243950-DEL-58 | chr18-69243990-DEL-56 |
| chr18-69292468-DEL-389 | chr18-69292469-DEL-389 |
| chr18-69299205-DEL-88 | chr18-69299206-DEL-88 |
| chr18-69324327-DEL-54 | chr18-69324332-DEL-62 |
| chr18-69409409-DEL-2105 | chr18-69409410-DEL-2105 |

| | |
|---|---|
| chr18-69426857-DEL-281 | chr18-69426858-DEL-281 |
| chr18-69484197-DEL-58 | chr18-69484198-DEL-58 |
| chr18-69539873-DEL-59 | chr18-69539847-DEL-56 |
| chr18-69547211-DEL-177 | chr18-69547210-DEL-178 |
| chr18-69657124-DEL-60 | chr18-69657125-DEL-60 |
| chr18-69665283-DEL-51 | chr18-69665284-DEL-51 |
| chr18-69690246-DEL-114 | chr18-69690247-DEL-114 |
| chr18-69744317-DEL-116 | chr18-69744318-DEL-116 |
| chr18-69831019-DEL-72 | chr18-69831022-DEL-70 |
| chr18-69858669-DEL-102 | chr18-69858663-DEL-80 |
| chr18-69871967-DEL-51 | chr18-69871968-DEL-51 |
| chr18-69888226-DEL-59 | chr18-69888226-DEL-52 |
| chr18-69888731-DEL-317 | chr18-69888732-DEL-317 |
| chr18-69892414-DEL-1893 | chr18-69892415-DEL-1893 |
| chr18-69928881-DEL-56 | chr18-69928884-DEL-56 |
| chr18-69941322-DEL-73 | chr18-69941326-DEL-71 |
| chr18-69957489-DEL-59 | chr18-69957484-DEL-64 |
| chr18-69992819-DEL-117 | chr18-69992820-DEL-117 |
| chr18-69996632-DEL-170 | chr18-69996634-DEL-169 |
| chr18-70036149-DEL-78 | chr18-70036150-DEL-78 |
| chr18-70045960-DEL-76 | chr18-70045960-DEL-76 |
| chr18-70049739-DEL-157 | chr18-70049740-DEL-157 |
| chr18-70060735-DEL-666 | chr18-70060736-DEL-666 |
| chr18-70064599-DEL-113 | chr18-70064600-DEL-113 |
| chr18-70090451-DEL-123 | chr18-70090450-DEL-150 |
| chr18-70106929-DEL-275 | chr18-70106930-DEL-275 |
| chr18-70138301-DEL-4499 | chr18-70138301-DEL-4499 |
| chr18-70140306-DEL-85 | chr18-70140305-DEL-86 |
| chr18-70140635-DEL-6342 | chr18-70140636-DEL-6342 |
| chr18-70157269-DEL-206 | chr18-70157270-DEL-206 |
| chr18-70164487-DEL-4150 | chr18-70164488-DEL-4150 |
| chr18-70178359-DEL-67 | chr18-70178312-DEL-88 |
| chr18-70179590-DEL-66 | chr18-70179595-DEL-60 |
| chr18-70209352-DEL-121 | chr18-70209353-DEL-121 |
| chr18-70266297-DEL-485 | chr18-70266298-DEL-485 |
| chr18-70280847-DEL-1883 | chr18-70280848-DEL-1883 |
| chr18-70284841-DEL-126 | chr18-70284842-DEL-126 |
| chr18-70310853-DEL-107 | chr18-70310832-DEL-123 |
| chr18-70327395-DEL-367 | chr18-70327397-DEL-366 |
| chr18-70336965-DEL-459 | chr18-70336964-DEL-460 |
| chr18-70347494-DEL-1479 | chr18-70347495-DEL-1479 |
| chr18-70349909-DEL-406 | chr18-70349940-DEL-379 |
| chr18-70366487-DEL-121 | chr18-70366488-DEL-121 |
| chr18-70370993-DEL-365 | chr18-70370994-DEL-365 |
| chr18-70373210-DEL-68 | chr18-70373211-DEL-67 |
| chr18-70405279-DEL-700 | chr18-70405276-DEL-704 |
| chr18-70444398-DEL-6377 | chr18-70444382-DEL-6393 |
| chr18-70520300-DEL-94 | chr18-70520298-DEL-72 |
| chr18-70547468-DEL-80 | chr18-70547472-DEL-69 |
| chr18-70551845-DEL-253 | chr18-70551838-DEL-261 |
| chr18-70557127-DEL-1433 | chr18-70557128-DEL-1433 |
| chr18-70571531-DEL-1073 | chr18-70571532-DEL-1073 |
| chr18-70607921-DEL-248 | chr18-70607922-DEL-248 |
| chr18-70619483-DEL-1447 | chr18-70619484-DEL-1447 |
| chr18-70621111-DEL-6443 | chr18-70621112-DEL-6443 |
| chr18-70627032-DEL-334 | chr18-70627033-DEL-334 |
| chr18-70636571-DEL-64 | chr18-70636572-DEL-60 |
| chr18-70640962-DEL-66 | chr18-70640990-DEL-65 |
| chr18-70643200-DEL-6186 | chr18-70643188-DEL-6187 |
| chr18-70690750-DEL-98 | chr18-70690751-DEL-98 |
| chr18-70691631-DEL-78 | chr18-70691632-DEL-78 |
| chr18-70708751-DEL-198 | chr18-70708752-DEL-198 |
| chr18-70719438-DEL-53 | chr18-70719438-DEL-53 |
| chr18-70729800-DEL-75 | chr18-70729801-DEL-75 |
| chr18-70744626-DEL-66 | chr18-70744627-DEL-66 |
| chr18-70756617-DEL-54 | chr18-70756618-DEL-54 |
| chr18-70766175-DEL-64 | chr18-70766176-DEL-64 |
| chr18-70801161-DEL-52 | chr18-70801162-DEL-52 |
| chr18-70832755-DEL-6619 | chr18-70832756-DEL-6619 |
| chr18-70854583-DEL-1355 | chr18-70854584-DEL-1355 |
| chr18-70862065-DEL-52 | chr18-70862076-DEL-54 |
| chr18-70871687-DEL-805 | chr18-70871688-DEL-805 |
| chr18-70882786-DEL-86 | chr18-70882787-DEL-86 |
| chr18-70902527-DEL-237 | chr18-70902528-DEL-237 |
| chr18-70903788-DEL-69 | chr18-70903789-DEL-69 |
| chr18-70930622-DEL-104 | chr18-70930617-DEL-107 |
| chr18-70938999-DEL-527 | chr18-70939000-DEL-527 |
| chr18-70971726-DEL-504 | chr18-70971727-DEL-504 |
| chr18-70975311-DEL-7052 | chr18-70975312-DEL-7052 |
| chr18-71006465-DEL-91 | chr18-71006466-DEL-91 |

| | |
|---|---|
| chr18-71012040-DEL-98 | chr18-71012040-DEL-98 |
| chr18-71014799-DEL-184 | chr18-71014800-DEL-184 |
| chr18-71016331-DEL-170 | chr18-71016332-DEL-170 |
| chr18-71038132-DEL-197 | chr18-71038144-DEL-195 |
| chr18-71043814-DEL-55 | chr18-71043815-DEL-50 |
| chr18-71082543-DEL-68 | chr18-71082548-DEL-64 |
| chr18-71088034-DEL-1824 | chr18-71088046-DEL-1824 |
| chr18-71101408-DEL-83 | chr18-71101409-DEL-71 |
| chr18-71165554-DEL-6376 | chr18-71165555-DEL-6376 |
| chr18-71176702-DEL-77 | chr18-71176703-DEL-77 |
| chr18-71177037-DEL-369 | chr18-71177038-DEL-369 |
| chr18-71180753-DEL-1972 | chr18-71180754-DEL-1972 |
| chr18-71214611-DEL-4490 | chr18-71214612-DEL-4490 |
| chr18-71222167-DEL-2983 | chr18-71222168-DEL-2983 |
| chr18-71225459-DEL-180 | chr18-71225460-DEL-180 |
| chr18-71261433-DEL-132 | chr18-71261434-DEL-132 |
| chr18-71272075-DEL-6359 | chr18-71272076-DEL-6359 |
| chr18-71284334-DEL-58 | chr18-71284336-DEL-57 |
| chr18-71310315-DEL-633 | chr18-71310316-DEL-633 |
| chr18-71328409-DEL-497 | chr18-71328410-DEL-497 |
| chr18-71334856-DEL-173 | chr18-71334857-DEL-173 |
| chr18-71341209-DEL-3200 | chr18-71341210-DEL-3200 |
| chr18-71400027-DEL-6401 | chr18-71400027-DEL-6401 |
| chr18-71435579-DEL-217 | chr18-71435580-DEL-217 |
| chr18-71452003-DEL-51 | chr18-71452003-DEL-51 |
| chr18-71485250-DEL-501 | chr18-71485250-DEL-499 |
| chr18-71486567-DEL-576 | chr18-71486567-DEL-574 |
| chr18-71508676-DEL-73 | chr18-71508677-DEL-73 |
| chr18-71521318-DEL-159 | chr18-71521319-DEL-161 |
| chr18-71525534-DEL-827 | chr18-71525534-DEL-802 |
| chr18-71542854-DEL-57 | chr18-71542855-DEL-57 |
| chr18-71633649-DEL-54 | chr18-71633650-DEL-56 |
| chr18-71660231-DEL-154 | chr18-71660231-DEL-154 |
| chr18-71662266-DEL-58 | chr18-71662267-DEL-58 |
| chr18-71704539-DEL-179 | chr18-71704540-DEL-179 |
| chr18-71719233-DEL-52 | chr18-71719240-DEL-60 |
| chr18-71732447-DEL-212 | chr18-71732448-DEL-212 |
| chr18-71748155-DEL-139 | chr18-71748155-DEL-139 |
| chr18-71763494-DEL-10555 | chr18-71763495-DEL-10555 |
| chr18-71764002-DEL-212 | chr18-71764003-DEL-212 |
| chr18-71776763-DEL-66 | chr18-71776726-DEL-54 |
| chr18-71806368-DEL-65 | chr18-71806365-DEL-69 |
| chr18-71807382-DEL-76 | chr18-71807383-DEL-76 |
| chr18-71819824-DEL-572 | chr18-71819825-DEL-572 |
| chr18-71825815-DEL-54 | chr18-71825820-DEL-62 |
| chr18-71906940-DEL-699 | chr18-71906941-DEL-699 |
| chr18-71934566-DEL-400 | chr18-71934567-DEL-400 |
| chr18-72008678-DEL-337 | chr18-72008679-DEL-337 |
| chr18-72025758-DEL-1016 | chr18-72025751-DEL-1017 |
| chr18-72040229-DEL-190 | chr18-72040230-DEL-190 |
| chr18-72049631-DEL-452 | chr18-72049632-DEL-452 |
| chr18-72053244-DEL-409 | chr18-72053245-DEL-409 |
| chr18-72130398-DEL-80 | chr18-72130399-DEL-80 |
| chr18-72147282-DEL-83 | chr18-72147287-DEL-96 |
| chr18-72163710-DEL-1325 | chr18-72163711-DEL-1325 |
| chr18-72189023-DEL-77 | chr18-72189021-DEL-84 |
| chr18-72200314-DEL-3164 | chr18-72200315-DEL-3164 |
| chr18-72253291-DEL-52 | chr18-72253292-DEL-52 |
| chr18-72258217-DEL-1892 | chr18-72258218-DEL-1892 |
| chr18-72262434-DEL-80 | chr18-72262435-DEL-80 |
| chr18-72275698-DEL-83 | chr18-72275683-DEL-50 |
| chr18-72285002-DEL-584 | chr18-72285003-DEL-584 |
| chr18-72306351-DEL-62 | chr18-72306342-DEL-63 |
| chr18-72335456-DEL-156 | chr18-72335457-DEL-156 |
| chr18-72385550-DEL-75 | chr18-72385551-DEL-75 |
| chr18-72387537-DEL-55 | chr18-72387538-DEL-55 |
| chr18-72402334-DEL-1369 | chr18-72402335-DEL-1369 |
| chr18-72471827-DEL-155 | chr18-72471828-DEL-155 |
| chr18-72485366-DEL-60 | chr18-72485359-DEL-68 |
| chr18-72492430-DEL-179 | chr18-72492431-DEL-179 |
| chr18-72513911-DEL-61 | chr18-72513909-DEL-61 |
| chr18-72515447-DEL-1587 | chr18-72515448-DEL-1587 |
| chr18-72526903-DEL-892 | chr18-72526904-DEL-892 |
| chr18-72562555-DEL-4552 | chr18-72562556-DEL-4552 |
| chr18-72575174-DEL-54 | chr18-72575152-DEL-60 |
| chr18-72583859-DEL-51 | chr18-72583860-DEL-51 |
| chr18-72585056-DEL-90 | chr18-72585056-DEL-102 |
| chr18-72590215-DEL-2990 | chr18-72590219-DEL-2985 |
| chr18-72595165-DEL-2276 | chr18-72595165-DEL-2276 |
| chr18-72599633-DEL-1324 | chr18-72599611-DEL-1309 |

| | |
|---|---|
| chr18-72603047-DEL-63 | chr18-72603048-DEL-56 |
| chr18-72626772-DEL-2858 | chr18-72626773-DEL-2858 |
| chr18-72639481-DEL-6694 | chr18-72639482-DEL-6694 |
| chr18-72672321-DEL-58 | chr18-72672329-DEL-53 |
| chr18-72678317-DEL-166 | chr18-72678317-DEL-166 |
| chr18-72698927-DEL-1809 | chr18-72698928-DEL-1809 |
| chr18-72709893-DEL-6824 | chr18-72709894-DEL-6824 |
| chr18-72745771-DEL-592 | chr18-72745772-DEL-592 |
| chr18-72755437-DEL-5200 | chr18-72755428-DEL-5201 |
| chr18-72768726-DEL-79 | chr18-72768691-DEL-125 |
| chr18-72775861-DEL-72 | chr18-72775862-DEL-72 |
| chr18-72793675-DEL-1372 | chr18-72793676-DEL-1373 |
| chr18-72799891-DEL-75 | chr18-72799892-DEL-75 |
| chr18-72831459-DEL-74 | chr18-72831460-DEL-74 |
| chr18-72868962-DEL-751 | chr18-72868963-DEL-751 |
| chr18-72886970-DEL-5766 | chr18-72886971-DEL-5766 |
| chr18-72894807-DEL-1011 | chr18-72894808-DEL-1011 |
| chr18-72905038-DEL-83 | chr18-72905036-DEL-84 |
| chr18-72908854-DEL-569 | chr18-72908855-DEL-569 |
| chr18-72917499-DEL-64 | chr18-72917507-DEL-58 |
| chr18-72920070-DEL-98 | chr18-72920068-DEL-103 |
| chr18-72922007-DEL-6108 | chr18-72922008-DEL-6108 |
| chr18-72956482-DEL-414 | chr18-72956483-DEL-414 |
| chr18-72972360-DEL-388 | chr18-72972350-DEL-385 |
| chr18-73004280-DEL-80 | chr18-73004272-DEL-68 |
| chr18-73032916-DEL-105 | chr18-73032904-DEL-99 |
| chr18-73056004-DEL-52 | chr18-73056005-DEL-52 |
| chr18-73067051-DEL-6428 | chr18-73067052-DEL-6428 |
| chr18-73078035-DEL-2905 | chr18-73078036-DEL-2905 |
| chr18-73095031-DEL-2051 | chr18-73095032-DEL-2051 |
| chr18-73117151-DEL-102 | chr18-73117154-DEL-92 |
| chr18-73137224-DEL-56 | chr18-73137237-DEL-60 |
| chr18-73144068-DEL-51 | chr18-73144069-DEL-51 |
| chr18-73145214-DEL-95 | chr18-73145215-DEL-95 |
| chr18-73149822-DEL-207 | chr18-73149800-DEL-208 |
| chr18-73157198-DEL-94 | chr18-73157199-DEL-94 |
| chr18-73183731-DEL-110 | chr18-73183760-DEL-102 |
| chr18-73215494-DEL-72 | chr18-73215495-DEL-73 |
| chr18-73222908-DEL-70 | chr18-73222909-DEL-70 |
| chr18-73225024-DEL-87 | chr18-73225068-DEL-55 |
| chr18-73243542-DEL-6218 | chr18-73243543-DEL-6218 |
| chr18-73334766-DEL-3141 | chr18-73334767-DEL-3141 |
| chr18-73334883-DEL-204 | chr18-73334884-DEL-204 |
| chr18-73343309-DEL-58 | chr18-73343271-DEL-79 |
| chr18-73352855-DEL-84 | chr18-73352885-DEL-53 |
| chr18-73370266-DEL-131 | chr18-73370266-DEL-131 |
| chr18-73377646-DEL-78 | chr18-73377647-DEL-78 |
| chr18-73392761-DEL-4322 | chr18-73392762-DEL-4322 |
| chr18-73402566-DEL-277 | chr18-73402567-DEL-277 |
| chr18-73460260-DEL-2555 | chr18-73460261-DEL-2555 |
| chr18-73472531-DEL-82 | chr18-73472532-DEL-73 |
| chr18-73476082-DEL-6632 | chr18-73476083-DEL-6632 |
| chr18-73482263-DEL-212 | chr18-73482264-DEL-212 |
| chr18-73516613-DEL-397 | chr18-73516614-DEL-397 |
| chr18-73531138-DEL-2394 | chr18-73531139-DEL-2394 |
| chr18-73539088-DEL-424 | chr18-73539089-DEL-424 |
| chr18-73540711-DEL-54 | chr18-73540712-DEL-54 |
| chr18-73548311-DEL-60 | chr18-73548288-DEL-60 |
| chr18-73580109-DEL-566 | chr18-73580110-DEL-566 |
| chr18-73613368-DEL-1604 | chr18-73613369-DEL-1604 |
| chr18-73613617-DEL-55 | chr18-73613615-DEL-55 |
| chr18-73620959-DEL-70 | chr18-73620959-DEL-70 |
| chr18-73624623-DEL-216 | chr18-73624624-DEL-216 |
| chr18-73628019-DEL-869 | chr18-73628018-DEL-869 |
| chr18-73632983-DEL-55 | chr18-73632966-DEL-73 |
| chr18-73633961-DEL-464 | chr18-73633962-DEL-464 |
| chr18-73653734-DEL-65 | chr18-73653735-DEL-65 |
| chr18-73662555-DEL-72 | chr18-73662556-DEL-72 |
| chr18-73663963-DEL-102 | chr18-73663977-DEL-72 |
| chr18-73682226-DEL-216 | chr18-73682227-DEL-216 |
| chr18-73691074-DEL-123 | chr18-73691075-DEL-123 |
| chr18-73715292-DEL-270 | chr18-73715293-DEL-270 |
| chr18-73746681-DEL-431 | chr18-73746682-DEL-431 |
| chr18-73757782-DEL-526 | chr18-73757783-DEL-526 |
| chr18-73762359-DEL-535 | chr18-73762360-DEL-535 |
| chr18-73814372-DEL-6250 | chr18-73814373-DEL-6249 |
| chr18-73822616-DEL-3900 | chr18-73822617-DEL-3900 |
| chr18-73824644-DEL-834 | chr18-73824643-DEL-834 |
| chr18-73847282-DEL-140 | chr18-73847283-DEL-140 |
| chr18-73849228-DEL-95 | chr18-73849229-DEL-95 |

| | |
|---|---|
| chr18-73856573-DEL-144 | chr18-73856581-DEL-143 |
| chr18-73857126-DEL-362 | chr18-73857127-DEL-362 |
| chr18-73905725-DEL-100 | chr18-73905722-DEL-104 |
| chr18-73930083-DEL-518 | chr18-73930084-DEL-518 |
| chr18-73931459-DEL-202 | chr18-73931460-DEL-202 |
| chr18-73933039-DEL-501 | chr18-73933039-DEL-500 |
| chr18-73941483-DEL-73 | chr18-73941484-DEL-73 |
| chr18-73955160-DEL-666 | chr18-73955161-DEL-666 |
| chr18-73964421-DEL-51 | chr18-73964422-DEL-51 |
| chr18-73975762-DEL-185 | chr18-73975759-DEL-169 |
| chr18-73984809-DEL-352 | chr18-73984810-DEL-351 |
| chr18-73990631-DEL-114 | chr18-73990632-DEL-114 |
| chr18-74004305-DEL-141 | chr18-74004306-DEL-141 |
| chr18-74006578-DEL-77 | chr18-74006579-DEL-93 |
| chr18-74042453-DEL-57 | chr18-74042454-DEL-57 |
| chr18-74046707-DEL-6356 | chr18-74046708-DEL-6356 |
| chr18-74057561-DEL-88 | chr18-74057565-DEL-79 |
| chr18-74072291-DEL-3683 | chr18-74072292-DEL-3683 |
| chr18-74075693-DEL-61 | chr18-74075694-DEL-61 |
| chr18-74080544-DEL-130 | chr18-74080545-DEL-130 |
| chr18-74083052-DEL-54 | chr18-74083053-DEL-55 |
| chr18-74092309-DEL-56 | chr18-74092272-DEL-57 |
| chr18-74125494-DEL-249 | chr18-74125495-DEL-249 |
| chr18-74168313-DEL-51 | chr18-74168282-DEL-61 |
| chr18-74180367-DEL-92 | chr18-74180368-DEL-92 |
| chr18-74183847-DEL-76 | chr18-74183851-DEL-73 |
| chr18-74203046-DEL-53 | chr18-74203039-DEL-61 |
| chr18-74206358-DEL-313 | chr18-74206359-DEL-313 |
| chr18-74225070-DEL-62 | chr18-74225071-DEL-62 |
| chr18-74236355-DEL-85 | chr18-74236356-DEL-75 |
| chr18-74237435-DEL-58 | chr18-74237436-DEL-82 |
| chr18-74240066-DEL-62 | chr18-74240067-DEL-62 |
| chr18-74289153-DEL-74 | chr18-74289154-DEL-74 |
| chr18-74297193-DEL-56 | chr18-74297194-DEL-56 |
| chr18-74299538-DEL-208 | chr18-74299539-DEL-207 |
| chr18-74317459-DEL-473 | chr18-74317464-DEL-470 |
| chr18-74332368-DEL-70 | chr18-74332370-DEL-68 |
| chr18-74368191-DEL-202 | chr18-74368192-DEL-202 |
| chr18-74370945-DEL-55 | chr18-74370946-DEL-55 |
| chr18-74433878-DEL-77 | chr18-74433879-DEL-77 |
| chr18-74450225-DEL-2264 | chr18-74450226-DEL-2264 |
| chr18-74457587-DEL-112 | chr18-74457588-DEL-112 |
| chr18-74461928-DEL-98 | chr18-74461929-DEL-104 |
| chr18-74465391-DEL-70 | chr18-74465392-DEL-70 |
| chr18-74469094-DEL-87 | chr18-74469095-DEL-87 |
| chr18-74509481-DEL-1077 | chr18-74509491-DEL-1063 |
| chr18-74516765-DEL-208 | chr18-74516766-DEL-208 |
| chr18-74566705-DEL-195 | chr18-74566706-DEL-195 |
| chr18-74567294-DEL-2579 | chr18-74567295-DEL-2579 |
| chr18-74596506-DEL-103 | chr18-74596507-DEL-103 |
| chr18-74598666-DEL-974 | chr18-74598667-DEL-974 |
| chr18-74639608-DEL-119 | chr18-74639609-DEL-119 |
| chr18-74650721-DEL-208 | chr18-74650727-DEL-209 |
| chr18-74668887-DEL-59 | chr18-74668888-DEL-59 |
| chr18-74696074-DEL-3443 | chr18-74696074-DEL-3443 |
| chr18-74701760-DEL-912 | chr18-74701761-DEL-912 |
| chr18-74709530-DEL-67 | chr18-74709527-DEL-68 |
| chr18-74711005-DEL-197 | chr18-74711019-DEL-179 |
| chr18-74714887-DEL-82 | chr18-74714888-DEL-82 |
| chr18-74746802-DEL-138 | chr18-74746803-DEL-138 |
| chr18-74806458-DEL-165 | chr18-74806459-DEL-165 |
| chr18-74811461-DEL-72 | chr18-74811462-DEL-72 |
| chr18-74815441-DEL-63 | chr18-74815441-DEL-69 |
| chr18-74819415-DEL-150 | chr18-74819416-DEL-150 |
| chr18-74824523-DEL-1819 | chr18-74824524-DEL-1819 |
| chr18-74849226-DEL-185 | chr18-74849227-DEL-185 |
| chr18-74856216-DEL-63 | chr18-74856217-DEL-58 |
| chr18-74878658-DEL-12839 | chr18-74878659-DEL-12839 |
| chr18-74915387-DEL-74 | chr18-74915388-DEL-74 |
| chr18-74926625-DEL-169 | chr18-74926610-DEL-170 |
| chr18-74930214-DEL-72 | chr18-74930165-DEL-74 |
| chr18-74944085-DEL-194 | chr18-74944081-DEL-195 |
| chr18-74954287-DEL-116 | chr18-74954287-DEL-116 |
| chr18-74955826-DEL-254 | chr18-74955827-DEL-254 |
| chr18-74960743-DEL-1079 | chr18-74960744-DEL-1079 |
| chr18-74965758-DEL-386 | chr18-74965759-DEL-386 |
| chr18-74969776-DEL-75 | chr18-74969777-DEL-75 |
| chr18-74972379-DEL-71 | chr18-74972380-DEL-71 |
| chr18-74994252-DEL-108 | chr18-74994253-DEL-91 |
| chr18-75028256-DEL-232 | chr18-75028257-DEL-232 |

| | |
|---|---|
| chr18-75058392-DEL-2011 | chr18-75058393-DEL-2008 |
| chr18-75096622-DEL-221 | chr18-75096621-DEL-206 |
| chr18-75110503-DEL-234 | chr18-75110502-DEL-235 |
| chr18-75118083-DEL-185 | chr18-75118084-DEL-185 |
| chr18-75128146-DEL-260 | chr18-75128129-DEL-253 |
| chr18-75146040-DEL-173 | chr18-75146041-DEL-173 |
| chr18-75158513-DEL-65 | chr18-75158514-DEL-65 |
| chr18-75164058-DEL-199 | chr18-75164059-DEL-199 |
| chr18-75165968-DEL-75 | chr18-75165969-DEL-75 |
| chr18-75189982-DEL-453 | chr18-75189975-DEL-461 |
| chr18-75191582-DEL-65 | chr18-75191583-DEL-65 |
| chr18-75199278-DEL-232 | chr18-75199279-DEL-232 |
| chr18-75208800-DEL-357 | chr18-75208801-DEL-357 |
| chr18-75218289-DEL-270 | chr18-75218290-DEL-269 |
| chr18-75233498-DEL-556 | chr18-75233499-DEL-556 |
| chr18-75240162-DEL-589 | chr18-75240163-DEL-589 |
| chr18-75269123-DEL-114 | chr18-75269124-DEL-114 |
| chr18-75291631-DEL-498 | chr18-75291632-DEL-498 |
| chr18-75344839-DEL-54 | chr18-75344840-DEL-54 |
| chr18-75349345-DEL-64 | chr18-75349346-DEL-64 |
| chr18-75366376-DEL-255 | chr18-75366391-DEL-268 |
| chr18-75379330-DEL-144 | chr18-75379329-DEL-145 |
| chr18-75400241-DEL-176 | chr18-75400244-DEL-177 |
| chr18-75404532-DEL-54 | chr18-75404533-DEL-54 |
| chr18-75415380-DEL-309 | chr18-75415381-DEL-309 |
| chr18-75418535-DEL-216 | chr18-75418536-DEL-216 |
| chr18-75433474-DEL-58 | chr18-75433475-DEL-87 |
| chr18-75434616-DEL-1695 | chr18-75434617-DEL-1695 |
| chr18-75436384-DEL-212 | chr18-75436385-DEL-213 |
| chr18-75446588-DEL-76 | chr18-75446610-DEL-55 |
| chr18-75449478-DEL-124 | chr18-75449507-DEL-96 |
| chr18-75466461-DEL-218 | chr18-75466462-DEL-218 |
| chr18-75539062-DEL-71 | chr18-75539037-DEL-57 |
| chr18-75540411-DEL-2390 | chr18-75540412-DEL-2390 |
| chr18-75591263-DEL-125 | chr18-75591251-DEL-139 |
| chr18-75682486-DEL-1889 | chr18-75682487-DEL-1889 |
| chr18-75700411-DEL-104 | chr18-75700412-DEL-104 |
| chr18-75701248-DEL-100 | chr18-75701249-DEL-100 |
| chr18-75732976-DEL-58 | chr18-75732977-DEL-58 |
| chr18-75746608-DEL-72 | chr18-75746619-DEL-64 |
| chr18-75748845-DEL-97 | chr18-75748846-DEL-97 |
| chr18-75764000-DEL-54 | chr18-75764001-DEL-54 |
| chr18-75778076-DEL-2431 | chr18-75778077-DEL-2431 |
| chr18-75787067-DEL-121 | chr18-75787068-DEL-121 |
| chr18-75836544-DEL-92 | chr18-75836536-DEL-76 |
| chr18-75847234-DEL-67 | chr18-75847234-DEL-67 |
| chr18-75876308-DEL-708 | chr18-75876309-DEL-708 |
| chr18-75889965-DEL-60 | chr18-75889943-DEL-78 |
| chr18-75913471-DEL-175 | chr18-75913472-DEL-175 |
| chr18-75925573-DEL-78 | chr18-75925574-DEL-78 |
| chr18-75998328-DEL-170 | chr18-75998329-DEL-170 |
| chr18-76008576-DEL-215 | chr18-76008577-DEL-215 |
| chr18-76015295-DEL-72 | chr18-76015254-DEL-110 |
| chr18-76017198-DEL-6584 | chr18-76017199-DEL-6584 |
| chr18-76023520-DEL-214 | chr18-76023521-DEL-214 |
| chr18-76065585-DEL-60 | chr18-76065586-DEL-50 |
| chr18-76086893-DEL-150 | chr18-76086894-DEL-150 |
| chr18-76100414-DEL-58 | chr18-76100414-DEL-58 |
| chr18-76134617-DEL-98 | chr18-76134595-DEL-99 |
| chr18-76143583-DEL-60 | chr18-76143536-DEL-78 |
| chr18-76154282-DEL-211 | chr18-76154280-DEL-211 |
| chr18-76155155-DEL-157 | chr18-76155156-DEL-156 |
| chr18-76160495-DEL-63 | chr18-76160496-DEL-63 |
| chr18-76235251-DEL-2080 | chr18-76235252-DEL-2080 |
| chr18-76310933-DEL-6403 | chr18-76310934-DEL-6403 |
| chr18-76320154-DEL-202 | chr18-76320155-DEL-202 |
| chr18-76339281-DEL-218 | chr18-76339282-DEL-218 |
| chr18-76345667-DEL-502 | chr18-76345668-DEL-502 |
| chr18-76352237-DEL-154 | chr18-76352238-DEL-154 |
| chr18-76359620-DEL-72 | chr18-76359629-DEL-63 |
| chr18-76440875-DEL-601 | chr18-76440876-DEL-601 |
| chr18-76453914-DEL-204 | chr18-76453915-DEL-204 |
| chr18-76455446-DEL-68 | chr18-76455447-DEL-66 |
| chr18-76468002-DEL-102 | chr18-76467974-DEL-124 |
| chr18-76475513-DEL-72 | chr18-76475481-DEL-122 |
| chr18-76487401-DEL-64 | chr18-76487434-DEL-53 |
| chr18-76496686-DEL-7040 | chr18-76496687-DEL-7040 |
| chr18-76496784-DEL-600 | chr18-76496785-DEL-600 |
| chr18-76497289-DEL-402 | chr18-76497290-DEL-402 |
| chr18-76559735-DEL-158 | chr18-76559736-DEL-158 |

| | |
|---|---|
| chr18-76562543-DEL-293 | chr18-76562544-DEL-293 |
| chr18-76585732-DEL-164 | chr18-76585712-DEL-158 |
| chr18-76587002-DEL-258 | chr18-76587003-DEL-258 |
| chr18-76593949-DEL-2091 | chr18-76593950-DEL-2091 |
| chr18-76627892-DEL-350 | chr18-76627892-DEL-350 |
| chr18-76656172-DEL-193 | chr18-76656205-DEL-189 |
| chr18-76660327-DEL-54 | chr18-76660327-DEL-54 |
| chr18-76663522-DEL-217 | chr18-76663523-DEL-217 |
| chr18-76667382-DEL-6550 | chr18-76667383-DEL-6550 |
| chr18-76710234-DEL-79 | chr18-76710235-DEL-79 |
| chr18-76711063-DEL-494 | chr18-76711064-DEL-494 |
| chr18-76721704-DEL-51 | chr18-76721705-DEL-51 |
| chr18-76722997-DEL-3608 | chr18-76722998-DEL-3608 |
| chr18-76735598-DEL-118 | chr18-76735599-DEL-118 |
| chr18-76741160-DEL-74 | chr18-76741161-DEL-71 |
| chr18-76745533-DEL-58 | chr18-76745515-DEL-85 |
| chr18-76747222-DEL-73 | chr18-76747223-DEL-73 |
| chr18-76753315-DEL-211 | chr18-76753316-DEL-211 |
| chr18-76768028-DEL-381 | chr18-76768029-DEL-381 |
| chr18-76782407-DEL-63 | chr18-76782408-DEL-63 |
| chr18-76789440-DEL-70 | chr18-76789425-DEL-110 |
| chr18-76790252-DEL-62 | chr18-76790294-DEL-58 |
| chr18-76809789-DEL-1049 | chr18-76809777-DEL-1041 |
| chr18-76812355-DEL-498 | chr18-76812356-DEL-498 |
| chr18-76834516-DEL-86 | chr18-76834517-DEL-86 |
| chr18-76874155-DEL-6225 | chr18-76874156-DEL-6225 |
| chr18-76897834-DEL-78 | chr18-76897835-DEL-78 |
| chr18-76899953-DEL-59 | chr18-76899954-DEL-59 |
| chr18-76917566-DEL-82 | chr18-76917579-DEL-61 |
| chr18-76921317-DEL-211 | chr18-76921318-DEL-211 |
| chr18-76987708-DEL-63 | chr18-76987675-DEL-66 |
| chr18-76996036-DEL-52 | chr18-76996037-DEL-52 |
| chr18-77012102-DEL-58 | chr18-77012103-DEL-58 |
| chr18-77034198-DEL-1117 | chr18-77034199-DEL-1117 |
| chr18-77055267-DEL-167 | chr18-77055263-DEL-136 |
| chr18-77055862-DEL-102 | chr18-77055863-DEL-102 |
| chr18-77064613-DEL-56 | chr18-77064603-DEL-53 |
| chr18-77066731-DEL-63 | chr18-77066732-DEL-63 |
| chr18-77089400-DEL-106 | chr18-77089401-DEL-106 |
| chr18-77111423-DEL-82 | chr18-77111426-DEL-82 |
| chr18-77140122-DEL-70 | chr18-77140123-DEL-70 |
| chr18-77167830-DEL-210 | chr18-77167831-DEL-210 |
| chr18-77170537-DEL-206 | chr18-77170538-DEL-206 |
| chr18-77183634-DEL-57 | chr18-77183635-DEL-57 |
| chr18-77192840-DEL-59 | chr18-77192841-DEL-59 |
| chr18-77225056-DEL-101 | chr18-77225057-DEL-101 |
| chr18-77254633-DEL-198 | chr18-77254634-DEL-198 |
| chr18-77302326-DEL-249 | chr18-77302327-DEL-249 |
| chr18-77322014-DEL-134 | chr18-77322012-DEL-132 |
| chr18-77323854-DEL-165 | chr18-77323849-DEL-171 |
| chr18-77324522-DEL-62 | chr18-77324491-DEL-80 |
| chr18-77343635-DEL-57 | chr18-77343636-DEL-57 |
| chr18-77352112-DEL-6400 | chr18-77352113-DEL-6400 |
| chr18-77382136-DEL-160 | chr18-77382137-DEL-160 |
| chr18-77398074-DEL-93 | chr18-77398075-DEL-93 |
| chr18-77399256-DEL-1168 | chr18-77399257-DEL-1168 |
| chr18-77401672-DEL-77 | chr18-77401673-DEL-77 |
| chr18-77410345-DEL-124 | chr18-77410340-DEL-124 |
| chr18-77411188-DEL-94 | chr18-77411188-DEL-94 |
| chr18-77417298-DEL-80 | chr18-77417302-DEL-76 |
| chr18-77425821-DEL-128 | chr18-77425816-DEL-118 |
| chr18-77454809-DEL-860 | chr18-77454810-DEL-860 |
| chr18-77456039-DEL-196 | chr18-77456040-DEL-196 |
| chr18-77475445-DEL-53 | chr18-77475446-DEL-53 |
| chr18-77479803-DEL-130 | chr18-77479804-DEL-130 |
| chr18-77494005-DEL-104 | chr18-77494006-DEL-104 |
| chr18-77513731-DEL-154 | chr18-77513732-DEL-154 |
| chr18-77585392-DEL-188 | chr18-77585393-DEL-212 |
| chr18-77631860-DEL-68 | chr18-77631861-DEL-68 |
| chr18-77662844-DEL-182 | chr18-77662845-DEL-182 |
| chr18-77667439-DEL-69 | chr18-77667440-DEL-69 |
| chr18-77672177-DEL-216 | chr18-77672178-DEL-216 |
| chr18-77677993-DEL-64 | chr18-77677994-DEL-64 |
| chr18-77696772-DEL-54 | chr18-77696752-DEL-54 |
| chr18-77702663-DEL-74 | chr18-77702656-DEL-72 |
| chr18-77725735-DEL-211 | chr18-77725736-DEL-211 |
| chr18-77736265-DEL-179 | chr18-77736266-DEL-179 |
| chr18-77756326-DEL-89 | chr18-77756328-DEL-88 |
| chr18-77797990-DEL-218 | chr18-77797991-DEL-213 |
| chr18-77813533-DEL-199 | chr18-77813534-DEL-199 |

| | |
|---|---|
| chr18-77823414-DEL-206 | chr18-77823415-DEL-206 |
| chr18-77826450-DEL-199 | chr18-77826451-DEL-198 |
| chr18-77835339-DEL-214 | chr18-77835340-DEL-214 |
| chr18-77844960-DEL-211 | chr18-77844961-DEL-211 |
| chr18-77845985-DEL-106 | chr18-77845986-DEL-106 |
| chr18-77846584-DEL-173 | chr18-77846587-DEL-158 |
| chr18-77850742-DEL-222 | chr18-77850743-DEL-222 |
| chr18-77875170-DEL-231 | chr18-77875171-DEL-231 |
| chr18-77885933-DEL-240 | chr18-77885934-DEL-240 |
| chr18-77900395-DEL-175 | chr18-77900396-DEL-175 |
| chr18-77921336-DEL-421 | chr18-77921337-DEL-421 |
| chr18-77928830-DEL-199 | chr18-77928831-DEL-200 |
| chr18-77939511-DEL-98 | chr18-77939517-DEL-56 |
| chr18-77947390-DEL-52 | chr18-77947390-DEL-52 |
| chr18-77951918-DEL-1122 | chr18-77951919-DEL-1122 |
| chr18-77952727-DEL-181 | chr18-77952728-DEL-181 |
| chr18-77957815-DEL-63 | chr18-77957807-DEL-81 |
| chr18-77962842-DEL-208 | chr18-77962843-DEL-208 |
| chr18-77965827-DEL-108 | chr18-77965828-DEL-108 |
| chr18-77971902-DEL-332 | chr18-77971894-DEL-322 |
| chr18-77989199-DEL-1504 | chr18-77989200-DEL-1504 |
| chr18-77990226-DEL-184 | chr18-77990227-DEL-184 |
| chr18-77990922-DEL-192 | chr18-77990923-DEL-192 |
| chr18-77993180-DEL-98 | chr18-77993181-DEL-98 |
| chr18-78019903-DEL-58 | chr18-78019880-DEL-58 |
| chr18-78020438-DEL-535 | chr18-78020395-DEL-536 |
| chr18-78043008-DEL-707 | chr18-78043009-DEL-670 |
| chr18-78044045-DEL-115 | chr18-78044046-DEL-115 |
| chr18-78045492-DEL-210 | chr18-78045504-DEL-165 |
| chr18-78080934-DEL-65 | chr18-78080941-DEL-65 |
| chr18-78101658-DEL-107 | chr18-78101659-DEL-107 |
| chr18-78123515-DEL-119 | chr18-78123516-DEL-119 |
| chr18-78161428-DEL-100 | chr18-78161429-DEL-100 |
| chr18-78170746-DEL-96 | chr18-78170747-DEL-90 |
| chr18-78172079-DEL-53 | chr18-78172080-DEL-53 |
| chr18-78173553-DEL-243 | chr18-78173582-DEL-246 |
| chr18-78181868-DEL-601 | chr18-78181868-DEL-601 |
| chr18-78243790-DEL-486 | chr18-78243791-DEL-485 |
| chr18-78270815-DEL-61 | chr18-78270816-DEL-61 |
| chr18-78398977-DEL-66 | chr18-78398978-DEL-66 |
| chr18-78445989-DEL-1803 | chr18-78445989-DEL-1802 |
| chr18-78451040-DEL-1041 | chr18-78451041-DEL-1041 |
| chr18-78465302-DEL-6402 | chr18-78465304-DEL-6401 |
| chr18-78547940-DEL-454 | chr18-78547965-DEL-430 |
| chr18-78548777-DEL-235 | chr18-78548778-DEL-235 |
| chr18-78559432-DEL-52 | chr18-78559433-DEL-52 |
| chr18-78561915-DEL-80 | chr18-78561920-DEL-78 |
| chr18-78629713-DEL-105 | chr18-78629714-DEL-105 |
| chr18-78630688-DEL-460 | chr18-78630689-DEL-461 |
| chr18-78691183-DEL-82 | chr18-78691198-DEL-68 |
| chr18-78709059-DEL-108 | chr18-78709060-DEL-72 |
| chr18-78736938-DEL-52 | chr18-78736938-DEL-52 |
| chr18-78739586-DEL-7542 | chr18-78739587-DEL-7542 |
| chr18-78739642-DEL-212 | chr18-78739643-DEL-212 |
| chr18-78754350-DEL-908 | chr18-78754351-DEL-908 |
| chr18-78765665-DEL-97 | chr18-78765666-DEL-97 |
| chr18-78802879-DEL-64 | chr18-78802880-DEL-62 |
| chr18-78812618-DEL-116 | chr18-78812611-DEL-124 |
| chr18-78819651-DEL-119 | chr18-78819652-DEL-119 |
| chr18-78850321-DEL-101 | chr18-78850322-DEL-75 |
| chr18-78904637-DEL-6526 | chr18-78904638-DEL-6526 |
| chr18-78992817-DEL-64 | chr18-78992818-DEL-64 |
| chr18-78993665-DEL-1200 | chr18-78993662-DEL-1200 |
| chr18-79003950-DEL-97 | chr18-79003951-DEL-97 |
| chr18-79008366-DEL-108 | chr18-79008381-DEL-64 |
| chr18-79014740-DEL-389 | chr18-79014741-DEL-389 |
| chr18-79015625-DEL-80 | chr18-79015623-DEL-80 |
| chr18-79071433-DEL-243 | chr18-79071421-DEL-256 |
| chr18-79122587-DEL-65 | chr18-79122634-DEL-52 |
| chr18-79158462-DEL-132 | chr18-79158463-DEL-132 |
| chr18-79170104-DEL-93 | chr18-79170105-DEL-93 |
| chr18-79176854-DEL-72 | chr18-79176861-DEL-54 |
| chr18-79184826-DEL-117 | chr18-79184848-DEL-91 |
| chr18-79223340-DEL-204 | chr18-79223336-DEL-205 |
| chr18-79227042-DEL-116 | chr18-79227043-DEL-116 |
| chr18-79234621-DEL-57 | chr18-79234622-DEL-57 |
| chr18-79326326-DEL-759 | chr18-79326327-DEL-759 |
| chr18-79357333-DEL-88 | chr18-79357335-DEL-87 |
| chr18-79411043-DEL-336 | chr18-79411044-DEL-336 |
| chr18-79423361-DEL-905 | chr18-79423362-DEL-906 |

| | |
|---|---|
| chr18-79426738-DEL-72 | chr18-79426739-DEL-72 |
| chr18-79463721-DEL-6569 | chr18-79463712-DEL-6570 |
| chr18-79469680-DEL-201 | chr18-79469681-DEL-201 |
| chr18-79470898-DEL-1563 | chr18-79470899-DEL-1563 |
| chr18-79473220-DEL-472 | chr18-79473221-DEL-472 |
| chr18-79550596-DEL-6864 | chr18-79550597-DEL-6864 |
| chr18-79646610-DEL-1162 | chr18-79646611-DEL-1162 |
| chr18-79648762-DEL-4247 | chr18-79648763-DEL-4247 |
| chr18-79649438-DEL-79 | chr18-79649436-DEL-79 |
| chr18-79701625-DEL-52 | chr18-79701626-DEL-52 |
| chr18-79710448-DEL-1337 | chr18-79710449-DEL-1337 |
| chr18-79743835-DEL-56 | chr18-79743833-DEL-56 |
| chr18-79747246-DEL-7130 | chr18-79747247-DEL-7130 |
| chr18-79766923-DEL-8232 | chr18-79766924-DEL-8232 |
| chr18-79812561-DEL-54 | chr18-79812562-DEL-66 |
| chr18-79816180-DEL-1098 | chr18-79816181-DEL-1098 |
| chr18-79838309-DEL-91 | chr18-79838336-DEL-56 |
| chr18-79865593-DEL-194 | chr18-79865594-DEL-194 |
| chr18-79871248-DEL-434 | chr18-79871249-DEL-434 |
| chr18-79902769-DEL-96 | chr18-79902760-DEL-93 |
| chr18-79907131-DEL-70 | chr18-79907132-DEL-70 |
| chr18-79915925-DEL-326 | chr18-79915922-DEL-326 |
| chr18-79921911-DEL-56 | chr18-79921912-DEL-56 |
| chr18-79929075-DEL-226 | chr18-79929081-DEL-221 |
| chr18-79942360-DEL-65 | chr18-79942361-DEL-65 |
| chr18-79965567-DEL-254 | chr18-79965567-DEL-254 |
| chr18-79975723-DEL-836 | chr18-79975724-DEL-836 |
| chr18-79990182-DEL-190 | chr18-79990178-DEL-191 |
| chr18-79990285-DEL-959 | chr18-79990286-DEL-959 |
| chr18-79998421-DEL-81 | chr18-79998422-DEL-67 |
| chr18-79999598-DEL-432 | chr18-79999599-DEL-432 |
| chr18-80008846-DEL-53 | chr18-80008840-DEL-54 |
| chr18-80010708-DEL-58 | chr18-80010709-DEL-58 |
| chr18-80025184-DEL-67 | chr18-80025185-DEL-67 |
| chr18-80026009-DEL-426 | chr18-80026010-DEL-426 |
| chr18-80035815-DEL-52 | chr18-80035809-DEL-67 |
| chr18-80042226-DEL-54 | chr18-80042229-DEL-52 |
| chr18-80110245-DEL-93 | chr18-80110197-DEL-129 |
| chr18-80151767-DEL-63 | chr18-80151768-DEL-63 |
| chr18-80201772-DEL-58 | chr18-80201773-DEL-58 |
| chr18-80241864-DEL-85 | chr18-80241865-DEL-85 |
| chr18-80245705-DEL-212 | chr18-80245706-DEL-212 |
| chr18-80247520-DEL-54 | chr18-80247521-DEL-54 |
| chr18-80259192-DEL-188 | chr18-80259191-DEL-188 |
| chr18-80266886-DEL-169 | chr18-80266867-DEL-169 |
| chr18-80277347-DEL-62 | chr18-80277348-DEL-62 |
| chr18-80282860-DEL-54 | chr18-80282860-DEL-54 |
| chr18-80313732-DEL-533 | chr18-80313733-DEL-500 |
| chr18-80315328-DEL-55 | chr18-80315329-DEL-55 |
| chr18-80317492-DEL-60 | chr18-80317493-DEL-60 |
| chr18-80320321-DEL-557 | chr18-80320322-DEL-557 |
| chr18-80344631-DEL-1413 | chr18-80344666-DEL-1375 |
| chr18-80350832-DEL-1893 | chr18-80350833-DEL-1893 |
| chr18-80362792-DEL-52 | chr18-80362793-DEL-52 |
| chr18-80377992-DEL-134 | chr18-80377977-DEL-134 |
| chr18-80388793-DEL-103 | chr18-80388751-DEL-102 |
| chr18-80392133-DEL-78 | chr18-80392132-DEL-73 |
| chr18-80402237-DEL-74 | chr18-80402238-DEL-74 |
| chr18-80420205-DEL-72 | chr18-80420206-DEL-72 |
| chr18-80421698-DEL-486 | chr18-80421699-DEL-486 |
| chr18-80428398-DEL-318 | chr18-80428399-DEL-321 |
| chr18-80432480-DEL-167 | chr18-80432481-DEL-167 |
| chr18-80435633-DEL-55 | chr18-80435634-DEL-55 |
| chr18-80448414-DEL-498 | chr18-80448415-DEL-498 |
| chr18-80450846-DEL-59 | chr18-80450847-DEL-59 |
| chr18-80470307-DEL-236 | chr18-80470308-DEL-236 |
| chr18-80479623-DEL-58 | chr18-80479624-DEL-58 |
| chr18-80496706-DEL-119 | chr18-80496707-DEL-119 |
| chr18-80507295-DEL-55 | chr18-80507296-DEL-55 |
| chr18-80513991-DEL-89 | chr18-80513992-DEL-89 |
| chr18-80524283-DEL-83 | chr18-80524284-DEL-83 |
| chr18-80537985-DEL-57 | chr18-80537986-DEL-57 |
| chr18-80548923-DEL-12859 | chr18-80548924-DEL-12859 |
| chr18-80568771-DEL-252 | chr18-80568772-DEL-233 |
| chr18-80571072-DEL-52 | chr18-80571073-DEL-52 |
| chr18-80590860-DEL-128 | chr18-80590860-DEL-128 |
| chr18-80622786-DEL-201 | chr18-80622787-DEL-201 |
| chr18-80701925-DEL-105 | chr18-80701946-DEL-100 |
| chr18-80716683-DEL-65 | chr18-80716723-DEL-54 |
| chr18-80765908-DEL-249 | chr18-80765909-DEL-249 |

| | |
|---|---|
| chr18-80770659-DEL-55 | chr18-80770674-DEL-55 |
| chr18-80771438-DEL-143 | chr18-80771439-DEL-143 |
| chr18-80804916-DEL-91 | chr18-80804917-DEL-91 |
| chr18-80816290-DEL-117 | chr18-80816244-DEL-117 |
| chr18-80844352-DEL-188 | chr18-80844352-DEL-187 |
| chr18-80899329-DEL-63 | chr18-80899330-DEL-63 |
| chr18-80900927-DEL-405 | chr18-80900927-DEL-405 |
| chr18-80904439-DEL-51 | chr18-80904439-DEL-51 |
| chr18-80923405-DEL-6739 | chr18-80923406-DEL-6739 |
| chr18-80929554-DEL-204 | chr18-80929555-DEL-204 |
| chr18-80953937-DEL-135 | chr18-80953935-DEL-136 |
| chr18-80963321-DEL-64 | chr18-80963322-DEL-64 |
| chr18-80985858-DEL-60 | chr18-80985863-DEL-60 |
| chr18-80992145-DEL-70 | chr18-80992143-DEL-63 |
| chr18-80997401-DEL-174 | chr18-80997402-DEL-174 |
| chr18-80998386-DEL-143 | chr18-80998387-DEL-143 |
| chr18-81046072-DEL-218 | chr18-81046057-DEL-219 |
| chr18-81073271-DEL-119 | chr18-81073272-DEL-119 |
| chr18-81074828-DEL-56 | chr18-81074829-DEL-56 |
| chr18-81085510-DEL-57 | chr18-81085511-DEL-57 |
| chr18-81100312-DEL-96 | chr18-81100298-DEL-97 |
| chr18-81122344-DEL-59 | chr18-81122333-DEL-73 |
| chr18-81165490-DEL-78 | chr18-81165487-DEL-82 |
| chr18-81207661-DEL-58 | chr18-81207632-DEL-50 |
| chr18-81242775-DEL-361 | chr18-81242776-DEL-361 |
| chr18-81257024-DEL-128 | chr18-81257043-DEL-110 |
| chr18-81297071-DEL-797 | chr18-81297072-DEL-797 |
| chr18-81298582-DEL-1160 | chr18-81298583-DEL-1160 |
| chr18-81311635-DEL-278 | chr18-81311627-DEL-276 |
| chr18-81342950-DEL-6442 | chr18-81342951-DEL-6442 |
| chr18-81361854-DEL-54 | chr18-81361853-DEL-55 |
| chr18-81362608-DEL-67 | chr18-81362609-DEL-67 |
| chr18-81365440-DEL-54 | chr18-81365415-DEL-52 |
| chr18-81378834-DEL-62 | chr18-81378835-DEL-62 |
| chr18-81403760-DEL-201 | chr18-81403755-DEL-174 |
| chr18-81418553-DEL-53 | chr18-81418543-DEL-51 |
| chr18-81420014-DEL-211 | chr18-81420015-DEL-211 |
| chr18-81445542-DEL-250 | chr18-81445570-DEL-203 |
| chr18-81521886-DEL-1064 | chr18-81521887-DEL-1064 |
| chr18-81583057-DEL-443 | chr18-81583058-DEL-443 |
| chr18-81588114-DEL-169 | chr18-81588109-DEL-167 |
| chr18-81618504-DEL-120 | chr18-81618505-DEL-120 |
| chr18-81619222-DEL-64 | chr18-81619223-DEL-64 |
| chr18-81648004-DEL-63 | chr18-81648005-DEL-63 |
| chr18-81660528-DEL-193 | chr18-81660529-DEL-193 |
| chr18-81662773-DEL-169 | chr18-81662774-DEL-169 |
| chr18-81669377-DEL-104 | chr18-81669378-DEL-124 |
| chr18-81706108-DEL-618 | chr18-81706109-DEL-618 |
| chr18-81711439-DEL-54 | chr18-81711440-DEL-54 |
| chr18-81735528-DEL-642 | chr18-81735529-DEL-642 |
| chr18-81737767-DEL-53 | chr18-81737768-DEL-53 |
| chr18-81754488-DEL-71 | chr18-81754489-DEL-71 |
| chr18-81765040-DEL-230 | chr18-81765057-DEL-212 |
| chr18-81781430-DEL-253 | chr18-81781431-DEL-253 |
| chr18-81811705-DEL-56 | chr18-81811702-DEL-60 |
| chr18-81856218-DEL-55 | chr18-81856210-DEL-65 |
| chr18-81888326-DEL-80 | chr18-81888327-DEL-80 |
| chr18-81901209-DEL-62 | chr18-81901174-DEL-64 |
| chr18-81908502-DEL-192 | chr18-81908503-DEL-192 |
| chr18-81948012-DEL-288 | chr18-81948013-DEL-288 |
| chr18-81968688-DEL-224 | chr18-81968689-DEL-224 |
| chr18-81983387-DEL-54 | chr18-81983378-DEL-55 |
| chr18-81991548-DEL-66 | chr18-81991535-DEL-60 |
| chr18-81998761-DEL-214 | chr18-81998731-DEL-234 |
| chr18-82020766-DEL-167 | chr18-82020767-DEL-167 |
| chr18-82031463-DEL-573 | chr18-82031464-DEL-573 |
| chr18-82039534-DEL-123 | chr18-82039535-DEL-123 |
| chr18-82057550-DEL-145 | chr18-82057551-DEL-145 |
| chr18-82105593-DEL-60 | chr18-82105588-DEL-50 |
| chr18-82115855-DEL-211 | chr18-82115861-DEL-208 |
| chr18-82162670-DEL-56 | chr18-82162671-DEL-56 |
| chr18-82172712-DEL-246 | chr18-82172713-DEL-246 |
| chr18-82193779-DEL-954 | chr18-82193780-DEL-954 |
| chr18-82198906-DEL-349 | chr18-82198907-DEL-349 |
| chr18-82201193-DEL-145 | chr18-82201194-DEL-145 |
| chr18-82203905-DEL-369 | chr18-82203923-DEL-328 |
| chr18-82267252-DEL-61 | chr18-82267252-DEL-61 |
| chr18-82283106-DEL-6285 | chr18-82283104-DEL-6286 |
| chr18-82302403-DEL-109 | chr18-82302431-DEL-109 |
| chr18-82304803-DEL-96 | chr18-82304828-DEL-72 |

| | |
|---|---|
| chr18-82316404-DEL-1670 | chr18-82316405-DEL-1670 |
| chr18-82328818-DEL-3001 | chr18-82328819-DEL-3001 |
| chr18-82337663-DEL-1199 | chr18-82337676-DEL-1187 |
| chr18-82347269-DEL-67 | chr18-82347229-DEL-88 |
| chr18-82354037-DEL-64 | chr18-82354038-DEL-64 |
| chr18-82356468-DEL-1644 | chr18-82356469-DEL-1644 |
| chr18-82362392-DEL-1659 | chr18-82362393-DEL-1659 |
| chr18-82367341-DEL-55 | chr18-82367342-DEL-55 |
| chr18-82382493-DEL-54 | chr18-82382482-DEL-54 |
| chr18-82388257-DEL-788 | chr18-82388258-DEL-788 |
| chr18-82398835-DEL-251 | chr18-82398836-DEL-251 |
| chr18-82469344-DEL-448 | chr18-82469345-DEL-448 |
| chr18-82475113-DEL-376 | chr18-82475114-DEL-376 |
| chr18-82476661-DEL-434 | chr18-82476662-DEL-434 |
| chr18-82483273-DEL-56 | chr18-82483291-DEL-57 |
| chr18-82488661-DEL-70 | chr18-82488662-DEL-70 |
| chr18-82510266-DEL-55 | chr18-82510267-DEL-55 |
| chr18-82529497-DEL-58 | chr18-82529498-DEL-58 |
| chr18-82599820-DEL-177 | chr18-82599821-DEL-177 |
| chr18-82609784-DEL-56 | chr18-82609764-DEL-54 |
| chr18-82684861-DEL-65 | chr18-82684862-DEL-65 |
| chr18-82699885-DEL-200 | chr18-82699886-DEL-200 |
| chr18-82711670-DEL-8884 | chr18-82711671-DEL-8884 |
| chr18-82728559-DEL-84 | chr18-82728560-DEL-84 |
| chr18-82729319-DEL-787 | chr18-82729319-DEL-787 |
| chr18-82747749-DEL-53 | chr18-82747750-DEL-70 |
| chr18-82754154-DEL-107 | chr18-82754155-DEL-107 |
| chr18-82772559-DEL-98 | chr18-82772559-DEL-98 |
| chr18-82788900-DEL-104 | chr18-82788903-DEL-108 |
| chr18-82808697-DEL-53 | chr18-82808700-DEL-50 |
| chr18-82820583-DEL-787 | chr18-82820584-DEL-787 |
| chr18-82851052-DEL-85 | chr18-82851053-DEL-85 |
| chr18-82896124-DEL-1392 | chr18-82896125-DEL-1392 |
| chr18-82901157-DEL-200 | chr18-82901158-DEL-200 |
| chr18-82952447-DEL-68 | chr18-82952414-DEL-102 |
| chr18-82989284-DEL-202 | chr18-82989285-DEL-202 |
| chr18-82993503-DEL-79 | chr18-82993504-DEL-79 |
| chr18-83015323-DEL-63 | chr18-83015324-DEL-63 |
| chr18-83016220-DEL-714 | chr18-83016221-DEL-714 |
| chr18-83046828-DEL-638 | chr18-83046829-DEL-638 |
| chr18-83062280-DEL-328 | chr18-83062281-DEL-328 |
| chr18-83074703-DEL-56 | chr18-83074704-DEL-56 |
| chr18-83080033-DEL-228 | chr18-83080034-DEL-228 |
| chr18-83102745-DEL-86 | chr18-83102785-DEL-50 |
| chr18-83104083-DEL-88 | chr18-83104084-DEL-88 |
| chr18-83107907-DEL-187 | chr18-83107908-DEL-187 |
| chr18-83108498-DEL-72 | chr18-83108499-DEL-72 |
| chr18-83114121-DEL-109 | chr18-83114122-DEL-109 |
| chr18-83117266-DEL-129 | chr18-83117279-DEL-117 |
| chr18-83129325-DEL-58 | chr18-83129303-DEL-53 |
| chr18-83152938-DEL-82 | chr18-83152939-DEL-82 |
| chr18-83166584-DEL-2022 | chr18-83166567-DEL-2027 |
| chr18-83189311-DEL-7104 | chr18-83189312-DEL-7104 |
| chr18-83197816-DEL-88 | chr18-83197817-DEL-88 |
| chr18-83215006-DEL-122 | chr18-83215007-DEL-122 |
| chr18-83254881-DEL-369 | chr18-83254882-DEL-369 |
| chr18-83268388-DEL-133 | chr18-83268389-DEL-133 |
| chr18-83271949-DEL-70 | chr18-83271950-DEL-70 |
| chr18-83276706-DEL-12737 | chr18-83276707-DEL-12737 |
| chr18-83290214-DEL-57 | chr18-83290215-DEL-57 |
| chr18-83294952-DEL-69 | chr18-83294953-DEL-69 |
| chr18-83319119-DEL-51 | chr18-83319098-DEL-82 |
| chr18-83325660-DEL-2055 | chr18-83325652-DEL-2056 |
| chr18-83376869-DEL-232 | chr18-83376850-DEL-203 |
| chr18-83430698-DEL-2157 | chr18-83430684-DEL-2158 |
| chr18-83434689-DEL-68 | chr18-83434690-DEL-68 |
| chr18-83455400-DEL-125 | chr18-83455367-DEL-114 |
| chr18-83463446-DEL-64 | chr18-83463441-DEL-68 |
| chr18-83501900-DEL-115 | chr18-83501898-DEL-116 |
| chr18-83513110-DEL-51 | chr18-83513111-DEL-51 |
| chr18-83517244-DEL-745 | chr18-83517236-DEL-746 |
| chr18-83559805-DEL-433 | chr18-83559809-DEL-434 |
| chr18-83572925-DEL-76 | chr18-83572926-DEL-76 |
| chr18-83586282-DEL-1744 | chr18-83586283-DEL-1744 |
| chr18-83588724-DEL-531 | chr18-83588725-DEL-531 |
| chr18-83598358-DEL-66 | chr18-83598359-DEL-66 |
| chr18-83599407-DEL-75 | chr18-83599414-DEL-72 |
| chr18-83609686-DEL-54 | chr18-83609687-DEL-54 |
| chr18-83642028-DEL-92 | chr18-83642016-DEL-54 |
| chr18-83664882-DEL-75 | chr18-83664871-DEL-114 |

| | |
|---|---|
| chr18-83671686-DEL-336 | chr18-83671687-DEL-336 |
| chr18-83698775-DEL-91 | chr18-83698776-DEL-91 |
| chr18-83708932-DEL-1878 | chr18-83708933-DEL-1878 |
| chr18-83711499-DEL-100 | chr18-83711500-DEL-100 |
| chr18-83712940-DEL-81 | chr18-83712953-DEL-52 |
| chr18-83715596-DEL-136 | chr18-83715601-DEL-127 |
| chr18-83718764-DEL-7139 | chr18-83718765-DEL-7139 |
| chr18-83747654-DEL-440 | chr18-83747655-DEL-440 |
| chr18-83758014-DEL-551 | chr18-83758015-DEL-551 |
| chr18-83796218-DEL-61 | chr18-83796219-DEL-61 |
| chr18-83854780-DEL-64 | chr18-83854780-DEL-64 |
| chr18-83870426-DEL-3829 | chr18-83870430-DEL-3826 |
| chr18-83892143-DEL-180 | chr18-83892144-DEL-180 |
| chr18-83906654-DEL-58 | chr18-83906655-DEL-58 |
| chr18-83944659-DEL-63 | chr18-83944660-DEL-63 |
| chr18-83958896-DEL-58 | chr18-83958897-DEL-58 |
| chr18-83992921-DEL-60 | chr18-83992922-DEL-60 |
| chr18-83999032-DEL-115 | chr18-83999033-DEL-91 |
| chr18-84020407-DEL-96 | chr18-84020408-DEL-96 |
| chr18-84065108-DEL-120 | chr18-84065098-DEL-150 |
| chr18-84108344-DEL-60 | chr18-84108345-DEL-60 |
| chr18-84167836-DEL-75 | chr18-84167837-DEL-75 |
| chr18-84170353-DEL-63 | chr18-84170353-DEL-56 |
| chr18-84176933-DEL-256 | chr18-84176982-DEL-240 |
| chr18-84185156-DEL-168 | chr18-84185157-DEL-168 |
| chr18-84196836-DEL-177 | chr18-84196788-DEL-180 |
| chr18-84198627-DEL-57 | chr18-84198624-DEL-60 |
| chr18-84200166-DEL-2737 | chr18-84200167-DEL-2737 |
| chr18-84213934-DEL-404 | chr18-84213933-DEL-405 |
| chr18-84231119-DEL-318 | chr18-84231120-DEL-318 |
| chr18-84291672-DEL-121 | chr18-84291673-DEL-121 |
| chr18-84326319-DEL-113 | chr18-84326362-DEL-115 |
| chr18-84379439-DEL-52 | chr18-84379440-DEL-52 |
| chr18-84384748-DEL-274 | chr18-84384749-DEL-274 |
| chr18-84436994-DEL-88 | chr18-84436995-DEL-88 |
| chr18-84499965-DEL-192 | chr18-84499975-DEL-160 |
| chr18-84521777-DEL-179 | chr18-84521766-DEL-170 |
| chr18-84536942-DEL-152 | chr18-84536943-DEL-152 |
| chr18-84537592-DEL-131 | chr18-84537592-DEL-131 |
| chr18-84544545-DEL-80 | chr18-84544532-DEL-81 |
| chr18-84588079-DEL-57 | chr18-84588081-DEL-50 |
| chr18-84609702-DEL-76 | chr18-84609703-DEL-76 |
| chr18-84621951-DEL-6599 | chr18-84621952-DEL-6599 |
| chr18-84643682-DEL-215 | chr18-84643646-DEL-230 |
| chr18-84650788-DEL-98 | chr18-84650804-DEL-83 |
| chr18-84653709-DEL-96 | chr18-84653738-DEL-68 |
| chr18-84659437-DEL-100 | chr18-84659434-DEL-97 |
| chr18-84674564-DEL-65 | chr18-84674565-DEL-65 |
| chr18-84677964-DEL-2975 | chr18-84677959-DEL-2980 |
| chr18-84698206-DEL-205 | chr18-84698207-DEL-204 |
| chr18-84716686-DEL-749 | chr18-84716687-DEL-749 |
| chr18-84722415-DEL-56 | chr18-84722416-DEL-56 |
| chr18-84727568-DEL-122 | chr18-84727569-DEL-122 |
| chr18-84732233-DEL-1623 | chr18-84732234-DEL-1623 |
| chr18-84750291-DEL-84 | chr18-84750290-DEL-83 |
| chr18-84751696-DEL-111 | chr18-84751697-DEL-111 |
| chr18-84753961-DEL-53 | chr18-84753961-DEL-53 |
| chr18-84770589-DEL-588 | chr18-84770590-DEL-588 |
| chr18-84774711-DEL-437 | chr18-84774716-DEL-435 |
| chr18-84776034-DEL-164 | chr18-84776035-DEL-164 |
| chr18-84785109-DEL-139 | chr18-84785110-DEL-139 |
| chr18-84810592-DEL-201 | chr18-84810593-DEL-201 |
| chr18-84816016-DEL-268 | chr18-84816017-DEL-268 |
| chr18-84838198-DEL-1239 | chr18-84838199-DEL-1239 |
| chr18-84843535-DEL-182 | chr18-84843535-DEL-183 |
| chr18-84858916-DEL-87 | chr18-84858917-DEL-87 |
| chr18-84861311-DEL-209 | chr18-84861312-DEL-209 |
| chr18-84867380-DEL-1377 | chr18-84867392-DEL-1372 |
| chr18-84874886-DEL-94 | chr18-84874878-DEL-94 |
| chr18-84881239-DEL-186 | chr18-84881288-DEL-178 |
| chr18-84883187-DEL-93 | chr18-84883188-DEL-93 |
| chr18-84904589-DEL-88 | chr18-84904590-DEL-88 |
| chr18-84916766-DEL-208 | chr18-84916767-DEL-208 |
| chr18-84918690-DEL-260 | chr18-84918691-DEL-260 |
| chr18-84919375-DEL-165 | chr18-84919376-DEL-165 |
| chr18-84943481-DEL-220 | chr18-84943482-DEL-220 |
| chr18-84963684-DEL-810 | chr18-84963685-DEL-810 |
| chr18-84977549-DEL-213 | chr18-84977551-DEL-211 |
| chr18-84989197-DEL-84 | chr18-84989198-DEL-84 |
| chr18-85000487-DEL-204 | chr18-85000488-DEL-204 |

| | |
|---|---|
| chr18-85027901-DEL-867 | chr18-85027902-DEL-867 |
| chr18-85028729-DEL-298 | chr18-85028730-DEL-290 |
| chr18-85030912-DEL-74 | chr18-85030913-DEL-74 |
| chr18-85034867-DEL-10516 | chr18-85034868-DEL-10516 |
| chr18-85060996-DEL-309 | chr18-85060997-DEL-309 |
| chr18-85065871-DEL-309 | chr18-85065872-DEL-309 |
| chr18-85073648-DEL-94 | chr18-85073649-DEL-94 |
| chr18-85074621-DEL-90 | chr18-85074614-DEL-98 |
| chr18-85083443-DEL-449 | chr18-85083444-DEL-449 |
| chr18-85087709-DEL-183 | chr18-85087710-DEL-183 |
| chr18-85092519-DEL-6178 | chr18-85092520-DEL-6178 |
| chr18-85101795-DEL-63 | chr18-85101771-DEL-87 |
| chr18-85114352-DEL-148 | chr18-85114353-DEL-148 |
| chr18-85123380-DEL-9901 | chr18-85123381-DEL-9901 |
| chr18-85127037-DEL-3683 | chr18-85127037-DEL-3683 |
| chr18-85146352-DEL-1652 | chr18-85146352-DEL-1652 |
| chr18-85158147-DEL-54 | chr18-85158101-DEL-86 |
| chr18-85184369-DEL-430 | chr18-85184370-DEL-430 |
| chr18-85187966-DEL-6748 | chr18-85187964-DEL-6749 |
| chr18-85214213-DEL-74 | chr18-85214214-DEL-74 |
| chr18-85317743-DEL-70 | chr18-85317744-DEL-70 |
| chr18-85323834-DEL-82 | chr18-85323836-DEL-70 |
| chr18-85337134-DEL-6439 | chr18-85337135-DEL-6439 |
| chr18-85343223-DEL-212 | chr18-85343224-DEL-212 |
| chr18-85396032-DEL-4882 | chr18-85396033-DEL-4882 |
| chr18-85423342-DEL-431 | chr18-85423343-DEL-431 |
| chr18-85482166-DEL-72 | chr18-85482167-DEL-70 |
| chr18-85523029-DEL-1695 | chr18-85523030-DEL-1695 |
| chr18-85655892-DEL-52 | chr18-85655893-DEL-70 |
| chr18-85669432-DEL-70 | chr18-85669433-DEL-70 |
| chr18-85704174-DEL-76 | chr18-85704162-DEL-74 |
| chr18-85726087-DEL-54 | chr18-85726088-DEL-54 |
| chr18-85728936-DEL-55 | chr18-85728936-DEL-71 |
| chr18-85729996-DEL-206 | chr18-85729997-DEL-206 |
| chr18-85774611-DEL-55 | chr18-85774612-DEL-51 |
| chr18-85788037-DEL-103 | chr18-85788038-DEL-103 |
| chr18-85836165-DEL-64 | chr18-85836147-DEL-112 |
| chr18-85838936-DEL-6573 | chr18-85838937-DEL-6573 |
| chr18-85852703-DEL-779 | chr18-85852704-DEL-779 |
| chr18-85882191-DEL-1007 | chr18-85882192-DEL-1007 |
| chr18-85890610-DEL-495 | chr18-85890605-DEL-499 |
| chr18-85897290-DEL-63 | chr18-85897291-DEL-63 |
| chr18-85910817-DEL-61 | chr18-85910818-DEL-61 |
| chr18-85915552-DEL-208 | chr18-85915553-DEL-208 |
| chr18-85970189-DEL-9179 | chr18-85970190-DEL-9179 |
| chr18-85987296-DEL-243 | chr18-85987297-DEL-243 |
| chr18-86030969-DEL-72 | chr18-86030970-DEL-72 |
| chr18-86034295-DEL-87 | chr18-86034296-DEL-87 |
| chr18-86036081-DEL-79 | chr18-86036105-DEL-83 |
| chr18-86062210-DEL-7131 | chr18-86062211-DEL-7131 |
| chr18-86073485-DEL-144 | chr18-86073509-DEL-166 |
| chr18-86083164-DEL-485 | chr18-86083164-DEL-485 |
| chr18-86103166-DEL-366 | chr18-86103173-DEL-362 |
| chr18-86143980-DEL-907 | chr18-86143981-DEL-907 |
| chr18-86145738-DEL-1036 | chr18-86145739-DEL-1036 |
| chr18-86151991-DEL-288 | chr18-86151992-DEL-288 |
| chr18-86161403-DEL-67 | chr18-86161404-DEL-67 |
| chr18-86183052-DEL-51 | chr18-86183053-DEL-51 |
| chr18-86190673-DEL-165 | chr18-86190674-DEL-165 |
| chr18-86200373-DEL-72 | chr18-86200374-DEL-72 |
| chr18-86210151-DEL-8843 | chr18-86210152-DEL-8843 |
| chr18-86219505-DEL-6800 | chr18-86219506-DEL-6800 |
| chr18-86225628-DEL-424 | chr18-86225628-DEL-424 |
| chr18-86254563-DEL-1084 | chr18-86254564-DEL-1084 |
| chr18-86259197-DEL-2576 | chr18-86259197-DEL-2576 |
| chr18-86267916-DEL-1384 | chr18-86267917-DEL-1384 |
| chr18-86275781-DEL-1734 | chr18-86275782-DEL-1735 |
| chr18-86285394-DEL-420 | chr18-86285395-DEL-420 |
| chr18-86316438-DEL-959 | chr18-86316439-DEL-959 |
| chr18-86344262-DEL-55 | chr18-86344263-DEL-55 |
| chr18-86348313-DEL-401 | chr18-86348314-DEL-401 |
| chr18-86385598-DEL-301 | chr18-86385599-DEL-301 |
| chr18-86389814-DEL-1985 | chr18-86389814-DEL-1985 |
| chr18-86397156-DEL-52 | chr18-86397157-DEL-50 |
| chr18-86482647-DEL-123 | chr18-86482687-DEL-75 |
| chr18-86530794-DEL-130 | chr18-86530795-DEL-130 |
| chr18-86536398-DEL-1000 | chr18-86536399-DEL-1000 |
| chr18-86566911-DEL-2430 | chr18-86566912-DEL-2430 |
| chr18-86573190-DEL-212 | chr18-86573191-DEL-212 |
| chr18-86578681-DEL-6859 | chr18-86578678-DEL-6860 |

| | |
|---|---|
| chr18-86578789-DEL-212 | chr18-86578790-DEL-212 |
| chr18-86600840-DEL-68 | chr18-86600850-DEL-69 |
| chr18-86601283-DEL-2995 | chr18-86601284-DEL-2995 |
| chr18-86674100-DEL-82 | chr18-86674101-DEL-82 |
| chr18-86687367-DEL-257 | chr18-86687368-DEL-257 |
| chr18-86688484-DEL-353 | chr18-86688485-DEL-353 |
| chr18-86833476-DEL-120 | chr18-86833477-DEL-120 |
| chr18-86863829-DEL-426 | chr18-86863830-DEL-426 |
| chr18-86886976-DEL-924 | chr18-86886983-DEL-918 |
| chr18-86893048-DEL-64 | chr18-86893049-DEL-64 |
| chr18-86913195-DEL-60 | chr18-86913187-DEL-58 |
| chr18-86917800-DEL-2869 | chr18-86917801-DEL-2869 |
| chr18-86950761-DEL-134 | chr18-86950765-DEL-121 |
| chr18-86951302-DEL-567 | chr18-86951303-DEL-567 |
| chr18-86989752-DEL-81 | chr18-86989745-DEL-89 |
| chr18-86998405-DEL-590 | chr18-86998406-DEL-590 |
| chr18-87003081-DEL-68 | chr18-87003082-DEL-68 |
| chr18-87047355-DEL-441 | chr18-87047356-DEL-436 |
| chr18-87062933-DEL-327 | chr18-87062930-DEL-328 |
| chr18-87073022-DEL-55 | chr18-87073022-DEL-70 |
| chr18-87089209-DEL-153 | chr18-87089212-DEL-118 |
| chr18-87103034-DEL-75 | chr18-87103035-DEL-75 |
| chr18-87205221-DEL-51 | chr18-87205222-DEL-51 |
| chr18-87224101-DEL-139 | chr18-87224102-DEL-139 |
| chr18-87240337-DEL-885 | chr18-87240338-DEL-885 |
| chr18-87245677-DEL-1102 | chr18-87245678-DEL-1102 |
| chr18-87267381-DEL-241 | chr18-87267387-DEL-239 |
| chr18-87306953-DEL-54 | chr18-87306943-DEL-52 |
| chr18-87328547-DEL-369 | chr18-87328548-DEL-369 |
| chr18-87340863-DEL-214 | chr18-87340864-DEL-214 |
| chr18-87363111-DEL-6367 | chr18-87363112-DEL-6366 |
| chr18-87381902-DEL-322 | chr18-87381903-DEL-322 |
| chr18-87401240-DEL-174 | chr18-87401241-DEL-174 |
| chr18-87430949-DEL-98 | chr18-87430950-DEL-98 |
| chr18-87435889-DEL-226 | chr18-87435890-DEL-226 |
| chr18-87473384-DEL-116 | chr18-87473384-DEL-116 |
| chr18-87474815-DEL-1896 | chr18-87474817-DEL-1895 |
| chr18-87511821-DEL-3030 | chr18-87511822-DEL-3030 |
| chr18-87550049-DEL-1676 | chr18-87550038-DEL-1677 |
| chr18-87569014-DEL-86 | chr18-87568999-DEL-102 |
| chr18-87586165-DEL-6383 | chr18-87586168-DEL-6384 |
| chr18-87592305-DEL-201 | chr18-87592306-DEL-201 |
| chr18-87602733-DEL-273 | chr18-87602735-DEL-275 |
| chr18-87615785-DEL-197 | chr18-87615786-DEL-197 |
| chr18-87616482-DEL-1198 | chr18-87616482-DEL-1198 |
| chr18-87661726-DEL-233 | chr18-87661727-DEL-233 |
| chr18-87670233-DEL-97 | chr18-87670256-DEL-54 |
| chr18-87674785-DEL-306 | chr18-87674786-DEL-306 |
| chr18-87676248-DEL-186 | chr18-87676249-DEL-186 |
| chr18-87707951-DEL-6248 | chr18-87707952-DEL-6248 |
| chr18-87713921-DEL-199 | chr18-87713922-DEL-199 |
| chr18-87774400-DEL-1892 | chr18-87774399-DEL-1891 |
| chr18-87837362-DEL-52 | chr18-87837363-DEL-52 |
| chr18-87860096-DEL-94 | chr18-87860118-DEL-60 |
| chr18-87869570-DEL-592 | chr18-87869572-DEL-590 |
| chr18-87921170-DEL-56 | chr18-87921171-DEL-56 |
| chr18-87922672-DEL-87 | chr18-87922673-DEL-87 |
| chr18-87983559-DEL-2765 | chr18-87983560-DEL-2765 |
| chr18-88045163-DEL-75 | chr18-88045164-DEL-75 |
| chr18-88112887-DEL-121 | chr18-88112888-DEL-121 |
| chr18-88143912-DEL-144 | chr18-88143912-DEL-145 |
| chr18-88145379-DEL-60 | chr18-88145378-DEL-82 |
| chr18-88146089-DEL-443 | chr18-88146090-DEL-443 |
| chr18-88159239-DEL-77 | chr18-88159240-DEL-77 |
| chr18-88163509-DEL-128 | chr18-88163510-DEL-128 |
| chr18-88167288-DEL-6207 | chr18-88167289-DEL-6207 |
| chr18-88211202-DEL-235 | chr18-88211203-DEL-235 |
| chr18-88241442-DEL-397 | chr18-88241429-DEL-398 |
| chr18-88276351-DEL-402 | chr18-88276352-DEL-402 |
| chr18-88282305-DEL-83 | chr18-88282304-DEL-83 |
| chr18-88302339-DEL-223 | chr18-88302340-DEL-223 |
| chr18-88304776-DEL-59 | chr18-88304777-DEL-59 |
| chr18-88315365-DEL-6510 | chr18-88315366-DEL-6510 |
| chr18-88382996-DEL-58 | chr18-88382997-DEL-58 |
| chr18-88384087-DEL-118 | chr18-88384126-DEL-74 |
| chr18-88411769-DEL-3032 | chr18-88411770-DEL-3032 |
| chr18-88459770-DEL-80 | chr18-88459771-DEL-80 |
| chr18-88467262-DEL-766 | chr18-88467264-DEL-765 |
| chr18-88489295-DEL-374 | chr18-88489296-DEL-374 |
| chr18-88496562-DEL-57 | chr18-88496563-DEL-57 |

| | |
|---|---|
| chr18-88499658-DEL-121 | chr18-88499659-DEL-121 |
| chr18-88509220-DEL-55 | chr18-88509221-DEL-55 |
| chr18-88521363-DEL-434 | chr18-88521364-DEL-434 |
| chr18-88531897-DEL-104 | chr18-88531902-DEL-100 |
| chr18-88540621-DEL-64 | chr18-88540623-DEL-65 |
| chr18-88542190-DEL-76 | chr18-88542194-DEL-75 |
| chr18-88545581-DEL-2757 | chr18-88545582-DEL-2757 |
| chr18-88547352-DEL-2436 | chr18-88547352-DEL-2435 |
| chr18-88554959-DEL-63 | chr18-88554960-DEL-63 |
| chr18-88567139-DEL-1103 | chr18-88567140-DEL-1103 |
| chr18-88570618-DEL-468 | chr18-88570619-DEL-468 |
| chr18-88581508-DEL-185 | chr18-88581509-DEL-185 |
| chr18-88637712-DEL-214 | chr18-88637713-DEL-214 |
| chr18-88658095-DEL-190 | chr18-88658096-DEL-190 |
| chr18-88707967-DEL-510 | chr18-88707968-DEL-510 |
| chr18-88709662-DEL-78 | chr18-88709663-DEL-78 |
| chr18-88746095-DEL-58 | chr18-88746097-DEL-72 |
| chr18-88758029-DEL-177 | chr18-88758030-DEL-177 |
| chr18-88773416-DEL-1147 | chr18-88773417-DEL-1147 |
| chr18-88782775-DEL-73 | chr18-88782782-DEL-67 |
| chr18-88799660-DEL-64 | chr18-88799661-DEL-64 |
| chr18-88849603-DEL-261 | chr18-88849604-DEL-261 |
| chr18-88893787-DEL-358 | chr18-88893788-DEL-358 |
| chr18-88919105-DEL-374 | chr18-88919106-DEL-374 |
| chr18-88925152-DEL-108 | chr18-88925165-DEL-72 |
| chr18-88953201-DEL-625 | chr18-88953202-DEL-625 |
| chr18-88957929-DEL-6444 | chr18-88957930-DEL-6444 |
| chr18-89090541-DEL-335 | chr18-89090542-DEL-335 |
| chr18-89110445-DEL-60 | chr18-89110446-DEL-60 |
| chr18-89122558-DEL-60 | chr18-89122541-DEL-86 |
| chr18-89135451-DEL-52 | chr18-89135452-DEL-52 |
| chr18-89154409-DEL-7078 | chr18-89154409-DEL-7078 |
| chr18-89154849-DEL-209 | chr18-89154850-DEL-209 |
| chr18-89178653-DEL-218 | chr18-89178654-DEL-218 |
| chr18-89179556-DEL-426 | chr18-89179557-DEL-426 |
| chr18-89183706-DEL-1667 | chr18-89183697-DEL-1678 |
| chr18-89189577-DEL-315 | chr18-89189579-DEL-324 |
| chr18-89193895-DEL-591 | chr18-89193912-DEL-579 |
| chr18-89196234-DEL-66 | chr18-89196235-DEL-66 |
| chr18-89212136-DEL-124 | chr18-89212137-DEL-124 |
| chr18-89230984-DEL-351 | chr18-89230985-DEL-351 |
| chr18-89243744-DEL-126 | chr18-89243745-DEL-126 |
| chr18-89257563-DEL-451 | chr18-89257564-DEL-451 |
| chr18-89269199-DEL-81 | chr18-89269200-DEL-81 |
| chr18-89295541-DEL-403 | chr18-89295542-DEL-403 |
| chr18-89330401-DEL-3552 | chr18-89330402-DEL-3552 |
| chr18-89333019-DEL-81 | chr18-89333020-DEL-81 |
| chr18-89377725-DEL-62 | chr18-89377726-DEL-62 |
| chr18-89386995-DEL-51 | chr18-89386989-DEL-59 |
| chr18-89388129-DEL-7397 | chr18-89388130-DEL-7397 |
| chr18-89396450-DEL-113 | chr18-89396450-DEL-113 |
| chr18-89400353-DEL-99 | chr18-89400401-DEL-51 |
| chr18-89403652-DEL-2366 | chr18-89403653-DEL-2366 |
| chr18-89412935-DEL-1722 | chr18-89412936-DEL-1722 |
| chr18-89455814-DEL-410 | chr18-89455815-DEL-410 |
| chr18-89497245-DEL-536 | chr18-89497246-DEL-536 |
| chr18-89498137-DEL-505 | chr18-89498138-DEL-505 |
| chr18-89604105-DEL-580 | chr18-89604106-DEL-580 |
| chr18-89616066-DEL-72 | chr18-89616059-DEL-64 |
| chr18-89633935-DEL-5060 | chr18-89633936-DEL-5060 |
| chr18-89644705-DEL-1962 | chr18-89644706-DEL-1962 |
| chr18-89659513-DEL-490 | chr18-89659514-DEL-490 |
| chr18-89660342-DEL-256 | chr18-89660343-DEL-256 |
| chr18-89666688-DEL-1890 | chr18-89666689-DEL-1890 |
| chr18-89700796-DEL-6357 | chr18-89700797-DEL-6357 |
| chr18-89734251-DEL-64 | chr18-89734263-DEL-52 |
| chr18-89735789-DEL-127 | chr18-89735790-DEL-127 |
| chr18-89736847-DEL-3373 | chr18-89736848-DEL-3373 |
| chr18-89836901-DEL-63 | chr18-89836902-DEL-61 |
| chr18-89853648-DEL-5046 | chr18-89853649-DEL-5046 |
| chr18-89869532-DEL-62 | chr18-89869532-DEL-58 |
| chr18-89928645-DEL-7678 | chr18-89928646-DEL-7678 |
| chr18-89935404-DEL-208 | chr18-89935405-DEL-208 |
| chr18-89937284-DEL-58 | chr18-89937285-DEL-58 |
| chr18-89939373-DEL-412 | chr18-89939374-DEL-412 |
| chr18-89956376-DEL-1212 | chr18-89956377-DEL-1210 |
| chr18-89970219-DEL-337 | chr18-89970220-DEL-337 |
| chr18-90001300-DEL-6485 | chr18-90001301-DEL-6485 |
| chr18-90057054-DEL-60 | chr18-90057054-DEL-60 |
| chr18-90119552-DEL-169 | chr18-90119553-DEL-169 |

| | |
|---|---|
| chr18-90128159-DEL-11121 | chr18-90128156-DEL-11100 |
| chr18-90131628-DEL-4970 | chr18-90131629-DEL-4970 |
| chr18-90141689-DEL-56 | chr18-90141678-DEL-62 |
| chr18-90221943-DEL-335 | chr18-90221944-DEL-335 |
| chr18-90222868-DEL-1711 | chr18-90222869-DEL-1711 |
| chr18-90248371-DEL-10582 | chr18-90248372-DEL-10582 |
| chr18-90274128-DEL-72 | chr18-90274087-DEL-102 |
| chr18-90288786-DEL-9419 | chr18-90288787-DEL-9419 |
| chr18-90288850-DEL-204 | chr18-90288851-DEL-204 |
| chr18-90316458-DEL-327 | chr18-90316459-DEL-327 |
| chr18-90322806-DEL-1558 | chr18-90322807-DEL-1558 |
| chr18-90328991-DEL-3150 | chr18-90328992-DEL-3150 |
| chr18-90391842-DEL-8001 | chr18-90391843-DEL-8001 |
| chr18-90481684-DEL-67 | chr18-90481685-DEL-67 |
| chr18-90482246-DEL-8265 | chr18-90482247-DEL-8265 |
| chr18-90513047-DEL-3467 | chr18-90513048-DEL-3467 |
| chr18-90596172-DEL-209 | chr18-90596173-DEL-209 |
| chr19-3129941-DEL-214 | chr19-3129942-DEL-214 |
| chr19-3211363-DEL-6400 | chr19-3211364-DEL-6400 |
| chr19-3273698-DEL-5098 | chr19-3273699-DEL-5098 |
| chr19-3297579-DEL-565 | chr19-3297580-DEL-565 |
| chr19-3349734-DEL-86 | chr19-3349735-DEL-86 |
| chr19-3352660-DEL-84 | chr19-3352661-DEL-72 |
| chr19-3365062-DEL-122 | chr19-3365093-DEL-129 |
| chr19-3377392-DEL-174 | chr19-3377433-DEL-134 |
| chr19-3430559-DEL-60 | chr19-3430560-DEL-60 |
| chr19-3559830-DEL-677 | chr19-3559818-DEL-693 |
| chr19-3567019-DEL-217 | chr19-3567020-DEL-217 |
| chr19-3601126-DEL-489 | chr19-3601127-DEL-489 |
| chr19-3621846-DEL-165 | chr19-3621847-DEL-165 |
| chr19-3650538-DEL-70 | chr19-3650539-DEL-70 |
| chr19-3657268-DEL-245 | chr19-3657269-DEL-245 |
| chr19-3682803-DEL-59 | chr19-3682784-DEL-60 |
| chr19-3707476-DEL-203 | chr19-3707477-DEL-203 |
| chr19-3725880-DEL-82 | chr19-3725889-DEL-50 |
| chr19-3728613-DEL-204 | chr19-3728614-DEL-204 |
| chr19-3747140-DEL-191 | chr19-3747140-DEL-192 |
| chr19-3789261-DEL-207 | chr19-3789262-DEL-207 |
| chr19-3799028-DEL-440 | chr19-3799023-DEL-439 |
| chr19-3801866-DEL-244 | chr19-3801867-DEL-244 |
| chr19-3811855-DEL-212 | chr19-3811856-DEL-212 |
| chr19-3821200-DEL-203 | chr19-3821201-DEL-203 |
| chr19-3851719-DEL-72 | chr19-3851720-DEL-72 |
| chr19-3873903-DEL-551 | chr19-3873904-DEL-512 |
| chr19-3877405-DEL-204 | chr19-3877420-DEL-202 |
| chr19-3881554-DEL-202 | chr19-3881555-DEL-202 |
| chr19-3888314-DEL-544 | chr19-3888316-DEL-542 |
| chr19-3904556-DEL-210 | chr19-3904570-DEL-214 |
| chr19-3907765-DEL-60 | chr19-3907759-DEL-54 |
| chr19-3917756-DEL-222 | chr19-3917757-DEL-222 |
| chr19-3950926-DEL-218 | chr19-3950927-DEL-218 |
| chr19-3977937-DEL-59 | chr19-3977938-DEL-59 |
| chr19-3990071-DEL-62 | chr19-3990072-DEL-62 |
| chr19-4004231-DEL-58 | chr19-4004231-DEL-52 |
| chr19-4027101-DEL-84 | chr19-4027102-DEL-84 |
| chr19-4072676-DEL-460 | chr19-4072677-DEL-460 |
| chr19-4082039-DEL-178 | chr19-4082040-DEL-178 |
| chr19-4093364-DEL-179 | chr19-4093365-DEL-184 |
| chr19-4095064-DEL-567 | chr19-4095065-DEL-567 |
| chr19-4099091-DEL-388 | chr19-4099092-DEL-388 |
| chr19-4111376-DEL-4993 | chr19-4111377-DEL-4993 |
| chr19-4138041-DEL-122 | chr19-4138042-DEL-122 |
| chr19-4142086-DEL-196 | chr19-4142087-DEL-196 |
| chr19-4227410-DEL-155 | chr19-4227421-DEL-145 |
| chr19-4258454-DEL-78 | chr19-4258454-DEL-78 |
| chr19-4297433-DEL-196 | chr19-4297440-DEL-197 |
| chr19-4338230-DEL-175 | chr19-4338231-DEL-175 |
| chr19-4360188-DEL-208 | chr19-4360190-DEL-233 |
| chr19-4395886-DEL-194 | chr19-4395887-DEL-194 |
| chr19-4408338-DEL-198 | chr19-4408339-DEL-198 |
| chr19-4438397-DEL-235 | chr19-4438398-DEL-235 |
| chr19-4457888-DEL-210 | chr19-4457889-DEL-210 |
| chr19-4478228-DEL-191 | chr19-4478229-DEL-191 |
| chr19-4481674-DEL-148 | chr19-4481675-DEL-148 |
| chr19-4500199-DEL-224 | chr19-4500200-DEL-224 |
| chr19-4518299-DEL-72 | chr19-4518300-DEL-72 |
| chr19-4528518-DEL-254 | chr19-4528519-DEL-254 |
| chr19-4533825-DEL-54 | chr19-4533825-DEL-54 |
| chr19-4535635-DEL-99 | chr19-4535636-DEL-99 |
| chr19-4536164-DEL-325 | chr19-4536165-DEL-325 |

| | |
|---|---|
| chr19-4537078-DEL-58 | chr19-4537081-DEL-56 |
| chr19-4558336-DEL-98 | chr19-4558337-DEL-98 |
| chr19-4572713-DEL-217 | chr19-4572708-DEL-217 |
| chr19-4635202-DEL-131 | chr19-4635203-DEL-131 |
| chr19-4635807-DEL-349 | chr19-4635808-DEL-349 |
| chr19-4643269-DEL-313 | chr19-4643270-DEL-313 |
| chr19-4680359-DEL-159 | chr19-4680360-DEL-159 |
| chr19-4689455-DEL-147 | chr19-4689454-DEL-150 |
| chr19-4690640-DEL-165 | chr19-4690641-DEL-165 |
| chr19-4705139-DEL-113 | chr19-4705140-DEL-113 |
| chr19-4707593-DEL-73 | chr19-4707593-DEL-73 |
| chr19-4714053-DEL-6146 | chr19-4714053-DEL-6146 |
| chr19-4737230-DEL-170 | chr19-4737228-DEL-169 |
| chr19-4759291-DEL-63 | chr19-4759292-DEL-63 |
| chr19-4767067-DEL-84 | chr19-4767040-DEL-112 |
| chr19-4778709-DEL-147 | chr19-4778710-DEL-147 |
| chr19-4866180-DEL-94 | chr19-4866180-DEL-75 |
| chr19-4895080-DEL-290 | chr19-4895081-DEL-290 |
| chr19-4918280-DEL-92 | chr19-4918307-DEL-58 |
| chr19-4918940-DEL-193 | chr19-4918951-DEL-183 |
| chr19-4923832-DEL-216 | chr19-4923833-DEL-216 |
| chr19-4945608-DEL-182 | chr19-4945609-DEL-182 |
| chr19-4975003-DEL-227 | chr19-4975004-DEL-227 |
| chr19-4989328-DEL-126 | chr19-4989329-DEL-126 |
| chr19-4996672-DEL-593 | chr19-4996673-DEL-593 |
| chr19-4999040-DEL-70 | chr19-4999041-DEL-70 |
| chr19-5004142-DEL-169 | chr19-5004143-DEL-169 |
| chr19-5006929-DEL-480 | chr19-5006930-DEL-480 |
| chr19-5007707-DEL-543 | chr19-5007708-DEL-543 |
| chr19-5017188-DEL-197 | chr19-5017189-DEL-197 |
| chr19-5018549-DEL-459 | chr19-5018547-DEL-461 |
| chr19-5023765-DEL-217 | chr19-5023769-DEL-216 |
| chr19-5029220-DEL-646 | chr19-5029221-DEL-646 |
| chr19-5059957-DEL-374 | chr19-5059958-DEL-374 |
| chr19-5127732-DEL-171 | chr19-5127733-DEL-171 |
| chr19-5148676-DEL-151 | chr19-5148677-DEL-151 |
| chr19-5199081-DEL-155 | chr19-5199096-DEL-140 |
| chr19-5209085-DEL-68 | chr19-5209086-DEL-68 |
| chr19-5234626-DEL-1778 | chr19-5234627-DEL-1778 |
| chr19-5236587-DEL-59 | chr19-5236579-DEL-65 |
| chr19-5255045-DEL-201 | chr19-5255046-DEL-201 |
| chr19-5271421-DEL-139 | chr19-5271422-DEL-139 |
| chr19-5280835-DEL-57 | chr19-5280819-DEL-76 |
| chr19-5293824-DEL-179 | chr19-5293825-DEL-179 |
| chr19-5308885-DEL-397 | chr19-5308885-DEL-394 |
| chr19-5359205-DEL-87 | chr19-5359167-DEL-89 |
| chr19-5371170-DEL-58 | chr19-5371171-DEL-58 |
| chr19-5379705-DEL-225 | chr19-5379704-DEL-227 |
| chr19-5410619-DEL-202 | chr19-5410621-DEL-181 |
| chr19-5413200-DEL-365 | chr19-5413202-DEL-366 |
| chr19-5436181-DEL-222 | chr19-5436182-DEL-222 |
| chr19-5449498-DEL-195 | chr19-5449499-DEL-195 |
| chr19-5456384-DEL-466 | chr19-5456385-DEL-466 |
| chr19-5467847-DEL-707 | chr19-5467848-DEL-707 |
| chr19-5485954-DEL-911 | chr19-5485955-DEL-911 |
| chr19-5493323-DEL-210 | chr19-5493324-DEL-210 |
| chr19-5495500-DEL-70 | chr19-5495475-DEL-96 |
| chr19-5512636-DEL-74 | chr19-5512637-DEL-74 |
| chr19-5545017-DEL-226 | chr19-5545018-DEL-226 |
| chr19-5557101-DEL-1843 | chr19-5557102-DEL-1843 |
| chr19-5557582-DEL-212 | chr19-5557589-DEL-166 |
| chr19-5564509-DEL-215 | chr19-5564510-DEL-194 |
| chr19-5624175-DEL-199 | chr19-5624176-DEL-199 |
| chr19-5640362-DEL-339 | chr19-5640363-DEL-339 |
| chr19-5643914-DEL-378 | chr19-5643915-DEL-378 |
| chr19-5647552-DEL-144 | chr19-5647553-DEL-144 |
| chr19-5672567-DEL-223 | chr19-5672568-DEL-223 |
| chr19-5700111-DEL-128 | chr19-5700074-DEL-174 |
| chr19-5720135-DEL-59 | chr19-5720136-DEL-59 |
| chr19-5724410-DEL-144 | chr19-5724380-DEL-144 |
| chr19-5762018-DEL-59 | chr19-5762019-DEL-59 |
| chr19-5783246-DEL-160 | chr19-5783246-DEL-160 |
| chr19-5784524-DEL-521 | chr19-5784525-DEL-521 |
| chr19-5790188-DEL-122 | chr19-5790189-DEL-122 |
| chr19-5827178-DEL-410 | chr19-5827179-DEL-410 |
| chr19-5831218-DEL-202 | chr19-5831219-DEL-202 |
| chr19-5842252-DEL-217 | chr19-5842269-DEL-216 |
| chr19-5859278-DEL-147 | chr19-5859275-DEL-163 |
| chr19-5871181-DEL-213 | chr19-5871182-DEL-213 |
| chr19-5872095-DEL-178 | chr19-5872068-DEL-206 |

| | |
|---|---|
| chr19-5890861-DEL-58 | chr19-5890826-DEL-57 |
| chr19-5904090-DEL-207 | chr19-5904090-DEL-208 |
| chr19-5908113-DEL-84 | chr19-5908126-DEL-72 |
| chr19-5909618-DEL-163 | chr19-5909619-DEL-163 |
| chr19-5911739-DEL-533 | chr19-5911689-DEL-582 |
| chr19-5938692-DEL-81 | chr19-5938646-DEL-81 |
| chr19-5949121-DEL-210 | chr19-5949123-DEL-207 |
| chr19-5959320-DEL-197 | chr19-5959321-DEL-197 |
| chr19-5971350-DEL-76 | chr19-5971350-DEL-76 |
| chr19-5981196-DEL-197 | chr19-5981196-DEL-197 |
| chr19-5986178-DEL-74 | chr19-5986179-DEL-74 |
| chr19-5986711-DEL-222 | chr19-5986712-DEL-222 |
| chr19-5991092-DEL-182 | chr19-5991093-DEL-182 |
| chr19-6019039-DEL-134 | chr19-6019040-DEL-134 |
| chr19-6024217-DEL-421 | chr19-6024218-DEL-421 |
| chr19-6032194-DEL-76 | chr19-6032195-DEL-76 |
| chr19-6055286-DEL-54 | chr19-6055287-DEL-54 |
| chr19-6056115-DEL-195 | chr19-6056116-DEL-195 |
| chr19-6057840-DEL-169 | chr19-6057831-DEL-170 |
| chr19-6072721-DEL-185 | chr19-6072722-DEL-185 |
| chr19-6076444-DEL-52 | chr19-6076445-DEL-52 |
| chr19-6113092-DEL-56 | chr19-6113093-DEL-56 |
| chr19-6138930-DEL-89 | chr19-6138931-DEL-89 |
| chr19-6161165-DEL-69 | chr19-6161166-DEL-69 |
| chr19-6175561-DEL-163 | chr19-6175562-DEL-163 |
| chr19-6186507-DEL-110 | chr19-6186508-DEL-110 |
| chr19-6188346-DEL-200 | chr19-6188347-DEL-199 |
| chr19-6198231-DEL-207 | chr19-6198232-DEL-207 |
| chr19-6199818-DEL-506 | chr19-6199819-DEL-506 |
| chr19-6210442-DEL-191 | chr19-6210422-DEL-181 |
| chr19-6240227-DEL-395 | chr19-6240231-DEL-419 |
| chr19-6243573-DEL-180 | chr19-6243574-DEL-180 |
| chr19-6298464-DEL-113 | chr19-6298464-DEL-111 |
| chr19-6321170-DEL-63 | chr19-6321171-DEL-63 |
| chr19-6337487-DEL-82 | chr19-6337514-DEL-82 |
| chr19-6340677-DEL-210 | chr19-6340678-DEL-210 |
| chr19-6344698-DEL-73 | chr19-6344699-DEL-73 |
| chr19-6406631-DEL-147 | chr19-6406632-DEL-147 |
| chr19-6407273-DEL-112 | chr19-6407274-DEL-112 |
| chr19-6430213-DEL-208 | chr19-6430214-DEL-208 |
| chr19-6441070-DEL-104 | chr19-6441071-DEL-104 |
| chr19-6441845-DEL-212 | chr19-6441842-DEL-211 |
| chr19-6445379-DEL-216 | chr19-6445380-DEL-216 |
| chr19-6484541-DEL-208 | chr19-6484542-DEL-208 |
| chr19-6501506-DEL-185 | chr19-6501507-DEL-185 |
| chr19-6528952-DEL-54 | chr19-6528939-DEL-64 |
| chr19-6612136-DEL-62 | chr19-6612136-DEL-62 |
| chr19-6615974-DEL-153 | chr19-6616019-DEL-152 |
| chr19-6616905-DEL-144 | chr19-6616892-DEL-145 |
| chr19-6640984-DEL-3171 | chr19-6640985-DEL-3171 |
| chr19-6653051-DEL-218 | chr19-6653052-DEL-218 |
| chr19-6676016-DEL-436 | chr19-6676017-DEL-436 |
| chr19-6679407-DEL-117 | chr19-6679392-DEL-118 |
| chr19-6693305-DEL-66 | chr19-6693319-DEL-67 |
| chr19-6708101-DEL-346 | chr19-6708092-DEL-351 |
| chr19-6711354-DEL-456 | chr19-6711355-DEL-456 |
| chr19-6717657-DEL-315 | chr19-6717704-DEL-315 |
| chr19-6729827-DEL-227 | chr19-6729828-DEL-227 |
| chr19-6811538-DEL-197 | chr19-6811537-DEL-198 |
| chr19-6843232-DEL-1073 | chr19-6843233-DEL-1073 |
| chr19-6880789-DEL-504 | chr19-6880790-DEL-504 |
| chr19-6924935-DEL-199 | chr19-6924936-DEL-199 |
| chr19-6978096-DEL-205 | chr19-6978097-DEL-205 |
| chr19-6984849-DEL-201 | chr19-6984850-DEL-201 |
| chr19-6986583-DEL-210 | chr19-6986584-DEL-210 |
| chr19-7004248-DEL-194 | chr19-7004249-DEL-194 |
| chr19-7011781-DEL-184 | chr19-7011782-DEL-184 |
| chr19-7019799-DEL-795 | chr19-7019800-DEL-795 |
| chr19-7128000-DEL-427 | chr19-7128001-DEL-427 |
| chr19-7141887-DEL-363 | chr19-7141888-DEL-363 |
| chr19-7185675-DEL-215 | chr19-7185676-DEL-215 |
| chr19-7196626-DEL-214 | chr19-7196627-DEL-215 |
| chr19-7210951-DEL-118 | chr19-7210973-DEL-99 |
| chr19-7230068-DEL-223 | chr19-7230069-DEL-223 |
| chr19-7328696-DEL-191 | chr19-7328697-DEL-191 |
| chr19-7331389-DEL-219 | chr19-7331389-DEL-219 |
| chr19-7370163-DEL-639 | chr19-7370164-DEL-639 |
| chr19-7383549-DEL-52 | chr19-7383558-DEL-72 |
| chr19-7409085-DEL-176 | chr19-7409086-DEL-176 |
| chr19-7431370-DEL-195 | chr19-7431372-DEL-193 |

| | |
|---|---|
| chr19-7458347-DEL-203 | chr19-7458348-DEL-203 |
| chr19-7473685-DEL-212 | chr19-7473686-DEL-212 |
| chr19-7481092-DEL-57 | chr19-7481091-DEL-59 |
| chr19-7496479-DEL-1019 | chr19-7496480-DEL-1019 |
| chr19-7554033-DEL-51 | chr19-7554035-DEL-52 |
| chr19-7560733-DEL-59 | chr19-7560734-DEL-59 |
| chr19-7566552-DEL-203 | chr19-7566553-DEL-203 |
| chr19-7570087-DEL-585 | chr19-7570087-DEL-585 |
| chr19-7570749-DEL-2056 | chr19-7570749-DEL-2056 |
| chr19-7590326-DEL-106 | chr19-7590327-DEL-106 |
| chr19-7602426-DEL-83 | chr19-7602464-DEL-84 |
| chr19-7635657-DEL-483 | chr19-7635669-DEL-473 |
| chr19-7645609-DEL-905 | chr19-7645610-DEL-904 |
| chr19-7678987-DEL-582 | chr19-7678988-DEL-582 |
| chr19-7690877-DEL-447 | chr19-7690878-DEL-447 |
| chr19-7693663-DEL-3748 | chr19-7693663-DEL-3748 |
| chr19-7703147-DEL-1281 | chr19-7703146-DEL-1284 |
| chr19-7705319-DEL-1114 | chr19-7705319-DEL-1114 |
| chr19-7745377-DEL-6799 | chr19-7745378-DEL-6799 |
| chr19-7796428-DEL-5828 | chr19-7796427-DEL-5829 |
| chr19-7854872-DEL-644 | chr19-7854873-DEL-644 |
| chr19-7872126-DEL-182 | chr19-7872127-DEL-182 |
| chr19-7875838-DEL-52 | chr19-7875839-DEL-50 |
| chr19-7890543-DEL-6980 | chr19-7890544-DEL-6980 |
| chr19-7966779-DEL-64 | chr19-7966780-DEL-64 |
| chr19-7969536-DEL-175 | chr19-7969536-DEL-176 |
| chr19-8000273-DEL-125 | chr19-8000274-DEL-125 |
| chr19-8001856-DEL-6212 | chr19-8001857-DEL-6212 |
| chr19-8015730-DEL-211 | chr19-8015731-DEL-211 |
| chr19-8051436-DEL-430 | chr19-8051437-DEL-430 |
| chr19-8065956-DEL-53 | chr19-8065957-DEL-53 |
| chr19-8066234-DEL-611 | chr19-8066235-DEL-611 |
| chr19-8192231-DEL-59 | chr19-8192230-DEL-60 |
| chr19-8212174-DEL-1888 | chr19-8212175-DEL-1888 |
| chr19-8238770-DEL-128 | chr19-8238768-DEL-128 |
| chr19-8289463-DEL-200 | chr19-8289464-DEL-200 |
| chr19-8323663-DEL-387 | chr19-8323664-DEL-387 |
| chr19-8427463-DEL-1851 | chr19-8427464-DEL-1851 |
| chr19-8462757-DEL-2871 | chr19-8462758-DEL-2871 |
| chr19-8488465-DEL-660 | chr19-8488466-DEL-660 |
| chr19-8505609-DEL-6448 | chr19-8505610-DEL-6448 |
| chr19-8569464-DEL-122 | chr19-8569513-DEL-74 |
| chr19-8611426-DEL-3354 | chr19-8611433-DEL-3307 |
| chr19-8619938-DEL-666 | chr19-8619940-DEL-669 |
| chr19-8629889-DEL-109 | chr19-8629854-DEL-125 |
| chr19-8647349-DEL-62 | chr19-8647353-DEL-50 |
| chr19-8678401-DEL-207 | chr19-8678402-DEL-207 |
| chr19-8682184-DEL-62 | chr19-8682185-DEL-57 |
| chr19-8683504-DEL-53 | chr19-8683503-DEL-51 |
| chr19-8684121-DEL-57 | chr19-8684122-DEL-57 |
| chr19-8684627-DEL-187 | chr19-8684627-DEL-187 |
| chr19-8694037-DEL-227 | chr19-8694038-DEL-227 |
| chr19-8703879-DEL-138 | chr19-8703880-DEL-138 |
| chr19-8707882-DEL-190 | chr19-8707883-DEL-190 |
| chr19-8708884-DEL-70 | chr19-8708885-DEL-70 |
| chr19-8709656-DEL-184 | chr19-8709657-DEL-184 |
| chr19-8713896-DEL-209 | chr19-8713897-DEL-209 |
| chr19-8756793-DEL-206 | chr19-8756794-DEL-206 |
| chr19-8773030-DEL-206 | chr19-8773031-DEL-206 |
| chr19-8778874-DEL-86 | chr19-8778874-DEL-86 |
| chr19-8825527-DEL-114 | chr19-8825528-DEL-114 |
| chr19-8842163-DEL-218 | chr19-8842164-DEL-189 |
| chr19-8853758-DEL-371 | chr19-8853775-DEL-362 |
| chr19-8854586-DEL-173 | chr19-8854586-DEL-173 |
| chr19-8926932-DEL-144 | chr19-8926933-DEL-144 |
| chr19-8934567-DEL-202 | chr19-8934528-DEL-203 |
| chr19-8941818-DEL-54 | chr19-8941820-DEL-55 |
| chr19-8999241-DEL-210 | chr19-8999242-DEL-210 |
| chr19-9013037-DEL-96 | chr19-9013038-DEL-96 |
| chr19-9023369-DEL-182 | chr19-9023370-DEL-182 |
| chr19-9041866-DEL-103 | chr19-9041867-DEL-103 |
| chr19-9045168-DEL-263 | chr19-9045169-DEL-263 |
| chr19-9055828-DEL-168 | chr19-9055806-DEL-165 |
| chr19-9069963-DEL-89 | chr19-9069965-DEL-74 |
| chr19-9139278-DEL-184 | chr19-9139279-DEL-184 |
| chr19-9193692-DEL-696 | chr19-9193692-DEL-696 |
| chr19-9398521-DEL-439 | chr19-9398522-DEL-439 |
| chr19-9564382-DEL-1291 | chr19-9564383-DEL-1291 |
| chr19-9575670-DEL-69 | chr19-9575674-DEL-71 |
| chr19-9586804-DEL-614 | chr19-9586802-DEL-617 |

| | |
|---|---|
| chr19-9617265-DEL-483 | chr19-9617267-DEL-484 |
| chr19-9731554-DEL-212 | chr19-9731555-DEL-212 |
| chr19-9755676-DEL-24506 | chr19-9755672-DEL-24509 |
| chr19-9802637-DEL-319 | chr19-9802638-DEL-319 |
| chr19-9838462-DEL-327 | chr19-9838463-DEL-327 |
| chr19-9883486-DEL-1623 | chr19-9883487-DEL-1618 |
| chr19-9885321-DEL-439 | chr19-9885322-DEL-439 |
| chr19-9895490-DEL-54 | chr19-9895490-DEL-55 |
| chr19-9926309-DEL-119 | chr19-9926310-DEL-117 |
| chr19-9930810-DEL-320 | chr19-9930806-DEL-321 |
| chr19-9945346-DEL-207 | chr19-9945339-DEL-222 |
| chr19-9972120-DEL-1261 | chr19-9972155-DEL-1223 |
| chr19-9984340-DEL-186 | chr19-9984350-DEL-177 |
| chr19-9997768-DEL-203 | chr19-9997763-DEL-204 |
| chr19-10028154-DEL-52 | chr19-10028125-DEL-80 |
| chr19-10051614-DEL-144 | chr19-10051615-DEL-144 |
| chr19-10058039-DEL-64 | chr19-10058040-DEL-64 |
| chr19-10083506-DEL-209 | chr19-10083507-DEL-209 |
| chr19-10087663-DEL-270 | chr19-10087663-DEL-269 |
| chr19-10119963-DEL-232 | chr19-10119964-DEL-232 |
| chr19-10122263-DEL-72 | chr19-10122264-DEL-72 |
| chr19-10134503-DEL-58 | chr19-10134504-DEL-58 |
| chr19-10135646-DEL-209 | chr19-10135647-DEL-209 |
| chr19-10142364-DEL-214 | chr19-10142365-DEL-214 |
| chr19-10143554-DEL-198 | chr19-10143555-DEL-198 |
| chr19-10156365-DEL-324 | chr19-10156366-DEL-324 |
| chr19-10157604-DEL-204 | chr19-10157605-DEL-204 |
| chr19-10179795-DEL-216 | chr19-10179796-DEL-216 |
| chr19-10191837-DEL-234 | chr19-10191838-DEL-234 |
| chr19-10243299-DEL-203 | chr19-10243314-DEL-207 |
| chr19-10305279-DEL-79 | chr19-10305280-DEL-79 |
| chr19-10323486-DEL-52 | chr19-10323487-DEL-52 |
| chr19-10324655-DEL-58 | chr19-10324656-DEL-62 |
| chr19-10325833-DEL-66 | chr19-10325834-DEL-66 |
| chr19-10335281-DEL-67 | chr19-10335282-DEL-67 |
| chr19-10338624-DEL-378 | chr19-10338625-DEL-379 |
| chr19-10339202-DEL-246 | chr19-10339203-DEL-246 |
| chr19-10358320-DEL-88 | chr19-10358317-DEL-100 |
| chr19-10448448-DEL-708 | chr19-10448440-DEL-744 |
| chr19-10448456-DEL-60 | chr19-10448459-DEL-58 |
| chr19-10449091-DEL-66 | chr19-10449091-DEL-66 |
| chr19-10463092-DEL-56 | chr19-10463093-DEL-56 |
| chr19-10463177-DEL-735 | chr19-10463179-DEL-733 |
| chr19-10467865-DEL-98 | chr19-10467908-DEL-55 |
| chr19-10501300-DEL-430 | chr19-10501301-DEL-430 |
| chr19-10508580-DEL-202 | chr19-10508581-DEL-202 |
| chr19-10517201-DEL-74 | chr19-10517202-DEL-74 |
| chr19-10535412-DEL-173 | chr19-10535413-DEL-173 |
| chr19-10536123-DEL-881 | chr19-10536124-DEL-881 |
| chr19-10544341-DEL-363 | chr19-10544342-DEL-363 |
| chr19-10545519-DEL-261 | chr19-10545520-DEL-261 |
| chr19-10572471-DEL-52 | chr19-10572472-DEL-52 |
| chr19-10616336-DEL-222 | chr19-10616337-DEL-222 |
| chr19-10625084-DEL-74 | chr19-10625085-DEL-74 |
| chr19-10629465-DEL-257 | chr19-10629463-DEL-260 |
| chr19-10638142-DEL-240 | chr19-10638143-DEL-240 |
| chr19-10648325-DEL-218 | chr19-10648326-DEL-218 |
| chr19-10650236-DEL-90 | chr19-10650237-DEL-84 |
| chr19-10692310-DEL-256 | chr19-10692311-DEL-256 |
| chr19-10717691-DEL-100 | chr19-10717692-DEL-108 |
| chr19-10722890-DEL-58 | chr19-10722889-DEL-60 |
| chr19-10726049-DEL-53 | chr19-10726050-DEL-53 |
| chr19-10727299-DEL-72 | chr19-10727326-DEL-62 |
| chr19-10729409-DEL-174 | chr19-10729405-DEL-172 |
| chr19-10779612-DEL-204 | chr19-10779613-DEL-204 |
| chr19-10784999-DEL-409 | chr19-10785000-DEL-409 |
| chr19-10785581-DEL-75 | chr19-10785582-DEL-75 |
| chr19-10792663-DEL-2284 | chr19-10792664-DEL-2284 |
| chr19-10817433-DEL-68 | chr19-10817430-DEL-72 |
| chr19-10821458-DEL-206 | chr19-10821459-DEL-206 |
| chr19-10827006-DEL-106 | chr19-10827013-DEL-100 |
| chr19-10838169-DEL-469 | chr19-10838170-DEL-469 |
| chr19-10843394-DEL-75 | chr19-10843400-DEL-91 |
| chr19-10865696-DEL-68 | chr19-10865691-DEL-72 |
| chr19-10875394-DEL-62 | chr19-10875360-DEL-62 |
| chr19-10898802-DEL-138 | chr19-10898804-DEL-124 |
| chr19-10903386-DEL-98 | chr19-10903387-DEL-98 |
| chr19-10905513-DEL-58 | chr19-10905518-DEL-70 |
| chr19-10931862-DEL-1904 | chr19-10931863-DEL-1904 |
| chr19-10938486-DEL-597 | chr19-10938487-DEL-597 |

| | |
|---|---|
| chr19-10955046-DEL-132 | chr19-10955047-DEL-132 |
| chr19-10997780-DEL-120 | chr19-10997787-DEL-70 |
| chr19-11050536-DEL-184 | chr19-11050537-DEL-136 |
| chr19-11059046-DEL-190 | chr19-11059043-DEL-191 |
| chr19-11064661-DEL-2165 | chr19-11064662-DEL-2165 |
| chr19-11098365-DEL-3384 | chr19-11098379-DEL-3390 |
| chr19-11108377-DEL-135 | chr19-11108387-DEL-97 |
| chr19-11110245-DEL-205 | chr19-11110246-DEL-205 |
| chr19-11123123-DEL-669 | chr19-11123124-DEL-669 |
| chr19-11127958-DEL-1609 | chr19-11127958-DEL-1609 |
| chr19-11135434-DEL-275 | chr19-11135435-DEL-275 |
| chr19-11137015-DEL-220 | chr19-11137016-DEL-220 |
| chr19-11176257-DEL-58 | chr19-11176258-DEL-58 |
| chr19-11194396-DEL-6924 | chr19-11194397-DEL-6924 |
| chr19-11206353-DEL-118 | chr19-11206356-DEL-103 |
| chr19-11213864-DEL-3404 | chr19-11213852-DEL-3405 |
| chr19-11269688-DEL-7302 | chr19-11269688-DEL-7302 |
| chr19-11295188-DEL-64 | chr19-11295183-DEL-61 |
| chr19-11321263-DEL-60 | chr19-11321264-DEL-60 |
| chr19-11334831-DEL-643 | chr19-11334832-DEL-643 |
| chr19-11387192-DEL-236 | chr19-11387180-DEL-239 |
| chr19-11399517-DEL-3213 | chr19-11399518-DEL-3213 |
| chr19-11405143-DEL-3283 | chr19-11405144-DEL-3283 |
| chr19-11450242-DEL-558 | chr19-11450243-DEL-558 |
| chr19-11453628-DEL-616 | chr19-11453629-DEL-616 |
| chr19-11466139-DEL-55 | chr19-11466128-DEL-50 |
| chr19-11467531-DEL-56 | chr19-11467526-DEL-70 |
| chr19-11497383-DEL-58 | chr19-11497384-DEL-58 |
| chr19-11554148-DEL-889 | chr19-11554149-DEL-889 |
| chr19-11570392-DEL-6807 | chr19-11570392-DEL-6807 |
| chr19-11586543-DEL-92 | chr19-11586544-DEL-92 |
| chr19-11591808-DEL-89 | chr19-11591809-DEL-89 |
| chr19-11598790-DEL-72 | chr19-11598788-DEL-72 |
| chr19-11602945-DEL-584 | chr19-11602944-DEL-582 |
| chr19-11627208-DEL-138 | chr19-11627209-DEL-138 |
| chr19-11671639-DEL-534 | chr19-11671648-DEL-520 |
| chr19-11711409-DEL-170 | chr19-11711406-DEL-171 |
| chr19-11740265-DEL-86 | chr19-11740266-DEL-86 |
| chr19-11744768-DEL-52 | chr19-11744769-DEL-52 |
| chr19-11758626-DEL-94 | chr19-11758627-DEL-94 |
| chr19-11763226-DEL-99 | chr19-11763227-DEL-99 |
| chr19-11768630-DEL-161 | chr19-11768637-DEL-155 |
| chr19-11777533-DEL-378 | chr19-11777530-DEL-356 |
| chr19-11793006-DEL-91 | chr19-11793007-DEL-91 |
| chr19-11820444-DEL-436 | chr19-11820445-DEL-436 |
| chr19-11821420-DEL-66 | chr19-11821420-DEL-80 |
| chr19-11826020-DEL-260 | chr19-11826021-DEL-260 |
| chr19-11871304-DEL-72 | chr19-11871305-DEL-72 |
| chr19-11882391-DEL-2031 | chr19-11882392-DEL-2031 |
| chr19-11883409-DEL-443 | chr19-11883410-DEL-443 |
| chr19-11896226-DEL-72 | chr19-11896227-DEL-68 |
| chr19-11908365-DEL-138 | chr19-11908326-DEL-138 |
| chr19-11915893-DEL-216 | chr19-11915894-DEL-216 |
| chr19-11919735-DEL-220 | chr19-11919736-DEL-220 |
| chr19-11926270-DEL-149 | chr19-11926244-DEL-149 |
| chr19-11939129-DEL-62 | chr19-11939130-DEL-62 |
| chr19-11962205-DEL-254 | chr19-11962206-DEL-254 |
| chr19-11994549-DEL-331 | chr19-11994549-DEL-331 |
| chr19-11997234-DEL-108 | chr19-11997237-DEL-100 |
| chr19-12046224-DEL-497 | chr19-12046225-DEL-497 |
| chr19-12059692-DEL-261 | chr19-12059693-DEL-261 |
| chr19-12060286-DEL-51 | chr19-12060301-DEL-58 |
| chr19-12061343-DEL-336 | chr19-12061344-DEL-335 |
| chr19-12062248-DEL-604 | chr19-12062249-DEL-604 |
| chr19-12063585-DEL-484 | chr19-12063586-DEL-484 |
| chr19-12072337-DEL-584 | chr19-12072330-DEL-592 |
| chr19-12087439-DEL-440 | chr19-12087440-DEL-440 |
| chr19-12089381-DEL-76 | chr19-12089379-DEL-76 |
| chr19-12110725-DEL-126 | chr19-12110730-DEL-124 |
| chr19-12117845-DEL-52 | chr19-12117817-DEL-70 |
| chr19-12121705-DEL-743 | chr19-12121706-DEL-743 |
| chr19-12123857-DEL-6641 | chr19-12123858-DEL-6641 |
| chr19-12129891-DEL-65 | chr19-12129892-DEL-64 |
| chr19-12135454-DEL-3388 | chr19-12135455-DEL-3388 |
| chr19-12167187-DEL-125 | chr19-12167188-DEL-125 |
| chr19-12181770-DEL-90 | chr19-12181771-DEL-90 |
| chr19-12189970-DEL-199 | chr19-12189943-DEL-227 |
| chr19-12224855-DEL-797 | chr19-12224853-DEL-794 |
| chr19-12250458-DEL-420 | chr19-12250459-DEL-420 |
| chr19-12261512-DEL-102 | chr19-12261512-DEL-102 |

| | |
|---|---|
| chr19-12274552-DEL-222 | chr19-12274553-DEL-222 |
| chr19-12292730-DEL-78 | chr19-12292731-DEL-84 |
| chr19-12307185-DEL-143 | chr19-12307184-DEL-177 |
| chr19-12338346-DEL-11214 | chr19-12338347-DEL-11214 |
| chr19-12339345-DEL-4883 | chr19-12339346-DEL-4883 |
| chr19-12378990-DEL-1367 | chr19-12378991-DEL-1367 |
| chr19-12382269-DEL-133 | chr19-12382267-DEL-134 |
| chr19-12424781-DEL-7156 | chr19-12424782-DEL-7156 |
| chr19-12442400-DEL-78 | chr19-12442401-DEL-78 |
| chr19-12480444-DEL-6243 | chr19-12480445-DEL-6243 |
| chr19-12493433-DEL-914 | chr19-12493436-DEL-912 |
| chr19-12509195-DEL-56 | chr19-12509188-DEL-56 |
| chr19-12533876-DEL-970 | chr19-12533877-DEL-970 |
| chr19-12556405-DEL-165 | chr19-12556405-DEL-165 |
| chr19-12556882-DEL-3425 | chr19-12556882-DEL-3425 |
| chr19-12570636-DEL-1844 | chr19-12570637-DEL-1844 |
| chr19-12640945-DEL-59 | chr19-12640946-DEL-59 |
| chr19-12641958-DEL-381 | chr19-12641959-DEL-381 |
| chr19-12643899-DEL-66 | chr19-12643900-DEL-66 |
| chr19-12660754-DEL-457 | chr19-12660755-DEL-457 |
| chr19-12668115-DEL-1570 | chr19-12668116-DEL-1570 |
| chr19-12712331-DEL-1671 | chr19-12712331-DEL-1669 |
| chr19-12717904-DEL-54 | chr19-12717887-DEL-54 |
| chr19-12721740-DEL-63 | chr19-12721742-DEL-62 |
| chr19-12723192-DEL-60 | chr19-12723194-DEL-58 |
| chr19-12727264-DEL-249 | chr19-12727265-DEL-249 |
| chr19-12736621-DEL-83 | chr19-12736589-DEL-83 |
| chr19-12784096-DEL-73 | chr19-12784097-DEL-89 |
| chr19-12792286-DEL-206 | chr19-12792287-DEL-206 |
| chr19-12799100-DEL-1312 | chr19-12799101-DEL-1312 |
| chr19-12881457-DEL-2906 | chr19-12881458-DEL-2906 |
| chr19-12885035-DEL-616 | chr19-12885036-DEL-616 |
| chr19-12889596-DEL-152 | chr19-12889597-DEL-152 |
| chr19-12894959-DEL-1582 | chr19-12894960-DEL-1582 |
| chr19-12903614-DEL-538 | chr19-12903633-DEL-551 |
| chr19-12905215-DEL-1271 | chr19-12905214-DEL-1272 |
| chr19-12916159-DEL-846 | chr19-12916159-DEL-846 |
| chr19-12925171-DEL-1220 | chr19-12925171-DEL-1220 |
| chr19-12967737-DEL-120 | chr19-12967738-DEL-120 |
| chr19-12978372-DEL-110 | chr19-12978376-DEL-109 |
| chr19-12999805-DEL-4380 | chr19-12999806-DEL-4379 |
| chr19-13006039-DEL-66 | chr19-13006046-DEL-64 |
| chr19-13014974-DEL-18620 | chr19-13014968-DEL-18614 |
| chr19-13053395-DEL-2492 | chr19-13053396-DEL-2492 |
| chr19-13071040-DEL-219 | chr19-13071041-DEL-219 |
| chr19-13097272-DEL-676 | chr19-13097271-DEL-676 |
| chr19-13101771-DEL-822 | chr19-13101771-DEL-822 |
| chr19-13130437-DEL-197 | chr19-13130438-DEL-196 |
| chr19-13133418-DEL-52 | chr19-13133420-DEL-74 |
| chr19-13137158-DEL-574 | chr19-13137159-DEL-574 |
| chr19-13172114-DEL-529 | chr19-13172115-DEL-530 |
| chr19-13184203-DEL-1180 | chr19-13184193-DEL-1194 |
| chr19-13202760-DEL-98 | chr19-13202766-DEL-92 |
| chr19-13217731-DEL-400 | chr19-13217732-DEL-400 |
| chr19-13234476-DEL-64 | chr19-13234477-DEL-64 |
| chr19-13238589-DEL-69 | chr19-13238556-DEL-58 |
| chr19-13243411-DEL-70 | chr19-13243410-DEL-72 |
| chr19-13243897-DEL-3671 | chr19-13243898-DEL-3671 |
| chr19-13357651-DEL-434 | chr19-13357652-DEL-434 |
| chr19-13365788-DEL-155 | chr19-13365789-DEL-155 |
| chr19-13410553-DEL-300 | chr19-13410557-DEL-303 |
| chr19-13433137-DEL-166 | chr19-13433138-DEL-166 |
| chr19-13457997-DEL-1888 | chr19-13457998-DEL-1888 |
| chr19-13515007-DEL-2755 | chr19-13515008-DEL-2755 |
| chr19-13526513-DEL-69 | chr19-13526515-DEL-68 |
| chr19-13528774-DEL-129 | chr19-13528775-DEL-129 |
| chr19-13579860-DEL-104 | chr19-13579846-DEL-87 |
| chr19-13587031-DEL-2554 | chr19-13587030-DEL-2555 |
| chr19-13642990-DEL-208 | chr19-13642991-DEL-208 |
| chr19-13747857-DEL-6677 | chr19-13747858-DEL-6677 |
| chr19-13790254-DEL-54 | chr19-13790255-DEL-54 |
| chr19-13844042-DEL-347 | chr19-13844042-DEL-347 |
| chr19-13849192-DEL-433 | chr19-13849192-DEL-433 |
| chr19-13876819-DEL-6853 | chr19-13876820-DEL-6853 |
| chr19-13884749-DEL-327 | chr19-13884750-DEL-327 |
| chr19-13887142-DEL-55 | chr19-13887143-DEL-55 |
| chr19-13900967-DEL-6764 | chr19-13900968-DEL-6764 |
| chr19-13908022-DEL-2757 | chr19-13908023-DEL-2757 |
| chr19-13908162-DEL-56 | chr19-13908163-DEL-56 |
| chr19-13916576-DEL-53 | chr19-13916574-DEL-54 |

| | |
|---|---|
| chr19-13919213-DEL-336 | chr19-13919208-DEL-337 |
| chr19-13924693-DEL-2749 | chr19-13924694-DEL-2749 |
| chr19-13949425-DEL-1558 | chr19-13949413-DEL-1558 |
| chr19-13975874-DEL-12931 | chr19-13975875-DEL-12931 |
| chr19-14001890-DEL-498 | chr19-14001891-DEL-498 |
| chr19-14011703-DEL-141 | chr19-14011704-DEL-141 |
| chr19-14024399-DEL-1680 | chr19-14024393-DEL-1681 |
| chr19-14048386-DEL-285 | chr19-14048387-DEL-284 |
| chr19-14054156-DEL-169 | chr19-14054157-DEL-169 |
| chr19-14061747-DEL-202 | chr19-14061748-DEL-202 |
| chr19-14076733-DEL-67 | chr19-14076715-DEL-70 |
| chr19-14077323-DEL-401 | chr19-14077327-DEL-403 |
| chr19-14077834-DEL-80 | chr19-14077835-DEL-80 |
| chr19-14079804-DEL-58 | chr19-14079805-DEL-56 |
| chr19-14084405-DEL-57 | chr19-14084406-DEL-57 |
| chr19-14088117-DEL-6138 | chr19-14088118-DEL-6138 |
| chr19-14124217-DEL-2573 | chr19-14124218-DEL-2573 |
| chr19-14126217-DEL-348 | chr19-14126218-DEL-348 |
| chr19-14139110-DEL-185 | chr19-14139111-DEL-193 |
| chr19-14149603-DEL-935 | chr19-14149614-DEL-936 |
| chr19-14180757-DEL-70 | chr19-14180758-DEL-70 |
| chr19-14183610-DEL-64 | chr19-14183621-DEL-54 |
| chr19-14188573-DEL-468 | chr19-14188574-DEL-468 |
| chr19-14205965-DEL-5063 | chr19-14205966-DEL-5064 |
| chr19-14212792-DEL-334 | chr19-14212793-DEL-334 |
| chr19-14217323-DEL-76 | chr19-14217310-DEL-81 |
| chr19-14228257-DEL-89 | chr19-14228258-DEL-89 |
| chr19-14234137-DEL-491 | chr19-14234138-DEL-491 |
| chr19-14238872-DEL-6468 | chr19-14238873-DEL-6468 |
| chr19-14247912-DEL-136 | chr19-14247913-DEL-136 |
| chr19-14288341-DEL-134 | chr19-14288338-DEL-126 |
| chr19-14291209-DEL-225 | chr19-14291173-DEL-262 |
| chr19-14305470-DEL-76 | chr19-14305464-DEL-90 |
| chr19-14341712-DEL-84 | chr19-14341713-DEL-84 |
| chr19-14342958-DEL-60 | chr19-14342959-DEL-65 |
| chr19-14349759-DEL-64 | chr19-14349772-DEL-52 |
| chr19-14370720-DEL-84 | chr19-14370721-DEL-66 |
| chr19-14405525-DEL-81 | chr19-14405516-DEL-109 |
| chr19-14467625-DEL-188 | chr19-14467626-DEL-188 |
| chr19-14546806-DEL-209 | chr19-14546807-DEL-209 |
| chr19-14606487-DEL-189 | chr19-14606485-DEL-190 |
| chr19-14618508-DEL-139 | chr19-14618509-DEL-139 |
| chr19-14627047-DEL-76 | chr19-14627048-DEL-76 |
| chr19-14631011-DEL-653 | chr19-14631012-DEL-653 |
| chr19-14674439-DEL-401 | chr19-14674431-DEL-403 |
| chr19-14679342-DEL-197 | chr19-14679343-DEL-197 |
| chr19-14716777-DEL-875 | chr19-14716773-DEL-876 |
| chr19-14725236-DEL-52 | chr19-14725208-DEL-80 |
| chr19-14743862-DEL-360 | chr19-14743863-DEL-360 |
| chr19-14768645-DEL-1364 | chr19-14768646-DEL-1364 |
| chr19-14859669-DEL-79 | chr19-14859699-DEL-60 |
| chr19-14875901-DEL-1372 | chr19-14875902-DEL-1372 |
| chr19-14906803-DEL-193 | chr19-14906804-DEL-193 |
| chr19-14909741-DEL-60 | chr19-14909703-DEL-56 |
| chr19-14914606-DEL-646 | chr19-14914607-DEL-646 |
| chr19-14960264-DEL-203 | chr19-14960235-DEL-204 |
| chr19-14981564-DEL-76 | chr19-14981564-DEL-76 |
| chr19-14990182-DEL-67 | chr19-14990183-DEL-67 |
| chr19-14993345-DEL-401 | chr19-14993346-DEL-401 |
| chr19-15023951-DEL-53 | chr19-15023951-DEL-51 |
| chr19-15053298-DEL-329 | chr19-15053299-DEL-329 |
| chr19-15067309-DEL-724 | chr19-15067310-DEL-724 |
| chr19-15151485-DEL-144 | chr19-15151486-DEL-144 |
| chr19-15170865-DEL-732 | chr19-15170866-DEL-732 |
| chr19-15197258-DEL-202 | chr19-15197235-DEL-226 |
| chr19-15233943-DEL-6525 | chr19-15233944-DEL-6525 |
| chr19-15259378-DEL-72 | chr19-15259379-DEL-72 |
| chr19-15277219-DEL-139 | chr19-15277220-DEL-139 |
| chr19-15285066-DEL-70 | chr19-15285067-DEL-70 |
| chr19-15289753-DEL-2150 | chr19-15289752-DEL-2150 |
| chr19-15323590-DEL-514 | chr19-15323591-DEL-514 |
| chr19-15359425-DEL-64 | chr19-15359471-DEL-68 |
| chr19-15410679-DEL-6232 | chr19-15410680-DEL-6232 |
| chr19-15443968-DEL-58 | chr19-15443969-DEL-58 |
| chr19-15448224-DEL-108 | chr19-15448225-DEL-108 |
| chr19-15462538-DEL-64 | chr19-15462539-DEL-64 |
| chr19-15504395-DEL-472 | chr19-15504399-DEL-469 |
| chr19-15538528-DEL-162 | chr19-15538529-DEL-162 |
| chr19-15558884-DEL-1647 | chr19-15558885-DEL-1647 |
| chr19-15604775-DEL-64 | chr19-15604776-DEL-64 |

| | |
|---|---|
| chr19-15607892-DEL-54 | chr19-15607889-DEL-56 |
| chr19-15647685-DEL-216 | chr19-15647686-DEL-216 |
| chr19-15648097-DEL-1154 | chr19-15648103-DEL-1155 |
| chr19-15650185-DEL-397 | chr19-15650186-DEL-397 |
| chr19-15654833-DEL-114 | chr19-15654834-DEL-114 |
| chr19-15689113-DEL-8189 | chr19-15689114-DEL-8189 |
| chr19-15713148-DEL-1488 | chr19-15713149-DEL-1488 |
| chr19-15734900-DEL-1258 | chr19-15734902-DEL-1268 |
| chr19-15749893-DEL-1038 | chr19-15749899-DEL-1022 |
| chr19-15758317-DEL-1789 | chr19-15758318-DEL-1789 |
| chr19-15771449-DEL-114 | chr19-15771456-DEL-106 |
| chr19-15789274-DEL-744 | chr19-15789266-DEL-745 |
| chr19-15790649-DEL-468 | chr19-15790644-DEL-469 |
| chr19-15816753-DEL-7206 | chr19-15816746-DEL-7207 |
| chr19-15827572-DEL-321 | chr19-15827573-DEL-321 |
| chr19-15854897-DEL-500 | chr19-15854901-DEL-496 |
| chr19-15863217-DEL-1130 | chr19-15863218-DEL-1130 |
| chr19-15865505-DEL-1563 | chr19-15865506-DEL-1563 |
| chr19-15877959-DEL-63 | chr19-15877960-DEL-63 |
| chr19-15878381-DEL-2110 | chr19-15878382-DEL-2110 |
| chr19-15881903-DEL-171 | chr19-15881904-DEL-171 |
| chr19-15904218-DEL-55 | chr19-15904219-DEL-60 |
| chr19-15904817-DEL-201 | chr19-15904818-DEL-201 |
| chr19-15907369-DEL-197 | chr19-15907370-DEL-196 |
| chr19-15910744-DEL-146 | chr19-15910745-DEL-150 |
| chr19-15922633-DEL-203 | chr19-15922634-DEL-203 |
| chr19-15928960-DEL-73 | chr19-15928961-DEL-73 |
| chr19-15933894-DEL-1885 | chr19-15933885-DEL-1906 |
| chr19-15947468-DEL-78 | chr19-15947463-DEL-76 |
| chr19-15970032-DEL-66 | chr19-15969989-DEL-67 |
| chr19-15978856-DEL-95 | chr19-15978857-DEL-95 |
| chr19-15989927-DEL-430 | chr19-15989928-DEL-430 |
| chr19-16049907-DEL-59 | chr19-16049908-DEL-53 |
| chr19-16098298-DEL-213 | chr19-16098299-DEL-213 |
| chr19-16141452-DEL-761 | chr19-16141453-DEL-761 |
| chr19-16148900-DEL-1165 | chr19-16148901-DEL-1165 |
| chr19-16229316-DEL-64 | chr19-16229334-DEL-62 |
| chr19-16260026-DEL-55 | chr19-16260021-DEL-59 |
| chr19-16291656-DEL-98 | chr19-16291657-DEL-98 |
| chr19-16327169-DEL-3632 | chr19-16327170-DEL-3632 |
| chr19-16331037-DEL-56 | chr19-16331038-DEL-56 |
| chr19-16345495-DEL-164 | chr19-16345496-DEL-164 |
| chr19-16374546-DEL-58 | chr19-16374547-DEL-58 |
| chr19-16378233-DEL-104 | chr19-16378234-DEL-104 |
| chr19-16385024-DEL-96 | chr19-16385025-DEL-96 |
| chr19-16389094-DEL-1689 | chr19-16389095-DEL-1689 |
| chr19-16397184-DEL-1947 | chr19-16397185-DEL-1948 |
| chr19-16409678-DEL-198 | chr19-16409679-DEL-198 |
| chr19-16427285-DEL-86 | chr19-16427284-DEL-87 |
| chr19-16429719-DEL-60 | chr19-16429720-DEL-60 |
| chr19-16446913-DEL-192 | chr19-16446914-DEL-192 |
| chr19-16447554-DEL-64 | chr19-16447555-DEL-64 |
| chr19-16450224-DEL-56 | chr19-16450225-DEL-56 |
| chr19-16469700-DEL-5115 | chr19-16469701-DEL-5115 |
| chr19-16484788-DEL-485 | chr19-16484788-DEL-483 |
| chr19-16521376-DEL-1154 | chr19-16521377-DEL-1154 |
| chr19-16530967-DEL-431 | chr19-16530968-DEL-431 |
| chr19-16539293-DEL-176 | chr19-16539294-DEL-176 |
| chr19-16552076-DEL-1037 | chr19-16552077-DEL-1037 |
| chr19-16626439-DEL-96 | chr19-16626423-DEL-112 |
| chr19-16644522-DEL-84 | chr19-16644523-DEL-84 |
| chr19-16647886-DEL-2811 | chr19-16647887-DEL-2811 |
| chr19-16671993-DEL-64 | chr19-16671994-DEL-64 |
| chr19-16684133-DEL-220 | chr19-16684134-DEL-220 |
| chr19-16704080-DEL-64 | chr19-16704081-DEL-64 |
| chr19-16707314-DEL-635 | chr19-16707315-DEL-635 |
| chr19-16735071-DEL-681 | chr19-16735072-DEL-681 |
| chr19-16744338-DEL-232 | chr19-16744334-DEL-236 |
| chr19-16775741-DEL-1498 | chr19-16775742-DEL-1498 |
| chr19-16777425-DEL-57 | chr19-16777425-DEL-57 |
| chr19-16782230-DEL-51 | chr19-16782230-DEL-56 |
| chr19-16783054-DEL-534 | chr19-16783055-DEL-534 |
| chr19-16799359-DEL-62 | chr19-16799360-DEL-62 |
| chr19-16811287-DEL-920 | chr19-16811288-DEL-920 |
| chr19-16830295-DEL-215 | chr19-16830296-DEL-215 |
| chr19-16862465-DEL-80 | chr19-16862466-DEL-80 |
| chr19-16865759-DEL-1742 | chr19-16865760-DEL-1742 |
| chr19-16874402-DEL-53 | chr19-16874364-DEL-63 |
| chr19-16897896-DEL-175 | chr19-16897900-DEL-174 |
| chr19-16950029-DEL-74 | chr19-16950030-DEL-74 |

| | |
|---|---|
| chr19-16951841-DEL-148 | chr19-16951842-DEL-148 |
| chr19-16995203-DEL-2647 | chr19-16995204-DEL-2647 |
| chr19-17038710-DEL-179 | chr19-17038711-DEL-179 |
| chr19-17050044-DEL-84 | chr19-17050045-DEL-84 |
| chr19-17052400-DEL-520 | chr19-17052401-DEL-522 |
| chr19-17053686-DEL-220 | chr19-17053687-DEL-220 |
| chr19-17060948-DEL-186 | chr19-17060955-DEL-180 |
| chr19-17146174-DEL-99 | chr19-17146191-DEL-78 |
| chr19-17151088-DEL-903 | chr19-17151089-DEL-903 |
| chr19-17178805-DEL-57 | chr19-17178806-DEL-56 |
| chr19-17206654-DEL-58 | chr19-17206655-DEL-58 |
| chr19-17209228-DEL-116 | chr19-17209229-DEL-116 |
| chr19-17222906-DEL-54 | chr19-17222907-DEL-54 |
| chr19-17223565-DEL-1251 | chr19-17223566-DEL-1249 |
| chr19-17224248-DEL-196 | chr19-17224244-DEL-196 |
| chr19-17235483-DEL-188 | chr19-17235484-DEL-168 |
| chr19-17238387-DEL-971 | chr19-17238388-DEL-971 |
| chr19-17257742-DEL-1232 | chr19-17257743-DEL-1232 |
| chr19-17264337-DEL-102 | chr19-17264338-DEL-102 |
| chr19-17279521-DEL-1074 | chr19-17279521-DEL-1074 |
| chr19-17280889-DEL-6051 | chr19-17280890-DEL-6050 |
| chr19-17302456-DEL-150 | chr19-17302456-DEL-151 |
| chr19-17313351-DEL-97 | chr19-17313352-DEL-97 |
| chr19-17320883-DEL-61 | chr19-17320884-DEL-61 |
| chr19-17340219-DEL-65 | chr19-17340225-DEL-75 |
| chr19-17348159-DEL-113 | chr19-17348160-DEL-113 |
| chr19-17364360-DEL-166 | chr19-17364361-DEL-166 |
| chr19-17368201-DEL-92 | chr19-17368202-DEL-92 |
| chr19-17392841-DEL-483 | chr19-17392840-DEL-484 |
| chr19-17410101-DEL-73 | chr19-17410097-DEL-78 |
| chr19-17416856-DEL-63 | chr19-17416857-DEL-63 |
| chr19-17431910-DEL-70 | chr19-17431893-DEL-67 |
| chr19-17444529-DEL-66 | chr19-17444530-DEL-66 |
| chr19-17460763-DEL-193 | chr19-17460764-DEL-193 |
| chr19-17466228-DEL-60 | chr19-17466241-DEL-56 |
| chr19-17486494-DEL-51 | chr19-17486492-DEL-53 |
| chr19-17496238-DEL-107 | chr19-17496239-DEL-107 |
| chr19-17517479-DEL-62 | chr19-17517484-DEL-58 |
| chr19-17541788-DEL-58 | chr19-17541791-DEL-56 |
| chr19-17543096-DEL-66 | chr19-17543097-DEL-66 |
| chr19-17557494-DEL-53 | chr19-17557495-DEL-53 |
| chr19-17569665-DEL-71 | chr19-17569661-DEL-72 |
| chr19-17616352-DEL-789 | chr19-17616352-DEL-786 |
| chr19-17633275-DEL-156 | chr19-17633274-DEL-156 |
| chr19-17645850-DEL-256 | chr19-17645851-DEL-256 |
| chr19-17729279-DEL-54 | chr19-17729280-DEL-54 |
| chr19-17734264-DEL-54 | chr19-17734265-DEL-54 |
| chr19-17783226-DEL-52 | chr19-17783227-DEL-52 |
| chr19-17791660-DEL-153 | chr19-17791661-DEL-153 |
| chr19-17821244-DEL-195 | chr19-17821245-DEL-195 |
| chr19-17834646-DEL-356 | chr19-17834647-DEL-356 |
| chr19-17845597-DEL-436 | chr19-17845598-DEL-436 |
| chr19-17848112-DEL-82 | chr19-17848112-DEL-68 |
| chr19-17892092-DEL-367 | chr19-17892093-DEL-367 |
| chr19-17905584-DEL-312 | chr19-17905585-DEL-312 |
| chr19-17913205-DEL-6444 | chr19-17913220-DEL-6443 |
| chr19-17919324-DEL-214 | chr19-17919324-DEL-214 |
| chr19-17932188-DEL-68 | chr19-17932177-DEL-92 |
| chr19-17935045-DEL-91 | chr19-17935021-DEL-68 |
| chr19-17968727-DEL-62 | chr19-17968728-DEL-62 |
| chr19-18004099-DEL-56 | chr19-18004099-DEL-56 |
| chr19-18036273-DEL-1883 | chr19-18036274-DEL-1883 |
| chr19-18045112-DEL-53 | chr19-18045088-DEL-54 |
| chr19-18056856-DEL-65 | chr19-18056857-DEL-70 |
| chr19-18066454-DEL-101 | chr19-18066497-DEL-62 |
| chr19-18074084-DEL-3877 | chr19-18074082-DEL-3878 |
| chr19-18099047-DEL-313 | chr19-18099048-DEL-313 |
| chr19-18121915-DEL-112 | chr19-18121916-DEL-112 |
| chr19-18172460-DEL-2183 | chr19-18172461-DEL-2183 |
| chr19-18187819-DEL-58 | chr19-18187820-DEL-54 |
| chr19-18203972-DEL-6230 | chr19-18203973-DEL-6230 |
| chr19-18220954-DEL-6145 | chr19-18220955-DEL-6145 |
| chr19-18241037-DEL-3272 | chr19-18241038-DEL-3272 |
| chr19-18245194-DEL-148 | chr19-18245195-DEL-148 |
| chr19-18263472-DEL-6345 | chr19-18263473-DEL-6345 |
| chr19-18282352-DEL-53 | chr19-18282353-DEL-53 |
| chr19-18303471-DEL-73 | chr19-18303472-DEL-73 |
| chr19-18335145-DEL-6395 | chr19-18335146-DEL-6395 |
| chr19-18341551-DEL-159 | chr19-18341550-DEL-159 |
| chr19-18360764-DEL-78 | chr19-18360767-DEL-94 |

| | |
|---|---|
| chr19-18374003-DEL-633 | chr19-18374004-DEL-633 |
| chr19-18377818-DEL-2432 | chr19-18377821-DEL-2441 |
| chr19-18394118-DEL-768 | chr19-18394119-DEL-768 |
| chr19-18422898-DEL-201 | chr19-18422899-DEL-201 |
| chr19-18436438-DEL-64 | chr19-18436438-DEL-64 |
| chr19-18451005-DEL-159 | chr19-18451009-DEL-156 |
| chr19-18451606-DEL-93 | chr19-18451607-DEL-93 |
| chr19-18460862-DEL-6765 | chr19-18460864-DEL-6766 |
| chr19-18469076-DEL-159 | chr19-18469077-DEL-158 |
| chr19-18477945-DEL-60 | chr19-18477956-DEL-82 |
| chr19-18509523-DEL-3323 | chr19-18509524-DEL-3323 |
| chr19-18520919-DEL-72 | chr19-18520920-DEL-72 |
| chr19-18538075-DEL-53 | chr19-18538076-DEL-53 |
| chr19-18548851-DEL-291 | chr19-18548852-DEL-291 |
| chr19-18555547-DEL-57 | chr19-18555548-DEL-57 |
| chr19-18590421-DEL-82 | chr19-18590382-DEL-78 |
| chr19-18594687-DEL-191 | chr19-18594688-DEL-191 |
| chr19-18611632-DEL-113 | chr19-18611674-DEL-77 |
| chr19-18623699-DEL-274 | chr19-18623700-DEL-274 |
| chr19-18642247-DEL-4042 | chr19-18642248-DEL-4042 |
| chr19-18642624-DEL-217 | chr19-18642625-DEL-217 |
| chr19-18676169-DEL-764 | chr19-18676170-DEL-764 |
| chr19-18689728-DEL-320 | chr19-18689729-DEL-320 |
| chr19-18701738-DEL-79 | chr19-18701737-DEL-77 |
| chr19-18707523-DEL-59 | chr19-18707523-DEL-60 |
| chr19-18786429-DEL-200 | chr19-18786430-DEL-200 |
| chr19-18828212-DEL-1602 | chr19-18828213-DEL-1600 |
| chr19-18840978-DEL-114 | chr19-18840979-DEL-114 |
| chr19-18876366-DEL-129 | chr19-18876367-DEL-129 |
| chr19-18897511-DEL-4155 | chr19-18897512-DEL-4155 |
| chr19-18903825-DEL-346 | chr19-18903826-DEL-346 |
| chr19-18935559-DEL-221 | chr19-18935560-DEL-221 |
| chr19-18966012-DEL-58 | chr19-18966025-DEL-50 |
| chr19-19016809-DEL-98 | chr19-19016809-DEL-90 |
| chr19-19043896-DEL-164 | chr19-19043902-DEL-159 |
| chr19-19100622-DEL-780 | chr19-19100622-DEL-780 |
| chr19-19114941-DEL-201 | chr19-19114942-DEL-201 |
| chr19-19130522-DEL-3065 | chr19-19130523-DEL-3065 |
| chr19-19195980-DEL-80 | chr19-19195979-DEL-81 |
| chr19-19221935-DEL-942 | chr19-19221935-DEL-942 |
| chr19-19225289-DEL-76 | chr19-19225290-DEL-76 |
| chr19-19232110-DEL-350 | chr19-19232114-DEL-347 |
| chr19-19232517-DEL-1051 | chr19-19232518-DEL-1051 |
| chr19-19258819-DEL-2709 | chr19-19258820-DEL-2709 |
| chr19-19276735-DEL-904 | chr19-19276749-DEL-900 |
| chr19-19279747-DEL-514 | chr19-19279748-DEL-514 |
| chr19-19282082-DEL-52 | chr19-19282085-DEL-52 |
| chr19-19292085-DEL-115 | chr19-19292097-DEL-107 |
| chr19-19293106-DEL-91 | chr19-19293103-DEL-89 |
| chr19-19299371-DEL-750 | chr19-19299372-DEL-750 |
| chr19-19319090-DEL-325 | chr19-19319079-DEL-326 |
| chr19-19344275-DEL-1841 | chr19-19344276-DEL-1841 |
| chr19-19367130-DEL-60 | chr19-19367131-DEL-60 |
| chr19-19372386-DEL-626 | chr19-19372384-DEL-628 |
| chr19-19391899-DEL-1667 | chr19-19391900-DEL-1667 |
| chr19-19401143-DEL-104 | chr19-19401156-DEL-103 |
| chr19-19456187-DEL-96 | chr19-19456188-DEL-96 |
| chr19-19463825-DEL-148 | chr19-19463826-DEL-148 |
| chr19-19476684-DEL-590 | chr19-19476685-DEL-590 |
| chr19-19486177-DEL-6177 | chr19-19486178-DEL-6177 |
| chr19-19552462-DEL-332 | chr19-19552463-DEL-332 |
| chr19-19572259-DEL-765 | chr19-19572260-DEL-765 |
| chr19-19575623-DEL-425 | chr19-19575622-DEL-426 |
| chr19-19578354-DEL-196 | chr19-19578355-DEL-196 |
| chr19-19585596-DEL-175 | chr19-19585595-DEL-176 |
| chr19-19640623-DEL-95 | chr19-19640648-DEL-71 |
| chr19-19644657-DEL-398 | chr19-19644656-DEL-396 |
| chr19-19647310-DEL-3368 | chr19-19647311-DEL-3368 |
| chr19-19672719-DEL-5739 | chr19-19672719-DEL-5740 |
| chr19-19678856-DEL-81 | chr19-19678857-DEL-81 |
| chr19-19691416-DEL-417 | chr19-19691417-DEL-417 |
| chr19-19705291-DEL-51 | chr19-19705291-DEL-51 |
| chr19-19718245-DEL-947 | chr19-19718246-DEL-948 |
| chr19-19740840-DEL-3689 | chr19-19740841-DEL-3688 |
| chr19-19772953-DEL-395 | chr19-19772954-DEL-395 |
| chr19-19805779-DEL-212 | chr19-19805779-DEL-212 |
| chr19-19816345-DEL-66 | chr19-19816346-DEL-64 |
| chr19-19831497-DEL-58 | chr19-19831500-DEL-56 |
| chr19-19847449-DEL-66 | chr19-19847449-DEL-66 |
| chr19-19847785-DEL-6440 | chr19-19847787-DEL-6440 |

| | |
|---|---|
| chr19-19880148-DEL-53 | chr19-19880149-DEL-52 |
| chr19-19890646-DEL-7350 | chr19-19890647-DEL-7350 |
| chr19-19893776-DEL-401 | chr19-19893777-DEL-401 |
| chr19-19932402-DEL-54 | chr19-19932403-DEL-54 |
| chr19-19935346-DEL-1933 | chr19-19935346-DEL-1933 |
| chr19-19941371-DEL-107 | chr19-19941378-DEL-103 |
| chr19-19944095-DEL-56 | chr19-19944096-DEL-51 |
| chr19-19961873-DEL-932 | chr19-19961874-DEL-932 |
| chr19-19972783-DEL-284 | chr19-19972784-DEL-284 |
| chr19-19995796-DEL-55 | chr19-19995797-DEL-55 |
| chr19-20006740-DEL-56 | chr19-20006741-DEL-56 |
| chr19-20020879-DEL-795 | chr19-20020878-DEL-790 |
| chr19-20040735-DEL-385 | chr19-20040736-DEL-385 |
| chr19-20060209-DEL-4965 | chr19-20060210-DEL-4965 |
| chr19-20062664-DEL-211 | chr19-20062660-DEL-215 |
| chr19-20063147-DEL-948 | chr19-20063130-DEL-965 |
| chr19-20072662-DEL-435 | chr19-20072664-DEL-434 |
| chr19-20117188-DEL-4548 | chr19-20117189-DEL-4548 |
| chr19-20169236-DEL-115 | chr19-20169239-DEL-113 |
| chr19-20178671-DEL-1934 | chr19-20178672-DEL-1934 |
| chr19-20182798-DEL-3571 | chr19-20182805-DEL-3569 |
| chr19-20184297-DEL-200 | chr19-20184298-DEL-200 |
| chr19-20263962-DEL-370 | chr19-20263963-DEL-370 |
| chr19-20265610-DEL-1892 | chr19-20265611-DEL-1892 |
| chr19-20355366-DEL-67 | chr19-20355366-DEL-68 |
| chr19-20368373-DEL-120 | chr19-20368374-DEL-102 |
| chr19-20375436-DEL-193 | chr19-20375437-DEL-193 |
| chr19-20405035-DEL-52 | chr19-20405034-DEL-56 |
| chr19-20407730-DEL-155 | chr19-20407696-DEL-144 |
| chr19-20431132-DEL-79 | chr19-20431133-DEL-79 |
| chr19-20432828-DEL-4964 | chr19-20432829-DEL-4964 |
| chr19-20451421-DEL-579 | chr19-20451422-DEL-579 |
| chr19-20493728-DEL-164 | chr19-20493729-DEL-163 |
| chr19-20498078-DEL-6352 | chr19-20498079-DEL-6352 |
| chr19-20517743-DEL-82 | chr19-20517744-DEL-123 |
| chr19-20538449-DEL-380 | chr19-20538450-DEL-380 |
| chr19-20540848-DEL-171 | chr19-20540849-DEL-171 |
| chr19-20544683-DEL-6473 | chr19-20544684-DEL-6473 |
| chr19-20551348-DEL-2316 | chr19-20551346-DEL-2317 |
| chr19-20562904-DEL-60 | chr19-20562909-DEL-56 |
| chr19-20563110-DEL-1285 | chr19-20563115-DEL-1282 |
| chr19-20589546-DEL-76 | chr19-20589547-DEL-72 |
| chr19-20593006-DEL-51 | chr19-20592962-DEL-96 |
| chr19-20636014-DEL-623 | chr19-20636015-DEL-623 |
| chr19-20640415-DEL-1384 | chr19-20640416-DEL-1384 |
| chr19-20689693-DEL-758 | chr19-20689693-DEL-758 |
| chr19-20721171-DEL-373 | chr19-20721172-DEL-373 |
| chr19-20726275-DEL-6468 | chr19-20726276-DEL-6468 |
| chr19-20745184-DEL-4244 | chr19-20745185-DEL-4244 |
| chr19-20753822-DEL-529 | chr19-20753821-DEL-529 |
| chr19-20755434-DEL-117 | chr19-20755420-DEL-86 |
| chr19-20764534-DEL-1982 | chr19-20764535-DEL-1982 |
| chr19-20768248-DEL-153 | chr19-20768249-DEL-153 |
| chr19-20774062-DEL-56 | chr19-20774062-DEL-57 |
| chr19-20785669-DEL-113 | chr19-20785670-DEL-113 |
| chr19-20793868-DEL-60 | chr19-20793869-DEL-60 |
| chr19-20796889-DEL-96 | chr19-20796890-DEL-60 |
| chr19-20817047-DEL-4049 | chr19-20817048-DEL-4049 |
| chr19-20833078-DEL-252 | chr19-20833079-DEL-231 |
| chr19-20839638-DEL-147 | chr19-20839639-DEL-147 |
| chr19-20860927-DEL-68 | chr19-20860928-DEL-68 |
| chr19-20883265-DEL-63 | chr19-20883266-DEL-61 |
| chr19-20894995-DEL-392 | chr19-20894995-DEL-392 |
| chr19-20900385-DEL-251 | chr19-20900386-DEL-251 |
| chr19-20916460-DEL-132 | chr19-20916461-DEL-132 |
| chr19-20923225-DEL-190 | chr19-20923225-DEL-190 |
| chr19-20958943-DEL-475 | chr19-20958944-DEL-475 |
| chr19-20968148-DEL-60 | chr19-20968143-DEL-81 |
| chr19-20989537-DEL-354 | chr19-20989538-DEL-354 |
| chr19-21003265-DEL-61 | chr19-21003303-DEL-66 |
| chr19-21035680-DEL-6325 | chr19-21035681-DEL-6325 |
| chr19-21046448-DEL-266 | chr19-21046449-DEL-266 |
| chr19-21067499-DEL-68 | chr19-21067500-DEL-68 |
| chr19-21079524-DEL-251 | chr19-21079525-DEL-251 |
| chr19-21166037-DEL-74 | chr19-21166037-DEL-74 |
| chr19-21184811-DEL-164 | chr19-21184812-DEL-164 |
| chr19-21195527-DEL-6648 | chr19-21195528-DEL-6648 |
| chr19-21214698-DEL-6524 | chr19-21214699-DEL-6524 |
| chr19-21225710-DEL-507 | chr19-21225711-DEL-507 |
| chr19-21252822-DEL-57 | chr19-21252823-DEL-57 |

| | |
|---|---|
| chr19-21269007-DEL-980 | chr19-21269008-DEL-980 |
| chr19-21270962-DEL-228 | chr19-21270963-DEL-228 |
| chr19-21274048-DEL-161 | chr19-21274047-DEL-162 |
| chr19-21282347-DEL-118 | chr19-21282336-DEL-130 |
| chr19-21304158-DEL-79 | chr19-21304162-DEL-77 |
| chr19-21362366-DEL-51 | chr19-21362373-DEL-58 |
| chr19-21365683-DEL-214 | chr19-21365684-DEL-214 |
| chr19-21367088-DEL-67 | chr19-21367089-DEL-67 |
| chr19-21369848-DEL-182 | chr19-21369844-DEL-182 |
| chr19-21381947-DEL-81 | chr19-21381948-DEL-79 |
| chr19-21411295-DEL-104 | chr19-21411304-DEL-96 |
| chr19-21417457-DEL-229 | chr19-21417464-DEL-233 |
| chr19-21421508-DEL-714 | chr19-21421509-DEL-714 |
| chr19-21425911-DEL-1158 | chr19-21425912-DEL-1158 |
| chr19-21432226-DEL-1253 | chr19-21432227-DEL-1253 |
| chr19-21433582-DEL-163 | chr19-21433583-DEL-163 |
| chr19-21451737-DEL-121 | chr19-21451738-DEL-121 |
| chr19-21453111-DEL-150 | chr19-21453115-DEL-158 |
| chr19-21492673-DEL-248 | chr19-21492676-DEL-249 |
| chr19-21506407-DEL-941 | chr19-21506408-DEL-941 |
| chr19-21509902-DEL-1888 | chr19-21509903-DEL-1888 |
| chr19-21533910-DEL-62 | chr19-21533911-DEL-62 |
| chr19-21538310-DEL-146 | chr19-21538311-DEL-146 |
| chr19-21546870-DEL-64 | chr19-21546879-DEL-52 |
| chr19-21568085-DEL-66 | chr19-21568086-DEL-66 |
| chr19-21583890-DEL-81 | chr19-21583878-DEL-56 |
| chr19-21591355-DEL-127 | chr19-21591356-DEL-127 |
| chr19-21609435-DEL-406 | chr19-21609436-DEL-406 |
| chr19-21631443-DEL-52 | chr19-21631444-DEL-52 |
| chr19-21696337-DEL-153 | chr19-21696338-DEL-153 |
| chr19-21702973-DEL-530 | chr19-21702974-DEL-530 |
| chr19-21725519-DEL-182 | chr19-21725520-DEL-182 |
| chr19-21726750-DEL-120 | chr19-21726783-DEL-88 |
| chr19-21727464-DEL-54 | chr19-21727465-DEL-54 |
| chr19-21730946-DEL-435 | chr19-21730947-DEL-435 |
| chr19-21800914-DEL-201 | chr19-21800915-DEL-201 |
| chr19-21813004-DEL-1418 | chr19-21813005-DEL-1418 |
| chr19-21814533-DEL-73 | chr19-21814534-DEL-73 |
| chr19-21816606-DEL-105 | chr19-21816580-DEL-108 |
| chr19-21817577-DEL-53 | chr19-21817573-DEL-50 |
| chr19-21858959-DEL-244 | chr19-21858960-DEL-244 |
| chr19-21868549-DEL-3055 | chr19-21868550-DEL-3053 |
| chr19-21878912-DEL-156 | chr19-21878913-DEL-156 |
| chr19-21887538-DEL-62 | chr19-21887539-DEL-62 |
| chr19-21888872-DEL-227 | chr19-21888873-DEL-227 |
| chr19-21895342-DEL-129 | chr19-21895343-DEL-129 |
| chr19-21897253-DEL-282 | chr19-21897253-DEL-282 |
| chr19-21911366-DEL-267 | chr19-21911351-DEL-268 |
| chr19-21914636-DEL-1326 | chr19-21914646-DEL-1333 |
| chr19-21918431-DEL-119 | chr19-21918393-DEL-159 |
| chr19-21926506-DEL-182 | chr19-21926507-DEL-182 |
| chr19-21927696-DEL-3138 | chr19-21927697-DEL-3138 |
| chr19-21942453-DEL-1494 | chr19-21942454-DEL-1494 |
| chr19-21949780-DEL-173 | chr19-21949781-DEL-173 |
| chr19-21953538-DEL-6227 | chr19-21953539-DEL-6227 |
| chr19-21978632-DEL-136 | chr19-21978633-DEL-136 |
| chr19-22037838-DEL-86 | chr19-22037839-DEL-79 |
| chr19-22067716-DEL-51 | chr19-22067717-DEL-51 |
| chr19-22075006-DEL-1986 | chr19-22075020-DEL-1983 |
| chr19-22085347-DEL-437 | chr19-22085348-DEL-437 |
| chr19-22095967-DEL-211 | chr19-22095968-DEL-211 |
| chr19-22102773-DEL-703 | chr19-22102774-DEL-703 |
| chr19-22109095-DEL-126 | chr19-22109096-DEL-126 |
| chr19-22113453-DEL-507 | chr19-22113454-DEL-507 |
| chr19-22122350-DEL-64 | chr19-22122345-DEL-97 |
| chr19-22132635-DEL-146 | chr19-22132658-DEL-104 |
| chr19-22135839-DEL-784 | chr19-22135840-DEL-784 |
| chr19-22154605-DEL-1383 | chr19-22154606-DEL-1383 |
| chr19-22169929-DEL-158 | chr19-22169930-DEL-158 |
| chr19-22182020-DEL-54 | chr19-22182021-DEL-51 |
| chr19-22187019-DEL-80 | chr19-22187020-DEL-80 |
| chr19-22231332-DEL-209 | chr19-22231333-DEL-209 |
| chr19-22240078-DEL-65 | chr19-22240074-DEL-66 |
| chr19-22251775-DEL-152 | chr19-22251776-DEL-152 |
| chr19-22255061-DEL-81 | chr19-22255062-DEL-81 |
| chr19-22257790-DEL-51 | chr19-22257791-DEL-51 |
| chr19-22262677-DEL-355 | chr19-22262678-DEL-355 |
| chr19-22264442-DEL-1219 | chr19-22264443-DEL-1219 |
| chr19-22269945-DEL-72 | chr19-22269994-DEL-52 |
| chr19-22337119-DEL-86 | chr19-22337119-DEL-86 |

| | |
|---|---|
| chr19-22383487-DEL-3565 | chr19-22383488-DEL-3565 |
| chr19-22391043-DEL-67 | chr19-22391044-DEL-67 |
| chr19-22402466-DEL-121 | chr19-22402467-DEL-121 |
| chr19-22408823-DEL-82 | chr19-22408824-DEL-82 |
| chr19-22415546-DEL-70 | chr19-22415546-DEL-71 |
| chr19-22418361-DEL-2666 | chr19-22418361-DEL-2666 |
| chr19-22437546-DEL-251 | chr19-22437547-DEL-251 |
| chr19-22452404-DEL-487 | chr19-22452405-DEL-487 |
| chr19-22482109-DEL-1762 | chr19-22482110-DEL-1762 |
| chr19-22497643-DEL-214 | chr19-22497644-DEL-214 |
| chr19-22518017-DEL-97 | chr19-22518018-DEL-97 |
| chr19-22554973-DEL-84 | chr19-22554974-DEL-84 |
| chr19-22565469-DEL-168 | chr19-22565470-DEL-168 |
| chr19-22567247-DEL-84 | chr19-22567246-DEL-86 |
| chr19-22574071-DEL-54 | chr19-22574072-DEL-50 |
| chr19-22606156-DEL-56 | chr19-22606150-DEL-60 |
| chr19-22617938-DEL-66 | chr19-22617939-DEL-66 |
| chr19-22621596-DEL-7448 | chr19-22621597-DEL-7448 |
| chr19-22698513-DEL-181 | chr19-22698514-DEL-164 |
| chr19-22699749-DEL-5849 | chr19-22699751-DEL-5850 |
| chr19-22716980-DEL-52 | chr19-22716981-DEL-52 |
| chr19-22753648-DEL-67 | chr19-22753648-DEL-68 |
| chr19-22774808-DEL-164 | chr19-22774809-DEL-164 |
| chr19-22785310-DEL-207 | chr19-22785311-DEL-207 |
| chr19-22810574-DEL-7371 | chr19-22810575-DEL-7371 |
| chr19-22861692-DEL-55 | chr19-22861697-DEL-53 |
| chr19-22885051-DEL-847 | chr19-22885047-DEL-848 |
| chr19-22885883-DEL-62 | chr19-22885884-DEL-58 |
| chr19-22911274-DEL-330 | chr19-22911275-DEL-330 |
| chr19-22911853-DEL-108 | chr19-22911826-DEL-114 |
| chr19-22937488-DEL-56 | chr19-22937489-DEL-66 |
| chr19-22987162-DEL-400 | chr19-22987163-DEL-400 |
| chr19-22991293-DEL-57 | chr19-22991266-DEL-87 |
| chr19-22995233-DEL-310 | chr19-22995234-DEL-310 |
| chr19-23002651-DEL-54 | chr19-23002647-DEL-60 |
| chr19-23017379-DEL-62 | chr19-23017380-DEL-62 |
| chr19-23036415-DEL-92 | chr19-23036416-DEL-92 |
| chr19-23036704-DEL-363 | chr19-23036705-DEL-363 |
| chr19-23037476-DEL-119 | chr19-23037476-DEL-103 |
| chr19-23053565-DEL-205 | chr19-23053566-DEL-205 |
| chr19-23060284-DEL-351 | chr19-23060285-DEL-351 |
| chr19-23061076-DEL-194 | chr19-23061077-DEL-194 |
| chr19-23070528-DEL-75 | chr19-23070501-DEL-72 |
| chr19-23075568-DEL-195 | chr19-23075569-DEL-195 |
| chr19-23091548-DEL-76 | chr19-23091549-DEL-76 |
| chr19-23135724-DEL-347 | chr19-23135725-DEL-347 |
| chr19-23156593-DEL-98 | chr19-23156594-DEL-98 |
| chr19-23197878-DEL-67 | chr19-23197879-DEL-67 |
| chr19-23210848-DEL-230 | chr19-23210849-DEL-230 |
| chr19-23235566-DEL-56 | chr19-23235567-DEL-56 |
| chr19-23241283-DEL-498 | chr19-23241284-DEL-498 |
| chr19-23247925-DEL-154 | chr19-23247925-DEL-132 |
| chr19-23257784-DEL-52 | chr19-23257785-DEL-52 |
| chr19-23261163-DEL-348 | chr19-23261164-DEL-348 |
| chr19-23319258-DEL-75 | chr19-23319251-DEL-83 |
| chr19-23335794-DEL-76 | chr19-23335794-DEL-76 |
| chr19-23347361-DEL-53 | chr19-23347361-DEL-53 |
| chr19-23426590-DEL-154 | chr19-23426591-DEL-154 |
| chr19-23436530-DEL-2191 | chr19-23436523-DEL-2192 |
| chr19-23455491-DEL-842 | chr19-23455492-DEL-842 |
| chr19-23461979-DEL-441 | chr19-23461980-DEL-441 |
| chr19-23488022-DEL-88 | chr19-23488023-DEL-88 |
| chr19-23503580-DEL-94 | chr19-23503595-DEL-88 |
| chr19-23523885-DEL-1896 | chr19-23523886-DEL-1894 |
| chr19-23545789-DEL-146 | chr19-23545792-DEL-142 |
| chr19-23575432-DEL-5468 | chr19-23575434-DEL-5467 |
| chr19-23578076-DEL-138 | chr19-23578068-DEL-147 |
| chr19-23596674-DEL-1053 | chr19-23596675-DEL-1053 |
| chr19-23596737-DEL-374 | chr19-23596738-DEL-375 |
| chr19-23599180-DEL-507 | chr19-23599181-DEL-507 |
| chr19-23653170-DEL-941 | chr19-23653171-DEL-942 |
| chr19-23676221-DEL-60 | chr19-23676222-DEL-60 |
| chr19-23710568-DEL-67 | chr19-23710569-DEL-67 |
| chr19-23723020-DEL-115 | chr19-23723021-DEL-115 |
| chr19-23728027-DEL-3331 | chr19-23728028-DEL-3331 |
| chr19-23751279-DEL-96 | chr19-23751280-DEL-96 |
| chr19-23760109-DEL-1977 | chr19-23760110-DEL-1977 |
| chr19-23770586-DEL-204 | chr19-23770587-DEL-204 |
| chr19-23784236-DEL-56 | chr19-23784237-DEL-56 |
| chr19-23816706-DEL-219 | chr19-23816707-DEL-219 |

| | |
|---|---|
| chr19-23832081-DEL-71 | chr19-23832082-DEL-71 |
| chr19-23902406-DEL-808 | chr19-23902407-DEL-808 |
| chr19-23921619-DEL-448 | chr19-23921620-DEL-448 |
| chr19-23934263-DEL-616 | chr19-23934264-DEL-616 |
| chr19-23962404-DEL-591 | chr19-23962373-DEL-592 |
| chr19-23963354-DEL-294 | chr19-23963354-DEL-341 |
| chr19-23983645-DEL-84 | chr19-23983646-DEL-77 |
| chr19-24017717-DEL-1897 | chr19-24017718-DEL-1897 |
| chr19-24020217-DEL-63 | chr19-24020218-DEL-63 |
| chr19-24027547-DEL-65 | chr19-24027548-DEL-65 |
| chr19-24028551-DEL-54 | chr19-24028548-DEL-62 |
| chr19-24029432-DEL-1211 | chr19-24029433-DEL-1211 |
| chr19-24055080-DEL-305 | chr19-24055081-DEL-305 |
| chr19-24094742-DEL-204 | chr19-24094743-DEL-204 |
| chr19-24124761-DEL-194 | chr19-24124759-DEL-195 |
| chr19-24139728-DEL-80 | chr19-24139717-DEL-74 |
| chr19-24144579-DEL-56 | chr19-24144581-DEL-57 |
| chr19-24151070-DEL-62 | chr19-24151071-DEL-62 |
| chr19-24160999-DEL-58 | chr19-24161004-DEL-54 |
| chr19-24163103-DEL-192 | chr19-24163104-DEL-192 |
| chr19-24166880-DEL-256 | chr19-24166881-DEL-256 |
| chr19-24169001-DEL-58 | chr19-24169002-DEL-50 |
| chr19-24179326-DEL-121 | chr19-24179326-DEL-122 |
| chr19-24204529-DEL-219 | chr19-24204540-DEL-208 |
| chr19-24207730-DEL-446 | chr19-24207731-DEL-446 |
| chr19-24234293-DEL-922 | chr19-24234293-DEL-920 |
| chr19-24247371-DEL-143 | chr19-24247385-DEL-130 |
| chr19-24250408-DEL-394 | chr19-24250402-DEL-395 |
| chr19-24250910-DEL-66 | chr19-24250911-DEL-66 |
| chr19-24260559-DEL-448 | chr19-24260560-DEL-448 |
| chr19-24264439-DEL-416 | chr19-24264440-DEL-416 |
| chr19-24273551-DEL-392 | chr19-24273550-DEL-392 |
| chr19-24277177-DEL-203 | chr19-24277178-DEL-203 |
| chr19-24286079-DEL-209 | chr19-24286080-DEL-209 |
| chr19-24300233-DEL-58 | chr19-24300235-DEL-50 |
| chr19-24302847-DEL-93 | chr19-24302815-DEL-93 |
| chr19-24304974-DEL-64 | chr19-24304975-DEL-64 |
| chr19-24323115-DEL-104 | chr19-24323116-DEL-104 |
| chr19-24326528-DEL-197 | chr19-24326529-DEL-197 |
| chr19-24327888-DEL-66 | chr19-24327889-DEL-72 |
| chr19-24340675-DEL-51 | chr19-24340676-DEL-51 |
| chr19-24342044-DEL-406 | chr19-24342045-DEL-406 |
| chr19-24369898-DEL-62 | chr19-24369899-DEL-62 |
| chr19-24417783-DEL-424 | chr19-24417784-DEL-424 |
| chr19-24425016-DEL-1610 | chr19-24425017-DEL-1610 |
| chr19-24429356-DEL-215 | chr19-24429357-DEL-215 |
| chr19-24448091-DEL-200 | chr19-24448092-DEL-200 |
| chr19-24471258-DEL-285 | chr19-24471259-DEL-285 |
| chr19-24479373-DEL-64 | chr19-24479387-DEL-51 |
| chr19-24503187-DEL-200 | chr19-24503196-DEL-192 |
| chr19-24511268-DEL-183 | chr19-24511261-DEL-191 |
| chr19-24522008-DEL-176 | chr19-24521994-DEL-191 |
| chr19-24568030-DEL-68 | chr19-24568035-DEL-68 |
| chr19-24585820-DEL-238 | chr19-24585821-DEL-238 |
| chr19-24590345-DEL-459 | chr19-24590346-DEL-459 |
| chr19-24600047-DEL-159 | chr19-24600046-DEL-158 |
| chr19-24601628-DEL-2948 | chr19-24601629-DEL-2948 |
| chr19-24636282-DEL-145 | chr19-24636283-DEL-145 |
| chr19-24639474-DEL-299 | chr19-24639475-DEL-299 |
| chr19-24640382-DEL-105 | chr19-24640404-DEL-84 |
| chr19-24653234-DEL-60 | chr19-24653223-DEL-56 |
| chr19-24672564-DEL-1782 | chr19-24672565-DEL-1782 |
| chr19-24680809-DEL-95 | chr19-24680833-DEL-70 |
| chr19-24747012-DEL-52 | chr19-24747013-DEL-56 |
| chr19-24752067-DEL-996 | chr19-24752068-DEL-996 |
| chr19-24780242-DEL-72 | chr19-24780243-DEL-69 |
| chr19-24796659-DEL-106 | chr19-24796657-DEL-104 |
| chr19-24798447-DEL-93 | chr19-24798450-DEL-109 |
| chr19-24804387-DEL-6384 | chr19-24804388-DEL-6384 |
| chr19-24814300-DEL-60 | chr19-24814301-DEL-60 |
| chr19-24820423-DEL-52 | chr19-24820424-DEL-52 |
| chr19-24823707-DEL-52 | chr19-24823681-DEL-50 |
| chr19-24852972-DEL-1271 | chr19-24852973-DEL-1271 |
| chr19-24856347-DEL-1412 | chr19-24856347-DEL-1412 |
| chr19-24858144-DEL-146 | chr19-24858148-DEL-118 |
| chr19-24861110-DEL-59 | chr19-24861111-DEL-59 |
| chr19-24885317-DEL-540 | chr19-24885318-DEL-540 |
| chr19-24887693-DEL-184 | chr19-24887694-DEL-184 |
| chr19-24947595-DEL-334 | chr19-24947596-DEL-334 |
| chr19-24996586-DEL-58 | chr19-24996587-DEL-57 |

| | |
|---|---|
| chr19-25007328-DEL-266 | chr19-25007329-DEL-266 |
| chr19-25010476-DEL-484 | chr19-25010477-DEL-484 |
| chr19-25034895-DEL-64 | chr19-25034895-DEL-64 |
| chr19-25045394-DEL-108 | chr19-25045381-DEL-122 |
| chr19-25054739-DEL-53 | chr19-25054740-DEL-53 |
| chr19-25057527-DEL-65 | chr19-25057528-DEL-65 |
| chr19-25061977-DEL-404 | chr19-25061978-DEL-404 |
| chr19-25070078-DEL-67 | chr19-25070079-DEL-67 |
| chr19-25098769-DEL-317 | chr19-25098770-DEL-317 |
| chr19-25102855-DEL-203 | chr19-25102848-DEL-204 |
| chr19-25126276-DEL-130 | chr19-25126262-DEL-132 |
| chr19-25138720-DEL-92 | chr19-25138721-DEL-92 |
| chr19-25153006-DEL-60 | chr19-25153007-DEL-50 |
| chr19-25180870-DEL-197 | chr19-25180871-DEL-197 |
| chr19-25187993-DEL-222 | chr19-25187994-DEL-222 |
| chr19-25205147-DEL-54 | chr19-25205148-DEL-54 |
| chr19-25207128-DEL-66 | chr19-25207128-DEL-66 |
| chr19-25210295-DEL-92 | chr19-25210296-DEL-92 |
| chr19-25239377-DEL-379 | chr19-25239378-DEL-379 |
| chr19-25242191-DEL-195 | chr19-25242192-DEL-195 |
| chr19-25251270-DEL-449 | chr19-25251271-DEL-449 |
| chr19-25260629-DEL-56 | chr19-25260630-DEL-56 |
| chr19-25261520-DEL-59 | chr19-25261525-DEL-57 |
| chr19-25265702-DEL-831 | chr19-25265703-DEL-831 |
| chr19-25272480-DEL-58 | chr19-25272480-DEL-58 |
| chr19-25277033-DEL-115 | chr19-25277034-DEL-115 |
| chr19-25290878-DEL-182 | chr19-25290881-DEL-180 |
| chr19-25292323-DEL-190 | chr19-25292324-DEL-190 |
| chr19-25299225-DEL-104 | chr19-25299242-DEL-77 |
| chr19-25312106-DEL-401 | chr19-25312107-DEL-401 |
| chr19-25323719-DEL-879 | chr19-25323720-DEL-875 |
| chr19-25392004-DEL-63 | chr19-25392005-DEL-57 |
| chr19-25402694-DEL-54 | chr19-25402695-DEL-54 |
| chr19-25458043-DEL-75 | chr19-25458044-DEL-75 |
| chr19-25467447-DEL-55 | chr19-25467452-DEL-73 |
| chr19-25488573-DEL-146 | chr19-25488573-DEL-146 |
| chr19-25518817-DEL-108 | chr19-25518818-DEL-108 |
| chr19-25532597-DEL-193 | chr19-25532598-DEL-193 |
| chr19-25534030-DEL-720 | chr19-25534008-DEL-746 |
| chr19-25545028-DEL-97 | chr19-25545029-DEL-97 |
| chr19-25558588-DEL-204 | chr19-25558589-DEL-203 |
| chr19-25567957-DEL-98 | chr19-25567958-DEL-98 |
| chr19-25574866-DEL-76 | chr19-25574866-DEL-52 |
| chr19-25609969-DEL-160 | chr19-25609948-DEL-165 |
| chr19-25620763-DEL-56 | chr19-25620764-DEL-56 |
| chr19-25646011-DEL-1441 | chr19-25646012-DEL-1441 |
| chr19-25651879-DEL-65 | chr19-25651879-DEL-75 |
| chr19-25656605-DEL-111 | chr19-25656606-DEL-111 |
| chr19-25677886-DEL-70 | chr19-25677887-DEL-70 |
| chr19-25695336-DEL-162 | chr19-25695362-DEL-137 |
| chr19-25720408-DEL-2499 | chr19-25720408-DEL-2499 |
| chr19-25741706-DEL-6249 | chr19-25741698-DEL-6250 |
| chr19-25763021-DEL-256 | chr19-25763024-DEL-254 |
| chr19-25775134-DEL-68 | chr19-25775134-DEL-68 |
| chr19-25789031-DEL-71 | chr19-25789016-DEL-90 |
| chr19-25789680-DEL-55 | chr19-25789681-DEL-55 |
| chr19-25795782-DEL-438 | chr19-25795783-DEL-438 |
| chr19-25796612-DEL-6151 | chr19-25796605-DEL-6152 |
| chr19-25805412-DEL-2262 | chr19-25805413-DEL-2262 |
| chr19-25807993-DEL-481 | chr19-25807994-DEL-481 |
| chr19-25816756-DEL-985 | chr19-25816754-DEL-986 |
| chr19-25821799-DEL-525 | chr19-25821803-DEL-524 |
| chr19-25831624-DEL-389 | chr19-25831609-DEL-389 |
| chr19-25841553-DEL-6525 | chr19-25841551-DEL-6526 |
| chr19-25852538-DEL-64 | chr19-25852537-DEL-66 |
| chr19-25872193-DEL-263 | chr19-25872194-DEL-263 |
| chr19-25888365-DEL-182 | chr19-25888373-DEL-176 |
| chr19-25894749-DEL-2700 | chr19-25894750-DEL-2700 |
| chr19-25909333-DEL-724 | chr19-25909334-DEL-724 |
| chr19-25929442-DEL-82 | chr19-25929443-DEL-82 |
| chr19-25933551-DEL-6720 | chr19-25933552-DEL-6720 |
| chr19-25987424-DEL-73 | chr19-25987425-DEL-73 |
| chr19-26027409-DEL-63 | chr19-26027409-DEL-63 |
| chr19-26031242-DEL-72 | chr19-26031241-DEL-58 |
| chr19-26051668-DEL-834 | chr19-26051654-DEL-827 |
| chr19-26055573-DEL-6530 | chr19-26055574-DEL-6530 |
| chr19-26067549-DEL-103 | chr19-26067550-DEL-103 |
| chr19-26072882-DEL-1085 | chr19-26072883-DEL-1085 |
| chr19-26144681-DEL-58 | chr19-26144684-DEL-52 |
| chr19-26167464-DEL-246 | chr19-26167470-DEL-241 |

| | |
|---|---|
| chr19-26187340-DEL-634 | chr19-26187341-DEL-634 |
| chr19-26194645-DEL-524 | chr19-26194646-DEL-524 |
| chr19-26214772-DEL-6438 | chr19-26214773-DEL-6438 |
| chr19-26247294-DEL-514 | chr19-26247295-DEL-514 |
| chr19-26267654-DEL-386 | chr19-26267655-DEL-386 |
| chr19-26297281-DEL-70 | chr19-26297271-DEL-74 |
| chr19-26317436-DEL-68 | chr19-26317436-DEL-68 |
| chr19-26321291-DEL-167 | chr19-26321293-DEL-164 |
| chr19-26353934-DEL-147 | chr19-26353935-DEL-147 |
| chr19-26357806-DEL-77 | chr19-26357830-DEL-53 |
| chr19-26369915-DEL-3218 | chr19-26369915-DEL-3218 |
| chr19-26389028-DEL-61 | chr19-26389028-DEL-58 |
| chr19-26423677-DEL-429 | chr19-26423678-DEL-429 |
| chr19-26457153-DEL-420 | chr19-26457154-DEL-420 |
| chr19-26486474-DEL-52 | chr19-26486440-DEL-55 |
| chr19-26514257-DEL-90 | chr19-26514258-DEL-90 |
| chr19-26568254-DEL-79 | chr19-26568254-DEL-79 |
| chr19-26579755-DEL-641 | chr19-26579758-DEL-635 |
| chr19-26616250-DEL-60 | chr19-26616255-DEL-88 |
| chr19-26648829-DEL-84 | chr19-26648830-DEL-84 |
| chr19-26669302-DEL-82 | chr19-26669303-DEL-82 |
| chr19-26680371-DEL-51 | chr19-26680372-DEL-51 |
| chr19-26684762-DEL-66 | chr19-26684760-DEL-64 |
| chr19-26691018-DEL-183 | chr19-26691018-DEL-182 |
| chr19-26702510-DEL-1168 | chr19-26702511-DEL-1168 |
| chr19-26707060-DEL-92 | chr19-26707061-DEL-86 |
| chr19-26735339-DEL-86 | chr19-26735340-DEL-80 |
| chr19-26760753-DEL-54 | chr19-26760754-DEL-54 |
| chr19-26774339-DEL-615 | chr19-26774340-DEL-615 |
| chr19-26927738-DEL-114 | chr19-26927733-DEL-120 |
| chr19-26942333-DEL-62 | chr19-26942334-DEL-68 |
| chr19-26970156-DEL-60 | chr19-26970165-DEL-52 |
| chr19-27003980-DEL-191 | chr19-27003991-DEL-190 |
| chr19-27005021-DEL-173 | chr19-27005025-DEL-170 |
| chr19-27009559-DEL-122 | chr19-27009559-DEL-122 |
| chr19-27108796-DEL-2080 | chr19-27108797-DEL-2080 |
| chr19-27178194-DEL-110 | chr19-27178195-DEL-110 |
| chr19-27183431-DEL-396 | chr19-27183432-DEL-396 |
| chr19-27251057-DEL-210 | chr19-27251059-DEL-182 |
| chr19-27257909-DEL-77 | chr19-27257917-DEL-81 |
| chr19-27278069-DEL-66 | chr19-27278070-DEL-66 |
| chr19-27313242-DEL-80 | chr19-27313243-DEL-80 |
| chr19-27331502-DEL-947 | chr19-27331503-DEL-947 |
| chr19-27333294-DEL-63 | chr19-27333295-DEL-63 |
| chr19-27335894-DEL-2070 | chr19-27335895-DEL-2070 |
| chr19-27379388-DEL-6176 | chr19-27379389-DEL-6176 |
| chr19-27403698-DEL-135 | chr19-27403699-DEL-136 |
| chr19-27416058-DEL-54 | chr19-27416057-DEL-62 |
| chr19-27459144-DEL-57 | chr19-27459119-DEL-84 |
| chr19-27470229-DEL-4937 | chr19-27470230-DEL-4937 |
| chr19-27479682-DEL-95 | chr19-27479684-DEL-92 |
| chr19-27501586-DEL-340 | chr19-27501587-DEL-340 |
| chr19-27546389-DEL-98 | chr19-27546390-DEL-98 |
| chr19-27561389-DEL-72 | chr19-27561390-DEL-72 |
| chr19-27588159-DEL-59 | chr19-27588156-DEL-57 |
| chr19-27601646-DEL-51 | chr19-27601646-DEL-51 |
| chr19-27615993-DEL-170 | chr19-27615994-DEL-170 |
| chr19-27625571-DEL-184 | chr19-27625572-DEL-184 |
| chr19-27680230-DEL-331 | chr19-27680231-DEL-331 |
| chr19-27691977-DEL-193 | chr19-27691978-DEL-193 |
| chr19-27728763-DEL-6800 | chr19-27728764-DEL-6800 |
| chr19-27761583-DEL-55 | chr19-27761584-DEL-55 |
| chr19-27763704-DEL-52 | chr19-27763702-DEL-52 |
| chr19-27769222-DEL-84 | chr19-27769241-DEL-60 |
| chr19-27881359-DEL-62 | chr19-27881360-DEL-62 |
| chr19-27997524-DEL-98 | chr19-27997524-DEL-107 |
| chr19-28000727-DEL-3683 | chr19-28000728-DEL-3683 |
| chr19-28004620-DEL-70 | chr19-28004621-DEL-70 |
| chr19-28045352-DEL-180 | chr19-28045353-DEL-180 |
| chr19-28092714-DEL-475 | chr19-28092715-DEL-475 |
| chr19-28101845-DEL-79 | chr19-28101856-DEL-69 |
| chr19-28112581-DEL-54 | chr19-28112582-DEL-54 |
| chr19-28115721-DEL-80 | chr19-28115716-DEL-81 |
| chr19-28126302-DEL-62 | chr19-28126303-DEL-62 |
| chr19-28126864-DEL-6668 | chr19-28126865-DEL-6668 |
| chr19-28171852-DEL-387 | chr19-28171853-DEL-387 |
| chr19-28185489-DEL-2118 | chr19-28185490-DEL-2119 |
| chr19-28299596-DEL-142 | chr19-28299597-DEL-142 |
| chr19-28320443-DEL-53 | chr19-28320452-DEL-64 |
| chr19-28433784-DEL-113 | chr19-28433785-DEL-113 |

| | |
|---|---|
| chr19-28447857-DEL-621 | chr19-28447858-DEL-621 |
| chr19-28456898-DEL-1109 | chr19-28456899-DEL-1109 |
| chr19-28467606-DEL-196 | chr19-28467607-DEL-196 |
| chr19-28487823-DEL-138 | chr19-28487849-DEL-123 |
| chr19-28523834-DEL-81 | chr19-28523794-DEL-105 |
| chr19-28540076-DEL-1349 | chr19-28540087-DEL-1347 |
| chr19-28552524-DEL-680 | chr19-28552525-DEL-680 |
| chr19-28630355-DEL-279 | chr19-28630356-DEL-279 |
| chr19-28662634-DEL-181 | chr19-28662635-DEL-181 |
| chr19-28676742-DEL-207 | chr19-28676729-DEL-207 |
| chr19-28725133-DEL-197 | chr19-28725135-DEL-179 |
| chr19-28727695-DEL-92 | chr19-28727696-DEL-93 |
| chr19-28731464-DEL-1215 | chr19-28731465-DEL-1215 |
| chr19-28739120-DEL-70 | chr19-28739121-DEL-70 |
| chr19-28740997-DEL-327 | chr19-28740998-DEL-327 |
| chr19-28751455-DEL-80 | chr19-28751456-DEL-76 |
| chr19-28756932-DEL-97 | chr19-28756933-DEL-96 |
| chr19-28766084-DEL-196 | chr19-28766076-DEL-197 |
| chr19-28778802-DEL-93 | chr19-28778801-DEL-62 |
| chr19-28787007-DEL-378 | chr19-28787006-DEL-376 |
| chr19-28794713-DEL-350 | chr19-28794714-DEL-350 |
| chr19-28824854-DEL-132 | chr19-28824855-DEL-132 |
| chr19-28825994-DEL-184 | chr19-28825995-DEL-184 |
| chr19-28827510-DEL-220 | chr19-28827511-DEL-220 |
| chr19-28834593-DEL-2292 | chr19-28834594-DEL-2292 |
| chr19-28866589-DEL-138 | chr19-28866589-DEL-139 |
| chr19-28907343-DEL-336 | chr19-28907357-DEL-332 |
| chr19-28967200-DEL-91 | chr19-28967201-DEL-91 |
| chr19-29021294-DEL-216 | chr19-29021298-DEL-216 |
| chr19-29043043-DEL-709 | chr19-29043044-DEL-709 |
| chr19-29052676-DEL-486 | chr19-29052677-DEL-486 |
| chr19-29058842-DEL-5206 | chr19-29058843-DEL-5206 |
| chr19-29073191-DEL-160 | chr19-29073191-DEL-161 |
| chr19-29082699-DEL-2740 | chr19-29082700-DEL-2740 |
| chr19-29092875-DEL-90 | chr19-29092831-DEL-135 |
| chr19-29137379-DEL-1302 | chr19-29137380-DEL-1302 |
| chr19-29139876-DEL-770 | chr19-29139877-DEL-770 |
| chr19-29166203-DEL-137 | chr19-29166204-DEL-137 |
| chr19-29194153-DEL-76 | chr19-29194154-DEL-68 |
| chr19-29194604-DEL-201 | chr19-29194605-DEL-201 |
| chr19-29196081-DEL-72 | chr19-29196086-DEL-52 |
| chr19-29203036-DEL-504 | chr19-29203037-DEL-504 |
| chr19-29235318-DEL-333 | chr19-29235319-DEL-333 |
| chr19-29242313-DEL-55 | chr19-29242314-DEL-54 |
| chr19-29279707-DEL-63 | chr19-29279708-DEL-63 |
| chr19-29300677-DEL-66 | chr19-29300679-DEL-71 |
| chr19-29302991-DEL-92 | chr19-29302992-DEL-92 |
| chr19-29316478-DEL-72 | chr19-29316477-DEL-73 |
| chr19-29322440-DEL-63 | chr19-29322441-DEL-63 |
| chr19-29419715-DEL-89 | chr19-29419716-DEL-89 |
| chr19-29428509-DEL-2433 | chr19-29428510-DEL-2433 |
| chr19-29443720-DEL-77 | chr19-29443721-DEL-77 |
| chr19-29451420-DEL-66 | chr19-29451421-DEL-66 |
| chr19-29454942-DEL-224 | chr19-29454943-DEL-224 |
| chr19-29502804-DEL-409 | chr19-29502805-DEL-409 |
| chr19-29565885-DEL-401 | chr19-29565886-DEL-401 |
| chr19-29571593-DEL-80 | chr19-29571588-DEL-81 |
| chr19-29596675-DEL-156 | chr19-29596666-DEL-157 |
| chr19-29603642-DEL-181 | chr19-29603643-DEL-152 |
| chr19-29606523-DEL-84 | chr19-29606524-DEL-84 |
| chr19-29629897-DEL-207 | chr19-29629898-DEL-207 |
| chr19-29637938-DEL-226 | chr19-29637939-DEL-226 |
| chr19-29648946-DEL-277 | chr19-29648948-DEL-276 |
| chr19-29658150-DEL-118 | chr19-29658151-DEL-118 |
| chr19-29659012-DEL-176 | chr19-29659029-DEL-138 |
| chr19-29663011-DEL-8750 | chr19-29663012-DEL-8750 |
| chr19-29717130-DEL-67 | chr19-29717131-DEL-67 |
| chr19-29792578-DEL-126 | chr19-29792584-DEL-111 |
| chr19-29803562-DEL-58 | chr19-29803549-DEL-74 |
| chr19-29862522-DEL-59 | chr19-29862524-DEL-58 |
| chr19-29896079-DEL-55 | chr19-29896080-DEL-55 |
| chr19-29911918-DEL-83 | chr19-29911919-DEL-83 |
| chr19-29912587-DEL-65 | chr19-29912588-DEL-65 |
| chr19-29933994-DEL-202 | chr19-29933995-DEL-202 |
| chr19-29961400-DEL-106 | chr19-29961401-DEL-106 |
| chr19-29968092-DEL-6477 | chr19-29968093-DEL-6477 |
| chr19-29987086-DEL-78 | chr19-29987087-DEL-78 |
| chr19-29991724-DEL-67 | chr19-29991730-DEL-87 |
| chr19-30005898-DEL-211 | chr19-30005899-DEL-211 |
| chr19-30051955-DEL-197 | chr19-30051956-DEL-197 |

| | |
|---|---|
| chr19-30080803-DEL-52 | chr19-30080804-DEL-52 |
| chr19-30081984-DEL-52 | chr19-30081985-DEL-52 |
| chr19-30103726-DEL-54 | chr19-30103731-DEL-50 |
| chr19-30108202-DEL-58 | chr19-30108203-DEL-58 |
| chr19-30119035-DEL-203 | chr19-30119036-DEL-203 |
| chr19-30130230-DEL-183 | chr19-30130224-DEL-199 |
| chr19-30136734-DEL-101 | chr19-30136735-DEL-101 |
| chr19-30158160-DEL-58 | chr19-30158161-DEL-58 |
| chr19-30190524-DEL-780 | chr19-30190525-DEL-780 |
| chr19-30193232-DEL-75 | chr19-30193233-DEL-75 |
| chr19-30203458-DEL-263 | chr19-30203459-DEL-263 |
| chr19-30208159-DEL-103 | chr19-30208160-DEL-101 |
| chr19-30208703-DEL-101 | chr19-30208693-DEL-102 |
| chr19-30219590-DEL-64 | chr19-30219591-DEL-76 |
| chr19-30230189-DEL-732 | chr19-30230190-DEL-732 |
| chr19-30238537-DEL-163 | chr19-30238544-DEL-147 |
| chr19-30240154-DEL-60 | chr19-30240155-DEL-60 |
| chr19-30241251-DEL-847 | chr19-30241252-DEL-847 |
| chr19-30247209-DEL-117 | chr19-30247210-DEL-117 |
| chr19-30248904-DEL-327 | chr19-30248907-DEL-328 |
| chr19-30257282-DEL-79 | chr19-30257283-DEL-79 |
| chr19-30271389-DEL-308 | chr19-30271390-DEL-307 |
| chr19-30273895-DEL-113 | chr19-30273896-DEL-113 |
| chr19-30276331-DEL-98 | chr19-30276332-DEL-98 |
| chr19-30295576-DEL-235 | chr19-30295577-DEL-235 |
| chr19-30298396-DEL-184 | chr19-30298397-DEL-184 |
| chr19-30316307-DEL-54 | chr19-30316307-DEL-54 |
| chr19-30318373-DEL-55 | chr19-30318372-DEL-53 |
| chr19-30319347-DEL-181 | chr19-30319337-DEL-182 |
| chr19-30377976-DEL-1746 | chr19-30377976-DEL-1746 |
| chr19-30394225-DEL-116 | chr19-30394226-DEL-110 |
| chr19-30459904-DEL-67 | chr19-30459905-DEL-63 |
| chr19-30481679-DEL-358 | chr19-30481680-DEL-358 |
| chr19-30520705-DEL-56 | chr19-30520706-DEL-56 |
| chr19-30543978-DEL-116 | chr19-30543983-DEL-112 |
| chr19-30571609-DEL-67 | chr19-30571623-DEL-73 |
| chr19-30574842-DEL-203 | chr19-30574841-DEL-204 |
| chr19-30582724-DEL-6394 | chr19-30582725-DEL-6394 |
| chr19-30605862-DEL-1635 | chr19-30605863-DEL-1635 |
| chr19-30622877-DEL-130 | chr19-30622878-DEL-130 |
| chr19-30632216-DEL-7440 | chr19-30632217-DEL-7440 |
| chr19-30638508-DEL-424 | chr19-30638509-DEL-424 |
| chr19-30648033-DEL-384 | chr19-30648034-DEL-384 |
| chr19-30660835-DEL-1908 | chr19-30660836-DEL-1908 |
| chr19-30671821-DEL-1592 | chr19-30671822-DEL-1592 |
| chr19-30677573-DEL-376 | chr19-30677557-DEL-377 |
| chr19-30730051-DEL-54 | chr19-30730052-DEL-54 |
| chr19-30734640-DEL-6491 | chr19-30734641-DEL-6491 |
| chr19-30752060-DEL-62 | chr19-30752061-DEL-62 |
| chr19-30799035-DEL-158 | chr19-30799036-DEL-158 |
| chr19-30803879-DEL-65 | chr19-30803880-DEL-65 |
| chr19-30828883-DEL-447 | chr19-30828883-DEL-447 |
| chr19-30914194-DEL-508 | chr19-30914195-DEL-508 |
| chr19-30941039-DEL-4186 | chr19-30941040-DEL-4186 |
| chr19-30966647-DEL-205 | chr19-30966648-DEL-205 |
| chr19-30975485-DEL-171 | chr19-30975486-DEL-171 |
| chr19-31033259-DEL-395 | chr19-31033260-DEL-395 |
| chr19-31110756-DEL-219 | chr19-31110757-DEL-219 |
| chr19-31124728-DEL-58 | chr19-31124729-DEL-58 |
| chr19-31141896-DEL-68 | chr19-31141897-DEL-68 |
| chr19-31161097-DEL-56 | chr19-31161098-DEL-54 |
| chr19-31165792-DEL-78 | chr19-31165796-DEL-74 |
| chr19-31211424-DEL-3940 | chr19-31211422-DEL-3941 |
| chr19-31254442-DEL-64 | chr19-31254445-DEL-62 |
| chr19-31268258-DEL-180 | chr19-31268259-DEL-180 |
| chr19-31325429-DEL-98 | chr19-31325430-DEL-98 |
| chr19-31336681-DEL-67 | chr19-31336672-DEL-77 |
| chr19-31406710-DEL-476 | chr19-31406711-DEL-476 |
| chr19-31414031-DEL-1819 | chr19-31414032-DEL-1819 |
| chr19-31463766-DEL-55 | chr19-31463767-DEL-55 |
| chr19-31480146-DEL-86 | chr19-31480143-DEL-90 |
| chr19-31506902-DEL-78 | chr19-31506898-DEL-69 |
| chr19-31509896-DEL-180 | chr19-31509897-DEL-180 |
| chr19-31510438-DEL-82 | chr19-31510439-DEL-82 |
| chr19-31511559-DEL-154 | chr19-31511560-DEL-154 |
| chr19-31518695-DEL-70 | chr19-31518698-DEL-72 |
| chr19-31526325-DEL-88 | chr19-31526326-DEL-88 |
| chr19-31530722-DEL-78 | chr19-31530723-DEL-78 |
| chr19-31615077-DEL-1030 | chr19-31615078-DEL-1030 |
| chr19-31685009-DEL-126 | chr19-31685003-DEL-85 |

| | |
|---|---|
| chr19-31685499-DEL-200 | chr19-31685500-DEL-200 |
| chr19-31700148-DEL-7374 | chr19-31700149-DEL-7374 |
| chr19-31712005-DEL-1218 | chr19-31712006-DEL-1218 |
| chr19-31761246-DEL-70 | chr19-31761247-DEL-70 |
| chr19-31802848-DEL-63 | chr19-31802842-DEL-63 |
| chr19-31806700-DEL-116 | chr19-31806701-DEL-116 |
| chr19-31813291-DEL-472 | chr19-31813292-DEL-472 |
| chr19-31815019-DEL-4323 | chr19-31815020-DEL-4323 |
| chr19-31820252-DEL-489 | chr19-31820253-DEL-489 |
| chr19-31826577-DEL-486 | chr19-31826578-DEL-486 |
| chr19-31833537-DEL-4674 | chr19-31833537-DEL-4674 |
| chr19-31843797-DEL-139 | chr19-31843798-DEL-139 |
| chr19-31849504-DEL-78 | chr19-31849505-DEL-78 |
| chr19-31854056-DEL-51 | chr19-31854057-DEL-51 |
| chr19-31881561-DEL-780 | chr19-31881555-DEL-781 |
| chr19-31902808-DEL-540 | chr19-31902809-DEL-541 |
| chr19-31914534-DEL-6675 | chr19-31914535-DEL-6675 |
| chr19-31921739-DEL-92 | chr19-31921723-DEL-74 |
| chr19-31966820-DEL-132 | chr19-31966821-DEL-132 |
| chr19-31974824-DEL-530 | chr19-31974824-DEL-530 |
| chr19-31983252-DEL-76 | chr19-31983223-DEL-108 |
| chr19-32000035-DEL-77 | chr19-32000036-DEL-77 |
| chr19-32010777-DEL-6372 | chr19-32010778-DEL-6372 |
| chr19-32062884-DEL-1728 | chr19-32062885-DEL-1728 |
| chr19-32066948-DEL-96 | chr19-32066909-DEL-96 |
| chr19-32138671-DEL-181 | chr19-32138672-DEL-181 |
| chr19-32147800-DEL-683 | chr19-32147801-DEL-683 |
| chr19-32170445-DEL-297 | chr19-32170438-DEL-270 |
| chr19-32180571-DEL-6400 | chr19-32180572-DEL-6400 |
| chr19-32189884-DEL-99 | chr19-32189895-DEL-109 |
| chr19-32191727-DEL-122 | chr19-32191728-DEL-116 |
| chr19-32205378-DEL-1206 | chr19-32205379-DEL-1206 |
| chr19-32287102-DEL-246 | chr19-32287107-DEL-242 |
| chr19-32354235-DEL-204 | chr19-32354236-DEL-204 |
| chr19-32378686-DEL-1341 | chr19-32378664-DEL-1371 |
| chr19-32409693-DEL-95 | chr19-32409695-DEL-94 |
| chr19-32410698-DEL-86 | chr19-32410699-DEL-86 |
| chr19-32416320-DEL-204 | chr19-32416321-DEL-204 |
| chr19-32419257-DEL-7103 | chr19-32419258-DEL-7103 |
| chr19-32434928-DEL-482 | chr19-32434929-DEL-482 |
| chr19-32439588-DEL-357 | chr19-32439589-DEL-357 |
| chr19-32443254-DEL-722 | chr19-32443255-DEL-722 |
| chr19-32465835-DEL-1969 | chr19-32465836-DEL-1969 |
| chr19-32467228-DEL-575 | chr19-32467229-DEL-575 |
| chr19-32504952-DEL-53 | chr19-32504953-DEL-53 |
| chr19-32505698-DEL-354 | chr19-32505699-DEL-354 |
| chr19-32536870-DEL-58 | chr19-32536871-DEL-58 |
| chr19-32553331-DEL-504 | chr19-32553332-DEL-504 |
| chr19-32555751-DEL-78 | chr19-32555764-DEL-95 |
| chr19-32562420-DEL-203 | chr19-32562421-DEL-203 |
| chr19-32640748-DEL-162 | chr19-32640750-DEL-161 |
| chr19-32666614-DEL-209 | chr19-32666615-DEL-209 |
| chr19-32707677-DEL-164 | chr19-32707654-DEL-161 |
| chr19-32708518-DEL-54 | chr19-32708519-DEL-54 |
| chr19-32715821-DEL-214 | chr19-32715822-DEL-214 |
| chr19-32720660-DEL-199 | chr19-32720661-DEL-199 |
| chr19-32727403-DEL-204 | chr19-32727404-DEL-204 |
| chr19-32728497-DEL-70 | chr19-32728515-DEL-52 |
| chr19-32744160-DEL-129 | chr19-32744157-DEL-129 |
| chr19-32765540-DEL-155 | chr19-32765541-DEL-155 |
| chr19-32807050-DEL-89 | chr19-32807051-DEL-89 |
| chr19-32827732-DEL-170 | chr19-32827738-DEL-163 |
| chr19-32828646-DEL-2875 | chr19-32828647-DEL-2875 |
| chr19-32832478-DEL-1139 | chr19-32832479-DEL-1139 |
| chr19-32845413-DEL-240 | chr19-32845414-DEL-240 |
| chr19-32853923-DEL-2919 | chr19-32853924-DEL-2919 |
| chr19-32877269-DEL-129 | chr19-32877272-DEL-95 |
| chr19-32962139-DEL-209 | chr19-32962140-DEL-209 |
| chr19-32967430-DEL-72 | chr19-32967431-DEL-72 |
| chr19-32980414-DEL-63 | chr19-32980393-DEL-109 |
| chr19-33009920-DEL-103 | chr19-33009921-DEL-103 |
| chr19-33044830-DEL-198 | chr19-33044831-DEL-198 |
| chr19-33050850-DEL-87 | chr19-33050857-DEL-90 |
| chr19-33055315-DEL-320 | chr19-33055316-DEL-320 |
| chr19-33062655-DEL-506 | chr19-33062656-DEL-506 |
| chr19-33108611-DEL-80 | chr19-33108612-DEL-80 |
| chr19-33157699-DEL-205 | chr19-33157700-DEL-205 |
| chr19-33164900-DEL-54 | chr19-33164901-DEL-54 |
| chr19-33195811-DEL-712 | chr19-33195812-DEL-712 |
| chr19-33196523-DEL-236 | chr19-33196523-DEL-237 |

| | |
|---|---|
| chr19-33225346-DEL-265 | chr19-33225347-DEL-265 |
| chr19-33263672-DEL-6469 | chr19-33263673-DEL-6469 |
| chr19-33292197-DEL-68 | chr19-33292191-DEL-84 |
| chr19-33303570-DEL-92 | chr19-33303571-DEL-92 |
| chr19-33317916-DEL-66 | chr19-33317917-DEL-66 |
| chr19-33324358-DEL-695 | chr19-33324359-DEL-695 |
| chr19-33350627-DEL-555 | chr19-33350628-DEL-555 |
| chr19-33403830-DEL-348 | chr19-33403831-DEL-348 |
| chr19-33404278-DEL-820 | chr19-33404279-DEL-820 |
| chr19-33405696-DEL-440 | chr19-33405697-DEL-440 |
| chr19-33455687-DEL-86 | chr19-33455688-DEL-86 |
| chr19-33479833-DEL-341 | chr19-33479834-DEL-341 |
| chr19-33572239-DEL-6943 | chr19-33572240-DEL-6943 |
| chr19-33591932-DEL-270 | chr19-33591933-DEL-270 |
| chr19-33656884-DEL-108 | chr19-33656885-DEL-156 |
| chr19-33707417-DEL-214 | chr19-33707410-DEL-214 |
| chr19-33780145-DEL-116 | chr19-33780147-DEL-86 |
| chr19-33782999-DEL-1592 | chr19-33782999-DEL-1592 |
| chr19-33791863-DEL-81 | chr19-33791864-DEL-81 |
| chr19-33827417-DEL-1347 | chr19-33827416-DEL-1345 |
| chr19-33834734-DEL-78 | chr19-33834735-DEL-78 |
| chr19-33852962-DEL-240 | chr19-33852963-DEL-240 |
| chr19-33856211-DEL-1347 | chr19-33856212-DEL-1347 |
| chr19-33858920-DEL-54 | chr19-33858921-DEL-54 |
| chr19-33946636-DEL-104 | chr19-33946641-DEL-96 |
| chr19-33992010-DEL-108 | chr19-33992050-DEL-80 |
| chr19-34010106-DEL-153 | chr19-34010107-DEL-153 |
| chr19-34063685-DEL-62 | chr19-34063666-DEL-70 |
| chr19-34068985-DEL-185 | chr19-34068986-DEL-185 |
| chr19-34082681-DEL-3608 | chr19-34082682-DEL-3608 |
| chr19-34090523-DEL-600 | chr19-34090524-DEL-600 |
| chr19-34108547-DEL-7110 | chr19-34108548-DEL-7110 |
| chr19-34132130-DEL-6440 | chr19-34132131-DEL-6440 |
| chr19-34142353-DEL-76 | chr19-34142339-DEL-87 |
| chr19-34183768-DEL-222 | chr19-34183771-DEL-220 |
| chr19-34200617-DEL-773 | chr19-34200618-DEL-773 |
| chr19-34276522-DEL-68 | chr19-34276522-DEL-68 |
| chr19-34343796-DEL-346 | chr19-34343800-DEL-347 |
| chr19-34345307-DEL-65 | chr19-34345288-DEL-66 |
| chr19-34356944-DEL-1039 | chr19-34356945-DEL-1039 |
| chr19-34362025-DEL-67 | chr19-34362017-DEL-68 |
| chr19-34369637-DEL-113 | chr19-34369638-DEL-113 |
| chr19-34396219-DEL-167 | chr19-34396219-DEL-166 |
| chr19-34434724-DEL-60 | chr19-34434725-DEL-62 |
| chr19-34442439-DEL-535 | chr19-34442440-DEL-535 |
| chr19-34518177-DEL-67 | chr19-34518192-DEL-54 |
| chr19-34535284-DEL-852 | chr19-34535285-DEL-852 |
| chr19-34609690-DEL-220 | chr19-34609691-DEL-220 |
| chr19-34615535-DEL-88 | chr19-34615537-DEL-87 |
| chr19-34653207-DEL-54 | chr19-34653199-DEL-82 |
| chr19-34739681-DEL-223 | chr19-34739682-DEL-225 |
| chr19-34757108-DEL-1723 | chr19-34757109-DEL-1723 |
| chr19-34760480-DEL-200 | chr19-34760481-DEL-200 |
| chr19-34775846-DEL-561 | chr19-34775846-DEL-561 |
| chr19-34781206-DEL-192 | chr19-34781207-DEL-192 |
| chr19-34791289-DEL-88 | chr19-34791290-DEL-88 |
| chr19-34795151-DEL-51 | chr19-34795148-DEL-55 |
| chr19-34800529-DEL-192 | chr19-34800530-DEL-192 |
| chr19-34842096-DEL-1793 | chr19-34842097-DEL-1793 |
| chr19-34852847-DEL-657 | chr19-34852846-DEL-655 |
| chr19-34860839-DEL-76 | chr19-34860867-DEL-50 |
| chr19-34873662-DEL-162 | chr19-34873663-DEL-162 |
| chr19-34879716-DEL-199 | chr19-34879714-DEL-199 |
| chr19-34888909-DEL-232 | chr19-34888910-DEL-232 |
| chr19-34892191-DEL-4078 | chr19-34892192-DEL-4078 |
| chr19-34900652-DEL-198 | chr19-34900653-DEL-198 |
| chr19-34903338-DEL-76 | chr19-34903339-DEL-73 |
| chr19-34924407-DEL-205 | chr19-34924408-DEL-205 |
| chr19-34946278-DEL-103 | chr19-34946279-DEL-103 |
| chr19-34956373-DEL-271 | chr19-34956374-DEL-270 |
| chr19-34959347-DEL-212 | chr19-34959348-DEL-212 |
| chr19-35007486-DEL-183 | chr19-35007493-DEL-155 |
| chr19-35035368-DEL-272 | chr19-35035369-DEL-272 |
| chr19-35053749-DEL-3462 | chr19-35053750-DEL-3462 |
| chr19-35069439-DEL-430 | chr19-35069440-DEL-430 |
| chr19-35087819-DEL-75 | chr19-35087793-DEL-51 |
| chr19-35105087-DEL-192 | chr19-35105088-DEL-192 |
| chr19-35107404-DEL-58 | chr19-35107404-DEL-59 |
| chr19-35134020-DEL-388 | chr19-35134021-DEL-388 |
| chr19-35157058-DEL-448 | chr19-35157059-DEL-447 |

| | |
|---|---|
| chr19-35166607-DEL-57 | chr19-35166608-DEL-57 |
| chr19-35178183-DEL-83 | chr19-35178184-DEL-67 |
| chr19-35185543-DEL-541 | chr19-35185544-DEL-541 |
| chr19-35187052-DEL-927 | chr19-35187053-DEL-927 |
| chr19-35188803-DEL-188 | chr19-35188802-DEL-189 |
| chr19-35199928-DEL-3239 | chr19-35199929-DEL-3239 |
| chr19-35211851-DEL-6517 | chr19-35211852-DEL-6517 |
| chr19-35212056-DEL-214 | chr19-35212057-DEL-214 |
| chr19-35222170-DEL-207 | chr19-35222171-DEL-208 |
| chr19-35249534-DEL-228 | chr19-35249535-DEL-230 |
| chr19-35251171-DEL-521 | chr19-35251170-DEL-518 |
| chr19-35253774-DEL-145 | chr19-35253785-DEL-100 |
| chr19-35271409-DEL-52 | chr19-35271400-DEL-50 |
| chr19-35278877-DEL-741 | chr19-35278878-DEL-741 |
| chr19-35285264-DEL-159 | chr19-35285265-DEL-159 |
| chr19-35320490-DEL-170 | chr19-35320491-DEL-170 |
| chr19-35386113-DEL-344 | chr19-35386115-DEL-343 |
| chr19-35399306-DEL-439 | chr19-35399307-DEL-439 |
| chr19-35404030-DEL-436 | chr19-35404031-DEL-436 |
| chr19-35421527-DEL-1361 | chr19-35421528-DEL-1361 |
| chr19-35443727-DEL-80 | chr19-35443728-DEL-80 |
| chr19-35445282-DEL-127 | chr19-35445283-DEL-127 |
| chr19-35446731-DEL-897 | chr19-35446719-DEL-900 |
| chr19-35452715-DEL-259 | chr19-35452716-DEL-259 |
| chr19-35493161-DEL-4795 | chr19-35493162-DEL-4795 |
| chr19-35534361-DEL-97 | chr19-35534343-DEL-89 |
| chr19-35536928-DEL-217 | chr19-35536929-DEL-217 |
| chr19-35537875-DEL-63 | chr19-35537876-DEL-63 |
| chr19-35588832-DEL-55 | chr19-35588823-DEL-65 |
| chr19-35590564-DEL-56 | chr19-35590570-DEL-63 |
| chr19-35603652-DEL-69 | chr19-35603665-DEL-50 |
| chr19-35605860-DEL-195 | chr19-35605861-DEL-195 |
| chr19-35627578-DEL-6813 | chr19-35627579-DEL-6813 |
| chr19-35657705-DEL-114 | chr19-35657706-DEL-114 |
| chr19-35668650-DEL-337 | chr19-35668651-DEL-338 |
| chr19-35675182-DEL-499 | chr19-35675183-DEL-499 |
| chr19-35685035-DEL-84 | chr19-35685036-DEL-70 |
| chr19-35727831-DEL-179 | chr19-35727832-DEL-179 |
| chr19-35740360-DEL-1938 | chr19-35740361-DEL-1938 |
| chr19-35783666-DEL-209 | chr19-35783667-DEL-209 |
| chr19-35785319-DEL-96 | chr19-35785320-DEL-96 |
| chr19-35788792-DEL-54 | chr19-35788793-DEL-54 |
| chr19-35802213-DEL-223 | chr19-35802214-DEL-223 |
| chr19-35810281-DEL-294 | chr19-35810282-DEL-294 |
| chr19-35815433-DEL-99 | chr19-35815434-DEL-99 |
| chr19-35819535-DEL-3579 | chr19-35819536-DEL-3579 |
| chr19-35821788-DEL-166 | chr19-35821789-DEL-166 |
| chr19-35831169-DEL-195 | chr19-35831177-DEL-195 |
| chr19-35852377-DEL-82 | chr19-35852389-DEL-82 |
| chr19-35881388-DEL-7223 | chr19-35881389-DEL-7223 |
| chr19-35896076-DEL-224 | chr19-35896110-DEL-200 |
| chr19-35910792-DEL-951 | chr19-35910793-DEL-951 |
| chr19-35920909-DEL-431 | chr19-35920910-DEL-431 |
| chr19-35923549-DEL-507 | chr19-35923550-DEL-507 |
| chr19-35926717-DEL-203 | chr19-35926718-DEL-203 |
| chr19-35977582-DEL-55 | chr19-35977583-DEL-55 |
| chr19-36035486-DEL-60 | chr19-36035487-DEL-56 |
| chr19-36083931-DEL-235 | chr19-36083932-DEL-235 |
| chr19-36091391-DEL-55 | chr19-36091392-DEL-52 |
| chr19-36112189-DEL-199 | chr19-36112190-DEL-199 |
| chr19-36116469-DEL-266 | chr19-36116470-DEL-266 |
| chr19-36132356-DEL-326 | chr19-36132357-DEL-326 |
| chr19-36135161-DEL-87 | chr19-36135168-DEL-112 |
| chr19-36146195-DEL-56 | chr19-36146196-DEL-56 |
| chr19-36175283-DEL-56 | chr19-36175284-DEL-56 |
| chr19-36302955-DEL-103 | chr19-36302957-DEL-104 |
| chr19-36311114-DEL-1519 | chr19-36311098-DEL-1520 |
| chr19-36332458-DEL-185 | chr19-36332459-DEL-185 |
| chr19-36334693-DEL-78 | chr19-36334694-DEL-78 |
| chr19-36360139-DEL-60 | chr19-36360140-DEL-59 |
| chr19-36363341-DEL-221 | chr19-36363342-DEL-221 |
| chr19-36374364-DEL-78 | chr19-36374366-DEL-75 |
| chr19-36377346-DEL-96 | chr19-36377347-DEL-96 |
| chr19-36394186-DEL-205 | chr19-36394187-DEL-205 |
| chr19-36446766-DEL-206 | chr19-36446767-DEL-206 |
| chr19-36451484-DEL-544 | chr19-36451483-DEL-541 |
| chr19-36456511-DEL-198 | chr19-36456512-DEL-198 |
| chr19-36468979-DEL-4700 | chr19-36468982-DEL-4698 |
| chr19-36519022-DEL-97 | chr19-36519027-DEL-121 |
| chr19-36550441-DEL-64 | chr19-36550442-DEL-64 |

| | |
|---|---|
| chr19-36573060-DEL-249 | chr19-36573061-DEL-249 |
| chr19-36573722-DEL-75 | chr19-36573723-DEL-75 |
| chr19-36625171-DEL-650 | chr19-36625172-DEL-650 |
| chr19-36627567-DEL-905 | chr19-36627568-DEL-905 |
| chr19-36635226-DEL-181 | chr19-36635227-DEL-181 |
| chr19-36641120-DEL-64 | chr19-36641121-DEL-64 |
| chr19-36737120-DEL-66 | chr19-36737121-DEL-66 |
| chr19-36739940-DEL-794 | chr19-36739941-DEL-794 |
| chr19-36740825-DEL-423 | chr19-36740823-DEL-427 |
| chr19-36754699-DEL-610 | chr19-36754700-DEL-610 |
| chr19-36766114-DEL-86 | chr19-36766115-DEL-86 |
| chr19-36780332-DEL-211 | chr19-36780333-DEL-211 |
| chr19-36797315-DEL-75 | chr19-36797316-DEL-75 |
| chr19-36841792-DEL-215 | chr19-36841793-DEL-215 |
| chr19-36864624-DEL-203 | chr19-36864625-DEL-203 |
| chr19-36887015-DEL-59 | chr19-36887011-DEL-59 |
| chr19-36898516-DEL-207 | chr19-36898517-DEL-207 |
| chr19-36952805-DEL-204 | chr19-36952806-DEL-204 |
| chr19-36979685-DEL-54 | chr19-36979685-DEL-54 |
| chr19-36991647-DEL-55 | chr19-36991648-DEL-55 |
| chr19-37009603-DEL-240 | chr19-37009604-DEL-240 |
| chr19-37035396-DEL-407 | chr19-37035397-DEL-408 |
| chr19-37083454-DEL-68 | chr19-37083455-DEL-68 |
| chr19-37092424-DEL-178 | chr19-37092397-DEL-166 |
| chr19-37095219-DEL-64 | chr19-37095220-DEL-64 |
| chr19-37097914-DEL-351 | chr19-37097915-DEL-351 |
| chr19-37102973-DEL-72 | chr19-37102974-DEL-72 |
| chr19-37112117-DEL-164 | chr19-37112122-DEL-160 |
| chr19-37171470-DEL-52 | chr19-37171470-DEL-50 |
| chr19-37190393-DEL-181 | chr19-37190394-DEL-181 |
| chr19-37206715-DEL-92 | chr19-37206716-DEL-104 |
| chr19-37234926-DEL-81 | chr19-37234927-DEL-81 |
| chr19-37242621-DEL-199 | chr19-37242622-DEL-199 |
| chr19-37272463-DEL-771 | chr19-37272464-DEL-771 |
| chr19-37309170-DEL-898 | chr19-37309170-DEL-898 |
| chr19-37335358-DEL-192 | chr19-37335359-DEL-192 |
| chr19-37336927-DEL-61 | chr19-37336930-DEL-59 |
| chr19-37341677-DEL-204 | chr19-37341675-DEL-207 |
| chr19-37382687-DEL-192 | chr19-37382688-DEL-192 |
| chr19-37383675-DEL-1007 | chr19-37383679-DEL-1008 |
| chr19-37403879-DEL-98 | chr19-37403880-DEL-98 |
| chr19-37407881-DEL-99 | chr19-37407878-DEL-95 |
| chr19-37420915-DEL-163 | chr19-37420916-DEL-163 |
| chr19-37434489-DEL-83 | chr19-37434503-DEL-78 |
| chr19-37443312-DEL-77 | chr19-37443313-DEL-77 |
| chr19-37466764-DEL-178 | chr19-37466764-DEL-178 |
| chr19-37469608-DEL-182 | chr19-37469609-DEL-182 |
| chr19-37513739-DEL-200 | chr19-37513747-DEL-192 |
| chr19-37545968-DEL-86 | chr19-37545924-DEL-85 |
| chr19-37553781-DEL-54 | chr19-37553785-DEL-50 |
| chr19-37557249-DEL-208 | chr19-37557250-DEL-208 |
| chr19-37575149-DEL-169 | chr19-37575150-DEL-169 |
| chr19-37578487-DEL-62 | chr19-37578486-DEL-60 |
| chr19-37583292-DEL-446 | chr19-37583293-DEL-446 |
| chr19-37584876-DEL-73 | chr19-37584877-DEL-73 |
| chr19-37592201-DEL-56 | chr19-37592202-DEL-57 |
| chr19-37605877-DEL-7056 | chr19-37605878-DEL-7056 |
| chr19-37623894-DEL-87 | chr19-37623917-DEL-82 |
| chr19-37632824-DEL-161 | chr19-37632825-DEL-161 |
| chr19-37636953-DEL-77 | chr19-37636954-DEL-77 |
| chr19-37651541-DEL-147 | chr19-37651542-DEL-147 |
| chr19-37724538-DEL-106 | chr19-37724539-DEL-106 |
| chr19-37726415-DEL-58 | chr19-37726414-DEL-58 |
| chr19-37735556-DEL-224 | chr19-37735543-DEL-225 |
| chr19-37774482-DEL-102 | chr19-37774483-DEL-102 |
| chr19-37787200-DEL-441 | chr19-37787201-DEL-441 |
| chr19-37803709-DEL-182 | chr19-37803710-DEL-206 |
| chr19-37832313-DEL-585 | chr19-37832314-DEL-585 |
| chr19-37861338-DEL-77 | chr19-37861325-DEL-91 |
| chr19-37874763-DEL-69 | chr19-37874731-DEL-101 |
| chr19-37884267-DEL-64 | chr19-37884267-DEL-64 |
| chr19-37926552-DEL-199 | chr19-37926553-DEL-199 |
| chr19-37953497-DEL-91 | chr19-37953498-DEL-91 |
| chr19-37972084-DEL-74 | chr19-37972085-DEL-74 |
| chr19-37991650-DEL-70 | chr19-37991653-DEL-56 |
| chr19-37999356-DEL-72 | chr19-37999357-DEL-72 |
| chr19-38007379-DEL-400 | chr19-38007380-DEL-400 |
| chr19-38009016-DEL-652 | chr19-38009017-DEL-652 |
| chr19-38010289-DEL-101 | chr19-38010290-DEL-101 |
| chr19-38012452-DEL-74 | chr19-38012488-DEL-84 |

| | |
|---|---|
| chr19-38036576-DEL-213 | chr19-38036577-DEL-213 |
| chr19-38055045-DEL-237 | chr19-38055046-DEL-237 |
| chr19-38067409-DEL-1347 | chr19-38067410-DEL-1347 |
| chr19-38068336-DEL-378 | chr19-38068337-DEL-378 |
| chr19-38107926-DEL-54 | chr19-38107927-DEL-54 |
| chr19-38108660-DEL-58 | chr19-38108661-DEL-58 |
| chr19-38162455-DEL-55 | chr19-38162456-DEL-55 |
| chr19-38203645-DEL-226 | chr19-38203639-DEL-227 |
| chr19-38218805-DEL-74 | chr19-38218806-DEL-74 |
| chr19-38224763-DEL-52 | chr19-38224764-DEL-52 |
| chr19-38232587-DEL-189 | chr19-38232588-DEL-189 |
| chr19-38262763-DEL-196 | chr19-38262757-DEL-197 |
| chr19-38312950-DEL-132 | chr19-38312951-DEL-132 |
| chr19-38321676-DEL-440 | chr19-38321677-DEL-440 |
| chr19-38322697-DEL-54 | chr19-38322698-DEL-54 |
| chr19-38326711-DEL-206 | chr19-38326699-DEL-208 |
| chr19-38363465-DEL-7066 | chr19-38363466-DEL-7066 |
| chr19-38378787-DEL-58 | chr19-38378770-DEL-76 |
| chr19-38380051-DEL-206 | chr19-38380052-DEL-206 |
| chr19-38383352-DEL-248 | chr19-38383353-DEL-248 |
| chr19-38390202-DEL-218 | chr19-38390203-DEL-218 |
| chr19-38409726-DEL-64 | chr19-38409755-DEL-60 |
| chr19-38413357-DEL-178 | chr19-38413358-DEL-178 |
| chr19-38419391-DEL-60 | chr19-38419392-DEL-60 |
| chr19-38432364-DEL-86 | chr19-38432365-DEL-84 |
| chr19-38452245-DEL-399 | chr19-38452246-DEL-387 |
| chr19-38477649-DEL-79 | chr19-38477650-DEL-79 |
| chr19-38487976-DEL-182 | chr19-38487977-DEL-182 |
| chr19-38504568-DEL-199 | chr19-38504569-DEL-199 |
| chr19-38518269-DEL-200 | chr19-38518270-DEL-200 |
| chr19-38522965-DEL-502 | chr19-38522966-DEL-502 |
| chr19-38525439-DEL-109 | chr19-38525425-DEL-107 |
| chr19-38550665-DEL-62 | chr19-38550667-DEL-61 |
| chr19-38560320-DEL-71 | chr19-38560321-DEL-67 |
| chr19-38593250-DEL-172 | chr19-38593252-DEL-135 |
| chr19-38735967-DEL-64 | chr19-38735968-DEL-64 |
| chr19-38744309-DEL-60 | chr19-38744312-DEL-58 |
| chr19-38780708-DEL-106 | chr19-38780705-DEL-106 |
| chr19-38793912-DEL-66 | chr19-38793877-DEL-58 |
| chr19-38816971-DEL-199 | chr19-38816972-DEL-199 |
| chr19-38836011-DEL-180 | chr19-38836012-DEL-180 |
| chr19-38858117-DEL-5322 | chr19-38858112-DEL-5323 |
| chr19-38872762-DEL-67 | chr19-38872763-DEL-67 |
| chr19-38911625-DEL-200 | chr19-38911626-DEL-200 |
| chr19-38912693-DEL-756 | chr19-38912703-DEL-731 |
| chr19-38940702-DEL-75 | chr19-38940703-DEL-75 |
| chr19-38951856-DEL-205 | chr19-38951857-DEL-205 |
| chr19-38961580-DEL-69 | chr19-38961581-DEL-63 |
| chr19-38967904-DEL-1627 | chr19-38967905-DEL-1627 |
| chr19-38976768-DEL-215 | chr19-38976755-DEL-216 |
| chr19-38980323-DEL-184 | chr19-38980324-DEL-184 |
| chr19-38984950-DEL-116 | chr19-38984951-DEL-116 |
| chr19-39002631-DEL-326 | chr19-39002632-DEL-326 |
| chr19-39030113-DEL-218 | chr19-39030114-DEL-218 |
| chr19-39032204-DEL-402 | chr19-39032204-DEL-402 |
| chr19-39045525-DEL-528 | chr19-39045523-DEL-528 |
| chr19-39051768-DEL-111 | chr19-39051768-DEL-130 |
| chr19-39107022-DEL-124 | chr19-39107023-DEL-124 |
| chr19-39111499-DEL-59809 | chr19-39111499-DEL-59809 |
| chr19-39119883-DEL-3873 | chr19-39119884-DEL-3873 |
| chr19-39127201-DEL-185 | chr19-39127202-DEL-185 |
| chr19-39127818-DEL-272 | chr19-39127819-DEL-272 |
| chr19-39131232-DEL-208 | chr19-39131233-DEL-208 |
| chr19-39134089-DEL-10696 | chr19-39134090-DEL-10696 |
| chr19-39206929-DEL-339 | chr19-39206930-DEL-339 |
| chr19-39237062-DEL-398 | chr19-39237063-DEL-398 |
| chr19-39323304-DEL-78 | chr19-39323275-DEL-75 |
| chr19-39340856-DEL-332 | chr19-39340857-DEL-332 |
| chr19-39363907-DEL-183 | chr19-39363908-DEL-183 |
| chr19-39369578-DEL-7184 | chr19-39369579-DEL-7184 |
| chr19-39397030-DEL-703 | chr19-39397031-DEL-703 |
| chr19-39400872-DEL-7395 | chr19-39400873-DEL-7395 |
| chr19-39409024-DEL-6265 | chr19-39409025-DEL-6265 |
| chr19-39414435-DEL-65 | chr19-39414436-DEL-65 |
| chr19-39486539-DEL-150 | chr19-39486540-DEL-150 |
| chr19-39494379-DEL-80 | chr19-39494380-DEL-80 |
| chr19-39504967-DEL-258 | chr19-39504968-DEL-258 |
| chr19-39506126-DEL-6577 | chr19-39506127-DEL-6577 |
| chr19-39507417-DEL-200 | chr19-39507418-DEL-200 |
| chr19-39512367-DEL-201 | chr19-39512367-DEL-201 |

| | |
|---|---|
| chr19-39525634-DEL-611 | chr19-39525635-DEL-611 |
| chr19-39535399-DEL-165 | chr19-39535400-DEL-165 |
| chr19-39574251-DEL-155 | chr19-39574252-DEL-155 |
| chr19-39588355-DEL-7183 | chr19-39588356-DEL-7183 |
| chr19-39649176-DEL-1377 | chr19-39649177-DEL-1377 |
| chr19-39666496-DEL-6401 | chr19-39666497-DEL-6401 |
| chr19-39676097-DEL-100 | chr19-39676089-DEL-95 |
| chr19-39680063-DEL-3663 | chr19-39680064-DEL-3663 |
| chr19-39712351-DEL-128 | chr19-39712371-DEL-109 |
| chr19-39735497-DEL-463 | chr19-39735498-DEL-463 |
| chr19-39852767-DEL-1423 | chr19-39852768-DEL-1421 |
| chr19-39904320-DEL-465 | chr19-39904321-DEL-465 |
| chr19-39935630-DEL-2112 | chr19-39935631-DEL-2112 |
| chr19-39939731-DEL-53 | chr19-39939732-DEL-53 |
| chr19-39947734-DEL-455 | chr19-39947754-DEL-437 |
| chr19-40003191-DEL-3663 | chr19-40003191-DEL-3663 |
| chr19-40008850-DEL-6295 | chr19-40008851-DEL-6295 |
| chr19-40028800-DEL-76 | chr19-40028798-DEL-76 |
| chr19-40054120-DEL-2954 | chr19-40054121-DEL-2954 |
| chr19-40054766-DEL-81 | chr19-40054767-DEL-81 |
| chr19-40084273-DEL-512 | chr19-40084274-DEL-512 |
| chr19-40090006-DEL-638 | chr19-40090007-DEL-638 |
| chr19-40094330-DEL-220 | chr19-40094331-DEL-220 |
| chr19-40098573-DEL-197 | chr19-40098540-DEL-198 |
| chr19-40125217-DEL-58 | chr19-40125228-DEL-62 |
| chr19-40135342-DEL-6429 | chr19-40135328-DEL-6441 |
| chr19-40146100-DEL-55 | chr19-40146056-DEL-95 |
| chr19-40147399-DEL-333 | chr19-40147400-DEL-333 |
| chr19-40181821-DEL-59 | chr19-40181821-DEL-58 |
| chr19-40192763-DEL-1051 | chr19-40192764-DEL-1051 |
| chr19-40194081-DEL-104 | chr19-40194082-DEL-104 |
| chr19-40196269-DEL-363 | chr19-40196270-DEL-363 |
| chr19-40197387-DEL-51 | chr19-40197387-DEL-51 |
| chr19-40203723-DEL-631 | chr19-40203724-DEL-631 |
| chr19-40222202-DEL-72 | chr19-40222203-DEL-72 |
| chr19-40225773-DEL-588 | chr19-40225774-DEL-587 |
| chr19-40234523-DEL-56 | chr19-40234524-DEL-53 |
| chr19-40262435-DEL-54 | chr19-40262435-DEL-54 |
| chr19-40277963-DEL-188 | chr19-40277963-DEL-189 |
| chr19-40333822-DEL-214 | chr19-40333823-DEL-214 |
| chr19-40346035-DEL-135 | chr19-40346036-DEL-135 |
| chr19-40407314-DEL-179 | chr19-40407315-DEL-178 |
| chr19-40410545-DEL-173 | chr19-40410546-DEL-172 |
| chr19-40412835-DEL-76 | chr19-40412839-DEL-80 |
| chr19-40432130-DEL-55 | chr19-40432132-DEL-55 |
| chr19-40438754-DEL-98 | chr19-40438753-DEL-72 |
| chr19-40442692-DEL-100 | chr19-40442693-DEL-100 |
| chr19-40450446-DEL-64 | chr19-40450448-DEL-54 |
| chr19-40462997-DEL-1607 | chr19-40462998-DEL-1607 |
| chr19-40467534-DEL-98 | chr19-40467541-DEL-92 |
| chr19-40491227-DEL-1137 | chr19-40491228-DEL-1137 |
| chr19-40496848-DEL-439 | chr19-40496849-DEL-432 |
| chr19-40505123-DEL-60 | chr19-40505124-DEL-50 |
| chr19-40531590-DEL-217 | chr19-40531591-DEL-217 |
| chr19-40535721-DEL-114 | chr19-40535722-DEL-114 |
| chr19-40547185-DEL-56 | chr19-40547186-DEL-56 |
| chr19-40547322-DEL-2876 | chr19-40547323-DEL-2876 |
| chr19-40556381-DEL-312 | chr19-40556383-DEL-311 |
| chr19-40560770-DEL-139 | chr19-40560771-DEL-139 |
| chr19-40580653-DEL-198 | chr19-40580654-DEL-198 |
| chr19-40581955-DEL-441 | chr19-40581954-DEL-443 |
| chr19-40582904-DEL-900 | chr19-40582894-DEL-898 |
| chr19-40586957-DEL-54 | chr19-40586958-DEL-54 |
| chr19-40603104-DEL-230 | chr19-40603105-DEL-230 |
| chr19-40605650-DEL-1328 | chr19-40605651-DEL-1328 |
| chr19-40609873-DEL-76 | chr19-40609874-DEL-76 |
| chr19-40610204-DEL-2846 | chr19-40610205-DEL-2846 |
| chr19-40614509-DEL-53 | chr19-40614510-DEL-53 |
| chr19-40644018-DEL-527 | chr19-40644019-DEL-527 |
| chr19-40658051-DEL-74 | chr19-40658052-DEL-74 |
| chr19-40666413-DEL-116 | chr19-40666414-DEL-116 |
| chr19-40672141-DEL-118 | chr19-40672142-DEL-118 |
| chr19-40675502-DEL-70 | chr19-40675508-DEL-63 |
| chr19-40697350-DEL-1252 | chr19-40697351-DEL-1252 |
| chr19-40702688-DEL-101 | chr19-40702688-DEL-101 |
| chr19-40728337-DEL-74 | chr19-40728338-DEL-74 |
| chr19-40734572-DEL-66 | chr19-40734573-DEL-66 |
| chr19-40743188-DEL-739 | chr19-40743189-DEL-739 |
| chr19-40761819-DEL-123 | chr19-40761797-DEL-122 |
| chr19-40765014-DEL-72 | chr19-40765015-DEL-70 |

| | |
|---|---|
| chr19-40831762-DEL-94 | chr19-40831763-DEL-94 |
| chr19-40839992-DEL-305 | chr19-40839993-DEL-305 |
| chr19-40843191-DEL-90 | chr19-40843192-DEL-90 |
| chr19-40845735-DEL-78 | chr19-40845738-DEL-93 |
| chr19-40854468-DEL-109 | chr19-40854467-DEL-110 |
| chr19-40857268-DEL-87 | chr19-40857271-DEL-107 |
| chr19-40872115-DEL-62 | chr19-40872131-DEL-89 |
| chr19-40890897-DEL-53 | chr19-40890898-DEL-53 |
| chr19-40910393-DEL-51 | chr19-40910405-DEL-52 |
| chr19-40911457-DEL-227 | chr19-40911458-DEL-227 |
| chr19-40923327-DEL-123 | chr19-40923328-DEL-123 |
| chr19-40933029-DEL-177 | chr19-40933036-DEL-171 |
| chr19-40940099-DEL-79 | chr19-40940083-DEL-80 |
| chr19-40942792-DEL-206 | chr19-40942793-DEL-206 |
| chr19-40944306-DEL-1780 | chr19-40944327-DEL-1760 |
| chr19-40963770-DEL-937 | chr19-40963771-DEL-937 |
| chr19-40965821-DEL-53 | chr19-40965822-DEL-53 |
| chr19-40972591-DEL-64 | chr19-40972592-DEL-60 |
| chr19-40990773-DEL-436 | chr19-40990774-DEL-436 |
| chr19-40999040-DEL-80 | chr19-40999041-DEL-80 |
| chr19-41022598-DEL-78 | chr19-41022593-DEL-73 |
| chr19-41029198-DEL-440 | chr19-41029199-DEL-440 |
| chr19-41032129-DEL-224 | chr19-41032123-DEL-222 |
| chr19-41038013-DEL-356 | chr19-41038014-DEL-356 |
| chr19-41044789-DEL-2680 | chr19-41044790-DEL-2680 |
| chr19-41054078-DEL-51 | chr19-41054079-DEL-51 |
| chr19-41082620-DEL-514 | chr19-41082621-DEL-514 |
| chr19-41131248-DEL-7089 | chr19-41131255-DEL-7085 |
| chr19-41212689-DEL-74 | chr19-41212698-DEL-72 |
| chr19-41297075-DEL-73 | chr19-41297076-DEL-73 |
| chr19-41325783-DEL-556 | chr19-41325784-DEL-555 |
| chr19-41346295-DEL-246 | chr19-41346295-DEL-244 |
| chr19-41359413-DEL-66 | chr19-41359414-DEL-66 |
| chr19-41382157-DEL-1255 | chr19-41382158-DEL-1255 |
| chr19-41388329-DEL-85 | chr19-41388330-DEL-85 |
| chr19-41411560-DEL-201 | chr19-41411561-DEL-201 |
| chr19-41418062-DEL-5395 | chr19-41418055-DEL-5401 |
| chr19-41443141-DEL-59 | chr19-41443155-DEL-78 |
| chr19-41456548-DEL-4852 | chr19-41456549-DEL-4852 |
| chr19-41477837-DEL-232 | chr19-41477838-DEL-232 |
| chr19-41513687-DEL-1723 | chr19-41513688-DEL-1723 |
| chr19-41602146-DEL-69 | chr19-41602147-DEL-53 |
| chr19-41616346-DEL-67 | chr19-41616347-DEL-67 |
| chr19-41636387-DEL-201 | chr19-41636388-DEL-201 |
| chr19-41655574-DEL-64 | chr19-41655575-DEL-64 |
| chr19-41698741-DEL-56 | chr19-41698740-DEL-58 |
| chr19-41746478-DEL-83 | chr19-41746477-DEL-84 |
| chr19-41747000-DEL-66 | chr19-41747001-DEL-66 |
| chr19-41751099-DEL-187 | chr19-41751100-DEL-187 |
| chr19-41771157-DEL-214 | chr19-41771158-DEL-214 |
| chr19-41777266-DEL-201 | chr19-41777267-DEL-201 |
| chr19-41800430-DEL-198 | chr19-41800431-DEL-198 |
| chr19-41810248-DEL-636 | chr19-41810248-DEL-635 |
| chr19-41864112-DEL-219 | chr19-41864118-DEL-212 |
| chr19-41872282-DEL-77 | chr19-41872279-DEL-78 |
| chr19-41879922-DEL-197 | chr19-41879923-DEL-197 |
| chr19-41891052-DEL-214 | chr19-41891053-DEL-179 |
| chr19-41892903-DEL-163 | chr19-41892902-DEL-163 |
| chr19-41910491-DEL-606 | chr19-41910492-DEL-606 |
| chr19-41913688-DEL-234 | chr19-41913689-DEL-234 |
| chr19-41917821-DEL-209 | chr19-41917822-DEL-209 |
| chr19-41924859-DEL-207 | chr19-41924860-DEL-207 |
| chr19-41935989-DEL-97 | chr19-41935990-DEL-97 |
| chr19-41950416-DEL-87 | chr19-41950417-DEL-87 |
| chr19-41953745-DEL-209 | chr19-41953746-DEL-209 |
| chr19-41996386-DEL-69 | chr19-41996376-DEL-79 |
| chr19-42043625-DEL-372 | chr19-42043628-DEL-367 |
| chr19-42047977-DEL-196 | chr19-42047978-DEL-196 |
| chr19-42053858-DEL-205 | chr19-42053859-DEL-205 |
| chr19-42054649-DEL-52 | chr19-42054645-DEL-51 |
| chr19-42073298-DEL-272 | chr19-42073299-DEL-272 |
| chr19-42082560-DEL-197 | chr19-42082558-DEL-198 |
| chr19-42086189-DEL-191 | chr19-42086190-DEL-191 |
| chr19-42089954-DEL-648 | chr19-42089955-DEL-648 |
| chr19-42094390-DEL-109 | chr19-42094413-DEL-95 |
| chr19-42098034-DEL-203 | chr19-42098035-DEL-203 |
| chr19-42100869-DEL-92 | chr19-42100870-DEL-92 |
| chr19-42102954-DEL-216 | chr19-42102955-DEL-216 |
| chr19-42111096-DEL-80 | chr19-42111097-DEL-80 |
| chr19-42126403-DEL-172 | chr19-42126404-DEL-172 |

| | |
|---|---|
| chr19-42141477-DEL-1512 | chr19-42141478-DEL-1512 |
| chr19-42145654-DEL-202 | chr19-42145655-DEL-202 |
| chr19-42148575-DEL-440 | chr19-42148576-DEL-440 |
| chr19-42150390-DEL-78 | chr19-42150402-DEL-50 |
| chr19-42158566-DEL-52 | chr19-42158567-DEL-52 |
| chr19-42170979-DEL-82 | chr19-42170980-DEL-82 |
| chr19-42206299-DEL-52 | chr19-42206300-DEL-52 |
| chr19-42210012-DEL-118 | chr19-42210013-DEL-118 |
| chr19-42249232-DEL-177 | chr19-42249238-DEL-172 |
| chr19-42256574-DEL-86 | chr19-42256575-DEL-86 |
| chr19-42263784-DEL-74 | chr19-42263785-DEL-74 |
| chr19-42313337-DEL-104 | chr19-42313338-DEL-104 |
| chr19-42334407-DEL-4341 | chr19-42334408-DEL-4341 |
| chr19-42341320-DEL-53 | chr19-42341321-DEL-53 |
| chr19-42343617-DEL-124 | chr19-42343618-DEL-124 |
| chr19-42348786-DEL-634 | chr19-42348787-DEL-634 |
| chr19-42367028-DEL-507 | chr19-42367029-DEL-507 |
| chr19-42385460-DEL-83 | chr19-42385459-DEL-83 |
| chr19-42389886-DEL-60 | chr19-42389878-DEL-68 |
| chr19-42429789-DEL-216 | chr19-42429794-DEL-212 |
| chr19-42451857-DEL-290 | chr19-42451858-DEL-290 |
| chr19-42493944-DEL-80 | chr19-42493974-DEL-64 |
| chr19-42508858-DEL-380 | chr19-42508856-DEL-381 |
| chr19-42519976-DEL-63 | chr19-42519977-DEL-63 |
| chr19-42528819-DEL-59 | chr19-42528820-DEL-59 |
| chr19-42537044-DEL-120 | chr19-42537032-DEL-133 |
| chr19-42543714-DEL-3608 | chr19-42543715-DEL-3608 |
| chr19-42545870-DEL-66 | chr19-42545871-DEL-66 |
| chr19-42553674-DEL-228 | chr19-42553672-DEL-229 |
| chr19-42597810-DEL-73 | chr19-42597797-DEL-86 |
| chr19-42613331-DEL-54 | chr19-42613320-DEL-66 |
| chr19-42636195-DEL-1888 | chr19-42636196-DEL-1888 |
| chr19-42657902-DEL-88 | chr19-42657923-DEL-66 |
| chr19-42662684-DEL-1539 | chr19-42662685-DEL-1539 |
| chr19-42675969-DEL-339 | chr19-42675971-DEL-339 |
| chr19-42688936-DEL-196 | chr19-42688923-DEL-197 |
| chr19-42694931-DEL-94 | chr19-42694935-DEL-95 |
| chr19-42757397-DEL-52 | chr19-42757397-DEL-52 |
| chr19-42771422-DEL-498 | chr19-42771423-DEL-498 |
| chr19-42772357-DEL-55 | chr19-42772358-DEL-55 |
| chr19-42773960-DEL-56 | chr19-42773961-DEL-56 |
| chr19-42817037-DEL-66 | chr19-42817038-DEL-65 |
| chr19-42869231-DEL-76 | chr19-42869232-DEL-76 |
| chr19-42875744-DEL-408 | chr19-42875745-DEL-408 |
| chr19-42877143-DEL-89 | chr19-42877161-DEL-50 |
| chr19-42884806-DEL-75 | chr19-42884807-DEL-75 |
| chr19-42927869-DEL-393 | chr19-42927870-DEL-393 |
| chr19-42950849-DEL-73 | chr19-42950850-DEL-73 |
| chr19-42951442-DEL-662 | chr19-42951443-DEL-661 |
| chr19-42986845-DEL-6653 | chr19-42986846-DEL-6653 |
| chr19-43007439-DEL-1049 | chr19-43007439-DEL-1034 |
| chr19-43035472-DEL-476 | chr19-43035473-DEL-476 |
| chr19-43063746-DEL-6252 | chr19-43063747-DEL-6252 |
| chr19-43084005-DEL-54 | chr19-43084006-DEL-54 |
| chr19-43106750-DEL-6380 | chr19-43106751-DEL-6380 |
| chr19-43120283-DEL-101 | chr19-43120309-DEL-99 |
| chr19-43133433-DEL-202 | chr19-43133434-DEL-202 |
| chr19-43220838-DEL-199 | chr19-43220839-DEL-199 |
| chr19-43227522-DEL-806 | chr19-43227523-DEL-806 |
| chr19-43288456-DEL-78 | chr19-43288505-DEL-50 |
| chr19-43335123-DEL-60 | chr19-43335124-DEL-59 |
| chr19-43372180-DEL-74 | chr19-43372164-DEL-75 |
| chr19-43394313-DEL-918 | chr19-43394314-DEL-918 |
| chr19-43408367-DEL-84 | chr19-43408367-DEL-84 |
| chr19-43414080-DEL-98 | chr19-43414080-DEL-110 |
| chr19-43419662-DEL-91 | chr19-43419663-DEL-91 |
| chr19-43475517-DEL-66 | chr19-43475512-DEL-50 |
| chr19-43490791-DEL-194 | chr19-43490798-DEL-190 |
| chr19-43500373-DEL-62 | chr19-43500374-DEL-62 |
| chr19-43511340-DEL-597 | chr19-43511345-DEL-593 |
| chr19-43524075-DEL-72 | chr19-43524076-DEL-72 |
| chr19-43526436-DEL-989 | chr19-43526437-DEL-989 |
| chr19-43531784-DEL-56 | chr19-43531784-DEL-56 |
| chr19-43557036-DEL-361 | chr19-43557036-DEL-362 |
| chr19-43566748-DEL-210 | chr19-43566749-DEL-210 |
| chr19-43622356-DEL-208 | chr19-43622357-DEL-208 |
| chr19-43701549-DEL-214 | chr19-43701550-DEL-214 |
| chr19-43738258-DEL-134 | chr19-43738260-DEL-126 |
| chr19-43751485-DEL-168 | chr19-43751472-DEL-166 |
| chr19-43796617-DEL-112 | chr19-43796645-DEL-84 |

| | |
|---|---|
| chr19-43846365-DEL-1197 | chr19-43846366-DEL-1197 |
| chr19-43853255-DEL-203 | chr19-43853256-DEL-203 |
| chr19-43882064-DEL-78 | chr19-43882056-DEL-53 |
| chr19-43883079-DEL-55 | chr19-43883080-DEL-55 |
| chr19-43940243-DEL-208 | chr19-43940244-DEL-208 |
| chr19-43941056-DEL-83 | chr19-43941057-DEL-83 |
| chr19-43942918-DEL-51 | chr19-43942919-DEL-51 |
| chr19-43953229-DEL-200 | chr19-43953230-DEL-200 |
| chr19-43959378-DEL-325 | chr19-43959379-DEL-325 |
| chr19-43977947-DEL-88 | chr19-43977948-DEL-88 |
| chr19-43981550-DEL-109 | chr19-43981569-DEL-75 |
| chr19-43983305-DEL-241 | chr19-43983308-DEL-239 |
| chr19-43991133-DEL-1445 | chr19-43991152-DEL-1434 |
| chr19-43998735-DEL-52 | chr19-43998736-DEL-52 |
| chr19-44007196-DEL-56 | chr19-44007197-DEL-56 |
| chr19-44038760-DEL-81 | chr19-44038805-DEL-61 |
| chr19-44041222-DEL-207 | chr19-44041223-DEL-207 |
| chr19-44089736-DEL-177 | chr19-44089737-DEL-177 |
| chr19-44108903-DEL-162 | chr19-44108904-DEL-162 |
| chr19-44112841-DEL-88 | chr19-44112842-DEL-88 |
| chr19-44124507-DEL-96 | chr19-44124508-DEL-96 |
| chr19-44125774-DEL-62 | chr19-44125787-DEL-66 |
| chr19-44129363-DEL-214 | chr19-44129354-DEL-215 |
| chr19-44150380-DEL-60 | chr19-44150377-DEL-64 |
| chr19-44153698-DEL-6401 | chr19-44153699-DEL-6401 |
| chr19-44188278-DEL-228 | chr19-44188279-DEL-228 |
| chr19-44206939-DEL-6442 | chr19-44206940-DEL-6442 |
| chr19-44216350-DEL-1769 | chr19-44216351-DEL-1769 |
| chr19-44221269-DEL-111 | chr19-44221268-DEL-97 |
| chr19-44242701-DEL-513 | chr19-44242702-DEL-513 |
| chr19-44248613-DEL-56 | chr19-44248614-DEL-56 |
| chr19-44258304-DEL-199 | chr19-44258305-DEL-198 |
| chr19-44288357-DEL-54 | chr19-44288358-DEL-54 |
| chr19-44319764-DEL-56 | chr19-44319764-DEL-56 |
| chr19-44328646-DEL-150 | chr19-44328695-DEL-101 |
| chr19-44333626-DEL-460 | chr19-44333627-DEL-460 |
| chr19-44352200-DEL-325 | chr19-44352201-DEL-325 |
| chr19-44353283-DEL-162 | chr19-44353284-DEL-162 |
| chr19-44377805-DEL-67 | chr19-44377804-DEL-69 |
| chr19-44408658-DEL-66 | chr19-44408661-DEL-61 |
| chr19-44422217-DEL-120 | chr19-44422218-DEL-120 |
| chr19-44430613-DEL-66 | chr19-44430607-DEL-55 |
| chr19-44437444-DEL-114 | chr19-44437445-DEL-114 |
| chr19-44456474-DEL-698 | chr19-44456475-DEL-696 |
| chr19-44467505-DEL-104 | chr19-44467506-DEL-104 |
| chr19-44471454-DEL-56 | chr19-44471453-DEL-61 |
| chr19-44472228-DEL-405 | chr19-44472229-DEL-405 |
| chr19-44479907-DEL-549 | chr19-44479908-DEL-549 |
| chr19-44495894-DEL-69 | chr19-44495895-DEL-69 |
| chr19-44498313-DEL-65 | chr19-44498314-DEL-65 |
| chr19-44504850-DEL-58 | chr19-44504851-DEL-58 |
| chr19-44511050-DEL-61 | chr19-44511051-DEL-61 |
| chr19-44517581-DEL-207 | chr19-44517582-DEL-207 |
| chr19-44521897-DEL-65 | chr19-44521899-DEL-64 |
| chr19-44547860-DEL-167 | chr19-44547877-DEL-151 |
| chr19-44574813-DEL-62 | chr19-44574814-DEL-62 |
| chr19-44582230-DEL-435 | chr19-44582234-DEL-434 |
| chr19-44600840-DEL-664 | chr19-44600841-DEL-664 |
| chr19-44610533-DEL-145 | chr19-44610533-DEL-144 |
| chr19-44614223-DEL-78 | chr19-44614224-DEL-84 |
| chr19-44614492-DEL-654 | chr19-44614493-DEL-654 |
| chr19-44701579-DEL-318 | chr19-44701537-DEL-339 |
| chr19-44754562-DEL-56 | chr19-44754586-DEL-58 |
| chr19-44834517-DEL-113 | chr19-44834512-DEL-111 |
| chr19-44938775-DEL-505 | chr19-44938750-DEL-506 |
| chr19-44939564-DEL-57 | chr19-44939565-DEL-57 |
| chr19-44966494-DEL-572 | chr19-44966495-DEL-572 |
| chr19-44993344-DEL-217 | chr19-44993345-DEL-217 |
| chr19-44997974-DEL-209 | chr19-44997975-DEL-209 |
| chr19-45029414-DEL-54 | chr19-45029415-DEL-54 |
| chr19-45048334-DEL-180 | chr19-45048335-DEL-180 |
| chr19-45058248-DEL-295 | chr19-45058249-DEL-295 |
| chr19-45076857-DEL-55 | chr19-45076857-DEL-56 |
| chr19-45077747-DEL-100 | chr19-45077748-DEL-100 |
| chr19-45078522-DEL-6399 | chr19-45078522-DEL-6399 |
| chr19-45095668-DEL-68 | chr19-45095667-DEL-54 |
| chr19-45107959-DEL-55 | chr19-45107959-DEL-55 |
| chr19-45117288-DEL-79 | chr19-45117267-DEL-88 |
| chr19-45119014-DEL-78 | chr19-45119015-DEL-60 |
| chr19-45119705-DEL-631 | chr19-45119706-DEL-631 |

| | |
|---|---|
| chr19-45129878-DEL-79 | chr19-45129879-DEL-79 |
| chr19-45146422-DEL-65 | chr19-45146422-DEL-66 |
| chr19-45152395-DEL-60 | chr19-45152394-DEL-62 |
| chr19-45163878-DEL-277 | chr19-45163879-DEL-277 |
| chr19-45169924-DEL-10577 | chr19-45169918-DEL-10578 |
| chr19-45179944-DEL-117 | chr19-45179945-DEL-117 |
| chr19-45185607-DEL-872 | chr19-45185608-DEL-872 |
| chr19-45202869-DEL-172 | chr19-45202870-DEL-172 |
| chr19-45208063-DEL-200 | chr19-45208055-DEL-201 |
| chr19-45228432-DEL-51 | chr19-45228421-DEL-50 |
| chr19-45242881-DEL-175 | chr19-45242882-DEL-175 |
| chr19-45273533-DEL-205 | chr19-45273534-DEL-205 |
| chr19-45276657-DEL-218 | chr19-45276658-DEL-200 |
| chr19-45284345-DEL-94 | chr19-45284346-DEL-94 |
| chr19-45294509-DEL-104 | chr19-45294510-DEL-104 |
| chr19-45366591-DEL-67 | chr19-45366590-DEL-64 |
| chr19-45370694-DEL-128 | chr19-45370695-DEL-128 |
| chr19-45376290-DEL-352 | chr19-45376283-DEL-353 |
| chr19-45401442-DEL-353 | chr19-45401443-DEL-353 |
| chr19-45424492-DEL-58 | chr19-45424493-DEL-60 |
| chr19-45425910-DEL-170 | chr19-45425911-DEL-170 |
| chr19-45535100-DEL-459 | chr19-45535101-DEL-459 |
| chr19-45544742-DEL-116 | chr19-45544783-DEL-74 |
| chr19-45577526-DEL-211 | chr19-45577527-DEL-211 |
| chr19-45608149-DEL-54 | chr19-45608150-DEL-54 |
| chr19-45637549-DEL-55 | chr19-45637550-DEL-55 |
| chr19-45654908-DEL-72 | chr19-45654909-DEL-72 |
| chr19-45676728-DEL-204 | chr19-45676729-DEL-204 |
| chr19-45687557-DEL-7109 | chr19-45687558-DEL-7109 |
| chr19-45702752-DEL-157 | chr19-45702753-DEL-157 |
| chr19-45747506-DEL-232 | chr19-45747507-DEL-232 |
| chr19-45797346-DEL-193 | chr19-45797347-DEL-193 |
| chr19-45799040-DEL-275 | chr19-45799041-DEL-275 |
| chr19-45812364-DEL-60 | chr19-45812365-DEL-60 |
| chr19-45818303-DEL-208 | chr19-45818304-DEL-208 |
| chr19-45837289-DEL-95 | chr19-45837290-DEL-95 |
| chr19-45839080-DEL-190 | chr19-45839081-DEL-190 |
| chr19-45841037-DEL-232 | chr19-45841038-DEL-232 |
| chr19-45842363-DEL-88 | chr19-45842364-DEL-88 |
| chr19-45850378-DEL-64 | chr19-45850397-DEL-66 |
| chr19-45855028-DEL-207 | chr19-45855029-DEL-207 |
| chr19-45869375-DEL-209 | chr19-45869379-DEL-206 |
| chr19-45881518-DEL-507 | chr19-45881517-DEL-502 |
| chr19-45883589-DEL-202 | chr19-45883590-DEL-202 |
| chr19-45903299-DEL-206 | chr19-45903300-DEL-206 |
| chr19-45918774-DEL-207 | chr19-45918775-DEL-206 |
| chr19-45930140-DEL-943 | chr19-45930141-DEL-943 |
| chr19-45944449-DEL-128 | chr19-45944450-DEL-116 |
| chr19-45950993-DEL-96 | chr19-45950994-DEL-96 |
| chr19-45956689-DEL-981 | chr19-45956690-DEL-981 |
| chr19-45973251-DEL-307 | chr19-45973252-DEL-307 |
| chr19-45980235-DEL-192 | chr19-45980236-DEL-192 |
| chr19-46001790-DEL-121 | chr19-46001791-DEL-121 |
| chr19-46003610-DEL-4687 | chr19-46003611-DEL-4687 |
| chr19-46006431-DEL-224 | chr19-46006432-DEL-224 |
| chr19-46037141-DEL-190 | chr19-46037142-DEL-190 |
| chr19-46057127-DEL-149 | chr19-46057128-DEL-149 |
| chr19-46074457-DEL-58 | chr19-46074458-DEL-58 |
| chr19-46110478-DEL-176 | chr19-46110479-DEL-176 |
| chr19-46116438-DEL-112 | chr19-46116439-DEL-112 |
| chr19-46139783-DEL-144 | chr19-46139784-DEL-141 |
| chr19-46154145-DEL-1070 | chr19-46154146-DEL-1070 |
| chr19-46155824-DEL-232 | chr19-46155825-DEL-232 |
| chr19-46169549-DEL-204 | chr19-46169550-DEL-204 |
| chr19-46175436-DEL-54 | chr19-46175437-DEL-54 |
| chr19-46198793-DEL-203 | chr19-46198794-DEL-203 |
| chr19-46206528-DEL-223 | chr19-46206529-DEL-223 |
| chr19-46215513-DEL-200 | chr19-46215514-DEL-200 |
| chr19-46217640-DEL-374 | chr19-46217672-DEL-343 |
| chr19-46242028-DEL-84 | chr19-46242029-DEL-66 |
| chr19-46295019-DEL-69 | chr19-46295020-DEL-69 |
| chr19-46325720-DEL-76 | chr19-46325721-DEL-76 |
| chr19-46330805-DEL-64 | chr19-46330812-DEL-60 |
| chr19-46343100-DEL-84 | chr19-46343108-DEL-76 |
| chr19-46348355-DEL-211 | chr19-46348356-DEL-211 |
| chr19-46352774-DEL-206 | chr19-46352775-DEL-206 |
| chr19-46355144-DEL-56 | chr19-46355145-DEL-56 |
| chr19-46409191-DEL-171 | chr19-46409192-DEL-167 |
| chr19-46481080-DEL-203 | chr19-46481081-DEL-200 |
| chr19-46511326-DEL-176 | chr19-46511327-DEL-176 |

| | |
|---|---|
| chr19-46522720-DEL-170 | chr19-46522721-DEL-170 |
| chr19-46524128-DEL-56 | chr19-46524155-DEL-54 |
| chr19-46525113-DEL-208 | chr19-46525114-DEL-208 |
| chr19-46527053-DEL-179 | chr19-46527053-DEL-179 |
| chr19-46533879-DEL-213 | chr19-46533880-DEL-213 |
| chr19-46540338-DEL-55 | chr19-46540308-DEL-63 |
| chr19-46555294-DEL-215 | chr19-46555295-DEL-215 |
| chr19-46568460-DEL-572 | chr19-46568461-DEL-572 |
| chr19-46572092-DEL-206 | chr19-46572088-DEL-217 |
| chr19-46593672-DEL-243 | chr19-46593673-DEL-243 |
| chr19-46616192-DEL-68 | chr19-46616193-DEL-68 |
| chr19-46624993-DEL-208 | chr19-46624994-DEL-208 |
| chr19-46639885-DEL-211 | chr19-46639888-DEL-210 |
| chr19-46641223-DEL-120 | chr19-46641214-DEL-108 |
| chr19-46651743-DEL-901 | chr19-46651744-DEL-901 |
| chr19-46693067-DEL-52 | chr19-46693047-DEL-78 |
| chr19-46700007-DEL-52 | chr19-46700008-DEL-52 |
| chr19-46702172-DEL-148 | chr19-46702186-DEL-135 |
| chr19-46707249-DEL-285 | chr19-46707250-DEL-285 |
| chr19-46712992-DEL-108 | chr19-46712993-DEL-108 |
| chr19-46713923-DEL-95 | chr19-46713924-DEL-95 |
| chr19-46720653-DEL-59 | chr19-46720654-DEL-59 |
| chr19-46721132-DEL-204 | chr19-46721133-DEL-204 |
| chr19-46741088-DEL-944 | chr19-46741089-DEL-944 |
| chr19-46744997-DEL-91 | chr19-46744998-DEL-91 |
| chr19-46746035-DEL-54 | chr19-46746036-DEL-54 |
| chr19-46804252-DEL-118 | chr19-46804267-DEL-118 |
| chr19-46821351-DEL-138 | chr19-46821311-DEL-179 |
| chr19-46838235-DEL-116 | chr19-46838236-DEL-116 |
| chr19-46844226-DEL-61 | chr19-46844219-DEL-69 |
| chr19-46860666-DEL-95 | chr19-46860681-DEL-81 |
| chr19-46884410-DEL-115 | chr19-46884411-DEL-80 |
| chr19-46926863-DEL-62 | chr19-46926862-DEL-63 |
| chr19-46935464-DEL-52 | chr19-46935465-DEL-52 |
| chr19-46943889-DEL-202 | chr19-46943890-DEL-202 |
| chr19-46969445-DEL-83 | chr19-46969446-DEL-83 |
| chr19-46974700-DEL-139 | chr19-46974694-DEL-138 |
| chr19-46975294-DEL-86 | chr19-46975298-DEL-86 |
| chr19-46986181-DEL-60 | chr19-46986182-DEL-60 |
| chr19-46991432-DEL-70 | chr19-46991433-DEL-70 |
| chr19-47014549-DEL-64 | chr19-47014550-DEL-63 |
| chr19-47015235-DEL-70 | chr19-47015235-DEL-70 |
| chr19-47018870-DEL-361 | chr19-47018870-DEL-361 |
| chr19-47045191-DEL-313 | chr19-47045214-DEL-290 |
| chr19-47046581-DEL-1493 | chr19-47046582-DEL-1493 |
| chr19-47073658-DEL-171 | chr19-47073659-DEL-171 |
| chr19-47110394-DEL-750 | chr19-47110395-DEL-750 |
| chr19-47114882-DEL-55 | chr19-47114883-DEL-55 |
| chr19-47135977-DEL-145 | chr19-47135978-DEL-145 |
| chr19-47144763-DEL-234 | chr19-47144793-DEL-233 |
| chr19-47146640-DEL-715 | chr19-47146641-DEL-715 |
| chr19-47151640-DEL-1275 | chr19-47151641-DEL-1275 |
| chr19-47160396-DEL-187 | chr19-47160398-DEL-187 |
| chr19-47183015-DEL-1871 | chr19-47183016-DEL-1871 |
| chr19-47197494-DEL-78 | chr19-47197495-DEL-78 |
| chr19-47202869-DEL-57 | chr19-47202870-DEL-57 |
| chr19-47204132-DEL-61 | chr19-47204133-DEL-61 |
| chr19-47204933-DEL-202 | chr19-47204927-DEL-203 |
| chr19-47312504-DEL-779 | chr19-47312505-DEL-779 |
| chr19-47316828-DEL-88 | chr19-47316830-DEL-118 |
| chr19-47347728-DEL-118 | chr19-47347729-DEL-118 |
| chr19-47358959-DEL-420 | chr19-47358960-DEL-420 |
| chr19-47378252-DEL-75 | chr19-47378253-DEL-75 |
| chr19-47409511-DEL-75 | chr19-47409512-DEL-83 |
| chr19-47440649-DEL-52 | chr19-47440650-DEL-52 |
| chr19-47441606-DEL-107 | chr19-47441607-DEL-97 |
| chr19-47450765-DEL-66 | chr19-47450766-DEL-66 |
| chr19-47466891-DEL-54 | chr19-47466892-DEL-54 |
| chr19-47474169-DEL-177 | chr19-47474170-DEL-177 |
| chr19-47478767-DEL-221 | chr19-47478767-DEL-215 |
| chr19-47492522-DEL-315 | chr19-47492520-DEL-318 |
| chr19-47511328-DEL-60 | chr19-47511329-DEL-60 |
| chr19-47534122-DEL-66 | chr19-47534122-DEL-66 |
| chr19-47536626-DEL-80 | chr19-47536626-DEL-81 |
| chr19-47538548-DEL-195 | chr19-47538549-DEL-195 |
| chr19-47554980-DEL-60 | chr19-47554980-DEL-61 |
| chr19-47561346-DEL-234 | chr19-47561354-DEL-226 |
| chr19-47578813-DEL-2941 | chr19-47578814-DEL-2941 |
| chr19-47581960-DEL-66 | chr19-47581959-DEL-56 |
| chr19-47584666-DEL-6399 | chr19-47584667-DEL-6399 |

| | |
|---|---|
| chr19-47595044-DEL-227 | chr19-47595044-DEL-227 |
| chr19-47681158-DEL-784 | chr19-47681159-DEL-784 |
| chr19-47682719-DEL-934 | chr19-47682718-DEL-934 |
| chr19-47692280-DEL-63 | chr19-47692281-DEL-63 |
| chr19-47721140-DEL-96 | chr19-47721146-DEL-96 |
| chr19-47742898-DEL-68 | chr19-47742899-DEL-68 |
| chr19-47774152-DEL-73 | chr19-47774157-DEL-53 |
| chr19-47775654-DEL-99 | chr19-47775655-DEL-99 |
| chr19-47777112-DEL-237 | chr19-47777113-DEL-237 |
| chr19-47782074-DEL-164 | chr19-47782077-DEL-156 |
| chr19-47792182-DEL-121 | chr19-47792187-DEL-117 |
| chr19-47793140-DEL-6834 | chr19-47793141-DEL-6834 |
| chr19-47799059-DEL-340 | chr19-47799060-DEL-340 |
| chr19-47850219-DEL-89 | chr19-47850220-DEL-89 |
| chr19-47912527-DEL-199 | chr19-47912528-DEL-199 |
| chr19-47937987-DEL-477 | chr19-47937988-DEL-477 |
| chr19-47961889-DEL-91 | chr19-47961890-DEL-91 |
| chr19-47962011-DEL-652 | chr19-47962012-DEL-652 |
| chr19-47970713-DEL-76 | chr19-47970714-DEL-76 |
| chr19-47979907-DEL-96 | chr19-47979908-DEL-96 |
| chr19-47999365-DEL-68 | chr19-47999366-DEL-64 |
| chr19-48042585-DEL-560 | chr19-48042586-DEL-559 |
| chr19-48084177-DEL-174 | chr19-48084178-DEL-174 |
| chr19-48092328-DEL-70 | chr19-48092296-DEL-80 |
| chr19-48094068-DEL-400 | chr19-48094069-DEL-400 |
| chr19-48100799-DEL-102 | chr19-48100798-DEL-103 |
| chr19-48113544-DEL-218 | chr19-48113545-DEL-218 |
| chr19-48118045-DEL-93 | chr19-48118042-DEL-94 |
| chr19-48133182-DEL-127 | chr19-48133183-DEL-126 |
| chr19-48154500-DEL-58 | chr19-48154507-DEL-50 |
| chr19-48162414-DEL-80 | chr19-48162417-DEL-81 |
| chr19-48199866-DEL-62 | chr19-48199867-DEL-62 |
| chr19-48212750-DEL-114 | chr19-48212750-DEL-115 |
| chr19-48214244-DEL-6483 | chr19-48214243-DEL-6482 |
| chr19-48259267-DEL-171 | chr19-48259279-DEL-154 |
| chr19-48260032-DEL-77 | chr19-48260054-DEL-56 |
| chr19-48344967-DEL-118 | chr19-48344968-DEL-118 |
| chr19-48381643-DEL-319 | chr19-48381644-DEL-319 |
| chr19-48390883-DEL-54 | chr19-48390885-DEL-59 |
| chr19-48416904-DEL-488 | chr19-48416905-DEL-488 |
| chr19-48421448-DEL-55 | chr19-48421449-DEL-55 |
| chr19-48422191-DEL-221 | chr19-48422192-DEL-222 |
| chr19-48483064-DEL-2037 | chr19-48483068-DEL-2037 |
| chr19-48503961-DEL-123 | chr19-48503962-DEL-123 |
| chr19-48505152-DEL-426 | chr19-48505153-DEL-426 |
| chr19-48572250-DEL-931 | chr19-48572251-DEL-931 |
| chr19-48582372-DEL-52 | chr19-48582373-DEL-50 |
| chr19-48651133-DEL-64 | chr19-48651134-DEL-64 |
| chr19-48697159-DEL-289 | chr19-48697160-DEL-289 |
| chr19-48708382-DEL-76 | chr19-48708383-DEL-81 |
| chr19-48716901-DEL-58 | chr19-48716902-DEL-60 |
| chr19-48730229-DEL-169 | chr19-48730262-DEL-137 |
| chr19-48732317-DEL-2963 | chr19-48732318-DEL-2963 |
| chr19-48736864-DEL-362 | chr19-48736865-DEL-362 |
| chr19-48744854-DEL-56 | chr19-48744845-DEL-97 |
| chr19-48757530-DEL-103 | chr19-48757487-DEL-102 |
| chr19-48805879-DEL-1888 | chr19-48805880-DEL-1888 |
| chr19-48810948-DEL-52 | chr19-48810950-DEL-51 |
| chr19-48875957-DEL-424 | chr19-48875958-DEL-424 |
| chr19-48890943-DEL-355 | chr19-48890944-DEL-355 |
| chr19-48910371-DEL-98 | chr19-48910364-DEL-106 |
| chr19-48920341-DEL-94 | chr19-48920337-DEL-92 |
| chr19-48920790-DEL-6644 | chr19-48920790-DEL-6644 |
| chr19-48920812-DEL-424 | chr19-48920813-DEL-424 |
| chr19-48964089-DEL-98 | chr19-48964089-DEL-98 |
| chr19-48965442-DEL-6536 | chr19-48965431-DEL-6537 |
| chr19-48996741-DEL-72 | chr19-48996732-DEL-81 |
| chr19-49000959-DEL-105 | chr19-49000960-DEL-135 |
| chr19-49002621-DEL-69 | chr19-49002622-DEL-69 |
| chr19-49010899-DEL-104 | chr19-49010900-DEL-69 |
| chr19-49058479-DEL-4796 | chr19-49058480-DEL-4796 |
| chr19-49067101-DEL-54 | chr19-49067102-DEL-54 |
| chr19-49125736-DEL-68 | chr19-49125753-DEL-65 |
| chr19-49135494-DEL-381 | chr19-49135495-DEL-381 |
| chr19-49139328-DEL-222 | chr19-49139331-DEL-220 |
| chr19-49159648-DEL-52 | chr19-49159638-DEL-58 |
| chr19-49165108-DEL-6420 | chr19-49165109-DEL-6420 |
| chr19-49189204-DEL-56 | chr19-49189205-DEL-56 |
| chr19-49272290-DEL-60 | chr19-49272295-DEL-56 |
| chr19-49292021-DEL-124 | chr19-49292022-DEL-124 |

| | |
|---|---|
| chr19-49328726-DEL-7211 | chr19-49328727-DEL-7211 |
| chr19-49361359-DEL-583 | chr19-49361380-DEL-580 |
| chr19-49372316-DEL-198 | chr19-49372317-DEL-198 |
| chr19-49379918-DEL-97 | chr19-49379925-DEL-115 |
| chr19-49397931-DEL-678 | chr19-49397932-DEL-678 |
| chr19-49414883-DEL-445 | chr19-49414884-DEL-445 |
| chr19-49430190-DEL-68 | chr19-49430189-DEL-68 |
| chr19-49434450-DEL-2239 | chr19-49434448-DEL-2242 |
| chr19-49448825-DEL-6865 | chr19-49448835-DEL-6856 |
| chr19-49495511-DEL-1297 | chr19-49495512-DEL-1297 |
| chr19-49504955-DEL-57 | chr19-49504948-DEL-56 |
| chr19-49511714-DEL-6471 | chr19-49511715-DEL-6471 |
| chr19-49530553-DEL-94 | chr19-49530554-DEL-94 |
| chr19-49549008-DEL-96 | chr19-49549009-DEL-96 |
| chr19-49569123-DEL-229 | chr19-49569133-DEL-215 |
| chr19-49637372-DEL-84 | chr19-49637373-DEL-84 |
| chr19-49647513-DEL-283 | chr19-49647514-DEL-283 |
| chr19-49683951-DEL-193 | chr19-49683952-DEL-193 |
| chr19-49731211-DEL-246 | chr19-49731212-DEL-246 |
| chr19-49745824-DEL-69 | chr19-49745825-DEL-69 |
| chr19-49800251-DEL-605 | chr19-49800253-DEL-606 |
| chr19-49832709-DEL-59 | chr19-49832664-DEL-69 |
| chr19-49850943-DEL-68 | chr19-49850932-DEL-96 |
| chr19-49859308-DEL-229 | chr19-49859309-DEL-229 |
| chr19-49879780-DEL-1380 | chr19-49879781-DEL-1380 |
| chr19-49898396-DEL-352 | chr19-49898397-DEL-352 |
| chr19-49908489-DEL-805 | chr19-49908490-DEL-805 |
| chr19-49918822-DEL-109 | chr19-49918805-DEL-109 |
| chr19-49983104-DEL-55 | chr19-49983105-DEL-55 |
| chr19-49988387-DEL-1385 | chr19-49988388-DEL-1385 |
| chr19-50000250-DEL-62 | chr19-50000206-DEL-81 |
| chr19-50008092-DEL-181 | chr19-50008091-DEL-133 |
| chr19-50008567-DEL-1011 | chr19-50008568-DEL-1011 |
| chr19-50008568-DEL-3769 | chr19-50008568-DEL-3769 |
| chr19-50045171-DEL-211 | chr19-50045174-DEL-211 |
| chr19-50047873-DEL-283 | chr19-50047875-DEL-282 |
| chr19-50051632-DEL-430 | chr19-50051633-DEL-430 |
| chr19-50056911-DEL-68 | chr19-50056912-DEL-68 |
| chr19-50096315-DEL-6558 | chr19-50096316-DEL-6558 |
| chr19-50107028-DEL-133 | chr19-50107029-DEL-133 |
| chr19-50162595-DEL-374 | chr19-50162593-DEL-374 |
| chr19-50166056-DEL-281 | chr19-50166057-DEL-281 |
| chr19-50183012-DEL-296 | chr19-50183013-DEL-296 |
| chr19-50183021-DEL-3619 | chr19-50183022-DEL-3618 |
| chr19-50226602-DEL-58 | chr19-50226603-DEL-58 |
| chr19-50261840-DEL-1474 | chr19-50261841-DEL-1474 |
| chr19-50278826-DEL-138 | chr19-50278827-DEL-138 |
| chr19-50317186-DEL-226 | chr19-50317187-DEL-226 |
| chr19-50341114-DEL-522 | chr19-50341115-DEL-522 |
| chr19-50348816-DEL-74 | chr19-50348817-DEL-74 |
| chr19-50351828-DEL-69 | chr19-50351817-DEL-69 |
| chr19-50356060-DEL-65 | chr19-50356061-DEL-65 |
| chr19-50374425-DEL-1007 | chr19-50374426-DEL-1007 |
| chr19-50388783-DEL-3015 | chr19-50388784-DEL-3015 |
| chr19-50411700-DEL-116 | chr19-50411693-DEL-124 |
| chr19-50412717-DEL-69 | chr19-50412718-DEL-69 |
| chr19-50414696-DEL-6860 | chr19-50414697-DEL-6860 |
| chr19-50423718-DEL-94 | chr19-50423731-DEL-60 |
| chr19-50431632-DEL-1228 | chr19-50431633-DEL-1228 |
| chr19-50448007-DEL-64 | chr19-50448008-DEL-64 |
| chr19-50456906-DEL-55 | chr19-50456907-DEL-55 |
| chr19-50459601-DEL-54 | chr19-50459601-DEL-54 |
| chr19-50460296-DEL-98 | chr19-50460297-DEL-98 |
| chr19-50517066-DEL-64 | chr19-50517066-DEL-65 |
| chr19-50529231-DEL-268 | chr19-50529232-DEL-268 |
| chr19-50532622-DEL-182 | chr19-50532626-DEL-183 |
| chr19-50539506-DEL-6809 | chr19-50539507-DEL-6809 |
| chr19-50549173-DEL-60 | chr19-50549178-DEL-72 |
| chr19-50586750-DEL-3644 | chr19-50586751-DEL-3644 |
| chr19-50592938-DEL-119 | chr19-50592939-DEL-119 |
| chr19-50602380-DEL-64 | chr19-50602381-DEL-64 |
| chr19-50615709-DEL-1997 | chr19-50615710-DEL-1997 |
| chr19-50623679-DEL-88 | chr19-50623675-DEL-96 |
| chr19-50636387-DEL-79 | chr19-50636388-DEL-79 |
| chr19-50662405-DEL-79 | chr19-50662402-DEL-79 |
| chr19-50677033-DEL-76 | chr19-50677034-DEL-76 |
| chr19-50699884-DEL-91 | chr19-50699885-DEL-91 |
| chr19-50729398-DEL-1807 | chr19-50729399-DEL-1807 |
| chr19-50742609-DEL-1992 | chr19-50742607-DEL-1993 |
| chr19-50760986-DEL-1676 | chr19-50760988-DEL-1675 |

| | |
|---|---|
| chr19-50784177-DEL-415 | chr19-50784178-DEL-415 |
| chr19-50793081-DEL-74 | chr19-50793076-DEL-85 |
| chr19-50817320-DEL-6185 | chr19-50817321-DEL-6185 |
| chr19-50828726-DEL-138 | chr19-50828726-DEL-138 |
| chr19-50836729-DEL-233 | chr19-50836729-DEL-233 |
| chr19-50851896-DEL-2529 | chr19-50851897-DEL-2529 |
| chr19-50857003-DEL-132 | chr19-50857004-DEL-132 |
| chr19-50863566-DEL-110 | chr19-50863567-DEL-110 |
| chr19-50872640-DEL-79 | chr19-50872641-DEL-79 |
| chr19-50875129-DEL-111 | chr19-50875129-DEL-111 |
| chr19-50879737-DEL-1094 | chr19-50879738-DEL-1094 |
| chr19-50896130-DEL-920 | chr19-50896130-DEL-920 |
| chr19-50911969-DEL-103 | chr19-50911990-DEL-94 |
| chr19-50985907-DEL-79 | chr19-50985904-DEL-79 |
| chr19-50990489-DEL-80 | chr19-50990490-DEL-86 |
| chr19-50999428-DEL-97 | chr19-50999437-DEL-101 |
| chr19-51036809-DEL-74 | chr19-51036810-DEL-74 |
| chr19-51058268-DEL-484 | chr19-51058271-DEL-485 |
| chr19-51074321-DEL-6479 | chr19-51074322-DEL-6479 |
| chr19-51094829-DEL-55 | chr19-51094827-DEL-57 |
| chr19-51096945-DEL-6645 | chr19-51096946-DEL-6645 |
| chr19-51103238-DEL-212 | chr19-51103239-DEL-212 |
| chr19-51108806-DEL-66 | chr19-51108821-DEL-54 |
| chr19-51124812-DEL-85 | chr19-51124813-DEL-85 |
| chr19-51129118-DEL-204 | chr19-51129119-DEL-204 |
| chr19-51142726-DEL-78 | chr19-51142711-DEL-90 |
| chr19-51151603-DEL-1416 | chr19-51151593-DEL-1417 |
| chr19-51156258-DEL-1610 | chr19-51156259-DEL-1610 |
| chr19-51188815-DEL-510 | chr19-51188816-DEL-510 |
| chr19-51200560-DEL-94 | chr19-51200561-DEL-94 |
| chr19-51214530-DEL-477 | chr19-51214531-DEL-476 |
| chr19-51238183-DEL-1025 | chr19-51238184-DEL-1025 |
| chr19-51297667-DEL-81 | chr19-51297687-DEL-62 |
| chr19-51300786-DEL-64 | chr19-51300770-DEL-92 |
| chr19-51324471-DEL-237 | chr19-51324471-DEL-237 |
| chr19-51334196-DEL-501 | chr19-51334197-DEL-501 |
| chr19-51376990-DEL-342 | chr19-51376990-DEL-340 |
| chr19-51408000-DEL-105 | chr19-51408000-DEL-105 |
| chr19-51411649-DEL-78 | chr19-51411658-DEL-62 |
| chr19-51419539-DEL-59 | chr19-51419540-DEL-59 |
| chr19-51424792-DEL-1805 | chr19-51424793-DEL-1805 |
| chr19-51430679-DEL-6116 | chr19-51430680-DEL-6114 |
| chr19-51449209-DEL-66 | chr19-51449210-DEL-66 |
| chr19-51493710-DEL-839 | chr19-51493711-DEL-839 |
| chr19-51502090-DEL-542 | chr19-51502086-DEL-545 |
| chr19-51507297-DEL-120 | chr19-51507312-DEL-93 |
| chr19-51512784-DEL-70 | chr19-51512785-DEL-70 |
| chr19-51532298-DEL-6174 | chr19-51532287-DEL-6171 |
| chr19-51539642-DEL-2121 | chr19-51539643-DEL-2121 |
| chr19-51542510-DEL-101 | chr19-51542507-DEL-102 |
| chr19-51549358-DEL-1878 | chr19-51549359-DEL-1878 |
| chr19-51558215-DEL-387 | chr19-51558216-DEL-387 |
| chr19-51560000-DEL-70 | chr19-51560000-DEL-71 |
| chr19-51570142-DEL-84 | chr19-51570139-DEL-68 |
| chr19-51579229-DEL-63 | chr19-51579219-DEL-64 |
| chr19-51582653-DEL-1261 | chr19-51582654-DEL-1261 |
| chr19-51619413-DEL-538 | chr19-51619414-DEL-538 |
| chr19-51625316-DEL-221 | chr19-51625318-DEL-220 |
| chr19-51638154-DEL-6213 | chr19-51638155-DEL-6213 |
| chr19-51664663-DEL-54 | chr19-51664664-DEL-54 |
| chr19-51728011-DEL-3043 | chr19-51728012-DEL-3043 |
| chr19-51729142-DEL-223 | chr19-51729143-DEL-223 |
| chr19-51730284-DEL-537 | chr19-51730282-DEL-537 |
| chr19-51751955-DEL-1872 | chr19-51751955-DEL-1872 |
| chr19-51761323-DEL-54 | chr19-51761323-DEL-54 |
| chr19-51772761-DEL-113 | chr19-51772762-DEL-125 |
| chr19-51787570-DEL-379 | chr19-51787571-DEL-379 |
| chr19-51808038-DEL-424 | chr19-51808039-DEL-424 |
| chr19-51821061-DEL-76 | chr19-51821062-DEL-76 |
| chr19-51848480-DEL-66 | chr19-51848479-DEL-68 |
| chr19-51874758-DEL-83 | chr19-51874759-DEL-83 |
| chr19-51882312-DEL-101 | chr19-51882313-DEL-93 |
| chr19-51888130-DEL-6641 | chr19-51888131-DEL-6641 |
| chr19-51921459-DEL-64 | chr19-51921459-DEL-64 |
| chr19-51930219-DEL-485 | chr19-51930220-DEL-485 |
| chr19-51937557-DEL-6623 | chr19-51937558-DEL-6623 |
| chr19-51946497-DEL-957 | chr19-51946498-DEL-957 |
| chr19-51955684-DEL-58 | chr19-51955685-DEL-62 |
| chr19-51980818-DEL-52 | chr19-51980819-DEL-52 |
| chr19-51983975-DEL-154 | chr19-51983976-DEL-154 |

| | |
|---|---|
| chr19-52022171-DEL-1541 | chr19-52022165-DEL-1542 |
| chr19-52028003-DEL-206 | chr19-52028005-DEL-205 |
| chr19-52038480-DEL-94 | chr19-52038481-DEL-92 |
| chr19-52039592-DEL-63 | chr19-52039593-DEL-63 |
| chr19-52042127-DEL-382 | chr19-52042127-DEL-382 |
| chr19-52069119-DEL-54 | chr19-52069120-DEL-54 |
| chr19-52107658-DEL-1975 | chr19-52107659-DEL-1975 |
| chr19-52140736-DEL-362 | chr19-52140719-DEL-363 |
| chr19-52141474-DEL-3648 | chr19-52141475-DEL-3648 |
| chr19-52147350-DEL-62 | chr19-52147351-DEL-62 |
| chr19-52203950-DEL-797 | chr19-52203951-DEL-797 |
| chr19-52206357-DEL-199 | chr19-52206360-DEL-197 |
| chr19-52243179-DEL-328 | chr19-52243182-DEL-327 |
| chr19-52256495-DEL-68 | chr19-52256516-DEL-58 |
| chr19-52282282-DEL-435 | chr19-52282283-DEL-435 |
| chr19-52291649-DEL-6445 | chr19-52291646-DEL-6449 |
| chr19-52305433-DEL-1350 | chr19-52305440-DEL-1344 |
| chr19-52316693-DEL-380 | chr19-52316694-DEL-380 |
| chr19-52338448-DEL-65 | chr19-52338449-DEL-65 |
| chr19-52341517-DEL-478 | chr19-52341518-DEL-478 |
| chr19-52344684-DEL-506 | chr19-52344685-DEL-506 |
| chr19-52357679-DEL-477 | chr19-52357663-DEL-455 |
| chr19-52368304-DEL-66 | chr19-52368280-DEL-86 |
| chr19-52370089-DEL-98 | chr19-52370090-DEL-98 |
| chr19-52370364-DEL-1876 | chr19-52370364-DEL-1876 |
| chr19-52372998-DEL-1070 | chr19-52372990-DEL-1064 |
| chr19-52392517-DEL-538 | chr19-52392518-DEL-538 |
| chr19-52400106-DEL-209 | chr19-52400107-DEL-209 |
| chr19-52409009-DEL-54 | chr19-52409010-DEL-54 |
| chr19-52420477-DEL-103 | chr19-52420478-DEL-103 |
| chr19-52444695-DEL-65 | chr19-52444675-DEL-54 |
| chr19-52449118-DEL-51 | chr19-52449143-DEL-55 |
| chr19-52473736-DEL-659 | chr19-52473737-DEL-659 |
| chr19-52506823-DEL-55 | chr19-52506812-DEL-67 |
| chr19-52512492-DEL-77 | chr19-52512527-DEL-91 |
| chr19-52517588-DEL-54 | chr19-52517589-DEL-56 |
| chr19-52521938-DEL-83 | chr19-52521939-DEL-83 |
| chr19-52534603-DEL-136 | chr19-52534604-DEL-136 |
| chr19-52552337-DEL-63 | chr19-52552338-DEL-63 |
| chr19-52635096-DEL-58 | chr19-52635106-DEL-95 |
| chr19-52640487-DEL-162 | chr19-52640488-DEL-162 |
| chr19-52644240-DEL-588 | chr19-52644241-DEL-588 |
| chr19-52652626-DEL-93 | chr19-52652611-DEL-71 |
| chr19-52662480-DEL-996 | chr19-52662478-DEL-995 |
| chr19-52713004-DEL-1221 | chr19-52713005-DEL-1221 |
| chr19-52730168-DEL-66 | chr19-52730169-DEL-68 |
| chr19-52739205-DEL-259 | chr19-52739205-DEL-257 |
| chr19-52740300-DEL-238 | chr19-52740296-DEL-239 |
| chr19-52757892-DEL-73 | chr19-52757893-DEL-73 |
| chr19-52766827-DEL-2786 | chr19-52766827-DEL-2786 |
| chr19-52786094-DEL-1826 | chr19-52786095-DEL-1826 |
| chr19-52822414-DEL-329 | chr19-52822422-DEL-327 |
| chr19-52823857-DEL-61 | chr19-52823858-DEL-61 |
| chr19-52827441-DEL-401 | chr19-52827442-DEL-401 |
| chr19-52829138-DEL-6398 | chr19-52829139-DEL-6398 |
| chr19-52883554-DEL-1174 | chr19-52883555-DEL-1169 |
| chr19-52892345-DEL-181 | chr19-52892346-DEL-181 |
| chr19-52916183-DEL-98 | chr19-52916178-DEL-110 |
| chr19-52916476-DEL-4164 | chr19-52916477-DEL-4164 |
| chr19-52934351-DEL-78 | chr19-52934388-DEL-54 |
| chr19-52935168-DEL-66 | chr19-52935169-DEL-66 |
| chr19-52955464-DEL-400 | chr19-52955469-DEL-404 |
| chr19-52956360-DEL-3377 | chr19-52956361-DEL-3377 |
| chr19-52973975-DEL-5402 | chr19-52973976-DEL-5402 |
| chr19-53012988-DEL-62 | chr19-53012989-DEL-62 |
| chr19-53024203-DEL-81 | chr19-53024207-DEL-79 |
| chr19-53025583-DEL-102 | chr19-53025584-DEL-102 |
| chr19-53033229-DEL-497 | chr19-53033230-DEL-497 |
| chr19-53044791-DEL-113 | chr19-53044791-DEL-114 |
| chr19-53070613-DEL-266 | chr19-53070614-DEL-266 |
| chr19-53093158-DEL-66 | chr19-53093159-DEL-66 |
| chr19-53106152-DEL-53 | chr19-53106153-DEL-53 |
| chr19-53150183-DEL-181 | chr19-53150184-DEL-181 |
| chr19-53188084-DEL-258 | chr19-53188085-DEL-258 |
| chr19-53193876-DEL-77 | chr19-53193877-DEL-77 |
| chr19-53200045-DEL-66 | chr19-53200046-DEL-66 |
| chr19-53215708-DEL-59 | chr19-53215709-DEL-59 |
| chr19-53275126-DEL-64 | chr19-53275127-DEL-64 |
| chr19-53336169-DEL-211 | chr19-53336169-DEL-211 |
| chr19-53337343-DEL-438 | chr19-53337296-DEL-474 |

| | |
|---|---|
| chr19-53356482-DEL-308 | chr19-53356483-DEL-308 |
| chr19-53387314-DEL-87 | chr19-53387339-DEL-56 |
| chr19-53395244-DEL-198 | chr19-53395245-DEL-198 |
| chr19-53401933-DEL-193 | chr19-53401934-DEL-193 |
| chr19-53408628-DEL-74 | chr19-53408619-DEL-84 |
| chr19-53410558-DEL-105 | chr19-53410548-DEL-92 |
| chr19-53430406-DEL-83 | chr19-53430364-DEL-84 |
| chr19-53434645-DEL-311 | chr19-53434646-DEL-311 |
| chr19-53462599-DEL-61 | chr19-53462581-DEL-79 |
| chr19-53468683-DEL-472 | chr19-53468684-DEL-471 |
| chr19-53544945-DEL-188 | chr19-53544946-DEL-188 |
| chr19-53561520-DEL-52 | chr19-53561507-DEL-54 |
| chr19-53565111-DEL-177 | chr19-53565123-DEL-178 |
| chr19-53576560-DEL-932 | chr19-53576561-DEL-932 |
| chr19-53597968-DEL-202 | chr19-53597968-DEL-203 |
| chr19-53632532-DEL-233 | chr19-53632524-DEL-231 |
| chr19-53644108-DEL-64 | chr19-53644067-DEL-102 |
| chr19-53674494-DEL-205 | chr19-53674495-DEL-205 |
| chr19-53687759-DEL-180 | chr19-53687760-DEL-180 |
| chr19-53697783-DEL-414 | chr19-53697784-DEL-414 |
| chr19-53712046-DEL-209 | chr19-53712047-DEL-183 |
| chr19-53724711-DEL-644 | chr19-53724712-DEL-644 |
| chr19-53752089-DEL-213 | chr19-53752090-DEL-213 |
| chr19-53754486-DEL-76 | chr19-53754497-DEL-56 |
| chr19-53765269-DEL-179 | chr19-53765268-DEL-179 |
| chr19-53813549-DEL-54 | chr19-53813550-DEL-54 |
| chr19-53830301-DEL-56 | chr19-53830302-DEL-56 |
| chr19-53867847-DEL-1370 | chr19-53867848-DEL-1370 |
| chr19-53871236-DEL-173 | chr19-53871237-DEL-173 |
| chr19-53872283-DEL-706 | chr19-53872278-DEL-714 |
| chr19-53874781-DEL-166 | chr19-53874782-DEL-166 |
| chr19-53878638-DEL-57 | chr19-53878639-DEL-57 |
| chr19-53928521-DEL-192 | chr19-53928518-DEL-196 |
| chr19-53951620-DEL-235 | chr19-53951621-DEL-235 |
| chr19-53966607-DEL-159 | chr19-53966608-DEL-159 |
| chr19-53992543-DEL-983 | chr19-53992544-DEL-983 |
| chr19-53996734-DEL-103 | chr19-53996735-DEL-134 |
| chr19-53997620-DEL-560 | chr19-53997634-DEL-558 |
| chr19-54006695-DEL-1600 | chr19-54006696-DEL-1600 |
| chr19-54074859-DEL-511 | chr19-54074854-DEL-512 |
| chr19-54080880-DEL-194 | chr19-54080881-DEL-194 |
| chr19-54095043-DEL-63 | chr19-54095044-DEL-63 |
| chr19-54120983-DEL-56 | chr19-54120977-DEL-59 |
| chr19-54152897-DEL-146 | chr19-54152898-DEL-150 |
| chr19-54201207-DEL-110 | chr19-54201208-DEL-110 |
| chr19-54222860-DEL-459 | chr19-54222861-DEL-459 |
| chr19-54234992-DEL-61 | chr19-54234998-DEL-56 |
| chr19-54252609-DEL-249 | chr19-54252609-DEL-249 |
| chr19-54270199-DEL-209 | chr19-54270200-DEL-209 |
| chr19-54274049-DEL-63 | chr19-54274050-DEL-63 |
| chr19-54290970-DEL-54 | chr19-54290936-DEL-51 |
| chr19-54297193-DEL-500 | chr19-54297194-DEL-500 |
| chr19-54298701-DEL-514 | chr19-54298702-DEL-514 |
| chr19-54330975-DEL-2493 | chr19-54330976-DEL-2493 |
| chr19-54338439-DEL-2048 | chr19-54338440-DEL-2048 |
| chr19-54356292-DEL-68 | chr19-54356293-DEL-52 |
| chr19-54361453-DEL-736 | chr19-54361454-DEL-736 |
| chr19-54375210-DEL-115 | chr19-54375211-DEL-115 |
| chr19-54406068-DEL-95 | chr19-54406068-DEL-96 |
| chr19-54471714-DEL-417 | chr19-54471715-DEL-417 |
| chr19-54474327-DEL-1274 | chr19-54474328-DEL-1274 |
| chr19-54477100-DEL-1489 | chr19-54477122-DEL-1470 |
| chr19-54493033-DEL-59 | chr19-54493033-DEL-59 |
| chr19-54495863-DEL-173 | chr19-54495864-DEL-173 |
| chr19-54496743-DEL-57 | chr19-54496744-DEL-56 |
| chr19-54520725-DEL-598 | chr19-54520735-DEL-586 |
| chr19-54528616-DEL-1995 | chr19-54528617-DEL-1995 |
| chr19-54563300-DEL-4109 | chr19-54563292-DEL-4107 |
| chr19-54579603-DEL-56 | chr19-54579588-DEL-59 |
| chr19-54584593-DEL-569 | chr19-54584592-DEL-570 |
| chr19-54629052-DEL-420 | chr19-54629053-DEL-420 |
| chr19-54631756-DEL-1417 | chr19-54631757-DEL-1417 |
| chr19-54654916-DEL-93 | chr19-54654912-DEL-67 |
| chr19-54675482-DEL-98 | chr19-54675493-DEL-88 |
| chr19-54682857-DEL-75 | chr19-54682858-DEL-75 |
| chr19-54687709-DEL-1092 | chr19-54687710-DEL-1092 |
| chr19-54691697-DEL-77 | chr19-54691700-DEL-88 |
| chr19-54726442-DEL-475 | chr19-54726406-DEL-476 |
| chr19-54838319-DEL-433 | chr19-54838320-DEL-433 |
| chr19-54866486-DEL-54 | chr19-54866487-DEL-54 |

| | |
|---|---|
| chr19-54878619-DEL-2828 | chr19-54878620-DEL-2828 |
| chr19-54887516-DEL-62 | chr19-54887516-DEL-62 |
| chr19-54900154-DEL-6215 | chr19-54900155-DEL-6215 |
| chr19-54912708-DEL-201 | chr19-54912709-DEL-201 |
| chr19-54958885-DEL-141 | chr19-54958886-DEL-141 |
| chr19-54960509-DEL-272 | chr19-54960500-DEL-233 |
| chr19-54983392-DEL-6578 | chr19-54983393-DEL-6578 |
| chr19-55013636-DEL-80 | chr19-55013641-DEL-78 |
| chr19-55024851-DEL-57 | chr19-55024846-DEL-57 |
| chr19-55030903-DEL-127 | chr19-55030903-DEL-128 |
| chr19-55041494-DEL-58 | chr19-55041507-DEL-59 |
| chr19-55148409-DEL-357 | chr19-55148410-DEL-357 |
| chr19-55179422-DEL-160 | chr19-55179422-DEL-160 |
| chr19-55193899-DEL-196 | chr19-55193916-DEL-197 |
| chr19-55222881-DEL-221 | chr19-55222882-DEL-221 |
| chr19-55227353-DEL-53 | chr19-55227354-DEL-53 |
| chr19-55228879-DEL-55 | chr19-55228879-DEL-55 |
| chr19-55267028-DEL-85 | chr19-55267029-DEL-85 |
| chr19-55323069-DEL-69 | chr19-55323070-DEL-69 |
| chr19-55344442-DEL-232 | chr19-55344443-DEL-232 |
| chr19-55409470-DEL-117 | chr19-55409471-DEL-117 |
| chr19-55410034-DEL-86 | chr19-55410019-DEL-58 |
| chr19-55415810-DEL-90 | chr19-55415811-DEL-90 |
| chr19-55423976-DEL-115 | chr19-55423977-DEL-115 |
| chr19-55432951-DEL-823 | chr19-55432952-DEL-823 |
| chr19-55441579-DEL-60 | chr19-55441580-DEL-60 |
| chr19-55443250-DEL-238 | chr19-55443251-DEL-238 |
| chr19-55455077-DEL-1970 | chr19-55455078-DEL-1970 |
| chr19-55460307-DEL-63 | chr19-55460308-DEL-63 |
| chr19-55473859-DEL-81 | chr19-55473843-DEL-54 |
| chr19-55503631-DEL-81 | chr19-55503634-DEL-79 |
| chr19-55506766-DEL-243 | chr19-55506767-DEL-243 |
| chr19-55507199-DEL-335 | chr19-55507200-DEL-335 |
| chr19-55517007-DEL-521 | chr19-55517017-DEL-519 |
| chr19-55520359-DEL-60 | chr19-55520360-DEL-60 |
| chr19-55522453-DEL-55 | chr19-55522454-DEL-55 |
| chr19-55534368-DEL-67 | chr19-55534369-DEL-67 |
| chr19-55538185-DEL-90 | chr19-55538190-DEL-62 |
| chr19-55546589-DEL-1187 | chr19-55546595-DEL-1183 |
| chr19-55575469-DEL-866 | chr19-55575469-DEL-866 |
| chr19-55582070-DEL-55 | chr19-55582071-DEL-55 |
| chr19-55640332-DEL-58 | chr19-55640335-DEL-56 |
| chr19-55663814-DEL-208 | chr19-55663815-DEL-208 |
| chr19-55697520-DEL-188 | chr19-55697521-DEL-175 |
| chr19-55707366-DEL-113 | chr19-55707367-DEL-113 |
| chr19-55713087-DEL-195 | chr19-55713084-DEL-196 |
| chr19-55727576-DEL-70 | chr19-55727577-DEL-70 |
| chr19-55756628-DEL-67 | chr19-55756628-DEL-65 |
| chr19-55776602-DEL-100 | chr19-55776598-DEL-105 |
| chr19-55821608-DEL-116 | chr19-55821578-DEL-122 |
| chr19-55830373-DEL-57 | chr19-55830374-DEL-57 |
| chr19-55847118-DEL-186 | chr19-55847119-DEL-186 |
| chr19-55928633-DEL-85 | chr19-55928631-DEL-79 |
| chr19-55965195-DEL-7067 | chr19-55965202-DEL-7063 |
| chr19-55986260-DEL-64 | chr19-55986245-DEL-65 |
| chr19-55997668-DEL-610 | chr19-55997668-DEL-614 |
| chr19-56028780-DEL-76 | chr19-56028784-DEL-116 |
| chr19-56029495-DEL-124 | chr19-56029496-DEL-124 |
| chr19-56030978-DEL-64 | chr19-56030981-DEL-62 |
| chr19-56040704-DEL-1159 | chr19-56040705-DEL-1159 |
| chr19-56042087-DEL-66 | chr19-56042088-DEL-66 |
| chr19-56058068-DEL-171 | chr19-56058069-DEL-171 |
| chr19-56105425-DEL-916 | chr19-56105425-DEL-916 |
| chr19-56107924-DEL-59 | chr19-56107935-DEL-50 |
| chr19-56115483-DEL-84 | chr19-56115484-DEL-84 |
| chr19-56147561-DEL-92 | chr19-56147522-DEL-112 |
| chr19-56172509-DEL-411 | chr19-56172507-DEL-411 |
| chr19-56202949-DEL-1731 | chr19-56202950-DEL-1731 |
| chr19-56204056-DEL-208 | chr19-56204057-DEL-208 |
| chr19-56214278-DEL-54 | chr19-56214279-DEL-54 |
| chr19-56228410-DEL-75 | chr19-56228408-DEL-60 |
| chr19-56245460-DEL-1901 | chr19-56245461-DEL-1901 |
| chr19-56254690-DEL-79 | chr19-56254691-DEL-79 |
| chr19-56255297-DEL-398 | chr19-56255298-DEL-398 |
| chr19-56281624-DEL-54 | chr19-56281625-DEL-52 |
| chr19-56283784-DEL-530 | chr19-56283785-DEL-530 |
| chr19-56285401-DEL-58 | chr19-56285362-DEL-58 |
| chr19-56298779-DEL-144 | chr19-56298780-DEL-144 |
| chr19-56328128-DEL-100 | chr19-56328129-DEL-100 |
| chr19-56341573-DEL-77 | chr19-56341559-DEL-98 |

| | |
|---|---|
| chr19-56369098-DEL-70 | chr19-56369098-DEL-56 |
| chr19-56371342-DEL-84 | chr19-56371344-DEL-83 |
| chr19-56394040-DEL-209 | chr19-56394041-DEL-209 |
| chr19-56409195-DEL-169 | chr19-56409198-DEL-166 |
| chr19-56423696-DEL-60 | chr19-56423697-DEL-60 |
| chr19-56433625-DEL-1191 | chr19-56433626-DEL-1191 |
| chr19-56496175-DEL-86 | chr19-56496176-DEL-86 |
| chr19-56499410-DEL-62 | chr19-56499382-DEL-93 |
| chr19-56516756-DEL-136 | chr19-56516757-DEL-136 |
| chr19-56578600-DEL-624 | chr19-56578603-DEL-623 |
| chr19-56588124-DEL-54 | chr19-56588125-DEL-54 |
| chr19-56600201-DEL-104 | chr19-56600188-DEL-68 |
| chr19-56622823-DEL-205 | chr19-56622835-DEL-187 |
| chr19-56629742-DEL-400 | chr19-56629743-DEL-400 |
| chr19-56647794-DEL-80 | chr19-56647795-DEL-80 |
| chr19-56680180-DEL-77 | chr19-56680184-DEL-73 |
| chr19-56687166-DEL-91 | chr19-56687167-DEL-91 |
| chr19-56723837-DEL-111 | chr19-56723875-DEL-77 |
| chr19-56731181-DEL-317 | chr19-56731138-DEL-333 |
| chr19-56740067-DEL-618 | chr19-56740068-DEL-618 |
| chr19-56745890-DEL-97 | chr19-56745866-DEL-121 |
| chr19-56786744-DEL-59 | chr19-56786745-DEL-59 |
| chr19-56792015-DEL-82 | chr19-56792016-DEL-82 |
| chr19-56805168-DEL-86 | chr19-56805169-DEL-86 |
| chr19-56847607-DEL-51 | chr19-56847608-DEL-57 |
| chr19-56867758-DEL-106 | chr19-56867798-DEL-108 |
| chr19-56875626-DEL-58 | chr19-56875627-DEL-58 |
| chr19-56910978-DEL-170 | chr19-56910971-DEL-159 |
| chr19-56939723-DEL-164 | chr19-56939752-DEL-133 |
| chr19-56941869-DEL-115 | chr19-56941870-DEL-114 |
| chr19-56956721-DEL-77 | chr19-56956704-DEL-87 |
| chr19-56959139-DEL-195 | chr19-56959140-DEL-195 |
| chr19-56969660-DEL-51 | chr19-56969661-DEL-51 |
| chr19-56977539-DEL-116 | chr19-56977540-DEL-116 |
| chr19-56979925-DEL-111 | chr19-56979937-DEL-111 |
| chr19-57006665-DEL-92 | chr19-57006666-DEL-114 |
| chr19-57059133-DEL-699 | chr19-57059134-DEL-699 |
| chr19-57062602-DEL-401 | chr19-57062603-DEL-401 |
| chr19-57070456-DEL-207 | chr19-57070457-DEL-207 |
| chr19-57084832-DEL-112 | chr19-57084833-DEL-112 |
| chr19-57085410-DEL-101 | chr19-57085442-DEL-61 |
| chr19-57094728-DEL-118 | chr19-57094729-DEL-118 |
| chr19-57101878-DEL-152 | chr19-57101861-DEL-170 |
| chr19-57109102-DEL-58 | chr19-57109105-DEL-84 |
| chr19-57114512-DEL-85 | chr19-57114513-DEL-85 |
| chr19-57123745-DEL-498 | chr19-57123746-DEL-498 |
| chr19-57133267-DEL-64 | chr19-57133292-DEL-56 |
| chr19-57147026-DEL-748 | chr19-57147027-DEL-748 |
| chr19-57160382-DEL-102 | chr19-57160383-DEL-100 |
| chr19-57202449-DEL-397 | chr19-57202450-DEL-397 |
| chr19-57208868-DEL-64 | chr19-57208869-DEL-64 |
| chr19-57214203-DEL-1100 | chr19-57214204-DEL-1100 |
| chr19-57218511-DEL-226 | chr19-57218529-DEL-205 |
| chr19-57221823-DEL-212 | chr19-57221852-DEL-198 |
| chr19-57259087-DEL-57 | chr19-57259088-DEL-57 |
| chr19-57269581-DEL-98 | chr19-57269623-DEL-88 |
| chr19-57309647-DEL-1174 | chr19-57309648-DEL-1174 |
| chr19-57326267-DEL-300 | chr19-57326268-DEL-300 |
| chr19-57339877-DEL-71 | chr19-57339891-DEL-74 |
| chr19-57347850-DEL-57 | chr19-57347851-DEL-57 |
| chr19-57368020-DEL-179 | chr19-57368020-DEL-179 |
| chr19-57392494-DEL-210 | chr19-57392495-DEL-210 |
| chr19-57418218-DEL-1516 | chr19-57418219-DEL-1516 |
| chr19-57426142-DEL-418 | chr19-57426143-DEL-418 |
| chr19-57436932-DEL-171 | chr19-57436931-DEL-175 |
| chr19-57444131-DEL-106 | chr19-57444132-DEL-106 |
| chr19-57447925-DEL-288 | chr19-57447926-DEL-288 |
| chr19-57449471-DEL-943 | chr19-57449472-DEL-943 |
| chr19-57451837-DEL-203 | chr19-57451838-DEL-202 |
| chr19-57502662-DEL-6401 | chr19-57502663-DEL-6401 |
| chr19-57525605-DEL-54 | chr19-57525606-DEL-54 |
| chr19-57549522-DEL-3413 | chr19-57549523-DEL-3413 |
| chr19-57570804-DEL-181 | chr19-57570805-DEL-181 |
| chr19-57590406-DEL-711 | chr19-57590407-DEL-711 |
| chr19-57608092-DEL-56 | chr19-57608097-DEL-52 |
| chr19-57616741-DEL-54 | chr19-57616742-DEL-54 |
| chr19-57622592-DEL-1282 | chr19-57622593-DEL-1282 |
| chr19-57622937-DEL-609 | chr19-57622938-DEL-609 |
| chr19-57627946-DEL-83 | chr19-57627947-DEL-83 |
| chr19-57645915-DEL-394 | chr19-57645916-DEL-394 |

| | |
|---|---|
| chr19-57650685-DEL-214 | chr19-57650685-DEL-214 |
| chr19-57667926-DEL-62 | chr19-57667927-DEL-62 |
| chr19-57672397-DEL-263 | chr19-57672406-DEL-227 |
| chr19-57696228-DEL-660 | chr19-57696229-DEL-660 |
| chr19-57728784-DEL-207 | chr19-57728791-DEL-206 |
| chr19-57729467-DEL-192 | chr19-57729468-DEL-192 |
| chr19-57762790-DEL-215 | chr19-57762791-DEL-215 |
| chr19-57766457-DEL-56 | chr19-57766458-DEL-56 |
| chr19-57775809-DEL-70 | chr19-57775810-DEL-70 |
| chr19-57812287-DEL-72 | chr19-57812317-DEL-70 |
| chr19-57815534-DEL-5402 | chr19-57815535-DEL-5402 |
| chr19-57824577-DEL-104 | chr19-57824584-DEL-100 |
| chr19-57838061-DEL-188 | chr19-57838027-DEL-189 |
| chr19-57880084-DEL-94 | chr19-57880087-DEL-94 |
| chr19-57931165-DEL-82 | chr19-57931166-DEL-82 |
| chr19-57935401-DEL-454 | chr19-57935402-DEL-454 |
| chr19-57963770-DEL-78 | chr19-57963771-DEL-78 |
| chr19-57968115-DEL-325 | chr19-57968116-DEL-325 |
| chr19-57971182-DEL-76 | chr19-57971183-DEL-62 |
| chr19-58001018-DEL-1891 | chr19-58001019-DEL-1891 |
| chr19-58039101-DEL-284 | chr19-58039102-DEL-284 |
| chr19-58103526-DEL-67 | chr19-58103533-DEL-68 |
| chr19-58115473-DEL-62 | chr19-58115474-DEL-62 |
| chr19-58138508-DEL-78 | chr19-58138511-DEL-82 |
| chr19-58142232-DEL-84 | chr19-58142233-DEL-84 |
| chr19-58157678-DEL-57 | chr19-58157706-DEL-60 |
| chr19-58177020-DEL-258 | chr19-58177021-DEL-258 |
| chr19-58178120-DEL-228 | chr19-58178121-DEL-228 |
| chr19-58185241-DEL-58 | chr19-58185212-DEL-68 |
| chr19-58306915-DEL-594 | chr19-58306920-DEL-578 |
| chr19-58318575-DEL-126 | chr19-58318576-DEL-126 |
| chr19-58348281-DEL-293 | chr19-58348281-DEL-293 |
| chr19-58349836-DEL-82 | chr19-58349841-DEL-78 |
| chr19-58351448-DEL-65 | chr19-58351449-DEL-65 |
| chr19-58452642-DEL-271 | chr19-58452643-DEL-271 |
| chr19-58482276-DEL-70 | chr19-58482269-DEL-68 |
| chr19-58494691-DEL-54 | chr19-58494684-DEL-79 |
| chr19-58506901-DEL-112 | chr19-58506904-DEL-74 |
| chr19-58532778-DEL-1816 | chr19-58532779-DEL-1816 |
| chr19-58549417-DEL-179 | chr19-58549418-DEL-179 |
| chr19-58607547-DEL-440 | chr19-58607548-DEL-440 |
| chr19-58641052-DEL-385 | chr19-58641053-DEL-385 |
| chr19-58644580-DEL-244 | chr19-58644581-DEL-244 |
| chr19-58654753-DEL-118 | chr19-58654738-DEL-134 |
| chr19-58660530-DEL-60 | chr19-58660529-DEL-58 |
| chr19-58679843-DEL-54 | chr19-58679844-DEL-54 |
| chr19-58695249-DEL-581 | chr19-58695250-DEL-581 |
| chr19-58725839-DEL-674 | chr19-58725840-DEL-674 |
| chr19-58726971-DEL-57 | chr19-58726972-DEL-57 |
| chr19-58758761-DEL-134 | chr19-58758762-DEL-134 |
| chr19-58764141-DEL-144 | chr19-58764142-DEL-144 |
| chr19-58850551-DEL-1833 | chr19-58850552-DEL-1833 |
| chr19-58874339-DEL-91 | chr19-58874332-DEL-89 |
| chr19-58897749-DEL-112 | chr19-58897750-DEL-112 |
| chr19-58899372-DEL-81 | chr19-58899373-DEL-81 |
| chr19-58903228-DEL-71 | chr19-58903229-DEL-71 |
| chr19-58914159-DEL-83 | chr19-58914160-DEL-83 |
| chr19-58954097-DEL-509 | chr19-58954082-DEL-510 |
| chr19-58955148-DEL-646 | chr19-58955149-DEL-646 |
| chr19-58971334-DEL-69 | chr19-58971335-DEL-69 |
| chr19-58976007-DEL-54 | chr19-58976007-DEL-54 |
| chr19-59095272-DEL-197 | chr19-59095273-DEL-197 |
| chr19-59110417-DEL-63 | chr19-59110423-DEL-62 |
| chr19-59141389-DEL-55 | chr19-59141390-DEL-55 |
| chr19-59210308-DEL-2907 | chr19-59210309-DEL-2907 |
| chr19-59266285-DEL-85 | chr19-59266288-DEL-83 |
| chr19-59274541-DEL-89 | chr19-59274519-DEL-90 |
| chr19-59310704-DEL-525 | chr19-59310705-DEL-525 |
| chr19-59313309-DEL-194 | chr19-59313310-DEL-194 |
| chr19-59342913-DEL-81 | chr19-59342912-DEL-78 |
| chr19-59367798-DEL-76 | chr19-59367799-DEL-76 |
| chr19-59379659-DEL-71 | chr19-59379639-DEL-111 |
| chr19-59405428-DEL-88 | chr19-59405428-DEL-88 |
| chr19-59410202-DEL-146 | chr19-59410202-DEL-146 |
| chr19-59422685-DEL-52 | chr19-59422670-DEL-56 |
| chr19-59468877-DEL-105 | chr19-59468877-DEL-100 |
| chr19-59469674-DEL-468 | chr19-59469675-DEL-468 |
| chr19-59475217-DEL-63 | chr19-59475206-DEL-75 |
| chr19-59489980-DEL-80 | chr19-59489964-DEL-104 |
| chr19-59493921-DEL-80 | chr19-59493964-DEL-70 |

| | |
|---|---|
| chr19-59581723-DEL-87 | chr19-59581724-DEL-87 |
| chr19-59590455-DEL-51 | chr19-59590456-DEL-51 |
| chr19-59591737-DEL-7201 | chr19-59591739-DEL-7197 |
| chr19-59650582-DEL-2881 | chr19-59650583-DEL-2881 |
| chr19-59663738-DEL-53 | chr19-59663711-DEL-80 |
| chr19-59730501-DEL-74 | chr19-59730502-DEL-74 |
| chr19-59735789-DEL-70 | chr19-59735795-DEL-65 |
| chr19-59748806-DEL-78 | chr19-59748807-DEL-57 |
| chr19-59755139-DEL-592 | chr19-59755140-DEL-592 |
| chr19-59762312-DEL-6005 | chr19-59762313-DEL-6005 |
| chr19-59777417-DEL-54 | chr19-59777373-DEL-98 |
| chr19-59802469-DEL-6186 | chr19-59802470-DEL-6186 |
| chr19-59813499-DEL-1611 | chr19-59813500-DEL-1609 |
| chr19-59838350-DEL-58 | chr19-59838359-DEL-52 |
| chr19-59845155-DEL-63 | chr19-59845158-DEL-64 |
| chr19-59860444-DEL-2873 | chr19-59860445-DEL-2873 |
| chr19-59940567-DEL-102 | chr19-59940568-DEL-102 |
| chr19-59941199-DEL-231 | chr19-59941200-DEL-231 |
| chr19-59965988-DEL-130 | chr19-59965989-DEL-130 |
| chr19-59971790-DEL-135 | chr19-59971791-DEL-135 |
| chr19-59986751-DEL-1527 | chr19-59986752-DEL-1527 |
| chr19-60007132-DEL-6245 | chr19-60007133-DEL-6245 |
| chr19-60033096-DEL-78 | chr19-60033097-DEL-78 |
| chr19-60044620-DEL-636 | chr19-60044621-DEL-636 |
| chr19-60058537-DEL-212 | chr19-60058538-DEL-212 |
| chr19-60097108-DEL-61 | chr19-60097109-DEL-63 |
| chr19-60160526-DEL-52 | chr19-60160529-DEL-50 |
| chr19-60175337-DEL-106 | chr19-60175336-DEL-106 |
| chr19-60220414-DEL-162 | chr19-60220415-DEL-162 |
| chr19-60223436-DEL-78 | chr19-60223437-DEL-78 |
| chr19-60229780-DEL-102 | chr19-60229781-DEL-102 |
| chr19-60254994-DEL-83 | chr19-60254961-DEL-117 |
| chr19-60279389-DEL-201 | chr19-60279390-DEL-201 |
| chr19-60283096-DEL-60 | chr19-60283093-DEL-68 |
| chr19-60297710-DEL-344 | chr19-60297711-DEL-344 |
| chr19-60303963-DEL-60 | chr19-60303964-DEL-60 |
| chr19-60308083-DEL-52 | chr19-60308084-DEL-52 |
| chr19-60335991-DEL-7227 | chr19-60335992-DEL-7227 |
| chr19-60365837-DEL-55 | chr19-60365838-DEL-55 |
| chr19-60401252-DEL-80 | chr19-60401253-DEL-76 |
| chr19-60412976-DEL-172 | chr19-60412977-DEL-172 |
| chr19-60453559-DEL-139 | chr19-60453554-DEL-143 |
| chr19-60458508-DEL-94 | chr19-60458524-DEL-78 |
| chr19-60468133-DEL-95 | chr19-60468137-DEL-107 |
| chr19-60541012-DEL-201 | chr19-60541013-DEL-200 |
| chr19-60561163-DEL-221 | chr19-60561164-DEL-221 |
| chr19-60574282-DEL-215 | chr19-60574283-DEL-215 |
| chr19-60605376-DEL-122 | chr19-60605377-DEL-122 |
| chr19-60615850-DEL-181 | chr19-60615851-DEL-181 |
| chr19-60629670-DEL-1106 | chr19-60629671-DEL-1106 |
| chr19-60632192-DEL-6477 | chr19-60632182-DEL-6480 |
| chr19-60641512-DEL-117 | chr19-60641517-DEL-107 |
| chr19-60641942-DEL-512 | chr19-60641943-DEL-512 |
| chr19-60645519-DEL-540 | chr19-60645520-DEL-540 |
| chr19-60657099-DEL-55 | chr19-60657079-DEL-52 |
| chr19-60658712-DEL-51 | chr19-60658713-DEL-51 |
| chr19-60679180-DEL-96 | chr19-60679181-DEL-96 |
| chr19-60680103-DEL-568 | chr19-60680104-DEL-568 |
| chr19-60692080-DEL-1132 | chr19-60692071-DEL-1133 |
| chr19-60713046-DEL-59 | chr19-60713046-DEL-59 |
| chr19-60732648-DEL-127 | chr19-60732649-DEL-127 |
| chr19-60735995-DEL-103 | chr19-60736002-DEL-124 |
| chr19-60789412-DEL-146 | chr19-60789413-DEL-110 |
| chr19-60789942-DEL-180 | chr19-60789943-DEL-180 |
| chr19-60801898-DEL-173 | chr19-60801899-DEL-173 |
| chr19-60817294-DEL-651 | chr19-60817254-DEL-692 |
| chr19-60819766-DEL-208 | chr19-60819761-DEL-198 |
| chr19-60822819-DEL-169 | chr19-60822820-DEL-169 |
| chr19-60826682-DEL-61 | chr19-60826683-DEL-61 |
| chr19-60867327-DEL-1569 | chr19-60867328-DEL-1569 |
| chr19-60879563-DEL-52 | chr19-60879564-DEL-52 |
| chr19-60899395-DEL-89 | chr19-60899396-DEL-89 |
| chr19-60910811-DEL-228 | chr19-60910812-DEL-228 |
| chr19-60913740-DEL-8731 | chr19-60913741-DEL-8731 |
| chr19-60937067-DEL-164 | chr19-60937042-DEL-156 |
| chr19-61009892-DEL-54 | chr19-61009893-DEL-54 |
| chr19-61014015-DEL-202 | chr19-61014016-DEL-202 |
| chr19-61015096-DEL-205 | chr19-61015099-DEL-203 |
| chr19-61101586-DEL-224 | chr19-61101557-DEL-219 |
| chr19-61102626-DEL-609 | chr19-61102627-DEL-609 |

| | |
|---|---|
| chr19-61107078-DEL-69 | chr19-61107079-DEL-69 |
| chr19-61120662-DEL-69 | chr19-61120663-DEL-69 |
| chr19-61123696-DEL-232 | chr19-61123702-DEL-227 |
| chr19-61157210-DEL-6469 | chr19-61157208-DEL-6468 |
| chr19-61172203-DEL-252 | chr19-61172204-DEL-252 |
| chr19-61180661-DEL-121 | chr19-61180662-DEL-121 |
| chr19-61202324-DEL-52 | chr19-61202325-DEL-52 |
| chr19-61226257-DEL-56 | chr19-61226258-DEL-56 |
| chr2-3051886-DEL-62 | chr2-3051899-DEL-56 |
| chr2-3105002-DEL-62 | chr2-3105001-DEL-70 |
| chr2-3174179-DEL-86 | chr2-3174177-DEL-52 |
| chr2-3218742-DEL-122 | chr2-3218743-DEL-122 |
| chr2-3251334-DEL-818 | chr2-3251335-DEL-818 |
| chr2-3282184-DEL-190 | chr2-3282185-DEL-190 |
| chr2-3315394-DEL-147 | chr2-3315395-DEL-147 |
| chr2-3397297-DEL-3916 | chr2-3397298-DEL-3916 |
| chr2-3433943-DEL-95 | chr2-3433944-DEL-95 |
| chr2-3440555-DEL-90 | chr2-3440556-DEL-90 |
| chr2-3469421-DEL-241 | chr2-3469422-DEL-241 |
| chr2-3484544-DEL-5266 | chr2-3484545-DEL-5266 |
| chr2-3502770-DEL-239 | chr2-3502771-DEL-239 |
| chr2-3525291-DEL-158 | chr2-3525292-DEL-158 |
| chr2-3548943-DEL-2148 | chr2-3548944-DEL-2148 |
| chr2-3567404-DEL-60 | chr2-3567405-DEL-60 |
| chr2-3568827-DEL-417 | chr2-3568828-DEL-417 |
| chr2-3606618-DEL-56 | chr2-3606618-DEL-56 |
| chr2-3615566-DEL-124 | chr2-3615567-DEL-124 |
| chr2-3632782-DEL-63 | chr2-3632783-DEL-63 |
| chr2-3742083-DEL-59 | chr2-3742072-DEL-50 |
| chr2-3747792-DEL-52 | chr2-3747793-DEL-52 |
| chr2-3768839-DEL-1418 | chr2-3768840-DEL-1418 |
| chr2-3779743-DEL-66 | chr2-3779744-DEL-66 |
| chr2-3796821-DEL-52 | chr2-3796844-DEL-54 |
| chr2-3801922-DEL-1957 | chr2-3801923-DEL-1957 |
| chr2-3822853-DEL-59 | chr2-3822854-DEL-59 |
| chr2-3843301-DEL-8371 | chr2-3843302-DEL-8371 |
| chr2-3848705-DEL-1912 | chr2-3848704-DEL-1912 |
| chr2-3922138-DEL-82 | chr2-3922139-DEL-82 |
| chr2-3924372-DEL-556 | chr2-3924373-DEL-556 |
| chr2-3933769-DEL-6440 | chr2-3933770-DEL-6440 |
| chr2-3970817-DEL-4874 | chr2-3970818-DEL-4874 |
| chr2-3988974-DEL-2244 | chr2-3988974-DEL-2244 |
| chr2-4040568-DEL-87 | chr2-4040585-DEL-83 |
| chr2-4063195-DEL-57 | chr2-4063194-DEL-53 |
| chr2-4071142-DEL-132 | chr2-4071151-DEL-124 |
| chr2-4099392-DEL-1266 | chr2-4099393-DEL-1264 |
| chr2-4130766-DEL-50 | chr2-4130791-DEL-54 |
| chr2-4161902-DEL-156 | chr2-4161903-DEL-156 |
| chr2-4170798-DEL-122 | chr2-4170799-DEL-122 |
| chr2-4191011-DEL-118 | chr2-4191012-DEL-118 |
| chr2-4194557-DEL-437 | chr2-4194555-DEL-437 |
| chr2-4202146-DEL-6964 | chr2-4202147-DEL-6964 |
| chr2-4258635-DEL-60 | chr2-4258643-DEL-54 |
| chr2-4263604-DEL-320 | chr2-4263605-DEL-320 |
| chr2-4270691-DEL-210 | chr2-4270692-DEL-210 |
| chr2-4276678-DEL-4892 | chr2-4276679-DEL-4892 |
| chr2-4286762-DEL-200 | chr2-4286763-DEL-200 |
| chr2-4298391-DEL-249 | chr2-4298392-DEL-249 |
| chr2-4305990-DEL-80 | chr2-4305999-DEL-72 |
| chr2-4308770-DEL-68 | chr2-4308784-DEL-66 |
| chr2-4322126-DEL-160 | chr2-4322127-DEL-160 |
| chr2-4326485-DEL-2975 | chr2-4326501-DEL-2962 |
| chr2-4332666-DEL-1140 | chr2-4332667-DEL-1140 |
| chr2-4341287-DEL-74 | chr2-4341288-DEL-74 |
| chr2-4347584-DEL-831 | chr2-4347585-DEL-831 |
| chr2-4458750-DEL-151 | chr2-4458751-DEL-151 |
| chr2-4460365-DEL-66 | chr2-4460367-DEL-65 |
| chr2-4470262-DEL-77 | chr2-4470258-DEL-115 |
| chr2-4476263-DEL-403 | chr2-4476264-DEL-403 |
| chr2-4485412-DEL-64 | chr2-4485412-DEL-64 |
| chr2-4512219-DEL-86 | chr2-4512220-DEL-86 |
| chr2-4520862-DEL-66 | chr2-4520863-DEL-66 |
| chr2-4521920-DEL-51 | chr2-4521948-DEL-56 |
| chr2-4527285-DEL-105 | chr2-4527300-DEL-64 |
| chr2-4535888-DEL-204 | chr2-4535889-DEL-204 |
| chr2-4597488-DEL-51 | chr2-4597489-DEL-51 |
| chr2-4609769-DEL-351 | chr2-4609770-DEL-351 |
| chr2-4619777-DEL-134 | chr2-4619778-DEL-138 |
| chr2-4644720-DEL-70 | chr2-4644721-DEL-70 |
| chr2-4655208-DEL-84 | chr2-4655235-DEL-78 |

| | |
|---|---|
| chr2-4657813-DEL-220 | chr2-4657814-DEL-201 |
| chr2-4661611-DEL-66 | chr2-4661612-DEL-66 |
| chr2-4689262-DEL-326 | chr2-4689263-DEL-326 |
| chr2-4717182-DEL-195 | chr2-4717185-DEL-193 |
| chr2-4742950-DEL-216 | chr2-4742951-DEL-216 |
| chr2-4747202-DEL-87 | chr2-4747203-DEL-87 |
| chr2-4749465-DEL-1353 | chr2-4749490-DEL-1353 |
| chr2-4768386-DEL-73 | chr2-4768381-DEL-74 |
| chr2-4783004-DEL-105 | chr2-4783005-DEL-105 |
| chr2-4787811-DEL-108 | chr2-4787812-DEL-108 |
| chr2-4812100-DEL-108 | chr2-4812101-DEL-108 |
| chr2-4841626-DEL-57 | chr2-4841626-DEL-58 |
| chr2-4852984-DEL-207 | chr2-4852985-DEL-159 |
| chr2-4869134-DEL-108 | chr2-4869135-DEL-90 |
| chr2-4891118-DEL-85 | chr2-4891119-DEL-85 |
| chr2-4899242-DEL-215 | chr2-4899256-DEL-224 |
| chr2-4920206-DEL-144 | chr2-4920220-DEL-96 |
| chr2-4938402-DEL-440 | chr2-4938403-DEL-440 |
| chr2-4947272-DEL-165 | chr2-4947273-DEL-165 |
| chr2-4956004-DEL-91 | chr2-4956004-DEL-91 |
| chr2-4958145-DEL-187 | chr2-4958146-DEL-187 |
| chr2-4958889-DEL-79 | chr2-4958879-DEL-78 |
| chr2-4964270-DEL-320 | chr2-4964247-DEL-348 |
| chr2-4974200-DEL-173 | chr2-4974201-DEL-173 |
| chr2-4987594-DEL-109 | chr2-4987595-DEL-109 |
| chr2-4996614-DEL-202 | chr2-4996615-DEL-202 |
| chr2-5004051-DEL-62 | chr2-5004052-DEL-62 |
| chr2-5020853-DEL-978 | chr2-5020854-DEL-978 |
| chr2-5033908-DEL-196 | chr2-5033909-DEL-196 |
| chr2-5043042-DEL-168 | chr2-5043043-DEL-168 |
| chr2-5055690-DEL-61 | chr2-5055691-DEL-59 |
| chr2-5056333-DEL-200 | chr2-5056334-DEL-199 |
| chr2-5056712-DEL-839 | chr2-5056713-DEL-839 |
| chr2-5070322-DEL-1882 | chr2-5070323-DEL-1882 |
| chr2-5078710-DEL-168 | chr2-5078711-DEL-168 |
| chr2-5086644-DEL-155 | chr2-5086643-DEL-146 |
| chr2-5091268-DEL-171 | chr2-5091269-DEL-171 |
| chr2-5108362-DEL-79 | chr2-5108327-DEL-124 |
| chr2-5114331-DEL-54 | chr2-5114343-DEL-52 |
| chr2-5115708-DEL-891 | chr2-5115740-DEL-880 |
| chr2-5131242-DEL-212 | chr2-5131243-DEL-212 |
| chr2-5136819-DEL-65 | chr2-5136820-DEL-65 |
| chr2-5140607-DEL-55 | chr2-5140608-DEL-55 |
| chr2-5153633-DEL-68 | chr2-5153636-DEL-66 |
| chr2-5168895-DEL-106 | chr2-5168937-DEL-88 |
| chr2-5193648-DEL-74 | chr2-5193687-DEL-74 |
| chr2-5228250-DEL-134 | chr2-5228251-DEL-134 |
| chr2-5243967-DEL-16548 | chr2-5243968-DEL-16548 |
| chr2-5247455-DEL-1802 | chr2-5247456-DEL-1802 |
| chr2-5254215-DEL-351 | chr2-5254216-DEL-351 |
| chr2-5258281-DEL-367 | chr2-5258282-DEL-367 |
| chr2-5266481-DEL-82 | chr2-5266438-DEL-89 |
| chr2-5281635-DEL-216 | chr2-5281636-DEL-216 |
| chr2-5288086-DEL-165 | chr2-5288087-DEL-165 |
| chr2-5311159-DEL-58 | chr2-5311149-DEL-68 |
| chr2-5316816-DEL-60 | chr2-5316817-DEL-60 |
| chr2-5321157-DEL-308 | chr2-5321158-DEL-308 |
| chr2-5382745-DEL-86 | chr2-5382746-DEL-86 |
| chr2-5390084-DEL-820 | chr2-5390085-DEL-820 |
| chr2-5398402-DEL-440 | chr2-5398403-DEL-440 |
| chr2-5445491-DEL-65 | chr2-5445492-DEL-65 |
| chr2-5450863-DEL-62 | chr2-5450851-DEL-60 |
| chr2-5470426-DEL-650 | chr2-5470427-DEL-650 |
| chr2-5484141-DEL-503 | chr2-5484142-DEL-503 |
| chr2-5488172-DEL-364 | chr2-5488166-DEL-344 |
| chr2-5507285-DEL-64 | chr2-5507286-DEL-64 |
| chr2-5527296-DEL-64 | chr2-5527297-DEL-64 |
| chr2-5531722-DEL-54 | chr2-5531715-DEL-90 |
| chr2-5559140-DEL-7103 | chr2-5559141-DEL-7103 |
| chr2-5579053-DEL-116 | chr2-5579034-DEL-110 |
| chr2-5587161-DEL-430 | chr2-5587162-DEL-430 |
| chr2-5626027-DEL-53 | chr2-5626027-DEL-53 |
| chr2-5659442-DEL-1503 | chr2-5659443-DEL-1500 |
| chr2-5672230-DEL-101 | chr2-5672230-DEL-79 |
| chr2-5676434-DEL-95 | chr2-5676435-DEL-94 |
| chr2-5688828-DEL-7109 | chr2-5688829-DEL-7109 |
| chr2-5707870-DEL-80 | chr2-5707871-DEL-80 |
| chr2-5716762-DEL-339 | chr2-5716763-DEL-339 |
| chr2-5728169-DEL-66 | chr2-5728188-DEL-57 |
| chr2-5736701-DEL-80 | chr2-5736700-DEL-80 |

| | |
|---|---|
| chr2-5746137-DEL-86 | chr2-5746156-DEL-64 |
| chr2-5764472-DEL-52 | chr2-5764473-DEL-52 |
| chr2-5767221-DEL-97 | chr2-5767212-DEL-74 |
| chr2-5778111-DEL-523 | chr2-5778106-DEL-528 |
| chr2-5779248-DEL-7095 | chr2-5779258-DEL-7094 |
| chr2-5785703-DEL-78 | chr2-5785704-DEL-78 |
| chr2-5840549-DEL-164 | chr2-5840550-DEL-164 |
| chr2-5844072-DEL-64 | chr2-5844073-DEL-64 |
| chr2-5848381-DEL-202 | chr2-5848382-DEL-202 |
| chr2-5853856-DEL-56 | chr2-5853857-DEL-56 |
| chr2-5868550-DEL-81 | chr2-5868551-DEL-81 |
| chr2-5870206-DEL-184 | chr2-5870207-DEL-184 |
| chr2-5871758-DEL-67 | chr2-5871759-DEL-67 |
| chr2-5898792-DEL-211 | chr2-5898793-DEL-211 |
| chr2-5900928-DEL-89 | chr2-5900929-DEL-89 |
| chr2-5906352-DEL-172 | chr2-5906353-DEL-172 |
| chr2-5912712-DEL-53 | chr2-5912713-DEL-53 |
| chr2-5928339-DEL-196 | chr2-5928340-DEL-196 |
| chr2-5929405-DEL-556 | chr2-5929406-DEL-556 |
| chr2-5961423-DEL-117 | chr2-5961424-DEL-117 |
| chr2-5963767-DEL-51 | chr2-5963743-DEL-52 |
| chr2-5994977-DEL-203 | chr2-5994978-DEL-203 |
| chr2-6016392-DEL-212 | chr2-6016393-DEL-212 |
| chr2-6019955-DEL-204 | chr2-6019956-DEL-204 |
| chr2-6041755-DEL-223 | chr2-6041756-DEL-223 |
| chr2-6074820-DEL-69 | chr2-6074795-DEL-69 |
| chr2-6083606-DEL-93 | chr2-6083607-DEL-93 |
| chr2-6090194-DEL-212 | chr2-6090192-DEL-211 |
| chr2-6115882-DEL-54 | chr2-6115898-DEL-55 |
| chr2-6141027-DEL-443 | chr2-6141041-DEL-430 |
| chr2-6202167-DEL-2871 | chr2-6202176-DEL-2886 |
| chr2-6212868-DEL-58 | chr2-6212869-DEL-58 |
| chr2-6229389-DEL-53 | chr2-6229390-DEL-53 |
| chr2-6241625-DEL-1205 | chr2-6241628-DEL-1203 |
| chr2-6245589-DEL-62 | chr2-6245590-DEL-62 |
| chr2-6267304-DEL-87 | chr2-6267306-DEL-86 |
| chr2-6286892-DEL-348 | chr2-6286893-DEL-348 |
| chr2-6291235-DEL-55 | chr2-6291236-DEL-55 |
| chr2-6300192-DEL-110 | chr2-6300193-DEL-110 |
| chr2-6336816-DEL-492 | chr2-6336817-DEL-492 |
| chr2-6349885-DEL-524 | chr2-6349886-DEL-524 |
| chr2-6369083-DEL-52 | chr2-6369084-DEL-52 |
| chr2-6408371-DEL-72 | chr2-6408405-DEL-71 |
| chr2-6424386-DEL-87 | chr2-6424387-DEL-116 |
| chr2-6427724-DEL-70 | chr2-6427701-DEL-68 |
| chr2-6514107-DEL-1465 | chr2-6514103-DEL-1469 |
| chr2-6534003-DEL-252 | chr2-6534004-DEL-252 |
| chr2-6534641-DEL-70 | chr2-6534627-DEL-84 |
| chr2-6588039-DEL-257 | chr2-6588033-DEL-250 |
| chr2-6591368-DEL-538 | chr2-6591369-DEL-538 |
| chr2-6634246-DEL-2326 | chr2-6634247-DEL-2326 |
| chr2-6675801-DEL-413 | chr2-6675799-DEL-407 |
| chr2-6695468-DEL-56 | chr2-6695469-DEL-56 |
| chr2-6704740-DEL-181 | chr2-6704741-DEL-181 |
| chr2-6705443-DEL-1554 | chr2-6705444-DEL-1554 |
| chr2-6751891-DEL-208 | chr2-6751892-DEL-208 |
| chr2-6776904-DEL-443 | chr2-6776905-DEL-443 |
| chr2-6812572-DEL-121 | chr2-6812573-DEL-110 |
| chr2-6844836-DEL-6641 | chr2-6844837-DEL-6641 |
| chr2-6855907-DEL-850 | chr2-6855908-DEL-850 |
| chr2-6953054-DEL-252 | chr2-6953055-DEL-252 |
| chr2-6990195-DEL-129 | chr2-6990196-DEL-129 |
| chr2-7009924-DEL-80 | chr2-7009925-DEL-80 |
| chr2-7037334-DEL-88 | chr2-7037332-DEL-89 |
| chr2-7040082-DEL-75 | chr2-7040083-DEL-75 |
| chr2-7079424-DEL-107 | chr2-7079438-DEL-75 |
| chr2-7080973-DEL-385 | chr2-7080974-DEL-385 |
| chr2-7082825-DEL-64 | chr2-7082826-DEL-96 |
| chr2-7110247-DEL-731 | chr2-7110248-DEL-731 |
| chr2-7139128-DEL-56 | chr2-7139125-DEL-52 |
| chr2-7142372-DEL-2178 | chr2-7142373-DEL-2178 |
| chr2-7190978-DEL-87 | chr2-7190980-DEL-86 |
| chr2-7194165-DEL-107 | chr2-7194178-DEL-83 |
| chr2-7199966-DEL-189 | chr2-7199967-DEL-189 |
| chr2-7204242-DEL-55 | chr2-7204243-DEL-55 |
| chr2-7210898-DEL-72 | chr2-7210864-DEL-76 |
| chr2-7279077-DEL-1799 | chr2-7279078-DEL-1799 |
| chr2-7311347-DEL-1137 | chr2-7311347-DEL-1137 |
| chr2-7332596-DEL-70 | chr2-7332595-DEL-70 |
| chr2-7345259-DEL-70 | chr2-7345260-DEL-70 |

| | |
|---|---|
| chr2-7352713-DEL-55 | chr2-7352697-DEL-56 |
| chr2-7360386-DEL-68 | chr2-7360387-DEL-68 |
| chr2-7378168-DEL-63 | chr2-7378169-DEL-63 |
| chr2-7393394-DEL-131 | chr2-7393392-DEL-132 |
| chr2-7408875-DEL-5286 | chr2-7408876-DEL-5286 |
| chr2-7433931-DEL-107 | chr2-7433932-DEL-107 |
| chr2-7451521-DEL-143 | chr2-7451522-DEL-143 |
| chr2-7452471-DEL-1142 | chr2-7452430-DEL-1142 |
| chr2-7456826-DEL-6247 | chr2-7456807-DEL-6266 |
| chr2-7488804-DEL-1045 | chr2-7488805-DEL-1045 |
| chr2-7497821-DEL-69 | chr2-7497815-DEL-63 |
| chr2-7507093-DEL-79 | chr2-7507094-DEL-78 |
| chr2-7508249-DEL-199 | chr2-7508250-DEL-197 |
| chr2-7513296-DEL-67 | chr2-7513297-DEL-67 |
| chr2-7546189-DEL-1600 | chr2-7546190-DEL-1600 |
| chr2-7591659-DEL-204 | chr2-7591660-DEL-204 |
| chr2-7607502-DEL-60 | chr2-7607513-DEL-50 |
| chr2-7632615-DEL-252 | chr2-7632637-DEL-230 |
| chr2-7661204-DEL-1862 | chr2-7661205-DEL-1862 |
| chr2-7716673-DEL-65 | chr2-7716676-DEL-70 |
| chr2-7744351-DEL-164 | chr2-7744346-DEL-161 |
| chr2-7750060-DEL-90 | chr2-7750061-DEL-90 |
| chr2-7766748-DEL-76 | chr2-7766760-DEL-78 |
| chr2-7822973-DEL-84 | chr2-7822974-DEL-84 |
| chr2-7857518-DEL-98 | chr2-7857519-DEL-98 |
| chr2-7858786-DEL-198 | chr2-7858787-DEL-198 |
| chr2-7897277-DEL-105 | chr2-7897281-DEL-104 |
| chr2-7920620-DEL-6449 | chr2-7920618-DEL-6450 |
| chr2-7954282-DEL-17307 | chr2-7954274-DEL-17302 |
| chr2-8001394-DEL-2175 | chr2-8001395-DEL-2175 |
| chr2-8042214-DEL-6453 | chr2-8042215-DEL-6453 |
| chr2-8057326-DEL-81 | chr2-8057327-DEL-81 |
| chr2-8065745-DEL-332 | chr2-8065746-DEL-332 |
| chr2-8072036-DEL-300 | chr2-8072036-DEL-301 |
| chr2-8090726-DEL-195 | chr2-8090727-DEL-195 |
| chr2-8106870-DEL-430 | chr2-8106871-DEL-430 |
| chr2-8111630-DEL-648 | chr2-8111631-DEL-648 |
| chr2-8132867-DEL-6221 | chr2-8132868-DEL-6221 |
| chr2-8159152-DEL-134 | chr2-8159153-DEL-134 |
| chr2-8188093-DEL-7388 | chr2-8188094-DEL-7388 |
| chr2-8196598-DEL-6680 | chr2-8196582-DEL-6681 |
| chr2-8227126-DEL-76 | chr2-8227127-DEL-76 |
| chr2-8229329-DEL-2814 | chr2-8229329-DEL-2814 |
| chr2-8244685-DEL-276 | chr2-8244686-DEL-274 |
| chr2-8247871-DEL-86 | chr2-8247872-DEL-86 |
| chr2-8248831-DEL-96 | chr2-8248832-DEL-96 |
| chr2-8263655-DEL-941 | chr2-8263659-DEL-942 |
| chr2-8286037-DEL-502 | chr2-8286038-DEL-497 |
| chr2-8364789-DEL-2201 | chr2-8364784-DEL-2197 |
| chr2-8438131-DEL-124 | chr2-8438140-DEL-118 |
| chr2-8444293-DEL-821 | chr2-8444294-DEL-823 |
| chr2-8486469-DEL-64 | chr2-8486455-DEL-54 |
| chr2-8500314-DEL-1395 | chr2-8500315-DEL-1395 |
| chr2-8520349-DEL-116 | chr2-8520361-DEL-112 |
| chr2-8533258-DEL-7157 | chr2-8533259-DEL-7157 |
| chr2-8546216-DEL-53 | chr2-8546217-DEL-53 |
| chr2-8565975-DEL-1201 | chr2-8565976-DEL-1201 |
| chr2-8673636-DEL-745 | chr2-8673637-DEL-745 |
| chr2-8688831-DEL-58 | chr2-8688824-DEL-54 |
| chr2-8692379-DEL-56 | chr2-8692380-DEL-56 |
| chr2-8713707-DEL-206 | chr2-8713708-DEL-206 |
| chr2-8724654-DEL-443 | chr2-8724655-DEL-443 |
| chr2-8728195-DEL-2096 | chr2-8728196-DEL-2096 |
| chr2-8779892-DEL-156 | chr2-8779893-DEL-156 |
| chr2-8852585-DEL-57 | chr2-8852567-DEL-56 |
| chr2-8864209-DEL-205 | chr2-8864210-DEL-205 |
| chr2-8876866-DEL-54 | chr2-8876867-DEL-54 |
| chr2-8889024-DEL-70 | chr2-8889024-DEL-70 |
| chr2-8891447-DEL-156 | chr2-8891446-DEL-145 |
| chr2-8937817-DEL-116 | chr2-8937821-DEL-115 |
| chr2-8939121-DEL-56 | chr2-8939122-DEL-74 |
| chr2-8953730-DEL-228 | chr2-8953731-DEL-228 |
| chr2-8983608-DEL-1089 | chr2-8983610-DEL-1088 |
| chr2-9045436-DEL-2046 | chr2-9045437-DEL-2046 |
| chr2-9072968-DEL-1015 | chr2-9072969-DEL-1015 |
| chr2-9074554-DEL-627 | chr2-9074555-DEL-627 |
| chr2-9079513-DEL-78 | chr2-9079537-DEL-74 |
| chr2-9088470-DEL-1852 | chr2-9088471-DEL-1852 |
| chr2-9090499-DEL-210 | chr2-9090500-DEL-210 |
| chr2-9091197-DEL-478 | chr2-9091198-DEL-477 |

| | |
|---|---|
| chr2-9102665-DEL-52 | chr2-9102666-DEL-52 |
| chr2-9145733-DEL-125 | chr2-9145744-DEL-99 |
| chr2-9159362-DEL-323 | chr2-9159363-DEL-323 |
| chr2-9168459-DEL-76 | chr2-9168441-DEL-68 |
| chr2-9171826-DEL-90 | chr2-9171870-DEL-50 |
| chr2-9175062-DEL-70 | chr2-9175052-DEL-69 |
| chr2-9205394-DEL-65 | chr2-9205395-DEL-65 |
| chr2-9211143-DEL-653 | chr2-9211147-DEL-650 |
| chr2-9217387-DEL-6721 | chr2-9217398-DEL-6717 |
| chr2-9269575-DEL-53 | chr2-9269576-DEL-53 |
| chr2-9275916-DEL-2266 | chr2-9275917-DEL-2265 |
| chr2-9298942-DEL-131 | chr2-9298943-DEL-131 |
| chr2-9316502-DEL-91 | chr2-9316501-DEL-92 |
| chr2-9319001-DEL-62 | chr2-9319002-DEL-62 |
| chr2-9325169-DEL-201 | chr2-9325170-DEL-201 |
| chr2-9351721-DEL-66 | chr2-9351723-DEL-58 |
| chr2-9386154-DEL-62 | chr2-9386155-DEL-62 |
| chr2-9421509-DEL-158 | chr2-9421510-DEL-158 |
| chr2-9463787-DEL-122 | chr2-9463775-DEL-156 |
| chr2-9501987-DEL-183 | chr2-9501988-DEL-173 |
| chr2-9502456-DEL-608 | chr2-9502467-DEL-598 |
| chr2-9510664-DEL-1098 | chr2-9510659-DEL-1099 |
| chr2-9513017-DEL-6778 | chr2-9513018-DEL-6778 |
| chr2-9520008-DEL-56 | chr2-9520008-DEL-56 |
| chr2-9527555-DEL-151 | chr2-9527556-DEL-151 |
| chr2-9532893-DEL-181 | chr2-9532892-DEL-182 |
| chr2-9548398-DEL-99 | chr2-9548399-DEL-99 |
| chr2-9551197-DEL-945 | chr2-9551198-DEL-945 |
| chr2-9557321-DEL-56 | chr2-9557322-DEL-56 |
| chr2-9613758-DEL-78 | chr2-9613743-DEL-59 |
| chr2-9618476-DEL-163 | chr2-9618477-DEL-163 |
| chr2-9622085-DEL-108 | chr2-9622083-DEL-111 |
| chr2-9727018-DEL-74 | chr2-9727019-DEL-74 |
| chr2-9730181-DEL-56 | chr2-9730182-DEL-54 |
| chr2-9735582-DEL-98 | chr2-9735583-DEL-98 |
| chr2-9743315-DEL-64 | chr2-9743316-DEL-64 |
| chr2-9746227-DEL-335 | chr2-9746228-DEL-335 |
| chr2-9763273-DEL-141 | chr2-9763274-DEL-141 |
| chr2-9805391-DEL-194 | chr2-9805392-DEL-194 |
| chr2-9821592-DEL-174 | chr2-9821593-DEL-174 |
| chr2-9824362-DEL-52 | chr2-9824363-DEL-52 |
| chr2-9859418-DEL-52 | chr2-9859419-DEL-52 |
| chr2-9875730-DEL-115 | chr2-9875731-DEL-115 |
| chr2-9899184-DEL-82 | chr2-9899199-DEL-68 |
| chr2-9915840-DEL-52 | chr2-9915841-DEL-52 |
| chr2-9931971-DEL-54 | chr2-9931972-DEL-54 |
| chr2-9952877-DEL-198 | chr2-9952878-DEL-198 |
| chr2-9991680-DEL-191 | chr2-9991681-DEL-191 |
| chr2-10058460-DEL-56 | chr2-10058461-DEL-58 |
| chr2-10059402-DEL-67 | chr2-10059402-DEL-68 |
| chr2-10080735-DEL-92 | chr2-10080775-DEL-76 |
| chr2-10086595-DEL-1076 | chr2-10086596-DEL-1076 |
| chr2-10087907-DEL-199 | chr2-10087908-DEL-199 |
| chr2-10093820-DEL-52 | chr2-10093821-DEL-52 |
| chr2-10109570-DEL-58 | chr2-10109568-DEL-63 |
| chr2-10116899-DEL-479 | chr2-10116900-DEL-479 |
| chr2-10118634-DEL-405 | chr2-10118656-DEL-411 |
| chr2-10160051-DEL-114 | chr2-10160059-DEL-93 |
| chr2-10178541-DEL-65 | chr2-10178542-DEL-66 |
| chr2-10181979-DEL-6224 | chr2-10181980-DEL-6224 |
| chr2-10192446-DEL-81 | chr2-10192447-DEL-80 |
| chr2-10200343-DEL-148 | chr2-10200374-DEL-118 |
| chr2-10201000-DEL-73 | chr2-10201004-DEL-72 |
| chr2-10218705-DEL-52 | chr2-10218706-DEL-52 |
| chr2-10223284-DEL-996 | chr2-10223285-DEL-996 |
| chr2-10273293-DEL-52 | chr2-10273293-DEL-53 |
| chr2-10299112-DEL-533 | chr2-10299113-DEL-533 |
| chr2-10329290-DEL-56 | chr2-10329305-DEL-88 |
| chr2-10330640-DEL-66 | chr2-10330660-DEL-56 |
| chr2-10366426-DEL-60 | chr2-10366427-DEL-76 |
| chr2-10390362-DEL-85 | chr2-10390363-DEL-51 |
| chr2-10408731-DEL-58 | chr2-10408732-DEL-58 |
| chr2-10415912-DEL-1560 | chr2-10415914-DEL-1561 |
| chr2-10424115-DEL-238 | chr2-10424116-DEL-238 |
| chr2-10451819-DEL-51 | chr2-10451820-DEL-51 |
| chr2-10540504-DEL-935 | chr2-10540505-DEL-935 |
| chr2-10542829-DEL-1072 | chr2-10542821-DEL-1074 |
| chr2-10544503-DEL-264 | chr2-10544504-DEL-264 |
| chr2-10552654-DEL-7010 | chr2-10552655-DEL-7010 |
| chr2-10564077-DEL-208 | chr2-10564078-DEL-208 |

| | |
|---|---|
| chr2-10594678-DEL-64 | chr2-10594679-DEL-64 |
| chr2-10656541-DEL-359 | chr2-10656543-DEL-358 |
| chr2-10660477-DEL-371 | chr2-10660478-DEL-371 |
| chr2-10665605-DEL-1571 | chr2-10665606-DEL-1571 |
| chr2-10678356-DEL-54 | chr2-10678357-DEL-54 |
| chr2-10696386-DEL-6209 | chr2-10696387-DEL-6209 |
| chr2-10736556-DEL-64 | chr2-10736531-DEL-72 |
| chr2-10760954-DEL-132 | chr2-10760969-DEL-118 |
| chr2-10769385-DEL-550 | chr2-10769385-DEL-550 |
| chr2-10772850-DEL-61 | chr2-10772821-DEL-87 |
| chr2-10781721-DEL-202 | chr2-10781722-DEL-198 |
| chr2-10813356-DEL-269 | chr2-10813357-DEL-269 |
| chr2-10822799-DEL-5986 | chr2-10822800-DEL-5986 |
| chr2-10841771-DEL-116 | chr2-10841782-DEL-70 |
| chr2-10880062-DEL-51 | chr2-10880062-DEL-51 |
| chr2-10888459-DEL-109 | chr2-10888452-DEL-99 |
| chr2-10891846-DEL-2880 | chr2-10891847-DEL-2880 |
| chr2-10923475-DEL-952 | chr2-10923476-DEL-952 |
| chr2-10951705-DEL-6224 | chr2-10951706-DEL-6224 |
| chr2-10972063-DEL-73 | chr2-10972069-DEL-69 |
| chr2-10975021-DEL-296 | chr2-10975013-DEL-297 |
| chr2-11000571-DEL-88 | chr2-11000572-DEL-88 |
| chr2-11013404-DEL-355 | chr2-11013405-DEL-355 |
| chr2-11018561-DEL-299 | chr2-11018562-DEL-299 |
| chr2-11022844-DEL-69 | chr2-11022886-DEL-67 |
| chr2-11024819-DEL-53 | chr2-11024820-DEL-53 |
| chr2-11026151-DEL-489 | chr2-11026152-DEL-489 |
| chr2-11046190-DEL-106 | chr2-11046191-DEL-110 |
| chr2-11065399-DEL-180 | chr2-11065400-DEL-180 |
| chr2-11066149-DEL-73 | chr2-11066150-DEL-73 |
| chr2-11074163-DEL-448 | chr2-11074164-DEL-448 |
| chr2-11079177-DEL-76 | chr2-11079175-DEL-79 |
| chr2-11109626-DEL-3053 | chr2-11109626-DEL-3053 |
| chr2-11116931-DEL-60 | chr2-11116932-DEL-60 |
| chr2-11126016-DEL-7300 | chr2-11126017-DEL-7300 |
| chr2-11126032-DEL-1060 | chr2-11126033-DEL-1060 |
| chr2-11136630-DEL-69 | chr2-11136627-DEL-73 |
| chr2-11136920-DEL-1065 | chr2-11136921-DEL-1065 |
| chr2-11165099-DEL-198 | chr2-11165100-DEL-206 |
| chr2-11168179-DEL-54 | chr2-11168179-DEL-54 |
| chr2-11184782-DEL-102 | chr2-11184750-DEL-102 |
| chr2-11193812-DEL-116 | chr2-11193774-DEL-120 |
| chr2-11224966-DEL-61 | chr2-11224967-DEL-61 |
| chr2-11244854-DEL-163 | chr2-11244851-DEL-172 |
| chr2-11266232-DEL-443 | chr2-11266233-DEL-443 |
| chr2-11267990-DEL-65 | chr2-11267990-DEL-71 |
| chr2-11303279-DEL-61 | chr2-11303280-DEL-61 |
| chr2-11307964-DEL-130 | chr2-11307973-DEL-122 |
| chr2-11337667-DEL-355 | chr2-11337668-DEL-355 |
| chr2-11371084-DEL-939 | chr2-11371049-DEL-937 |
| chr2-11373276-DEL-56 | chr2-11373277-DEL-56 |
| chr2-11384920-DEL-62 | chr2-11384921-DEL-62 |
| chr2-11387120-DEL-52 | chr2-11387117-DEL-56 |
| chr2-11412593-DEL-220 | chr2-11412594-DEL-220 |
| chr2-11413622-DEL-1293 | chr2-11413623-DEL-1293 |
| chr2-11432879-DEL-55 | chr2-11432880-DEL-55 |
| chr2-11456781-DEL-159 | chr2-11456781-DEL-155 |
| chr2-11459387-DEL-111 | chr2-11459424-DEL-68 |
| chr2-11462415-DEL-60 | chr2-11462422-DEL-54 |
| chr2-11467251-DEL-1109 | chr2-11467247-DEL-1112 |
| chr2-11477683-DEL-51 | chr2-11477680-DEL-55 |
| chr2-11510863-DEL-163 | chr2-11510864-DEL-163 |
| chr2-11518806-DEL-125 | chr2-11518807-DEL-125 |
| chr2-11524316-DEL-298 | chr2-11524317-DEL-298 |
| chr2-11538796-DEL-196 | chr2-11538797-DEL-196 |
| chr2-11547507-DEL-52 | chr2-11547508-DEL-52 |
| chr2-11580887-DEL-201 | chr2-11580888-DEL-201 |
| chr2-11610294-DEL-175 | chr2-11610303-DEL-177 |
| chr2-11636518-DEL-63 | chr2-11636526-DEL-64 |
| chr2-11652967-DEL-1602 | chr2-11652968-DEL-1602 |
| chr2-11663546-DEL-138 | chr2-11663550-DEL-134 |
| chr2-11664037-DEL-100 | chr2-11664038-DEL-100 |
| chr2-11672584-DEL-51 | chr2-11672596-DEL-52 |
| chr2-11682982-DEL-589 | chr2-11682983-DEL-589 |
| chr2-11704975-DEL-87 | chr2-11704983-DEL-60 |
| chr2-11715990-DEL-56 | chr2-11715981-DEL-66 |
| chr2-11722065-DEL-72 | chr2-11722066-DEL-78 |
| chr2-11886470-DEL-2348 | chr2-11886471-DEL-2348 |
| chr2-11902208-DEL-56 | chr2-11902205-DEL-52 |
| chr2-11923012-DEL-762 | chr2-11923027-DEL-744 |

| | |
|---|---|
| chr2-11956520-DEL-1643 | chr2-11956521-DEL-1643 |
| chr2-11960738-DEL-3254 | chr2-11960737-DEL-3255 |
| chr2-12001069-DEL-2838 | chr2-12001070-DEL-2838 |
| chr2-12031395-DEL-515 | chr2-12031396-DEL-515 |
| chr2-12035034-DEL-1833 | chr2-12035035-DEL-1833 |
| chr2-12045609-DEL-59 | chr2-12045610-DEL-59 |
| chr2-12048005-DEL-3233 | chr2-12048006-DEL-3233 |
| chr2-12053412-DEL-54 | chr2-12053413-DEL-54 |
| chr2-12065860-DEL-54 | chr2-12065853-DEL-78 |
| chr2-12077742-DEL-82 | chr2-12077743-DEL-82 |
| chr2-12079435-DEL-663 | chr2-12079435-DEL-663 |
| chr2-12085151-DEL-202 | chr2-12085152-DEL-202 |
| chr2-12133688-DEL-1906 | chr2-12133689-DEL-1905 |
| chr2-12140780-DEL-821 | chr2-12140781-DEL-821 |
| chr2-12166421-DEL-6164 | chr2-12166422-DEL-6164 |
| chr2-12177529-DEL-99 | chr2-12177530-DEL-99 |
| chr2-12188187-DEL-218 | chr2-12188188-DEL-218 |
| chr2-12191017-DEL-1417 | chr2-12191018-DEL-1417 |
| chr2-12208926-DEL-59 | chr2-12208927-DEL-59 |
| chr2-12240968-DEL-111 | chr2-12240971-DEL-109 |
| chr2-12285423-DEL-58 | chr2-12285424-DEL-52 |
| chr2-12286027-DEL-142 | chr2-12286028-DEL-142 |
| chr2-12293869-DEL-74 | chr2-12293870-DEL-74 |
| chr2-12359647-DEL-74 | chr2-12359649-DEL-73 |
| chr2-12438191-DEL-98 | chr2-12438184-DEL-83 |
| chr2-12454184-DEL-342 | chr2-12454185-DEL-342 |
| chr2-12483741-DEL-120 | chr2-12483742-DEL-120 |
| chr2-12502123-DEL-6237 | chr2-12502127-DEL-6234 |
| chr2-12502341-DEL-64 | chr2-12502342-DEL-64 |
| chr2-12510950-DEL-58 | chr2-12510951-DEL-52 |
| chr2-12513827-DEL-267 | chr2-12513828-DEL-265 |
| chr2-12523780-DEL-54 | chr2-12523787-DEL-50 |
| chr2-12578415-DEL-3283 | chr2-12578416-DEL-3283 |
| chr2-12619820-DEL-6199 | chr2-12619821-DEL-6199 |
| chr2-12645623-DEL-9774 | chr2-12645623-DEL-9774 |
| chr2-12655580-DEL-792 | chr2-12655539-DEL-793 |
| chr2-12667744-DEL-594 | chr2-12667745-DEL-593 |
| chr2-12671950-DEL-1487 | chr2-12671950-DEL-1487 |
| chr2-12718093-DEL-540 | chr2-12718094-DEL-540 |
| chr2-12730462-DEL-263 | chr2-12730458-DEL-267 |
| chr2-12742836-DEL-490 | chr2-12742836-DEL-490 |
| chr2-12747527-DEL-666 | chr2-12747528-DEL-666 |
| chr2-12789583-DEL-1341 | chr2-12789583-DEL-1341 |
| chr2-12822105-DEL-83 | chr2-12822085-DEL-107 |
| chr2-12861475-DEL-162 | chr2-12861476-DEL-182 |
| chr2-12880970-DEL-187 | chr2-12880971-DEL-187 |
| chr2-12943893-DEL-6615 | chr2-12943893-DEL-6615 |
| chr2-12955345-DEL-3942 | chr2-12955345-DEL-3942 |
| chr2-12987892-DEL-64 | chr2-12987891-DEL-68 |
| chr2-12993706-DEL-156 | chr2-12993707-DEL-156 |
| chr2-12995829-DEL-78 | chr2-12995844-DEL-60 |
| chr2-13028020-DEL-190 | chr2-13028021-DEL-190 |
| chr2-13077506-DEL-149 | chr2-13077506-DEL-166 |
| chr2-13109111-DEL-124 | chr2-13109070-DEL-148 |
| chr2-13141007-DEL-797 | chr2-13140994-DEL-798 |
| chr2-13162145-DEL-3904 | chr2-13162146-DEL-3904 |
| chr2-13176564-DEL-953 | chr2-13176565-DEL-953 |
| chr2-13185223-DEL-2025 | chr2-13185224-DEL-2025 |
| chr2-13230598-DEL-52 | chr2-13230599-DEL-52 |
| chr2-13251156-DEL-88 | chr2-13251157-DEL-88 |
| chr2-13254481-DEL-2816 | chr2-13254482-DEL-2816 |
| chr2-13277241-DEL-176 | chr2-13277242-DEL-176 |
| chr2-13292925-DEL-879 | chr2-13292926-DEL-879 |
| chr2-13301578-DEL-773 | chr2-13301579-DEL-773 |
| chr2-13331942-DEL-401 | chr2-13331943-DEL-401 |
| chr2-13337868-DEL-265 | chr2-13337869-DEL-265 |
| chr2-13369078-DEL-960 | chr2-13369079-DEL-960 |
| chr2-13380606-DEL-234 | chr2-13380607-DEL-234 |
| chr2-13382484-DEL-90 | chr2-13382488-DEL-111 |
| chr2-13397673-DEL-55 | chr2-13397674-DEL-55 |
| chr2-13413018-DEL-809 | chr2-13413019-DEL-809 |
| chr2-13419135-DEL-192 | chr2-13419136-DEL-192 |
| chr2-13512538-DEL-212 | chr2-13512541-DEL-211 |
| chr2-13516106-DEL-67 | chr2-13516102-DEL-68 |
| chr2-13536062-DEL-776 | chr2-13536095-DEL-779 |
| chr2-13616065-DEL-6579 | chr2-13616066-DEL-6579 |
| chr2-13622100-DEL-208 | chr2-13622101-DEL-208 |
| chr2-13623321-DEL-70 | chr2-13623292-DEL-71 |
| chr2-13632142-DEL-378 | chr2-13632143-DEL-378 |
| chr2-13643258-DEL-815 | chr2-13643259-DEL-815 |

| | |
|---|---|
| chr2-13662814-DEL-6873 | chr2-13662815-DEL-6873 |
| chr2-13681818-DEL-75 | chr2-13681808-DEL-60 |
| chr2-13756667-DEL-84 | chr2-13756667-DEL-84 |
| chr2-13770748-DEL-1319 | chr2-13770749-DEL-1319 |
| chr2-13772048-DEL-196 | chr2-13772049-DEL-196 |
| chr2-13791877-DEL-62 | chr2-13791878-DEL-60 |
| chr2-13792936-DEL-77 | chr2-13792922-DEL-70 |
| chr2-13793081-DEL-560 | chr2-13793082-DEL-560 |
| chr2-13814284-DEL-717 | chr2-13814285-DEL-717 |
| chr2-13824614-DEL-129 | chr2-13824615-DEL-129 |
| chr2-13826576-DEL-87 | chr2-13826579-DEL-85 |
| chr2-13828697-DEL-9577 | chr2-13828698-DEL-9577 |
| chr2-13853002-DEL-1894 | chr2-13853003-DEL-1894 |
| chr2-13880350-DEL-214 | chr2-13880351-DEL-214 |
| chr2-13889454-DEL-322 | chr2-13889455-DEL-322 |
| chr2-13948950-DEL-6427 | chr2-13948950-DEL-6427 |
| chr2-13976134-DEL-201 | chr2-13976118-DEL-202 |
| chr2-13991666-DEL-58 | chr2-13991665-DEL-60 |
| chr2-14007714-DEL-80 | chr2-14007715-DEL-80 |
| chr2-14015010-DEL-326 | chr2-14015011-DEL-326 |
| chr2-14028385-DEL-56 | chr2-14028386-DEL-82 |
| chr2-14030312-DEL-92 | chr2-14030287-DEL-100 |
| chr2-14033925-DEL-6217 | chr2-14033929-DEL-6218 |
| chr2-14046305-DEL-80 | chr2-14046306-DEL-80 |
| chr2-14081988-DEL-9316 | chr2-14081988-DEL-9316 |
| chr2-14123617-DEL-6655 | chr2-14123618-DEL-6655 |
| chr2-14194320-DEL-429 | chr2-14194321-DEL-429 |
| chr2-14208734-DEL-253 | chr2-14208758-DEL-207 |
| chr2-14248304-DEL-583 | chr2-14248305-DEL-583 |
| chr2-14252413-DEL-59 | chr2-14252402-DEL-68 |
| chr2-14270281-DEL-203 | chr2-14270282-DEL-203 |
| chr2-14274490-DEL-63 | chr2-14274491-DEL-63 |
| chr2-14278822-DEL-516 | chr2-14278822-DEL-516 |
| chr2-14293419-DEL-212 | chr2-14293420-DEL-212 |
| chr2-14314174-DEL-99 | chr2-14314175-DEL-99 |
| chr2-14338904-DEL-172 | chr2-14338905-DEL-172 |
| chr2-14344897-DEL-247 | chr2-14344893-DEL-247 |
| chr2-14346050-DEL-63 | chr2-14346055-DEL-58 |
| chr2-14410653-DEL-122 | chr2-14410651-DEL-123 |
| chr2-14425243-DEL-611 | chr2-14425247-DEL-605 |
| chr2-14475318-DEL-62 | chr2-14475319-DEL-62 |
| chr2-14496102-DEL-1693 | chr2-14496103-DEL-1693 |
| chr2-14506213-DEL-89 | chr2-14506214-DEL-89 |
| chr2-14511946-DEL-2413 | chr2-14511943-DEL-2414 |
| chr2-14525978-DEL-2905 | chr2-14525979-DEL-2905 |
| chr2-14596511-DEL-65 | chr2-14596512-DEL-65 |
| chr2-14631623-DEL-553 | chr2-14631624-DEL-553 |
| chr2-14756325-DEL-60 | chr2-14756320-DEL-51 |
| chr2-14771744-DEL-357 | chr2-14771784-DEL-322 |
| chr2-14816550-DEL-803 | chr2-14816550-DEL-773 |
| chr2-14895810-DEL-406 | chr2-14895811-DEL-406 |
| chr2-14923063-DEL-6221 | chr2-14923064-DEL-6221 |
| chr2-14983300-DEL-2472 | chr2-14983301-DEL-2472 |
| chr2-15009422-DEL-217 | chr2-15009423-DEL-217 |
| chr2-15031057-DEL-82 | chr2-15031058-DEL-64 |
| chr2-15101313-DEL-138 | chr2-15101314-DEL-138 |
| chr2-15119386-DEL-6216 | chr2-15119387-DEL-6216 |
| chr2-15137200-DEL-53 | chr2-15137201-DEL-53 |
| chr2-15139648-DEL-253 | chr2-15139649-DEL-253 |
| chr2-15154577-DEL-183 | chr2-15154578-DEL-183 |
| chr2-15156049-DEL-96 | chr2-15156049-DEL-97 |
| chr2-15160090-DEL-98 | chr2-15160091-DEL-98 |
| chr2-15193036-DEL-130 | chr2-15193038-DEL-125 |
| chr2-15198352-DEL-7496 | chr2-15198353-DEL-7496 |
| chr2-15235521-DEL-2056 | chr2-15235522-DEL-2056 |
| chr2-15235565-DEL-426 | chr2-15235566-DEL-426 |
| chr2-15238738-DEL-3160 | chr2-15238739-DEL-3160 |
| chr2-15258625-DEL-2342 | chr2-15258626-DEL-2342 |
| chr2-15271380-DEL-413 | chr2-15271381-DEL-413 |
| chr2-15284775-DEL-558 | chr2-15284776-DEL-558 |
| chr2-15302745-DEL-66 | chr2-15302753-DEL-62 |
| chr2-15381932-DEL-6575 | chr2-15381933-DEL-6575 |
| chr2-15410307-DEL-341 | chr2-15410308-DEL-341 |
| chr2-15435151-DEL-4679 | chr2-15435152-DEL-4679 |
| chr2-15457536-DEL-520 | chr2-15457537-DEL-520 |
| chr2-15490252-DEL-104 | chr2-15490267-DEL-90 |
| chr2-15501674-DEL-2723 | chr2-15501687-DEL-2711 |
| chr2-15544466-DEL-58 | chr2-15544466-DEL-53 |
| chr2-15550498-DEL-328 | chr2-15550499-DEL-328 |
| chr2-15573141-DEL-365 | chr2-15573141-DEL-365 |

| | |
|---|---|
| chr2-15580731-DEL-7808 | chr2-15580724-DEL-7809 |
| chr2-15580740-DEL-216 | chr2-15580736-DEL-214 |
| chr2-15601704-DEL-6009 | chr2-15601705-DEL-6009 |
| chr2-15608154-DEL-63 | chr2-15608155-DEL-63 |
| chr2-15639423-DEL-600 | chr2-15639424-DEL-599 |
| chr2-15649265-DEL-247 | chr2-15649266-DEL-247 |
| chr2-15660323-DEL-306 | chr2-15660324-DEL-306 |
| chr2-15662640-DEL-6297 | chr2-15662641-DEL-6297 |
| chr2-15700969-DEL-445 | chr2-15700970-DEL-445 |
| chr2-15715461-DEL-60 | chr2-15715462-DEL-60 |
| chr2-15721367-DEL-6183 | chr2-15721368-DEL-6183 |
| chr2-15769930-DEL-965 | chr2-15769931-DEL-965 |
| chr2-15772959-DEL-2270 | chr2-15772960-DEL-2270 |
| chr2-15795898-DEL-70 | chr2-15795899-DEL-70 |
| chr2-15800718-DEL-56 | chr2-15800719-DEL-56 |
| chr2-15846034-DEL-117 | chr2-15846042-DEL-109 |
| chr2-15856656-DEL-103 | chr2-15856657-DEL-103 |
| chr2-15857094-DEL-1129 | chr2-15857095-DEL-1129 |
| chr2-15895145-DEL-639 | chr2-15895146-DEL-639 |
| chr2-15909460-DEL-344 | chr2-15909461-DEL-331 |
| chr2-15919820-DEL-6625 | chr2-15919821-DEL-6625 |
| chr2-16025306-DEL-9046 | chr2-16025301-DEL-9046 |
| chr2-16083697-DEL-170 | chr2-16083698-DEL-170 |
| chr2-16107227-DEL-6815 | chr2-16107228-DEL-6815 |
| chr2-16123773-DEL-363 | chr2-16123774-DEL-363 |
| chr2-16215792-DEL-471 | chr2-16215793-DEL-471 |
| chr2-16216681-DEL-894 | chr2-16216682-DEL-894 |
| chr2-16225643-DEL-1655 | chr2-16225644-DEL-1655 |
| chr2-16261898-DEL-221 | chr2-16261899-DEL-221 |
| chr2-16282440-DEL-494 | chr2-16282441-DEL-494 |
| chr2-16313260-DEL-207 | chr2-16313261-DEL-207 |
| chr2-16314454-DEL-163 | chr2-16314455-DEL-163 |
| chr2-16322177-DEL-1612 | chr2-16322178-DEL-1612 |
| chr2-16325066-DEL-6401 | chr2-16325067-DEL-6401 |
| chr2-16334803-DEL-63 | chr2-16334804-DEL-54 |
| chr2-16348734-DEL-4654 | chr2-16348735-DEL-4654 |
| chr2-16350951-DEL-79 | chr2-16350952-DEL-79 |
| chr2-16380015-DEL-7104 | chr2-16380016-DEL-7104 |
| chr2-16413768-DEL-110 | chr2-16413770-DEL-108 |
| chr2-16415395-DEL-324 | chr2-16415397-DEL-322 |
| chr2-16427761-DEL-70 | chr2-16427757-DEL-55 |
| chr2-16436086-DEL-52 | chr2-16436087-DEL-52 |
| chr2-16529533-DEL-105 | chr2-16529537-DEL-68 |
| chr2-16541166-DEL-93 | chr2-16541167-DEL-92 |
| chr2-16585885-DEL-4987 | chr2-16585886-DEL-4987 |
| chr2-16609757-DEL-227 | chr2-16609758-DEL-227 |
| chr2-16625765-DEL-6174 | chr2-16625766-DEL-6174 |
| chr2-16752998-DEL-180 | chr2-16752999-DEL-180 |
| chr2-16767466-DEL-6645 | chr2-16767467-DEL-6645 |
| chr2-16804455-DEL-110 | chr2-16804457-DEL-110 |
| chr2-16824263-DEL-130 | chr2-16824264-DEL-130 |
| chr2-16824456-DEL-853 | chr2-16824457-DEL-853 |
| chr2-16856204-DEL-5123 | chr2-16856201-DEL-5120 |
| chr2-16876807-DEL-70 | chr2-16876812-DEL-66 |
| chr2-16906327-DEL-107 | chr2-16906298-DEL-108 |
| chr2-16908947-DEL-74 | chr2-16908924-DEL-76 |
| chr2-16950265-DEL-5247 | chr2-16950257-DEL-5248 |
| chr2-16965780-DEL-95 | chr2-16965751-DEL-119 |
| chr2-16996004-DEL-475 | chr2-16996004-DEL-475 |
| chr2-17000698-DEL-56 | chr2-17000687-DEL-60 |
| chr2-17002346-DEL-80 | chr2-17002353-DEL-64 |
| chr2-17083923-DEL-786 | chr2-17083923-DEL-786 |
| chr2-17089326-DEL-4684 | chr2-17089327-DEL-4684 |
| chr2-17101090-DEL-66 | chr2-17101091-DEL-86 |
| chr2-17128953-DEL-7286 | chr2-17128954-DEL-7286 |
| chr2-17136836-DEL-54 | chr2-17136836-DEL-50 |
| chr2-17139804-DEL-118 | chr2-17139805-DEL-114 |
| chr2-17185464-DEL-57 | chr2-17185444-DEL-58 |
| chr2-17213074-DEL-1842 | chr2-17213075-DEL-1842 |
| chr2-17226538-DEL-1189 | chr2-17226539-DEL-1189 |
| chr2-17241755-DEL-2730 | chr2-17241756-DEL-2730 |
| chr2-17244720-DEL-183 | chr2-17244721-DEL-183 |
| chr2-17264123-DEL-54 | chr2-17264123-DEL-54 |
| chr2-17275935-DEL-6539 | chr2-17275936-DEL-6539 |
| chr2-17282062-DEL-204 | chr2-17282062-DEL-204 |
| chr2-17309234-DEL-82 | chr2-17309265-DEL-52 |
| chr2-17322601-DEL-6727 | chr2-17322596-DEL-6733 |
| chr2-17343355-DEL-390 | chr2-17343356-DEL-390 |
| chr2-17344544-DEL-64 | chr2-17344557-DEL-54 |
| chr2-17408516-DEL-83 | chr2-17408515-DEL-85 |

| | |
|---|---|
| chr2-17426929-DEL-87 | chr2-17426918-DEL-79 |
| chr2-17490858-DEL-98 | chr2-17490859-DEL-98 |
| chr2-17525488-DEL-2336 | chr2-17525489-DEL-2336 |
| chr2-17542154-DEL-66 | chr2-17542154-DEL-66 |
| chr2-17606559-DEL-66 | chr2-17606560-DEL-66 |
| chr2-17617346-DEL-5392 | chr2-17617347-DEL-5392 |
| chr2-17627061-DEL-943 | chr2-17627062-DEL-943 |
| chr2-17637536-DEL-1888 | chr2-17637537-DEL-1888 |
| chr2-17656458-DEL-2806 | chr2-17656459-DEL-2806 |
| chr2-17670897-DEL-80 | chr2-17670902-DEL-51 |
| chr2-17723830-DEL-482 | chr2-17723831-DEL-481 |
| chr2-17736775-DEL-250 | chr2-17736775-DEL-251 |
| chr2-17750716-DEL-103 | chr2-17750717-DEL-103 |
| chr2-17752142-DEL-5423 | chr2-17752143-DEL-5423 |
| chr2-17780369-DEL-78 | chr2-17780365-DEL-79 |
| chr2-17823736-DEL-213 | chr2-17823737-DEL-213 |
| chr2-17850471-DEL-228 | chr2-17850472-DEL-228 |
| chr2-17856772-DEL-194 | chr2-17856773-DEL-194 |
| chr2-17890120-DEL-64 | chr2-17890121-DEL-64 |
| chr2-17897250-DEL-354 | chr2-17897251-DEL-354 |
| chr2-17909299-DEL-113 | chr2-17909300-DEL-113 |
| chr2-17955216-DEL-259 | chr2-17955208-DEL-260 |
| chr2-17969128-DEL-214 | chr2-17969129-DEL-214 |
| chr2-17970970-DEL-148 | chr2-17970971-DEL-148 |
| chr2-17978586-DEL-120 | chr2-17978587-DEL-120 |
| chr2-17982868-DEL-72 | chr2-17982870-DEL-64 |
| chr2-18007827-DEL-56 | chr2-18007832-DEL-52 |
| chr2-18024394-DEL-209 | chr2-18024395-DEL-209 |
| chr2-18073386-DEL-176 | chr2-18073387-DEL-176 |
| chr2-18082022-DEL-457 | chr2-18082022-DEL-459 |
| chr2-18093000-DEL-188 | chr2-18093000-DEL-189 |
| chr2-18110820-DEL-80 | chr2-18110851-DEL-58 |
| chr2-18114077-DEL-206 | chr2-18114078-DEL-206 |
| chr2-18140106-DEL-370 | chr2-18140107-DEL-370 |
| chr2-18145789-DEL-227 | chr2-18145790-DEL-226 |
| chr2-18195137-DEL-2130 | chr2-18195137-DEL-2130 |
| chr2-18287500-DEL-203 | chr2-18287501-DEL-203 |
| chr2-18327820-DEL-544 | chr2-18327821-DEL-544 |
| chr2-18333725-DEL-477 | chr2-18333726-DEL-477 |
| chr2-18384161-DEL-56 | chr2-18384162-DEL-56 |
| chr2-18404545-DEL-365 | chr2-18404546-DEL-365 |
| chr2-18406546-DEL-54 | chr2-18406534-DEL-56 |
| chr2-18407351-DEL-113 | chr2-18407350-DEL-72 |
| chr2-18417387-DEL-191 | chr2-18417388-DEL-190 |
| chr2-18450373-DEL-76 | chr2-18450374-DEL-76 |
| chr2-18466708-DEL-61 | chr2-18466709-DEL-61 |
| chr2-18467682-DEL-66 | chr2-18467683-DEL-66 |
| chr2-18481746-DEL-10810 | chr2-18481747-DEL-10810 |
| chr2-18481904-DEL-214 | chr2-18481904-DEL-214 |
| chr2-18493167-DEL-109 | chr2-18493154-DEL-121 |
| chr2-18503751-DEL-63 | chr2-18503752-DEL-63 |
| chr2-18508636-DEL-2236 | chr2-18508636-DEL-2236 |
| chr2-18545632-DEL-462 | chr2-18545633-DEL-462 |
| chr2-18554890-DEL-6496 | chr2-18554891-DEL-6496 |
| chr2-18582306-DEL-2844 | chr2-18582307-DEL-2844 |
| chr2-18602895-DEL-109 | chr2-18602896-DEL-109 |
| chr2-18626149-DEL-1553 | chr2-18626150-DEL-1553 |
| chr2-18633181-DEL-96 | chr2-18633190-DEL-104 |
| chr2-18642525-DEL-204 | chr2-18642526-DEL-204 |
| chr2-18646202-DEL-71 | chr2-18646200-DEL-76 |
| chr2-18742260-DEL-2215 | chr2-18742261-DEL-2215 |
| chr2-18770578-DEL-72 | chr2-18770578-DEL-72 |
| chr2-18774331-DEL-236 | chr2-18774332-DEL-236 |
| chr2-18777632-DEL-58 | chr2-18777633-DEL-56 |
| chr2-18817411-DEL-91 | chr2-18817412-DEL-91 |
| chr2-18825404-DEL-70 | chr2-18825405-DEL-70 |
| chr2-18840492-DEL-53 | chr2-18840493-DEL-53 |
| chr2-18962603-DEL-565 | chr2-18962604-DEL-533 |
| chr2-18970825-DEL-75 | chr2-18970825-DEL-75 |
| chr2-19037421-DEL-905 | chr2-19037422-DEL-905 |
| chr2-19047050-DEL-3804 | chr2-19047051-DEL-3804 |
| chr2-19075371-DEL-6407 | chr2-19075372-DEL-6407 |
| chr2-19104262-DEL-2962 | chr2-19104261-DEL-2963 |
| chr2-19108188-DEL-54 | chr2-19108202-DEL-52 |
| chr2-19109258-DEL-63 | chr2-19109259-DEL-63 |
| chr2-19116205-DEL-1819 | chr2-19116206-DEL-1819 |
| chr2-19126741-DEL-52 | chr2-19126742-DEL-52 |
| chr2-19171537-DEL-63 | chr2-19171535-DEL-64 |
| chr2-19176198-DEL-640 | chr2-19176203-DEL-641 |
| chr2-19199865-DEL-84 | chr2-19199866-DEL-84 |

| | |
|---|---|
| chr2-19217099-DEL-1343 | chr2-19217099-DEL-1343 |
| chr2-19225787-DEL-4734 | chr2-19225788-DEL-4734 |
| chr2-19231081-DEL-4536 | chr2-19231081-DEL-4536 |
| chr2-19250803-DEL-349 | chr2-19250804-DEL-349 |
| chr2-19253938-DEL-195 | chr2-19253939-DEL-195 |
| chr2-19256456-DEL-60 | chr2-19256454-DEL-61 |
| chr2-19278451-DEL-64 | chr2-19278452-DEL-64 |
| chr2-19308105-DEL-206 | chr2-19308106-DEL-206 |
| chr2-19334672-DEL-138 | chr2-19334673-DEL-140 |
| chr2-19337889-DEL-3089 | chr2-19337890-DEL-3089 |
| chr2-19359005-DEL-7279 | chr2-19359006-DEL-7279 |
| chr2-19380380-DEL-66 | chr2-19380381-DEL-66 |
| chr2-19405122-DEL-55 | chr2-19405077-DEL-100 |
| chr2-19418803-DEL-96 | chr2-19418804-DEL-96 |
| chr2-19421761-DEL-54 | chr2-19421762-DEL-54 |
| chr2-19423406-DEL-76 | chr2-19423405-DEL-77 |
| chr2-19425966-DEL-72 | chr2-19425967-DEL-72 |
| chr2-19513489-DEL-81 | chr2-19513490-DEL-81 |
| chr2-19537313-DEL-649 | chr2-19537315-DEL-650 |
| chr2-19539477-DEL-548 | chr2-19539478-DEL-548 |
| chr2-19541140-DEL-202 | chr2-19541141-DEL-202 |
| chr2-19576436-DEL-70 | chr2-19576437-DEL-70 |
| chr2-19582565-DEL-85 | chr2-19582566-DEL-85 |
| chr2-19586068-DEL-6999 | chr2-19586069-DEL-6999 |
| chr2-19586435-DEL-424 | chr2-19586436-DEL-424 |
| chr2-19643636-DEL-200 | chr2-19643637-DEL-200 |
| chr2-19707634-DEL-64 | chr2-19707635-DEL-64 |
| chr2-19757250-DEL-70 | chr2-19757253-DEL-68 |
| chr2-19765332-DEL-65 | chr2-19765333-DEL-65 |
| chr2-19794318-DEL-59 | chr2-19794318-DEL-59 |
| chr2-19804414-DEL-60 | chr2-19804419-DEL-57 |
| chr2-19843199-DEL-259 | chr2-19843200-DEL-259 |
| chr2-19859887-DEL-654 | chr2-19859888-DEL-654 |
| chr2-19903029-DEL-117 | chr2-19903041-DEL-114 |
| chr2-19915546-DEL-52 | chr2-19915547-DEL-52 |
| chr2-19919625-DEL-372 | chr2-19919626-DEL-372 |
| chr2-19926532-DEL-1812 | chr2-19926533-DEL-1812 |
| chr2-19960262-DEL-6338 | chr2-19960262-DEL-6338 |
| chr2-19988008-DEL-514 | chr2-19988009-DEL-514 |
| chr2-19997065-DEL-2815 | chr2-19997066-DEL-2815 |
| chr2-20001844-DEL-105 | chr2-20001845-DEL-105 |
| chr2-20003614-DEL-77 | chr2-20003635-DEL-72 |
| chr2-20007684-DEL-3699 | chr2-20007685-DEL-3699 |
| chr2-20036500-DEL-649 | chr2-20036501-DEL-649 |
| chr2-20041240-DEL-58 | chr2-20041246-DEL-52 |
| chr2-20043373-DEL-6440 | chr2-20043374-DEL-6440 |
| chr2-20096794-DEL-877 | chr2-20096795-DEL-877 |
| chr2-20110255-DEL-140 | chr2-20110256-DEL-112 |
| chr2-20127675-DEL-232 | chr2-20127676-DEL-232 |
| chr2-20151558-DEL-6266 | chr2-20151559-DEL-6266 |
| chr2-20159364-DEL-55 | chr2-20159365-DEL-55 |
| chr2-20162139-DEL-195 | chr2-20162140-DEL-195 |
| chr2-20162824-DEL-342 | chr2-20162821-DEL-342 |
| chr2-20189116-DEL-3070 | chr2-20189117-DEL-3070 |
| chr2-20206283-DEL-66 | chr2-20206284-DEL-66 |
| chr2-20224964-DEL-64 | chr2-20224984-DEL-50 |
| chr2-20233294-DEL-87 | chr2-20233295-DEL-87 |
| chr2-20239682-DEL-1620 | chr2-20239683-DEL-1620 |
| chr2-20251773-DEL-534 | chr2-20251774-DEL-534 |
| chr2-20258752-DEL-4983 | chr2-20258753-DEL-4983 |
| chr2-20265872-DEL-201 | chr2-20265873-DEL-201 |
| chr2-20270634-DEL-245 | chr2-20270634-DEL-246 |
| chr2-20343086-DEL-85 | chr2-20343087-DEL-85 |
| chr2-20347067-DEL-82 | chr2-20347107-DEL-78 |
| chr2-20349590-DEL-207 | chr2-20349591-DEL-207 |
| chr2-20372638-DEL-93 | chr2-20372639-DEL-93 |
| chr2-20383096-DEL-1002 | chr2-20383097-DEL-1002 |
| chr2-20385935-DEL-8409 | chr2-20385945-DEL-8395 |
| chr2-20415448-DEL-70 | chr2-20415449-DEL-69 |
| chr2-20431631-DEL-6442 | chr2-20431632-DEL-6442 |
| chr2-20462715-DEL-200 | chr2-20462716-DEL-200 |
| chr2-20480047-DEL-157 | chr2-20480037-DEL-149 |
| chr2-20496043-DEL-175 | chr2-20496044-DEL-175 |
| chr2-20502563-DEL-2801 | chr2-20502564-DEL-2801 |
| chr2-20540669-DEL-88 | chr2-20540695-DEL-58 |
| chr2-20567621-DEL-80 | chr2-20567622-DEL-80 |
| chr2-20584169-DEL-204 | chr2-20584170-DEL-204 |
| chr2-20586085-DEL-56 | chr2-20586069-DEL-64 |
| chr2-20588793-DEL-69 | chr2-20588792-DEL-70 |
| chr2-20600699-DEL-328 | chr2-20600700-DEL-328 |

| | |
|---|---|
| chr2-20610102-DEL-435 | chr2-20610132-DEL-386 |
| chr2-20626889-DEL-238 | chr2-20626890-DEL-238 |
| chr2-20661184-DEL-802 | chr2-20661185-DEL-802 |
| chr2-20668480-DEL-230 | chr2-20668481-DEL-231 |
| chr2-20682640-DEL-218 | chr2-20682641-DEL-218 |
| chr2-20688460-DEL-2276 | chr2-20688460-DEL-2276 |
| chr2-20691711-DEL-197 | chr2-20691701-DEL-198 |
| chr2-20711335-DEL-196 | chr2-20711336-DEL-196 |
| chr2-20734140-DEL-226 | chr2-20734141-DEL-226 |
| chr2-20781456-DEL-235 | chr2-20781457-DEL-235 |
| chr2-20804109-DEL-171 | chr2-20804110-DEL-171 |
| chr2-20828225-DEL-292 | chr2-20828226-DEL-292 |
| chr2-20835294-DEL-202 | chr2-20835295-DEL-202 |
| chr2-20861708-DEL-195 | chr2-20861709-DEL-195 |
| chr2-20867329-DEL-80 | chr2-20867330-DEL-80 |
| chr2-20896157-DEL-205 | chr2-20896158-DEL-205 |
| chr2-20902492-DEL-245 | chr2-20902493-DEL-245 |
| chr2-20904652-DEL-809 | chr2-20904651-DEL-810 |
| chr2-20920503-DEL-102 | chr2-20920504-DEL-102 |
| chr2-20929237-DEL-853 | chr2-20929238-DEL-853 |
| chr2-20936539-DEL-200 | chr2-20936540-DEL-200 |
| chr2-20982736-DEL-210 | chr2-20982737-DEL-210 |
| chr2-20996701-DEL-169 | chr2-20996702-DEL-169 |
| chr2-21011309-DEL-59 | chr2-21011310-DEL-59 |
| chr2-21024029-DEL-207 | chr2-21024030-DEL-207 |
| chr2-21033966-DEL-64 | chr2-21033967-DEL-64 |
| chr2-21072675-DEL-62 | chr2-21072685-DEL-56 |
| chr2-21079746-DEL-7084 | chr2-21079747-DEL-7084 |
| chr2-21094713-DEL-601 | chr2-21094709-DEL-602 |
| chr2-21135573-DEL-195 | chr2-21135574-DEL-195 |
| chr2-21137984-DEL-101 | chr2-21137983-DEL-98 |
| chr2-21140380-DEL-81 | chr2-21140384-DEL-54 |
| chr2-21145002-DEL-763 | chr2-21145003-DEL-763 |
| chr2-21149522-DEL-118 | chr2-21149523-DEL-118 |
| chr2-21152465-DEL-205 | chr2-21152466-DEL-205 |
| chr2-21154473-DEL-544 | chr2-21154474-DEL-544 |
| chr2-21155223-DEL-374 | chr2-21155224-DEL-374 |
| chr2-21166330-DEL-60 | chr2-21166331-DEL-60 |
| chr2-21175144-DEL-2247 | chr2-21175145-DEL-2243 |
| chr2-21198046-DEL-55 | chr2-21197996-DEL-60 |
| chr2-21248919-DEL-339 | chr2-21248913-DEL-339 |
| chr2-21251257-DEL-6394 | chr2-21251258-DEL-6394 |
| chr2-21304814-DEL-6588 | chr2-21304815-DEL-6588 |
| chr2-21311821-DEL-6822 | chr2-21311822-DEL-6822 |
| chr2-21329216-DEL-474 | chr2-21329208-DEL-472 |
| chr2-21331219-DEL-272 | chr2-21331219-DEL-272 |
| chr2-21437341-DEL-174 | chr2-21437342-DEL-174 |
| chr2-21448365-DEL-387 | chr2-21448366-DEL-387 |
| chr2-21457508-DEL-182 | chr2-21457509-DEL-182 |
| chr2-21464798-DEL-232 | chr2-21464799-DEL-232 |
| chr2-21484933-DEL-353 | chr2-21484934-DEL-353 |
| chr2-21489344-DEL-396 | chr2-21489344-DEL-392 |
| chr2-21491716-DEL-6405 | chr2-21491703-DEL-6416 |
| chr2-21515395-DEL-76 | chr2-21515396-DEL-76 |
| chr2-21584005-DEL-6632 | chr2-21584005-DEL-6632 |
| chr2-21609676-DEL-164 | chr2-21609677-DEL-164 |
| chr2-21629856-DEL-6446 | chr2-21629857-DEL-6446 |
| chr2-21640694-DEL-76 | chr2-21640738-DEL-57 |
| chr2-21641412-DEL-742 | chr2-21641413-DEL-748 |
| chr2-21659018-DEL-340 | chr2-21659019-DEL-340 |
| chr2-21691178-DEL-52 | chr2-21691184-DEL-52 |
| chr2-21755432-DEL-56 | chr2-21755433-DEL-56 |
| chr2-21761149-DEL-289 | chr2-21761142-DEL-291 |
| chr2-21761438-DEL-6944 | chr2-21761424-DEL-6958 |
| chr2-21769166-DEL-422 | chr2-21769167-DEL-422 |
| chr2-21772021-DEL-200 | chr2-21772022-DEL-200 |
| chr2-21806223-DEL-1699 | chr2-21806224-DEL-1699 |
| chr2-21812548-DEL-94 | chr2-21812506-DEL-95 |
| chr2-21813679-DEL-116 | chr2-21813680-DEL-124 |
| chr2-21837483-DEL-4483 | chr2-21837484-DEL-4483 |
| chr2-21866453-DEL-143 | chr2-21866454-DEL-143 |
| chr2-21877497-DEL-689 | chr2-21877498-DEL-689 |
| chr2-21901292-DEL-148 | chr2-21901293-DEL-148 |
| chr2-21907229-DEL-617 | chr2-21907230-DEL-617 |
| chr2-21920478-DEL-103 | chr2-21920479-DEL-103 |
| chr2-21937530-DEL-4961 | chr2-21937531-DEL-4961 |
| chr2-21956020-DEL-3811 | chr2-21956021-DEL-3811 |
| chr2-21957210-DEL-969 | chr2-21957211-DEL-969 |
| chr2-21987928-DEL-342 | chr2-21987927-DEL-343 |
| chr2-22000723-DEL-11412 | chr2-22000724-DEL-11412 |

| | |
|---|---|
| chr2-22006683-DEL-208 | chr2-22006684-DEL-208 |
| chr2-22043063-DEL-350 | chr2-22043064-DEL-350 |
| chr2-22048573-DEL-606 | chr2-22048574-DEL-606 |
| chr2-22054494-DEL-6254 | chr2-22054483-DEL-6255 |
| chr2-22062812-DEL-58 | chr2-22062813-DEL-58 |
| chr2-22070430-DEL-52 | chr2-22070431-DEL-52 |
| chr2-22125759-DEL-140 | chr2-22125760-DEL-140 |
| chr2-22141780-DEL-6280 | chr2-22141783-DEL-6277 |
| chr2-22149244-DEL-247 | chr2-22149247-DEL-242 |
| chr2-22163254-DEL-7080 | chr2-22163255-DEL-7080 |
| chr2-22224540-DEL-2360 | chr2-22224541-DEL-2360 |
| chr2-22228970-DEL-74 | chr2-22228971-DEL-74 |
| chr2-22279988-DEL-354 | chr2-22279989-DEL-354 |
| chr2-22303775-DEL-6216 | chr2-22303776-DEL-6216 |
| chr2-22313758-DEL-1075 | chr2-22313759-DEL-1075 |
| chr2-22343374-DEL-88 | chr2-22343375-DEL-88 |
| chr2-22345157-DEL-430 | chr2-22345158-DEL-430 |
| chr2-22368057-DEL-6194 | chr2-22368058-DEL-6194 |
| chr2-22377128-DEL-752 | chr2-22377129-DEL-752 |
| chr2-22378987-DEL-60 | chr2-22378988-DEL-60 |
| chr2-22379446-DEL-1604 | chr2-22379446-DEL-1604 |
| chr2-22386842-DEL-480 | chr2-22386843-DEL-480 |
| chr2-22403148-DEL-155 | chr2-22403134-DEL-148 |
| chr2-22405446-DEL-4934 | chr2-22405447-DEL-4934 |
| chr2-22415088-DEL-479 | chr2-22415089-DEL-479 |
| chr2-22417064-DEL-59 | chr2-22417065-DEL-59 |
| chr2-22419992-DEL-713 | chr2-22419993-DEL-713 |
| chr2-22449173-DEL-5044 | chr2-22449174-DEL-5044 |
| chr2-22460061-DEL-2073 | chr2-22460061-DEL-2073 |
| chr2-22495238-DEL-3424 | chr2-22495239-DEL-3424 |
| chr2-22539802-DEL-347 | chr2-22539790-DEL-339 |
| chr2-22562565-DEL-286 | chr2-22562566-DEL-285 |
| chr2-22568303-DEL-1456 | chr2-22568304-DEL-1456 |
| chr2-22576112-DEL-146 | chr2-22576113-DEL-146 |
| chr2-22606517-DEL-427 | chr2-22606518-DEL-427 |
| chr2-22628731-DEL-516 | chr2-22628730-DEL-515 |
| chr2-22661899-DEL-283 | chr2-22661900-DEL-283 |
| chr2-22672770-DEL-517 | chr2-22672771-DEL-517 |
| chr2-22682276-DEL-145 | chr2-22682277-DEL-145 |
| chr2-22696950-DEL-51 | chr2-22696951-DEL-51 |
| chr2-22738438-DEL-161 | chr2-22738439-DEL-161 |
| chr2-22746084-DEL-68 | chr2-22746085-DEL-68 |
| chr2-22814131-DEL-227 | chr2-22814132-DEL-227 |
| chr2-22816144-DEL-251 | chr2-22816145-DEL-251 |
| chr2-22865667-DEL-208 | chr2-22865668-DEL-208 |
| chr2-22899736-DEL-5320 | chr2-22899737-DEL-5320 |
| chr2-22939796-DEL-368 | chr2-22939797-DEL-368 |
| chr2-22944312-DEL-90 | chr2-22944340-DEL-64 |
| chr2-22954785-DEL-216 | chr2-22954786-DEL-214 |
| chr2-23024302-DEL-171 | chr2-23024303-DEL-171 |
| chr2-23034733-DEL-57 | chr2-23034731-DEL-58 |
| chr2-23044247-DEL-199 | chr2-23044248-DEL-199 |
| chr2-23058145-DEL-56 | chr2-23058151-DEL-56 |
| chr2-23074505-DEL-693 | chr2-23074506-DEL-693 |
| chr2-23081570-DEL-193 | chr2-23081571-DEL-193 |
| chr2-23086589-DEL-212 | chr2-23086590-DEL-212 |
| chr2-23186024-DEL-196 | chr2-23186025-DEL-196 |
| chr2-23213428-DEL-78 | chr2-23213429-DEL-78 |
| chr2-23229549-DEL-1268 | chr2-23229549-DEL-1268 |
| chr2-23270678-DEL-1776 | chr2-23270679-DEL-1776 |
| chr2-23310834-DEL-6134 | chr2-23310834-DEL-6134 |
| chr2-23317503-DEL-94 | chr2-23317503-DEL-94 |
| chr2-23345023-DEL-8170 | chr2-23345024-DEL-8170 |
| chr2-23376232-DEL-318 | chr2-23376233-DEL-318 |
| chr2-23390286-DEL-925 | chr2-23390287-DEL-925 |
| chr2-23394522-DEL-61 | chr2-23394523-DEL-61 |
| chr2-23412480-DEL-1552 | chr2-23412481-DEL-1552 |
| chr2-23458878-DEL-174 | chr2-23458879-DEL-174 |
| chr2-23462299-DEL-112 | chr2-23462328-DEL-73 |
| chr2-23512664-DEL-241 | chr2-23512673-DEL-241 |
| chr2-23528822-DEL-89 | chr2-23528823-DEL-89 |
| chr2-23551788-DEL-85 | chr2-23551815-DEL-56 |
| chr2-23659950-DEL-79 | chr2-23659930-DEL-94 |
| chr2-23664551-DEL-311 | chr2-23664552-DEL-311 |
| chr2-23675400-DEL-3123 | chr2-23675401-DEL-3123 |
| chr2-23702940-DEL-52 | chr2-23702949-DEL-54 |
| chr2-23712649-DEL-84 | chr2-23712650-DEL-84 |
| chr2-23717608-DEL-57 | chr2-23717609-DEL-64 |
| chr2-23760291-DEL-60 | chr2-23760298-DEL-54 |
| chr2-23763933-DEL-61 | chr2-23763934-DEL-61 |

| | |
|---|---|
| chr2-23786901-DEL-84 | chr2-23786902-DEL-84 |
| chr2-23798449-DEL-434 | chr2-23798448-DEL-435 |
| chr2-23851428-DEL-718 | chr2-23851428-DEL-718 |
| chr2-23871382-DEL-54 | chr2-23871383-DEL-54 |
| chr2-23907875-DEL-498 | chr2-23907879-DEL-498 |
| chr2-23983986-DEL-68 | chr2-23983987-DEL-68 |
| chr2-23996550-DEL-192 | chr2-23996551-DEL-192 |
| chr2-24006625-DEL-7309 | chr2-24006626-DEL-7309 |
| chr2-24013494-DEL-204 | chr2-24013495-DEL-204 |
| chr2-24053960-DEL-75 | chr2-24053965-DEL-71 |
| chr2-24055018-DEL-3424 | chr2-24055019-DEL-3424 |
| chr2-24059976-DEL-802 | chr2-24059977-DEL-802 |
| chr2-24066602-DEL-204 | chr2-24066603-DEL-204 |
| chr2-24072565-DEL-127 | chr2-24072566-DEL-127 |
| chr2-24083289-DEL-5735 | chr2-24083290-DEL-5735 |
| chr2-24118008-DEL-176 | chr2-24118009-DEL-176 |
| chr2-24118918-DEL-297 | chr2-24118919-DEL-297 |
| chr2-24127829-DEL-6194 | chr2-24127830-DEL-6194 |
| chr2-24138364-DEL-135 | chr2-24138365-DEL-135 |
| chr2-24188268-DEL-1097 | chr2-24188269-DEL-1097 |
| chr2-24214796-DEL-874 | chr2-24214797-DEL-874 |
| chr2-24220962-DEL-476 | chr2-24220963-DEL-476 |
| chr2-24278154-DEL-1556 | chr2-24278154-DEL-1556 |
| chr2-24282951-DEL-147 | chr2-24282952-DEL-147 |
| chr2-24285408-DEL-199 | chr2-24285408-DEL-199 |
| chr2-24314281-DEL-73 | chr2-24314282-DEL-73 |
| chr2-24380284-DEL-52 | chr2-24380278-DEL-70 |
| chr2-24423002-DEL-287 | chr2-24423050-DEL-240 |
| chr2-24441830-DEL-285 | chr2-24441823-DEL-286 |
| chr2-24447539-DEL-101 | chr2-24447572-DEL-69 |
| chr2-24465148-DEL-65 | chr2-24465144-DEL-70 |
| chr2-24468516-DEL-54 | chr2-24468517-DEL-54 |
| chr2-24470164-DEL-103 | chr2-24470169-DEL-99 |
| chr2-24485798-DEL-743 | chr2-24485799-DEL-743 |
| chr2-24542462-DEL-74 | chr2-24542463-DEL-74 |
| chr2-24553131-DEL-78 | chr2-24553132-DEL-78 |
| chr2-24622689-DEL-79 | chr2-24622690-DEL-79 |
| chr2-24631525-DEL-252 | chr2-24631526-DEL-252 |
| chr2-24654803-DEL-206 | chr2-24654804-DEL-206 |
| chr2-24750443-DEL-246 | chr2-24750444-DEL-246 |
| chr2-24772773-DEL-149 | chr2-24772774-DEL-149 |
| chr2-24775330-DEL-206 | chr2-24775331-DEL-206 |
| chr2-24784913-DEL-101 | chr2-24784914-DEL-103 |
| chr2-24803651-DEL-220 | chr2-24803652-DEL-220 |
| chr2-24825962-DEL-212 | chr2-24825963-DEL-212 |
| chr2-24831642-DEL-66 | chr2-24831643-DEL-66 |
| chr2-24841619-DEL-214 | chr2-24841620-DEL-214 |
| chr2-24849929-DEL-71 | chr2-24849930-DEL-71 |
| chr2-24875946-DEL-580 | chr2-24875947-DEL-580 |
| chr2-24884350-DEL-78 | chr2-24884350-DEL-78 |
| chr2-24895077-DEL-1278 | chr2-24895078-DEL-1278 |
| chr2-24895251-DEL-311 | chr2-24895252-DEL-311 |
| chr2-24914821-DEL-189 | chr2-24914822-DEL-189 |
| chr2-24918027-DEL-109 | chr2-24918028-DEL-107 |
| chr2-24962788-DEL-205 | chr2-24962789-DEL-205 |
| chr2-24967142-DEL-52 | chr2-24967100-DEL-59 |
| chr2-24980617-DEL-54 | chr2-24980606-DEL-55 |
| chr2-24995987-DEL-122 | chr2-24995988-DEL-122 |
| chr2-25013295-DEL-182 | chr2-25013296-DEL-182 |
| chr2-25038032-DEL-86 | chr2-25038033-DEL-87 |
| chr2-25043192-DEL-1703 | chr2-25043197-DEL-1699 |
| chr2-25079339-DEL-208 | chr2-25079340-DEL-208 |
| chr2-25082699-DEL-210 | chr2-25082700-DEL-210 |
| chr2-25146676-DEL-238 | chr2-25146677-DEL-238 |
| chr2-25149909-DEL-203 | chr2-25149910-DEL-203 |
| chr2-25199602-DEL-270 | chr2-25199603-DEL-264 |
| chr2-25220586-DEL-205 | chr2-25220587-DEL-205 |
| chr2-25229079-DEL-208 | chr2-25229080-DEL-208 |
| chr2-25230360-DEL-116 | chr2-25230339-DEL-117 |
| chr2-25231193-DEL-66 | chr2-25231190-DEL-62 |
| chr2-25232007-DEL-220 | chr2-25232008-DEL-220 |
| chr2-25248216-DEL-266 | chr2-25248217-DEL-266 |
| chr2-25255955-DEL-213 | chr2-25255956-DEL-213 |
| chr2-25257213-DEL-165 | chr2-25257214-DEL-165 |
| chr2-25278266-DEL-209 | chr2-25278267-DEL-209 |
| chr2-25305611-DEL-194 | chr2-25305612-DEL-195 |
| chr2-25341687-DEL-189 | chr2-25341688-DEL-186 |
| chr2-25344492-DEL-78 | chr2-25344493-DEL-78 |
| chr2-25363888-DEL-58 | chr2-25363883-DEL-59 |
| chr2-25371120-DEL-98 | chr2-25371121-DEL-98 |

| | |
|---|---|
| chr2-25379870-DEL-57 | chr2-25379871-DEL-57 |
| chr2-25399740-DEL-155 | chr2-25399741-DEL-155 |
| chr2-25404992-DEL-82 | chr2-25405004-DEL-70 |
| chr2-25405502-DEL-138 | chr2-25405539-DEL-102 |
| chr2-25518855-DEL-54 | chr2-25518829-DEL-81 |
| chr2-25530913-DEL-73 | chr2-25530941-DEL-69 |
| chr2-25535578-DEL-1586 | chr2-25535579-DEL-1586 |
| chr2-25550369-DEL-62 | chr2-25550368-DEL-63 |
| chr2-25562081-DEL-3271 | chr2-25562082-DEL-3271 |
| chr2-25576606-DEL-78 | chr2-25576606-DEL-75 |
| chr2-25607489-DEL-857 | chr2-25607488-DEL-858 |
| chr2-25608950-DEL-8656 | chr2-25608951-DEL-8656 |
| chr2-25625033-DEL-1114 | chr2-25625034-DEL-1114 |
| chr2-25633013-DEL-5189 | chr2-25633014-DEL-5189 |
| chr2-25633228-DEL-66 | chr2-25633229-DEL-66 |
| chr2-25643351-DEL-303 | chr2-25643352-DEL-303 |
| chr2-25661067-DEL-64 | chr2-25661068-DEL-64 |
| chr2-25689595-DEL-5662 | chr2-25689596-DEL-5662 |
| chr2-25707814-DEL-216 | chr2-25707815-DEL-216 |
| chr2-25763247-DEL-638 | chr2-25763248-DEL-638 |
| chr2-25784197-DEL-60 | chr2-25784198-DEL-60 |
| chr2-25785232-DEL-1150 | chr2-25785233-DEL-1150 |
| chr2-25879757-DEL-53 | chr2-25879758-DEL-53 |
| chr2-25880873-DEL-156 | chr2-25880889-DEL-146 |
| chr2-25889761-DEL-211 | chr2-25889762-DEL-211 |
| chr2-25891660-DEL-339 | chr2-25891661-DEL-340 |
| chr2-25920852-DEL-199 | chr2-25920853-DEL-199 |
| chr2-25940011-DEL-1498 | chr2-25940012-DEL-1498 |
| chr2-25940821-DEL-206 | chr2-25940822-DEL-206 |
| chr2-25959878-DEL-203 | chr2-25959879-DEL-203 |
| chr2-25993527-DEL-52 | chr2-25993506-DEL-79 |
| chr2-26015028-DEL-198 | chr2-26015014-DEL-190 |
| chr2-26017401-DEL-200 | chr2-26017402-DEL-200 |
| chr2-26026128-DEL-176 | chr2-26026113-DEL-177 |
| chr2-26035573-DEL-70 | chr2-26035574-DEL-70 |
| chr2-26039768-DEL-148 | chr2-26039765-DEL-152 |
| chr2-26042768-DEL-62 | chr2-26042781-DEL-50 |
| chr2-26051747-DEL-52 | chr2-26051748-DEL-52 |
| chr2-26066803-DEL-102 | chr2-26066804-DEL-102 |
| chr2-26069742-DEL-201 | chr2-26069743-DEL-201 |
| chr2-26071291-DEL-199 | chr2-26071292-DEL-199 |
| chr2-26123501-DEL-207 | chr2-26123502-DEL-207 |
| chr2-26223224-DEL-5259 | chr2-26223225-DEL-5259 |
| chr2-26261440-DEL-84 | chr2-26261441-DEL-84 |
| chr2-26265517-DEL-57 | chr2-26265518-DEL-57 |
| chr2-26342504-DEL-2503 | chr2-26342505-DEL-2503 |
| chr2-26444442-DEL-57 | chr2-26444443-DEL-57 |
| chr2-26455211-DEL-168 | chr2-26455228-DEL-152 |
| chr2-26509134-DEL-98 | chr2-26509136-DEL-97 |
| chr2-26510617-DEL-213 | chr2-26510618-DEL-213 |
| chr2-26514999-DEL-188 | chr2-26515000-DEL-188 |
| chr2-26521409-DEL-181 | chr2-26521410-DEL-181 |
| chr2-26529518-DEL-54 | chr2-26529519-DEL-54 |
| chr2-26559886-DEL-72 | chr2-26559887-DEL-72 |
| chr2-26567286-DEL-679 | chr2-26567287-DEL-679 |
| chr2-26647351-DEL-6622 | chr2-26647352-DEL-6622 |
| chr2-26678217-DEL-59 | chr2-26678218-DEL-59 |
| chr2-26683754-DEL-4673 | chr2-26683755-DEL-4673 |
| chr2-26684126-DEL-1674 | chr2-26684126-DEL-1674 |
| chr2-26697384-DEL-82 | chr2-26697385-DEL-82 |
| chr2-26707219-DEL-547 | chr2-26707220-DEL-547 |
| chr2-26711072-DEL-652 | chr2-26711072-DEL-652 |
| chr2-26744263-DEL-74 | chr2-26744235-DEL-74 |
| chr2-26745811-DEL-176 | chr2-26745805-DEL-177 |
| chr2-26748138-DEL-2227 | chr2-26748139-DEL-2227 |
| chr2-26749064-DEL-208 | chr2-26749065-DEL-208 |
| chr2-26758343-DEL-173 | chr2-26758344-DEL-173 |
| chr2-26766442-DEL-211 | chr2-26766443-DEL-211 |
| chr2-26770165-DEL-527 | chr2-26770166-DEL-527 |
| chr2-26858854-DEL-217 | chr2-26858855-DEL-198 |
| chr2-26888303-DEL-6026 | chr2-26888304-DEL-6026 |
| chr2-26943851-DEL-318 | chr2-26943851-DEL-318 |
| chr2-26954606-DEL-593 | chr2-26954605-DEL-598 |
| chr2-26964352-DEL-232 | chr2-26964352-DEL-232 |
| chr2-27001619-DEL-133 | chr2-27001620-DEL-86 |
| chr2-27036100-DEL-440 | chr2-27036101-DEL-440 |
| chr2-27040326-DEL-152 | chr2-27040327-DEL-152 |
| chr2-27048252-DEL-66 | chr2-27048253-DEL-64 |
| chr2-27058348-DEL-116 | chr2-27058349-DEL-116 |
| chr2-27077474-DEL-78 | chr2-27077475-DEL-78 |

| | |
|---|---|
| chr2-27080865-DEL-128 | chr2-27080864-DEL-129 |
| chr2-27091157-DEL-52 | chr2-27091158-DEL-52 |
| chr2-27106358-DEL-188 | chr2-27106359-DEL-188 |
| chr2-27107565-DEL-79 | chr2-27107566-DEL-79 |
| chr2-27130643-DEL-209 | chr2-27130644-DEL-209 |
| chr2-27135794-DEL-95 | chr2-27135808-DEL-104 |
| chr2-27140283-DEL-76 | chr2-27140284-DEL-76 |
| chr2-27147499-DEL-59 | chr2-27147498-DEL-61 |
| chr2-27163208-DEL-72 | chr2-27163209-DEL-72 |
| chr2-27196148-DEL-50 | chr2-27196149-DEL-50 |
| chr2-27196987-DEL-64 | chr2-27196988-DEL-64 |
| chr2-27201306-DEL-185 | chr2-27201307-DEL-185 |
| chr2-27222249-DEL-104 | chr2-27222250-DEL-104 |
| chr2-27239947-DEL-757 | chr2-27239948-DEL-757 |
| chr2-27348037-DEL-220 | chr2-27348038-DEL-220 |
| chr2-27379623-DEL-142 | chr2-27379624-DEL-134 |
| chr2-27382785-DEL-51 | chr2-27382786-DEL-51 |
| chr2-27386552-DEL-54 | chr2-27386553-DEL-54 |
| chr2-27396081-DEL-736 | chr2-27396082-DEL-736 |
| chr2-27408322-DEL-83 | chr2-27408323-DEL-83 |
| chr2-27411951-DEL-201 | chr2-27411952-DEL-201 |
| chr2-27422102-DEL-572 | chr2-27422103-DEL-572 |
| chr2-27438259-DEL-414 | chr2-27438260-DEL-414 |
| chr2-27445137-DEL-4314 | chr2-27445138-DEL-4314 |
| chr2-27447815-DEL-235 | chr2-27447816-DEL-235 |
| chr2-27465468-DEL-52 | chr2-27465464-DEL-53 |
| chr2-27495228-DEL-65 | chr2-27495228-DEL-60 |
| chr2-27503394-DEL-58 | chr2-27503395-DEL-58 |
| chr2-27505947-DEL-431 | chr2-27505927-DEL-468 |
| chr2-27548643-DEL-1356 | chr2-27548644-DEL-1356 |
| chr2-27559350-DEL-95 | chr2-27559351-DEL-95 |
| chr2-27597859-DEL-215 | chr2-27597861-DEL-214 |
| chr2-27639050-DEL-259 | chr2-27639051-DEL-259 |
| chr2-27682361-DEL-1344 | chr2-27682362-DEL-1344 |
| chr2-27693586-DEL-86 | chr2-27693587-DEL-86 |
| chr2-27712880-DEL-66 | chr2-27712881-DEL-66 |
| chr2-27731445-DEL-66 | chr2-27731446-DEL-66 |
| chr2-27800344-DEL-796 | chr2-27800347-DEL-797 |
| chr2-27819959-DEL-62 | chr2-27819973-DEL-56 |
| chr2-27838350-DEL-80 | chr2-27838371-DEL-60 |
| chr2-27847443-DEL-69 | chr2-27847444-DEL-69 |
| chr2-27868912-DEL-75 | chr2-27868941-DEL-73 |
| chr2-27945677-DEL-1874 | chr2-27945674-DEL-1875 |
| chr2-27959996-DEL-1005 | chr2-27959997-DEL-1005 |
| chr2-27990899-DEL-790 | chr2-27990900-DEL-790 |
| chr2-28029669-DEL-54 | chr2-28029670-DEL-78 |
| chr2-28041161-DEL-1687 | chr2-28041162-DEL-1687 |
| chr2-28065283-DEL-94 | chr2-28065268-DEL-110 |
| chr2-28074016-DEL-292 | chr2-28074047-DEL-292 |
| chr2-28081007-DEL-66 | chr2-28081008-DEL-66 |
| chr2-28114478-DEL-81 | chr2-28114479-DEL-101 |
| chr2-28124293-DEL-314 | chr2-28124289-DEL-312 |
| chr2-28171279-DEL-504 | chr2-28171280-DEL-504 |
| chr2-28182821-DEL-580 | chr2-28182816-DEL-581 |
| chr2-28195815-DEL-54 | chr2-28195816-DEL-54 |
| chr2-28231862-DEL-290 | chr2-28231863-DEL-290 |
| chr2-28297834-DEL-53 | chr2-28297835-DEL-53 |
| chr2-28300553-DEL-684 | chr2-28300554-DEL-684 |
| chr2-28313117-DEL-69 | chr2-28313115-DEL-62 |
| chr2-28316941-DEL-296 | chr2-28316988-DEL-284 |
| chr2-28362513-DEL-863 | chr2-28362514-DEL-863 |
| chr2-28364315-DEL-1402 | chr2-28364316-DEL-1402 |
| chr2-28365958-DEL-188 | chr2-28365959-DEL-188 |
| chr2-28367111-DEL-1052 | chr2-28367114-DEL-1043 |
| chr2-28376558-DEL-7055 | chr2-28376559-DEL-7055 |
| chr2-28465000-DEL-174 | chr2-28464999-DEL-174 |
| chr2-28475998-DEL-60 | chr2-28475999-DEL-60 |
| chr2-28495248-DEL-244 | chr2-28495249-DEL-244 |
| chr2-28498228-DEL-76 | chr2-28498227-DEL-70 |
| chr2-28502653-DEL-68 | chr2-28502654-DEL-68 |
| chr2-28518887-DEL-581 | chr2-28518888-DEL-581 |
| chr2-28523391-DEL-199 | chr2-28523392-DEL-199 |
| chr2-28537963-DEL-252 | chr2-28537964-DEL-252 |
| chr2-28572342-DEL-200 | chr2-28572343-DEL-200 |
| chr2-28647170-DEL-51 | chr2-28647171-DEL-51 |
| chr2-28680655-DEL-176 | chr2-28680656-DEL-176 |
| chr2-28685856-DEL-205 | chr2-28685857-DEL-205 |
| chr2-28694122-DEL-198 | chr2-28694123-DEL-198 |
| chr2-28700552-DEL-103 | chr2-28700580-DEL-67 |
| chr2-28762021-DEL-1957 | chr2-28762022-DEL-1957 |

| | |
|---|---|
| chr2-28814259-DEL-168 | chr2-28814261-DEL-160 |
| chr2-28823404-DEL-1290 | chr2-28823405-DEL-1290 |
| chr2-28831641-DEL-53 | chr2-28831642-DEL-53 |
| chr2-28832517-DEL-207 | chr2-28832518-DEL-207 |
| chr2-28857492-DEL-58 | chr2-28857493-DEL-58 |
| chr2-28869534-DEL-78 | chr2-28869549-DEL-50 |
| chr2-28890467-DEL-273 | chr2-28890436-DEL-293 |
| chr2-28909499-DEL-59 | chr2-28909500-DEL-59 |
| chr2-28918976-DEL-212 | chr2-28918977-DEL-212 |
| chr2-28925422-DEL-228 | chr2-28925423-DEL-228 |
| chr2-28934207-DEL-221 | chr2-28934208-DEL-221 |
| chr2-28949247-DEL-202 | chr2-28949248-DEL-202 |
| chr2-28963456-DEL-203 | chr2-28963457-DEL-203 |
| chr2-28967386-DEL-401 | chr2-28967387-DEL-401 |
| chr2-28972941-DEL-167 | chr2-28972942-DEL-167 |
| chr2-28976912-DEL-212 | chr2-28976913-DEL-213 |
| chr2-28998256-DEL-216 | chr2-28998257-DEL-216 |
| chr2-29000427-DEL-186 | chr2-29000428-DEL-186 |
| chr2-29018443-DEL-181 | chr2-29018429-DEL-179 |
| chr2-29027313-DEL-155 | chr2-29027314-DEL-155 |
| chr2-29072800-DEL-181 | chr2-29072801-DEL-181 |
| chr2-29078674-DEL-523 | chr2-29078675-DEL-523 |
| chr2-29111044-DEL-72 | chr2-29111044-DEL-72 |
| chr2-29153432-DEL-72 | chr2-29153436-DEL-51 |
| chr2-29154635-DEL-94 | chr2-29154636-DEL-94 |
| chr2-29178194-DEL-66 | chr2-29178194-DEL-54 |
| chr2-29188355-DEL-6991 | chr2-29188356-DEL-6991 |
| chr2-29221582-DEL-3243 | chr2-29221583-DEL-3243 |
| chr2-29232167-DEL-74 | chr2-29232131-DEL-98 |
| chr2-29247702-DEL-122 | chr2-29247703-DEL-122 |
| chr2-29266603-DEL-222 | chr2-29266604-DEL-222 |
| chr2-29273476-DEL-823 | chr2-29273475-DEL-824 |
| chr2-29279931-DEL-205 | chr2-29279932-DEL-205 |
| chr2-29323358-DEL-207 | chr2-29323359-DEL-207 |
| chr2-29332019-DEL-204 | chr2-29332020-DEL-204 |
| chr2-29333073-DEL-195 | chr2-29333074-DEL-195 |
| chr2-29380901-DEL-370 | chr2-29380900-DEL-371 |
| chr2-29424179-DEL-266 | chr2-29424187-DEL-263 |
| chr2-29446862-DEL-212 | chr2-29446863-DEL-212 |
| chr2-29471651-DEL-195 | chr2-29471666-DEL-220 |
| chr2-29478997-DEL-96 | chr2-29478989-DEL-104 |
| chr2-29487063-DEL-430 | chr2-29487064-DEL-430 |
| chr2-29487710-DEL-203 | chr2-29487723-DEL-212 |
| chr2-29528574-DEL-289 | chr2-29528575-DEL-289 |
| chr2-29547451-DEL-54 | chr2-29547452-DEL-54 |
| chr2-29572791-DEL-234 | chr2-29572792-DEL-234 |
| chr2-29596392-DEL-196 | chr2-29596393-DEL-196 |
| chr2-29598902-DEL-211 | chr2-29598903-DEL-211 |
| chr2-29631805-DEL-305 | chr2-29631806-DEL-305 |
| chr2-29648029-DEL-456 | chr2-29648058-DEL-458 |
| chr2-29698791-DEL-218 | chr2-29698792-DEL-220 |
| chr2-29711484-DEL-75 | chr2-29711458-DEL-103 |
| chr2-29712464-DEL-72 | chr2-29712465-DEL-72 |
| chr2-29724135-DEL-224 | chr2-29724136-DEL-224 |
| chr2-29749801-DEL-131 | chr2-29749801-DEL-130 |
| chr2-29751848-DEL-326 | chr2-29751849-DEL-320 |
| chr2-29768654-DEL-212 | chr2-29768651-DEL-213 |
| chr2-29798454-DEL-79 | chr2-29798417-DEL-80 |
| chr2-29822939-DEL-96 | chr2-29822941-DEL-54 |
| chr2-29841843-DEL-57 | chr2-29841843-DEL-57 |
| chr2-29926954-DEL-54 | chr2-29926955-DEL-54 |
| chr2-29945831-DEL-207 | chr2-29945829-DEL-210 |
| chr2-29966239-DEL-245 | chr2-29966242-DEL-238 |
| chr2-29966760-DEL-179 | chr2-29966761-DEL-179 |
| chr2-29977880-DEL-215 | chr2-29977881-DEL-215 |
| chr2-30091496-DEL-222 | chr2-30091497-DEL-222 |
| chr2-30108501-DEL-7101 | chr2-30108502-DEL-7101 |
| chr2-30139010-DEL-75 | chr2-30139011-DEL-75 |
| chr2-30161964-DEL-182 | chr2-30161964-DEL-182 |
| chr2-30168149-DEL-72 | chr2-30168150-DEL-72 |
| chr2-30171358-DEL-215 | chr2-30171358-DEL-215 |
| chr2-30201920-DEL-224 | chr2-30201921-DEL-224 |
| chr2-30267386-DEL-72 | chr2-30267363-DEL-96 |
| chr2-30279859-DEL-211 | chr2-30279860-DEL-211 |
| chr2-30328620-DEL-97 | chr2-30328621-DEL-97 |
| chr2-30352676-DEL-205 | chr2-30352677-DEL-207 |
| chr2-30379389-DEL-182 | chr2-30379390-DEL-182 |
| chr2-30414343-DEL-51 | chr2-30414344-DEL-51 |
| chr2-30427597-DEL-52 | chr2-30427598-DEL-60 |
| chr2-30433018-DEL-65 | chr2-30433019-DEL-65 |

| | |
|---|---|
| chr2-30436665-DEL-76 | chr2-30436657-DEL-59 |
| chr2-30449293-DEL-98 | chr2-30449294-DEL-98 |
| chr2-30525463-DEL-88 | chr2-30525462-DEL-85 |
| chr2-30538799-DEL-194 | chr2-30538800-DEL-194 |
| chr2-30584328-DEL-208 | chr2-30584329-DEL-208 |
| chr2-30586765-DEL-124 | chr2-30586747-DEL-136 |
| chr2-30591857-DEL-210 | chr2-30591858-DEL-210 |
| chr2-30616691-DEL-299 | chr2-30616692-DEL-299 |
| chr2-30617411-DEL-952 | chr2-30617412-DEL-951 |
| chr2-30618400-DEL-172 | chr2-30618401-DEL-172 |
| chr2-30623336-DEL-193 | chr2-30623337-DEL-193 |
| chr2-30625136-DEL-223 | chr2-30625137-DEL-223 |
| chr2-30629745-DEL-214 | chr2-30629746-DEL-214 |
| chr2-30640771-DEL-206 | chr2-30640772-DEL-206 |
| chr2-30722476-DEL-198 | chr2-30722477-DEL-198 |
| chr2-30767932-DEL-96 | chr2-30767933-DEL-92 |
| chr2-30772880-DEL-205 | chr2-30772881-DEL-205 |
| chr2-30807715-DEL-57 | chr2-30807715-DEL-57 |
| chr2-30809617-DEL-165 | chr2-30809618-DEL-165 |
| chr2-30811684-DEL-224 | chr2-30811687-DEL-222 |
| chr2-30820587-DEL-51 | chr2-30820588-DEL-51 |
| chr2-30831326-DEL-129 | chr2-30831327-DEL-129 |
| chr2-30834389-DEL-216 | chr2-30834390-DEL-216 |
| chr2-30840714-DEL-175 | chr2-30840707-DEL-176 |
| chr2-30869662-DEL-62 | chr2-30869661-DEL-63 |
| chr2-30935735-DEL-133 | chr2-30935736-DEL-133 |
| chr2-30949114-DEL-235 | chr2-30949115-DEL-235 |
| chr2-30960373-DEL-183 | chr2-30960374-DEL-183 |
| chr2-30974796-DEL-230 | chr2-30974796-DEL-230 |
| chr2-30999498-DEL-53 | chr2-30999499-DEL-55 |
| chr2-31010705-DEL-223 | chr2-31010706-DEL-223 |
| chr2-31035766-DEL-260 | chr2-31035767-DEL-260 |
| chr2-31040478-DEL-164 | chr2-31040479-DEL-164 |
| chr2-31048516-DEL-72 | chr2-31048517-DEL-72 |
| chr2-31085832-DEL-109 | chr2-31085834-DEL-107 |
| chr2-31092524-DEL-1875 | chr2-31092525-DEL-1875 |
| chr2-31109276-DEL-184 | chr2-31109277-DEL-184 |
| chr2-31128841-DEL-51 | chr2-31128841-DEL-51 |
| chr2-31162505-DEL-101 | chr2-31162506-DEL-101 |
| chr2-31164779-DEL-148 | chr2-31164780-DEL-148 |
| chr2-31189737-DEL-311 | chr2-31189738-DEL-311 |
| chr2-31198642-DEL-172 | chr2-31198643-DEL-172 |
| chr2-31216340-DEL-399 | chr2-31216341-DEL-399 |
| chr2-31220068-DEL-332 | chr2-31220069-DEL-332 |
| chr2-31266053-DEL-62 | chr2-31266054-DEL-62 |
| chr2-31294208-DEL-133 | chr2-31294203-DEL-98 |
| chr2-31306380-DEL-199 | chr2-31306381-DEL-199 |
| chr2-31351408-DEL-56 | chr2-31351409-DEL-57 |
| chr2-31352497-DEL-320 | chr2-31352459-DEL-320 |
| chr2-31368171-DEL-399 | chr2-31368172-DEL-399 |
| chr2-31421580-DEL-221 | chr2-31421581-DEL-221 |
| chr2-31425902-DEL-133 | chr2-31425922-DEL-83 |
| chr2-31446091-DEL-54 | chr2-31446092-DEL-54 |
| chr2-31447683-DEL-404 | chr2-31447684-DEL-404 |
| chr2-31515702-DEL-182 | chr2-31515703-DEL-182 |
| chr2-31520127-DEL-220 | chr2-31520128-DEL-220 |
| chr2-31558403-DEL-199 | chr2-31558404-DEL-199 |
| chr2-31558922-DEL-90 | chr2-31558929-DEL-60 |
| chr2-31571130-DEL-207 | chr2-31571131-DEL-207 |
| chr2-31588460-DEL-62 | chr2-31588461-DEL-62 |
| chr2-31589100-DEL-190 | chr2-31589101-DEL-190 |
| chr2-31597559-DEL-870 | chr2-31597560-DEL-870 |
| chr2-31621667-DEL-199 | chr2-31621668-DEL-199 |
| chr2-31622333-DEL-202 | chr2-31622334-DEL-202 |
| chr2-31623886-DEL-210 | chr2-31623887-DEL-210 |
| chr2-31685574-DEL-284 | chr2-31685585-DEL-264 |
| chr2-31786659-DEL-243 | chr2-31786660-DEL-243 |
| chr2-31801115-DEL-107 | chr2-31801123-DEL-84 |
| chr2-31828837-DEL-65 | chr2-31828836-DEL-67 |
| chr2-31848145-DEL-58 | chr2-31848153-DEL-50 |
| chr2-31916049-DEL-220 | chr2-31916050-DEL-220 |
| chr2-31936718-DEL-260 | chr2-31936718-DEL-260 |
| chr2-31966390-DEL-211 | chr2-31966391-DEL-211 |
| chr2-31991872-DEL-208 | chr2-31991873-DEL-208 |
| chr2-32002330-DEL-120 | chr2-32002373-DEL-119 |
| chr2-32007118-DEL-2883 | chr2-32007118-DEL-2887 |
| chr2-32015895-DEL-212 | chr2-32015897-DEL-213 |
| chr2-32025000-DEL-117 | chr2-32024992-DEL-92 |
| chr2-32034365-DEL-210 | chr2-32034366-DEL-210 |
| chr2-32039719-DEL-58 | chr2-32039719-DEL-55 |

| | |
|---|---|
| chr2-32060055-DEL-2018 | chr2-32060056-DEL-2016 |
| chr2-32060978-DEL-51 | chr2-32060979-DEL-51 |
| chr2-32062371-DEL-209 | chr2-32062372-DEL-209 |
| chr2-32076925-DEL-81 | chr2-32076926-DEL-81 |
| chr2-32082141-DEL-183 | chr2-32082155-DEL-176 |
| chr2-32087547-DEL-406 | chr2-32087548-DEL-406 |
| chr2-32094911-DEL-96 | chr2-32094912-DEL-96 |
| chr2-32115284-DEL-58 | chr2-32115285-DEL-58 |
| chr2-32117326-DEL-188 | chr2-32117327-DEL-188 |
| chr2-32125786-DEL-188 | chr2-32125787-DEL-188 |
| chr2-32130168-DEL-205 | chr2-32130169-DEL-205 |
| chr2-32164163-DEL-521 | chr2-32164164-DEL-521 |
| chr2-32245375-DEL-200 | chr2-32245364-DEL-224 |
| chr2-32298107-DEL-72 | chr2-32298114-DEL-62 |
| chr2-32304894-DEL-106 | chr2-32304895-DEL-106 |
| chr2-32357022-DEL-1069 | chr2-32357021-DEL-1070 |
| chr2-32388122-DEL-328 | chr2-32388123-DEL-328 |
| chr2-32396942-DEL-84 | chr2-32396943-DEL-84 |
| chr2-32450723-DEL-83 | chr2-32450724-DEL-83 |
| chr2-32531282-DEL-219 | chr2-32531283-DEL-219 |
| chr2-32547201-DEL-185 | chr2-32547202-DEL-185 |
| chr2-32556188-DEL-407 | chr2-32556189-DEL-407 |
| chr2-32556805-DEL-54 | chr2-32556806-DEL-54 |
| chr2-32578612-DEL-64 | chr2-32578620-DEL-64 |
| chr2-32650979-DEL-79 | chr2-32650980-DEL-78 |
| chr2-32697191-DEL-196 | chr2-32697192-DEL-196 |
| chr2-32727371-DEL-202 | chr2-32727372-DEL-202 |
| chr2-32819970-DEL-81 | chr2-32819971-DEL-81 |
| chr2-32842448-DEL-60 | chr2-32842449-DEL-60 |
| chr2-32847564-DEL-122 | chr2-32847565-DEL-122 |
| chr2-32852384-DEL-58 | chr2-32852385-DEL-58 |
| chr2-32926717-DEL-203 | chr2-32926718-DEL-203 |
| chr2-32984167-DEL-837 | chr2-32984167-DEL-838 |
| chr2-33203937-DEL-233 | chr2-33203938-DEL-233 |
| chr2-33216528-DEL-173 | chr2-33216529-DEL-173 |
| chr2-33282433-DEL-205 | chr2-33282434-DEL-205 |
| chr2-33295516-DEL-207 | chr2-33295530-DEL-207 |
| chr2-33306508-DEL-231 | chr2-33306509-DEL-231 |
| chr2-33311039-DEL-400 | chr2-33311040-DEL-400 |
| chr2-33348692-DEL-229 | chr2-33348693-DEL-229 |
| chr2-33369845-DEL-205 | chr2-33369846-DEL-205 |
| chr2-33391312-DEL-681 | chr2-33391313-DEL-681 |
| chr2-33398622-DEL-218 | chr2-33398623-DEL-218 |
| chr2-33400973-DEL-60 | chr2-33400974-DEL-60 |
| chr2-33416088-DEL-129 | chr2-33416089-DEL-129 |
| chr2-33434894-DEL-58 | chr2-33434895-DEL-58 |
| chr2-33435568-DEL-226 | chr2-33435569-DEL-226 |
| chr2-33489979-DEL-54 | chr2-33489983-DEL-50 |
| chr2-33543875-DEL-210 | chr2-33543876-DEL-210 |
| chr2-33544570-DEL-221 | chr2-33544573-DEL-223 |
| chr2-33546211-DEL-232 | chr2-33546212-DEL-232 |
| chr2-33550446-DEL-84 | chr2-33550447-DEL-90 |
| chr2-33570750-DEL-238 | chr2-33570751-DEL-238 |
| chr2-33575352-DEL-202 | chr2-33575353-DEL-202 |
| chr2-33615458-DEL-69 | chr2-33615456-DEL-70 |
| chr2-33821045-DEL-2239 | chr2-33821047-DEL-2238 |
| chr2-33910695-DEL-213 | chr2-33910696-DEL-213 |
| chr2-33919281-DEL-63 | chr2-33919263-DEL-60 |
| chr2-33963429-DEL-54 | chr2-33963430-DEL-54 |
| chr2-34079183-DEL-227 | chr2-34079184-DEL-227 |
| chr2-34088920-DEL-2783 | chr2-34088921-DEL-2784 |
| chr2-34155642-DEL-228 | chr2-34155643-DEL-228 |
| chr2-34196576-DEL-77 | chr2-34196559-DEL-86 |
| chr2-34265293-DEL-56 | chr2-34265294-DEL-56 |
| chr2-34272073-DEL-217 | chr2-34272074-DEL-217 |
| chr2-34278033-DEL-55 | chr2-34278034-DEL-55 |
| chr2-34307031-DEL-239 | chr2-34307032-DEL-239 |
| chr2-34376200-DEL-51 | chr2-34376200-DEL-51 |
| chr2-34394852-DEL-184 | chr2-34394853-DEL-184 |
| chr2-34411886-DEL-394 | chr2-34411887-DEL-394 |
| chr2-34415317-DEL-246 | chr2-34415318-DEL-246 |
| chr2-34441498-DEL-172 | chr2-34441499-DEL-172 |
| chr2-34480367-DEL-225 | chr2-34480368-DEL-225 |
| chr2-34497082-DEL-199 | chr2-34497083-DEL-199 |
| chr2-34538865-DEL-1525 | chr2-34538858-DEL-1527 |
| chr2-34549602-DEL-90 | chr2-34549603-DEL-90 |
| chr2-34560869-DEL-206 | chr2-34560870-DEL-206 |
| chr2-34578200-DEL-54 | chr2-34578201-DEL-58 |
| chr2-34605708-DEL-211 | chr2-34605708-DEL-211 |
| chr2-34613786-DEL-210 | chr2-34613774-DEL-205 |

| | |
|---|---|
| chr2-34640665-DEL-55 | chr2-34640666-DEL-55 |
| chr2-34644633-DEL-57 | chr2-34644634-DEL-57 |
| chr2-34649368-DEL-1890 | chr2-34649369-DEL-1890 |
| chr2-34677208-DEL-1168 | chr2-34677209-DEL-1168 |
| chr2-34679579-DEL-213 | chr2-34679580-DEL-213 |
| chr2-34682319-DEL-199 | chr2-34682320-DEL-199 |
| chr2-34691139-DEL-202 | chr2-34691140-DEL-202 |
| chr2-34693032-DEL-64 | chr2-34693033-DEL-64 |
| chr2-34707122-DEL-68 | chr2-34707123-DEL-68 |
| chr2-34710477-DEL-215 | chr2-34710478-DEL-215 |
| chr2-34717079-DEL-174 | chr2-34717082-DEL-172 |
| chr2-34721317-DEL-212 | chr2-34721306-DEL-213 |
| chr2-34734104-DEL-52 | chr2-34734076-DEL-62 |
| chr2-34735112-DEL-208 | chr2-34735113-DEL-208 |
| chr2-34735926-DEL-108 | chr2-34735927-DEL-108 |
| chr2-34736624-DEL-2851 | chr2-34736624-DEL-2852 |
| chr2-34751425-DEL-541 | chr2-34751426-DEL-535 |
| chr2-34765592-DEL-431 | chr2-34765593-DEL-431 |
| chr2-34778515-DEL-1500 | chr2-34778516-DEL-1500 |
| chr2-34819302-DEL-6474 | chr2-34819303-DEL-6474 |
| chr2-34932389-DEL-399 | chr2-34932390-DEL-399 |
| chr2-34936019-DEL-112 | chr2-34936020-DEL-112 |
| chr2-34946380-DEL-56 | chr2-34946381-DEL-55 |
| chr2-34951852-DEL-117 | chr2-34951853-DEL-122 |
| chr2-34960762-DEL-1370 | chr2-34960765-DEL-1368 |
| chr2-34980662-DEL-2728 | chr2-34980663-DEL-2728 |
| chr2-34983559-DEL-290 | chr2-34983560-DEL-290 |
| chr2-35004133-DEL-60 | chr2-35004134-DEL-60 |
| chr2-35016869-DEL-223 | chr2-35016834-DEL-224 |
| chr2-35055108-DEL-173 | chr2-35055109-DEL-173 |
| chr2-35135512-DEL-120 | chr2-35135513-DEL-120 |
| chr2-35230326-DEL-1274 | chr2-35230328-DEL-1273 |
| chr2-35260682-DEL-110 | chr2-35260683-DEL-110 |
| chr2-35274934-DEL-191 | chr2-35274935-DEL-191 |
| chr2-35295273-DEL-204 | chr2-35295274-DEL-204 |
| chr2-35368777-DEL-62 | chr2-35368778-DEL-52 |
| chr2-35381604-DEL-439 | chr2-35381603-DEL-439 |
| chr2-35393075-DEL-197 | chr2-35393076-DEL-197 |
| chr2-35401308-DEL-58 | chr2-35401286-DEL-80 |
| chr2-35405552-DEL-435 | chr2-35405553-DEL-435 |
| chr2-35409530-DEL-7079 | chr2-35409531-DEL-7079 |
| chr2-35440945-DEL-77 | chr2-35440913-DEL-78 |
| chr2-35466102-DEL-69 | chr2-35466143-DEL-61 |
| chr2-35489372-DEL-54 | chr2-35489373-DEL-54 |
| chr2-35515097-DEL-132 | chr2-35515093-DEL-133 |
| chr2-35520749-DEL-398 | chr2-35520750-DEL-398 |
| chr2-35528765-DEL-72 | chr2-35528766-DEL-72 |
| chr2-35531309-DEL-380 | chr2-35531314-DEL-352 |
| chr2-35567360-DEL-114 | chr2-35567361-DEL-114 |
| chr2-35598987-DEL-168 | chr2-35599002-DEL-167 |
| chr2-35630004-DEL-257 | chr2-35630005-DEL-257 |
| chr2-35638885-DEL-76 | chr2-35638886-DEL-76 |
| chr2-35690354-DEL-56 | chr2-35690354-DEL-59 |
| chr2-35692104-DEL-72 | chr2-35692104-DEL-84 |
| chr2-35730078-DEL-95 | chr2-35730094-DEL-92 |
| chr2-35755629-DEL-77 | chr2-35755630-DEL-77 |
| chr2-35817681-DEL-124 | chr2-35817728-DEL-120 |
| chr2-35821262-DEL-248 | chr2-35821263-DEL-248 |
| chr2-35824462-DEL-52 | chr2-35824463-DEL-52 |
| chr2-35834587-DEL-58 | chr2-35834600-DEL-58 |
| chr2-35861666-DEL-549 | chr2-35861683-DEL-578 |
| chr2-35904558-DEL-4693 | chr2-35904559-DEL-4693 |
| chr2-35945205-DEL-393 | chr2-35945164-DEL-393 |
| chr2-36000603-DEL-207 | chr2-36000604-DEL-207 |
| chr2-36048456-DEL-56 | chr2-36048457-DEL-56 |
| chr2-36082795-DEL-578 | chr2-36082795-DEL-578 |
| chr2-36139949-DEL-620 | chr2-36139950-DEL-620 |
| chr2-36177203-DEL-104 | chr2-36177204-DEL-104 |
| chr2-36180459-DEL-52 | chr2-36180460-DEL-52 |
| chr2-36183492-DEL-6753 | chr2-36183493-DEL-6753 |
| chr2-36189834-DEL-204 | chr2-36189834-DEL-204 |
| chr2-36236357-DEL-53 | chr2-36236357-DEL-53 |
| chr2-36263742-DEL-75 | chr2-36263743-DEL-75 |
| chr2-36270997-DEL-166 | chr2-36270998-DEL-166 |
| chr2-36401191-DEL-2689 | chr2-36401191-DEL-2689 |
| chr2-36432806-DEL-6616 | chr2-36432807-DEL-6616 |
| chr2-36471483-DEL-1666 | chr2-36471484-DEL-1666 |
| chr2-36517287-DEL-204 | chr2-36517288-DEL-204 |
| chr2-36525800-DEL-96 | chr2-36525794-DEL-93 |
| chr2-36534121-DEL-54 | chr2-36534122-DEL-54 |

| | |
|---|---|
| chr2-36552176-DEL-3432 | chr2-36552177-DEL-3432 |
| chr2-36569030-DEL-4251 | chr2-36569031-DEL-4251 |
| chr2-36573616-DEL-199 | chr2-36573617-DEL-199 |
| chr2-36580525-DEL-87 | chr2-36580526-DEL-87 |
| chr2-36581454-DEL-65 | chr2-36581455-DEL-65 |
| chr2-36592747-DEL-779 | chr2-36592747-DEL-779 |
| chr2-36650182-DEL-6361 | chr2-36650182-DEL-6361 |
| chr2-36754264-DEL-96 | chr2-36754281-DEL-84 |
| chr2-36848535-DEL-881 | chr2-36848542-DEL-879 |
| chr2-36884033-DEL-1659 | chr2-36884034-DEL-1659 |
| chr2-36900474-DEL-401 | chr2-36900475-DEL-401 |
| chr2-36915301-DEL-202 | chr2-36915302-DEL-202 |
| chr2-36998162-DEL-6862 | chr2-36998163-DEL-6862 |
| chr2-37015384-DEL-101 | chr2-37015385-DEL-101 |
| chr2-37035103-DEL-195 | chr2-37035104-DEL-195 |
| chr2-37046797-DEL-4871 | chr2-37046797-DEL-4871 |
| chr2-37106395-DEL-51 | chr2-37106398-DEL-55 |
| chr2-37135753-DEL-276 | chr2-37135754-DEL-276 |
| chr2-37220577-DEL-853 | chr2-37220578-DEL-853 |
| chr2-37229747-DEL-7095 | chr2-37229748-DEL-7096 |
| chr2-37250308-DEL-434 | chr2-37250309-DEL-434 |
| chr2-37253667-DEL-199 | chr2-37253671-DEL-195 |
| chr2-37296232-DEL-238 | chr2-37296233-DEL-238 |
| chr2-37310075-DEL-206 | chr2-37310076-DEL-206 |
| chr2-37361417-DEL-212 | chr2-37361441-DEL-203 |
| chr2-37543730-DEL-203 | chr2-37543731-DEL-203 |
| chr2-37560416-DEL-228 | chr2-37560417-DEL-228 |
| chr2-37578040-DEL-243 | chr2-37578041-DEL-200 |
| chr2-37648131-DEL-131 | chr2-37648132-DEL-131 |
| chr2-37652374-DEL-82 | chr2-37652375-DEL-82 |
| chr2-37791382-DEL-118 | chr2-37791383-DEL-118 |
| chr2-37806424-DEL-414 | chr2-37806424-DEL-414 |
| chr2-37826869-DEL-201 | chr2-37826869-DEL-202 |
| chr2-37852029-DEL-596 | chr2-37852030-DEL-596 |
| chr2-37867131-DEL-73 | chr2-37867123-DEL-74 |
| chr2-37943719-DEL-7016 | chr2-37943720-DEL-7016 |
| chr2-38018825-DEL-792 | chr2-38018826-DEL-792 |
| chr2-38120064-DEL-55 | chr2-38120033-DEL-57 |
| chr2-38154710-DEL-102 | chr2-38154711-DEL-102 |
| chr2-38224561-DEL-227 | chr2-38224562-DEL-227 |
| chr2-38255178-DEL-211 | chr2-38255179-DEL-211 |
| chr2-38277421-DEL-114 | chr2-38277422-DEL-114 |
| chr2-38290538-DEL-856 | chr2-38290539-DEL-856 |
| chr2-38337507-DEL-76 | chr2-38337516-DEL-68 |
| chr2-38341663-DEL-92 | chr2-38341677-DEL-52 |
| chr2-38369373-DEL-164 | chr2-38369375-DEL-166 |
| chr2-38387272-DEL-221 | chr2-38387273-DEL-221 |
| chr2-38401080-DEL-116 | chr2-38401081-DEL-108 |
| chr2-38454679-DEL-1888 | chr2-38454680-DEL-1888 |
| chr2-38471798-DEL-172 | chr2-38471799-DEL-172 |
| chr2-38472782-DEL-325 | chr2-38472783-DEL-325 |
| chr2-38473544-DEL-87 | chr2-38473545-DEL-87 |
| chr2-38498535-DEL-109 | chr2-38498533-DEL-98 |
| chr2-38549508-DEL-60 | chr2-38549518-DEL-53 |
| chr2-38565738-DEL-117 | chr2-38565785-DEL-83 |
| chr2-38605581-DEL-133 | chr2-38605582-DEL-95 |
| chr2-38740385-DEL-200 | chr2-38740386-DEL-200 |
| chr2-38758035-DEL-200 | chr2-38758036-DEL-200 |
| chr2-38777992-DEL-201 | chr2-38777993-DEL-201 |
| chr2-38803874-DEL-72 | chr2-38803875-DEL-72 |
| chr2-38854088-DEL-64 | chr2-38854089-DEL-64 |
| chr2-38858427-DEL-52 | chr2-38858428-DEL-52 |
| chr2-38864935-DEL-85 | chr2-38864976-DEL-82 |
| chr2-38900071-DEL-214 | chr2-38900072-DEL-214 |
| chr2-38919584-DEL-234 | chr2-38919585-DEL-234 |
| chr2-38925481-DEL-750 | chr2-38925482-DEL-750 |
| chr2-39130476-DEL-6554 | chr2-39130477-DEL-6554 |
| chr2-39212276-DEL-870 | chr2-39212266-DEL-871 |
| chr2-39229210-DEL-438 | chr2-39229210-DEL-438 |
| chr2-39250688-DEL-7102 | chr2-39250689-DEL-7102 |
| chr2-39275728-DEL-309 | chr2-39275729-DEL-309 |
| chr2-39283177-DEL-1611 | chr2-39283177-DEL-1611 |
| chr2-39415956-DEL-4995 | chr2-39415957-DEL-4995 |
| chr2-39449644-DEL-94 | chr2-39449645-DEL-94 |
| chr2-39492052-DEL-280 | chr2-39492053-DEL-280 |
| chr2-39519026-DEL-3689 | chr2-39519026-DEL-3689 |
| chr2-39529388-DEL-6455 | chr2-39529374-DEL-6469 |
| chr2-39542444-DEL-3766 | chr2-39542445-DEL-3766 |
| chr2-39547265-DEL-75 | chr2-39547266-DEL-75 |
| chr2-39575581-DEL-6490 | chr2-39575582-DEL-6490 |

| | |
|---|---|
| chr2-39634101-DEL-1117 | chr2-39634102-DEL-1117 |
| chr2-39653398-DEL-4327 | chr2-39653399-DEL-4327 |
| chr2-39664924-DEL-313 | chr2-39664924-DEL-313 |
| chr2-39668581-DEL-7081 | chr2-39668581-DEL-7081 |
| chr2-39693202-DEL-354 | chr2-39693203-DEL-354 |
| chr2-39720807-DEL-158 | chr2-39720808-DEL-158 |
| chr2-39724951-DEL-619 | chr2-39724952-DEL-619 |
| chr2-39782655-DEL-6372 | chr2-39782656-DEL-6372 |
| chr2-39801302-DEL-4786 | chr2-39801303-DEL-4786 |
| chr2-39837776-DEL-53161 | chr2-39837776-DEL-53161 |
| chr2-39858232-DEL-75 | chr2-39858233-DEL-75 |
| chr2-39871984-DEL-537 | chr2-39871985-DEL-537 |
| chr2-39878952-DEL-2760 | chr2-39878953-DEL-2760 |
| chr2-39883094-DEL-5222 | chr2-39883095-DEL-5222 |
| chr2-39907710-DEL-3490 | chr2-39907710-DEL-3490 |
| chr2-39917948-DEL-2748 | chr2-39917949-DEL-2748 |
| chr2-39967890-DEL-80 | chr2-39967891-DEL-80 |
| chr2-39979793-DEL-133 | chr2-39979794-DEL-133 |
| chr2-40008042-DEL-52 | chr2-40008032-DEL-68 |
| chr2-40059948-DEL-6198 | chr2-40059949-DEL-6198 |
| chr2-40122852-DEL-135 | chr2-40122833-DEL-155 |
| chr2-40125608-DEL-5060 | chr2-40125609-DEL-5060 |
| chr2-40135860-DEL-148 | chr2-40135861-DEL-147 |
| chr2-40185871-DEL-1369 | chr2-40185870-DEL-1369 |
| chr2-40198609-DEL-103 | chr2-40198654-DEL-100 |
| chr2-40203840-DEL-96 | chr2-40203838-DEL-96 |
| chr2-40219861-DEL-911 | chr2-40219862-DEL-911 |
| chr2-40220942-DEL-1192 | chr2-40220943-DEL-1192 |
| chr2-40230094-DEL-55 | chr2-40230092-DEL-51 |
| chr2-40245840-DEL-6724 | chr2-40245841-DEL-6724 |
| chr2-40271025-DEL-140 | chr2-40271036-DEL-106 |
| chr2-40316805-DEL-3195 | chr2-40316818-DEL-3173 |
| chr2-40326910-DEL-249 | chr2-40326909-DEL-232 |
| chr2-40327120-DEL-783 | chr2-40327121-DEL-783 |
| chr2-40328188-DEL-333 | chr2-40328189-DEL-333 |
| chr2-40361657-DEL-168 | chr2-40361658-DEL-168 |
| chr2-40365870-DEL-189 | chr2-40365871-DEL-189 |
| chr2-40376971-DEL-358 | chr2-40376972-DEL-358 |
| chr2-40380231-DEL-359 | chr2-40380232-DEL-359 |
| chr2-40381447-DEL-399 | chr2-40381448-DEL-399 |
| chr2-40386693-DEL-114 | chr2-40386694-DEL-114 |
| chr2-40410643-DEL-862 | chr2-40410643-DEL-862 |
| chr2-40412594-DEL-482 | chr2-40412594-DEL-482 |
| chr2-40423248-DEL-542 | chr2-40423249-DEL-542 |
| chr2-40428765-DEL-229 | chr2-40428765-DEL-229 |
| chr2-40442622-DEL-68 | chr2-40442623-DEL-68 |
| chr2-40452557-DEL-417 | chr2-40452558-DEL-415 |
| chr2-40524023-DEL-334 | chr2-40524022-DEL-329 |
| chr2-40547908-DEL-90 | chr2-40547922-DEL-89 |
| chr2-40563319-DEL-403 | chr2-40563320-DEL-402 |
| chr2-40571936-DEL-233 | chr2-40571937-DEL-232 |
| chr2-40573997-DEL-1375 | chr2-40574002-DEL-1371 |
| chr2-40707409-DEL-254 | chr2-40707410-DEL-254 |
| chr2-40725732-DEL-59 | chr2-40725733-DEL-59 |
| chr2-40799002-DEL-62 | chr2-40799007-DEL-58 |
| chr2-40819519-DEL-6580 | chr2-40819520-DEL-6580 |
| chr2-40839943-DEL-56 | chr2-40839944-DEL-56 |
| chr2-40861243-DEL-133 | chr2-40861244-DEL-133 |
| chr2-40880956-DEL-56 | chr2-40880957-DEL-56 |
| chr2-40892352-DEL-94 | chr2-40892353-DEL-94 |
| chr2-40944723-DEL-78 | chr2-40944724-DEL-78 |
| chr2-40949394-DEL-1366 | chr2-40949395-DEL-1366 |
| chr2-40953636-DEL-6487 | chr2-40953635-DEL-6488 |
| chr2-40974234-DEL-51 | chr2-40974235-DEL-51 |
| chr2-40980108-DEL-1586 | chr2-40980109-DEL-1635 |
| chr2-40995434-DEL-151 | chr2-40995427-DEL-148 |
| chr2-41038704-DEL-63 | chr2-41038704-DEL-63 |
| chr2-41055557-DEL-196 | chr2-41055551-DEL-195 |
| chr2-41056584-DEL-475 | chr2-41056585-DEL-475 |
| chr2-41058695-DEL-2774 | chr2-41058696-DEL-2774 |
| chr2-41102457-DEL-8809 | chr2-41102458-DEL-8809 |
| chr2-41121569-DEL-130 | chr2-41121567-DEL-131 |
| chr2-41176434-DEL-57 | chr2-41176435-DEL-57 |
| chr2-41190871-DEL-410 | chr2-41190879-DEL-405 |
| chr2-41249893-DEL-3572 | chr2-41249894-DEL-3572 |
| chr2-41256833-DEL-104 | chr2-41256834-DEL-104 |
| chr2-41371209-DEL-90 | chr2-41371210-DEL-89 |
| chr2-41373385-DEL-198 | chr2-41373384-DEL-199 |
| chr2-41389191-DEL-60 | chr2-41389192-DEL-55 |
| chr2-41391560-DEL-213 | chr2-41391561-DEL-213 |

| | |
|---|---|
| chr2-41393265-DEL-107 | chr2-41393300-DEL-95 |
| chr2-41430716-DEL-1364 | chr2-41430717-DEL-1361 |
| chr2-41443568-DEL-881 | chr2-41443569-DEL-881 |
| chr2-41465308-DEL-402 | chr2-41465309-DEL-402 |
| chr2-41467659-DEL-92 | chr2-41467675-DEL-77 |
| chr2-41487805-DEL-727 | chr2-41487806-DEL-727 |
| chr2-41513772-DEL-52 | chr2-41513773-DEL-52 |
| chr2-41554697-DEL-341 | chr2-41554698-DEL-341 |
| chr2-41578417-DEL-300 | chr2-41578418-DEL-300 |
| chr2-41582692-DEL-2175 | chr2-41582692-DEL-2175 |
| chr2-41588224-DEL-1126 | chr2-41588234-DEL-1109 |
| chr2-41590776-DEL-76 | chr2-41590784-DEL-68 |
| chr2-41593734-DEL-113 | chr2-41593735-DEL-113 |
| chr2-41609113-DEL-89 | chr2-41609115-DEL-88 |
| chr2-41634375-DEL-7090 | chr2-41634376-DEL-7090 |
| chr2-41664484-DEL-7188 | chr2-41664485-DEL-7184 |
| chr2-41665107-DEL-208 | chr2-41665107-DEL-208 |
| chr2-41733192-DEL-2783 | chr2-41733192-DEL-2784 |
| chr2-41745311-DEL-164 | chr2-41745312-DEL-164 |
| chr2-41757783-DEL-107 | chr2-41757784-DEL-107 |
| chr2-41776731-DEL-51 | chr2-41776732-DEL-51 |
| chr2-41777098-DEL-1735 | chr2-41777099-DEL-1735 |
| chr2-41818480-DEL-75 | chr2-41818491-DEL-68 |
| chr2-41820692-DEL-378 | chr2-41820678-DEL-379 |
| chr2-41875976-DEL-2912 | chr2-41875966-DEL-2913 |
| chr2-41884921-DEL-584 | chr2-41884922-DEL-584 |
| chr2-41896542-DEL-588 | chr2-41896535-DEL-547 |
| chr2-41934471-DEL-7212 | chr2-41934472-DEL-7212 |
| chr2-41982944-DEL-77 | chr2-41982945-DEL-77 |
| chr2-41998313-DEL-62 | chr2-41998314-DEL-56 |
| chr2-42002041-DEL-352 | chr2-42002042-DEL-352 |
| chr2-42013944-DEL-51 | chr2-42013942-DEL-52 |
| chr2-42016823-DEL-249 | chr2-42016824-DEL-249 |
| chr2-42044110-DEL-76 | chr2-42044123-DEL-64 |
| chr2-42090169-DEL-2819 | chr2-42090164-DEL-2840 |
| chr2-42098307-DEL-6645 | chr2-42098308-DEL-6645 |
| chr2-42124761-DEL-256 | chr2-42124762-DEL-256 |
| chr2-42129395-DEL-1332 | chr2-42129396-DEL-1329 |
| chr2-42139622-DEL-88 | chr2-42139636-DEL-75 |
| chr2-42166238-DEL-3537 | chr2-42166239-DEL-3537 |
| chr2-42188023-DEL-314 | chr2-42188024-DEL-314 |
| chr2-42194960-DEL-106 | chr2-42194961-DEL-106 |
| chr2-42203323-DEL-58 | chr2-42203320-DEL-50 |
| chr2-42227940-DEL-2191 | chr2-42227940-DEL-2190 |
| chr2-42236197-DEL-158 | chr2-42236198-DEL-158 |
| chr2-42236988-DEL-60 | chr2-42236989-DEL-60 |
| chr2-42244532-DEL-54 | chr2-42244532-DEL-54 |
| chr2-42251296-DEL-52 | chr2-42251295-DEL-54 |
| chr2-42263401-DEL-120 | chr2-42263397-DEL-155 |
| chr2-42298058-DEL-192 | chr2-42298059-DEL-192 |
| chr2-42335976-DEL-57 | chr2-42335977-DEL-57 |
| chr2-42448023-DEL-1657 | chr2-42448024-DEL-1657 |
| chr2-42459701-DEL-6289 | chr2-42459702-DEL-6289 |
| chr2-42485822-DEL-62 | chr2-42485792-DEL-74 |
| chr2-42514784-DEL-303 | chr2-42514785-DEL-303 |
| chr2-42550805-DEL-53 | chr2-42550801-DEL-57 |
| chr2-42585342-DEL-76 | chr2-42585343-DEL-66 |
| chr2-42610574-DEL-1072 | chr2-42610575-DEL-1072 |
| chr2-42612891-DEL-53 | chr2-42612882-DEL-50 |
| chr2-42617065-DEL-54 | chr2-42617066-DEL-54 |
| chr2-42640714-DEL-343 | chr2-42640715-DEL-343 |
| chr2-42685569-DEL-200 | chr2-42685572-DEL-204 |
| chr2-42700211-DEL-71 | chr2-42700193-DEL-63 |
| chr2-42737502-DEL-3032 | chr2-42737503-DEL-3032 |
| chr2-42757571-DEL-80 | chr2-42757572-DEL-80 |
| chr2-42819118-DEL-434 | chr2-42819119-DEL-434 |
| chr2-42831858-DEL-436 | chr2-42831824-DEL-441 |
| chr2-42834138-DEL-131 | chr2-42834139-DEL-131 |
| chr2-42840146-DEL-6265 | chr2-42840147-DEL-6265 |
| chr2-42858284-DEL-234 | chr2-42858285-DEL-234 |
| chr2-42883048-DEL-55 | chr2-42883046-DEL-56 |
| chr2-42886694-DEL-446 | chr2-42886695-DEL-445 |
| chr2-42903972-DEL-1443 | chr2-42903973-DEL-1443 |
| chr2-42912525-DEL-78 | chr2-42912524-DEL-61 |
| chr2-42961389-DEL-410 | chr2-42961386-DEL-414 |
| chr2-42968714-DEL-69 | chr2-42968715-DEL-69 |
| chr2-43002311-DEL-85 | chr2-43002337-DEL-55 |
| chr2-43009673-DEL-165 | chr2-43009674-DEL-165 |
| chr2-43013409-DEL-62 | chr2-43013410-DEL-62 |
| chr2-43040132-DEL-3195 | chr2-43040133-DEL-3194 |

| | |
|---|---|
| chr2-43061045-DEL-84 | chr2-43061080-DEL-50 |
| chr2-43085675-DEL-899 | chr2-43085676-DEL-899 |
| chr2-43089666-DEL-1579 | chr2-43089667-DEL-1579 |
| chr2-43134011-DEL-53 | chr2-43134001-DEL-54 |
| chr2-43145857-DEL-78 | chr2-43145858-DEL-78 |
| chr2-43147059-DEL-126 | chr2-43147063-DEL-123 |
| chr2-43151361-DEL-422 | chr2-43151362-DEL-422 |
| chr2-43167144-DEL-207 | chr2-43167145-DEL-207 |
| chr2-43187706-DEL-65 | chr2-43187706-DEL-65 |
| chr2-43190117-DEL-253 | chr2-43190118-DEL-253 |
| chr2-43191007-DEL-6301 | chr2-43191008-DEL-6301 |
| chr2-43210047-DEL-183 | chr2-43210048-DEL-182 |
| chr2-43298597-DEL-6480 | chr2-43298598-DEL-6480 |
| chr2-43298837-DEL-212 | chr2-43298838-DEL-212 |
| chr2-43311850-DEL-326 | chr2-43311851-DEL-326 |
| chr2-43372155-DEL-287 | chr2-43372123-DEL-295 |
| chr2-43377366-DEL-227 | chr2-43377377-DEL-228 |
| chr2-43408255-DEL-1226 | chr2-43408256-DEL-1226 |
| chr2-43457085-DEL-769 | chr2-43457086-DEL-769 |
| chr2-43519074-DEL-1372 | chr2-43519075-DEL-1372 |
| chr2-43552363-DEL-822 | chr2-43552364-DEL-822 |
| chr2-43555422-DEL-783 | chr2-43555423-DEL-783 |
| chr2-43568383-DEL-192 | chr2-43568384-DEL-192 |
| chr2-43616012-DEL-649 | chr2-43616013-DEL-649 |
| chr2-43622281-DEL-1302 | chr2-43622279-DEL-1299 |
| chr2-43624484-DEL-144 | chr2-43624480-DEL-146 |
| chr2-43658659-DEL-202 | chr2-43658660-DEL-201 |
| chr2-43680729-DEL-55 | chr2-43680724-DEL-61 |
| chr2-43759117-DEL-826 | chr2-43759118-DEL-826 |
| chr2-43773435-DEL-5664 | chr2-43773436-DEL-5664 |
| chr2-43785352-DEL-1905 | chr2-43785353-DEL-1905 |
| chr2-43790374-DEL-206 | chr2-43790402-DEL-188 |
| chr2-43822085-DEL-73 | chr2-43822086-DEL-56 |
| chr2-43880958-DEL-85 | chr2-43880959-DEL-80 |
| chr2-43951762-DEL-325 | chr2-43951763-DEL-325 |
| chr2-43996021-DEL-55 | chr2-43996043-DEL-54 |
| chr2-43998923-DEL-540 | chr2-43998924-DEL-540 |
| chr2-44029711-DEL-859 | chr2-44029711-DEL-860 |
| chr2-44057998-DEL-6658 | chr2-44057999-DEL-6658 |
| chr2-44076963-DEL-261 | chr2-44076964-DEL-261 |
| chr2-44103057-DEL-154 | chr2-44103061-DEL-151 |
| chr2-44103562-DEL-1418 | chr2-44103563-DEL-1418 |
| chr2-44118618-DEL-290 | chr2-44118619-DEL-290 |
| chr2-44135541-DEL-175 | chr2-44135533-DEL-180 |
| chr2-44136238-DEL-249 | chr2-44136239-DEL-249 |
| chr2-44142212-DEL-134 | chr2-44142213-DEL-134 |
| chr2-44166289-DEL-6367 | chr2-44166292-DEL-6369 |
| chr2-44209263-DEL-117 | chr2-44209265-DEL-123 |
| chr2-44217148-DEL-67 | chr2-44217149-DEL-51 |
| chr2-44224406-DEL-53 | chr2-44224405-DEL-54 |
| chr2-44249052-DEL-6454 | chr2-44249053-DEL-6454 |
| chr2-44270907-DEL-92 | chr2-44270908-DEL-92 |
| chr2-44369154-DEL-51 | chr2-44369155-DEL-51 |
| chr2-44384027-DEL-52 | chr2-44384028-DEL-52 |
| chr2-44420747-DEL-1237 | chr2-44420748-DEL-1237 |
| chr2-44422549-DEL-1743 | chr2-44422550-DEL-1743 |
| chr2-44432643-DEL-64 | chr2-44432644-DEL-64 |
| chr2-44444058-DEL-272 | chr2-44444059-DEL-272 |
| chr2-44493794-DEL-198 | chr2-44493795-DEL-198 |
| chr2-44501443-DEL-6466 | chr2-44501444-DEL-6466 |
| chr2-44565397-DEL-3724 | chr2-44565398-DEL-3724 |
| chr2-44599621-DEL-65 | chr2-44599600-DEL-50 |
| chr2-44654105-DEL-194 | chr2-44654106-DEL-194 |
| chr2-44685048-DEL-433 | chr2-44685049-DEL-433 |
| chr2-44718289-DEL-223 | chr2-44718290-DEL-223 |
| chr2-44748899-DEL-95 | chr2-44748898-DEL-96 |
| chr2-44793186-DEL-58 | chr2-44793187-DEL-58 |
| chr2-44826225-DEL-204 | chr2-44826232-DEL-204 |
| chr2-44833768-DEL-390 | chr2-44833769-DEL-390 |
| chr2-44836442-DEL-1068 | chr2-44836425-DEL-1077 |
| chr2-44854342-DEL-93 | chr2-44854327-DEL-109 |
| chr2-44894829-DEL-66 | chr2-44894814-DEL-62 |
| chr2-44906435-DEL-864 | chr2-44906436-DEL-864 |
| chr2-45017714-DEL-157 | chr2-45017715-DEL-157 |
| chr2-45026115-DEL-246 | chr2-45026116-DEL-246 |
| chr2-45047894-DEL-53 | chr2-45047895-DEL-53 |
| chr2-45076678-DEL-52 | chr2-45076679-DEL-52 |
| chr2-45114327-DEL-94 | chr2-45114317-DEL-93 |
| chr2-45161889-DEL-95 | chr2-45161894-DEL-88 |
| chr2-45191386-DEL-225 | chr2-45191387-DEL-225 |

| | |
|---|---|
| chr2-45210305-DEL-285 | chr2-45210306-DEL-285 |
| chr2-45243786-DEL-83 | chr2-45243784-DEL-86 |
| chr2-45357077-DEL-176 | chr2-45357079-DEL-174 |
| chr2-45361393-DEL-68 | chr2-45361386-DEL-78 |
| chr2-45373839-DEL-206 | chr2-45373840-DEL-205 |
| chr2-45424251-DEL-119 | chr2-45424222-DEL-123 |
| chr2-45447768-DEL-170 | chr2-45447769-DEL-170 |
| chr2-45571706-DEL-2437 | chr2-45571707-DEL-2437 |
| chr2-45709500-DEL-101 | chr2-45709501-DEL-101 |
| chr2-45729941-DEL-2825 | chr2-45729942-DEL-2825 |
| chr2-45746039-DEL-498 | chr2-45746040-DEL-498 |
| chr2-45771036-DEL-435 | chr2-45771037-DEL-435 |
| chr2-45793996-DEL-202 | chr2-45793997-DEL-202 |
| chr2-45805577-DEL-105 | chr2-45805578-DEL-83 |
| chr2-45820362-DEL-452 | chr2-45820363-DEL-452 |
| chr2-45829719-DEL-264 | chr2-45829720-DEL-264 |
| chr2-45869195-DEL-76 | chr2-45869194-DEL-78 |
| chr2-45872384-DEL-612 | chr2-45872385-DEL-612 |
| chr2-45921844-DEL-52 | chr2-45921852-DEL-51 |
| chr2-45964810-DEL-67 | chr2-45964811-DEL-67 |
| chr2-45994587-DEL-139 | chr2-45994588-DEL-139 |
| chr2-46027600-DEL-1012 | chr2-46027599-DEL-1013 |
| chr2-46054907-DEL-690 | chr2-46054908-DEL-690 |
| chr2-46063407-DEL-516 | chr2-46063408-DEL-516 |
| chr2-46066102-DEL-1593 | chr2-46066103-DEL-1593 |
| chr2-46079708-DEL-1707 | chr2-46079709-DEL-1707 |
| chr2-46099100-DEL-52 | chr2-46099103-DEL-50 |
| chr2-46155277-DEL-6859 | chr2-46155278-DEL-6859 |
| chr2-46169645-DEL-410 | chr2-46169644-DEL-411 |
| chr2-46170882-DEL-207 | chr2-46170881-DEL-208 |
| chr2-46180793-DEL-6279 | chr2-46180794-DEL-6279 |
| chr2-46199364-DEL-1887 | chr2-46199365-DEL-1887 |
| chr2-46271662-DEL-7166 | chr2-46271663-DEL-7166 |
| chr2-46295825-DEL-399 | chr2-46295826-DEL-399 |
| chr2-46354081-DEL-56 | chr2-46354082-DEL-84 |
| chr2-46440128-DEL-487 | chr2-46440129-DEL-487 |
| chr2-46464325-DEL-5237 | chr2-46464326-DEL-5237 |
| chr2-46482606-DEL-3315 | chr2-46482607-DEL-3315 |
| chr2-46507826-DEL-60 | chr2-46507829-DEL-59 |
| chr2-46511920-DEL-59 | chr2-46511921-DEL-59 |
| chr2-46525000-DEL-886 | chr2-46525001-DEL-886 |
| chr2-46570550-DEL-12627 | chr2-46570551-DEL-12627 |
| chr2-46575711-DEL-5846 | chr2-46575711-DEL-5846 |
| chr2-46586308-DEL-2908 | chr2-46586309-DEL-2908 |
| chr2-46613409-DEL-7218 | chr2-46613410-DEL-7218 |
| chr2-46640431-DEL-89 | chr2-46640421-DEL-88 |
| chr2-46665066-DEL-64 | chr2-46665067-DEL-64 |
| chr2-46707603-DEL-64 | chr2-46707611-DEL-53 |
| chr2-46723023-DEL-1106 | chr2-46723023-DEL-1106 |
| chr2-46757956-DEL-69 | chr2-46757974-DEL-77 |
| chr2-46760189-DEL-60 | chr2-46760190-DEL-60 |
| chr2-46762997-DEL-576 | chr2-46762998-DEL-576 |
| chr2-46804493-DEL-60 | chr2-46804494-DEL-62 |
| chr2-46817402-DEL-336 | chr2-46817411-DEL-328 |
| chr2-46818032-DEL-57 | chr2-46818022-DEL-99 |
| chr2-46828099-DEL-594 | chr2-46828100-DEL-594 |
| chr2-46843782-DEL-1336 | chr2-46843781-DEL-1337 |
| chr2-46849727-DEL-192 | chr2-46849725-DEL-193 |
| chr2-46866822-DEL-2248 | chr2-46866823-DEL-2248 |
| chr2-46898665-DEL-7398 | chr2-46898666-DEL-7398 |
| chr2-46907293-DEL-6235 | chr2-46907294-DEL-6235 |
| chr2-46922404-DEL-148 | chr2-46922406-DEL-145 |
| chr2-46923572-DEL-244 | chr2-46923573-DEL-244 |
| chr2-47012559-DEL-6955 | chr2-47012560-DEL-6955 |
| chr2-47062776-DEL-63 | chr2-47062764-DEL-64 |
| chr2-47068946-DEL-145 | chr2-47068947-DEL-143 |
| chr2-47069461-DEL-51 | chr2-47069462-DEL-51 |
| chr2-47081738-DEL-1174 | chr2-47081736-DEL-1207 |
| chr2-47085669-DEL-2646 | chr2-47085670-DEL-2646 |
| chr2-47107395-DEL-2484 | chr2-47107387-DEL-2476 |
| chr2-47112889-DEL-92 | chr2-47112891-DEL-91 |
| chr2-47119311-DEL-942 | chr2-47119312-DEL-942 |
| chr2-47182011-DEL-168 | chr2-47182012-DEL-168 |
| chr2-47197433-DEL-279 | chr2-47197431-DEL-280 |
| chr2-47211632-DEL-398 | chr2-47211633-DEL-398 |
| chr2-47278626-DEL-55 | chr2-47278622-DEL-54 |
| chr2-47278774-DEL-1723 | chr2-47278775-DEL-1723 |
| chr2-47295193-DEL-321 | chr2-47295194-DEL-321 |
| chr2-47301213-DEL-1890 | chr2-47301214-DEL-1890 |
| chr2-47316473-DEL-122 | chr2-47316471-DEL-123 |

| | |
|---|---|
| chr2-47353781-DEL-4482 | chr2-47353782-DEL-4482 |
| chr2-47382604-DEL-888 | chr2-47382604-DEL-889 |
| chr2-47413862-DEL-56 | chr2-47413863-DEL-56 |
| chr2-47446970-DEL-62 | chr2-47446948-DEL-84 |
| chr2-47461588-DEL-3365 | chr2-47461576-DEL-3364 |
| chr2-47484892-DEL-804 | chr2-47484893-DEL-804 |
| chr2-47490688-DEL-1035 | chr2-47490689-DEL-1035 |
| chr2-47491972-DEL-1919 | chr2-47491972-DEL-1919 |
| chr2-47505689-DEL-3021 | chr2-47505690-DEL-3021 |
| chr2-47509460-DEL-60 | chr2-47509461-DEL-60 |
| chr2-47525753-DEL-7267 | chr2-47525743-DEL-7268 |
| chr2-47531860-DEL-212 | chr2-47531860-DEL-212 |
| chr2-47546221-DEL-104 | chr2-47546222-DEL-104 |
| chr2-47547513-DEL-266 | chr2-47547514-DEL-266 |
| chr2-47553433-DEL-6629 | chr2-47553435-DEL-6630 |
| chr2-47563486-DEL-1090 | chr2-47563487-DEL-1090 |
| chr2-47593279-DEL-105 | chr2-47593246-DEL-106 |
| chr2-47596872-DEL-421 | chr2-47596873-DEL-421 |
| chr2-47617462-DEL-3356 | chr2-47617462-DEL-3356 |
| chr2-47648135-DEL-1258 | chr2-47648134-DEL-1260 |
| chr2-47703279-DEL-106 | chr2-47703280-DEL-70 |
| chr2-47710194-DEL-1258 | chr2-47710195-DEL-1258 |
| chr2-47721023-DEL-308 | chr2-47721024-DEL-308 |
| chr2-47742219-DEL-386 | chr2-47742216-DEL-382 |
| chr2-47763416-DEL-1786 | chr2-47763417-DEL-1786 |
| chr2-47768960-DEL-177 | chr2-47768961-DEL-177 |
| chr2-47774767-DEL-2436 | chr2-47774768-DEL-2436 |
| chr2-47795904-DEL-1188 | chr2-47795904-DEL-1188 |
| chr2-47807734-DEL-1244 | chr2-47807735-DEL-1244 |
| chr2-47825119-DEL-570 | chr2-47825120-DEL-570 |
| chr2-47831304-DEL-451 | chr2-47831305-DEL-451 |
| chr2-47838139-DEL-69 | chr2-47838140-DEL-69 |
| chr2-47841600-DEL-58 | chr2-47841606-DEL-54 |
| chr2-47854125-DEL-203 | chr2-47854126-DEL-203 |
| chr2-47858697-DEL-3750 | chr2-47858698-DEL-3750 |
| chr2-47910683-DEL-50 | chr2-47910683-DEL-56 |
| chr2-47914291-DEL-53 | chr2-47914295-DEL-51 |
| chr2-47919109-DEL-477 | chr2-47919110-DEL-477 |
| chr2-47924810-DEL-66 | chr2-47924823-DEL-57 |
| chr2-47944125-DEL-2570 | chr2-47944126-DEL-2570 |
| chr2-47959590-DEL-6187 | chr2-47959591-DEL-6187 |
| chr2-48047077-DEL-704 | chr2-48047078-DEL-703 |
| chr2-48047835-DEL-3238 | chr2-48047835-DEL-3238 |
| chr2-48052148-DEL-111 | chr2-48052149-DEL-111 |
| chr2-48073426-DEL-57 | chr2-48073426-DEL-57 |
| chr2-48126861-DEL-406 | chr2-48126862-DEL-406 |
| chr2-48165503-DEL-68 | chr2-48165504-DEL-68 |
| chr2-48261631-DEL-60 | chr2-48261632-DEL-60 |
| chr2-48285717-DEL-54 | chr2-48285718-DEL-54 |
| chr2-48319994-DEL-189 | chr2-48319995-DEL-188 |
| chr2-48356623-DEL-58 | chr2-48356644-DEL-54 |
| chr2-48361105-DEL-102 | chr2-48361106-DEL-102 |
| chr2-48373513-DEL-7123 | chr2-48373514-DEL-7123 |
| chr2-48385103-DEL-153 | chr2-48385108-DEL-149 |
| chr2-48386442-DEL-121 | chr2-48386443-DEL-113 |
| chr2-48403191-DEL-191 | chr2-48403192-DEL-190 |
| chr2-48468857-DEL-6813 | chr2-48468858-DEL-6813 |
| chr2-48469135-DEL-212 | chr2-48469136-DEL-212 |
| chr2-48478448-DEL-54 | chr2-48478449-DEL-54 |
| chr2-48497049-DEL-508 | chr2-48497050-DEL-508 |
| chr2-48500316-DEL-75 | chr2-48500317-DEL-75 |
| chr2-48508758-DEL-51 | chr2-48508759-DEL-51 |
| chr2-48617220-DEL-101 | chr2-48617221-DEL-101 |
| chr2-48618859-DEL-220 | chr2-48618860-DEL-220 |
| chr2-48674242-DEL-3258 | chr2-48674243-DEL-3258 |
| chr2-48697618-DEL-1075 | chr2-48697619-DEL-1075 |
| chr2-48796602-DEL-192 | chr2-48796601-DEL-192 |
| chr2-48829927-DEL-80 | chr2-48829928-DEL-80 |
| chr2-48900636-DEL-3829 | chr2-48900637-DEL-3830 |
| chr2-48921692-DEL-638 | chr2-48921693-DEL-638 |
| chr2-48942091-DEL-7068 | chr2-48942092-DEL-7068 |
| chr2-48985020-DEL-315 | chr2-48985021-DEL-315 |
| chr2-49029850-DEL-724 | chr2-49029851-DEL-724 |
| chr2-49065408-DEL-468 | chr2-49065409-DEL-466 |
| chr2-49099033-DEL-52 | chr2-49099032-DEL-52 |
| chr2-49113706-DEL-319 | chr2-49113707-DEL-319 |
| chr2-49120925-DEL-508 | chr2-49120926-DEL-507 |
| chr2-49124827-DEL-88 | chr2-49124826-DEL-94 |
| chr2-49197577-DEL-81 | chr2-49197578-DEL-78 |
| chr2-49198019-DEL-1924 | chr2-49198020-DEL-1924 |

| | |
|---|---|
| chr2-49215936-DEL-52 | chr2-49215937-DEL-52 |
| chr2-49219535-DEL-98 | chr2-49219536-DEL-98 |
| chr2-49222877-DEL-66 | chr2-49222874-DEL-70 |
| chr2-49224246-DEL-63 | chr2-49224241-DEL-96 |
| chr2-49299868-DEL-171 | chr2-49299869-DEL-171 |
| chr2-49343655-DEL-171 | chr2-49343655-DEL-176 |
| chr2-49365321-DEL-6271 | chr2-49365322-DEL-6271 |
| chr2-49442137-DEL-52 | chr2-49442138-DEL-56 |
| chr2-49468042-DEL-200 | chr2-49468043-DEL-200 |
| chr2-49479645-DEL-171 | chr2-49479632-DEL-183 |
| chr2-49493748-DEL-2176 | chr2-49493748-DEL-2177 |
| chr2-49498000-DEL-66 | chr2-49498001-DEL-66 |
| chr2-49502641-DEL-58 | chr2-49502642-DEL-58 |
| chr2-49506181-DEL-70 | chr2-49506175-DEL-76 |
| chr2-49538131-DEL-77 | chr2-49538087-DEL-78 |
| chr2-49607864-DEL-75 | chr2-49607865-DEL-75 |
| chr2-49618069-DEL-119 | chr2-49618070-DEL-119 |
| chr2-49658050-DEL-55 | chr2-49658051-DEL-55 |
| chr2-49666197-DEL-66 | chr2-49666197-DEL-67 |
| chr2-49683734-DEL-96 | chr2-49683738-DEL-92 |
| chr2-49752160-DEL-92 | chr2-49752161-DEL-91 |
| chr2-49763148-DEL-857 | chr2-49763149-DEL-857 |
| chr2-49774374-DEL-101 | chr2-49774375-DEL-101 |
| chr2-49788679-DEL-125 | chr2-49788680-DEL-125 |
| chr2-49794073-DEL-57 | chr2-49794045-DEL-58 |
| chr2-49803718-DEL-401 | chr2-49803719-DEL-401 |
| chr2-49810166-DEL-232 | chr2-49810167-DEL-232 |
| chr2-49886852-DEL-401 | chr2-49886853-DEL-401 |
| chr2-49899883-DEL-231 | chr2-49899884-DEL-231 |
| chr2-49907443-DEL-212 | chr2-49907444-DEL-212 |
| chr2-49964907-DEL-81 | chr2-49964920-DEL-60 |
| chr2-49972074-DEL-76 | chr2-49972091-DEL-60 |
| chr2-49973711-DEL-54 | chr2-49973713-DEL-53 |
| chr2-49993607-DEL-62 | chr2-49993598-DEL-58 |
| chr2-50004802-DEL-51 | chr2-50004803-DEL-52 |
| chr2-50046244-DEL-87 | chr2-50046245-DEL-65 |
| chr2-50048426-DEL-381 | chr2-50048427-DEL-381 |
| chr2-50056339-DEL-60 | chr2-50056339-DEL-60 |
| chr2-50057503-DEL-169 | chr2-50057538-DEL-135 |
| chr2-50066530-DEL-548 | chr2-50066565-DEL-514 |
| chr2-50141093-DEL-68 | chr2-50141098-DEL-64 |
| chr2-50219281-DEL-60 | chr2-50219282-DEL-60 |
| chr2-50240354-DEL-6740 | chr2-50240355-DEL-6740 |
| chr2-50241344-DEL-62 | chr2-50241341-DEL-51 |
| chr2-50309784-DEL-52 | chr2-50309785-DEL-52 |
| chr2-50329224-DEL-123 | chr2-50329225-DEL-123 |
| chr2-50350589-DEL-291 | chr2-50350585-DEL-288 |
| chr2-50362705-DEL-520 | chr2-50362706-DEL-520 |
| chr2-50377411-DEL-2286 | chr2-50377412-DEL-2286 |
| chr2-50381432-DEL-57 | chr2-50381417-DEL-58 |
| chr2-50395832-DEL-2130 | chr2-50395826-DEL-2131 |
| chr2-50417645-DEL-3048 | chr2-50417646-DEL-3048 |
| chr2-50419608-DEL-138 | chr2-50419628-DEL-102 |
| chr2-50430423-DEL-112 | chr2-50430428-DEL-92 |
| chr2-50432928-DEL-1304 | chr2-50432929-DEL-1304 |
| chr2-50478385-DEL-66 | chr2-50478386-DEL-66 |
| chr2-50490308-DEL-8191 | chr2-50490309-DEL-8191 |
| chr2-50544039-DEL-335 | chr2-50544040-DEL-335 |
| chr2-50545989-DEL-207 | chr2-50545990-DEL-207 |
| chr2-50569620-DEL-7095 | chr2-50569621-DEL-7095 |
| chr2-50579954-DEL-59 | chr2-50579962-DEL-64 |
| chr2-50603106-DEL-54 | chr2-50603107-DEL-60 |
| chr2-50631475-DEL-66 | chr2-50631476-DEL-66 |
| chr2-50637261-DEL-2172 | chr2-50637262-DEL-2172 |
| chr2-50654190-DEL-6483 | chr2-50654191-DEL-6483 |
| chr2-50690696-DEL-396 | chr2-50690696-DEL-396 |
| chr2-50721404-DEL-411 | chr2-50721405-DEL-411 |
| chr2-50727019-DEL-119 | chr2-50727020-DEL-119 |
| chr2-50742864-DEL-99 | chr2-50742875-DEL-87 |
| chr2-50755924-DEL-92 | chr2-50755968-DEL-52 |
| chr2-50783635-DEL-387 | chr2-50783636-DEL-387 |
| chr2-50786877-DEL-96 | chr2-50786878-DEL-96 |
| chr2-50823619-DEL-78 | chr2-50823628-DEL-66 |
| chr2-50828839-DEL-961 | chr2-50828840-DEL-961 |
| chr2-50880473-DEL-10261 | chr2-50880474-DEL-10259 |
| chr2-50902010-DEL-76 | chr2-50902011-DEL-77 |
| chr2-50910952-DEL-61 | chr2-50910953-DEL-61 |
| chr2-50916071-DEL-6469 | chr2-50916072-DEL-6469 |
| chr2-50940802-DEL-1668 | chr2-50940803-DEL-1668 |
| chr2-50970650-DEL-191 | chr2-50970651-DEL-191 |

| | |
|---|---|
| chr2-50982382-DEL-208 | chr2-50982383-DEL-191 |
| chr2-50985088-DEL-125 | chr2-50985089-DEL-125 |
| chr2-50996470-DEL-1793 | chr2-50996470-DEL-1794 |
| chr2-50997776-DEL-140 | chr2-50997781-DEL-130 |
| chr2-50999875-DEL-58 | chr2-50999876-DEL-58 |
| chr2-51012712-DEL-1484 | chr2-51012713-DEL-1484 |
| chr2-51039629-DEL-94 | chr2-51039630-DEL-51 |
| chr2-51044919-DEL-325 | chr2-51044920-DEL-325 |
| chr2-51077308-DEL-207 | chr2-51077309-DEL-207 |
| chr2-51096802-DEL-171 | chr2-51096813-DEL-161 |
| chr2-51140405-DEL-62 | chr2-51140394-DEL-64 |
| chr2-51147102-DEL-92 | chr2-51147118-DEL-68 |
| chr2-51163782-DEL-115 | chr2-51163763-DEL-113 |
| chr2-51164833-DEL-1545 | chr2-51164834-DEL-1545 |
| chr2-51171104-DEL-80 | chr2-51171105-DEL-80 |
| chr2-51230212-DEL-103 | chr2-51230213-DEL-103 |
| chr2-51233200-DEL-129 | chr2-51233201-DEL-129 |
| chr2-51248133-DEL-66 | chr2-51248134-DEL-66 |
| chr2-51289265-DEL-88 | chr2-51289247-DEL-104 |
| chr2-51330082-DEL-68 | chr2-51330083-DEL-68 |
| chr2-51361315-DEL-186 | chr2-51361316-DEL-182 |
| chr2-51369956-DEL-359 | chr2-51369957-DEL-359 |
| chr2-51371621-DEL-187 | chr2-51371622-DEL-187 |
| chr2-51373834-DEL-6281 | chr2-51373835-DEL-6281 |
| chr2-51386988-DEL-81 | chr2-51386985-DEL-76 |
| chr2-51395219-DEL-75 | chr2-51395220-DEL-75 |
| chr2-51416634-DEL-322 | chr2-51416636-DEL-315 |
| chr2-51417470-DEL-429 | chr2-51417471-DEL-429 |
| chr2-51439684-DEL-382 | chr2-51439685-DEL-382 |
| chr2-51442867-DEL-564 | chr2-51442868-DEL-564 |
| chr2-51444190-DEL-120 | chr2-51444191-DEL-114 |
| chr2-51468052-DEL-4407 | chr2-51468053-DEL-4407 |
| chr2-51469469-DEL-245 | chr2-51469470-DEL-245 |
| chr2-51472516-DEL-1596 | chr2-51472516-DEL-1596 |
| chr2-51484806-DEL-53 | chr2-51484780-DEL-51 |
| chr2-51496537-DEL-56 | chr2-51496538-DEL-56 |
| chr2-51505187-DEL-6349 | chr2-51505188-DEL-6349 |
| chr2-51514292-DEL-69 | chr2-51514293-DEL-69 |
| chr2-51529475-DEL-70 | chr2-51529476-DEL-70 |
| chr2-51538439-DEL-94 | chr2-51538446-DEL-54 |
| chr2-51543832-DEL-51 | chr2-51543833-DEL-51 |
| chr2-51552233-DEL-490 | chr2-51552234-DEL-490 |
| chr2-51563193-DEL-328 | chr2-51563194-DEL-328 |
| chr2-51584556-DEL-1012 | chr2-51584557-DEL-1012 |
| chr2-51612409-DEL-120 | chr2-51612410-DEL-120 |
| chr2-51614739-DEL-185 | chr2-51614738-DEL-186 |
| chr2-51617557-DEL-69 | chr2-51617558-DEL-69 |
| chr2-51639719-DEL-52 | chr2-51639720-DEL-52 |
| chr2-51649379-DEL-1384 | chr2-51649380-DEL-1384 |
| chr2-51668364-DEL-771 | chr2-51668365-DEL-771 |
| chr2-51671244-DEL-60 | chr2-51671245-DEL-60 |
| chr2-51678260-DEL-53 | chr2-51678225-DEL-52 |
| chr2-51760905-DEL-768 | chr2-51760906-DEL-768 |
| chr2-51764516-DEL-68 | chr2-51764517-DEL-68 |
| chr2-51765185-DEL-60 | chr2-51765186-DEL-54 |
| chr2-51766327-DEL-301 | chr2-51766328-DEL-301 |
| chr2-51771597-DEL-1456 | chr2-51771598-DEL-1456 |
| chr2-51776210-DEL-4390 | chr2-51776210-DEL-4390 |
| chr2-51804020-DEL-53 | chr2-51804021-DEL-50 |
| chr2-51810340-DEL-120 | chr2-51810341-DEL-120 |
| chr2-51835372-DEL-319 | chr2-51835351-DEL-339 |
| chr2-51851867-DEL-173 | chr2-51851882-DEL-162 |
| chr2-51853175-DEL-64 | chr2-51853176-DEL-64 |
| chr2-51867538-DEL-433 | chr2-51867539-DEL-433 |
| chr2-51889624-DEL-399 | chr2-51889625-DEL-399 |
| chr2-51893344-DEL-55 | chr2-51893361-DEL-52 |
| chr2-51894743-DEL-80 | chr2-51894744-DEL-80 |
| chr2-51901398-DEL-117 | chr2-51901403-DEL-113 |
| chr2-51905657-DEL-205 | chr2-51905683-DEL-173 |
| chr2-51922089-DEL-438 | chr2-51922090-DEL-438 |
| chr2-51944232-DEL-68 | chr2-51944233-DEL-68 |
| chr2-51949426-DEL-434 | chr2-51949427-DEL-434 |
| chr2-51951160-DEL-206 | chr2-51951161-DEL-206 |
| chr2-51953340-DEL-204 | chr2-51953341-DEL-204 |
| chr2-51989397-DEL-478 | chr2-51989392-DEL-484 |
| chr2-52060902-DEL-66 | chr2-52060895-DEL-58 |
| chr2-52065975-DEL-215 | chr2-52065976-DEL-215 |
| chr2-52083811-DEL-63 | chr2-52083812-DEL-63 |
| chr2-52106563-DEL-169 | chr2-52106564-DEL-169 |
| chr2-52121648-DEL-98 | chr2-52121656-DEL-95 |

| | |
|---|---|
| chr2-52142282-DEL-96 | chr2-52142283-DEL-96 |
| chr2-52153245-DEL-82 | chr2-52153250-DEL-78 |
| chr2-52159385-DEL-520 | chr2-52159384-DEL-516 |
| chr2-52160509-DEL-76 | chr2-52160510-DEL-76 |
| chr2-52164027-DEL-582 | chr2-52164028-DEL-582 |
| chr2-52213119-DEL-181 | chr2-52213120-DEL-181 |
| chr2-52263682-DEL-1038 | chr2-52263665-DEL-1053 |
| chr2-52266398-DEL-65 | chr2-52266397-DEL-65 |
| chr2-52270892-DEL-579 | chr2-52270893-DEL-579 |
| chr2-52271573-DEL-2364 | chr2-52271574-DEL-2364 |
| chr2-52286138-DEL-74 | chr2-52286139-DEL-74 |
| chr2-52339547-DEL-294 | chr2-52339516-DEL-285 |
| chr2-52347630-DEL-55 | chr2-52347631-DEL-55 |
| chr2-52350124-DEL-64 | chr2-52350125-DEL-64 |
| chr2-52363448-DEL-696 | chr2-52363449-DEL-696 |
| chr2-52371863-DEL-993 | chr2-52371864-DEL-993 |
| chr2-52392770-DEL-60 | chr2-52392771-DEL-51 |
| chr2-52422639-DEL-434 | chr2-52422640-DEL-434 |
| chr2-52431935-DEL-237 | chr2-52431936-DEL-237 |
| chr2-52463180-DEL-67 | chr2-52463180-DEL-63 |
| chr2-52490528-DEL-93 | chr2-52490529-DEL-93 |
| chr2-52550297-DEL-934 | chr2-52550298-DEL-934 |
| chr2-52554660-DEL-321 | chr2-52554661-DEL-321 |
| chr2-52555161-DEL-1048 | chr2-52555162-DEL-1048 |
| chr2-52567570-DEL-52 | chr2-52567571-DEL-52 |
| chr2-52572391-DEL-827 | chr2-52572392-DEL-827 |
| chr2-52586442-DEL-2580 | chr2-52586417-DEL-2572 |
| chr2-52607441-DEL-1278 | chr2-52607454-DEL-1266 |
| chr2-52618302-DEL-987 | chr2-52618303-DEL-987 |
| chr2-52649539-DEL-195 | chr2-52649497-DEL-195 |
| chr2-52650829-DEL-536 | chr2-52650830-DEL-536 |
| chr2-52671941-DEL-198 | chr2-52671942-DEL-198 |
| chr2-52673935-DEL-603 | chr2-52673936-DEL-603 |
| chr2-52674711-DEL-498 | chr2-52674712-DEL-498 |
| chr2-52688599-DEL-224 | chr2-52688600-DEL-224 |
| chr2-52693350-DEL-215 | chr2-52693351-DEL-215 |
| chr2-52695782-DEL-612 | chr2-52695767-DEL-627 |
| chr2-52704082-DEL-138 | chr2-52704063-DEL-168 |
| chr2-52721482-DEL-64 | chr2-52721483-DEL-64 |
| chr2-52732651-DEL-539 | chr2-52732652-DEL-539 |
| chr2-52733609-DEL-692 | chr2-52733610-DEL-692 |
| chr2-52782739-DEL-379 | chr2-52782740-DEL-379 |
| chr2-52803942-DEL-69 | chr2-52803943-DEL-69 |
| chr2-52842825-DEL-65 | chr2-52842826-DEL-65 |
| chr2-52879906-DEL-443 | chr2-52879907-DEL-443 |
| chr2-52882605-DEL-50 | chr2-52882577-DEL-75 |
| chr2-52886688-DEL-493 | chr2-52886683-DEL-494 |
| chr2-52951523-DEL-247 | chr2-52951524-DEL-247 |
| chr2-52959828-DEL-52 | chr2-52959829-DEL-52 |
| chr2-52975705-DEL-71 | chr2-52975706-DEL-71 |
| chr2-52984587-DEL-206 | chr2-52984588-DEL-206 |
| chr2-53044825-DEL-218 | chr2-53044826-DEL-218 |
| chr2-53062090-DEL-2858 | chr2-53062091-DEL-2858 |
| chr2-53096033-DEL-66 | chr2-53096035-DEL-65 |
| chr2-53118941-DEL-1887 | chr2-53118942-DEL-1887 |
| chr2-53138784-DEL-270 | chr2-53138785-DEL-248 |
| chr2-53146548-DEL-84 | chr2-53146555-DEL-78 |
| chr2-53151667-DEL-1717 | chr2-53151668-DEL-1717 |
| chr2-53173420-DEL-170 | chr2-53173421-DEL-170 |
| chr2-53223118-DEL-101 | chr2-53223117-DEL-102 |
| chr2-53299409-DEL-991 | chr2-53299410-DEL-991 |
| chr2-53307818-DEL-789 | chr2-53307819-DEL-789 |
| chr2-53339151-DEL-7038 | chr2-53339152-DEL-7038 |
| chr2-53379214-DEL-72 | chr2-53379215-DEL-72 |
| chr2-53385461-DEL-1877 | chr2-53385462-DEL-1877 |
| chr2-53393345-DEL-6260 | chr2-53393346-DEL-6260 |
| chr2-53400362-DEL-537 | chr2-53400359-DEL-540 |
| chr2-53413641-DEL-689 | chr2-53413642-DEL-689 |
| chr2-53423506-DEL-216 | chr2-53423507-DEL-216 |
| chr2-53424499-DEL-480 | chr2-53424500-DEL-480 |
| chr2-53438322-DEL-216 | chr2-53438323-DEL-216 |
| chr2-53451115-DEL-744 | chr2-53451116-DEL-744 |
| chr2-53458384-DEL-174 | chr2-53458373-DEL-186 |
| chr2-53500351-DEL-107 | chr2-53500352-DEL-107 |
| chr2-53505165-DEL-60 | chr2-53505166-DEL-60 |
| chr2-53513899-DEL-456 | chr2-53513896-DEL-457 |
| chr2-53522304-DEL-111 | chr2-53522305-DEL-111 |
| chr2-53522839-DEL-3516 | chr2-53522840-DEL-3516 |
| chr2-53522854-DEL-64 | chr2-53522855-DEL-64 |
| chr2-53528268-DEL-6193 | chr2-53528269-DEL-6193 |

| | |
|---|---|
| chr2-53539056-DEL-1478 | chr2-53539057-DEL-1478 |
| chr2-53544473-DEL-428 | chr2-53544472-DEL-429 |
| chr2-53559406-DEL-51 | chr2-53559407-DEL-51 |
| chr2-53562650-DEL-176 | chr2-53562651-DEL-176 |
| chr2-53581440-DEL-51 | chr2-53581438-DEL-63 |
| chr2-53584341-DEL-2530 | chr2-53584342-DEL-2530 |
| chr2-53597001-DEL-62 | chr2-53597002-DEL-62 |
| chr2-53607788-DEL-789 | chr2-53607789-DEL-789 |
| chr2-53613458-DEL-325 | chr2-53613459-DEL-325 |
| chr2-53626913-DEL-54 | chr2-53626914-DEL-54 |
| chr2-53629863-DEL-353 | chr2-53629864-DEL-353 |
| chr2-53645925-DEL-6997 | chr2-53645926-DEL-6997 |
| chr2-53700372-DEL-1226 | chr2-53700373-DEL-1226 |
| chr2-53704256-DEL-72 | chr2-53704268-DEL-54 |
| chr2-53710471-DEL-1913 | chr2-53710470-DEL-1914 |
| chr2-53735497-DEL-206 | chr2-53735498-DEL-206 |
| chr2-53804278-DEL-142 | chr2-53804279-DEL-142 |
| chr2-53809712-DEL-6304 | chr2-53809713-DEL-6304 |
| chr2-53830457-DEL-81 | chr2-53830458-DEL-81 |
| chr2-53845008-DEL-495 | chr2-53845009-DEL-495 |
| chr2-53896590-DEL-74 | chr2-53896595-DEL-70 |
| chr2-53910581-DEL-202 | chr2-53910582-DEL-202 |
| chr2-53915494-DEL-874 | chr2-53915455-DEL-913 |
| chr2-53923907-DEL-6208 | chr2-53923908-DEL-6208 |
| chr2-53954902-DEL-6451 | chr2-53954903-DEL-6451 |
| chr2-53965260-DEL-386 | chr2-53965261-DEL-386 |
| chr2-53984259-DEL-113 | chr2-53984260-DEL-113 |
| chr2-53987894-DEL-75 | chr2-53987893-DEL-77 |
| chr2-54027021-DEL-53 | chr2-54026979-DEL-51 |
| chr2-54053387-DEL-51 | chr2-54053340-DEL-61 |
| chr2-54114873-DEL-94 | chr2-54114874-DEL-94 |
| chr2-54127136-DEL-88 | chr2-54127139-DEL-86 |
| chr2-54142562-DEL-6210 | chr2-54142563-DEL-6210 |
| chr2-54144851-DEL-82 | chr2-54144852-DEL-82 |
| chr2-54169003-DEL-107 | chr2-54169004-DEL-107 |
| chr2-54187219-DEL-119 | chr2-54187220-DEL-119 |
| chr2-54199379-DEL-64 | chr2-54199380-DEL-64 |
| chr2-54231197-DEL-1321 | chr2-54231198-DEL-1321 |
| chr2-54240608-DEL-57 | chr2-54240609-DEL-57 |
| chr2-54268385-DEL-214 | chr2-54268386-DEL-214 |
| chr2-54275026-DEL-6937 | chr2-54275027-DEL-6937 |
| chr2-54275042-DEL-424 | chr2-54275042-DEL-424 |
| chr2-54287325-DEL-59 | chr2-54287326-DEL-59 |
| chr2-54289958-DEL-1780 | chr2-54289959-DEL-1780 |
| chr2-54302342-DEL-291 | chr2-54302343-DEL-291 |
| chr2-54309089-DEL-1234 | chr2-54309090-DEL-1234 |
| chr2-54351275-DEL-58 | chr2-54351276-DEL-58 |
| chr2-54358651-DEL-217 | chr2-54358652-DEL-217 |
| chr2-54364413-DEL-651 | chr2-54364414-DEL-651 |
| chr2-54461868-DEL-2123 | chr2-54461869-DEL-2123 |
| chr2-54480755-DEL-171 | chr2-54480756-DEL-172 |
| chr2-54483574-DEL-7064 | chr2-54483575-DEL-7064 |
| chr2-54528246-DEL-94 | chr2-54528247-DEL-98 |
| chr2-54536309-DEL-1244 | chr2-54536310-DEL-1244 |
| chr2-54539976-DEL-1621 | chr2-54539977-DEL-1621 |
| chr2-54556514-DEL-99 | chr2-54556515-DEL-99 |
| chr2-54581188-DEL-203 | chr2-54581189-DEL-203 |
| chr2-54581958-DEL-988 | chr2-54581959-DEL-988 |
| chr2-54631517-DEL-6433 | chr2-54631518-DEL-6433 |
| chr2-54641885-DEL-52 | chr2-54641919-DEL-54 |
| chr2-54684995-DEL-53 | chr2-54684995-DEL-53 |
| chr2-54693731-DEL-399 | chr2-54693732-DEL-399 |
| chr2-54696688-DEL-67 | chr2-54696689-DEL-67 |
| chr2-54707391-DEL-334 | chr2-54707392-DEL-334 |
| chr2-54711125-DEL-1253 | chr2-54711126-DEL-1253 |
| chr2-54713805-DEL-397 | chr2-54713806-DEL-397 |
| chr2-54752566-DEL-3770 | chr2-54752567-DEL-3770 |
| chr2-54777727-DEL-2696 | chr2-54777728-DEL-2696 |
| chr2-54810246-DEL-622 | chr2-54810247-DEL-622 |
| chr2-54814794-DEL-2840 | chr2-54814795-DEL-2840 |
| chr2-54827081-DEL-64 | chr2-54827082-DEL-64 |
| chr2-54841214-DEL-62 | chr2-54841215-DEL-62 |
| chr2-54868707-DEL-64 | chr2-54868708-DEL-64 |
| chr2-54890945-DEL-1710 | chr2-54890946-DEL-1710 |
| chr2-54914074-DEL-146 | chr2-54914060-DEL-146 |
| chr2-54916458-DEL-6594 | chr2-54916459-DEL-6594 |
| chr2-54933829-DEL-661 | chr2-54933830-DEL-661 |
| chr2-54973703-DEL-189 | chr2-54973704-DEL-189 |
| chr2-54976918-DEL-425 | chr2-54976919-DEL-425 |
| chr2-55017303-DEL-75 | chr2-55017264-DEL-76 |

| | |
|---|---|
| chr2-55019993-DEL-54 | chr2-55019992-DEL-54 |
| chr2-55059826-DEL-1579 | chr2-55059829-DEL-1579 |
| chr2-55093468-DEL-214 | chr2-55093469-DEL-214 |
| chr2-55093987-DEL-922 | chr2-55093988-DEL-922 |
| chr2-55132859-DEL-1343 | chr2-55132860-DEL-1343 |
| chr2-55143063-DEL-82 | chr2-55143060-DEL-87 |
| chr2-55152513-DEL-124 | chr2-55152514-DEL-124 |
| chr2-55158898-DEL-55 | chr2-55158899-DEL-55 |
| chr2-55195512-DEL-202 | chr2-55195513-DEL-202 |
| chr2-55198297-DEL-1064 | chr2-55198298-DEL-1064 |
| chr2-55207247-DEL-3441 | chr2-55207248-DEL-3437 |
| chr2-55207899-DEL-6352 | chr2-55207900-DEL-6352 |
| chr2-55220782-DEL-1207 | chr2-55220779-DEL-1203 |
| chr2-55236914-DEL-6813 | chr2-55236915-DEL-6813 |
| chr2-55275936-DEL-837 | chr2-55275937-DEL-837 |
| chr2-55287135-DEL-4431 | chr2-55287136-DEL-4431 |
| chr2-55338682-DEL-479 | chr2-55338683-DEL-479 |
| chr2-55356898-DEL-767 | chr2-55356897-DEL-769 |
| chr2-55382225-DEL-6225 | chr2-55382226-DEL-6225 |
| chr2-55405130-DEL-62 | chr2-55405113-DEL-76 |
| chr2-55434060-DEL-1486 | chr2-55434061-DEL-1484 |
| chr2-55490166-DEL-1320 | chr2-55490167-DEL-1320 |
| chr2-55518909-DEL-214 | chr2-55518884-DEL-214 |
| chr2-55548863-DEL-1411 | chr2-55548864-DEL-1411 |
| chr2-55561388-DEL-6875 | chr2-55561389-DEL-6875 |
| chr2-55589323-DEL-364 | chr2-55589324-DEL-364 |
| chr2-55594690-DEL-558 | chr2-55594697-DEL-558 |
| chr2-55616772-DEL-356 | chr2-55616773-DEL-356 |
| chr2-55671361-DEL-60 | chr2-55671362-DEL-60 |
| chr2-55671951-DEL-8700 | chr2-55671952-DEL-8700 |
| chr2-55713765-DEL-6320 | chr2-55713746-DEL-6314 |
| chr2-55749191-DEL-2034 | chr2-55749192-DEL-2034 |
| chr2-55755914-DEL-147 | chr2-55755915-DEL-147 |
| chr2-55757187-DEL-7084 | chr2-55757188-DEL-7084 |
| chr2-55802814-DEL-235 | chr2-55802815-DEL-235 |
| chr2-55814231-DEL-405 | chr2-55814232-DEL-405 |
| chr2-55817674-DEL-311 | chr2-55817675-DEL-308 |
| chr2-55821947-DEL-144 | chr2-55821948-DEL-144 |
| chr2-55831384-DEL-6497 | chr2-55831385-DEL-6497 |
| chr2-55841623-DEL-56 | chr2-55841599-DEL-92 |
| chr2-55855456-DEL-380 | chr2-55855457-DEL-380 |
| chr2-55920117-DEL-74 | chr2-55920118-DEL-74 |
| chr2-55928156-DEL-69 | chr2-55928162-DEL-54 |
| chr2-55939734-DEL-1331 | chr2-55939734-DEL-1331 |
| chr2-55954633-DEL-4633 | chr2-55954629-DEL-4637 |
| chr2-55967972-DEL-1092 | chr2-55967973-DEL-1092 |
| chr2-55970499-DEL-88 | chr2-55970500-DEL-88 |
| chr2-55976468-DEL-61 | chr2-55976469-DEL-61 |
| chr2-56030958-DEL-91 | chr2-56030961-DEL-57 |
| chr2-56035081-DEL-172 | chr2-56035082-DEL-172 |
| chr2-56081939-DEL-69 | chr2-56081985-DEL-59 |
| chr2-56086855-DEL-92 | chr2-56086853-DEL-92 |
| chr2-56106056-DEL-6393 | chr2-56106057-DEL-6393 |
| chr2-56188584-DEL-6395 | chr2-56188585-DEL-6395 |
| chr2-56248016-DEL-90 | chr2-56248017-DEL-90 |
| chr2-56299027-DEL-54 | chr2-56299011-DEL-76 |
| chr2-56306267-DEL-69 | chr2-56306252-DEL-50 |
| chr2-56355011-DEL-6919 | chr2-56355012-DEL-6919 |
| chr2-56419170-DEL-325 | chr2-56419160-DEL-291 |
| chr2-56443767-DEL-140 | chr2-56443768-DEL-140 |
| chr2-56505162-DEL-1092 | chr2-56505163-DEL-1092 |
| chr2-56633174-DEL-244 | chr2-56633175-DEL-244 |
| chr2-56654412-DEL-871 | chr2-56654413-DEL-871 |
| chr2-56655741-DEL-8178 | chr2-56655742-DEL-8176 |
| chr2-56656645-DEL-1893 | chr2-56656646-DEL-1893 |
| chr2-56682762-DEL-8561 | chr2-56682763-DEL-8561 |
| chr2-56706226-DEL-142 | chr2-56706227-DEL-142 |
| chr2-56715664-DEL-64 | chr2-56715665-DEL-64 |
| chr2-56722840-DEL-52 | chr2-56722841-DEL-54 |
| chr2-56728025-DEL-1176 | chr2-56728023-DEL-1174 |
| chr2-56742734-DEL-2556 | chr2-56742735-DEL-2556 |
| chr2-56762814-DEL-6357 | chr2-56762815-DEL-6357 |
| chr2-56804695-DEL-78 | chr2-56804696-DEL-78 |
| chr2-56821682-DEL-1329 | chr2-56821683-DEL-1330 |
| chr2-56830471-DEL-6422 | chr2-56830472-DEL-6422 |
| chr2-56851763-DEL-58 | chr2-56851764-DEL-58 |
| chr2-56878688-DEL-1490 | chr2-56878689-DEL-1490 |
| chr2-56895572-DEL-431 | chr2-56895573-DEL-431 |
| chr2-56948268-DEL-685 | chr2-56948269-DEL-685 |
| chr2-56962052-DEL-56 | chr2-56962052-DEL-56 |

| | |
|---|---|
| chr2-57025090-DEL-51 | chr2-57025090-DEL-51 |
| chr2-57033629-DEL-64 | chr2-57033630-DEL-64 |
| chr2-57038107-DEL-60 | chr2-57038108-DEL-60 |
| chr2-57044703-DEL-1954 | chr2-57044704-DEL-1954 |
| chr2-57056438-DEL-6438 | chr2-57056439-DEL-6438 |
| chr2-57073451-DEL-2624 | chr2-57073452-DEL-2624 |
| chr2-57074621-DEL-64 | chr2-57074620-DEL-54 |
| chr2-57104419-DEL-8985 | chr2-57104420-DEL-8985 |
| chr2-57119482-DEL-195 | chr2-57119483-DEL-195 |
| chr2-57125969-DEL-261 | chr2-57125970-DEL-261 |
| chr2-57163868-DEL-226 | chr2-57163859-DEL-227 |
| chr2-57167386-DEL-124 | chr2-57167387-DEL-116 |
| chr2-57181929-DEL-88 | chr2-57181930-DEL-88 |
| chr2-57190926-DEL-72 | chr2-57190947-DEL-56 |
| chr2-57198262-DEL-52 | chr2-57198274-DEL-68 |
| chr2-57207142-DEL-107 | chr2-57207142-DEL-107 |
| chr2-57210610-DEL-117 | chr2-57210611-DEL-117 |
| chr2-57220371-DEL-75 | chr2-57220374-DEL-64 |
| chr2-57294035-DEL-71 | chr2-57294053-DEL-82 |
| chr2-57307424-DEL-2739 | chr2-57307425-DEL-2739 |
| chr2-57334869-DEL-1738 | chr2-57334870-DEL-1738 |
| chr2-57443114-DEL-6379 | chr2-57443115-DEL-6379 |
| chr2-57463793-DEL-1807 | chr2-57463793-DEL-1807 |
| chr2-57508123-DEL-88 | chr2-57508122-DEL-82 |
| chr2-57519500-DEL-232 | chr2-57519501-DEL-232 |
| chr2-57610330-DEL-54 | chr2-57610331-DEL-54 |
| chr2-57625277-DEL-71 | chr2-57625278-DEL-71 |
| chr2-57664718-DEL-124 | chr2-57664719-DEL-124 |
| chr2-57729752-DEL-67 | chr2-57729753-DEL-67 |
| chr2-57767344-DEL-57 | chr2-57767347-DEL-55 |
| chr2-57797529-DEL-56 | chr2-57797530-DEL-56 |
| chr2-57827176-DEL-100 | chr2-57827177-DEL-100 |
| chr2-57853547-DEL-2712 | chr2-57853547-DEL-2712 |
| chr2-57861390-DEL-62 | chr2-57861383-DEL-62 |
| chr2-57877794-DEL-64 | chr2-57877801-DEL-56 |
| chr2-57878544-DEL-72 | chr2-57878545-DEL-72 |
| chr2-57898344-DEL-52 | chr2-57898345-DEL-52 |
| chr2-57900196-DEL-2333 | chr2-57900197-DEL-2333 |
| chr2-57926705-DEL-52 | chr2-57926686-DEL-81 |
| chr2-57939548-DEL-54 | chr2-57939551-DEL-51 |
| chr2-57963621-DEL-199 | chr2-57963622-DEL-199 |
| chr2-57998036-DEL-3207 | chr2-57998037-DEL-3207 |
| chr2-58018947-DEL-100 | chr2-58018956-DEL-92 |
| chr2-58055420-DEL-3786 | chr2-58055421-DEL-3786 |
| chr2-58100434-DEL-417 | chr2-58100435-DEL-417 |
| chr2-58119450-DEL-6515 | chr2-58119451-DEL-6515 |
| chr2-58119607-DEL-214 | chr2-58119608-DEL-214 |
| chr2-58140202-DEL-92 | chr2-58140201-DEL-106 |
| chr2-58146775-DEL-6435 | chr2-58146776-DEL-6435 |
| chr2-58151738-DEL-2044 | chr2-58151739-DEL-2044 |
| chr2-58191185-DEL-52 | chr2-58191137-DEL-80 |
| chr2-58194509-DEL-327 | chr2-58194510-DEL-327 |
| chr2-58231395-DEL-2764 | chr2-58231396-DEL-2764 |
| chr2-58251241-DEL-63 | chr2-58251242-DEL-63 |
| chr2-58286576-DEL-54 | chr2-58286577-DEL-54 |
| chr2-58327790-DEL-284 | chr2-58327788-DEL-277 |
| chr2-58488235-DEL-477 | chr2-58488236-DEL-477 |
| chr2-58517526-DEL-56 | chr2-58517527-DEL-56 |
| chr2-58564108-DEL-4095 | chr2-58564108-DEL-4095 |
| chr2-58575505-DEL-6317 | chr2-58575506-DEL-6317 |
| chr2-58582490-DEL-145 | chr2-58582488-DEL-141 |
| chr2-58612314-DEL-5922 | chr2-58612315-DEL-5922 |
| chr2-58630275-DEL-506 | chr2-58630276-DEL-506 |
| chr2-58636825-DEL-136 | chr2-58636826-DEL-136 |
| chr2-58646594-DEL-112 | chr2-58646583-DEL-92 |
| chr2-58650085-DEL-67 | chr2-58650073-DEL-99 |
| chr2-58651888-DEL-84 | chr2-58651888-DEL-91 |
| chr2-58697510-DEL-72 | chr2-58697511-DEL-72 |
| chr2-58723133-DEL-963 | chr2-58723134-DEL-963 |
| chr2-58745686-DEL-68 | chr2-58745687-DEL-68 |
| chr2-58748043-DEL-3848 | chr2-58748044-DEL-3848 |
| chr2-58748267-DEL-1531 | chr2-58748268-DEL-1531 |
| chr2-58834333-DEL-404 | chr2-58834334-DEL-404 |
| chr2-58836179-DEL-57 | chr2-58836183-DEL-54 |
| chr2-58849358-DEL-60 | chr2-58849382-DEL-63 |
| chr2-58883479-DEL-53 | chr2-58883478-DEL-52 |
| chr2-58889912-DEL-399 | chr2-58889913-DEL-399 |
| chr2-58896989-DEL-54 | chr2-58896988-DEL-56 |
| chr2-58922555-DEL-59 | chr2-58922540-DEL-58 |
| chr2-58927304-DEL-76 | chr2-58927351-DEL-52 |

| | |
|---|---|
| chr2-58955038-DEL-196 | chr2-58955039-DEL-196 |
| chr2-58995493-DEL-110 | chr2-58995494-DEL-110 |
| chr2-59164775-DEL-60 | chr2-59164776-DEL-60 |
| chr2-59178427-DEL-54 | chr2-59178428-DEL-84 |
| chr2-59197011-DEL-732 | chr2-59197012-DEL-732 |
| chr2-59198467-DEL-188 | chr2-59198503-DEL-181 |
| chr2-59200746-DEL-72 | chr2-59200741-DEL-72 |
| chr2-59205731-DEL-215 | chr2-59205732-DEL-215 |
| chr2-59209430-DEL-386 | chr2-59209431-DEL-386 |
| chr2-59259673-DEL-601 | chr2-59259671-DEL-602 |
| chr2-59277759-DEL-52 | chr2-59277715-DEL-52 |
| chr2-59296458-DEL-452 | chr2-59296459-DEL-452 |
| chr2-59298303-DEL-579 | chr2-59298304-DEL-579 |
| chr2-59302275-DEL-164 | chr2-59302313-DEL-127 |
| chr2-59323264-DEL-70 | chr2-59323261-DEL-74 |
| chr2-59326582-DEL-269 | chr2-59326583-DEL-269 |
| chr2-59333723-DEL-96 | chr2-59333724-DEL-96 |
| chr2-59347291-DEL-229 | chr2-59347292-DEL-229 |
| chr2-59366388-DEL-621 | chr2-59366391-DEL-620 |
| chr2-59370485-DEL-56 | chr2-59370486-DEL-56 |
| chr2-59376119-DEL-69 | chr2-59376115-DEL-81 |
| chr2-59380222-DEL-62 | chr2-59380223-DEL-62 |
| chr2-59394324-DEL-944 | chr2-59394315-DEL-953 |
| chr2-59395751-DEL-199 | chr2-59395752-DEL-199 |
| chr2-59414947-DEL-205 | chr2-59414948-DEL-205 |
| chr2-59421138-DEL-748 | chr2-59421139-DEL-748 |
| chr2-59466130-DEL-80 | chr2-59466131-DEL-80 |
| chr2-59467197-DEL-104 | chr2-59467191-DEL-102 |
| chr2-59476524-DEL-126 | chr2-59476525-DEL-126 |
| chr2-59477235-DEL-602 | chr2-59477269-DEL-569 |
| chr2-59487814-DEL-262 | chr2-59487815-DEL-262 |
| chr2-59494129-DEL-66 | chr2-59494128-DEL-66 |
| chr2-59499486-DEL-491 | chr2-59499487-DEL-491 |
| chr2-59519354-DEL-64 | chr2-59519355-DEL-64 |
| chr2-59526277-DEL-58 | chr2-59526278-DEL-58 |
| chr2-59532682-DEL-211 | chr2-59532683-DEL-211 |
| chr2-59537675-DEL-85 | chr2-59537676-DEL-85 |
| chr2-59560013-DEL-136 | chr2-59560014-DEL-136 |
| chr2-59560412-DEL-180 | chr2-59560413-DEL-180 |
| chr2-59571715-DEL-224 | chr2-59571716-DEL-224 |
| chr2-59579893-DEL-54 | chr2-59579894-DEL-54 |
| chr2-59584028-DEL-194 | chr2-59584027-DEL-194 |
| chr2-59584836-DEL-52 | chr2-59584837-DEL-52 |
| chr2-59626675-DEL-78 | chr2-59626674-DEL-79 |
| chr2-59645036-DEL-218 | chr2-59645037-DEL-218 |
| chr2-59653959-DEL-180 | chr2-59653960-DEL-180 |
| chr2-59667666-DEL-56 | chr2-59667667-DEL-56 |
| chr2-59678792-DEL-58 | chr2-59678754-DEL-74 |
| chr2-59682211-DEL-191 | chr2-59682212-DEL-191 |
| chr2-59695656-DEL-107 | chr2-59695657-DEL-107 |
| chr2-59709615-DEL-194 | chr2-59709617-DEL-190 |
| chr2-59717404-DEL-57 | chr2-59717408-DEL-55 |
| chr2-59773979-DEL-2045 | chr2-59773980-DEL-2045 |
| chr2-59777243-DEL-205 | chr2-59777244-DEL-205 |
| chr2-59801856-DEL-953 | chr2-59801827-DEL-954 |
| chr2-59860642-DEL-570 | chr2-59860643-DEL-570 |
| chr2-59896583-DEL-211 | chr2-59896584-DEL-211 |
| chr2-59993355-DEL-202 | chr2-59993356-DEL-202 |
| chr2-59995744-DEL-206 | chr2-59995745-DEL-206 |
| chr2-60005349-DEL-180 | chr2-60005350-DEL-180 |
| chr2-60012660-DEL-57 | chr2-60012661-DEL-57 |
| chr2-60020115-DEL-418 | chr2-60020116-DEL-418 |
| chr2-60028048-DEL-215 | chr2-60028049-DEL-215 |
| chr2-60043209-DEL-181 | chr2-60043210-DEL-181 |
| chr2-60053079-DEL-52 | chr2-60053079-DEL-50 |
| chr2-60084303-DEL-201 | chr2-60084304-DEL-201 |
| chr2-60098930-DEL-434 | chr2-60098931-DEL-434 |
| chr2-60103159-DEL-1553 | chr2-60103164-DEL-1549 |
| chr2-60120718-DEL-73 | chr2-60120719-DEL-73 |
| chr2-60126748-DEL-1264 | chr2-60126748-DEL-1264 |
| chr2-60148854-DEL-51 | chr2-60148855-DEL-51 |
| chr2-60150900-DEL-52 | chr2-60150904-DEL-53 |
| chr2-60175566-DEL-1189 | chr2-60175567-DEL-1189 |
| chr2-60202001-DEL-1689 | chr2-60202002-DEL-1689 |
| chr2-60203488-DEL-119 | chr2-60203490-DEL-115 |
| chr2-60253465-DEL-132 | chr2-60253461-DEL-119 |
| chr2-60256761-DEL-64 | chr2-60256762-DEL-73 |
| chr2-60305517-DEL-114 | chr2-60305518-DEL-114 |
| chr2-60318361-DEL-1691 | chr2-60318362-DEL-1691 |
| chr2-60325229-DEL-210 | chr2-60325230-DEL-210 |

| | |
|---|---|
| chr2-60326775-DEL-98 | chr2-60326786-DEL-72 |
| chr2-60342056-DEL-129 | chr2-60342057-DEL-129 |
| chr2-60371411-DEL-64 | chr2-60371412-DEL-62 |
| chr2-60388090-DEL-154 | chr2-60388091-DEL-154 |
| chr2-60389032-DEL-1685 | chr2-60389033-DEL-1685 |
| chr2-60404858-DEL-1296 | chr2-60404859-DEL-1296 |
| chr2-60411004-DEL-60 | chr2-60411005-DEL-60 |
| chr2-60426051-DEL-217 | chr2-60426043-DEL-175 |
| chr2-60461908-DEL-465 | chr2-60461909-DEL-465 |
| chr2-60496450-DEL-67 | chr2-60496451-DEL-67 |
| chr2-60505837-DEL-2016 | chr2-60505854-DEL-1996 |
| chr2-60537627-DEL-2264 | chr2-60537628-DEL-2264 |
| chr2-60560923-DEL-63 | chr2-60560924-DEL-63 |
| chr2-60574351-DEL-92 | chr2-60574352-DEL-92 |
| chr2-60621246-DEL-422 | chr2-60621247-DEL-422 |
| chr2-60644205-DEL-81 | chr2-60644203-DEL-84 |
| chr2-60656112-DEL-61 | chr2-60656113-DEL-61 |
| chr2-60665314-DEL-504 | chr2-60665315-DEL-504 |
| chr2-60692895-DEL-468 | chr2-60692896-DEL-468 |
| chr2-60756687-DEL-88 | chr2-60756676-DEL-80 |
| chr2-60768031-DEL-60 | chr2-60768032-DEL-60 |
| chr2-60818369-DEL-56 | chr2-60818374-DEL-54 |
| chr2-60821303-DEL-85 | chr2-60821337-DEL-52 |
| chr2-60840106-DEL-66 | chr2-60840107-DEL-66 |
| chr2-60867127-DEL-5979 | chr2-60867127-DEL-5979 |
| chr2-60885974-DEL-63 | chr2-60885974-DEL-64 |
| chr2-60923498-DEL-995 | chr2-60923499-DEL-995 |
| chr2-60952099-DEL-61 | chr2-60952100-DEL-65 |
| chr2-60971282-DEL-92 | chr2-60971285-DEL-84 |
| chr2-60982832-DEL-52 | chr2-60982822-DEL-56 |
| chr2-61008664-DEL-399 | chr2-61008665-DEL-399 |
| chr2-61038822-DEL-94 | chr2-61038823-DEL-94 |
| chr2-61051308-DEL-680 | chr2-61051308-DEL-680 |
| chr2-61064830-DEL-90 | chr2-61064843-DEL-78 |
| chr2-61088124-DEL-51 | chr2-61088124-DEL-51 |
| chr2-61114062-DEL-52 | chr2-61114063-DEL-52 |
| chr2-61124363-DEL-449 | chr2-61124364-DEL-449 |
| chr2-61130673-DEL-81 | chr2-61130650-DEL-82 |
| chr2-61202558-DEL-833 | chr2-61202559-DEL-833 |
| chr2-61206465-DEL-1085 | chr2-61206466-DEL-1085 |
| chr2-61212476-DEL-59 | chr2-61212477-DEL-59 |
| chr2-61269176-DEL-6206 | chr2-61269178-DEL-6205 |
| chr2-61275601-DEL-704 | chr2-61275602-DEL-704 |
| chr2-61281640-DEL-91 | chr2-61281641-DEL-91 |
| chr2-61296492-DEL-55 | chr2-61296467-DEL-50 |
| chr2-61355388-DEL-446 | chr2-61355389-DEL-446 |
| chr2-61356505-DEL-420 | chr2-61356514-DEL-414 |
| chr2-61398736-DEL-280 | chr2-61398737-DEL-280 |
| chr2-61415535-DEL-401 | chr2-61415536-DEL-401 |
| chr2-61427228-DEL-144 | chr2-61427229-DEL-144 |
| chr2-61492436-DEL-88 | chr2-61492437-DEL-88 |
| chr2-61505956-DEL-84 | chr2-61505957-DEL-84 |
| chr2-61513078-DEL-203 | chr2-61513079-DEL-203 |
| chr2-61536062-DEL-1309 | chr2-61536060-DEL-1312 |
| chr2-61538082-DEL-1618 | chr2-61538083-DEL-1618 |
| chr2-61541342-DEL-91 | chr2-61541343-DEL-91 |
| chr2-61661589-DEL-198 | chr2-61661590-DEL-198 |
| chr2-61683479-DEL-214 | chr2-61683480-DEL-214 |
| chr2-61706729-DEL-864 | chr2-61706730-DEL-864 |
| chr2-61744424-DEL-2099 | chr2-61744418-DEL-2099 |
| chr2-61747172-DEL-776 | chr2-61747173-DEL-776 |
| chr2-61748612-DEL-1132 | chr2-61748613-DEL-1132 |
| chr2-61776015-DEL-112 | chr2-61776004-DEL-142 |
| chr2-61818669-DEL-549 | chr2-61818670-DEL-549 |
| chr2-61838637-DEL-214 | chr2-61838638-DEL-214 |
| chr2-61851784-DEL-1495 | chr2-61851785-DEL-1495 |
| chr2-61864526-DEL-6404 | chr2-61864527-DEL-6404 |
| chr2-61891939-DEL-68 | chr2-61891945-DEL-60 |
| chr2-61969671-DEL-179 | chr2-61969671-DEL-175 |
| chr2-61975874-DEL-228 | chr2-61975875-DEL-228 |
| chr2-61991400-DEL-94 | chr2-61991401-DEL-94 |
| chr2-61999418-DEL-6548 | chr2-61999419-DEL-6548 |
| chr2-62024379-DEL-6572 | chr2-62024380-DEL-6572 |
| chr2-62043457-DEL-1740 | chr2-62043458-DEL-1740 |
| chr2-62048439-DEL-406 | chr2-62048440-DEL-406 |
| chr2-62163957-DEL-52 | chr2-62163958-DEL-52 |
| chr2-62165708-DEL-78 | chr2-62165709-DEL-78 |
| chr2-62176447-DEL-90 | chr2-62176448-DEL-90 |
| chr2-62188535-DEL-237 | chr2-62188536-DEL-237 |
| chr2-62252139-DEL-417 | chr2-62252140-DEL-417 |

| | |
|---|---|
| chr2-62335295-DEL-55 | chr2-62335280-DEL-71 |
| chr2-62346782-DEL-80 | chr2-62346775-DEL-88 |
| chr2-62357084-DEL-186 | chr2-62357085-DEL-186 |
| chr2-62391142-DEL-525 | chr2-62391143-DEL-525 |
| chr2-62472709-DEL-99 | chr2-62472719-DEL-90 |
| chr2-62478535-DEL-51 | chr2-62478536-DEL-51 |
| chr2-62529546-DEL-104 | chr2-62529547-DEL-104 |
| chr2-62543371-DEL-661 | chr2-62543372-DEL-661 |
| chr2-62620612-DEL-168 | chr2-62620613-DEL-168 |
| chr2-62623420-DEL-781 | chr2-62623416-DEL-777 |
| chr2-62650942-DEL-6537 | chr2-62650943-DEL-6537 |
| chr2-62661859-DEL-2459 | chr2-62661860-DEL-2459 |
| chr2-62665205-DEL-67 | chr2-62665206-DEL-67 |
| chr2-62672665-DEL-72 | chr2-62672668-DEL-70 |
| chr2-62691380-DEL-6257 | chr2-62691381-DEL-6257 |
| chr2-62703363-DEL-96 | chr2-62703366-DEL-126 |
| chr2-62714689-DEL-133 | chr2-62714690-DEL-133 |
| chr2-62720192-DEL-400 | chr2-62720193-DEL-400 |
| chr2-62721717-DEL-122 | chr2-62721697-DEL-143 |
| chr2-62725965-DEL-59 | chr2-62725966-DEL-59 |
| chr2-62738748-DEL-1483 | chr2-62738749-DEL-1482 |
| chr2-62751047-DEL-2064 | chr2-62751048-DEL-2064 |
| chr2-62759976-DEL-771 | chr2-62759977-DEL-771 |
| chr2-62803093-DEL-54 | chr2-62803094-DEL-54 |
| chr2-62808997-DEL-1164 | chr2-62808998-DEL-1164 |
| chr2-62817256-DEL-112 | chr2-62817257-DEL-84 |
| chr2-62825073-DEL-54 | chr2-62825073-DEL-54 |
| chr2-62862532-DEL-91 | chr2-62862533-DEL-91 |
| chr2-62867171-DEL-1014 | chr2-62867172-DEL-1014 |
| chr2-62870926-DEL-479 | chr2-62870917-DEL-480 |
| chr2-62877587-DEL-222 | chr2-62877588-DEL-222 |
| chr2-62881304-DEL-126 | chr2-62881305-DEL-126 |
| chr2-62904917-DEL-4729 | chr2-62904918-DEL-4729 |
| chr2-62941499-DEL-4135 | chr2-62941500-DEL-4135 |
| chr2-62947223-DEL-64 | chr2-62947224-DEL-64 |
| chr2-62950372-DEL-80 | chr2-62950370-DEL-62 |
| chr2-62956784-DEL-80 | chr2-62956750-DEL-112 |
| chr2-62972171-DEL-64 | chr2-62972190-DEL-68 |
| chr2-62987838-DEL-70 | chr2-62987837-DEL-68 |
| chr2-63036791-DEL-109 | chr2-63036792-DEL-109 |
| chr2-63038280-DEL-136 | chr2-63038277-DEL-164 |
| chr2-63053524-DEL-939 | chr2-63053525-DEL-939 |
| chr2-63075855-DEL-100 | chr2-63075856-DEL-100 |
| chr2-63089751-DEL-753 | chr2-63089752-DEL-753 |
| chr2-63119257-DEL-606 | chr2-63119258-DEL-606 |
| chr2-63127712-DEL-5380 | chr2-63127713-DEL-5380 |
| chr2-63188865-DEL-52 | chr2-63188866-DEL-50 |
| chr2-63198394-DEL-6496 | chr2-63198395-DEL-6496 |
| chr2-63208565-DEL-94 | chr2-63208566-DEL-94 |
| chr2-63210867-DEL-179 | chr2-63210868-DEL-179 |
| chr2-63227599-DEL-6656 | chr2-63227600-DEL-6656 |
| chr2-63234238-DEL-141 | chr2-63234242-DEL-138 |
| chr2-63263468-DEL-588 | chr2-63263467-DEL-588 |
| chr2-63270916-DEL-3543 | chr2-63270917-DEL-3543 |
| chr2-63290616-DEL-443 | chr2-63290617-DEL-443 |
| chr2-63292712-DEL-1047 | chr2-63292713-DEL-1047 |
| chr2-63296925-DEL-228 | chr2-63296921-DEL-228 |
| chr2-63303314-DEL-98 | chr2-63303297-DEL-99 |
| chr2-63309203-DEL-6182 | chr2-63309204-DEL-6182 |
| chr2-63315093-DEL-199 | chr2-63315094-DEL-199 |
| chr2-63318251-DEL-62 | chr2-63318217-DEL-63 |
| chr2-63366040-DEL-998 | chr2-63366041-DEL-998 |
| chr2-63429133-DEL-155 | chr2-63429134-DEL-155 |
| chr2-63436792-DEL-7108 | chr2-63436793-DEL-7108 |
| chr2-63452203-DEL-61 | chr2-63452204-DEL-61 |
| chr2-63460417-DEL-825 | chr2-63460418-DEL-825 |
| chr2-63469118-DEL-59 | chr2-63469120-DEL-54 |
| chr2-63512871-DEL-248 | chr2-63512860-DEL-249 |
| chr2-63518247-DEL-92 | chr2-63518252-DEL-60 |
| chr2-63545522-DEL-54 | chr2-63545523-DEL-54 |
| chr2-63548512-DEL-7219 | chr2-63548513-DEL-7219 |
| chr2-63592899-DEL-1030 | chr2-63592900-DEL-1030 |
| chr2-63645808-DEL-1665 | chr2-63645809-DEL-1665 |
| chr2-63668452-DEL-7177 | chr2-63668453-DEL-7177 |
| chr2-63698458-DEL-203 | chr2-63698459-DEL-203 |
| chr2-63772091-DEL-63 | chr2-63772092-DEL-63 |
| chr2-63784310-DEL-220 | chr2-63784311-DEL-220 |
| chr2-63834250-DEL-57 | chr2-63834278-DEL-54 |
| chr2-63836964-DEL-587 | chr2-63836965-DEL-587 |
| chr2-63937694-DEL-121 | chr2-63937690-DEL-123 |

| | |
|---|---|
| chr2-63944535-DEL-237 | chr2-63944536-DEL-237 |
| chr2-63949448-DEL-70 | chr2-63949448-DEL-70 |
| chr2-63975049-DEL-6794 | chr2-63975050-DEL-6794 |
| chr2-64024360-DEL-6416 | chr2-64024361-DEL-6416 |
| chr2-64038055-DEL-103 | chr2-64038056-DEL-103 |
| chr2-64142923-DEL-7027 | chr2-64142924-DEL-7027 |
| chr2-64175548-DEL-302 | chr2-64175549-DEL-302 |
| chr2-64182675-DEL-2169 | chr2-64182676-DEL-2169 |
| chr2-64198264-DEL-87 | chr2-64198258-DEL-114 |
| chr2-64208686-DEL-124 | chr2-64208687-DEL-124 |
| chr2-64214541-DEL-379 | chr2-64214542-DEL-379 |
| chr2-64245126-DEL-1351 | chr2-64245127-DEL-1351 |
| chr2-64304559-DEL-729 | chr2-64304560-DEL-729 |
| chr2-64311170-DEL-1183 | chr2-64311179-DEL-1183 |
| chr2-64366189-DEL-54 | chr2-64366189-DEL-55 |
| chr2-64433528-DEL-54 | chr2-64433529-DEL-54 |
| chr2-64438711-DEL-1173 | chr2-64438712-DEL-1173 |
| chr2-64450921-DEL-1688 | chr2-64450922-DEL-1688 |
| chr2-64455189-DEL-8455 | chr2-64455190-DEL-8455 |
| chr2-64497564-DEL-3106 | chr2-64497565-DEL-3106 |
| chr2-64529227-DEL-4110 | chr2-64529228-DEL-4110 |
| chr2-64560730-DEL-498 | chr2-64560731-DEL-498 |
| chr2-64579704-DEL-201 | chr2-64579705-DEL-201 |
| chr2-64603871-DEL-297 | chr2-64603872-DEL-297 |
| chr2-64619242-DEL-7322 | chr2-64619243-DEL-7322 |
| chr2-64754111-DEL-185 | chr2-64754112-DEL-185 |
| chr2-64756879-DEL-344 | chr2-64756883-DEL-339 |
| chr2-64780434-DEL-225 | chr2-64780435-DEL-225 |
| chr2-64827661-DEL-123 | chr2-64827667-DEL-106 |
| chr2-64846014-DEL-56 | chr2-64846003-DEL-50 |
| chr2-64859504-DEL-100 | chr2-64859505-DEL-100 |
| chr2-64894932-DEL-58 | chr2-64894933-DEL-58 |
| chr2-64910125-DEL-61 | chr2-64910126-DEL-61 |
| chr2-64915322-DEL-72 | chr2-64915323-DEL-72 |
| chr2-64973768-DEL-129 | chr2-64973769-DEL-129 |
| chr2-65007667-DEL-70 | chr2-65007668-DEL-70 |
| chr2-65082822-DEL-995 | chr2-65082826-DEL-995 |
| chr2-65090489-DEL-234 | chr2-65090489-DEL-223 |
| chr2-65098688-DEL-67 | chr2-65098689-DEL-67 |
| chr2-65103283-DEL-58 | chr2-65103284-DEL-58 |
| chr2-65143642-DEL-64 | chr2-65143621-DEL-78 |
| chr2-65144912-DEL-184 | chr2-65144913-DEL-184 |
| chr2-65149157-DEL-78 | chr2-65149134-DEL-79 |
| chr2-65150269-DEL-386 | chr2-65150270-DEL-386 |
| chr2-65154691-DEL-59 | chr2-65154692-DEL-59 |
| chr2-65183599-DEL-811 | chr2-65183600-DEL-811 |
| chr2-65225329-DEL-55 | chr2-65225330-DEL-55 |
| chr2-65229436-DEL-201 | chr2-65229448-DEL-202 |
| chr2-65235406-DEL-792 | chr2-65235407-DEL-792 |
| chr2-65247959-DEL-448 | chr2-65247960-DEL-448 |
| chr2-65254306-DEL-437 | chr2-65254307-DEL-437 |
| chr2-65258501-DEL-830 | chr2-65258503-DEL-825 |
| chr2-65266326-DEL-57 | chr2-65266326-DEL-66 |
| chr2-65282022-DEL-479 | chr2-65282024-DEL-480 |
| chr2-65295410-DEL-687 | chr2-65295411-DEL-687 |
| chr2-65303130-DEL-929 | chr2-65303131-DEL-929 |
| chr2-65306385-DEL-98 | chr2-65306385-DEL-99 |
| chr2-65310779-DEL-187 | chr2-65310780-DEL-187 |
| chr2-65344330-DEL-181 | chr2-65344331-DEL-181 |
| chr2-65410181-DEL-3721 | chr2-65410187-DEL-3707 |
| chr2-65415592-DEL-401 | chr2-65415593-DEL-401 |
| chr2-65422766-DEL-5926 | chr2-65422767-DEL-5926 |
| chr2-65443667-DEL-680 | chr2-65443668-DEL-680 |
| chr2-65454950-DEL-224 | chr2-65454951-DEL-224 |
| chr2-65456976-DEL-98 | chr2-65456977-DEL-98 |
| chr2-65483161-DEL-204 | chr2-65483162-DEL-204 |
| chr2-65526291-DEL-1296 | chr2-65526292-DEL-1296 |
| chr2-65557809-DEL-149 | chr2-65557810-DEL-149 |
| chr2-65562690-DEL-54 | chr2-65562704-DEL-72 |
| chr2-65563296-DEL-57 | chr2-65563297-DEL-57 |
| chr2-65585203-DEL-1882 | chr2-65585201-DEL-1880 |
| chr2-65599524-DEL-393 | chr2-65599519-DEL-390 |
| chr2-65603331-DEL-75 | chr2-65603332-DEL-75 |
| chr2-65605424-DEL-6437 | chr2-65605425-DEL-6437 |
| chr2-65616410-DEL-63 | chr2-65616400-DEL-64 |
| chr2-65661006-DEL-56 | chr2-65661015-DEL-58 |
| chr2-65703284-DEL-57 | chr2-65703284-DEL-58 |
| chr2-65728220-DEL-351 | chr2-65728221-DEL-351 |
| chr2-65776671-DEL-53 | chr2-65776672-DEL-53 |
| chr2-65881588-DEL-51 | chr2-65881589-DEL-51 |

| | |
|---|---|
| chr2-65903644-DEL-68 | chr2-65903641-DEL-84 |
| chr2-65918694-DEL-97 | chr2-65918695-DEL-97 |
| chr2-65923515-DEL-77 | chr2-65923514-DEL-79 |
| chr2-65942565-DEL-75 | chr2-65942566-DEL-75 |
| chr2-65973042-DEL-81 | chr2-65973043-DEL-81 |
| chr2-65983542-DEL-54 | chr2-65983543-DEL-54 |
| chr2-65996934-DEL-58 | chr2-65996940-DEL-56 |
| chr2-65999654-DEL-82 | chr2-65999655-DEL-82 |
| chr2-66005964-DEL-785 | chr2-66005965-DEL-785 |
| chr2-66011037-DEL-61 | chr2-66011038-DEL-61 |
| chr2-66019578-DEL-233 | chr2-66019570-DEL-212 |
| chr2-66060456-DEL-189 | chr2-66060452-DEL-190 |
| chr2-66100146-DEL-341 | chr2-66100147-DEL-341 |
| chr2-66163710-DEL-63 | chr2-66163712-DEL-62 |
| chr2-66188308-DEL-1932 | chr2-66188309-DEL-1932 |
| chr2-66190519-DEL-63 | chr2-66190507-DEL-66 |
| chr2-66192750-DEL-318 | chr2-66192725-DEL-305 |
| chr2-66201382-DEL-436 | chr2-66201383-DEL-436 |
| chr2-66205935-DEL-91 | chr2-66205936-DEL-91 |
| chr2-66206549-DEL-59 | chr2-66206530-DEL-60 |
| chr2-66259898-DEL-6456 | chr2-66259899-DEL-6456 |
| chr2-66260097-DEL-214 | chr2-66260098-DEL-214 |
| chr2-66275539-DEL-200 | chr2-66275540-DEL-200 |
| chr2-66307030-DEL-230 | chr2-66307031-DEL-230 |
| chr2-66310231-DEL-77 | chr2-66310230-DEL-79 |
| chr2-66360508-DEL-90 | chr2-66360496-DEL-123 |
| chr2-66365008-DEL-56 | chr2-66365009-DEL-56 |
| chr2-66409200-DEL-51 | chr2-66409201-DEL-51 |
| chr2-66414058-DEL-74 | chr2-66414073-DEL-50 |
| chr2-66462177-DEL-78 | chr2-66462155-DEL-76 |
| chr2-66478026-DEL-164 | chr2-66478027-DEL-164 |
| chr2-66480082-DEL-134 | chr2-66480083-DEL-134 |
| chr2-66508826-DEL-74 | chr2-66508827-DEL-74 |
| chr2-66536471-DEL-2072 | chr2-66536472-DEL-2072 |
| chr2-66607961-DEL-663 | chr2-66607964-DEL-662 |
| chr2-66609773-DEL-282 | chr2-66609769-DEL-282 |
| chr2-66618569-DEL-197 | chr2-66618570-DEL-197 |
| chr2-66666982-DEL-112 | chr2-66667009-DEL-86 |
| chr2-66675442-DEL-7164 | chr2-66675441-DEL-7165 |
| chr2-66715406-DEL-386 | chr2-66715407-DEL-387 |
| chr2-66718675-DEL-671 | chr2-66718676-DEL-671 |
| chr2-66722616-DEL-3339 | chr2-66722617-DEL-3339 |
| chr2-66736277-DEL-1464 | chr2-66736278-DEL-1464 |
| chr2-66783022-DEL-750 | chr2-66783023-DEL-750 |
| chr2-66790488-DEL-135 | chr2-66790489-DEL-135 |
| chr2-66815466-DEL-80 | chr2-66815467-DEL-72 |
| chr2-66844381-DEL-120 | chr2-66844382-DEL-96 |
| chr2-66853728-DEL-56 | chr2-66853727-DEL-57 |
| chr2-66884030-DEL-5231 | chr2-66884031-DEL-5231 |
| chr2-66890991-DEL-731 | chr2-66890992-DEL-731 |
| chr2-66904089-DEL-113 | chr2-66904077-DEL-98 |
| chr2-66909829-DEL-88 | chr2-66909829-DEL-89 |
| chr2-66913919-DEL-7129 | chr2-66913920-DEL-7129 |
| chr2-66970377-DEL-439 | chr2-66970378-DEL-439 |
| chr2-66974328-DEL-72 | chr2-66974329-DEL-74 |
| chr2-66977593-DEL-441 | chr2-66977594-DEL-441 |
| chr2-66994078-DEL-664 | chr2-66994076-DEL-665 |
| chr2-67000101-DEL-70 | chr2-67000102-DEL-70 |
| chr2-67047335-DEL-1577 | chr2-67047336-DEL-1577 |
| chr2-67070009-DEL-771 | chr2-67070010-DEL-771 |
| chr2-67080210-DEL-2064 | chr2-67080211-DEL-2064 |
| chr2-67094829-DEL-315 | chr2-67094830-DEL-315 |
| chr2-67099589-DEL-6174 | chr2-67099590-DEL-6174 |
| chr2-67111015-DEL-456 | chr2-67111016-DEL-456 |
| chr2-67116812-DEL-116 | chr2-67116813-DEL-116 |
| chr2-67136054-DEL-101 | chr2-67136063-DEL-109 |
| chr2-67145862-DEL-1697 | chr2-67145863-DEL-1697 |
| chr2-67149571-DEL-6397 | chr2-67149571-DEL-6397 |
| chr2-67157398-DEL-632 | chr2-67157386-DEL-644 |
| chr2-67184090-DEL-204 | chr2-67184091-DEL-204 |
| chr2-67211869-DEL-1106 | chr2-67211870-DEL-1106 |
| chr2-67218142-DEL-168 | chr2-67218143-DEL-168 |
| chr2-67270891-DEL-101 | chr2-67270892-DEL-101 |
| chr2-67279660-DEL-2262 | chr2-67279661-DEL-2262 |
| chr2-67297956-DEL-56 | chr2-67297911-DEL-56 |
| chr2-67299393-DEL-131 | chr2-67299394-DEL-131 |
| chr2-67300106-DEL-431 | chr2-67300105-DEL-430 |
| chr2-67320679-DEL-663 | chr2-67320683-DEL-662 |
| chr2-67327649-DEL-333 | chr2-67327650-DEL-333 |
| chr2-67364444-DEL-710 | chr2-67364446-DEL-702 |

| | |
|---|---|
| chr2-67367498-DEL-259 | chr2-67367498-DEL-259 |
| chr2-67409218-DEL-1922 | chr2-67409219-DEL-1922 |
| chr2-67414988-DEL-1596 | chr2-67414989-DEL-1596 |
| chr2-67415811-DEL-86 | chr2-67415812-DEL-86 |
| chr2-67438775-DEL-64 | chr2-67438769-DEL-56 |
| chr2-67454435-DEL-87 | chr2-67454436-DEL-58 |
| chr2-67464789-DEL-3192 | chr2-67464790-DEL-3192 |
| chr2-67467256-DEL-257 | chr2-67467256-DEL-257 |
| chr2-67489120-DEL-53 | chr2-67489121-DEL-53 |
| chr2-67495767-DEL-119 | chr2-67495768-DEL-119 |
| chr2-67513812-DEL-87 | chr2-67513810-DEL-88 |
| chr2-67519093-DEL-1487 | chr2-67519115-DEL-1444 |
| chr2-67570747-DEL-55 | chr2-67570738-DEL-57 |
| chr2-67585350-DEL-52 | chr2-67585351-DEL-52 |
| chr2-67724384-DEL-57 | chr2-67724389-DEL-53 |
| chr2-67737927-DEL-352 | chr2-67737928-DEL-352 |
| chr2-67765949-DEL-300 | chr2-67765950-DEL-300 |
| chr2-67769452-DEL-6900 | chr2-67769453-DEL-6900 |
| chr2-67769778-DEL-428 | chr2-67769779-DEL-428 |
| chr2-67795893-DEL-65 | chr2-67795894-DEL-65 |
| chr2-67811399-DEL-368 | chr2-67811398-DEL-368 |
| chr2-67827347-DEL-80 | chr2-67827348-DEL-80 |
| chr2-67936010-DEL-2093 | chr2-67936011-DEL-2093 |
| chr2-67954593-DEL-56 | chr2-67954594-DEL-60 |
| chr2-67957023-DEL-329 | chr2-67957024-DEL-328 |
| chr2-67960080-DEL-8727 | chr2-67960081-DEL-8727 |
| chr2-67972892-DEL-518 | chr2-67972893-DEL-518 |
| chr2-67981343-DEL-63 | chr2-67981343-DEL-63 |
| chr2-67982962-DEL-2087 | chr2-67982963-DEL-2087 |
| chr2-68016233-DEL-1077 | chr2-68016233-DEL-1077 |
| chr2-68065936-DEL-401 | chr2-68065937-DEL-401 |
| chr2-68074215-DEL-73 | chr2-68074219-DEL-62 |
| chr2-68120111-DEL-94 | chr2-68120112-DEL-94 |
| chr2-68148110-DEL-81 | chr2-68148111-DEL-69 |
| chr2-68172891-DEL-130 | chr2-68172892-DEL-121 |
| chr2-68177234-DEL-66 | chr2-68177235-DEL-66 |
| chr2-68180579-DEL-75 | chr2-68180580-DEL-75 |
| chr2-68199115-DEL-166 | chr2-68199116-DEL-166 |
| chr2-68201869-DEL-85 | chr2-68201871-DEL-84 |
| chr2-68257025-DEL-86 | chr2-68257028-DEL-114 |
| chr2-68279383-DEL-130 | chr2-68279384-DEL-130 |
| chr2-68283183-DEL-60 | chr2-68283157-DEL-108 |
| chr2-68289738-DEL-63 | chr2-68289740-DEL-62 |
| chr2-68309106-DEL-122 | chr2-68309112-DEL-113 |
| chr2-68322804-DEL-210 | chr2-68322805-DEL-210 |
| chr2-68331871-DEL-69 | chr2-68331869-DEL-69 |
| chr2-68337061-DEL-66 | chr2-68337044-DEL-72 |
| chr2-68341779-DEL-58 | chr2-68341764-DEL-70 |
| chr2-68347395-DEL-2173 | chr2-68347395-DEL-2174 |
| chr2-68350860-DEL-1188 | chr2-68350867-DEL-1182 |
| chr2-68373585-DEL-358 | chr2-68373586-DEL-358 |
| chr2-68376812-DEL-5761 | chr2-68376813-DEL-5757 |
| chr2-68384831-DEL-1292 | chr2-68384832-DEL-1292 |
| chr2-68548774-DEL-6437 | chr2-68548776-DEL-6435 |
| chr2-68600685-DEL-6422 | chr2-68600686-DEL-6422 |
| chr2-68629686-DEL-445 | chr2-68629687-DEL-444 |
| chr2-68632794-DEL-56 | chr2-68632795-DEL-56 |
| chr2-68664522-DEL-170 | chr2-68664523-DEL-170 |
| chr2-68664972-DEL-2540 | chr2-68664973-DEL-2540 |
| chr2-68680790-DEL-1539 | chr2-68680791-DEL-1539 |
| chr2-68683999-DEL-1378 | chr2-68684000-DEL-1378 |
| chr2-68711130-DEL-92 | chr2-68711131-DEL-92 |
| chr2-68711788-DEL-121 | chr2-68711789-DEL-121 |
| chr2-68729321-DEL-152 | chr2-68729364-DEL-146 |
| chr2-68748140-DEL-6297 | chr2-68748141-DEL-6297 |
| chr2-68762103-DEL-91 | chr2-68762104-DEL-96 |
| chr2-68783140-DEL-52 | chr2-68783141-DEL-52 |
| chr2-68809828-DEL-170 | chr2-68809829-DEL-170 |
| chr2-68821205-DEL-1097 | chr2-68821206-DEL-1097 |
| chr2-68831455-DEL-68 | chr2-68831461-DEL-85 |
| chr2-68859613-DEL-61 | chr2-68859614-DEL-61 |
| chr2-68904886-DEL-1124 | chr2-68904887-DEL-1124 |
| chr2-68925985-DEL-118 | chr2-68925986-DEL-118 |
| chr2-68936802-DEL-135 | chr2-68936803-DEL-135 |
| chr2-68972004-DEL-538 | chr2-68972005-DEL-536 |
| chr2-68973064-DEL-53 | chr2-68973065-DEL-53 |
| chr2-68982434-DEL-61 | chr2-68982433-DEL-62 |
| chr2-68985394-DEL-64 | chr2-68985395-DEL-64 |
| chr2-68988425-DEL-70 | chr2-68988426-DEL-70 |
| chr2-68991754-DEL-398 | chr2-68991755-DEL-398 |

| | |
|---|---|
| chr2-69029068-DEL-277 | chr2-69029069-DEL-277 |
| chr2-69038374-DEL-211 | chr2-69038375-DEL-211 |
| chr2-69067877-DEL-173 | chr2-69067878-DEL-173 |
| chr2-69082732-DEL-89 | chr2-69082733-DEL-89 |
| chr2-69086063-DEL-151 | chr2-69086064-DEL-152 |
| chr2-69098723-DEL-340 | chr2-69098724-DEL-340 |
| chr2-69119043-DEL-62 | chr2-69119044-DEL-62 |
| chr2-69161551-DEL-446 | chr2-69161552-DEL-446 |
| chr2-69163520-DEL-119 | chr2-69163523-DEL-116 |
| chr2-69164997-DEL-658 | chr2-69164998-DEL-658 |
| chr2-69166449-DEL-59 | chr2-69166452-DEL-55 |
| chr2-69170253-DEL-62 | chr2-69170252-DEL-88 |
| chr2-69179004-DEL-76 | chr2-69179031-DEL-52 |
| chr2-69185882-DEL-127 | chr2-69185882-DEL-127 |
| chr2-69188850-DEL-86 | chr2-69188851-DEL-86 |
| chr2-69204617-DEL-400 | chr2-69204618-DEL-400 |
| chr2-69226959-DEL-85 | chr2-69226942-DEL-103 |
| chr2-69233867-DEL-54 | chr2-69233868-DEL-50 |
| chr2-69239694-DEL-1596 | chr2-69239694-DEL-1596 |
| chr2-69246235-DEL-56 | chr2-69246236-DEL-56 |
| chr2-69274546-DEL-78 | chr2-69274557-DEL-68 |
| chr2-69293730-DEL-90 | chr2-69293721-DEL-66 |
| chr2-69295267-DEL-108 | chr2-69295268-DEL-108 |
| chr2-69298114-DEL-355 | chr2-69298115-DEL-355 |
| chr2-69304221-DEL-729 | chr2-69304222-DEL-729 |
| chr2-69326956-DEL-165 | chr2-69326957-DEL-165 |
| chr2-69372381-DEL-79 | chr2-69372338-DEL-116 |
| chr2-69374180-DEL-71 | chr2-69374181-DEL-71 |
| chr2-69374418-DEL-580 | chr2-69374421-DEL-568 |
| chr2-69398504-DEL-238 | chr2-69398505-DEL-238 |
| chr2-69401434-DEL-388 | chr2-69401435-DEL-388 |
| chr2-69404877-DEL-77 | chr2-69404876-DEL-75 |
| chr2-69424607-DEL-70 | chr2-69424608-DEL-70 |
| chr2-69428709-DEL-61 | chr2-69428710-DEL-61 |
| chr2-69470334-DEL-75 | chr2-69470335-DEL-75 |
| chr2-69476041-DEL-218 | chr2-69476042-DEL-218 |
| chr2-69479516-DEL-1424 | chr2-69479517-DEL-1424 |
| chr2-69489239-DEL-212 | chr2-69489240-DEL-212 |
| chr2-69505266-DEL-66 | chr2-69505256-DEL-75 |
| chr2-69514146-DEL-174 | chr2-69514147-DEL-174 |
| chr2-69561246-DEL-52 | chr2-69561262-DEL-74 |
| chr2-69570101-DEL-157 | chr2-69570102-DEL-157 |
| chr2-69573040-DEL-198 | chr2-69573041-DEL-198 |
| chr2-69602747-DEL-209 | chr2-69602748-DEL-209 |
| chr2-69629697-DEL-1298 | chr2-69629698-DEL-1298 |
| chr2-69633090-DEL-121 | chr2-69633091-DEL-121 |
| chr2-69638470-DEL-2202 | chr2-69638471-DEL-2202 |
| chr2-69647654-DEL-78 | chr2-69647655-DEL-78 |
| chr2-69659791-DEL-190 | chr2-69659792-DEL-190 |
| chr2-69660939-DEL-127 | chr2-69660940-DEL-127 |
| chr2-69675735-DEL-977 | chr2-69675736-DEL-977 |
| chr2-69682710-DEL-60 | chr2-69682719-DEL-58 |
| chr2-69705006-DEL-204 | chr2-69704999-DEL-205 |
| chr2-69706469-DEL-214 | chr2-69706470-DEL-214 |
| chr2-69737408-DEL-210 | chr2-69737409-DEL-210 |
| chr2-69737990-DEL-213 | chr2-69737991-DEL-213 |
| chr2-69751933-DEL-221 | chr2-69751934-DEL-221 |
| chr2-69757181-DEL-222 | chr2-69757182-DEL-222 |
| chr2-69771699-DEL-136 | chr2-69771700-DEL-136 |
| chr2-69792839-DEL-2591 | chr2-69792841-DEL-2590 |
| chr2-69795388-DEL-228 | chr2-69795389-DEL-228 |
| chr2-69796487-DEL-372 | chr2-69796492-DEL-369 |
| chr2-69804107-DEL-113 | chr2-69804103-DEL-82 |
| chr2-69817294-DEL-56 | chr2-69817295-DEL-56 |
| chr2-69824660-DEL-234 | chr2-69824661-DEL-232 |
| chr2-69826075-DEL-70 | chr2-69826075-DEL-71 |
| chr2-69873182-DEL-342 | chr2-69873183-DEL-342 |
| chr2-69886817-DEL-118 | chr2-69886816-DEL-115 |
| chr2-69892189-DEL-66 | chr2-69892190-DEL-66 |
| chr2-69902884-DEL-6188 | chr2-69902885-DEL-6188 |
| chr2-69954352-DEL-110 | chr2-69954353-DEL-92 |
| chr2-69976583-DEL-166 | chr2-69976584-DEL-166 |
| chr2-69979712-DEL-578 | chr2-69979697-DEL-576 |
| chr2-69982115-DEL-64 | chr2-69982116-DEL-64 |
| chr2-69985533-DEL-415 | chr2-69985533-DEL-415 |
| chr2-70030115-DEL-52 | chr2-70030114-DEL-54 |
| chr2-70057829-DEL-1265 | chr2-70057830-DEL-1282 |
| chr2-70093409-DEL-7728 | chr2-70093410-DEL-7727 |
| chr2-70104543-DEL-53 | chr2-70104544-DEL-53 |
| chr2-70108550-DEL-69 | chr2-70108551-DEL-69 |

| | |
|---|---|
| chr2-70157196-DEL-54 | chr2-70157194-DEL-58 |
| chr2-70236556-DEL-376 | chr2-70236542-DEL-367 |
| chr2-70262313-DEL-315 | chr2-70262316-DEL-315 |
| chr2-70271904-DEL-94 | chr2-70271905-DEL-94 |
| chr2-70289272-DEL-768 | chr2-70289273-DEL-768 |
| chr2-70295950-DEL-58 | chr2-70295923-DEL-50 |
| chr2-70329329-DEL-358 | chr2-70329330-DEL-358 |
| chr2-70348531-DEL-197 | chr2-70348532-DEL-197 |
| chr2-70365609-DEL-142 | chr2-70365608-DEL-139 |
| chr2-70385414-DEL-549 | chr2-70385415-DEL-549 |
| chr2-70430627-DEL-81 | chr2-70430607-DEL-78 |
| chr2-70441446-DEL-222 | chr2-70441447-DEL-222 |
| chr2-70446755-DEL-96 | chr2-70446756-DEL-94 |
| chr2-70471526-DEL-124 | chr2-70471527-DEL-124 |
| chr2-70540195-DEL-151 | chr2-70540219-DEL-108 |
| chr2-70609515-DEL-210 | chr2-70609516-DEL-210 |
| chr2-70612109-DEL-258 | chr2-70612110-DEL-258 |
| chr2-70630854-DEL-82 | chr2-70630855-DEL-82 |
| chr2-70664283-DEL-53 | chr2-70664284-DEL-53 |
| chr2-70701210-DEL-124 | chr2-70701211-DEL-124 |
| chr2-70710510-DEL-4568 | chr2-70710511-DEL-4568 |
| chr2-70718667-DEL-99 | chr2-70718662-DEL-111 |
| chr2-70783260-DEL-90 | chr2-70783258-DEL-93 |
| chr2-70793332-DEL-62 | chr2-70793333-DEL-62 |
| chr2-70803957-DEL-462 | chr2-70803971-DEL-495 |
| chr2-70812720-DEL-73 | chr2-70812721-DEL-73 |
| chr2-70817242-DEL-78 | chr2-70817243-DEL-78 |
| chr2-70828312-DEL-228 | chr2-70828313-DEL-228 |
| chr2-70832937-DEL-87 | chr2-70832938-DEL-87 |
| chr2-70840589-DEL-207 | chr2-70840581-DEL-226 |
| chr2-70851457-DEL-64 | chr2-70851448-DEL-64 |
| chr2-70869018-DEL-75 | chr2-70869019-DEL-75 |
| chr2-70878265-DEL-93 | chr2-70878266-DEL-93 |
| chr2-70879076-DEL-103 | chr2-70879101-DEL-79 |
| chr2-70895149-DEL-6440 | chr2-70895150-DEL-6440 |
| chr2-70909074-DEL-121 | chr2-70909071-DEL-113 |
| chr2-70956570-DEL-75 | chr2-70956571-DEL-75 |
| chr2-70959539-DEL-62 | chr2-70959540-DEL-62 |
| chr2-70964918-DEL-68 | chr2-70964919-DEL-68 |
| chr2-70988425-DEL-173 | chr2-70988426-DEL-173 |
| chr2-70988738-DEL-481 | chr2-70988739-DEL-481 |
| chr2-70997921-DEL-220 | chr2-70997929-DEL-220 |
| chr2-71000935-DEL-1549 | chr2-71000936-DEL-1549 |
| chr2-71038995-DEL-148 | chr2-71038996-DEL-148 |
| chr2-71060893-DEL-115 | chr2-71060894-DEL-115 |
| chr2-71083649-DEL-66 | chr2-71083650-DEL-66 |
| chr2-71089437-DEL-351 | chr2-71089434-DEL-352 |
| chr2-71099274-DEL-91 | chr2-71099275-DEL-91 |
| chr2-71103322-DEL-646 | chr2-71103323-DEL-646 |
| chr2-71127206-DEL-207 | chr2-71127207-DEL-207 |
| chr2-71134684-DEL-117 | chr2-71134676-DEL-122 |
| chr2-71138838-DEL-188 | chr2-71138863-DEL-204 |
| chr2-71140020-DEL-106 | chr2-71140021-DEL-106 |
| chr2-71150076-DEL-568 | chr2-71150077-DEL-568 |
| chr2-71182448-DEL-190 | chr2-71182449-DEL-190 |
| chr2-71186816-DEL-163 | chr2-71186817-DEL-162 |
| chr2-71187689-DEL-309 | chr2-71187690-DEL-309 |
| chr2-71191689-DEL-68 | chr2-71191690-DEL-72 |
| chr2-71194206-DEL-248 | chr2-71194207-DEL-248 |
| chr2-71207844-DEL-270 | chr2-71207845-DEL-270 |
| chr2-71228597-DEL-4687 | chr2-71228598-DEL-4687 |
| chr2-71229063-DEL-228 | chr2-71229064-DEL-228 |
| chr2-71230877-DEL-204 | chr2-71230878-DEL-204 |
| chr2-71232911-DEL-104 | chr2-71232892-DEL-125 |
| chr2-71234599-DEL-1786 | chr2-71234595-DEL-1787 |
| chr2-71247478-DEL-177 | chr2-71247479-DEL-177 |
| chr2-71261769-DEL-263 | chr2-71261770-DEL-263 |
| chr2-71273662-DEL-580 | chr2-71273663-DEL-580 |
| chr2-71281858-DEL-96 | chr2-71281839-DEL-116 |
| chr2-71295399-DEL-58 | chr2-71295400-DEL-58 |
| chr2-71299831-DEL-82 | chr2-71299831-DEL-82 |
| chr2-71316671-DEL-247 | chr2-71316668-DEL-248 |
| chr2-71317761-DEL-226 | chr2-71317762-DEL-226 |
| chr2-71319620-DEL-108 | chr2-71319621-DEL-108 |
| chr2-71380383-DEL-52 | chr2-71380384-DEL-52 |
| chr2-71394886-DEL-66 | chr2-71394893-DEL-60 |
| chr2-71412410-DEL-51 | chr2-71412410-DEL-51 |
| chr2-71434238-DEL-76 | chr2-71434239-DEL-76 |
| chr2-71441610-DEL-140 | chr2-71441601-DEL-144 |
| chr2-71447249-DEL-104 | chr2-71447250-DEL-104 |

| | |
|---|---|
| chr2-71479939-DEL-294 | chr2-71479940-DEL-294 |
| chr2-71537467-DEL-176 | chr2-71537468-DEL-177 |
| chr2-71596747-DEL-196 | chr2-71596748-DEL-196 |
| chr2-71604039-DEL-72 | chr2-71604040-DEL-72 |
| chr2-71607548-DEL-708 | chr2-71607549-DEL-708 |
| chr2-71645491-DEL-71 | chr2-71645492-DEL-71 |
| chr2-71682700-DEL-332 | chr2-71682701-DEL-332 |
| chr2-71712449-DEL-370 | chr2-71712450-DEL-370 |
| chr2-71736187-DEL-401 | chr2-71736188-DEL-401 |
| chr2-71752566-DEL-204 | chr2-71752567-DEL-204 |
| chr2-71755553-DEL-201 | chr2-71755554-DEL-201 |
| chr2-71772466-DEL-7170 | chr2-71772475-DEL-7165 |
| chr2-71783262-DEL-78 | chr2-71783265-DEL-76 |
| chr2-71792329-DEL-400 | chr2-71792330-DEL-400 |
| chr2-71793601-DEL-199 | chr2-71793592-DEL-200 |
| chr2-71803595-DEL-124 | chr2-71803596-DEL-124 |
| chr2-71846746-DEL-86 | chr2-71846747-DEL-86 |
| chr2-71851266-DEL-58 | chr2-71851267-DEL-58 |
| chr2-71887872-DEL-120 | chr2-71887873-DEL-120 |
| chr2-71905800-DEL-935 | chr2-71905801-DEL-935 |
| chr2-71927656-DEL-785 | chr2-71927657-DEL-785 |
| chr2-71937471-DEL-58 | chr2-71937474-DEL-60 |
| chr2-71954838-DEL-119 | chr2-71954815-DEL-120 |
| chr2-72023558-DEL-184 | chr2-72023559-DEL-184 |
| chr2-72058343-DEL-62 | chr2-72058344-DEL-62 |
| chr2-72079131-DEL-60 | chr2-72079093-DEL-60 |
| chr2-72099831-DEL-56 | chr2-72099832-DEL-56 |
| chr2-72100699-DEL-5244 | chr2-72100700-DEL-5244 |
| chr2-72120000-DEL-204 | chr2-72120001-DEL-204 |
| chr2-72149804-DEL-875 | chr2-72149805-DEL-875 |
| chr2-72156252-DEL-75 | chr2-72156253-DEL-75 |
| chr2-72159456-DEL-104 | chr2-72159457-DEL-104 |
| chr2-72274734-DEL-77 | chr2-72274735-DEL-76 |
| chr2-72283329-DEL-92 | chr2-72283330-DEL-92 |
| chr2-72290655-DEL-1313 | chr2-72290656-DEL-1313 |
| chr2-72293919-DEL-134 | chr2-72293920-DEL-128 |
| chr2-72300025-DEL-101 | chr2-72300026-DEL-101 |
| chr2-72340470-DEL-246 | chr2-72340471-DEL-246 |
| chr2-72369591-DEL-69 | chr2-72369567-DEL-70 |
| chr2-72382964-DEL-82 | chr2-72382965-DEL-82 |
| chr2-72389444-DEL-103 | chr2-72389445-DEL-103 |
| chr2-72396964-DEL-56 | chr2-72396965-DEL-56 |
| chr2-72438946-DEL-350 | chr2-72438947-DEL-350 |
| chr2-72464175-DEL-1651 | chr2-72464176-DEL-1651 |
| chr2-72466359-DEL-55 | chr2-72466360-DEL-55 |
| chr2-72469300-DEL-80 | chr2-72469301-DEL-80 |
| chr2-72472857-DEL-287 | chr2-72472858-DEL-287 |
| chr2-72485499-DEL-63 | chr2-72485491-DEL-65 |
| chr2-72489222-DEL-56 | chr2-72489229-DEL-50 |
| chr2-72495781-DEL-194 | chr2-72495782-DEL-194 |
| chr2-72540771-DEL-248 | chr2-72540772-DEL-248 |
| chr2-72572844-DEL-149 | chr2-72572845-DEL-149 |
| chr2-72611171-DEL-59 | chr2-72611170-DEL-60 |
| chr2-72613526-DEL-535 | chr2-72613527-DEL-535 |
| chr2-72633852-DEL-279 | chr2-72633857-DEL-275 |
| chr2-72639947-DEL-339 | chr2-72639954-DEL-345 |
| chr2-72641846-DEL-216 | chr2-72641847-DEL-216 |
| chr2-72651814-DEL-59 | chr2-72651800-DEL-52 |
| chr2-72693201-DEL-75 | chr2-72693202-DEL-75 |
| chr2-72707924-DEL-56 | chr2-72707917-DEL-64 |
| chr2-72722982-DEL-631 | chr2-72722983-DEL-631 |
| chr2-72752137-DEL-1585 | chr2-72752132-DEL-1586 |
| chr2-72784203-DEL-367 | chr2-72784198-DEL-368 |
| chr2-72784752-DEL-2298 | chr2-72784753-DEL-2298 |
| chr2-72812469-DEL-99 | chr2-72812470-DEL-99 |
| chr2-72829561-DEL-333 | chr2-72829562-DEL-333 |
| chr2-72859816-DEL-192 | chr2-72859817-DEL-192 |
| chr2-72900486-DEL-205 | chr2-72900487-DEL-205 |
| chr2-72978151-DEL-52 | chr2-72978152-DEL-52 |
| chr2-73032428-DEL-188 | chr2-73032429-DEL-187 |
| chr2-73055556-DEL-58 | chr2-73055557-DEL-58 |
| chr2-73069011-DEL-457 | chr2-73069012-DEL-457 |
| chr2-73074344-DEL-52 | chr2-73074346-DEL-50 |
| chr2-73080557-DEL-66 | chr2-73080558-DEL-61 |
| chr2-73084891-DEL-215 | chr2-73084892-DEL-215 |
| chr2-73088540-DEL-529 | chr2-73088538-DEL-530 |
| chr2-73124381-DEL-327 | chr2-73124382-DEL-327 |
| chr2-73132710-DEL-119 | chr2-73132711-DEL-119 |
| chr2-73191544-DEL-453 | chr2-73191545-DEL-453 |
| chr2-73257462-DEL-67 | chr2-73257499-DEL-67 |

| | |
|---|---|
| chr2-73266686-DEL-97 | chr2-73266687-DEL-95 |
| chr2-73277152-DEL-309 | chr2-73277153-DEL-309 |
| chr2-73279144-DEL-102 | chr2-73279145-DEL-54 |
| chr2-73281495-DEL-182 | chr2-73281496-DEL-182 |
| chr2-73296115-DEL-60 | chr2-73296116-DEL-55 |
| chr2-73303394-DEL-101 | chr2-73303395-DEL-101 |
| chr2-73343098-DEL-130 | chr2-73343099-DEL-130 |
| chr2-73347581-DEL-312 | chr2-73347582-DEL-313 |
| chr2-73362430-DEL-78 | chr2-73362431-DEL-78 |
| chr2-73386573-DEL-997 | chr2-73386574-DEL-997 |
| chr2-73394669-DEL-186 | chr2-73394670-DEL-186 |
| chr2-73429851-DEL-400 | chr2-73429852-DEL-400 |
| chr2-73444530-DEL-65 | chr2-73444531-DEL-65 |
| chr2-73578134-DEL-7133 | chr2-73578135-DEL-7133 |
| chr2-73592837-DEL-214 | chr2-73592838-DEL-214 |
| chr2-73602153-DEL-60 | chr2-73602154-DEL-60 |
| chr2-73617209-DEL-354 | chr2-73617210-DEL-354 |
| chr2-73624984-DEL-325 | chr2-73624985-DEL-325 |
| chr2-73630260-DEL-520 | chr2-73630261-DEL-520 |
| chr2-73641844-DEL-516 | chr2-73641845-DEL-516 |
| chr2-73668439-DEL-215 | chr2-73668440-DEL-215 |
| chr2-73674005-DEL-217 | chr2-73674006-DEL-217 |
| chr2-73700063-DEL-231 | chr2-73700064-DEL-231 |
| chr2-73736821-DEL-210 | chr2-73736822-DEL-210 |
| chr2-73744638-DEL-211 | chr2-73744633-DEL-212 |
| chr2-73752136-DEL-96 | chr2-73752137-DEL-96 |
| chr2-73768657-DEL-484 | chr2-73768658-DEL-484 |
| chr2-73778826-DEL-169 | chr2-73778827-DEL-169 |
| chr2-73805558-DEL-100 | chr2-73805559-DEL-100 |
| chr2-73816651-DEL-99 | chr2-73816650-DEL-100 |
| chr2-73817571-DEL-208 | chr2-73817572-DEL-208 |
| chr2-73831169-DEL-1581 | chr2-73831170-DEL-1581 |
| chr2-73833376-DEL-56 | chr2-73833383-DEL-50 |
| chr2-73837483-DEL-61 | chr2-73837488-DEL-57 |
| chr2-73839770-DEL-763 | chr2-73839782-DEL-756 |
| chr2-73842192-DEL-171 | chr2-73842193-DEL-171 |
| chr2-73871119-DEL-57 | chr2-73871120-DEL-57 |
| chr2-73874483-DEL-100 | chr2-73874484-DEL-100 |
| chr2-73898608-DEL-81 | chr2-73898609-DEL-81 |
| chr2-73907055-DEL-52 | chr2-73907024-DEL-88 |
| chr2-73920712-DEL-109 | chr2-73920713-DEL-109 |
| chr2-73943134-DEL-2515 | chr2-73943135-DEL-2515 |
| chr2-73973134-DEL-54 | chr2-73973123-DEL-66 |
| chr2-73989570-DEL-66 | chr2-73989571-DEL-65 |
| chr2-74056546-DEL-59 | chr2-74056547-DEL-59 |
| chr2-74068131-DEL-55 | chr2-74068120-DEL-67 |
| chr2-74175753-DEL-698 | chr2-74175754-DEL-698 |
| chr2-74194788-DEL-7129 | chr2-74194789-DEL-7129 |
| chr2-74205940-DEL-1962 | chr2-74205941-DEL-1962 |
| chr2-74226668-DEL-6268 | chr2-74226669-DEL-6268 |
| chr2-74268017-DEL-7162 | chr2-74268018-DEL-7162 |
| chr2-74286146-DEL-176 | chr2-74286147-DEL-164 |
| chr2-74332007-DEL-142 | chr2-74332008-DEL-142 |
| chr2-74390125-DEL-56 | chr2-74390126-DEL-56 |
| chr2-74405073-DEL-111 | chr2-74405058-DEL-127 |
| chr2-74421661-DEL-591 | chr2-74421662-DEL-591 |
| chr2-74430230-DEL-57 | chr2-74430242-DEL-53 |
| chr2-74515203-DEL-58 | chr2-74515200-DEL-56 |
| chr2-74541124-DEL-156 | chr2-74541125-DEL-155 |
| chr2-74548746-DEL-50 | chr2-74548728-DEL-68 |
| chr2-74589303-DEL-70 | chr2-74589304-DEL-70 |
| chr2-74612403-DEL-212 | chr2-74612404-DEL-212 |
| chr2-74616515-DEL-222 | chr2-74616516-DEL-222 |
| chr2-74634960-DEL-119 | chr2-74634961-DEL-119 |
| chr2-74733432-DEL-1576 | chr2-74733433-DEL-1576 |
| chr2-74819501-DEL-52 | chr2-74819501-DEL-52 |
| chr2-74821522-DEL-139 | chr2-74821521-DEL-141 |
| chr2-74842018-DEL-90 | chr2-74842019-DEL-90 |
| chr2-74874490-DEL-146 | chr2-74874479-DEL-158 |
| chr2-74880267-DEL-468 | chr2-74880268-DEL-468 |
| chr2-74883292-DEL-128 | chr2-74883278-DEL-129 |
| chr2-74888604-DEL-64 | chr2-74888605-DEL-64 |
| chr2-74898815-DEL-160 | chr2-74898816-DEL-160 |
| chr2-74916399-DEL-148 | chr2-74916400-DEL-148 |
| chr2-75021335-DEL-864 | chr2-75021336-DEL-864 |
| chr2-75057196-DEL-6739 | chr2-75057197-DEL-6739 |
| chr2-75081638-DEL-67 | chr2-75081639-DEL-67 |
| chr2-75104123-DEL-480 | chr2-75104124-DEL-480 |
| chr2-75106390-DEL-283 | chr2-75106379-DEL-284 |
| chr2-75165996-DEL-1211 | chr2-75165984-DEL-1212 |

| | |
|---|---|
| chr2-75170955-DEL-341 | chr2-75170956-DEL-341 |
| chr2-75247622-DEL-141 | chr2-75247623-DEL-141 |
| chr2-75263633-DEL-176 | chr2-75263634-DEL-176 |
| chr2-75271104-DEL-148 | chr2-75271105-DEL-148 |
| chr2-75290149-DEL-63 | chr2-75290150-DEL-63 |
| chr2-75300046-DEL-203 | chr2-75300047-DEL-203 |
| chr2-75301499-DEL-52 | chr2-75301500-DEL-52 |
| chr2-75317539-DEL-60 | chr2-75317519-DEL-106 |
| chr2-75329745-DEL-152 | chr2-75329746-DEL-152 |
| chr2-75346672-DEL-55 | chr2-75346673-DEL-55 |
| chr2-75367615-DEL-53 | chr2-75367614-DEL-54 |
| chr2-75382464-DEL-60 | chr2-75382467-DEL-84 |
| chr2-75399830-DEL-209 | chr2-75399831-DEL-209 |
| chr2-75405823-DEL-65 | chr2-75405824-DEL-65 |
| chr2-75406736-DEL-216 | chr2-75406731-DEL-217 |
| chr2-75421006-DEL-318 | chr2-75421007-DEL-318 |
| chr2-75421816-DEL-89 | chr2-75421840-DEL-65 |
| chr2-75427123-DEL-285 | chr2-75427123-DEL-284 |
| chr2-75447326-DEL-1422 | chr2-75447327-DEL-1422 |
| chr2-75465179-DEL-206 | chr2-75465180-DEL-206 |
| chr2-75468441-DEL-4848 | chr2-75468442-DEL-4848 |
| chr2-75474039-DEL-56 | chr2-75474039-DEL-56 |
| chr2-75477843-DEL-171 | chr2-75477826-DEL-171 |
| chr2-75520722-DEL-257 | chr2-75520723-DEL-257 |
| chr2-75553044-DEL-54 | chr2-75553037-DEL-62 |
| chr2-75557217-DEL-198 | chr2-75557218-DEL-198 |
| chr2-75571849-DEL-86 | chr2-75571850-DEL-86 |
| chr2-75591674-DEL-93 | chr2-75591675-DEL-91 |
| chr2-75613782-DEL-199 | chr2-75613783-DEL-199 |
| chr2-75621638-DEL-204 | chr2-75621654-DEL-189 |
| chr2-75622666-DEL-95 | chr2-75622682-DEL-66 |
| chr2-75630625-DEL-152 | chr2-75630626-DEL-152 |
| chr2-75641537-DEL-58 | chr2-75641536-DEL-59 |
| chr2-75691175-DEL-241 | chr2-75691176-DEL-241 |
| chr2-75711204-DEL-153 | chr2-75711205-DEL-153 |
| chr2-75750687-DEL-86 | chr2-75750688-DEL-86 |
| chr2-75771048-DEL-500 | chr2-75771050-DEL-499 |
| chr2-75777572-DEL-53 | chr2-75777573-DEL-53 |
| chr2-75875498-DEL-1474 | chr2-75875499-DEL-1475 |
| chr2-75879912-DEL-78 | chr2-75879913-DEL-78 |
| chr2-75891885-DEL-81 | chr2-75891913-DEL-60 |
| chr2-75914860-DEL-68 | chr2-75914861-DEL-68 |
| chr2-75936638-DEL-500 | chr2-75936639-DEL-500 |
| chr2-75961699-DEL-61 | chr2-75961700-DEL-61 |
| chr2-75985890-DEL-268 | chr2-75985889-DEL-269 |
| chr2-76024259-DEL-90 | chr2-76024261-DEL-89 |
| chr2-76059480-DEL-1102 | chr2-76059481-DEL-1102 |
| chr2-76068769-DEL-66 | chr2-76068770-DEL-66 |
| chr2-76071781-DEL-183 | chr2-76071782-DEL-183 |
| chr2-76081179-DEL-140 | chr2-76081180-DEL-140 |
| chr2-76096556-DEL-54 | chr2-76096553-DEL-54 |
| chr2-76104452-DEL-6257 | chr2-76104453-DEL-6257 |
| chr2-76116246-DEL-61 | chr2-76116253-DEL-56 |
| chr2-76138145-DEL-297 | chr2-76138146-DEL-297 |
| chr2-76158831-DEL-55 | chr2-76158835-DEL-53 |
| chr2-76194200-DEL-2042 | chr2-76194201-DEL-2042 |
| chr2-76230083-DEL-1595 | chr2-76230084-DEL-1595 |
| chr2-76252344-DEL-202 | chr2-76252345-DEL-202 |
| chr2-76263347-DEL-198 | chr2-76263348-DEL-198 |
| chr2-76266012-DEL-53 | chr2-76266029-DEL-60 |
| chr2-76280907-DEL-6594 | chr2-76280902-DEL-6595 |
| chr2-76309929-DEL-172 | chr2-76309930-DEL-172 |
| chr2-76310301-DEL-629 | chr2-76310302-DEL-629 |
| chr2-76324128-DEL-78 | chr2-76324129-DEL-78 |
| chr2-76332668-DEL-70 | chr2-76332669-DEL-72 |
| chr2-76336874-DEL-78 | chr2-76336875-DEL-78 |
| chr2-76345195-DEL-180 | chr2-76345196-DEL-180 |
| chr2-76346293-DEL-400 | chr2-76346294-DEL-400 |
| chr2-76348106-DEL-140 | chr2-76348107-DEL-140 |
| chr2-76365583-DEL-77 | chr2-76365584-DEL-72 |
| chr2-76389793-DEL-172 | chr2-76389794-DEL-172 |
| chr2-76404029-DEL-52 | chr2-76404030-DEL-52 |
| chr2-76409106-DEL-53 | chr2-76409107-DEL-53 |
| chr2-76445776-DEL-60 | chr2-76445777-DEL-60 |
| chr2-76450096-DEL-284 | chr2-76450097-DEL-284 |
| chr2-76498776-DEL-57 | chr2-76498776-DEL-57 |
| chr2-76509395-DEL-196 | chr2-76509396-DEL-197 |
| chr2-76511263-DEL-1518 | chr2-76511264-DEL-1517 |
| chr2-76516697-DEL-60 | chr2-76516698-DEL-60 |
| chr2-76650178-DEL-203 | chr2-76650179-DEL-203 |

| | |
|---|---|
| chr2-76741903-DEL-123 | chr2-76741895-DEL-124 |
| chr2-76785633-DEL-6224 | chr2-76785634-DEL-6224 |
| chr2-76815863-DEL-190 | chr2-76815864-DEL-190 |
| chr2-76825650-DEL-1914 | chr2-76825636-DEL-1915 |
| chr2-76869327-DEL-362 | chr2-76869288-DEL-396 |
| chr2-76906179-DEL-623 | chr2-76906180-DEL-623 |
| chr2-76922859-DEL-61 | chr2-76922860-DEL-56 |
| chr2-76928425-DEL-61 | chr2-76928426-DEL-61 |
| chr2-76931651-DEL-203 | chr2-76931652-DEL-203 |
| chr2-76946364-DEL-117 | chr2-76946365-DEL-117 |
| chr2-76951459-DEL-1332 | chr2-76951441-DEL-1316 |
| chr2-77007281-DEL-69 | chr2-77007270-DEL-52 |
| chr2-77030921-DEL-64 | chr2-77030922-DEL-64 |
| chr2-77054336-DEL-6415 | chr2-77054337-DEL-6415 |
| chr2-77121358-DEL-1919 | chr2-77121359-DEL-1919 |
| chr2-77125549-DEL-606 | chr2-77125550-DEL-606 |
| chr2-77139267-DEL-181 | chr2-77139268-DEL-181 |
| chr2-77145952-DEL-210 | chr2-77145953-DEL-210 |
| chr2-77151435-DEL-771 | chr2-77151436-DEL-771 |
| chr2-77185836-DEL-195 | chr2-77185837-DEL-195 |
| chr2-77282440-DEL-116 | chr2-77282441-DEL-116 |
| chr2-77296238-DEL-311 | chr2-77296243-DEL-307 |
| chr2-77321539-DEL-64 | chr2-77321540-DEL-64 |
| chr2-77358851-DEL-82 | chr2-77358852-DEL-82 |
| chr2-77368071-DEL-107 | chr2-77368072-DEL-107 |
| chr2-77376349-DEL-99 | chr2-77376350-DEL-99 |
| chr2-77380849-DEL-59 | chr2-77380850-DEL-59 |
| chr2-77418266-DEL-704 | chr2-77418267-DEL-704 |
| chr2-77429894-DEL-6468 | chr2-77429895-DEL-6468 |
| chr2-77437119-DEL-83 | chr2-77437120-DEL-83 |
| chr2-77450550-DEL-90 | chr2-77450551-DEL-60 |
| chr2-77462933-DEL-69 | chr2-77462930-DEL-73 |
| chr2-77477060-DEL-80 | chr2-77477061-DEL-104 |
| chr2-77488530-DEL-1208 | chr2-77488531-DEL-1208 |
| chr2-77505177-DEL-288 | chr2-77505178-DEL-288 |
| chr2-77507948-DEL-52 | chr2-77507949-DEL-52 |
| chr2-77536144-DEL-78 | chr2-77536161-DEL-78 |
| chr2-77541181-DEL-92 | chr2-77541177-DEL-92 |
| chr2-77557296-DEL-52 | chr2-77557297-DEL-52 |
| chr2-77590199-DEL-382 | chr2-77590201-DEL-381 |
| chr2-77659945-DEL-81 | chr2-77659991-DEL-53 |
| chr2-77663102-DEL-52 | chr2-77663099-DEL-56 |
| chr2-77674897-DEL-172 | chr2-77674898-DEL-172 |
| chr2-77701894-DEL-1177 | chr2-77701895-DEL-1177 |
| chr2-77732280-DEL-58 | chr2-77732292-DEL-84 |
| chr2-77734773-DEL-86 | chr2-77734778-DEL-82 |
| chr2-77773370-DEL-527 | chr2-77773371-DEL-527 |
| chr2-77779848-DEL-53 | chr2-77779841-DEL-54 |
| chr2-77788348-DEL-208 | chr2-77788349-DEL-208 |
| chr2-77843107-DEL-7004 | chr2-77843108-DEL-7004 |
| chr2-77857703-DEL-100 | chr2-77857703-DEL-98 |
| chr2-77897553-DEL-56 | chr2-77897548-DEL-62 |
| chr2-77933039-DEL-609 | chr2-77933049-DEL-601 |
| chr2-77936510-DEL-128 | chr2-77936511-DEL-128 |
| chr2-77955957-DEL-164 | chr2-77955974-DEL-168 |
| chr2-77970520-DEL-6550 | chr2-77970521-DEL-6550 |
| chr2-77990872-DEL-60 | chr2-77990873-DEL-60 |
| chr2-78032337-DEL-2344 | chr2-78032332-DEL-2346 |
| chr2-78067989-DEL-76 | chr2-78067993-DEL-75 |
| chr2-78074213-DEL-801 | chr2-78074214-DEL-801 |
| chr2-78093302-DEL-60 | chr2-78093287-DEL-63 |
| chr2-78146490-DEL-475 | chr2-78146500-DEL-471 |
| chr2-78147153-DEL-2801 | chr2-78147154-DEL-2801 |
| chr2-78160812-DEL-66 | chr2-78160813-DEL-66 |
| chr2-78163639-DEL-6277 | chr2-78163640-DEL-6277 |
| chr2-78214069-DEL-683 | chr2-78214070-DEL-683 |
| chr2-78221843-DEL-65 | chr2-78221844-DEL-65 |
| chr2-78250188-DEL-84 | chr2-78250189-DEL-84 |
| chr2-78263441-DEL-56 | chr2-78263430-DEL-76 |
| chr2-78264672-DEL-836 | chr2-78264673-DEL-836 |
| chr2-78296377-DEL-127 | chr2-78296377-DEL-128 |
| chr2-78335222-DEL-6566 | chr2-78335220-DEL-6567 |
| chr2-78425595-DEL-106 | chr2-78425597-DEL-104 |
| chr2-78447177-DEL-6605 | chr2-78447178-DEL-6605 |
| chr2-78456897-DEL-924 | chr2-78456898-DEL-924 |
| chr2-78468984-DEL-84 | chr2-78468974-DEL-92 |
| chr2-78472866-DEL-632 | chr2-78472866-DEL-632 |
| chr2-78491003-DEL-1481 | chr2-78491003-DEL-1481 |
| chr2-78492505-DEL-6582 | chr2-78492505-DEL-6582 |
| chr2-78502154-DEL-626 | chr2-78502158-DEL-627 |

| | |
|---|---|
| chr2-78515013-DEL-332 | chr2-78515014-DEL-332 |
| chr2-78542191-DEL-172 | chr2-78542192-DEL-172 |
| chr2-78622488-DEL-74 | chr2-78622489-DEL-74 |
| chr2-78626608-DEL-1284 | chr2-78626609-DEL-1283 |
| chr2-78636932-DEL-97 | chr2-78636935-DEL-104 |
| chr2-78639995-DEL-528 | chr2-78639996-DEL-528 |
| chr2-78643085-DEL-140 | chr2-78643078-DEL-141 |
| chr2-78664144-DEL-55 | chr2-78664145-DEL-55 |
| chr2-78705844-DEL-88 | chr2-78705845-DEL-88 |
| chr2-78718088-DEL-66 | chr2-78718069-DEL-62 |
| chr2-78778479-DEL-6176 | chr2-78778480-DEL-6176 |
| chr2-78827759-DEL-56 | chr2-78827760-DEL-56 |
| chr2-78836640-DEL-60 | chr2-78836626-DEL-58 |
| chr2-78852419-DEL-210 | chr2-78852410-DEL-191 |
| chr2-78858160-DEL-54 | chr2-78858159-DEL-54 |
| chr2-78875382-DEL-165 | chr2-78875383-DEL-165 |
| chr2-78886276-DEL-54 | chr2-78886277-DEL-54 |
| chr2-78921286-DEL-409 | chr2-78921287-DEL-409 |
| chr2-78947179-DEL-1383 | chr2-78947180-DEL-1383 |
| chr2-78954903-DEL-208 | chr2-78954894-DEL-209 |
| chr2-78964249-DEL-2128 | chr2-78964250-DEL-2128 |
| chr2-78990803-DEL-2545 | chr2-78990804-DEL-2545 |
| chr2-79046482-DEL-81 | chr2-79046483-DEL-81 |
| chr2-79047128-DEL-181 | chr2-79047129-DEL-181 |
| chr2-79053390-DEL-120 | chr2-79053391-DEL-120 |
| chr2-79071463-DEL-70 | chr2-79071474-DEL-60 |
| chr2-79120476-DEL-52 | chr2-79120477-DEL-52 |
| chr2-79123980-DEL-799 | chr2-79123981-DEL-799 |
| chr2-79130679-DEL-541 | chr2-79130684-DEL-535 |
| chr2-79135664-DEL-598 | chr2-79135665-DEL-598 |
| chr2-79137289-DEL-52 | chr2-79137290-DEL-50 |
| chr2-79242452-DEL-100 | chr2-79242453-DEL-100 |
| chr2-79267595-DEL-4800 | chr2-79267612-DEL-4800 |
| chr2-79272957-DEL-75 | chr2-79272961-DEL-98 |
| chr2-79300536-DEL-294 | chr2-79300539-DEL-293 |
| chr2-79318182-DEL-1892 | chr2-79318183-DEL-1892 |
| chr2-79340516-DEL-103 | chr2-79340517-DEL-103 |
| chr2-79344577-DEL-87 | chr2-79344578-DEL-87 |
| chr2-79384528-DEL-52 | chr2-79384529-DEL-52 |
| chr2-79394555-DEL-84 | chr2-79394556-DEL-84 |
| chr2-79404766-DEL-179 | chr2-79404767-DEL-179 |
| chr2-79507286-DEL-204 | chr2-79507287-DEL-204 |
| chr2-79511004-DEL-52 | chr2-79511025-DEL-68 |
| chr2-79518867-DEL-74 | chr2-79518867-DEL-72 |
| chr2-79542549-DEL-53 | chr2-79542551-DEL-52 |
| chr2-79546811-DEL-178 | chr2-79546812-DEL-178 |
| chr2-79567681-DEL-1467 | chr2-79567674-DEL-1462 |
| chr2-79577082-DEL-115 | chr2-79577083-DEL-115 |
| chr2-79585844-DEL-345 | chr2-79585845-DEL-345 |
| chr2-79597036-DEL-408 | chr2-79597037-DEL-408 |
| chr2-79604616-DEL-65 | chr2-79604620-DEL-64 |
| chr2-79608065-DEL-78 | chr2-79608066-DEL-78 |
| chr2-79608322-DEL-1224 | chr2-79608345-DEL-1212 |
| chr2-79635323-DEL-85 | chr2-79635324-DEL-85 |
| chr2-79641558-DEL-53 | chr2-79641559-DEL-53 |
| chr2-79670982-DEL-62 | chr2-79670979-DEL-54 |
| chr2-79690728-DEL-1474 | chr2-79690729-DEL-1474 |
| chr2-79702245-DEL-52 | chr2-79702246-DEL-52 |
| chr2-79708942-DEL-52 | chr2-79708943-DEL-50 |
| chr2-79752056-DEL-64 | chr2-79752042-DEL-86 |
| chr2-79766987-DEL-7328 | chr2-79766988-DEL-7328 |
| chr2-79767123-DEL-211 | chr2-79767124-DEL-211 |
| chr2-79780982-DEL-106 | chr2-79780983-DEL-104 |
| chr2-79799686-DEL-2386 | chr2-79799687-DEL-2386 |
| chr2-79810543-DEL-406 | chr2-79810543-DEL-406 |
| chr2-79826500-DEL-1533 | chr2-79826501-DEL-1533 |
| chr2-79846454-DEL-352 | chr2-79846455-DEL-352 |
| chr2-79881011-DEL-61 | chr2-79881014-DEL-58 |
| chr2-79978909-DEL-227 | chr2-79978910-DEL-227 |
| chr2-79981595-DEL-52 | chr2-79981596-DEL-54 |
| chr2-80044966-DEL-1009 | chr2-80044967-DEL-1009 |
| chr2-80056413-DEL-4757 | chr2-80056414-DEL-4757 |
| chr2-80061662-DEL-69 | chr2-80061663-DEL-69 |
| chr2-80064730-DEL-183 | chr2-80064731-DEL-183 |
| chr2-80081419-DEL-6748 | chr2-80081420-DEL-6748 |
| chr2-80110285-DEL-59 | chr2-80110286-DEL-59 |
| chr2-80117984-DEL-90 | chr2-80117985-DEL-90 |
| chr2-80118718-DEL-753 | chr2-80118719-DEL-753 |
| chr2-80127119-DEL-797 | chr2-80127120-DEL-797 |
| chr2-80133122-DEL-70 | chr2-80133123-DEL-70 |

| | |
|---|---|
| chr2-80137055-DEL-248 | chr2-80137056-DEL-248 |
| chr2-80139428-DEL-63 | chr2-80139474-DEL-54 |
| chr2-80201182-DEL-112 | chr2-80201183-DEL-112 |
| chr2-80203655-DEL-164 | chr2-80203647-DEL-173 |
| chr2-80209477-DEL-65 | chr2-80209480-DEL-66 |
| chr2-80213823-DEL-64 | chr2-80213823-DEL-64 |
| chr2-80238236-DEL-187 | chr2-80238237-DEL-187 |
| chr2-80250236-DEL-134 | chr2-80250237-DEL-134 |
| chr2-80253038-DEL-343 | chr2-80253039-DEL-343 |
| chr2-80284142-DEL-3792 | chr2-80284143-DEL-3792 |
| chr2-80284998-DEL-132 | chr2-80284999-DEL-132 |
| chr2-80298189-DEL-71 | chr2-80298187-DEL-68 |
| chr2-80306685-DEL-685 | chr2-80306686-DEL-685 |
| chr2-80352400-DEL-217 | chr2-80352401-DEL-217 |
| chr2-80387833-DEL-204 | chr2-80387834-DEL-204 |
| chr2-80437638-DEL-62 | chr2-80437640-DEL-60 |
| chr2-80456667-DEL-207 | chr2-80456668-DEL-207 |
| chr2-80458969-DEL-1014 | chr2-80458970-DEL-1014 |
| chr2-80478696-DEL-512 | chr2-80478697-DEL-512 |
| chr2-80482334-DEL-247 | chr2-80482335-DEL-247 |
| chr2-80492452-DEL-2000 | chr2-80492453-DEL-2000 |
| chr2-80507026-DEL-103 | chr2-80507016-DEL-122 |
| chr2-80528931-DEL-427 | chr2-80528932-DEL-427 |
| chr2-80562681-DEL-464 | chr2-80562682-DEL-464 |
| chr2-80642336-DEL-4368 | chr2-80642335-DEL-4368 |
| chr2-80667423-DEL-1689 | chr2-80667424-DEL-1689 |
| chr2-80673698-DEL-221 | chr2-80673699-DEL-221 |
| chr2-80686742-DEL-108 | chr2-80686743-DEL-109 |
| chr2-80707047-DEL-120 | chr2-80707048-DEL-116 |
| chr2-80709280-DEL-1335 | chr2-80709281-DEL-1334 |
| chr2-80786847-DEL-7421 | chr2-80786848-DEL-7421 |
| chr2-80806564-DEL-3355 | chr2-80806581-DEL-3338 |
| chr2-80829715-DEL-6720 | chr2-80829716-DEL-6720 |
| chr2-80835959-DEL-424 | chr2-80835960-DEL-424 |
| chr2-80879533-DEL-1536 | chr2-80879534-DEL-1536 |
| chr2-80884226-DEL-943 | chr2-80884227-DEL-943 |
| chr2-80987832-DEL-201 | chr2-80987833-DEL-201 |
| chr2-81013261-DEL-361 | chr2-81013262-DEL-361 |
| chr2-81086662-DEL-487 | chr2-81086663-DEL-487 |
| chr2-81122289-DEL-579 | chr2-81122290-DEL-579 |
| chr2-81142265-DEL-214 | chr2-81142266-DEL-214 |
| chr2-81150365-DEL-461 | chr2-81150366-DEL-461 |
| chr2-81168791-DEL-81 | chr2-81168792-DEL-81 |
| chr2-81171783-DEL-52 | chr2-81171784-DEL-60 |
| chr2-81197089-DEL-390 | chr2-81197089-DEL-390 |
| chr2-81204453-DEL-87 | chr2-81204481-DEL-77 |
| chr2-81288938-DEL-600 | chr2-81288941-DEL-599 |
| chr2-81305490-DEL-63 | chr2-81305491-DEL-63 |
| chr2-81337840-DEL-4306 | chr2-81337841-DEL-4306 |
| chr2-81350780-DEL-68 | chr2-81350777-DEL-58 |
| chr2-81394862-DEL-351 | chr2-81394863-DEL-351 |
| chr2-81445900-DEL-367 | chr2-81445895-DEL-368 |
| chr2-81469644-DEL-5731 | chr2-81469645-DEL-5731 |
| chr2-81475082-DEL-250 | chr2-81475034-DEL-249 |
| chr2-81530390-DEL-888 | chr2-81530388-DEL-889 |
| chr2-81620375-DEL-5770 | chr2-81620376-DEL-5770 |
| chr2-81680655-DEL-81 | chr2-81680656-DEL-81 |
| chr2-81683399-DEL-2097 | chr2-81683400-DEL-2097 |
| chr2-81694366-DEL-57 | chr2-81694367-DEL-57 |
| chr2-81741880-DEL-94 | chr2-81741925-DEL-75 |
| chr2-81752513-DEL-178 | chr2-81752515-DEL-177 |
| chr2-81753511-DEL-75 | chr2-81753509-DEL-80 |
| chr2-81769199-DEL-356 | chr2-81769232-DEL-331 |
| chr2-81771894-DEL-2617 | chr2-81771895-DEL-2617 |
| chr2-81783475-DEL-85 | chr2-81783465-DEL-84 |
| chr2-81790721-DEL-394 | chr2-81790722-DEL-394 |
| chr2-81819458-DEL-98 | chr2-81819459-DEL-98 |
| chr2-81842587-DEL-66 | chr2-81842588-DEL-74 |
| chr2-81955459-DEL-566 | chr2-81955460-DEL-566 |
| chr2-81985036-DEL-59 | chr2-81985027-DEL-70 |
| chr2-82009947-DEL-578 | chr2-82009948-DEL-578 |
| chr2-82033697-DEL-6351 | chr2-82033698-DEL-6351 |
| chr2-82155629-DEL-91 | chr2-82155634-DEL-85 |
| chr2-82160276-DEL-10200 | chr2-82160277-DEL-10200 |
| chr2-82165652-DEL-4062 | chr2-82165651-DEL-4062 |
| chr2-82175451-DEL-68 | chr2-82175439-DEL-59 |
| chr2-82191050-DEL-56 | chr2-82191051-DEL-56 |
| chr2-82218830-DEL-162 | chr2-82218831-DEL-162 |
| chr2-82230474-DEL-216 | chr2-82230475-DEL-216 |
| chr2-82276878-DEL-5947 | chr2-82276880-DEL-5946 |

| | |
|---|---|
| chr2-82316048-DEL-10342 | chr2-82316064-DEL-10343 |
| chr2-82326008-DEL-212 | chr2-82326009-DEL-212 |
| chr2-82433840-DEL-204 | chr2-82433840-DEL-204 |
| chr2-82493641-DEL-102 | chr2-82493652-DEL-92 |
| chr2-82504116-DEL-6001 | chr2-82504117-DEL-6001 |
| chr2-82539832-DEL-349 | chr2-82539833-DEL-349 |
| chr2-82552317-DEL-6478 | chr2-82552318-DEL-6478 |
| chr2-82560866-DEL-58 | chr2-82560867-DEL-58 |
| chr2-82616141-DEL-513 | chr2-82616142-DEL-513 |
| chr2-82649474-DEL-77 | chr2-82649475-DEL-77 |
| chr2-82668108-DEL-68 | chr2-82668121-DEL-56 |
| chr2-82680529-DEL-433 | chr2-82680530-DEL-433 |
| chr2-82712884-DEL-6483 | chr2-82712883-DEL-6482 |
| chr2-82719116-DEL-214 | chr2-82719117-DEL-214 |
| chr2-82746546-DEL-6696 | chr2-82746547-DEL-6696 |
| chr2-82776618-DEL-164 | chr2-82776619-DEL-164 |
| chr2-82789802-DEL-1642 | chr2-82789803-DEL-1642 |
| chr2-82858761-DEL-56 | chr2-82858762-DEL-56 |
| chr2-82859744-DEL-1182 | chr2-82859744-DEL-1182 |
| chr2-82864090-DEL-1738 | chr2-82864091-DEL-1738 |
| chr2-82878601-DEL-2739 | chr2-82878602-DEL-2739 |
| chr2-82948375-DEL-4573 | chr2-82948376-DEL-4573 |
| chr2-82952236-DEL-81 | chr2-82952237-DEL-81 |
| chr2-82964939-DEL-1606 | chr2-82964940-DEL-1606 |
| chr2-82977831-DEL-531 | chr2-82977832-DEL-531 |
| chr2-83004128-DEL-765 | chr2-83004129-DEL-765 |
| chr2-83015524-DEL-6401 | chr2-83015525-DEL-6401 |
| chr2-83058052-DEL-3209 | chr2-83058050-DEL-3196 |
| chr2-83063735-DEL-2763 | chr2-83063735-DEL-2762 |
| chr2-83138728-DEL-586 | chr2-83138729-DEL-586 |
| chr2-83158755-DEL-277 | chr2-83158756-DEL-277 |
| chr2-83169971-DEL-509 | chr2-83169960-DEL-510 |
| chr2-83202809-DEL-1732 | chr2-83202810-DEL-1731 |
| chr2-83244125-DEL-424 | chr2-83244126-DEL-424 |
| chr2-83260379-DEL-636 | chr2-83260379-DEL-636 |
| chr2-83268996-DEL-68 | chr2-83268997-DEL-68 |
| chr2-83323453-DEL-91 | chr2-83323452-DEL-92 |
| chr2-83324025-DEL-1498 | chr2-83324024-DEL-1500 |
| chr2-83338256-DEL-341 | chr2-83338257-DEL-339 |
| chr2-83342839-DEL-796 | chr2-83342846-DEL-788 |
| chr2-83343624-DEL-3271 | chr2-83343625-DEL-3271 |
| chr2-83351077-DEL-51 | chr2-83351078-DEL-51 |
| chr2-83361043-DEL-1901 | chr2-83361044-DEL-1901 |
| chr2-83410637-DEL-7142 | chr2-83410637-DEL-7142 |
| chr2-83493589-DEL-224 | chr2-83493590-DEL-224 |
| chr2-83499428-DEL-90 | chr2-83499429-DEL-90 |
| chr2-83527563-DEL-307 | chr2-83527571-DEL-308 |
| chr2-83529512-DEL-140 | chr2-83529524-DEL-134 |
| chr2-83541540-DEL-77 | chr2-83541541-DEL-77 |
| chr2-83548145-DEL-299 | chr2-83548146-DEL-299 |
| chr2-83550000-DEL-110 | chr2-83550003-DEL-128 |
| chr2-83565025-DEL-173 | chr2-83565026-DEL-173 |
| chr2-83572513-DEL-56 | chr2-83572514-DEL-56 |
| chr2-83579962-DEL-213 | chr2-83579964-DEL-194 |
| chr2-83615108-DEL-173 | chr2-83615109-DEL-173 |
| chr2-83658118-DEL-200 | chr2-83658119-DEL-200 |
| chr2-83682562-DEL-1088 | chr2-83682563-DEL-1088 |
| chr2-83696017-DEL-6546 | chr2-83696018-DEL-6546 |
| chr2-83715152-DEL-169 | chr2-83715153-DEL-169 |
| chr2-83728336-DEL-769 | chr2-83728337-DEL-769 |
| chr2-83737085-DEL-72 | chr2-83737085-DEL-71 |
| chr2-83901880-DEL-117 | chr2-83901883-DEL-123 |
| chr2-83909378-DEL-189 | chr2-83909379-DEL-189 |
| chr2-83911208-DEL-6464 | chr2-83911209-DEL-6464 |
| chr2-83932661-DEL-1102 | chr2-83932659-DEL-1103 |
| chr2-83941737-DEL-425 | chr2-83941732-DEL-426 |
| chr2-83952696-DEL-383 | chr2-83952697-DEL-383 |
| chr2-83994579-DEL-6604 | chr2-83994579-DEL-6604 |
| chr2-84007023-DEL-1019 | chr2-84007014-DEL-1020 |
| chr2-84008626-DEL-430 | chr2-84008627-DEL-430 |
| chr2-84020715-DEL-966 | chr2-84020716-DEL-966 |
| chr2-84066383-DEL-340 | chr2-84066381-DEL-340 |
| chr2-84105766-DEL-72 | chr2-84105767-DEL-72 |
| chr2-84172762-DEL-92 | chr2-84172759-DEL-74 |
| chr2-84215056-DEL-52 | chr2-84215059-DEL-52 |
| chr2-84222177-DEL-491 | chr2-84222178-DEL-491 |
| chr2-84230153-DEL-796 | chr2-84230153-DEL-796 |
| chr2-84250196-DEL-193 | chr2-84250197-DEL-193 |
| chr2-84266089-DEL-159 | chr2-84266090-DEL-159 |
| chr2-84332954-DEL-190 | chr2-84332955-DEL-190 |

| | |
|---|---|
| chr2-84335547-DEL-5218 | chr2-84335548-DEL-5218 |
| chr2-84353702-DEL-123 | chr2-84353700-DEL-126 |
| chr2-84362373-DEL-72 | chr2-84362380-DEL-66 |
| chr2-84362989-DEL-1885 | chr2-84362990-DEL-1885 |
| chr2-84367686-DEL-56 | chr2-84367686-DEL-56 |
| chr2-84377277-DEL-354 | chr2-84377278-DEL-354 |
| chr2-84395339-DEL-7125 | chr2-84395340-DEL-7125 |
| chr2-84407312-DEL-328 | chr2-84407289-DEL-327 |
| chr2-84411853-DEL-209 | chr2-84411854-DEL-209 |
| chr2-84419505-DEL-74 | chr2-84419506-DEL-74 |
| chr2-84434524-DEL-80 | chr2-84434525-DEL-80 |
| chr2-84475774-DEL-185 | chr2-84475775-DEL-185 |
| chr2-84484302-DEL-214 | chr2-84484303-DEL-214 |
| chr2-84490055-DEL-266 | chr2-84490056-DEL-266 |
| chr2-84512916-DEL-52 | chr2-84512897-DEL-96 |
| chr2-84538574-DEL-337 | chr2-84538575-DEL-337 |
| chr2-84553093-DEL-258 | chr2-84553094-DEL-258 |
| chr2-84575354-DEL-70 | chr2-84575354-DEL-70 |
| chr2-84615652-DEL-52 | chr2-84615653-DEL-52 |
| chr2-84616604-DEL-72 | chr2-84616604-DEL-72 |
| chr2-84623280-DEL-99 | chr2-84623281-DEL-87 |
| chr2-84629357-DEL-151 | chr2-84629358-DEL-151 |
| chr2-84663810-DEL-236 | chr2-84663798-DEL-237 |
| chr2-84728876-DEL-8873 | chr2-84728877-DEL-8873 |
| chr2-84756928-DEL-1490 | chr2-84756929-DEL-1490 |
| chr2-84795446-DEL-127 | chr2-84795444-DEL-128 |
| chr2-84908864-DEL-6728 | chr2-84908865-DEL-6728 |
| chr2-84938597-DEL-441 | chr2-84938598-DEL-441 |
| chr2-84942661-DEL-343 | chr2-84942662-DEL-341 |
| chr2-84942965-DEL-3355 | chr2-84942975-DEL-3352 |
| chr2-84991878-DEL-85 | chr2-84991868-DEL-76 |
| chr2-85014720-DEL-52 | chr2-85014721-DEL-52 |
| chr2-85036324-DEL-314 | chr2-85036325-DEL-314 |
| chr2-85063577-DEL-400 | chr2-85063560-DEL-409 |
| chr2-85084369-DEL-166 | chr2-85084363-DEL-173 |
| chr2-85111833-DEL-194 | chr2-85111830-DEL-198 |
| chr2-85112439-DEL-441 | chr2-85112440-DEL-441 |
| chr2-85124551-DEL-6519 | chr2-85124552-DEL-6519 |
| chr2-85147952-DEL-80 | chr2-85147952-DEL-80 |
| chr2-85186763-DEL-413 | chr2-85186764-DEL-413 |
| chr2-85195478-DEL-1509 | chr2-85195479-DEL-1509 |
| chr2-85248254-DEL-129 | chr2-85248267-DEL-128 |
| chr2-85273775-DEL-85 | chr2-85273776-DEL-85 |
| chr2-85313393-DEL-91 | chr2-85313394-DEL-91 |
| chr2-85322836-DEL-52 | chr2-85322871-DEL-53 |
| chr2-85341678-DEL-52 | chr2-85341668-DEL-50 |
| chr2-85386999-DEL-1748 | chr2-85387000-DEL-1748 |
| chr2-85416565-DEL-64 | chr2-85416566-DEL-64 |
| chr2-85436111-DEL-70 | chr2-85436112-DEL-70 |
| chr2-85437997-DEL-62 | chr2-85437998-DEL-62 |
| chr2-85453314-DEL-5703 | chr2-85453315-DEL-5703 |
| chr2-85471457-DEL-860 | chr2-85471458-DEL-860 |
| chr2-85472735-DEL-3522 | chr2-85472736-DEL-3522 |
| chr2-85511198-DEL-1874 | chr2-85511200-DEL-1872 |
| chr2-85512446-DEL-301 | chr2-85512447-DEL-301 |
| chr2-85524916-DEL-7116 | chr2-85524917-DEL-7116 |
| chr2-85587642-DEL-353 | chr2-85587643-DEL-353 |
| chr2-85592111-DEL-490 | chr2-85592112-DEL-490 |
| chr2-85620928-DEL-178 | chr2-85620929-DEL-178 |
| chr2-85635661-DEL-62 | chr2-85635661-DEL-60 |
| chr2-85643122-DEL-2757 | chr2-85643123-DEL-2757 |
| chr2-85648207-DEL-144 | chr2-85648208-DEL-144 |
| chr2-85651119-DEL-51 | chr2-85651120-DEL-51 |
| chr2-85707166-DEL-1259 | chr2-85707162-DEL-1260 |
| chr2-85732116-DEL-561 | chr2-85732117-DEL-561 |
| chr2-85774913-DEL-76 | chr2-85774888-DEL-108 |
| chr2-85788803-DEL-124 | chr2-85788804-DEL-124 |
| chr2-85823815-DEL-939 | chr2-85823816-DEL-939 |
| chr2-85841972-DEL-189 | chr2-85841973-DEL-189 |
| chr2-85845131-DEL-56 | chr2-85845131-DEL-56 |
| chr2-85848705-DEL-5223 | chr2-85848706-DEL-5223 |
| chr2-85861448-DEL-192 | chr2-85861459-DEL-194 |
| chr2-85880317-DEL-1289 | chr2-85880318-DEL-1289 |
| chr2-85933609-DEL-61 | chr2-85933608-DEL-63 |
| chr2-85961768-DEL-15071 | chr2-85961767-DEL-15071 |
| chr2-86019386-DEL-1697 | chr2-86019387-DEL-1697 |
| chr2-86024683-DEL-107 | chr2-86024657-DEL-84 |
| chr2-86052386-DEL-255 | chr2-86052387-DEL-255 |
| chr2-86085856-DEL-17790 | chr2-86085856-DEL-17790 |
| chr2-86121972-DEL-56 | chr2-86121977-DEL-54 |

| | |
|---|---|
| chr2-86153623-DEL-52 | chr2-86153624-DEL-52 |
| chr2-86170084-DEL-7085 | chr2-86170085-DEL-7085 |
| chr2-86182945-DEL-57 | chr2-86182917-DEL-58 |
| chr2-86196518-DEL-2065 | chr2-86196521-DEL-2063 |
| chr2-86261196-DEL-58 | chr2-86261197-DEL-58 |
| chr2-86286203-DEL-59 | chr2-86286204-DEL-59 |
| chr2-86290651-DEL-138 | chr2-86290650-DEL-137 |
| chr2-86294951-DEL-332 | chr2-86294952-DEL-332 |
| chr2-86299744-DEL-930 | chr2-86299745-DEL-930 |
| chr2-86322015-DEL-5262 | chr2-86322016-DEL-5262 |
| chr2-86329424-DEL-3113 | chr2-86329425-DEL-3113 |
| chr2-86389434-DEL-68 | chr2-86389443-DEL-60 |
| chr2-86389601-DEL-1107 | chr2-86389602-DEL-1107 |
| chr2-86391508-DEL-53 | chr2-86391513-DEL-55 |
| chr2-86392131-DEL-2229 | chr2-86392132-DEL-2229 |
| chr2-86403256-DEL-2784 | chr2-86403257-DEL-2784 |
| chr2-86446638-DEL-56 | chr2-86446636-DEL-57 |
| chr2-86454352-DEL-54 | chr2-86454353-DEL-54 |
| chr2-86479881-DEL-374 | chr2-86479882-DEL-374 |
| chr2-86527799-DEL-1405 | chr2-86527800-DEL-1405 |
| chr2-86557561-DEL-817 | chr2-86557561-DEL-817 |
| chr2-86572904-DEL-726 | chr2-86572905-DEL-726 |
| chr2-86580429-DEL-1039 | chr2-86580430-DEL-1039 |
| chr2-86672859-DEL-7515 | chr2-86672860-DEL-7515 |
| chr2-86672916-DEL-208 | chr2-86672917-DEL-208 |
| chr2-86727766-DEL-78 | chr2-86727767-DEL-78 |
| chr2-86763221-DEL-73 | chr2-86763231-DEL-52 |
| chr2-86824728-DEL-843 | chr2-86824729-DEL-843 |
| chr2-86891191-DEL-400 | chr2-86891192-DEL-400 |
| chr2-86893496-DEL-152 | chr2-86893497-DEL-152 |
| chr2-86894921-DEL-148 | chr2-86894921-DEL-131 |
| chr2-86906500-DEL-192 | chr2-86906499-DEL-193 |
| chr2-86915209-DEL-316 | chr2-86915210-DEL-316 |
| chr2-86929811-DEL-193 | chr2-86929812-DEL-193 |
| chr2-86936249-DEL-192 | chr2-86936253-DEL-192 |
| chr2-86942975-DEL-61 | chr2-86942972-DEL-55 |
| chr2-86976437-DEL-74 | chr2-86976438-DEL-74 |
| chr2-86989112-DEL-59 | chr2-86989113-DEL-59 |
| chr2-86991303-DEL-4999 | chr2-86991303-DEL-4999 |
| chr2-87015869-DEL-490 | chr2-87015869-DEL-490 |
| chr2-87017586-DEL-84 | chr2-87017596-DEL-69 |
| chr2-87156067-DEL-51 | chr2-87156063-DEL-50 |
| chr2-87166591-DEL-1725 | chr2-87166592-DEL-1725 |
| chr2-87169225-DEL-135 | chr2-87169210-DEL-147 |
| chr2-87199812-DEL-1895 | chr2-87199813-DEL-1895 |
| chr2-87207658-DEL-397 | chr2-87207659-DEL-397 |
| chr2-87211333-DEL-1907 | chr2-87211318-DEL-1908 |
| chr2-87221153-DEL-184 | chr2-87221154-DEL-184 |
| chr2-87243518-DEL-1300 | chr2-87243517-DEL-1301 |
| chr2-87253130-DEL-60 | chr2-87253131-DEL-50 |
| chr2-87265779-DEL-175 | chr2-87265780-DEL-175 |
| chr2-87266589-DEL-85 | chr2-87266590-DEL-85 |
| chr2-87268057-DEL-300 | chr2-87268056-DEL-300 |
| chr2-87269870-DEL-393 | chr2-87269871-DEL-393 |
| chr2-87325169-DEL-191 | chr2-87325170-DEL-191 |
| chr2-87347686-DEL-57 | chr2-87347693-DEL-59 |
| chr2-87348517-DEL-307 | chr2-87348518-DEL-307 |
| chr2-87356336-DEL-784 | chr2-87356337-DEL-784 |
| chr2-87368151-DEL-68 | chr2-87368152-DEL-68 |
| chr2-87371760-DEL-205 | chr2-87371761-DEL-205 |
| chr2-87373246-DEL-247 | chr2-87373247-DEL-247 |
| chr2-87384494-DEL-909 | chr2-87384486-DEL-913 |
| chr2-87424499-DEL-2487 | chr2-87424500-DEL-2487 |
| chr2-87548249-DEL-133 | chr2-87548250-DEL-133 |
| chr2-87551950-DEL-212 | chr2-87551951-DEL-212 |
| chr2-87599684-DEL-4212 | chr2-87599677-DEL-4217 |
| chr2-87620585-DEL-58 | chr2-87620586-DEL-58 |
| chr2-87678883-DEL-1693 | chr2-87678884-DEL-1693 |
| chr2-87696666-DEL-395 | chr2-87696667-DEL-395 |
| chr2-87702899-DEL-2669 | chr2-87702896-DEL-2672 |
| chr2-87710154-DEL-487 | chr2-87710155-DEL-487 |
| chr2-87711226-DEL-1087 | chr2-87711227-DEL-1087 |
| chr2-87720103-DEL-660 | chr2-87720103-DEL-660 |
| chr2-87736715-DEL-6458 | chr2-87736716-DEL-6458 |
| chr2-87764554-DEL-7086 | chr2-87764555-DEL-7086 |
| chr2-87848102-DEL-1063 | chr2-87848103-DEL-1063 |
| chr2-87859380-DEL-2576 | chr2-87859381-DEL-2576 |
| chr2-87869158-DEL-4646 | chr2-87869156-DEL-4649 |
| chr2-87883192-DEL-857 | chr2-87883193-DEL-857 |
| chr2-87940576-DEL-158 | chr2-87940577-DEL-159 |

| | |
|---|---|
| chr2-87947109-DEL-6528 | chr2-87947108-DEL-6529 |
| chr2-88002297-DEL-7159 | chr2-88002292-DEL-7160 |
| chr2-88036838-DEL-6203 | chr2-88036837-DEL-6203 |
| chr2-88071540-DEL-6446 | chr2-88071540-DEL-6446 |
| chr2-88093252-DEL-505 | chr2-88093253-DEL-505 |
| chr2-88190770-DEL-1353 | chr2-88190771-DEL-1353 |
| chr2-88193779-DEL-5715 | chr2-88193780-DEL-5715 |
| chr2-88235264-DEL-113 | chr2-88235301-DEL-72 |
| chr2-88240161-DEL-323 | chr2-88240162-DEL-323 |
| chr2-88285073-DEL-60 | chr2-88285076-DEL-68 |
| chr2-88303503-DEL-390 | chr2-88303504-DEL-390 |
| chr2-88306996-DEL-87 | chr2-88306997-DEL-87 |
| chr2-88307029-DEL-6181 | chr2-88307030-DEL-6181 |
| chr2-88338580-DEL-5505 | chr2-88338581-DEL-5505 |
| chr2-88362561-DEL-335 | chr2-88362561-DEL-335 |
| chr2-88386710-DEL-55 | chr2-88386696-DEL-65 |
| chr2-88400923-DEL-275 | chr2-88400924-DEL-275 |
| chr2-88415989-DEL-149 | chr2-88415984-DEL-150 |
| chr2-88417730-DEL-198 | chr2-88417731-DEL-198 |
| chr2-88423360-DEL-78 | chr2-88423359-DEL-79 |
| chr2-88426450-DEL-5881 | chr2-88426451-DEL-5881 |
| chr2-88435480-DEL-67 | chr2-88435480-DEL-67 |
| chr2-88447265-DEL-6277 | chr2-88447266-DEL-6277 |
| chr2-88452279-DEL-348 | chr2-88452280-DEL-348 |
| chr2-88457539-DEL-221 | chr2-88457540-DEL-221 |
| chr2-88464837-DEL-255 | chr2-88464842-DEL-250 |
| chr2-88469937-DEL-57 | chr2-88469946-DEL-54 |
| chr2-88535393-DEL-6533 | chr2-88535394-DEL-6533 |
| chr2-88614458-DEL-1779 | chr2-88614458-DEL-1779 |
| chr2-88637830-DEL-1236 | chr2-88637831-DEL-1236 |
| chr2-88669347-DEL-2565 | chr2-88669348-DEL-2565 |
| chr2-88687723-DEL-72 | chr2-88687724-DEL-72 |
| chr2-88689233-DEL-934 | chr2-88689234-DEL-934 |
| chr2-88701545-DEL-191 | chr2-88701546-DEL-191 |
| chr2-88707863-DEL-1626 | chr2-88707863-DEL-1626 |
| chr2-88709391-DEL-200 | chr2-88709392-DEL-200 |
| chr2-88731505-DEL-97 | chr2-88731505-DEL-92 |
| chr2-88734186-DEL-55 | chr2-88734187-DEL-55 |
| chr2-88763089-DEL-81 | chr2-88763090-DEL-81 |
| chr2-88799619-DEL-199 | chr2-88799620-DEL-199 |
| chr2-88856177-DEL-110 | chr2-88856213-DEL-85 |
| chr2-88892138-DEL-52 | chr2-88892137-DEL-52 |
| chr2-88916769-DEL-72 | chr2-88916770-DEL-72 |
| chr2-88929520-DEL-2979 | chr2-88929521-DEL-2979 |
| chr2-88938527-DEL-56 | chr2-88938524-DEL-60 |
| chr2-88978805-DEL-1969 | chr2-88978806-DEL-1969 |
| chr2-89094921-DEL-6342 | chr2-89094919-DEL-6349 |
| chr2-89095722-DEL-1888 | chr2-89095723-DEL-1888 |
| chr2-89104902-DEL-2584 | chr2-89104905-DEL-2585 |
| chr2-89110107-DEL-466 | chr2-89110108-DEL-466 |
| chr2-89198284-DEL-66 | chr2-89198285-DEL-64 |
| chr2-89210468-DEL-57 | chr2-89210469-DEL-57 |
| chr2-89218849-DEL-52 | chr2-89218850-DEL-52 |
| chr2-89295847-DEL-134 | chr2-89295848-DEL-134 |
| chr2-89344295-DEL-6202 | chr2-89344296-DEL-6202 |
| chr2-89356548-DEL-55 | chr2-89356549-DEL-55 |
| chr2-89362231-DEL-55 | chr2-89362232-DEL-55 |
| chr2-89363163-DEL-167 | chr2-89363164-DEL-167 |
| chr2-89377968-DEL-4139 | chr2-89377969-DEL-4139 |
| chr2-89397590-DEL-87 | chr2-89397591-DEL-81 |
| chr2-89399963-DEL-548 | chr2-89399964-DEL-548 |
| chr2-89412902-DEL-1525 | chr2-89412903-DEL-1525 |
| chr2-89428928-DEL-2247 | chr2-89428929-DEL-2247 |
| chr2-89541250-DEL-276 | chr2-89541251-DEL-276 |
| chr2-89546251-DEL-413 | chr2-89546252-DEL-413 |
| chr2-89546942-DEL-335 | chr2-89546943-DEL-335 |
| chr2-89590602-DEL-524 | chr2-89590601-DEL-524 |
| chr2-89629284-DEL-148 | chr2-89629286-DEL-145 |
| chr2-89669091-DEL-477 | chr2-89669092-DEL-477 |
| chr2-89686706-DEL-1886 | chr2-89686707-DEL-1886 |
| chr2-89691167-DEL-159 | chr2-89691168-DEL-159 |
| chr2-89715460-DEL-1084 | chr2-89715460-DEL-1084 |
| chr2-89723402-DEL-6684 | chr2-89723403-DEL-6684 |
| chr2-89734229-DEL-202 | chr2-89734230-DEL-202 |
| chr2-89739560-DEL-428 | chr2-89739561-DEL-428 |
| chr2-89748481-DEL-2025 | chr2-89748482-DEL-2025 |
| chr2-89776797-DEL-304 | chr2-89776798-DEL-304 |
| chr2-89803960-DEL-516 | chr2-89803961-DEL-515 |
| chr2-89819082-DEL-964 | chr2-89819083-DEL-964 |
| chr2-89850334-DEL-98 | chr2-89850335-DEL-98 |

| | |
|---|---|
| chr2-89895162-DEL-56 | chr2-89895163-DEL-56 |
| chr2-89905598-DEL-1130 | chr2-89905599-DEL-1130 |
| chr2-89913878-DEL-592 | chr2-89913879-DEL-592 |
| chr2-89917721-DEL-7003 | chr2-89917722-DEL-7003 |
| chr2-89918244-DEL-424 | chr2-89918245-DEL-424 |
| chr2-89929629-DEL-2604 | chr2-89929630-DEL-2604 |
| chr2-89965848-DEL-518 | chr2-89965852-DEL-515 |
| chr2-89978744-DEL-576 | chr2-89978745-DEL-576 |
| chr2-90102964-DEL-201 | chr2-90102965-DEL-201 |
| chr2-90128356-DEL-292 | chr2-90128342-DEL-342 |
| chr2-90133100-DEL-60 | chr2-90133100-DEL-61 |
| chr2-90149904-DEL-760 | chr2-90149905-DEL-760 |
| chr2-90203817-DEL-212 | chr2-90203818-DEL-212 |
| chr2-90247069-DEL-1494 | chr2-90247067-DEL-1495 |
| chr2-90248775-DEL-872 | chr2-90248778-DEL-856 |
| chr2-90269690-DEL-304 | chr2-90269678-DEL-305 |
| chr2-90286241-DEL-77 | chr2-90286242-DEL-77 |
| chr2-90453924-DEL-330 | chr2-90453924-DEL-294 |
| chr2-90517259-DEL-177 | chr2-90517260-DEL-177 |
| chr2-90582185-DEL-209 | chr2-90582186-DEL-209 |
| chr2-90587231-DEL-171 | chr2-90587232-DEL-171 |
| chr2-90603855-DEL-51 | chr2-90603834-DEL-51 |
| chr2-90612122-DEL-184 | chr2-90612123-DEL-184 |
| chr2-90622072-DEL-160 | chr2-90622041-DEL-174 |
| chr2-90652998-DEL-217 | chr2-90652999-DEL-217 |
| chr2-90656613-DEL-153 | chr2-90656613-DEL-153 |
| chr2-90671913-DEL-260 | chr2-90671927-DEL-261 |
| chr2-90680875-DEL-525 | chr2-90680876-DEL-525 |
| chr2-90684365-DEL-200 | chr2-90684366-DEL-200 |
| chr2-90730733-DEL-213 | chr2-90730734-DEL-213 |
| chr2-90750074-DEL-65 | chr2-90750075-DEL-65 |
| chr2-90794279-DEL-263 | chr2-90794280-DEL-263 |
| chr2-90807091-DEL-707 | chr2-90807092-DEL-707 |
| chr2-90837576-DEL-228 | chr2-90837577-DEL-228 |
| chr2-90876187-DEL-396 | chr2-90876180-DEL-397 |
| chr2-90883877-DEL-80 | chr2-90883872-DEL-82 |
| chr2-90904306-DEL-382 | chr2-90904307-DEL-382 |
| chr2-90913696-DEL-52 | chr2-90913697-DEL-52 |
| chr2-90946506-DEL-437 | chr2-90946503-DEL-439 |
| chr2-90996444-DEL-263 | chr2-90996445-DEL-263 |
| chr2-91018401-DEL-64 | chr2-91018405-DEL-60 |
| chr2-91021395-DEL-352 | chr2-91021396-DEL-352 |
| chr2-91029873-DEL-206 | chr2-91029884-DEL-205 |
| chr2-91056784-DEL-126 | chr2-91056785-DEL-126 |
| chr2-91079670-DEL-58 | chr2-91079671-DEL-52 |
| chr2-91110632-DEL-64 | chr2-91110633-DEL-64 |
| chr2-91112281-DEL-72 | chr2-91112296-DEL-71 |
| chr2-91116652-DEL-2026 | chr2-91116653-DEL-2026 |
| chr2-91123043-DEL-53 | chr2-91123045-DEL-54 |
| chr2-91123696-DEL-134 | chr2-91123697-DEL-134 |
| chr2-91125223-DEL-339 | chr2-91125224-DEL-339 |
| chr2-91138656-DEL-53 | chr2-91138656-DEL-52 |
| chr2-91171601-DEL-97 | chr2-91171617-DEL-76 |
| chr2-91200289-DEL-214 | chr2-91200290-DEL-214 |
| chr2-91323553-DEL-317 | chr2-91323550-DEL-316 |
| chr2-91324920-DEL-84 | chr2-91324921-DEL-84 |
| chr2-91326129-DEL-62 | chr2-91326130-DEL-62 |
| chr2-91332586-DEL-182 | chr2-91332586-DEL-181 |
| chr2-91363670-DEL-121 | chr2-91363673-DEL-99 |
| chr2-91374961-DEL-1027 | chr2-91374962-DEL-1027 |
| chr2-91468557-DEL-161 | chr2-91468558-DEL-161 |
| chr2-91470154-DEL-180 | chr2-91470155-DEL-180 |
| chr2-91471201-DEL-157 | chr2-91471202-DEL-157 |
| chr2-91471881-DEL-1195 | chr2-91471882-DEL-1195 |
| chr2-91490689-DEL-86 | chr2-91490654-DEL-84 |
| chr2-91494864-DEL-348 | chr2-91494865-DEL-348 |
| chr2-91496185-DEL-274 | chr2-91496186-DEL-274 |
| chr2-91504668-DEL-61 | chr2-91504669-DEL-61 |
| chr2-91511633-DEL-56 | chr2-91511634-DEL-56 |
| chr2-91522167-DEL-183 | chr2-91522168-DEL-183 |
| chr2-91532510-DEL-56 | chr2-91532507-DEL-54 |
| chr2-91554719-DEL-175 | chr2-91554736-DEL-130 |
| chr2-91564767-DEL-208 | chr2-91564768-DEL-208 |
| chr2-91573400-DEL-184 | chr2-91573401-DEL-184 |
| chr2-91577881-DEL-204 | chr2-91577882-DEL-204 |
| chr2-91578905-DEL-212 | chr2-91578906-DEL-212 |
| chr2-91582692-DEL-202 | chr2-91582693-DEL-202 |
| chr2-91590554-DEL-90 | chr2-91590565-DEL-61 |
| chr2-91613029-DEL-67 | chr2-91613030-DEL-67 |
| chr2-91625981-DEL-210 | chr2-91625982-DEL-210 |

| | |
|---|---|
| chr2-91667434-DEL-1731 | chr2-91667442-DEL-1725 |
| chr2-91668462-DEL-714 | chr2-91668463-DEL-716 |
| chr2-91676474-DEL-3792 | chr2-91676475-DEL-3792 |
| chr2-91693551-DEL-98 | chr2-91693552-DEL-98 |
| chr2-91738199-DEL-218 | chr2-91738200-DEL-218 |
| chr2-91756678-DEL-315 | chr2-91756679-DEL-315 |
| chr2-91811048-DEL-152 | chr2-91811025-DEL-151 |
| chr2-91828611-DEL-354 | chr2-91828612-DEL-354 |
| chr2-91832682-DEL-74 | chr2-91832679-DEL-70 |
| chr2-91846007-DEL-60 | chr2-91845969-DEL-78 |
| chr2-91855447-DEL-82 | chr2-91855444-DEL-86 |
| chr2-91878400-DEL-52 | chr2-91878401-DEL-52 |
| chr2-91896826-DEL-80 | chr2-91896827-DEL-80 |
| chr2-91905913-DEL-63 | chr2-91905906-DEL-71 |
| chr2-91936002-DEL-65 | chr2-91936008-DEL-66 |
| chr2-91940924-DEL-68 | chr2-91940925-DEL-56 |
| chr2-91946844-DEL-53 | chr2-91946823-DEL-55 |
| chr2-91951814-DEL-54 | chr2-91951815-DEL-54 |
| chr2-91952174-DEL-206 | chr2-91952175-DEL-206 |
| chr2-91965302-DEL-203 | chr2-91965303-DEL-202 |
| chr2-91968422-DEL-221 | chr2-91968423-DEL-221 |
| chr2-91972660-DEL-252 | chr2-91972616-DEL-251 |
| chr2-91977576-DEL-90 | chr2-91977577-DEL-90 |
| chr2-91980037-DEL-166 | chr2-91980038-DEL-166 |
| chr2-92010114-DEL-285 | chr2-92010115-DEL-285 |
| chr2-92025532-DEL-212 | chr2-92025533-DEL-212 |
| chr2-92034157-DEL-61 | chr2-92034157-DEL-71 |
| chr2-92037634-DEL-236 | chr2-92037635-DEL-236 |
| chr2-92045385-DEL-162 | chr2-92045386-DEL-162 |
| chr2-92080727-DEL-71 | chr2-92080728-DEL-71 |
| chr2-92111983-DEL-213 | chr2-92111984-DEL-213 |
| chr2-92115197-DEL-120 | chr2-92115198-DEL-120 |
| chr2-92118685-DEL-2781 | chr2-92118686-DEL-2781 |
| chr2-92126459-DEL-113 | chr2-92126460-DEL-113 |
| chr2-92141913-DEL-364 | chr2-92141914-DEL-364 |
| chr2-92176127-DEL-74 | chr2-92176142-DEL-60 |
| chr2-92275215-DEL-107 | chr2-92275215-DEL-122 |
| chr2-92277236-DEL-192 | chr2-92277237-DEL-192 |
| chr2-92307637-DEL-52 | chr2-92307630-DEL-60 |
| chr2-92312663-DEL-62 | chr2-92312663-DEL-62 |
| chr2-92315188-DEL-693 | chr2-92315189-DEL-693 |
| chr2-92350951-DEL-238 | chr2-92350952-DEL-238 |
| chr2-92361864-DEL-83 | chr2-92361865-DEL-83 |
| chr2-92449326-DEL-183 | chr2-92449328-DEL-180 |
| chr2-92457739-DEL-56 | chr2-92457740-DEL-56 |
| chr2-92467425-DEL-56 | chr2-92467426-DEL-56 |
| chr2-92470945-DEL-67 | chr2-92470946-DEL-67 |
| chr2-92474671-DEL-86 | chr2-92474672-DEL-86 |
| chr2-92477452-DEL-55 | chr2-92477453-DEL-55 |
| chr2-92488468-DEL-559 | chr2-92488467-DEL-560 |
| chr2-92568935-DEL-60 | chr2-92568935-DEL-60 |
| chr2-92575045-DEL-52 | chr2-92575046-DEL-52 |
| chr2-92595449-DEL-127 | chr2-92595450-DEL-127 |
| chr2-92623401-DEL-394 | chr2-92623402-DEL-394 |
| chr2-92641578-DEL-483 | chr2-92641579-DEL-483 |
| chr2-92661501-DEL-63 | chr2-92661502-DEL-63 |
| chr2-92682478-DEL-115 | chr2-92682481-DEL-114 |
| chr2-92708298-DEL-97 | chr2-92708274-DEL-98 |
| chr2-92710503-DEL-830 | chr2-92710505-DEL-832 |
| chr2-92714401-DEL-188 | chr2-92714402-DEL-188 |
| chr2-92719204-DEL-1798 | chr2-92719205-DEL-1798 |
| chr2-92731126-DEL-167 | chr2-92731127-DEL-167 |
| chr2-92768870-DEL-404 | chr2-92768871-DEL-404 |
| chr2-92825481-DEL-994 | chr2-92825482-DEL-994 |
| chr2-92829379-DEL-164 | chr2-92829375-DEL-160 |
| chr2-92837072-DEL-84 | chr2-92837065-DEL-86 |
| chr2-92840330-DEL-606 | chr2-92840331-DEL-606 |
| chr2-92922781-DEL-61 | chr2-92922782-DEL-61 |
| chr2-92961006-DEL-82 | chr2-92961019-DEL-70 |
| chr2-92964541-DEL-82 | chr2-92964539-DEL-83 |
| chr2-92968199-DEL-88 | chr2-92968200-DEL-88 |
| chr2-93008222-DEL-93 | chr2-93008207-DEL-85 |
| chr2-93033489-DEL-64 | chr2-93033472-DEL-92 |
| chr2-93036675-DEL-169 | chr2-93036676-DEL-169 |
| chr2-93091573-DEL-59 | chr2-93091574-DEL-59 |
| chr2-93101329-DEL-209 | chr2-93101330-DEL-209 |
| chr2-93125020-DEL-2862 | chr2-93125021-DEL-2862 |
| chr2-93130488-DEL-68 | chr2-93130485-DEL-72 |
| chr2-93155598-DEL-139 | chr2-93155599-DEL-139 |
| chr2-93156376-DEL-155 | chr2-93156376-DEL-155 |

| | |
|---|---|
| chr2-93182928-DEL-171 | chr2-93182929-DEL-171 |
| chr2-93186685-DEL-112 | chr2-93186686-DEL-112 |
| chr2-93187880-DEL-64 | chr2-93187881-DEL-64 |
| chr2-93189687-DEL-102 | chr2-93189690-DEL-100 |
| chr2-93207234-DEL-54 | chr2-93207267-DEL-52 |
| chr2-93231723-DEL-52 | chr2-93231720-DEL-59 |
| chr2-93238619-DEL-55 | chr2-93238619-DEL-60 |
| chr2-93240212-DEL-54 | chr2-93240213-DEL-65 |
| chr2-93332498-DEL-81 | chr2-93332499-DEL-81 |
| chr2-93358327-DEL-116 | chr2-93358338-DEL-122 |
| chr2-93389309-DEL-441 | chr2-93389310-DEL-441 |
| chr2-93391100-DEL-1311 | chr2-93391101-DEL-1311 |
| chr2-93412373-DEL-82 | chr2-93412374-DEL-82 |
| chr2-93478667-DEL-1557 | chr2-93478670-DEL-1558 |
| chr2-93505450-DEL-58 | chr2-93505456-DEL-72 |
| chr2-93509991-DEL-225 | chr2-93509992-DEL-225 |
| chr2-93553039-DEL-369 | chr2-93553040-DEL-371 |
| chr2-93564643-DEL-80 | chr2-93564610-DEL-120 |
| chr2-93579114-DEL-81 | chr2-93579115-DEL-81 |
| chr2-93617019-DEL-280 | chr2-93617020-DEL-280 |
| chr2-93649288-DEL-327 | chr2-93649289-DEL-327 |
| chr2-93650027-DEL-69 | chr2-93650005-DEL-92 |
| chr2-93655118-DEL-7082 | chr2-93655119-DEL-7082 |
| chr2-93664699-DEL-951 | chr2-93664700-DEL-951 |
| chr2-93675955-DEL-72 | chr2-93675956-DEL-72 |
| chr2-93707681-DEL-75 | chr2-93707692-DEL-55 |
| chr2-93726468-DEL-392 | chr2-93726466-DEL-395 |
| chr2-93763364-DEL-84 | chr2-93763365-DEL-84 |
| chr2-93765443-DEL-82 | chr2-93765435-DEL-92 |
| chr2-93789634-DEL-82 | chr2-93789635-DEL-82 |
| chr2-93796728-DEL-4429 | chr2-93796729-DEL-4429 |
| chr2-93814538-DEL-79 | chr2-93814542-DEL-76 |
| chr2-93854023-DEL-68 | chr2-93854024-DEL-64 |
| chr2-93857239-DEL-59 | chr2-93857240-DEL-59 |
| chr2-93858462-DEL-870 | chr2-93858481-DEL-861 |
| chr2-93861833-DEL-112 | chr2-93861834-DEL-112 |
| chr2-93866746-DEL-180 | chr2-93866747-DEL-180 |
| chr2-93900969-DEL-69 | chr2-93900970-DEL-69 |
| chr2-93910950-DEL-62 | chr2-93910949-DEL-64 |
| chr2-93928540-DEL-51 | chr2-93928541-DEL-51 |
| chr2-93936661-DEL-1691 | chr2-93936662-DEL-1691 |
| chr2-93969106-DEL-61 | chr2-93969103-DEL-62 |
| chr2-94016651-DEL-6455 | chr2-94016652-DEL-6455 |
| chr2-94016726-DEL-64 | chr2-94016727-DEL-64 |
| chr2-94042068-DEL-200 | chr2-94042069-DEL-199 |
| chr2-94099780-DEL-56 | chr2-94099781-DEL-56 |
| chr2-94104705-DEL-55 | chr2-94104699-DEL-62 |
| chr2-94107070-DEL-76 | chr2-94107071-DEL-76 |
| chr2-94109135-DEL-92 | chr2-94109134-DEL-92 |
| chr2-94143123-DEL-218 | chr2-94143124-DEL-218 |
| chr2-94155227-DEL-61 | chr2-94155228-DEL-61 |
| chr2-94179113-DEL-765 | chr2-94179116-DEL-775 |
| chr2-94189954-DEL-206 | chr2-94189955-DEL-205 |
| chr2-94209367-DEL-70 | chr2-94209368-DEL-70 |
| chr2-94214299-DEL-1171 | chr2-94214300-DEL-1171 |
| chr2-94220505-DEL-132 | chr2-94220506-DEL-132 |
| chr2-94221155-DEL-506 | chr2-94221156-DEL-506 |
| chr2-94230174-DEL-148 | chr2-94230175-DEL-148 |
| chr2-94277879-DEL-11362 | chr2-94277880-DEL-11362 |
| chr2-94296157-DEL-109 | chr2-94296158-DEL-109 |
| chr2-94324652-DEL-76 | chr2-94324616-DEL-76 |
| chr2-94327343-DEL-58 | chr2-94327348-DEL-50 |
| chr2-94332905-DEL-53 | chr2-94332870-DEL-60 |
| chr2-94350567-DEL-180 | chr2-94350564-DEL-204 |
| chr2-94433396-DEL-4633 | chr2-94433397-DEL-4633 |
| chr2-94449757-DEL-51 | chr2-94449754-DEL-51 |
| chr2-94464462-DEL-888 | chr2-94464463-DEL-888 |
| chr2-94492992-DEL-59 | chr2-94492993-DEL-59 |
| chr2-94507315-DEL-141 | chr2-94507316-DEL-141 |
| chr2-94515658-DEL-436 | chr2-94515659-DEL-436 |
| chr2-94524429-DEL-120 | chr2-94524417-DEL-130 |
| chr2-94541351-DEL-199 | chr2-94541395-DEL-200 |
| chr2-94581439-DEL-1398 | chr2-94581440-DEL-1398 |
| chr2-94594586-DEL-68 | chr2-94594583-DEL-67 |
| chr2-94631404-DEL-8418 | chr2-94631405-DEL-8418 |
| chr2-94650040-DEL-466 | chr2-94650041-DEL-466 |
| chr2-94675123-DEL-160 | chr2-94675131-DEL-150 |
| chr2-94679697-DEL-356 | chr2-94679697-DEL-356 |
| chr2-94714865-DEL-6872 | chr2-94714866-DEL-6872 |
| chr2-94735435-DEL-84 | chr2-94735436-DEL-84 |

| | |
|---|---|
| chr2-94742499-DEL-107 | chr2-94742500-DEL-107 |
| chr2-94759418-DEL-349 | chr2-94759419-DEL-349 |
| chr2-94774485-DEL-218 | chr2-94774484-DEL-216 |
| chr2-94781897-DEL-416 | chr2-94781898-DEL-416 |
| chr2-94794173-DEL-3112 | chr2-94794175-DEL-3111 |
| chr2-94826928-DEL-1661 | chr2-94826929-DEL-1661 |
| chr2-94895106-DEL-5352 | chr2-94895107-DEL-5352 |
| chr2-94905621-DEL-82 | chr2-94905645-DEL-56 |
| chr2-94935730-DEL-66 | chr2-94935727-DEL-67 |
| chr2-94986527-DEL-363 | chr2-94986528-DEL-363 |
| chr2-94995911-DEL-2304 | chr2-94995912-DEL-2304 |
| chr2-95016278-DEL-111 | chr2-95016279-DEL-111 |
| chr2-95017652-DEL-249 | chr2-95017649-DEL-249 |
| chr2-95046177-DEL-8068 | chr2-95046178-DEL-8068 |
| chr2-95047612-DEL-1677 | chr2-95047612-DEL-1677 |
| chr2-95053934-DEL-201 | chr2-95053935-DEL-201 |
| chr2-95067942-DEL-59 | chr2-95067947-DEL-55 |
| chr2-95080914-DEL-10530 | chr2-95080915-DEL-10530 |
| chr2-95093015-DEL-1888 | chr2-95093016-DEL-1888 |
| chr2-95095523-DEL-219 | chr2-95095524-DEL-219 |
| chr2-95147473-DEL-961 | chr2-95147474-DEL-961 |
| chr2-95149783-DEL-1213 | chr2-95149784-DEL-1213 |
| chr2-95182625-DEL-7317 | chr2-95182626-DEL-7317 |
| chr2-95232808-DEL-52 | chr2-95232818-DEL-62 |
| chr2-95244409-DEL-6679 | chr2-95244410-DEL-6679 |
| chr2-95244528-DEL-212 | chr2-95244529-DEL-212 |
| chr2-95255369-DEL-352 | chr2-95255370-DEL-352 |
| chr2-95257285-DEL-51 | chr2-95257267-DEL-70 |
| chr2-95257527-DEL-560 | chr2-95257528-DEL-560 |
| chr2-95264373-DEL-2280 | chr2-95264373-DEL-2280 |
| chr2-95291444-DEL-1435 | chr2-95291445-DEL-1435 |
| chr2-95311938-DEL-80 | chr2-95311939-DEL-80 |
| chr2-95312437-DEL-824 | chr2-95312438-DEL-824 |
| chr2-95322512-DEL-490 | chr2-95322513-DEL-490 |
| chr2-95337620-DEL-773 | chr2-95337621-DEL-773 |
| chr2-95343651-DEL-55 | chr2-95343652-DEL-55 |
| chr2-95354293-DEL-157 | chr2-95354294-DEL-157 |
| chr2-95366840-DEL-1888 | chr2-95366841-DEL-1888 |
| chr2-95439555-DEL-75 | chr2-95439556-DEL-75 |
| chr2-95453147-DEL-55 | chr2-95453148-DEL-53 |
| chr2-95453781-DEL-105 | chr2-95453782-DEL-105 |
| chr2-95478266-DEL-6476 | chr2-95478263-DEL-6477 |
| chr2-95478369-DEL-214 | chr2-95478370-DEL-214 |
| chr2-95500019-DEL-143 | chr2-95500020-DEL-143 |
| chr2-95500586-DEL-438 | chr2-95500587-DEL-438 |
| chr2-95598704-DEL-307 | chr2-95598705-DEL-307 |
| chr2-95600272-DEL-439 | chr2-95600268-DEL-440 |
| chr2-95618979-DEL-1004 | chr2-95618980-DEL-1004 |
| chr2-95619524-DEL-3523 | chr2-95619525-DEL-3523 |
| chr2-95623613-DEL-6605 | chr2-95623614-DEL-6605 |
| chr2-95688884-DEL-136 | chr2-95688917-DEL-92 |
| chr2-95693907-DEL-505 | chr2-95693908-DEL-505 |
| chr2-95695289-DEL-500 | chr2-95695290-DEL-500 |
| chr2-95728425-DEL-53 | chr2-95728395-DEL-68 |
| chr2-95758530-DEL-6634 | chr2-95758531-DEL-6634 |
| chr2-95764830-DEL-214 | chr2-95764831-DEL-214 |
| chr2-95810851-DEL-51 | chr2-95810812-DEL-81 |
| chr2-95844894-DEL-72 | chr2-95844899-DEL-56 |
| chr2-95871276-DEL-72 | chr2-95871277-DEL-72 |
| chr2-95900257-DEL-3405 | chr2-95900258-DEL-3405 |
| chr2-95926162-DEL-6227 | chr2-95926163-DEL-6227 |
| chr2-95935827-DEL-98 | chr2-95935828-DEL-98 |
| chr2-95946003-DEL-1222 | chr2-95946004-DEL-1222 |
| chr2-95947665-DEL-253 | chr2-95947666-DEL-253 |
| chr2-96021849-DEL-6984 | chr2-96021850-DEL-6984 |
| chr2-96037490-DEL-1529 | chr2-96037491-DEL-1528 |
| chr2-96038868-DEL-201 | chr2-96038869-DEL-201 |
| chr2-96041209-DEL-1398 | chr2-96041210-DEL-1398 |
| chr2-96061028-DEL-1066 | chr2-96061022-DEL-1068 |
| chr2-96103327-DEL-81 | chr2-96103308-DEL-101 |
| chr2-96111493-DEL-219 | chr2-96111494-DEL-219 |
| chr2-96188500-DEL-263 | chr2-96188501-DEL-263 |
| chr2-96261192-DEL-6774 | chr2-96261193-DEL-6774 |
| chr2-96269613-DEL-2894 | chr2-96269614-DEL-2894 |
| chr2-96353806-DEL-2209 | chr2-96353807-DEL-2209 |
| chr2-96376613-DEL-55 | chr2-96376614-DEL-55 |
| chr2-96378334-DEL-1407 | chr2-96378335-DEL-1407 |
| chr2-96429632-DEL-3507 | chr2-96429633-DEL-3507 |
| chr2-96447803-DEL-541 | chr2-96447804-DEL-541 |
| chr2-96489950-DEL-4608 | chr2-96489951-DEL-4608 |

| | |
|---|---|
| chr2-96504776-DEL-58 | chr2-96504777-DEL-58 |
| chr2-96507517-DEL-104 | chr2-96507518-DEL-104 |
| chr2-96585919-DEL-961 | chr2-96585928-DEL-961 |
| chr2-96590185-DEL-91 | chr2-96590180-DEL-89 |
| chr2-96659550-DEL-628 | chr2-96659551-DEL-628 |
| chr2-96675134-DEL-64 | chr2-96675141-DEL-58 |
| chr2-96695972-DEL-54 | chr2-96695969-DEL-58 |
| chr2-96699507-DEL-655 | chr2-96699508-DEL-655 |
| chr2-96730113-DEL-343 | chr2-96730114-DEL-343 |
| chr2-96735417-DEL-354 | chr2-96735418-DEL-354 |
| chr2-96741774-DEL-579 | chr2-96741775-DEL-579 |
| chr2-96747011-DEL-199 | chr2-96747012-DEL-199 |
| chr2-96766943-DEL-52 | chr2-96766944-DEL-52 |
| chr2-96782495-DEL-651 | chr2-96782495-DEL-652 |
| chr2-96785027-DEL-6321 | chr2-96785027-DEL-6321 |
| chr2-96825592-DEL-64 | chr2-96825593-DEL-64 |
| chr2-96826925-DEL-108 | chr2-96826926-DEL-116 |
| chr2-96860017-DEL-6146 | chr2-96860018-DEL-6146 |
| chr2-96865778-DEL-201 | chr2-96865779-DEL-201 |
| chr2-96872217-DEL-378 | chr2-96872218-DEL-379 |
| chr2-96909941-DEL-68 | chr2-96909938-DEL-72 |
| chr2-96927393-DEL-417 | chr2-96927404-DEL-407 |
| chr2-96947906-DEL-98 | chr2-96947907-DEL-98 |
| chr2-96984520-DEL-63 | chr2-96984519-DEL-74 |
| chr2-96990923-DEL-174 | chr2-96990924-DEL-174 |
| chr2-97010158-DEL-9395 | chr2-97010159-DEL-9395 |
| chr2-97050622-DEL-1082 | chr2-97050622-DEL-1082 |
| chr2-97060475-DEL-412 | chr2-97060475-DEL-412 |
| chr2-97070440-DEL-6463 | chr2-97070441-DEL-6463 |
| chr2-97083781-DEL-340 | chr2-97083783-DEL-341 |
| chr2-97091007-DEL-996 | chr2-97091008-DEL-996 |
| chr2-97101423-DEL-117 | chr2-97101409-DEL-132 |
| chr2-97109144-DEL-149 | chr2-97109149-DEL-126 |
| chr2-97110925-DEL-59 | chr2-97110932-DEL-53 |
| chr2-97120386-DEL-81 | chr2-97120387-DEL-81 |
| chr2-97151385-DEL-6601 | chr2-97151385-DEL-6602 |
| chr2-97160460-DEL-71 | chr2-97160461-DEL-71 |
| chr2-97187944-DEL-105 | chr2-97187945-DEL-104 |
| chr2-97222986-DEL-426 | chr2-97222986-DEL-424 |
| chr2-97224949-DEL-665 | chr2-97224949-DEL-665 |
| chr2-97231203-DEL-1714 | chr2-97231200-DEL-1715 |
| chr2-97267936-DEL-68 | chr2-97267939-DEL-66 |
| chr2-97268949-DEL-498 | chr2-97268950-DEL-498 |
| chr2-97288730-DEL-2756 | chr2-97288731-DEL-2756 |
| chr2-97306778-DEL-53 | chr2-97306779-DEL-53 |
| chr2-97310228-DEL-220 | chr2-97310229-DEL-220 |
| chr2-97319553-DEL-50 | chr2-97319568-DEL-56 |
| chr2-97322594-DEL-81 | chr2-97322590-DEL-71 |
| chr2-97405667-DEL-210 | chr2-97405668-DEL-210 |
| chr2-97424696-DEL-189 | chr2-97424701-DEL-185 |
| chr2-97453006-DEL-74 | chr2-97453007-DEL-78 |
| chr2-97455829-DEL-66 | chr2-97455828-DEL-68 |
| chr2-97500655-DEL-6657 | chr2-97500656-DEL-6657 |
| chr2-97515317-DEL-1427 | chr2-97515318-DEL-1427 |
| chr2-97522449-DEL-54 | chr2-97522450-DEL-54 |
| chr2-97531271-DEL-79 | chr2-97531272-DEL-79 |
| chr2-97532747-DEL-59 | chr2-97532733-DEL-60 |
| chr2-97542690-DEL-259 | chr2-97542691-DEL-259 |
| chr2-97547482-DEL-91 | chr2-97547483-DEL-91 |
| chr2-97551163-DEL-92 | chr2-97551164-DEL-92 |
| chr2-97557369-DEL-6363 | chr2-97557370-DEL-6363 |
| chr2-97571547-DEL-215 | chr2-97571545-DEL-218 |
| chr2-97590958-DEL-6125 | chr2-97590959-DEL-6125 |
| chr2-97631887-DEL-4023 | chr2-97631881-DEL-4025 |
| chr2-97668629-DEL-6455 | chr2-97668630-DEL-6455 |
| chr2-97715336-DEL-2005 | chr2-97715337-DEL-2005 |
| chr2-97720481-DEL-73 | chr2-97720470-DEL-71 |
| chr2-97725556-DEL-6607 | chr2-97725557-DEL-6607 |
| chr2-97746148-DEL-114 | chr2-97746149-DEL-114 |
| chr2-97750793-DEL-54 | chr2-97750793-DEL-62 |
| chr2-97751136-DEL-4585 | chr2-97751137-DEL-4585 |
| chr2-97761143-DEL-1568 | chr2-97761144-DEL-1568 |
| chr2-97769307-DEL-427 | chr2-97769308-DEL-427 |
| chr2-97773127-DEL-1893 | chr2-97773128-DEL-1893 |
| chr2-97776066-DEL-102 | chr2-97776063-DEL-103 |
| chr2-97780925-DEL-693 | chr2-97780929-DEL-690 |
| chr2-97786579-DEL-2123 | chr2-97786580-DEL-2123 |
| chr2-97804372-DEL-507 | chr2-97804373-DEL-507 |
| chr2-97813336-DEL-245 | chr2-97813337-DEL-245 |
| chr2-97814833-DEL-1297 | chr2-97814833-DEL-1297 |

| | |
|---|---|
| chr2-97848505-DEL-75 | chr2-97848506-DEL-75 |
| chr2-97889042-DEL-52 | chr2-97889041-DEL-54 |
| chr2-97939934-DEL-136 | chr2-97939935-DEL-136 |
| chr2-97946574-DEL-2824 | chr2-97946575-DEL-2824 |
| chr2-97959653-DEL-4041 | chr2-97959655-DEL-4040 |
| chr2-98020485-DEL-6251 | chr2-98020486-DEL-6251 |
| chr2-98030230-DEL-232 | chr2-98030231-DEL-232 |
| chr2-98036039-DEL-350 | chr2-98036040-DEL-350 |
| chr2-98052628-DEL-56 | chr2-98052629-DEL-56 |
| chr2-98058610-DEL-246 | chr2-98058609-DEL-247 |
| chr2-98068827-DEL-163 | chr2-98068828-DEL-163 |
| chr2-98084246-DEL-63 | chr2-98084247-DEL-63 |
| chr2-98101240-DEL-254 | chr2-98101259-DEL-236 |
| chr2-98103525-DEL-1436 | chr2-98103526-DEL-1436 |
| chr2-98137740-DEL-6659 | chr2-98137741-DEL-6660 |
| chr2-98148493-DEL-120 | chr2-98148494-DEL-120 |
| chr2-98162023-DEL-2043 | chr2-98162024-DEL-2043 |
| chr2-98172281-DEL-544 | chr2-98172281-DEL-544 |
| chr2-98180277-DEL-1428 | chr2-98180278-DEL-1428 |
| chr2-98192103-DEL-1888 | chr2-98192103-DEL-1888 |
| chr2-98201560-DEL-2614 | chr2-98201561-DEL-2614 |
| chr2-98225750-DEL-519 | chr2-98225751-DEL-519 |
| chr2-98259233-DEL-6902 | chr2-98259234-DEL-6902 |
| chr2-98272880-DEL-81 | chr2-98272902-DEL-52 |
| chr2-98296084-DEL-395 | chr2-98296085-DEL-395 |
| chr2-98337228-DEL-80 | chr2-98337236-DEL-62 |
| chr2-98362666-DEL-55 | chr2-98362657-DEL-56 |
| chr2-98374588-DEL-1547 | chr2-98374589-DEL-1547 |
| chr2-98401459-DEL-60 | chr2-98401460-DEL-60 |
| chr2-98401918-DEL-238 | chr2-98401919-DEL-238 |
| chr2-98414013-DEL-64 | chr2-98413988-DEL-54 |
| chr2-98415447-DEL-264 | chr2-98415448-DEL-264 |
| chr2-98416434-DEL-63 | chr2-98416434-DEL-62 |
| chr2-98423605-DEL-1344 | chr2-98423606-DEL-1344 |
| chr2-98462408-DEL-6792 | chr2-98462409-DEL-6792 |
| chr2-98491449-DEL-6233 | chr2-98491468-DEL-6236 |
| chr2-98511143-DEL-123 | chr2-98511147-DEL-117 |
| chr2-98557226-DEL-2795 | chr2-98557227-DEL-2795 |
| chr2-98576391-DEL-51 | chr2-98576390-DEL-51 |
| chr2-98582424-DEL-5095 | chr2-98582425-DEL-5095 |
| chr2-98604534-DEL-1094 | chr2-98604535-DEL-1095 |
| chr2-98610580-DEL-1103 | chr2-98610581-DEL-1103 |
| chr2-98626969-DEL-5269 | chr2-98626970-DEL-5269 |
| chr2-98632645-DEL-224 | chr2-98632646-DEL-223 |
| chr2-98660207-DEL-111 | chr2-98660207-DEL-107 |
| chr2-98667872-DEL-494 | chr2-98667873-DEL-494 |
| chr2-98684350-DEL-1317 | chr2-98684351-DEL-1317 |
| chr2-98693025-DEL-292 | chr2-98693026-DEL-310 |
| chr2-98703506-DEL-493 | chr2-98703506-DEL-491 |
| chr2-98736989-DEL-88 | chr2-98736990-DEL-88 |
| chr2-98766500-DEL-391 | chr2-98766500-DEL-391 |
| chr2-98771232-DEL-151 | chr2-98771233-DEL-151 |
| chr2-98774080-DEL-2603 | chr2-98774081-DEL-2603 |
| chr2-98782790-DEL-1599 | chr2-98782788-DEL-1554 |
| chr2-98802501-DEL-561 | chr2-98802502-DEL-561 |
| chr2-98803763-DEL-972 | chr2-98803764-DEL-972 |
| chr2-98810690-DEL-333 | chr2-98810691-DEL-333 |
| chr2-98848230-DEL-214 | chr2-98848235-DEL-213 |
| chr2-98850252-DEL-1102 | chr2-98850253-DEL-1102 |
| chr2-98860050-DEL-1912 | chr2-98860051-DEL-1912 |
| chr2-98881312-DEL-6487 | chr2-98881313-DEL-6487 |
| chr2-98894961-DEL-328 | chr2-98894962-DEL-328 |
| chr2-98900441-DEL-638 | chr2-98900442-DEL-638 |
| chr2-98909809-DEL-178 | chr2-98909810-DEL-178 |
| chr2-98949817-DEL-1388 | chr2-98949818-DEL-1388 |
| chr2-98974488-DEL-168 | chr2-98974489-DEL-168 |
| chr2-98982567-DEL-143 | chr2-98982568-DEL-143 |
| chr2-98991297-DEL-6658 | chr2-98991298-DEL-6658 |
| chr2-98997689-DEL-201 | chr2-98997690-DEL-201 |
| chr2-99013607-DEL-71 | chr2-99013585-DEL-51 |
| chr2-99023339-DEL-2435 | chr2-99023340-DEL-2435 |
| chr2-99025697-DEL-1099 | chr2-99025698-DEL-1099 |
| chr2-99056296-DEL-2942 | chr2-99056297-DEL-2942 |
| chr2-99079542-DEL-77 | chr2-99079543-DEL-77 |
| chr2-99082577-DEL-1286 | chr2-99082578-DEL-1286 |
| chr2-99086303-DEL-60 | chr2-99086306-DEL-50 |
| chr2-99098455-DEL-430 | chr2-99098456-DEL-430 |
| chr2-99100037-DEL-70 | chr2-99100037-DEL-70 |
| chr2-99114878-DEL-54 | chr2-99114875-DEL-58 |
| chr2-99126745-DEL-51 | chr2-99126746-DEL-51 |

| | |
|---|---|
| chr2-99127744-DEL-54 | chr2-99127745-DEL-98 |
| chr2-99129345-DEL-304 | chr2-99129346-DEL-304 |
| chr2-99130231-DEL-6545 | chr2-99130232-DEL-6545 |
| chr2-99130842-DEL-216 | chr2-99130843-DEL-216 |
| chr2-99198908-DEL-3217 | chr2-99198909-DEL-3217 |
| chr2-99231423-DEL-186 | chr2-99231424-DEL-186 |
| chr2-99259022-DEL-84 | chr2-99259017-DEL-85 |
| chr2-99265962-DEL-3106 | chr2-99265963-DEL-3106 |
| chr2-99291081-DEL-5129 | chr2-99291081-DEL-5131 |
| chr2-99323163-DEL-578 | chr2-99323164-DEL-578 |
| chr2-99359812-DEL-1408 | chr2-99359813-DEL-1408 |
| chr2-99374632-DEL-2349 | chr2-99374632-DEL-2349 |
| chr2-99375563-DEL-7431 | chr2-99375564-DEL-7431 |
| chr2-99413587-DEL-335 | chr2-99413588-DEL-335 |
| chr2-99427436-DEL-64 | chr2-99427437-DEL-64 |
| chr2-99429964-DEL-53 | chr2-99429965-DEL-53 |
| chr2-99434672-DEL-256 | chr2-99434673-DEL-256 |
| chr2-99455263-DEL-350 | chr2-99455255-DEL-358 |
| chr2-99483952-DEL-162 | chr2-99483953-DEL-161 |
| chr2-99504821-DEL-184 | chr2-99504822-DEL-184 |
| chr2-99506699-DEL-342 | chr2-99506700-DEL-342 |
| chr2-99509161-DEL-53 | chr2-99509162-DEL-53 |
| chr2-99513062-DEL-160 | chr2-99513062-DEL-161 |
| chr2-99513211-DEL-6227 | chr2-99513212-DEL-6227 |
| chr2-99548997-DEL-200 | chr2-99549003-DEL-194 |
| chr2-99551547-DEL-76 | chr2-99551548-DEL-64 |
| chr2-99563862-DEL-706 | chr2-99563863-DEL-706 |
| chr2-99588350-DEL-70 | chr2-99588351-DEL-70 |
| chr2-99596521-DEL-302 | chr2-99596522-DEL-302 |
| chr2-99611333-DEL-2743 | chr2-99611334-DEL-2743 |
| chr2-99637919-DEL-132 | chr2-99637918-DEL-133 |
| chr2-99676159-DEL-4507 | chr2-99676159-DEL-4507 |
| chr2-99680706-DEL-1040 | chr2-99680706-DEL-1040 |
| chr2-99692711-DEL-2527 | chr2-99692712-DEL-2527 |
| chr2-99707593-DEL-331 | chr2-99707594-DEL-331 |
| chr2-99711884-DEL-1717 | chr2-99711885-DEL-1717 |
| chr2-99729736-DEL-352 | chr2-99729737-DEL-352 |
| chr2-99740512-DEL-333 | chr2-99740513-DEL-333 |
| chr2-99752415-DEL-82 | chr2-99752416-DEL-82 |
| chr2-99776436-DEL-1727 | chr2-99776437-DEL-1727 |
| chr2-99781931-DEL-4245 | chr2-99781932-DEL-4245 |
| chr2-99787122-DEL-807 | chr2-99787123-DEL-807 |
| chr2-99804474-DEL-110 | chr2-99804473-DEL-110 |
| chr2-99818238-DEL-2178 | chr2-99818239-DEL-2178 |
| chr2-99828831-DEL-226 | chr2-99828832-DEL-226 |
| chr2-99844818-DEL-286 | chr2-99844819-DEL-286 |
| chr2-99855505-DEL-222 | chr2-99855506-DEL-222 |
| chr2-99865151-DEL-5244 | chr2-99865152-DEL-5244 |
| chr2-99871248-DEL-1185 | chr2-99871249-DEL-1185 |
| chr2-99874893-DEL-200 | chr2-99874894-DEL-200 |
| chr2-99875665-DEL-53 | chr2-99875666-DEL-53 |
| chr2-99903685-DEL-66 | chr2-99903686-DEL-66 |
| chr2-99921696-DEL-3808 | chr2-99921697-DEL-3808 |
| chr2-99931062-DEL-55 | chr2-99931063-DEL-55 |
| chr2-99933524-DEL-614 | chr2-99933525-DEL-614 |
| chr2-99965873-DEL-1295 | chr2-99965874-DEL-1295 |
| chr2-99974777-DEL-481 | chr2-99974778-DEL-481 |
| chr2-99990269-DEL-650 | chr2-99990270-DEL-650 |
| chr2-100001510-DEL-4439 | chr2-100001511-DEL-4439 |
| chr2-100007745-DEL-57 | chr2-100007747-DEL-56 |
| chr2-100009668-DEL-51 | chr2-100009658-DEL-52 |
| chr2-100014627-DEL-1546 | chr2-100014628-DEL-1546 |
| chr2-100021372-DEL-1922 | chr2-100021373-DEL-1922 |
| chr2-100047311-DEL-659 | chr2-100047312-DEL-659 |
| chr2-100062067-DEL-103 | chr2-100062068-DEL-103 |
| chr2-100097891-DEL-54 | chr2-100097892-DEL-54 |
| chr2-100106727-DEL-314 | chr2-100106729-DEL-314 |
| chr2-100108801-DEL-518 | chr2-100108802-DEL-518 |
| chr2-100132645-DEL-3602 | chr2-100132646-DEL-3602 |
| chr2-100134465-DEL-455 | chr2-100134466-DEL-455 |
| chr2-100137806-DEL-4614 | chr2-100137807-DEL-4614 |
| chr2-100141874-DEL-70 | chr2-100141875-DEL-70 |
| chr2-100163984-DEL-270 | chr2-100163985-DEL-270 |
| chr2-100188346-DEL-83 | chr2-100188347-DEL-83 |
| chr2-100230696-DEL-51 | chr2-100230697-DEL-51 |
| chr2-100243220-DEL-56 | chr2-100243221-DEL-56 |
| chr2-100248723-DEL-4899 | chr2-100248723-DEL-4899 |
| chr2-100289971-DEL-4678 | chr2-100289971-DEL-4679 |
| chr2-100302571-DEL-1534 | chr2-100302571-DEL-1535 |
| chr2-100305062-DEL-9491 | chr2-100305062-DEL-9491 |

| | |
|---|---|
| chr2-100376952-DEL-206 | chr2-100376953-DEL-206 |
| chr2-100377472-DEL-412 | chr2-100377472-DEL-412 |
| chr2-100414600-DEL-294 | chr2-100414598-DEL-297 |
| chr2-100435379-DEL-4286 | chr2-100435380-DEL-4287 |
| chr2-100611766-DEL-60 | chr2-100611769-DEL-58 |
| chr2-100617574-DEL-69 | chr2-100617575-DEL-69 |
| chr2-100630879-DEL-316 | chr2-100630880-DEL-316 |
| chr2-100653003-DEL-929 | chr2-100653004-DEL-929 |
| chr2-100676094-DEL-74 | chr2-100676097-DEL-71 |
| chr2-100681097-DEL-69 | chr2-100681098-DEL-69 |
| chr2-100695986-DEL-69 | chr2-100695987-DEL-69 |
| chr2-100700082-DEL-120 | chr2-100700083-DEL-120 |
| chr2-100706946-DEL-131 | chr2-100706947-DEL-131 |
| chr2-100713412-DEL-51 | chr2-100713413-DEL-51 |
| chr2-100723330-DEL-128 | chr2-100723331-DEL-125 |
| chr2-100757197-DEL-462 | chr2-100757197-DEL-462 |
| chr2-100761744-DEL-6521 | chr2-100761745-DEL-6521 |
| chr2-100768846-DEL-5984 | chr2-100768841-DEL-5985 |
| chr2-100775266-DEL-75 | chr2-100775276-DEL-50 |
| chr2-100800199-DEL-53 | chr2-100800196-DEL-60 |
| chr2-100802546-DEL-70 | chr2-100802549-DEL-68 |
| chr2-100826650-DEL-281 | chr2-100826651-DEL-281 |
| chr2-100837837-DEL-998 | chr2-100837840-DEL-1002 |
| chr2-100860687-DEL-58 | chr2-100860688-DEL-58 |
| chr2-100861260-DEL-385 | chr2-100861261-DEL-385 |
| chr2-100876775-DEL-107 | chr2-100876776-DEL-107 |
| chr2-100884095-DEL-1690 | chr2-100884096-DEL-1690 |
| chr2-100901798-DEL-251 | chr2-100901799-DEL-251 |
| chr2-100928861-DEL-182 | chr2-100928862-DEL-182 |
| chr2-100939047-DEL-242 | chr2-100939046-DEL-244 |
| chr2-100944736-DEL-86 | chr2-100944737-DEL-84 |
| chr2-100956753-DEL-7210 | chr2-100956754-DEL-7210 |
| chr2-100976682-DEL-7116 | chr2-100976683-DEL-7116 |
| chr2-101016616-DEL-2261 | chr2-101016617-DEL-2261 |
| chr2-101022506-DEL-92 | chr2-101022503-DEL-96 |
| chr2-101046464-DEL-118 | chr2-101046465-DEL-118 |
| chr2-101046753-DEL-2228 | chr2-101046753-DEL-2229 |
| chr2-101097822-DEL-54 | chr2-101097823-DEL-54 |
| chr2-101106538-DEL-269 | chr2-101106539-DEL-269 |
| chr2-101143507-DEL-1893 | chr2-101143508-DEL-1893 |
| chr2-101155640-DEL-6366 | chr2-101155641-DEL-6366 |
| chr2-101174574-DEL-6789 | chr2-101174575-DEL-6789 |
| chr2-101218798-DEL-1362 | chr2-101218799-DEL-1362 |
| chr2-101233450-DEL-6360 | chr2-101233451-DEL-6360 |
| chr2-101243179-DEL-69 | chr2-101243180-DEL-69 |
| chr2-101267119-DEL-69 | chr2-101267120-DEL-69 |
| chr2-101267415-DEL-441 | chr2-101267416-DEL-441 |
| chr2-101290345-DEL-67 | chr2-101290345-DEL-67 |
| chr2-101297642-DEL-54 | chr2-101297643-DEL-54 |
| chr2-101326748-DEL-138 | chr2-101326749-DEL-138 |
| chr2-101332323-DEL-2094 | chr2-101332324-DEL-2094 |
| chr2-101342968-DEL-3599 | chr2-101342969-DEL-3598 |
| chr2-101354216-DEL-914 | chr2-101354218-DEL-913 |
| chr2-101365646-DEL-398 | chr2-101365647-DEL-398 |
| chr2-101455705-DEL-187 | chr2-101455706-DEL-187 |
| chr2-101524738-DEL-68 | chr2-101524739-DEL-68 |
| chr2-101560759-DEL-158 | chr2-101560760-DEL-158 |
| chr2-101575910-DEL-196 | chr2-101575924-DEL-181 |
| chr2-101587052-DEL-62 | chr2-101587055-DEL-60 |
| chr2-101619464-DEL-84 | chr2-101619451-DEL-58 |
| chr2-101650793-DEL-124 | chr2-101650794-DEL-124 |
| chr2-101703232-DEL-68 | chr2-101703235-DEL-66 |
| chr2-101710696-DEL-58 | chr2-101710697-DEL-54 |
| chr2-101737105-DEL-92 | chr2-101737104-DEL-93 |
| chr2-101740847-DEL-1759 | chr2-101740848-DEL-1759 |
| chr2-101800178-DEL-64 | chr2-101800179-DEL-64 |
| chr2-101804877-DEL-52 | chr2-101804878-DEL-50 |
| chr2-101810453-DEL-68 | chr2-101810453-DEL-68 |
| chr2-101831120-DEL-79 | chr2-101831140-DEL-61 |
| chr2-101841082-DEL-64 | chr2-101841083-DEL-64 |
| chr2-101855520-DEL-122 | chr2-101855521-DEL-122 |
| chr2-101858666-DEL-55 | chr2-101858665-DEL-55 |
| chr2-101868001-DEL-64 | chr2-101868002-DEL-64 |
| chr2-101876289-DEL-52 | chr2-101876290-DEL-52 |
| chr2-101896703-DEL-120 | chr2-101896702-DEL-120 |
| chr2-101906868-DEL-620 | chr2-101906869-DEL-620 |
| chr2-101911229-DEL-51 | chr2-101911230-DEL-51 |
| chr2-101948811-DEL-90 | chr2-101948812-DEL-90 |
| chr2-101959002-DEL-101 | chr2-101959003-DEL-101 |
| chr2-102002657-DEL-878 | chr2-102002658-DEL-878 |

| | |
|---|---|
| chr2-102031488-DEL-198 | chr2-102031490-DEL-197 |
| chr2-102051402-DEL-149 | chr2-102051406-DEL-145 |
| chr2-102056689-DEL-450 | chr2-102056692-DEL-443 |
| chr2-102057201-DEL-52 | chr2-102057201-DEL-52 |
| chr2-102057775-DEL-152 | chr2-102057778-DEL-154 |
| chr2-102093344-DEL-239 | chr2-102093345-DEL-239 |
| chr2-102112259-DEL-1112 | chr2-102112260-DEL-1112 |
| chr2-102120109-DEL-61 | chr2-102120106-DEL-56 |
| chr2-102150320-DEL-87 | chr2-102150321-DEL-87 |
| chr2-102183847-DEL-64 | chr2-102183848-DEL-64 |
| chr2-102195598-DEL-154 | chr2-102195599-DEL-154 |
| chr2-102202255-DEL-559 | chr2-102202256-DEL-559 |
| chr2-102248884-DEL-537 | chr2-102248885-DEL-537 |
| chr2-102291013-DEL-201 | chr2-102291014-DEL-201 |
| chr2-102362132-DEL-92 | chr2-102362132-DEL-92 |
| chr2-102371770-DEL-73 | chr2-102371740-DEL-67 |
| chr2-102375594-DEL-247 | chr2-102375595-DEL-247 |
| chr2-102377233-DEL-392 | chr2-102377234-DEL-392 |
| chr2-102386445-DEL-2827 | chr2-102386446-DEL-2827 |
| chr2-102402604-DEL-70 | chr2-102402605-DEL-70 |
| chr2-102417361-DEL-184 | chr2-102417362-DEL-184 |
| chr2-102429041-DEL-86 | chr2-102429042-DEL-86 |
| chr2-102433171-DEL-6876 | chr2-102433172-DEL-6876 |
| chr2-102439504-DEL-201 | chr2-102439505-DEL-201 |
| chr2-102446469-DEL-77 | chr2-102446470-DEL-77 |
| chr2-102458445-DEL-64 | chr2-102458446-DEL-64 |
| chr2-102495626-DEL-150 | chr2-102495627-DEL-150 |
| chr2-102503019-DEL-86 | chr2-102503020-DEL-86 |
| chr2-102508115-DEL-957 | chr2-102508116-DEL-957 |
| chr2-102509132-DEL-903 | chr2-102509133-DEL-903 |
| chr2-102544959-DEL-90 | chr2-102544960-DEL-90 |
| chr2-102549211-DEL-73 | chr2-102549196-DEL-89 |
| chr2-102555464-DEL-1165 | chr2-102555465-DEL-1165 |
| chr2-102580902-DEL-1918 | chr2-102580903-DEL-1918 |
| chr2-102612544-DEL-56 | chr2-102612545-DEL-56 |
| chr2-102624835-DEL-64 | chr2-102624843-DEL-62 |
| chr2-102630913-DEL-80 | chr2-102630914-DEL-80 |
| chr2-102653644-DEL-85 | chr2-102653645-DEL-85 |
| chr2-102689804-DEL-99 | chr2-102689805-DEL-99 |
| chr2-102698437-DEL-60 | chr2-102698442-DEL-56 |
| chr2-102720374-DEL-349 | chr2-102720375-DEL-349 |
| chr2-102734541-DEL-68 | chr2-102734542-DEL-68 |
| chr2-102735450-DEL-159 | chr2-102735451-DEL-159 |
| chr2-102744255-DEL-397 | chr2-102744256-DEL-397 |
| chr2-102759661-DEL-250 | chr2-102759662-DEL-250 |
| chr2-102810433-DEL-127 | chr2-102810425-DEL-90 |
| chr2-102832460-DEL-81 | chr2-102832461-DEL-81 |
| chr2-102833092-DEL-852 | chr2-102833093-DEL-852 |
| chr2-102894649-DEL-2433 | chr2-102894650-DEL-2433 |
| chr2-102914662-DEL-198 | chr2-102914663-DEL-198 |
| chr2-102948343-DEL-60 | chr2-102948322-DEL-94 |
| chr2-102962370-DEL-95 | chr2-102962382-DEL-62 |
| chr2-102970166-DEL-104 | chr2-102970168-DEL-60 |
| chr2-103019183-DEL-63 | chr2-103019177-DEL-52 |
| chr2-103162646-DEL-1522 | chr2-103162647-DEL-1522 |
| chr2-103177675-DEL-56 | chr2-103177638-DEL-102 |
| chr2-103201575-DEL-62 | chr2-103201576-DEL-59 |
| chr2-103212363-DEL-60 | chr2-103212363-DEL-60 |
| chr2-103233563-DEL-107 | chr2-103233564-DEL-107 |
| chr2-103263556-DEL-61 | chr2-103263557-DEL-63 |
| chr2-103271396-DEL-56 | chr2-103271397-DEL-56 |
| chr2-103278326-DEL-507 | chr2-103278331-DEL-502 |
| chr2-103306113-DEL-853 | chr2-103306114-DEL-853 |
| chr2-103310873-DEL-90 | chr2-103310831-DEL-90 |
| chr2-103316889-DEL-430 | chr2-103316878-DEL-396 |
| chr2-103357202-DEL-56 | chr2-103357203-DEL-56 |
| chr2-103366179-DEL-196 | chr2-103366177-DEL-198 |
| chr2-103402412-DEL-75 | chr2-103402413-DEL-75 |
| chr2-103414911-DEL-635 | chr2-103414912-DEL-635 |
| chr2-103447709-DEL-208 | chr2-103447710-DEL-208 |
| chr2-103449499-DEL-190 | chr2-103449500-DEL-190 |
| chr2-103462357-DEL-208 | chr2-103462358-DEL-208 |
| chr2-103477754-DEL-72 | chr2-103477758-DEL-66 |
| chr2-103499404-DEL-907 | chr2-103499404-DEL-904 |
| chr2-103500504-DEL-1045 | chr2-103500505-DEL-1045 |
| chr2-103523697-DEL-62 | chr2-103523698-DEL-62 |
| chr2-103543417-DEL-52 | chr2-103543418-DEL-52 |
| chr2-103549915-DEL-52 | chr2-103549921-DEL-60 |
| chr2-103583324-DEL-108 | chr2-103583325-DEL-108 |
| chr2-103611525-DEL-217 | chr2-103611526-DEL-217 |

| | |
|---|---|
| chr2-103691854-DEL-1869 | chr2-103691855-DEL-1869 |
| chr2-103703570-DEL-192 | chr2-103703571-DEL-192 |
| chr2-103718220-DEL-211 | chr2-103718221-DEL-211 |
| chr2-103815603-DEL-209 | chr2-103815604-DEL-209 |
| chr2-103818160-DEL-328 | chr2-103818161-DEL-328 |
| chr2-103818742-DEL-260 | chr2-103818743-DEL-260 |
| chr2-103830901-DEL-132 | chr2-103830922-DEL-123 |
| chr2-103836813-DEL-53 | chr2-103836814-DEL-53 |
| chr2-103839129-DEL-175 | chr2-103839130-DEL-175 |
| chr2-103839544-DEL-7079 | chr2-103839561-DEL-7063 |
| chr2-103849584-DEL-177 | chr2-103849585-DEL-177 |
| chr2-103861420-DEL-383 | chr2-103861421-DEL-382 |
| chr2-103877056-DEL-722 | chr2-103877056-DEL-722 |
| chr2-103942712-DEL-193 | chr2-103942713-DEL-193 |
| chr2-103943202-DEL-372 | chr2-103943203-DEL-372 |
| chr2-103968365-DEL-56 | chr2-103968354-DEL-60 |
| chr2-103970257-DEL-939 | chr2-103970258-DEL-939 |
| chr2-103989351-DEL-613 | chr2-103989352-DEL-613 |
| chr2-103990665-DEL-1051 | chr2-103990666-DEL-1051 |
| chr2-103999863-DEL-823 | chr2-103999864-DEL-823 |
| chr2-104032694-DEL-179 | chr2-104032688-DEL-164 |
| chr2-104050548-DEL-56 | chr2-104050549-DEL-84 |
| chr2-104070474-DEL-54 | chr2-104070475-DEL-54 |
| chr2-104094699-DEL-601 | chr2-104094700-DEL-601 |
| chr2-104100170-DEL-149 | chr2-104100171-DEL-149 |
| chr2-104103123-DEL-327 | chr2-104103124-DEL-327 |
| chr2-104110979-DEL-328 | chr2-104110980-DEL-328 |
| chr2-104118108-DEL-72 | chr2-104118109-DEL-72 |
| chr2-104124537-DEL-135 | chr2-104124538-DEL-135 |
| chr2-104141220-DEL-1482 | chr2-104141221-DEL-1482 |
| chr2-104160440-DEL-74 | chr2-104160441-DEL-74 |
| chr2-104201924-DEL-185 | chr2-104201923-DEL-185 |
| chr2-104249914-DEL-99 | chr2-104249915-DEL-99 |
| chr2-104267716-DEL-191 | chr2-104267717-DEL-189 |
| chr2-104351622-DEL-170 | chr2-104351612-DEL-171 |
| chr2-104357990-DEL-71 | chr2-104357971-DEL-70 |
| chr2-104374707-DEL-99 | chr2-104374709-DEL-98 |
| chr2-104375526-DEL-144 | chr2-104375527-DEL-144 |
| chr2-104396928-DEL-305 | chr2-104396915-DEL-319 |
| chr2-104415852-DEL-59 | chr2-104415851-DEL-60 |
| chr2-104436960-DEL-110 | chr2-104436961-DEL-110 |
| chr2-104438880-DEL-214 | chr2-104438881-DEL-214 |
| chr2-104471303-DEL-141 | chr2-104471309-DEL-124 |
| chr2-104486482-DEL-153 | chr2-104486483-DEL-153 |
| chr2-104501090-DEL-4104 | chr2-104501091-DEL-4104 |
| chr2-104501672-DEL-104 | chr2-104501673-DEL-104 |
| chr2-104503182-DEL-2339 | chr2-104503183-DEL-2341 |
| chr2-104524671-DEL-2996 | chr2-104524655-DEL-3046 |
| chr2-104530956-DEL-271 | chr2-104530957-DEL-271 |
| chr2-104601488-DEL-195 | chr2-104601489-DEL-195 |
| chr2-104605659-DEL-399 | chr2-104605660-DEL-399 |
| chr2-104607958-DEL-58 | chr2-104607959-DEL-56 |
| chr2-104625136-DEL-60 | chr2-104625137-DEL-60 |
| chr2-104661839-DEL-94 | chr2-104661840-DEL-94 |
| chr2-104664931-DEL-2574 | chr2-104664932-DEL-2574 |
| chr2-104712102-DEL-209 | chr2-104712103-DEL-209 |
| chr2-104729850-DEL-247 | chr2-104729851-DEL-247 |
| chr2-104785811-DEL-1076 | chr2-104785812-DEL-1076 |
| chr2-104790801-DEL-235 | chr2-104790802-DEL-235 |
| chr2-104815641-DEL-171 | chr2-104815642-DEL-141 |
| chr2-104828788-DEL-104 | chr2-104828789-DEL-104 |
| chr2-104858874-DEL-214 | chr2-104858875-DEL-214 |
| chr2-104934673-DEL-84 | chr2-104934684-DEL-74 |
| chr2-104937951-DEL-443 | chr2-104937952-DEL-443 |
| chr2-104941205-DEL-76 | chr2-104941206-DEL-76 |
| chr2-104941459-DEL-845 | chr2-104941460-DEL-844 |
| chr2-105028097-DEL-400 | chr2-105028098-DEL-400 |
| chr2-105061261-DEL-315 | chr2-105061262-DEL-315 |
| chr2-105088612-DEL-741 | chr2-105088613-DEL-741 |
| chr2-105104204-DEL-52 | chr2-105104205-DEL-52 |
| chr2-105118559-DEL-350 | chr2-105118560-DEL-350 |
| chr2-105121997-DEL-126 | chr2-105122008-DEL-116 |
| chr2-105125484-DEL-73 | chr2-105125485-DEL-73 |
| chr2-105146426-DEL-1128 | chr2-105146429-DEL-1131 |
| chr2-105156887-DEL-1166 | chr2-105156888-DEL-1166 |
| chr2-105193220-DEL-110 | chr2-105193218-DEL-119 |
| chr2-105268869-DEL-55 | chr2-105268855-DEL-56 |
| chr2-105292978-DEL-156 | chr2-105292979-DEL-156 |
| chr2-105298406-DEL-90 | chr2-105298407-DEL-90 |
| chr2-105302509-DEL-64 | chr2-105302499-DEL-65 |

| | |
|---|---|
| chr2-105324467-DEL-138 | chr2-105324468-DEL-138 |
| chr2-105333807-DEL-231 | chr2-105333808-DEL-231 |
| chr2-105341496-DEL-99 | chr2-105341497-DEL-99 |
| chr2-105368270-DEL-6024 | chr2-105368256-DEL-6025 |
| chr2-105384448-DEL-783 | chr2-105384449-DEL-783 |
| chr2-105388244-DEL-5132 | chr2-105388245-DEL-5132 |
| chr2-105399555-DEL-361 | chr2-105399556-DEL-361 |
| chr2-105401968-DEL-54 | chr2-105401969-DEL-54 |
| chr2-105417693-DEL-102 | chr2-105417694-DEL-102 |
| chr2-105445704-DEL-211 | chr2-105445705-DEL-211 |
| chr2-105460257-DEL-1966 | chr2-105460258-DEL-1966 |
| chr2-105461460-DEL-66 | chr2-105461461-DEL-60 |
| chr2-105462694-DEL-58 | chr2-105462694-DEL-68 |
| chr2-105555509-DEL-683 | chr2-105555510-DEL-683 |
| chr2-105599379-DEL-137 | chr2-105599380-DEL-137 |
| chr2-105675702-DEL-212 | chr2-105675703-DEL-212 |
| chr2-105688922-DEL-398 | chr2-105688923-DEL-398 |
| chr2-105715557-DEL-1668 | chr2-105715558-DEL-1668 |
| chr2-105772741-DEL-6597 | chr2-105772742-DEL-6597 |
| chr2-105921101-DEL-761 | chr2-105921102-DEL-761 |
| chr2-105959620-DEL-686 | chr2-105959621-DEL-686 |
| chr2-105972023-DEL-110 | chr2-105972024-DEL-110 |
| chr2-106001428-DEL-78 | chr2-106001448-DEL-90 |
| chr2-106071034-DEL-1939 | chr2-106071036-DEL-1937 |
| chr2-106071864-DEL-272 | chr2-106071865-DEL-272 |
| chr2-106106558-DEL-834 | chr2-106106559-DEL-834 |
| chr2-106118283-DEL-1516 | chr2-106118284-DEL-1516 |
| chr2-106123577-DEL-520 | chr2-106123578-DEL-520 |
| chr2-106144045-DEL-84 | chr2-106144046-DEL-84 |
| chr2-106173043-DEL-467 | chr2-106173047-DEL-464 |
| chr2-106174848-DEL-107 | chr2-106174849-DEL-107 |
| chr2-106183598-DEL-1262 | chr2-106183599-DEL-1262 |
| chr2-106190674-DEL-851 | chr2-106190675-DEL-851 |
| chr2-106208266-DEL-169 | chr2-106208267-DEL-169 |
| chr2-106247001-DEL-95 | chr2-106247002-DEL-95 |
| chr2-106251378-DEL-2272 | chr2-106251379-DEL-2272 |
| chr2-106270982-DEL-274 | chr2-106270991-DEL-266 |
| chr2-106290383-DEL-56 | chr2-106290384-DEL-56 |
| chr2-106324474-DEL-103 | chr2-106324459-DEL-119 |
| chr2-106348566-DEL-91 | chr2-106348566-DEL-91 |
| chr2-106365646-DEL-88 | chr2-106365626-DEL-87 |
| chr2-106379393-DEL-195 | chr2-106379394-DEL-195 |
| chr2-106395929-DEL-329 | chr2-106395933-DEL-326 |
| chr2-106409516-DEL-178 | chr2-106409517-DEL-178 |
| chr2-106423171-DEL-960 | chr2-106423172-DEL-960 |
| chr2-106435938-DEL-68 | chr2-106435941-DEL-56 |
| chr2-106466358-DEL-56 | chr2-106466359-DEL-56 |
| chr2-106466938-DEL-3283 | chr2-106466939-DEL-3283 |
| chr2-106470282-DEL-205 | chr2-106470283-DEL-205 |
| chr2-106516003-DEL-200 | chr2-106516004-DEL-200 |
| chr2-106539620-DEL-60 | chr2-106539620-DEL-60 |
| chr2-106583051-DEL-76 | chr2-106583063-DEL-71 |
| chr2-106608022-DEL-490 | chr2-106608023-DEL-490 |
| chr2-106668817-DEL-60 | chr2-106668818-DEL-60 |
| chr2-106702171-DEL-54 | chr2-106702172-DEL-54 |
| chr2-106711743-DEL-75 | chr2-106711744-DEL-75 |
| chr2-106713128-DEL-90 | chr2-106713129-DEL-90 |
| chr2-106715742-DEL-57 | chr2-106715743-DEL-57 |
| chr2-106728506-DEL-401 | chr2-106728507-DEL-401 |
| chr2-106741259-DEL-56 | chr2-106741258-DEL-60 |
| chr2-106766109-DEL-70 | chr2-106766110-DEL-70 |
| chr2-106769447-DEL-54 | chr2-106769447-DEL-55 |
| chr2-106774939-DEL-498 | chr2-106774940-DEL-498 |
| chr2-106802373-DEL-213 | chr2-106802374-DEL-213 |
| chr2-106892130-DEL-102 | chr2-106892120-DEL-104 |
| chr2-106904917-DEL-74 | chr2-106904919-DEL-94 |
| chr2-106914385-DEL-144 | chr2-106914386-DEL-144 |
| chr2-106931163-DEL-6543 | chr2-106931164-DEL-6543 |
| chr2-106947252-DEL-98 | chr2-106947255-DEL-94 |
| chr2-106952326-DEL-2632 | chr2-106952327-DEL-2632 |
| chr2-106955196-DEL-74 | chr2-106955197-DEL-74 |
| chr2-106955972-DEL-732 | chr2-106955973-DEL-732 |
| chr2-106956009-DEL-143 | chr2-106956010-DEL-143 |
| chr2-106956579-DEL-54 | chr2-106956580-DEL-54 |
| chr2-106964418-DEL-6808 | chr2-106964419-DEL-6808 |
| chr2-106972625-DEL-204 | chr2-106972644-DEL-174 |
| chr2-106995871-DEL-6267 | chr2-106995872-DEL-6267 |
| chr2-107015106-DEL-60 | chr2-107015107-DEL-60 |
| chr2-107026189-DEL-63 | chr2-107026190-DEL-57 |
| chr2-107029721-DEL-195 | chr2-107029745-DEL-194 |

| | |
|---|---|
| chr2-107034449-DEL-55 | chr2-107034449-DEL-55 |
| chr2-107077244-DEL-623 | chr2-107077260-DEL-615 |
| chr2-107090692-DEL-156 | chr2-107090693-DEL-156 |
| chr2-107141097-DEL-55 | chr2-107141088-DEL-56 |
| chr2-107174377-DEL-54 | chr2-107174382-DEL-80 |
| chr2-107193915-DEL-58 | chr2-107193916-DEL-58 |
| chr2-107224605-DEL-2040 | chr2-107224605-DEL-2040 |
| chr2-107226680-DEL-1339 | chr2-107226680-DEL-1339 |
| chr2-107242681-DEL-491 | chr2-107242682-DEL-491 |
| chr2-107278555-DEL-70 | chr2-107278556-DEL-70 |
| chr2-107283520-DEL-202 | chr2-107283521-DEL-202 |
| chr2-107289175-DEL-414 | chr2-107289176-DEL-414 |
| chr2-107292446-DEL-67 | chr2-107292452-DEL-62 |
| chr2-107380559-DEL-1008 | chr2-107380560-DEL-1008 |
| chr2-107409916-DEL-429 | chr2-107409917-DEL-429 |
| chr2-107420838-DEL-160 | chr2-107420838-DEL-160 |
| chr2-107424391-DEL-7099 | chr2-107424392-DEL-7099 |
| chr2-107443089-DEL-99 | chr2-107443075-DEL-70 |
| chr2-107448094-DEL-62 | chr2-107448074-DEL-62 |
| chr2-107449562-DEL-918 | chr2-107449563-DEL-917 |
| chr2-107478886-DEL-3878 | chr2-107478887-DEL-3878 |
| chr2-107536748-DEL-586 | chr2-107536749-DEL-586 |
| chr2-107567560-DEL-341 | chr2-107567561-DEL-341 |
| chr2-107572112-DEL-6389 | chr2-107572113-DEL-6389 |
| chr2-107584856-DEL-2075 | chr2-107584857-DEL-2075 |
| chr2-107602395-DEL-70 | chr2-107602398-DEL-68 |
| chr2-107628706-DEL-3235 | chr2-107628708-DEL-3234 |
| chr2-107633953-DEL-410 | chr2-107633954-DEL-410 |
| chr2-107647310-DEL-81 | chr2-107647311-DEL-81 |
| chr2-107647439-DEL-473 | chr2-107647440-DEL-473 |
| chr2-107668929-DEL-167 | chr2-107668930-DEL-165 |
| chr2-107682450-DEL-1418 | chr2-107682451-DEL-1418 |
| chr2-107698529-DEL-210 | chr2-107698530-DEL-210 |
| chr2-107714373-DEL-280 | chr2-107714374-DEL-280 |
| chr2-107726383-DEL-146 | chr2-107726384-DEL-146 |
| chr2-107733119-DEL-73 | chr2-107733100-DEL-71 |
| chr2-107750876-DEL-201 | chr2-107750877-DEL-201 |
| chr2-107762764-DEL-399 | chr2-107762765-DEL-399 |
| chr2-107782710-DEL-3385 | chr2-107782711-DEL-3385 |
| chr2-107784534-DEL-527 | chr2-107784535-DEL-527 |
| chr2-107797365-DEL-72 | chr2-107797366-DEL-60 |
| chr2-107807157-DEL-6619 | chr2-107807158-DEL-6619 |
| chr2-107845832-DEL-1066 | chr2-107845833-DEL-1066 |
| chr2-107871235-DEL-260 | chr2-107871236-DEL-260 |
| chr2-107877179-DEL-2901 | chr2-107877179-DEL-2901 |
| chr2-107877902-DEL-56 | chr2-107877903-DEL-56 |
| chr2-107891547-DEL-1330 | chr2-107891548-DEL-1330 |
| chr2-107907101-DEL-62 | chr2-107907102-DEL-62 |
| chr2-107920925-DEL-151 | chr2-107920926-DEL-151 |
| chr2-107937193-DEL-70 | chr2-107937191-DEL-73 |
| chr2-107949365-DEL-6387 | chr2-107949366-DEL-6387 |
| chr2-107956512-DEL-64 | chr2-107956505-DEL-72 |
| chr2-107996560-DEL-80 | chr2-107996531-DEL-110 |
| chr2-108028515-DEL-1022 | chr2-108028513-DEL-1023 |
| chr2-108067309-DEL-195 | chr2-108067310-DEL-195 |
| chr2-108068424-DEL-928 | chr2-108068425-DEL-928 |
| chr2-108112069-DEL-7048 | chr2-108112070-DEL-7048 |
| chr2-108114434-DEL-204 | chr2-108114436-DEL-202 |
| chr2-108114850-DEL-517 | chr2-108114851-DEL-517 |
| chr2-108117293-DEL-51 | chr2-108117261-DEL-51 |
| chr2-108157646-DEL-6833 | chr2-108157647-DEL-6833 |
| chr2-108173820-DEL-2443 | chr2-108173821-DEL-2443 |
| chr2-108185093-DEL-1115 | chr2-108185094-DEL-1115 |
| chr2-108190137-DEL-598 | chr2-108190138-DEL-598 |
| chr2-108193582-DEL-455 | chr2-108193583-DEL-455 |
| chr2-108212164-DEL-735 | chr2-108212167-DEL-739 |
| chr2-108233380-DEL-1191 | chr2-108233381-DEL-1191 |
| chr2-108245137-DEL-226 | chr2-108245138-DEL-226 |
| chr2-108247846-DEL-150 | chr2-108247847-DEL-150 |
| chr2-108251153-DEL-2787 | chr2-108251153-DEL-2787 |
| chr2-108266206-DEL-85 | chr2-108266225-DEL-78 |
| chr2-108267114-DEL-2449 | chr2-108267115-DEL-2449 |
| chr2-108272854-DEL-57 | chr2-108272855-DEL-57 |
| chr2-108319826-DEL-203 | chr2-108319827-DEL-203 |
| chr2-108338883-DEL-77 | chr2-108338891-DEL-53 |
| chr2-108383176-DEL-6391 | chr2-108383177-DEL-6391 |
| chr2-108410493-DEL-229 | chr2-108410494-DEL-229 |
| chr2-108424818-DEL-55 | chr2-108424818-DEL-55 |
| chr2-108457852-DEL-785 | chr2-108457849-DEL-797 |
| chr2-108468180-DEL-484 | chr2-108468181-DEL-484 |

| | |
|---|---|
| chr2-108548323-DEL-64 | chr2-108548326-DEL-54 |
| chr2-108549974-DEL-130 | chr2-108549964-DEL-141 |
| chr2-108588203-DEL-54 | chr2-108588204-DEL-54 |
| chr2-108640363-DEL-79 | chr2-108640347-DEL-78 |
| chr2-108705208-DEL-450 | chr2-108705209-DEL-450 |
| chr2-108723416-DEL-58 | chr2-108723417-DEL-58 |
| chr2-108729925-DEL-61 | chr2-108729926-DEL-61 |
| chr2-108738030-DEL-84 | chr2-108738031-DEL-84 |
| chr2-108769320-DEL-103 | chr2-108769331-DEL-105 |
| chr2-108779098-DEL-1988 | chr2-108779099-DEL-1988 |
| chr2-108802164-DEL-1100 | chr2-108802165-DEL-1100 |
| chr2-108824851-DEL-1555 | chr2-108824852-DEL-1555 |
| chr2-108835150-DEL-1778 | chr2-108835151-DEL-1778 |
| chr2-108928126-DEL-6380 | chr2-108928127-DEL-6380 |
| chr2-108954064-DEL-118 | chr2-108954065-DEL-118 |
| chr2-109046815-DEL-280 | chr2-109046815-DEL-280 |
| chr2-109132004-DEL-181 | chr2-109132022-DEL-166 |
| chr2-109164919-DEL-1119 | chr2-109164907-DEL-1118 |
| chr2-109176748-DEL-60 | chr2-109176783-DEL-50 |
| chr2-109185653-DEL-90 | chr2-109185656-DEL-88 |
| chr2-109189931-DEL-1699 | chr2-109189932-DEL-1700 |
| chr2-109198310-DEL-1515 | chr2-109198311-DEL-1514 |
| chr2-109198317-DEL-83 | chr2-109198316-DEL-50 |
| chr2-109202055-DEL-152 | chr2-109202056-DEL-151 |
| chr2-109210081-DEL-753 | chr2-109210082-DEL-753 |
| chr2-109254795-DEL-85 | chr2-109254796-DEL-85 |
| chr2-109258533-DEL-57 | chr2-109258510-DEL-60 |
| chr2-109290151-DEL-683 | chr2-109290152-DEL-683 |
| chr2-109346003-DEL-134 | chr2-109346004-DEL-134 |
| chr2-109416310-DEL-503 | chr2-109416311-DEL-503 |
| chr2-109431339-DEL-91 | chr2-109431340-DEL-91 |
| chr2-109459694-DEL-6576 | chr2-109459695-DEL-6576 |
| chr2-109479232-DEL-74 | chr2-109479233-DEL-74 |
| chr2-109484062-DEL-118 | chr2-109484103-DEL-120 |
| chr2-109497674-DEL-1054 | chr2-109497675-DEL-1054 |
| chr2-109499683-DEL-307 | chr2-109499684-DEL-307 |
| chr2-109559472-DEL-146 | chr2-109559473-DEL-146 |
| chr2-109591415-DEL-201 | chr2-109591416-DEL-201 |
| chr2-109620817-DEL-462 | chr2-109620816-DEL-463 |
| chr2-109651605-DEL-56 | chr2-109651606-DEL-54 |
| chr2-109654748-DEL-1067 | chr2-109654740-DEL-1066 |
| chr2-109656677-DEL-1453 | chr2-109656678-DEL-1453 |
| chr2-109682341-DEL-593 | chr2-109682334-DEL-594 |
| chr2-109691689-DEL-357 | chr2-109691690-DEL-357 |
| chr2-109701950-DEL-1353 | chr2-109701951-DEL-1353 |
| chr2-109734506-DEL-61 | chr2-109734507-DEL-61 |
| chr2-109754794-DEL-208 | chr2-109754795-DEL-208 |
| chr2-109780279-DEL-55 | chr2-109780280-DEL-55 |
| chr2-109829495-DEL-209 | chr2-109829496-DEL-209 |
| chr2-109862017-DEL-207 | chr2-109862018-DEL-207 |
| chr2-109906570-DEL-753 | chr2-109906571-DEL-753 |
| chr2-109924861-DEL-137 | chr2-109924862-DEL-137 |
| chr2-109987477-DEL-6108 | chr2-109987478-DEL-6108 |
| chr2-110004873-DEL-475 | chr2-110004868-DEL-476 |
| chr2-110067549-DEL-6418 | chr2-110067545-DEL-6419 |
| chr2-110123957-DEL-57 | chr2-110123958-DEL-57 |
| chr2-110139659-DEL-605 | chr2-110139652-DEL-606 |
| chr2-110165212-DEL-52 | chr2-110165213-DEL-52 |
| chr2-110170422-DEL-6632 | chr2-110170420-DEL-6633 |
| chr2-110189586-DEL-74 | chr2-110189575-DEL-84 |
| chr2-110218128-DEL-548 | chr2-110218129-DEL-548 |
| chr2-110259389-DEL-6868 | chr2-110259378-DEL-6867 |
| chr2-110284854-DEL-55 | chr2-110284855-DEL-55 |
| chr2-110288728-DEL-331 | chr2-110288729-DEL-331 |
| chr2-110328039-DEL-496 | chr2-110328037-DEL-497 |
| chr2-110365583-DEL-87 | chr2-110365584-DEL-87 |
| chr2-110369981-DEL-489 | chr2-110369982-DEL-489 |
| chr2-110375986-DEL-6221 | chr2-110375987-DEL-6221 |
| chr2-110458591-DEL-51 | chr2-110458604-DEL-60 |
| chr2-110460089-DEL-206 | chr2-110460090-DEL-206 |
| chr2-110467926-DEL-727 | chr2-110467927-DEL-727 |
| chr2-110480865-DEL-1541 | chr2-110480866-DEL-1541 |
| chr2-110515455-DEL-63 | chr2-110515456-DEL-63 |
| chr2-110518033-DEL-86 | chr2-110518020-DEL-55 |
| chr2-110519953-DEL-1298 | chr2-110519954-DEL-1298 |
| chr2-110530088-DEL-668 | chr2-110530089-DEL-662 |
| chr2-110539144-DEL-522 | chr2-110539145-DEL-522 |
| chr2-110541664-DEL-86 | chr2-110541665-DEL-82 |
| chr2-110556075-DEL-385 | chr2-110556076-DEL-385 |
| chr2-110582382-DEL-2806 | chr2-110582383-DEL-2806 |

| | |
|---|---|
| chr2-110588210-DEL-71 | chr2-110588208-DEL-72 |
| chr2-110592818-DEL-605 | chr2-110592818-DEL-606 |
| chr2-110596112-DEL-660 | chr2-110596113-DEL-660 |
| chr2-110597052-DEL-529 | chr2-110597053-DEL-529 |
| chr2-110649655-DEL-442 | chr2-110649656-DEL-442 |
| chr2-110652470-DEL-88 | chr2-110652466-DEL-92 |
| chr2-110653660-DEL-52 | chr2-110653661-DEL-52 |
| chr2-110655663-DEL-95 | chr2-110655653-DEL-98 |
| chr2-110663471-DEL-206 | chr2-110663472-DEL-206 |
| chr2-110695560-DEL-59 | chr2-110695561-DEL-74 |
| chr2-110697691-DEL-1482 | chr2-110697692-DEL-1482 |
| chr2-110725188-DEL-69 | chr2-110725189-DEL-69 |
| chr2-110733641-DEL-195 | chr2-110733642-DEL-195 |
| chr2-110776833-DEL-134 | chr2-110776834-DEL-134 |
| chr2-110781447-DEL-58 | chr2-110781448-DEL-58 |
| chr2-110791440-DEL-390 | chr2-110791441-DEL-390 |
| chr2-110830275-DEL-256 | chr2-110830276-DEL-256 |
| chr2-110832532-DEL-181 | chr2-110832546-DEL-173 |
| chr2-110879128-DEL-80 | chr2-110879129-DEL-80 |
| chr2-110890220-DEL-85 | chr2-110890221-DEL-85 |
| chr2-110910879-DEL-55 | chr2-110910880-DEL-55 |
| chr2-110914957-DEL-485 | chr2-110914958-DEL-485 |
| chr2-110993557-DEL-66 | chr2-110993560-DEL-52 |
| chr2-111004340-DEL-6582 | chr2-111004340-DEL-6582 |
| chr2-111046389-DEL-1780 | chr2-111046390-DEL-1780 |
| chr2-111086703-DEL-110 | chr2-111086716-DEL-92 |
| chr2-111105664-DEL-1875 | chr2-111105665-DEL-1875 |
| chr2-111111833-DEL-6478 | chr2-111111834-DEL-6478 |
| chr2-111163514-DEL-93 | chr2-111163514-DEL-93 |
| chr2-111175578-DEL-6167 | chr2-111175579-DEL-6167 |
| chr2-111192043-DEL-160 | chr2-111192044-DEL-160 |
| chr2-111201439-DEL-64 | chr2-111201436-DEL-64 |
| chr2-111305759-DEL-73 | chr2-111305760-DEL-73 |
| chr2-111318661-DEL-283 | chr2-111318658-DEL-284 |
| chr2-111340651-DEL-160 | chr2-111340656-DEL-159 |
| chr2-111341342-DEL-3044 | chr2-111341343-DEL-3044 |
| chr2-111363590-DEL-399 | chr2-111363591-DEL-399 |
| chr2-111378556-DEL-5066 | chr2-111378557-DEL-5066 |
| chr2-111412253-DEL-200 | chr2-111412252-DEL-201 |
| chr2-111427259-DEL-403 | chr2-111427260-DEL-403 |
| chr2-111428560-DEL-128 | chr2-111428561-DEL-128 |
| chr2-111435646-DEL-51 | chr2-111435638-DEL-52 |
| chr2-111480107-DEL-6374 | chr2-111480108-DEL-6374 |
| chr2-111520667-DEL-52 | chr2-111520668-DEL-52 |
| chr2-111547083-DEL-6544 | chr2-111547084-DEL-6544 |
| chr2-111553278-DEL-201 | chr2-111553292-DEL-201 |
| chr2-111603904-DEL-107 | chr2-111603897-DEL-96 |
| chr2-111614346-DEL-70 | chr2-111614347-DEL-70 |
| chr2-111622150-DEL-101 | chr2-111622141-DEL-61 |
| chr2-111687672-DEL-1888 | chr2-111687673-DEL-1888 |
| chr2-111691332-DEL-5416 | chr2-111691333-DEL-5416 |
| chr2-111698213-DEL-139 | chr2-111698214-DEL-139 |
| chr2-111717960-DEL-14546 | chr2-111717961-DEL-14546 |
| chr2-111754696-DEL-98 | chr2-111754685-DEL-99 |
| chr2-111803347-DEL-602 | chr2-111803348-DEL-602 |
| chr2-111884061-DEL-214 | chr2-111884062-DEL-214 |
| chr2-111912081-DEL-7534 | chr2-111912082-DEL-7534 |
| chr2-111913486-DEL-961 | chr2-111913487-DEL-961 |
| chr2-112005714-DEL-436 | chr2-112005715-DEL-436 |
| chr2-112030974-DEL-58 | chr2-112030975-DEL-58 |
| chr2-112031975-DEL-461 | chr2-112031976-DEL-461 |
| chr2-112038394-DEL-63 | chr2-112038396-DEL-55 |
| chr2-112053892-DEL-62 | chr2-112053893-DEL-62 |
| chr2-112066782-DEL-219 | chr2-112066783-DEL-219 |
| chr2-112067599-DEL-201 | chr2-112067600-DEL-201 |
| chr2-112118505-DEL-64 | chr2-112118506-DEL-64 |
| chr2-112138514-DEL-239 | chr2-112138515-DEL-239 |
| chr2-112143628-DEL-350 | chr2-112143629-DEL-350 |
| chr2-112150102-DEL-6442 | chr2-112150103-DEL-6442 |
| chr2-112159220-DEL-122 | chr2-112159218-DEL-122 |
| chr2-112166658-DEL-87 | chr2-112166659-DEL-87 |
| chr2-112184802-DEL-103 | chr2-112184803-DEL-103 |
| chr2-112190145-DEL-91 | chr2-112190146-DEL-91 |
| chr2-112230964-DEL-101 | chr2-112230968-DEL-97 |
| chr2-112231674-DEL-74 | chr2-112231675-DEL-74 |
| chr2-112247018-DEL-6411 | chr2-112247019-DEL-6411 |
| chr2-112324443-DEL-148 | chr2-112324444-DEL-154 |
| chr2-112325514-DEL-206 | chr2-112325515-DEL-206 |
| chr2-112341275-DEL-701 | chr2-112341276-DEL-701 |
| chr2-112357677-DEL-184 | chr2-112357678-DEL-184 |

| | |
|---|---|
| chr2-112379291-DEL-4835 | chr2-112379292-DEL-4835 |
| chr2-112398674-DEL-1890 | chr2-112398675-DEL-1890 |
| chr2-112414560-DEL-60 | chr2-112414561-DEL-54 |
| chr2-112417395-DEL-78 | chr2-112417364-DEL-96 |
| chr2-112450614-DEL-203 | chr2-112450615-DEL-203 |
| chr2-112519953-DEL-113 | chr2-112519954-DEL-113 |
| chr2-112521217-DEL-1085 | chr2-112521218-DEL-1085 |
| chr2-112528888-DEL-961 | chr2-112528889-DEL-961 |
| chr2-112531745-DEL-6432 | chr2-112531752-DEL-6434 |
| chr2-112568312-DEL-3777 | chr2-112568312-DEL-3777 |
| chr2-112601248-DEL-52 | chr2-112601249-DEL-52 |
| chr2-112632649-DEL-172 | chr2-112632671-DEL-188 |
| chr2-112651921-DEL-271 | chr2-112651922-DEL-271 |
| chr2-112654579-DEL-138 | chr2-112654580-DEL-138 |
| chr2-112656026-DEL-60 | chr2-112656026-DEL-60 |
| chr2-112656768-DEL-67 | chr2-112656769-DEL-66 |
| chr2-112657581-DEL-163 | chr2-112657582-DEL-163 |
| chr2-112700890-DEL-289 | chr2-112700891-DEL-289 |
| chr2-112793737-DEL-113 | chr2-112793765-DEL-78 |
| chr2-112805976-DEL-68 | chr2-112805977-DEL-68 |
| chr2-112833984-DEL-129 | chr2-112833983-DEL-130 |
| chr2-112926127-DEL-6475 | chr2-112926128-DEL-6475 |
| chr2-112937716-DEL-1130 | chr2-112937717-DEL-1130 |
| chr2-112952606-DEL-75 | chr2-112952579-DEL-75 |
| chr2-112955301-DEL-55 | chr2-112955302-DEL-55 |
| chr2-112958427-DEL-6626 | chr2-112958428-DEL-6626 |
| chr2-112971215-DEL-58 | chr2-112971216-DEL-58 |
| chr2-112979970-DEL-78 | chr2-112979971-DEL-78 |
| chr2-113010852-DEL-241 | chr2-113010853-DEL-241 |
| chr2-113034199-DEL-471 | chr2-113034200-DEL-471 |
| chr2-113034807-DEL-421 | chr2-113034808-DEL-421 |
| chr2-113057994-DEL-154 | chr2-113057995-DEL-154 |
| chr2-113065360-DEL-55 | chr2-113065360-DEL-55 |
| chr2-113072721-DEL-3332 | chr2-113072722-DEL-3332 |
| chr2-113092284-DEL-60 | chr2-113092284-DEL-60 |
| chr2-113103468-DEL-2234 | chr2-113103469-DEL-2234 |
| chr2-113114351-DEL-498 | chr2-113114352-DEL-498 |
| chr2-113152533-DEL-72 | chr2-113152534-DEL-72 |
| chr2-113158601-DEL-67 | chr2-113158601-DEL-68 |
| chr2-113161463-DEL-1891 | chr2-113161464-DEL-1891 |
| chr2-113168367-DEL-58 | chr2-113168369-DEL-58 |
| chr2-113183204-DEL-100 | chr2-113183190-DEL-93 |
| chr2-113218254-DEL-100 | chr2-113218255-DEL-100 |
| chr2-113220085-DEL-1052 | chr2-113220086-DEL-1052 |
| chr2-113244837-DEL-58 | chr2-113244831-DEL-59 |
| chr2-113278761-DEL-80 | chr2-113278762-DEL-80 |
| chr2-113290377-DEL-53 | chr2-113290378-DEL-53 |
| chr2-113298926-DEL-51 | chr2-113298927-DEL-51 |
| chr2-113348807-DEL-6401 | chr2-113348808-DEL-6401 |
| chr2-113395420-DEL-66 | chr2-113395421-DEL-66 |
| chr2-113405312-DEL-58 | chr2-113405270-DEL-60 |
| chr2-113502152-DEL-1170 | chr2-113502153-DEL-1170 |
| chr2-113559267-DEL-6990 | chr2-113559268-DEL-6990 |
| chr2-113568278-DEL-54 | chr2-113568278-DEL-54 |
| chr2-113617488-DEL-72 | chr2-113617489-DEL-72 |
| chr2-113626460-DEL-327 | chr2-113626461-DEL-327 |
| chr2-113640280-DEL-7071 | chr2-113640281-DEL-7071 |
| chr2-113650770-DEL-153 | chr2-113650771-DEL-153 |
| chr2-113654271-DEL-56 | chr2-113654268-DEL-60 |
| chr2-113657159-DEL-136 | chr2-113657160-DEL-136 |
| chr2-113662677-DEL-510 | chr2-113662633-DEL-510 |
| chr2-113715179-DEL-195 | chr2-113715191-DEL-184 |
| chr2-113721623-DEL-1249 | chr2-113721624-DEL-1249 |
| chr2-113723656-DEL-76 | chr2-113723657-DEL-50 |
| chr2-113735368-DEL-55 | chr2-113735369-DEL-55 |
| chr2-113750129-DEL-1357 | chr2-113750130-DEL-1357 |
| chr2-113770536-DEL-1885 | chr2-113770537-DEL-1885 |
| chr2-113777776-DEL-66 | chr2-113777777-DEL-66 |
| chr2-113798321-DEL-228 | chr2-113798322-DEL-228 |
| chr2-113902827-DEL-52 | chr2-113902828-DEL-52 |
| chr2-113906782-DEL-73 | chr2-113906783-DEL-73 |
| chr2-113907708-DEL-54 | chr2-113907746-DEL-52 |
| chr2-113988200-DEL-79 | chr2-113988212-DEL-70 |
| chr2-113990559-DEL-58 | chr2-113990560-DEL-58 |
| chr2-114009807-DEL-434 | chr2-114009808-DEL-434 |
| chr2-114011346-DEL-6473 | chr2-114011347-DEL-6473 |
| chr2-114040931-DEL-82 | chr2-114040932-DEL-81 |
| chr2-114066205-DEL-87 | chr2-114066206-DEL-87 |
| chr2-114108747-DEL-197 | chr2-114108740-DEL-198 |
| chr2-114114694-DEL-193 | chr2-114114695-DEL-193 |

| | |
|---|---|
| chr2-114126878-DEL-479 | chr2-114126879-DEL-479 |
| chr2-114127787-DEL-1550 | chr2-114127788-DEL-1550 |
| chr2-114143872-DEL-390 | chr2-114143873-DEL-390 |
| chr2-114172382-DEL-118 | chr2-114172383-DEL-118 |
| chr2-114184627-DEL-127 | chr2-114184628-DEL-127 |
| chr2-114186294-DEL-503 | chr2-114186295-DEL-503 |
| chr2-114213959-DEL-408 | chr2-114213959-DEL-408 |
| chr2-114230964-DEL-177 | chr2-114230965-DEL-177 |
| chr2-114256270-DEL-331 | chr2-114256271-DEL-331 |
| chr2-114310470-DEL-6437 | chr2-114310471-DEL-6437 |
| chr2-114327969-DEL-6577 | chr2-114327970-DEL-6577 |
| chr2-114344750-DEL-172 | chr2-114344751-DEL-172 |
| chr2-114357207-DEL-639 | chr2-114357208-DEL-638 |
| chr2-114385879-DEL-204 | chr2-114385880-DEL-204 |
| chr2-114395230-DEL-84 | chr2-114395231-DEL-84 |
| chr2-114398705-DEL-80 | chr2-114398709-DEL-80 |
| chr2-114403096-DEL-53 | chr2-114403094-DEL-51 |
| chr2-114412681-DEL-445 | chr2-114412682-DEL-445 |
| chr2-114486512-DEL-52 | chr2-114486513-DEL-52 |
| chr2-114530932-DEL-58 | chr2-114530915-DEL-98 |
| chr2-114539847-DEL-392 | chr2-114539848-DEL-392 |
| chr2-114542834-DEL-169 | chr2-114542806-DEL-207 |
| chr2-114581565-DEL-58 | chr2-114581566-DEL-58 |
| chr2-114625536-DEL-125 | chr2-114625537-DEL-125 |
| chr2-114690780-DEL-428 | chr2-114690781-DEL-428 |
| chr2-114695436-DEL-189 | chr2-114695396-DEL-195 |
| chr2-114701558-DEL-56 | chr2-114701558-DEL-56 |
| chr2-114727484-DEL-64 | chr2-114727485-DEL-64 |
| chr2-114739428-DEL-813 | chr2-114739429-DEL-813 |
| chr2-114757552-DEL-691 | chr2-114757553-DEL-691 |
| chr2-114766486-DEL-59 | chr2-114766487-DEL-59 |
| chr2-114797472-DEL-180 | chr2-114797473-DEL-181 |
| chr2-114828499-DEL-188 | chr2-114828500-DEL-188 |
| chr2-114856102-DEL-55 | chr2-114856103-DEL-55 |
| chr2-114867817-DEL-183 | chr2-114867816-DEL-183 |
| chr2-114869128-DEL-81 | chr2-114869127-DEL-82 |
| chr2-114884199-DEL-66 | chr2-114884200-DEL-66 |
| chr2-114890953-DEL-171 | chr2-114890952-DEL-172 |
| chr2-114892191-DEL-103 | chr2-114892192-DEL-103 |
| chr2-114900007-DEL-92 | chr2-114900008-DEL-88 |
| chr2-114939067-DEL-661 | chr2-114939068-DEL-661 |
| chr2-114952023-DEL-51 | chr2-114952024-DEL-51 |
| chr2-114980778-DEL-12858 | chr2-114980779-DEL-12858 |
| chr2-114986299-DEL-6386 | chr2-114986300-DEL-6386 |
| chr2-114999010-DEL-54 | chr2-114999023-DEL-57 |
| chr2-115023324-DEL-80 | chr2-115023325-DEL-80 |
| chr2-115028435-DEL-71 | chr2-115028435-DEL-71 |
| chr2-115047533-DEL-58 | chr2-115047532-DEL-59 |
| chr2-115088428-DEL-1213 | chr2-115088429-DEL-1213 |
| chr2-115113043-DEL-68 | chr2-115113044-DEL-68 |
| chr2-115122132-DEL-63 | chr2-115122124-DEL-71 |
| chr2-115136981-DEL-51 | chr2-115136952-DEL-81 |
| chr2-115156504-DEL-91 | chr2-115156505-DEL-91 |
| chr2-115161626-DEL-3972 | chr2-115161627-DEL-3972 |
| chr2-115192272-DEL-176 | chr2-115192274-DEL-174 |
| chr2-115219261-DEL-1169 | chr2-115219262-DEL-1169 |
| chr2-115281597-DEL-62 | chr2-115281608-DEL-51 |
| chr2-115306120-DEL-116 | chr2-115306121-DEL-116 |
| chr2-115308115-DEL-948 | chr2-115308116-DEL-945 |
| chr2-115321689-DEL-156 | chr2-115321690-DEL-156 |
| chr2-115326649-DEL-140 | chr2-115326650-DEL-140 |
| chr2-115485434-DEL-840 | chr2-115485461-DEL-814 |
| chr2-115578117-DEL-53 | chr2-115578118-DEL-53 |
| chr2-115622188-DEL-2464 | chr2-115622189-DEL-2464 |
| chr2-115634961-DEL-3390 | chr2-115634962-DEL-3390 |
| chr2-115674979-DEL-195 | chr2-115674980-DEL-195 |
| chr2-115678823-DEL-75 | chr2-115678828-DEL-71 |
| chr2-115708720-DEL-65 | chr2-115708721-DEL-65 |
| chr2-115776987-DEL-158 | chr2-115776988-DEL-158 |
| chr2-115811481-DEL-97 | chr2-115811457-DEL-101 |
| chr2-115842781-DEL-117 | chr2-115842782-DEL-117 |
| chr2-115871767-DEL-350 | chr2-115871768-DEL-350 |
| chr2-115894815-DEL-156 | chr2-115894816-DEL-142 |
| chr2-115912080-DEL-67 | chr2-115912081-DEL-67 |
| chr2-115925133-DEL-509 | chr2-115925118-DEL-524 |
| chr2-115939762-DEL-1250 | chr2-115939749-DEL-1251 |
| chr2-115942510-DEL-231 | chr2-115942511-DEL-231 |
| chr2-115963070-DEL-104 | chr2-115963080-DEL-70 |
| chr2-115964896-DEL-132 | chr2-115964901-DEL-128 |
| chr2-115980313-DEL-6628 | chr2-115980317-DEL-6629 |

| | |
|---|---|
| chr2-115998215-DEL-192 | chr2-115998204-DEL-193 |
| chr2-116002941-DEL-244 | chr2-116002942-DEL-244 |
| chr2-116021222-DEL-401 | chr2-116021223-DEL-401 |
| chr2-116054843-DEL-96 | chr2-116054844-DEL-96 |
| chr2-116059931-DEL-402 | chr2-116059932-DEL-402 |
| chr2-116066474-DEL-84 | chr2-116066479-DEL-80 |
| chr2-116069999-DEL-174 | chr2-116070000-DEL-174 |
| chr2-116098783-DEL-6249 | chr2-116098784-DEL-6249 |
| chr2-116165155-DEL-186 | chr2-116165140-DEL-178 |
| chr2-116280580-DEL-369 | chr2-116280573-DEL-371 |
| chr2-116284768-DEL-6229 | chr2-116284769-DEL-6229 |
| chr2-116314659-DEL-471 | chr2-116314660-DEL-471 |
| chr2-116362903-DEL-78 | chr2-116362904-DEL-78 |
| chr2-116376721-DEL-227 | chr2-116376722-DEL-227 |
| chr2-116379504-DEL-117 | chr2-116379505-DEL-117 |
| chr2-116381304-DEL-58 | chr2-116381304-DEL-58 |
| chr2-116382501-DEL-202 | chr2-116382502-DEL-202 |
| chr2-116383791-DEL-147 | chr2-116383792-DEL-147 |
| chr2-116386467-DEL-430 | chr2-116386468-DEL-430 |
| chr2-116387951-DEL-301 | chr2-116387952-DEL-301 |
| chr2-116389808-DEL-58 | chr2-116389809-DEL-58 |
| chr2-116392199-DEL-58 | chr2-116392200-DEL-58 |
| chr2-116392730-DEL-58 | chr2-116392731-DEL-58 |
| chr2-116414856-DEL-58 | chr2-116414847-DEL-85 |
| chr2-116417584-DEL-6702 | chr2-116417585-DEL-6702 |
| chr2-116424286-DEL-6011 | chr2-116424286-DEL-6011 |
| chr2-116452414-DEL-55 | chr2-116452415-DEL-55 |
| chr2-116483013-DEL-300 | chr2-116483014-DEL-300 |
| chr2-116525938-DEL-652 | chr2-116525939-DEL-652 |
| chr2-116544021-DEL-58 | chr2-116544022-DEL-58 |
| chr2-116548732-DEL-358 | chr2-116548733-DEL-355 |
| chr2-116568376-DEL-2722 | chr2-116568377-DEL-2722 |
| chr2-116573436-DEL-52 | chr2-116573437-DEL-52 |
| chr2-116601632-DEL-67 | chr2-116601633-DEL-67 |
| chr2-116634485-DEL-472 | chr2-116634486-DEL-471 |
| chr2-116641713-DEL-58 | chr2-116641714-DEL-58 |
| chr2-116685695-DEL-64 | chr2-116685675-DEL-99 |
| chr2-116726065-DEL-246 | chr2-116726073-DEL-242 |
| chr2-116727909-DEL-96 | chr2-116727910-DEL-96 |
| chr2-116731417-DEL-454 | chr2-116731418-DEL-454 |
| chr2-116739041-DEL-69 | chr2-116739042-DEL-69 |
| chr2-116741443-DEL-62 | chr2-116741419-DEL-58 |
| chr2-116743930-DEL-126 | chr2-116743927-DEL-124 |
| chr2-116753386-DEL-55 | chr2-116753387-DEL-55 |
| chr2-116756685-DEL-60 | chr2-116756686-DEL-60 |
| chr2-116763572-DEL-74 | chr2-116763573-DEL-74 |
| chr2-116766554-DEL-155 | chr2-116766555-DEL-155 |
| chr2-116786252-DEL-181 | chr2-116786255-DEL-179 |
| chr2-116833038-DEL-52 | chr2-116833000-DEL-61 |
| chr2-116844123-DEL-66 | chr2-116844124-DEL-72 |
| chr2-116854442-DEL-97 | chr2-116854443-DEL-95 |
| chr2-116877954-DEL-537 | chr2-116877955-DEL-537 |
| chr2-116886348-DEL-2296 | chr2-116886348-DEL-2296 |
| chr2-116888715-DEL-4537 | chr2-116888716-DEL-4537 |
| chr2-116922243-DEL-73 | chr2-116922244-DEL-65 |
| chr2-116948144-DEL-58 | chr2-116948142-DEL-59 |
| chr2-116973088-DEL-228 | chr2-116973089-DEL-228 |
| chr2-116988766-DEL-58 | chr2-116988767-DEL-58 |
| chr2-117004333-DEL-60 | chr2-117004338-DEL-52 |
| chr2-117063950-DEL-76 | chr2-117063951-DEL-76 |
| chr2-117135601-DEL-52 | chr2-117135602-DEL-52 |
| chr2-117137456-DEL-326 | chr2-117137459-DEL-311 |
| chr2-117148498-DEL-52 | chr2-117148501-DEL-50 |
| chr2-117161810-DEL-53 | chr2-117161813-DEL-51 |
| chr2-117202944-DEL-87 | chr2-117202946-DEL-89 |
| chr2-117206508-DEL-85 | chr2-117206514-DEL-109 |
| chr2-117209396-DEL-51 | chr2-117209357-DEL-76 |
| chr2-117213703-DEL-68 | chr2-117213704-DEL-68 |
| chr2-117224888-DEL-91 | chr2-117224891-DEL-92 |
| chr2-117265959-DEL-143 | chr2-117265960-DEL-143 |
| chr2-117273671-DEL-54 | chr2-117273668-DEL-54 |
| chr2-117279342-DEL-56 | chr2-117279343-DEL-56 |
| chr2-117293705-DEL-2013 | chr2-117293706-DEL-2013 |
| chr2-117302460-DEL-417 | chr2-117302470-DEL-413 |
| chr2-117355638-DEL-54 | chr2-117355639-DEL-54 |
| chr2-117359817-DEL-101 | chr2-117359818-DEL-100 |
| chr2-117496065-DEL-193 | chr2-117496066-DEL-193 |
| chr2-117497409-DEL-128 | chr2-117497410-DEL-128 |
| chr2-117508522-DEL-259 | chr2-117508532-DEL-263 |
| chr2-117554526-DEL-125 | chr2-117554527-DEL-125 |

| | |
|---|---|
| chr2-117565315-DEL-83 | chr2-117565316-DEL-80 |
| chr2-117601084-DEL-2018 | chr2-117601085-DEL-2018 |
| chr2-117635632-DEL-163 | chr2-117635633-DEL-163 |
| chr2-117674947-DEL-164 | chr2-117674948-DEL-164 |
| chr2-117688283-DEL-1887 | chr2-117688278-DEL-1888 |
| chr2-117700596-DEL-56 | chr2-117700597-DEL-56 |
| chr2-117704915-DEL-116 | chr2-117704916-DEL-116 |
| chr2-117709345-DEL-62 | chr2-117709346-DEL-62 |
| chr2-117715726-DEL-702 | chr2-117715727-DEL-702 |
| chr2-117723664-DEL-2378 | chr2-117723665-DEL-2378 |
| chr2-117761891-DEL-73 | chr2-117761892-DEL-73 |
| chr2-117804230-DEL-241 | chr2-117804234-DEL-238 |
| chr2-117806788-DEL-3187 | chr2-117806789-DEL-3187 |
| chr2-117825948-DEL-196 | chr2-117825954-DEL-195 |
| chr2-117848091-DEL-2627 | chr2-117848092-DEL-2627 |
| chr2-117851722-DEL-470 | chr2-117851709-DEL-439 |
| chr2-117855757-DEL-67 | chr2-117855758-DEL-67 |
| chr2-117902042-DEL-60 | chr2-117902043-DEL-60 |
| chr2-117962479-DEL-89 | chr2-117962480-DEL-89 |
| chr2-117963197-DEL-2330 | chr2-117963198-DEL-2330 |
| chr2-118002779-DEL-315 | chr2-118002780-DEL-315 |
| chr2-118024576-DEL-54 | chr2-118024577-DEL-54 |
| chr2-118061737-DEL-72 | chr2-118061726-DEL-84 |
| chr2-118072539-DEL-135 | chr2-118072528-DEL-122 |
| chr2-118073717-DEL-212 | chr2-118073718-DEL-212 |
| chr2-118087513-DEL-174 | chr2-118087514-DEL-174 |
| chr2-118089782-DEL-74 | chr2-118089783-DEL-74 |
| chr2-118095475-DEL-478 | chr2-118095476-DEL-478 |
| chr2-118154616-DEL-1061 | chr2-118154617-DEL-1061 |
| chr2-118188943-DEL-60 | chr2-118188944-DEL-60 |
| chr2-118239514-DEL-106 | chr2-118239514-DEL-107 |
| chr2-118253913-DEL-210 | chr2-118253917-DEL-213 |
| chr2-118281220-DEL-89 | chr2-118281213-DEL-97 |
| chr2-118308038-DEL-786 | chr2-118308036-DEL-784 |
| chr2-118337046-DEL-60 | chr2-118337047-DEL-60 |
| chr2-118348338-DEL-72 | chr2-118348316-DEL-56 |
| chr2-118385240-DEL-3621 | chr2-118385241-DEL-3621 |
| chr2-118393903-DEL-267 | chr2-118393904-DEL-267 |
| chr2-118399081-DEL-84 | chr2-118399093-DEL-86 |
| chr2-118405665-DEL-120 | chr2-118405666-DEL-120 |
| chr2-118425722-DEL-103 | chr2-118425721-DEL-104 |
| chr2-118442792-DEL-215 | chr2-118442793-DEL-215 |
| chr2-118461946-DEL-60 | chr2-118461987-DEL-57 |
| chr2-118565479-DEL-177 | chr2-118565480-DEL-177 |
| chr2-118570262-DEL-125 | chr2-118570248-DEL-140 |
| chr2-118571434-DEL-59 | chr2-118571421-DEL-60 |
| chr2-118584402-DEL-206 | chr2-118584403-DEL-206 |
| chr2-118608387-DEL-63 | chr2-118608388-DEL-63 |
| chr2-118629075-DEL-163 | chr2-118629073-DEL-166 |
| chr2-118636739-DEL-82 | chr2-118636750-DEL-54 |
| chr2-118651964-DEL-554 | chr2-118651965-DEL-554 |
| chr2-118677659-DEL-84 | chr2-118677617-DEL-84 |
| chr2-118683659-DEL-67 | chr2-118683660-DEL-67 |
| chr2-118688895-DEL-78 | chr2-118688900-DEL-74 |
| chr2-118690042-DEL-169 | chr2-118690039-DEL-163 |
| chr2-118693595-DEL-72 | chr2-118693600-DEL-68 |
| chr2-118698084-DEL-1445 | chr2-118698085-DEL-1445 |
| chr2-118739909-DEL-52 | chr2-118739910-DEL-52 |
| chr2-118761797-DEL-623 | chr2-118761797-DEL-623 |
| chr2-118764012-DEL-54 | chr2-118764015-DEL-52 |
| chr2-118822148-DEL-52 | chr2-118822149-DEL-52 |
| chr2-118831361-DEL-66 | chr2-118831362-DEL-58 |
| chr2-118883410-DEL-207 | chr2-118883407-DEL-211 |
| chr2-118916664-DEL-56 | chr2-118916665-DEL-56 |
| chr2-118920022-DEL-186 | chr2-118920013-DEL-187 |
| chr2-118929522-DEL-53 | chr2-118929523-DEL-53 |
| chr2-118949313-DEL-2114 | chr2-118949313-DEL-2115 |
| chr2-118950648-DEL-76 | chr2-118950649-DEL-76 |
| chr2-118996954-DEL-80 | chr2-118996921-DEL-96 |
| chr2-119001444-DEL-281 | chr2-119001445-DEL-281 |
| chr2-119012407-DEL-199 | chr2-119012408-DEL-198 |
| chr2-119023153-DEL-396 | chr2-119023154-DEL-396 |
| chr2-119036929-DEL-220 | chr2-119036930-DEL-220 |
| chr2-119050741-DEL-72 | chr2-119050756-DEL-56 |
| chr2-119061235-DEL-59 | chr2-119061231-DEL-56 |
| chr2-119063213-DEL-77 | chr2-119063213-DEL-73 |
| chr2-119097993-DEL-275 | chr2-119097994-DEL-271 |
| chr2-119102767-DEL-232 | chr2-119102768-DEL-232 |
| chr2-119109587-DEL-68 | chr2-119109573-DEL-56 |
| chr2-119110125-DEL-176 | chr2-119110126-DEL-176 |

| | |
|---|---|
| chr2-119126941-DEL-195 | chr2-119126942-DEL-195 |
| chr2-119139726-DEL-155 | chr2-119139727-DEL-155 |
| chr2-119154987-DEL-58 | chr2-119154988-DEL-58 |
| chr2-119169201-DEL-203 | chr2-119169202-DEL-203 |
| chr2-119174498-DEL-55 | chr2-119174499-DEL-55 |
| chr2-119187183-DEL-104 | chr2-119187184-DEL-104 |
| chr2-119190719-DEL-95 | chr2-119190720-DEL-95 |
| chr2-119202042-DEL-154 | chr2-119202043-DEL-154 |
| chr2-119209114-DEL-84 | chr2-119209115-DEL-84 |
| chr2-119213175-DEL-55 | chr2-119213176-DEL-55 |
| chr2-119217124-DEL-56 | chr2-119217109-DEL-72 |
| chr2-119222636-DEL-180 | chr2-119222637-DEL-180 |
| chr2-119234004-DEL-127 | chr2-119234005-DEL-127 |
| chr2-119235879-DEL-169 | chr2-119235879-DEL-170 |
| chr2-119273327-DEL-232 | chr2-119273328-DEL-232 |
| chr2-119274578-DEL-60 | chr2-119274579-DEL-58 |
| chr2-119285353-DEL-197 | chr2-119285354-DEL-197 |
| chr2-119320229-DEL-225 | chr2-119320230-DEL-225 |
| chr2-119324938-DEL-68 | chr2-119324931-DEL-66 |
| chr2-119336172-DEL-134 | chr2-119336167-DEL-144 |
| chr2-119341259-DEL-96 | chr2-119341260-DEL-96 |
| chr2-119341871-DEL-135 | chr2-119341872-DEL-130 |
| chr2-119343865-DEL-71 | chr2-119343875-DEL-70 |
| chr2-119357838-DEL-182 | chr2-119357839-DEL-182 |
| chr2-119374449-DEL-244 | chr2-119374450-DEL-244 |
| chr2-119396323-DEL-206 | chr2-119396324-DEL-206 |
| chr2-119401873-DEL-216 | chr2-119401873-DEL-216 |
| chr2-119406618-DEL-197 | chr2-119406619-DEL-197 |
| chr2-119408747-DEL-228 | chr2-119408748-DEL-228 |
| chr2-119410617-DEL-64 | chr2-119410616-DEL-66 |
| chr2-119420785-DEL-87 | chr2-119420768-DEL-105 |
| chr2-119451286-DEL-209 | chr2-119451287-DEL-209 |
| chr2-119453501-DEL-201 | chr2-119453502-DEL-201 |
| chr2-119492854-DEL-130 | chr2-119492855-DEL-130 |
| chr2-119518223-DEL-244 | chr2-119518225-DEL-240 |
| chr2-119525592-DEL-187 | chr2-119525593-DEL-187 |
| chr2-119541600-DEL-78 | chr2-119541601-DEL-78 |
| chr2-119554945-DEL-52 | chr2-119554946-DEL-52 |
| chr2-119590380-DEL-81 | chr2-119590385-DEL-79 |
| chr2-119609568-DEL-196 | chr2-119609560-DEL-208 |
| chr2-119610904-DEL-100 | chr2-119610905-DEL-100 |
| chr2-119616488-DEL-210 | chr2-119616489-DEL-210 |
| chr2-119619355-DEL-218 | chr2-119619356-DEL-218 |
| chr2-119624168-DEL-356 | chr2-119624169-DEL-356 |
| chr2-119625525-DEL-101 | chr2-119625535-DEL-92 |
| chr2-119659936-DEL-205 | chr2-119659937-DEL-205 |
| chr2-119670091-DEL-95 | chr2-119670092-DEL-94 |
| chr2-119679775-DEL-63 | chr2-119679751-DEL-64 |
| chr2-119687493-DEL-74 | chr2-119687486-DEL-82 |
| chr2-119697514-DEL-268 | chr2-119697515-DEL-268 |
| chr2-119699871-DEL-51 | chr2-119699869-DEL-54 |
| chr2-119712406-DEL-111 | chr2-119712400-DEL-117 |
| chr2-119726730-DEL-75 | chr2-119726731-DEL-75 |
| chr2-119736226-DEL-114 | chr2-119736236-DEL-107 |
| chr2-119770461-DEL-210 | chr2-119770462-DEL-210 |
| chr2-119793885-DEL-54 | chr2-119793904-DEL-52 |
| chr2-119821419-DEL-330 | chr2-119821420-DEL-331 |
| chr2-119830839-DEL-203 | chr2-119830840-DEL-203 |
| chr2-119837522-DEL-63 | chr2-119837521-DEL-64 |
| chr2-119858585-DEL-94 | chr2-119858586-DEL-94 |
| chr2-119878006-DEL-63 | chr2-119878007-DEL-57 |
| chr2-119883549-DEL-1235 | chr2-119883550-DEL-1235 |
| chr2-119990622-DEL-62 | chr2-119990623-DEL-62 |
| chr2-120016242-DEL-96 | chr2-120016243-DEL-96 |
| chr2-120018576-DEL-130 | chr2-120018577-DEL-128 |
| chr2-120041512-DEL-62 | chr2-120041513-DEL-62 |
| chr2-120042842-DEL-213 | chr2-120042843-DEL-213 |
| chr2-120046010-DEL-876 | chr2-120046011-DEL-876 |
| chr2-120047975-DEL-7095 | chr2-120047976-DEL-7095 |
| chr2-120078191-DEL-234 | chr2-120078190-DEL-234 |
| chr2-120104230-DEL-210 | chr2-120104231-DEL-208 |
| chr2-120125471-DEL-74 | chr2-120125472-DEL-74 |
| chr2-120131891-DEL-69 | chr2-120131865-DEL-70 |
| chr2-120160045-DEL-231 | chr2-120160046-DEL-231 |
| chr2-120172225-DEL-54 | chr2-120172226-DEL-54 |
| chr2-120200201-DEL-93 | chr2-120200202-DEL-93 |
| chr2-120208961-DEL-219 | chr2-120208962-DEL-219 |
| chr2-120242242-DEL-208 | chr2-120242243-DEL-208 |
| chr2-120260223-DEL-59 | chr2-120260221-DEL-62 |
| chr2-120268882-DEL-128 | chr2-120268883-DEL-126 |

| | |
|---|---|
| chr2-120320564-DEL-141 | chr2-120320565-DEL-145 |
| chr2-120323658-DEL-52 | chr2-120323653-DEL-58 |
| chr2-120363140-DEL-183 | chr2-120363141-DEL-183 |
| chr2-120367715-DEL-211 | chr2-120367716-DEL-210 |
| chr2-120372890-DEL-56 | chr2-120372891-DEL-56 |
| chr2-120374988-DEL-59 | chr2-120374989-DEL-59 |
| chr2-120379648-DEL-2762 | chr2-120379649-DEL-2762 |
| chr2-120395597-DEL-231 | chr2-120395598-DEL-231 |
| chr2-120400489-DEL-80 | chr2-120400494-DEL-54 |
| chr2-120410166-DEL-247 | chr2-120410167-DEL-247 |
| chr2-120445749-DEL-67 | chr2-120445750-DEL-67 |
| chr2-120456588-DEL-60 | chr2-120456589-DEL-60 |
| chr2-120458764-DEL-253 | chr2-120458765-DEL-253 |
| chr2-120466294-DEL-3458 | chr2-120466295-DEL-3458 |
| chr2-120474944-DEL-382 | chr2-120474945-DEL-382 |
| chr2-120478628-DEL-272 | chr2-120478635-DEL-268 |
| chr2-120481469-DEL-99 | chr2-120481448-DEL-100 |
| chr2-120486745-DEL-141 | chr2-120486746-DEL-141 |
| chr2-120489495-DEL-185 | chr2-120489496-DEL-186 |
| chr2-120496169-DEL-227 | chr2-120496170-DEL-227 |
| chr2-120502762-DEL-62 | chr2-120502763-DEL-62 |
| chr2-120507888-DEL-186 | chr2-120507889-DEL-186 |
| chr2-120512709-DEL-222 | chr2-120512710-DEL-222 |
| chr2-120524855-DEL-203 | chr2-120524856-DEL-203 |
| chr2-120579846-DEL-147 | chr2-120579851-DEL-143 |
| chr2-120583846-DEL-489 | chr2-120583847-DEL-489 |
| chr2-120605138-DEL-218 | chr2-120605139-DEL-218 |
| chr2-120613648-DEL-82 | chr2-120613647-DEL-88 |
| chr2-120639501-DEL-203 | chr2-120639502-DEL-203 |
| chr2-120640758-DEL-1490 | chr2-120640759-DEL-1490 |
| chr2-120664682-DEL-143 | chr2-120664671-DEL-151 |
| chr2-120678058-DEL-68 | chr2-120678047-DEL-80 |
| chr2-120720004-DEL-200 | chr2-120720005-DEL-200 |
| chr2-120737814-DEL-202 | chr2-120737815-DEL-202 |
| chr2-120743187-DEL-978 | chr2-120743188-DEL-978 |
| chr2-120783157-DEL-69 | chr2-120783165-DEL-70 |
| chr2-120809079-DEL-76 | chr2-120809080-DEL-70 |
| chr2-120827396-DEL-52 | chr2-120827397-DEL-52 |
| chr2-120841317-DEL-96 | chr2-120841318-DEL-96 |
| chr2-120868691-DEL-202 | chr2-120868692-DEL-202 |
| chr2-120870981-DEL-78 | chr2-120870982-DEL-78 |
| chr2-120877960-DEL-220 | chr2-120877961-DEL-220 |
| chr2-120895695-DEL-892 | chr2-120895696-DEL-892 |
| chr2-120899998-DEL-51 | chr2-120899999-DEL-51 |
| chr2-120913638-DEL-69 | chr2-120913620-DEL-69 |
| chr2-120931944-DEL-56 | chr2-120931945-DEL-56 |
| chr2-120936241-DEL-192 | chr2-120936242-DEL-192 |
| chr2-120937607-DEL-62 | chr2-120937608-DEL-62 |
| chr2-120962130-DEL-72 | chr2-120962131-DEL-60 |
| chr2-120971363-DEL-354 | chr2-120971364-DEL-354 |
| chr2-120997333-DEL-81 | chr2-120997334-DEL-81 |
| chr2-121032606-DEL-65 | chr2-121032608-DEL-66 |
| chr2-121057682-DEL-207 | chr2-121057683-DEL-207 |
| chr2-121062106-DEL-80 | chr2-121062107-DEL-80 |
| chr2-121090257-DEL-215 | chr2-121090258-DEL-215 |
| chr2-121096542-DEL-93 | chr2-121096543-DEL-92 |
| chr2-121108921-DEL-55 | chr2-121108927-DEL-50 |
| chr2-121159797-DEL-102 | chr2-121159798-DEL-102 |
| chr2-121165416-DEL-203 | chr2-121165417-DEL-203 |
| chr2-121217533-DEL-341 | chr2-121217534-DEL-341 |
| chr2-121267022-DEL-203 | chr2-121267025-DEL-201 |
| chr2-121319903-DEL-55 | chr2-121319903-DEL-55 |
| chr2-121351048-DEL-684 | chr2-121351049-DEL-684 |
| chr2-121352794-DEL-603 | chr2-121352795-DEL-603 |
| chr2-121353807-DEL-202 | chr2-121353808-DEL-202 |
| chr2-121390877-DEL-224 | chr2-121390878-DEL-221 |
| chr2-121403024-DEL-203 | chr2-121403025-DEL-203 |
| chr2-121405034-DEL-109 | chr2-121405035-DEL-109 |
| chr2-121423397-DEL-195 | chr2-121423389-DEL-204 |
| chr2-121452824-DEL-156 | chr2-121452825-DEL-156 |
| chr2-121476234-DEL-66 | chr2-121476245-DEL-56 |
| chr2-121493970-DEL-591 | chr2-121493971-DEL-591 |
| chr2-121497237-DEL-331 | chr2-121497238-DEL-333 |
| chr2-121505174-DEL-74 | chr2-121505175-DEL-72 |
| chr2-121515645-DEL-60 | chr2-121515646-DEL-68 |
| chr2-121558768-DEL-76 | chr2-121558755-DEL-78 |
| chr2-121569503-DEL-254 | chr2-121569504-DEL-254 |
| chr2-121573567-DEL-54 | chr2-121573568-DEL-54 |
| chr2-121584732-DEL-175 | chr2-121584733-DEL-175 |
| chr2-121615786-DEL-58 | chr2-121615779-DEL-56 |

| | |
|---|---|
| chr2-121625932-DEL-354 | chr2-121625933-DEL-354 |
| chr2-121629221-DEL-52 | chr2-121629197-DEL-96 |
| chr2-121651474-DEL-1582 | chr2-121651475-DEL-1583 |
| chr2-121655024-DEL-60 | chr2-121655025-DEL-60 |
| chr2-121679422-DEL-132 | chr2-121679423-DEL-132 |
| chr2-121686606-DEL-80 | chr2-121686607-DEL-80 |
| chr2-121719991-DEL-201 | chr2-121719992-DEL-201 |
| chr2-121723281-DEL-70 | chr2-121723281-DEL-70 |
| chr2-121735628-DEL-176 | chr2-121735618-DEL-161 |
| chr2-121766494-DEL-203 | chr2-121766495-DEL-203 |
| chr2-121769342-DEL-1893 | chr2-121769343-DEL-1893 |
| chr2-121777427-DEL-207 | chr2-121777428-DEL-207 |
| chr2-121795189-DEL-361 | chr2-121795190-DEL-361 |
| chr2-121800618-DEL-201 | chr2-121800619-DEL-201 |
| chr2-121817003-DEL-207 | chr2-121817004-DEL-207 |
| chr2-121828832-DEL-147 | chr2-121828833-DEL-147 |
| chr2-121862003-DEL-718 | chr2-121862004-DEL-712 |
| chr2-121873208-DEL-83 | chr2-121873176-DEL-80 |
| chr2-121881172-DEL-62 | chr2-121881177-DEL-55 |
| chr2-121887952-DEL-56 | chr2-121887953-DEL-56 |
| chr2-121889343-DEL-202 | chr2-121889344-DEL-202 |
| chr2-121897040-DEL-52 | chr2-121897041-DEL-52 |
| chr2-121898328-DEL-65 | chr2-121898329-DEL-65 |
| chr2-121922117-DEL-76 | chr2-121922118-DEL-76 |
| chr2-121932921-DEL-52 | chr2-121932921-DEL-52 |
| chr2-121936827-DEL-67 | chr2-121936828-DEL-67 |
| chr2-121943727-DEL-127 | chr2-121943728-DEL-127 |
| chr2-121950241-DEL-56 | chr2-121950242-DEL-56 |
| chr2-121972716-DEL-848 | chr2-121972717-DEL-848 |
| chr2-121993215-DEL-126 | chr2-121993215-DEL-127 |
| chr2-122022823-DEL-168 | chr2-122022824-DEL-168 |
| chr2-122026928-DEL-1081 | chr2-122026929-DEL-1081 |
| chr2-122029154-DEL-110 | chr2-122029155-DEL-110 |
| chr2-122029965-DEL-1343 | chr2-122029966-DEL-1343 |
| chr2-122031172-DEL-150 | chr2-122031173-DEL-148 |
| chr2-122039119-DEL-2248 | chr2-122039120-DEL-2248 |
| chr2-122044577-DEL-1284 | chr2-122044571-DEL-1285 |
| chr2-122046081-DEL-205 | chr2-122046082-DEL-205 |
| chr2-122049871-DEL-165 | chr2-122049859-DEL-166 |
| chr2-122050391-DEL-504 | chr2-122050391-DEL-465 |
| chr2-122058381-DEL-348 | chr2-122058382-DEL-348 |
| chr2-122098507-DEL-158 | chr2-122098526-DEL-161 |
| chr2-122138811-DEL-113 | chr2-122138812-DEL-116 |
| chr2-122207231-DEL-263 | chr2-122207229-DEL-266 |
| chr2-122221134-DEL-474 | chr2-122221135-DEL-474 |
| chr2-122254711-DEL-4120 | chr2-122254711-DEL-4120 |
| chr2-122281764-DEL-95 | chr2-122281765-DEL-95 |
| chr2-122298855-DEL-56 | chr2-122298855-DEL-56 |
| chr2-122308131-DEL-748 | chr2-122308132-DEL-748 |
| chr2-122310144-DEL-59 | chr2-122310144-DEL-51 |
| chr2-122316312-DEL-485 | chr2-122316313-DEL-484 |
| chr2-122318581-DEL-1160 | chr2-122318587-DEL-1149 |
| chr2-122319859-DEL-671 | chr2-122319860-DEL-671 |
| chr2-122338512-DEL-308 | chr2-122338515-DEL-309 |
| chr2-122350747-DEL-1391 | chr2-122350748-DEL-1391 |
| chr2-122396517-DEL-3393 | chr2-122396518-DEL-3393 |
| chr2-122407681-DEL-81 | chr2-122407655-DEL-80 |
| chr2-122425383-DEL-75 | chr2-122425393-DEL-75 |
| chr2-122440949-DEL-96 | chr2-122440950-DEL-84 |
| chr2-122450417-DEL-91 | chr2-122450422-DEL-87 |
| chr2-122452016-DEL-215 | chr2-122452017-DEL-214 |
| chr2-122452931-DEL-211 | chr2-122452932-DEL-211 |
| chr2-122459878-DEL-52 | chr2-122459878-DEL-54 |
| chr2-122468557-DEL-494 | chr2-122468572-DEL-480 |
| chr2-122485260-DEL-6885 | chr2-122485261-DEL-6885 |
| chr2-122514041-DEL-56 | chr2-122514042-DEL-56 |
| chr2-122539300-DEL-55 | chr2-122539302-DEL-53 |
| chr2-122557522-DEL-232 | chr2-122557520-DEL-232 |
| chr2-122637665-DEL-1072 | chr2-122637666-DEL-1072 |
| chr2-122640931-DEL-52 | chr2-122640932-DEL-58 |
| chr2-122657276-DEL-766 | chr2-122657277-DEL-766 |
| chr2-122658417-DEL-298 | chr2-122658418-DEL-298 |
| chr2-122661635-DEL-228 | chr2-122661598-DEL-266 |
| chr2-122671641-DEL-78 | chr2-122671642-DEL-78 |
| chr2-122697260-DEL-58 | chr2-122697281-DEL-50 |
| chr2-122720010-DEL-540 | chr2-122720011-DEL-539 |
| chr2-122753493-DEL-838 | chr2-122753494-DEL-838 |
| chr2-122784739-DEL-56 | chr2-122784740-DEL-56 |
| chr2-122792283-DEL-6433 | chr2-122792284-DEL-6433 |
| chr2-122801632-DEL-72 | chr2-122801629-DEL-80 |

| | |
|---|---|
| chr2-122822150-DEL-55 | chr2-122822132-DEL-58 |
| chr2-122835246-DEL-6219 | chr2-122835247-DEL-6219 |
| chr2-122843062-DEL-62 | chr2-122843063-DEL-62 |
| chr2-122858063-DEL-200 | chr2-122858063-DEL-200 |
| chr2-122867898-DEL-328 | chr2-122867899-DEL-328 |
| chr2-122879146-DEL-167 | chr2-122879147-DEL-167 |
| chr2-122933482-DEL-130 | chr2-122933483-DEL-130 |
| chr2-122937197-DEL-209 | chr2-122937198-DEL-209 |
| chr2-123066980-DEL-4440 | chr2-123066981-DEL-4440 |
| chr2-123073662-DEL-2002 | chr2-123073663-DEL-2002 |
| chr2-123094791-DEL-1263 | chr2-123094792-DEL-1263 |
| chr2-123105838-DEL-97 | chr2-123105839-DEL-97 |
| chr2-123129846-DEL-6370 | chr2-123129847-DEL-6370 |
| chr2-123151879-DEL-1160 | chr2-123151880-DEL-1160 |
| chr2-123201217-DEL-275 | chr2-123201216-DEL-272 |
| chr2-123205881-DEL-187 | chr2-123205882-DEL-187 |
| chr2-123230716-DEL-480 | chr2-123230715-DEL-481 |
| chr2-123261255-DEL-84 | chr2-123261260-DEL-80 |
| chr2-123268035-DEL-155 | chr2-123268036-DEL-155 |
| chr2-123272153-DEL-60 | chr2-123272154-DEL-60 |
| chr2-123321786-DEL-56 | chr2-123321787-DEL-54 |
| chr2-123338471-DEL-214 | chr2-123338472-DEL-214 |
| chr2-123341777-DEL-3668 | chr2-123341776-DEL-3669 |
| chr2-123355540-DEL-11657 | chr2-123355541-DEL-11657 |
| chr2-123368316-DEL-52 | chr2-123368317-DEL-52 |
| chr2-123421222-DEL-60 | chr2-123421223-DEL-60 |
| chr2-123428178-DEL-1016 | chr2-123428179-DEL-1016 |
| chr2-123432920-DEL-54 | chr2-123432921-DEL-54 |
| chr2-123444565-DEL-3611 | chr2-123444566-DEL-3611 |
| chr2-123465369-DEL-288 | chr2-123465370-DEL-288 |
| chr2-123484777-DEL-604 | chr2-123484776-DEL-606 |
| chr2-123488454-DEL-785 | chr2-123488455-DEL-785 |
| chr2-123522794-DEL-50 | chr2-123522795-DEL-50 |
| chr2-123532798-DEL-103 | chr2-123532798-DEL-104 |
| chr2-123561074-DEL-69 | chr2-123561075-DEL-69 |
| chr2-123576471-DEL-131 | chr2-123576472-DEL-131 |
| chr2-123586509-DEL-3881 | chr2-123586510-DEL-3881 |
| chr2-123596434-DEL-478 | chr2-123596435-DEL-478 |
| chr2-123613347-DEL-59 | chr2-123613348-DEL-59 |
| chr2-123621916-DEL-89 | chr2-123621936-DEL-62 |
| chr2-123626386-DEL-6257 | chr2-123626387-DEL-6257 |
| chr2-123643269-DEL-1300 | chr2-123643270-DEL-1300 |
| chr2-123648384-DEL-173 | chr2-123648385-DEL-173 |
| chr2-123651461-DEL-2181 | chr2-123651462-DEL-2181 |
| chr2-123652444-DEL-76 | chr2-123652445-DEL-52 |
| chr2-123722087-DEL-6462 | chr2-123722088-DEL-6462 |
| chr2-123800987-DEL-70 | chr2-123800984-DEL-74 |
| chr2-123875183-DEL-54 | chr2-123875184-DEL-52 |
| chr2-123925400-DEL-702 | chr2-123925400-DEL-702 |
| chr2-123950133-DEL-66 | chr2-123950134-DEL-66 |
| chr2-123967878-DEL-266 | chr2-123967879-DEL-266 |
| chr2-124028969-DEL-529 | chr2-124028970-DEL-529 |
| chr2-124051464-DEL-2756 | chr2-124051465-DEL-2756 |
| chr2-124058581-DEL-449 | chr2-124058582-DEL-449 |
| chr2-124069676-DEL-396 | chr2-124069677-DEL-396 |
| chr2-124077981-DEL-96 | chr2-124077982-DEL-96 |
| chr2-124091026-DEL-6870 | chr2-124091027-DEL-6870 |
| chr2-124097059-DEL-209 | chr2-124097060-DEL-209 |
| chr2-124103752-DEL-506 | chr2-124103753-DEL-506 |
| chr2-124107925-DEL-600 | chr2-124107926-DEL-600 |
| chr2-124110749-DEL-232 | chr2-124110749-DEL-232 |
| chr2-124112122-DEL-338 | chr2-124112112-DEL-346 |
| chr2-124139415-DEL-52 | chr2-124139416-DEL-52 |
| chr2-124160716-DEL-466 | chr2-124160717-DEL-466 |
| chr2-124218111-DEL-182 | chr2-124218112-DEL-182 |
| chr2-124237976-DEL-57 | chr2-124237977-DEL-57 |
| chr2-124257575-DEL-56 | chr2-124257572-DEL-60 |
| chr2-124266404-DEL-53 | chr2-124266406-DEL-54 |
| chr2-124304621-DEL-179 | chr2-124304622-DEL-179 |
| chr2-124347202-DEL-52 | chr2-124347203-DEL-69 |
| chr2-124358420-DEL-984 | chr2-124358421-DEL-984 |
| chr2-124393175-DEL-505 | chr2-124393176-DEL-505 |
| chr2-124401049-DEL-167 | chr2-124401050-DEL-167 |
| chr2-124416204-DEL-127 | chr2-124416205-DEL-127 |
| chr2-124430007-DEL-78 | chr2-124430008-DEL-90 |
| chr2-124483774-DEL-77 | chr2-124483775-DEL-77 |
| chr2-124523283-DEL-60 | chr2-124523284-DEL-60 |
| chr2-124536390-DEL-124 | chr2-124536391-DEL-124 |
| chr2-124539862-DEL-68 | chr2-124539863-DEL-56 |
| chr2-124556739-DEL-90 | chr2-124556749-DEL-83 |

| | |
|---|---|
| chr2-124577679-DEL-205 | chr2-124577680-DEL-205 |
| chr2-124640830-DEL-7172 | chr2-124640831-DEL-7172 |
| chr2-124666974-DEL-10821 | chr2-124666961-DEL-10822 |
| chr2-124668501-DEL-4420 | chr2-124668502-DEL-4420 |
| chr2-124700045-DEL-67 | chr2-124700054-DEL-61 |
| chr2-124703769-DEL-126 | chr2-124703770-DEL-126 |
| chr2-124704336-DEL-348 | chr2-124704337-DEL-348 |
| chr2-124717535-DEL-2800 | chr2-124717536-DEL-2800 |
| chr2-124723045-DEL-6237 | chr2-124723046-DEL-6237 |
| chr2-124733132-DEL-204 | chr2-124733133-DEL-204 |
| chr2-124734308-DEL-136 | chr2-124734290-DEL-144 |
| chr2-124831443-DEL-316 | chr2-124831440-DEL-320 |
| chr2-124832618-DEL-93 | chr2-124832596-DEL-97 |
| chr2-124839653-DEL-106 | chr2-124839652-DEL-108 |
| chr2-124840573-DEL-2280 | chr2-124840574-DEL-2280 |
| chr2-124848488-DEL-1705 | chr2-124848489-DEL-1705 |
| chr2-124868150-DEL-109 | chr2-124868151-DEL-109 |
| chr2-124871224-DEL-73 | chr2-124871188-DEL-63 |
| chr2-124876950-DEL-256 | chr2-124876967-DEL-253 |
| chr2-124891069-DEL-218 | chr2-124891070-DEL-218 |
| chr2-124901746-DEL-68 | chr2-124901747-DEL-68 |
| chr2-124909703-DEL-211 | chr2-124909704-DEL-211 |
| chr2-124926823-DEL-84 | chr2-124926823-DEL-84 |
| chr2-124933953-DEL-215 | chr2-124933954-DEL-215 |
| chr2-124936304-DEL-139 | chr2-124936305-DEL-139 |
| chr2-124939686-DEL-63 | chr2-124939687-DEL-63 |
| chr2-124975342-DEL-336 | chr2-124975377-DEL-295 |
| chr2-124991725-DEL-66 | chr2-124991726-DEL-66 |
| chr2-125002747-DEL-70 | chr2-125002748-DEL-70 |
| chr2-125009449-DEL-136 | chr2-125009450-DEL-136 |
| chr2-125026886-DEL-65 | chr2-125026904-DEL-62 |
| chr2-125072417-DEL-1647 | chr2-125072418-DEL-1647 |
| chr2-125084757-DEL-212 | chr2-125084758-DEL-212 |
| chr2-125086605-DEL-88 | chr2-125086605-DEL-89 |
| chr2-125092614-DEL-149 | chr2-125092615-DEL-149 |
| chr2-125094424-DEL-54 | chr2-125094425-DEL-54 |
| chr2-125103889-DEL-66 | chr2-125103890-DEL-66 |
| chr2-125122001-DEL-100 | chr2-125121981-DEL-100 |
| chr2-125125916-DEL-842 | chr2-125125917-DEL-842 |
| chr2-125146417-DEL-53 | chr2-125146418-DEL-53 |
| chr2-125168077-DEL-677 | chr2-125168082-DEL-673 |
| chr2-125239093-DEL-77 | chr2-125239094-DEL-77 |
| chr2-125241901-DEL-182 | chr2-125241902-DEL-182 |
| chr2-125247105-DEL-6450 | chr2-125247106-DEL-6450 |
| chr2-125261199-DEL-176 | chr2-125261200-DEL-176 |
| chr2-125262938-DEL-96 | chr2-125262943-DEL-100 |
| chr2-125273181-DEL-132 | chr2-125273184-DEL-130 |
| chr2-125283216-DEL-51 | chr2-125283217-DEL-51 |
| chr2-125329614-DEL-425 | chr2-125329615-DEL-425 |
| chr2-125341614-DEL-74 | chr2-125341602-DEL-75 |
| chr2-125368960-DEL-1337 | chr2-125368961-DEL-1337 |
| chr2-125371125-DEL-94 | chr2-125371124-DEL-110 |
| chr2-125377513-DEL-275 | chr2-125377514-DEL-275 |
| chr2-125383880-DEL-1221 | chr2-125383881-DEL-1221 |
| chr2-125423392-DEL-60 | chr2-125423393-DEL-60 |
| chr2-125437487-DEL-69 | chr2-125437488-DEL-69 |
| chr2-125450090-DEL-1914 | chr2-125450091-DEL-1914 |
| chr2-125451125-DEL-528 | chr2-125451126-DEL-528 |
| chr2-125472743-DEL-5044 | chr2-125472742-DEL-5045 |
| chr2-125483198-DEL-448 | chr2-125483196-DEL-450 |
| chr2-125486163-DEL-54 | chr2-125486188-DEL-60 |
| chr2-125502723-DEL-64 | chr2-125502724-DEL-64 |
| chr2-125600389-DEL-64 | chr2-125600390-DEL-64 |
| chr2-125603804-DEL-62 | chr2-125603815-DEL-53 |
| chr2-125608174-DEL-153 | chr2-125608173-DEL-153 |
| chr2-125618037-DEL-210 | chr2-125618038-DEL-210 |
| chr2-125680349-DEL-147 | chr2-125680350-DEL-147 |
| chr2-125680594-DEL-462 | chr2-125680595-DEL-462 |
| chr2-125688421-DEL-258 | chr2-125688422-DEL-260 |
| chr2-125715231-DEL-185 | chr2-125715232-DEL-185 |
| chr2-125716489-DEL-1289 | chr2-125716489-DEL-1289 |
| chr2-125721275-DEL-336 | chr2-125721276-DEL-336 |
| chr2-125735954-DEL-92 | chr2-125735955-DEL-92 |
| chr2-125760186-DEL-65 | chr2-125760187-DEL-65 |
| chr2-125806914-DEL-54 | chr2-125806915-DEL-51 |
| chr2-125813798-DEL-976 | chr2-125813799-DEL-976 |
| chr2-125834700-DEL-191 | chr2-125834701-DEL-191 |
| chr2-125836819-DEL-6401 | chr2-125836820-DEL-6401 |
| chr2-125850465-DEL-61 | chr2-125850462-DEL-51 |
| chr2-125855254-DEL-162 | chr2-125855255-DEL-162 |

| | |
|---|---|
| chr2-125979256-DEL-2056 | chr2-125979257-DEL-2056 |
| chr2-125997328-DEL-1130 | chr2-125997329-DEL-1130 |
| chr2-126021234-DEL-204 | chr2-126021235-DEL-204 |
| chr2-126031839-DEL-1121 | chr2-126031840-DEL-1121 |
| chr2-126050650-DEL-258 | chr2-126050651-DEL-258 |
| chr2-126055090-DEL-152 | chr2-126055100-DEL-168 |
| chr2-126059619-DEL-2145 | chr2-126059619-DEL-2145 |
| chr2-126066210-DEL-442 | chr2-126066211-DEL-442 |
| chr2-126080799-DEL-6427 | chr2-126080792-DEL-6421 |
| chr2-126100372-DEL-6167 | chr2-126100373-DEL-6167 |
| chr2-126125710-DEL-54 | chr2-126125710-DEL-62 |
| chr2-126130610-DEL-1584 | chr2-126130611-DEL-1584 |
| chr2-126136239-DEL-198 | chr2-126136236-DEL-199 |
| chr2-126165850-DEL-608 | chr2-126165851-DEL-608 |
| chr2-126195205-DEL-251 | chr2-126195206-DEL-251 |
| chr2-126222752-DEL-128 | chr2-126222749-DEL-132 |
| chr2-126235249-DEL-89 | chr2-126235250-DEL-89 |
| chr2-126304953-DEL-352 | chr2-126304954-DEL-352 |
| chr2-126311985-DEL-55 | chr2-126311985-DEL-55 |
| chr2-126336488-DEL-196 | chr2-126336489-DEL-195 |
| chr2-126350438-DEL-198 | chr2-126350439-DEL-198 |
| chr2-126359226-DEL-586 | chr2-126359227-DEL-586 |
| chr2-126374332-DEL-2501 | chr2-126374333-DEL-2501 |
| chr2-126379092-DEL-195 | chr2-126379093-DEL-195 |
| chr2-126382770-DEL-548 | chr2-126382771-DEL-548 |
| chr2-126445209-DEL-62 | chr2-126445219-DEL-65 |
| chr2-126454649-DEL-51 | chr2-126454650-DEL-51 |
| chr2-126458814-DEL-130 | chr2-126458815-DEL-130 |
| chr2-126463788-DEL-201 | chr2-126463789-DEL-201 |
| chr2-126496644-DEL-195 | chr2-126496643-DEL-197 |
| chr2-126505867-DEL-182 | chr2-126505868-DEL-182 |
| chr2-126520887-DEL-206 | chr2-126520888-DEL-206 |
| chr2-126527651-DEL-1649 | chr2-126527651-DEL-1649 |
| chr2-126538861-DEL-171 | chr2-126538862-DEL-171 |
| chr2-126543497-DEL-63 | chr2-126543499-DEL-68 |
| chr2-126551062-DEL-427 | chr2-126551063-DEL-427 |
| chr2-126606879-DEL-198 | chr2-126606880-DEL-198 |
| chr2-126650392-DEL-68 | chr2-126650393-DEL-68 |
| chr2-126688913-DEL-210 | chr2-126688914-DEL-210 |
| chr2-126753842-DEL-52 | chr2-126753843-DEL-52 |
| chr2-126788665-DEL-1415 | chr2-126788666-DEL-1415 |
| chr2-126800244-DEL-515 | chr2-126800245-DEL-515 |
| chr2-126800942-DEL-205 | chr2-126800943-DEL-205 |
| chr2-126806646-DEL-220 | chr2-126806646-DEL-220 |
| chr2-126806915-DEL-7109 | chr2-126806916-DEL-7109 |
| chr2-126818888-DEL-110 | chr2-126818917-DEL-64 |
| chr2-126827475-DEL-99 | chr2-126827470-DEL-100 |
| chr2-126828937-DEL-2174 | chr2-126828938-DEL-2174 |
| chr2-126830454-DEL-130 | chr2-126830455-DEL-130 |
| chr2-126837848-DEL-173 | chr2-126837849-DEL-173 |
| chr2-126840760-DEL-204 | chr2-126840761-DEL-204 |
| chr2-126841879-DEL-170 | chr2-126841880-DEL-170 |
| chr2-126842483-DEL-132 | chr2-126842484-DEL-132 |
| chr2-126844054-DEL-221 | chr2-126844055-DEL-221 |
| chr2-126849270-DEL-110 | chr2-126849271-DEL-110 |
| chr2-126853936-DEL-51 | chr2-126853937-DEL-51 |
| chr2-126859520-DEL-205 | chr2-126859521-DEL-205 |
| chr2-126864179-DEL-633 | chr2-126864178-DEL-633 |
| chr2-126866298-DEL-221 | chr2-126866299-DEL-221 |
| chr2-126872037-DEL-680 | chr2-126872038-DEL-680 |
| chr2-126891962-DEL-91 | chr2-126891963-DEL-91 |
| chr2-126919278-DEL-217 | chr2-126919279-DEL-217 |
| chr2-126925679-DEL-67 | chr2-126925678-DEL-84 |
| chr2-126946446-DEL-786 | chr2-126946447-DEL-786 |
| chr2-126948461-DEL-612 | chr2-126948456-DEL-605 |
| chr2-126962108-DEL-56 | chr2-126962109-DEL-56 |
| chr2-126977399-DEL-386 | chr2-126977412-DEL-377 |
| chr2-126988008-DEL-194 | chr2-126988009-DEL-194 |
| chr2-126993637-DEL-3311 | chr2-126993638-DEL-3311 |
| chr2-127011919-DEL-563 | chr2-127011920-DEL-563 |
| chr2-127024171-DEL-1854 | chr2-127024172-DEL-1854 |
| chr2-127061779-DEL-222 | chr2-127061786-DEL-220 |
| chr2-127108527-DEL-219 | chr2-127108528-DEL-219 |
| chr2-127144665-DEL-1482 | chr2-127144670-DEL-1478 |
| chr2-127149908-DEL-546 | chr2-127149909-DEL-546 |
| chr2-127159816-DEL-57 | chr2-127159817-DEL-57 |
| chr2-127167396-DEL-182 | chr2-127167395-DEL-183 |
| chr2-127204407-DEL-54 | chr2-127204408-DEL-54 |
| chr2-127222889-DEL-897 | chr2-127222890-DEL-897 |
| chr2-127257140-DEL-172 | chr2-127257134-DEL-171 |

| | |
|---|---|
| chr2-127260354-DEL-428 | chr2-127260355-DEL-426 |
| chr2-127260878-DEL-53 | chr2-127260874-DEL-58 |
| chr2-127284078-DEL-61 | chr2-127284078-DEL-61 |
| chr2-127290386-DEL-373 | chr2-127290392-DEL-367 |
| chr2-127291931-DEL-3612 | chr2-127291932-DEL-3612 |
| chr2-127297131-DEL-82 | chr2-127297131-DEL-82 |
| chr2-127300466-DEL-147 | chr2-127300467-DEL-147 |
| chr2-127315680-DEL-101 | chr2-127315681-DEL-101 |
| chr2-127332752-DEL-73 | chr2-127332751-DEL-78 |
| chr2-127355920-DEL-98 | chr2-127355921-DEL-98 |
| chr2-127368102-DEL-100 | chr2-127368071-DEL-98 |
| chr2-127412765-DEL-54 | chr2-127412766-DEL-50 |
| chr2-127420271-DEL-168 | chr2-127420272-DEL-168 |
| chr2-127421042-DEL-676 | chr2-127421043-DEL-676 |
| chr2-127421769-DEL-1040 | chr2-127421770-DEL-1040 |
| chr2-127505859-DEL-344 | chr2-127505860-DEL-348 |
| chr2-127509178-DEL-60 | chr2-127509186-DEL-53 |
| chr2-127512827-DEL-6278 | chr2-127512828-DEL-6278 |
| chr2-127523186-DEL-79 | chr2-127523187-DEL-79 |
| chr2-127551403-DEL-68 | chr2-127551396-DEL-76 |
| chr2-127552601-DEL-112 | chr2-127552602-DEL-112 |
| chr2-127569867-DEL-94 | chr2-127569868-DEL-94 |
| chr2-127573872-DEL-1894 | chr2-127573873-DEL-1894 |
| chr2-127580499-DEL-58 | chr2-127580507-DEL-59 |
| chr2-127589227-DEL-443 | chr2-127589228-DEL-443 |
| chr2-127594401-DEL-1354 | chr2-127594402-DEL-1354 |
| chr2-127611751-DEL-148 | chr2-127611752-DEL-148 |
| chr2-127628694-DEL-367 | chr2-127628695-DEL-367 |
| chr2-127635066-DEL-6921 | chr2-127635067-DEL-6921 |
| chr2-127647701-DEL-99 | chr2-127647702-DEL-99 |
| chr2-127651352-DEL-175 | chr2-127651353-DEL-175 |
| chr2-127667384-DEL-217 | chr2-127667385-DEL-217 |
| chr2-127689990-DEL-147 | chr2-127689991-DEL-147 |
| chr2-127711744-DEL-56 | chr2-127711745-DEL-56 |
| chr2-127739386-DEL-152 | chr2-127739387-DEL-152 |
| chr2-127765871-DEL-188 | chr2-127765871-DEL-189 |
| chr2-127782946-DEL-51 | chr2-127782947-DEL-51 |
| chr2-127790145-DEL-76 | chr2-127790146-DEL-60 |
| chr2-127806969-DEL-2059 | chr2-127806970-DEL-2059 |
| chr2-127836562-DEL-51 | chr2-127836558-DEL-52 |
| chr2-127863417-DEL-651 | chr2-127863418-DEL-651 |
| chr2-127943542-DEL-70 | chr2-127943543-DEL-70 |
| chr2-127987085-DEL-65 | chr2-127987084-DEL-66 |
| chr2-128055844-DEL-63 | chr2-128055845-DEL-63 |
| chr2-128066119-DEL-158 | chr2-128066120-DEL-158 |
| chr2-128094766-DEL-145 | chr2-128094763-DEL-146 |
| chr2-128100804-DEL-82 | chr2-128100805-DEL-82 |
| chr2-128138558-DEL-207 | chr2-128138559-DEL-207 |
| chr2-128168180-DEL-52 | chr2-128168181-DEL-52 |
| chr2-128247846-DEL-60 | chr2-128247847-DEL-60 |
| chr2-128284644-DEL-76 | chr2-128284673-DEL-57 |
| chr2-128284772-DEL-7190 | chr2-128284767-DEL-7196 |
| chr2-128305290-DEL-54 | chr2-128305291-DEL-54 |
| chr2-128320169-DEL-537 | chr2-128320170-DEL-537 |
| chr2-128322396-DEL-6442 | chr2-128322397-DEL-6442 |
| chr2-128332011-DEL-327 | chr2-128332012-DEL-327 |
| chr2-128343977-DEL-356 | chr2-128343978-DEL-356 |
| chr2-128357625-DEL-63 | chr2-128357626-DEL-63 |
| chr2-128374310-DEL-76 | chr2-128374289-DEL-51 |
| chr2-128404060-DEL-147 | chr2-128404084-DEL-97 |
| chr2-128439578-DEL-170 | chr2-128439579-DEL-170 |
| chr2-128448660-DEL-1221 | chr2-128448661-DEL-1221 |
| chr2-128482743-DEL-204 | chr2-128482744-DEL-204 |
| chr2-128488164-DEL-170 | chr2-128488165-DEL-170 |
| chr2-128524625-DEL-1774 | chr2-128524626-DEL-1774 |
| chr2-128534081-DEL-262 | chr2-128534069-DEL-275 |
| chr2-128535790-DEL-53 | chr2-128535792-DEL-52 |
| chr2-128562543-DEL-103 | chr2-128562549-DEL-98 |
| chr2-128582164-DEL-199 | chr2-128582165-DEL-199 |
| chr2-128601395-DEL-210 | chr2-128601396-DEL-210 |
| chr2-128613511-DEL-102 | chr2-128613512-DEL-102 |
| chr2-128614533-DEL-1893 | chr2-128614534-DEL-1893 |
| chr2-128628822-DEL-119 | chr2-128628823-DEL-119 |
| chr2-128631775-DEL-184 | chr2-128631778-DEL-182 |
| chr2-128649352-DEL-425 | chr2-128649353-DEL-425 |
| chr2-128653511-DEL-147 | chr2-128653512-DEL-147 |
| chr2-128658638-DEL-236 | chr2-128658639-DEL-236 |
| chr2-128666915-DEL-385 | chr2-128666916-DEL-385 |
| chr2-128673584-DEL-189 | chr2-128673585-DEL-189 |
| chr2-128703163-DEL-153 | chr2-128703164-DEL-153 |

| | |
|---|---|
| chr2-128719655-DEL-714 | chr2-128719689-DEL-718 |
| chr2-128732307-DEL-111 | chr2-128732308-DEL-111 |
| chr2-128761102-DEL-171 | chr2-128761103-DEL-171 |
| chr2-128833301-DEL-1349 | chr2-128833302-DEL-1349 |
| chr2-128861310-DEL-200 | chr2-128861311-DEL-200 |
| chr2-128865498-DEL-618 | chr2-128865499-DEL-618 |
| chr2-128892297-DEL-318 | chr2-128892298-DEL-318 |
| chr2-128895096-DEL-118 | chr2-128895093-DEL-113 |
| chr2-128900656-DEL-3346 | chr2-128900657-DEL-3346 |
| chr2-128902313-DEL-120 | chr2-128902344-DEL-90 |
| chr2-128924948-DEL-218 | chr2-128924949-DEL-218 |
| chr2-128931443-DEL-90 | chr2-128931444-DEL-90 |
| chr2-128939196-DEL-84 | chr2-128939197-DEL-84 |
| chr2-128946533-DEL-72 | chr2-128946534-DEL-72 |
| chr2-128976187-DEL-3243 | chr2-128976188-DEL-3243 |
| chr2-128996401-DEL-270 | chr2-128996399-DEL-281 |
| chr2-129005312-DEL-56 | chr2-129005304-DEL-54 |
| chr2-129030852-DEL-206 | chr2-129030853-DEL-206 |
| chr2-129096778-DEL-202 | chr2-129096779-DEL-202 |
| chr2-129097384-DEL-110 | chr2-129097385-DEL-110 |
| chr2-129104530-DEL-328 | chr2-129104531-DEL-328 |
| chr2-129106609-DEL-155 | chr2-129106610-DEL-155 |
| chr2-129113424-DEL-55 | chr2-129113419-DEL-55 |
| chr2-129130759-DEL-126 | chr2-129130760-DEL-126 |
| chr2-129146191-DEL-87 | chr2-129146192-DEL-87 |
| chr2-129154196-DEL-66 | chr2-129154176-DEL-75 |
| chr2-129246446-DEL-258 | chr2-129246447-DEL-255 |
| chr2-129262161-DEL-174 | chr2-129262162-DEL-174 |
| chr2-129272751-DEL-52 | chr2-129272752-DEL-52 |
| chr2-129290052-DEL-6963 | chr2-129290060-DEL-6960 |
| chr2-129296754-DEL-214 | chr2-129296755-DEL-214 |
| chr2-129318772-DEL-880 | chr2-129318773-DEL-880 |
| chr2-129323185-DEL-88 | chr2-129323186-DEL-88 |
| chr2-129333520-DEL-59 | chr2-129333517-DEL-60 |
| chr2-129341220-DEL-1289 | chr2-129341221-DEL-1289 |
| chr2-129363189-DEL-3875 | chr2-129363190-DEL-3875 |
| chr2-129395774-DEL-152 | chr2-129395775-DEL-152 |
| chr2-129427354-DEL-103 | chr2-129427350-DEL-100 |
| chr2-129453628-DEL-1858 | chr2-129453622-DEL-1860 |
| chr2-129500114-DEL-85 | chr2-129500098-DEL-102 |
| chr2-129517797-DEL-80 | chr2-129517798-DEL-80 |
| chr2-129518783-DEL-272 | chr2-129518784-DEL-272 |
| chr2-129548053-DEL-82 | chr2-129548054-DEL-82 |
| chr2-129591150-DEL-65 | chr2-129591158-DEL-64 |
| chr2-129605224-DEL-1188 | chr2-129605225-DEL-1188 |
| chr2-129610868-DEL-64 | chr2-129610869-DEL-64 |
| chr2-129612990-DEL-255 | chr2-129612991-DEL-255 |
| chr2-129614359-DEL-2942 | chr2-129614360-DEL-2942 |
| chr2-129636020-DEL-188 | chr2-129636021-DEL-173 |
| chr2-129639564-DEL-180 | chr2-129639565-DEL-180 |
| chr2-129684153-DEL-4631 | chr2-129684164-DEL-4621 |
| chr2-129725865-DEL-88 | chr2-129725869-DEL-85 |
| chr2-129733583-DEL-77 | chr2-129733584-DEL-77 |
| chr2-129748175-DEL-453 | chr2-129748176-DEL-453 |
| chr2-129811963-DEL-142 | chr2-129811964-DEL-142 |
| chr2-129824557-DEL-185 | chr2-129824558-DEL-185 |
| chr2-129826122-DEL-70 | chr2-129826099-DEL-65 |
| chr2-129861897-DEL-87 | chr2-129861880-DEL-105 |
| chr2-129893105-DEL-508 | chr2-129893106-DEL-508 |
| chr2-129893784-DEL-59 | chr2-129893785-DEL-59 |
| chr2-129904318-DEL-1444 | chr2-129904319-DEL-1444 |
| chr2-129955544-DEL-54 | chr2-129955545-DEL-54 |
| chr2-129991856-DEL-93 | chr2-129991835-DEL-81 |
| chr2-130010357-DEL-109 | chr2-130010358-DEL-109 |
| chr2-130011384-DEL-82 | chr2-130011385-DEL-82 |
| chr2-130057942-DEL-108 | chr2-130057943-DEL-108 |
| chr2-130059838-DEL-175 | chr2-130059839-DEL-175 |
| chr2-130067549-DEL-52 | chr2-130067550-DEL-52 |
| chr2-130068873-DEL-442 | chr2-130068874-DEL-442 |
| chr2-130072510-DEL-62 | chr2-130072511-DEL-62 |
| chr2-130107052-DEL-69 | chr2-130107053-DEL-67 |
| chr2-130162538-DEL-1199 | chr2-130162539-DEL-1199 |
| chr2-130163838-DEL-631 | chr2-130163839-DEL-631 |
| chr2-130167152-DEL-208 | chr2-130167153-DEL-208 |
| chr2-130171103-DEL-84 | chr2-130171128-DEL-58 |
| chr2-130180204-DEL-650 | chr2-130180252-DEL-603 |
| chr2-130185577-DEL-70 | chr2-130185578-DEL-70 |
| chr2-130206178-DEL-54 | chr2-130206179-DEL-54 |
| chr2-130226721-DEL-551 | chr2-130226722-DEL-551 |
| chr2-130229499-DEL-51 | chr2-130229500-DEL-51 |

| | |
|---|---|
| chr2-130231418-DEL-51 | chr2-130231419-DEL-51 |
| chr2-130249324-DEL-1249 | chr2-130249325-DEL-1249 |
| chr2-130251127-DEL-79 | chr2-130251128-DEL-79 |
| chr2-130263337-DEL-76 | chr2-130263338-DEL-76 |
| chr2-130270215-DEL-171 | chr2-130270216-DEL-171 |
| chr2-130286641-DEL-208 | chr2-130286642-DEL-208 |
| chr2-130304700-DEL-832 | chr2-130304701-DEL-830 |
| chr2-130316524-DEL-213 | chr2-130316525-DEL-213 |
| chr2-130351839-DEL-150 | chr2-130351840-DEL-127 |
| chr2-130353479-DEL-210 | chr2-130353480-DEL-210 |
| chr2-130354424-DEL-266 | chr2-130354425-DEL-266 |
| chr2-130398999-DEL-118 | chr2-130399000-DEL-118 |
| chr2-130409295-DEL-216 | chr2-130409296-DEL-216 |
| chr2-130436107-DEL-95 | chr2-130436100-DEL-80 |
| chr2-130465148-DEL-56 | chr2-130465149-DEL-60 |
| chr2-130466408-DEL-173 | chr2-130466409-DEL-173 |
| chr2-130472604-DEL-63 | chr2-130472600-DEL-58 |
| chr2-130514759-DEL-213 | chr2-130514760-DEL-213 |
| chr2-130530798-DEL-222 | chr2-130530830-DEL-191 |
| chr2-130556853-DEL-67 | chr2-130556854-DEL-61 |
| chr2-130621000-DEL-261 | chr2-130621001-DEL-261 |
| chr2-130663965-DEL-222 | chr2-130663966-DEL-222 |
| chr2-130676327-DEL-62 | chr2-130676328-DEL-62 |
| chr2-130678360-DEL-213 | chr2-130678361-DEL-213 |
| chr2-130689817-DEL-364 | chr2-130689815-DEL-365 |
| chr2-130705776-DEL-544 | chr2-130705773-DEL-545 |
| chr2-130717319-DEL-95 | chr2-130717293-DEL-95 |
| chr2-130733046-DEL-102 | chr2-130733047-DEL-102 |
| chr2-130758142-DEL-196 | chr2-130758143-DEL-196 |
| chr2-130762007-DEL-58 | chr2-130762010-DEL-56 |
| chr2-130795216-DEL-57 | chr2-130795189-DEL-58 |
| chr2-130859437-DEL-221 | chr2-130859438-DEL-221 |
| chr2-130922091-DEL-65 | chr2-130922084-DEL-78 |
| chr2-131006762-DEL-287 | chr2-131006763-DEL-287 |
| chr2-131011849-DEL-123 | chr2-131011850-DEL-123 |
| chr2-131049806-DEL-208 | chr2-131049807-DEL-208 |
| chr2-131061468-DEL-171 | chr2-131061461-DEL-172 |
| chr2-131066307-DEL-105 | chr2-131066304-DEL-109 |
| chr2-131115572-DEL-93 | chr2-131115573-DEL-93 |
| chr2-131152263-DEL-516 | chr2-131152264-DEL-516 |
| chr2-131164944-DEL-121 | chr2-131164945-DEL-121 |
| chr2-131178943-DEL-3510 | chr2-131178944-DEL-3510 |
| chr2-131186890-DEL-201 | chr2-131186891-DEL-201 |
| chr2-131187391-DEL-2224 | chr2-131187392-DEL-2224 |
| chr2-131222057-DEL-2034 | chr2-131222058-DEL-2034 |
| chr2-131224149-DEL-434 | chr2-131224150-DEL-434 |
| chr2-131240631-DEL-65 | chr2-131240585-DEL-66 |
| chr2-131247264-DEL-354 | chr2-131247265-DEL-332 |
| chr2-131275235-DEL-212 | chr2-131275236-DEL-212 |
| chr2-131280592-DEL-756 | chr2-131280593-DEL-756 |
| chr2-131293674-DEL-211 | chr2-131293675-DEL-211 |
| chr2-131295573-DEL-68 | chr2-131295584-DEL-72 |
| chr2-131323510-DEL-64 | chr2-131323558-DEL-52 |
| chr2-131330568-DEL-89 | chr2-131330569-DEL-89 |
| chr2-131342105-DEL-132 | chr2-131342106-DEL-132 |
| chr2-131408541-DEL-224 | chr2-131408542-DEL-224 |
| chr2-131415540-DEL-364 | chr2-131415541-DEL-365 |
| chr2-131423301-DEL-69 | chr2-131423302-DEL-69 |
| chr2-131478451-DEL-68 | chr2-131478454-DEL-66 |
| chr2-131487400-DEL-2812 | chr2-131487401-DEL-2812 |
| chr2-131590254-DEL-56 | chr2-131590255-DEL-56 |
| chr2-131601090-DEL-57 | chr2-131601091-DEL-51 |
| chr2-131604718-DEL-1377 | chr2-131604719-DEL-1377 |
| chr2-131621815-DEL-148 | chr2-131621816-DEL-147 |
| chr2-131631011-DEL-6159 | chr2-131631012-DEL-6159 |
| chr2-131648042-DEL-218 | chr2-131648043-DEL-218 |
| chr2-131690625-DEL-123 | chr2-131690626-DEL-123 |
| chr2-131694465-DEL-57 | chr2-131694466-DEL-57 |
| chr2-131709114-DEL-145 | chr2-131709114-DEL-145 |
| chr2-131714894-DEL-200 | chr2-131714895-DEL-200 |
| chr2-131717442-DEL-287 | chr2-131717442-DEL-285 |
| chr2-131737395-DEL-446 | chr2-131737394-DEL-442 |
| chr2-131759945-DEL-12992 | chr2-131759946-DEL-12992 |
| chr2-131762388-DEL-6399 | chr2-131762389-DEL-6399 |
| chr2-131797243-DEL-295 | chr2-131797244-DEL-295 |
| chr2-131805007-DEL-7381 | chr2-131805008-DEL-7381 |
| chr2-131819829-DEL-60 | chr2-131819830-DEL-60 |
| chr2-131820479-DEL-374 | chr2-131820480-DEL-374 |
| chr2-131821145-DEL-85 | chr2-131821146-DEL-85 |
| chr2-131822351-DEL-90 | chr2-131822352-DEL-90 |

| | |
|---|---|
| chr2-131833374-DEL-66 | chr2-131833376-DEL-64 |
| chr2-131840594-DEL-73 | chr2-131840594-DEL-74 |
| chr2-131857237-DEL-56 | chr2-131857237-DEL-56 |
| chr2-131879129-DEL-56 | chr2-131879130-DEL-56 |
| chr2-131909131-DEL-178 | chr2-131909132-DEL-178 |
| chr2-131927294-DEL-58 | chr2-131927295-DEL-58 |
| chr2-131959681-DEL-52 | chr2-131959682-DEL-52 |
| chr2-131961861-DEL-59 | chr2-131961825-DEL-68 |
| chr2-131969023-DEL-65 | chr2-131969024-DEL-65 |
| chr2-131992383-DEL-150 | chr2-131992384-DEL-150 |
| chr2-132005898-DEL-1393 | chr2-132005899-DEL-1393 |
| chr2-132007868-DEL-1222 | chr2-132007869-DEL-1222 |
| chr2-132044158-DEL-212 | chr2-132044159-DEL-212 |
| chr2-132070520-DEL-92 | chr2-132070521-DEL-92 |
| chr2-132073088-DEL-588 | chr2-132073089-DEL-588 |
| chr2-132097489-DEL-221 | chr2-132097491-DEL-220 |
| chr2-132100438-DEL-180 | chr2-132100439-DEL-180 |
| chr2-132121016-DEL-202 | chr2-132121017-DEL-202 |
| chr2-132131932-DEL-213 | chr2-132131933-DEL-213 |
| chr2-132147673-DEL-134 | chr2-132147674-DEL-134 |
| chr2-132155335-DEL-52 | chr2-132155335-DEL-52 |
| chr2-132159577-DEL-2184 | chr2-132159578-DEL-2184 |
| chr2-132183231-DEL-999 | chr2-132183232-DEL-999 |
| chr2-132218747-DEL-54 | chr2-132218748-DEL-54 |
| chr2-132234668-DEL-179 | chr2-132234669-DEL-179 |
| chr2-132268051-DEL-114 | chr2-132268052-DEL-84 |
| chr2-132272845-DEL-58 | chr2-132272845-DEL-58 |
| chr2-132277057-DEL-1286 | chr2-132277058-DEL-1286 |
| chr2-132287891-DEL-142 | chr2-132287892-DEL-142 |
| chr2-132321028-DEL-1817 | chr2-132321028-DEL-1817 |
| chr2-132323578-DEL-146 | chr2-132323579-DEL-146 |
| chr2-132365819-DEL-76 | chr2-132365820-DEL-76 |
| chr2-132408096-DEL-172 | chr2-132408097-DEL-172 |
| chr2-132450575-DEL-193 | chr2-132450576-DEL-193 |
| chr2-132473385-DEL-180 | chr2-132473376-DEL-181 |
| chr2-132486609-DEL-75 | chr2-132486603-DEL-76 |
| chr2-132487557-DEL-498 | chr2-132487558-DEL-498 |
| chr2-132494845-DEL-87 | chr2-132494846-DEL-88 |
| chr2-132501979-DEL-51 | chr2-132501968-DEL-63 |
| chr2-132516966-DEL-193 | chr2-132516959-DEL-193 |
| chr2-132527453-DEL-103 | chr2-132527435-DEL-102 |
| chr2-132534744-DEL-64 | chr2-132534745-DEL-64 |
| chr2-132596046-DEL-195 | chr2-132596047-DEL-195 |
| chr2-132614654-DEL-80 | chr2-132614655-DEL-80 |
| chr2-132723584-DEL-1810 | chr2-132723585-DEL-1810 |
| chr2-132740597-DEL-52 | chr2-132740598-DEL-52 |
| chr2-132743426-DEL-56 | chr2-132743433-DEL-55 |
| chr2-132746665-DEL-282 | chr2-132746666-DEL-282 |
| chr2-132759774-DEL-52 | chr2-132759779-DEL-56 |
| chr2-132768816-DEL-51 | chr2-132768783-DEL-88 |
| chr2-132775407-DEL-88 | chr2-132775410-DEL-110 |
| chr2-132782987-DEL-125 | chr2-132783005-DEL-123 |
| chr2-132794435-DEL-1891 | chr2-132794436-DEL-1891 |
| chr2-132814895-DEL-283 | chr2-132814896-DEL-283 |
| chr2-132830895-DEL-144 | chr2-132830892-DEL-176 |
| chr2-132837044-DEL-3921 | chr2-132837045-DEL-3921 |
| chr2-132844592-DEL-432 | chr2-132844593-DEL-432 |
| chr2-132851125-DEL-476 | chr2-132851126-DEL-476 |
| chr2-132874922-DEL-63 | chr2-132874923-DEL-63 |
| chr2-132879958-DEL-431 | chr2-132879959-DEL-431 |
| chr2-132900255-DEL-800 | chr2-132900256-DEL-800 |
| chr2-132909643-DEL-7042 | chr2-132909644-DEL-7042 |
| chr2-132922203-DEL-3575 | chr2-132922204-DEL-3575 |
| chr2-132956629-DEL-88 | chr2-132956643-DEL-78 |
| chr2-132958700-DEL-196 | chr2-132958701-DEL-196 |
| chr2-132968295-DEL-360 | chr2-132968296-DEL-360 |
| chr2-133006396-DEL-574 | chr2-133006397-DEL-574 |
| chr2-133070548-DEL-66 | chr2-133070549-DEL-66 |
| chr2-133081137-DEL-6038 | chr2-133081126-DEL-6048 |
| chr2-133095029-DEL-54 | chr2-133095030-DEL-54 |
| chr2-133096497-DEL-1012 | chr2-133096499-DEL-1013 |
| chr2-133097480-DEL-6416 | chr2-133097481-DEL-6416 |
| chr2-133103607-DEL-201 | chr2-133103608-DEL-201 |
| chr2-133114004-DEL-774 | chr2-133114005-DEL-774 |
| chr2-133157673-DEL-54 | chr2-133157670-DEL-58 |
| chr2-133163974-DEL-94 | chr2-133164000-DEL-56 |
| chr2-133195278-DEL-625 | chr2-133195279-DEL-625 |
| chr2-133259741-DEL-108 | chr2-133259745-DEL-88 |
| chr2-133283546-DEL-986 | chr2-133283536-DEL-987 |
| chr2-133315517-DEL-58 | chr2-133315526-DEL-50 |

| | |
|---|---|
| chr2-133317586-DEL-691 | chr2-133317587-DEL-691 |
| chr2-133328872-DEL-369 | chr2-133328868-DEL-370 |
| chr2-133355868-DEL-98 | chr2-133355869-DEL-98 |
| chr2-133370089-DEL-73 | chr2-133370096-DEL-52 |
| chr2-133381608-DEL-116 | chr2-133381609-DEL-116 |
| chr2-133400292-DEL-141 | chr2-133400289-DEL-145 |
| chr2-133407648-DEL-78 | chr2-133407637-DEL-74 |
| chr2-133415461-DEL-126 | chr2-133415462-DEL-126 |
| chr2-133510894-DEL-69 | chr2-133510894-DEL-80 |
| chr2-133533314-DEL-882 | chr2-133533315-DEL-882 |
| chr2-133537335-DEL-2137 | chr2-133537336-DEL-2137 |
| chr2-133542885-DEL-1332 | chr2-133542886-DEL-1331 |
| chr2-133555129-DEL-82 | chr2-133555134-DEL-78 |
| chr2-133574170-DEL-69 | chr2-133574171-DEL-63 |
| chr2-133619973-DEL-1595 | chr2-133619974-DEL-1595 |
| chr2-133649711-DEL-6378 | chr2-133649712-DEL-6378 |
| chr2-133720244-DEL-65 | chr2-133720259-DEL-66 |
| chr2-133727794-DEL-401 | chr2-133727795-DEL-401 |
| chr2-133739559-DEL-206 | chr2-133739560-DEL-206 |
| chr2-133769065-DEL-566 | chr2-133769066-DEL-563 |
| chr2-133774063-DEL-6395 | chr2-133774064-DEL-6395 |
| chr2-133803141-DEL-76 | chr2-133803157-DEL-65 |
| chr2-133804945-DEL-339 | chr2-133804946-DEL-339 |
| chr2-133805859-DEL-343 | chr2-133805864-DEL-345 |
| chr2-133807001-DEL-6798 | chr2-133807002-DEL-6798 |
| chr2-133816400-DEL-51 | chr2-133816383-DEL-80 |
| chr2-133833001-DEL-591 | chr2-133833002-DEL-591 |
| chr2-133855325-DEL-306 | chr2-133855326-DEL-306 |
| chr2-133865676-DEL-788 | chr2-133865677-DEL-789 |
| chr2-133879794-DEL-65 | chr2-133879795-DEL-65 |
| chr2-133883288-DEL-116 | chr2-133883337-DEL-108 |
| chr2-133926991-DEL-57 | chr2-133926991-DEL-58 |
| chr2-133930578-DEL-371 | chr2-133930576-DEL-370 |
| chr2-133934418-DEL-66 | chr2-133934419-DEL-66 |
| chr2-133957313-DEL-266 | chr2-133957314-DEL-266 |
| chr2-134025045-DEL-108 | chr2-134025046-DEL-108 |
| chr2-134059492-DEL-730 | chr2-134059490-DEL-731 |
| chr2-134072537-DEL-62 | chr2-134072538-DEL-60 |
| chr2-134076449-DEL-932 | chr2-134076450-DEL-932 |
| chr2-134081000-DEL-1751 | chr2-134081001-DEL-1751 |
| chr2-134083134-DEL-135 | chr2-134083125-DEL-136 |
| chr2-134118609-DEL-125 | chr2-134118610-DEL-125 |
| chr2-134119591-DEL-52 | chr2-134119592-DEL-54 |
| chr2-134155537-DEL-3485 | chr2-134155538-DEL-3485 |
| chr2-134164627-DEL-2845 | chr2-134164627-DEL-2844 |
| chr2-134243142-DEL-99 | chr2-134243167-DEL-80 |
| chr2-134256170-DEL-79 | chr2-134256171-DEL-79 |
| chr2-134317465-DEL-944 | chr2-134317466-DEL-944 |
| chr2-134347819-DEL-6797 | chr2-134347820-DEL-6797 |
| chr2-134358029-DEL-919 | chr2-134358030-DEL-919 |
| chr2-134373445-DEL-333 | chr2-134373450-DEL-327 |
| chr2-134393504-DEL-513 | chr2-134393504-DEL-508 |
| chr2-134413435-DEL-161 | chr2-134413436-DEL-161 |
| chr2-134416967-DEL-2848 | chr2-134416968-DEL-2848 |
| chr2-134444680-DEL-1052 | chr2-134444684-DEL-1049 |
| chr2-134449204-DEL-600 | chr2-134449205-DEL-600 |
| chr2-134455409-DEL-6275 | chr2-134455410-DEL-6275 |
| chr2-134473401-DEL-2700 | chr2-134473402-DEL-2700 |
| chr2-134476405-DEL-2005 | chr2-134476406-DEL-2005 |
| chr2-134488511-DEL-171 | chr2-134488512-DEL-171 |
| chr2-134502508-DEL-12467 | chr2-134502509-DEL-12467 |
| chr2-134520216-DEL-1389 | chr2-134520217-DEL-1389 |
| chr2-134532194-DEL-116 | chr2-134532195-DEL-116 |
| chr2-134535609-DEL-606 | chr2-134535623-DEL-579 |
| chr2-134540980-DEL-60 | chr2-134540981-DEL-60 |
| chr2-134588571-DEL-4457 | chr2-134588572-DEL-4457 |
| chr2-134603961-DEL-1610 | chr2-134603960-DEL-1610 |
| chr2-134631681-DEL-171 | chr2-134631682-DEL-171 |
| chr2-134645799-DEL-686 | chr2-134645799-DEL-686 |
| chr2-134741626-DEL-76 | chr2-134741662-DEL-56 |
| chr2-134748412-DEL-422 | chr2-134748413-DEL-422 |
| chr2-134782682-DEL-88 | chr2-134782691-DEL-56 |
| chr2-134785542-DEL-485 | chr2-134785543-DEL-485 |
| chr2-134801603-DEL-108 | chr2-134801624-DEL-88 |
| chr2-134841066-DEL-91 | chr2-134841067-DEL-91 |
| chr2-134845872-DEL-106 | chr2-134845852-DEL-138 |
| chr2-134848660-DEL-114 | chr2-134848660-DEL-115 |
| chr2-134874286-DEL-574 | chr2-134874288-DEL-573 |
| chr2-134900688-DEL-1097 | chr2-134900689-DEL-1097 |
| chr2-134901046-DEL-516 | chr2-134901047-DEL-516 |

| | |
|---|---|
| chr2-134907900-DEL-400 | chr2-134907901-DEL-400 |
| chr2-134947960-DEL-618 | chr2-134947961-DEL-618 |
| chr2-134988597-DEL-2395 | chr2-134988598-DEL-2395 |
| chr2-135031613-DEL-104 | chr2-135031607-DEL-100 |
| chr2-135034662-DEL-498 | chr2-135034663-DEL-496 |
| chr2-135064286-DEL-77 | chr2-135064287-DEL-77 |
| chr2-135081589-DEL-157 | chr2-135081590-DEL-157 |
| chr2-135114181-DEL-497 | chr2-135114182-DEL-497 |
| chr2-135202379-DEL-155 | chr2-135202378-DEL-157 |
| chr2-135215019-DEL-133 | chr2-135215022-DEL-131 |
| chr2-135258421-DEL-80 | chr2-135258422-DEL-80 |
| chr2-135275044-DEL-208 | chr2-135275045-DEL-208 |
| chr2-135283459-DEL-113 | chr2-135283458-DEL-128 |
| chr2-135284522-DEL-3033 | chr2-135284523-DEL-3033 |
| chr2-135288114-DEL-297 | chr2-135288115-DEL-297 |
| chr2-135289735-DEL-1408 | chr2-135289736-DEL-1408 |
| chr2-135332920-DEL-69 | chr2-135332921-DEL-69 |
| chr2-135354071-DEL-6446 | chr2-135354072-DEL-6446 |
| chr2-135387158-DEL-110 | chr2-135387155-DEL-114 |
| chr2-135398787-DEL-72 | chr2-135398788-DEL-72 |
| chr2-135400080-DEL-59 | chr2-135400081-DEL-59 |
| chr2-135421067-DEL-67 | chr2-135421068-DEL-67 |
| chr2-135424398-DEL-82 | chr2-135424399-DEL-82 |
| chr2-135434581-DEL-865 | chr2-135434582-DEL-865 |
| chr2-135437901-DEL-6262 | chr2-135437949-DEL-6214 |
| chr2-135487339-DEL-58 | chr2-135487332-DEL-50 |
| chr2-135538996-DEL-82 | chr2-135538992-DEL-85 |
| chr2-135600620-DEL-318 | chr2-135600621-DEL-318 |
| chr2-135683378-DEL-1795 | chr2-135683379-DEL-1795 |
| chr2-135693438-DEL-240 | chr2-135693439-DEL-240 |
| chr2-135793813-DEL-83 | chr2-135793814-DEL-83 |
| chr2-135819820-DEL-54 | chr2-135819811-DEL-56 |
| chr2-135846858-DEL-58 | chr2-135846859-DEL-58 |
| chr2-135850299-DEL-58 | chr2-135850300-DEL-58 |
| chr2-135862314-DEL-3012 | chr2-135862315-DEL-3012 |
| chr2-135873558-DEL-1073 | chr2-135873559-DEL-1073 |
| chr2-135888932-DEL-89 | chr2-135888933-DEL-89 |
| chr2-135936808-DEL-490 | chr2-135936799-DEL-500 |
| chr2-135944654-DEL-907 | chr2-135944655-DEL-907 |
| chr2-135946281-DEL-1295 | chr2-135946282-DEL-1295 |
| chr2-135977818-DEL-59 | chr2-135977821-DEL-57 |
| chr2-136014728-DEL-456 | chr2-136014729-DEL-456 |
| chr2-136034744-DEL-395 | chr2-136034745-DEL-395 |
| chr2-136036933-DEL-252 | chr2-136036934-DEL-252 |
| chr2-136046655-DEL-920 | chr2-136046656-DEL-920 |
| chr2-136076134-DEL-6169 | chr2-136076135-DEL-6169 |
| chr2-136091784-DEL-75 | chr2-136091785-DEL-75 |
| chr2-136103114-DEL-2455 | chr2-136103115-DEL-2455 |
| chr2-136116357-DEL-4957 | chr2-136116358-DEL-4957 |
| chr2-136122929-DEL-72 | chr2-136122930-DEL-92 |
| chr2-136123531-DEL-585 | chr2-136123531-DEL-586 |
| chr2-136126616-DEL-2268 | chr2-136126617-DEL-2268 |
| chr2-136143999-DEL-80 | chr2-136143998-DEL-70 |
| chr2-136153341-DEL-543 | chr2-136153342-DEL-543 |
| chr2-136251029-DEL-70 | chr2-136251030-DEL-70 |
| chr2-136320681-DEL-59 | chr2-136320684-DEL-55 |
| chr2-136336571-DEL-3568 | chr2-136336572-DEL-3568 |
| chr2-136409725-DEL-350 | chr2-136409726-DEL-350 |
| chr2-136476090-DEL-419 | chr2-136476091-DEL-419 |
| chr2-136487495-DEL-74 | chr2-136487496-DEL-74 |
| chr2-136498046-DEL-9700 | chr2-136498047-DEL-9700 |
| chr2-136548303-DEL-655 | chr2-136548304-DEL-655 |
| chr2-136563717-DEL-213 | chr2-136563721-DEL-214 |
| chr2-136627059-DEL-78 | chr2-136627066-DEL-72 |
| chr2-136680405-DEL-58 | chr2-136680406-DEL-58 |
| chr2-136682915-DEL-76 | chr2-136682916-DEL-76 |
| chr2-136702307-DEL-177 | chr2-136702307-DEL-184 |
| chr2-136703151-DEL-82 | chr2-136703191-DEL-85 |
| chr2-136705614-DEL-6698 | chr2-136705615-DEL-6698 |
| chr2-136720737-DEL-566 | chr2-136720738-DEL-566 |
| chr2-136724700-DEL-74 | chr2-136724705-DEL-68 |
| chr2-136743371-DEL-215 | chr2-136743372-DEL-215 |
| chr2-136770800-DEL-90 | chr2-136770816-DEL-60 |
| chr2-136824984-DEL-81 | chr2-136824988-DEL-77 |
| chr2-136843306-DEL-65 | chr2-136843318-DEL-53 |
| chr2-136864863-DEL-1295 | chr2-136864864-DEL-1295 |
| chr2-136871328-DEL-7130 | chr2-136871328-DEL-7130 |
| chr2-136877200-DEL-1773 | chr2-136877201-DEL-1773 |
| chr2-137048686-DEL-141 | chr2-137048687-DEL-141 |
| chr2-137049401-DEL-284 | chr2-137049402-DEL-284 |

| | |
|---|---|
| chr2-137068779-DEL-162 | chr2-137068780-DEL-162 |
| chr2-137076225-DEL-853 | chr2-137076226-DEL-853 |
| chr2-137093503-DEL-51 | chr2-137093504-DEL-51 |
| chr2-137097565-DEL-63 | chr2-137097566-DEL-63 |
| chr2-137102820-DEL-727 | chr2-137102821-DEL-727 |
| chr2-137108023-DEL-544 | chr2-137108024-DEL-544 |
| chr2-137116000-DEL-2299 | chr2-137116001-DEL-2299 |
| chr2-137119220-DEL-827 | chr2-137119221-DEL-827 |
| chr2-137122093-DEL-1374 | chr2-137122093-DEL-1374 |
| chr2-137144935-DEL-77 | chr2-137144936-DEL-83 |
| chr2-137236587-DEL-3425 | chr2-137236588-DEL-3425 |
| chr2-137241132-DEL-6440 | chr2-137241133-DEL-6440 |
| chr2-137249024-DEL-105 | chr2-137249038-DEL-91 |
| chr2-137268376-DEL-130 | chr2-137268401-DEL-130 |
| chr2-137274171-DEL-145 | chr2-137274169-DEL-146 |
| chr2-137292442-DEL-82 | chr2-137292446-DEL-50 |
| chr2-137329147-DEL-58 | chr2-137329148-DEL-58 |
| chr2-137382687-DEL-641 | chr2-137382683-DEL-642 |
| chr2-137521685-DEL-613 | chr2-137521686-DEL-612 |
| chr2-137526915-DEL-198 | chr2-137526916-DEL-198 |
| chr2-137549787-DEL-131 | chr2-137549788-DEL-131 |
| chr2-137552937-DEL-1586 | chr2-137552938-DEL-1586 |
| chr2-137589436-DEL-1519 | chr2-137589438-DEL-1520 |
| chr2-137590650-DEL-56 | chr2-137590655-DEL-52 |
| chr2-137611437-DEL-1155 | chr2-137611438-DEL-1155 |
| chr2-137645602-DEL-432 | chr2-137645603-DEL-432 |
| chr2-137731368-DEL-534 | chr2-137731369-DEL-534 |
| chr2-137753829-DEL-660 | chr2-137753830-DEL-660 |
| chr2-137756818-DEL-2400 | chr2-137756819-DEL-2400 |
| chr2-137762124-DEL-87 | chr2-137762124-DEL-87 |
| chr2-137776653-DEL-362 | chr2-137776654-DEL-362 |
| chr2-137791552-DEL-991 | chr2-137791553-DEL-990 |
| chr2-137804241-DEL-401 | chr2-137804242-DEL-401 |
| chr2-137832578-DEL-82 | chr2-137832584-DEL-52 |
| chr2-137876999-DEL-5209 | chr2-137877000-DEL-5209 |
| chr2-137891024-DEL-55 | chr2-137891025-DEL-55 |
| chr2-137898453-DEL-57 | chr2-137898454-DEL-57 |
| chr2-137954597-DEL-536 | chr2-137954598-DEL-536 |
| chr2-137974271-DEL-55 | chr2-137974275-DEL-52 |
| chr2-137978405-DEL-57 | chr2-137978424-DEL-78 |
| chr2-138008826-DEL-1668 | chr2-138008827-DEL-1668 |
| chr2-138021809-DEL-63 | chr2-138021810-DEL-63 |
| chr2-138024277-DEL-56 | chr2-138024278-DEL-54 |
| chr2-138029893-DEL-139 | chr2-138029899-DEL-131 |
| chr2-138032835-DEL-84 | chr2-138032836-DEL-90 |
| chr2-138059611-DEL-58 | chr2-138059623-DEL-50 |
| chr2-138102873-DEL-58 | chr2-138102872-DEL-59 |
| chr2-138106343-DEL-60 | chr2-138106342-DEL-66 |
| chr2-138140337-DEL-330 | chr2-138140338-DEL-330 |
| chr2-138167555-DEL-64 | chr2-138167556-DEL-64 |
| chr2-138230960-DEL-6479 | chr2-138230960-DEL-6479 |
| chr2-138269117-DEL-56 | chr2-138269118-DEL-56 |
| chr2-138270506-DEL-68 | chr2-138270509-DEL-75 |
| chr2-138294668-DEL-2993 | chr2-138294669-DEL-2993 |
| chr2-138303671-DEL-5789 | chr2-138303672-DEL-5789 |
| chr2-138335981-DEL-51 | chr2-138335982-DEL-51 |
| chr2-138338815-DEL-131 | chr2-138338816-DEL-131 |
| chr2-138342387-DEL-1968 | chr2-138342388-DEL-1968 |
| chr2-138358918-DEL-8460 | chr2-138358919-DEL-8460 |
| chr2-138459922-DEL-56 | chr2-138459923-DEL-56 |
| chr2-138463387-DEL-2812 | chr2-138463388-DEL-2812 |
| chr2-138479560-DEL-549 | chr2-138479561-DEL-549 |
| chr2-138533767-DEL-1581 | chr2-138533768-DEL-1581 |
| chr2-138538002-DEL-457 | chr2-138538004-DEL-456 |
| chr2-138546857-DEL-69 | chr2-138546815-DEL-116 |
| chr2-138624078-DEL-403 | chr2-138624079-DEL-403 |
| chr2-138668990-DEL-129 | chr2-138668991-DEL-129 |
| chr2-138675557-DEL-4396 | chr2-138675558-DEL-4396 |
| chr2-138686924-DEL-8162 | chr2-138686925-DEL-8162 |
| chr2-138691263-DEL-1888 | chr2-138691264-DEL-1888 |
| chr2-138695922-DEL-101 | chr2-138695903-DEL-102 |
| chr2-138696542-DEL-66 | chr2-138696535-DEL-66 |
| chr2-138698422-DEL-189 | chr2-138698423-DEL-189 |
| chr2-138737483-DEL-351 | chr2-138737488-DEL-347 |
| chr2-138757216-DEL-1146 | chr2-138757217-DEL-1146 |
| chr2-138775025-DEL-2738 | chr2-138775026-DEL-2738 |
| chr2-138788895-DEL-90 | chr2-138788896-DEL-90 |
| chr2-138800716-DEL-1640 | chr2-138800701-DEL-1653 |
| chr2-138815138-DEL-55 | chr2-138815139-DEL-55 |
| chr2-138820428-DEL-2192 | chr2-138820429-DEL-2192 |

| | |
|---|---|
| chr2-138829817-DEL-95 | chr2-138829812-DEL-93 |
| chr2-138835716-DEL-2967 | chr2-138835724-DEL-2963 |
| chr2-138863460-DEL-55 | chr2-138863465-DEL-50 |
| chr2-138907050-DEL-2902 | chr2-138907051-DEL-2902 |
| chr2-138938159-DEL-3211 | chr2-138938160-DEL-3211 |
| chr2-138952370-DEL-60 | chr2-138952371-DEL-60 |
| chr2-138967541-DEL-84 | chr2-138967541-DEL-84 |
| chr2-138983978-DEL-1200 | chr2-138983979-DEL-1200 |
| chr2-139004632-DEL-696 | chr2-139004633-DEL-696 |
| chr2-139033321-DEL-1207 | chr2-139033322-DEL-1207 |
| chr2-139080104-DEL-4097 | chr2-139080115-DEL-4081 |
| chr2-139169659-DEL-56 | chr2-139169635-DEL-84 |
| chr2-139171881-DEL-571 | chr2-139171897-DEL-575 |
| chr2-139180853-DEL-559 | chr2-139180854-DEL-559 |
| chr2-139186159-DEL-1644 | chr2-139186146-DEL-1642 |
| chr2-139233077-DEL-198 | chr2-139233078-DEL-198 |
| chr2-139243519-DEL-6419 | chr2-139243520-DEL-6419 |
| chr2-139276363-DEL-325 | chr2-139276364-DEL-325 |
| chr2-139281474-DEL-64 | chr2-139281475-DEL-64 |
| chr2-139286110-DEL-106 | chr2-139286110-DEL-107 |
| chr2-139312201-DEL-6219 | chr2-139312202-DEL-6219 |
| chr2-139318973-DEL-66 | chr2-139318974-DEL-50 |
| chr2-139364032-DEL-6671 | chr2-139364033-DEL-6671 |
| chr2-139377284-DEL-351 | chr2-139377284-DEL-351 |
| chr2-139378196-DEL-509 | chr2-139378197-DEL-509 |
| chr2-139381383-DEL-206 | chr2-139381384-DEL-206 |
| chr2-139473581-DEL-612 | chr2-139473582-DEL-612 |
| chr2-139476069-DEL-64 | chr2-139476063-DEL-106 |
| chr2-139477666-DEL-248 | chr2-139477686-DEL-229 |
| chr2-139484798-DEL-86 | chr2-139484799-DEL-86 |
| chr2-139578587-DEL-248 | chr2-139578592-DEL-244 |
| chr2-139593293-DEL-93 | chr2-139593291-DEL-96 |
| chr2-139621915-DEL-313 | chr2-139621916-DEL-313 |
| chr2-139625380-DEL-65 | chr2-139625381-DEL-59 |
| chr2-139650133-DEL-505 | chr2-139650134-DEL-505 |
| chr2-139704556-DEL-187 | chr2-139704557-DEL-187 |
| chr2-139706418-DEL-212 | chr2-139706431-DEL-213 |
| chr2-139734110-DEL-103 | chr2-139734097-DEL-116 |
| chr2-139756231-DEL-4140 | chr2-139756232-DEL-4140 |
| chr2-139767364-DEL-57 | chr2-139767365-DEL-57 |
| chr2-139802031-DEL-134 | chr2-139802030-DEL-134 |
| chr2-139820172-DEL-682 | chr2-139820196-DEL-681 |
| chr2-139825008-DEL-57 | chr2-139825009-DEL-57 |
| chr2-139829927-DEL-54 | chr2-139829928-DEL-54 |
| chr2-139833864-DEL-1393 | chr2-139833865-DEL-1393 |
| chr2-139838511-DEL-98 | chr2-139838518-DEL-94 |
| chr2-139838732-DEL-2953 | chr2-139838733-DEL-2953 |
| chr2-139841338-DEL-96 | chr2-139841339-DEL-96 |
| chr2-139845578-DEL-51 | chr2-139845585-DEL-57 |
| chr2-139860508-DEL-759 | chr2-139860510-DEL-764 |
| chr2-139875792-DEL-4210 | chr2-139875793-DEL-4210 |
| chr2-139877669-DEL-123 | chr2-139877670-DEL-120 |
| chr2-139893924-DEL-1974 | chr2-139893925-DEL-1974 |
| chr2-139911609-DEL-111 | chr2-139911648-DEL-65 |
| chr2-139920877-DEL-58 | chr2-139920878-DEL-56 |
| chr2-139933369-DEL-60 | chr2-139933370-DEL-60 |
| chr2-139941842-DEL-50 | chr2-139941847-DEL-68 |
| chr2-139947427-DEL-215 | chr2-139947428-DEL-215 |
| chr2-139969004-DEL-205 | chr2-139969005-DEL-205 |
| chr2-139996917-DEL-60 | chr2-139996918-DEL-60 |
| chr2-140026254-DEL-52 | chr2-140026257-DEL-51 |
| chr2-140028515-DEL-335 | chr2-140028516-DEL-335 |
| chr2-140037840-DEL-396 | chr2-140037839-DEL-393 |
| chr2-140049946-DEL-206 | chr2-140049946-DEL-206 |
| chr2-140114933-DEL-68 | chr2-140114934-DEL-68 |
| chr2-140129668-DEL-183 | chr2-140129669-DEL-183 |
| chr2-140130176-DEL-486 | chr2-140130177-DEL-486 |
| chr2-140140638-DEL-59 | chr2-140140634-DEL-60 |
| chr2-140148221-DEL-2688 | chr2-140148222-DEL-2688 |
| chr2-140151009-DEL-60 | chr2-140151017-DEL-77 |
| chr2-140161823-DEL-4158 | chr2-140161824-DEL-4158 |
| chr2-140187794-DEL-6475 | chr2-140187795-DEL-6475 |
| chr2-140226151-DEL-116 | chr2-140226152-DEL-117 |
| chr2-140247122-DEL-5680 | chr2-140247123-DEL-5680 |
| chr2-140253083-DEL-440 | chr2-140253084-DEL-440 |
| chr2-140256165-DEL-94 | chr2-140256166-DEL-89 |
| chr2-140270253-DEL-909 | chr2-140270254-DEL-909 |
| chr2-140278969-DEL-127 | chr2-140278969-DEL-127 |
| chr2-140294882-DEL-239 | chr2-140294883-DEL-239 |
| chr2-140304111-DEL-421 | chr2-140304118-DEL-456 |

| | |
|---|---|
| chr2-140306596-DEL-56 | chr2-140306600-DEL-51 |
| chr2-140342298-DEL-1421 | chr2-140342299-DEL-1421 |
| chr2-140432289-DEL-63 | chr2-140432290-DEL-61 |
| chr2-140433421-DEL-545 | chr2-140433422-DEL-545 |
| chr2-140436160-DEL-112 | chr2-140436157-DEL-108 |
| chr2-140485795-DEL-120 | chr2-140485801-DEL-115 |
| chr2-140536291-DEL-69 | chr2-140536292-DEL-69 |
| chr2-140562022-DEL-54 | chr2-140562023-DEL-54 |
| chr2-140575197-DEL-6776 | chr2-140575198-DEL-6776 |
| chr2-140592464-DEL-6599 | chr2-140592465-DEL-6599 |
| chr2-140637858-DEL-52 | chr2-140637859-DEL-52 |
| chr2-140652117-DEL-72 | chr2-140652118-DEL-72 |
| chr2-140658422-DEL-174 | chr2-140658423-DEL-168 |
| chr2-140687760-DEL-58 | chr2-140687761-DEL-58 |
| chr2-140701646-DEL-59 | chr2-140701631-DEL-51 |
| chr2-140722547-DEL-1040 | chr2-140722567-DEL-1021 |
| chr2-140733344-DEL-280 | chr2-140733346-DEL-278 |
| chr2-140745959-DEL-72 | chr2-140745960-DEL-72 |
| chr2-140765073-DEL-2814 | chr2-140765074-DEL-2814 |
| chr2-140789229-DEL-66 | chr2-140789230-DEL-66 |
| chr2-140824550-DEL-520 | chr2-140824551-DEL-520 |
| chr2-140865666-DEL-57 | chr2-140865649-DEL-94 |
| chr2-140895854-DEL-931 | chr2-140895855-DEL-931 |
| chr2-140917006-DEL-1351 | chr2-140917007-DEL-1351 |
| chr2-140933380-DEL-1158 | chr2-140933381-DEL-1158 |
| chr2-140945033-DEL-6768 | chr2-140945034-DEL-6768 |
| chr2-140969146-DEL-62 | chr2-140969147-DEL-62 |
| chr2-140972841-DEL-58 | chr2-140972842-DEL-62 |
| chr2-140991495-DEL-192 | chr2-140991510-DEL-226 |
| chr2-140996199-DEL-744 | chr2-140996200-DEL-744 |
| chr2-141019208-DEL-207 | chr2-141019209-DEL-207 |
| chr2-141075314-DEL-137 | chr2-141075315-DEL-137 |
| chr2-141274846-DEL-56 | chr2-141274839-DEL-58 |
| chr2-141294062-DEL-296 | chr2-141294063-DEL-296 |
| chr2-141295752-DEL-584 | chr2-141295753-DEL-584 |
| chr2-141377277-DEL-6288 | chr2-141377278-DEL-6287 |
| chr2-141412414-DEL-199 | chr2-141412415-DEL-199 |
| chr2-141415482-DEL-53 | chr2-141415483-DEL-53 |
| chr2-141437337-DEL-124 | chr2-141437337-DEL-136 |
| chr2-141443664-DEL-603 | chr2-141443665-DEL-603 |
| chr2-141462152-DEL-161 | chr2-141462152-DEL-161 |
| chr2-141471162-DEL-120 | chr2-141471159-DEL-124 |
| chr2-141482395-DEL-2917 | chr2-141482396-DEL-2917 |
| chr2-141488218-DEL-68 | chr2-141488225-DEL-62 |
| chr2-141559886-DEL-1252 | chr2-141559887-DEL-1252 |
| chr2-141591692-DEL-62 | chr2-141591679-DEL-76 |
| chr2-141603826-DEL-1194 | chr2-141603827-DEL-1194 |
| chr2-141617198-DEL-1038 | chr2-141617199-DEL-1038 |
| chr2-141651708-DEL-70 | chr2-141651711-DEL-68 |
| chr2-141663587-DEL-75 | chr2-141663588-DEL-75 |
| chr2-141673308-DEL-54 | chr2-141673309-DEL-54 |
| chr2-141680942-DEL-82 | chr2-141680977-DEL-57 |
| chr2-141684425-DEL-35408 | chr2-141684423-DEL-35412 |
| chr2-141697048-DEL-58 | chr2-141697052-DEL-54 |
| chr2-141730258-DEL-83 | chr2-141730259-DEL-83 |
| chr2-141740982-DEL-79 | chr2-141740978-DEL-112 |
| chr2-141741231-DEL-6389 | chr2-141741232-DEL-6389 |
| chr2-141751187-DEL-924 | chr2-141751188-DEL-924 |
| chr2-141764562-DEL-60 | chr2-141764563-DEL-58 |
| chr2-141814130-DEL-661 | chr2-141814131-DEL-667 |
| chr2-141865656-DEL-56 | chr2-141865655-DEL-57 |
| chr2-141898627-DEL-199 | chr2-141898628-DEL-199 |
| chr2-141901794-DEL-87 | chr2-141901795-DEL-87 |
| chr2-141905371-DEL-550 | chr2-141905372-DEL-550 |
| chr2-141943631-DEL-149 | chr2-141943632-DEL-149 |
| chr2-141966420-DEL-99 | chr2-141966421-DEL-99 |
| chr2-141983793-DEL-60 | chr2-141983781-DEL-72 |
| chr2-142039057-DEL-7125 | chr2-142039058-DEL-7125 |
| chr2-142055266-DEL-290 | chr2-142055267-DEL-290 |
| chr2-142069503-DEL-98 | chr2-142069504-DEL-98 |
| chr2-142126955-DEL-116 | chr2-142126936-DEL-118 |
| chr2-142147949-DEL-5459 | chr2-142147950-DEL-5459 |
| chr2-142197321-DEL-204 | chr2-142197322-DEL-204 |
| chr2-142338071-DEL-55 | chr2-142338072-DEL-54 |
| chr2-142353588-DEL-520 | chr2-142353589-DEL-520 |
| chr2-142396619-DEL-55 | chr2-142396620-DEL-55 |
| chr2-142404167-DEL-370 | chr2-142404168-DEL-370 |
| chr2-142444308-DEL-264 | chr2-142444308-DEL-265 |
| chr2-142496266-DEL-60 | chr2-142496267-DEL-60 |
| chr2-142497410-DEL-370 | chr2-142497397-DEL-370 |

| | |
|---|---|
| chr2-142588780-DEL-434 | chr2-142588781-DEL-434 |
| chr2-142631063-DEL-6452 | chr2-142631065-DEL-6452 |
| chr2-142677821-DEL-82 | chr2-142677815-DEL-52 |
| chr2-142684037-DEL-3051 | chr2-142684037-DEL-3051 |
| chr2-142697596-DEL-203 | chr2-142697597-DEL-203 |
| chr2-142708923-DEL-123 | chr2-142708924-DEL-123 |
| chr2-142711345-DEL-180 | chr2-142711346-DEL-180 |
| chr2-142754608-DEL-528 | chr2-142754609-DEL-528 |
| chr2-142778148-DEL-202 | chr2-142778149-DEL-202 |
| chr2-142788364-DEL-8049 | chr2-142788365-DEL-8048 |
| chr2-142789099-DEL-90 | chr2-142789100-DEL-90 |
| chr2-142793389-DEL-1745 | chr2-142793390-DEL-1745 |
| chr2-142825814-DEL-808 | chr2-142825815-DEL-808 |
| chr2-142879129-DEL-201 | chr2-142879124-DEL-201 |
| chr2-142884555-DEL-6616 | chr2-142884556-DEL-6616 |
| chr2-142916437-DEL-55 | chr2-142916438-DEL-55 |
| chr2-142948698-DEL-1056 | chr2-142948698-DEL-1056 |
| chr2-142955548-DEL-208 | chr2-142955549-DEL-209 |
| chr2-142963311-DEL-770 | chr2-142963312-DEL-770 |
| chr2-142964133-DEL-61 | chr2-142964134-DEL-61 |
| chr2-143011639-DEL-64 | chr2-143011612-DEL-72 |
| chr2-143029081-DEL-202 | chr2-143029082-DEL-202 |
| chr2-143029892-DEL-87 | chr2-143029893-DEL-87 |
| chr2-143044243-DEL-539 | chr2-143044193-DEL-547 |
| chr2-143052899-DEL-652 | chr2-143052900-DEL-652 |
| chr2-143095025-DEL-157 | chr2-143095026-DEL-157 |
| chr2-143096256-DEL-204 | chr2-143096257-DEL-204 |
| chr2-143101305-DEL-1579 | chr2-143101306-DEL-1579 |
| chr2-143116694-DEL-251 | chr2-143116695-DEL-251 |
| chr2-143125268-DEL-946 | chr2-143125269-DEL-946 |
| chr2-143157852-DEL-63 | chr2-143157853-DEL-63 |
| chr2-143232026-DEL-101 | chr2-143232033-DEL-75 |
| chr2-143238438-DEL-54 | chr2-143238439-DEL-54 |
| chr2-143238975-DEL-1112 | chr2-143238976-DEL-1112 |
| chr2-143272092-DEL-465 | chr2-143272093-DEL-465 |
| chr2-143287388-DEL-681 | chr2-143287389-DEL-681 |
| chr2-143299785-DEL-65 | chr2-143299786-DEL-65 |
| chr2-143302107-DEL-67 | chr2-143302108-DEL-67 |
| chr2-143316245-DEL-375 | chr2-143316240-DEL-373 |
| chr2-143328708-DEL-292 | chr2-143328709-DEL-292 |
| chr2-143345542-DEL-154 | chr2-143345543-DEL-154 |
| chr2-143354316-DEL-405 | chr2-143354317-DEL-405 |
| chr2-143370458-DEL-57 | chr2-143370459-DEL-57 |
| chr2-143372145-DEL-51 | chr2-143372145-DEL-51 |
| chr2-143405383-DEL-152 | chr2-143405384-DEL-152 |
| chr2-143426175-DEL-1012 | chr2-143426176-DEL-1012 |
| chr2-143428287-DEL-85 | chr2-143428288-DEL-85 |
| chr2-143462536-DEL-87 | chr2-143462537-DEL-87 |
| chr2-143464266-DEL-150 | chr2-143464226-DEL-176 |
| chr2-143467145-DEL-6610 | chr2-143467146-DEL-6610 |
| chr2-143473485-DEL-200 | chr2-143473486-DEL-200 |
| chr2-143475344-DEL-93 | chr2-143475350-DEL-94 |
| chr2-143479048-DEL-220 | chr2-143479049-DEL-220 |
| chr2-143491459-DEL-98 | chr2-143491460-DEL-64 |
| chr2-143510878-DEL-117 | chr2-143510881-DEL-122 |
| chr2-143546543-DEL-100 | chr2-143546544-DEL-100 |
| chr2-143548636-DEL-841 | chr2-143548637-DEL-841 |
| chr2-143565432-DEL-136 | chr2-143565446-DEL-137 |
| chr2-143567795-DEL-6640 | chr2-143567796-DEL-6640 |
| chr2-143586638-DEL-1409 | chr2-143586639-DEL-1409 |
| chr2-143586808-DEL-197 | chr2-143586809-DEL-197 |
| chr2-143600200-DEL-6491 | chr2-143600201-DEL-6491 |
| chr2-143636752-DEL-306 | chr2-143636745-DEL-331 |
| chr2-143671513-DEL-331 | chr2-143671514-DEL-331 |
| chr2-143702792-DEL-66 | chr2-143702793-DEL-66 |
| chr2-143708646-DEL-378 | chr2-143708647-DEL-378 |
| chr2-143714314-DEL-104 | chr2-143714312-DEL-104 |
| chr2-143722087-DEL-851 | chr2-143722089-DEL-850 |
| chr2-143723974-DEL-58 | chr2-143723975-DEL-58 |
| chr2-143728517-DEL-102 | chr2-143728518-DEL-102 |
| chr2-143733978-DEL-432 | chr2-143733981-DEL-428 |
| chr2-143769733-DEL-545 | chr2-143769734-DEL-545 |
| chr2-143812618-DEL-75 | chr2-143812619-DEL-67 |
| chr2-143835833-DEL-858 | chr2-143835834-DEL-858 |
| chr2-143841298-DEL-58 | chr2-143841299-DEL-58 |
| chr2-143870134-DEL-73 | chr2-143870134-DEL-73 |
| chr2-143899437-DEL-55 | chr2-143899438-DEL-55 |
| chr2-143923599-DEL-105 | chr2-143923600-DEL-131 |
| chr2-143931751-DEL-276 | chr2-143931748-DEL-280 |
| chr2-143946526-DEL-168 | chr2-143946527-DEL-168 |

| | |
|---|---|
| chr2-143967086-DEL-79 | chr2-143967099-DEL-67 |
| chr2-143990893-DEL-7483 | chr2-143990894-DEL-7483 |
| chr2-144020452-DEL-55 | chr2-144020453-DEL-55 |
| chr2-144040012-DEL-178 | chr2-144040013-DEL-178 |
| chr2-144067909-DEL-1643 | chr2-144067910-DEL-1643 |
| chr2-144071185-DEL-251 | chr2-144071185-DEL-251 |
| chr2-144094972-DEL-349 | chr2-144094973-DEL-349 |
| chr2-144131831-DEL-123 | chr2-144131832-DEL-123 |
| chr2-144135508-DEL-245 | chr2-144135509-DEL-245 |
| chr2-144180564-DEL-1419 | chr2-144180565-DEL-1419 |
| chr2-144186077-DEL-54 | chr2-144186060-DEL-50 |
| chr2-144186493-DEL-1246 | chr2-144186494-DEL-1246 |
| chr2-144188379-DEL-1596 | chr2-144188380-DEL-1596 |
| chr2-144242237-DEL-98 | chr2-144242238-DEL-95 |
| chr2-144257749-DEL-180 | chr2-144257750-DEL-180 |
| chr2-144271056-DEL-71 | chr2-144271057-DEL-71 |
| chr2-144278368-DEL-989 | chr2-144278369-DEL-989 |
| chr2-144281231-DEL-241 | chr2-144281232-DEL-241 |
| chr2-144321784-DEL-158 | chr2-144321795-DEL-160 |
| chr2-144347221-DEL-103 | chr2-144347222-DEL-103 |
| chr2-144377332-DEL-334 | chr2-144377330-DEL-335 |
| chr2-144391155-DEL-51 | chr2-144391156-DEL-51 |
| chr2-144400700-DEL-76 | chr2-144400701-DEL-76 |
| chr2-144402783-DEL-225 | chr2-144402783-DEL-225 |
| chr2-144418957-DEL-211 | chr2-144418944-DEL-212 |
| chr2-144424318-DEL-52 | chr2-144424299-DEL-53 |
| chr2-144457298-DEL-71 | chr2-144457299-DEL-71 |
| chr2-144469809-DEL-4464 | chr2-144469810-DEL-4464 |
| chr2-144494268-DEL-2632 | chr2-144494269-DEL-2632 |
| chr2-144510909-DEL-580 | chr2-144510910-DEL-580 |
| chr2-144527870-DEL-1698 | chr2-144527871-DEL-1698 |
| chr2-144528368-DEL-151 | chr2-144528369-DEL-151 |
| chr2-144561336-DEL-299 | chr2-144561337-DEL-299 |
| chr2-144592803-DEL-420 | chr2-144592804-DEL-420 |
| chr2-144603283-DEL-2080 | chr2-144603284-DEL-2080 |
| chr2-144607809-DEL-199 | chr2-144607810-DEL-199 |
| chr2-144613917-DEL-59 | chr2-144613918-DEL-59 |
| chr2-144621874-DEL-60 | chr2-144621875-DEL-60 |
| chr2-144658203-DEL-2648 | chr2-144658204-DEL-2648 |
| chr2-144691347-DEL-1971 | chr2-144691348-DEL-1971 |
| chr2-144699296-DEL-80 | chr2-144699264-DEL-70 |
| chr2-144718191-DEL-54 | chr2-144718192-DEL-54 |
| chr2-144733494-DEL-2387 | chr2-144733494-DEL-2387 |
| chr2-144747488-DEL-226 | chr2-144747489-DEL-226 |
| chr2-144804905-DEL-54 | chr2-144804906-DEL-54 |
| chr2-144829876-DEL-1267 | chr2-144829877-DEL-1267 |
| chr2-144868906-DEL-7341 | chr2-144868907-DEL-7341 |
| chr2-144881658-DEL-56 | chr2-144881659-DEL-56 |
| chr2-144889196-DEL-1501 | chr2-144889197-DEL-1501 |
| chr2-144896520-DEL-395 | chr2-144896521-DEL-395 |
| chr2-144959735-DEL-125 | chr2-144959737-DEL-124 |
| chr2-144960317-DEL-5321 | chr2-144960318-DEL-5321 |
| chr2-144976799-DEL-72 | chr2-144976801-DEL-73 |
| chr2-144981953-DEL-189 | chr2-144981954-DEL-189 |
| chr2-144988943-DEL-947 | chr2-144988944-DEL-947 |
| chr2-145078201-DEL-6394 | chr2-145078202-DEL-6394 |
| chr2-145176294-DEL-80 | chr2-145176295-DEL-80 |
| chr2-145183717-DEL-53 | chr2-145183716-DEL-65 |
| chr2-145195124-DEL-84 | chr2-145195125-DEL-84 |
| chr2-145209713-DEL-52 | chr2-145209714-DEL-52 |
| chr2-145221649-DEL-84 | chr2-145221650-DEL-82 |
| chr2-145264251-DEL-51 | chr2-145264244-DEL-61 |
| chr2-145293010-DEL-703 | chr2-145293012-DEL-702 |
| chr2-145368737-DEL-87 | chr2-145368738-DEL-87 |
| chr2-145382165-DEL-54 | chr2-145382130-DEL-54 |
| chr2-145409766-DEL-54 | chr2-145409767-DEL-54 |
| chr2-145444260-DEL-88 | chr2-145444260-DEL-88 |
| chr2-145446610-DEL-119 | chr2-145446625-DEL-113 |
| chr2-145461339-DEL-110 | chr2-145461340-DEL-108 |
| chr2-145467152-DEL-5402 | chr2-145467153-DEL-5402 |
| chr2-145505593-DEL-216 | chr2-145505594-DEL-216 |
| chr2-145508518-DEL-100 | chr2-145508519-DEL-100 |
| chr2-145528705-DEL-369 | chr2-145528706-DEL-365 |
| chr2-145533158-DEL-357 | chr2-145533159-DEL-357 |
| chr2-145553292-DEL-56 | chr2-145553293-DEL-56 |
| chr2-145559598-DEL-97 | chr2-145559596-DEL-98 |
| chr2-145574587-DEL-1898 | chr2-145574588-DEL-1898 |
| chr2-145579963-DEL-403 | chr2-145579995-DEL-379 |
| chr2-145604600-DEL-70 | chr2-145604605-DEL-68 |
| chr2-145618665-DEL-115 | chr2-145618666-DEL-115 |

| | |
|---|---|
| chr2-145632041-DEL-60 | chr2-145632044-DEL-58 |
| chr2-145633824-DEL-475 | chr2-145633825-DEL-475 |
| chr2-145645156-DEL-169 | chr2-145645149-DEL-170 |
| chr2-145650974-DEL-54 | chr2-145650975-DEL-54 |
| chr2-145652113-DEL-195 | chr2-145652114-DEL-195 |
| chr2-145656420-DEL-94 | chr2-145656442-DEL-80 |
| chr2-145679835-DEL-166 | chr2-145679836-DEL-166 |
| chr2-145737743-DEL-199 | chr2-145737744-DEL-199 |
| chr2-145765129-DEL-198 | chr2-145765130-DEL-198 |
| chr2-145808419-DEL-93 | chr2-145808422-DEL-90 |
| chr2-145809593-DEL-286 | chr2-145809594-DEL-286 |
| chr2-145813109-DEL-119 | chr2-145813110-DEL-119 |
| chr2-145835154-DEL-3346 | chr2-145835155-DEL-3347 |
| chr2-145854432-DEL-181 | chr2-145854433-DEL-181 |
| chr2-145860210-DEL-791 | chr2-145860211-DEL-790 |
| chr2-145867910-DEL-345 | chr2-145867887-DEL-351 |
| chr2-145881119-DEL-2291 | chr2-145881120-DEL-2291 |
| chr2-145897553-DEL-180 | chr2-145897554-DEL-173 |
| chr2-145902025-DEL-144 | chr2-145902026-DEL-144 |
| chr2-145915866-DEL-458 | chr2-145915867-DEL-458 |
| chr2-145926565-DEL-986 | chr2-145926566-DEL-986 |
| chr2-145930144-DEL-60 | chr2-145930145-DEL-60 |
| chr2-145934140-DEL-81 | chr2-145934147-DEL-52 |
| chr2-145938053-DEL-199 | chr2-145938054-DEL-199 |
| chr2-145944312-DEL-97 | chr2-145944313-DEL-97 |
| chr2-145946184-DEL-205 | chr2-145946185-DEL-205 |
| chr2-145949514-DEL-56 | chr2-145949527-DEL-50 |
| chr2-145955057-DEL-274 | chr2-145955050-DEL-269 |
| chr2-145977684-DEL-4519 | chr2-145977685-DEL-4519 |
| chr2-145999341-DEL-6286 | chr2-145999342-DEL-6286 |
| chr2-146035433-DEL-76 | chr2-146035428-DEL-82 |
| chr2-146038393-DEL-1948 | chr2-146038394-DEL-1948 |
| chr2-146058882-DEL-52 | chr2-146058883-DEL-52 |
| chr2-146118060-DEL-218 | chr2-146118061-DEL-218 |
| chr2-146127076-DEL-279 | chr2-146127078-DEL-277 |
| chr2-146138791-DEL-143 | chr2-146138792-DEL-143 |
| chr2-146147449-DEL-1584 | chr2-146147450-DEL-1584 |
| chr2-146226047-DEL-73 | chr2-146226048-DEL-73 |
| chr2-146274006-DEL-1787 | chr2-146274006-DEL-1785 |
| chr2-146374345-DEL-96 | chr2-146374345-DEL-95 |
| chr2-146379013-DEL-72 | chr2-146379014-DEL-78 |
| chr2-146381519-DEL-205 | chr2-146381520-DEL-205 |
| chr2-146384250-DEL-82 | chr2-146384251-DEL-72 |
| chr2-146409441-DEL-54 | chr2-146409442-DEL-54 |
| chr2-146414153-DEL-2155 | chr2-146414154-DEL-2155 |
| chr2-146426283-DEL-51 | chr2-146426282-DEL-54 |
| chr2-146448338-DEL-97 | chr2-146448350-DEL-91 |
| chr2-146481862-DEL-1677 | chr2-146481863-DEL-1677 |
| chr2-146486162-DEL-57 | chr2-146486163-DEL-57 |
| chr2-146493897-DEL-78 | chr2-146493898-DEL-78 |
| chr2-146519005-DEL-78 | chr2-146518985-DEL-63 |
| chr2-146538655-DEL-119 | chr2-146538656-DEL-119 |
| chr2-146573548-DEL-216 | chr2-146573546-DEL-213 |
| chr2-146580054-DEL-1086 | chr2-146580055-DEL-1086 |
| chr2-146600277-DEL-520 | chr2-146600278-DEL-520 |
| chr2-146612300-DEL-210 | chr2-146612294-DEL-211 |
| chr2-146663443-DEL-88 | chr2-146663448-DEL-80 |
| chr2-146666710-DEL-1115 | chr2-146666711-DEL-1115 |
| chr2-146687835-DEL-248 | chr2-146687848-DEL-236 |
| chr2-146707296-DEL-51 | chr2-146707297-DEL-51 |
| chr2-146731855-DEL-539 | chr2-146731856-DEL-539 |
| chr2-146760475-DEL-825 | chr2-146760476-DEL-825 |
| chr2-146768290-DEL-474 | chr2-146768291-DEL-474 |
| chr2-146819417-DEL-64 | chr2-146819418-DEL-64 |
| chr2-146830274-DEL-347 | chr2-146830275-DEL-347 |
| chr2-146833467-DEL-206 | chr2-146833468-DEL-206 |
| chr2-146837591-DEL-9511 | chr2-146837592-DEL-9511 |
| chr2-146887538-DEL-86 | chr2-146887539-DEL-86 |
| chr2-146963988-DEL-1561 | chr2-146964003-DEL-1560 |
| chr2-146979551-DEL-6652 | chr2-146979552-DEL-6652 |
| chr2-146981539-DEL-2154 | chr2-146981539-DEL-2154 |
| chr2-147004323-DEL-64 | chr2-147004324-DEL-64 |
| chr2-147006226-DEL-78 | chr2-147006227-DEL-78 |
| chr2-147012133-DEL-92 | chr2-147012146-DEL-100 |
| chr2-147012643-DEL-506 | chr2-147012644-DEL-506 |
| chr2-147022096-DEL-54 | chr2-147022095-DEL-55 |
| chr2-147027016-DEL-177 | chr2-147026979-DEL-210 |
| chr2-147048789-DEL-317 | chr2-147048790-DEL-318 |
| chr2-147062463-DEL-255 | chr2-147062464-DEL-255 |
| chr2-147069719-DEL-178 | chr2-147069720-DEL-178 |

| | |
|---|---|
| chr2-147085444-DEL-174 | chr2-147085445-DEL-170 |
| chr2-147087345-DEL-661 | chr2-147087346-DEL-661 |
| chr2-147092268-DEL-54 | chr2-147092269-DEL-54 |
| chr2-147097049-DEL-59 | chr2-147097044-DEL-59 |
| chr2-147099360-DEL-69 | chr2-147099361-DEL-69 |
| chr2-147102778-DEL-109 | chr2-147102789-DEL-99 |
| chr2-147131359-DEL-99 | chr2-147131348-DEL-66 |
| chr2-147134078-DEL-60 | chr2-147134079-DEL-60 |
| chr2-147136719-DEL-59 | chr2-147136720-DEL-59 |
| chr2-147137401-DEL-158 | chr2-147137400-DEL-156 |
| chr2-147189948-DEL-1857 | chr2-147189949-DEL-1857 |
| chr2-147210686-DEL-665 | chr2-147210687-DEL-665 |
| chr2-147239000-DEL-72 | chr2-147239001-DEL-70 |
| chr2-147239806-DEL-78 | chr2-147239828-DEL-60 |
| chr2-147272462-DEL-104 | chr2-147272436-DEL-88 |
| chr2-147285420-DEL-3500 | chr2-147285421-DEL-3500 |
| chr2-147304512-DEL-7775 | chr2-147304513-DEL-7775 |
| chr2-147318723-DEL-66 | chr2-147318724-DEL-66 |
| chr2-147323707-DEL-68 | chr2-147323708-DEL-64 |
| chr2-147345012-DEL-200 | chr2-147345013-DEL-200 |
| chr2-147356358-DEL-61 | chr2-147356359-DEL-61 |
| chr2-147364710-DEL-88 | chr2-147364706-DEL-93 |
| chr2-147385136-DEL-72 | chr2-147385137-DEL-72 |
| chr2-147430777-DEL-351 | chr2-147430778-DEL-351 |
| chr2-147491101-DEL-51 | chr2-147491102-DEL-51 |
| chr2-147514181-DEL-6204 | chr2-147514182-DEL-6204 |
| chr2-147536388-DEL-136 | chr2-147536389-DEL-136 |
| chr2-147543717-DEL-390 | chr2-147543718-DEL-390 |
| chr2-147576234-DEL-87 | chr2-147576235-DEL-87 |
| chr2-147585621-DEL-81 | chr2-147585621-DEL-81 |
| chr2-147596038-DEL-86 | chr2-147596069-DEL-56 |
| chr2-147611617-DEL-2110 | chr2-147611618-DEL-2110 |
| chr2-147640410-DEL-70 | chr2-147640420-DEL-80 |
| chr2-147652442-DEL-64 | chr2-147652442-DEL-69 |
| chr2-147662775-DEL-55 | chr2-147662776-DEL-55 |
| chr2-147663549-DEL-67 | chr2-147663550-DEL-68 |
| chr2-147684741-DEL-970 | chr2-147684742-DEL-970 |
| chr2-147697007-DEL-58 | chr2-147697006-DEL-58 |
| chr2-147760692-DEL-6225 | chr2-147760693-DEL-6225 |
| chr2-147785764-DEL-159 | chr2-147785780-DEL-144 |
| chr2-147817380-DEL-60 | chr2-147817381-DEL-60 |
| chr2-147821672-DEL-51 | chr2-147821673-DEL-51 |
| chr2-147904918-DEL-211 | chr2-147904919-DEL-211 |
| chr2-147919190-DEL-455 | chr2-147919194-DEL-453 |
| chr2-147932761-DEL-434 | chr2-147932762-DEL-434 |
| chr2-147946414-DEL-205 | chr2-147946415-DEL-205 |
| chr2-147946991-DEL-629 | chr2-147946992-DEL-629 |
| chr2-147950313-DEL-51 | chr2-147950314-DEL-51 |
| chr2-147964330-DEL-72 | chr2-147964331-DEL-64 |
| chr2-147966998-DEL-429 | chr2-147966986-DEL-430 |
| chr2-147985388-DEL-108 | chr2-147985398-DEL-138 |
| chr2-148024067-DEL-83 | chr2-148024068-DEL-83 |
| chr2-148086809-DEL-64 | chr2-148086810-DEL-64 |
| chr2-148089227-DEL-1365 | chr2-148089228-DEL-1365 |
| chr2-148145289-DEL-1486 | chr2-148145290-DEL-1486 |
| chr2-148201193-DEL-323 | chr2-148201194-DEL-323 |
| chr2-148202430-DEL-52 | chr2-148202431-DEL-52 |
| chr2-148243120-DEL-197 | chr2-148243121-DEL-197 |
| chr2-148252520-DEL-5272 | chr2-148252521-DEL-5272 |
| chr2-148293288-DEL-55 | chr2-148293286-DEL-56 |
| chr2-148299378-DEL-335 | chr2-148299379-DEL-335 |
| chr2-148300106-DEL-58 | chr2-148300109-DEL-56 |
| chr2-148302255-DEL-6720 | chr2-148302256-DEL-6720 |
| chr2-148336063-DEL-712 | chr2-148336064-DEL-712 |
| chr2-148337144-DEL-1070 | chr2-148337145-DEL-1070 |
| chr2-148342865-DEL-122 | chr2-148342866-DEL-122 |
| chr2-148354927-DEL-350 | chr2-148354928-DEL-350 |
| chr2-148372035-DEL-86 | chr2-148372036-DEL-86 |
| chr2-148385398-DEL-64 | chr2-148385399-DEL-64 |
| chr2-148430282-DEL-92 | chr2-148430283-DEL-92 |
| chr2-148432058-DEL-1836 | chr2-148432059-DEL-1836 |
| chr2-148454572-DEL-2988 | chr2-148454573-DEL-2988 |
| chr2-148458243-DEL-3674 | chr2-148458244-DEL-3674 |
| chr2-148469651-DEL-53 | chr2-148469651-DEL-51 |
| chr2-148477070-DEL-126 | chr2-148477114-DEL-80 |
| chr2-148481722-DEL-1116 | chr2-148481723-DEL-1116 |
| chr2-148486204-DEL-52 | chr2-148486201-DEL-56 |
| chr2-148494413-DEL-1662 | chr2-148494413-DEL-1662 |
| chr2-148495108-DEL-433 | chr2-148495109-DEL-433 |
| chr2-148542671-DEL-64 | chr2-148542665-DEL-60 |

| | |
|---|---|
| chr2-148569012-DEL-61 | chr2-148568997-DEL-65 |
| chr2-148577232-DEL-524 | chr2-148577233-DEL-524 |
| chr2-148586772-DEL-2838 | chr2-148586773-DEL-2838 |
| chr2-148599041-DEL-6457 | chr2-148599042-DEL-6457 |
| chr2-148599231-DEL-214 | chr2-148599232-DEL-214 |
| chr2-148606780-DEL-1215 | chr2-148606781-DEL-1215 |
| chr2-148617239-DEL-436 | chr2-148617240-DEL-436 |
| chr2-148676318-DEL-94 | chr2-148676340-DEL-70 |
| chr2-148701210-DEL-59 | chr2-148701210-DEL-59 |
| chr2-148706412-DEL-352 | chr2-148706373-DEL-358 |
| chr2-148712222-DEL-114 | chr2-148712237-DEL-72 |
| chr2-148713467-DEL-100 | chr2-148713468-DEL-100 |
| chr2-148743798-DEL-90 | chr2-148743799-DEL-90 |
| chr2-148793559-DEL-573 | chr2-148793560-DEL-573 |
| chr2-148814150-DEL-6653 | chr2-148814152-DEL-6652 |
| chr2-148856936-DEL-2361 | chr2-148856937-DEL-2361 |
| chr2-148879427-DEL-593 | chr2-148879428-DEL-593 |
| chr2-148895516-DEL-1084 | chr2-148895523-DEL-1078 |
| chr2-148941473-DEL-68 | chr2-148941494-DEL-70 |
| chr2-148943244-DEL-3024 | chr2-148943245-DEL-3024 |
| chr2-148946758-DEL-4964 | chr2-148946791-DEL-4965 |
| chr2-148960308-DEL-56 | chr2-148960309-DEL-56 |
| chr2-148972533-DEL-230 | chr2-148972534-DEL-230 |
| chr2-149000179-DEL-408 | chr2-149000185-DEL-402 |
| chr2-149011473-DEL-4984 | chr2-149011474-DEL-4984 |
| chr2-149014258-DEL-951 | chr2-149014258-DEL-951 |
| chr2-149022608-DEL-69 | chr2-149022601-DEL-69 |
| chr2-149024213-DEL-52 | chr2-149024213-DEL-52 |
| chr2-149026955-DEL-6215 | chr2-149026956-DEL-6215 |
| chr2-149034693-DEL-64 | chr2-149034694-DEL-64 |
| chr2-149060259-DEL-782 | chr2-149060260-DEL-782 |
| chr2-149082797-DEL-157 | chr2-149082796-DEL-158 |
| chr2-149090418-DEL-390 | chr2-149090419-DEL-390 |
| chr2-149159786-DEL-251 | chr2-149159787-DEL-251 |
| chr2-149161442-DEL-13085 | chr2-149161443-DEL-13085 |
| chr2-149190182-DEL-482 | chr2-149190183-DEL-482 |
| chr2-149197885-DEL-705 | chr2-149197887-DEL-704 |
| chr2-149202687-DEL-2010 | chr2-149202688-DEL-2010 |
| chr2-149206846-DEL-167 | chr2-149206847-DEL-159 |
| chr2-149207919-DEL-62 | chr2-149207920-DEL-62 |
| chr2-149230387-DEL-1942 | chr2-149230388-DEL-1942 |
| chr2-149236572-DEL-2032 | chr2-149236573-DEL-2032 |
| chr2-149238689-DEL-153 | chr2-149238688-DEL-154 |
| chr2-149250086-DEL-1735 | chr2-149250087-DEL-1735 |
| chr2-149254592-DEL-73 | chr2-149254593-DEL-73 |
| chr2-149280065-DEL-2876 | chr2-149280066-DEL-2876 |
| chr2-149319566-DEL-131 | chr2-149319567-DEL-131 |
| chr2-149325482-DEL-6562 | chr2-149325483-DEL-6562 |
| chr2-149404718-DEL-60 | chr2-149404718-DEL-56 |
| chr2-149412919-DEL-251 | chr2-149412920-DEL-251 |
| chr2-149422490-DEL-238 | chr2-149422491-DEL-238 |
| chr2-149430363-DEL-7183 | chr2-149430364-DEL-7183 |
| chr2-149449284-DEL-554 | chr2-149449285-DEL-554 |
| chr2-149473158-DEL-606 | chr2-149473159-DEL-606 |
| chr2-149476361-DEL-635 | chr2-149476369-DEL-639 |
| chr2-149501608-DEL-64 | chr2-149501609-DEL-64 |
| chr2-149515177-DEL-125 | chr2-149515178-DEL-125 |
| chr2-149522574-DEL-493 | chr2-149522575-DEL-493 |
| chr2-149530181-DEL-2893 | chr2-149530182-DEL-2893 |
| chr2-149605433-DEL-205 | chr2-149605434-DEL-205 |
| chr2-149635756-DEL-165 | chr2-149635757-DEL-165 |
| chr2-149657827-DEL-51 | chr2-149657828-DEL-51 |
| chr2-149663970-DEL-59 | chr2-149663971-DEL-59 |
| chr2-149694048-DEL-62 | chr2-149694031-DEL-80 |
| chr2-149704884-DEL-600 | chr2-149704885-DEL-600 |
| chr2-149711743-DEL-75 | chr2-149711744-DEL-75 |
| chr2-149720535-DEL-54 | chr2-149720534-DEL-56 |
| chr2-149780974-DEL-96 | chr2-149780975-DEL-96 |
| chr2-149793123-DEL-77 | chr2-149793124-DEL-77 |
| chr2-149797413-DEL-2314 | chr2-149797414-DEL-2314 |
| chr2-149801131-DEL-5482 | chr2-149801132-DEL-5482 |
| chr2-149808794-DEL-474 | chr2-149808795-DEL-474 |
| chr2-149810911-DEL-1474 | chr2-149810912-DEL-1474 |
| chr2-149827628-DEL-674 | chr2-149827629-DEL-674 |
| chr2-149852473-DEL-60 | chr2-149852474-DEL-60 |
| chr2-149863510-DEL-103 | chr2-149863511-DEL-103 |
| chr2-149870804-DEL-376 | chr2-149870805-DEL-376 |
| chr2-149875158-DEL-290 | chr2-149875159-DEL-290 |
| chr2-149877987-DEL-602 | chr2-149877989-DEL-594 |
| chr2-149878885-DEL-5438 | chr2-149878886-DEL-5438 |

| | |
|---|---|
| chr2-149907231-DEL-93 | chr2-149907232-DEL-93 |
| chr2-149929819-DEL-4271 | chr2-149929820-DEL-4271 |
| chr2-149997640-DEL-891 | chr2-149997641-DEL-891 |
| chr2-150003329-DEL-168 | chr2-150003330-DEL-168 |
| chr2-150051284-DEL-42595 | chr2-150051284-DEL-42595 |
| chr2-150080900-DEL-84 | chr2-150080901-DEL-84 |
| chr2-150131191-DEL-107 | chr2-150131198-DEL-101 |
| chr2-150139256-DEL-4010 | chr2-150139257-DEL-4010 |
| chr2-150150084-DEL-68 | chr2-150150085-DEL-80 |
| chr2-150184793-DEL-84 | chr2-150184794-DEL-84 |
| chr2-150208400-DEL-101 | chr2-150208401-DEL-101 |
| chr2-150209871-DEL-71 | chr2-150209918-DEL-56 |
| chr2-150224221-DEL-3801 | chr2-150224222-DEL-3801 |
| chr2-150240679-DEL-172 | chr2-150240680-DEL-172 |
| chr2-150282448-DEL-6834 | chr2-150282449-DEL-6834 |
| chr2-150306980-DEL-427 | chr2-150306981-DEL-427 |
| chr2-150307712-DEL-120 | chr2-150307713-DEL-120 |
| chr2-150318232-DEL-132 | chr2-150318233-DEL-132 |
| chr2-150331186-DEL-83 | chr2-150331187-DEL-83 |
| chr2-150336740-DEL-107 | chr2-150336736-DEL-108 |
| chr2-150345444-DEL-5353 | chr2-150345443-DEL-5353 |
| chr2-150402514-DEL-450 | chr2-150402514-DEL-450 |
| chr2-150404542-DEL-202 | chr2-150404543-DEL-202 |
| chr2-150439223-DEL-274 | chr2-150439224-DEL-271 |
| chr2-150462075-DEL-204 | chr2-150462076-DEL-204 |
| chr2-150474637-DEL-362 | chr2-150474658-DEL-342 |
| chr2-150492446-DEL-172 | chr2-150492447-DEL-172 |
| chr2-150517783-DEL-91 | chr2-150517805-DEL-69 |
| chr2-150539349-DEL-2450 | chr2-150539350-DEL-2450 |
| chr2-150543603-DEL-5226 | chr2-150543604-DEL-5226 |
| chr2-150551792-DEL-80 | chr2-150551793-DEL-80 |
| chr2-150561716-DEL-76 | chr2-150561717-DEL-88 |
| chr2-150570408-DEL-6337 | chr2-150570409-DEL-6337 |
| chr2-150597271-DEL-197 | chr2-150597272-DEL-197 |
| chr2-150621630-DEL-172 | chr2-150621636-DEL-164 |
| chr2-150663000-DEL-318 | chr2-150663001-DEL-318 |
| chr2-150718191-DEL-51 | chr2-150718187-DEL-52 |
| chr2-150730567-DEL-207 | chr2-150730568-DEL-207 |
| chr2-150752745-DEL-121 | chr2-150752752-DEL-73 |
| chr2-150767119-DEL-139 | chr2-150767120-DEL-95 |
| chr2-150780324-DEL-1246 | chr2-150780325-DEL-1246 |
| chr2-150825878-DEL-314 | chr2-150825879-DEL-314 |
| chr2-150836925-DEL-66 | chr2-150836927-DEL-65 |
| chr2-150845140-DEL-2041 | chr2-150845141-DEL-2041 |
| chr2-150852148-DEL-67 | chr2-150852149-DEL-67 |
| chr2-150906667-DEL-4542 | chr2-150906667-DEL-4542 |
| chr2-150972968-DEL-781 | chr2-150972968-DEL-781 |
| chr2-150977838-DEL-3172 | chr2-150977838-DEL-3172 |
| chr2-150990275-DEL-951 | chr2-150990276-DEL-951 |
| chr2-151036105-DEL-1130 | chr2-151036105-DEL-1130 |
| chr2-151046881-DEL-7080 | chr2-151046882-DEL-7080 |
| chr2-151079158-DEL-214 | chr2-151079159-DEL-214 |
| chr2-151100439-DEL-898 | chr2-151100440-DEL-898 |
| chr2-151123375-DEL-715 | chr2-151123376-DEL-715 |
| chr2-151186862-DEL-10456 | chr2-151186861-DEL-10456 |
| chr2-151254006-DEL-899 | chr2-151254007-DEL-899 |
| chr2-151287121-DEL-364 | chr2-151287138-DEL-365 |
| chr2-151304219-DEL-6136 | chr2-151304220-DEL-6136 |
| chr2-151369605-DEL-328 | chr2-151369606-DEL-328 |
| chr2-151375594-DEL-58 | chr2-151375595-DEL-58 |
| chr2-151404609-DEL-218 | chr2-151404610-DEL-218 |
| chr2-151411962-DEL-1277 | chr2-151411963-DEL-1277 |
| chr2-151452800-DEL-92 | chr2-151452801-DEL-92 |
| chr2-151471546-DEL-67 | chr2-151471547-DEL-67 |
| chr2-151480618-DEL-173 | chr2-151480619-DEL-172 |
| chr2-151492170-DEL-1215 | chr2-151492171-DEL-1216 |
| chr2-151501719-DEL-191 | chr2-151501720-DEL-191 |
| chr2-151516331-DEL-480 | chr2-151516332-DEL-480 |
| chr2-151540995-DEL-74 | chr2-151540997-DEL-77 |
| chr2-151563963-DEL-51 | chr2-151563964-DEL-51 |
| chr2-151570057-DEL-124 | chr2-151570060-DEL-93 |
| chr2-151572777-DEL-114 | chr2-151572778-DEL-76 |
| chr2-151620642-DEL-74 | chr2-151620643-DEL-74 |
| chr2-151636669-DEL-68 | chr2-151636670-DEL-68 |
| chr2-151652928-DEL-92 | chr2-151652929-DEL-92 |
| chr2-151722337-DEL-6186 | chr2-151722338-DEL-6186 |
| chr2-151738894-DEL-498 | chr2-151738895-DEL-498 |
| chr2-151774698-DEL-1035 | chr2-151774699-DEL-1033 |
| chr2-151785447-DEL-53 | chr2-151785448-DEL-53 |
| chr2-151787416-DEL-976 | chr2-151787417-DEL-975 |

| | |
|---|---|
| chr2-151789537-DEL-88 | chr2-151789538-DEL-88 |
| chr2-151791835-DEL-211 | chr2-151791836-DEL-211 |
| chr2-151831775-DEL-184 | chr2-151831776-DEL-184 |
| chr2-151904750-DEL-198 | chr2-151904732-DEL-175 |
| chr2-151948582-DEL-80 | chr2-151948583-DEL-80 |
| chr2-151958635-DEL-168 | chr2-151958636-DEL-173 |
| chr2-151998943-DEL-1887 | chr2-151998950-DEL-1879 |
| chr2-152012934-DEL-56 | chr2-152012929-DEL-71 |
| chr2-152013968-DEL-67 | chr2-152013975-DEL-56 |
| chr2-152016566-DEL-183 | chr2-152016567-DEL-183 |
| chr2-152026388-DEL-155 | chr2-152026389-DEL-155 |
| chr2-152027153-DEL-475 | chr2-152027154-DEL-475 |
| chr2-152027946-DEL-173 | chr2-152027947-DEL-173 |
| chr2-152033493-DEL-436 | chr2-152033494-DEL-436 |
| chr2-152040624-DEL-160 | chr2-152040625-DEL-160 |
| chr2-152041948-DEL-1406 | chr2-152041949-DEL-1407 |
| chr2-152048309-DEL-4964 | chr2-152048310-DEL-4964 |
| chr2-152059491-DEL-337 | chr2-152059492-DEL-337 |
| chr2-152105276-DEL-6442 | chr2-152105277-DEL-6442 |
| chr2-152159896-DEL-202 | chr2-152159897-DEL-202 |
| chr2-152196670-DEL-225 | chr2-152196658-DEL-225 |
| chr2-152201138-DEL-106 | chr2-152201102-DEL-102 |
| chr2-152219912-DEL-514 | chr2-152219907-DEL-509 |
| chr2-152223448-DEL-173 | chr2-152223449-DEL-173 |
| chr2-152228141-DEL-275 | chr2-152228142-DEL-275 |
| chr2-152233172-DEL-119 | chr2-152233165-DEL-120 |
| chr2-152243151-DEL-93 | chr2-152243152-DEL-93 |
| chr2-152249287-DEL-1033 | chr2-152249288-DEL-1033 |
| chr2-152268386-DEL-194 | chr2-152268382-DEL-186 |
| chr2-152270167-DEL-406 | chr2-152270168-DEL-406 |
| chr2-152288239-DEL-54 | chr2-152288254-DEL-61 |
| chr2-152293326-DEL-129 | chr2-152293327-DEL-129 |
| chr2-152359004-DEL-60 | chr2-152359005-DEL-60 |
| chr2-152373265-DEL-103 | chr2-152373266-DEL-103 |
| chr2-152394073-DEL-89 | chr2-152394068-DEL-90 |
| chr2-152427107-DEL-62 | chr2-152427108-DEL-62 |
| chr2-152455714-DEL-209 | chr2-152455715-DEL-209 |
| chr2-152472535-DEL-56 | chr2-152472545-DEL-55 |
| chr2-152493306-DEL-62 | chr2-152493309-DEL-60 |
| chr2-152507893-DEL-217 | chr2-152507894-DEL-217 |
| chr2-152522713-DEL-69 | chr2-152522714-DEL-69 |
| chr2-152573540-DEL-74 | chr2-152573540-DEL-80 |
| chr2-152719293-DEL-235 | chr2-152719294-DEL-235 |
| chr2-152731494-DEL-213 | chr2-152731495-DEL-213 |
| chr2-152738841-DEL-333 | chr2-152738840-DEL-334 |
| chr2-152771238-DEL-799 | chr2-152771258-DEL-786 |
| chr2-152806109-DEL-408 | chr2-152806109-DEL-408 |
| chr2-152819970-DEL-54 | chr2-152819971-DEL-54 |
| chr2-152860267-DEL-178 | chr2-152860264-DEL-173 |
| chr2-152886967-DEL-66 | chr2-152886967-DEL-66 |
| chr2-152926736-DEL-1640 | chr2-152926737-DEL-1640 |
| chr2-152961144-DEL-197 | chr2-152961145-DEL-197 |
| chr2-152983732-DEL-464 | chr2-152983733-DEL-464 |
| chr2-152999063-DEL-126 | chr2-152999064-DEL-126 |
| chr2-153005543-DEL-337 | chr2-153005544-DEL-337 |
| chr2-153015264-DEL-227 | chr2-153015280-DEL-226 |
| chr2-153025239-DEL-53 | chr2-153025226-DEL-55 |
| chr2-153042887-DEL-498 | chr2-153042888-DEL-498 |
| chr2-153047597-DEL-243 | chr2-153047596-DEL-245 |
| chr2-153121834-DEL-249 | chr2-153121835-DEL-249 |
| chr2-153127461-DEL-112 | chr2-153127467-DEL-106 |
| chr2-153142633-DEL-207 | chr2-153142634-DEL-207 |
| chr2-153150226-DEL-204 | chr2-153150227-DEL-204 |
| chr2-153153280-DEL-88 | chr2-153153281-DEL-88 |
| chr2-153178684-DEL-57 | chr2-153178680-DEL-51 |
| chr2-153215621-DEL-79 | chr2-153215622-DEL-79 |
| chr2-153236104-DEL-61 | chr2-153236105-DEL-61 |
| chr2-153281534-DEL-423 | chr2-153281535-DEL-423 |
| chr2-153322741-DEL-213 | chr2-153322742-DEL-213 |
| chr2-153400946-DEL-59 | chr2-153400948-DEL-57 |
| chr2-153430870-DEL-204 | chr2-153430871-DEL-204 |
| chr2-153436126-DEL-79 | chr2-153436127-DEL-79 |
| chr2-153437919-DEL-213 | chr2-153437920-DEL-213 |
| chr2-153456117-DEL-75 | chr2-153456118-DEL-75 |
| chr2-153460231-DEL-104 | chr2-153460232-DEL-104 |
| chr2-153523392-DEL-142 | chr2-153523393-DEL-142 |
| chr2-153524033-DEL-207 | chr2-153524034-DEL-207 |
| chr2-153532946-DEL-2161 | chr2-153532947-DEL-2161 |
| chr2-153542389-DEL-213 | chr2-153542384-DEL-214 |
| chr2-153561697-DEL-100 | chr2-153561698-DEL-100 |

| | |
|---|---|
| chr2-153566177-DEL-88 | chr2-153566174-DEL-89 |
| chr2-153619813-DEL-408 | chr2-153619814-DEL-408 |
| chr2-153635618-DEL-54 | chr2-153635576-DEL-54 |
| chr2-153647767-DEL-1737 | chr2-153647776-DEL-1728 |
| chr2-153653198-DEL-84 | chr2-153653199-DEL-84 |
| chr2-153667955-DEL-1052 | chr2-153667956-DEL-1052 |
| chr2-153675212-DEL-88 | chr2-153675206-DEL-89 |
| chr2-153677407-DEL-60 | chr2-153677408-DEL-60 |
| chr2-153704551-DEL-54 | chr2-153704552-DEL-54 |
| chr2-153725642-DEL-1158 | chr2-153725643-DEL-1158 |
| chr2-153727134-DEL-2443 | chr2-153727135-DEL-2443 |
| chr2-153727154-DEL-128 | chr2-153727155-DEL-126 |
| chr2-153797106-DEL-96 | chr2-153797106-DEL-96 |
| chr2-153818826-DEL-55 | chr2-153818827-DEL-55 |
| chr2-153827770-DEL-163 | chr2-153827768-DEL-164 |
| chr2-153838332-DEL-3514 | chr2-153838333-DEL-3514 |
| chr2-153858029-DEL-168 | chr2-153858030-DEL-168 |
| chr2-153873565-DEL-6233 | chr2-153873566-DEL-6233 |
| chr2-153899518-DEL-64 | chr2-153899514-DEL-68 |
| chr2-153905631-DEL-255 | chr2-153905629-DEL-255 |
| chr2-153940413-DEL-73 | chr2-153940414-DEL-73 |
| chr2-153969245-DEL-801 | chr2-153969246-DEL-801 |
| chr2-153973200-DEL-75 | chr2-153973201-DEL-75 |
| chr2-153982452-DEL-64 | chr2-153982453-DEL-64 |
| chr2-154014697-DEL-190 | chr2-154014698-DEL-190 |
| chr2-154015685-DEL-6264 | chr2-154015686-DEL-6264 |
| chr2-154015859-DEL-128 | chr2-154015860-DEL-128 |
| chr2-154025252-DEL-126 | chr2-154025253-DEL-126 |
| chr2-154026836-DEL-102 | chr2-154026837-DEL-102 |
| chr2-154038328-DEL-54 | chr2-154038329-DEL-54 |
| chr2-154041322-DEL-615 | chr2-154041323-DEL-615 |
| chr2-154046492-DEL-61 | chr2-154046492-DEL-62 |
| chr2-154049550-DEL-155 | chr2-154049557-DEL-152 |
| chr2-154077273-DEL-514 | chr2-154077274-DEL-512 |
| chr2-154122658-DEL-199 | chr2-154122659-DEL-199 |
| chr2-154125227-DEL-161 | chr2-154125228-DEL-161 |
| chr2-154125934-DEL-157 | chr2-154125935-DEL-157 |
| chr2-154163386-DEL-86 | chr2-154163387-DEL-86 |
| chr2-154196176-DEL-5336 | chr2-154196177-DEL-5336 |
| chr2-154227999-DEL-59 | chr2-154227981-DEL-60 |
| chr2-154257191-DEL-233 | chr2-154257192-DEL-217 |
| chr2-154272299-DEL-220 | chr2-154272300-DEL-220 |
| chr2-154281045-DEL-60 | chr2-154281046-DEL-60 |
| chr2-154332321-DEL-81 | chr2-154332322-DEL-81 |
| chr2-154344036-DEL-96 | chr2-154344037-DEL-96 |
| chr2-154346985-DEL-70 | chr2-154346986-DEL-70 |
| chr2-154369761-DEL-206 | chr2-154369762-DEL-206 |
| chr2-154385226-DEL-107 | chr2-154385227-DEL-107 |
| chr2-154402285-DEL-56 | chr2-154402269-DEL-54 |
| chr2-154417341-DEL-211 | chr2-154417342-DEL-211 |
| chr2-154470402-DEL-81 | chr2-154470394-DEL-77 |
| chr2-154495678-DEL-212 | chr2-154495679-DEL-212 |
| chr2-154502082-DEL-51 | chr2-154502083-DEL-51 |
| chr2-154506849-DEL-60 | chr2-154506850-DEL-60 |
| chr2-154509065-DEL-1096 | chr2-154509066-DEL-1096 |
| chr2-154513601-DEL-253 | chr2-154513597-DEL-248 |
| chr2-154514164-DEL-1290 | chr2-154514164-DEL-1290 |
| chr2-154517926-DEL-165 | chr2-154517927-DEL-165 |
| chr2-154538029-DEL-59 | chr2-154538030-DEL-59 |
| chr2-154550433-DEL-60 | chr2-154550434-DEL-60 |
| chr2-154562658-DEL-54 | chr2-154562658-DEL-80 |
| chr2-154566904-DEL-97 | chr2-154566905-DEL-97 |
| chr2-154568679-DEL-200 | chr2-154568684-DEL-196 |
| chr2-154593221-DEL-54 | chr2-154593221-DEL-54 |
| chr2-154598239-DEL-215 | chr2-154598227-DEL-201 |
| chr2-154608963-DEL-930 | chr2-154608964-DEL-930 |
| chr2-154616941-DEL-171 | chr2-154616942-DEL-171 |
| chr2-154622627-DEL-80 | chr2-154622623-DEL-79 |
| chr2-154643584-DEL-234 | chr2-154643587-DEL-232 |
| chr2-154665998-DEL-1024 | chr2-154665999-DEL-1024 |
| chr2-154667357-DEL-217 | chr2-154667358-DEL-217 |
| chr2-154680399-DEL-206 | chr2-154680400-DEL-206 |
| chr2-154711977-DEL-74 | chr2-154711978-DEL-74 |
| chr2-154714088-DEL-69 | chr2-154714089-DEL-69 |
| chr2-154725361-DEL-140 | chr2-154725362-DEL-140 |
| chr2-154736307-DEL-829 | chr2-154736308-DEL-829 |
| chr2-154739908-DEL-3290 | chr2-154739909-DEL-3290 |
| chr2-154747673-DEL-115 | chr2-154747674-DEL-115 |
| chr2-154750077-DEL-181 | chr2-154750078-DEL-181 |
| chr2-154750812-DEL-52 | chr2-154750827-DEL-60 |

| | |
|---|---|
| chr2-154760761-DEL-320 | chr2-154760774-DEL-323 |
| chr2-154761671-DEL-54 | chr2-154761672-DEL-54 |
| chr2-154764270-DEL-452 | chr2-154764271-DEL-452 |
| chr2-154768300-DEL-64 | chr2-154768301-DEL-64 |
| chr2-154770780-DEL-204 | chr2-154770781-DEL-204 |
| chr2-154782723-DEL-223 | chr2-154782724-DEL-223 |
| chr2-154812351-DEL-55 | chr2-154812352-DEL-50 |
| chr2-154836149-DEL-195 | chr2-154836150-DEL-195 |
| chr2-154848250-DEL-299 | chr2-154848251-DEL-299 |
| chr2-154856866-DEL-14705 | chr2-154856867-DEL-14705 |
| chr2-154874740-DEL-50 | chr2-154874723-DEL-68 |
| chr2-154896369-DEL-205 | chr2-154896370-DEL-205 |
| chr2-154908938-DEL-130 | chr2-154908939-DEL-130 |
| chr2-154913312-DEL-633 | chr2-154913313-DEL-633 |
| chr2-154932892-DEL-62 | chr2-154932893-DEL-62 |
| chr2-154934141-DEL-209 | chr2-154934142-DEL-209 |
| chr2-154954353-DEL-523 | chr2-154954354-DEL-523 |
| chr2-154962987-DEL-7192 | chr2-154962988-DEL-7192 |
| chr2-154979015-DEL-52 | chr2-154979016-DEL-52 |
| chr2-155018851-DEL-204 | chr2-155018852-DEL-204 |
| chr2-155051533-DEL-79 | chr2-155051517-DEL-92 |
| chr2-155063983-DEL-60 | chr2-155063987-DEL-56 |
| chr2-155065223-DEL-785 | chr2-155065224-DEL-785 |
| chr2-155077368-DEL-156 | chr2-155077369-DEL-156 |
| chr2-155097501-DEL-170 | chr2-155097502-DEL-170 |
| chr2-155103038-DEL-66 | chr2-155103017-DEL-67 |
| chr2-155106864-DEL-344 | chr2-155106865-DEL-344 |
| chr2-155109406-DEL-250 | chr2-155109450-DEL-223 |
| chr2-155130229-DEL-97 | chr2-155130230-DEL-65 |
| chr2-155153237-DEL-148 | chr2-155153238-DEL-148 |
| chr2-155154695-DEL-54 | chr2-155154690-DEL-60 |
| chr2-155186601-DEL-807 | chr2-155186602-DEL-807 |
| chr2-155202443-DEL-117 | chr2-155202444-DEL-117 |
| chr2-155217163-DEL-75 | chr2-155217174-DEL-65 |
| chr2-155228141-DEL-75 | chr2-155228142-DEL-75 |
| chr2-155235934-DEL-106 | chr2-155235939-DEL-102 |
| chr2-155242522-DEL-76 | chr2-155242510-DEL-57 |
| chr2-155308785-DEL-78 | chr2-155308786-DEL-78 |
| chr2-155361961-DEL-322 | chr2-155361962-DEL-322 |
| chr2-155376868-DEL-102 | chr2-155376831-DEL-103 |
| chr2-155380365-DEL-394 | chr2-155380352-DEL-382 |
| chr2-155385964-DEL-867 | chr2-155385965-DEL-867 |
| chr2-155421926-DEL-1154 | chr2-155421927-DEL-1154 |
| chr2-155500942-DEL-62 | chr2-155500943-DEL-62 |
| chr2-155507721-DEL-60 | chr2-155507722-DEL-60 |
| chr2-155539920-DEL-82 | chr2-155539921-DEL-80 |
| chr2-155567725-DEL-247 | chr2-155567726-DEL-247 |
| chr2-155574428-DEL-203 | chr2-155574429-DEL-203 |
| chr2-155583007-DEL-77 | chr2-155583008-DEL-77 |
| chr2-155600092-DEL-72 | chr2-155600093-DEL-60 |
| chr2-155607624-DEL-171 | chr2-155607625-DEL-171 |
| chr2-155608519-DEL-2072 | chr2-155608520-DEL-2072 |
| chr2-155613044-DEL-64 | chr2-155613045-DEL-64 |
| chr2-155638838-DEL-52 | chr2-155638839-DEL-52 |
| chr2-155665543-DEL-96 | chr2-155665544-DEL-96 |
| chr2-155685867-DEL-216 | chr2-155685868-DEL-216 |
| chr2-155700646-DEL-207 | chr2-155700648-DEL-205 |
| chr2-155732423-DEL-83 | chr2-155732424-DEL-83 |
| chr2-155762057-DEL-982 | chr2-155762041-DEL-989 |
| chr2-155767341-DEL-149 | chr2-155767342-DEL-149 |
| chr2-155784958-DEL-53 | chr2-155784959-DEL-53 |
| chr2-155795962-DEL-271 | chr2-155795956-DEL-277 |
| chr2-155812023-DEL-63 | chr2-155812023-DEL-63 |
| chr2-155853789-DEL-102 | chr2-155853790-DEL-102 |
| chr2-155867954-DEL-292 | chr2-155867959-DEL-288 |
| chr2-155872808-DEL-202 | chr2-155872809-DEL-202 |
| chr2-155892595-DEL-97 | chr2-155892590-DEL-94 |
| chr2-155897485-DEL-257 | chr2-155897492-DEL-263 |
| chr2-155899768-DEL-96 | chr2-155899766-DEL-98 |
| chr2-155954358-DEL-151 | chr2-155954340-DEL-132 |
| chr2-155956241-DEL-4996 | chr2-155956239-DEL-4995 |
| chr2-155972010-DEL-164 | chr2-155972015-DEL-162 |
| chr2-155978613-DEL-714 | chr2-155978612-DEL-714 |
| chr2-155984311-DEL-124 | chr2-155984312-DEL-124 |
| chr2-156011676-DEL-206 | chr2-156011677-DEL-206 |
| chr2-156012718-DEL-471 | chr2-156012719-DEL-471 |
| chr2-156029979-DEL-228 | chr2-156029980-DEL-228 |
| chr2-156052896-DEL-3421 | chr2-156052897-DEL-3421 |
| chr2-156063795-DEL-90 | chr2-156063788-DEL-98 |
| chr2-156064653-DEL-211 | chr2-156064654-DEL-211 |

| | |
|---|---|
| chr2-156080906-DEL-141 | chr2-156080906-DEL-142 |
| chr2-156081174-DEL-1341 | chr2-156081175-DEL-1341 |
| chr2-156084325-DEL-86 | chr2-156084326-DEL-86 |
| chr2-156109833-DEL-61 | chr2-156109834-DEL-61 |
| chr2-156142114-DEL-234 | chr2-156142113-DEL-235 |
| chr2-156156074-DEL-81 | chr2-156156056-DEL-85 |
| chr2-156160884-DEL-148 | chr2-156160885-DEL-148 |
| chr2-156162836-DEL-207 | chr2-156162837-DEL-207 |
| chr2-156168901-DEL-87 | chr2-156168902-DEL-87 |
| chr2-156198471-DEL-7054 | chr2-156198472-DEL-7054 |
| chr2-156211115-DEL-92 | chr2-156211116-DEL-92 |
| chr2-156245290-DEL-689 | chr2-156245291-DEL-689 |
| chr2-156281251-DEL-162 | chr2-156281269-DEL-145 |
| chr2-156286388-DEL-167 | chr2-156286389-DEL-167 |
| chr2-156314026-DEL-199 | chr2-156314027-DEL-199 |
| chr2-156371453-DEL-92 | chr2-156371478-DEL-68 |
| chr2-156400099-DEL-690 | chr2-156400100-DEL-690 |
| chr2-156402185-DEL-1804 | chr2-156402186-DEL-1804 |
| chr2-156416816-DEL-90 | chr2-156416833-DEL-50 |
| chr2-156424848-DEL-60 | chr2-156424848-DEL-60 |
| chr2-156573229-DEL-170 | chr2-156573230-DEL-170 |
| chr2-156593548-DEL-221 | chr2-156593542-DEL-222 |
| chr2-156593847-DEL-1256 | chr2-156593848-DEL-1256 |
| chr2-156594976-DEL-242 | chr2-156594977-DEL-242 |
| chr2-156654479-DEL-192 | chr2-156654480-DEL-192 |
| chr2-156663396-DEL-194 | chr2-156663396-DEL-195 |
| chr2-156722823-DEL-104 | chr2-156722824-DEL-56 |
| chr2-156744654-DEL-241 | chr2-156744655-DEL-241 |
| chr2-156749703-DEL-431 | chr2-156749704-DEL-431 |
| chr2-156753724-DEL-358 | chr2-156753723-DEL-359 |
| chr2-156754643-DEL-188 | chr2-156754644-DEL-188 |
| chr2-156760383-DEL-56 | chr2-156760388-DEL-52 |
| chr2-156765595-DEL-401 | chr2-156765596-DEL-401 |
| chr2-156773250-DEL-51 | chr2-156773241-DEL-52 |
| chr2-156778496-DEL-200 | chr2-156778497-DEL-200 |
| chr2-156801301-DEL-167 | chr2-156801301-DEL-174 |
| chr2-156809898-DEL-94 | chr2-156809899-DEL-74 |
| chr2-156825378-DEL-238 | chr2-156825379-DEL-238 |
| chr2-156866031-DEL-609 | chr2-156866032-DEL-608 |
| chr2-156866818-DEL-225 | chr2-156866819-DEL-225 |
| chr2-156874046-DEL-65 | chr2-156874065-DEL-66 |
| chr2-156876570-DEL-498 | chr2-156876571-DEL-498 |
| chr2-156901867-DEL-118 | chr2-156901868-DEL-118 |
| chr2-156943859-DEL-228 | chr2-156943860-DEL-228 |
| chr2-156950600-DEL-54 | chr2-156950601-DEL-54 |
| chr2-156965060-DEL-56 | chr2-156965061-DEL-54 |
| chr2-157010063-DEL-221 | chr2-157010064-DEL-221 |
| chr2-157027530-DEL-66 | chr2-157027531-DEL-66 |
| chr2-157028555-DEL-83 | chr2-157028556-DEL-83 |
| chr2-157039959-DEL-111 | chr2-157039960-DEL-111 |
| chr2-157079621-DEL-277 | chr2-157079660-DEL-234 |
| chr2-157081975-DEL-181 | chr2-157081957-DEL-181 |
| chr2-157084623-DEL-498 | chr2-157084624-DEL-498 |
| chr2-157116569-DEL-150 | chr2-157116592-DEL-137 |
| chr2-157123847-DEL-193 | chr2-157123883-DEL-176 |
| chr2-157153817-DEL-185 | chr2-157153847-DEL-185 |
| chr2-157161550-DEL-171 | chr2-157161551-DEL-171 |
| chr2-157222745-DEL-206 | chr2-157222745-DEL-206 |
| chr2-157230682-DEL-445 | chr2-157230681-DEL-447 |
| chr2-157243705-DEL-178 | chr2-157243692-DEL-220 |
| chr2-157361279-DEL-51 | chr2-157361280-DEL-51 |
| chr2-157363531-DEL-478 | chr2-157363532-DEL-478 |
| chr2-157369559-DEL-1335 | chr2-157369560-DEL-1335 |
| chr2-157425224-DEL-58 | chr2-157425219-DEL-64 |
| chr2-157434588-DEL-400 | chr2-157434589-DEL-400 |
| chr2-157439378-DEL-164 | chr2-157439379-DEL-164 |
| chr2-157468982-DEL-69 | chr2-157468980-DEL-72 |
| chr2-157482451-DEL-75 | chr2-157482452-DEL-75 |
| chr2-157489475-DEL-55 | chr2-157489517-DEL-62 |
| chr2-157493314-DEL-107 | chr2-157493309-DEL-149 |
| chr2-157494162-DEL-4173 | chr2-157494163-DEL-4173 |
| chr2-157508775-DEL-174 | chr2-157508776-DEL-174 |
| chr2-157529295-DEL-66 | chr2-157529295-DEL-66 |
| chr2-157539927-DEL-205 | chr2-157539928-DEL-205 |
| chr2-157582788-DEL-52 | chr2-157582789-DEL-52 |
| chr2-157613358-DEL-237 | chr2-157613359-DEL-237 |
| chr2-157620965-DEL-79 | chr2-157620961-DEL-80 |
| chr2-157662369-DEL-234 | chr2-157662370-DEL-234 |
| chr2-157697683-DEL-205 | chr2-157697684-DEL-205 |
| chr2-157739744-DEL-52 | chr2-157739745-DEL-52 |

| | |
|---|---|
| chr2-157743357-DEL-1113 | chr2-157743358-DEL-1113 |
| chr2-157746979-DEL-103 | chr2-157746980-DEL-103 |
| chr2-157775263-DEL-185 | chr2-157775264-DEL-185 |
| chr2-157787366-DEL-181 | chr2-157787387-DEL-161 |
| chr2-157794528-DEL-211 | chr2-157794529-DEL-211 |
| chr2-157800849-DEL-51 | chr2-157800839-DEL-64 |
| chr2-157811680-DEL-323 | chr2-157811681-DEL-323 |
| chr2-157818590-DEL-204 | chr2-157818595-DEL-195 |
| chr2-157841117-DEL-102 | chr2-157841137-DEL-77 |
| chr2-157853842-DEL-109 | chr2-157853843-DEL-109 |
| chr2-157896166-DEL-216 | chr2-157896167-DEL-216 |
| chr2-157921337-DEL-79 | chr2-157921338-DEL-79 |
| chr2-157927450-DEL-117 | chr2-157927452-DEL-110 |
| chr2-157939453-DEL-52 | chr2-157939454-DEL-52 |
| chr2-157968367-DEL-56 | chr2-157968378-DEL-76 |
| chr2-157992504-DEL-69 | chr2-157992508-DEL-51 |
| chr2-158041697-DEL-54 | chr2-158041698-DEL-54 |
| chr2-158043494-DEL-58 | chr2-158043496-DEL-56 |
| chr2-158045435-DEL-96 | chr2-158045444-DEL-95 |
| chr2-158055803-DEL-490 | chr2-158055804-DEL-490 |
| chr2-158075684-DEL-466 | chr2-158075685-DEL-466 |
| chr2-158078173-DEL-51 | chr2-158078174-DEL-51 |
| chr2-158086871-DEL-227 | chr2-158086872-DEL-227 |
| chr2-158089590-DEL-86 | chr2-158089590-DEL-86 |
| chr2-158093463-DEL-54 | chr2-158093464-DEL-54 |
| chr2-158113042-DEL-66 | chr2-158113048-DEL-57 |
| chr2-158114327-DEL-52 | chr2-158114350-DEL-52 |
| chr2-158128190-DEL-6442 | chr2-158128191-DEL-6442 |
| chr2-158135724-DEL-225 | chr2-158135725-DEL-225 |
| chr2-158137076-DEL-481 | chr2-158137077-DEL-481 |
| chr2-158140673-DEL-455 | chr2-158140674-DEL-455 |
| chr2-158168751-DEL-78 | chr2-158168752-DEL-78 |
| chr2-158180544-DEL-919 | chr2-158180545-DEL-919 |
| chr2-158204534-DEL-164 | chr2-158204535-DEL-164 |
| chr2-158211034-DEL-215 | chr2-158211035-DEL-215 |
| chr2-158220146-DEL-938 | chr2-158220147-DEL-938 |
| chr2-158226535-DEL-91 | chr2-158226536-DEL-91 |
| chr2-158242637-DEL-398 | chr2-158242634-DEL-402 |
| chr2-158243853-DEL-185 | chr2-158243860-DEL-148 |
| chr2-158270512-DEL-185 | chr2-158270513-DEL-185 |
| chr2-158330215-DEL-197 | chr2-158330231-DEL-197 |
| chr2-158330884-DEL-908 | chr2-158330883-DEL-909 |
| chr2-158362095-DEL-420 | chr2-158362095-DEL-419 |
| chr2-158383609-DEL-52 | chr2-158383610-DEL-60 |
| chr2-158389204-DEL-103 | chr2-158389205-DEL-103 |
| chr2-158390200-DEL-403 | chr2-158390201-DEL-403 |
| chr2-158427605-DEL-56 | chr2-158427606-DEL-52 |
| chr2-158450410-DEL-56 | chr2-158450413-DEL-56 |
| chr2-158479088-DEL-664 | chr2-158479089-DEL-664 |
| chr2-158495248-DEL-72 | chr2-158495254-DEL-59 |
| chr2-158538906-DEL-576 | chr2-158538907-DEL-576 |
| chr2-158546309-DEL-706 | chr2-158546310-DEL-706 |
| chr2-158576236-DEL-56 | chr2-158576197-DEL-57 |
| chr2-158606726-DEL-226 | chr2-158606727-DEL-226 |
| chr2-158620196-DEL-53 | chr2-158620197-DEL-53 |
| chr2-158644130-DEL-198 | chr2-158644155-DEL-198 |
| chr2-158650627-DEL-131 | chr2-158650599-DEL-130 |
| chr2-158655472-DEL-56 | chr2-158655473-DEL-56 |
| chr2-158693912-DEL-371 | chr2-158693913-DEL-371 |
| chr2-158728116-DEL-181 | chr2-158728117-DEL-181 |
| chr2-158731838-DEL-89 | chr2-158731839-DEL-89 |
| chr2-158742329-DEL-221 | chr2-158742330-DEL-221 |
| chr2-158810796-DEL-490 | chr2-158810797-DEL-490 |
| chr2-158845705-DEL-185 | chr2-158845702-DEL-186 |
| chr2-158871865-DEL-56 | chr2-158871878-DEL-62 |
| chr2-158902749-DEL-54 | chr2-158902750-DEL-54 |
| chr2-158982596-DEL-56 | chr2-158982595-DEL-58 |
| chr2-158986631-DEL-782 | chr2-158986632-DEL-782 |
| chr2-159014479-DEL-631 | chr2-159014469-DEL-632 |
| chr2-159040800-DEL-835 | chr2-159040800-DEL-835 |
| chr2-159061536-DEL-1293 | chr2-159061537-DEL-1293 |
| chr2-159067432-DEL-54 | chr2-159067433-DEL-54 |
| chr2-159068623-DEL-60 | chr2-159068606-DEL-86 |
| chr2-159069967-DEL-75 | chr2-159069966-DEL-75 |
| chr2-159114084-DEL-267 | chr2-159114085-DEL-267 |
| chr2-159138384-DEL-101 | chr2-159138385-DEL-101 |
| chr2-159164579-DEL-93 | chr2-159164574-DEL-52 |
| chr2-159188030-DEL-6519 | chr2-159188031-DEL-6519 |
| chr2-159195898-DEL-60 | chr2-159195899-DEL-60 |
| chr2-159208639-DEL-966 | chr2-159208640-DEL-966 |

| | |
|---|---|
| chr2-159221384-DEL-54 | chr2-159221422-DEL-55 |
| chr2-159276872-DEL-64 | chr2-159276873-DEL-56 |
| chr2-159280483-DEL-63 | chr2-159280492-DEL-55 |
| chr2-159285318-DEL-64 | chr2-159285319-DEL-64 |
| chr2-159294981-DEL-404 | chr2-159294981-DEL-403 |
| chr2-159297252-DEL-7105 | chr2-159297253-DEL-7105 |
| chr2-159307193-DEL-6744 | chr2-159307194-DEL-6744 |
| chr2-159311705-DEL-552 | chr2-159311706-DEL-552 |
| chr2-159339212-DEL-92 | chr2-159339213-DEL-92 |
| chr2-159370965-DEL-54 | chr2-159370966-DEL-54 |
| chr2-159371776-DEL-3083 | chr2-159371777-DEL-3083 |
| chr2-159434560-DEL-88 | chr2-159434554-DEL-94 |
| chr2-159443074-DEL-70 | chr2-159443089-DEL-68 |
| chr2-159463019-DEL-95 | chr2-159463017-DEL-95 |
| chr2-159479839-DEL-7011 | chr2-159479839-DEL-7011 |
| chr2-159488972-DEL-73 | chr2-159488973-DEL-74 |
| chr2-159498151-DEL-6491 | chr2-159498152-DEL-6491 |
| chr2-159542186-DEL-74 | chr2-159542187-DEL-74 |
| chr2-159554276-DEL-59 | chr2-159554276-DEL-59 |
| chr2-159593990-DEL-742 | chr2-159593991-DEL-742 |
| chr2-159615072-DEL-54 | chr2-159615073-DEL-54 |
| chr2-159620994-DEL-636 | chr2-159620995-DEL-636 |
| chr2-159630504-DEL-6381 | chr2-159630505-DEL-6381 |
| chr2-159643118-DEL-130 | chr2-159643112-DEL-131 |
| chr2-159646585-DEL-211 | chr2-159646586-DEL-222 |
| chr2-159649018-DEL-71 | chr2-159649023-DEL-67 |
| chr2-159684154-DEL-386 | chr2-159684155-DEL-386 |
| chr2-159723353-DEL-70 | chr2-159723354-DEL-70 |
| chr2-159750226-DEL-103 | chr2-159750227-DEL-103 |
| chr2-159755518-DEL-328 | chr2-159755519-DEL-328 |
| chr2-159759407-DEL-52 | chr2-159759408-DEL-52 |
| chr2-159771125-DEL-72 | chr2-159771126-DEL-72 |
| chr2-159774496-DEL-6393 | chr2-159774497-DEL-6393 |
| chr2-159804595-DEL-100 | chr2-159804611-DEL-92 |
| chr2-159807989-DEL-54 | chr2-159807990-DEL-54 |
| chr2-159840756-DEL-66 | chr2-159840757-DEL-78 |
| chr2-159915615-DEL-102 | chr2-159915616-DEL-102 |
| chr2-159918815-DEL-81 | chr2-159918816-DEL-81 |
| chr2-159929274-DEL-1038 | chr2-159929275-DEL-1038 |
| chr2-159937051-DEL-205 | chr2-159937052-DEL-205 |
| chr2-159952249-DEL-131 | chr2-159952250-DEL-131 |
| chr2-159952523-DEL-2480 | chr2-159952524-DEL-2480 |
| chr2-159959416-DEL-78 | chr2-159959415-DEL-80 |
| chr2-159965089-DEL-60 | chr2-159965090-DEL-58 |
| chr2-159975823-DEL-85 | chr2-159975830-DEL-79 |
| chr2-159977053-DEL-440 | chr2-159977054-DEL-440 |
| chr2-159992694-DEL-67 | chr2-159992731-DEL-75 |
| chr2-160061045-DEL-74 | chr2-160061046-DEL-77 |
| chr2-160069966-DEL-119 | chr2-160069967-DEL-119 |
| chr2-160072032-DEL-164 | chr2-160072034-DEL-163 |
| chr2-160078296-DEL-54 | chr2-160078297-DEL-50 |
| chr2-160113131-DEL-164 | chr2-160113128-DEL-161 |
| chr2-160215054-DEL-120 | chr2-160215055-DEL-120 |
| chr2-160286511-DEL-72 | chr2-160286512-DEL-72 |
| chr2-160298046-DEL-56 | chr2-160298049-DEL-54 |
| chr2-160306644-DEL-231 | chr2-160306645-DEL-231 |
| chr2-160323986-DEL-93 | chr2-160323987-DEL-93 |
| chr2-160348325-DEL-68 | chr2-160348326-DEL-68 |
| chr2-160373638-DEL-69 | chr2-160373592-DEL-88 |
| chr2-160376324-DEL-58 | chr2-160376327-DEL-56 |
| chr2-160378151-DEL-57 | chr2-160378136-DEL-72 |
| chr2-160405624-DEL-192 | chr2-160405625-DEL-192 |
| chr2-160412702-DEL-242 | chr2-160412702-DEL-245 |
| chr2-160447419-DEL-678 | chr2-160447406-DEL-674 |
| chr2-160471951-DEL-199 | chr2-160471939-DEL-232 |
| chr2-160473719-DEL-6637 | chr2-160473720-DEL-6637 |
| chr2-160499980-DEL-180 | chr2-160499981-DEL-180 |
| chr2-160509038-DEL-213 | chr2-160509039-DEL-213 |
| chr2-160541123-DEL-218 | chr2-160541124-DEL-218 |
| chr2-160630779-DEL-79 | chr2-160630780-DEL-79 |
| chr2-160650776-DEL-838 | chr2-160650777-DEL-838 |
| chr2-160659649-DEL-7865 | chr2-160659650-DEL-7865 |
| chr2-160671234-DEL-64 | chr2-160671235-DEL-64 |
| chr2-160675205-DEL-2356 | chr2-160675206-DEL-2356 |
| chr2-160709755-DEL-202 | chr2-160709757-DEL-201 |
| chr2-160736513-DEL-112 | chr2-160736513-DEL-108 |
| chr2-160762350-DEL-54 | chr2-160762354-DEL-52 |
| chr2-160765337-DEL-93 | chr2-160765348-DEL-108 |
| chr2-160771849-DEL-1081 | chr2-160771850-DEL-1081 |
| chr2-160779673-DEL-389 | chr2-160779674-DEL-389 |

| | |
|---|---|
| chr2-160784084-DEL-79 | chr2-160784035-DEL-85 |
| chr2-160800300-DEL-247 | chr2-160800253-DEL-248 |
| chr2-160906131-DEL-2796 | chr2-160906132-DEL-2796 |
| chr2-160919451-DEL-187 | chr2-160919452-DEL-187 |
| chr2-161006727-DEL-83 | chr2-161006710-DEL-52 |
| chr2-161030253-DEL-1109 | chr2-161030253-DEL-1109 |
| chr2-161054123-DEL-220 | chr2-161054124-DEL-220 |
| chr2-161067622-DEL-54 | chr2-161067623-DEL-50 |
| chr2-161113297-DEL-606 | chr2-161113298-DEL-606 |
| chr2-161124312-DEL-111 | chr2-161124311-DEL-112 |
| chr2-161127453-DEL-114 | chr2-161127454-DEL-114 |
| chr2-161149294-DEL-82 | chr2-161149294-DEL-83 |
| chr2-161158727-DEL-65 | chr2-161158728-DEL-60 |
| chr2-161174324-DEL-60 | chr2-161174325-DEL-60 |
| chr2-161178815-DEL-71 | chr2-161178816-DEL-71 |
| chr2-161210741-DEL-7242 | chr2-161210742-DEL-7242 |
| chr2-161230440-DEL-122 | chr2-161230445-DEL-120 |
| chr2-161243968-DEL-99 | chr2-161243969-DEL-99 |
| chr2-161283897-DEL-1191 | chr2-161283898-DEL-1191 |
| chr2-161292132-DEL-82 | chr2-161292121-DEL-94 |
| chr2-161326125-DEL-1071 | chr2-161326126-DEL-1071 |
| chr2-161343906-DEL-560 | chr2-161343907-DEL-560 |
| chr2-161348053-DEL-60 | chr2-161348051-DEL-50 |
| chr2-161353559-DEL-484 | chr2-161353560-DEL-484 |
| chr2-161362277-DEL-70 | chr2-161362278-DEL-94 |
| chr2-161427035-DEL-242 | chr2-161427036-DEL-242 |
| chr2-161483735-DEL-568 | chr2-161483736-DEL-568 |
| chr2-161490141-DEL-7536 | chr2-161490142-DEL-7536 |
| chr2-161513777-DEL-87 | chr2-161513777-DEL-88 |
| chr2-161573711-DEL-52 | chr2-161573712-DEL-52 |
| chr2-161589350-DEL-105 | chr2-161589348-DEL-108 |
| chr2-161589679-DEL-548 | chr2-161589682-DEL-542 |
| chr2-161597357-DEL-207 | chr2-161597357-DEL-207 |
| chr2-161604559-DEL-83 | chr2-161604560-DEL-83 |
| chr2-161645631-DEL-1948 | chr2-161645632-DEL-1946 |
| chr2-161658338-DEL-107 | chr2-161658336-DEL-108 |
| chr2-161663284-DEL-53 | chr2-161663285-DEL-53 |
| chr2-161712215-DEL-67 | chr2-161712216-DEL-67 |
| chr2-161738559-DEL-60 | chr2-161738559-DEL-60 |
| chr2-161763899-DEL-104 | chr2-161763900-DEL-104 |
| chr2-161767586-DEL-179 | chr2-161767582-DEL-180 |
| chr2-161808154-DEL-199 | chr2-161808155-DEL-199 |
| chr2-161814721-DEL-96 | chr2-161814722-DEL-101 |
| chr2-161826483-DEL-6843 | chr2-161826484-DEL-6843 |
| chr2-161833090-DEL-214 | chr2-161833091-DEL-214 |
| chr2-161839549-DEL-62 | chr2-161839550-DEL-60 |
| chr2-161881029-DEL-62 | chr2-161881030-DEL-58 |
| chr2-161938328-DEL-77 | chr2-161938329-DEL-77 |
| chr2-161956698-DEL-797 | chr2-161956700-DEL-796 |
| chr2-161984830-DEL-728 | chr2-161984831-DEL-728 |
| chr2-162021438-DEL-356 | chr2-162021439-DEL-356 |
| chr2-162036772-DEL-198 | chr2-162036777-DEL-187 |
| chr2-162081688-DEL-57 | chr2-162081689-DEL-57 |
| chr2-162087763-DEL-60 | chr2-162087764-DEL-60 |
| chr2-162162835-DEL-66 | chr2-162162842-DEL-53 |
| chr2-162173575-DEL-68 | chr2-162173572-DEL-66 |
| chr2-162182873-DEL-70 | chr2-162182874-DEL-70 |
| chr2-162198862-DEL-369 | chr2-162198863-DEL-369 |
| chr2-162200287-DEL-102 | chr2-162200287-DEL-102 |
| chr2-162210959-DEL-2257 | chr2-162210960-DEL-2257 |
| chr2-162215433-DEL-3277 | chr2-162215434-DEL-3277 |
| chr2-162238337-DEL-59 | chr2-162238330-DEL-50 |
| chr2-162245197-DEL-78 | chr2-162245216-DEL-74 |
| chr2-162278858-DEL-2771 | chr2-162278860-DEL-2771 |
| chr2-162285746-DEL-66 | chr2-162285738-DEL-54 |
| chr2-162305071-DEL-55 | chr2-162305072-DEL-55 |
| chr2-162312706-DEL-147 | chr2-162312705-DEL-153 |
| chr2-162318229-DEL-71 | chr2-162318230-DEL-71 |
| chr2-162327617-DEL-101 | chr2-162327660-DEL-104 |
| chr2-162354386-DEL-52 | chr2-162354387-DEL-52 |
| chr2-162357684-DEL-162 | chr2-162357666-DEL-136 |
| chr2-162367068-DEL-81 | chr2-162367069-DEL-81 |
| chr2-162436397-DEL-8249 | chr2-162436398-DEL-8249 |
| chr2-162533737-DEL-68 | chr2-162533739-DEL-69 |
| chr2-162544347-DEL-90 | chr2-162544348-DEL-90 |
| chr2-162546014-DEL-61 | chr2-162546013-DEL-62 |
| chr2-162567876-DEL-168 | chr2-162567877-DEL-168 |
| chr2-162570093-DEL-1267 | chr2-162570094-DEL-1267 |
| chr2-162574460-DEL-1329 | chr2-162574461-DEL-1329 |
| chr2-162631355-DEL-152 | chr2-162631354-DEL-148 |

| | |
|---|---|
| chr2-162649313-DEL-94 | chr2-162649312-DEL-92 |
| chr2-162653450-DEL-54 | chr2-162653451-DEL-54 |
| chr2-162664640-DEL-393 | chr2-162664641-DEL-393 |
| chr2-162734460-DEL-174 | chr2-162734461-DEL-174 |
| chr2-162740682-DEL-768 | chr2-162740683-DEL-768 |
| chr2-162760977-DEL-770 | chr2-162760978-DEL-770 |
| chr2-162779807-DEL-76 | chr2-162779810-DEL-74 |
| chr2-162815067-DEL-56 | chr2-162815068-DEL-56 |
| chr2-162830701-DEL-224 | chr2-162830700-DEL-225 |
| chr2-162832595-DEL-64 | chr2-162832592-DEL-88 |
| chr2-162846365-DEL-164 | chr2-162846366-DEL-164 |
| chr2-162850490-DEL-1199 | chr2-162850491-DEL-1199 |
| chr2-162852418-DEL-897 | chr2-162852424-DEL-894 |
| chr2-162872681-DEL-68 | chr2-162872682-DEL-68 |
| chr2-162879823-DEL-222 | chr2-162879824-DEL-222 |
| chr2-162885257-DEL-205 | chr2-162885258-DEL-205 |
| chr2-162922107-DEL-248 | chr2-162922097-DEL-244 |
| chr2-162923401-DEL-237 | chr2-162923402-DEL-237 |
| chr2-162935766-DEL-211 | chr2-162935767-DEL-211 |
| chr2-162937248-DEL-63 | chr2-162937248-DEL-63 |
| chr2-162944576-DEL-74 | chr2-162944577-DEL-74 |
| chr2-162967410-DEL-78 | chr2-162967411-DEL-78 |
| chr2-163018027-DEL-205 | chr2-163018028-DEL-205 |
| chr2-163030042-DEL-118 | chr2-163030043-DEL-118 |
| chr2-163107646-DEL-205 | chr2-163107647-DEL-205 |
| chr2-163121655-DEL-66 | chr2-163121656-DEL-66 |
| chr2-163137037-DEL-52 | chr2-163137038-DEL-52 |
| chr2-163143747-DEL-911 | chr2-163143748-DEL-911 |
| chr2-163144067-DEL-76 | chr2-163144068-DEL-76 |
| chr2-163146805-DEL-62 | chr2-163146806-DEL-62 |
| chr2-163155154-DEL-173 | chr2-163155155-DEL-173 |
| chr2-163158058-DEL-162 | chr2-163158059-DEL-162 |
| chr2-163188049-DEL-150 | chr2-163188053-DEL-132 |
| chr2-163263742-DEL-884 | chr2-163263743-DEL-884 |
| chr2-163303518-DEL-54 | chr2-163303502-DEL-55 |
| chr2-163324575-DEL-146 | chr2-163324576-DEL-146 |
| chr2-163325123-DEL-160 | chr2-163325124-DEL-160 |
| chr2-163361038-DEL-6440 | chr2-163361039-DEL-6440 |
| chr2-163392452-DEL-68 | chr2-163392454-DEL-64 |
| chr2-163450466-DEL-66 | chr2-163450466-DEL-64 |
| chr2-163450854-DEL-228 | chr2-163450855-DEL-228 |
| chr2-163492804-DEL-4984 | chr2-163492805-DEL-4984 |
| chr2-163508515-DEL-203 | chr2-163508516-DEL-203 |
| chr2-163510656-DEL-111 | chr2-163510657-DEL-111 |
| chr2-163514846-DEL-56 | chr2-163514847-DEL-56 |
| chr2-163519326-DEL-198 | chr2-163519327-DEL-198 |
| chr2-163549330-DEL-62 | chr2-163549330-DEL-52 |
| chr2-163554637-DEL-278 | chr2-163554638-DEL-277 |
| chr2-163555757-DEL-90 | chr2-163555745-DEL-92 |
| chr2-163562180-DEL-87 | chr2-163562181-DEL-87 |
| chr2-163601055-DEL-70 | chr2-163601056-DEL-70 |
| chr2-163603906-DEL-89 | chr2-163603907-DEL-89 |
| chr2-163622190-DEL-63 | chr2-163622188-DEL-63 |
| chr2-163627948-DEL-400 | chr2-163627949-DEL-400 |
| chr2-163630939-DEL-7087 | chr2-163630940-DEL-7087 |
| chr2-163675734-DEL-2737 | chr2-163675735-DEL-2737 |
| chr2-163684711-DEL-6442 | chr2-163684712-DEL-6442 |
| chr2-163715949-DEL-52 | chr2-163715939-DEL-54 |
| chr2-163734345-DEL-202 | chr2-163734346-DEL-202 |
| chr2-163752877-DEL-302 | chr2-163752878-DEL-302 |
| chr2-163800127-DEL-4634 | chr2-163800128-DEL-4634 |
| chr2-163826017-DEL-763 | chr2-163826018-DEL-763 |
| chr2-163832961-DEL-60 | chr2-163832962-DEL-60 |
| chr2-163871144-DEL-138 | chr2-163871177-DEL-106 |
| chr2-163882482-DEL-102 | chr2-163882515-DEL-85 |
| chr2-163889742-DEL-80 | chr2-163889743-DEL-72 |
| chr2-163898674-DEL-108 | chr2-163898675-DEL-108 |
| chr2-163915645-DEL-52 | chr2-163915646-DEL-52 |
| chr2-163926895-DEL-174 | chr2-163926899-DEL-173 |
| chr2-163932245-DEL-1044 | chr2-163932246-DEL-1044 |
| chr2-163937313-DEL-224 | chr2-163937314-DEL-224 |
| chr2-163938981-DEL-197 | chr2-163938982-DEL-197 |
| chr2-164004666-DEL-101 | chr2-164004667-DEL-101 |
| chr2-164027476-DEL-54 | chr2-164027479-DEL-52 |
| chr2-164028765-DEL-84 | chr2-164028766-DEL-84 |
| chr2-164054936-DEL-290 | chr2-164054937-DEL-290 |
| chr2-164055351-DEL-776 | chr2-164055354-DEL-774 |
| chr2-164058524-DEL-463 | chr2-164058525-DEL-463 |
| chr2-164067367-DEL-62 | chr2-164067370-DEL-60 |
| chr2-164067982-DEL-2000 | chr2-164067991-DEL-2001 |

| | |
|---|---|
| chr2-164071374-DEL-1202 | chr2-164071345-DEL-1200 |
| chr2-164075808-DEL-95 | chr2-164075809-DEL-95 |
| chr2-164095185-DEL-52 | chr2-164095186-DEL-52 |
| chr2-164106397-DEL-5064 | chr2-164106398-DEL-5064 |
| chr2-164111609-DEL-58 | chr2-164111610-DEL-58 |
| chr2-164128627-DEL-96 | chr2-164128627-DEL-96 |
| chr2-164144987-DEL-242 | chr2-164145001-DEL-233 |
| chr2-164155608-DEL-2194 | chr2-164155609-DEL-2194 |
| chr2-164171205-DEL-73 | chr2-164171202-DEL-57 |
| chr2-164183634-DEL-171 | chr2-164183635-DEL-171 |
| chr2-164185897-DEL-2348 | chr2-164185898-DEL-2348 |
| chr2-164190623-DEL-59 | chr2-164190612-DEL-71 |
| chr2-164192174-DEL-129 | chr2-164192175-DEL-129 |
| chr2-164204586-DEL-167 | chr2-164204593-DEL-161 |
| chr2-164227333-DEL-207 | chr2-164227334-DEL-207 |
| chr2-164239272-DEL-72 | chr2-164239273-DEL-72 |
| chr2-164255065-DEL-124 | chr2-164255090-DEL-100 |
| chr2-164288600-DEL-199 | chr2-164288600-DEL-198 |
| chr2-164323021-DEL-223 | chr2-164322971-DEL-226 |
| chr2-164333152-DEL-60 | chr2-164333153-DEL-60 |
| chr2-164353648-DEL-181 | chr2-164353621-DEL-212 |
| chr2-164382102-DEL-202 | chr2-164382103-DEL-202 |
| chr2-164398561-DEL-102 | chr2-164398562-DEL-102 |
| chr2-164400045-DEL-66 | chr2-164400046-DEL-66 |
| chr2-164425272-DEL-66 | chr2-164425273-DEL-66 |
| chr2-164442488-DEL-960 | chr2-164442489-DEL-960 |
| chr2-164464426-DEL-329 | chr2-164464427-DEL-329 |
| chr2-164490154-DEL-69 | chr2-164490156-DEL-68 |
| chr2-164524143-DEL-201 | chr2-164524144-DEL-201 |
| chr2-164547138-DEL-308 | chr2-164547109-DEL-338 |
| chr2-164548390-DEL-56 | chr2-164548391-DEL-56 |
| chr2-164558143-DEL-144 | chr2-164558144-DEL-144 |
| chr2-164566392-DEL-56 | chr2-164566393-DEL-56 |
| chr2-164570766-DEL-72 | chr2-164570763-DEL-72 |
| chr2-164610715-DEL-217 | chr2-164610722-DEL-211 |
| chr2-164642984-DEL-442 | chr2-164642985-DEL-442 |
| chr2-164677955-DEL-212 | chr2-164677956-DEL-212 |
| chr2-164691615-DEL-73 | chr2-164691616-DEL-73 |
| chr2-164705478-DEL-437 | chr2-164705479-DEL-437 |
| chr2-164708088-DEL-201 | chr2-164708088-DEL-201 |
| chr2-164719690-DEL-206 | chr2-164719690-DEL-206 |
| chr2-164787235-DEL-70 | chr2-164787236-DEL-70 |
| chr2-164816059-DEL-54 | chr2-164816060-DEL-54 |
| chr2-164821018-DEL-78 | chr2-164821019-DEL-78 |
| chr2-164871912-DEL-217 | chr2-164871913-DEL-217 |
| chr2-164889738-DEL-172 | chr2-164889739-DEL-172 |
| chr2-164909888-DEL-80 | chr2-164909854-DEL-80 |
| chr2-164949266-DEL-65 | chr2-164949255-DEL-91 |
| chr2-164981065-DEL-222 | chr2-164981066-DEL-222 |
| chr2-165030543-DEL-502 | chr2-165030543-DEL-501 |
| chr2-165038986-DEL-171 | chr2-165038987-DEL-171 |
| chr2-165112204-DEL-52 | chr2-165112205-DEL-52 |
| chr2-165134578-DEL-111 | chr2-165134579-DEL-111 |
| chr2-165171253-DEL-57 | chr2-165171254-DEL-56 |
| chr2-165178062-DEL-80 | chr2-165178065-DEL-78 |
| chr2-165192041-DEL-77 | chr2-165192042-DEL-77 |
| chr2-165218369-DEL-59 | chr2-165218370-DEL-59 |
| chr2-165272772-DEL-242 | chr2-165272773-DEL-242 |
| chr2-165277217-DEL-72 | chr2-165277200-DEL-90 |
| chr2-165281578-DEL-2074 | chr2-165281579-DEL-2074 |
| chr2-165286682-DEL-92 | chr2-165286683-DEL-92 |
| chr2-165291311-DEL-52 | chr2-165291312-DEL-52 |
| chr2-165319337-DEL-1864 | chr2-165319338-DEL-1864 |
| chr2-165345603-DEL-54 | chr2-165345604-DEL-52 |
| chr2-165348794-DEL-1890 | chr2-165348795-DEL-1890 |
| chr2-165351169-DEL-55 | chr2-165351168-DEL-55 |
| chr2-165372479-DEL-907 | chr2-165372483-DEL-903 |
| chr2-165398208-DEL-51 | chr2-165398158-DEL-51 |
| chr2-165401397-DEL-118 | chr2-165401398-DEL-118 |
| chr2-165441045-DEL-78 | chr2-165441036-DEL-87 |
| chr2-165461688-DEL-78 | chr2-165461689-DEL-72 |
| chr2-165482237-DEL-162 | chr2-165482275-DEL-134 |
| chr2-165486679-DEL-2686 | chr2-165486680-DEL-2686 |
| chr2-165486878-DEL-76 | chr2-165486871-DEL-52 |
| chr2-165513548-DEL-309 | chr2-165513549-DEL-309 |
| chr2-165517101-DEL-4412 | chr2-165517102-DEL-4412 |
| chr2-165524988-DEL-1744 | chr2-165524989-DEL-1744 |
| chr2-165528989-DEL-255 | chr2-165529036-DEL-210 |
| chr2-165533792-DEL-62 | chr2-165533793-DEL-60 |
| chr2-165543295-DEL-320 | chr2-165543295-DEL-320 |

| | |
|---|---|
| chr2-165556676-DEL-187 | chr2-165556677-DEL-187 |
| chr2-165577653-DEL-175 | chr2-165577654-DEL-140 |
| chr2-165602420-DEL-182 | chr2-165602421-DEL-182 |
| chr2-165610281-DEL-162 | chr2-165610282-DEL-162 |
| chr2-165618959-DEL-87 | chr2-165618960-DEL-87 |
| chr2-165620539-DEL-196 | chr2-165620540-DEL-196 |
| chr2-165622850-DEL-440 | chr2-165622851-DEL-440 |
| chr2-165634411-DEL-165 | chr2-165634412-DEL-165 |
| chr2-165642398-DEL-207 | chr2-165642399-DEL-207 |
| chr2-165651667-DEL-63 | chr2-165651658-DEL-63 |
| chr2-165657980-DEL-321 | chr2-165657981-DEL-321 |
| chr2-165669632-DEL-214 | chr2-165669633-DEL-214 |
| chr2-165682111-DEL-97 | chr2-165682112-DEL-97 |
| chr2-165683430-DEL-103 | chr2-165683450-DEL-103 |
| chr2-165692340-DEL-55 | chr2-165692341-DEL-55 |
| chr2-165745154-DEL-212 | chr2-165745155-DEL-212 |
| chr2-165751145-DEL-207 | chr2-165751146-DEL-206 |
| chr2-165754171-DEL-267 | chr2-165754164-DEL-275 |
| chr2-165758175-DEL-250 | chr2-165758176-DEL-236 |
| chr2-165762763-DEL-197 | chr2-165762749-DEL-203 |
| chr2-165770354-DEL-5240 | chr2-165770355-DEL-5240 |
| chr2-165776998-DEL-83 | chr2-165776999-DEL-83 |
| chr2-165785537-DEL-190 | chr2-165785538-DEL-190 |
| chr2-165788999-DEL-120 | chr2-165788966-DEL-126 |
| chr2-165792213-DEL-56 | chr2-165792238-DEL-64 |
| chr2-165793171-DEL-170 | chr2-165793187-DEL-167 |
| chr2-165797703-DEL-88 | chr2-165797709-DEL-98 |
| chr2-165801445-DEL-201 | chr2-165801446-DEL-201 |
| chr2-165813723-DEL-112 | chr2-165813726-DEL-106 |
| chr2-165821797-DEL-207 | chr2-165821797-DEL-208 |
| chr2-165837584-DEL-56 | chr2-165837585-DEL-56 |
| chr2-165845817-DEL-178 | chr2-165845820-DEL-167 |
| chr2-165857809-DEL-219 | chr2-165857810-DEL-219 |
| chr2-165870612-DEL-240 | chr2-165870613-DEL-240 |
| chr2-165874064-DEL-57 | chr2-165874065-DEL-57 |
| chr2-165876410-DEL-207 | chr2-165876411-DEL-207 |
| chr2-165884326-DEL-274 | chr2-165884334-DEL-272 |
| chr2-165899333-DEL-256 | chr2-165899334-DEL-256 |
| chr2-165918841-DEL-57 | chr2-165918842-DEL-57 |
| chr2-165938554-DEL-86 | chr2-165938555-DEL-86 |
| chr2-165951611-DEL-85 | chr2-165951612-DEL-85 |
| chr2-165957319-DEL-231 | chr2-165957304-DEL-232 |
| chr2-165972299-DEL-168 | chr2-165972298-DEL-168 |
| chr2-165981950-DEL-86 | chr2-165981942-DEL-87 |
| chr2-166006326-DEL-166 | chr2-166006327-DEL-166 |
| chr2-166035335-DEL-62 | chr2-166035324-DEL-57 |
| chr2-166042003-DEL-57 | chr2-166042002-DEL-58 |
| chr2-166048617-DEL-216 | chr2-166048618-DEL-215 |
| chr2-166073486-DEL-400 | chr2-166073487-DEL-400 |
| chr2-166086042-DEL-365 | chr2-166086031-DEL-365 |
| chr2-166086627-DEL-96 | chr2-166086642-DEL-97 |
| chr2-166088333-DEL-67 | chr2-166088341-DEL-50 |
| chr2-166099377-DEL-88 | chr2-166099376-DEL-89 |
| chr2-166131415-DEL-53 | chr2-166131435-DEL-69 |
| chr2-166135446-DEL-56 | chr2-166135447-DEL-56 |
| chr2-166168066-DEL-76 | chr2-166168067-DEL-76 |
| chr2-166206648-DEL-82 | chr2-166206649-DEL-82 |
| chr2-166207986-DEL-54 | chr2-166207989-DEL-68 |
| chr2-166246327-DEL-399 | chr2-166246328-DEL-399 |
| chr2-166262534-DEL-111 | chr2-166262535-DEL-111 |
| chr2-166266231-DEL-74 | chr2-166266232-DEL-56 |
| chr2-166272488-DEL-449 | chr2-166272489-DEL-449 |
| chr2-166276398-DEL-74 | chr2-166276399-DEL-74 |
| chr2-166283852-DEL-62 | chr2-166283852-DEL-62 |
| chr2-166309617-DEL-92 | chr2-166309617-DEL-92 |
| chr2-166311907-DEL-136 | chr2-166311906-DEL-136 |
| chr2-166313839-DEL-104 | chr2-166313833-DEL-105 |
| chr2-166321852-DEL-625 | chr2-166321853-DEL-623 |
| chr2-166324630-DEL-58 | chr2-166324631-DEL-58 |
| chr2-166335011-DEL-65 | chr2-166335012-DEL-65 |
| chr2-166348622-DEL-552 | chr2-166348623-DEL-552 |
| chr2-166361792-DEL-400 | chr2-166361793-DEL-400 |
| chr2-166365785-DEL-62 | chr2-166365786-DEL-62 |
| chr2-166372567-DEL-4443 | chr2-166372566-DEL-4444 |
| chr2-166397218-DEL-184 | chr2-166397219-DEL-184 |
| chr2-166399586-DEL-178 | chr2-166399587-DEL-178 |
| chr2-166402091-DEL-132 | chr2-166402092-DEL-132 |
| chr2-166442341-DEL-76 | chr2-166442342-DEL-76 |
| chr2-166462114-DEL-120 | chr2-166462115-DEL-120 |
| chr2-166468062-DEL-629 | chr2-166468063-DEL-629 |

| | |
|---|---|
| chr2-166573153-DEL-72 | chr2-166573154-DEL-54 |
| chr2-166590999-DEL-202 | chr2-166591000-DEL-202 |
| chr2-166595903-DEL-207 | chr2-166595904-DEL-207 |
| chr2-166602164-DEL-494 | chr2-166602165-DEL-494 |
| chr2-166623962-DEL-241 | chr2-166623950-DEL-242 |
| chr2-166654112-DEL-5277 | chr2-166654113-DEL-5277 |
| chr2-166695958-DEL-135 | chr2-166695959-DEL-135 |
| chr2-166714520-DEL-198 | chr2-166714521-DEL-198 |
| chr2-166723523-DEL-185 | chr2-166723524-DEL-185 |
| chr2-166726991-DEL-61 | chr2-166726992-DEL-61 |
| chr2-166735701-DEL-118 | chr2-166735702-DEL-118 |
| chr2-166745969-DEL-707 | chr2-166745970-DEL-707 |
| chr2-166749910-DEL-274 | chr2-166749911-DEL-274 |
| chr2-166758649-DEL-494 | chr2-166758650-DEL-494 |
| chr2-166791445-DEL-87 | chr2-166791479-DEL-57 |
| chr2-166808111-DEL-182 | chr2-166808112-DEL-182 |
| chr2-166822996-DEL-211 | chr2-166822986-DEL-205 |
| chr2-166829211-DEL-237 | chr2-166829212-DEL-237 |
| chr2-166849875-DEL-434 | chr2-166849876-DEL-434 |
| chr2-166853232-DEL-206 | chr2-166853233-DEL-206 |
| chr2-166853747-DEL-193 | chr2-166853752-DEL-189 |
| chr2-166855787-DEL-633 | chr2-166855788-DEL-633 |
| chr2-166863082-DEL-186 | chr2-166863074-DEL-188 |
| chr2-166875039-DEL-69 | chr2-166875040-DEL-69 |
| chr2-166906354-DEL-206 | chr2-166906355-DEL-206 |
| chr2-166908498-DEL-139 | chr2-166908498-DEL-136 |
| chr2-166916793-DEL-66 | chr2-166916793-DEL-66 |
| chr2-166925774-DEL-554 | chr2-166925775-DEL-554 |
| chr2-166932631-DEL-134 | chr2-166932634-DEL-113 |
| chr2-166934122-DEL-66 | chr2-166934123-DEL-66 |
| chr2-166956374-DEL-240 | chr2-166956375-DEL-240 |
| chr2-166990764-DEL-95 | chr2-166990765-DEL-95 |
| chr2-167037751-DEL-223 | chr2-167037752-DEL-223 |
| chr2-167108068-DEL-172 | chr2-167108069-DEL-172 |
| chr2-167109181-DEL-197 | chr2-167109182-DEL-197 |
| chr2-167138794-DEL-207 | chr2-167138795-DEL-207 |
| chr2-167161713-DEL-70 | chr2-167161703-DEL-60 |
| chr2-167239771-DEL-301 | chr2-167239772-DEL-301 |
| chr2-167242536-DEL-227 | chr2-167242537-DEL-227 |
| chr2-167249957-DEL-5309 | chr2-167249958-DEL-5309 |
| chr2-167276082-DEL-142 | chr2-167276083-DEL-142 |
| chr2-167289634-DEL-58 | chr2-167289635-DEL-58 |
| chr2-167293691-DEL-53 | chr2-167293691-DEL-53 |
| chr2-167294385-DEL-377 | chr2-167294397-DEL-365 |
| chr2-167297610-DEL-208 | chr2-167297611-DEL-208 |
| chr2-167360132-DEL-176 | chr2-167360133-DEL-176 |
| chr2-167422600-DEL-59 | chr2-167422601-DEL-59 |
| chr2-167426586-DEL-55 | chr2-167426598-DEL-60 |
| chr2-167431961-DEL-53 | chr2-167431962-DEL-53 |
| chr2-167434341-DEL-71 | chr2-167434342-DEL-70 |
| chr2-167466217-DEL-218 | chr2-167466218-DEL-218 |
| chr2-167506330-DEL-174 | chr2-167506310-DEL-166 |
| chr2-167545266-DEL-62 | chr2-167545224-DEL-57 |
| chr2-167554615-DEL-216 | chr2-167554616-DEL-216 |
| chr2-167635188-DEL-105 | chr2-167635219-DEL-75 |
| chr2-167726369-DEL-170 | chr2-167726353-DEL-187 |
| chr2-167734532-DEL-84 | chr2-167734533-DEL-84 |
| chr2-167737338-DEL-68 | chr2-167737317-DEL-66 |
| chr2-167740431-DEL-198 | chr2-167740432-DEL-198 |
| chr2-167755942-DEL-1471 | chr2-167755943-DEL-1471 |
| chr2-167761550-DEL-96 | chr2-167761555-DEL-110 |
| chr2-167829606-DEL-72 | chr2-167829607-DEL-72 |
| chr2-167836258-DEL-81 | chr2-167836259-DEL-82 |
| chr2-167857029-DEL-152 | chr2-167857066-DEL-131 |
| chr2-167875160-DEL-180 | chr2-167875131-DEL-190 |
| chr2-167906218-DEL-217 | chr2-167906216-DEL-218 |
| chr2-167913303-DEL-213 | chr2-167913301-DEL-214 |
| chr2-167914381-DEL-54 | chr2-167914382-DEL-54 |
| chr2-167919343-DEL-68 | chr2-167919344-DEL-68 |
| chr2-167948459-DEL-93 | chr2-167948460-DEL-93 |
| chr2-167962586-DEL-946 | chr2-167962586-DEL-947 |
| chr2-167973515-DEL-88 | chr2-167973516-DEL-88 |
| chr2-167981845-DEL-206 | chr2-167981846-DEL-206 |
| chr2-167998664-DEL-203 | chr2-167998665-DEL-203 |
| chr2-167999296-DEL-63 | chr2-167999297-DEL-67 |
| chr2-167999786-DEL-122 | chr2-167999790-DEL-119 |
| chr2-168002876-DEL-98 | chr2-168002877-DEL-98 |
| chr2-168061233-DEL-600 | chr2-168061234-DEL-600 |
| chr2-168079630-DEL-602 | chr2-168079631-DEL-602 |
| chr2-168082120-DEL-205 | chr2-168082121-DEL-205 |

| | |
|---|---|
| chr2-168100149-DEL-169 | chr2-168100150-DEL-169 |
| chr2-168120551-DEL-63 | chr2-168120532-DEL-83 |
| chr2-168128156-DEL-70 | chr2-168128157-DEL-70 |
| chr2-168165784-DEL-59 | chr2-168165785-DEL-59 |
| chr2-168190640-DEL-540 | chr2-168190641-DEL-540 |
| chr2-168207311-DEL-895 | chr2-168207312-DEL-895 |
| chr2-168212491-DEL-70 | chr2-168212492-DEL-70 |
| chr2-168226961-DEL-130 | chr2-168226962-DEL-130 |
| chr2-168284773-DEL-214 | chr2-168284774-DEL-214 |
| chr2-168308476-DEL-537 | chr2-168308491-DEL-511 |
| chr2-168350290-DEL-74 | chr2-168350285-DEL-50 |
| chr2-168377192-DEL-137 | chr2-168377195-DEL-138 |
| chr2-168388664-DEL-56 | chr2-168388665-DEL-56 |
| chr2-168453439-DEL-215 | chr2-168453442-DEL-172 |
| chr2-168494045-DEL-204 | chr2-168494030-DEL-205 |
| chr2-168513392-DEL-200 | chr2-168513392-DEL-200 |
| chr2-168515968-DEL-138 | chr2-168515969-DEL-138 |
| chr2-168543005-DEL-214 | chr2-168543005-DEL-214 |
| chr2-168547536-DEL-207 | chr2-168547536-DEL-207 |
| chr2-168556954-DEL-231 | chr2-168556955-DEL-231 |
| chr2-168557813-DEL-479 | chr2-168557814-DEL-479 |
| chr2-168562946-DEL-202 | chr2-168562946-DEL-202 |
| chr2-168567231-DEL-453 | chr2-168567232-DEL-453 |
| chr2-168570662-DEL-199 | chr2-168570647-DEL-196 |
| chr2-168586577-DEL-260 | chr2-168586617-DEL-234 |
| chr2-168589534-DEL-195 | chr2-168589504-DEL-198 |
| chr2-168602564-DEL-82 | chr2-168602564-DEL-83 |
| chr2-168631288-DEL-222 | chr2-168631289-DEL-222 |
| chr2-168675987-DEL-206 | chr2-168675988-DEL-206 |
| chr2-168678218-DEL-192 | chr2-168678219-DEL-192 |
| chr2-168691144-DEL-58 | chr2-168691145-DEL-58 |
| chr2-168695884-DEL-132 | chr2-168695885-DEL-132 |
| chr2-168703468-DEL-51 | chr2-168703468-DEL-51 |
| chr2-168705968-DEL-267 | chr2-168705970-DEL-266 |
| chr2-168726294-DEL-106 | chr2-168726295-DEL-106 |
| chr2-168734382-DEL-53 | chr2-168734383-DEL-53 |
| chr2-168743017-DEL-1157 | chr2-168743018-DEL-1157 |
| chr2-168743197-DEL-153 | chr2-168743198-DEL-153 |
| chr2-168756178-DEL-222 | chr2-168756179-DEL-222 |
| chr2-168759534-DEL-86 | chr2-168759535-DEL-86 |
| chr2-168763651-DEL-165 | chr2-168763652-DEL-165 |
| chr2-168778123-DEL-104 | chr2-168778124-DEL-104 |
| chr2-168783322-DEL-212 | chr2-168783323-DEL-212 |
| chr2-168788274-DEL-744 | chr2-168788275-DEL-744 |
| chr2-168823862-DEL-81 | chr2-168823863-DEL-84 |
| chr2-168838895-DEL-197 | chr2-168838896-DEL-197 |
| chr2-168868850-DEL-440 | chr2-168868851-DEL-440 |
| chr2-168874438-DEL-108 | chr2-168874481-DEL-64 |
| chr2-168880525-DEL-149 | chr2-168880530-DEL-144 |
| chr2-168881509-DEL-138 | chr2-168881537-DEL-123 |
| chr2-168884629-DEL-1862 | chr2-168884636-DEL-1872 |
| chr2-168893241-DEL-154 | chr2-168893242-DEL-140 |
| chr2-168895519-DEL-204 | chr2-168895520-DEL-204 |
| chr2-168910695-DEL-179 | chr2-168910696-DEL-179 |
| chr2-168962078-DEL-62 | chr2-168962079-DEL-62 |
| chr2-169025667-DEL-59 | chr2-169025668-DEL-54 |
| chr2-169030451-DEL-98 | chr2-169030490-DEL-97 |
| chr2-169041559-DEL-150 | chr2-169041560-DEL-150 |
| chr2-169060056-DEL-117 | chr2-169060057-DEL-117 |
| chr2-169080145-DEL-66 | chr2-169080146-DEL-66 |
| chr2-169081238-DEL-71 | chr2-169081229-DEL-50 |
| chr2-169091596-DEL-1383 | chr2-169091597-DEL-1383 |
| chr2-169139484-DEL-100 | chr2-169139461-DEL-150 |
| chr2-169165927-DEL-242 | chr2-169165926-DEL-243 |
| chr2-169201028-DEL-59 | chr2-169201029-DEL-59 |
| chr2-169226104-DEL-432 | chr2-169226105-DEL-432 |
| chr2-169243002-DEL-114 | chr2-169243040-DEL-104 |
| chr2-169257059-DEL-113 | chr2-169257060-DEL-113 |
| chr2-169269026-DEL-68 | chr2-169269027-DEL-68 |
| chr2-169293988-DEL-84 | chr2-169293989-DEL-84 |
| chr2-169302241-DEL-1335 | chr2-169302242-DEL-1335 |
| chr2-169327445-DEL-75 | chr2-169327445-DEL-75 |
| chr2-169329124-DEL-73 | chr2-169329125-DEL-73 |
| chr2-169354803-DEL-226 | chr2-169354804-DEL-226 |
| chr2-169377044-DEL-80 | chr2-169377045-DEL-80 |
| chr2-169397591-DEL-254 | chr2-169397592-DEL-254 |
| chr2-169417051-DEL-216 | chr2-169417052-DEL-216 |
| chr2-169459738-DEL-144 | chr2-169459739-DEL-144 |
| chr2-169460374-DEL-90 | chr2-169460375-DEL-90 |
| chr2-169490977-DEL-78 | chr2-169490978-DEL-78 |

| | |
|---|---|
| chr2-169510383-DEL-52 | chr2-169510382-DEL-52 |
| chr2-169548078-DEL-69 | chr2-169548055-DEL-93 |
| chr2-169565149-DEL-68 | chr2-169565122-DEL-80 |
| chr2-169582462-DEL-57 | chr2-169582463-DEL-57 |
| chr2-169630497-DEL-249 | chr2-169630535-DEL-212 |
| chr2-169642863-DEL-1864 | chr2-169642864-DEL-1864 |
| chr2-169646770-DEL-146 | chr2-169646771-DEL-146 |
| chr2-169659090-DEL-177 | chr2-169659091-DEL-177 |
| chr2-169680752-DEL-113 | chr2-169680753-DEL-113 |
| chr2-169717121-DEL-72 | chr2-169717122-DEL-72 |
| chr2-169749495-DEL-70 | chr2-169749500-DEL-54 |
| chr2-169777874-DEL-114 | chr2-169777873-DEL-116 |
| chr2-169795584-DEL-56 | chr2-169795571-DEL-57 |
| chr2-169846760-DEL-81 | chr2-169846759-DEL-83 |
| chr2-169847680-DEL-69 | chr2-169847681-DEL-69 |
| chr2-169917981-DEL-103 | chr2-169917981-DEL-78 |
| chr2-169926100-DEL-55 | chr2-169926105-DEL-51 |
| chr2-169930824-DEL-54 | chr2-169930824-DEL-50 |
| chr2-169998015-DEL-55 | chr2-169997981-DEL-72 |
| chr2-170012977-DEL-70 | chr2-170012978-DEL-68 |
| chr2-170014843-DEL-414 | chr2-170014844-DEL-414 |
| chr2-170023030-DEL-1317 | chr2-170023031-DEL-1317 |
| chr2-170034033-DEL-185 | chr2-170034030-DEL-185 |
| chr2-170039448-DEL-218 | chr2-170039449-DEL-218 |
| chr2-170041398-DEL-209 | chr2-170041399-DEL-209 |
| chr2-170056214-DEL-99 | chr2-170056215-DEL-99 |
| chr2-170092196-DEL-437 | chr2-170092197-DEL-437 |
| chr2-170097887-DEL-85 | chr2-170097888-DEL-105 |
| chr2-170126887-DEL-1113 | chr2-170126883-DEL-1114 |
| chr2-170135761-DEL-200 | chr2-170135762-DEL-200 |
| chr2-170140149-DEL-439 | chr2-170140150-DEL-439 |
| chr2-170157852-DEL-497 | chr2-170157862-DEL-489 |
| chr2-170171702-DEL-71 | chr2-170171666-DEL-75 |
| chr2-170194921-DEL-11003 | chr2-170194922-DEL-11003 |
| chr2-170204990-DEL-204 | chr2-170204974-DEL-205 |
| chr2-170210736-DEL-80 | chr2-170210737-DEL-80 |
| chr2-170234687-DEL-63 | chr2-170234688-DEL-63 |
| chr2-170284341-DEL-197 | chr2-170284341-DEL-198 |
| chr2-170290639-DEL-202 | chr2-170290640-DEL-202 |
| chr2-170304654-DEL-1157 | chr2-170304655-DEL-1157 |
| chr2-170321997-DEL-204 | chr2-170321985-DEL-205 |
| chr2-170327153-DEL-92 | chr2-170327153-DEL-68 |
| chr2-170334899-DEL-77 | chr2-170334900-DEL-77 |
| chr2-170359535-DEL-61 | chr2-170359536-DEL-61 |
| chr2-170400764-DEL-613 | chr2-170400765-DEL-613 |
| chr2-170405301-DEL-57 | chr2-170405302-DEL-57 |
| chr2-170413437-DEL-117 | chr2-170413438-DEL-117 |
| chr2-170423789-DEL-75 | chr2-170423790-DEL-75 |
| chr2-170456640-DEL-79 | chr2-170456632-DEL-117 |
| chr2-170468699-DEL-607 | chr2-170468708-DEL-586 |
| chr2-170509793-DEL-78 | chr2-170509793-DEL-78 |
| chr2-170514414-DEL-1016 | chr2-170514415-DEL-1016 |
| chr2-170520111-DEL-116 | chr2-170520102-DEL-126 |
| chr2-170523945-DEL-77 | chr2-170523945-DEL-77 |
| chr2-170535591-DEL-452 | chr2-170535592-DEL-452 |
| chr2-170537417-DEL-60 | chr2-170537407-DEL-50 |
| chr2-170539103-DEL-1260 | chr2-170539104-DEL-1260 |
| chr2-170543235-DEL-1889 | chr2-170543235-DEL-1889 |
| chr2-170554106-DEL-212 | chr2-170554107-DEL-212 |
| chr2-170569659-DEL-52 | chr2-170569660-DEL-52 |
| chr2-170588416-DEL-90 | chr2-170588423-DEL-84 |
| chr2-170623795-DEL-220 | chr2-170623796-DEL-234 |
| chr2-170642512-DEL-172 | chr2-170642513-DEL-172 |
| chr2-170647165-DEL-123 | chr2-170647173-DEL-108 |
| chr2-170670079-DEL-79 | chr2-170670080-DEL-79 |
| chr2-170705260-DEL-57 | chr2-170705254-DEL-58 |
| chr2-170708427-DEL-123 | chr2-170708428-DEL-123 |
| chr2-170714170-DEL-71 | chr2-170714190-DEL-51 |
| chr2-170719872-DEL-68 | chr2-170719873-DEL-68 |
| chr2-170753289-DEL-198 | chr2-170753290-DEL-198 |
| chr2-170768611-DEL-536 | chr2-170768612-DEL-536 |
| chr2-170769389-DEL-59 | chr2-170769390-DEL-75 |
| chr2-170784427-DEL-72 | chr2-170784428-DEL-66 |
| chr2-170786538-DEL-58 | chr2-170786539-DEL-58 |
| chr2-170824039-DEL-57 | chr2-170824035-DEL-58 |
| chr2-170834093-DEL-74 | chr2-170834087-DEL-80 |
| chr2-170844533-DEL-79 | chr2-170844534-DEL-79 |
| chr2-170862891-DEL-66 | chr2-170862892-DEL-66 |
| chr2-170882807-DEL-395 | chr2-170882808-DEL-395 |
| chr2-170892796-DEL-873 | chr2-170892797-DEL-873 |

| | |
|---|---|
| chr2-170925565-DEL-6034 | chr2-170925566-DEL-6034 |
| chr2-170962193-DEL-93 | chr2-170962194-DEL-93 |
| chr2-170982770-DEL-67 | chr2-170982771-DEL-67 |
| chr2-171016800-DEL-2712 | chr2-171016801-DEL-2712 |
| chr2-171017819-DEL-86 | chr2-171017819-DEL-56 |
| chr2-171069729-DEL-75 | chr2-171069725-DEL-75 |
| chr2-171090985-DEL-405 | chr2-171090984-DEL-405 |
| chr2-171116028-DEL-86 | chr2-171116048-DEL-64 |
| chr2-171131171-DEL-248 | chr2-171131164-DEL-243 |
| chr2-171159233-DEL-59 | chr2-171159225-DEL-65 |
| chr2-171164572-DEL-61 | chr2-171164573-DEL-61 |
| chr2-171168386-DEL-51 | chr2-171168376-DEL-52 |
| chr2-171187654-DEL-100 | chr2-171187654-DEL-50 |
| chr2-171188707-DEL-90 | chr2-171188717-DEL-80 |
| chr2-171193449-DEL-52 | chr2-171193450-DEL-52 |
| chr2-171194140-DEL-302 | chr2-171194141-DEL-302 |
| chr2-171222030-DEL-55 | chr2-171222031-DEL-55 |
| chr2-171250820-DEL-63 | chr2-171250811-DEL-65 |
| chr2-171253206-DEL-127 | chr2-171253207-DEL-127 |
| chr2-171260624-DEL-56 | chr2-171260580-DEL-59 |
| chr2-171286313-DEL-52 | chr2-171286302-DEL-53 |
| chr2-171287752-DEL-71 | chr2-171287756-DEL-69 |
| chr2-171290660-DEL-54 | chr2-171290661-DEL-56 |
| chr2-171338261-DEL-52 | chr2-171338262-DEL-52 |
| chr2-171341857-DEL-176 | chr2-171341866-DEL-168 |
| chr2-171368332-DEL-124 | chr2-171368333-DEL-124 |
| chr2-171405128-DEL-307 | chr2-171405129-DEL-307 |
| chr2-171406440-DEL-68 | chr2-171406441-DEL-68 |
| chr2-171418979-DEL-6909 | chr2-171418980-DEL-6909 |
| chr2-171441245-DEL-67 | chr2-171441246-DEL-67 |
| chr2-171449323-DEL-62 | chr2-171449324-DEL-62 |
| chr2-171505996-DEL-135 | chr2-171505982-DEL-136 |
| chr2-171516623-DEL-972 | chr2-171516624-DEL-972 |
| chr2-171607892-DEL-128 | chr2-171607893-DEL-128 |
| chr2-171631723-DEL-74 | chr2-171631736-DEL-76 |
| chr2-171638434-DEL-114 | chr2-171638435-DEL-114 |
| chr2-171659949-DEL-507 | chr2-171659950-DEL-496 |
| chr2-171680422-DEL-167 | chr2-171680423-DEL-167 |
| chr2-171680891-DEL-470 | chr2-171680892-DEL-470 |
| chr2-171686983-DEL-107 | chr2-171686984-DEL-107 |
| chr2-171728789-DEL-87 | chr2-171728790-DEL-87 |
| chr2-171743109-DEL-51 | chr2-171743109-DEL-52 |
| chr2-171754075-DEL-2369 | chr2-171754076-DEL-2369 |
| chr2-171767640-DEL-6454 | chr2-171767633-DEL-6447 |
| chr2-171800145-DEL-52 | chr2-171800146-DEL-52 |
| chr2-171843069-DEL-321 | chr2-171843070-DEL-321 |
| chr2-171847315-DEL-56 | chr2-171847316-DEL-56 |
| chr2-171848961-DEL-7303 | chr2-171848960-DEL-7303 |
| chr2-171886669-DEL-64 | chr2-171886670-DEL-64 |
| chr2-171921433-DEL-72 | chr2-171921434-DEL-72 |
| chr2-171932162-DEL-94 | chr2-171932167-DEL-90 |
| chr2-171948169-DEL-64 | chr2-171948170-DEL-64 |
| chr2-171964430-DEL-68 | chr2-171964423-DEL-80 |
| chr2-171968243-DEL-62 | chr2-171968229-DEL-76 |
| chr2-171991453-DEL-91 | chr2-171991483-DEL-57 |
| chr2-172018681-DEL-135 | chr2-172018682-DEL-135 |
| chr2-172024367-DEL-65 | chr2-172024375-DEL-53 |
| chr2-172026672-DEL-654 | chr2-172026668-DEL-656 |
| chr2-172036981-DEL-56 | chr2-172036982-DEL-58 |
| chr2-172040695-DEL-308 | chr2-172040696-DEL-308 |
| chr2-172041262-DEL-614 | chr2-172041263-DEL-614 |
| chr2-172049764-DEL-193 | chr2-172049777-DEL-210 |
| chr2-172051200-DEL-318 | chr2-172051200-DEL-318 |
| chr2-172058126-DEL-58 | chr2-172058111-DEL-74 |
| chr2-172081426-DEL-58 | chr2-172081427-DEL-60 |
| chr2-172109417-DEL-204 | chr2-172109418-DEL-204 |
| chr2-172119786-DEL-70 | chr2-172119787-DEL-70 |
| chr2-172131092-DEL-858 | chr2-172131093-DEL-858 |
| chr2-172136557-DEL-76 | chr2-172136558-DEL-57 |
| chr2-172143295-DEL-77 | chr2-172143295-DEL-61 |
| chr2-172147997-DEL-398 | chr2-172147998-DEL-398 |
| chr2-172154864-DEL-207 | chr2-172154865-DEL-207 |
| chr2-172159014-DEL-332 | chr2-172159015-DEL-332 |
| chr2-172161861-DEL-179 | chr2-172161860-DEL-181 |
| chr2-172178485-DEL-1070 | chr2-172178486-DEL-1070 |
| chr2-172193707-DEL-80 | chr2-172193708-DEL-80 |
| chr2-172225877-DEL-421 | chr2-172225878-DEL-421 |
| chr2-172249223-DEL-91 | chr2-172249246-DEL-89 |
| chr2-172251162-DEL-181 | chr2-172251163-DEL-181 |
| chr2-172266117-DEL-52 | chr2-172266118-DEL-52 |

| | |
|---|---|
| chr2-172312555-DEL-214 | chr2-172312556-DEL-214 |
| chr2-172321192-DEL-256 | chr2-172321193-DEL-256 |
| chr2-172322846-DEL-204 | chr2-172322847-DEL-204 |
| chr2-172326524-DEL-160 | chr2-172326529-DEL-166 |
| chr2-172351311-DEL-346 | chr2-172351312-DEL-346 |
| chr2-172398635-DEL-52 | chr2-172398636-DEL-52 |
| chr2-172416776-DEL-132 | chr2-172416777-DEL-132 |
| chr2-172421824-DEL-621 | chr2-172421825-DEL-621 |
| chr2-172461302-DEL-647 | chr2-172461302-DEL-647 |
| chr2-172478107-DEL-78 | chr2-172478108-DEL-66 |
| chr2-172479561-DEL-205 | chr2-172479562-DEL-205 |
| chr2-172481458-DEL-400 | chr2-172481464-DEL-394 |
| chr2-172492299-DEL-91 | chr2-172492285-DEL-94 |
| chr2-172493562-DEL-197 | chr2-172493563-DEL-197 |
| chr2-172514119-DEL-88 | chr2-172514120-DEL-88 |
| chr2-172526406-DEL-81 | chr2-172526407-DEL-81 |
| chr2-172536546-DEL-698 | chr2-172536547-DEL-698 |
| chr2-172551588-DEL-178 | chr2-172551589-DEL-178 |
| chr2-172552241-DEL-222 | chr2-172552242-DEL-222 |
| chr2-172572956-DEL-219 | chr2-172572957-DEL-219 |
| chr2-172585528-DEL-7264 | chr2-172585529-DEL-7264 |
| chr2-172595617-DEL-330 | chr2-172595617-DEL-330 |
| chr2-172601852-DEL-68 | chr2-172601853-DEL-68 |
| chr2-172607865-DEL-91 | chr2-172607866-DEL-91 |
| chr2-172630200-DEL-149 | chr2-172630205-DEL-155 |
| chr2-172651492-DEL-180 | chr2-172651493-DEL-180 |
| chr2-172657515-DEL-306 | chr2-172657517-DEL-295 |
| chr2-172681836-DEL-164 | chr2-172681840-DEL-163 |
| chr2-172686080-DEL-112 | chr2-172686104-DEL-112 |
| chr2-172710878-DEL-264 | chr2-172710911-DEL-228 |
| chr2-172728071-DEL-76 | chr2-172728072-DEL-76 |
| chr2-172755150-DEL-66 | chr2-172755141-DEL-64 |
| chr2-172785903-DEL-55 | chr2-172785931-DEL-54 |
| chr2-172786650-DEL-137 | chr2-172786651-DEL-137 |
| chr2-172788536-DEL-112 | chr2-172788537-DEL-112 |
| chr2-172789366-DEL-347 | chr2-172789367-DEL-347 |
| chr2-172794706-DEL-62 | chr2-172794702-DEL-51 |
| chr2-172797180-DEL-72 | chr2-172797180-DEL-72 |
| chr2-172810177-DEL-218 | chr2-172810178-DEL-218 |
| chr2-172826206-DEL-201 | chr2-172826207-DEL-201 |
| chr2-172851543-DEL-448 | chr2-172851544-DEL-448 |
| chr2-172858394-DEL-67 | chr2-172858395-DEL-67 |
| chr2-172893613-DEL-56 | chr2-172893614-DEL-56 |
| chr2-172919456-DEL-63 | chr2-172919457-DEL-63 |
| chr2-172920323-DEL-239 | chr2-172920324-DEL-237 |
| chr2-172931116-DEL-63 | chr2-172931117-DEL-63 |
| chr2-172964435-DEL-58 | chr2-172964436-DEL-58 |
| chr2-172967278-DEL-66 | chr2-172967279-DEL-66 |
| chr2-172969906-DEL-60 | chr2-172969877-DEL-60 |
| chr2-172987667-DEL-177 | chr2-172987668-DEL-177 |
| chr2-172999647-DEL-63 | chr2-172999597-DEL-63 |
| chr2-173040330-DEL-91 | chr2-173040331-DEL-92 |
| chr2-173046661-DEL-67 | chr2-173046660-DEL-68 |
| chr2-173055007-DEL-306 | chr2-173055002-DEL-294 |
| chr2-173056201-DEL-72 | chr2-173056202-DEL-72 |
| chr2-173094587-DEL-51 | chr2-173094586-DEL-51 |
| chr2-173096358-DEL-56 | chr2-173096359-DEL-56 |
| chr2-173097300-DEL-105 | chr2-173097301-DEL-105 |
| chr2-173101887-DEL-86 | chr2-173101888-DEL-86 |
| chr2-173109167-DEL-58 | chr2-173109167-DEL-58 |
| chr2-173132755-DEL-1718 | chr2-173132756-DEL-1719 |
| chr2-173154314-DEL-54 | chr2-173154315-DEL-54 |
| chr2-173162122-DEL-208 | chr2-173162138-DEL-220 |
| chr2-173175563-DEL-54 | chr2-173175563-DEL-54 |
| chr2-173190172-DEL-54 | chr2-173190173-DEL-54 |
| chr2-173190852-DEL-237 | chr2-173190846-DEL-238 |
| chr2-173192588-DEL-53 | chr2-173192589-DEL-53 |
| chr2-173193152-DEL-66 | chr2-173193153-DEL-66 |
| chr2-173199122-DEL-116 | chr2-173199123-DEL-116 |
| chr2-173209683-DEL-55 | chr2-173209684-DEL-55 |
| chr2-173212914-DEL-1149 | chr2-173212915-DEL-1149 |
| chr2-173216860-DEL-301 | chr2-173216861-DEL-301 |
| chr2-173219209-DEL-54 | chr2-173219210-DEL-54 |
| chr2-173225730-DEL-250 | chr2-173225731-DEL-250 |
| chr2-173231877-DEL-71 | chr2-173231878-DEL-71 |
| chr2-173238431-DEL-65 | chr2-173238432-DEL-65 |
| chr2-173248066-DEL-83 | chr2-173248105-DEL-53 |
| chr2-173260084-DEL-75 | chr2-173260094-DEL-105 |
| chr2-173260606-DEL-56 | chr2-173260611-DEL-52 |
| chr2-173263525-DEL-1886 | chr2-173263526-DEL-1886 |

| | |
|---|---|
| chr2-173279849-DEL-200 | chr2-173279861-DEL-199 |
| chr2-173295131-DEL-73 | chr2-173295132-DEL-73 |
| chr2-173308415-DEL-70 | chr2-173308411-DEL-50 |
| chr2-173320246-DEL-70 | chr2-173320247-DEL-70 |
| chr2-173328087-DEL-120 | chr2-173328087-DEL-115 |
| chr2-173339045-DEL-73 | chr2-173339043-DEL-57 |
| chr2-173475366-DEL-624 | chr2-173475377-DEL-623 |
| chr2-173488313-DEL-63 | chr2-173488292-DEL-86 |
| chr2-173493460-DEL-53 | chr2-173493461-DEL-53 |
| chr2-173542068-DEL-105 | chr2-173542104-DEL-70 |
| chr2-173596106-DEL-155 | chr2-173596107-DEL-155 |
| chr2-173603126-DEL-102 | chr2-173603127-DEL-68 |
| chr2-173647345-DEL-83 | chr2-173647345-DEL-83 |
| chr2-173678088-DEL-342 | chr2-173678090-DEL-341 |
| chr2-173684950-DEL-113 | chr2-173684951-DEL-113 |
| chr2-173688093-DEL-96 | chr2-173688089-DEL-97 |
| chr2-173689644-DEL-57 | chr2-173689645-DEL-57 |
| chr2-173690418-DEL-64 | chr2-173690419-DEL-65 |
| chr2-173695579-DEL-66 | chr2-173695578-DEL-88 |
| chr2-173703164-DEL-51 | chr2-173703165-DEL-51 |
| chr2-173705278-DEL-415 | chr2-173705279-DEL-415 |
| chr2-173706561-DEL-114 | chr2-173706561-DEL-115 |
| chr2-173708930-DEL-93 | chr2-173708929-DEL-91 |
| chr2-173768411-DEL-52 | chr2-173768412-DEL-52 |
| chr2-173795523-DEL-58 | chr2-173795524-DEL-58 |
| chr2-173797084-DEL-129 | chr2-173797085-DEL-129 |
| chr2-173798767-DEL-2066 | chr2-173798768-DEL-2066 |
| chr2-173837135-DEL-180 | chr2-173837098-DEL-201 |
| chr2-173840677-DEL-116 | chr2-173840678-DEL-116 |
| chr2-173906654-DEL-460 | chr2-173906655-DEL-460 |
| chr2-174007605-DEL-56 | chr2-174007606-DEL-56 |
| chr2-174031721-DEL-60 | chr2-174031706-DEL-60 |
| chr2-174054124-DEL-79 | chr2-174054125-DEL-79 |
| chr2-174066375-DEL-668 | chr2-174066376-DEL-667 |
| chr2-174070819-DEL-60 | chr2-174070820-DEL-60 |
| chr2-174150600-DEL-481 | chr2-174150601-DEL-481 |
| chr2-174157307-DEL-129 | chr2-174157329-DEL-129 |
| chr2-174163244-DEL-202 | chr2-174163245-DEL-202 |
| chr2-174181934-DEL-184 | chr2-174181973-DEL-146 |
| chr2-174189472-DEL-162 | chr2-174189473-DEL-162 |
| chr2-174196548-DEL-72 | chr2-174196548-DEL-72 |
| chr2-174209893-DEL-57 | chr2-174209889-DEL-66 |
| chr2-174229932-DEL-3000 | chr2-174229933-DEL-2999 |
| chr2-174236864-DEL-80 | chr2-174236865-DEL-80 |
| chr2-174237242-DEL-242 | chr2-174237243-DEL-244 |
| chr2-174239112-DEL-72 | chr2-174239113-DEL-72 |
| chr2-174240946-DEL-1838 | chr2-174240947-DEL-1838 |
| chr2-174248394-DEL-72 | chr2-174248443-DEL-55 |
| chr2-174252080-DEL-55 | chr2-174252081-DEL-55 |
| chr2-174254331-DEL-62 | chr2-174254334-DEL-62 |
| chr2-174262522-DEL-92 | chr2-174262523-DEL-92 |
| chr2-174272711-DEL-91 | chr2-174272711-DEL-91 |
| chr2-174282602-DEL-70 | chr2-174282593-DEL-80 |
| chr2-174286403-DEL-56 | chr2-174286404-DEL-56 |
| chr2-174320245-DEL-161 | chr2-174320246-DEL-159 |
| chr2-174347337-DEL-2016 | chr2-174347338-DEL-2016 |
| chr2-174365881-DEL-130 | chr2-174365882-DEL-130 |
| chr2-174369401-DEL-63 | chr2-174369430-DEL-70 |
| chr2-174370995-DEL-935 | chr2-174370995-DEL-935 |
| chr2-174386722-DEL-60 | chr2-174386709-DEL-57 |
| chr2-174410287-DEL-85 | chr2-174410328-DEL-57 |
| chr2-174415335-DEL-120 | chr2-174415360-DEL-120 |
| chr2-174416564-DEL-87 | chr2-174416565-DEL-81 |
| chr2-174428477-DEL-147 | chr2-174428478-DEL-147 |
| chr2-174446223-DEL-162 | chr2-174446225-DEL-161 |
| chr2-174467662-DEL-465 | chr2-174467663-DEL-465 |
| chr2-174470337-DEL-57 | chr2-174470316-DEL-57 |
| chr2-174532647-DEL-80 | chr2-174532648-DEL-80 |
| chr2-174533153-DEL-88 | chr2-174533154-DEL-88 |
| chr2-174533900-DEL-112 | chr2-174533854-DEL-140 |
| chr2-174573120-DEL-245 | chr2-174573121-DEL-245 |
| chr2-174628632-DEL-308 | chr2-174628633-DEL-308 |
| chr2-174633311-DEL-74 | chr2-174633312-DEL-74 |
| chr2-174661518-DEL-211 | chr2-174661506-DEL-212 |
| chr2-174667194-DEL-72 | chr2-174667207-DEL-60 |
| chr2-174672172-DEL-166 | chr2-174672173-DEL-166 |
| chr2-176769731-DEL-676 | chr2-176769731-DEL-676 |
| chr2-176772609-DEL-217 | chr2-176772609-DEL-217 |
| chr2-176890023-DEL-1894 | chr2-176890024-DEL-1894 |
| chr2-177005486-DEL-148 | chr2-177005486-DEL-148 |

| | |
|---|---|
| chr2-177802959-DEL-61 | chr2-177802960-DEL-61 |
| chr2-177820636-DEL-189 | chr2-177820608-DEL-194 |
| chr2-177875637-DEL-65 | chr2-177875638-DEL-65 |
| chr2-178084043-DEL-6802 | chr2-178084044-DEL-6802 |
| chr2-178090100-DEL-603 | chr2-178090100-DEL-603 |
| chr2-178094997-DEL-155 | chr2-178094998-DEL-158 |
| chr2-178106602-DEL-112 | chr2-178106593-DEL-100 |
| chr2-178107294-DEL-81 | chr2-178107295-DEL-81 |
| chr2-178112575-DEL-80 | chr2-178112586-DEL-80 |
| chr2-178121282-DEL-176 | chr2-178121283-DEL-176 |
| chr2-178174487-DEL-101 | chr2-178174488-DEL-101 |
| chr2-178175166-DEL-169 | chr2-178175167-DEL-169 |
| chr2-178193944-DEL-77 | chr2-178193899-DEL-91 |
| chr2-178218814-DEL-349 | chr2-178218815-DEL-349 |
| chr2-178225623-DEL-122 | chr2-178225624-DEL-122 |
| chr2-178228593-DEL-154 | chr2-178228594-DEL-154 |
| chr2-178245809-DEL-645 | chr2-178245810-DEL-645 |
| chr2-178253303-DEL-56 | chr2-178253304-DEL-56 |
| chr2-178281114-DEL-318 | chr2-178281099-DEL-318 |
| chr2-178307833-DEL-105 | chr2-178307834-DEL-105 |
| chr2-178357652-DEL-96 | chr2-178357635-DEL-64 |
| chr2-178377382-DEL-57 | chr2-178377381-DEL-59 |
| chr2-178383459-DEL-529 | chr2-178383460-DEL-529 |
| chr2-178386027-DEL-56 | chr2-178386028-DEL-56 |
| chr2-178388072-DEL-155 | chr2-178388073-DEL-124 |
| chr2-178405070-DEL-154 | chr2-178405071-DEL-154 |
| chr2-178415986-DEL-6439 | chr2-178415987-DEL-6439 |
| chr2-178430131-DEL-108 | chr2-178430114-DEL-106 |
| chr2-178438173-DEL-306 | chr2-178438174-DEL-306 |
| chr2-178456456-DEL-74 | chr2-178456457-DEL-74 |
| chr2-178461203-DEL-677 | chr2-178461204-DEL-677 |
| chr2-178474480-DEL-54 | chr2-178474481-DEL-52 |
| chr2-178481548-DEL-218 | chr2-178481549-DEL-218 |
| chr2-178506618-DEL-175 | chr2-178506620-DEL-177 |
| chr2-178516670-DEL-2260 | chr2-178516671-DEL-2260 |
| chr2-178533400-DEL-124 | chr2-178533401-DEL-124 |
| chr2-178607824-DEL-187 | chr2-178607825-DEL-187 |
| chr2-178614449-DEL-51 | chr2-178614450-DEL-51 |
| chr2-178614991-DEL-56 | chr2-178614992-DEL-58 |
| chr2-178628459-DEL-51 | chr2-178628460-DEL-51 |
| chr2-178636041-DEL-3144 | chr2-178636042-DEL-3144 |
| chr2-178643941-DEL-144 | chr2-178643942-DEL-144 |
| chr2-178659968-DEL-58 | chr2-178659972-DEL-70 |
| chr2-178674250-DEL-91 | chr2-178674254-DEL-95 |
| chr2-178674761-DEL-82 | chr2-178674762-DEL-82 |
| chr2-178698492-DEL-63 | chr2-178698490-DEL-64 |
| chr2-178725208-DEL-76 | chr2-178725212-DEL-71 |
| chr2-178737792-DEL-62 | chr2-178737796-DEL-63 |
| chr2-178771318-DEL-111 | chr2-178771319-DEL-111 |
| chr2-178772176-DEL-104 | chr2-178772177-DEL-104 |
| chr2-178819015-DEL-1690 | chr2-178819016-DEL-1690 |
| chr2-178858129-DEL-62 | chr2-178858154-DEL-61 |
| chr2-178866228-DEL-98 | chr2-178866229-DEL-98 |
| chr2-178869103-DEL-412 | chr2-178869104-DEL-412 |
| chr2-178885363-DEL-102 | chr2-178885364-DEL-102 |
| chr2-178898492-DEL-94 | chr2-178898492-DEL-92 |
| chr2-178907921-DEL-51 | chr2-178907902-DEL-51 |
| chr2-178937466-DEL-596 | chr2-178937467-DEL-596 |
| chr2-178940301-DEL-2862 | chr2-178940302-DEL-2862 |
| chr2-178951896-DEL-760 | chr2-178951898-DEL-759 |
| chr2-178974450-DEL-54 | chr2-178974451-DEL-54 |
| chr2-178985614-DEL-86 | chr2-178985614-DEL-86 |
| chr2-179007418-DEL-76 | chr2-179007431-DEL-113 |
| chr2-179064557-DEL-79 | chr2-179064559-DEL-78 |
| chr2-179075005-DEL-182 | chr2-179075005-DEL-212 |
| chr2-179075390-DEL-898 | chr2-179075391-DEL-898 |
| chr2-179077376-DEL-132 | chr2-179077393-DEL-125 |
| chr2-179080063-DEL-52 | chr2-179080064-DEL-52 |
| chr2-179087952-DEL-220 | chr2-179087953-DEL-220 |
| chr2-179121993-DEL-55 | chr2-179121994-DEL-53 |
| chr2-179127569-DEL-72 | chr2-179127570-DEL-72 |
| chr2-179144921-DEL-70 | chr2-179144922-DEL-70 |
| chr2-179177987-DEL-527 | chr2-179177987-DEL-512 |
| chr2-179218386-DEL-158 | chr2-179218376-DEL-158 |
| chr2-179231617-DEL-546 | chr2-179231617-DEL-546 |
| chr2-179247823-DEL-216 | chr2-179247824-DEL-216 |
| chr2-179254250-DEL-2570 | chr2-179254251-DEL-2570 |
| chr2-179260811-DEL-52 | chr2-179260812-DEL-52 |
| chr2-179280448-DEL-170 | chr2-179280449-DEL-170 |
| chr2-179293855-DEL-198 | chr2-179293887-DEL-195 |

| | |
|---|---|
| chr2-179300527-DEL-52 | chr2-179300540-DEL-76 |
| chr2-179339233-DEL-106 | chr2-179339234-DEL-106 |
| chr2-179406611-DEL-226 | chr2-179406612-DEL-226 |
| chr2-179419128-DEL-75 | chr2-179419128-DEL-75 |
| chr2-179419517-DEL-398 | chr2-179419515-DEL-399 |
| chr2-179433918-DEL-56 | chr2-179433919-DEL-56 |
| chr2-179456345-DEL-74 | chr2-179456346-DEL-74 |
| chr2-179456942-DEL-1424 | chr2-179456943-DEL-1424 |
| chr2-179483005-DEL-175 | chr2-179483003-DEL-155 |
| chr2-179486147-DEL-244 | chr2-179486114-DEL-268 |
| chr2-179548162-DEL-216 | chr2-179548163-DEL-216 |
| chr2-179639028-DEL-75 | chr2-179639003-DEL-52 |
| chr2-179645640-DEL-208 | chr2-179645641-DEL-208 |
| chr2-179702077-DEL-175 | chr2-179702078-DEL-175 |
| chr2-179722221-DEL-202 | chr2-179722222-DEL-202 |
| chr2-179733206-DEL-1893 | chr2-179733211-DEL-1893 |
| chr2-179776943-DEL-190 | chr2-179776944-DEL-190 |
| chr2-179826769-DEL-90 | chr2-179826814-DEL-90 |
| chr2-179841444-DEL-835 | chr2-179841445-DEL-835 |
| chr2-179854490-DEL-202 | chr2-179854491-DEL-202 |
| chr2-179857734-DEL-105 | chr2-179857734-DEL-105 |
| chr2-179874129-DEL-121 | chr2-179874124-DEL-136 |
| chr2-179885798-DEL-889 | chr2-179885799-DEL-889 |
| chr2-179891792-DEL-81 | chr2-179891793-DEL-81 |
| chr2-179892585-DEL-202 | chr2-179892586-DEL-202 |
| chr2-179895169-DEL-185 | chr2-179895149-DEL-194 |
| chr2-179941266-DEL-78 | chr2-179941295-DEL-53 |
| chr2-179955478-DEL-52 | chr2-179955457-DEL-62 |
| chr2-179958317-DEL-408 | chr2-179958318-DEL-408 |
| chr2-179995344-DEL-62 | chr2-179995367-DEL-60 |
| chr2-180012255-DEL-189 | chr2-180012255-DEL-189 |
| chr2-180039192-DEL-54 | chr2-180039191-DEL-56 |
| chr2-180048897-DEL-92 | chr2-180048898-DEL-92 |
| chr2-180117163-DEL-280 | chr2-180117164-DEL-280 |
| chr2-180183104-DEL-56 | chr2-180183105-DEL-56 |
| chr2-180193103-DEL-276 | chr2-180193103-DEL-276 |
| chr2-180194123-DEL-66 | chr2-180194138-DEL-84 |
| chr2-180237939-DEL-78 | chr2-180237940-DEL-78 |
| chr2-180312406-DEL-52 | chr2-180312407-DEL-52 |
| chr2-180357077-DEL-213 | chr2-180357078-DEL-213 |
| chr2-180393420-DEL-207 | chr2-180393421-DEL-207 |
| chr2-180399420-DEL-181 | chr2-180399421-DEL-181 |
| chr2-180425851-DEL-84 | chr2-180425860-DEL-100 |
| chr2-180427318-DEL-186 | chr2-180427319-DEL-186 |
| chr2-180450077-DEL-124 | chr2-180450082-DEL-120 |
| chr2-180451530-DEL-107 | chr2-180451531-DEL-107 |
| chr2-180476497-DEL-170 | chr2-180476498-DEL-170 |
| chr2-180493369-DEL-479 | chr2-180493370-DEL-479 |
| chr2-180506827-DEL-1889 | chr2-180506828-DEL-1889 |
| chr2-180513480-DEL-67 | chr2-180513481-DEL-67 |
| chr2-180514843-DEL-286 | chr2-180514844-DEL-286 |
| chr2-180563589-DEL-586 | chr2-180563598-DEL-584 |
| chr2-180582801-DEL-195 | chr2-180582802-DEL-195 |
| chr2-180704503-DEL-203 | chr2-180704504-DEL-203 |
| chr2-180811740-DEL-3990 | chr2-180811746-DEL-3983 |
| chr2-180818892-DEL-162 | chr2-180818893-DEL-162 |
| chr2-180833256-DEL-516 | chr2-180833257-DEL-516 |
| chr2-180880622-DEL-246 | chr2-180880623-DEL-246 |
| chr2-180986188-DEL-273 | chr2-180986196-DEL-272 |
| chr2-181004798-DEL-207 | chr2-181004799-DEL-207 |
| chr2-181069778-DEL-164 | chr2-181069790-DEL-187 |
| chr2-181081168-DEL-52 | chr2-181081167-DEL-54 |
| chr2-181095334-DEL-108 | chr2-181095341-DEL-58 |
| chr2-181116329-DEL-217 | chr2-181116321-DEL-220 |
| chr2-181119061-DEL-211 | chr2-181119062-DEL-211 |
| chr2-181124669-DEL-84 | chr2-181124670-DEL-51 |
| chr2-181137837-DEL-205 | chr2-181137838-DEL-205 |
| chr2-181186984-DEL-244 | chr2-181186985-DEL-244 |
| chr2-181197374-DEL-68 | chr2-181197381-DEL-63 |
| chr2-181202320-DEL-222 | chr2-181202321-DEL-222 |
| chr2-181205628-DEL-210 | chr2-181205629-DEL-210 |
| chr2-181206688-DEL-55 | chr2-181206689-DEL-55 |
| chr2-181245978-DEL-74 | chr2-181246006-DEL-90 |
| chr2-181269760-DEL-175 | chr2-181269761-DEL-175 |
| chr2-181280502-DEL-508 | chr2-181280502-DEL-506 |
| chr2-181288284-DEL-207 | chr2-181288285-DEL-207 |
| chr2-181341740-DEL-218 | chr2-181341741-DEL-218 |
| chr2-181346602-DEL-187 | chr2-181346616-DEL-185 |
| chr2-181381564-DEL-6389 | chr2-181381565-DEL-6389 |
| chr2-181392689-DEL-597 | chr2-181392686-DEL-598 |

| | |
|---|---|
| chr2-181440493-DEL-104 | chr2-181440505-DEL-76 |
| chr2-181521401-DEL-448 | chr2-181521402-DEL-448 |
| chr2-181530724-DEL-189 | chr2-181530725-DEL-189 |
| chr2-181558664-DEL-400 | chr2-181558665-DEL-400 |
| chr2-181561079-DEL-8810 | chr2-181561080-DEL-8810 |
| chr2-181599554-DEL-216 | chr2-181599554-DEL-217 |
| chr2-181607783-DEL-559 | chr2-181607784-DEL-559 |
| chr3-3008197-DEL-117 | chr3-3008195-DEL-117 |
| chr3-3038667-DEL-63 | chr3-3038668-DEL-63 |
| chr3-3046333-DEL-3119 | chr3-3046333-DEL-3119 |
| chr3-3048970-DEL-51 | chr3-3048971-DEL-51 |
| chr3-3052258-DEL-83 | chr3-3052259-DEL-83 |
| chr3-3054105-DEL-1583 | chr3-3054105-DEL-1583 |
| chr3-3062400-DEL-104 | chr3-3062401-DEL-104 |
| chr3-3091150-DEL-399 | chr3-3091151-DEL-399 |
| chr3-3101915-DEL-201 | chr3-3101916-DEL-201 |
| chr3-3181454-DEL-12302 | chr3-3181456-DEL-12303 |
| chr3-3232331-DEL-94 | chr3-3232332-DEL-94 |
| chr3-3242185-DEL-573 | chr3-3242186-DEL-573 |
| chr3-3291571-DEL-415 | chr3-3291572-DEL-415 |
| chr3-3297346-DEL-345 | chr3-3297347-DEL-345 |
| chr3-3378607-DEL-6405 | chr3-3378608-DEL-6405 |
| chr3-3398931-DEL-73 | chr3-3398896-DEL-63 |
| chr3-3428266-DEL-814 | chr3-3428267-DEL-814 |
| chr3-3471605-DEL-399 | chr3-3471594-DEL-411 |
| chr3-3506473-DEL-219 | chr3-3506474-DEL-219 |
| chr3-3523100-DEL-686 | chr3-3523099-DEL-687 |
| chr3-3529831-DEL-498 | chr3-3529832-DEL-498 |
| chr3-3599486-DEL-72 | chr3-3599487-DEL-72 |
| chr3-3628684-DEL-54 | chr3-3628684-DEL-54 |
| chr3-3701067-DEL-184 | chr3-3701068-DEL-184 |
| chr3-3723509-DEL-56 | chr3-3723508-DEL-56 |
| chr3-3748251-DEL-938 | chr3-3748252-DEL-938 |
| chr3-3752239-DEL-7226 | chr3-3752240-DEL-7226 |
| chr3-3786291-DEL-5233 | chr3-3786292-DEL-5233 |
| chr3-3819413-DEL-167 | chr3-3819365-DEL-168 |
| chr3-3897770-DEL-316 | chr3-3897771-DEL-316 |
| chr3-3914072-DEL-258 | chr3-3914073-DEL-258 |
| chr3-3930290-DEL-4417 | chr3-3930291-DEL-4417 |
| chr3-3975350-DEL-3317 | chr3-3975351-DEL-3317 |
| chr3-3978315-DEL-58 | chr3-3978308-DEL-58 |
| chr3-3980898-DEL-164 | chr3-3980899-DEL-164 |
| chr3-3986287-DEL-2038 | chr3-3986288-DEL-2038 |
| chr3-3994026-DEL-6692 | chr3-3994018-DEL-6693 |
| chr3-4036204-DEL-142 | chr3-4036205-DEL-142 |
| chr3-4069084-DEL-2930 | chr3-4069085-DEL-2930 |
| chr3-4101314-DEL-256 | chr3-4101315-DEL-255 |
| chr3-4123064-DEL-146 | chr3-4123065-DEL-146 |
| chr3-4127873-DEL-4425 | chr3-4127874-DEL-4425 |
| chr3-4220903-DEL-7436 | chr3-4220903-DEL-7436 |
| chr3-4229240-DEL-6674 | chr3-4229241-DEL-6674 |
| chr3-4249668-DEL-478 | chr3-4249669-DEL-478 |
| chr3-4255112-DEL-6609 | chr3-4255113-DEL-6609 |
| chr3-4267305-DEL-203 | chr3-4267306-DEL-201 |
| chr3-4276553-DEL-2358 | chr3-4276554-DEL-2359 |
| chr3-4288487-DEL-969 | chr3-4288488-DEL-969 |
| chr3-4313429-DEL-69 | chr3-4313429-DEL-61 |
| chr3-4353316-DEL-371 | chr3-4353317-DEL-371 |
| chr3-4356723-DEL-326 | chr3-4356724-DEL-326 |
| chr3-4374280-DEL-2842 | chr3-4374281-DEL-2842 |
| chr3-4406556-DEL-403 | chr3-4406556-DEL-403 |
| chr3-4420062-DEL-90 | chr3-4420067-DEL-86 |
| chr3-4431578-DEL-774 | chr3-4431579-DEL-774 |
| chr3-4491733-DEL-498 | chr3-4491734-DEL-498 |
| chr3-4501977-DEL-6216 | chr3-4501978-DEL-6216 |
| chr3-4520493-DEL-179 | chr3-4520494-DEL-179 |
| chr3-4605170-DEL-51 | chr3-4605159-DEL-52 |
| chr3-4649563-DEL-6880 | chr3-4649564-DEL-6880 |
| chr3-4674424-DEL-382 | chr3-4674425-DEL-382 |
| chr3-4676200-DEL-399 | chr3-4676201-DEL-399 |
| chr3-4698690-DEL-3679 | chr3-4698691-DEL-3679 |
| chr3-4762691-DEL-119 | chr3-4762727-DEL-72 |
| chr3-4783364-DEL-7205 | chr3-4783365-DEL-7205 |
| chr3-4806717-DEL-173 | chr3-4806718-DEL-173 |
| chr3-4883173-DEL-6734 | chr3-4883169-DEL-6735 |
| chr3-4889819-DEL-54 | chr3-4889818-DEL-56 |
| chr3-4896992-DEL-89 | chr3-4896993-DEL-89 |
| chr3-4926516-DEL-325 | chr3-4926513-DEL-326 |
| chr3-5018714-DEL-51 | chr3-5018715-DEL-51 |
| chr3-5030761-DEL-76 | chr3-5030782-DEL-56 |

| | |
|---|---|
| chr3-5045682-DEL-69 | chr3-5045683-DEL-69 |
| chr3-5071128-DEL-1566 | chr3-5071129-DEL-1566 |
| chr3-5074452-DEL-422 | chr3-5074445-DEL-395 |
| chr3-5114062-DEL-6843 | chr3-5114063-DEL-6843 |
| chr3-5134936-DEL-6433 | chr3-5134937-DEL-6433 |
| chr3-5134992-DEL-201 | chr3-5134993-DEL-201 |
| chr3-5146314-DEL-81 | chr3-5146315-DEL-81 |
| chr3-5254849-DEL-71 | chr3-5254850-DEL-71 |
| chr3-5368076-DEL-1547 | chr3-5368077-DEL-1547 |
| chr3-5447066-DEL-332 | chr3-5447067-DEL-332 |
| chr3-5484527-DEL-52 | chr3-5484528-DEL-52 |
| chr3-5500469-DEL-7053 | chr3-5500469-DEL-7053 |
| chr3-5500482-DEL-411 | chr3-5500483-DEL-411 |
| chr3-5529411-DEL-579 | chr3-5529412-DEL-579 |
| chr3-5530949-DEL-233 | chr3-5530950-DEL-233 |
| chr3-5563829-DEL-1331 | chr3-5563830-DEL-1331 |
| chr3-5582804-DEL-7187 | chr3-5582805-DEL-7187 |
| chr3-5605516-DEL-52 | chr3-5605517-DEL-52 |
| chr3-5617646-DEL-71 | chr3-5617653-DEL-66 |
| chr3-5656521-DEL-1108 | chr3-5656522-DEL-1108 |
| chr3-5668055-DEL-1220 | chr3-5668053-DEL-1221 |
| chr3-5677423-DEL-62 | chr3-5677424-DEL-62 |
| chr3-5684599-DEL-773 | chr3-5684600-DEL-773 |
| chr3-5706264-DEL-1172 | chr3-5706265-DEL-1172 |
| chr3-5713008-DEL-6412 | chr3-5713009-DEL-6412 |
| chr3-5732425-DEL-56 | chr3-5732426-DEL-51 |
| chr3-5749399-DEL-110 | chr3-5749400-DEL-110 |
| chr3-5752790-DEL-212 | chr3-5752791-DEL-211 |
| chr3-5755429-DEL-591 | chr3-5755430-DEL-591 |
| chr3-5757895-DEL-1524 | chr3-5757896-DEL-1524 |
| chr3-5770192-DEL-944 | chr3-5770193-DEL-944 |
| chr3-5798322-DEL-56 | chr3-5798323-DEL-56 |
| chr3-5810326-DEL-398 | chr3-5810327-DEL-398 |
| chr3-5829014-DEL-461 | chr3-5829015-DEL-461 |
| chr3-5847575-DEL-58 | chr3-5847578-DEL-56 |
| chr3-5852086-DEL-103 | chr3-5852084-DEL-104 |
| chr3-5905626-DEL-63 | chr3-5905627-DEL-63 |
| chr3-5925383-DEL-3209 | chr3-5925384-DEL-3209 |
| chr3-5945888-DEL-5077 | chr3-5945889-DEL-5077 |
| chr3-5956605-DEL-63 | chr3-5956606-DEL-63 |
| chr3-5963365-DEL-79 | chr3-5963366-DEL-79 |
| chr3-5970163-DEL-1835 | chr3-5970164-DEL-1835 |
| chr3-5973189-DEL-220 | chr3-5973190-DEL-220 |
| chr3-5989172-DEL-56 | chr3-5989173-DEL-56 |
| chr3-5991220-DEL-138 | chr3-5991197-DEL-135 |
| chr3-6009003-DEL-6534 | chr3-6009004-DEL-6534 |
| chr3-6018710-DEL-1286 | chr3-6018711-DEL-1286 |
| chr3-6034478-DEL-58 | chr3-6034472-DEL-71 |
| chr3-6036240-DEL-1541 | chr3-6036241-DEL-1541 |
| chr3-6045956-DEL-144 | chr3-6045969-DEL-132 |
| chr3-6048472-DEL-204 | chr3-6048473-DEL-204 |
| chr3-6060007-DEL-90 | chr3-6060038-DEL-51 |
| chr3-6119427-DEL-4828 | chr3-6119428-DEL-4828 |
| chr3-6138844-DEL-3642 | chr3-6138845-DEL-3642 |
| chr3-6143473-DEL-103 | chr3-6143474-DEL-103 |
| chr3-6152093-DEL-358 | chr3-6152094-DEL-358 |
| chr3-6180673-DEL-57 | chr3-6180674-DEL-57 |
| chr3-6187861-DEL-69 | chr3-6187862-DEL-69 |
| chr3-6193143-DEL-350 | chr3-6193144-DEL-350 |
| chr3-6222599-DEL-7077 | chr3-6222601-DEL-7076 |
| chr3-6256800-DEL-733 | chr3-6256805-DEL-729 |
| chr3-6362454-DEL-18334 | chr3-6362455-DEL-18334 |
| chr3-6376450-DEL-2182 | chr3-6376451-DEL-2182 |
| chr3-6401615-DEL-199 | chr3-6401616-DEL-199 |
| chr3-6403734-DEL-5095 | chr3-6403735-DEL-5095 |
| chr3-6412405-DEL-474 | chr3-6412406-DEL-474 |
| chr3-6426655-DEL-179 | chr3-6426656-DEL-179 |
| chr3-6456991-DEL-409 | chr3-6456992-DEL-409 |
| chr3-6489200-DEL-1819 | chr3-6489200-DEL-1819 |
| chr3-6520913-DEL-6219 | chr3-6520914-DEL-6219 |
| chr3-6545773-DEL-357 | chr3-6545773-DEL-357 |
| chr3-6547677-DEL-55 | chr3-6547664-DEL-52 |
| chr3-6639194-DEL-55 | chr3-6639195-DEL-55 |
| chr3-6641268-DEL-208 | chr3-6641269-DEL-208 |
| chr3-6666860-DEL-6825 | chr3-6666861-DEL-6825 |
| chr3-6686404-DEL-6112 | chr3-6686405-DEL-6112 |
| chr3-6705953-DEL-72 | chr3-6705954-DEL-72 |
| chr3-6732072-DEL-495 | chr3-6732073-DEL-495 |
| chr3-6744009-DEL-1638 | chr3-6744010-DEL-1638 |
| chr3-6764065-DEL-3050 | chr3-6764066-DEL-3050 |

| | |
|---|---|
| chr3-6769454-DEL-66 | chr3-6769455-DEL-66 |
| chr3-6801809-DEL-65 | chr3-6801807-DEL-58 |
| chr3-6803461-DEL-6400 | chr3-6803444-DEL-6401 |
| chr3-6803744-DEL-199 | chr3-6803745-DEL-199 |
| chr3-6822110-DEL-5013 | chr3-6822111-DEL-5013 |
| chr3-6848867-DEL-1725 | chr3-6848868-DEL-1725 |
| chr3-6865583-DEL-71 | chr3-6865584-DEL-71 |
| chr3-6879391-DEL-962 | chr3-6879392-DEL-962 |
| chr3-6915012-DEL-6512 | chr3-6915013-DEL-6512 |
| chr3-6934901-DEL-919 | chr3-6934902-DEL-919 |
| chr3-6960038-DEL-372 | chr3-6960039-DEL-372 |
| chr3-6974986-DEL-6437 | chr3-6974987-DEL-6437 |
| chr3-6985956-DEL-75 | chr3-6985957-DEL-75 |
| chr3-7004891-DEL-1221 | chr3-7004892-DEL-1221 |
| chr3-7099210-DEL-197 | chr3-7099178-DEL-196 |
| chr3-7134739-DEL-680 | chr3-7134740-DEL-680 |
| chr3-7184707-DEL-116 | chr3-7184708-DEL-116 |
| chr3-7201098-DEL-488 | chr3-7201099-DEL-492 |
| chr3-7238054-DEL-55 | chr3-7238055-DEL-55 |
| chr3-7254653-DEL-10371 | chr3-7254654-DEL-10371 |
| chr3-7282362-DEL-6851 | chr3-7282363-DEL-6851 |
| chr3-7315362-DEL-168 | chr3-7315353-DEL-184 |
| chr3-7320631-DEL-327 | chr3-7320632-DEL-327 |
| chr3-7331560-DEL-1114 | chr3-7331561-DEL-1114 |
| chr3-7332813-DEL-324 | chr3-7332814-DEL-324 |
| chr3-7344180-DEL-69 | chr3-7344187-DEL-65 |
| chr3-7347130-DEL-1016 | chr3-7347132-DEL-1016 |
| chr3-7366543-DEL-3186 | chr3-7366544-DEL-3186 |
| chr3-7376188-DEL-6453 | chr3-7376189-DEL-6453 |
| chr3-7382630-DEL-303 | chr3-7382640-DEL-294 |
| chr3-7391513-DEL-1500 | chr3-7391513-DEL-1500 |
| chr3-7401781-DEL-231 | chr3-7401782-DEL-231 |
| chr3-7405823-DEL-92 | chr3-7405820-DEL-95 |
| chr3-7433651-DEL-58 | chr3-7433653-DEL-56 |
| chr3-7434794-DEL-52 | chr3-7434795-DEL-52 |
| chr3-7463367-DEL-1384 | chr3-7463368-DEL-1384 |
| chr3-7504948-DEL-139 | chr3-7504949-DEL-139 |
| chr3-7548622-DEL-64 | chr3-7548623-DEL-64 |
| chr3-7571483-DEL-100 | chr3-7571484-DEL-100 |
| chr3-7582427-DEL-72 | chr3-7582393-DEL-67 |
| chr3-7585453-DEL-74 | chr3-7585454-DEL-74 |
| chr3-7598247-DEL-110 | chr3-7598248-DEL-110 |
| chr3-7603599-DEL-303 | chr3-7603594-DEL-304 |
| chr3-7613505-DEL-523 | chr3-7613506-DEL-523 |
| chr3-7628094-DEL-5523 | chr3-7628095-DEL-5523 |
| chr3-7661387-DEL-60 | chr3-7661388-DEL-75 |
| chr3-7730611-DEL-435 | chr3-7730612-DEL-435 |
| chr3-7738399-DEL-2299 | chr3-7738395-DEL-2296 |
| chr3-7794271-DEL-462 | chr3-7794272-DEL-462 |
| chr3-7795946-DEL-51 | chr3-7795947-DEL-51 |
| chr3-7797765-DEL-6424 | chr3-7797766-DEL-6424 |
| chr3-7805741-DEL-2997 | chr3-7805742-DEL-2997 |
| chr3-7840959-DEL-2920 | chr3-7840960-DEL-2920 |
| chr3-7853993-DEL-94 | chr3-7853994-DEL-94 |
| chr3-7864828-DEL-56 | chr3-7864829-DEL-56 |
| chr3-7899248-DEL-2368 | chr3-7899249-DEL-2368 |
| chr3-7914970-DEL-2085 | chr3-7914971-DEL-2085 |
| chr3-7921522-DEL-6821 | chr3-7921523-DEL-6821 |
| chr3-7975398-DEL-6244 | chr3-7975399-DEL-6244 |
| chr3-8026126-DEL-1535 | chr3-8026127-DEL-1535 |
| chr3-8107513-DEL-119 | chr3-8107479-DEL-151 |
| chr3-8128338-DEL-1478 | chr3-8128338-DEL-1478 |
| chr3-8134907-DEL-1347 | chr3-8134908-DEL-1347 |
| chr3-8136522-DEL-710 | chr3-8136522-DEL-710 |
| chr3-8168227-DEL-2883 | chr3-8168228-DEL-2879 |
| chr3-8194659-DEL-6848 | chr3-8194651-DEL-6856 |
| chr3-8249671-DEL-6061 | chr3-8249672-DEL-6061 |
| chr3-8302861-DEL-530 | chr3-8302864-DEL-528 |
| chr3-8314688-DEL-54 | chr3-8314689-DEL-54 |
| chr3-8327570-DEL-472 | chr3-8327571-DEL-472 |
| chr3-8332921-DEL-111 | chr3-8332922-DEL-136 |
| chr3-8340622-DEL-52 | chr3-8340623-DEL-52 |
| chr3-8362797-DEL-64 | chr3-8362798-DEL-64 |
| chr3-8364668-DEL-2266 | chr3-8364669-DEL-2266 |
| chr3-8395054-DEL-89 | chr3-8395054-DEL-89 |
| chr3-8409648-DEL-376 | chr3-8409649-DEL-376 |
| chr3-8412511-DEL-85 | chr3-8412512-DEL-85 |
| chr3-8425042-DEL-180 | chr3-8425040-DEL-179 |
| chr3-8458130-DEL-208 | chr3-8458131-DEL-208 |
| chr3-8462919-DEL-177 | chr3-8462920-DEL-177 |

| | |
|---|---|
| chr3-8468423-DEL-633 | chr3-8468423-DEL-633 |
| chr3-8473384-DEL-381 | chr3-8473384-DEL-381 |
| chr3-8494202-DEL-4188 | chr3-8494203-DEL-4188 |
| chr3-8499524-DEL-1350 | chr3-8499525-DEL-1350 |
| chr3-8510294-DEL-6869 | chr3-8510295-DEL-6869 |
| chr3-8519349-DEL-80 | chr3-8519362-DEL-72 |
| chr3-8519743-DEL-610 | chr3-8519744-DEL-610 |
| chr3-8529169-DEL-107 | chr3-8529167-DEL-108 |
| chr3-8530296-DEL-363 | chr3-8530297-DEL-363 |
| chr3-8553231-DEL-66 | chr3-8553213-DEL-57 |
| chr3-8561209-DEL-1128 | chr3-8561210-DEL-1128 |
| chr3-8587560-DEL-213 | chr3-8587579-DEL-195 |
| chr3-8592790-DEL-1166 | chr3-8592791-DEL-1166 |
| chr3-8595998-DEL-445 | chr3-8595998-DEL-443 |
| chr3-8600295-DEL-2529 | chr3-8600296-DEL-2529 |
| chr3-8614190-DEL-138 | chr3-8614191-DEL-138 |
| chr3-8639126-DEL-2756 | chr3-8639127-DEL-2756 |
| chr3-8647218-DEL-1852 | chr3-8647219-DEL-1852 |
| chr3-8662340-DEL-84 | chr3-8662352-DEL-89 |
| chr3-8712882-DEL-183 | chr3-8712883-DEL-183 |
| chr3-8726436-DEL-147 | chr3-8726437-DEL-147 |
| chr3-8738270-DEL-248 | chr3-8738271-DEL-248 |
| chr3-8748164-DEL-64 | chr3-8748165-DEL-64 |
| chr3-8751162-DEL-1212 | chr3-8751163-DEL-1212 |
| chr3-8764267-DEL-92 | chr3-8764268-DEL-92 |
| chr3-8788029-DEL-82 | chr3-8788030-DEL-82 |
| chr3-8800598-DEL-280 | chr3-8800599-DEL-280 |
| chr3-8830838-DEL-64 | chr3-8830838-DEL-64 |
| chr3-8839417-DEL-676 | chr3-8839418-DEL-676 |
| chr3-8868794-DEL-2049 | chr3-8868795-DEL-2049 |
| chr3-8874263-DEL-182 | chr3-8874263-DEL-180 |
| chr3-8911302-DEL-67 | chr3-8911301-DEL-68 |
| chr3-8946834-DEL-339 | chr3-8946835-DEL-339 |
| chr3-8955224-DEL-112 | chr3-8955192-DEL-104 |
| chr3-8960458-DEL-59 | chr3-8960432-DEL-91 |
| chr3-9002430-DEL-180 | chr3-9002431-DEL-180 |
| chr3-9018066-DEL-51 | chr3-9018067-DEL-51 |
| chr3-9018598-DEL-76 | chr3-9018598-DEL-76 |
| chr3-9019615-DEL-230 | chr3-9019616-DEL-230 |
| chr3-9025304-DEL-208 | chr3-9025305-DEL-208 |
| chr3-9043867-DEL-193 | chr3-9043868-DEL-193 |
| chr3-9045709-DEL-184 | chr3-9045710-DEL-184 |
| chr3-9050055-DEL-512 | chr3-9050056-DEL-512 |
| chr3-9056612-DEL-78 | chr3-9056613-DEL-78 |
| chr3-9058208-DEL-54 | chr3-9058224-DEL-54 |
| chr3-9063219-DEL-200 | chr3-9063220-DEL-200 |
| chr3-9070053-DEL-139 | chr3-9070075-DEL-114 |
| chr3-9102323-DEL-106 | chr3-9102320-DEL-110 |
| chr3-9105330-DEL-177 | chr3-9105330-DEL-177 |
| chr3-9108187-DEL-190 | chr3-9108187-DEL-190 |
| chr3-9110219-DEL-194 | chr3-9110219-DEL-194 |
| chr3-9113869-DEL-74 | chr3-9113897-DEL-74 |
| chr3-9115037-DEL-411 | chr3-9115037-DEL-411 |
| chr3-9119246-DEL-710 | chr3-9119247-DEL-710 |
| chr3-9133604-DEL-57 | chr3-9133603-DEL-99 |
| chr3-9138236-DEL-115 | chr3-9138236-DEL-111 |
| chr3-9151076-DEL-149 | chr3-9151068-DEL-148 |
| chr3-9156509-DEL-5314 | chr3-9156510-DEL-5314 |
| chr3-9204517-DEL-57 | chr3-9204506-DEL-58 |
| chr3-9232856-DEL-400 | chr3-9232857-DEL-400 |
| chr3-9237102-DEL-52 | chr3-9237103-DEL-52 |
| chr3-9250223-DEL-343 | chr3-9250224-DEL-343 |
| chr3-9255938-DEL-178 | chr3-9255939-DEL-178 |
| chr3-9278161-DEL-205 | chr3-9278162-DEL-205 |
| chr3-9295269-DEL-218 | chr3-9295270-DEL-218 |
| chr3-9301702-DEL-64 | chr3-9301703-DEL-66 |
| chr3-9386310-DEL-87 | chr3-9386279-DEL-99 |
| chr3-9447541-DEL-2019 | chr3-9447542-DEL-2019 |
| chr3-9522389-DEL-211 | chr3-9522389-DEL-211 |
| chr3-9528276-DEL-401 | chr3-9528278-DEL-400 |
| chr3-9551505-DEL-920 | chr3-9551506-DEL-920 |
| chr3-9553323-DEL-201 | chr3-9553324-DEL-201 |
| chr3-9572963-DEL-249 | chr3-9572964-DEL-249 |
| chr3-9581471-DEL-59 | chr3-9581472-DEL-52 |
| chr3-9592843-DEL-790 | chr3-9592869-DEL-756 |
| chr3-9601168-DEL-179 | chr3-9601169-DEL-179 |
| chr3-9610382-DEL-112 | chr3-9610387-DEL-108 |
| chr3-9625937-DEL-124 | chr3-9625938-DEL-124 |
| chr3-9658704-DEL-126 | chr3-9658705-DEL-84 |
| chr3-9699521-DEL-196 | chr3-9699522-DEL-196 |

| | |
|---|---|
| chr3-9707480-DEL-57 | chr3-9707481-DEL-57 |
| chr3-9720621-DEL-63 | chr3-9720622-DEL-63 |
| chr3-9735612-DEL-135 | chr3-9735613-DEL-135 |
| chr3-9748081-DEL-108 | chr3-9748082-DEL-108 |
| chr3-9753506-DEL-53 | chr3-9753507-DEL-53 |
| chr3-9779496-DEL-366 | chr3-9779499-DEL-364 |
| chr3-9781766-DEL-60 | chr3-9781767-DEL-60 |
| chr3-9796223-DEL-7158 | chr3-9796224-DEL-7158 |
| chr3-9842629-DEL-816 | chr3-9842654-DEL-816 |
| chr3-9843181-DEL-192 | chr3-9843182-DEL-192 |
| chr3-9867392-DEL-246 | chr3-9867393-DEL-246 |
| chr3-9903202-DEL-440 | chr3-9903203-DEL-440 |
| chr3-9947501-DEL-140 | chr3-9947520-DEL-136 |
| chr3-9969124-DEL-6659 | chr3-9969125-DEL-6659 |
| chr3-9973814-DEL-76 | chr3-9973815-DEL-76 |
| chr3-9986039-DEL-234 | chr3-9986040-DEL-234 |
| chr3-9993690-DEL-90 | chr3-9993680-DEL-91 |
| chr3-10013617-DEL-7546 | chr3-10013618-DEL-7546 |
| chr3-10026322-DEL-6333 | chr3-10026331-DEL-6325 |
| chr3-10032395-DEL-198 | chr3-10032374-DEL-199 |
| chr3-10049686-DEL-81 | chr3-10049686-DEL-81 |
| chr3-10053516-DEL-7510 | chr3-10053517-DEL-7510 |
| chr3-10054597-DEL-1023 | chr3-10054598-DEL-1023 |
| chr3-10061916-DEL-89 | chr3-10061917-DEL-89 |
| chr3-10085060-DEL-68 | chr3-10085060-DEL-68 |
| chr3-10089445-DEL-236 | chr3-10089446-DEL-236 |
| chr3-10099384-DEL-63 | chr3-10099385-DEL-63 |
| chr3-10103496-DEL-419 | chr3-10103496-DEL-386 |
| chr3-10118761-DEL-128 | chr3-10118762-DEL-128 |
| chr3-10122296-DEL-363 | chr3-10122297-DEL-363 |
| chr3-10125493-DEL-1386 | chr3-10125483-DEL-1387 |
| chr3-10148758-DEL-64 | chr3-10148759-DEL-64 |
| chr3-10151562-DEL-3108 | chr3-10151563-DEL-3108 |
| chr3-10153154-DEL-2131 | chr3-10153155-DEL-2131 |
| chr3-10162938-DEL-4792 | chr3-10162939-DEL-4792 |
| chr3-10167302-DEL-60 | chr3-10167255-DEL-60 |
| chr3-10169543-DEL-91 | chr3-10169544-DEL-91 |
| chr3-10177321-DEL-53 | chr3-10177322-DEL-53 |
| chr3-10177986-DEL-1898 | chr3-10177987-DEL-1898 |
| chr3-10202769-DEL-80 | chr3-10202770-DEL-80 |
| chr3-10207302-DEL-204 | chr3-10207303-DEL-204 |
| chr3-10243167-DEL-60 | chr3-10243168-DEL-60 |
| chr3-10276377-DEL-123 | chr3-10276378-DEL-123 |
| chr3-10277466-DEL-76 | chr3-10277466-DEL-106 |
| chr3-10285372-DEL-245 | chr3-10285373-DEL-245 |
| chr3-10314329-DEL-332 | chr3-10314330-DEL-332 |
| chr3-10338006-DEL-58 | chr3-10338007-DEL-56 |
| chr3-10341305-DEL-60 | chr3-10341306-DEL-60 |
| chr3-10361683-DEL-180 | chr3-10361684-DEL-180 |
| chr3-10372450-DEL-194 | chr3-10372451-DEL-194 |
| chr3-10373269-DEL-283 | chr3-10373276-DEL-277 |
| chr3-10380651-DEL-76 | chr3-10380640-DEL-77 |
| chr3-10390239-DEL-63 | chr3-10390240-DEL-63 |
| chr3-10408210-DEL-77 | chr3-10408211-DEL-77 |
| chr3-10412973-DEL-526 | chr3-10412974-DEL-526 |
| chr3-10418026-DEL-291 | chr3-10418027-DEL-291 |
| chr3-10429771-DEL-94 | chr3-10429772-DEL-94 |
| chr3-10445952-DEL-192 | chr3-10445953-DEL-192 |
| chr3-10449301-DEL-204 | chr3-10449302-DEL-204 |
| chr3-10462181-DEL-71 | chr3-10462182-DEL-71 |
| chr3-10481635-DEL-165 | chr3-10481636-DEL-165 |
| chr3-10507227-DEL-268 | chr3-10507228-DEL-268 |
| chr3-10530746-DEL-199 | chr3-10530747-DEL-199 |
| chr3-10558845-DEL-58 | chr3-10558848-DEL-57 |
| chr3-10562397-DEL-519 | chr3-10562398-DEL-519 |
| chr3-10570264-DEL-1288 | chr3-10570265-DEL-1288 |
| chr3-10570924-DEL-145 | chr3-10570925-DEL-145 |
| chr3-10601970-DEL-2978 | chr3-10601971-DEL-2978 |
| chr3-10617288-DEL-221 | chr3-10617289-DEL-221 |
| chr3-10617303-DEL-868 | chr3-10617302-DEL-870 |
| chr3-10656416-DEL-2908 | chr3-10656416-DEL-2908 |
| chr3-10658673-DEL-1101 | chr3-10658673-DEL-1101 |
| chr3-10660092-DEL-102 | chr3-10660077-DEL-98 |
| chr3-10717136-DEL-65 | chr3-10717135-DEL-66 |
| chr3-10741314-DEL-2344 | chr3-10741315-DEL-2344 |
| chr3-10744409-DEL-1667 | chr3-10744411-DEL-1668 |
| chr3-10747151-DEL-104 | chr3-10747132-DEL-84 |
| chr3-10776607-DEL-61 | chr3-10776608-DEL-61 |
| chr3-10792375-DEL-338 | chr3-10792376-DEL-338 |
| chr3-10795357-DEL-192 | chr3-10795358-DEL-192 |

| | |
|---|---|
| chr3-10809149-DEL-60 | chr3-10809150-DEL-58 |
| chr3-10813510-DEL-1273 | chr3-10813511-DEL-1273 |
| chr3-10859308-DEL-691 | chr3-10859311-DEL-692 |
| chr3-10862525-DEL-63 | chr3-10862514-DEL-59 |
| chr3-10869215-DEL-6785 | chr3-10869216-DEL-6785 |
| chr3-10878579-DEL-155 | chr3-10878622-DEL-113 |
| chr3-10883165-DEL-102 | chr3-10883162-DEL-100 |
| chr3-10890992-DEL-75 | chr3-10890990-DEL-60 |
| chr3-10891752-DEL-489 | chr3-10891753-DEL-489 |
| chr3-10905679-DEL-6212 | chr3-10905680-DEL-6212 |
| chr3-10940249-DEL-8772 | chr3-10940250-DEL-8772 |
| chr3-10940961-DEL-1530 | chr3-10940962-DEL-1530 |
| chr3-11018032-DEL-97 | chr3-11018033-DEL-97 |
| chr3-11026776-DEL-6817 | chr3-11026777-DEL-6817 |
| chr3-11048135-DEL-60 | chr3-11048135-DEL-60 |
| chr3-11067929-DEL-299 | chr3-11067930-DEL-299 |
| chr3-11068508-DEL-57 | chr3-11068509-DEL-57 |
| chr3-11075483-DEL-1374 | chr3-11075484-DEL-1374 |
| chr3-11078524-DEL-1729 | chr3-11078525-DEL-1729 |
| chr3-11081232-DEL-200 | chr3-11081233-DEL-200 |
| chr3-11082072-DEL-585 | chr3-11082073-DEL-585 |
| chr3-11115176-DEL-9663 | chr3-11115170-DEL-9654 |
| chr3-11120011-DEL-3617 | chr3-11120011-DEL-3616 |
| chr3-11131154-DEL-4806 | chr3-11131155-DEL-4806 |
| chr3-11146116-DEL-119 | chr3-11146117-DEL-119 |
| chr3-11163275-DEL-1386 | chr3-11163275-DEL-1386 |
| chr3-11189069-DEL-7248 | chr3-11189070-DEL-7248 |
| chr3-11199320-DEL-201 | chr3-11199321-DEL-201 |
| chr3-11219328-DEL-399 | chr3-11219324-DEL-400 |
| chr3-11224038-DEL-109 | chr3-11224039-DEL-109 |
| chr3-11225675-DEL-51 | chr3-11225688-DEL-55 |
| chr3-11240536-DEL-218 | chr3-11240570-DEL-188 |
| chr3-11249411-DEL-179 | chr3-11249412-DEL-179 |
| chr3-11254197-DEL-479 | chr3-11254198-DEL-479 |
| chr3-11263437-DEL-358 | chr3-11263438-DEL-358 |
| chr3-11270496-DEL-1573 | chr3-11270497-DEL-1573 |
| chr3-11284288-DEL-217 | chr3-11284289-DEL-217 |
| chr3-11290906-DEL-64 | chr3-11290907-DEL-64 |
| chr3-11301540-DEL-160 | chr3-11301541-DEL-160 |
| chr3-11305682-DEL-568 | chr3-11305683-DEL-568 |
| chr3-11334879-DEL-63 | chr3-11334880-DEL-63 |
| chr3-11376898-DEL-285 | chr3-11376899-DEL-285 |
| chr3-11382409-DEL-62 | chr3-11382392-DEL-64 |
| chr3-11410654-DEL-72 | chr3-11410654-DEL-76 |
| chr3-11421205-DEL-51 | chr3-11421190-DEL-67 |
| chr3-11446850-DEL-6427 | chr3-11446851-DEL-6427 |
| chr3-11510166-DEL-76 | chr3-11510167-DEL-68 |
| chr3-11603516-DEL-60 | chr3-11603517-DEL-60 |
| chr3-11611776-DEL-7382 | chr3-11611777-DEL-7382 |
| chr3-11645811-DEL-58 | chr3-11645811-DEL-58 |
| chr3-11655501-DEL-7020 | chr3-11655502-DEL-7020 |
| chr3-11690792-DEL-75 | chr3-11690793-DEL-75 |
| chr3-11695917-DEL-1137 | chr3-11695918-DEL-1137 |
| chr3-11750441-DEL-301 | chr3-11750440-DEL-303 |
| chr3-11755753-DEL-1892 | chr3-11755754-DEL-1892 |
| chr3-11780801-DEL-1892 | chr3-11780802-DEL-1892 |
| chr3-11783126-DEL-58 | chr3-11783127-DEL-58 |
| chr3-11789272-DEL-284 | chr3-11789273-DEL-284 |
| chr3-11797717-DEL-7065 | chr3-11797718-DEL-7065 |
| chr3-11835874-DEL-56 | chr3-11835875-DEL-58 |
| chr3-11843904-DEL-1911 | chr3-11843900-DEL-1923 |
| chr3-11852389-DEL-51 | chr3-11852401-DEL-63 |
| chr3-11856337-DEL-6458 | chr3-11856335-DEL-6459 |
| chr3-11856397-DEL-214 | chr3-11856398-DEL-214 |
| chr3-11868537-DEL-2591 | chr3-11868537-DEL-2591 |
| chr3-11875469-DEL-202 | chr3-11875470-DEL-202 |
| chr3-11898966-DEL-70 | chr3-11898967-DEL-70 |
| chr3-11916989-DEL-6318 | chr3-11916990-DEL-6318 |
| chr3-11917019-DEL-214 | chr3-11917020-DEL-214 |
| chr3-12023391-DEL-70 | chr3-12023423-DEL-70 |
| chr3-12031138-DEL-54 | chr3-12031139-DEL-50 |
| chr3-12039224-DEL-264 | chr3-12039225-DEL-264 |
| chr3-12043348-DEL-1953 | chr3-12043350-DEL-1952 |
| chr3-12053001-DEL-80 | chr3-12053002-DEL-80 |
| chr3-12060720-DEL-6394 | chr3-12060721-DEL-6394 |
| chr3-12085568-DEL-313 | chr3-12085569-DEL-313 |
| chr3-12096444-DEL-624 | chr3-12096445-DEL-624 |
| chr3-12101001-DEL-567 | chr3-12101002-DEL-567 |
| chr3-12172242-DEL-5297 | chr3-12172242-DEL-5297 |
| chr3-12199233-DEL-381 | chr3-12199234-DEL-381 |

| | |
|---|---|
| chr3-12200730-DEL-970 | chr3-12200731-DEL-970 |
| chr3-12201952-DEL-51 | chr3-12201950-DEL-51 |
| chr3-12219502-DEL-452 | chr3-12219497-DEL-453 |
| chr3-12225644-DEL-106 | chr3-12225645-DEL-106 |
| chr3-12270328-DEL-62 | chr3-12270323-DEL-63 |
| chr3-12286162-DEL-3124 | chr3-12286163-DEL-3124 |
| chr3-12290560-DEL-919 | chr3-12290561-DEL-919 |
| chr3-12295492-DEL-214 | chr3-12295493-DEL-214 |
| chr3-12337980-DEL-544 | chr3-12337981-DEL-544 |
| chr3-12394194-DEL-53 | chr3-12394195-DEL-53 |
| chr3-12425521-DEL-6444 | chr3-12425522-DEL-6444 |
| chr3-12473995-DEL-3839 | chr3-12473996-DEL-3839 |
| chr3-12516401-DEL-168 | chr3-12516402-DEL-168 |
| chr3-12533745-DEL-7092 | chr3-12533746-DEL-7092 |
| chr3-12546628-DEL-68 | chr3-12546629-DEL-68 |
| chr3-12565539-DEL-136 | chr3-12565540-DEL-136 |
| chr3-12566048-DEL-94 | chr3-12566035-DEL-108 |
| chr3-12579610-DEL-6228 | chr3-12579611-DEL-6228 |
| chr3-12586966-DEL-92 | chr3-12586967-DEL-92 |
| chr3-12588040-DEL-76 | chr3-12587998-DEL-70 |
| chr3-12644971-DEL-253 | chr3-12644971-DEL-253 |
| chr3-12649980-DEL-4964 | chr3-12649980-DEL-4964 |
| chr3-12671863-DEL-400 | chr3-12671864-DEL-400 |
| chr3-12675591-DEL-1186 | chr3-12675592-DEL-1186 |
| chr3-12682308-DEL-52 | chr3-12682309-DEL-52 |
| chr3-12716279-DEL-401 | chr3-12716276-DEL-405 |
| chr3-12727940-DEL-496 | chr3-12727941-DEL-496 |
| chr3-12740646-DEL-162 | chr3-12740647-DEL-162 |
| chr3-12744387-DEL-1577 | chr3-12744387-DEL-1588 |
| chr3-12757985-DEL-80 | chr3-12757980-DEL-73 |
| chr3-12762132-DEL-74 | chr3-12762139-DEL-68 |
| chr3-12765592-DEL-215 | chr3-12765593-DEL-213 |
| chr3-12806531-DEL-485 | chr3-12806532-DEL-485 |
| chr3-12809913-DEL-104 | chr3-12809914-DEL-104 |
| chr3-12811583-DEL-60 | chr3-12811583-DEL-60 |
| chr3-12831291-DEL-83 | chr3-12831271-DEL-83 |
| chr3-12838440-DEL-89 | chr3-12838441-DEL-89 |
| chr3-12868115-DEL-64 | chr3-12868115-DEL-64 |
| chr3-12870938-DEL-56 | chr3-12870939-DEL-56 |
| chr3-12879455-DEL-5253 | chr3-12879455-DEL-5253 |
| chr3-12890605-DEL-52 | chr3-12890606-DEL-50 |
| chr3-12901960-DEL-3113 | chr3-12901961-DEL-3113 |
| chr3-12920847-DEL-2861 | chr3-12920848-DEL-2861 |
| chr3-12947675-DEL-122 | chr3-12947676-DEL-122 |
| chr3-12950140-DEL-76 | chr3-12950149-DEL-68 |
| chr3-12955972-DEL-107 | chr3-12955971-DEL-107 |
| chr3-12973615-DEL-214 | chr3-12973616-DEL-214 |
| chr3-12983361-DEL-358 | chr3-12983362-DEL-358 |
| chr3-12989811-DEL-809 | chr3-12989812-DEL-809 |
| chr3-12994231-DEL-7436 | chr3-12994232-DEL-7436 |
| chr3-13004536-DEL-172 | chr3-13004537-DEL-172 |
| chr3-13036471-DEL-65 | chr3-13036480-DEL-57 |
| chr3-13040416-DEL-345 | chr3-13040417-DEL-345 |
| chr3-13046149-DEL-520 | chr3-13046150-DEL-522 |
| chr3-13052714-DEL-287 | chr3-13052715-DEL-287 |
| chr3-13058135-DEL-6314 | chr3-13058136-DEL-6314 |
| chr3-13079650-DEL-5339 | chr3-13079651-DEL-5339 |
| chr3-13088582-DEL-2421 | chr3-13088583-DEL-2421 |
| chr3-13102882-DEL-63 | chr3-13102883-DEL-63 |
| chr3-13120736-DEL-2144 | chr3-13120737-DEL-2144 |
| chr3-13131273-DEL-953 | chr3-13131274-DEL-953 |
| chr3-13165270-DEL-56 | chr3-13165271-DEL-60 |
| chr3-13167207-DEL-98 | chr3-13167208-DEL-98 |
| chr3-13208593-DEL-52 | chr3-13208594-DEL-62 |
| chr3-13211075-DEL-427 | chr3-13211072-DEL-428 |
| chr3-13241138-DEL-54 | chr3-13241133-DEL-60 |
| chr3-13249353-DEL-52 | chr3-13249353-DEL-52 |
| chr3-13260617-DEL-53 | chr3-13260618-DEL-53 |
| chr3-13276377-DEL-844 | chr3-13276378-DEL-844 |
| chr3-13284399-DEL-7030 | chr3-13284400-DEL-7030 |
| chr3-13303503-DEL-172 | chr3-13303502-DEL-173 |
| chr3-13325213-DEL-117 | chr3-13325168-DEL-117 |
| chr3-13326460-DEL-55 | chr3-13326460-DEL-55 |
| chr3-13337740-DEL-1197 | chr3-13337741-DEL-1197 |
| chr3-13345105-DEL-66 | chr3-13345106-DEL-66 |
| chr3-13356802-DEL-4436 | chr3-13356803-DEL-4436 |
| chr3-13363666-DEL-74 | chr3-13363667-DEL-74 |
| chr3-13455403-DEL-228 | chr3-13455409-DEL-214 |
| chr3-13509941-DEL-54 | chr3-13509937-DEL-58 |
| chr3-13539712-DEL-63 | chr3-13539713-DEL-63 |

| | |
|---|---|
| chr3-13578160-DEL-51 | chr3-13578161-DEL-51 |
| chr3-13580959-DEL-430 | chr3-13580960-DEL-430 |
| chr3-13652427-DEL-60 | chr3-13652428-DEL-60 |
| chr3-13673756-DEL-969 | chr3-13673757-DEL-969 |
| chr3-13712624-DEL-515 | chr3-13712627-DEL-516 |
| chr3-13734063-DEL-1302 | chr3-13734055-DEL-1303 |
| chr3-13783886-DEL-172 | chr3-13783887-DEL-172 |
| chr3-13793573-DEL-81 | chr3-13793574-DEL-81 |
| chr3-13823475-DEL-86 | chr3-13823494-DEL-62 |
| chr3-13830351-DEL-758 | chr3-13830352-DEL-758 |
| chr3-13848228-DEL-114 | chr3-13848229-DEL-114 |
| chr3-13853222-DEL-382 | chr3-13853223-DEL-382 |
| chr3-13855808-DEL-4463 | chr3-13855809-DEL-4463 |
| chr3-13885989-DEL-120 | chr3-13885990-DEL-117 |
| chr3-13886317-DEL-5279 | chr3-13886318-DEL-5279 |
| chr3-13894501-DEL-51 | chr3-13894458-DEL-66 |
| chr3-13926619-DEL-1500 | chr3-13926620-DEL-1500 |
| chr3-13939538-DEL-365 | chr3-13939539-DEL-365 |
| chr3-13946087-DEL-230 | chr3-13946088-DEL-230 |
| chr3-13948935-DEL-6014 | chr3-13948936-DEL-6014 |
| chr3-13981845-DEL-966 | chr3-13981846-DEL-966 |
| chr3-13994113-DEL-802 | chr3-13994114-DEL-802 |
| chr3-14030729-DEL-6412 | chr3-14030744-DEL-6411 |
| chr3-14165889-DEL-439 | chr3-14165890-DEL-439 |
| chr3-14175288-DEL-66 | chr3-14175319-DEL-52 |
| chr3-14185073-DEL-82 | chr3-14185074-DEL-82 |
| chr3-14188787-DEL-852 | chr3-14188788-DEL-852 |
| chr3-14201575-DEL-855 | chr3-14201576-DEL-855 |
| chr3-14226786-DEL-1109 | chr3-14226791-DEL-1108 |
| chr3-14231351-DEL-453 | chr3-14231352-DEL-452 |
| chr3-14237467-DEL-111 | chr3-14237477-DEL-102 |
| chr3-14252928-DEL-5727 | chr3-14252929-DEL-5727 |
| chr3-14263144-DEL-93 | chr3-14263143-DEL-94 |
| chr3-14277134-DEL-200 | chr3-14277135-DEL-200 |
| chr3-14299790-DEL-7581 | chr3-14299791-DEL-7581 |
| chr3-14308004-DEL-342 | chr3-14308010-DEL-357 |
| chr3-14400114-DEL-1491 | chr3-14400115-DEL-1491 |
| chr3-14418206-DEL-99 | chr3-14418207-DEL-99 |
| chr3-14433391-DEL-54 | chr3-14433392-DEL-54 |
| chr3-14450530-DEL-1807 | chr3-14450531-DEL-1807 |
| chr3-14458651-DEL-3556 | chr3-14458652-DEL-3556 |
| chr3-14469901-DEL-1982 | chr3-14469902-DEL-1982 |
| chr3-14512608-DEL-199 | chr3-14512609-DEL-199 |
| chr3-14607050-DEL-140 | chr3-14607024-DEL-141 |
| chr3-14685505-DEL-283 | chr3-14685506-DEL-283 |
| chr3-14704118-DEL-2080 | chr3-14704119-DEL-2080 |
| chr3-14704362-DEL-219 | chr3-14704363-DEL-219 |
| chr3-14733518-DEL-5498 | chr3-14733519-DEL-5498 |
| chr3-14749791-DEL-91 | chr3-14749792-DEL-91 |
| chr3-14753069-DEL-195 | chr3-14753070-DEL-195 |
| chr3-14781392-DEL-3477 | chr3-14781393-DEL-3477 |
| chr3-14786968-DEL-214 | chr3-14786969-DEL-214 |
| chr3-14790688-DEL-305 | chr3-14790689-DEL-304 |
| chr3-14835985-DEL-529 | chr3-14835986-DEL-529 |
| chr3-14855228-DEL-55 | chr3-14855214-DEL-90 |
| chr3-14860385-DEL-114 | chr3-14860386-DEL-114 |
| chr3-14882320-DEL-65 | chr3-14882321-DEL-65 |
| chr3-14896401-DEL-65 | chr3-14896402-DEL-65 |
| chr3-14898172-DEL-3796 | chr3-14898173-DEL-3796 |
| chr3-14907154-DEL-51 | chr3-14907155-DEL-51 |
| chr3-14921824-DEL-491 | chr3-14921825-DEL-491 |
| chr3-14932210-DEL-138 | chr3-14932211-DEL-138 |
| chr3-14949192-DEL-100 | chr3-14949193-DEL-100 |
| chr3-14958365-DEL-400 | chr3-14958366-DEL-400 |
| chr3-14969194-DEL-69 | chr3-14969200-DEL-69 |
| chr3-15005376-DEL-56 | chr3-15005399-DEL-55 |
| chr3-15035446-DEL-78 | chr3-15035447-DEL-78 |
| chr3-15037475-DEL-135 | chr3-15037476-DEL-138 |
| chr3-15048753-DEL-189 | chr3-15048754-DEL-189 |
| chr3-15052381-DEL-201 | chr3-15052382-DEL-201 |
| chr3-15053915-DEL-187 | chr3-15053916-DEL-187 |
| chr3-15091764-DEL-202 | chr3-15091765-DEL-202 |
| chr3-15093503-DEL-6481 | chr3-15093504-DEL-6481 |
| chr3-15102770-DEL-971 | chr3-15102769-DEL-972 |
| chr3-15111448-DEL-505 | chr3-15111449-DEL-505 |
| chr3-15120117-DEL-362 | chr3-15120122-DEL-363 |
| chr3-15144525-DEL-6234 | chr3-15144526-DEL-6234 |
| chr3-15180789-DEL-63 | chr3-15180832-DEL-63 |
| chr3-15188030-DEL-96 | chr3-15188031-DEL-96 |
| chr3-15194449-DEL-480 | chr3-15194450-DEL-480 |

| | |
|---|---|
| chr3-15197683-DEL-564 | chr3-15197684-DEL-563 |
| chr3-15215340-DEL-506 | chr3-15215341-DEL-506 |
| chr3-15227080-DEL-297 | chr3-15227081-DEL-297 |
| chr3-15302368-DEL-55 | chr3-15302348-DEL-64 |
| chr3-15313241-DEL-6451 | chr3-15313242-DEL-6451 |
| chr3-15332242-DEL-561 | chr3-15332243-DEL-561 |
| chr3-15333732-DEL-351 | chr3-15333738-DEL-343 |
| chr3-15339148-DEL-952 | chr3-15339149-DEL-952 |
| chr3-15343085-DEL-59 | chr3-15343086-DEL-59 |
| chr3-15371743-DEL-6437 | chr3-15371744-DEL-6437 |
| chr3-15371773-DEL-208 | chr3-15371774-DEL-208 |
| chr3-15381729-DEL-83 | chr3-15381730-DEL-83 |
| chr3-15386729-DEL-2498 | chr3-15386730-DEL-2498 |
| chr3-15445525-DEL-105 | chr3-15445488-DEL-106 |
| chr3-15451706-DEL-1186 | chr3-15451706-DEL-1186 |
| chr3-15483504-DEL-204 | chr3-15483480-DEL-239 |
| chr3-15578787-DEL-2950 | chr3-15578788-DEL-2950 |
| chr3-15606785-DEL-53 | chr3-15606788-DEL-51 |
| chr3-15615096-DEL-231 | chr3-15615091-DEL-237 |
| chr3-15621107-DEL-3982 | chr3-15621106-DEL-3982 |
| chr3-15633223-DEL-110 | chr3-15633224-DEL-110 |
| chr3-15640250-DEL-158 | chr3-15640251-DEL-158 |
| chr3-15655127-DEL-4055 | chr3-15655128-DEL-4055 |
| chr3-15666397-DEL-92 | chr3-15666433-DEL-59 |
| chr3-15671901-DEL-87 | chr3-15671902-DEL-87 |
| chr3-15678337-DEL-61 | chr3-15678338-DEL-61 |
| chr3-15685313-DEL-1463 | chr3-15685313-DEL-1463 |
| chr3-15890155-DEL-537 | chr3-15890156-DEL-537 |
| chr3-15933444-DEL-329 | chr3-15933445-DEL-329 |
| chr3-15941185-DEL-105 | chr3-15941186-DEL-105 |
| chr3-15951776-DEL-1452 | chr3-15951777-DEL-1452 |
| chr3-15967100-DEL-2164 | chr3-15967101-DEL-2164 |
| chr3-15968573-DEL-132 | chr3-15968571-DEL-138 |
| chr3-15976698-DEL-6531 | chr3-15976699-DEL-6531 |
| chr3-16004500-DEL-69 | chr3-16004502-DEL-67 |
| chr3-16008457-DEL-152 | chr3-16008449-DEL-120 |
| chr3-16009035-DEL-843 | chr3-16009036-DEL-843 |
| chr3-16010780-DEL-376 | chr3-16010781-DEL-376 |
| chr3-16013014-DEL-803 | chr3-16013015-DEL-803 |
| chr3-16028790-DEL-97 | chr3-16028790-DEL-97 |
| chr3-16032608-DEL-218 | chr3-16032608-DEL-218 |
| chr3-16038266-DEL-4870 | chr3-16038266-DEL-4870 |
| chr3-16055859-DEL-1292 | chr3-16055860-DEL-1292 |
| chr3-16063773-DEL-2990 | chr3-16063774-DEL-2990 |
| chr3-16079337-DEL-332 | chr3-16079338-DEL-332 |
| chr3-16132125-DEL-154 | chr3-16132125-DEL-154 |
| chr3-16138056-DEL-210 | chr3-16138056-DEL-211 |
| chr3-16139014-DEL-99 | chr3-16139019-DEL-95 |
| chr3-16142749-DEL-6236 | chr3-16142750-DEL-6236 |
| chr3-16158086-DEL-74 | chr3-16158086-DEL-74 |
| chr3-16164710-DEL-166 | chr3-16164710-DEL-166 |
| chr3-16180736-DEL-1039 | chr3-16180737-DEL-1039 |
| chr3-16188587-DEL-130 | chr3-16188588-DEL-130 |
| chr3-16195424-DEL-651 | chr3-16195414-DEL-652 |
| chr3-16207820-DEL-96 | chr3-16207821-DEL-96 |
| chr3-16211546-DEL-621 | chr3-16211545-DEL-621 |
| chr3-16218626-DEL-266 | chr3-16218626-DEL-266 |
| chr3-16285790-DEL-8675 | chr3-16285791-DEL-8675 |
| chr3-16329959-DEL-293 | chr3-16329956-DEL-293 |
| chr3-16335556-DEL-54 | chr3-16335530-DEL-60 |
| chr3-16339058-DEL-203 | chr3-16339059-DEL-203 |
| chr3-16349009-DEL-88 | chr3-16349010-DEL-56 |
| chr3-16361611-DEL-90 | chr3-16361638-DEL-56 |
| chr3-16362256-DEL-65 | chr3-16362257-DEL-65 |
| chr3-16373349-DEL-201 | chr3-16373350-DEL-201 |
| chr3-16375309-DEL-2644 | chr3-16375310-DEL-2644 |
| chr3-16400827-DEL-73 | chr3-16400828-DEL-56 |
| chr3-16405568-DEL-6372 | chr3-16405569-DEL-6372 |
| chr3-16422963-DEL-928 | chr3-16422964-DEL-928 |
| chr3-16456529-DEL-2061 | chr3-16456539-DEL-2064 |
| chr3-16462546-DEL-271 | chr3-16462560-DEL-250 |
| chr3-16465623-DEL-510 | chr3-16465624-DEL-509 |
| chr3-16475266-DEL-138 | chr3-16475267-DEL-138 |
| chr3-16477494-DEL-1039 | chr3-16477495-DEL-1039 |
| chr3-16503958-DEL-410 | chr3-16503958-DEL-410 |
| chr3-16507432-DEL-3320 | chr3-16507432-DEL-3320 |
| chr3-16521955-DEL-1190 | chr3-16521956-DEL-1190 |
| chr3-16545572-DEL-6591 | chr3-16545573-DEL-6591 |
| chr3-16567948-DEL-436 | chr3-16567949-DEL-436 |
| chr3-16579920-DEL-1917 | chr3-16579921-DEL-1917 |

| | |
|---|---|
| chr3-16591962-DEL-52 | chr3-16591963-DEL-52 |
| chr3-16604697-DEL-108 | chr3-16604698-DEL-108 |
| chr3-16632464-DEL-6827 | chr3-16632465-DEL-6827 |
| chr3-16667110-DEL-1456 | chr3-16667111-DEL-1456 |
| chr3-16670243-DEL-178 | chr3-16670251-DEL-129 |
| chr3-16679616-DEL-10113 | chr3-16679617-DEL-10113 |
| chr3-16698841-DEL-180 | chr3-16698842-DEL-180 |
| chr3-16710716-DEL-533 | chr3-16710717-DEL-533 |
| chr3-16715886-DEL-5055 | chr3-16715886-DEL-5055 |
| chr3-16737474-DEL-3688 | chr3-16737475-DEL-3688 |
| chr3-16740979-DEL-121 | chr3-16740978-DEL-121 |
| chr3-16753492-DEL-576 | chr3-16753493-DEL-576 |
| chr3-16766047-DEL-264 | chr3-16766048-DEL-263 |
| chr3-16767704-DEL-206 | chr3-16767712-DEL-200 |
| chr3-16785796-DEL-277 | chr3-16785797-DEL-278 |
| chr3-16846689-DEL-8418 | chr3-16846690-DEL-8418 |
| chr3-16883914-DEL-99 | chr3-16883915-DEL-99 |
| chr3-16932282-DEL-420 | chr3-16932283-DEL-420 |
| chr3-17024293-DEL-64 | chr3-17024294-DEL-64 |
| chr3-17062786-DEL-200 | chr3-17062789-DEL-198 |
| chr3-17073197-DEL-2453 | chr3-17073198-DEL-2453 |
| chr3-17077216-DEL-13672 | chr3-17077218-DEL-13670 |
| chr3-17121225-DEL-103 | chr3-17121225-DEL-103 |
| chr3-17150834-DEL-2369 | chr3-17150834-DEL-2369 |
| chr3-17182671-DEL-1100 | chr3-17182659-DEL-1107 |
| chr3-17193402-DEL-2157 | chr3-17193403-DEL-2154 |
| chr3-17212052-DEL-1651 | chr3-17212053-DEL-1651 |
| chr3-17220332-DEL-739 | chr3-17220332-DEL-737 |
| chr3-17240600-DEL-1764 | chr3-17240601-DEL-1764 |
| chr3-17243577-DEL-75 | chr3-17243578-DEL-75 |
| chr3-17274562-DEL-6362 | chr3-17274563-DEL-6362 |
| chr3-17290411-DEL-63 | chr3-17290413-DEL-64 |
| chr3-17295808-DEL-165 | chr3-17295809-DEL-165 |
| chr3-17315111-DEL-1159 | chr3-17315112-DEL-1159 |
| chr3-17320959-DEL-83 | chr3-17320972-DEL-71 |
| chr3-17331483-DEL-1773 | chr3-17331484-DEL-1773 |
| chr3-17333778-DEL-72 | chr3-17333742-DEL-109 |
| chr3-17347948-DEL-6247 | chr3-17347949-DEL-6247 |
| chr3-17388681-DEL-53 | chr3-17388681-DEL-54 |
| chr3-17392067-DEL-358 | chr3-17392068-DEL-358 |
| chr3-17436401-DEL-916 | chr3-17436402-DEL-916 |
| chr3-17437247-DEL-133 | chr3-17437248-DEL-133 |
| chr3-17437721-DEL-6287 | chr3-17437722-DEL-6287 |
| chr3-17449855-DEL-57 | chr3-17449851-DEL-62 |
| chr3-17454300-DEL-78 | chr3-17454295-DEL-80 |
| chr3-17485505-DEL-81 | chr3-17485506-DEL-81 |
| chr3-17492964-DEL-66 | chr3-17492968-DEL-63 |
| chr3-17495431-DEL-6430 | chr3-17495432-DEL-6430 |
| chr3-17520112-DEL-586 | chr3-17520113-DEL-586 |
| chr3-17532135-DEL-73 | chr3-17532127-DEL-74 |
| chr3-17558370-DEL-79 | chr3-17558367-DEL-66 |
| chr3-17568115-DEL-535 | chr3-17568116-DEL-536 |
| chr3-17592423-DEL-52 | chr3-17592424-DEL-52 |
| chr3-17642537-DEL-52 | chr3-17642505-DEL-68 |
| chr3-17651353-DEL-551 | chr3-17651354-DEL-551 |
| chr3-17685686-DEL-397 | chr3-17685687-DEL-395 |
| chr3-17695400-DEL-1015 | chr3-17695401-DEL-1015 |
| chr3-17703427-DEL-6920 | chr3-17703428-DEL-6920 |
| chr3-17733669-DEL-6188 | chr3-17733670-DEL-6188 |
| chr3-17765450-DEL-246 | chr3-17765451-DEL-246 |
| chr3-17781968-DEL-7512 | chr3-17781969-DEL-7512 |
| chr3-17803385-DEL-60 | chr3-17803386-DEL-60 |
| chr3-17822878-DEL-394 | chr3-17822879-DEL-394 |
| chr3-17823581-DEL-103 | chr3-17823582-DEL-103 |
| chr3-17875246-DEL-142 | chr3-17875251-DEL-128 |
| chr3-17935636-DEL-425 | chr3-17935637-DEL-425 |
| chr3-17956795-DEL-7173 | chr3-17956796-DEL-7173 |
| chr3-17983760-DEL-66 | chr3-17983761-DEL-70 |
| chr3-17984143-DEL-641 | chr3-17984164-DEL-625 |
| chr3-17986084-DEL-508 | chr3-17986085-DEL-508 |
| chr3-17991159-DEL-52 | chr3-17991160-DEL-52 |
| chr3-17992214-DEL-74 | chr3-17992215-DEL-74 |
| chr3-18008300-DEL-4984 | chr3-18008301-DEL-4984 |
| chr3-18023798-DEL-326 | chr3-18023799-DEL-326 |
| chr3-18058069-DEL-3538 | chr3-18058070-DEL-3538 |
| chr3-18062904-DEL-2775 | chr3-18062905-DEL-2775 |
| chr3-18086429-DEL-6010 | chr3-18086430-DEL-6010 |
| chr3-18129585-DEL-96 | chr3-18129586-DEL-96 |
| chr3-18148734-DEL-995 | chr3-18148735-DEL-995 |
| chr3-18170882-DEL-77 | chr3-18170883-DEL-77 |

| | |
|---|---|
| chr3-18181704-DEL-184 | chr3-18181705-DEL-184 |
| chr3-18189840-DEL-478 | chr3-18189848-DEL-468 |
| chr3-18225951-DEL-1178 | chr3-18225952-DEL-1178 |
| chr3-18244845-DEL-66 | chr3-18244843-DEL-66 |
| chr3-18294653-DEL-66 | chr3-18294656-DEL-64 |
| chr3-18416683-DEL-196 | chr3-18416684-DEL-196 |
| chr3-18421557-DEL-1036 | chr3-18421558-DEL-1036 |
| chr3-18433884-DEL-3553 | chr3-18433885-DEL-3553 |
| chr3-18485416-DEL-6259 | chr3-18485417-DEL-6259 |
| chr3-18505846-DEL-440 | chr3-18505847-DEL-440 |
| chr3-18509011-DEL-265 | chr3-18509012-DEL-265 |
| chr3-18564613-DEL-127 | chr3-18564614-DEL-127 |
| chr3-18577076-DEL-1529 | chr3-18577077-DEL-1529 |
| chr3-18584642-DEL-231 | chr3-18584672-DEL-227 |
| chr3-18598136-DEL-153 | chr3-18598122-DEL-154 |
| chr3-18603609-DEL-7632 | chr3-18603610-DEL-7632 |
| chr3-18624531-DEL-66 | chr3-18624532-DEL-66 |
| chr3-18626756-DEL-113 | chr3-18626757-DEL-113 |
| chr3-18638793-DEL-6186 | chr3-18638794-DEL-6186 |
| chr3-18645605-DEL-82 | chr3-18645606-DEL-82 |
| chr3-18676827-DEL-780 | chr3-18676828-DEL-777 |
| chr3-18681667-DEL-308 | chr3-18681668-DEL-308 |
| chr3-18687104-DEL-56 | chr3-18687103-DEL-57 |
| chr3-18704188-DEL-144 | chr3-18704189-DEL-144 |
| chr3-18713345-DEL-472 | chr3-18713346-DEL-472 |
| chr3-18715740-DEL-608 | chr3-18715741-DEL-608 |
| chr3-18741444-DEL-63 | chr3-18741445-DEL-63 |
| chr3-18750680-DEL-549 | chr3-18750681-DEL-549 |
| chr3-18765564-DEL-52 | chr3-18765559-DEL-58 |
| chr3-18771117-DEL-6380 | chr3-18771117-DEL-6380 |
| chr3-18805681-DEL-53 | chr3-18805682-DEL-53 |
| chr3-18815078-DEL-275 | chr3-18815079-DEL-275 |
| chr3-18835243-DEL-112 | chr3-18835244-DEL-112 |
| chr3-18878398-DEL-500 | chr3-18878399-DEL-500 |
| chr3-18886537-DEL-527 | chr3-18886538-DEL-527 |
| chr3-18895277-DEL-892 | chr3-18895278-DEL-893 |
| chr3-18906891-DEL-6655 | chr3-18906892-DEL-6655 |
| chr3-18906996-DEL-201 | chr3-18906996-DEL-201 |
| chr3-18928120-DEL-52 | chr3-18928121-DEL-52 |
| chr3-18976269-DEL-1610 | chr3-18976270-DEL-1610 |
| chr3-19019375-DEL-7571 | chr3-19019369-DEL-7569 |
| chr3-19026679-DEL-214 | chr3-19026680-DEL-214 |
| chr3-19036366-DEL-1573 | chr3-19036366-DEL-1573 |
| chr3-19038933-DEL-1037 | chr3-19038934-DEL-1037 |
| chr3-19069618-DEL-53 | chr3-19069584-DEL-53 |
| chr3-19070056-DEL-6243 | chr3-19070057-DEL-6243 |
| chr3-19124510-DEL-177 | chr3-19124511-DEL-177 |
| chr3-19200165-DEL-128 | chr3-19200166-DEL-128 |
| chr3-19220651-DEL-163 | chr3-19220652-DEL-163 |
| chr3-19256884-DEL-213 | chr3-19256885-DEL-213 |
| chr3-19289412-DEL-73 | chr3-19289413-DEL-73 |
| chr3-19321470-DEL-519 | chr3-19321471-DEL-519 |
| chr3-19348318-DEL-62 | chr3-19348316-DEL-60 |
| chr3-19360997-DEL-99 | chr3-19360998-DEL-99 |
| chr3-19371734-DEL-100 | chr3-19371735-DEL-100 |
| chr3-19374931-DEL-423 | chr3-19374932-DEL-422 |
| chr3-19382823-DEL-133 | chr3-19382821-DEL-134 |
| chr3-19395582-DEL-130 | chr3-19395581-DEL-130 |
| chr3-19410029-DEL-3554 | chr3-19410045-DEL-3552 |
| chr3-19429046-DEL-78 | chr3-19429047-DEL-76 |
| chr3-19500048-DEL-67 | chr3-19500002-DEL-68 |
| chr3-19506391-DEL-81 | chr3-19506393-DEL-80 |
| chr3-19516304-DEL-6365 | chr3-19516305-DEL-6365 |
| chr3-19525836-DEL-70 | chr3-19525822-DEL-67 |
| chr3-19542852-DEL-1937 | chr3-19542850-DEL-1939 |
| chr3-19557613-DEL-2546 | chr3-19557614-DEL-2546 |
| chr3-19571486-DEL-1672 | chr3-19571487-DEL-1672 |
| chr3-19596351-DEL-78 | chr3-19596352-DEL-78 |
| chr3-19625136-DEL-104 | chr3-19625137-DEL-104 |
| chr3-19637793-DEL-1699 | chr3-19637794-DEL-1699 |
| chr3-19658625-DEL-82 | chr3-19658626-DEL-82 |
| chr3-19720868-DEL-1027 | chr3-19720868-DEL-1027 |
| chr3-19798626-DEL-215 | chr3-19798627-DEL-215 |
| chr3-19826779-DEL-263 | chr3-19826804-DEL-257 |
| chr3-19894456-DEL-131 | chr3-19894454-DEL-132 |
| chr3-19900613-DEL-762 | chr3-19900614-DEL-762 |
| chr3-19903935-DEL-4836 | chr3-19903936-DEL-4836 |
| chr3-19913382-DEL-253 | chr3-19913377-DEL-254 |
| chr3-19921207-DEL-135 | chr3-19921210-DEL-105 |
| chr3-19925185-DEL-689 | chr3-19925186-DEL-689 |

| | |
|---|---|
| chr3-19959246-DEL-60 | chr3-19959246-DEL-60 |
| chr3-19961462-DEL-316 | chr3-19961461-DEL-314 |
| chr3-19964797-DEL-2529 | chr3-19964798-DEL-2529 |
| chr3-20003630-DEL-65 | chr3-20003631-DEL-65 |
| chr3-20010602-DEL-202 | chr3-20010603-DEL-202 |
| chr3-20044732-DEL-84 | chr3-20044733-DEL-81 |
| chr3-20050011-DEL-233 | chr3-20050012-DEL-233 |
| chr3-20059181-DEL-330 | chr3-20059180-DEL-331 |
| chr3-20064086-DEL-219 | chr3-20064081-DEL-219 |
| chr3-20065233-DEL-95 | chr3-20065234-DEL-95 |
| chr3-20087579-DEL-95 | chr3-20087580-DEL-95 |
| chr3-20089697-DEL-160 | chr3-20089698-DEL-160 |
| chr3-20111182-DEL-148 | chr3-20111183-DEL-148 |
| chr3-20114837-DEL-268 | chr3-20114838-DEL-268 |
| chr3-20121643-DEL-66 | chr3-20121645-DEL-66 |
| chr3-20124591-DEL-227 | chr3-20124608-DEL-211 |
| chr3-20125734-DEL-79 | chr3-20125725-DEL-84 |
| chr3-20126696-DEL-66 | chr3-20126665-DEL-98 |
| chr3-20129686-DEL-205 | chr3-20129687-DEL-205 |
| chr3-20139270-DEL-79 | chr3-20139271-DEL-79 |
| chr3-20147854-DEL-1856 | chr3-20147854-DEL-1854 |
| chr3-20154260-DEL-100 | chr3-20154261-DEL-100 |
| chr3-20157260-DEL-934 | chr3-20157261-DEL-934 |
| chr3-20162694-DEL-51 | chr3-20162695-DEL-51 |
| chr3-20183008-DEL-1097 | chr3-20183009-DEL-1097 |
| chr3-20186678-DEL-57 | chr3-20186679-DEL-58 |
| chr3-20188434-DEL-62 | chr3-20188433-DEL-62 |
| chr3-20224454-DEL-186 | chr3-20224455-DEL-186 |
| chr3-20233869-DEL-497 | chr3-20233870-DEL-497 |
| chr3-20238586-DEL-275 | chr3-20238538-DEL-276 |
| chr3-20244374-DEL-76 | chr3-20244376-DEL-74 |
| chr3-20250642-DEL-150 | chr3-20250643-DEL-150 |
| chr3-20252689-DEL-59 | chr3-20252689-DEL-59 |
| chr3-20260860-DEL-1410 | chr3-20260861-DEL-1410 |
| chr3-20266930-DEL-54 | chr3-20266930-DEL-54 |
| chr3-20291138-DEL-101 | chr3-20291139-DEL-101 |
| chr3-20297251-DEL-57 | chr3-20297279-DEL-60 |
| chr3-20298985-DEL-126 | chr3-20298986-DEL-126 |
| chr3-20321777-DEL-2036 | chr3-20321778-DEL-2036 |
| chr3-20342264-DEL-105 | chr3-20342267-DEL-103 |
| chr3-20351682-DEL-400 | chr3-20351683-DEL-400 |
| chr3-20355042-DEL-400 | chr3-20355044-DEL-401 |
| chr3-20363901-DEL-251 | chr3-20363901-DEL-251 |
| chr3-20364514-DEL-166 | chr3-20364515-DEL-165 |
| chr3-20366815-DEL-1709 | chr3-20366816-DEL-1709 |
| chr3-20376983-DEL-158 | chr3-20377025-DEL-162 |
| chr3-20414858-DEL-654 | chr3-20414859-DEL-654 |
| chr3-20424507-DEL-1299 | chr3-20424508-DEL-1299 |
| chr3-20438972-DEL-128 | chr3-20438973-DEL-128 |
| chr3-20441637-DEL-908 | chr3-20441638-DEL-908 |
| chr3-20442756-DEL-1806 | chr3-20442756-DEL-1806 |
| chr3-20455965-DEL-3727 | chr3-20455966-DEL-3727 |
| chr3-20467504-DEL-146 | chr3-20467505-DEL-146 |
| chr3-20485972-DEL-96 | chr3-20485973-DEL-131 |
| chr3-20493917-DEL-474 | chr3-20493918-DEL-474 |
| chr3-20528935-DEL-687 | chr3-20528937-DEL-686 |
| chr3-20539139-DEL-2937 | chr3-20539140-DEL-2937 |
| chr3-20544925-DEL-1409 | chr3-20544926-DEL-1409 |
| chr3-20546679-DEL-176 | chr3-20546680-DEL-176 |
| chr3-20565183-DEL-92 | chr3-20565157-DEL-100 |
| chr3-20568955-DEL-155 | chr3-20568955-DEL-155 |
| chr3-20581097-DEL-137 | chr3-20581098-DEL-137 |
| chr3-20588695-DEL-667 | chr3-20588696-DEL-667 |
| chr3-20596324-DEL-371 | chr3-20596327-DEL-379 |
| chr3-20627427-DEL-2631 | chr3-20627428-DEL-2631 |
| chr3-20677636-DEL-416 | chr3-20677637-DEL-416 |
| chr3-20692203-DEL-857 | chr3-20692204-DEL-857 |
| chr3-20725713-DEL-141 | chr3-20725714-DEL-141 |
| chr3-20742147-DEL-7250 | chr3-20742148-DEL-7250 |
| chr3-20742648-DEL-81 | chr3-20742649-DEL-81 |
| chr3-20778791-DEL-169 | chr3-20778792-DEL-169 |
| chr3-20785531-DEL-267 | chr3-20785532-DEL-267 |
| chr3-20794762-DEL-8207 | chr3-20794763-DEL-8207 |
| chr3-20797175-DEL-1328 | chr3-20797175-DEL-1328 |
| chr3-20827937-DEL-456 | chr3-20827944-DEL-451 |
| chr3-20832679-DEL-71 | chr3-20832682-DEL-72 |
| chr3-20842407-DEL-544 | chr3-20842408-DEL-544 |
| chr3-20868626-DEL-95 | chr3-20868627-DEL-95 |
| chr3-20937858-DEL-2198 | chr3-20937857-DEL-2199 |
| chr3-20951401-DEL-6238 | chr3-20951392-DEL-6238 |

| | |
|---|---|
| chr3-21003227-DEL-54 | chr3-21003228-DEL-52 |
| chr3-21014658-DEL-64 | chr3-21014659-DEL-62 |
| chr3-21061847-DEL-64 | chr3-21061848-DEL-64 |
| chr3-21067712-DEL-4543 | chr3-21067713-DEL-4543 |
| chr3-21084532-DEL-52 | chr3-21084532-DEL-52 |
| chr3-21085873-DEL-76 | chr3-21085859-DEL-98 |
| chr3-21099371-DEL-72 | chr3-21099390-DEL-54 |
| chr3-21102925-DEL-116 | chr3-21102926-DEL-116 |
| chr3-21103596-DEL-5020 | chr3-21103596-DEL-5020 |
| chr3-21111276-DEL-1244 | chr3-21111275-DEL-1244 |
| chr3-21117399-DEL-3530 | chr3-21117399-DEL-3530 |
| chr3-21145862-DEL-98 | chr3-21145863-DEL-98 |
| chr3-21147717-DEL-63 | chr3-21147717-DEL-62 |
| chr3-21176755-DEL-1551 | chr3-21176756-DEL-1551 |
| chr3-21210833-DEL-3753 | chr3-21210827-DEL-3754 |
| chr3-21246065-DEL-6850 | chr3-21246066-DEL-6850 |
| chr3-21286036-DEL-827 | chr3-21286036-DEL-828 |
| chr3-21295021-DEL-237 | chr3-21294992-DEL-230 |
| chr3-21298478-DEL-237 | chr3-21298479-DEL-237 |
| chr3-21301610-DEL-2762 | chr3-21301611-DEL-2762 |
| chr3-21312883-DEL-1995 | chr3-21312883-DEL-1993 |
| chr3-21316485-DEL-1932 | chr3-21316486-DEL-1931 |
| chr3-21337941-DEL-1767 | chr3-21337942-DEL-1767 |
| chr3-21348841-DEL-156 | chr3-21348842-DEL-156 |
| chr3-21371943-DEL-6835 | chr3-21371952-DEL-6836 |
| chr3-21408913-DEL-914 | chr3-21408913-DEL-912 |
| chr3-21437825-DEL-5279 | chr3-21437849-DEL-5279 |
| chr3-21453243-DEL-84 | chr3-21453244-DEL-84 |
| chr3-21458366-DEL-337 | chr3-21458367-DEL-337 |
| chr3-21469918-DEL-799 | chr3-21469919-DEL-799 |
| chr3-21526747-DEL-538 | chr3-21526748-DEL-538 |
| chr3-21530957-DEL-278 | chr3-21530958-DEL-278 |
| chr3-21577111-DEL-377 | chr3-21577112-DEL-377 |
| chr3-21591584-DEL-191 | chr3-21591586-DEL-189 |
| chr3-21630515-DEL-264 | chr3-21630516-DEL-232 |
| chr3-21655365-DEL-209 | chr3-21655366-DEL-209 |
| chr3-21663694-DEL-84 | chr3-21663719-DEL-60 |
| chr3-21668115-DEL-372 | chr3-21668116-DEL-372 |
| chr3-21677364-DEL-60 | chr3-21677365-DEL-60 |
| chr3-21715905-DEL-1810 | chr3-21715905-DEL-1810 |
| chr3-21719526-DEL-549 | chr3-21719527-DEL-549 |
| chr3-21725059-DEL-60 | chr3-21725075-DEL-60 |
| chr3-21739520-DEL-6671 | chr3-21739521-DEL-6671 |
| chr3-21755794-DEL-56 | chr3-21755795-DEL-56 |
| chr3-21761609-DEL-4731 | chr3-21761610-DEL-4731 |
| chr3-21773716-DEL-945 | chr3-21773716-DEL-942 |
| chr3-21800717-DEL-62 | chr3-21800718-DEL-62 |
| chr3-21805084-DEL-59 | chr3-21805085-DEL-59 |
| chr3-21818109-DEL-95 | chr3-21818091-DEL-96 |
| chr3-21820097-DEL-218 | chr3-21820098-DEL-218 |
| chr3-21829875-DEL-740 | chr3-21829876-DEL-740 |
| chr3-21857736-DEL-96 | chr3-21857737-DEL-96 |
| chr3-21875334-DEL-88 | chr3-21875359-DEL-83 |
| chr3-21880222-DEL-70 | chr3-21880211-DEL-74 |
| chr3-21902431-DEL-1642 | chr3-21902432-DEL-1642 |
| chr3-21903797-DEL-217 | chr3-21903798-DEL-217 |
| chr3-21954522-DEL-3374 | chr3-21954523-DEL-3374 |
| chr3-22024564-DEL-151 | chr3-22024565-DEL-151 |
| chr3-22033117-DEL-127 | chr3-22033141-DEL-87 |
| chr3-22052529-DEL-411 | chr3-22052530-DEL-411 |
| chr3-22061028-DEL-60 | chr3-22061029-DEL-60 |
| chr3-22065000-DEL-1891 | chr3-22065001-DEL-1891 |
| chr3-22074341-DEL-220 | chr3-22074343-DEL-219 |
| chr3-22079060-DEL-75 | chr3-22079061-DEL-75 |
| chr3-22086237-DEL-169 | chr3-22086238-DEL-169 |
| chr3-22087622-DEL-95 | chr3-22087587-DEL-96 |
| chr3-22092626-DEL-245 | chr3-22092627-DEL-245 |
| chr3-22108250-DEL-67 | chr3-22108253-DEL-64 |
| chr3-22121239-DEL-616 | chr3-22121239-DEL-617 |
| chr3-22192342-DEL-86 | chr3-22192344-DEL-87 |
| chr3-22219219-DEL-105 | chr3-22219219-DEL-107 |
| chr3-22271102-DEL-301 | chr3-22271103-DEL-301 |
| chr3-22274796-DEL-142 | chr3-22274797-DEL-142 |
| chr3-22279469-DEL-80 | chr3-22279470-DEL-80 |
| chr3-22324832-DEL-6585 | chr3-22324833-DEL-6585 |
| chr3-22333101-DEL-6173 | chr3-22333102-DEL-6173 |
| chr3-22340638-DEL-62 | chr3-22340639-DEL-62 |
| chr3-22347479-DEL-1239 | chr3-22347480-DEL-1239 |
| chr3-22350951-DEL-3670 | chr3-22350952-DEL-3670 |
| chr3-22374066-DEL-126 | chr3-22374067-DEL-126 |

| | |
|---|---|
| chr3-22376603-DEL-176 | chr3-22376604-DEL-176 |
| chr3-22389832-DEL-69 | chr3-22389833-DEL-69 |
| chr3-22390224-DEL-6406 | chr3-22390225-DEL-6406 |
| chr3-22428139-DEL-3005 | chr3-22428139-DEL-3005 |
| chr3-22466787-DEL-405 | chr3-22466788-DEL-405 |
| chr3-22467603-DEL-9358 | chr3-22467589-DEL-9346 |
| chr3-22477316-DEL-1272 | chr3-22477309-DEL-1273 |
| chr3-22489390-DEL-54 | chr3-22489389-DEL-56 |
| chr3-22490780-DEL-168 | chr3-22490800-DEL-148 |
| chr3-22493925-DEL-491 | chr3-22493926-DEL-491 |
| chr3-22507840-DEL-6912 | chr3-22507841-DEL-6912 |
| chr3-22526537-DEL-886 | chr3-22526538-DEL-886 |
| chr3-22543047-DEL-54 | chr3-22543048-DEL-54 |
| chr3-22566992-DEL-297 | chr3-22566993-DEL-297 |
| chr3-22621010-DEL-6194 | chr3-22621011-DEL-6194 |
| chr3-22628055-DEL-342 | chr3-22628056-DEL-342 |
| chr3-22633907-DEL-63 | chr3-22633908-DEL-63 |
| chr3-22650469-DEL-67 | chr3-22650470-DEL-67 |
| chr3-22655949-DEL-269 | chr3-22655950-DEL-269 |
| chr3-22663651-DEL-110 | chr3-22663653-DEL-96 |
| chr3-22680078-DEL-193 | chr3-22680078-DEL-193 |
| chr3-22756786-DEL-2745 | chr3-22756787-DEL-2745 |
| chr3-22807780-DEL-6265 | chr3-22807782-DEL-6264 |
| chr3-22837642-DEL-1904 | chr3-22837642-DEL-1904 |
| chr3-22851650-DEL-1220 | chr3-22851642-DEL-1218 |
| chr3-22858504-DEL-55 | chr3-22858505-DEL-55 |
| chr3-22865942-DEL-106 | chr3-22865943-DEL-106 |
| chr3-22920919-DEL-3110 | chr3-22920920-DEL-3110 |
| chr3-22928163-DEL-58 | chr3-22928164-DEL-58 |
| chr3-22935734-DEL-66 | chr3-22935741-DEL-60 |
| chr3-22954804-DEL-72 | chr3-22954781-DEL-96 |
| chr3-22955703-DEL-13535 | chr3-22955704-DEL-13535 |
| chr3-22986861-DEL-75 | chr3-22986854-DEL-76 |
| chr3-22994692-DEL-749 | chr3-22994692-DEL-749 |
| chr3-23041775-DEL-884 | chr3-23041776-DEL-884 |
| chr3-23059154-DEL-1965 | chr3-23059155-DEL-1965 |
| chr3-23064142-DEL-9710 | chr3-23064142-DEL-9710 |
| chr3-23080976-DEL-187 | chr3-23080977-DEL-187 |
| chr3-23081819-DEL-56 | chr3-23081819-DEL-56 |
| chr3-23131314-DEL-6116 | chr3-23131315-DEL-6116 |
| chr3-23194398-DEL-237 | chr3-23194399-DEL-237 |
| chr3-23227860-DEL-7087 | chr3-23227861-DEL-7087 |
| chr3-23267566-DEL-54 | chr3-23267577-DEL-61 |
| chr3-23273026-DEL-82 | chr3-23273026-DEL-82 |
| chr3-23279783-DEL-5625 | chr3-23279784-DEL-5625 |
| chr3-23294162-DEL-456 | chr3-23294162-DEL-455 |
| chr3-23295897-DEL-61 | chr3-23295898-DEL-61 |
| chr3-23312231-DEL-6186 | chr3-23312247-DEL-6186 |
| chr3-23361872-DEL-6595 | chr3-23361873-DEL-6595 |
| chr3-23379024-DEL-203 | chr3-23379025-DEL-203 |
| chr3-23386828-DEL-539 | chr3-23386813-DEL-554 |
| chr3-23392297-DEL-232 | chr3-23392298-DEL-232 |
| chr3-23407705-DEL-6427 | chr3-23407706-DEL-6427 |
| chr3-23414930-DEL-254 | chr3-23414931-DEL-254 |
| chr3-23415363-DEL-847 | chr3-23415364-DEL-847 |
| chr3-23434350-DEL-191 | chr3-23434351-DEL-191 |
| chr3-23439692-DEL-325 | chr3-23439693-DEL-325 |
| chr3-23442037-DEL-70 | chr3-23442058-DEL-60 |
| chr3-23447194-DEL-256 | chr3-23447195-DEL-256 |
| chr3-23468922-DEL-104 | chr3-23468894-DEL-105 |
| chr3-23494913-DEL-737 | chr3-23494912-DEL-755 |
| chr3-23539404-DEL-225 | chr3-23539404-DEL-225 |
| chr3-23557443-DEL-55 | chr3-23557444-DEL-55 |
| chr3-23588066-DEL-496 | chr3-23588067-DEL-496 |
| chr3-23591494-DEL-4150 | chr3-23591494-DEL-4150 |
| chr3-23600970-DEL-136 | chr3-23600969-DEL-134 |
| chr3-23621948-DEL-247 | chr3-23621948-DEL-247 |
| chr3-23632534-DEL-78 | chr3-23632534-DEL-78 |
| chr3-23643552-DEL-594 | chr3-23643553-DEL-594 |
| chr3-23644423-DEL-147 | chr3-23644423-DEL-148 |
| chr3-23660812-DEL-12472 | chr3-23660813-DEL-12472 |
| chr3-23689129-DEL-445 | chr3-23689130-DEL-445 |
| chr3-23725635-DEL-205 | chr3-23725636-DEL-205 |
| chr3-23727067-DEL-56 | chr3-23727074-DEL-54 |
| chr3-23738227-DEL-9020 | chr3-23738228-DEL-9020 |
| chr3-23758889-DEL-74 | chr3-23758908-DEL-56 |
| chr3-23761647-DEL-620 | chr3-23761648-DEL-620 |
| chr3-23777551-DEL-56 | chr3-23777552-DEL-56 |
| chr3-23821305-DEL-401 | chr3-23821306-DEL-401 |
| chr3-23824869-DEL-2101 | chr3-23824870-DEL-2101 |

| | |
|---|---|
| chr3-23828442-DEL-523 | chr3-23828443-DEL-523 |
| chr3-23833006-DEL-555 | chr3-23833007-DEL-555 |
| chr3-23873905-DEL-1155 | chr3-23873906-DEL-1155 |
| chr3-23895769-DEL-1717 | chr3-23895770-DEL-1717 |
| chr3-23912514-DEL-1792 | chr3-23912515-DEL-1791 |
| chr3-23941878-DEL-72 | chr3-23941880-DEL-70 |
| chr3-23948342-DEL-2593 | chr3-23948341-DEL-2593 |
| chr3-23955097-DEL-55 | chr3-23955096-DEL-57 |
| chr3-23965839-DEL-4908 | chr3-23965840-DEL-4908 |
| chr3-24005518-DEL-143 | chr3-24005519-DEL-143 |
| chr3-24018586-DEL-60 | chr3-24018587-DEL-60 |
| chr3-24065665-DEL-104 | chr3-24065666-DEL-105 |
| chr3-24067169-DEL-245 | chr3-24067168-DEL-242 |
| chr3-24072294-DEL-6563 | chr3-24072291-DEL-6562 |
| chr3-24078491-DEL-204 | chr3-24078492-DEL-204 |
| chr3-24080639-DEL-96 | chr3-24080654-DEL-66 |
| chr3-24095870-DEL-1034 | chr3-24095868-DEL-1036 |
| chr3-24123320-DEL-116 | chr3-24123321-DEL-116 |
| chr3-24142918-DEL-6200 | chr3-24142919-DEL-6200 |
| chr3-24161316-DEL-117 | chr3-24161317-DEL-117 |
| chr3-24173746-DEL-149 | chr3-24173747-DEL-140 |
| chr3-24189649-DEL-72 | chr3-24189662-DEL-56 |
| chr3-24191282-DEL-1754 | chr3-24191283-DEL-1754 |
| chr3-24193142-DEL-581 | chr3-24193143-DEL-581 |
| chr3-24208829-DEL-51 | chr3-24208830-DEL-51 |
| chr3-24214066-DEL-54 | chr3-24214066-DEL-53 |
| chr3-24241325-DEL-6381 | chr3-24241326-DEL-6381 |
| chr3-24281360-DEL-2360 | chr3-24281361-DEL-2360 |
| chr3-24286422-DEL-14007 | chr3-24286423-DEL-14007 |
| chr3-24293387-DEL-6456 | chr3-24293388-DEL-6456 |
| chr3-24306983-DEL-172 | chr3-24306984-DEL-172 |
| chr3-24327722-DEL-110224 | chr3-24327722-DEL-110224 |
| chr3-24371355-DEL-62 | chr3-24371356-DEL-62 |
| chr3-24387199-DEL-548 | chr3-24387200-DEL-548 |
| chr3-24399305-DEL-67 | chr3-24399330-DEL-55 |
| chr3-24443317-DEL-54 | chr3-24443330-DEL-50 |
| chr3-24453353-DEL-6404 | chr3-24453354-DEL-6404 |
| chr3-24508135-DEL-246 | chr3-24508130-DEL-252 |
| chr3-24524794-DEL-1516 | chr3-24524795-DEL-1516 |
| chr3-24527325-DEL-170 | chr3-24527326-DEL-170 |
| chr3-24537892-DEL-58 | chr3-24537893-DEL-58 |
| chr3-24548388-DEL-1509 | chr3-24548389-DEL-1509 |
| chr3-24553916-DEL-227 | chr3-24553917-DEL-227 |
| chr3-24581736-DEL-3680 | chr3-24581737-DEL-3678 |
| chr3-24591632-DEL-87 | chr3-24591633-DEL-85 |
| chr3-24616894-DEL-127 | chr3-24616893-DEL-123 |
| chr3-24620518-DEL-205 | chr3-24620519-DEL-205 |
| chr3-24636861-DEL-86 | chr3-24636862-DEL-86 |
| chr3-24664360-DEL-60 | chr3-24664361-DEL-60 |
| chr3-24689284-DEL-615 | chr3-24689285-DEL-615 |
| chr3-24704646-DEL-624 | chr3-24704647-DEL-624 |
| chr3-24718476-DEL-1039 | chr3-24718477-DEL-1039 |
| chr3-24770528-DEL-491 | chr3-24770529-DEL-491 |
| chr3-24771801-DEL-97 | chr3-24771802-DEL-92 |
| chr3-24812815-DEL-6542 | chr3-24812816-DEL-6542 |
| chr3-24835342-DEL-100 | chr3-24835313-DEL-82 |
| chr3-24852167-DEL-255 | chr3-24852168-DEL-255 |
| chr3-24864277-DEL-461 | chr3-24864278-DEL-461 |
| chr3-24865441-DEL-128 | chr3-24865442-DEL-102 |
| chr3-24928838-DEL-2014 | chr3-24928839-DEL-2014 |
| chr3-24950161-DEL-786 | chr3-24950162-DEL-786 |
| chr3-24953994-DEL-929 | chr3-24953995-DEL-929 |
| chr3-25016611-DEL-6651 | chr3-25016612-DEL-6651 |
| chr3-25025051-DEL-68 | chr3-25025052-DEL-68 |
| chr3-25038835-DEL-6307 | chr3-25038822-DEL-6308 |
| chr3-25088303-DEL-6443 | chr3-25088300-DEL-6444 |
| chr3-25088430-DEL-214 | chr3-25088430-DEL-214 |
| chr3-25144354-DEL-51 | chr3-25144355-DEL-51 |
| chr3-25155111-DEL-6422 | chr3-25155112-DEL-6422 |
| chr3-25175947-DEL-545 | chr3-25175948-DEL-544 |
| chr3-25180217-DEL-645 | chr3-25180218-DEL-645 |
| chr3-25192413-DEL-1483 | chr3-25192414-DEL-1483 |
| chr3-25209276-DEL-7248 | chr3-25209277-DEL-7248 |
| chr3-25221057-DEL-84 | chr3-25221053-DEL-95 |
| chr3-25222873-DEL-55 | chr3-25222870-DEL-58 |
| chr3-25229899-DEL-91 | chr3-25229910-DEL-93 |
| chr3-25239675-DEL-60 | chr3-25239676-DEL-60 |
| chr3-25249402-DEL-51 | chr3-25249403-DEL-51 |
| chr3-25252394-DEL-450 | chr3-25252395-DEL-450 |
| chr3-25264903-DEL-1993 | chr3-25264904-DEL-1993 |

| | |
|---|---|
| chr3-25301287-DEL-216 | chr3-25301288-DEL-216 |
| chr3-25344398-DEL-6065 | chr3-25344399-DEL-6065 |
| chr3-25461257-DEL-203 | chr3-25461242-DEL-204 |
| chr3-25471005-DEL-4606 | chr3-25471006-DEL-4606 |
| chr3-25527042-DEL-6383 | chr3-25527042-DEL-6383 |
| chr3-25544035-DEL-534 | chr3-25544036-DEL-533 |
| chr3-25654745-DEL-486 | chr3-25654746-DEL-486 |
| chr3-25662805-DEL-267 | chr3-25662804-DEL-268 |
| chr3-25699907-DEL-230 | chr3-25699908-DEL-229 |
| chr3-25712143-DEL-80 | chr3-25712144-DEL-80 |
| chr3-25785346-DEL-1469 | chr3-25785334-DEL-1469 |
| chr3-25818981-DEL-836 | chr3-25818982-DEL-836 |
| chr3-25821118-DEL-256 | chr3-25821119-DEL-256 |
| chr3-25873926-DEL-54 | chr3-25873916-DEL-85 |
| chr3-25900822-DEL-470 | chr3-25900831-DEL-465 |
| chr3-25934319-DEL-62 | chr3-25934320-DEL-62 |
| chr3-25975737-DEL-953 | chr3-25975738-DEL-953 |
| chr3-25989861-DEL-105 | chr3-25989839-DEL-127 |
| chr3-26001218-DEL-631 | chr3-26001219-DEL-631 |
| chr3-26016794-DEL-89 | chr3-26016795-DEL-89 |
| chr3-26018526-DEL-59 | chr3-26018528-DEL-60 |
| chr3-26043439-DEL-1032 | chr3-26043440-DEL-1032 |
| chr3-26110322-DEL-82 | chr3-26110309-DEL-89 |
| chr3-26114220-DEL-465 | chr3-26114221-DEL-465 |
| chr3-26143300-DEL-2556 | chr3-26143301-DEL-2556 |
| chr3-26151131-DEL-1951 | chr3-26151132-DEL-1951 |
| chr3-26160884-DEL-6680 | chr3-26160885-DEL-6680 |
| chr3-26173811-DEL-52 | chr3-26173795-DEL-70 |
| chr3-26180266-DEL-84 | chr3-26180286-DEL-58 |
| chr3-26205344-DEL-140 | chr3-26205345-DEL-140 |
| chr3-26217886-DEL-2380 | chr3-26217887-DEL-2380 |
| chr3-26241864-DEL-254 | chr3-26241865-DEL-254 |
| chr3-26244126-DEL-261 | chr3-26244127-DEL-261 |
| chr3-26261443-DEL-75 | chr3-26261444-DEL-75 |
| chr3-26278317-DEL-648 | chr3-26278318-DEL-648 |
| chr3-26291171-DEL-954 | chr3-26291172-DEL-954 |
| chr3-26295046-DEL-74 | chr3-26295047-DEL-74 |
| chr3-26306054-DEL-68 | chr3-26306055-DEL-68 |
| chr3-26313898-DEL-1978 | chr3-26313899-DEL-1978 |
| chr3-26342667-DEL-1718 | chr3-26342668-DEL-1718 |
| chr3-26354752-DEL-72 | chr3-26354753-DEL-72 |
| chr3-26361005-DEL-69 | chr3-26361006-DEL-71 |
| chr3-26368260-DEL-182 | chr3-26368261-DEL-182 |
| chr3-26384907-DEL-58 | chr3-26384908-DEL-58 |
| chr3-26401000-DEL-7266 | chr3-26401001-DEL-7266 |
| chr3-26445419-DEL-6378 | chr3-26445420-DEL-6378 |
| chr3-26458415-DEL-56 | chr3-26458418-DEL-54 |
| chr3-26459073-DEL-145 | chr3-26459074-DEL-145 |
| chr3-26470027-DEL-1398 | chr3-26470028-DEL-1398 |
| chr3-26498181-DEL-6590 | chr3-26498182-DEL-6589 |
| chr3-26502238-DEL-188 | chr3-26502238-DEL-188 |
| chr3-26504409-DEL-208 | chr3-26504409-DEL-208 |
| chr3-26508561-DEL-163 | chr3-26508562-DEL-169 |
| chr3-26515264-DEL-55 | chr3-26515265-DEL-55 |
| chr3-26616095-DEL-67 | chr3-26616065-DEL-67 |
| chr3-26625956-DEL-7166 | chr3-26625957-DEL-7166 |
| chr3-26646295-DEL-1526 | chr3-26646296-DEL-1526 |
| chr3-26654878-DEL-177 | chr3-26654878-DEL-177 |
| chr3-26672150-DEL-72 | chr3-26672162-DEL-60 |
| chr3-26690441-DEL-719 | chr3-26690438-DEL-721 |
| chr3-26699599-DEL-1018 | chr3-26699606-DEL-1013 |
| chr3-26720462-DEL-792 | chr3-26720463-DEL-792 |
| chr3-26723214-DEL-60 | chr3-26723205-DEL-60 |
| chr3-26728689-DEL-66 | chr3-26728690-DEL-66 |
| chr3-26734976-DEL-323 | chr3-26734977-DEL-323 |
| chr3-26740008-DEL-74 | chr3-26740009-DEL-74 |
| chr3-26747710-DEL-60 | chr3-26747663-DEL-72 |
| chr3-26806915-DEL-404 | chr3-26806916-DEL-404 |
| chr3-26816086-DEL-6272 | chr3-26816087-DEL-6272 |
| chr3-26838139-DEL-6291 | chr3-26838140-DEL-6291 |
| chr3-26854442-DEL-113 | chr3-26854443-DEL-113 |
| chr3-26856967-DEL-60 | chr3-26856973-DEL-55 |
| chr3-26884245-DEL-70 | chr3-26884246-DEL-70 |
| chr3-26912468-DEL-201 | chr3-26912469-DEL-201 |
| chr3-26921297-DEL-135 | chr3-26921298-DEL-136 |
| chr3-26928506-DEL-88 | chr3-26928497-DEL-66 |
| chr3-26936796-DEL-61 | chr3-26936798-DEL-60 |
| chr3-26940486-DEL-901 | chr3-26940486-DEL-901 |
| chr3-26954392-DEL-6211 | chr3-26954393-DEL-6211 |
| chr3-26961730-DEL-51 | chr3-26961698-DEL-89 |

| | |
|---|---|
| chr3-26998595-DEL-2188 | chr3-26998595-DEL-2188 |
| chr3-26998927-DEL-199 | chr3-26998928-DEL-199 |
| chr3-27001669-DEL-4617 | chr3-27001669-DEL-4617 |
| chr3-27009712-DEL-393 | chr3-27009713-DEL-393 |
| chr3-27028400-DEL-2364 | chr3-27028401-DEL-2364 |
| chr3-27042306-DEL-50 | chr3-27042307-DEL-50 |
| chr3-27069495-DEL-8845 | chr3-27069488-DEL-8846 |
| chr3-27105749-DEL-152 | chr3-27105759-DEL-143 |
| chr3-27122192-DEL-811 | chr3-27122191-DEL-811 |
| chr3-27130519-DEL-212 | chr3-27130520-DEL-212 |
| chr3-27136479-DEL-179 | chr3-27136480-DEL-179 |
| chr3-27142635-DEL-1714 | chr3-27142636-DEL-1714 |
| chr3-27167255-DEL-203 | chr3-27167301-DEL-204 |
| chr3-27212050-DEL-384 | chr3-27212051-DEL-384 |
| chr3-27213590-DEL-431 | chr3-27213591-DEL-431 |
| chr3-27227198-DEL-1138 | chr3-27227221-DEL-1140 |
| chr3-27257265-DEL-358 | chr3-27257266-DEL-358 |
| chr3-27278414-DEL-108 | chr3-27278413-DEL-109 |
| chr3-27340278-DEL-69 | chr3-27340279-DEL-69 |
| chr3-27342339-DEL-115 | chr3-27342340-DEL-115 |
| chr3-27372038-DEL-1865 | chr3-27372039-DEL-1865 |
| chr3-27386396-DEL-97 | chr3-27386397-DEL-97 |
| chr3-27396548-DEL-75 | chr3-27396534-DEL-60 |
| chr3-27404466-DEL-148 | chr3-27404468-DEL-145 |
| chr3-27408928-DEL-1865 | chr3-27408929-DEL-1865 |
| chr3-27418617-DEL-52 | chr3-27418617-DEL-52 |
| chr3-27419560-DEL-2726 | chr3-27419560-DEL-2724 |
| chr3-27441197-DEL-1325 | chr3-27441198-DEL-1325 |
| chr3-27446821-DEL-774 | chr3-27446822-DEL-774 |
| chr3-27532163-DEL-68 | chr3-27532164-DEL-68 |
| chr3-27534894-DEL-58 | chr3-27534868-DEL-59 |
| chr3-27538555-DEL-164 | chr3-27538568-DEL-157 |
| chr3-27540390-DEL-736 | chr3-27540391-DEL-736 |
| chr3-27571535-DEL-108 | chr3-27571536-DEL-108 |
| chr3-27582181-DEL-337 | chr3-27582196-DEL-334 |
| chr3-27649633-DEL-53 | chr3-27649635-DEL-51 |
| chr3-27677399-DEL-1132 | chr3-27677406-DEL-1129 |
| chr3-27677805-DEL-401 | chr3-27677806-DEL-401 |
| chr3-27733312-DEL-181 | chr3-27733310-DEL-184 |
| chr3-27761389-DEL-58 | chr3-27761390-DEL-76 |
| chr3-27768709-DEL-138 | chr3-27768688-DEL-140 |
| chr3-27779473-DEL-80 | chr3-27779474-DEL-80 |
| chr3-27807087-DEL-2371 | chr3-27807088-DEL-2371 |
| chr3-27839923-DEL-76 | chr3-27839913-DEL-78 |
| chr3-27873458-DEL-509 | chr3-27873460-DEL-510 |
| chr3-27879083-DEL-89 | chr3-27879084-DEL-89 |
| chr3-27891829-DEL-84 | chr3-27891823-DEL-87 |
| chr3-27897526-DEL-102 | chr3-27897526-DEL-102 |
| chr3-27898301-DEL-379 | chr3-27898302-DEL-379 |
| chr3-27911836-DEL-168 | chr3-27911837-DEL-168 |
| chr3-27914572-DEL-51 | chr3-27914536-DEL-52 |
| chr3-27926081-DEL-500 | chr3-27926082-DEL-500 |
| chr3-27927809-DEL-1033 | chr3-27927810-DEL-1033 |
| chr3-28035044-DEL-625 | chr3-28035045-DEL-625 |
| chr3-28105540-DEL-257 | chr3-28105541-DEL-257 |
| chr3-28106485-DEL-90 | chr3-28106486-DEL-90 |
| chr3-28108506-DEL-801 | chr3-28108507-DEL-801 |
| chr3-28116884-DEL-7234 | chr3-28116885-DEL-7234 |
| chr3-28122805-DEL-69 | chr3-28122806-DEL-69 |
| chr3-28125509-DEL-1996 | chr3-28125510-DEL-1996 |
| chr3-28170673-DEL-427 | chr3-28170661-DEL-410 |
| chr3-28202080-DEL-105 | chr3-28202081-DEL-105 |
| chr3-28203639-DEL-93 | chr3-28203632-DEL-95 |
| chr3-28206587-DEL-203 | chr3-28206588-DEL-203 |
| chr3-28210085-DEL-58 | chr3-28210085-DEL-58 |
| chr3-28252344-DEL-3096 | chr3-28252345-DEL-3096 |
| chr3-28268627-DEL-448 | chr3-28268628-DEL-448 |
| chr3-28295452-DEL-68 | chr3-28295453-DEL-68 |
| chr3-28333481-DEL-68 | chr3-28333469-DEL-80 |
| chr3-28422870-DEL-94 | chr3-28422871-DEL-94 |
| chr3-28440462-DEL-279 | chr3-28440463-DEL-279 |
| chr3-28454290-DEL-93 | chr3-28454242-DEL-132 |
| chr3-28535827-DEL-158 | chr3-28535830-DEL-156 |
| chr3-28567970-DEL-8092 | chr3-28567971-DEL-8092 |
| chr3-28588374-DEL-1569 | chr3-28588375-DEL-1569 |
| chr3-28591514-DEL-256 | chr3-28591515-DEL-256 |
| chr3-28596626-DEL-72 | chr3-28596627-DEL-72 |
| chr3-28617552-DEL-55 | chr3-28617553-DEL-55 |
| chr3-28619406-DEL-118 | chr3-28619407-DEL-118 |
| chr3-28622343-DEL-1424 | chr3-28622344-DEL-1424 |

| | |
|---|---|
| chr3-28642914-DEL-178 | chr3-28642915-DEL-178 |
| chr3-28647240-DEL-65 | chr3-28647241-DEL-65 |
| chr3-28656390-DEL-957 | chr3-28656390-DEL-954 |
| chr3-28668667-DEL-509 | chr3-28668668-DEL-509 |
| chr3-28747716-DEL-55 | chr3-28747717-DEL-55 |
| chr3-28748499-DEL-454 | chr3-28748500-DEL-454 |
| chr3-28758559-DEL-96 | chr3-28758559-DEL-96 |
| chr3-28780109-DEL-218 | chr3-28780110-DEL-218 |
| chr3-28785957-DEL-839 | chr3-28785958-DEL-839 |
| chr3-28792631-DEL-633 | chr3-28792632-DEL-633 |
| chr3-28817644-DEL-1904 | chr3-28817645-DEL-1904 |
| chr3-28820307-DEL-93 | chr3-28820308-DEL-93 |
| chr3-28859090-DEL-188 | chr3-28859091-DEL-187 |
| chr3-28866104-DEL-52 | chr3-28866105-DEL-52 |
| chr3-28867622-DEL-86 | chr3-28867616-DEL-83 |
| chr3-28868205-DEL-130 | chr3-28868205-DEL-131 |
| chr3-28872358-DEL-65 | chr3-28872359-DEL-65 |
| chr3-28880697-DEL-898 | chr3-28880698-DEL-898 |
| chr3-28897018-DEL-614 | chr3-28897010-DEL-623 |
| chr3-28898161-DEL-98 | chr3-28898162-DEL-98 |
| chr3-28902528-DEL-56 | chr3-28902555-DEL-52 |
| chr3-28920946-DEL-6503 | chr3-28920947-DEL-6503 |
| chr3-28946404-DEL-55 | chr3-28946420-DEL-83 |
| chr3-28962729-DEL-136 | chr3-28962732-DEL-92 |
| chr3-28966074-DEL-552 | chr3-28966075-DEL-552 |
| chr3-28968011-DEL-63 | chr3-28968028-DEL-60 |
| chr3-28977325-DEL-1040 | chr3-28977326-DEL-1040 |
| chr3-29007785-DEL-988 | chr3-29007786-DEL-988 |
| chr3-29025841-DEL-106 | chr3-29025842-DEL-106 |
| chr3-29027078-DEL-369 | chr3-29027091-DEL-360 |
| chr3-29039165-DEL-641 | chr3-29039166-DEL-641 |
| chr3-29040377-DEL-223 | chr3-29040378-DEL-223 |
| chr3-29072850-DEL-307 | chr3-29072851-DEL-307 |
| chr3-29090654-DEL-108 | chr3-29090655-DEL-99 |
| chr3-29098968-DEL-61 | chr3-29098969-DEL-61 |
| chr3-29106082-DEL-1927 | chr3-29106083-DEL-1927 |
| chr3-29107039-DEL-440 | chr3-29107040-DEL-440 |
| chr3-29118048-DEL-1403 | chr3-29118049-DEL-1403 |
| chr3-29134704-DEL-74 | chr3-29134705-DEL-74 |
| chr3-29137527-DEL-1049 | chr3-29137528-DEL-1049 |
| chr3-29142013-DEL-6550 | chr3-29142014-DEL-6550 |
| chr3-29158946-DEL-211 | chr3-29158937-DEL-212 |
| chr3-29168503-DEL-102 | chr3-29168504-DEL-102 |
| chr3-29177851-DEL-774 | chr3-29177852-DEL-774 |
| chr3-29186166-DEL-74 | chr3-29186158-DEL-80 |
| chr3-29195268-DEL-712 | chr3-29195269-DEL-712 |
| chr3-29243034-DEL-63 | chr3-29243034-DEL-69 |
| chr3-29271721-DEL-53 | chr3-29271732-DEL-67 |
| chr3-29286256-DEL-64 | chr3-29286257-DEL-64 |
| chr3-29292660-DEL-100 | chr3-29292661-DEL-100 |
| chr3-29294672-DEL-84 | chr3-29294677-DEL-80 |
| chr3-29303032-DEL-72 | chr3-29303033-DEL-72 |
| chr3-29329442-DEL-60 | chr3-29329443-DEL-66 |
| chr3-29348677-DEL-536 | chr3-29348675-DEL-534 |
| chr3-29369506-DEL-6210 | chr3-29369507-DEL-6210 |
| chr3-29381447-DEL-354 | chr3-29381487-DEL-346 |
| chr3-29391265-DEL-51 | chr3-29391266-DEL-51 |
| chr3-29425005-DEL-350 | chr3-29425004-DEL-351 |
| chr3-29441473-DEL-538 | chr3-29441474-DEL-538 |
| chr3-29476935-DEL-1635 | chr3-29476943-DEL-1630 |
| chr3-29507001-DEL-225 | chr3-29507002-DEL-225 |
| chr3-29515602-DEL-6417 | chr3-29515603-DEL-6417 |
| chr3-29522650-DEL-70 | chr3-29522650-DEL-60 |
| chr3-29547690-DEL-199 | chr3-29547697-DEL-199 |
| chr3-29573137-DEL-6227 | chr3-29573138-DEL-6227 |
| chr3-29592569-DEL-216 | chr3-29592570-DEL-216 |
| chr3-29596628-DEL-1582 | chr3-29596629-DEL-1582 |
| chr3-29598390-DEL-202 | chr3-29598391-DEL-202 |
| chr3-29604598-DEL-425 | chr3-29604603-DEL-423 |
| chr3-29619579-DEL-60 | chr3-29619580-DEL-60 |
| chr3-29632098-DEL-64 | chr3-29632099-DEL-64 |
| chr3-29698633-DEL-64 | chr3-29698652-DEL-62 |
| chr3-29706900-DEL-73 | chr3-29706901-DEL-77 |
| chr3-29708457-DEL-54 | chr3-29708458-DEL-54 |
| chr3-29732800-DEL-1976 | chr3-29732801-DEL-1976 |
| chr3-29739592-DEL-951 | chr3-29739593-DEL-916 |
| chr3-29745472-DEL-102 | chr3-29745463-DEL-102 |
| chr3-29748088-DEL-187 | chr3-29748088-DEL-187 |
| chr3-29749447-DEL-990 | chr3-29749448-DEL-990 |
| chr3-29773613-DEL-6397 | chr3-29773614-DEL-6397 |

| | |
|---|---|
| chr3-29783028-DEL-69 | chr3-29783029-DEL-68 |
| chr3-29784180-DEL-55 | chr3-29784181-DEL-55 |
| chr3-29784970-DEL-64 | chr3-29784995-DEL-65 |
| chr3-29790562-DEL-2870 | chr3-29790563-DEL-2870 |
| chr3-29804316-DEL-61 | chr3-29804317-DEL-61 |
| chr3-29805251-DEL-102 | chr3-29805240-DEL-86 |
| chr3-29820933-DEL-201 | chr3-29820934-DEL-201 |
| chr3-29823810-DEL-104 | chr3-29823813-DEL-82 |
| chr3-29834697-DEL-277 | chr3-29834699-DEL-276 |
| chr3-29837512-DEL-389 | chr3-29837512-DEL-389 |
| chr3-29838129-DEL-1482 | chr3-29838130-DEL-1482 |
| chr3-29869399-DEL-1143 | chr3-29869400-DEL-1143 |
| chr3-29869932-DEL-197 | chr3-29869931-DEL-162 |
| chr3-29872967-DEL-1359 | chr3-29872969-DEL-1360 |
| chr3-29878816-DEL-53 | chr3-29878810-DEL-50 |
| chr3-29895154-DEL-65 | chr3-29895155-DEL-65 |
| chr3-29920270-DEL-6670 | chr3-29920271-DEL-6670 |
| chr3-29938932-DEL-700 | chr3-29938933-DEL-700 |
| chr3-29974855-DEL-327 | chr3-29974856-DEL-327 |
| chr3-29990569-DEL-70 | chr3-29990570-DEL-70 |
| chr3-30049748-DEL-65 | chr3-30049749-DEL-65 |
| chr3-30050364-DEL-61 | chr3-30050377-DEL-84 |
| chr3-30078079-DEL-7580 | chr3-30078080-DEL-7580 |
| chr3-30084881-DEL-206 | chr3-30084880-DEL-206 |
| chr3-30138435-DEL-51 | chr3-30138407-DEL-84 |
| chr3-30160487-DEL-53 | chr3-30160487-DEL-54 |
| chr3-30179865-DEL-79 | chr3-30179865-DEL-80 |
| chr3-30191513-DEL-102 | chr3-30191514-DEL-102 |
| chr3-30224462-DEL-1955 | chr3-30224463-DEL-1955 |
| chr3-30238084-DEL-104 | chr3-30238083-DEL-106 |
| chr3-30239430-DEL-196 | chr3-30239431-DEL-196 |
| chr3-30279676-DEL-52 | chr3-30279677-DEL-52 |
| chr3-30291733-DEL-52 | chr3-30291734-DEL-52 |
| chr3-30335775-DEL-64 | chr3-30335772-DEL-68 |
| chr3-30341846-DEL-64 | chr3-30341847-DEL-64 |
| chr3-30342315-DEL-126 | chr3-30342316-DEL-126 |
| chr3-30352935-DEL-53 | chr3-30352935-DEL-53 |
| chr3-30372679-DEL-51 | chr3-30372674-DEL-51 |
| chr3-30385750-DEL-72 | chr3-30385791-DEL-54 |
| chr3-30399494-DEL-58 | chr3-30399495-DEL-58 |
| chr3-30406074-DEL-130 | chr3-30406075-DEL-130 |
| chr3-30410247-DEL-1220 | chr3-30410248-DEL-1220 |
| chr3-30421049-DEL-141 | chr3-30421050-DEL-141 |
| chr3-30453542-DEL-1543 | chr3-30453543-DEL-1543 |
| chr3-30473450-DEL-184 | chr3-30473434-DEL-188 |
| chr3-30501513-DEL-145 | chr3-30501520-DEL-146 |
| chr3-30514598-DEL-54 | chr3-30514599-DEL-54 |
| chr3-30580563-DEL-95 | chr3-30580564-DEL-95 |
| chr3-30581755-DEL-123 | chr3-30581762-DEL-123 |
| chr3-30587694-DEL-65 | chr3-30587681-DEL-66 |
| chr3-30609579-DEL-1816 | chr3-30609580-DEL-1816 |
| chr3-30614348-DEL-627 | chr3-30614349-DEL-628 |
| chr3-30618199-DEL-3561 | chr3-30618200-DEL-3561 |
| chr3-30624201-DEL-1319 | chr3-30624202-DEL-1319 |
| chr3-30625730-DEL-214 | chr3-30625731-DEL-214 |
| chr3-30630374-DEL-214 | chr3-30630375-DEL-214 |
| chr3-30633196-DEL-92 | chr3-30633191-DEL-93 |
| chr3-30736435-DEL-300 | chr3-30736434-DEL-301 |
| chr3-30784805-DEL-123 | chr3-30784806-DEL-123 |
| chr3-30796139-DEL-90 | chr3-30796137-DEL-100 |
| chr3-30810103-DEL-55 | chr3-30810127-DEL-51 |
| chr3-30818778-DEL-62 | chr3-30818778-DEL-62 |
| chr3-30823922-DEL-91 | chr3-30823922-DEL-91 |
| chr3-30838602-DEL-149 | chr3-30838603-DEL-149 |
| chr3-30842259-DEL-200 | chr3-30842263-DEL-196 |
| chr3-30881429-DEL-113 | chr3-30881432-DEL-107 |
| chr3-30890235-DEL-3978 | chr3-30890236-DEL-3978 |
| chr3-30893282-DEL-73 | chr3-30893283-DEL-75 |
| chr3-30905425-DEL-61 | chr3-30905404-DEL-68 |
| chr3-30911583-DEL-52 | chr3-30911584-DEL-60 |
| chr3-30919959-DEL-74 | chr3-30919959-DEL-60 |
| chr3-30937711-DEL-910 | chr3-30937730-DEL-891 |
| chr3-30943859-DEL-193 | chr3-30943860-DEL-193 |
| chr3-30964790-DEL-1166 | chr3-30964773-DEL-1168 |
| chr3-30969721-DEL-62 | chr3-30969722-DEL-62 |
| chr3-30973699-DEL-519 | chr3-30973700-DEL-519 |
| chr3-31010334-DEL-242 | chr3-31010335-DEL-242 |
| chr3-31033057-DEL-207 | chr3-31033058-DEL-207 |
| chr3-31063152-DEL-566 | chr3-31063153-DEL-566 |
| chr3-31120668-DEL-239 | chr3-31120669-DEL-239 |

| | |
|---|---|
| chr3-31121673-DEL-436 | chr3-31121674-DEL-436 |
| chr3-31123102-DEL-231 | chr3-31123103-DEL-231 |
| chr3-31194166-DEL-78 | chr3-31194134-DEL-78 |
| chr3-31196241-DEL-215 | chr3-31196242-DEL-215 |
| chr3-31196666-DEL-399 | chr3-31196677-DEL-359 |
| chr3-31213799-DEL-208 | chr3-31213799-DEL-203 |
| chr3-31214393-DEL-905 | chr3-31214394-DEL-905 |
| chr3-31222480-DEL-276 | chr3-31222481-DEL-276 |
| chr3-31230471-DEL-2873 | chr3-31230471-DEL-2872 |
| chr3-31235898-DEL-7077 | chr3-31235896-DEL-7086 |
| chr3-31254023-DEL-137 | chr3-31254024-DEL-125 |
| chr3-31260362-DEL-201 | chr3-31260361-DEL-202 |
| chr3-31270725-DEL-202 | chr3-31270715-DEL-203 |
| chr3-31278971-DEL-375 | chr3-31278972-DEL-375 |
| chr3-31287366-DEL-54 | chr3-31287345-DEL-55 |
| chr3-31302278-DEL-62 | chr3-31302279-DEL-62 |
| chr3-31342887-DEL-54 | chr3-31342870-DEL-55 |
| chr3-31355971-DEL-131 | chr3-31355970-DEL-131 |
| chr3-31356576-DEL-74 | chr3-31356585-DEL-66 |
| chr3-31394955-DEL-1890 | chr3-31394956-DEL-1890 |
| chr3-31404780-DEL-6563 | chr3-31404781-DEL-6563 |
| chr3-31418375-DEL-60 | chr3-31418376-DEL-60 |
| chr3-31429923-DEL-1198 | chr3-31429924-DEL-1198 |
| chr3-31435010-DEL-187 | chr3-31435059-DEL-165 |
| chr3-31438151-DEL-320 | chr3-31438153-DEL-319 |
| chr3-31450284-DEL-60 | chr3-31450285-DEL-60 |
| chr3-31465942-DEL-202 | chr3-31465943-DEL-202 |
| chr3-31475784-DEL-58 | chr3-31475785-DEL-58 |
| chr3-31476297-DEL-1476 | chr3-31476298-DEL-1476 |
| chr3-31487687-DEL-366 | chr3-31487687-DEL-367 |
| chr3-31490708-DEL-693 | chr3-31490709-DEL-693 |
| chr3-31497046-DEL-94 | chr3-31497047-DEL-94 |
| chr3-31499054-DEL-60 | chr3-31499055-DEL-60 |
| chr3-31502370-DEL-3808 | chr3-31502371-DEL-3808 |
| chr3-31524076-DEL-7789 | chr3-31524077-DEL-7789 |
| chr3-31539295-DEL-1906 | chr3-31539296-DEL-1906 |
| chr3-31545063-DEL-60 | chr3-31545063-DEL-60 |
| chr3-31552311-DEL-223 | chr3-31552312-DEL-223 |
| chr3-31570146-DEL-54 | chr3-31570147-DEL-54 |
| chr3-31582104-DEL-433 | chr3-31582105-DEL-433 |
| chr3-31621633-DEL-1316 | chr3-31621628-DEL-1319 |
| chr3-31633150-DEL-90 | chr3-31633169-DEL-72 |
| chr3-31635153-DEL-1602 | chr3-31635154-DEL-1602 |
| chr3-31636954-DEL-3005 | chr3-31636955-DEL-3005 |
| chr3-31662684-DEL-198 | chr3-31662685-DEL-198 |
| chr3-31693681-DEL-1228 | chr3-31693682-DEL-1228 |
| chr3-31694013-DEL-4175 | chr3-31694014-DEL-4175 |
| chr3-31702694-DEL-147 | chr3-31702723-DEL-119 |
| chr3-31740344-DEL-200 | chr3-31740345-DEL-214 |
| chr3-31745384-DEL-197 | chr3-31745389-DEL-195 |
| chr3-31755489-DEL-6207 | chr3-31755490-DEL-6207 |
| chr3-31763060-DEL-194 | chr3-31763061-DEL-194 |
| chr3-31801648-DEL-341 | chr3-31801649-DEL-341 |
| chr3-31802822-DEL-1834 | chr3-31802822-DEL-1834 |
| chr3-31816285-DEL-2952 | chr3-31816286-DEL-2952 |
| chr3-31821192-DEL-70 | chr3-31821182-DEL-80 |
| chr3-31826027-DEL-52 | chr3-31826025-DEL-50 |
| chr3-31854176-DEL-445 | chr3-31854177-DEL-445 |
| chr3-31873463-DEL-7777 | chr3-31873463-DEL-7777 |
| chr3-31877308-DEL-1071 | chr3-31877309-DEL-1071 |
| chr3-31882191-DEL-427 | chr3-31882191-DEL-427 |
| chr3-31896473-DEL-2513 | chr3-31896474-DEL-2513 |
| chr3-31899618-DEL-176 | chr3-31899631-DEL-173 |
| chr3-31902821-DEL-619 | chr3-31902822-DEL-619 |
| chr3-31910847-DEL-723 | chr3-31910848-DEL-723 |
| chr3-31929698-DEL-54 | chr3-31929658-DEL-52 |
| chr3-31936179-DEL-345 | chr3-31936180-DEL-345 |
| chr3-31973077-DEL-110 | chr3-31973074-DEL-111 |
| chr3-31990826-DEL-70 | chr3-31990827-DEL-70 |
| chr3-32000538-DEL-399 | chr3-32000539-DEL-399 |
| chr3-32014593-DEL-411 | chr3-32014584-DEL-413 |
| chr3-32041599-DEL-58 | chr3-32041599-DEL-59 |
| chr3-32058063-DEL-53 | chr3-32058060-DEL-51 |
| chr3-32061087-DEL-178 | chr3-32061088-DEL-178 |
| chr3-32071548-DEL-51 | chr3-32071543-DEL-57 |
| chr3-32101320-DEL-104 | chr3-32101336-DEL-65 |
| chr3-32115766-DEL-56 | chr3-32115767-DEL-56 |
| chr3-32129701-DEL-7123 | chr3-32129702-DEL-7123 |
| chr3-32185913-DEL-64 | chr3-32185914-DEL-64 |
| chr3-32216068-DEL-6666 | chr3-32216069-DEL-6666 |

| | |
|---|---|
| chr3-32222310-DEL-201 | chr3-32222311-DEL-201 |
| chr3-32223376-DEL-57 | chr3-32223377-DEL-57 |
| chr3-32273919-DEL-212 | chr3-32273920-DEL-211 |
| chr3-32286960-DEL-177 | chr3-32286960-DEL-178 |
| chr3-32294687-DEL-6381 | chr3-32294688-DEL-6381 |
| chr3-32301976-DEL-204 | chr3-32301977-DEL-204 |
| chr3-32317131-DEL-1027 | chr3-32317132-DEL-1027 |
| chr3-32319793-DEL-1808 | chr3-32319794-DEL-1808 |
| chr3-32320566-DEL-147 | chr3-32320574-DEL-139 |
| chr3-32330423-DEL-201 | chr3-32330424-DEL-201 |
| chr3-32332015-DEL-64 | chr3-32332036-DEL-91 |
| chr3-32332667-DEL-52 | chr3-32332668-DEL-52 |
| chr3-32339455-DEL-135 | chr3-32339456-DEL-135 |
| chr3-32350432-DEL-270 | chr3-32350433-DEL-270 |
| chr3-32362215-DEL-6195 | chr3-32362216-DEL-6195 |
| chr3-32382949-DEL-64 | chr3-32382944-DEL-68 |
| chr3-32414922-DEL-72 | chr3-32414917-DEL-71 |
| chr3-32424747-DEL-5688 | chr3-32424735-DEL-5698 |
| chr3-32428014-DEL-96 | chr3-32428022-DEL-94 |
| chr3-32429536-DEL-435 | chr3-32429537-DEL-435 |
| chr3-32472327-DEL-5098 | chr3-32472328-DEL-5098 |
| chr3-32474574-DEL-199 | chr3-32474582-DEL-198 |
| chr3-32478510-DEL-84 | chr3-32478512-DEL-85 |
| chr3-32509135-DEL-154 | chr3-32509172-DEL-110 |
| chr3-32555395-DEL-268 | chr3-32555396-DEL-268 |
| chr3-32561898-DEL-200 | chr3-32561899-DEL-170 |
| chr3-32579696-DEL-100 | chr3-32579697-DEL-100 |
| chr3-32585821-DEL-190 | chr3-32585822-DEL-190 |
| chr3-32593196-DEL-66 | chr3-32593197-DEL-66 |
| chr3-32635810-DEL-388 | chr3-32635811-DEL-388 |
| chr3-32645994-DEL-90 | chr3-32645995-DEL-90 |
| chr3-32656624-DEL-486 | chr3-32656625-DEL-486 |
| chr3-32686052-DEL-116 | chr3-32686048-DEL-114 |
| chr3-32713410-DEL-185 | chr3-32713460-DEL-136 |
| chr3-32736750-DEL-446 | chr3-32736751-DEL-446 |
| chr3-32748812-DEL-59 | chr3-32748839-DEL-72 |
| chr3-32771254-DEL-412 | chr3-32771255-DEL-412 |
| chr3-32783128-DEL-187 | chr3-32783129-DEL-187 |
| chr3-32783933-DEL-199 | chr3-32783934-DEL-199 |
| chr3-32786347-DEL-184 | chr3-32786348-DEL-184 |
| chr3-32813153-DEL-154 | chr3-32813154-DEL-154 |
| chr3-32821373-DEL-1493 | chr3-32821374-DEL-1493 |
| chr3-32825746-DEL-1637 | chr3-32825739-DEL-1637 |
| chr3-32838631-DEL-6222 | chr3-32838632-DEL-6222 |
| chr3-32890980-DEL-135 | chr3-32890979-DEL-142 |
| chr3-32892449-DEL-171 | chr3-32892450-DEL-171 |
| chr3-32896876-DEL-195 | chr3-32896862-DEL-224 |
| chr3-32898218-DEL-74 | chr3-32898219-DEL-74 |
| chr3-32917096-DEL-764 | chr3-32917097-DEL-764 |
| chr3-32923553-DEL-102 | chr3-32923559-DEL-58 |
| chr3-32927193-DEL-702 | chr3-32927194-DEL-702 |
| chr3-32932313-DEL-3004 | chr3-32932312-DEL-2998 |
| chr3-32940886-DEL-58 | chr3-32940887-DEL-58 |
| chr3-32944761-DEL-97 | chr3-32944762-DEL-97 |
| chr3-32950753-DEL-51 | chr3-32950764-DEL-51 |
| chr3-32954191-DEL-72 | chr3-32954191-DEL-70 |
| chr3-33000348-DEL-180 | chr3-33000349-DEL-180 |
| chr3-33028437-DEL-718 | chr3-33028438-DEL-718 |
| chr3-33041230-DEL-130 | chr3-33041196-DEL-130 |
| chr3-33088213-DEL-1502 | chr3-33088214-DEL-1502 |
| chr3-33123773-DEL-554 | chr3-33123774-DEL-554 |
| chr3-33139675-DEL-100 | chr3-33139676-DEL-100 |
| chr3-33159822-DEL-226 | chr3-33159823-DEL-226 |
| chr3-33163551-DEL-273 | chr3-33163517-DEL-309 |
| chr3-33167516-DEL-64 | chr3-33167517-DEL-64 |
| chr3-33180086-DEL-245 | chr3-33180088-DEL-242 |
| chr3-33186188-DEL-491 | chr3-33186189-DEL-491 |
| chr3-33200096-DEL-558 | chr3-33200097-DEL-558 |
| chr3-33210063-DEL-3103 | chr3-33210064-DEL-3103 |
| chr3-33221655-DEL-6186 | chr3-33221656-DEL-6186 |
| chr3-33255115-DEL-6390 | chr3-33255116-DEL-6390 |
| chr3-33291866-DEL-3910 | chr3-33291867-DEL-3910 |
| chr3-33293587-DEL-572 | chr3-33293588-DEL-572 |
| chr3-33297572-DEL-6871 | chr3-33297573-DEL-6871 |
| chr3-33305641-DEL-84 | chr3-33305642-DEL-84 |
| chr3-33313586-DEL-64 | chr3-33313565-DEL-62 |
| chr3-33323381-DEL-735 | chr3-33323382-DEL-735 |
| chr3-33325980-DEL-587 | chr3-33325981-DEL-587 |
| chr3-33332144-DEL-324 | chr3-33332145-DEL-324 |
| chr3-33335857-DEL-531 | chr3-33335857-DEL-531 |

| | |
|---|---|
| chr3-33339925-DEL-2247 | chr3-33339926-DEL-2247 |
| chr3-33341289-DEL-54 | chr3-33341290-DEL-54 |
| chr3-33342416-DEL-87 | chr3-33342417-DEL-87 |
| chr3-33353055-DEL-70 | chr3-33353056-DEL-70 |
| chr3-33375784-DEL-3156 | chr3-33375776-DEL-3164 |
| chr3-33378508-DEL-74 | chr3-33378509-DEL-74 |
| chr3-33385486-DEL-391 | chr3-33385486-DEL-391 |
| chr3-33388807-DEL-51 | chr3-33388808-DEL-51 |
| chr3-33390283-DEL-1100 | chr3-33390284-DEL-1100 |
| chr3-33420312-DEL-824 | chr3-33420313-DEL-824 |
| chr3-33441278-DEL-457 | chr3-33441279-DEL-457 |
| chr3-33480487-DEL-2710 | chr3-33480488-DEL-2710 |
| chr3-33502652-DEL-56 | chr3-33502653-DEL-56 |
| chr3-33563544-DEL-1439 | chr3-33563542-DEL-1440 |
| chr3-33636885-DEL-216 | chr3-33636861-DEL-217 |
| chr3-33640808-DEL-1423 | chr3-33640809-DEL-1423 |
| chr3-33650262-DEL-105 | chr3-33650261-DEL-106 |
| chr3-33697239-DEL-1002 | chr3-33697240-DEL-1002 |
| chr3-33704602-DEL-85 | chr3-33704603-DEL-85 |
| chr3-33717043-DEL-3089 | chr3-33717032-DEL-3090 |
| chr3-33731233-DEL-275 | chr3-33731234-DEL-275 |
| chr3-33738035-DEL-317 | chr3-33738036-DEL-317 |
| chr3-33747424-DEL-156 | chr3-33747421-DEL-178 |
| chr3-33807290-DEL-565 | chr3-33807291-DEL-565 |
| chr3-33822262-DEL-6873 | chr3-33822263-DEL-6873 |
| chr3-33835390-DEL-4471 | chr3-33835391-DEL-4471 |
| chr3-33876310-DEL-473 | chr3-33876311-DEL-473 |
| chr3-33893540-DEL-1110 | chr3-33893541-DEL-1110 |
| chr3-33910407-DEL-337 | chr3-33910408-DEL-337 |
| chr3-33928381-DEL-201 | chr3-33928381-DEL-201 |
| chr3-33932550-DEL-110 | chr3-33932565-DEL-74 |
| chr3-33953336-DEL-90 | chr3-33953337-DEL-90 |
| chr3-33974393-DEL-1601 | chr3-33974394-DEL-1599 |
| chr3-33983051-DEL-375 | chr3-33983053-DEL-374 |
| chr3-33996514-DEL-6440 | chr3-33996515-DEL-6440 |
| chr3-34004609-DEL-98 | chr3-34004609-DEL-98 |
| chr3-34006931-DEL-72 | chr3-34006927-DEL-64 |
| chr3-34017981-DEL-209 | chr3-34017982-DEL-209 |
| chr3-34024243-DEL-84 | chr3-34024244-DEL-84 |
| chr3-34055097-DEL-214 | chr3-34055098-DEL-214 |
| chr3-34058919-DEL-780 | chr3-34058891-DEL-784 |
| chr3-34087668-DEL-70 | chr3-34087682-DEL-55 |
| chr3-34140180-DEL-218 | chr3-34140181-DEL-218 |
| chr3-34141900-DEL-62 | chr3-34141913-DEL-52 |
| chr3-34156771-DEL-124 | chr3-34156793-DEL-82 |
| chr3-34159904-DEL-72 | chr3-34159898-DEL-74 |
| chr3-34183656-DEL-76 | chr3-34183657-DEL-76 |
| chr3-34202584-DEL-129 | chr3-34202591-DEL-128 |
| chr3-34208747-DEL-111 | chr3-34208748-DEL-111 |
| chr3-34236294-DEL-1034 | chr3-34236295-DEL-1034 |
| chr3-34242978-DEL-1997 | chr3-34242979-DEL-1997 |
| chr3-34247776-DEL-52 | chr3-34247819-DEL-58 |
| chr3-34278266-DEL-203 | chr3-34278267-DEL-203 |
| chr3-34305794-DEL-81 | chr3-34305786-DEL-71 |
| chr3-34312518-DEL-698 | chr3-34312519-DEL-698 |
| chr3-34334585-DEL-5346 | chr3-34334583-DEL-5347 |
| chr3-34370514-DEL-51 | chr3-34370515-DEL-51 |
| chr3-34417354-DEL-6111 | chr3-34417380-DEL-6083 |
| chr3-34438052-DEL-51 | chr3-34438020-DEL-93 |
| chr3-34493092-DEL-72 | chr3-34493098-DEL-67 |
| chr3-34566620-DEL-52 | chr3-34566621-DEL-52 |
| chr3-34570552-DEL-1895 | chr3-34570553-DEL-1895 |
| chr3-34588478-DEL-93 | chr3-34588479-DEL-93 |
| chr3-34593702-DEL-73 | chr3-34593706-DEL-71 |
| chr3-34595081-DEL-121 | chr3-34595087-DEL-88 |
| chr3-34622787-DEL-166 | chr3-34622788-DEL-166 |
| chr3-34637790-DEL-54 | chr3-34637791-DEL-54 |
| chr3-34639905-DEL-127 | chr3-34639906-DEL-127 |
| chr3-34647721-DEL-59 | chr3-34647721-DEL-53 |
| chr3-34659491-DEL-62 | chr3-34659492-DEL-62 |
| chr3-34674955-DEL-65 | chr3-34674928-DEL-95 |
| chr3-34675122-DEL-741 | chr3-34675123-DEL-741 |
| chr3-34678172-DEL-70 | chr3-34678173-DEL-70 |
| chr3-34685865-DEL-571 | chr3-34685866-DEL-570 |
| chr3-34699706-DEL-147 | chr3-34699707-DEL-147 |
| chr3-34735046-DEL-51 | chr3-34735043-DEL-54 |
| chr3-34758583-DEL-219 | chr3-34758584-DEL-219 |
| chr3-34765528-DEL-91 | chr3-34765531-DEL-101 |
| chr3-34774991-DEL-194 | chr3-34774989-DEL-195 |
| chr3-34797218-DEL-186 | chr3-34797219-DEL-192 |

| | |
|---|---|
| chr3-34811684-DEL-85 | chr3-34811685-DEL-85 |
| chr3-34817878-DEL-208 | chr3-34817879-DEL-208 |
| chr3-34827495-DEL-68 | chr3-34827509-DEL-73 |
| chr3-34831445-DEL-58 | chr3-34831446-DEL-58 |
| chr3-34843535-DEL-222 | chr3-34843536-DEL-222 |
| chr3-34845188-DEL-81 | chr3-34845189-DEL-81 |
| chr3-34866415-DEL-521 | chr3-34866415-DEL-522 |
| chr3-34872619-DEL-362 | chr3-34872620-DEL-362 |
| chr3-34874242-DEL-79 | chr3-34874245-DEL-52 |
| chr3-34900571-DEL-7107 | chr3-34900572-DEL-7107 |
| chr3-34912095-DEL-69 | chr3-34912097-DEL-67 |
| chr3-34939089-DEL-54 | chr3-34939090-DEL-54 |
| chr3-34941021-DEL-185 | chr3-34941022-DEL-185 |
| chr3-34946181-DEL-474 | chr3-34946182-DEL-474 |
| chr3-34992626-DEL-1364 | chr3-34992627-DEL-1364 |
| chr3-34997454-DEL-74 | chr3-34997442-DEL-86 |
| chr3-35053838-DEL-88 | chr3-35053839-DEL-88 |
| chr3-35080667-DEL-249 | chr3-35080668-DEL-249 |
| chr3-35112567-DEL-699 | chr3-35112565-DEL-699 |
| chr3-35119922-DEL-63 | chr3-35119923-DEL-64 |
| chr3-35127653-DEL-1768 | chr3-35127654-DEL-1768 |
| chr3-35128450-DEL-95 | chr3-35128451-DEL-95 |
| chr3-35139467-DEL-60 | chr3-35139468-DEL-60 |
| chr3-35178733-DEL-6886 | chr3-35178734-DEL-6886 |
| chr3-35213685-DEL-133 | chr3-35213656-DEL-134 |
| chr3-35219988-DEL-711 | chr3-35219989-DEL-711 |
| chr3-35223672-DEL-430 | chr3-35223673-DEL-430 |
| chr3-35224416-DEL-67 | chr3-35224405-DEL-81 |
| chr3-35225661-DEL-507 | chr3-35225662-DEL-507 |
| chr3-35226779-DEL-100 | chr3-35226786-DEL-50 |
| chr3-35228822-DEL-58 | chr3-35228823-DEL-58 |
| chr3-35286575-DEL-80 | chr3-35286576-DEL-80 |
| chr3-35306416-DEL-136 | chr3-35306417-DEL-136 |
| chr3-35321497-DEL-98 | chr3-35321466-DEL-84 |
| chr3-35340894-DEL-2253 | chr3-35340894-DEL-2253 |
| chr3-35349641-DEL-75 | chr3-35349641-DEL-75 |
| chr3-35366188-DEL-78 | chr3-35366171-DEL-110 |
| chr3-35398607-DEL-52 | chr3-35398588-DEL-56 |
| chr3-35404239-DEL-80 | chr3-35404240-DEL-83 |
| chr3-35434156-DEL-5994 | chr3-35434157-DEL-5994 |
| chr3-35444468-DEL-63 | chr3-35444466-DEL-64 |
| chr3-35454173-DEL-58 | chr3-35454174-DEL-57 |
| chr3-35456027-DEL-282 | chr3-35456028-DEL-329 |
| chr3-35458361-DEL-116 | chr3-35458361-DEL-116 |
| chr3-35480091-DEL-6412 | chr3-35480090-DEL-6413 |
| chr3-35511287-DEL-683 | chr3-35511288-DEL-683 |
| chr3-35511874-DEL-52 | chr3-35511874-DEL-64 |
| chr3-35545573-DEL-72 | chr3-35545577-DEL-60 |
| chr3-35548447-DEL-85 | chr3-35548447-DEL-85 |
| chr3-35554048-DEL-129 | chr3-35554049-DEL-129 |
| chr3-35582372-DEL-62 | chr3-35582373-DEL-60 |
| chr3-35593102-DEL-124 | chr3-35593103-DEL-124 |
| chr3-35595249-DEL-63 | chr3-35595250-DEL-57 |
| chr3-35604353-DEL-110 | chr3-35604354-DEL-110 |
| chr3-35604837-DEL-401 | chr3-35604838-DEL-401 |
| chr3-35630455-DEL-332 | chr3-35630456-DEL-332 |
| chr3-35630958-DEL-119 | chr3-35630959-DEL-119 |
| chr3-35635055-DEL-2449 | chr3-35635055-DEL-2449 |
| chr3-35647771-DEL-379 | chr3-35647772-DEL-379 |
| chr3-35648519-DEL-3051 | chr3-35648520-DEL-3051 |
| chr3-35657584-DEL-73 | chr3-35657593-DEL-66 |
| chr3-35658161-DEL-1781 | chr3-35658162-DEL-1781 |
| chr3-35661421-DEL-6179 | chr3-35661422-DEL-6179 |
| chr3-35690122-DEL-386 | chr3-35690123-DEL-386 |
| chr3-35704333-DEL-114 | chr3-35704324-DEL-121 |
| chr3-35718091-DEL-400 | chr3-35718092-DEL-400 |
| chr3-35721707-DEL-89 | chr3-35721708-DEL-89 |
| chr3-35761168-DEL-6445 | chr3-35761169-DEL-6445 |
| chr3-35771732-DEL-299 | chr3-35771733-DEL-299 |
| chr3-35776464-DEL-312 | chr3-35776465-DEL-263 |
| chr3-35781486-DEL-92 | chr3-35781487-DEL-92 |
| chr3-35786211-DEL-216 | chr3-35786211-DEL-209 |
| chr3-35800377-DEL-80 | chr3-35800378-DEL-80 |
| chr3-35890565-DEL-292 | chr3-35890566-DEL-292 |
| chr3-35925032-DEL-3780 | chr3-35925033-DEL-3780 |
| chr3-35926645-DEL-79 | chr3-35926645-DEL-79 |
| chr3-35960434-DEL-473 | chr3-35960435-DEL-473 |
| chr3-35979451-DEL-122 | chr3-35979452-DEL-122 |
| chr3-36032672-DEL-169 | chr3-36032671-DEL-170 |
| chr3-36043397-DEL-122 | chr3-36043398-DEL-122 |

| | |
|---|---|
| chr3-36047251-DEL-175 | chr3-36047252-DEL-175 |
| chr3-36056166-DEL-267 | chr3-36056167-DEL-267 |
| chr3-36066248-DEL-64 | chr3-36066259-DEL-54 |
| chr3-36066959-DEL-860 | chr3-36066955-DEL-861 |
| chr3-36091298-DEL-411 | chr3-36091299-DEL-411 |
| chr3-36092741-DEL-97 | chr3-36092742-DEL-97 |
| chr3-36097204-DEL-62 | chr3-36097223-DEL-52 |
| chr3-36139772-DEL-84 | chr3-36139773-DEL-84 |
| chr3-36152570-DEL-830 | chr3-36152571-DEL-830 |
| chr3-36153927-DEL-173 | chr3-36153927-DEL-173 |
| chr3-36160580-DEL-2775 | chr3-36160581-DEL-2775 |
| chr3-36169691-DEL-54 | chr3-36169692-DEL-52 |
| chr3-36199201-DEL-328 | chr3-36199202-DEL-328 |
| chr3-36204156-DEL-58 | chr3-36204157-DEL-56 |
| chr3-36220606-DEL-194 | chr3-36220607-DEL-194 |
| chr3-36231634-DEL-60 | chr3-36231635-DEL-60 |
| chr3-36271852-DEL-52 | chr3-36271853-DEL-52 |
| chr3-36281267-DEL-63 | chr3-36281268-DEL-63 |
| chr3-36287109-DEL-51 | chr3-36287110-DEL-51 |
| chr3-36297386-DEL-51 | chr3-36297387-DEL-51 |
| chr3-36297687-DEL-328 | chr3-36297688-DEL-328 |
| chr3-36304227-DEL-263 | chr3-36304228-DEL-263 |
| chr3-36309924-DEL-2926 | chr3-36309924-DEL-2926 |
| chr3-36335424-DEL-120 | chr3-36335419-DEL-114 |
| chr3-36339491-DEL-6449 | chr3-36339492-DEL-6449 |
| chr3-36345530-DEL-398 | chr3-36345530-DEL-398 |
| chr3-36350491-DEL-111 | chr3-36350492-DEL-111 |
| chr3-36354636-DEL-52 | chr3-36354616-DEL-73 |
| chr3-36372044-DEL-62 | chr3-36372045-DEL-62 |
| chr3-36380459-DEL-2549 | chr3-36380460-DEL-2549 |
| chr3-36391298-DEL-56 | chr3-36391299-DEL-56 |
| chr3-36397860-DEL-84 | chr3-36397864-DEL-81 |
| chr3-36420443-DEL-936 | chr3-36420443-DEL-935 |
| chr3-36422591-DEL-122 | chr3-36422595-DEL-123 |
| chr3-36432352-DEL-73 | chr3-36432350-DEL-56 |
| chr3-36499740-DEL-56 | chr3-36499741-DEL-62 |
| chr3-36522660-DEL-56 | chr3-36522661-DEL-56 |
| chr3-36550209-DEL-105 | chr3-36550210-DEL-105 |
| chr3-36562387-DEL-1890 | chr3-36562388-DEL-1890 |
| chr3-36576224-DEL-72 | chr3-36576241-DEL-64 |
| chr3-36580070-DEL-268 | chr3-36580070-DEL-268 |
| chr3-36588120-DEL-265 | chr3-36588121-DEL-265 |
| chr3-36616326-DEL-205 | chr3-36616327-DEL-205 |
| chr3-36647192-DEL-66 | chr3-36647193-DEL-66 |
| chr3-36654662-DEL-751 | chr3-36654663-DEL-751 |
| chr3-36686272-DEL-146 | chr3-36686264-DEL-147 |
| chr3-36698570-DEL-82 | chr3-36698575-DEL-74 |
| chr3-36756506-DEL-55 | chr3-36756507-DEL-54 |
| chr3-36781983-DEL-56 | chr3-36782003-DEL-52 |
| chr3-36792726-DEL-188 | chr3-36792734-DEL-190 |
| chr3-36793998-DEL-1677 | chr3-36793999-DEL-1677 |
| chr3-36812045-DEL-1860 | chr3-36812046-DEL-1860 |
| chr3-36815438-DEL-62 | chr3-36815434-DEL-63 |
| chr3-36840039-DEL-61 | chr3-36840041-DEL-60 |
| chr3-36843799-DEL-341 | chr3-36843791-DEL-342 |
| chr3-36853590-DEL-282 | chr3-36853591-DEL-282 |
| chr3-36854307-DEL-332 | chr3-36854314-DEL-323 |
| chr3-36893404-DEL-94 | chr3-36893399-DEL-97 |
| chr3-36913178-DEL-52 | chr3-36913179-DEL-52 |
| chr3-36930729-DEL-1010 | chr3-36930730-DEL-1010 |
| chr3-36953400-DEL-6187 | chr3-36953401-DEL-6187 |
| chr3-36968028-DEL-504 | chr3-36968029-DEL-504 |
| chr3-37016629-DEL-1042 | chr3-37016630-DEL-1042 |
| chr3-37019584-DEL-57 | chr3-37019584-DEL-58 |
| chr3-37037016-DEL-54 | chr3-37036988-DEL-57 |
| chr3-37083794-DEL-68 | chr3-37083795-DEL-60 |
| chr3-37136691-DEL-60 | chr3-37136692-DEL-60 |
| chr3-37152746-DEL-318 | chr3-37152742-DEL-319 |
| chr3-37160737-DEL-54 | chr3-37160738-DEL-54 |
| chr3-37204935-DEL-56 | chr3-37204935-DEL-56 |
| chr3-37205809-DEL-137 | chr3-37205810-DEL-137 |
| chr3-37211611-DEL-52 | chr3-37211612-DEL-52 |
| chr3-37235948-DEL-870 | chr3-37235949-DEL-870 |
| chr3-37268770-DEL-2814 | chr3-37268771-DEL-2814 |
| chr3-37277228-DEL-66 | chr3-37277211-DEL-62 |
| chr3-37283226-DEL-56 | chr3-37283227-DEL-56 |
| chr3-37299364-DEL-3168 | chr3-37299364-DEL-3168 |
| chr3-37310678-DEL-572 | chr3-37310677-DEL-572 |
| chr3-37320671-DEL-812 | chr3-37320672-DEL-812 |
| chr3-37346455-DEL-62 | chr3-37346448-DEL-70 |

| | |
|---|---|
| chr3-37348598-DEL-1370 | chr3-37348599-DEL-1370 |
| chr3-37365936-DEL-92 | chr3-37365937-DEL-91 |
| chr3-37377732-DEL-125 | chr3-37377733-DEL-126 |
| chr3-37387319-DEL-164 | chr3-37387320-DEL-164 |
| chr3-37388736-DEL-116 | chr3-37388737-DEL-116 |
| chr3-37396904-DEL-53 | chr3-37396902-DEL-52 |
| chr3-37399781-DEL-86 | chr3-37399782-DEL-86 |
| chr3-37410118-DEL-678 | chr3-37410119-DEL-678 |
| chr3-37436110-DEL-102 | chr3-37436111-DEL-102 |
| chr3-37440830-DEL-55 | chr3-37440831-DEL-51 |
| chr3-37444733-DEL-4859 | chr3-37444734-DEL-4859 |
| chr3-37445241-DEL-263 | chr3-37445242-DEL-263 |
| chr3-37457958-DEL-297 | chr3-37457959-DEL-297 |
| chr3-37494969-DEL-183 | chr3-37494970-DEL-183 |
| chr3-37516698-DEL-176 | chr3-37516699-DEL-176 |
| chr3-37522965-DEL-233 | chr3-37522966-DEL-233 |
| chr3-37542523-DEL-610 | chr3-37542524-DEL-610 |
| chr3-37599543-DEL-3602 | chr3-37599544-DEL-3602 |
| chr3-37613224-DEL-205 | chr3-37613225-DEL-205 |
| chr3-37615358-DEL-6503 | chr3-37615359-DEL-6503 |
| chr3-37628949-DEL-58 | chr3-37628950-DEL-58 |
| chr3-37653497-DEL-54 | chr3-37653498-DEL-54 |
| chr3-37672821-DEL-203 | chr3-37672822-DEL-201 |
| chr3-37684208-DEL-66 | chr3-37684209-DEL-66 |
| chr3-37689242-DEL-64 | chr3-37689255-DEL-52 |
| chr3-37746547-DEL-370 | chr3-37746548-DEL-370 |
| chr3-37764284-DEL-88 | chr3-37764286-DEL-87 |
| chr3-37768843-DEL-532 | chr3-37768844-DEL-532 |
| chr3-37776428-DEL-174 | chr3-37776429-DEL-174 |
| chr3-37798765-DEL-208 | chr3-37798750-DEL-197 |
| chr3-37806708-DEL-52 | chr3-37806709-DEL-52 |
| chr3-37811661-DEL-58 | chr3-37811660-DEL-59 |
| chr3-37839216-DEL-72 | chr3-37839220-DEL-71 |
| chr3-37840553-DEL-56 | chr3-37840554-DEL-56 |
| chr3-37841244-DEL-194 | chr3-37841245-DEL-194 |
| chr3-37879027-DEL-446 | chr3-37879028-DEL-446 |
| chr3-37888585-DEL-54 | chr3-37888586-DEL-54 |
| chr3-37932953-DEL-1376 | chr3-37932954-DEL-1376 |
| chr3-37970876-DEL-60 | chr3-37970876-DEL-72 |
| chr3-37974837-DEL-67 | chr3-37974849-DEL-50 |
| chr3-37979819-DEL-71 | chr3-37979820-DEL-71 |
| chr3-37987396-DEL-171 | chr3-37987396-DEL-171 |
| chr3-38002883-DEL-142 | chr3-38002883-DEL-143 |
| chr3-38004719-DEL-197 | chr3-38004720-DEL-197 |
| chr3-38030441-DEL-62 | chr3-38030442-DEL-62 |
| chr3-38043620-DEL-551 | chr3-38043621-DEL-551 |
| chr3-38068686-DEL-52 | chr3-38068687-DEL-52 |
| chr3-38071130-DEL-102 | chr3-38071131-DEL-84 |
| chr3-38092394-DEL-971 | chr3-38092395-DEL-971 |
| chr3-38098399-DEL-77 | chr3-38098397-DEL-77 |
| chr3-38112109-DEL-91 | chr3-38112110-DEL-91 |
| chr3-38114017-DEL-790 | chr3-38114018-DEL-790 |
| chr3-38116916-DEL-416 | chr3-38116917-DEL-416 |
| chr3-38117974-DEL-1000 | chr3-38117978-DEL-987 |
| chr3-38120153-DEL-532 | chr3-38120154-DEL-532 |
| chr3-38140334-DEL-1314 | chr3-38140335-DEL-1314 |
| chr3-38154528-DEL-797 | chr3-38154529-DEL-797 |
| chr3-38160272-DEL-115 | chr3-38160273-DEL-115 |
| chr3-38173144-DEL-6623 | chr3-38173145-DEL-6623 |
| chr3-38218940-DEL-239 | chr3-38218941-DEL-239 |
| chr3-38250448-DEL-140 | chr3-38250449-DEL-140 |
| chr3-38251638-DEL-203 | chr3-38251639-DEL-203 |
| chr3-38253397-DEL-923 | chr3-38253398-DEL-923 |
| chr3-38254559-DEL-75 | chr3-38254560-DEL-80 |
| chr3-38255742-DEL-220 | chr3-38255727-DEL-236 |
| chr3-38271898-DEL-1015 | chr3-38271899-DEL-1015 |
| chr3-38283636-DEL-143 | chr3-38283637-DEL-143 |
| chr3-38289110-DEL-70 | chr3-38289111-DEL-54 |
| chr3-38293683-DEL-552 | chr3-38293684-DEL-552 |
| chr3-38296296-DEL-201 | chr3-38296297-DEL-201 |
| chr3-38312529-DEL-257 | chr3-38312534-DEL-258 |
| chr3-38321964-DEL-90 | chr3-38321965-DEL-90 |
| chr3-38328264-DEL-559 | chr3-38328265-DEL-559 |
| chr3-38348369-DEL-76 | chr3-38348370-DEL-76 |
| chr3-38355496-DEL-165 | chr3-38355493-DEL-163 |
| chr3-38369668-DEL-52 | chr3-38369669-DEL-52 |
| chr3-38421806-DEL-54 | chr3-38421804-DEL-55 |
| chr3-38423539-DEL-71 | chr3-38423538-DEL-72 |
| chr3-38445720-DEL-373 | chr3-38445721-DEL-373 |
| chr3-38472579-DEL-108 | chr3-38472580-DEL-108 |

| | |
|---|---|
| chr3-38486419-DEL-167 | chr3-38486423-DEL-165 |
| chr3-38560867-DEL-53 | chr3-38560868-DEL-53 |
| chr3-38618411-DEL-70 | chr3-38618412-DEL-70 |
| chr3-38630280-DEL-86 | chr3-38630235-DEL-87 |
| chr3-38633879-DEL-96 | chr3-38633883-DEL-91 |
| chr3-38637346-DEL-98 | chr3-38637348-DEL-97 |
| chr3-38640480-DEL-881 | chr3-38640481-DEL-881 |
| chr3-38647964-DEL-86 | chr3-38647965-DEL-86 |
| chr3-38654829-DEL-51 | chr3-38654830-DEL-51 |
| chr3-38656811-DEL-4589 | chr3-38656812-DEL-4589 |
| chr3-38681007-DEL-6145 | chr3-38681008-DEL-6142 |
| chr3-38693047-DEL-61 | chr3-38693049-DEL-59 |
| chr3-38698723-DEL-396 | chr3-38698724-DEL-396 |
| chr3-38709416-DEL-190 | chr3-38709417-DEL-190 |
| chr3-38713324-DEL-1278 | chr3-38713325-DEL-1278 |
| chr3-38726235-DEL-436 | chr3-38726236-DEL-436 |
| chr3-38765912-DEL-55 | chr3-38765906-DEL-62 |
| chr3-38773253-DEL-62 | chr3-38773254-DEL-66 |
| chr3-38780505-DEL-406 | chr3-38780506-DEL-405 |
| chr3-38782237-DEL-2490 | chr3-38782238-DEL-2489 |
| chr3-38786636-DEL-138 | chr3-38786637-DEL-138 |
| chr3-38788558-DEL-959 | chr3-38788559-DEL-959 |
| chr3-38814794-DEL-64 | chr3-38814795-DEL-64 |
| chr3-38816545-DEL-6476 | chr3-38816546-DEL-6476 |
| chr3-38825780-DEL-67 | chr3-38825781-DEL-67 |
| chr3-38831651-DEL-840 | chr3-38831651-DEL-840 |
| chr3-38864065-DEL-602 | chr3-38864066-DEL-602 |
| chr3-38870677-DEL-419 | chr3-38870678-DEL-419 |
| chr3-38873052-DEL-106 | chr3-38873088-DEL-78 |
| chr3-38874188-DEL-59 | chr3-38874189-DEL-59 |
| chr3-38895893-DEL-69 | chr3-38895894-DEL-69 |
| chr3-38898423-DEL-54 | chr3-38898424-DEL-54 |
| chr3-38913883-DEL-836 | chr3-38913884-DEL-836 |
| chr3-38915760-DEL-420 | chr3-38915759-DEL-420 |
| chr3-39031835-DEL-64 | chr3-39031836-DEL-64 |
| chr3-39092434-DEL-88 | chr3-39092418-DEL-89 |
| chr3-39094081-DEL-1302 | chr3-39094082-DEL-1302 |
| chr3-39114745-DEL-2095 | chr3-39114746-DEL-2095 |
| chr3-39162870-DEL-69 | chr3-39162871-DEL-69 |
| chr3-39174912-DEL-180 | chr3-39174913-DEL-180 |
| chr3-39191489-DEL-88 | chr3-39191494-DEL-84 |
| chr3-39198501-DEL-70 | chr3-39198475-DEL-96 |
| chr3-39199804-DEL-1117 | chr3-39199814-DEL-1110 |
| chr3-39212542-DEL-60 | chr3-39212543-DEL-60 |
| chr3-39237406-DEL-6575 | chr3-39237407-DEL-6575 |
| chr3-39272328-DEL-52 | chr3-39272329-DEL-52 |
| chr3-39272394-DEL-1395 | chr3-39272407-DEL-1379 |
| chr3-39291872-DEL-6279 | chr3-39291873-DEL-6279 |
| chr3-39313484-DEL-52 | chr3-39313485-DEL-52 |
| chr3-39333517-DEL-66 | chr3-39333507-DEL-80 |
| chr3-39352890-DEL-2807 | chr3-39352891-DEL-2807 |
| chr3-39404823-DEL-58 | chr3-39404818-DEL-59 |
| chr3-39411506-DEL-1858 | chr3-39411504-DEL-1848 |
| chr3-39452355-DEL-6126 | chr3-39452310-DEL-6150 |
| chr3-39458325-DEL-64 | chr3-39458326-DEL-64 |
| chr3-39500159-DEL-3685 | chr3-39500160-DEL-3685 |
| chr3-39532275-DEL-74 | chr3-39532265-DEL-74 |
| chr3-39535568-DEL-90 | chr3-39535574-DEL-64 |
| chr3-39547840-DEL-480 | chr3-39547841-DEL-480 |
| chr3-39557083-DEL-94 | chr3-39557084-DEL-94 |
| chr3-39583662-DEL-846 | chr3-39583663-DEL-846 |
| chr3-39600610-DEL-6440 | chr3-39600611-DEL-6440 |
| chr3-39620713-DEL-269 | chr3-39620714-DEL-269 |
| chr3-39628262-DEL-188 | chr3-39628263-DEL-188 |
| chr3-39630768-DEL-269 | chr3-39630769-DEL-269 |
| chr3-39634742-DEL-213 | chr3-39634743-DEL-213 |
| chr3-39635298-DEL-299 | chr3-39635299-DEL-287 |
| chr3-39637048-DEL-74 | chr3-39637041-DEL-75 |
| chr3-39656114-DEL-6660 | chr3-39656115-DEL-6660 |
| chr3-39656298-DEL-212 | chr3-39656299-DEL-212 |
| chr3-39678560-DEL-153 | chr3-39678594-DEL-150 |
| chr3-39688788-DEL-122 | chr3-39688789-DEL-122 |
| chr3-39700653-DEL-500 | chr3-39700654-DEL-500 |
| chr3-39722988-DEL-1976 | chr3-39722989-DEL-1976 |
| chr3-39725069-DEL-2303 | chr3-39725070-DEL-2303 |
| chr3-39737408-DEL-52 | chr3-39737407-DEL-50 |
| chr3-39742070-DEL-88 | chr3-39742071-DEL-88 |
| chr3-39757113-DEL-264 | chr3-39757114-DEL-264 |
| chr3-39758262-DEL-55 | chr3-39758263-DEL-55 |
| chr3-39785719-DEL-1910 | chr3-39785720-DEL-1909 |

| | |
|---|---|
| chr3-39821733-DEL-76 | chr3-39821722-DEL-88 |
| chr3-39841486-DEL-329 | chr3-39841487-DEL-327 |
| chr3-39846528-DEL-56 | chr3-39846529-DEL-56 |
| chr3-39854102-DEL-136 | chr3-39854103-DEL-136 |
| chr3-39875876-DEL-6366 | chr3-39875877-DEL-6366 |
| chr3-39884451-DEL-212 | chr3-39884450-DEL-213 |
| chr3-39887827-DEL-552 | chr3-39887826-DEL-550 |
| chr3-39897961-DEL-510 | chr3-39897962-DEL-510 |
| chr3-39908741-DEL-56 | chr3-39908742-DEL-56 |
| chr3-39974046-DEL-224 | chr3-39974047-DEL-224 |
| chr3-40003777-DEL-89 | chr3-40003778-DEL-89 |
| chr3-40010126-DEL-233 | chr3-40010127-DEL-233 |
| chr3-40021136-DEL-61 | chr3-40021137-DEL-61 |
| chr3-40024291-DEL-877 | chr3-40024292-DEL-877 |
| chr3-40072472-DEL-328 | chr3-40072473-DEL-328 |
| chr3-40092189-DEL-1282 | chr3-40092225-DEL-1232 |
| chr3-40098931-DEL-234 | chr3-40098932-DEL-234 |
| chr3-40107122-DEL-184 | chr3-40107122-DEL-184 |
| chr3-40125007-DEL-59 | chr3-40124989-DEL-56 |
| chr3-40125884-DEL-2449 | chr3-40125884-DEL-2449 |
| chr3-40135910-DEL-99 | chr3-40135895-DEL-100 |
| chr3-40141643-DEL-94 | chr3-40141644-DEL-94 |
| chr3-40146596-DEL-54 | chr3-40146579-DEL-72 |
| chr3-40181987-DEL-216 | chr3-40181976-DEL-225 |
| chr3-40182937-DEL-64 | chr3-40182938-DEL-64 |
| chr3-40190886-DEL-1239 | chr3-40190887-DEL-1239 |
| chr3-40195372-DEL-102 | chr3-40195373-DEL-102 |
| chr3-40205408-DEL-6379 | chr3-40205409-DEL-6379 |
| chr3-40238024-DEL-77 | chr3-40238022-DEL-78 |
| chr3-40251621-DEL-573 | chr3-40251622-DEL-573 |
| chr3-40277046-DEL-144 | chr3-40277047-DEL-144 |
| chr3-40334109-DEL-58 | chr3-40334110-DEL-58 |
| chr3-40359193-DEL-70 | chr3-40359206-DEL-58 |
| chr3-40396906-DEL-106 | chr3-40396916-DEL-97 |
| chr3-40400400-DEL-6378 | chr3-40400401-DEL-6378 |
| chr3-40406370-DEL-201 | chr3-40406371-DEL-201 |
| chr3-40441362-DEL-97 | chr3-40441363-DEL-97 |
| chr3-40475286-DEL-54 | chr3-40475291-DEL-50 |
| chr3-40501947-DEL-1152 | chr3-40501948-DEL-1152 |
| chr3-40519150-DEL-2652 | chr3-40519151-DEL-2652 |
| chr3-40567878-DEL-82 | chr3-40567879-DEL-82 |
| chr3-40580509-DEL-72 | chr3-40580510-DEL-72 |
| chr3-40598128-DEL-60 | chr3-40598126-DEL-61 |
| chr3-40631170-DEL-63 | chr3-40631167-DEL-64 |
| chr3-40644827-DEL-223 | chr3-40644828-DEL-223 |
| chr3-40657075-DEL-399 | chr3-40657076-DEL-399 |
| chr3-40667701-DEL-159 | chr3-40667702-DEL-142 |
| chr3-40673878-DEL-62 | chr3-40673879-DEL-62 |
| chr3-40762657-DEL-208 | chr3-40762658-DEL-208 |
| chr3-40772435-DEL-69 | chr3-40772436-DEL-69 |
| chr3-40807817-DEL-121 | chr3-40807818-DEL-120 |
| chr3-40809944-DEL-54 | chr3-40809945-DEL-54 |
| chr3-40814829-DEL-3833 | chr3-40814829-DEL-3833 |
| chr3-40874492-DEL-293 | chr3-40874493-DEL-293 |
| chr3-40878690-DEL-670 | chr3-40878711-DEL-670 |
| chr3-40893316-DEL-158 | chr3-40893317-DEL-142 |
| chr3-40907483-DEL-372 | chr3-40907484-DEL-372 |
| chr3-40947445-DEL-122 | chr3-40947446-DEL-122 |
| chr3-40981401-DEL-138 | chr3-40981407-DEL-92 |
| chr3-40985388-DEL-313 | chr3-40985389-DEL-313 |
| chr3-40986177-DEL-97 | chr3-40986178-DEL-97 |
| chr3-40994229-DEL-65 | chr3-40994230-DEL-65 |
| chr3-41050196-DEL-61 | chr3-41050213-DEL-58 |
| chr3-41113816-DEL-397 | chr3-41113817-DEL-397 |
| chr3-41128861-DEL-873 | chr3-41128863-DEL-872 |
| chr3-41146164-DEL-66 | chr3-41146165-DEL-66 |
| chr3-41170065-DEL-203 | chr3-41170066-DEL-203 |
| chr3-41182253-DEL-69 | chr3-41182253-DEL-69 |
| chr3-41184483-DEL-222 | chr3-41184484-DEL-222 |
| chr3-41198013-DEL-128 | chr3-41198014-DEL-128 |
| chr3-41205740-DEL-895 | chr3-41205727-DEL-895 |
| chr3-41234340-DEL-687 | chr3-41234341-DEL-687 |
| chr3-41290409-DEL-77 | chr3-41290406-DEL-81 |
| chr3-41291071-DEL-6798 | chr3-41291072-DEL-6798 |
| chr3-41299751-DEL-209 | chr3-41299752-DEL-209 |
| chr3-41333702-DEL-330 | chr3-41333703-DEL-330 |
| chr3-41336741-DEL-240 | chr3-41336742-DEL-240 |
| chr3-41356846-DEL-56 | chr3-41356847-DEL-56 |
| chr3-41379267-DEL-104 | chr3-41379267-DEL-105 |
| chr3-41390011-DEL-118 | chr3-41390012-DEL-118 |

| | |
|---|---|
| chr3-41428397-DEL-647 | chr3-41428398-DEL-647 |
| chr3-41438527-DEL-54 | chr3-41438528-DEL-54 |
| chr3-41449466-DEL-63 | chr3-41449467-DEL-63 |
| chr3-41459440-DEL-135 | chr3-41459441-DEL-135 |
| chr3-41472241-DEL-92 | chr3-41472242-DEL-92 |
| chr3-41482135-DEL-72 | chr3-41482130-DEL-78 |
| chr3-41494402-DEL-55 | chr3-41494381-DEL-55 |
| chr3-41582986-DEL-110 | chr3-41582965-DEL-104 |
| chr3-41596694-DEL-60 | chr3-41596692-DEL-56 |
| chr3-41602574-DEL-212 | chr3-41602575-DEL-212 |
| chr3-41638069-DEL-299 | chr3-41638069-DEL-299 |
| chr3-41644610-DEL-458 | chr3-41644610-DEL-458 |
| chr3-41692298-DEL-833 | chr3-41692299-DEL-833 |
| chr3-41714258-DEL-931 | chr3-41714259-DEL-931 |
| chr3-41718272-DEL-6534 | chr3-41718273-DEL-6534 |
| chr3-41725695-DEL-432 | chr3-41725696-DEL-432 |
| chr3-41753693-DEL-1130 | chr3-41753686-DEL-1130 |
| chr3-41761541-DEL-109 | chr3-41761542-DEL-109 |
| chr3-41769413-DEL-60 | chr3-41769414-DEL-60 |
| chr3-41804707-DEL-380 | chr3-41804708-DEL-380 |
| chr3-41826255-DEL-7732 | chr3-41826255-DEL-7732 |
| chr3-41827377-DEL-1132 | chr3-41827377-DEL-1132 |
| chr3-41833514-DEL-201 | chr3-41833515-DEL-201 |
| chr3-41858528-DEL-889 | chr3-41858529-DEL-889 |
| chr3-41861291-DEL-373 | chr3-41861288-DEL-374 |
| chr3-41887675-DEL-148 | chr3-41887676-DEL-148 |
| chr3-41896731-DEL-412 | chr3-41896732-DEL-412 |
| chr3-41906461-DEL-206 | chr3-41906462-DEL-206 |
| chr3-41907377-DEL-136 | chr3-41907391-DEL-114 |
| chr3-41967031-DEL-594 | chr3-41967032-DEL-594 |
| chr3-41979066-DEL-6207 | chr3-41979067-DEL-6207 |
| chr3-42026925-DEL-310 | chr3-42026926-DEL-310 |
| chr3-42036332-DEL-1436 | chr3-42036332-DEL-1436 |
| chr3-42061943-DEL-130 | chr3-42061943-DEL-130 |
| chr3-42070475-DEL-6842 | chr3-42070476-DEL-6842 |
| chr3-42098482-DEL-200 | chr3-42098483-DEL-200 |
| chr3-42114814-DEL-51 | chr3-42114815-DEL-51 |
| chr3-42131242-DEL-66 | chr3-42131243-DEL-66 |
| chr3-42131934-DEL-200 | chr3-42131924-DEL-200 |
| chr3-42137274-DEL-4526 | chr3-42137275-DEL-4526 |
| chr3-42164027-DEL-108 | chr3-42164028-DEL-96 |
| chr3-42183954-DEL-2625 | chr3-42183955-DEL-2625 |
| chr3-42192680-DEL-438 | chr3-42192681-DEL-438 |
| chr3-42201337-DEL-1579 | chr3-42201338-DEL-1579 |
| chr3-42229256-DEL-654 | chr3-42229257-DEL-654 |
| chr3-42233882-DEL-802 | chr3-42233883-DEL-802 |
| chr3-42250810-DEL-2045 | chr3-42250822-DEL-2044 |
| chr3-42260231-DEL-123 | chr3-42260232-DEL-121 |
| chr3-42272642-DEL-6912 | chr3-42272643-DEL-6912 |
| chr3-42336092-DEL-127 | chr3-42336093-DEL-127 |
| chr3-42356560-DEL-52 | chr3-42356561-DEL-52 |
| chr3-42361239-DEL-524 | chr3-42361240-DEL-524 |
| chr3-42384209-DEL-129 | chr3-42384210-DEL-129 |
| chr3-42401202-DEL-76 | chr3-42401216-DEL-60 |
| chr3-42408423-DEL-127 | chr3-42408424-DEL-127 |
| chr3-42457082-DEL-3039 | chr3-42457083-DEL-3039 |
| chr3-42500759-DEL-2175 | chr3-42500760-DEL-2175 |
| chr3-42506789-DEL-70 | chr3-42506766-DEL-96 |
| chr3-42541354-DEL-59 | chr3-42541332-DEL-81 |
| chr3-42559982-DEL-52 | chr3-42559977-DEL-64 |
| chr3-42560929-DEL-313 | chr3-42560930-DEL-313 |
| chr3-42565640-DEL-102 | chr3-42565661-DEL-82 |
| chr3-42600767-DEL-849 | chr3-42600773-DEL-850 |
| chr3-42624655-DEL-135 | chr3-42624655-DEL-135 |
| chr3-42670588-DEL-6731 | chr3-42670589-DEL-6731 |
| chr3-42677825-DEL-58 | chr3-42677827-DEL-57 |
| chr3-42733346-DEL-2849 | chr3-42733347-DEL-2849 |
| chr3-42733415-DEL-57 | chr3-42733416-DEL-57 |
| chr3-42769861-DEL-81 | chr3-42769860-DEL-58 |
| chr3-42774661-DEL-7189 | chr3-42774662-DEL-7189 |
| chr3-42789181-DEL-150 | chr3-42789175-DEL-151 |
| chr3-42812131-DEL-426 | chr3-42812132-DEL-426 |
| chr3-42864329-DEL-56 | chr3-42864330-DEL-52 |
| chr3-42868090-DEL-138 | chr3-42868114-DEL-126 |
| chr3-42915498-DEL-912 | chr3-42915499-DEL-912 |
| chr3-42941684-DEL-127 | chr3-42941685-DEL-127 |
| chr3-42951203-DEL-339 | chr3-42951204-DEL-339 |
| chr3-42960884-DEL-6207 | chr3-42960885-DEL-6207 |
| chr3-43015492-DEL-135 | chr3-43015493-DEL-135 |
| chr3-43042150-DEL-4630 | chr3-43042156-DEL-4625 |

| | |
|---|---|
| chr3-43053884-DEL-1171 | chr3-43053885-DEL-1171 |
| chr3-43062455-DEL-3336 | chr3-43062455-DEL-3334 |
| chr3-43078247-DEL-114 | chr3-43078248-DEL-114 |
| chr3-43087370-DEL-80 | chr3-43087371-DEL-84 |
| chr3-43099475-DEL-535 | chr3-43099476-DEL-535 |
| chr3-43101479-DEL-3048 | chr3-43101480-DEL-3048 |
| chr3-43114442-DEL-127 | chr3-43114443-DEL-127 |
| chr3-43116547-DEL-256 | chr3-43116548-DEL-256 |
| chr3-43126390-DEL-57 | chr3-43126391-DEL-57 |
| chr3-43128784-DEL-1843 | chr3-43128785-DEL-1843 |
| chr3-43139266-DEL-9215 | chr3-43139267-DEL-9215 |
| chr3-43165479-DEL-1522 | chr3-43165480-DEL-1522 |
| chr3-43190045-DEL-1089 | chr3-43190046-DEL-1089 |
| chr3-43254364-DEL-6569 | chr3-43254365-DEL-6569 |
| chr3-43268347-DEL-238 | chr3-43268348-DEL-237 |
| chr3-43328069-DEL-2986 | chr3-43328070-DEL-2986 |
| chr3-43332409-DEL-1639 | chr3-43332410-DEL-1638 |
| chr3-43334197-DEL-626 | chr3-43334198-DEL-626 |
| chr3-43355749-DEL-667 | chr3-43355750-DEL-667 |
| chr3-43364814-DEL-418 | chr3-43364815-DEL-418 |
| chr3-43365547-DEL-200 | chr3-43365548-DEL-200 |
| chr3-43408448-DEL-346 | chr3-43408449-DEL-346 |
| chr3-43430200-DEL-62 | chr3-43430201-DEL-104 |
| chr3-43440379-DEL-6220 | chr3-43440380-DEL-6220 |
| chr3-43493224-DEL-3086 | chr3-43493225-DEL-3086 |
| chr3-43501400-DEL-1894 | chr3-43501401-DEL-1894 |
| chr3-43549811-DEL-86 | chr3-43549816-DEL-81 |
| chr3-43608426-DEL-369 | chr3-43608427-DEL-369 |
| chr3-43612308-DEL-1406 | chr3-43612265-DEL-1447 |
| chr3-43647318-DEL-6667 | chr3-43647319-DEL-6667 |
| chr3-43684433-DEL-3766 | chr3-43684433-DEL-3765 |
| chr3-43708310-DEL-434 | chr3-43708311-DEL-434 |
| chr3-43728753-DEL-311 | chr3-43728754-DEL-311 |
| chr3-43786851-DEL-80 | chr3-43786847-DEL-74 |
| chr3-43806476-DEL-63 | chr3-43806477-DEL-63 |
| chr3-43807352-DEL-13080 | chr3-43807353-DEL-13080 |
| chr3-43901414-DEL-270 | chr3-43901415-DEL-270 |
| chr3-43901415-DEL-1097 | chr3-43901415-DEL-1097 |
| chr3-43929204-DEL-437 | chr3-43929205-DEL-437 |
| chr3-43931597-DEL-163 | chr3-43931598-DEL-163 |
| chr3-43933803-DEL-52 | chr3-43933801-DEL-60 |
| chr3-43938017-DEL-1066 | chr3-43938018-DEL-1066 |
| chr3-43951272-DEL-227 | chr3-43951273-DEL-227 |
| chr3-43953663-DEL-160 | chr3-43953639-DEL-160 |
| chr3-43959041-DEL-6180 | chr3-43959042-DEL-6179 |
| chr3-43975603-DEL-2047 | chr3-43975604-DEL-2047 |
| chr3-43978837-DEL-199 | chr3-43978838-DEL-199 |
| chr3-44034675-DEL-6668 | chr3-44034676-DEL-6668 |
| chr3-44042424-DEL-537 | chr3-44042425-DEL-537 |
| chr3-44053692-DEL-7592 | chr3-44053684-DEL-7590 |
| chr3-44067982-DEL-512 | chr3-44067983-DEL-512 |
| chr3-44075642-DEL-393 | chr3-44075643-DEL-393 |
| chr3-44105548-DEL-73 | chr3-44105549-DEL-76 |
| chr3-44140268-DEL-6948 | chr3-44140269-DEL-6948 |
| chr3-44173665-DEL-190 | chr3-44173666-DEL-190 |
| chr3-44178150-DEL-67 | chr3-44178151-DEL-67 |
| chr3-44179997-DEL-1135 | chr3-44179998-DEL-1135 |
| chr3-44195626-DEL-6787 | chr3-44195627-DEL-6787 |
| chr3-44281394-DEL-119 | chr3-44281392-DEL-120 |
| chr3-44284117-DEL-179 | chr3-44284118-DEL-179 |
| chr3-44288935-DEL-68 | chr3-44288936-DEL-68 |
| chr3-44290583-DEL-979 | chr3-44290584-DEL-978 |
| chr3-44294235-DEL-2223 | chr3-44294236-DEL-2223 |
| chr3-44294625-DEL-378 | chr3-44294626-DEL-378 |
| chr3-44341263-DEL-144 | chr3-44341264-DEL-144 |
| chr3-44355127-DEL-6855 | chr3-44355128-DEL-6855 |
| chr3-44355150-DEL-200 | chr3-44355151-DEL-200 |
| chr3-44359770-DEL-447 | chr3-44359771-DEL-447 |
| chr3-44363676-DEL-110 | chr3-44363677-DEL-110 |
| chr3-44377578-DEL-944 | chr3-44377579-DEL-944 |
| chr3-44384585-DEL-56 | chr3-44384586-DEL-56 |
| chr3-44392219-DEL-66 | chr3-44392236-DEL-50 |
| chr3-44393883-DEL-121 | chr3-44393884-DEL-121 |
| chr3-44398853-DEL-101 | chr3-44398854-DEL-101 |
| chr3-44408832-DEL-52 | chr3-44408837-DEL-50 |
| chr3-44418655-DEL-90 | chr3-44418642-DEL-98 |
| chr3-44425337-DEL-3915 | chr3-44425338-DEL-3915 |
| chr3-44426808-DEL-96 | chr3-44426809-DEL-96 |
| chr3-44434389-DEL-61 | chr3-44434402-DEL-55 |
| chr3-44448558-DEL-88 | chr3-44448557-DEL-84 |

| | |
|---|---|
| chr3-44459898-DEL-2167 | chr3-44459899-DEL-2167 |
| chr3-44466534-DEL-1851 | chr3-44466535-DEL-1851 |
| chr3-44482421-DEL-2023 | chr3-44482422-DEL-2023 |
| chr3-44513209-DEL-7813 | chr3-44513210-DEL-7813 |
| chr3-44525052-DEL-65 | chr3-44525062-DEL-54 |
| chr3-44540619-DEL-722 | chr3-44540620-DEL-722 |
| chr3-44557124-DEL-64 | chr3-44557125-DEL-64 |
| chr3-44568448-DEL-1815 | chr3-44568448-DEL-1815 |
| chr3-44576968-DEL-6487 | chr3-44576969-DEL-6487 |
| chr3-44583616-DEL-494 | chr3-44583617-DEL-494 |
| chr3-44585129-DEL-575 | chr3-44585130-DEL-575 |
| chr3-44604949-DEL-1648 | chr3-44604950-DEL-1648 |
| chr3-44620169-DEL-6674 | chr3-44620170-DEL-6674 |
| chr3-44650417-DEL-53 | chr3-44650416-DEL-52 |
| chr3-44673279-DEL-1223 | chr3-44673280-DEL-1223 |
| chr3-44718051-DEL-2124 | chr3-44718045-DEL-2115 |
| chr3-44738612-DEL-66 | chr3-44738587-DEL-90 |
| chr3-44792960-DEL-226 | chr3-44792961-DEL-226 |
| chr3-44846018-DEL-424 | chr3-44846019-DEL-424 |
| chr3-44850226-DEL-6255 | chr3-44850227-DEL-6255 |
| chr3-44858982-DEL-87 | chr3-44858969-DEL-88 |
| chr3-44862020-DEL-638 | chr3-44862021-DEL-636 |
| chr3-44869133-DEL-981 | chr3-44869138-DEL-986 |
| chr3-44885997-DEL-3029 | chr3-44886002-DEL-3030 |
| chr3-44891820-DEL-4148 | chr3-44891825-DEL-4144 |
| chr3-44935409-DEL-96 | chr3-44935410-DEL-96 |
| chr3-44950185-DEL-111 | chr3-44950186-DEL-115 |
| chr3-44951361-DEL-3919 | chr3-44951362-DEL-3919 |
| chr3-44973349-DEL-199 | chr3-44973350-DEL-199 |
| chr3-44978382-DEL-2048 | chr3-44978383-DEL-2048 |
| chr3-45028793-DEL-454 | chr3-45028794-DEL-454 |
| chr3-45031402-DEL-633 | chr3-45031403-DEL-633 |
| chr3-45037742-DEL-466 | chr3-45037743-DEL-466 |
| chr3-45041351-DEL-659 | chr3-45041352-DEL-659 |
| chr3-45065898-DEL-314 | chr3-45065899-DEL-314 |
| chr3-45076791-DEL-63 | chr3-45076805-DEL-50 |
| chr3-45114928-DEL-1001 | chr3-45114893-DEL-1000 |
| chr3-45117301-DEL-6091 | chr3-45117302-DEL-6091 |
| chr3-45167251-DEL-148 | chr3-45167225-DEL-130 |
| chr3-45215845-DEL-6857 | chr3-45215846-DEL-6857 |
| chr3-45270006-DEL-229 | chr3-45270007-DEL-229 |
| chr3-45278563-DEL-74 | chr3-45278564-DEL-74 |
| chr3-45284052-DEL-292 | chr3-45284051-DEL-293 |
| chr3-45302609-DEL-6255 | chr3-45302610-DEL-6255 |
| chr3-45312304-DEL-405 | chr3-45312305-DEL-405 |
| chr3-45449038-DEL-320 | chr3-45449039-DEL-320 |
| chr3-45451014-DEL-4009 | chr3-45451015-DEL-4009 |
| chr3-45464022-DEL-1470 | chr3-45464023-DEL-1470 |
| chr3-45468233-DEL-1019 | chr3-45468234-DEL-1019 |
| chr3-45486874-DEL-846 | chr3-45486875-DEL-846 |
| chr3-45502941-DEL-1689 | chr3-45502941-DEL-1689 |
| chr3-45532640-DEL-7338 | chr3-45532641-DEL-7338 |
| chr3-45562817-DEL-484 | chr3-45562816-DEL-484 |
| chr3-45564712-DEL-3337 | chr3-45564713-DEL-3337 |
| chr3-45568463-DEL-202 | chr3-45568464-DEL-202 |
| chr3-45606747-DEL-1916 | chr3-45606748-DEL-1916 |
| chr3-45627354-DEL-62 | chr3-45627355-DEL-58 |
| chr3-45668692-DEL-2101 | chr3-45668690-DEL-2101 |
| chr3-45669505-DEL-60 | chr3-45669511-DEL-52 |
| chr3-45669700-DEL-1050 | chr3-45669701-DEL-1047 |
| chr3-45678695-DEL-399 | chr3-45678696-DEL-399 |
| chr3-45701387-DEL-654 | chr3-45701388-DEL-654 |
| chr3-45712071-DEL-62 | chr3-45712075-DEL-58 |
| chr3-45726558-DEL-120 | chr3-45726559-DEL-120 |
| chr3-45740733-DEL-274 | chr3-45740734-DEL-274 |
| chr3-45746406-DEL-942 | chr3-45746407-DEL-942 |
| chr3-45815853-DEL-1893 | chr3-45815854-DEL-1893 |
| chr3-45832085-DEL-92 | chr3-45832070-DEL-83 |
| chr3-45845594-DEL-282 | chr3-45845595-DEL-282 |
| chr3-45847246-DEL-6223 | chr3-45847247-DEL-6223 |
| chr3-45895648-DEL-96 | chr3-45895648-DEL-96 |
| chr3-45938082-DEL-60 | chr3-45938080-DEL-60 |
| chr3-45940313-DEL-6245 | chr3-45940332-DEL-6227 |
| chr3-45946264-DEL-64 | chr3-45946265-DEL-64 |
| chr3-45986042-DEL-97 | chr3-45986040-DEL-97 |
| chr3-45997802-DEL-52 | chr3-45997819-DEL-80 |
| chr3-46006200-DEL-86 | chr3-46006193-DEL-86 |
| chr3-46049676-DEL-6537 | chr3-46049677-DEL-6537 |
| chr3-46055804-DEL-214 | chr3-46055805-DEL-214 |
| chr3-46095346-DEL-7224 | chr3-46095347-DEL-7224 |

| | |
|---|---|
| chr3-46124912-DEL-2885 | chr3-46124913-DEL-2885 |
| chr3-46147459-DEL-95 | chr3-46147460-DEL-95 |
| chr3-46152999-DEL-6656 | chr3-46153000-DEL-6656 |
| chr3-46162502-DEL-572 | chr3-46162503-DEL-572 |
| chr3-46165920-DEL-111 | chr3-46165894-DEL-138 |
| chr3-46192331-DEL-200 | chr3-46192332-DEL-200 |
| chr3-46222946-DEL-430 | chr3-46222947-DEL-429 |
| chr3-46228666-DEL-1558 | chr3-46228667-DEL-1558 |
| chr3-46228735-DEL-50 | chr3-46228733-DEL-52 |
| chr3-46234037-DEL-209 | chr3-46234038-DEL-209 |
| chr3-46260854-DEL-1004 | chr3-46260855-DEL-1004 |
| chr3-46263189-DEL-2063 | chr3-46263190-DEL-2063 |
| chr3-46292722-DEL-1893 | chr3-46292723-DEL-1893 |
| chr3-46324286-DEL-15741 | chr3-46324287-DEL-15741 |
| chr3-46328073-DEL-942 | chr3-46328074-DEL-942 |
| chr3-46331935-DEL-381 | chr3-46331932-DEL-380 |
| chr3-46358343-DEL-52 | chr3-46358340-DEL-88 |
| chr3-46380842-DEL-61 | chr3-46380843-DEL-61 |
| chr3-46407318-DEL-66 | chr3-46407363-DEL-53 |
| chr3-46415585-DEL-4463 | chr3-46415586-DEL-4463 |
| chr3-46434688-DEL-115 | chr3-46434689-DEL-115 |
| chr3-46456539-DEL-1636 | chr3-46456540-DEL-1636 |
| chr3-46484102-DEL-87 | chr3-46484103-DEL-87 |
| chr3-46549855-DEL-64 | chr3-46549856-DEL-64 |
| chr3-46550701-DEL-129 | chr3-46550702-DEL-124 |
| chr3-46576316-DEL-54 | chr3-46576317-DEL-54 |
| chr3-46590752-DEL-56 | chr3-46590759-DEL-52 |
| chr3-46603524-DEL-6665 | chr3-46603524-DEL-6666 |
| chr3-46681893-DEL-62 | chr3-46681914-DEL-82 |
| chr3-46689332-DEL-7028 | chr3-46689333-DEL-7028 |
| chr3-46699062-DEL-138 | chr3-46699063-DEL-138 |
| chr3-46722943-DEL-108 | chr3-46722958-DEL-94 |
| chr3-46740792-DEL-1001 | chr3-46740793-DEL-1001 |
| chr3-46749570-DEL-10565 | chr3-46749570-DEL-10565 |
| chr3-46822321-DEL-53 | chr3-46822322-DEL-53 |
| chr3-46875563-DEL-204 | chr3-46875564-DEL-204 |
| chr3-46890455-DEL-208 | chr3-46890456-DEL-208 |
| chr3-46904910-DEL-163 | chr3-46904911-DEL-163 |
| chr3-46928652-DEL-6532 | chr3-46928653-DEL-6532 |
| chr3-47003523-DEL-2716 | chr3-47003524-DEL-2716 |
| chr3-47008928-DEL-60 | chr3-47008929-DEL-60 |
| chr3-47011692-DEL-448 | chr3-47011693-DEL-448 |
| chr3-47051380-DEL-135 | chr3-47051364-DEL-119 |
| chr3-47063607-DEL-110 | chr3-47063610-DEL-108 |
| chr3-47072585-DEL-625 | chr3-47072585-DEL-626 |
| chr3-47073900-DEL-153 | chr3-47073901-DEL-153 |
| chr3-47075189-DEL-626 | chr3-47075190-DEL-626 |
| chr3-47093100-DEL-402 | chr3-47093101-DEL-402 |
| chr3-47102380-DEL-66 | chr3-47102396-DEL-66 |
| chr3-47133788-DEL-439 | chr3-47133789-DEL-439 |
| chr3-47136750-DEL-53 | chr3-47136744-DEL-53 |
| chr3-47152724-DEL-80 | chr3-47152727-DEL-78 |
| chr3-47168146-DEL-355 | chr3-47168147-DEL-355 |
| chr3-47187743-DEL-55 | chr3-47187745-DEL-54 |
| chr3-47197936-DEL-337 | chr3-47197936-DEL-334 |
| chr3-47245647-DEL-1527 | chr3-47245648-DEL-1527 |
| chr3-47267883-DEL-2039 | chr3-47267884-DEL-2039 |
| chr3-47269823-DEL-98 | chr3-47269826-DEL-101 |
| chr3-47326167-DEL-85 | chr3-47326169-DEL-86 |
| chr3-47361836-DEL-193 | chr3-47361837-DEL-193 |
| chr3-47405117-DEL-485 | chr3-47405118-DEL-485 |
| chr3-47405694-DEL-1168 | chr3-47405699-DEL-1128 |
| chr3-47407646-DEL-401 | chr3-47407647-DEL-400 |
| chr3-47435717-DEL-4431 | chr3-47435722-DEL-4435 |
| chr3-47450408-DEL-56 | chr3-47450409-DEL-56 |
| chr3-47461692-DEL-60 | chr3-47461698-DEL-54 |
| chr3-47468330-DEL-95 | chr3-47468331-DEL-95 |
| chr3-47480686-DEL-548 | chr3-47480687-DEL-548 |
| chr3-47487169-DEL-173 | chr3-47487170-DEL-173 |
| chr3-47518666-DEL-634 | chr3-47518667-DEL-634 |
| chr3-47537880-DEL-614 | chr3-47537876-DEL-614 |
| chr3-47546202-DEL-6404 | chr3-47546203-DEL-6404 |
| chr3-47561898-DEL-2985 | chr3-47561899-DEL-2985 |
| chr3-47576991-DEL-3964 | chr3-47576991-DEL-3964 |
| chr3-47593529-DEL-440 | chr3-47593530-DEL-440 |
| chr3-47600404-DEL-3433 | chr3-47600380-DEL-3433 |
| chr3-47690624-DEL-509 | chr3-47690623-DEL-509 |
| chr3-47695696-DEL-802 | chr3-47695697-DEL-802 |
| chr3-47708007-DEL-98 | chr3-47708008-DEL-98 |
| chr3-47721965-DEL-2633 | chr3-47721965-DEL-2633 |

| | |
|---|---|
| chr3-47736813-DEL-420 | chr3-47736822-DEL-416 |
| chr3-47798574-DEL-3569 | chr3-47798575-DEL-3569 |
| chr3-47800783-DEL-56 | chr3-47800784-DEL-56 |
| chr3-47804018-DEL-73 | chr3-47804021-DEL-74 |
| chr3-47806704-DEL-903 | chr3-47806705-DEL-903 |
| chr3-47809701-DEL-775 | chr3-47809701-DEL-775 |
| chr3-47826952-DEL-106 | chr3-47826954-DEL-107 |
| chr3-47836486-DEL-59 | chr3-47836487-DEL-59 |
| chr3-47844846-DEL-54 | chr3-47844847-DEL-54 |
| chr3-47865594-DEL-346 | chr3-47865594-DEL-346 |
| chr3-47919337-DEL-4281 | chr3-47919338-DEL-4281 |
| chr3-47933443-DEL-54 | chr3-47933444-DEL-54 |
| chr3-47940106-DEL-515 | chr3-47940107-DEL-515 |
| chr3-47954363-DEL-446 | chr3-47954364-DEL-446 |
| chr3-47958757-DEL-6171 | chr3-47958757-DEL-6171 |
| chr3-48000698-DEL-62 | chr3-48000692-DEL-72 |
| chr3-48062903-DEL-63 | chr3-48062903-DEL-55 |
| chr3-48070560-DEL-53 | chr3-48070561-DEL-53 |
| chr3-48109728-DEL-2584 | chr3-48109729-DEL-2584 |
| chr3-48121015-DEL-6356 | chr3-48121016-DEL-6356 |
| chr3-48129225-DEL-511 | chr3-48129225-DEL-511 |
| chr3-48133473-DEL-1227 | chr3-48133474-DEL-1227 |
| chr3-48214916-DEL-81 | chr3-48214928-DEL-81 |
| chr3-48230227-DEL-389 | chr3-48230228-DEL-389 |
| chr3-48240969-DEL-120 | chr3-48240967-DEL-123 |
| chr3-48241842-DEL-721 | chr3-48241843-DEL-721 |
| chr3-48247259-DEL-75 | chr3-48247260-DEL-75 |
| chr3-48252285-DEL-334 | chr3-48252285-DEL-334 |
| chr3-48275571-DEL-214 | chr3-48275572-DEL-214 |
| chr3-48284345-DEL-99 | chr3-48284349-DEL-94 |
| chr3-48298036-DEL-295 | chr3-48298036-DEL-295 |
| chr3-48307451-DEL-263 | chr3-48307452-DEL-263 |
| chr3-48310458-DEL-359 | chr3-48310459-DEL-359 |
| chr3-48314422-DEL-1596 | chr3-48314405-DEL-1613 |
| chr3-48339592-DEL-1989 | chr3-48339592-DEL-1989 |
| chr3-48345305-DEL-89 | chr3-48345314-DEL-89 |
| chr3-48357084-DEL-159 | chr3-48357085-DEL-159 |
| chr3-48364186-DEL-80 | chr3-48364187-DEL-80 |
| chr3-48386598-DEL-56 | chr3-48386599-DEL-56 |
| chr3-48429639-DEL-78 | chr3-48429640-DEL-78 |
| chr3-48431960-DEL-190 | chr3-48431965-DEL-185 |
| chr3-48459518-DEL-156 | chr3-48459506-DEL-174 |
| chr3-48483984-DEL-142 | chr3-48483983-DEL-143 |
| chr3-48513038-DEL-1808 | chr3-48513039-DEL-1807 |
| chr3-48521655-DEL-1612 | chr3-48521656-DEL-1612 |
| chr3-48523770-DEL-141 | chr3-48523771-DEL-101 |
| chr3-48531879-DEL-655 | chr3-48531880-DEL-655 |
| chr3-48552425-DEL-364 | chr3-48552426-DEL-364 |
| chr3-48552967-DEL-86 | chr3-48552968-DEL-89 |
| chr3-48569672-DEL-122 | chr3-48569686-DEL-125 |
| chr3-48572750-DEL-658 | chr3-48572751-DEL-658 |
| chr3-48592833-DEL-100 | chr3-48592834-DEL-100 |
| chr3-48627518-DEL-2408 | chr3-48627519-DEL-2408 |
| chr3-48634258-DEL-51 | chr3-48634259-DEL-51 |
| chr3-48640154-DEL-203 | chr3-48640155-DEL-203 |
| chr3-48661100-DEL-52 | chr3-48661101-DEL-52 |
| chr3-48688174-DEL-3135 | chr3-48688175-DEL-3135 |
| chr3-48753941-DEL-1595 | chr3-48753942-DEL-1595 |
| chr3-48764966-DEL-51 | chr3-48764967-DEL-51 |
| chr3-48776073-DEL-58 | chr3-48776074-DEL-58 |
| chr3-48804171-DEL-109 | chr3-48804175-DEL-118 |
| chr3-48808104-DEL-2575 | chr3-48808105-DEL-2575 |
| chr3-48839786-DEL-70 | chr3-48839792-DEL-80 |
| chr3-48848137-DEL-72 | chr3-48848142-DEL-74 |
| chr3-48850553-DEL-2263 | chr3-48850537-DEL-2278 |
| chr3-48861092-DEL-64 | chr3-48861091-DEL-60 |
| chr3-48884817-DEL-4682 | chr3-48884818-DEL-4682 |
| chr3-48895737-DEL-287 | chr3-48895737-DEL-287 |
| chr3-48911561-DEL-2944 | chr3-48911562-DEL-2945 |
| chr3-48950779-DEL-76 | chr3-48950748-DEL-82 |
| chr3-48952888-DEL-103 | chr3-48952866-DEL-108 |
| chr3-48969359-DEL-708 | chr3-48969360-DEL-708 |
| chr3-48974708-DEL-60 | chr3-48974713-DEL-56 |
| chr3-48990588-DEL-538 | chr3-48990589-DEL-538 |
| chr3-48992414-DEL-69 | chr3-48992415-DEL-69 |
| chr3-48997171-DEL-221 | chr3-48997169-DEL-216 |
| chr3-49025452-DEL-6303 | chr3-49025453-DEL-6303 |
| chr3-49077299-DEL-5291 | chr3-49077300-DEL-5291 |
| chr3-49106362-DEL-72 | chr3-49106359-DEL-73 |
| chr3-49114821-DEL-4950 | chr3-49114822-DEL-4950 |

| | |
|---|---|
| chr3-49153640-DEL-302 | chr3-49153641-DEL-302 |
| chr3-49176781-DEL-107 | chr3-49176782-DEL-107 |
| chr3-49179291-DEL-1475 | chr3-49179291-DEL-1475 |
| chr3-49209626-DEL-2292 | chr3-49209623-DEL-2296 |
| chr3-49220242-DEL-106 | chr3-49220242-DEL-105 |
| chr3-49221274-DEL-6182 | chr3-49221275-DEL-6182 |
| chr3-49229552-DEL-54 | chr3-49229564-DEL-50 |
| chr3-49241362-DEL-1716 | chr3-49241363-DEL-1716 |
| chr3-49246396-DEL-4339 | chr3-49246391-DEL-4343 |
| chr3-49269260-DEL-60 | chr3-49269257-DEL-57 |
| chr3-49277497-DEL-60 | chr3-49277498-DEL-60 |
| chr3-49280384-DEL-1844 | chr3-49280385-DEL-1844 |
| chr3-49283907-DEL-791 | chr3-49283906-DEL-792 |
| chr3-49286181-DEL-65 | chr3-49286176-DEL-61 |
| chr3-49292465-DEL-574 | chr3-49292466-DEL-574 |
| chr3-49331820-DEL-5472 | chr3-49331820-DEL-5472 |
| chr3-49342477-DEL-62 | chr3-49342482-DEL-54 |
| chr3-49352716-DEL-68 | chr3-49352717-DEL-68 |
| chr3-49359479-DEL-437 | chr3-49359480-DEL-437 |
| chr3-49365391-DEL-358 | chr3-49365392-DEL-358 |
| chr3-49395624-DEL-199 | chr3-49395625-DEL-199 |
| chr3-49397999-DEL-1554 | chr3-49397999-DEL-1554 |
| chr3-49455855-DEL-95 | chr3-49455848-DEL-96 |
| chr3-49465692-DEL-113 | chr3-49465691-DEL-109 |
| chr3-49484300-DEL-68 | chr3-49484304-DEL-54 |
| chr3-49488930-DEL-323 | chr3-49488930-DEL-324 |
| chr3-49491398-DEL-61 | chr3-49491399-DEL-61 |
| chr3-49533073-DEL-790 | chr3-49533074-DEL-790 |
| chr3-49541952-DEL-76 | chr3-49541953-DEL-77 |
| chr3-49549018-DEL-837 | chr3-49549019-DEL-837 |
| chr3-49562859-DEL-65 | chr3-49562846-DEL-54 |
| chr3-49566767-DEL-87 | chr3-49566770-DEL-80 |
| chr3-49566804-DEL-1970 | chr3-49566805-DEL-1970 |
| chr3-49569688-DEL-528 | chr3-49569689-DEL-528 |
| chr3-49589182-DEL-150 | chr3-49589215-DEL-118 |
| chr3-49594193-DEL-1478 | chr3-49594194-DEL-1478 |
| chr3-49605265-DEL-1673 | chr3-49605266-DEL-1673 |
| chr3-49611181-DEL-370 | chr3-49611188-DEL-363 |
| chr3-49615328-DEL-52 | chr3-49615329-DEL-52 |
| chr3-49635110-DEL-52 | chr3-49635111-DEL-52 |
| chr3-49645044-DEL-110 | chr3-49645071-DEL-84 |
| chr3-49652011-DEL-66 | chr3-49652012-DEL-66 |
| chr3-49692025-DEL-358 | chr3-49692026-DEL-358 |
| chr3-49700281-DEL-58 | chr3-49700278-DEL-62 |
| chr3-49739782-DEL-53 | chr3-49739781-DEL-54 |
| chr3-49740588-DEL-101 | chr3-49740589-DEL-101 |
| chr3-49751462-DEL-6301 | chr3-49751463-DEL-6301 |
| chr3-49771303-DEL-5774 | chr3-49771304-DEL-5774 |
| chr3-49777261-DEL-6396 | chr3-49777262-DEL-6396 |
| chr3-49802178-DEL-302 | chr3-49802179-DEL-302 |
| chr3-49845123-DEL-57 | chr3-49845124-DEL-57 |
| chr3-49902362-DEL-371 | chr3-49902362-DEL-371 |
| chr3-49921232-DEL-79 | chr3-49921228-DEL-80 |
| chr3-49929191-DEL-85 | chr3-49929191-DEL-86 |
| chr3-49934733-DEL-2447 | chr3-49934733-DEL-2448 |
| chr3-49941603-DEL-55 | chr3-49941604-DEL-55 |
| chr3-49943604-DEL-935 | chr3-49943605-DEL-935 |
| chr3-49963783-DEL-463 | chr3-49963784-DEL-463 |
| chr3-49971449-DEL-69 | chr3-49971449-DEL-69 |
| chr3-49989390-DEL-138 | chr3-49989377-DEL-141 |
| chr3-49998746-DEL-101 | chr3-49998747-DEL-99 |
| chr3-50008132-DEL-6340 | chr3-50008133-DEL-6340 |
| chr3-50015580-DEL-54 | chr3-50015581-DEL-66 |
| chr3-50019942-DEL-157 | chr3-50019943-DEL-157 |
| chr3-50025372-DEL-162 | chr3-50025373-DEL-162 |
| chr3-50035222-DEL-94 | chr3-50035223-DEL-94 |
| chr3-50044370-DEL-2318 | chr3-50044371-DEL-2318 |
| chr3-50095905-DEL-1967 | chr3-50095906-DEL-1967 |
| chr3-50114606-DEL-63 | chr3-50114607-DEL-67 |
| chr3-50136654-DEL-71 | chr3-50136655-DEL-71 |
| chr3-50191755-DEL-918 | chr3-50191756-DEL-918 |
| chr3-50200683-DEL-633 | chr3-50200684-DEL-633 |
| chr3-50201918-DEL-302 | chr3-50201919-DEL-302 |
| chr3-50209217-DEL-502 | chr3-50209218-DEL-502 |
| chr3-50227022-DEL-98 | chr3-50227023-DEL-98 |
| chr3-50245555-DEL-102 | chr3-50245556-DEL-102 |
| chr3-50251901-DEL-52 | chr3-50251902-DEL-52 |
| chr3-50264917-DEL-345 | chr3-50264927-DEL-342 |
| chr3-50306529-DEL-519 | chr3-50306530-DEL-519 |
| chr3-50316275-DEL-432 | chr3-50316276-DEL-432 |

| | |
|---|---|
| chr3-50342787-DEL-74 | chr3-50342788-DEL-74 |
| chr3-50368994-DEL-61 | chr3-50369016-DEL-62 |
| chr3-50407047-DEL-167 | chr3-50407048-DEL-167 |
| chr3-50419279-DEL-6773 | chr3-50419280-DEL-6773 |
| chr3-50427002-DEL-1490 | chr3-50427003-DEL-1490 |
| chr3-50444745-DEL-55 | chr3-50444746-DEL-55 |
| chr3-50469435-DEL-112 | chr3-50469436-DEL-112 |
| chr3-50476360-DEL-476 | chr3-50476360-DEL-477 |
| chr3-50491811-DEL-72 | chr3-50491812-DEL-75 |
| chr3-50495328-DEL-6221 | chr3-50495329-DEL-6221 |
| chr3-50538723-DEL-80 | chr3-50538685-DEL-80 |
| chr3-50558784-DEL-372 | chr3-50558785-DEL-372 |
| chr3-50583451-DEL-317 | chr3-50583444-DEL-315 |
| chr3-50593894-DEL-270 | chr3-50593895-DEL-294 |
| chr3-50621685-DEL-6198 | chr3-50621686-DEL-6198 |
| chr3-50638242-DEL-93 | chr3-50638243-DEL-93 |
| chr3-50650857-DEL-6189 | chr3-50650858-DEL-6189 |
| chr3-50663848-DEL-68 | chr3-50663848-DEL-68 |
| chr3-50680120-DEL-52 | chr3-50680121-DEL-52 |
| chr3-50692144-DEL-54 | chr3-50692145-DEL-54 |
| chr3-50692446-DEL-343 | chr3-50692453-DEL-337 |
| chr3-50719802-DEL-274 | chr3-50719803-DEL-274 |
| chr3-50737002-DEL-719 | chr3-50737003-DEL-719 |
| chr3-50760806-DEL-90 | chr3-50760807-DEL-90 |
| chr3-50768055-DEL-282 | chr3-50768056-DEL-282 |
| chr3-50769115-DEL-199 | chr3-50769116-DEL-199 |
| chr3-50770187-DEL-85 | chr3-50770188-DEL-85 |
| chr3-50786017-DEL-68 | chr3-50786018-DEL-68 |
| chr3-50820827-DEL-153 | chr3-50820817-DEL-151 |
| chr3-50822979-DEL-71 | chr3-50822976-DEL-62 |
| chr3-50852310-DEL-67 | chr3-50852311-DEL-67 |
| chr3-50853682-DEL-56 | chr3-50853683-DEL-56 |
| chr3-50859107-DEL-152 | chr3-50859108-DEL-152 |
| chr3-50864693-DEL-6544 | chr3-50864682-DEL-6531 |
| chr3-50917654-DEL-183 | chr3-50917655-DEL-183 |
| chr3-50927928-DEL-241 | chr3-50927926-DEL-244 |
| chr3-50939658-DEL-98 | chr3-50939678-DEL-88 |
| chr3-50943465-DEL-395 | chr3-50943466-DEL-395 |
| chr3-50947449-DEL-221 | chr3-50947450-DEL-221 |
| chr3-50964413-DEL-267 | chr3-50964414-DEL-267 |
| chr3-50965216-DEL-124 | chr3-50965217-DEL-124 |
| chr3-50978871-DEL-59 | chr3-50978872-DEL-59 |
| chr3-50986669-DEL-52 | chr3-50986670-DEL-60 |
| chr3-50989722-DEL-54 | chr3-50989723-DEL-54 |
| chr3-50992630-DEL-70 | chr3-50992631-DEL-64 |
| chr3-51000190-DEL-201 | chr3-51000190-DEL-202 |
| chr3-51008474-DEL-4201 | chr3-51008469-DEL-4172 |
| chr3-51041842-DEL-88 | chr3-51041839-DEL-92 |
| chr3-51087072-DEL-94 | chr3-51087073-DEL-94 |
| chr3-51087922-DEL-54 | chr3-51087923-DEL-52 |
| chr3-51091517-DEL-85 | chr3-51091516-DEL-84 |
| chr3-51101152-DEL-53 | chr3-51101145-DEL-53 |
| chr3-51111043-DEL-88 | chr3-51111044-DEL-88 |
| chr3-51115130-DEL-453 | chr3-51115131-DEL-452 |
| chr3-51120989-DEL-78 | chr3-51121000-DEL-68 |
| chr3-51122128-DEL-53 | chr3-51122126-DEL-56 |
| chr3-51123412-DEL-481 | chr3-51123413-DEL-481 |
| chr3-51140727-DEL-112 | chr3-51140728-DEL-112 |
| chr3-51147382-DEL-5340 | chr3-51147383-DEL-5340 |
| chr3-51203413-DEL-238 | chr3-51203414-DEL-238 |
| chr3-51212127-DEL-52 | chr3-51212128-DEL-52 |
| chr3-51253032-DEL-70 | chr3-51253025-DEL-102 |
| chr3-51254251-DEL-352 | chr3-51254252-DEL-352 |
| chr3-51261688-DEL-65 | chr3-51261689-DEL-65 |
| chr3-51263750-DEL-60 | chr3-51263746-DEL-52 |
| chr3-51271821-DEL-97 | chr3-51271822-DEL-97 |
| chr3-51277290-DEL-65 | chr3-51277291-DEL-65 |
| chr3-51278907-DEL-153 | chr3-51278897-DEL-162 |
| chr3-51287969-DEL-212 | chr3-51287970-DEL-212 |
| chr3-51302667-DEL-59 | chr3-51302684-DEL-57 |
| chr3-51307379-DEL-441 | chr3-51307380-DEL-441 |
| chr3-51326218-DEL-124 | chr3-51326219-DEL-124 |
| chr3-51334330-DEL-115 | chr3-51334331-DEL-115 |
| chr3-51347357-DEL-201 | chr3-51347358-DEL-201 |
| chr3-51410023-DEL-181 | chr3-51410024-DEL-181 |
| chr3-51418113-DEL-121 | chr3-51418115-DEL-92 |
| chr3-51435619-DEL-70 | chr3-51435612-DEL-74 |
| chr3-51442486-DEL-199 | chr3-51442487-DEL-199 |
| chr3-51448699-DEL-59 | chr3-51448700-DEL-59 |
| chr3-51477643-DEL-672 | chr3-51477645-DEL-662 |

| | |
|---|---|
| chr3-51479012-DEL-174 | chr3-51479013-DEL-173 |
| chr3-51479347-DEL-467 | chr3-51479348-DEL-467 |
| chr3-51509805-DEL-206 | chr3-51509806-DEL-206 |
| chr3-51515766-DEL-712 | chr3-51515765-DEL-712 |
| chr3-51520369-DEL-127 | chr3-51520370-DEL-127 |
| chr3-51523597-DEL-7382 | chr3-51523598-DEL-7382 |
| chr3-51556072-DEL-499 | chr3-51556075-DEL-497 |
| chr3-51566971-DEL-275 | chr3-51566972-DEL-275 |
| chr3-51572501-DEL-96 | chr3-51572522-DEL-68 |
| chr3-51586870-DEL-1588 | chr3-51586871-DEL-1588 |
| chr3-51587187-DEL-752 | chr3-51587188-DEL-752 |
| chr3-51603882-DEL-76 | chr3-51603859-DEL-75 |
| chr3-51623423-DEL-55 | chr3-51623426-DEL-55 |
| chr3-51638988-DEL-95 | chr3-51638989-DEL-95 |
| chr3-51647383-DEL-92 | chr3-51647377-DEL-56 |
| chr3-51654046-DEL-80 | chr3-51654047-DEL-80 |
| chr3-51659910-DEL-205 | chr3-51659909-DEL-205 |
| chr3-51677937-DEL-238 | chr3-51677936-DEL-237 |
| chr3-51678617-DEL-174 | chr3-51678618-DEL-174 |
| chr3-51685565-DEL-56 | chr3-51685566-DEL-56 |
| chr3-51786458-DEL-247 | chr3-51786459-DEL-247 |
| chr3-51810357-DEL-57 | chr3-51810358-DEL-57 |
| chr3-51818998-DEL-52 | chr3-51818997-DEL-50 |
| chr3-51837519-DEL-4391 | chr3-51837520-DEL-4391 |
| chr3-51860050-DEL-213 | chr3-51860051-DEL-213 |
| chr3-51907308-DEL-200 | chr3-51907309-DEL-197 |
| chr3-51914818-DEL-65 | chr3-51914811-DEL-66 |
| chr3-51921466-DEL-127 | chr3-51921467-DEL-127 |
| chr3-51963753-DEL-57 | chr3-51963754-DEL-57 |
| chr3-51977049-DEL-53 | chr3-51977047-DEL-54 |
| chr3-52000320-DEL-84 | chr3-52000321-DEL-84 |
| chr3-52027398-DEL-151 | chr3-52027399-DEL-151 |
| chr3-52029119-DEL-69 | chr3-52029120-DEL-70 |
| chr3-52087590-DEL-64 | chr3-52087591-DEL-64 |
| chr3-52091448-DEL-75 | chr3-52091449-DEL-75 |
| chr3-52110005-DEL-2828 | chr3-52110006-DEL-2828 |
| chr3-52111572-DEL-102 | chr3-52111617-DEL-57 |
| chr3-52115581-DEL-508 | chr3-52115582-DEL-508 |
| chr3-52135911-DEL-239 | chr3-52135912-DEL-239 |
| chr3-52137355-DEL-3192 | chr3-52137356-DEL-3192 |
| chr3-52147674-DEL-344 | chr3-52147675-DEL-344 |
| chr3-52159416-DEL-305 | chr3-52159442-DEL-298 |
| chr3-52184408-DEL-244 | chr3-52184411-DEL-242 |
| chr3-52199914-DEL-4937 | chr3-52199915-DEL-4937 |
| chr3-52292498-DEL-117 | chr3-52292499-DEL-117 |
| chr3-52322230-DEL-86 | chr3-52322257-DEL-60 |
| chr3-52333766-DEL-494 | chr3-52333767-DEL-494 |
| chr3-52338084-DEL-6641 | chr3-52338085-DEL-6641 |
| chr3-52345853-DEL-597 | chr3-52345854-DEL-597 |
| chr3-52348455-DEL-1327 | chr3-52348456-DEL-1327 |
| chr3-52348851-DEL-82 | chr3-52348856-DEL-80 |
| chr3-52361749-DEL-72 | chr3-52361750-DEL-72 |
| chr3-52364023-DEL-82 | chr3-52364021-DEL-86 |
| chr3-52373543-DEL-159 | chr3-52373544-DEL-159 |
| chr3-52379551-DEL-81 | chr3-52379579-DEL-78 |
| chr3-52382906-DEL-218 | chr3-52382907-DEL-218 |
| chr3-52393324-DEL-189 | chr3-52393325-DEL-189 |
| chr3-52399583-DEL-80 | chr3-52399584-DEL-80 |
| chr3-52412713-DEL-132 | chr3-52412714-DEL-132 |
| chr3-52419409-DEL-628 | chr3-52419410-DEL-628 |
| chr3-52440975-DEL-65 | chr3-52440938-DEL-64 |
| chr3-52454445-DEL-151 | chr3-52454446-DEL-151 |
| chr3-52490289-DEL-136 | chr3-52490334-DEL-132 |
| chr3-52520155-DEL-1296 | chr3-52520156-DEL-1297 |
| chr3-52542760-DEL-57 | chr3-52542755-DEL-58 |
| chr3-52561106-DEL-350 | chr3-52561106-DEL-350 |
| chr3-52576043-DEL-519 | chr3-52576044-DEL-519 |
| chr3-52597987-DEL-210 | chr3-52597987-DEL-212 |
| chr3-52625742-DEL-722 | chr3-52625743-DEL-722 |
| chr3-52651275-DEL-83 | chr3-52651276-DEL-83 |
| chr3-52714813-DEL-179 | chr3-52714814-DEL-179 |
| chr3-52721135-DEL-68 | chr3-52721136-DEL-68 |
| chr3-52788159-DEL-103 | chr3-52788172-DEL-91 |
| chr3-52790467-DEL-1510 | chr3-52790468-DEL-1510 |
| chr3-52812519-DEL-90 | chr3-52812520-DEL-90 |
| chr3-52818278-DEL-7616 | chr3-52818279-DEL-7616 |
| chr3-52881951-DEL-84 | chr3-52881952-DEL-84 |
| chr3-52893018-DEL-60 | chr3-52893019-DEL-60 |
| chr3-52895713-DEL-156 | chr3-52895675-DEL-156 |
| chr3-52899119-DEL-87 | chr3-52899120-DEL-87 |

| | |
|---|---|
| chr3-52901016-DEL-275 | chr3-52901017-DEL-275 |
| chr3-52901939-DEL-60 | chr3-52901940-DEL-60 |
| chr3-52954532-DEL-72 | chr3-52954545-DEL-60 |
| chr3-52959065-DEL-152 | chr3-52959066-DEL-152 |
| chr3-52968987-DEL-53 | chr3-52968988-DEL-68 |
| chr3-52974288-DEL-897 | chr3-52974295-DEL-890 |
| chr3-53006126-DEL-66 | chr3-53006127-DEL-66 |
| chr3-53020833-DEL-5486 | chr3-53020834-DEL-5486 |
| chr3-53028732-DEL-1691 | chr3-53028733-DEL-1691 |
| chr3-53078873-DEL-355 | chr3-53078874-DEL-355 |
| chr3-53083941-DEL-52 | chr3-53083942-DEL-50 |
| chr3-53093809-DEL-6182 | chr3-53093811-DEL-6198 |
| chr3-53120336-DEL-91 | chr3-53120352-DEL-61 |
| chr3-53127824-DEL-56 | chr3-53127858-DEL-55 |
| chr3-53135935-DEL-187 | chr3-53135936-DEL-187 |
| chr3-53137559-DEL-71 | chr3-53137560-DEL-71 |
| chr3-53158255-DEL-399 | chr3-53158256-DEL-399 |
| chr3-53164909-DEL-415 | chr3-53164910-DEL-415 |
| chr3-53186047-DEL-238 | chr3-53186046-DEL-238 |
| chr3-53210990-DEL-360 | chr3-53210991-DEL-360 |
| chr3-53245377-DEL-204 | chr3-53245378-DEL-204 |
| chr3-53253715-DEL-115 | chr3-53253716-DEL-115 |
| chr3-53257103-DEL-1458 | chr3-53257104-DEL-1458 |
| chr3-53259566-DEL-642 | chr3-53259567-DEL-642 |
| chr3-53271426-DEL-5262 | chr3-53271425-DEL-5262 |
| chr3-53275940-DEL-77 | chr3-53275941-DEL-77 |
| chr3-53276893-DEL-60 | chr3-53276894-DEL-60 |
| chr3-53314831-DEL-161 | chr3-53314832-DEL-155 |
| chr3-53320831-DEL-183 | chr3-53320832-DEL-183 |
| chr3-53343115-DEL-127 | chr3-53343113-DEL-155 |
| chr3-53373216-DEL-75 | chr3-53373217-DEL-75 |
| chr3-53401646-DEL-102 | chr3-53401612-DEL-137 |
| chr3-53407374-DEL-100 | chr3-53407374-DEL-100 |
| chr3-53413954-DEL-86 | chr3-53413968-DEL-68 |
| chr3-53416100-DEL-78 | chr3-53416135-DEL-79 |
| chr3-53428319-DEL-76 | chr3-53428315-DEL-77 |
| chr3-53435172-DEL-304 | chr3-53435173-DEL-304 |
| chr3-53435759-DEL-65 | chr3-53435754-DEL-63 |
| chr3-53447263-DEL-61 | chr3-53447264-DEL-61 |
| chr3-53448976-DEL-463 | chr3-53448977-DEL-463 |
| chr3-53478271-DEL-116 | chr3-53478272-DEL-116 |
| chr3-53499993-DEL-81 | chr3-53499998-DEL-77 |
| chr3-53502555-DEL-92 | chr3-53502602-DEL-88 |
| chr3-53505541-DEL-60 | chr3-53505522-DEL-57 |
| chr3-53511666-DEL-78 | chr3-53511667-DEL-78 |
| chr3-53545214-DEL-1475 | chr3-53545210-DEL-1476 |
| chr3-53599759-DEL-578 | chr3-53599760-DEL-578 |
| chr3-53601365-DEL-1118 | chr3-53601366-DEL-1118 |
| chr3-53602907-DEL-56 | chr3-53602904-DEL-60 |
| chr3-53617228-DEL-69 | chr3-53617229-DEL-69 |
| chr3-53622502-DEL-6879 | chr3-53622503-DEL-6879 |
| chr3-53630362-DEL-508 | chr3-53630363-DEL-508 |
| chr3-53634946-DEL-78 | chr3-53634937-DEL-103 |
| chr3-53643126-DEL-78 | chr3-53643127-DEL-78 |
| chr3-53650029-DEL-878 | chr3-53650030-DEL-878 |
| chr3-53654830-DEL-90 | chr3-53654831-DEL-84 |
| chr3-53656756-DEL-580 | chr3-53656745-DEL-581 |
| chr3-53667739-DEL-206 | chr3-53667740-DEL-206 |
| chr3-53749688-DEL-201 | chr3-53749688-DEL-201 |
| chr3-53755493-DEL-4369 | chr3-53755494-DEL-4369 |
| chr3-53759559-DEL-66 | chr3-53759580-DEL-50 |
| chr3-53798998-DEL-1619 | chr3-53798999-DEL-1619 |
| chr3-53801436-DEL-242 | chr3-53801437-DEL-242 |
| chr3-53850449-DEL-327 | chr3-53850450-DEL-327 |
| chr3-53867668-DEL-72 | chr3-53867681-DEL-60 |
| chr3-53899342-DEL-51 | chr3-53899337-DEL-57 |
| chr3-53946551-DEL-64 | chr3-53946552-DEL-64 |
| chr3-53951589-DEL-1294 | chr3-53951590-DEL-1294 |
| chr3-53955528-DEL-60 | chr3-53955529-DEL-60 |
| chr3-53962412-DEL-78 | chr3-53962413-DEL-76 |
| chr3-53971524-DEL-62 | chr3-53971525-DEL-62 |
| chr3-53979357-DEL-143 | chr3-53979358-DEL-143 |
| chr3-53995186-DEL-412 | chr3-53995186-DEL-409 |
| chr3-54008304-DEL-533 | chr3-54008305-DEL-533 |
| chr3-54011998-DEL-60 | chr3-54011998-DEL-60 |
| chr3-54027184-DEL-263 | chr3-54027188-DEL-260 |
| chr3-54056378-DEL-2445 | chr3-54056379-DEL-2445 |
| chr3-54076444-DEL-425 | chr3-54076450-DEL-419 |
| chr3-54085046-DEL-81 | chr3-54085047-DEL-81 |
| chr3-54168979-DEL-1957 | chr3-54168980-DEL-1957 |

| | |
|---|---|
| chr3-54189111-DEL-59 | chr3-54189112-DEL-59 |
| chr3-54203746-DEL-563 | chr3-54203742-DEL-564 |
| chr3-54205723-DEL-358 | chr3-54205724-DEL-358 |
| chr3-54209847-DEL-54 | chr3-54209848-DEL-54 |
| chr3-54227770-DEL-68 | chr3-54227771-DEL-68 |
| chr3-54253100-DEL-1900 | chr3-54253101-DEL-1900 |
| chr3-54309661-DEL-56 | chr3-54309662-DEL-56 |
| chr3-54316002-DEL-166 | chr3-54316003-DEL-166 |
| chr3-54316893-DEL-1009 | chr3-54316894-DEL-1009 |
| chr3-54322551-DEL-60 | chr3-54322552-DEL-60 |
| chr3-54334194-DEL-68 | chr3-54334189-DEL-62 |
| chr3-54352414-DEL-67 | chr3-54352411-DEL-79 |
| chr3-54368534-DEL-92 | chr3-54368496-DEL-119 |
| chr3-54382384-DEL-93 | chr3-54382385-DEL-93 |
| chr3-54388822-DEL-1535 | chr3-54388823-DEL-1535 |
| chr3-54395552-DEL-170 | chr3-54395553-DEL-170 |
| chr3-54408475-DEL-282 | chr3-54408476-DEL-282 |
| chr3-54422208-DEL-289 | chr3-54422209-DEL-289 |
| chr3-54424553-DEL-118 | chr3-54424554-DEL-118 |
| chr3-54437681-DEL-81 | chr3-54437685-DEL-79 |
| chr3-54444671-DEL-3779 | chr3-54444676-DEL-3776 |
| chr3-54462447-DEL-86 | chr3-54462469-DEL-52 |
| chr3-54463298-DEL-147 | chr3-54463299-DEL-147 |
| chr3-54470241-DEL-66 | chr3-54470239-DEL-69 |
| chr3-54470555-DEL-378 | chr3-54470556-DEL-378 |
| chr3-54476678-DEL-494 | chr3-54476679-DEL-494 |
| chr3-54481437-DEL-86 | chr3-54481446-DEL-72 |
| chr3-54482769-DEL-58 | chr3-54482770-DEL-58 |
| chr3-54489347-DEL-485 | chr3-54489356-DEL-479 |
| chr3-54496357-DEL-1176 | chr3-54496358-DEL-1176 |
| chr3-54512666-DEL-405 | chr3-54512667-DEL-405 |
| chr3-54518308-DEL-62 | chr3-54518309-DEL-62 |
| chr3-54532267-DEL-313 | chr3-54532268-DEL-313 |
| chr3-54533988-DEL-169 | chr3-54533989-DEL-169 |
| chr3-54556880-DEL-88 | chr3-54556881-DEL-88 |
| chr3-54593236-DEL-152 | chr3-54593237-DEL-152 |
| chr3-54667217-DEL-123 | chr3-54667217-DEL-124 |
| chr3-54678808-DEL-551 | chr3-54678809-DEL-551 |
| chr3-54712508-DEL-111 | chr3-54712509-DEL-111 |
| chr3-54717390-DEL-1435 | chr3-54717391-DEL-1435 |
| chr3-54729221-DEL-76 | chr3-54729224-DEL-78 |
| chr3-54731498-DEL-2818 | chr3-54731499-DEL-2818 |
| chr3-54756594-DEL-7160 | chr3-54756595-DEL-7160 |
| chr3-54764535-DEL-120 | chr3-54764536-DEL-120 |
| chr3-54778595-DEL-67 | chr3-54778596-DEL-67 |
| chr3-54779253-DEL-2538 | chr3-54779254-DEL-2538 |
| chr3-54810211-DEL-241 | chr3-54810212-DEL-241 |
| chr3-54814283-DEL-70 | chr3-54814284-DEL-68 |
| chr3-54817706-DEL-54 | chr3-54817700-DEL-54 |
| chr3-54830677-DEL-80 | chr3-54830678-DEL-80 |
| chr3-54847872-DEL-55 | chr3-54847847-DEL-70 |
| chr3-54881807-DEL-270 | chr3-54881802-DEL-271 |
| chr3-54886884-DEL-81 | chr3-54886885-DEL-81 |
| chr3-54888185-DEL-142 | chr3-54888195-DEL-138 |
| chr3-54892096-DEL-68 | chr3-54892089-DEL-69 |
| chr3-54900212-DEL-99 | chr3-54900186-DEL-100 |
| chr3-54903768-DEL-122 | chr3-54903768-DEL-123 |
| chr3-54905730-DEL-73 | chr3-54905731-DEL-73 |
| chr3-54913931-DEL-187 | chr3-54913932-DEL-187 |
| chr3-54914552-DEL-104 | chr3-54914565-DEL-92 |
| chr3-54957670-DEL-72 | chr3-54957665-DEL-61 |
| chr3-54968745-DEL-184 | chr3-54968708-DEL-190 |
| chr3-54986487-DEL-61 | chr3-54986484-DEL-62 |
| chr3-55021722-DEL-66 | chr3-55021723-DEL-66 |
| chr3-55055530-DEL-55 | chr3-55055528-DEL-58 |
| chr3-55127113-DEL-878 | chr3-55127114-DEL-878 |
| chr3-55131192-DEL-107 | chr3-55131193-DEL-107 |
| chr3-55144642-DEL-153 | chr3-55144653-DEL-143 |
| chr3-55162246-DEL-130 | chr3-55162241-DEL-131 |
| chr3-55204902-DEL-67 | chr3-55204903-DEL-67 |
| chr3-55209585-DEL-2559 | chr3-55209586-DEL-2559 |
| chr3-55226679-DEL-72 | chr3-55226680-DEL-72 |
| chr3-55235221-DEL-75 | chr3-55235222-DEL-75 |
| chr3-55235911-DEL-112 | chr3-55235912-DEL-112 |
| chr3-55274476-DEL-88 | chr3-55274456-DEL-64 |
| chr3-55326363-DEL-94 | chr3-55326360-DEL-94 |
| chr3-55329793-DEL-117 | chr3-55329794-DEL-117 |
| chr3-55355328-DEL-54 | chr3-55355329-DEL-54 |
| chr3-55390582-DEL-65 | chr3-55390562-DEL-87 |
| chr3-55408688-DEL-474 | chr3-55408691-DEL-473 |

| | |
|---|---|
| chr3-55412003-DEL-61 | chr3-55411998-DEL-63 |
| chr3-55413196-DEL-465 | chr3-55413197-DEL-465 |
| chr3-55419485-DEL-117 | chr3-55419486-DEL-117 |
| chr3-55470983-DEL-1041 | chr3-55470984-DEL-1041 |
| chr3-55485412-DEL-8407 | chr3-55485413-DEL-8407 |
| chr3-55505316-DEL-218 | chr3-55505319-DEL-216 |
| chr3-55509587-DEL-58 | chr3-55509588-DEL-58 |
| chr3-55510182-DEL-7550 | chr3-55510183-DEL-7550 |
| chr3-55521308-DEL-81 | chr3-55521260-DEL-120 |
| chr3-55529647-DEL-606 | chr3-55529648-DEL-607 |
| chr3-55544558-DEL-68 | chr3-55544555-DEL-76 |
| chr3-55548161-DEL-60 | chr3-55548134-DEL-58 |
| chr3-55568942-DEL-196 | chr3-55568978-DEL-164 |
| chr3-55580628-DEL-54 | chr3-55580629-DEL-54 |
| chr3-55594439-DEL-104 | chr3-55594442-DEL-68 |
| chr3-55740328-DEL-196 | chr3-55740329-DEL-196 |
| chr3-55745273-DEL-52 | chr3-55745266-DEL-56 |
| chr3-55746679-DEL-303 | chr3-55746680-DEL-302 |
| chr3-55758862-DEL-1078 | chr3-55758863-DEL-1078 |
| chr3-55779455-DEL-202 | chr3-55779456-DEL-202 |
| chr3-55823666-DEL-6243 | chr3-55823667-DEL-6243 |
| chr3-55843720-DEL-53 | chr3-55843721-DEL-53 |
| chr3-55928509-DEL-5304 | chr3-55928510-DEL-5304 |
| chr3-55960044-DEL-193 | chr3-55960045-DEL-193 |
| chr3-55963091-DEL-181 | chr3-55963092-DEL-181 |
| chr3-55965153-DEL-56 | chr3-55965148-DEL-62 |
| chr3-55977057-DEL-6289 | chr3-55977058-DEL-6289 |
| chr3-56009047-DEL-663 | chr3-56009048-DEL-663 |
| chr3-56016638-DEL-127 | chr3-56016639-DEL-131 |
| chr3-56026633-DEL-2019 | chr3-56026633-DEL-2019 |
| chr3-56043654-DEL-6438 | chr3-56043654-DEL-6438 |
| chr3-56055305-DEL-93 | chr3-56055305-DEL-93 |
| chr3-56116685-DEL-6554 | chr3-56116686-DEL-6554 |
| chr3-56132128-DEL-105 | chr3-56132114-DEL-106 |
| chr3-56141053-DEL-90 | chr3-56141045-DEL-94 |
| chr3-56142606-DEL-399 | chr3-56142601-DEL-398 |
| chr3-56149501-DEL-99 | chr3-56149502-DEL-87 |
| chr3-56154307-DEL-6344 | chr3-56154308-DEL-6344 |
| chr3-56176173-DEL-498 | chr3-56176174-DEL-498 |
| chr3-56189718-DEL-1894 | chr3-56189719-DEL-1894 |
| chr3-56190582-DEL-142 | chr3-56190583-DEL-142 |
| chr3-56196894-DEL-403 | chr3-56196898-DEL-401 |
| chr3-56200090-DEL-96 | chr3-56200091-DEL-96 |
| chr3-56231537-DEL-758 | chr3-56231538-DEL-759 |
| chr3-56264280-DEL-66 | chr3-56264281-DEL-66 |
| chr3-56278755-DEL-6804 | chr3-56278756-DEL-6804 |
| chr3-56286702-DEL-64 | chr3-56286703-DEL-64 |
| chr3-56311793-DEL-70 | chr3-56311794-DEL-70 |
| chr3-56333057-DEL-1058 | chr3-56333059-DEL-1056 |
| chr3-56337196-DEL-417 | chr3-56337197-DEL-417 |
| chr3-56353369-DEL-58 | chr3-56353355-DEL-54 |
| chr3-56398735-DEL-85 | chr3-56398738-DEL-83 |
| chr3-56421009-DEL-6668 | chr3-56421010-DEL-6668 |
| chr3-56430548-DEL-421 | chr3-56430549-DEL-421 |
| chr3-56436613-DEL-499 | chr3-56436614-DEL-499 |
| chr3-56446260-DEL-6421 | chr3-56446261-DEL-6421 |
| chr3-56490068-DEL-579 | chr3-56490069-DEL-579 |
| chr3-56505189-DEL-6398 | chr3-56505190-DEL-6398 |
| chr3-56530789-DEL-60 | chr3-56530790-DEL-50 |
| chr3-56551569-DEL-1308 | chr3-56551570-DEL-1308 |
| chr3-56574883-DEL-108 | chr3-56574882-DEL-107 |
| chr3-56635745-DEL-141 | chr3-56635746-DEL-141 |
| chr3-56639496-DEL-52 | chr3-56639477-DEL-84 |
| chr3-56641012-DEL-127 | chr3-56641010-DEL-128 |
| chr3-56666638-DEL-2506 | chr3-56666639-DEL-2506 |
| chr3-56698725-DEL-289 | chr3-56698726-DEL-289 |
| chr3-56712166-DEL-3399 | chr3-56712167-DEL-3399 |
| chr3-56716505-DEL-101 | chr3-56716506-DEL-101 |
| chr3-56718751-DEL-83 | chr3-56718752-DEL-83 |
| chr3-56725849-DEL-2197 | chr3-56725850-DEL-2197 |
| chr3-56733686-DEL-62 | chr3-56733687-DEL-62 |
| chr3-56747039-DEL-92 | chr3-56747026-DEL-77 |
| chr3-56747782-DEL-487 | chr3-56747783-DEL-487 |
| chr3-56755523-DEL-6140 | chr3-56755524-DEL-6140 |
| chr3-56762337-DEL-107 | chr3-56762338-DEL-107 |
| chr3-56765342-DEL-58 | chr3-56765343-DEL-58 |
| chr3-56778960-DEL-898 | chr3-56778961-DEL-898 |
| chr3-56786504-DEL-72 | chr3-56786505-DEL-72 |
| chr3-56855919-DEL-782 | chr3-56855920-DEL-782 |
| chr3-56857814-DEL-170 | chr3-56857815-DEL-170 |

| | |
|---|---|
| chr3-56907175-DEL-341 | chr3-56907176-DEL-341 |
| chr3-56919845-DEL-302 | chr3-56919846-DEL-302 |
| chr3-56920960-DEL-2308 | chr3-56920953-DEL-2282 |
| chr3-56932614-DEL-163 | chr3-56932610-DEL-152 |
| chr3-56935904-DEL-52 | chr3-56935905-DEL-52 |
| chr3-56939287-DEL-52 | chr3-56939288-DEL-52 |
| chr3-56950315-DEL-53 | chr3-56950316-DEL-53 |
| chr3-56953646-DEL-67 | chr3-56953622-DEL-91 |
| chr3-56968232-DEL-7010 | chr3-56968233-DEL-7010 |
| chr3-56992808-DEL-507 | chr3-56992809-DEL-507 |
| chr3-56995675-DEL-62 | chr3-56995676-DEL-62 |
| chr3-56997658-DEL-54 | chr3-56997659-DEL-54 |
| chr3-57014940-DEL-612 | chr3-57014941-DEL-612 |
| chr3-57039083-DEL-411 | chr3-57039084-DEL-412 |
| chr3-57044118-DEL-308 | chr3-57044121-DEL-307 |
| chr3-57052977-DEL-163 | chr3-57052974-DEL-158 |
| chr3-57054250-DEL-1345 | chr3-57054251-DEL-1345 |
| chr3-57066801-DEL-62 | chr3-57066802-DEL-62 |
| chr3-57079259-DEL-52 | chr3-57079230-DEL-80 |
| chr3-57089978-DEL-6160 | chr3-57089979-DEL-6160 |
| chr3-57127678-DEL-64 | chr3-57127669-DEL-66 |
| chr3-57141548-DEL-51 | chr3-57141537-DEL-58 |
| chr3-57143301-DEL-195 | chr3-57143302-DEL-195 |
| chr3-57143980-DEL-3295 | chr3-57143981-DEL-3295 |
| chr3-57180057-DEL-364 | chr3-57180058-DEL-364 |
| chr3-57191859-DEL-757 | chr3-57191860-DEL-757 |
| chr3-57204737-DEL-201 | chr3-57204738-DEL-201 |
| chr3-57211275-DEL-128 | chr3-57211276-DEL-126 |
| chr3-57229285-DEL-6199 | chr3-57229286-DEL-6199 |
| chr3-57257218-DEL-1459 | chr3-57257219-DEL-1459 |
| chr3-57259431-DEL-61 | chr3-57259431-DEL-61 |
| chr3-57289619-DEL-76 | chr3-57289622-DEL-74 |
| chr3-57290483-DEL-519 | chr3-57290484-DEL-519 |
| chr3-57319785-DEL-365 | chr3-57319786-DEL-365 |
| chr3-57334900-DEL-111 | chr3-57334901-DEL-111 |
| chr3-57335654-DEL-2038 | chr3-57335673-DEL-2019 |
| chr3-57345379-DEL-58 | chr3-57345380-DEL-52 |
| chr3-57349219-DEL-59 | chr3-57349220-DEL-59 |
| chr3-57350845-DEL-71 | chr3-57350861-DEL-50 |
| chr3-57356447-DEL-2629 | chr3-57356448-DEL-2629 |
| chr3-57361299-DEL-59 | chr3-57361300-DEL-59 |
| chr3-57391589-DEL-1105 | chr3-57391586-DEL-1093 |
| chr3-57406588-DEL-112 | chr3-57406590-DEL-110 |
| chr3-57468957-DEL-1016 | chr3-57468958-DEL-1016 |
| chr3-57502187-DEL-83 | chr3-57502188-DEL-79 |
| chr3-57559476-DEL-68 | chr3-57559477-DEL-68 |
| chr3-57573889-DEL-59 | chr3-57573890-DEL-59 |
| chr3-57589629-DEL-5510 | chr3-57589624-DEL-5504 |
| chr3-57602198-DEL-2000 | chr3-57602199-DEL-2000 |
| chr3-57640911-DEL-116 | chr3-57640912-DEL-110 |
| chr3-57646623-DEL-259 | chr3-57646632-DEL-260 |
| chr3-57653193-DEL-60 | chr3-57653193-DEL-56 |
| chr3-57687249-DEL-78 | chr3-57687236-DEL-72 |
| chr3-57721768-DEL-68 | chr3-57721769-DEL-68 |
| chr3-57725508-DEL-55 | chr3-57725505-DEL-59 |
| chr3-57756415-DEL-147 | chr3-57756416-DEL-147 |
| chr3-57776963-DEL-53 | chr3-57776964-DEL-53 |
| chr3-57788865-DEL-154 | chr3-57788868-DEL-155 |
| chr3-57791791-DEL-54 | chr3-57791792-DEL-54 |
| chr3-57812475-DEL-60 | chr3-57812476-DEL-60 |
| chr3-57814832-DEL-785 | chr3-57814833-DEL-785 |
| chr3-57835664-DEL-817 | chr3-57835664-DEL-817 |
| chr3-57841991-DEL-131 | chr3-57842012-DEL-92 |
| chr3-57871515-DEL-1718 | chr3-57871516-DEL-1718 |
| chr3-57878310-DEL-188 | chr3-57878311-DEL-188 |
| chr3-57913957-DEL-6404 | chr3-57913948-DEL-6405 |
| chr3-57923297-DEL-169 | chr3-57923298-DEL-169 |
| chr3-57939059-DEL-105 | chr3-57939060-DEL-112 |
| chr3-57941981-DEL-336 | chr3-57941977-DEL-337 |
| chr3-57953963-DEL-287 | chr3-57953972-DEL-283 |
| chr3-57962816-DEL-292 | chr3-57962817-DEL-292 |
| chr3-57976064-DEL-114 | chr3-57976051-DEL-96 |
| chr3-58009970-DEL-1742 | chr3-58009971-DEL-1742 |
| chr3-58060589-DEL-5744 | chr3-58060590-DEL-5744 |
| chr3-58081756-DEL-1004 | chr3-58081757-DEL-1004 |
| chr3-58094108-DEL-6566 | chr3-58094110-DEL-6565 |
| chr3-58105132-DEL-57 | chr3-58105133-DEL-57 |
| chr3-58167194-DEL-88 | chr3-58167197-DEL-62 |
| chr3-58178953-DEL-6664 | chr3-58178954-DEL-6664 |
| chr3-58203762-DEL-280 | chr3-58203771-DEL-270 |

| | |
|---|---|
| chr3-58300719-DEL-101 | chr3-58300720-DEL-101 |
| chr3-58383818-DEL-61 | chr3-58383819-DEL-52 |
| chr3-58400432-DEL-119 | chr3-58400433-DEL-119 |
| chr3-58413024-DEL-1409 | chr3-58413025-DEL-1409 |
| chr3-58450752-DEL-113 | chr3-58450748-DEL-87 |
| chr3-58454274-DEL-305 | chr3-58454275-DEL-305 |
| chr3-58465596-DEL-200 | chr3-58465597-DEL-200 |
| chr3-58480334-DEL-555 | chr3-58480329-DEL-553 |
| chr3-58492905-DEL-207 | chr3-58492906-DEL-207 |
| chr3-58519991-DEL-406 | chr3-58519992-DEL-406 |
| chr3-58553218-DEL-162 | chr3-58553221-DEL-160 |
| chr3-58573262-DEL-210 | chr3-58573266-DEL-186 |
| chr3-58653985-DEL-62 | chr3-58653986-DEL-62 |
| chr3-58670105-DEL-6423 | chr3-58670106-DEL-6423 |
| chr3-58687676-DEL-324 | chr3-58687676-DEL-324 |
| chr3-58735863-DEL-83 | chr3-58735864-DEL-83 |
| chr3-58759567-DEL-6810 | chr3-58759568-DEL-6810 |
| chr3-58779414-DEL-95 | chr3-58779364-DEL-96 |
| chr3-58803265-DEL-74 | chr3-58803266-DEL-73 |
| chr3-58806288-DEL-88 | chr3-58806313-DEL-60 |
| chr3-58840612-DEL-571 | chr3-58840613-DEL-571 |
| chr3-58846347-DEL-1052 | chr3-58846348-DEL-1052 |
| chr3-58855378-DEL-199 | chr3-58855379-DEL-199 |
| chr3-58894541-DEL-740 | chr3-58894553-DEL-709 |
| chr3-58948615-DEL-82 | chr3-58948622-DEL-70 |
| chr3-58952766-DEL-99 | chr3-58952767-DEL-51 |
| chr3-58976567-DEL-80 | chr3-58976568-DEL-80 |
| chr3-58995623-DEL-300 | chr3-58995624-DEL-300 |
| chr3-59033989-DEL-106 | chr3-59033990-DEL-106 |
| chr3-59036136-DEL-56 | chr3-59036126-DEL-84 |
| chr3-59062671-DEL-210 | chr3-59062672-DEL-210 |
| chr3-59144971-DEL-467 | chr3-59144972-DEL-467 |
| chr3-59158459-DEL-740 | chr3-59158460-DEL-740 |
| chr3-59163512-DEL-497 | chr3-59163513-DEL-497 |
| chr3-59187283-DEL-7016 | chr3-59187284-DEL-7016 |
| chr3-59209154-DEL-2057 | chr3-59209155-DEL-2057 |
| chr3-59214154-DEL-341 | chr3-59214155-DEL-341 |
| chr3-59234015-DEL-88 | chr3-59234009-DEL-89 |
| chr3-59234574-DEL-81 | chr3-59234575-DEL-80 |
| chr3-59252440-DEL-368 | chr3-59252441-DEL-368 |
| chr3-59278223-DEL-7086 | chr3-59278224-DEL-7086 |
| chr3-59346103-DEL-327 | chr3-59346104-DEL-328 |
| chr3-59346705-DEL-86 | chr3-59346706-DEL-87 |
| chr3-59354888-DEL-96 | chr3-59354889-DEL-96 |
| chr3-59366598-DEL-62 | chr3-59366599-DEL-56 |
| chr3-59368186-DEL-1698 | chr3-59368187-DEL-1698 |
| chr3-59372142-DEL-82 | chr3-59372161-DEL-80 |
| chr3-59426141-DEL-208 | chr3-59426142-DEL-208 |
| chr3-59430712-DEL-68 | chr3-59430713-DEL-68 |
| chr3-59443004-DEL-450 | chr3-59443005-DEL-450 |
| chr3-59458526-DEL-56 | chr3-59458527-DEL-56 |
| chr3-59472699-DEL-5768 | chr3-59472700-DEL-5768 |
| chr3-59475270-DEL-66 | chr3-59475271-DEL-66 |
| chr3-59532811-DEL-7354 | chr3-59532812-DEL-7354 |
| chr3-59545374-DEL-58 | chr3-59545375-DEL-58 |
| chr3-59550389-DEL-306 | chr3-59550390-DEL-306 |
| chr3-59555771-DEL-88 | chr3-59555772-DEL-86 |
| chr3-59577440-DEL-178 | chr3-59577441-DEL-178 |
| chr3-59612000-DEL-464 | chr3-59611963-DEL-502 |
| chr3-59694492-DEL-83 | chr3-59694489-DEL-65 |
| chr3-59699157-DEL-1554 | chr3-59699163-DEL-1553 |
| chr3-59704443-DEL-7114 | chr3-59704444-DEL-7114 |
| chr3-59718760-DEL-2642 | chr3-59718761-DEL-2642 |
| chr3-59721462-DEL-1396 | chr3-59721463-DEL-1397 |
| chr3-59726766-DEL-437 | chr3-59726767-DEL-437 |
| chr3-59732299-DEL-56 | chr3-59732309-DEL-72 |
| chr3-59744936-DEL-52 | chr3-59744937-DEL-52 |
| chr3-59791356-DEL-70 | chr3-59791366-DEL-64 |
| chr3-59922609-DEL-741 | chr3-59922610-DEL-741 |
| chr3-59952639-DEL-215 | chr3-59952640-DEL-186 |
| chr3-59978101-DEL-851 | chr3-59978102-DEL-851 |
| chr3-59990576-DEL-62 | chr3-59990575-DEL-78 |
| chr3-60029721-DEL-81 | chr3-60029709-DEL-82 |
| chr3-60060479-DEL-170 | chr3-60060480-DEL-170 |
| chr3-60068583-DEL-89 | chr3-60068598-DEL-54 |
| chr3-60103999-DEL-69 | chr3-60103996-DEL-57 |
| chr3-60138958-DEL-6358 | chr3-60138959-DEL-6358 |
| chr3-60162253-DEL-208 | chr3-60162254-DEL-208 |
| chr3-60202967-DEL-69 | chr3-60202968-DEL-69 |
| chr3-60240201-DEL-377 | chr3-60240202-DEL-377 |

| | |
|---|---|
| chr3-60281847-DEL-2320 | chr3-60281848-DEL-2320 |
| chr3-60347282-DEL-193 | chr3-60347283-DEL-193 |
| chr3-60365799-DEL-614 | chr3-60365800-DEL-614 |
| chr3-60397286-DEL-5218 | chr3-60397287-DEL-5218 |
| chr3-60519106-DEL-185 | chr3-60519107-DEL-185 |
| chr3-60585287-DEL-56 | chr3-60585287-DEL-60 |
| chr3-60589085-DEL-2643 | chr3-60589086-DEL-2643 |
| chr3-60604052-DEL-1288 | chr3-60604053-DEL-1288 |
| chr3-60661343-DEL-53 | chr3-60661344-DEL-53 |
| chr3-60700044-DEL-128 | chr3-60700045-DEL-134 |
| chr3-60712844-DEL-142 | chr3-60712845-DEL-142 |
| chr3-60736948-DEL-132 | chr3-60736949-DEL-132 |
| chr3-60757881-DEL-847 | chr3-60757873-DEL-850 |
| chr3-60779140-DEL-62 | chr3-60779145-DEL-62 |
| chr3-60784406-DEL-1866 | chr3-60784407-DEL-1866 |
| chr3-60807554-DEL-454 | chr3-60807555-DEL-454 |
| chr3-60870189-DEL-325 | chr3-60870190-DEL-325 |
| chr3-60875767-DEL-6446 | chr3-60875768-DEL-6446 |
| chr3-60883507-DEL-54 | chr3-60883514-DEL-64 |
| chr3-60952281-DEL-260 | chr3-60952282-DEL-260 |
| chr3-60966194-DEL-1762 | chr3-60966195-DEL-1762 |
| chr3-60985460-DEL-6534 | chr3-60985461-DEL-6534 |
| chr3-60991401-DEL-1623 | chr3-60991402-DEL-1623 |
| chr3-60993219-DEL-62 | chr3-60993219-DEL-62 |
| chr3-61007591-DEL-1604 | chr3-61007591-DEL-1604 |
| chr3-61039957-DEL-52 | chr3-61039958-DEL-52 |
| chr3-61079127-DEL-6249 | chr3-61079128-DEL-6249 |
| chr3-61102502-DEL-193 | chr3-61102503-DEL-193 |
| chr3-61114366-DEL-196 | chr3-61114367-DEL-196 |
| chr3-61135380-DEL-2377 | chr3-61135381-DEL-2377 |
| chr3-61157150-DEL-87 | chr3-61157150-DEL-87 |
| chr3-61159356-DEL-525 | chr3-61159357-DEL-525 |
| chr3-61163877-DEL-57 | chr3-61163887-DEL-65 |
| chr3-61176808-DEL-5462 | chr3-61176809-DEL-5462 |
| chr3-61189210-DEL-69 | chr3-61189186-DEL-63 |
| chr3-61195142-DEL-110 | chr3-61195143-DEL-105 |
| chr3-61239694-DEL-269 | chr3-61239695-DEL-269 |
| chr3-61308881-DEL-119 | chr3-61308927-DEL-76 |
| chr3-61319371-DEL-53 | chr3-61319409-DEL-52 |
| chr3-61329297-DEL-218 | chr3-61329298-DEL-218 |
| chr3-61351997-DEL-6974 | chr3-61351981-DEL-6990 |
| chr3-61369073-DEL-1268 | chr3-61369075-DEL-1268 |
| chr3-61396477-DEL-199 | chr3-61396478-DEL-199 |
| chr3-61402758-DEL-6191 | chr3-61402759-DEL-6191 |
| chr3-61411380-DEL-65 | chr3-61411380-DEL-65 |
| chr3-61494731-DEL-507 | chr3-61494732-DEL-507 |
| chr3-61543961-DEL-6481 | chr3-61543962-DEL-6481 |
| chr3-61585801-DEL-167 | chr3-61585802-DEL-167 |
| chr3-61591743-DEL-1041 | chr3-61591744-DEL-1039 |
| chr3-61667256-DEL-1202 | chr3-61667257-DEL-1202 |
| chr3-61704424-DEL-61 | chr3-61704425-DEL-61 |
| chr3-61732874-DEL-4740 | chr3-61732875-DEL-4740 |
| chr3-61760360-DEL-6179 | chr3-61760361-DEL-6179 |
| chr3-61838212-DEL-87 | chr3-61838227-DEL-79 |
| chr3-61865894-DEL-319 | chr3-61865895-DEL-319 |
| chr3-61877714-DEL-67 | chr3-61877715-DEL-67 |
| chr3-61878872-DEL-54 | chr3-61878873-DEL-54 |
| chr3-61906303-DEL-104 | chr3-61906282-DEL-120 |
| chr3-61931885-DEL-126 | chr3-61931886-DEL-126 |
| chr3-61933365-DEL-54 | chr3-61933347-DEL-70 |
| chr3-61936972-DEL-114 | chr3-61936973-DEL-114 |
| chr3-62007354-DEL-346 | chr3-62007355-DEL-346 |
| chr3-62023867-DEL-24689 | chr3-62023866-DEL-24689 |
| chr3-62095821-DEL-3065 | chr3-62095822-DEL-3065 |
| chr3-62110603-DEL-411 | chr3-62110604-DEL-425 |
| chr3-62113318-DEL-6926 | chr3-62113319-DEL-6926 |
| chr3-62196244-DEL-368 | chr3-62196245-DEL-368 |
| chr3-62224724-DEL-331 | chr3-62224725-DEL-331 |
| chr3-62264317-DEL-6474 | chr3-62264315-DEL-6475 |
| chr3-62271295-DEL-3811 | chr3-62271296-DEL-3811 |
| chr3-62279530-DEL-172 | chr3-62279531-DEL-172 |
| chr3-62324102-DEL-57 | chr3-62324081-DEL-54 |
| chr3-62329925-DEL-175 | chr3-62329946-DEL-170 |
| chr3-62387051-DEL-146 | chr3-62387074-DEL-124 |
| chr3-62390682-DEL-97 | chr3-62390683-DEL-97 |
| chr3-62410358-DEL-813 | chr3-62410359-DEL-813 |
| chr3-62429216-DEL-378 | chr3-62429217-DEL-378 |
| chr3-62446674-DEL-57 | chr3-62446674-DEL-57 |
| chr3-62450376-DEL-6238 | chr3-62450377-DEL-6238 |
| chr3-62460886-DEL-102 | chr3-62460887-DEL-102 |

| | |
|---|---|
| chr3-62474299-DEL-76 | chr3-62474316-DEL-60 |
| chr3-62477189-DEL-175 | chr3-62477188-DEL-175 |
| chr3-62480430-DEL-62 | chr3-62480430-DEL-62 |
| chr3-62480812-DEL-6434 | chr3-62480813-DEL-6434 |
| chr3-62558654-DEL-552 | chr3-62558666-DEL-545 |
| chr3-62631894-DEL-390 | chr3-62631895-DEL-390 |
| chr3-62643671-DEL-6656 | chr3-62643672-DEL-6656 |
| chr3-62671810-DEL-5983 | chr3-62671811-DEL-5983 |
| chr3-62780416-DEL-6681 | chr3-62780401-DEL-6696 |
| chr3-62796332-DEL-373 | chr3-62796333-DEL-373 |
| chr3-62855202-DEL-195 | chr3-62855189-DEL-205 |
| chr3-62862169-DEL-6154 | chr3-62862169-DEL-6154 |
| chr3-62879420-DEL-85 | chr3-62879421-DEL-85 |
| chr3-62890397-DEL-74 | chr3-62890398-DEL-74 |
| chr3-62892931-DEL-6424 | chr3-62892932-DEL-6424 |
| chr3-62915678-DEL-1374 | chr3-62915678-DEL-1374 |
| chr3-62915810-DEL-203 | chr3-62915811-DEL-203 |
| chr3-62918697-DEL-331 | chr3-62918697-DEL-331 |
| chr3-62937388-DEL-1439 | chr3-62937387-DEL-1438 |
| chr3-62941673-DEL-68 | chr3-62941678-DEL-69 |
| chr3-62949360-DEL-50 | chr3-62949321-DEL-69 |
| chr3-62950796-DEL-247 | chr3-62950797-DEL-247 |
| chr3-63011344-DEL-4049 | chr3-63011345-DEL-4049 |
| chr3-63021095-DEL-227 | chr3-63021096-DEL-227 |
| chr3-63040697-DEL-195 | chr3-63040710-DEL-186 |
| chr3-63125878-DEL-66 | chr3-63125879-DEL-63 |
| chr3-63138280-DEL-219 | chr3-63138266-DEL-221 |
| chr3-63142089-DEL-109 | chr3-63142088-DEL-109 |
| chr3-63336977-DEL-773 | chr3-63336978-DEL-773 |
| chr3-63377731-DEL-871 | chr3-63377732-DEL-871 |
| chr3-63399377-DEL-440 | chr3-63399378-DEL-440 |
| chr3-63406228-DEL-893 | chr3-63406229-DEL-893 |
| chr3-63417466-DEL-1005 | chr3-63417426-DEL-1035 |
| chr3-63427801-DEL-1778 | chr3-63427802-DEL-1778 |
| chr3-63431171-DEL-6414 | chr3-63431172-DEL-6414 |
| chr3-63443548-DEL-84 | chr3-63443569-DEL-64 |
| chr3-63473834-DEL-95 | chr3-63473835-DEL-95 |
| chr3-63476787-DEL-66 | chr3-63476768-DEL-62 |
| chr3-63497442-DEL-1476 | chr3-63497443-DEL-1476 |
| chr3-63505459-DEL-56 | chr3-63505460-DEL-56 |
| chr3-63526464-DEL-77 | chr3-63526448-DEL-78 |
| chr3-63558741-DEL-54 | chr3-63558709-DEL-64 |
| chr3-63563076-DEL-58 | chr3-63563080-DEL-54 |
| chr3-63577124-DEL-7290 | chr3-63577125-DEL-7290 |
| chr3-63583921-DEL-211 | chr3-63583908-DEL-212 |
| chr3-63641994-DEL-451 | chr3-63641995-DEL-451 |
| chr3-63663632-DEL-4186 | chr3-63663633-DEL-4186 |
| chr3-63684002-DEL-186 | chr3-63683999-DEL-182 |
| chr3-63820622-DEL-184 | chr3-63820623-DEL-184 |
| chr3-63823366-DEL-79 | chr3-63823367-DEL-79 |
| chr3-63839663-DEL-313 | chr3-63839664-DEL-313 |
| chr3-63948108-DEL-58 | chr3-63948109-DEL-58 |
| chr3-64078668-DEL-935 | chr3-64078669-DEL-935 |
| chr3-64138308-DEL-211 | chr3-64138309-DEL-211 |
| chr3-64140866-DEL-328 | chr3-64140871-DEL-329 |
| chr3-64144886-DEL-8806 | chr3-64144887-DEL-8806 |
| chr3-64153265-DEL-208 | chr3-64153266-DEL-208 |
| chr3-64157217-DEL-203 | chr3-64157218-DEL-203 |
| chr3-64179634-DEL-92 | chr3-64179635-DEL-91 |
| chr3-64480345-DEL-6790 | chr3-64480346-DEL-6790 |
| chr3-64577373-DEL-382 | chr3-64577374-DEL-382 |
| chr3-64631570-DEL-60 | chr3-64631561-DEL-60 |
| chr3-64721197-DEL-7210 | chr3-64721198-DEL-7210 |
| chr3-64781082-DEL-76 | chr3-64781083-DEL-76 |
| chr3-64793911-DEL-51 | chr3-64793912-DEL-51 |
| chr3-64830288-DEL-6166 | chr3-64830289-DEL-6166 |
| chr3-64862277-DEL-6189 | chr3-64862278-DEL-6189 |
| chr3-64876813-DEL-2751 | chr3-64876814-DEL-2751 |
| chr3-64887764-DEL-339 | chr3-64887766-DEL-340 |
| chr3-64888482-DEL-799 | chr3-64888484-DEL-797 |
| chr3-64893665-DEL-738 | chr3-64893666-DEL-738 |
| chr3-64917182-DEL-6113 | chr3-64917183-DEL-6113 |
| chr3-64926904-DEL-142 | chr3-64926905-DEL-142 |
| chr3-64933660-DEL-83 | chr3-64933656-DEL-84 |
| chr3-64967770-DEL-185 | chr3-64967777-DEL-179 |
| chr3-64970372-DEL-180 | chr3-64970373-DEL-180 |
| chr3-64971223-DEL-80 | chr3-64971225-DEL-51 |
| chr3-64980595-DEL-341 | chr3-64980596-DEL-341 |
| chr3-65043111-DEL-391 | chr3-65043160-DEL-366 |
| chr3-65044741-DEL-946 | chr3-65044742-DEL-947 |

| | |
|---|---|
| chr3-65055085-DEL-696 | chr3-65055071-DEL-697 |
| chr3-65062849-DEL-274 | chr3-65062850-DEL-274 |
| chr3-65066202-DEL-164 | chr3-65066203-DEL-164 |
| chr3-65069059-DEL-109 | chr3-65069060-DEL-109 |
| chr3-65085355-DEL-829 | chr3-65085356-DEL-829 |
| chr3-65096783-DEL-8157 | chr3-65096784-DEL-8157 |
| chr3-65106673-DEL-62 | chr3-65106674-DEL-60 |
| chr3-65107568-DEL-436 | chr3-65107563-DEL-432 |
| chr3-65111681-DEL-52 | chr3-65111682-DEL-52 |
| chr3-65123705-DEL-51 | chr3-65123706-DEL-51 |
| chr3-65127338-DEL-103 | chr3-65127339-DEL-103 |
| chr3-65147545-DEL-51 | chr3-65147546-DEL-51 |
| chr3-65168305-DEL-54 | chr3-65168306-DEL-60 |
| chr3-65171657-DEL-99 | chr3-65171657-DEL-100 |
| chr3-65187789-DEL-940 | chr3-65187790-DEL-940 |
| chr3-65214353-DEL-152 | chr3-65214362-DEL-149 |
| chr3-65230727-DEL-493 | chr3-65230728-DEL-493 |
| chr3-65257877-DEL-86 | chr3-65257878-DEL-84 |
| chr3-65258770-DEL-55 | chr3-65258781-DEL-64 |
| chr3-65300702-DEL-3014 | chr3-65300703-DEL-3014 |
| chr3-65330549-DEL-6453 | chr3-65330549-DEL-6453 |
| chr3-65359508-DEL-129 | chr3-65359509-DEL-129 |
| chr3-65369687-DEL-948 | chr3-65369688-DEL-948 |
| chr3-65370745-DEL-199 | chr3-65370746-DEL-199 |
| chr3-65379543-DEL-187 | chr3-65379539-DEL-188 |
| chr3-65390155-DEL-1992 | chr3-65390156-DEL-1992 |
| chr3-65423095-DEL-54 | chr3-65423096-DEL-78 |
| chr3-65432703-DEL-66 | chr3-65432704-DEL-66 |
| chr3-65446346-DEL-53 | chr3-65446347-DEL-53 |
| chr3-65484440-DEL-92 | chr3-65484441-DEL-92 |
| chr3-65498980-DEL-70 | chr3-65498980-DEL-70 |
| chr3-65642312-DEL-724 | chr3-65642313-DEL-724 |
| chr3-65663225-DEL-6389 | chr3-65663226-DEL-6389 |
| chr3-65663369-DEL-201 | chr3-65663370-DEL-201 |
| chr3-65675978-DEL-68 | chr3-65675979-DEL-68 |
| chr3-65677666-DEL-264 | chr3-65677667-DEL-264 |
| chr3-65704168-DEL-55 | chr3-65704169-DEL-55 |
| chr3-65749725-DEL-104 | chr3-65749726-DEL-88 |
| chr3-65763245-DEL-140 | chr3-65763288-DEL-94 |
| chr3-65794152-DEL-176 | chr3-65794155-DEL-174 |
| chr3-65808045-DEL-101 | chr3-65808046-DEL-101 |
| chr3-65814429-DEL-132 | chr3-65814444-DEL-117 |
| chr3-65815524-DEL-97 | chr3-65815525-DEL-97 |
| chr3-65831087-DEL-63 | chr3-65831088-DEL-63 |
| chr3-65842984-DEL-56 | chr3-65842985-DEL-56 |
| chr3-65871941-DEL-75 | chr3-65871940-DEL-77 |
| chr3-65908989-DEL-1890 | chr3-65908990-DEL-1890 |
| chr3-65909008-DEL-703 | chr3-65909009-DEL-703 |
| chr3-65930573-DEL-116 | chr3-65930574-DEL-116 |
| chr3-65937379-DEL-1211 | chr3-65937380-DEL-1211 |
| chr3-65969127-DEL-80 | chr3-65969128-DEL-81 |
| chr3-65994257-DEL-55 | chr3-65994258-DEL-55 |
| chr3-66008329-DEL-119 | chr3-66008330-DEL-119 |
| chr3-66019816-DEL-255 | chr3-66019817-DEL-255 |
| chr3-66022022-DEL-193 | chr3-66022023-DEL-193 |
| chr3-66027105-DEL-608 | chr3-66027107-DEL-607 |
| chr3-66046049-DEL-532 | chr3-66046050-DEL-532 |
| chr3-66053389-DEL-197 | chr3-66053390-DEL-197 |
| chr3-66056273-DEL-71 | chr3-66056273-DEL-71 |
| chr3-66061626-DEL-277 | chr3-66061626-DEL-277 |
| chr3-66071597-DEL-120 | chr3-66071598-DEL-120 |
| chr3-66072993-DEL-139 | chr3-66072994-DEL-139 |
| chr3-66074179-DEL-1349 | chr3-66074179-DEL-1349 |
| chr3-66081933-DEL-464 | chr3-66081932-DEL-461 |
| chr3-66089436-DEL-71 | chr3-66089449-DEL-71 |
| chr3-66120190-DEL-515 | chr3-66120191-DEL-515 |
| chr3-66133692-DEL-7073 | chr3-66133693-DEL-7073 |
| chr3-66139889-DEL-212 | chr3-66139888-DEL-212 |
| chr3-66153857-DEL-64 | chr3-66153858-DEL-52 |
| chr3-66165861-DEL-54 | chr3-66165863-DEL-51 |
| chr3-66191512-DEL-1369 | chr3-66191513-DEL-1369 |
| chr3-66219005-DEL-2644 | chr3-66219006-DEL-2644 |
| chr3-66288010-DEL-55 | chr3-66288011-DEL-55 |
| chr3-66328704-DEL-52 | chr3-66328705-DEL-52 |
| chr3-66341139-DEL-2065 | chr3-66341140-DEL-2065 |
| chr3-66356368-DEL-64 | chr3-66356369-DEL-64 |
| chr3-66357902-DEL-6549 | chr3-66357903-DEL-6549 |
| chr3-66381222-DEL-153 | chr3-66381223-DEL-153 |
| chr3-66381990-DEL-89 | chr3-66381991-DEL-89 |
| chr3-66392817-DEL-53 | chr3-66392814-DEL-51 |

| | |
|---|---|
| chr3-66429908-DEL-82 | chr3-66429907-DEL-72 |
| chr3-66440831-DEL-987 | chr3-66440832-DEL-987 |
| chr3-66461958-DEL-60 | chr3-66461961-DEL-58 |
| chr3-66462873-DEL-95 | chr3-66462889-DEL-108 |
| chr3-66504923-DEL-64 | chr3-66504924-DEL-90 |
| chr3-66535421-DEL-8199 | chr3-66535422-DEL-8199 |
| chr3-66577355-DEL-7415 | chr3-66577356-DEL-7415 |
| chr3-66615499-DEL-3752 | chr3-66615500-DEL-3752 |
| chr3-66635927-DEL-209 | chr3-66635884-DEL-237 |
| chr3-66644846-DEL-52 | chr3-66644810-DEL-68 |
| chr3-66661379-DEL-94 | chr3-66661382-DEL-92 |
| chr3-66681396-DEL-205 | chr3-66681397-DEL-205 |
| chr3-66687837-DEL-71 | chr3-66687838-DEL-71 |
| chr3-66725814-DEL-676 | chr3-66725815-DEL-676 |
| chr3-66729453-DEL-743 | chr3-66729454-DEL-743 |
| chr3-66742376-DEL-331 | chr3-66742377-DEL-331 |
| chr3-66743068-DEL-6746 | chr3-66743052-DEL-6748 |
| chr3-66762652-DEL-6413 | chr3-66762653-DEL-6413 |
| chr3-66774348-DEL-87 | chr3-66774349-DEL-87 |
| chr3-66798325-DEL-220 | chr3-66798326-DEL-174 |
| chr3-66935268-DEL-2245 | chr3-66935269-DEL-2245 |
| chr3-66956853-DEL-63 | chr3-66956854-DEL-63 |
| chr3-66958496-DEL-3064 | chr3-66958497-DEL-3064 |
| chr3-66963813-DEL-5863 | chr3-66963814-DEL-5863 |
| chr3-66980981-DEL-68 | chr3-66980982-DEL-112 |
| chr3-67002500-DEL-439 | chr3-67002501-DEL-439 |
| chr3-67003406-DEL-7721 | chr3-67003407-DEL-7721 |
| chr3-67061269-DEL-8510 | chr3-67061270-DEL-8510 |
| chr3-67086622-DEL-780 | chr3-67086623-DEL-780 |
| chr3-67118002-DEL-204 | chr3-67118003-DEL-204 |
| chr3-67135230-DEL-150 | chr3-67135236-DEL-150 |
| chr3-67169915-DEL-351 | chr3-67169931-DEL-351 |
| chr3-67170416-DEL-81 | chr3-67170417-DEL-81 |
| chr3-67200641-DEL-4215 | chr3-67200642-DEL-4215 |
| chr3-67315197-DEL-232 | chr3-67315198-DEL-232 |
| chr3-67323938-DEL-8696 | chr3-67323939-DEL-8696 |
| chr3-67338574-DEL-121 | chr3-67338575-DEL-121 |
| chr3-67388275-DEL-124 | chr3-67388310-DEL-120 |
| chr3-67394677-DEL-64 | chr3-67394678-DEL-63 |
| chr3-67397363-DEL-80 | chr3-67397364-DEL-80 |
| chr3-67401974-DEL-80 | chr3-67401994-DEL-64 |
| chr3-67412097-DEL-152 | chr3-67412070-DEL-175 |
| chr3-67427283-DEL-232 | chr3-67427284-DEL-231 |
| chr3-67438753-DEL-192 | chr3-67438754-DEL-192 |
| chr3-67442611-DEL-6457 | chr3-67442612-DEL-6457 |
| chr3-67472575-DEL-68 | chr3-67472576-DEL-56 |
| chr3-67481449-DEL-61 | chr3-67481415-DEL-85 |
| chr3-67502769-DEL-458 | chr3-67502769-DEL-458 |
| chr3-67512889-DEL-195 | chr3-67512889-DEL-196 |
| chr3-67525468-DEL-70 | chr3-67525463-DEL-68 |
| chr3-67529407-DEL-1643 | chr3-67529408-DEL-1643 |
| chr3-67541336-DEL-1112 | chr3-67541337-DEL-1112 |
| chr3-67541416-DEL-172 | chr3-67541417-DEL-172 |
| chr3-67548473-DEL-256 | chr3-67548477-DEL-257 |
| chr3-67554978-DEL-186 | chr3-67554979-DEL-186 |
| chr3-67571257-DEL-72 | chr3-67571258-DEL-72 |
| chr3-67592625-DEL-2818 | chr3-67592625-DEL-2818 |
| chr3-67596771-DEL-54 | chr3-67596753-DEL-70 |
| chr3-67601664-DEL-62 | chr3-67601669-DEL-58 |
| chr3-67626875-DEL-768 | chr3-67626876-DEL-768 |
| chr3-67633918-DEL-838 | chr3-67633918-DEL-839 |
| chr3-67654567-DEL-6390 | chr3-67654568-DEL-6390 |
| chr3-67694033-DEL-6882 | chr3-67694034-DEL-6882 |
| chr3-67702662-DEL-859 | chr3-67702663-DEL-859 |
| chr3-67728325-DEL-2613 | chr3-67728326-DEL-2613 |
| chr3-67732595-DEL-859 | chr3-67732596-DEL-859 |
| chr3-67737049-DEL-68 | chr3-67737068-DEL-67 |
| chr3-67758023-DEL-1300 | chr3-67758024-DEL-1300 |
| chr3-67759060-DEL-60 | chr3-67759051-DEL-56 |
| chr3-67809705-DEL-210 | chr3-67809706-DEL-210 |
| chr3-67822344-DEL-82 | chr3-67822302-DEL-118 |
| chr3-67833371-DEL-1039 | chr3-67833372-DEL-1039 |
| chr3-67848753-DEL-310 | chr3-67848754-DEL-310 |
| chr3-67861284-DEL-121 | chr3-67861285-DEL-121 |
| chr3-67885523-DEL-60 | chr3-67885541-DEL-54 |
| chr3-67897719-DEL-224 | chr3-67897723-DEL-221 |
| chr3-67906020-DEL-190 | chr3-67906021-DEL-190 |
| chr3-67932522-DEL-670 | chr3-67932519-DEL-674 |
| chr3-67949205-DEL-264 | chr3-67949206-DEL-264 |
| chr3-67949753-DEL-183 | chr3-67949754-DEL-183 |

| | |
|---|---|
| chr3-67983553-DEL-185 | chr3-67983554-DEL-185 |
| chr3-67984600-DEL-57 | chr3-67984600-DEL-58 |
| chr3-67985594-DEL-3688 | chr3-67985595-DEL-3688 |
| chr3-67994870-DEL-152 | chr3-67994914-DEL-128 |
| chr3-68003992-DEL-76 | chr3-68003993-DEL-76 |
| chr3-68014532-DEL-445 | chr3-68014533-DEL-445 |
| chr3-68021783-DEL-5146 | chr3-68021784-DEL-5146 |
| chr3-68088137-DEL-54 | chr3-68088138-DEL-54 |
| chr3-68105388-DEL-54 | chr3-68105389-DEL-54 |
| chr3-68106148-DEL-101 | chr3-68106149-DEL-101 |
| chr3-68131944-DEL-502 | chr3-68131952-DEL-496 |
| chr3-68133348-DEL-431 | chr3-68133349-DEL-431 |
| chr3-68156047-DEL-64 | chr3-68156054-DEL-58 |
| chr3-68164797-DEL-58 | chr3-68164798-DEL-58 |
| chr3-68206931-DEL-108 | chr3-68206932-DEL-108 |
| chr3-68209730-DEL-1372 | chr3-68209731-DEL-1372 |
| chr3-68243908-DEL-328 | chr3-68243909-DEL-328 |
| chr3-68261191-DEL-1649 | chr3-68261192-DEL-1649 |
| chr3-68268571-DEL-64 | chr3-68268571-DEL-64 |
| chr3-68280144-DEL-203 | chr3-68280145-DEL-203 |
| chr3-68327308-DEL-90 | chr3-68327309-DEL-90 |
| chr3-68347183-DEL-72 | chr3-68347183-DEL-72 |
| chr3-68358485-DEL-209 | chr3-68358454-DEL-221 |
| chr3-68394222-DEL-205 | chr3-68394223-DEL-205 |
| chr3-68407460-DEL-100 | chr3-68407477-DEL-86 |
| chr3-68426283-DEL-6434 | chr3-68426284-DEL-6434 |
| chr3-68461214-DEL-78 | chr3-68461186-DEL-56 |
| chr3-68495440-DEL-60 | chr3-68495441-DEL-60 |
| chr3-68517816-DEL-189 | chr3-68517818-DEL-188 |
| chr3-68576259-DEL-54 | chr3-68576259-DEL-54 |
| chr3-68581569-DEL-607 | chr3-68581567-DEL-608 |
| chr3-68611419-DEL-229 | chr3-68611444-DEL-185 |
| chr3-68629596-DEL-295 | chr3-68629597-DEL-295 |
| chr3-68632139-DEL-7126 | chr3-68632140-DEL-7126 |
| chr3-68706567-DEL-213 | chr3-68706568-DEL-213 |
| chr3-68754138-DEL-235 | chr3-68754128-DEL-211 |
| chr3-68807449-DEL-4755 | chr3-68807451-DEL-4754 |
| chr3-68822557-DEL-77 | chr3-68822558-DEL-77 |
| chr3-68857113-DEL-181 | chr3-68857114-DEL-181 |
| chr3-68859338-DEL-553 | chr3-68859339-DEL-553 |
| chr3-68862198-DEL-56 | chr3-68862199-DEL-56 |
| chr3-68894066-DEL-1114 | chr3-68894067-DEL-1114 |
| chr3-68984717-DEL-218 | chr3-68984718-DEL-218 |
| chr3-69002321-DEL-137 | chr3-69002322-DEL-137 |
| chr3-69036471-DEL-61 | chr3-69036472-DEL-61 |
| chr3-69051910-DEL-2165 | chr3-69051911-DEL-2165 |
| chr3-69124137-DEL-1135 | chr3-69124164-DEL-1109 |
| chr3-69135965-DEL-1864 | chr3-69135966-DEL-1864 |
| chr3-69139429-DEL-564 | chr3-69139430-DEL-564 |
| chr3-69145501-DEL-68 | chr3-69145502-DEL-68 |
| chr3-69155862-DEL-483 | chr3-69155863-DEL-483 |
| chr3-69163140-DEL-538 | chr3-69163141-DEL-539 |
| chr3-69165433-DEL-143 | chr3-69165450-DEL-110 |
| chr3-69180102-DEL-831 | chr3-69180103-DEL-831 |
| chr3-69217044-DEL-486 | chr3-69217045-DEL-486 |
| chr3-69235794-DEL-206 | chr3-69235786-DEL-207 |
| chr3-69262348-DEL-198 | chr3-69262349-DEL-196 |
| chr3-69299369-DEL-54 | chr3-69299370-DEL-54 |
| chr3-69301135-DEL-189 | chr3-69301151-DEL-174 |
| chr3-69316614-DEL-321 | chr3-69316615-DEL-321 |
| chr3-69349884-DEL-210 | chr3-69349885-DEL-210 |
| chr3-69360956-DEL-1339 | chr3-69360957-DEL-1339 |
| chr3-69371032-DEL-88 | chr3-69371033-DEL-88 |
| chr3-69386159-DEL-183 | chr3-69386147-DEL-186 |
| chr3-69420402-DEL-186 | chr3-69420403-DEL-186 |
| chr3-69436517-DEL-54 | chr3-69436518-DEL-54 |
| chr3-69437893-DEL-215 | chr3-69437894-DEL-215 |
| chr3-69470939-DEL-6568 | chr3-69470940-DEL-6568 |
| chr3-69495590-DEL-55 | chr3-69495591-DEL-55 |
| chr3-69496385-DEL-74 | chr3-69496384-DEL-70 |
| chr3-69496573-DEL-2388 | chr3-69496574-DEL-2388 |
| chr3-69498862-DEL-70 | chr3-69498863-DEL-70 |
| chr3-69525573-DEL-4567 | chr3-69525574-DEL-4567 |
| chr3-69536285-DEL-227 | chr3-69536270-DEL-242 |
| chr3-69545266-DEL-7142 | chr3-69545267-DEL-7142 |
| chr3-69590307-DEL-211 | chr3-69590325-DEL-216 |
| chr3-69605472-DEL-60 | chr3-69605471-DEL-60 |
| chr3-69608151-DEL-84 | chr3-69608152-DEL-84 |
| chr3-69619668-DEL-181 | chr3-69619669-DEL-181 |
| chr3-69623511-DEL-336 | chr3-69623512-DEL-336 |

| | |
|---|---|
| chr3-69624026-DEL-60 | chr3-69624027-DEL-60 |
| chr3-69631783-DEL-183 | chr3-69631788-DEL-182 |
| chr3-69640615-DEL-76 | chr3-69640591-DEL-53 |
| chr3-69646512-DEL-212 | chr3-69646513-DEL-212 |
| chr3-69675539-DEL-128 | chr3-69675540-DEL-84 |
| chr3-69702066-DEL-56 | chr3-69702035-DEL-68 |
| chr3-69735260-DEL-90 | chr3-69735267-DEL-86 |
| chr3-69742971-DEL-52 | chr3-69742972-DEL-52 |
| chr3-69745520-DEL-1489 | chr3-69745521-DEL-1489 |
| chr3-69754610-DEL-193 | chr3-69754611-DEL-193 |
| chr3-69755711-DEL-6413 | chr3-69755712-DEL-6413 |
| chr3-69771555-DEL-134 | chr3-69771556-DEL-134 |
| chr3-69803171-DEL-2212 | chr3-69803172-DEL-2212 |
| chr3-69813319-DEL-72 | chr3-69813320-DEL-72 |
| chr3-69820869-DEL-990 | chr3-69820870-DEL-990 |
| chr3-69823411-DEL-2865 | chr3-69823412-DEL-2866 |
| chr3-69834757-DEL-3865 | chr3-69834758-DEL-3865 |
| chr3-69837986-DEL-85 | chr3-69837968-DEL-85 |
| chr3-69840429-DEL-61 | chr3-69840430-DEL-61 |
| chr3-69848902-DEL-504 | chr3-69848903-DEL-504 |
| chr3-69865870-DEL-58 | chr3-69865871-DEL-58 |
| chr3-69870503-DEL-190 | chr3-69870504-DEL-190 |
| chr3-69879357-DEL-711 | chr3-69879358-DEL-711 |
| chr3-69885649-DEL-115 | chr3-69885650-DEL-115 |
| chr3-69903730-DEL-96 | chr3-69903727-DEL-100 |
| chr3-69907089-DEL-1620 | chr3-69907090-DEL-1620 |
| chr3-69947933-DEL-53 | chr3-69947934-DEL-53 |
| chr3-69988820-DEL-941 | chr3-69988821-DEL-941 |
| chr3-70039189-DEL-90 | chr3-70039190-DEL-90 |
| chr3-70072426-DEL-6120 | chr3-70072427-DEL-6120 |
| chr3-70081217-DEL-159 | chr3-70081218-DEL-159 |
| chr3-70096246-DEL-359 | chr3-70096247-DEL-359 |
| chr3-70115099-DEL-54 | chr3-70115100-DEL-74 |
| chr3-70135136-DEL-118 | chr3-70135137-DEL-118 |
| chr3-70197466-DEL-58 | chr3-70197467-DEL-56 |
| chr3-70201405-DEL-90 | chr3-70201406-DEL-99 |
| chr3-70207572-DEL-2819 | chr3-70207574-DEL-2818 |
| chr3-70208098-DEL-349 | chr3-70208099-DEL-349 |
| chr3-70253840-DEL-3486 | chr3-70253841-DEL-3486 |
| chr3-70268053-DEL-614 | chr3-70268054-DEL-614 |
| chr3-70286118-DEL-546 | chr3-70286119-DEL-545 |
| chr3-70294120-DEL-71 | chr3-70294127-DEL-63 |
| chr3-70299423-DEL-91 | chr3-70299424-DEL-91 |
| chr3-70322233-DEL-886 | chr3-70322234-DEL-886 |
| chr3-70332446-DEL-522 | chr3-70332447-DEL-522 |
| chr3-70362685-DEL-784 | chr3-70362686-DEL-784 |
| chr3-70402900-DEL-84 | chr3-70402905-DEL-80 |
| chr3-70418388-DEL-955 | chr3-70418389-DEL-955 |
| chr3-70432524-DEL-358 | chr3-70432525-DEL-358 |
| chr3-70441961-DEL-58 | chr3-70441971-DEL-79 |
| chr3-70479038-DEL-4165 | chr3-70479039-DEL-4165 |
| chr3-70504700-DEL-52 | chr3-70504699-DEL-52 |
| chr3-70526403-DEL-96 | chr3-70526414-DEL-86 |
| chr3-70534177-DEL-84 | chr3-70534178-DEL-84 |
| chr3-70571350-DEL-1198 | chr3-70571351-DEL-1198 |
| chr3-70620844-DEL-1888 | chr3-70620837-DEL-1929 |
| chr3-70625080-DEL-64 | chr3-70625043-DEL-102 |
| chr3-70625949-DEL-6443 | chr3-70625950-DEL-6443 |
| chr3-70651600-DEL-3132 | chr3-70651601-DEL-3132 |
| chr3-70666532-DEL-204 | chr3-70666533-DEL-204 |
| chr3-70670006-DEL-75 | chr3-70670006-DEL-85 |
| chr3-70682028-DEL-1486 | chr3-70682029-DEL-1481 |
| chr3-70703000-DEL-492 | chr3-70703001-DEL-492 |
| chr3-70704557-DEL-57 | chr3-70704570-DEL-54 |
| chr3-70710645-DEL-78 | chr3-70710646-DEL-78 |
| chr3-70716435-DEL-1920 | chr3-70716436-DEL-1920 |
| chr3-70762030-DEL-198 | chr3-70762015-DEL-199 |
| chr3-70814051-DEL-103 | chr3-70814062-DEL-93 |
| chr3-70835951-DEL-212 | chr3-70835949-DEL-213 |
| chr3-70847464-DEL-246 | chr3-70847465-DEL-246 |
| chr3-70848682-DEL-64 | chr3-70848683-DEL-64 |
| chr3-70849289-DEL-374 | chr3-70849288-DEL-375 |
| chr3-70872855-DEL-154 | chr3-70872855-DEL-153 |
| chr3-70894465-DEL-78 | chr3-70894466-DEL-78 |
| chr3-70947472-DEL-394 | chr3-70947473-DEL-394 |
| chr3-70965799-DEL-6839 | chr3-70965781-DEL-6857 |
| chr3-70972142-DEL-212 | chr3-70972143-DEL-212 |
| chr3-70984083-DEL-582 | chr3-70984084-DEL-582 |
| chr3-70995341-DEL-1948 | chr3-70995342-DEL-1948 |
| chr3-71056573-DEL-322 | chr3-71056574-DEL-322 |

| | |
|---|---|
| chr3-71086428-DEL-1855 | chr3-71086429-DEL-1856 |
| chr3-71088351-DEL-75 | chr3-71088352-DEL-90 |
| chr3-71096195-DEL-57 | chr3-71096196-DEL-57 |
| chr3-71097141-DEL-426 | chr3-71097145-DEL-408 |
| chr3-71192599-DEL-62 | chr3-71192600-DEL-62 |
| chr3-71194863-DEL-79 | chr3-71194864-DEL-79 |
| chr3-71197198-DEL-228 | chr3-71197199-DEL-228 |
| chr3-71244494-DEL-80 | chr3-71244495-DEL-80 |
| chr3-71265118-DEL-102 | chr3-71265110-DEL-79 |
| chr3-71265619-DEL-95 | chr3-71265620-DEL-95 |
| chr3-71276778-DEL-5738 | chr3-71276778-DEL-5738 |
| chr3-71297122-DEL-6225 | chr3-71297105-DEL-6242 |
| chr3-71317722-DEL-1593 | chr3-71317723-DEL-1593 |
| chr3-71335453-DEL-4557 | chr3-71335454-DEL-4557 |
| chr3-71341046-DEL-1392 | chr3-71341047-DEL-1392 |
| chr3-71344735-DEL-3693 | chr3-71344736-DEL-3693 |
| chr3-71348768-DEL-6392 | chr3-71348754-DEL-6390 |
| chr3-71364915-DEL-403 | chr3-71364916-DEL-403 |
| chr3-71374488-DEL-144 | chr3-71374480-DEL-150 |
| chr3-71380417-DEL-77 | chr3-71380416-DEL-77 |
| chr3-71385682-DEL-787 | chr3-71385673-DEL-788 |
| chr3-71386298-DEL-125 | chr3-71386299-DEL-125 |
| chr3-71390779-DEL-57 | chr3-71390780-DEL-57 |
| chr3-71417551-DEL-66 | chr3-71417552-DEL-64 |
| chr3-71421553-DEL-254 | chr3-71421554-DEL-254 |
| chr3-71426184-DEL-68 | chr3-71426185-DEL-68 |
| chr3-71432354-DEL-5723 | chr3-71432355-DEL-5723 |
| chr3-71449881-DEL-82 | chr3-71449882-DEL-82 |
| chr3-71492928-DEL-1756 | chr3-71492929-DEL-1756 |
| chr3-71502960-DEL-127 | chr3-71502960-DEL-125 |
| chr3-71538854-DEL-402 | chr3-71538855-DEL-402 |
| chr3-71556172-DEL-1439 | chr3-71556173-DEL-1439 |
| chr3-71644606-DEL-1149 | chr3-71644607-DEL-1149 |
| chr3-71648658-DEL-59 | chr3-71648659-DEL-59 |
| chr3-71656463-DEL-1145 | chr3-71656464-DEL-1145 |
| chr3-71678553-DEL-2663 | chr3-71678554-DEL-2663 |
| chr3-71703938-DEL-54 | chr3-71703940-DEL-52 |
| chr3-71719180-DEL-144 | chr3-71719190-DEL-126 |
| chr3-71727286-DEL-56 | chr3-71727287-DEL-56 |
| chr3-71729429-DEL-6385 | chr3-71729425-DEL-6386 |
| chr3-71745005-DEL-85 | chr3-71745006-DEL-85 |
| chr3-71760634-DEL-222 | chr3-71760635-DEL-222 |
| chr3-71761592-DEL-3902 | chr3-71761593-DEL-3905 |
| chr3-71767973-DEL-3736 | chr3-71767974-DEL-3736 |
| chr3-71796102-DEL-59 | chr3-71796104-DEL-57 |
| chr3-71798169-DEL-628 | chr3-71798170-DEL-628 |
| chr3-71799326-DEL-786 | chr3-71799320-DEL-787 |
| chr3-71810755-DEL-62 | chr3-71810756-DEL-62 |
| chr3-71817592-DEL-51 | chr3-71817579-DEL-91 |
| chr3-71837388-DEL-751 | chr3-71837386-DEL-751 |
| chr3-71849356-DEL-821 | chr3-71849357-DEL-821 |
| chr3-71853022-DEL-2760 | chr3-71853023-DEL-2760 |
| chr3-71875964-DEL-1289 | chr3-71875965-DEL-1289 |
| chr3-71877050-DEL-83 | chr3-71877078-DEL-84 |
| chr3-71880277-DEL-2465 | chr3-71880278-DEL-2465 |
| chr3-71905289-DEL-84 | chr3-71905296-DEL-78 |
| chr3-71922462-DEL-691 | chr3-71922463-DEL-691 |
| chr3-71932489-DEL-5653 | chr3-71932490-DEL-5653 |
| chr3-71944015-DEL-76 | chr3-71944016-DEL-76 |
| chr3-72001428-DEL-102 | chr3-72001429-DEL-102 |
| chr3-72042399-DEL-1300 | chr3-72042399-DEL-1300 |
| chr3-72043576-DEL-370 | chr3-72043577-DEL-370 |
| chr3-72052740-DEL-3542 | chr3-72052741-DEL-3542 |
| chr3-72058308-DEL-6626 | chr3-72058309-DEL-6626 |
| chr3-72064380-DEL-409 | chr3-72064380-DEL-409 |
| chr3-72074049-DEL-94 | chr3-72074050-DEL-94 |
| chr3-72083176-DEL-110 | chr3-72083177-DEL-110 |
| chr3-72097739-DEL-1130 | chr3-72097740-DEL-1130 |
| chr3-72103307-DEL-158 | chr3-72103308-DEL-158 |
| chr3-72112491-DEL-821 | chr3-72112492-DEL-821 |
| chr3-72123727-DEL-68 | chr3-72123728-DEL-68 |
| chr3-72125658-DEL-86 | chr3-72125680-DEL-63 |
| chr3-72146148-DEL-58 | chr3-72146150-DEL-56 |
| chr3-72180757-DEL-54 | chr3-72180760-DEL-52 |
| chr3-72194242-DEL-6800 | chr3-72194240-DEL-6801 |
| chr3-72218067-DEL-11614 | chr3-72218068-DEL-11614 |
| chr3-72229182-DEL-401 | chr3-72229183-DEL-401 |
| chr3-72291926-DEL-1519 | chr3-72291927-DEL-1519 |
| chr3-72316137-DEL-74 | chr3-72316137-DEL-122 |
| chr3-72356339-DEL-324 | chr3-72356340-DEL-324 |

| | |
|---|---|
| chr3-72400079-DEL-134 | chr3-72400105-DEL-92 |
| chr3-72540280-DEL-131 | chr3-72540281-DEL-131 |
| chr3-72594166-DEL-84 | chr3-72594167-DEL-78 |
| chr3-72595762-DEL-81 | chr3-72595766-DEL-83 |
| chr3-72600583-DEL-942 | chr3-72600584-DEL-942 |
| chr3-72605448-DEL-7089 | chr3-72605449-DEL-7089 |
| chr3-72722264-DEL-56 | chr3-72722265-DEL-56 |
| chr3-72725317-DEL-204 | chr3-72725318-DEL-205 |
| chr3-72783987-DEL-578 | chr3-72783988-DEL-578 |
| chr3-72785006-DEL-1079 | chr3-72785007-DEL-1079 |
| chr3-72800590-DEL-6590 | chr3-72800591-DEL-6590 |
| chr3-72815201-DEL-104 | chr3-72815198-DEL-62 |
| chr3-72822493-DEL-1212 | chr3-72822494-DEL-1212 |
| chr3-72880801-DEL-64 | chr3-72880803-DEL-63 |
| chr3-72921436-DEL-2001 | chr3-72921436-DEL-2001 |
| chr3-72925836-DEL-183 | chr3-72925837-DEL-183 |
| chr3-72927254-DEL-158 | chr3-72927251-DEL-158 |
| chr3-72987909-DEL-281 | chr3-72987910-DEL-281 |
| chr3-72989600-DEL-66 | chr3-72989601-DEL-66 |
| chr3-72995431-DEL-6520 | chr3-72995432-DEL-6520 |
| chr3-73018822-DEL-72 | chr3-73018826-DEL-70 |
| chr3-73082799-DEL-530 | chr3-73082788-DEL-531 |
| chr3-73110768-DEL-70 | chr3-73110771-DEL-66 |
| chr3-73110869-DEL-874 | chr3-73110870-DEL-874 |
| chr3-73112503-DEL-116 | chr3-73112504-DEL-116 |
| chr3-73118429-DEL-68 | chr3-73118430-DEL-68 |
| chr3-73143488-DEL-774 | chr3-73143489-DEL-774 |
| chr3-73150099-DEL-3335 | chr3-73150100-DEL-3335 |
| chr3-73159049-DEL-6401 | chr3-73159050-DEL-6401 |
| chr3-73170528-DEL-1831 | chr3-73170529-DEL-1831 |
| chr3-73175246-DEL-66 | chr3-73175248-DEL-65 |
| chr3-73206981-DEL-1080 | chr3-73206982-DEL-1080 |
| chr3-73209792-DEL-198 | chr3-73209793-DEL-198 |
| chr3-73222424-DEL-131 | chr3-73222425-DEL-131 |
| chr3-73225038-DEL-495 | chr3-73225039-DEL-495 |
| chr3-73248476-DEL-1611 | chr3-73248477-DEL-1611 |
| chr3-73272311-DEL-411 | chr3-73272312-DEL-411 |
| chr3-73272943-DEL-54 | chr3-73272947-DEL-60 |
| chr3-73286784-DEL-486 | chr3-73286785-DEL-486 |
| chr3-73305611-DEL-207 | chr3-73305612-DEL-207 |
| chr3-73321831-DEL-858 | chr3-73321832-DEL-858 |
| chr3-73325820-DEL-171 | chr3-73325821-DEL-171 |
| chr3-73330046-DEL-1928 | chr3-73330047-DEL-1928 |
| chr3-73335452-DEL-6211 | chr3-73335452-DEL-6211 |
| chr3-73354856-DEL-438 | chr3-73354857-DEL-438 |
| chr3-73357822-DEL-6373 | chr3-73357823-DEL-6373 |
| chr3-73370422-DEL-5678 | chr3-73370423-DEL-5678 |
| chr3-73387011-DEL-80 | chr3-73387012-DEL-78 |
| chr3-73395586-DEL-95 | chr3-73395587-DEL-95 |
| chr3-73418109-DEL-100 | chr3-73418110-DEL-100 |
| chr3-73431239-DEL-88 | chr3-73431239-DEL-88 |
| chr3-73433567-DEL-96 | chr3-73433562-DEL-94 |
| chr3-73436357-DEL-65 | chr3-73436358-DEL-65 |
| chr3-73458912-DEL-80 | chr3-73458906-DEL-63 |
| chr3-73462763-DEL-58 | chr3-73462726-DEL-59 |
| chr3-73469026-DEL-222 | chr3-73469027-DEL-222 |
| chr3-73470705-DEL-135 | chr3-73470706-DEL-135 |
| chr3-73495320-DEL-57 | chr3-73495321-DEL-57 |
| chr3-73511017-DEL-6070 | chr3-73511018-DEL-6070 |
| chr3-73519983-DEL-1055 | chr3-73519984-DEL-1055 |
| chr3-73553839-DEL-64 | chr3-73553820-DEL-58 |
| chr3-73583144-DEL-4315 | chr3-73583145-DEL-4315 |
| chr3-73607267-DEL-88 | chr3-73607272-DEL-84 |
| chr3-73640518-DEL-5260 | chr3-73640511-DEL-5261 |
| chr3-73665142-DEL-552 | chr3-73665138-DEL-553 |
| chr3-73671867-DEL-1062 | chr3-73671868-DEL-1062 |
| chr3-73673687-DEL-180 | chr3-73673688-DEL-180 |
| chr3-73684430-DEL-2185 | chr3-73684431-DEL-2185 |
| chr3-73715351-DEL-11002 | chr3-73715350-DEL-11003 |
| chr3-73760201-DEL-291 | chr3-73760202-DEL-291 |
| chr3-73771366-DEL-6664 | chr3-73771367-DEL-6664 |
| chr3-73771586-DEL-212 | chr3-73771587-DEL-212 |
| chr3-73781753-DEL-2321 | chr3-73781754-DEL-2321 |
| chr3-73798408-DEL-1102 | chr3-73798400-DEL-1103 |
| chr3-73823936-DEL-208 | chr3-73823936-DEL-208 |
| chr3-73834936-DEL-144 | chr3-73834937-DEL-144 |
| chr3-73857159-DEL-226 | chr3-73857160-DEL-226 |
| chr3-73866137-DEL-427 | chr3-73866138-DEL-427 |
| chr3-73897592-DEL-213 | chr3-73897593-DEL-213 |
| chr3-73933497-DEL-1050 | chr3-73933498-DEL-1050 |

| | |
|---|---|
| chr3-73934742-DEL-251 | chr3-73934743-DEL-251 |
| chr3-73938334-DEL-1337 | chr3-73938332-DEL-1336 |
| chr3-73975799-DEL-1605 | chr3-73975800-DEL-1605 |
| chr3-73988252-DEL-509 | chr3-73988253-DEL-509 |
| chr3-74044432-DEL-63 | chr3-74044433-DEL-63 |
| chr3-74050900-DEL-55 | chr3-74050901-DEL-55 |
| chr3-74053033-DEL-76 | chr3-74052999-DEL-80 |
| chr3-74055955-DEL-175 | chr3-74055951-DEL-174 |
| chr3-74068515-DEL-199 | chr3-74068514-DEL-199 |
| chr3-74082689-DEL-57 | chr3-74082690-DEL-57 |
| chr3-74093740-DEL-574 | chr3-74093739-DEL-574 |
| chr3-74095126-DEL-1915 | chr3-74095125-DEL-1915 |
| chr3-74118536-DEL-6241 | chr3-74118537-DEL-6241 |
| chr3-74128526-DEL-209 | chr3-74128527-DEL-209 |
| chr3-74137849-DEL-4511 | chr3-74137850-DEL-4511 |
| chr3-74138041-DEL-1233 | chr3-74138041-DEL-1233 |
| chr3-74140626-DEL-231 | chr3-74140611-DEL-232 |
| chr3-74143422-DEL-1008 | chr3-74143423-DEL-1008 |
| chr3-74147957-DEL-940 | chr3-74147955-DEL-930 |
| chr3-74182817-DEL-231 | chr3-74182817-DEL-231 |
| chr3-74258451-DEL-164 | chr3-74258451-DEL-164 |
| chr3-74288444-DEL-371 | chr3-74288445-DEL-371 |
| chr3-74309967-DEL-1178 | chr3-74309968-DEL-1178 |
| chr3-74324749-DEL-169 | chr3-74324750-DEL-169 |
| chr3-74343866-DEL-6099 | chr3-74343867-DEL-6099 |
| chr3-74356138-DEL-101 | chr3-74356097-DEL-103 |
| chr3-74365648-DEL-204 | chr3-74365649-DEL-204 |
| chr3-74399402-DEL-228 | chr3-74399403-DEL-228 |
| chr3-74419779-DEL-216 | chr3-74419779-DEL-217 |
| chr3-74422640-DEL-356 | chr3-74422641-DEL-356 |
| chr3-74448875-DEL-423 | chr3-74448876-DEL-423 |
| chr3-74498426-DEL-7151 | chr3-74498427-DEL-7151 |
| chr3-74515575-DEL-830 | chr3-74515576-DEL-830 |
| chr3-74521641-DEL-903 | chr3-74521650-DEL-890 |
| chr3-74526186-DEL-4592 | chr3-74526186-DEL-4592 |
| chr3-74597305-DEL-51 | chr3-74597288-DEL-69 |
| chr3-74653841-DEL-95 | chr3-74653842-DEL-95 |
| chr3-74658760-DEL-214 | chr3-74658761-DEL-214 |
| chr3-74666584-DEL-6419 | chr3-74666572-DEL-6431 |
| chr3-74707404-DEL-1000 | chr3-74707393-DEL-999 |
| chr3-74730690-DEL-56 | chr3-74730691-DEL-55 |
| chr3-74733287-DEL-52 | chr3-74733290-DEL-50 |
| chr3-74741634-DEL-6642 | chr3-74741635-DEL-6642 |
| chr3-74863479-DEL-731 | chr3-74863480-DEL-731 |
| chr3-74867016-DEL-1174 | chr3-74867017-DEL-1173 |
| chr3-74887651-DEL-94 | chr3-74887652-DEL-94 |
| chr3-74937285-DEL-161 | chr3-74937286-DEL-161 |
| chr3-75001700-DEL-857 | chr3-75001701-DEL-857 |
| chr3-75002738-DEL-1191 | chr3-75002733-DEL-1180 |
| chr3-75017154-DEL-427 | chr3-75017155-DEL-427 |
| chr3-75029236-DEL-130 | chr3-75029237-DEL-130 |
| chr3-75038168-DEL-1977 | chr3-75038169-DEL-1977 |
| chr3-75104925-DEL-54 | chr3-75104910-DEL-69 |
| chr3-75196040-DEL-489 | chr3-75196041-DEL-489 |
| chr3-75207081-DEL-92 | chr3-75207069-DEL-80 |
| chr3-75235195-DEL-4154 | chr3-75235196-DEL-4154 |
| chr3-75253577-DEL-112 | chr3-75253578-DEL-112 |
| chr3-75255758-DEL-4816 | chr3-75255771-DEL-4802 |
| chr3-75264432-DEL-758 | chr3-75264433-DEL-758 |
| chr3-75276093-DEL-406 | chr3-75276094-DEL-406 |
| chr3-75279632-DEL-6448 | chr3-75279633-DEL-6448 |
| chr3-75316141-DEL-6880 | chr3-75316142-DEL-6880 |
| chr3-75324635-DEL-6689 | chr3-75324636-DEL-6689 |
| chr3-75348687-DEL-104 | chr3-75348686-DEL-64 |
| chr3-75367844-DEL-6173 | chr3-75367845-DEL-6173 |
| chr3-75398387-DEL-219 | chr3-75398388-DEL-219 |
| chr3-75421757-DEL-627 | chr3-75421758-DEL-625 |
| chr3-75468539-DEL-146 | chr3-75468540-DEL-144 |
| chr3-75474499-DEL-54 | chr3-75474494-DEL-62 |
| chr3-75496921-DEL-1593 | chr3-75496922-DEL-1593 |
| chr3-75551430-DEL-56 | chr3-75551433-DEL-56 |
| chr3-75557639-DEL-90 | chr3-75557640-DEL-90 |
| chr3-75570124-DEL-1490 | chr3-75570125-DEL-1490 |
| chr3-75581305-DEL-201 | chr3-75581306-DEL-201 |
| chr3-75607030-DEL-60 | chr3-75607031-DEL-60 |
| chr3-75614302-DEL-59 | chr3-75614308-DEL-55 |
| chr3-75621678-DEL-51 | chr3-75621679-DEL-51 |
| chr3-75622246-DEL-352 | chr3-75622247-DEL-352 |
| chr3-75658043-DEL-79 | chr3-75658036-DEL-79 |
| chr3-75672876-DEL-66 | chr3-75672871-DEL-70 |

| | |
|---|---|
| chr3-75686960-DEL-6376 | chr3-75686942-DEL-6369 |
| chr3-75711703-DEL-158 | chr3-75711706-DEL-157 |
| chr3-75715825-DEL-657 | chr3-75715826-DEL-657 |
| chr3-75737965-DEL-538 | chr3-75737953-DEL-561 |
| chr3-75748340-DEL-132 | chr3-75748339-DEL-138 |
| chr3-75748742-DEL-6024 | chr3-75748743-DEL-6024 |
| chr3-75758934-DEL-138 | chr3-75758935-DEL-138 |
| chr3-75789174-DEL-66 | chr3-75789175-DEL-99 |
| chr3-75797279-DEL-94 | chr3-75797272-DEL-102 |
| chr3-75803183-DEL-136 | chr3-75803184-DEL-136 |
| chr3-75838654-DEL-411 | chr3-75838655-DEL-411 |
| chr3-75857562-DEL-66 | chr3-75857514-DEL-102 |
| chr3-75875775-DEL-2561 | chr3-75875776-DEL-2561 |
| chr3-75903530-DEL-299 | chr3-75903530-DEL-299 |
| chr3-75904535-DEL-492 | chr3-75904535-DEL-482 |
| chr3-75911173-DEL-52 | chr3-75911173-DEL-52 |
| chr3-75919279-DEL-100 | chr3-75919280-DEL-100 |
| chr3-75961234-DEL-92 | chr3-75961235-DEL-92 |
| chr3-75992098-DEL-52 | chr3-75992099-DEL-52 |
| chr3-75994171-DEL-6407 | chr3-75994172-DEL-6407 |
| chr3-76026949-DEL-78 | chr3-76026982-DEL-50 |
| chr3-76033015-DEL-8518 | chr3-76033016-DEL-8518 |
| chr3-76042876-DEL-51 | chr3-76042836-DEL-64 |
| chr3-76085251-DEL-57 | chr3-76085252-DEL-57 |
| chr3-76090196-DEL-54 | chr3-76090199-DEL-52 |
| chr3-76114900-DEL-125 | chr3-76114900-DEL-125 |
| chr3-76117317-DEL-206 | chr3-76117323-DEL-206 |
| chr3-76123162-DEL-364 | chr3-76123163-DEL-364 |
| chr3-76123912-DEL-79 | chr3-76123917-DEL-75 |
| chr3-76127408-DEL-64 | chr3-76127409-DEL-64 |
| chr3-76206056-DEL-196 | chr3-76206057-DEL-197 |
| chr3-76224929-DEL-89 | chr3-76224928-DEL-90 |
| chr3-76227217-DEL-51 | chr3-76227217-DEL-51 |
| chr3-76267583-DEL-55 | chr3-76267584-DEL-55 |
| chr3-76280381-DEL-54 | chr3-76280382-DEL-54 |
| chr3-76319403-DEL-651 | chr3-76319404-DEL-651 |
| chr3-76339030-DEL-201 | chr3-76339031-DEL-201 |
| chr3-76346164-DEL-1394 | chr3-76346165-DEL-1394 |
| chr3-76366075-DEL-69 | chr3-76366075-DEL-68 |
| chr3-76425429-DEL-947 | chr3-76425430-DEL-947 |
| chr3-76462097-DEL-3020 | chr3-76462098-DEL-3020 |
| chr3-76489285-DEL-54 | chr3-76489286-DEL-54 |
| chr3-76498357-DEL-58 | chr3-76498360-DEL-56 |
| chr3-76505999-DEL-6274 | chr3-76505998-DEL-6274 |
| chr3-76523328-DEL-598 | chr3-76523331-DEL-599 |
| chr3-76535225-DEL-90 | chr3-76535226-DEL-90 |
| chr3-76542209-DEL-358 | chr3-76542209-DEL-358 |
| chr3-76546032-DEL-448 | chr3-76546037-DEL-445 |
| chr3-76553329-DEL-67 | chr3-76553330-DEL-67 |
| chr3-76596897-DEL-78 | chr3-76596901-DEL-72 |
| chr3-76602434-DEL-56 | chr3-76602435-DEL-56 |
| chr3-76603798-DEL-163 | chr3-76603799-DEL-163 |
| chr3-76624313-DEL-669 | chr3-76624314-DEL-669 |
| chr3-76625774-DEL-1207 | chr3-76625774-DEL-1207 |
| chr3-76653822-DEL-58 | chr3-76653823-DEL-58 |
| chr3-76680101-DEL-72 | chr3-76680104-DEL-52 |
| chr3-76740760-DEL-750 | chr3-76740761-DEL-750 |
| chr3-76757594-DEL-2716 | chr3-76757595-DEL-2716 |
| chr3-76766209-DEL-56 | chr3-76766216-DEL-50 |
| chr3-76783248-DEL-636 | chr3-76783249-DEL-636 |
| chr3-76793850-DEL-6433 | chr3-76793851-DEL-6433 |
| chr3-76811196-DEL-57 | chr3-76811197-DEL-57 |
| chr3-76828728-DEL-2212 | chr3-76828729-DEL-2212 |
| chr3-76857158-DEL-60 | chr3-76857127-DEL-50 |
| chr3-76862326-DEL-105 | chr3-76862321-DEL-111 |
| chr3-76903748-DEL-269 | chr3-76903750-DEL-268 |
| chr3-76918909-DEL-129 | chr3-76918910-DEL-131 |
| chr3-76928133-DEL-356 | chr3-76928134-DEL-356 |
| chr3-76930238-DEL-443 | chr3-76930240-DEL-442 |
| chr3-76959752-DEL-1913 | chr3-76959752-DEL-1913 |
| chr3-76961983-DEL-911 | chr3-76961983-DEL-911 |
| chr3-76979398-DEL-1869 | chr3-76979400-DEL-1865 |
| chr3-76990279-DEL-52 | chr3-76990280-DEL-52 |
| chr3-77002816-DEL-943 | chr3-77002817-DEL-943 |
| chr3-77005676-DEL-517 | chr3-77005677-DEL-516 |
| chr3-77047083-DEL-72 | chr3-77047062-DEL-66 |
| chr3-77052209-DEL-1876 | chr3-77052210-DEL-1876 |
| chr3-77084381-DEL-1290 | chr3-77084382-DEL-1290 |
| chr3-77087663-DEL-398 | chr3-77087664-DEL-398 |
| chr3-77091325-DEL-6423 | chr3-77091326-DEL-6423 |

| | |
|---|---|
| chr3-77097419-DEL-214 | chr3-77097420-DEL-214 |
| chr3-77108917-DEL-52 | chr3-77108917-DEL-80 |
| chr3-77134308-DEL-286 | chr3-77134309-DEL-286 |
| chr3-77140936-DEL-6443 | chr3-77140937-DEL-6443 |
| chr3-77160261-DEL-6283 | chr3-77160262-DEL-6283 |
| chr3-77170413-DEL-215 | chr3-77170414-DEL-215 |
| chr3-77176401-DEL-787 | chr3-77176402-DEL-787 |
| chr3-77180108-DEL-1844 | chr3-77180109-DEL-1844 |
| chr3-77192530-DEL-83 | chr3-77192531-DEL-83 |
| chr3-77204881-DEL-5167 | chr3-77204882-DEL-5167 |
| chr3-77219152-DEL-1299 | chr3-77219153-DEL-1299 |
| chr3-77231462-DEL-740 | chr3-77231463-DEL-739 |
| chr3-77284662-DEL-1712 | chr3-77284663-DEL-1712 |
| chr3-77295820-DEL-481 | chr3-77295820-DEL-482 |
| chr3-77318132-DEL-256 | chr3-77318133-DEL-256 |
| chr3-77331857-DEL-65 | chr3-77331858-DEL-65 |
| chr3-77341570-DEL-149 | chr3-77341571-DEL-149 |
| chr3-77349437-DEL-2263 | chr3-77349423-DEL-2277 |
| chr3-77353211-DEL-1001 | chr3-77353212-DEL-1001 |
| chr3-77363950-DEL-57 | chr3-77363951-DEL-57 |
| chr3-77389885-DEL-3430 | chr3-77389887-DEL-3424 |
| chr3-77457156-DEL-53 | chr3-77457157-DEL-53 |
| chr3-77469642-DEL-325 | chr3-77469643-DEL-325 |
| chr3-77480447-DEL-1035 | chr3-77480448-DEL-1035 |
| chr3-77490282-DEL-58 | chr3-77490283-DEL-58 |
| chr3-77495612-DEL-65 | chr3-77495608-DEL-50 |
| chr3-77501481-DEL-57 | chr3-77501482-DEL-57 |
| chr3-77504675-DEL-6811 | chr3-77504676-DEL-6811 |
| chr3-77523969-DEL-592 | chr3-77523970-DEL-592 |
| chr3-77535267-DEL-56 | chr3-77535266-DEL-64 |
| chr3-77538224-DEL-349 | chr3-77538225-DEL-349 |
| chr3-77569814-DEL-869 | chr3-77569820-DEL-866 |
| chr3-77586160-DEL-2651 | chr3-77586161-DEL-2651 |
| chr3-77589417-DEL-4090 | chr3-77589418-DEL-4090 |
| chr3-77606491-DEL-325 | chr3-77606492-DEL-325 |
| chr3-77612475-DEL-214 | chr3-77612476-DEL-214 |
| chr3-77643672-DEL-71 | chr3-77643671-DEL-71 |
| chr3-77648296-DEL-994 | chr3-77648294-DEL-995 |
| chr3-77649975-DEL-226 | chr3-77649976-DEL-226 |
| chr3-77656630-DEL-76 | chr3-77656631-DEL-76 |
| chr3-77682114-DEL-67 | chr3-77682115-DEL-67 |
| chr3-77755370-DEL-118 | chr3-77755371-DEL-118 |
| chr3-77860920-DEL-1748 | chr3-77860921-DEL-1748 |
| chr3-77863878-DEL-1987 | chr3-77863879-DEL-1987 |
| chr3-77888509-DEL-51 | chr3-77888510-DEL-51 |
| chr3-77914737-DEL-1309 | chr3-77914738-DEL-1309 |
| chr3-77960406-DEL-358 | chr3-77960407-DEL-358 |
| chr3-77972961-DEL-112 | chr3-77972962-DEL-112 |
| chr3-78004869-DEL-90 | chr3-78004870-DEL-90 |
| chr3-78018288-DEL-255 | chr3-78018289-DEL-255 |
| chr3-78049476-DEL-103 | chr3-78049476-DEL-104 |
| chr3-78050216-DEL-3149 | chr3-78050217-DEL-3149 |
| chr3-78055825-DEL-78 | chr3-78055821-DEL-77 |
| chr3-78063272-DEL-62 | chr3-78063272-DEL-62 |
| chr3-78077385-DEL-681 | chr3-78077384-DEL-681 |
| chr3-78084068-DEL-1021 | chr3-78084069-DEL-1021 |
| chr3-78101946-DEL-57 | chr3-78101947-DEL-57 |
| chr3-78102972-DEL-85 | chr3-78102973-DEL-85 |
| chr3-78111558-DEL-1412 | chr3-78111559-DEL-1412 |
| chr3-78123735-DEL-117 | chr3-78123736-DEL-117 |
| chr3-78124742-DEL-401 | chr3-78124743-DEL-401 |
| chr3-78130536-DEL-298 | chr3-78130537-DEL-298 |
| chr3-78147216-DEL-746 | chr3-78147218-DEL-744 |
| chr3-78177451-DEL-961 | chr3-78177452-DEL-961 |
| chr3-78188261-DEL-53 | chr3-78188261-DEL-53 |
| chr3-78189224-DEL-1894 | chr3-78189225-DEL-1894 |
| chr3-78206986-DEL-58 | chr3-78206987-DEL-58 |
| chr3-78239791-DEL-7380 | chr3-78239792-DEL-7380 |
| chr3-78255407-DEL-1050 | chr3-78255408-DEL-1050 |
| chr3-78269394-DEL-170 | chr3-78269395-DEL-170 |
| chr3-78282844-DEL-623 | chr3-78282845-DEL-623 |
| chr3-78283925-DEL-213 | chr3-78283926-DEL-213 |
| chr3-78289159-DEL-390 | chr3-78289160-DEL-390 |
| chr3-78291110-DEL-67 | chr3-78291109-DEL-67 |
| chr3-78304174-DEL-240 | chr3-78304175-DEL-240 |
| chr3-78308430-DEL-1237 | chr3-78308430-DEL-1237 |
| chr3-78343302-DEL-493 | chr3-78343303-DEL-493 |
| chr3-78358529-DEL-6394 | chr3-78358530-DEL-6394 |
| chr3-78378332-DEL-677 | chr3-78378332-DEL-683 |
| chr3-78384395-DEL-69 | chr3-78384396-DEL-69 |

| | |
|---|---|
| chr3-78390191-DEL-652 | chr3-78390192-DEL-652 |
| chr3-78418899-DEL-211 | chr3-78418900-DEL-211 |
| chr3-78436701-DEL-100 | chr3-78436729-DEL-58 |
| chr3-78459611-DEL-7478 | chr3-78459612-DEL-7478 |
| chr3-78495227-DEL-140 | chr3-78495228-DEL-140 |
| chr3-78504929-DEL-62 | chr3-78504930-DEL-62 |
| chr3-78511274-DEL-6283 | chr3-78511275-DEL-6283 |
| chr3-78517712-DEL-711 | chr3-78517713-DEL-711 |
| chr3-78523191-DEL-362 | chr3-78523180-DEL-366 |
| chr3-78535369-DEL-2267 | chr3-78535370-DEL-2267 |
| chr3-78557889-DEL-55 | chr3-78557890-DEL-75 |
| chr3-78604516-DEL-1212 | chr3-78604517-DEL-1212 |
| chr3-78615523-DEL-85 | chr3-78615524-DEL-85 |
| chr3-78626748-DEL-1661 | chr3-78626749-DEL-1661 |
| chr3-78677535-DEL-7009 | chr3-78677536-DEL-7009 |
| chr3-78692434-DEL-74 | chr3-78692434-DEL-74 |
| chr3-78698011-DEL-98 | chr3-78698012-DEL-94 |
| chr3-78701023-DEL-80 | chr3-78701038-DEL-66 |
| chr3-78720469-DEL-3870 | chr3-78720470-DEL-3870 |
| chr3-78724577-DEL-7941 | chr3-78724578-DEL-7941 |
| chr3-78759179-DEL-67 | chr3-78759180-DEL-67 |
| chr3-78768058-DEL-57 | chr3-78768057-DEL-58 |
| chr3-78847767-DEL-69 | chr3-78847768-DEL-69 |
| chr3-78852018-DEL-61 | chr3-78852019-DEL-61 |
| chr3-78872755-DEL-1768 | chr3-78872756-DEL-1768 |
| chr3-78915925-DEL-435 | chr3-78915926-DEL-435 |
| chr3-78920678-DEL-849 | chr3-78920659-DEL-850 |
| chr3-78964679-DEL-60 | chr3-78964682-DEL-61 |
| chr3-78974391-DEL-228 | chr3-78974392-DEL-228 |
| chr3-79017387-DEL-59 | chr3-79017378-DEL-57 |
| chr3-79047092-DEL-181 | chr3-79047093-DEL-181 |
| chr3-79115177-DEL-83 | chr3-79115178-DEL-83 |
| chr3-79170258-DEL-241 | chr3-79170259-DEL-241 |
| chr3-79211071-DEL-116 | chr3-79211072-DEL-116 |
| chr3-79294817-DEL-66 | chr3-79294813-DEL-71 |
| chr3-79306580-DEL-148 | chr3-79306581-DEL-148 |
| chr3-79320204-DEL-210 | chr3-79320205-DEL-210 |
| chr3-79390242-DEL-100 | chr3-79390264-DEL-82 |
| chr3-79397108-DEL-112 | chr3-79397157-DEL-64 |
| chr3-79404163-DEL-68 | chr3-79404132-DEL-100 |
| chr3-79414650-DEL-60 | chr3-79414651-DEL-58 |
| chr3-79416483-DEL-89 | chr3-79416487-DEL-85 |
| chr3-79429910-DEL-60 | chr3-79429907-DEL-52 |
| chr3-79451891-DEL-184 | chr3-79451892-DEL-184 |
| chr3-79468208-DEL-229 | chr3-79468208-DEL-229 |
| chr3-79477378-DEL-91 | chr3-79477395-DEL-54 |
| chr3-79477587-DEL-632 | chr3-79477588-DEL-632 |
| chr3-79491352-DEL-58 | chr3-79491362-DEL-58 |
| chr3-79508107-DEL-111 | chr3-79508108-DEL-111 |
| chr3-79514835-DEL-203 | chr3-79514833-DEL-194 |
| chr3-79525521-DEL-80 | chr3-79525522-DEL-80 |
| chr3-79532732-DEL-201 | chr3-79532729-DEL-199 |
| chr3-79534564-DEL-168 | chr3-79534565-DEL-168 |
| chr3-79541454-DEL-6657 | chr3-79541455-DEL-6657 |
| chr3-79541860-DEL-212 | chr3-79541861-DEL-212 |
| chr3-79558229-DEL-201 | chr3-79558230-DEL-201 |
| chr3-79566033-DEL-523 | chr3-79566034-DEL-523 |
| chr3-79570131-DEL-52 | chr3-79570132-DEL-52 |
| chr3-79598264-DEL-101 | chr3-79598269-DEL-107 |
| chr3-79620069-DEL-54 | chr3-79620045-DEL-54 |
| chr3-79626377-DEL-52 | chr3-79626378-DEL-52 |
| chr3-79643377-DEL-86 | chr3-79643378-DEL-86 |
| chr3-79668816-DEL-58 | chr3-79668821-DEL-68 |
| chr3-79669630-DEL-1425 | chr3-79669632-DEL-1424 |
| chr3-79685817-DEL-1588 | chr3-79685818-DEL-1588 |
| chr3-79698181-DEL-60 | chr3-79698182-DEL-52 |
| chr3-79698869-DEL-68 | chr3-79698866-DEL-72 |
| chr3-79712881-DEL-672 | chr3-79712882-DEL-672 |
| chr3-79726452-DEL-94 | chr3-79726451-DEL-98 |
| chr3-79735962-DEL-76 | chr3-79735963-DEL-76 |
| chr3-79760675-DEL-65 | chr3-79760676-DEL-65 |
| chr3-79799742-DEL-1893 | chr3-79799743-DEL-1893 |
| chr3-79807646-DEL-773 | chr3-79807647-DEL-774 |
| chr3-79866787-DEL-6489 | chr3-79866788-DEL-6489 |
| chr3-79872921-DEL-212 | chr3-79872922-DEL-212 |
| chr3-79907538-DEL-68 | chr3-79907518-DEL-75 |
| chr3-79918188-DEL-8709 | chr3-79918199-DEL-8703 |
| chr3-79944384-DEL-934 | chr3-79944385-DEL-934 |
| chr3-79977377-DEL-64 | chr3-79977387-DEL-80 |
| chr3-79981837-DEL-1059 | chr3-79981840-DEL-1058 |

| | |
|---|---|
| chr3-80004576-DEL-420 | chr3-80004576-DEL-421 |
| chr3-80020019-DEL-69 | chr3-80020020-DEL-69 |
| chr3-80036686-DEL-3542 | chr3-80036687-DEL-3542 |
| chr3-80093770-DEL-88 | chr3-80093776-DEL-87 |
| chr3-80108764-DEL-875 | chr3-80108765-DEL-875 |
| chr3-80113855-DEL-67 | chr3-80113856-DEL-55 |
| chr3-80127932-DEL-192 | chr3-80127933-DEL-192 |
| chr3-80130320-DEL-2593 | chr3-80130321-DEL-2593 |
| chr3-80138914-DEL-152 | chr3-80138915-DEL-152 |
| chr3-80144556-DEL-6770 | chr3-80144557-DEL-6770 |
| chr3-80152239-DEL-63 | chr3-80152240-DEL-63 |
| chr3-80186185-DEL-115 | chr3-80186186-DEL-115 |
| chr3-80209689-DEL-96 | chr3-80209691-DEL-81 |
| chr3-80243315-DEL-145 | chr3-80243322-DEL-139 |
| chr3-80250764-DEL-189 | chr3-80250765-DEL-189 |
| chr3-80256551-DEL-6521 | chr3-80256552-DEL-6521 |
| chr3-80262654-DEL-212 | chr3-80262655-DEL-212 |
| chr3-80309341-DEL-68 | chr3-80309342-DEL-68 |
| chr3-80314517-DEL-322 | chr3-80314518-DEL-322 |
| chr3-80329331-DEL-53 | chr3-80329332-DEL-53 |
| chr3-80345626-DEL-6397 | chr3-80345627-DEL-6397 |
| chr3-80358252-DEL-149 | chr3-80358253-DEL-149 |
| chr3-80364757-DEL-51 | chr3-80364758-DEL-51 |
| chr3-80368763-DEL-73 | chr3-80368761-DEL-64 |
| chr3-80372684-DEL-6665 | chr3-80372685-DEL-6665 |
| chr3-80383095-DEL-1880 | chr3-80383096-DEL-1880 |
| chr3-80406234-DEL-202 | chr3-80406235-DEL-202 |
| chr3-80419423-DEL-54 | chr3-80419437-DEL-54 |
| chr3-80428841-DEL-1890 | chr3-80428842-DEL-1890 |
| chr3-80444515-DEL-989 | chr3-80444549-DEL-978 |
| chr3-80445834-DEL-1328 | chr3-80445835-DEL-1328 |
| chr3-80449650-DEL-851 | chr3-80449655-DEL-851 |
| chr3-80529657-DEL-204 | chr3-80529658-DEL-204 |
| chr3-80550467-DEL-746 | chr3-80550468-DEL-746 |
| chr3-80555511-DEL-149 | chr3-80555512-DEL-149 |
| chr3-80564551-DEL-50 | chr3-80564550-DEL-52 |
| chr3-80570195-DEL-98 | chr3-80570196-DEL-104 |
| chr3-80618521-DEL-66 | chr3-80618522-DEL-66 |
| chr3-80620951-DEL-82 | chr3-80620952-DEL-82 |
| chr3-80647077-DEL-536 | chr3-80647078-DEL-533 |
| chr3-80649715-DEL-6684 | chr3-80649716-DEL-6684 |
| chr3-80668721-DEL-51 | chr3-80668722-DEL-51 |
| chr3-80675465-DEL-565 | chr3-80675466-DEL-565 |
| chr3-80696598-DEL-110 | chr3-80696599-DEL-110 |
| chr3-80718044-DEL-76 | chr3-80718053-DEL-84 |
| chr3-80729443-DEL-1755 | chr3-80729444-DEL-1755 |
| chr3-80734305-DEL-94 | chr3-80734312-DEL-88 |
| chr3-80755777-DEL-82 | chr3-80755778-DEL-82 |
| chr3-80763277-DEL-516 | chr3-80763278-DEL-516 |
| chr3-80816365-DEL-165 | chr3-80816355-DEL-176 |
| chr3-80824687-DEL-879 | chr3-80824688-DEL-879 |
| chr3-80850259-DEL-221 | chr3-80850260-DEL-221 |
| chr3-80856615-DEL-366 | chr3-80856616-DEL-366 |
| chr3-80872759-DEL-489 | chr3-80872760-DEL-488 |
| chr3-80877287-DEL-64 | chr3-80877288-DEL-64 |
| chr3-80878130-DEL-165 | chr3-80878131-DEL-165 |
| chr3-80886802-DEL-221 | chr3-80886803-DEL-221 |
| chr3-80893637-DEL-2131 | chr3-80893638-DEL-2131 |
| chr3-80977516-DEL-7405 | chr3-80977517-DEL-7405 |
| chr3-81000669-DEL-156 | chr3-81000670-DEL-156 |
| chr3-81057329-DEL-76 | chr3-81057330-DEL-76 |
| chr3-81082992-DEL-54 | chr3-81082993-DEL-54 |
| chr3-81091492-DEL-50 | chr3-81091504-DEL-52 |
| chr3-81103104-DEL-235 | chr3-81103105-DEL-235 |
| chr3-81146847-DEL-1475 | chr3-81146848-DEL-1475 |
| chr3-81149092-DEL-490 | chr3-81149094-DEL-491 |
| chr3-81156538-DEL-302 | chr3-81156527-DEL-311 |
| chr3-81159514-DEL-845 | chr3-81159515-DEL-846 |
| chr3-81160342-DEL-6217 | chr3-81160357-DEL-6199 |
| chr3-81183155-DEL-89 | chr3-81183156-DEL-89 |
| chr3-81194161-DEL-293 | chr3-81194170-DEL-292 |
| chr3-81200036-DEL-64 | chr3-81200037-DEL-64 |
| chr3-81205770-DEL-2581 | chr3-81205771-DEL-2581 |
| chr3-81215960-DEL-6375 | chr3-81215961-DEL-6375 |
| chr3-81224118-DEL-1296 | chr3-81224119-DEL-1296 |
| chr3-81258509-DEL-306 | chr3-81258510-DEL-305 |
| chr3-81268479-DEL-6512 | chr3-81268480-DEL-6512 |
| chr3-81292550-DEL-66 | chr3-81292551-DEL-66 |
| chr3-81307842-DEL-54 | chr3-81307843-DEL-54 |
| chr3-81324802-DEL-54 | chr3-81324803-DEL-54 |

| | |
|---|---|
| chr3-81363147-DEL-118 | chr3-81363148-DEL-118 |
| chr3-81376568-DEL-1832 | chr3-81376569-DEL-1832 |
| chr3-81394164-DEL-388 | chr3-81394165-DEL-388 |
| chr3-81397082-DEL-615 | chr3-81397083-DEL-615 |
| chr3-81409626-DEL-123 | chr3-81409627-DEL-123 |
| chr3-81423143-DEL-54 | chr3-81423144-DEL-54 |
| chr3-81429890-DEL-2421 | chr3-81429891-DEL-2421 |
| chr3-81434843-DEL-74 | chr3-81434846-DEL-83 |
| chr3-81443974-DEL-194 | chr3-81443975-DEL-194 |
| chr3-81447867-DEL-1962 | chr3-81447868-DEL-1962 |
| chr3-81451443-DEL-60 | chr3-81451444-DEL-60 |
| chr3-81452846-DEL-52 | chr3-81452847-DEL-52 |
| chr3-81484830-DEL-124 | chr3-81484831-DEL-124 |
| chr3-81538489-DEL-81 | chr3-81538490-DEL-81 |
| chr3-81544354-DEL-52 | chr3-81544355-DEL-52 |
| chr3-81544614-DEL-843 | chr3-81544615-DEL-843 |
| chr3-81549967-DEL-60 | chr3-81549968-DEL-60 |
| chr3-81551728-DEL-1784 | chr3-81551729-DEL-1784 |
| chr3-81552311-DEL-206 | chr3-81552312-DEL-206 |
| chr3-81553216-DEL-51 | chr3-81553217-DEL-52 |
| chr3-81557517-DEL-3255 | chr3-81557518-DEL-3255 |
| chr3-81561873-DEL-175 | chr3-81561852-DEL-170 |
| chr3-81562389-DEL-130 | chr3-81562389-DEL-129 |
| chr3-81586068-DEL-338 | chr3-81586068-DEL-338 |
| chr3-81588378-DEL-558 | chr3-81588379-DEL-558 |
| chr3-81589608-DEL-393 | chr3-81589609-DEL-393 |
| chr3-81596852-DEL-8348 | chr3-81596853-DEL-8348 |
| chr3-81634362-DEL-102 | chr3-81634369-DEL-90 |
| chr3-81635211-DEL-1001 | chr3-81635212-DEL-1001 |
| chr3-81669063-DEL-283 | chr3-81669064-DEL-283 |
| chr3-81670768-DEL-393 | chr3-81670769-DEL-393 |
| chr3-81730583-DEL-60 | chr3-81730584-DEL-60 |
| chr3-81736727-DEL-205 | chr3-81736728-DEL-205 |
| chr3-81750241-DEL-60 | chr3-81750241-DEL-60 |
| chr3-81771295-DEL-402 | chr3-81771296-DEL-402 |
| chr3-81786838-DEL-305 | chr3-81786839-DEL-305 |
| chr3-81832292-DEL-68 | chr3-81832293-DEL-68 |
| chr3-81836070-DEL-52 | chr3-81836067-DEL-56 |
| chr3-81850247-DEL-180 | chr3-81850248-DEL-180 |
| chr3-81917777-DEL-582 | chr3-81917778-DEL-582 |
| chr3-81943064-DEL-54 | chr3-81943065-DEL-52 |
| chr3-82006450-DEL-52 | chr3-82006451-DEL-52 |
| chr3-82013766-DEL-51 | chr3-82013767-DEL-51 |
| chr3-82044819-DEL-99 | chr3-82044819-DEL-99 |
| chr3-82048333-DEL-126 | chr3-82048334-DEL-126 |
| chr3-82084866-DEL-64 | chr3-82084875-DEL-56 |
| chr3-82114928-DEL-849 | chr3-82114919-DEL-844 |
| chr3-82139253-DEL-65 | chr3-82139254-DEL-65 |
| chr3-82145627-DEL-77 | chr3-82145623-DEL-80 |
| chr3-82151748-DEL-1488 | chr3-82151749-DEL-1488 |
| chr3-82179162-DEL-128 | chr3-82179194-DEL-86 |
| chr3-82180911-DEL-101 | chr3-82180912-DEL-101 |
| chr3-82190667-DEL-5293 | chr3-82190668-DEL-5293 |
| chr3-82207152-DEL-96 | chr3-82207153-DEL-96 |
| chr3-82254209-DEL-65 | chr3-82254208-DEL-66 |
| chr3-82288052-DEL-200 | chr3-82288053-DEL-200 |
| chr3-82289665-DEL-66 | chr3-82289666-DEL-66 |
| chr3-82294728-DEL-676 | chr3-82294729-DEL-676 |
| chr3-82296156-DEL-66 | chr3-82296153-DEL-70 |
| chr3-82331218-DEL-134 | chr3-82331229-DEL-92 |
| chr3-82331226-DEL-1783 | chr3-82331227-DEL-1783 |
| chr3-82359796-DEL-903 | chr3-82359797-DEL-903 |
| chr3-82369376-DEL-1109 | chr3-82369377-DEL-1109 |
| chr3-82372741-DEL-399 | chr3-82372742-DEL-399 |
| chr3-82388094-DEL-7217 | chr3-82388095-DEL-7217 |
| chr3-82409653-DEL-52 | chr3-82409615-DEL-88 |
| chr3-82414710-DEL-1856 | chr3-82414709-DEL-1856 |
| chr3-82431295-DEL-259 | chr3-82431296-DEL-259 |
| chr3-82436792-DEL-1996 | chr3-82436793-DEL-1996 |
| chr3-82449600-DEL-69 | chr3-82449591-DEL-79 |
| chr3-82459683-DEL-213 | chr3-82459684-DEL-213 |
| chr3-82467282-DEL-74 | chr3-82467310-DEL-71 |
| chr3-82480474-DEL-327 | chr3-82480499-DEL-314 |
| chr3-82493865-DEL-170 | chr3-82493866-DEL-170 |
| chr3-82496539-DEL-5056 | chr3-82496539-DEL-5056 |
| chr3-82517151-DEL-6555 | chr3-82517152-DEL-6555 |
| chr3-82555205-DEL-1889 | chr3-82555206-DEL-1889 |
| chr3-82566863-DEL-498 | chr3-82566864-DEL-498 |
| chr3-82574272-DEL-135 | chr3-82574273-DEL-135 |
| chr3-82576288-DEL-485 | chr3-82576289-DEL-482 |

| | |
|---|---|
| chr3-82615792-DEL-450 | chr3-82615793-DEL-450 |
| chr3-82636320-DEL-65 | chr3-82636352-DEL-52 |
| chr3-82648556-DEL-68 | chr3-82648557-DEL-66 |
| chr3-82665880-DEL-73 | chr3-82665884-DEL-99 |
| chr3-82671753-DEL-2232 | chr3-82671754-DEL-2232 |
| chr3-82673503-DEL-103 | chr3-82673504-DEL-103 |
| chr3-82686878-DEL-1810 | chr3-82686879-DEL-1810 |
| chr3-82694268-DEL-51 | chr3-82694240-DEL-52 |
| chr3-82709776-DEL-350 | chr3-82709768-DEL-351 |
| chr3-82720579-DEL-55 | chr3-82720593-DEL-56 |
| chr3-82776361-DEL-110 | chr3-82776366-DEL-106 |
| chr3-82799386-DEL-119 | chr3-82799387-DEL-119 |
| chr3-82867667-DEL-106 | chr3-82867668-DEL-106 |
| chr3-82879826-DEL-7157 | chr3-82879827-DEL-7157 |
| chr3-82905513-DEL-90 | chr3-82905464-DEL-91 |
| chr3-83042930-DEL-72 | chr3-83042962-DEL-72 |
| chr3-83057064-DEL-56 | chr3-83057065-DEL-56 |
| chr3-83093271-DEL-378 | chr3-83093256-DEL-377 |
| chr3-83096834-DEL-55 | chr3-83096835-DEL-55 |
| chr3-83111927-DEL-1857 | chr3-83111928-DEL-1857 |
| chr3-83162915-DEL-1251 | chr3-83162916-DEL-1251 |
| chr3-83200083-DEL-6427 | chr3-83200084-DEL-6427 |
| chr3-83214717-DEL-432 | chr3-83214711-DEL-433 |
| chr3-83218746-DEL-272 | chr3-83218747-DEL-272 |
| chr3-83238154-DEL-2131 | chr3-83238155-DEL-2131 |
| chr3-83271736-DEL-391 | chr3-83271737-DEL-391 |
| chr3-83282402-DEL-193 | chr3-83282388-DEL-196 |
| chr3-83342810-DEL-120 | chr3-83342811-DEL-121 |
| chr3-83362240-DEL-1357 | chr3-83362241-DEL-1357 |
| chr3-83368503-DEL-62 | chr3-83368504-DEL-62 |
| chr3-83422912-DEL-78 | chr3-83422913-DEL-78 |
| chr3-83432871-DEL-567 | chr3-83432872-DEL-567 |
| chr3-83439280-DEL-52 | chr3-83439261-DEL-72 |
| chr3-83450695-DEL-5181 | chr3-83450728-DEL-5181 |
| chr3-83511778-DEL-64 | chr3-83511779-DEL-64 |
| chr3-83563648-DEL-66 | chr3-83563649-DEL-66 |
| chr3-83572106-DEL-226 | chr3-83572107-DEL-226 |
| chr3-83576886-DEL-107 | chr3-83576887-DEL-107 |
| chr3-83595233-DEL-217 | chr3-83595234-DEL-217 |
| chr3-83627752-DEL-113 | chr3-83627753-DEL-113 |
| chr3-83694393-DEL-110 | chr3-83694397-DEL-107 |
| chr3-83725846-DEL-228 | chr3-83725845-DEL-229 |
| chr3-83728413-DEL-197 | chr3-83728414-DEL-197 |
| chr3-83732880-DEL-209 | chr3-83732884-DEL-208 |
| chr3-83737601-DEL-208 | chr3-83737603-DEL-206 |
| chr3-83739890-DEL-211 | chr3-83739878-DEL-211 |
| chr3-83742865-DEL-193 | chr3-83742861-DEL-192 |
| chr3-83765017-DEL-68 | chr3-83765018-DEL-68 |
| chr3-83773238-DEL-874 | chr3-83773236-DEL-877 |
| chr3-83775868-DEL-328 | chr3-83775869-DEL-328 |
| chr3-83778290-DEL-281 | chr3-83778291-DEL-280 |
| chr3-83819825-DEL-89 | chr3-83819826-DEL-89 |
| chr3-83830163-DEL-79 | chr3-83830164-DEL-79 |
| chr3-83840803-DEL-177 | chr3-83840804-DEL-177 |
| chr3-83859845-DEL-52 | chr3-83859825-DEL-56 |
| chr3-83863157-DEL-102 | chr3-83863192-DEL-68 |
| chr3-83889605-DEL-321 | chr3-83889606-DEL-321 |
| chr3-83899042-DEL-1241 | chr3-83899043-DEL-1241 |
| chr3-83905924-DEL-223 | chr3-83905925-DEL-223 |
| chr3-83924497-DEL-414 | chr3-83924534-DEL-381 |
| chr3-83965182-DEL-1432 | chr3-83965183-DEL-1432 |
| chr3-83979467-DEL-144 | chr3-83979453-DEL-158 |
| chr3-84021826-DEL-550 | chr3-84021827-DEL-550 |
| chr3-84025162-DEL-4220 | chr3-84025163-DEL-4220 |
| chr3-84042777-DEL-108 | chr3-84042812-DEL-75 |
| chr3-84049687-DEL-7232 | chr3-84049682-DEL-7230 |
| chr3-84086096-DEL-60 | chr3-84086097-DEL-60 |
| chr3-84096528-DEL-62 | chr3-84096513-DEL-62 |
| chr3-84100891-DEL-57 | chr3-84100892-DEL-57 |
| chr3-84172817-DEL-75 | chr3-84172818-DEL-75 |
| chr3-84186814-DEL-208 | chr3-84186818-DEL-210 |
| chr3-84187424-DEL-777 | chr3-84187425-DEL-777 |
| chr3-84194881-DEL-158 | chr3-84194880-DEL-123 |
| chr3-84209853-DEL-144 | chr3-84209854-DEL-144 |
| chr3-84266462-DEL-7161 | chr3-84266463-DEL-7161 |
| chr3-84277623-DEL-409 | chr3-84277624-DEL-409 |
| chr3-84290751-DEL-1174 | chr3-84290752-DEL-1174 |
| chr3-84304725-DEL-92 | chr3-84304726-DEL-92 |
| chr3-84308618-DEL-52 | chr3-84308615-DEL-57 |
| chr3-84322595-DEL-183 | chr3-84322617-DEL-181 |

| | |
|---|---|
| chr3-84348500-DEL-123 | chr3-84348498-DEL-120 |
| chr3-84357313-DEL-65 | chr3-84357314-DEL-65 |
| chr3-84365754-DEL-476 | chr3-84365755-DEL-476 |
| chr3-84372225-DEL-214 | chr3-84372226-DEL-214 |
| chr3-84386052-DEL-59 | chr3-84386053-DEL-59 |
| chr3-84442728-DEL-255 | chr3-84442729-DEL-255 |
| chr3-84449748-DEL-142 | chr3-84449749-DEL-142 |
| chr3-84457896-DEL-63 | chr3-84457857-DEL-64 |
| chr3-84496559-DEL-212 | chr3-84496560-DEL-212 |
| chr3-84515958-DEL-702 | chr3-84515957-DEL-704 |
| chr3-84517958-DEL-60 | chr3-84517958-DEL-60 |
| chr3-84538603-DEL-72 | chr3-84538630-DEL-50 |
| chr3-84540418-DEL-70 | chr3-84540419-DEL-78 |
| chr3-84557686-DEL-475 | chr3-84557687-DEL-475 |
| chr3-84563409-DEL-98 | chr3-84563411-DEL-97 |
| chr3-84579140-DEL-58 | chr3-84579141-DEL-58 |
| chr3-84590813-DEL-61 | chr3-84590822-DEL-53 |
| chr3-84606259-DEL-414 | chr3-84606260-DEL-414 |
| chr3-84613730-DEL-483 | chr3-84613731-DEL-483 |
| chr3-84619462-DEL-99 | chr3-84619445-DEL-117 |
| chr3-84620590-DEL-1227 | chr3-84620591-DEL-1227 |
| chr3-84635684-DEL-300 | chr3-84635685-DEL-300 |
| chr3-84650471-DEL-174 | chr3-84650472-DEL-174 |
| chr3-84658010-DEL-1896 | chr3-84658011-DEL-1896 |
| chr3-84669439-DEL-5315 | chr3-84669440-DEL-5315 |
| chr3-84675906-DEL-76 | chr3-84675907-DEL-76 |
| chr3-84692799-DEL-60 | chr3-84692804-DEL-56 |
| chr3-84695508-DEL-58 | chr3-84695509-DEL-58 |
| chr3-84748968-DEL-382 | chr3-84748969-DEL-382 |
| chr3-84766519-DEL-76 | chr3-84766533-DEL-79 |
| chr3-84769368-DEL-214 | chr3-84769369-DEL-214 |
| chr3-84905754-DEL-1000 | chr3-84905755-DEL-1000 |
| chr3-84940074-DEL-156 | chr3-84940075-DEL-172 |
| chr3-84942247-DEL-815 | chr3-84942247-DEL-815 |
| chr3-84942496-DEL-2077 | chr3-84942497-DEL-2077 |
| chr3-85009001-DEL-152 | chr3-85009002-DEL-152 |
| chr3-85012273-DEL-72 | chr3-85012274-DEL-72 |
| chr3-85019346-DEL-59 | chr3-85019347-DEL-59 |
| chr3-85027876-DEL-88 | chr3-85027877-DEL-88 |
| chr3-85054380-DEL-125 | chr3-85054381-DEL-124 |
| chr3-85073832-DEL-714 | chr3-85073833-DEL-714 |
| chr3-85110282-DEL-702 | chr3-85110281-DEL-703 |
| chr3-85167743-DEL-492 | chr3-85167744-DEL-492 |
| chr3-85174413-DEL-183 | chr3-85174414-DEL-183 |
| chr3-85186088-DEL-82 | chr3-85186089-DEL-82 |
| chr3-85205310-DEL-56 | chr3-85205314-DEL-61 |
| chr3-85212526-DEL-2872 | chr3-85212527-DEL-2872 |
| chr3-85231497-DEL-91 | chr3-85231530-DEL-59 |
| chr3-85232421-DEL-1594 | chr3-85232422-DEL-1594 |
| chr3-85241953-DEL-60 | chr3-85241929-DEL-84 |
| chr3-85254218-DEL-72 | chr3-85254219-DEL-72 |
| chr3-85257284-DEL-173 | chr3-85257285-DEL-173 |
| chr3-85308573-DEL-142 | chr3-85308571-DEL-145 |
| chr3-85309897-DEL-336 | chr3-85309898-DEL-336 |
| chr3-85346603-DEL-57 | chr3-85346595-DEL-70 |
| chr3-85352562-DEL-80 | chr3-85352551-DEL-50 |
| chr3-85370950-DEL-683 | chr3-85370951-DEL-683 |
| chr3-85408382-DEL-72 | chr3-85408382-DEL-72 |
| chr3-85409840-DEL-143 | chr3-85409841-DEL-143 |
| chr3-85529581-DEL-7116 | chr3-85529582-DEL-7116 |
| chr3-85633712-DEL-58 | chr3-85633713-DEL-58 |
| chr3-85652212-DEL-100 | chr3-85652213-DEL-100 |
| chr3-85671293-DEL-125 | chr3-85671294-DEL-125 |
| chr3-85675559-DEL-58 | chr3-85675560-DEL-58 |
| chr3-85685229-DEL-1558 | chr3-85685230-DEL-1558 |
| chr3-85703642-DEL-490 | chr3-85703643-DEL-490 |
| chr3-85706841-DEL-270 | chr3-85706842-DEL-270 |
| chr3-85741107-DEL-99 | chr3-85741108-DEL-99 |
| chr3-85759564-DEL-89 | chr3-85759558-DEL-90 |
| chr3-85836123-DEL-58 | chr3-85836124-DEL-54 |
| chr3-85840953-DEL-183 | chr3-85840954-DEL-183 |
| chr3-85865170-DEL-341 | chr3-85865173-DEL-310 |
| chr3-85871091-DEL-568 | chr3-85871090-DEL-568 |
| chr3-85872279-DEL-178 | chr3-85872280-DEL-178 |
| chr3-85874280-DEL-74 | chr3-85874277-DEL-50 |
| chr3-85884014-DEL-215 | chr3-85884015-DEL-215 |
| chr3-85913715-DEL-71 | chr3-85913716-DEL-71 |
| chr3-85918165-DEL-151 | chr3-85918166-DEL-152 |
| chr3-85919907-DEL-60 | chr3-85919907-DEL-60 |
| chr3-85936436-DEL-727 | chr3-85936437-DEL-727 |

| | |
|---|---|
| chr3-85964898-DEL-160 | chr3-85964915-DEL-144 |
| chr3-85989660-DEL-157 | chr3-85989661-DEL-157 |
| chr3-86003513-DEL-528 | chr3-86003514-DEL-528 |
| chr3-86011226-DEL-86 | chr3-86011227-DEL-86 |
| chr3-86012705-DEL-403 | chr3-86012706-DEL-403 |
| chr3-86024561-DEL-211 | chr3-86024562-DEL-211 |
| chr3-86024892-DEL-690 | chr3-86024884-DEL-708 |
| chr3-86055622-DEL-486 | chr3-86055623-DEL-486 |
| chr3-86060480-DEL-6379 | chr3-86060481-DEL-6379 |
| chr3-86086264-DEL-323 | chr3-86086265-DEL-323 |
| chr3-86128498-DEL-407 | chr3-86128499-DEL-450 |
| chr3-86142356-DEL-158 | chr3-86142357-DEL-158 |
| chr3-86143463-DEL-62 | chr3-86143463-DEL-62 |
| chr3-86145981-DEL-2804 | chr3-86145981-DEL-2804 |
| chr3-86149621-DEL-117 | chr3-86149622-DEL-112 |
| chr3-86159462-DEL-173 | chr3-86159463-DEL-173 |
| chr3-86173451-DEL-875 | chr3-86173453-DEL-876 |
| chr3-86209696-DEL-51 | chr3-86209697-DEL-68 |
| chr3-86229393-DEL-377 | chr3-86229394-DEL-377 |
| chr3-86231928-DEL-88 | chr3-86231929-DEL-88 |
| chr3-86236600-DEL-7123 | chr3-86236601-DEL-7123 |
| chr3-86287539-DEL-68 | chr3-86287541-DEL-64 |
| chr3-86307997-DEL-106 | chr3-86308002-DEL-63 |
| chr3-86388059-DEL-2067 | chr3-86388060-DEL-2067 |
| chr3-86398949-DEL-73 | chr3-86398950-DEL-73 |
| chr3-86399907-DEL-6436 | chr3-86399908-DEL-6436 |
| chr3-86464974-DEL-220 | chr3-86464975-DEL-220 |
| chr3-86556825-DEL-174 | chr3-86556826-DEL-173 |
| chr3-86564745-DEL-71 | chr3-86564746-DEL-71 |
| chr3-86571275-DEL-189 | chr3-86571276-DEL-189 |
| chr3-86591158-DEL-1976 | chr3-86591159-DEL-1976 |
| chr3-86629339-DEL-117 | chr3-86629340-DEL-117 |
| chr3-86732509-DEL-114 | chr3-86732510-DEL-114 |
| chr3-86734397-DEL-84 | chr3-86734398-DEL-84 |
| chr3-86747705-DEL-218 | chr3-86747706-DEL-218 |
| chr3-86750627-DEL-73 | chr3-86750628-DEL-72 |
| chr3-86786575-DEL-199 | chr3-86786576-DEL-199 |
| chr3-86800377-DEL-204 | chr3-86800378-DEL-204 |
| chr3-86808210-DEL-54 | chr3-86808211-DEL-54 |
| chr3-86822662-DEL-81 | chr3-86822663-DEL-81 |
| chr3-86837774-DEL-98 | chr3-86837775-DEL-98 |
| chr3-86842227-DEL-337 | chr3-86842228-DEL-337 |
| chr3-86849453-DEL-4898 | chr3-86849454-DEL-4898 |
| chr3-86858322-DEL-194 | chr3-86858323-DEL-194 |
| chr3-86917281-DEL-103 | chr3-86917244-DEL-147 |
| chr3-86947730-DEL-53 | chr3-86947731-DEL-53 |
| chr3-86964343-DEL-66 | chr3-86964343-DEL-66 |
| chr3-87016014-DEL-681 | chr3-87016015-DEL-681 |
| chr3-87035837-DEL-51 | chr3-87035838-DEL-51 |
| chr3-87046461-DEL-80 | chr3-87046459-DEL-73 |
| chr3-87054333-DEL-94 | chr3-87054346-DEL-50 |
| chr3-87088834-DEL-186 | chr3-87088835-DEL-186 |
| chr3-87095812-DEL-197 | chr3-87095813-DEL-197 |
| chr3-87107043-DEL-582 | chr3-87107044-DEL-582 |
| chr3-87117465-DEL-88 | chr3-87117466-DEL-88 |
| chr3-87119429-DEL-161 | chr3-87119458-DEL-133 |
| chr3-87120613-DEL-204 | chr3-87120614-DEL-204 |
| chr3-87124938-DEL-60 | chr3-87124939-DEL-60 |
| chr3-87174339-DEL-52 | chr3-87174340-DEL-52 |
| chr3-87176296-DEL-61 | chr3-87176293-DEL-62 |
| chr3-87186400-DEL-2099 | chr3-87186401-DEL-2099 |
| chr3-87192551-DEL-2279 | chr3-87192550-DEL-2280 |
| chr3-87209137-DEL-201 | chr3-87209137-DEL-201 |
| chr3-87238967-DEL-1555 | chr3-87238968-DEL-1555 |
| chr3-87244878-DEL-54 | chr3-87244879-DEL-60 |
| chr3-87261021-DEL-142 | chr3-87261022-DEL-142 |
| chr3-87266945-DEL-202 | chr3-87266946-DEL-202 |
| chr3-87284882-DEL-55 | chr3-87284883-DEL-55 |
| chr3-87304348-DEL-77 | chr3-87304349-DEL-77 |
| chr3-87311880-DEL-779 | chr3-87311877-DEL-814 |
| chr3-87315084-DEL-109 | chr3-87315085-DEL-109 |
| chr3-87332431-DEL-60 | chr3-87332432-DEL-60 |
| chr3-87348470-DEL-862 | chr3-87348471-DEL-862 |
| chr3-87362695-DEL-105 | chr3-87362696-DEL-105 |
| chr3-87371505-DEL-76 | chr3-87371506-DEL-76 |
| chr3-87376620-DEL-5486 | chr3-87376621-DEL-5486 |
| chr3-87395979-DEL-58 | chr3-87395980-DEL-58 |
| chr3-87398163-DEL-225 | chr3-87398164-DEL-225 |
| chr3-87404469-DEL-532 | chr3-87404470-DEL-532 |
| chr3-87442633-DEL-51 | chr3-87442634-DEL-51 |

| | |
|---|---|
| chr3-87474641-DEL-98 | chr3-87474642-DEL-98 |
| chr3-87478162-DEL-88 | chr3-87478141-DEL-110 |
| chr3-87502738-DEL-782 | chr3-87502740-DEL-781 |
| chr3-87561768-DEL-228 | chr3-87561779-DEL-215 |
| chr3-87669730-DEL-58 | chr3-87669731-DEL-58 |
| chr3-87721949-DEL-102 | chr3-87721949-DEL-99 |
| chr3-87733162-DEL-150 | chr3-87733163-DEL-150 |
| chr3-87742160-DEL-441 | chr3-87742161-DEL-441 |
| chr3-87743151-DEL-392 | chr3-87743152-DEL-392 |
| chr3-87770187-DEL-58 | chr3-87770183-DEL-74 |
| chr3-87789275-DEL-215 | chr3-87789276-DEL-215 |
| chr3-87797004-DEL-852 | chr3-87797004-DEL-852 |
| chr3-87798609-DEL-220 | chr3-87798609-DEL-220 |
| chr3-87803744-DEL-203 | chr3-87803744-DEL-203 |
| chr3-87804862-DEL-404 | chr3-87804863-DEL-404 |
| chr3-87827833-DEL-200 | chr3-87827834-DEL-200 |
| chr3-87833570-DEL-213 | chr3-87833571-DEL-213 |
| chr3-87840389-DEL-219 | chr3-87840390-DEL-219 |
| chr3-87883875-DEL-132 | chr3-87883876-DEL-132 |
| chr3-87921531-DEL-183 | chr3-87921532-DEL-183 |
| chr3-87925970-DEL-205 | chr3-87925971-DEL-205 |
| chr3-87951322-DEL-628 | chr3-87951324-DEL-628 |
| chr3-87967605-DEL-180 | chr3-87967603-DEL-181 |
| chr3-87970858-DEL-2693 | chr3-87970859-DEL-2693 |
| chr3-88010303-DEL-331 | chr3-88010304-DEL-331 |
| chr3-88036316-DEL-51 | chr3-88036320-DEL-64 |
| chr3-88061331-DEL-263 | chr3-88061344-DEL-231 |
| chr3-88104733-DEL-141 | chr3-88104734-DEL-141 |
| chr3-88160719-DEL-95 | chr3-88160720-DEL-95 |
| chr3-88167214-DEL-210 | chr3-88167215-DEL-209 |
| chr3-88170454-DEL-467 | chr3-88170455-DEL-467 |
| chr3-88171186-DEL-193 | chr3-88171190-DEL-194 |
| chr3-88186357-DEL-213 | chr3-88186358-DEL-213 |
| chr3-88193037-DEL-217 | chr3-88193048-DEL-207 |
| chr3-88198656-DEL-465 | chr3-88198657-DEL-465 |
| chr3-88200361-DEL-402 | chr3-88200362-DEL-402 |
| chr3-88202371-DEL-170 | chr3-88202419-DEL-171 |
| chr3-88214707-DEL-191 | chr3-88214709-DEL-190 |
| chr3-88294800-DEL-606 | chr3-88294799-DEL-603 |
| chr3-88338019-DEL-180 | chr3-88338020-DEL-180 |
| chr3-88386224-DEL-60 | chr3-88386225-DEL-59 |
| chr3-88406570-DEL-306 | chr3-88406571-DEL-308 |
| chr3-88481419-DEL-3240 | chr3-88481420-DEL-3240 |
| chr3-88484715-DEL-1147 | chr3-88484716-DEL-1147 |
| chr3-88489832-DEL-176 | chr3-88489833-DEL-176 |
| chr3-88491235-DEL-110 | chr3-88491236-DEL-110 |
| chr3-88499304-DEL-52 | chr3-88499305-DEL-52 |
| chr3-88520780-DEL-228 | chr3-88520787-DEL-222 |
| chr3-88544794-DEL-214 | chr3-88544795-DEL-214 |
| chr3-88573711-DEL-7082 | chr3-88573712-DEL-7082 |
| chr3-88588553-DEL-204 | chr3-88588554-DEL-204 |
| chr3-88589713-DEL-250 | chr3-88589755-DEL-254 |
| chr3-88591919-DEL-198 | chr3-88591920-DEL-198 |
| chr3-88599437-DEL-228 | chr3-88599437-DEL-228 |
| chr3-88623391-DEL-198 | chr3-88623392-DEL-198 |
| chr3-88629872-DEL-361 | chr3-88629871-DEL-398 |
| chr3-88634225-DEL-178 | chr3-88634226-DEL-178 |
| chr3-88641287-DEL-340 | chr3-88641288-DEL-340 |
| chr3-88649026-DEL-546 | chr3-88649027-DEL-546 |
| chr3-88670406-DEL-207 | chr3-88670407-DEL-207 |
| chr3-88676258-DEL-145 | chr3-88676273-DEL-110 |
| chr3-88676768-DEL-186 | chr3-88676781-DEL-184 |
| chr3-88689754-DEL-54 | chr3-88689733-DEL-52 |
| chr3-88708987-DEL-210 | chr3-88708988-DEL-210 |
| chr3-88718894-DEL-187 | chr3-88718895-DEL-187 |
| chr3-88727568-DEL-7661 | chr3-88727573-DEL-7656 |
| chr3-88736059-DEL-88 | chr3-88736060-DEL-88 |
| chr3-88778832-DEL-200 | chr3-88778836-DEL-202 |
| chr3-88860288-DEL-68 | chr3-88860283-DEL-76 |
| chr3-88868737-DEL-201 | chr3-88868738-DEL-201 |
| chr3-88904735-DEL-674 | chr3-88904736-DEL-674 |
| chr3-88920188-DEL-169 | chr3-88920189-DEL-169 |
| chr3-88924723-DEL-3885 | chr3-88924724-DEL-3885 |
| chr3-88981825-DEL-111 | chr3-88981826-DEL-111 |
| chr3-89012940-DEL-2258 | chr3-89012941-DEL-2258 |
| chr3-89013035-DEL-53 | chr3-89013036-DEL-52 |
| chr3-89017559-DEL-58 | chr3-89017556-DEL-66 |
| chr3-89019276-DEL-81 | chr3-89019271-DEL-104 |
| chr3-89026269-DEL-208 | chr3-89026279-DEL-206 |
| chr3-89031939-DEL-226 | chr3-89031940-DEL-225 |

| | |
|---|---|
| chr3-89032974-DEL-329 | chr3-89032975-DEL-329 |
| chr3-89034399-DEL-211 | chr3-89034400-DEL-210 |
| chr3-89062926-DEL-201 | chr3-89062927-DEL-201 |
| chr3-89069764-DEL-554 | chr3-89069765-DEL-554 |
| chr3-89128039-DEL-207 | chr3-89128040-DEL-207 |
| chr3-89137753-DEL-126 | chr3-89137754-DEL-126 |
| chr3-89207794-DEL-59 | chr3-89207795-DEL-59 |
| chr3-89373660-DEL-211 | chr3-89373661-DEL-211 |
| chr3-89391457-DEL-203 | chr3-89391459-DEL-202 |
| chr3-89407718-DEL-179 | chr3-89407706-DEL-180 |
| chr3-89440891-DEL-54 | chr3-89440902-DEL-64 |
| chr3-89591543-DEL-84 | chr3-89591542-DEL-86 |
| chr3-89596320-DEL-56 | chr3-89596356-DEL-57 |
| chr3-89704750-DEL-228 | chr3-89704751-DEL-228 |
| chr3-89743720-DEL-700 | chr3-89743721-DEL-700 |
| chr3-89764483-DEL-119 | chr3-89764484-DEL-119 |
| chr3-89783737-DEL-63 | chr3-89783753-DEL-60 |
| chr3-89791843-DEL-203 | chr3-89791844-DEL-203 |
| chr3-89798190-DEL-55 | chr3-89798190-DEL-55 |
| chr3-89814144-DEL-856 | chr3-89814145-DEL-856 |
| chr3-89834840-DEL-342 | chr3-89834841-DEL-342 |
| chr3-89839667-DEL-397 | chr3-89839668-DEL-397 |
| chr3-89881795-DEL-69 | chr3-89881775-DEL-69 |
| chr3-89885938-DEL-231 | chr3-89885939-DEL-231 |
| chr3-89897307-DEL-86 | chr3-89897308-DEL-86 |
| chr3-89898473-DEL-231 | chr3-89898474-DEL-231 |
| chr3-89982369-DEL-167 | chr3-89982370-DEL-167 |
| chr3-90015482-DEL-197 | chr3-90015489-DEL-191 |
| chr3-90025968-DEL-53 | chr3-90025969-DEL-53 |
| chr3-90027829-DEL-431 | chr3-90027830-DEL-431 |
| chr3-90033749-DEL-1165 | chr3-90033750-DEL-1165 |
| chr3-90044531-DEL-1861 | chr3-90044532-DEL-1861 |
| chr3-90055357-DEL-150 | chr3-90055358-DEL-145 |
| chr3-90062441-DEL-210 | chr3-90062442-DEL-210 |
| chr3-90084139-DEL-197 | chr3-90084139-DEL-197 |
| chr3-90084885-DEL-1696 | chr3-90084886-DEL-1687 |
| chr3-90090677-DEL-396 | chr3-90090678-DEL-396 |
| chr3-90093652-DEL-54 | chr3-90093653-DEL-54 |
| chr3-90098714-DEL-299 | chr3-90098716-DEL-297 |
| chr3-90123746-DEL-176 | chr3-90123747-DEL-176 |
| chr3-90134258-DEL-97 | chr3-90134259-DEL-97 |
| chr3-90188327-DEL-525 | chr3-90188328-DEL-524 |
| chr3-90194839-DEL-56 | chr3-90194840-DEL-73 |
| chr3-90212358-DEL-203 | chr3-90212359-DEL-203 |
| chr3-90215016-DEL-201 | chr3-90215017-DEL-201 |
| chr3-90217271-DEL-825 | chr3-90217272-DEL-825 |
| chr3-90224730-DEL-199 | chr3-90224731-DEL-199 |
| chr3-90295154-DEL-190 | chr3-90295155-DEL-190 |
| chr3-90327029-DEL-129 | chr3-90327072-DEL-123 |
| chr3-90349201-DEL-100 | chr3-90349202-DEL-100 |
| chr3-90367489-DEL-104 | chr3-90367491-DEL-93 |
| chr3-90389022-DEL-60 | chr3-90389023-DEL-60 |
| chr3-90390380-DEL-198 | chr3-90390381-DEL-198 |
| chr3-90452067-DEL-56 | chr3-90452068-DEL-52 |
| chr3-90459316-DEL-51 | chr3-90459289-DEL-62 |
| chr3-90483671-DEL-1556 | chr3-90483672-DEL-1556 |
| chr3-90488239-DEL-950 | chr3-90488240-DEL-950 |
| chr3-90490723-DEL-64 | chr3-90490718-DEL-61 |
| chr3-90591999-DEL-456 | chr3-90592000-DEL-456 |
| chr3-90636702-DEL-596 | chr3-90636703-DEL-596 |
| chr3-90649999-DEL-1511 | chr3-90650000-DEL-1511 |
| chr3-90766081-DEL-3227 | chr3-90766082-DEL-3227 |
| chr3-90766302-DEL-694 | chr3-90766303-DEL-694 |
| chr3-90840827-DEL-2292 | chr3-90840828-DEL-2292 |
| chr3-90885437-DEL-580 | chr3-90885438-DEL-580 |
| chr3-90895645-DEL-63 | chr3-90895646-DEL-63 |
| chr3-90898508-DEL-5561 | chr3-90898509-DEL-5561 |
| chr3-90902705-DEL-133 | chr3-90902705-DEL-133 |
| chr3-91002296-DEL-11160 | chr3-91002285-DEL-11161 |
| chr3-91002568-DEL-1040 | chr3-91002567-DEL-1040 |
| chr3-91003771-DEL-624 | chr3-91003772-DEL-624 |
| chr3-91004941-DEL-1871 | chr3-91004942-DEL-1871 |
| chr3-91024343-DEL-68 | chr3-91024342-DEL-60 |
| chr3-91034777-DEL-212 | chr3-91034778-DEL-212 |
| chr3-91056440-DEL-85 | chr3-91056441-DEL-85 |
| chr3-91076104-DEL-51 | chr3-91076103-DEL-52 |
| chr3-91084446-DEL-495 | chr3-91084447-DEL-495 |
| chr3-91395302-DEL-55 | chr3-91395302-DEL-55 |
| chr3-91416377-DEL-1155 | chr3-91416377-DEL-1155 |
| chr3-91490412-DEL-60 | chr3-91490414-DEL-58 |

| | |
|---|---|
| chr3-91512826-DEL-497 | chr3-91512826-DEL-497 |
| chr3-91550877-DEL-248 | chr3-91550878-DEL-248 |
| chr3-91648332-DEL-85 | chr3-91648333-DEL-85 |
| chr3-91662746-DEL-203 | chr3-91662742-DEL-204 |
| chr3-91714208-DEL-54 | chr3-91714202-DEL-60 |
| chr3-91773219-DEL-152 | chr3-91773219-DEL-152 |
| chr3-91780534-DEL-55 | chr3-91780534-DEL-55 |
| chr3-91783223-DEL-74 | chr3-91783223-DEL-74 |
| chr3-91788849-DEL-6630 | chr3-91788850-DEL-6630 |
| chr3-91820970-DEL-9052 | chr3-91820971-DEL-9052 |
| chr3-91870035-DEL-242 | chr3-91870036-DEL-242 |
| chr3-91870635-DEL-3486 | chr3-91870636-DEL-3486 |
| chr3-91893642-DEL-56 | chr3-91893643-DEL-56 |
| chr3-91909352-DEL-124 | chr3-91909353-DEL-124 |
| chr3-91930869-DEL-204 | chr3-91930863-DEL-205 |
| chr3-91935246-DEL-3524 | chr3-91935247-DEL-3524 |
| chr3-91970562-DEL-54 | chr3-91970562-DEL-55 |
| chr3-92003592-DEL-74 | chr3-92003593-DEL-74 |
| chr3-92007685-DEL-153 | chr3-92007686-DEL-153 |
| chr3-92033944-DEL-73 | chr3-92033922-DEL-115 |
| chr3-92069476-DEL-2337 | chr3-92069477-DEL-2337 |
| chr3-92092700-DEL-7154 | chr3-92092701-DEL-7154 |
| chr3-92218354-DEL-155 | chr3-92218351-DEL-156 |
| chr3-92224837-DEL-53 | chr3-92224827-DEL-54 |
| chr3-92253925-DEL-396 | chr3-92253926-DEL-395 |
| chr3-92255067-DEL-1024 | chr3-92255068-DEL-1024 |
| chr3-92256059-DEL-682 | chr3-92256060-DEL-678 |
| chr3-92269962-DEL-70 | chr3-92269979-DEL-62 |
| chr3-92300560-DEL-86 | chr3-92300563-DEL-90 |
| chr3-92335324-DEL-1406 | chr3-92335325-DEL-1406 |
| chr3-92387081-DEL-363 | chr3-92387082-DEL-363 |
| chr3-92396718-DEL-71 | chr3-92396719-DEL-71 |
| chr3-92454185-DEL-7415 | chr3-92454186-DEL-7415 |
| chr3-92460452-DEL-832 | chr3-92460452-DEL-832 |
| chr3-92469625-DEL-1337 | chr3-92469626-DEL-1337 |
| chr3-92479211-DEL-87 | chr3-92479196-DEL-87 |
| chr3-92519273-DEL-56 | chr3-92519271-DEL-54 |
| chr3-92520856-DEL-921 | chr3-92520857-DEL-921 |
| chr3-92549963-DEL-70 | chr3-92549964-DEL-70 |
| chr3-92560486-DEL-148 | chr3-92560484-DEL-148 |
| chr3-92596587-DEL-1905 | chr3-92596588-DEL-1905 |
| chr3-92631490-DEL-126 | chr3-92631491-DEL-126 |
| chr3-92645179-DEL-60 | chr3-92645180-DEL-60 |
| chr3-92652617-DEL-416 | chr3-92652618-DEL-416 |
| chr3-92684750-DEL-60 | chr3-92684750-DEL-60 |
| chr3-92694176-DEL-120 | chr3-92694177-DEL-120 |
| chr3-92705162-DEL-604 | chr3-92705163-DEL-604 |
| chr3-92740225-DEL-369 | chr3-92740226-DEL-369 |
| chr3-92743552-DEL-320 | chr3-92743555-DEL-318 |
| chr3-92796410-DEL-52 | chr3-92796411-DEL-52 |
| chr3-92800213-DEL-53 | chr3-92800214-DEL-55 |
| chr3-92806273-DEL-81 | chr3-92806295-DEL-60 |
| chr3-92821277-DEL-1884 | chr3-92821278-DEL-1890 |
| chr3-92849915-DEL-1628 | chr3-92849914-DEL-1628 |
| chr3-92884832-DEL-4694 | chr3-92884833-DEL-4693 |
| chr3-92896721-DEL-1612 | chr3-92896722-DEL-1612 |
| chr3-92902825-DEL-55 | chr3-92902833-DEL-66 |
| chr3-92939171-DEL-609 | chr3-92939172-DEL-609 |
| chr3-92979843-DEL-401 | chr3-92979844-DEL-401 |
| chr3-92985970-DEL-7132 | chr3-92985971-DEL-7132 |
| chr3-93003698-DEL-52 | chr3-93003699-DEL-51 |
| chr3-93060677-DEL-70 | chr3-93060678-DEL-70 |
| chr3-93061970-DEL-83 | chr3-93061971-DEL-83 |
| chr3-93073810-DEL-384 | chr3-93073811-DEL-397 |
| chr3-93075813-DEL-4368 | chr3-93075814-DEL-4368 |
| chr3-93092218-DEL-2157 | chr3-93092219-DEL-2157 |
| chr3-93133951-DEL-66 | chr3-93133911-DEL-66 |
| chr3-93154064-DEL-51 | chr3-93154057-DEL-51 |
| chr3-93174185-DEL-266 | chr3-93174190-DEL-260 |
| chr3-93208380-DEL-93 | chr3-93208366-DEL-108 |
| chr3-93298196-DEL-385 | chr3-93298194-DEL-367 |
| chr3-93312990-DEL-160 | chr3-93312991-DEL-158 |
| chr3-93353126-DEL-120 | chr3-93353127-DEL-120 |
| chr3-93353997-DEL-54 | chr3-93353998-DEL-72 |
| chr3-93364266-DEL-214 | chr3-93364267-DEL-214 |
| chr3-93365482-DEL-2900 | chr3-93365483-DEL-2900 |
| chr3-93369185-DEL-538 | chr3-93369186-DEL-538 |
| chr3-93387875-DEL-53 | chr3-93387876-DEL-53 |
| chr3-93395787-DEL-60 | chr3-93395788-DEL-60 |
| chr3-93429795-DEL-52 | chr3-93429790-DEL-52 |

| | |
|---|---|
| chr3-93437750-DEL-1218 | chr3-93437751-DEL-1218 |
| chr3-93480671-DEL-184 | chr3-93480672-DEL-184 |
| chr3-93483590-DEL-60 | chr3-93483587-DEL-64 |
| chr3-93521491-DEL-6326 | chr3-93521492-DEL-6326 |
| chr3-93569201-DEL-52 | chr3-93569202-DEL-52 |
| chr3-93609570-DEL-560 | chr3-93609560-DEL-548 |
| chr3-93653298-DEL-210 | chr3-93653299-DEL-210 |
| chr3-93670055-DEL-172 | chr3-93670055-DEL-172 |
| chr3-93680139-DEL-57 | chr3-93680120-DEL-56 |
| chr3-93694119-DEL-54 | chr3-93694118-DEL-54 |
| chr3-93706593-DEL-1928 | chr3-93706593-DEL-1928 |
| chr3-93716084-DEL-1083 | chr3-93716087-DEL-1080 |
| chr3-93733069-DEL-74 | chr3-93733073-DEL-70 |
| chr3-93745376-DEL-668 | chr3-93745377-DEL-668 |
| chr3-93752733-DEL-78 | chr3-93752733-DEL-78 |
| chr3-93776656-DEL-54 | chr3-93776656-DEL-54 |
| chr3-93806020-DEL-72 | chr3-93806019-DEL-72 |
| chr3-93905259-DEL-498 | chr3-93905264-DEL-497 |
| chr3-93922158-DEL-62 | chr3-93922159-DEL-62 |
| chr3-93942168-DEL-268 | chr3-93942169-DEL-268 |
| chr3-93945217-DEL-205 | chr3-93945218-DEL-205 |
| chr3-94175680-DEL-6684 | chr3-94175681-DEL-6684 |
| chr3-94204913-DEL-178 | chr3-94204914-DEL-178 |
| chr3-94363113-DEL-93 | chr3-94363111-DEL-94 |
| chr3-94370358-DEL-194 | chr3-94370359-DEL-194 |
| chr3-94376066-DEL-56 | chr3-94376066-DEL-56 |
| chr3-94382657-DEL-99 | chr3-94382655-DEL-84 |
| chr3-94456869-DEL-6804 | chr3-94456870-DEL-6804 |
| chr3-94508042-DEL-141 | chr3-94508043-DEL-141 |
| chr3-94516518-DEL-213 | chr3-94516519-DEL-213 |
| chr3-94524727-DEL-215 | chr3-94524736-DEL-184 |
| chr3-94550441-DEL-1297 | chr3-94550442-DEL-1297 |
| chr3-94553574-DEL-52 | chr3-94553575-DEL-52 |
| chr3-94574147-DEL-126 | chr3-94574148-DEL-126 |
| chr3-94612577-DEL-213 | chr3-94612579-DEL-212 |
| chr3-94615126-DEL-162 | chr3-94615123-DEL-158 |
| chr3-94622811-DEL-153 | chr3-94622796-DEL-158 |
| chr3-94632958-DEL-6595 | chr3-94632955-DEL-6598 |
| chr3-94645213-DEL-208 | chr3-94645253-DEL-168 |
| chr3-94659304-DEL-63 | chr3-94659312-DEL-55 |
| chr3-94661626-DEL-208 | chr3-94661626-DEL-208 |
| chr3-94672060-DEL-67 | chr3-94672061-DEL-67 |
| chr3-94673573-DEL-327 | chr3-94673574-DEL-309 |
| chr3-94702285-DEL-76 | chr3-94702286-DEL-76 |
| chr3-94727620-DEL-192 | chr3-94727621-DEL-192 |
| chr3-94755635-DEL-102 | chr3-94755648-DEL-93 |
| chr3-94765335-DEL-113 | chr3-94765336-DEL-113 |
| chr3-94773616-DEL-54 | chr3-94773621-DEL-50 |
| chr3-94774630-DEL-98 | chr3-94774631-DEL-98 |
| chr3-94819833-DEL-188 | chr3-94819836-DEL-181 |
| chr3-94850421-DEL-205 | chr3-94850422-DEL-205 |
| chr3-94948033-DEL-217 | chr3-94948034-DEL-217 |
| chr3-94974481-DEL-60 | chr3-94974479-DEL-74 |
| chr3-94982755-DEL-319 | chr3-94982756-DEL-319 |
| chr3-95001770-DEL-124 | chr3-95001771-DEL-124 |
| chr3-95011648-DEL-181 | chr3-95011649-DEL-181 |
| chr3-95014862-DEL-205 | chr3-95014863-DEL-205 |
| chr3-95023806-DEL-205 | chr3-95023807-DEL-205 |
| chr3-95027991-DEL-204 | chr3-95027992-DEL-204 |
| chr3-95034083-DEL-213 | chr3-95034084-DEL-211 |
| chr3-95037964-DEL-209 | chr3-95037965-DEL-209 |
| chr3-95120439-DEL-215 | chr3-95120440-DEL-215 |
| chr3-95160403-DEL-67 | chr3-95160404-DEL-67 |
| chr3-95177912-DEL-219 | chr3-95177914-DEL-210 |
| chr3-95255739-DEL-203 | chr3-95255740-DEL-203 |
| chr3-95270889-DEL-6171 | chr3-95270886-DEL-6180 |
| chr3-95277840-DEL-205 | chr3-95277841-DEL-205 |
| chr3-95284120-DEL-168 | chr3-95284121-DEL-168 |
| chr3-95290073-DEL-69 | chr3-95290074-DEL-69 |
| chr3-95291152-DEL-270 | chr3-95291138-DEL-284 |
| chr3-95292582-DEL-245 | chr3-95292582-DEL-245 |
| chr3-95294796-DEL-1281 | chr3-95294797-DEL-1281 |
| chr3-95297015-DEL-162 | chr3-95296972-DEL-166 |
| chr3-95298735-DEL-2489 | chr3-95298736-DEL-2489 |
| chr3-95310934-DEL-139 | chr3-95310935-DEL-135 |
| chr3-95338668-DEL-495 | chr3-95338676-DEL-498 |
| chr3-95376297-DEL-1019 | chr3-95376296-DEL-1016 |
| chr3-95376416-DEL-127 | chr3-95376418-DEL-123 |
| chr3-95414905-DEL-66 | chr3-95414906-DEL-72 |
| chr3-95425139-DEL-146 | chr3-95425144-DEL-108 |

| | |
|---|---|
| chr3-95449675-DEL-106 | chr3-95449678-DEL-124 |
| chr3-95470824-DEL-116 | chr3-95470824-DEL-116 |
| chr3-95474270-DEL-175 | chr3-95474271-DEL-175 |
| chr3-95477244-DEL-221 | chr3-95477245-DEL-221 |
| chr3-95495182-DEL-197 | chr3-95495183-DEL-197 |
| chr3-95499865-DEL-204 | chr3-95499866-DEL-204 |
| chr3-95563749-DEL-62 | chr3-95563750-DEL-62 |
| chr3-95573115-DEL-207 | chr3-95573116-DEL-207 |
| chr3-95619750-DEL-201 | chr3-95619751-DEL-201 |
| chr3-95632778-DEL-213 | chr3-95632778-DEL-213 |
| chr3-95675007-DEL-54 | chr3-95675008-DEL-54 |
| chr3-95684113-DEL-80 | chr3-95684114-DEL-72 |
| chr3-95696008-DEL-132 | chr3-95696009-DEL-132 |
| chr3-95703295-DEL-203 | chr3-95703296-DEL-203 |
| chr3-95715036-DEL-202 | chr3-95715037-DEL-202 |
| chr3-95724336-DEL-69 | chr3-95724337-DEL-69 |
| chr3-95737286-DEL-189 | chr3-95737286-DEL-189 |
| chr3-95743161-DEL-58 | chr3-95743162-DEL-58 |
| chr3-95749252-DEL-204 | chr3-95749253-DEL-204 |
| chr3-95754194-DEL-163 | chr3-95754195-DEL-163 |
| chr3-95785589-DEL-210 | chr3-95785593-DEL-208 |
| chr3-95818817-DEL-59 | chr3-95818818-DEL-59 |
| chr3-95819629-DEL-361 | chr3-95819637-DEL-362 |
| chr3-95821813-DEL-179 | chr3-95821826-DEL-176 |
| chr3-95832211-DEL-218 | chr3-95832212-DEL-218 |
| chr3-95862450-DEL-102 | chr3-95862451-DEL-102 |
| chr3-95877937-DEL-209 | chr3-95877948-DEL-208 |
| chr3-95895293-DEL-114 | chr3-95895294-DEL-114 |
| chr3-95933578-DEL-120 | chr3-95933579-DEL-120 |
| chr3-95984857-DEL-202 | chr3-95984858-DEL-202 |
| chr3-96000387-DEL-113 | chr3-96000388-DEL-113 |
| chr3-96042951-DEL-215 | chr3-96042952-DEL-215 |
| chr3-96082934-DEL-223 | chr3-96082935-DEL-223 |
| chr3-96105789-DEL-328 | chr3-96105790-DEL-328 |
| chr3-96107554-DEL-6442 | chr3-96107555-DEL-6442 |
| chr3-96123303-DEL-208 | chr3-96123304-DEL-208 |
| chr3-96136562-DEL-57 | chr3-96136562-DEL-58 |
| chr3-96141787-DEL-579 | chr3-96141788-DEL-579 |
| chr3-96167243-DEL-77 | chr3-96167257-DEL-76 |
| chr3-96169918-DEL-218 | chr3-96169919-DEL-218 |
| chr3-96177623-DEL-3188 | chr3-96177624-DEL-3188 |
| chr3-96196592-DEL-437 | chr3-96196594-DEL-436 |
| chr3-96197123-DEL-141 | chr3-96197154-DEL-138 |
| chr3-96198906-DEL-168 | chr3-96198907-DEL-166 |
| chr3-96211764-DEL-60 | chr3-96211734-DEL-60 |
| chr3-96217634-DEL-375 | chr3-96217635-DEL-375 |
| chr3-96220772-DEL-7120 | chr3-96220773-DEL-7120 |
| chr3-96249630-DEL-201 | chr3-96249631-DEL-202 |
| chr3-96250547-DEL-120 | chr3-96250569-DEL-118 |
| chr3-96304093-DEL-88 | chr3-96304112-DEL-114 |
| chr3-96312603-DEL-181 | chr3-96312604-DEL-181 |
| chr3-96374804-DEL-97 | chr3-96374802-DEL-96 |
| chr3-96384127-DEL-331 | chr3-96384128-DEL-331 |
| chr3-96415516-DEL-7110 | chr3-96415517-DEL-7110 |
| chr3-96460953-DEL-69 | chr3-96460954-DEL-69 |
| chr3-96527754-DEL-269 | chr3-96527755-DEL-267 |
| chr3-96534382-DEL-215 | chr3-96534383-DEL-215 |
| chr3-96573255-DEL-217 | chr3-96573287-DEL-206 |
| chr3-96625424-DEL-199 | chr3-96625425-DEL-199 |
| chr3-96638370-DEL-89 | chr3-96638342-DEL-87 |
| chr3-96643107-DEL-211 | chr3-96643108-DEL-211 |
| chr3-96660875-DEL-65 | chr3-96660876-DEL-65 |
| chr3-96703842-DEL-177 | chr3-96703843-DEL-177 |
| chr3-96715229-DEL-64 | chr3-96715274-DEL-60 |
| chr3-96746216-DEL-171 | chr3-96746218-DEL-165 |
| chr3-96747137-DEL-51 | chr3-96747139-DEL-60 |
| chr3-96810075-DEL-424 | chr3-96810080-DEL-421 |
| chr3-96864561-DEL-85 | chr3-96864562-DEL-85 |
| chr3-96869857-DEL-75 | chr3-96869855-DEL-64 |
| chr3-96892236-DEL-199 | chr3-96892237-DEL-199 |
| chr3-96898731-DEL-430 | chr3-96898732-DEL-430 |
| chr3-96933684-DEL-72 | chr3-96933685-DEL-72 |
| chr3-96956017-DEL-58 | chr3-96956019-DEL-56 |
| chr3-96979122-DEL-116 | chr3-96979123-DEL-116 |
| chr3-96989392-DEL-349 | chr3-96989393-DEL-349 |
| chr3-96993254-DEL-220 | chr3-96993255-DEL-220 |
| chr3-96994099-DEL-2711 | chr3-96994100-DEL-2711 |
| chr3-97004391-DEL-400 | chr3-97004392-DEL-400 |
| chr3-97011015-DEL-402 | chr3-97011014-DEL-403 |
| chr3-97035143-DEL-635 | chr3-97035137-DEL-638 |

| | |
|---|---|
| chr3-97050642-DEL-6179 | chr3-97050643-DEL-6179 |
| chr3-97063633-DEL-158 | chr3-97063634-DEL-158 |
| chr3-97081433-DEL-204 | chr3-97081434-DEL-204 |
| chr3-97085646-DEL-194 | chr3-97085635-DEL-195 |
| chr3-97097493-DEL-468 | chr3-97097494-DEL-468 |
| chr3-97098803-DEL-118 | chr3-97098772-DEL-150 |
| chr3-97145556-DEL-71 | chr3-97145553-DEL-92 |
| chr3-97152461-DEL-6413 | chr3-97152461-DEL-6413 |
| chr3-97187090-DEL-55 | chr3-97187091-DEL-55 |
| chr3-97222528-DEL-59 | chr3-97222542-DEL-57 |
| chr3-97227167-DEL-165 | chr3-97227168-DEL-165 |
| chr3-97236984-DEL-54 | chr3-97236985-DEL-54 |
| chr3-97273494-DEL-206 | chr3-97273495-DEL-206 |
| chr3-97275192-DEL-760 | chr3-97275193-DEL-760 |
| chr3-97284969-DEL-4756 | chr3-97284970-DEL-4756 |
| chr3-97303955-DEL-631 | chr3-97303956-DEL-631 |
| chr3-97327524-DEL-52 | chr3-97327527-DEL-50 |
| chr3-97330369-DEL-51 | chr3-97330370-DEL-51 |
| chr3-97334200-DEL-985 | chr3-97334204-DEL-984 |
| chr3-97336081-DEL-382 | chr3-97336082-DEL-382 |
| chr3-97348850-DEL-6241 | chr3-97348851-DEL-6241 |
| chr3-97365926-DEL-129 | chr3-97365927-DEL-129 |
| chr3-97371146-DEL-88 | chr3-97371147-DEL-88 |
| chr3-97397179-DEL-67 | chr3-97397179-DEL-68 |
| chr3-97398316-DEL-54 | chr3-97398317-DEL-55 |
| chr3-97455634-DEL-62 | chr3-97455635-DEL-62 |
| chr3-97465991-DEL-55 | chr3-97466009-DEL-52 |
| chr3-97478483-DEL-51 | chr3-97478484-DEL-51 |
| chr3-97507713-DEL-459 | chr3-97507739-DEL-428 |
| chr3-97518420-DEL-113 | chr3-97518421-DEL-113 |
| chr3-97539336-DEL-180 | chr3-97539383-DEL-134 |
| chr3-97619005-DEL-57 | chr3-97619006-DEL-57 |
| chr3-97651382-DEL-64 | chr3-97651383-DEL-64 |
| chr3-97662531-DEL-51 | chr3-97662532-DEL-51 |
| chr3-97692800-DEL-108 | chr3-97692803-DEL-96 |
| chr3-97707705-DEL-80 | chr3-97707706-DEL-80 |
| chr3-97711968-DEL-126 | chr3-97711972-DEL-113 |
| chr3-97713865-DEL-82 | chr3-97713898-DEL-50 |
| chr3-97726733-DEL-96 | chr3-97726708-DEL-90 |
| chr3-97729008-DEL-211 | chr3-97729009-DEL-211 |
| chr3-97739960-DEL-296 | chr3-97739961-DEL-296 |
| chr3-97755547-DEL-54 | chr3-97755548-DEL-54 |
| chr3-97763517-DEL-1310 | chr3-97763518-DEL-1310 |
| chr3-97788057-DEL-929 | chr3-97788058-DEL-929 |
| chr3-97789298-DEL-163 | chr3-97789299-DEL-163 |
| chr3-97793711-DEL-54 | chr3-97793712-DEL-54 |
| chr3-97824506-DEL-240 | chr3-97824507-DEL-240 |
| chr3-97846686-DEL-72 | chr3-97846689-DEL-60 |
| chr3-97854314-DEL-381 | chr3-97854315-DEL-381 |
| chr3-97866420-DEL-289 | chr3-97866421-DEL-289 |
| chr3-97868001-DEL-442 | chr3-97868007-DEL-436 |
| chr3-97910214-DEL-190 | chr3-97910215-DEL-179 |
| chr3-97921938-DEL-123 | chr3-97921939-DEL-123 |
| chr3-97980629-DEL-64 | chr3-97980627-DEL-65 |
| chr3-97981599-DEL-106 | chr3-97981600-DEL-106 |
| chr3-98018437-DEL-101 | chr3-98018403-DEL-101 |
| chr3-98032846-DEL-51 | chr3-98032847-DEL-51 |
| chr3-98048343-DEL-73 | chr3-98048344-DEL-73 |
| chr3-98109401-DEL-73 | chr3-98109402-DEL-73 |
| chr3-98123942-DEL-52 | chr3-98123943-DEL-52 |
| chr3-98153703-DEL-69 | chr3-98153718-DEL-66 |
| chr3-98163631-DEL-481 | chr3-98163632-DEL-481 |
| chr3-98191015-DEL-185 | chr3-98191016-DEL-185 |
| chr3-98210332-DEL-76 | chr3-98210333-DEL-76 |
| chr3-98211058-DEL-4100 | chr3-98211059-DEL-4100 |
| chr3-98229107-DEL-452 | chr3-98229108-DEL-452 |
| chr3-98264871-DEL-147 | chr3-98264872-DEL-147 |
| chr3-98293902-DEL-70 | chr3-98293903-DEL-70 |
| chr3-98309576-DEL-9085 | chr3-98309573-DEL-9086 |
| chr3-98466509-DEL-65 | chr3-98466549-DEL-65 |
| chr3-98512978-DEL-76 | chr3-98512972-DEL-72 |
| chr3-98519488-DEL-103 | chr3-98519489-DEL-103 |
| chr3-98525296-DEL-51 | chr3-98525276-DEL-52 |
| chr3-98544122-DEL-6770 | chr3-98544123-DEL-6770 |
| chr3-98567063-DEL-1577 | chr3-98567063-DEL-1577 |
| chr3-98583824-DEL-402 | chr3-98583825-DEL-402 |
| chr3-98611704-DEL-1304 | chr3-98611705-DEL-1304 |
| chr3-98624118-DEL-142 | chr3-98624119-DEL-142 |
| chr3-98636719-DEL-339 | chr3-98636720-DEL-339 |
| chr3-98644256-DEL-55 | chr3-98644257-DEL-83 |

| | |
|---|---|
| chr3-98670809-DEL-58 | chr3-98670800-DEL-70 |
| chr3-98680446-DEL-6246 | chr3-98680447-DEL-6246 |
| chr3-98742520-DEL-5643 | chr3-98742521-DEL-5643 |
| chr3-98751759-DEL-102 | chr3-98751780-DEL-82 |
| chr3-98754087-DEL-668 | chr3-98754088-DEL-668 |
| chr3-98770679-DEL-3342 | chr3-98770680-DEL-3342 |
| chr3-98782908-DEL-82 | chr3-98782909-DEL-84 |
| chr3-98798245-DEL-955 | chr3-98798246-DEL-955 |
| chr3-98832420-DEL-93 | chr3-98832421-DEL-93 |
| chr3-98841976-DEL-173 | chr3-98841977-DEL-173 |
| chr3-98849180-DEL-273 | chr3-98849181-DEL-273 |
| chr3-98851529-DEL-112 | chr3-98851529-DEL-112 |
| chr3-98860178-DEL-61 | chr3-98860183-DEL-55 |
| chr3-98885233-DEL-2624 | chr3-98885235-DEL-2621 |
| chr3-98906908-DEL-63 | chr3-98906909-DEL-63 |
| chr3-98911048-DEL-10834 | chr3-98911073-DEL-10809 |
| chr3-98913840-DEL-4634 | chr3-98913840-DEL-4634 |
| chr3-98923005-DEL-54 | chr3-98923006-DEL-54 |
| chr3-98936948-DEL-396 | chr3-98936962-DEL-368 |
| chr3-98969784-DEL-2413 | chr3-98969785-DEL-2413 |
| chr3-98978761-DEL-53 | chr3-98978762-DEL-53 |
| chr3-98983634-DEL-3422 | chr3-98983635-DEL-3422 |
| chr3-98993199-DEL-278 | chr3-98993200-DEL-278 |
| chr3-99002318-DEL-218 | chr3-99002319-DEL-218 |
| chr3-99024949-DEL-192 | chr3-99024950-DEL-192 |
| chr3-99034858-DEL-87 | chr3-99034838-DEL-58 |
| chr3-99074309-DEL-843 | chr3-99074310-DEL-843 |
| chr3-99104790-DEL-132 | chr3-99104788-DEL-133 |
| chr3-99106176-DEL-1351 | chr3-99106177-DEL-1351 |
| chr3-99126568-DEL-80 | chr3-99126566-DEL-81 |
| chr3-99194565-DEL-56 | chr3-99194566-DEL-56 |
| chr3-99222275-DEL-225 | chr3-99222276-DEL-225 |
| chr3-99237326-DEL-58 | chr3-99237327-DEL-58 |
| chr3-99264520-DEL-52 | chr3-99264521-DEL-52 |
| chr3-99265308-DEL-6842 | chr3-99265308-DEL-6842 |
| chr3-99271854-DEL-214 | chr3-99271900-DEL-214 |
| chr3-99278862-DEL-151 | chr3-99278865-DEL-152 |
| chr3-99294753-DEL-184 | chr3-99294754-DEL-184 |
| chr3-99304996-DEL-389 | chr3-99304997-DEL-389 |
| chr3-99305656-DEL-53 | chr3-99305657-DEL-53 |
| chr3-99322688-DEL-7077 | chr3-99322689-DEL-7077 |
| chr3-99364357-DEL-175 | chr3-99364381-DEL-156 |
| chr3-99389928-DEL-316 | chr3-99389929-DEL-316 |
| chr3-99407020-DEL-316 | chr3-99407021-DEL-316 |
| chr3-99412067-DEL-125 | chr3-99412068-DEL-125 |
| chr3-99417464-DEL-1222 | chr3-99417465-DEL-1222 |
| chr3-99427474-DEL-453 | chr3-99427474-DEL-454 |
| chr3-99448742-DEL-240 | chr3-99448743-DEL-240 |
| chr3-99449468-DEL-264 | chr3-99449480-DEL-255 |
| chr3-99475226-DEL-337 | chr3-99475225-DEL-335 |
| chr3-99496714-DEL-74 | chr3-99496715-DEL-74 |
| chr3-99519980-DEL-6399 | chr3-99519980-DEL-6399 |
| chr3-99547162-DEL-126 | chr3-99547163-DEL-126 |
| chr3-99548268-DEL-56 | chr3-99548269-DEL-56 |
| chr3-99569615-DEL-54 | chr3-99569616-DEL-54 |
| chr3-99591311-DEL-123 | chr3-99591312-DEL-123 |
| chr3-99600068-DEL-667 | chr3-99600069-DEL-667 |
| chr3-99605681-DEL-65 | chr3-99605683-DEL-61 |
| chr3-99606726-DEL-1017 | chr3-99606727-DEL-1017 |
| chr3-99610492-DEL-4671 | chr3-99610493-DEL-4671 |
| chr3-99653648-DEL-194 | chr3-99653649-DEL-194 |
| chr3-99661103-DEL-74 | chr3-99661110-DEL-53 |
| chr3-99663390-DEL-2699 | chr3-99663391-DEL-2699 |
| chr3-99668208-DEL-544 | chr3-99668209-DEL-544 |
| chr3-99790224-DEL-1145 | chr3-99790225-DEL-1145 |
| chr3-99791487-DEL-64 | chr3-99791488-DEL-64 |
| chr3-99793265-DEL-70 | chr3-99793266-DEL-70 |
| chr3-99800362-DEL-1723 | chr3-99800363-DEL-1720 |
| chr3-99808958-DEL-6397 | chr3-99808959-DEL-6397 |
| chr3-99879048-DEL-57 | chr3-99879049-DEL-57 |
| chr3-99881425-DEL-158 | chr3-99881426-DEL-158 |
| chr3-99885177-DEL-96 | chr3-99885178-DEL-96 |
| chr3-99888410-DEL-292 | chr3-99888410-DEL-292 |
| chr3-99951454-DEL-417 | chr3-99951455-DEL-417 |
| chr3-99952539-DEL-859 | chr3-99952521-DEL-877 |
| chr3-99987974-DEL-126 | chr3-99987975-DEL-128 |
| chr3-99996933-DEL-1188 | chr3-99996934-DEL-1188 |
| chr3-100012685-DEL-469 | chr3-100012686-DEL-469 |
| chr3-100014297-DEL-1089 | chr3-100014299-DEL-1090 |
| chr3-100025613-DEL-1028 | chr3-100025614-DEL-1028 |

| | |
|---|---|
| chr3-100026947-DEL-393 | chr3-100026957-DEL-378 |
| chr3-100032854-DEL-6914 | chr3-100032855-DEL-6914 |
| chr3-100038993-DEL-212 | chr3-100038994-DEL-212 |
| chr3-100070587-DEL-66 | chr3-100070588-DEL-66 |
| chr3-100074137-DEL-63 | chr3-100074138-DEL-63 |
| chr3-100083706-DEL-60 | chr3-100083710-DEL-86 |
| chr3-100086043-DEL-87 | chr3-100086044-DEL-87 |
| chr3-100124736-DEL-83 | chr3-100124718-DEL-66 |
| chr3-100130430-DEL-70 | chr3-100130380-DEL-117 |
| chr3-100143629-DEL-69 | chr3-100143630-DEL-69 |
| chr3-100154541-DEL-1702 | chr3-100154542-DEL-1702 |
| chr3-100216675-DEL-4306 | chr3-100216676-DEL-4306 |
| chr3-100223760-DEL-95 | chr3-100223759-DEL-96 |
| chr3-100285981-DEL-1144 | chr3-100285982-DEL-1144 |
| chr3-100298278-DEL-495 | chr3-100298279-DEL-495 |
| chr3-100311854-DEL-92 | chr3-100311874-DEL-72 |
| chr3-100342707-DEL-56 | chr3-100342712-DEL-55 |
| chr3-100343010-DEL-898 | chr3-100343011-DEL-898 |
| chr3-100354270-DEL-322 | chr3-100354271-DEL-322 |
| chr3-100439692-DEL-186 | chr3-100439694-DEL-175 |
| chr3-100449035-DEL-70 | chr3-100449036-DEL-70 |
| chr3-100452008-DEL-133 | chr3-100452009-DEL-133 |
| chr3-100465215-DEL-65 | chr3-100465215-DEL-68 |
| chr3-100475600-DEL-66 | chr3-100475601-DEL-66 |
| chr3-100499958-DEL-1463 | chr3-100499961-DEL-1464 |
| chr3-100531972-DEL-68 | chr3-100531941-DEL-92 |
| chr3-100538048-DEL-75 | chr3-100538049-DEL-75 |
| chr3-100568126-DEL-276 | chr3-100568127-DEL-276 |
| chr3-100575607-DEL-206 | chr3-100575608-DEL-206 |
| chr3-100598996-DEL-60 | chr3-100598995-DEL-56 |
| chr3-100634461-DEL-56 | chr3-100634462-DEL-56 |
| chr3-100647586-DEL-64 | chr3-100647590-DEL-50 |
| chr3-100648473-DEL-308 | chr3-100648467-DEL-309 |
| chr3-100650529-DEL-137 | chr3-100650530-DEL-137 |
| chr3-100651514-DEL-99 | chr3-100651515-DEL-99 |
| chr3-100671370-DEL-137 | chr3-100671385-DEL-120 |
| chr3-100705715-DEL-237 | chr3-100705716-DEL-237 |
| chr3-100721953-DEL-60 | chr3-100721917-DEL-76 |
| chr3-100724530-DEL-181 | chr3-100724531-DEL-181 |
| chr3-100727138-DEL-3310 | chr3-100727139-DEL-3310 |
| chr3-100745483-DEL-207 | chr3-100745484-DEL-207 |
| chr3-100750522-DEL-190 | chr3-100750523-DEL-190 |
| chr3-100754583-DEL-211 | chr3-100754584-DEL-211 |
| chr3-100794216-DEL-79 | chr3-100794217-DEL-79 |
| chr3-100806377-DEL-93 | chr3-100806378-DEL-93 |
| chr3-100819660-DEL-56 | chr3-100819661-DEL-56 |
| chr3-100825117-DEL-96 | chr3-100825118-DEL-96 |
| chr3-100835031-DEL-61 | chr3-100835008-DEL-66 |
| chr3-100847683-DEL-153 | chr3-100847684-DEL-159 |
| chr3-100908365-DEL-665 | chr3-100908366-DEL-665 |
| chr3-100968863-DEL-52 | chr3-100968877-DEL-72 |
| chr3-100992449-DEL-328 | chr3-100992450-DEL-328 |
| chr3-100994666-DEL-387 | chr3-100994667-DEL-387 |
| chr3-101022112-DEL-603 | chr3-101022113-DEL-639 |
| chr3-101025961-DEL-2563 | chr3-101025962-DEL-2563 |
| chr3-101038607-DEL-116 | chr3-101038608-DEL-116 |
| chr3-101060424-DEL-56 | chr3-101060423-DEL-58 |
| chr3-101099810-DEL-1711 | chr3-101099811-DEL-1711 |
| chr3-101123227-DEL-78 | chr3-101123228-DEL-78 |
| chr3-101130852-DEL-62 | chr3-101130853-DEL-62 |
| chr3-101145533-DEL-200 | chr3-101145534-DEL-200 |
| chr3-101198047-DEL-113 | chr3-101198048-DEL-113 |
| chr3-101199519-DEL-198 | chr3-101199510-DEL-204 |
| chr3-101213704-DEL-804 | chr3-101213705-DEL-804 |
| chr3-101216452-DEL-131 | chr3-101216472-DEL-160 |
| chr3-101224276-DEL-55 | chr3-101224276-DEL-56 |
| chr3-101384128-DEL-240 | chr3-101384129-DEL-240 |
| chr3-101396040-DEL-163 | chr3-101396041-DEL-163 |
| chr3-101412521-DEL-454 | chr3-101412522-DEL-454 |
| chr3-101432406-DEL-274 | chr3-101432407-DEL-274 |
| chr3-101444991-DEL-119 | chr3-101445041-DEL-69 |
| chr3-101455824-DEL-66 | chr3-101455825-DEL-66 |
| chr3-101467038-DEL-399 | chr3-101467039-DEL-399 |
| chr3-101471367-DEL-66 | chr3-101471368-DEL-82 |
| chr3-101530595-DEL-55 | chr3-101530593-DEL-51 |
| chr3-101582799-DEL-78 | chr3-101582800-DEL-78 |
| chr3-101584084-DEL-92 | chr3-101584100-DEL-76 |
| chr3-101584631-DEL-55 | chr3-101584632-DEL-55 |
| chr3-101606392-DEL-177 | chr3-101606393-DEL-177 |
| chr3-101613372-DEL-116 | chr3-101613373-DEL-116 |

| | |
|---|---|
| chr3-101623679-DEL-90 | chr3-101623680-DEL-90 |
| chr3-101626660-DEL-63 | chr3-101626642-DEL-102 |
| chr3-101638959-DEL-70 | chr3-101638960-DEL-70 |
| chr3-101661534-DEL-1113 | chr3-101661535-DEL-1113 |
| chr3-101674593-DEL-541 | chr3-101674597-DEL-537 |
| chr3-101679506-DEL-143 | chr3-101679544-DEL-141 |
| chr3-101745006-DEL-217 | chr3-101745007-DEL-217 |
| chr3-101752187-DEL-361 | chr3-101752185-DEL-362 |
| chr3-101778529-DEL-115 | chr3-101778530-DEL-115 |
| chr3-101795945-DEL-6393 | chr3-101795946-DEL-6393 |
| chr3-101815188-DEL-166 | chr3-101815203-DEL-152 |
| chr3-101828307-DEL-172 | chr3-101828308-DEL-172 |
| chr3-101842540-DEL-1417 | chr3-101842541-DEL-1417 |
| chr3-101883018-DEL-206 | chr3-101883019-DEL-205 |
| chr3-101903263-DEL-222 | chr3-101903264-DEL-222 |
| chr3-101959894-DEL-561 | chr3-101959895-DEL-561 |
| chr3-101962066-DEL-221 | chr3-101962067-DEL-221 |
| chr3-101987670-DEL-670 | chr3-101987671-DEL-670 |
| chr3-102010219-DEL-586 | chr3-102010220-DEL-586 |
| chr3-102010854-DEL-67 | chr3-102010853-DEL-67 |
| chr3-102012952-DEL-1286 | chr3-102012953-DEL-1286 |
| chr3-102016358-DEL-99 | chr3-102016357-DEL-100 |
| chr3-102017745-DEL-205 | chr3-102017746-DEL-205 |
| chr3-102070129-DEL-132 | chr3-102070130-DEL-132 |
| chr3-102112203-DEL-119 | chr3-102112203-DEL-119 |
| chr3-102114795-DEL-198 | chr3-102114795-DEL-197 |
| chr3-102122220-DEL-80 | chr3-102122221-DEL-80 |
| chr3-102127919-DEL-165 | chr3-102127920-DEL-165 |
| chr3-102162917-DEL-260 | chr3-102162878-DEL-280 |
| chr3-102186029-DEL-60 | chr3-102186030-DEL-60 |
| chr3-102243303-DEL-86 | chr3-102243304-DEL-86 |
| chr3-102300662-DEL-6410 | chr3-102300663-DEL-6410 |
| chr3-102310936-DEL-94 | chr3-102310937-DEL-94 |
| chr3-102336978-DEL-6714 | chr3-102336979-DEL-6714 |
| chr3-102412981-DEL-63 | chr3-102412981-DEL-63 |
| chr3-102445886-DEL-111 | chr3-102445893-DEL-114 |
| chr3-102449269-DEL-93 | chr3-102449270-DEL-93 |
| chr3-102465978-DEL-63 | chr3-102465979-DEL-63 |
| chr3-102487750-DEL-1278 | chr3-102487751-DEL-1278 |
| chr3-102503555-DEL-51 | chr3-102503556-DEL-57 |
| chr3-102509392-DEL-114 | chr3-102509431-DEL-76 |
| chr3-102532680-DEL-2178 | chr3-102532691-DEL-2176 |
| chr3-102553359-DEL-126 | chr3-102553360-DEL-126 |
| chr3-102569193-DEL-63 | chr3-102569194-DEL-63 |
| chr3-102586460-DEL-180 | chr3-102586461-DEL-180 |
| chr3-102688500-DEL-361 | chr3-102688501-DEL-361 |
| chr3-102739954-DEL-6688 | chr3-102739955-DEL-6689 |
| chr3-102787910-DEL-12298 | chr3-102787910-DEL-12298 |
| chr3-102793596-DEL-55 | chr3-102793597-DEL-55 |
| chr3-102849573-DEL-208 | chr3-102849574-DEL-208 |
| chr3-102861234-DEL-91 | chr3-102861235-DEL-91 |
| chr3-102890835-DEL-6644 | chr3-102890836-DEL-6644 |
| chr3-102930527-DEL-541 | chr3-102930522-DEL-540 |
| chr3-102953578-DEL-212 | chr3-102953579-DEL-212 |
| chr3-102977383-DEL-68 | chr3-102977384-DEL-68 |
| chr3-102980750-DEL-52 | chr3-102980780-DEL-54 |
| chr3-102990975-DEL-189 | chr3-102990975-DEL-189 |
| chr3-102993865-DEL-280 | chr3-102993862-DEL-281 |
| chr3-102995393-DEL-88 | chr3-102995394-DEL-88 |
| chr3-103000290-DEL-64 | chr3-103000291-DEL-64 |
| chr3-103002708-DEL-54 | chr3-103002709-DEL-54 |
| chr3-103022321-DEL-73 | chr3-103022346-DEL-98 |
| chr3-103042392-DEL-329 | chr3-103042373-DEL-310 |
| chr3-103086278-DEL-199 | chr3-103086279-DEL-199 |
| chr3-103094552-DEL-255 | chr3-103094551-DEL-253 |
| chr3-103096834-DEL-207 | chr3-103096835-DEL-207 |
| chr3-103123725-DEL-240 | chr3-103123726-DEL-238 |
| chr3-103145060-DEL-401 | chr3-103145061-DEL-401 |
| chr3-103146292-DEL-470 | chr3-103146293-DEL-470 |
| chr3-103148179-DEL-214 | chr3-103148180-DEL-214 |
| chr3-103154209-DEL-56 | chr3-103154210-DEL-56 |
| chr3-103164215-DEL-86 | chr3-103164216-DEL-84 |
| chr3-103204023-DEL-67 | chr3-103204028-DEL-65 |
| chr3-103249155-DEL-215 | chr3-103249156-DEL-215 |
| chr3-103287713-DEL-192 | chr3-103287714-DEL-192 |
| chr3-103289661-DEL-420 | chr3-103289662-DEL-420 |
| chr3-103317234-DEL-66 | chr3-103317235-DEL-66 |
| chr3-103321228-DEL-629 | chr3-103321229-DEL-629 |
| chr3-103323398-DEL-72 | chr3-103323399-DEL-72 |
| chr3-103334735-DEL-1895 | chr3-103334736-DEL-1895 |

| | |
|---|---|
| chr3-103338044-DEL-351 | chr3-103338045-DEL-351 |
| chr3-103395419-DEL-89 | chr3-103395420-DEL-89 |
| chr3-103400696-DEL-164 | chr3-103400697-DEL-164 |
| chr3-103401495-DEL-492 | chr3-103401496-DEL-492 |
| chr3-103432594-DEL-643 | chr3-103432595-DEL-641 |
| chr3-103435243-DEL-60 | chr3-103435254-DEL-50 |
| chr3-103435634-DEL-181 | chr3-103435635-DEL-181 |
| chr3-103462336-DEL-1491 | chr3-103462337-DEL-1491 |
| chr3-103465281-DEL-54 | chr3-103465262-DEL-52 |
| chr3-103485545-DEL-62 | chr3-103485546-DEL-62 |
| chr3-103498826-DEL-3230 | chr3-103498827-DEL-3230 |
| chr3-103515573-DEL-1771 | chr3-103515574-DEL-1771 |
| chr3-103525502-DEL-93 | chr3-103525503-DEL-93 |
| chr3-103551237-DEL-2148 | chr3-103551238-DEL-2148 |
| chr3-103564194-DEL-450 | chr3-103564195-DEL-450 |
| chr3-103583035-DEL-67 | chr3-103583035-DEL-68 |
| chr3-103670896-DEL-57 | chr3-103670869-DEL-74 |
| chr3-103681641-DEL-65 | chr3-103681642-DEL-65 |
| chr3-103695450-DEL-205 | chr3-103695451-DEL-205 |
| chr3-103757576-DEL-109 | chr3-103757577-DEL-109 |
| chr3-103774520-DEL-136 | chr3-103774521-DEL-136 |
| chr3-103810935-DEL-87 | chr3-103810936-DEL-85 |
| chr3-103824903-DEL-214 | chr3-103824904-DEL-214 |
| chr3-103842233-DEL-187 | chr3-103842234-DEL-187 |
| chr3-104034235-DEL-206 | chr3-104034236-DEL-206 |
| chr3-104046560-DEL-572 | chr3-104046562-DEL-570 |
| chr3-104181766-DEL-204 | chr3-104181767-DEL-204 |
| chr3-104188792-DEL-51 | chr3-104188793-DEL-51 |
| chr3-104192146-DEL-56 | chr3-104192147-DEL-56 |
| chr3-104346660-DEL-63 | chr3-104346666-DEL-84 |
| chr3-104357104-DEL-210 | chr3-104357105-DEL-210 |
| chr3-104414559-DEL-1088 | chr3-104414559-DEL-1089 |
| chr3-104450743-DEL-268 | chr3-104450744-DEL-268 |
| chr3-104454886-DEL-440 | chr3-104454887-DEL-440 |
| chr3-104495499-DEL-7214 | chr3-104495500-DEL-7214 |
| chr3-104529756-DEL-112 | chr3-104529757-DEL-112 |
| chr3-104532242-DEL-1892 | chr3-104532243-DEL-1892 |
| chr3-104573180-DEL-198 | chr3-104573181-DEL-198 |
| chr3-104601622-DEL-1885 | chr3-104601623-DEL-1885 |
| chr3-104626591-DEL-206 | chr3-104626592-DEL-206 |
| chr3-104646698-DEL-1490 | chr3-104646699-DEL-1490 |
| chr3-104648762-DEL-64 | chr3-104648771-DEL-56 |
| chr3-104652045-DEL-1228 | chr3-104652046-DEL-1228 |
| chr3-104658855-DEL-1529 | chr3-104658856-DEL-1524 |
| chr3-104660457-DEL-377 | chr3-104660458-DEL-377 |
| chr3-104669676-DEL-262 | chr3-104669677-DEL-262 |
| chr3-104674062-DEL-198 | chr3-104674062-DEL-198 |
| chr3-104682691-DEL-156 | chr3-104682700-DEL-144 |
| chr3-104705878-DEL-444 | chr3-104705879-DEL-444 |
| chr3-104710477-DEL-212 | chr3-104710478-DEL-212 |
| chr3-104716292-DEL-1894 | chr3-104716293-DEL-1894 |
| chr3-104729625-DEL-215 | chr3-104729626-DEL-215 |
| chr3-104733650-DEL-54 | chr3-104733650-DEL-54 |
| chr3-104749271-DEL-246 | chr3-104749274-DEL-244 |
| chr3-104774218-DEL-225 | chr3-104774219-DEL-225 |
| chr3-104790778-DEL-206 | chr3-104790779-DEL-206 |
| chr3-104813411-DEL-64 | chr3-104813412-DEL-64 |
| chr3-104823096-DEL-54 | chr3-104823097-DEL-54 |
| chr3-104864964-DEL-54 | chr3-104864965-DEL-54 |
| chr3-104866748-DEL-119 | chr3-104866750-DEL-120 |
| chr3-104874492-DEL-52 | chr3-104874493-DEL-52 |
| chr3-104880895-DEL-203 | chr3-104880896-DEL-203 |
| chr3-104882860-DEL-3844 | chr3-104882861-DEL-3844 |
| chr3-104888757-DEL-164 | chr3-104888760-DEL-165 |
| chr3-104937635-DEL-54 | chr3-104937636-DEL-54 |
| chr3-104965074-DEL-202 | chr3-104965075-DEL-202 |
| chr3-104976724-DEL-189 | chr3-104976725-DEL-189 |
| chr3-104990859-DEL-283 | chr3-104990860-DEL-283 |
| chr3-105012878-DEL-103 | chr3-105012879-DEL-103 |
| chr3-105016345-DEL-615 | chr3-105016346-DEL-615 |
| chr3-105019362-DEL-227 | chr3-105019363-DEL-227 |
| chr3-105039264-DEL-370 | chr3-105039265-DEL-370 |
| chr3-105043704-DEL-66 | chr3-105043705-DEL-66 |
| chr3-105076022-DEL-108 | chr3-105076023-DEL-103 |
| chr3-105126622-DEL-60 | chr3-105126623-DEL-60 |
| chr3-105128800-DEL-56 | chr3-105128801-DEL-52 |
| chr3-105132792-DEL-1863 | chr3-105132793-DEL-1863 |
| chr3-105133892-DEL-109 | chr3-105133880-DEL-114 |
| chr3-105143217-DEL-433 | chr3-105143217-DEL-434 |
| chr3-105153865-DEL-6181 | chr3-105153866-DEL-6181 |

| | |
|---|---|
| chr3-105191221-DEL-64 | chr3-105191207-DEL-91 |
| chr3-105192018-DEL-567 | chr3-105192019-DEL-567 |
| chr3-105239094-DEL-60 | chr3-105239052-DEL-69 |
| chr3-105240808-DEL-225 | chr3-105240809-DEL-225 |
| chr3-105244847-DEL-6555 | chr3-105244849-DEL-6554 |
| chr3-105252623-DEL-2070 | chr3-105252624-DEL-2069 |
| chr3-105265685-DEL-857 | chr3-105265686-DEL-857 |
| chr3-105283325-DEL-4947 | chr3-105283326-DEL-4947 |
| chr3-105330245-DEL-260 | chr3-105330244-DEL-261 |
| chr3-105334630-DEL-69 | chr3-105334631-DEL-69 |
| chr3-105370024-DEL-56 | chr3-105370025-DEL-56 |
| chr3-105381628-DEL-90 | chr3-105381661-DEL-76 |
| chr3-105430197-DEL-222 | chr3-105430198-DEL-222 |
| chr3-105491452-DEL-777 | chr3-105491453-DEL-777 |
| chr3-105515908-DEL-85 | chr3-105515909-DEL-85 |
| chr3-105518013-DEL-195 | chr3-105518014-DEL-195 |
| chr3-105531499-DEL-52 | chr3-105531500-DEL-56 |
| chr3-105582752-DEL-345 | chr3-105582752-DEL-333 |
| chr3-105592205-DEL-213 | chr3-105592206-DEL-213 |
| chr3-105691450-DEL-206 | chr3-105691451-DEL-206 |
| chr3-105743954-DEL-180 | chr3-105743955-DEL-180 |
| chr3-105755549-DEL-130 | chr3-105755549-DEL-130 |
| chr3-105763711-DEL-228 | chr3-105763712-DEL-228 |
| chr3-105776331-DEL-163 | chr3-105776332-DEL-163 |
| chr3-105820488-DEL-69 | chr3-105820488-DEL-70 |
| chr3-105837203-DEL-66 | chr3-105837204-DEL-66 |
| chr3-105855609-DEL-368 | chr3-105855610-DEL-368 |
| chr3-105891214-DEL-62 | chr3-105891215-DEL-62 |
| chr3-105901040-DEL-312 | chr3-105901041-DEL-312 |
| chr3-105910911-DEL-372 | chr3-105910912-DEL-372 |
| chr3-105940800-DEL-511 | chr3-105940801-DEL-511 |
| chr3-105984811-DEL-54 | chr3-105984814-DEL-52 |
| chr3-105999377-DEL-603 | chr3-105999378-DEL-603 |
| chr3-106022819-DEL-115 | chr3-106022832-DEL-103 |
| chr3-106049245-DEL-199 | chr3-106049246-DEL-199 |
| chr3-106147806-DEL-576 | chr3-106147807-DEL-576 |
| chr3-106173514-DEL-6790 | chr3-106173514-DEL-6790 |
| chr3-106192461-DEL-6605 | chr3-106192462-DEL-6605 |
| chr3-106231963-DEL-374 | chr3-106231964-DEL-374 |
| chr3-106313027-DEL-56 | chr3-106313027-DEL-56 |
| chr3-106321755-DEL-6477 | chr3-106321755-DEL-6477 |
| chr3-106344687-DEL-53 | chr3-106344688-DEL-53 |
| chr3-106345555-DEL-397 | chr3-106345556-DEL-397 |
| chr3-106361308-DEL-143 | chr3-106361309-DEL-143 |
| chr3-106382012-DEL-115 | chr3-106382006-DEL-108 |
| chr3-106418318-DEL-76 | chr3-106418319-DEL-76 |
| chr3-106496493-DEL-537 | chr3-106496494-DEL-537 |
| chr3-106497973-DEL-2015 | chr3-106497974-DEL-2015 |
| chr3-106504788-DEL-65 | chr3-106504789-DEL-65 |
| chr3-106530111-DEL-533 | chr3-106530112-DEL-533 |
| chr3-106542173-DEL-174 | chr3-106542171-DEL-174 |
| chr3-106576725-DEL-58 | chr3-106576726-DEL-58 |
| chr3-106610290-DEL-1581 | chr3-106610290-DEL-1581 |
| chr3-106620362-DEL-1244 | chr3-106620363-DEL-1244 |
| chr3-106657152-DEL-72 | chr3-106657153-DEL-72 |
| chr3-106800826-DEL-1812 | chr3-106800827-DEL-1812 |
| chr3-106829845-DEL-10382 | chr3-106829846-DEL-10382 |
| chr3-106844271-DEL-57 | chr3-106844272-DEL-57 |
| chr3-106844528-DEL-203 | chr3-106844529-DEL-203 |
| chr3-106852581-DEL-395 | chr3-106852582-DEL-395 |
| chr3-106882776-DEL-110 | chr3-106882777-DEL-110 |
| chr3-106889261-DEL-58 | chr3-106889258-DEL-62 |
| chr3-106901765-DEL-104 | chr3-106901766-DEL-104 |
| chr3-106918019-DEL-2172 | chr3-106918019-DEL-2172 |
| chr3-106986771-DEL-137 | chr3-106986772-DEL-137 |
| chr3-107027427-DEL-178 | chr3-107027428-DEL-178 |
| chr3-107069095-DEL-419 | chr3-107069096-DEL-419 |
| chr3-107085078-DEL-655 | chr3-107085079-DEL-655 |
| chr3-107141634-DEL-699 | chr3-107141635-DEL-699 |
| chr3-107184919-DEL-101 | chr3-107184920-DEL-101 |
| chr3-107229131-DEL-53 | chr3-107229132-DEL-53 |
| chr3-107253689-DEL-210 | chr3-107253690-DEL-210 |
| chr3-107254621-DEL-247 | chr3-107254622-DEL-247 |
| chr3-107255967-DEL-105 | chr3-107255968-DEL-105 |
| chr3-107263131-DEL-51 | chr3-107263132-DEL-51 |
| chr3-107268268-DEL-80 | chr3-107268269-DEL-81 |
| chr3-107284231-DEL-365 | chr3-107284232-DEL-365 |
| chr3-107299713-DEL-139 | chr3-107299714-DEL-139 |
| chr3-107330625-DEL-278 | chr3-107330626-DEL-278 |
| chr3-107388446-DEL-1248 | chr3-107388447-DEL-1248 |

| | |
|---|---|
| chr3-107442820-DEL-66 | chr3-107442821-DEL-66 |
| chr3-107447326-DEL-51 | chr3-107447327-DEL-51 |
| chr3-107458472-DEL-76 | chr3-107458472-DEL-76 |
| chr3-107479522-DEL-757 | chr3-107479523-DEL-757 |
| chr3-107485976-DEL-550 | chr3-107485977-DEL-550 |
| chr3-107496085-DEL-460 | chr3-107496086-DEL-460 |
| chr3-107510464-DEL-210 | chr3-107510465-DEL-208 |
| chr3-107532232-DEL-521 | chr3-107532233-DEL-521 |
| chr3-107546760-DEL-197 | chr3-107546761-DEL-197 |
| chr3-107585663-DEL-407 | chr3-107585664-DEL-407 |
| chr3-107606862-DEL-1529 | chr3-107606863-DEL-1529 |
| chr3-107790523-DEL-52 | chr3-107790524-DEL-50 |
| chr3-107829578-DEL-231 | chr3-107829579-DEL-231 |
| chr3-107855973-DEL-54 | chr3-107855974-DEL-54 |
| chr3-107872848-DEL-66 | chr3-107872849-DEL-66 |
| chr3-107878857-DEL-445 | chr3-107878858-DEL-445 |
| chr3-107884332-DEL-559 | chr3-107884332-DEL-559 |
| chr3-107894946-DEL-5993 | chr3-107894947-DEL-5993 |
| chr3-107902127-DEL-267 | chr3-107902128-DEL-267 |
| chr3-107911147-DEL-1353 | chr3-107911149-DEL-1352 |
| chr3-107934383-DEL-99 | chr3-107934383-DEL-99 |
| chr3-107963484-DEL-170 | chr3-107963485-DEL-170 |
| chr3-107970786-DEL-231 | chr3-107970787-DEL-231 |
| chr3-107975417-DEL-148 | chr3-107975422-DEL-144 |
| chr3-108003066-DEL-288 | chr3-108003067-DEL-288 |
| chr3-108006249-DEL-81 | chr3-108006250-DEL-81 |
| chr3-108144190-DEL-74 | chr3-108144191-DEL-74 |
| chr3-108145300-DEL-197 | chr3-108145301-DEL-197 |
| chr3-108149291-DEL-4818 | chr3-108149292-DEL-4818 |
| chr3-108167758-DEL-436 | chr3-108167755-DEL-443 |
| chr3-108183934-DEL-809 | chr3-108183938-DEL-806 |
| chr3-108201395-DEL-54 | chr3-108201396-DEL-54 |
| chr3-108233375-DEL-215 | chr3-108233376-DEL-215 |
| chr3-108273614-DEL-192 | chr3-108273615-DEL-192 |
| chr3-108277867-DEL-203 | chr3-108277868-DEL-203 |
| chr3-108357294-DEL-82 | chr3-108357296-DEL-64 |
| chr3-108376762-DEL-68 | chr3-108376760-DEL-54 |
| chr3-108380734-DEL-69 | chr3-108380735-DEL-69 |
| chr3-108411584-DEL-395 | chr3-108411585-DEL-395 |
| chr3-108415158-DEL-63 | chr3-108415159-DEL-81 |
| chr3-108416835-DEL-68 | chr3-108416833-DEL-69 |
| chr3-108427444-DEL-914 | chr3-108427445-DEL-914 |
| chr3-108434333-DEL-198 | chr3-108434334-DEL-198 |
| chr3-108513368-DEL-2477 | chr3-108513369-DEL-2477 |
| chr3-108523508-DEL-467 | chr3-108523499-DEL-468 |
| chr3-108524556-DEL-298 | chr3-108524557-DEL-298 |
| chr3-108548345-DEL-725 | chr3-108548346-DEL-725 |
| chr3-108560937-DEL-67 | chr3-108560944-DEL-74 |
| chr3-108563597-DEL-181 | chr3-108563598-DEL-181 |
| chr3-108582580-DEL-573 | chr3-108582581-DEL-573 |
| chr3-108597169-DEL-103 | chr3-108597166-DEL-103 |
| chr3-108598510-DEL-2688 | chr3-108598511-DEL-2688 |
| chr3-108652413-DEL-66 | chr3-108652414-DEL-66 |
| chr3-108660233-DEL-201 | chr3-108660234-DEL-202 |
| chr3-108694445-DEL-191 | chr3-108694446-DEL-189 |
| chr3-108697564-DEL-203 | chr3-108697567-DEL-201 |
| chr3-108699415-DEL-62 | chr3-108699416-DEL-62 |
| chr3-108716680-DEL-60 | chr3-108716681-DEL-60 |
| chr3-108742225-DEL-359 | chr3-108742225-DEL-360 |
| chr3-108749966-DEL-165 | chr3-108749967-DEL-165 |
| chr3-108753603-DEL-1672 | chr3-108753604-DEL-1672 |
| chr3-108755447-DEL-1166 | chr3-108755448-DEL-1166 |
| chr3-108771036-DEL-57 | chr3-108771038-DEL-55 |
| chr3-108787780-DEL-78 | chr3-108787781-DEL-78 |
| chr3-108792465-DEL-63 | chr3-108792466-DEL-63 |
| chr3-108799730-DEL-345 | chr3-108799743-DEL-328 |
| chr3-108803212-DEL-1191 | chr3-108803213-DEL-1191 |
| chr3-108824002-DEL-174 | chr3-108824003-DEL-174 |
| chr3-108842962-DEL-2909 | chr3-108842963-DEL-2909 |
| chr3-108862914-DEL-234 | chr3-108862960-DEL-193 |
| chr3-108874537-DEL-286 | chr3-108874537-DEL-286 |
| chr3-108880838-DEL-694 | chr3-108880837-DEL-690 |
| chr3-108884380-DEL-61 | chr3-108884383-DEL-59 |
| chr3-108887226-DEL-62 | chr3-108887226-DEL-63 |
| chr3-108888902-DEL-57 | chr3-108888883-DEL-77 |
| chr3-108906750-DEL-230 | chr3-108906756-DEL-242 |
| chr3-108945680-DEL-1056 | chr3-108945681-DEL-1056 |
| chr3-108962999-DEL-54 | chr3-108963000-DEL-54 |
| chr3-108981055-DEL-1002 | chr3-108981056-DEL-1002 |
| chr3-108992020-DEL-162 | chr3-108992021-DEL-162 |

| | |
|---|---|
| chr3-109011433-DEL-184 | chr3-109011434-DEL-182 |
| chr3-109017752-DEL-59 | chr3-109017753-DEL-59 |
| chr3-109042432-DEL-196 | chr3-109042433-DEL-196 |
| chr3-109046488-DEL-6445 | chr3-109046489-DEL-6445 |
| chr3-109060485-DEL-203 | chr3-109060486-DEL-203 |
| chr3-109062835-DEL-83 | chr3-109062836-DEL-83 |
| chr3-109115333-DEL-60 | chr3-109115306-DEL-68 |
| chr3-109124893-DEL-684 | chr3-109124892-DEL-685 |
| chr3-109127869-DEL-58 | chr3-109127870-DEL-58 |
| chr3-109133732-DEL-60 | chr3-109133733-DEL-60 |
| chr3-109134303-DEL-62 | chr3-109134304-DEL-58 |
| chr3-109138113-DEL-237 | chr3-109138114-DEL-237 |
| chr3-109145891-DEL-148 | chr3-109145893-DEL-149 |
| chr3-109146935-DEL-60 | chr3-109146927-DEL-53 |
| chr3-109148041-DEL-710 | chr3-109148040-DEL-710 |
| chr3-109165411-DEL-6285 | chr3-109165412-DEL-6285 |
| chr3-109182750-DEL-1878 | chr3-109182746-DEL-1883 |
| chr3-109199940-DEL-1237 | chr3-109199941-DEL-1237 |
| chr3-109211724-DEL-108 | chr3-109211732-DEL-86 |
| chr3-109227699-DEL-357 | chr3-109227700-DEL-357 |
| chr3-109254772-DEL-70 | chr3-109254805-DEL-50 |
| chr3-109261772-DEL-492 | chr3-109261773-DEL-492 |
| chr3-109272876-DEL-168 | chr3-109272874-DEL-170 |
| chr3-109280746-DEL-114 | chr3-109280755-DEL-122 |
| chr3-109301320-DEL-426 | chr3-109301321-DEL-426 |
| chr3-109320052-DEL-509 | chr3-109320053-DEL-509 |
| chr3-109348893-DEL-283 | chr3-109348894-DEL-283 |
| chr3-109372818-DEL-54 | chr3-109372819-DEL-54 |
| chr3-109373305-DEL-328 | chr3-109373306-DEL-330 |
| chr3-109378012-DEL-59 | chr3-109378013-DEL-59 |
| chr3-109417818-DEL-168 | chr3-109417819-DEL-168 |
| chr3-109421149-DEL-887 | chr3-109421150-DEL-887 |
| chr3-109458412-DEL-78 | chr3-109458413-DEL-78 |
| chr3-109520095-DEL-6806 | chr3-109520096-DEL-6806 |
| chr3-109576310-DEL-2112 | chr3-109576311-DEL-2112 |
| chr3-109605212-DEL-934 | chr3-109605213-DEL-934 |
| chr3-109606603-DEL-722 | chr3-109606604-DEL-722 |
| chr3-109633362-DEL-159 | chr3-109633363-DEL-159 |
| chr3-109633683-DEL-1661 | chr3-109633684-DEL-1661 |
| chr3-109653359-DEL-169 | chr3-109653360-DEL-169 |
| chr3-109654622-DEL-639 | chr3-109654619-DEL-642 |
| chr3-109716335-DEL-76 | chr3-109716335-DEL-63 |
| chr3-109763818-DEL-157 | chr3-109763820-DEL-158 |
| chr3-109770397-DEL-499 | chr3-109770398-DEL-499 |
| chr3-109805937-DEL-70 | chr3-109805974-DEL-80 |
| chr3-109820222-DEL-52 | chr3-109820223-DEL-52 |
| chr3-109844499-DEL-80 | chr3-109844500-DEL-80 |
| chr3-109853772-DEL-82 | chr3-109853773-DEL-82 |
| chr3-109872006-DEL-54 | chr3-109872007-DEL-54 |
| chr3-109882318-DEL-54 | chr3-109882319-DEL-54 |
| chr3-109919772-DEL-1316 | chr3-109919773-DEL-1316 |
| chr3-109964589-DEL-6769 | chr3-109964590-DEL-6769 |
| chr3-109977195-DEL-80 | chr3-109977196-DEL-80 |
| chr3-109999860-DEL-58 | chr3-109999857-DEL-51 |
| chr3-110014503-DEL-6241 | chr3-110014504-DEL-6241 |
| chr3-110067735-DEL-1895 | chr3-110067736-DEL-1895 |
| chr3-110078289-DEL-90 | chr3-110078292-DEL-88 |
| chr3-110085632-DEL-96 | chr3-110085633-DEL-96 |
| chr3-110099355-DEL-157 | chr3-110099367-DEL-113 |
| chr3-110116617-DEL-6389 | chr3-110116618-DEL-6389 |
| chr3-110138709-DEL-54 | chr3-110138710-DEL-54 |
| chr3-110198826-DEL-3693 | chr3-110198823-DEL-3694 |
| chr3-110227127-DEL-6691 | chr3-110227128-DEL-6691 |
| chr3-110263020-DEL-470 | chr3-110263021-DEL-470 |
| chr3-110322941-DEL-6223 | chr3-110322942-DEL-6223 |
| chr3-110322963-DEL-181 | chr3-110322964-DEL-181 |
| chr3-110330168-DEL-96 | chr3-110330168-DEL-96 |
| chr3-110339072-DEL-2643 | chr3-110339073-DEL-2644 |
| chr3-110356367-DEL-1895 | chr3-110356368-DEL-1895 |
| chr3-110374341-DEL-120 | chr3-110374326-DEL-136 |
| chr3-110386326-DEL-56 | chr3-110386326-DEL-70 |
| chr3-110394565-DEL-66 | chr3-110394579-DEL-52 |
| chr3-110397216-DEL-2000 | chr3-110397217-DEL-2000 |
| chr3-110433167-DEL-56 | chr3-110433164-DEL-60 |
| chr3-110454287-DEL-80 | chr3-110454286-DEL-84 |
| chr3-110484067-DEL-202 | chr3-110484068-DEL-202 |
| chr3-110514965-DEL-163 | chr3-110514964-DEL-161 |
| chr3-110528352-DEL-213 | chr3-110528353-DEL-213 |
| chr3-110564064-DEL-53 | chr3-110564066-DEL-51 |
| chr3-110584118-DEL-6378 | chr3-110584119-DEL-6378 |

| | |
|---|---|
| chr3-110629693-DEL-141 | chr3-110629694-DEL-141 |
| chr3-110654980-DEL-72 | chr3-110654987-DEL-70 |
| chr3-110677299-DEL-5145 | chr3-110677300-DEL-5145 |
| chr3-110692369-DEL-353 | chr3-110692374-DEL-350 |
| chr3-110698475-DEL-1841 | chr3-110698488-DEL-1830 |
| chr3-110727943-DEL-54 | chr3-110727944-DEL-54 |
| chr3-110797982-DEL-717 | chr3-110797983-DEL-717 |
| chr3-110798726-DEL-67 | chr3-110798727-DEL-67 |
| chr3-110806495-DEL-58 | chr3-110806486-DEL-108 |
| chr3-110840705-DEL-98 | chr3-110840714-DEL-110 |
| chr3-110849054-DEL-529 | chr3-110849055-DEL-529 |
| chr3-110918499-DEL-81 | chr3-110918480-DEL-76 |
| chr3-110947067-DEL-6270 | chr3-110947068-DEL-6270 |
| chr3-110984786-DEL-150 | chr3-110984787-DEL-150 |
| chr3-111018258-DEL-123 | chr3-111018262-DEL-87 |
| chr3-111043153-DEL-442 | chr3-111043154-DEL-442 |
| chr3-111072343-DEL-712 | chr3-111072344-DEL-712 |
| chr3-111074793-DEL-90 | chr3-111074794-DEL-90 |
| chr3-111081212-DEL-52 | chr3-111081215-DEL-50 |
| chr3-111094674-DEL-105 | chr3-111094672-DEL-108 |
| chr3-111114330-DEL-65 | chr3-111114331-DEL-65 |
| chr3-111118850-DEL-54 | chr3-111118851-DEL-54 |
| chr3-111131887-DEL-287 | chr3-111131888-DEL-287 |
| chr3-111169573-DEL-6413 | chr3-111169574-DEL-6413 |
| chr3-111209222-DEL-84 | chr3-111209223-DEL-84 |
| chr3-111235048-DEL-110 | chr3-111235047-DEL-110 |
| chr3-111249268-DEL-60 | chr3-111249269-DEL-60 |
| chr3-111298566-DEL-652 | chr3-111298567-DEL-651 |
| chr3-111345368-DEL-1102 | chr3-111345360-DEL-1125 |
| chr3-111362586-DEL-435 | chr3-111362587-DEL-435 |
| chr3-111366512-DEL-1548 | chr3-111366513-DEL-1548 |
| chr3-111379689-DEL-64 | chr3-111379691-DEL-50 |
| chr3-111407568-DEL-64 | chr3-111407561-DEL-62 |
| chr3-111416539-DEL-1206 | chr3-111416540-DEL-1206 |
| chr3-111418492-DEL-57 | chr3-111418493-DEL-57 |
| chr3-111426310-DEL-4298 | chr3-111426311-DEL-4298 |
| chr3-111432833-DEL-3748 | chr3-111432834-DEL-3748 |
| chr3-111465690-DEL-94 | chr3-111465691-DEL-94 |
| chr3-111484569-DEL-2605 | chr3-111484570-DEL-2605 |
| chr3-111505494-DEL-7003 | chr3-111505495-DEL-7003 |
| chr3-111529717-DEL-76 | chr3-111529722-DEL-72 |
| chr3-111563129-DEL-52 | chr3-111563128-DEL-54 |
| chr3-111593464-DEL-6841 | chr3-111593465-DEL-6841 |
| chr3-111598736-DEL-262 | chr3-111598737-DEL-260 |
| chr3-111619387-DEL-79 | chr3-111619388-DEL-79 |
| chr3-111632125-DEL-4841 | chr3-111632126-DEL-4841 |
| chr3-111639971-DEL-217 | chr3-111639972-DEL-217 |
| chr3-111646577-DEL-1276 | chr3-111646570-DEL-1277 |
| chr3-111655821-DEL-1501 | chr3-111655821-DEL-1501 |
| chr3-111668961-DEL-6355 | chr3-111668962-DEL-6355 |
| chr3-111700314-DEL-72 | chr3-111700324-DEL-62 |
| chr3-111706344-DEL-53 | chr3-111706340-DEL-52 |
| chr3-111711477-DEL-1133 | chr3-111711477-DEL-1133 |
| chr3-111737538-DEL-6172 | chr3-111737539-DEL-6172 |
| chr3-111774369-DEL-282 | chr3-111774370-DEL-282 |
| chr3-111783364-DEL-833 | chr3-111783371-DEL-825 |
| chr3-111806079-DEL-200 | chr3-111806080-DEL-200 |
| chr3-111891189-DEL-52 | chr3-111891190-DEL-52 |
| chr3-111896823-DEL-391 | chr3-111896824-DEL-391 |
| chr3-111911752-DEL-688 | chr3-111911753-DEL-688 |
| chr3-111913667-DEL-3450 | chr3-111913668-DEL-3450 |
| chr3-111971955-DEL-4723 | chr3-111971956-DEL-4723 |
| chr3-112031377-DEL-330 | chr3-112031378-DEL-330 |
| chr3-112032482-DEL-201 | chr3-112032479-DEL-204 |
| chr3-112050441-DEL-63 | chr3-112050442-DEL-63 |
| chr3-112053070-DEL-397 | chr3-112053068-DEL-398 |
| chr3-112058932-DEL-6383 | chr3-112058932-DEL-6383 |
| chr3-112092731-DEL-7427 | chr3-112092732-DEL-7427 |
| chr3-112127537-DEL-96 | chr3-112127538-DEL-96 |
| chr3-112173707-DEL-498 | chr3-112173708-DEL-498 |
| chr3-112239167-DEL-6404 | chr3-112239168-DEL-6404 |
| chr3-112256599-DEL-88 | chr3-112256597-DEL-89 |
| chr3-112267429-DEL-978 | chr3-112267424-DEL-979 |
| chr3-112269850-DEL-60 | chr3-112269847-DEL-63 |
| chr3-112281235-DEL-72 | chr3-112281215-DEL-98 |
| chr3-112285654-DEL-100 | chr3-112285657-DEL-98 |
| chr3-112299403-DEL-232 | chr3-112299404-DEL-232 |
| chr3-112324212-DEL-605 | chr3-112324213-DEL-605 |
| chr3-112355642-DEL-66 | chr3-112355651-DEL-54 |
| chr3-112356427-DEL-104 | chr3-112356428-DEL-104 |

| | |
|---|---|
| chr3-112359727-DEL-54 | chr3-112359728-DEL-78 |
| chr3-112397226-DEL-56 | chr3-112397227-DEL-56 |
| chr3-112400333-DEL-424 | chr3-112400334-DEL-424 |
| chr3-112412836-DEL-63 | chr3-112412837-DEL-63 |
| chr3-112431842-DEL-82 | chr3-112431843-DEL-82 |
| chr3-112435287-DEL-471 | chr3-112435288-DEL-471 |
| chr3-112467756-DEL-6160 | chr3-112467757-DEL-6160 |
| chr3-112478542-DEL-84 | chr3-112478543-DEL-84 |
| chr3-112491368-DEL-2691 | chr3-112491369-DEL-2691 |
| chr3-112495250-DEL-419 | chr3-112495251-DEL-419 |
| chr3-112500391-DEL-598 | chr3-112500392-DEL-598 |
| chr3-112514227-DEL-279 | chr3-112514228-DEL-279 |
| chr3-112517350-DEL-74 | chr3-112517322-DEL-86 |
| chr3-112536798-DEL-125 | chr3-112536799-DEL-125 |
| chr3-112568810-DEL-533 | chr3-112568811-DEL-533 |
| chr3-112604308-DEL-326 | chr3-112604309-DEL-326 |
| chr3-112618968-DEL-54 | chr3-112618969-DEL-54 |
| chr3-112636871-DEL-196 | chr3-112636851-DEL-176 |
| chr3-112673779-DEL-532 | chr3-112673780-DEL-529 |
| chr3-112700528-DEL-1360 | chr3-112700531-DEL-1359 |
| chr3-112726868-DEL-79 | chr3-112726866-DEL-82 |
| chr3-112738798-DEL-54 | chr3-112738799-DEL-54 |
| chr3-112741308-DEL-6489 | chr3-112741309-DEL-6489 |
| chr3-112771538-DEL-502 | chr3-112771539-DEL-502 |
| chr3-112862105-DEL-196 | chr3-112862061-DEL-241 |
| chr3-112865312-DEL-107 | chr3-112865313-DEL-107 |
| chr3-112871894-DEL-1839 | chr3-112871895-DEL-1839 |
| chr3-112880138-DEL-154 | chr3-112880139-DEL-154 |
| chr3-112881310-DEL-272 | chr3-112881311-DEL-272 |
| chr3-112911761-DEL-741 | chr3-112911762-DEL-740 |
| chr3-112932478-DEL-1006 | chr3-112932519-DEL-977 |
| chr3-112975123-DEL-573 | chr3-112975124-DEL-573 |
| chr3-113019333-DEL-200 | chr3-113019334-DEL-200 |
| chr3-113027174-DEL-72 | chr3-113027175-DEL-72 |
| chr3-113045813-DEL-975 | chr3-113045821-DEL-975 |
| chr3-113088805-DEL-246 | chr3-113088806-DEL-246 |
| chr3-113092017-DEL-5367 | chr3-113092018-DEL-5367 |
| chr3-113102910-DEL-576 | chr3-113102910-DEL-576 |
| chr3-113108004-DEL-1712 | chr3-113108004-DEL-1712 |
| chr3-113116853-DEL-58 | chr3-113116859-DEL-53 |
| chr3-113129314-DEL-124 | chr3-113129313-DEL-122 |
| chr3-113156834-DEL-375 | chr3-113156834-DEL-375 |
| chr3-113157981-DEL-95 | chr3-113157980-DEL-95 |
| chr3-113314322-DEL-58 | chr3-113314323-DEL-58 |
| chr3-113352614-DEL-1320 | chr3-113352615-DEL-1320 |
| chr3-113359935-DEL-136 | chr3-113359936-DEL-136 |
| chr3-113406436-DEL-285 | chr3-113406461-DEL-285 |
| chr3-113429206-DEL-1488 | chr3-113429207-DEL-1488 |
| chr3-113434539-DEL-6031 | chr3-113434536-DEL-6032 |
| chr3-113455275-DEL-337 | chr3-113455271-DEL-338 |
| chr3-113474021-DEL-52 | chr3-113474022-DEL-58 |
| chr3-113510670-DEL-196 | chr3-113510670-DEL-195 |
| chr3-113514919-DEL-612 | chr3-113514920-DEL-612 |
| chr3-113557527-DEL-385 | chr3-113557552-DEL-385 |
| chr3-113565982-DEL-848 | chr3-113565983-DEL-848 |
| chr3-113584577-DEL-134 | chr3-113584577-DEL-131 |
| chr3-113627813-DEL-6217 | chr3-113627814-DEL-6217 |
| chr3-113636594-DEL-156 | chr3-113636595-DEL-156 |
| chr3-113658498-DEL-3255 | chr3-113658499-DEL-3255 |
| chr3-113675322-DEL-52 | chr3-113675325-DEL-50 |
| chr3-113683211-DEL-76 | chr3-113683163-DEL-91 |
| chr3-113684710-DEL-3191 | chr3-113684708-DEL-3191 |
| chr3-113702121-DEL-133 | chr3-113702121-DEL-133 |
| chr3-113712740-DEL-81 | chr3-113712745-DEL-76 |
| chr3-113718937-DEL-2167 | chr3-113718938-DEL-2167 |
| chr3-113729657-DEL-58 | chr3-113729662-DEL-54 |
| chr3-113757434-DEL-1627 | chr3-113757435-DEL-1627 |
| chr3-113808929-DEL-64 | chr3-113808924-DEL-70 |
| chr3-113841024-DEL-534 | chr3-113841025-DEL-534 |
| chr3-113895186-DEL-293 | chr3-113895187-DEL-293 |
| chr3-113909899-DEL-99 | chr3-113909887-DEL-100 |
| chr3-113985033-DEL-370 | chr3-113985034-DEL-370 |
| chr3-114002096-DEL-71 | chr3-114002097-DEL-71 |
| chr3-114031823-DEL-1031 | chr3-114031824-DEL-1031 |
| chr3-114036406-DEL-201 | chr3-114036406-DEL-201 |
| chr3-114061021-DEL-199 | chr3-114061022-DEL-199 |
| chr3-114063054-DEL-632 | chr3-114063055-DEL-632 |
| chr3-114063928-DEL-1317 | chr3-114063929-DEL-1317 |
| chr3-114066124-DEL-6587 | chr3-114066114-DEL-6588 |
| chr3-114077225-DEL-90 | chr3-114077226-DEL-90 |

| | |
|---|---|
| chr3-114080326-DEL-61 | chr3-114080327-DEL-61 |
| chr3-114096280-DEL-199 | chr3-114096274-DEL-200 |
| chr3-114101574-DEL-359 | chr3-114101575-DEL-359 |
| chr3-114136741-DEL-194 | chr3-114136742-DEL-194 |
| chr3-114143292-DEL-1137 | chr3-114143293-DEL-1137 |
| chr3-114201758-DEL-727 | chr3-114201759-DEL-727 |
| chr3-114206706-DEL-1928 | chr3-114206707-DEL-1928 |
| chr3-114210500-DEL-4824 | chr3-114210501-DEL-4824 |
| chr3-114224165-DEL-435 | chr3-114224166-DEL-435 |
| chr3-114259077-DEL-162 | chr3-114259078-DEL-162 |
| chr3-114265741-DEL-7509 | chr3-114265742-DEL-7509 |
| chr3-114287401-DEL-6249 | chr3-114287402-DEL-6248 |
| chr3-114329322-DEL-440 | chr3-114329323-DEL-440 |
| chr3-114354088-DEL-594 | chr3-114354089-DEL-594 |
| chr3-114369508-DEL-96 | chr3-114369513-DEL-92 |
| chr3-114381476-DEL-2801 | chr3-114381477-DEL-2801 |
| chr3-114384746-DEL-871 | chr3-114384750-DEL-869 |
| chr3-114386761-DEL-2824 | chr3-114386762-DEL-2824 |
| chr3-114400967-DEL-71 | chr3-114400968-DEL-71 |
| chr3-114414213-DEL-4638 | chr3-114414214-DEL-4638 |
| chr3-114442187-DEL-56 | chr3-114442188-DEL-56 |
| chr3-114482268-DEL-296 | chr3-114482269-DEL-296 |
| chr3-114483086-DEL-390 | chr3-114483087-DEL-390 |
| chr3-114488824-DEL-98 | chr3-114488826-DEL-114 |
| chr3-114514459-DEL-60 | chr3-114514464-DEL-56 |
| chr3-114534553-DEL-295 | chr3-114534554-DEL-295 |
| chr3-114536091-DEL-59 | chr3-114536092-DEL-59 |
| chr3-114540130-DEL-204 | chr3-114540131-DEL-204 |
| chr3-114556146-DEL-307 | chr3-114556147-DEL-307 |
| chr3-114559169-DEL-854 | chr3-114559170-DEL-854 |
| chr3-114569974-DEL-68 | chr3-114569977-DEL-66 |
| chr3-114574522-DEL-4243 | chr3-114574519-DEL-4243 |
| chr3-114579008-DEL-125 | chr3-114579008-DEL-125 |
| chr3-114599248-DEL-238 | chr3-114599225-DEL-232 |
| chr3-114604276-DEL-1897 | chr3-114604277-DEL-1897 |
| chr3-114634391-DEL-1329 | chr3-114634376-DEL-1344 |
| chr3-114642311-DEL-543 | chr3-114642312-DEL-543 |
| chr3-114657238-DEL-3507 | chr3-114657239-DEL-3507 |
| chr3-114668247-DEL-545 | chr3-114668248-DEL-545 |
| chr3-114714001-DEL-529 | chr3-114714002-DEL-529 |
| chr3-114735408-DEL-81 | chr3-114735404-DEL-77 |
| chr3-114744407-DEL-55 | chr3-114744408-DEL-55 |
| chr3-114770301-DEL-6204 | chr3-114770302-DEL-6204 |
| chr3-114787374-DEL-1063 | chr3-114787375-DEL-1063 |
| chr3-114814380-DEL-6295 | chr3-114814381-DEL-6295 |
| chr3-114842107-DEL-243 | chr3-114842108-DEL-243 |
| chr3-114945012-DEL-217 | chr3-114945013-DEL-217 |
| chr3-114954775-DEL-52 | chr3-114954776-DEL-52 |
| chr3-114958574-DEL-148 | chr3-114958575-DEL-148 |
| chr3-114966769-DEL-326 | chr3-114966767-DEL-325 |
| chr3-115000412-DEL-90 | chr3-115000411-DEL-91 |
| chr3-115014673-DEL-864 | chr3-115014674-DEL-864 |
| chr3-115032626-DEL-60 | chr3-115032627-DEL-60 |
| chr3-115035089-DEL-106 | chr3-115035090-DEL-106 |
| chr3-115038057-DEL-5365 | chr3-115038058-DEL-5365 |
| chr3-115065628-DEL-1184 | chr3-115065629-DEL-1184 |
| chr3-115079598-DEL-6044 | chr3-115079598-DEL-6044 |
| chr3-115085812-DEL-112 | chr3-115085791-DEL-125 |
| chr3-115124319-DEL-6250 | chr3-115124359-DEL-6210 |
| chr3-115138897-DEL-1379 | chr3-115138898-DEL-1379 |
| chr3-115161407-DEL-4324 | chr3-115161407-DEL-4325 |
| chr3-115187433-DEL-6204 | chr3-115187434-DEL-6204 |
| chr3-115194932-DEL-194 | chr3-115194933-DEL-195 |
| chr3-115214379-DEL-64 | chr3-115214380-DEL-64 |
| chr3-115221668-DEL-2005 | chr3-115221669-DEL-2005 |
| chr3-115245913-DEL-444 | chr3-115245914-DEL-444 |
| chr3-115272106-DEL-113 | chr3-115272107-DEL-113 |
| chr3-115292629-DEL-1062 | chr3-115292630-DEL-1062 |
| chr3-115294164-DEL-623 | chr3-115294165-DEL-623 |
| chr3-115301868-DEL-62 | chr3-115301869-DEL-58 |
| chr3-115313818-DEL-107 | chr3-115313819-DEL-107 |
| chr3-115338664-DEL-76 | chr3-115338674-DEL-77 |
| chr3-115380594-DEL-90 | chr3-115380593-DEL-78 |
| chr3-115412015-DEL-58 | chr3-115412024-DEL-50 |
| chr3-115460614-DEL-62 | chr3-115460631-DEL-54 |
| chr3-115470509-DEL-1613 | chr3-115470528-DEL-1595 |
| chr3-115499637-DEL-201 | chr3-115499638-DEL-201 |
| chr3-115511623-DEL-65 | chr3-115511624-DEL-65 |
| chr3-115517288-DEL-91 | chr3-115517292-DEL-50 |
| chr3-115522306-DEL-195 | chr3-115522309-DEL-193 |

| | |
|---|---|
| chr3-115530903-DEL-64 | chr3-115530904-DEL-75 |
| chr3-115545292-DEL-92 | chr3-115545292-DEL-84 |
| chr3-115549891-DEL-52 | chr3-115549872-DEL-51 |
| chr3-115621809-DEL-96 | chr3-115621817-DEL-69 |
| chr3-115622506-DEL-521 | chr3-115622507-DEL-521 |
| chr3-115646226-DEL-54 | chr3-115646227-DEL-60 |
| chr3-115658223-DEL-1856 | chr3-115658224-DEL-1856 |
| chr3-115695773-DEL-669 | chr3-115695776-DEL-670 |
| chr3-115699912-DEL-1250 | chr3-115699913-DEL-1250 |
| chr3-115701741-DEL-979 | chr3-115701741-DEL-979 |
| chr3-115702769-DEL-5319 | chr3-115702770-DEL-5319 |
| chr3-115709543-DEL-7064 | chr3-115709544-DEL-7064 |
| chr3-115722403-DEL-75 | chr3-115722404-DEL-75 |
| chr3-115736193-DEL-151 | chr3-115736194-DEL-151 |
| chr3-115753615-DEL-1232 | chr3-115753616-DEL-1233 |
| chr3-115815852-DEL-1083 | chr3-115815853-DEL-1083 |
| chr3-115823902-DEL-68 | chr3-115823903-DEL-68 |
| chr3-115825251-DEL-97 | chr3-115825252-DEL-97 |
| chr3-115828201-DEL-1548 | chr3-115828202-DEL-1548 |
| chr3-115853020-DEL-216 | chr3-115853021-DEL-216 |
| chr3-115861085-DEL-162 | chr3-115861086-DEL-162 |
| chr3-115901956-DEL-213 | chr3-115901957-DEL-213 |
| chr3-115953665-DEL-168 | chr3-115953647-DEL-174 |
| chr3-115971574-DEL-55 | chr3-115971574-DEL-53 |
| chr3-116021619-DEL-56 | chr3-116021625-DEL-60 |
| chr3-116037256-DEL-52 | chr3-116037257-DEL-52 |
| chr3-116038804-DEL-88 | chr3-116038823-DEL-84 |
| chr3-116059621-DEL-171 | chr3-116059622-DEL-171 |
| chr3-116060572-DEL-200 | chr3-116060541-DEL-200 |
| chr3-116060969-DEL-377 | chr3-116060970-DEL-377 |
| chr3-116064335-DEL-304 | chr3-116064336-DEL-304 |
| chr3-116076026-DEL-389 | chr3-116076027-DEL-389 |
| chr3-116079672-DEL-57 | chr3-116079672-DEL-72 |
| chr3-116106677-DEL-162 | chr3-116106726-DEL-163 |
| chr3-116115102-DEL-489 | chr3-116115103-DEL-489 |
| chr3-116116656-DEL-141 | chr3-116116667-DEL-126 |
| chr3-116125430-DEL-78 | chr3-116125398-DEL-107 |
| chr3-116131893-DEL-66 | chr3-116131885-DEL-78 |
| chr3-116133346-DEL-340 | chr3-116133346-DEL-346 |
| chr3-116139757-DEL-1845 | chr3-116139758-DEL-1845 |
| chr3-116141392-DEL-165 | chr3-116141392-DEL-165 |
| chr3-116176218-DEL-66 | chr3-116176209-DEL-75 |
| chr3-116186940-DEL-73 | chr3-116186941-DEL-103 |
| chr3-116245104-DEL-74 | chr3-116245092-DEL-55 |
| chr3-116288779-DEL-229 | chr3-116288780-DEL-229 |
| chr3-116293730-DEL-308 | chr3-116293731-DEL-308 |
| chr3-116295985-DEL-417 | chr3-116295984-DEL-418 |
| chr3-116311061-DEL-189 | chr3-116311062-DEL-189 |
| chr3-116318786-DEL-206 | chr3-116318787-DEL-204 |
| chr3-116324379-DEL-100 | chr3-116324380-DEL-100 |
| chr3-116335789-DEL-670 | chr3-116335790-DEL-670 |
| chr3-116342789-DEL-82 | chr3-116342790-DEL-83 |
| chr3-116351127-DEL-64 | chr3-116351126-DEL-65 |
| chr3-116351834-DEL-647 | chr3-116351835-DEL-647 |
| chr3-116354235-DEL-957 | chr3-116354236-DEL-957 |
| chr3-116356381-DEL-135 | chr3-116356372-DEL-135 |
| chr3-116374047-DEL-201 | chr3-116374060-DEL-204 |
| chr3-116394538-DEL-211 | chr3-116394539-DEL-211 |
| chr3-116400067-DEL-208 | chr3-116400068-DEL-208 |
| chr3-116417049-DEL-107 | chr3-116417039-DEL-99 |
| chr3-116418304-DEL-393 | chr3-116418299-DEL-394 |
| chr3-116477874-DEL-176 | chr3-116477875-DEL-176 |
| chr3-116481376-DEL-176 | chr3-116481377-DEL-171 |
| chr3-116486131-DEL-378 | chr3-116486132-DEL-378 |
| chr3-116514268-DEL-1410 | chr3-116514269-DEL-1410 |
| chr3-116516956-DEL-238 | chr3-116516957-DEL-238 |
| chr3-116520991-DEL-162 | chr3-116520992-DEL-162 |
| chr3-116529717-DEL-60 | chr3-116529718-DEL-60 |
| chr3-116534057-DEL-196 | chr3-116534058-DEL-196 |
| chr3-116542023-DEL-1125 | chr3-116542023-DEL-1125 |
| chr3-116543756-DEL-60 | chr3-116543762-DEL-50 |
| chr3-116546109-DEL-209 | chr3-116546110-DEL-209 |
| chr3-116553539-DEL-213 | chr3-116553540-DEL-213 |
| chr3-116554057-DEL-5075 | chr3-116554058-DEL-5075 |
| chr3-116561521-DEL-168 | chr3-116561522-DEL-168 |
| chr3-116586067-DEL-211 | chr3-116586068-DEL-209 |
| chr3-116589045-DEL-208 | chr3-116589046-DEL-208 |
| chr3-116598182-DEL-53 | chr3-116598183-DEL-53 |
| chr3-116600559-DEL-53 | chr3-116600560-DEL-53 |
| chr3-116611178-DEL-5485 | chr3-116611179-DEL-5485 |

| | |
|---|---|
| chr3-116618043-DEL-134 | chr3-116618036-DEL-142 |
| chr3-116621101-DEL-69 | chr3-116621101-DEL-77 |
| chr3-116624570-DEL-211 | chr3-116624571-DEL-211 |
| chr3-116635275-DEL-69 | chr3-116635276-DEL-68 |
| chr3-116662795-DEL-60 | chr3-116662795-DEL-60 |
| chr3-116698133-DEL-70 | chr3-116698142-DEL-62 |
| chr3-116714943-DEL-200 | chr3-116714944-DEL-200 |
| chr3-116743611-DEL-63 | chr3-116743611-DEL-63 |
| chr3-116753668-DEL-123 | chr3-116753669-DEL-123 |
| chr3-116767558-DEL-655 | chr3-116767559-DEL-655 |
| chr3-116770891-DEL-54 | chr3-116770875-DEL-52 |
| chr3-116786426-DEL-50 | chr3-116786403-DEL-74 |
| chr3-116803242-DEL-51 | chr3-116803243-DEL-51 |
| chr3-116811783-DEL-2123 | chr3-116811784-DEL-2123 |
| chr3-116823604-DEL-6506 | chr3-116823605-DEL-6507 |
| chr3-116830132-DEL-378 | chr3-116830137-DEL-374 |
| chr3-116836028-DEL-55 | chr3-116836027-DEL-56 |
| chr3-116856988-DEL-57 | chr3-116856991-DEL-57 |
| chr3-116858138-DEL-195 | chr3-116858128-DEL-190 |
| chr3-116860439-DEL-414 | chr3-116860439-DEL-414 |
| chr3-116884656-DEL-1727 | chr3-116884657-DEL-1727 |
| chr3-116927514-DEL-171 | chr3-116927515-DEL-171 |
| chr3-116951454-DEL-404 | chr3-116951454-DEL-404 |
| chr3-116962324-DEL-440 | chr3-116962325-DEL-440 |
| chr3-116964384-DEL-1132 | chr3-116964385-DEL-1132 |
| chr3-116974166-DEL-80 | chr3-116974167-DEL-80 |
| chr3-116993894-DEL-10327 | chr3-116993895-DEL-10330 |
| chr3-116995864-DEL-2929 | chr3-116995865-DEL-2929 |
| chr3-117016740-DEL-265 | chr3-117016741-DEL-265 |
| chr3-117046293-DEL-101 | chr3-117046294-DEL-101 |
| chr3-117050275-DEL-2509 | chr3-117050276-DEL-2509 |
| chr3-117056935-DEL-64 | chr3-117056936-DEL-64 |
| chr3-117060728-DEL-1876 | chr3-117060729-DEL-1876 |
| chr3-117079213-DEL-6926 | chr3-117079200-DEL-6927 |
| chr3-117097419-DEL-97 | chr3-117097420-DEL-97 |
| chr3-117107974-DEL-64 | chr3-117107975-DEL-64 |
| chr3-117134009-DEL-432 | chr3-117134010-DEL-432 |
| chr3-117143271-DEL-74 | chr3-117143272-DEL-74 |
| chr3-117144672-DEL-970 | chr3-117144673-DEL-969 |
| chr3-117146127-DEL-85 | chr3-117146128-DEL-83 |
| chr3-117157276-DEL-54 | chr3-117157277-DEL-54 |
| chr3-117162450-DEL-773 | chr3-117162451-DEL-773 |
| chr3-117172811-DEL-7088 | chr3-117172812-DEL-7088 |
| chr3-117180593-DEL-97 | chr3-117180594-DEL-97 |
| chr3-117216884-DEL-229 | chr3-117216875-DEL-230 |
| chr3-117233609-DEL-60 | chr3-117233610-DEL-60 |
| chr3-117234171-DEL-91 | chr3-117234171-DEL-91 |
| chr3-117282213-DEL-158 | chr3-117282215-DEL-176 |
| chr3-117322962-DEL-6184 | chr3-117322963-DEL-6184 |
| chr3-117342036-DEL-76 | chr3-117342037-DEL-76 |
| chr3-117382437-DEL-1431 | chr3-117382438-DEL-1431 |
| chr3-117403396-DEL-294 | chr3-117403397-DEL-294 |
| chr3-117406376-DEL-52 | chr3-117406363-DEL-63 |
| chr3-117420000-DEL-382 | chr3-117420001-DEL-382 |
| chr3-117429826-DEL-599 | chr3-117429826-DEL-598 |
| chr3-117472425-DEL-70 | chr3-117472426-DEL-70 |
| chr3-117506996-DEL-52 | chr3-117506998-DEL-51 |
| chr3-117534252-DEL-126 | chr3-117534262-DEL-81 |
| chr3-117547008-DEL-132 | chr3-117547000-DEL-94 |
| chr3-117556846-DEL-94 | chr3-117556895-DEL-56 |
| chr3-117560122-DEL-76 | chr3-117560125-DEL-58 |
| chr3-117564981-DEL-54 | chr3-117564982-DEL-54 |
| chr3-117565057-DEL-1806 | chr3-117565058-DEL-1806 |
| chr3-117573606-DEL-74 | chr3-117573607-DEL-74 |
| chr3-117613654-DEL-81 | chr3-117613655-DEL-84 |
| chr3-117618869-DEL-321 | chr3-117618870-DEL-321 |
| chr3-117644377-DEL-137 | chr3-117644378-DEL-153 |
| chr3-117656373-DEL-391 | chr3-117656374-DEL-391 |
| chr3-117699915-DEL-330 | chr3-117699916-DEL-330 |
| chr3-117718676-DEL-51 | chr3-117718677-DEL-51 |
| chr3-117750808-DEL-86 | chr3-117750809-DEL-86 |
| chr3-117755468-DEL-466 | chr3-117755483-DEL-446 |
| chr3-117774616-DEL-1639 | chr3-117774617-DEL-1639 |
| chr3-117781699-DEL-171 | chr3-117781700-DEL-171 |
| chr3-117848693-DEL-4981 | chr3-117848694-DEL-4981 |
| chr3-117858845-DEL-252 | chr3-117858846-DEL-252 |
| chr3-117898484-DEL-6395 | chr3-117898485-DEL-6395 |
| chr3-117913150-DEL-51 | chr3-117913151-DEL-51 |
| chr3-117972835-DEL-6355 | chr3-117972836-DEL-6355 |
| chr3-117997920-DEL-103 | chr3-117997919-DEL-97 |

| | |
|---|---|
| chr3-118003088-DEL-52 | chr3-118003088-DEL-52 |
| chr3-118010622-DEL-72 | chr3-118010610-DEL-50 |
| chr3-118074234-DEL-439 | chr3-118074235-DEL-439 |
| chr3-118152070-DEL-2804 | chr3-118152071-DEL-2804 |
| chr3-118156386-DEL-293 | chr3-118156386-DEL-293 |
| chr3-118170346-DEL-92 | chr3-118170349-DEL-94 |
| chr3-118195414-DEL-980 | chr3-118195415-DEL-980 |
| chr3-118264500-DEL-504 | chr3-118264501-DEL-504 |
| chr3-118273391-DEL-65 | chr3-118273392-DEL-64 |
| chr3-118275438-DEL-532 | chr3-118275439-DEL-532 |
| chr3-118290836-DEL-198 | chr3-118290831-DEL-199 |
| chr3-118291755-DEL-65 | chr3-118291761-DEL-77 |
| chr3-118313846-DEL-319 | chr3-118313847-DEL-319 |
| chr3-118330607-DEL-58 | chr3-118330608-DEL-58 |
| chr3-118361787-DEL-581 | chr3-118361796-DEL-571 |
| chr3-118362536-DEL-148 | chr3-118362537-DEL-148 |
| chr3-118365507-DEL-6386 | chr3-118365508-DEL-6386 |
| chr3-118382358-DEL-79 | chr3-118382362-DEL-68 |
| chr3-118389918-DEL-72 | chr3-118389919-DEL-72 |
| chr3-118403421-DEL-262 | chr3-118403422-DEL-262 |
| chr3-118408300-DEL-548 | chr3-118408301-DEL-548 |
| chr3-118427633-DEL-81 | chr3-118427634-DEL-81 |
| chr3-118440835-DEL-54 | chr3-118440836-DEL-54 |
| chr3-118450982-DEL-66 | chr3-118450983-DEL-66 |
| chr3-118475700-DEL-350 | chr3-118475710-DEL-351 |
| chr3-118501076-DEL-57 | chr3-118501076-DEL-57 |
| chr3-118507358-DEL-6480 | chr3-118507359-DEL-6480 |
| chr3-118568979-DEL-68 | chr3-118568984-DEL-64 |
| chr3-118577152-DEL-96 | chr3-118577177-DEL-72 |
| chr3-118584546-DEL-70 | chr3-118584548-DEL-57 |
| chr3-118587257-DEL-244 | chr3-118587258-DEL-244 |
| chr3-118645683-DEL-52 | chr3-118645686-DEL-50 |
| chr3-118646868-DEL-123 | chr3-118646869-DEL-123 |
| chr3-118693446-DEL-106 | chr3-118693481-DEL-70 |
| chr3-118698557-DEL-396 | chr3-118698558-DEL-396 |
| chr3-118713088-DEL-1467 | chr3-118713089-DEL-1467 |
| chr3-118714628-DEL-574 | chr3-118714629-DEL-574 |
| chr3-118718458-DEL-1996 | chr3-118718459-DEL-1996 |
| chr3-118739103-DEL-92 | chr3-118739115-DEL-56 |
| chr3-118743436-DEL-98 | chr3-118743428-DEL-146 |
| chr3-118752224-DEL-519 | chr3-118752225-DEL-519 |
| chr3-118759933-DEL-103 | chr3-118759926-DEL-102 |
| chr3-118770425-DEL-74 | chr3-118770419-DEL-87 |
| chr3-118803369-DEL-1848 | chr3-118803370-DEL-1848 |
| chr3-118850179-DEL-64 | chr3-118850182-DEL-52 |
| chr3-118889603-DEL-615 | chr3-118889604-DEL-615 |
| chr3-118904627-DEL-2735 | chr3-118904628-DEL-2735 |
| chr3-118910885-DEL-224 | chr3-118910888-DEL-220 |
| chr3-118914760-DEL-67 | chr3-118914738-DEL-77 |
| chr3-118931607-DEL-56 | chr3-118931614-DEL-50 |
| chr3-118964461-DEL-109 | chr3-118964462-DEL-109 |
| chr3-118969143-DEL-64 | chr3-118969142-DEL-70 |
| chr3-118985787-DEL-191 | chr3-118985788-DEL-191 |
| chr3-119028905-DEL-262 | chr3-119028906-DEL-262 |
| chr3-119053234-DEL-1234 | chr3-119053235-DEL-1234 |
| chr3-119148497-DEL-6191 | chr3-119148498-DEL-6191 |
| chr3-119188697-DEL-237 | chr3-119188698-DEL-237 |
| chr3-119196233-DEL-76 | chr3-119196240-DEL-56 |
| chr3-119206759-DEL-81 | chr3-119206760-DEL-91 |
| chr3-119225365-DEL-52 | chr3-119225366-DEL-52 |
| chr3-119237959-DEL-72 | chr3-119237966-DEL-54 |
| chr3-119246448-DEL-66 | chr3-119246447-DEL-52 |
| chr3-119259962-DEL-77 | chr3-119259965-DEL-82 |
| chr3-119273017-DEL-65 | chr3-119273018-DEL-65 |
| chr3-119276864-DEL-419 | chr3-119276848-DEL-436 |
| chr3-119338244-DEL-1166 | chr3-119338245-DEL-1166 |
| chr3-119342922-DEL-4101 | chr3-119342923-DEL-4101 |
| chr3-119374530-DEL-56 | chr3-119374531-DEL-84 |
| chr3-119390508-DEL-74 | chr3-119390483-DEL-64 |
| chr3-119403157-DEL-498 | chr3-119403158-DEL-498 |
| chr3-119449520-DEL-4145 | chr3-119449521-DEL-4145 |
| chr3-119459228-DEL-2351 | chr3-119459229-DEL-2351 |
| chr3-119503155-DEL-419 | chr3-119503156-DEL-419 |
| chr3-119567742-DEL-1355 | chr3-119567743-DEL-1355 |
| chr3-119573593-DEL-65 | chr3-119573598-DEL-61 |
| chr3-119625377-DEL-6408 | chr3-119625378-DEL-6408 |
| chr3-119673617-DEL-194 | chr3-119673618-DEL-194 |
| chr3-119678818-DEL-101 | chr3-119678819-DEL-101 |
| chr3-119685494-DEL-744 | chr3-119685511-DEL-728 |
| chr3-119693302-DEL-2075 | chr3-119693303-DEL-2075 |

| | |
|---|---|
| chr3-119698540-DEL-74 | chr3-119698517-DEL-56 |
| chr3-119771279-DEL-6352 | chr3-119771280-DEL-6352 |
| chr3-119803294-DEL-58 | chr3-119803294-DEL-58 |
| chr3-119803718-DEL-254 | chr3-119803719-DEL-254 |
| chr3-119833591-DEL-50 | chr3-119833592-DEL-75 |
| chr3-119839774-DEL-93 | chr3-119839770-DEL-68 |
| chr3-119869945-DEL-859 | chr3-119869946-DEL-859 |
| chr3-119879380-DEL-60 | chr3-119879381-DEL-60 |
| chr3-119894460-DEL-84 | chr3-119894505-DEL-58 |
| chr3-119898309-DEL-6936 | chr3-119898310-DEL-6936 |
| chr3-119913398-DEL-1394 | chr3-119913399-DEL-1394 |
| chr3-119956549-DEL-93 | chr3-119956550-DEL-93 |
| chr3-119989047-DEL-56 | chr3-119989048-DEL-54 |
| chr3-120016571-DEL-67 | chr3-120016572-DEL-70 |
| chr3-120097055-DEL-68 | chr3-120097056-DEL-68 |
| chr3-120128107-DEL-69 | chr3-120128109-DEL-66 |
| chr3-120148942-DEL-127 | chr3-120148941-DEL-107 |
| chr3-120170443-DEL-127 | chr3-120170444-DEL-97 |
| chr3-120183546-DEL-6244 | chr3-120183547-DEL-6244 |
| chr3-120240774-DEL-481 | chr3-120240775-DEL-481 |
| chr3-120341434-DEL-74 | chr3-120341437-DEL-72 |
| chr3-120351117-DEL-302 | chr3-120351141-DEL-259 |
| chr3-120486461-DEL-822 | chr3-120486462-DEL-822 |
| chr3-120494508-DEL-149 | chr3-120494509-DEL-149 |
| chr3-120502040-DEL-166 | chr3-120502041-DEL-166 |
| chr3-120508860-DEL-1185 | chr3-120508862-DEL-1181 |
| chr3-120510151-DEL-2155 | chr3-120510152-DEL-2155 |
| chr3-120510788-DEL-388 | chr3-120510789-DEL-388 |
| chr3-120528683-DEL-7467 | chr3-120528682-DEL-7468 |
| chr3-120544503-DEL-276 | chr3-120544504-DEL-276 |
| chr3-120567199-DEL-56 | chr3-120567189-DEL-58 |
| chr3-120594514-DEL-491 | chr3-120594512-DEL-494 |
| chr3-120607765-DEL-341 | chr3-120607766-DEL-341 |
| chr3-120618846-DEL-7507 | chr3-120618846-DEL-7507 |
| chr3-120640450-DEL-4809 | chr3-120640451-DEL-4809 |
| chr3-120686472-DEL-56 | chr3-120686473-DEL-56 |
| chr3-120697194-DEL-53 | chr3-120697195-DEL-53 |
| chr3-120701392-DEL-111 | chr3-120701393-DEL-111 |
| chr3-120745289-DEL-218 | chr3-120745282-DEL-226 |
| chr3-120774300-DEL-71 | chr3-120774290-DEL-84 |
| chr3-120786583-DEL-76 | chr3-120786584-DEL-76 |
| chr3-120787853-DEL-202 | chr3-120787854-DEL-202 |
| chr3-120805656-DEL-58 | chr3-120805659-DEL-56 |
| chr3-120808118-DEL-54 | chr3-120808118-DEL-104 |
| chr3-120810272-DEL-1157 | chr3-120810273-DEL-1157 |
| chr3-120814111-DEL-312 | chr3-120814112-DEL-312 |
| chr3-120837353-DEL-186 | chr3-120837353-DEL-186 |
| chr3-120840989-DEL-7225 | chr3-120840990-DEL-7225 |
| chr3-120856486-DEL-3046 | chr3-120856487-DEL-3046 |
| chr3-120879365-DEL-3879 | chr3-120879366-DEL-3879 |
| chr3-120923782-DEL-2524 | chr3-120923783-DEL-2524 |
| chr3-120931019-DEL-2007 | chr3-120931020-DEL-2007 |
| chr3-120939925-DEL-63 | chr3-120939926-DEL-63 |
| chr3-120943479-DEL-1883 | chr3-120943480-DEL-1883 |
| chr3-120954790-DEL-86 | chr3-120954783-DEL-77 |
| chr3-120970902-DEL-52 | chr3-120970903-DEL-52 |
| chr3-120976142-DEL-218 | chr3-120976155-DEL-232 |
| chr3-120990495-DEL-400 | chr3-120990496-DEL-400 |
| chr3-121003789-DEL-311 | chr3-121003790-DEL-311 |
| chr3-121016295-DEL-51 | chr3-121016276-DEL-71 |
| chr3-121029882-DEL-153 | chr3-121029873-DEL-166 |
| chr3-121033386-DEL-52 | chr3-121033381-DEL-66 |
| chr3-121040738-DEL-216 | chr3-121040739-DEL-216 |
| chr3-121082646-DEL-514 | chr3-121082647-DEL-514 |
| chr3-121104264-DEL-153 | chr3-121104265-DEL-150 |
| chr3-121110006-DEL-1141 | chr3-121110008-DEL-1143 |
| chr3-121117695-DEL-133 | chr3-121117696-DEL-133 |
| chr3-121129977-DEL-81 | chr3-121129978-DEL-81 |
| chr3-121136210-DEL-97 | chr3-121136210-DEL-94 |
| chr3-121150234-DEL-253 | chr3-121150234-DEL-250 |
| chr3-121152745-DEL-93 | chr3-121152746-DEL-93 |
| chr3-121162637-DEL-84 | chr3-121162638-DEL-84 |
| chr3-121177662-DEL-156 | chr3-121177666-DEL-153 |
| chr3-121186395-DEL-280 | chr3-121186396-DEL-280 |
| chr3-121187100-DEL-57 | chr3-121187101-DEL-51 |
| chr3-121192941-DEL-68 | chr3-121192917-DEL-57 |
| chr3-121199545-DEL-160 | chr3-121199550-DEL-156 |
| chr3-121207049-DEL-196 | chr3-121207041-DEL-197 |
| chr3-121214282-DEL-485 | chr3-121214283-DEL-485 |
| chr3-121244077-DEL-68 | chr3-121244078-DEL-68 |

| | |
|---|---|
| chr3-121260405-DEL-60 | chr3-121260406-DEL-60 |
| chr3-121287852-DEL-162 | chr3-121287853-DEL-162 |
| chr3-121316861-DEL-1371 | chr3-121316862-DEL-1371 |
| chr3-121321419-DEL-56 | chr3-121321420-DEL-56 |
| chr3-121335411-DEL-291 | chr3-121335419-DEL-282 |
| chr3-121344332-DEL-77 | chr3-121344333-DEL-77 |
| chr3-121358079-DEL-223 | chr3-121358080-DEL-223 |
| chr3-121390308-DEL-53 | chr3-121390309-DEL-53 |
| chr3-121408531-DEL-160 | chr3-121408532-DEL-160 |
| chr3-121423197-DEL-119 | chr3-121423196-DEL-119 |
| chr3-121447733-DEL-104 | chr3-121447742-DEL-96 |
| chr3-121497962-DEL-215 | chr3-121497963-DEL-215 |
| chr3-121524151-DEL-247 | chr3-121524152-DEL-247 |
| chr3-121539800-DEL-486 | chr3-121539801-DEL-486 |
| chr3-121541835-DEL-3530 | chr3-121541835-DEL-3530 |
| chr3-121555938-DEL-2666 | chr3-121555939-DEL-2666 |
| chr3-121564423-DEL-113 | chr3-121564466-DEL-113 |
| chr3-121578968-DEL-198 | chr3-121578969-DEL-198 |
| chr3-121586611-DEL-372 | chr3-121586612-DEL-372 |
| chr3-121591214-DEL-137 | chr3-121591215-DEL-137 |
| chr3-121626878-DEL-111 | chr3-121626879-DEL-111 |
| chr3-121653787-DEL-346 | chr3-121653788-DEL-346 |
| chr3-121657990-DEL-227 | chr3-121657991-DEL-227 |
| chr3-121704885-DEL-51 | chr3-121704886-DEL-51 |
| chr3-121711873-DEL-60 | chr3-121711874-DEL-60 |
| chr3-121792913-DEL-73 | chr3-121792936-DEL-59 |
| chr3-121803953-DEL-110 | chr3-121803954-DEL-110 |
| chr3-121822703-DEL-418 | chr3-121822704-DEL-418 |
| chr3-121839203-DEL-71 | chr3-121839205-DEL-72 |
| chr3-121857575-DEL-440 | chr3-121857576-DEL-440 |
| chr3-121912836-DEL-445 | chr3-121912834-DEL-442 |
| chr3-121938683-DEL-1507 | chr3-121938684-DEL-1507 |
| chr3-121939166-DEL-104 | chr3-121939167-DEL-104 |
| chr3-121943638-DEL-289 | chr3-121943639-DEL-289 |
| chr3-121958789-DEL-149 | chr3-121958790-DEL-149 |
| chr3-121962209-DEL-373 | chr3-121962192-DEL-374 |
| chr3-121972117-DEL-406 | chr3-121972118-DEL-406 |
| chr3-121984598-DEL-166 | chr3-121984599-DEL-162 |
| chr3-122023632-DEL-1289 | chr3-122023629-DEL-1290 |
| chr3-122026356-DEL-210 | chr3-122026357-DEL-210 |
| chr3-122043102-DEL-220 | chr3-122043103-DEL-219 |
| chr3-122080634-DEL-200 | chr3-122080635-DEL-200 |
| chr3-122082968-DEL-200 | chr3-122082969-DEL-200 |
| chr3-122092228-DEL-795 | chr3-122092229-DEL-795 |
| chr3-122099382-DEL-360 | chr3-122099383-DEL-360 |
| chr3-122109385-DEL-209 | chr3-122109386-DEL-209 |
| chr3-122111358-DEL-787 | chr3-122111359-DEL-787 |
| chr3-122120468-DEL-1280 | chr3-122120469-DEL-1280 |
| chr3-122122545-DEL-340 | chr3-122122506-DEL-380 |
| chr3-122145668-DEL-572 | chr3-122145669-DEL-572 |
| chr3-122184366-DEL-221 | chr3-122184367-DEL-221 |
| chr3-122187008-DEL-72 | chr3-122187009-DEL-72 |
| chr3-122232854-DEL-54 | chr3-122232855-DEL-54 |
| chr3-122238471-DEL-77 | chr3-122238449-DEL-50 |
| chr3-122250457-DEL-66 | chr3-122250458-DEL-66 |
| chr3-122266610-DEL-201 | chr3-122266611-DEL-201 |
| chr3-122273495-DEL-82 | chr3-122273500-DEL-58 |
| chr3-122292012-DEL-105 | chr3-122292013-DEL-105 |
| chr3-122315800-DEL-161 | chr3-122315801-DEL-161 |
| chr3-122318319-DEL-79 | chr3-122318301-DEL-80 |
| chr3-122425061-DEL-60 | chr3-122425062-DEL-60 |
| chr3-122431813-DEL-207 | chr3-122431814-DEL-207 |
| chr3-122447865-DEL-2977 | chr3-122447866-DEL-2977 |
| chr3-122459308-DEL-336 | chr3-122459309-DEL-336 |
| chr3-122460202-DEL-53 | chr3-122460203-DEL-53 |
| chr3-122474869-DEL-902 | chr3-122474870-DEL-899 |
| chr3-122496077-DEL-210 | chr3-122496100-DEL-188 |
| chr3-122509011-DEL-83 | chr3-122509010-DEL-87 |
| chr3-122521389-DEL-207 | chr3-122521390-DEL-207 |
| chr3-122532772-DEL-53 | chr3-122532771-DEL-57 |
| chr3-122559072-DEL-52 | chr3-122559073-DEL-52 |
| chr3-122581108-DEL-59 | chr3-122581109-DEL-58 |
| chr3-122610224-DEL-162 | chr3-122610225-DEL-171 |
| chr3-122616159-DEL-115 | chr3-122616158-DEL-104 |
| chr3-122639427-DEL-718 | chr3-122639428-DEL-718 |
| chr3-122641206-DEL-144 | chr3-122641207-DEL-144 |
| chr3-122698949-DEL-94 | chr3-122698936-DEL-96 |
| chr3-122736795-DEL-51 | chr3-122736796-DEL-51 |
| chr3-122745056-DEL-200 | chr3-122745057-DEL-200 |
| chr3-122764435-DEL-253 | chr3-122764436-DEL-230 |

| | |
|---|---|
| chr3-122796422-DEL-54 | chr3-122796423-DEL-54 |
| chr3-122801719-DEL-60 | chr3-122801720-DEL-60 |
| chr3-122808715-DEL-508 | chr3-122808716-DEL-508 |
| chr3-122811690-DEL-402 | chr3-122811691-DEL-402 |
| chr3-122838643-DEL-52 | chr3-122838644-DEL-52 |
| chr3-122849394-DEL-52 | chr3-122849395-DEL-52 |
| chr3-122889786-DEL-92 | chr3-122889787-DEL-92 |
| chr3-122896472-DEL-52 | chr3-122896473-DEL-52 |
| chr3-122898915-DEL-105 | chr3-122898916-DEL-105 |
| chr3-122905266-DEL-79 | chr3-122905267-DEL-79 |
| chr3-122926124-DEL-58 | chr3-122926125-DEL-58 |
| chr3-122960347-DEL-315 | chr3-122960348-DEL-315 |
| chr3-122978690-DEL-51 | chr3-122978682-DEL-60 |
| chr3-122980749-DEL-106 | chr3-122980744-DEL-112 |
| chr3-122988438-DEL-6293 | chr3-122988439-DEL-6293 |
| chr3-123013725-DEL-1074 | chr3-123013726-DEL-1074 |
| chr3-123019739-DEL-72 | chr3-123019740-DEL-72 |
| chr3-123025883-DEL-204 | chr3-123025884-DEL-204 |
| chr3-123055789-DEL-126 | chr3-123055790-DEL-132 |
| chr3-123058337-DEL-102 | chr3-123058337-DEL-102 |
| chr3-123097460-DEL-202 | chr3-123097461-DEL-202 |
| chr3-123119864-DEL-136 | chr3-123119865-DEL-117 |
| chr3-123124453-DEL-931 | chr3-123124454-DEL-931 |
| chr3-123125994-DEL-58 | chr3-123125990-DEL-62 |
| chr3-123135059-DEL-736 | chr3-123135060-DEL-736 |
| chr3-123158263-DEL-56 | chr3-123158263-DEL-56 |
| chr3-123189683-DEL-62 | chr3-123189684-DEL-62 |
| chr3-123192528-DEL-87 | chr3-123192491-DEL-79 |
| chr3-123203766-DEL-1433 | chr3-123203767-DEL-1433 |
| chr3-123214159-DEL-505 | chr3-123214160-DEL-505 |
| chr3-123216494-DEL-77 | chr3-123216495-DEL-77 |
| chr3-123245168-DEL-577 | chr3-123245167-DEL-579 |
| chr3-123265402-DEL-4379 | chr3-123265403-DEL-4379 |
| chr3-123294298-DEL-1680 | chr3-123294299-DEL-1680 |
| chr3-123296167-DEL-386 | chr3-123296168-DEL-386 |
| chr3-123305845-DEL-81 | chr3-123305846-DEL-81 |
| chr3-123309761-DEL-59 | chr3-123309750-DEL-71 |
| chr3-123311101-DEL-404 | chr3-123311090-DEL-413 |
| chr3-123317133-DEL-57 | chr3-123317121-DEL-58 |
| chr3-123347060-DEL-196 | chr3-123347061-DEL-196 |
| chr3-123356012-DEL-550 | chr3-123356013-DEL-550 |
| chr3-123374039-DEL-402 | chr3-123374040-DEL-402 |
| chr3-123377397-DEL-123 | chr3-123377398-DEL-103 |
| chr3-123397536-DEL-87 | chr3-123397537-DEL-87 |
| chr3-123398566-DEL-273 | chr3-123398567-DEL-273 |
| chr3-123416751-DEL-1914 | chr3-123416752-DEL-1914 |
| chr3-123421894-DEL-72 | chr3-123421923-DEL-84 |
| chr3-123485650-DEL-80 | chr3-123485652-DEL-78 |
| chr3-123497654-DEL-219 | chr3-123497642-DEL-217 |
| chr3-123499980-DEL-73 | chr3-123499979-DEL-73 |
| chr3-123523305-DEL-92 | chr3-123523306-DEL-92 |
| chr3-123532403-DEL-4649 | chr3-123532404-DEL-4649 |
| chr3-123557221-DEL-97 | chr3-123557222-DEL-96 |
| chr3-123566902-DEL-247 | chr3-123566906-DEL-243 |
| chr3-123567699-DEL-293 | chr3-123567700-DEL-293 |
| chr3-123606181-DEL-62 | chr3-123606183-DEL-63 |
| chr3-123611165-DEL-6204 | chr3-123611166-DEL-6204 |
| chr3-123616669-DEL-105 | chr3-123616670-DEL-105 |
| chr3-123619641-DEL-276 | chr3-123619642-DEL-276 |
| chr3-123627082-DEL-57 | chr3-123627083-DEL-57 |
| chr3-123659741-DEL-122 | chr3-123659738-DEL-123 |
| chr3-123662175-DEL-64 | chr3-123662176-DEL-60 |
| chr3-123664181-DEL-74 | chr3-123664194-DEL-74 |
| chr3-123670382-DEL-420 | chr3-123670383-DEL-420 |
| chr3-123672869-DEL-6205 | chr3-123672870-DEL-6205 |
| chr3-123706507-DEL-64 | chr3-123706508-DEL-64 |
| chr3-123720699-DEL-7146 | chr3-123720700-DEL-7146 |
| chr3-123739351-DEL-61 | chr3-123739352-DEL-61 |
| chr3-123744657-DEL-955 | chr3-123744658-DEL-955 |
| chr3-123765379-DEL-65 | chr3-123765376-DEL-66 |
| chr3-123770056-DEL-71 | chr3-123770057-DEL-71 |
| chr3-123784581-DEL-313 | chr3-123784586-DEL-309 |
| chr3-123814687-DEL-2504 | chr3-123814688-DEL-2504 |
| chr3-123823542-DEL-51 | chr3-123823544-DEL-52 |
| chr3-123825883-DEL-13513 | chr3-123825884-DEL-13512 |
| chr3-123849821-DEL-222 | chr3-123849822-DEL-220 |
| chr3-123852471-DEL-2492 | chr3-123852472-DEL-2492 |
| chr3-123855929-DEL-2327 | chr3-123855930-DEL-2327 |
| chr3-123866610-DEL-401 | chr3-123866611-DEL-401 |
| chr3-123904877-DEL-183 | chr3-123904878-DEL-183 |

| | |
|---|---|
| chr3-123911077-DEL-387 | chr3-123911078-DEL-387 |
| chr3-123911792-DEL-410 | chr3-123911794-DEL-409 |
| chr3-123916344-DEL-55 | chr3-123916345-DEL-55 |
| chr3-123929712-DEL-70 | chr3-123929713-DEL-70 |
| chr3-123939537-DEL-90 | chr3-123939538-DEL-90 |
| chr3-123947300-DEL-177 | chr3-123947301-DEL-177 |
| chr3-123954187-DEL-6169 | chr3-123954188-DEL-6169 |
| chr3-123982505-DEL-514 | chr3-123982506-DEL-514 |
| chr3-123998364-DEL-340 | chr3-123998365-DEL-340 |
| chr3-124031487-DEL-2126 | chr3-124031487-DEL-2126 |
| chr3-124033785-DEL-73 | chr3-124033785-DEL-73 |
| chr3-124040554-DEL-138 | chr3-124040547-DEL-146 |
| chr3-124075843-DEL-7749 | chr3-124075829-DEL-7741 |
| chr3-124087153-DEL-112 | chr3-124087182-DEL-84 |
| chr3-124102168-DEL-1653 | chr3-124102169-DEL-1653 |
| chr3-124124030-DEL-99 | chr3-124124030-DEL-100 |
| chr3-124130627-DEL-154 | chr3-124130628-DEL-154 |
| chr3-124133191-DEL-221 | chr3-124133192-DEL-221 |
| chr3-124136637-DEL-149 | chr3-124136638-DEL-149 |
| chr3-124143715-DEL-1016 | chr3-124143716-DEL-1016 |
| chr3-124145999-DEL-1937 | chr3-124146000-DEL-1937 |
| chr3-124152613-DEL-1437 | chr3-124152614-DEL-1437 |
| chr3-124179145-DEL-1201 | chr3-124179146-DEL-1201 |
| chr3-124190148-DEL-1516 | chr3-124190149-DEL-1516 |
| chr3-124203262-DEL-140 | chr3-124203263-DEL-140 |
| chr3-124238943-DEL-176 | chr3-124238944-DEL-176 |
| chr3-124243123-DEL-859 | chr3-124243124-DEL-859 |
| chr3-124251442-DEL-90 | chr3-124251442-DEL-89 |
| chr3-124301598-DEL-3069 | chr3-124301599-DEL-3069 |
| chr3-124360241-DEL-140 | chr3-124360242-DEL-140 |
| chr3-124361813-DEL-2671 | chr3-124361813-DEL-2671 |
| chr3-124408255-DEL-184 | chr3-124408256-DEL-184 |
| chr3-124411110-DEL-54 | chr3-124411090-DEL-71 |
| chr3-124440001-DEL-1000 | chr3-124440002-DEL-1000 |
| chr3-124441115-DEL-198 | chr3-124441116-DEL-198 |
| chr3-124442283-DEL-239 | chr3-124442284-DEL-239 |
| chr3-124457118-DEL-66 | chr3-124457111-DEL-74 |
| chr3-124458324-DEL-401 | chr3-124458325-DEL-401 |
| chr3-124459090-DEL-194 | chr3-124459091-DEL-194 |
| chr3-124471804-DEL-3117 | chr3-124471805-DEL-3117 |
| chr3-124509444-DEL-300 | chr3-124509445-DEL-300 |
| chr3-124511112-DEL-940 | chr3-124511113-DEL-940 |
| chr3-124517089-DEL-1002 | chr3-124517090-DEL-1002 |
| chr3-124540703-DEL-391 | chr3-124540704-DEL-391 |
| chr3-124557388-DEL-2968 | chr3-124557389-DEL-2968 |
| chr3-124569692-DEL-92 | chr3-124569687-DEL-84 |
| chr3-124591834-DEL-109 | chr3-124591835-DEL-109 |
| chr3-124637436-DEL-273 | chr3-124637433-DEL-274 |
| chr3-124643867-DEL-186 | chr3-124643868-DEL-176 |
| chr3-124663949-DEL-6076 | chr3-124663950-DEL-6076 |
| chr3-124684139-DEL-58 | chr3-124684140-DEL-58 |
| chr3-124700373-DEL-60 | chr3-124700374-DEL-60 |
| chr3-124706779-DEL-60 | chr3-124706782-DEL-56 |
| chr3-124715690-DEL-76 | chr3-124715687-DEL-78 |
| chr3-124736740-DEL-3163 | chr3-124736741-DEL-3163 |
| chr3-124754456-DEL-80 | chr3-124754457-DEL-82 |
| chr3-124765694-DEL-1966 | chr3-124765695-DEL-1966 |
| chr3-124805252-DEL-634 | chr3-124805253-DEL-634 |
| chr3-124823351-DEL-6160 | chr3-124823352-DEL-6160 |
| chr3-124848870-DEL-74 | chr3-124848871-DEL-74 |
| chr3-124855488-DEL-791 | chr3-124855466-DEL-814 |
| chr3-124857053-DEL-199 | chr3-124857054-DEL-199 |
| chr3-124857936-DEL-66 | chr3-124857939-DEL-64 |
| chr3-124869502-DEL-6584 | chr3-124869503-DEL-6584 |
| chr3-124891317-DEL-3426 | chr3-124891318-DEL-3426 |
| chr3-124903756-DEL-72 | chr3-124903754-DEL-73 |
| chr3-124926636-DEL-558 | chr3-124926637-DEL-558 |
| chr3-124928988-DEL-320 | chr3-124928989-DEL-319 |
| chr3-124952981-DEL-345 | chr3-124952982-DEL-345 |
| chr3-124962924-DEL-124 | chr3-124962925-DEL-124 |
| chr3-124967141-DEL-105 | chr3-124967142-DEL-105 |
| chr3-125002042-DEL-7241 | chr3-125002043-DEL-7241 |
| chr3-125012605-DEL-65 | chr3-125012606-DEL-65 |
| chr3-125019304-DEL-3109 | chr3-125019304-DEL-3110 |
| chr3-125027167-DEL-214 | chr3-125027168-DEL-214 |
| chr3-125030964-DEL-1335 | chr3-125030962-DEL-1333 |
| chr3-125038074-DEL-2272 | chr3-125038075-DEL-2272 |
| chr3-125068238-DEL-103 | chr3-125068235-DEL-60 |
| chr3-125107613-DEL-54 | chr3-125107609-DEL-52 |
| chr3-125118895-DEL-985 | chr3-125118896-DEL-985 |

| | |
|---|---|
| chr3-125165942-DEL-184 | chr3-125165945-DEL-182 |
| chr3-125183746-DEL-4682 | chr3-125183747-DEL-4682 |
| chr3-125211965-DEL-206 | chr3-125211966-DEL-206 |
| chr3-125248698-DEL-317 | chr3-125248684-DEL-332 |
| chr3-125272091-DEL-63 | chr3-125272092-DEL-63 |
| chr3-125301663-DEL-1506 | chr3-125301674-DEL-1514 |
| chr3-125329996-DEL-105 | chr3-125329997-DEL-105 |
| chr3-125352245-DEL-196 | chr3-125352246-DEL-196 |
| chr3-125360017-DEL-51 | chr3-125360016-DEL-51 |
| chr3-125364578-DEL-2573 | chr3-125364578-DEL-2573 |
| chr3-125368053-DEL-246 | chr3-125368054-DEL-246 |
| chr3-125453487-DEL-262 | chr3-125453490-DEL-250 |
| chr3-125455710-DEL-120 | chr3-125455709-DEL-97 |
| chr3-125493418-DEL-287 | chr3-125493419-DEL-287 |
| chr3-125540631-DEL-1677 | chr3-125540630-DEL-1678 |
| chr3-125549657-DEL-467 | chr3-125549670-DEL-455 |
| chr3-125598170-DEL-58 | chr3-125598171-DEL-60 |
| chr3-125600789-DEL-199 | chr3-125600790-DEL-199 |
| chr3-125610774-DEL-51 | chr3-125610774-DEL-51 |
| chr3-125623842-DEL-78 | chr3-125623841-DEL-52 |
| chr3-125624977-DEL-110 | chr3-125624978-DEL-110 |
| chr3-125650472-DEL-83 | chr3-125650477-DEL-88 |
| chr3-125662891-DEL-99 | chr3-125662892-DEL-99 |
| chr3-125669786-DEL-1042 | chr3-125669787-DEL-1042 |
| chr3-125673119-DEL-133 | chr3-125673120-DEL-133 |
| chr3-125696437-DEL-6180 | chr3-125696438-DEL-6180 |
| chr3-125713028-DEL-348 | chr3-125713065-DEL-320 |
| chr3-125718850-DEL-53 | chr3-125718850-DEL-54 |
| chr3-125752828-DEL-6861 | chr3-125752829-DEL-6861 |
| chr3-125758938-DEL-416 | chr3-125758939-DEL-416 |
| chr3-125786128-DEL-75 | chr3-125786113-DEL-75 |
| chr3-125797846-DEL-1854 | chr3-125797847-DEL-1854 |
| chr3-125834477-DEL-153 | chr3-125834478-DEL-153 |
| chr3-125873359-DEL-406 | chr3-125873360-DEL-406 |
| chr3-125878570-DEL-184 | chr3-125878588-DEL-173 |
| chr3-125880506-DEL-107 | chr3-125880507-DEL-107 |
| chr3-125905011-DEL-81 | chr3-125905012-DEL-80 |
| chr3-125907776-DEL-762 | chr3-125907777-DEL-762 |
| chr3-125913232-DEL-887 | chr3-125913233-DEL-887 |
| chr3-125921020-DEL-8532 | chr3-125921021-DEL-8532 |
| chr3-125952088-DEL-231 | chr3-125952091-DEL-226 |
| chr3-125952857-DEL-6401 | chr3-125952841-DEL-6417 |
| chr3-125971372-DEL-629 | chr3-125971373-DEL-629 |
| chr3-125974524-DEL-160 | chr3-125974527-DEL-158 |
| chr3-125998791-DEL-76 | chr3-125998792-DEL-76 |
| chr3-126036710-DEL-51 | chr3-126036690-DEL-72 |
| chr3-126047668-DEL-2872 | chr3-126047669-DEL-2872 |
| chr3-126054420-DEL-105 | chr3-126054423-DEL-104 |
| chr3-126056625-DEL-68 | chr3-126056637-DEL-50 |
| chr3-126057024-DEL-4149 | chr3-126057025-DEL-4149 |
| chr3-126068346-DEL-324 | chr3-126068347-DEL-324 |
| chr3-126114655-DEL-99 | chr3-126114656-DEL-94 |
| chr3-126125913-DEL-106 | chr3-126125914-DEL-106 |
| chr3-126128519-DEL-155 | chr3-126128520-DEL-155 |
| chr3-126165234-DEL-201 | chr3-126165235-DEL-201 |
| chr3-126170170-DEL-378 | chr3-126170171-DEL-378 |
| chr3-126174641-DEL-111 | chr3-126174642-DEL-114 |
| chr3-126188299-DEL-407 | chr3-126188300-DEL-407 |
| chr3-126216562-DEL-631 | chr3-126216562-DEL-631 |
| chr3-126218449-DEL-230 | chr3-126218450-DEL-230 |
| chr3-126226331-DEL-52 | chr3-126226329-DEL-54 |
| chr3-126241357-DEL-335 | chr3-126241358-DEL-335 |
| chr3-126273675-DEL-64 | chr3-126273682-DEL-67 |
| chr3-126279847-DEL-1112 | chr3-126279848-DEL-1112 |
| chr3-126300012-DEL-338 | chr3-126300013-DEL-338 |
| chr3-126302488-DEL-89 | chr3-126302489-DEL-89 |
| chr3-126314241-DEL-796 | chr3-126314225-DEL-797 |
| chr3-126371919-DEL-75 | chr3-126371920-DEL-75 |
| chr3-126373416-DEL-68 | chr3-126373413-DEL-72 |
| chr3-126384343-DEL-1617 | chr3-126384344-DEL-1617 |
| chr3-126405184-DEL-51 | chr3-126405185-DEL-51 |
| chr3-126408179-DEL-54 | chr3-126408180-DEL-54 |
| chr3-126419844-DEL-403 | chr3-126419854-DEL-394 |
| chr3-126429267-DEL-59 | chr3-126429265-DEL-61 |
| chr3-126462572-DEL-105 | chr3-126462573-DEL-105 |
| chr3-126499743-DEL-612 | chr3-126499744-DEL-612 |
| chr3-126536888-DEL-166 | chr3-126536889-DEL-166 |
| chr3-126547711-DEL-580 | chr3-126547712-DEL-580 |
| chr3-126554122-DEL-60 | chr3-126554123-DEL-60 |
| chr3-126569199-DEL-171 | chr3-126569199-DEL-169 |

| | |
|---|---|
| chr3-126654417-DEL-4669 | chr3-126654418-DEL-4669 |
| chr3-126660016-DEL-55 | chr3-126660017-DEL-55 |
| chr3-126664042-DEL-199 | chr3-126664043-DEL-199 |
| chr3-126696905-DEL-196 | chr3-126696906-DEL-196 |
| chr3-126729506-DEL-231 | chr3-126729507-DEL-231 |
| chr3-126777531-DEL-200 | chr3-126777532-DEL-201 |
| chr3-126806028-DEL-52 | chr3-126806027-DEL-56 |
| chr3-126820924-DEL-139 | chr3-126820956-DEL-108 |
| chr3-126833761-DEL-86 | chr3-126833762-DEL-86 |
| chr3-126899300-DEL-765 | chr3-126899301-DEL-765 |
| chr3-126912386-DEL-116 | chr3-126912387-DEL-116 |
| chr3-126922121-DEL-99 | chr3-126922122-DEL-99 |
| chr3-126928932-DEL-64 | chr3-126928935-DEL-64 |
| chr3-126934507-DEL-6370 | chr3-126934508-DEL-6370 |
| chr3-126942083-DEL-95 | chr3-126942084-DEL-95 |
| chr3-126943263-DEL-247 | chr3-126943264-DEL-221 |
| chr3-127012286-DEL-111 | chr3-127012287-DEL-111 |
| chr3-127013346-DEL-1441 | chr3-127013347-DEL-1441 |
| chr3-127029203-DEL-77 | chr3-127029192-DEL-73 |
| chr3-127047964-DEL-60 | chr3-127047965-DEL-60 |
| chr3-127073099-DEL-60 | chr3-127073100-DEL-60 |
| chr3-127078654-DEL-1576 | chr3-127078655-DEL-1576 |
| chr3-127081434-DEL-110 | chr3-127081435-DEL-62 |
| chr3-127092097-DEL-326 | chr3-127092098-DEL-326 |
| chr3-127100129-DEL-52 | chr3-127100130-DEL-52 |
| chr3-127165543-DEL-1033 | chr3-127165544-DEL-1032 |
| chr3-127173547-DEL-84 | chr3-127173548-DEL-84 |
| chr3-127179265-DEL-66 | chr3-127179269-DEL-63 |
| chr3-127186770-DEL-424 | chr3-127186773-DEL-422 |
| chr3-127187540-DEL-168 | chr3-127187541-DEL-168 |
| chr3-127194301-DEL-112 | chr3-127194294-DEL-120 |
| chr3-127215212-DEL-2413 | chr3-127215213-DEL-2413 |
| chr3-127226616-DEL-72 | chr3-127226617-DEL-72 |
| chr3-127228888-DEL-191 | chr3-127228887-DEL-192 |
| chr3-127232896-DEL-193 | chr3-127232897-DEL-191 |
| chr3-127236317-DEL-231 | chr3-127236318-DEL-231 |
| chr3-127237527-DEL-1241 | chr3-127237528-DEL-1241 |
| chr3-127242377-DEL-61 | chr3-127242407-DEL-53 |
| chr3-127243259-DEL-212 | chr3-127243260-DEL-212 |
| chr3-127250430-DEL-220 | chr3-127250431-DEL-220 |
| chr3-127251475-DEL-220 | chr3-127251476-DEL-216 |
| chr3-127255475-DEL-71 | chr3-127255485-DEL-51 |
| chr3-127258823-DEL-447 | chr3-127258824-DEL-447 |
| chr3-127273488-DEL-222 | chr3-127273482-DEL-223 |
| chr3-127276011-DEL-197 | chr3-127276012-DEL-197 |
| chr3-127294280-DEL-74 | chr3-127294281-DEL-74 |
| chr3-127298695-DEL-60 | chr3-127298704-DEL-65 |
| chr3-127316254-DEL-1177 | chr3-127316250-DEL-1175 |
| chr3-127320801-DEL-115 | chr3-127320802-DEL-115 |
| chr3-127344158-DEL-68 | chr3-127344159-DEL-68 |
| chr3-127374397-DEL-415 | chr3-127374391-DEL-416 |
| chr3-127388346-DEL-1042 | chr3-127388346-DEL-1042 |
| chr3-127442029-DEL-206 | chr3-127442030-DEL-206 |
| chr3-127469850-DEL-54 | chr3-127469851-DEL-54 |
| chr3-127515988-DEL-222 | chr3-127515989-DEL-222 |
| chr3-127530967-DEL-2452 | chr3-127530968-DEL-2451 |
| chr3-127565230-DEL-7291 | chr3-127565231-DEL-7291 |
| chr3-127565256-DEL-416 | chr3-127565257-DEL-416 |
| chr3-127581280-DEL-63 | chr3-127581280-DEL-51 |
| chr3-127694281-DEL-54 | chr3-127694280-DEL-60 |
| chr3-127701542-DEL-539 | chr3-127701543-DEL-539 |
| chr3-127717770-DEL-51 | chr3-127717771-DEL-51 |
| chr3-127720210-DEL-59 | chr3-127720211-DEL-59 |
| chr3-127723367-DEL-185 | chr3-127723359-DEL-186 |
| chr3-127735799-DEL-223 | chr3-127735800-DEL-185 |
| chr3-127741521-DEL-63 | chr3-127741524-DEL-61 |
| chr3-127750693-DEL-62 | chr3-127750691-DEL-62 |
| chr3-127757195-DEL-74 | chr3-127757196-DEL-74 |
| chr3-127776913-DEL-326 | chr3-127776914-DEL-326 |
| chr3-127794341-DEL-212 | chr3-127794359-DEL-210 |
| chr3-127799531-DEL-86 | chr3-127799532-DEL-86 |
| chr3-127866922-DEL-513 | chr3-127866918-DEL-518 |
| chr3-127879752-DEL-64 | chr3-127879752-DEL-64 |
| chr3-127895021-DEL-53 | chr3-127895012-DEL-60 |
| chr3-127910340-DEL-122 | chr3-127910341-DEL-122 |
| chr3-127927506-DEL-55 | chr3-127927507-DEL-55 |
| chr3-127931626-DEL-6671 | chr3-127931627-DEL-6671 |
| chr3-127940730-DEL-184 | chr3-127940731-DEL-184 |
| chr3-127962310-DEL-55 | chr3-127962262-DEL-68 |
| chr3-127975828-DEL-60 | chr3-127975829-DEL-60 |

| | |
|---|---|
| chr3-128019483-DEL-298 | chr3-128019484-DEL-294 |
| chr3-128052718-DEL-439 | chr3-128052719-DEL-439 |
| chr3-128090464-DEL-128 | chr3-128090467-DEL-126 |
| chr3-128106800-DEL-216 | chr3-128106801-DEL-214 |
| chr3-128134968-DEL-53 | chr3-128134969-DEL-53 |
| chr3-128146361-DEL-72 | chr3-128146364-DEL-71 |
| chr3-128154199-DEL-218 | chr3-128154200-DEL-217 |
| chr3-128154647-DEL-511 | chr3-128154648-DEL-511 |
| chr3-128182469-DEL-1413 | chr3-128182469-DEL-1398 |
| chr3-128233775-DEL-6354 | chr3-128233776-DEL-6354 |
| chr3-128260401-DEL-979 | chr3-128260401-DEL-979 |
| chr3-128276343-DEL-70 | chr3-128276344-DEL-66 |
| chr3-128295934-DEL-971 | chr3-128295935-DEL-971 |
| chr3-128301642-DEL-432 | chr3-128301643-DEL-432 |
| chr3-128327099-DEL-78 | chr3-128327092-DEL-68 |
| chr3-128332945-DEL-668 | chr3-128332946-DEL-666 |
| chr3-128336748-DEL-843 | chr3-128336749-DEL-843 |
| chr3-128338187-DEL-68 | chr3-128338188-DEL-68 |
| chr3-128357304-DEL-397 | chr3-128357305-DEL-397 |
| chr3-128360014-DEL-480 | chr3-128360035-DEL-460 |
| chr3-128385479-DEL-432 | chr3-128385480-DEL-432 |
| chr3-128392734-DEL-57 | chr3-128392735-DEL-57 |
| chr3-128396137-DEL-165 | chr3-128396138-DEL-165 |
| chr3-128398055-DEL-651 | chr3-128398056-DEL-651 |
| chr3-128415780-DEL-4286 | chr3-128415781-DEL-4286 |
| chr3-128424524-DEL-57 | chr3-128424549-DEL-52 |
| chr3-128432524-DEL-54 | chr3-128432523-DEL-55 |
| chr3-128448500-DEL-56 | chr3-128448501-DEL-56 |
| chr3-128461379-DEL-51 | chr3-128461380-DEL-51 |
| chr3-128484398-DEL-6906 | chr3-128484399-DEL-6906 |
| chr3-128501461-DEL-223 | chr3-128501462-DEL-223 |
| chr3-128501904-DEL-528 | chr3-128501908-DEL-525 |
| chr3-128528264-DEL-61 | chr3-128528268-DEL-51 |
| chr3-128529417-DEL-58 | chr3-128529418-DEL-58 |
| chr3-128530486-DEL-4430 | chr3-128530487-DEL-4430 |
| chr3-128537910-DEL-106 | chr3-128537911-DEL-106 |
| chr3-128539476-DEL-95 | chr3-128539477-DEL-95 |
| chr3-128552051-DEL-502 | chr3-128552052-DEL-500 |
| chr3-128554799-DEL-59 | chr3-128554797-DEL-65 |
| chr3-128575168-DEL-1466 | chr3-128575169-DEL-1466 |
| chr3-128595848-DEL-6187 | chr3-128595848-DEL-6187 |
| chr3-128602619-DEL-56 | chr3-128602592-DEL-54 |
| chr3-128610026-DEL-311 | chr3-128610027-DEL-311 |
| chr3-128615794-DEL-361 | chr3-128615795-DEL-361 |
| chr3-128626464-DEL-353 | chr3-128626465-DEL-353 |
| chr3-128687414-DEL-244 | chr3-128687415-DEL-244 |
| chr3-128691136-DEL-58 | chr3-128691137-DEL-56 |
| chr3-128692745-DEL-174 | chr3-128692746-DEL-174 |
| chr3-128698952-DEL-106 | chr3-128698952-DEL-106 |
| chr3-128708565-DEL-436 | chr3-128708565-DEL-436 |
| chr3-128724393-DEL-56 | chr3-128724394-DEL-56 |
| chr3-128730363-DEL-54 | chr3-128730364-DEL-54 |
| chr3-128736165-DEL-63 | chr3-128736154-DEL-69 |
| chr3-128754260-DEL-6180 | chr3-128754260-DEL-6180 |
| chr3-128771073-DEL-788 | chr3-128771074-DEL-788 |
| chr3-128791246-DEL-856 | chr3-128791247-DEL-856 |
| chr3-128811235-DEL-1063 | chr3-128811236-DEL-1063 |
| chr3-128828508-DEL-64 | chr3-128828509-DEL-64 |
| chr3-128834965-DEL-251 | chr3-128834965-DEL-248 |
| chr3-128839037-DEL-62 | chr3-128839037-DEL-63 |
| chr3-128845839-DEL-82 | chr3-128845840-DEL-82 |
| chr3-128854247-DEL-132 | chr3-128854248-DEL-128 |
| chr3-128865258-DEL-55 | chr3-128865259-DEL-60 |
| chr3-128894849-DEL-550 | chr3-128894850-DEL-550 |
| chr3-128903917-DEL-132 | chr3-128903914-DEL-136 |
| chr3-128917817-DEL-341 | chr3-128917818-DEL-341 |
| chr3-128919355-DEL-1017 | chr3-128919355-DEL-1017 |
| chr3-128933568-DEL-56 | chr3-128933569-DEL-56 |
| chr3-128956595-DEL-66 | chr3-128956596-DEL-66 |
| chr3-128986778-DEL-247 | chr3-128986779-DEL-247 |
| chr3-128987434-DEL-142 | chr3-128987432-DEL-165 |
| chr3-129094228-DEL-1755 | chr3-129094226-DEL-1756 |
| chr3-129105190-DEL-106 | chr3-129105199-DEL-98 |
| chr3-129127823-DEL-276 | chr3-129127824-DEL-276 |
| chr3-129176423-DEL-485 | chr3-129176392-DEL-477 |
| chr3-129184123-DEL-64 | chr3-129184124-DEL-64 |
| chr3-129188339-DEL-1150 | chr3-129188340-DEL-1150 |
| chr3-129192594-DEL-451 | chr3-129192595-DEL-451 |
| chr3-129212970-DEL-196 | chr3-129212971-DEL-196 |
| chr3-129215278-DEL-192 | chr3-129215279-DEL-192 |

| | |
|---|---|
| chr3-129224081-DEL-54 | chr3-129224046-DEL-54 |
| chr3-129230519-DEL-91 | chr3-129230523-DEL-88 |
| chr3-129247754-DEL-76 | chr3-129247745-DEL-73 |
| chr3-129255585-DEL-874 | chr3-129255582-DEL-875 |
| chr3-129288970-DEL-142 | chr3-129288971-DEL-142 |
| chr3-129300534-DEL-52 | chr3-129300535-DEL-50 |
| chr3-129305151-DEL-78 | chr3-129305150-DEL-80 |
| chr3-129318229-DEL-326 | chr3-129318230-DEL-326 |
| chr3-129355453-DEL-73 | chr3-129355454-DEL-61 |
| chr3-129370631-DEL-201 | chr3-129370632-DEL-201 |
| chr3-129378274-DEL-53 | chr3-129378275-DEL-53 |
| chr3-129393266-DEL-194 | chr3-129393271-DEL-220 |
| chr3-129400619-DEL-246 | chr3-129400620-DEL-246 |
| chr3-129411333-DEL-80 | chr3-129411334-DEL-81 |
| chr3-129453528-DEL-60 | chr3-129453529-DEL-60 |
| chr3-129457905-DEL-106 | chr3-129457905-DEL-106 |
| chr3-129459725-DEL-7447 | chr3-129459726-DEL-7447 |
| chr3-129480251-DEL-157 | chr3-129480252-DEL-157 |
| chr3-129507687-DEL-168 | chr3-129507688-DEL-168 |
| chr3-129519397-DEL-149 | chr3-129519398-DEL-149 |
| chr3-129540395-DEL-320 | chr3-129540394-DEL-320 |
| chr3-129560033-DEL-197 | chr3-129560034-DEL-197 |
| chr3-129583336-DEL-201 | chr3-129583337-DEL-201 |
| chr3-129599246-DEL-58 | chr3-129599247-DEL-58 |
| chr3-129610782-DEL-1075 | chr3-129610783-DEL-1075 |
| chr3-129613572-DEL-601 | chr3-129613573-DEL-595 |
| chr3-129614117-DEL-60 | chr3-129614118-DEL-60 |
| chr3-129650014-DEL-51 | chr3-129650015-DEL-51 |
| chr3-129656157-DEL-97 | chr3-129656158-DEL-97 |
| chr3-129702558-DEL-53 | chr3-129702565-DEL-53 |
| chr3-129718552-DEL-52 | chr3-129718553-DEL-51 |
| chr3-129723554-DEL-110 | chr3-129723547-DEL-114 |
| chr3-129763988-DEL-225 | chr3-129763989-DEL-225 |
| chr3-129778370-DEL-295 | chr3-129778371-DEL-294 |
| chr3-129779773-DEL-186 | chr3-129779774-DEL-186 |
| chr3-129841503-DEL-211 | chr3-129841504-DEL-211 |
| chr3-129844483-DEL-98 | chr3-129844497-DEL-80 |
| chr3-129862026-DEL-663 | chr3-129862027-DEL-663 |
| chr3-129894181-DEL-713 | chr3-129894182-DEL-713 |
| chr3-129907944-DEL-2123 | chr3-129907945-DEL-2123 |
| chr3-129918163-DEL-152 | chr3-129918181-DEL-114 |
| chr3-129934440-DEL-1070 | chr3-129934441-DEL-1070 |
| chr3-129950226-DEL-399 | chr3-129950227-DEL-399 |
| chr3-129951980-DEL-78 | chr3-129951955-DEL-58 |
| chr3-129956548-DEL-135 | chr3-129956568-DEL-137 |
| chr3-129999099-DEL-4633 | chr3-129999100-DEL-4633 |
| chr3-130017430-DEL-177 | chr3-130017431-DEL-177 |
| chr3-130042501-DEL-74 | chr3-130042501-DEL-73 |
| chr3-130046654-DEL-659 | chr3-130046655-DEL-659 |
| chr3-130119149-DEL-959 | chr3-130119150-DEL-959 |
| chr3-130146018-DEL-383 | chr3-130146019-DEL-383 |
| chr3-130192182-DEL-187 | chr3-130192183-DEL-187 |
| chr3-130207811-DEL-56 | chr3-130207812-DEL-56 |
| chr3-130225226-DEL-67 | chr3-130225227-DEL-67 |
| chr3-130225988-DEL-60 | chr3-130225989-DEL-65 |
| chr3-130240722-DEL-1078 | chr3-130240721-DEL-1079 |
| chr3-130248593-DEL-1408 | chr3-130248594-DEL-1408 |
| chr3-130297830-DEL-2204 | chr3-130297831-DEL-2204 |
| chr3-130299056-DEL-486 | chr3-130299057-DEL-486 |
| chr3-130302429-DEL-6398 | chr3-130302417-DEL-6399 |
| chr3-130339071-DEL-169 | chr3-130339071-DEL-169 |
| chr3-130432569-DEL-1970 | chr3-130432570-DEL-1970 |
| chr3-130436243-DEL-401 | chr3-130436244-DEL-401 |
| chr3-130441057-DEL-113 | chr3-130441058-DEL-113 |
| chr3-130481401-DEL-1203 | chr3-130481402-DEL-1203 |
| chr3-130511643-DEL-226 | chr3-130511643-DEL-226 |
| chr3-130527346-DEL-236 | chr3-130527347-DEL-236 |
| chr3-130549543-DEL-780 | chr3-130549544-DEL-780 |
| chr3-130567942-DEL-6446 | chr3-130567943-DEL-6446 |
| chr3-130587831-DEL-53 | chr3-130587834-DEL-51 |
| chr3-130594267-DEL-214 | chr3-130594258-DEL-206 |
| chr3-130615215-DEL-278 | chr3-130615216-DEL-278 |
| chr3-130618896-DEL-51 | chr3-130618896-DEL-51 |
| chr3-130641111-DEL-68 | chr3-130641112-DEL-67 |
| chr3-130651525-DEL-725 | chr3-130651526-DEL-725 |
| chr3-130661552-DEL-75 | chr3-130661553-DEL-75 |
| chr3-130690201-DEL-116 | chr3-130690202-DEL-116 |
| chr3-130692559-DEL-62 | chr3-130692560-DEL-60 |
| chr3-130701541-DEL-69 | chr3-130701542-DEL-69 |
| chr3-130711748-DEL-2488 | chr3-130711749-DEL-2488 |

| | |
|---|---|
| chr3-130733011-DEL-69 | chr3-130733009-DEL-72 |
| chr3-130775710-DEL-61 | chr3-130775711-DEL-62 |
| chr3-130809212-DEL-59 | chr3-130809185-DEL-60 |
| chr3-130827128-DEL-95 | chr3-130827156-DEL-68 |
| chr3-130860714-DEL-79 | chr3-130860745-DEL-87 |
| chr3-130864934-DEL-933 | chr3-130864932-DEL-936 |
| chr3-130898203-DEL-212 | chr3-130898211-DEL-205 |
| chr3-130925186-DEL-97 | chr3-130925167-DEL-117 |
| chr3-130928144-DEL-57 | chr3-130928145-DEL-57 |
| chr3-130942023-DEL-109 | chr3-130942023-DEL-109 |
| chr3-130943465-DEL-1154 | chr3-130943466-DEL-1154 |
| chr3-130944320-DEL-144 | chr3-130944321-DEL-144 |
| chr3-130945132-DEL-184 | chr3-130945133-DEL-184 |
| chr3-130946990-DEL-62 | chr3-130946991-DEL-62 |
| chr3-130985458-DEL-90 | chr3-130985447-DEL-95 |
| chr3-131003176-DEL-1424 | chr3-131003161-DEL-1440 |
| chr3-131015653-DEL-66 | chr3-131015654-DEL-66 |
| chr3-131060149-DEL-206 | chr3-131060150-DEL-204 |
| chr3-131077382-DEL-57 | chr3-131077368-DEL-62 |
| chr3-131082644-DEL-338 | chr3-131082645-DEL-338 |
| chr3-131131478-DEL-515 | chr3-131131479-DEL-515 |
| chr3-131150782-DEL-70 | chr3-131150783-DEL-70 |
| chr3-131155206-DEL-120 | chr3-131155210-DEL-81 |
| chr3-131166740-DEL-202 | chr3-131166741-DEL-202 |
| chr3-131167581-DEL-479 | chr3-131167562-DEL-474 |
| chr3-131180681-DEL-1136 | chr3-131180682-DEL-1136 |
| chr3-131218690-DEL-2647 | chr3-131218691-DEL-2647 |
| chr3-131227274-DEL-172 | chr3-131227275-DEL-172 |
| chr3-131239072-DEL-126 | chr3-131239069-DEL-123 |
| chr3-131249997-DEL-3718 | chr3-131249998-DEL-3718 |
| chr3-131253774-DEL-444 | chr3-131253774-DEL-444 |
| chr3-131273066-DEL-1382 | chr3-131273078-DEL-1379 |
| chr3-131277428-DEL-87 | chr3-131277429-DEL-87 |
| chr3-131286147-DEL-312 | chr3-131286148-DEL-312 |
| chr3-131325344-DEL-2885 | chr3-131325345-DEL-2885 |
| chr3-131345668-DEL-1574 | chr3-131345669-DEL-1574 |
| chr3-131412402-DEL-6314 | chr3-131412403-DEL-6314 |
| chr3-131421879-DEL-2712 | chr3-131421880-DEL-2712 |
| chr3-131506622-DEL-143 | chr3-131506615-DEL-142 |
| chr3-131531882-DEL-408 | chr3-131531925-DEL-366 |
| chr3-131534786-DEL-145 | chr3-131534777-DEL-155 |
| chr3-131542531-DEL-774 | chr3-131542532-DEL-774 |
| chr3-131580740-DEL-417 | chr3-131580741-DEL-417 |
| chr3-131596186-DEL-77 | chr3-131596186-DEL-77 |
| chr3-131602021-DEL-72 | chr3-131602022-DEL-92 |
| chr3-131604648-DEL-751 | chr3-131604649-DEL-751 |
| chr3-131650999-DEL-6615 | chr3-131651000-DEL-6615 |
| chr3-131681321-DEL-152 | chr3-131681294-DEL-153 |
| chr3-131718263-DEL-58 | chr3-131718264-DEL-62 |
| chr3-131722494-DEL-96 | chr3-131722495-DEL-96 |
| chr3-131735280-DEL-206 | chr3-131735277-DEL-207 |
| chr3-131750918-DEL-890 | chr3-131750919-DEL-890 |
| chr3-131754850-DEL-370 | chr3-131754851-DEL-370 |
| chr3-131778866-DEL-948 | chr3-131778880-DEL-946 |
| chr3-131797996-DEL-57 | chr3-131798017-DEL-52 |
| chr3-131814564-DEL-72 | chr3-131814564-DEL-66 |
| chr3-131830292-DEL-290 | chr3-131830305-DEL-292 |
| chr3-131868407-DEL-156 | chr3-131868408-DEL-156 |
| chr3-131912336-DEL-81 | chr3-131912344-DEL-79 |
| chr3-131931483-DEL-809 | chr3-131931484-DEL-809 |
| chr3-131998250-DEL-328 | chr3-131998250-DEL-328 |
| chr3-131999568-DEL-443 | chr3-131999569-DEL-443 |
| chr3-132021318-DEL-385 | chr3-132021319-DEL-385 |
| chr3-132022680-DEL-462 | chr3-132022681-DEL-462 |
| chr3-132025349-DEL-451 | chr3-132025350-DEL-451 |
| chr3-132035118-DEL-94 | chr3-132035119-DEL-94 |
| chr3-132050588-DEL-54 | chr3-132050589-DEL-54 |
| chr3-132071533-DEL-197 | chr3-132071534-DEL-197 |
| chr3-132077621-DEL-142 | chr3-132077622-DEL-142 |
| chr3-132086070-DEL-331 | chr3-132086071-DEL-331 |
| chr3-132097571-DEL-92 | chr3-132097575-DEL-91 |
| chr3-132105181-DEL-250 | chr3-132105182-DEL-250 |
| chr3-132111726-DEL-6918 | chr3-132111727-DEL-6918 |
| chr3-132119709-DEL-381 | chr3-132119710-DEL-381 |
| chr3-132127638-DEL-72 | chr3-132127639-DEL-72 |
| chr3-132160343-DEL-1016 | chr3-132160344-DEL-1016 |
| chr3-132162401-DEL-400 | chr3-132162402-DEL-400 |
| chr3-132166013-DEL-168 | chr3-132166013-DEL-168 |
| chr3-132170558-DEL-76 | chr3-132170563-DEL-95 |
| chr3-132172501-DEL-74 | chr3-132172483-DEL-74 |

| | |
|---|---|
| chr3-132174266-DEL-235 | chr3-132174267-DEL-235 |
| chr3-132175845-DEL-102 | chr3-132175846-DEL-98 |
| chr3-132178655-DEL-294 | chr3-132178663-DEL-291 |
| chr3-132186647-DEL-53 | chr3-132186648-DEL-53 |
| chr3-132197577-DEL-1200 | chr3-132197578-DEL-1200 |
| chr3-132209037-DEL-410 | chr3-132209037-DEL-412 |
| chr3-132215368-DEL-60 | chr3-132215369-DEL-60 |
| chr3-132222141-DEL-60 | chr3-132222142-DEL-68 |
| chr3-132225323-DEL-3776 | chr3-132225324-DEL-3776 |
| chr3-132234382-DEL-83 | chr3-132234378-DEL-63 |
| chr3-132249266-DEL-408 | chr3-132249267-DEL-408 |
| chr3-132250650-DEL-60 | chr3-132250651-DEL-60 |
| chr3-132262515-DEL-58 | chr3-132262491-DEL-85 |
| chr3-132273101-DEL-52 | chr3-132273102-DEL-52 |
| chr3-132283987-DEL-85 | chr3-132283962-DEL-105 |
| chr3-132324076-DEL-155 | chr3-132324080-DEL-152 |
| chr3-132327513-DEL-363 | chr3-132327475-DEL-402 |
| chr3-132331366-DEL-861 | chr3-132331367-DEL-861 |
| chr3-132360751-DEL-154 | chr3-132360775-DEL-116 |
| chr3-132363377-DEL-987 | chr3-132363378-DEL-987 |
| chr3-132382210-DEL-224 | chr3-132382211-DEL-224 |
| chr3-132412100-DEL-246 | chr3-132412101-DEL-246 |
| chr3-132446185-DEL-470 | chr3-132446186-DEL-470 |
| chr3-132447955-DEL-64 | chr3-132447956-DEL-64 |
| chr3-132465822-DEL-953 | chr3-132465823-DEL-953 |
| chr3-132481315-DEL-257 | chr3-132481316-DEL-257 |
| chr3-132505267-DEL-100 | chr3-132505268-DEL-100 |
| chr3-132515252-DEL-101 | chr3-132515252-DEL-99 |
| chr3-132528122-DEL-6678 | chr3-132528110-DEL-6677 |
| chr3-132534256-DEL-212 | chr3-132534257-DEL-212 |
| chr3-132555141-DEL-270 | chr3-132555142-DEL-270 |
| chr3-132557774-DEL-77 | chr3-132557773-DEL-78 |
| chr3-132564131-DEL-210 | chr3-132564126-DEL-205 |
| chr3-132599444-DEL-91 | chr3-132599443-DEL-89 |
| chr3-132609292-DEL-90 | chr3-132609293-DEL-90 |
| chr3-132615052-DEL-100 | chr3-132615046-DEL-104 |
| chr3-132653006-DEL-90 | chr3-132653022-DEL-84 |
| chr3-132660220-DEL-254 | chr3-132660221-DEL-254 |
| chr3-132694817-DEL-127 | chr3-132694818-DEL-127 |
| chr3-132697309-DEL-443 | chr3-132697306-DEL-444 |
| chr3-132734425-DEL-325 | chr3-132734426-DEL-325 |
| chr3-132736526-DEL-961 | chr3-132736527-DEL-961 |
| chr3-132813154-DEL-123 | chr3-132813155-DEL-123 |
| chr3-132829417-DEL-7070 | chr3-132829418-DEL-7070 |
| chr3-132860897-DEL-100 | chr3-132860908-DEL-98 |
| chr3-132876832-DEL-574 | chr3-132876833-DEL-574 |
| chr3-132898083-DEL-507 | chr3-132898084-DEL-507 |
| chr3-132908017-DEL-372 | chr3-132908018-DEL-372 |
| chr3-132919058-DEL-57 | chr3-132919059-DEL-57 |
| chr3-132954788-DEL-3821 | chr3-132954789-DEL-3821 |
| chr3-132964636-DEL-96 | chr3-132964637-DEL-96 |
| chr3-132977569-DEL-100 | chr3-132977570-DEL-100 |
| chr3-132986549-DEL-97 | chr3-132986532-DEL-99 |
| chr3-133003479-DEL-4390 | chr3-133003480-DEL-4390 |
| chr3-133024467-DEL-76 | chr3-133024467-DEL-76 |
| chr3-133026053-DEL-58 | chr3-133026054-DEL-58 |
| chr3-133048896-DEL-59 | chr3-133048896-DEL-59 |
| chr3-133055352-DEL-1348 | chr3-133055353-DEL-1348 |
| chr3-133060086-DEL-106 | chr3-133060089-DEL-100 |
| chr3-133068203-DEL-243 | chr3-133068204-DEL-243 |
| chr3-133069203-DEL-250 | chr3-133069205-DEL-251 |
| chr3-133071729-DEL-177 | chr3-133071723-DEL-178 |
| chr3-133094115-DEL-179 | chr3-133094116-DEL-179 |
| chr3-133104355-DEL-333 | chr3-133104356-DEL-333 |
| chr3-133135434-DEL-58 | chr3-133135435-DEL-58 |
| chr3-133137345-DEL-492 | chr3-133137346-DEL-492 |
| chr3-133165396-DEL-219 | chr3-133165397-DEL-219 |
| chr3-133166425-DEL-59 | chr3-133166424-DEL-60 |
| chr3-133181898-DEL-90 | chr3-133181899-DEL-90 |
| chr3-133187312-DEL-197 | chr3-133187305-DEL-191 |
| chr3-133197678-DEL-1259 | chr3-133197679-DEL-1259 |
| chr3-133202003-DEL-231 | chr3-133202004-DEL-231 |
| chr3-133207572-DEL-2706 | chr3-133207573-DEL-2706 |
| chr3-133231212-DEL-51 | chr3-133231213-DEL-51 |
| chr3-133265804-DEL-71 | chr3-133265803-DEL-73 |
| chr3-133277522-DEL-326 | chr3-133277522-DEL-322 |
| chr3-133291656-DEL-760 | chr3-133291657-DEL-760 |
| chr3-133294370-DEL-51 | chr3-133294361-DEL-74 |
| chr3-133334768-DEL-61 | chr3-133334766-DEL-61 |
| chr3-133341678-DEL-6872 | chr3-133341679-DEL-6872 |

| | |
|---|---|
| chr3-133368564-DEL-64 | chr3-133368560-DEL-93 |
| chr3-133373241-DEL-57 | chr3-133373242-DEL-57 |
| chr3-133375920-DEL-266 | chr3-133375921-DEL-266 |
| chr3-133376723-DEL-129 | chr3-133376724-DEL-129 |
| chr3-133434935-DEL-420 | chr3-133434936-DEL-420 |
| chr3-133447965-DEL-60 | chr3-133447966-DEL-60 |
| chr3-133468849-DEL-177 | chr3-133468850-DEL-177 |
| chr3-133472245-DEL-96 | chr3-133472246-DEL-96 |
| chr3-133476302-DEL-63 | chr3-133476288-DEL-64 |
| chr3-133476899-DEL-333 | chr3-133476900-DEL-333 |
| chr3-133486031-DEL-338 | chr3-133486032-DEL-338 |
| chr3-133491757-DEL-1826 | chr3-133491755-DEL-1825 |
| chr3-133511437-DEL-88 | chr3-133511438-DEL-88 |
| chr3-133512408-DEL-1307 | chr3-133512409-DEL-1307 |
| chr3-133542702-DEL-2980 | chr3-133542703-DEL-2980 |
| chr3-133549182-DEL-1840 | chr3-133549183-DEL-1840 |
| chr3-133575447-DEL-337 | chr3-133575451-DEL-334 |
| chr3-133590789-DEL-164 | chr3-133590790-DEL-164 |
| chr3-133599578-DEL-1921 | chr3-133599579-DEL-1921 |
| chr3-133636650-DEL-453 | chr3-133636650-DEL-453 |
| chr3-133653465-DEL-5659 | chr3-133653464-DEL-5659 |
| chr3-133695277-DEL-58 | chr3-133695278-DEL-58 |
| chr3-133706926-DEL-231 | chr3-133706927-DEL-231 |
| chr3-133713195-DEL-6436 | chr3-133713196-DEL-6436 |
| chr3-133766078-DEL-350 | chr3-133766079-DEL-350 |
| chr3-133767400-DEL-839 | chr3-133767401-DEL-839 |
| chr3-133812762-DEL-93 | chr3-133812757-DEL-54 |
| chr3-133829445-DEL-172 | chr3-133829446-DEL-172 |
| chr3-133871105-DEL-726 | chr3-133871106-DEL-726 |
| chr3-133889210-DEL-61 | chr3-133889211-DEL-61 |
| chr3-133957998-DEL-54 | chr3-133957975-DEL-78 |
| chr3-133993509-DEL-61 | chr3-133993510-DEL-61 |
| chr3-134005828-DEL-56 | chr3-134005829-DEL-56 |
| chr3-134018480-DEL-358 | chr3-134018481-DEL-358 |
| chr3-134032098-DEL-62 | chr3-134032053-DEL-102 |
| chr3-134063044-DEL-326 | chr3-134063045-DEL-327 |
| chr3-134094484-DEL-1143 | chr3-134094485-DEL-1143 |
| chr3-134126860-DEL-725 | chr3-134126861-DEL-725 |
| chr3-134155076-DEL-6533 | chr3-134155077-DEL-6533 |
| chr3-134163321-DEL-299 | chr3-134163322-DEL-299 |
| chr3-134181148-DEL-232 | chr3-134181149-DEL-232 |
| chr3-134203737-DEL-253 | chr3-134203738-DEL-253 |
| chr3-134208127-DEL-798 | chr3-134208128-DEL-798 |
| chr3-134220547-DEL-122 | chr3-134220548-DEL-146 |
| chr3-134262630-DEL-2076 | chr3-134262631-DEL-2076 |
| chr3-134323305-DEL-73 | chr3-134323306-DEL-73 |
| chr3-134344685-DEL-173 | chr3-134344684-DEL-177 |
| chr3-134361901-DEL-69 | chr3-134361866-DEL-82 |
| chr3-134366665-DEL-421 | chr3-134366666-DEL-421 |
| chr3-134389342-DEL-961 | chr3-134389343-DEL-961 |
| chr3-134404947-DEL-205 | chr3-134404948-DEL-205 |
| chr3-134407547-DEL-218 | chr3-134407527-DEL-225 |
| chr3-134417213-DEL-328 | chr3-134417214-DEL-328 |
| chr3-134420208-DEL-1453 | chr3-134420209-DEL-1453 |
| chr3-134442260-DEL-60 | chr3-134442261-DEL-54 |
| chr3-134454438-DEL-923 | chr3-134454440-DEL-922 |
| chr3-134456727-DEL-839 | chr3-134456727-DEL-840 |
| chr3-134479482-DEL-60 | chr3-134479483-DEL-60 |
| chr3-134483834-DEL-51 | chr3-134483835-DEL-57 |
| chr3-134514383-DEL-139 | chr3-134514384-DEL-139 |
| chr3-134521050-DEL-6495 | chr3-134521051-DEL-6495 |
| chr3-134533139-DEL-71 | chr3-134533140-DEL-71 |
| chr3-134536166-DEL-107 | chr3-134536167-DEL-107 |
| chr3-134540144-DEL-101 | chr3-134540149-DEL-97 |
| chr3-134564784-DEL-52 | chr3-134564785-DEL-52 |
| chr3-134586685-DEL-7511 | chr3-134586686-DEL-7511 |
| chr3-134587514-DEL-636 | chr3-134587514-DEL-636 |
| chr3-134602273-DEL-1531 | chr3-134602274-DEL-1531 |
| chr3-134607237-DEL-498 | chr3-134607238-DEL-498 |
| chr3-134626562-DEL-87 | chr3-134626563-DEL-87 |
| chr3-134626896-DEL-1748 | chr3-134626897-DEL-1748 |
| chr3-134627137-DEL-55 | chr3-134627129-DEL-63 |
| chr3-134645958-DEL-386 | chr3-134645959-DEL-385 |
| chr3-134678323-DEL-1009 | chr3-134678322-DEL-1010 |
| chr3-134678948-DEL-3339 | chr3-134678949-DEL-3340 |
| chr3-134680205-DEL-855 | chr3-134680206-DEL-855 |
| chr3-134708313-DEL-1444 | chr3-134708305-DEL-1445 |
| chr3-134713948-DEL-84 | chr3-134713949-DEL-84 |
| chr3-134762775-DEL-517 | chr3-134762776-DEL-517 |
| chr3-134770077-DEL-209 | chr3-134770078-DEL-209 |

| | |
|---|---|
| chr3-134793714-DEL-1884 | chr3-134793715-DEL-1884 |
| chr3-134802229-DEL-3410 | chr3-134802230-DEL-3410 |
| chr3-134821652-DEL-563 | chr3-134821653-DEL-562 |
| chr3-134853952-DEL-1163 | chr3-134853953-DEL-1163 |
| chr3-134862479-DEL-6397 | chr3-134862480-DEL-6397 |
| chr3-134868698-DEL-128 | chr3-134868699-DEL-128 |
| chr3-134870162-DEL-68 | chr3-134870163-DEL-68 |
| chr3-134890964-DEL-1340 | chr3-134890965-DEL-1339 |
| chr3-134893161-DEL-156 | chr3-134893162-DEL-156 |
| chr3-134898752-DEL-498 | chr3-134898753-DEL-498 |
| chr3-134931988-DEL-51 | chr3-134931989-DEL-51 |
| chr3-134949660-DEL-2044 | chr3-134949661-DEL-2043 |
| chr3-134955012-DEL-78 | chr3-134955010-DEL-79 |
| chr3-134955931-DEL-128 | chr3-134955949-DEL-111 |
| chr3-134956380-DEL-259 | chr3-134956381-DEL-296 |
| chr3-135035747-DEL-69 | chr3-135035747-DEL-70 |
| chr3-135083440-DEL-186 | chr3-135083441-DEL-186 |
| chr3-135087070-DEL-93 | chr3-135087071-DEL-93 |
| chr3-135106612-DEL-131 | chr3-135106610-DEL-130 |
| chr3-135121166-DEL-781 | chr3-135121167-DEL-781 |
| chr3-135187722-DEL-176 | chr3-135187723-DEL-176 |
| chr3-135190616-DEL-166 | chr3-135190617-DEL-166 |
| chr3-135194437-DEL-53 | chr3-135194423-DEL-52 |
| chr3-135203509-DEL-215 | chr3-135203510-DEL-215 |
| chr3-135209782-DEL-211 | chr3-135209781-DEL-211 |
| chr3-135216572-DEL-59 | chr3-135216573-DEL-59 |
| chr3-135223357-DEL-211 | chr3-135223358-DEL-211 |
| chr3-135229206-DEL-437 | chr3-135229206-DEL-437 |
| chr3-135230603-DEL-1842 | chr3-135230604-DEL-1842 |
| chr3-135253394-DEL-690 | chr3-135253395-DEL-690 |
| chr3-135255443-DEL-57 | chr3-135255436-DEL-81 |
| chr3-135256636-DEL-158 | chr3-135256637-DEL-158 |
| chr3-135272108-DEL-62 | chr3-135272109-DEL-62 |
| chr3-135293085-DEL-512 | chr3-135293087-DEL-509 |
| chr3-135310725-DEL-54 | chr3-135310726-DEL-54 |
| chr3-135335356-DEL-68 | chr3-135335356-DEL-68 |
| chr3-135344432-DEL-60 | chr3-135344433-DEL-60 |
| chr3-135380076-DEL-53 | chr3-135380062-DEL-54 |
| chr3-135390111-DEL-66 | chr3-135390112-DEL-66 |
| chr3-135413656-DEL-1879 | chr3-135413657-DEL-1879 |
| chr3-135420936-DEL-106 | chr3-135420901-DEL-107 |
| chr3-135500531-DEL-103 | chr3-135500532-DEL-103 |
| chr3-135505856-DEL-2930 | chr3-135505857-DEL-2930 |
| chr3-135507289-DEL-1453 | chr3-135507289-DEL-1453 |
| chr3-135543510-DEL-537 | chr3-135543511-DEL-537 |
| chr3-135571319-DEL-95 | chr3-135571316-DEL-88 |
| chr3-135595190-DEL-136 | chr3-135595191-DEL-136 |
| chr3-135635678-DEL-57 | chr3-135635675-DEL-57 |
| chr3-135641717-DEL-390 | chr3-135641718-DEL-390 |
| chr3-135661669-DEL-2729 | chr3-135661670-DEL-2729 |
| chr3-135670314-DEL-315 | chr3-135670315-DEL-313 |
| chr3-135681990-DEL-90 | chr3-135681976-DEL-80 |
| chr3-135688053-DEL-263 | chr3-135688054-DEL-263 |
| chr3-135699159-DEL-76 | chr3-135699160-DEL-76 |
| chr3-135710910-DEL-94 | chr3-135710909-DEL-94 |
| chr3-135719979-DEL-53 | chr3-135719979-DEL-53 |
| chr3-135724942-DEL-2245 | chr3-135724943-DEL-2245 |
| chr3-135736537-DEL-62 | chr3-135736570-DEL-54 |
| chr3-135740268-DEL-120 | chr3-135740269-DEL-120 |
| chr3-135744025-DEL-54 | chr3-135744026-DEL-54 |
| chr3-135767796-DEL-1722 | chr3-135767797-DEL-1722 |
| chr3-135787957-DEL-455 | chr3-135787958-DEL-455 |
| chr3-135808939-DEL-142 | chr3-135808942-DEL-145 |
| chr3-135823007-DEL-435 | chr3-135823008-DEL-435 |
| chr3-135823984-DEL-4385 | chr3-135823979-DEL-4377 |
| chr3-135879533-DEL-7111 | chr3-135879534-DEL-7111 |
| chr3-135907480-DEL-3285 | chr3-135907481-DEL-3285 |
| chr3-135910602-DEL-56 | chr3-135910603-DEL-56 |
| chr3-135925297-DEL-317 | chr3-135925298-DEL-317 |
| chr3-135941769-DEL-56 | chr3-135941770-DEL-56 |
| chr3-135949053-DEL-72 | chr3-135949054-DEL-72 |
| chr3-135978362-DEL-118 | chr3-135978363-DEL-118 |
| chr3-135979748-DEL-54 | chr3-135979749-DEL-54 |
| chr3-135991277-DEL-1293 | chr3-135991278-DEL-1289 |
| chr3-136005812-DEL-70 | chr3-136005817-DEL-65 |
| chr3-136075192-DEL-80 | chr3-136075190-DEL-81 |
| chr3-136093915-DEL-669 | chr3-136093916-DEL-669 |
| chr3-136097477-DEL-62 | chr3-136097478-DEL-62 |
| chr3-136098588-DEL-51 | chr3-136098593-DEL-52 |
| chr3-136099938-DEL-1832 | chr3-136099939-DEL-1832 |

| | |
|---|---|
| chr3-136130141-DEL-771 | chr3-136130142-DEL-771 |
| chr3-136140938-DEL-62 | chr3-136140939-DEL-62 |
| chr3-136154494-DEL-60 | chr3-136154511-DEL-50 |
| chr3-136163857-DEL-108 | chr3-136163863-DEL-126 |
| chr3-136216495-DEL-6214 | chr3-136216496-DEL-6214 |
| chr3-136220774-DEL-222 | chr3-136220775-DEL-222 |
| chr3-136247333-DEL-440 | chr3-136247334-DEL-438 |
| chr3-136252062-DEL-64 | chr3-136252071-DEL-62 |
| chr3-136270334-DEL-56 | chr3-136270292-DEL-57 |
| chr3-136278744-DEL-204 | chr3-136278745-DEL-204 |
| chr3-136309892-DEL-123 | chr3-136309893-DEL-123 |
| chr3-136320290-DEL-6133 | chr3-136320291-DEL-6133 |
| chr3-136357848-DEL-767 | chr3-136357849-DEL-767 |
| chr3-136389571-DEL-77 | chr3-136389572-DEL-77 |
| chr3-136393598-DEL-55 | chr3-136393599-DEL-55 |
| chr3-136395379-DEL-101 | chr3-136395380-DEL-101 |
| chr3-136405556-DEL-188 | chr3-136405574-DEL-187 |
| chr3-136408001-DEL-253 | chr3-136408002-DEL-253 |
| chr3-136436720-DEL-792 | chr3-136436721-DEL-792 |
| chr3-136437918-DEL-58 | chr3-136437912-DEL-58 |
| chr3-136454355-DEL-76 | chr3-136454356-DEL-76 |
| chr3-136459335-DEL-180 | chr3-136459336-DEL-180 |
| chr3-136463336-DEL-84 | chr3-136463337-DEL-84 |
| chr3-136470564-DEL-2193 | chr3-136470565-DEL-2193 |
| chr3-136552575-DEL-58 | chr3-136552575-DEL-59 |
| chr3-136636348-DEL-149 | chr3-136636347-DEL-150 |
| chr3-136652063-DEL-69 | chr3-136652064-DEL-69 |
| chr3-136661544-DEL-963 | chr3-136661545-DEL-963 |
| chr3-136727040-DEL-658 | chr3-136727041-DEL-657 |
| chr3-136765209-DEL-194 | chr3-136765208-DEL-188 |
| chr3-136766721-DEL-606 | chr3-136766722-DEL-606 |
| chr3-136768795-DEL-566 | chr3-136768796-DEL-566 |
| chr3-136772061-DEL-173 | chr3-136772060-DEL-174 |
| chr3-136778872-DEL-412 | chr3-136778873-DEL-412 |
| chr3-136779539-DEL-147 | chr3-136779540-DEL-147 |
| chr3-136795404-DEL-1014 | chr3-136795405-DEL-1014 |
| chr3-136811443-DEL-60 | chr3-136811444-DEL-60 |
| chr3-136817052-DEL-125 | chr3-136817061-DEL-117 |
| chr3-136825376-DEL-98 | chr3-136825376-DEL-99 |
| chr3-136833690-DEL-1348 | chr3-136833691-DEL-1348 |
| chr3-136837071-DEL-101 | chr3-136837072-DEL-101 |
| chr3-136846113-DEL-345 | chr3-136846114-DEL-345 |
| chr3-136873434-DEL-626 | chr3-136873436-DEL-627 |
| chr3-136881206-DEL-72 | chr3-136881207-DEL-81 |
| chr3-136883070-DEL-105 | chr3-136883071-DEL-105 |
| chr3-136913799-DEL-68 | chr3-136913800-DEL-68 |
| chr3-136916706-DEL-67 | chr3-136916723-DEL-52 |
| chr3-136930308-DEL-88 | chr3-136930309-DEL-68 |
| chr3-136934679-DEL-3572 | chr3-136934679-DEL-3572 |
| chr3-136960268-DEL-250 | chr3-136960269-DEL-250 |
| chr3-136987351-DEL-4324 | chr3-136987352-DEL-4324 |
| chr3-136998148-DEL-102 | chr3-136998159-DEL-123 |
| chr3-137005776-DEL-185 | chr3-137005777-DEL-185 |
| chr3-137023991-DEL-300 | chr3-137023992-DEL-300 |
| chr3-137031195-DEL-427 | chr3-137031196-DEL-427 |
| chr3-137049217-DEL-56 | chr3-137049218-DEL-56 |
| chr3-137050058-DEL-83 | chr3-137050059-DEL-83 |
| chr3-137056695-DEL-76 | chr3-137056696-DEL-76 |
| chr3-137064070-DEL-2033 | chr3-137064071-DEL-2033 |
| chr3-137175001-DEL-1165 | chr3-137175001-DEL-1166 |
| chr3-137227638-DEL-58 | chr3-137227639-DEL-58 |
| chr3-137292982-DEL-1235 | chr3-137292983-DEL-1235 |
| chr3-137298417-DEL-82 | chr3-137298418-DEL-92 |
| chr3-137299087-DEL-304 | chr3-137299088-DEL-304 |
| chr3-137306165-DEL-6506 | chr3-137306166-DEL-6506 |
| chr3-137351543-DEL-440 | chr3-137351544-DEL-440 |
| chr3-137374054-DEL-7415 | chr3-137374055-DEL-7415 |
| chr3-137407533-DEL-190 | chr3-137407534-DEL-190 |
| chr3-137424959-DEL-3250 | chr3-137424960-DEL-3250 |
| chr3-137429172-DEL-79 | chr3-137429173-DEL-79 |
| chr3-137444407-DEL-86 | chr3-137444408-DEL-86 |
| chr3-137453615-DEL-633 | chr3-137453611-DEL-632 |
| chr3-137459724-DEL-102 | chr3-137459725-DEL-102 |
| chr3-137478025-DEL-209 | chr3-137478026-DEL-209 |
| chr3-137480319-DEL-156 | chr3-137480320-DEL-156 |
| chr3-137494085-DEL-79 | chr3-137494098-DEL-80 |
| chr3-137495072-DEL-68 | chr3-137495073-DEL-68 |
| chr3-137495758-DEL-200 | chr3-137495759-DEL-200 |
| chr3-137506399-DEL-180 | chr3-137506400-DEL-180 |
| chr3-137528663-DEL-87 | chr3-137528664-DEL-87 |

| | |
|---|---|
| chr3-137534171-DEL-194 | chr3-137534172-DEL-194 |
| chr3-137543561-DEL-977 | chr3-137543562-DEL-977 |
| chr3-137558906-DEL-88 | chr3-137558907-DEL-88 |
| chr3-137581006-DEL-221 | chr3-137581007-DEL-221 |
| chr3-137582972-DEL-56 | chr3-137582973-DEL-56 |
| chr3-137593178-DEL-58 | chr3-137593183-DEL-54 |
| chr3-137608181-DEL-1011 | chr3-137608182-DEL-1011 |
| chr3-137639739-DEL-73 | chr3-137639737-DEL-69 |
| chr3-137666765-DEL-69 | chr3-137666761-DEL-74 |
| chr3-137682270-DEL-52 | chr3-137682271-DEL-52 |
| chr3-137688052-DEL-68 | chr3-137688052-DEL-68 |
| chr3-137690726-DEL-92 | chr3-137690727-DEL-92 |
| chr3-137704079-DEL-96 | chr3-137704080-DEL-96 |
| chr3-137718302-DEL-59 | chr3-137718303-DEL-59 |
| chr3-137741467-DEL-194 | chr3-137741468-DEL-194 |
| chr3-137741987-DEL-381 | chr3-137741988-DEL-381 |
| chr3-137751600-DEL-2104 | chr3-137751601-DEL-2104 |
| chr3-137763221-DEL-145 | chr3-137763222-DEL-140 |
| chr3-137779863-DEL-202 | chr3-137779897-DEL-177 |
| chr3-137787608-DEL-261 | chr3-137787609-DEL-261 |
| chr3-137805480-DEL-2307 | chr3-137805481-DEL-2307 |
| chr3-137831933-DEL-64 | chr3-137831934-DEL-64 |
| chr3-137835212-DEL-77 | chr3-137835250-DEL-50 |
| chr3-137846882-DEL-92 | chr3-137846881-DEL-87 |
| chr3-137872435-DEL-6002 | chr3-137872436-DEL-6002 |
| chr3-137905865-DEL-2496 | chr3-137905866-DEL-2496 |
| chr3-137911517-DEL-2414 | chr3-137911518-DEL-2414 |
| chr3-137972571-DEL-2441 | chr3-137972570-DEL-2442 |
| chr3-137985189-DEL-128 | chr3-137985174-DEL-122 |
| chr3-137994307-DEL-57 | chr3-137994308-DEL-57 |
| chr3-138023275-DEL-149 | chr3-138023276-DEL-149 |
| chr3-138025778-DEL-74 | chr3-138025779-DEL-74 |
| chr3-138028924-DEL-190 | chr3-138028925-DEL-190 |
| chr3-138036560-DEL-117 | chr3-138036557-DEL-119 |
| chr3-138038376-DEL-67 | chr3-138038369-DEL-68 |
| chr3-138051450-DEL-70 | chr3-138051451-DEL-70 |
| chr3-138078698-DEL-1640 | chr3-138078699-DEL-1640 |
| chr3-138090773-DEL-108 | chr3-138090774-DEL-108 |
| chr3-138091843-DEL-1026 | chr3-138091844-DEL-1026 |
| chr3-138096238-DEL-108 | chr3-138096238-DEL-108 |
| chr3-138119843-DEL-60 | chr3-138119844-DEL-60 |
| chr3-138129202-DEL-139 | chr3-138129200-DEL-134 |
| chr3-138131008-DEL-717 | chr3-138131009-DEL-717 |
| chr3-138139551-DEL-2500 | chr3-138139552-DEL-2500 |
| chr3-138154812-DEL-252 | chr3-138154830-DEL-253 |
| chr3-138169420-DEL-948 | chr3-138169420-DEL-948 |
| chr3-138175569-DEL-68 | chr3-138175570-DEL-68 |
| chr3-138184769-DEL-105 | chr3-138184770-DEL-105 |
| chr3-138199405-DEL-78 | chr3-138199397-DEL-78 |
| chr3-138225311-DEL-271 | chr3-138225312-DEL-271 |
| chr3-138272865-DEL-115 | chr3-138272850-DEL-116 |
| chr3-138291730-DEL-145 | chr3-138291731-DEL-145 |
| chr3-138301220-DEL-63 | chr3-138301221-DEL-74 |
| chr3-138323152-DEL-111 | chr3-138323153-DEL-111 |
| chr3-138329387-DEL-61 | chr3-138329387-DEL-61 |
| chr3-138330621-DEL-506 | chr3-138330622-DEL-506 |
| chr3-138337647-DEL-201 | chr3-138337648-DEL-201 |
| chr3-138350796-DEL-60 | chr3-138350797-DEL-60 |
| chr3-138374660-DEL-68 | chr3-138374660-DEL-68 |
| chr3-138388851-DEL-2880 | chr3-138388852-DEL-2880 |
| chr3-138403800-DEL-347 | chr3-138403801-DEL-347 |
| chr3-138432679-DEL-100 | chr3-138432679-DEL-100 |
| chr3-138460607-DEL-221 | chr3-138460608-DEL-221 |
| chr3-138464380-DEL-86 | chr3-138464381-DEL-86 |
| chr3-138469247-DEL-150 | chr3-138469246-DEL-151 |
| chr3-138500246-DEL-71 | chr3-138500247-DEL-71 |
| chr3-138514000-DEL-725 | chr3-138514001-DEL-725 |
| chr3-138529265-DEL-51 | chr3-138529267-DEL-52 |
| chr3-138550479-DEL-76 | chr3-138550480-DEL-76 |
| chr3-138585991-DEL-61 | chr3-138585992-DEL-61 |
| chr3-138612057-DEL-94 | chr3-138612058-DEL-94 |
| chr3-138640578-DEL-513 | chr3-138640579-DEL-513 |
| chr3-138684917-DEL-162 | chr3-138684918-DEL-162 |
| chr3-138701104-DEL-249 | chr3-138701110-DEL-240 |
| chr3-138712117-DEL-102 | chr3-138712132-DEL-85 |
| chr3-138721970-DEL-51 | chr3-138721971-DEL-51 |
| chr3-138743880-DEL-108 | chr3-138743881-DEL-105 |
| chr3-138759668-DEL-89 | chr3-138759669-DEL-89 |
| chr3-138760576-DEL-833 | chr3-138760577-DEL-833 |
| chr3-138787033-DEL-473 | chr3-138787034-DEL-473 |

| | |
|---|---|
| chr3-138802414-DEL-74 | chr3-138802409-DEL-92 |
| chr3-138880796-DEL-550 | chr3-138880797-DEL-550 |
| chr3-138922179-DEL-74 | chr3-138922180-DEL-74 |
| chr3-138979391-DEL-54 | chr3-138979392-DEL-54 |
| chr3-138982860-DEL-112 | chr3-138982861-DEL-156 |
| chr3-138990980-DEL-83 | chr3-138990980-DEL-83 |
| chr3-139025203-DEL-492 | chr3-139025204-DEL-492 |
| chr3-139032711-DEL-110 | chr3-139032712-DEL-110 |
| chr3-139051490-DEL-204 | chr3-139051491-DEL-204 |
| chr3-139054524-DEL-1118 | chr3-139054524-DEL-1118 |
| chr3-139054543-DEL-183 | chr3-139054544-DEL-183 |
| chr3-139069473-DEL-55 | chr3-139069474-DEL-55 |
| chr3-139089230-DEL-54 | chr3-139089231-DEL-54 |
| chr3-139105066-DEL-191 | chr3-139105067-DEL-191 |
| chr3-139108532-DEL-67 | chr3-139108559-DEL-52 |
| chr3-139119329-DEL-4293 | chr3-139119330-DEL-4293 |
| chr3-139129912-DEL-65 | chr3-139129882-DEL-66 |
| chr3-139179756-DEL-54 | chr3-139179743-DEL-68 |
| chr3-139189197-DEL-400 | chr3-139189198-DEL-400 |
| chr3-139206085-DEL-487 | chr3-139206083-DEL-488 |
| chr3-139236446-DEL-6874 | chr3-139236447-DEL-6874 |
| chr3-139236555-DEL-208 | chr3-139236556-DEL-208 |
| chr3-139248148-DEL-235 | chr3-139248149-DEL-235 |
| chr3-139265775-DEL-4165 | chr3-139265776-DEL-4165 |
| chr3-139270477-DEL-7057 | chr3-139270478-DEL-7057 |
| chr3-139331582-DEL-84 | chr3-139331583-DEL-84 |
| chr3-139359755-DEL-177 | chr3-139359756-DEL-212 |
| chr3-139397743-DEL-56 | chr3-139397744-DEL-56 |
| chr3-139410285-DEL-2997 | chr3-139410286-DEL-2997 |
| chr3-139462867-DEL-142 | chr3-139462868-DEL-142 |
| chr3-139476827-DEL-476 | chr3-139476828-DEL-476 |
| chr3-139479378-DEL-467 | chr3-139479379-DEL-467 |
| chr3-139480594-DEL-521 | chr3-139480595-DEL-521 |
| chr3-139492820-DEL-54 | chr3-139492821-DEL-54 |
| chr3-139495261-DEL-714 | chr3-139495262-DEL-714 |
| chr3-139499553-DEL-90 | chr3-139499554-DEL-90 |
| chr3-139510123-DEL-60 | chr3-139510115-DEL-68 |
| chr3-139533643-DEL-194 | chr3-139533644-DEL-194 |
| chr3-139535395-DEL-169 | chr3-139535396-DEL-169 |
| chr3-139540438-DEL-370 | chr3-139540439-DEL-370 |
| chr3-139554695-DEL-64 | chr3-139554696-DEL-50 |
| chr3-139622314-DEL-76 | chr3-139622310-DEL-80 |
| chr3-139653706-DEL-54 | chr3-139653707-DEL-52 |
| chr3-139673330-DEL-53 | chr3-139673331-DEL-53 |
| chr3-139706084-DEL-54 | chr3-139706095-DEL-90 |
| chr3-139723230-DEL-54 | chr3-139723231-DEL-52 |
| chr3-139725692-DEL-7101 | chr3-139725693-DEL-7101 |
| chr3-139745470-DEL-69 | chr3-139745462-DEL-70 |
| chr3-139747941-DEL-100 | chr3-139747942-DEL-104 |
| chr3-139825764-DEL-5291 | chr3-139825765-DEL-5291 |
| chr3-139838647-DEL-72 | chr3-139838648-DEL-72 |
| chr3-139845492-DEL-475 | chr3-139845493-DEL-475 |
| chr3-139846447-DEL-798 | chr3-139846428-DEL-798 |
| chr3-139858288-DEL-116 | chr3-139858289-DEL-116 |
| chr3-139862372-DEL-6908 | chr3-139862373-DEL-6908 |
| chr3-139893812-DEL-96 | chr3-139893821-DEL-88 |
| chr3-139899650-DEL-573 | chr3-139899651-DEL-573 |
| chr3-139902154-DEL-296 | chr3-139902183-DEL-264 |
| chr3-139910438-DEL-112 | chr3-139910439-DEL-112 |
| chr3-139920642-DEL-54 | chr3-139920643-DEL-54 |
| chr3-139922232-DEL-665 | chr3-139922233-DEL-665 |
| chr3-139927689-DEL-92 | chr3-139927688-DEL-90 |
| chr3-139981401-DEL-274 | chr3-139981402-DEL-274 |
| chr3-139985896-DEL-871 | chr3-139985897-DEL-871 |
| chr3-140018201-DEL-66 | chr3-140018199-DEL-66 |
| chr3-140020312-DEL-94 | chr3-140020295-DEL-112 |
| chr3-140052285-DEL-58 | chr3-140052286-DEL-58 |
| chr3-140082786-DEL-7091 | chr3-140082787-DEL-7091 |
| chr3-140098975-DEL-60 | chr3-140098978-DEL-60 |
| chr3-140112189-DEL-794 | chr3-140112193-DEL-793 |
| chr3-140136605-DEL-56 | chr3-140136599-DEL-58 |
| chr3-140137483-DEL-58 | chr3-140137482-DEL-64 |
| chr3-140137833-DEL-269 | chr3-140137843-DEL-269 |
| chr3-140145486-DEL-68 | chr3-140145487-DEL-66 |
| chr3-140207853-DEL-350 | chr3-140207854-DEL-350 |
| chr3-140209051-DEL-1771 | chr3-140209054-DEL-1768 |
| chr3-140218198-DEL-6497 | chr3-140218199-DEL-6497 |
| chr3-140253501-DEL-62 | chr3-140253502-DEL-62 |
| chr3-140261975-DEL-183 | chr3-140261976-DEL-183 |
| chr3-140268240-DEL-52 | chr3-140268241-DEL-52 |

| | |
|---|---|
| chr3-140276789-DEL-110 | chr3-140276794-DEL-92 |
| chr3-140280278-DEL-211 | chr3-140280279-DEL-211 |
| chr3-140286009-DEL-54 | chr3-140285994-DEL-54 |
| chr3-140291796-DEL-1209 | chr3-140291796-DEL-1210 |
| chr3-140295048-DEL-876 | chr3-140295049-DEL-876 |
| chr3-140308514-DEL-137 | chr3-140308515-DEL-137 |
| chr3-140318941-DEL-60 | chr3-140318945-DEL-57 |
| chr3-140383178-DEL-60 | chr3-140383179-DEL-60 |
| chr3-140390632-DEL-1890 | chr3-140390633-DEL-1890 |
| chr3-140401954-DEL-105 | chr3-140401949-DEL-106 |
| chr3-140404021-DEL-149 | chr3-140404056-DEL-116 |
| chr3-140416772-DEL-213 | chr3-140416773-DEL-213 |
| chr3-140431957-DEL-6236 | chr3-140431958-DEL-6236 |
| chr3-140464951-DEL-205 | chr3-140464952-DEL-205 |
| chr3-140481685-DEL-58 | chr3-140481686-DEL-58 |
| chr3-140496755-DEL-1230 | chr3-140496756-DEL-1230 |
| chr3-140509766-DEL-982 | chr3-140509767-DEL-982 |
| chr3-140519958-DEL-184 | chr3-140519959-DEL-184 |
| chr3-140522289-DEL-53 | chr3-140522290-DEL-53 |
| chr3-140529118-DEL-220 | chr3-140529119-DEL-220 |
| chr3-140530285-DEL-53 | chr3-140530285-DEL-53 |
| chr3-140548290-DEL-118 | chr3-140548297-DEL-124 |
| chr3-140570740-DEL-6156 | chr3-140570741-DEL-6156 |
| chr3-140584109-DEL-120 | chr3-140584110-DEL-120 |
| chr3-140591512-DEL-386 | chr3-140591513-DEL-386 |
| chr3-140604138-DEL-189 | chr3-140604139-DEL-233 |
| chr3-140607933-DEL-90 | chr3-140607932-DEL-74 |
| chr3-140645955-DEL-73 | chr3-140645943-DEL-76 |
| chr3-140660626-DEL-1908 | chr3-140660627-DEL-1908 |
| chr3-140671546-DEL-54 | chr3-140671564-DEL-53 |
| chr3-140691359-DEL-219 | chr3-140691360-DEL-219 |
| chr3-140714220-DEL-162 | chr3-140714221-DEL-162 |
| chr3-140720451-DEL-425 | chr3-140720452-DEL-425 |
| chr3-140742274-DEL-1232 | chr3-140742275-DEL-1232 |
| chr3-140758622-DEL-6178 | chr3-140758626-DEL-6175 |
| chr3-140765131-DEL-83 | chr3-140765132-DEL-83 |
| chr3-140775522-DEL-362 | chr3-140775523-DEL-362 |
| chr3-140782586-DEL-537 | chr3-140782592-DEL-532 |
| chr3-140787669-DEL-74 | chr3-140787676-DEL-70 |
| chr3-140798700-DEL-389 | chr3-140798697-DEL-390 |
| chr3-140826594-DEL-84 | chr3-140826595-DEL-84 |
| chr3-140829732-DEL-96 | chr3-140829734-DEL-92 |
| chr3-140833746-DEL-1012 | chr3-140833747-DEL-1012 |
| chr3-140843143-DEL-62 | chr3-140843144-DEL-62 |
| chr3-140851111-DEL-53 | chr3-140851111-DEL-53 |
| chr3-140853519-DEL-7158 | chr3-140853529-DEL-7155 |
| chr3-140862297-DEL-57 | chr3-140862296-DEL-57 |
| chr3-140905140-DEL-97 | chr3-140905141-DEL-97 |
| chr3-140914743-DEL-56 | chr3-140914744-DEL-54 |
| chr3-140918385-DEL-917 | chr3-140918386-DEL-917 |
| chr3-140934058-DEL-66 | chr3-140934059-DEL-66 |
| chr3-140938489-DEL-928 | chr3-140938489-DEL-928 |
| chr3-140974066-DEL-474 | chr3-140974059-DEL-475 |
| chr3-140980502-DEL-181 | chr3-140980503-DEL-181 |
| chr3-140995013-DEL-739 | chr3-140995014-DEL-739 |
| chr3-140999441-DEL-2659 | chr3-140999442-DEL-2659 |
| chr3-141000948-DEL-60 | chr3-141000949-DEL-60 |
| chr3-141047375-DEL-64 | chr3-141047376-DEL-64 |
| chr3-141053423-DEL-268 | chr3-141053423-DEL-268 |
| chr3-141084425-DEL-6428 | chr3-141084426-DEL-6428 |
| chr3-141093457-DEL-64 | chr3-141093458-DEL-64 |
| chr3-141124733-DEL-108 | chr3-141124730-DEL-112 |
| chr3-141195743-DEL-358 | chr3-141195744-DEL-358 |
| chr3-141200441-DEL-282 | chr3-141200442-DEL-282 |
| chr3-141203056-DEL-70 | chr3-141203057-DEL-70 |
| chr3-141250817-DEL-66 | chr3-141250818-DEL-66 |
| chr3-141261181-DEL-68 | chr3-141261176-DEL-74 |
| chr3-141289600-DEL-857 | chr3-141289601-DEL-857 |
| chr3-141297110-DEL-56 | chr3-141297127-DEL-56 |
| chr3-141309401-DEL-6987 | chr3-141309402-DEL-6987 |
| chr3-141338052-DEL-2281 | chr3-141338053-DEL-2281 |
| chr3-141362445-DEL-61 | chr3-141362443-DEL-86 |
| chr3-141364049-DEL-69 | chr3-141364052-DEL-67 |
| chr3-141370972-DEL-1468 | chr3-141370973-DEL-1468 |
| chr3-141413446-DEL-497 | chr3-141413447-DEL-497 |
| chr3-141465713-DEL-6670 | chr3-141465714-DEL-6669 |
| chr3-141473359-DEL-82 | chr3-141473353-DEL-53 |
| chr3-141550383-DEL-58 | chr3-141550383-DEL-61 |
| chr3-141591414-DEL-522 | chr3-141591415-DEL-522 |
| chr3-141605685-DEL-86 | chr3-141605686-DEL-96 |

| | |
|---|---|
| chr3-141616628-DEL-1592 | chr3-141616624-DEL-1593 |
| chr3-141629104-DEL-66 | chr3-141629073-DEL-52 |
| chr3-141633195-DEL-99 | chr3-141633204-DEL-98 |
| chr3-141643808-DEL-60 | chr3-141643809-DEL-52 |
| chr3-141669488-DEL-145 | chr3-141669489-DEL-145 |
| chr3-141670098-DEL-262 | chr3-141670099-DEL-262 |
| chr3-141671527-DEL-57 | chr3-141671527-DEL-57 |
| chr3-141708682-DEL-7042 | chr3-141708683-DEL-7042 |
| chr3-141723729-DEL-52 | chr3-141723730-DEL-52 |
| chr3-141760632-DEL-611 | chr3-141760633-DEL-611 |
| chr3-141796924-DEL-470 | chr3-141796925-DEL-470 |
| chr3-141850133-DEL-124 | chr3-141850143-DEL-156 |
| chr3-141975659-DEL-332 | chr3-141975660-DEL-332 |
| chr3-141999984-DEL-327 | chr3-141999985-DEL-327 |
| chr3-142030989-DEL-67 | chr3-142031022-DEL-51 |
| chr3-142032737-DEL-607 | chr3-142032738-DEL-607 |
| chr3-142034782-DEL-138 | chr3-142034810-DEL-140 |
| chr3-142070235-DEL-106 | chr3-142070236-DEL-106 |
| chr3-142074758-DEL-123 | chr3-142074759-DEL-123 |
| chr3-142092935-DEL-62 | chr3-142092936-DEL-50 |
| chr3-142108988-DEL-205 | chr3-142108989-DEL-205 |
| chr3-142138432-DEL-51 | chr3-142138398-DEL-54 |
| chr3-142142995-DEL-201 | chr3-142142996-DEL-201 |
| chr3-142144602-DEL-214 | chr3-142144641-DEL-190 |
| chr3-142155534-DEL-1255 | chr3-142155535-DEL-1255 |
| chr3-142156311-DEL-51 | chr3-142156311-DEL-51 |
| chr3-142161678-DEL-57 | chr3-142161677-DEL-58 |
| chr3-142173106-DEL-154 | chr3-142173098-DEL-146 |
| chr3-142190332-DEL-1223 | chr3-142190333-DEL-1223 |
| chr3-142205984-DEL-200 | chr3-142205985-DEL-200 |
| chr3-142217978-DEL-6661 | chr3-142217979-DEL-6661 |
| chr3-142224160-DEL-203 | chr3-142224160-DEL-203 |
| chr3-142225046-DEL-456 | chr3-142225047-DEL-456 |
| chr3-142256934-DEL-64 | chr3-142256935-DEL-64 |
| chr3-142257734-DEL-90 | chr3-142257735-DEL-86 |
| chr3-142293511-DEL-67 | chr3-142293510-DEL-67 |
| chr3-142298002-DEL-345 | chr3-142298003-DEL-345 |
| chr3-142298527-DEL-306 | chr3-142298525-DEL-305 |
| chr3-142304300-DEL-170 | chr3-142304299-DEL-170 |
| chr3-142305492-DEL-82 | chr3-142305493-DEL-68 |
| chr3-142336840-DEL-101 | chr3-142336841-DEL-101 |
| chr3-142351993-DEL-129 | chr3-142351986-DEL-110 |
| chr3-142356814-DEL-70 | chr3-142356814-DEL-70 |
| chr3-142358106-DEL-561 | chr3-142358099-DEL-570 |
| chr3-142365446-DEL-52 | chr3-142365447-DEL-52 |
| chr3-142374208-DEL-60 | chr3-142374209-DEL-56 |
| chr3-142395631-DEL-121 | chr3-142395632-DEL-121 |
| chr3-142404552-DEL-76 | chr3-142404553-DEL-76 |
| chr3-142420663-DEL-67 | chr3-142420658-DEL-68 |
| chr3-142466559-DEL-200 | chr3-142466560-DEL-200 |
| chr3-142475345-DEL-60 | chr3-142475346-DEL-60 |
| chr3-142477649-DEL-657 | chr3-142477650-DEL-657 |
| chr3-142501811-DEL-52 | chr3-142501808-DEL-56 |
| chr3-142528707-DEL-149 | chr3-142528708-DEL-149 |
| chr3-142539846-DEL-54 | chr3-142539847-DEL-54 |
| chr3-142565863-DEL-93 | chr3-142565865-DEL-92 |
| chr3-142612553-DEL-51 | chr3-142612554-DEL-51 |
| chr3-142661454-DEL-3682 | chr3-142661455-DEL-3682 |
| chr3-142668283-DEL-1294 | chr3-142668284-DEL-1294 |
| chr3-142674130-DEL-66 | chr3-142674131-DEL-66 |
| chr3-142677570-DEL-86 | chr3-142677571-DEL-86 |
| chr3-142723178-DEL-115 | chr3-142723218-DEL-69 |
| chr3-142765395-DEL-118 | chr3-142765396-DEL-118 |
| chr3-142836044-DEL-2657 | chr3-142836045-DEL-2657 |
| chr3-142874837-DEL-80 | chr3-142874836-DEL-82 |
| chr3-142879197-DEL-61 | chr3-142879191-DEL-62 |
| chr3-142891275-DEL-70 | chr3-142891276-DEL-70 |
| chr3-142921843-DEL-2653 | chr3-142921844-DEL-2653 |
| chr3-142936882-DEL-2418 | chr3-142936883-DEL-2418 |
| chr3-142961444-DEL-665 | chr3-142961445-DEL-665 |
| chr3-142985547-DEL-181 | chr3-142985534-DEL-197 |
| chr3-142992161-DEL-70 | chr3-142992168-DEL-98 |
| chr3-143010451-DEL-740 | chr3-143010452-DEL-740 |
| chr3-143015259-DEL-59 | chr3-143015250-DEL-60 |
| chr3-143025373-DEL-64 | chr3-143025374-DEL-64 |
| chr3-143210724-DEL-205 | chr3-143210727-DEL-204 |
| chr3-143245858-DEL-431 | chr3-143245856-DEL-432 |
| chr3-143261539-DEL-179 | chr3-143261588-DEL-131 |
| chr3-143273959-DEL-278 | chr3-143273960-DEL-278 |
| chr3-143311435-DEL-180 | chr3-143311436-DEL-175 |

| | |
|---|---|
| chr3-143323334-DEL-67 | chr3-143323335-DEL-67 |
| chr3-143392549-DEL-452 | chr3-143392550-DEL-452 |
| chr3-143404482-DEL-964 | chr3-143404503-DEL-943 |
| chr3-143414314-DEL-743 | chr3-143414315-DEL-743 |
| chr3-143440952-DEL-94 | chr3-143440950-DEL-83 |
| chr3-143511858-DEL-276 | chr3-143511859-DEL-276 |
| chr3-143520712-DEL-94 | chr3-143520741-DEL-60 |
| chr3-143546507-DEL-6252 | chr3-143546508-DEL-6252 |
| chr3-143557170-DEL-1068 | chr3-143557171-DEL-1068 |
| chr3-143589263-DEL-5215 | chr3-143589264-DEL-5215 |
| chr3-143626388-DEL-66 | chr3-143626389-DEL-66 |
| chr3-143640412-DEL-190 | chr3-143640382-DEL-225 |
| chr3-143645047-DEL-192 | chr3-143645048-DEL-192 |
| chr3-143646019-DEL-61 | chr3-143646020-DEL-61 |
| chr3-143681636-DEL-207 | chr3-143681667-DEL-203 |
| chr3-143704928-DEL-74 | chr3-143704931-DEL-72 |
| chr3-143725892-DEL-85 | chr3-143725842-DEL-131 |
| chr3-143727874-DEL-64 | chr3-143727875-DEL-64 |
| chr3-143761071-DEL-114 | chr3-143761072-DEL-114 |
| chr3-143787577-DEL-1567 | chr3-143787578-DEL-1567 |
| chr3-143808752-DEL-112 | chr3-143808753-DEL-82 |
| chr3-143811222-DEL-58 | chr3-143811189-DEL-86 |
| chr3-143850269-DEL-2732 | chr3-143850270-DEL-2732 |
| chr3-143871869-DEL-74 | chr3-143871870-DEL-72 |
| chr3-143888035-DEL-51 | chr3-143888036-DEL-51 |
| chr3-143923098-DEL-229 | chr3-143923096-DEL-230 |
| chr3-143955972-DEL-266 | chr3-143955932-DEL-289 |
| chr3-143976531-DEL-56 | chr3-143976532-DEL-56 |
| chr3-143981217-DEL-64 | chr3-143981212-DEL-61 |
| chr3-144006561-DEL-80 | chr3-144006562-DEL-110 |
| chr3-144016283-DEL-54 | chr3-144016284-DEL-54 |
| chr3-144047466-DEL-230 | chr3-144047467-DEL-230 |
| chr3-144048626-DEL-1890 | chr3-144048627-DEL-1890 |
| chr3-144048960-DEL-54 | chr3-144048961-DEL-54 |
| chr3-144052043-DEL-1234 | chr3-144052044-DEL-1234 |
| chr3-144112641-DEL-210 | chr3-144112645-DEL-202 |
| chr3-144118044-DEL-58 | chr3-144118045-DEL-54 |
| chr3-144125372-DEL-159 | chr3-144125373-DEL-159 |
| chr3-144164015-DEL-65 | chr3-144164019-DEL-61 |
| chr3-144179797-DEL-363 | chr3-144179798-DEL-363 |
| chr3-144189940-DEL-217 | chr3-144189941-DEL-217 |
| chr3-144196440-DEL-418 | chr3-144196441-DEL-418 |
| chr3-144213065-DEL-63 | chr3-144213066-DEL-63 |
| chr3-144243359-DEL-82 | chr3-144243360-DEL-82 |
| chr3-144258657-DEL-113 | chr3-144258658-DEL-113 |
| chr3-144293583-DEL-374 | chr3-144293584-DEL-374 |
| chr3-144299814-DEL-66 | chr3-144299815-DEL-66 |
| chr3-144304508-DEL-396 | chr3-144304509-DEL-396 |
| chr3-144305206-DEL-86 | chr3-144305207-DEL-86 |
| chr3-144314317-DEL-109 | chr3-144314318-DEL-109 |
| chr3-144324737-DEL-3594 | chr3-144324738-DEL-3594 |
| chr3-144337351-DEL-642 | chr3-144337349-DEL-643 |
| chr3-144351994-DEL-361 | chr3-144351995-DEL-361 |
| chr3-144360166-DEL-6394 | chr3-144360168-DEL-6395 |
| chr3-144386541-DEL-87 | chr3-144386542-DEL-87 |
| chr3-144394420-DEL-56 | chr3-144394414-DEL-55 |
| chr3-144456742-DEL-174 | chr3-144456748-DEL-169 |
| chr3-144482582-DEL-56 | chr3-144482583-DEL-56 |
| chr3-144534136-DEL-7449 | chr3-144534137-DEL-7449 |
| chr3-144534660-DEL-212 | chr3-144534661-DEL-212 |
| chr3-144546545-DEL-135 | chr3-144546573-DEL-108 |
| chr3-144563680-DEL-414 | chr3-144563681-DEL-414 |
| chr3-144574325-DEL-3595 | chr3-144574326-DEL-3595 |
| chr3-144618702-DEL-67 | chr3-144618687-DEL-68 |
| chr3-144625198-DEL-1397 | chr3-144625198-DEL-1397 |
| chr3-144628068-DEL-329 | chr3-144628069-DEL-328 |
| chr3-144630600-DEL-230 | chr3-144630601-DEL-230 |
| chr3-144641649-DEL-3153 | chr3-144641642-DEL-3158 |
| chr3-144662557-DEL-208 | chr3-144662558-DEL-208 |
| chr3-144689223-DEL-78 | chr3-144689218-DEL-68 |
| chr3-144696859-DEL-178 | chr3-144696860-DEL-178 |
| chr3-144698075-DEL-293 | chr3-144698076-DEL-293 |
| chr3-144731323-DEL-271 | chr3-144731308-DEL-272 |
| chr3-144761071-DEL-6455 | chr3-144761072-DEL-6455 |
| chr3-144808498-DEL-67 | chr3-144808499-DEL-67 |
| chr3-144814785-DEL-5178 | chr3-144814786-DEL-5178 |
| chr3-144832536-DEL-173 | chr3-144832537-DEL-173 |
| chr3-144846790-DEL-719 | chr3-144846792-DEL-718 |
| chr3-144897126-DEL-61 | chr3-144897127-DEL-61 |
| chr3-144989377-DEL-811 | chr3-144989378-DEL-811 |

| | |
|---|---|
| chr3-145007336-DEL-1794 | chr3-145007337-DEL-1794 |
| chr3-145022502-DEL-59 | chr3-145022485-DEL-77 |
| chr3-145024410-DEL-358 | chr3-145024411-DEL-359 |
| chr3-145038973-DEL-126 | chr3-145038974-DEL-126 |
| chr3-145060222-DEL-581 | chr3-145060223-DEL-581 |
| chr3-145062179-DEL-57 | chr3-145062180-DEL-64 |
| chr3-145115221-DEL-682 | chr3-145115222-DEL-682 |
| chr3-145129101-DEL-225 | chr3-145129102-DEL-225 |
| chr3-145136579-DEL-54 | chr3-145136580-DEL-54 |
| chr3-145138539-DEL-173 | chr3-145138524-DEL-174 |
| chr3-145146436-DEL-56 | chr3-145146436-DEL-56 |
| chr3-145159687-DEL-72 | chr3-145159690-DEL-70 |
| chr3-145166642-DEL-64 | chr3-145166643-DEL-52 |
| chr3-145173454-DEL-223 | chr3-145173455-DEL-223 |
| chr3-145180838-DEL-68 | chr3-145180824-DEL-63 |
| chr3-145199767-DEL-116 | chr3-145199768-DEL-116 |
| chr3-145202698-DEL-345 | chr3-145202698-DEL-345 |
| chr3-145237977-DEL-228 | chr3-145237978-DEL-228 |
| chr3-145263773-DEL-61 | chr3-145263774-DEL-61 |
| chr3-145267739-DEL-127 | chr3-145267740-DEL-127 |
| chr3-145270262-DEL-53 | chr3-145270263-DEL-53 |
| chr3-145272641-DEL-348 | chr3-145272642-DEL-348 |
| chr3-145279291-DEL-878 | chr3-145279292-DEL-878 |
| chr3-145316096-DEL-67 | chr3-145316097-DEL-67 |
| chr3-145322586-DEL-107 | chr3-145322587-DEL-107 |
| chr3-145333286-DEL-139 | chr3-145333287-DEL-139 |
| chr3-145372962-DEL-110 | chr3-145372963-DEL-110 |
| chr3-145377887-DEL-873 | chr3-145377888-DEL-873 |
| chr3-145383226-DEL-76 | chr3-145383227-DEL-76 |
| chr3-145395096-DEL-399 | chr3-145395097-DEL-399 |
| chr3-145401817-DEL-60 | chr3-145401818-DEL-60 |
| chr3-145414226-DEL-195 | chr3-145414227-DEL-195 |
| chr3-145424347-DEL-55 | chr3-145424348-DEL-55 |
| chr3-145427464-DEL-199 | chr3-145427463-DEL-198 |
| chr3-145439404-DEL-52 | chr3-145439393-DEL-64 |
| chr3-145471439-DEL-165 | chr3-145471440-DEL-165 |
| chr3-145479956-DEL-179 | chr3-145479957-DEL-177 |
| chr3-145483426-DEL-58 | chr3-145483461-DEL-58 |
| chr3-145513165-DEL-134 | chr3-145513173-DEL-104 |
| chr3-145515643-DEL-72 | chr3-145515644-DEL-72 |
| chr3-145570841-DEL-6783 | chr3-145570842-DEL-6783 |
| chr3-145584727-DEL-422 | chr3-145584728-DEL-422 |
| chr3-145589339-DEL-54 | chr3-145589373-DEL-54 |
| chr3-145595355-DEL-82 | chr3-145595372-DEL-66 |
| chr3-145613773-DEL-58 | chr3-145613774-DEL-58 |
| chr3-145616228-DEL-252 | chr3-145616235-DEL-246 |
| chr3-145654911-DEL-53 | chr3-145654912-DEL-53 |
| chr3-145666376-DEL-61 | chr3-145666367-DEL-68 |
| chr3-145676783-DEL-162 | chr3-145676784-DEL-162 |
| chr3-145679535-DEL-213 | chr3-145679536-DEL-213 |
| chr3-145685041-DEL-324 | chr3-145685002-DEL-324 |
| chr3-145696644-DEL-226 | chr3-145696645-DEL-226 |
| chr3-145700189-DEL-189 | chr3-145700187-DEL-190 |
| chr3-145702210-DEL-185 | chr3-145702211-DEL-185 |
| chr3-145705778-DEL-59 | chr3-145705774-DEL-57 |
| chr3-145709713-DEL-300 | chr3-145709707-DEL-298 |
| chr3-145714307-DEL-440 | chr3-145714308-DEL-440 |
| chr3-145718827-DEL-82 | chr3-145718828-DEL-82 |
| chr3-145731644-DEL-373 | chr3-145731644-DEL-373 |
| chr3-145740807-DEL-101 | chr3-145740808-DEL-101 |
| chr3-145758394-DEL-54 | chr3-145758375-DEL-68 |
| chr3-145789335-DEL-6817 | chr3-145789336-DEL-6817 |
| chr3-145831908-DEL-97 | chr3-145831936-DEL-111 |
| chr3-145851498-DEL-1587 | chr3-145851499-DEL-1587 |
| chr3-145864116-DEL-87 | chr3-145864114-DEL-83 |
| chr3-145885056-DEL-55 | chr3-145885056-DEL-55 |
| chr3-145938177-DEL-111 | chr3-145938178-DEL-111 |
| chr3-145948167-DEL-64 | chr3-145948168-DEL-64 |
| chr3-145963038-DEL-198 | chr3-145963039-DEL-198 |
| chr3-145999479-DEL-6174 | chr3-145999480-DEL-6174 |
| chr3-146012494-DEL-403 | chr3-146012489-DEL-404 |
| chr3-146029190-DEL-56 | chr3-146029189-DEL-57 |
| chr3-146032812-DEL-53 | chr3-146032813-DEL-53 |
| chr3-146062076-DEL-6423 | chr3-146062074-DEL-6424 |
| chr3-146062101-DEL-199 | chr3-146062102-DEL-199 |
| chr3-146074829-DEL-78 | chr3-146074830-DEL-78 |
| chr3-146097612-DEL-678 | chr3-146097613-DEL-678 |
| chr3-146106562-DEL-62 | chr3-146106563-DEL-62 |
| chr3-146147544-DEL-61 | chr3-146147547-DEL-59 |
| chr3-146152281-DEL-225 | chr3-146152282-DEL-225 |

| | |
|---|---|
| chr3-146184074-DEL-218 | chr3-146184075-DEL-218 |
| chr3-146195954-DEL-144 | chr3-146195955-DEL-145 |
| chr3-146240863-DEL-90 | chr3-146240864-DEL-89 |
| chr3-146244796-DEL-533 | chr3-146244797-DEL-533 |
| chr3-146290579-DEL-51 | chr3-146290576-DEL-51 |
| chr3-146309963-DEL-1601 | chr3-146309964-DEL-1601 |
| chr3-146311722-DEL-84 | chr3-146311723-DEL-84 |
| chr3-146324472-DEL-856 | chr3-146324475-DEL-847 |
| chr3-146361552-DEL-57 | chr3-146361589-DEL-69 |
| chr3-146405099-DEL-78 | chr3-146405100-DEL-78 |
| chr3-146432662-DEL-153 | chr3-146432663-DEL-153 |
| chr3-146470358-DEL-456 | chr3-146470359-DEL-456 |
| chr3-146474014-DEL-86 | chr3-146474015-DEL-86 |
| chr3-146488882-DEL-202 | chr3-146488883-DEL-202 |
| chr3-146508739-DEL-216 | chr3-146508740-DEL-216 |
| chr3-146535496-DEL-2922 | chr3-146535497-DEL-2922 |
| chr3-146544501-DEL-617 | chr3-146544502-DEL-617 |
| chr3-146574387-DEL-1438 | chr3-146574388-DEL-1438 |
| chr3-146617465-DEL-498 | chr3-146617465-DEL-498 |
| chr3-146624557-DEL-6216 | chr3-146624516-DEL-6241 |
| chr3-146643635-DEL-110 | chr3-146643636-DEL-110 |
| chr3-146653611-DEL-177 | chr3-146653612-DEL-177 |
| chr3-146698306-DEL-93 | chr3-146698308-DEL-95 |
| chr3-146758643-DEL-116 | chr3-146758656-DEL-112 |
| chr3-146771442-DEL-259 | chr3-146771446-DEL-258 |
| chr3-146812905-DEL-6191 | chr3-146812906-DEL-6191 |
| chr3-146857979-DEL-212 | chr3-146857989-DEL-204 |
| chr3-146890703-DEL-54 | chr3-146890704-DEL-54 |
| chr3-146891542-DEL-59 | chr3-146891549-DEL-53 |
| chr3-146893108-DEL-67 | chr3-146893108-DEL-67 |
| chr3-146900641-DEL-100 | chr3-146900642-DEL-100 |
| chr3-146908228-DEL-78 | chr3-146908229-DEL-67 |
| chr3-146918010-DEL-82 | chr3-146918011-DEL-82 |
| chr3-146918989-DEL-60 | chr3-146918990-DEL-60 |
| chr3-146924688-DEL-6482 | chr3-146924689-DEL-6482 |
| chr3-146974822-DEL-60 | chr3-146974823-DEL-58 |
| chr3-146989128-DEL-731 | chr3-146989129-DEL-731 |
| chr3-147005816-DEL-6261 | chr3-147005817-DEL-6261 |
| chr3-147025643-DEL-141 | chr3-147025644-DEL-141 |
| chr3-147058528-DEL-94 | chr3-147058532-DEL-92 |
| chr3-147062021-DEL-6168 | chr3-147062022-DEL-6168 |
| chr3-147068954-DEL-851 | chr3-147068955-DEL-851 |
| chr3-147092351-DEL-61 | chr3-147092352-DEL-53 |
| chr3-147101055-DEL-57 | chr3-147101056-DEL-57 |
| chr3-147110411-DEL-54 | chr3-147110412-DEL-54 |
| chr3-147159433-DEL-282 | chr3-147159434-DEL-282 |
| chr3-147170451-DEL-52 | chr3-147170436-DEL-62 |
| chr3-147203922-DEL-298 | chr3-147203921-DEL-297 |
| chr3-147205631-DEL-535 | chr3-147205631-DEL-536 |
| chr3-147213229-DEL-105 | chr3-147213230-DEL-105 |
| chr3-147261526-DEL-6442 | chr3-147261527-DEL-6442 |
| chr3-147290672-DEL-193 | chr3-147290673-DEL-193 |
| chr3-147295306-DEL-52 | chr3-147295307-DEL-52 |
| chr3-147320442-DEL-124 | chr3-147320443-DEL-124 |
| chr3-147324981-DEL-73 | chr3-147324979-DEL-69 |
| chr3-147341145-DEL-370 | chr3-147341146-DEL-370 |
| chr3-147346726-DEL-56 | chr3-147346726-DEL-56 |
| chr3-147353492-DEL-82 | chr3-147353493-DEL-123 |
| chr3-147456800-DEL-120 | chr3-147456801-DEL-120 |
| chr3-147466254-DEL-52 | chr3-147466261-DEL-50 |
| chr3-147474645-DEL-96 | chr3-147474646-DEL-96 |
| chr3-147474851-DEL-1831 | chr3-147474848-DEL-1831 |
| chr3-147477003-DEL-1051 | chr3-147477004-DEL-1051 |
| chr3-147510054-DEL-400 | chr3-147510055-DEL-400 |
| chr3-147519149-DEL-237 | chr3-147519150-DEL-237 |
| chr3-147576536-DEL-6904 | chr3-147576532-DEL-6905 |
| chr3-147620575-DEL-96 | chr3-147620575-DEL-104 |
| chr3-147641925-DEL-569 | chr3-147641926-DEL-569 |
| chr3-147677962-DEL-343 | chr3-147677989-DEL-334 |
| chr3-147680055-DEL-52 | chr3-147680056-DEL-52 |
| chr3-147733031-DEL-825 | chr3-147733032-DEL-825 |
| chr3-147750025-DEL-375 | chr3-147750026-DEL-375 |
| chr3-147750787-DEL-507 | chr3-147750789-DEL-504 |
| chr3-147771320-DEL-89 | chr3-147771321-DEL-89 |
| chr3-147792745-DEL-1104 | chr3-147792746-DEL-1104 |
| chr3-147804092-DEL-66 | chr3-147804093-DEL-58 |
| chr3-147824269-DEL-75 | chr3-147824270-DEL-75 |
| chr3-147846103-DEL-61 | chr3-147846104-DEL-61 |
| chr3-147851889-DEL-82 | chr3-147851893-DEL-79 |
| chr3-147876505-DEL-380 | chr3-147876506-DEL-380 |

| | |
|---|---|
| chr3-147926933-DEL-79 | chr3-147926932-DEL-81 |
| chr3-147953668-DEL-83 | chr3-147953644-DEL-72 |
| chr3-147973431-DEL-1038 | chr3-147973430-DEL-1039 |
| chr3-147999412-DEL-86 | chr3-147999409-DEL-60 |
| chr3-148023579-DEL-2850 | chr3-148023580-DEL-2850 |
| chr3-148051258-DEL-170 | chr3-148051259-DEL-170 |
| chr3-148052965-DEL-70 | chr3-148052966-DEL-70 |
| chr3-148061261-DEL-54 | chr3-148061262-DEL-52 |
| chr3-148077475-DEL-502 | chr3-148077476-DEL-502 |
| chr3-148081253-DEL-64 | chr3-148081254-DEL-64 |
| chr3-148103235-DEL-108 | chr3-148103236-DEL-99 |
| chr3-148114116-DEL-378 | chr3-148114117-DEL-378 |
| chr3-148126339-DEL-336 | chr3-148126340-DEL-336 |
| chr3-148136690-DEL-264 | chr3-148136733-DEL-228 |
| chr3-148153447-DEL-54 | chr3-148153471-DEL-58 |
| chr3-148188940-DEL-163 | chr3-148188949-DEL-162 |
| chr3-148312435-DEL-129 | chr3-148312436-DEL-129 |
| chr3-148313448-DEL-58 | chr3-148313441-DEL-70 |
| chr3-148321814-DEL-1989 | chr3-148321815-DEL-1989 |
| chr3-148323761-DEL-498 | chr3-148323762-DEL-498 |
| chr3-148324320-DEL-396 | chr3-148324321-DEL-396 |
| chr3-148367760-DEL-206 | chr3-148367761-DEL-206 |
| chr3-148391496-DEL-57 | chr3-148391493-DEL-81 |
| chr3-148397492-DEL-455 | chr3-148397493-DEL-449 |
| chr3-148416157-DEL-6440 | chr3-148416157-DEL-6440 |
| chr3-148448100-DEL-119 | chr3-148448101-DEL-119 |
| chr3-148453515-DEL-78 | chr3-148453516-DEL-78 |
| chr3-148455726-DEL-7222 | chr3-148455727-DEL-7222 |
| chr3-148464716-DEL-94 | chr3-148464762-DEL-92 |
| chr3-148503967-DEL-78 | chr3-148503966-DEL-78 |
| chr3-148556205-DEL-77 | chr3-148556206-DEL-57 |
| chr3-148589813-DEL-55 | chr3-148589814-DEL-55 |
| chr3-148596926-DEL-162 | chr3-148596927-DEL-162 |
| chr3-148615243-DEL-380 | chr3-148615241-DEL-379 |
| chr3-148628667-DEL-81 | chr3-148628668-DEL-81 |
| chr3-148724178-DEL-514 | chr3-148724179-DEL-514 |
| chr3-148734551-DEL-622 | chr3-148734552-DEL-622 |
| chr3-148736824-DEL-142 | chr3-148736777-DEL-156 |
| chr3-148756369-DEL-226 | chr3-148756370-DEL-226 |
| chr3-148758135-DEL-276 | chr3-148758136-DEL-270 |
| chr3-148771236-DEL-1926 | chr3-148771237-DEL-1926 |
| chr3-148828588-DEL-59 | chr3-148828604-DEL-60 |
| chr3-148829450-DEL-1893 | chr3-148829451-DEL-1893 |
| chr3-148849207-DEL-59 | chr3-148849208-DEL-59 |
| chr3-148899027-DEL-110 | chr3-148899028-DEL-110 |
| chr3-148916766-DEL-330 | chr3-148916767-DEL-330 |
| chr3-148966718-DEL-85 | chr3-148966719-DEL-85 |
| chr3-148968885-DEL-60 | chr3-148968886-DEL-60 |
| chr3-148979225-DEL-165 | chr3-148979230-DEL-162 |
| chr3-148994041-DEL-551 | chr3-148994042-DEL-551 |
| chr3-149007444-DEL-115 | chr3-149007469-DEL-91 |
| chr3-149039865-DEL-3373 | chr3-149039866-DEL-3370 |
| chr3-149047109-DEL-144 | chr3-149047110-DEL-144 |
| chr3-149075234-DEL-75 | chr3-149075196-DEL-76 |
| chr3-149093618-DEL-64 | chr3-149093619-DEL-64 |
| chr3-149095336-DEL-103 | chr3-149095341-DEL-71 |
| chr3-149104036-DEL-108 | chr3-149104036-DEL-108 |
| chr3-149135302-DEL-187 | chr3-149135303-DEL-187 |
| chr3-149170757-DEL-68 | chr3-149170758-DEL-68 |
| chr3-149173736-DEL-1477 | chr3-149173737-DEL-1477 |
| chr3-149209807-DEL-529 | chr3-149209808-DEL-529 |
| chr3-149228645-DEL-114 | chr3-149228646-DEL-106 |
| chr3-149234519-DEL-70 | chr3-149234520-DEL-70 |
| chr3-149245934-DEL-185 | chr3-149245935-DEL-185 |
| chr3-149275077-DEL-398 | chr3-149275078-DEL-398 |
| chr3-149278984-DEL-386 | chr3-149278985-DEL-386 |
| chr3-149316224-DEL-214 | chr3-149316225-DEL-214 |
| chr3-149401373-DEL-486 | chr3-149401372-DEL-479 |
| chr3-149413496-DEL-3284 | chr3-149413497-DEL-3284 |
| chr3-149420662-DEL-64 | chr3-149420663-DEL-64 |
| chr3-149462020-DEL-81 | chr3-149462021-DEL-81 |
| chr3-149477139-DEL-1047 | chr3-149477140-DEL-1047 |
| chr3-149565551-DEL-674 | chr3-149565552-DEL-674 |
| chr3-149585240-DEL-157 | chr3-149585274-DEL-158 |
| chr3-149662272-DEL-58 | chr3-149662272-DEL-58 |
| chr3-149675327-DEL-205 | chr3-149675328-DEL-205 |
| chr3-149676746-DEL-6117 | chr3-149676747-DEL-6117 |
| chr3-149688578-DEL-85 | chr3-149688579-DEL-85 |
| chr3-149713958-DEL-68 | chr3-149713965-DEL-58 |
| chr3-149773785-DEL-1498 | chr3-149773780-DEL-1470 |

| | |
|---|---|
| chr3-149774415-DEL-59 | chr3-149774416-DEL-59 |
| chr3-149776847-DEL-532 | chr3-149776845-DEL-533 |
| chr3-149778192-DEL-1065 | chr3-149778195-DEL-1063 |
| chr3-149782628-DEL-9257 | chr3-149782629-DEL-9257 |
| chr3-149791020-DEL-422 | chr3-149791021-DEL-422 |
| chr3-149826667-DEL-2842 | chr3-149826670-DEL-2842 |
| chr3-149850715-DEL-1094 | chr3-149850716-DEL-1094 |
| chr3-149852601-DEL-1686 | chr3-149852602-DEL-1686 |
| chr3-149860641-DEL-361 | chr3-149860642-DEL-361 |
| chr3-149864292-DEL-6345 | chr3-149864293-DEL-6345 |
| chr3-149907189-DEL-92 | chr3-149907186-DEL-88 |
| chr3-149923585-DEL-58 | chr3-149923586-DEL-62 |
| chr3-149929899-DEL-281 | chr3-149929900-DEL-281 |
| chr3-149948491-DEL-68 | chr3-149948536-DEL-64 |
| chr3-149975382-DEL-51 | chr3-149975384-DEL-52 |
| chr3-149986848-DEL-188 | chr3-149986849-DEL-185 |
| chr3-150029875-DEL-86 | chr3-150029862-DEL-86 |
| chr3-150042483-DEL-4482 | chr3-150042484-DEL-4482 |
| chr3-150061029-DEL-173 | chr3-150061030-DEL-173 |
| chr3-150120055-DEL-97 | chr3-150120056-DEL-97 |
| chr3-150125988-DEL-405 | chr3-150125989-DEL-405 |
| chr3-150169776-DEL-107 | chr3-150169750-DEL-106 |
| chr3-150174070-DEL-52 | chr3-150174071-DEL-50 |
| chr3-150215608-DEL-111 | chr3-150215602-DEL-117 |
| chr3-150247940-DEL-182 | chr3-150247927-DEL-208 |
| chr3-150253560-DEL-118 | chr3-150253553-DEL-118 |
| chr3-150254700-DEL-118 | chr3-150254701-DEL-118 |
| chr3-150263444-DEL-64 | chr3-150263439-DEL-70 |
| chr3-150269869-DEL-55 | chr3-150269831-DEL-60 |
| chr3-150276183-DEL-263 | chr3-150276184-DEL-263 |
| chr3-150289411-DEL-2002 | chr3-150289412-DEL-2002 |
| chr3-150300857-DEL-101 | chr3-150300858-DEL-74 |
| chr3-150329802-DEL-59 | chr3-150329803-DEL-59 |
| chr3-150338491-DEL-122 | chr3-150338492-DEL-122 |
| chr3-150420434-DEL-90 | chr3-150420439-DEL-86 |
| chr3-150429720-DEL-305 | chr3-150429721-DEL-305 |
| chr3-150456771-DEL-115 | chr3-150456772-DEL-115 |
| chr3-150499637-DEL-832 | chr3-150499638-DEL-832 |
| chr3-150588859-DEL-525 | chr3-150588860-DEL-525 |
| chr3-150594862-DEL-828 | chr3-150594863-DEL-828 |
| chr3-150607927-DEL-54 | chr3-150607928-DEL-54 |
| chr3-150615586-DEL-60 | chr3-150615586-DEL-60 |
| chr3-150626612-DEL-1161 | chr3-150626612-DEL-1161 |
| chr3-150630824-DEL-78 | chr3-150630831-DEL-72 |
| chr3-150636485-DEL-2262 | chr3-150636486-DEL-2262 |
| chr3-150644490-DEL-50 | chr3-150644491-DEL-54 |
| chr3-150658796-DEL-78 | chr3-150658799-DEL-84 |
| chr3-150665059-DEL-1002 | chr3-150665060-DEL-1002 |
| chr3-150691122-DEL-986 | chr3-150691123-DEL-986 |
| chr3-150732265-DEL-58 | chr3-150732282-DEL-50 |
| chr3-150734505-DEL-2615 | chr3-150734506-DEL-2615 |
| chr3-150752844-DEL-58 | chr3-150752845-DEL-58 |
| chr3-150770300-DEL-787 | chr3-150770301-DEL-787 |
| chr3-150783827-DEL-791 | chr3-150783828-DEL-791 |
| chr3-150795506-DEL-106 | chr3-150795507-DEL-106 |
| chr3-150802068-DEL-507 | chr3-150802067-DEL-507 |
| chr3-150814944-DEL-198 | chr3-150814945-DEL-198 |
| chr3-150816811-DEL-63 | chr3-150816815-DEL-51 |
| chr3-150854151-DEL-83 | chr3-150854152-DEL-83 |
| chr3-150856032-DEL-364 | chr3-150856033-DEL-364 |
| chr3-150935462-DEL-73 | chr3-150935463-DEL-73 |
| chr3-150939094-DEL-3050 | chr3-150939095-DEL-3050 |
| chr3-150947863-DEL-52 | chr3-150947864-DEL-52 |
| chr3-150993057-DEL-240 | chr3-150993063-DEL-228 |
| chr3-151027998-DEL-1072 | chr3-151027997-DEL-1073 |
| chr3-151050276-DEL-351 | chr3-151050277-DEL-351 |
| chr3-151069202-DEL-95 | chr3-151069196-DEL-96 |
| chr3-151116552-DEL-533 | chr3-151116553-DEL-533 |
| chr3-151119213-DEL-366 | chr3-151119214-DEL-366 |
| chr3-151152658-DEL-85 | chr3-151152658-DEL-67 |
| chr3-151163145-DEL-142 | chr3-151163164-DEL-116 |
| chr3-151198836-DEL-1887 | chr3-151198837-DEL-1887 |
| chr3-151255662-DEL-60 | chr3-151255667-DEL-56 |
| chr3-151267237-DEL-54 | chr3-151267238-DEL-51 |
| chr3-151286386-DEL-268 | chr3-151286412-DEL-243 |
| chr3-151288216-DEL-101 | chr3-151288206-DEL-66 |
| chr3-151313965-DEL-516 | chr3-151313966-DEL-516 |
| chr3-151317343-DEL-428 | chr3-151317338-DEL-430 |
| chr3-151329530-DEL-6802 | chr3-151329531-DEL-6802 |
| chr3-151338687-DEL-74 | chr3-151338688-DEL-74 |

| | |
|---|---|
| chr3-151344164-DEL-181 | chr3-151344166-DEL-176 |
| chr3-151378190-DEL-7288 | chr3-151378191-DEL-7288 |
| chr3-151384560-DEL-424 | chr3-151384560-DEL-424 |
| chr3-151425138-DEL-92 | chr3-151425139-DEL-92 |
| chr3-151446286-DEL-472 | chr3-151446287-DEL-472 |
| chr3-151449801-DEL-536 | chr3-151449802-DEL-536 |
| chr3-151459765-DEL-313 | chr3-151459766-DEL-313 |
| chr3-151469046-DEL-677 | chr3-151469078-DEL-641 |
| chr3-151485193-DEL-1657 | chr3-151485194-DEL-1657 |
| chr3-151520438-DEL-162 | chr3-151520403-DEL-154 |
| chr3-151568873-DEL-709 | chr3-151568873-DEL-709 |
| chr3-151571020-DEL-246 | chr3-151571026-DEL-241 |
| chr3-151592546-DEL-498 | chr3-151592547-DEL-498 |
| chr3-151656665-DEL-7201 | chr3-151656666-DEL-7201 |
| chr3-151665640-DEL-7131 | chr3-151665641-DEL-7131 |
| chr3-151674607-DEL-117 | chr3-151674608-DEL-117 |
| chr3-151677444-DEL-63 | chr3-151677440-DEL-64 |
| chr3-151690435-DEL-65 | chr3-151690436-DEL-65 |
| chr3-151720219-DEL-2582 | chr3-151720220-DEL-2582 |
| chr3-151742065-DEL-3613 | chr3-151742068-DEL-3612 |
| chr3-151779812-DEL-52 | chr3-151779772-DEL-56 |
| chr3-151788950-DEL-80 | chr3-151788951-DEL-80 |
| chr3-151802478-DEL-131 | chr3-151802479-DEL-129 |
| chr3-151814030-DEL-150 | chr3-151814031-DEL-150 |
| chr3-151834914-DEL-55 | chr3-151834901-DEL-104 |
| chr3-151850059-DEL-123 | chr3-151850044-DEL-124 |
| chr3-151880019-DEL-54 | chr3-151880020-DEL-54 |
| chr3-151887164-DEL-238 | chr3-151887165-DEL-238 |
| chr3-151922050-DEL-119 | chr3-151922051-DEL-119 |
| chr3-151932501-DEL-199 | chr3-151932502-DEL-199 |
| chr3-151960359-DEL-199 | chr3-151960360-DEL-199 |
| chr3-151965006-DEL-189 | chr3-151965007-DEL-189 |
| chr3-151991341-DEL-218 | chr3-151991342-DEL-218 |
| chr3-152028772-DEL-9047 | chr3-152028773-DEL-9047 |
| chr3-152042391-DEL-159 | chr3-152042392-DEL-159 |
| chr3-152044101-DEL-207 | chr3-152044085-DEL-212 |
| chr3-152044856-DEL-230 | chr3-152044857-DEL-230 |
| chr3-152049480-DEL-415 | chr3-152049473-DEL-422 |
| chr3-152060841-DEL-161 | chr3-152060842-DEL-158 |
| chr3-152066824-DEL-148 | chr3-152066825-DEL-148 |
| chr3-152144994-DEL-82 | chr3-152144995-DEL-82 |
| chr3-152148255-DEL-66 | chr3-152148255-DEL-66 |
| chr3-152159161-DEL-62 | chr3-152159162-DEL-60 |
| chr3-152186762-DEL-60 | chr3-152186763-DEL-60 |
| chr3-152188825-DEL-383 | chr3-152188817-DEL-347 |
| chr3-152216774-DEL-177 | chr3-152216805-DEL-145 |
| chr3-152217744-DEL-58 | chr3-152217745-DEL-58 |
| chr3-152241416-DEL-6442 | chr3-152241417-DEL-6442 |
| chr3-152251432-DEL-206 | chr3-152251433-DEL-206 |
| chr3-152254702-DEL-352 | chr3-152254703-DEL-352 |
| chr3-152276132-DEL-212 | chr3-152276132-DEL-212 |
| chr3-152289323-DEL-115 | chr3-152289306-DEL-138 |
| chr3-152313461-DEL-98 | chr3-152313462-DEL-98 |
| chr3-152323014-DEL-131 | chr3-152323015-DEL-131 |
| chr3-152331140-DEL-67 | chr3-152331141-DEL-67 |
| chr3-152338243-DEL-86 | chr3-152338228-DEL-102 |
| chr3-152355190-DEL-94 | chr3-152355191-DEL-94 |
| chr3-152374909-DEL-197 | chr3-152374910-DEL-197 |
| chr3-152391035-DEL-70 | chr3-152391036-DEL-69 |
| chr3-152393105-DEL-436 | chr3-152393106-DEL-436 |
| chr3-152415972-DEL-1267 | chr3-152415973-DEL-1267 |
| chr3-152420432-DEL-67 | chr3-152420433-DEL-67 |
| chr3-152431450-DEL-253 | chr3-152431451-DEL-253 |
| chr3-152436830-DEL-697 | chr3-152436826-DEL-698 |
| chr3-152439013-DEL-348 | chr3-152439012-DEL-332 |
| chr3-152454252-DEL-126 | chr3-152454253-DEL-126 |
| chr3-152465133-DEL-80 | chr3-152465134-DEL-80 |
| chr3-152466517-DEL-67 | chr3-152466518-DEL-67 |
| chr3-152471366-DEL-62 | chr3-152471367-DEL-62 |
| chr3-152473228-DEL-59 | chr3-152473229-DEL-59 |
| chr3-152491866-DEL-240 | chr3-152491867-DEL-240 |
| chr3-152507102-DEL-7417 | chr3-152507103-DEL-7417 |
| chr3-152514824-DEL-344 | chr3-152514817-DEL-360 |
| chr3-152524987-DEL-351 | chr3-152524988-DEL-351 |
| chr3-152534995-DEL-1284 | chr3-152534996-DEL-1284 |
| chr3-152600294-DEL-2858 | chr3-152600295-DEL-2858 |
| chr3-152608038-DEL-150 | chr3-152608039-DEL-150 |
| chr3-152644474-DEL-126 | chr3-152644524-DEL-80 |
| chr3-152658844-DEL-72 | chr3-152658845-DEL-72 |
| chr3-152664627-DEL-456 | chr3-152664628-DEL-456 |

| | |
|---|---|
| chr3-152670666-DEL-140 | chr3-152670667-DEL-140 |
| chr3-152684518-DEL-59 | chr3-152684506-DEL-60 |
| chr3-152690855-DEL-320 | chr3-152690856-DEL-320 |
| chr3-152693693-DEL-1320 | chr3-152693694-DEL-1320 |
| chr3-152701122-DEL-80 | chr3-152701123-DEL-80 |
| chr3-152726467-DEL-57 | chr3-152726468-DEL-57 |
| chr3-152730111-DEL-697 | chr3-152730111-DEL-697 |
| chr3-152749459-DEL-92 | chr3-152749455-DEL-58 |
| chr3-152751413-DEL-82 | chr3-152751414-DEL-82 |
| chr3-152770017-DEL-52 | chr3-152770012-DEL-58 |
| chr3-152771073-DEL-68 | chr3-152771074-DEL-68 |
| chr3-152774553-DEL-118 | chr3-152774554-DEL-118 |
| chr3-152793038-DEL-239 | chr3-152793039-DEL-239 |
| chr3-152796269-DEL-3099 | chr3-152796270-DEL-3099 |
| chr3-152802071-DEL-480 | chr3-152802072-DEL-480 |
| chr3-152817062-DEL-135 | chr3-152817065-DEL-133 |
| chr3-152829438-DEL-95 | chr3-152829439-DEL-95 |
| chr3-152833081-DEL-104 | chr3-152833082-DEL-104 |
| chr3-152890132-DEL-74 | chr3-152890133-DEL-74 |
| chr3-152914289-DEL-92 | chr3-152914294-DEL-88 |
| chr3-152915155-DEL-807 | chr3-152915156-DEL-807 |
| chr3-152925473-DEL-215 | chr3-152925474-DEL-215 |
| chr3-152928113-DEL-123 | chr3-152928112-DEL-124 |
| chr3-152939741-DEL-82 | chr3-152939742-DEL-82 |
| chr3-152957729-DEL-144 | chr3-152957730-DEL-144 |
| chr3-152961258-DEL-338 | chr3-152961259-DEL-338 |
| chr3-153019372-DEL-59 | chr3-153019373-DEL-59 |
| chr3-153054802-DEL-67 | chr3-153054803-DEL-67 |
| chr3-153060093-DEL-57 | chr3-153060080-DEL-57 |
| chr3-153077450-DEL-519 | chr3-153077451-DEL-519 |
| chr3-153102766-DEL-464 | chr3-153102767-DEL-464 |
| chr3-153116112-DEL-260 | chr3-153116113-DEL-260 |
| chr3-153126383-DEL-65 | chr3-153126384-DEL-65 |
| chr3-153141234-DEL-76 | chr3-153141268-DEL-76 |
| chr3-153144407-DEL-1894 | chr3-153144407-DEL-1894 |
| chr3-153154063-DEL-156 | chr3-153154021-DEL-171 |
| chr3-153168199-DEL-145 | chr3-153168200-DEL-145 |
| chr3-153246324-DEL-157 | chr3-153246325-DEL-157 |
| chr3-153252953-DEL-54 | chr3-153252958-DEL-55 |
| chr3-153275294-DEL-573 | chr3-153275295-DEL-573 |
| chr3-153280552-DEL-67 | chr3-153280553-DEL-67 |
| chr3-153287221-DEL-74 | chr3-153287252-DEL-68 |
| chr3-153309955-DEL-584 | chr3-153309956-DEL-584 |
| chr3-153317336-DEL-242 | chr3-153317338-DEL-240 |
| chr3-153334250-DEL-70 | chr3-153334226-DEL-68 |
| chr3-153351067-DEL-60 | chr3-153351068-DEL-60 |
| chr3-153352488-DEL-64 | chr3-153352489-DEL-64 |
| chr3-153353444-DEL-59 | chr3-153353436-DEL-60 |
| chr3-153391329-DEL-168 | chr3-153391330-DEL-168 |
| chr3-153392566-DEL-1017 | chr3-153392528-DEL-1017 |
| chr3-153393509-DEL-53 | chr3-153393510-DEL-53 |
| chr3-153411978-DEL-3071 | chr3-153411977-DEL-3072 |
| chr3-153419383-DEL-697 | chr3-153419384-DEL-698 |
| chr3-153424475-DEL-928 | chr3-153424476-DEL-928 |
| chr3-153427753-DEL-934 | chr3-153427756-DEL-933 |
| chr3-153438959-DEL-281 | chr3-153438960-DEL-281 |
| chr3-153450607-DEL-58 | chr3-153450608-DEL-58 |
| chr3-153453117-DEL-725 | chr3-153453118-DEL-725 |
| chr3-153453711-DEL-62 | chr3-153453712-DEL-62 |
| chr3-153457867-DEL-113 | chr3-153457868-DEL-113 |
| chr3-153458125-DEL-398 | chr3-153458126-DEL-398 |
| chr3-153459177-DEL-94 | chr3-153459144-DEL-95 |
| chr3-153469992-DEL-146 | chr3-153469993-DEL-146 |
| chr3-153481018-DEL-498 | chr3-153481019-DEL-498 |
| chr3-153486261-DEL-51 | chr3-153486262-DEL-51 |
| chr3-153487015-DEL-1240 | chr3-153487016-DEL-1240 |
| chr3-153491021-DEL-56 | chr3-153490979-DEL-76 |
| chr3-153493717-DEL-842 | chr3-153493718-DEL-842 |
| chr3-153498907-DEL-53 | chr3-153498876-DEL-56 |
| chr3-153500163-DEL-298 | chr3-153500164-DEL-298 |
| chr3-153514513-DEL-229 | chr3-153514514-DEL-229 |
| chr3-153530358-DEL-60 | chr3-153530359-DEL-60 |
| chr3-153536244-DEL-406 | chr3-153536245-DEL-406 |
| chr3-153538298-DEL-450 | chr3-153538299-DEL-450 |
| chr3-153544813-DEL-233 | chr3-153544814-DEL-233 |
| chr3-153548769-DEL-479 | chr3-153548774-DEL-475 |
| chr3-153557370-DEL-164 | chr3-153557371-DEL-164 |
| chr3-153564612-DEL-230 | chr3-153564613-DEL-230 |
| chr3-153589922-DEL-204 | chr3-153589923-DEL-204 |
| chr3-153591796-DEL-226 | chr3-153591797-DEL-226 |

| | |
|---|---|
| chr3-153595234-DEL-208 | chr3-153595235-DEL-208 |
| chr3-153636204-DEL-210 | chr3-153636205-DEL-210 |
| chr3-153686686-DEL-1341 | chr3-153686687-DEL-1341 |
| chr3-153694739-DEL-189 | chr3-153694740-DEL-216 |
| chr3-153702386-DEL-118 | chr3-153702387-DEL-118 |
| chr3-153709127-DEL-1341 | chr3-153709128-DEL-1341 |
| chr3-153721723-DEL-106 | chr3-153721728-DEL-102 |
| chr3-153726835-DEL-804 | chr3-153726836-DEL-804 |
| chr3-153729077-DEL-6802 | chr3-153729078-DEL-6802 |
| chr3-153749330-DEL-114 | chr3-153749331-DEL-112 |
| chr3-153767331-DEL-291 | chr3-153767330-DEL-290 |
| chr3-153777539-DEL-613 | chr3-153777540-DEL-614 |
| chr3-153782287-DEL-57 | chr3-153782288-DEL-57 |
| chr3-153789927-DEL-113 | chr3-153789937-DEL-109 |
| chr3-153806026-DEL-152 | chr3-153806027-DEL-149 |
| chr3-153826165-DEL-348 | chr3-153826166-DEL-348 |
| chr3-153828967-DEL-97 | chr3-153829004-DEL-50 |
| chr3-153848799-DEL-53 | chr3-153848800-DEL-53 |
| chr3-153876928-DEL-192 | chr3-153876929-DEL-192 |
| chr3-153881579-DEL-512 | chr3-153881580-DEL-512 |
| chr3-153884689-DEL-263 | chr3-153884693-DEL-247 |
| chr3-153885270-DEL-225 | chr3-153885268-DEL-233 |
| chr3-153917265-DEL-544 | chr3-153917266-DEL-544 |
| chr3-153921746-DEL-210 | chr3-153921759-DEL-193 |
| chr3-153923160-DEL-58 | chr3-153923160-DEL-58 |
| chr3-153925198-DEL-57 | chr3-153925199-DEL-57 |
| chr3-153926172-DEL-147 | chr3-153926173-DEL-147 |
| chr3-153935217-DEL-54 | chr3-153935190-DEL-58 |
| chr3-153959469-DEL-92 | chr3-153959467-DEL-56 |
| chr3-153999160-DEL-68 | chr3-153999168-DEL-63 |
| chr3-154015100-DEL-112 | chr3-154015073-DEL-122 |
| chr3-154019398-DEL-64 | chr3-154019399-DEL-76 |
| chr3-154042553-DEL-69 | chr3-154042559-DEL-52 |
| chr3-154048242-DEL-287 | chr3-154048227-DEL-288 |
| chr3-154052135-DEL-1215 | chr3-154052136-DEL-1215 |
| chr3-154061354-DEL-127 | chr3-154061355-DEL-127 |
| chr3-154076415-DEL-450 | chr3-154076416-DEL-450 |
| chr3-154098656-DEL-56 | chr3-154098657-DEL-56 |
| chr3-154128820-DEL-524 | chr3-154128818-DEL-525 |
| chr3-154152646-DEL-66 | chr3-154152647-DEL-66 |
| chr3-154159037-DEL-107 | chr3-154159035-DEL-112 |
| chr3-154192411-DEL-106 | chr3-154192412-DEL-106 |
| chr3-154202216-DEL-6413 | chr3-154202217-DEL-6413 |
| chr3-154225329-DEL-234 | chr3-154225330-DEL-234 |
| chr3-154232074-DEL-81 | chr3-154232075-DEL-81 |
| chr3-154244370-DEL-52 | chr3-154244367-DEL-53 |
| chr3-154287371-DEL-81 | chr3-154287372-DEL-81 |
| chr3-154289117-DEL-238 | chr3-154289113-DEL-239 |
| chr3-154328549-DEL-52 | chr3-154328550-DEL-52 |
| chr3-154336587-DEL-881 | chr3-154336588-DEL-881 |
| chr3-154343765-DEL-59 | chr3-154343766-DEL-59 |
| chr3-154345178-DEL-7170 | chr3-154345198-DEL-7143 |
| chr3-154367746-DEL-59 | chr3-154367730-DEL-87 |
| chr3-154385829-DEL-76 | chr3-154385832-DEL-78 |
| chr3-154403387-DEL-76 | chr3-154403389-DEL-60 |
| chr3-154435564-DEL-53 | chr3-154435565-DEL-53 |
| chr3-154468053-DEL-69 | chr3-154468054-DEL-69 |
| chr3-154560609-DEL-230 | chr3-154560614-DEL-226 |
| chr3-154584234-DEL-56 | chr3-154584235-DEL-56 |
| chr3-154594022-DEL-110 | chr3-154594023-DEL-110 |
| chr3-154597830-DEL-56 | chr3-154597833-DEL-54 |
| chr3-154615287-DEL-88 | chr3-154615288-DEL-88 |
| chr3-154624767-DEL-456 | chr3-154624768-DEL-456 |
| chr3-154651125-DEL-137 | chr3-154651126-DEL-137 |
| chr3-154693504-DEL-920 | chr3-154693505-DEL-920 |
| chr3-154710304-DEL-138 | chr3-154710305-DEL-138 |
| chr3-154734950-DEL-56 | chr3-154734950-DEL-56 |
| chr3-154737259-DEL-83 | chr3-154737260-DEL-83 |
| chr3-154786203-DEL-1812 | chr3-154786200-DEL-1813 |
| chr3-154786902-DEL-58 | chr3-154786903-DEL-58 |
| chr3-154787374-DEL-1187 | chr3-154787375-DEL-1187 |
| chr3-154825241-DEL-52 | chr3-154825225-DEL-59 |
| chr3-154921128-DEL-4676 | chr3-154921126-DEL-4677 |
| chr3-154957725-DEL-202 | chr3-154957695-DEL-203 |
| chr3-154971729-DEL-63 | chr3-154971702-DEL-90 |
| chr3-154980083-DEL-330 | chr3-154980076-DEL-330 |
| chr3-154991601-DEL-68 | chr3-154991598-DEL-84 |
| chr3-155000265-DEL-87 | chr3-155000226-DEL-88 |
| chr3-155006826-DEL-83 | chr3-155006827-DEL-83 |
| chr3-155009300-DEL-6615 | chr3-155009301-DEL-6615 |

| | |
|---|---|
| chr3-155040033-DEL-1894 | chr3-155040034-DEL-1894 |
| chr3-155044362-DEL-78 | chr3-155044363-DEL-86 |
| chr3-155052893-DEL-666 | chr3-155052894-DEL-666 |
| chr3-155060795-DEL-99 | chr3-155060796-DEL-99 |
| chr3-155064584-DEL-73 | chr3-155064565-DEL-74 |
| chr3-155157503-DEL-1039 | chr3-155157497-DEL-1040 |
| chr3-155219723-DEL-105 | chr3-155219724-DEL-105 |
| chr3-155221302-DEL-2249 | chr3-155221303-DEL-2249 |
| chr3-155290386-DEL-1425 | chr3-155290387-DEL-1425 |
| chr3-155303024-DEL-63 | chr3-155303004-DEL-64 |
| chr3-155319258-DEL-123 | chr3-155319267-DEL-115 |
| chr3-155341511-DEL-1510 | chr3-155341512-DEL-1510 |
| chr3-155358963-DEL-118 | chr3-155358964-DEL-118 |
| chr3-155387001-DEL-70 | chr3-155387002-DEL-70 |
| chr3-155398605-DEL-1310 | chr3-155398606-DEL-1310 |
| chr3-155425053-DEL-56 | chr3-155425054-DEL-56 |
| chr3-155440421-DEL-75 | chr3-155440422-DEL-74 |
| chr3-155451254-DEL-536 | chr3-155451255-DEL-536 |
| chr3-155471402-DEL-6733 | chr3-155471403-DEL-6733 |
| chr3-155477694-DEL-205 | chr3-155477695-DEL-205 |
| chr3-155496616-DEL-985 | chr3-155496617-DEL-985 |
| chr3-155512731-DEL-1656 | chr3-155512732-DEL-1656 |
| chr3-155530794-DEL-267 | chr3-155530795-DEL-267 |
| chr3-155547221-DEL-2491 | chr3-155547223-DEL-2490 |
| chr3-155567247-DEL-803 | chr3-155567248-DEL-803 |
| chr3-155568598-DEL-58 | chr3-155568599-DEL-84 |
| chr3-155573368-DEL-80 | chr3-155573369-DEL-76 |
| chr3-155618847-DEL-69 | chr3-155618848-DEL-69 |
| chr3-155625658-DEL-74 | chr3-155625664-DEL-66 |
| chr3-155634854-DEL-54 | chr3-155634855-DEL-54 |
| chr3-155655606-DEL-57 | chr3-155655607-DEL-57 |
| chr3-155720641-DEL-1887 | chr3-155720642-DEL-1887 |
| chr3-155753388-DEL-389 | chr3-155753392-DEL-386 |
| chr3-155802498-DEL-95 | chr3-155802499-DEL-95 |
| chr3-155828017-DEL-87 | chr3-155828018-DEL-86 |
| chr3-155833032-DEL-64 | chr3-155833033-DEL-64 |
| chr3-155842493-DEL-52 | chr3-155842494-DEL-52 |
| chr3-155883617-DEL-4581 | chr3-155883618-DEL-4581 |
| chr3-155892431-DEL-109 | chr3-155892468-DEL-73 |
| chr3-155908035-DEL-51 | chr3-155908036-DEL-51 |
| chr3-155940983-DEL-100 | chr3-155941007-DEL-124 |
| chr3-155990385-DEL-6292 | chr3-155990372-DEL-6293 |
| chr3-156003283-DEL-76 | chr3-156003284-DEL-88 |
| chr3-156015198-DEL-3309 | chr3-156015199-DEL-3309 |
| chr3-156022049-DEL-6250 | chr3-156022050-DEL-6250 |
| chr3-156063993-DEL-204 | chr3-156063994-DEL-204 |
| chr3-156097996-DEL-881 | chr3-156097997-DEL-881 |
| chr3-156104315-DEL-223 | chr3-156104316-DEL-223 |
| chr3-156132576-DEL-89 | chr3-156132577-DEL-89 |
| chr3-156140293-DEL-120 | chr3-156140293-DEL-120 |
| chr3-156143621-DEL-6431 | chr3-156143622-DEL-6431 |
| chr3-156163033-DEL-51 | chr3-156163034-DEL-51 |
| chr3-156168445-DEL-340 | chr3-156168446-DEL-340 |
| chr3-156195037-DEL-492 | chr3-156195038-DEL-492 |
| chr3-156206886-DEL-356 | chr3-156206887-DEL-356 |
| chr3-156242655-DEL-103 | chr3-156242654-DEL-103 |
| chr3-156256975-DEL-66 | chr3-156257000-DEL-84 |
| chr3-156324632-DEL-140 | chr3-156324633-DEL-140 |
| chr3-156327074-DEL-109 | chr3-156327063-DEL-110 |
| chr3-156355572-DEL-410 | chr3-156355569-DEL-411 |
| chr3-156364443-DEL-72 | chr3-156364444-DEL-72 |
| chr3-156411487-DEL-909 | chr3-156411488-DEL-909 |
| chr3-156425408-DEL-68 | chr3-156425409-DEL-68 |
| chr3-156430584-DEL-6866 | chr3-156430585-DEL-6866 |
| chr3-156440816-DEL-52 | chr3-156440817-DEL-52 |
| chr3-156468432-DEL-450 | chr3-156468433-DEL-450 |
| chr3-156472745-DEL-73 | chr3-156472771-DEL-64 |
| chr3-156505173-DEL-5225 | chr3-156505174-DEL-5225 |
| chr3-156546724-DEL-221 | chr3-156546725-DEL-221 |
| chr3-156598163-DEL-53 | chr3-156598164-DEL-53 |
| chr3-156627729-DEL-221 | chr3-156627731-DEL-220 |
| chr3-156649159-DEL-2154 | chr3-156649160-DEL-2154 |
| chr3-156666556-DEL-6429 | chr3-156666557-DEL-6429 |
| chr3-156672574-DEL-71 | chr3-156672576-DEL-70 |
| chr3-156681186-DEL-185 | chr3-156681187-DEL-185 |
| chr3-156694498-DEL-208 | chr3-156694520-DEL-183 |
| chr3-156712368-DEL-6385 | chr3-156712369-DEL-6385 |
| chr3-156719033-DEL-142 | chr3-156719070-DEL-138 |
| chr3-156740059-DEL-58 | chr3-156740060-DEL-58 |
| chr3-156793710-DEL-63 | chr3-156793711-DEL-63 |

| | |
|---|---|
| chr3-156819284-DEL-52 | chr3-156819285-DEL-52 |
| chr3-156822839-DEL-1754 | chr3-156822840-DEL-1754 |
| chr3-156826901-DEL-140 | chr3-156826887-DEL-136 |
| chr3-156906786-DEL-63 | chr3-156906785-DEL-64 |
| chr3-156925868-DEL-426 | chr3-156925869-DEL-426 |
| chr3-157003858-DEL-187 | chr3-157003859-DEL-187 |
| chr3-157040883-DEL-368 | chr3-157040884-DEL-368 |
| chr3-157050251-DEL-1297 | chr3-157050252-DEL-1297 |
| chr3-157053513-DEL-113 | chr3-157053511-DEL-104 |
| chr3-157055999-DEL-316 | chr3-157056000-DEL-316 |
| chr3-157074429-DEL-63 | chr3-157074404-DEL-57 |
| chr3-157084896-DEL-71 | chr3-157084897-DEL-71 |
| chr3-157088012-DEL-590 | chr3-157088013-DEL-590 |
| chr3-157116730-DEL-11492 | chr3-157116730-DEL-11491 |
| chr3-157122008-DEL-5285 | chr3-157122009-DEL-5285 |
| chr3-157151165-DEL-140 | chr3-157151164-DEL-140 |
| chr3-157223059-DEL-82 | chr3-157223058-DEL-116 |
| chr3-157244030-DEL-62 | chr3-157244031-DEL-62 |
| chr3-157325653-DEL-52 | chr3-157325654-DEL-52 |
| chr3-157340774-DEL-237 | chr3-157340775-DEL-234 |
| chr3-157343589-DEL-111 | chr3-157343590-DEL-111 |
| chr3-157352400-DEL-6196 | chr3-157352401-DEL-6196 |
| chr3-157364559-DEL-59 | chr3-157364568-DEL-74 |
| chr3-157373475-DEL-122 | chr3-157373476-DEL-127 |
| chr3-157388023-DEL-498 | chr3-157388024-DEL-498 |
| chr3-157393329-DEL-70 | chr3-157393330-DEL-70 |
| chr3-157396325-DEL-514 | chr3-157396326-DEL-514 |
| chr3-157402403-DEL-106 | chr3-157402404-DEL-106 |
| chr3-157408006-DEL-1994 | chr3-157408007-DEL-1994 |
| chr3-157420260-DEL-67 | chr3-157420265-DEL-68 |
| chr3-157422258-DEL-191 | chr3-157422259-DEL-191 |
| chr3-157430098-DEL-52 | chr3-157430099-DEL-52 |
| chr3-157430667-DEL-203 | chr3-157430668-DEL-203 |
| chr3-157431846-DEL-498 | chr3-157431847-DEL-498 |
| chr3-157483218-DEL-92 | chr3-157483223-DEL-88 |
| chr3-157497753-DEL-265 | chr3-157497749-DEL-247 |
| chr3-157499927-DEL-66 | chr3-157499928-DEL-56 |
| chr3-157505760-DEL-123 | chr3-157505770-DEL-104 |
| chr3-157536201-DEL-497 | chr3-157536202-DEL-497 |
| chr3-157548873-DEL-82 | chr3-157548874-DEL-82 |
| chr3-157563278-DEL-2040 | chr3-157563279-DEL-2040 |
| chr3-157596184-DEL-174 | chr3-157596185-DEL-174 |
| chr3-157596747-DEL-54 | chr3-157596748-DEL-54 |
| chr3-157599054-DEL-128 | chr3-157599055-DEL-128 |
| chr3-157617278-DEL-174 | chr3-157617307-DEL-160 |
| chr3-157619642-DEL-1120 | chr3-157619643-DEL-1119 |
| chr3-157623408-DEL-134 | chr3-157623409-DEL-134 |
| chr3-157631159-DEL-225 | chr3-157631160-DEL-225 |
| chr3-157634413-DEL-51 | chr3-157634414-DEL-51 |
| chr3-157642037-DEL-143 | chr3-157642038-DEL-143 |
| chr3-157684559-DEL-94 | chr3-157684560-DEL-94 |
| chr3-157806925-DEL-212 | chr3-157806926-DEL-212 |
| chr3-157887913-DEL-130 | chr3-157887916-DEL-128 |
| chr3-157901630-DEL-430 | chr3-157901631-DEL-430 |
| chr3-157928587-DEL-60 | chr3-157928588-DEL-60 |
| chr3-157938203-DEL-98 | chr3-157938204-DEL-98 |
| chr3-157941194-DEL-449 | chr3-157941195-DEL-449 |
| chr3-157955539-DEL-54 | chr3-157955540-DEL-58 |
| chr3-157969614-DEL-121 | chr3-157969658-DEL-78 |
| chr3-157979204-DEL-53 | chr3-157979205-DEL-53 |
| chr3-158017026-DEL-163 | chr3-158017027-DEL-163 |
| chr3-158055181-DEL-279 | chr3-158055182-DEL-280 |
| chr3-158083390-DEL-59 | chr3-158083391-DEL-59 |
| chr3-158147371-DEL-6485 | chr3-158147372-DEL-6485 |
| chr3-158185702-DEL-60 | chr3-158185703-DEL-60 |
| chr3-158188057-DEL-221 | chr3-158188058-DEL-221 |
| chr3-158333543-DEL-7200 | chr3-158333544-DEL-7200 |
| chr3-158376946-DEL-62 | chr3-158376947-DEL-62 |
| chr3-158398650-DEL-387 | chr3-158398651-DEL-387 |
| chr3-158402914-DEL-6986 | chr3-158402915-DEL-6986 |
| chr3-158414675-DEL-1447 | chr3-158414676-DEL-1447 |
| chr3-158426159-DEL-1476 | chr3-158426160-DEL-1476 |
| chr3-158435690-DEL-9592 | chr3-158435691-DEL-9592 |
| chr3-158442816-DEL-208 | chr3-158442817-DEL-208 |
| chr3-158447904-DEL-206 | chr3-158447899-DEL-207 |
| chr3-158452185-DEL-472 | chr3-158452186-DEL-472 |
| chr3-158467512-DEL-930 | chr3-158467510-DEL-924 |
| chr3-158495118-DEL-89 | chr3-158495119-DEL-89 |
| chr3-158511710-DEL-764 | chr3-158511711-DEL-764 |
| chr3-158518630-DEL-215 | chr3-158518631-DEL-215 |

| | |
|---|---|
| chr3-158534837-DEL-4666 | chr3-158534844-DEL-4660 |
| chr3-158553761-DEL-719 | chr3-158553762-DEL-719 |
| chr3-158580989-DEL-5478 | chr3-158580990-DEL-5478 |
| chr3-158606528-DEL-1727 | chr3-158606529-DEL-1727 |
| chr3-158665123-DEL-52 | chr3-158665112-DEL-64 |
| chr3-158674790-DEL-117 | chr3-158674791-DEL-117 |
| chr3-158683582-DEL-1662 | chr3-158683582-DEL-1662 |
| chr3-158693491-DEL-6464 | chr3-158693508-DEL-6464 |
| chr3-158717807-DEL-677 | chr3-158717808-DEL-677 |
| chr3-158725928-DEL-55 | chr3-158725924-DEL-60 |
| chr3-158742565-DEL-490 | chr3-158742566-DEL-490 |
| chr3-158780254-DEL-134 | chr3-158780255-DEL-134 |
| chr3-158815235-DEL-394 | chr3-158815236-DEL-394 |
| chr3-158836383-DEL-714 | chr3-158836384-DEL-714 |
| chr3-158877782-DEL-201 | chr3-158877783-DEL-201 |
| chr3-158894335-DEL-61 | chr3-158894336-DEL-61 |
| chr3-158916227-DEL-498 | chr3-158916224-DEL-499 |
| chr3-158931536-DEL-5192 | chr3-158931537-DEL-5192 |
| chr3-158938511-DEL-6407 | chr3-158938512-DEL-6407 |
| chr3-159104399-DEL-273 | chr3-159104388-DEL-280 |
| chr3-159105960-DEL-61 | chr3-159105960-DEL-62 |
| chr3-159151799-DEL-216 | chr3-159151787-DEL-214 |
| chr3-159162821-DEL-709 | chr3-159162822-DEL-709 |
| chr3-159185781-DEL-53 | chr3-159185782-DEL-54 |
| chr3-159192174-DEL-246 | chr3-159192174-DEL-247 |
| chr3-159216225-DEL-511 | chr3-159216226-DEL-511 |
| chr3-159285468-DEL-1296 | chr3-159285469-DEL-1296 |
| chr3-159319527-DEL-94 | chr3-159319528-DEL-91 |
| chr3-159332020-DEL-57 | chr3-159332021-DEL-57 |
| chr3-159359867-DEL-6459 | chr3-159359870-DEL-6455 |
| chr3-159380393-DEL-210 | chr3-159380394-DEL-210 |
| chr3-159401031-DEL-3334 | chr3-159401032-DEL-3334 |
| chr3-159406349-DEL-2688 | chr3-159406350-DEL-2688 |
| chr3-159420005-DEL-8322 | chr3-159420005-DEL-8322 |
| chr3-159472700-DEL-7324 | chr3-159472701-DEL-7324 |
| chr3-159515017-DEL-1591 | chr3-159515018-DEL-1591 |
| chr3-159611174-DEL-55 | chr3-159611176-DEL-53 |
| chr4-3056582-DEL-1886 | chr4-3056583-DEL-1886 |
| chr4-3066347-DEL-72 | chr4-3066347-DEL-72 |
| chr4-3066961-DEL-84 | chr4-3067011-DEL-84 |
| chr4-3094207-DEL-542 | chr4-3094208-DEL-542 |
| chr4-3181655-DEL-308 | chr4-3181656-DEL-308 |
| chr4-3192204-DEL-8325 | chr4-3192205-DEL-8325 |
| chr4-3236871-DEL-69 | chr4-3236890-DEL-86 |
| chr4-3312199-DEL-5573 | chr4-3312198-DEL-5573 |
| chr4-3352580-DEL-204 | chr4-3352581-DEL-204 |
| chr4-3370018-DEL-210 | chr4-3370019-DEL-210 |
| chr4-3373980-DEL-6401 | chr4-3373981-DEL-6401 |
| chr4-3412166-DEL-73 | chr4-3412179-DEL-61 |
| chr4-3431640-DEL-60 | chr4-3431636-DEL-62 |
| chr4-3436648-DEL-226 | chr4-3436646-DEL-224 |
| chr4-3443390-DEL-6209 | chr4-3443391-DEL-6209 |
| chr4-3457742-DEL-54 | chr4-3457743-DEL-54 |
| chr4-3463337-DEL-1650 | chr4-3463337-DEL-1647 |
| chr4-3471048-DEL-54 | chr4-3471049-DEL-54 |
| chr4-3479383-DEL-407 | chr4-3479384-DEL-407 |
| chr4-3510568-DEL-203 | chr4-3510569-DEL-203 |
| chr4-3554000-DEL-54 | chr4-3554022-DEL-50 |
| chr4-3672675-DEL-83 | chr4-3672661-DEL-98 |
| chr4-3763680-DEL-80 | chr4-3763680-DEL-80 |
| chr4-3781774-DEL-381 | chr4-3781775-DEL-381 |
| chr4-3808818-DEL-67 | chr4-3808800-DEL-80 |
| chr4-3819594-DEL-61 | chr4-3819594-DEL-58 |
| chr4-3822170-DEL-139 | chr4-3822171-DEL-138 |
| chr4-3847781-DEL-53 | chr4-3847782-DEL-53 |
| chr4-3853384-DEL-684 | chr4-3853385-DEL-682 |
| chr4-3910792-DEL-176 | chr4-3910793-DEL-176 |
| chr4-3919807-DEL-63 | chr4-3919808-DEL-63 |
| chr4-3973081-DEL-514 | chr4-3973082-DEL-514 |
| chr4-3994608-DEL-2509 | chr4-3994609-DEL-2509 |
| chr4-4006738-DEL-52 | chr4-4006739-DEL-52 |
| chr4-4090553-DEL-251 | chr4-4090554-DEL-251 |
| chr4-4094294-DEL-8300 | chr4-4094295-DEL-8300 |
| chr4-4171384-DEL-1512 | chr4-4171385-DEL-1512 |
| chr4-4208219-DEL-52 | chr4-4208220-DEL-52 |
| chr4-4220405-DEL-54 | chr4-4220406-DEL-50 |
| chr4-4236955-DEL-5496 | chr4-4236956-DEL-5496 |
| chr4-4265609-DEL-1283 | chr4-4265610-DEL-1283 |
| chr4-4268743-DEL-6534 | chr4-4268744-DEL-6534 |
| chr4-4334532-DEL-384 | chr4-4334533-DEL-384 |

| | |
|---|---|
| chr4-4340342-DEL-4099 | chr4-4340336-DEL-4104 |
| chr4-4360763-DEL-173 | chr4-4360764-DEL-173 |
| chr4-4382428-DEL-611 | chr4-4382429-DEL-611 |
| chr4-4390221-DEL-1789 | chr4-4390222-DEL-1789 |
| chr4-4409031-DEL-160 | chr4-4409032-DEL-160 |
| chr4-4432726-DEL-82 | chr4-4432726-DEL-82 |
| chr4-4442437-DEL-66 | chr4-4442438-DEL-66 |
| chr4-4537606-DEL-122 | chr4-4537607-DEL-122 |
| chr4-4541889-DEL-57 | chr4-4541890-DEL-57 |
| chr4-4549352-DEL-811 | chr4-4549353-DEL-811 |
| chr4-4553228-DEL-515 | chr4-4553229-DEL-515 |
| chr4-4608595-DEL-1298 | chr4-4608596-DEL-1298 |
| chr4-4616551-DEL-191 | chr4-4616552-DEL-191 |
| chr4-4628693-DEL-57 | chr4-4628693-DEL-58 |
| chr4-4630569-DEL-197 | chr4-4630570-DEL-197 |
| chr4-4652114-DEL-7163 | chr4-4652115-DEL-7163 |
| chr4-4665794-DEL-69 | chr4-4665794-DEL-69 |
| chr4-4682941-DEL-131 | chr4-4682942-DEL-131 |
| chr4-4694200-DEL-548 | chr4-4694201-DEL-548 |
| chr4-4723166-DEL-69 | chr4-4723167-DEL-69 |
| chr4-4724608-DEL-58 | chr4-4724609-DEL-58 |
| chr4-4728440-DEL-974 | chr4-4728441-DEL-974 |
| chr4-4737326-DEL-610 | chr4-4737327-DEL-610 |
| chr4-4819118-DEL-311 | chr4-4819119-DEL-311 |
| chr4-4824474-DEL-1456 | chr4-4824475-DEL-1456 |
| chr4-4846518-DEL-183 | chr4-4846519-DEL-183 |
| chr4-4848328-DEL-249 | chr4-4848329-DEL-249 |
| chr4-4850441-DEL-323 | chr4-4850442-DEL-323 |
| chr4-4856313-DEL-3201 | chr4-4856314-DEL-3201 |
| chr4-4883507-DEL-7178 | chr4-4883508-DEL-7178 |
| chr4-4906846-DEL-71 | chr4-4906847-DEL-71 |
| chr4-4921133-DEL-51 | chr4-4921134-DEL-51 |
| chr4-4933344-DEL-6581 | chr4-4933345-DEL-6581 |
| chr4-4964308-DEL-147 | chr4-4964309-DEL-147 |
| chr4-4966655-DEL-220 | chr4-4966656-DEL-220 |
| chr4-4969635-DEL-245 | chr4-4969636-DEL-245 |
| chr4-4974497-DEL-2282 | chr4-4974498-DEL-2282 |
| chr4-4984058-DEL-112 | chr4-4984059-DEL-112 |
| chr4-4987192-DEL-52 | chr4-4987193-DEL-52 |
| chr4-5000567-DEL-6331 | chr4-5000549-DEL-6346 |
| chr4-5048218-DEL-239 | chr4-5048219-DEL-239 |
| chr4-5091942-DEL-1082 | chr4-5091943-DEL-1082 |
| chr4-5096886-DEL-214 | chr4-5096887-DEL-214 |
| chr4-5143191-DEL-4222 | chr4-5143192-DEL-4222 |
| chr4-5162570-DEL-170 | chr4-5162571-DEL-170 |
| chr4-5163890-DEL-386 | chr4-5163890-DEL-386 |
| chr4-5184633-DEL-7124 | chr4-5184634-DEL-7124 |
| chr4-5228835-DEL-460 | chr4-5228836-DEL-460 |
| chr4-5238358-DEL-82 | chr4-5238360-DEL-81 |
| chr4-5254513-DEL-216 | chr4-5254514-DEL-216 |
| chr4-5259466-DEL-4220 | chr4-5259467-DEL-4220 |
| chr4-5278944-DEL-114 | chr4-5278975-DEL-82 |
| chr4-5305665-DEL-6185 | chr4-5305666-DEL-6185 |
| chr4-5321718-DEL-542 | chr4-5321719-DEL-542 |
| chr4-5326884-DEL-58 | chr4-5326884-DEL-58 |
| chr4-5345816-DEL-389 | chr4-5345817-DEL-389 |
| chr4-5350508-DEL-5995 | chr4-5350509-DEL-5995 |
| chr4-5363178-DEL-180 | chr4-5363177-DEL-181 |
| chr4-5371723-DEL-231 | chr4-5371724-DEL-231 |
| chr4-5412850-DEL-6158 | chr4-5412851-DEL-6158 |
| chr4-5460810-DEL-4563 | chr4-5460811-DEL-4563 |
| chr4-5506833-DEL-73 | chr4-5506834-DEL-73 |
| chr4-5515367-DEL-67 | chr4-5515368-DEL-67 |
| chr4-5557240-DEL-759 | chr4-5557241-DEL-759 |
| chr4-5563972-DEL-110 | chr4-5563973-DEL-110 |
| chr4-5576804-DEL-6150 | chr4-5576805-DEL-6150 |
| chr4-5607771-DEL-2586 | chr4-5607772-DEL-2586 |
| chr4-5660928-DEL-95 | chr4-5660962-DEL-95 |
| chr4-5708644-DEL-6671 | chr4-5708645-DEL-6671 |
| chr4-5714669-DEL-212 | chr4-5714670-DEL-212 |
| chr4-5716450-DEL-107 | chr4-5716451-DEL-107 |
| chr4-5756458-DEL-6583 | chr4-5756459-DEL-6583 |
| chr4-5776604-DEL-184 | chr4-5776593-DEL-148 |
| chr4-5815261-DEL-327 | chr4-5815262-DEL-327 |
| chr4-5819396-DEL-6796 | chr4-5819397-DEL-6796 |
| chr4-5902585-DEL-1661 | chr4-5902586-DEL-1661 |
| chr4-5915793-DEL-2835 | chr4-5915794-DEL-2835 |
| chr4-5927567-DEL-214 | chr4-5927568-DEL-214 |
| chr4-5997885-DEL-211 | chr4-5997876-DEL-212 |
| chr4-6054492-DEL-510 | chr4-6054493-DEL-510 |

| | |
|---|---|
| chr4-6090450-DEL-2264 | chr4-6090451-DEL-2264 |
| chr4-6110408-DEL-1116 | chr4-6110409-DEL-1116 |
| chr4-6111956-DEL-206 | chr4-6111956-DEL-202 |
| chr4-6112307-DEL-698 | chr4-6112308-DEL-698 |
| chr4-6151277-DEL-1289 | chr4-6151324-DEL-1289 |
| chr4-6159022-DEL-92 | chr4-6159023-DEL-92 |
| chr4-6164743-DEL-71 | chr4-6164745-DEL-76 |
| chr4-6226075-DEL-199 | chr4-6226076-DEL-199 |
| chr4-6241957-DEL-144 | chr4-6241958-DEL-124 |
| chr4-6287025-DEL-716 | chr4-6287026-DEL-716 |
| chr4-6316629-DEL-146 | chr4-6316629-DEL-147 |
| chr4-6320153-DEL-860 | chr4-6320154-DEL-860 |
| chr4-6326351-DEL-340 | chr4-6326352-DEL-340 |
| chr4-6338672-DEL-83 | chr4-6338673-DEL-101 |
| chr4-6439499-DEL-94 | chr4-6439543-DEL-91 |
| chr4-6450564-DEL-67 | chr4-6450565-DEL-67 |
| chr4-6516872-DEL-198 | chr4-6516873-DEL-198 |
| chr4-6524919-DEL-190 | chr4-6524920-DEL-189 |
| chr4-6535204-DEL-924 | chr4-6535205-DEL-924 |
| chr4-6540810-DEL-58 | chr4-6540811-DEL-58 |
| chr4-6541025-DEL-482 | chr4-6541026-DEL-482 |
| chr4-6542315-DEL-861 | chr4-6542316-DEL-861 |
| chr4-6550997-DEL-1495 | chr4-6550998-DEL-1495 |
| chr4-6566674-DEL-458 | chr4-6566675-DEL-458 |
| chr4-6586823-DEL-2406 | chr4-6586824-DEL-2406 |
| chr4-6609384-DEL-199 | chr4-6609385-DEL-199 |
| chr4-6614862-DEL-194 | chr4-6614863-DEL-194 |
| chr4-6623258-DEL-443 | chr4-6623259-DEL-443 |
| chr4-6632737-DEL-395 | chr4-6632738-DEL-395 |
| chr4-6649173-DEL-6816 | chr4-6649174-DEL-6816 |
| chr4-6657508-DEL-70 | chr4-6657509-DEL-70 |
| chr4-6663393-DEL-103 | chr4-6663417-DEL-80 |
| chr4-6664739-DEL-1044 | chr4-6664739-DEL-1044 |
| chr4-6738492-DEL-94 | chr4-6738493-DEL-94 |
| chr4-6795036-DEL-327 | chr4-6795038-DEL-328 |
| chr4-6834934-DEL-136 | chr4-6834935-DEL-136 |
| chr4-6867561-DEL-399 | chr4-6867562-DEL-399 |
| chr4-6927977-DEL-278 | chr4-6927978-DEL-278 |
| chr4-7056054-DEL-60 | chr4-7056055-DEL-60 |
| chr4-7086888-DEL-1133 | chr4-7086889-DEL-1133 |
| chr4-7096749-DEL-5343 | chr4-7096750-DEL-5343 |
| chr4-7109903-DEL-51 | chr4-7109904-DEL-51 |
| chr4-7116732-DEL-1072 | chr4-7116733-DEL-1072 |
| chr4-7120163-DEL-3685 | chr4-7120162-DEL-3685 |
| chr4-7139069-DEL-84 | chr4-7139056-DEL-112 |
| chr4-7151154-DEL-459 | chr4-7151155-DEL-459 |
| chr4-7165353-DEL-4874 | chr4-7165354-DEL-4874 |
| chr4-7213227-DEL-53 | chr4-7213228-DEL-53 |
| chr4-7234514-DEL-97 | chr4-7234519-DEL-50 |
| chr4-7273144-DEL-6515 | chr4-7273145-DEL-6515 |
| chr4-7279361-DEL-201 | chr4-7279362-DEL-201 |
| chr4-7297677-DEL-5135 | chr4-7297678-DEL-5135 |
| chr4-7311536-DEL-333 | chr4-7311537-DEL-333 |
| chr4-7324843-DEL-58 | chr4-7324844-DEL-58 |
| chr4-7349171-DEL-6484 | chr4-7349169-DEL-6485 |
| chr4-7369842-DEL-139 | chr4-7369843-DEL-139 |
| chr4-7446636-DEL-68 | chr4-7446637-DEL-67 |
| chr4-7447428-DEL-126 | chr4-7447437-DEL-118 |
| chr4-7463075-DEL-1369 | chr4-7463076-DEL-1369 |
| chr4-7470857-DEL-2968 | chr4-7470843-DEL-2969 |
| chr4-7483687-DEL-429 | chr4-7483680-DEL-431 |
| chr4-7542334-DEL-62 | chr4-7542340-DEL-54 |
| chr4-7552242-DEL-2978 | chr4-7552243-DEL-2978 |
| chr4-7576419-DEL-759 | chr4-7576419-DEL-760 |
| chr4-7595686-DEL-86 | chr4-7595656-DEL-81 |
| chr4-7604853-DEL-955 | chr4-7604854-DEL-955 |
| chr4-7616692-DEL-244 | chr4-7616715-DEL-203 |
| chr4-7618467-DEL-7203 | chr4-7618468-DEL-7203 |
| chr4-7628316-DEL-408 | chr4-7628317-DEL-408 |
| chr4-7651984-DEL-409 | chr4-7651985-DEL-409 |
| chr4-7681016-DEL-453 | chr4-7681018-DEL-452 |
| chr4-7702641-DEL-4622 | chr4-7702641-DEL-4622 |
| chr4-7714645-DEL-131 | chr4-7714646-DEL-131 |
| chr4-7717321-DEL-71 | chr4-7717341-DEL-69 |
| chr4-7733843-DEL-213 | chr4-7733845-DEL-214 |
| chr4-7758542-DEL-938 | chr4-7758543-DEL-938 |
| chr4-7773219-DEL-84 | chr4-7773219-DEL-85 |
| chr4-7782816-DEL-117 | chr4-7782817-DEL-117 |
| chr4-7823535-DEL-540 | chr4-7823536-DEL-540 |
| chr4-7830486-DEL-51 | chr4-7830487-DEL-51 |

| | |
|---|---|
| chr4-7830666-DEL-7198 | chr4-7830667-DEL-7198 |
| chr4-7880481-DEL-5109 | chr4-7880482-DEL-5109 |
| chr4-7924800-DEL-1975 | chr4-7924801-DEL-1975 |
| chr4-7940900-DEL-135 | chr4-7940901-DEL-135 |
| chr4-7961022-DEL-158 | chr4-7961021-DEL-170 |
| chr4-7991237-DEL-68 | chr4-7991244-DEL-65 |
| chr4-8021754-DEL-204 | chr4-8021755-DEL-204 |
| chr4-8037528-DEL-325 | chr4-8037529-DEL-325 |
| chr4-8043897-DEL-1424 | chr4-8043898-DEL-1424 |
| chr4-8081732-DEL-1442 | chr4-8081733-DEL-1442 |
| chr4-8172659-DEL-781 | chr4-8172660-DEL-781 |
| chr4-8181100-DEL-202 | chr4-8181115-DEL-201 |
| chr4-8191732-DEL-6238 | chr4-8191733-DEL-6238 |
| chr4-8227743-DEL-497 | chr4-8227744-DEL-497 |
| chr4-8235341-DEL-72 | chr4-8235292-DEL-102 |
| chr4-8240376-DEL-1467 | chr4-8240392-DEL-1460 |
| chr4-8258996-DEL-1193 | chr4-8258997-DEL-1193 |
| chr4-8261164-DEL-1316 | chr4-8261165-DEL-1316 |
| chr4-8286193-DEL-79 | chr4-8286190-DEL-83 |
| chr4-8287416-DEL-299 | chr4-8287417-DEL-299 |
| chr4-8297738-DEL-8146 | chr4-8297739-DEL-8146 |
| chr4-8313409-DEL-3028 | chr4-8313410-DEL-3028 |
| chr4-8314299-DEL-75 | chr4-8314300-DEL-75 |
| chr4-8329722-DEL-180 | chr4-8329724-DEL-185 |
| chr4-8341214-DEL-68 | chr4-8341215-DEL-68 |
| chr4-8385607-DEL-13183 | chr4-8385608-DEL-13183 |
| chr4-8392442-DEL-5700 | chr4-8392442-DEL-5700 |
| chr4-8405508-DEL-2762 | chr4-8405509-DEL-2762 |
| chr4-8409919-DEL-1095 | chr4-8409924-DEL-1056 |
| chr4-8433893-DEL-401 | chr4-8433894-DEL-401 |
| chr4-8443028-DEL-6558 | chr4-8443029-DEL-6558 |
| chr4-8472528-DEL-400 | chr4-8472529-DEL-400 |
| chr4-8477047-DEL-460 | chr4-8477048-DEL-460 |
| chr4-8488242-DEL-6401 | chr4-8488243-DEL-6401 |
| chr4-8501046-DEL-954 | chr4-8501047-DEL-954 |
| chr4-8523859-DEL-1844 | chr4-8523859-DEL-1829 |
| chr4-8527471-DEL-596 | chr4-8527472-DEL-596 |
| chr4-8590572-DEL-156 | chr4-8590573-DEL-156 |
| chr4-8593323-DEL-97 | chr4-8593322-DEL-93 |
| chr4-8594570-DEL-869 | chr4-8594571-DEL-869 |
| chr4-8629829-DEL-168 | chr4-8629830-DEL-168 |
| chr4-8631947-DEL-119 | chr4-8631940-DEL-84 |
| chr4-8642129-DEL-58 | chr4-8642115-DEL-90 |
| chr4-8654339-DEL-6410 | chr4-8654340-DEL-6410 |
| chr4-8661546-DEL-396 | chr4-8661547-DEL-396 |
| chr4-8677187-DEL-438 | chr4-8677188-DEL-438 |
| chr4-8713109-DEL-53 | chr4-8713131-DEL-53 |
| chr4-8917938-DEL-120 | chr4-8917930-DEL-140 |
| chr4-8964944-DEL-57 | chr4-8964937-DEL-58 |
| chr4-8978068-DEL-126 | chr4-8978097-DEL-104 |
| chr4-8982983-DEL-52 | chr4-8982984-DEL-50 |
| chr4-9047092-DEL-60 | chr4-9047094-DEL-52 |
| chr4-9067668-DEL-52 | chr4-9067668-DEL-52 |
| chr4-9084495-DEL-146 | chr4-9084489-DEL-147 |
| chr4-9085519-DEL-107 | chr4-9085494-DEL-108 |
| chr4-9102482-DEL-177 | chr4-9102483-DEL-177 |
| chr4-9144663-DEL-78 | chr4-9144664-DEL-78 |
| chr4-9150429-DEL-6192 | chr4-9150430-DEL-6192 |
| chr4-9158705-DEL-343 | chr4-9158706-DEL-343 |
| chr4-9171239-DEL-68 | chr4-9171240-DEL-68 |
| chr4-9173578-DEL-56 | chr4-9173566-DEL-52 |
| chr4-9179516-DEL-836 | chr4-9179517-DEL-836 |
| chr4-9206981-DEL-51 | chr4-9206982-DEL-51 |
| chr4-9207793-DEL-78 | chr4-9207794-DEL-76 |
| chr4-9238337-DEL-122 | chr4-9238326-DEL-105 |
| chr4-9253644-DEL-481 | chr4-9253644-DEL-481 |
| chr4-9257387-DEL-331 | chr4-9257388-DEL-331 |
| chr4-9257961-DEL-194 | chr4-9257962-DEL-194 |
| chr4-9263289-DEL-54 | chr4-9263290-DEL-54 |
| chr4-9308843-DEL-151 | chr4-9308859-DEL-116 |
| chr4-9343390-DEL-66 | chr4-9343391-DEL-66 |
| chr4-9360521-DEL-199 | chr4-9360522-DEL-199 |
| chr4-9385276-DEL-127 | chr4-9385277-DEL-125 |
| chr4-9398588-DEL-51 | chr4-9398589-DEL-51 |
| chr4-9401176-DEL-442 | chr4-9401177-DEL-442 |
| chr4-9414684-DEL-273 | chr4-9414687-DEL-270 |
| chr4-9454506-DEL-100 | chr4-9454507-DEL-100 |
| chr4-9465499-DEL-265 | chr4-9465500-DEL-265 |
| chr4-9478375-DEL-68 | chr4-9478376-DEL-68 |
| chr4-9488606-DEL-159 | chr4-9488609-DEL-157 |

| | |
|---|---|
| chr4-9491262-DEL-758 | chr4-9491263-DEL-758 |
| chr4-9493808-DEL-6469 | chr4-9493809-DEL-6469 |
| chr4-9531604-DEL-112 | chr4-9531605-DEL-112 |
| chr4-9544489-DEL-55 | chr4-9544490-DEL-55 |
| chr4-9547253-DEL-65 | chr4-9547253-DEL-65 |
| chr4-9549901-DEL-94 | chr4-9549902-DEL-94 |
| chr4-9578761-DEL-89 | chr4-9578762-DEL-89 |
| chr4-9580028-DEL-94 | chr4-9580066-DEL-94 |
| chr4-9585449-DEL-7268 | chr4-9585450-DEL-7268 |
| chr4-9593892-DEL-511 | chr4-9593892-DEL-512 |
| chr4-9600184-DEL-1011 | chr4-9600185-DEL-1011 |
| chr4-9636392-DEL-59 | chr4-9636393-DEL-59 |
| chr4-9648440-DEL-93 | chr4-9648438-DEL-96 |
| chr4-9661532-DEL-399 | chr4-9661533-DEL-399 |
| chr4-9671862-DEL-196 | chr4-9671863-DEL-196 |
| chr4-9673239-DEL-281 | chr4-9673240-DEL-281 |
| chr4-9699562-DEL-99 | chr4-9699563-DEL-99 |
| chr4-9710998-DEL-109 | chr4-9711001-DEL-98 |
| chr4-9737238-DEL-54 | chr4-9737235-DEL-95 |
| chr4-9781091-DEL-109 | chr4-9781092-DEL-111 |
| chr4-9784126-DEL-87 | chr4-9784127-DEL-87 |
| chr4-9801179-DEL-1969 | chr4-9801180-DEL-1969 |
| chr4-9817506-DEL-191 | chr4-9817512-DEL-192 |
| chr4-9827972-DEL-570 | chr4-9827968-DEL-563 |
| chr4-9864455-DEL-322 | chr4-9864456-DEL-322 |
| chr4-9865290-DEL-793 | chr4-9865302-DEL-787 |
| chr4-9870681-DEL-1473 | chr4-9870682-DEL-1473 |
| chr4-9877751-DEL-225 | chr4-9877755-DEL-228 |
| chr4-9919172-DEL-350 | chr4-9919173-DEL-350 |
| chr4-9921873-DEL-6603 | chr4-9921874-DEL-6603 |
| chr4-9954861-DEL-131 | chr4-9954883-DEL-110 |
| chr4-9958019-DEL-56 | chr4-9958000-DEL-56 |
| chr4-9997486-DEL-479 | chr4-9997487-DEL-479 |
| chr4-10002723-DEL-6710 | chr4-10002724-DEL-6710 |
| chr4-10014317-DEL-100 | chr4-10014318-DEL-100 |
| chr4-10050179-DEL-54 | chr4-10050180-DEL-60 |
| chr4-10101754-DEL-9946 | chr4-10101755-DEL-9946 |
| chr4-10107430-DEL-2966 | chr4-10107431-DEL-2966 |
| chr4-10111224-DEL-208 | chr4-10111225-DEL-208 |
| chr4-10117071-DEL-69 | chr4-10117072-DEL-69 |
| chr4-10137064-DEL-999 | chr4-10137106-DEL-958 |
| chr4-10156557-DEL-800 | chr4-10156558-DEL-800 |
| chr4-10157872-DEL-68 | chr4-10157862-DEL-76 |
| chr4-10165303-DEL-192 | chr4-10165304-DEL-192 |
| chr4-10176598-DEL-800 | chr4-10176599-DEL-800 |
| chr4-10188447-DEL-203 | chr4-10188448-DEL-203 |
| chr4-10200032-DEL-54 | chr4-10200031-DEL-56 |
| chr4-10220288-DEL-1172 | chr4-10220289-DEL-1172 |
| chr4-10252718-DEL-81 | chr4-10252713-DEL-82 |
| chr4-10310405-DEL-559 | chr4-10310406-DEL-559 |
| chr4-10354235-DEL-325 | chr4-10354232-DEL-326 |
| chr4-10376754-DEL-2782 | chr4-10376755-DEL-2782 |
| chr4-10401513-DEL-202 | chr4-10401506-DEL-210 |
| chr4-10405759-DEL-228 | chr4-10405760-DEL-228 |
| chr4-10428872-DEL-5503 | chr4-10428873-DEL-5503 |
| chr4-10448741-DEL-5124 | chr4-10448741-DEL-5124 |
| chr4-10458232-DEL-917 | chr4-10458233-DEL-917 |
| chr4-10466355-DEL-1118 | chr4-10466356-DEL-1117 |
| chr4-10478139-DEL-64 | chr4-10478140-DEL-64 |
| chr4-10486890-DEL-2795 | chr4-10486891-DEL-2795 |
| chr4-10491976-DEL-6665 | chr4-10491977-DEL-6665 |
| chr4-10518741-DEL-67 | chr4-10518740-DEL-69 |
| chr4-10555721-DEL-75 | chr4-10555722-DEL-75 |
| chr4-10592186-DEL-145 | chr4-10592187-DEL-143 |
| chr4-10593661-DEL-177 | chr4-10593662-DEL-177 |
| chr4-10644698-DEL-1831 | chr4-10644698-DEL-1831 |
| chr4-10672462-DEL-6536 | chr4-10672456-DEL-6537 |
| chr4-10694304-DEL-54 | chr4-10694305-DEL-54 |
| chr4-10709154-DEL-207 | chr4-10709155-DEL-207 |
| chr4-10713795-DEL-828 | chr4-10713795-DEL-828 |
| chr4-10728713-DEL-90 | chr4-10728714-DEL-82 |
| chr4-10734560-DEL-6690 | chr4-10734561-DEL-6690 |
| chr4-10747671-DEL-486 | chr4-10747672-DEL-486 |
| chr4-10762587-DEL-189 | chr4-10762588-DEL-189 |
| chr4-10786432-DEL-295 | chr4-10786433-DEL-295 |
| chr4-10798553-DEL-54 | chr4-10798554-DEL-54 |
| chr4-10838243-DEL-527 | chr4-10838243-DEL-527 |
| chr4-10848378-DEL-728 | chr4-10848379-DEL-728 |
| chr4-10852166-DEL-200 | chr4-10852167-DEL-200 |
| chr4-10865326-DEL-6045 | chr4-10865327-DEL-6045 |

| | |
|---|---|
| chr4-10904183-DEL-66 | chr4-10904167-DEL-74 |
| chr4-10907306-DEL-560 | chr4-10907296-DEL-560 |
| chr4-10910539-DEL-240 | chr4-10910538-DEL-241 |
| chr4-10914300-DEL-226 | chr4-10914301-DEL-226 |
| chr4-10926969-DEL-71 | chr4-10926970-DEL-71 |
| chr4-10938491-DEL-86 | chr4-10938492-DEL-86 |
| chr4-10957200-DEL-86 | chr4-10957202-DEL-87 |
| chr4-10957864-DEL-1898 | chr4-10957865-DEL-1898 |
| chr4-10961832-DEL-240 | chr4-10961833-DEL-240 |
| chr4-10974147-DEL-58 | chr4-10974148-DEL-58 |
| chr4-10983757-DEL-337 | chr4-10983755-DEL-347 |
| chr4-11000930-DEL-58 | chr4-11000943-DEL-58 |
| chr4-11006020-DEL-155 | chr4-11006021-DEL-155 |
| chr4-11021419-DEL-99 | chr4-11021420-DEL-99 |
| chr4-11036188-DEL-488 | chr4-11036189-DEL-488 |
| chr4-11040373-DEL-74 | chr4-11040373-DEL-78 |
| chr4-11044762-DEL-550 | chr4-11044763-DEL-550 |
| chr4-11054214-DEL-90 | chr4-11054215-DEL-90 |
| chr4-11057127-DEL-54 | chr4-11057128-DEL-54 |
| chr4-11067368-DEL-182 | chr4-11067369-DEL-182 |
| chr4-11078735-DEL-179 | chr4-11078743-DEL-172 |
| chr4-11079927-DEL-363 | chr4-11079928-DEL-363 |
| chr4-11093134-DEL-58 | chr4-11093135-DEL-58 |
| chr4-11114440-DEL-243 | chr4-11114441-DEL-243 |
| chr4-11121823-DEL-221 | chr4-11121824-DEL-221 |
| chr4-11127664-DEL-144 | chr4-11127665-DEL-144 |
| chr4-11128903-DEL-189 | chr4-11128904-DEL-189 |
| chr4-11171007-DEL-78 | chr4-11170995-DEL-52 |
| chr4-11174510-DEL-257 | chr4-11174511-DEL-257 |
| chr4-11185377-DEL-55 | chr4-11185378-DEL-55 |
| chr4-11210188-DEL-204 | chr4-11210189-DEL-204 |
| chr4-11224232-DEL-456 | chr4-11224233-DEL-456 |
| chr4-11251495-DEL-209 | chr4-11251496-DEL-209 |
| chr4-11262863-DEL-55 | chr4-11262864-DEL-55 |
| chr4-11284183-DEL-62 | chr4-11284184-DEL-62 |
| chr4-11288277-DEL-110 | chr4-11288280-DEL-108 |
| chr4-11300068-DEL-73 | chr4-11300060-DEL-73 |
| chr4-11309057-DEL-204 | chr4-11309058-DEL-204 |
| chr4-11341887-DEL-70 | chr4-11341888-DEL-70 |
| chr4-11364070-DEL-54 | chr4-11364071-DEL-54 |
| chr4-11400127-DEL-89 | chr4-11400123-DEL-87 |
| chr4-11402542-DEL-213 | chr4-11402543-DEL-213 |
| chr4-11450607-DEL-203 | chr4-11450608-DEL-203 |
| chr4-11454187-DEL-99 | chr4-11454205-DEL-50 |
| chr4-11478284-DEL-168 | chr4-11478280-DEL-167 |
| chr4-11484583-DEL-222 | chr4-11484584-DEL-222 |
| chr4-11552121-DEL-418 | chr4-11552122-DEL-418 |
| chr4-11558695-DEL-68 | chr4-11558696-DEL-68 |
| chr4-11608397-DEL-92 | chr4-11608398-DEL-92 |
| chr4-11618514-DEL-647 | chr4-11618515-DEL-647 |
| chr4-11674875-DEL-57 | chr4-11674876-DEL-57 |
| chr4-11692792-DEL-61 | chr4-11692793-DEL-61 |
| chr4-11698904-DEL-185 | chr4-11698880-DEL-204 |
| chr4-11702269-DEL-76 | chr4-11702270-DEL-52 |
| chr4-11718156-DEL-108 | chr4-11718138-DEL-112 |
| chr4-11751063-DEL-362 | chr4-11751064-DEL-362 |
| chr4-11829718-DEL-89 | chr4-11829718-DEL-86 |
| chr4-11849737-DEL-56 | chr4-11849740-DEL-55 |
| chr4-11856178-DEL-60 | chr4-11856142-DEL-104 |
| chr4-11921852-DEL-590 | chr4-11921853-DEL-590 |
| chr4-11943980-DEL-519 | chr4-11943980-DEL-520 |
| chr4-11948866-DEL-56 | chr4-11948867-DEL-56 |
| chr4-11995071-DEL-177 | chr4-11995097-DEL-175 |
| chr4-12043833-DEL-1548 | chr4-12043834-DEL-1548 |
| chr4-12099235-DEL-233 | chr4-12099210-DEL-234 |
| chr4-12117043-DEL-190 | chr4-12117043-DEL-190 |
| chr4-12124462-DEL-56 | chr4-12124461-DEL-50 |
| chr4-12128655-DEL-52 | chr4-12128663-DEL-52 |
| chr4-12133148-DEL-5005 | chr4-12133149-DEL-5005 |
| chr4-12139405-DEL-101 | chr4-12139412-DEL-84 |
| chr4-12154522-DEL-208 | chr4-12154523-DEL-208 |
| chr4-12212338-DEL-7007 | chr4-12212339-DEL-7007 |
| chr4-12232780-DEL-5275 | chr4-12232767-DEL-5276 |
| chr4-12292841-DEL-161 | chr4-12292842-DEL-161 |
| chr4-12319638-DEL-6441 | chr4-12319639-DEL-6441 |
| chr4-12341314-DEL-366 | chr4-12341324-DEL-356 |
| chr4-12341374-DEL-1916 | chr4-12341366-DEL-1924 |
| chr4-12370742-DEL-333 | chr4-12370743-DEL-333 |
| chr4-12398247-DEL-5356 | chr4-12398248-DEL-5356 |
| chr4-12431137-DEL-6619 | chr4-12431138-DEL-6619 |

| | |
|---|---|
| chr4-12464357-DEL-1475 | chr4-12464358-DEL-1475 |
| chr4-12467393-DEL-6725 | chr4-12467394-DEL-6725 |
| chr4-12484969-DEL-75 | chr4-12484932-DEL-75 |
| chr4-12495953-DEL-393 | chr4-12495954-DEL-393 |
| chr4-12528944-DEL-193 | chr4-12528945-DEL-193 |
| chr4-12608028-DEL-2209 | chr4-12608029-DEL-2209 |
| chr4-12647322-DEL-233 | chr4-12647323-DEL-233 |
| chr4-12651788-DEL-86 | chr4-12651789-DEL-86 |
| chr4-12706919-DEL-6590 | chr4-12706920-DEL-6590 |
| chr4-12712902-DEL-208 | chr4-12712903-DEL-208 |
| chr4-12721641-DEL-87 | chr4-12721630-DEL-78 |
| chr4-12749351-DEL-59 | chr4-12749352-DEL-59 |
| chr4-12756067-DEL-549 | chr4-12756068-DEL-549 |
| chr4-12765941-DEL-109 | chr4-12765942-DEL-109 |
| chr4-12769984-DEL-53 | chr4-12769978-DEL-50 |
| chr4-12778949-DEL-51 | chr4-12778950-DEL-51 |
| chr4-12806918-DEL-133 | chr4-12806945-DEL-102 |
| chr4-12809061-DEL-107 | chr4-12809062-DEL-107 |
| chr4-12812388-DEL-54 | chr4-12812377-DEL-60 |
| chr4-12813930-DEL-291 | chr4-12813925-DEL-289 |
| chr4-12840129-DEL-11354 | chr4-12840130-DEL-11354 |
| chr4-12845433-DEL-5144 | chr4-12845432-DEL-5144 |
| chr4-12921719-DEL-816 | chr4-12921720-DEL-816 |
| chr4-12934175-DEL-6662 | chr4-12934168-DEL-6663 |
| chr4-12953488-DEL-6476 | chr4-12953489-DEL-6476 |
| chr4-12960490-DEL-60 | chr4-12960491-DEL-60 |
| chr4-12966264-DEL-90 | chr4-12966265-DEL-90 |
| chr4-12979769-DEL-62 | chr4-12979770-DEL-62 |
| chr4-12981637-DEL-56 | chr4-12981638-DEL-56 |
| chr4-12988529-DEL-1065 | chr4-12988530-DEL-1065 |
| chr4-13015269-DEL-3186 | chr4-13015270-DEL-3186 |
| chr4-13024493-DEL-86 | chr4-13024493-DEL-84 |
| chr4-13044165-DEL-6178 | chr4-13044164-DEL-6177 |
| chr4-13044181-DEL-197 | chr4-13044165-DEL-197 |
| chr4-13084669-DEL-177 | chr4-13084670-DEL-177 |
| chr4-13094722-DEL-293 | chr4-13094723-DEL-293 |
| chr4-13102390-DEL-82 | chr4-13102391-DEL-56 |
| chr4-13118645-DEL-6435 | chr4-13118646-DEL-6435 |
| chr4-13139872-DEL-2331 | chr4-13139869-DEL-2332 |
| chr4-13161858-DEL-140 | chr4-13161859-DEL-140 |
| chr4-13180876-DEL-51 | chr4-13180877-DEL-51 |
| chr4-13191096-DEL-140 | chr4-13191097-DEL-140 |
| chr4-13200396-DEL-54 | chr4-13200397-DEL-54 |
| chr4-13209355-DEL-1464 | chr4-13209356-DEL-1464 |
| chr4-13217481-DEL-991 | chr4-13217488-DEL-985 |
| chr4-13266969-DEL-144 | chr4-13266970-DEL-144 |
| chr4-13277848-DEL-78 | chr4-13277849-DEL-78 |
| chr4-13289022-DEL-7549 | chr4-13289023-DEL-7549 |
| chr4-13296077-DEL-424 | chr4-13296078-DEL-424 |
| chr4-13337129-DEL-51 | chr4-13337130-DEL-51 |
| chr4-13355196-DEL-400 | chr4-13355195-DEL-401 |
| chr4-13360369-DEL-171 | chr4-13360377-DEL-163 |
| chr4-13372937-DEL-346 | chr4-13372938-DEL-346 |
| chr4-13377434-DEL-4301 | chr4-13377435-DEL-4301 |
| chr4-13380783-DEL-58 | chr4-13380784-DEL-57 |
| chr4-13425210-DEL-7308 | chr4-13425211-DEL-7308 |
| chr4-13473516-DEL-196 | chr4-13473523-DEL-195 |
| chr4-13490297-DEL-56 | chr4-13490298-DEL-56 |
| chr4-13651445-DEL-511 | chr4-13651444-DEL-512 |
| chr4-13664437-DEL-87 | chr4-13664428-DEL-96 |
| chr4-13668555-DEL-399 | chr4-13668556-DEL-399 |
| chr4-13726778-DEL-89 | chr4-13726763-DEL-73 |
| chr4-13762574-DEL-60 | chr4-13762575-DEL-60 |
| chr4-13763372-DEL-91 | chr4-13763412-DEL-92 |
| chr4-13828175-DEL-52 | chr4-13828176-DEL-52 |
| chr4-13838718-DEL-72 | chr4-13838719-DEL-72 |
| chr4-13871853-DEL-52 | chr4-13871854-DEL-52 |
| chr4-13874154-DEL-464 | chr4-13874155-DEL-464 |
| chr4-13899686-DEL-199 | chr4-13899687-DEL-199 |
| chr4-13913503-DEL-526 | chr4-13913504-DEL-526 |
| chr4-13927088-DEL-914 | chr4-13927089-DEL-914 |
| chr4-13930318-DEL-201 | chr4-13930319-DEL-201 |
| chr4-13936680-DEL-6807 | chr4-13936680-DEL-6807 |
| chr4-13939124-DEL-131 | chr4-13939125-DEL-131 |
| chr4-13952180-DEL-57 | chr4-13952136-DEL-76 |
| chr4-13983570-DEL-125 | chr4-13983571-DEL-125 |
| chr4-13984182-DEL-263 | chr4-13984183-DEL-263 |
| chr4-14000530-DEL-472 | chr4-14000531-DEL-472 |
| chr4-14005205-DEL-54 | chr4-14005206-DEL-54 |
| chr4-14026979-DEL-1288 | chr4-14026967-DEL-1289 |

| | |
|---|---|
| chr4-14036919-DEL-875 | chr4-14036920-DEL-875 |
| chr4-14041922-DEL-7202 | chr4-14041923-DEL-7202 |
| chr4-14042410-DEL-141 | chr4-14042411-DEL-141 |
| chr4-14048284-DEL-3234 | chr4-14048285-DEL-3235 |
| chr4-14053395-DEL-95 | chr4-14053396-DEL-95 |
| chr4-14119852-DEL-56 | chr4-14119855-DEL-54 |
| chr4-14126537-DEL-1506 | chr4-14126537-DEL-1506 |
| chr4-14137244-DEL-214 | chr4-14137214-DEL-204 |
| chr4-14158933-DEL-71 | chr4-14158906-DEL-72 |
| chr4-14171013-DEL-155 | chr4-14171014-DEL-155 |
| chr4-14192755-DEL-6160 | chr4-14192756-DEL-6160 |
| chr4-14207250-DEL-62 | chr4-14207202-DEL-63 |
| chr4-14219374-DEL-146 | chr4-14219375-DEL-146 |
| chr4-14220774-DEL-6321 | chr4-14220774-DEL-6321 |
| chr4-14231161-DEL-910 | chr4-14231164-DEL-909 |
| chr4-14249151-DEL-149 | chr4-14249140-DEL-150 |
| chr4-14250052-DEL-89 | chr4-14250053-DEL-89 |
| chr4-14252857-DEL-3495 | chr4-14252855-DEL-3496 |
| chr4-14276625-DEL-1610 | chr4-14276626-DEL-1610 |
| chr4-14340918-DEL-88 | chr4-14340919-DEL-88 |
| chr4-14366907-DEL-2648 | chr4-14366909-DEL-2649 |
| chr4-14377877-DEL-510 | chr4-14377883-DEL-509 |
| chr4-14424070-DEL-5995 | chr4-14424057-DEL-5996 |
| chr4-14446810-DEL-1354 | chr4-14446810-DEL-1354 |
| chr4-14448004-DEL-905 | chr4-14448005-DEL-899 |
| chr4-14450809-DEL-201 | chr4-14450810-DEL-201 |
| chr4-14482617-DEL-470 | chr4-14482618-DEL-470 |
| chr4-14494200-DEL-1109 | chr4-14494201-DEL-1109 |
| chr4-14543707-DEL-95 | chr4-14543707-DEL-122 |
| chr4-14582399-DEL-6410 | chr4-14582400-DEL-6410 |
| chr4-14592153-DEL-566 | chr4-14592154-DEL-566 |
| chr4-14592781-DEL-834 | chr4-14592782-DEL-834 |
| chr4-14593946-DEL-66 | chr4-14593947-DEL-66 |
| chr4-14602361-DEL-215 | chr4-14602353-DEL-252 |
| chr4-14626605-DEL-6849 | chr4-14626603-DEL-6850 |
| chr4-14638321-DEL-3985 | chr4-14638322-DEL-3985 |
| chr4-14649688-DEL-1249 | chr4-14649705-DEL-1233 |
| chr4-14660918-DEL-1778 | chr4-14660919-DEL-1778 |
| chr4-14673216-DEL-6863 | chr4-14673217-DEL-6863 |
| chr4-14679566-DEL-212 | chr4-14679567-DEL-212 |
| chr4-14700131-DEL-58 | chr4-14700114-DEL-74 |
| chr4-14710620-DEL-148 | chr4-14710610-DEL-138 |
| chr4-14729125-DEL-63 | chr4-14729150-DEL-73 |
| chr4-14729533-DEL-1165 | chr4-14729540-DEL-1173 |
| chr4-14739052-DEL-84 | chr4-14739053-DEL-81 |
| chr4-14744772-DEL-176 | chr4-14744774-DEL-137 |
| chr4-14767583-DEL-109 | chr4-14767584-DEL-109 |
| chr4-14786212-DEL-87 | chr4-14786213-DEL-87 |
| chr4-14801234-DEL-224 | chr4-14801235-DEL-224 |
| chr4-14808260-DEL-87 | chr4-14808261-DEL-87 |
| chr4-14816013-DEL-93 | chr4-14816014-DEL-93 |
| chr4-14816243-DEL-695 | chr4-14816244-DEL-695 |
| chr4-14821681-DEL-213 | chr4-14821687-DEL-211 |
| chr4-14832430-DEL-66 | chr4-14832431-DEL-66 |
| chr4-14835781-DEL-200 | chr4-14835768-DEL-201 |
| chr4-14839686-DEL-167 | chr4-14839687-DEL-167 |
| chr4-14848860-DEL-422 | chr4-14848861-DEL-422 |
| chr4-14852026-DEL-1005 | chr4-14852026-DEL-1006 |
| chr4-14888014-DEL-304 | chr4-14888000-DEL-314 |
| chr4-14905767-DEL-118 | chr4-14905768-DEL-118 |
| chr4-14905777-DEL-3378 | chr4-14905788-DEL-3368 |
| chr4-14908825-DEL-80 | chr4-14908826-DEL-80 |
| chr4-14915841-DEL-507 | chr4-14915841-DEL-508 |
| chr4-14921578-DEL-337 | chr4-14921579-DEL-337 |
| chr4-14929350-DEL-63 | chr4-14929336-DEL-52 |
| chr4-14934099-DEL-72 | chr4-14934100-DEL-72 |
| chr4-14942084-DEL-58 | chr4-14942085-DEL-58 |
| chr4-15009298-DEL-92 | chr4-15009299-DEL-92 |
| chr4-15014215-DEL-6371 | chr4-15014216-DEL-6371 |
| chr4-15024011-DEL-55 | chr4-15024015-DEL-56 |
| chr4-15054429-DEL-6667 | chr4-15054430-DEL-6667 |
| chr4-15054616-DEL-424 | chr4-15054616-DEL-424 |
| chr4-15070759-DEL-182 | chr4-15070760-DEL-182 |
| chr4-15071502-DEL-1746 | chr4-15071503-DEL-1746 |
| chr4-15128911-DEL-589 | chr4-15128912-DEL-589 |
| chr4-15132974-DEL-54 | chr4-15132977-DEL-52 |
| chr4-15155687-DEL-52 | chr4-15155720-DEL-52 |
| chr4-15173913-DEL-588 | chr4-15173914-DEL-587 |
| chr4-15189106-DEL-198 | chr4-15189107-DEL-198 |
| chr4-15223795-DEL-217 | chr4-15223796-DEL-217 |

| | |
|---|---|
| chr4-15229519-DEL-185 | chr4-15229520-DEL-185 |
| chr4-15236200-DEL-6610 | chr4-15236201-DEL-6610 |
| chr4-15269499-DEL-54 | chr4-15269483-DEL-82 |
| chr4-15297660-DEL-1430 | chr4-15297642-DEL-1424 |
| chr4-15307089-DEL-68 | chr4-15307090-DEL-68 |
| chr4-15326924-DEL-66 | chr4-15326925-DEL-66 |
| chr4-15341319-DEL-2858 | chr4-15341320-DEL-2858 |
| chr4-15367628-DEL-643 | chr4-15367629-DEL-643 |
| chr4-15390745-DEL-59 | chr4-15390746-DEL-59 |
| chr4-15422788-DEL-315 | chr4-15422788-DEL-314 |
| chr4-15444562-DEL-86 | chr4-15444563-DEL-86 |
| chr4-15480720-DEL-62 | chr4-15480720-DEL-54 |
| chr4-15517048-DEL-2120 | chr4-15517049-DEL-2120 |
| chr4-15529000-DEL-248 | chr4-15529001-DEL-248 |
| chr4-15530706-DEL-76 | chr4-15530711-DEL-72 |
| chr4-15546032-DEL-69 | chr4-15546033-DEL-69 |
| chr4-15577949-DEL-6802 | chr4-15577949-DEL-6802 |
| chr4-15600341-DEL-1269 | chr4-15600342-DEL-1269 |
| chr4-15626352-DEL-78 | chr4-15626368-DEL-52 |
| chr4-15651203-DEL-469 | chr4-15651203-DEL-464 |
| chr4-15670927-DEL-220 | chr4-15670928-DEL-220 |
| chr4-15674661-DEL-60 | chr4-15674662-DEL-60 |
| chr4-15678293-DEL-115 | chr4-15678290-DEL-115 |
| chr4-15700899-DEL-135 | chr4-15700900-DEL-135 |
| chr4-15723217-DEL-60 | chr4-15723218-DEL-60 |
| chr4-15731087-DEL-56 | chr4-15731087-DEL-57 |
| chr4-15755731-DEL-584 | chr4-15755732-DEL-584 |
| chr4-15759215-DEL-739 | chr4-15759216-DEL-739 |
| chr4-15801921-DEL-1291 | chr4-15801922-DEL-1291 |
| chr4-15811165-DEL-462 | chr4-15811166-DEL-462 |
| chr4-15811735-DEL-1599 | chr4-15811736-DEL-1599 |
| chr4-15831962-DEL-91 | chr4-15831950-DEL-98 |
| chr4-15853140-DEL-60 | chr4-15853149-DEL-52 |
| chr4-15853513-DEL-6143 | chr4-15853514-DEL-6143 |
| chr4-15862143-DEL-203 | chr4-15862143-DEL-203 |
| chr4-15871576-DEL-173 | chr4-15871576-DEL-173 |
| chr4-15877769-DEL-82 | chr4-15877770-DEL-82 |
| chr4-15919369-DEL-167 | chr4-15919370-DEL-167 |
| chr4-15921254-DEL-98 | chr4-15921255-DEL-98 |
| chr4-15981115-DEL-55 | chr4-15981118-DEL-55 |
| chr4-16000464-DEL-56 | chr4-16000476-DEL-54 |
| chr4-16030401-DEL-900 | chr4-16030396-DEL-902 |
| chr4-16057261-DEL-1204 | chr4-16057262-DEL-1204 |
| chr4-16058698-DEL-6333 | chr4-16058699-DEL-6333 |
| chr4-16067762-DEL-66 | chr4-16067761-DEL-59 |
| chr4-16077636-DEL-119 | chr4-16077637-DEL-119 |
| chr4-16084047-DEL-442 | chr4-16084092-DEL-404 |
| chr4-16095904-DEL-3480 | chr4-16095905-DEL-3480 |
| chr4-16108844-DEL-2890 | chr4-16108845-DEL-2890 |
| chr4-16133690-DEL-51 | chr4-16133691-DEL-51 |
| chr4-16134463-DEL-1978 | chr4-16134464-DEL-1978 |
| chr4-16144911-DEL-2107 | chr4-16144912-DEL-2107 |
| chr4-16171497-DEL-1542 | chr4-16171498-DEL-1542 |
| chr4-16172749-DEL-108 | chr4-16172750-DEL-108 |
| chr4-16174663-DEL-64 | chr4-16174664-DEL-68 |
| chr4-16175117-DEL-911 | chr4-16175118-DEL-911 |
| chr4-16237073-DEL-173 | chr4-16237072-DEL-174 |
| chr4-16240095-DEL-397 | chr4-16240096-DEL-397 |
| chr4-16267314-DEL-129 | chr4-16267318-DEL-135 |
| chr4-16268780-DEL-149 | chr4-16268783-DEL-147 |
| chr4-16309775-DEL-208 | chr4-16309774-DEL-208 |
| chr4-16333261-DEL-1167 | chr4-16333262-DEL-1167 |
| chr4-16333446-DEL-3126 | chr4-16333446-DEL-3126 |
| chr4-16335705-DEL-474 | chr4-16335706-DEL-474 |
| chr4-16341017-DEL-61 | chr4-16341017-DEL-61 |
| chr4-16361791-DEL-3075 | chr4-16361792-DEL-3075 |
| chr4-16367735-DEL-66 | chr4-16367708-DEL-67 |
| chr4-16368940-DEL-96 | chr4-16368941-DEL-97 |
| chr4-16396913-DEL-810 | chr4-16396914-DEL-810 |
| chr4-16413701-DEL-59 | chr4-16413688-DEL-60 |
| chr4-16423126-DEL-56 | chr4-16423125-DEL-56 |
| chr4-16433271-DEL-2506 | chr4-16433272-DEL-2506 |
| chr4-16442829-DEL-73 | chr4-16442827-DEL-74 |
| chr4-16444527-DEL-523 | chr4-16444528-DEL-523 |
| chr4-16474227-DEL-1549 | chr4-16474228-DEL-1549 |
| chr4-16485015-DEL-141 | chr4-16485016-DEL-141 |
| chr4-16515028-DEL-279 | chr4-16515029-DEL-279 |
| chr4-16530672-DEL-183 | chr4-16530673-DEL-181 |
| chr4-16545261-DEL-6965 | chr4-16545262-DEL-6965 |
| chr4-16566905-DEL-50 | chr4-16566914-DEL-51 |

| | |
|---|---|
| chr4-16570580-DEL-1869 | chr4-16570584-DEL-1867 |
| chr4-16586198-DEL-193 | chr4-16586199-DEL-193 |
| chr4-16588926-DEL-3567 | chr4-16588927-DEL-3567 |
| chr4-16610540-DEL-260 | chr4-16610541-DEL-260 |
| chr4-16612704-DEL-51 | chr4-16612700-DEL-52 |
| chr4-16618003-DEL-298 | chr4-16618004-DEL-298 |
| chr4-16631669-DEL-538 | chr4-16631669-DEL-538 |
| chr4-16633432-DEL-94 | chr4-16633433-DEL-94 |
| chr4-16641275-DEL-4169 | chr4-16641276-DEL-4169 |
| chr4-16675906-DEL-5178 | chr4-16675907-DEL-5178 |
| chr4-16706323-DEL-1961 | chr4-16706324-DEL-1961 |
| chr4-16720686-DEL-105 | chr4-16720710-DEL-58 |
| chr4-16730589-DEL-185 | chr4-16730590-DEL-185 |
| chr4-16777126-DEL-6591 | chr4-16777140-DEL-6584 |
| chr4-16784850-DEL-5062 | chr4-16784851-DEL-5062 |
| chr4-16804722-DEL-6266 | chr4-16804723-DEL-6266 |
| chr4-16824922-DEL-58 | chr4-16824923-DEL-54 |
| chr4-16850538-DEL-192 | chr4-16850539-DEL-192 |
| chr4-16858021-DEL-52 | chr4-16858022-DEL-52 |
| chr4-16870990-DEL-56 | chr4-16870991-DEL-72 |
| chr4-16878812-DEL-6400 | chr4-16878813-DEL-6400 |
| chr4-16885644-DEL-6046 | chr4-16885645-DEL-6046 |
| chr4-16888070-DEL-866 | chr4-16888071-DEL-866 |
| chr4-16894534-DEL-68 | chr4-16894523-DEL-92 |
| chr4-16926610-DEL-390 | chr4-16926605-DEL-391 |
| chr4-16935765-DEL-225 | chr4-16935765-DEL-225 |
| chr4-16956934-DEL-1328 | chr4-16956935-DEL-1328 |
| chr4-16969564-DEL-109 | chr4-16969565-DEL-109 |
| chr4-16995040-DEL-53 | chr4-16995073-DEL-51 |
| chr4-17024311-DEL-64 | chr4-17024312-DEL-64 |
| chr4-17062952-DEL-144 | chr4-17062953-DEL-144 |
| chr4-17064922-DEL-461 | chr4-17064923-DEL-461 |
| chr4-17065919-DEL-211 | chr4-17065920-DEL-211 |
| chr4-17077182-DEL-1540 | chr4-17077183-DEL-1540 |
| chr4-17080308-DEL-61 | chr4-17080309-DEL-61 |
| chr4-17097852-DEL-1622 | chr4-17097853-DEL-1622 |
| chr4-17128036-DEL-2304 | chr4-17128036-DEL-2304 |
| chr4-17128384-DEL-6366 | chr4-17128385-DEL-6366 |
| chr4-17134824-DEL-1027 | chr4-17134825-DEL-1027 |
| chr4-17142288-DEL-850 | chr4-17142289-DEL-850 |
| chr4-17148505-DEL-1594 | chr4-17148506-DEL-1594 |
| chr4-17150146-DEL-2141 | chr4-17150147-DEL-2141 |
| chr4-17166028-DEL-1080 | chr4-17166029-DEL-1080 |
| chr4-17167195-DEL-72 | chr4-17167196-DEL-72 |
| chr4-17199152-DEL-445 | chr4-17199153-DEL-445 |
| chr4-17217236-DEL-6686 | chr4-17217237-DEL-6686 |
| chr4-17217669-DEL-212 | chr4-17217670-DEL-212 |
| chr4-17236775-DEL-234 | chr4-17236790-DEL-212 |
| chr4-17243196-DEL-1932 | chr4-17243197-DEL-1932 |
| chr4-17245984-DEL-87 | chr4-17245985-DEL-87 |
| chr4-17281822-DEL-214 | chr4-17281823-DEL-214 |
| chr4-17309374-DEL-214 | chr4-17309375-DEL-214 |
| chr4-17379233-DEL-351 | chr4-17379233-DEL-364 |
| chr4-17408771-DEL-6850 | chr4-17408772-DEL-6850 |
| chr4-17409060-DEL-212 | chr4-17409061-DEL-212 |
| chr4-17420935-DEL-305 | chr4-17420942-DEL-304 |
| chr4-17429708-DEL-55 | chr4-17429705-DEL-59 |
| chr4-17444671-DEL-6662 | chr4-17444672-DEL-6662 |
| chr4-17467323-DEL-153 | chr4-17467323-DEL-153 |
| chr4-17470103-DEL-486 | chr4-17470104-DEL-486 |
| chr4-17494994-DEL-1213 | chr4-17494995-DEL-1213 |
| chr4-17511164-DEL-634 | chr4-17511165-DEL-634 |
| chr4-17512982-DEL-358 | chr4-17512983-DEL-358 |
| chr4-17561447-DEL-579 | chr4-17561436-DEL-585 |
| chr4-17563241-DEL-2627 | chr4-17563242-DEL-2627 |
| chr4-17571105-DEL-6702 | chr4-17571106-DEL-6702 |
| chr4-17603729-DEL-105 | chr4-17603730-DEL-105 |
| chr4-17629947-DEL-3106 | chr4-17629941-DEL-3102 |
| chr4-17655501-DEL-2719 | chr4-17655501-DEL-2719 |
| chr4-17687693-DEL-2720 | chr4-17687694-DEL-2720 |
| chr4-17702881-DEL-934 | chr4-17702882-DEL-935 |
| chr4-17711408-DEL-853 | chr4-17711408-DEL-854 |
| chr4-17744581-DEL-199 | chr4-17744582-DEL-199 |
| chr4-17775753-DEL-401 | chr4-17775754-DEL-400 |
| chr4-17777347-DEL-6010 | chr4-17777348-DEL-6010 |
| chr4-17817621-DEL-306 | chr4-17817622-DEL-306 |
| chr4-17826919-DEL-477 | chr4-17826920-DEL-477 |
| chr4-17844800-DEL-1261 | chr4-17844801-DEL-1261 |
| chr4-17872651-DEL-119 | chr4-17872652-DEL-119 |
| chr4-17900860-DEL-5205 | chr4-17900861-DEL-5205 |

| | |
|---|---|
| chr4-17910577-DEL-50 | chr4-17910578-DEL-52 |
| chr4-18021572-DEL-6462 | chr4-18021573-DEL-6462 |
| chr4-18044231-DEL-681 | chr4-18044232-DEL-682 |
| chr4-18071215-DEL-220 | chr4-18071216-DEL-220 |
| chr4-18076230-DEL-72 | chr4-18076195-DEL-111 |
| chr4-18127801-DEL-57 | chr4-18127804-DEL-56 |
| chr4-18136133-DEL-911 | chr4-18136134-DEL-911 |
| chr4-18138927-DEL-52 | chr4-18138917-DEL-62 |
| chr4-18193519-DEL-231 | chr4-18193520-DEL-230 |
| chr4-18209742-DEL-2886 | chr4-18209743-DEL-2887 |
| chr4-18212831-DEL-7209 | chr4-18212832-DEL-7209 |
| chr4-18221606-DEL-6433 | chr4-18221607-DEL-6433 |
| chr4-18231421-DEL-81 | chr4-18231422-DEL-81 |
| chr4-18239930-DEL-72 | chr4-18239931-DEL-72 |
| chr4-18244579-DEL-358 | chr4-18244580-DEL-358 |
| chr4-18272662-DEL-1962 | chr4-18272662-DEL-1962 |
| chr4-18281489-DEL-1632 | chr4-18281484-DEL-1637 |
| chr4-18289801-DEL-64 | chr4-18289787-DEL-89 |
| chr4-18301368-DEL-82 | chr4-18301387-DEL-66 |
| chr4-18378878-DEL-6549 | chr4-18378879-DEL-6549 |
| chr4-18385009-DEL-204 | chr4-18385010-DEL-204 |
| chr4-18411963-DEL-226 | chr4-18411963-DEL-226 |
| chr4-18421926-DEL-6497 | chr4-18421927-DEL-6497 |
| chr4-18428118-DEL-212 | chr4-18428119-DEL-212 |
| chr4-18447803-DEL-64 | chr4-18447795-DEL-72 |
| chr4-18456961-DEL-54 | chr4-18456960-DEL-52 |
| chr4-18474320-DEL-7154 | chr4-18474310-DEL-7155 |
| chr4-18512963-DEL-6198 | chr4-18512963-DEL-6198 |
| chr4-18535852-DEL-831 | chr4-18535853-DEL-831 |
| chr4-18550846-DEL-67 | chr4-18550850-DEL-63 |
| chr4-18553531-DEL-136 | chr4-18553574-DEL-112 |
| chr4-18565674-DEL-729 | chr4-18565671-DEL-727 |
| chr4-18606830-DEL-6942 | chr4-18606831-DEL-6942 |
| chr4-18619866-DEL-207 | chr4-18619867-DEL-207 |
| chr4-18677970-DEL-211 | chr4-18677971-DEL-211 |
| chr4-18710378-DEL-6250 | chr4-18710379-DEL-6250 |
| chr4-18717932-DEL-2742 | chr4-18717933-DEL-2742 |
| chr4-18727438-DEL-2030 | chr4-18727434-DEL-2052 |
| chr4-18729789-DEL-466 | chr4-18729774-DEL-466 |
| chr4-18737779-DEL-93 | chr4-18737780-DEL-93 |
| chr4-18741254-DEL-882 | chr4-18741255-DEL-882 |
| chr4-18744058-DEL-78 | chr4-18744059-DEL-77 |
| chr4-18747602-DEL-588 | chr4-18747603-DEL-588 |
| chr4-18763540-DEL-696 | chr4-18763541-DEL-696 |
| chr4-18774166-DEL-671 | chr4-18774166-DEL-672 |
| chr4-18807971-DEL-6403 | chr4-18807971-DEL-6403 |
| chr4-18830481-DEL-858 | chr4-18830482-DEL-858 |
| chr4-18912181-DEL-64 | chr4-18912133-DEL-64 |
| chr4-18948146-DEL-498 | chr4-18948147-DEL-498 |
| chr4-18975367-DEL-71 | chr4-18975389-DEL-77 |
| chr4-18979512-DEL-648 | chr4-18979513-DEL-648 |
| chr4-19006316-DEL-198 | chr4-19006317-DEL-198 |
| chr4-19009246-DEL-440 | chr4-19009247-DEL-440 |
| chr4-19072458-DEL-70 | chr4-19072461-DEL-56 |
| chr4-19072720-DEL-6359 | chr4-19072721-DEL-6359 |
| chr4-19166776-DEL-241 | chr4-19166777-DEL-241 |
| chr4-19190043-DEL-6602 | chr4-19190044-DEL-6602 |
| chr4-19196658-DEL-2494 | chr4-19196659-DEL-2493 |
| chr4-19246404-DEL-443 | chr4-19246405-DEL-443 |
| chr4-19313096-DEL-181 | chr4-19313097-DEL-181 |
| chr4-19325156-DEL-60 | chr4-19325161-DEL-50 |
| chr4-19364594-DEL-1004 | chr4-19364603-DEL-1005 |
| chr4-19372321-DEL-911 | chr4-19372322-DEL-911 |
| chr4-19375576-DEL-173 | chr4-19375577-DEL-173 |
| chr4-19416955-DEL-1107 | chr4-19416955-DEL-1107 |
| chr4-19422882-DEL-380 | chr4-19422883-DEL-380 |
| chr4-19432700-DEL-1782 | chr4-19432699-DEL-1784 |
| chr4-19466422-DEL-176 | chr4-19466423-DEL-176 |
| chr4-19476227-DEL-53 | chr4-19476228-DEL-53 |
| chr4-19527360-DEL-53 | chr4-19527361-DEL-53 |
| chr4-19614847-DEL-282 | chr4-19614848-DEL-282 |
| chr4-19663726-DEL-6600 | chr4-19663727-DEL-6600 |
| chr4-19706753-DEL-79 | chr4-19706752-DEL-78 |
| chr4-19715696-DEL-181 | chr4-19715699-DEL-179 |
| chr4-19744058-DEL-6665 | chr4-19744059-DEL-6665 |
| chr4-19750218-DEL-214 | chr4-19750219-DEL-214 |
| chr4-19759706-DEL-1241 | chr4-19759710-DEL-1238 |
| chr4-19774564-DEL-2361 | chr4-19774565-DEL-2361 |
| chr4-19806343-DEL-70 | chr4-19806344-DEL-70 |
| chr4-19854662-DEL-85 | chr4-19854663-DEL-85 |

| | |
|---|---|
| chr4-19863769-DEL-52 | chr4-19863770-DEL-52 |
| chr4-19896807-DEL-52 | chr4-19896829-DEL-58 |
| chr4-19925247-DEL-200 | chr4-19925248-DEL-195 |
| chr4-19959543-DEL-810 | chr4-19959544-DEL-810 |
| chr4-19969460-DEL-674 | chr4-19969461-DEL-674 |
| chr4-19970679-DEL-57 | chr4-19970642-DEL-58 |
| chr4-19978240-DEL-341 | chr4-19978241-DEL-341 |
| chr4-19981690-DEL-407 | chr4-19981691-DEL-407 |
| chr4-19985721-DEL-2173 | chr4-19985722-DEL-2173 |
| chr4-20006943-DEL-56 | chr4-20006915-DEL-56 |
| chr4-20050042-DEL-237 | chr4-20050043-DEL-237 |
| chr4-20069551-DEL-1557 | chr4-20069552-DEL-1557 |
| chr4-20076302-DEL-8724 | chr4-20076303-DEL-8724 |
| chr4-20081381-DEL-3047 | chr4-20081382-DEL-3047 |
| chr4-20125826-DEL-70 | chr4-20125827-DEL-70 |
| chr4-20183145-DEL-60 | chr4-20183098-DEL-92 |
| chr4-20184489-DEL-73 | chr4-20184501-DEL-70 |
| chr4-20188473-DEL-52 | chr4-20188474-DEL-52 |
| chr4-20207262-DEL-6158 | chr4-20207263-DEL-6158 |
| chr4-20207309-DEL-199 | chr4-20207310-DEL-199 |
| chr4-20215135-DEL-62 | chr4-20215136-DEL-62 |
| chr4-20220499-DEL-155 | chr4-20220500-DEL-154 |
| chr4-20243830-DEL-163 | chr4-20243831-DEL-163 |
| chr4-20249533-DEL-715 | chr4-20249534-DEL-715 |
| chr4-20253770-DEL-84 | chr4-20253769-DEL-75 |
| chr4-20268052-DEL-104 | chr4-20268052-DEL-104 |
| chr4-20289146-DEL-6430 | chr4-20289147-DEL-6430 |
| chr4-20295667-DEL-382 | chr4-20295671-DEL-383 |
| chr4-20299631-DEL-720 | chr4-20299632-DEL-720 |
| chr4-20307758-DEL-1276 | chr4-20307759-DEL-1276 |
| chr4-20330095-DEL-294 | chr4-20330092-DEL-289 |
| chr4-20354552-DEL-6282 | chr4-20354553-DEL-6282 |
| chr4-20380609-DEL-133 | chr4-20380610-DEL-133 |
| chr4-20393557-DEL-110 | chr4-20393572-DEL-96 |
| chr4-20409117-DEL-3411 | chr4-20409113-DEL-3413 |
| chr4-20420619-DEL-6700 | chr4-20420627-DEL-6699 |
| chr4-20446374-DEL-504 | chr4-20446375-DEL-504 |
| chr4-20498046-DEL-6912 | chr4-20498047-DEL-6912 |
| chr4-20558423-DEL-519 | chr4-20558424-DEL-519 |
| chr4-20567873-DEL-536 | chr4-20567874-DEL-536 |
| chr4-20575530-DEL-53 | chr4-20575531-DEL-53 |
| chr4-20600343-DEL-573 | chr4-20600344-DEL-573 |
| chr4-20602780-DEL-70 | chr4-20602803-DEL-50 |
| chr4-20608266-DEL-3070 | chr4-20608267-DEL-3070 |
| chr4-20608891-DEL-56 | chr4-20608891-DEL-58 |
| chr4-20608998-DEL-1499 | chr4-20608995-DEL-1499 |
| chr4-20620335-DEL-56 | chr4-20620336-DEL-56 |
| chr4-20647938-DEL-399 | chr4-20647939-DEL-399 |
| chr4-20714203-DEL-273 | chr4-20714210-DEL-270 |
| chr4-20738494-DEL-51 | chr4-20738495-DEL-51 |
| chr4-20770485-DEL-1447 | chr4-20770514-DEL-1419 |
| chr4-20781829-DEL-140 | chr4-20781830-DEL-139 |
| chr4-20790470-DEL-95 | chr4-20790471-DEL-95 |
| chr4-20810096-DEL-201 | chr4-20810096-DEL-201 |
| chr4-20818679-DEL-58 | chr4-20818680-DEL-58 |
| chr4-20868807-DEL-1456 | chr4-20868808-DEL-1456 |
| chr4-20875743-DEL-454 | chr4-20875744-DEL-454 |
| chr4-20886108-DEL-952 | chr4-20886109-DEL-952 |
| chr4-20889435-DEL-528 | chr4-20889434-DEL-529 |
| chr4-20894768-DEL-82 | chr4-20894769-DEL-82 |
| chr4-20896707-DEL-588 | chr4-20896704-DEL-592 |
| chr4-20899266-DEL-332 | chr4-20899267-DEL-332 |
| chr4-20905800-DEL-364 | chr4-20905801-DEL-364 |
| chr4-20919891-DEL-199 | chr4-20919892-DEL-199 |
| chr4-20938801-DEL-6466 | chr4-20938802-DEL-6466 |
| chr4-20961421-DEL-568 | chr4-20961422-DEL-568 |
| chr4-20965954-DEL-177 | chr4-20965955-DEL-177 |
| chr4-20971791-DEL-648 | chr4-20971792-DEL-648 |
| chr4-20990444-DEL-58 | chr4-20990445-DEL-58 |
| chr4-21004087-DEL-359 | chr4-21004088-DEL-359 |
| chr4-21036949-DEL-198 | chr4-21036950-DEL-198 |
| chr4-21054878-DEL-690 | chr4-21054879-DEL-690 |
| chr4-21073822-DEL-7197 | chr4-21073823-DEL-7197 |
| chr4-21093066-DEL-1864 | chr4-21093067-DEL-1864 |
| chr4-21129222-DEL-2417 | chr4-21129223-DEL-2417 |
| chr4-21194145-DEL-427 | chr4-21194145-DEL-426 |
| chr4-21205976-DEL-1627 | chr4-21205977-DEL-1627 |
| chr4-21222944-DEL-578 | chr4-21222945-DEL-578 |
| chr4-21262723-DEL-310 | chr4-21262724-DEL-310 |
| chr4-21263305-DEL-1695 | chr4-21263309-DEL-1690 |

| | |
|---|---|
| chr4-21281250-DEL-6727 | chr4-21281251-DEL-6727 |
| chr4-21293062-DEL-58 | chr4-21293063-DEL-58 |
| chr4-21296853-DEL-87 | chr4-21296854-DEL-87 |
| chr4-21302611-DEL-928 | chr4-21302612-DEL-928 |
| chr4-21312402-DEL-6235 | chr4-21312405-DEL-6235 |
| chr4-21340641-DEL-201 | chr4-21340641-DEL-201 |
| chr4-21363861-DEL-65 | chr4-21363870-DEL-57 |
| chr4-21381483-DEL-416 | chr4-21381484-DEL-416 |
| chr4-21453142-DEL-1577 | chr4-21453143-DEL-1577 |
| chr4-21465837-DEL-1362 | chr4-21465838-DEL-1362 |
| chr4-21524474-DEL-144 | chr4-21524469-DEL-139 |
| chr4-21550945-DEL-1147 | chr4-21550946-DEL-1147 |
| chr4-21552414-DEL-1700 | chr4-21552415-DEL-1700 |
| chr4-21555259-DEL-1206 | chr4-21555250-DEL-1207 |
| chr4-21562983-DEL-491 | chr4-21562984-DEL-491 |
| chr4-21570792-DEL-833 | chr4-21570793-DEL-833 |
| chr4-21572273-DEL-88 | chr4-21572274-DEL-88 |
| chr4-21573575-DEL-1000 | chr4-21573576-DEL-1000 |
| chr4-21577627-DEL-93 | chr4-21577630-DEL-94 |
| chr4-21593874-DEL-1894 | chr4-21593874-DEL-1894 |
| chr4-21598534-DEL-72 | chr4-21598500-DEL-75 |
| chr4-21627196-DEL-748 | chr4-21627197-DEL-748 |
| chr4-21647242-DEL-84 | chr4-21647243-DEL-84 |
| chr4-21650299-DEL-7252 | chr4-21650283-DEL-7251 |
| chr4-21651225-DEL-212 | chr4-21651226-DEL-212 |
| chr4-21698422-DEL-3546 | chr4-21698423-DEL-3546 |
| chr4-21722104-DEL-95 | chr4-21722113-DEL-97 |
| chr4-21724969-DEL-213 | chr4-21724970-DEL-213 |
| chr4-21730945-DEL-629 | chr4-21730958-DEL-627 |
| chr4-21767598-DEL-168 | chr4-21767598-DEL-168 |
| chr4-21770908-DEL-214 | chr4-21770903-DEL-217 |
| chr4-21773020-DEL-773 | chr4-21773021-DEL-773 |
| chr4-21786755-DEL-341 | chr4-21786741-DEL-337 |
| chr4-21797890-DEL-79 | chr4-21797891-DEL-79 |
| chr4-21801960-DEL-256 | chr4-21801961-DEL-256 |
| chr4-21805098-DEL-51 | chr4-21805099-DEL-51 |
| chr4-21833145-DEL-446 | chr4-21833146-DEL-446 |
| chr4-21889754-DEL-60 | chr4-21889755-DEL-60 |
| chr4-21916602-DEL-93 | chr4-21916557-DEL-96 |
| chr4-21940122-DEL-77 | chr4-21940123-DEL-77 |
| chr4-21948143-DEL-526 | chr4-21948132-DEL-522 |
| chr4-21985684-DEL-341 | chr4-21985686-DEL-338 |
| chr4-21988199-DEL-532 | chr4-21988200-DEL-532 |
| chr4-22006203-DEL-126 | chr4-22006204-DEL-126 |
| chr4-22031783-DEL-390 | chr4-22031784-DEL-390 |
| chr4-22036881-DEL-1250 | chr4-22036882-DEL-1250 |
| chr4-22115933-DEL-161 | chr4-22115934-DEL-161 |
| chr4-22163242-DEL-60 | chr4-22163243-DEL-60 |
| chr4-22209802-DEL-268 | chr4-22209803-DEL-268 |
| chr4-22235958-DEL-56 | chr4-22235959-DEL-56 |
| chr4-22250553-DEL-60 | chr4-22250554-DEL-58 |
| chr4-22251827-DEL-74 | chr4-22251828-DEL-77 |
| chr4-22372623-DEL-271 | chr4-22372615-DEL-278 |
| chr4-22424130-DEL-218 | chr4-22424131-DEL-218 |
| chr4-22439591-DEL-618 | chr4-22439582-DEL-619 |
| chr4-22442076-DEL-237 | chr4-22442077-DEL-237 |
| chr4-22450382-DEL-358 | chr4-22450383-DEL-358 |
| chr4-22453250-DEL-441 | chr4-22453251-DEL-441 |
| chr4-22463230-DEL-5939 | chr4-22463231-DEL-5939 |
| chr4-22518725-DEL-837 | chr4-22518726-DEL-837 |
| chr4-22535850-DEL-6489 | chr4-22535851-DEL-6489 |
| chr4-22548660-DEL-226 | chr4-22548661-DEL-226 |
| chr4-22549485-DEL-143 | chr4-22549486-DEL-143 |
| chr4-22554016-DEL-393 | chr4-22554037-DEL-374 |
| chr4-22603273-DEL-138 | chr4-22603274-DEL-138 |
| chr4-22618231-DEL-443 | chr4-22618232-DEL-443 |
| chr4-22619408-DEL-87 | chr4-22619419-DEL-80 |
| chr4-22625063-DEL-529 | chr4-22625064-DEL-529 |
| chr4-22628066-DEL-1831 | chr4-22628067-DEL-1831 |
| chr4-22646198-DEL-93 | chr4-22646199-DEL-93 |
| chr4-22679205-DEL-498 | chr4-22679206-DEL-498 |
| chr4-22705330-DEL-928 | chr4-22705329-DEL-929 |
| chr4-22723149-DEL-472 | chr4-22723150-DEL-472 |
| chr4-22739084-DEL-2356 | chr4-22739084-DEL-2356 |
| chr4-22761205-DEL-10564 | chr4-22761206-DEL-10564 |
| chr4-22763855-DEL-114 | chr4-22763856-DEL-114 |
| chr4-22775289-DEL-52 | chr4-22775298-DEL-50 |
| chr4-22781361-DEL-401 | chr4-22781362-DEL-401 |
| chr4-22808157-DEL-6199 | chr4-22808158-DEL-6199 |
| chr4-22824148-DEL-191 | chr4-22824149-DEL-191 |

| | |
|---|---|
| chr4-22840652-DEL-57 | chr4-22840652-DEL-57 |
| chr4-22849677-DEL-173 | chr4-22849678-DEL-173 |
| chr4-22851450-DEL-3532 | chr4-22851451-DEL-3532 |
| chr4-22899323-DEL-64 | chr4-22899335-DEL-59 |
| chr4-22900985-DEL-7955 | chr4-22900986-DEL-7955 |
| chr4-22909714-DEL-297 | chr4-22909715-DEL-297 |
| chr4-22922110-DEL-52 | chr4-22922113-DEL-50 |
| chr4-22963476-DEL-58 | chr4-22963447-DEL-90 |
| chr4-23008753-DEL-56 | chr4-23008739-DEL-50 |
| chr4-23021197-DEL-232 | chr4-23021198-DEL-232 |
| chr4-23034365-DEL-92 | chr4-23034366-DEL-92 |
| chr4-23060277-DEL-1177 | chr4-23060278-DEL-1177 |
| chr4-23071519-DEL-1511 | chr4-23071520-DEL-1511 |
| chr4-23079892-DEL-4534 | chr4-23079893-DEL-4534 |
| chr4-23100613-DEL-423 | chr4-23100576-DEL-424 |
| chr4-23140705-DEL-509 | chr4-23140705-DEL-509 |
| chr4-23180946-DEL-57 | chr4-23180947-DEL-57 |
| chr4-23195117-DEL-15256 | chr4-23195118-DEL-15256 |
| chr4-23213929-DEL-2790 | chr4-23213930-DEL-2790 |
| chr4-23374603-DEL-2151 | chr4-23374604-DEL-2151 |
| chr4-23399901-DEL-132 | chr4-23399934-DEL-89 |
| chr4-23421293-DEL-2712 | chr4-23421294-DEL-2712 |
| chr4-23438540-DEL-2914 | chr4-23438536-DEL-2916 |
| chr4-23451787-DEL-565 | chr4-23451787-DEL-565 |
| chr4-23469549-DEL-183 | chr4-23469550-DEL-183 |
| chr4-23543181-DEL-6206 | chr4-23543166-DEL-6207 |
| chr4-23585151-DEL-358 | chr4-23585152-DEL-358 |
| chr4-23604680-DEL-811 | chr4-23604681-DEL-811 |
| chr4-23616663-DEL-107 | chr4-23616672-DEL-83 |
| chr4-23646394-DEL-7091 | chr4-23646395-DEL-7091 |
| chr4-23652522-DEL-212 | chr4-23652523-DEL-212 |
| chr4-23656960-DEL-670 | chr4-23656961-DEL-670 |
| chr4-23675162-DEL-455 | chr4-23675163-DEL-455 |
| chr4-23675645-DEL-85 | chr4-23675659-DEL-50 |
| chr4-23717480-DEL-652 | chr4-23717481-DEL-652 |
| chr4-23749611-DEL-102 | chr4-23749624-DEL-99 |
| chr4-23793378-DEL-66 | chr4-23793397-DEL-84 |
| chr4-23849562-DEL-574 | chr4-23849563-DEL-574 |
| chr4-23851110-DEL-6878 | chr4-23851111-DEL-6878 |
| chr4-23911135-DEL-148 | chr4-23911136-DEL-148 |
| chr4-23942407-DEL-6222 | chr4-23942408-DEL-6222 |
| chr4-23956383-DEL-1845 | chr4-23956384-DEL-1845 |
| chr4-23995257-DEL-4595 | chr4-23995258-DEL-4595 |
| chr4-23995728-DEL-791 | chr4-23995729-DEL-791 |
| chr4-24000781-DEL-559 | chr4-24000782-DEL-558 |
| chr4-24011349-DEL-72 | chr4-24011349-DEL-72 |
| chr4-24021941-DEL-100 | chr4-24021942-DEL-100 |
| chr4-24041264-DEL-136 | chr4-24041261-DEL-140 |
| chr4-24054050-DEL-1017 | chr4-24054051-DEL-1017 |
| chr4-24084326-DEL-6394 | chr4-24084327-DEL-6394 |
| chr4-24105341-DEL-73 | chr4-24105340-DEL-73 |
| chr4-24116954-DEL-5982 | chr4-24116955-DEL-5982 |
| chr4-24193687-DEL-7536 | chr4-24193688-DEL-7536 |
| chr4-24324093-DEL-553 | chr4-24324094-DEL-553 |
| chr4-24331745-DEL-392 | chr4-24331746-DEL-392 |
| chr4-24334254-DEL-191 | chr4-24334255-DEL-191 |
| chr4-24360396-DEL-191 | chr4-24360397-DEL-189 |
| chr4-24362480-DEL-779 | chr4-24362481-DEL-779 |
| chr4-24392525-DEL-498 | chr4-24392526-DEL-498 |
| chr4-24416111-DEL-80 | chr4-24416115-DEL-73 |
| chr4-24417517-DEL-63 | chr4-24417518-DEL-63 |
| chr4-24459668-DEL-2331 | chr4-24459653-DEL-2300 |
| chr4-24469695-DEL-1309 | chr4-24469696-DEL-1309 |
| chr4-24472258-DEL-6973 | chr4-24472259-DEL-6973 |
| chr4-24490053-DEL-71 | chr4-24490054-DEL-71 |
| chr4-24491797-DEL-160 | chr4-24491795-DEL-162 |
| chr4-24493976-DEL-1313 | chr4-24493983-DEL-1311 |
| chr4-24499410-DEL-321 | chr4-24499411-DEL-321 |
| chr4-24500676-DEL-75 | chr4-24500677-DEL-75 |
| chr4-24503267-DEL-99 | chr4-24503268-DEL-99 |
| chr4-24619581-DEL-161 | chr4-24619582-DEL-161 |
| chr4-24638487-DEL-63 | chr4-24638487-DEL-64 |
| chr4-24643501-DEL-2700 | chr4-24643502-DEL-2700 |
| chr4-24652755-DEL-86 | chr4-24652756-DEL-86 |
| chr4-24655179-DEL-1188 | chr4-24655204-DEL-1172 |
| chr4-24657191-DEL-704 | chr4-24657192-DEL-704 |
| chr4-24666964-DEL-1176 | chr4-24666965-DEL-1176 |
| chr4-24722717-DEL-61 | chr4-24722718-DEL-61 |
| chr4-24730235-DEL-393 | chr4-24730229-DEL-392 |
| chr4-24731964-DEL-1183 | chr4-24731965-DEL-1178 |

| | |
|---|---|
| chr4-24740036-DEL-526 | chr4-24740030-DEL-532 |
| chr4-24741101-DEL-108 | chr4-24741102-DEL-108 |
| chr4-24772457-DEL-4166 | chr4-24772458-DEL-4166 |
| chr4-24809019-DEL-62 | chr4-24809019-DEL-60 |
| chr4-24844165-DEL-920 | chr4-24844166-DEL-920 |
| chr4-24904810-DEL-144 | chr4-24904811-DEL-144 |
| chr4-24911058-DEL-582 | chr4-24911059-DEL-582 |
| chr4-24922410-DEL-55 | chr4-24922407-DEL-56 |
| chr4-24934867-DEL-56 | chr4-24934868-DEL-56 |
| chr4-24939615-DEL-439 | chr4-24939616-DEL-439 |
| chr4-24951935-DEL-1024 | chr4-24951935-DEL-1024 |
| chr4-24960555-DEL-890 | chr4-24960555-DEL-866 |
| chr4-24963953-DEL-67 | chr4-24963956-DEL-68 |
| chr4-25026450-DEL-140 | chr4-25026450-DEL-140 |
| chr4-25063772-DEL-66 | chr4-25063773-DEL-60 |
| chr4-25159422-DEL-6222 | chr4-25159423-DEL-6222 |
| chr4-25182138-DEL-7229 | chr4-25182139-DEL-7229 |
| chr4-25188365-DEL-424 | chr4-25188366-DEL-424 |
| chr4-25212079-DEL-79 | chr4-25212080-DEL-79 |
| chr4-25225918-DEL-116 | chr4-25225927-DEL-128 |
| chr4-25234790-DEL-258 | chr4-25234791-DEL-258 |
| chr4-25262682-DEL-878 | chr4-25262683-DEL-878 |
| chr4-25274723-DEL-64 | chr4-25274724-DEL-64 |
| chr4-25286821-DEL-13883 | chr4-25286822-DEL-13883 |
| chr4-25287705-DEL-427 | chr4-25287717-DEL-425 |
| chr4-25293792-DEL-391 | chr4-25293793-DEL-391 |
| chr4-25304353-DEL-321 | chr4-25304359-DEL-301 |
| chr4-25306638-DEL-96 | chr4-25306645-DEL-90 |
| chr4-25322283-DEL-86 | chr4-25322274-DEL-113 |
| chr4-25323362-DEL-938 | chr4-25323363-DEL-938 |
| chr4-25330898-DEL-103 | chr4-25330899-DEL-103 |
| chr4-25356043-DEL-159 | chr4-25356043-DEL-156 |
| chr4-25357567-DEL-6171 | chr4-25357568-DEL-6171 |
| chr4-25372404-DEL-156 | chr4-25372418-DEL-158 |
| chr4-25374607-DEL-4001 | chr4-25374608-DEL-4001 |
| chr4-25387902-DEL-335 | chr4-25387903-DEL-335 |
| chr4-25394119-DEL-1936 | chr4-25394119-DEL-1937 |
| chr4-25396306-DEL-193 | chr4-25396307-DEL-193 |
| chr4-25400042-DEL-315 | chr4-25400032-DEL-315 |
| chr4-25431853-DEL-503 | chr4-25431854-DEL-503 |
| chr4-25438126-DEL-86 | chr4-25438127-DEL-86 |
| chr4-25439066-DEL-1321 | chr4-25439068-DEL-1319 |
| chr4-25493039-DEL-16390 | chr4-25493040-DEL-16390 |
| chr4-25494804-DEL-2195 | chr4-25494804-DEL-2195 |
| chr4-25581740-DEL-75 | chr4-25581741-DEL-75 |
| chr4-25604452-DEL-442 | chr4-25604453-DEL-442 |
| chr4-25631784-DEL-113 | chr4-25631785-DEL-113 |
| chr4-25651118-DEL-54 | chr4-25651119-DEL-54 |
| chr4-25651376-DEL-1832 | chr4-25651377-DEL-1832 |
| chr4-25658057-DEL-72 | chr4-25658058-DEL-72 |
| chr4-25670444-DEL-1269 | chr4-25670445-DEL-1269 |
| chr4-25671092-DEL-441 | chr4-25671093-DEL-441 |
| chr4-25701418-DEL-1238 | chr4-25701419-DEL-1238 |
| chr4-25720070-DEL-71 | chr4-25720071-DEL-50 |
| chr4-25727671-DEL-7234 | chr4-25727672-DEL-7234 |
| chr4-25790688-DEL-813 | chr4-25790689-DEL-813 |
| chr4-25861032-DEL-504 | chr4-25861032-DEL-504 |
| chr4-25866960-DEL-924 | chr4-25866961-DEL-925 |
| chr4-25879529-DEL-87 | chr4-25879530-DEL-87 |
| chr4-25888335-DEL-463 | chr4-25888336-DEL-463 |
| chr4-25889407-DEL-65 | chr4-25889406-DEL-66 |
| chr4-25906361-DEL-54 | chr4-25906362-DEL-54 |
| chr4-25918801-DEL-124 | chr4-25918802-DEL-124 |
| chr4-25920276-DEL-53 | chr4-25920276-DEL-53 |
| chr4-25921815-DEL-3809 | chr4-25921816-DEL-3809 |
| chr4-25941149-DEL-53 | chr4-25941115-DEL-52 |
| chr4-25966695-DEL-227 | chr4-25966696-DEL-227 |
| chr4-26134790-DEL-344 | chr4-26134791-DEL-344 |
| chr4-26138224-DEL-6712 | chr4-26138225-DEL-6712 |
| chr4-26146903-DEL-1525 | chr4-26146904-DEL-1525 |
| chr4-26160502-DEL-86 | chr4-26160503-DEL-86 |
| chr4-26182621-DEL-201 | chr4-26182622-DEL-201 |
| chr4-26236488-DEL-134 | chr4-26236489-DEL-134 |
| chr4-26244817-DEL-1383 | chr4-26244817-DEL-1383 |
| chr4-26268351-DEL-190 | chr4-26268324-DEL-218 |
| chr4-26277795-DEL-234 | chr4-26277796-DEL-234 |
| chr4-26294231-DEL-1031 | chr4-26294243-DEL-1019 |
| chr4-26295724-DEL-4177 | chr4-26295725-DEL-4176 |
| chr4-26303313-DEL-277 | chr4-26303314-DEL-276 |
| chr4-26314815-DEL-2126 | chr4-26314816-DEL-2124 |

| | |
|---|---|
| chr4-26330382-DEL-629 | chr4-26330383-DEL-629 |
| chr4-26332079-DEL-54 | chr4-26332082-DEL-56 |
| chr4-26350133-DEL-157 | chr4-26350134-DEL-157 |
| chr4-26368381-DEL-1895 | chr4-26368382-DEL-1895 |
| chr4-26407881-DEL-434 | chr4-26407881-DEL-434 |
| chr4-26416496-DEL-53 | chr4-26416497-DEL-53 |
| chr4-26438700-DEL-862 | chr4-26438701-DEL-862 |
| chr4-26441064-DEL-65 | chr4-26441064-DEL-65 |
| chr4-26448636-DEL-532 | chr4-26448636-DEL-532 |
| chr4-26457718-DEL-885 | chr4-26457719-DEL-881 |
| chr4-26525522-DEL-1753 | chr4-26525523-DEL-1753 |
| chr4-26538186-DEL-70 | chr4-26538186-DEL-70 |
| chr4-26562229-DEL-6551 | chr4-26562230-DEL-6551 |
| chr4-26576671-DEL-732 | chr4-26576671-DEL-732 |
| chr4-26587758-DEL-870 | chr4-26587759-DEL-870 |
| chr4-26601121-DEL-60 | chr4-26601122-DEL-60 |
| chr4-26608253-DEL-123 | chr4-26608254-DEL-123 |
| chr4-26611021-DEL-204 | chr4-26611022-DEL-204 |
| chr4-26646356-DEL-270 | chr4-26646357-DEL-270 |
| chr4-26656436-DEL-7209 | chr4-26656437-DEL-7209 |
| chr4-26701779-DEL-65 | chr4-26701770-DEL-62 |
| chr4-26702026-DEL-924 | chr4-26702027-DEL-924 |
| chr4-26752784-DEL-106 | chr4-26752785-DEL-106 |
| chr4-26794462-DEL-88 | chr4-26794463-DEL-88 |
| chr4-26806906-DEL-1064 | chr4-26806907-DEL-1064 |
| chr4-26839020-DEL-65 | chr4-26839021-DEL-65 |
| chr4-26863805-DEL-637 | chr4-26863806-DEL-637 |
| chr4-26873498-DEL-72 | chr4-26873499-DEL-72 |
| chr4-26899121-DEL-428 | chr4-26899122-DEL-428 |
| chr4-26901978-DEL-1019 | chr4-26901979-DEL-1019 |
| chr4-26905670-DEL-1580 | chr4-26905671-DEL-1580 |
| chr4-26954844-DEL-839 | chr4-26954845-DEL-839 |
| chr4-26962139-DEL-78 | chr4-26962140-DEL-78 |
| chr4-27007431-DEL-6225 | chr4-27007423-DEL-6231 |
| chr4-27018447-DEL-1137 | chr4-27018448-DEL-1137 |
| chr4-27028799-DEL-60 | chr4-27028798-DEL-61 |
| chr4-27096101-DEL-89 | chr4-27096090-DEL-108 |
| chr4-27132767-DEL-826 | chr4-27132768-DEL-826 |
| chr4-27134365-DEL-52 | chr4-27134366-DEL-52 |
| chr4-27184685-DEL-56 | chr4-27184686-DEL-56 |
| chr4-27254038-DEL-195 | chr4-27254039-DEL-195 |
| chr4-27262802-DEL-74 | chr4-27262805-DEL-77 |
| chr4-27267779-DEL-173 | chr4-27267773-DEL-173 |
| chr4-27270882-DEL-389 | chr4-27270883-DEL-389 |
| chr4-27290330-DEL-1012 | chr4-27290331-DEL-1012 |
| chr4-27303120-DEL-56 | chr4-27303121-DEL-56 |
| chr4-27329097-DEL-1871 | chr4-27329098-DEL-1871 |
| chr4-27353733-DEL-54 | chr4-27353734-DEL-54 |
| chr4-27354280-DEL-56 | chr4-27354281-DEL-56 |
| chr4-27368365-DEL-2708 | chr4-27368363-DEL-2711 |
| chr4-27378523-DEL-51 | chr4-27378524-DEL-51 |
| chr4-27381238-DEL-344 | chr4-27381238-DEL-342 |
| chr4-27410044-DEL-7590 | chr4-27410044-DEL-7590 |
| chr4-27475096-DEL-59 | chr4-27475097-DEL-59 |
| chr4-27511285-DEL-203 | chr4-27511286-DEL-203 |
| chr4-27514467-DEL-83 | chr4-27514468-DEL-83 |
| chr4-27553748-DEL-474 | chr4-27553749-DEL-474 |
| chr4-27556093-DEL-793 | chr4-27556094-DEL-793 |
| chr4-27567644-DEL-68 | chr4-27567650-DEL-67 |
| chr4-27575055-DEL-62 | chr4-27575056-DEL-62 |
| chr4-27576157-DEL-69 | chr4-27576158-DEL-69 |
| chr4-27586356-DEL-61 | chr4-27586357-DEL-61 |
| chr4-27587608-DEL-146 | chr4-27587636-DEL-127 |
| chr4-27587742-DEL-1631 | chr4-27587743-DEL-1631 |
| chr4-27593202-DEL-757 | chr4-27593203-DEL-757 |
| chr4-27604229-DEL-237 | chr4-27604227-DEL-238 |
| chr4-27611924-DEL-95 | chr4-27611926-DEL-94 |
| chr4-27640412-DEL-72 | chr4-27640413-DEL-72 |
| chr4-27700655-DEL-3738 | chr4-27700656-DEL-3738 |
| chr4-27729927-DEL-1890 | chr4-27729928-DEL-1890 |
| chr4-27741749-DEL-400 | chr4-27741750-DEL-400 |
| chr4-27759175-DEL-199 | chr4-27759174-DEL-200 |
| chr4-27770085-DEL-656 | chr4-27770086-DEL-656 |
| chr4-27788531-DEL-72 | chr4-27788532-DEL-72 |
| chr4-27841636-DEL-51 | chr4-27841637-DEL-51 |
| chr4-27857759-DEL-425 | chr4-27857759-DEL-425 |
| chr4-27861101-DEL-4432 | chr4-27861102-DEL-4432 |
| chr4-27888075-DEL-6449 | chr4-27888069-DEL-6450 |
| chr4-27895144-DEL-101 | chr4-27895145-DEL-101 |
| chr4-27917791-DEL-6182 | chr4-27917792-DEL-6182 |

| | |
|---|---|
| chr4-27934651-DEL-1213 | chr4-27934652-DEL-1213 |
| chr4-27979063-DEL-54 | chr4-27979064-DEL-54 |
| chr4-27988844-DEL-66 | chr4-27988845-DEL-50 |
| chr4-28022484-DEL-3789 | chr4-28022484-DEL-3789 |
| chr4-28048886-DEL-113 | chr4-28048887-DEL-113 |
| chr4-28054835-DEL-146 | chr4-28054836-DEL-146 |
| chr4-28079822-DEL-60 | chr4-28079798-DEL-84 |
| chr4-28080303-DEL-2174 | chr4-28080304-DEL-2174 |
| chr4-28100977-DEL-80 | chr4-28100978-DEL-80 |
| chr4-28107701-DEL-183 | chr4-28107702-DEL-183 |
| chr4-28182847-DEL-55 | chr4-28182847-DEL-56 |
| chr4-28186855-DEL-54 | chr4-28186855-DEL-54 |
| chr4-28194495-DEL-806 | chr4-28194496-DEL-806 |
| chr4-28207558-DEL-54 | chr4-28207571-DEL-54 |
| chr4-28235721-DEL-1296 | chr4-28235722-DEL-1296 |
| chr4-28235733-DEL-127 | chr4-28235734-DEL-127 |
| chr4-28280971-DEL-62 | chr4-28280969-DEL-63 |
| chr4-28336301-DEL-4156 | chr4-28336284-DEL-4155 |
| chr4-28356052-DEL-100 | chr4-28356050-DEL-54 |
| chr4-28356525-DEL-253 | chr4-28356526-DEL-253 |
| chr4-28360867-DEL-138 | chr4-28360868-DEL-138 |
| chr4-28373503-DEL-1096 | chr4-28373503-DEL-1096 |
| chr4-28412103-DEL-7051 | chr4-28412143-DEL-7012 |
| chr4-28454342-DEL-79 | chr4-28454342-DEL-72 |
| chr4-28457770-DEL-564 | chr4-28457771-DEL-564 |
| chr4-28461491-DEL-6409 | chr4-28461491-DEL-6409 |
| chr4-28496424-DEL-60 | chr4-28496425-DEL-60 |
| chr4-28515989-DEL-214 | chr4-28515990-DEL-214 |
| chr4-28587178-DEL-1235 | chr4-28587179-DEL-1235 |
| chr4-28599798-DEL-1456 | chr4-28599799-DEL-1456 |
| chr4-28605737-DEL-1664 | chr4-28605738-DEL-1664 |
| chr4-28614207-DEL-12749 | chr4-28614219-DEL-12750 |
| chr4-28637876-DEL-84 | chr4-28637876-DEL-84 |
| chr4-28647393-DEL-74 | chr4-28647394-DEL-74 |
| chr4-28653904-DEL-931 | chr4-28653909-DEL-930 |
| chr4-28657170-DEL-538 | chr4-28657171-DEL-538 |
| chr4-28660586-DEL-425 | chr4-28660586-DEL-425 |
| chr4-28661837-DEL-6263 | chr4-28661837-DEL-6263 |
| chr4-28710168-DEL-72 | chr4-28710169-DEL-72 |
| chr4-28710495-DEL-448 | chr4-28710496-DEL-448 |
| chr4-28723821-DEL-61 | chr4-28723822-DEL-61 |
| chr4-28734431-DEL-83 | chr4-28734432-DEL-83 |
| chr4-28740700-DEL-3780 | chr4-28740701-DEL-3780 |
| chr4-28746628-DEL-6312 | chr4-28746628-DEL-6312 |
| chr4-28754737-DEL-114 | chr4-28754737-DEL-114 |
| chr4-28800492-DEL-62 | chr4-28800493-DEL-61 |
| chr4-28823026-DEL-53 | chr4-28823027-DEL-53 |
| chr4-28930764-DEL-144 | chr4-28930765-DEL-144 |
| chr4-29012781-DEL-2934 | chr4-29012782-DEL-2934 |
| chr4-29049445-DEL-6243 | chr4-29049442-DEL-6244 |
| chr4-29057288-DEL-101 | chr4-29057309-DEL-52 |
| chr4-29083496-DEL-92 | chr4-29083497-DEL-92 |
| chr4-29089009-DEL-1247 | chr4-29089010-DEL-1247 |
| chr4-29130253-DEL-191 | chr4-29130253-DEL-191 |
| chr4-29152913-DEL-84 | chr4-29152879-DEL-102 |
| chr4-29214952-DEL-2192 | chr4-29214951-DEL-2193 |
| chr4-29240888-DEL-4183 | chr4-29240889-DEL-4183 |
| chr4-29246915-DEL-6587 | chr4-29246916-DEL-6586 |
| chr4-29365133-DEL-1026 | chr4-29365134-DEL-1026 |
| chr4-29378569-DEL-61 | chr4-29378535-DEL-102 |
| chr4-29383231-DEL-59 | chr4-29383227-DEL-58 |
| chr4-29406859-DEL-516 | chr4-29406860-DEL-514 |
| chr4-29409232-DEL-73 | chr4-29409246-DEL-61 |
| chr4-29412923-DEL-87 | chr4-29412924-DEL-87 |
| chr4-29440882-DEL-2361 | chr4-29440888-DEL-2357 |
| chr4-29446477-DEL-237 | chr4-29446478-DEL-237 |
| chr4-29458704-DEL-413 | chr4-29458686-DEL-423 |
| chr4-29489756-DEL-92 | chr4-29489751-DEL-106 |
| chr4-29505250-DEL-707 | chr4-29505238-DEL-717 |
| chr4-29514516-DEL-1513 | chr4-29514517-DEL-1513 |
| chr4-29536123-DEL-704 | chr4-29536124-DEL-704 |
| chr4-29567297-DEL-391 | chr4-29567298-DEL-391 |
| chr4-29574668-DEL-121 | chr4-29574669-DEL-121 |
| chr4-29577061-DEL-51 | chr4-29577098-DEL-63 |
| chr4-29587812-DEL-67 | chr4-29587813-DEL-66 |
| chr4-29592796-DEL-642 | chr4-29592797-DEL-642 |
| chr4-29619394-DEL-1043 | chr4-29619395-DEL-1043 |
| chr4-29623922-DEL-805 | chr4-29623923-DEL-805 |
| chr4-29640693-DEL-962 | chr4-29640694-DEL-962 |
| chr4-29649392-DEL-198 | chr4-29649393-DEL-198 |

| | |
|---|---|
| chr4-29659256-DEL-463 | chr4-29659255-DEL-464 |
| chr4-29701981-DEL-4297 | chr4-29701982-DEL-4297 |
| chr4-29710983-DEL-6830 | chr4-29710984-DEL-6830 |
| chr4-29721455-DEL-64 | chr4-29721456-DEL-64 |
| chr4-29723983-DEL-68 | chr4-29723998-DEL-54 |
| chr4-29743209-DEL-56 | chr4-29743208-DEL-56 |
| chr4-29775880-DEL-63 | chr4-29775890-DEL-54 |
| chr4-29788569-DEL-102 | chr4-29788570-DEL-106 |
| chr4-29815376-DEL-797 | chr4-29815377-DEL-797 |
| chr4-29817819-DEL-106 | chr4-29817838-DEL-58 |
| chr4-29871797-DEL-221 | chr4-29871798-DEL-221 |
| chr4-29905633-DEL-267 | chr4-29905632-DEL-267 |
| chr4-29936157-DEL-237 | chr4-29936158-DEL-237 |
| chr4-29936980-DEL-613 | chr4-29936981-DEL-613 |
| chr4-29938759-DEL-236 | chr4-29938764-DEL-234 |
| chr4-29991636-DEL-532 | chr4-29991637-DEL-532 |
| chr4-30030348-DEL-56 | chr4-30030349-DEL-62 |
| chr4-30034612-DEL-54 | chr4-30034592-DEL-74 |
| chr4-30079803-DEL-727 | chr4-30079804-DEL-727 |
| chr4-30151278-DEL-55 | chr4-30151279-DEL-55 |
| chr4-30183307-DEL-5679 | chr4-30183307-DEL-5679 |
| chr4-30228547-DEL-7425 | chr4-30228573-DEL-7383 |
| chr4-30257640-DEL-677 | chr4-30257640-DEL-677 |
| chr4-30296213-DEL-111 | chr4-30296219-DEL-110 |
| chr4-30326838-DEL-6833 | chr4-30326839-DEL-6833 |
| chr4-30335101-DEL-957 | chr4-30335102-DEL-956 |
| chr4-30350396-DEL-5843 | chr4-30350397-DEL-5843 |
| chr4-30350444-DEL-208 | chr4-30350445-DEL-208 |
| chr4-30357638-DEL-114 | chr4-30357638-DEL-111 |
| chr4-30359854-DEL-2128 | chr4-30359854-DEL-2128 |
| chr4-30394575-DEL-301 | chr4-30394576-DEL-301 |
| chr4-30435942-DEL-520 | chr4-30435943-DEL-520 |
| chr4-30487407-DEL-197 | chr4-30487408-DEL-197 |
| chr4-30516809-DEL-2162 | chr4-30516810-DEL-2162 |
| chr4-30525590-DEL-430 | chr4-30525591-DEL-430 |
| chr4-30544387-DEL-58 | chr4-30544360-DEL-66 |
| chr4-30549260-DEL-1192 | chr4-30549261-DEL-1192 |
| chr4-30575860-DEL-3117 | chr4-30575861-DEL-3117 |
| chr4-30623274-DEL-2765 | chr4-30623275-DEL-2765 |
| chr4-30634942-DEL-55 | chr4-30634938-DEL-83 |
| chr4-30639827-DEL-1584 | chr4-30639817-DEL-1592 |
| chr4-30644759-DEL-6341 | chr4-30644760-DEL-6340 |
| chr4-30657169-DEL-159 | chr4-30657170-DEL-159 |
| chr4-30670202-DEL-1295 | chr4-30670203-DEL-1295 |
| chr4-30718831-DEL-168 | chr4-30718832-DEL-168 |
| chr4-30720017-DEL-108 | chr4-30720021-DEL-93 |
| chr4-30720681-DEL-1210 | chr4-30720682-DEL-1208 |
| chr4-30726441-DEL-135 | chr4-30726441-DEL-135 |
| chr4-30733083-DEL-126 | chr4-30733084-DEL-126 |
| chr4-30737959-DEL-1891 | chr4-30737959-DEL-1891 |
| chr4-30765541-DEL-1800 | chr4-30765542-DEL-1800 |
| chr4-30768881-DEL-949 | chr4-30768882-DEL-949 |
| chr4-30772264-DEL-135 | chr4-30772261-DEL-139 |
| chr4-30822292-DEL-606 | chr4-30822293-DEL-606 |
| chr4-30852195-DEL-113 | chr4-30852196-DEL-113 |
| chr4-30854208-DEL-148 | chr4-30854207-DEL-150 |
| chr4-30859217-DEL-65 | chr4-30859218-DEL-65 |
| chr4-30867193-DEL-94 | chr4-30867194-DEL-94 |
| chr4-30905872-DEL-3626 | chr4-30905873-DEL-3626 |
| chr4-30923663-DEL-6584 | chr4-30923664-DEL-6584 |
| chr4-30937985-DEL-82 | chr4-30937986-DEL-82 |
| chr4-30946191-DEL-155 | chr4-30946192-DEL-152 |
| chr4-30989936-DEL-8701 | chr4-30989937-DEL-8701 |
| chr4-30999407-DEL-81 | chr4-30999408-DEL-81 |
| chr4-31061996-DEL-54 | chr4-31061957-DEL-72 |
| chr4-31099712-DEL-567 | chr4-31099713-DEL-565 |
| chr4-31138797-DEL-187 | chr4-31138807-DEL-190 |
| chr4-31144030-DEL-98 | chr4-31144031-DEL-98 |
| chr4-31149515-DEL-54 | chr4-31149516-DEL-54 |
| chr4-31158646-DEL-247 | chr4-31158647-DEL-247 |
| chr4-31191127-DEL-400 | chr4-31191124-DEL-401 |
| chr4-31199769-DEL-491 | chr4-31199770-DEL-491 |
| chr4-31236952-DEL-65 | chr4-31236953-DEL-65 |
| chr4-31244805-DEL-1055 | chr4-31244806-DEL-1055 |
| chr4-31253948-DEL-72 | chr4-31253948-DEL-73 |
| chr4-31269041-DEL-778 | chr4-31269042-DEL-778 |
| chr4-31306987-DEL-604 | chr4-31306988-DEL-604 |
| chr4-31309234-DEL-614 | chr4-31309235-DEL-614 |
| chr4-31320255-DEL-91 | chr4-31320256-DEL-91 |
| chr4-31354253-DEL-686 | chr4-31354294-DEL-646 |

| | |
|---|---|
| chr4-31374639-DEL-486 | chr4-31374640-DEL-486 |
| chr4-31394667-DEL-431 | chr4-31394668-DEL-431 |
| chr4-31467148-DEL-612 | chr4-31467123-DEL-657 |
| chr4-31495463-DEL-1307 | chr4-31495464-DEL-1308 |
| chr4-31527771-DEL-6568 | chr4-31527772-DEL-6568 |
| chr4-31534723-DEL-1667 | chr4-31534724-DEL-1667 |
| chr4-31542385-DEL-58 | chr4-31542386-DEL-58 |
| chr4-31567601-DEL-88 | chr4-31567602-DEL-88 |
| chr4-31571269-DEL-1367 | chr4-31571270-DEL-1367 |
| chr4-31577906-DEL-350 | chr4-31577917-DEL-338 |
| chr4-31616143-DEL-52 | chr4-31616139-DEL-56 |
| chr4-31632370-DEL-4355 | chr4-31632371-DEL-4355 |
| chr4-31691008-DEL-273 | chr4-31691009-DEL-273 |
| chr4-31697654-DEL-118 | chr4-31697679-DEL-72 |
| chr4-31759939-DEL-57 | chr4-31759938-DEL-55 |
| chr4-31775667-DEL-9082 | chr4-31775668-DEL-9082 |
| chr4-31827942-DEL-139 | chr4-31827943-DEL-139 |
| chr4-31833289-DEL-98 | chr4-31833318-DEL-70 |
| chr4-31885602-DEL-343 | chr4-31885603-DEL-343 |
| chr4-31892070-DEL-6675 | chr4-31892071-DEL-6675 |
| chr4-31932393-DEL-55 | chr4-31932398-DEL-51 |
| chr4-31950246-DEL-2996 | chr4-31950247-DEL-2996 |
| chr4-31953988-DEL-105 | chr4-31953983-DEL-101 |
| chr4-31958961-DEL-938 | chr4-31958962-DEL-938 |
| chr4-31961733-DEL-104 | chr4-31961729-DEL-100 |
| chr4-31970895-DEL-998 | chr4-31970896-DEL-998 |
| chr4-31983449-DEL-201 | chr4-31983450-DEL-201 |
| chr4-31983876-DEL-1271 | chr4-31983877-DEL-1269 |
| chr4-31991119-DEL-116 | chr4-31991120-DEL-116 |
| chr4-32029663-DEL-7247 | chr4-32029660-DEL-7248 |
| chr4-32043399-DEL-1367 | chr4-32043400-DEL-1367 |
| chr4-32043975-DEL-440 | chr4-32043975-DEL-456 |
| chr4-32046507-DEL-2931 | chr4-32046508-DEL-2931 |
| chr4-32063032-DEL-62 | chr4-32063033-DEL-62 |
| chr4-32064490-DEL-54 | chr4-32064491-DEL-54 |
| chr4-32070470-DEL-56 | chr4-32070471-DEL-56 |
| chr4-32071313-DEL-192 | chr4-32071314-DEL-192 |
| chr4-32072150-DEL-519 | chr4-32072151-DEL-519 |
| chr4-32076038-DEL-113 | chr4-32076039-DEL-113 |
| chr4-32091719-DEL-193 | chr4-32091720-DEL-193 |
| chr4-32116646-DEL-204 | chr4-32116647-DEL-204 |
| chr4-32120139-DEL-51 | chr4-32120098-DEL-60 |
| chr4-32132116-DEL-207 | chr4-32132117-DEL-207 |
| chr4-32134917-DEL-65 | chr4-32134918-DEL-65 |
| chr4-32143534-DEL-90 | chr4-32143535-DEL-88 |
| chr4-32190007-DEL-167 | chr4-32190027-DEL-139 |
| chr4-32194494-DEL-54 | chr4-32194495-DEL-54 |
| chr4-32228442-DEL-53 | chr4-32228442-DEL-53 |
| chr4-32233776-DEL-52 | chr4-32233788-DEL-53 |
| chr4-32321471-DEL-87 | chr4-32321472-DEL-87 |
| chr4-32354538-DEL-55 | chr4-32354539-DEL-55 |
| chr4-32408313-DEL-141 | chr4-32408314-DEL-141 |
| chr4-32448459-DEL-127 | chr4-32448460-DEL-127 |
| chr4-32514403-DEL-86 | chr4-32514404-DEL-86 |
| chr4-32529285-DEL-99 | chr4-32529307-DEL-99 |
| chr4-32559350-DEL-127 | chr4-32559349-DEL-129 |
| chr4-32703966-DEL-373 | chr4-32703967-DEL-373 |
| chr4-32721308-DEL-64 | chr4-32721309-DEL-50 |
| chr4-32747156-DEL-202 | chr4-32747157-DEL-202 |
| chr4-32753427-DEL-205 | chr4-32753428-DEL-205 |
| chr4-32763998-DEL-214 | chr4-32763999-DEL-214 |
| chr4-32778589-DEL-204 | chr4-32778590-DEL-204 |
| chr4-32790350-DEL-7366 | chr4-32790351-DEL-7366 |
| chr4-32819727-DEL-74 | chr4-32819728-DEL-74 |
| chr4-32820384-DEL-84 | chr4-32820385-DEL-84 |
| chr4-32829874-DEL-1545 | chr4-32829875-DEL-1545 |
| chr4-32838913-DEL-57 | chr4-32838914-DEL-57 |
| chr4-32865886-DEL-181 | chr4-32865897-DEL-192 |
| chr4-32870590-DEL-58 | chr4-32870591-DEL-58 |
| chr4-32876511-DEL-1888 | chr4-32876512-DEL-1888 |
| chr4-32885362-DEL-202 | chr4-32885363-DEL-202 |
| chr4-32887622-DEL-204 | chr4-32887610-DEL-204 |
| chr4-32900909-DEL-195 | chr4-32900909-DEL-195 |
| chr4-32910004-DEL-937 | chr4-32910005-DEL-937 |
| chr4-32915072-DEL-1337 | chr4-32915073-DEL-1337 |
| chr4-32916581-DEL-1983 | chr4-32916582-DEL-1983 |
| chr4-32933864-DEL-113 | chr4-32933863-DEL-113 |
| chr4-32935378-DEL-121 | chr4-32935376-DEL-124 |
| chr4-32947186-DEL-269 | chr4-32947187-DEL-269 |
| chr4-32953429-DEL-51 | chr4-32953430-DEL-51 |

| | |
|---|---|
| chr4-32953884-DEL-118 | chr4-32953885-DEL-118 |
| chr4-32959317-DEL-93 | chr4-32959318-DEL-93 |
| chr4-32959854-DEL-196 | chr4-32959855-DEL-196 |
| chr4-32964758-DEL-538 | chr4-32964759-DEL-538 |
| chr4-32973231-DEL-410 | chr4-32973260-DEL-365 |
| chr4-32976649-DEL-200 | chr4-32976649-DEL-200 |
| chr4-32987287-DEL-80 | chr4-32987288-DEL-80 |
| chr4-32997072-DEL-71 | chr4-32997073-DEL-71 |
| chr4-33005996-DEL-226 | chr4-33005997-DEL-226 |
| chr4-33066665-DEL-1086 | chr4-33066666-DEL-1086 |
| chr4-33087371-DEL-232 | chr4-33087367-DEL-234 |
| chr4-33096250-DEL-122 | chr4-33096255-DEL-118 |
| chr4-33103227-DEL-1337 | chr4-33103227-DEL-1333 |
| chr4-33109196-DEL-323 | chr4-33109203-DEL-321 |
| chr4-33119456-DEL-59 | chr4-33119457-DEL-59 |
| chr4-33122973-DEL-860 | chr4-33122974-DEL-860 |
| chr4-33141417-DEL-202 | chr4-33141418-DEL-202 |
| chr4-33143449-DEL-2097 | chr4-33143450-DEL-2097 |
| chr4-33145025-DEL-99 | chr4-33145026-DEL-99 |
| chr4-33156688-DEL-122 | chr4-33156689-DEL-122 |
| chr4-33158902-DEL-432 | chr4-33158903-DEL-432 |
| chr4-33169878-DEL-205 | chr4-33169879-DEL-205 |
| chr4-33174176-DEL-60 | chr4-33174177-DEL-60 |
| chr4-33176457-DEL-92 | chr4-33176437-DEL-93 |
| chr4-33183450-DEL-160 | chr4-33183459-DEL-152 |
| chr4-33195712-DEL-76 | chr4-33195713-DEL-76 |
| chr4-33203647-DEL-151 | chr4-33203648-DEL-151 |
| chr4-33206133-DEL-58 | chr4-33206105-DEL-87 |
| chr4-33217890-DEL-1219 | chr4-33217891-DEL-1219 |
| chr4-33240308-DEL-70 | chr4-33240309-DEL-70 |
| chr4-33269870-DEL-538 | chr4-33269871-DEL-538 |
| chr4-33277790-DEL-137 | chr4-33277791-DEL-137 |
| chr4-33279753-DEL-72 | chr4-33279754-DEL-84 |
| chr4-33301668-DEL-309 | chr4-33301669-DEL-309 |
| chr4-33304795-DEL-198 | chr4-33304796-DEL-198 |
| chr4-33314384-DEL-182 | chr4-33314383-DEL-182 |
| chr4-33358491-DEL-235 | chr4-33358481-DEL-233 |
| chr4-33397093-DEL-1589 | chr4-33397094-DEL-1589 |
| chr4-33425916-DEL-915 | chr4-33425917-DEL-915 |
| chr4-33432687-DEL-63 | chr4-33432688-DEL-63 |
| chr4-33483785-DEL-60 | chr4-33483801-DEL-52 |
| chr4-33495114-DEL-134 | chr4-33495086-DEL-150 |
| chr4-33532235-DEL-60 | chr4-33532209-DEL-87 |
| chr4-33533109-DEL-69 | chr4-33533110-DEL-69 |
| chr4-33546208-DEL-3394 | chr4-33546209-DEL-3394 |
| chr4-33578403-DEL-56 | chr4-33578404-DEL-56 |
| chr4-33617683-DEL-72 | chr4-33617676-DEL-53 |
| chr4-33618705-DEL-523 | chr4-33618727-DEL-531 |
| chr4-33643449-DEL-923 | chr4-33643450-DEL-923 |
| chr4-33653481-DEL-60 | chr4-33653484-DEL-58 |
| chr4-33654990-DEL-53 | chr4-33654991-DEL-51 |
| chr4-33674499-DEL-72 | chr4-33674500-DEL-72 |
| chr4-33676375-DEL-1665 | chr4-33676359-DEL-1679 |
| chr4-33710517-DEL-80 | chr4-33710512-DEL-70 |
| chr4-33724348-DEL-2139 | chr4-33724349-DEL-2139 |
| chr4-33750272-DEL-139 | chr4-33750273-DEL-139 |
| chr4-33750817-DEL-87 | chr4-33750824-DEL-68 |
| chr4-33751869-DEL-82 | chr4-33751870-DEL-82 |
| chr4-33763088-DEL-67 | chr4-33763089-DEL-67 |
| chr4-33793823-DEL-86 | chr4-33793824-DEL-86 |
| chr4-33801166-DEL-109 | chr4-33801169-DEL-107 |
| chr4-33837287-DEL-77 | chr4-33837288-DEL-77 |
| chr4-33837721-DEL-395 | chr4-33837722-DEL-395 |
| chr4-33955166-DEL-68 | chr4-33955167-DEL-68 |
| chr4-33979516-DEL-248 | chr4-33979517-DEL-248 |
| chr4-34006887-DEL-1880 | chr4-34006888-DEL-1880 |
| chr4-34017605-DEL-156 | chr4-34017586-DEL-157 |
| chr4-34094443-DEL-99 | chr4-34094485-DEL-57 |
| chr4-34176103-DEL-56 | chr4-34176104-DEL-56 |
| chr4-34193528-DEL-2715 | chr4-34193529-DEL-2715 |
| chr4-34250147-DEL-9086 | chr4-34250148-DEL-9086 |
| chr4-34256607-DEL-1865 | chr4-34256606-DEL-1865 |
| chr4-34298492-DEL-93 | chr4-34298493-DEL-93 |
| chr4-34341153-DEL-8421 | chr4-34341154-DEL-8421 |
| chr4-34350806-DEL-4611 | chr4-34350807-DEL-4611 |
| chr4-34364175-DEL-211 | chr4-34364176-DEL-211 |
| chr4-34366090-DEL-100 | chr4-34366091-DEL-50 |
| chr4-34383740-DEL-101 | chr4-34383741-DEL-101 |
| chr4-34388442-DEL-164 | chr4-34388443-DEL-164 |
| chr4-34392053-DEL-1891 | chr4-34392054-DEL-1892 |

| | |
|---|---|
| chr4-34394526-DEL-96 | chr4-34394527-DEL-96 |
| chr4-34396003-DEL-620 | chr4-34396004-DEL-620 |
| chr4-34398982-DEL-254 | chr4-34398983-DEL-254 |
| chr4-34399859-DEL-87 | chr4-34399864-DEL-83 |
| chr4-34426179-DEL-196 | chr4-34426180-DEL-196 |
| chr4-34442299-DEL-54 | chr4-34442299-DEL-54 |
| chr4-34442986-DEL-6586 | chr4-34442987-DEL-6586 |
| chr4-34456405-DEL-64 | chr4-34456401-DEL-65 |
| chr4-34478770-DEL-309 | chr4-34478771-DEL-309 |
| chr4-34485710-DEL-803 | chr4-34485711-DEL-803 |
| chr4-34512858-DEL-6401 | chr4-34512859-DEL-6401 |
| chr4-34520033-DEL-464 | chr4-34520034-DEL-464 |
| chr4-34530636-DEL-178 | chr4-34530636-DEL-177 |
| chr4-34534858-DEL-217 | chr4-34534858-DEL-217 |
| chr4-34535139-DEL-7055 | chr4-34535140-DEL-7055 |
| chr4-34553610-DEL-352 | chr4-34553608-DEL-326 |
| chr4-34568373-DEL-52 | chr4-34568374-DEL-52 |
| chr4-34585719-DEL-115 | chr4-34585720-DEL-114 |
| chr4-34593902-DEL-297 | chr4-34593898-DEL-293 |
| chr4-34677863-DEL-225 | chr4-34677864-DEL-225 |
| chr4-34695325-DEL-842 | chr4-34695331-DEL-836 |
| chr4-34700866-DEL-114 | chr4-34700867-DEL-114 |
| chr4-34702111-DEL-175 | chr4-34702112-DEL-175 |
| chr4-34704976-DEL-1598 | chr4-34704977-DEL-1598 |
| chr4-34707028-DEL-149 | chr4-34707029-DEL-149 |
| chr4-34718830-DEL-59 | chr4-34718830-DEL-60 |
| chr4-34733076-DEL-364 | chr4-34733071-DEL-375 |
| chr4-34737241-DEL-486 | chr4-34737242-DEL-486 |
| chr4-34746307-DEL-76 | chr4-34746308-DEL-76 |
| chr4-34756774-DEL-3859 | chr4-34756775-DEL-3859 |
| chr4-34764390-DEL-213 | chr4-34764368-DEL-213 |
| chr4-34765340-DEL-67 | chr4-34765341-DEL-67 |
| chr4-34780735-DEL-216 | chr4-34780736-DEL-216 |
| chr4-34783972-DEL-578 | chr4-34783973-DEL-578 |
| chr4-34816878-DEL-145 | chr4-34816879-DEL-145 |
| chr4-34834852-DEL-62 | chr4-34834853-DEL-62 |
| chr4-34842481-DEL-92 | chr4-34842480-DEL-91 |
| chr4-34859711-DEL-95 | chr4-34859703-DEL-90 |
| chr4-34892260-DEL-179 | chr4-34892261-DEL-180 |
| chr4-34898158-DEL-65 | chr4-34898173-DEL-50 |
| chr4-34908741-DEL-519 | chr4-34908742-DEL-519 |
| chr4-34921556-DEL-53 | chr4-34921599-DEL-57 |
| chr4-34923405-DEL-970 | chr4-34923405-DEL-971 |
| chr4-34926244-DEL-561 | chr4-34926245-DEL-561 |
| chr4-34928767-DEL-93 | chr4-34928762-DEL-84 |
| chr4-34929780-DEL-96 | chr4-34929764-DEL-87 |
| chr4-34931128-DEL-180 | chr4-34931129-DEL-180 |
| chr4-34935241-DEL-1172 | chr4-34935242-DEL-1172 |
| chr4-34935758-DEL-241 | chr4-34935755-DEL-242 |
| chr4-34944837-DEL-201 | chr4-34944838-DEL-200 |
| chr4-34950693-DEL-66 | chr4-34950694-DEL-66 |
| chr4-34959046-DEL-238 | chr4-34959047-DEL-238 |
| chr4-34977531-DEL-755 | chr4-34977532-DEL-755 |
| chr4-34996946-DEL-191 | chr4-34996947-DEL-167 |
| chr4-35042614-DEL-6220 | chr4-35042615-DEL-6220 |
| chr4-35070177-DEL-7612 | chr4-35070178-DEL-7612 |
| chr4-35078994-DEL-1760 | chr4-35078995-DEL-1760 |
| chr4-35088936-DEL-57 | chr4-35088938-DEL-58 |
| chr4-35097751-DEL-60 | chr4-35097764-DEL-61 |
| chr4-35107432-DEL-88 | chr4-35107433-DEL-88 |
| chr4-35135863-DEL-89 | chr4-35135864-DEL-89 |
| chr4-35138779-DEL-771 | chr4-35138780-DEL-771 |
| chr4-35151385-DEL-96 | chr4-35151386-DEL-50 |
| chr4-35153684-DEL-79 | chr4-35153685-DEL-79 |
| chr4-35166493-DEL-222 | chr4-35166494-DEL-222 |
| chr4-35175615-DEL-87 | chr4-35175616-DEL-87 |
| chr4-35176061-DEL-227 | chr4-35176062-DEL-227 |
| chr4-35180406-DEL-6789 | chr4-35180407-DEL-6789 |
| chr4-35186804-DEL-65 | chr4-35186805-DEL-65 |
| chr4-35198503-DEL-72 | chr4-35198506-DEL-58 |
| chr4-35232069-DEL-193 | chr4-35232070-DEL-193 |
| chr4-35240057-DEL-320 | chr4-35240057-DEL-321 |
| chr4-35242505-DEL-144 | chr4-35242507-DEL-145 |
| chr4-35258230-DEL-478 | chr4-35258231-DEL-478 |
| chr4-35260807-DEL-138 | chr4-35260808-DEL-138 |
| chr4-35285125-DEL-149 | chr4-35285126-DEL-149 |
| chr4-35288044-DEL-3825 | chr4-35288045-DEL-3825 |
| chr4-35293198-DEL-341 | chr4-35293199-DEL-341 |
| chr4-35317766-DEL-223 | chr4-35317767-DEL-223 |
| chr4-35377356-DEL-7104 | chr4-35377357-DEL-7104 |

| | |
|---|---|
| chr4-35407209-DEL-275 | chr4-35407204-DEL-276 |
| chr4-35413863-DEL-170 | chr4-35413864-DEL-170 |
| chr4-35467847-DEL-206 | chr4-35467848-DEL-206 |
| chr4-35492457-DEL-384 | chr4-35492458-DEL-384 |
| chr4-35536267-DEL-6676 | chr4-35536268-DEL-6676 |
| chr4-35589475-DEL-52 | chr4-35589482-DEL-50 |
| chr4-35639731-DEL-101 | chr4-35639704-DEL-104 |
| chr4-35642517-DEL-4136 | chr4-35642518-DEL-4136 |
| chr4-35671698-DEL-191 | chr4-35671699-DEL-191 |
| chr4-35712004-DEL-331 | chr4-35712005-DEL-331 |
| chr4-35719299-DEL-300 | chr4-35719300-DEL-300 |
| chr4-35732327-DEL-652 | chr4-35732328-DEL-652 |
| chr4-35775479-DEL-6208 | chr4-35775480-DEL-6208 |
| chr4-35785280-DEL-844 | chr4-35785283-DEL-845 |
| chr4-35798981-DEL-1103 | chr4-35798982-DEL-1103 |
| chr4-35816668-DEL-51 | chr4-35816669-DEL-51 |
| chr4-35824631-DEL-1562 | chr4-35824614-DEL-1562 |
| chr4-35866775-DEL-7012 | chr4-35866776-DEL-7012 |
| chr4-35873779-DEL-59 | chr4-35873780-DEL-59 |
| chr4-35875562-DEL-87 | chr4-35875515-DEL-87 |
| chr4-35885186-DEL-148 | chr4-35885187-DEL-148 |
| chr4-35914698-DEL-1386 | chr4-35914699-DEL-1386 |
| chr4-35955806-DEL-104 | chr4-35955796-DEL-113 |
| chr4-35977317-DEL-52 | chr4-35977318-DEL-52 |
| chr4-35990694-DEL-549 | chr4-35990694-DEL-549 |
| chr4-35995049-DEL-7529 | chr4-35995053-DEL-7526 |
| chr4-36049546-DEL-823 | chr4-36049538-DEL-824 |
| chr4-36083229-DEL-68 | chr4-36083236-DEL-62 |
| chr4-36135612-DEL-121 | chr4-36135613-DEL-121 |
| chr4-36193839-DEL-67 | chr4-36193838-DEL-67 |
| chr4-36205770-DEL-78 | chr4-36205771-DEL-78 |
| chr4-36270567-DEL-6356 | chr4-36270568-DEL-6356 |
| chr4-36302275-DEL-143 | chr4-36302257-DEL-191 |
| chr4-36336474-DEL-1821 | chr4-36336475-DEL-1821 |
| chr4-36393294-DEL-81 | chr4-36393300-DEL-69 |
| chr4-36401991-DEL-2136 | chr4-36401992-DEL-2136 |
| chr4-36413451-DEL-829 | chr4-36413452-DEL-829 |
| chr4-36432033-DEL-995 | chr4-36432034-DEL-994 |
| chr4-36440896-DEL-337 | chr4-36440899-DEL-330 |
| chr4-36472127-DEL-2210 | chr4-36472128-DEL-2210 |
| chr4-36478482-DEL-55 | chr4-36478483-DEL-55 |
| chr4-36503432-DEL-71 | chr4-36503433-DEL-71 |
| chr4-36517805-DEL-461 | chr4-36517818-DEL-461 |
| chr4-36547119-DEL-72 | chr4-36547120-DEL-72 |
| chr4-36563086-DEL-598 | chr4-36563087-DEL-598 |
| chr4-36579960-DEL-188 | chr4-36579958-DEL-189 |
| chr4-36584961-DEL-570 | chr4-36584962-DEL-570 |
| chr4-36635705-DEL-5406 | chr4-36635706-DEL-5406 |
| chr4-36645013-DEL-54 | chr4-36645018-DEL-68 |
| chr4-36647109-DEL-261 | chr4-36647059-DEL-271 |
| chr4-36650897-DEL-231 | chr4-36650884-DEL-238 |
| chr4-36654305-DEL-59 | chr4-36654272-DEL-59 |
| chr4-36660434-DEL-122 | chr4-36660435-DEL-122 |
| chr4-36678622-DEL-240 | chr4-36678623-DEL-240 |
| chr4-36738433-DEL-572 | chr4-36738434-DEL-572 |
| chr4-36769538-DEL-3716 | chr4-36769539-DEL-3716 |
| chr4-36860607-DEL-60 | chr4-36860600-DEL-68 |
| chr4-36862923-DEL-1103 | chr4-36862924-DEL-1103 |
| chr4-36876344-DEL-72 | chr4-36876367-DEL-50 |
| chr4-36884672-DEL-1250 | chr4-36884673-DEL-1250 |
| chr4-36891505-DEL-1093 | chr4-36891504-DEL-1093 |
| chr4-36906016-DEL-308 | chr4-36906017-DEL-308 |
| chr4-36955044-DEL-195 | chr4-36955045-DEL-195 |
| chr4-36988183-DEL-67 | chr4-36988169-DEL-68 |
| chr4-37001933-DEL-5986 | chr4-37001934-DEL-5986 |
| chr4-37022302-DEL-748 | chr4-37022303-DEL-748 |
| chr4-37061651-DEL-200 | chr4-37061652-DEL-200 |
| chr4-37131262-DEL-610 | chr4-37131263-DEL-610 |
| chr4-37166354-DEL-456 | chr4-37166355-DEL-456 |
| chr4-37177212-DEL-123 | chr4-37177213-DEL-123 |
| chr4-37201615-DEL-1675 | chr4-37201616-DEL-1675 |
| chr4-37208521-DEL-131 | chr4-37208495-DEL-108 |
| chr4-37218058-DEL-7091 | chr4-37218059-DEL-7091 |
| chr4-37243507-DEL-249 | chr4-37243508-DEL-249 |
| chr4-37251355-DEL-96 | chr4-37251333-DEL-77 |
| chr4-37252911-DEL-1896 | chr4-37252911-DEL-1896 |
| chr4-37262229-DEL-826 | chr4-37262222-DEL-826 |
| chr4-37315710-DEL-154 | chr4-37315747-DEL-118 |
| chr4-37343160-DEL-6245 | chr4-37343161-DEL-6245 |
| chr4-37366827-DEL-1303 | chr4-37366828-DEL-1303 |

| | |
|---|---|
| chr4-37420936-DEL-1083 | chr4-37420938-DEL-1083 |
| chr4-37442107-DEL-691 | chr4-37442108-DEL-691 |
| chr4-37469061-DEL-439 | chr4-37469062-DEL-439 |
| chr4-37471253-DEL-374 | chr4-37471254-DEL-386 |
| chr4-37479200-DEL-1665 | chr4-37479200-DEL-1665 |
| chr4-37481155-DEL-61 | chr4-37481155-DEL-61 |
| chr4-37481445-DEL-521 | chr4-37481446-DEL-521 |
| chr4-37489192-DEL-67 | chr4-37489192-DEL-67 |
| chr4-37490800-DEL-145 | chr4-37490800-DEL-145 |
| chr4-37500930-DEL-205 | chr4-37500931-DEL-205 |
| chr4-37502104-DEL-94 | chr4-37502105-DEL-94 |
| chr4-37520969-DEL-59 | chr4-37520970-DEL-59 |
| chr4-37532338-DEL-88 | chr4-37532340-DEL-86 |
| chr4-37534229-DEL-198 | chr4-37534230-DEL-198 |
| chr4-37562767-DEL-63 | chr4-37562768-DEL-63 |
| chr4-37601325-DEL-256 | chr4-37601326-DEL-249 |
| chr4-37610970-DEL-6966 | chr4-37610971-DEL-6966 |
| chr4-37618066-DEL-1420 | chr4-37618067-DEL-1420 |
| chr4-37675884-DEL-392 | chr4-37675885-DEL-392 |
| chr4-37679833-DEL-176 | chr4-37679819-DEL-167 |
| chr4-37703102-DEL-4622 | chr4-37703103-DEL-4622 |
| chr4-37712389-DEL-873 | chr4-37712390-DEL-873 |
| chr4-37713739-DEL-155 | chr4-37713740-DEL-155 |
| chr4-37723732-DEL-699 | chr4-37723733-DEL-699 |
| chr4-37801770-DEL-1529 | chr4-37801771-DEL-1529 |
| chr4-37804637-DEL-225 | chr4-37804638-DEL-225 |
| chr4-37830343-DEL-93 | chr4-37830344-DEL-93 |
| chr4-37847258-DEL-1699 | chr4-37847259-DEL-1699 |
| chr4-37859895-DEL-867 | chr4-37859896-DEL-867 |
| chr4-37863414-DEL-321 | chr4-37863415-DEL-321 |
| chr4-37916944-DEL-6221 | chr4-37916945-DEL-6221 |
| chr4-37955280-DEL-3800 | chr4-37955281-DEL-3800 |
| chr4-37956225-DEL-1880 | chr4-37956226-DEL-1880 |
| chr4-37983026-DEL-4254 | chr4-37983028-DEL-4255 |
| chr4-38061188-DEL-6679 | chr4-38061189-DEL-6679 |
| chr4-38080226-DEL-906 | chr4-38080229-DEL-907 |
| chr4-38112149-DEL-336 | chr4-38112149-DEL-336 |
| chr4-38215047-DEL-15325 | chr4-38215047-DEL-15325 |
| chr4-38293314-DEL-361 | chr4-38293315-DEL-362 |
| chr4-38313048-DEL-416 | chr4-38313049-DEL-416 |
| chr4-38315759-DEL-6460 | chr4-38315760-DEL-6460 |
| chr4-38333053-DEL-66 | chr4-38333054-DEL-66 |
| chr4-38368229-DEL-195 | chr4-38368231-DEL-194 |
| chr4-38383885-DEL-142 | chr4-38383886-DEL-142 |
| chr4-38402031-DEL-6369 | chr4-38402032-DEL-6369 |
| chr4-38427592-DEL-263 | chr4-38427593-DEL-263 |
| chr4-38436293-DEL-359 | chr4-38436294-DEL-359 |
| chr4-38508286-DEL-3793 | chr4-38508280-DEL-3799 |
| chr4-38513902-DEL-52 | chr4-38513903-DEL-52 |
| chr4-38549408-DEL-6456 | chr4-38549408-DEL-6456 |
| chr4-38559426-DEL-447 | chr4-38559426-DEL-447 |
| chr4-38577105-DEL-6268 | chr4-38577093-DEL-6280 |
| chr4-38641642-DEL-244 | chr4-38641643-DEL-244 |
| chr4-38656495-DEL-122 | chr4-38656493-DEL-112 |
| chr4-38660220-DEL-542 | chr4-38660221-DEL-542 |
| chr4-38671522-DEL-127 | chr4-38671525-DEL-125 |
| chr4-38677803-DEL-1979 | chr4-38677755-DEL-2020 |
| chr4-38713156-DEL-546 | chr4-38713157-DEL-546 |
| chr4-38750222-DEL-6222 | chr4-38750223-DEL-6221 |
| chr4-38757669-DEL-354 | chr4-38757671-DEL-355 |
| chr4-38761964-DEL-894 | chr4-38761964-DEL-894 |
| chr4-38762012-DEL-194 | chr4-38762017-DEL-187 |
| chr4-38798340-DEL-634 | chr4-38798340-DEL-634 |
| chr4-38840968-DEL-52 | chr4-38840969-DEL-52 |
| chr4-38870088-DEL-7106 | chr4-38870089-DEL-7106 |
| chr4-38905900-DEL-7066 | chr4-38905901-DEL-7066 |
| chr4-38928845-DEL-424 | chr4-38928846-DEL-424 |
| chr4-38968339-DEL-78 | chr4-38968322-DEL-58 |
| chr4-38980093-DEL-1121 | chr4-38980094-DEL-1121 |
| chr4-38983259-DEL-219 | chr4-38983260-DEL-219 |
| chr4-38994628-DEL-60 | chr4-38994629-DEL-60 |
| chr4-39040460-DEL-7715 | chr4-39040454-DEL-7719 |
| chr4-39049772-DEL-6374 | chr4-39049773-DEL-6374 |
| chr4-39080299-DEL-5109 | chr4-39080300-DEL-5109 |
| chr4-39080309-DEL-564 | chr4-39080310-DEL-564 |
| chr4-39106784-DEL-70 | chr4-39106789-DEL-68 |
| chr4-39111385-DEL-116 | chr4-39111386-DEL-116 |
| chr4-39120349-DEL-784 | chr4-39120350-DEL-785 |
| chr4-39139160-DEL-318 | chr4-39139161-DEL-318 |
| chr4-39151326-DEL-99 | chr4-39151364-DEL-69 |

| | |
|---|---|
| chr4-39175623-DEL-3887 | chr4-39175624-DEL-3887 |
| chr4-39180706-DEL-150 | chr4-39180707-DEL-150 |
| chr4-39182658-DEL-1145 | chr4-39182659-DEL-1145 |
| chr4-39219976-DEL-660 | chr4-39219977-DEL-660 |
| chr4-39228103-DEL-9990 | chr4-39228103-DEL-9991 |
| chr4-39238923-DEL-1891 | chr4-39238924-DEL-1891 |
| chr4-39256445-DEL-1352 | chr4-39256446-DEL-1352 |
| chr4-39266352-DEL-6298 | chr4-39266355-DEL-6300 |
| chr4-39276863-DEL-88 | chr4-39276864-DEL-88 |
| chr4-39277545-DEL-1929 | chr4-39277546-DEL-1929 |
| chr4-39280933-DEL-184 | chr4-39280934-DEL-184 |
| chr4-39282938-DEL-955 | chr4-39282939-DEL-955 |
| chr4-39284226-DEL-60 | chr4-39284227-DEL-60 |
| chr4-39288215-DEL-59 | chr4-39288216-DEL-59 |
| chr4-39293000-DEL-483 | chr4-39293001-DEL-483 |
| chr4-39336207-DEL-521 | chr4-39336208-DEL-521 |
| chr4-39374043-DEL-75 | chr4-39374044-DEL-75 |
| chr4-39382284-DEL-8797 | chr4-39382285-DEL-8797 |
| chr4-39385131-DEL-400 | chr4-39385132-DEL-400 |
| chr4-39410694-DEL-60 | chr4-39410695-DEL-60 |
| chr4-39426910-DEL-66 | chr4-39426905-DEL-76 |
| chr4-39430663-DEL-1875 | chr4-39430664-DEL-1874 |
| chr4-39433094-DEL-520 | chr4-39433063-DEL-520 |
| chr4-39434571-DEL-634 | chr4-39434572-DEL-634 |
| chr4-39435917-DEL-62 | chr4-39435912-DEL-56 |
| chr4-39441655-DEL-72 | chr4-39441656-DEL-70 |
| chr4-39508438-DEL-384 | chr4-39508439-DEL-384 |
| chr4-39534077-DEL-60 | chr4-39534078-DEL-59 |
| chr4-39558542-DEL-59 | chr4-39558543-DEL-59 |
| chr4-39561959-DEL-395 | chr4-39561960-DEL-395 |
| chr4-39596652-DEL-131 | chr4-39596652-DEL-132 |
| chr4-39598186-DEL-1127 | chr4-39598187-DEL-1127 |
| chr4-39647466-DEL-6401 | chr4-39647467-DEL-6401 |
| chr4-39658381-DEL-986 | chr4-39658379-DEL-985 |
| chr4-39723214-DEL-171 | chr4-39723209-DEL-173 |
| chr4-39755080-DEL-6455 | chr4-39755081-DEL-6455 |
| chr4-39774260-DEL-6591 | chr4-39774261-DEL-6591 |
| chr4-39815346-DEL-76 | chr4-39815347-DEL-76 |
| chr4-39866095-DEL-52 | chr4-39866092-DEL-56 |
| chr4-39918601-DEL-487 | chr4-39918602-DEL-487 |
| chr4-39952751-DEL-5615 | chr4-39952752-DEL-5615 |
| chr4-40014572-DEL-66 | chr4-40014567-DEL-68 |
| chr4-40026253-DEL-2518 | chr4-40026254-DEL-2518 |
| chr4-40053806-DEL-86 | chr4-40053807-DEL-86 |
| chr4-40054163-DEL-7241 | chr4-40054164-DEL-7241 |
| chr4-40076143-DEL-554 | chr4-40076144-DEL-554 |
| chr4-40083091-DEL-6259 | chr4-40083091-DEL-6258 |
| chr4-40127359-DEL-820 | chr4-40127360-DEL-820 |
| chr4-40132592-DEL-1135 | chr4-40132594-DEL-1134 |
| chr4-40212080-DEL-212 | chr4-40212081-DEL-212 |
| chr4-40214103-DEL-206 | chr4-40214104-DEL-206 |
| chr4-40224506-DEL-206 | chr4-40224505-DEL-207 |
| chr4-40235625-DEL-525 | chr4-40235626-DEL-525 |
| chr4-40258712-DEL-400 | chr4-40258713-DEL-400 |
| chr4-40271676-DEL-203 | chr4-40271677-DEL-203 |
| chr4-40274317-DEL-205 | chr4-40274318-DEL-205 |
| chr4-40296780-DEL-1376 | chr4-40296779-DEL-1374 |
| chr4-40318827-DEL-94 | chr4-40318828-DEL-94 |
| chr4-40331436-DEL-1196 | chr4-40331437-DEL-1196 |
| chr4-40369335-DEL-397 | chr4-40369336-DEL-397 |
| chr4-40377085-DEL-69 | chr4-40377085-DEL-70 |
| chr4-40402275-DEL-413 | chr4-40402276-DEL-411 |
| chr4-40418097-DEL-54 | chr4-40418073-DEL-51 |
| chr4-40438375-DEL-3431 | chr4-40438376-DEL-3431 |
| chr4-40452434-DEL-223 | chr4-40452432-DEL-224 |
| chr4-40460628-DEL-6480 | chr4-40460629-DEL-6481 |
| chr4-40499286-DEL-56 | chr4-40499287-DEL-56 |
| chr4-40504230-DEL-740 | chr4-40504231-DEL-740 |
| chr4-40520770-DEL-54 | chr4-40520782-DEL-50 |
| chr4-40527318-DEL-737 | chr4-40527318-DEL-737 |
| chr4-40539594-DEL-527 | chr4-40539595-DEL-527 |
| chr4-40577482-DEL-2159 | chr4-40577482-DEL-2159 |
| chr4-40586785-DEL-326 | chr4-40586786-DEL-326 |
| chr4-40592270-DEL-1525 | chr4-40592271-DEL-1525 |
| chr4-40594750-DEL-13769 | chr4-40594751-DEL-13769 |
| chr4-40641815-DEL-62 | chr4-40641818-DEL-63 |
| chr4-40642761-DEL-181 | chr4-40642762-DEL-181 |
| chr4-40644327-DEL-84 | chr4-40644327-DEL-84 |
| chr4-40646965-DEL-628 | chr4-40646965-DEL-628 |
| chr4-40651443-DEL-8817 | chr4-40651444-DEL-8817 |

| | |
|---|---|
| chr4-40674394-DEL-66 | chr4-40674383-DEL-78 |
| chr4-40679627-DEL-186 | chr4-40679628-DEL-186 |
| chr4-40680261-DEL-162 | chr4-40680261-DEL-162 |
| chr4-40689413-DEL-207 | chr4-40689414-DEL-208 |
| chr4-40689976-DEL-71 | chr4-40689963-DEL-72 |
| chr4-40691893-DEL-242 | chr4-40691894-DEL-242 |
| chr4-40743539-DEL-186 | chr4-40743540-DEL-186 |
| chr4-40756914-DEL-55 | chr4-40756919-DEL-51 |
| chr4-40766523-DEL-51 | chr4-40766524-DEL-51 |
| chr4-40780448-DEL-231 | chr4-40780448-DEL-205 |
| chr4-40782876-DEL-202 | chr4-40782877-DEL-202 |
| chr4-40838593-DEL-192 | chr4-40838571-DEL-176 |
| chr4-40869575-DEL-252 | chr4-40869576-DEL-252 |
| chr4-40870167-DEL-78 | chr4-40870168-DEL-78 |
| chr4-40909675-DEL-204 | chr4-40909675-DEL-205 |
| chr4-40911691-DEL-1803 | chr4-40911694-DEL-1812 |
| chr4-40964054-DEL-145 | chr4-40964020-DEL-145 |
| chr4-41019815-DEL-57 | chr4-41019843-DEL-58 |
| chr4-41022716-DEL-387 | chr4-41022717-DEL-387 |
| chr4-41028766-DEL-205 | chr4-41028767-DEL-205 |
| chr4-41037878-DEL-61 | chr4-41037879-DEL-60 |
| chr4-41039049-DEL-853 | chr4-41039050-DEL-853 |
| chr4-41042885-DEL-65 | chr4-41042886-DEL-65 |
| chr4-41049251-DEL-54 | chr4-41049252-DEL-54 |
| chr4-41059258-DEL-114 | chr4-41059258-DEL-114 |
| chr4-41079428-DEL-67 | chr4-41079428-DEL-66 |
| chr4-41103687-DEL-211 | chr4-41103688-DEL-211 |
| chr4-41105733-DEL-156 | chr4-41105734-DEL-160 |
| chr4-41142490-DEL-210 | chr4-41142485-DEL-205 |
| chr4-41177185-DEL-201 | chr4-41177186-DEL-201 |
| chr4-41234780-DEL-74 | chr4-41234781-DEL-74 |
| chr4-41259914-DEL-93 | chr4-41259915-DEL-93 |
| chr4-41263627-DEL-174 | chr4-41263628-DEL-174 |
| chr4-41287296-DEL-113 | chr4-41287292-DEL-114 |
| chr4-41323969-DEL-70 | chr4-41323970-DEL-70 |
| chr4-41365340-DEL-204 | chr4-41365332-DEL-205 |
| chr4-41409185-DEL-180 | chr4-41409186-DEL-180 |
| chr4-41436080-DEL-106 | chr4-41436081-DEL-106 |
| chr4-41454728-DEL-5272 | chr4-41454729-DEL-5272 |
| chr4-41487739-DEL-338 | chr4-41487740-DEL-338 |
| chr4-41534569-DEL-70 | chr4-41534570-DEL-54 |
| chr4-41541363-DEL-67 | chr4-41541364-DEL-67 |
| chr4-41561015-DEL-208 | chr4-41561016-DEL-207 |
| chr4-41565872-DEL-70 | chr4-41565873-DEL-70 |
| chr4-41571756-DEL-75 | chr4-41571757-DEL-75 |
| chr4-41644645-DEL-141 | chr4-41644646-DEL-141 |
| chr4-41666099-DEL-131 | chr4-41666050-DEL-135 |
| chr4-41681627-DEL-240 | chr4-41681604-DEL-271 |
| chr4-41786011-DEL-70 | chr4-41786011-DEL-69 |
| chr4-42479739-DEL-274 | chr4-42479713-DEL-315 |
| chr4-42776008-DEL-60 | chr4-42776009-DEL-60 |
| chr4-42807363-DEL-68 | chr4-42807364-DEL-68 |
| chr4-42825752-DEL-83 | chr4-42825766-DEL-63 |
| chr4-42858315-DEL-70 | chr4-42858316-DEL-70 |
| chr4-42881861-DEL-63 | chr4-42881862-DEL-63 |
| chr4-42907398-DEL-94 | chr4-42907398-DEL-92 |
| chr4-42930597-DEL-92 | chr4-42930594-DEL-96 |
| chr4-42992514-DEL-212 | chr4-42992514-DEL-212 |
| chr4-43008293-DEL-53 | chr4-43008294-DEL-53 |
| chr4-43015099-DEL-1196 | chr4-43015100-DEL-1196 |
| chr4-43053715-DEL-397 | chr4-43053716-DEL-397 |
| chr4-43070436-DEL-99 | chr4-43070437-DEL-99 |
| chr4-43071309-DEL-55 | chr4-43071311-DEL-53 |
| chr4-43079545-DEL-675 | chr4-43079546-DEL-675 |
| chr4-43081660-DEL-276 | chr4-43081661-DEL-276 |
| chr4-43086169-DEL-87 | chr4-43086170-DEL-87 |
| chr4-43123356-DEL-6273 | chr4-43123357-DEL-6273 |
| chr4-43132201-DEL-99 | chr4-43132202-DEL-99 |
| chr4-43142283-DEL-53 | chr4-43142284-DEL-53 |
| chr4-43146288-DEL-328 | chr4-43146289-DEL-328 |
| chr4-43196364-DEL-460 | chr4-43196365-DEL-460 |
| chr4-43203760-DEL-119 | chr4-43203761-DEL-119 |
| chr4-43223516-DEL-103 | chr4-43223517-DEL-103 |
| chr4-43297717-DEL-4405 | chr4-43297718-DEL-4405 |
| chr4-43302434-DEL-69 | chr4-43302435-DEL-69 |
| chr4-43310615-DEL-52 | chr4-43310616-DEL-52 |
| chr4-43319973-DEL-68 | chr4-43319980-DEL-96 |
| chr4-43344237-DEL-125 | chr4-43344238-DEL-125 |
| chr4-43348491-DEL-1181 | chr4-43348492-DEL-1181 |
| chr4-43351469-DEL-603 | chr4-43351470-DEL-603 |

| | |
|---|---|
| chr4-43365008-DEL-58 | chr4-43365009-DEL-58 |
| chr4-43371711-DEL-270 | chr4-43371696-DEL-270 |
| chr4-43387452-DEL-245 | chr4-43387453-DEL-245 |
| chr4-43409788-DEL-84 | chr4-43409789-DEL-84 |
| chr4-43431329-DEL-115 | chr4-43431330-DEL-115 |
| chr4-43432656-DEL-202 | chr4-43432657-DEL-202 |
| chr4-43434986-DEL-74 | chr4-43434983-DEL-51 |
| chr4-43440238-DEL-71 | chr4-43440239-DEL-71 |
| chr4-43456305-DEL-214 | chr4-43456306-DEL-214 |
| chr4-43459943-DEL-94 | chr4-43459934-DEL-93 |
| chr4-43464076-DEL-206 | chr4-43464077-DEL-206 |
| chr4-43472099-DEL-52 | chr4-43472099-DEL-52 |
| chr4-43477413-DEL-195 | chr4-43477414-DEL-195 |
| chr4-43491377-DEL-670 | chr4-43491378-DEL-670 |
| chr4-43511402-DEL-84 | chr4-43511399-DEL-88 |
| chr4-43527278-DEL-1858 | chr4-43527279-DEL-1858 |
| chr4-43541107-DEL-123 | chr4-43541067-DEL-123 |
| chr4-43596527-DEL-110 | chr4-43596526-DEL-132 |
| chr4-43623290-DEL-211 | chr4-43623291-DEL-211 |
| chr4-43663903-DEL-521 | chr4-43663904-DEL-521 |
| chr4-43666001-DEL-69 | chr4-43666002-DEL-69 |
| chr4-43703231-DEL-421 | chr4-43703232-DEL-421 |
| chr4-43714199-DEL-136 | chr4-43714217-DEL-128 |
| chr4-43715637-DEL-67 | chr4-43715638-DEL-67 |
| chr4-43717334-DEL-60 | chr4-43717322-DEL-94 |
| chr4-43723278-DEL-400 | chr4-43723279-DEL-400 |
| chr4-43751458-DEL-4606 | chr4-43751457-DEL-4605 |
| chr4-43758138-DEL-164 | chr4-43758138-DEL-164 |
| chr4-43775638-DEL-183 | chr4-43775637-DEL-184 |
| chr4-43782993-DEL-80 | chr4-43783009-DEL-60 |
| chr4-43785920-DEL-5655 | chr4-43785921-DEL-5655 |
| chr4-43802786-DEL-544 | chr4-43802793-DEL-538 |
| chr4-43804213-DEL-93 | chr4-43804214-DEL-93 |
| chr4-43811506-DEL-116 | chr4-43811507-DEL-117 |
| chr4-43812286-DEL-6465 | chr4-43812287-DEL-6465 |
| chr4-43824797-DEL-62 | chr4-43824798-DEL-62 |
| chr4-43830146-DEL-60 | chr4-43830147-DEL-60 |
| chr4-43842912-DEL-1476 | chr4-43842913-DEL-1476 |
| chr4-43843027-DEL-430 | chr4-43843028-DEL-430 |
| chr4-43880844-DEL-288 | chr4-43880845-DEL-240 |
| chr4-43891671-DEL-255 | chr4-43891672-DEL-255 |
| chr4-43920291-DEL-415 | chr4-43920292-DEL-413 |
| chr4-43937930-DEL-75 | chr4-43937931-DEL-75 |
| chr4-43961250-DEL-204 | chr4-43961250-DEL-204 |
| chr4-43971321-DEL-220 | chr4-43971322-DEL-220 |
| chr4-43975505-DEL-434 | chr4-43975505-DEL-434 |
| chr4-44001313-DEL-195 | chr4-44001314-DEL-195 |
| chr4-44006331-DEL-214 | chr4-44006332-DEL-214 |
| chr4-44009034-DEL-514 | chr4-44009035-DEL-514 |
| chr4-44033283-DEL-1175 | chr4-44033284-DEL-1175 |
| chr4-44056010-DEL-134 | chr4-44056011-DEL-134 |
| chr4-44061069-DEL-204 | chr4-44061070-DEL-204 |
| chr4-44089568-DEL-64 | chr4-44089578-DEL-64 |
| chr4-44093493-DEL-200 | chr4-44093493-DEL-200 |
| chr4-44109471-DEL-210 | chr4-44109472-DEL-210 |
| chr4-44118644-DEL-192 | chr4-44118644-DEL-192 |
| chr4-44123605-DEL-137 | chr4-44123606-DEL-137 |
| chr4-44139385-DEL-1867 | chr4-44139386-DEL-1867 |
| chr4-44168114-DEL-65 | chr4-44168115-DEL-75 |
| chr4-44190809-DEL-207 | chr4-44190810-DEL-207 |
| chr4-44192248-DEL-1378 | chr4-44192249-DEL-1378 |
| chr4-44193809-DEL-405 | chr4-44193810-DEL-405 |
| chr4-44198716-DEL-244 | chr4-44198717-DEL-244 |
| chr4-44221687-DEL-1701 | chr4-44221687-DEL-1701 |
| chr4-44256802-DEL-428 | chr4-44256796-DEL-428 |
| chr4-44264085-DEL-52 | chr4-44264086-DEL-52 |
| chr4-44268797-DEL-85 | chr4-44268798-DEL-85 |
| chr4-44272557-DEL-64 | chr4-44272558-DEL-62 |
| chr4-44276338-DEL-948 | chr4-44276339-DEL-948 |
| chr4-44284643-DEL-221 | chr4-44284644-DEL-220 |
| chr4-44286309-DEL-83 | chr4-44286337-DEL-84 |
| chr4-44293839-DEL-226 | chr4-44293840-DEL-226 |
| chr4-44294540-DEL-1040 | chr4-44294541-DEL-1040 |
| chr4-44328972-DEL-105 | chr4-44328973-DEL-105 |
| chr4-44333767-DEL-103 | chr4-44333788-DEL-84 |
| chr4-44339951-DEL-56 | chr4-44339952-DEL-56 |
| chr4-44345703-DEL-86 | chr4-44345687-DEL-56 |
| chr4-44351800-DEL-6619 | chr4-44351801-DEL-6619 |
| chr4-44367945-DEL-230 | chr4-44367946-DEL-230 |
| chr4-44371462-DEL-358 | chr4-44371463-DEL-358 |

| | |
|---|---|
| chr4-44396241-DEL-56 | chr4-44396230-DEL-62 |
| chr4-44432244-DEL-1183 | chr4-44432232-DEL-1184 |
| chr4-44435251-DEL-79 | chr4-44435252-DEL-79 |
| chr4-44439528-DEL-134 | chr4-44439529-DEL-134 |
| chr4-44460183-DEL-540 | chr4-44460184-DEL-540 |
| chr4-44464090-DEL-1249 | chr4-44464092-DEL-1248 |
| chr4-44485471-DEL-64 | chr4-44485463-DEL-60 |
| chr4-44492375-DEL-68 | chr4-44492380-DEL-64 |
| chr4-44507222-DEL-94 | chr4-44507223-DEL-94 |
| chr4-44552360-DEL-1750 | chr4-44552378-DEL-1747 |
| chr4-44569160-DEL-62 | chr4-44569161-DEL-62 |
| chr4-44594481-DEL-133 | chr4-44594482-DEL-133 |
| chr4-44627185-DEL-84 | chr4-44627178-DEL-92 |
| chr4-44627693-DEL-174 | chr4-44627694-DEL-174 |
| chr4-44652199-DEL-60 | chr4-44652204-DEL-56 |
| chr4-44752116-DEL-1083 | chr4-44752117-DEL-1083 |
| chr4-44779958-DEL-57 | chr4-44779959-DEL-57 |
| chr4-44787237-DEL-165 | chr4-44787238-DEL-165 |
| chr4-44955187-DEL-196 | chr4-44955188-DEL-196 |
| chr4-44956270-DEL-53 | chr4-44956270-DEL-54 |
| chr4-45000554-DEL-191 | chr4-45000555-DEL-190 |
| chr4-45021494-DEL-58 | chr4-45021495-DEL-58 |
| chr4-45061713-DEL-121 | chr4-45061714-DEL-121 |
| chr4-45076246-DEL-60 | chr4-45076247-DEL-60 |
| chr4-45078368-DEL-179 | chr4-45078369-DEL-179 |
| chr4-45083907-DEL-65 | chr4-45083908-DEL-65 |
| chr4-45098126-DEL-152 | chr4-45098127-DEL-152 |
| chr4-45127988-DEL-58 | chr4-45127994-DEL-57 |
| chr4-45133689-DEL-177 | chr4-45133690-DEL-177 |
| chr4-45160140-DEL-89 | chr4-45160140-DEL-70 |
| chr4-45178509-DEL-714 | chr4-45178508-DEL-715 |
| chr4-45193331-DEL-550 | chr4-45193290-DEL-552 |
| chr4-45195104-DEL-58 | chr4-45195105-DEL-58 |
| chr4-45217261-DEL-69 | chr4-45217262-DEL-69 |
| chr4-45221867-DEL-304 | chr4-45221891-DEL-287 |
| chr4-45245519-DEL-76 | chr4-45245520-DEL-76 |
| chr4-45247620-DEL-185 | chr4-45247625-DEL-181 |
| chr4-45249051-DEL-51 | chr4-45249052-DEL-51 |
| chr4-45249790-DEL-161 | chr4-45249791-DEL-161 |
| chr4-45282492-DEL-143 | chr4-45282493-DEL-143 |
| chr4-45289981-DEL-6460 | chr4-45289982-DEL-6460 |
| chr4-45306627-DEL-175 | chr4-45306628-DEL-175 |
| chr4-45309971-DEL-98 | chr4-45309986-DEL-80 |
| chr4-45317211-DEL-66 | chr4-45317224-DEL-56 |
| chr4-45328950-DEL-759 | chr4-45328951-DEL-759 |
| chr4-45329889-DEL-1652 | chr4-45329889-DEL-1653 |
| chr4-45360980-DEL-168 | chr4-45360976-DEL-168 |
| chr4-45405984-DEL-217 | chr4-45405977-DEL-217 |
| chr4-45414404-DEL-80 | chr4-45414421-DEL-64 |
| chr4-45451483-DEL-51 | chr4-45451484-DEL-51 |
| chr4-45481385-DEL-53 | chr4-45481386-DEL-53 |
| chr4-45505052-DEL-179 | chr4-45505053-DEL-179 |
| chr4-45545287-DEL-56 | chr4-45545288-DEL-56 |
| chr4-45549965-DEL-577 | chr4-45549965-DEL-577 |
| chr4-45563837-DEL-220 | chr4-45563833-DEL-219 |
| chr4-45592178-DEL-128 | chr4-45592179-DEL-128 |
| chr4-45593886-DEL-436 | chr4-45593887-DEL-442 |
| chr4-45597371-DEL-79 | chr4-45597372-DEL-79 |
| chr4-45612635-DEL-72 | chr4-45612636-DEL-72 |
| chr4-45620543-DEL-953 | chr4-45620541-DEL-951 |
| chr4-45633261-DEL-78 | chr4-45633252-DEL-54 |
| chr4-45633970-DEL-232 | chr4-45633967-DEL-225 |
| chr4-45648859-DEL-146 | chr4-45648859-DEL-143 |
| chr4-45658030-DEL-53 | chr4-45658030-DEL-53 |
| chr4-45664360-DEL-651 | chr4-45664361-DEL-651 |
| chr4-45665135-DEL-6412 | chr4-45665136-DEL-6412 |
| chr4-45725747-DEL-65 | chr4-45725748-DEL-65 |
| chr4-45744960-DEL-60 | chr4-45744961-DEL-60 |
| chr4-45801866-DEL-202 | chr4-45801867-DEL-202 |
| chr4-45812225-DEL-199 | chr4-45812226-DEL-199 |
| chr4-45835768-DEL-10889 | chr4-45835769-DEL-10889 |
| chr4-45836850-DEL-4925 | chr4-45836851-DEL-4925 |
| chr4-45846411-DEL-54 | chr4-45846412-DEL-54 |
| chr4-45852650-DEL-184 | chr4-45852665-DEL-185 |
| chr4-45857525-DEL-67 | chr4-45857526-DEL-67 |
| chr4-45859755-DEL-121 | chr4-45859756-DEL-121 |
| chr4-45863845-DEL-213 | chr4-45863846-DEL-213 |
| chr4-45871568-DEL-213 | chr4-45871569-DEL-213 |
| chr4-45952642-DEL-1550 | chr4-45952643-DEL-1550 |
| chr4-46014403-DEL-607 | chr4-46014404-DEL-607 |

| | |
|---|---|
| chr4-46024140-DEL-253 | chr4-46024141-DEL-253 |
| chr4-46031917-DEL-773 | chr4-46031918-DEL-773 |
| chr4-46046348-DEL-1657 | chr4-46046345-DEL-1661 |
| chr4-46082385-DEL-82 | chr4-46082385-DEL-82 |
| chr4-46104307-DEL-80 | chr4-46104313-DEL-75 |
| chr4-46108741-DEL-68 | chr4-46108741-DEL-68 |
| chr4-46129643-DEL-102 | chr4-46129644-DEL-102 |
| chr4-46160501-DEL-311 | chr4-46160502-DEL-311 |
| chr4-46219232-DEL-390 | chr4-46219253-DEL-404 |
| chr4-46241354-DEL-547 | chr4-46241354-DEL-547 |
| chr4-46247814-DEL-1765 | chr4-46247815-DEL-1765 |
| chr4-46274447-DEL-65 | chr4-46274448-DEL-65 |
| chr4-46275367-DEL-81 | chr4-46275365-DEL-82 |
| chr4-46301092-DEL-199 | chr4-46301093-DEL-199 |
| chr4-46305594-DEL-54 | chr4-46305584-DEL-56 |
| chr4-46314049-DEL-82 | chr4-46314050-DEL-82 |
| chr4-46319852-DEL-56 | chr4-46319869-DEL-52 |
| chr4-46322618-DEL-1273 | chr4-46322621-DEL-1271 |
| chr4-46324457-DEL-58 | chr4-46324429-DEL-51 |
| chr4-46341205-DEL-366 | chr4-46341208-DEL-364 |
| chr4-46391357-DEL-133 | chr4-46391358-DEL-133 |
| chr4-46395324-DEL-56 | chr4-46395342-DEL-64 |
| chr4-46418190-DEL-60 | chr4-46418191-DEL-60 |
| chr4-46448056-DEL-79 | chr4-46448057-DEL-79 |
| chr4-46511078-DEL-6559 | chr4-46511079-DEL-6559 |
| chr4-46521240-DEL-61 | chr4-46521240-DEL-61 |
| chr4-46533426-DEL-182 | chr4-46533427-DEL-182 |
| chr4-46549211-DEL-1414 | chr4-46549212-DEL-1414 |
| chr4-46561312-DEL-340 | chr4-46561327-DEL-306 |
| chr4-46562661-DEL-129 | chr4-46562662-DEL-129 |
| chr4-46573762-DEL-65 | chr4-46573763-DEL-65 |
| chr4-46575137-DEL-9907 | chr4-46575138-DEL-9907 |
| chr4-46602586-DEL-463 | chr4-46602587-DEL-461 |
| chr4-46636364-DEL-131 | chr4-46636365-DEL-131 |
| chr4-46640700-DEL-192 | chr4-46640701-DEL-192 |
| chr4-46645156-DEL-220 | chr4-46645157-DEL-220 |
| chr4-46647300-DEL-202 | chr4-46647301-DEL-202 |
| chr4-46658571-DEL-191 | chr4-46658572-DEL-191 |
| chr4-46692497-DEL-233 | chr4-46692496-DEL-233 |
| chr4-46743521-DEL-54 | chr4-46743521-DEL-55 |
| chr4-46754692-DEL-55 | chr4-46754693-DEL-55 |
| chr4-46760150-DEL-152 | chr4-46760143-DEL-150 |
| chr4-46793065-DEL-191 | chr4-46793061-DEL-192 |
| chr4-46860104-DEL-177 | chr4-46860105-DEL-177 |
| chr4-46916353-DEL-63 | chr4-46916354-DEL-51 |
| chr4-46936849-DEL-86 | chr4-46936850-DEL-86 |
| chr4-46941744-DEL-789 | chr4-46941745-DEL-789 |
| chr4-46947899-DEL-105 | chr4-46947900-DEL-105 |
| chr4-46998059-DEL-318 | chr4-46998060-DEL-318 |
| chr4-47001966-DEL-989 | chr4-47001967-DEL-987 |
| chr4-47007792-DEL-54 | chr4-47007793-DEL-54 |
| chr4-47035131-DEL-296 | chr4-47035132-DEL-295 |
| chr4-47067993-DEL-3594 | chr4-47067993-DEL-3594 |
| chr4-47099494-DEL-3333 | chr4-47099495-DEL-3333 |
| chr4-47114096-DEL-52 | chr4-47114096-DEL-52 |
| chr4-47152870-DEL-51 | chr4-47152859-DEL-52 |
| chr4-47161723-DEL-104 | chr4-47161724-DEL-104 |
| chr4-47164939-DEL-289 | chr4-47164940-DEL-295 |
| chr4-47166400-DEL-191 | chr4-47166402-DEL-190 |
| chr4-47174117-DEL-74 | chr4-47174118-DEL-74 |
| chr4-47180221-DEL-165 | chr4-47180218-DEL-165 |
| chr4-47185792-DEL-57 | chr4-47185793-DEL-57 |
| chr4-47218537-DEL-56 | chr4-47218538-DEL-56 |
| chr4-47246831-DEL-73 | chr4-47246833-DEL-70 |
| chr4-47305940-DEL-177 | chr4-47305938-DEL-171 |
| chr4-47311630-DEL-318 | chr4-47311631-DEL-318 |
| chr4-47329823-DEL-64 | chr4-47329812-DEL-76 |
| chr4-47335212-DEL-77 | chr4-47335209-DEL-75 |
| chr4-47464668-DEL-80 | chr4-47464661-DEL-75 |
| chr4-47497343-DEL-399 | chr4-47497344-DEL-399 |
| chr4-47522147-DEL-553 | chr4-47522148-DEL-553 |
| chr4-47530129-DEL-65 | chr4-47530118-DEL-66 |
| chr4-47555055-DEL-2781 | chr4-47555056-DEL-2781 |
| chr4-47559709-DEL-115 | chr4-47559694-DEL-115 |
| chr4-47632994-DEL-113 | chr4-47632995-DEL-113 |
| chr4-47673160-DEL-5066 | chr4-47673153-DEL-5074 |
| chr4-47681937-DEL-1718 | chr4-47681938-DEL-1718 |
| chr4-47717326-DEL-411 | chr4-47717340-DEL-381 |
| chr4-47746062-DEL-141 | chr4-47746063-DEL-141 |
| chr4-47751260-DEL-636 | chr4-47751261-DEL-636 |

| | |
|---|---|
| chr4-47752586-DEL-80 | chr4-47752616-DEL-57 |
| chr4-47767240-DEL-2010 | chr4-47767241-DEL-2010 |
| chr4-47772117-DEL-54 | chr4-47772118-DEL-54 |
| chr4-47775850-DEL-4038 | chr4-47775850-DEL-4030 |
| chr4-47787356-DEL-263 | chr4-47787357-DEL-263 |
| chr4-47801362-DEL-172 | chr4-47801363-DEL-172 |
| chr4-47813348-DEL-59 | chr4-47813344-DEL-60 |
| chr4-47839853-DEL-808 | chr4-47839852-DEL-801 |
| chr4-47842814-DEL-51 | chr4-47842785-DEL-81 |
| chr4-47879415-DEL-92 | chr4-47879416-DEL-92 |
| chr4-47896574-DEL-61 | chr4-47896575-DEL-61 |
| chr4-47898157-DEL-160 | chr4-47898170-DEL-148 |
| chr4-47899494-DEL-74 | chr4-47899495-DEL-70 |
| chr4-47918528-DEL-152 | chr4-47918529-DEL-152 |
| chr4-47941464-DEL-88 | chr4-47941471-DEL-90 |
| chr4-47974443-DEL-872 | chr4-47974444-DEL-872 |
| chr4-47992385-DEL-51 | chr4-47992386-DEL-51 |
| chr4-48005576-DEL-70 | chr4-48005577-DEL-66 |
| chr4-48026523-DEL-1204 | chr4-48026514-DEL-1205 |
| chr4-48093621-DEL-346 | chr4-48093622-DEL-346 |
| chr4-48146907-DEL-1980 | chr4-48146908-DEL-1980 |
| chr4-48168729-DEL-138 | chr4-48168777-DEL-95 |
| chr4-48228739-DEL-62 | chr4-48228716-DEL-64 |
| chr4-48280271-DEL-710 | chr4-48280272-DEL-710 |
| chr4-48313979-DEL-75 | chr4-48313962-DEL-57 |
| chr4-48343576-DEL-1708 | chr4-48343577-DEL-1708 |
| chr4-48351632-DEL-64 | chr4-48351633-DEL-68 |
| chr4-48360493-DEL-211 | chr4-48360494-DEL-211 |
| chr4-48366612-DEL-54 | chr4-48366613-DEL-54 |
| chr4-48391492-DEL-55 | chr4-48391493-DEL-55 |
| chr4-48393915-DEL-196 | chr4-48393916-DEL-196 |
| chr4-48405954-DEL-251 | chr4-48405955-DEL-251 |
| chr4-48439977-DEL-1119 | chr4-48439978-DEL-1119 |
| chr4-48491301-DEL-369 | chr4-48491300-DEL-370 |
| chr4-48495417-DEL-68 | chr4-48495427-DEL-63 |
| chr4-48512946-DEL-221 | chr4-48512947-DEL-221 |
| chr4-48528126-DEL-487 | chr4-48528127-DEL-487 |
| chr4-48528842-DEL-176 | chr4-48528844-DEL-175 |
| chr4-48529887-DEL-222 | chr4-48529888-DEL-222 |
| chr4-48562217-DEL-135 | chr4-48562218-DEL-135 |
| chr4-48564470-DEL-71 | chr4-48564471-DEL-71 |
| chr4-48590646-DEL-196 | chr4-48590647-DEL-196 |
| chr4-48602148-DEL-332 | chr4-48602146-DEL-329 |
| chr4-48619199-DEL-433 | chr4-48619200-DEL-433 |
| chr4-48666349-DEL-210 | chr4-48666337-DEL-212 |
| chr4-48683653-DEL-345 | chr4-48683653-DEL-345 |
| chr4-48685191-DEL-317 | chr4-48685192-DEL-317 |
| chr4-48686611-DEL-505 | chr4-48686612-DEL-505 |
| chr4-48697044-DEL-204 | chr4-48697045-DEL-204 |
| chr4-48702515-DEL-443 | chr4-48702524-DEL-435 |
| chr4-48711560-DEL-168 | chr4-48711561-DEL-170 |
| chr4-48740881-DEL-343 | chr4-48740882-DEL-343 |
| chr4-48742399-DEL-2816 | chr4-48742400-DEL-2816 |
| chr4-48743685-DEL-498 | chr4-48743686-DEL-498 |
| chr4-48766443-DEL-170 | chr4-48766444-DEL-170 |
| chr4-48786193-DEL-336 | chr4-48786212-DEL-318 |
| chr4-48792964-DEL-430 | chr4-48792965-DEL-430 |
| chr4-48795135-DEL-6726 | chr4-48795136-DEL-6726 |
| chr4-48867963-DEL-1025 | chr4-48867964-DEL-1025 |
| chr4-48872714-DEL-575 | chr4-48872715-DEL-575 |
| chr4-48890041-DEL-572 | chr4-48890042-DEL-572 |
| chr4-48895673-DEL-197 | chr4-48895669-DEL-192 |
| chr4-48934367-DEL-52 | chr4-48934368-DEL-52 |
| chr4-48940333-DEL-106 | chr4-48940334-DEL-106 |
| chr4-48942871-DEL-54 | chr4-48942850-DEL-76 |
| chr4-48958206-DEL-187 | chr4-48958205-DEL-186 |
| chr4-48965563-DEL-1059 | chr4-48965564-DEL-1059 |
| chr4-48970075-DEL-89 | chr4-48970057-DEL-80 |
| chr4-48986104-DEL-649 | chr4-48986106-DEL-648 |
| chr4-49008618-DEL-4292 | chr4-49008619-DEL-4292 |
| chr4-49063717-DEL-426 | chr4-49063718-DEL-427 |
| chr4-49065147-DEL-3008 | chr4-49065146-DEL-3008 |
| chr4-49085977-DEL-1243 | chr4-49085978-DEL-1243 |
| chr4-49086078-DEL-71 | chr4-49086079-DEL-63 |
| chr4-49096772-DEL-86 | chr4-49096773-DEL-86 |
| chr4-49165478-DEL-467 | chr4-49165471-DEL-471 |
| chr4-49184082-DEL-78 | chr4-49184083-DEL-78 |
| chr4-49212299-DEL-56 | chr4-49212299-DEL-56 |
| chr4-49221222-DEL-106 | chr4-49221233-DEL-96 |
| chr4-49241811-DEL-809 | chr4-49241812-DEL-809 |

| | |
|---|---|
| chr4-49265741-DEL-6586 | chr4-49265742-DEL-6586 |
| chr4-49277263-DEL-224 | chr4-49277264-DEL-224 |
| chr4-49328507-DEL-7298 | chr4-49328508-DEL-7298 |
| chr4-49342685-DEL-138 | chr4-49342686-DEL-138 |
| chr4-49347151-DEL-240 | chr4-49347151-DEL-240 |
| chr4-49384191-DEL-2306 | chr4-49384192-DEL-2306 |
| chr4-49390750-DEL-50 | chr4-49390740-DEL-58 |
| chr4-49406955-DEL-420 | chr4-49406955-DEL-421 |
| chr4-49419070-DEL-62 | chr4-49419070-DEL-62 |
| chr4-49466362-DEL-212 | chr4-49466363-DEL-212 |
| chr4-49561383-DEL-1876 | chr4-49561384-DEL-1876 |
| chr4-49604748-DEL-54 | chr4-49604747-DEL-53 |
| chr4-49622278-DEL-56 | chr4-49622254-DEL-59 |
| chr4-49628648-DEL-75 | chr4-49628649-DEL-75 |
| chr4-49642594-DEL-95 | chr4-49642595-DEL-95 |
| chr4-49643469-DEL-160 | chr4-49643470-DEL-160 |
| chr4-49643988-DEL-108 | chr4-49643989-DEL-108 |
| chr4-49686463-DEL-596 | chr4-49686477-DEL-593 |
| chr4-49698554-DEL-1488 | chr4-49698555-DEL-1488 |
| chr4-49731430-DEL-7046 | chr4-49731431-DEL-7046 |
| chr4-49731997-DEL-208 | chr4-49731998-DEL-208 |
| chr4-49753620-DEL-87 | chr4-49753621-DEL-87 |
| chr4-49791516-DEL-90 | chr4-49791525-DEL-82 |
| chr4-49802852-DEL-54 | chr4-49802815-DEL-58 |
| chr4-49808561-DEL-1424 | chr4-49808562-DEL-1424 |
| chr4-49813530-DEL-249 | chr4-49813531-DEL-249 |
| chr4-49817047-DEL-68 | chr4-49817048-DEL-68 |
| chr4-49818161-DEL-510 | chr4-49818162-DEL-510 |
| chr4-49818769-DEL-96 | chr4-49818770-DEL-96 |
| chr4-49824925-DEL-77 | chr4-49824926-DEL-77 |
| chr4-49847655-DEL-57 | chr4-49847656-DEL-57 |
| chr4-49848274-DEL-135 | chr4-49848277-DEL-133 |
| chr4-49888446-DEL-87 | chr4-49888449-DEL-123 |
| chr4-49993709-DEL-546 | chr4-49993710-DEL-546 |
| chr4-50056958-DEL-6232 | chr4-50056959-DEL-6232 |
| chr4-50104451-DEL-191 | chr4-50104449-DEL-192 |
| chr4-50114421-DEL-371 | chr4-50114422-DEL-371 |
| chr4-50120188-DEL-53 | chr4-50120189-DEL-53 |
| chr4-50124393-DEL-736 | chr4-50124394-DEL-736 |
| chr4-50134021-DEL-67 | chr4-50134022-DEL-67 |
| chr4-50136888-DEL-1074 | chr4-50136889-DEL-1074 |
| chr4-50155652-DEL-2460 | chr4-50155653-DEL-2460 |
| chr4-50164953-DEL-594 | chr4-50164954-DEL-594 |
| chr4-50165768-DEL-6303 | chr4-50165769-DEL-6303 |
| chr4-50177779-DEL-4511 | chr4-50177780-DEL-4511 |
| chr4-50184173-DEL-69 | chr4-50184174-DEL-69 |
| chr4-50191019-DEL-6380 | chr4-50191013-DEL-6379 |
| chr4-50196967-DEL-208 | chr4-50196968-DEL-208 |
| chr4-50252544-DEL-56 | chr4-50252545-DEL-56 |
| chr4-50261753-DEL-120 | chr4-50261754-DEL-120 |
| chr4-50266968-DEL-84 | chr4-50266969-DEL-85 |
| chr4-50316035-DEL-887 | chr4-50316036-DEL-887 |
| chr4-50319793-DEL-473 | chr4-50319794-DEL-473 |
| chr4-50327564-DEL-2117 | chr4-50327563-DEL-2114 |
| chr4-50346238-DEL-2895 | chr4-50346239-DEL-2895 |
| chr4-50358491-DEL-247 | chr4-50358488-DEL-233 |
| chr4-50380355-DEL-6589 | chr4-50380356-DEL-6589 |
| chr4-50406252-DEL-2762 | chr4-50406253-DEL-2762 |
| chr4-50409530-DEL-65 | chr4-50409531-DEL-65 |
| chr4-50411004-DEL-1545 | chr4-50411005-DEL-1545 |
| chr4-50417916-DEL-130 | chr4-50417917-DEL-130 |
| chr4-50441365-DEL-64 | chr4-50441357-DEL-57 |
| chr4-50454330-DEL-58 | chr4-50454331-DEL-58 |
| chr4-50524741-DEL-201 | chr4-50524741-DEL-201 |
| chr4-50528012-DEL-7725 | chr4-50528013-DEL-7725 |
| chr4-50535885-DEL-352 | chr4-50535885-DEL-352 |
| chr4-50552003-DEL-54 | chr4-50552004-DEL-54 |
| chr4-50562392-DEL-839 | chr4-50562393-DEL-839 |
| chr4-50565163-DEL-1688 | chr4-50565163-DEL-1682 |
| chr4-50568074-DEL-514 | chr4-50568075-DEL-514 |
| chr4-50571352-DEL-63 | chr4-50571353-DEL-63 |
| chr4-50574394-DEL-58 | chr4-50574389-DEL-64 |
| chr4-50622015-DEL-401 | chr4-50622016-DEL-401 |
| chr4-50623567-DEL-7584 | chr4-50623568-DEL-7584 |
| chr4-50624905-DEL-1340 | chr4-50624905-DEL-1340 |
| chr4-50654811-DEL-115 | chr4-50654814-DEL-116 |
| chr4-50659503-DEL-1995 | chr4-50659502-DEL-1996 |
| chr4-50687480-DEL-7110 | chr4-50687481-DEL-7110 |
| chr4-50736182-DEL-320 | chr4-50736183-DEL-320 |
| chr4-50770113-DEL-401 | chr4-50770083-DEL-432 |

| | |
|---|---|
| chr4-50780550-DEL-88 | chr4-50780551-DEL-88 |
| chr4-50785777-DEL-113 | chr4-50785778-DEL-113 |
| chr4-50791303-DEL-1230 | chr4-50791304-DEL-1230 |
| chr4-50797976-DEL-108 | chr4-50797965-DEL-92 |
| chr4-50802632-DEL-51 | chr4-50802633-DEL-51 |
| chr4-50815780-DEL-236 | chr4-50815781-DEL-236 |
| chr4-50826758-DEL-52 | chr4-50826758-DEL-52 |
| chr4-50845874-DEL-3251 | chr4-50845873-DEL-3251 |
| chr4-50848822-DEL-549 | chr4-50848823-DEL-549 |
| chr4-50855199-DEL-7484 | chr4-50855200-DEL-7484 |
| chr4-50891267-DEL-99 | chr4-50891268-DEL-99 |
| chr4-50892848-DEL-363 | chr4-50892849-DEL-363 |
| chr4-50908598-DEL-3665 | chr4-50908599-DEL-3665 |
| chr4-50915311-DEL-84 | chr4-50915312-DEL-85 |
| chr4-50917357-DEL-821 | chr4-50917358-DEL-821 |
| chr4-50921623-DEL-433 | chr4-50921611-DEL-432 |
| chr4-50938935-DEL-67 | chr4-50938936-DEL-67 |
| chr4-50973906-DEL-760 | chr4-50973907-DEL-760 |
| chr4-51012283-DEL-6423 | chr4-51012287-DEL-6422 |
| chr4-51018307-DEL-208 | chr4-51018308-DEL-208 |
| chr4-51027390-DEL-389 | chr4-51027391-DEL-389 |
| chr4-51038188-DEL-3927 | chr4-51038189-DEL-3927 |
| chr4-51067272-DEL-6275 | chr4-51067273-DEL-6275 |
| chr4-51098255-DEL-7146 | chr4-51098256-DEL-7146 |
| chr4-51162310-DEL-2293 | chr4-51162311-DEL-2293 |
| chr4-51200879-DEL-82 | chr4-51200874-DEL-81 |
| chr4-51228990-DEL-81 | chr4-51228991-DEL-81 |
| chr4-51252457-DEL-1906 | chr4-51252472-DEL-1892 |
| chr4-51261652-DEL-532 | chr4-51261653-DEL-533 |
| chr4-51267182-DEL-310 | chr4-51267183-DEL-310 |
| chr4-51289785-DEL-1041 | chr4-51289786-DEL-1041 |
| chr4-51298865-DEL-7000 | chr4-51298866-DEL-7000 |
| chr4-51341247-DEL-1060 | chr4-51341252-DEL-1061 |
| chr4-51352264-DEL-80 | chr4-51352265-DEL-80 |
| chr4-51371085-DEL-81 | chr4-51371086-DEL-80 |
| chr4-51373090-DEL-84 | chr4-51373089-DEL-78 |
| chr4-51536395-DEL-81 | chr4-51536396-DEL-81 |
| chr4-51581570-DEL-103 | chr4-51581549-DEL-101 |
| chr4-51587563-DEL-2404 | chr4-51587563-DEL-2404 |
| chr4-51614282-DEL-57 | chr4-51614283-DEL-57 |
| chr4-51625770-DEL-95 | chr4-51625771-DEL-95 |
| chr4-51626689-DEL-116 | chr4-51626670-DEL-117 |
| chr4-51648217-DEL-6784 | chr4-51648218-DEL-6784 |
| chr4-51654598-DEL-212 | chr4-51654599-DEL-212 |
| chr4-51678606-DEL-66 | chr4-51678607-DEL-66 |
| chr4-51691245-DEL-124 | chr4-51691246-DEL-124 |
| chr4-51778769-DEL-527 | chr4-51778770-DEL-527 |
| chr4-51782138-DEL-213 | chr4-51782139-DEL-213 |
| chr4-51803696-DEL-65 | chr4-51803697-DEL-65 |
| chr4-51805639-DEL-456 | chr4-51805640-DEL-456 |
| chr4-51810039-DEL-2423 | chr4-51810040-DEL-2423 |
| chr4-51812420-DEL-226 | chr4-51812421-DEL-226 |
| chr4-51818784-DEL-66 | chr4-51818785-DEL-66 |
| chr4-51825162-DEL-6622 | chr4-51825163-DEL-6622 |
| chr4-51831594-DEL-263 | chr4-51831595-DEL-263 |
| chr4-51832676-DEL-172 | chr4-51832685-DEL-164 |
| chr4-51849770-DEL-494 | chr4-51849771-DEL-494 |
| chr4-51871445-DEL-8424 | chr4-51871445-DEL-8420 |
| chr4-51885831-DEL-1118 | chr4-51885832-DEL-1118 |
| chr4-51894688-DEL-5391 | chr4-51894689-DEL-5391 |
| chr4-51924689-DEL-71 | chr4-51924690-DEL-71 |
| chr4-51967251-DEL-182 | chr4-51967252-DEL-182 |
| chr4-51974328-DEL-1251 | chr4-51974329-DEL-1251 |
| chr4-52142188-DEL-2361 | chr4-52142189-DEL-2361 |
| chr4-52145422-DEL-108 | chr4-52145423-DEL-108 |
| chr4-52150424-DEL-172 | chr4-52150425-DEL-173 |
| chr4-52161602-DEL-382 | chr4-52161603-DEL-381 |
| chr4-52177663-DEL-86 | chr4-52177696-DEL-56 |
| chr4-52186175-DEL-638 | chr4-52186176-DEL-638 |
| chr4-52186998-DEL-4456 | chr4-52186999-DEL-4456 |
| chr4-52202767-DEL-788 | chr4-52202766-DEL-788 |
| chr4-52223553-DEL-1450 | chr4-52223554-DEL-1450 |
| chr4-52228568-DEL-6274 | chr4-52228559-DEL-6275 |
| chr4-52248483-DEL-64 | chr4-52248484-DEL-64 |
| chr4-52257967-DEL-418 | chr4-52257968-DEL-418 |
| chr4-52266745-DEL-2136 | chr4-52266746-DEL-2136 |
| chr4-52330326-DEL-149 | chr4-52330328-DEL-148 |
| chr4-52378169-DEL-478 | chr4-52378170-DEL-478 |
| chr4-52379286-DEL-6471 | chr4-52379287-DEL-6471 |
| chr4-52405004-DEL-57 | chr4-52405005-DEL-57 |

| | |
|---|---|
| chr4-52407418-DEL-3691 | chr4-52407419-DEL-3691 |
| chr4-52429851-DEL-97 | chr4-52429867-DEL-87 |
| chr4-52432082-DEL-87 | chr4-52432083-DEL-71 |
| chr4-52433026-DEL-206 | chr4-52433027-DEL-206 |
| chr4-52459325-DEL-53 | chr4-52459326-DEL-53 |
| chr4-52494155-DEL-112 | chr4-52494161-DEL-68 |
| chr4-52517948-DEL-199 | chr4-52517949-DEL-199 |
| chr4-52529676-DEL-325 | chr4-52529677-DEL-325 |
| chr4-52535836-DEL-201 | chr4-52535837-DEL-201 |
| chr4-52545140-DEL-90 | chr4-52545141-DEL-90 |
| chr4-52570264-DEL-941 | chr4-52570265-DEL-941 |
| chr4-52572788-DEL-56 | chr4-52572789-DEL-56 |
| chr4-52606391-DEL-52 | chr4-52606392-DEL-52 |
| chr4-52615010-DEL-489 | chr4-52615011-DEL-489 |
| chr4-52640555-DEL-215 | chr4-52640556-DEL-215 |
| chr4-52674168-DEL-6345 | chr4-52674169-DEL-6345 |
| chr4-52683159-DEL-564 | chr4-52683160-DEL-564 |
| chr4-52708696-DEL-434 | chr4-52708697-DEL-434 |
| chr4-52749728-DEL-285 | chr4-52749729-DEL-285 |
| chr4-52760597-DEL-768 | chr4-52760615-DEL-732 |
| chr4-52764374-DEL-56 | chr4-52764375-DEL-57 |
| chr4-52769680-DEL-94 | chr4-52769678-DEL-91 |
| chr4-52774955-DEL-382 | chr4-52774956-DEL-382 |
| chr4-52775446-DEL-587 | chr4-52775462-DEL-587 |
| chr4-52788822-DEL-558 | chr4-52788823-DEL-558 |
| chr4-52819878-DEL-65 | chr4-52819881-DEL-63 |
| chr4-52844805-DEL-104 | chr4-52844806-DEL-104 |
| chr4-52845592-DEL-691 | chr4-52845594-DEL-690 |
| chr4-52860235-DEL-239 | chr4-52860216-DEL-233 |
| chr4-52887216-DEL-5126 | chr4-52887217-DEL-5126 |
| chr4-52924823-DEL-802 | chr4-52924823-DEL-802 |
| chr4-52937050-DEL-216 | chr4-52937051-DEL-216 |
| chr4-52942485-DEL-69 | chr4-52942482-DEL-65 |
| chr4-52963849-DEL-151 | chr4-52963850-DEL-151 |
| chr4-52998662-DEL-695 | chr4-52998663-DEL-695 |
| chr4-53003081-DEL-1194 | chr4-53003082-DEL-1194 |
| chr4-53005004-DEL-71 | chr4-53004963-DEL-71 |
| chr4-53052066-DEL-7148 | chr4-53052067-DEL-7148 |
| chr4-53126490-DEL-70 | chr4-53126491-DEL-68 |
| chr4-53160635-DEL-51 | chr4-53160610-DEL-89 |
| chr4-53164144-DEL-85 | chr4-53164126-DEL-88 |
| chr4-53165747-DEL-90 | chr4-53165751-DEL-86 |
| chr4-53173689-DEL-128 | chr4-53173735-DEL-91 |
| chr4-53225209-DEL-76 | chr4-53225208-DEL-82 |
| chr4-53229033-DEL-400 | chr4-53229034-DEL-400 |
| chr4-53262835-DEL-139 | chr4-53262859-DEL-107 |
| chr4-53271310-DEL-52 | chr4-53271311-DEL-52 |
| chr4-53272922-DEL-200 | chr4-53272923-DEL-200 |
| chr4-53286760-DEL-650 | chr4-53286736-DEL-674 |
| chr4-53289648-DEL-2335 | chr4-53289649-DEL-2335 |
| chr4-53293092-DEL-81 | chr4-53293093-DEL-81 |
| chr4-53293479-DEL-400 | chr4-53293480-DEL-400 |
| chr4-53301952-DEL-198 | chr4-53301953-DEL-198 |
| chr4-53320895-DEL-51 | chr4-53320896-DEL-51 |
| chr4-53328468-DEL-6403 | chr4-53328469-DEL-6403 |
| chr4-53341566-DEL-593 | chr4-53341567-DEL-592 |
| chr4-53383853-DEL-52 | chr4-53383854-DEL-52 |
| chr4-53386784-DEL-6318 | chr4-53386785-DEL-6318 |
| chr4-53394004-DEL-104 | chr4-53394019-DEL-100 |
| chr4-53399015-DEL-60 | chr4-53399016-DEL-60 |
| chr4-53403770-DEL-493 | chr4-53403771-DEL-493 |
| chr4-53422371-DEL-552 | chr4-53422372-DEL-552 |
| chr4-53476472-DEL-74 | chr4-53476473-DEL-62 |
| chr4-53494281-DEL-212 | chr4-53494282-DEL-212 |
| chr4-53539397-DEL-62 | chr4-53539405-DEL-62 |
| chr4-53559089-DEL-418 | chr4-53559078-DEL-399 |
| chr4-53608058-DEL-6440 | chr4-53608059-DEL-6440 |
| chr4-53623688-DEL-55 | chr4-53623684-DEL-55 |
| chr4-53668386-DEL-7529 | chr4-53668387-DEL-7529 |
| chr4-53669598-DEL-212 | chr4-53669597-DEL-212 |
| chr4-53678458-DEL-64 | chr4-53678458-DEL-65 |
| chr4-53719631-DEL-68 | chr4-53719632-DEL-66 |
| chr4-53728571-DEL-88 | chr4-53728573-DEL-87 |
| chr4-53752702-DEL-101 | chr4-53752733-DEL-102 |
| chr4-53759567-DEL-138 | chr4-53759568-DEL-138 |
| chr4-53778357-DEL-98 | chr4-53778368-DEL-50 |
| chr4-53783300-DEL-62 | chr4-53783301-DEL-62 |
| chr4-53784163-DEL-193 | chr4-53784164-DEL-193 |
| chr4-53807585-DEL-75 | chr4-53807537-DEL-80 |
| chr4-53837880-DEL-78 | chr4-53837881-DEL-78 |

| | |
|---|---|
| chr4-53864340-DEL-316 | chr4-53864341-DEL-316 |
| chr4-53930666-DEL-397 | chr4-53930667-DEL-397 |
| chr4-53937100-DEL-674 | chr4-53937101-DEL-674 |
| chr4-53970900-DEL-52 | chr4-53970901-DEL-52 |
| chr4-53995796-DEL-385 | chr4-53995797-DEL-385 |
| chr4-53997471-DEL-68 | chr4-53997460-DEL-71 |
| chr4-54011343-DEL-54 | chr4-54011344-DEL-54 |
| chr4-54019426-DEL-164 | chr4-54019427-DEL-164 |
| chr4-54040127-DEL-78 | chr4-54040118-DEL-74 |
| chr4-54042559-DEL-160 | chr4-54042560-DEL-185 |
| chr4-54042940-DEL-423 | chr4-54042951-DEL-413 |
| chr4-54072413-DEL-104 | chr4-54072434-DEL-58 |
| chr4-54109309-DEL-54 | chr4-54109310-DEL-54 |
| chr4-54133466-DEL-512 | chr4-54133467-DEL-515 |
| chr4-54138048-DEL-180 | chr4-54138054-DEL-139 |
| chr4-54140469-DEL-61 | chr4-54140474-DEL-64 |
| chr4-54149456-DEL-6389 | chr4-54149452-DEL-6387 |
| chr4-54171547-DEL-126 | chr4-54171544-DEL-130 |
| chr4-54214671-DEL-6390 | chr4-54214672-DEL-6390 |
| chr4-54220646-DEL-201 | chr4-54220646-DEL-201 |
| chr4-54235389-DEL-110 | chr4-54235390-DEL-110 |
| chr4-54239512-DEL-100 | chr4-54239525-DEL-88 |
| chr4-54243564-DEL-6396 | chr4-54243565-DEL-6396 |
| chr4-54256547-DEL-213 | chr4-54256548-DEL-219 |
| chr4-54287653-DEL-60 | chr4-54287654-DEL-60 |
| chr4-54301802-DEL-737 | chr4-54301803-DEL-737 |
| chr4-54394731-DEL-99 | chr4-54394731-DEL-99 |
| chr4-54440996-DEL-6826 | chr4-54440988-DEL-6827 |
| chr4-54447472-DEL-1542 | chr4-54447472-DEL-1542 |
| chr4-54484672-DEL-6375 | chr4-54484673-DEL-6375 |
| chr4-54495946-DEL-194 | chr4-54495947-DEL-194 |
| chr4-54508982-DEL-135 | chr4-54509027-DEL-107 |
| chr4-54516334-DEL-74 | chr4-54516322-DEL-62 |
| chr4-54524983-DEL-52 | chr4-54524984-DEL-51 |
| chr4-54553167-DEL-60 | chr4-54553167-DEL-60 |
| chr4-54564099-DEL-60 | chr4-54564100-DEL-60 |
| chr4-54564672-DEL-315 | chr4-54564673-DEL-315 |
| chr4-54572350-DEL-60 | chr4-54572351-DEL-60 |
| chr4-54586786-DEL-52 | chr4-54586779-DEL-60 |
| chr4-54626291-DEL-72 | chr4-54626292-DEL-60 |
| chr4-54626670-DEL-6485 | chr4-54626671-DEL-6485 |
| chr4-54634487-DEL-64 | chr4-54634488-DEL-64 |
| chr4-54640166-DEL-58 | chr4-54640167-DEL-58 |
| chr4-54651566-DEL-60 | chr4-54651567-DEL-72 |
| chr4-54655397-DEL-6669 | chr4-54655398-DEL-6668 |
| chr4-54670180-DEL-6704 | chr4-54670181-DEL-6704 |
| chr4-54685633-DEL-70 | chr4-54685634-DEL-70 |
| chr4-54718756-DEL-86 | chr4-54718757-DEL-84 |
| chr4-54729787-DEL-56 | chr4-54729791-DEL-55 |
| chr4-54738859-DEL-52 | chr4-54738862-DEL-50 |
| chr4-54750721-DEL-80 | chr4-54750720-DEL-80 |
| chr4-54755024-DEL-572 | chr4-54755025-DEL-572 |
| chr4-54793332-DEL-94 | chr4-54793337-DEL-90 |
| chr4-54794747-DEL-60 | chr4-54794748-DEL-60 |
| chr4-54806250-DEL-542 | chr4-54806251-DEL-542 |
| chr4-54817317-DEL-60 | chr4-54817317-DEL-60 |
| chr4-54821028-DEL-6286 | chr4-54821029-DEL-6286 |
| chr4-54827111-DEL-65 | chr4-54827112-DEL-65 |
| chr4-54828323-DEL-90 | chr4-54828332-DEL-78 |
| chr4-54841637-DEL-395 | chr4-54841635-DEL-395 |
| chr4-54846401-DEL-96 | chr4-54846402-DEL-96 |
| chr4-54870282-DEL-83 | chr4-54870289-DEL-80 |
| chr4-54879713-DEL-2104 | chr4-54879714-DEL-2104 |
| chr4-54884824-DEL-52 | chr4-54884825-DEL-52 |
| chr4-54908456-DEL-221 | chr4-54908457-DEL-221 |
| chr4-54917383-DEL-68 | chr4-54917384-DEL-66 |
| chr4-54970662-DEL-52 | chr4-54970663-DEL-72 |
| chr4-54983434-DEL-240 | chr4-54983435-DEL-241 |
| chr4-54998981-DEL-52 | chr4-54998985-DEL-55 |
| chr4-55064885-DEL-66 | chr4-55064890-DEL-58 |
| chr4-55067820-DEL-52 | chr4-55067821-DEL-52 |
| chr4-55085120-DEL-51 | chr4-55085086-DEL-74 |
| chr4-55096643-DEL-126 | chr4-55096628-DEL-128 |
| chr4-55109870-DEL-65 | chr4-55109876-DEL-68 |
| chr4-55129497-DEL-136 | chr4-55129500-DEL-132 |
| chr4-55150892-DEL-74 | chr4-55150893-DEL-74 |
| chr4-55159673-DEL-228 | chr4-55159674-DEL-228 |
| chr4-55167659-DEL-56 | chr4-55167660-DEL-56 |
| chr4-55177411-DEL-92 | chr4-55177412-DEL-91 |
| chr4-55197028-DEL-182 | chr4-55197029-DEL-182 |

| | |
|---|---|
| chr4-55207040-DEL-93 | chr4-55207041-DEL-93 |
| chr4-55219529-DEL-92 | chr4-55219530-DEL-92 |
| chr4-55238858-DEL-209 | chr4-55238859-DEL-209 |
| chr4-55246160-DEL-1254 | chr4-55246161-DEL-1254 |
| chr4-55266203-DEL-264 | chr4-55266204-DEL-264 |
| chr4-55267986-DEL-102 | chr4-55267987-DEL-60 |
| chr4-55278469-DEL-55 | chr4-55278470-DEL-55 |
| chr4-55287911-DEL-77 | chr4-55287912-DEL-77 |
| chr4-55289543-DEL-210 | chr4-55289544-DEL-210 |
| chr4-55299841-DEL-144 | chr4-55299842-DEL-144 |
| chr4-55303824-DEL-191 | chr4-55303825-DEL-191 |
| chr4-55308698-DEL-7477 | chr4-55308699-DEL-7477 |
| chr4-55324478-DEL-377 | chr4-55324480-DEL-376 |
| chr4-55329138-DEL-205 | chr4-55329139-DEL-205 |
| chr4-55337724-DEL-54 | chr4-55337727-DEL-52 |
| chr4-55354761-DEL-56 | chr4-55354760-DEL-50 |
| chr4-55364696-DEL-489 | chr4-55364696-DEL-488 |
| chr4-55397473-DEL-66 | chr4-55397474-DEL-66 |
| chr4-55404718-DEL-96 | chr4-55404719-DEL-96 |
| chr4-55409369-DEL-410 | chr4-55409370-DEL-410 |
| chr4-55417565-DEL-165 | chr4-55417566-DEL-165 |
| chr4-55418203-DEL-5125 | chr4-55418204-DEL-5125 |
| chr4-55429593-DEL-178 | chr4-55429594-DEL-178 |
| chr4-55430666-DEL-75 | chr4-55430667-DEL-75 |
| chr4-55433018-DEL-208 | chr4-55433019-DEL-208 |
| chr4-55435160-DEL-82 | chr4-55435171-DEL-52 |
| chr4-55436064-DEL-202 | chr4-55436065-DEL-202 |
| chr4-55446382-DEL-104 | chr4-55446384-DEL-102 |
| chr4-55450211-DEL-203 | chr4-55450212-DEL-203 |
| chr4-55455661-DEL-255 | chr4-55455662-DEL-255 |
| chr4-55459288-DEL-68 | chr4-55459289-DEL-68 |
| chr4-55466215-DEL-78 | chr4-55466216-DEL-78 |
| chr4-55467855-DEL-222 | chr4-55467856-DEL-222 |
| chr4-55469686-DEL-64 | chr4-55469689-DEL-72 |
| chr4-55473417-DEL-58 | chr4-55473418-DEL-58 |
| chr4-55482495-DEL-98 | chr4-55482496-DEL-98 |
| chr4-55483236-DEL-2667 | chr4-55483237-DEL-2667 |
| chr4-55483910-DEL-733 | chr4-55483911-DEL-733 |
| chr4-55491895-DEL-252 | chr4-55491896-DEL-252 |
| chr4-55507032-DEL-160 | chr4-55507020-DEL-174 |
| chr4-55517346-DEL-51 | chr4-55517347-DEL-51 |
| chr4-55528675-DEL-305 | chr4-55528674-DEL-306 |
| chr4-55537115-DEL-365 | chr4-55537110-DEL-390 |
| chr4-55590355-DEL-80 | chr4-55590340-DEL-96 |
| chr4-55636106-DEL-65 | chr4-55636114-DEL-77 |
| chr4-55654783-DEL-60 | chr4-55654776-DEL-72 |
| chr4-55666446-DEL-107 | chr4-55666446-DEL-107 |
| chr4-55688536-DEL-53 | chr4-55688537-DEL-53 |
| chr4-55767414-DEL-93 | chr4-55767415-DEL-93 |
| chr4-55773811-DEL-996 | chr4-55773812-DEL-996 |
| chr4-55798688-DEL-60 | chr4-55798688-DEL-68 |
| chr4-55833326-DEL-52 | chr4-55833327-DEL-53 |
| chr4-55846931-DEL-53 | chr4-55846919-DEL-50 |
| chr4-55852955-DEL-1061 | chr4-55852956-DEL-1061 |
| chr4-55861383-DEL-6265 | chr4-55861384-DEL-6265 |
| chr4-55894975-DEL-83 | chr4-55894988-DEL-67 |
| chr4-55901369-DEL-762 | chr4-55901370-DEL-762 |
| chr4-55917197-DEL-70 | chr4-55917198-DEL-70 |
| chr4-55939795-DEL-55 | chr4-55939795-DEL-56 |
| chr4-55991079-DEL-545 | chr4-55991080-DEL-545 |
| chr4-55994504-DEL-61 | chr4-55994505-DEL-61 |
| chr4-56008368-DEL-108 | chr4-56008374-DEL-109 |
| chr4-56039829-DEL-6199 | chr4-56039832-DEL-6186 |
| chr4-56047943-DEL-106 | chr4-56047944-DEL-106 |
| chr4-56075448-DEL-58 | chr4-56075449-DEL-58 |
| chr4-56077560-DEL-56 | chr4-56077565-DEL-55 |
| chr4-56096550-DEL-157 | chr4-56096551-DEL-157 |
| chr4-56131876-DEL-1405 | chr4-56131877-DEL-1405 |
| chr4-56141672-DEL-69 | chr4-56141670-DEL-60 |
| chr4-56189025-DEL-54 | chr4-56189028-DEL-52 |
| chr4-56209899-DEL-878 | chr4-56209900-DEL-878 |
| chr4-56246701-DEL-91 | chr4-56246669-DEL-78 |
| chr4-56268697-DEL-54 | chr4-56268698-DEL-54 |
| chr4-56279227-DEL-56 | chr4-56279227-DEL-56 |
| chr4-56300016-DEL-1603 | chr4-56300017-DEL-1603 |
| chr4-56303768-DEL-76 | chr4-56303809-DEL-82 |
| chr4-56329175-DEL-89 | chr4-56329172-DEL-100 |
| chr4-56330141-DEL-52 | chr4-56330132-DEL-58 |
| chr4-56391169-DEL-166 | chr4-56391170-DEL-165 |
| chr4-56392026-DEL-538 | chr4-56392036-DEL-540 |

| | |
|---|---|
| chr4-56408300-DEL-397 | chr4-56408301-DEL-397 |
| chr4-56425668-DEL-288 | chr4-56425669-DEL-288 |
| chr4-56444280-DEL-982 | chr4-56444320-DEL-945 |
| chr4-56445242-DEL-89 | chr4-56445243-DEL-89 |
| chr4-56447732-DEL-134 | chr4-56447683-DEL-140 |
| chr4-56452067-DEL-162 | chr4-56452068-DEL-160 |
| chr4-56461286-DEL-62 | chr4-56461287-DEL-62 |
| chr4-56462979-DEL-92 | chr4-56462980-DEL-92 |
| chr4-56470270-DEL-276 | chr4-56470271-DEL-276 |
| chr4-56497655-DEL-87 | chr4-56497656-DEL-87 |
| chr4-56508686-DEL-544 | chr4-56508687-DEL-544 |
| chr4-56511059-DEL-1013 | chr4-56511060-DEL-1013 |
| chr4-56515825-DEL-2946 | chr4-56515826-DEL-2946 |
| chr4-56559876-DEL-199 | chr4-56559838-DEL-199 |
| chr4-56585604-DEL-1215 | chr4-56585605-DEL-1215 |
| chr4-56627790-DEL-66 | chr4-56627791-DEL-70 |
| chr4-56690737-DEL-426 | chr4-56690746-DEL-442 |
| chr4-56700023-DEL-108 | chr4-56700060-DEL-72 |
| chr4-56729030-DEL-6419 | chr4-56729031-DEL-6419 |
| chr4-56737136-DEL-339 | chr4-56737137-DEL-339 |
| chr4-56764101-DEL-199 | chr4-56764102-DEL-199 |
| chr4-56821474-DEL-189 | chr4-56821475-DEL-189 |
| chr4-56828728-DEL-100 | chr4-56828729-DEL-100 |
| chr4-56845276-DEL-211 | chr4-56845277-DEL-211 |
| chr4-56849989-DEL-222 | chr4-56849989-DEL-220 |
| chr4-56855253-DEL-174 | chr4-56855254-DEL-174 |
| chr4-56857447-DEL-211 | chr4-56857448-DEL-211 |
| chr4-56862696-DEL-72 | chr4-56862697-DEL-72 |
| chr4-56875663-DEL-212 | chr4-56875664-DEL-212 |
| chr4-56897205-DEL-157 | chr4-56897206-DEL-157 |
| chr4-56919287-DEL-94 | chr4-56919288-DEL-94 |
| chr4-56930937-DEL-2163 | chr4-56930938-DEL-2163 |
| chr4-56982009-DEL-54 | chr4-56982010-DEL-54 |
| chr4-57000897-DEL-245 | chr4-57000898-DEL-245 |
| chr4-57017638-DEL-400 | chr4-57017639-DEL-400 |
| chr4-57104942-DEL-138 | chr4-57104943-DEL-138 |
| chr4-57110242-DEL-71 | chr4-57110243-DEL-71 |
| chr4-57111156-DEL-198 | chr4-57111157-DEL-198 |
| chr4-57161370-DEL-56 | chr4-57161360-DEL-59 |
| chr4-57161885-DEL-189 | chr4-57161886-DEL-189 |
| chr4-57188746-DEL-79 | chr4-57188747-DEL-79 |
| chr4-57189511-DEL-201 | chr4-57189512-DEL-201 |
| chr4-57233038-DEL-58 | chr4-57233039-DEL-58 |
| chr4-57291575-DEL-373 | chr4-57291576-DEL-373 |
| chr4-57294341-DEL-204 | chr4-57294342-DEL-204 |
| chr4-57324426-DEL-174 | chr4-57324427-DEL-174 |
| chr4-57327660-DEL-72 | chr4-57327661-DEL-72 |
| chr4-57340081-DEL-148 | chr4-57340081-DEL-148 |
| chr4-57344482-DEL-59 | chr4-57344483-DEL-59 |
| chr4-57348021-DEL-213 | chr4-57348017-DEL-175 |
| chr4-57370730-DEL-1296 | chr4-57370731-DEL-1296 |
| chr4-57409130-DEL-340 | chr4-57409128-DEL-341 |
| chr4-57431834-DEL-100 | chr4-57431831-DEL-104 |
| chr4-57448833-DEL-2639 | chr4-57448834-DEL-2639 |
| chr4-57461985-DEL-504 | chr4-57461986-DEL-504 |
| chr4-57476787-DEL-1322 | chr4-57476788-DEL-1322 |
| chr4-57501653-DEL-196 | chr4-57501654-DEL-196 |
| chr4-57510500-DEL-60 | chr4-57510501-DEL-60 |
| chr4-57527154-DEL-1193 | chr4-57527155-DEL-1193 |
| chr4-57546646-DEL-88 | chr4-57546651-DEL-86 |
| chr4-57578776-DEL-59 | chr4-57578777-DEL-50 |
| chr4-57609041-DEL-299 | chr4-57609040-DEL-299 |
| chr4-57626742-DEL-113 | chr4-57626743-DEL-110 |
| chr4-57652680-DEL-91 | chr4-57652664-DEL-91 |
| chr4-57663182-DEL-308 | chr4-57663183-DEL-308 |
| chr4-57683642-DEL-411 | chr4-57683643-DEL-411 |
| chr4-57692537-DEL-55 | chr4-57692538-DEL-55 |
| chr4-57694671-DEL-89 | chr4-57694671-DEL-89 |
| chr4-57695730-DEL-115 | chr4-57695722-DEL-95 |
| chr4-57747449-DEL-62 | chr4-57747472-DEL-56 |
| chr4-57767754-DEL-454 | chr4-57767755-DEL-454 |
| chr4-57772193-DEL-307 | chr4-57772196-DEL-308 |
| chr4-57779644-DEL-65 | chr4-57779644-DEL-65 |
| chr4-57781778-DEL-58 | chr4-57781779-DEL-58 |
| chr4-57788041-DEL-108 | chr4-57788042-DEL-108 |
| chr4-57791671-DEL-171 | chr4-57791678-DEL-165 |
| chr4-57793597-DEL-54 | chr4-57793598-DEL-54 |
| chr4-57826162-DEL-1587 | chr4-57826163-DEL-1587 |
| chr4-57837331-DEL-114 | chr4-57837332-DEL-114 |
| chr4-57849268-DEL-57 | chr4-57849269-DEL-57 |

| | |
|---|---|
| chr4-57854016-DEL-204 | chr4-57854020-DEL-201 |
| chr4-57872963-DEL-4533 | chr4-57872964-DEL-4533 |
| chr4-57887254-DEL-53 | chr4-57887255-DEL-53 |
| chr4-57898742-DEL-77 | chr4-57898743-DEL-77 |
| chr4-57933169-DEL-52 | chr4-57933169-DEL-52 |
| chr4-57944602-DEL-56 | chr4-57944615-DEL-64 |
| chr4-57979552-DEL-2427 | chr4-57979553-DEL-2427 |
| chr4-57990539-DEL-56 | chr4-57990540-DEL-56 |
| chr4-58011955-DEL-6243 | chr4-58011956-DEL-6243 |
| chr4-58027816-DEL-1362 | chr4-58027817-DEL-1362 |
| chr4-58091008-DEL-84 | chr4-58091009-DEL-84 |
| chr4-58136297-DEL-1701 | chr4-58136298-DEL-1701 |
| chr4-58159132-DEL-90 | chr4-58159133-DEL-90 |
| chr4-58160215-DEL-480 | chr4-58160216-DEL-480 |
| chr4-58164222-DEL-223 | chr4-58164223-DEL-223 |
| chr4-58181585-DEL-200 | chr4-58181586-DEL-200 |
| chr4-58224789-DEL-6242 | chr4-58224790-DEL-6242 |
| chr4-58236600-DEL-91 | chr4-58236601-DEL-91 |
| chr4-58254031-DEL-54 | chr4-58254018-DEL-72 |
| chr4-58260910-DEL-196 | chr4-58260911-DEL-196 |
| chr4-58312055-DEL-99 | chr4-58312081-DEL-104 |
| chr4-58321473-DEL-562 | chr4-58321474-DEL-562 |
| chr4-58347420-DEL-60 | chr4-58347412-DEL-53 |
| chr4-58360771-DEL-87 | chr4-58360780-DEL-68 |
| chr4-58385628-DEL-7066 | chr4-58385635-DEL-7065 |
| chr4-58427589-DEL-500 | chr4-58427590-DEL-500 |
| chr4-58429047-DEL-73 | chr4-58429048-DEL-73 |
| chr4-58429233-DEL-1623 | chr4-58429234-DEL-1623 |
| chr4-58432297-DEL-54 | chr4-58432288-DEL-55 |
| chr4-58440262-DEL-67 | chr4-58440310-DEL-61 |
| chr4-58443889-DEL-60 | chr4-58443890-DEL-60 |
| chr4-58470998-DEL-70 | chr4-58471000-DEL-71 |
| chr4-58473873-DEL-64 | chr4-58473874-DEL-64 |
| chr4-58494552-DEL-231 | chr4-58494553-DEL-231 |
| chr4-58505408-DEL-285 | chr4-58505409-DEL-285 |
| chr4-58514598-DEL-398 | chr4-58514599-DEL-398 |
| chr4-58519271-DEL-63 | chr4-58519248-DEL-64 |
| chr4-58527408-DEL-218 | chr4-58527409-DEL-217 |
| chr4-58532793-DEL-68 | chr4-58532761-DEL-73 |
| chr4-58560830-DEL-338 | chr4-58560831-DEL-338 |
| chr4-58567272-DEL-879 | chr4-58567273-DEL-879 |
| chr4-58569478-DEL-427 | chr4-58569479-DEL-427 |
| chr4-58578366-DEL-140 | chr4-58578354-DEL-141 |
| chr4-58589555-DEL-80 | chr4-58589550-DEL-92 |
| chr4-58597635-DEL-67 | chr4-58597636-DEL-67 |
| chr4-58607028-DEL-2233 | chr4-58607029-DEL-2233 |
| chr4-58608598-DEL-276 | chr4-58608599-DEL-276 |
| chr4-58637114-DEL-408 | chr4-58637115-DEL-408 |
| chr4-58648344-DEL-734 | chr4-58648345-DEL-735 |
| chr4-58681763-DEL-58 | chr4-58681772-DEL-50 |
| chr4-58682501-DEL-6057 | chr4-58682501-DEL-6057 |
| chr4-58694594-DEL-289 | chr4-58694595-DEL-289 |
| chr4-58724389-DEL-204 | chr4-58724390-DEL-202 |
| chr4-58753002-DEL-2808 | chr4-58753003-DEL-2808 |
| chr4-58756823-DEL-133 | chr4-58756824-DEL-133 |
| chr4-58758928-DEL-52 | chr4-58758927-DEL-54 |
| chr4-58769382-DEL-3960 | chr4-58769383-DEL-3960 |
| chr4-58788500-DEL-196 | chr4-58788503-DEL-193 |
| chr4-58818364-DEL-54 | chr4-58818393-DEL-62 |
| chr4-58938284-DEL-70 | chr4-58938285-DEL-70 |
| chr4-58988086-DEL-67 | chr4-58988087-DEL-67 |
| chr4-59001592-DEL-59 | chr4-59001593-DEL-55 |
| chr4-59030284-DEL-2752 | chr4-59030285-DEL-2752 |
| chr4-59033715-DEL-69 | chr4-59033716-DEL-69 |
| chr4-59037183-DEL-218 | chr4-59037184-DEL-218 |
| chr4-59057653-DEL-78 | chr4-59057673-DEL-79 |
| chr4-59070978-DEL-182 | chr4-59070979-DEL-182 |
| chr4-59076799-DEL-114 | chr4-59076796-DEL-122 |
| chr4-59086315-DEL-201 | chr4-59086316-DEL-201 |
| chr4-59101747-DEL-120 | chr4-59101748-DEL-120 |
| chr4-59109295-DEL-119 | chr4-59109296-DEL-119 |
| chr4-59123845-DEL-62 | chr4-59123846-DEL-62 |
| chr4-59138809-DEL-503 | chr4-59138810-DEL-503 |
| chr4-59139481-DEL-95 | chr4-59139477-DEL-131 |
| chr4-59144377-DEL-189 | chr4-59144382-DEL-188 |
| chr4-59148325-DEL-218 | chr4-59148326-DEL-218 |
| chr4-59171115-DEL-190 | chr4-59171116-DEL-190 |
| chr4-59185473-DEL-98 | chr4-59185501-DEL-64 |
| chr4-59197770-DEL-255 | chr4-59197771-DEL-255 |
| chr4-59276495-DEL-72 | chr4-59276493-DEL-74 |

| | |
|---|---|
| chr4-59278987-DEL-210 | chr4-59278988-DEL-210 |
| chr4-59280551-DEL-188 | chr4-59280552-DEL-188 |
| chr4-59291671-DEL-135 | chr4-59291663-DEL-136 |
| chr4-59316702-DEL-72 | chr4-59316703-DEL-72 |
| chr4-59331045-DEL-60 | chr4-59331046-DEL-60 |
| chr4-59375425-DEL-56 | chr4-59375425-DEL-56 |
| chr4-59433113-DEL-51 | chr4-59433104-DEL-61 |
| chr4-59498654-DEL-482 | chr4-59498647-DEL-483 |
| chr4-59502064-DEL-450 | chr4-59502065-DEL-450 |
| chr4-59527326-DEL-196 | chr4-59527327-DEL-195 |
| chr4-59556046-DEL-1403 | chr4-59556018-DEL-1446 |
| chr4-59570732-DEL-203 | chr4-59570733-DEL-203 |
| chr4-59636461-DEL-116 | chr4-59636462-DEL-116 |
| chr4-59654054-DEL-347 | chr4-59654054-DEL-347 |
| chr4-59681925-DEL-62 | chr4-59681926-DEL-62 |
| chr4-59687876-DEL-58 | chr4-59687877-DEL-58 |
| chr4-59776148-DEL-81 | chr4-59776154-DEL-75 |
| chr4-59778061-DEL-546 | chr4-59778062-DEL-546 |
| chr4-59796509-DEL-235 | chr4-59796513-DEL-234 |
| chr4-59802893-DEL-72 | chr4-59802910-DEL-72 |
| chr4-59813294-DEL-330 | chr4-59813295-DEL-330 |
| chr4-59817372-DEL-463 | chr4-59817373-DEL-463 |
| chr4-59837365-DEL-56 | chr4-59837365-DEL-56 |
| chr4-59875700-DEL-52 | chr4-59875701-DEL-52 |
| chr4-59987359-DEL-150 | chr4-59987360-DEL-150 |
| chr4-59997739-DEL-85 | chr4-59997740-DEL-85 |
| chr4-59999927-DEL-298 | chr4-59999946-DEL-276 |
| chr4-61399515-DEL-100 | chr4-61399516-DEL-100 |
| chr4-61460399-DEL-363 | chr4-61460399-DEL-363 |
| chr4-61483563-DEL-411 | chr4-61483564-DEL-411 |
| chr4-61655348-DEL-921 | chr4-61655349-DEL-921 |
| chr4-61674661-DEL-808 | chr4-61674662-DEL-808 |
| chr4-61707661-DEL-788 | chr4-61707661-DEL-787 |
| chr4-61715723-DEL-51 | chr4-61715724-DEL-51 |
| chr4-61717383-DEL-660 | chr4-61717384-DEL-660 |
| chr4-61721379-DEL-95 | chr4-61721380-DEL-95 |
| chr4-61735456-DEL-2804 | chr4-61735457-DEL-2804 |
| chr4-61749392-DEL-155 | chr4-61749391-DEL-154 |
| chr4-61813498-DEL-2052 | chr4-61813499-DEL-2052 |
| chr4-61841606-DEL-1887 | chr4-61841607-DEL-1887 |
| chr4-61867816-DEL-7101 | chr4-61867817-DEL-7101 |
| chr4-61910775-DEL-1881 | chr4-61910776-DEL-1881 |
| chr4-61915064-DEL-1573 | chr4-61915038-DEL-1575 |
| chr4-61956073-DEL-109 | chr4-61956074-DEL-109 |
| chr4-61993158-DEL-496 | chr4-61993159-DEL-496 |
| chr4-61997119-DEL-172 | chr4-61997119-DEL-172 |
| chr4-62002433-DEL-2852 | chr4-62002432-DEL-2852 |
| chr4-62043005-DEL-3140 | chr4-62043006-DEL-3140 |
| chr4-62070137-DEL-197 | chr4-62070138-DEL-197 |
| chr4-62080285-DEL-151 | chr4-62080285-DEL-151 |
| chr4-62101008-DEL-397 | chr4-62101009-DEL-397 |
| chr4-62180084-DEL-220 | chr4-62180085-DEL-220 |
| chr4-62213911-DEL-127 | chr4-62213912-DEL-127 |
| chr4-62227197-DEL-1891 | chr4-62227198-DEL-1891 |
| chr4-62255958-DEL-242 | chr4-62255959-DEL-242 |
| chr4-62270125-DEL-63 | chr4-62270117-DEL-92 |
| chr4-62305739-DEL-130 | chr4-62305740-DEL-130 |
| chr4-62323359-DEL-2793 | chr4-62323360-DEL-2793 |
| chr4-62351748-DEL-228 | chr4-62351749-DEL-228 |
| chr4-62361040-DEL-67 | chr4-62361021-DEL-83 |
| chr4-62362039-DEL-5692 | chr4-62362040-DEL-5692 |
| chr4-62397877-DEL-250 | chr4-62397878-DEL-250 |
| chr4-62432742-DEL-233 | chr4-62432731-DEL-245 |
| chr4-62450239-DEL-61 | chr4-62450216-DEL-63 |
| chr4-62471367-DEL-72 | chr4-62471365-DEL-79 |
| chr4-62472819-DEL-240 | chr4-62472829-DEL-234 |
| chr4-62473216-DEL-509 | chr4-62473217-DEL-509 |
| chr4-62479297-DEL-72 | chr4-62479296-DEL-73 |
| chr4-62484002-DEL-55 | chr4-62484003-DEL-55 |
| chr4-62491853-DEL-5477 | chr4-62491854-DEL-5477 |
| chr4-62512076-DEL-194 | chr4-62512056-DEL-195 |
| chr4-62515789-DEL-88 | chr4-62515786-DEL-92 |
| chr4-62526556-DEL-51 | chr4-62526573-DEL-73 |
| chr4-62529794-DEL-204 | chr4-62529795-DEL-204 |
| chr4-62534354-DEL-56 | chr4-62534366-DEL-88 |
| chr4-62607858-DEL-54 | chr4-62607859-DEL-54 |
| chr4-62630501-DEL-6604 | chr4-62630502-DEL-6604 |
| chr4-62668487-DEL-82 | chr4-62668488-DEL-82 |
| chr4-62689751-DEL-191 | chr4-62689752-DEL-191 |
| chr4-62723580-DEL-54 | chr4-62723581-DEL-54 |

| | |
|---|---|
| chr4-62726017-DEL-174 | chr4-62726018-DEL-174 |
| chr4-62755970-DEL-65 | chr4-62755973-DEL-63 |
| chr4-62782096-DEL-5034 | chr4-62782097-DEL-5034 |
| chr4-62800333-DEL-204 | chr4-62800333-DEL-204 |
| chr4-62803466-DEL-61 | chr4-62803467-DEL-61 |
| chr4-62821757-DEL-77 | chr4-62821788-DEL-56 |
| chr4-62826497-DEL-128 | chr4-62826498-DEL-128 |
| chr4-62827145-DEL-55 | chr4-62827146-DEL-55 |
| chr4-62843383-DEL-6706 | chr4-62843384-DEL-6706 |
| chr4-62934664-DEL-1901 | chr4-62934660-DEL-1890 |
| chr4-63078467-DEL-52 | chr4-63078468-DEL-52 |
| chr4-63098615-DEL-589 | chr4-63098615-DEL-595 |
| chr4-63211046-DEL-60 | chr4-63211047-DEL-60 |
| chr4-63331445-DEL-608 | chr4-63331446-DEL-608 |
| chr4-63367016-DEL-134 | chr4-63367015-DEL-136 |
| chr4-63390169-DEL-66 | chr4-63390170-DEL-72 |
| chr4-63396713-DEL-215 | chr4-63396711-DEL-216 |
| chr4-63420642-DEL-136 | chr4-63420640-DEL-136 |
| chr4-63440886-DEL-103 | chr4-63440887-DEL-103 |
| chr4-63492200-DEL-3589 | chr4-63492204-DEL-3579 |
| chr4-63507965-DEL-95 | chr4-63507966-DEL-95 |
| chr4-63528515-DEL-451 | chr4-63528516-DEL-451 |
| chr4-63564048-DEL-6713 | chr4-63564049-DEL-6713 |
| chr4-63627768-DEL-80 | chr4-63627793-DEL-57 |
| chr4-63654368-DEL-694 | chr4-63654369-DEL-694 |
| chr4-63705320-DEL-2163 | chr4-63705309-DEL-2149 |
| chr4-63713365-DEL-277 | chr4-63713366-DEL-277 |
| chr4-63722683-DEL-200 | chr4-63722684-DEL-200 |
| chr4-63734469-DEL-852 | chr4-63734470-DEL-852 |
| chr4-63778399-DEL-78 | chr4-63778418-DEL-50 |
| chr4-63783568-DEL-664 | chr4-63783569-DEL-664 |
| chr4-63803085-DEL-105 | chr4-63803086-DEL-105 |
| chr4-63837285-DEL-3239 | chr4-63837286-DEL-3239 |
| chr4-63840081-DEL-204 | chr4-63840082-DEL-204 |
| chr4-63841442-DEL-53 | chr4-63841443-DEL-53 |
| chr4-63871864-DEL-71 | chr4-63871863-DEL-73 |
| chr4-63874871-DEL-76 | chr4-63874872-DEL-76 |
| chr4-63876056-DEL-278 | chr4-63876056-DEL-278 |
| chr4-64026582-DEL-6296 | chr4-64026583-DEL-6296 |
| chr4-64052170-DEL-104 | chr4-64052175-DEL-100 |
| chr4-64076886-DEL-62 | chr4-64076906-DEL-64 |
| chr4-64104349-DEL-1796 | chr4-64104372-DEL-1774 |
| chr4-64160409-DEL-6266 | chr4-64160410-DEL-6266 |
| chr4-64177242-DEL-116 | chr4-64177223-DEL-98 |
| chr4-64205573-DEL-6656 | chr4-64205574-DEL-6656 |
| chr4-64221080-DEL-144 | chr4-64221104-DEL-136 |
| chr4-64260785-DEL-337 | chr4-64260786-DEL-337 |
| chr4-64264627-DEL-95 | chr4-64264628-DEL-95 |
| chr4-64266006-DEL-406 | chr4-64266007-DEL-394 |
| chr4-64279062-DEL-128 | chr4-64279062-DEL-128 |
| chr4-64280461-DEL-6382 | chr4-64280462-DEL-6382 |
| chr4-64310611-DEL-1462 | chr4-64310613-DEL-1461 |
| chr4-64330354-DEL-1657 | chr4-64330355-DEL-1657 |
| chr4-64338699-DEL-6456 | chr4-64338700-DEL-6456 |
| chr4-64361477-DEL-52 | chr4-64361480-DEL-50 |
| chr4-64362525-DEL-64 | chr4-64362526-DEL-64 |
| chr4-64372797-DEL-7307 | chr4-64372798-DEL-7307 |
| chr4-64372811-DEL-1060 | chr4-64372812-DEL-1060 |
| chr4-64406380-DEL-60 | chr4-64406393-DEL-76 |
| chr4-64429564-DEL-143 | chr4-64429611-DEL-106 |
| chr4-64430950-DEL-225 | chr4-64430952-DEL-224 |
| chr4-64432055-DEL-706 | chr4-64432056-DEL-706 |
| chr4-64448020-DEL-63 | chr4-64447996-DEL-50 |
| chr4-64481443-DEL-82 | chr4-64481444-DEL-82 |
| chr4-64492155-DEL-78 | chr4-64492156-DEL-75 |
| chr4-64503669-DEL-1217 | chr4-64503670-DEL-1217 |
| chr4-64506352-DEL-68 | chr4-64506352-DEL-68 |
| chr4-64522548-DEL-102 | chr4-64522548-DEL-102 |
| chr4-64606687-DEL-845 | chr4-64606687-DEL-845 |
| chr4-64612558-DEL-1092 | chr4-64612559-DEL-1092 |
| chr4-64615262-DEL-637 | chr4-64615263-DEL-637 |
| chr4-64623102-DEL-3764 | chr4-64623103-DEL-3764 |
| chr4-64645766-DEL-81 | chr4-64645767-DEL-81 |
| chr4-64664110-DEL-124 | chr4-64664111-DEL-126 |
| chr4-64665356-DEL-246 | chr4-64665357-DEL-246 |
| chr4-64689429-DEL-380 | chr4-64689430-DEL-380 |
| chr4-64714248-DEL-95 | chr4-64714249-DEL-95 |
| chr4-64722712-DEL-130 | chr4-64722713-DEL-130 |
| chr4-64728144-DEL-383 | chr4-64728145-DEL-383 |
| chr4-64749664-DEL-98 | chr4-64749683-DEL-101 |

| | |
|---|---|
| chr4-64779715-DEL-7129 | chr4-64779716-DEL-7128 |
| chr4-64802309-DEL-731 | chr4-64802308-DEL-731 |
| chr4-64861525-DEL-308 | chr4-64861526-DEL-306 |
| chr4-64899130-DEL-340 | chr4-64899131-DEL-340 |
| chr4-64902801-DEL-6463 | chr4-64902802-DEL-6463 |
| chr4-64914647-DEL-183 | chr4-64914648-DEL-183 |
| chr4-64923023-DEL-552 | chr4-64923024-DEL-552 |
| chr4-64926046-DEL-1200 | chr4-64926032-DEL-1201 |
| chr4-64935542-DEL-993 | chr4-64935555-DEL-982 |
| chr4-64959054-DEL-2537 | chr4-64959055-DEL-2537 |
| chr4-64974492-DEL-2551 | chr4-64974493-DEL-2551 |
| chr4-65009551-DEL-201 | chr4-65009553-DEL-202 |
| chr4-65023243-DEL-1094 | chr4-65023244-DEL-1094 |
| chr4-65047134-DEL-3347 | chr4-65047135-DEL-3347 |
| chr4-65086232-DEL-273 | chr4-65086233-DEL-273 |
| chr4-65095818-DEL-72 | chr4-65095822-DEL-58 |
| chr4-65101857-DEL-116 | chr4-65101858-DEL-114 |
| chr4-65127076-DEL-423 | chr4-65127077-DEL-423 |
| chr4-65169889-DEL-88 | chr4-65169904-DEL-69 |
| chr4-65187067-DEL-1186 | chr4-65187068-DEL-1186 |
| chr4-65187299-DEL-58 | chr4-65187300-DEL-58 |
| chr4-65203563-DEL-3722 | chr4-65203564-DEL-3722 |
| chr4-65328469-DEL-115 | chr4-65328470-DEL-115 |
| chr4-65337029-DEL-670 | chr4-65337032-DEL-669 |
| chr4-65342516-DEL-1888 | chr4-65342517-DEL-1888 |
| chr4-65347370-DEL-66 | chr4-65347359-DEL-84 |
| chr4-65350525-DEL-67 | chr4-65350529-DEL-64 |
| chr4-65397473-DEL-265 | chr4-65397474-DEL-265 |
| chr4-65415963-DEL-807 | chr4-65415963-DEL-792 |
| chr4-65438548-DEL-957 | chr4-65438551-DEL-955 |
| chr4-65442301-DEL-3615 | chr4-65442302-DEL-3615 |
| chr4-65445041-DEL-97 | chr4-65445042-DEL-97 |
| chr4-65457447-DEL-381 | chr4-65457438-DEL-377 |
| chr4-65465399-DEL-84 | chr4-65465400-DEL-84 |
| chr4-65493433-DEL-66 | chr4-65493436-DEL-76 |
| chr4-65506937-DEL-68 | chr4-65506940-DEL-66 |
| chr4-65517334-DEL-629 | chr4-65517372-DEL-629 |
| chr4-65541270-DEL-905 | chr4-65541271-DEL-905 |
| chr4-65564770-DEL-130 | chr4-65564771-DEL-130 |
| chr4-65596292-DEL-319 | chr4-65596293-DEL-319 |
| chr4-65602683-DEL-89 | chr4-65602684-DEL-89 |
| chr4-65611404-DEL-96 | chr4-65611404-DEL-96 |
| chr4-65641774-DEL-58 | chr4-65641773-DEL-58 |
| chr4-65676136-DEL-60 | chr4-65676146-DEL-58 |
| chr4-65693748-DEL-6744 | chr4-65693782-DEL-6712 |
| chr4-65697396-DEL-476 | chr4-65697397-DEL-476 |
| chr4-65726149-DEL-217 | chr4-65726150-DEL-217 |
| chr4-65741518-DEL-6249 | chr4-65741519-DEL-6249 |
| chr4-65760584-DEL-203 | chr4-65760585-DEL-203 |
| chr4-65838980-DEL-246 | chr4-65838981-DEL-246 |
| chr4-65847149-DEL-107 | chr4-65847150-DEL-107 |
| chr4-65881786-DEL-56 | chr4-65881815-DEL-64 |
| chr4-65902390-DEL-6575 | chr4-65902391-DEL-6575 |
| chr4-65928168-DEL-783 | chr4-65928169-DEL-783 |
| chr4-65961031-DEL-52 | chr4-65961012-DEL-88 |
| chr4-65962671-DEL-6358 | chr4-65962672-DEL-6358 |
| chr4-66006784-DEL-165 | chr4-66006789-DEL-166 |
| chr4-66047738-DEL-402 | chr4-66047739-DEL-402 |
| chr4-66060317-DEL-1516 | chr4-66060318-DEL-1517 |
| chr4-66147282-DEL-68 | chr4-66147263-DEL-76 |
| chr4-66208314-DEL-2751 | chr4-66208315-DEL-2751 |
| chr4-66273245-DEL-7112 | chr4-66273246-DEL-7112 |
| chr4-66326265-DEL-2242 | chr4-66326266-DEL-2242 |
| chr4-66336919-DEL-124 | chr4-66336920-DEL-124 |
| chr4-66350885-DEL-200 | chr4-66350886-DEL-200 |
| chr4-66364727-DEL-56 | chr4-66364730-DEL-64 |
| chr4-66404800-DEL-292 | chr4-66404801-DEL-292 |
| chr4-66409157-DEL-587 | chr4-66409158-DEL-587 |
| chr4-66415659-DEL-2562 | chr4-66415657-DEL-2558 |
| chr4-66450866-DEL-95 | chr4-66450867-DEL-95 |
| chr4-66488532-DEL-2405 | chr4-66488528-DEL-2406 |
| chr4-66492572-DEL-1592 | chr4-66492573-DEL-1592 |
| chr4-66494992-DEL-6178 | chr4-66494993-DEL-6178 |
| chr4-66502179-DEL-70 | chr4-66502185-DEL-54 |
| chr4-66516270-DEL-52 | chr4-66516271-DEL-52 |
| chr4-66519346-DEL-57 | chr4-66519349-DEL-54 |
| chr4-66529280-DEL-358 | chr4-66529281-DEL-358 |
| chr4-66548218-DEL-71 | chr4-66548219-DEL-71 |
| chr4-66549548-DEL-6952 | chr4-66549549-DEL-6952 |
| chr4-66565680-DEL-117 | chr4-66565681-DEL-117 |

| | |
|---|---|
| chr4-66594040-DEL-6631 | chr4-66594037-DEL-6632 |
| chr4-66600338-DEL-214 | chr4-66600339-DEL-214 |
| chr4-66636529-DEL-6390 | chr4-66636530-DEL-6390 |
| chr4-66652205-DEL-92 | chr4-66652206-DEL-92 |
| chr4-66660275-DEL-66 | chr4-66660276-DEL-66 |
| chr4-66693500-DEL-214 | chr4-66693501-DEL-214 |
| chr4-66741537-DEL-103 | chr4-66741538-DEL-97 |
| chr4-66771285-DEL-661 | chr4-66771286-DEL-661 |
| chr4-66802597-DEL-57 | chr4-66802598-DEL-57 |
| chr4-66803112-DEL-146 | chr4-66803113-DEL-146 |
| chr4-66876598-DEL-52 | chr4-66876591-DEL-68 |
| chr4-66883934-DEL-6987 | chr4-66883918-DEL-7001 |
| chr4-66909120-DEL-175 | chr4-66909119-DEL-177 |
| chr4-66940348-DEL-62 | chr4-66940349-DEL-62 |
| chr4-66957571-DEL-51 | chr4-66957572-DEL-51 |
| chr4-66958506-DEL-199 | chr4-66958507-DEL-199 |
| chr4-66966374-DEL-135 | chr4-66966374-DEL-136 |
| chr4-67013641-DEL-2676 | chr4-67013642-DEL-2676 |
| chr4-67040076-DEL-223 | chr4-67040077-DEL-223 |
| chr4-67046004-DEL-95 | chr4-67046013-DEL-87 |
| chr4-67051338-DEL-392 | chr4-67051344-DEL-387 |
| chr4-67075159-DEL-793 | chr4-67075153-DEL-796 |
| chr4-67079525-DEL-98 | chr4-67079526-DEL-98 |
| chr4-67129534-DEL-54 | chr4-67129535-DEL-54 |
| chr4-67144710-DEL-63 | chr4-67144711-DEL-63 |
| chr4-67151647-DEL-6513 | chr4-67151648-DEL-6513 |
| chr4-67205880-DEL-413 | chr4-67205881-DEL-413 |
| chr4-67236043-DEL-485 | chr4-67236044-DEL-485 |
| chr4-67241721-DEL-6212 | chr4-67241722-DEL-6212 |
| chr4-67268135-DEL-397 | chr4-67268136-DEL-397 |
| chr4-67280118-DEL-4153 | chr4-67280119-DEL-4153 |
| chr4-67293932-DEL-904 | chr4-67293933-DEL-904 |
| chr4-67318177-DEL-52 | chr4-67318178-DEL-52 |
| chr4-67324736-DEL-3962 | chr4-67324737-DEL-3962 |
| chr4-67345525-DEL-489 | chr4-67345526-DEL-489 |
| chr4-67367759-DEL-195 | chr4-67367762-DEL-191 |
| chr4-67381219-DEL-77 | chr4-67381209-DEL-60 |
| chr4-67384010-DEL-471 | chr4-67384011-DEL-471 |
| chr4-67494397-DEL-2185 | chr4-67494398-DEL-2185 |
| chr4-67524482-DEL-774 | chr4-67524483-DEL-774 |
| chr4-67541326-DEL-673 | chr4-67541327-DEL-673 |
| chr4-67577812-DEL-65 | chr4-67577813-DEL-65 |
| chr4-67631967-DEL-56 | chr4-67631966-DEL-58 |
| chr4-67650944-DEL-55 | chr4-67650944-DEL-55 |
| chr4-67662498-DEL-58 | chr4-67662499-DEL-58 |
| chr4-67664390-DEL-180 | chr4-67664391-DEL-179 |
| chr4-67668663-DEL-304 | chr4-67668664-DEL-304 |
| chr4-67681174-DEL-58 | chr4-67681175-DEL-58 |
| chr4-67716680-DEL-88 | chr4-67716692-DEL-84 |
| chr4-67738461-DEL-659 | chr4-67738462-DEL-659 |
| chr4-67757555-DEL-283 | chr4-67757556-DEL-283 |
| chr4-67759219-DEL-6376 | chr4-67759220-DEL-6376 |
| chr4-67801408-DEL-149 | chr4-67801409-DEL-149 |
| chr4-67832595-DEL-61 | chr4-67832596-DEL-61 |
| chr4-67834217-DEL-6943 | chr4-67834216-DEL-6944 |
| chr4-67849087-DEL-943 | chr4-67849077-DEL-940 |
| chr4-67855931-DEL-8058 | chr4-67855932-DEL-8058 |
| chr4-67891649-DEL-75 | chr4-67891650-DEL-75 |
| chr4-67957924-DEL-5076 | chr4-67957925-DEL-5076 |
| chr4-67978915-DEL-50 | chr4-67978907-DEL-58 |
| chr4-68033630-DEL-5364 | chr4-68033631-DEL-5364 |
| chr4-68073761-DEL-332 | chr4-68073762-DEL-332 |
| chr4-68090524-DEL-280 | chr4-68090525-DEL-280 |
| chr4-68095446-DEL-570 | chr4-68095447-DEL-570 |
| chr4-68111670-DEL-56 | chr4-68111671-DEL-56 |
| chr4-68173123-DEL-437 | chr4-68173124-DEL-437 |
| chr4-68208308-DEL-7032 | chr4-68208309-DEL-7032 |
| chr4-68355222-DEL-208 | chr4-68355223-DEL-208 |
| chr4-68391130-DEL-136 | chr4-68391137-DEL-134 |
| chr4-68420201-DEL-369 | chr4-68420204-DEL-367 |
| chr4-68490296-DEL-60 | chr4-68490297-DEL-60 |
| chr4-68497401-DEL-197 | chr4-68497402-DEL-197 |
| chr4-68545809-DEL-246 | chr4-68545810-DEL-246 |
| chr4-68573217-DEL-92 | chr4-68573223-DEL-95 |
| chr4-68578669-DEL-87 | chr4-68578670-DEL-87 |
| chr4-68586126-DEL-6924 | chr4-68586127-DEL-6924 |
| chr4-68586232-DEL-402 | chr4-68586214-DEL-402 |
| chr4-68592672-DEL-312 | chr4-68592718-DEL-313 |
| chr4-68615642-DEL-801 | chr4-68615642-DEL-801 |
| chr4-68652581-DEL-110 | chr4-68652580-DEL-110 |

| | |
|---|---|
| chr4-68703072-DEL-100 | chr4-68703068-DEL-114 |
| chr4-68722333-DEL-66 | chr4-68722311-DEL-50 |
| chr4-68735076-DEL-835 | chr4-68735077-DEL-835 |
| chr4-68780132-DEL-737 | chr4-68780133-DEL-737 |
| chr4-68797110-DEL-194 | chr4-68797111-DEL-193 |
| chr4-68855037-DEL-60 | chr4-68855038-DEL-54 |
| chr4-68879027-DEL-7104 | chr4-68879028-DEL-7104 |
| chr4-68895674-DEL-2045 | chr4-68895674-DEL-2045 |
| chr4-68924794-DEL-834 | chr4-68924794-DEL-834 |
| chr4-68972374-DEL-204 | chr4-68972375-DEL-204 |
| chr4-68990295-DEL-154 | chr4-68990295-DEL-154 |
| chr4-69076288-DEL-6467 | chr4-69076289-DEL-6467 |
| chr4-69098203-DEL-2547 | chr4-69098204-DEL-2547 |
| chr4-69130765-DEL-409 | chr4-69130766-DEL-409 |
| chr4-69145901-DEL-208 | chr4-69145902-DEL-208 |
| chr4-69188886-DEL-211 | chr4-69188887-DEL-211 |
| chr4-69193472-DEL-167 | chr4-69193473-DEL-167 |
| chr4-69207172-DEL-199 | chr4-69207173-DEL-199 |
| chr4-69210125-DEL-64 | chr4-69210134-DEL-56 |
| chr4-69253955-DEL-163 | chr4-69253956-DEL-163 |
| chr4-69280225-DEL-75 | chr4-69280217-DEL-79 |
| chr4-69286814-DEL-5799 | chr4-69286814-DEL-5799 |
| chr4-69290337-DEL-127 | chr4-69290338-DEL-125 |
| chr4-69322099-DEL-57 | chr4-69322100-DEL-57 |
| chr4-69328427-DEL-4926 | chr4-69328427-DEL-4926 |
| chr4-69347001-DEL-754 | chr4-69347002-DEL-754 |
| chr4-69428151-DEL-6392 | chr4-69428152-DEL-6392 |
| chr4-69524758-DEL-56 | chr4-69524759-DEL-56 |
| chr4-69528479-DEL-3018 | chr4-69528480-DEL-3018 |
| chr4-69564637-DEL-3313 | chr4-69564638-DEL-3312 |
| chr4-69577610-DEL-72 | chr4-69577569-DEL-79 |
| chr4-69613729-DEL-66 | chr4-69613730-DEL-66 |
| chr4-69625853-DEL-6262 | chr4-69625854-DEL-6262 |
| chr4-69632934-DEL-7313 | chr4-69632935-DEL-7313 |
| chr4-69705115-DEL-6407 | chr4-69705112-DEL-6408 |
| chr4-69717307-DEL-2230 | chr4-69717309-DEL-2229 |
| chr4-69720117-DEL-3817 | chr4-69720118-DEL-3817 |
| chr4-69725685-DEL-344 | chr4-69725686-DEL-343 |
| chr4-69737063-DEL-390 | chr4-69737064-DEL-390 |
| chr4-69746078-DEL-260 | chr4-69746079-DEL-260 |
| chr4-69792333-DEL-71 | chr4-69792334-DEL-71 |
| chr4-69831703-DEL-6112 | chr4-69831704-DEL-6112 |
| chr4-69843400-DEL-1006 | chr4-69843401-DEL-1006 |
| chr4-69868559-DEL-56 | chr4-69868560-DEL-56 |
| chr4-69873155-DEL-132 | chr4-69873156-DEL-132 |
| chr4-69949315-DEL-69 | chr4-69949293-DEL-69 |
| chr4-69964521-DEL-74 | chr4-69964522-DEL-64 |
| chr4-70035933-DEL-2182 | chr4-70035934-DEL-2182 |
| chr4-70036030-DEL-157 | chr4-70036031-DEL-157 |
| chr4-70044545-DEL-115 | chr4-70044546-DEL-115 |
| chr4-70046821-DEL-118 | chr4-70046822-DEL-118 |
| chr4-70062473-DEL-56 | chr4-70062474-DEL-56 |
| chr4-70067668-DEL-2195 | chr4-70067669-DEL-2195 |
| chr4-70113851-DEL-1004 | chr4-70113852-DEL-1004 |
| chr4-70118888-DEL-64 | chr4-70118889-DEL-64 |
| chr4-70158616-DEL-59 | chr4-70158617-DEL-59 |
| chr4-70166159-DEL-1521 | chr4-70166160-DEL-1521 |
| chr4-70176424-DEL-144 | chr4-70176425-DEL-144 |
| chr4-70187271-DEL-527 | chr4-70187272-DEL-527 |
| chr4-70209955-DEL-52 | chr4-70209938-DEL-53 |
| chr4-70216905-DEL-62 | chr4-70216904-DEL-72 |
| chr4-70228966-DEL-105 | chr4-70228953-DEL-105 |
| chr4-70258365-DEL-6211 | chr4-70258366-DEL-6211 |
| chr4-70267506-DEL-2360 | chr4-70267507-DEL-2360 |
| chr4-70289095-DEL-1892 | chr4-70289096-DEL-1892 |
| chr4-70293897-DEL-321 | chr4-70293895-DEL-318 |
| chr4-70305371-DEL-61 | chr4-70305372-DEL-61 |
| chr4-70322475-DEL-333 | chr4-70322477-DEL-332 |
| chr4-70334843-DEL-1399 | chr4-70334844-DEL-1399 |
| chr4-70375332-DEL-1391 | chr4-70375333-DEL-1391 |
| chr4-70376953-DEL-2333 | chr4-70376954-DEL-2333 |
| chr4-70386223-DEL-100 | chr4-70386224-DEL-100 |
| chr4-70390550-DEL-6452 | chr4-70390551-DEL-6452 |
| chr4-70545691-DEL-1630 | chr4-70545692-DEL-1630 |
| chr4-70554376-DEL-60 | chr4-70554377-DEL-60 |
| chr4-70557256-DEL-512 | chr4-70557257-DEL-512 |
| chr4-70597916-DEL-56 | chr4-70597917-DEL-56 |
| chr4-70600602-DEL-70 | chr4-70600624-DEL-62 |
| chr4-70601134-DEL-443 | chr4-70601140-DEL-443 |
| chr4-70611052-DEL-71 | chr4-70611040-DEL-71 |

| | |
|---|---|
| chr4-70612057-DEL-344 | chr4-70612054-DEL-338 |
| chr4-70616227-DEL-73 | chr4-70616228-DEL-73 |
| chr4-70617968-DEL-188 | chr4-70617969-DEL-188 |
| chr4-70619940-DEL-1368 | chr4-70619941-DEL-1368 |
| chr4-70621561-DEL-139 | chr4-70621562-DEL-136 |
| chr4-70657448-DEL-72 | chr4-70657407-DEL-64 |
| chr4-70662163-DEL-113 | chr4-70662140-DEL-137 |
| chr4-70709180-DEL-108 | chr4-70709181-DEL-108 |
| chr4-70732306-DEL-71 | chr4-70732307-DEL-71 |
| chr4-70739593-DEL-235 | chr4-70739596-DEL-233 |
| chr4-70748158-DEL-3492 | chr4-70748159-DEL-3492 |
| chr4-70764719-DEL-3070 | chr4-70764720-DEL-3070 |
| chr4-70783566-DEL-6450 | chr4-70783566-DEL-6450 |
| chr4-70794818-DEL-1522 | chr4-70794819-DEL-1522 |
| chr4-70802589-DEL-107 | chr4-70802584-DEL-102 |
| chr4-70812517-DEL-458 | chr4-70812518-DEL-458 |
| chr4-70813903-DEL-137 | chr4-70813904-DEL-137 |
| chr4-70837252-DEL-856 | chr4-70837253-DEL-856 |
| chr4-70839011-DEL-56 | chr4-70839012-DEL-56 |
| chr4-70861691-DEL-74 | chr4-70861692-DEL-74 |
| chr4-70869546-DEL-55 | chr4-70869543-DEL-56 |
| chr4-70887573-DEL-403 | chr4-70887573-DEL-403 |
| chr4-70891136-DEL-1407 | chr4-70891137-DEL-1407 |
| chr4-70904544-DEL-116 | chr4-70904545-DEL-116 |
| chr4-70910056-DEL-372 | chr4-70910055-DEL-372 |
| chr4-70917643-DEL-3207 | chr4-70917643-DEL-3207 |
| chr4-70921560-DEL-1137 | chr4-70921560-DEL-1137 |
| chr4-70938201-DEL-2788 | chr4-70938202-DEL-2788 |
| chr4-70957128-DEL-708 | chr4-70957129-DEL-708 |
| chr4-70971002-DEL-197 | chr4-70971003-DEL-197 |
| chr4-70991018-DEL-367 | chr4-70991019-DEL-367 |
| chr4-70998223-DEL-52 | chr4-70998209-DEL-50 |
| chr4-71005474-DEL-413 | chr4-71005475-DEL-413 |
| chr4-71012159-DEL-89 | chr4-71012159-DEL-94 |
| chr4-71031967-DEL-88 | chr4-71031968-DEL-88 |
| chr4-71041003-DEL-3140 | chr4-71041004-DEL-3140 |
| chr4-71073980-DEL-234 | chr4-71073976-DEL-235 |
| chr4-71078928-DEL-1015 | chr4-71078929-DEL-1020 |
| chr4-71140411-DEL-1243 | chr4-71140412-DEL-1243 |
| chr4-71149246-DEL-108 | chr4-71149245-DEL-103 |
| chr4-71152818-DEL-1695 | chr4-71152819-DEL-1695 |
| chr4-71158190-DEL-245 | chr4-71158190-DEL-245 |
| chr4-71197214-DEL-68 | chr4-71197211-DEL-84 |
| chr4-71206941-DEL-87 | chr4-71206942-DEL-87 |
| chr4-71207862-DEL-77 | chr4-71207867-DEL-73 |
| chr4-71221215-DEL-156 | chr4-71221203-DEL-188 |
| chr4-71262519-DEL-60 | chr4-71262520-DEL-61 |
| chr4-71288911-DEL-564 | chr4-71288912-DEL-564 |
| chr4-71299889-DEL-388 | chr4-71299890-DEL-388 |
| chr4-71302234-DEL-68 | chr4-71302213-DEL-90 |
| chr4-71323890-DEL-54 | chr4-71323891-DEL-54 |
| chr4-71336985-DEL-4879 | chr4-71336986-DEL-4879 |
| chr4-71352485-DEL-6484 | chr4-71352475-DEL-6485 |
| chr4-71377980-DEL-55 | chr4-71377980-DEL-56 |
| chr4-71426810-DEL-5382 | chr4-71426811-DEL-5382 |
| chr4-71448463-DEL-129 | chr4-71448464-DEL-129 |
| chr4-71615428-DEL-109 | chr4-71615425-DEL-109 |
| chr4-71631450-DEL-93 | chr4-71631451-DEL-93 |
| chr4-71637564-DEL-56 | chr4-71637565-DEL-56 |
| chr4-71647396-DEL-304 | chr4-71647397-DEL-304 |
| chr4-71650419-DEL-201 | chr4-71650420-DEL-201 |
| chr4-71653907-DEL-71 | chr4-71653901-DEL-75 |
| chr4-71656776-DEL-3040 | chr4-71656777-DEL-3040 |
| chr4-71701213-DEL-5966 | chr4-71701213-DEL-5966 |
| chr4-71701468-DEL-53 | chr4-71701469-DEL-53 |
| chr4-71705445-DEL-15552 | chr4-71705446-DEL-15552 |
| chr4-71745036-DEL-1606 | chr4-71745037-DEL-1606 |
| chr4-71774604-DEL-3189 | chr4-71774605-DEL-3189 |
| chr4-71782832-DEL-657 | chr4-71782833-DEL-657 |
| chr4-71801557-DEL-1132 | chr4-71801557-DEL-1132 |
| chr4-71858309-DEL-2351 | chr4-71858310-DEL-2351 |
| chr4-71863465-DEL-202 | chr4-71863466-DEL-202 |
| chr4-71881416-DEL-4551 | chr4-71881417-DEL-4551 |
| chr4-71917692-DEL-69 | chr4-71917693-DEL-69 |
| chr4-71943021-DEL-488 | chr4-71943022-DEL-488 |
| chr4-71946172-DEL-227 | chr4-71946171-DEL-227 |
| chr4-71963848-DEL-674 | chr4-71963849-DEL-674 |
| chr4-71972127-DEL-3423 | chr4-71972128-DEL-3423 |
| chr4-71974161-DEL-2652 | chr4-71974162-DEL-2652 |
| chr4-71992138-DEL-6900 | chr4-71992139-DEL-6900 |

| | |
|---|---|
| chr4-72006393-DEL-204 | chr4-72006394-DEL-204 |
| chr4-72018583-DEL-60 | chr4-72018584-DEL-60 |
| chr4-72041545-DEL-66 | chr4-72041540-DEL-72 |
| chr4-72131130-DEL-78 | chr4-72131131-DEL-78 |
| chr4-72142647-DEL-187 | chr4-72142648-DEL-187 |
| chr4-72154413-DEL-59 | chr4-72154375-DEL-71 |
| chr4-72184937-DEL-2258 | chr4-72184938-DEL-2258 |
| chr4-72231814-DEL-152 | chr4-72231815-DEL-152 |
| chr4-72244993-DEL-247 | chr4-72244994-DEL-247 |
| chr4-72252225-DEL-6646 | chr4-72252225-DEL-6646 |
| chr4-72266246-DEL-608 | chr4-72266247-DEL-608 |
| chr4-72315125-DEL-60 | chr4-72315108-DEL-78 |
| chr4-72321551-DEL-6540 | chr4-72321552-DEL-6540 |
| chr4-72352037-DEL-4316 | chr4-72352038-DEL-4316 |
| chr4-72353755-DEL-202 | chr4-72353756-DEL-202 |
| chr4-72371946-DEL-3605 | chr4-72371947-DEL-3605 |
| chr4-72384196-DEL-1347 | chr4-72384197-DEL-1347 |
| chr4-72394570-DEL-1306 | chr4-72394571-DEL-1306 |
| chr4-72396989-DEL-963 | chr4-72396990-DEL-963 |
| chr4-72406521-DEL-1326 | chr4-72406522-DEL-1326 |
| chr4-72419051-DEL-1338 | chr4-72419052-DEL-1337 |
| chr4-72427846-DEL-58 | chr4-72427847-DEL-58 |
| chr4-72444091-DEL-654 | chr4-72444092-DEL-654 |
| chr4-72461306-DEL-6422 | chr4-72461307-DEL-6422 |
| chr4-72482377-DEL-1499 | chr4-72482378-DEL-1499 |
| chr4-72492584-DEL-160 | chr4-72492583-DEL-156 |
| chr4-72497318-DEL-6993 | chr4-72497319-DEL-6993 |
| chr4-72554202-DEL-713 | chr4-72554203-DEL-713 |
| chr4-72557777-DEL-3389 | chr4-72557778-DEL-3389 |
| chr4-72560764-DEL-834 | chr4-72560765-DEL-832 |
| chr4-72583411-DEL-96 | chr4-72583428-DEL-76 |
| chr4-72588680-DEL-113 | chr4-72588680-DEL-113 |
| chr4-72593893-DEL-1166 | chr4-72593893-DEL-1166 |
| chr4-72631958-DEL-116 | chr4-72631953-DEL-108 |
| chr4-72671682-DEL-61 | chr4-72671692-DEL-52 |
| chr4-72683231-DEL-54 | chr4-72683218-DEL-68 |
| chr4-72695883-DEL-248 | chr4-72695859-DEL-249 |
| chr4-72743774-DEL-60 | chr4-72743777-DEL-50 |
| chr4-72757832-DEL-225 | chr4-72757829-DEL-226 |
| chr4-72758524-DEL-784 | chr4-72758529-DEL-787 |
| chr4-72790423-DEL-332 | chr4-72790424-DEL-332 |
| chr4-72807451-DEL-253 | chr4-72807451-DEL-254 |
| chr4-72808171-DEL-835 | chr4-72808169-DEL-836 |
| chr4-72828263-DEL-54 | chr4-72828258-DEL-60 |
| chr4-72830544-DEL-1057 | chr4-72830544-DEL-1035 |
| chr4-72870861-DEL-62 | chr4-72870862-DEL-62 |
| chr4-72887848-DEL-58 | chr4-72887849-DEL-58 |
| chr4-72918670-DEL-376 | chr4-72918671-DEL-376 |
| chr4-72922309-DEL-395 | chr4-72922290-DEL-409 |
| chr4-72930214-DEL-399 | chr4-72930215-DEL-399 |
| chr4-72957166-DEL-81 | chr4-72957167-DEL-81 |
| chr4-72960310-DEL-90 | chr4-72960308-DEL-75 |
| chr4-72983743-DEL-309 | chr4-72983744-DEL-309 |
| chr4-72997038-DEL-80 | chr4-72996999-DEL-78 |
| chr4-73013853-DEL-74 | chr4-73013854-DEL-74 |
| chr4-73031821-DEL-266 | chr4-73031822-DEL-266 |
| chr4-73036953-DEL-70 | chr4-73036919-DEL-65 |
| chr4-73042503-DEL-276 | chr4-73042504-DEL-276 |
| chr4-73059924-DEL-62 | chr4-73059972-DEL-54 |
| chr4-73064143-DEL-5305 | chr4-73064144-DEL-5303 |
| chr4-73181335-DEL-59 | chr4-73181329-DEL-60 |
| chr4-73188585-DEL-66 | chr4-73188586-DEL-66 |
| chr4-73212748-DEL-7971 | chr4-73212749-DEL-7971 |
| chr4-73250858-DEL-63 | chr4-73250859-DEL-63 |
| chr4-73279570-DEL-1734 | chr4-73279571-DEL-1734 |
| chr4-73305809-DEL-56 | chr4-73305809-DEL-56 |
| chr4-73351805-DEL-1076 | chr4-73351806-DEL-1076 |
| chr4-73375018-DEL-61 | chr4-73375019-DEL-61 |
| chr4-73449276-DEL-86 | chr4-73449277-DEL-86 |
| chr4-73466468-DEL-1825 | chr4-73466468-DEL-1825 |
| chr4-73479734-DEL-5422 | chr4-73479734-DEL-5417 |
| chr4-73491629-DEL-650 | chr4-73491630-DEL-649 |
| chr4-73494769-DEL-14298 | chr4-73494770-DEL-14298 |
| chr4-73521858-DEL-130 | chr4-73521859-DEL-130 |
| chr4-73659986-DEL-54 | chr4-73659987-DEL-54 |
| chr4-73668728-DEL-214 | chr4-73668729-DEL-214 |
| chr4-73732809-DEL-5934 | chr4-73732792-DEL-5929 |
| chr4-73764995-DEL-6443 | chr4-73764996-DEL-6443 |
| chr4-73912215-DEL-4215 | chr4-73912216-DEL-4215 |
| chr4-73921903-DEL-68 | chr4-73921904-DEL-68 |

| | |
|---|---|
| chr4-73951782-DEL-618 | chr4-73951783-DEL-618 |
| chr4-73965183-DEL-1168 | chr4-73965184-DEL-1168 |
| chr4-73970884-DEL-294 | chr4-73970887-DEL-294 |
| chr4-73972212-DEL-6444 | chr4-73972213-DEL-6444 |
| chr4-73998506-DEL-2189 | chr4-73998507-DEL-2189 |
| chr4-74017576-DEL-439 | chr4-74017577-DEL-439 |
| chr4-74038555-DEL-68 | chr4-74038556-DEL-68 |
| chr4-74043965-DEL-342 | chr4-74043966-DEL-334 |
| chr4-74046818-DEL-810 | chr4-74046823-DEL-809 |
| chr4-74053280-DEL-230 | chr4-74053281-DEL-230 |
| chr4-74070561-DEL-103 | chr4-74070562-DEL-103 |
| chr4-74073415-DEL-51 | chr4-74073416-DEL-51 |
| chr4-74136892-DEL-240 | chr4-74136892-DEL-240 |
| chr4-74139135-DEL-498 | chr4-74139136-DEL-498 |
| chr4-74140186-DEL-138 | chr4-74140184-DEL-139 |
| chr4-74143491-DEL-53 | chr4-74143492-DEL-53 |
| chr4-74157905-DEL-550 | chr4-74157906-DEL-550 |
| chr4-74201572-DEL-175 | chr4-74201573-DEL-175 |
| chr4-74204003-DEL-153 | chr4-74204004-DEL-153 |
| chr4-74205252-DEL-209 | chr4-74205253-DEL-209 |
| chr4-74219186-DEL-345 | chr4-74219187-DEL-345 |
| chr4-74264486-DEL-80 | chr4-74264487-DEL-80 |
| chr4-74284850-DEL-1529 | chr4-74284848-DEL-1530 |
| chr4-74289552-DEL-2007 | chr4-74289556-DEL-2006 |
| chr4-74300918-DEL-60 | chr4-74300919-DEL-60 |
| chr4-74301613-DEL-224 | chr4-74301609-DEL-231 |
| chr4-74304989-DEL-324 | chr4-74304990-DEL-324 |
| chr4-74305596-DEL-75 | chr4-74305597-DEL-75 |
| chr4-74313187-DEL-58 | chr4-74313190-DEL-56 |
| chr4-74315008-DEL-908 | chr4-74315009-DEL-908 |
| chr4-74332554-DEL-1434 | chr4-74332555-DEL-1434 |
| chr4-74352861-DEL-6433 | chr4-74352862-DEL-6433 |
| chr4-74364142-DEL-106 | chr4-74364142-DEL-103 |
| chr4-74365091-DEL-2889 | chr4-74365091-DEL-2889 |
| chr4-74370260-DEL-64 | chr4-74370261-DEL-64 |
| chr4-74376205-DEL-451 | chr4-74376206-DEL-451 |
| chr4-74387774-DEL-56 | chr4-74387776-DEL-60 |
| chr4-74441430-DEL-24180 | chr4-74441419-DEL-24191 |
| chr4-74458648-DEL-742 | chr4-74458636-DEL-750 |
| chr4-74491167-DEL-2948 | chr4-74491168-DEL-2948 |
| chr4-74512818-DEL-86 | chr4-74512824-DEL-51 |
| chr4-74517693-DEL-6159 | chr4-74517694-DEL-6159 |
| chr4-74543786-DEL-90 | chr4-74543787-DEL-90 |
| chr4-74555828-DEL-6373 | chr4-74555829-DEL-6373 |
| chr4-74566503-DEL-1890 | chr4-74566504-DEL-1890 |
| chr4-74592302-DEL-66 | chr4-74592319-DEL-61 |
| chr4-74593769-DEL-6238 | chr4-74593770-DEL-6238 |
| chr4-74597680-DEL-2878 | chr4-74597681-DEL-2878 |
| chr4-74600197-DEL-484 | chr4-74600198-DEL-484 |
| chr4-74611471-DEL-64 | chr4-74611471-DEL-64 |
| chr4-74616794-DEL-6145 | chr4-74616801-DEL-6144 |
| chr4-74626428-DEL-2053 | chr4-74626428-DEL-2053 |
| chr4-74626745-DEL-73 | chr4-74626746-DEL-74 |
| chr4-74638429-DEL-1674 | chr4-74638429-DEL-1674 |
| chr4-74643924-DEL-113 | chr4-74643924-DEL-111 |
| chr4-74661301-DEL-1637 | chr4-74661302-DEL-1637 |
| chr4-74782396-DEL-6609 | chr4-74782396-DEL-6609 |
| chr4-74845389-DEL-5454 | chr4-74845390-DEL-5454 |
| chr4-74851955-DEL-6592 | chr4-74851956-DEL-6592 |
| chr4-74858091-DEL-208 | chr4-74858092-DEL-208 |
| chr4-74860292-DEL-1242 | chr4-74860293-DEL-1242 |
| chr4-74866811-DEL-65 | chr4-74866812-DEL-65 |
| chr4-74934693-DEL-953 | chr4-74934694-DEL-953 |
| chr4-74935530-DEL-200 | chr4-74935531-DEL-200 |
| chr4-74951134-DEL-56 | chr4-74951129-DEL-62 |
| chr4-74985289-DEL-8424 | chr4-74985290-DEL-8424 |
| chr4-75017148-DEL-5763 | chr4-75017149-DEL-5763 |
| chr4-75028410-DEL-203 | chr4-75028411-DEL-203 |
| chr4-75030618-DEL-76 | chr4-75030619-DEL-76 |
| chr4-75049608-DEL-6489 | chr4-75049599-DEL-6490 |
| chr4-75064774-DEL-214 | chr4-75064775-DEL-214 |
| chr4-75080770-DEL-50 | chr4-75080771-DEL-50 |
| chr4-75135730-DEL-322 | chr4-75135731-DEL-322 |
| chr4-75145518-DEL-104 | chr4-75145525-DEL-102 |
| chr4-75159096-DEL-606 | chr4-75159097-DEL-606 |
| chr4-75161658-DEL-621 | chr4-75161658-DEL-621 |
| chr4-75164810-DEL-558 | chr4-75164806-DEL-557 |
| chr4-75221440-DEL-7002 | chr4-75221441-DEL-7002 |
| chr4-75252098-DEL-71 | chr4-75252104-DEL-72 |
| chr4-75266967-DEL-6999 | chr4-75266968-DEL-6999 |

| | |
|---|---|
| chr4-75351994-DEL-740 | chr4-75351995-DEL-740 |
| chr4-75369744-DEL-1682 | chr4-75369745-DEL-1682 |
| chr4-75403952-DEL-203 | chr4-75403953-DEL-203 |
| chr4-75429013-DEL-212 | chr4-75429014-DEL-212 |
| chr4-75431720-DEL-139 | chr4-75431721-DEL-139 |
| chr4-75439787-DEL-52 | chr4-75439788-DEL-64 |
| chr4-75441464-DEL-393 | chr4-75441465-DEL-393 |
| chr4-75461124-DEL-495 | chr4-75461125-DEL-495 |
| chr4-75498563-DEL-712 | chr4-75498564-DEL-712 |
| chr4-75535068-DEL-7125 | chr4-75535069-DEL-7125 |
| chr4-75586687-DEL-424 | chr4-75586688-DEL-424 |
| chr4-75653597-DEL-3732 | chr4-75653598-DEL-3732 |
| chr4-75663740-DEL-56 | chr4-75663735-DEL-62 |
| chr4-75713566-DEL-62 | chr4-75713566-DEL-62 |
| chr4-75717723-DEL-881 | chr4-75717724-DEL-881 |
| chr4-75725225-DEL-59 | chr4-75725218-DEL-67 |
| chr4-75725375-DEL-509 | chr4-75725392-DEL-500 |
| chr4-75749729-DEL-510 | chr4-75749730-DEL-510 |
| chr4-75849238-DEL-212 | chr4-75849239-DEL-212 |
| chr4-75855178-DEL-597 | chr4-75855179-DEL-597 |
| chr4-75882836-DEL-69 | chr4-75882837-DEL-69 |
| chr4-76029039-DEL-1117 | chr4-76029040-DEL-1117 |
| chr4-76033149-DEL-1628 | chr4-76033150-DEL-1628 |
| chr4-76069074-DEL-80 | chr4-76069075-DEL-80 |
| chr4-76075813-DEL-215 | chr4-76075814-DEL-215 |
| chr4-76089493-DEL-66 | chr4-76089492-DEL-63 |
| chr4-76125970-DEL-7404 | chr4-76125971-DEL-7404 |
| chr4-76132911-DEL-208 | chr4-76132910-DEL-208 |
| chr4-76168447-DEL-96 | chr4-76168448-DEL-96 |
| chr4-76245106-DEL-93 | chr4-76245107-DEL-93 |
| chr4-76249496-DEL-124 | chr4-76249496-DEL-122 |
| chr4-76258547-DEL-673 | chr4-76258548-DEL-673 |
| chr4-76261243-DEL-1023 | chr4-76261244-DEL-1023 |
| chr4-76278071-DEL-52 | chr4-76278072-DEL-52 |
| chr4-76322382-DEL-610 | chr4-76322383-DEL-610 |
| chr4-76329731-DEL-1568 | chr4-76329732-DEL-1568 |
| chr4-76335915-DEL-129 | chr4-76335916-DEL-129 |
| chr4-76344083-DEL-705 | chr4-76344081-DEL-706 |
| chr4-76462045-DEL-899 | chr4-76462046-DEL-899 |
| chr4-76476857-DEL-68 | chr4-76476844-DEL-82 |
| chr4-76489904-DEL-272 | chr4-76489905-DEL-272 |
| chr4-76514929-DEL-74 | chr4-76514930-DEL-74 |
| chr4-76516325-DEL-70 | chr4-76516326-DEL-70 |
| chr4-76521656-DEL-498 | chr4-76521652-DEL-499 |
| chr4-76523943-DEL-1055 | chr4-76523944-DEL-1055 |
| chr4-76532700-DEL-543 | chr4-76532701-DEL-543 |
| chr4-76560182-DEL-861 | chr4-76560183-DEL-861 |
| chr4-76563901-DEL-145 | chr4-76563902-DEL-145 |
| chr4-76568660-DEL-53 | chr4-76568657-DEL-77 |
| chr4-76619817-DEL-68 | chr4-76619811-DEL-66 |
| chr4-76641530-DEL-1199 | chr4-76641531-DEL-1199 |
| chr4-76643838-DEL-136 | chr4-76643850-DEL-125 |
| chr4-76646916-DEL-6623 | chr4-76646917-DEL-6623 |
| chr4-76659894-DEL-974 | chr4-76659895-DEL-974 |
| chr4-76721615-DEL-92 | chr4-76721610-DEL-96 |
| chr4-76737449-DEL-170 | chr4-76737450-DEL-170 |
| chr4-76771055-DEL-167 | chr4-76771057-DEL-168 |
| chr4-76779374-DEL-143 | chr4-76779375-DEL-143 |
| chr4-76794450-DEL-96 | chr4-76794417-DEL-96 |
| chr4-76821378-DEL-52 | chr4-76821379-DEL-52 |
| chr4-76832341-DEL-574 | chr4-76832342-DEL-574 |
| chr4-76852192-DEL-125 | chr4-76852193-DEL-125 |
| chr4-76853331-DEL-126 | chr4-76853332-DEL-126 |
| chr4-76864945-DEL-95 | chr4-76864946-DEL-95 |
| chr4-76881982-DEL-738 | chr4-76881978-DEL-740 |
| chr4-76888217-DEL-58 | chr4-76888218-DEL-58 |
| chr4-76922602-DEL-442 | chr4-76922603-DEL-442 |
| chr4-76930676-DEL-148 | chr4-76930670-DEL-149 |
| chr4-76933533-DEL-1082 | chr4-76933534-DEL-1082 |
| chr4-76943591-DEL-109 | chr4-76943594-DEL-103 |
| chr4-76945747-DEL-183 | chr4-76945747-DEL-183 |
| chr4-76954007-DEL-389 | chr4-76954008-DEL-389 |
| chr4-76959086-DEL-486 | chr4-76959087-DEL-486 |
| chr4-76980937-DEL-6648 | chr4-76980938-DEL-6648 |
| chr4-76991191-DEL-71 | chr4-76991192-DEL-71 |
| chr4-77001702-DEL-51 | chr4-77001703-DEL-51 |
| chr4-77004003-DEL-2007 | chr4-77004004-DEL-2007 |
| chr4-77031084-DEL-5093 | chr4-77031084-DEL-5093 |
| chr4-77053385-DEL-116 | chr4-77053430-DEL-113 |
| chr4-77056985-DEL-678 | chr4-77056986-DEL-678 |

| | |
|---|---|
| chr4-77066792-DEL-90 | chr4-77066793-DEL-90 |
| chr4-77067562-DEL-528 | chr4-77067563-DEL-528 |
| chr4-77078022-DEL-64 | chr4-77078006-DEL-50 |
| chr4-77089346-DEL-53 | chr4-77089347-DEL-53 |
| chr4-77113767-DEL-56 | chr4-77113768-DEL-56 |
| chr4-77120979-DEL-91 | chr4-77120973-DEL-92 |
| chr4-77133753-DEL-72 | chr4-77133754-DEL-72 |
| chr4-77166374-DEL-1426 | chr4-77166375-DEL-1425 |
| chr4-77195163-DEL-52 | chr4-77195164-DEL-52 |
| chr4-77201536-DEL-69 | chr4-77201532-DEL-69 |
| chr4-77217817-DEL-342 | chr4-77217818-DEL-342 |
| chr4-77223459-DEL-471 | chr4-77223460-DEL-471 |
| chr4-77260289-DEL-72 | chr4-77260288-DEL-63 |
| chr4-77278894-DEL-65 | chr4-77278895-DEL-65 |
| chr4-77283001-DEL-283 | chr4-77283002-DEL-283 |
| chr4-77286416-DEL-410 | chr4-77286418-DEL-410 |
| chr4-77312320-DEL-205 | chr4-77312321-DEL-205 |
| chr4-77328679-DEL-151 | chr4-77328680-DEL-151 |
| chr4-77333458-DEL-59 | chr4-77333459-DEL-59 |
| chr4-77352300-DEL-764 | chr4-77352301-DEL-764 |
| chr4-77353346-DEL-162 | chr4-77353344-DEL-160 |
| chr4-77355531-DEL-61 | chr4-77355532-DEL-61 |
| chr4-77368292-DEL-87 | chr4-77368321-DEL-63 |
| chr4-77375698-DEL-96 | chr4-77375739-DEL-50 |
| chr4-77405614-DEL-427 | chr4-77405615-DEL-427 |
| chr4-77443919-DEL-76 | chr4-77443945-DEL-50 |
| chr4-77469870-DEL-54 | chr4-77469852-DEL-55 |
| chr4-77470247-DEL-1176 | chr4-77470248-DEL-1176 |
| chr4-77587238-DEL-670 | chr4-77587239-DEL-670 |
| chr4-77604457-DEL-75 | chr4-77604458-DEL-75 |
| chr4-77625014-DEL-86 | chr4-77625012-DEL-87 |
| chr4-77630367-DEL-648 | chr4-77630368-DEL-648 |
| chr4-77638534-DEL-557 | chr4-77638536-DEL-556 |
| chr4-77654411-DEL-133 | chr4-77654424-DEL-128 |
| chr4-77670278-DEL-1358 | chr4-77670279-DEL-1358 |
| chr4-77710814-DEL-1251 | chr4-77710816-DEL-1250 |
| chr4-77719450-DEL-528 | chr4-77719451-DEL-528 |
| chr4-77735173-DEL-455 | chr4-77735174-DEL-455 |
| chr4-77736361-DEL-686 | chr4-77736362-DEL-686 |
| chr4-77829851-DEL-198 | chr4-77829852-DEL-198 |
| chr4-77849567-DEL-1638 | chr4-77849567-DEL-1638 |
| chr4-77891383-DEL-6893 | chr4-77891384-DEL-6893 |
| chr4-77899744-DEL-56 | chr4-77899745-DEL-56 |
| chr4-77902599-DEL-497 | chr4-77902600-DEL-497 |
| chr4-77932416-DEL-6402 | chr4-77932417-DEL-6402 |
| chr4-77962183-DEL-59 | chr4-77962184-DEL-57 |
| chr4-77966037-DEL-161 | chr4-77966037-DEL-161 |
| chr4-78004626-DEL-368 | chr4-78004619-DEL-365 |
| chr4-78011442-DEL-130 | chr4-78011443-DEL-131 |
| chr4-78011926-DEL-241 | chr4-78011933-DEL-235 |
| chr4-78041964-DEL-6220 | chr4-78041965-DEL-6220 |
| chr4-78055204-DEL-1358 | chr4-78055205-DEL-1358 |
| chr4-78082703-DEL-390 | chr4-78082704-DEL-390 |
| chr4-78088407-DEL-392 | chr4-78088421-DEL-376 |
| chr4-78093393-DEL-493 | chr4-78093394-DEL-493 |
| chr4-78115565-DEL-67 | chr4-78115570-DEL-64 |
| chr4-78141943-DEL-1315 | chr4-78141944-DEL-1315 |
| chr4-78155709-DEL-3133 | chr4-78155710-DEL-3133 |
| chr4-78165528-DEL-70 | chr4-78165529-DEL-70 |
| chr4-78168537-DEL-161 | chr4-78168537-DEL-161 |
| chr4-78188449-DEL-56 | chr4-78188450-DEL-58 |
| chr4-78206221-DEL-568 | chr4-78206220-DEL-569 |
| chr4-78225846-DEL-109 | chr4-78225847-DEL-109 |
| chr4-78226742-DEL-476 | chr4-78226742-DEL-476 |
| chr4-78230448-DEL-5569 | chr4-78230449-DEL-5569 |
| chr4-78238356-DEL-654 | chr4-78238385-DEL-650 |
| chr4-78241221-DEL-59 | chr4-78241224-DEL-55 |
| chr4-78243853-DEL-84 | chr4-78243854-DEL-84 |
| chr4-78245102-DEL-2543 | chr4-78245103-DEL-2544 |
| chr4-78271598-DEL-132 | chr4-78271599-DEL-130 |
| chr4-78288453-DEL-871 | chr4-78288454-DEL-871 |
| chr4-78290437-DEL-6240 | chr4-78290437-DEL-6242 |
| chr4-78361295-DEL-111 | chr4-78361298-DEL-109 |
| chr4-78395453-DEL-1862 | chr4-78395453-DEL-1862 |
| chr4-78430391-DEL-5200 | chr4-78430392-DEL-5200 |
| chr4-78451371-DEL-1832 | chr4-78451372-DEL-1832 |
| chr4-78458875-DEL-118 | chr4-78458879-DEL-116 |
| chr4-78459479-DEL-1481 | chr4-78459479-DEL-1481 |
| chr4-78463162-DEL-610 | chr4-78463162-DEL-610 |
| chr4-78464228-DEL-342 | chr4-78464228-DEL-342 |

| | |
|---|---|
| chr4-78471687-DEL-168 | chr4-78471687-DEL-169 |
| chr4-78478676-DEL-480 | chr4-78478671-DEL-480 |
| chr4-78480844-DEL-89 | chr4-78480844-DEL-89 |
| chr4-78503540-DEL-54 | chr4-78503546-DEL-51 |
| chr4-78505091-DEL-60 | chr4-78505092-DEL-60 |
| chr4-78553065-DEL-194 | chr4-78553066-DEL-192 |
| chr4-78576501-DEL-329 | chr4-78576501-DEL-329 |
| chr4-78576997-DEL-3074 | chr4-78576998-DEL-3074 |
| chr4-78598672-DEL-1134 | chr4-78598673-DEL-1134 |
| chr4-78618643-DEL-1456 | chr4-78618644-DEL-1456 |
| chr4-78641430-DEL-439 | chr4-78641431-DEL-439 |
| chr4-78644725-DEL-916 | chr4-78644732-DEL-908 |
| chr4-78683237-DEL-59 | chr4-78683226-DEL-71 |
| chr4-78687266-DEL-139 | chr4-78687267-DEL-139 |
| chr4-78716716-DEL-96 | chr4-78716717-DEL-96 |
| chr4-78721959-DEL-234 | chr4-78721929-DEL-264 |
| chr4-78750005-DEL-3705 | chr4-78749994-DEL-3699 |
| chr4-78764454-DEL-52 | chr4-78764429-DEL-78 |
| chr4-78767253-DEL-1148 | chr4-78767254-DEL-1148 |
| chr4-78772184-DEL-6379 | chr4-78772185-DEL-6379 |
| chr4-78776404-DEL-1988 | chr4-78776405-DEL-1988 |
| chr4-78782402-DEL-665 | chr4-78782403-DEL-665 |
| chr4-78789827-DEL-1174 | chr4-78789828-DEL-1174 |
| chr4-78798195-DEL-1926 | chr4-78798196-DEL-1926 |
| chr4-78804557-DEL-138 | chr4-78804558-DEL-136 |
| chr4-78815024-DEL-624 | chr4-78815025-DEL-624 |
| chr4-78833303-DEL-114 | chr4-78833303-DEL-114 |
| chr4-78908821-DEL-119 | chr4-78908825-DEL-116 |
| chr4-78918197-DEL-91 | chr4-78918200-DEL-89 |
| chr4-78928965-DEL-57 | chr4-78928966-DEL-57 |
| chr4-78956805-DEL-288 | chr4-78956806-DEL-288 |
| chr4-78958231-DEL-54 | chr4-78958236-DEL-50 |
| chr4-78985617-DEL-480 | chr4-78985616-DEL-481 |
| chr4-79038394-DEL-2813 | chr4-79038395-DEL-2813 |
| chr4-79061014-DEL-6669 | chr4-79061015-DEL-6669 |
| chr4-79145909-DEL-2296 | chr4-79145910-DEL-2296 |
| chr4-79159787-DEL-66 | chr4-79159804-DEL-50 |
| chr4-79170007-DEL-764 | chr4-79170008-DEL-764 |
| chr4-79185374-DEL-54 | chr4-79185375-DEL-54 |
| chr4-79210875-DEL-65 | chr4-79210876-DEL-65 |
| chr4-79215299-DEL-9856 | chr4-79215300-DEL-9856 |
| chr4-79234255-DEL-84 | chr4-79234256-DEL-84 |
| chr4-79241101-DEL-477 | chr4-79241105-DEL-467 |
| chr4-79245892-DEL-866 | chr4-79245887-DEL-867 |
| chr4-79247264-DEL-273 | chr4-79247265-DEL-273 |
| chr4-79263517-DEL-560 | chr4-79263518-DEL-560 |
| chr4-79269520-DEL-968 | chr4-79269521-DEL-966 |
| chr4-79276294-DEL-68 | chr4-79276295-DEL-68 |
| chr4-79287673-DEL-140 | chr4-79287674-DEL-140 |
| chr4-79308720-DEL-423 | chr4-79308721-DEL-423 |
| chr4-79313250-DEL-178 | chr4-79313251-DEL-178 |
| chr4-79384729-DEL-2337 | chr4-79384732-DEL-2332 |
| chr4-79397317-DEL-96 | chr4-79397309-DEL-92 |
| chr4-79411835-DEL-92 | chr4-79411872-DEL-68 |
| chr4-79427895-DEL-97 | chr4-79427900-DEL-52 |
| chr4-79472516-DEL-13446 | chr4-79472517-DEL-13446 |
| chr4-79493227-DEL-2750 | chr4-79493228-DEL-2749 |
| chr4-79496820-DEL-1425 | chr4-79496821-DEL-1425 |
| chr4-79512095-DEL-60 | chr4-79512096-DEL-60 |
| chr4-79546142-DEL-6380 | chr4-79546143-DEL-6380 |
| chr4-79550557-DEL-317 | chr4-79550558-DEL-315 |
| chr4-79553749-DEL-244 | chr4-79553750-DEL-244 |
| chr4-79563469-DEL-462 | chr4-79563470-DEL-461 |
| chr4-79589318-DEL-59 | chr4-79589350-DEL-60 |
| chr4-79598447-DEL-62 | chr4-79598448-DEL-62 |
| chr4-79607944-DEL-475 | chr4-79607945-DEL-475 |
| chr4-79632526-DEL-6944 | chr4-79632527-DEL-6944 |
| chr4-79638586-DEL-201 | chr4-79638587-DEL-201 |
| chr4-79653044-DEL-74 | chr4-79653045-DEL-67 |
| chr4-79687729-DEL-3624 | chr4-79687730-DEL-3624 |
| chr4-79744619-DEL-52 | chr4-79744619-DEL-52 |
| chr4-79760482-DEL-137 | chr4-79760483-DEL-137 |
| chr4-79773032-DEL-404 | chr4-79773033-DEL-404 |
| chr4-79811340-DEL-70 | chr4-79811341-DEL-70 |
| chr4-79825829-DEL-60 | chr4-79825829-DEL-58 |
| chr4-79857999-DEL-1924 | chr4-79858000-DEL-1923 |
| chr4-79869084-DEL-1209 | chr4-79869086-DEL-1208 |
| chr4-79881274-DEL-85 | chr4-79881275-DEL-85 |
| chr4-79886698-DEL-296 | chr4-79886699-DEL-296 |
| chr4-79910566-DEL-345 | chr4-79910567-DEL-345 |

| | |
|---|---|
| chr4-79917465-DEL-2260 | chr4-79917466-DEL-2260 |
| chr4-79920566-DEL-2881 | chr4-79920573-DEL-2882 |
| chr4-79926848-DEL-208 | chr4-79926858-DEL-192 |
| chr4-79949394-DEL-349 | chr4-79949395-DEL-349 |
| chr4-79960477-DEL-2576 | chr4-79960478-DEL-2576 |
| chr4-79975089-DEL-65 | chr4-79975090-DEL-65 |
| chr4-79979379-DEL-501 | chr4-79979380-DEL-501 |
| chr4-80000424-DEL-74 | chr4-80000425-DEL-74 |
| chr4-80005169-DEL-104 | chr4-80005169-DEL-104 |
| chr4-80005971-DEL-529 | chr4-80005971-DEL-529 |
| chr4-80006724-DEL-247 | chr4-80006724-DEL-247 |
| chr4-80017101-DEL-56 | chr4-80017106-DEL-52 |
| chr4-80027637-DEL-133 | chr4-80027631-DEL-135 |
| chr4-80048548-DEL-945 | chr4-80048549-DEL-945 |
| chr4-80053566-DEL-1184 | chr4-80053566-DEL-1184 |
| chr4-80077469-DEL-140 | chr4-80077470-DEL-140 |
| chr4-80084202-DEL-188 | chr4-80084204-DEL-187 |
| chr4-80107855-DEL-2009 | chr4-80107856-DEL-2010 |
| chr4-80110324-DEL-256 | chr4-80110325-DEL-256 |
| chr4-80165410-DEL-214 | chr4-80165411-DEL-214 |
| chr4-80175053-DEL-8806 | chr4-80175041-DEL-8818 |
| chr4-80206917-DEL-8358 | chr4-80206918-DEL-8358 |
| chr4-80219694-DEL-1428 | chr4-80219695-DEL-1428 |
| chr4-80226074-DEL-2670 | chr4-80226074-DEL-2671 |
| chr4-80233391-DEL-1302 | chr4-80233392-DEL-1302 |
| chr4-80254535-DEL-74 | chr4-80254556-DEL-99 |
| chr4-80257360-DEL-77 | chr4-80257361-DEL-77 |
| chr4-80314636-DEL-68 | chr4-80314637-DEL-68 |
| chr4-80320972-DEL-522 | chr4-80320973-DEL-522 |
| chr4-80327137-DEL-5578 | chr4-80327138-DEL-5578 |
| chr4-80333000-DEL-1028 | chr4-80333001-DEL-1028 |
| chr4-80366286-DEL-2252 | chr4-80366287-DEL-2252 |
| chr4-80375084-DEL-162 | chr4-80375085-DEL-162 |
| chr4-80399545-DEL-3296 | chr4-80399546-DEL-3296 |
| chr4-80481212-DEL-6475 | chr4-80481213-DEL-6475 |
| chr4-80500835-DEL-3290 | chr4-80500836-DEL-3290 |
| chr4-80515300-DEL-6173 | chr4-80515301-DEL-6173 |
| chr4-80526374-DEL-84 | chr4-80526372-DEL-85 |
| chr4-80541784-DEL-452 | chr4-80541785-DEL-452 |
| chr4-80551483-DEL-669 | chr4-80551484-DEL-669 |
| chr4-80552509-DEL-144 | chr4-80552510-DEL-144 |
| chr4-80587777-DEL-1233 | chr4-80587778-DEL-1233 |
| chr4-80591881-DEL-403 | chr4-80591882-DEL-403 |
| chr4-80649242-DEL-4466 | chr4-80649243-DEL-4466 |
| chr4-80701241-DEL-113 | chr4-80701240-DEL-110 |
| chr4-80710860-DEL-694 | chr4-80710861-DEL-694 |
| chr4-80731238-DEL-6242 | chr4-80731244-DEL-6241 |
| chr4-80760126-DEL-52 | chr4-80760127-DEL-52 |
| chr4-80777885-DEL-131 | chr4-80777882-DEL-135 |
| chr4-80782556-DEL-86 | chr4-80782557-DEL-86 |
| chr4-80790871-DEL-65 | chr4-80790835-DEL-81 |
| chr4-80798374-DEL-150 | chr4-80798369-DEL-140 |
| chr4-80800934-DEL-365 | chr4-80800936-DEL-364 |
| chr4-80817789-DEL-89 | chr4-80817790-DEL-89 |
| chr4-80818033-DEL-860 | chr4-80818034-DEL-860 |
| chr4-80845913-DEL-2882 | chr4-80845914-DEL-2882 |
| chr4-80870403-DEL-165 | chr4-80870404-DEL-165 |
| chr4-80879429-DEL-6211 | chr4-80879430-DEL-6211 |
| chr4-80894242-DEL-960 | chr4-80894243-DEL-960 |
| chr4-80897416-DEL-136 | chr4-80897421-DEL-135 |
| chr4-80929627-DEL-338 | chr4-80929628-DEL-333 |
| chr4-80932336-DEL-75 | chr4-80932337-DEL-75 |
| chr4-80959428-DEL-624 | chr4-80959440-DEL-623 |
| chr4-80963447-DEL-52 | chr4-80963448-DEL-52 |
| chr4-80984975-DEL-6410 | chr4-80984976-DEL-6410 |
| chr4-81026922-DEL-1128 | chr4-81026923-DEL-1128 |
| chr4-81033389-DEL-108 | chr4-81033390-DEL-108 |
| chr4-81053605-DEL-1007 | chr4-81053606-DEL-1007 |
| chr4-81060610-DEL-320 | chr4-81060611-DEL-320 |
| chr4-81107101-DEL-141 | chr4-81107102-DEL-135 |
| chr4-81116446-DEL-66 | chr4-81116442-DEL-84 |
| chr4-81117751-DEL-5621 | chr4-81117741-DEL-5586 |
| chr4-81138632-DEL-81 | chr4-81138589-DEL-82 |
| chr4-81147279-DEL-6077 | chr4-81147280-DEL-6093 |
| chr4-81154256-DEL-128 | chr4-81154257-DEL-128 |
| chr4-81172043-DEL-90 | chr4-81172044-DEL-90 |
| chr4-81182251-DEL-497 | chr4-81182252-DEL-497 |
| chr4-81224712-DEL-154 | chr4-81224713-DEL-152 |
| chr4-81260970-DEL-60 | chr4-81260971-DEL-60 |
| chr4-81270356-DEL-92 | chr4-81270357-DEL-90 |

| | |
|---|---|
| chr4-81270774-DEL-194 | chr4-81270775-DEL-195 |
| chr4-81276153-DEL-182 | chr4-81276186-DEL-150 |
| chr4-81284536-DEL-157 | chr4-81284537-DEL-157 |
| chr4-81289409-DEL-136 | chr4-81289410-DEL-136 |
| chr4-81317940-DEL-52 | chr4-81317941-DEL-50 |
| chr4-81323639-DEL-6337 | chr4-81323640-DEL-6337 |
| chr4-81339520-DEL-203 | chr4-81339521-DEL-203 |
| chr4-81341362-DEL-205 | chr4-81341363-DEL-205 |
| chr4-81345451-DEL-209 | chr4-81345452-DEL-209 |
| chr4-81366306-DEL-201 | chr4-81366307-DEL-201 |
| chr4-81368615-DEL-182 | chr4-81368616-DEL-182 |
| chr4-81382982-DEL-166 | chr4-81382983-DEL-166 |
| chr4-81384328-DEL-66 | chr4-81384329-DEL-66 |
| chr4-81417244-DEL-1438 | chr4-81417245-DEL-1438 |
| chr4-81424866-DEL-205 | chr4-81424867-DEL-205 |
| chr4-81437671-DEL-74 | chr4-81437672-DEL-92 |
| chr4-81438346-DEL-163 | chr4-81438347-DEL-163 |
| chr4-81441151-DEL-1155 | chr4-81441152-DEL-1155 |
| chr4-81441179-DEL-383 | chr4-81441180-DEL-383 |
| chr4-81450658-DEL-632 | chr4-81450659-DEL-633 |
| chr4-81503750-DEL-190 | chr4-81503750-DEL-190 |
| chr4-81605420-DEL-101 | chr4-81605421-DEL-100 |
| chr4-81617215-DEL-7922 | chr4-81617216-DEL-7922 |
| chr4-81623524-DEL-208 | chr4-81623525-DEL-208 |
| chr4-81634530-DEL-3565 | chr4-81634531-DEL-3565 |
| chr4-81669148-DEL-818 | chr4-81669149-DEL-818 |
| chr4-81690686-DEL-8404 | chr4-81690687-DEL-8404 |
| chr4-81719771-DEL-384 | chr4-81719772-DEL-384 |
| chr4-81761512-DEL-349 | chr4-81761513-DEL-349 |
| chr4-81772946-DEL-662 | chr4-81772946-DEL-662 |
| chr4-81779884-DEL-140 | chr4-81779885-DEL-141 |
| chr4-81783344-DEL-737 | chr4-81783347-DEL-740 |
| chr4-81785616-DEL-196 | chr4-81785617-DEL-196 |
| chr4-81792437-DEL-488 | chr4-81792438-DEL-488 |
| chr4-81805906-DEL-3814 | chr4-81805907-DEL-3814 |
| chr4-81820396-DEL-6755 | chr4-81820397-DEL-6755 |
| chr4-81837885-DEL-1273 | chr4-81837886-DEL-1273 |
| chr4-81851464-DEL-440 | chr4-81851465-DEL-440 |
| chr4-81887431-DEL-79 | chr4-81887459-DEL-51 |
| chr4-81902271-DEL-106 | chr4-81902272-DEL-106 |
| chr4-81959395-DEL-6888 | chr4-81959396-DEL-6888 |
| chr4-81979332-DEL-66 | chr4-81979333-DEL-66 |
| chr4-81982042-DEL-1139 | chr4-81982043-DEL-1139 |
| chr4-81996444-DEL-158 | chr4-81996445-DEL-158 |
| chr4-82020828-DEL-329 | chr4-82020829-DEL-329 |
| chr4-82031250-DEL-88 | chr4-82031251-DEL-88 |
| chr4-82044998-DEL-52 | chr4-82044999-DEL-52 |
| chr4-82105420-DEL-6484 | chr4-82105421-DEL-6484 |
| chr4-82120514-DEL-257 | chr4-82120515-DEL-257 |
| chr4-82170541-DEL-417 | chr4-82170542-DEL-417 |
| chr4-82172414-DEL-127 | chr4-82172417-DEL-134 |
| chr4-82233480-DEL-174 | chr4-82233481-DEL-174 |
| chr4-82249304-DEL-180 | chr4-82249305-DEL-180 |
| chr4-82286542-DEL-66 | chr4-82286544-DEL-65 |
| chr4-82345121-DEL-235 | chr4-82345122-DEL-235 |
| chr4-82380976-DEL-79 | chr4-82380997-DEL-59 |
| chr4-82405513-DEL-188 | chr4-82405511-DEL-188 |
| chr4-82520664-DEL-192 | chr4-82520665-DEL-192 |
| chr4-82565226-DEL-711 | chr4-82565227-DEL-711 |
| chr4-82604097-DEL-112 | chr4-82604095-DEL-108 |
| chr4-82630654-DEL-208 | chr4-82630655-DEL-208 |
| chr4-82632934-DEL-191 | chr4-82632934-DEL-192 |
| chr4-82640001-DEL-432 | chr4-82640002-DEL-432 |
| chr4-82647358-DEL-472 | chr4-82647359-DEL-472 |
| chr4-82656310-DEL-6450 | chr4-82656307-DEL-6451 |
| chr4-82662335-DEL-201 | chr4-82662336-DEL-201 |
| chr4-82671644-DEL-377 | chr4-82671639-DEL-378 |
| chr4-82682623-DEL-197 | chr4-82682624-DEL-197 |
| chr4-82688846-DEL-96 | chr4-82688847-DEL-97 |
| chr4-82702885-DEL-191 | chr4-82702886-DEL-191 |
| chr4-82703699-DEL-69 | chr4-82703714-DEL-67 |
| chr4-82704360-DEL-6364 | chr4-82704361-DEL-6364 |
| chr4-82728338-DEL-65 | chr4-82728339-DEL-65 |
| chr4-82738982-DEL-82 | chr4-82738983-DEL-78 |
| chr4-82746389-DEL-344 | chr4-82746390-DEL-344 |
| chr4-82766451-DEL-82 | chr4-82766452-DEL-82 |
| chr4-82778532-DEL-56 | chr4-82778537-DEL-55 |
| chr4-82782999-DEL-182 | chr4-82783000-DEL-182 |
| chr4-82844401-DEL-1227 | chr4-82844402-DEL-1227 |
| chr4-82872873-DEL-167 | chr4-82872874-DEL-167 |

| | |
|---|---|
| chr4-82884785-DEL-54 | chr4-82884786-DEL-54 |
| chr4-82935368-DEL-66 | chr4-82935369-DEL-66 |
| chr4-82981301-DEL-62 | chr4-82981304-DEL-56 |
| chr4-82983973-DEL-7432 | chr4-82983974-DEL-7432 |
| chr4-82999641-DEL-169 | chr4-82999642-DEL-169 |
| chr4-83012290-DEL-86 | chr4-83012290-DEL-83 |
| chr4-83032218-DEL-333 | chr4-83032219-DEL-328 |
| chr4-83040149-DEL-303 | chr4-83040150-DEL-303 |
| chr4-83055116-DEL-2284 | chr4-83055117-DEL-2284 |
| chr4-83059748-DEL-122 | chr4-83059761-DEL-93 |
| chr4-83060605-DEL-202 | chr4-83060606-DEL-202 |
| chr4-83074517-DEL-6485 | chr4-83074518-DEL-6485 |
| chr4-83129659-DEL-87 | chr4-83129652-DEL-63 |
| chr4-83132020-DEL-165 | chr4-83132025-DEL-154 |
| chr4-83155336-DEL-58 | chr4-83155337-DEL-58 |
| chr4-83157559-DEL-66 | chr4-83157560-DEL-66 |
| chr4-83196273-DEL-59 | chr4-83196274-DEL-59 |
| chr4-83199680-DEL-64 | chr4-83199681-DEL-64 |
| chr4-83217717-DEL-215 | chr4-83217718-DEL-215 |
| chr4-83219347-DEL-154 | chr4-83219348-DEL-154 |
| chr4-83222327-DEL-207 | chr4-83222328-DEL-207 |
| chr4-83231895-DEL-236 | chr4-83231896-DEL-236 |
| chr4-83234134-DEL-80 | chr4-83234135-DEL-80 |
| chr4-83257984-DEL-54 | chr4-83257985-DEL-54 |
| chr4-83259309-DEL-228 | chr4-83259310-DEL-228 |
| chr4-83280754-DEL-79 | chr4-83280755-DEL-79 |
| chr4-83284480-DEL-6306 | chr4-83284481-DEL-6306 |
| chr4-83293596-DEL-194 | chr4-83293597-DEL-194 |
| chr4-83339708-DEL-919 | chr4-83339709-DEL-919 |
| chr4-83361038-DEL-6439 | chr4-83361039-DEL-6439 |
| chr4-83377461-DEL-60 | chr4-83377462-DEL-60 |
| chr4-83399643-DEL-220 | chr4-83399644-DEL-220 |
| chr4-83400551-DEL-63 | chr4-83400552-DEL-63 |
| chr4-83408974-DEL-1864 | chr4-83408975-DEL-1864 |
| chr4-83417980-DEL-132 | chr4-83417981-DEL-132 |
| chr4-83423351-DEL-1633 | chr4-83423360-DEL-1634 |
| chr4-83425163-DEL-54 | chr4-83425164-DEL-54 |
| chr4-83434361-DEL-211 | chr4-83434362-DEL-211 |
| chr4-83438170-DEL-647 | chr4-83438171-DEL-646 |
| chr4-83455102-DEL-5086 | chr4-83455103-DEL-5086 |
| chr4-83497857-DEL-53 | chr4-83497858-DEL-53 |
| chr4-83555310-DEL-7089 | chr4-83555311-DEL-7089 |
| chr4-83581985-DEL-78 | chr4-83581986-DEL-78 |
| chr4-83584359-DEL-142 | chr4-83584360-DEL-142 |
| chr4-83585287-DEL-1056 | chr4-83585288-DEL-1056 |
| chr4-83630545-DEL-846 | chr4-83630543-DEL-846 |
| chr4-83644653-DEL-439 | chr4-83644654-DEL-439 |
| chr4-83662385-DEL-6603 | chr4-83662386-DEL-6603 |
| chr4-83706529-DEL-60 | chr4-83706527-DEL-56 |
| chr4-83742912-DEL-121 | chr4-83742913-DEL-88 |
| chr4-83757165-DEL-52 | chr4-83757166-DEL-52 |
| chr4-83776325-DEL-54 | chr4-83776326-DEL-54 |
| chr4-83845222-DEL-1775 | chr4-83845223-DEL-1775 |
| chr4-83854730-DEL-134 | chr4-83854731-DEL-134 |
| chr4-83855263-DEL-83 | chr4-83855264-DEL-83 |
| chr4-83860919-DEL-76 | chr4-83860920-DEL-77 |
| chr4-83876109-DEL-121 | chr4-83876110-DEL-121 |
| chr4-83877010-DEL-6887 | chr4-83877011-DEL-6887 |
| chr4-84002759-DEL-72 | chr4-84002759-DEL-62 |
| chr4-84059686-DEL-351 | chr4-84059687-DEL-351 |
| chr4-84077273-DEL-434 | chr4-84077274-DEL-434 |
| chr4-84079417-DEL-70 | chr4-84079418-DEL-70 |
| chr4-84096388-DEL-1890 | chr4-84096389-DEL-1890 |
| chr4-84111392-DEL-70 | chr4-84111436-DEL-52 |
| chr4-84116217-DEL-76 | chr4-84116218-DEL-52 |
| chr4-84156873-DEL-91 | chr4-84156874-DEL-91 |
| chr4-84166432-DEL-691 | chr4-84166433-DEL-691 |
| chr4-84240433-DEL-703 | chr4-84240434-DEL-703 |
| chr4-84253191-DEL-80 | chr4-84253192-DEL-80 |
| chr4-84256036-DEL-58 | chr4-84256037-DEL-56 |
| chr4-84327932-DEL-176 | chr4-84327933-DEL-176 |
| chr4-84351780-DEL-85 | chr4-84351781-DEL-85 |
| chr4-84449812-DEL-198 | chr4-84449813-DEL-198 |
| chr4-84471737-DEL-56 | chr4-84471738-DEL-56 |
| chr4-84504946-DEL-58 | chr4-84504947-DEL-58 |
| chr4-84507273-DEL-220 | chr4-84507271-DEL-221 |
| chr4-84510371-DEL-207 | chr4-84510372-DEL-207 |
| chr4-84529153-DEL-1126 | chr4-84529152-DEL-1127 |
| chr4-84620241-DEL-187 | chr4-84620242-DEL-187 |
| chr4-84638469-DEL-544 | chr4-84638470-DEL-544 |

| | |
|---|---|
| chr4-84689984-DEL-259 | chr4-84689985-DEL-259 |
| chr4-84718588-DEL-355 | chr4-84718589-DEL-355 |
| chr4-84749014-DEL-182 | chr4-84749015-DEL-182 |
| chr4-84772885-DEL-60 | chr4-84772886-DEL-60 |
| chr4-84780666-DEL-209 | chr4-84780667-DEL-209 |
| chr4-84781355-DEL-445 | chr4-84781356-DEL-445 |
| chr4-84800486-DEL-192 | chr4-84800460-DEL-233 |
| chr4-84817081-DEL-4681 | chr4-84817082-DEL-4681 |
| chr4-84834056-DEL-1964 | chr4-84834057-DEL-1964 |
| chr4-84854897-DEL-149 | chr4-84854898-DEL-149 |
| chr4-84953407-DEL-6434 | chr4-84953407-DEL-6434 |
| chr4-84960902-DEL-166 | chr4-84960903-DEL-166 |
| chr4-84962055-DEL-237 | chr4-84962056-DEL-237 |
| chr4-84984089-DEL-155 | chr4-84984093-DEL-154 |
| chr4-84997490-DEL-1422 | chr4-84997491-DEL-1422 |
| chr4-85000978-DEL-80 | chr4-85000979-DEL-80 |
| chr4-85004111-DEL-60 | chr4-85004094-DEL-58 |
| chr4-85026767-DEL-608 | chr4-85026768-DEL-608 |
| chr4-85083773-DEL-1248 | chr4-85083773-DEL-1248 |
| chr4-85089738-DEL-55 | chr4-85089726-DEL-68 |
| chr4-85103136-DEL-54 | chr4-85103137-DEL-54 |
| chr4-85132407-DEL-117 | chr4-85132408-DEL-117 |
| chr4-85132766-DEL-618 | chr4-85132768-DEL-617 |
| chr4-85134480-DEL-70 | chr4-85134481-DEL-70 |
| chr4-85147500-DEL-626 | chr4-85147499-DEL-623 |
| chr4-85152510-DEL-6441 | chr4-85152511-DEL-6441 |
| chr4-85186714-DEL-56 | chr4-85186711-DEL-58 |
| chr4-85219129-DEL-58 | chr4-85219130-DEL-56 |
| chr4-85244681-DEL-72 | chr4-85244682-DEL-72 |
| chr4-85250668-DEL-75 | chr4-85250668-DEL-75 |
| chr4-85254207-DEL-198 | chr4-85254208-DEL-198 |
| chr4-85300408-DEL-72 | chr4-85300409-DEL-72 |
| chr4-85315793-DEL-52 | chr4-85315789-DEL-51 |
| chr4-85321808-DEL-132 | chr4-85321821-DEL-122 |
| chr4-85325597-DEL-6272 | chr4-85325598-DEL-6272 |
| chr4-85331612-DEL-214 | chr4-85331613-DEL-214 |
| chr4-85340944-DEL-93 | chr4-85340945-DEL-93 |
| chr4-85348247-DEL-132 | chr4-85348251-DEL-125 |
| chr4-85378316-DEL-490 | chr4-85378317-DEL-490 |
| chr4-85391348-DEL-2295 | chr4-85391349-DEL-2295 |
| chr4-85395412-DEL-6457 | chr4-85395413-DEL-6457 |
| chr4-85402640-DEL-559 | chr4-85402641-DEL-559 |
| chr4-85427097-DEL-216 | chr4-85427098-DEL-216 |
| chr4-85486820-DEL-133 | chr4-85486819-DEL-135 |
| chr4-85552359-DEL-2269 | chr4-85552360-DEL-2269 |
| chr4-85597994-DEL-171 | chr4-85597994-DEL-169 |
| chr4-85619315-DEL-79 | chr4-85619324-DEL-77 |
| chr4-85633360-DEL-472 | chr4-85633361-DEL-472 |
| chr4-85681775-DEL-198 | chr4-85681776-DEL-198 |
| chr4-85714433-DEL-192 | chr4-85714434-DEL-191 |
| chr4-85730805-DEL-159 | chr4-85730806-DEL-159 |
| chr4-85735065-DEL-72 | chr4-85735066-DEL-72 |
| chr4-85750900-DEL-270 | chr4-85750859-DEL-288 |
| chr4-85767123-DEL-218 | chr4-85767124-DEL-218 |
| chr4-85809861-DEL-6465 | chr4-85809862-DEL-6465 |
| chr4-85822859-DEL-56 | chr4-85822860-DEL-54 |
| chr4-85841622-DEL-189 | chr4-85841623-DEL-189 |
| chr4-85843602-DEL-6288 | chr4-85843603-DEL-6288 |
| chr4-85859433-DEL-81 | chr4-85859434-DEL-81 |
| chr4-85883277-DEL-750 | chr4-85883278-DEL-750 |
| chr4-85894206-DEL-60 | chr4-85894207-DEL-60 |
| chr4-85917243-DEL-4009 | chr4-85917244-DEL-4009 |
| chr4-86019263-DEL-594 | chr4-86019264-DEL-593 |
| chr4-86051789-DEL-111 | chr4-86051789-DEL-96 |
| chr4-86061388-DEL-3032 | chr4-86061389-DEL-3032 |
| chr4-86061632-DEL-202 | chr4-86061633-DEL-202 |
| chr4-86071954-DEL-79 | chr4-86071955-DEL-77 |
| chr4-86073201-DEL-380 | chr4-86073216-DEL-365 |
| chr4-86078635-DEL-117 | chr4-86078660-DEL-93 |
| chr4-86084439-DEL-372 | chr4-86084440-DEL-372 |
| chr4-86088435-DEL-266 | chr4-86088436-DEL-266 |
| chr4-86105842-DEL-128 | chr4-86105848-DEL-125 |
| chr4-86117862-DEL-112 | chr4-86117857-DEL-111 |
| chr4-86123085-DEL-467 | chr4-86123086-DEL-467 |
| chr4-86131380-DEL-294 | chr4-86131381-DEL-294 |
| chr4-86132045-DEL-147 | chr4-86132046-DEL-147 |
| chr4-86173836-DEL-5699 | chr4-86173837-DEL-5699 |
| chr4-86203828-DEL-191 | chr4-86203829-DEL-191 |
| chr4-86215348-DEL-1010 | chr4-86215349-DEL-1010 |
| chr4-86253229-DEL-60 | chr4-86253230-DEL-96 |

| | |
|---|---|
| chr4-86283262-DEL-63 | chr4-86283263-DEL-63 |
| chr4-86293699-DEL-189 | chr4-86293700-DEL-189 |
| chr4-86361621-DEL-70 | chr4-86361621-DEL-70 |
| chr4-86392474-DEL-1162 | chr4-86392474-DEL-1162 |
| chr4-86426965-DEL-162 | chr4-86426964-DEL-162 |
| chr4-86429880-DEL-145 | chr4-86429881-DEL-143 |
| chr4-86433436-DEL-1266 | chr4-86433437-DEL-1265 |
| chr4-86454933-DEL-1015 | chr4-86454934-DEL-1016 |
| chr4-86466271-DEL-56 | chr4-86466273-DEL-52 |
| chr4-86533441-DEL-5343 | chr4-86533442-DEL-5343 |
| chr4-86553522-DEL-55 | chr4-86553523-DEL-50 |
| chr4-86557148-DEL-205 | chr4-86557149-DEL-205 |
| chr4-86569336-DEL-55 | chr4-86569330-DEL-61 |
| chr4-86577785-DEL-417 | chr4-86577786-DEL-417 |
| chr4-86578644-DEL-83 | chr4-86578678-DEL-55 |
| chr4-86579254-DEL-203 | chr4-86579255-DEL-203 |
| chr4-86585365-DEL-400 | chr4-86585366-DEL-400 |
| chr4-86598511-DEL-66 | chr4-86598524-DEL-54 |
| chr4-86627084-DEL-68 | chr4-86627090-DEL-56 |
| chr4-86638457-DEL-248 | chr4-86638458-DEL-248 |
| chr4-86642900-DEL-221 | chr4-86642901-DEL-221 |
| chr4-86647518-DEL-67 | chr4-86647519-DEL-67 |
| chr4-86651962-DEL-204 | chr4-86651963-DEL-204 |
| chr4-86653838-DEL-158 | chr4-86653839-DEL-158 |
| chr4-86673099-DEL-6383 | chr4-86673100-DEL-6383 |
| chr4-86691717-DEL-210 | chr4-86691718-DEL-210 |
| chr4-86693551-DEL-57 | chr4-86693565-DEL-58 |
| chr4-86721714-DEL-210 | chr4-86721700-DEL-238 |
| chr4-86733354-DEL-205 | chr4-86733355-DEL-205 |
| chr4-86741332-DEL-54 | chr4-86741333-DEL-52 |
| chr4-86752316-DEL-61 | chr4-86752317-DEL-61 |
| chr4-86784849-DEL-144 | chr4-86784846-DEL-146 |
| chr4-86791087-DEL-60 | chr4-86791087-DEL-60 |
| chr4-86810876-DEL-3580 | chr4-86810877-DEL-3580 |
| chr4-86816574-DEL-51 | chr4-86816574-DEL-51 |
| chr4-86844756-DEL-230 | chr4-86844755-DEL-230 |
| chr4-86845010-DEL-859 | chr4-86845010-DEL-859 |
| chr4-86855883-DEL-400 | chr4-86855884-DEL-400 |
| chr4-86872767-DEL-68 | chr4-86872768-DEL-71 |
| chr4-86877257-DEL-267 | chr4-86877258-DEL-267 |
| chr4-86877671-DEL-1008 | chr4-86877672-DEL-1008 |
| chr4-86913211-DEL-56 | chr4-86913231-DEL-54 |
| chr4-86947262-DEL-441 | chr4-86947263-DEL-441 |
| chr4-86961682-DEL-256 | chr4-86961683-DEL-256 |
| chr4-86983211-DEL-66 | chr4-86983212-DEL-66 |
| chr4-87021586-DEL-1726 | chr4-87021587-DEL-1726 |
| chr4-87051988-DEL-716 | chr4-87051986-DEL-707 |
| chr4-87055398-DEL-416 | chr4-87055399-DEL-416 |
| chr4-87087528-DEL-1677 | chr4-87087529-DEL-1677 |
| chr4-87089475-DEL-265 | chr4-87089476-DEL-265 |
| chr4-87090903-DEL-121 | chr4-87090904-DEL-121 |
| chr4-87095847-DEL-100 | chr4-87095845-DEL-98 |
| chr4-87105319-DEL-953 | chr4-87105324-DEL-952 |
| chr4-87110274-DEL-72 | chr4-87110276-DEL-61 |
| chr4-87158996-DEL-193 | chr4-87158995-DEL-194 |
| chr4-87174105-DEL-582 | chr4-87174106-DEL-582 |
| chr4-87175188-DEL-86 | chr4-87175189-DEL-92 |
| chr4-87229531-DEL-5541 | chr4-87229532-DEL-5541 |
| chr4-87262093-DEL-65 | chr4-87262094-DEL-65 |
| chr4-87284115-DEL-181 | chr4-87284121-DEL-176 |
| chr4-87324876-DEL-207 | chr4-87324877-DEL-207 |
| chr4-87331077-DEL-824 | chr4-87331078-DEL-824 |
| chr4-87349720-DEL-131 | chr4-87349699-DEL-109 |
| chr4-87371092-DEL-400 | chr4-87371093-DEL-400 |
| chr4-87392613-DEL-64 | chr4-87392607-DEL-59 |
| chr4-87421213-DEL-6458 | chr4-87421214-DEL-6458 |
| chr4-87430035-DEL-64 | chr4-87430018-DEL-74 |
| chr4-87442069-DEL-1863 | chr4-87442118-DEL-1818 |
| chr4-87473179-DEL-66 | chr4-87473182-DEL-64 |
| chr4-87475895-DEL-390 | chr4-87475896-DEL-390 |
| chr4-87490115-DEL-920 | chr4-87490116-DEL-920 |
| chr4-87499144-DEL-199 | chr4-87499145-DEL-199 |
| chr4-87547571-DEL-92 | chr4-87547568-DEL-93 |
| chr4-87548477-DEL-1364 | chr4-87548478-DEL-1364 |
| chr4-87578301-DEL-108 | chr4-87578302-DEL-108 |
| chr4-87592701-DEL-6475 | chr4-87592702-DEL-6475 |
| chr4-87599590-DEL-894 | chr4-87599591-DEL-894 |
| chr4-87606840-DEL-2423 | chr4-87606841-DEL-2423 |
| chr4-87668840-DEL-2418 | chr4-87668841-DEL-2418 |
| chr4-87697908-DEL-56 | chr4-87697913-DEL-52 |

| | |
|---|---|
| chr4-87745007-DEL-1834 | chr4-87745008-DEL-1834 |
| chr4-87779401-DEL-64 | chr4-87779390-DEL-76 |
| chr4-87808745-DEL-101 | chr4-87808795-DEL-52 |
| chr4-87837229-DEL-1037 | chr4-87837230-DEL-1037 |
| chr4-87840407-DEL-150 | chr4-87840408-DEL-125 |
| chr4-87869072-DEL-1552 | chr4-87869073-DEL-1552 |
| chr4-87882254-DEL-1017 | chr4-87882255-DEL-1018 |
| chr4-87892583-DEL-160 | chr4-87892552-DEL-159 |
| chr4-87899333-DEL-494 | chr4-87899334-DEL-494 |
| chr4-87900351-DEL-52 | chr4-87900352-DEL-52 |
| chr4-87942936-DEL-631 | chr4-87942937-DEL-631 |
| chr4-87971861-DEL-6138 | chr4-87971835-DEL-6136 |
| chr4-88013709-DEL-81 | chr4-88013710-DEL-79 |
| chr4-88042108-DEL-62 | chr4-88042109-DEL-62 |
| chr4-88062617-DEL-213 | chr4-88062618-DEL-213 |
| chr4-88109617-DEL-340 | chr4-88109618-DEL-340 |
| chr4-88110925-DEL-59 | chr4-88110916-DEL-60 |
| chr4-88151530-DEL-1938 | chr4-88151531-DEL-1938 |
| chr4-88172612-DEL-173 | chr4-88172613-DEL-173 |
| chr4-88172827-DEL-2848 | chr4-88172827-DEL-2848 |
| chr4-88176380-DEL-1641 | chr4-88176381-DEL-1639 |
| chr4-88199380-DEL-6935 | chr4-88199381-DEL-6935 |
| chr4-88205535-DEL-636 | chr4-88205536-DEL-636 |
| chr4-88212727-DEL-383 | chr4-88212728-DEL-383 |
| chr4-88218162-DEL-61 | chr4-88218163-DEL-61 |
| chr4-88271248-DEL-115 | chr4-88271255-DEL-87 |
| chr4-88276988-DEL-70 | chr4-88277028-DEL-70 |
| chr4-88294421-DEL-6989 | chr4-88294422-DEL-6989 |
| chr4-88323354-DEL-121 | chr4-88323355-DEL-121 |
| chr4-88334420-DEL-3382 | chr4-88334421-DEL-3381 |
| chr4-88340144-DEL-102 | chr4-88340145-DEL-102 |
| chr4-88409448-DEL-133 | chr4-88409449-DEL-133 |
| chr4-88415386-DEL-3353 | chr4-88415387-DEL-3353 |
| chr4-88503063-DEL-3230 | chr4-88503064-DEL-3230 |
| chr4-88517202-DEL-961 | chr4-88517192-DEL-922 |
| chr4-88586524-DEL-6462 | chr4-88586525-DEL-6462 |
| chr4-88633904-DEL-278 | chr4-88633905-DEL-278 |
| chr4-88648257-DEL-2923 | chr4-88648258-DEL-2923 |
| chr4-88657506-DEL-396 | chr4-88657507-DEL-396 |
| chr4-88664535-DEL-52 | chr4-88664536-DEL-52 |
| chr4-88665399-DEL-64 | chr4-88665398-DEL-65 |
| chr4-88729471-DEL-60 | chr4-88729472-DEL-60 |
| chr4-88771696-DEL-54 | chr4-88771698-DEL-56 |
| chr4-88779173-DEL-112 | chr4-88779173-DEL-112 |
| chr4-88785406-DEL-330 | chr4-88785407-DEL-330 |
| chr4-88883340-DEL-128 | chr4-88883366-DEL-119 |
| chr4-88885430-DEL-199 | chr4-88885431-DEL-199 |
| chr4-88890646-DEL-526 | chr4-88890640-DEL-527 |
| chr4-88901179-DEL-56 | chr4-88901185-DEL-54 |
| chr4-88916893-DEL-80 | chr4-88916893-DEL-80 |
| chr4-88922656-DEL-1582 | chr4-88922657-DEL-1582 |
| chr4-88932799-DEL-597 | chr4-88932800-DEL-597 |
| chr4-88958673-DEL-53 | chr4-88958674-DEL-53 |
| chr4-88964884-DEL-343 | chr4-88964885-DEL-343 |
| chr4-88999730-DEL-204 | chr4-88999731-DEL-204 |
| chr4-89005684-DEL-162 | chr4-89005686-DEL-161 |
| chr4-89009438-DEL-105 | chr4-89009439-DEL-105 |
| chr4-89028724-DEL-1333 | chr4-89028723-DEL-1334 |
| chr4-89062120-DEL-203 | chr4-89062115-DEL-204 |
| chr4-89073490-DEL-71 | chr4-89073491-DEL-71 |
| chr4-89076778-DEL-697 | chr4-89076779-DEL-697 |
| chr4-89077896-DEL-189 | chr4-89077897-DEL-189 |
| chr4-89083235-DEL-391 | chr4-89083236-DEL-390 |
| chr4-89097207-DEL-55 | chr4-89097211-DEL-51 |
| chr4-89113260-DEL-834 | chr4-89113261-DEL-834 |
| chr4-89119954-DEL-3146 | chr4-89119955-DEL-3146 |
| chr4-89136807-DEL-1151 | chr4-89136800-DEL-1149 |
| chr4-89144197-DEL-6447 | chr4-89144198-DEL-6447 |
| chr4-89182651-DEL-437 | chr4-89182658-DEL-443 |
| chr4-89222453-DEL-55 | chr4-89222454-DEL-55 |
| chr4-89230447-DEL-106 | chr4-89230448-DEL-106 |
| chr4-89243962-DEL-272 | chr4-89243945-DEL-293 |
| chr4-89258612-DEL-1729 | chr4-89258613-DEL-1729 |
| chr4-89311813-DEL-59 | chr4-89311814-DEL-59 |
| chr4-89324655-DEL-150 | chr4-89324655-DEL-154 |
| chr4-89337755-DEL-235 | chr4-89337756-DEL-235 |
| chr4-89344800-DEL-68 | chr4-89344801-DEL-68 |
| chr4-89362705-DEL-206 | chr4-89362706-DEL-206 |
| chr4-89413721-DEL-221 | chr4-89413722-DEL-221 |
| chr4-89443705-DEL-71 | chr4-89443712-DEL-65 |

| | |
|---|---|
| chr4-89456991-DEL-63 | chr4-89457005-DEL-62 |
| chr4-89460903-DEL-175 | chr4-89460901-DEL-176 |
| chr4-89472108-DEL-81 | chr4-89472109-DEL-77 |
| chr4-89482648-DEL-91 | chr4-89482649-DEL-91 |
| chr4-89500125-DEL-7042 | chr4-89500126-DEL-7042 |
| chr4-89513020-DEL-191 | chr4-89513021-DEL-191 |
| chr4-89524195-DEL-58 | chr4-89524196-DEL-58 |
| chr4-89548578-DEL-53 | chr4-89548579-DEL-53 |
| chr4-89590843-DEL-6264 | chr4-89590844-DEL-6264 |
| chr4-89602619-DEL-198 | chr4-89602620-DEL-198 |
| chr4-89606946-DEL-11513 | chr4-89606947-DEL-11512 |
| chr4-89607173-DEL-1628 | chr4-89607174-DEL-1628 |
| chr4-89627570-DEL-83 | chr4-89627571-DEL-83 |
| chr4-89646153-DEL-788 | chr4-89646154-DEL-788 |
| chr4-89648904-DEL-68 | chr4-89648905-DEL-68 |
| chr4-89654036-DEL-3319 | chr4-89654037-DEL-3319 |
| chr4-89680873-DEL-6685 | chr4-89680873-DEL-6685 |
| chr4-89691275-DEL-416 | chr4-89691276-DEL-416 |
| chr4-89692109-DEL-432 | chr4-89692109-DEL-432 |
| chr4-89697817-DEL-285 | chr4-89697820-DEL-288 |
| chr4-89715142-DEL-3804 | chr4-89715143-DEL-3804 |
| chr4-89733435-DEL-2697 | chr4-89733435-DEL-2697 |
| chr4-89740758-DEL-817 | chr4-89740759-DEL-817 |
| chr4-89755221-DEL-292 | chr4-89755222-DEL-292 |
| chr4-89764057-DEL-484 | chr4-89764058-DEL-484 |
| chr4-89770833-DEL-221 | chr4-89770834-DEL-221 |
| chr4-89772832-DEL-76 | chr4-89772833-DEL-76 |
| chr4-89785123-DEL-326 | chr4-89785124-DEL-326 |
| chr4-89787149-DEL-98 | chr4-89787150-DEL-98 |
| chr4-89790015-DEL-58 | chr4-89790015-DEL-76 |
| chr4-89920246-DEL-192 | chr4-89920199-DEL-192 |
| chr4-89974132-DEL-419 | chr4-89974133-DEL-419 |
| chr4-89999080-DEL-6557 | chr4-89999081-DEL-6557 |
| chr4-90005146-DEL-64 | chr4-90005147-DEL-64 |
| chr4-90006326-DEL-249 | chr4-90006327-DEL-249 |
| chr4-90102329-DEL-1968 | chr4-90102330-DEL-1968 |
| chr4-90105165-DEL-104 | chr4-90105190-DEL-105 |
| chr4-90122645-DEL-61 | chr4-90122644-DEL-62 |
| chr4-90134380-DEL-72 | chr4-90134381-DEL-78 |
| chr4-90148721-DEL-331 | chr4-90148722-DEL-331 |
| chr4-90197722-DEL-76 | chr4-90197723-DEL-76 |
| chr4-90204965-DEL-5235 | chr4-90204966-DEL-5235 |
| chr4-90222599-DEL-69 | chr4-90222613-DEL-66 |
| chr4-90270662-DEL-2264 | chr4-90270663-DEL-2264 |
| chr4-90273973-DEL-175 | chr4-90273974-DEL-174 |
| chr4-90280771-DEL-4487 | chr4-90280772-DEL-4487 |
| chr4-90293867-DEL-96 | chr4-90293868-DEL-96 |
| chr4-90302568-DEL-62 | chr4-90302569-DEL-62 |
| chr4-90304791-DEL-2710 | chr4-90304792-DEL-2710 |
| chr4-90319589-DEL-577 | chr4-90319590-DEL-577 |
| chr4-90321438-DEL-85 | chr4-90321438-DEL-85 |
| chr4-90323294-DEL-123 | chr4-90323295-DEL-123 |
| chr4-90431487-DEL-113 | chr4-90431488-DEL-113 |
| chr4-90440510-DEL-61 | chr4-90440511-DEL-61 |
| chr4-90441989-DEL-6496 | chr4-90441990-DEL-6496 |
| chr4-90488278-DEL-68 | chr4-90488265-DEL-69 |
| chr4-90557265-DEL-83 | chr4-90557276-DEL-73 |
| chr4-90638360-DEL-80 | chr4-90638352-DEL-81 |
| chr4-90669295-DEL-75 | chr4-90669296-DEL-75 |
| chr4-90671559-DEL-90 | chr4-90671560-DEL-90 |
| chr4-90698988-DEL-597 | chr4-90698989-DEL-597 |
| chr4-90701448-DEL-501 | chr4-90701449-DEL-501 |
| chr4-90713810-DEL-285 | chr4-90713812-DEL-284 |
| chr4-90715689-DEL-214 | chr4-90715699-DEL-196 |
| chr4-90759521-DEL-218 | chr4-90759522-DEL-218 |
| chr4-90763122-DEL-8369 | chr4-90763123-DEL-8369 |
| chr4-90781617-DEL-531 | chr4-90781618-DEL-531 |
| chr4-90782257-DEL-322 | chr4-90782258-DEL-321 |
| chr4-90798219-DEL-1746 | chr4-90798220-DEL-1746 |
| chr4-90803741-DEL-1287 | chr4-90803742-DEL-1287 |
| chr4-90824788-DEL-83 | chr4-90824789-DEL-83 |
| chr4-90840482-DEL-393 | chr4-90840483-DEL-393 |
| chr4-90846674-DEL-246 | chr4-90846681-DEL-240 |
| chr4-90850075-DEL-1214 | chr4-90850076-DEL-1214 |
| chr4-90929314-DEL-52 | chr4-90929310-DEL-56 |
| chr4-90949651-DEL-1598 | chr4-90949647-DEL-1597 |
| chr4-90965744-DEL-271 | chr4-90965745-DEL-271 |
| chr4-90967735-DEL-84 | chr4-90967736-DEL-84 |
| chr4-91042268-DEL-3052 | chr4-91042269-DEL-3052 |
| chr4-91071873-DEL-6414 | chr4-91071874-DEL-6414 |

| | |
|---|---|
| chr4-91084368-DEL-92 | chr4-91084367-DEL-94 |
| chr4-91088373-DEL-1398 | chr4-91088374-DEL-1398 |
| chr4-91133289-DEL-228 | chr4-91133290-DEL-228 |
| chr4-91138318-DEL-451 | chr4-91138319-DEL-451 |
| chr4-91188434-DEL-59 | chr4-91188435-DEL-65 |
| chr4-91191071-DEL-72 | chr4-91191072-DEL-68 |
| chr4-91205299-DEL-419 | chr4-91205300-DEL-419 |
| chr4-91285295-DEL-84 | chr4-91285296-DEL-84 |
| chr4-91303992-DEL-99 | chr4-91303995-DEL-97 |
| chr4-91322436-DEL-388 | chr4-91322437-DEL-388 |
| chr4-91325332-DEL-1102 | chr4-91325349-DEL-1086 |
| chr4-91352133-DEL-59 | chr4-91352134-DEL-59 |
| chr4-91367339-DEL-1025 | chr4-91367340-DEL-1025 |
| chr4-91375685-DEL-6239 | chr4-91375686-DEL-6239 |
| chr4-91395346-DEL-342 | chr4-91395347-DEL-342 |
| chr4-91396538-DEL-62 | chr4-91396540-DEL-58 |
| chr4-91400688-DEL-303 | chr4-91400689-DEL-303 |
| chr4-91402698-DEL-170 | chr4-91402698-DEL-170 |
| chr4-91426419-DEL-393 | chr4-91426429-DEL-393 |
| chr4-91438528-DEL-222 | chr4-91438529-DEL-222 |
| chr4-91458747-DEL-6250 | chr4-91458748-DEL-6250 |
| chr4-91468935-DEL-52 | chr4-91468936-DEL-52 |
| chr4-91514686-DEL-443 | chr4-91514687-DEL-443 |
| chr4-91520579-DEL-8354 | chr4-91520592-DEL-8353 |
| chr4-91534408-DEL-205 | chr4-91534409-DEL-205 |
| chr4-91542842-DEL-7123 | chr4-91542841-DEL-7123 |
| chr4-91612210-DEL-247 | chr4-91612211-DEL-247 |
| chr4-91622363-DEL-2235 | chr4-91622364-DEL-2235 |
| chr4-91651453-DEL-54 | chr4-91651446-DEL-61 |
| chr4-91670920-DEL-6359 | chr4-91670924-DEL-6360 |
| chr4-91686939-DEL-402 | chr4-91686939-DEL-402 |
| chr4-91687963-DEL-1287 | chr4-91687963-DEL-1287 |
| chr4-91739690-DEL-1288 | chr4-91739691-DEL-1288 |
| chr4-91747670-DEL-285 | chr4-91747671-DEL-285 |
| chr4-91773000-DEL-1512 | chr4-91773001-DEL-1512 |
| chr4-91782658-DEL-6441 | chr4-91782659-DEL-6441 |
| chr4-91802861-DEL-1743 | chr4-91802862-DEL-1743 |
| chr4-91837371-DEL-293 | chr4-91837372-DEL-293 |
| chr4-91844790-DEL-448 | chr4-91844791-DEL-448 |
| chr4-91897281-DEL-1323 | chr4-91897282-DEL-1323 |
| chr4-91900573-DEL-803 | chr4-91900574-DEL-803 |
| chr4-91925802-DEL-2508 | chr4-91925803-DEL-2508 |
| chr4-91931442-DEL-1225 | chr4-91931442-DEL-1225 |
| chr4-91939638-DEL-211 | chr4-91939639-DEL-211 |
| chr4-92025596-DEL-56 | chr4-92025611-DEL-74 |
| chr4-92070453-DEL-79 | chr4-92070454-DEL-79 |
| chr4-92078948-DEL-1142 | chr4-92078941-DEL-1141 |
| chr4-92097490-DEL-456 | chr4-92097491-DEL-456 |
| chr4-92108924-DEL-367 | chr4-92108930-DEL-364 |
| chr4-92119039-DEL-6593 | chr4-92119040-DEL-6593 |
| chr4-92168894-DEL-2595 | chr4-92168895-DEL-2595 |
| chr4-92182846-DEL-66 | chr4-92182847-DEL-66 |
| chr4-92186916-DEL-732 | chr4-92186917-DEL-732 |
| chr4-92213110-DEL-1097 | chr4-92213116-DEL-1086 |
| chr4-92214557-DEL-1255 | chr4-92214558-DEL-1255 |
| chr4-92395626-DEL-6309 | chr4-92395627-DEL-6309 |
| chr4-92406549-DEL-128 | chr4-92406550-DEL-128 |
| chr4-92411770-DEL-390 | chr4-92411771-DEL-390 |
| chr4-92476511-DEL-56 | chr4-92476474-DEL-94 |
| chr4-92483536-DEL-201 | chr4-92483536-DEL-202 |
| chr4-92489080-DEL-220 | chr4-92489081-DEL-219 |
| chr4-92496165-DEL-1318 | chr4-92496166-DEL-1318 |
| chr4-92505199-DEL-580 | chr4-92505200-DEL-580 |
| chr4-92541323-DEL-61 | chr4-92541324-DEL-62 |
| chr4-92541895-DEL-72 | chr4-92541896-DEL-72 |
| chr4-92545445-DEL-59 | chr4-92545448-DEL-57 |
| chr4-92551625-DEL-444 | chr4-92551626-DEL-444 |
| chr4-92571974-DEL-205 | chr4-92571975-DEL-205 |
| chr4-92592722-DEL-6268 | chr4-92592716-DEL-6269 |
| chr4-92619335-DEL-59 | chr4-92619331-DEL-60 |
| chr4-92638802-DEL-1894 | chr4-92638803-DEL-1894 |
| chr4-92646259-DEL-6147 | chr4-92646260-DEL-6147 |
| chr4-92687600-DEL-412 | chr4-92687601-DEL-412 |
| chr4-92695987-DEL-338 | chr4-92695988-DEL-338 |
| chr4-92699109-DEL-6498 | chr4-92699110-DEL-6498 |
| chr4-92758483-DEL-65 | chr4-92758484-DEL-65 |
| chr4-92760334-DEL-4856 | chr4-92760335-DEL-4856 |
| chr4-92775441-DEL-7389 | chr4-92775441-DEL-7389 |
| chr4-92789406-DEL-51 | chr4-92789403-DEL-57 |
| chr4-92813136-DEL-6186 | chr4-92813136-DEL-6186 |

| | |
|---|---|
| chr4-92852777-DEL-54 | chr4-92852778-DEL-54 |
| chr4-92854520-DEL-2459 | chr4-92854521-DEL-2459 |
| chr4-92885911-DEL-7110 | chr4-92885912-DEL-7110 |
| chr4-92909339-DEL-200 | chr4-92909340-DEL-200 |
| chr4-92945416-DEL-51 | chr4-92945416-DEL-51 |
| chr4-92947152-DEL-57 | chr4-92947153-DEL-57 |
| chr4-92947724-DEL-60 | chr4-92947725-DEL-60 |
| chr4-92969990-DEL-59 | chr4-92969955-DEL-94 |
| chr4-92991322-DEL-507 | chr4-92991328-DEL-504 |
| chr4-92992276-DEL-1991 | chr4-92992275-DEL-1992 |
| chr4-93005492-DEL-4480 | chr4-93005493-DEL-4480 |
| chr4-93021967-DEL-188 | chr4-93021984-DEL-172 |
| chr4-93043433-DEL-2998 | chr4-93043434-DEL-2998 |
| chr4-93055443-DEL-1428 | chr4-93055445-DEL-1429 |
| chr4-93063834-DEL-443 | chr4-93063838-DEL-441 |
| chr4-93068309-DEL-90 | chr4-93068273-DEL-76 |
| chr4-93074685-DEL-612 | chr4-93074686-DEL-612 |
| chr4-93076123-DEL-124 | chr4-93076134-DEL-113 |
| chr4-93194324-DEL-75 | chr4-93194325-DEL-75 |
| chr4-93217192-DEL-381 | chr4-93217193-DEL-381 |
| chr4-93219219-DEL-51 | chr4-93219220-DEL-51 |
| chr4-93244986-DEL-133 | chr4-93244997-DEL-123 |
| chr4-93252188-DEL-56 | chr4-93252189-DEL-56 |
| chr4-93289981-DEL-87 | chr4-93289982-DEL-87 |
| chr4-93343933-DEL-363 | chr4-93343934-DEL-362 |
| chr4-93371031-DEL-51 | chr4-93371023-DEL-60 |
| chr4-93389673-DEL-66 | chr4-93389656-DEL-51 |
| chr4-93390516-DEL-122 | chr4-93390517-DEL-122 |
| chr4-93412108-DEL-146 | chr4-93412109-DEL-146 |
| chr4-93418912-DEL-77 | chr4-93418925-DEL-65 |
| chr4-93419881-DEL-1879 | chr4-93419882-DEL-1879 |
| chr4-93424774-DEL-887 | chr4-93424774-DEL-888 |
| chr4-93430959-DEL-423 | chr4-93430960-DEL-423 |
| chr4-93457933-DEL-52 | chr4-93457934-DEL-60 |
| chr4-93460314-DEL-669 | chr4-93460315-DEL-669 |
| chr4-93495834-DEL-254 | chr4-93495835-DEL-254 |
| chr4-93510729-DEL-2500 | chr4-93510730-DEL-2500 |
| chr4-93514693-DEL-5585 | chr4-93514694-DEL-5585 |
| chr4-93526886-DEL-89 | chr4-93526887-DEL-89 |
| chr4-93560057-DEL-325 | chr4-93560057-DEL-324 |
| chr4-93570679-DEL-340 | chr4-93570680-DEL-340 |
| chr4-93591951-DEL-7332 | chr4-93591952-DEL-7332 |
| chr4-93633045-DEL-331 | chr4-93633046-DEL-331 |
| chr4-93677755-DEL-599 | chr4-93677756-DEL-599 |
| chr4-93699096-DEL-232 | chr4-93699097-DEL-232 |
| chr4-93723139-DEL-56 | chr4-93723140-DEL-56 |
| chr4-93731185-DEL-68 | chr4-93731166-DEL-86 |
| chr4-93732120-DEL-438 | chr4-93732121-DEL-438 |
| chr4-93755542-DEL-51 | chr4-93755543-DEL-51 |
| chr4-93777408-DEL-6260 | chr4-93777409-DEL-6260 |
| chr4-93791050-DEL-3115 | chr4-93791051-DEL-3115 |
| chr4-93798231-DEL-80 | chr4-93798233-DEL-85 |
| chr4-93807905-DEL-51 | chr4-93807906-DEL-52 |
| chr4-93839894-DEL-5183 | chr4-93839895-DEL-5183 |
| chr4-93854480-DEL-121 | chr4-93854469-DEL-120 |
| chr4-93887590-DEL-80 | chr4-93887605-DEL-66 |
| chr4-93892961-DEL-63 | chr4-93892955-DEL-64 |
| chr4-93987157-DEL-272 | chr4-93987158-DEL-272 |
| chr4-94012525-DEL-323 | chr4-94012526-DEL-323 |
| chr4-94045538-DEL-206 | chr4-94045539-DEL-206 |
| chr4-94074548-DEL-380 | chr4-94074549-DEL-380 |
| chr4-94084726-DEL-6627 | chr4-94084727-DEL-6627 |
| chr4-94102587-DEL-54 | chr4-94102588-DEL-54 |
| chr4-94125865-DEL-2478 | chr4-94125862-DEL-2479 |
| chr4-94176935-DEL-329 | chr4-94176936-DEL-329 |
| chr4-94270895-DEL-55 | chr4-94270896-DEL-55 |
| chr4-94322046-DEL-54 | chr4-94322046-DEL-54 |
| chr4-94345148-DEL-394 | chr4-94345149-DEL-394 |
| chr4-94354805-DEL-161 | chr4-94354806-DEL-161 |
| chr4-94367266-DEL-71 | chr4-94367241-DEL-72 |
| chr4-94368247-DEL-54 | chr4-94368246-DEL-56 |
| chr4-94369862-DEL-161 | chr4-94369853-DEL-162 |
| chr4-94376242-DEL-66 | chr4-94376228-DEL-69 |
| chr4-94395521-DEL-79 | chr4-94395522-DEL-79 |
| chr4-94403783-DEL-56 | chr4-94403784-DEL-56 |
| chr4-94427774-DEL-162 | chr4-94427775-DEL-162 |
| chr4-94451040-DEL-3057 | chr4-94451041-DEL-3057 |
| chr4-94484443-DEL-68 | chr4-94484444-DEL-68 |
| chr4-94510478-DEL-185 | chr4-94510479-DEL-185 |
| chr4-94541847-DEL-216 | chr4-94541848-DEL-216 |

| | |
|---|---|
| chr4-94554955-DEL-131 | chr4-94554958-DEL-127 |
| chr4-94562679-DEL-816 | chr4-94562665-DEL-817 |
| chr4-94578246-DEL-207 | chr4-94578247-DEL-207 |
| chr4-94587536-DEL-6452 | chr4-94587537-DEL-6452 |
| chr4-94616544-DEL-68 | chr4-94616529-DEL-62 |
| chr4-94640259-DEL-55 | chr4-94640260-DEL-56 |
| chr4-94658208-DEL-1743 | chr4-94658209-DEL-1743 |
| chr4-94661212-DEL-1119 | chr4-94661213-DEL-1119 |
| chr4-94665502-DEL-125 | chr4-94665503-DEL-125 |
| chr4-94679811-DEL-6607 | chr4-94679812-DEL-6607 |
| chr4-94701473-DEL-6276 | chr4-94701474-DEL-6276 |
| chr4-94727552-DEL-6290 | chr4-94727553-DEL-6290 |
| chr4-94738525-DEL-55 | chr4-94738488-DEL-50 |
| chr4-94779390-DEL-7013 | chr4-94779391-DEL-7013 |
| chr4-94800838-DEL-6987 | chr4-94800839-DEL-6987 |
| chr4-94801170-DEL-423 | chr4-94801170-DEL-423 |
| chr4-94812405-DEL-96 | chr4-94812406-DEL-96 |
| chr4-94818413-DEL-74 | chr4-94818414-DEL-74 |
| chr4-94823882-DEL-922 | chr4-94823883-DEL-922 |
| chr4-94838637-DEL-358 | chr4-94838638-DEL-358 |
| chr4-94869290-DEL-54 | chr4-94869291-DEL-54 |
| chr4-94905357-DEL-127 | chr4-94905363-DEL-128 |
| chr4-94973695-DEL-1464 | chr4-94973696-DEL-1464 |
| chr4-94985652-DEL-59 | chr4-94985653-DEL-59 |
| chr4-95019193-DEL-73 | chr4-95019193-DEL-73 |
| chr4-95028365-DEL-125 | chr4-95028366-DEL-125 |
| chr4-95047670-DEL-199 | chr4-95047671-DEL-199 |
| chr4-95082885-DEL-73 | chr4-95082868-DEL-91 |
| chr4-95119190-DEL-1134 | chr4-95119191-DEL-1134 |
| chr4-95134325-DEL-467 | chr4-95134326-DEL-467 |
| chr4-95142487-DEL-8415 | chr4-95142488-DEL-8415 |
| chr4-95155516-DEL-392 | chr4-95155517-DEL-392 |
| chr4-95169491-DEL-98 | chr4-95169492-DEL-98 |
| chr4-95186136-DEL-177 | chr4-95186137-DEL-177 |
| chr4-95251631-DEL-51 | chr4-95251620-DEL-52 |
| chr4-95270575-DEL-58 | chr4-95270576-DEL-58 |
| chr4-95292832-DEL-328 | chr4-95292833-DEL-328 |
| chr4-95296285-DEL-102 | chr4-95296286-DEL-102 |
| chr4-95302573-DEL-576 | chr4-95302574-DEL-576 |
| chr4-95331355-DEL-92 | chr4-95331340-DEL-108 |
| chr4-95371783-DEL-52 | chr4-95371752-DEL-61 |
| chr4-95385538-DEL-1581 | chr4-95385539-DEL-1581 |
| chr4-95397968-DEL-7236 | chr4-95397969-DEL-7236 |
| chr4-95432350-DEL-1474 | chr4-95432351-DEL-1474 |
| chr4-95449696-DEL-317 | chr4-95449697-DEL-317 |
| chr4-95543455-DEL-1157 | chr4-95543456-DEL-1157 |
| chr4-95554173-DEL-80 | chr4-95554172-DEL-78 |
| chr4-95555751-DEL-120 | chr4-95555758-DEL-118 |
| chr4-95580086-DEL-77 | chr4-95580087-DEL-77 |
| chr4-95599042-DEL-160 | chr4-95599043-DEL-160 |
| chr4-95636885-DEL-290 | chr4-95636886-DEL-290 |
| chr4-95641555-DEL-80 | chr4-95641558-DEL-52 |
| chr4-95661378-DEL-141 | chr4-95661379-DEL-141 |
| chr4-95662356-DEL-78 | chr4-95662354-DEL-79 |
| chr4-95689566-DEL-239 | chr4-95689567-DEL-239 |
| chr4-95717312-DEL-55 | chr4-95717360-DEL-53 |
| chr4-95722012-DEL-2292 | chr4-95722013-DEL-2292 |
| chr4-95722141-DEL-1001 | chr4-95722142-DEL-1001 |
| chr4-95738927-DEL-62 | chr4-95738928-DEL-54 |
| chr4-95766663-DEL-244 | chr4-95766664-DEL-244 |
| chr4-95830270-DEL-828 | chr4-95830271-DEL-828 |
| chr4-95845258-DEL-84 | chr4-95845259-DEL-84 |
| chr4-95917910-DEL-172 | chr4-95917911-DEL-172 |
| chr4-95936968-DEL-7066 | chr4-95936969-DEL-7066 |
| chr4-95957981-DEL-87 | chr4-95957982-DEL-86 |
| chr4-95960938-DEL-607 | chr4-95960939-DEL-607 |
| chr4-96009186-DEL-149 | chr4-96009187-DEL-149 |
| chr4-96040995-DEL-4194 | chr4-96040996-DEL-4194 |
| chr4-96052253-DEL-5944 | chr4-96052254-DEL-5944 |
| chr4-96055913-DEL-601 | chr4-96055914-DEL-601 |
| chr4-96059000-DEL-81 | chr4-96059001-DEL-81 |
| chr4-96067069-DEL-1685 | chr4-96067070-DEL-1685 |
| chr4-96074641-DEL-1070 | chr4-96074642-DEL-1070 |
| chr4-96087065-DEL-440 | chr4-96087066-DEL-440 |
| chr4-96101351-DEL-231 | chr4-96101352-DEL-231 |
| chr4-96114908-DEL-659 | chr4-96114909-DEL-659 |
| chr4-96119281-DEL-511 | chr4-96119282-DEL-511 |
| chr4-96150115-DEL-400 | chr4-96150116-DEL-400 |
| chr4-96151464-DEL-6237 | chr4-96151465-DEL-6237 |
| chr4-96239757-DEL-972 | chr4-96239758-DEL-971 |

| | |
|---|---|
| chr4-96246772-DEL-72 | chr4-96246773-DEL-72 |
| chr4-96272864-DEL-1551 | chr4-96272865-DEL-1551 |
| chr4-96277505-DEL-3216 | chr4-96277506-DEL-3216 |
| chr4-96283053-DEL-2787 | chr4-96283054-DEL-2787 |
| chr4-96318786-DEL-73 | chr4-96318787-DEL-73 |
| chr4-96320279-DEL-6324 | chr4-96320280-DEL-6324 |
| chr4-96375780-DEL-1258 | chr4-96375781-DEL-1258 |
| chr4-96379676-DEL-161 | chr4-96379677-DEL-161 |
| chr4-96385920-DEL-1059 | chr4-96385921-DEL-1059 |
| chr4-96400426-DEL-2423 | chr4-96400426-DEL-2423 |
| chr4-96411396-DEL-246 | chr4-96411397-DEL-246 |
| chr4-96413481-DEL-90 | chr4-96413482-DEL-60 |
| chr4-96463992-DEL-205 | chr4-96463993-DEL-205 |
| chr4-96474259-DEL-58 | chr4-96474255-DEL-56 |
| chr4-96596597-DEL-829 | chr4-96596598-DEL-829 |
| chr4-96621468-DEL-69 | chr4-96621445-DEL-105 |
| chr4-96636785-DEL-541 | chr4-96636786-DEL-541 |
| chr4-96642100-DEL-52 | chr4-96642101-DEL-52 |
| chr4-96679891-DEL-158 | chr4-96679891-DEL-158 |
| chr4-96694607-DEL-51 | chr4-96694561-DEL-50 |
| chr4-96720117-DEL-51 | chr4-96720118-DEL-51 |
| chr4-96736410-DEL-4489 | chr4-96736413-DEL-4487 |
| chr4-96776014-DEL-591 | chr4-96776015-DEL-591 |
| chr4-96781249-DEL-105 | chr4-96781260-DEL-55 |
| chr4-96796189-DEL-1244 | chr4-96796190-DEL-1244 |
| chr4-96814215-DEL-159 | chr4-96814210-DEL-159 |
| chr4-96814944-DEL-69 | chr4-96814954-DEL-61 |
| chr4-96827576-DEL-3883 | chr4-96827577-DEL-3883 |
| chr4-96834846-DEL-7272 | chr4-96834846-DEL-7273 |
| chr4-96845378-DEL-731 | chr4-96845379-DEL-731 |
| chr4-96882768-DEL-6773 | chr4-96882769-DEL-6773 |
| chr4-96907898-DEL-117 | chr4-96907899-DEL-117 |
| chr4-96967059-DEL-174 | chr4-96967060-DEL-174 |
| chr4-96968359-DEL-6288 | chr4-96968360-DEL-6288 |
| chr4-96983310-DEL-1997 | chr4-96983311-DEL-1997 |
| chr4-96985183-DEL-694 | chr4-96985184-DEL-694 |
| chr4-96999591-DEL-310 | chr4-96999592-DEL-310 |
| chr4-97042605-DEL-51 | chr4-97042569-DEL-85 |
| chr4-97086990-DEL-498 | chr4-97086991-DEL-498 |
| chr4-97117045-DEL-284 | chr4-97117046-DEL-284 |
| chr4-97128811-DEL-94 | chr4-97128801-DEL-94 |
| chr4-97165569-DEL-142 | chr4-97165570-DEL-142 |
| chr4-97176991-DEL-1627 | chr4-97176992-DEL-1627 |
| chr4-97179743-DEL-339 | chr4-97179739-DEL-340 |
| chr4-97203509-DEL-6083 | chr4-97203510-DEL-6083 |
| chr4-97218543-DEL-162 | chr4-97218544-DEL-162 |
| chr4-97222444-DEL-1288 | chr4-97222445-DEL-1288 |
| chr4-97238404-DEL-2838 | chr4-97238405-DEL-2838 |
| chr4-97247666-DEL-58 | chr4-97247671-DEL-54 |
| chr4-97292155-DEL-354 | chr4-97292156-DEL-354 |
| chr4-97297688-DEL-297 | chr4-97297689-DEL-297 |
| chr4-97298394-DEL-3100 | chr4-97298395-DEL-3100 |
| chr4-97309805-DEL-79 | chr4-97309806-DEL-79 |
| chr4-97345595-DEL-899 | chr4-97345596-DEL-899 |
| chr4-97348112-DEL-199 | chr4-97348113-DEL-199 |
| chr4-97352362-DEL-688 | chr4-97352358-DEL-688 |
| chr4-97377207-DEL-122 | chr4-97377208-DEL-122 |
| chr4-97382104-DEL-1416 | chr4-97382105-DEL-1416 |
| chr4-97388916-DEL-78 | chr4-97388917-DEL-78 |
| chr4-97413091-DEL-232 | chr4-97413090-DEL-233 |
| chr4-97416561-DEL-53 | chr4-97416564-DEL-54 |
| chr4-97422047-DEL-495 | chr4-97422048-DEL-495 |
| chr4-97507960-DEL-633 | chr4-97507961-DEL-633 |
| chr4-97563264-DEL-67 | chr4-97563260-DEL-72 |
| chr4-97583014-DEL-74 | chr4-97582989-DEL-75 |
| chr4-97680882-DEL-174 | chr4-97680883-DEL-174 |
| chr4-97909446-DEL-66 | chr4-97909447-DEL-65 |
| chr4-97925933-DEL-451 | chr4-97925933-DEL-451 |
| chr4-97947320-DEL-137 | chr4-97947321-DEL-137 |
| chr4-97984719-DEL-86 | chr4-97984720-DEL-86 |
| chr4-98030126-DEL-165 | chr4-98030127-DEL-165 |
| chr4-98036054-DEL-52 | chr4-98036051-DEL-56 |
| chr4-98041706-DEL-139 | chr4-98041708-DEL-137 |
| chr4-98043204-DEL-1643 | chr4-98043205-DEL-1643 |
| chr4-98073459-DEL-76 | chr4-98073476-DEL-60 |
| chr4-98085985-DEL-65 | chr4-98085982-DEL-66 |
| chr4-98125205-DEL-62 | chr4-98125184-DEL-83 |
| chr4-98161308-DEL-1875 | chr4-98161309-DEL-1875 |
| chr4-98173579-DEL-501 | chr4-98173580-DEL-501 |
| chr4-98213728-DEL-193 | chr4-98213729-DEL-193 |

| | |
|---|---|
| chr4-98237565-DEL-327 | chr4-98237566-DEL-327 |
| chr4-98247140-DEL-64 | chr4-98247141-DEL-64 |
| chr4-98247479-DEL-331 | chr4-98247480-DEL-331 |
| chr4-98255520-DEL-246 | chr4-98255521-DEL-247 |
| chr4-98266562-DEL-1081 | chr4-98266563-DEL-1081 |
| chr4-98288516-DEL-193 | chr4-98288504-DEL-194 |
| chr4-98300906-DEL-248 | chr4-98300907-DEL-248 |
| chr4-98319202-DEL-181 | chr4-98319203-DEL-181 |
| chr4-98324891-DEL-154 | chr4-98324869-DEL-152 |
| chr4-98342640-DEL-358 | chr4-98342641-DEL-358 |
| chr4-98351109-DEL-317 | chr4-98351110-DEL-317 |
| chr4-98355044-DEL-60 | chr4-98355060-DEL-61 |
| chr4-98368276-DEL-1338 | chr4-98368277-DEL-1338 |
| chr4-98405345-DEL-162 | chr4-98405346-DEL-162 |
| chr4-98413098-DEL-7095 | chr4-98413099-DEL-7095 |
| chr4-98446629-DEL-2706 | chr4-98446630-DEL-2706 |
| chr4-98482812-DEL-216 | chr4-98482813-DEL-216 |
| chr4-98525795-DEL-1099 | chr4-98525796-DEL-1099 |
| chr4-98529415-DEL-7957 | chr4-98529416-DEL-7957 |
| chr4-98538811-DEL-55 | chr4-98538812-DEL-55 |
| chr4-98553169-DEL-399 | chr4-98553170-DEL-399 |
| chr4-98600239-DEL-1183 | chr4-98600244-DEL-1184 |
| chr4-98603438-DEL-56 | chr4-98603438-DEL-56 |
| chr4-98620907-DEL-117 | chr4-98620908-DEL-117 |
| chr4-98632265-DEL-55 | chr4-98632266-DEL-55 |
| chr4-98636892-DEL-76 | chr4-98636893-DEL-76 |
| chr4-98638404-DEL-213 | chr4-98638405-DEL-213 |
| chr4-98658087-DEL-510 | chr4-98658088-DEL-510 |
| chr4-98672513-DEL-503 | chr4-98672516-DEL-504 |
| chr4-98680261-DEL-60 | chr4-98680252-DEL-70 |
| chr4-98712455-DEL-84 | chr4-98712445-DEL-88 |
| chr4-98740461-DEL-1549 | chr4-98740462-DEL-1549 |
| chr4-98740621-DEL-378 | chr4-98740622-DEL-378 |
| chr4-98743672-DEL-510 | chr4-98743673-DEL-510 |
| chr4-98752281-DEL-79 | chr4-98752284-DEL-51 |
| chr4-98753479-DEL-220 | chr4-98753481-DEL-206 |
| chr4-98754494-DEL-68 | chr4-98754477-DEL-78 |
| chr4-98756676-DEL-64 | chr4-98756645-DEL-59 |
| chr4-98757718-DEL-201 | chr4-98757719-DEL-201 |
| chr4-98763285-DEL-98 | chr4-98763286-DEL-98 |
| chr4-98766392-DEL-514 | chr4-98766393-DEL-514 |
| chr4-98773837-DEL-84 | chr4-98773838-DEL-108 |
| chr4-98796233-DEL-1893 | chr4-98796234-DEL-1893 |
| chr4-98809027-DEL-131 | chr4-98809028-DEL-131 |
| chr4-98831472-DEL-321 | chr4-98831473-DEL-321 |
| chr4-98936631-DEL-54 | chr4-98936632-DEL-54 |
| chr4-98973128-DEL-66 | chr4-98973129-DEL-66 |
| chr4-98984053-DEL-1626 | chr4-98984043-DEL-1636 |
| chr4-98987594-DEL-200 | chr4-98987595-DEL-198 |
| chr4-98995401-DEL-143 | chr4-98995402-DEL-143 |
| chr4-99012506-DEL-52 | chr4-99012507-DEL-52 |
| chr4-99028379-DEL-376 | chr4-99028356-DEL-383 |
| chr4-99036740-DEL-58 | chr4-99036741-DEL-58 |
| chr4-99040017-DEL-96 | chr4-99040018-DEL-78 |
| chr4-99049107-DEL-154 | chr4-99049108-DEL-154 |
| chr4-99070638-DEL-74 | chr4-99070639-DEL-74 |
| chr4-99075537-DEL-199 | chr4-99075537-DEL-199 |
| chr4-99083703-DEL-401 | chr4-99083704-DEL-401 |
| chr4-99118159-DEL-66 | chr4-99118160-DEL-66 |
| chr4-99131097-DEL-3818 | chr4-99131098-DEL-3818 |
| chr4-99147441-DEL-68 | chr4-99147442-DEL-68 |
| chr4-99175968-DEL-1256 | chr4-99175969-DEL-1256 |
| chr4-99229832-DEL-339 | chr4-99229833-DEL-339 |
| chr4-99267133-DEL-78 | chr4-99267134-DEL-78 |
| chr4-99282348-DEL-79 | chr4-99282349-DEL-79 |
| chr4-99285941-DEL-64 | chr4-99285944-DEL-62 |
| chr4-99291338-DEL-245 | chr4-99291356-DEL-253 |
| chr4-99321672-DEL-6777 | chr4-99321673-DEL-6777 |
| chr4-99350718-DEL-217 | chr4-99350719-DEL-217 |
| chr4-99396359-DEL-709 | chr4-99396359-DEL-709 |
| chr4-99404200-DEL-1732 | chr4-99404201-DEL-1732 |
| chr4-99408525-DEL-1459 | chr4-99408526-DEL-1459 |
| chr4-99425475-DEL-200 | chr4-99425476-DEL-200 |
| chr4-99441748-DEL-210 | chr4-99441749-DEL-210 |
| chr4-99446810-DEL-124 | chr4-99446811-DEL-100 |
| chr4-99470809-DEL-52 | chr4-99470810-DEL-52 |
| chr4-99474799-DEL-123 | chr4-99474790-DEL-123 |
| chr4-99481108-DEL-212 | chr4-99481109-DEL-212 |
| chr4-99482668-DEL-204 | chr4-99482668-DEL-204 |
| chr4-99491976-DEL-193 | chr4-99491977-DEL-193 |

| | |
|---|---|
| chr4-99513669-DEL-714 | chr4-99513670-DEL-714 |
| chr4-99519247-DEL-100 | chr4-99519222-DEL-132 |
| chr4-99520761-DEL-214 | chr4-99520762-DEL-214 |
| chr4-99525913-DEL-150 | chr4-99525907-DEL-151 |
| chr4-99534844-DEL-519 | chr4-99534845-DEL-519 |
| chr4-99561395-DEL-96 | chr4-99561396-DEL-96 |
| chr4-99563841-DEL-70 | chr4-99563875-DEL-69 |
| chr4-99566836-DEL-89 | chr4-99566837-DEL-89 |
| chr4-99575447-DEL-209 | chr4-99575448-DEL-209 |
| chr4-99619386-DEL-6702 | chr4-99619387-DEL-6702 |
| chr4-99652162-DEL-392 | chr4-99652163-DEL-392 |
| chr4-99682018-DEL-201 | chr4-99682019-DEL-201 |
| chr4-99702422-DEL-1364 | chr4-99702423-DEL-1364 |
| chr4-99706609-DEL-495 | chr4-99706610-DEL-495 |
| chr4-99728096-DEL-191 | chr4-99728097-DEL-191 |
| chr4-99731099-DEL-177 | chr4-99731100-DEL-177 |
| chr4-99734250-DEL-118 | chr4-99734251-DEL-118 |
| chr4-99755851-DEL-59 | chr4-99755852-DEL-59 |
| chr4-99763401-DEL-479 | chr4-99763402-DEL-479 |
| chr4-99775666-DEL-106 | chr4-99775667-DEL-106 |
| chr4-99778523-DEL-68 | chr4-99778524-DEL-68 |
| chr4-99801204-DEL-2863 | chr4-99801201-DEL-2864 |
| chr4-99851696-DEL-6210 | chr4-99851697-DEL-6210 |
| chr4-99863790-DEL-1563 | chr4-99863791-DEL-1563 |
| chr4-99879972-DEL-199 | chr4-99879973-DEL-199 |
| chr4-99881368-DEL-64 | chr4-99881369-DEL-56 |
| chr4-99883199-DEL-52 | chr4-99883198-DEL-52 |
| chr4-99886956-DEL-217 | chr4-99886956-DEL-217 |
| chr4-99892547-DEL-376 | chr4-99892548-DEL-376 |
| chr4-99900761-DEL-69 | chr4-99900717-DEL-66 |
| chr4-99903891-DEL-163 | chr4-99903889-DEL-136 |
| chr4-99906263-DEL-54 | chr4-99906263-DEL-54 |
| chr4-99966217-DEL-63 | chr4-99966218-DEL-63 |
| chr4-100126810-DEL-88 | chr4-100126811-DEL-64 |
| chr4-100213929-DEL-390 | chr4-100213930-DEL-390 |
| chr4-100225011-DEL-63 | chr4-100225041-DEL-54 |
| chr4-100226758-DEL-1146 | chr4-100226759-DEL-1146 |
| chr4-100255110-DEL-128 | chr4-100255111-DEL-128 |
| chr4-100295925-DEL-627 | chr4-100295926-DEL-627 |
| chr4-100296759-DEL-941 | chr4-100296760-DEL-941 |
| chr4-100327239-DEL-82 | chr4-100327262-DEL-87 |
| chr4-100332682-DEL-2452 | chr4-100332683-DEL-2452 |
| chr4-100342462-DEL-680 | chr4-100342463-DEL-680 |
| chr4-100343050-DEL-376 | chr4-100343061-DEL-370 |
| chr4-100352893-DEL-817 | chr4-100352894-DEL-817 |
| chr4-100358944-DEL-58 | chr4-100358945-DEL-58 |
| chr4-100376335-DEL-958 | chr4-100376336-DEL-958 |
| chr4-100395972-DEL-151 | chr4-100396003-DEL-121 |
| chr4-100413875-DEL-2232 | chr4-100413875-DEL-2232 |
| chr4-100451686-DEL-89 | chr4-100451687-DEL-90 |
| chr4-100463794-DEL-230 | chr4-100463795-DEL-230 |
| chr4-100464645-DEL-6406 | chr4-100464646-DEL-6406 |
| chr4-100478162-DEL-71 | chr4-100478163-DEL-71 |
| chr4-100507206-DEL-6791 | chr4-100507207-DEL-6791 |
| chr4-100522605-DEL-54 | chr4-100522607-DEL-52 |
| chr4-100526287-DEL-318 | chr4-100526288-DEL-318 |
| chr4-100533935-DEL-6254 | chr4-100533936-DEL-6254 |
| chr4-100554993-DEL-66 | chr4-100554994-DEL-65 |
| chr4-100558101-DEL-58 | chr4-100558116-DEL-76 |
| chr4-100565320-DEL-54 | chr4-100565320-DEL-55 |
| chr4-100597476-DEL-79 | chr4-100597477-DEL-67 |
| chr4-100621268-DEL-268 | chr4-100621269-DEL-268 |
| chr4-100625168-DEL-251 | chr4-100625205-DEL-210 |
| chr4-100630386-DEL-213 | chr4-100630387-DEL-213 |
| chr4-100646795-DEL-90 | chr4-100646796-DEL-90 |
| chr4-100659739-DEL-202 | chr4-100659740-DEL-202 |
| chr4-100674744-DEL-509 | chr4-100674745-DEL-509 |
| chr4-100698218-DEL-272 | chr4-100698219-DEL-272 |
| chr4-100741004-DEL-413 | chr4-100741005-DEL-413 |
| chr4-100756127-DEL-1235 | chr4-100756128-DEL-1235 |
| chr4-100770220-DEL-300 | chr4-100770221-DEL-300 |
| chr4-100771809-DEL-61 | chr4-100771809-DEL-61 |
| chr4-100773816-DEL-118 | chr4-100773817-DEL-118 |
| chr4-100790309-DEL-4868 | chr4-100790310-DEL-4868 |
| chr4-100797167-DEL-295 | chr4-100797168-DEL-295 |
| chr4-100813961-DEL-86 | chr4-100813960-DEL-88 |
| chr4-100828689-DEL-148 | chr4-100828690-DEL-148 |
| chr4-100850382-DEL-103 | chr4-100850383-DEL-103 |
| chr4-100853851-DEL-93 | chr4-100853852-DEL-93 |
| chr4-100866233-DEL-6943 | chr4-100866234-DEL-6943 |

| | |
|---|---|
| chr4-100872287-DEL-3473 | chr4-100872288-DEL-3473 |
| chr4-100907306-DEL-166 | chr4-100907307-DEL-166 |
| chr4-100910159-DEL-203 | chr4-100910160-DEL-203 |
| chr4-100911826-DEL-6619 | chr4-100911827-DEL-6618 |
| chr4-100934111-DEL-68 | chr4-100934112-DEL-68 |
| chr4-100948085-DEL-81 | chr4-100948086-DEL-81 |
| chr4-100967050-DEL-541 | chr4-100967051-DEL-541 |
| chr4-100976762-DEL-72 | chr4-100976763-DEL-72 |
| chr4-100981244-DEL-5173 | chr4-100981245-DEL-5173 |
| chr4-100987415-DEL-208 | chr4-100987416-DEL-208 |
| chr4-101002641-DEL-127 | chr4-101002642-DEL-127 |
| chr4-101025687-DEL-107 | chr4-101025686-DEL-107 |
| chr4-101163161-DEL-76 | chr4-101163162-DEL-76 |
| chr4-101189273-DEL-163 | chr4-101189258-DEL-204 |
| chr4-101198772-DEL-309 | chr4-101198773-DEL-309 |
| chr4-101201837-DEL-872 | chr4-101201838-DEL-872 |
| chr4-101218221-DEL-284 | chr4-101218222-DEL-284 |
| chr4-101243836-DEL-240 | chr4-101243837-DEL-240 |
| chr4-101251498-DEL-210 | chr4-101251499-DEL-210 |
| chr4-101264860-DEL-6440 | chr4-101264865-DEL-6437 |
| chr4-101280080-DEL-188 | chr4-101280081-DEL-188 |
| chr4-101332515-DEL-61 | chr4-101332526-DEL-59 |
| chr4-101413760-DEL-539 | chr4-101413761-DEL-539 |
| chr4-101439590-DEL-95 | chr4-101439570-DEL-133 |
| chr4-101457437-DEL-512 | chr4-101457418-DEL-518 |
| chr4-101465430-DEL-693 | chr4-101465431-DEL-691 |
| chr4-101466282-DEL-63 | chr4-101466270-DEL-76 |
| chr4-101534375-DEL-52 | chr4-101534372-DEL-56 |
| chr4-101604311-DEL-128 | chr4-101604312-DEL-128 |
| chr4-101610761-DEL-6363 | chr4-101610762-DEL-6363 |
| chr4-101624067-DEL-66 | chr4-101624068-DEL-66 |
| chr4-101626502-DEL-887 | chr4-101626504-DEL-888 |
| chr4-101655022-DEL-6399 | chr4-101655023-DEL-6399 |
| chr4-101677070-DEL-197 | chr4-101677060-DEL-166 |
| chr4-101678617-DEL-205 | chr4-101678618-DEL-205 |
| chr4-101684190-DEL-108 | chr4-101684191-DEL-108 |
| chr4-101691670-DEL-91 | chr4-101691671-DEL-91 |
| chr4-101720349-DEL-8986 | chr4-101720350-DEL-8986 |
| chr4-101729609-DEL-53 | chr4-101729609-DEL-53 |
| chr4-101778775-DEL-6430 | chr4-101778776-DEL-6430 |
| chr4-101822288-DEL-206 | chr4-101822289-DEL-206 |
| chr4-101830925-DEL-51 | chr4-101830923-DEL-54 |
| chr4-101832106-DEL-6442 | chr4-101832107-DEL-6442 |
| chr4-101863805-DEL-331 | chr4-101863806-DEL-331 |
| chr4-101887576-DEL-892 | chr4-101887577-DEL-892 |
| chr4-101892075-DEL-311 | chr4-101892076-DEL-309 |
| chr4-101901090-DEL-231 | chr4-101901091-DEL-231 |
| chr4-101906646-DEL-475 | chr4-101906647-DEL-475 |
| chr4-101914411-DEL-210 | chr4-101914406-DEL-216 |
| chr4-101916670-DEL-1888 | chr4-101916671-DEL-1888 |
| chr4-101948587-DEL-387 | chr4-101948588-DEL-344 |
| chr4-101962723-DEL-57 | chr4-101962700-DEL-84 |
| chr4-101977427-DEL-751 | chr4-101977428-DEL-751 |
| chr4-102027322-DEL-101 | chr4-102027323-DEL-101 |
| chr4-102041013-DEL-839 | chr4-102041014-DEL-839 |
| chr4-102062032-DEL-307 | chr4-102062033-DEL-307 |
| chr4-102089091-DEL-84 | chr4-102089058-DEL-98 |
| chr4-102119921-DEL-68 | chr4-102119922-DEL-68 |
| chr4-102129010-DEL-420 | chr4-102129011-DEL-420 |
| chr4-102134728-DEL-108 | chr4-102134729-DEL-108 |
| chr4-102157866-DEL-501 | chr4-102157867-DEL-501 |
| chr4-102160886-DEL-597 | chr4-102160887-DEL-597 |
| chr4-102174749-DEL-56 | chr4-102174750-DEL-56 |
| chr4-102201689-DEL-57 | chr4-102201690-DEL-57 |
| chr4-102204958-DEL-520 | chr4-102204959-DEL-520 |
| chr4-102275961-DEL-61 | chr4-102275953-DEL-99 |
| chr4-102287573-DEL-66 | chr4-102287559-DEL-70 |
| chr4-102305203-DEL-135 | chr4-102305201-DEL-131 |
| chr4-102322896-DEL-63 | chr4-102322904-DEL-51 |
| chr4-102406132-DEL-64 | chr4-102406123-DEL-52 |
| chr4-102448430-DEL-79 | chr4-102448433-DEL-77 |
| chr4-102479945-DEL-54 | chr4-102479946-DEL-54 |
| chr4-102482775-DEL-125 | chr4-102482756-DEL-145 |
| chr4-102549591-DEL-96 | chr4-102549592-DEL-96 |
| chr4-102552854-DEL-1415 | chr4-102552855-DEL-1415 |
| chr4-102562845-DEL-403 | chr4-102562846-DEL-403 |
| chr4-102584974-DEL-82 | chr4-102584973-DEL-110 |
| chr4-102602191-DEL-6225 | chr4-102602192-DEL-6225 |
| chr4-102654057-DEL-168 | chr4-102654058-DEL-168 |
| chr4-102672034-DEL-56 | chr4-102672035-DEL-54 |

| | |
|---|---|
| chr4-102682883-DEL-91 | chr4-102682884-DEL-91 |
| chr4-102704410-DEL-1354 | chr4-102704411-DEL-1354 |
| chr4-102716639-DEL-84 | chr4-102716640-DEL-84 |
| chr4-102727267-DEL-151 | chr4-102727268-DEL-151 |
| chr4-102728218-DEL-70 | chr4-102728217-DEL-72 |
| chr4-102732852-DEL-7179 | chr4-102732853-DEL-7179 |
| chr4-102757988-DEL-176 | chr4-102757989-DEL-176 |
| chr4-102825849-DEL-270 | chr4-102825850-DEL-270 |
| chr4-102876154-DEL-201 | chr4-102876155-DEL-201 |
| chr4-102883328-DEL-203 | chr4-102883330-DEL-202 |
| chr4-102888120-DEL-575 | chr4-102888121-DEL-575 |
| chr4-102890450-DEL-628 | chr4-102890451-DEL-628 |
| chr4-102891337-DEL-115 | chr4-102891341-DEL-114 |
| chr4-102946454-DEL-159 | chr4-102946455-DEL-159 |
| chr4-102950963-DEL-68 | chr4-102950964-DEL-68 |
| chr4-102970556-DEL-68 | chr4-102970557-DEL-68 |
| chr4-103015611-DEL-239 | chr4-103015612-DEL-239 |
| chr4-103024088-DEL-1371 | chr4-103024088-DEL-1371 |
| chr4-103052902-DEL-64 | chr4-103052903-DEL-64 |
| chr4-103060124-DEL-88 | chr4-103060152-DEL-59 |
| chr4-103083715-DEL-431 | chr4-103083716-DEL-431 |
| chr4-103112995-DEL-242 | chr4-103112996-DEL-242 |
| chr4-103130497-DEL-6438 | chr4-103130498-DEL-6438 |
| chr4-103141833-DEL-454 | chr4-103141834-DEL-453 |
| chr4-103162123-DEL-958 | chr4-103162130-DEL-942 |
| chr4-103175816-DEL-96 | chr4-103175822-DEL-87 |
| chr4-103220689-DEL-1332 | chr4-103220690-DEL-1332 |
| chr4-103243581-DEL-87 | chr4-103243582-DEL-58 |
| chr4-103261160-DEL-76 | chr4-103261161-DEL-76 |
| chr4-103269509-DEL-551 | chr4-103269510-DEL-551 |
| chr4-103273937-DEL-2890 | chr4-103273938-DEL-2890 |
| chr4-103295173-DEL-202 | chr4-103295174-DEL-202 |
| chr4-103314014-DEL-1954 | chr4-103314015-DEL-1954 |
| chr4-103326676-DEL-60 | chr4-103326675-DEL-61 |
| chr4-103327112-DEL-132 | chr4-103327109-DEL-128 |
| chr4-103328982-DEL-1890 | chr4-103328983-DEL-1890 |
| chr4-103336115-DEL-212 | chr4-103336116-DEL-212 |
| chr4-103346565-DEL-904 | chr4-103346566-DEL-904 |
| chr4-103362682-DEL-526 | chr4-103362683-DEL-526 |
| chr4-103366126-DEL-97 | chr4-103366134-DEL-91 |
| chr4-103381648-DEL-458 | chr4-103381649-DEL-458 |
| chr4-103384400-DEL-60 | chr4-103384405-DEL-56 |
| chr4-103387958-DEL-1043 | chr4-103387961-DEL-1041 |
| chr4-103396395-DEL-878 | chr4-103396392-DEL-886 |
| chr4-103420505-DEL-1945 | chr4-103420506-DEL-1945 |
| chr4-103441902-DEL-55 | chr4-103441852-DEL-56 |
| chr4-103465059-DEL-1712 | chr4-103465059-DEL-1713 |
| chr4-103522277-DEL-1767 | chr4-103522278-DEL-1767 |
| chr4-103533457-DEL-1166 | chr4-103533458-DEL-1166 |
| chr4-103576716-DEL-140 | chr4-103576717-DEL-140 |
| chr4-103590289-DEL-144 | chr4-103590290-DEL-144 |
| chr4-103606794-DEL-55 | chr4-103606747-DEL-59 |
| chr4-103641306-DEL-612 | chr4-103641307-DEL-612 |
| chr4-103660127-DEL-332 | chr4-103660128-DEL-332 |
| chr4-103681193-DEL-104 | chr4-103681194-DEL-104 |
| chr4-103687587-DEL-6333 | chr4-103687588-DEL-6333 |
| chr4-103693596-DEL-96 | chr4-103693613-DEL-64 |
| chr4-103697646-DEL-86 | chr4-103697665-DEL-68 |
| chr4-103700922-DEL-100 | chr4-103700923-DEL-100 |
| chr4-103716297-DEL-55 | chr4-103716298-DEL-55 |
| chr4-103761662-DEL-96 | chr4-103761663-DEL-96 |
| chr4-103765932-DEL-134 | chr4-103765934-DEL-130 |
| chr4-103806178-DEL-64 | chr4-103806179-DEL-64 |
| chr4-103806707-DEL-286 | chr4-103806708-DEL-286 |
| chr4-103822067-DEL-206 | chr4-103822068-DEL-206 |
| chr4-103904358-DEL-419 | chr4-103904359-DEL-417 |
| chr4-103905568-DEL-2306 | chr4-103905569-DEL-2306 |
| chr4-103946898-DEL-6706 | chr4-103946899-DEL-6706 |
| chr4-104080960-DEL-129 | chr4-104080942-DEL-100 |
| chr4-104086601-DEL-102 | chr4-104086602-DEL-102 |
| chr4-104119958-DEL-6572 | chr4-104119959-DEL-6572 |
| chr4-104145007-DEL-132 | chr4-104145001-DEL-133 |
| chr4-104151193-DEL-1750 | chr4-104151194-DEL-1750 |
| chr4-104173019-DEL-1082 | chr4-104173020-DEL-1082 |
| chr4-104177504-DEL-89 | chr4-104177505-DEL-89 |
| chr4-104179800-DEL-52 | chr4-104179793-DEL-60 |
| chr4-104187612-DEL-164 | chr4-104187613-DEL-164 |
| chr4-104192991-DEL-143 | chr4-104192992-DEL-143 |
| chr4-104207278-DEL-74 | chr4-104207272-DEL-85 |
| chr4-104213211-DEL-548 | chr4-104213212-DEL-548 |

| | |
|---|---|
| chr4-104220166-DEL-189 | chr4-104220167-DEL-189 |
| chr4-104317698-DEL-202 | chr4-104317699-DEL-202 |
| chr4-104343706-DEL-658 | chr4-104343707-DEL-658 |
| chr4-104436435-DEL-126 | chr4-104436452-DEL-96 |
| chr4-104459085-DEL-101 | chr4-104459085-DEL-101 |
| chr4-104484110-DEL-51 | chr4-104484110-DEL-51 |
| chr4-104484809-DEL-903 | chr4-104484810-DEL-896 |
| chr4-104519756-DEL-75 | chr4-104519757-DEL-75 |
| chr4-104528965-DEL-135 | chr4-104528966-DEL-135 |
| chr4-104550968-DEL-54 | chr4-104550966-DEL-56 |
| chr4-104561352-DEL-152 | chr4-104561353-DEL-152 |
| chr4-104610140-DEL-110 | chr4-104610141-DEL-110 |
| chr4-104620630-DEL-103 | chr4-104620631-DEL-103 |
| chr4-104629462-DEL-113 | chr4-104629463-DEL-108 |
| chr4-104664106-DEL-245 | chr4-104664107-DEL-245 |
| chr4-104703816-DEL-6969 | chr4-104703817-DEL-6969 |
| chr4-104727841-DEL-190 | chr4-104727842-DEL-190 |
| chr4-104763437-DEL-155 | chr4-104763438-DEL-115 |
| chr4-104781541-DEL-197 | chr4-104781542-DEL-197 |
| chr4-104800798-DEL-92 | chr4-104800839-DEL-52 |
| chr4-104841145-DEL-77 | chr4-104841147-DEL-76 |
| chr4-104849179-DEL-1109 | chr4-104849180-DEL-1109 |
| chr4-104861912-DEL-135 | chr4-104861913-DEL-162 |
| chr4-104879660-DEL-246 | chr4-104879660-DEL-246 |
| chr4-104880967-DEL-581 | chr4-104880968-DEL-581 |
| chr4-104894462-DEL-420 | chr4-104894463-DEL-420 |
| chr4-104920454-DEL-6985 | chr4-104920455-DEL-6985 |
| chr4-104954523-DEL-1008 | chr4-104954524-DEL-1008 |
| chr4-104959843-DEL-57 | chr4-104959844-DEL-57 |
| chr4-104980631-DEL-275 | chr4-104980632-DEL-275 |
| chr4-104995155-DEL-214 | chr4-104995156-DEL-214 |
| chr4-105051156-DEL-80 | chr4-105051157-DEL-80 |
| chr4-105101713-DEL-475 | chr4-105101713-DEL-475 |
| chr4-105125195-DEL-64 | chr4-105125196-DEL-64 |
| chr4-105126572-DEL-193 | chr4-105126573-DEL-193 |
| chr4-105126951-DEL-376 | chr4-105126955-DEL-388 |
| chr4-105149694-DEL-6906 | chr4-105149708-DEL-6905 |
| chr4-105164320-DEL-51 | chr4-105164320-DEL-55 |
| chr4-105200486-DEL-134 | chr4-105200508-DEL-135 |
| chr4-105230972-DEL-53 | chr4-105230973-DEL-53 |
| chr4-105233572-DEL-66 | chr4-105233558-DEL-78 |
| chr4-105270707-DEL-252 | chr4-105270708-DEL-252 |
| chr4-105295258-DEL-6068 | chr4-105295259-DEL-6068 |
| chr4-105311111-DEL-308 | chr4-105311112-DEL-308 |
| chr4-105319623-DEL-53 | chr4-105319624-DEL-53 |
| chr4-105329303-DEL-109 | chr4-105329304-DEL-109 |
| chr4-105357241-DEL-82 | chr4-105357242-DEL-82 |
| chr4-105359768-DEL-498 | chr4-105359769-DEL-498 |
| chr4-105369059-DEL-159 | chr4-105369062-DEL-157 |
| chr4-105416083-DEL-59 | chr4-105416083-DEL-69 |
| chr4-105449470-DEL-53 | chr4-105449471-DEL-53 |
| chr4-105451156-DEL-6466 | chr4-105451156-DEL-6466 |
| chr4-105467888-DEL-54 | chr4-105467889-DEL-54 |
| chr4-105476323-DEL-6659 | chr4-105476324-DEL-6659 |
| chr4-105492376-DEL-1944 | chr4-105492376-DEL-1944 |
| chr4-105537129-DEL-124 | chr4-105537130-DEL-124 |
| chr4-105551381-DEL-375 | chr4-105551382-DEL-376 |
| chr4-105592576-DEL-6680 | chr4-105592577-DEL-6680 |
| chr4-105662198-DEL-498 | chr4-105662199-DEL-498 |
| chr4-105679407-DEL-6172 | chr4-105679408-DEL-6172 |
| chr4-105703314-DEL-54 | chr4-105703315-DEL-54 |
| chr4-105707306-DEL-258 | chr4-105707307-DEL-258 |
| chr4-105720193-DEL-506 | chr4-105720194-DEL-506 |
| chr4-105740748-DEL-1036 | chr4-105740749-DEL-1036 |
| chr4-105776723-DEL-1400 | chr4-105776724-DEL-1400 |
| chr4-105796586-DEL-79 | chr4-105796595-DEL-71 |
| chr4-105841752-DEL-518 | chr4-105841752-DEL-492 |
| chr4-105847354-DEL-1049 | chr4-105847355-DEL-1049 |
| chr4-105853999-DEL-56 | chr4-105853996-DEL-57 |
| chr4-105896911-DEL-105 | chr4-105896912-DEL-105 |
| chr4-105923279-DEL-108 | chr4-105923260-DEL-104 |
| chr4-105928196-DEL-102 | chr4-105928197-DEL-102 |
| chr4-105937085-DEL-814 | chr4-105937086-DEL-814 |
| chr4-105943170-DEL-52 | chr4-105943171-DEL-52 |
| chr4-105946917-DEL-54 | chr4-105946918-DEL-54 |
| chr4-105989902-DEL-188 | chr4-105989935-DEL-156 |
| chr4-105994166-DEL-1502 | chr4-105994167-DEL-1502 |
| chr4-106003308-DEL-151 | chr4-106003309-DEL-151 |
| chr4-106034157-DEL-6622 | chr4-106034158-DEL-6622 |
| chr4-106082549-DEL-100 | chr4-106082550-DEL-100 |

| | |
|---|---|
| chr4-106091899-DEL-81 | chr4-106091900-DEL-81 |
| chr4-106094011-DEL-4591 | chr4-106094021-DEL-4585 |
| chr4-106107193-DEL-82 | chr4-106107194-DEL-82 |
| chr4-106110951-DEL-162 | chr4-106110952-DEL-162 |
| chr4-106127260-DEL-1894 | chr4-106127261-DEL-1894 |
| chr4-106159729-DEL-63 | chr4-106159730-DEL-63 |
| chr4-106185695-DEL-459 | chr4-106185696-DEL-459 |
| chr4-106186410-DEL-263 | chr4-106186411-DEL-263 |
| chr4-106235599-DEL-510 | chr4-106235600-DEL-510 |
| chr4-106236747-DEL-127 | chr4-106236790-DEL-80 |
| chr4-106249958-DEL-3666 | chr4-106249959-DEL-3666 |
| chr4-106278749-DEL-1080 | chr4-106278750-DEL-1080 |
| chr4-106308927-DEL-91 | chr4-106308922-DEL-55 |
| chr4-106322972-DEL-114 | chr4-106322956-DEL-137 |
| chr4-106333225-DEL-54 | chr4-106333226-DEL-54 |
| chr4-106342941-DEL-113 | chr4-106342940-DEL-118 |
| chr4-106388883-DEL-66 | chr4-106388884-DEL-66 |
| chr4-106392773-DEL-179 | chr4-106392774-DEL-179 |
| chr4-106395198-DEL-62 | chr4-106395190-DEL-63 |
| chr4-106473777-DEL-52 | chr4-106473744-DEL-71 |
| chr4-106489794-DEL-56 | chr4-106489775-DEL-56 |
| chr4-106497955-DEL-125 | chr4-106497956-DEL-125 |
| chr4-106555242-DEL-86 | chr4-106555243-DEL-86 |
| chr4-106557587-DEL-108 | chr4-106557587-DEL-109 |
| chr4-106588630-DEL-311 | chr4-106588636-DEL-312 |
| chr4-106602216-DEL-85 | chr4-106602217-DEL-85 |
| chr4-106607915-DEL-69 | chr4-106607916-DEL-69 |
| chr4-106622090-DEL-219 | chr4-106622100-DEL-220 |
| chr4-106623727-DEL-1868 | chr4-106623728-DEL-1868 |
| chr4-106639570-DEL-193 | chr4-106639571-DEL-193 |
| chr4-106687721-DEL-56 | chr4-106687721-DEL-56 |
| chr4-106692565-DEL-181 | chr4-106692566-DEL-181 |
| chr4-106717724-DEL-92 | chr4-106717714-DEL-106 |
| chr4-106733244-DEL-200 | chr4-106733245-DEL-200 |
| chr4-106736938-DEL-220 | chr4-106736939-DEL-220 |
| chr4-106748257-DEL-180 | chr4-106748258-DEL-180 |
| chr4-106809676-DEL-404 | chr4-106809677-DEL-404 |
| chr4-106902480-DEL-78 | chr4-106902497-DEL-97 |
| chr4-106955446-DEL-207 | chr4-106955447-DEL-207 |
| chr4-106980359-DEL-405 | chr4-106980360-DEL-405 |
| chr4-107013470-DEL-58 | chr4-107013470-DEL-59 |
| chr4-107044076-DEL-64 | chr4-107044074-DEL-62 |
| chr4-107067527-DEL-226 | chr4-107067529-DEL-224 |
| chr4-107076864-DEL-99 | chr4-107076884-DEL-61 |
| chr4-107078286-DEL-209 | chr4-107078287-DEL-209 |
| chr4-107084777-DEL-118 | chr4-107084778-DEL-118 |
| chr4-107099814-DEL-157 | chr4-107099815-DEL-161 |
| chr4-107158011-DEL-103 | chr4-107158007-DEL-104 |
| chr4-107175839-DEL-135 | chr4-107175841-DEL-131 |
| chr4-107183749-DEL-60 | chr4-107183750-DEL-60 |
| chr4-107190023-DEL-207 | chr4-107190010-DEL-208 |
| chr4-107204109-DEL-206 | chr4-107204110-DEL-206 |
| chr4-107222072-DEL-216 | chr4-107222058-DEL-214 |
| chr4-107223772-DEL-56 | chr4-107223763-DEL-55 |
| chr4-107253711-DEL-119 | chr4-107253712-DEL-119 |
| chr4-107261824-DEL-327 | chr4-107261825-DEL-375 |
| chr4-107273739-DEL-58 | chr4-107273750-DEL-72 |
| chr4-107286326-DEL-134 | chr4-107286327-DEL-134 |
| chr4-107289014-DEL-219 | chr4-107289015-DEL-219 |
| chr4-107305356-DEL-120 | chr4-107305357-DEL-120 |
| chr4-107337631-DEL-206 | chr4-107337632-DEL-206 |
| chr4-107342858-DEL-105 | chr4-107342859-DEL-105 |
| chr4-107363661-DEL-6452 | chr4-107363662-DEL-6452 |
| chr4-107371540-DEL-516 | chr4-107371541-DEL-516 |
| chr4-107391044-DEL-72 | chr4-107391045-DEL-72 |
| chr4-107393026-DEL-58 | chr4-107393027-DEL-58 |
| chr4-107450394-DEL-98 | chr4-107450395-DEL-98 |
| chr4-107502325-DEL-3882 | chr4-107502326-DEL-3882 |
| chr4-107518419-DEL-6327 | chr4-107518420-DEL-6327 |
| chr4-107534913-DEL-92 | chr4-107534914-DEL-69 |
| chr4-107626802-DEL-199 | chr4-107626803-DEL-199 |
| chr4-107629174-DEL-56 | chr4-107629134-DEL-65 |
| chr4-107639471-DEL-61 | chr4-107639461-DEL-86 |
| chr4-107701536-DEL-174 | chr4-107701537-DEL-174 |
| chr4-107722301-DEL-447 | chr4-107722300-DEL-447 |
| chr4-107734733-DEL-87 | chr4-107734716-DEL-87 |
| chr4-107789336-DEL-201 | chr4-107789337-DEL-201 |
| chr4-107791446-DEL-183 | chr4-107791452-DEL-188 |
| chr4-107801421-DEL-81 | chr4-107801402-DEL-101 |
| chr4-107802527-DEL-195 | chr4-107802524-DEL-194 |

| | |
|---|---|
| chr4-107808550-DEL-105 | chr4-107808569-DEL-75 |
| chr4-107821425-DEL-313 | chr4-107821423-DEL-311 |
| chr4-107834867-DEL-99 | chr4-107834868-DEL-99 |
| chr4-107846944-DEL-75 | chr4-107846960-DEL-60 |
| chr4-107869020-DEL-200 | chr4-107869021-DEL-200 |
| chr4-107920517-DEL-164 | chr4-107920518-DEL-164 |
| chr4-107941398-DEL-89 | chr4-107941399-DEL-89 |
| chr4-107965985-DEL-208 | chr4-107965986-DEL-208 |
| chr4-107975161-DEL-215 | chr4-107975162-DEL-215 |
| chr4-107976530-DEL-246 | chr4-107976531-DEL-246 |
| chr4-107983060-DEL-461 | chr4-107983061-DEL-461 |
| chr4-107986889-DEL-65 | chr4-107986888-DEL-67 |
| chr4-107994115-DEL-2479 | chr4-107994116-DEL-2479 |
| chr4-108003223-DEL-8991 | chr4-108003223-DEL-8991 |
| chr4-108048020-DEL-614 | chr4-108048021-DEL-614 |
| chr4-108053095-DEL-53 | chr4-108053096-DEL-52 |
| chr4-108063136-DEL-215 | chr4-108063137-DEL-215 |
| chr4-108064092-DEL-73 | chr4-108064083-DEL-83 |
| chr4-108080501-DEL-53 | chr4-108080502-DEL-53 |
| chr4-108085753-DEL-444 | chr4-108085754-DEL-444 |
| chr4-108101490-DEL-205 | chr4-108101491-DEL-205 |
| chr4-108103479-DEL-1894 | chr4-108103480-DEL-1894 |
| chr4-108120571-DEL-194 | chr4-108120590-DEL-165 |
| chr4-108141263-DEL-7086 | chr4-108141271-DEL-7087 |
| chr4-108181653-DEL-195 | chr4-108181654-DEL-195 |
| chr4-108182673-DEL-203 | chr4-108182674-DEL-203 |
| chr4-108190904-DEL-128 | chr4-108190905-DEL-128 |
| chr4-108209796-DEL-202 | chr4-108209798-DEL-204 |
| chr4-108230781-DEL-55 | chr4-108230782-DEL-55 |
| chr4-108247089-DEL-71 | chr4-108247090-DEL-71 |
| chr4-108250782-DEL-197 | chr4-108250777-DEL-198 |
| chr4-108252385-DEL-54 | chr4-108252382-DEL-55 |
| chr4-108273890-DEL-215 | chr4-108273891-DEL-215 |
| chr4-108275988-DEL-51 | chr4-108275991-DEL-70 |
| chr4-108278440-DEL-1478 | chr4-108278440-DEL-1478 |
| chr4-108281368-DEL-85 | chr4-108281369-DEL-85 |
| chr4-108284094-DEL-787 | chr4-108284095-DEL-787 |
| chr4-108330378-DEL-183 | chr4-108330379-DEL-183 |
| chr4-108331225-DEL-236 | chr4-108331226-DEL-236 |
| chr4-108364612-DEL-182 | chr4-108364613-DEL-182 |
| chr4-108393541-DEL-203 | chr4-108393542-DEL-203 |
| chr4-108409371-DEL-1145 | chr4-108409372-DEL-1145 |
| chr4-108451096-DEL-58 | chr4-108451097-DEL-58 |
| chr4-108499678-DEL-90 | chr4-108499679-DEL-90 |
| chr4-108502364-DEL-216 | chr4-108502365-DEL-216 |
| chr4-108532027-DEL-200 | chr4-108532041-DEL-181 |
| chr4-108557734-DEL-7336 | chr4-108557735-DEL-7336 |
| chr4-108568911-DEL-51 | chr4-108568912-DEL-51 |
| chr4-108581793-DEL-420 | chr4-108581795-DEL-419 |
| chr4-108600337-DEL-197 | chr4-108600338-DEL-197 |
| chr4-108626517-DEL-100 | chr4-108626552-DEL-61 |
| chr4-108636769-DEL-53 | chr4-108636770-DEL-53 |
| chr4-108639197-DEL-52 | chr4-108639198-DEL-52 |
| chr4-108656251-DEL-103 | chr4-108656252-DEL-103 |
| chr4-108659323-DEL-600 | chr4-108659324-DEL-600 |
| chr4-108686635-DEL-664 | chr4-108686637-DEL-656 |
| chr4-108703745-DEL-199 | chr4-108703746-DEL-199 |
| chr4-108707948-DEL-201 | chr4-108707934-DEL-202 |
| chr4-108710030-DEL-196 | chr4-108710031-DEL-196 |
| chr4-108722532-DEL-170 | chr4-108722532-DEL-170 |
| chr4-108723825-DEL-51 | chr4-108723825-DEL-51 |
| chr4-108738134-DEL-1149 | chr4-108738135-DEL-1149 |
| chr4-108760793-DEL-197 | chr4-108760794-DEL-197 |
| chr4-108763554-DEL-52 | chr4-108763555-DEL-52 |
| chr4-108787598-DEL-57 | chr4-108787599-DEL-57 |
| chr4-108794888-DEL-175 | chr4-108794889-DEL-175 |
| chr4-108796506-DEL-58 | chr4-108796507-DEL-58 |
| chr4-108803903-DEL-430 | chr4-108803904-DEL-430 |
| chr4-108838591-DEL-251 | chr4-108838592-DEL-251 |
| chr4-109015305-DEL-970 | chr4-109015306-DEL-970 |
| chr4-109033748-DEL-457 | chr4-109033749-DEL-457 |
| chr4-109051011-DEL-124 | chr4-109051012-DEL-124 |
| chr4-109104562-DEL-228 | chr4-109104563-DEL-228 |
| chr4-109158105-DEL-1232 | chr4-109158105-DEL-1232 |
| chr4-109205315-DEL-6310 | chr4-109205316-DEL-6310 |
| chr4-109229802-DEL-73 | chr4-109229803-DEL-73 |
| chr4-109265790-DEL-188 | chr4-109265791-DEL-188 |
| chr4-109292754-DEL-977 | chr4-109292753-DEL-977 |
| chr4-109302088-DEL-178 | chr4-109302089-DEL-178 |
| chr4-109313352-DEL-204 | chr4-109313353-DEL-204 |

| | |
|---|---|
| chr4-109346544-DEL-170 | chr4-109346545-DEL-170 |
| chr4-109376589-DEL-574 | chr4-109376590-DEL-574 |
| chr4-109390283-DEL-51 | chr4-109390268-DEL-75 |
| chr4-109410230-DEL-250 | chr4-109410231-DEL-250 |
| chr4-109459460-DEL-6278 | chr4-109459461-DEL-6278 |
| chr4-109465936-DEL-76 | chr4-109465936-DEL-80 |
| chr4-109508258-DEL-340 | chr4-109508259-DEL-340 |
| chr4-109583221-DEL-209 | chr4-109583230-DEL-203 |
| chr4-109591575-DEL-229 | chr4-109591576-DEL-229 |
| chr4-109665944-DEL-271 | chr4-109665945-DEL-271 |
| chr4-109708654-DEL-1827 | chr4-109708655-DEL-1827 |
| chr4-109763017-DEL-1464 | chr4-109763018-DEL-1464 |
| chr4-109763058-DEL-218 | chr4-109763051-DEL-219 |
| chr4-109771035-DEL-220 | chr4-109771036-DEL-220 |
| chr4-109778647-DEL-64 | chr4-109778597-DEL-83 |
| chr4-109781303-DEL-76 | chr4-109781304-DEL-76 |
| chr4-109786198-DEL-69 | chr4-109786199-DEL-69 |
| chr4-109788019-DEL-76 | chr4-109788020-DEL-72 |
| chr4-109815434-DEL-176 | chr4-109815435-DEL-176 |
| chr4-109824043-DEL-452 | chr4-109824044-DEL-452 |
| chr4-109858290-DEL-1902 | chr4-109858291-DEL-1902 |
| chr4-109893705-DEL-608 | chr4-109893706-DEL-608 |
| chr4-109896293-DEL-578 | chr4-109896294-DEL-578 |
| chr4-109901824-DEL-138 | chr4-109901825-DEL-132 |
| chr4-109904180-DEL-1016 | chr4-109904181-DEL-1016 |
| chr4-109936027-DEL-718 | chr4-109936028-DEL-718 |
| chr4-109947242-DEL-726 | chr4-109947243-DEL-726 |
| chr4-109954769-DEL-6683 | chr4-109954770-DEL-6683 |
| chr4-110020862-DEL-1078 | chr4-110020863-DEL-1078 |
| chr4-110038638-DEL-663 | chr4-110038639-DEL-663 |
| chr4-110043904-DEL-6693 | chr4-110043905-DEL-6695 |
| chr4-110043939-DEL-430 | chr4-110043940-DEL-430 |
| chr4-110160038-DEL-518 | chr4-110160039-DEL-518 |
| chr4-110172688-DEL-6677 | chr4-110172689-DEL-6677 |
| chr4-110196047-DEL-77 | chr4-110196039-DEL-84 |
| chr4-110303256-DEL-383 | chr4-110303257-DEL-383 |
| chr4-110305724-DEL-1159 | chr4-110305725-DEL-1159 |
| chr4-110308493-DEL-639 | chr4-110308494-DEL-639 |
| chr4-110310346-DEL-861 | chr4-110310347-DEL-861 |
| chr4-110311738-DEL-61 | chr4-110311738-DEL-62 |
| chr4-110332908-DEL-416 | chr4-110332909-DEL-416 |
| chr4-110350501-DEL-207 | chr4-110350502-DEL-207 |
| chr4-110410179-DEL-6188 | chr4-110410171-DEL-6196 |
| chr4-110452746-DEL-56 | chr4-110452747-DEL-68 |
| chr4-110465353-DEL-851 | chr4-110465352-DEL-852 |
| chr4-110467741-DEL-192 | chr4-110467742-DEL-162 |
| chr4-110507719-DEL-601 | chr4-110507720-DEL-601 |
| chr4-110511196-DEL-296 | chr4-110511197-DEL-296 |
| chr4-110530004-DEL-107 | chr4-110530006-DEL-102 |
| chr4-110533801-DEL-71 | chr4-110533802-DEL-73 |
| chr4-110538231-DEL-763 | chr4-110538232-DEL-763 |
| chr4-110545518-DEL-6371 | chr4-110545519-DEL-6371 |
| chr4-110568956-DEL-6475 | chr4-110568957-DEL-6475 |
| chr4-110585248-DEL-1056 | chr4-110585249-DEL-1056 |
| chr4-110587312-DEL-1375 | chr4-110587313-DEL-1375 |
| chr4-110603475-DEL-1327 | chr4-110603476-DEL-1327 |
| chr4-110618450-DEL-6334 | chr4-110618499-DEL-6284 |
| chr4-110688580-DEL-587 | chr4-110688581-DEL-587 |
| chr4-110705068-DEL-930 | chr4-110705071-DEL-929 |
| chr4-110720768-DEL-357 | chr4-110720769-DEL-357 |
| chr4-110731175-DEL-623 | chr4-110731176-DEL-623 |
| chr4-110733740-DEL-401 | chr4-110733763-DEL-386 |
| chr4-110764353-DEL-514 | chr4-110764354-DEL-514 |
| chr4-110796732-DEL-52 | chr4-110796733-DEL-50 |
| chr4-110805065-DEL-247 | chr4-110805066-DEL-247 |
| chr4-110823313-DEL-921 | chr4-110823313-DEL-921 |
| chr4-110826962-DEL-87 | chr4-110826970-DEL-52 |
| chr4-110886386-DEL-109 | chr4-110886387-DEL-109 |
| chr4-110920597-DEL-78 | chr4-110920594-DEL-70 |
| chr4-110942686-DEL-2018 | chr4-110942687-DEL-2018 |
| chr4-110950384-DEL-890 | chr4-110950386-DEL-892 |
| chr4-110990933-DEL-61 | chr4-110990908-DEL-105 |
| chr4-110992753-DEL-361 | chr4-110992754-DEL-361 |
| chr4-111025533-DEL-5995 | chr4-111025534-DEL-5995 |
| chr4-111039115-DEL-3975 | chr4-111039116-DEL-3975 |
| chr4-111050279-DEL-143 | chr4-111050280-DEL-144 |
| chr4-111063663-DEL-2285 | chr4-111063664-DEL-2285 |
| chr4-111065911-DEL-174 | chr4-111065925-DEL-177 |
| chr4-111094583-DEL-12816 | chr4-111094582-DEL-12816 |
| chr4-111117514-DEL-56 | chr4-111117512-DEL-56 |

| | |
|---|---|
| chr4-111128134-DEL-3074 | chr4-111128135-DEL-3074 |
| chr4-111145217-DEL-703 | chr4-111145218-DEL-703 |
| chr4-111149404-DEL-121 | chr4-111149396-DEL-124 |
| chr4-111179825-DEL-408 | chr4-111179826-DEL-408 |
| chr4-111192328-DEL-516 | chr4-111192328-DEL-516 |
| chr4-111252199-DEL-1147 | chr4-111252200-DEL-1147 |
| chr4-111296496-DEL-220 | chr4-111296484-DEL-262 |
| chr4-111341165-DEL-62 | chr4-111341172-DEL-66 |
| chr4-111369726-DEL-396 | chr4-111369731-DEL-392 |
| chr4-111392670-DEL-3584 | chr4-111392671-DEL-3584 |
| chr4-111407955-DEL-108 | chr4-111407971-DEL-92 |
| chr4-111428939-DEL-5414 | chr4-111428940-DEL-5414 |
| chr4-111439895-DEL-871 | chr4-111439898-DEL-869 |
| chr4-111498821-DEL-308 | chr4-111498822-DEL-308 |
| chr4-111519490-DEL-94 | chr4-111519506-DEL-88 |
| chr4-111551892-DEL-328 | chr4-111551893-DEL-328 |
| chr4-111599432-DEL-840 | chr4-111599433-DEL-840 |
| chr4-111661438-DEL-1452 | chr4-111661439-DEL-1452 |
| chr4-111664164-DEL-1499 | chr4-111664165-DEL-1499 |
| chr4-111683753-DEL-123 | chr4-111683754-DEL-123 |
| chr4-111691029-DEL-5098 | chr4-111691030-DEL-5098 |
| chr4-111707404-DEL-1274 | chr4-111707405-DEL-1274 |
| chr4-111712959-DEL-6221 | chr4-111712960-DEL-6221 |
| chr4-111738826-DEL-195 | chr4-111738827-DEL-195 |
| chr4-111745395-DEL-80 | chr4-111745396-DEL-65 |
| chr4-111749770-DEL-78 | chr4-111749749-DEL-100 |
| chr4-111768113-DEL-705 | chr4-111768114-DEL-669 |
| chr4-111786813-DEL-6679 | chr4-111786814-DEL-6679 |
| chr4-111816463-DEL-1989 | chr4-111816464-DEL-1989 |
| chr4-111835714-DEL-1419 | chr4-111835715-DEL-1420 |
| chr4-111843648-DEL-73 | chr4-111843624-DEL-72 |
| chr4-111861055-DEL-633 | chr4-111861057-DEL-646 |
| chr4-111862016-DEL-141 | chr4-111862017-DEL-141 |
| chr4-111873994-DEL-59 | chr4-111873995-DEL-59 |
| chr4-111876907-DEL-572 | chr4-111876908-DEL-571 |
| chr4-111881778-DEL-474 | chr4-111881779-DEL-474 |
| chr4-112472970-DEL-59 | chr4-112472970-DEL-59 |
| chr4-112489060-DEL-2371 | chr4-112489061-DEL-2371 |
| chr4-112501813-DEL-70 | chr4-112501814-DEL-70 |
| chr4-112502873-DEL-1382 | chr4-112502874-DEL-1382 |
| chr4-112525395-DEL-818 | chr4-112525396-DEL-818 |
| chr4-112539592-DEL-70 | chr4-112539583-DEL-70 |
| chr4-112545253-DEL-1353 | chr4-112545254-DEL-1353 |
| chr4-112547038-DEL-595 | chr4-112547039-DEL-595 |
| chr4-112548565-DEL-3841 | chr4-112548566-DEL-3841 |
| chr4-112560212-DEL-56 | chr4-112560208-DEL-72 |
| chr4-112597025-DEL-1898 | chr4-112597026-DEL-1898 |
| chr4-112612516-DEL-2407 | chr4-112612517-DEL-2407 |
| chr4-112623170-DEL-180 | chr4-112623171-DEL-180 |
| chr4-113303144-DEL-485 | chr4-113303144-DEL-485 |
| chr4-113305685-DEL-1052 | chr4-113305685-DEL-1052 |
| chr4-113314001-DEL-81 | chr4-113314001-DEL-81 |
| chr4-113488328-DEL-225 | chr4-113488327-DEL-187 |
| chr4-113504374-DEL-108 | chr4-113504373-DEL-108 |
| chr4-113523576-DEL-54 | chr4-113523576-DEL-54 |
| chr4-113526099-DEL-239 | chr4-113526098-DEL-239 |
| chr4-113555387-DEL-1420 | chr4-113555387-DEL-1420 |
| chr4-113623138-DEL-535 | chr4-113623137-DEL-535 |
| chr4-113670596-DEL-104 | chr4-113670596-DEL-104 |
| chr4-113672345-DEL-449 | chr4-113672345-DEL-449 |
| chr4-113683226-DEL-1477 | chr4-113683223-DEL-1479 |
| chr4-113759604-DEL-166 | chr4-113759604-DEL-166 |
| chr4-113772679-DEL-215 | chr4-113772679-DEL-215 |
| chr4-113790121-DEL-1888 | chr4-113790121-DEL-1888 |
| chr4-113795699-DEL-105 | chr4-113795700-DEL-105 |
| chr4-113800804-DEL-219 | chr4-113800815-DEL-203 |
| chr4-113808846-DEL-90 | chr4-113808888-DEL-50 |
| chr4-113817864-DEL-92 | chr4-113817868-DEL-60 |
| chr4-113820893-DEL-6301 | chr4-113820894-DEL-6301 |
| chr4-113842603-DEL-230 | chr4-113842603-DEL-230 |
| chr4-113845029-DEL-3653 | chr4-113845030-DEL-3653 |
| chr4-113884144-DEL-82 | chr4-113884152-DEL-79 |
| chr4-113886579-DEL-244 | chr4-113886576-DEL-244 |
| chr4-113920534-DEL-291 | chr4-113920534-DEL-291 |
| chr4-113930074-DEL-4908 | chr4-113930039-DEL-4908 |
| chr4-114002158-DEL-69 | chr4-114002130-DEL-66 |
| chr4-114009412-DEL-506 | chr4-114009412-DEL-506 |
| chr4-114022118-DEL-1524 | chr4-114022117-DEL-1524 |
| chr4-114026067-DEL-616 | chr4-114026063-DEL-632 |
| chr4-114029993-DEL-332 | chr4-114029992-DEL-332 |

| | |
|---|---|
| chr4-114031706-DEL-8517 | chr4-114031705-DEL-8517 |
| chr4-114042997-DEL-2380 | chr4-114042998-DEL-2380 |
| chr4-114056668-DEL-1217 | chr4-114056667-DEL-1217 |
| chr4-114153820-DEL-64 | chr4-114153821-DEL-63 |
| chr4-114154123-DEL-279 | chr4-114154124-DEL-279 |
| chr4-114164077-DEL-116 | chr4-114164078-DEL-116 |
| chr4-114186485-DEL-65 | chr4-114186507-DEL-50 |
| chr4-114204305-DEL-1162 | chr4-114204306-DEL-1162 |
| chr4-114213033-DEL-72 | chr4-114213034-DEL-72 |
| chr4-114223482-DEL-74 | chr4-114223484-DEL-73 |
| chr4-114237627-DEL-52 | chr4-114237630-DEL-50 |
| chr4-114282669-DEL-72 | chr4-114282670-DEL-72 |
| chr4-114352093-DEL-69 | chr4-114352136-DEL-57 |
| chr4-114377695-DEL-91 | chr4-114377696-DEL-91 |
| chr4-114382902-DEL-177 | chr4-114382903-DEL-174 |
| chr4-114384498-DEL-480 | chr4-114384499-DEL-480 |
| chr4-114385716-DEL-244 | chr4-114385717-DEL-244 |
| chr4-114387373-DEL-55 | chr4-114387378-DEL-55 |
| chr4-114393051-DEL-97 | chr4-114393052-DEL-97 |
| chr4-114427842-DEL-54 | chr4-114427843-DEL-54 |
| chr4-114431033-DEL-171 | chr4-114431034-DEL-171 |
| chr4-114454787-DEL-59 | chr4-114454780-DEL-60 |
| chr4-114455989-DEL-56 | chr4-114455990-DEL-56 |
| chr4-114470633-DEL-112 | chr4-114470634-DEL-112 |
| chr4-114510759-DEL-2751 | chr4-114510760-DEL-2751 |
| chr4-114557038-DEL-60 | chr4-114557039-DEL-60 |
| chr4-114584854-DEL-153 | chr4-114584855-DEL-153 |
| chr4-114602648-DEL-328 | chr4-114602649-DEL-328 |
| chr4-114635835-DEL-165 | chr4-114635836-DEL-165 |
| chr4-114677021-DEL-1324 | chr4-114677022-DEL-1324 |
| chr4-114694792-DEL-55 | chr4-114694793-DEL-55 |
| chr4-114717519-DEL-225 | chr4-114717520-DEL-225 |
| chr4-114747139-DEL-55 | chr4-114747140-DEL-55 |
| chr4-114768473-DEL-209 | chr4-114768474-DEL-209 |
| chr4-114778373-DEL-178 | chr4-114778372-DEL-210 |
| chr4-114819182-DEL-208 | chr4-114819183-DEL-208 |
| chr4-114821121-DEL-236 | chr4-114821122-DEL-236 |
| chr4-114831514-DEL-268 | chr4-114831515-DEL-268 |
| chr4-114854388-DEL-84 | chr4-114854389-DEL-84 |
| chr4-114876830-DEL-187 | chr4-114876831-DEL-187 |
| chr4-114882503-DEL-410 | chr4-114882504-DEL-410 |
| chr4-114896965-DEL-208 | chr4-114896966-DEL-208 |
| chr4-114943186-DEL-266 | chr4-114943187-DEL-266 |
| chr4-114960678-DEL-66 | chr4-114960647-DEL-102 |
| chr4-114963837-DEL-311 | chr4-114963840-DEL-310 |
| chr4-114964250-DEL-746 | chr4-114964251-DEL-744 |
| chr4-115063395-DEL-452 | chr4-115063396-DEL-452 |
| chr4-115108959-DEL-205 | chr4-115108960-DEL-205 |
| chr4-115111913-DEL-4412 | chr4-115111914-DEL-4412 |
| chr4-115138259-DEL-470 | chr4-115138260-DEL-470 |
| chr4-115165312-DEL-348 | chr4-115165312-DEL-348 |
| chr4-115177868-DEL-682 | chr4-115177869-DEL-682 |
| chr4-115194429-DEL-6399 | chr4-115194430-DEL-6399 |
| chr4-115201643-DEL-7470 | chr4-115201644-DEL-7470 |
| chr4-115228669-DEL-7086 | chr4-115228670-DEL-7086 |
| chr4-115278203-DEL-332 | chr4-115278204-DEL-332 |
| chr4-115334482-DEL-309 | chr4-115334483-DEL-309 |
| chr4-115396976-DEL-2498 | chr4-115396977-DEL-2498 |
| chr4-115423000-DEL-178 | chr4-115423001-DEL-178 |
| chr4-115460474-DEL-3097 | chr4-115460475-DEL-3097 |
| chr4-115467171-DEL-178 | chr4-115467160-DEL-179 |
| chr4-115504544-DEL-59 | chr4-115504508-DEL-60 |
| chr4-115508921-DEL-193 | chr4-115508922-DEL-193 |
| chr4-115529835-DEL-620 | chr4-115529835-DEL-606 |
| chr4-115569431-DEL-554 | chr4-115569432-DEL-554 |
| chr4-115586355-DEL-85 | chr4-115586377-DEL-64 |
| chr4-115625959-DEL-718 | chr4-115625960-DEL-718 |
| chr4-115649671-DEL-208 | chr4-115649672-DEL-208 |
| chr4-115673488-DEL-711 | chr4-115673489-DEL-711 |
| chr4-115674202-DEL-129 | chr4-115674208-DEL-124 |
| chr4-115718074-DEL-60 | chr4-115718075-DEL-54 |
| chr4-115724538-DEL-91 | chr4-115724538-DEL-86 |
| chr4-115729139-DEL-533 | chr4-115729140-DEL-533 |
| chr4-115767942-DEL-62 | chr4-115767953-DEL-70 |
| chr4-115783071-DEL-52 | chr4-115783073-DEL-54 |
| chr4-115783470-DEL-265 | chr4-115783471-DEL-261 |
| chr4-115787745-DEL-98 | chr4-115787746-DEL-98 |
| chr4-115845937-DEL-98 | chr4-115845938-DEL-98 |
| chr4-115870231-DEL-100 | chr4-115870232-DEL-100 |
| chr4-115877937-DEL-98 | chr4-115877938-DEL-98 |

| | |
|---|---|
| chr4-115878767-DEL-198 | chr4-115878809-DEL-148 |
| chr4-115882063-DEL-180 | chr4-115882064-DEL-180 |
| chr4-115882689-DEL-262 | chr4-115882690-DEL-262 |
| chr4-115899064-DEL-205 | chr4-115899066-DEL-204 |
| chr4-115919578-DEL-187 | chr4-115919579-DEL-187 |
| chr4-115922348-DEL-217 | chr4-115922349-DEL-217 |
| chr4-115973083-DEL-132 | chr4-115973084-DEL-132 |
| chr4-115974147-DEL-96 | chr4-115974142-DEL-102 |
| chr4-115976394-DEL-285 | chr4-115976395-DEL-285 |
| chr4-115979486-DEL-195 | chr4-115979487-DEL-195 |
| chr4-116025207-DEL-191 | chr4-116025208-DEL-191 |
| chr4-116041555-DEL-101 | chr4-116041556-DEL-101 |
| chr4-116044479-DEL-191 | chr4-116044480-DEL-191 |
| chr4-116053010-DEL-200 | chr4-116053011-DEL-200 |
| chr4-116054475-DEL-211 | chr4-116054466-DEL-209 |
| chr4-116074856-DEL-76 | chr4-116074860-DEL-73 |
| chr4-116093145-DEL-104 | chr4-116093148-DEL-104 |
| chr4-116099128-DEL-73 | chr4-116099129-DEL-73 |
| chr4-116106354-DEL-185 | chr4-116106340-DEL-186 |
| chr4-116118831-DEL-52 | chr4-116118806-DEL-60 |
| chr4-116129347-DEL-64 | chr4-116129348-DEL-64 |
| chr4-116132102-DEL-204 | chr4-116132103-DEL-204 |
| chr4-116145007-DEL-266 | chr4-116145008-DEL-266 |
| chr4-116182996-DEL-172 | chr4-116182997-DEL-172 |
| chr4-116209230-DEL-92 | chr4-116209231-DEL-92 |
| chr4-116235988-DEL-400 | chr4-116235989-DEL-400 |
| chr4-116249763-DEL-75 | chr4-116249764-DEL-75 |
| chr4-116264751-DEL-763 | chr4-116264756-DEL-757 |
| chr4-116344303-DEL-561 | chr4-116344305-DEL-562 |
| chr4-116375467-DEL-81 | chr4-116375439-DEL-82 |
| chr4-116377065-DEL-208 | chr4-116377065-DEL-208 |
| chr4-116399680-DEL-6547 | chr4-116399681-DEL-6547 |
| chr4-116409307-DEL-198 | chr4-116409308-DEL-198 |
| chr4-116424190-DEL-201 | chr4-116424191-DEL-201 |
| chr4-116439898-DEL-81 | chr4-116439899-DEL-81 |
| chr4-116449359-DEL-254 | chr4-116449360-DEL-254 |
| chr4-116460871-DEL-199 | chr4-116460872-DEL-199 |
| chr4-116480901-DEL-215 | chr4-116480902-DEL-215 |
| chr4-116485937-DEL-198 | chr4-116485938-DEL-197 |
| chr4-116520516-DEL-209 | chr4-116520517-DEL-209 |
| chr4-116523564-DEL-257 | chr4-116523565-DEL-252 |
| chr4-116528845-DEL-184 | chr4-116528846-DEL-184 |
| chr4-116538186-DEL-242 | chr4-116538187-DEL-242 |
| chr4-116538934-DEL-196 | chr4-116538934-DEL-196 |
| chr4-116553469-DEL-206 | chr4-116553499-DEL-178 |
| chr4-116557362-DEL-62 | chr4-116557362-DEL-62 |
| chr4-116559061-DEL-328 | chr4-116559060-DEL-329 |
| chr4-116562779-DEL-187 | chr4-116562779-DEL-187 |
| chr4-116567093-DEL-55 | chr4-116567094-DEL-55 |
| chr4-116570416-DEL-202 | chr4-116570417-DEL-202 |
| chr4-116602228-DEL-209 | chr4-116602229-DEL-209 |
| chr4-116602896-DEL-172 | chr4-116602897-DEL-172 |
| chr4-116607443-DEL-65 | chr4-116607428-DEL-63 |
| chr4-116611131-DEL-52 | chr4-116611132-DEL-52 |
| chr4-116612861-DEL-66 | chr4-116612888-DEL-84 |
| chr4-116672282-DEL-92 | chr4-116672277-DEL-60 |
| chr4-116694968-DEL-305 | chr4-116694964-DEL-310 |
| chr4-116718442-DEL-203 | chr4-116718443-DEL-203 |
| chr4-116731059-DEL-193 | chr4-116731060-DEL-193 |
| chr4-116737496-DEL-785 | chr4-116737497-DEL-785 |
| chr4-116738029-DEL-72 | chr4-116738002-DEL-103 |
| chr4-116775058-DEL-4629 | chr4-116775059-DEL-4629 |
| chr4-116783479-DEL-210 | chr4-116783480-DEL-210 |
| chr4-116833494-DEL-80 | chr4-116833502-DEL-84 |
| chr4-116842412-DEL-52 | chr4-116842413-DEL-52 |
| chr4-116864051-DEL-160 | chr4-116864052-DEL-160 |
| chr4-116871134-DEL-201 | chr4-116871135-DEL-201 |
| chr4-116884788-DEL-229 | chr4-116884788-DEL-229 |
| chr4-116887410-DEL-229 | chr4-116887411-DEL-229 |
| chr4-116889346-DEL-184 | chr4-116889347-DEL-184 |
| chr4-116911878-DEL-200 | chr4-116911879-DEL-200 |
| chr4-116916959-DEL-69 | chr4-116916944-DEL-84 |
| chr4-116937858-DEL-266 | chr4-116937860-DEL-287 |
| chr4-116941858-DEL-388 | chr4-116941859-DEL-388 |
| chr4-116949611-DEL-298 | chr4-116949612-DEL-297 |
| chr4-116963460-DEL-195 | chr4-116963461-DEL-195 |
| chr4-116964144-DEL-60 | chr4-116964145-DEL-74 |
| chr4-116970942-DEL-141 | chr4-116970933-DEL-136 |
| chr4-117000770-DEL-60 | chr4-117000761-DEL-61 |
| chr4-117006093-DEL-290 | chr4-117006094-DEL-290 |

| | |
|---|---|
| chr4-117009179-DEL-220 | chr4-117009180-DEL-220 |
| chr4-117014416-DEL-76 | chr4-117014417-DEL-76 |
| chr4-117015481-DEL-169 | chr4-117015482-DEL-168 |
| chr4-117049315-DEL-7093 | chr4-117049316-DEL-7093 |
| chr4-117057763-DEL-412 | chr4-117057753-DEL-413 |
| chr4-117060408-DEL-198 | chr4-117060409-DEL-198 |
| chr4-117062090-DEL-83 | chr4-117062090-DEL-83 |
| chr4-117071820-DEL-70 | chr4-117071821-DEL-70 |
| chr4-117089265-DEL-94 | chr4-117089268-DEL-95 |
| chr4-117092136-DEL-908 | chr4-117092137-DEL-908 |
| chr4-117122274-DEL-148 | chr4-117122275-DEL-148 |
| chr4-117186577-DEL-58 | chr4-117186578-DEL-60 |
| chr4-117197184-DEL-99 | chr4-117197142-DEL-126 |
| chr4-117203997-DEL-230 | chr4-117203998-DEL-230 |
| chr4-117292698-DEL-192 | chr4-117292699-DEL-192 |
| chr4-117294360-DEL-67 | chr4-117294361-DEL-67 |
| chr4-117300030-DEL-93 | chr4-117300020-DEL-94 |
| chr4-117362784-DEL-195 | chr4-117362787-DEL-190 |
| chr4-117378213-DEL-77 | chr4-117378214-DEL-77 |
| chr4-117379278-DEL-63 | chr4-117379281-DEL-54 |
| chr4-117414891-DEL-72 | chr4-117414896-DEL-68 |
| chr4-117467766-DEL-54 | chr4-117467767-DEL-54 |
| chr4-117496378-DEL-82 | chr4-117496388-DEL-86 |
| chr4-117513241-DEL-229 | chr4-117513242-DEL-229 |
| chr4-117545444-DEL-84 | chr4-117545445-DEL-84 |
| chr4-117546613-DEL-7116 | chr4-117546614-DEL-7116 |
| chr4-117555900-DEL-3625 | chr4-117555891-DEL-3635 |
| chr4-117561704-DEL-459 | chr4-117561705-DEL-459 |
| chr4-117564754-DEL-574 | chr4-117564746-DEL-574 |
| chr4-117588226-DEL-544 | chr4-117588227-DEL-542 |
| chr4-117595050-DEL-219 | chr4-117595051-DEL-219 |
| chr4-117598338-DEL-371 | chr4-117598339-DEL-371 |
| chr4-117600286-DEL-295 | chr4-117600287-DEL-295 |
| chr4-117609509-DEL-59 | chr4-117609510-DEL-59 |
| chr4-117622957-DEL-108 | chr4-117622958-DEL-108 |
| chr4-117626328-DEL-88 | chr4-117626329-DEL-92 |
| chr4-117654771-DEL-162 | chr4-117654772-DEL-162 |
| chr4-117658968-DEL-234 | chr4-117658969-DEL-234 |
| chr4-117711033-DEL-218 | chr4-117711034-DEL-218 |
| chr4-117750211-DEL-196 | chr4-117750212-DEL-196 |
| chr4-117764253-DEL-384 | chr4-117764254-DEL-384 |
| chr4-117774772-DEL-252 | chr4-117774773-DEL-252 |
| chr4-117777692-DEL-54 | chr4-117777693-DEL-54 |
| chr4-117792090-DEL-60 | chr4-117792092-DEL-58 |
| chr4-117793328-DEL-193 | chr4-117793329-DEL-192 |
| chr4-117801403-DEL-107 | chr4-117801404-DEL-106 |
| chr4-117875938-DEL-294 | chr4-117875938-DEL-294 |
| chr4-117879918-DEL-65 | chr4-117879920-DEL-64 |
| chr4-117895516-DEL-62 | chr4-117895517-DEL-62 |
| chr4-117914535-DEL-146 | chr4-117914536-DEL-146 |
| chr4-117920468-DEL-205 | chr4-117920469-DEL-205 |
| chr4-117931709-DEL-214 | chr4-117931708-DEL-197 |
| chr4-117932373-DEL-275 | chr4-117932374-DEL-275 |
| chr4-117934001-DEL-88 | chr4-117934002-DEL-88 |
| chr4-117935809-DEL-63 | chr4-117935810-DEL-63 |
| chr4-118038339-DEL-414 | chr4-118038322-DEL-415 |
| chr4-118038737-DEL-751 | chr4-118038727-DEL-752 |
| chr4-118059165-DEL-154 | chr4-118059166-DEL-154 |
| chr4-118060365-DEL-109 | chr4-118060366-DEL-109 |
| chr4-118094294-DEL-271 | chr4-118094295-DEL-271 |
| chr4-118110135-DEL-52 | chr4-118110136-DEL-52 |
| chr4-118157926-DEL-203 | chr4-118157927-DEL-203 |
| chr4-118187609-DEL-56 | chr4-118187610-DEL-56 |
| chr4-118190504-DEL-64 | chr4-118190505-DEL-56 |
| chr4-118191296-DEL-203 | chr4-118191297-DEL-203 |
| chr4-118203739-DEL-115 | chr4-118203740-DEL-115 |
| chr4-118224241-DEL-185 | chr4-118224270-DEL-183 |
| chr4-118228193-DEL-76 | chr4-118228202-DEL-58 |
| chr4-118261704-DEL-54 | chr4-118261678-DEL-56 |
| chr4-118296019-DEL-250 | chr4-118296020-DEL-250 |
| chr4-118298746-DEL-195 | chr4-118298747-DEL-195 |
| chr4-118308608-DEL-180 | chr4-118308609-DEL-192 |
| chr4-118326721-DEL-619 | chr4-118326722-DEL-619 |
| chr4-118348236-DEL-6434 | chr4-118348237-DEL-6434 |
| chr4-118362210-DEL-56 | chr4-118362211-DEL-56 |
| chr4-118364890-DEL-156 | chr4-118364902-DEL-145 |
| chr4-118402844-DEL-155 | chr4-118402848-DEL-150 |
| chr4-118403501-DEL-212 | chr4-118403502-DEL-212 |
| chr4-118406953-DEL-213 | chr4-118406954-DEL-213 |
| chr4-118410801-DEL-139 | chr4-118410802-DEL-139 |

| | |
|---|---|
| chr4-118416248-DEL-105 | chr4-118416260-DEL-104 |
| chr4-118431171-DEL-169 | chr4-118431172-DEL-169 |
| chr4-118453630-DEL-209 | chr4-118453631-DEL-209 |
| chr4-118459739-DEL-2546 | chr4-118459740-DEL-2546 |
| chr4-118496024-DEL-132 | chr4-118496025-DEL-132 |
| chr4-118501048-DEL-454 | chr4-118501045-DEL-455 |
| chr4-118511169-DEL-450 | chr4-118511170-DEL-450 |
| chr4-118528982-DEL-196 | chr4-118528983-DEL-196 |
| chr4-118530852-DEL-4115 | chr4-118530852-DEL-4115 |
| chr4-118538635-DEL-67 | chr4-118538630-DEL-61 |
| chr4-118563921-DEL-55 | chr4-118563917-DEL-56 |
| chr4-118575839-DEL-278 | chr4-118575840-DEL-278 |
| chr4-118580055-DEL-1892 | chr4-118580056-DEL-1892 |
| chr4-118595618-DEL-259 | chr4-118595626-DEL-252 |
| chr4-118602326-DEL-328 | chr4-118602327-DEL-328 |
| chr4-118624428-DEL-185 | chr4-118624429-DEL-185 |
| chr4-118640487-DEL-60 | chr4-118640490-DEL-58 |
| chr4-118646134-DEL-55 | chr4-118646134-DEL-55 |
| chr4-118651203-DEL-401 | chr4-118651204-DEL-401 |
| chr4-118652414-DEL-80 | chr4-118652414-DEL-64 |
| chr4-118656399-DEL-125 | chr4-118656390-DEL-134 |
| chr4-118661326-DEL-266 | chr4-118661327-DEL-266 |
| chr4-118703785-DEL-6485 | chr4-118703787-DEL-6485 |
| chr4-118709479-DEL-206 | chr4-118709480-DEL-206 |
| chr4-118713613-DEL-599 | chr4-118713614-DEL-599 |
| chr4-118733371-DEL-507 | chr4-118733372-DEL-507 |
| chr4-118759156-DEL-78 | chr4-118759119-DEL-119 |
| chr4-118779953-DEL-168 | chr4-118779954-DEL-168 |
| chr4-118790200-DEL-491 | chr4-118790200-DEL-491 |
| chr4-118810121-DEL-131 | chr4-118810128-DEL-125 |
| chr4-118816558-DEL-820 | chr4-118816559-DEL-820 |
| chr4-118823049-DEL-54 | chr4-118823050-DEL-54 |
| chr4-118829036-DEL-198 | chr4-118829037-DEL-198 |
| chr4-118832643-DEL-63 | chr4-118832644-DEL-63 |
| chr4-118851202-DEL-224 | chr4-118851203-DEL-224 |
| chr4-118873523-DEL-461 | chr4-118873524-DEL-461 |
| chr4-118874076-DEL-1385 | chr4-118874077-DEL-1385 |
| chr4-118894519-DEL-100 | chr4-118894520-DEL-100 |
| chr4-118905963-DEL-66 | chr4-118905963-DEL-54 |
| chr4-118931703-DEL-136 | chr4-118931703-DEL-136 |
| chr4-118959143-DEL-73 | chr4-118959167-DEL-50 |
| chr4-119006303-DEL-185 | chr4-119006304-DEL-185 |
| chr4-119010819-DEL-173 | chr4-119010820-DEL-173 |
| chr4-119014671-DEL-205 | chr4-119014672-DEL-205 |
| chr4-119016324-DEL-227 | chr4-119016325-DEL-227 |
| chr4-119043853-DEL-189 | chr4-119043854-DEL-189 |
| chr4-119057485-DEL-203 | chr4-119057486-DEL-203 |
| chr4-119079092-DEL-203 | chr4-119079093-DEL-203 |
| chr4-119126825-DEL-204 | chr4-119126826-DEL-204 |
| chr4-119137702-DEL-217 | chr4-119137702-DEL-217 |
| chr4-119205684-DEL-907 | chr4-119205685-DEL-907 |
| chr4-119217195-DEL-1296 | chr4-119217196-DEL-1296 |
| chr4-119227042-DEL-199 | chr4-119227043-DEL-199 |
| chr4-119252474-DEL-58 | chr4-119252474-DEL-58 |
| chr4-119367298-DEL-203 | chr4-119367299-DEL-203 |
| chr4-119371256-DEL-65 | chr4-119371227-DEL-55 |
| chr4-119390209-DEL-209 | chr4-119390210-DEL-209 |
| chr4-119399478-DEL-169 | chr4-119399479-DEL-169 |
| chr4-119422666-DEL-1042 | chr4-119422667-DEL-1042 |
| chr4-119451624-DEL-56 | chr4-119451625-DEL-56 |
| chr4-119452496-DEL-4068 | chr4-119452496-DEL-4068 |
| chr4-119457086-DEL-80 | chr4-119457087-DEL-81 |
| chr4-119470793-DEL-90 | chr4-119470794-DEL-90 |
| chr4-119482698-DEL-80 | chr4-119482699-DEL-80 |
| chr4-119500953-DEL-9168 | chr4-119500954-DEL-9168 |
| chr4-119517627-DEL-112 | chr4-119517621-DEL-111 |
| chr4-119520882-DEL-1232 | chr4-119520882-DEL-1232 |
| chr4-119524020-DEL-314 | chr4-119524023-DEL-314 |
| chr4-119535313-DEL-1887 | chr4-119535314-DEL-1887 |
| chr4-119543859-DEL-2869 | chr4-119543860-DEL-2869 |
| chr4-119579334-DEL-52 | chr4-119579335-DEL-52 |
| chr4-119585380-DEL-266 | chr4-119585381-DEL-266 |
| chr4-119651291-DEL-1580 | chr4-119651292-DEL-1580 |
| chr4-119664595-DEL-65 | chr4-119664593-DEL-66 |
| chr4-119670925-DEL-80 | chr4-119670926-DEL-80 |
| chr4-119675226-DEL-1757 | chr4-119675227-DEL-1757 |
| chr4-119691119-DEL-1701 | chr4-119691120-DEL-1701 |
| chr4-119697216-DEL-75 | chr4-119697217-DEL-75 |
| chr4-119759874-DEL-81 | chr4-119759875-DEL-81 |
| chr4-119764836-DEL-513 | chr4-119764837-DEL-513 |

| | |
|---|---|
| chr4-119783624-DEL-4171 | chr4-119783625-DEL-4171 |
| chr4-119796697-DEL-68 | chr4-119796692-DEL-69 |
| chr4-119799629-DEL-71 | chr4-119799593-DEL-66 |
| chr4-119821432-DEL-52 | chr4-119821433-DEL-52 |
| chr4-119836037-DEL-68 | chr4-119836061-DEL-64 |
| chr4-119854020-DEL-52 | chr4-119854027-DEL-51 |
| chr4-119858719-DEL-115 | chr4-119858720-DEL-115 |
| chr4-119878848-DEL-59 | chr4-119878839-DEL-55 |
| chr4-119906572-DEL-245 | chr4-119906573-DEL-245 |
| chr4-119923595-DEL-73 | chr4-119923602-DEL-75 |
| chr4-119966662-DEL-3362 | chr4-119966663-DEL-3361 |
| chr4-120149127-DEL-56 | chr4-120149128-DEL-50 |
| chr4-120151355-DEL-240 | chr4-120151356-DEL-240 |
| chr4-120176133-DEL-202 | chr4-120176134-DEL-202 |
| chr4-120186611-DEL-1893 | chr4-120186612-DEL-1893 |
| chr4-120200803-DEL-95 | chr4-120200805-DEL-94 |
| chr4-120206126-DEL-52 | chr4-120206127-DEL-51 |
| chr4-120214001-DEL-96 | chr4-120214002-DEL-92 |
| chr4-120233642-DEL-130 | chr4-120233646-DEL-120 |
| chr4-120248320-DEL-171 | chr4-120248321-DEL-171 |
| chr4-120251653-DEL-210 | chr4-120251654-DEL-210 |
| chr4-120270932-DEL-350 | chr4-120270933-DEL-350 |
| chr4-120287596-DEL-122 | chr4-120287597-DEL-122 |
| chr4-120295287-DEL-4001 | chr4-120295288-DEL-4001 |
| chr4-120296058-DEL-610 | chr4-120296059-DEL-610 |
| chr4-120345999-DEL-206 | chr4-120346000-DEL-206 |
| chr4-120395338-DEL-446 | chr4-120395340-DEL-445 |
| chr4-120421622-DEL-51 | chr4-120421623-DEL-51 |
| chr4-120447448-DEL-74 | chr4-120447449-DEL-74 |
| chr4-120453711-DEL-237 | chr4-120453712-DEL-237 |
| chr4-120457338-DEL-195 | chr4-120457339-DEL-195 |
| chr4-120458074-DEL-81 | chr4-120458105-DEL-89 |
| chr4-120532677-DEL-56 | chr4-120532688-DEL-52 |
| chr4-120542088-DEL-205 | chr4-120542089-DEL-205 |
| chr4-120542733-DEL-208 | chr4-120542734-DEL-208 |
| chr4-120633952-DEL-73 | chr4-120633953-DEL-73 |
| chr4-120640049-DEL-422 | chr4-120640046-DEL-425 |
| chr4-120649064-DEL-114 | chr4-120649069-DEL-98 |
| chr4-120692321-DEL-202 | chr4-120692322-DEL-202 |
| chr4-120694474-DEL-2452 | chr4-120694475-DEL-2452 |
| chr4-120745829-DEL-206 | chr4-120745830-DEL-206 |
| chr4-120761897-DEL-1281 | chr4-120761898-DEL-1281 |
| chr4-120768080-DEL-60 | chr4-120768082-DEL-62 |
| chr4-120774506-DEL-53 | chr4-120774509-DEL-51 |
| chr4-120793347-DEL-7111 | chr4-120793348-DEL-7111 |
| chr4-120805067-DEL-81 | chr4-120805068-DEL-81 |
| chr4-120815160-DEL-80 | chr4-120815161-DEL-80 |
| chr4-120855849-DEL-209 | chr4-120855850-DEL-209 |
| chr4-120875188-DEL-68 | chr4-120875223-DEL-50 |
| chr4-120887560-DEL-54 | chr4-120887553-DEL-50 |
| chr4-120893919-DEL-416 | chr4-120893920-DEL-416 |
| chr4-120894496-DEL-481 | chr4-120894497-DEL-481 |
| chr4-120918646-DEL-224 | chr4-120918647-DEL-224 |
| chr4-120924217-DEL-210 | chr4-120924218-DEL-210 |
| chr4-120925296-DEL-103 | chr4-120925283-DEL-103 |
| chr4-120928835-DEL-224 | chr4-120928835-DEL-224 |
| chr4-120951011-DEL-544 | chr4-120951012-DEL-544 |
| chr4-120971212-DEL-254 | chr4-120971212-DEL-254 |
| chr4-120972749-DEL-174 | chr4-120972750-DEL-174 |
| chr4-120975160-DEL-4158 | chr4-120975159-DEL-4158 |
| chr4-121023989-DEL-535 | chr4-121023990-DEL-535 |
| chr4-121029120-DEL-6546 | chr4-121029121-DEL-6546 |
| chr4-121041072-DEL-233 | chr4-121041073-DEL-233 |
| chr4-121061661-DEL-88 | chr4-121061662-DEL-88 |
| chr4-121101161-DEL-3533 | chr4-121101162-DEL-3533 |
| chr4-121105723-DEL-7354 | chr4-121105724-DEL-7354 |
| chr4-121166617-DEL-489 | chr4-121166618-DEL-489 |
| chr4-121217140-DEL-199 | chr4-121217141-DEL-199 |
| chr4-121235894-DEL-365 | chr4-121235895-DEL-365 |
| chr4-121245089-DEL-877 | chr4-121245090-DEL-877 |
| chr4-121258955-DEL-56 | chr4-121258928-DEL-52 |
| chr4-121293936-DEL-76 | chr4-121293936-DEL-76 |
| chr4-121299599-DEL-54 | chr4-121299599-DEL-54 |
| chr4-121303889-DEL-62 | chr4-121303889-DEL-62 |
| chr4-121312120-DEL-1350 | chr4-121312120-DEL-1350 |
| chr4-121364139-DEL-293 | chr4-121364140-DEL-293 |
| chr4-121371248-DEL-55 | chr4-121371248-DEL-55 |
| chr4-121376695-DEL-158 | chr4-121376695-DEL-158 |
| chr4-121396018-DEL-63 | chr4-121396018-DEL-63 |
| chr4-121415375-DEL-4383 | chr4-121415376-DEL-4383 |

| | |
|---|---|
| chr4-121426418-DEL-400 | chr4-121426418-DEL-400 |
| chr4-121441333-DEL-844 | chr4-121441333-DEL-844 |
| chr4-121463476-DEL-219 | chr4-121463470-DEL-223 |
| chr4-121484501-DEL-2098 | chr4-121484502-DEL-2098 |
| chr4-121497074-DEL-65 | chr4-121497073-DEL-65 |
| chr4-121499408-DEL-4774 | chr4-121499409-DEL-4774 |
| chr4-121511991-DEL-76 | chr4-121511991-DEL-76 |
| chr4-121521838-DEL-1703 | chr4-121521836-DEL-1704 |
| chr4-121524392-DEL-1765 | chr4-121524391-DEL-1765 |
| chr4-121572977-DEL-402 | chr4-121572977-DEL-402 |
| chr4-121598586-DEL-1693 | chr4-121598587-DEL-1693 |
| chr4-121631142-DEL-68 | chr4-121631129-DEL-60 |
| chr4-122425313-DEL-1461 | chr4-122425313-DEL-1461 |
| chr4-122456140-DEL-134 | chr4-122456153-DEL-130 |
| chr4-122463675-DEL-1303 | chr4-122463673-DEL-1305 |
| chr4-122487349-DEL-181 | chr4-122487350-DEL-181 |
| chr4-122632731-DEL-614 | chr4-122632732-DEL-610 |
| chr4-122667900-DEL-3134 | chr4-122667901-DEL-3134 |
| chr4-122676516-DEL-189 | chr4-122676466-DEL-235 |
| chr4-122677721-DEL-115 | chr4-122677722-DEL-115 |
| chr4-122679715-DEL-74 | chr4-122679685-DEL-77 |
| chr4-122691147-DEL-275 | chr4-122691148-DEL-275 |
| chr4-122696483-DEL-70 | chr4-122696488-DEL-80 |
| chr4-122719170-DEL-483 | chr4-122719171-DEL-483 |
| chr4-122738930-DEL-221 | chr4-122738931-DEL-207 |
| chr4-122760907-DEL-201 | chr4-122760908-DEL-201 |
| chr4-122761616-DEL-171 | chr4-122761617-DEL-171 |
| chr4-122813745-DEL-80 | chr4-122813746-DEL-80 |
| chr4-122814507-DEL-54 | chr4-122814510-DEL-52 |
| chr4-122821430-DEL-100 | chr4-122821430-DEL-100 |
| chr4-122871344-DEL-108 | chr4-122871321-DEL-132 |
| chr4-122872170-DEL-160 | chr4-122872171-DEL-160 |
| chr4-122918494-DEL-211 | chr4-122918495-DEL-211 |
| chr4-122923404-DEL-168 | chr4-122923405-DEL-168 |
| chr4-122934746-DEL-177 | chr4-122934749-DEL-175 |
| chr4-122952506-DEL-416 | chr4-122952507-DEL-416 |
| chr4-122964321-DEL-51 | chr4-122964278-DEL-95 |
| chr4-122966677-DEL-54 | chr4-122966669-DEL-86 |
| chr4-122979202-DEL-77 | chr4-122979203-DEL-84 |
| chr4-122984041-DEL-52 | chr4-122984040-DEL-54 |
| chr4-122996341-DEL-1133 | chr4-122996342-DEL-1133 |
| chr4-123003816-DEL-129 | chr4-123003817-DEL-90 |
| chr4-123018881-DEL-72 | chr4-123018882-DEL-72 |
| chr4-123039939-DEL-54 | chr4-123039917-DEL-55 |
| chr4-123049153-DEL-388 | chr4-123049153-DEL-389 |
| chr4-123071663-DEL-116 | chr4-123071661-DEL-76 |
| chr4-123116726-DEL-322 | chr4-123116727-DEL-320 |
| chr4-123132578-DEL-68 | chr4-123132579-DEL-68 |
| chr4-123144103-DEL-116 | chr4-123144104-DEL-116 |
| chr4-123148109-DEL-174 | chr4-123148110-DEL-174 |
| chr4-123171972-DEL-166 | chr4-123171962-DEL-167 |
| chr4-123184207-DEL-209 | chr4-123184208-DEL-209 |
| chr4-123198995-DEL-147 | chr4-123198996-DEL-147 |
| chr4-123214813-DEL-93 | chr4-123214809-DEL-65 |
| chr4-123230726-DEL-95 | chr4-123230763-DEL-63 |
| chr4-123231908-DEL-58 | chr4-123231909-DEL-58 |
| chr4-123253072-DEL-221 | chr4-123253073-DEL-220 |
| chr4-123337267-DEL-510 | chr4-123337268-DEL-510 |
| chr4-123376598-DEL-201 | chr4-123376599-DEL-201 |
| chr4-123397725-DEL-216 | chr4-123397726-DEL-216 |
| chr4-123424023-DEL-73 | chr4-123424024-DEL-73 |
| chr4-123460900-DEL-401 | chr4-123460890-DEL-402 |
| chr4-123466030-DEL-76 | chr4-123466031-DEL-70 |
| chr4-123470245-DEL-51 | chr4-123470267-DEL-50 |
| chr4-123554571-DEL-169 | chr4-123554572-DEL-169 |
| chr4-123582654-DEL-222 | chr4-123582655-DEL-222 |
| chr4-123590106-DEL-206 | chr4-123590107-DEL-205 |
| chr4-123590747-DEL-385 | chr4-123590748-DEL-385 |
| chr4-123616985-DEL-7107 | chr4-123616986-DEL-7107 |
| chr4-123631306-DEL-207 | chr4-123631307-DEL-207 |
| chr4-123637713-DEL-60 | chr4-123637714-DEL-66 |
| chr4-123640791-DEL-66 | chr4-123640792-DEL-66 |
| chr4-123649114-DEL-224 | chr4-123649115-DEL-202 |
| chr4-123651314-DEL-198 | chr4-123651315-DEL-198 |
| chr4-123655325-DEL-196 | chr4-123655322-DEL-200 |
| chr4-123662594-DEL-129 | chr4-123662595-DEL-129 |
| chr4-123663209-DEL-102 | chr4-123663197-DEL-103 |
| chr4-123664563-DEL-7103 | chr4-123664564-DEL-7103 |
| chr4-123676921-DEL-1562 | chr4-123676922-DEL-1562 |
| chr4-123710031-DEL-411 | chr4-123710032-DEL-411 |

| | |
|---|---|
| chr4-123712238-DEL-180 | chr4-123712252-DEL-166 |
| chr4-123718511-DEL-64 | chr4-123718496-DEL-80 |
| chr4-123726628-DEL-75 | chr4-123726629-DEL-75 |
| chr4-123731591-DEL-304 | chr4-123731592-DEL-304 |
| chr4-123755786-DEL-1366 | chr4-123755787-DEL-1366 |
| chr4-123769690-DEL-197 | chr4-123769690-DEL-197 |
| chr4-123831643-DEL-75 | chr4-123831644-DEL-75 |
| chr4-123837162-DEL-194 | chr4-123837163-DEL-194 |
| chr4-123865224-DEL-386 | chr4-123865225-DEL-386 |
| chr4-123952278-DEL-284 | chr4-123952279-DEL-284 |
| chr4-123962386-DEL-1820 | chr4-123962387-DEL-1820 |
| chr4-123972532-DEL-287 | chr4-123972537-DEL-278 |
| chr4-123973839-DEL-383 | chr4-123973840-DEL-383 |
| chr4-123982730-DEL-68 | chr4-123982731-DEL-68 |
| chr4-123984279-DEL-54 | chr4-123984280-DEL-54 |
| chr4-124015149-DEL-120 | chr4-124015108-DEL-124 |
| chr4-124026577-DEL-385 | chr4-124026572-DEL-361 |
| chr4-124048144-DEL-341 | chr4-124048145-DEL-341 |
| chr4-124058959-DEL-120 | chr4-124058960-DEL-120 |
| chr4-124088207-DEL-312 | chr4-124088208-DEL-312 |
| chr4-124091669-DEL-70 | chr4-124091678-DEL-62 |
| chr4-124099406-DEL-52 | chr4-124099407-DEL-52 |
| chr4-124109243-DEL-59 | chr4-124109244-DEL-58 |
| chr4-124128797-DEL-66 | chr4-124128802-DEL-56 |
| chr4-124172300-DEL-90 | chr4-124172279-DEL-98 |
| chr4-124184064-DEL-172 | chr4-124184071-DEL-164 |
| chr4-124246846-DEL-5560 | chr4-124246847-DEL-5560 |
| chr4-124318623-DEL-81 | chr4-124318638-DEL-64 |
| chr4-124347462-DEL-65 | chr4-124347463-DEL-65 |
| chr4-124357556-DEL-176 | chr4-124357544-DEL-177 |
| chr4-124357921-DEL-847 | chr4-124357924-DEL-848 |
| chr4-124378860-DEL-67 | chr4-124378861-DEL-67 |
| chr4-124425606-DEL-257 | chr4-124425601-DEL-275 |
| chr4-124460997-DEL-60 | chr4-124461000-DEL-58 |
| chr4-124485107-DEL-132 | chr4-124485108-DEL-132 |
| chr4-124515943-DEL-203 | chr4-124515944-DEL-203 |
| chr4-124542928-DEL-293 | chr4-124542929-DEL-293 |
| chr4-124549730-DEL-111 | chr4-124549727-DEL-119 |
| chr4-124563506-DEL-207 | chr4-124563507-DEL-207 |
| chr4-124575638-DEL-241 | chr4-124575639-DEL-241 |
| chr4-124584227-DEL-207 | chr4-124584228-DEL-207 |
| chr4-124591111-DEL-174 | chr4-124591112-DEL-174 |
| chr4-124600072-DEL-59 | chr4-124600073-DEL-59 |
| chr4-124603579-DEL-113 | chr4-124603580-DEL-113 |
| chr4-124605708-DEL-378 | chr4-124605714-DEL-379 |
| chr4-124623713-DEL-120 | chr4-124623714-DEL-120 |
| chr4-124650632-DEL-204 | chr4-124650633-DEL-204 |
| chr4-124652889-DEL-66 | chr4-124652890-DEL-66 |
| chr4-124670233-DEL-67 | chr4-124670234-DEL-67 |
| chr4-124674634-DEL-136 | chr4-124674630-DEL-136 |
| chr4-124681958-DEL-237 | chr4-124681963-DEL-233 |
| chr4-124708487-DEL-219 | chr4-124708488-DEL-219 |
| chr4-124758563-DEL-118 | chr4-124758564-DEL-118 |
| chr4-124759113-DEL-54 | chr4-124759118-DEL-53 |
| chr4-124761221-DEL-231 | chr4-124761222-DEL-231 |
| chr4-124762614-DEL-944 | chr4-124762615-DEL-944 |
| chr4-124766001-DEL-210 | chr4-124766002-DEL-210 |
| chr4-124784496-DEL-56 | chr4-124784501-DEL-52 |
| chr4-124801457-DEL-114 | chr4-124801458-DEL-114 |
| chr4-124804435-DEL-192 | chr4-124804436-DEL-192 |
| chr4-124825430-DEL-1725 | chr4-124825429-DEL-1725 |
| chr4-124839471-DEL-1842 | chr4-124839472-DEL-1842 |
| chr4-124858444-DEL-52 | chr4-124858445-DEL-52 |
| chr4-124877583-DEL-65 | chr4-124877584-DEL-65 |
| chr4-124933330-DEL-138 | chr4-124933333-DEL-136 |
| chr4-124952229-DEL-91 | chr4-124952230-DEL-91 |
| chr4-125034031-DEL-136 | chr4-125034032-DEL-136 |
| chr4-125034537-DEL-52 | chr4-125034538-DEL-52 |
| chr4-125035513-DEL-202 | chr4-125035522-DEL-160 |
| chr4-125056101-DEL-70 | chr4-125056100-DEL-51 |
| chr4-125057157-DEL-117 | chr4-125057158-DEL-117 |
| chr4-125057931-DEL-668 | chr4-125057932-DEL-668 |
| chr4-125071324-DEL-84 | chr4-125071282-DEL-82 |
| chr4-125073012-DEL-60 | chr4-125073005-DEL-72 |
| chr4-125086351-DEL-71 | chr4-125086341-DEL-75 |
| chr4-125104257-DEL-56 | chr4-125104258-DEL-56 |
| chr4-125127243-DEL-359 | chr4-125127244-DEL-359 |
| chr4-125141296-DEL-700 | chr4-125141297-DEL-700 |
| chr4-125160384-DEL-78 | chr4-125160385-DEL-70 |
| chr4-125184830-DEL-192 | chr4-125184831-DEL-192 |

| | |
|---|---|
| chr4-125215876-DEL-152 | chr4-125215877-DEL-152 |
| chr4-125237381-DEL-66 | chr4-125237382-DEL-66 |
| chr4-125260354-DEL-2641 | chr4-125260355-DEL-2641 |
| chr4-125263493-DEL-1579 | chr4-125263494-DEL-1579 |
| chr4-125273751-DEL-72 | chr4-125273731-DEL-98 |
| chr4-125285851-DEL-53 | chr4-125285852-DEL-53 |
| chr4-125325704-DEL-80 | chr4-125325727-DEL-62 |
| chr4-125332872-DEL-62 | chr4-125332879-DEL-54 |
| chr4-125362093-DEL-208 | chr4-125362117-DEL-185 |
| chr4-125377354-DEL-91 | chr4-125377365-DEL-81 |
| chr4-125381660-DEL-90 | chr4-125381661-DEL-90 |
| chr4-125397873-DEL-957 | chr4-125397874-DEL-957 |
| chr4-125401194-DEL-88 | chr4-125401185-DEL-87 |
| chr4-125449758-DEL-214 | chr4-125449759-DEL-214 |
| chr4-125499736-DEL-252 | chr4-125499737-DEL-252 |
| chr4-125502287-DEL-549 | chr4-125502288-DEL-549 |
| chr4-125544899-DEL-54 | chr4-125544900-DEL-54 |
| chr4-125582654-DEL-68 | chr4-125582683-DEL-52 |
| chr4-125680298-DEL-846 | chr4-125680299-DEL-846 |
| chr4-125681499-DEL-51 | chr4-125681500-DEL-51 |
| chr4-125719757-DEL-98 | chr4-125719758-DEL-98 |
| chr4-125733780-DEL-74 | chr4-125733781-DEL-74 |
| chr4-125757421-DEL-328 | chr4-125757422-DEL-328 |
| chr4-125762217-DEL-69 | chr4-125762218-DEL-69 |
| chr4-125785591-DEL-75 | chr4-125785592-DEL-75 |
| chr4-125804417-DEL-234 | chr4-125804418-DEL-234 |
| chr4-125808288-DEL-52 | chr4-125808282-DEL-53 |
| chr4-125848242-DEL-54 | chr4-125848235-DEL-61 |
| chr4-125860629-DEL-192 | chr4-125860630-DEL-192 |
| chr4-125872217-DEL-118 | chr4-125872218-DEL-116 |
| chr4-125874501-DEL-175 | chr4-125874489-DEL-176 |
| chr4-125883919-DEL-64 | chr4-125883920-DEL-60 |
| chr4-125886525-DEL-1801 | chr4-125886526-DEL-1800 |
| chr4-125888727-DEL-194 | chr4-125888728-DEL-194 |
| chr4-125906601-DEL-203 | chr4-125906622-DEL-176 |
| chr4-125963573-DEL-539 | chr4-125963574-DEL-539 |
| chr4-125973854-DEL-207 | chr4-125973855-DEL-207 |
| chr4-125990120-DEL-56 | chr4-125990121-DEL-56 |
| chr4-126106251-DEL-219 | chr4-126106252-DEL-219 |
| chr4-126112274-DEL-667 | chr4-126112276-DEL-665 |
| chr4-126118921-DEL-200 | chr4-126118922-DEL-200 |
| chr4-126165422-DEL-192 | chr4-126165423-DEL-192 |
| chr4-126224502-DEL-102 | chr4-126224503-DEL-102 |
| chr4-126263476-DEL-115 | chr4-126263444-DEL-113 |
| chr4-126271819-DEL-3438 | chr4-126271820-DEL-3438 |
| chr4-126272313-DEL-201 | chr4-126272314-DEL-201 |
| chr4-126283358-DEL-175 | chr4-126283362-DEL-173 |
| chr4-126306969-DEL-277 | chr4-126306970-DEL-277 |
| chr4-126306985-DEL-779 | chr4-126306985-DEL-775 |
| chr4-126309760-DEL-56 | chr4-126309761-DEL-56 |
| chr4-126314397-DEL-1462 | chr4-126314398-DEL-1462 |
| chr4-126318818-DEL-205 | chr4-126318819-DEL-205 |
| chr4-126339359-DEL-200 | chr4-126339360-DEL-200 |
| chr4-126350912-DEL-65 | chr4-126350916-DEL-61 |
| chr4-126367909-DEL-193 | chr4-126367910-DEL-193 |
| chr4-126391857-DEL-54 | chr4-126391858-DEL-54 |
| chr4-126393448-DEL-154 | chr4-126393445-DEL-158 |
| chr4-126440741-DEL-421 | chr4-126440741-DEL-422 |
| chr4-126477052-DEL-52 | chr4-126477054-DEL-51 |
| chr4-126485184-DEL-63 | chr4-126485185-DEL-63 |
| chr4-126489577-DEL-335 | chr4-126489601-DEL-302 |
| chr4-126498666-DEL-216 | chr4-126498681-DEL-240 |
| chr4-126501535-DEL-112 | chr4-126501532-DEL-116 |
| chr4-126503089-DEL-193 | chr4-126503091-DEL-192 |
| chr4-126510409-DEL-1932 | chr4-126510410-DEL-1932 |
| chr4-126517021-DEL-2479 | chr4-126517022-DEL-2479 |
| chr4-126525002-DEL-193 | chr4-126525003-DEL-193 |
| chr4-126533978-DEL-213 | chr4-126533979-DEL-213 |
| chr4-126546437-DEL-77 | chr4-126546439-DEL-84 |
| chr4-126567166-DEL-241 | chr4-126567167-DEL-241 |
| chr4-126621104-DEL-207 | chr4-126621105-DEL-207 |
| chr4-126651143-DEL-214 | chr4-126651144-DEL-214 |
| chr4-126665648-DEL-1946 | chr4-126665649-DEL-1946 |
| chr4-126698567-DEL-256 | chr4-126698568-DEL-256 |
| chr4-126713337-DEL-360 | chr4-126713338-DEL-360 |
| chr4-126725252-DEL-207 | chr4-126725255-DEL-201 |
| chr4-126743995-DEL-76 | chr4-126743996-DEL-76 |
| chr4-126792991-DEL-214 | chr4-126792992-DEL-214 |
| chr4-126805197-DEL-210 | chr4-126805198-DEL-210 |
| chr4-126822883-DEL-205 | chr4-126822884-DEL-205 |

| | |
|---|---|
| chr4-126837275-DEL-3847 | chr4-126837276-DEL-3847 |
| chr4-126850590-DEL-71 | chr4-126850591-DEL-71 |
| chr4-126856019-DEL-194 | chr4-126856020-DEL-194 |
| chr4-126873622-DEL-178 | chr4-126873619-DEL-179 |
| chr4-126877820-DEL-217 | chr4-126877811-DEL-231 |
| chr4-126891144-DEL-207 | chr4-126891145-DEL-207 |
| chr4-126893066-DEL-128 | chr4-126893067-DEL-128 |
| chr4-126902306-DEL-7178 | chr4-126902307-DEL-7178 |
| chr4-126932689-DEL-173 | chr4-126932690-DEL-173 |
| chr4-126949933-DEL-234 | chr4-126949934-DEL-234 |
| chr4-126955817-DEL-65 | chr4-126955818-DEL-65 |
| chr4-126956519-DEL-716 | chr4-126956520-DEL-716 |
| chr4-126966017-DEL-755 | chr4-126966018-DEL-755 |
| chr4-126982703-DEL-60 | chr4-126982703-DEL-60 |
| chr4-126985279-DEL-130 | chr4-126985280-DEL-130 |
| chr4-126992612-DEL-212 | chr4-126992613-DEL-212 |
| chr4-126993272-DEL-220 | chr4-126993273-DEL-219 |
| chr4-127005014-DEL-430 | chr4-127005015-DEL-430 |
| chr4-127041563-DEL-226 | chr4-127041564-DEL-226 |
| chr4-127096672-DEL-70 | chr4-127096673-DEL-100 |
| chr4-127107289-DEL-136 | chr4-127107290-DEL-136 |
| chr4-127142397-DEL-172 | chr4-127142398-DEL-172 |
| chr4-127188434-DEL-214 | chr4-127188429-DEL-212 |
| chr4-127197288-DEL-176 | chr4-127197293-DEL-178 |
| chr4-127212610-DEL-222 | chr4-127212611-DEL-222 |
| chr4-127218013-DEL-85 | chr4-127218014-DEL-85 |
| chr4-127260175-DEL-136 | chr4-127260176-DEL-136 |
| chr4-127260781-DEL-266 | chr4-127260782-DEL-266 |
| chr4-127261542-DEL-68 | chr4-127261543-DEL-68 |
| chr4-127273745-DEL-186 | chr4-127273746-DEL-186 |
| chr4-127296943-DEL-62 | chr4-127296946-DEL-58 |
| chr4-127301507-DEL-525 | chr4-127301508-DEL-525 |
| chr4-127311532-DEL-77 | chr4-127311533-DEL-77 |
| chr4-127332678-DEL-98 | chr4-127332679-DEL-98 |
| chr4-127338487-DEL-251 | chr4-127338488-DEL-251 |
| chr4-127343284-DEL-220 | chr4-127343285-DEL-220 |
| chr4-127366533-DEL-416 | chr4-127366534-DEL-416 |
| chr4-127367590-DEL-823 | chr4-127367591-DEL-823 |
| chr4-127379408-DEL-177 | chr4-127379408-DEL-178 |
| chr4-127402096-DEL-629 | chr4-127402117-DEL-600 |
| chr4-127413845-DEL-1542 | chr4-127413845-DEL-1542 |
| chr4-127424124-DEL-81 | chr4-127424125-DEL-81 |
| chr4-127455334-DEL-153 | chr4-127455335-DEL-153 |
| chr4-127468733-DEL-703 | chr4-127468733-DEL-704 |
| chr4-127508676-DEL-640 | chr4-127508677-DEL-640 |
| chr4-127510095-DEL-62 | chr4-127510079-DEL-100 |
| chr4-127531732-DEL-72 | chr4-127531732-DEL-58 |
| chr4-127572867-DEL-104 | chr4-127572868-DEL-104 |
| chr4-127620911-DEL-2319 | chr4-127620912-DEL-2319 |
| chr4-127687638-DEL-71 | chr4-127687639-DEL-71 |
| chr4-127705953-DEL-99 | chr4-127705954-DEL-66 |
| chr4-127729257-DEL-60 | chr4-127729257-DEL-51 |
| chr4-127768179-DEL-3720 | chr4-127768180-DEL-3720 |
| chr4-127778051-DEL-84 | chr4-127778056-DEL-80 |
| chr4-127803141-DEL-1888 | chr4-127803143-DEL-1889 |
| chr4-127867737-DEL-1061 | chr4-127867738-DEL-1061 |
| chr4-127968842-DEL-64 | chr4-127968888-DEL-62 |
| chr4-127982820-DEL-91 | chr4-127982821-DEL-91 |
| chr4-127989755-DEL-96 | chr4-127989766-DEL-78 |
| chr4-127996309-DEL-54 | chr4-127996302-DEL-50 |
| chr4-128006091-DEL-207 | chr4-128006092-DEL-207 |
| chr4-128040715-DEL-54 | chr4-128040716-DEL-54 |
| chr4-128059617-DEL-92 | chr4-128059618-DEL-92 |
| chr4-128059906-DEL-323 | chr4-128059907-DEL-323 |
| chr4-128061293-DEL-52 | chr4-128061293-DEL-52 |
| chr4-128073943-DEL-96 | chr4-128073944-DEL-96 |
| chr4-128092311-DEL-196 | chr4-128092312-DEL-196 |
| chr4-128100654-DEL-132 | chr4-128100653-DEL-134 |
| chr4-128141486-DEL-493 | chr4-128141486-DEL-493 |
| chr4-128159564-DEL-96 | chr4-128159565-DEL-96 |
| chr4-128175761-DEL-58 | chr4-128175760-DEL-60 |
| chr4-128227107-DEL-76 | chr4-128227135-DEL-71 |
| chr4-128234231-DEL-293 | chr4-128234230-DEL-288 |
| chr4-128235422-DEL-52 | chr4-128235423-DEL-52 |
| chr4-128245694-DEL-4513 | chr4-128245695-DEL-4513 |
| chr4-128260420-DEL-59 | chr4-128260416-DEL-60 |
| chr4-128272036-DEL-79 | chr4-128272040-DEL-78 |
| chr4-128286053-DEL-97 | chr4-128286054-DEL-97 |
| chr4-128294356-DEL-54 | chr4-128294357-DEL-54 |
| chr4-128317507-DEL-86 | chr4-128317508-DEL-86 |

| | |
|---|---|
| chr4-128326377-DEL-7137 | chr4-128326378-DEL-7137 |
| chr4-128343616-DEL-68 | chr4-128343617-DEL-68 |
| chr4-128350233-DEL-103 | chr4-128350234-DEL-103 |
| chr4-128375236-DEL-64 | chr4-128375205-DEL-60 |
| chr4-128375749-DEL-889 | chr4-128375750-DEL-889 |
| chr4-128388223-DEL-113 | chr4-128388224-DEL-101 |
| chr4-128395096-DEL-66 | chr4-128395097-DEL-78 |
| chr4-128396324-DEL-138 | chr4-128396325-DEL-138 |
| chr4-128414809-DEL-122 | chr4-128414842-DEL-81 |
| chr4-128422509-DEL-76 | chr4-128422510-DEL-76 |
| chr4-128441490-DEL-119 | chr4-128441488-DEL-116 |
| chr4-128475487-DEL-479 | chr4-128475488-DEL-479 |
| chr4-128496554-DEL-69 | chr4-128496555-DEL-69 |
| chr4-128523064-DEL-160 | chr4-128523033-DEL-176 |
| chr4-128529647-DEL-594 | chr4-128529648-DEL-594 |
| chr4-128534042-DEL-57 | chr4-128534043-DEL-57 |
| chr4-128540915-DEL-1319 | chr4-128540916-DEL-1294 |
| chr4-128542870-DEL-58 | chr4-128542871-DEL-58 |
| chr4-128556341-DEL-70 | chr4-128556339-DEL-67 |
| chr4-128575288-DEL-55 | chr4-128575284-DEL-54 |
| chr4-128588941-DEL-1330 | chr4-128588942-DEL-1330 |
| chr4-128591018-DEL-116 | chr4-128591019-DEL-116 |
| chr4-128646934-DEL-94 | chr4-128646935-DEL-94 |
| chr4-128648550-DEL-80 | chr4-128648569-DEL-66 |
| chr4-128719887-DEL-132 | chr4-128719850-DEL-149 |
| chr4-128741415-DEL-218 | chr4-128741416-DEL-218 |
| chr4-128752026-DEL-93 | chr4-128752033-DEL-87 |
| chr4-128787787-DEL-64 | chr4-128787788-DEL-64 |
| chr4-128791193-DEL-154 | chr4-128791194-DEL-154 |
| chr4-128806476-DEL-386 | chr4-128806477-DEL-386 |
| chr4-128809801-DEL-177 | chr4-128809802-DEL-177 |
| chr4-128813402-DEL-844 | chr4-128813400-DEL-820 |
| chr4-128815432-DEL-88 | chr4-128815433-DEL-88 |
| chr4-128824819-DEL-121 | chr4-128824819-DEL-121 |
| chr4-128839452-DEL-164 | chr4-128839453-DEL-164 |
| chr4-128848678-DEL-58 | chr4-128848679-DEL-58 |
| chr4-128871494-DEL-259 | chr4-128871495-DEL-259 |
| chr4-128908853-DEL-97 | chr4-128908854-DEL-97 |
| chr4-128920275-DEL-174 | chr4-128920276-DEL-174 |
| chr4-128921832-DEL-507 | chr4-128921833-DEL-507 |
| chr4-128921895-DEL-2506 | chr4-128921944-DEL-2506 |
| chr4-128939173-DEL-54 | chr4-128939174-DEL-52 |
| chr4-128943393-DEL-66 | chr4-128943394-DEL-66 |
| chr4-128958012-DEL-887 | chr4-128958013-DEL-887 |
| chr4-128964007-DEL-211 | chr4-128964008-DEL-211 |
| chr4-128985051-DEL-270 | chr4-128985052-DEL-270 |
| chr4-128987351-DEL-184 | chr4-128987352-DEL-184 |
| chr4-128991141-DEL-74 | chr4-128991142-DEL-74 |
| chr4-128992189-DEL-57 | chr4-128992190-DEL-57 |
| chr4-128992896-DEL-52 | chr4-128992890-DEL-62 |
| chr4-128995562-DEL-208 | chr4-128995563-DEL-208 |
| chr4-128996668-DEL-217 | chr4-128996669-DEL-217 |
| chr4-129025761-DEL-60 | chr4-129025769-DEL-50 |
| chr4-129047578-DEL-189 | chr4-129047579-DEL-189 |
| chr4-129072962-DEL-244 | chr4-129072963-DEL-244 |
| chr4-129092848-DEL-164 | chr4-129092864-DEL-165 |
| chr4-129093503-DEL-469 | chr4-129093504-DEL-468 |
| chr4-129110912-DEL-64 | chr4-129110913-DEL-64 |
| chr4-129162096-DEL-52 | chr4-129162097-DEL-52 |
| chr4-129190725-DEL-206 | chr4-129190714-DEL-207 |
| chr4-129205491-DEL-157 | chr4-129205492-DEL-157 |
| chr4-129206809-DEL-703 | chr4-129206810-DEL-703 |
| chr4-129213341-DEL-221 | chr4-129213342-DEL-221 |
| chr4-129250510-DEL-213 | chr4-129250510-DEL-214 |
| chr4-129256157-DEL-63 | chr4-129256158-DEL-63 |
| chr4-129260469-DEL-217 | chr4-129260470-DEL-217 |
| chr4-129267853-DEL-204 | chr4-129267854-DEL-204 |
| chr4-129277693-DEL-206 | chr4-129277694-DEL-206 |
| chr4-129281620-DEL-51 | chr4-129281619-DEL-52 |
| chr4-129287026-DEL-165 | chr4-129287027-DEL-165 |
| chr4-129309326-DEL-227 | chr4-129309327-DEL-227 |
| chr4-129310910-DEL-177 | chr4-129310901-DEL-175 |
| chr4-129312613-DEL-243 | chr4-129312593-DEL-249 |
| chr4-129314287-DEL-207 | chr4-129314288-DEL-207 |
| chr4-129314412-DEL-1131 | chr4-129314412-DEL-1131 |
| chr4-129315413-DEL-200 | chr4-129315414-DEL-200 |
| chr4-129318596-DEL-114 | chr4-129318597-DEL-114 |
| chr4-129329000-DEL-81 | chr4-129329001-DEL-81 |
| chr4-129343725-DEL-311 | chr4-129343734-DEL-306 |
| chr4-129352358-DEL-172 | chr4-129352359-DEL-172 |

| | |
|---|---|
| chr4-129366980-DEL-199 | chr4-129366981-DEL-199 |
| chr4-129368752-DEL-264 | chr4-129368753-DEL-264 |
| chr4-129413255-DEL-210 | chr4-129413256-DEL-210 |
| chr4-129421028-DEL-1352 | chr4-129421029-DEL-1352 |
| chr4-129428877-DEL-223 | chr4-129428878-DEL-223 |
| chr4-129439731-DEL-113 | chr4-129439723-DEL-90 |
| chr4-129447947-DEL-52 | chr4-129447947-DEL-52 |
| chr4-129455317-DEL-202 | chr4-129455318-DEL-202 |
| chr4-129458722-DEL-122 | chr4-129458723-DEL-122 |
| chr4-129469608-DEL-252 | chr4-129469609-DEL-252 |
| chr4-129473989-DEL-111 | chr4-129473990-DEL-109 |
| chr4-129489832-DEL-62 | chr4-129489853-DEL-56 |
| chr4-129499506-DEL-225 | chr4-129499507-DEL-225 |
| chr4-129504604-DEL-51 | chr4-129504605-DEL-51 |
| chr4-129508628-DEL-161 | chr4-129508629-DEL-161 |
| chr4-129513353-DEL-202 | chr4-129513354-DEL-202 |
| chr4-129531501-DEL-215 | chr4-129531502-DEL-215 |
| chr4-129582107-DEL-1895 | chr4-129582108-DEL-1895 |
| chr4-129585126-DEL-175 | chr4-129585127-DEL-175 |
| chr4-129601513-DEL-680 | chr4-129601514-DEL-680 |
| chr4-129637998-DEL-218 | chr4-129637999-DEL-218 |
| chr4-129641513-DEL-182 | chr4-129641514-DEL-182 |
| chr4-129642186-DEL-328 | chr4-129642187-DEL-328 |
| chr4-129656296-DEL-63 | chr4-129656303-DEL-54 |
| chr4-129658813-DEL-2094 | chr4-129658814-DEL-2094 |
| chr4-129666803-DEL-111 | chr4-129666804-DEL-111 |
| chr4-129667369-DEL-193 | chr4-129667370-DEL-193 |
| chr4-129668316-DEL-54 | chr4-129668317-DEL-54 |
| chr4-129673543-DEL-199 | chr4-129673544-DEL-199 |
| chr4-129684630-DEL-68 | chr4-129684630-DEL-68 |
| chr4-129685905-DEL-221 | chr4-129685906-DEL-221 |
| chr4-129696022-DEL-219 | chr4-129696023-DEL-219 |
| chr4-129696688-DEL-75 | chr4-129696689-DEL-75 |
| chr4-129709349-DEL-138 | chr4-129709350-DEL-138 |
| chr4-129756854-DEL-1003 | chr4-129756827-DEL-1005 |
| chr4-129763233-DEL-103 | chr4-129763246-DEL-87 |
| chr4-129803772-DEL-81 | chr4-129803773-DEL-81 |
| chr4-129896587-DEL-72 | chr4-129896607-DEL-53 |
| chr4-129920228-DEL-56 | chr4-129920229-DEL-56 |
| chr4-129973796-DEL-87 | chr4-129973808-DEL-59 |
| chr4-130031605-DEL-61 | chr4-130031605-DEL-68 |
| chr4-130055784-DEL-51 | chr4-130055779-DEL-57 |
| chr4-130087571-DEL-56 | chr4-130087590-DEL-62 |
| chr4-130105829-DEL-1263 | chr4-130105830-DEL-1263 |
| chr4-130128535-DEL-75 | chr4-130128536-DEL-75 |
| chr4-130134809-DEL-94 | chr4-130134810-DEL-52 |
| chr4-130138375-DEL-72 | chr4-130138376-DEL-72 |
| chr4-130145554-DEL-57 | chr4-130145555-DEL-57 |
| chr4-130173615-DEL-73 | chr4-130173592-DEL-78 |
| chr4-130183943-DEL-200 | chr4-130183944-DEL-200 |
| chr4-130191175-DEL-136 | chr4-130191176-DEL-136 |
| chr4-130200097-DEL-57 | chr4-130200100-DEL-56 |
| chr4-130234462-DEL-209 | chr4-130234463-DEL-209 |
| chr4-130235056-DEL-222 | chr4-130235057-DEL-222 |
| chr4-130259255-DEL-230 | chr4-130259256-DEL-230 |
| chr4-130259806-DEL-315 | chr4-130259807-DEL-313 |
| chr4-130264945-DEL-1095 | chr4-130264946-DEL-1095 |
| chr4-130287279-DEL-53 | chr4-130287294-DEL-51 |
| chr4-130291368-DEL-265 | chr4-130291369-DEL-265 |
| chr4-130317634-DEL-203 | chr4-130317635-DEL-203 |
| chr4-130325732-DEL-52 | chr4-130325733-DEL-52 |
| chr4-130354129-DEL-208 | chr4-130354138-DEL-199 |
| chr4-130354868-DEL-136 | chr4-130354862-DEL-127 |
| chr4-130360114-DEL-201 | chr4-130360115-DEL-202 |
| chr4-130364544-DEL-222 | chr4-130364545-DEL-222 |
| chr4-130368745-DEL-171 | chr4-130368746-DEL-171 |
| chr4-130373462-DEL-182 | chr4-130373457-DEL-168 |
| chr4-130387779-DEL-52 | chr4-130387781-DEL-51 |
| chr4-130410591-DEL-210 | chr4-130410581-DEL-222 |
| chr4-130419406-DEL-120 | chr4-130419429-DEL-96 |
| chr4-130426955-DEL-208 | chr4-130426956-DEL-208 |
| chr4-130511754-DEL-188 | chr4-130511761-DEL-182 |
| chr4-130562650-DEL-80 | chr4-130562651-DEL-80 |
| chr4-130579724-DEL-400 | chr4-130579725-DEL-400 |
| chr4-130595053-DEL-88 | chr4-130595070-DEL-72 |
| chr4-130618069-DEL-132 | chr4-130618070-DEL-98 |
| chr4-130685109-DEL-113 | chr4-130685110-DEL-113 |
| chr4-130689657-DEL-569 | chr4-130689658-DEL-569 |
| chr4-130690461-DEL-97 | chr4-130690462-DEL-97 |
| chr4-130691174-DEL-150 | chr4-130691175-DEL-150 |

| | |
|---|---|
| chr4-130701998-DEL-58 | chr4-130701999-DEL-58 |
| chr4-130721020-DEL-208 | chr4-130721021-DEL-208 |
| chr4-130722724-DEL-82 | chr4-130722725-DEL-82 |
| chr4-130735035-DEL-56 | chr4-130735046-DEL-58 |
| chr4-130743989-DEL-64 | chr4-130743990-DEL-64 |
| chr4-130745301-DEL-153 | chr4-130745280-DEL-175 |
| chr4-130758197-DEL-132 | chr4-130758183-DEL-103 |
| chr4-130768143-DEL-1898 | chr4-130768144-DEL-1898 |
| chr4-130768449-DEL-56 | chr4-130768449-DEL-54 |
| chr4-130776838-DEL-1192 | chr4-130776839-DEL-1192 |
| chr4-130777750-DEL-156 | chr4-130777751-DEL-156 |
| chr4-130812094-DEL-83 | chr4-130812095-DEL-83 |
| chr4-130865798-DEL-270 | chr4-130865799-DEL-270 |
| chr4-130874492-DEL-183 | chr4-130874493-DEL-183 |
| chr4-130903523-DEL-55 | chr4-130903524-DEL-55 |
| chr4-130914231-DEL-54 | chr4-130914232-DEL-50 |
| chr4-130922441-DEL-154 | chr4-130922442-DEL-154 |
| chr4-130965303-DEL-59 | chr4-130965304-DEL-55 |
| chr4-130969320-DEL-209 | chr4-130969321-DEL-209 |
| chr4-130978139-DEL-285 | chr4-130978139-DEL-285 |
| chr4-130993048-DEL-314 | chr4-130993049-DEL-314 |
| chr4-130996584-DEL-60 | chr4-130996585-DEL-60 |
| chr4-131025329-DEL-682 | chr4-131025328-DEL-682 |
| chr4-131050966-DEL-62 | chr4-131050967-DEL-66 |
| chr4-131054582-DEL-54 | chr4-131054583-DEL-54 |
| chr4-131120329-DEL-88 | chr4-131120365-DEL-64 |
| chr4-131132369-DEL-64 | chr4-131132366-DEL-56 |
| chr4-131133685-DEL-108 | chr4-131133683-DEL-109 |
| chr4-131157934-DEL-194 | chr4-131157932-DEL-195 |
| chr4-131210976-DEL-81 | chr4-131210977-DEL-81 |
| chr4-131213146-DEL-109 | chr4-131213147-DEL-109 |
| chr4-131232347-DEL-110 | chr4-131232348-DEL-110 |
| chr4-131237742-DEL-75 | chr4-131237743-DEL-75 |
| chr4-131247060-DEL-401 | chr4-131247061-DEL-401 |
| chr4-131259031-DEL-118 | chr4-131259031-DEL-118 |
| chr4-131294176-DEL-104 | chr4-131294177-DEL-104 |
| chr4-131301983-DEL-464 | chr4-131301995-DEL-462 |
| chr4-131308423-DEL-53 | chr4-131308424-DEL-53 |
| chr4-131332785-DEL-62 | chr4-131332785-DEL-62 |
| chr4-131349876-DEL-156 | chr4-131349877-DEL-156 |
| chr4-131361511-DEL-101 | chr4-131361481-DEL-129 |
| chr4-131363615-DEL-70 | chr4-131363616-DEL-70 |
| chr4-131394770-DEL-152 | chr4-131394765-DEL-153 |
| chr4-131402220-DEL-60 | chr4-131402221-DEL-60 |
| chr4-131436849-DEL-176 | chr4-131436850-DEL-176 |
| chr4-131452201-DEL-199 | chr4-131452202-DEL-199 |
| chr4-131461911-DEL-111 | chr4-131461929-DEL-91 |
| chr4-131472723-DEL-636 | chr4-131472724-DEL-636 |
| chr4-131474641-DEL-117 | chr4-131474642-DEL-117 |
| chr4-131475617-DEL-61 | chr4-131475618-DEL-61 |
| chr4-131526377-DEL-59 | chr4-131526378-DEL-59 |
| chr4-131537568-DEL-64 | chr4-131537569-DEL-62 |
| chr4-131540994-DEL-203 | chr4-131540995-DEL-203 |
| chr4-131580191-DEL-74 | chr4-131580192-DEL-74 |
| chr4-131637990-DEL-53 | chr4-131637991-DEL-53 |
| chr4-131661932-DEL-64 | chr4-131661933-DEL-64 |
| chr4-131702364-DEL-95 | chr4-131702365-DEL-95 |
| chr4-131714193-DEL-636 | chr4-131714193-DEL-636 |
| chr4-131722021-DEL-70 | chr4-131722022-DEL-66 |
| chr4-131747927-DEL-107 | chr4-131747928-DEL-107 |
| chr4-131757272-DEL-141 | chr4-131757273-DEL-141 |
| chr4-131791137-DEL-125 | chr4-131791138-DEL-125 |
| chr4-131805251-DEL-78 | chr4-131805251-DEL-79 |
| chr4-131816063-DEL-90 | chr4-131816053-DEL-88 |
| chr4-131817349-DEL-668 | chr4-131817350-DEL-668 |
| chr4-131818449-DEL-54 | chr4-131818450-DEL-60 |
| chr4-131846816-DEL-69 | chr4-131846793-DEL-81 |
| chr4-131873103-DEL-200 | chr4-131873104-DEL-200 |
| chr4-131875298-DEL-114 | chr4-131875299-DEL-114 |
| chr4-131890695-DEL-205 | chr4-131890696-DEL-205 |
| chr4-131906199-DEL-72 | chr4-131906200-DEL-72 |
| chr4-131908022-DEL-128 | chr4-131908023-DEL-128 |
| chr4-131918329-DEL-217 | chr4-131918330-DEL-217 |
| chr4-131934231-DEL-96 | chr4-131934232-DEL-96 |
| chr4-131948175-DEL-59 | chr4-131948176-DEL-59 |
| chr4-131978158-DEL-171 | chr4-131978159-DEL-171 |
| chr4-131981803-DEL-82 | chr4-131981803-DEL-82 |
| chr4-131993068-DEL-389 | chr4-131993069-DEL-389 |
| chr4-132016395-DEL-111 | chr4-132016395-DEL-111 |
| chr4-132033659-DEL-216 | chr4-132033660-DEL-216 |

| | |
|---|---|
| chr4-132042694-DEL-212 | chr4-132042695-DEL-212 |
| chr4-132089850-DEL-218 | chr4-132089851-DEL-218 |
| chr4-132091453-DEL-198 | chr4-132091454-DEL-198 |
| chr4-132119127-DEL-171 | chr4-132119128-DEL-171 |
| chr4-132141311-DEL-264 | chr4-132141312-DEL-264 |
| chr4-132142619-DEL-96 | chr4-132142620-DEL-96 |
| chr4-132155946-DEL-202 | chr4-132155946-DEL-202 |
| chr4-132158390-DEL-51 | chr4-132158391-DEL-51 |
| chr4-132176289-DEL-469 | chr4-132176286-DEL-457 |
| chr4-132214245-DEL-209 | chr4-132214246-DEL-209 |
| chr4-132227544-DEL-191 | chr4-132227545-DEL-191 |
| chr4-132229536-DEL-172 | chr4-132229534-DEL-169 |
| chr4-132244335-DEL-291 | chr4-132244334-DEL-292 |
| chr4-132249117-DEL-70 | chr4-132249118-DEL-68 |
| chr4-132255957-DEL-157 | chr4-132255957-DEL-157 |
| chr4-132267896-DEL-220 | chr4-132267896-DEL-220 |
| chr4-132276016-DEL-147 | chr4-132276017-DEL-147 |
| chr4-132278485-DEL-168 | chr4-132278478-DEL-169 |
| chr4-132318722-DEL-383 | chr4-132318723-DEL-383 |
| chr4-132325889-DEL-390 | chr4-132325878-DEL-414 |
| chr4-132390104-DEL-198 | chr4-132390105-DEL-198 |
| chr4-132403669-DEL-70 | chr4-132403680-DEL-76 |
| chr4-132409685-DEL-234 | chr4-132409686-DEL-234 |
| chr4-132418218-DEL-1878 | chr4-132418219-DEL-1878 |
| chr4-132429298-DEL-111 | chr4-132429299-DEL-111 |
| chr4-132432314-DEL-1418 | chr4-132432315-DEL-1418 |
| chr4-132443281-DEL-230 | chr4-132443282-DEL-230 |
| chr4-132443555-DEL-1120 | chr4-132443574-DEL-1112 |
| chr4-132469413-DEL-220 | chr4-132469414-DEL-220 |
| chr4-132475643-DEL-233 | chr4-132475644-DEL-233 |
| chr4-132480915-DEL-72 | chr4-132480916-DEL-72 |
| chr4-132481453-DEL-178 | chr4-132481454-DEL-178 |
| chr4-132500651-DEL-197 | chr4-132500652-DEL-197 |
| chr4-132507380-DEL-188 | chr4-132507381-DEL-188 |
| chr4-132518793-DEL-200 | chr4-132518795-DEL-199 |
| chr4-132558247-DEL-194 | chr4-132558248-DEL-194 |
| chr4-132578785-DEL-207 | chr4-132578786-DEL-207 |
| chr4-132605776-DEL-246 | chr4-132605777-DEL-246 |
| chr4-132617174-DEL-91 | chr4-132617125-DEL-98 |
| chr4-132630049-DEL-608 | chr4-132630050-DEL-608 |
| chr4-132630870-DEL-206 | chr4-132630870-DEL-207 |
| chr4-132661009-DEL-207 | chr4-132661010-DEL-207 |
| chr4-132669760-DEL-334 | chr4-132669761-DEL-334 |
| chr4-132692954-DEL-1102 | chr4-132692955-DEL-1102 |
| chr4-132711851-DEL-51 | chr4-132711853-DEL-52 |
| chr4-132733605-DEL-177 | chr4-132733585-DEL-203 |
| chr4-132784296-DEL-202 | chr4-132784297-DEL-202 |
| chr4-132810644-DEL-146 | chr4-132810645-DEL-146 |
| chr4-132815712-DEL-472 | chr4-132815713-DEL-472 |
| chr4-132823263-DEL-5671 | chr4-132823264-DEL-5671 |
| chr4-132831987-DEL-60 | chr4-132831988-DEL-72 |
| chr4-132852215-DEL-180 | chr4-132852216-DEL-180 |
| chr4-132860790-DEL-192 | chr4-132860790-DEL-192 |
| chr4-132865300-DEL-52 | chr4-132865300-DEL-58 |
| chr4-132875402-DEL-114 | chr4-132875402-DEL-114 |
| chr4-132879143-DEL-187 | chr4-132879143-DEL-187 |
| chr4-132879964-DEL-109 | chr4-132879965-DEL-109 |
| chr4-132881727-DEL-235 | chr4-132881720-DEL-228 |
| chr4-132882334-DEL-326 | chr4-132882332-DEL-323 |
| chr4-132893045-DEL-217 | chr4-132893046-DEL-217 |
| chr4-132895613-DEL-186 | chr4-132895615-DEL-185 |
| chr4-132896172-DEL-88 | chr4-132896189-DEL-84 |
| chr4-132914605-DEL-109 | chr4-132914606-DEL-109 |
| chr4-132921332-DEL-53 | chr4-132921333-DEL-53 |
| chr4-132930014-DEL-56 | chr4-132930015-DEL-56 |
| chr4-132933813-DEL-335 | chr4-132933814-DEL-335 |
| chr4-132941416-DEL-2022 | chr4-132941393-DEL-2045 |
| chr4-132999441-DEL-62 | chr4-132999441-DEL-62 |
| chr4-133007725-DEL-189 | chr4-133007725-DEL-189 |
| chr4-133011281-DEL-97 | chr4-133011295-DEL-84 |
| chr4-133033431-DEL-203 | chr4-133033431-DEL-204 |
| chr4-133039943-DEL-202 | chr4-133039944-DEL-202 |
| chr4-133044949-DEL-81 | chr4-133044961-DEL-90 |
| chr4-133046089-DEL-604 | chr4-133046075-DEL-602 |
| chr4-133062245-DEL-68 | chr4-133062246-DEL-68 |
| chr4-133062379-DEL-2149 | chr4-133062380-DEL-2149 |
| chr4-133083858-DEL-99 | chr4-133083852-DEL-92 |
| chr4-133085565-DEL-65 | chr4-133085566-DEL-65 |
| chr4-133088037-DEL-3926 | chr4-133088038-DEL-3926 |
| chr4-133092419-DEL-62 | chr4-133092413-DEL-60 |

| | |
|---|---|
| chr4-133095105-DEL-87 | chr4-133095105-DEL-84 |
| chr4-133100229-DEL-316 | chr4-133100230-DEL-316 |
| chr4-133106816-DEL-52 | chr4-133106816-DEL-56 |
| chr4-133115474-DEL-74 | chr4-133115475-DEL-74 |
| chr4-133125998-DEL-898 | chr4-133125999-DEL-898 |
| chr4-133138468-DEL-190 | chr4-133138469-DEL-190 |
| chr4-133161722-DEL-52 | chr4-133161723-DEL-50 |
| chr4-133164950-DEL-92 | chr4-133164951-DEL-94 |
| chr4-133179603-DEL-88 | chr4-133179650-DEL-88 |
| chr4-133245048-DEL-168 | chr4-133245034-DEL-166 |
| chr4-133265097-DEL-83 | chr4-133265098-DEL-83 |
| chr4-133274705-DEL-218 | chr4-133274713-DEL-213 |
| chr4-133282415-DEL-98 | chr4-133282442-DEL-86 |
| chr4-133293324-DEL-80 | chr4-133293325-DEL-80 |
| chr4-133298009-DEL-575 | chr4-133298010-DEL-575 |
| chr4-133323320-DEL-122 | chr4-133323321-DEL-122 |
| chr4-133331881-DEL-186 | chr4-133331882-DEL-186 |
| chr4-133339317-DEL-568 | chr4-133339312-DEL-569 |
| chr4-133354734-DEL-216 | chr4-133354735-DEL-216 |
| chr4-133356128-DEL-56 | chr4-133356129-DEL-56 |
| chr4-133362675-DEL-604 | chr4-133362676-DEL-604 |
| chr4-133372731-DEL-152 | chr4-133372732-DEL-152 |
| chr4-133373157-DEL-822 | chr4-133373158-DEL-822 |
| chr4-133375205-DEL-187 | chr4-133375206-DEL-187 |
| chr4-133379868-DEL-54 | chr4-133379869-DEL-54 |
| chr4-133382174-DEL-228 | chr4-133382175-DEL-228 |
| chr4-133417384-DEL-58 | chr4-133417385-DEL-58 |
| chr4-133425125-DEL-399 | chr4-133425126-DEL-399 |
| chr4-133464081-DEL-200 | chr4-133464082-DEL-186 |
| chr4-133490008-DEL-228 | chr4-133490017-DEL-225 |
| chr4-133492604-DEL-55 | chr4-133492598-DEL-56 |
| chr4-133500364-DEL-175 | chr4-133500365-DEL-175 |
| chr4-133505207-DEL-103 | chr4-133505208-DEL-100 |
| chr4-133508704-DEL-258 | chr4-133508704-DEL-259 |
| chr4-133518516-DEL-189 | chr4-133518517-DEL-189 |
| chr4-133525262-DEL-5154 | chr4-133525263-DEL-5154 |
| chr4-133527802-DEL-54 | chr4-133527800-DEL-55 |
| chr4-133530636-DEL-78 | chr4-133530639-DEL-81 |
| chr4-133535954-DEL-229 | chr4-133535955-DEL-229 |
| chr4-133539547-DEL-200 | chr4-133539548-DEL-200 |
| chr4-133551971-DEL-303 | chr4-133551972-DEL-303 |
| chr4-133558163-DEL-124 | chr4-133558175-DEL-118 |
| chr4-133558442-DEL-789 | chr4-133558443-DEL-789 |
| chr4-133560483-DEL-1443 | chr4-133560484-DEL-1443 |
| chr4-133562853-DEL-217 | chr4-133562852-DEL-171 |
| chr4-133566037-DEL-98 | chr4-133566037-DEL-53 |
| chr4-133569712-DEL-191 | chr4-133569713-DEL-191 |
| chr4-133576415-DEL-198 | chr4-133576416-DEL-197 |
| chr4-133579067-DEL-199 | chr4-133579065-DEL-202 |
| chr4-133585687-DEL-178 | chr4-133585688-DEL-156 |
| chr4-133590286-DEL-201 | chr4-133590287-DEL-201 |
| chr4-133627181-DEL-431 | chr4-133627182-DEL-431 |
| chr4-133630136-DEL-165 | chr4-133630137-DEL-165 |
| chr4-133642657-DEL-92 | chr4-133642658-DEL-92 |
| chr4-133644887-DEL-3241 | chr4-133644888-DEL-3241 |
| chr4-133654219-DEL-184 | chr4-133654220-DEL-184 |
| chr4-133655183-DEL-96 | chr4-133655184-DEL-106 |
| chr4-133657200-DEL-842 | chr4-133657201-DEL-842 |
| chr4-133663580-DEL-206 | chr4-133663581-DEL-172 |
| chr4-133664557-DEL-304 | chr4-133664558-DEL-304 |
| chr4-133670725-DEL-189 | chr4-133670726-DEL-181 |
| chr4-133677174-DEL-437 | chr4-133677175-DEL-437 |
| chr4-133711116-DEL-215 | chr4-133711117-DEL-215 |
| chr4-133724857-DEL-213 | chr4-133724858-DEL-213 |
| chr4-133734593-DEL-84 | chr4-133734596-DEL-82 |
| chr4-133775830-DEL-78 | chr4-133775831-DEL-78 |
| chr4-133778663-DEL-64 | chr4-133778672-DEL-56 |
| chr4-133793465-DEL-58 | chr4-133793446-DEL-68 |
| chr4-133811795-DEL-201 | chr4-133811796-DEL-201 |
| chr4-133827262-DEL-104 | chr4-133827263-DEL-104 |
| chr4-133839541-DEL-66 | chr4-133839542-DEL-66 |
| chr4-133876805-DEL-204 | chr4-133876806-DEL-204 |
| chr4-133886696-DEL-5316 | chr4-133886697-DEL-5316 |
| chr4-133917246-DEL-189 | chr4-133917283-DEL-174 |
| chr4-133923680-DEL-52 | chr4-133923681-DEL-52 |
| chr4-133966613-DEL-222 | chr4-133966617-DEL-221 |
| chr4-133967568-DEL-213 | chr4-133967569-DEL-213 |
| chr4-133994864-DEL-195 | chr4-133994879-DEL-182 |
| chr4-134024868-DEL-61 | chr4-134024869-DEL-61 |
| chr4-134030382-DEL-685 | chr4-134030383-DEL-685 |

| | |
|---|---|
| chr4-134032818-DEL-91 | chr4-134032817-DEL-56 |
| chr4-134040872-DEL-199 | chr4-134040873-DEL-199 |
| chr4-134047982-DEL-153 | chr4-134047978-DEL-158 |
| chr4-134052378-DEL-136 | chr4-134052379-DEL-136 |
| chr4-134057941-DEL-1321 | chr4-134057942-DEL-1321 |
| chr4-134065329-DEL-204 | chr4-134065330-DEL-204 |
| chr4-134067408-DEL-171 | chr4-134067399-DEL-172 |
| chr4-134069453-DEL-84 | chr4-134069447-DEL-70 |
| chr4-134102751-DEL-80 | chr4-134102751-DEL-80 |
| chr4-134111227-DEL-3111 | chr4-134111228-DEL-3110 |
| chr4-134118564-DEL-477 | chr4-134118565-DEL-477 |
| chr4-134120192-DEL-51 | chr4-134120193-DEL-51 |
| chr4-134120910-DEL-60 | chr4-134120911-DEL-60 |
| chr4-134121224-DEL-1318 | chr4-134121225-DEL-1318 |
| chr4-134125394-DEL-154 | chr4-134125395-DEL-154 |
| chr4-134137243-DEL-306 | chr4-134137244-DEL-306 |
| chr4-134172620-DEL-65 | chr4-134172620-DEL-50 |
| chr4-134175901-DEL-97 | chr4-134175902-DEL-97 |
| chr4-134209418-DEL-63 | chr4-134209418-DEL-63 |
| chr4-134226356-DEL-52 | chr4-134226357-DEL-52 |
| chr4-134246298-DEL-207 | chr4-134246299-DEL-207 |
| chr4-134247496-DEL-252 | chr4-134247497-DEL-252 |
| chr4-134281692-DEL-212 | chr4-134281691-DEL-214 |
| chr4-134294634-DEL-112 | chr4-134294625-DEL-113 |
| chr4-134300948-DEL-191 | chr4-134300949-DEL-191 |
| chr4-134322922-DEL-203 | chr4-134322923-DEL-203 |
| chr4-134350847-DEL-156 | chr4-134350850-DEL-154 |
| chr4-134408745-DEL-232 | chr4-134408739-DEL-233 |
| chr4-134418095-DEL-196 | chr4-134418096-DEL-196 |
| chr4-134438439-DEL-63 | chr4-134438399-DEL-60 |
| chr4-134441901-DEL-223 | chr4-134441902-DEL-226 |
| chr4-134446374-DEL-220 | chr4-134446375-DEL-220 |
| chr4-134521534-DEL-201 | chr4-134521536-DEL-200 |
| chr4-134523531-DEL-59 | chr4-134523538-DEL-53 |
| chr4-134526241-DEL-66 | chr4-134526241-DEL-66 |
| chr4-134569787-DEL-209 | chr4-134569788-DEL-209 |
| chr4-134598598-DEL-196 | chr4-134598607-DEL-191 |
| chr4-134602612-DEL-195 | chr4-134602595-DEL-200 |
| chr4-134605012-DEL-202 | chr4-134605009-DEL-199 |
| chr4-134612899-DEL-102 | chr4-134612891-DEL-109 |
| chr4-134638421-DEL-507 | chr4-134638413-DEL-508 |
| chr4-134666561-DEL-214 | chr4-134666562-DEL-214 |
| chr4-134678908-DEL-189 | chr4-134678879-DEL-190 |
| chr4-134686619-DEL-85 | chr4-134686645-DEL-66 |
| chr4-134755119-DEL-61 | chr4-134755120-DEL-59 |
| chr4-134760157-DEL-69 | chr4-134760181-DEL-52 |
| chr4-134770280-DEL-54 | chr4-134770281-DEL-54 |
| chr4-134780739-DEL-66 | chr4-134780740-DEL-66 |
| chr4-134787593-DEL-65 | chr4-134787592-DEL-67 |
| chr4-134791299-DEL-91 | chr4-134791300-DEL-91 |
| chr4-134813941-DEL-71 | chr4-134813942-DEL-71 |
| chr4-134821350-DEL-819 | chr4-134821351-DEL-819 |
| chr4-134837598-DEL-71 | chr4-134837599-DEL-71 |
| chr4-134897310-DEL-166 | chr4-134897311-DEL-167 |
| chr4-134911799-DEL-4291 | chr4-134911800-DEL-4291 |
| chr4-134917777-DEL-101 | chr4-134917746-DEL-86 |
| chr4-134983556-DEL-201 | chr4-134983557-DEL-201 |
| chr4-134987082-DEL-565 | chr4-134987083-DEL-565 |
| chr4-135008783-DEL-74 | chr4-135008784-DEL-74 |
| chr4-135011131-DEL-160 | chr4-135011133-DEL-152 |
| chr4-135020367-DEL-107 | chr4-135020368-DEL-107 |
| chr4-135075367-DEL-223 | chr4-135075368-DEL-223 |
| chr4-135094555-DEL-473 | chr4-135094556-DEL-473 |
| chr4-135099564-DEL-50 | chr4-135099565-DEL-50 |
| chr4-135101194-DEL-108 | chr4-135101222-DEL-81 |
| chr4-135104564-DEL-641 | chr4-135104565-DEL-641 |
| chr4-135105986-DEL-86 | chr4-135105987-DEL-86 |
| chr4-135126646-DEL-201 | chr4-135126647-DEL-201 |
| chr4-135134329-DEL-72 | chr4-135134330-DEL-72 |
| chr4-135171758-DEL-176 | chr4-135171759-DEL-176 |
| chr4-135174764-DEL-192 | chr4-135174765-DEL-192 |
| chr4-135177429-DEL-116 | chr4-135177430-DEL-116 |
| chr4-135183108-DEL-7080 | chr4-135183109-DEL-7080 |
| chr4-135192417-DEL-482 | chr4-135192418-DEL-482 |
| chr4-135202653-DEL-57 | chr4-135202654-DEL-57 |
| chr4-135213367-DEL-135 | chr4-135213368-DEL-135 |
| chr4-135282543-DEL-66 | chr4-135282544-DEL-66 |
| chr4-135284035-DEL-51 | chr4-135284035-DEL-51 |
| chr4-135342359-DEL-57 | chr4-135342377-DEL-52 |
| chr4-135343108-DEL-79 | chr4-135343109-DEL-79 |

| | |
|---|---|
| chr4-135360415-DEL-6306 | chr4-135360416-DEL-6306 |
| chr4-135364276-DEL-122 | chr4-135364277-DEL-122 |
| chr4-135366929-DEL-203 | chr4-135366930-DEL-203 |
| chr4-135381632-DEL-2064 | chr4-135381633-DEL-2064 |
| chr4-135388108-DEL-55 | chr4-135388109-DEL-55 |
| chr4-135391360-DEL-92 | chr4-135391361-DEL-87 |
| chr4-135443003-DEL-1005 | chr4-135443004-DEL-1005 |
| chr4-135450252-DEL-790 | chr4-135450253-DEL-790 |
| chr4-135495701-DEL-56 | chr4-135495702-DEL-56 |
| chr4-135513648-DEL-121 | chr4-135513634-DEL-122 |
| chr4-135540375-DEL-99 | chr4-135540376-DEL-99 |
| chr4-135547043-DEL-77 | chr4-135547044-DEL-77 |
| chr4-135562292-DEL-207 | chr4-135562293-DEL-207 |
| chr4-135563475-DEL-1684 | chr4-135563485-DEL-1675 |
| chr4-135570173-DEL-179 | chr4-135570174-DEL-179 |
| chr4-135597517-DEL-79 | chr4-135597518-DEL-87 |
| chr4-135602643-DEL-52 | chr4-135602643-DEL-52 |
| chr4-135606817-DEL-460 | chr4-135606818-DEL-460 |
| chr4-135614527-DEL-416 | chr4-135614528-DEL-416 |
| chr4-135668696-DEL-150 | chr4-135668697-DEL-150 |
| chr4-135680852-DEL-235 | chr4-135680853-DEL-235 |
| chr4-135704943-DEL-92 | chr4-135704944-DEL-92 |
| chr4-135707104-DEL-172 | chr4-135707105-DEL-172 |
| chr4-135721156-DEL-96 | chr4-135721157-DEL-96 |
| chr4-135735784-DEL-203 | chr4-135735785-DEL-202 |
| chr4-135739464-DEL-209 | chr4-135739465-DEL-209 |
| chr4-135753984-DEL-260 | chr4-135753985-DEL-260 |
| chr4-135763403-DEL-215 | chr4-135763404-DEL-215 |
| chr4-135769135-DEL-53 | chr4-135769136-DEL-53 |
| chr4-135785015-DEL-1475 | chr4-135785016-DEL-1475 |
| chr4-135787701-DEL-159 | chr4-135787704-DEL-157 |
| chr4-135791819-DEL-214 | chr4-135791820-DEL-214 |
| chr4-135799367-DEL-599 | chr4-135799368-DEL-599 |
| chr4-135800217-DEL-1607 | chr4-135800218-DEL-1607 |
| chr4-135807615-DEL-72 | chr4-135807616-DEL-72 |
| chr4-135815320-DEL-132 | chr4-135815317-DEL-136 |
| chr4-135831533-DEL-56 | chr4-135831534-DEL-56 |
| chr4-135834583-DEL-221 | chr4-135834584-DEL-221 |
| chr4-135835405-DEL-490 | chr4-135835402-DEL-488 |
| chr4-135840632-DEL-197 | chr4-135840633-DEL-196 |
| chr4-135841385-DEL-173 | chr4-135841386-DEL-173 |
| chr4-135855423-DEL-926 | chr4-135855424-DEL-926 |
| chr4-135875535-DEL-128 | chr4-135875536-DEL-128 |
| chr4-135877107-DEL-73 | chr4-135877108-DEL-73 |
| chr4-135931725-DEL-205 | chr4-135931726-DEL-205 |
| chr4-135949917-DEL-94 | chr4-135949918-DEL-94 |
| chr4-135954376-DEL-230 | chr4-135954377-DEL-230 |
| chr4-135958180-DEL-73 | chr4-135958180-DEL-73 |
| chr4-135960170-DEL-234 | chr4-135960171-DEL-234 |
| chr4-135962897-DEL-194 | chr4-135962898-DEL-194 |
| chr4-135978224-DEL-131 | chr4-135978225-DEL-131 |
| chr4-135991984-DEL-51 | chr4-135991985-DEL-51 |
| chr4-136007098-DEL-406 | chr4-136007099-DEL-406 |
| chr4-136023967-DEL-4561 | chr4-136023968-DEL-4561 |
| chr4-136027878-DEL-153 | chr4-136027886-DEL-146 |
| chr4-136061016-DEL-202 | chr4-136061017-DEL-202 |
| chr4-136092939-DEL-77 | chr4-136092940-DEL-87 |
| chr4-136099645-DEL-124 | chr4-136099647-DEL-127 |
| chr4-136101830-DEL-56 | chr4-136101830-DEL-56 |
| chr4-136102530-DEL-987 | chr4-136102531-DEL-987 |
| chr4-136112269-DEL-208 | chr4-136112270-DEL-208 |
| chr4-136115602-DEL-106 | chr4-136115606-DEL-102 |
| chr4-136120564-DEL-85 | chr4-136120565-DEL-85 |
| chr4-136164481-DEL-76 | chr4-136164478-DEL-60 |
| chr4-136180008-DEL-177 | chr4-136180006-DEL-180 |
| chr4-136187728-DEL-214 | chr4-136187729-DEL-214 |
| chr4-136194120-DEL-52 | chr4-136194121-DEL-57 |
| chr4-136207671-DEL-98 | chr4-136207666-DEL-104 |
| chr4-136208497-DEL-185 | chr4-136208489-DEL-186 |
| chr4-136220237-DEL-203 | chr4-136220238-DEL-203 |
| chr4-136223726-DEL-172 | chr4-136223726-DEL-173 |
| chr4-136225083-DEL-964 | chr4-136225084-DEL-964 |
| chr4-136250011-DEL-68 | chr4-136250011-DEL-68 |
| chr4-136282720-DEL-231 | chr4-136282721-DEL-231 |
| chr4-136307680-DEL-74 | chr4-136307675-DEL-85 |
| chr4-136323531-DEL-76 | chr4-136323536-DEL-72 |
| chr4-136332920-DEL-157 | chr4-136332921-DEL-157 |
| chr4-136335523-DEL-178 | chr4-136335524-DEL-178 |
| chr4-136338505-DEL-128 | chr4-136338506-DEL-128 |
| chr4-136347202-DEL-259 | chr4-136347212-DEL-251 |

| | |
|---|---|
| chr4-136356045-DEL-62 | chr4-136356046-DEL-62 |
| chr4-136359214-DEL-163 | chr4-136359215-DEL-163 |
| chr4-136366700-DEL-119 | chr4-136366701-DEL-119 |
| chr4-136396853-DEL-60 | chr4-136396854-DEL-60 |
| chr4-136425522-DEL-129 | chr4-136425506-DEL-104 |
| chr4-136450020-DEL-61 | chr4-136450021-DEL-61 |
| chr4-136450854-DEL-108 | chr4-136450864-DEL-83 |
| chr4-136466350-DEL-388 | chr4-136466351-DEL-388 |
| chr4-136487296-DEL-191 | chr4-136487297-DEL-191 |
| chr4-136489796-DEL-62 | chr4-136489797-DEL-62 |
| chr4-136512462-DEL-213 | chr4-136512463-DEL-213 |
| chr4-136539517-DEL-666 | chr4-136539518-DEL-666 |
| chr4-136598722-DEL-51 | chr4-136598716-DEL-57 |
| chr4-136615782-DEL-184 | chr4-136615783-DEL-184 |
| chr4-136625901-DEL-94 | chr4-136625900-DEL-93 |
| chr4-136691811-DEL-755 | chr4-136691812-DEL-755 |
| chr4-136715516-DEL-213 | chr4-136715517-DEL-213 |
| chr4-136739522-DEL-250 | chr4-136739523-DEL-250 |
| chr4-136748493-DEL-79 | chr4-136748494-DEL-67 |
| chr4-136782009-DEL-242 | chr4-136782010-DEL-242 |
| chr4-136796695-DEL-62 | chr4-136796696-DEL-62 |
| chr4-136802531-DEL-472 | chr4-136802532-DEL-472 |
| chr4-136819882-DEL-132 | chr4-136819883-DEL-132 |
| chr4-136820319-DEL-401 | chr4-136820320-DEL-401 |
| chr4-136847729-DEL-70 | chr4-136847732-DEL-64 |
| chr4-136866072-DEL-79 | chr4-136866073-DEL-79 |
| chr4-136873460-DEL-71 | chr4-136873465-DEL-67 |
| chr4-136876627-DEL-112 | chr4-136876624-DEL-100 |
| chr4-136878727-DEL-68 | chr4-136878705-DEL-69 |
| chr4-136897430-DEL-52 | chr4-136897431-DEL-52 |
| chr4-136906976-DEL-54 | chr4-136906977-DEL-50 |
| chr4-136911342-DEL-72 | chr4-136911331-DEL-66 |
| chr4-136923514-DEL-107 | chr4-136923511-DEL-111 |
| chr4-136943078-DEL-78 | chr4-136943073-DEL-94 |
| chr4-136999691-DEL-66 | chr4-136999692-DEL-66 |
| chr4-137012634-DEL-145 | chr4-137012635-DEL-145 |
| chr4-137029712-DEL-815 | chr4-137029713-DEL-815 |
| chr4-137052528-DEL-71 | chr4-137052529-DEL-71 |
| chr4-137066497-DEL-65 | chr4-137066462-DEL-60 |
| chr4-137073042-DEL-98 | chr4-137073043-DEL-98 |
| chr4-137090849-DEL-168 | chr4-137090850-DEL-165 |
| chr4-137108788-DEL-5512 | chr4-137108789-DEL-5512 |
| chr4-137125571-DEL-190 | chr4-137125572-DEL-178 |
| chr4-137132524-DEL-56 | chr4-137132525-DEL-56 |
| chr4-137134205-DEL-91 | chr4-137134209-DEL-92 |
| chr4-137135781-DEL-78 | chr4-137135782-DEL-78 |
| chr4-137145319-DEL-90 | chr4-137145320-DEL-90 |
| chr4-137153050-DEL-56 | chr4-137153035-DEL-72 |
| chr4-137161031-DEL-51 | chr4-137161031-DEL-78 |
| chr4-137168945-DEL-77 | chr4-137168963-DEL-77 |
| chr4-137170457-DEL-103 | chr4-137170461-DEL-101 |
| chr4-137188467-DEL-68 | chr4-137188468-DEL-68 |
| chr4-137194039-DEL-52 | chr4-137194040-DEL-52 |
| chr4-137205309-DEL-290 | chr4-137205310-DEL-290 |
| chr4-137236554-DEL-66 | chr4-137236555-DEL-66 |
| chr4-137250647-DEL-164 | chr4-137250649-DEL-160 |
| chr4-137288095-DEL-147 | chr4-137288088-DEL-132 |
| chr4-137313329-DEL-171 | chr4-137313330-DEL-171 |
| chr4-137325518-DEL-445 | chr4-137325519-DEL-445 |
| chr4-137328902-DEL-212 | chr4-137328903-DEL-212 |
| chr4-137333429-DEL-174 | chr4-137333430-DEL-174 |
| chr4-137388807-DEL-74 | chr4-137388808-DEL-74 |
| chr4-137396264-DEL-214 | chr4-137396265-DEL-214 |
| chr4-137397421-DEL-248 | chr4-137397425-DEL-245 |
| chr4-137398651-DEL-85 | chr4-137398652-DEL-85 |
| chr4-137421664-DEL-164 | chr4-137421665-DEL-164 |
| chr4-137422383-DEL-59 | chr4-137422399-DEL-66 |
| chr4-137429204-DEL-57 | chr4-137429202-DEL-63 |
| chr4-137442116-DEL-70 | chr4-137442117-DEL-70 |
| chr4-137461385-DEL-89 | chr4-137461386-DEL-90 |
| chr4-137487420-DEL-195 | chr4-137487424-DEL-179 |
| chr4-137488088-DEL-57 | chr4-137488107-DEL-51 |
| chr4-137503313-DEL-78 | chr4-137503299-DEL-97 |
| chr4-137515024-DEL-53 | chr4-137515025-DEL-52 |
| chr4-137517153-DEL-60 | chr4-137517133-DEL-84 |
| chr4-137550255-DEL-68 | chr4-137550256-DEL-68 |
| chr4-137564603-DEL-55 | chr4-137564604-DEL-55 |
| chr4-137565681-DEL-428 | chr4-137565682-DEL-428 |
| chr4-137568239-DEL-57 | chr4-137568201-DEL-85 |
| chr4-137569228-DEL-153 | chr4-137569256-DEL-137 |

| | |
|---|---|
| chr4-137582850-DEL-53 | chr4-137582822-DEL-66 |
| chr4-137586173-DEL-109 | chr4-137586172-DEL-110 |
| chr4-137606981-DEL-53 | chr4-137606980-DEL-53 |
| chr4-137607600-DEL-188 | chr4-137607601-DEL-188 |
| chr4-137637207-DEL-284 | chr4-137637208-DEL-284 |
| chr4-137646515-DEL-55 | chr4-137646516-DEL-55 |
| chr4-137668934-DEL-62 | chr4-137668935-DEL-62 |
| chr4-137676359-DEL-140 | chr4-137676359-DEL-136 |
| chr4-137679976-DEL-68 | chr4-137679977-DEL-68 |
| chr4-137684626-DEL-98 | chr4-137684627-DEL-98 |
| chr4-137712535-DEL-734 | chr4-137712536-DEL-738 |
| chr4-137713848-DEL-2345 | chr4-137713849-DEL-2345 |
| chr4-137724808-DEL-208 | chr4-137724798-DEL-209 |
| chr4-137735275-DEL-332 | chr4-137735241-DEL-335 |
| chr4-137741284-DEL-231 | chr4-137741285-DEL-231 |
| chr4-137746834-DEL-53 | chr4-137746837-DEL-51 |
| chr4-137749620-DEL-785 | chr4-137749621-DEL-785 |
| chr4-137751661-DEL-100 | chr4-137751649-DEL-108 |
| chr4-137757009-DEL-259 | chr4-137757010-DEL-259 |
| chr4-137770696-DEL-1013 | chr4-137770697-DEL-1013 |
| chr4-137786028-DEL-1313 | chr4-137786029-DEL-1313 |
| chr4-137787594-DEL-208 | chr4-137787595-DEL-208 |
| chr4-137790026-DEL-84 | chr4-137790027-DEL-84 |
| chr4-137849086-DEL-419 | chr4-137849087-DEL-418 |
| chr4-137855491-DEL-216 | chr4-137855492-DEL-216 |
| chr4-137858163-DEL-51 | chr4-137858164-DEL-51 |
| chr4-137861217-DEL-58 | chr4-137861218-DEL-58 |
| chr4-137917108-DEL-138 | chr4-137917108-DEL-132 |
| chr4-137924875-DEL-78 | chr4-137924876-DEL-78 |
| chr4-137928688-DEL-209 | chr4-137928689-DEL-209 |
| chr4-137930171-DEL-151 | chr4-137930172-DEL-151 |
| chr4-137958391-DEL-96 | chr4-137958392-DEL-96 |
| chr4-137966034-DEL-166 | chr4-137966035-DEL-166 |
| chr4-138003267-DEL-202 | chr4-138003268-DEL-202 |
| chr4-138009101-DEL-64 | chr4-138009102-DEL-64 |
| chr4-138024933-DEL-153 | chr4-138024943-DEL-118 |
| chr4-138037259-DEL-100 | chr4-138037264-DEL-96 |
| chr4-138045690-DEL-69 | chr4-138045691-DEL-69 |
| chr4-138054690-DEL-479 | chr4-138054691-DEL-479 |
| chr4-138085141-DEL-51 | chr4-138085142-DEL-51 |
| chr4-138103456-DEL-56 | chr4-138103456-DEL-56 |
| chr4-138110783-DEL-154 | chr4-138110784-DEL-152 |
| chr4-138128300-DEL-200 | chr4-138128301-DEL-200 |
| chr4-138130595-DEL-200 | chr4-138130596-DEL-200 |
| chr4-138135073-DEL-179 | chr4-138135074-DEL-179 |
| chr4-138144792-DEL-62 | chr4-138144793-DEL-59 |
| chr4-138147706-DEL-171 | chr4-138147749-DEL-159 |
| chr4-138153036-DEL-369 | chr4-138153037-DEL-369 |
| chr4-138153976-DEL-70 | chr4-138153977-DEL-70 |
| chr4-138167369-DEL-63 | chr4-138167370-DEL-63 |
| chr4-138170642-DEL-58 | chr4-138170643-DEL-58 |
| chr4-138205132-DEL-72 | chr4-138205145-DEL-56 |
| chr4-138214066-DEL-77 | chr4-138214073-DEL-71 |
| chr4-138227947-DEL-92 | chr4-138227948-DEL-84 |
| chr4-138229881-DEL-203 | chr4-138229876-DEL-243 |
| chr4-138229891-DEL-713 | chr4-138229876-DEL-741 |
| chr4-138242499-DEL-92 | chr4-138242500-DEL-92 |
| chr4-138268955-DEL-137 | chr4-138268950-DEL-143 |
| chr4-138272613-DEL-53 | chr4-138272614-DEL-53 |
| chr4-138273509-DEL-54 | chr4-138273510-DEL-50 |
| chr4-138284553-DEL-362 | chr4-138284555-DEL-360 |
| chr4-138314653-DEL-52 | chr4-138314654-DEL-54 |
| chr4-138322919-DEL-80 | chr4-138322917-DEL-82 |
| chr4-138343490-DEL-71 | chr4-138343490-DEL-71 |
| chr4-138348423-DEL-201 | chr4-138348424-DEL-201 |
| chr4-138428812-DEL-538 | chr4-138428813-DEL-538 |
| chr4-138441266-DEL-58 | chr4-138441263-DEL-62 |
| chr4-138486859-DEL-70 | chr4-138486853-DEL-76 |
| chr4-138512916-DEL-56 | chr4-138512919-DEL-54 |
| chr4-138570080-DEL-384 | chr4-138570081-DEL-384 |
| chr4-138600622-DEL-1480 | chr4-138600623-DEL-1480 |
| chr4-138616386-DEL-289 | chr4-138616386-DEL-287 |
| chr4-138639785-DEL-364 | chr4-138639786-DEL-364 |
| chr4-138651867-DEL-1226 | chr4-138651846-DEL-1214 |
| chr4-138656398-DEL-72 | chr4-138656402-DEL-72 |
| chr4-138679489-DEL-399 | chr4-138679490-DEL-399 |
| chr4-138716519-DEL-108 | chr4-138716537-DEL-90 |
| chr4-138730385-DEL-108 | chr4-138730386-DEL-108 |
| chr4-138739029-DEL-121 | chr4-138739003-DEL-119 |
| chr4-138744529-DEL-51 | chr4-138744536-DEL-55 |

| | |
|---|---|
| chr4-138777295-DEL-232 | chr4-138777296-DEL-232 |
| chr4-138784493-DEL-52 | chr4-138784496-DEL-50 |
| chr4-138790435-DEL-75 | chr4-138790399-DEL-76 |
| chr4-138798246-DEL-78 | chr4-138798251-DEL-79 |
| chr4-138807375-DEL-124 | chr4-138807381-DEL-121 |
| chr4-138840567-DEL-2513 | chr4-138840568-DEL-2513 |
| chr4-138852641-DEL-181 | chr4-138852642-DEL-181 |
| chr4-138860697-DEL-196 | chr4-138860698-DEL-196 |
| chr4-138861316-DEL-215 | chr4-138861317-DEL-219 |
| chr4-138866675-DEL-188 | chr4-138866660-DEL-204 |
| chr4-138879263-DEL-197 | chr4-138879251-DEL-198 |
| chr4-138884361-DEL-173 | chr4-138884362-DEL-173 |
| chr4-138899066-DEL-66 | chr4-138899067-DEL-66 |
| chr4-138899420-DEL-1269 | chr4-138899421-DEL-1269 |
| chr4-139031062-DEL-65 | chr4-139031063-DEL-65 |
| chr4-139042115-DEL-1586 | chr4-139042115-DEL-1586 |
| chr4-139046005-DEL-194 | chr4-139046006-DEL-194 |
| chr4-139049602-DEL-59 | chr4-139049599-DEL-58 |
| chr4-139052004-DEL-1398 | chr4-139051955-DEL-1399 |
| chr4-139057669-DEL-225 | chr4-139057670-DEL-225 |
| chr4-139060348-DEL-874 | chr4-139060349-DEL-874 |
| chr4-139065413-DEL-342 | chr4-139065414-DEL-342 |
| chr4-139080883-DEL-167 | chr4-139080882-DEL-168 |
| chr4-139082711-DEL-199 | chr4-139082712-DEL-199 |
| chr4-139104557-DEL-70 | chr4-139104558-DEL-70 |
| chr4-139123198-DEL-52 | chr4-139123198-DEL-52 |
| chr4-139251904-DEL-52 | chr4-139251905-DEL-52 |
| chr4-139262162-DEL-56 | chr4-139262184-DEL-56 |
| chr4-139267650-DEL-213 | chr4-139267651-DEL-213 |
| chr4-139268994-DEL-115 | chr4-139268995-DEL-115 |
| chr4-139286648-DEL-52 | chr4-139286649-DEL-52 |
| chr4-139288786-DEL-67 | chr4-139288824-DEL-57 |
| chr4-139297981-DEL-55 | chr4-139297979-DEL-56 |
| chr4-139298642-DEL-208 | chr4-139298643-DEL-208 |
| chr4-139354680-DEL-83 | chr4-139354681-DEL-83 |
| chr4-139384961-DEL-400 | chr4-139384962-DEL-400 |
| chr4-139390573-DEL-71 | chr4-139390611-DEL-55 |
| chr4-139411421-DEL-68 | chr4-139411422-DEL-68 |
| chr4-139416147-DEL-62 | chr4-139416148-DEL-62 |
| chr4-139424626-DEL-64 | chr4-139424579-DEL-112 |
| chr4-139432302-DEL-90 | chr4-139432303-DEL-90 |
| chr4-139444698-DEL-96 | chr4-139444698-DEL-96 |
| chr4-139462126-DEL-397 | chr4-139462127-DEL-397 |
| chr4-139508488-DEL-76 | chr4-139508489-DEL-76 |
| chr4-139539808-DEL-132 | chr4-139539809-DEL-132 |
| chr4-139564511-DEL-68 | chr4-139564502-DEL-68 |
| chr4-139598196-DEL-57 | chr4-139598183-DEL-58 |
| chr4-139631284-DEL-171 | chr4-139631285-DEL-171 |
| chr4-139631680-DEL-507 | chr4-139631639-DEL-511 |
| chr4-139641525-DEL-58 | chr4-139641520-DEL-61 |
| chr4-139651141-DEL-98 | chr4-139651142-DEL-98 |
| chr4-139662526-DEL-4218 | chr4-139662527-DEL-4218 |
| chr4-139662625-DEL-165 | chr4-139662626-DEL-165 |
| chr4-139668595-DEL-145 | chr4-139668628-DEL-113 |
| chr4-139673791-DEL-268 | chr4-139673792-DEL-268 |
| chr4-139678155-DEL-76 | chr4-139678156-DEL-76 |
| chr4-139680153-DEL-5501 | chr4-139680154-DEL-5501 |
| chr4-139688684-DEL-438 | chr4-139688685-DEL-438 |
| chr4-139696197-DEL-58 | chr4-139696198-DEL-58 |
| chr4-139711428-DEL-78 | chr4-139711427-DEL-54 |
| chr4-139728959-DEL-85 | chr4-139728960-DEL-85 |
| chr4-139768852-DEL-72 | chr4-139768849-DEL-64 |
| chr4-139814376-DEL-85 | chr4-139814397-DEL-105 |
| chr4-139833863-DEL-56 | chr4-139833864-DEL-56 |
| chr4-139839497-DEL-74 | chr4-139839498-DEL-74 |
| chr4-139849407-DEL-389 | chr4-139849403-DEL-390 |
| chr4-139869348-DEL-98 | chr4-139869356-DEL-91 |
| chr4-139874075-DEL-82 | chr4-139874076-DEL-82 |
| chr4-139883622-DEL-237 | chr4-139883623-DEL-237 |
| chr4-139885848-DEL-56 | chr4-139885860-DEL-52 |
| chr4-139892359-DEL-98 | chr4-139892360-DEL-98 |
| chr4-139897604-DEL-687 | chr4-139897605-DEL-687 |
| chr4-139989426-DEL-123 | chr4-139989427-DEL-123 |
| chr4-139994164-DEL-52 | chr4-139994165-DEL-62 |
| chr4-140037697-DEL-68 | chr4-140037698-DEL-68 |
| chr4-140045309-DEL-63 | chr4-140045310-DEL-63 |
| chr4-140064769-DEL-328 | chr4-140064770-DEL-328 |
| chr4-140119331-DEL-72 | chr4-140119332-DEL-72 |
| chr4-140138468-DEL-335 | chr4-140138467-DEL-336 |
| chr4-140144893-DEL-59 | chr4-140144873-DEL-52 |

| | |
|---|---|
| chr4-140164270-DEL-381 | chr4-140164271-DEL-364 |
| chr4-140181049-DEL-148 | chr4-140181046-DEL-152 |
| chr4-140207953-DEL-63 | chr4-140207954-DEL-63 |
| chr4-140208966-DEL-170 | chr4-140208967-DEL-170 |
| chr4-140210286-DEL-2066 | chr4-140210287-DEL-2063 |
| chr4-140214595-DEL-64 | chr4-140214596-DEL-64 |
| chr4-140220139-DEL-57 | chr4-140220140-DEL-57 |
| chr4-140222582-DEL-132 | chr4-140222582-DEL-132 |
| chr4-140226861-DEL-53 | chr4-140226867-DEL-60 |
| chr4-140232186-DEL-52 | chr4-140232187-DEL-52 |
| chr4-140235967-DEL-1182 | chr4-140236000-DEL-1150 |
| chr4-140244579-DEL-100 | chr4-140244610-DEL-100 |
| chr4-140252847-DEL-89 | chr4-140252848-DEL-89 |
| chr4-140258608-DEL-53 | chr4-140258609-DEL-53 |
| chr4-140273552-DEL-51 | chr4-140273535-DEL-56 |
| chr4-140282017-DEL-54 | chr4-140282018-DEL-54 |
| chr4-140286234-DEL-188 | chr4-140286235-DEL-188 |
| chr4-140287756-DEL-128 | chr4-140287755-DEL-128 |
| chr4-140293631-DEL-66 | chr4-140293647-DEL-65 |
| chr4-140302351-DEL-156 | chr4-140302352-DEL-156 |
| chr4-140333546-DEL-55 | chr4-140333547-DEL-55 |
| chr4-140345702-DEL-92 | chr4-140345696-DEL-99 |
| chr4-140346701-DEL-195 | chr4-140346702-DEL-195 |
| chr4-140355182-DEL-88 | chr4-140355172-DEL-126 |
| chr4-140372445-DEL-61 | chr4-140372445-DEL-61 |
| chr4-140376923-DEL-92 | chr4-140376920-DEL-91 |
| chr4-140378987-DEL-92 | chr4-140378985-DEL-78 |
| chr4-140380255-DEL-177 | chr4-140380256-DEL-177 |
| chr4-140381288-DEL-75 | chr4-140381289-DEL-75 |
| chr4-140394589-DEL-97 | chr4-140394589-DEL-97 |
| chr4-140396421-DEL-51 | chr4-140396470-DEL-50 |
| chr4-140399107-DEL-532 | chr4-140399114-DEL-526 |
| chr4-140400347-DEL-202 | chr4-140400370-DEL-202 |
| chr4-140402695-DEL-172 | chr4-140402694-DEL-168 |
| chr4-140405676-DEL-660 | chr4-140405677-DEL-660 |
| chr4-140405724-DEL-51 | chr4-140405725-DEL-51 |
| chr4-140410715-DEL-201 | chr4-140410716-DEL-201 |
| chr4-140419621-DEL-275 | chr4-140419622-DEL-275 |
| chr4-140424416-DEL-59 | chr4-140424434-DEL-53 |
| chr4-140433822-DEL-263 | chr4-140433823-DEL-263 |
| chr4-140442245-DEL-208 | chr4-140442246-DEL-208 |
| chr4-140444667-DEL-208 | chr4-140444668-DEL-208 |
| chr4-140562158-DEL-54 | chr4-140562159-DEL-52 |
| chr4-140588881-DEL-464 | chr4-140588882-DEL-464 |
| chr4-140593005-DEL-57 | chr4-140593006-DEL-57 |
| chr4-140594989-DEL-54 | chr4-140594989-DEL-54 |
| chr4-140615591-DEL-52 | chr4-140615592-DEL-52 |
| chr4-140615991-DEL-2792 | chr4-140615992-DEL-2792 |
| chr4-140624723-DEL-755 | chr4-140624718-DEL-756 |
| chr4-140659726-DEL-147 | chr4-140659727-DEL-147 |
| chr4-140666319-DEL-4828 | chr4-140666320-DEL-4828 |
| chr4-140674733-DEL-65 | chr4-140674705-DEL-111 |
| chr4-140682598-DEL-92 | chr4-140682599-DEL-92 |
| chr4-140690916-DEL-209 | chr4-140690917-DEL-209 |
| chr4-140701409-DEL-64 | chr4-140701410-DEL-64 |
| chr4-140705895-DEL-112 | chr4-140705896-DEL-106 |
| chr4-140725697-DEL-202 | chr4-140725706-DEL-203 |
| chr4-140734654-DEL-440 | chr4-140734655-DEL-440 |
| chr4-140771667-DEL-72 | chr4-140771666-DEL-62 |
| chr4-140772344-DEL-61 | chr4-140772345-DEL-61 |
| chr4-140785315-DEL-73 | chr4-140785266-DEL-73 |
| chr4-140813238-DEL-255 | chr4-140813239-DEL-255 |
| chr4-140820588-DEL-60 | chr4-140820595-DEL-54 |
| chr4-140848827-DEL-52 | chr4-140848828-DEL-52 |
| chr4-140856178-DEL-360 | chr4-140856190-DEL-359 |
| chr4-140857455-DEL-239 | chr4-140857456-DEL-239 |
| chr4-140867457-DEL-263 | chr4-140867458-DEL-263 |
| chr4-140872491-DEL-280 | chr4-140872492-DEL-280 |
| chr4-140882209-DEL-1495 | chr4-140882210-DEL-1495 |
| chr4-140887123-DEL-370 | chr4-140887124-DEL-370 |
| chr4-140904943-DEL-68 | chr4-140904948-DEL-60 |
| chr4-140925031-DEL-58 | chr4-140925018-DEL-76 |
| chr4-140933439-DEL-175 | chr4-140933463-DEL-140 |
| chr4-140934906-DEL-2996 | chr4-140934907-DEL-2996 |
| chr4-140944105-DEL-93 | chr4-140944106-DEL-93 |
| chr4-140944780-DEL-211 | chr4-140944770-DEL-212 |
| chr4-140956729-DEL-584 | chr4-140956730-DEL-584 |
| chr4-140994010-DEL-75 | chr4-140994011-DEL-75 |
| chr4-140996120-DEL-90 | chr4-140996121-DEL-70 |
| chr4-140998344-DEL-164 | chr4-140998344-DEL-164 |

| | |
|---|---|
| chr4-141007016-DEL-87 | chr4-141007016-DEL-87 |
| chr4-141010386-DEL-173 | chr4-141010387-DEL-173 |
| chr4-141013669-DEL-157 | chr4-141013652-DEL-178 |
| chr4-141069521-DEL-72 | chr4-141069538-DEL-56 |
| chr4-141073192-DEL-232 | chr4-141073193-DEL-232 |
| chr4-141105577-DEL-654 | chr4-141105578-DEL-654 |
| chr4-141114177-DEL-71 | chr4-141114197-DEL-70 |
| chr4-141119663-DEL-112 | chr4-141119665-DEL-111 |
| chr4-141129666-DEL-58 | chr4-141129667-DEL-58 |
| chr4-141137320-DEL-201 | chr4-141137321-DEL-201 |
| chr4-141140371-DEL-58 | chr4-141140371-DEL-59 |
| chr4-141144095-DEL-52 | chr4-141144095-DEL-52 |
| chr4-141163816-DEL-1659 | chr4-141163817-DEL-1659 |
| chr4-141169454-DEL-68 | chr4-141169455-DEL-68 |
| chr4-141184226-DEL-190 | chr4-141184233-DEL-192 |
| chr4-141188071-DEL-67 | chr4-141188072-DEL-67 |
| chr4-141211745-DEL-473 | chr4-141211746-DEL-473 |
| chr4-141220430-DEL-100 | chr4-141220430-DEL-100 |
| chr4-141222572-DEL-163 | chr4-141222573-DEL-163 |
| chr4-141227955-DEL-58 | chr4-141227954-DEL-56 |
| chr4-141232248-DEL-422 | chr4-141232249-DEL-422 |
| chr4-141239609-DEL-105 | chr4-141239611-DEL-104 |
| chr4-141246610-DEL-66 | chr4-141246611-DEL-62 |
| chr4-141281806-DEL-188 | chr4-141281807-DEL-188 |
| chr4-141313451-DEL-75 | chr4-141313452-DEL-75 |
| chr4-141315003-DEL-64 | chr4-141315015-DEL-52 |
| chr4-141317852-DEL-181 | chr4-141317853-DEL-181 |
| chr4-141323892-DEL-51 | chr4-141323893-DEL-51 |
| chr4-141334603-DEL-138 | chr4-141334604-DEL-138 |
| chr4-141360110-DEL-581 | chr4-141360111-DEL-575 |
| chr4-141365304-DEL-249 | chr4-141365305-DEL-249 |
| chr4-141365851-DEL-59 | chr4-141365854-DEL-56 |
| chr4-141372082-DEL-197 | chr4-141372067-DEL-194 |
| chr4-141385420-DEL-204 | chr4-141385422-DEL-216 |
| chr4-141399226-DEL-91 | chr4-141399226-DEL-90 |
| chr4-141414841-DEL-405 | chr4-141414842-DEL-405 |
| chr4-141416735-DEL-51 | chr4-141416736-DEL-51 |
| chr4-141417291-DEL-166 | chr4-141417293-DEL-165 |
| chr4-141435161-DEL-66 | chr4-141435162-DEL-66 |
| chr4-141439174-DEL-75 | chr4-141439175-DEL-75 |
| chr4-141466656-DEL-242 | chr4-141466657-DEL-242 |
| chr4-141472881-DEL-818 | chr4-141472882-DEL-818 |
| chr4-141480755-DEL-310 | chr4-141480756-DEL-310 |
| chr4-141484937-DEL-152 | chr4-141484938-DEL-152 |
| chr4-141492132-DEL-1628 | chr4-141492133-DEL-1628 |
| chr4-141492700-DEL-175 | chr4-141492701-DEL-175 |
| chr4-141494123-DEL-68 | chr4-141494073-DEL-69 |
| chr4-141525773-DEL-4762 | chr4-141525774-DEL-4760 |
| chr4-141550976-DEL-823 | chr4-141550976-DEL-823 |
| chr4-141556381-DEL-78 | chr4-141556382-DEL-78 |
| chr4-141559905-DEL-526 | chr4-141559906-DEL-526 |
| chr4-141563089-DEL-182 | chr4-141563090-DEL-182 |
| chr4-141565728-DEL-64 | chr4-141565729-DEL-64 |
| chr4-141568187-DEL-175 | chr4-141568187-DEL-175 |
| chr4-141576932-DEL-3622 | chr4-141576927-DEL-3623 |
| chr4-141582010-DEL-208 | chr4-141582010-DEL-208 |
| chr4-141590139-DEL-59 | chr4-141590139-DEL-57 |
| chr4-141612426-DEL-117 | chr4-141612427-DEL-116 |
| chr4-141614905-DEL-90 | chr4-141614882-DEL-104 |
| chr4-141640026-DEL-512 | chr4-141640027-DEL-512 |
| chr4-141658522-DEL-62 | chr4-141658538-DEL-71 |
| chr4-141660929-DEL-202 | chr4-141660930-DEL-202 |
| chr4-141661943-DEL-1596 | chr4-141661944-DEL-1596 |
| chr4-141663778-DEL-186 | chr4-141663792-DEL-186 |
| chr4-141665235-DEL-113 | chr4-141665257-DEL-98 |
| chr4-141667602-DEL-172 | chr4-141667603-DEL-172 |
| chr4-141698620-DEL-199 | chr4-141698621-DEL-199 |
| chr4-141705253-DEL-66 | chr4-141705224-DEL-58 |
| chr4-141752227-DEL-195 | chr4-141752219-DEL-196 |
| chr4-141752800-DEL-60 | chr4-141752801-DEL-60 |
| chr4-141769895-DEL-329 | chr4-141769896-DEL-329 |
| chr4-141782427-DEL-206 | chr4-141782428-DEL-206 |
| chr4-141797187-DEL-129 | chr4-141797207-DEL-139 |
| chr4-141806781-DEL-55 | chr4-141806781-DEL-55 |
| chr4-141818937-DEL-121 | chr4-141818938-DEL-121 |
| chr4-141821015-DEL-230 | chr4-141821016-DEL-230 |
| chr4-141850122-DEL-85 | chr4-141850122-DEL-85 |
| chr4-141853737-DEL-144 | chr4-141853737-DEL-144 |
| chr4-141883620-DEL-71 | chr4-141883621-DEL-71 |
| chr4-141887751-DEL-92 | chr4-141887752-DEL-92 |

| | |
|---|---|
| chr4-141958145-DEL-64 | chr4-141958148-DEL-62 |
| chr4-141985783-DEL-73 | chr4-141985788-DEL-69 |
| chr4-141997557-DEL-213 | chr4-141997556-DEL-213 |
| chr4-141998672-DEL-60 | chr4-141998658-DEL-76 |
| chr4-142001738-DEL-175 | chr4-142001782-DEL-175 |
| chr4-142017424-DEL-51 | chr4-142017412-DEL-52 |
| chr4-142049817-DEL-882 | chr4-142049816-DEL-882 |
| chr4-142051111-DEL-306 | chr4-142051112-DEL-306 |
| chr4-142057443-DEL-427 | chr4-142057444-DEL-427 |
| chr4-142076271-DEL-2614 | chr4-142076276-DEL-2612 |
| chr4-142084066-DEL-1610 | chr4-142084067-DEL-1610 |
| chr4-142087680-DEL-52 | chr4-142087709-DEL-50 |
| chr4-142121077-DEL-189 | chr4-142121078-DEL-189 |
| chr4-142150916-DEL-75 | chr4-142150917-DEL-75 |
| chr4-142159776-DEL-58 | chr4-142159735-DEL-97 |
| chr4-142160804-DEL-66 | chr4-142160805-DEL-66 |
| chr4-142177372-DEL-242 | chr4-142177369-DEL-246 |
| chr4-142178785-DEL-2088 | chr4-142178786-DEL-2088 |
| chr4-142181192-DEL-58 | chr4-142181187-DEL-64 |
| chr4-142241808-DEL-193 | chr4-142241809-DEL-193 |
| chr4-142258790-DEL-95 | chr4-142258774-DEL-94 |
| chr4-142282694-DEL-3232 | chr4-142282695-DEL-3232 |
| chr4-142288395-DEL-797 | chr4-142288396-DEL-797 |
| chr4-142289865-DEL-55 | chr4-142289866-DEL-55 |
| chr4-142320601-DEL-872 | chr4-142320602-DEL-872 |
| chr4-142327768-DEL-84 | chr4-142327818-DEL-64 |
| chr4-142338567-DEL-509 | chr4-142338568-DEL-509 |
| chr4-142341123-DEL-51 | chr4-142341124-DEL-51 |
| chr4-142344105-DEL-52 | chr4-142344106-DEL-52 |
| chr4-142363796-DEL-485 | chr4-142363798-DEL-477 |
| chr4-142384348-DEL-58 | chr4-142384349-DEL-58 |
| chr4-142417640-DEL-199 | chr4-142417641-DEL-190 |
| chr4-142436355-DEL-60 | chr4-142436356-DEL-60 |
| chr4-142460452-DEL-358 | chr4-142460453-DEL-358 |
| chr4-142487835-DEL-84 | chr4-142487836-DEL-75 |
| chr4-142506052-DEL-1384 | chr4-142506053-DEL-1384 |
| chr4-142508197-DEL-143 | chr4-142508198-DEL-143 |
| chr4-142512722-DEL-469 | chr4-142512723-DEL-469 |
| chr4-142527953-DEL-2027 | chr4-142527954-DEL-2027 |
| chr4-142532840-DEL-1583 | chr4-142532841-DEL-1583 |
| chr4-142584970-DEL-75 | chr4-142584971-DEL-75 |
| chr4-142636476-DEL-812 | chr4-142636477-DEL-812 |
| chr4-142647994-DEL-216 | chr4-142647993-DEL-208 |
| chr4-142653765-DEL-52 | chr4-142653766-DEL-52 |
| chr4-142662962-DEL-6380 | chr4-142662963-DEL-6380 |
| chr4-142681993-DEL-113 | chr4-142681994-DEL-113 |
| chr4-142697264-DEL-63 | chr4-142697265-DEL-63 |
| chr4-142704014-DEL-2559 | chr4-142704015-DEL-2559 |
| chr4-142709558-DEL-66 | chr4-142709559-DEL-66 |
| chr4-142719087-DEL-115 | chr4-142719115-DEL-80 |
| chr4-142736509-DEL-76 | chr4-142736511-DEL-75 |
| chr4-142745626-DEL-57 | chr4-142745620-DEL-63 |
| chr4-142760939-DEL-146 | chr4-142760939-DEL-145 |
| chr4-142761639-DEL-66 | chr4-142761623-DEL-58 |
| chr4-142766198-DEL-1644 | chr4-142766199-DEL-1644 |
| chr4-142770066-DEL-56 | chr4-142770067-DEL-56 |
| chr4-142774546-DEL-198 | chr4-142774547-DEL-198 |
| chr4-142788867-DEL-93 | chr4-142788868-DEL-91 |
| chr4-142806565-DEL-76 | chr4-142806566-DEL-74 |
| chr4-142809917-DEL-292 | chr4-142809914-DEL-309 |
| chr4-142811136-DEL-980 | chr4-142811137-DEL-979 |
| chr4-142865927-DEL-123 | chr4-142865928-DEL-123 |
| chr4-142899587-DEL-180 | chr4-142899588-DEL-180 |
| chr4-142922446-DEL-151 | chr4-142922441-DEL-127 |
| chr4-142923169-DEL-179 | chr4-142923170-DEL-179 |
| chr4-142943122-DEL-191 | chr4-142943121-DEL-186 |
| chr4-142944395-DEL-488 | chr4-142944396-DEL-488 |
| chr4-142944989-DEL-55 | chr4-142944989-DEL-55 |
| chr4-142948134-DEL-54 | chr4-142948132-DEL-57 |
| chr4-142968129-DEL-1832 | chr4-142968130-DEL-1832 |
| chr4-142971182-DEL-809 | chr4-142971183-DEL-809 |
| chr4-142983981-DEL-56 | chr4-142983982-DEL-56 |
| chr4-142984310-DEL-274 | chr4-142984311-DEL-274 |
| chr4-142992315-DEL-171 | chr4-142992316-DEL-171 |
| chr4-143019443-DEL-218 | chr4-143019444-DEL-218 |
| chr4-143030346-DEL-216 | chr4-143030347-DEL-216 |
| chr4-143036628-DEL-955 | chr4-143036629-DEL-955 |
| chr4-143038328-DEL-777 | chr4-143038329-DEL-777 |
| chr4-143048348-DEL-94 | chr4-143048349-DEL-94 |
| chr4-143089847-DEL-192 | chr4-143089848-DEL-192 |

| | |
|---|---|
| chr4-143107672-DEL-60 | chr4-143107669-DEL-60 |
| chr4-143113920-DEL-229 | chr4-143113917-DEL-240 |
| chr4-143118499-DEL-102 | chr4-143118500-DEL-102 |
| chr4-143126893-DEL-77 | chr4-143126886-DEL-77 |
| chr4-143150400-DEL-102 | chr4-143150401-DEL-102 |
| chr4-143175703-DEL-67 | chr4-143175717-DEL-56 |
| chr4-143185195-DEL-135 | chr4-143185196-DEL-115 |
| chr4-143189608-DEL-53 | chr4-143189609-DEL-53 |
| chr4-143201268-DEL-6390 | chr4-143201269-DEL-6390 |
| chr4-143238891-DEL-365 | chr4-143238893-DEL-363 |
| chr4-143256673-DEL-56 | chr4-143256673-DEL-72 |
| chr4-143267780-DEL-8892 | chr4-143267779-DEL-8892 |
| chr4-143331853-DEL-55 | chr4-143331853-DEL-55 |
| chr4-143337299-DEL-1433 | chr4-143337299-DEL-1433 |
| chr4-143349332-DEL-379 | chr4-143349333-DEL-379 |
| chr4-143371443-DEL-7194 | chr4-143371444-DEL-7194 |
| chr4-143433924-DEL-74 | chr4-143433926-DEL-101 |
| chr4-143459548-DEL-57 | chr4-143459549-DEL-57 |
| chr4-143488792-DEL-58 | chr4-143488798-DEL-51 |
| chr4-143493397-DEL-911 | chr4-143493398-DEL-911 |
| chr4-143517795-DEL-64 | chr4-143517794-DEL-66 |
| chr4-143574796-DEL-13360 | chr4-143574797-DEL-13360 |
| chr4-143657186-DEL-3495 | chr4-143657187-DEL-3495 |
| chr4-143657956-DEL-53 | chr4-143657957-DEL-53 |
| chr4-143679499-DEL-76 | chr4-143679510-DEL-81 |
| chr4-143688679-DEL-91 | chr4-143688679-DEL-91 |
| chr4-143716567-DEL-60 | chr4-143716568-DEL-60 |
| chr4-143732777-DEL-6442 | chr4-143732777-DEL-6442 |
| chr4-143755187-DEL-179 | chr4-143755188-DEL-179 |
| chr4-143766823-DEL-177 | chr4-143766824-DEL-177 |
| chr4-143770874-DEL-207 | chr4-143770875-DEL-207 |
| chr4-143780445-DEL-468 | chr4-143780446-DEL-468 |
| chr4-143784262-DEL-177 | chr4-143784263-DEL-177 |
| chr4-143898639-DEL-6285 | chr4-143898640-DEL-6285 |
| chr4-143950009-DEL-301 | chr4-143950009-DEL-298 |
| chr4-143954654-DEL-196 | chr4-143954650-DEL-197 |
| chr4-143955879-DEL-6347 | chr4-143955879-DEL-6347 |
| chr4-143974332-DEL-1019 | chr4-143974333-DEL-1019 |
| chr4-143993331-DEL-248 | chr4-143993332-DEL-246 |
| chr4-144016918-DEL-53 | chr4-144016938-DEL-54 |
| chr4-144040081-DEL-88 | chr4-144040082-DEL-80 |
| chr4-144155916-DEL-470 | chr4-144155917-DEL-470 |
| chr4-144207718-DEL-328 | chr4-144207719-DEL-328 |
| chr4-144215570-DEL-65 | chr4-144215553-DEL-96 |
| chr4-144220887-DEL-389 | chr4-144220888-DEL-390 |
| chr4-144230993-DEL-337 | chr4-144230994-DEL-337 |
| chr4-144262089-DEL-635 | chr4-144262090-DEL-635 |
| chr4-144265136-DEL-252 | chr4-144265137-DEL-252 |
| chr4-144271771-DEL-67 | chr4-144271771-DEL-67 |
| chr4-144278690-DEL-65 | chr4-144278684-DEL-71 |
| chr4-144307685-DEL-89 | chr4-144307701-DEL-59 |
| chr4-144312776-DEL-2765 | chr4-144312793-DEL-2765 |
| chr4-144348957-DEL-179 | chr4-144348958-DEL-179 |
| chr4-144350413-DEL-96 | chr4-144350429-DEL-81 |
| chr4-144359826-DEL-1893 | chr4-144359827-DEL-1893 |
| chr4-144384709-DEL-63 | chr4-144384701-DEL-72 |
| chr4-144434443-DEL-683 | chr4-144434444-DEL-683 |
| chr4-144448821-DEL-1031 | chr4-144448820-DEL-1033 |
| chr4-144451734-DEL-2514 | chr4-144451735-DEL-2514 |
| chr4-144455767-DEL-67 | chr4-144455768-DEL-67 |
| chr4-144464453-DEL-595 | chr4-144464454-DEL-592 |
| chr4-144465880-DEL-220 | chr4-144465881-DEL-220 |
| chr4-144493042-DEL-105 | chr4-144493041-DEL-66 |
| chr4-144518755-DEL-233 | chr4-144518756-DEL-233 |
| chr4-144526119-DEL-612 | chr4-144526120-DEL-612 |
| chr4-144542100-DEL-1073 | chr4-144542101-DEL-1073 |
| chr4-144543248-DEL-6541 | chr4-144543249-DEL-6541 |
| chr4-144586271-DEL-165 | chr4-144586272-DEL-165 |
| chr4-144601086-DEL-60 | chr4-144601087-DEL-60 |
| chr4-144622356-DEL-258 | chr4-144622357-DEL-258 |
| chr4-144639137-DEL-72 | chr4-144639154-DEL-60 |
| chr4-144652105-DEL-52 | chr4-144652101-DEL-52 |
| chr4-144662227-DEL-55 | chr4-144662227-DEL-52 |
| chr4-144673895-DEL-68 | chr4-144673898-DEL-66 |
| chr4-144689994-DEL-67 | chr4-144689951-DEL-111 |
| chr4-144692373-DEL-215 | chr4-144692374-DEL-215 |
| chr4-144693011-DEL-493 | chr4-144693012-DEL-493 |
| chr4-144775748-DEL-152 | chr4-144775749-DEL-152 |
| chr4-144843559-DEL-64 | chr4-144843560-DEL-60 |
| chr4-144856082-DEL-159 | chr4-144856088-DEL-149 |

| | |
|---|---|
| chr4-144869526-DEL-181 | chr4-144869526-DEL-134 |
| chr4-144880491-DEL-106 | chr4-144880510-DEL-88 |
| chr4-144892411-DEL-56 | chr4-144892412-DEL-56 |
| chr4-144893405-DEL-91 | chr4-144893416-DEL-98 |
| chr4-144958830-DEL-645 | chr4-144958831-DEL-645 |
| chr4-144988956-DEL-408 | chr4-144988957-DEL-408 |
| chr4-145010689-DEL-60 | chr4-145010689-DEL-60 |
| chr4-145016612-DEL-104 | chr4-145016613-DEL-104 |
| chr4-145032393-DEL-71 | chr4-145032394-DEL-71 |
| chr4-145052841-DEL-191 | chr4-145052842-DEL-191 |
| chr4-145060566-DEL-62 | chr4-145060567-DEL-62 |
| chr4-145072445-DEL-54 | chr4-145072438-DEL-62 |
| chr4-145075442-DEL-62 | chr4-145075443-DEL-62 |
| chr4-145088467-DEL-76 | chr4-145088476-DEL-68 |
| chr4-145099526-DEL-139 | chr4-145099527-DEL-139 |
| chr4-145104914-DEL-53 | chr4-145104915-DEL-53 |
| chr4-145109597-DEL-99 | chr4-145109616-DEL-81 |
| chr4-145124817-DEL-2869 | chr4-145124817-DEL-2869 |
| chr4-145151111-DEL-1815 | chr4-145151110-DEL-1815 |
| chr4-145208053-DEL-205 | chr4-145208053-DEL-205 |
| chr4-145218628-DEL-363 | chr4-145218628-DEL-363 |
| chr4-145252163-DEL-87 | chr4-145252169-DEL-69 |
| chr4-145265575-DEL-2913 | chr4-145265574-DEL-2913 |
| chr4-145311952-DEL-178 | chr4-145311952-DEL-178 |
| chr4-145317800-DEL-411 | chr4-145317799-DEL-411 |
| chr4-145321294-DEL-58 | chr4-145321278-DEL-65 |
| chr4-145396091-DEL-56 | chr4-145396091-DEL-56 |
| chr4-145455769-DEL-1172 | chr4-145455780-DEL-1171 |
| chr4-145468412-DEL-214 | chr4-145468411-DEL-214 |
| chr4-145576669-DEL-169 | chr4-145576669-DEL-169 |
| chr4-145998460-DEL-338 | chr4-145998459-DEL-338 |
| chr4-146115686-DEL-210 | chr4-146115680-DEL-211 |
| chr4-146176003-DEL-73 | chr4-146176002-DEL-73 |
| chr4-146177025-DEL-222 | chr4-146177025-DEL-222 |
| chr4-146210395-DEL-169 | chr4-146210396-DEL-169 |
| chr4-146275586-DEL-169 | chr4-146275586-DEL-169 |
| chr4-146279205-DEL-831 | chr4-146279210-DEL-839 |
| chr4-146308561-DEL-233 | chr4-146308562-DEL-233 |
| chr4-146426129-DEL-197 | chr4-146426116-DEL-209 |
| chr4-146469444-DEL-519 | chr4-146469444-DEL-519 |
| chr4-146474237-DEL-58 | chr4-146474237-DEL-64 |
| chr4-146478583-DEL-222 | chr4-146478583-DEL-222 |
| chr4-146486020-DEL-272 | chr4-146486020-DEL-272 |
| chr4-146489441-DEL-204 | chr4-146489442-DEL-204 |
| chr4-146493349-DEL-212 | chr4-146493350-DEL-212 |
| chr4-146515758-DEL-183 | chr4-146515759-DEL-183 |
| chr4-146535738-DEL-76 | chr4-146535761-DEL-52 |
| chr4-146586021-DEL-225 | chr4-146586020-DEL-225 |
| chr4-146598996-DEL-132 | chr4-146598996-DEL-132 |
| chr4-146617253-DEL-209 | chr4-146617254-DEL-209 |
| chr4-146631013-DEL-575 | chr4-146631021-DEL-566 |
| chr4-146983765-DEL-1209 | chr4-146983764-DEL-1209 |
| chr4-146988796-DEL-209 | chr4-146988787-DEL-207 |
| chr4-146990762-DEL-869 | chr4-146990780-DEL-870 |
| chr4-147014460-DEL-944 | chr4-147014460-DEL-944 |
| chr4-147019883-DEL-161 | chr4-147019884-DEL-161 |
| chr4-147052199-DEL-428 | chr4-147052199-DEL-428 |
| chr4-147090187-DEL-327 | chr4-147090195-DEL-328 |
| chr4-147105868-DEL-198 | chr4-147105883-DEL-186 |
| chr4-147170428-DEL-1175 | chr4-147170427-DEL-1175 |
| chr4-147190279-DEL-55 | chr4-147190252-DEL-55 |
| chr4-147204552-DEL-181 | chr4-147204552-DEL-181 |
| chr4-147209091-DEL-96 | chr4-147209091-DEL-96 |
| chr4-147219971-DEL-170 | chr4-147219972-DEL-170 |
| chr4-147386968-DEL-123 | chr4-147386957-DEL-150 |
| chr4-147409052-DEL-70 | chr4-147409042-DEL-68 |
| chr4-147434342-DEL-55 | chr4-147434328-DEL-55 |
| chr4-147443747-DEL-687 | chr4-147443764-DEL-671 |
| chr4-147445321-DEL-202 | chr4-147445302-DEL-210 |
| chr4-147473229-DEL-216 | chr4-147473230-DEL-216 |
| chr4-147490771-DEL-1281 | chr4-147490771-DEL-1281 |
| chr4-147503242-DEL-509 | chr4-147503242-DEL-509 |
| chr4-147605875-DEL-61 | chr4-147605874-DEL-61 |
| chr4-147635263-DEL-159 | chr4-147635264-DEL-158 |
| chr4-147651932-DEL-201 | chr4-147651933-DEL-201 |
| chr4-147657604-DEL-210 | chr4-147657604-DEL-210 |
| chr4-147672007-DEL-6298 | chr4-147672008-DEL-6298 |
| chr4-147727208-DEL-199 | chr4-147727197-DEL-211 |
| chr4-147728244-DEL-707 | chr4-147728249-DEL-671 |
| chr4-147835890-DEL-229 | chr4-147835891-DEL-229 |

| | |
|---|---|
| chr4-147878375-DEL-211 | chr4-147878374-DEL-211 |
| chr4-147891923-DEL-168 | chr4-147891893-DEL-213 |
| chr4-147899283-DEL-104 | chr4-147899284-DEL-68 |
| chr4-147900554-DEL-237 | chr4-147900555-DEL-237 |
| chr4-147901189-DEL-115 | chr4-147901189-DEL-115 |
| chr4-147909369-DEL-175 | chr4-147909370-DEL-175 |
| chr4-147915043-DEL-63 | chr4-147915044-DEL-63 |
| chr4-147917951-DEL-7163 | chr4-147917952-DEL-7163 |
| chr4-147938167-DEL-226 | chr4-147938168-DEL-226 |
| chr4-147939758-DEL-204 | chr4-147939759-DEL-204 |
| chr4-147940086-DEL-609 | chr4-147940084-DEL-612 |
| chr4-147944118-DEL-290 | chr4-147944119-DEL-290 |
| chr4-147945013-DEL-206 | chr4-147945014-DEL-205 |
| chr4-147954745-DEL-1342 | chr4-147954746-DEL-1342 |
| chr4-147965188-DEL-311 | chr4-147965191-DEL-313 |
| chr4-147973903-DEL-214 | chr4-147973904-DEL-214 |
| chr4-147977089-DEL-119 | chr4-147977090-DEL-111 |
| chr4-147996709-DEL-202 | chr4-147996710-DEL-202 |
| chr4-148006858-DEL-71 | chr4-148006859-DEL-71 |
| chr4-148026822-DEL-87 | chr4-148026823-DEL-87 |
| chr4-148034970-DEL-184 | chr4-148034971-DEL-184 |
| chr4-148038654-DEL-171 | chr4-148038655-DEL-171 |
| chr4-148039547-DEL-170 | chr4-148039571-DEL-185 |
| chr4-148075775-DEL-124 | chr4-148075774-DEL-124 |
| chr4-148080302-DEL-232 | chr4-148080303-DEL-226 |
| chr4-148088002-DEL-62 | chr4-148088003-DEL-62 |
| chr4-148110177-DEL-212 | chr4-148110178-DEL-212 |
| chr4-148113414-DEL-78 | chr4-148113417-DEL-76 |
| chr4-148129785-DEL-57 | chr4-148129786-DEL-57 |
| chr4-148140072-DEL-52 | chr4-148140073-DEL-52 |
| chr4-148174896-DEL-201 | chr4-148174897-DEL-201 |
| chr4-148177937-DEL-223 | chr4-148177923-DEL-225 |
| chr4-148187918-DEL-202 | chr4-148187919-DEL-202 |
| chr4-148192049-DEL-112 | chr4-148192050-DEL-112 |
| chr4-148204612-DEL-79 | chr4-148204613-DEL-79 |
| chr4-148215872-DEL-56 | chr4-148215873-DEL-62 |
| chr4-148223610-DEL-52 | chr4-148223611-DEL-52 |
| chr4-148251326-DEL-52 | chr4-148251329-DEL-50 |
| chr4-148267941-DEL-70 | chr4-148267942-DEL-60 |
| chr4-148300948-DEL-86 | chr4-148300949-DEL-70 |
| chr4-148308686-DEL-85 | chr4-148308687-DEL-73 |
| chr4-148310295-DEL-85 | chr4-148310296-DEL-85 |
| chr4-148333861-DEL-97 | chr4-148333862-DEL-97 |
| chr4-148346052-DEL-63 | chr4-148346053-DEL-63 |
| chr4-148352193-DEL-347 | chr4-148352202-DEL-339 |
| chr4-148379113-DEL-327 | chr4-148379114-DEL-327 |
| chr4-148400960-DEL-493 | chr4-148400961-DEL-493 |
| chr4-148404453-DEL-52 | chr4-148404454-DEL-52 |
| chr4-148407367-DEL-56 | chr4-148407368-DEL-52 |
| chr4-148436777-DEL-112 | chr4-148436778-DEL-112 |
| chr4-148439957-DEL-199 | chr4-148439958-DEL-199 |
| chr4-148444341-DEL-67 | chr4-148444342-DEL-67 |
| chr4-148466677-DEL-55 | chr4-148466697-DEL-50 |
| chr4-148487171-DEL-105 | chr4-148487130-DEL-122 |
| chr4-148491237-DEL-124 | chr4-148491238-DEL-124 |
| chr4-148508466-DEL-94 | chr4-148508467-DEL-94 |
| chr4-148591369-DEL-52 | chr4-148591370-DEL-52 |
| chr4-148606950-DEL-85 | chr4-148606951-DEL-85 |
| chr4-148609161-DEL-50 | chr4-148609134-DEL-90 |
| chr4-148625767-DEL-937 | chr4-148625768-DEL-937 |
| chr4-148626379-DEL-214 | chr4-148626380-DEL-214 |
| chr4-148629518-DEL-74 | chr4-148629519-DEL-74 |
| chr4-148644376-DEL-137 | chr4-148644390-DEL-124 |
| chr4-148672694-DEL-205 | chr4-148672695-DEL-205 |
| chr4-148681602-DEL-199 | chr4-148681603-DEL-199 |
| chr4-148684286-DEL-73 | chr4-148684257-DEL-65 |
| chr4-148691132-DEL-938 | chr4-148691133-DEL-938 |
| chr4-148714189-DEL-69 | chr4-148714157-DEL-70 |
| chr4-148714882-DEL-62 | chr4-148714883-DEL-62 |
| chr4-148719154-DEL-103 | chr4-148719154-DEL-103 |
| chr4-148753929-DEL-256 | chr4-148753930-DEL-256 |
| chr4-148756482-DEL-338 | chr4-148756483-DEL-332 |
| chr4-148758523-DEL-159 | chr4-148758524-DEL-159 |
| chr4-148802115-DEL-297 | chr4-148802134-DEL-276 |
| chr4-148815414-DEL-66 | chr4-148815411-DEL-69 |
| chr4-148826037-DEL-254 | chr4-148826030-DEL-252 |
| chr4-148828372-DEL-65 | chr4-148828372-DEL-75 |
| chr4-148839272-DEL-192 | chr4-148839273-DEL-192 |
| chr4-148842514-DEL-88 | chr4-148842519-DEL-76 |
| chr4-148880023-DEL-365 | chr4-148880024-DEL-365 |

| | |
|---|---|
| chr4-148948335-DEL-52 | chr4-148948336-DEL-52 |
| chr4-148967597-DEL-424 | chr4-148967598-DEL-424 |
| chr4-148992298-DEL-63 | chr4-148992299-DEL-63 |
| chr4-149012286-DEL-74 | chr4-149012287-DEL-74 |
| chr4-149014140-DEL-101 | chr4-149014141-DEL-101 |
| chr4-149077161-DEL-205 | chr4-149077162-DEL-205 |
| chr4-149099468-DEL-121 | chr4-149099450-DEL-122 |
| chr4-149113629-DEL-52 | chr4-149113630-DEL-52 |
| chr4-149115952-DEL-63 | chr4-149115953-DEL-63 |
| chr4-149165692-DEL-66 | chr4-149165666-DEL-66 |
| chr4-149172990-DEL-51 | chr4-149172972-DEL-70 |
| chr4-149181497-DEL-88 | chr4-149181494-DEL-92 |
| chr4-149194132-DEL-168 | chr4-149194133-DEL-168 |
| chr4-149197037-DEL-209 | chr4-149197043-DEL-204 |
| chr4-149197672-DEL-682 | chr4-149197673-DEL-682 |
| chr4-149206973-DEL-204 | chr4-149206974-DEL-204 |
| chr4-149212801-DEL-82 | chr4-149212801-DEL-82 |
| chr4-149220543-DEL-206 | chr4-149220543-DEL-206 |
| chr4-149236014-DEL-58 | chr4-149236015-DEL-58 |
| chr4-149254289-DEL-217 | chr4-149254290-DEL-217 |
| chr4-149255617-DEL-203 | chr4-149255618-DEL-200 |
| chr4-149273328-DEL-212 | chr4-149273329-DEL-212 |
| chr4-149279702-DEL-206 | chr4-149279703-DEL-206 |
| chr4-149284338-DEL-261 | chr4-149284351-DEL-262 |
| chr4-149309440-DEL-100 | chr4-149309441-DEL-100 |
| chr4-149372951-DEL-222 | chr4-149372952-DEL-222 |
| chr4-149378347-DEL-214 | chr4-149378348-DEL-214 |
| chr4-149404841-DEL-600 | chr4-149404845-DEL-597 |
| chr4-149423978-DEL-53 | chr4-149423979-DEL-53 |
| chr4-149432918-DEL-83 | chr4-149432914-DEL-84 |
| chr4-149438867-DEL-134 | chr4-149438868-DEL-134 |
| chr4-149454899-DEL-236 | chr4-149454900-DEL-236 |
| chr4-149485040-DEL-196 | chr4-149485041-DEL-195 |
| chr4-149491858-DEL-84 | chr4-149491859-DEL-84 |
| chr4-149507935-DEL-209 | chr4-149507936-DEL-209 |
| chr4-149518434-DEL-5220 | chr4-149518435-DEL-5220 |
| chr4-149532100-DEL-70 | chr4-149532101-DEL-70 |
| chr4-149541730-DEL-619 | chr4-149541731-DEL-619 |
| chr4-149550950-DEL-630 | chr4-149550980-DEL-600 |
| chr4-149558568-DEL-56 | chr4-149558569-DEL-56 |
| chr4-149579871-DEL-219 | chr4-149579872-DEL-219 |
| chr4-149586645-DEL-175 | chr4-149586647-DEL-174 |
| chr4-149615177-DEL-67 | chr4-149615178-DEL-67 |
| chr4-149628743-DEL-211 | chr4-149628744-DEL-211 |
| chr4-149644993-DEL-439 | chr4-149644994-DEL-439 |
| chr4-149685148-DEL-422 | chr4-149685149-DEL-422 |
| chr4-149686540-DEL-340 | chr4-149686541-DEL-340 |
| chr4-149704027-DEL-207 | chr4-149704028-DEL-207 |
| chr4-149713610-DEL-116 | chr4-149713606-DEL-121 |
| chr4-149714891-DEL-85 | chr4-149714892-DEL-85 |
| chr4-149762010-DEL-56 | chr4-149762011-DEL-63 |
| chr4-149781765-DEL-100 | chr4-149781766-DEL-100 |
| chr4-149788775-DEL-855 | chr4-149788775-DEL-851 |
| chr4-149789362-DEL-54 | chr4-149789361-DEL-52 |
| chr4-149794592-DEL-191 | chr4-149794598-DEL-186 |
| chr4-149839569-DEL-189 | chr4-149839570-DEL-189 |
| chr4-149854926-DEL-263 | chr4-149854927-DEL-263 |
| chr4-149882561-DEL-100 | chr4-149882532-DEL-80 |
| chr4-149887800-DEL-115 | chr4-149887801-DEL-115 |
| chr4-149890770-DEL-54 | chr4-149890770-DEL-54 |
| chr4-149902174-DEL-100 | chr4-149902175-DEL-100 |
| chr4-149924490-DEL-69 | chr4-149924491-DEL-69 |
| chr4-149947664-DEL-160 | chr4-149947665-DEL-160 |
| chr4-149963465-DEL-451 | chr4-149963466-DEL-451 |
| chr4-149965663-DEL-238 | chr4-149965664-DEL-240 |
| chr4-149971961-DEL-166 | chr4-149971964-DEL-186 |
| chr4-150001652-DEL-87 | chr4-150001653-DEL-87 |
| chr4-150034006-DEL-195 | chr4-150034007-DEL-195 |
| chr4-150037966-DEL-68 | chr4-150037944-DEL-50 |
| chr4-150044460-DEL-60 | chr4-150044461-DEL-60 |
| chr4-150045890-DEL-486 | chr4-150045891-DEL-486 |
| chr4-150052774-DEL-64 | chr4-150052775-DEL-64 |
| chr4-150124136-DEL-214 | chr4-150124122-DEL-196 |
| chr4-150131313-DEL-146 | chr4-150131309-DEL-153 |
| chr4-150151493-DEL-900 | chr4-150151492-DEL-901 |
| chr4-150154053-DEL-178 | chr4-150154050-DEL-182 |
| chr4-150159302-DEL-95 | chr4-150159303-DEL-95 |
| chr4-150160930-DEL-221 | chr4-150160931-DEL-221 |
| chr4-150163474-DEL-136 | chr4-150163463-DEL-148 |
| chr4-150164488-DEL-240 | chr4-150164489-DEL-240 |

| | |
|---|---|
| chr4-150168022-DEL-533 | chr4-150168022-DEL-532 |
| chr4-150225687-DEL-189 | chr4-150225688-DEL-189 |
| chr4-150237461-DEL-51 | chr4-150237462-DEL-51 |
| chr4-150240966-DEL-131 | chr4-150240970-DEL-128 |
| chr4-150241419-DEL-223 | chr4-150241402-DEL-213 |
| chr4-150245205-DEL-813 | chr4-150245208-DEL-812 |
| chr4-150245980-DEL-65 | chr4-150245981-DEL-65 |
| chr4-150257725-DEL-79 | chr4-150257726-DEL-79 |
| chr4-150259691-DEL-747 | chr4-150259692-DEL-747 |
| chr4-150262083-DEL-125 | chr4-150262084-DEL-125 |
| chr4-150266431-DEL-438 | chr4-150266432-DEL-438 |
| chr4-150283490-DEL-87 | chr4-150283479-DEL-88 |
| chr4-150299207-DEL-149 | chr4-150299208-DEL-149 |
| chr4-150301696-DEL-115 | chr4-150301697-DEL-114 |
| chr4-150304206-DEL-62 | chr4-150304207-DEL-62 |
| chr4-150337923-DEL-220 | chr4-150337924-DEL-220 |
| chr4-150359697-DEL-64 | chr4-150359698-DEL-64 |
| chr4-150363699-DEL-220 | chr4-150363692-DEL-221 |
| chr4-150369235-DEL-214 | chr4-150369236-DEL-212 |
| chr4-150422872-DEL-200 | chr4-150422873-DEL-200 |
| chr4-150425349-DEL-1605 | chr4-150425350-DEL-1605 |
| chr4-150438439-DEL-160 | chr4-150438440-DEL-160 |
| chr4-150465150-DEL-3697 | chr4-150465151-DEL-3697 |
| chr4-150510675-DEL-58 | chr4-150510676-DEL-58 |
| chr4-150513460-DEL-405 | chr4-150513461-DEL-405 |
| chr4-150514445-DEL-51 | chr4-150514443-DEL-51 |
| chr4-150570288-DEL-151 | chr4-150570289-DEL-151 |
| chr4-150576095-DEL-382 | chr4-150576096-DEL-382 |
| chr4-150578534-DEL-206 | chr4-150578535-DEL-206 |
| chr4-150588969-DEL-184 | chr4-150588975-DEL-178 |
| chr4-150589516-DEL-101 | chr4-150589517-DEL-101 |
| chr4-150590555-DEL-56 | chr4-150590551-DEL-61 |
| chr4-150596892-DEL-771 | chr4-150596893-DEL-771 |
| chr4-150599754-DEL-240 | chr4-150599755-DEL-240 |
| chr4-150622997-DEL-199 | chr4-150622998-DEL-199 |
| chr4-150637666-DEL-170 | chr4-150637667-DEL-170 |
| chr4-150641925-DEL-84 | chr4-150641926-DEL-84 |
| chr4-150657769-DEL-187 | chr4-150657770-DEL-184 |
| chr4-150658333-DEL-135 | chr4-150658334-DEL-135 |
| chr4-150665598-DEL-168 | chr4-150665599-DEL-192 |
| chr4-150688201-DEL-60 | chr4-150688201-DEL-60 |
| chr4-150707551-DEL-59 | chr4-150707552-DEL-59 |
| chr4-150743779-DEL-414 | chr4-150743780-DEL-382 |
| chr4-150751583-DEL-1906 | chr4-150751584-DEL-1906 |
| chr4-150756568-DEL-56 | chr4-150756569-DEL-56 |
| chr4-150760558-DEL-80 | chr4-150760559-DEL-80 |
| chr4-150764197-DEL-202 | chr4-150764198-DEL-202 |
| chr4-150779239-DEL-2205 | chr4-150779240-DEL-2205 |
| chr4-150795014-DEL-77 | chr4-150795042-DEL-71 |
| chr4-150796930-DEL-189 | chr4-150796931-DEL-189 |
| chr4-150821466-DEL-86 | chr4-150821467-DEL-86 |
| chr4-150824026-DEL-83 | chr4-150823981-DEL-84 |
| chr4-150825024-DEL-2031 | chr4-150825025-DEL-2031 |
| chr4-150829288-DEL-68 | chr4-150829289-DEL-68 |
| chr4-150844314-DEL-71 | chr4-150844315-DEL-71 |
| chr4-150859362-DEL-54 | chr4-150859353-DEL-63 |
| chr4-150931533-DEL-479 | chr4-150931534-DEL-479 |
| chr4-150946129-DEL-211 | chr4-150946130-DEL-211 |
| chr4-150959095-DEL-100 | chr4-150959096-DEL-60 |
| chr4-150970936-DEL-95 | chr4-150970963-DEL-79 |
| chr4-150978474-DEL-178 | chr4-150978460-DEL-179 |
| chr4-151032660-DEL-203 | chr4-151032661-DEL-203 |
| chr4-151034524-DEL-95 | chr4-151034525-DEL-95 |
| chr4-151042279-DEL-222 | chr4-151042276-DEL-223 |
| chr4-151068338-DEL-3100 | chr4-151068339-DEL-3100 |
| chr4-151071174-DEL-92 | chr4-151071175-DEL-92 |
| chr4-151131355-DEL-108 | chr4-151131356-DEL-108 |
| chr4-151135961-DEL-75 | chr4-151135962-DEL-75 |
| chr4-151189830-DEL-73 | chr4-151189793-DEL-74 |
| chr4-151210783-DEL-54 | chr4-151210779-DEL-57 |
| chr4-151220670-DEL-63 | chr4-151220671-DEL-63 |
| chr4-151231502-DEL-617 | chr4-151231503-DEL-617 |
| chr4-151313445-DEL-102 | chr4-151313402-DEL-139 |
| chr4-151402556-DEL-118 | chr4-151402556-DEL-98 |
| chr4-151433938-DEL-149 | chr4-151433924-DEL-148 |
| chr4-151443336-DEL-124 | chr4-151443337-DEL-124 |
| chr4-151444684-DEL-64 | chr4-151444685-DEL-64 |
| chr4-151478478-DEL-76 | chr4-151478479-DEL-76 |
| chr4-151505713-DEL-89 | chr4-151505714-DEL-89 |
| chr4-151545305-DEL-83 | chr4-151545306-DEL-83 |

| | |
|---|---|
| chr4-151608235-DEL-368 | chr4-151608236-DEL-368 |
| chr4-151616195-DEL-176 | chr4-151616196-DEL-176 |
| chr4-151631517-DEL-122 | chr4-151631514-DEL-112 |
| chr4-151638225-DEL-53 | chr4-151638226-DEL-53 |
| chr4-151656194-DEL-67 | chr4-151656195-DEL-67 |
| chr4-151728235-DEL-76 | chr4-151728236-DEL-76 |
| chr4-151738086-DEL-94 | chr4-151738087-DEL-94 |
| chr4-151748027-DEL-80 | chr4-151748028-DEL-80 |
| chr4-151771366-DEL-121 | chr4-151771367-DEL-121 |
| chr4-151793174-DEL-52 | chr4-151793175-DEL-56 |
| chr4-151861462-DEL-225 | chr4-151861463-DEL-225 |
| chr4-151866947-DEL-69 | chr4-151866948-DEL-69 |
| chr4-151912701-DEL-86 | chr4-151912697-DEL-68 |
| chr4-151956827-DEL-111 | chr4-151956828-DEL-111 |
| chr4-151982542-DEL-95 | chr4-151982545-DEL-93 |
| chr4-151998255-DEL-210 | chr4-151998256-DEL-210 |
| chr4-152006632-DEL-122 | chr4-152006633-DEL-122 |
| chr4-152010229-DEL-208 | chr4-152010231-DEL-207 |
| chr4-152024587-DEL-189 | chr4-152024588-DEL-189 |
| chr4-152035242-DEL-100 | chr4-152035243-DEL-100 |
| chr4-152099325-DEL-64 | chr4-152099329-DEL-52 |
| chr4-152101426-DEL-105 | chr4-152101432-DEL-93 |
| chr4-152120800-DEL-196 | chr4-152120801-DEL-196 |
| chr4-152127388-DEL-54 | chr4-152127389-DEL-54 |
| chr4-152138627-DEL-705 | chr4-152138625-DEL-706 |
| chr4-152139195-DEL-93 | chr4-152139193-DEL-97 |
| chr4-152141795-DEL-191 | chr4-152141796-DEL-188 |
| chr4-152150646-DEL-185 | chr4-152150647-DEL-185 |
| chr4-152157632-DEL-122 | chr4-152157615-DEL-143 |
| chr4-152165196-DEL-1299 | chr4-152165194-DEL-1299 |
| chr4-152165445-DEL-218 | chr4-152165446-DEL-218 |
| chr4-152166729-DEL-56 | chr4-152166705-DEL-56 |
| chr4-152170185-DEL-207 | chr4-152170186-DEL-207 |
| chr4-152196437-DEL-53 | chr4-152196438-DEL-53 |
| chr4-152206988-DEL-437 | chr4-152206983-DEL-433 |
| chr4-152222082-DEL-223 | chr4-152222092-DEL-222 |
| chr4-152227527-DEL-110 | chr4-152227550-DEL-86 |
| chr4-152240577-DEL-236 | chr4-152240578-DEL-236 |
| chr4-152250838-DEL-199 | chr4-152250839-DEL-198 |
| chr4-152252163-DEL-203 | chr4-152252164-DEL-203 |
| chr4-152267322-DEL-65 | chr4-152267317-DEL-53 |
| chr4-152292863-DEL-229 | chr4-152292864-DEL-229 |
| chr4-152295238-DEL-169 | chr4-152295239-DEL-169 |
| chr4-152318551-DEL-1524 | chr4-152318552-DEL-1524 |
| chr4-152322687-DEL-122 | chr4-152322688-DEL-122 |
| chr4-152323882-DEL-98 | chr4-152323883-DEL-98 |
| chr4-152327392-DEL-56 | chr4-152327365-DEL-56 |
| chr4-152332546-DEL-96 | chr4-152332547-DEL-96 |
| chr4-152342243-DEL-59 | chr4-152342244-DEL-59 |
| chr4-152351237-DEL-573 | chr4-152351238-DEL-573 |
| chr4-152392788-DEL-85 | chr4-152392789-DEL-85 |
| chr4-152404599-DEL-174 | chr4-152404600-DEL-174 |
| chr4-152441490-DEL-915 | chr4-152441493-DEL-912 |
| chr4-152482182-DEL-210 | chr4-152482183-DEL-210 |
| chr4-152494495-DEL-110 | chr4-152494524-DEL-83 |
| chr4-152523830-DEL-1676 | chr4-152523831-DEL-1676 |
| chr4-152524205-DEL-131 | chr4-152524206-DEL-131 |
| chr4-152525760-DEL-230 | chr4-152525761-DEL-230 |
| chr4-152527256-DEL-191 | chr4-152527257-DEL-191 |
| chr4-152548636-DEL-52 | chr4-152548628-DEL-50 |
| chr4-152552409-DEL-1166 | chr4-152552410-DEL-1166 |
| chr4-152554009-DEL-251 | chr4-152554010-DEL-251 |
| chr4-152578284-DEL-80 | chr4-152578285-DEL-80 |
| chr4-152588872-DEL-80 | chr4-152588873-DEL-80 |
| chr4-152591490-DEL-61 | chr4-152591489-DEL-63 |
| chr4-152593079-DEL-62 | chr4-152593050-DEL-72 |
| chr4-152596186-DEL-120 | chr4-152596187-DEL-120 |
| chr4-152597402-DEL-74 | chr4-152597402-DEL-74 |
| chr4-152599028-DEL-64 | chr4-152598998-DEL-54 |
| chr4-152600658-DEL-1826 | chr4-152600659-DEL-1826 |
| chr4-152605826-DEL-2072 | chr4-152605826-DEL-2072 |
| chr4-152627008-DEL-57 | chr4-152627028-DEL-51 |
| chr4-152634250-DEL-61 | chr4-152634251-DEL-61 |
| chr4-152635708-DEL-364 | chr4-152635701-DEL-365 |
| chr4-152636874-DEL-84 | chr4-152636875-DEL-112 |
| chr4-152677261-DEL-80 | chr4-152677262-DEL-80 |
| chr4-152679393-DEL-132 | chr4-152679394-DEL-132 |
| chr4-152681899-DEL-81 | chr4-152681900-DEL-81 |
| chr4-152686019-DEL-196 | chr4-152686020-DEL-196 |
| chr4-152689202-DEL-273 | chr4-152689203-DEL-273 |

| | |
|---|---|
| chr4-152728613-DEL-66 | chr4-152728567-DEL-96 |
| chr4-152737466-DEL-542 | chr4-152737424-DEL-582 |
| chr4-152739188-DEL-75 | chr4-152739191-DEL-51 |
| chr4-152754908-DEL-224 | chr4-152754909-DEL-228 |
| chr4-152768253-DEL-52 | chr4-152768254-DEL-52 |
| chr4-152769627-DEL-556 | chr4-152769628-DEL-556 |
| chr4-152773216-DEL-54 | chr4-152773217-DEL-54 |
| chr4-152773890-DEL-216 | chr4-152773891-DEL-216 |
| chr4-152779793-DEL-60 | chr4-152779794-DEL-58 |
| chr4-152790555-DEL-101 | chr4-152790581-DEL-75 |
| chr4-152802672-DEL-53 | chr4-152802667-DEL-54 |
| chr4-152813518-DEL-213 | chr4-152813519-DEL-213 |
| chr4-152823977-DEL-275 | chr4-152823959-DEL-294 |
| chr4-152829845-DEL-72 | chr4-152829846-DEL-72 |
| chr4-152837242-DEL-612 | chr4-152837243-DEL-612 |
| chr4-152839317-DEL-108 | chr4-152839318-DEL-108 |
| chr4-152865834-DEL-60 | chr4-152865837-DEL-56 |
| chr4-152886010-DEL-113 | chr4-152886014-DEL-112 |
| chr4-152888136-DEL-69 | chr4-152888137-DEL-69 |
| chr4-152899519-DEL-90 | chr4-152899519-DEL-90 |
| chr4-152902398-DEL-59 | chr4-152902369-DEL-66 |
| chr4-152912969-DEL-62 | chr4-152912970-DEL-62 |
| chr4-152913248-DEL-464 | chr4-152913249-DEL-464 |
| chr4-152931242-DEL-1876 | chr4-152931243-DEL-1876 |
| chr4-152933171-DEL-74 | chr4-152933172-DEL-74 |
| chr4-152968310-DEL-52 | chr4-152968311-DEL-52 |
| chr4-153020952-DEL-69 | chr4-153020953-DEL-69 |
| chr4-153027264-DEL-2974 | chr4-153027265-DEL-2975 |
| chr4-153064140-DEL-80 | chr4-153064116-DEL-64 |
| chr4-153085175-DEL-129 | chr4-153085176-DEL-129 |
| chr4-153085187-DEL-1309 | chr4-153085188-DEL-1309 |
| chr4-153109791-DEL-105 | chr4-153109792-DEL-105 |
| chr4-153110534-DEL-63 | chr4-153110529-DEL-66 |
| chr4-153141258-DEL-116 | chr4-153141289-DEL-116 |
| chr4-153141886-DEL-54 | chr4-153141887-DEL-54 |
| chr4-153146137-DEL-65 | chr4-153146138-DEL-87 |
| chr4-153161170-DEL-183 | chr4-153161171-DEL-183 |
| chr4-153190624-DEL-64 | chr4-153190625-DEL-64 |
| chr4-153235388-DEL-1539 | chr4-153235389-DEL-1539 |
| chr4-153245682-DEL-316 | chr4-153245683-DEL-316 |
| chr4-153249513-DEL-53 | chr4-153249514-DEL-53 |
| chr4-153258880-DEL-100 | chr4-153258881-DEL-100 |
| chr4-153275856-DEL-64 | chr4-153275857-DEL-64 |
| chr4-153284461-DEL-186 | chr4-153284461-DEL-183 |
| chr4-153288717-DEL-83 | chr4-153288717-DEL-83 |
| chr4-153311757-DEL-414 | chr4-153311758-DEL-414 |
| chr4-153319385-DEL-105 | chr4-153319418-DEL-113 |
| chr4-153337871-DEL-766 | chr4-153337872-DEL-766 |
| chr4-153346473-DEL-91 | chr4-153346475-DEL-96 |
| chr4-153362048-DEL-84 | chr4-153362023-DEL-96 |
| chr4-153389612-DEL-1203 | chr4-153389613-DEL-1203 |
| chr4-153390876-DEL-1165 | chr4-153390877-DEL-1165 |
| chr4-153392241-DEL-114 | chr4-153392242-DEL-114 |
| chr4-153402448-DEL-60 | chr4-153402449-DEL-60 |
| chr4-153427411-DEL-396 | chr4-153427412-DEL-396 |
| chr4-153433167-DEL-302 | chr4-153433168-DEL-302 |
| chr4-153443301-DEL-67 | chr4-153443302-DEL-67 |
| chr4-153444130-DEL-197 | chr4-153444131-DEL-197 |
| chr4-153453784-DEL-66 | chr4-153453785-DEL-66 |
| chr4-153494246-DEL-156 | chr4-153494255-DEL-148 |
| chr4-153519051-DEL-56 | chr4-153519052-DEL-56 |
| chr4-153524858-DEL-101 | chr4-153524859-DEL-101 |
| chr4-153552934-DEL-123 | chr4-153552935-DEL-123 |
| chr4-153578228-DEL-59 | chr4-153578229-DEL-59 |
| chr4-153593186-DEL-75 | chr4-153593186-DEL-125 |
| chr4-153612669-DEL-81 | chr4-153612670-DEL-81 |
| chr4-153625577-DEL-93 | chr4-153625578-DEL-93 |
| chr4-153628010-DEL-157 | chr4-153627983-DEL-138 |
| chr4-153652457-DEL-123 | chr4-153652458-DEL-123 |
| chr4-153704657-DEL-184 | chr4-153704655-DEL-172 |
| chr4-153763151-DEL-72 | chr4-153763170-DEL-54 |
| chr4-153767839-DEL-216 | chr4-153767840-DEL-216 |
| chr4-153776565-DEL-381 | chr4-153776566-DEL-381 |
| chr4-153777693-DEL-113 | chr4-153777694-DEL-113 |
| chr4-153793492-DEL-57 | chr4-153793493-DEL-57 |
| chr4-153804610-DEL-196 | chr4-153804611-DEL-196 |
| chr4-153827020-DEL-7103 | chr4-153827021-DEL-7103 |
| chr4-153837310-DEL-117 | chr4-153837311-DEL-117 |
| chr4-153844472-DEL-59 | chr4-153844473-DEL-59 |
| chr4-153846276-DEL-54 | chr4-153846277-DEL-54 |

| | |
|---|---|
| chr4-153851202-DEL-56 | chr4-153851203-DEL-56 |
| chr4-153885347-DEL-361 | chr4-153885348-DEL-361 |
| chr4-153890106-DEL-90 | chr4-153890106-DEL-90 |
| chr4-153896742-DEL-168 | chr4-153896743-DEL-168 |
| chr4-153898864-DEL-72 | chr4-153898861-DEL-76 |
| chr4-153900779-DEL-120 | chr4-153900780-DEL-120 |
| chr4-153910635-DEL-126 | chr4-153910636-DEL-126 |
| chr4-153946723-DEL-61 | chr4-153946706-DEL-84 |
| chr4-153948327-DEL-119 | chr4-153948326-DEL-119 |
| chr4-153953527-DEL-168 | chr4-153953528-DEL-216 |
| chr4-153955408-DEL-68 | chr4-153955409-DEL-68 |
| chr4-153971718-DEL-145 | chr4-153971719-DEL-145 |
| chr4-153977323-DEL-2394 | chr4-153977328-DEL-2392 |
| chr4-153981267-DEL-89 | chr4-153981282-DEL-75 |
| chr4-154001269-DEL-1868 | chr4-154001270-DEL-1868 |
| chr4-154007427-DEL-219 | chr4-154007428-DEL-217 |
| chr4-154025620-DEL-120 | chr4-154025621-DEL-120 |
| chr4-154052539-DEL-57 | chr4-154052498-DEL-55 |
| chr4-154053102-DEL-99 | chr4-154053103-DEL-102 |
| chr4-154058622-DEL-168 | chr4-154058613-DEL-200 |
| chr4-154069128-DEL-57 | chr4-154069129-DEL-57 |
| chr4-154075190-DEL-75 | chr4-154075191-DEL-75 |
| chr4-154085529-DEL-240 | chr4-154085530-DEL-240 |
| chr4-154101317-DEL-76 | chr4-154101318-DEL-76 |
| chr4-154106246-DEL-209 | chr4-154106247-DEL-209 |
| chr4-154137832-DEL-212 | chr4-154137833-DEL-212 |
| chr4-154160503-DEL-315 | chr4-154160504-DEL-315 |
| chr4-154194121-DEL-80 | chr4-154194122-DEL-80 |
| chr4-154217211-DEL-211 | chr4-154217212-DEL-211 |
| chr4-154219692-DEL-873 | chr4-154219693-DEL-873 |
| chr4-154230435-DEL-209 | chr4-154230451-DEL-210 |
| chr4-154231361-DEL-58 | chr4-154231362-DEL-58 |
| chr4-154234232-DEL-65 | chr4-154234233-DEL-65 |
| chr4-154237939-DEL-82 | chr4-154237899-DEL-123 |
| chr4-154246372-DEL-395 | chr4-154246373-DEL-395 |
| chr4-154246986-DEL-169 | chr4-154246987-DEL-169 |
| chr4-154248224-DEL-201 | chr4-154248225-DEL-201 |
| chr4-154275749-DEL-251 | chr4-154275758-DEL-259 |
| chr4-154277898-DEL-375 | chr4-154277884-DEL-376 |
| chr4-154284756-DEL-95 | chr4-154284757-DEL-95 |
| chr4-154298772-DEL-120 | chr4-154298758-DEL-85 |
| chr4-154339608-DEL-68 | chr4-154339609-DEL-68 |
| chr4-154345537-DEL-349 | chr4-154345538-DEL-349 |
| chr4-154374497-DEL-177 | chr4-154374498-DEL-177 |
| chr4-154392742-DEL-435 | chr4-154392743-DEL-436 |
| chr4-154411911-DEL-207 | chr4-154411899-DEL-215 |
| chr4-154457031-DEL-77 | chr4-154457016-DEL-71 |
| chr4-154462465-DEL-286 | chr4-154462466-DEL-286 |
| chr4-154554378-DEL-200 | chr4-154554379-DEL-200 |
| chr4-154652030-DEL-54 | chr4-154652031-DEL-54 |
| chr4-154686748-DEL-78 | chr4-154686749-DEL-78 |
| chr4-154705932-DEL-52 | chr4-154705933-DEL-52 |
| chr4-154712924-DEL-68 | chr4-154712925-DEL-68 |
| chr4-154729402-DEL-105 | chr4-154729429-DEL-62 |
| chr4-154739998-DEL-96 | chr4-154739999-DEL-95 |
| chr4-154753971-DEL-89 | chr4-154753936-DEL-90 |
| chr4-154770042-DEL-2078 | chr4-154770049-DEL-2069 |
| chr4-154772647-DEL-178 | chr4-154772648-DEL-178 |
| chr4-154797526-DEL-64 | chr4-154797491-DEL-66 |
| chr4-154808020-DEL-1029 | chr4-154808021-DEL-1029 |
| chr4-154823151-DEL-52 | chr4-154823152-DEL-52 |
| chr4-154843238-DEL-78 | chr4-154843239-DEL-78 |
| chr4-154854788-DEL-250 | chr4-154854789-DEL-250 |
| chr4-154882271-DEL-72 | chr4-154882272-DEL-72 |
| chr4-154888009-DEL-342 | chr4-154888010-DEL-342 |
| chr4-154892298-DEL-68 | chr4-154892299-DEL-68 |
| chr4-154902882-DEL-126 | chr4-154902881-DEL-105 |
| chr4-154914799-DEL-894 | chr4-154914800-DEL-894 |
| chr4-154916032-DEL-60 | chr4-154916033-DEL-82 |
| chr4-154917278-DEL-67 | chr4-154917266-DEL-84 |
| chr4-154918533-DEL-317 | chr4-154918534-DEL-317 |
| chr4-154919670-DEL-77 | chr4-154919668-DEL-74 |
| chr4-154937207-DEL-120 | chr4-154937207-DEL-120 |
| chr4-154968899-DEL-223 | chr4-154968900-DEL-223 |
| chr4-154973146-DEL-102 | chr4-154973147-DEL-102 |
| chr4-154982643-DEL-2056 | chr4-154982644-DEL-2056 |
| chr4-154992725-DEL-62 | chr4-154992726-DEL-62 |
| chr4-154994728-DEL-57 | chr4-154994729-DEL-57 |
| chr4-155001621-DEL-169 | chr4-155001622-DEL-169 |
| chr4-155017846-DEL-56 | chr4-155017847-DEL-56 |

| | |
|---|---|
| chr4-155019739-DEL-245 | chr4-155019740-DEL-245 |
| chr4-155026638-DEL-357 | chr4-155026639-DEL-357 |
| chr4-155149781-DEL-173 | chr4-155149782-DEL-173 |
| chr4-155163518-DEL-212 | chr4-155163519-DEL-212 |
| chr4-155197831-DEL-100 | chr4-155197823-DEL-64 |
| chr4-155204038-DEL-210 | chr4-155204039-DEL-210 |
| chr4-155226026-DEL-65 | chr4-155226029-DEL-53 |
| chr4-155233783-DEL-282 | chr4-155233784-DEL-282 |
| chr4-155323421-DEL-79 | chr4-155323445-DEL-89 |
| chr4-155325024-DEL-120 | chr4-155325025-DEL-105 |
| chr4-155325639-DEL-184 | chr4-155325640-DEL-184 |
| chr4-155344059-DEL-302 | chr4-155344065-DEL-296 |
| chr4-155358254-DEL-460 | chr4-155358255-DEL-460 |
| chr4-155362274-DEL-68 | chr4-155362324-DEL-52 |
| chr4-155403679-DEL-404 | chr4-155403680-DEL-404 |
| chr4-155412861-DEL-51 | chr4-155412850-DEL-65 |
| chr4-155413671-DEL-52 | chr4-155413672-DEL-52 |
| chr4-155448273-DEL-53 | chr4-155448274-DEL-53 |
| chr4-155454861-DEL-192 | chr4-155454900-DEL-150 |
| chr4-155457425-DEL-241 | chr4-155457426-DEL-241 |
| chr4-155462170-DEL-56 | chr4-155462150-DEL-83 |
| chr4-155476041-DEL-200 | chr4-155476042-DEL-200 |
| chr4-155487371-DEL-326 | chr4-155487372-DEL-326 |
| chr4-155490151-DEL-110 | chr4-155490155-DEL-100 |
| chr4-155500616-DEL-600 | chr4-155500568-DEL-598 |
| chr4-155506132-DEL-373 | chr4-155506133-DEL-373 |
| chr4-155535684-DEL-206 | chr4-155535685-DEL-206 |
| chr4-155548890-DEL-124 | chr4-155548884-DEL-114 |
| chr4-155558673-DEL-191 | chr4-155558674-DEL-191 |
| chr4-155581452-DEL-132 | chr4-155581447-DEL-137 |
| chr4-155582155-DEL-222 | chr4-155582156-DEL-222 |
| chr4-155586710-DEL-488 | chr4-155586716-DEL-481 |
| chr4-155604729-DEL-221 | chr4-155604730-DEL-221 |
| chr4-155606136-DEL-64 | chr4-155606137-DEL-64 |
| chr4-155616934-DEL-202 | chr4-155616928-DEL-203 |
| chr4-155625834-DEL-208 | chr4-155625835-DEL-208 |
| chr4-155645854-DEL-71 | chr4-155645855-DEL-71 |
| chr4-155650312-DEL-51 | chr4-155650312-DEL-51 |
| chr4-155677653-DEL-56 | chr4-155677654-DEL-50 |
| chr4-155683198-DEL-58 | chr4-155683199-DEL-58 |
| chr4-155695116-DEL-218 | chr4-155695117-DEL-218 |
| chr4-155700955-DEL-208 | chr4-155700956-DEL-208 |
| chr4-155705803-DEL-52 | chr4-155705802-DEL-53 |
| chr4-155714379-DEL-153 | chr4-155714380-DEL-153 |
| chr4-155716985-DEL-169 | chr4-155716986-DEL-169 |
| chr4-155719765-DEL-572 | chr4-155719766-DEL-572 |
| chr4-155721152-DEL-68 | chr4-155721153-DEL-68 |
| chr4-155752154-DEL-214 | chr4-155752155-DEL-196 |
| chr4-155757098-DEL-187 | chr4-155757097-DEL-187 |
| chr4-155757593-DEL-1203 | chr4-155757586-DEL-1202 |
| chr4-155759657-DEL-175 | chr4-155759658-DEL-175 |
| chr4-155760855-DEL-61 | chr4-155760856-DEL-61 |
| chr4-155761061-DEL-7205 | chr4-155761062-DEL-7205 |
| chr4-155768556-DEL-205 | chr4-155768557-DEL-204 |
| chr4-155772621-DEL-79 | chr4-155772622-DEL-76 |
| chr4-155792295-DEL-498 | chr4-155792296-DEL-498 |
| chr4-155803163-DEL-217 | chr4-155803164-DEL-217 |
| chr4-155847535-DEL-183 | chr4-155847531-DEL-184 |
| chr4-155847593-DEL-1775 | chr4-155847594-DEL-1775 |
| chr4-155848177-DEL-192 | chr4-155848178-DEL-192 |
| chr4-155853114-DEL-61 | chr4-155853115-DEL-61 |
| chr4-155862743-DEL-260 | chr4-155862744-DEL-260 |
| chr4-155863842-DEL-159 | chr4-155863843-DEL-141 |
| chr4-155871597-DEL-151 | chr4-155871595-DEL-152 |
| chr4-155872191-DEL-157 | chr4-155872178-DEL-157 |
| chr4-155879490-DEL-159 | chr4-155879483-DEL-159 |
| chr4-155881475-DEL-74 | chr4-155881476-DEL-74 |
| chr4-155885680-DEL-196 | chr4-155885681-DEL-196 |
| chr4-155887024-DEL-128 | chr4-155887025-DEL-128 |
| chr4-155913322-DEL-199 | chr4-155913323-DEL-199 |
| chr4-155913869-DEL-125 | chr4-155913839-DEL-150 |
| chr4-155915389-DEL-205 | chr4-155915390-DEL-205 |
| chr4-155926707-DEL-58 | chr4-155926704-DEL-62 |
| chr4-155929315-DEL-186 | chr4-155929316-DEL-186 |
| chr4-155958636-DEL-145 | chr4-155958640-DEL-116 |
| chr4-155961638-DEL-108 | chr4-155961631-DEL-106 |
| chr4-155962595-DEL-58 | chr4-155962598-DEL-56 |
| chr4-155981900-DEL-191 | chr4-155981864-DEL-192 |
| chr4-155995882-DEL-16310 | chr4-155995883-DEL-16310 |
| chr4-155999632-DEL-7457 | chr4-155999633-DEL-7457 |

| | |
|---|---|
| chr4-156013800-DEL-461 | chr4-156013800-DEL-462 |
| chr4-156014396-DEL-218 | chr4-156014397-DEL-218 |
| chr4-156015464-DEL-198 | chr4-156015465-DEL-198 |
| chr4-156060094-DEL-181 | chr4-156060095-DEL-181 |
| chr4-156065813-DEL-179 | chr4-156065814-DEL-179 |
| chr4-156091167-DEL-206 | chr4-156091168-DEL-206 |
| chr4-156109165-DEL-233 | chr4-156109166-DEL-233 |
| chr4-156114457-DEL-114 | chr4-156114457-DEL-114 |
| chr4-156135485-DEL-124 | chr4-156135481-DEL-123 |
| chr4-156163550-DEL-238 | chr4-156163551-DEL-238 |
| chr4-156179226-DEL-833 | chr4-156179221-DEL-834 |
| chr4-156180307-DEL-63 | chr4-156180305-DEL-81 |
| chr4-156186748-DEL-162 | chr4-156186749-DEL-162 |
| chr4-156190903-DEL-214 | chr4-156190904-DEL-214 |
| chr4-156193374-DEL-200 | chr4-156193375-DEL-200 |
| chr4-156220825-DEL-314 | chr4-156220826-DEL-312 |
| chr4-156221770-DEL-780 | chr4-156221799-DEL-762 |
| chr4-156223676-DEL-199 | chr4-156223677-DEL-199 |
| chr4-156224419-DEL-6740 | chr4-156224420-DEL-6740 |
| chr4-156237619-DEL-932 | chr4-156237620-DEL-932 |
| chr4-156240864-DEL-51 | chr4-156240865-DEL-51 |
| chr4-156290943-DEL-183 | chr4-156290943-DEL-183 |
| chr4-156298811-DEL-322 | chr4-156298818-DEL-316 |
| chr4-156318981-DEL-100 | chr4-156318982-DEL-77 |
| chr4-156338532-DEL-200 | chr4-156338532-DEL-187 |
| chr4-156341098-DEL-199 | chr4-156341098-DEL-199 |
| chr4-156374207-DEL-203 | chr4-156374207-DEL-203 |
| chr4-156378543-DEL-198 | chr4-156378543-DEL-198 |
| chr4-156389359-DEL-435 | chr4-156389345-DEL-427 |
| chr4-156394195-DEL-82 | chr4-156394196-DEL-82 |
| chr4-156394788-DEL-60 | chr4-156394789-DEL-60 |
| chr4-156396641-DEL-207 | chr4-156396642-DEL-207 |
| chr4-156408090-DEL-148 | chr4-156408091-DEL-148 |
| chr4-156412777-DEL-75 | chr4-156412778-DEL-75 |
| chr4-156418435-DEL-75 | chr4-156418435-DEL-75 |
| chr4-156421660-DEL-88 | chr4-156421662-DEL-87 |
| chr4-156422377-DEL-571 | chr4-156422338-DEL-572 |
| chr4-156451827-DEL-248 | chr4-156451828-DEL-248 |
| chr4-156463417-DEL-327 | chr4-156463418-DEL-327 |
| chr4-156464349-DEL-171 | chr4-156464350-DEL-169 |
| chr4-156502195-DEL-144 | chr4-156502243-DEL-96 |
| chr4-156563401-DEL-189 | chr4-156563402-DEL-189 |
| chr4-156564341-DEL-1812 | chr4-156564342-DEL-1812 |
| chr4-156568061-DEL-93 | chr4-156568058-DEL-94 |
| chr4-156568702-DEL-145 | chr4-156568703-DEL-144 |
| chr5-3141688-DEL-70 | chr5-3141689-DEL-70 |
| chr5-3194444-DEL-54 | chr5-3194445-DEL-54 |
| chr5-3197024-DEL-207 | chr5-3197025-DEL-207 |
| chr5-3225335-DEL-11666 | chr5-3225332-DEL-11666 |
| chr5-3244080-DEL-178 | chr5-3244077-DEL-174 |
| chr5-3286373-DEL-566 | chr5-3286374-DEL-566 |
| chr5-3301608-DEL-5517 | chr5-3301609-DEL-5517 |
| chr5-3316570-DEL-119 | chr5-3316532-DEL-120 |
| chr5-3361110-DEL-203 | chr5-3361111-DEL-203 |
| chr5-3453268-DEL-64 | chr5-3453277-DEL-56 |
| chr5-3477230-DEL-68 | chr5-3477231-DEL-68 |
| chr5-3507313-DEL-54 | chr5-3507313-DEL-60 |
| chr5-3530239-DEL-162 | chr5-3530200-DEL-190 |
| chr5-3543596-DEL-52 | chr5-3543597-DEL-52 |
| chr5-3558019-DEL-71 | chr5-3558020-DEL-70 |
| chr5-3593187-DEL-96 | chr5-3593188-DEL-96 |
| chr5-3600883-DEL-192 | chr5-3600884-DEL-192 |
| chr5-3613290-DEL-121 | chr5-3613291-DEL-121 |
| chr5-3649823-DEL-69 | chr5-3649824-DEL-69 |
| chr5-3663377-DEL-1100 | chr5-3663378-DEL-1100 |
| chr5-3703117-DEL-343 | chr5-3703118-DEL-343 |
| chr5-3736568-DEL-66 | chr5-3736568-DEL-66 |
| chr5-3771729-DEL-241 | chr5-3771701-DEL-242 |
| chr5-3781059-DEL-768 | chr5-3781060-DEL-768 |
| chr5-3791070-DEL-1795 | chr5-3791071-DEL-1795 |
| chr5-3902394-DEL-102 | chr5-3902434-DEL-63 |
| chr5-3916859-DEL-330 | chr5-3916860-DEL-330 |
| chr5-3927369-DEL-64 | chr5-3927378-DEL-56 |
| chr5-3946524-DEL-6744 | chr5-3946524-DEL-6744 |
| chr5-3954342-DEL-163 | chr5-3954343-DEL-163 |
| chr5-3955811-DEL-382 | chr5-3955812-DEL-382 |
| chr5-3960862-DEL-65 | chr5-3960863-DEL-65 |
| chr5-3968463-DEL-384 | chr5-3968470-DEL-385 |
| chr5-4029571-DEL-111 | chr5-4029572-DEL-111 |
| chr5-4031522-DEL-7038 | chr5-4031523-DEL-7038 |

| | |
|---|---|
| chr5-4083262-DEL-658 | chr5-4083263-DEL-658 |
| chr5-4087114-DEL-64 | chr5-4087115-DEL-64 |
| chr5-4131126-DEL-162 | chr5-4131160-DEL-166 |
| chr5-4138362-DEL-62 | chr5-4138362-DEL-62 |
| chr5-4142990-DEL-579 | chr5-4142991-DEL-578 |
| chr5-4144595-DEL-197 | chr5-4144596-DEL-197 |
| chr5-4183712-DEL-333 | chr5-4183713-DEL-333 |
| chr5-4196389-DEL-794 | chr5-4196384-DEL-795 |
| chr5-4198261-DEL-275 | chr5-4198262-DEL-276 |
| chr5-4203666-DEL-70 | chr5-4203667-DEL-75 |
| chr5-4219959-DEL-58 | chr5-4219960-DEL-58 |
| chr5-4221239-DEL-9537 | chr5-4221239-DEL-9537 |
| chr5-4262802-DEL-192 | chr5-4262803-DEL-192 |
| chr5-4364178-DEL-89 | chr5-4364192-DEL-86 |
| chr5-4390186-DEL-70 | chr5-4390197-DEL-60 |
| chr5-4400838-DEL-78 | chr5-4400833-DEL-78 |
| chr5-4418517-DEL-6285 | chr5-4418518-DEL-6285 |
| chr5-4424470-DEL-203 | chr5-4424471-DEL-203 |
| chr5-4428541-DEL-1284 | chr5-4428542-DEL-1280 |
| chr5-4430265-DEL-3212 | chr5-4430266-DEL-3212 |
| chr5-4442704-DEL-400 | chr5-4442705-DEL-400 |
| chr5-4455150-DEL-603 | chr5-4455151-DEL-603 |
| chr5-4467859-DEL-153 | chr5-4467859-DEL-152 |
| chr5-4472695-DEL-55 | chr5-4472716-DEL-71 |
| chr5-4473209-DEL-65 | chr5-4473210-DEL-65 |
| chr5-4492912-DEL-154 | chr5-4492913-DEL-154 |
| chr5-4508301-DEL-5073 | chr5-4508302-DEL-5073 |
| chr5-4527868-DEL-92 | chr5-4527869-DEL-92 |
| chr5-4564490-DEL-4064 | chr5-4564491-DEL-4064 |
| chr5-4631067-DEL-139 | chr5-4631070-DEL-131 |
| chr5-4647447-DEL-51 | chr5-4647450-DEL-55 |
| chr5-4651262-DEL-6438 | chr5-4651263-DEL-6438 |
| chr5-4664386-DEL-84 | chr5-4664387-DEL-78 |
| chr5-4711088-DEL-378 | chr5-4711089-DEL-378 |
| chr5-4725934-DEL-74 | chr5-4725933-DEL-76 |
| chr5-4733166-DEL-65 | chr5-4733161-DEL-67 |
| chr5-4773672-DEL-94 | chr5-4773683-DEL-84 |
| chr5-4788453-DEL-1197 | chr5-4788454-DEL-1197 |
| chr5-4790221-DEL-64 | chr5-4790222-DEL-62 |
| chr5-4809180-DEL-68 | chr5-4809181-DEL-68 |
| chr5-4821223-DEL-58 | chr5-4821224-DEL-58 |
| chr5-4852472-DEL-54 | chr5-4852473-DEL-54 |
| chr5-4894462-DEL-1290 | chr5-4894463-DEL-1290 |
| chr5-4904220-DEL-62 | chr5-4904221-DEL-62 |
| chr5-4921079-DEL-65 | chr5-4921056-DEL-66 |
| chr5-4978585-DEL-78 | chr5-4978586-DEL-76 |
| chr5-4987500-DEL-6450 | chr5-4987501-DEL-6450 |
| chr5-5007805-DEL-52 | chr5-5007804-DEL-53 |
| chr5-5026151-DEL-52 | chr5-5026152-DEL-52 |
| chr5-5029074-DEL-51 | chr5-5029075-DEL-51 |
| chr5-5036193-DEL-56 | chr5-5036194-DEL-56 |
| chr5-5042551-DEL-390 | chr5-5042552-DEL-390 |
| chr5-5056442-DEL-267 | chr5-5056431-DEL-260 |
| chr5-5078668-DEL-281 | chr5-5078669-DEL-281 |
| chr5-5100056-DEL-211 | chr5-5100057-DEL-211 |
| chr5-5126717-DEL-3664 | chr5-5126718-DEL-3664 |
| chr5-5143242-DEL-527 | chr5-5143243-DEL-519 |
| chr5-5163137-DEL-2242 | chr5-5163138-DEL-2242 |
| chr5-5168683-DEL-463 | chr5-5168684-DEL-463 |
| chr5-5182099-DEL-68 | chr5-5182100-DEL-68 |
| chr5-5183972-DEL-92 | chr5-5183973-DEL-92 |
| chr5-5191397-DEL-76 | chr5-5191398-DEL-76 |
| chr5-5209231-DEL-85 | chr5-5209232-DEL-85 |
| chr5-5282029-DEL-168 | chr5-5282030-DEL-168 |
| chr5-5300780-DEL-112 | chr5-5300757-DEL-110 |
| chr5-5322023-DEL-333 | chr5-5322027-DEL-329 |
| chr5-5328554-DEL-1309 | chr5-5328543-DEL-1310 |
| chr5-5361967-DEL-137 | chr5-5361968-DEL-137 |
| chr5-5378507-DEL-91 | chr5-5378469-DEL-92 |
| chr5-5382221-DEL-58 | chr5-5382221-DEL-58 |
| chr5-5432033-DEL-310 | chr5-5432034-DEL-310 |
| chr5-5435231-DEL-51 | chr5-5435232-DEL-51 |
| chr5-5447031-DEL-247 | chr5-5447032-DEL-247 |
| chr5-5453382-DEL-6199 | chr5-5453386-DEL-6196 |
| chr5-5482661-DEL-1270 | chr5-5482658-DEL-1271 |
| chr5-5503450-DEL-102 | chr5-5503451-DEL-102 |
| chr5-5517339-DEL-6367 | chr5-5517340-DEL-6367 |
| chr5-5528405-DEL-205 | chr5-5528404-DEL-206 |
| chr5-5543651-DEL-465 | chr5-5543652-DEL-465 |
| chr5-5586915-DEL-83 | chr5-5586916-DEL-83 |

| | |
|---|---|
| chr5-5592592-DEL-66 | chr5-5592593-DEL-66 |
| chr5-5596017-DEL-203 | chr5-5596018-DEL-203 |
| chr5-5603393-DEL-429 | chr5-5603394-DEL-429 |
| chr5-5647282-DEL-1650 | chr5-5647283-DEL-1650 |
| chr5-5660278-DEL-191 | chr5-5660279-DEL-191 |
| chr5-5680454-DEL-6435 | chr5-5680455-DEL-6435 |
| chr5-5751583-DEL-212 | chr5-5751584-DEL-212 |
| chr5-5754364-DEL-1336 | chr5-5754365-DEL-1336 |
| chr5-5762615-DEL-719 | chr5-5762597-DEL-720 |
| chr5-5774054-DEL-60 | chr5-5774059-DEL-54 |
| chr5-5788968-DEL-193 | chr5-5788966-DEL-194 |
| chr5-5808673-DEL-307 | chr5-5808674-DEL-307 |
| chr5-5833877-DEL-2627 | chr5-5833877-DEL-2618 |
| chr5-5835727-DEL-75 | chr5-5835728-DEL-75 |
| chr5-5853294-DEL-222 | chr5-5853295-DEL-222 |
| chr5-5855779-DEL-557 | chr5-5855780-DEL-557 |
| chr5-5897997-DEL-621 | chr5-5897998-DEL-621 |
| chr5-5914463-DEL-400 | chr5-5914464-DEL-400 |
| chr5-5946670-DEL-608 | chr5-5946671-DEL-608 |
| chr5-5977529-DEL-221 | chr5-5977530-DEL-221 |
| chr5-5978915-DEL-569 | chr5-5978916-DEL-569 |
| chr5-5994813-DEL-408 | chr5-5994814-DEL-408 |
| chr5-6050963-DEL-54 | chr5-6050964-DEL-54 |
| chr5-6077035-DEL-52 | chr5-6077036-DEL-52 |
| chr5-6131454-DEL-54 | chr5-6131453-DEL-70 |
| chr5-6133091-DEL-775 | chr5-6133091-DEL-775 |
| chr5-6140587-DEL-822 | chr5-6140588-DEL-822 |
| chr5-6145680-DEL-80 | chr5-6145707-DEL-80 |
| chr5-6149039-DEL-135 | chr5-6149039-DEL-135 |
| chr5-6153704-DEL-206 | chr5-6153705-DEL-203 |
| chr5-6201651-DEL-498 | chr5-6201652-DEL-498 |
| chr5-6234066-DEL-7324 | chr5-6234067-DEL-7324 |
| chr5-6255178-DEL-173 | chr5-6255179-DEL-173 |
| chr5-6262520-DEL-10848 | chr5-6262516-DEL-10852 |
| chr5-6307327-DEL-4987 | chr5-6307328-DEL-4987 |
| chr5-6340143-DEL-91 | chr5-6340137-DEL-98 |
| chr5-6353940-DEL-404 | chr5-6353941-DEL-404 |
| chr5-6360145-DEL-1661 | chr5-6360146-DEL-1661 |
| chr5-6365393-DEL-400 | chr5-6365394-DEL-400 |
| chr5-6381891-DEL-932 | chr5-6381892-DEL-932 |
| chr5-6383050-DEL-152 | chr5-6383051-DEL-152 |
| chr5-6411767-DEL-307 | chr5-6411768-DEL-307 |
| chr5-6432168-DEL-135 | chr5-6432169-DEL-135 |
| chr5-6440509-DEL-148 | chr5-6440510-DEL-148 |
| chr5-6446430-DEL-6236 | chr5-6446431-DEL-6236 |
| chr5-6521680-DEL-64 | chr5-6521681-DEL-64 |
| chr5-6533427-DEL-8851 | chr5-6533428-DEL-8851 |
| chr5-6559424-DEL-3608 | chr5-6559425-DEL-3608 |
| chr5-6606585-DEL-10699 | chr5-6606586-DEL-10699 |
| chr5-6640842-DEL-2740 | chr5-6640843-DEL-2740 |
| chr5-6652668-DEL-983 | chr5-6652669-DEL-983 |
| chr5-6657080-DEL-846 | chr5-6657081-DEL-846 |
| chr5-6669916-DEL-358 | chr5-6669917-DEL-358 |
| chr5-6676067-DEL-201 | chr5-6676068-DEL-201 |
| chr5-6678865-DEL-215 | chr5-6678866-DEL-215 |
| chr5-6679624-DEL-1350 | chr5-6679621-DEL-1350 |
| chr5-6696018-DEL-517 | chr5-6696019-DEL-517 |
| chr5-6714002-DEL-71 | chr5-6714012-DEL-61 |
| chr5-6715144-DEL-79 | chr5-6715145-DEL-79 |
| chr5-6717238-DEL-63 | chr5-6717238-DEL-63 |
| chr5-6738403-DEL-157 | chr5-6738404-DEL-157 |
| chr5-6744600-DEL-535 | chr5-6744601-DEL-534 |
| chr5-6754965-DEL-51 | chr5-6754933-DEL-99 |
| chr5-6768863-DEL-53 | chr5-6768864-DEL-53 |
| chr5-6774344-DEL-89 | chr5-6774345-DEL-89 |
| chr5-6786005-DEL-6460 | chr5-6786006-DEL-6460 |
| chr5-6803749-DEL-361 | chr5-6803750-DEL-361 |
| chr5-6844971-DEL-281 | chr5-6844972-DEL-281 |
| chr5-6860831-DEL-64 | chr5-6860832-DEL-64 |
| chr5-6917568-DEL-156 | chr5-6917569-DEL-156 |
| chr5-6919191-DEL-62 | chr5-6919192-DEL-64 |
| chr5-6924172-DEL-1607 | chr5-6924173-DEL-1607 |
| chr5-6938958-DEL-2770 | chr5-6938959-DEL-2770 |
| chr5-6969379-DEL-456 | chr5-6969380-DEL-456 |
| chr5-6979292-DEL-1531 | chr5-6979293-DEL-1531 |
| chr5-7002318-DEL-56 | chr5-7002319-DEL-56 |
| chr5-7005417-DEL-237 | chr5-7005418-DEL-238 |
| chr5-7008101-DEL-956 | chr5-7008103-DEL-956 |
| chr5-7015229-DEL-6170 | chr5-7015230-DEL-6170 |
| chr5-7022326-DEL-1053 | chr5-7022327-DEL-1053 |

| | |
|---|---|
| chr5-7029073-DEL-52 | chr5-7029062-DEL-64 |
| chr5-7061315-DEL-898 | chr5-7061315-DEL-899 |
| chr5-7105349-DEL-86 | chr5-7105350-DEL-86 |
| chr5-7112822-DEL-4269 | chr5-7112823-DEL-4270 |
| chr5-7120072-DEL-489 | chr5-7120073-DEL-489 |
| chr5-7134543-DEL-212 | chr5-7134544-DEL-212 |
| chr5-7135456-DEL-6667 | chr5-7135457-DEL-6667 |
| chr5-7162125-DEL-327 | chr5-7162126-DEL-327 |
| chr5-7180292-DEL-1303 | chr5-7180290-DEL-1312 |
| chr5-7194654-DEL-99 | chr5-7194655-DEL-99 |
| chr5-7200365-DEL-5612 | chr5-7200364-DEL-5613 |
| chr5-7236420-DEL-53 | chr5-7236421-DEL-53 |
| chr5-7237319-DEL-1064 | chr5-7237318-DEL-1063 |
| chr5-7241570-DEL-54 | chr5-7241571-DEL-54 |
| chr5-7242538-DEL-113 | chr5-7242535-DEL-114 |
| chr5-7244789-DEL-1772 | chr5-7244790-DEL-1772 |
| chr5-7250212-DEL-6227 | chr5-7250213-DEL-6227 |
| chr5-7277037-DEL-473 | chr5-7277038-DEL-473 |
| chr5-7304853-DEL-773 | chr5-7304854-DEL-773 |
| chr5-7306063-DEL-429 | chr5-7306064-DEL-428 |
| chr5-7307086-DEL-567 | chr5-7307087-DEL-565 |
| chr5-7318306-DEL-558 | chr5-7318307-DEL-558 |
| chr5-7319237-DEL-261 | chr5-7319229-DEL-291 |
| chr5-7342401-DEL-2035 | chr5-7342402-DEL-2035 |
| chr5-7366230-DEL-99 | chr5-7366226-DEL-104 |
| chr5-7368605-DEL-59 | chr5-7368606-DEL-59 |
| chr5-7383041-DEL-54 | chr5-7383042-DEL-54 |
| chr5-7403628-DEL-2353 | chr5-7403631-DEL-2351 |
| chr5-7409513-DEL-542 | chr5-7409514-DEL-542 |
| chr5-7447432-DEL-109 | chr5-7447433-DEL-109 |
| chr5-7455520-DEL-7674 | chr5-7455518-DEL-7671 |
| chr5-7459431-DEL-759 | chr5-7459431-DEL-759 |
| chr5-7464100-DEL-1283 | chr5-7464101-DEL-1283 |
| chr5-7469768-DEL-1652 | chr5-7469769-DEL-1652 |
| chr5-7478873-DEL-6902 | chr5-7478874-DEL-6902 |
| chr5-7499394-DEL-55 | chr5-7499395-DEL-55 |
| chr5-7542591-DEL-2183 | chr5-7542592-DEL-2183 |
| chr5-7566616-DEL-131 | chr5-7566617-DEL-131 |
| chr5-7600390-DEL-85 | chr5-7600391-DEL-85 |
| chr5-7626268-DEL-59 | chr5-7626269-DEL-58 |
| chr5-7627212-DEL-54 | chr5-7627213-DEL-54 |
| chr5-7665357-DEL-7236 | chr5-7665358-DEL-7236 |
| chr5-7688664-DEL-430 | chr5-7688665-DEL-430 |
| chr5-7708839-DEL-2244 | chr5-7708840-DEL-2244 |
| chr5-7717252-DEL-5965 | chr5-7717253-DEL-5965 |
| chr5-7790105-DEL-53 | chr5-7790109-DEL-50 |
| chr5-7801568-DEL-2925 | chr5-7801569-DEL-2925 |
| chr5-7806140-DEL-1451 | chr5-7806141-DEL-1452 |
| chr5-7826169-DEL-82 | chr5-7826170-DEL-82 |
| chr5-7850095-DEL-207 | chr5-7850096-DEL-199 |
| chr5-7854922-DEL-6428 | chr5-7854923-DEL-6428 |
| chr5-7868320-DEL-2047 | chr5-7868321-DEL-2047 |
| chr5-7872550-DEL-2665 | chr5-7872551-DEL-2665 |
| chr5-7912043-DEL-202 | chr5-7912044-DEL-202 |
| chr5-7921829-DEL-327 | chr5-7921830-DEL-327 |
| chr5-7931676-DEL-66 | chr5-7931677-DEL-62 |
| chr5-7945110-DEL-314 | chr5-7945111-DEL-314 |
| chr5-7970317-DEL-61 | chr5-7970359-DEL-56 |
| chr5-7970984-DEL-56 | chr5-7970985-DEL-60 |
| chr5-7978604-DEL-53 | chr5-7978605-DEL-53 |
| chr5-7983609-DEL-84 | chr5-7983622-DEL-83 |
| chr5-7987274-DEL-201 | chr5-7987300-DEL-201 |
| chr5-7988029-DEL-7205 | chr5-7988030-DEL-7205 |
| chr5-8003179-DEL-106 | chr5-8003184-DEL-104 |
| chr5-8020988-DEL-1680 | chr5-8020989-DEL-1680 |
| chr5-8029458-DEL-160 | chr5-8029459-DEL-160 |
| chr5-8033559-DEL-339 | chr5-8033560-DEL-339 |
| chr5-8054405-DEL-82 | chr5-8054405-DEL-82 |
| chr5-8057604-DEL-226 | chr5-8057605-DEL-226 |
| chr5-8066866-DEL-74 | chr5-8066884-DEL-100 |
| chr5-8088933-DEL-95 | chr5-8088905-DEL-96 |
| chr5-8089542-DEL-1234 | chr5-8089543-DEL-1234 |
| chr5-8103024-DEL-352 | chr5-8103026-DEL-351 |
| chr5-8190399-DEL-686 | chr5-8190400-DEL-686 |
| chr5-8219153-DEL-6335 | chr5-8219154-DEL-6335 |
| chr5-8243792-DEL-54 | chr5-8243792-DEL-54 |
| chr5-8262185-DEL-53 | chr5-8262186-DEL-53 |
| chr5-8290659-DEL-78 | chr5-8290660-DEL-78 |
| chr5-8307001-DEL-141 | chr5-8307002-DEL-141 |
| chr5-8308738-DEL-6622 | chr5-8308739-DEL-6622 |

| | |
|---|---|
| chr5-8316583-DEL-6224 | chr5-8316584-DEL-6224 |
| chr5-8331783-DEL-1785 | chr5-8331782-DEL-1786 |
| chr5-8376033-DEL-57 | chr5-8376034-DEL-57 |
| chr5-8422167-DEL-317 | chr5-8422168-DEL-317 |
| chr5-8426298-DEL-1480 | chr5-8426299-DEL-1480 |
| chr5-8432146-DEL-6351 | chr5-8432147-DEL-6351 |
| chr5-8491280-DEL-431 | chr5-8491282-DEL-432 |
| chr5-8511120-DEL-209 | chr5-8511121-DEL-209 |
| chr5-8514636-DEL-4177 | chr5-8514637-DEL-4177 |
| chr5-8529397-DEL-73 | chr5-8529398-DEL-73 |
| chr5-8533077-DEL-118 | chr5-8533078-DEL-118 |
| chr5-8560123-DEL-569 | chr5-8560124-DEL-569 |
| chr5-8580036-DEL-6164 | chr5-8580037-DEL-6164 |
| chr5-8599867-DEL-154 | chr5-8599866-DEL-156 |
| chr5-8601588-DEL-6270 | chr5-8601589-DEL-6270 |
| chr5-8611093-DEL-379 | chr5-8611094-DEL-379 |
| chr5-8620304-DEL-134 | chr5-8620305-DEL-134 |
| chr5-8646128-DEL-2484 | chr5-8646129-DEL-2484 |
| chr5-8681998-DEL-1749 | chr5-8681999-DEL-1749 |
| chr5-8702974-DEL-6421 | chr5-8702975-DEL-6421 |
| chr5-8713738-DEL-176 | chr5-8713739-DEL-176 |
| chr5-8740569-DEL-66 | chr5-8740564-DEL-72 |
| chr5-8747371-DEL-76 | chr5-8747372-DEL-76 |
| chr5-8754388-DEL-7813 | chr5-8754388-DEL-7814 |
| chr5-8767978-DEL-80 | chr5-8767979-DEL-80 |
| chr5-8804242-DEL-610 | chr5-8804243-DEL-610 |
| chr5-8827245-DEL-112 | chr5-8827246-DEL-112 |
| chr5-8829392-DEL-54 | chr5-8829396-DEL-51 |
| chr5-8838219-DEL-249 | chr5-8838213-DEL-250 |
| chr5-8881422-DEL-60 | chr5-8881423-DEL-60 |
| chr5-8916823-DEL-154 | chr5-8916856-DEL-124 |
| chr5-8919100-DEL-976 | chr5-8919101-DEL-976 |
| chr5-8930732-DEL-552 | chr5-8930760-DEL-552 |
| chr5-8985897-DEL-93 | chr5-8985898-DEL-93 |
| chr5-8995185-DEL-150 | chr5-8995140-DEL-150 |
| chr5-9015303-DEL-92 | chr5-9015298-DEL-87 |
| chr5-9023164-DEL-58 | chr5-9023165-DEL-58 |
| chr5-9083222-DEL-229 | chr5-9083223-DEL-229 |
| chr5-9086162-DEL-222 | chr5-9086163-DEL-222 |
| chr5-9091078-DEL-106 | chr5-9091061-DEL-120 |
| chr5-9096407-DEL-69 | chr5-9096427-DEL-52 |
| chr5-9099256-DEL-6675 | chr5-9099257-DEL-6675 |
| chr5-9099671-DEL-212 | chr5-9099672-DEL-212 |
| chr5-9145808-DEL-200 | chr5-9145809-DEL-200 |
| chr5-9153096-DEL-207 | chr5-9153097-DEL-207 |
| chr5-9190525-DEL-76 | chr5-9190526-DEL-76 |
| chr5-9204003-DEL-165 | chr5-9204004-DEL-165 |
| chr5-9238450-DEL-190 | chr5-9238457-DEL-194 |
| chr5-9240688-DEL-6187 | chr5-9240689-DEL-6187 |
| chr5-9310631-DEL-764 | chr5-9310632-DEL-764 |
| chr5-9317953-DEL-62 | chr5-9317954-DEL-62 |
| chr5-9353641-DEL-7403 | chr5-9353642-DEL-7403 |
| chr5-9363206-DEL-89 | chr5-9363207-DEL-89 |
| chr5-9371270-DEL-1775 | chr5-9371270-DEL-1775 |
| chr5-9371642-DEL-159 | chr5-9371641-DEL-160 |
| chr5-9380377-DEL-6404 | chr5-9380378-DEL-6404 |
| chr5-9392451-DEL-312 | chr5-9392451-DEL-312 |
| chr5-9401846-DEL-76 | chr5-9401847-DEL-76 |
| chr5-9415106-DEL-118 | chr5-9415105-DEL-119 |
| chr5-9429544-DEL-70 | chr5-9429545-DEL-70 |
| chr5-9446436-DEL-536 | chr5-9446439-DEL-534 |
| chr5-9483024-DEL-888 | chr5-9483025-DEL-888 |
| chr5-9502492-DEL-148 | chr5-9502493-DEL-147 |
| chr5-9553278-DEL-252 | chr5-9553279-DEL-252 |
| chr5-9574990-DEL-322 | chr5-9574991-DEL-322 |
| chr5-9626051-DEL-312 | chr5-9626052-DEL-312 |
| chr5-9645909-DEL-315 | chr5-9645910-DEL-315 |
| chr5-9695880-DEL-52 | chr5-9695873-DEL-60 |
| chr5-9709519-DEL-1017 | chr5-9709494-DEL-1018 |
| chr5-9710362-DEL-257 | chr5-9710363-DEL-257 |
| chr5-9855646-DEL-2324 | chr5-9855647-DEL-2324 |
| chr5-9860872-DEL-171 | chr5-9860902-DEL-150 |
| chr5-9864566-DEL-6236 | chr5-9864567-DEL-6236 |
| chr5-9875279-DEL-207 | chr5-9875280-DEL-207 |
| chr5-9876323-DEL-1650 | chr5-9876324-DEL-1650 |
| chr5-9882633-DEL-82 | chr5-9882634-DEL-82 |
| chr5-9888934-DEL-192 | chr5-9888935-DEL-190 |
| chr5-9890324-DEL-138 | chr5-9890349-DEL-114 |
| chr5-9893519-DEL-56 | chr5-9893520-DEL-60 |
| chr5-9917129-DEL-191 | chr5-9917130-DEL-191 |

| | |
|---|---|
| chr5-9938978-DEL-142 | chr5-9938999-DEL-157 |
| chr5-9942033-DEL-174 | chr5-9942034-DEL-174 |
| chr5-9981136-DEL-4769 | chr5-9981137-DEL-4769 |
| chr5-10001510-DEL-404 | chr5-10001510-DEL-404 |
| chr5-10028054-DEL-74 | chr5-10028055-DEL-74 |
| chr5-10045954-DEL-444 | chr5-10045955-DEL-444 |
| chr5-10085448-DEL-102 | chr5-10085449-DEL-102 |
| chr5-10094678-DEL-1794 | chr5-10094679-DEL-1794 |
| chr5-10120653-DEL-7073 | chr5-10120654-DEL-7073 |
| chr5-10151067-DEL-244 | chr5-10151068-DEL-244 |
| chr5-10154294-DEL-108 | chr5-10154283-DEL-70 |
| chr5-10162308-DEL-666 | chr5-10162309-DEL-666 |
| chr5-10163791-DEL-660 | chr5-10163792-DEL-660 |
| chr5-10190383-DEL-889 | chr5-10190383-DEL-889 |
| chr5-10212333-DEL-52 | chr5-10212334-DEL-52 |
| chr5-10224269-DEL-218 | chr5-10224270-DEL-218 |
| chr5-10228343-DEL-745 | chr5-10228340-DEL-746 |
| chr5-10277788-DEL-145 | chr5-10277789-DEL-145 |
| chr5-10280019-DEL-52 | chr5-10280036-DEL-60 |
| chr5-10339672-DEL-4428 | chr5-10339673-DEL-4428 |
| chr5-10349133-DEL-136 | chr5-10349134-DEL-136 |
| chr5-10417939-DEL-52 | chr5-10417940-DEL-53 |
| chr5-10432014-DEL-109 | chr5-10432015-DEL-109 |
| chr5-10455789-DEL-342 | chr5-10455786-DEL-343 |
| chr5-10460714-DEL-105 | chr5-10460715-DEL-105 |
| chr5-10502090-DEL-6247 | chr5-10502091-DEL-6247 |
| chr5-10523688-DEL-1584 | chr5-10523689-DEL-1584 |
| chr5-10536102-DEL-964 | chr5-10536103-DEL-964 |
| chr5-10586955-DEL-70 | chr5-10586956-DEL-70 |
| chr5-10615681-DEL-6710 | chr5-10615682-DEL-6709 |
| chr5-10649183-DEL-219 | chr5-10649187-DEL-213 |
| chr5-10653849-DEL-504 | chr5-10653849-DEL-505 |
| chr5-10667250-DEL-403 | chr5-10667251-DEL-403 |
| chr5-10671345-DEL-8662 | chr5-10671345-DEL-8662 |
| chr5-10686910-DEL-1862 | chr5-10686910-DEL-1862 |
| chr5-10715616-DEL-321 | chr5-10715617-DEL-321 |
| chr5-10743731-DEL-692 | chr5-10743730-DEL-698 |
| chr5-10776220-DEL-83 | chr5-10776219-DEL-81 |
| chr5-10870762-DEL-6356 | chr5-10870763-DEL-6356 |
| chr5-11220049-DEL-288 | chr5-11220048-DEL-288 |
| chr5-11484506-DEL-64 | chr5-11484507-DEL-64 |
| chr5-11981835-DEL-68 | chr5-11981836-DEL-68 |
| chr5-12003467-DEL-399 | chr5-12003468-DEL-399 |
| chr5-12007255-DEL-6421 | chr5-12007256-DEL-6421 |
| chr5-12013428-DEL-214 | chr5-12013429-DEL-214 |
| chr5-12039044-DEL-6781 | chr5-12039045-DEL-6781 |
| chr5-12039581-DEL-2781 | chr5-12039582-DEL-2781 |
| chr5-12049336-DEL-233 | chr5-12049337-DEL-233 |
| chr5-12059376-DEL-67 | chr5-12059378-DEL-66 |
| chr5-12073256-DEL-899 | chr5-12073257-DEL-899 |
| chr5-12120525-DEL-1212 | chr5-12120526-DEL-1212 |
| chr5-12137788-DEL-90 | chr5-12137788-DEL-88 |
| chr5-12139302-DEL-379 | chr5-12139303-DEL-379 |
| chr5-12144551-DEL-358 | chr5-12144552-DEL-358 |
| chr5-12171223-DEL-623 | chr5-12171224-DEL-621 |
| chr5-12171983-DEL-70 | chr5-12171984-DEL-70 |
| chr5-12187284-DEL-89 | chr5-12187278-DEL-50 |
| chr5-12190267-DEL-1584 | chr5-12190268-DEL-1584 |
| chr5-12192222-DEL-705 | chr5-12192224-DEL-703 |
| chr5-12193787-DEL-403 | chr5-12193788-DEL-403 |
| chr5-12195124-DEL-170 | chr5-12195125-DEL-170 |
| chr5-12200475-DEL-87 | chr5-12200476-DEL-87 |
| chr5-12205058-DEL-110 | chr5-12205072-DEL-118 |
| chr5-12214265-DEL-5194 | chr5-12214266-DEL-5194 |
| chr5-12236755-DEL-499 | chr5-12236756-DEL-499 |
| chr5-12243578-DEL-67 | chr5-12243554-DEL-62 |
| chr5-12267975-DEL-67 | chr5-12267934-DEL-66 |
| chr5-12290618-DEL-179 | chr5-12290615-DEL-180 |
| chr5-12296612-DEL-1742 | chr5-12296608-DEL-1740 |
| chr5-12298822-DEL-62 | chr5-12298823-DEL-60 |
| chr5-12299606-DEL-4350 | chr5-12299607-DEL-4350 |
| chr5-12315144-DEL-108 | chr5-12315145-DEL-108 |
| chr5-12321520-DEL-87 | chr5-12321521-DEL-87 |
| chr5-12328783-DEL-6381 | chr5-12328784-DEL-6381 |
| chr5-12341363-DEL-774 | chr5-12341355-DEL-775 |
| chr5-12369949-DEL-995 | chr5-12369940-DEL-993 |
| chr5-12386988-DEL-214 | chr5-12386989-DEL-214 |
| chr5-12483349-DEL-1668 | chr5-12483350-DEL-1668 |
| chr5-12487806-DEL-239 | chr5-12487807-DEL-239 |
| chr5-12496935-DEL-313 | chr5-12496936-DEL-313 |

| | |
|---|---|
| chr5-12503909-DEL-82 | chr5-12503910-DEL-82 |
| chr5-12520272-DEL-3921 | chr5-12520269-DEL-3923 |
| chr5-12528779-DEL-824 | chr5-12528780-DEL-824 |
| chr5-12539544-DEL-1153 | chr5-12539533-DEL-1151 |
| chr5-12567051-DEL-4063 | chr5-12567052-DEL-4063 |
| chr5-12581866-DEL-65 | chr5-12581867-DEL-65 |
| chr5-12591799-DEL-95 | chr5-12591798-DEL-96 |
| chr5-12592628-DEL-6441 | chr5-12592629-DEL-6441 |
| chr5-12599407-DEL-51 | chr5-12599408-DEL-51 |
| chr5-12639231-DEL-541 | chr5-12639232-DEL-540 |
| chr5-12667837-DEL-111 | chr5-12667838-DEL-111 |
| chr5-12710087-DEL-66 | chr5-12710042-DEL-104 |
| chr5-12728553-DEL-1736 | chr5-12728553-DEL-1737 |
| chr5-12758500-DEL-732 | chr5-12758501-DEL-732 |
| chr5-12767094-DEL-740 | chr5-12767095-DEL-740 |
| chr5-12778356-DEL-220 | chr5-12778357-DEL-220 |
| chr5-12788668-DEL-5117 | chr5-12788669-DEL-5117 |
| chr5-12810317-DEL-154 | chr5-12810318-DEL-154 |
| chr5-12811465-DEL-1266 | chr5-12811466-DEL-1266 |
| chr5-12826341-DEL-1846 | chr5-12826341-DEL-1847 |
| chr5-12840039-DEL-6251 | chr5-12840039-DEL-6251 |
| chr5-12850484-DEL-1908 | chr5-12850484-DEL-1908 |
| chr5-12855392-DEL-1541 | chr5-12855393-DEL-1541 |
| chr5-12895650-DEL-212 | chr5-12895651-DEL-212 |
| chr5-12909123-DEL-206 | chr5-12909124-DEL-206 |
| chr5-12926162-DEL-187 | chr5-12926145-DEL-189 |
| chr5-12952404-DEL-334 | chr5-12952404-DEL-334 |
| chr5-12975189-DEL-217 | chr5-12975190-DEL-217 |
| chr5-12978815-DEL-96 | chr5-12978816-DEL-96 |
| chr5-12990119-DEL-527 | chr5-12990120-DEL-527 |
| chr5-13001922-DEL-1470 | chr5-13001923-DEL-1470 |
| chr5-13030156-DEL-508 | chr5-13030157-DEL-508 |
| chr5-13037986-DEL-99 | chr5-13037987-DEL-62 |
| chr5-13045830-DEL-52 | chr5-13045831-DEL-52 |
| chr5-13064547-DEL-637 | chr5-13064548-DEL-637 |
| chr5-13074696-DEL-408 | chr5-13074697-DEL-408 |
| chr5-13130851-DEL-394 | chr5-13130852-DEL-394 |
| chr5-13170830-DEL-968 | chr5-13170831-DEL-968 |
| chr5-13191637-DEL-710 | chr5-13191643-DEL-709 |
| chr5-13236378-DEL-129 | chr5-13236379-DEL-129 |
| chr5-13249077-DEL-66 | chr5-13249078-DEL-66 |
| chr5-13259489-DEL-900 | chr5-13259490-DEL-900 |
| chr5-13265360-DEL-940 | chr5-13265361-DEL-940 |
| chr5-13267678-DEL-759 | chr5-13267679-DEL-759 |
| chr5-13276009-DEL-1176 | chr5-13276010-DEL-1176 |
| chr5-13285851-DEL-93 | chr5-13285865-DEL-69 |
| chr5-13301517-DEL-171 | chr5-13301518-DEL-171 |
| chr5-13328172-DEL-74 | chr5-13328173-DEL-74 |
| chr5-13336538-DEL-535 | chr5-13336539-DEL-535 |
| chr5-13342626-DEL-2303 | chr5-13342627-DEL-2303 |
| chr5-13413233-DEL-71 | chr5-13413227-DEL-81 |
| chr5-13414650-DEL-62 | chr5-13414647-DEL-62 |
| chr5-13417005-DEL-445 | chr5-13417006-DEL-445 |
| chr5-13425936-DEL-58 | chr5-13425949-DEL-62 |
| chr5-13432220-DEL-676 | chr5-13432221-DEL-676 |
| chr5-13460514-DEL-69 | chr5-13460505-DEL-79 |
| chr5-13462765-DEL-53 | chr5-13462783-DEL-52 |
| chr5-13463671-DEL-82 | chr5-13463674-DEL-80 |
| chr5-13568661-DEL-111 | chr5-13568662-DEL-111 |
| chr5-13579908-DEL-565 | chr5-13579909-DEL-565 |
| chr5-13586452-DEL-1889 | chr5-13586453-DEL-1889 |
| chr5-13621323-DEL-79 | chr5-13621324-DEL-79 |
| chr5-13624430-DEL-453 | chr5-13624431-DEL-453 |
| chr5-13631413-DEL-54 | chr5-13631413-DEL-54 |
| chr5-13636182-DEL-723 | chr5-13636231-DEL-675 |
| chr5-13645294-DEL-64 | chr5-13645276-DEL-65 |
| chr5-13695034-DEL-61 | chr5-13695035-DEL-61 |
| chr5-13724875-DEL-98 | chr5-13724876-DEL-98 |
| chr5-13725831-DEL-3770 | chr5-13725832-DEL-3770 |
| chr5-13746188-DEL-1641 | chr5-13746189-DEL-1641 |
| chr5-13746400-DEL-232 | chr5-13746401-DEL-232 |
| chr5-13747238-DEL-1673 | chr5-13747232-DEL-1674 |
| chr5-13764938-DEL-445 | chr5-13764939-DEL-445 |
| chr5-13766940-DEL-120 | chr5-13766988-DEL-96 |
| chr5-13774084-DEL-6289 | chr5-13774085-DEL-6289 |
| chr5-13787488-DEL-362 | chr5-13787489-DEL-360 |
| chr5-13818006-DEL-246 | chr5-13818007-DEL-246 |
| chr5-13889899-DEL-915 | chr5-13889900-DEL-915 |
| chr5-13898143-DEL-1241 | chr5-13898144-DEL-1241 |
| chr5-13907007-DEL-2549 | chr5-13907008-DEL-2549 |

| | |
|---|---|
| chr5-13909792-DEL-4316 | chr5-13909793-DEL-4316 |
| chr5-13914944-DEL-6851 | chr5-13914945-DEL-6851 |
| chr5-13934034-DEL-2651 | chr5-13934026-DEL-2652 |
| chr5-13936795-DEL-55 | chr5-13936796-DEL-55 |
| chr5-13968155-DEL-767 | chr5-13968156-DEL-767 |
| chr5-13985262-DEL-130 | chr5-13985261-DEL-130 |
| chr5-14010016-DEL-59 | chr5-14010017-DEL-59 |
| chr5-14014570-DEL-202 | chr5-14014571-DEL-202 |
| chr5-14015541-DEL-100 | chr5-14015542-DEL-100 |
| chr5-14022293-DEL-85 | chr5-14022294-DEL-85 |
| chr5-14056895-DEL-52 | chr5-14056896-DEL-52 |
| chr5-14111305-DEL-61 | chr5-14111306-DEL-61 |
| chr5-14119273-DEL-589 | chr5-14119274-DEL-589 |
| chr5-14136718-DEL-165 | chr5-14136719-DEL-165 |
| chr5-14149185-DEL-220 | chr5-14149186-DEL-220 |
| chr5-14190954-DEL-198 | chr5-14190955-DEL-198 |
| chr5-14195327-DEL-72 | chr5-14195328-DEL-72 |
| chr5-14198006-DEL-2205 | chr5-14198008-DEL-2206 |
| chr5-14217244-DEL-675 | chr5-14217246-DEL-674 |
| chr5-14219448-DEL-555 | chr5-14219449-DEL-555 |
| chr5-14224092-DEL-6538 | chr5-14224093-DEL-6538 |
| chr5-14230102-DEL-220 | chr5-14230103-DEL-220 |
| chr5-14235312-DEL-55 | chr5-14235313-DEL-55 |
| chr5-14238847-DEL-1202 | chr5-14238848-DEL-1202 |
| chr5-14252968-DEL-416 | chr5-14252976-DEL-415 |
| chr5-14261051-DEL-702 | chr5-14261052-DEL-702 |
| chr5-14330205-DEL-108 | chr5-14330206-DEL-108 |
| chr5-14395496-DEL-254 | chr5-14395495-DEL-255 |
| chr5-14401468-DEL-185 | chr5-14401462-DEL-181 |
| chr5-14405913-DEL-66 | chr5-14405914-DEL-66 |
| chr5-14412364-DEL-80 | chr5-14412365-DEL-80 |
| chr5-14422412-DEL-203 | chr5-14422413-DEL-203 |
| chr5-14427064-DEL-6042 | chr5-14427065-DEL-6042 |
| chr5-14445474-DEL-6472 | chr5-14445475-DEL-6472 |
| chr5-14453323-DEL-543 | chr5-14453324-DEL-543 |
| chr5-14467237-DEL-59 | chr5-14467267-DEL-52 |
| chr5-14481583-DEL-2545 | chr5-14481584-DEL-2545 |
| chr5-14493594-DEL-400 | chr5-14493593-DEL-400 |
| chr5-14517215-DEL-96 | chr5-14517216-DEL-96 |
| chr5-14531798-DEL-52 | chr5-14531798-DEL-60 |
| chr5-14552550-DEL-71 | chr5-14552555-DEL-67 |
| chr5-14571966-DEL-90 | chr5-14571958-DEL-60 |
| chr5-14586697-DEL-212 | chr5-14586698-DEL-212 |
| chr5-14596488-DEL-6270 | chr5-14596489-DEL-6270 |
| chr5-14602497-DEL-206 | chr5-14602498-DEL-206 |
| chr5-14606191-DEL-560 | chr5-14606192-DEL-560 |
| chr5-14627241-DEL-851 | chr5-14627242-DEL-851 |
| chr5-14648369-DEL-2850 | chr5-14648370-DEL-2850 |
| chr5-14655502-DEL-475 | chr5-14655503-DEL-475 |
| chr5-14682812-DEL-1932 | chr5-14682813-DEL-1932 |
| chr5-14684869-DEL-1876 | chr5-14684870-DEL-1876 |
| chr5-14692663-DEL-1861 | chr5-14692663-DEL-1861 |
| chr5-14713149-DEL-3420 | chr5-14713141-DEL-3408 |
| chr5-14718620-DEL-201 | chr5-14718609-DEL-186 |
| chr5-14776031-DEL-145 | chr5-14776032-DEL-145 |
| chr5-14781240-DEL-78 | chr5-14781241-DEL-78 |
| chr5-14783048-DEL-261 | chr5-14783049-DEL-261 |
| chr5-14788096-DEL-201 | chr5-14788097-DEL-201 |
| chr5-14822589-DEL-1018 | chr5-14822590-DEL-1018 |
| chr5-14858146-DEL-71 | chr5-14858181-DEL-65 |
| chr5-14885839-DEL-2555 | chr5-14885840-DEL-2555 |
| chr5-14890794-DEL-1718 | chr5-14890795-DEL-1718 |
| chr5-14894861-DEL-566 | chr5-14894862-DEL-566 |
| chr5-14898770-DEL-351 | chr5-14898769-DEL-352 |
| chr5-14900602-DEL-1181 | chr5-14900603-DEL-1181 |
| chr5-14928829-DEL-1523 | chr5-14928830-DEL-1523 |
| chr5-14995762-DEL-924 | chr5-14995763-DEL-925 |
| chr5-15644388-DEL-66 | chr5-15644389-DEL-66 |
| chr5-15695761-DEL-934 | chr5-15695760-DEL-934 |
| chr5-15701300-DEL-123 | chr5-15701299-DEL-123 |
| chr5-15726156-DEL-106 | chr5-15726156-DEL-106 |
| chr5-15796634-DEL-270 | chr5-15796633-DEL-270 |
| chr5-15809735-DEL-72 | chr5-15809713-DEL-64 |
| chr5-15875521-DEL-214 | chr5-15875521-DEL-214 |
| chr5-15887098-DEL-66 | chr5-15887098-DEL-66 |
| chr5-15925965-DEL-721 | chr5-15925966-DEL-721 |
| chr5-15929755-DEL-2175 | chr5-15929755-DEL-2175 |
| chr5-15944482-DEL-1973 | chr5-15944483-DEL-1973 |
| chr5-15946822-DEL-401 | chr5-15946823-DEL-401 |
| chr5-15976235-DEL-82 | chr5-15976236-DEL-82 |

| | |
|---|---|
| chr5-15986571-DEL-6255 | chr5-15986572-DEL-6255 |
| chr5-16011947-DEL-7625 | chr5-16011948-DEL-7625 |
| chr5-16036698-DEL-75 | chr5-16036693-DEL-67 |
| chr5-16042954-DEL-84 | chr5-16042955-DEL-74 |
| chr5-16049124-DEL-679 | chr5-16049125-DEL-679 |
| chr5-16071600-DEL-158 | chr5-16071598-DEL-159 |
| chr5-16090741-DEL-448 | chr5-16090742-DEL-448 |
| chr5-16092037-DEL-68 | chr5-16092038-DEL-68 |
| chr5-16105123-DEL-597 | chr5-16105124-DEL-592 |
| chr5-16109027-DEL-809 | chr5-16109028-DEL-809 |
| chr5-16151309-DEL-1789 | chr5-16151310-DEL-1789 |
| chr5-16158121-DEL-1392 | chr5-16158122-DEL-1392 |
| chr5-16227345-DEL-183 | chr5-16227311-DEL-192 |
| chr5-16247448-DEL-2974 | chr5-16247449-DEL-2974 |
| chr5-16255560-DEL-493 | chr5-16255561-DEL-493 |
| chr5-16262814-DEL-778 | chr5-16262815-DEL-778 |
| chr5-16271294-DEL-6884 | chr5-16271294-DEL-6884 |
| chr5-16297187-DEL-219 | chr5-16297188-DEL-219 |
| chr5-16327851-DEL-96 | chr5-16327846-DEL-95 |
| chr5-16373967-DEL-6744 | chr5-16373968-DEL-6744 |
| chr5-16402587-DEL-576 | chr5-16402593-DEL-565 |
| chr5-16409799-DEL-52 | chr5-16409802-DEL-54 |
| chr5-16412327-DEL-70 | chr5-16412328-DEL-56 |
| chr5-16418017-DEL-72 | chr5-16418018-DEL-78 |
| chr5-16428158-DEL-81 | chr5-16428159-DEL-81 |
| chr5-16446248-DEL-167 | chr5-16446249-DEL-167 |
| chr5-16479032-DEL-842 | chr5-16479033-DEL-842 |
| chr5-16489902-DEL-7076 | chr5-16489903-DEL-7076 |
| chr5-16516713-DEL-54 | chr5-16516687-DEL-63 |
| chr5-16518523-DEL-60 | chr5-16518523-DEL-60 |
| chr5-16551537-DEL-68 | chr5-16551538-DEL-68 |
| chr5-16573858-DEL-73 | chr5-16573856-DEL-76 |
| chr5-16591910-DEL-103 | chr5-16591911-DEL-103 |
| chr5-16596274-DEL-239 | chr5-16596275-DEL-239 |
| chr5-16605142-DEL-337 | chr5-16605143-DEL-337 |
| chr5-16631296-DEL-66 | chr5-16631330-DEL-50 |
| chr5-16632096-DEL-51 | chr5-16632092-DEL-52 |
| chr5-16633818-DEL-3273 | chr5-16633819-DEL-3273 |
| chr5-16654425-DEL-1491 | chr5-16654441-DEL-1491 |
| chr5-16658630-DEL-2796 | chr5-16658631-DEL-2796 |
| chr5-16660651-DEL-81 | chr5-16660652-DEL-81 |
| chr5-16669861-DEL-83 | chr5-16669837-DEL-60 |
| chr5-16686502-DEL-60 | chr5-16686503-DEL-60 |
| chr5-16697565-DEL-62 | chr5-16697566-DEL-62 |
| chr5-16706250-DEL-1026 | chr5-16706251-DEL-1026 |
| chr5-16723237-DEL-70 | chr5-16723228-DEL-55 |
| chr5-16776919-DEL-76 | chr5-16776920-DEL-76 |
| chr5-16793610-DEL-1657 | chr5-16793611-DEL-1657 |
| chr5-16798123-DEL-414 | chr5-16798124-DEL-414 |
| chr5-16819001-DEL-53 | chr5-16818980-DEL-52 |
| chr5-16864778-DEL-3329 | chr5-16864779-DEL-3329 |
| chr5-16868493-DEL-89 | chr5-16868494-DEL-87 |
| chr5-16869408-DEL-114 | chr5-16869409-DEL-114 |
| chr5-16872418-DEL-7301 | chr5-16872419-DEL-7301 |
| chr5-16909559-DEL-2820 | chr5-16909567-DEL-2819 |
| chr5-16934563-DEL-1546 | chr5-16934564-DEL-1546 |
| chr5-16954690-DEL-6602 | chr5-16954691-DEL-6602 |
| chr5-16973295-DEL-325 | chr5-16973296-DEL-325 |
| chr5-17026074-DEL-60 | chr5-17026075-DEL-84 |
| chr5-17062606-DEL-707 | chr5-17062607-DEL-707 |
| chr5-17081201-DEL-78 | chr5-17081202-DEL-80 |
| chr5-17093844-DEL-3757 | chr5-17093844-DEL-3757 |
| chr5-17108401-DEL-7073 | chr5-17108402-DEL-7073 |
| chr5-17145842-DEL-6497 | chr5-17145843-DEL-6497 |
| chr5-17168609-DEL-2680 | chr5-17168610-DEL-2680 |
| chr5-17191025-DEL-256 | chr5-17191026-DEL-256 |
| chr5-17207258-DEL-161 | chr5-17207259-DEL-161 |
| chr5-17213717-DEL-70 | chr5-17213718-DEL-70 |
| chr5-17215986-DEL-688 | chr5-17215987-DEL-688 |
| chr5-17237832-DEL-6810 | chr5-17237833-DEL-6826 |
| chr5-17262195-DEL-421 | chr5-17262196-DEL-421 |
| chr5-17301807-DEL-6520 | chr5-17301808-DEL-6520 |
| chr5-17307951-DEL-214 | chr5-17307952-DEL-214 |
| chr5-17324558-DEL-267 | chr5-17324559-DEL-267 |
| chr5-17326442-DEL-295 | chr5-17326429-DEL-296 |
| chr5-17371549-DEL-231 | chr5-17371550-DEL-231 |
| chr5-17403297-DEL-180 | chr5-17403266-DEL-212 |
| chr5-17411406-DEL-94 | chr5-17411407-DEL-94 |
| chr5-17417888-DEL-6382 | chr5-17417889-DEL-6382 |
| chr5-17443239-DEL-9014 | chr5-17443240-DEL-9014 |

| | |
|---|---|
| chr5-17468304-DEL-112 | chr5-17468305-DEL-112 |
| chr5-17470892-DEL-4159 | chr5-17470893-DEL-4159 |
| chr5-17503874-DEL-678 | chr5-17503875-DEL-679 |
| chr5-17507205-DEL-66 | chr5-17507206-DEL-66 |
| chr5-17509343-DEL-92 | chr5-17509350-DEL-78 |
| chr5-17526850-DEL-324 | chr5-17526861-DEL-314 |
| chr5-17559447-DEL-3024 | chr5-17559448-DEL-3024 |
| chr5-17569615-DEL-1138 | chr5-17569616-DEL-1138 |
| chr5-17621553-DEL-55 | chr5-17621554-DEL-55 |
| chr5-17668297-DEL-3084 | chr5-17668298-DEL-3084 |
| chr5-17678929-DEL-595 | chr5-17678931-DEL-594 |
| chr5-17685391-DEL-90 | chr5-17685407-DEL-90 |
| chr5-17716005-DEL-87 | chr5-17716008-DEL-84 |
| chr5-17752556-DEL-82 | chr5-17752551-DEL-80 |
| chr5-17783229-DEL-64 | chr5-17783249-DEL-71 |
| chr5-17805404-DEL-86 | chr5-17805399-DEL-72 |
| chr5-17813430-DEL-7282 | chr5-17813431-DEL-7282 |
| chr5-17813445-DEL-212 | chr5-17813446-DEL-212 |
| chr5-17838238-DEL-322 | chr5-17838238-DEL-323 |
| chr5-17868289-DEL-1136 | chr5-17868290-DEL-1136 |
| chr5-17887142-DEL-3298 | chr5-17887143-DEL-3298 |
| chr5-17904747-DEL-1647 | chr5-17904760-DEL-1636 |
| chr5-17952702-DEL-141 | chr5-17952703-DEL-141 |
| chr5-17958171-DEL-65 | chr5-17958172-DEL-65 |
| chr5-17962668-DEL-233 | chr5-17962669-DEL-233 |
| chr5-17966683-DEL-56 | chr5-17966682-DEL-58 |
| chr5-18004076-DEL-6926 | chr5-18004077-DEL-6926 |
| chr5-18023122-DEL-60 | chr5-18023123-DEL-60 |
| chr5-18032352-DEL-176 | chr5-18032353-DEL-176 |
| chr5-18106331-DEL-64 | chr5-18106332-DEL-64 |
| chr5-18107976-DEL-80 | chr5-18107979-DEL-78 |
| chr5-18114673-DEL-6385 | chr5-18114674-DEL-6385 |
| chr5-18127126-DEL-412 | chr5-18127127-DEL-412 |
| chr5-18135571-DEL-80 | chr5-18135576-DEL-66 |
| chr5-18136896-DEL-67 | chr5-18136911-DEL-67 |
| chr5-18137771-DEL-56 | chr5-18137772-DEL-56 |
| chr5-18142756-DEL-104 | chr5-18142717-DEL-143 |
| chr5-18168986-DEL-52 | chr5-18168989-DEL-50 |
| chr5-18179525-DEL-84 | chr5-18179533-DEL-85 |
| chr5-18181679-DEL-649 | chr5-18181680-DEL-649 |
| chr5-18184067-DEL-574 | chr5-18184068-DEL-574 |
| chr5-18215893-DEL-90 | chr5-18215919-DEL-50 |
| chr5-18235017-DEL-407 | chr5-18235018-DEL-407 |
| chr5-18251872-DEL-205 | chr5-18251872-DEL-200 |
| chr5-18298428-DEL-341 | chr5-18298429-DEL-341 |
| chr5-18325999-DEL-3865 | chr5-18326000-DEL-3865 |
| chr5-18343926-DEL-193 | chr5-18343927-DEL-193 |
| chr5-18365518-DEL-147 | chr5-18365519-DEL-147 |
| chr5-18386706-DEL-394 | chr5-18386707-DEL-394 |
| chr5-18419847-DEL-444 | chr5-18419848-DEL-444 |
| chr5-18426926-DEL-88 | chr5-18426928-DEL-87 |
| chr5-18448218-DEL-64 | chr5-18448231-DEL-90 |
| chr5-18452244-DEL-64 | chr5-18452245-DEL-64 |
| chr5-18463352-DEL-241 | chr5-18463348-DEL-222 |
| chr5-18473253-DEL-58 | chr5-18473254-DEL-52 |
| chr5-18479917-DEL-6736 | chr5-18479925-DEL-6738 |
| chr5-18503233-DEL-6387 | chr5-18503234-DEL-6387 |
| chr5-18518671-DEL-450 | chr5-18518672-DEL-450 |
| chr5-18522227-DEL-465 | chr5-18522228-DEL-465 |
| chr5-18553729-DEL-101 | chr5-18553734-DEL-101 |
| chr5-18554732-DEL-185 | chr5-18554732-DEL-185 |
| chr5-18574136-DEL-91 | chr5-18574137-DEL-91 |
| chr5-18578264-DEL-209 | chr5-18578266-DEL-208 |
| chr5-18592017-DEL-56 | chr5-18592028-DEL-50 |
| chr5-18610322-DEL-129 | chr5-18610323-DEL-129 |
| chr5-18633435-DEL-52 | chr5-18633435-DEL-52 |
| chr5-18651605-DEL-220 | chr5-18651606-DEL-220 |
| chr5-18664989-DEL-122 | chr5-18664990-DEL-122 |
| chr5-18669021-DEL-180 | chr5-18669034-DEL-177 |
| chr5-18703521-DEL-351 | chr5-18703522-DEL-351 |
| chr5-18712456-DEL-388 | chr5-18712457-DEL-388 |
| chr5-18723300-DEL-1238 | chr5-18723284-DEL-1252 |
| chr5-18729878-DEL-324 | chr5-18729876-DEL-321 |
| chr5-18740470-DEL-488 | chr5-18740471-DEL-488 |
| chr5-18741668-DEL-6216 | chr5-18741669-DEL-6216 |
| chr5-18753121-DEL-58 | chr5-18753137-DEL-59 |
| chr5-18755853-DEL-3911 | chr5-18755854-DEL-3912 |
| chr5-18801949-DEL-351 | chr5-18801950-DEL-351 |
| chr5-18808939-DEL-190 | chr5-18808940-DEL-190 |
| chr5-18821474-DEL-178 | chr5-18821475-DEL-178 |

| | |
|---|---|
| chr5-18831985-DEL-2425 | chr5-18831986-DEL-2425 |
| chr5-18845228-DEL-125 | chr5-18845229-DEL-125 |
| chr5-18850315-DEL-1216 | chr5-18850316-DEL-1216 |
| chr5-18861257-DEL-4396 | chr5-18861258-DEL-4396 |
| chr5-18869921-DEL-1374 | chr5-18869922-DEL-1374 |
| chr5-18882741-DEL-895 | chr5-18882741-DEL-895 |
| chr5-18888667-DEL-63 | chr5-18888672-DEL-56 |
| chr5-18902148-DEL-1706 | chr5-18902146-DEL-1707 |
| chr5-18909453-DEL-2191 | chr5-18909454-DEL-2191 |
| chr5-18921555-DEL-758 | chr5-18921556-DEL-757 |
| chr5-18933160-DEL-78 | chr5-18933146-DEL-87 |
| chr5-18937540-DEL-2137 | chr5-18937541-DEL-2137 |
| chr5-18939873-DEL-6204 | chr5-18939874-DEL-6204 |
| chr5-18959678-DEL-1981 | chr5-18959679-DEL-1981 |
| chr5-18968398-DEL-266 | chr5-18968399-DEL-266 |
| chr5-18997147-DEL-1397 | chr5-18997148-DEL-1397 |
| chr5-19008529-DEL-187 | chr5-19008530-DEL-187 |
| chr5-19013339-DEL-51 | chr5-19013340-DEL-63 |
| chr5-19023389-DEL-74 | chr5-19023417-DEL-74 |
| chr5-19028369-DEL-6959 | chr5-19028370-DEL-6959 |
| chr5-19028644-DEL-208 | chr5-19028645-DEL-208 |
| chr5-19038499-DEL-207 | chr5-19038501-DEL-201 |
| chr5-19057340-DEL-6711 | chr5-19057352-DEL-6715 |
| chr5-19057696-DEL-214 | chr5-19057696-DEL-214 |
| chr5-19065375-DEL-2135 | chr5-19065381-DEL-2135 |
| chr5-19112421-DEL-293 | chr5-19112425-DEL-292 |
| chr5-19133130-DEL-6368 | chr5-19133131-DEL-6368 |
| chr5-19178061-DEL-1864 | chr5-19178062-DEL-1864 |
| chr5-19185972-DEL-6308 | chr5-19185973-DEL-6308 |
| chr5-19198955-DEL-2774 | chr5-19198956-DEL-2774 |
| chr5-19221360-DEL-742 | chr5-19221351-DEL-743 |
| chr5-19262297-DEL-6117 | chr5-19262288-DEL-6117 |
| chr5-19279466-DEL-984 | chr5-19279467-DEL-984 |
| chr5-19287702-DEL-284 | chr5-19287703-DEL-284 |
| chr5-19307445-DEL-828 | chr5-19307446-DEL-828 |
| chr5-19361616-DEL-60 | chr5-19361627-DEL-50 |
| chr5-19372942-DEL-1462 | chr5-19372943-DEL-1462 |
| chr5-19388561-DEL-447 | chr5-19388562-DEL-447 |
| chr5-19515002-DEL-70 | chr5-19515003-DEL-70 |
| chr5-19516508-DEL-850 | chr5-19516509-DEL-850 |
| chr5-19546644-DEL-6356 | chr5-19546645-DEL-6356 |
| chr5-19553960-DEL-54 | chr5-19553951-DEL-64 |
| chr5-19570593-DEL-95 | chr5-19570592-DEL-96 |
| chr5-19607520-DEL-65 | chr5-19607509-DEL-66 |
| chr5-19622582-DEL-691 | chr5-19622583-DEL-691 |
| chr5-19626196-DEL-437 | chr5-19626197-DEL-437 |
| chr5-19638880-DEL-61 | chr5-19638881-DEL-61 |
| chr5-19639752-DEL-78 | chr5-19639753-DEL-78 |
| chr5-19641090-DEL-70 | chr5-19641100-DEL-73 |
| chr5-19643858-DEL-388 | chr5-19643859-DEL-388 |
| chr5-19660098-DEL-449 | chr5-19660099-DEL-449 |
| chr5-19661835-DEL-672 | chr5-19661836-DEL-672 |
| chr5-19662976-DEL-60 | chr5-19662977-DEL-60 |
| chr5-19681570-DEL-6241 | chr5-19681568-DEL-6242 |
| chr5-19740668-DEL-1367 | chr5-19740669-DEL-1366 |
| chr5-19772983-DEL-56 | chr5-19772984-DEL-56 |
| chr5-19776573-DEL-201 | chr5-19776583-DEL-200 |
| chr5-19815635-DEL-94 | chr5-19815661-DEL-62 |
| chr5-19829378-DEL-70 | chr5-19829379-DEL-70 |
| chr5-19837115-DEL-1836 | chr5-19837116-DEL-1836 |
| chr5-19854112-DEL-6525 | chr5-19854113-DEL-6525 |
| chr5-19870265-DEL-139 | chr5-19870266-DEL-137 |
| chr5-19920705-DEL-168 | chr5-19920706-DEL-168 |
| chr5-19947987-DEL-553 | chr5-19947988-DEL-553 |
| chr5-19969759-DEL-56 | chr5-19969760-DEL-56 |
| chr5-19987310-DEL-601 | chr5-19987311-DEL-601 |
| chr5-19989728-DEL-63 | chr5-19989729-DEL-63 |
| chr5-19996746-DEL-70 | chr5-19996747-DEL-70 |
| chr5-20045799-DEL-63 | chr5-20045800-DEL-63 |
| chr5-20071927-DEL-244 | chr5-20071928-DEL-244 |
| chr5-20078171-DEL-67 | chr5-20078179-DEL-64 |
| chr5-20122341-DEL-228 | chr5-20122351-DEL-219 |
| chr5-20134893-DEL-81 | chr5-20134893-DEL-81 |
| chr5-20159572-DEL-61 | chr5-20159573-DEL-61 |
| chr5-20161461-DEL-6546 | chr5-20161462-DEL-6546 |
| chr5-20180195-DEL-3858 | chr5-20180196-DEL-3858 |
| chr5-20196483-DEL-84 | chr5-20196517-DEL-68 |
| chr5-20247194-DEL-266 | chr5-20247184-DEL-267 |
| chr5-20332822-DEL-237 | chr5-20332823-DEL-237 |
| chr5-20343488-DEL-656 | chr5-20343486-DEL-651 |

| | |
|---|---|
| chr5-20354879-DEL-55 | chr5-20354880-DEL-55 |
| chr5-20381328-DEL-83 | chr5-20381312-DEL-58 |
| chr5-20388737-DEL-7216 | chr5-20388738-DEL-7216 |
| chr5-20412068-DEL-374 | chr5-20412054-DEL-375 |
| chr5-20451445-DEL-65 | chr5-20451434-DEL-65 |
| chr5-20466780-DEL-224 | chr5-20466787-DEL-195 |
| chr5-20485772-DEL-80 | chr5-20485796-DEL-50 |
| chr5-20544543-DEL-2411 | chr5-20544544-DEL-2411 |
| chr5-20569032-DEL-6464 | chr5-20569033-DEL-6464 |
| chr5-20575131-DEL-212 | chr5-20575132-DEL-212 |
| chr5-20579766-DEL-63 | chr5-20579767-DEL-63 |
| chr5-20580104-DEL-308 | chr5-20580105-DEL-305 |
| chr5-20606852-DEL-748 | chr5-20606853-DEL-748 |
| chr5-20611262-DEL-51 | chr5-20611263-DEL-51 |
| chr5-20618095-DEL-149 | chr5-20618093-DEL-151 |
| chr5-20619900-DEL-170 | chr5-20619902-DEL-169 |
| chr5-20629432-DEL-325 | chr5-20629440-DEL-318 |
| chr5-20630610-DEL-1389 | chr5-20630611-DEL-1389 |
| chr5-20648431-DEL-667 | chr5-20648432-DEL-667 |
| chr5-20654161-DEL-363 | chr5-20654164-DEL-361 |
| chr5-20693537-DEL-53 | chr5-20693538-DEL-53 |
| chr5-20700351-DEL-532 | chr5-20700352-DEL-532 |
| chr5-20701092-DEL-153 | chr5-20701093-DEL-153 |
| chr5-20716540-DEL-1510 | chr5-20716541-DEL-1510 |
| chr5-20732683-DEL-265 | chr5-20732684-DEL-264 |
| chr5-20758990-DEL-96 | chr5-20758991-DEL-96 |
| chr5-20775930-DEL-63 | chr5-20775881-DEL-96 |
| chr5-20791322-DEL-372 | chr5-20791323-DEL-372 |
| chr5-20812389-DEL-324 | chr5-20812390-DEL-324 |
| chr5-20818620-DEL-56 | chr5-20818621-DEL-56 |
| chr5-20861494-DEL-229 | chr5-20861495-DEL-229 |
| chr5-20878502-DEL-67 | chr5-20878509-DEL-60 |
| chr5-20923267-DEL-523 | chr5-20923268-DEL-523 |
| chr5-20937378-DEL-1840 | chr5-20937379-DEL-1840 |
| chr5-20963410-DEL-89 | chr5-20963411-DEL-89 |
| chr5-20974907-DEL-946 | chr5-20974908-DEL-946 |
| chr5-21028577-DEL-6441 | chr5-21028583-DEL-6436 |
| chr5-21035066-DEL-481 | chr5-21035066-DEL-481 |
| chr5-21052583-DEL-210 | chr5-21052584-DEL-210 |
| chr5-21092293-DEL-277 | chr5-21092294-DEL-277 |
| chr5-21111432-DEL-211 | chr5-21111433-DEL-211 |
| chr5-21130702-DEL-84 | chr5-21130703-DEL-92 |
| chr5-21142546-DEL-205 | chr5-21142547-DEL-205 |
| chr5-21149323-DEL-194 | chr5-21149315-DEL-195 |
| chr5-21160842-DEL-774 | chr5-21160843-DEL-774 |
| chr5-21181519-DEL-203 | chr5-21181520-DEL-203 |
| chr5-21189557-DEL-213 | chr5-21189558-DEL-213 |
| chr5-21198621-DEL-113 | chr5-21198621-DEL-114 |
| chr5-21204356-DEL-106 | chr5-21204357-DEL-106 |
| chr5-21225401-DEL-198 | chr5-21225402-DEL-198 |
| chr5-21230286-DEL-184 | chr5-21230287-DEL-184 |
| chr5-21232774-DEL-201 | chr5-21232775-DEL-201 |
| chr5-21244104-DEL-1887 | chr5-21244105-DEL-1885 |
| chr5-21253230-DEL-56 | chr5-21253231-DEL-56 |
| chr5-21265631-DEL-292 | chr5-21265631-DEL-292 |
| chr5-21268758-DEL-535 | chr5-21268759-DEL-528 |
| chr5-21301060-DEL-2376 | chr5-21301060-DEL-2376 |
| chr5-21305142-DEL-169 | chr5-21305143-DEL-169 |
| chr5-21305730-DEL-7073 | chr5-21305731-DEL-7073 |
| chr5-21321207-DEL-78 | chr5-21321208-DEL-52 |
| chr5-21346102-DEL-60 | chr5-21346103-DEL-60 |
| chr5-21356729-DEL-244 | chr5-21356730-DEL-244 |
| chr5-21360150-DEL-1285 | chr5-21360151-DEL-1285 |
| chr5-21370810-DEL-59 | chr5-21370810-DEL-60 |
| chr5-21385744-DEL-59 | chr5-21385728-DEL-56 |
| chr5-21388332-DEL-118 | chr5-21388333-DEL-118 |
| chr5-21406063-DEL-996 | chr5-21406064-DEL-996 |
| chr5-21432772-DEL-55 | chr5-21432783-DEL-53 |
| chr5-21437546-DEL-1785 | chr5-21437547-DEL-1783 |
| chr5-21440409-DEL-7191 | chr5-21440410-DEL-7191 |
| chr5-21448623-DEL-67 | chr5-21448634-DEL-51 |
| chr5-21469933-DEL-86 | chr5-21469934-DEL-76 |
| chr5-21473931-DEL-287 | chr5-21473932-DEL-287 |
| chr5-21479668-DEL-72 | chr5-21479669-DEL-72 |
| chr5-21505459-DEL-473 | chr5-21505460-DEL-473 |
| chr5-21527047-DEL-154 | chr5-21527030-DEL-157 |
| chr5-21533383-DEL-54 | chr5-21533384-DEL-54 |
| chr5-21539814-DEL-739 | chr5-21539814-DEL-739 |
| chr5-21548248-DEL-1888 | chr5-21548249-DEL-1888 |
| chr5-21556215-DEL-1379 | chr5-21556216-DEL-1379 |

| | |
|---|---|
| chr5-21562420-DEL-70 | chr5-21562424-DEL-64 |
| chr5-21586283-DEL-58 | chr5-21586283-DEL-58 |
| chr5-21592779-DEL-224 | chr5-21592780-DEL-226 |
| chr5-21611688-DEL-2019 | chr5-21611689-DEL-2019 |
| chr5-21636225-DEL-549 | chr5-21636226-DEL-549 |
| chr5-21658193-DEL-439 | chr5-21658194-DEL-439 |
| chr5-21678043-DEL-82 | chr5-21678044-DEL-82 |
| chr5-21716119-DEL-54 | chr5-21716120-DEL-54 |
| chr5-21720402-DEL-165 | chr5-21720402-DEL-166 |
| chr5-21730049-DEL-54 | chr5-21730050-DEL-54 |
| chr5-21741241-DEL-52 | chr5-21741242-DEL-52 |
| chr5-21762934-DEL-337 | chr5-21762935-DEL-337 |
| chr5-21770273-DEL-61 | chr5-21770274-DEL-61 |
| chr5-21773930-DEL-60 | chr5-21773931-DEL-60 |
| chr5-21781233-DEL-55 | chr5-21781234-DEL-55 |
| chr5-21782427-DEL-4373 | chr5-21782427-DEL-4373 |
| chr5-21785004-DEL-78 | chr5-21785005-DEL-78 |
| chr5-21808552-DEL-351 | chr5-21808553-DEL-351 |
| chr5-21814654-DEL-58 | chr5-21814655-DEL-58 |
| chr5-21832049-DEL-75 | chr5-21832049-DEL-76 |
| chr5-21832365-DEL-646 | chr5-21832366-DEL-646 |
| chr5-21839117-DEL-68 | chr5-21839118-DEL-68 |
| chr5-21857871-DEL-146 | chr5-21857872-DEL-146 |
| chr5-21898205-DEL-168 | chr5-21898206-DEL-168 |
| chr5-21910664-DEL-196 | chr5-21910665-DEL-196 |
| chr5-21934987-DEL-69 | chr5-21934988-DEL-69 |
| chr5-21935748-DEL-271 | chr5-21935721-DEL-262 |
| chr5-21942541-DEL-110 | chr5-21942542-DEL-110 |
| chr5-21944532-DEL-357 | chr5-21944533-DEL-357 |
| chr5-21951238-DEL-162 | chr5-21951258-DEL-155 |
| chr5-21955349-DEL-75 | chr5-21955346-DEL-76 |
| chr5-21959006-DEL-167 | chr5-21959007-DEL-167 |
| chr5-21972525-DEL-89 | chr5-21972482-DEL-121 |
| chr5-21983960-DEL-794 | chr5-21983961-DEL-794 |
| chr5-21985666-DEL-138 | chr5-21985701-DEL-104 |
| chr5-22011638-DEL-110 | chr5-22011639-DEL-91 |
| chr5-22013108-DEL-178 | chr5-22013109-DEL-178 |
| chr5-22016966-DEL-88 | chr5-22016990-DEL-61 |
| chr5-22026772-DEL-169 | chr5-22026773-DEL-169 |
| chr5-22048365-DEL-715 | chr5-22048366-DEL-715 |
| chr5-22082643-DEL-78 | chr5-22082644-DEL-78 |
| chr5-22126002-DEL-96 | chr5-22126003-DEL-96 |
| chr5-22166433-DEL-2689 | chr5-22166432-DEL-2689 |
| chr5-22171441-DEL-222 | chr5-22171434-DEL-220 |
| chr5-22205176-DEL-1533 | chr5-22205177-DEL-1533 |
| chr5-22235293-DEL-202 | chr5-22235294-DEL-202 |
| chr5-22289769-DEL-6445 | chr5-22289770-DEL-6445 |
| chr5-22341563-DEL-352 | chr5-22341564-DEL-352 |
| chr5-22355671-DEL-150 | chr5-22355716-DEL-120 |
| chr5-22369083-DEL-57 | chr5-22369084-DEL-60 |
| chr5-22377500-DEL-83 | chr5-22377502-DEL-80 |
| chr5-22385796-DEL-445 | chr5-22385797-DEL-445 |
| chr5-22387221-DEL-210 | chr5-22387222-DEL-210 |
| chr5-22395213-DEL-67 | chr5-22395214-DEL-67 |
| chr5-22406027-DEL-5403 | chr5-22406028-DEL-5403 |
| chr5-22436424-DEL-645 | chr5-22436425-DEL-645 |
| chr5-22480201-DEL-479 | chr5-22480202-DEL-479 |
| chr5-22567223-DEL-11113 | chr5-22567224-DEL-11113 |
| chr5-22569494-DEL-214 | chr5-22569494-DEL-214 |
| chr5-22609076-DEL-62 | chr5-22609077-DEL-62 |
| chr5-22624311-DEL-5926 | chr5-22624312-DEL-5926 |
| chr5-22636292-DEL-418 | chr5-22636293-DEL-418 |
| chr5-22640108-DEL-300 | chr5-22640068-DEL-334 |
| chr5-22659187-DEL-144 | chr5-22659188-DEL-144 |
| chr5-22661273-DEL-7766 | chr5-22661274-DEL-7766 |
| chr5-22676752-DEL-1240 | chr5-22676753-DEL-1240 |
| chr5-22683904-DEL-58 | chr5-22683905-DEL-58 |
| chr5-22709862-DEL-140 | chr5-22709861-DEL-100 |
| chr5-22717271-DEL-1572 | chr5-22717272-DEL-1572 |
| chr5-22720589-DEL-245 | chr5-22720590-DEL-245 |
| chr5-22722566-DEL-4229 | chr5-22722567-DEL-4229 |
| chr5-22733210-DEL-506 | chr5-22733211-DEL-506 |
| chr5-22734715-DEL-68 | chr5-22734716-DEL-68 |
| chr5-22736591-DEL-211 | chr5-22736592-DEL-211 |
| chr5-22771109-DEL-6403 | chr5-22771110-DEL-6403 |
| chr5-22803669-DEL-7232 | chr5-22803667-DEL-7235 |
| chr5-22810408-DEL-204 | chr5-22810409-DEL-204 |
| chr5-22815325-DEL-1356 | chr5-22815326-DEL-1356 |
| chr5-22817541-DEL-712 | chr5-22817542-DEL-712 |
| chr5-22836737-DEL-157 | chr5-22836738-DEL-157 |

| | |
|---|---|
| chr5-22850618-DEL-513 | chr5-22850617-DEL-514 |
| chr5-22863461-DEL-97 | chr5-22863473-DEL-88 |
| chr5-22865659-DEL-502 | chr5-22865660-DEL-502 |
| chr5-22869424-DEL-1667 | chr5-22869425-DEL-1667 |
| chr5-22912391-DEL-2448 | chr5-22912392-DEL-2446 |
| chr5-22916181-DEL-144 | chr5-22916182-DEL-144 |
| chr5-22924863-DEL-82 | chr5-22924862-DEL-58 |
| chr5-22925749-DEL-68 | chr5-22925750-DEL-68 |
| chr5-22929037-DEL-2231 | chr5-22929063-DEL-2202 |
| chr5-22934792-DEL-1520 | chr5-22934793-DEL-1518 |
| chr5-22959847-DEL-54 | chr5-22959848-DEL-54 |
| chr5-22973912-DEL-973 | chr5-22973913-DEL-973 |
| chr5-22982523-DEL-1063 | chr5-22982532-DEL-1055 |
| chr5-22991217-DEL-306 | chr5-22991218-DEL-306 |
| chr5-22992103-DEL-363 | chr5-22992104-DEL-363 |
| chr5-23016982-DEL-233 | chr5-23016983-DEL-233 |
| chr5-23025241-DEL-891 | chr5-23025243-DEL-892 |
| chr5-23037667-DEL-280 | chr5-23037668-DEL-280 |
| chr5-23041618-DEL-89 | chr5-23041619-DEL-89 |
| chr5-23043055-DEL-64 | chr5-23043049-DEL-76 |
| chr5-23060838-DEL-79 | chr5-23060839-DEL-83 |
| chr5-23067650-DEL-85 | chr5-23067651-DEL-85 |
| chr5-23069693-DEL-2562 | chr5-23069694-DEL-2562 |
| chr5-23083818-DEL-162 | chr5-23083819-DEL-162 |
| chr5-23097585-DEL-94 | chr5-23097586-DEL-94 |
| chr5-23214219-DEL-126 | chr5-23214220-DEL-126 |
| chr5-23227189-DEL-86 | chr5-23227202-DEL-104 |
| chr5-23229989-DEL-6635 | chr5-23229990-DEL-6635 |
| chr5-23237625-DEL-204 | chr5-23237626-DEL-204 |
| chr5-23238885-DEL-1082 | chr5-23238886-DEL-1081 |
| chr5-23250472-DEL-687 | chr5-23250473-DEL-687 |
| chr5-23252651-DEL-56 | chr5-23252652-DEL-56 |
| chr5-23263946-DEL-64 | chr5-23263946-DEL-64 |
| chr5-23290961-DEL-80 | chr5-23290960-DEL-76 |
| chr5-23302412-DEL-70 | chr5-23302418-DEL-65 |
| chr5-23309493-DEL-305 | chr5-23309494-DEL-305 |
| chr5-23328016-DEL-659 | chr5-23328017-DEL-659 |
| chr5-23359086-DEL-52 | chr5-23359087-DEL-52 |
| chr5-23366021-DEL-151 | chr5-23366021-DEL-147 |
| chr5-23385725-DEL-763 | chr5-23385726-DEL-763 |
| chr5-23390031-DEL-3510 | chr5-23390032-DEL-3510 |
| chr5-23398781-DEL-1551 | chr5-23398782-DEL-1551 |
| chr5-23435446-DEL-4366 | chr5-23435447-DEL-4366 |
| chr5-23437363-DEL-620 | chr5-23437363-DEL-620 |
| chr5-23438005-DEL-1090 | chr5-23438005-DEL-1090 |
| chr5-23442613-DEL-55 | chr5-23442613-DEL-55 |
| chr5-23450119-DEL-191 | chr5-23450120-DEL-191 |
| chr5-23459751-DEL-119 | chr5-23459744-DEL-120 |
| chr5-23467831-DEL-69 | chr5-23467832-DEL-69 |
| chr5-23477868-DEL-269 | chr5-23477865-DEL-269 |
| chr5-23482158-DEL-84 | chr5-23482159-DEL-84 |
| chr5-23486061-DEL-1510 | chr5-23486062-DEL-1510 |
| chr5-23495911-DEL-4903 | chr5-23495912-DEL-4903 |
| chr5-23508419-DEL-59 | chr5-23508420-DEL-59 |
| chr5-23570165-DEL-5105 | chr5-23570170-DEL-5101 |
| chr5-23575442-DEL-349 | chr5-23575442-DEL-349 |
| chr5-23578139-DEL-196 | chr5-23578140-DEL-196 |
| chr5-23578874-DEL-60 | chr5-23578875-DEL-60 |
| chr5-23584556-DEL-485 | chr5-23584557-DEL-485 |
| chr5-23594598-DEL-59 | chr5-23594601-DEL-57 |
| chr5-23607314-DEL-56 | chr5-23607315-DEL-56 |
| chr5-23609474-DEL-52 | chr5-23609475-DEL-52 |
| chr5-23651411-DEL-52 | chr5-23651412-DEL-52 |
| chr5-23665066-DEL-200 | chr5-23665067-DEL-200 |
| chr5-23699279-DEL-449 | chr5-23699280-DEL-449 |
| chr5-23717322-DEL-136 | chr5-23717324-DEL-150 |
| chr5-23721422-DEL-207 | chr5-23721419-DEL-211 |
| chr5-23722722-DEL-86 | chr5-23722755-DEL-53 |
| chr5-23753976-DEL-532 | chr5-23753977-DEL-532 |
| chr5-23758459-DEL-103 | chr5-23758416-DEL-104 |
| chr5-23774975-DEL-3420 | chr5-23774976-DEL-3420 |
| chr5-23779641-DEL-590 | chr5-23779642-DEL-590 |
| chr5-23811683-DEL-180 | chr5-23811684-DEL-180 |
| chr5-23816710-DEL-287 | chr5-23816711-DEL-287 |
| chr5-23818668-DEL-268 | chr5-23818669-DEL-268 |
| chr5-23826508-DEL-70 | chr5-23826509-DEL-70 |
| chr5-23848085-DEL-1702 | chr5-23848072-DEL-1715 |
| chr5-23850145-DEL-56 | chr5-23850146-DEL-56 |
| chr5-23900901-DEL-118 | chr5-23900902-DEL-118 |
| chr5-23902500-DEL-1267 | chr5-23902501-DEL-1267 |

| | |
|---|---|
| chr5-23905572-DEL-8671 | chr5-23905573-DEL-8671 |
| chr5-23915431-DEL-282 | chr5-23915432-DEL-282 |
| chr5-23935246-DEL-119 | chr5-23935247-DEL-119 |
| chr5-23936626-DEL-159 | chr5-23936622-DEL-163 |
| chr5-23937339-DEL-189 | chr5-23937340-DEL-189 |
| chr5-23940160-DEL-73 | chr5-23940161-DEL-60 |
| chr5-23941012-DEL-157 | chr5-23941013-DEL-157 |
| chr5-23945114-DEL-70 | chr5-23945115-DEL-66 |
| chr5-23966355-DEL-68 | chr5-23966356-DEL-72 |
| chr5-23972949-DEL-55 | chr5-23972915-DEL-56 |
| chr5-23977890-DEL-196 | chr5-23977891-DEL-196 |
| chr5-24004469-DEL-979 | chr5-24004470-DEL-979 |
| chr5-24007717-DEL-203 | chr5-24007718-DEL-203 |
| chr5-24009449-DEL-288 | chr5-24009450-DEL-288 |
| chr5-24011667-DEL-168 | chr5-24011668-DEL-167 |
| chr5-24013099-DEL-173 | chr5-24013090-DEL-174 |
| chr5-24017236-DEL-84 | chr5-24017237-DEL-84 |
| chr5-24019932-DEL-84 | chr5-24019933-DEL-84 |
| chr5-24021250-DEL-210 | chr5-24021251-DEL-210 |
| chr5-24025680-DEL-212 | chr5-24025681-DEL-211 |
| chr5-24026391-DEL-71 | chr5-24026398-DEL-65 |
| chr5-24028640-DEL-207 | chr5-24028641-DEL-207 |
| chr5-24030464-DEL-56 | chr5-24030436-DEL-74 |
| chr5-24033551-DEL-4013 | chr5-24033552-DEL-4013 |
| chr5-24057382-DEL-89 | chr5-24057383-DEL-89 |
| chr5-24083198-DEL-3045 | chr5-24083199-DEL-3045 |
| chr5-24086940-DEL-196 | chr5-24086941-DEL-196 |
| chr5-24096887-DEL-51 | chr5-24096864-DEL-58 |
| chr5-24097700-DEL-194 | chr5-24097679-DEL-198 |
| chr5-24110208-DEL-190 | chr5-24110199-DEL-178 |
| chr5-24113592-DEL-209 | chr5-24113593-DEL-209 |
| chr5-24121503-DEL-60 | chr5-24121498-DEL-70 |
| chr5-24126372-DEL-57 | chr5-24126372-DEL-57 |
| chr5-24127187-DEL-57 | chr5-24127188-DEL-57 |
| chr5-24131811-DEL-400 | chr5-24131812-DEL-400 |
| chr5-24143260-DEL-486 | chr5-24143261-DEL-486 |
| chr5-24157536-DEL-227 | chr5-24157525-DEL-243 |
| chr5-24161315-DEL-90 | chr5-24161346-DEL-60 |
| chr5-24181563-DEL-91 | chr5-24181548-DEL-99 |
| chr5-24201647-DEL-179 | chr5-24201648-DEL-179 |
| chr5-24221273-DEL-214 | chr5-24221274-DEL-214 |
| chr5-24236089-DEL-56 | chr5-24236090-DEL-56 |
| chr5-24243749-DEL-63 | chr5-24243719-DEL-93 |
| chr5-24250903-DEL-61 | chr5-24250876-DEL-62 |
| chr5-24255852-DEL-70 | chr5-24255851-DEL-72 |
| chr5-24261171-DEL-65 | chr5-24261172-DEL-65 |
| chr5-24263769-DEL-252 | chr5-24263770-DEL-252 |
| chr5-24270557-DEL-598 | chr5-24270558-DEL-598 |
| chr5-24286793-DEL-147 | chr5-24286794-DEL-147 |
| chr5-24291106-DEL-150 | chr5-24291107-DEL-149 |
| chr5-24292650-DEL-73 | chr5-24292651-DEL-71 |
| chr5-24304176-DEL-207 | chr5-24304177-DEL-207 |
| chr5-24312850-DEL-58 | chr5-24312851-DEL-58 |
| chr5-24322766-DEL-962 | chr5-24322764-DEL-963 |
| chr5-24341296-DEL-697 | chr5-24341338-DEL-698 |
| chr5-24383665-DEL-63 | chr5-24383666-DEL-63 |
| chr5-24385395-DEL-211 | chr5-24385396-DEL-208 |
| chr5-24399027-DEL-109 | chr5-24399029-DEL-108 |
| chr5-24409408-DEL-449 | chr5-24409409-DEL-449 |
| chr5-24409927-DEL-164 | chr5-24409928-DEL-164 |
| chr5-24420287-DEL-6863 | chr5-24420301-DEL-6850 |
| chr5-24448599-DEL-110 | chr5-24448604-DEL-132 |
| chr5-24450081-DEL-75 | chr5-24450081-DEL-75 |
| chr5-24451027-DEL-512 | chr5-24451028-DEL-512 |
| chr5-24452631-DEL-779 | chr5-24452626-DEL-775 |
| chr5-24454451-DEL-446 | chr5-24454452-DEL-447 |
| chr5-24455407-DEL-2240 | chr5-24455408-DEL-2240 |
| chr5-24461695-DEL-962 | chr5-24461696-DEL-962 |
| chr5-24475189-DEL-150 | chr5-24475190-DEL-150 |
| chr5-24520117-DEL-193 | chr5-24520137-DEL-184 |
| chr5-24584252-DEL-171 | chr5-24584253-DEL-171 |
| chr5-24602416-DEL-202 | chr5-24602417-DEL-202 |
| chr5-24670344-DEL-117 | chr5-24670345-DEL-117 |
| chr5-24693783-DEL-340 | chr5-24693783-DEL-340 |
| chr5-24716297-DEL-52 | chr5-24716297-DEL-52 |
| chr5-24746139-DEL-70 | chr5-24746140-DEL-70 |
| chr5-24786383-DEL-520 | chr5-24786384-DEL-520 |
| chr5-24793426-DEL-198 | chr5-24793427-DEL-198 |
| chr5-24815604-DEL-87 | chr5-24815605-DEL-87 |
| chr5-24817669-DEL-73 | chr5-24817670-DEL-73 |

| | |
|---|---|
| chr5-24818064-DEL-589 | chr5-24818103-DEL-551 |
| chr5-24833718-DEL-52 | chr5-24833719-DEL-72 |
| chr5-24837470-DEL-51 | chr5-24837467-DEL-57 |
| chr5-24848660-DEL-86 | chr5-24848641-DEL-84 |
| chr5-24861758-DEL-72 | chr5-24861797-DEL-56 |
| chr5-24886175-DEL-203 | chr5-24886176-DEL-203 |
| chr5-24898136-DEL-208 | chr5-24898137-DEL-208 |
| chr5-24909103-DEL-1459 | chr5-24909104-DEL-1459 |
| chr5-24922527-DEL-60 | chr5-24922528-DEL-60 |
| chr5-24925210-DEL-418 | chr5-24925211-DEL-418 |
| chr5-24946075-DEL-703 | chr5-24946076-DEL-703 |
| chr5-24947612-DEL-227 | chr5-24947613-DEL-227 |
| chr5-24960193-DEL-383 | chr5-24960194-DEL-383 |
| chr5-24966007-DEL-6823 | chr5-24966008-DEL-6823 |
| chr5-24975554-DEL-1289 | chr5-24975555-DEL-1289 |
| chr5-24986754-DEL-182 | chr5-24986755-DEL-182 |
| chr5-24997018-DEL-191 | chr5-24997011-DEL-192 |
| chr5-25014302-DEL-103 | chr5-25014295-DEL-110 |
| chr5-25027291-DEL-481 | chr5-25027292-DEL-481 |
| chr5-25043456-DEL-90 | chr5-25043457-DEL-90 |
| chr5-25070760-DEL-111 | chr5-25070745-DEL-107 |
| chr5-25076152-DEL-101 | chr5-25076146-DEL-95 |
| chr5-25093788-DEL-238 | chr5-25093789-DEL-238 |
| chr5-25164259-DEL-58 | chr5-25164257-DEL-59 |
| chr5-25208886-DEL-94 | chr5-25208887-DEL-94 |
| chr5-25218581-DEL-155 | chr5-25218582-DEL-155 |
| chr5-25220423-DEL-317 | chr5-25220424-DEL-317 |
| chr5-25236136-DEL-135 | chr5-25236136-DEL-135 |
| chr5-25266615-DEL-191 | chr5-25266616-DEL-191 |
| chr5-25281061-DEL-757 | chr5-25281076-DEL-758 |
| chr5-25323506-DEL-354 | chr5-25323507-DEL-354 |
| chr5-25327657-DEL-2801 | chr5-25327658-DEL-2801 |
| chr5-25330618-DEL-188 | chr5-25330619-DEL-188 |
| chr5-25333261-DEL-348 | chr5-25333262-DEL-348 |
| chr5-25334972-DEL-239 | chr5-25334973-DEL-239 |
| chr5-25336184-DEL-6168 | chr5-25336184-DEL-6168 |
| chr5-25352961-DEL-175 | chr5-25352962-DEL-175 |
| chr5-25364775-DEL-400 | chr5-25364776-DEL-400 |
| chr5-25376873-DEL-115 | chr5-25376874-DEL-115 |
| chr5-25406469-DEL-227 | chr5-25406470-DEL-227 |
| chr5-25418308-DEL-78 | chr5-25418309-DEL-117 |
| chr5-25422717-DEL-369 | chr5-25422718-DEL-369 |
| chr5-25428824-DEL-93 | chr5-25428825-DEL-90 |
| chr5-25435691-DEL-8976 | chr5-25435692-DEL-8976 |
| chr5-25445346-DEL-604 | chr5-25445347-DEL-604 |
| chr5-25447705-DEL-96 | chr5-25447706-DEL-96 |
| chr5-25454326-DEL-83 | chr5-25454327-DEL-83 |
| chr5-25471301-DEL-2097 | chr5-25471300-DEL-2098 |
| chr5-25584739-DEL-142 | chr5-25584740-DEL-142 |
| chr5-25601966-DEL-1492 | chr5-25601967-DEL-1492 |
| chr5-25641126-DEL-111 | chr5-25641127-DEL-110 |
| chr5-25684402-DEL-89 | chr5-25684400-DEL-91 |
| chr5-25728500-DEL-80 | chr5-25728501-DEL-82 |
| chr5-25736463-DEL-180 | chr5-25736464-DEL-180 |
| chr5-25752402-DEL-371 | chr5-25752403-DEL-371 |
| chr5-25756461-DEL-112 | chr5-25756462-DEL-112 |
| chr5-25784237-DEL-3485 | chr5-25784265-DEL-3457 |
| chr5-25789296-DEL-66 | chr5-25789296-DEL-62 |
| chr5-25800697-DEL-200 | chr5-25800698-DEL-200 |
| chr5-25808418-DEL-104 | chr5-25808413-DEL-113 |
| chr5-25820759-DEL-435 | chr5-25820760-DEL-435 |
| chr5-25823554-DEL-128 | chr5-25823555-DEL-120 |
| chr5-25825820-DEL-116 | chr5-25825821-DEL-116 |
| chr5-25827848-DEL-78 | chr5-25827849-DEL-78 |
| chr5-25836519-DEL-91 | chr5-25836520-DEL-91 |
| chr5-25841744-DEL-671 | chr5-25841745-DEL-671 |
| chr5-25849157-DEL-704 | chr5-25849159-DEL-700 |
| chr5-25853305-DEL-104 | chr5-25853306-DEL-104 |
| chr5-25899288-DEL-56 | chr5-25899275-DEL-52 |
| chr5-25918379-DEL-96 | chr5-25918380-DEL-96 |
| chr5-25963588-DEL-185 | chr5-25963590-DEL-184 |
| chr5-25974723-DEL-92 | chr5-25974724-DEL-92 |
| chr5-25990877-DEL-858 | chr5-25990879-DEL-850 |
| chr5-25992452-DEL-548 | chr5-25992452-DEL-548 |
| chr5-25998749-DEL-90 | chr5-25998750-DEL-72 |
| chr5-26018459-DEL-86 | chr5-26018420-DEL-83 |
| chr5-26059060-DEL-287 | chr5-26059061-DEL-287 |
| chr5-26077156-DEL-171 | chr5-26077153-DEL-173 |
| chr5-26104744-DEL-170 | chr5-26104745-DEL-169 |
| chr5-26109246-DEL-906 | chr5-26109247-DEL-906 |

| | |
|---|---|
| chr5-26121113-DEL-336 | chr5-26121114-DEL-336 |
| chr5-26207768-DEL-126 | chr5-26207770-DEL-102 |
| chr5-26242515-DEL-140 | chr5-26242515-DEL-140 |
| chr5-26370415-DEL-52 | chr5-26370416-DEL-52 |
| chr5-26381582-DEL-1268 | chr5-26381583-DEL-1268 |
| chr5-26397241-DEL-697 | chr5-26397242-DEL-697 |
| chr5-26403772-DEL-206 | chr5-26403773-DEL-206 |
| chr5-26436829-DEL-125 | chr5-26436805-DEL-123 |
| chr5-26501311-DEL-133 | chr5-26501312-DEL-133 |
| chr5-26538116-DEL-1878 | chr5-26538117-DEL-1878 |
| chr5-26584431-DEL-1060 | chr5-26584432-DEL-1060 |
| chr5-26597493-DEL-2318 | chr5-26597494-DEL-2318 |
| chr5-26609055-DEL-1047 | chr5-26609056-DEL-1047 |
| chr5-26610763-DEL-1405 | chr5-26610764-DEL-1405 |
| chr5-26616136-DEL-928 | chr5-26616137-DEL-928 |
| chr5-26623976-DEL-85 | chr5-26623977-DEL-85 |
| chr5-26631091-DEL-334 | chr5-26631092-DEL-334 |
| chr5-26635602-DEL-51 | chr5-26635603-DEL-51 |
| chr5-26639428-DEL-6311 | chr5-26639429-DEL-6311 |
| chr5-26651354-DEL-306 | chr5-26651355-DEL-306 |
| chr5-26681121-DEL-138 | chr5-26681122-DEL-138 |
| chr5-26705276-DEL-5782 | chr5-26705277-DEL-5782 |
| chr5-26828884-DEL-7091 | chr5-26828885-DEL-7091 |
| chr5-26860394-DEL-9127 | chr5-26860395-DEL-9127 |
| chr5-26884333-DEL-507 | chr5-26884334-DEL-507 |
| chr5-26900415-DEL-103 | chr5-26900416-DEL-103 |
| chr5-26916756-DEL-1131 | chr5-26916757-DEL-1129 |
| chr5-26920526-DEL-67 | chr5-26920537-DEL-57 |
| chr5-26949856-DEL-92 | chr5-26949885-DEL-64 |
| chr5-26953464-DEL-448 | chr5-26953465-DEL-448 |
| chr5-26969070-DEL-135 | chr5-26969071-DEL-135 |
| chr5-26978372-DEL-166 | chr5-26978373-DEL-166 |
| chr5-27009351-DEL-171 | chr5-27009352-DEL-171 |
| chr5-27148414-DEL-106 | chr5-27148415-DEL-106 |
| chr5-27158234-DEL-204 | chr5-27158235-DEL-204 |
| chr5-27186051-DEL-137 | chr5-27186043-DEL-130 |
| chr5-27226648-DEL-975 | chr5-27226649-DEL-975 |
| chr5-27237336-DEL-68 | chr5-27237337-DEL-68 |
| chr5-27249690-DEL-490 | chr5-27249691-DEL-490 |
| chr5-27257463-DEL-99 | chr5-27257471-DEL-94 |
| chr5-27294423-DEL-3589 | chr5-27294424-DEL-3589 |
| chr5-27298985-DEL-53 | chr5-27298986-DEL-53 |
| chr5-27307649-DEL-548 | chr5-27307650-DEL-548 |
| chr5-27335515-DEL-73 | chr5-27335532-DEL-74 |
| chr5-27346942-DEL-1540 | chr5-27346943-DEL-1540 |
| chr5-27406712-DEL-92 | chr5-27406698-DEL-90 |
| chr5-27414565-DEL-54 | chr5-27414566-DEL-54 |
| chr5-27430179-DEL-6246 | chr5-27430180-DEL-6246 |
| chr5-27446249-DEL-71 | chr5-27446252-DEL-69 |
| chr5-27450229-DEL-90 | chr5-27450230-DEL-86 |
| chr5-27501122-DEL-6466 | chr5-27501123-DEL-6466 |
| chr5-27537641-DEL-570 | chr5-27537642-DEL-571 |
| chr5-27541769-DEL-78 | chr5-27541770-DEL-78 |
| chr5-27546051-DEL-164 | chr5-27546052-DEL-164 |
| chr5-27550605-DEL-587 | chr5-27550606-DEL-587 |
| chr5-27563884-DEL-6706 | chr5-27563885-DEL-6706 |
| chr5-27602894-DEL-79 | chr5-27602895-DEL-79 |
| chr5-27622948-DEL-284 | chr5-27622989-DEL-262 |
| chr5-27630591-DEL-56 | chr5-27630592-DEL-56 |
| chr5-27633264-DEL-51 | chr5-27633260-DEL-52 |
| chr5-27638931-DEL-1302 | chr5-27638932-DEL-1302 |
| chr5-27644579-DEL-164 | chr5-27644580-DEL-164 |
| chr5-27674626-DEL-121 | chr5-27674627-DEL-121 |
| chr5-27677807-DEL-683 | chr5-27677808-DEL-683 |
| chr5-27708830-DEL-561 | chr5-27708830-DEL-562 |
| chr5-27747709-DEL-52 | chr5-27747710-DEL-52 |
| chr5-27750798-DEL-69 | chr5-27750799-DEL-63 |
| chr5-27767253-DEL-2644 | chr5-27767254-DEL-2644 |
| chr5-27778376-DEL-286 | chr5-27778377-DEL-286 |
| chr5-27806038-DEL-406 | chr5-27806039-DEL-406 |
| chr5-27826418-DEL-67 | chr5-27826431-DEL-51 |
| chr5-27848664-DEL-1269 | chr5-27848665-DEL-1269 |
| chr5-27870989-DEL-75 | chr5-27870990-DEL-50 |
| chr5-27883340-DEL-161 | chr5-27883338-DEL-162 |
| chr5-27891671-DEL-52 | chr5-27891672-DEL-52 |
| chr5-27895491-DEL-145 | chr5-27895490-DEL-145 |
| chr5-27921803-DEL-54 | chr5-27921804-DEL-54 |
| chr5-27929897-DEL-90 | chr5-27929910-DEL-88 |
| chr5-28032591-DEL-53 | chr5-28032592-DEL-53 |
| chr5-28043814-DEL-196 | chr5-28043815-DEL-196 |

| | |
|---|---|
| chr5-28044371-DEL-100 | chr5-28044374-DEL-98 |
| chr5-28048701-DEL-147 | chr5-28048702-DEL-147 |
| chr5-28051539-DEL-194 | chr5-28051540-DEL-188 |
| chr5-28064481-DEL-1651 | chr5-28064482-DEL-1651 |
| chr5-28083793-DEL-95 | chr5-28083794-DEL-95 |
| chr5-28120785-DEL-309 | chr5-28120787-DEL-308 |
| chr5-28137991-DEL-62 | chr5-28137994-DEL-60 |
| chr5-28147335-DEL-1105 | chr5-28147336-DEL-1105 |
| chr5-28162759-DEL-68 | chr5-28162748-DEL-50 |
| chr5-28190322-DEL-131 | chr5-28190322-DEL-131 |
| chr5-28211473-DEL-156 | chr5-28211474-DEL-156 |
| chr5-28239555-DEL-158 | chr5-28239556-DEL-158 |
| chr5-28262813-DEL-2468 | chr5-28262814-DEL-2468 |
| chr5-28283939-DEL-212 | chr5-28283940-DEL-212 |
| chr5-28303343-DEL-194 | chr5-28303343-DEL-194 |
| chr5-28303640-DEL-513 | chr5-28303665-DEL-537 |
| chr5-28306782-DEL-54 | chr5-28306783-DEL-54 |
| chr5-28319860-DEL-1343 | chr5-28319861-DEL-1343 |
| chr5-28323986-DEL-134 | chr5-28324005-DEL-116 |
| chr5-28336587-DEL-516 | chr5-28336595-DEL-509 |
| chr5-28393167-DEL-113 | chr5-28393167-DEL-113 |
| chr5-28437045-DEL-165 | chr5-28437046-DEL-165 |
| chr5-28458645-DEL-388 | chr5-28458645-DEL-388 |
| chr5-28470844-DEL-288 | chr5-28470845-DEL-288 |
| chr5-28512434-DEL-201 | chr5-28512435-DEL-201 |
| chr5-28533428-DEL-80 | chr5-28533431-DEL-70 |
| chr5-28550767-DEL-208 | chr5-28550768-DEL-208 |
| chr5-28575214-DEL-2731 | chr5-28575213-DEL-2733 |
| chr5-28585961-DEL-868 | chr5-28585962-DEL-868 |
| chr5-28588227-DEL-215 | chr5-28588228-DEL-215 |
| chr5-28629730-DEL-76 | chr5-28629727-DEL-87 |
| chr5-28641399-DEL-1730 | chr5-28641400-DEL-1730 |
| chr5-28645888-DEL-6742 | chr5-28645889-DEL-6742 |
| chr5-28652338-DEL-204 | chr5-28652338-DEL-204 |
| chr5-28660244-DEL-209 | chr5-28660245-DEL-209 |
| chr5-28682818-DEL-58 | chr5-28682819-DEL-58 |
| chr5-28690006-DEL-6828 | chr5-28690007-DEL-6828 |
| chr5-28712336-DEL-6374 | chr5-28712337-DEL-6374 |
| chr5-28743485-DEL-1290 | chr5-28743486-DEL-1290 |
| chr5-28746466-DEL-887 | chr5-28746467-DEL-887 |
| chr5-28747047-DEL-1272 | chr5-28747048-DEL-1272 |
| chr5-28762823-DEL-160 | chr5-28762824-DEL-160 |
| chr5-28873742-DEL-64 | chr5-28873742-DEL-64 |
| chr5-28874125-DEL-223 | chr5-28874125-DEL-223 |
| chr5-28922319-DEL-546 | chr5-28922320-DEL-546 |
| chr5-28938169-DEL-1049 | chr5-28938170-DEL-1049 |
| chr5-28964255-DEL-90 | chr5-28964255-DEL-90 |
| chr5-28992809-DEL-73 | chr5-28992810-DEL-73 |
| chr5-28995382-DEL-212 | chr5-28995363-DEL-228 |
| chr5-29082706-DEL-3096 | chr5-29082707-DEL-3096 |
| chr5-29120116-DEL-185 | chr5-29120117-DEL-185 |
| chr5-29220280-DEL-188 | chr5-29220266-DEL-204 |
| chr5-29222522-DEL-242 | chr5-29222523-DEL-242 |
| chr5-29233940-DEL-53 | chr5-29233941-DEL-50 |
| chr5-29246172-DEL-752 | chr5-29246173-DEL-752 |
| chr5-29248518-DEL-282 | chr5-29248519-DEL-282 |
| chr5-29255675-DEL-1721 | chr5-29255676-DEL-1721 |
| chr5-29260391-DEL-7041 | chr5-29260392-DEL-7041 |
| chr5-29260970-DEL-583 | chr5-29260971-DEL-583 |
| chr5-29271664-DEL-1750 | chr5-29271665-DEL-1750 |
| chr5-29282628-DEL-1259 | chr5-29282629-DEL-1259 |
| chr5-29286000-DEL-70 | chr5-29285991-DEL-80 |
| chr5-29377410-DEL-1737 | chr5-29377411-DEL-1737 |
| chr5-29380821-DEL-81 | chr5-29380820-DEL-82 |
| chr5-29381718-DEL-6668 | chr5-29381719-DEL-6668 |
| chr5-29471478-DEL-1077 | chr5-29471479-DEL-1077 |
| chr5-29511035-DEL-815 | chr5-29511036-DEL-815 |
| chr5-29542289-DEL-203 | chr5-29542298-DEL-197 |
| chr5-29554467-DEL-7204 | chr5-29554469-DEL-7205 |
| chr5-29589092-DEL-63 | chr5-29589093-DEL-63 |
| chr5-29597969-DEL-436 | chr5-29597970-DEL-436 |
| chr5-29600918-DEL-1658 | chr5-29600919-DEL-1658 |
| chr5-29616848-DEL-207 | chr5-29616849-DEL-207 |
| chr5-29624213-DEL-338 | chr5-29624214-DEL-338 |
| chr5-29626469-DEL-243 | chr5-29626470-DEL-243 |
| chr5-29641602-DEL-80 | chr5-29641607-DEL-78 |
| chr5-29649848-DEL-68 | chr5-29649849-DEL-68 |
| chr5-29670177-DEL-58 | chr5-29670197-DEL-84 |
| chr5-29698002-DEL-463 | chr5-29698003-DEL-457 |
| chr5-29708575-DEL-191 | chr5-29708564-DEL-208 |

| | |
|---|---|
| chr5-29730529-DEL-203 | chr5-29730530-DEL-203 |
| chr5-29753527-DEL-215 | chr5-29753528-DEL-215 |
| chr5-29800424-DEL-103 | chr5-29800419-DEL-129 |
| chr5-29808287-DEL-87 | chr5-29808288-DEL-87 |
| chr5-29813552-DEL-122 | chr5-29813553-DEL-104 |
| chr5-29888315-DEL-222 | chr5-29888316-DEL-222 |
| chr5-29895677-DEL-161 | chr5-29895678-DEL-161 |
| chr5-29904012-DEL-204 | chr5-29904013-DEL-204 |
| chr5-29918615-DEL-189 | chr5-29918616-DEL-189 |
| chr5-29934117-DEL-90 | chr5-29934128-DEL-64 |
| chr5-29937701-DEL-528 | chr5-29937702-DEL-528 |
| chr5-29953909-DEL-206 | chr5-29953910-DEL-206 |
| chr5-29984699-DEL-181 | chr5-29984700-DEL-181 |
| chr5-29999148-DEL-1078 | chr5-29999149-DEL-1078 |
| chr5-30009022-DEL-2247 | chr5-30009023-DEL-2247 |
| chr5-30082055-DEL-2241 | chr5-30082056-DEL-2241 |
| chr5-30094861-DEL-56 | chr5-30094862-DEL-52 |
| chr5-30109869-DEL-414 | chr5-30109870-DEL-414 |
| chr5-30120815-DEL-60 | chr5-30120816-DEL-60 |
| chr5-30122555-DEL-226 | chr5-30122556-DEL-226 |
| chr5-30146378-DEL-80 | chr5-30146342-DEL-90 |
| chr5-30147462-DEL-539 | chr5-30147463-DEL-538 |
| chr5-30168856-DEL-100 | chr5-30168857-DEL-100 |
| chr5-30188142-DEL-603 | chr5-30188143-DEL-603 |
| chr5-30213195-DEL-52 | chr5-30213208-DEL-53 |
| chr5-30221840-DEL-410 | chr5-30221841-DEL-410 |
| chr5-30222392-DEL-213 | chr5-30222393-DEL-213 |
| chr5-30226063-DEL-70 | chr5-30226068-DEL-60 |
| chr5-30228056-DEL-249 | chr5-30228055-DEL-249 |
| chr5-30242188-DEL-92 | chr5-30242189-DEL-92 |
| chr5-30254214-DEL-2523 | chr5-30254215-DEL-2523 |
| chr5-30255725-DEL-5248 | chr5-30255726-DEL-5248 |
| chr5-30267691-DEL-227 | chr5-30267692-DEL-227 |
| chr5-30276868-DEL-58 | chr5-30276869-DEL-58 |
| chr5-30289673-DEL-77 | chr5-30289640-DEL-111 |
| chr5-30304844-DEL-68 | chr5-30304845-DEL-68 |
| chr5-30387016-DEL-500 | chr5-30387018-DEL-497 |
| chr5-30408959-DEL-166 | chr5-30408966-DEL-165 |
| chr5-30411271-DEL-624 | chr5-30411272-DEL-624 |
| chr5-30412935-DEL-1399 | chr5-30412936-DEL-1399 |
| chr5-30494914-DEL-734 | chr5-30494930-DEL-749 |
| chr5-30502599-DEL-65 | chr5-30502600-DEL-65 |
| chr5-30503609-DEL-57 | chr5-30503610-DEL-57 |
| chr5-30507096-DEL-64 | chr5-30507105-DEL-56 |
| chr5-30509958-DEL-173 | chr5-30509941-DEL-176 |
| chr5-30520737-DEL-188 | chr5-30520762-DEL-160 |
| chr5-30559264-DEL-99 | chr5-30559259-DEL-100 |
| chr5-30589245-DEL-86 | chr5-30589227-DEL-85 |
| chr5-30596180-DEL-143 | chr5-30596228-DEL-144 |
| chr5-30624556-DEL-64 | chr5-30624557-DEL-64 |
| chr5-30625113-DEL-61 | chr5-30625114-DEL-61 |
| chr5-30668240-DEL-508 | chr5-30668241-DEL-508 |
| chr5-30696195-DEL-53 | chr5-30696195-DEL-53 |
| chr5-30702410-DEL-568 | chr5-30702409-DEL-562 |
| chr5-30715622-DEL-79 | chr5-30715643-DEL-59 |
| chr5-30721247-DEL-88 | chr5-30721248-DEL-88 |
| chr5-30737328-DEL-188 | chr5-30737329-DEL-188 |
| chr5-30769516-DEL-497 | chr5-30769517-DEL-497 |
| chr5-30825478-DEL-164 | chr5-30825479-DEL-164 |
| chr5-30826250-DEL-342 | chr5-30826251-DEL-342 |
| chr5-30831837-DEL-1032 | chr5-30831838-DEL-1032 |
| chr5-30832929-DEL-1175 | chr5-30832930-DEL-1175 |
| chr5-30837823-DEL-56 | chr5-30837824-DEL-56 |
| chr5-30850766-DEL-2708 | chr5-30850767-DEL-2708 |
| chr5-30860224-DEL-64 | chr5-30860225-DEL-60 |
| chr5-30860786-DEL-199 | chr5-30860787-DEL-199 |
| chr5-30878359-DEL-469 | chr5-30878360-DEL-469 |
| chr5-30908148-DEL-54 | chr5-30908149-DEL-54 |
| chr5-30912183-DEL-218 | chr5-30912184-DEL-217 |
| chr5-30915786-DEL-3427 | chr5-30915787-DEL-3427 |
| chr5-30925210-DEL-196 | chr5-30925211-DEL-196 |
| chr5-30925883-DEL-2517 | chr5-30925884-DEL-2517 |
| chr5-30943681-DEL-191 | chr5-30943682-DEL-191 |
| chr5-30945051-DEL-52 | chr5-30945051-DEL-52 |
| chr5-30946903-DEL-74 | chr5-30946896-DEL-75 |
| chr5-31040934-DEL-124 | chr5-31040917-DEL-138 |
| chr5-31052911-DEL-67 | chr5-31052939-DEL-50 |
| chr5-31054720-DEL-68 | chr5-31054721-DEL-68 |
| chr5-31089123-DEL-74 | chr5-31089124-DEL-74 |
| chr5-31093488-DEL-985 | chr5-31093489-DEL-985 |

| | |
|---|---|
| chr5-31124936-DEL-60 | chr5-31124937-DEL-60 |
| chr5-31130170-DEL-64 | chr5-31130182-DEL-63 |
| chr5-31135319-DEL-135 | chr5-31135320-DEL-135 |
| chr5-31141064-DEL-58 | chr5-31141065-DEL-58 |
| chr5-31143425-DEL-333 | chr5-31143426-DEL-333 |
| chr5-31145371-DEL-324 | chr5-31145372-DEL-324 |
| chr5-31147735-DEL-85 | chr5-31147736-DEL-85 |
| chr5-31153997-DEL-509 | chr5-31153999-DEL-508 |
| chr5-31162368-DEL-966 | chr5-31162369-DEL-966 |
| chr5-31168219-DEL-54 | chr5-31168220-DEL-54 |
| chr5-31168944-DEL-52 | chr5-31168945-DEL-52 |
| chr5-31179350-DEL-210 | chr5-31179351-DEL-210 |
| chr5-31242806-DEL-79 | chr5-31242807-DEL-79 |
| chr5-31270688-DEL-1892 | chr5-31270689-DEL-1892 |
| chr5-31276078-DEL-108 | chr5-31276081-DEL-100 |
| chr5-31284923-DEL-190 | chr5-31284924-DEL-190 |
| chr5-31299636-DEL-216 | chr5-31299637-DEL-216 |
| chr5-31302255-DEL-82 | chr5-31302256-DEL-82 |
| chr5-31388958-DEL-82 | chr5-31388918-DEL-123 |
| chr5-31391440-DEL-2409 | chr5-31391441-DEL-2409 |
| chr5-31426077-DEL-179 | chr5-31426078-DEL-179 |
| chr5-31428106-DEL-218 | chr5-31428107-DEL-218 |
| chr5-31459176-DEL-173 | chr5-31459177-DEL-173 |
| chr5-31472797-DEL-337 | chr5-31472798-DEL-337 |
| chr5-31484965-DEL-701 | chr5-31484966-DEL-701 |
| chr5-31486772-DEL-4628 | chr5-31486773-DEL-4628 |
| chr5-31528152-DEL-240 | chr5-31528153-DEL-240 |
| chr5-31548510-DEL-969 | chr5-31548511-DEL-969 |
| chr5-31613658-DEL-1893 | chr5-31613659-DEL-1893 |
| chr5-31615401-DEL-197 | chr5-31615403-DEL-196 |
| chr5-31629422-DEL-102 | chr5-31629423-DEL-102 |
| chr5-31675092-DEL-207 | chr5-31675093-DEL-207 |
| chr5-31749960-DEL-497 | chr5-31749961-DEL-497 |
| chr5-31755273-DEL-78 | chr5-31755274-DEL-78 |
| chr5-31845816-DEL-677 | chr5-31845817-DEL-677 |
| chr5-31902597-DEL-211 | chr5-31902598-DEL-211 |
| chr5-31912348-DEL-74 | chr5-31912301-DEL-74 |
| chr5-31913233-DEL-420 | chr5-31913234-DEL-420 |
| chr5-31921047-DEL-178 | chr5-31921048-DEL-178 |
| chr5-31938379-DEL-88 | chr5-31938380-DEL-88 |
| chr5-31952670-DEL-56 | chr5-31952674-DEL-56 |
| chr5-31959845-DEL-292 | chr5-31959843-DEL-293 |
| chr5-31971922-DEL-183 | chr5-31971923-DEL-183 |
| chr5-31987712-DEL-72 | chr5-31987707-DEL-84 |
| chr5-32028969-DEL-56 | chr5-32028970-DEL-52 |
| chr5-32040298-DEL-924 | chr5-32040299-DEL-924 |
| chr5-32097232-DEL-113 | chr5-32097233-DEL-113 |
| chr5-32145952-DEL-6193 | chr5-32145953-DEL-6193 |
| chr5-32222952-DEL-79 | chr5-32222953-DEL-79 |
| chr5-32247759-DEL-131 | chr5-32247761-DEL-132 |
| chr5-32251818-DEL-55 | chr5-32251819-DEL-55 |
| chr5-32265486-DEL-141 | chr5-32265487-DEL-141 |
| chr5-32335756-DEL-464 | chr5-32335757-DEL-464 |
| chr5-32359361-DEL-70 | chr5-32359349-DEL-120 |
| chr5-32371597-DEL-63 | chr5-32371595-DEL-57 |
| chr5-32373182-DEL-152 | chr5-32373183-DEL-151 |
| chr5-32382271-DEL-763 | chr5-32382272-DEL-763 |
| chr5-32393750-DEL-73 | chr5-32393751-DEL-73 |
| chr5-32433126-DEL-90 | chr5-32433130-DEL-85 |
| chr5-32489477-DEL-72 | chr5-32489478-DEL-72 |
| chr5-32520298-DEL-111 | chr5-32520297-DEL-111 |
| chr5-32569567-DEL-53 | chr5-32569568-DEL-53 |
| chr5-32593219-DEL-1562 | chr5-32593220-DEL-1562 |
| chr5-32659865-DEL-598 | chr5-32659866-DEL-598 |
| chr5-32689827-DEL-440 | chr5-32689827-DEL-440 |
| chr5-32694387-DEL-215 | chr5-32694388-DEL-215 |
| chr5-32700757-DEL-2141 | chr5-32700757-DEL-2141 |
| chr5-32735625-DEL-389 | chr5-32735626-DEL-389 |
| chr5-32778764-DEL-201 | chr5-32778765-DEL-201 |
| chr5-32781422-DEL-265 | chr5-32781423-DEL-265 |
| chr5-32784118-DEL-193 | chr5-32784119-DEL-193 |
| chr5-32786975-DEL-1864 | chr5-32786976-DEL-1864 |
| chr5-32793875-DEL-353 | chr5-32793876-DEL-353 |
| chr5-32831738-DEL-6401 | chr5-32831739-DEL-6401 |
| chr5-32849193-DEL-196 | chr5-32849194-DEL-196 |
| chr5-32870946-DEL-76 | chr5-32870947-DEL-76 |
| chr5-32872259-DEL-234 | chr5-32872278-DEL-206 |
| chr5-32888815-DEL-1892 | chr5-32888816-DEL-1892 |
| chr5-32913506-DEL-1171 | chr5-32913507-DEL-1183 |
| chr5-32945720-DEL-180 | chr5-32945720-DEL-180 |

| | |
|---|---|
| chr5-32964092-DEL-243 | chr5-32964093-DEL-243 |
| chr5-32990855-DEL-203 | chr5-32990856-DEL-203 |
| chr5-33012833-DEL-205 | chr5-33012834-DEL-205 |
| chr5-33013779-DEL-60 | chr5-33013780-DEL-60 |
| chr5-33031285-DEL-67 | chr5-33031286-DEL-67 |
| chr5-33052224-DEL-195 | chr5-33052225-DEL-195 |
| chr5-33077586-DEL-859 | chr5-33077587-DEL-859 |
| chr5-33111349-DEL-636 | chr5-33111350-DEL-636 |
| chr5-33128713-DEL-212 | chr5-33128714-DEL-212 |
| chr5-33138630-DEL-202 | chr5-33138631-DEL-202 |
| chr5-33190960-DEL-199 | chr5-33190961-DEL-199 |
| chr5-33195292-DEL-853 | chr5-33195293-DEL-853 |
| chr5-33215041-DEL-132 | chr5-33215044-DEL-94 |
| chr5-33228735-DEL-215 | chr5-33228734-DEL-215 |
| chr5-33233308-DEL-459 | chr5-33233309-DEL-459 |
| chr5-33250421-DEL-56 | chr5-33250422-DEL-56 |
| chr5-33269018-DEL-229 | chr5-33269006-DEL-230 |
| chr5-33311597-DEL-205 | chr5-33311598-DEL-205 |
| chr5-33335810-DEL-210 | chr5-33335810-DEL-210 |
| chr5-33341794-DEL-3784 | chr5-33341795-DEL-3784 |
| chr5-33347797-DEL-861 | chr5-33347798-DEL-861 |
| chr5-33351431-DEL-60 | chr5-33351434-DEL-62 |
| chr5-33352027-DEL-163 | chr5-33352030-DEL-160 |
| chr5-33353225-DEL-118 | chr5-33353225-DEL-118 |
| chr5-33356364-DEL-232 | chr5-33356365-DEL-232 |
| chr5-33444176-DEL-186 | chr5-33444177-DEL-186 |
| chr5-33481187-DEL-445 | chr5-33481188-DEL-445 |
| chr5-33485122-DEL-4928 | chr5-33485123-DEL-4928 |
| chr5-33622158-DEL-51 | chr5-33622152-DEL-58 |
| chr5-33623680-DEL-85 | chr5-33623658-DEL-86 |
| chr5-33681882-DEL-53 | chr5-33681883-DEL-53 |
| chr5-33689945-DEL-69 | chr5-33689946-DEL-69 |
| chr5-33718166-DEL-498 | chr5-33718167-DEL-498 |
| chr5-33728326-DEL-333 | chr5-33728327-DEL-333 |
| chr5-33731804-DEL-79 | chr5-33731805-DEL-79 |
| chr5-33802357-DEL-206 | chr5-33802358-DEL-206 |
| chr5-33806258-DEL-207 | chr5-33806259-DEL-208 |
| chr5-33822305-DEL-203 | chr5-33822306-DEL-203 |
| chr5-33836937-DEL-387 | chr5-33836938-DEL-387 |
| chr5-33916676-DEL-804 | chr5-33916686-DEL-798 |
| chr5-33917721-DEL-209 | chr5-33917722-DEL-209 |
| chr5-33935837-DEL-54 | chr5-33935838-DEL-54 |
| chr5-33965409-DEL-67 | chr5-33965440-DEL-59 |
| chr5-34061466-DEL-8153 | chr5-34061467-DEL-8153 |
| chr5-34103407-DEL-393 | chr5-34103362-DEL-425 |
| chr5-34104878-DEL-223 | chr5-34104878-DEL-223 |
| chr5-34126532-DEL-334 | chr5-34126533-DEL-334 |
| chr5-34192817-DEL-1892 | chr5-34192818-DEL-1892 |
| chr5-34199426-DEL-6269 | chr5-34199427-DEL-6269 |
| chr5-34212474-DEL-185 | chr5-34212473-DEL-183 |
| chr5-34222445-DEL-107 | chr5-34222446-DEL-107 |
| chr5-34222938-DEL-1164 | chr5-34222939-DEL-1163 |
| chr5-34235452-DEL-152 | chr5-34235453-DEL-144 |
| chr5-34251451-DEL-190 | chr5-34251452-DEL-183 |
| chr5-34278564-DEL-867 | chr5-34278565-DEL-867 |
| chr5-34402026-DEL-60 | chr5-34402027-DEL-60 |
| chr5-34406261-DEL-54 | chr5-34406262-DEL-54 |
| chr5-34419464-DEL-3414 | chr5-34419465-DEL-3414 |
| chr5-34446313-DEL-55 | chr5-34446315-DEL-50 |
| chr5-34463475-DEL-207 | chr5-34463478-DEL-209 |
| chr5-34488937-DEL-201 | chr5-34488938-DEL-201 |
| chr5-34500498-DEL-214 | chr5-34500499-DEL-214 |
| chr5-34546300-DEL-213 | chr5-34546317-DEL-230 |
| chr5-34554513-DEL-80 | chr5-34554514-DEL-80 |
| chr5-34566713-DEL-332 | chr5-34566714-DEL-332 |
| chr5-34587837-DEL-59 | chr5-34587838-DEL-59 |
| chr5-34651738-DEL-210 | chr5-34651739-DEL-210 |
| chr5-34663197-DEL-1590 | chr5-34663198-DEL-1590 |
| chr5-34675919-DEL-236 | chr5-34675920-DEL-236 |
| chr5-34700225-DEL-274 | chr5-34700226-DEL-274 |
| chr5-34725826-DEL-88 | chr5-34725827-DEL-88 |
| chr5-34729616-DEL-429 | chr5-34729617-DEL-429 |
| chr5-34738580-DEL-213 | chr5-34738566-DEL-211 |
| chr5-34739227-DEL-133 | chr5-34739184-DEL-142 |
| chr5-34769241-DEL-195 | chr5-34769242-DEL-195 |
| chr5-34799640-DEL-544 | chr5-34799641-DEL-544 |
| chr5-34894995-DEL-1983 | chr5-34894996-DEL-1983 |
| chr5-34940553-DEL-234 | chr5-34940563-DEL-225 |
| chr5-34957258-DEL-139 | chr5-34957259-DEL-139 |
| chr5-35061074-DEL-102 | chr5-35061066-DEL-118 |

| | |
|---|---|
| chr5-35135237-DEL-63 | chr5-35135210-DEL-76 |
| chr5-35163480-DEL-102 | chr5-35163481-DEL-102 |
| chr5-35218023-DEL-935 | chr5-35218024-DEL-935 |
| chr5-35218323-DEL-124 | chr5-35218338-DEL-121 |
| chr5-35223216-DEL-51 | chr5-35223217-DEL-51 |
| chr5-35251594-DEL-217 | chr5-35251595-DEL-217 |
| chr5-35271809-DEL-58 | chr5-35271810-DEL-58 |
| chr5-35294428-DEL-271 | chr5-35294426-DEL-272 |
| chr5-35303554-DEL-79 | chr5-35303555-DEL-79 |
| chr5-35310584-DEL-3732 | chr5-35310585-DEL-3732 |
| chr5-35327012-DEL-134 | chr5-35326979-DEL-176 |
| chr5-35338866-DEL-71 | chr5-35338878-DEL-55 |
| chr5-35377777-DEL-52 | chr5-35377778-DEL-52 |
| chr5-35382207-DEL-74 | chr5-35382173-DEL-74 |
| chr5-35383405-DEL-252 | chr5-35383406-DEL-252 |
| chr5-35402393-DEL-300 | chr5-35402394-DEL-300 |
| chr5-35446039-DEL-686 | chr5-35446045-DEL-680 |
| chr5-35465064-DEL-202 | chr5-35465065-DEL-202 |
| chr5-35498030-DEL-178 | chr5-35498031-DEL-178 |
| chr5-35499383-DEL-495 | chr5-35499382-DEL-489 |
| chr5-35586820-DEL-998 | chr5-35586821-DEL-998 |
| chr5-35640391-DEL-56 | chr5-35640389-DEL-50 |
| chr5-35642438-DEL-399 | chr5-35642439-DEL-399 |
| chr5-35655645-DEL-84 | chr5-35655661-DEL-74 |
| chr5-35667790-DEL-60 | chr5-35667791-DEL-60 |
| chr5-35672110-DEL-400 | chr5-35672105-DEL-401 |
| chr5-35673438-DEL-82 | chr5-35673439-DEL-82 |
| chr5-35699577-DEL-204 | chr5-35699578-DEL-203 |
| chr5-35705635-DEL-2739 | chr5-35705636-DEL-2739 |
| chr5-35772115-DEL-152 | chr5-35772116-DEL-152 |
| chr5-35834072-DEL-52 | chr5-35834073-DEL-52 |
| chr5-35855168-DEL-844 | chr5-35855169-DEL-844 |
| chr5-35879026-DEL-56 | chr5-35879027-DEL-56 |
| chr5-35894016-DEL-310 | chr5-35894017-DEL-310 |
| chr5-35899962-DEL-72 | chr5-35899963-DEL-72 |
| chr5-35914711-DEL-56 | chr5-35914712-DEL-56 |
| chr5-35948001-DEL-82 | chr5-35948002-DEL-82 |
| chr5-35986828-DEL-150 | chr5-35986855-DEL-157 |
| chr5-36009686-DEL-66 | chr5-36009687-DEL-66 |
| chr5-36028352-DEL-252 | chr5-36028353-DEL-252 |
| chr5-36028936-DEL-60 | chr5-36028939-DEL-61 |
| chr5-36036298-DEL-63 | chr5-36036299-DEL-63 |
| chr5-36047916-DEL-210 | chr5-36047917-DEL-210 |
| chr5-36050021-DEL-188 | chr5-36050022-DEL-188 |
| chr5-36056680-DEL-56 | chr5-36056681-DEL-56 |
| chr5-36059195-DEL-592 | chr5-36059196-DEL-592 |
| chr5-36116591-DEL-567 | chr5-36116592-DEL-567 |
| chr5-36124074-DEL-60 | chr5-36124075-DEL-60 |
| chr5-36132242-DEL-64 | chr5-36132239-DEL-68 |
| chr5-36143528-DEL-186 | chr5-36143531-DEL-185 |
| chr5-36150088-DEL-78 | chr5-36150091-DEL-91 |
| chr5-36245051-DEL-62 | chr5-36245052-DEL-62 |
| chr5-36265402-DEL-169 | chr5-36265403-DEL-169 |
| chr5-36336531-DEL-68 | chr5-36336532-DEL-66 |
| chr5-36395119-DEL-8157 | chr5-36395120-DEL-8157 |
| chr5-36395316-DEL-1272 | chr5-36395317-DEL-1272 |
| chr5-36407088-DEL-54 | chr5-36407088-DEL-54 |
| chr5-36410164-DEL-89 | chr5-36410165-DEL-89 |
| chr5-36413133-DEL-79 | chr5-36413144-DEL-69 |
| chr5-36465846-DEL-53 | chr5-36465847-DEL-53 |
| chr5-36467643-DEL-102 | chr5-36467644-DEL-102 |
| chr5-36490072-DEL-193 | chr5-36490063-DEL-195 |
| chr5-36529551-DEL-76 | chr5-36529552-DEL-76 |
| chr5-36548001-DEL-61 | chr5-36547983-DEL-60 |
| chr5-36549927-DEL-92 | chr5-36549906-DEL-69 |
| chr5-36559232-DEL-70 | chr5-36559244-DEL-58 |
| chr5-36563550-DEL-58 | chr5-36563552-DEL-50 |
| chr5-36576934-DEL-59 | chr5-36576935-DEL-59 |
| chr5-36581919-DEL-59 | chr5-36581920-DEL-57 |
| chr5-36587948-DEL-775 | chr5-36587949-DEL-775 |
| chr5-36617407-DEL-97 | chr5-36617408-DEL-97 |
| chr5-36624670-DEL-78 | chr5-36624671-DEL-78 |
| chr5-36717266-DEL-965 | chr5-36717267-DEL-965 |
| chr5-36746375-DEL-200 | chr5-36746376-DEL-200 |
| chr5-36891911-DEL-779 | chr5-36891912-DEL-779 |
| chr5-36912047-DEL-223 | chr5-36912035-DEL-205 |
| chr5-36912925-DEL-116 | chr5-36912926-DEL-116 |
| chr5-36936043-DEL-129 | chr5-36936055-DEL-118 |
| chr5-36937071-DEL-137 | chr5-36937058-DEL-128 |
| chr5-36938375-DEL-2596 | chr5-36938376-DEL-2596 |

| | |
|---|---|
| chr5-36943603-DEL-220 | chr5-36943604-DEL-220 |
| chr5-36956087-DEL-640 | chr5-36956088-DEL-640 |
| chr5-36962935-DEL-62 | chr5-36962947-DEL-51 |
| chr5-36969633-DEL-209 | chr5-36969634-DEL-209 |
| chr5-36975494-DEL-157 | chr5-36975495-DEL-157 |
| chr5-37051635-DEL-1127 | chr5-37051636-DEL-1127 |
| chr5-37067691-DEL-326 | chr5-37067692-DEL-326 |
| chr5-37070758-DEL-60 | chr5-37070759-DEL-60 |
| chr5-37072841-DEL-64 | chr5-37072842-DEL-74 |
| chr5-37083172-DEL-173 | chr5-37083173-DEL-173 |
| chr5-37091005-DEL-117 | chr5-37091008-DEL-115 |
| chr5-37096224-DEL-333 | chr5-37096225-DEL-333 |
| chr5-37120065-DEL-52 | chr5-37120066-DEL-50 |
| chr5-37123342-DEL-52 | chr5-37123343-DEL-52 |
| chr5-37128032-DEL-538 | chr5-37128033-DEL-538 |
| chr5-37149930-DEL-58 | chr5-37149931-DEL-58 |
| chr5-37151879-DEL-342 | chr5-37151880-DEL-342 |
| chr5-37160349-DEL-118 | chr5-37160350-DEL-118 |
| chr5-37174143-DEL-205 | chr5-37174144-DEL-205 |
| chr5-37254475-DEL-201 | chr5-37254476-DEL-201 |
| chr5-37267248-DEL-1865 | chr5-37267249-DEL-1865 |
| chr5-37268599-DEL-265 | chr5-37268600-DEL-265 |
| chr5-37279169-DEL-67 | chr5-37279170-DEL-67 |
| chr5-37314732-DEL-60 | chr5-37314774-DEL-63 |
| chr5-37321851-DEL-58 | chr5-37321852-DEL-58 |
| chr5-37333341-DEL-64 | chr5-37333342-DEL-64 |
| chr5-37335885-DEL-655 | chr5-37335886-DEL-655 |
| chr5-37340600-DEL-858 | chr5-37340600-DEL-856 |
| chr5-37358915-DEL-116 | chr5-37358939-DEL-93 |
| chr5-37372950-DEL-3028 | chr5-37372951-DEL-3028 |
| chr5-37389895-DEL-53 | chr5-37389896-DEL-53 |
| chr5-37442480-DEL-126 | chr5-37442481-DEL-126 |
| chr5-37464268-DEL-51 | chr5-37464305-DEL-51 |
| chr5-37504130-DEL-74 | chr5-37504131-DEL-72 |
| chr5-37511400-DEL-648 | chr5-37511401-DEL-648 |
| chr5-37528471-DEL-176 | chr5-37528482-DEL-184 |
| chr5-37531115-DEL-298 | chr5-37531114-DEL-298 |
| chr5-37548186-DEL-939 | chr5-37548187-DEL-939 |
| chr5-37586179-DEL-220 | chr5-37586180-DEL-220 |
| chr5-37597256-DEL-80 | chr5-37597257-DEL-80 |
| chr5-37599290-DEL-278 | chr5-37599291-DEL-278 |
| chr5-37604840-DEL-62 | chr5-37604833-DEL-63 |
| chr5-37622510-DEL-215 | chr5-37622511-DEL-215 |
| chr5-37633876-DEL-204 | chr5-37633877-DEL-204 |
| chr5-37656619-DEL-6512 | chr5-37656617-DEL-6513 |
| chr5-37656700-DEL-214 | chr5-37656701-DEL-214 |
| chr5-37663780-DEL-127 | chr5-37663776-DEL-128 |
| chr5-37680801-DEL-98 | chr5-37680797-DEL-54 |
| chr5-37692488-DEL-109 | chr5-37692512-DEL-85 |
| chr5-37693520-DEL-464 | chr5-37693521-DEL-464 |
| chr5-37698970-DEL-53 | chr5-37698969-DEL-54 |
| chr5-37713662-DEL-207 | chr5-37713650-DEL-208 |
| chr5-37722154-DEL-1959 | chr5-37722155-DEL-1960 |
| chr5-37724515-DEL-53 | chr5-37724515-DEL-61 |
| chr5-37736604-DEL-6584 | chr5-37736605-DEL-6584 |
| chr5-37755596-DEL-73 | chr5-37755595-DEL-74 |
| chr5-37757423-DEL-358 | chr5-37757424-DEL-358 |
| chr5-37764426-DEL-3695 | chr5-37764427-DEL-3695 |
| chr5-37771530-DEL-640 | chr5-37771531-DEL-640 |
| chr5-37775939-DEL-1815 | chr5-37775941-DEL-1813 |
| chr5-37779818-DEL-61 | chr5-37779818-DEL-62 |
| chr5-37791032-DEL-78 | chr5-37791029-DEL-76 |
| chr5-37795294-DEL-52 | chr5-37795295-DEL-52 |
| chr5-37799548-DEL-1815 | chr5-37799549-DEL-1815 |
| chr5-37809219-DEL-408 | chr5-37809220-DEL-408 |
| chr5-37828052-DEL-61 | chr5-37828045-DEL-50 |
| chr5-37843788-DEL-3752 | chr5-37843789-DEL-3752 |
| chr5-37867417-DEL-263 | chr5-37867418-DEL-263 |
| chr5-37868215-DEL-56 | chr5-37868216-DEL-56 |
| chr5-37868993-DEL-608 | chr5-37868994-DEL-608 |
| chr5-37884518-DEL-1188 | chr5-37884519-DEL-1188 |
| chr5-37890624-DEL-70 | chr5-37890625-DEL-62 |
| chr5-37923743-DEL-72 | chr5-37923744-DEL-68 |
| chr5-37935763-DEL-130 | chr5-37935759-DEL-117 |
| chr5-37936325-DEL-77 | chr5-37936330-DEL-79 |
| chr5-37947007-DEL-209 | chr5-37946987-DEL-230 |
| chr5-37951758-DEL-52 | chr5-37951761-DEL-50 |
| chr5-37968702-DEL-64 | chr5-37968703-DEL-64 |
| chr5-37976810-DEL-108 | chr5-37976811-DEL-107 |
| chr5-37987691-DEL-210 | chr5-37987692-DEL-210 |

| | |
|---|---|
| chr5-38007139-DEL-94 | chr5-38007140-DEL-78 |
| chr5-38010855-DEL-76 | chr5-38010856-DEL-76 |
| chr5-38024401-DEL-92 | chr5-38024389-DEL-108 |
| chr5-38028710-DEL-111 | chr5-38028711-DEL-111 |
| chr5-38034547-DEL-211 | chr5-38034548-DEL-211 |
| chr5-38038353-DEL-149 | chr5-38038354-DEL-149 |
| chr5-38041534-DEL-714 | chr5-38041522-DEL-692 |
| chr5-38075952-DEL-221 | chr5-38075953-DEL-221 |
| chr5-38081690-DEL-4734 | chr5-38081691-DEL-4734 |
| chr5-38138926-DEL-177 | chr5-38138927-DEL-177 |
| chr5-38148264-DEL-57 | chr5-38148265-DEL-57 |
| chr5-38148963-DEL-55 | chr5-38148964-DEL-55 |
| chr5-38191172-DEL-796 | chr5-38191173-DEL-796 |
| chr5-38192389-DEL-194 | chr5-38192391-DEL-195 |
| chr5-38203678-DEL-904 | chr5-38203679-DEL-904 |
| chr5-38230288-DEL-74 | chr5-38230277-DEL-66 |
| chr5-38250015-DEL-2858 | chr5-38250016-DEL-2858 |
| chr5-38287681-DEL-4350 | chr5-38287682-DEL-4350 |
| chr5-38326198-DEL-58 | chr5-38326199-DEL-58 |
| chr5-38335558-DEL-352 | chr5-38335559-DEL-352 |
| chr5-38397784-DEL-112 | chr5-38397777-DEL-96 |
| chr5-38399944-DEL-139 | chr5-38399945-DEL-139 |
| chr5-38429686-DEL-55 | chr5-38429687-DEL-55 |
| chr5-38450031-DEL-56 | chr5-38450037-DEL-64 |
| chr5-38486450-DEL-99 | chr5-38486450-DEL-99 |
| chr5-38503256-DEL-84 | chr5-38503257-DEL-84 |
| chr5-38517758-DEL-77 | chr5-38517758-DEL-77 |
| chr5-38531012-DEL-164 | chr5-38531037-DEL-140 |
| chr5-38539939-DEL-504 | chr5-38539941-DEL-503 |
| chr5-38548061-DEL-383 | chr5-38548052-DEL-384 |
| chr5-38550513-DEL-52 | chr5-38550508-DEL-54 |
| chr5-38563026-DEL-7165 | chr5-38563027-DEL-7165 |
| chr5-38604390-DEL-639 | chr5-38604391-DEL-639 |
| chr5-38611219-DEL-5958 | chr5-38611220-DEL-5958 |
| chr5-38628805-DEL-4191 | chr5-38628806-DEL-4191 |
| chr5-38663993-DEL-99 | chr5-38663994-DEL-99 |
| chr5-38680914-DEL-54 | chr5-38680915-DEL-54 |
| chr5-38690517-DEL-161 | chr5-38690518-DEL-161 |
| chr5-38709585-DEL-77 | chr5-38709580-DEL-78 |
| chr5-38736972-DEL-198 | chr5-38736958-DEL-190 |
| chr5-38754239-DEL-102 | chr5-38754240-DEL-102 |
| chr5-38755714-DEL-498 | chr5-38755715-DEL-498 |
| chr5-38784472-DEL-58 | chr5-38784473-DEL-58 |
| chr5-38816934-DEL-63 | chr5-38816935-DEL-63 |
| chr5-38851920-DEL-104 | chr5-38851929-DEL-102 |
| chr5-38898068-DEL-121 | chr5-38898069-DEL-121 |
| chr5-38902076-DEL-342 | chr5-38902077-DEL-342 |
| chr5-38926757-DEL-140 | chr5-38926757-DEL-140 |
| chr5-38933917-DEL-55 | chr5-38933918-DEL-55 |
| chr5-38938546-DEL-63 | chr5-38938548-DEL-58 |
| chr5-38944752-DEL-218 | chr5-38944753-DEL-218 |
| chr5-38961432-DEL-445 | chr5-38961433-DEL-446 |
| chr5-38988547-DEL-6623 | chr5-38988548-DEL-6623 |
| chr5-38994692-DEL-212 | chr5-38994693-DEL-212 |
| chr5-39007460-DEL-133 | chr5-39007461-DEL-133 |
| chr5-39069484-DEL-6222 | chr5-39069485-DEL-6222 |
| chr5-39107320-DEL-51 | chr5-39107321-DEL-51 |
| chr5-39117877-DEL-2507 | chr5-39117878-DEL-2507 |
| chr5-39125368-DEL-284 | chr5-39125369-DEL-282 |
| chr5-39139633-DEL-1864 | chr5-39139634-DEL-1864 |
| chr5-39142021-DEL-73 | chr5-39142022-DEL-73 |
| chr5-39159356-DEL-1011 | chr5-39159357-DEL-1011 |
| chr5-39167002-DEL-162 | chr5-39167003-DEL-162 |
| chr5-39167538-DEL-53 | chr5-39167539-DEL-53 |
| chr5-39189017-DEL-6449 | chr5-39189018-DEL-6449 |
| chr5-39238802-DEL-70 | chr5-39238810-DEL-54 |
| chr5-39262684-DEL-58 | chr5-39262681-DEL-52 |
| chr5-39341736-DEL-111 | chr5-39341735-DEL-112 |
| chr5-39348301-DEL-6829 | chr5-39348302-DEL-6829 |
| chr5-39362044-DEL-1485 | chr5-39362045-DEL-1485 |
| chr5-39382172-DEL-7147 | chr5-39382168-DEL-7148 |
| chr5-39404166-DEL-198 | chr5-39404167-DEL-198 |
| chr5-39421901-DEL-90 | chr5-39421902-DEL-90 |
| chr5-39468624-DEL-94 | chr5-39468625-DEL-94 |
| chr5-39513872-DEL-158 | chr5-39513873-DEL-158 |
| chr5-39524667-DEL-342 | chr5-39524668-DEL-338 |
| chr5-39536910-DEL-60 | chr5-39536911-DEL-60 |
| chr5-39563440-DEL-64 | chr5-39563440-DEL-56 |
| chr5-39570292-DEL-102 | chr5-39570293-DEL-102 |
| chr5-39589427-DEL-53 | chr5-39589427-DEL-50 |

| | |
|---|---|
| chr5-39613697-DEL-62 | chr5-39613698-DEL-62 |
| chr5-39658422-DEL-6392 | chr5-39658423-DEL-6392 |
| chr5-39674542-DEL-331 | chr5-39674544-DEL-330 |
| chr5-39678550-DEL-120 | chr5-39678551-DEL-120 |
| chr5-39686135-DEL-292 | chr5-39686136-DEL-292 |
| chr5-39690064-DEL-866 | chr5-39690065-DEL-866 |
| chr5-39702659-DEL-686 | chr5-39702660-DEL-686 |
| chr5-39713273-DEL-122 | chr5-39713274-DEL-122 |
| chr5-39720617-DEL-1181 | chr5-39720618-DEL-1181 |
| chr5-39778379-DEL-56 | chr5-39778380-DEL-56 |
| chr5-39807364-DEL-55 | chr5-39807364-DEL-50 |
| chr5-39832775-DEL-64 | chr5-39832776-DEL-50 |
| chr5-39852490-DEL-149 | chr5-39852492-DEL-149 |
| chr5-39857573-DEL-210 | chr5-39857574-DEL-210 |
| chr5-39864019-DEL-790 | chr5-39864020-DEL-790 |
| chr5-39868472-DEL-248 | chr5-39868473-DEL-248 |
| chr5-39938292-DEL-99 | chr5-39938293-DEL-99 |
| chr5-39951986-DEL-67 | chr5-39951987-DEL-67 |
| chr5-39955433-DEL-1491 | chr5-39955434-DEL-1491 |
| chr5-39997107-DEL-68 | chr5-39997124-DEL-52 |
| chr5-40006709-DEL-1719 | chr5-40006706-DEL-1715 |
| chr5-40072921-DEL-1183 | chr5-40072922-DEL-1183 |
| chr5-40076473-DEL-84 | chr5-40076474-DEL-84 |
| chr5-40088145-DEL-401 | chr5-40088145-DEL-401 |
| chr5-40090817-DEL-277 | chr5-40090815-DEL-277 |
| chr5-40102999-DEL-83 | chr5-40102981-DEL-84 |
| chr5-40106840-DEL-2410 | chr5-40106841-DEL-2410 |
| chr5-40112223-DEL-196 | chr5-40112224-DEL-190 |
| chr5-40125114-DEL-92 | chr5-40125137-DEL-69 |
| chr5-40215211-DEL-200 | chr5-40215210-DEL-200 |
| chr5-40226715-DEL-7406 | chr5-40226716-DEL-7406 |
| chr5-40276560-DEL-52 | chr5-40276561-DEL-52 |
| chr5-40305519-DEL-96 | chr5-40305519-DEL-96 |
| chr5-40306928-DEL-3713 | chr5-40306929-DEL-3713 |
| chr5-40311352-DEL-87 | chr5-40311353-DEL-87 |
| chr5-40320004-DEL-1740 | chr5-40320005-DEL-1740 |
| chr5-40344560-DEL-76 | chr5-40344569-DEL-100 |
| chr5-40370681-DEL-442 | chr5-40370682-DEL-442 |
| chr5-40371901-DEL-1821 | chr5-40371902-DEL-1821 |
| chr5-40374680-DEL-345 | chr5-40374681-DEL-345 |
| chr5-40375998-DEL-426 | chr5-40375999-DEL-426 |
| chr5-40383613-DEL-74 | chr5-40383615-DEL-60 |
| chr5-40407481-DEL-102 | chr5-40407482-DEL-102 |
| chr5-40409322-DEL-64 | chr5-40409327-DEL-57 |
| chr5-40410922-DEL-94 | chr5-40410944-DEL-94 |
| chr5-40425854-DEL-54 | chr5-40425855-DEL-50 |
| chr5-40434831-DEL-77 | chr5-40434832-DEL-77 |
| chr5-40477770-DEL-61 | chr5-40477780-DEL-58 |
| chr5-40485389-DEL-2640 | chr5-40485390-DEL-2640 |
| chr5-40496716-DEL-883 | chr5-40496717-DEL-883 |
| chr5-40501462-DEL-97 | chr5-40501463-DEL-97 |
| chr5-40508102-DEL-1631 | chr5-40508103-DEL-1628 |
| chr5-40510650-DEL-1415 | chr5-40510650-DEL-1415 |
| chr5-40518419-DEL-288 | chr5-40518420-DEL-287 |
| chr5-40564556-DEL-6402 | chr5-40564557-DEL-6402 |
| chr5-40601476-DEL-88 | chr5-40601477-DEL-88 |
| chr5-40619736-DEL-77 | chr5-40619756-DEL-97 |
| chr5-40628603-DEL-7425 | chr5-40628604-DEL-7425 |
| chr5-40643809-DEL-600 | chr5-40643810-DEL-600 |
| chr5-40650421-DEL-1883 | chr5-40650422-DEL-1883 |
| chr5-40651720-DEL-322 | chr5-40651721-DEL-322 |
| chr5-40695052-DEL-1015 | chr5-40695053-DEL-1015 |
| chr5-40701566-DEL-6044 | chr5-40701559-DEL-6051 |
| chr5-40718271-DEL-892 | chr5-40718272-DEL-892 |
| chr5-40739348-DEL-1893 | chr5-40739349-DEL-1893 |
| chr5-40749052-DEL-924 | chr5-40749053-DEL-923 |
| chr5-40759942-DEL-2785 | chr5-40759940-DEL-2786 |
| chr5-40788826-DEL-160 | chr5-40788827-DEL-160 |
| chr5-40815218-DEL-930 | chr5-40815219-DEL-930 |
| chr5-40820040-DEL-2509 | chr5-40820041-DEL-2509 |
| chr5-40843642-DEL-358 | chr5-40843643-DEL-358 |
| chr5-40845342-DEL-390 | chr5-40845343-DEL-390 |
| chr5-40849563-DEL-572 | chr5-40849564-DEL-572 |
| chr5-40853677-DEL-771 | chr5-40853678-DEL-771 |
| chr5-40937884-DEL-91 | chr5-40937884-DEL-91 |
| chr5-40957962-DEL-3536 | chr5-40957963-DEL-3536 |
| chr5-40974908-DEL-4871 | chr5-40974909-DEL-4871 |
| chr5-41003822-DEL-250 | chr5-41003823-DEL-249 |
| chr5-41004786-DEL-296 | chr5-41004787-DEL-296 |
| chr5-41061904-DEL-116 | chr5-41061879-DEL-116 |

| | |
|---|---|
| chr5-41071993-DEL-64 | chr5-41071994-DEL-64 |
| chr5-41076084-DEL-352 | chr5-41076085-DEL-352 |
| chr5-41080420-DEL-3729 | chr5-41080421-DEL-3729 |
| chr5-41112431-DEL-398 | chr5-41112432-DEL-398 |
| chr5-41128492-DEL-452 | chr5-41128493-DEL-451 |
| chr5-41159991-DEL-72 | chr5-41159992-DEL-72 |
| chr5-41175051-DEL-1099 | chr5-41175052-DEL-1099 |
| chr5-41212965-DEL-62 | chr5-41212972-DEL-55 |
| chr5-41218268-DEL-52 | chr5-41218269-DEL-52 |
| chr5-41238757-DEL-62 | chr5-41238758-DEL-62 |
| chr5-41326979-DEL-494 | chr5-41326980-DEL-494 |
| chr5-41395307-DEL-67 | chr5-41395313-DEL-61 |
| chr5-41410225-DEL-89 | chr5-41410229-DEL-88 |
| chr5-41444127-DEL-1396 | chr5-41444127-DEL-1396 |
| chr5-41479459-DEL-55 | chr5-41479455-DEL-56 |
| chr5-41552619-DEL-7187 | chr5-41552620-DEL-7187 |
| chr5-41578257-DEL-97 | chr5-41578258-DEL-97 |
| chr5-41622631-DEL-208 | chr5-41622632-DEL-208 |
| chr5-41646835-DEL-206 | chr5-41646836-DEL-206 |
| chr5-41668782-DEL-568 | chr5-41668783-DEL-568 |
| chr5-41705261-DEL-241 | chr5-41705262-DEL-241 |
| chr5-41723497-DEL-6558 | chr5-41723498-DEL-6558 |
| chr5-41723554-DEL-204 | chr5-41723555-DEL-204 |
| chr5-41734021-DEL-241 | chr5-41734022-DEL-241 |
| chr5-41742298-DEL-765 | chr5-41742299-DEL-765 |
| chr5-41745190-DEL-412 | chr5-41745186-DEL-416 |
| chr5-41761721-DEL-94 | chr5-41761722-DEL-94 |
| chr5-41817241-DEL-1224 | chr5-41817242-DEL-1224 |
| chr5-41822819-DEL-570 | chr5-41822820-DEL-570 |
| chr5-41861346-DEL-128 | chr5-41861347-DEL-128 |
| chr5-41868293-DEL-72 | chr5-41868294-DEL-72 |
| chr5-41875953-DEL-81 | chr5-41875964-DEL-84 |
| chr5-41880335-DEL-1082 | chr5-41880336-DEL-1082 |
| chr5-41890861-DEL-185 | chr5-41890862-DEL-184 |
| chr5-41912882-DEL-62 | chr5-41912883-DEL-62 |
| chr5-41929531-DEL-1885 | chr5-41929532-DEL-1885 |
| chr5-41932437-DEL-52 | chr5-41932433-DEL-56 |
| chr5-41942252-DEL-93 | chr5-41942256-DEL-90 |
| chr5-41993626-DEL-700 | chr5-41993627-DEL-700 |
| chr5-42001949-DEL-96 | chr5-42001950-DEL-56 |
| chr5-42011626-DEL-84 | chr5-42011647-DEL-64 |
| chr5-42018856-DEL-2366 | chr5-42018857-DEL-2366 |
| chr5-42034611-DEL-61 | chr5-42034637-DEL-51 |
| chr5-42043070-DEL-87 | chr5-42043086-DEL-89 |
| chr5-42117633-DEL-255 | chr5-42117634-DEL-255 |
| chr5-42129519-DEL-478 | chr5-42129520-DEL-472 |
| chr5-42145606-DEL-133 | chr5-42145602-DEL-128 |
| chr5-42157838-DEL-13297 | chr5-42157839-DEL-13297 |
| chr5-42186914-DEL-6221 | chr5-42186915-DEL-6221 |
| chr5-42291816-DEL-2119 | chr5-42291816-DEL-2119 |
| chr5-42293486-DEL-6924 | chr5-42293487-DEL-6920 |
| chr5-42294152-DEL-212 | chr5-42294153-DEL-212 |
| chr5-42302428-DEL-138 | chr5-42302429-DEL-138 |
| chr5-42333813-DEL-242 | chr5-42333814-DEL-242 |
| chr5-42345642-DEL-7024 | chr5-42345643-DEL-7024 |
| chr5-42346276-DEL-212 | chr5-42346277-DEL-212 |
| chr5-42403543-DEL-66 | chr5-42403543-DEL-76 |
| chr5-42416101-DEL-1909 | chr5-42416102-DEL-1909 |
| chr5-42422607-DEL-412 | chr5-42422608-DEL-412 |
| chr5-42437562-DEL-6031 | chr5-42437563-DEL-6031 |
| chr5-42443830-DEL-708 | chr5-42443831-DEL-708 |
| chr5-42463110-DEL-84 | chr5-42463111-DEL-84 |
| chr5-42471437-DEL-5363 | chr5-42471437-DEL-5364 |
| chr5-42492078-DEL-52 | chr5-42492079-DEL-52 |
| chr5-42533214-DEL-6301 | chr5-42533215-DEL-6301 |
| chr5-42543681-DEL-52 | chr5-42543682-DEL-52 |
| chr5-42547334-DEL-59 | chr5-42547336-DEL-51 |
| chr5-42578455-DEL-588 | chr5-42578455-DEL-589 |
| chr5-42676573-DEL-426 | chr5-42676574-DEL-426 |
| chr5-42726137-DEL-243 | chr5-42726138-DEL-243 |
| chr5-42750658-DEL-191 | chr5-42750659-DEL-189 |
| chr5-42761295-DEL-56 | chr5-42761296-DEL-56 |
| chr5-42767985-DEL-218 | chr5-42767986-DEL-218 |
| chr5-42839458-DEL-188 | chr5-42839459-DEL-188 |
| chr5-42841250-DEL-309 | chr5-42841251-DEL-309 |
| chr5-42846413-DEL-990 | chr5-42846413-DEL-990 |
| chr5-42848017-DEL-623 | chr5-42848018-DEL-623 |
| chr5-42856837-DEL-1018 | chr5-42856838-DEL-1018 |
| chr5-42861177-DEL-6825 | chr5-42861178-DEL-6825 |
| chr5-42861527-DEL-200 | chr5-42861527-DEL-200 |

| | |
|---|---|
| chr5-42866956-DEL-1464 | chr5-42866957-DEL-1464 |
| chr5-42905789-DEL-6567 | chr5-42905790-DEL-6567 |
| chr5-42911746-DEL-208 | chr5-42911746-DEL-208 |
| chr5-42945506-DEL-51 | chr5-42945507-DEL-51 |
| chr5-42960673-DEL-79 | chr5-42960715-DEL-78 |
| chr5-42975829-DEL-182 | chr5-42975830-DEL-182 |
| chr5-42978304-DEL-7250 | chr5-42978305-DEL-7250 |
| chr5-42984688-DEL-208 | chr5-42984689-DEL-208 |
| chr5-43003979-DEL-485 | chr5-43003980-DEL-485 |
| chr5-43016805-DEL-192 | chr5-43016806-DEL-192 |
| chr5-43027080-DEL-58 | chr5-43027055-DEL-58 |
| chr5-43034365-DEL-1579 | chr5-43034366-DEL-1579 |
| chr5-43077982-DEL-800 | chr5-43077983-DEL-800 |
| chr5-43090810-DEL-1201 | chr5-43090811-DEL-1201 |
| chr5-43112830-DEL-343 | chr5-43112824-DEL-343 |
| chr5-43168946-DEL-470 | chr5-43168947-DEL-470 |
| chr5-43169565-DEL-88 | chr5-43169566-DEL-88 |
| chr5-43216389-DEL-1891 | chr5-43216390-DEL-1891 |
| chr5-43221748-DEL-6975 | chr5-43221749-DEL-6975 |
| chr5-43227926-DEL-611 | chr5-43227927-DEL-611 |
| chr5-43229358-DEL-65 | chr5-43229359-DEL-65 |
| chr5-43252475-DEL-6731 | chr5-43252476-DEL-6731 |
| chr5-43259723-DEL-126 | chr5-43259687-DEL-122 |
| chr5-43264968-DEL-207 | chr5-43264969-DEL-207 |
| chr5-43290130-DEL-158 | chr5-43290127-DEL-129 |
| chr5-43329442-DEL-70 | chr5-43329443-DEL-70 |
| chr5-43354104-DEL-142 | chr5-43354105-DEL-142 |
| chr5-43362092-DEL-317 | chr5-43362089-DEL-318 |
| chr5-43365280-DEL-9045 | chr5-43365281-DEL-9045 |
| chr5-43398164-DEL-81 | chr5-43398164-DEL-81 |
| chr5-43465076-DEL-72 | chr5-43465077-DEL-78 |
| chr5-43470931-DEL-5801 | chr5-43470933-DEL-5801 |
| chr5-43489225-DEL-236 | chr5-43489229-DEL-235 |
| chr5-43502309-DEL-4530 | chr5-43502310-DEL-4530 |
| chr5-43507903-DEL-54 | chr5-43507904-DEL-54 |
| chr5-43518226-DEL-782 | chr5-43518227-DEL-782 |
| chr5-43533525-DEL-440 | chr5-43533526-DEL-440 |
| chr5-43545407-DEL-183 | chr5-43545408-DEL-183 |
| chr5-43545440-DEL-1939 | chr5-43545453-DEL-1927 |
| chr5-43547798-DEL-593 | chr5-43547799-DEL-593 |
| chr5-43566030-DEL-5278 | chr5-43566031-DEL-5278 |
| chr5-43620394-DEL-51 | chr5-43620394-DEL-51 |
| chr5-43641311-DEL-204 | chr5-43641312-DEL-204 |
| chr5-43699141-DEL-100 | chr5-43699173-DEL-72 |
| chr5-43701027-DEL-56 | chr5-43701028-DEL-53 |
| chr5-43718432-DEL-70 | chr5-43718425-DEL-94 |
| chr5-43737191-DEL-67 | chr5-43737192-DEL-67 |
| chr5-43742710-DEL-124 | chr5-43742711-DEL-124 |
| chr5-43751281-DEL-107 | chr5-43751307-DEL-68 |
| chr5-43770058-DEL-557 | chr5-43770059-DEL-557 |
| chr5-43781685-DEL-56 | chr5-43781686-DEL-56 |
| chr5-43807060-DEL-197 | chr5-43807054-DEL-198 |
| chr5-43811747-DEL-3776 | chr5-43811749-DEL-3775 |
| chr5-43815019-DEL-377 | chr5-43815020-DEL-377 |
| chr5-43851128-DEL-68 | chr5-43851129-DEL-68 |
| chr5-43852827-DEL-524 | chr5-43852828-DEL-524 |
| chr5-43865197-DEL-100 | chr5-43865198-DEL-100 |
| chr5-43877839-DEL-487 | chr5-43877855-DEL-488 |
| chr5-43883803-DEL-82 | chr5-43883783-DEL-81 |
| chr5-43894660-DEL-72 | chr5-43894661-DEL-72 |
| chr5-43901165-DEL-58 | chr5-43901166-DEL-58 |
| chr5-43986165-DEL-7261 | chr5-43986166-DEL-7261 |
| chr5-44009392-DEL-486 | chr5-44009393-DEL-486 |
| chr5-44069650-DEL-1924 | chr5-44069651-DEL-1924 |
| chr5-44099295-DEL-244 | chr5-44099295-DEL-237 |
| chr5-44101388-DEL-616 | chr5-44101389-DEL-616 |
| chr5-44151233-DEL-76 | chr5-44151234-DEL-76 |
| chr5-44161480-DEL-125 | chr5-44161481-DEL-125 |
| chr5-44190933-DEL-386 | chr5-44190934-DEL-386 |
| chr5-44211225-DEL-1224 | chr5-44211226-DEL-1218 |
| chr5-44212919-DEL-62 | chr5-44212920-DEL-62 |
| chr5-44255238-DEL-158 | chr5-44255239-DEL-158 |
| chr5-44273827-DEL-254 | chr5-44273828-DEL-254 |
| chr5-44275684-DEL-59 | chr5-44275685-DEL-60 |
| chr5-44291697-DEL-8986 | chr5-44291698-DEL-8986 |
| chr5-44303518-DEL-75 | chr5-44303519-DEL-75 |
| chr5-44329549-DEL-109 | chr5-44329550-DEL-109 |
| chr5-44347102-DEL-1155 | chr5-44347103-DEL-1155 |
| chr5-44369877-DEL-505 | chr5-44369871-DEL-506 |
| chr5-44415793-DEL-108 | chr5-44415794-DEL-108 |

| | |
|---|---|
| chr5-44419551-DEL-569 | chr5-44419555-DEL-551 |
| chr5-44421997-DEL-740 | chr5-44421998-DEL-740 |
| chr5-44422974-DEL-84 | chr5-44422975-DEL-84 |
| chr5-44426748-DEL-62 | chr5-44426748-DEL-69 |
| chr5-44438115-DEL-82 | chr5-44438124-DEL-76 |
| chr5-44443702-DEL-72 | chr5-44443703-DEL-72 |
| chr5-44462471-DEL-405 | chr5-44462472-DEL-405 |
| chr5-44469208-DEL-2153 | chr5-44469209-DEL-2153 |
| chr5-44472276-DEL-1209 | chr5-44472277-DEL-1209 |
| chr5-44486609-DEL-63 | chr5-44486610-DEL-63 |
| chr5-44502382-DEL-56 | chr5-44502383-DEL-54 |
| chr5-44513467-DEL-115 | chr5-44513468-DEL-115 |
| chr5-44516207-DEL-804 | chr5-44516208-DEL-804 |
| chr5-44518363-DEL-58 | chr5-44518364-DEL-58 |
| chr5-44545653-DEL-54 | chr5-44545654-DEL-52 |
| chr5-44583512-DEL-172 | chr5-44583513-DEL-171 |
| chr5-44594285-DEL-66 | chr5-44594286-DEL-66 |
| chr5-44604908-DEL-6169 | chr5-44604909-DEL-6169 |
| chr5-44618482-DEL-56 | chr5-44618487-DEL-52 |
| chr5-44679713-DEL-234 | chr5-44679714-DEL-234 |
| chr5-44749891-DEL-81 | chr5-44749869-DEL-107 |
| chr5-44780487-DEL-75 | chr5-44780488-DEL-75 |
| chr5-44809892-DEL-70 | chr5-44809893-DEL-68 |
| chr5-44814874-DEL-79 | chr5-44814875-DEL-79 |
| chr5-44828293-DEL-62 | chr5-44828294-DEL-62 |
| chr5-44828837-DEL-681 | chr5-44828835-DEL-682 |
| chr5-44848540-DEL-5029 | chr5-44848541-DEL-5029 |
| chr5-44860636-DEL-148 | chr5-44860645-DEL-144 |
| chr5-44861542-DEL-110 | chr5-44861585-DEL-108 |
| chr5-44888687-DEL-684 | chr5-44888688-DEL-684 |
| chr5-44909486-DEL-70 | chr5-44909487-DEL-70 |
| chr5-44936089-DEL-52 | chr5-44936090-DEL-52 |
| chr5-44941223-DEL-58 | chr5-44941223-DEL-56 |
| chr5-44962340-DEL-81 | chr5-44962340-DEL-81 |
| chr5-44980752-DEL-1493 | chr5-44980752-DEL-1493 |
| chr5-45001632-DEL-67 | chr5-45001625-DEL-50 |
| chr5-45008306-DEL-65 | chr5-45008312-DEL-61 |
| chr5-45030720-DEL-137 | chr5-45030769-DEL-100 |
| chr5-45038254-DEL-105 | chr5-45038257-DEL-116 |
| chr5-45049568-DEL-1064 | chr5-45049569-DEL-1065 |
| chr5-45050288-DEL-739 | chr5-45050289-DEL-739 |
| chr5-45072061-DEL-60 | chr5-45072061-DEL-60 |
| chr5-45074232-DEL-81 | chr5-45074236-DEL-54 |
| chr5-45105716-DEL-257 | chr5-45105716-DEL-257 |
| chr5-45208157-DEL-212 | chr5-45208158-DEL-212 |
| chr5-45226345-DEL-147 | chr5-45226323-DEL-147 |
| chr5-45239155-DEL-56 | chr5-45239152-DEL-78 |
| chr5-45245924-DEL-484 | chr5-45245925-DEL-483 |
| chr5-45248573-DEL-610 | chr5-45248574-DEL-610 |
| chr5-45281240-DEL-259 | chr5-45281240-DEL-218 |
| chr5-45286792-DEL-62 | chr5-45286793-DEL-62 |
| chr5-45289281-DEL-1131 | chr5-45289282-DEL-1131 |
| chr5-45336550-DEL-1798 | chr5-45336551-DEL-1798 |
| chr5-45405813-DEL-1185 | chr5-45405814-DEL-1185 |
| chr5-45414642-DEL-195 | chr5-45414642-DEL-196 |
| chr5-45417823-DEL-621 | chr5-45417824-DEL-621 |
| chr5-45419090-DEL-765 | chr5-45419091-DEL-765 |
| chr5-45419893-DEL-130 | chr5-45419894-DEL-130 |
| chr5-45432433-DEL-183 | chr5-45432434-DEL-183 |
| chr5-45437974-DEL-346 | chr5-45437975-DEL-346 |
| chr5-45450606-DEL-67 | chr5-45450607-DEL-67 |
| chr5-45452804-DEL-55 | chr5-45452805-DEL-55 |
| chr5-45453617-DEL-69 | chr5-45453618-DEL-69 |
| chr5-45460204-DEL-232 | chr5-45460176-DEL-247 |
| chr5-45485144-DEL-214 | chr5-45485145-DEL-214 |
| chr5-45489959-DEL-97 | chr5-45489960-DEL-97 |
| chr5-45528349-DEL-142 | chr5-45528350-DEL-142 |
| chr5-45548557-DEL-74 | chr5-45548556-DEL-74 |
| chr5-45552762-DEL-4524 | chr5-45552763-DEL-4524 |
| chr5-45565847-DEL-6539 | chr5-45565848-DEL-6539 |
| chr5-45580120-DEL-339 | chr5-45580120-DEL-339 |
| chr5-45600265-DEL-88 | chr5-45600243-DEL-89 |
| chr5-45632481-DEL-110 | chr5-45632482-DEL-109 |
| chr5-45637867-DEL-58 | chr5-45637868-DEL-58 |
| chr5-45647718-DEL-114 | chr5-45647727-DEL-106 |
| chr5-45650257-DEL-86 | chr5-45650258-DEL-86 |
| chr5-45718794-DEL-70 | chr5-45718795-DEL-70 |
| chr5-45726683-DEL-2962 | chr5-45726684-DEL-2962 |
| chr5-45733045-DEL-807 | chr5-45733046-DEL-805 |
| chr5-45740416-DEL-311 | chr5-45740414-DEL-312 |

| | |
|---|---|
| chr5-45760974-DEL-4920 | chr5-45760975-DEL-4920 |
| chr5-45766888-DEL-477 | chr5-45766888-DEL-478 |
| chr5-45775328-DEL-514 | chr5-45775329-DEL-514 |
| chr5-45803421-DEL-4950 | chr5-45803422-DEL-4950 |
| chr5-45804070-DEL-651 | chr5-45804071-DEL-651 |
| chr5-45811663-DEL-2173 | chr5-45811664-DEL-2173 |
| chr5-45825123-DEL-66 | chr5-45825124-DEL-66 |
| chr5-45828230-DEL-102 | chr5-45828230-DEL-102 |
| chr5-45860715-DEL-1373 | chr5-45860716-DEL-1373 |
| chr5-45874162-DEL-53 | chr5-45874163-DEL-53 |
| chr5-45931083-DEL-338 | chr5-45931084-DEL-338 |
| chr5-45972540-DEL-7660 | chr5-45972540-DEL-7660 |
| chr5-46005089-DEL-6929 | chr5-46005090-DEL-6929 |
| chr5-46012649-DEL-214 | chr5-46012650-DEL-213 |
| chr5-46020680-DEL-364 | chr5-46020681-DEL-364 |
| chr5-46024427-DEL-103 | chr5-46024428-DEL-103 |
| chr5-46057606-DEL-130 | chr5-46057622-DEL-130 |
| chr5-46076963-DEL-331 | chr5-46076964-DEL-331 |
| chr5-46084837-DEL-270 | chr5-46084838-DEL-270 |
| chr5-46095748-DEL-117 | chr5-46095749-DEL-116 |
| chr5-46128288-DEL-2555 | chr5-46128289-DEL-2555 |
| chr5-46142401-DEL-1535 | chr5-46142402-DEL-1535 |
| chr5-46165135-DEL-350 | chr5-46165136-DEL-350 |
| chr5-46196960-DEL-177 | chr5-46196961-DEL-177 |
| chr5-46209084-DEL-813 | chr5-46209085-DEL-813 |
| chr5-46225420-DEL-3320 | chr5-46225421-DEL-3320 |
| chr5-46256369-DEL-3193 | chr5-46256370-DEL-3208 |
| chr5-46299022-DEL-210 | chr5-46299023-DEL-207 |
| chr5-46302707-DEL-94 | chr5-46302698-DEL-80 |
| chr5-46313460-DEL-789 | chr5-46313461-DEL-789 |
| chr5-46351013-DEL-448 | chr5-46351014-DEL-448 |
| chr5-46351897-DEL-469 | chr5-46351898-DEL-469 |
| chr5-46357533-DEL-54 | chr5-46357496-DEL-68 |
| chr5-46375185-DEL-72 | chr5-46375185-DEL-73 |
| chr5-46393854-DEL-60 | chr5-46393855-DEL-60 |
| chr5-46395447-DEL-132 | chr5-46395448-DEL-132 |
| chr5-46412889-DEL-134396 | chr5-46412887-DEL-134398 |
| chr5-46420182-DEL-6091 | chr5-46420183-DEL-6091 |
| chr5-46442538-DEL-3554 | chr5-46442534-DEL-3552 |
| chr5-46450571-DEL-2385 | chr5-46450572-DEL-2385 |
| chr5-46467553-DEL-1446 | chr5-46467554-DEL-1446 |
| chr5-46486386-DEL-52 | chr5-46486394-DEL-53 |
| chr5-46530706-DEL-457 | chr5-46530707-DEL-457 |
| chr5-46551688-DEL-6904 | chr5-46551689-DEL-6904 |
| chr5-46571583-DEL-1182 | chr5-46571584-DEL-1182 |
| chr5-46607375-DEL-84 | chr5-46607376-DEL-84 |
| chr5-46607481-DEL-718 | chr5-46607482-DEL-718 |
| chr5-46610772-DEL-497 | chr5-46610773-DEL-497 |
| chr5-46615804-DEL-7413 | chr5-46615805-DEL-7413 |
| chr5-46615926-DEL-208 | chr5-46615927-DEL-208 |
| chr5-46676974-DEL-80 | chr5-46676972-DEL-86 |
| chr5-46681323-DEL-64 | chr5-46681322-DEL-65 |
| chr5-46681994-DEL-576 | chr5-46681995-DEL-576 |
| chr5-46691735-DEL-2381 | chr5-46691736-DEL-2381 |
| chr5-46702502-DEL-2627 | chr5-46702503-DEL-2627 |
| chr5-46727321-DEL-142 | chr5-46727325-DEL-128 |
| chr5-46738646-DEL-2220 | chr5-46738647-DEL-2220 |
| chr5-46759286-DEL-360 | chr5-46759288-DEL-364 |
| chr5-46762162-DEL-63 | chr5-46762163-DEL-63 |
| chr5-46774671-DEL-139 | chr5-46774642-DEL-136 |
| chr5-46778498-DEL-100 | chr5-46778503-DEL-96 |
| chr5-46792011-DEL-277 | chr5-46792012-DEL-277 |
| chr5-46817415-DEL-50 | chr5-46817416-DEL-50 |
| chr5-46824297-DEL-68 | chr5-46824298-DEL-68 |
| chr5-46857495-DEL-2278 | chr5-46857495-DEL-2278 |
| chr5-46865538-DEL-175 | chr5-46865539-DEL-175 |
| chr5-46882217-DEL-384 | chr5-46882218-DEL-384 |
| chr5-46934599-DEL-1403 | chr5-46934600-DEL-1403 |
| chr5-46956673-DEL-1831 | chr5-46956670-DEL-1832 |
| chr5-46967047-DEL-589 | chr5-46967048-DEL-589 |
| chr5-46992130-DEL-58 | chr5-46992131-DEL-58 |
| chr5-47015571-DEL-103 | chr5-47015574-DEL-101 |
| chr5-47018974-DEL-72 | chr5-47018979-DEL-74 |
| chr5-47034314-DEL-2664 | chr5-47034315-DEL-2664 |
| chr5-47048181-DEL-190 | chr5-47048182-DEL-190 |
| chr5-47067442-DEL-61 | chr5-47067445-DEL-62 |
| chr5-47097855-DEL-7236 | chr5-47097856-DEL-7236 |
| chr5-47111159-DEL-1336 | chr5-47111160-DEL-1336 |
| chr5-47127605-DEL-65 | chr5-47127610-DEL-50 |
| chr5-47128815-DEL-433 | chr5-47128816-DEL-433 |

| | |
|---|---|
| chr5-47157909-DEL-57 | chr5-47157910-DEL-57 |
| chr5-47182655-DEL-222 | chr5-47182656-DEL-222 |
| chr5-47202905-DEL-52 | chr5-47202906-DEL-52 |
| chr5-47218686-DEL-278 | chr5-47218687-DEL-278 |
| chr5-47228541-DEL-58 | chr5-47228544-DEL-56 |
| chr5-47254411-DEL-51 | chr5-47254413-DEL-52 |
| chr5-47259923-DEL-377 | chr5-47259924-DEL-377 |
| chr5-47265995-DEL-1371 | chr5-47265996-DEL-1371 |
| chr5-47299209-DEL-2426 | chr5-47299210-DEL-2426 |
| chr5-47308278-DEL-13093 | chr5-47308279-DEL-13093 |
| chr5-47371896-DEL-188 | chr5-47371897-DEL-188 |
| chr5-47392620-DEL-503 | chr5-47392621-DEL-503 |
| chr5-47393780-DEL-58 | chr5-47393777-DEL-70 |
| chr5-47406703-DEL-1996 | chr5-47406704-DEL-1996 |
| chr5-47437497-DEL-703 | chr5-47437498-DEL-703 |
| chr5-47438615-DEL-6453 | chr5-47438616-DEL-6453 |
| chr5-47449211-DEL-2830 | chr5-47449212-DEL-2830 |
| chr5-47454844-DEL-69 | chr5-47454845-DEL-69 |
| chr5-47460468-DEL-507 | chr5-47460469-DEL-507 |
| chr5-47483948-DEL-52 | chr5-47483949-DEL-52 |
| chr5-47502008-DEL-152 | chr5-47502009-DEL-152 |
| chr5-47509881-DEL-396 | chr5-47509882-DEL-396 |
| chr5-47514898-DEL-91 | chr5-47514899-DEL-91 |
| chr5-47594805-DEL-173 | chr5-47594761-DEL-189 |
| chr5-47597546-DEL-2739 | chr5-47597547-DEL-2739 |
| chr5-47624020-DEL-91 | chr5-47624050-DEL-70 |
| chr5-47644393-DEL-58 | chr5-47644394-DEL-58 |
| chr5-47700430-DEL-53 | chr5-47700421-DEL-54 |
| chr5-47702213-DEL-120 | chr5-47702214-DEL-120 |
| chr5-47705851-DEL-62 | chr5-47705817-DEL-50 |
| chr5-47710026-DEL-596 | chr5-47710027-DEL-596 |
| chr5-47716126-DEL-120 | chr5-47716127-DEL-120 |
| chr5-47769111-DEL-264 | chr5-47769112-DEL-264 |
| chr5-47800692-DEL-466 | chr5-47800693-DEL-466 |
| chr5-47814692-DEL-84 | chr5-47814693-DEL-84 |
| chr5-47823874-DEL-6218 | chr5-47823875-DEL-6218 |
| chr5-47842987-DEL-419 | chr5-47842988-DEL-419 |
| chr5-47843812-DEL-117 | chr5-47843818-DEL-112 |
| chr5-47845431-DEL-122 | chr5-47845432-DEL-122 |
| chr5-47863537-DEL-1132 | chr5-47863538-DEL-1132 |
| chr5-47886209-DEL-114 | chr5-47886210-DEL-114 |
| chr5-47892205-DEL-189 | chr5-47892206-DEL-189 |
| chr5-47897358-DEL-486 | chr5-47897359-DEL-486 |
| chr5-47932091-DEL-101 | chr5-47932089-DEL-102 |
| chr5-47950869-DEL-525 | chr5-47950867-DEL-524 |
| chr5-47953605-DEL-73 | chr5-47953606-DEL-73 |
| chr5-47975143-DEL-1120 | chr5-47975144-DEL-1120 |
| chr5-47980764-DEL-82 | chr5-47980759-DEL-63 |
| chr5-48000928-DEL-1391 | chr5-48000928-DEL-1391 |
| chr5-48016496-DEL-709 | chr5-48016497-DEL-709 |
| chr5-48018590-DEL-1137 | chr5-48018591-DEL-1137 |
| chr5-48025537-DEL-85 | chr5-48025538-DEL-83 |
| chr5-48033221-DEL-212 | chr5-48033236-DEL-198 |
| chr5-48052923-DEL-4291 | chr5-48052925-DEL-4290 |
| chr5-48055213-DEL-712 | chr5-48055214-DEL-712 |
| chr5-48067546-DEL-76 | chr5-48067546-DEL-76 |
| chr5-48100515-DEL-186 | chr5-48100516-DEL-186 |
| chr5-48112235-DEL-60 | chr5-48112236-DEL-60 |
| chr5-48113774-DEL-1201 | chr5-48113775-DEL-1201 |
| chr5-48122999-DEL-66 | chr5-48123000-DEL-66 |
| chr5-48123503-DEL-5165 | chr5-48123504-DEL-5165 |
| chr5-48171975-DEL-88 | chr5-48171976-DEL-88 |
| chr5-48182023-DEL-738 | chr5-48182024-DEL-738 |
| chr5-48229928-DEL-73 | chr5-48229929-DEL-73 |
| chr5-48309559-DEL-176 | chr5-48309560-DEL-176 |
| chr5-48355928-DEL-78 | chr5-48355929-DEL-78 |
| chr5-48380695-DEL-70 | chr5-48380696-DEL-70 |
| chr5-48384304-DEL-95 | chr5-48384291-DEL-78 |
| chr5-48396268-DEL-64 | chr5-48396269-DEL-63 |
| chr5-48451779-DEL-146 | chr5-48451782-DEL-147 |
| chr5-48470023-DEL-108 | chr5-48470024-DEL-108 |
| chr5-48470873-DEL-921 | chr5-48470874-DEL-921 |
| chr5-48504203-DEL-100 | chr5-48504202-DEL-102 |
| chr5-48522225-DEL-56 | chr5-48522226-DEL-56 |
| chr5-48575063-DEL-532 | chr5-48575076-DEL-504 |
| chr5-48575789-DEL-99 | chr5-48575790-DEL-99 |
| chr5-48584211-DEL-2234 | chr5-48584212-DEL-2234 |
| chr5-48625549-DEL-84 | chr5-48625550-DEL-84 |
| chr5-48631026-DEL-56 | chr5-48631027-DEL-56 |
| chr5-48640486-DEL-232 | chr5-48640487-DEL-232 |

| | |
|---|---|
| chr5-48675554-DEL-61 | chr5-48675555-DEL-61 |
| chr5-48681484-DEL-4963 | chr5-48681485-DEL-4963 |
| chr5-48690972-DEL-2387 | chr5-48690973-DEL-2387 |
| chr5-48703670-DEL-81 | chr5-48703671-DEL-81 |
| chr5-48727519-DEL-1469 | chr5-48727520-DEL-1469 |
| chr5-48743475-DEL-674 | chr5-48743476-DEL-674 |
| chr5-48763272-DEL-298 | chr5-48763273-DEL-298 |
| chr5-48769235-DEL-54 | chr5-48769214-DEL-76 |
| chr5-48781552-DEL-54 | chr5-48781553-DEL-54 |
| chr5-48794635-DEL-512 | chr5-48794636-DEL-512 |
| chr5-48795290-DEL-6402 | chr5-48795280-DEL-6362 |
| chr5-48812168-DEL-306 | chr5-48812175-DEL-304 |
| chr5-48812567-DEL-1592 | chr5-48812568-DEL-1592 |
| chr5-48827095-DEL-4733 | chr5-48827096-DEL-4733 |
| chr5-48837908-DEL-429 | chr5-48837912-DEL-429 |
| chr5-48856129-DEL-87 | chr5-48856140-DEL-77 |
| chr5-48883984-DEL-327 | chr5-48883985-DEL-327 |
| chr5-48888964-DEL-1722 | chr5-48888965-DEL-1722 |
| chr5-48916606-DEL-52 | chr5-48916603-DEL-53 |
| chr5-48919048-DEL-84 | chr5-48919049-DEL-85 |
| chr5-48926276-DEL-88 | chr5-48926267-DEL-78 |
| chr5-48934105-DEL-56 | chr5-48934106-DEL-56 |
| chr5-48948223-DEL-796 | chr5-48948224-DEL-796 |
| chr5-48961276-DEL-72 | chr5-48961283-DEL-66 |
| chr5-48971646-DEL-189 | chr5-48971645-DEL-190 |
| chr5-48972023-DEL-616 | chr5-48972024-DEL-616 |
| chr5-48973008-DEL-54 | chr5-48972998-DEL-60 |
| chr5-48976672-DEL-548 | chr5-48976673-DEL-548 |
| chr5-48983559-DEL-205 | chr5-48983560-DEL-205 |
| chr5-48985020-DEL-459 | chr5-48985021-DEL-459 |
| chr5-48990615-DEL-148 | chr5-48990616-DEL-148 |
| chr5-49010949-DEL-90 | chr5-49010956-DEL-95 |
| chr5-49074879-DEL-203 | chr5-49074879-DEL-203 |
| chr5-49111095-DEL-1801 | chr5-49111096-DEL-1801 |
| chr5-49113594-DEL-74 | chr5-49113595-DEL-74 |
| chr5-49135200-DEL-136 | chr5-49135201-DEL-136 |
| chr5-49192256-DEL-57 | chr5-49192257-DEL-57 |
| chr5-49232560-DEL-2321 | chr5-49232561-DEL-2321 |
| chr5-49250585-DEL-337 | chr5-49250586-DEL-337 |
| chr5-49287418-DEL-52 | chr5-49287386-DEL-52 |
| chr5-49293967-DEL-134 | chr5-49293944-DEL-152 |
| chr5-49297104-DEL-88 | chr5-49297099-DEL-94 |
| chr5-49308516-DEL-84 | chr5-49308517-DEL-84 |
| chr5-49311986-DEL-384 | chr5-49311987-DEL-384 |
| chr5-49320241-DEL-60 | chr5-49320242-DEL-59 |
| chr5-49322162-DEL-71 | chr5-49322163-DEL-71 |
| chr5-49343260-DEL-1076 | chr5-49343262-DEL-1079 |
| chr5-49372070-DEL-187 | chr5-49372060-DEL-180 |
| chr5-49399280-DEL-74 | chr5-49399281-DEL-72 |
| chr5-49407290-DEL-53 | chr5-49407291-DEL-81 |
| chr5-49416282-DEL-175 | chr5-49416291-DEL-174 |
| chr5-49464651-DEL-197 | chr5-49464652-DEL-197 |
| chr5-49474517-DEL-98 | chr5-49474518-DEL-98 |
| chr5-49477036-DEL-383 | chr5-49477037-DEL-383 |
| chr5-49552774-DEL-3524 | chr5-49552775-DEL-3524 |
| chr5-49557553-DEL-83 | chr5-49557585-DEL-58 |
| chr5-49564158-DEL-262 | chr5-49564159-DEL-267 |
| chr5-49566097-DEL-102 | chr5-49566098-DEL-102 |
| chr5-49583167-DEL-991 | chr5-49583168-DEL-991 |
| chr5-49588126-DEL-68 | chr5-49588127-DEL-68 |
| chr5-49588702-DEL-508 | chr5-49588694-DEL-509 |
| chr5-49592585-DEL-52 | chr5-49592586-DEL-52 |
| chr5-49610906-DEL-3895 | chr5-49610907-DEL-3895 |
| chr5-49611538-DEL-749 | chr5-49611539-DEL-740 |
| chr5-49613236-DEL-396 | chr5-49613237-DEL-396 |
| chr5-49637755-DEL-114 | chr5-49637756-DEL-114 |
| chr5-49649189-DEL-104 | chr5-49649190-DEL-104 |
| chr5-49658413-DEL-1041 | chr5-49658415-DEL-1040 |
| chr5-49695304-DEL-6949 | chr5-49695305-DEL-6949 |
| chr5-49726538-DEL-84 | chr5-49726539-DEL-84 |
| chr5-49730934-DEL-126 | chr5-49730935-DEL-126 |
| chr5-49733822-DEL-98 | chr5-49733819-DEL-87 |
| chr5-49743758-DEL-100 | chr5-49743765-DEL-94 |
| chr5-49759924-DEL-54 | chr5-49759925-DEL-54 |
| chr5-49771703-DEL-508 | chr5-49771704-DEL-508 |
| chr5-49780822-DEL-78 | chr5-49780823-DEL-78 |
| chr5-49783796-DEL-242 | chr5-49783797-DEL-242 |
| chr5-49805953-DEL-54 | chr5-49805954-DEL-54 |
| chr5-49818671-DEL-56 | chr5-49818672-DEL-56 |
| chr5-49824211-DEL-7243 | chr5-49824212-DEL-7243 |

| | |
|---|---|
| chr5-49854464-DEL-4885 | chr5-49854464-DEL-4885 |
| chr5-49862409-DEL-196 | chr5-49862410-DEL-196 |
| chr5-49863272-DEL-119 | chr5-49863275-DEL-117 |
| chr5-49929977-DEL-769 | chr5-49929978-DEL-765 |
| chr5-49941771-DEL-66 | chr5-49941769-DEL-75 |
| chr5-49969597-DEL-217 | chr5-49969596-DEL-218 |
| chr5-49981630-DEL-1301 | chr5-49981630-DEL-1301 |
| chr5-50002698-DEL-51 | chr5-50002699-DEL-51 |
| chr5-50007513-DEL-91 | chr5-50007514-DEL-91 |
| chr5-50008234-DEL-1877 | chr5-50008235-DEL-1877 |
| chr5-50042407-DEL-202 | chr5-50042384-DEL-175 |
| chr5-50048473-DEL-88 | chr5-50048474-DEL-88 |
| chr5-50111752-DEL-58 | chr5-50111750-DEL-57 |
| chr5-50175839-DEL-992 | chr5-50175840-DEL-992 |
| chr5-50202918-DEL-1754 | chr5-50202919-DEL-1754 |
| chr5-50238753-DEL-1037 | chr5-50238754-DEL-1037 |
| chr5-50265086-DEL-6380 | chr5-50265087-DEL-6380 |
| chr5-50301109-DEL-1195 | chr5-50301110-DEL-1195 |
| chr5-50307868-DEL-113 | chr5-50307869-DEL-113 |
| chr5-50308125-DEL-1249 | chr5-50308126-DEL-1249 |
| chr5-50344106-DEL-64 | chr5-50344069-DEL-65 |
| chr5-50388060-DEL-90 | chr5-50388061-DEL-90 |
| chr5-50391467-DEL-278 | chr5-50391468-DEL-276 |
| chr5-50396066-DEL-62 | chr5-50396074-DEL-55 |
| chr5-50400719-DEL-59 | chr5-50400714-DEL-65 |
| chr5-50405472-DEL-5121 | chr5-50405473-DEL-5121 |
| chr5-50409977-DEL-201 | chr5-50409978-DEL-201 |
| chr5-50425474-DEL-2484 | chr5-50425475-DEL-2484 |
| chr5-50437768-DEL-196 | chr5-50437761-DEL-187 |
| chr5-50442498-DEL-326 | chr5-50442498-DEL-326 |
| chr5-50442960-DEL-4876 | chr5-50442961-DEL-4876 |
| chr5-50469473-DEL-7133 | chr5-50469474-DEL-7133 |
| chr5-50482883-DEL-76 | chr5-50482884-DEL-74 |
| chr5-50484431-DEL-420 | chr5-50484432-DEL-420 |
| chr5-50487555-DEL-625 | chr5-50487556-DEL-625 |
| chr5-50492336-DEL-92 | chr5-50492337-DEL-92 |
| chr5-50496456-DEL-58 | chr5-50496439-DEL-59 |
| chr5-50501834-DEL-115 | chr5-50501834-DEL-115 |
| chr5-50502610-DEL-1386 | chr5-50502611-DEL-1383 |
| chr5-50514948-DEL-66 | chr5-50514948-DEL-66 |
| chr5-50552765-DEL-88 | chr5-50552804-DEL-65 |
| chr5-50604403-DEL-56 | chr5-50604404-DEL-56 |
| chr5-50610486-DEL-249 | chr5-50610487-DEL-249 |
| chr5-50630745-DEL-608 | chr5-50630746-DEL-608 |
| chr5-50635326-DEL-98 | chr5-50635327-DEL-98 |
| chr5-50642348-DEL-70 | chr5-50642349-DEL-70 |
| chr5-50652722-DEL-262 | chr5-50652723-DEL-262 |
| chr5-50654740-DEL-176 | chr5-50654741-DEL-176 |
| chr5-50685712-DEL-56 | chr5-50685713-DEL-51 |
| chr5-50694707-DEL-223 | chr5-50694708-DEL-223 |
| chr5-50695177-DEL-2370 | chr5-50695178-DEL-2370 |
| chr5-50731435-DEL-118 | chr5-50731436-DEL-118 |
| chr5-50735704-DEL-1109 | chr5-50735713-DEL-1106 |
| chr5-50737947-DEL-2713 | chr5-50737946-DEL-2714 |
| chr5-50750583-DEL-70 | chr5-50750584-DEL-70 |
| chr5-50760101-DEL-68 | chr5-50760100-DEL-70 |
| chr5-50818117-DEL-137 | chr5-50818117-DEL-137 |
| chr5-50838909-DEL-82 | chr5-50838908-DEL-80 |
| chr5-50840693-DEL-66 | chr5-50840680-DEL-85 |
| chr5-50851418-DEL-354 | chr5-50851419-DEL-354 |
| chr5-50864141-DEL-129 | chr5-50864142-DEL-129 |
| chr5-50873070-DEL-59 | chr5-50873065-DEL-56 |
| chr5-50918098-DEL-1096 | chr5-50918099-DEL-1096 |
| chr5-50936309-DEL-55 | chr5-50936310-DEL-55 |
| chr5-50965862-DEL-59 | chr5-50965852-DEL-84 |
| chr5-50967137-DEL-940 | chr5-50967138-DEL-940 |
| chr5-50968869-DEL-1864 | chr5-50968869-DEL-1864 |
| chr5-50973877-DEL-524 | chr5-50973877-DEL-524 |
| chr5-50979079-DEL-911 | chr5-50979080-DEL-911 |
| chr5-51006284-DEL-6595 | chr5-51006285-DEL-6595 |
| chr5-51016331-DEL-60 | chr5-51016329-DEL-61 |
| chr5-51035090-DEL-76 | chr5-51035091-DEL-76 |
| chr5-51035977-DEL-204 | chr5-51035978-DEL-201 |
| chr5-51061890-DEL-52 | chr5-51061889-DEL-52 |
| chr5-51083381-DEL-373 | chr5-51083382-DEL-373 |
| chr5-51129145-DEL-76 | chr5-51129146-DEL-76 |
| chr5-51132838-DEL-820 | chr5-51132839-DEL-820 |
| chr5-51133091-DEL-58 | chr5-51133092-DEL-58 |
| chr5-51136940-DEL-1069 | chr5-51136941-DEL-1069 |
| chr5-51143327-DEL-151 | chr5-51143328-DEL-151 |

| | |
|---|---|
| chr5-51158201-DEL-7187 | chr5-51158200-DEL-7188 |
| chr5-51176940-DEL-932 | chr5-51176941-DEL-932 |
| chr5-51179701-DEL-2697 | chr5-51179705-DEL-2696 |
| chr5-51224408-DEL-730 | chr5-51224409-DEL-730 |
| chr5-51235246-DEL-53 | chr5-51235247-DEL-53 |
| chr5-51236266-DEL-132 | chr5-51236267-DEL-132 |
| chr5-51251307-DEL-96 | chr5-51251307-DEL-92 |
| chr5-51272780-DEL-1746 | chr5-51272780-DEL-1745 |
| chr5-51275927-DEL-136 | chr5-51275941-DEL-134 |
| chr5-51293303-DEL-6208 | chr5-51293304-DEL-6208 |
| chr5-51323084-DEL-81 | chr5-51323087-DEL-73 |
| chr5-51329770-DEL-88 | chr5-51329771-DEL-88 |
| chr5-51344013-DEL-94 | chr5-51343998-DEL-62 |
| chr5-51375403-DEL-4225 | chr5-51375404-DEL-4225 |
| chr5-51380444-DEL-56 | chr5-51380445-DEL-56 |
| chr5-51395267-DEL-2886 | chr5-51395268-DEL-2886 |
| chr5-51481778-DEL-187 | chr5-51481779-DEL-187 |
| chr5-51497451-DEL-207 | chr5-51497452-DEL-207 |
| chr5-51524055-DEL-1633 | chr5-51524056-DEL-1633 |
| chr5-51533129-DEL-6382 | chr5-51533130-DEL-6383 |
| chr5-51539512-DEL-62 | chr5-51539514-DEL-61 |
| chr5-51542571-DEL-677 | chr5-51542572-DEL-677 |
| chr5-51566187-DEL-60 | chr5-51566187-DEL-60 |
| chr5-51593928-DEL-70 | chr5-51593928-DEL-67 |
| chr5-51629815-DEL-70 | chr5-51629816-DEL-70 |
| chr5-51638010-DEL-1406 | chr5-51638011-DEL-1406 |
| chr5-51683680-DEL-63 | chr5-51683681-DEL-63 |
| chr5-51700631-DEL-67 | chr5-51700637-DEL-56 |
| chr5-51711804-DEL-52 | chr5-51711805-DEL-52 |
| chr5-51743282-DEL-56 | chr5-51743287-DEL-52 |
| chr5-51775389-DEL-422 | chr5-51775390-DEL-422 |
| chr5-51808479-DEL-54 | chr5-51808476-DEL-55 |
| chr5-51836140-DEL-104 | chr5-51836141-DEL-104 |
| chr5-51846796-DEL-106 | chr5-51846797-DEL-106 |
| chr5-51848745-DEL-59 | chr5-51848746-DEL-59 |
| chr5-51850673-DEL-64 | chr5-51850674-DEL-64 |
| chr5-51859784-DEL-52 | chr5-51859777-DEL-56 |
| chr5-51866477-DEL-54 | chr5-51866448-DEL-60 |
| chr5-51942381-DEL-58 | chr5-51942382-DEL-58 |
| chr5-51992251-DEL-164 | chr5-51992252-DEL-164 |
| chr5-52018625-DEL-126 | chr5-52018626-DEL-126 |
| chr5-52054544-DEL-55 | chr5-52054544-DEL-56 |
| chr5-52097515-DEL-396 | chr5-52097516-DEL-396 |
| chr5-52106403-DEL-76 | chr5-52106408-DEL-72 |
| chr5-52130879-DEL-6585 | chr5-52130871-DEL-6586 |
| chr5-52158931-DEL-576 | chr5-52158931-DEL-574 |
| chr5-52165062-DEL-779 | chr5-52165063-DEL-779 |
| chr5-52171288-DEL-53 | chr5-52171289-DEL-53 |
| chr5-52183450-DEL-78 | chr5-52183451-DEL-54 |
| chr5-52184285-DEL-244 | chr5-52184286-DEL-245 |
| chr5-52194787-DEL-105 | chr5-52194784-DEL-106 |
| chr5-52200958-DEL-73 | chr5-52200967-DEL-70 |
| chr5-52249084-DEL-134 | chr5-52249089-DEL-114 |
| chr5-52257093-DEL-166 | chr5-52257099-DEL-133 |
| chr5-52266298-DEL-114 | chr5-52266317-DEL-80 |
| chr5-52279864-DEL-84 | chr5-52279844-DEL-105 |
| chr5-52287557-DEL-94 | chr5-52287558-DEL-94 |
| chr5-52299451-DEL-111 | chr5-52299452-DEL-111 |
| chr5-52352857-DEL-100 | chr5-52352858-DEL-100 |
| chr5-52366542-DEL-113 | chr5-52366543-DEL-113 |
| chr5-52369104-DEL-207 | chr5-52369105-DEL-207 |
| chr5-52374381-DEL-114 | chr5-52374367-DEL-93 |
| chr5-52389512-DEL-54 | chr5-52389503-DEL-60 |
| chr5-52434286-DEL-201 | chr5-52434287-DEL-201 |
| chr5-52441835-DEL-68 | chr5-52441839-DEL-65 |
| chr5-52442238-DEL-1227 | chr5-52442239-DEL-1227 |
| chr5-52503864-DEL-172 | chr5-52503865-DEL-172 |
| chr5-52508288-DEL-72 | chr5-52508289-DEL-72 |
| chr5-52558302-DEL-201 | chr5-52558303-DEL-201 |
| chr5-52562254-DEL-60 | chr5-52562216-DEL-72 |
| chr5-52565395-DEL-59 | chr5-52565396-DEL-60 |
| chr5-52600049-DEL-625 | chr5-52600050-DEL-625 |
| chr5-52604505-DEL-309 | chr5-52604506-DEL-307 |
| chr5-52609297-DEL-820 | chr5-52609299-DEL-819 |
| chr5-52612593-DEL-82 | chr5-52612594-DEL-82 |
| chr5-52624339-DEL-68 | chr5-52624340-DEL-68 |
| chr5-52632816-DEL-64 | chr5-52632817-DEL-64 |
| chr5-52643845-DEL-84 | chr5-52643846-DEL-84 |
| chr5-52677539-DEL-78 | chr5-52677540-DEL-78 |
| chr5-52682703-DEL-2064 | chr5-52682704-DEL-2064 |

| | |
|---|---|
| chr5-52706405-DEL-95 | chr5-52706395-DEL-91 |
| chr5-52744597-DEL-307 | chr5-52744597-DEL-307 |
| chr5-52778154-DEL-722 | chr5-52778155-DEL-722 |
| chr5-52832317-DEL-96 | chr5-52832318-DEL-96 |
| chr5-52837742-DEL-56 | chr5-52837743-DEL-56 |
| chr5-52849065-DEL-81 | chr5-52849066-DEL-81 |
| chr5-52851295-DEL-83 | chr5-52851296-DEL-83 |
| chr5-52852893-DEL-173 | chr5-52852894-DEL-173 |
| chr5-52855763-DEL-70 | chr5-52855730-DEL-54 |
| chr5-52869189-DEL-60 | chr5-52869190-DEL-60 |
| chr5-52870527-DEL-206 | chr5-52870528-DEL-201 |
| chr5-52874878-DEL-90 | chr5-52874879-DEL-90 |
| chr5-52875816-DEL-1297 | chr5-52875816-DEL-1297 |
| chr5-52881010-DEL-498 | chr5-52881011-DEL-498 |
| chr5-52884925-DEL-64 | chr5-52884926-DEL-64 |
| chr5-52895188-DEL-52 | chr5-52895189-DEL-50 |
| chr5-52918013-DEL-134 | chr5-52918014-DEL-134 |
| chr5-52935548-DEL-197 | chr5-52935549-DEL-197 |
| chr5-52964940-DEL-99 | chr5-52964947-DEL-90 |
| chr5-52985554-DEL-1681 | chr5-52985555-DEL-1681 |
| chr5-53029474-DEL-54 | chr5-53029475-DEL-54 |
| chr5-53034171-DEL-217 | chr5-53034172-DEL-211 |
| chr5-53052638-DEL-7547 | chr5-53052639-DEL-7547 |
| chr5-53064989-DEL-64 | chr5-53064990-DEL-64 |
| chr5-53066588-DEL-236 | chr5-53066589-DEL-235 |
| chr5-53069837-DEL-134 | chr5-53069838-DEL-134 |
| chr5-53081810-DEL-324 | chr5-53081810-DEL-324 |
| chr5-53089900-DEL-106 | chr5-53089901-DEL-106 |
| chr5-53095406-DEL-114 | chr5-53095425-DEL-99 |
| chr5-53096239-DEL-140 | chr5-53096239-DEL-140 |
| chr5-53106190-DEL-66 | chr5-53106191-DEL-66 |
| chr5-53106705-DEL-61 | chr5-53106755-DEL-50 |
| chr5-53119325-DEL-1413 | chr5-53119326-DEL-1413 |
| chr5-53129984-DEL-107 | chr5-53129985-DEL-107 |
| chr5-53133330-DEL-179 | chr5-53133331-DEL-179 |
| chr5-53157472-DEL-91 | chr5-53157473-DEL-89 |
| chr5-53166052-DEL-129 | chr5-53166068-DEL-111 |
| chr5-53179632-DEL-122 | chr5-53179619-DEL-136 |
| chr5-53193242-DEL-113 | chr5-53193246-DEL-111 |
| chr5-53237848-DEL-401 | chr5-53237859-DEL-391 |
| chr5-53289453-DEL-52 | chr5-53289454-DEL-52 |
| chr5-53393246-DEL-59 | chr5-53393247-DEL-59 |
| chr5-53402396-DEL-65 | chr5-53402396-DEL-65 |
| chr5-53413338-DEL-64 | chr5-53413339-DEL-64 |
| chr5-53431160-DEL-98 | chr5-53431161-DEL-98 |
| chr5-53492328-DEL-57 | chr5-53492329-DEL-57 |
| chr5-53494572-DEL-97 | chr5-53494578-DEL-71 |
| chr5-53548096-DEL-74 | chr5-53548096-DEL-62 |
| chr5-53599902-DEL-92 | chr5-53599903-DEL-92 |
| chr5-53688362-DEL-498 | chr5-53688363-DEL-498 |
| chr5-53715736-DEL-177 | chr5-53715737-DEL-177 |
| chr5-53740121-DEL-876 | chr5-53740122-DEL-876 |
| chr5-53742621-DEL-80 | chr5-53742648-DEL-50 |
| chr5-53780550-DEL-2609 | chr5-53780551-DEL-2609 |
| chr5-53785062-DEL-2031 | chr5-53785063-DEL-2031 |
| chr5-53788978-DEL-597 | chr5-53788979-DEL-597 |
| chr5-53791815-DEL-96 | chr5-53791816-DEL-96 |
| chr5-53802109-DEL-106 | chr5-53802110-DEL-106 |
| chr5-53809170-DEL-104 | chr5-53809153-DEL-96 |
| chr5-53824676-DEL-267 | chr5-53824675-DEL-267 |
| chr5-53830110-DEL-54 | chr5-53830090-DEL-52 |
| chr5-53905320-DEL-6624 | chr5-53905321-DEL-6624 |
| chr5-53915072-DEL-144 | chr5-53915061-DEL-105 |
| chr5-53941709-DEL-160 | chr5-53941749-DEL-115 |
| chr5-53956615-DEL-83 | chr5-53956616-DEL-83 |
| chr5-53961032-DEL-1178 | chr5-53961037-DEL-1178 |
| chr5-53990527-DEL-1488 | chr5-53990535-DEL-1484 |
| chr5-54021966-DEL-459 | chr5-54021963-DEL-460 |
| chr5-54027327-DEL-774 | chr5-54027328-DEL-774 |
| chr5-54045768-DEL-65 | chr5-54045800-DEL-77 |
| chr5-54047682-DEL-136 | chr5-54047683-DEL-136 |
| chr5-54130751-DEL-2839 | chr5-54130752-DEL-2839 |
| chr5-54141904-DEL-199 | chr5-54141905-DEL-199 |
| chr5-54142919-DEL-93 | chr5-54142920-DEL-93 |
| chr5-54167952-DEL-80 | chr5-54167922-DEL-69 |
| chr5-54205952-DEL-256 | chr5-54205953-DEL-256 |
| chr5-54207968-DEL-291 | chr5-54207969-DEL-291 |
| chr5-54221304-DEL-140 | chr5-54221305-DEL-140 |
| chr5-54233041-DEL-320 | chr5-54233042-DEL-319 |
| chr5-54236025-DEL-2558 | chr5-54236026-DEL-2558 |

| | |
|---|---|
| chr5-54284583-DEL-122 | chr5-54284591-DEL-76 |
| chr5-54288469-DEL-58 | chr5-54288470-DEL-58 |
| chr5-54293405-DEL-11775 | chr5-54293406-DEL-11775 |
| chr5-54293652-DEL-214 | chr5-54293653-DEL-214 |
| chr5-54309875-DEL-546 | chr5-54309876-DEL-546 |
| chr5-54334541-DEL-717 | chr5-54334542-DEL-717 |
| chr5-54386573-DEL-80 | chr5-54386566-DEL-60 |
| chr5-54393231-DEL-748 | chr5-54393232-DEL-748 |
| chr5-54441217-DEL-166 | chr5-54441218-DEL-166 |
| chr5-54459524-DEL-1755 | chr5-54459525-DEL-1755 |
| chr5-54461353-DEL-2561 | chr5-54461352-DEL-2560 |
| chr5-54493297-DEL-344 | chr5-54493298-DEL-344 |
| chr5-54496610-DEL-66 | chr5-54496636-DEL-90 |
| chr5-54565214-DEL-56 | chr5-54565211-DEL-66 |
| chr5-54573822-DEL-508 | chr5-54573823-DEL-508 |
| chr5-54574957-DEL-57 | chr5-54574958-DEL-57 |
| chr5-54575222-DEL-344 | chr5-54575223-DEL-344 |
| chr5-54588780-DEL-240 | chr5-54588811-DEL-193 |
| chr5-54591546-DEL-190 | chr5-54591547-DEL-190 |
| chr5-54594987-DEL-364 | chr5-54594989-DEL-363 |
| chr5-54596215-DEL-292 | chr5-54596216-DEL-292 |
| chr5-54599416-DEL-85 | chr5-54599417-DEL-85 |
| chr5-54606083-DEL-4576 | chr5-54606084-DEL-4576 |
| chr5-54618629-DEL-193 | chr5-54618626-DEL-193 |
| chr5-54632653-DEL-701 | chr5-54632649-DEL-700 |
| chr5-54638187-DEL-86 | chr5-54638184-DEL-84 |
| chr5-54638446-DEL-438 | chr5-54638447-DEL-438 |
| chr5-54674804-DEL-122 | chr5-54674787-DEL-110 |
| chr5-54690763-DEL-474 | chr5-54690764-DEL-474 |
| chr5-54733630-DEL-65 | chr5-54733624-DEL-66 |
| chr5-54743887-DEL-58 | chr5-54743888-DEL-58 |
| chr5-54744655-DEL-120 | chr5-54744656-DEL-120 |
| chr5-54754443-DEL-67 | chr5-54754444-DEL-67 |
| chr5-54762510-DEL-1671 | chr5-54762511-DEL-1671 |
| chr5-54805776-DEL-72 | chr5-54805777-DEL-72 |
| chr5-54834534-DEL-52 | chr5-54834523-DEL-60 |
| chr5-54846247-DEL-5991 | chr5-54846248-DEL-5991 |
| chr5-54862873-DEL-150 | chr5-54862874-DEL-149 |
| chr5-54890012-DEL-51 | chr5-54890013-DEL-51 |
| chr5-54909494-DEL-206 | chr5-54909495-DEL-206 |
| chr5-54930613-DEL-6434 | chr5-54930614-DEL-6434 |
| chr5-54930682-DEL-201 | chr5-54930682-DEL-201 |
| chr5-54968184-DEL-188 | chr5-54968189-DEL-184 |
| chr5-54969065-DEL-687 | chr5-54969066-DEL-687 |
| chr5-55006798-DEL-330 | chr5-55006799-DEL-327 |
| chr5-55017470-DEL-118 | chr5-55017491-DEL-98 |
| chr5-55068724-DEL-391 | chr5-55068725-DEL-391 |
| chr5-55100544-DEL-817 | chr5-55100545-DEL-817 |
| chr5-55114391-DEL-771 | chr5-55114392-DEL-772 |
| chr5-55119766-DEL-671 | chr5-55119754-DEL-669 |
| chr5-55131732-DEL-214 | chr5-55131733-DEL-214 |
| chr5-55145209-DEL-55 | chr5-55145210-DEL-55 |
| chr5-55167398-DEL-3149 | chr5-55167399-DEL-3149 |
| chr5-55180414-DEL-69 | chr5-55180415-DEL-69 |
| chr5-55183294-DEL-62 | chr5-55183295-DEL-58 |
| chr5-55186061-DEL-2703 | chr5-55186050-DEL-2704 |
| chr5-55216390-DEL-258 | chr5-55216391-DEL-258 |
| chr5-55286322-DEL-1536 | chr5-55286323-DEL-1536 |
| chr5-55288539-DEL-390 | chr5-55288539-DEL-391 |
| chr5-55290916-DEL-94 | chr5-55290917-DEL-94 |
| chr5-55328113-DEL-639 | chr5-55328114-DEL-639 |
| chr5-55359235-DEL-6647 | chr5-55359236-DEL-6647 |
| chr5-55413290-DEL-179 | chr5-55413291-DEL-179 |
| chr5-55415399-DEL-1933 | chr5-55415400-DEL-1933 |
| chr5-55422201-DEL-60 | chr5-55422197-DEL-64 |
| chr5-55426158-DEL-53 | chr5-55426158-DEL-53 |
| chr5-55484758-DEL-54 | chr5-55484759-DEL-54 |
| chr5-55488801-DEL-145 | chr5-55488805-DEL-146 |
| chr5-55494971-DEL-3698 | chr5-55494970-DEL-3699 |
| chr5-55501427-DEL-190 | chr5-55501428-DEL-190 |
| chr5-55555341-DEL-816 | chr5-55555342-DEL-816 |
| chr5-55566187-DEL-74 | chr5-55566188-DEL-74 |
| chr5-55607405-DEL-1941 | chr5-55607406-DEL-1941 |
| chr5-55644282-DEL-768 | chr5-55644283-DEL-768 |
| chr5-55653075-DEL-1129 | chr5-55653076-DEL-1129 |
| chr5-55659962-DEL-58 | chr5-55659959-DEL-56 |
| chr5-55672980-DEL-68 | chr5-55672999-DEL-82 |
| chr5-55675718-DEL-52 | chr5-55675718-DEL-56 |
| chr5-55683215-DEL-63 | chr5-55683216-DEL-63 |
| chr5-55685770-DEL-392 | chr5-55685771-DEL-391 |

| | |
|---|---|
| chr5-55690808-DEL-717 | chr5-55690808-DEL-717 |
| chr5-55694842-DEL-70 | chr5-55694858-DEL-52 |
| chr5-55704509-DEL-98 | chr5-55704546-DEL-72 |
| chr5-55752789-DEL-146 | chr5-55752790-DEL-146 |
| chr5-55781312-DEL-101 | chr5-55781311-DEL-100 |
| chr5-55798104-DEL-80 | chr5-55798105-DEL-80 |
| chr5-55804346-DEL-200 | chr5-55804347-DEL-200 |
| chr5-55810060-DEL-4989 | chr5-55810061-DEL-4989 |
| chr5-55821581-DEL-880 | chr5-55821582-DEL-880 |
| chr5-55837139-DEL-686 | chr5-55837140-DEL-686 |
| chr5-55858931-DEL-56 | chr5-55858932-DEL-56 |
| chr5-55883225-DEL-6635 | chr5-55883226-DEL-6635 |
| chr5-55919527-DEL-6340 | chr5-55919517-DEL-6339 |
| chr5-55934328-DEL-196 | chr5-55934329-DEL-196 |
| chr5-55941210-DEL-52 | chr5-55941211-DEL-52 |
| chr5-55945057-DEL-71 | chr5-55945058-DEL-71 |
| chr5-55954468-DEL-55 | chr5-55954469-DEL-55 |
| chr5-55964873-DEL-63 | chr5-55964874-DEL-63 |
| chr5-55966848-DEL-106 | chr5-55966849-DEL-106 |
| chr5-55975799-DEL-51 | chr5-55975799-DEL-56 |
| chr5-55977171-DEL-869 | chr5-55977172-DEL-869 |
| chr5-55994615-DEL-6932 | chr5-55994616-DEL-6932 |
| chr5-56018452-DEL-184 | chr5-56018453-DEL-172 |
| chr5-56056171-DEL-2395 | chr5-56056172-DEL-2395 |
| chr5-56072533-DEL-234 | chr5-56072533-DEL-234 |
| chr5-56083401-DEL-171 | chr5-56083402-DEL-171 |
| chr5-56100170-DEL-55 | chr5-56100172-DEL-54 |
| chr5-56100943-DEL-144 | chr5-56100944-DEL-144 |
| chr5-56101288-DEL-1381 | chr5-56101289-DEL-1380 |
| chr5-56107495-DEL-2973 | chr5-56107496-DEL-2973 |
| chr5-56126312-DEL-104 | chr5-56126309-DEL-108 |
| chr5-56147722-DEL-9762 | chr5-56147723-DEL-9762 |
| chr5-56173176-DEL-3656 | chr5-56173177-DEL-3656 |
| chr5-56183702-DEL-63 | chr5-56183703-DEL-63 |
| chr5-56186284-DEL-1052 | chr5-56186283-DEL-1053 |
| chr5-56195000-DEL-76 | chr5-56195001-DEL-76 |
| chr5-56201603-DEL-187 | chr5-56201604-DEL-187 |
| chr5-56220581-DEL-346 | chr5-56220582-DEL-346 |
| chr5-56222236-DEL-73 | chr5-56222231-DEL-77 |
| chr5-56232392-DEL-83 | chr5-56232394-DEL-87 |
| chr5-56243551-DEL-143 | chr5-56243552-DEL-143 |
| chr5-56244768-DEL-310 | chr5-56244769-DEL-310 |
| chr5-56272617-DEL-430 | chr5-56272618-DEL-430 |
| chr5-56278083-DEL-66 | chr5-56278082-DEL-73 |
| chr5-56280176-DEL-6872 | chr5-56280177-DEL-6872 |
| chr5-56286542-DEL-213 | chr5-56286500-DEL-214 |
| chr5-56290164-DEL-488 | chr5-56290165-DEL-488 |
| chr5-56296454-DEL-102 | chr5-56296489-DEL-60 |
| chr5-56309048-DEL-81 | chr5-56309047-DEL-82 |
| chr5-56328421-DEL-785 | chr5-56328422-DEL-785 |
| chr5-56335794-DEL-80 | chr5-56335801-DEL-76 |
| chr5-56346358-DEL-6840 | chr5-56346359-DEL-6840 |
| chr5-56358666-DEL-152 | chr5-56358667-DEL-152 |
| chr5-56373687-DEL-659 | chr5-56373685-DEL-666 |
| chr5-56394845-DEL-585 | chr5-56394846-DEL-585 |
| chr5-56410748-DEL-6480 | chr5-56410749-DEL-6480 |
| chr5-56418748-DEL-1607 | chr5-56418749-DEL-1606 |
| chr5-56429951-DEL-208 | chr5-56429952-DEL-208 |
| chr5-56443391-DEL-1341 | chr5-56443392-DEL-1342 |
| chr5-56444201-DEL-445 | chr5-56444202-DEL-445 |
| chr5-56446967-DEL-16534 | chr5-56446968-DEL-16534 |
| chr5-56461877-DEL-59 | chr5-56461878-DEL-59 |
| chr5-56491539-DEL-214 | chr5-56491540-DEL-214 |
| chr5-56544537-DEL-77 | chr5-56544536-DEL-81 |
| chr5-56548908-DEL-119 | chr5-56548909-DEL-119 |
| chr5-56575261-DEL-459 | chr5-56575262-DEL-459 |
| chr5-56579139-DEL-411 | chr5-56579140-DEL-411 |
| chr5-56583651-DEL-266 | chr5-56583652-DEL-266 |
| chr5-56611429-DEL-72 | chr5-56611430-DEL-72 |
| chr5-56628832-DEL-6887 | chr5-56628832-DEL-6887 |
| chr5-56650499-DEL-6454 | chr5-56650500-DEL-6454 |
| chr5-56732368-DEL-6248 | chr5-56732369-DEL-6248 |
| chr5-56742977-DEL-3356 | chr5-56742977-DEL-3356 |
| chr5-56751338-DEL-1141 | chr5-56751339-DEL-1141 |
| chr5-56776120-DEL-165 | chr5-56776119-DEL-164 |
| chr5-56779734-DEL-412 | chr5-56779735-DEL-412 |
| chr5-56784513-DEL-1444 | chr5-56784514-DEL-1444 |
| chr5-56790497-DEL-174 | chr5-56790498-DEL-180 |
| chr5-56801076-DEL-74 | chr5-56801077-DEL-74 |
| chr5-56804758-DEL-1604 | chr5-56804759-DEL-1604 |

| | |
|---|---|
| chr5-56814292-DEL-1389 | chr5-56814293-DEL-1389 |
| chr5-56825877-DEL-1384 | chr5-56825878-DEL-1384 |
| chr5-56857213-DEL-1753 | chr5-56857214-DEL-1753 |
| chr5-56861862-DEL-1734 | chr5-56861863-DEL-1734 |
| chr5-56864493-DEL-892 | chr5-56864494-DEL-892 |
| chr5-56871759-DEL-255 | chr5-56871760-DEL-255 |
| chr5-56877444-DEL-766 | chr5-56877445-DEL-766 |
| chr5-56938679-DEL-4172 | chr5-56938680-DEL-4172 |
| chr5-56971743-DEL-3272 | chr5-56971733-DEL-3272 |
| chr5-57029428-DEL-1884 | chr5-57029429-DEL-1884 |
| chr5-57037435-DEL-389 | chr5-57037436-DEL-389 |
| chr5-57116928-DEL-104 | chr5-57116884-DEL-102 |
| chr5-57119971-DEL-57 | chr5-57119972-DEL-57 |
| chr5-57126105-DEL-2301 | chr5-57126106-DEL-2301 |
| chr5-57127118-DEL-6227 | chr5-57127122-DEL-6228 |
| chr5-57175046-DEL-7109 | chr5-57175047-DEL-7109 |
| chr5-57190411-DEL-57 | chr5-57190412-DEL-57 |
| chr5-57208650-DEL-70 | chr5-57208659-DEL-58 |
| chr5-57266292-DEL-58 | chr5-57266295-DEL-56 |
| chr5-57303841-DEL-6189 | chr5-57303830-DEL-6198 |
| chr5-57332957-DEL-531 | chr5-57332958-DEL-528 |
| chr5-57358846-DEL-263 | chr5-57358847-DEL-263 |
| chr5-57362705-DEL-427 | chr5-57362706-DEL-427 |
| chr5-57367266-DEL-659 | chr5-57367267-DEL-659 |
| chr5-57368267-DEL-183 | chr5-57368268-DEL-183 |
| chr5-57404240-DEL-208 | chr5-57404241-DEL-208 |
| chr5-57408815-DEL-54 | chr5-57408816-DEL-54 |
| chr5-57446930-DEL-102 | chr5-57446931-DEL-102 |
| chr5-57454236-DEL-271 | chr5-57454237-DEL-271 |
| chr5-57455277-DEL-64 | chr5-57455278-DEL-64 |
| chr5-57468401-DEL-54 | chr5-57468402-DEL-54 |
| chr5-57499145-DEL-106 | chr5-57499144-DEL-98 |
| chr5-57552517-DEL-104 | chr5-57552531-DEL-105 |
| chr5-57576130-DEL-62 | chr5-57576131-DEL-62 |
| chr5-57589704-DEL-6595 | chr5-57589705-DEL-6595 |
| chr5-57619757-DEL-390 | chr5-57619758-DEL-390 |
| chr5-57624562-DEL-54 | chr5-57624563-DEL-54 |
| chr5-57633667-DEL-60 | chr5-57633668-DEL-60 |
| chr5-57661303-DEL-287 | chr5-57661304-DEL-287 |
| chr5-57697337-DEL-214 | chr5-57697338-DEL-214 |
| chr5-57712943-DEL-6632 | chr5-57712944-DEL-6632 |
| chr5-57834236-DEL-400 | chr5-57834237-DEL-402 |
| chr5-57836566-DEL-399 | chr5-57836567-DEL-399 |
| chr5-57848703-DEL-5222 | chr5-57848704-DEL-5222 |
| chr5-57891258-DEL-86 | chr5-57891258-DEL-88 |
| chr5-57957805-DEL-1094 | chr5-57957806-DEL-1094 |
| chr5-57965897-DEL-64 | chr5-57965898-DEL-64 |
| chr5-57983044-DEL-139 | chr5-57983048-DEL-125 |
| chr5-57995776-DEL-70 | chr5-57995777-DEL-70 |
| chr5-58017340-DEL-3510 | chr5-58017341-DEL-3510 |
| chr5-58072157-DEL-60 | chr5-58072158-DEL-58 |
| chr5-58076159-DEL-56 | chr5-58076160-DEL-54 |
| chr5-58094381-DEL-64 | chr5-58094382-DEL-64 |
| chr5-58096479-DEL-120 | chr5-58096499-DEL-127 |
| chr5-58139580-DEL-2171 | chr5-58139581-DEL-2171 |
| chr5-58151848-DEL-6327 | chr5-58151851-DEL-6328 |
| chr5-58158523-DEL-52 | chr5-58158524-DEL-52 |
| chr5-58190434-DEL-60 | chr5-58190425-DEL-64 |
| chr5-58331937-DEL-287 | chr5-58331938-DEL-287 |
| chr5-58345788-DEL-66 | chr5-58345789-DEL-66 |
| chr5-58351771-DEL-55 | chr5-58351772-DEL-55 |
| chr5-58356037-DEL-342 | chr5-58356038-DEL-342 |
| chr5-58364869-DEL-163 | chr5-58364870-DEL-163 |
| chr5-58376006-DEL-57 | chr5-58375995-DEL-89 |
| chr5-58394344-DEL-75 | chr5-58394345-DEL-75 |
| chr5-58402695-DEL-6271 | chr5-58402696-DEL-6271 |
| chr5-58411210-DEL-52 | chr5-58411217-DEL-58 |
| chr5-58417268-DEL-88 | chr5-58417235-DEL-82 |
| chr5-58520962-DEL-64 | chr5-58520923-DEL-65 |
| chr5-58525831-DEL-62 | chr5-58525832-DEL-66 |
| chr5-58555991-DEL-707 | chr5-58555989-DEL-708 |
| chr5-58556926-DEL-6855 | chr5-58556927-DEL-6855 |
| chr5-58563871-DEL-407 | chr5-58563871-DEL-407 |
| chr5-58603893-DEL-1621 | chr5-58603894-DEL-1621 |
| chr5-58612232-DEL-117 | chr5-58612233-DEL-117 |
| chr5-58623333-DEL-51 | chr5-58623334-DEL-51 |
| chr5-58625313-DEL-402 | chr5-58625314-DEL-402 |
| chr5-58647087-DEL-451 | chr5-58647088-DEL-451 |
| chr5-58651960-DEL-76 | chr5-58651980-DEL-72 |
| chr5-58655879-DEL-176 | chr5-58655879-DEL-176 |

| | |
|---|---|
| chr5-58655929-DEL-1852 | chr5-58655930-DEL-1852 |
| chr5-58665354-DEL-70 | chr5-58665386-DEL-78 |
| chr5-58673639-DEL-4922 | chr5-58673640-DEL-4922 |
| chr5-58685808-DEL-80 | chr5-58685855-DEL-51 |
| chr5-58697133-DEL-230 | chr5-58697134-DEL-230 |
| chr5-58707335-DEL-463 | chr5-58707336-DEL-463 |
| chr5-58709106-DEL-134 | chr5-58709105-DEL-133 |
| chr5-58710395-DEL-54 | chr5-58710396-DEL-54 |
| chr5-58721110-DEL-85 | chr5-58721111-DEL-85 |
| chr5-58727909-DEL-3278 | chr5-58727910-DEL-3278 |
| chr5-58791781-DEL-3439 | chr5-58791781-DEL-3440 |
| chr5-58808544-DEL-52 | chr5-58808545-DEL-52 |
| chr5-58836276-DEL-66 | chr5-58836277-DEL-68 |
| chr5-58854374-DEL-2700 | chr5-58854375-DEL-2700 |
| chr5-58882248-DEL-7688 | chr5-58882249-DEL-7688 |
| chr5-58891127-DEL-88 | chr5-58891145-DEL-63 |
| chr5-58935024-DEL-6654 | chr5-58935025-DEL-6654 |
| chr5-58935184-DEL-214 | chr5-58935185-DEL-214 |
| chr5-58943849-DEL-1437 | chr5-58943850-DEL-1437 |
| chr5-58987465-DEL-1139 | chr5-58987463-DEL-1140 |
| chr5-58989093-DEL-51 | chr5-58989094-DEL-51 |
| chr5-59007775-DEL-448 | chr5-59007776-DEL-448 |
| chr5-59012929-DEL-58 | chr5-59012930-DEL-54 |
| chr5-59019745-DEL-2545 | chr5-59019746-DEL-2545 |
| chr5-59030888-DEL-6380 | chr5-59030929-DEL-6331 |
| chr5-59038084-DEL-92 | chr5-59038085-DEL-92 |
| chr5-59046945-DEL-172 | chr5-59046946-DEL-172 |
| chr5-59082272-DEL-7208 | chr5-59082273-DEL-7208 |
| chr5-59082797-DEL-208 | chr5-59082798-DEL-208 |
| chr5-59090731-DEL-6433 | chr5-59090732-DEL-6433 |
| chr5-59115918-DEL-64 | chr5-59115879-DEL-58 |
| chr5-59123639-DEL-6412 | chr5-59123640-DEL-6412 |
| chr5-59130613-DEL-68 | chr5-59130601-DEL-50 |
| chr5-59131149-DEL-284 | chr5-59131150-DEL-284 |
| chr5-59160523-DEL-7078 | chr5-59160524-DEL-7078 |
| chr5-59189912-DEL-204 | chr5-59189914-DEL-203 |
| chr5-59203758-DEL-295 | chr5-59203758-DEL-295 |
| chr5-59236486-DEL-92 | chr5-59236487-DEL-92 |
| chr5-59242340-DEL-100 | chr5-59242341-DEL-100 |
| chr5-59247103-DEL-1580 | chr5-59247104-DEL-1580 |
| chr5-59254090-DEL-390 | chr5-59254090-DEL-390 |
| chr5-59263944-DEL-179 | chr5-59263941-DEL-183 |
| chr5-59276801-DEL-7233 | chr5-59276802-DEL-7233 |
| chr5-59292072-DEL-400 | chr5-59292073-DEL-400 |
| chr5-59315479-DEL-298 | chr5-59315480-DEL-298 |
| chr5-59321359-DEL-56 | chr5-59321318-DEL-58 |
| chr5-59336517-DEL-213 | chr5-59336518-DEL-213 |
| chr5-59338726-DEL-4523 | chr5-59338727-DEL-4523 |
| chr5-59355335-DEL-734 | chr5-59355336-DEL-734 |
| chr5-59383883-DEL-64 | chr5-59383895-DEL-56 |
| chr5-59388048-DEL-2271 | chr5-59388049-DEL-2271 |
| chr5-59410555-DEL-1512 | chr5-59410556-DEL-1512 |
| chr5-59412456-DEL-6198 | chr5-59412457-DEL-6198 |
| chr5-59421657-DEL-517 | chr5-59421658-DEL-517 |
| chr5-59422811-DEL-752 | chr5-59422812-DEL-752 |
| chr5-59425926-DEL-53 | chr5-59425907-DEL-98 |
| chr5-59441889-DEL-66 | chr5-59441890-DEL-66 |
| chr5-59453606-DEL-233 | chr5-59453597-DEL-257 |
| chr5-59467971-DEL-6378 | chr5-59467972-DEL-6378 |
| chr5-59482883-DEL-52 | chr5-59482884-DEL-52 |
| chr5-59491902-DEL-560 | chr5-59491901-DEL-558 |
| chr5-59500324-DEL-2594 | chr5-59500325-DEL-2594 |
| chr5-59532245-DEL-945 | chr5-59532245-DEL-945 |
| chr5-59596920-DEL-4923 | chr5-59596921-DEL-4923 |
| chr5-59641692-DEL-72 | chr5-59641693-DEL-72 |
| chr5-59646202-DEL-73 | chr5-59646203-DEL-74 |
| chr5-59648518-DEL-969 | chr5-59648515-DEL-970 |
| chr5-59691427-DEL-147 | chr5-59691423-DEL-140 |
| chr5-59716711-DEL-383 | chr5-59716712-DEL-383 |
| chr5-59729907-DEL-1865 | chr5-59729908-DEL-1865 |
| chr5-59735652-DEL-1066 | chr5-59735652-DEL-1077 |
| chr5-59747424-DEL-60 | chr5-59747454-DEL-50 |
| chr5-59750704-DEL-1111 | chr5-59750704-DEL-1111 |
| chr5-59752670-DEL-51 | chr5-59752671-DEL-51 |
| chr5-59791469-DEL-70 | chr5-59791470-DEL-70 |
| chr5-59793654-DEL-70 | chr5-59793655-DEL-70 |
| chr5-59801164-DEL-6446 | chr5-59801189-DEL-6417 |
| chr5-59832305-DEL-398 | chr5-59832306-DEL-398 |
| chr5-59842032-DEL-1861 | chr5-59842033-DEL-1861 |
| chr5-59845609-DEL-82 | chr5-59845591-DEL-92 |

| | |
|---|---|
| chr5-59861833-DEL-1174 | chr5-59861846-DEL-1173 |
| chr5-59865820-DEL-74 | chr5-59865825-DEL-70 |
| chr5-59874301-DEL-6343 | chr5-59874302-DEL-6343 |
| chr5-59886752-DEL-67 | chr5-59886747-DEL-68 |
| chr5-59892486-DEL-80 | chr5-59892488-DEL-79 |
| chr5-59897750-DEL-72 | chr5-59897765-DEL-106 |
| chr5-59901982-DEL-65 | chr5-59901983-DEL-62 |
| chr5-59907345-DEL-6197 | chr5-59907334-DEL-6198 |
| chr5-59923806-DEL-60 | chr5-59923807-DEL-60 |
| chr5-59945389-DEL-204 | chr5-59945368-DEL-198 |
| chr5-59959406-DEL-135 | chr5-59959407-DEL-136 |
| chr5-59974316-DEL-56 | chr5-59974317-DEL-58 |
| chr5-59979191-DEL-734 | chr5-59979192-DEL-734 |
| chr5-60005583-DEL-60 | chr5-60005584-DEL-60 |
| chr5-60005922-DEL-854 | chr5-60005923-DEL-854 |
| chr5-60011457-DEL-236 | chr5-60011458-DEL-236 |
| chr5-60032537-DEL-76 | chr5-60032538-DEL-70 |
| chr5-60081200-DEL-199 | chr5-60081201-DEL-199 |
| chr5-60083223-DEL-357 | chr5-60083224-DEL-357 |
| chr5-60105909-DEL-78 | chr5-60105910-DEL-78 |
| chr5-60166044-DEL-81 | chr5-60166051-DEL-64 |
| chr5-60200054-DEL-392 | chr5-60200053-DEL-393 |
| chr5-60226421-DEL-3652 | chr5-60226419-DEL-3652 |
| chr5-60274195-DEL-1071 | chr5-60274196-DEL-1071 |
| chr5-60294961-DEL-344 | chr5-60294962-DEL-344 |
| chr5-60314000-DEL-6380 | chr5-60314001-DEL-6380 |
| chr5-60332676-DEL-85 | chr5-60332668-DEL-86 |
| chr5-60336217-DEL-143 | chr5-60336218-DEL-143 |
| chr5-60388722-DEL-815 | chr5-60388723-DEL-814 |
| chr5-60392291-DEL-2106 | chr5-60392292-DEL-2106 |
| chr5-60398535-DEL-78 | chr5-60398518-DEL-106 |
| chr5-60425041-DEL-71 | chr5-60425019-DEL-72 |
| chr5-60430313-DEL-54 | chr5-60430314-DEL-54 |
| chr5-60445343-DEL-53 | chr5-60445344-DEL-53 |
| chr5-60456913-DEL-1898 | chr5-60456914-DEL-1898 |
| chr5-60472896-DEL-92 | chr5-60472932-DEL-56 |
| chr5-60490282-DEL-418 | chr5-60490283-DEL-418 |
| chr5-60503309-DEL-6335 | chr5-60503310-DEL-6335 |
| chr5-60532467-DEL-62 | chr5-60532468-DEL-62 |
| chr5-60547318-DEL-288 | chr5-60547319-DEL-288 |
| chr5-60627823-DEL-3307 | chr5-60627824-DEL-3307 |
| chr5-60686297-DEL-519 | chr5-60686298-DEL-519 |
| chr5-60698826-DEL-993 | chr5-60698843-DEL-977 |
| chr5-60701752-DEL-15522 | chr5-60701742-DEL-15523 |
| chr5-60707846-DEL-6760 | chr5-60707847-DEL-6760 |
| chr5-60768021-DEL-52 | chr5-60768022-DEL-52 |
| chr5-60808988-DEL-6598 | chr5-60808989-DEL-6598 |
| chr5-60828095-DEL-626 | chr5-60828092-DEL-626 |
| chr5-60837972-DEL-1895 | chr5-60837964-DEL-1893 |
| chr5-60845075-DEL-83 | chr5-60845093-DEL-65 |
| chr5-60869830-DEL-338 | chr5-60869831-DEL-338 |
| chr5-60886089-DEL-7004 | chr5-60886090-DEL-7004 |
| chr5-60886602-DEL-413 | chr5-60886570-DEL-414 |
| chr5-60911815-DEL-4497 | chr5-60911816-DEL-4497 |
| chr5-60922827-DEL-62 | chr5-60922828-DEL-62 |
| chr5-60931189-DEL-401 | chr5-60931190-DEL-401 |
| chr5-60933578-DEL-203 | chr5-60933579-DEL-203 |
| chr5-60937316-DEL-62 | chr5-60937317-DEL-62 |
| chr5-60943055-DEL-2461 | chr5-60943056-DEL-2461 |
| chr5-60960975-DEL-5408 | chr5-60960976-DEL-5408 |
| chr5-60986557-DEL-188 | chr5-60986558-DEL-188 |
| chr5-61012373-DEL-504 | chr5-61012374-DEL-504 |
| chr5-61018014-DEL-187 | chr5-61018015-DEL-187 |
| chr5-61035023-DEL-63 | chr5-61035013-DEL-64 |
| chr5-61136211-DEL-144 | chr5-61136212-DEL-144 |
| chr5-61140979-DEL-2562 | chr5-61140980-DEL-2562 |
| chr5-61169703-DEL-5396 | chr5-61169704-DEL-5396 |
| chr5-61202179-DEL-104 | chr5-61202157-DEL-104 |
| chr5-61203351-DEL-63 | chr5-61203352-DEL-63 |
| chr5-61204087-DEL-359 | chr5-61204087-DEL-359 |
| chr5-61207467-DEL-93 | chr5-61207467-DEL-93 |
| chr5-61370289-DEL-6442 | chr5-61370277-DEL-6439 |
| chr5-61378471-DEL-75 | chr5-61378472-DEL-75 |
| chr5-61379109-DEL-70 | chr5-61379114-DEL-60 |
| chr5-61488841-DEL-3082 | chr5-61488841-DEL-3082 |
| chr5-61497950-DEL-434 | chr5-61497951-DEL-434 |
| chr5-61550530-DEL-277 | chr5-61550537-DEL-281 |
| chr5-61556856-DEL-279 | chr5-61556857-DEL-279 |
| chr5-61604597-DEL-123 | chr5-61604598-DEL-123 |
| chr5-61622541-DEL-2833 | chr5-61622541-DEL-2836 |

| | |
|---|---|
| chr5-61649871-DEL-116 | chr5-61649872-DEL-100 |
| chr5-61660155-DEL-480 | chr5-61660164-DEL-480 |
| chr5-61751852-DEL-7116 | chr5-61751852-DEL-7116 |
| chr5-61765873-DEL-69 | chr5-61765874-DEL-69 |
| chr5-61859031-DEL-4122 | chr5-61859032-DEL-4122 |
| chr5-61861881-DEL-486 | chr5-61861882-DEL-486 |
| chr5-61866710-DEL-224 | chr5-61866710-DEL-224 |
| chr5-61878176-DEL-552 | chr5-61878177-DEL-552 |
| chr5-61909508-DEL-425 | chr5-61909509-DEL-425 |
| chr5-61913090-DEL-55 | chr5-61913090-DEL-50 |
| chr5-61914433-DEL-161 | chr5-61914434-DEL-161 |
| chr5-61932561-DEL-228 | chr5-61932562-DEL-228 |
| chr5-61935639-DEL-6866 | chr5-61935640-DEL-6866 |
| chr5-61959736-DEL-1135 | chr5-61959737-DEL-1135 |
| chr5-61989370-DEL-84 | chr5-61989369-DEL-52 |
| chr5-62002005-DEL-3988 | chr5-62002006-DEL-3988 |
| chr5-62103786-DEL-487 | chr5-62103787-DEL-487 |
| chr5-62108525-DEL-527 | chr5-62108527-DEL-522 |
| chr5-62133173-DEL-53 | chr5-62133174-DEL-61 |
| chr5-62135671-DEL-74 | chr5-62135672-DEL-74 |
| chr5-62136215-DEL-3496 | chr5-62136216-DEL-3495 |
| chr5-62185390-DEL-952 | chr5-62185392-DEL-951 |
| chr5-62191692-DEL-1328 | chr5-62191691-DEL-1328 |
| chr5-62247402-DEL-55 | chr5-62247403-DEL-55 |
| chr5-62262999-DEL-1941 | chr5-62262999-DEL-1941 |
| chr5-62267291-DEL-457 | chr5-62267290-DEL-457 |
| chr5-62270108-DEL-58 | chr5-62270109-DEL-58 |
| chr5-62278771-DEL-758 | chr5-62278772-DEL-758 |
| chr5-62287210-DEL-207 | chr5-62287211-DEL-207 |
| chr5-62290367-DEL-688 | chr5-62290368-DEL-688 |
| chr5-62335618-DEL-278 | chr5-62335619-DEL-278 |
| chr5-62353253-DEL-401 | chr5-62353254-DEL-401 |
| chr5-62385146-DEL-424 | chr5-62385147-DEL-424 |
| chr5-62413004-DEL-2478 | chr5-62413005-DEL-2478 |
| chr5-62445181-DEL-228 | chr5-62445182-DEL-227 |
| chr5-62449142-DEL-425 | chr5-62449143-DEL-425 |
| chr5-62459634-DEL-1092 | chr5-62459635-DEL-1092 |
| chr5-62464950-DEL-6719 | chr5-62464951-DEL-6719 |
| chr5-62493274-DEL-1823 | chr5-62493275-DEL-1823 |
| chr5-62512168-DEL-56 | chr5-62512149-DEL-76 |
| chr5-62536120-DEL-1883 | chr5-62536121-DEL-1883 |
| chr5-62584842-DEL-6485 | chr5-62584843-DEL-6485 |
| chr5-62597231-DEL-213 | chr5-62597232-DEL-213 |
| chr5-62640101-DEL-85 | chr5-62640116-DEL-85 |
| chr5-62692712-DEL-1920 | chr5-62692713-DEL-1920 |
| chr5-62709649-DEL-66 | chr5-62709652-DEL-64 |
| chr5-62716053-DEL-234 | chr5-62716054-DEL-234 |
| chr5-62725009-DEL-938 | chr5-62725010-DEL-938 |
| chr5-62758389-DEL-51 | chr5-62758387-DEL-50 |
| chr5-62766551-DEL-64 | chr5-62766553-DEL-63 |
| chr5-62830923-DEL-416 | chr5-62830924-DEL-416 |
| chr5-62898849-DEL-146 | chr5-62898838-DEL-158 |
| chr5-62940431-DEL-56 | chr5-62940431-DEL-56 |
| chr5-62952022-DEL-61 | chr5-62952023-DEL-61 |
| chr5-62965336-DEL-89 | chr5-62965337-DEL-89 |
| chr5-63007445-DEL-51 | chr5-63007446-DEL-51 |
| chr5-63009417-DEL-365 | chr5-63009415-DEL-394 |
| chr5-63025832-DEL-6977 | chr5-63025830-DEL-6978 |
| chr5-63034664-DEL-55 | chr5-63034665-DEL-55 |
| chr5-63058091-DEL-186 | chr5-63058092-DEL-186 |
| chr5-63076676-DEL-861 | chr5-63076677-DEL-861 |
| chr5-63112434-DEL-6016 | chr5-63112435-DEL-6016 |
| chr5-63121047-DEL-1178 | chr5-63121048-DEL-1178 |
| chr5-63143450-DEL-62 | chr5-63143455-DEL-58 |
| chr5-63158806-DEL-78 | chr5-63158807-DEL-78 |
| chr5-63168950-DEL-2121 | chr5-63168951-DEL-2121 |
| chr5-63183579-DEL-52 | chr5-63183580-DEL-52 |
| chr5-63263388-DEL-523 | chr5-63263389-DEL-523 |
| chr5-63266245-DEL-57 | chr5-63266249-DEL-57 |
| chr5-63298065-DEL-808 | chr5-63298065-DEL-808 |
| chr5-63317963-DEL-283 | chr5-63317964-DEL-283 |
| chr5-63367774-DEL-1023 | chr5-63367775-DEL-1022 |
| chr5-63373049-DEL-1119 | chr5-63373050-DEL-1119 |
| chr5-63382301-DEL-3057 | chr5-63382302-DEL-3057 |
| chr5-63409671-DEL-877 | chr5-63409685-DEL-874 |
| chr5-63435897-DEL-175 | chr5-63435895-DEL-174 |
| chr5-63468463-DEL-2066 | chr5-63468463-DEL-2066 |
| chr5-63481024-DEL-322 | chr5-63481025-DEL-322 |
| chr5-63481604-DEL-155 | chr5-63481589-DEL-144 |
| chr5-63489336-DEL-1081 | chr5-63489337-DEL-1081 |

| | |
|---|---|
| chr5-63494006-DEL-702 | chr5-63494007-DEL-702 |
| chr5-63499336-DEL-999 | chr5-63499337-DEL-999 |
| chr5-63504316-DEL-400 | chr5-63504317-DEL-400 |
| chr5-63515013-DEL-240 | chr5-63515014-DEL-240 |
| chr5-63544472-DEL-2377 | chr5-63544460-DEL-2378 |
| chr5-63548256-DEL-362 | chr5-63548258-DEL-361 |
| chr5-63556785-DEL-864 | chr5-63556786-DEL-864 |
| chr5-63570133-DEL-684 | chr5-63570134-DEL-642 |
| chr5-63576806-DEL-82 | chr5-63576823-DEL-66 |
| chr5-63589950-DEL-386 | chr5-63589951-DEL-386 |
| chr5-63653623-DEL-73 | chr5-63653624-DEL-73 |
| chr5-63671471-DEL-1498 | chr5-63671472-DEL-1498 |
| chr5-63748072-DEL-6730 | chr5-63748057-DEL-6731 |
| chr5-63803289-DEL-205 | chr5-63803290-DEL-205 |
| chr5-63814724-DEL-6469 | chr5-63814725-DEL-6469 |
| chr5-63832724-DEL-101 | chr5-63832711-DEL-122 |
| chr5-63891313-DEL-85 | chr5-63891344-DEL-65 |
| chr5-63895557-DEL-171 | chr5-63895554-DEL-175 |
| chr5-63910345-DEL-57 | chr5-63910346-DEL-50 |
| chr5-63933092-DEL-77 | chr5-63933093-DEL-77 |
| chr5-63938754-DEL-220 | chr5-63938745-DEL-224 |
| chr5-63973126-DEL-135 | chr5-63973132-DEL-98 |
| chr5-63994010-DEL-76 | chr5-63994014-DEL-50 |
| chr5-63997762-DEL-246 | chr5-63997763-DEL-246 |
| chr5-64005560-DEL-6401 | chr5-64005561-DEL-6401 |
| chr5-64015017-DEL-472 | chr5-64015018-DEL-471 |
| chr5-64069728-DEL-101 | chr5-64069729-DEL-101 |
| chr5-64073258-DEL-98 | chr5-64073259-DEL-98 |
| chr5-64075363-DEL-54 | chr5-64075316-DEL-74 |
| chr5-64077185-DEL-62 | chr5-64077190-DEL-68 |
| chr5-64084845-DEL-284 | chr5-64084850-DEL-280 |
| chr5-64091614-DEL-1083 | chr5-64091613-DEL-1084 |
| chr5-64117809-DEL-90 | chr5-64117810-DEL-90 |
| chr5-64129959-DEL-52 | chr5-64129960-DEL-52 |
| chr5-64130498-DEL-125 | chr5-64130499-DEL-125 |
| chr5-64141694-DEL-178 | chr5-64141695-DEL-178 |
| chr5-64166079-DEL-160 | chr5-64166082-DEL-137 |
| chr5-64171972-DEL-172 | chr5-64171973-DEL-172 |
| chr5-64184047-DEL-116 | chr5-64184048-DEL-116 |
| chr5-64188170-DEL-7475 | chr5-64188171-DEL-7475 |
| chr5-64213274-DEL-341 | chr5-64213275-DEL-341 |
| chr5-64220158-DEL-70 | chr5-64220170-DEL-67 |
| chr5-64223443-DEL-126 | chr5-64223449-DEL-126 |
| chr5-64225266-DEL-224 | chr5-64225267-DEL-224 |
| chr5-64232836-DEL-164 | chr5-64232820-DEL-179 |
| chr5-64257869-DEL-54 | chr5-64257870-DEL-54 |
| chr5-64275077-DEL-6635 | chr5-64275078-DEL-6635 |
| chr5-64275239-DEL-212 | chr5-64275240-DEL-212 |
| chr5-64284409-DEL-169 | chr5-64284410-DEL-169 |
| chr5-64287986-DEL-59 | chr5-64287952-DEL-53 |
| chr5-64292059-DEL-2439 | chr5-64292060-DEL-2439 |
| chr5-64309968-DEL-202 | chr5-64309969-DEL-202 |
| chr5-64333731-DEL-95 | chr5-64333732-DEL-95 |
| chr5-64349963-DEL-148 | chr5-64349964-DEL-148 |
| chr5-64405585-DEL-64 | chr5-64405586-DEL-64 |
| chr5-64406262-DEL-60 | chr5-64406255-DEL-50 |
| chr5-64407216-DEL-126 | chr5-64407217-DEL-126 |
| chr5-64409630-DEL-223 | chr5-64409631-DEL-219 |
| chr5-64417218-DEL-385 | chr5-64417219-DEL-385 |
| chr5-64418046-DEL-618 | chr5-64418074-DEL-635 |
| chr5-64454355-DEL-99 | chr5-64454357-DEL-59 |
| chr5-64456124-DEL-54 | chr5-64456125-DEL-54 |
| chr5-64498047-DEL-179 | chr5-64498048-DEL-179 |
| chr5-64505267-DEL-208 | chr5-64505268-DEL-208 |
| chr5-64534112-DEL-116 | chr5-64534123-DEL-117 |
| chr5-64545154-DEL-72 | chr5-64545155-DEL-72 |
| chr5-64565411-DEL-156 | chr5-64565412-DEL-156 |
| chr5-64576585-DEL-215 | chr5-64576586-DEL-215 |
| chr5-64577305-DEL-90 | chr5-64577306-DEL-90 |
| chr5-64583392-DEL-2841 | chr5-64583393-DEL-2841 |
| chr5-64675761-DEL-428 | chr5-64675764-DEL-426 |
| chr5-64692224-DEL-77 | chr5-64692225-DEL-77 |
| chr5-64699964-DEL-68 | chr5-64699965-DEL-68 |
| chr5-64715297-DEL-7108 | chr5-64715298-DEL-7108 |
| chr5-64746408-DEL-212 | chr5-64746409-DEL-212 |
| chr5-64765542-DEL-98 | chr5-64765538-DEL-96 |
| chr5-64770601-DEL-93 | chr5-64770602-DEL-93 |
| chr5-64877210-DEL-222 | chr5-64877211-DEL-222 |
| chr5-64894741-DEL-55 | chr5-64894703-DEL-58 |
| chr5-64897886-DEL-51 | chr5-64897886-DEL-51 |

| | |
|---|---|
| chr5-64975710-DEL-95 | chr5-64975711-DEL-95 |
| chr5-64982649-DEL-140 | chr5-64982650-DEL-139 |
| chr5-65012119-DEL-62 | chr5-65012134-DEL-56 |
| chr5-65014763-DEL-96 | chr5-65014764-DEL-96 |
| chr5-65037103-DEL-97 | chr5-65037110-DEL-93 |
| chr5-65046767-DEL-179 | chr5-65046767-DEL-179 |
| chr5-65047040-DEL-6978 | chr5-65047041-DEL-6978 |
| chr5-65055881-DEL-135 | chr5-65055882-DEL-162 |
| chr5-65060712-DEL-193 | chr5-65060713-DEL-193 |
| chr5-65063647-DEL-72 | chr5-65063648-DEL-72 |
| chr5-65071673-DEL-498 | chr5-65071674-DEL-498 |
| chr5-65098058-DEL-853 | chr5-65098053-DEL-858 |
| chr5-65135530-DEL-51 | chr5-65135530-DEL-51 |
| chr5-65144199-DEL-605 | chr5-65144207-DEL-587 |
| chr5-65151340-DEL-85 | chr5-65151375-DEL-52 |
| chr5-65179975-DEL-98 | chr5-65179982-DEL-92 |
| chr5-65201628-DEL-62 | chr5-65201631-DEL-58 |
| chr5-65223960-DEL-1077 | chr5-65223961-DEL-1077 |
| chr5-65263258-DEL-188 | chr5-65263259-DEL-188 |
| chr5-65329808-DEL-1848 | chr5-65329809-DEL-1848 |
| chr5-65334043-DEL-111 | chr5-65334044-DEL-74 |
| chr5-65372733-DEL-204 | chr5-65372734-DEL-204 |
| chr5-65396094-DEL-82 | chr5-65396095-DEL-82 |
| chr5-65399822-DEL-98 | chr5-65399860-DEL-60 |
| chr5-65424264-DEL-86 | chr5-65424253-DEL-127 |
| chr5-65448854-DEL-1170 | chr5-65448855-DEL-1169 |
| chr5-65457245-DEL-129 | chr5-65457260-DEL-93 |
| chr5-65461360-DEL-70 | chr5-65461327-DEL-66 |
| chr5-65467293-DEL-105 | chr5-65467294-DEL-69 |
| chr5-65467578-DEL-71 | chr5-65467578-DEL-71 |
| chr5-65473031-DEL-272 | chr5-65473032-DEL-272 |
| chr5-65485803-DEL-184 | chr5-65485783-DEL-185 |
| chr5-65489896-DEL-211 | chr5-65489897-DEL-211 |
| chr5-65512013-DEL-196 | chr5-65512014-DEL-196 |
| chr5-65523451-DEL-76 | chr5-65523452-DEL-76 |
| chr5-65547015-DEL-107 | chr5-65547016-DEL-107 |
| chr5-65572342-DEL-204 | chr5-65572343-DEL-204 |
| chr5-65582951-DEL-206 | chr5-65582952-DEL-206 |
| chr5-65587086-DEL-70 | chr5-65587076-DEL-69 |
| chr5-65596956-DEL-209 | chr5-65596953-DEL-204 |
| chr5-65608448-DEL-177 | chr5-65608449-DEL-177 |
| chr5-65609396-DEL-197 | chr5-65609397-DEL-197 |
| chr5-65618843-DEL-53 | chr5-65618836-DEL-54 |
| chr5-65619472-DEL-53 | chr5-65619473-DEL-53 |
| chr5-65630784-DEL-78 | chr5-65630785-DEL-78 |
| chr5-65642815-DEL-226 | chr5-65642816-DEL-226 |
| chr5-65647183-DEL-52 | chr5-65647184-DEL-52 |
| chr5-65672620-DEL-64 | chr5-65672618-DEL-64 |
| chr5-65703225-DEL-1724 | chr5-65703226-DEL-1724 |
| chr5-65703324-DEL-208 | chr5-65703325-DEL-208 |
| chr5-65707503-DEL-95 | chr5-65707504-DEL-95 |
| chr5-65716971-DEL-51 | chr5-65716971-DEL-51 |
| chr5-65726690-DEL-227 | chr5-65726691-DEL-227 |
| chr5-65743117-DEL-206 | chr5-65743118-DEL-206 |
| chr5-65744553-DEL-57 | chr5-65744554-DEL-57 |
| chr5-65756476-DEL-70 | chr5-65756477-DEL-70 |
| chr5-65759938-DEL-2067 | chr5-65759937-DEL-2066 |
| chr5-65783950-DEL-190 | chr5-65783950-DEL-190 |
| chr5-65800297-DEL-105 | chr5-65800298-DEL-105 |
| chr5-65839475-DEL-215 | chr5-65839476-DEL-215 |
| chr5-65841805-DEL-212 | chr5-65841806-DEL-212 |
| chr5-65848158-DEL-57 | chr5-65848159-DEL-57 |
| chr5-65863287-DEL-62 | chr5-65863288-DEL-62 |
| chr5-65877873-DEL-53 | chr5-65877874-DEL-53 |
| chr5-65900226-DEL-209 | chr5-65900227-DEL-209 |
| chr5-65901452-DEL-206 | chr5-65901453-DEL-206 |
| chr5-65902247-DEL-169 | chr5-65902248-DEL-169 |
| chr5-65924492-DEL-194 | chr5-65924493-DEL-194 |
| chr5-65972660-DEL-150 | chr5-65972661-DEL-150 |
| chr5-65983068-DEL-206 | chr5-65983069-DEL-206 |
| chr5-66015651-DEL-108 | chr5-66015652-DEL-108 |
| chr5-66058826-DEL-68 | chr5-66058827-DEL-68 |
| chr5-66124139-DEL-133 | chr5-66124140-DEL-129 |
| chr5-66127076-DEL-807 | chr5-66127077-DEL-807 |
| chr5-66163601-DEL-51 | chr5-66163601-DEL-60 |
| chr5-66164824-DEL-199 | chr5-66164827-DEL-198 |
| chr5-66172613-DEL-126 | chr5-66172614-DEL-126 |
| chr5-66179177-DEL-84 | chr5-66179178-DEL-84 |
| chr5-66185410-DEL-1863 | chr5-66185411-DEL-1863 |
| chr5-66207329-DEL-173 | chr5-66207330-DEL-173 |

| | |
|---|---|
| chr5-66212870-DEL-209 | chr5-66212871-DEL-209 |
| chr5-66215264-DEL-77 | chr5-66215265-DEL-77 |
| chr5-66222658-DEL-183 | chr5-66222659-DEL-183 |
| chr5-66228420-DEL-86 | chr5-66228420-DEL-86 |
| chr5-66252294-DEL-161 | chr5-66252290-DEL-166 |
| chr5-66273143-DEL-317 | chr5-66273144-DEL-317 |
| chr5-66274003-DEL-143 | chr5-66274004-DEL-143 |
| chr5-66276293-DEL-1630 | chr5-66276296-DEL-1628 |
| chr5-66284177-DEL-117 | chr5-66284178-DEL-117 |
| chr5-66289090-DEL-52 | chr5-66289052-DEL-52 |
| chr5-66291261-DEL-70 | chr5-66291262-DEL-70 |
| chr5-66360223-DEL-80 | chr5-66360225-DEL-80 |
| chr5-66403266-DEL-181 | chr5-66403267-DEL-181 |
| chr5-66410262-DEL-202 | chr5-66410263-DEL-202 |
| chr5-66411264-DEL-123 | chr5-66411267-DEL-141 |
| chr5-66420435-DEL-214 | chr5-66420436-DEL-214 |
| chr5-66426340-DEL-60 | chr5-66426341-DEL-60 |
| chr5-66438604-DEL-301 | chr5-66438605-DEL-301 |
| chr5-66561588-DEL-122 | chr5-66561608-DEL-75 |
| chr5-66565309-DEL-53 | chr5-66565310-DEL-53 |
| chr5-66566575-DEL-211 | chr5-66566577-DEL-210 |
| chr5-66588194-DEL-86 | chr5-66588195-DEL-86 |
| chr5-66601679-DEL-628 | chr5-66601680-DEL-628 |
| chr5-66623700-DEL-212 | chr5-66623680-DEL-192 |
| chr5-66633423-DEL-55 | chr5-66633424-DEL-55 |
| chr5-66637617-DEL-107 | chr5-66637607-DEL-108 |
| chr5-66644712-DEL-146 | chr5-66644744-DEL-136 |
| chr5-66650125-DEL-7117 | chr5-66650126-DEL-7117 |
| chr5-66657498-DEL-79 | chr5-66657499-DEL-79 |
| chr5-66679061-DEL-52 | chr5-66679062-DEL-52 |
| chr5-66716178-DEL-52 | chr5-66716179-DEL-52 |
| chr5-66725902-DEL-945 | chr5-66725903-DEL-945 |
| chr5-66729160-DEL-65 | chr5-66729136-DEL-74 |
| chr5-66750601-DEL-107 | chr5-66750602-DEL-107 |
| chr5-66781214-DEL-568 | chr5-66781215-DEL-568 |
| chr5-66791522-DEL-108 | chr5-66791541-DEL-129 |
| chr5-66809742-DEL-183 | chr5-66809743-DEL-183 |
| chr5-66854450-DEL-98 | chr5-66854451-DEL-98 |
| chr5-66863123-DEL-54 | chr5-66863124-DEL-54 |
| chr5-66863984-DEL-65 | chr5-66863987-DEL-64 |
| chr5-66867107-DEL-59 | chr5-66867108-DEL-50 |
| chr5-66894135-DEL-1335 | chr5-66894137-DEL-1334 |
| chr5-66899350-DEL-110 | chr5-66899358-DEL-102 |
| chr5-66907126-DEL-153 | chr5-66907124-DEL-154 |
| chr5-66915548-DEL-82 | chr5-66915506-DEL-123 |
| chr5-66933500-DEL-64 | chr5-66933501-DEL-64 |
| chr5-66951397-DEL-367 | chr5-66951398-DEL-367 |
| chr5-66969264-DEL-57 | chr5-66969265-DEL-57 |
| chr5-66983944-DEL-73 | chr5-66983945-DEL-73 |
| chr5-66992594-DEL-54 | chr5-66992595-DEL-54 |
| chr5-67035981-DEL-66 | chr5-67035984-DEL-64 |
| chr5-67040047-DEL-58 | chr5-67040046-DEL-50 |
| chr5-67052041-DEL-94 | chr5-67052038-DEL-99 |
| chr5-67053996-DEL-213 | chr5-67053997-DEL-213 |
| chr5-67106399-DEL-62 | chr5-67106400-DEL-62 |
| chr5-67120160-DEL-56 | chr5-67120186-DEL-62 |
| chr5-67175568-DEL-90 | chr5-67175530-DEL-118 |
| chr5-67181384-DEL-52 | chr5-67181377-DEL-52 |
| chr5-67190895-DEL-58 | chr5-67190898-DEL-56 |
| chr5-67194946-DEL-6435 | chr5-67194947-DEL-6435 |
| chr5-67229844-DEL-236 | chr5-67229845-DEL-236 |
| chr5-67268576-DEL-158 | chr5-67268577-DEL-158 |
| chr5-67291660-DEL-59 | chr5-67291661-DEL-59 |
| chr5-67325936-DEL-76 | chr5-67325937-DEL-76 |
| chr5-67338074-DEL-141 | chr5-67338081-DEL-135 |
| chr5-67341630-DEL-84 | chr5-67341624-DEL-77 |
| chr5-67345648-DEL-145 | chr5-67345667-DEL-102 |
| chr5-67393245-DEL-199 | chr5-67393246-DEL-199 |
| chr5-67401204-DEL-283 | chr5-67401205-DEL-283 |
| chr5-67408205-DEL-1160 | chr5-67408206-DEL-1159 |
| chr5-67411154-DEL-253 | chr5-67411154-DEL-253 |
| chr5-67427464-DEL-6440 | chr5-67427465-DEL-6440 |
| chr5-67436297-DEL-52 | chr5-67436298-DEL-52 |
| chr5-67441629-DEL-201 | chr5-67441630-DEL-201 |
| chr5-67452672-DEL-75 | chr5-67452673-DEL-75 |
| chr5-67489447-DEL-236 | chr5-67489448-DEL-236 |
| chr5-67495923-DEL-212 | chr5-67495924-DEL-212 |
| chr5-67521954-DEL-220 | chr5-67521955-DEL-220 |
| chr5-67533823-DEL-462 | chr5-67533823-DEL-462 |
| chr5-67541614-DEL-6596 | chr5-67541615-DEL-6596 |

| | |
|---|---|
| chr5-67612343-DEL-209 | chr5-67612344-DEL-209 |
| chr5-67614657-DEL-226 | chr5-67614653-DEL-213 |
| chr5-67637589-DEL-59 | chr5-67637590-DEL-59 |
| chr5-67652351-DEL-84 | chr5-67652352-DEL-84 |
| chr5-67671705-DEL-933 | chr5-67671706-DEL-933 |
| chr5-67699746-DEL-91 | chr5-67699747-DEL-91 |
| chr5-67705755-DEL-555 | chr5-67705756-DEL-555 |
| chr5-67708052-DEL-74 | chr5-67708053-DEL-74 |
| chr5-67714453-DEL-52 | chr5-67714468-DEL-50 |
| chr5-67738352-DEL-150 | chr5-67738353-DEL-161 |
| chr5-67742117-DEL-4833 | chr5-67742118-DEL-4833 |
| chr5-67773742-DEL-238 | chr5-67773742-DEL-238 |
| chr5-67853131-DEL-52 | chr5-67853132-DEL-50 |
| chr5-67865731-DEL-6185 | chr5-67865732-DEL-6185 |
| chr5-67877494-DEL-1087 | chr5-67877478-DEL-1087 |
| chr5-67878711-DEL-659 | chr5-67878712-DEL-646 |
| chr5-67888137-DEL-80 | chr5-67888122-DEL-96 |
| chr5-67888959-DEL-520 | chr5-67888960-DEL-520 |
| chr5-67898913-DEL-60 | chr5-67898914-DEL-60 |
| chr5-67949480-DEL-64 | chr5-67949481-DEL-64 |
| chr5-67950684-DEL-452 | chr5-67950685-DEL-452 |
| chr5-67953730-DEL-379 | chr5-67953730-DEL-379 |
| chr5-67956244-DEL-96 | chr5-67956231-DEL-64 |
| chr5-67991474-DEL-103 | chr5-67991475-DEL-103 |
| chr5-67993530-DEL-55 | chr5-67993531-DEL-55 |
| chr5-68016700-DEL-81 | chr5-68016696-DEL-86 |
| chr5-68047562-DEL-66 | chr5-68047563-DEL-66 |
| chr5-68059687-DEL-66 | chr5-68059694-DEL-67 |
| chr5-68067289-DEL-6642 | chr5-68067279-DEL-6644 |
| chr5-68073381-DEL-212 | chr5-68073382-DEL-212 |
| chr5-68118645-DEL-244 | chr5-68118646-DEL-244 |
| chr5-68126893-DEL-2012 | chr5-68126894-DEL-2012 |
| chr5-68131701-DEL-118 | chr5-68131702-DEL-118 |
| chr5-68135131-DEL-2190 | chr5-68135132-DEL-2190 |
| chr5-68140995-DEL-77 | chr5-68140996-DEL-77 |
| chr5-68162583-DEL-53 | chr5-68162542-DEL-93 |
| chr5-68205281-DEL-94 | chr5-68205282-DEL-94 |
| chr5-68229605-DEL-73 | chr5-68229605-DEL-123 |
| chr5-68256870-DEL-6445 | chr5-68256871-DEL-6445 |
| chr5-68269734-DEL-106 | chr5-68269735-DEL-106 |
| chr5-68276661-DEL-145 | chr5-68276662-DEL-145 |
| chr5-68294533-DEL-6880 | chr5-68294534-DEL-6880 |
| chr5-68330050-DEL-197 | chr5-68330051-DEL-197 |
| chr5-68358524-DEL-647 | chr5-68358525-DEL-647 |
| chr5-68361221-DEL-1299 | chr5-68361222-DEL-1299 |
| chr5-68367724-DEL-498 | chr5-68367725-DEL-498 |
| chr5-68370511-DEL-9069 | chr5-68370512-DEL-9069 |
| chr5-68377360-DEL-208 | chr5-68377359-DEL-208 |
| chr5-68394572-DEL-56 | chr5-68394583-DEL-94 |
| chr5-68446339-DEL-51 | chr5-68446340-DEL-54 |
| chr5-68473564-DEL-399 | chr5-68473565-DEL-399 |
| chr5-68521835-DEL-384 | chr5-68521836-DEL-384 |
| chr5-68568162-DEL-313 | chr5-68568163-DEL-313 |
| chr5-68570329-DEL-4902 | chr5-68570330-DEL-4902 |
| chr5-68586926-DEL-8468 | chr5-68586927-DEL-8468 |
| chr5-68628945-DEL-353 | chr5-68628944-DEL-354 |
| chr5-68650272-DEL-4792 | chr5-68650262-DEL-4794 |
| chr5-68660954-DEL-160 | chr5-68660954-DEL-161 |
| chr5-68666304-DEL-557 | chr5-68666304-DEL-557 |
| chr5-68684347-DEL-196 | chr5-68684362-DEL-168 |
| chr5-68707321-DEL-418 | chr5-68707321-DEL-418 |
| chr5-68710447-DEL-6327 | chr5-68710448-DEL-6327 |
| chr5-68719842-DEL-267 | chr5-68719842-DEL-268 |
| chr5-68734214-DEL-548 | chr5-68734215-DEL-548 |
| chr5-68744386-DEL-614 | chr5-68744378-DEL-615 |
| chr5-68747408-DEL-189 | chr5-68747409-DEL-189 |
| chr5-68754354-DEL-458 | chr5-68754354-DEL-458 |
| chr5-68759066-DEL-1551 | chr5-68759068-DEL-1540 |
| chr5-68780792-DEL-66 | chr5-68780793-DEL-66 |
| chr5-68793321-DEL-213 | chr5-68793322-DEL-213 |
| chr5-68808291-DEL-70 | chr5-68808291-DEL-70 |
| chr5-68809731-DEL-400 | chr5-68809732-DEL-400 |
| chr5-68813940-DEL-383 | chr5-68813941-DEL-383 |
| chr5-68823352-DEL-597 | chr5-68823391-DEL-591 |
| chr5-68862121-DEL-698 | chr5-68862122-DEL-698 |
| chr5-68895094-DEL-6341 | chr5-68895095-DEL-6341 |
| chr5-68901181-DEL-53 | chr5-68901182-DEL-53 |
| chr5-68914811-DEL-70 | chr5-68914770-DEL-95 |
| chr5-68922570-DEL-2087 | chr5-68922571-DEL-2087 |
| chr5-68925176-DEL-56 | chr5-68925177-DEL-56 |

| | |
|---|---|
| chr5-68942029-DEL-241 | chr5-68942030-DEL-241 |
| chr5-69017073-DEL-340 | chr5-69017074-DEL-340 |
| chr5-69017937-DEL-188 | chr5-69017938-DEL-188 |
| chr5-69036097-DEL-1229 | chr5-69036098-DEL-1229 |
| chr5-69049762-DEL-66 | chr5-69049760-DEL-66 |
| chr5-69117338-DEL-238 | chr5-69117339-DEL-238 |
| chr5-69121515-DEL-62 | chr5-69121516-DEL-62 |
| chr5-69128645-DEL-619 | chr5-69128646-DEL-619 |
| chr5-69146839-DEL-393 | chr5-69146840-DEL-393 |
| chr5-69181149-DEL-490 | chr5-69181163-DEL-477 |
| chr5-69181926-DEL-400 | chr5-69181927-DEL-400 |
| chr5-69192166-DEL-1478 | chr5-69192167-DEL-1478 |
| chr5-69202785-DEL-3266 | chr5-69202786-DEL-3266 |
| chr5-69214465-DEL-75 | chr5-69214466-DEL-75 |
| chr5-69242990-DEL-103 | chr5-69242991-DEL-103 |
| chr5-69255587-DEL-51 | chr5-69255568-DEL-51 |
| chr5-69274544-DEL-7411 | chr5-69274545-DEL-7411 |
| chr5-69280923-DEL-208 | chr5-69280924-DEL-208 |
| chr5-69308466-DEL-362 | chr5-69308440-DEL-348 |
| chr5-69458228-DEL-74 | chr5-69458229-DEL-74 |
| chr5-69504636-DEL-200 | chr5-69504637-DEL-200 |
| chr5-69525629-DEL-6419 | chr5-69525630-DEL-6419 |
| chr5-69555857-DEL-113 | chr5-69555858-DEL-113 |
| chr5-69561823-DEL-88 | chr5-69561812-DEL-81 |
| chr5-69562638-DEL-85 | chr5-69562639-DEL-85 |
| chr5-69580695-DEL-422 | chr5-69580696-DEL-422 |
| chr5-69599011-DEL-652 | chr5-69599016-DEL-651 |
| chr5-69632861-DEL-6586 | chr5-69632862-DEL-6586 |
| chr5-69656397-DEL-417 | chr5-69656396-DEL-401 |
| chr5-69660643-DEL-1864 | chr5-69660644-DEL-1864 |
| chr5-69787961-DEL-89 | chr5-69787962-DEL-89 |
| chr5-69795850-DEL-3070 | chr5-69795851-DEL-3070 |
| chr5-69854660-DEL-2838 | chr5-69854661-DEL-2838 |
| chr5-69872216-DEL-165 | chr5-69872215-DEL-167 |
| chr5-69882837-DEL-10948 | chr5-69882838-DEL-10948 |
| chr5-69929149-DEL-6440 | chr5-69929150-DEL-6440 |
| chr5-69940543-DEL-50 | chr5-69940536-DEL-58 |
| chr5-69993136-DEL-425 | chr5-69993137-DEL-425 |
| chr5-69997123-DEL-530 | chr5-69997117-DEL-538 |
| chr5-70038122-DEL-1489 | chr5-70038118-DEL-1490 |
| chr5-70040130-DEL-1802 | chr5-70040131-DEL-1802 |
| chr5-70068869-DEL-6224 | chr5-70068870-DEL-6224 |
| chr5-70104168-DEL-144 | chr5-70104169-DEL-144 |
| chr5-70105331-DEL-227 | chr5-70105332-DEL-227 |
| chr5-70105864-DEL-122 | chr5-70105870-DEL-111 |
| chr5-70153000-DEL-439 | chr5-70153001-DEL-439 |
| chr5-70176314-DEL-438 | chr5-70176315-DEL-438 |
| chr5-70177277-DEL-1626 | chr5-70177278-DEL-1626 |
| chr5-70180370-DEL-1620 | chr5-70180371-DEL-1620 |
| chr5-70207640-DEL-106 | chr5-70207641-DEL-106 |
| chr5-70216801-DEL-74 | chr5-70216802-DEL-50 |
| chr5-70228391-DEL-6852 | chr5-70228392-DEL-6852 |
| chr5-70340057-DEL-3146 | chr5-70340058-DEL-3146 |
| chr5-70349956-DEL-327 | chr5-70349957-DEL-327 |
| chr5-70360344-DEL-7428 | chr5-70360345-DEL-7428 |
| chr5-70374332-DEL-630 | chr5-70374333-DEL-630 |
| chr5-70401461-DEL-67 | chr5-70401462-DEL-67 |
| chr5-70437914-DEL-71 | chr5-70437905-DEL-58 |
| chr5-70440287-DEL-175 | chr5-70440288-DEL-175 |
| chr5-70485427-DEL-62 | chr5-70485428-DEL-62 |
| chr5-70498461-DEL-203 | chr5-70498462-DEL-203 |
| chr5-70549958-DEL-6450 | chr5-70549959-DEL-6450 |
| chr5-70580879-DEL-6675 | chr5-70580880-DEL-6675 |
| chr5-70596684-DEL-248 | chr5-70596685-DEL-248 |
| chr5-70624482-DEL-3367 | chr5-70624483-DEL-3367 |
| chr5-70656094-DEL-1027 | chr5-70656096-DEL-1026 |
| chr5-70659876-DEL-53 | chr5-70659908-DEL-56 |
| chr5-70667506-DEL-290 | chr5-70667507-DEL-290 |
| chr5-70732661-DEL-266 | chr5-70732660-DEL-267 |
| chr5-70765489-DEL-1890 | chr5-70765490-DEL-1890 |
| chr5-70785476-DEL-69 | chr5-70785477-DEL-69 |
| chr5-70802955-DEL-52 | chr5-70802956-DEL-52 |
| chr5-70803964-DEL-3452 | chr5-70803965-DEL-3452 |
| chr5-70818394-DEL-3150 | chr5-70818394-DEL-3150 |
| chr5-70850483-DEL-937 | chr5-70850483-DEL-937 |
| chr5-70855026-DEL-68 | chr5-70855050-DEL-60 |
| chr5-70868883-DEL-297 | chr5-70868884-DEL-297 |
| chr5-70875163-DEL-52 | chr5-70875162-DEL-52 |
| chr5-70918826-DEL-7122 | chr5-70918827-DEL-7122 |
| chr5-70939887-DEL-698 | chr5-70939888-DEL-698 |

| | |
|---|---|
| chr5-71014164-DEL-6203 | chr5-71014165-DEL-6203 |
| chr5-71136962-DEL-196 | chr5-71136954-DEL-197 |
| chr5-71192330-DEL-627 | chr5-71192330-DEL-627 |
| chr5-71199179-DEL-177 | chr5-71199180-DEL-177 |
| chr5-71204838-DEL-963 | chr5-71204862-DEL-940 |
| chr5-71216725-DEL-476 | chr5-71216726-DEL-476 |
| chr5-71259441-DEL-176 | chr5-71259442-DEL-176 |
| chr5-71264398-DEL-85 | chr5-71264399-DEL-85 |
| chr5-71296422-DEL-55 | chr5-71296423-DEL-55 |
| chr5-71311741-DEL-63 | chr5-71311742-DEL-63 |
| chr5-71323636-DEL-2230 | chr5-71323637-DEL-2230 |
| chr5-71341151-DEL-6391 | chr5-71341152-DEL-6391 |
| chr5-71378547-DEL-190 | chr5-71378548-DEL-190 |
| chr5-71403778-DEL-142 | chr5-71403779-DEL-142 |
| chr5-71411199-DEL-6585 | chr5-71411202-DEL-6577 |
| chr5-71517862-DEL-230 | chr5-71517861-DEL-232 |
| chr5-71536097-DEL-2466 | chr5-71536098-DEL-2466 |
| chr5-71539899-DEL-6861 | chr5-71539899-DEL-6861 |
| chr5-71546098-DEL-212 | chr5-71546099-DEL-212 |
| chr5-71576799-DEL-200 | chr5-71576800-DEL-200 |
| chr5-71620754-DEL-553 | chr5-71620755-DEL-553 |
| chr5-71651989-DEL-80 | chr5-71651990-DEL-80 |
| chr5-71653406-DEL-89 | chr5-71653407-DEL-89 |
| chr5-71666497-DEL-93 | chr5-71666497-DEL-93 |
| chr5-71683073-DEL-8050 | chr5-71683074-DEL-8050 |
| chr5-71683088-DEL-206 | chr5-71683089-DEL-206 |
| chr5-71713288-DEL-7397 | chr5-71713289-DEL-7397 |
| chr5-71733843-DEL-358 | chr5-71733844-DEL-358 |
| chr5-71773329-DEL-356 | chr5-71773330-DEL-356 |
| chr5-71776401-DEL-1048 | chr5-71776441-DEL-1026 |
| chr5-71778750-DEL-657 | chr5-71778751-DEL-657 |
| chr5-71781771-DEL-186 | chr5-71781772-DEL-186 |
| chr5-71789123-DEL-168 | chr5-71789119-DEL-169 |
| chr5-71822893-DEL-56 | chr5-71822894-DEL-56 |
| chr5-71823559-DEL-121 | chr5-71823560-DEL-121 |
| chr5-71833315-DEL-266 | chr5-71833316-DEL-266 |
| chr5-71840071-DEL-96 | chr5-71840096-DEL-60 |
| chr5-71848307-DEL-164 | chr5-71848298-DEL-165 |
| chr5-71885881-DEL-232 | chr5-71885882-DEL-232 |
| chr5-71919220-DEL-845 | chr5-71919221-DEL-845 |
| chr5-71989165-DEL-7749 | chr5-71989166-DEL-7749 |
| chr5-72106512-DEL-495 | chr5-72106513-DEL-495 |
| chr5-72136174-DEL-6089 | chr5-72136175-DEL-6089 |
| chr5-72142327-DEL-60 | chr5-72142327-DEL-61 |
| chr5-72146145-DEL-100 | chr5-72146146-DEL-100 |
| chr5-72148109-DEL-811 | chr5-72148110-DEL-815 |
| chr5-72171416-DEL-456 | chr5-72171415-DEL-457 |
| chr5-72174883-DEL-243 | chr5-72174876-DEL-244 |
| chr5-72182197-DEL-793 | chr5-72182198-DEL-793 |
| chr5-72227009-DEL-7102 | chr5-72227010-DEL-7102 |
| chr5-72258091-DEL-54 | chr5-72258092-DEL-54 |
| chr5-72267461-DEL-9488 | chr5-72267459-DEL-9489 |
| chr5-72277278-DEL-1218 | chr5-72277279-DEL-1218 |
| chr5-72307936-DEL-136 | chr5-72307937-DEL-136 |
| chr5-72314507-DEL-287 | chr5-72314508-DEL-279 |
| chr5-72337246-DEL-206 | chr5-72337246-DEL-206 |
| chr5-72372695-DEL-406 | chr5-72372696-DEL-406 |
| chr5-72416619-DEL-884 | chr5-72416620-DEL-884 |
| chr5-72454965-DEL-116 | chr5-72454967-DEL-117 |
| chr5-72467838-DEL-55 | chr5-72467839-DEL-55 |
| chr5-72502076-DEL-1864 | chr5-72502077-DEL-1864 |
| chr5-72519975-DEL-1136 | chr5-72519976-DEL-1136 |
| chr5-72529727-DEL-835 | chr5-72529728-DEL-835 |
| chr5-72542764-DEL-66 | chr5-72542733-DEL-66 |
| chr5-72548449-DEL-76 | chr5-72548452-DEL-74 |
| chr5-72551537-DEL-568 | chr5-72551538-DEL-568 |
| chr5-72623320-DEL-313 | chr5-72623321-DEL-313 |
| chr5-72630495-DEL-1151 | chr5-72630496-DEL-1151 |
| chr5-72642911-DEL-324 | chr5-72642934-DEL-302 |
| chr5-72652242-DEL-73 | chr5-72652241-DEL-69 |
| chr5-72672619-DEL-920 | chr5-72672607-DEL-888 |
| chr5-72672625-DEL-71 | chr5-72672626-DEL-71 |
| chr5-72699660-DEL-6141 | chr5-72699661-DEL-6141 |
| chr5-72717912-DEL-723 | chr5-72717913-DEL-723 |
| chr5-72794825-DEL-98 | chr5-72794798-DEL-132 |
| chr5-72799294-DEL-61 | chr5-72799301-DEL-61 |
| chr5-72823842-DEL-54 | chr5-72823843-DEL-54 |
| chr5-72884962-DEL-163 | chr5-72884963-DEL-163 |
| chr5-72890937-DEL-289 | chr5-72890934-DEL-286 |
| chr5-72895293-DEL-293 | chr5-72895280-DEL-264 |

| | |
|---|---|
| chr5-72896766-DEL-403 | chr5-72896767-DEL-403 |
| chr5-72897938-DEL-94 | chr5-72897973-DEL-64 |
| chr5-72904584-DEL-188 | chr5-72904585-DEL-188 |
| chr5-72942574-DEL-264 | chr5-72942600-DEL-240 |
| chr5-72960764-DEL-304 | chr5-72960765-DEL-304 |
| chr5-72963685-DEL-136 | chr5-72963686-DEL-136 |
| chr5-72974565-DEL-271 | chr5-72974566-DEL-271 |
| chr5-72975714-DEL-92 | chr5-72975715-DEL-92 |
| chr5-72998601-DEL-128 | chr5-72998601-DEL-128 |
| chr5-73041905-DEL-173 | chr5-73041906-DEL-173 |
| chr5-73043711-DEL-181 | chr5-73043709-DEL-182 |
| chr5-73052466-DEL-570 | chr5-73052468-DEL-569 |
| chr5-73104276-DEL-1003 | chr5-73104276-DEL-1003 |
| chr5-73104643-DEL-477 | chr5-73104647-DEL-474 |
| chr5-73139020-DEL-168 | chr5-73139021-DEL-168 |
| chr5-73140725-DEL-71 | chr5-73140739-DEL-60 |
| chr5-73143951-DEL-59 | chr5-73143943-DEL-60 |
| chr5-73149766-DEL-60 | chr5-73149783-DEL-52 |
| chr5-73166990-DEL-138 | chr5-73166971-DEL-160 |
| chr5-73181031-DEL-108 | chr5-73181042-DEL-106 |
| chr5-73187831-DEL-1141 | chr5-73187832-DEL-1141 |
| chr5-73194219-DEL-4208 | chr5-73194220-DEL-4208 |
| chr5-73201561-DEL-67 | chr5-73201543-DEL-68 |
| chr5-73217845-DEL-74 | chr5-73217846-DEL-70 |
| chr5-73221091-DEL-110 | chr5-73221092-DEL-110 |
| chr5-73235722-DEL-257 | chr5-73235723-DEL-257 |
| chr5-73237639-DEL-401 | chr5-73237640-DEL-401 |
| chr5-73281602-DEL-118 | chr5-73281630-DEL-90 |
| chr5-73285245-DEL-357 | chr5-73285246-DEL-357 |
| chr5-73294128-DEL-62 | chr5-73294123-DEL-60 |
| chr5-73315441-DEL-288 | chr5-73315442-DEL-288 |
| chr5-73331169-DEL-74 | chr5-73331167-DEL-82 |
| chr5-73361858-DEL-250 | chr5-73361859-DEL-250 |
| chr5-73365872-DEL-55 | chr5-73365873-DEL-55 |
| chr5-73387599-DEL-64 | chr5-73387600-DEL-64 |
| chr5-73397217-DEL-213 | chr5-73397218-DEL-213 |
| chr5-73432051-DEL-63 | chr5-73432061-DEL-52 |
| chr5-73432685-DEL-79 | chr5-73432683-DEL-78 |
| chr5-73441950-DEL-117 | chr5-73441951-DEL-117 |
| chr5-73469194-DEL-204 | chr5-73469195-DEL-204 |
| chr5-73492905-DEL-221 | chr5-73492906-DEL-221 |
| chr5-73502368-DEL-1720 | chr5-73502369-DEL-1720 |
| chr5-73509584-DEL-81 | chr5-73509576-DEL-62 |
| chr5-73543352-DEL-400 | chr5-73543353-DEL-400 |
| chr5-73552007-DEL-358 | chr5-73552008-DEL-362 |
| chr5-73559289-DEL-67 | chr5-73559290-DEL-67 |
| chr5-73575586-DEL-78 | chr5-73575587-DEL-52 |
| chr5-73616464-DEL-7251 | chr5-73616465-DEL-7251 |
| chr5-73631116-DEL-58 | chr5-73631111-DEL-62 |
| chr5-73650814-DEL-52 | chr5-73650831-DEL-55 |
| chr5-73652873-DEL-172 | chr5-73652874-DEL-172 |
| chr5-73656747-DEL-55 | chr5-73656752-DEL-51 |
| chr5-73663695-DEL-117 | chr5-73663696-DEL-117 |
| chr5-73664472-DEL-206 | chr5-73664473-DEL-206 |
| chr5-73700567-DEL-51 | chr5-73700582-DEL-50 |
| chr5-73727781-DEL-146 | chr5-73727782-DEL-146 |
| chr5-73772300-DEL-56 | chr5-73772256-DEL-66 |
| chr5-73830868-DEL-126 | chr5-73830866-DEL-124 |
| chr5-73850223-DEL-606 | chr5-73850224-DEL-606 |
| chr5-73852628-DEL-1841 | chr5-73852629-DEL-1841 |
| chr5-73889239-DEL-52 | chr5-73889232-DEL-59 |
| chr5-73901308-DEL-59 | chr5-73901309-DEL-59 |
| chr5-73918307-DEL-116 | chr5-73918324-DEL-100 |
| chr5-73931545-DEL-313 | chr5-73931546-DEL-313 |
| chr5-73937656-DEL-67 | chr5-73937642-DEL-70 |
| chr5-73950493-DEL-75 | chr5-73950490-DEL-65 |
| chr5-73975107-DEL-62 | chr5-73975100-DEL-70 |
| chr5-74009496-DEL-98 | chr5-74009464-DEL-102 |
| chr5-74017090-DEL-332 | chr5-74017091-DEL-332 |
| chr5-74023642-DEL-1686 | chr5-74023643-DEL-1686 |
| chr5-74027170-DEL-238 | chr5-74027170-DEL-239 |
| chr5-74035294-DEL-451 | chr5-74035302-DEL-443 |
| chr5-74045372-DEL-186 | chr5-74045373-DEL-186 |
| chr5-74062158-DEL-2818 | chr5-74062159-DEL-2818 |
| chr5-74067253-DEL-151 | chr5-74067254-DEL-151 |
| chr5-74088938-DEL-831 | chr5-74088937-DEL-832 |
| chr5-74092007-DEL-67 | chr5-74091998-DEL-57 |
| chr5-74105223-DEL-774 | chr5-74105224-DEL-774 |
| chr5-74106350-DEL-1874 | chr5-74106351-DEL-1874 |
| chr5-74116003-DEL-130 | chr5-74116004-DEL-130 |

| | |
|---|---|
| chr5-74124105-DEL-2102 | chr5-74124106-DEL-2101 |
| chr5-74126600-DEL-165 | chr5-74126602-DEL-164 |
| chr5-74128596-DEL-124 | chr5-74128595-DEL-126 |
| chr5-74130045-DEL-502 | chr5-74130046-DEL-502 |
| chr5-74141730-DEL-6751 | chr5-74141731-DEL-6751 |
| chr5-74179049-DEL-875 | chr5-74179050-DEL-871 |
| chr5-74203869-DEL-202 | chr5-74203870-DEL-202 |
| chr5-74243302-DEL-739 | chr5-74243303-DEL-739 |
| chr5-74271365-DEL-56 | chr5-74271366-DEL-56 |
| chr5-74301753-DEL-142 | chr5-74301754-DEL-142 |
| chr5-74306131-DEL-58 | chr5-74306129-DEL-58 |
| chr5-74332789-DEL-53 | chr5-74332767-DEL-52 |
| chr5-74435112-DEL-78 | chr5-74435113-DEL-78 |
| chr5-74457768-DEL-650 | chr5-74457769-DEL-650 |
| chr5-74498622-DEL-2902 | chr5-74498623-DEL-2902 |
| chr5-74505238-DEL-199 | chr5-74505237-DEL-201 |
| chr5-74569532-DEL-60 | chr5-74569537-DEL-52 |
| chr5-74591993-DEL-150 | chr5-74591994-DEL-150 |
| chr5-74599588-DEL-81 | chr5-74599589-DEL-81 |
| chr5-74607172-DEL-81 | chr5-74607161-DEL-82 |
| chr5-74623796-DEL-951 | chr5-74623797-DEL-951 |
| chr5-74663326-DEL-2919 | chr5-74663327-DEL-2919 |
| chr5-74674875-DEL-380 | chr5-74674876-DEL-380 |
| chr5-74683486-DEL-172 | chr5-74683487-DEL-172 |
| chr5-74686758-DEL-595 | chr5-74686759-DEL-595 |
| chr5-74735192-DEL-1343 | chr5-74735193-DEL-1343 |
| chr5-74749700-DEL-114 | chr5-74749701-DEL-114 |
| chr5-74852587-DEL-396 | chr5-74852588-DEL-396 |
| chr5-74860038-DEL-178 | chr5-74860039-DEL-178 |
| chr5-74887364-DEL-71 | chr5-74887364-DEL-71 |
| chr5-74897035-DEL-83 | chr5-74897036-DEL-83 |
| chr5-74897573-DEL-97 | chr5-74897574-DEL-97 |
| chr5-74904656-DEL-991 | chr5-74904657-DEL-991 |
| chr5-74908937-DEL-415 | chr5-74908938-DEL-415 |
| chr5-74916916-DEL-3741 | chr5-74916917-DEL-3741 |
| chr5-74945781-DEL-2843 | chr5-74945782-DEL-2843 |
| chr5-74960178-DEL-52 | chr5-74960175-DEL-53 |
| chr5-75014101-DEL-459 | chr5-75014102-DEL-459 |
| chr5-75030707-DEL-73 | chr5-75030708-DEL-73 |
| chr5-75045241-DEL-61 | chr5-75045242-DEL-60 |
| chr5-75046883-DEL-938 | chr5-75046884-DEL-938 |
| chr5-75048936-DEL-80 | chr5-75048937-DEL-80 |
| chr5-75052892-DEL-60 | chr5-75052895-DEL-58 |
| chr5-75060393-DEL-1759 | chr5-75060394-DEL-1759 |
| chr5-75067684-DEL-144 | chr5-75067685-DEL-144 |
| chr5-75073649-DEL-58 | chr5-75073652-DEL-56 |
| chr5-75107377-DEL-436 | chr5-75107376-DEL-437 |
| chr5-75120586-DEL-76 | chr5-75120595-DEL-68 |
| chr5-75122164-DEL-168 | chr5-75122165-DEL-168 |
| chr5-75138713-DEL-423 | chr5-75138711-DEL-424 |
| chr5-75142370-DEL-1680 | chr5-75142371-DEL-1680 |
| chr5-75205681-DEL-76 | chr5-75205688-DEL-65 |
| chr5-75253430-DEL-65 | chr5-75253426-DEL-52 |
| chr5-75267565-DEL-56 | chr5-75267594-DEL-60 |
| chr5-75268310-DEL-208 | chr5-75268311-DEL-208 |
| chr5-75272871-DEL-52 | chr5-75272852-DEL-57 |
| chr5-75291075-DEL-3597 | chr5-75291076-DEL-3597 |
| chr5-75301581-DEL-1143 | chr5-75301582-DEL-1143 |
| chr5-75305664-DEL-61 | chr5-75305665-DEL-61 |
| chr5-75324704-DEL-52 | chr5-75324705-DEL-52 |
| chr5-75355198-DEL-75 | chr5-75355199-DEL-75 |
| chr5-75380893-DEL-91 | chr5-75380904-DEL-57 |
| chr5-75384329-DEL-122 | chr5-75384321-DEL-98 |
| chr5-75389158-DEL-1116 | chr5-75389159-DEL-1116 |
| chr5-75440294-DEL-4612 | chr5-75440295-DEL-4612 |
| chr5-75465140-DEL-80 | chr5-75465141-DEL-80 |
| chr5-75465924-DEL-4779 | chr5-75465925-DEL-4779 |
| chr5-75534896-DEL-55 | chr5-75534896-DEL-59 |
| chr5-75553638-DEL-81 | chr5-75553633-DEL-78 |
| chr5-75565946-DEL-58 | chr5-75565947-DEL-50 |
| chr5-75569556-DEL-71 | chr5-75569557-DEL-71 |
| chr5-75578187-DEL-1892 | chr5-75578188-DEL-1892 |
| chr5-75586440-DEL-400 | chr5-75586441-DEL-400 |
| chr5-75636723-DEL-60 | chr5-75636724-DEL-60 |
| chr5-75639939-DEL-1119 | chr5-75639940-DEL-1119 |
| chr5-75657073-DEL-62 | chr5-75657052-DEL-56 |
| chr5-75689094-DEL-79 | chr5-75689095-DEL-79 |
| chr5-75714200-DEL-98 | chr5-75714171-DEL-128 |
| chr5-75803049-DEL-61 | chr5-75803049-DEL-59 |
| chr5-75803227-DEL-2912 | chr5-75803228-DEL-2912 |

| | |
|---|---|
| chr5-75825146-DEL-60 | chr5-75825135-DEL-56 |
| chr5-75828433-DEL-66 | chr5-75828434-DEL-64 |
| chr5-75840252-DEL-158 | chr5-75840245-DEL-156 |
| chr5-75875931-DEL-54 | chr5-75875924-DEL-54 |
| chr5-75901255-DEL-106 | chr5-75901256-DEL-106 |
| chr5-75907680-DEL-62 | chr5-75907680-DEL-61 |
| chr5-75927081-DEL-91 | chr5-75927079-DEL-94 |
| chr5-75941445-DEL-98 | chr5-75941446-DEL-98 |
| chr5-75962605-DEL-62 | chr5-75962624-DEL-50 |
| chr5-76016488-DEL-115 | chr5-76016489-DEL-90 |
| chr5-76021862-DEL-77 | chr5-76021863-DEL-77 |
| chr5-76113850-DEL-445 | chr5-76113851-DEL-445 |
| chr5-76145390-DEL-61 | chr5-76145391-DEL-61 |
| chr5-76149765-DEL-157 | chr5-76149765-DEL-156 |
| chr5-76246717-DEL-504 | chr5-76246718-DEL-504 |
| chr5-76265704-DEL-211 | chr5-76265705-DEL-211 |
| chr5-76278736-DEL-168 | chr5-76278736-DEL-167 |
| chr5-76317024-DEL-402 | chr5-76317025-DEL-402 |
| chr5-76365408-DEL-6786 | chr5-76365409-DEL-6786 |
| chr5-76371556-DEL-204 | chr5-76371557-DEL-204 |
| chr5-76379729-DEL-1223 | chr5-76379730-DEL-1223 |
| chr5-76423690-DEL-4085 | chr5-76423691-DEL-4085 |
| chr5-76488062-DEL-202 | chr5-76488063-DEL-202 |
| chr5-76497270-DEL-172 | chr5-76497271-DEL-172 |
| chr5-76524425-DEL-172 | chr5-76524422-DEL-168 |
| chr5-76551033-DEL-228 | chr5-76551034-DEL-228 |
| chr5-76565493-DEL-60 | chr5-76565494-DEL-60 |
| chr5-76581600-DEL-87 | chr5-76581598-DEL-84 |
| chr5-76587851-DEL-401 | chr5-76587852-DEL-400 |
| chr5-76692238-DEL-93 | chr5-76692239-DEL-93 |
| chr5-76700934-DEL-218 | chr5-76700935-DEL-218 |
| chr5-76710426-DEL-92 | chr5-76710410-DEL-109 |
| chr5-76738165-DEL-552 | chr5-76738166-DEL-552 |
| chr5-76743622-DEL-53 | chr5-76743623-DEL-53 |
| chr5-76772241-DEL-178 | chr5-76772242-DEL-178 |
| chr5-76794536-DEL-249 | chr5-76794537-DEL-249 |
| chr5-76796647-DEL-54 | chr5-76796648-DEL-60 |
| chr5-76822654-DEL-270 | chr5-76822655-DEL-273 |
| chr5-76864903-DEL-188 | chr5-76864904-DEL-147 |
| chr5-76869502-DEL-114 | chr5-76869552-DEL-65 |
| chr5-76894504-DEL-1084 | chr5-76894505-DEL-1084 |
| chr5-76897662-DEL-81 | chr5-76897663-DEL-81 |
| chr5-76945345-DEL-91 | chr5-76945337-DEL-100 |
| chr5-76982672-DEL-212 | chr5-76982673-DEL-212 |
| chr5-77051657-DEL-194 | chr5-77051657-DEL-191 |
| chr5-77078357-DEL-8986 | chr5-77078357-DEL-8986 |
| chr5-77139507-DEL-118 | chr5-77139549-DEL-74 |
| chr5-77150162-DEL-64 | chr5-77150210-DEL-57 |
| chr5-77166817-DEL-4351 | chr5-77166818-DEL-4351 |
| chr5-77170596-DEL-78 | chr5-77170596-DEL-78 |
| chr5-77224268-DEL-441 | chr5-77224269-DEL-441 |
| chr5-77225304-DEL-68 | chr5-77225305-DEL-68 |
| chr5-77239159-DEL-57 | chr5-77239154-DEL-66 |
| chr5-77241475-DEL-210 | chr5-77241476-DEL-210 |
| chr5-77249494-DEL-80 | chr5-77249495-DEL-80 |
| chr5-77273115-DEL-538 | chr5-77273116-DEL-538 |
| chr5-77289716-DEL-203 | chr5-77289717-DEL-203 |
| chr5-77298431-DEL-77 | chr5-77298428-DEL-74 |
| chr5-77308480-DEL-90 | chr5-77308486-DEL-79 |
| chr5-77320106-DEL-1459 | chr5-77320107-DEL-1471 |
| chr5-77331055-DEL-54 | chr5-77331101-DEL-51 |
| chr5-77343737-DEL-206 | chr5-77343738-DEL-206 |
| chr5-77358533-DEL-208 | chr5-77358534-DEL-208 |
| chr5-77372697-DEL-218 | chr5-77372700-DEL-217 |
| chr5-77374098-DEL-121 | chr5-77374098-DEL-121 |
| chr5-77382491-DEL-146 | chr5-77382492-DEL-146 |
| chr5-77414725-DEL-78 | chr5-77414726-DEL-78 |
| chr5-77417047-DEL-217 | chr5-77417046-DEL-219 |
| chr5-77438189-DEL-171 | chr5-77438190-DEL-171 |
| chr5-77450081-DEL-260 | chr5-77450082-DEL-260 |
| chr5-77451203-DEL-196 | chr5-77451225-DEL-187 |
| chr5-77494273-DEL-202 | chr5-77494274-DEL-202 |
| chr5-77515087-DEL-217 | chr5-77515088-DEL-217 |
| chr5-77520555-DEL-54 | chr5-77520556-DEL-54 |
| chr5-77526526-DEL-162 | chr5-77526526-DEL-170 |
| chr5-77585194-DEL-142 | chr5-77585195-DEL-142 |
| chr5-77586778-DEL-51 | chr5-77586779-DEL-51 |
| chr5-77593808-DEL-225 | chr5-77593809-DEL-225 |
| chr5-77606869-DEL-137 | chr5-77606870-DEL-137 |
| chr5-77683087-DEL-123 | chr5-77683090-DEL-121 |

| | |
|---|---|
| chr5-77684328-DEL-6185 | chr5-77684329-DEL-6185 |
| chr5-77707469-DEL-7135 | chr5-77707470-DEL-7135 |
| chr5-77734900-DEL-1896 | chr5-77734920-DEL-1893 |
| chr5-77747143-DEL-85 | chr5-77747168-DEL-52 |
| chr5-77752791-DEL-546 | chr5-77752792-DEL-546 |
| chr5-77756504-DEL-78 | chr5-77756509-DEL-85 |
| chr5-77771417-DEL-9540 | chr5-77771418-DEL-9539 |
| chr5-77793135-DEL-274 | chr5-77793136-DEL-274 |
| chr5-77805730-DEL-211 | chr5-77805731-DEL-211 |
| chr5-77827569-DEL-53 | chr5-77827566-DEL-54 |
| chr5-77834378-DEL-191 | chr5-77834378-DEL-190 |
| chr5-77843237-DEL-6380 | chr5-77843237-DEL-6380 |
| chr5-77850333-DEL-84 | chr5-77850334-DEL-84 |
| chr5-77868165-DEL-192 | chr5-77868166-DEL-192 |
| chr5-77902688-DEL-1192 | chr5-77902687-DEL-1192 |
| chr5-77905928-DEL-2559 | chr5-77905929-DEL-2559 |
| chr5-77991196-DEL-6246 | chr5-77991197-DEL-6246 |
| chr5-78010823-DEL-61 | chr5-78010824-DEL-61 |
| chr5-78019623-DEL-115 | chr5-78019624-DEL-115 |
| chr5-78048377-DEL-1033 | chr5-78048378-DEL-1033 |
| chr5-78108224-DEL-6315 | chr5-78108225-DEL-6315 |
| chr5-78116041-DEL-6366 | chr5-78116042-DEL-6366 |
| chr5-78116299-DEL-198 | chr5-78116300-DEL-198 |
| chr5-78181483-DEL-101 | chr5-78181484-DEL-101 |
| chr5-78190377-DEL-78 | chr5-78190378-DEL-78 |
| chr5-78197056-DEL-70 | chr5-78197057-DEL-68 |
| chr5-78234624-DEL-4258 | chr5-78234625-DEL-4258 |
| chr5-78347004-DEL-6377 | chr5-78347005-DEL-6377 |
| chr5-78379318-DEL-520 | chr5-78379319-DEL-521 |
| chr5-78402779-DEL-2792 | chr5-78402780-DEL-2792 |
| chr5-78420070-DEL-1461 | chr5-78420071-DEL-1461 |
| chr5-78428413-DEL-2269 | chr5-78428414-DEL-2269 |
| chr5-78456545-DEL-1751 | chr5-78456546-DEL-1751 |
| chr5-78463698-DEL-286 | chr5-78463698-DEL-286 |
| chr5-78464392-DEL-2578 | chr5-78464392-DEL-2578 |
| chr5-78464691-DEL-717 | chr5-78464683-DEL-718 |
| chr5-78467258-DEL-517 | chr5-78467258-DEL-517 |
| chr5-78478583-DEL-6293 | chr5-78478584-DEL-6293 |
| chr5-78496774-DEL-827 | chr5-78496775-DEL-827 |
| chr5-78510329-DEL-66 | chr5-78510328-DEL-63 |
| chr5-78526989-DEL-262 | chr5-78526990-DEL-262 |
| chr5-78530902-DEL-4132 | chr5-78530903-DEL-4132 |
| chr5-78539735-DEL-89 | chr5-78539736-DEL-89 |
| chr5-78594489-DEL-52 | chr5-78594490-DEL-52 |
| chr5-78621707-DEL-5428 | chr5-78621708-DEL-5428 |
| chr5-78636672-DEL-433 | chr5-78636673-DEL-433 |
| chr5-78659489-DEL-52 | chr5-78659490-DEL-52 |
| chr5-78689553-DEL-51 | chr5-78689538-DEL-66 |
| chr5-78706234-DEL-950 | chr5-78706235-DEL-950 |
| chr5-78719932-DEL-151 | chr5-78719933-DEL-151 |
| chr5-78725670-DEL-148 | chr5-78725671-DEL-147 |
| chr5-78743845-DEL-180 | chr5-78743846-DEL-180 |
| chr5-78761079-DEL-130 | chr5-78761086-DEL-116 |
| chr5-78771062-DEL-1089 | chr5-78771062-DEL-1089 |
| chr5-78805913-DEL-52 | chr5-78805926-DEL-53 |
| chr5-78875733-DEL-74 | chr5-78875736-DEL-73 |
| chr5-78887423-DEL-76 | chr5-78887429-DEL-72 |
| chr5-78891056-DEL-114 | chr5-78891024-DEL-111 |
| chr5-78897826-DEL-54 | chr5-78897827-DEL-60 |
| chr5-78902093-DEL-52 | chr5-78902094-DEL-52 |
| chr5-78954760-DEL-90 | chr5-78954761-DEL-90 |
| chr5-78961796-DEL-479 | chr5-78961797-DEL-479 |
| chr5-78962509-DEL-54 | chr5-78962510-DEL-54 |
| chr5-78997313-DEL-66 | chr5-78997313-DEL-66 |
| chr5-79020687-DEL-52 | chr5-79020683-DEL-60 |
| chr5-79045634-DEL-70 | chr5-79045635-DEL-70 |
| chr5-79051189-DEL-154 | chr5-79051190-DEL-154 |
| chr5-79057922-DEL-150 | chr5-79057923-DEL-150 |
| chr5-79078033-DEL-800 | chr5-79078034-DEL-800 |
| chr5-79106681-DEL-108 | chr5-79106648-DEL-145 |
| chr5-79122044-DEL-144 | chr5-79122045-DEL-144 |
| chr5-79144031-DEL-227 | chr5-79144023-DEL-236 |
| chr5-79167622-DEL-4829 | chr5-79167619-DEL-4830 |
| chr5-79180706-DEL-7332 | chr5-79180707-DEL-7332 |
| chr5-79191306-DEL-78 | chr5-79191284-DEL-78 |
| chr5-79194693-DEL-2394 | chr5-79194694-DEL-2394 |
| chr5-79201778-DEL-100 | chr5-79201778-DEL-100 |
| chr5-79207218-DEL-539 | chr5-79207219-DEL-539 |
| chr5-79224955-DEL-1701 | chr5-79224948-DEL-1700 |
| chr5-79231346-DEL-127 | chr5-79231345-DEL-129 |

| | |
|---|---|
| chr5-79236108-DEL-78 | chr5-79236107-DEL-79 |
| chr5-79236283-DEL-648 | chr5-79236283-DEL-648 |
| chr5-79258931-DEL-6241 | chr5-79258931-DEL-6241 |
| chr5-79307702-DEL-53 | chr5-79307703-DEL-53 |
| chr5-79309240-DEL-1749 | chr5-79309241-DEL-1749 |
| chr5-79313513-DEL-211 | chr5-79313514-DEL-211 |
| chr5-79320592-DEL-6207 | chr5-79320593-DEL-6207 |
| chr5-79374543-DEL-53 | chr5-79374536-DEL-54 |
| chr5-79387706-DEL-1519 | chr5-79387707-DEL-1519 |
| chr5-79397309-DEL-6344 | chr5-79397299-DEL-6345 |
| chr5-79423036-DEL-157 | chr5-79423037-DEL-157 |
| chr5-79423598-DEL-401 | chr5-79423597-DEL-402 |
| chr5-79432934-DEL-1337 | chr5-79432935-DEL-1337 |
| chr5-79445515-DEL-908 | chr5-79445506-DEL-909 |
| chr5-79462284-DEL-6378 | chr5-79462268-DEL-6394 |
| chr5-79478834-DEL-51 | chr5-79478834-DEL-56 |
| chr5-79482714-DEL-511 | chr5-79482714-DEL-511 |
| chr5-79593053-DEL-1354 | chr5-79593054-DEL-1354 |
| chr5-79665359-DEL-312 | chr5-79665360-DEL-312 |
| chr5-79686302-DEL-115 | chr5-79686329-DEL-78 |
| chr5-79765465-DEL-59 | chr5-79765466-DEL-59 |
| chr5-79767817-DEL-505 | chr5-79767812-DEL-507 |
| chr5-79769238-DEL-95 | chr5-79769232-DEL-74 |
| chr5-79775411-DEL-74 | chr5-79775414-DEL-72 |
| chr5-79796645-DEL-186 | chr5-79796646-DEL-186 |
| chr5-79823690-DEL-836 | chr5-79823691-DEL-836 |
| chr5-79824452-DEL-448 | chr5-79824453-DEL-447 |
| chr5-79826851-DEL-66 | chr5-79826852-DEL-66 |
| chr5-79839213-DEL-57 | chr5-79839214-DEL-57 |
| chr5-79844334-DEL-402 | chr5-79844335-DEL-402 |
| chr5-79846751-DEL-222 | chr5-79846753-DEL-221 |
| chr5-79874522-DEL-2949 | chr5-79874522-DEL-2949 |
| chr5-79883169-DEL-67 | chr5-79883151-DEL-67 |
| chr5-79884535-DEL-6373 | chr5-79884536-DEL-6373 |
| chr5-79898116-DEL-189 | chr5-79898117-DEL-189 |
| chr5-79952430-DEL-2429 | chr5-79952431-DEL-2429 |
| chr5-79965143-DEL-613 | chr5-79965144-DEL-613 |
| chr5-79986401-DEL-1213 | chr5-79986402-DEL-1213 |
| chr5-79995926-DEL-6878 | chr5-79995927-DEL-6878 |
| chr5-80004328-DEL-454 | chr5-80004325-DEL-455 |
| chr5-80010436-DEL-55 | chr5-80010437-DEL-56 |
| chr5-80023216-DEL-597 | chr5-80023217-DEL-597 |
| chr5-80031821-DEL-1557 | chr5-80031822-DEL-1557 |
| chr5-80051845-DEL-449 | chr5-80051845-DEL-449 |
| chr5-80104532-DEL-399 | chr5-80104533-DEL-399 |
| chr5-80108746-DEL-203 | chr5-80108747-DEL-203 |
| chr5-80140986-DEL-200 | chr5-80140987-DEL-200 |
| chr5-80189296-DEL-62 | chr5-80189294-DEL-63 |
| chr5-80190358-DEL-85 | chr5-80190361-DEL-88 |
| chr5-80206054-DEL-4338 | chr5-80206055-DEL-4338 |
| chr5-80274493-DEL-726 | chr5-80274494-DEL-723 |
| chr5-80351966-DEL-260 | chr5-80351967-DEL-260 |
| chr5-80355638-DEL-311 | chr5-80355639-DEL-310 |
| chr5-80427583-DEL-6395 | chr5-80427584-DEL-6395 |
| chr5-80433676-DEL-201 | chr5-80433655-DEL-201 |
| chr5-80445449-DEL-4691 | chr5-80445450-DEL-4691 |
| chr5-80521793-DEL-58 | chr5-80521794-DEL-58 |
| chr5-80525099-DEL-70 | chr5-80525099-DEL-64 |
| chr5-80544595-DEL-81 | chr5-80544559-DEL-77 |
| chr5-80579945-DEL-83 | chr5-80579946-DEL-83 |
| chr5-80594163-DEL-82 | chr5-80594164-DEL-82 |
| chr5-80597512-DEL-110 | chr5-80597513-DEL-110 |
| chr5-80599402-DEL-6431 | chr5-80599403-DEL-6431 |
| chr5-80619009-DEL-6723 | chr5-80619010-DEL-6723 |
| chr5-80645277-DEL-74 | chr5-80645278-DEL-62 |
| chr5-80652786-DEL-54 | chr5-80652786-DEL-54 |
| chr5-80675936-DEL-82 | chr5-80675937-DEL-82 |
| chr5-80701895-DEL-63 | chr5-80701896-DEL-63 |
| chr5-80723742-DEL-2550 | chr5-80723742-DEL-2550 |
| chr5-80753948-DEL-2182 | chr5-80753949-DEL-2182 |
| chr5-80775564-DEL-63 | chr5-80775565-DEL-63 |
| chr5-80786348-DEL-278 | chr5-80786349-DEL-278 |
| chr5-80788036-DEL-794 | chr5-80788037-DEL-794 |
| chr5-80794833-DEL-647 | chr5-80794834-DEL-647 |
| chr5-80840024-DEL-480 | chr5-80840025-DEL-480 |
| chr5-80848105-DEL-63 | chr5-80848118-DEL-52 |
| chr5-80851008-DEL-209 | chr5-80851009-DEL-209 |
| chr5-80868518-DEL-66 | chr5-80868519-DEL-66 |
| chr5-80901763-DEL-558 | chr5-80901766-DEL-553 |
| chr5-80906690-DEL-67 | chr5-80906691-DEL-65 |

| | |
|---|---|
| chr5-80909158-DEL-1187 | chr5-80909158-DEL-1187 |
| chr5-80915136-DEL-112 | chr5-80915137-DEL-112 |
| chr5-80918344-DEL-78 | chr5-80918341-DEL-68 |
| chr5-80922149-DEL-2563 | chr5-80922150-DEL-2563 |
| chr5-80934065-DEL-1429 | chr5-80934066-DEL-1429 |
| chr5-80991873-DEL-6395 | chr5-80991874-DEL-6395 |
| chr5-81036596-DEL-57 | chr5-81036597-DEL-57 |
| chr5-81103355-DEL-173 | chr5-81103352-DEL-193 |
| chr5-81121152-DEL-390 | chr5-81121153-DEL-390 |
| chr5-81136474-DEL-1872 | chr5-81136475-DEL-1872 |
| chr5-81293788-DEL-69 | chr5-81293795-DEL-63 |
| chr5-81300525-DEL-66 | chr5-81300546-DEL-50 |
| chr5-81301261-DEL-529 | chr5-81301262-DEL-529 |
| chr5-81317372-DEL-829 | chr5-81317373-DEL-829 |
| chr5-81352755-DEL-6730 | chr5-81352756-DEL-6730 |
| chr5-81364339-DEL-179 | chr5-81364321-DEL-211 |
| chr5-81368252-DEL-70 | chr5-81368253-DEL-70 |
| chr5-81387827-DEL-60 | chr5-81387828-DEL-60 |
| chr5-81405114-DEL-230 | chr5-81405115-DEL-230 |
| chr5-81469684-DEL-134 | chr5-81469685-DEL-134 |
| chr5-81520782-DEL-177 | chr5-81520783-DEL-177 |
| chr5-81525540-DEL-7109 | chr5-81525541-DEL-7109 |
| chr5-81545373-DEL-2289 | chr5-81545374-DEL-2289 |
| chr5-81585526-DEL-688 | chr5-81585553-DEL-667 |
| chr5-81597551-DEL-4840 | chr5-81597552-DEL-4840 |
| chr5-81650641-DEL-202 | chr5-81650624-DEL-204 |
| chr5-81661714-DEL-198 | chr5-81661715-DEL-198 |
| chr5-81675820-DEL-163 | chr5-81675821-DEL-163 |
| chr5-81719460-DEL-6220 | chr5-81719445-DEL-6235 |
| chr5-81773471-DEL-8418 | chr5-81773472-DEL-8418 |
| chr5-81782271-DEL-276 | chr5-81782272-DEL-276 |
| chr5-81798654-DEL-72 | chr5-81798655-DEL-72 |
| chr5-81849754-DEL-6447 | chr5-81849755-DEL-6447 |
| chr5-81895237-DEL-51 | chr5-81895238-DEL-57 |
| chr5-81901227-DEL-311 | chr5-81901228-DEL-311 |
| chr5-81903016-DEL-506 | chr5-81903016-DEL-502 |
| chr5-81922784-DEL-56 | chr5-81922785-DEL-56 |
| chr5-82006675-DEL-414 | chr5-82006676-DEL-414 |
| chr5-82023819-DEL-1772 | chr5-82023812-DEL-1773 |
| chr5-82052375-DEL-768 | chr5-82052376-DEL-768 |
| chr5-82066806-DEL-562 | chr5-82066807-DEL-543 |
| chr5-82184937-DEL-6269 | chr5-82184938-DEL-6269 |
| chr5-82286542-DEL-172 | chr5-82286536-DEL-173 |
| chr5-82306402-DEL-75 | chr5-82306410-DEL-58 |
| chr5-82342141-DEL-6479 | chr5-82342142-DEL-6479 |
| chr5-82412357-DEL-56 | chr5-82412358-DEL-56 |
| chr5-82428963-DEL-2141 | chr5-82428964-DEL-2141 |
| chr5-82455286-DEL-201 | chr5-82455287-DEL-201 |
| chr5-82510934-DEL-100 | chr5-82510942-DEL-88 |
| chr5-82533041-DEL-699 | chr5-82533042-DEL-699 |
| chr5-82537859-DEL-120 | chr5-82537860-DEL-120 |
| chr5-82559975-DEL-6182 | chr5-82559976-DEL-6182 |
| chr5-82583299-DEL-80 | chr5-82583300-DEL-80 |
| chr5-82592951-DEL-1084 | chr5-82592952-DEL-1084 |
| chr5-82598875-DEL-62 | chr5-82598881-DEL-52 |
| chr5-82606760-DEL-390 | chr5-82606761-DEL-390 |
| chr5-82617566-DEL-1271 | chr5-82617556-DEL-1267 |
| chr5-82631360-DEL-67 | chr5-82631361-DEL-67 |
| chr5-82637277-DEL-173 | chr5-82637278-DEL-173 |
| chr5-82641726-DEL-1567 | chr5-82641727-DEL-1567 |
| chr5-82678743-DEL-193 | chr5-82678743-DEL-193 |
| chr5-82752872-DEL-665 | chr5-82752872-DEL-665 |
| chr5-82799113-DEL-52 | chr5-82799114-DEL-52 |
| chr5-82847801-DEL-63 | chr5-82847806-DEL-62 |
| chr5-82884827-DEL-57 | chr5-82884827-DEL-57 |
| chr5-82889884-DEL-2991 | chr5-82889885-DEL-2991 |
| chr5-82893943-DEL-491 | chr5-82893939-DEL-492 |
| chr5-82913622-DEL-53 | chr5-82913620-DEL-52 |
| chr5-82915916-DEL-72 | chr5-82915917-DEL-72 |
| chr5-82931460-DEL-264 | chr5-82931461-DEL-264 |
| chr5-82948662-DEL-96 | chr5-82948667-DEL-86 |
| chr5-82967583-DEL-390 | chr5-82967584-DEL-390 |
| chr5-82970526-DEL-56 | chr5-82970527-DEL-56 |
| chr5-83018878-DEL-2330 | chr5-83018872-DEL-2333 |
| chr5-83023778-DEL-7193 | chr5-83023779-DEL-7193 |
| chr5-83031353-DEL-615 | chr5-83031354-DEL-615 |
| chr5-83054696-DEL-6717 | chr5-83054697-DEL-6717 |
| chr5-83067348-DEL-55 | chr5-83067344-DEL-57 |
| chr5-83133118-DEL-51 | chr5-83133119-DEL-51 |
| chr5-83169000-DEL-7153 | chr5-83169001-DEL-7153 |

| | |
|---|---|
| chr5-83218171-DEL-1892 | chr5-83218172-DEL-1892 |
| chr5-83218340-DEL-429 | chr5-83218340-DEL-429 |
| chr5-83237661-DEL-83 | chr5-83237653-DEL-84 |
| chr5-83256565-DEL-6444 | chr5-83256566-DEL-6444 |
| chr5-83277882-DEL-64 | chr5-83277883-DEL-64 |
| chr5-83285812-DEL-4432 | chr5-83285813-DEL-4432 |
| chr5-83296398-DEL-473 | chr5-83296399-DEL-473 |
| chr5-83302388-DEL-1023 | chr5-83302389-DEL-1023 |
| chr5-83310093-DEL-2973 | chr5-83310094-DEL-2973 |
| chr5-83319380-DEL-875 | chr5-83319381-DEL-874 |
| chr5-83358320-DEL-117 | chr5-83358331-DEL-106 |
| chr5-83444321-DEL-624 | chr5-83444322-DEL-624 |
| chr5-83500992-DEL-82 | chr5-83500975-DEL-108 |
| chr5-83504203-DEL-117 | chr5-83504234-DEL-87 |
| chr5-83527201-DEL-1377 | chr5-83527202-DEL-1377 |
| chr5-83533153-DEL-4710 | chr5-83533154-DEL-4710 |
| chr5-83540118-DEL-93 | chr5-83540119-DEL-93 |
| chr5-83540467-DEL-2391 | chr5-83540468-DEL-2391 |
| chr5-83577843-DEL-113 | chr5-83577864-DEL-96 |
| chr5-83586223-DEL-62 | chr5-83586238-DEL-52 |
| chr5-83590638-DEL-52 | chr5-83590637-DEL-56 |
| chr5-83618125-DEL-215 | chr5-83618126-DEL-215 |
| chr5-83679637-DEL-72 | chr5-83679638-DEL-72 |
| chr5-83684619-DEL-828 | chr5-83684620-DEL-828 |
| chr5-83696844-DEL-4390 | chr5-83696845-DEL-4390 |
| chr5-83733966-DEL-5482 | chr5-83733967-DEL-5482 |
| chr5-83799939-DEL-54 | chr5-83799940-DEL-54 |
| chr5-83810744-DEL-52 | chr5-83810745-DEL-52 |
| chr5-83829522-DEL-73 | chr5-83829544-DEL-74 |
| chr5-83913032-DEL-52 | chr5-83913033-DEL-52 |
| chr5-83931356-DEL-6706 | chr5-83931357-DEL-6706 |
| chr5-83931516-DEL-212 | chr5-83931517-DEL-212 |
| chr5-83985008-DEL-170 | chr5-83985009-DEL-170 |
| chr5-83996849-DEL-51 | chr5-83996809-DEL-67 |
| chr5-84013457-DEL-62 | chr5-84013435-DEL-56 |
| chr5-84073220-DEL-52 | chr5-84073188-DEL-53 |
| chr5-84186473-DEL-68 | chr5-84186472-DEL-72 |
| chr5-84209502-DEL-60 | chr5-84209503-DEL-60 |
| chr5-84251151-DEL-348 | chr5-84251140-DEL-349 |
| chr5-84251696-DEL-209 | chr5-84251659-DEL-209 |
| chr5-84271059-DEL-1196 | chr5-84271060-DEL-1196 |
| chr5-84346407-DEL-1376 | chr5-84346423-DEL-1360 |
| chr5-84368079-DEL-1475 | chr5-84368080-DEL-1475 |
| chr5-84370895-DEL-115 | chr5-84370892-DEL-105 |
| chr5-84373643-DEL-200 | chr5-84373644-DEL-200 |
| chr5-84446306-DEL-78 | chr5-84446307-DEL-78 |
| chr5-84511061-DEL-342 | chr5-84511062-DEL-342 |
| chr5-84519551-DEL-7094 | chr5-84519552-DEL-7094 |
| chr5-84604813-DEL-106 | chr5-84604811-DEL-108 |
| chr5-84613284-DEL-56 | chr5-84613275-DEL-66 |
| chr5-84662182-DEL-1124 | chr5-84662183-DEL-1124 |
| chr5-84677466-DEL-1160 | chr5-84677467-DEL-1160 |
| chr5-84695029-DEL-226 | chr5-84695030-DEL-226 |
| chr5-84805308-DEL-2923 | chr5-84805309-DEL-2923 |
| chr5-84808960-DEL-92 | chr5-84808961-DEL-92 |
| chr5-84812361-DEL-155 | chr5-84812361-DEL-155 |
| chr5-84868350-DEL-7314 | chr5-84868351-DEL-7314 |
| chr5-84872671-DEL-242 | chr5-84872680-DEL-253 |
| chr5-84901775-DEL-213 | chr5-84901768-DEL-214 |
| chr5-84974803-DEL-55 | chr5-84974804-DEL-54 |
| chr5-85085246-DEL-9522 | chr5-85085249-DEL-9521 |
| chr5-85132402-DEL-6135 | chr5-85132402-DEL-6135 |
| chr5-85143400-DEL-476 | chr5-85143399-DEL-478 |
| chr5-85154923-DEL-2951 | chr5-85154924-DEL-2951 |
| chr5-85195196-DEL-208 | chr5-85195197-DEL-208 |
| chr5-85221727-DEL-7306 | chr5-85221734-DEL-7298 |
| chr5-85242132-DEL-360 | chr5-85242133-DEL-360 |
| chr5-85289567-DEL-2983 | chr5-85289551-DEL-2999 |
| chr5-85315146-DEL-6672 | chr5-85315147-DEL-6672 |
| chr5-85364300-DEL-1025 | chr5-85364301-DEL-1025 |
| chr5-85368979-DEL-781 | chr5-85368980-DEL-781 |
| chr5-85399488-DEL-4576 | chr5-85399492-DEL-4575 |
| chr5-85406250-DEL-85 | chr5-85406251-DEL-85 |
| chr5-85411713-DEL-6414 | chr5-85411714-DEL-6414 |
| chr5-85474546-DEL-2371 | chr5-85474547-DEL-2371 |
| chr5-85525483-DEL-205 | chr5-85525484-DEL-205 |
| chr5-85564552-DEL-1878 | chr5-85564553-DEL-1878 |
| chr5-85574254-DEL-60 | chr5-85574250-DEL-62 |
| chr5-85596520-DEL-5998 | chr5-85596521-DEL-5998 |
| chr5-85624944-DEL-7129 | chr5-85624945-DEL-7129 |

| | |
|---|---|
| chr5-85646045-DEL-112 | chr5-85646082-DEL-70 |
| chr5-85661852-DEL-3265 | chr5-85661853-DEL-3265 |
| chr5-85688500-DEL-6385 | chr5-85688501-DEL-6385 |
| chr5-85708079-DEL-6270 | chr5-85708080-DEL-6270 |
| chr5-85797505-DEL-106 | chr5-85797486-DEL-126 |
| chr5-85860633-DEL-579 | chr5-85860634-DEL-579 |
| chr5-85865568-DEL-339 | chr5-85865569-DEL-339 |
| chr5-85906594-DEL-207 | chr5-85906595-DEL-207 |
| chr5-86034776-DEL-213 | chr5-86034777-DEL-213 |
| chr5-86039137-DEL-1637 | chr5-86039139-DEL-1636 |
| chr5-86058075-DEL-84 | chr5-86058076-DEL-84 |
| chr5-86098869-DEL-1890 | chr5-86098870-DEL-1890 |
| chr5-86171630-DEL-70 | chr5-86171631-DEL-70 |
| chr5-86230922-DEL-210 | chr5-86230925-DEL-208 |
| chr5-86254682-DEL-407 | chr5-86254683-DEL-407 |
| chr5-86286436-DEL-226 | chr5-86286437-DEL-226 |
| chr5-86309449-DEL-531 | chr5-86309450-DEL-531 |
| chr5-86317159-DEL-129 | chr5-86317160-DEL-129 |
| chr5-86335802-DEL-682 | chr5-86335803-DEL-682 |
| chr5-86354344-DEL-546 | chr5-86354345-DEL-546 |
| chr5-86355320-DEL-246 | chr5-86355320-DEL-246 |
| chr5-86367794-DEL-892 | chr5-86367795-DEL-892 |
| chr5-86427772-DEL-64 | chr5-86427773-DEL-62 |
| chr5-86434666-DEL-181 | chr5-86434667-DEL-181 |
| chr5-86441827-DEL-68 | chr5-86441828-DEL-68 |
| chr5-86444643-DEL-1141 | chr5-86444644-DEL-1141 |
| chr5-86458104-DEL-508 | chr5-86458104-DEL-508 |
| chr5-86460833-DEL-53 | chr5-86460834-DEL-53 |
| chr5-86466125-DEL-482 | chr5-86466126-DEL-482 |
| chr5-86499904-DEL-101 | chr5-86499905-DEL-101 |
| chr5-86510763-DEL-6657 | chr5-86510764-DEL-6657 |
| chr5-86548799-DEL-247 | chr5-86548800-DEL-247 |
| chr5-86568920-DEL-127 | chr5-86568921-DEL-127 |
| chr5-86590548-DEL-78 | chr5-86590549-DEL-78 |
| chr5-86598172-DEL-6260 | chr5-86598173-DEL-6260 |
| chr5-86688201-DEL-173 | chr5-86688193-DEL-181 |
| chr5-86696011-DEL-6304 | chr5-86696012-DEL-6304 |
| chr5-86720690-DEL-105 | chr5-86720668-DEL-106 |
| chr5-86724987-DEL-126 | chr5-86724997-DEL-96 |
| chr5-86740510-DEL-263 | chr5-86740502-DEL-286 |
| chr5-86741979-DEL-6675 | chr5-86741980-DEL-6675 |
| chr5-86754321-DEL-6873 | chr5-86754322-DEL-6872 |
| chr5-86769147-DEL-62 | chr5-86769148-DEL-62 |
| chr5-86772559-DEL-116 | chr5-86772562-DEL-109 |
| chr5-86777858-DEL-60 | chr5-86777868-DEL-67 |
| chr5-86816760-DEL-262 | chr5-86816770-DEL-263 |
| chr5-86826327-DEL-340 | chr5-86826328-DEL-340 |
| chr5-86830664-DEL-890 | chr5-86830664-DEL-890 |
| chr5-86833576-DEL-6187 | chr5-86833577-DEL-6187 |
| chr5-86840198-DEL-1046 | chr5-86840199-DEL-1046 |
| chr5-86844132-DEL-4518 | chr5-86844133-DEL-4517 |
| chr5-86856093-DEL-620 | chr5-86856094-DEL-619 |
| chr5-86909302-DEL-59 | chr5-86909307-DEL-58 |
| chr5-86915861-DEL-205 | chr5-86915859-DEL-205 |
| chr5-86996560-DEL-389 | chr5-86996561-DEL-389 |
| chr5-87011770-DEL-561 | chr5-87011750-DEL-559 |
| chr5-87013021-DEL-397 | chr5-87013021-DEL-397 |
| chr5-87059884-DEL-4297 | chr5-87059884-DEL-4296 |
| chr5-87102463-DEL-172 | chr5-87102464-DEL-172 |
| chr5-87115602-DEL-2457 | chr5-87115603-DEL-2457 |
| chr5-87153830-DEL-770 | chr5-87153830-DEL-770 |
| chr5-87176020-DEL-204 | chr5-87176009-DEL-203 |
| chr5-87182721-DEL-7476 | chr5-87182722-DEL-7476 |
| chr5-87245686-DEL-2451 | chr5-87245687-DEL-2451 |
| chr5-87300412-DEL-4870 | chr5-87300413-DEL-4870 |
| chr5-87405433-DEL-95 | chr5-87405434-DEL-95 |
| chr5-87408945-DEL-2847 | chr5-87408946-DEL-2847 |
| chr5-87415566-DEL-248 | chr5-87415566-DEL-249 |
| chr5-87415938-DEL-4161 | chr5-87415939-DEL-4161 |
| chr5-87433032-DEL-111 | chr5-87433058-DEL-91 |
| chr5-87444071-DEL-598 | chr5-87444071-DEL-598 |
| chr5-87447171-DEL-70 | chr5-87447138-DEL-71 |
| chr5-87455982-DEL-281 | chr5-87455983-DEL-281 |
| chr5-87505535-DEL-142 | chr5-87505567-DEL-143 |
| chr5-87569387-DEL-4446 | chr5-87569388-DEL-4446 |
| chr5-87618239-DEL-60 | chr5-87618240-DEL-60 |
| chr5-87635675-DEL-90 | chr5-87635676-DEL-90 |
| chr5-87657995-DEL-2105 | chr5-87657996-DEL-2105 |
| chr5-87687616-DEL-6454 | chr5-87687598-DEL-6448 |
| chr5-87766399-DEL-473 | chr5-87766400-DEL-473 |

| | |
|---|---|
| chr5-87791297-DEL-696 | chr5-87791297-DEL-696 |
| chr5-87847621-DEL-62 | chr5-87847622-DEL-62 |
| chr5-87891868-DEL-68 | chr5-87891869-DEL-68 |
| chr5-87893270-DEL-6203 | chr5-87893271-DEL-6203 |
| chr5-87968370-DEL-78 | chr5-87968371-DEL-78 |
| chr5-87975252-DEL-66 | chr5-87975253-DEL-66 |
| chr5-87982216-DEL-502 | chr5-87982217-DEL-502 |
| chr5-88006398-DEL-1543 | chr5-88006399-DEL-1543 |
| chr5-88010806-DEL-66 | chr5-88010807-DEL-66 |
| chr5-88033844-DEL-175 | chr5-88033844-DEL-170 |
| chr5-88065757-DEL-1179 | chr5-88065758-DEL-1179 |
| chr5-88068698-DEL-786 | chr5-88068689-DEL-796 |
| chr5-88090479-DEL-55 | chr5-88090480-DEL-55 |
| chr5-88114323-DEL-410 | chr5-88114324-DEL-410 |
| chr5-88193006-DEL-193 | chr5-88193007-DEL-193 |
| chr5-88251465-DEL-768 | chr5-88251466-DEL-768 |
| chr5-88260365-DEL-117 | chr5-88260365-DEL-117 |
| chr5-88293708-DEL-83 | chr5-88293709-DEL-83 |
| chr5-88319261-DEL-598 | chr5-88319262-DEL-598 |
| chr5-88364075-DEL-97 | chr5-88364076-DEL-96 |
| chr5-88365271-DEL-10973 | chr5-88365272-DEL-10976 |
| chr5-88366122-DEL-3071 | chr5-88366123-DEL-3071 |
| chr5-88378027-DEL-6018 | chr5-88378031-DEL-6017 |
| chr5-88413357-DEL-305 | chr5-88413360-DEL-303 |
| chr5-88432835-DEL-489 | chr5-88432836-DEL-489 |
| chr5-88442897-DEL-2509 | chr5-88442899-DEL-2506 |
| chr5-88467797-DEL-54 | chr5-88467798-DEL-54 |
| chr5-88468949-DEL-154 | chr5-88468950-DEL-154 |
| chr5-88476261-DEL-54 | chr5-88476262-DEL-54 |
| chr5-88497464-DEL-150 | chr5-88497465-DEL-150 |
| chr5-88510522-DEL-304 | chr5-88510522-DEL-304 |
| chr5-88519900-DEL-194 | chr5-88519885-DEL-195 |
| chr5-88570152-DEL-199 | chr5-88570146-DEL-201 |
| chr5-88570969-DEL-2410 | chr5-88570970-DEL-2410 |
| chr5-88583998-DEL-1671 | chr5-88583999-DEL-1671 |
| chr5-88597028-DEL-187 | chr5-88597029-DEL-187 |
| chr5-88599277-DEL-154 | chr5-88599278-DEL-154 |
| chr5-88639581-DEL-163 | chr5-88639582-DEL-163 |
| chr5-88684210-DEL-66 | chr5-88684211-DEL-66 |
| chr5-88693731-DEL-7313 | chr5-88693732-DEL-7313 |
| chr5-88693743-DEL-213 | chr5-88693745-DEL-212 |
| chr5-88701411-DEL-61 | chr5-88701442-DEL-58 |
| chr5-88715724-DEL-179 | chr5-88715725-DEL-179 |
| chr5-88753427-DEL-6754 | chr5-88753428-DEL-6754 |
| chr5-88829540-DEL-365 | chr5-88829549-DEL-358 |
| chr5-88831514-DEL-66 | chr5-88831515-DEL-66 |
| chr5-88846257-DEL-197 | chr5-88846257-DEL-198 |
| chr5-88856069-DEL-423 | chr5-88856070-DEL-423 |
| chr5-88881297-DEL-5680 | chr5-88881298-DEL-5680 |
| chr5-88926551-DEL-86 | chr5-88926537-DEL-68 |
| chr5-88940213-DEL-63 | chr5-88940214-DEL-63 |
| chr5-88988738-DEL-495 | chr5-88988736-DEL-496 |
| chr5-88997213-DEL-1229 | chr5-88997213-DEL-1229 |
| chr5-89016846-DEL-6608 | chr5-89016847-DEL-6608 |
| chr5-89040075-DEL-414 | chr5-89040076-DEL-414 |
| chr5-89067092-DEL-77 | chr5-89067092-DEL-77 |
| chr5-89091885-DEL-156 | chr5-89091886-DEL-156 |
| chr5-89099592-DEL-74 | chr5-89099590-DEL-71 |
| chr5-89134545-DEL-2113 | chr5-89134546-DEL-2113 |
| chr5-89148762-DEL-7071 | chr5-89148763-DEL-7071 |
| chr5-89259772-DEL-2044 | chr5-89259773-DEL-2044 |
| chr5-89259838-DEL-6219 | chr5-89259839-DEL-6219 |
| chr5-89274096-DEL-198 | chr5-89274097-DEL-197 |
| chr5-89276673-DEL-69 | chr5-89276674-DEL-69 |
| chr5-89279691-DEL-231 | chr5-89279689-DEL-232 |
| chr5-89302209-DEL-51 | chr5-89302210-DEL-51 |
| chr5-89328788-DEL-64 | chr5-89328788-DEL-64 |
| chr5-89351629-DEL-5560 | chr5-89351630-DEL-5560 |
| chr5-89457660-DEL-208 | chr5-89457661-DEL-208 |
| chr5-89476057-DEL-214 | chr5-89476058-DEL-214 |
| chr5-89482466-DEL-1292 | chr5-89482467-DEL-1292 |
| chr5-89487032-DEL-396 | chr5-89487033-DEL-396 |
| chr5-89487599-DEL-68 | chr5-89487602-DEL-60 |
| chr5-89507205-DEL-111 | chr5-89507206-DEL-111 |
| chr5-89515427-DEL-106 | chr5-89515428-DEL-64 |
| chr5-89536794-DEL-962 | chr5-89536795-DEL-962 |
| chr5-89543323-DEL-6190 | chr5-89543325-DEL-6190 |
| chr5-89552001-DEL-427 | chr5-89552002-DEL-427 |
| chr5-89562360-DEL-269 | chr5-89562361-DEL-269 |
| chr5-89575425-DEL-6907 | chr5-89575426-DEL-6907 |

| | |
|---|---|
| chr5-89599879-DEL-62 | chr5-89599880-DEL-62 |
| chr5-89640383-DEL-89 | chr5-89640375-DEL-84 |
| chr5-89679761-DEL-2766 | chr5-89679762-DEL-2766 |
| chr5-89702932-DEL-210 | chr5-89702933-DEL-210 |
| chr5-89704541-DEL-1892 | chr5-89704542-DEL-1892 |
| chr5-89713556-DEL-62 | chr5-89713557-DEL-62 |
| chr5-89719087-DEL-99 | chr5-89719088-DEL-99 |
| chr5-89726211-DEL-61 | chr5-89726212-DEL-61 |
| chr5-89750319-DEL-354 | chr5-89750320-DEL-354 |
| chr5-89788707-DEL-2716 | chr5-89788708-DEL-2716 |
| chr5-89811097-DEL-114 | chr5-89811103-DEL-108 |
| chr5-89815483-DEL-2324 | chr5-89815483-DEL-2324 |
| chr5-89836999-DEL-1191 | chr5-89837000-DEL-1191 |
| chr5-89897945-DEL-133 | chr5-89897946-DEL-133 |
| chr5-89903646-DEL-91 | chr5-89903647-DEL-91 |
| chr5-89904488-DEL-90 | chr5-89904489-DEL-90 |
| chr5-89933181-DEL-469 | chr5-89933182-DEL-469 |
| chr5-89949535-DEL-2486 | chr5-89949536-DEL-2486 |
| chr5-89978574-DEL-428 | chr5-89978575-DEL-428 |
| chr5-89999665-DEL-3268 | chr5-89999666-DEL-3268 |
| chr5-90082413-DEL-54 | chr5-90082403-DEL-75 |
| chr5-90100796-DEL-408 | chr5-90100797-DEL-408 |
| chr5-90101948-DEL-184 | chr5-90101949-DEL-184 |
| chr5-90106374-DEL-1289 | chr5-90106375-DEL-1289 |
| chr5-90128526-DEL-60 | chr5-90128527-DEL-60 |
| chr5-90142085-DEL-929 | chr5-90142086-DEL-929 |
| chr5-90147595-DEL-247 | chr5-90147608-DEL-235 |
| chr5-90184843-DEL-181 | chr5-90184844-DEL-181 |
| chr5-90187655-DEL-694 | chr5-90187655-DEL-695 |
| chr5-90196203-DEL-501 | chr5-90196198-DEL-498 |
| chr5-90233539-DEL-230 | chr5-90233535-DEL-231 |
| chr5-90237717-DEL-58 | chr5-90237733-DEL-50 |
| chr5-90242687-DEL-220 | chr5-90242690-DEL-220 |
| chr5-90243243-DEL-6395 | chr5-90243242-DEL-6395 |
| chr5-90249839-DEL-126 | chr5-90249849-DEL-120 |
| chr5-90263196-DEL-290 | chr5-90263197-DEL-290 |
| chr5-90291478-DEL-132 | chr5-90291479-DEL-132 |
| chr5-90294191-DEL-1042 | chr5-90294192-DEL-1042 |
| chr5-90335940-DEL-212 | chr5-90335936-DEL-210 |
| chr5-90337856-DEL-177 | chr5-90337857-DEL-177 |
| chr5-90355774-DEL-1109 | chr5-90355776-DEL-1102 |
| chr5-90406496-DEL-872 | chr5-90406497-DEL-872 |
| chr5-90414361-DEL-178 | chr5-90414362-DEL-178 |
| chr5-90439931-DEL-70 | chr5-90439932-DEL-70 |
| chr5-90454655-DEL-202 | chr5-90454654-DEL-204 |
| chr5-90455885-DEL-1266 | chr5-90455885-DEL-1266 |
| chr5-90534782-DEL-151 | chr5-90534781-DEL-132 |
| chr5-90558186-DEL-239 | chr5-90558187-DEL-239 |
| chr5-90565216-DEL-72 | chr5-90565217-DEL-72 |
| chr5-90566385-DEL-733 | chr5-90566386-DEL-733 |
| chr5-90585135-DEL-198 | chr5-90585139-DEL-195 |
| chr5-90594935-DEL-69 | chr5-90594922-DEL-70 |
| chr5-90620899-DEL-877 | chr5-90620900-DEL-877 |
| chr5-90664599-DEL-87 | chr5-90664589-DEL-77 |
| chr5-90676752-DEL-426 | chr5-90676753-DEL-426 |
| chr5-90680777-DEL-9945 | chr5-90680778-DEL-9945 |
| chr5-90715303-DEL-241 | chr5-90715304-DEL-241 |
| chr5-90760622-DEL-90 | chr5-90760627-DEL-86 |
| chr5-90777969-DEL-167 | chr5-90777955-DEL-168 |
| chr5-90780298-DEL-66 | chr5-90780299-DEL-66 |
| chr5-90794147-DEL-53 | chr5-90794150-DEL-51 |
| chr5-90815812-DEL-88 | chr5-90815813-DEL-88 |
| chr5-90827431-DEL-372 | chr5-90827432-DEL-372 |
| chr5-90887906-DEL-92 | chr5-90887907-DEL-92 |
| chr5-90891581-DEL-252 | chr5-90891582-DEL-252 |
| chr5-90908110-DEL-68 | chr5-90908095-DEL-84 |
| chr5-90908715-DEL-53 | chr5-90908710-DEL-59 |
| chr5-90946277-DEL-177 | chr5-90946278-DEL-177 |
| chr5-90960317-DEL-6873 | chr5-90960318-DEL-6873 |
| chr5-90975009-DEL-1486 | chr5-90975010-DEL-1486 |
| chr5-91013030-DEL-374 | chr5-91013031-DEL-374 |
| chr5-91052522-DEL-174 | chr5-91052523-DEL-174 |
| chr5-91072775-DEL-5525 | chr5-91072776-DEL-5525 |
| chr5-91101490-DEL-3231 | chr5-91101491-DEL-3231 |
| chr5-91165532-DEL-52 | chr5-91165512-DEL-51 |
| chr5-91167618-DEL-183 | chr5-91167619-DEL-183 |
| chr5-91169248-DEL-581 | chr5-91169249-DEL-581 |
| chr5-91175564-DEL-84 | chr5-91175564-DEL-84 |
| chr5-91203273-DEL-6497 | chr5-91203274-DEL-6497 |
| chr5-91223365-DEL-53 | chr5-91223347-DEL-54 |

| | |
|---|---|
| chr5-91226409-DEL-874 | chr5-91226411-DEL-873 |
| chr5-91268457-DEL-712 | chr5-91268458-DEL-712 |
| chr5-91307091-DEL-78 | chr5-91307092-DEL-86 |
| chr5-91319857-DEL-59 | chr5-91319861-DEL-67 |
| chr5-91332218-DEL-107 | chr5-91332210-DEL-65 |
| chr5-91337885-DEL-63 | chr5-91337884-DEL-64 |
| chr5-91345165-DEL-77 | chr5-91345135-DEL-90 |
| chr5-91365380-DEL-483 | chr5-91365381-DEL-483 |
| chr5-91370427-DEL-162 | chr5-91370428-DEL-162 |
| chr5-91374625-DEL-7267 | chr5-91374626-DEL-7267 |
| chr5-91392231-DEL-106 | chr5-91392249-DEL-62 |
| chr5-91395405-DEL-4623 | chr5-91395406-DEL-4623 |
| chr5-91452571-DEL-85 | chr5-91452572-DEL-85 |
| chr5-91479341-DEL-604 | chr5-91479343-DEL-605 |
| chr5-91535640-DEL-1928 | chr5-91535641-DEL-1928 |
| chr5-91542105-DEL-819 | chr5-91542106-DEL-819 |
| chr5-91559533-DEL-1059 | chr5-91559534-DEL-1059 |
| chr5-91564812-DEL-7015 | chr5-91564830-DEL-7023 |
| chr5-91586496-DEL-137 | chr5-91586497-DEL-137 |
| chr5-91611080-DEL-489 | chr5-91611077-DEL-487 |
| chr5-91612023-DEL-87 | chr5-91612024-DEL-87 |
| chr5-91628001-DEL-61 | chr5-91627996-DEL-62 |
| chr5-91688323-DEL-97 | chr5-91688324-DEL-93 |
| chr5-91690354-DEL-2151 | chr5-91690354-DEL-2152 |
| chr5-91698871-DEL-1442 | chr5-91698851-DEL-1445 |
| chr5-91705880-DEL-79 | chr5-91705879-DEL-80 |
| chr5-91778810-DEL-6580 | chr5-91778811-DEL-6580 |
| chr5-91786905-DEL-151 | chr5-91786906-DEL-151 |
| chr5-91798719-DEL-54 | chr5-91798716-DEL-64 |
| chr5-91828371-DEL-6399 | chr5-91828372-DEL-6399 |
| chr5-91835015-DEL-3380 | chr5-91835016-DEL-3380 |
| chr5-91847020-DEL-66 | chr5-91847009-DEL-81 |
| chr5-91869439-DEL-117 | chr5-91869440-DEL-117 |
| chr5-91885378-DEL-624 | chr5-91885380-DEL-626 |
| chr5-91918970-DEL-206 | chr5-91918949-DEL-207 |
| chr5-91922316-DEL-53 | chr5-91922293-DEL-59 |
| chr5-91927691-DEL-236 | chr5-91927692-DEL-236 |
| chr5-91953868-DEL-98 | chr5-91953868-DEL-98 |
| chr5-91979941-DEL-6266 | chr5-91979942-DEL-6266 |
| chr5-91997554-DEL-57 | chr5-91997555-DEL-57 |
| chr5-91998694-DEL-400 | chr5-91998695-DEL-400 |
| chr5-92001910-DEL-422 | chr5-92001911-DEL-420 |
| chr5-92003397-DEL-52 | chr5-92003398-DEL-52 |
| chr5-92007059-DEL-232 | chr5-92007060-DEL-232 |
| chr5-92016389-DEL-62 | chr5-92016390-DEL-62 |
| chr5-92035277-DEL-247 | chr5-92035278-DEL-247 |
| chr5-92072676-DEL-6294 | chr5-92072677-DEL-6294 |
| chr5-92079728-DEL-205 | chr5-92079729-DEL-205 |
| chr5-92081179-DEL-1719 | chr5-92081179-DEL-1719 |
| chr5-92120738-DEL-399 | chr5-92120739-DEL-399 |
| chr5-92138964-DEL-71 | chr5-92138968-DEL-71 |
| chr5-92189042-DEL-76 | chr5-92189056-DEL-88 |
| chr5-92236616-DEL-96 | chr5-92236617-DEL-96 |
| chr5-92244615-DEL-108 | chr5-92244616-DEL-108 |
| chr5-92259454-DEL-177 | chr5-92259454-DEL-159 |
| chr5-92288074-DEL-751 | chr5-92288075-DEL-751 |
| chr5-92335353-DEL-76 | chr5-92335360-DEL-70 |
| chr5-92364064-DEL-206 | chr5-92364065-DEL-206 |
| chr5-92371889-DEL-528 | chr5-92371890-DEL-528 |
| chr5-92385075-DEL-55 | chr5-92385054-DEL-78 |
| chr5-92391515-DEL-423 | chr5-92391516-DEL-423 |
| chr5-92422935-DEL-117 | chr5-92422936-DEL-117 |
| chr5-92428126-DEL-203 | chr5-92428127-DEL-203 |
| chr5-92446220-DEL-240 | chr5-92446220-DEL-241 |
| chr5-92500432-DEL-212 | chr5-92500433-DEL-212 |
| chr5-92538426-DEL-72 | chr5-92538427-DEL-72 |
| chr5-92543368-DEL-203 | chr5-92543369-DEL-203 |
| chr5-92585256-DEL-400 | chr5-92585257-DEL-400 |
| chr5-92608963-DEL-60 | chr5-92608964-DEL-60 |
| chr5-92623958-DEL-5999 | chr5-92623959-DEL-5999 |
| chr5-92645104-DEL-183 | chr5-92645105-DEL-183 |
| chr5-92660472-DEL-7348 | chr5-92660473-DEL-7348 |
| chr5-92661200-DEL-220 | chr5-92661196-DEL-221 |
| chr5-92679512-DEL-383 | chr5-92679513-DEL-383 |
| chr5-92690262-DEL-190 | chr5-92690253-DEL-186 |
| chr5-92690808-DEL-8837 | chr5-92690809-DEL-8837 |
| chr5-92704492-DEL-206 | chr5-92704493-DEL-206 |
| chr5-92707353-DEL-3798 | chr5-92707354-DEL-3798 |
| chr5-92724368-DEL-61 | chr5-92724356-DEL-59 |
| chr5-92762396-DEL-7236 | chr5-92762397-DEL-7236 |

| | |
|---|---|
| chr5-92777670-DEL-72 | chr5-92777644-DEL-87 |
| chr5-92789172-DEL-197 | chr5-92789173-DEL-197 |
| chr5-92811244-DEL-96 | chr5-92811221-DEL-120 |
| chr5-92848671-DEL-334 | chr5-92848654-DEL-349 |
| chr5-92853012-DEL-143 | chr5-92853040-DEL-126 |
| chr5-92882561-DEL-162 | chr5-92882562-DEL-162 |
| chr5-92897996-DEL-179 | chr5-92897997-DEL-179 |
| chr5-92903499-DEL-160 | chr5-92903500-DEL-160 |
| chr5-92916830-DEL-66 | chr5-92916831-DEL-66 |
| chr5-92918202-DEL-73 | chr5-92918203-DEL-73 |
| chr5-92938020-DEL-1265 | chr5-92938021-DEL-1265 |
| chr5-92942464-DEL-120 | chr5-92942465-DEL-120 |
| chr5-92965688-DEL-176 | chr5-92965702-DEL-163 |
| chr5-92995108-DEL-96 | chr5-92995142-DEL-74 |
| chr5-93002001-DEL-173 | chr5-93002002-DEL-173 |
| chr5-93003408-DEL-246 | chr5-93003409-DEL-246 |
| chr5-93034562-DEL-96 | chr5-93034550-DEL-66 |
| chr5-93067645-DEL-69 | chr5-93067646-DEL-69 |
| chr5-93068079-DEL-574 | chr5-93068080-DEL-574 |
| chr5-93147133-DEL-66 | chr5-93147128-DEL-60 |
| chr5-93159591-DEL-172 | chr5-93159592-DEL-172 |
| chr5-93167296-DEL-97 | chr5-93167259-DEL-84 |
| chr5-93174517-DEL-280 | chr5-93174518-DEL-280 |
| chr5-93182561-DEL-222 | chr5-93182562-DEL-222 |
| chr5-93186374-DEL-126 | chr5-93186350-DEL-134 |
| chr5-93202460-DEL-87 | chr5-93202441-DEL-107 |
| chr5-93214657-DEL-3184 | chr5-93214658-DEL-3184 |
| chr5-93227948-DEL-226 | chr5-93227949-DEL-226 |
| chr5-93253112-DEL-233 | chr5-93253113-DEL-233 |
| chr5-93255967-DEL-74 | chr5-93255968-DEL-74 |
| chr5-93318296-DEL-217 | chr5-93318293-DEL-217 |
| chr5-93342142-DEL-275 | chr5-93342138-DEL-276 |
| chr5-93369162-DEL-237 | chr5-93369163-DEL-237 |
| chr5-93374490-DEL-56 | chr5-93374459-DEL-62 |
| chr5-93430537-DEL-53 | chr5-93430538-DEL-53 |
| chr5-93440651-DEL-244 | chr5-93440649-DEL-245 |
| chr5-93457618-DEL-6226 | chr5-93457619-DEL-6226 |
| chr5-93466947-DEL-264 | chr5-93466948-DEL-264 |
| chr5-93479965-DEL-6425 | chr5-93479965-DEL-6425 |
| chr5-93495657-DEL-206 | chr5-93495657-DEL-206 |
| chr5-93555210-DEL-6192 | chr5-93555211-DEL-6192 |
| chr5-93566730-DEL-387 | chr5-93566731-DEL-387 |
| chr5-93588031-DEL-6571 | chr5-93588032-DEL-6571 |
| chr5-93601541-DEL-206 | chr5-93601542-DEL-206 |
| chr5-93614813-DEL-152 | chr5-93614832-DEL-151 |
| chr5-93624818-DEL-59 | chr5-93624859-DEL-52 |
| chr5-93698366-DEL-6850 | chr5-93698366-DEL-6850 |
| chr5-93758701-DEL-114 | chr5-93758702-DEL-114 |
| chr5-96004451-DEL-7753 | chr5-96004452-DEL-7753 |
| chr5-96070581-DEL-866 | chr5-96070582-DEL-867 |
| chr5-96074834-DEL-6591 | chr5-96074835-DEL-6591 |
| chr5-96075089-DEL-208 | chr5-96075090-DEL-208 |
| chr5-96131599-DEL-52 | chr5-96131600-DEL-52 |
| chr5-96161549-DEL-7691 | chr5-96161550-DEL-7691 |
| chr5-96167660-DEL-624 | chr5-96167661-DEL-624 |
| chr5-96204429-DEL-66 | chr5-96204430-DEL-66 |
| chr5-96211239-DEL-66 | chr5-96211239-DEL-66 |
| chr5-96265368-DEL-104 | chr5-96265369-DEL-104 |
| chr5-96267431-DEL-6443 | chr5-96267432-DEL-6443 |
| chr5-96328156-DEL-2753 | chr5-96328157-DEL-2753 |
| chr5-96351527-DEL-229 | chr5-96351528-DEL-229 |
| chr5-96391349-DEL-64 | chr5-96391350-DEL-64 |
| chr5-96394809-DEL-194 | chr5-96394810-DEL-194 |
| chr5-96420594-DEL-3478 | chr5-96420595-DEL-3478 |
| chr5-96427354-DEL-92 | chr5-96427355-DEL-92 |
| chr5-96440761-DEL-208 | chr5-96440761-DEL-208 |
| chr5-96444601-DEL-3412 | chr5-96444601-DEL-3412 |
| chr5-96510356-DEL-59 | chr5-96510363-DEL-58 |
| chr5-96515669-DEL-90 | chr5-96515667-DEL-94 |
| chr5-96533317-DEL-270 | chr5-96533318-DEL-270 |
| chr5-96546587-DEL-488 | chr5-96546588-DEL-488 |
| chr5-96581993-DEL-54 | chr5-96581994-DEL-54 |
| chr5-96587106-DEL-54 | chr5-96587107-DEL-54 |
| chr5-96592538-DEL-1071 | chr5-96592539-DEL-1071 |
| chr5-96615735-DEL-1874 | chr5-96615735-DEL-1874 |
| chr5-96633356-DEL-5190 | chr5-96633357-DEL-5190 |
| chr5-96644685-DEL-54 | chr5-96644650-DEL-72 |
| chr5-96660341-DEL-154 | chr5-96660342-DEL-154 |
| chr5-96708888-DEL-6392 | chr5-96708873-DEL-6393 |
| chr5-96735681-DEL-62 | chr5-96735676-DEL-60 |

| | |
|---|---|
| chr5-96741977-DEL-1183 | chr5-96741978-DEL-1183 |
| chr5-96753764-DEL-491 | chr5-96753765-DEL-491 |
| chr5-96760458-DEL-165 | chr5-96760481-DEL-141 |
| chr5-96763485-DEL-273 | chr5-96763486-DEL-273 |
| chr5-96803030-DEL-79 | chr5-96803031-DEL-79 |
| chr5-96819998-DEL-358 | chr5-96819999-DEL-358 |
| chr5-96875963-DEL-52 | chr5-96875964-DEL-52 |
| chr5-96877458-DEL-105 | chr5-96877459-DEL-99 |
| chr5-96889495-DEL-162 | chr5-96889513-DEL-162 |
| chr5-96904297-DEL-59 | chr5-96904298-DEL-59 |
| chr5-96946731-DEL-1899 | chr5-96946732-DEL-1899 |
| chr5-96964844-DEL-643 | chr5-96964845-DEL-643 |
| chr5-96977835-DEL-59 | chr5-96977836-DEL-59 |
| chr5-96979169-DEL-183 | chr5-96979168-DEL-181 |
| chr5-96983095-DEL-618 | chr5-96983096-DEL-618 |
| chr5-96995029-DEL-288 | chr5-96995066-DEL-252 |
| chr5-97001067-DEL-57 | chr5-97001116-DEL-58 |
| chr5-97042279-DEL-815 | chr5-97042280-DEL-815 |
| chr5-97050449-DEL-51 | chr5-97050450-DEL-51 |
| chr5-97056503-DEL-188 | chr5-97056504-DEL-185 |
| chr5-97081958-DEL-72 | chr5-97081971-DEL-60 |
| chr5-97095880-DEL-68 | chr5-97095871-DEL-68 |
| chr5-97103367-DEL-54 | chr5-97103368-DEL-54 |
| chr5-97106748-DEL-202 | chr5-97106749-DEL-202 |
| chr5-97125855-DEL-85 | chr5-97125856-DEL-91 |
| chr5-97136460-DEL-229 | chr5-97136461-DEL-229 |
| chr5-97158731-DEL-52 | chr5-97158730-DEL-50 |
| chr5-97198441-DEL-694 | chr5-97198442-DEL-694 |
| chr5-97264802-DEL-86 | chr5-97264803-DEL-86 |
| chr5-97291036-DEL-76 | chr5-97291037-DEL-76 |
| chr5-97297723-DEL-880 | chr5-97297724-DEL-880 |
| chr5-97310048-DEL-59 | chr5-97310045-DEL-63 |
| chr5-97318916-DEL-505 | chr5-97318917-DEL-505 |
| chr5-97344515-DEL-58 | chr5-97344516-DEL-58 |
| chr5-97363200-DEL-83 | chr5-97363201-DEL-83 |
| chr5-97400892-DEL-445 | chr5-97400893-DEL-445 |
| chr5-97419953-DEL-1259 | chr5-97419954-DEL-1259 |
| chr5-97427748-DEL-204 | chr5-97427749-DEL-204 |
| chr5-97428614-DEL-54 | chr5-97428582-DEL-64 |
| chr5-97429331-DEL-1237 | chr5-97429332-DEL-1237 |
| chr5-97434946-DEL-66 | chr5-97434957-DEL-50 |
| chr5-97444586-DEL-400 | chr5-97444587-DEL-400 |
| chr5-97467455-DEL-75 | chr5-97467463-DEL-78 |
| chr5-97467763-DEL-2773 | chr5-97467764-DEL-2773 |
| chr5-97494408-DEL-124 | chr5-97494409-DEL-124 |
| chr5-97542595-DEL-495 | chr5-97542596-DEL-495 |
| chr5-97543363-DEL-447 | chr5-97543364-DEL-445 |
| chr5-97543876-DEL-1702 | chr5-97543877-DEL-1702 |
| chr5-97559739-DEL-57 | chr5-97559740-DEL-57 |
| chr5-97606674-DEL-315 | chr5-97606673-DEL-316 |
| chr5-97635960-DEL-3141 | chr5-97635961-DEL-3141 |
| chr5-97650425-DEL-54 | chr5-97650426-DEL-54 |
| chr5-97667557-DEL-6475 | chr5-97667558-DEL-6475 |
| chr5-97720901-DEL-98 | chr5-97720949-DEL-84 |
| chr5-97756275-DEL-2588 | chr5-97756276-DEL-2588 |
| chr5-97763503-DEL-53 | chr5-97763504-DEL-53 |
| chr5-97799132-DEL-105 | chr5-97799129-DEL-103 |
| chr5-97865434-DEL-222 | chr5-97865435-DEL-222 |
| chr5-97866518-DEL-120 | chr5-97866519-DEL-120 |
| chr5-97889433-DEL-107 | chr5-97889434-DEL-107 |
| chr5-97915307-DEL-58 | chr5-97915274-DEL-50 |
| chr5-97922670-DEL-130 | chr5-97922715-DEL-110 |
| chr5-97929789-DEL-102 | chr5-97929793-DEL-99 |
| chr5-97937788-DEL-70 | chr5-97937759-DEL-58 |
| chr5-97939587-DEL-91 | chr5-97939588-DEL-91 |
| chr5-97946398-DEL-52 | chr5-97946399-DEL-52 |
| chr5-97950686-DEL-3234 | chr5-97950685-DEL-3235 |
| chr5-97953777-DEL-67 | chr5-97953778-DEL-68 |
| chr5-97977829-DEL-150 | chr5-97977802-DEL-142 |
| chr5-98041760-DEL-52 | chr5-98041757-DEL-56 |
| chr5-98074220-DEL-58 | chr5-98074221-DEL-58 |
| chr5-98081024-DEL-58 | chr5-98081006-DEL-77 |
| chr5-98093810-DEL-82 | chr5-98093802-DEL-80 |
| chr5-98136571-DEL-7125 | chr5-98136572-DEL-7125 |
| chr5-98175755-DEL-302 | chr5-98175756-DEL-302 |
| chr5-98221089-DEL-148 | chr5-98221054-DEL-145 |
| chr5-98222048-DEL-504 | chr5-98222049-DEL-504 |
| chr5-98243106-DEL-643 | chr5-98243107-DEL-641 |
| chr5-98257781-DEL-67 | chr5-98257782-DEL-67 |
| chr5-98260890-DEL-115 | chr5-98260891-DEL-115 |

| | |
|---|---|
| chr5-98298087-DEL-68 | chr5-98298062-DEL-52 |
| chr5-98339211-DEL-168 | chr5-98339212-DEL-168 |
| chr5-98350551-DEL-62 | chr5-98350552-DEL-62 |
| chr5-98352573-DEL-7228 | chr5-98352574-DEL-7228 |
| chr5-98364549-DEL-206 | chr5-98364550-DEL-205 |
| chr5-98369269-DEL-147 | chr5-98369270-DEL-147 |
| chr5-98435737-DEL-201 | chr5-98435738-DEL-201 |
| chr5-98456890-DEL-216 | chr5-98456891-DEL-216 |
| chr5-98482666-DEL-653 | chr5-98482668-DEL-652 |
| chr5-98564206-DEL-57 | chr5-98564206-DEL-57 |
| chr5-98571353-DEL-197 | chr5-98571361-DEL-198 |
| chr5-98582654-DEL-210 | chr5-98582655-DEL-210 |
| chr5-98601465-DEL-791 | chr5-98601466-DEL-790 |
| chr5-98623319-DEL-252 | chr5-98623320-DEL-252 |
| chr5-98635166-DEL-4452 | chr5-98635167-DEL-4452 |
| chr5-98643553-DEL-124 | chr5-98643554-DEL-124 |
| chr5-98644351-DEL-337 | chr5-98644352-DEL-337 |
| chr5-98675883-DEL-892 | chr5-98675884-DEL-892 |
| chr5-98678874-DEL-198 | chr5-98678875-DEL-198 |
| chr5-98711887-DEL-66 | chr5-98711874-DEL-115 |
| chr5-98713232-DEL-223 | chr5-98713227-DEL-223 |
| chr5-98751867-DEL-110 | chr5-98751868-DEL-110 |
| chr5-98775404-DEL-240 | chr5-98775405-DEL-240 |
| chr5-98828528-DEL-52 | chr5-98828525-DEL-53 |
| chr5-98864583-DEL-75 | chr5-98864583-DEL-75 |
| chr5-98905061-DEL-322 | chr5-98905062-DEL-312 |
| chr5-98945387-DEL-62 | chr5-98945392-DEL-68 |
| chr5-98953810-DEL-119 | chr5-98953811-DEL-119 |
| chr5-98970861-DEL-88 | chr5-98970862-DEL-88 |
| chr5-98978591-DEL-53 | chr5-98978582-DEL-63 |
| chr5-98997984-DEL-202 | chr5-98997985-DEL-202 |
| chr5-99038856-DEL-62 | chr5-99038857-DEL-62 |
| chr5-99044361-DEL-412 | chr5-99044359-DEL-413 |
| chr5-99057221-DEL-6561 | chr5-99057222-DEL-6561 |
| chr5-99169499-DEL-574 | chr5-99169506-DEL-569 |
| chr5-99183167-DEL-159 | chr5-99183168-DEL-159 |
| chr5-99213264-DEL-417 | chr5-99213263-DEL-417 |
| chr5-99227146-DEL-187 | chr5-99227147-DEL-187 |
| chr5-99260299-DEL-5358 | chr5-99260299-DEL-5358 |
| chr5-99311017-DEL-508 | chr5-99311018-DEL-508 |
| chr5-99321862-DEL-172 | chr5-99321863-DEL-172 |
| chr5-99385750-DEL-77 | chr5-99385763-DEL-81 |
| chr5-99409859-DEL-131 | chr5-99409880-DEL-89 |
| chr5-99429512-DEL-57 | chr5-99429510-DEL-58 |
| chr5-99470918-DEL-219 | chr5-99470919-DEL-219 |
| chr5-99478555-DEL-360 | chr5-99478555-DEL-358 |
| chr5-99483178-DEL-71 | chr5-99483197-DEL-53 |
| chr5-99489365-DEL-62 | chr5-99489373-DEL-50 |
| chr5-99513837-DEL-70 | chr5-99513838-DEL-64 |
| chr5-99528748-DEL-79 | chr5-99528749-DEL-79 |
| chr5-99540990-DEL-61 | chr5-99540995-DEL-56 |
| chr5-99557707-DEL-212 | chr5-99557704-DEL-211 |
| chr5-99585236-DEL-672 | chr5-99585237-DEL-672 |
| chr5-99593692-DEL-60 | chr5-99593692-DEL-60 |
| chr5-99629679-DEL-74 | chr5-99629713-DEL-75 |
| chr5-99631116-DEL-111 | chr5-99631117-DEL-111 |
| chr5-99638506-DEL-52 | chr5-99638507-DEL-52 |
| chr5-99670191-DEL-75 | chr5-99670192-DEL-75 |
| chr5-99680718-DEL-215 | chr5-99680741-DEL-193 |
| chr5-99686106-DEL-148 | chr5-99686107-DEL-148 |
| chr5-99701543-DEL-52 | chr5-99701544-DEL-52 |
| chr5-99707469-DEL-82 | chr5-99707470-DEL-81 |
| chr5-99720237-DEL-806 | chr5-99720249-DEL-806 |
| chr5-99721440-DEL-78 | chr5-99721464-DEL-60 |
| chr5-99731338-DEL-78 | chr5-99731350-DEL-67 |
| chr5-99756672-DEL-81 | chr5-99756673-DEL-81 |
| chr5-99760972-DEL-82 | chr5-99760973-DEL-81 |
| chr5-99763927-DEL-99 | chr5-99763928-DEL-99 |
| chr5-99789659-DEL-52 | chr5-99789652-DEL-56 |
| chr5-99791804-DEL-192 | chr5-99791846-DEL-195 |
| chr5-99808248-DEL-172 | chr5-99808249-DEL-172 |
| chr5-99821894-DEL-52 | chr5-99821895-DEL-52 |
| chr5-99840218-DEL-58 | chr5-99840219-DEL-58 |
| chr5-99881688-DEL-142 | chr5-99881689-DEL-142 |
| chr5-99883019-DEL-99 | chr5-99883065-DEL-51 |
| chr5-99911487-DEL-60 | chr5-99911488-DEL-60 |
| chr5-99942321-DEL-1315 | chr5-99942313-DEL-1313 |
| chr5-99956929-DEL-218 | chr5-99956930-DEL-218 |
| chr5-99959506-DEL-2756 | chr5-99959507-DEL-2756 |
| chr5-99973171-DEL-404 | chr5-99973172-DEL-404 |

| | |
|---|---|
| chr5-99975447-DEL-355 | chr5-99975448-DEL-355 |
| chr5-99991899-DEL-77 | chr5-99991900-DEL-77 |
| chr5-100001992-DEL-51 | chr5-100001993-DEL-51 |
| chr5-100030803-DEL-106 | chr5-100030804-DEL-106 |
| chr5-100044398-DEL-215 | chr5-100044388-DEL-208 |
| chr5-100054895-DEL-7106 | chr5-100054896-DEL-7106 |
| chr5-100083945-DEL-1896 | chr5-100083946-DEL-1897 |
| chr5-100088795-DEL-211 | chr5-100088796-DEL-211 |
| chr5-100091746-DEL-489 | chr5-100091747-DEL-489 |
| chr5-100105964-DEL-102 | chr5-100105965-DEL-102 |
| chr5-100129241-DEL-179 | chr5-100129242-DEL-179 |
| chr5-100170555-DEL-214 | chr5-100170556-DEL-214 |
| chr5-100199111-DEL-505 | chr5-100199112-DEL-505 |
| chr5-100199736-DEL-204 | chr5-100199737-DEL-204 |
| chr5-100200973-DEL-1157 | chr5-100200974-DEL-1157 |
| chr5-100211832-DEL-51 | chr5-100211832-DEL-52 |
| chr5-100224956-DEL-209 | chr5-100224946-DEL-208 |
| chr5-100248672-DEL-66 | chr5-100248673-DEL-66 |
| chr5-100253098-DEL-64 | chr5-100253099-DEL-64 |
| chr5-100267956-DEL-83 | chr5-100267957-DEL-83 |
| chr5-100272810-DEL-84 | chr5-100272775-DEL-87 |
| chr5-100287863-DEL-56 | chr5-100287864-DEL-56 |
| chr5-100304461-DEL-62 | chr5-100304462-DEL-60 |
| chr5-100355543-DEL-114 | chr5-100355544-DEL-114 |
| chr5-100356635-DEL-148 | chr5-100356659-DEL-146 |
| chr5-100370565-DEL-1185 | chr5-100370566-DEL-1185 |
| chr5-100380858-DEL-262 | chr5-100380882-DEL-260 |
| chr5-100410961-DEL-2069 | chr5-100410981-DEL-2063 |
| chr5-100416780-DEL-193 | chr5-100416781-DEL-193 |
| chr5-100419688-DEL-550 | chr5-100419687-DEL-550 |
| chr5-100431739-DEL-222 | chr5-100431740-DEL-222 |
| chr5-100437799-DEL-65 | chr5-100437800-DEL-65 |
| chr5-100440410-DEL-126 | chr5-100440411-DEL-126 |
| chr5-100447831-DEL-200 | chr5-100447832-DEL-200 |
| chr5-100487760-DEL-194 | chr5-100487761-DEL-194 |
| chr5-100494661-DEL-245 | chr5-100494662-DEL-245 |
| chr5-100498000-DEL-57 | chr5-100498005-DEL-53 |
| chr5-100564722-DEL-80 | chr5-100564723-DEL-80 |
| chr5-100621310-DEL-229 | chr5-100621311-DEL-229 |
| chr5-100632169-DEL-96 | chr5-100632167-DEL-97 |
| chr5-100670051-DEL-879 | chr5-100670052-DEL-879 |
| chr5-100723523-DEL-100 | chr5-100723524-DEL-100 |
| chr5-100733373-DEL-259 | chr5-100733374-DEL-259 |
| chr5-100749123-DEL-52 | chr5-100749124-DEL-52 |
| chr5-100757193-DEL-184 | chr5-100757194-DEL-184 |
| chr5-100792641-DEL-54 | chr5-100792642-DEL-54 |
| chr5-100798725-DEL-97 | chr5-100798746-DEL-97 |
| chr5-100840356-DEL-202 | chr5-100840357-DEL-202 |
| chr5-100843110-DEL-65 | chr5-100843111-DEL-65 |
| chr5-100875787-DEL-280 | chr5-100875788-DEL-280 |
| chr5-100888482-DEL-7087 | chr5-100888483-DEL-7087 |
| chr5-100897534-DEL-148 | chr5-100897550-DEL-139 |
| chr5-100938440-DEL-195 | chr5-100938439-DEL-195 |
| chr5-100972967-DEL-239 | chr5-100972968-DEL-239 |
| chr5-100988411-DEL-56 | chr5-100988388-DEL-52 |
| chr5-101005133-DEL-227 | chr5-101005134-DEL-227 |
| chr5-101012977-DEL-384 | chr5-101012978-DEL-384 |
| chr5-101015512-DEL-296 | chr5-101015485-DEL-294 |
| chr5-101019259-DEL-511 | chr5-101019260-DEL-511 |
| chr5-101020214-DEL-192 | chr5-101020238-DEL-189 |
| chr5-101021098-DEL-82 | chr5-101021099-DEL-82 |
| chr5-101025631-DEL-107 | chr5-101025648-DEL-92 |
| chr5-101029363-DEL-246 | chr5-101029363-DEL-246 |
| chr5-101030285-DEL-85 | chr5-101030276-DEL-95 |
| chr5-101036007-DEL-70 | chr5-101036008-DEL-70 |
| chr5-101039779-DEL-51 | chr5-101039750-DEL-51 |
| chr5-101050103-DEL-89 | chr5-101050126-DEL-52 |
| chr5-101061868-DEL-52 | chr5-101061868-DEL-52 |
| chr5-101070386-DEL-135 | chr5-101070387-DEL-135 |
| chr5-101080155-DEL-63 | chr5-101080165-DEL-51 |
| chr5-101083845-DEL-55 | chr5-101083846-DEL-55 |
| chr5-101096852-DEL-165 | chr5-101096853-DEL-165 |
| chr5-101134697-DEL-105 | chr5-101134697-DEL-105 |
| chr5-101135549-DEL-577 | chr5-101135550-DEL-577 |
| chr5-101136483-DEL-221 | chr5-101136484-DEL-221 |
| chr5-101156732-DEL-3786 | chr5-101156733-DEL-3786 |
| chr5-101185976-DEL-922 | chr5-101185977-DEL-922 |
| chr5-101201187-DEL-4913 | chr5-101201188-DEL-4913 |
| chr5-101244163-DEL-474 | chr5-101244164-DEL-474 |
| chr5-101264877-DEL-80 | chr5-101264918-DEL-62 |

| | |
|---|---|
| chr5-101279940-DEL-3233 | chr5-101279941-DEL-3233 |
| chr5-101280604-DEL-714 | chr5-101280605-DEL-714 |
| chr5-101286953-DEL-918 | chr5-101286955-DEL-916 |
| chr5-101292587-DEL-915 | chr5-101292588-DEL-915 |
| chr5-101304123-DEL-224 | chr5-101304124-DEL-224 |
| chr5-101357916-DEL-817 | chr5-101357916-DEL-817 |
| chr5-101372866-DEL-77 | chr5-101372867-DEL-77 |
| chr5-101379017-DEL-1094 | chr5-101379018-DEL-1094 |
| chr5-101420057-DEL-328 | chr5-101420058-DEL-328 |
| chr5-101422309-DEL-476 | chr5-101422309-DEL-476 |
| chr5-101423106-DEL-3754 | chr5-101423107-DEL-3754 |
| chr5-101444475-DEL-111 | chr5-101444476-DEL-111 |
| chr5-101447179-DEL-60 | chr5-101447180-DEL-65 |
| chr5-101448950-DEL-52 | chr5-101448951-DEL-52 |
| chr5-101458529-DEL-304 | chr5-101458577-DEL-259 |
| chr5-101460196-DEL-6488 | chr5-101460197-DEL-6488 |
| chr5-101482622-DEL-1727 | chr5-101482623-DEL-1727 |
| chr5-101489515-DEL-459 | chr5-101489516-DEL-459 |
| chr5-101559368-DEL-58 | chr5-101559373-DEL-54 |
| chr5-101559641-DEL-481 | chr5-101559642-DEL-481 |
| chr5-101610042-DEL-77 | chr5-101610043-DEL-77 |
| chr5-101617578-DEL-67 | chr5-101617579-DEL-67 |
| chr5-101617791-DEL-1219 | chr5-101617792-DEL-1219 |
| chr5-101621180-DEL-62 | chr5-101621181-DEL-62 |
| chr5-101660543-DEL-234 | chr5-101660544-DEL-234 |
| chr5-101679190-DEL-196 | chr5-101679191-DEL-196 |
| chr5-101694278-DEL-260 | chr5-101694323-DEL-249 |
| chr5-101697324-DEL-620 | chr5-101697325-DEL-620 |
| chr5-101763341-DEL-3839 | chr5-101763342-DEL-3839 |
| chr5-101771502-DEL-136 | chr5-101771499-DEL-137 |
| chr5-101818934-DEL-265 | chr5-101818939-DEL-261 |
| chr5-101845374-DEL-69 | chr5-101845375-DEL-69 |
| chr5-101851218-DEL-70 | chr5-101851218-DEL-68 |
| chr5-101853917-DEL-1786 | chr5-101853928-DEL-1782 |
| chr5-101855181-DEL-60 | chr5-101855182-DEL-62 |
| chr5-101881530-DEL-645 | chr5-101881529-DEL-646 |
| chr5-101888738-DEL-61 | chr5-101888739-DEL-61 |
| chr5-101904614-DEL-401 | chr5-101904615-DEL-401 |
| chr5-101948365-DEL-84 | chr5-101948366-DEL-84 |
| chr5-101993562-DEL-1256 | chr5-101993563-DEL-1256 |
| chr5-102013245-DEL-219 | chr5-102013246-DEL-219 |
| chr5-102087409-DEL-64 | chr5-102087410-DEL-64 |
| chr5-102123864-DEL-1930 | chr5-102123865-DEL-1929 |
| chr5-102126513-DEL-78 | chr5-102126514-DEL-78 |
| chr5-102132990-DEL-93 | chr5-102132991-DEL-93 |
| chr5-102138873-DEL-254 | chr5-102138874-DEL-254 |
| chr5-102154870-DEL-54 | chr5-102154871-DEL-54 |
| chr5-102171372-DEL-61 | chr5-102171373-DEL-61 |
| chr5-102192358-DEL-202 | chr5-102192359-DEL-202 |
| chr5-102219032-DEL-52 | chr5-102219033-DEL-52 |
| chr5-102253568-DEL-1154 | chr5-102253569-DEL-1156 |
| chr5-102301791-DEL-58 | chr5-102301792-DEL-58 |
| chr5-102319583-DEL-72 | chr5-102319583-DEL-72 |
| chr5-102338633-DEL-6668 | chr5-102338634-DEL-6668 |
| chr5-102344945-DEL-212 | chr5-102344946-DEL-212 |
| chr5-102357756-DEL-68 | chr5-102357758-DEL-66 |
| chr5-102360786-DEL-75 | chr5-102360822-DEL-60 |
| chr5-102362585-DEL-119 | chr5-102362584-DEL-120 |
| chr5-102364811-DEL-6201 | chr5-102364812-DEL-6201 |
| chr5-102391437-DEL-933 | chr5-102391450-DEL-925 |
| chr5-102418208-DEL-68 | chr5-102418205-DEL-68 |
| chr5-102419570-DEL-3290 | chr5-102419571-DEL-3290 |
| chr5-102457424-DEL-103 | chr5-102457421-DEL-103 |
| chr5-102457950-DEL-932 | chr5-102457951-DEL-932 |
| chr5-102519907-DEL-57 | chr5-102519908-DEL-57 |
| chr5-102530061-DEL-6407 | chr5-102530062-DEL-6407 |
| chr5-102556654-DEL-7300 | chr5-102556655-DEL-7300 |
| chr5-102564115-DEL-106 | chr5-102564116-DEL-106 |
| chr5-102570513-DEL-783 | chr5-102570511-DEL-784 |
| chr5-102588651-DEL-6213 | chr5-102588652-DEL-6213 |
| chr5-102595903-DEL-51 | chr5-102595904-DEL-51 |
| chr5-102613632-DEL-973 | chr5-102613633-DEL-973 |
| chr5-102632102-DEL-74 | chr5-102632101-DEL-76 |
| chr5-102639125-DEL-60 | chr5-102639125-DEL-75 |
| chr5-102648420-DEL-60 | chr5-102648418-DEL-60 |
| chr5-102650007-DEL-57 | chr5-102650008-DEL-57 |
| chr5-102672007-DEL-1008 | chr5-102672008-DEL-1008 |
| chr5-102679009-DEL-576 | chr5-102679010-DEL-576 |
| chr5-102725649-DEL-94 | chr5-102725650-DEL-92 |
| chr5-102729697-DEL-197 | chr5-102729698-DEL-197 |

| | |
|---|---|
| chr5-102747292-DEL-6439 | chr5-102747293-DEL-6439 |
| chr5-102772433-DEL-212 | chr5-102772432-DEL-209 |
| chr5-102772939-DEL-915 | chr5-102772939-DEL-915 |
| chr5-102792997-DEL-6830 | chr5-102792998-DEL-6830 |
| chr5-102800328-DEL-168 | chr5-102800317-DEL-167 |
| chr5-102810277-DEL-69 | chr5-102810278-DEL-67 |
| chr5-102837285-DEL-78 | chr5-102837286-DEL-78 |
| chr5-102847096-DEL-3387 | chr5-102847096-DEL-3387 |
| chr5-102901648-DEL-576 | chr5-102901649-DEL-576 |
| chr5-102944994-DEL-457 | chr5-102944994-DEL-458 |
| chr5-102947157-DEL-92 | chr5-102947144-DEL-90 |
| chr5-102947513-DEL-901 | chr5-102947514-DEL-901 |
| chr5-102985713-DEL-110 | chr5-102985714-DEL-110 |
| chr5-102987206-DEL-86 | chr5-102987207-DEL-86 |
| chr5-103021233-DEL-1361 | chr5-103021234-DEL-1361 |
| chr5-103081539-DEL-6208 | chr5-103081543-DEL-6204 |
| chr5-103102953-DEL-1454 | chr5-103102954-DEL-1454 |
| chr5-103127299-DEL-52 | chr5-103127300-DEL-52 |
| chr5-103156320-DEL-178 | chr5-103156321-DEL-178 |
| chr5-103165129-DEL-158 | chr5-103165127-DEL-158 |
| chr5-103174853-DEL-79 | chr5-103174854-DEL-79 |
| chr5-103188813-DEL-67 | chr5-103188797-DEL-68 |
| chr5-103200370-DEL-399 | chr5-103200357-DEL-438 |
| chr5-103227418-DEL-605 | chr5-103227419-DEL-605 |
| chr5-103250599-DEL-181 | chr5-103250600-DEL-181 |
| chr5-103268443-DEL-70 | chr5-103268444-DEL-70 |
| chr5-103270539-DEL-6437 | chr5-103270585-DEL-6391 |
| chr5-103291975-DEL-6477 | chr5-103291978-DEL-6476 |
| chr5-103299216-DEL-1203 | chr5-103299217-DEL-1203 |
| chr5-103322064-DEL-86 | chr5-103322065-DEL-86 |
| chr5-103409732-DEL-124 | chr5-103409724-DEL-128 |
| chr5-103437741-DEL-437 | chr5-103437742-DEL-437 |
| chr5-103446060-DEL-696 | chr5-103446059-DEL-698 |
| chr5-103454719-DEL-52 | chr5-103454720-DEL-52 |
| chr5-103458651-DEL-140 | chr5-103458652-DEL-140 |
| chr5-103473414-DEL-681 | chr5-103473415-DEL-681 |
| chr5-103477095-DEL-667 | chr5-103477096-DEL-667 |
| chr5-103478356-DEL-129 | chr5-103478357-DEL-129 |
| chr5-103482552-DEL-239 | chr5-103482549-DEL-240 |
| chr5-103502130-DEL-79 | chr5-103502131-DEL-79 |
| chr5-103519757-DEL-100 | chr5-103519758-DEL-100 |
| chr5-103536546-DEL-209 | chr5-103536547-DEL-209 |
| chr5-103539206-DEL-79 | chr5-103539189-DEL-73 |
| chr5-103545660-DEL-129 | chr5-103545661-DEL-129 |
| chr5-103554551-DEL-2578 | chr5-103554552-DEL-2578 |
| chr5-103579745-DEL-106 | chr5-103579743-DEL-107 |
| chr5-103598260-DEL-197 | chr5-103598261-DEL-197 |
| chr5-103626617-DEL-704 | chr5-103626618-DEL-704 |
| chr5-103644099-DEL-78 | chr5-103644100-DEL-78 |
| chr5-103654721-DEL-225 | chr5-103654722-DEL-225 |
| chr5-103674597-DEL-54 | chr5-103674598-DEL-54 |
| chr5-103688200-DEL-203 | chr5-103688193-DEL-204 |
| chr5-103737818-DEL-58 | chr5-103737819-DEL-58 |
| chr5-103751884-DEL-203 | chr5-103751885-DEL-203 |
| chr5-103773908-DEL-66 | chr5-103773909-DEL-54 |
| chr5-103778455-DEL-6402 | chr5-103778456-DEL-6402 |
| chr5-103786791-DEL-1363 | chr5-103786792-DEL-1363 |
| chr5-103823421-DEL-191 | chr5-103823422-DEL-191 |
| chr5-103836005-DEL-238 | chr5-103836006-DEL-238 |
| chr5-103863790-DEL-161 | chr5-103863788-DEL-162 |
| chr5-103891327-DEL-180 | chr5-103891328-DEL-180 |
| chr5-103898187-DEL-52 | chr5-103898188-DEL-50 |
| chr5-103899772-DEL-211 | chr5-103899773-DEL-211 |
| chr5-104010754-DEL-98 | chr5-104010755-DEL-98 |
| chr5-104045441-DEL-162 | chr5-104045442-DEL-162 |
| chr5-104063365-DEL-199 | chr5-104063366-DEL-199 |
| chr5-104072616-DEL-250 | chr5-104072617-DEL-250 |
| chr5-104075962-DEL-216 | chr5-104075963-DEL-215 |
| chr5-104079960-DEL-345 | chr5-104079961-DEL-345 |
| chr5-104082156-DEL-637 | chr5-104082159-DEL-635 |
| chr5-104083673-DEL-358 | chr5-104083674-DEL-358 |
| chr5-104158896-DEL-201 | chr5-104158897-DEL-201 |
| chr5-104172275-DEL-222 | chr5-104172276-DEL-222 |
| chr5-104181329-DEL-446 | chr5-104181327-DEL-446 |
| chr5-104185614-DEL-298 | chr5-104185615-DEL-299 |
| chr5-104208960-DEL-88 | chr5-104208960-DEL-88 |
| chr5-104213316-DEL-3367 | chr5-104213316-DEL-3367 |
| chr5-104216277-DEL-89 | chr5-104216267-DEL-90 |
| chr5-104231364-DEL-68 | chr5-104231330-DEL-76 |
| chr5-104247946-DEL-231 | chr5-104247947-DEL-231 |

| | |
|---|---|
| chr5-104265515-DEL-56 | chr5-104265508-DEL-57 |
| chr5-104267253-DEL-96 | chr5-104267254-DEL-96 |
| chr5-104292339-DEL-209 | chr5-104292340-DEL-209 |
| chr5-104294298-DEL-201 | chr5-104294299-DEL-201 |
| chr5-104294946-DEL-1089 | chr5-104294947-DEL-1089 |
| chr5-104304350-DEL-53 | chr5-104304351-DEL-53 |
| chr5-104333918-DEL-115 | chr5-104333919-DEL-115 |
| chr5-104336172-DEL-4067 | chr5-104336173-DEL-4067 |
| chr5-104355873-DEL-377 | chr5-104355874-DEL-377 |
| chr5-104362365-DEL-748 | chr5-104362368-DEL-749 |
| chr5-104390822-DEL-142 | chr5-104390823-DEL-142 |
| chr5-104399412-DEL-87 | chr5-104399413-DEL-89 |
| chr5-104417483-DEL-54 | chr5-104417484-DEL-52 |
| chr5-104420565-DEL-184 | chr5-104420525-DEL-194 |
| chr5-104426606-DEL-6214 | chr5-104426606-DEL-6214 |
| chr5-104476123-DEL-183 | chr5-104476124-DEL-183 |
| chr5-104504854-DEL-59 | chr5-104504855-DEL-59 |
| chr5-104510315-DEL-1331 | chr5-104510316-DEL-1331 |
| chr5-104511108-DEL-1431 | chr5-104511107-DEL-1431 |
| chr5-104521580-DEL-150 | chr5-104521581-DEL-150 |
| chr5-104529332-DEL-328 | chr5-104529333-DEL-328 |
| chr5-104542056-DEL-103 | chr5-104542057-DEL-102 |
| chr5-104554101-DEL-144 | chr5-104554102-DEL-144 |
| chr5-104559614-DEL-181 | chr5-104559615-DEL-181 |
| chr5-104569779-DEL-168 | chr5-104569780-DEL-168 |
| chr5-104570622-DEL-248 | chr5-104570634-DEL-216 |
| chr5-104571249-DEL-74 | chr5-104571242-DEL-70 |
| chr5-104591816-DEL-516 | chr5-104591817-DEL-516 |
| chr5-104595618-DEL-54 | chr5-104595619-DEL-54 |
| chr5-104604863-DEL-338 | chr5-104604864-DEL-338 |
| chr5-104621484-DEL-3187 | chr5-104621485-DEL-3187 |
| chr5-104688110-DEL-207 | chr5-104688111-DEL-207 |
| chr5-104710367-DEL-180 | chr5-104710366-DEL-182 |
| chr5-104718607-DEL-102 | chr5-104718607-DEL-102 |
| chr5-104725116-DEL-212 | chr5-104725117-DEL-212 |
| chr5-104728822-DEL-375 | chr5-104728822-DEL-375 |
| chr5-104736617-DEL-304 | chr5-104736618-DEL-304 |
| chr5-104773512-DEL-196 | chr5-104773513-DEL-196 |
| chr5-104786405-DEL-84 | chr5-104786406-DEL-84 |
| chr5-104787043-DEL-83 | chr5-104787044-DEL-83 |
| chr5-104788020-DEL-1935 | chr5-104788020-DEL-1935 |
| chr5-104923636-DEL-3646 | chr5-104923637-DEL-3646 |
| chr5-104952648-DEL-69 | chr5-104952651-DEL-63 |
| chr5-104957517-DEL-2936 | chr5-104957514-DEL-2937 |
| chr5-104962344-DEL-86 | chr5-104962333-DEL-84 |
| chr5-104968293-DEL-264 | chr5-104968294-DEL-263 |
| chr5-104975296-DEL-75 | chr5-104975297-DEL-75 |
| chr5-104980241-DEL-60 | chr5-104980242-DEL-58 |
| chr5-104991117-DEL-624 | chr5-104991091-DEL-623 |
| chr5-105007538-DEL-272 | chr5-105007539-DEL-272 |
| chr5-105009452-DEL-96 | chr5-105009453-DEL-96 |
| chr5-105097185-DEL-81 | chr5-105097215-DEL-82 |
| chr5-105101852-DEL-65 | chr5-105101853-DEL-65 |
| chr5-105115075-DEL-201 | chr5-105115076-DEL-203 |
| chr5-105169949-DEL-6432 | chr5-105169950-DEL-6432 |
| chr5-105190972-DEL-7177 | chr5-105190973-DEL-7177 |
| chr5-105195535-DEL-215 | chr5-105195536-DEL-215 |
| chr5-105214508-DEL-223 | chr5-105214509-DEL-223 |
| chr5-105234955-DEL-6652 | chr5-105234956-DEL-6652 |
| chr5-105272885-DEL-70 | chr5-105272886-DEL-70 |
| chr5-105295213-DEL-203 | chr5-105295214-DEL-203 |
| chr5-105382768-DEL-52 | chr5-105382753-DEL-52 |
| chr5-105384805-DEL-729 | chr5-105384806-DEL-729 |
| chr5-105396016-DEL-53 | chr5-105396009-DEL-51 |
| chr5-105399174-DEL-3122 | chr5-105399175-DEL-3122 |
| chr5-105400088-DEL-120 | chr5-105400088-DEL-120 |
| chr5-105405540-DEL-483 | chr5-105405541-DEL-483 |
| chr5-105409002-DEL-67 | chr5-105409035-DEL-80 |
| chr5-105425039-DEL-433 | chr5-105425040-DEL-433 |
| chr5-105455993-DEL-131 | chr5-105455994-DEL-131 |
| chr5-105483441-DEL-127 | chr5-105483442-DEL-127 |
| chr5-105485390-DEL-1219 | chr5-105485391-DEL-1219 |
| chr5-105507919-DEL-7604 | chr5-105507913-DEL-7610 |
| chr5-105549838-DEL-116 | chr5-105549839-DEL-116 |
| chr5-105567195-DEL-5377 | chr5-105567196-DEL-5377 |
| chr5-105587974-DEL-429 | chr5-105587975-DEL-429 |
| chr5-105594781-DEL-1244 | chr5-105594787-DEL-1239 |
| chr5-105605911-DEL-280 | chr5-105605912-DEL-280 |
| chr5-105638941-DEL-60 | chr5-105638942-DEL-60 |
| chr5-105715900-DEL-54 | chr5-105715945-DEL-53 |

| | |
|---|---|
| chr5-105780647-DEL-51 | chr5-105780648-DEL-51 |
| chr5-105793575-DEL-106 | chr5-105793578-DEL-120 |
| chr5-105808913-DEL-78 | chr5-105808914-DEL-78 |
| chr5-105834279-DEL-458 | chr5-105834280-DEL-458 |
| chr5-105858916-DEL-236 | chr5-105858917-DEL-236 |
| chr5-105872013-DEL-5248 | chr5-105872014-DEL-5248 |
| chr5-105896966-DEL-341 | chr5-105896967-DEL-341 |
| chr5-105911282-DEL-102 | chr5-105911238-DEL-111 |
| chr5-105913519-DEL-202 | chr5-105913520-DEL-202 |
| chr5-105919352-DEL-103 | chr5-105919353-DEL-103 |
| chr5-105927892-DEL-442 | chr5-105927893-DEL-442 |
| chr5-105936345-DEL-536 | chr5-105936325-DEL-537 |
| chr5-105936862-DEL-107 | chr5-105936865-DEL-104 |
| chr5-105988485-DEL-2859 | chr5-105988486-DEL-2859 |
| chr5-105998680-DEL-1019 | chr5-105998680-DEL-986 |
| chr5-105999123-DEL-203 | chr5-105999124-DEL-203 |
| chr5-106010354-DEL-1287 | chr5-106010355-DEL-1287 |
| chr5-106067844-DEL-74 | chr5-106067845-DEL-74 |
| chr5-106079275-DEL-143 | chr5-106079276-DEL-142 |
| chr5-106090911-DEL-100 | chr5-106090912-DEL-100 |
| chr5-106092990-DEL-101 | chr5-106092989-DEL-94 |
| chr5-106096222-DEL-209 | chr5-106096223-DEL-209 |
| chr5-106188461-DEL-96 | chr5-106188456-DEL-102 |
| chr5-106268504-DEL-75 | chr5-106268505-DEL-75 |
| chr5-106275678-DEL-175 | chr5-106275675-DEL-176 |
| chr5-106288759-DEL-1929 | chr5-106288760-DEL-1929 |
| chr5-106299426-DEL-504 | chr5-106299420-DEL-531 |
| chr5-106305084-DEL-262 | chr5-106305085-DEL-250 |
| chr5-106317183-DEL-60 | chr5-106317186-DEL-58 |
| chr5-106358624-DEL-60 | chr5-106358625-DEL-60 |
| chr5-106359527-DEL-1331 | chr5-106359528-DEL-1331 |
| chr5-106362998-DEL-1842 | chr5-106362999-DEL-1842 |
| chr5-106381626-DEL-75 | chr5-106381661-DEL-50 |
| chr5-106400486-DEL-843 | chr5-106400487-DEL-843 |
| chr5-106404867-DEL-64 | chr5-106404868-DEL-64 |
| chr5-106406641-DEL-401 | chr5-106406642-DEL-401 |
| chr5-106436367-DEL-58 | chr5-106436370-DEL-56 |
| chr5-106476126-DEL-52 | chr5-106476115-DEL-64 |
| chr5-106484366-DEL-64 | chr5-106484367-DEL-60 |
| chr5-106485374-DEL-188 | chr5-106485375-DEL-188 |
| chr5-106486001-DEL-53 | chr5-106485993-DEL-50 |
| chr5-106490460-DEL-131 | chr5-106490461-DEL-131 |
| chr5-106544707-DEL-105 | chr5-106544708-DEL-105 |
| chr5-106566575-DEL-441 | chr5-106566576-DEL-441 |
| chr5-106593888-DEL-112 | chr5-106593889-DEL-70 |
| chr5-106641477-DEL-86 | chr5-106641488-DEL-52 |
| chr5-106643094-DEL-190 | chr5-106643095-DEL-190 |
| chr5-106671865-DEL-159 | chr5-106671866-DEL-159 |
| chr5-106827119-DEL-91 | chr5-106827118-DEL-90 |
| chr5-106849681-DEL-60 | chr5-106849675-DEL-67 |
| chr5-106870915-DEL-58 | chr5-106870916-DEL-58 |
| chr5-106902267-DEL-1211 | chr5-106902260-DEL-1212 |
| chr5-106912857-DEL-229 | chr5-106912858-DEL-229 |
| chr5-106916028-DEL-177 | chr5-106916013-DEL-152 |
| chr5-106920540-DEL-124 | chr5-106920541-DEL-124 |
| chr5-106928903-DEL-71 | chr5-106928904-DEL-71 |
| chr5-106930945-DEL-62 | chr5-106930946-DEL-62 |
| chr5-106937750-DEL-177 | chr5-106937751-DEL-177 |
| chr5-106939755-DEL-1203 | chr5-106939756-DEL-1203 |
| chr5-106944783-DEL-216 | chr5-106944785-DEL-214 |
| chr5-106946330-DEL-222 | chr5-106946331-DEL-222 |
| chr5-106957637-DEL-142 | chr5-106957642-DEL-138 |
| chr5-106985530-DEL-52 | chr5-106985489-DEL-94 |
| chr5-107043585-DEL-204 | chr5-107043586-DEL-204 |
| chr5-107051596-DEL-52 | chr5-107051597-DEL-52 |
| chr5-107068460-DEL-507 | chr5-107068461-DEL-507 |
| chr5-107079803-DEL-68 | chr5-107079804-DEL-68 |
| chr5-107083442-DEL-6036 | chr5-107083443-DEL-6036 |
| chr5-107097634-DEL-71 | chr5-107097635-DEL-71 |
| chr5-107103124-DEL-1640 | chr5-107103125-DEL-1640 |
| chr5-107135966-DEL-72 | chr5-107135967-DEL-72 |
| chr5-107150648-DEL-66 | chr5-107150649-DEL-66 |
| chr5-107166120-DEL-204 | chr5-107166121-DEL-204 |
| chr5-107171473-DEL-54 | chr5-107171474-DEL-54 |
| chr5-107189287-DEL-215 | chr5-107189285-DEL-216 |
| chr5-107193600-DEL-68 | chr5-107193601-DEL-68 |
| chr5-107195043-DEL-54 | chr5-107195044-DEL-54 |
| chr5-107239575-DEL-51 | chr5-107239576-DEL-51 |
| chr5-107260129-DEL-83 | chr5-107260130-DEL-83 |
| chr5-107263397-DEL-223 | chr5-107263398-DEL-223 |

| | |
|---|---|
| chr5-107305929-DEL-89 | chr5-107305912-DEL-95 |
| chr5-107336918-DEL-54 | chr5-107336919-DEL-54 |
| chr5-107345889-DEL-445 | chr5-107345884-DEL-446 |
| chr5-107350278-DEL-76 | chr5-107350279-DEL-76 |
| chr5-107371240-DEL-141 | chr5-107371241-DEL-141 |
| chr5-107379556-DEL-107 | chr5-107379565-DEL-95 |
| chr5-107403201-DEL-3263 | chr5-107403202-DEL-3263 |
| chr5-107422122-DEL-128 | chr5-107422123-DEL-127 |
| chr5-107425443-DEL-113 | chr5-107425444-DEL-113 |
| chr5-107520046-DEL-51 | chr5-107520008-DEL-51 |
| chr5-107596424-DEL-218 | chr5-107596425-DEL-218 |
| chr5-107602148-DEL-246 | chr5-107602190-DEL-205 |
| chr5-107624160-DEL-52 | chr5-107624161-DEL-52 |
| chr5-107628137-DEL-51 | chr5-107628138-DEL-51 |
| chr5-107677864-DEL-280 | chr5-107677860-DEL-278 |
| chr5-107840376-DEL-66 | chr5-107840379-DEL-64 |
| chr5-107858729-DEL-188 | chr5-107858771-DEL-138 |
| chr5-107903733-DEL-60 | chr5-107903734-DEL-60 |
| chr5-107945315-DEL-79 | chr5-107945333-DEL-68 |
| chr5-107968808-DEL-482 | chr5-107968809-DEL-482 |
| chr5-107970138-DEL-3831 | chr5-107970140-DEL-3829 |
| chr5-108005830-DEL-200 | chr5-108005831-DEL-200 |
| chr5-108007708-DEL-86 | chr5-108007709-DEL-86 |
| chr5-108030070-DEL-205 | chr5-108030071-DEL-205 |
| chr5-108033179-DEL-151 | chr5-108033180-DEL-151 |
| chr5-108042642-DEL-66 | chr5-108042597-DEL-66 |
| chr5-108043427-DEL-212 | chr5-108043428-DEL-212 |
| chr5-108058741-DEL-55 | chr5-108058742-DEL-55 |
| chr5-108061250-DEL-95 | chr5-108061251-DEL-95 |
| chr5-108067121-DEL-193 | chr5-108067122-DEL-193 |
| chr5-108074747-DEL-2309 | chr5-108074748-DEL-2309 |
| chr5-108077203-DEL-2621 | chr5-108077204-DEL-2621 |
| chr5-108113856-DEL-59 | chr5-108113857-DEL-58 |
| chr5-108136145-DEL-205 | chr5-108136138-DEL-203 |
| chr5-108161812-DEL-172 | chr5-108161813-DEL-172 |
| chr5-108168609-DEL-489 | chr5-108168611-DEL-488 |
| chr5-108238179-DEL-56 | chr5-108238179-DEL-64 |
| chr5-108266886-DEL-203 | chr5-108266887-DEL-203 |
| chr5-108282157-DEL-216 | chr5-108282158-DEL-216 |
| chr5-108307628-DEL-392 | chr5-108307629-DEL-392 |
| chr5-108328469-DEL-176 | chr5-108328470-DEL-176 |
| chr5-108358413-DEL-162 | chr5-108358414-DEL-162 |
| chr5-108360818-DEL-86 | chr5-108360819-DEL-86 |
| chr5-108381699-DEL-170 | chr5-108381700-DEL-170 |
| chr5-108386434-DEL-68 | chr5-108386435-DEL-68 |
| chr5-108388146-DEL-197 | chr5-108388147-DEL-197 |
| chr5-108402353-DEL-166 | chr5-108402354-DEL-166 |
| chr5-108406195-DEL-70 | chr5-108406229-DEL-70 |
| chr5-108431079-DEL-281 | chr5-108431080-DEL-281 |
| chr5-108432674-DEL-60 | chr5-108432671-DEL-64 |
| chr5-108450363-DEL-62 | chr5-108450360-DEL-54 |
| chr5-108455284-DEL-1375 | chr5-108455285-DEL-1375 |
| chr5-108479564-DEL-213 | chr5-108479564-DEL-213 |
| chr5-108501821-DEL-203 | chr5-108501822-DEL-203 |
| chr5-108542057-DEL-55 | chr5-108542059-DEL-51 |
| chr5-108548987-DEL-314 | chr5-108548988-DEL-312 |
| chr5-108555896-DEL-180 | chr5-108555897-DEL-180 |
| chr5-108561628-DEL-108 | chr5-108561646-DEL-90 |
| chr5-108563889-DEL-51 | chr5-108563870-DEL-52 |
| chr5-108595989-DEL-81 | chr5-108595990-DEL-81 |
| chr5-108606914-DEL-1172 | chr5-108606915-DEL-1172 |
| chr5-108634272-DEL-264 | chr5-108634273-DEL-264 |
| chr5-108639510-DEL-1886 | chr5-108639511-DEL-1886 |
| chr5-108655997-DEL-208 | chr5-108655998-DEL-208 |
| chr5-108659358-DEL-75 | chr5-108659359-DEL-75 |
| chr5-108683443-DEL-98 | chr5-108683444-DEL-97 |
| chr5-108689751-DEL-57 | chr5-108689752-DEL-57 |
| chr5-108710368-DEL-107 | chr5-108710387-DEL-88 |
| chr5-108716502-DEL-100 | chr5-108716503-DEL-100 |
| chr5-108734904-DEL-87 | chr5-108734912-DEL-64 |
| chr5-108756256-DEL-83 | chr5-108756248-DEL-85 |
| chr5-108768449-DEL-51 | chr5-108768449-DEL-54 |
| chr5-108781869-DEL-208 | chr5-108781871-DEL-207 |
| chr5-108811269-DEL-417 | chr5-108811264-DEL-418 |
| chr5-108883045-DEL-1891 | chr5-108883046-DEL-1891 |
| chr5-108895972-DEL-279 | chr5-108895973-DEL-279 |
| chr5-108909465-DEL-665 | chr5-108909449-DEL-665 |
| chr5-108953408-DEL-1063 | chr5-108953408-DEL-1063 |
| chr5-108976764-DEL-1259 | chr5-108976765-DEL-1259 |
| chr5-109031223-DEL-66 | chr5-109031224-DEL-66 |

| | |
|---|---|
| chr5-109051074-DEL-1133 | chr5-109051075-DEL-1133 |
| chr5-109071985-DEL-59 | chr5-109071980-DEL-60 |
| chr5-109085175-DEL-3171 | chr5-109085176-DEL-3171 |
| chr5-109197629-DEL-1724 | chr5-109197630-DEL-1724 |
| chr5-109212453-DEL-6474 | chr5-109212454-DEL-6474 |
| chr5-109231074-DEL-314 | chr5-109231104-DEL-282 |
| chr5-109242962-DEL-201 | chr5-109242963-DEL-201 |
| chr5-109270502-DEL-6913 | chr5-109270489-DEL-6914 |
| chr5-109294111-DEL-624 | chr5-109294112-DEL-624 |
| chr5-109308488-DEL-6265 | chr5-109308489-DEL-6265 |
| chr5-109363044-DEL-546 | chr5-109363045-DEL-546 |
| chr5-109393487-DEL-4528 | chr5-109393488-DEL-4528 |
| chr5-109429494-DEL-442 | chr5-109429495-DEL-442 |
| chr5-109516404-DEL-2161 | chr5-109516405-DEL-2161 |
| chr5-109557577-DEL-475 | chr5-109557578-DEL-475 |
| chr5-109589381-DEL-143 | chr5-109589382-DEL-143 |
| chr5-109647795-DEL-62 | chr5-109647796-DEL-66 |
| chr5-109674124-DEL-194 | chr5-109674125-DEL-194 |
| chr5-109697109-DEL-438 | chr5-109697110-DEL-438 |
| chr5-109709682-DEL-667 | chr5-109709682-DEL-667 |
| chr5-109712906-DEL-62 | chr5-109712907-DEL-62 |
| chr5-109740180-DEL-387 | chr5-109740181-DEL-387 |
| chr5-109742661-DEL-78 | chr5-109742702-DEL-71 |
| chr5-109752866-DEL-180 | chr5-109752867-DEL-180 |
| chr5-109792653-DEL-8327 | chr5-109792654-DEL-8327 |
| chr5-109823482-DEL-84 | chr5-109823483-DEL-84 |
| chr5-109873016-DEL-5087 | chr5-109873016-DEL-5087 |
| chr5-109906641-DEL-202 | chr5-109906641-DEL-202 |
| chr5-109917111-DEL-151 | chr5-109917112-DEL-151 |
| chr5-109988554-DEL-426 | chr5-109988555-DEL-426 |
| chr5-110037271-DEL-9953 | chr5-110037272-DEL-9953 |
| chr5-110039234-DEL-3207 | chr5-110039235-DEL-3207 |
| chr5-110046640-DEL-177 | chr5-110046641-DEL-177 |
| chr5-110048410-DEL-7470 | chr5-110048411-DEL-7470 |
| chr5-110064426-DEL-204 | chr5-110064423-DEL-204 |
| chr5-110065911-DEL-203 | chr5-110065933-DEL-198 |
| chr5-110077998-DEL-207 | chr5-110077999-DEL-207 |
| chr5-110090083-DEL-83 | chr5-110090065-DEL-84 |
| chr5-110095796-DEL-218 | chr5-110095796-DEL-210 |
| chr5-110121714-DEL-188 | chr5-110121715-DEL-188 |
| chr5-110134491-DEL-235 | chr5-110134476-DEL-232 |
| chr5-110136723-DEL-184 | chr5-110136724-DEL-184 |
| chr5-110179910-DEL-201 | chr5-110179911-DEL-201 |
| chr5-110181977-DEL-326 | chr5-110181978-DEL-326 |
| chr5-110193480-DEL-197 | chr5-110193481-DEL-197 |
| chr5-110208136-DEL-435 | chr5-110208136-DEL-436 |
| chr5-110209121-DEL-429 | chr5-110209122-DEL-429 |
| chr5-110226067-DEL-84 | chr5-110226068-DEL-84 |
| chr5-110229418-DEL-189 | chr5-110229418-DEL-189 |
| chr5-110238166-DEL-233 | chr5-110238164-DEL-234 |
| chr5-110239775-DEL-209 | chr5-110239763-DEL-209 |
| chr5-110262179-DEL-185 | chr5-110262180-DEL-185 |
| chr5-110280499-DEL-471 | chr5-110280500-DEL-471 |
| chr5-110328701-DEL-196 | chr5-110328703-DEL-195 |
| chr5-110342753-DEL-4963 | chr5-110342754-DEL-4963 |
| chr5-110363317-DEL-64 | chr5-110363318-DEL-64 |
| chr5-110376097-DEL-202 | chr5-110376078-DEL-201 |
| chr5-110394826-DEL-1369 | chr5-110394827-DEL-1369 |
| chr5-110398742-DEL-197 | chr5-110398743-DEL-197 |
| chr5-110424986-DEL-98 | chr5-110424987-DEL-98 |
| chr5-110434773-DEL-205 | chr5-110434774-DEL-205 |
| chr5-110449492-DEL-81 | chr5-110449479-DEL-65 |
| chr5-110468592-DEL-219 | chr5-110468611-DEL-201 |
| chr5-110481250-DEL-94 | chr5-110481264-DEL-58 |
| chr5-110582268-DEL-61 | chr5-110582269-DEL-61 |
| chr5-110612640-DEL-85 | chr5-110612641-DEL-85 |
| chr5-110623919-DEL-223 | chr5-110623917-DEL-226 |
| chr5-110650494-DEL-52 | chr5-110650495-DEL-52 |
| chr5-110668469-DEL-171 | chr5-110668470-DEL-171 |
| chr5-110672133-DEL-132 | chr5-110672134-DEL-126 |
| chr5-110677152-DEL-618 | chr5-110677153-DEL-618 |
| chr5-110739447-DEL-83 | chr5-110739448-DEL-83 |
| chr5-110759973-DEL-194 | chr5-110759974-DEL-194 |
| chr5-110774965-DEL-84 | chr5-110774966-DEL-84 |
| chr5-110899187-DEL-120 | chr5-110899188-DEL-120 |
| chr5-110911729-DEL-2417 | chr5-110911729-DEL-2418 |
| chr5-110972337-DEL-411 | chr5-110972338-DEL-411 |
| chr5-110973662-DEL-244 | chr5-110973663-DEL-244 |
| chr5-110991431-DEL-4418 | chr5-110991432-DEL-4418 |
| chr5-110998187-DEL-62 | chr5-110998188-DEL-62 |

| | |
|---|---|
| chr5-111054856-DEL-64 | chr5-111054857-DEL-64 |
| chr5-111075464-DEL-90 | chr5-111075465-DEL-90 |
| chr5-111097374-DEL-213 | chr5-111097375-DEL-213 |
| chr5-111110674-DEL-245 | chr5-111110675-DEL-245 |
| chr5-111111283-DEL-216 | chr5-111111284-DEL-216 |
| chr5-111143975-DEL-70 | chr5-111143976-DEL-70 |
| chr5-111235442-DEL-7077 | chr5-111235443-DEL-7077 |
| chr5-111269539-DEL-60 | chr5-111269540-DEL-60 |
| chr5-111275242-DEL-549 | chr5-111275243-DEL-549 |
| chr5-111294203-DEL-58 | chr5-111294204-DEL-58 |
| chr5-111331562-DEL-80 | chr5-111331565-DEL-78 |
| chr5-111349874-DEL-95 | chr5-111349916-DEL-54 |
| chr5-111353584-DEL-123 | chr5-111353574-DEL-118 |
| chr5-111379518-DEL-120 | chr5-111379518-DEL-120 |
| chr5-111394935-DEL-148 | chr5-111394938-DEL-98 |
| chr5-111396488-DEL-632 | chr5-111396489-DEL-632 |
| chr5-111438094-DEL-59 | chr5-111438095-DEL-59 |
| chr5-111441916-DEL-213 | chr5-111441917-DEL-213 |
| chr5-111448080-DEL-65 | chr5-111448081-DEL-65 |
| chr5-111454171-DEL-76 | chr5-111454172-DEL-76 |
| chr5-111454990-DEL-107 | chr5-111454991-DEL-106 |
| chr5-111506178-DEL-57 | chr5-111506179-DEL-57 |
| chr5-111536486-DEL-206 | chr5-111536487-DEL-206 |
| chr5-111550884-DEL-1062 | chr5-111550885-DEL-1062 |
| chr5-111594155-DEL-59 | chr5-111594156-DEL-59 |
| chr5-111616792-DEL-59 | chr5-111616792-DEL-59 |
| chr5-111646817-DEL-64 | chr5-111646818-DEL-64 |
| chr5-111679535-DEL-52 | chr5-111679536-DEL-52 |
| chr5-111681505-DEL-86 | chr5-111681506-DEL-74 |
| chr5-111715866-DEL-60 | chr5-111715867-DEL-60 |
| chr5-111736646-DEL-206 | chr5-111736643-DEL-210 |
| chr5-111740301-DEL-152 | chr5-111740313-DEL-141 |
| chr5-111761126-DEL-116 | chr5-111761127-DEL-115 |
| chr5-111762793-DEL-137 | chr5-111762794-DEL-137 |
| chr5-111810679-DEL-57 | chr5-111810680-DEL-56 |
| chr5-111834362-DEL-56 | chr5-111834354-DEL-60 |
| chr5-111844308-DEL-78 | chr5-111844309-DEL-78 |
| chr5-111869523-DEL-666 | chr5-111869524-DEL-666 |
| chr5-111886389-DEL-254 | chr5-111886394-DEL-282 |
| chr5-111904286-DEL-242 | chr5-111904275-DEL-240 |
| chr5-111951373-DEL-84 | chr5-111951374-DEL-84 |
| chr5-112004051-DEL-196 | chr5-112004052-DEL-196 |
| chr5-112006819-DEL-114 | chr5-112006817-DEL-112 |
| chr5-112037635-DEL-136 | chr5-112037636-DEL-136 |
| chr5-112050974-DEL-96 | chr5-112050975-DEL-96 |
| chr5-112062888-DEL-78 | chr5-112062889-DEL-78 |
| chr5-112080455-DEL-1959 | chr5-112080455-DEL-1959 |
| chr5-112101489-DEL-305 | chr5-112101490-DEL-305 |
| chr5-112103389-DEL-1721 | chr5-112103390-DEL-1721 |
| chr5-112124118-DEL-2971 | chr5-112124119-DEL-2971 |
| chr5-112128743-DEL-100 | chr5-112128744-DEL-100 |
| chr5-112156938-DEL-182 | chr5-112156935-DEL-186 |
| chr5-112170225-DEL-1364 | chr5-112170226-DEL-1364 |
| chr5-112174765-DEL-72 | chr5-112174766-DEL-96 |
| chr5-112184365-DEL-51 | chr5-112184366-DEL-77 |
| chr5-112190358-DEL-122 | chr5-112190359-DEL-122 |
| chr5-112204860-DEL-70 | chr5-112204861-DEL-70 |
| chr5-112207818-DEL-60 | chr5-112207818-DEL-60 |
| chr5-112244000-DEL-66 | chr5-112244001-DEL-58 |
| chr5-112334873-DEL-6452 | chr5-112334874-DEL-6452 |
| chr5-112348456-DEL-93 | chr5-112348457-DEL-89 |
| chr5-112351483-DEL-105 | chr5-112351483-DEL-102 |
| chr5-112441897-DEL-53 | chr5-112441898-DEL-53 |
| chr5-112496967-DEL-92 | chr5-112496974-DEL-86 |
| chr5-112515104-DEL-59 | chr5-112515105-DEL-59 |
| chr5-112530824-DEL-66 | chr5-112530822-DEL-88 |
| chr5-112546079-DEL-112 | chr5-112546081-DEL-109 |
| chr5-112555839-DEL-51 | chr5-112555839-DEL-51 |
| chr5-112558336-DEL-68 | chr5-112558337-DEL-68 |
| chr5-112587207-DEL-63 | chr5-112587206-DEL-61 |
| chr5-112604774-DEL-89 | chr5-112604775-DEL-89 |
| chr5-112619942-DEL-238 | chr5-112619943-DEL-238 |
| chr5-112646407-DEL-69 | chr5-112646407-DEL-69 |
| chr5-112660696-DEL-88 | chr5-112660671-DEL-90 |
| chr5-112671798-DEL-100 | chr5-112671831-DEL-99 |
| chr5-112689310-DEL-52 | chr5-112689306-DEL-60 |
| chr5-112691230-DEL-51 | chr5-112691208-DEL-79 |
| chr5-112716136-DEL-174 | chr5-112716136-DEL-175 |
| chr5-112726964-DEL-87 | chr5-112726974-DEL-81 |
| chr5-112727703-DEL-51 | chr5-112727704-DEL-50 |

| | |
|---|---|
| chr5-112769923-DEL-180 | chr5-112769924-DEL-180 |
| chr5-112774402-DEL-104 | chr5-112774388-DEL-102 |
| chr5-112793423-DEL-108 | chr5-112793424-DEL-108 |
| chr5-112825507-DEL-61 | chr5-112825508-DEL-61 |
| chr5-112830401-DEL-825 | chr5-112830402-DEL-825 |
| chr5-112843894-DEL-117 | chr5-112843895-DEL-117 |
| chr5-112851488-DEL-104 | chr5-112851493-DEL-100 |
| chr5-112860340-DEL-155 | chr5-112860341-DEL-155 |
| chr5-112886116-DEL-92 | chr5-112886127-DEL-82 |
| chr5-112921972-DEL-491 | chr5-112921985-DEL-493 |
| chr5-112935814-DEL-127 | chr5-112935860-DEL-80 |
| chr5-112958332-DEL-427 | chr5-112958333-DEL-427 |
| chr5-112969841-DEL-86 | chr5-112969854-DEL-84 |
| chr5-112976151-DEL-116 | chr5-112976152-DEL-116 |
| chr5-112998731-DEL-136 | chr5-112998732-DEL-136 |
| chr5-113028908-DEL-96 | chr5-113028917-DEL-89 |
| chr5-113047137-DEL-68 | chr5-113047138-DEL-68 |
| chr5-113047902-DEL-184 | chr5-113047910-DEL-183 |
| chr5-113055849-DEL-60 | chr5-113055850-DEL-60 |
| chr5-113064248-DEL-86 | chr5-113064249-DEL-86 |
| chr5-113074073-DEL-122 | chr5-113074074-DEL-122 |
| chr5-113079889-DEL-780 | chr5-113079890-DEL-780 |
| chr5-113138380-DEL-95 | chr5-113138375-DEL-96 |
| chr5-113176849-DEL-56 | chr5-113176860-DEL-62 |
| chr5-113208322-DEL-573 | chr5-113208323-DEL-573 |
| chr5-113209068-DEL-52 | chr5-113209069-DEL-52 |
| chr5-113292558-DEL-64 | chr5-113292559-DEL-64 |
| chr5-113333693-DEL-173 | chr5-113333694-DEL-173 |
| chr5-113370120-DEL-72 | chr5-113370122-DEL-73 |
| chr5-113372291-DEL-72 | chr5-113372292-DEL-72 |
| chr5-113388058-DEL-495 | chr5-113388059-DEL-495 |
| chr5-113389706-DEL-84 | chr5-113389707-DEL-84 |
| chr5-113406109-DEL-184 | chr5-113406110-DEL-184 |
| chr5-113415365-DEL-323 | chr5-113415366-DEL-323 |
| chr5-113417179-DEL-54 | chr5-113417178-DEL-56 |
| chr5-113420487-DEL-59 | chr5-113420488-DEL-59 |
| chr5-113421009-DEL-54 | chr5-113421010-DEL-56 |
| chr5-113459749-DEL-1494 | chr5-113459754-DEL-1494 |
| chr5-113461309-DEL-461 | chr5-113461311-DEL-459 |
| chr5-113507211-DEL-58 | chr5-113507212-DEL-58 |
| chr5-113531719-DEL-6444 | chr5-113531720-DEL-6444 |
| chr5-113554296-DEL-136 | chr5-113554307-DEL-126 |
| chr5-113563496-DEL-606 | chr5-113563497-DEL-606 |
| chr5-113565506-DEL-52 | chr5-113565506-DEL-52 |
| chr5-113566828-DEL-53 | chr5-113566828-DEL-53 |
| chr5-113595707-DEL-176 | chr5-113595708-DEL-176 |
| chr5-113626554-DEL-78 | chr5-113626550-DEL-58 |
| chr5-113628666-DEL-71 | chr5-113628659-DEL-79 |
| chr5-113711176-DEL-63 | chr5-113711177-DEL-63 |
| chr5-113742542-DEL-52 | chr5-113742543-DEL-52 |
| chr5-113780949-DEL-175 | chr5-113780950-DEL-175 |
| chr5-113796467-DEL-56 | chr5-113796470-DEL-60 |
| chr5-113805321-DEL-3289 | chr5-113805322-DEL-3289 |
| chr5-113841259-DEL-52 | chr5-113841259-DEL-50 |
| chr5-113847147-DEL-185 | chr5-113847135-DEL-161 |
| chr5-113863698-DEL-96 | chr5-113863699-DEL-96 |
| chr5-113905145-DEL-339 | chr5-113905146-DEL-339 |
| chr5-113915458-DEL-211 | chr5-113915459-DEL-211 |
| chr5-113949312-DEL-207 | chr5-113949296-DEL-207 |
| chr5-113950137-DEL-589 | chr5-113950138-DEL-589 |
| chr5-113957726-DEL-60 | chr5-113957727-DEL-60 |
| chr5-113959147-DEL-191 | chr5-113959144-DEL-188 |
| chr5-113991237-DEL-52 | chr5-113991232-DEL-58 |
| chr5-114033073-DEL-77 | chr5-114033074-DEL-77 |
| chr5-114058541-DEL-70 | chr5-114058547-DEL-64 |
| chr5-114062188-DEL-91 | chr5-114062189-DEL-91 |
| chr5-114074590-DEL-131 | chr5-114074591-DEL-131 |
| chr5-114084964-DEL-58 | chr5-114084965-DEL-58 |
| chr5-114123419-DEL-71 | chr5-114123421-DEL-67 |
| chr5-114126346-DEL-295 | chr5-114126347-DEL-295 |
| chr5-114136052-DEL-220 | chr5-114136052-DEL-214 |
| chr5-114146737-DEL-229 | chr5-114146735-DEL-230 |
| chr5-114149901-DEL-57 | chr5-114149901-DEL-57 |
| chr5-114156570-DEL-89 | chr5-114156574-DEL-89 |
| chr5-114177357-DEL-70 | chr5-114177395-DEL-57 |
| chr5-114210627-DEL-760 | chr5-114210628-DEL-760 |
| chr5-114218149-DEL-220 | chr5-114218150-DEL-220 |
| chr5-114256544-DEL-106 | chr5-114256548-DEL-101 |
| chr5-114273163-DEL-104 | chr5-114273164-DEL-104 |
| chr5-114274590-DEL-172 | chr5-114274602-DEL-161 |

| | |
|---|---|
| chr5-114281693-DEL-51 | chr5-114281694-DEL-51 |
| chr5-114297412-DEL-205 | chr5-114297413-DEL-205 |
| chr5-114305822-DEL-363 | chr5-114305838-DEL-333 |
| chr5-114311350-DEL-186 | chr5-114311352-DEL-185 |
| chr5-114324530-DEL-198 | chr5-114324531-DEL-198 |
| chr5-114343412-DEL-54 | chr5-114343413-DEL-54 |
| chr5-114420721-DEL-54 | chr5-114420716-DEL-60 |
| chr5-114455777-DEL-68 | chr5-114455778-DEL-68 |
| chr5-114467679-DEL-63 | chr5-114467680-DEL-63 |
| chr5-114471870-DEL-131 | chr5-114471871-DEL-131 |
| chr5-114492376-DEL-92 | chr5-114492376-DEL-93 |
| chr5-114514692-DEL-280 | chr5-114514693-DEL-280 |
| chr5-114516105-DEL-101 | chr5-114516106-DEL-101 |
| chr5-114534679-DEL-84 | chr5-114534680-DEL-72 |
| chr5-114577778-DEL-96 | chr5-114577779-DEL-96 |
| chr5-114593745-DEL-74 | chr5-114593731-DEL-72 |
| chr5-114600308-DEL-81 | chr5-114600309-DEL-81 |
| chr5-114602767-DEL-191 | chr5-114602768-DEL-190 |
| chr5-114630452-DEL-2062 | chr5-114630453-DEL-2062 |
| chr5-114649817-DEL-66 | chr5-114649818-DEL-66 |
| chr5-114656971-DEL-202 | chr5-114656972-DEL-202 |
| chr5-114662216-DEL-78 | chr5-114662217-DEL-78 |
| chr5-114671748-DEL-211 | chr5-114671749-DEL-211 |
| chr5-114684954-DEL-87 | chr5-114684955-DEL-87 |
| chr5-114754005-DEL-197 | chr5-114754006-DEL-197 |
| chr5-114766230-DEL-94 | chr5-114766231-DEL-94 |
| chr5-114771776-DEL-366 | chr5-114771777-DEL-366 |
| chr5-114793093-DEL-180 | chr5-114793094-DEL-180 |
| chr5-114809424-DEL-1378 | chr5-114809426-DEL-1375 |
| chr5-114822766-DEL-71 | chr5-114822767-DEL-71 |
| chr5-114873248-DEL-180 | chr5-114873249-DEL-180 |
| chr5-114879208-DEL-217 | chr5-114879209-DEL-217 |
| chr5-114918468-DEL-71 | chr5-114918469-DEL-71 |
| chr5-114957107-DEL-157 | chr5-114957108-DEL-156 |
| chr5-114971712-DEL-236 | chr5-114971713-DEL-236 |
| chr5-114974279-DEL-191 | chr5-114974280-DEL-191 |
| chr5-115033232-DEL-160 | chr5-115033232-DEL-160 |
| chr5-115069035-DEL-194 | chr5-115069037-DEL-193 |
| chr5-115078209-DEL-61 | chr5-115078211-DEL-60 |
| chr5-115095123-DEL-345 | chr5-115095124-DEL-345 |
| chr5-115099402-DEL-204 | chr5-115099403-DEL-203 |
| chr5-115133948-DEL-287 | chr5-115133947-DEL-289 |
| chr5-115136423-DEL-117 | chr5-115136424-DEL-117 |
| chr5-115150558-DEL-94 | chr5-115150559-DEL-94 |
| chr5-115179226-DEL-100 | chr5-115179227-DEL-100 |
| chr5-115241226-DEL-394 | chr5-115241227-DEL-394 |
| chr5-115244321-DEL-206 | chr5-115244322-DEL-206 |
| chr5-115268856-DEL-838 | chr5-115268857-DEL-838 |
| chr5-115273838-DEL-51 | chr5-115273829-DEL-60 |
| chr5-115287160-DEL-218 | chr5-115287161-DEL-218 |
| chr5-115309696-DEL-159 | chr5-115309694-DEL-163 |
| chr5-115312925-DEL-132 | chr5-115312926-DEL-126 |
| chr5-115317521-DEL-91 | chr5-115317522-DEL-91 |
| chr5-115344122-DEL-55 | chr5-115344092-DEL-56 |
| chr5-115359880-DEL-66 | chr5-115359881-DEL-66 |
| chr5-115374876-DEL-165 | chr5-115374877-DEL-165 |
| chr5-115381437-DEL-172 | chr5-115381438-DEL-172 |
| chr5-115390005-DEL-1888 | chr5-115390006-DEL-1888 |
| chr5-115405982-DEL-211 | chr5-115405983-DEL-211 |
| chr5-115420147-DEL-74 | chr5-115420148-DEL-74 |
| chr5-115423917-DEL-385 | chr5-115423918-DEL-385 |
| chr5-115425243-DEL-226 | chr5-115425244-DEL-226 |
| chr5-115444948-DEL-203 | chr5-115444949-DEL-203 |
| chr5-115460530-DEL-222 | chr5-115460531-DEL-222 |
| chr5-115491217-DEL-197 | chr5-115491218-DEL-196 |
| chr5-115500660-DEL-233 | chr5-115500660-DEL-233 |
| chr5-115501953-DEL-201 | chr5-115501954-DEL-201 |
| chr5-115549484-DEL-74 | chr5-115549485-DEL-74 |
| chr5-115555093-DEL-204 | chr5-115555094-DEL-204 |
| chr5-115606608-DEL-1047 | chr5-115606609-DEL-1047 |
| chr5-115607374-DEL-212 | chr5-115607375-DEL-212 |
| chr5-115637943-DEL-192 | chr5-115637944-DEL-192 |
| chr5-115639332-DEL-210 | chr5-115639333-DEL-210 |
| chr5-115644052-DEL-242 | chr5-115644053-DEL-242 |
| chr5-115656213-DEL-181 | chr5-115656214-DEL-181 |
| chr5-115668288-DEL-57 | chr5-115668255-DEL-58 |
| chr5-115677764-DEL-54 | chr5-115677765-DEL-54 |
| chr5-115699061-DEL-94 | chr5-115699062-DEL-94 |
| chr5-115699619-DEL-117 | chr5-115699620-DEL-117 |
| chr5-115713726-DEL-200 | chr5-115713727-DEL-200 |

| | |
|---|---|
| chr5-115719602-DEL-105 | chr5-115719603-DEL-105 |
| chr5-115739604-DEL-500 | chr5-115739605-DEL-500 |
| chr5-115745251-DEL-129 | chr5-115745252-DEL-128 |
| chr5-115748592-DEL-105 | chr5-115748593-DEL-105 |
| chr5-115763895-DEL-138 | chr5-115763896-DEL-138 |
| chr5-115768194-DEL-211 | chr5-115768195-DEL-211 |
| chr5-115774347-DEL-70 | chr5-115774347-DEL-96 |
| chr5-115872767-DEL-67 | chr5-115872768-DEL-67 |
| chr5-115879993-DEL-197 | chr5-115879994-DEL-196 |
| chr5-115897783-DEL-71 | chr5-115897788-DEL-72 |
| chr5-115902041-DEL-66 | chr5-115902042-DEL-66 |
| chr5-115902477-DEL-248 | chr5-115902478-DEL-248 |
| chr5-115914625-DEL-185 | chr5-115914642-DEL-189 |
| chr5-115919491-DEL-115 | chr5-115919492-DEL-115 |
| chr5-115924820-DEL-58 | chr5-115924821-DEL-58 |
| chr5-115926153-DEL-207 | chr5-115926154-DEL-207 |
| chr5-115939600-DEL-677 | chr5-115939590-DEL-679 |
| chr5-115947501-DEL-1008 | chr5-115947502-DEL-996 |
| chr5-115958187-DEL-84 | chr5-115958188-DEL-84 |
| chr5-115962675-DEL-80 | chr5-115962676-DEL-80 |
| chr5-115972312-DEL-205 | chr5-115972313-DEL-205 |
| chr5-115975339-DEL-171 | chr5-115975344-DEL-153 |
| chr5-115978868-DEL-80 | chr5-115978868-DEL-80 |
| chr5-115994264-DEL-235 | chr5-115994265-DEL-235 |
| chr5-116034736-DEL-127 | chr5-116034737-DEL-121 |
| chr5-116037687-DEL-67 | chr5-116037692-DEL-50 |
| chr5-116054114-DEL-217 | chr5-116054115-DEL-217 |
| chr5-116055518-DEL-115 | chr5-116055514-DEL-120 |
| chr5-116087594-DEL-60 | chr5-116087587-DEL-64 |
| chr5-116111564-DEL-63 | chr5-116111565-DEL-63 |
| chr5-116112303-DEL-57 | chr5-116112311-DEL-57 |
| chr5-116134726-DEL-84 | chr5-116134727-DEL-84 |
| chr5-116149420-DEL-124 | chr5-116149460-DEL-84 |
| chr5-116153317-DEL-143 | chr5-116153318-DEL-143 |
| chr5-116172263-DEL-502 | chr5-116172264-DEL-502 |
| chr5-116185044-DEL-281 | chr5-116185045-DEL-281 |
| chr5-116201044-DEL-1326 | chr5-116201054-DEL-1279 |
| chr5-116204055-DEL-183 | chr5-116204056-DEL-183 |
| chr5-116204795-DEL-292 | chr5-116204796-DEL-292 |
| chr5-116208765-DEL-60 | chr5-116208768-DEL-62 |
| chr5-116229840-DEL-179 | chr5-116229823-DEL-180 |
| chr5-116235154-DEL-60 | chr5-116235155-DEL-60 |
| chr5-116251463-DEL-155 | chr5-116251463-DEL-153 |
| chr5-116258848-DEL-133 | chr5-116258853-DEL-129 |
| chr5-116274273-DEL-68 | chr5-116274274-DEL-68 |
| chr5-116285103-DEL-6440 | chr5-116285104-DEL-6440 |
| chr5-116305526-DEL-245 | chr5-116305527-DEL-243 |
| chr5-116315771-DEL-203 | chr5-116315771-DEL-203 |
| chr5-116321619-DEL-52 | chr5-116321620-DEL-52 |
| chr5-116333037-DEL-1082 | chr5-116333038-DEL-1082 |
| chr5-116396309-DEL-62 | chr5-116396324-DEL-80 |
| chr5-116408758-DEL-2003 | chr5-116408759-DEL-2003 |
| chr5-116444949-DEL-137 | chr5-116444950-DEL-137 |
| chr5-116453852-DEL-296 | chr5-116453853-DEL-296 |
| chr5-116475523-DEL-400 | chr5-116475524-DEL-400 |
| chr5-116477270-DEL-66 | chr5-116477271-DEL-66 |
| chr5-116483123-DEL-295 | chr5-116483124-DEL-295 |
| chr5-116515490-DEL-55 | chr5-116515482-DEL-71 |
| chr5-116522560-DEL-53 | chr5-116522529-DEL-81 |
| chr5-116589939-DEL-743 | chr5-116589940-DEL-743 |
| chr5-116590854-DEL-74 | chr5-116590857-DEL-50 |
| chr5-116600722-DEL-71 | chr5-116600721-DEL-72 |
| chr5-116618792-DEL-71 | chr5-116618793-DEL-71 |
| chr5-116672538-DEL-444 | chr5-116672539-DEL-444 |
| chr5-116689082-DEL-140 | chr5-116689083-DEL-140 |
| chr5-116741149-DEL-72 | chr5-116741150-DEL-72 |
| chr5-116743222-DEL-6422 | chr5-116743223-DEL-6422 |
| chr5-116776068-DEL-781 | chr5-116776069-DEL-781 |
| chr5-116836312-DEL-80 | chr5-116836313-DEL-80 |
| chr5-116868456-DEL-52 | chr5-116868457-DEL-52 |
| chr5-116897254-DEL-340 | chr5-116897255-DEL-340 |
| chr5-116906148-DEL-1885 | chr5-116906149-DEL-1885 |
| chr5-116914798-DEL-400 | chr5-116914806-DEL-393 |
| chr5-116923130-DEL-102 | chr5-116923131-DEL-102 |
| chr5-116939700-DEL-86 | chr5-116939701-DEL-86 |
| chr5-116942863-DEL-607 | chr5-116942870-DEL-600 |
| chr5-116946693-DEL-62 | chr5-116946706-DEL-50 |
| chr5-116950129-DEL-61 | chr5-116950129-DEL-61 |
| chr5-116971893-DEL-75 | chr5-116971889-DEL-75 |
| chr5-116993860-DEL-262 | chr5-116993861-DEL-262 |

| | |
|---|---|
| chr5-117013658-DEL-74 | chr5-117013674-DEL-52 |
| chr5-117025439-DEL-100 | chr5-117025447-DEL-92 |
| chr5-117058161-DEL-299 | chr5-117058162-DEL-299 |
| chr5-117059061-DEL-59 | chr5-117059064-DEL-60 |
| chr5-117085000-DEL-78 | chr5-117085001-DEL-78 |
| chr5-117090403-DEL-369 | chr5-117090404-DEL-369 |
| chr5-117101034-DEL-351 | chr5-117101035-DEL-351 |
| chr5-117109404-DEL-1025 | chr5-117109405-DEL-1025 |
| chr5-117109634-DEL-60 | chr5-117109635-DEL-60 |
| chr5-117143473-DEL-537 | chr5-117143474-DEL-537 |
| chr5-117146304-DEL-1826 | chr5-117146305-DEL-1826 |
| chr5-117151635-DEL-384 | chr5-117151636-DEL-384 |
| chr5-117164621-DEL-54 | chr5-117164621-DEL-54 |
| chr5-117166006-DEL-498 | chr5-117166007-DEL-498 |
| chr5-117210268-DEL-72 | chr5-117210237-DEL-72 |
| chr5-117219097-DEL-100 | chr5-117219098-DEL-100 |
| chr5-117220463-DEL-92 | chr5-117220471-DEL-85 |
| chr5-117248960-DEL-78 | chr5-117248970-DEL-74 |
| chr5-117268597-DEL-68 | chr5-117268598-DEL-68 |
| chr5-117271067-DEL-279 | chr5-117271068-DEL-249 |
| chr5-117284737-DEL-59 | chr5-117284737-DEL-57 |
| chr5-117286753-DEL-132 | chr5-117286754-DEL-132 |
| chr5-117293602-DEL-832 | chr5-117293603-DEL-832 |
| chr5-117301480-DEL-163 | chr5-117301481-DEL-163 |
| chr5-117340384-DEL-57 | chr5-117340385-DEL-57 |
| chr5-117360789-DEL-152 | chr5-117360792-DEL-148 |
| chr5-117369552-DEL-228 | chr5-117369553-DEL-228 |
| chr5-117372895-DEL-67 | chr5-117372896-DEL-67 |
| chr5-117397490-DEL-61 | chr5-117397491-DEL-61 |
| chr5-117427593-DEL-198 | chr5-117427594-DEL-198 |
| chr5-117433511-DEL-7077 | chr5-117433512-DEL-7077 |
| chr5-117450699-DEL-198 | chr5-117450704-DEL-197 |
| chr5-117469717-DEL-99 | chr5-117469718-DEL-99 |
| chr5-117471455-DEL-86 | chr5-117471456-DEL-86 |
| chr5-117471592-DEL-943 | chr5-117471593-DEL-943 |
| chr5-117472300-DEL-86 | chr5-117472297-DEL-84 |
| chr5-117512451-DEL-127 | chr5-117512441-DEL-128 |
| chr5-117535159-DEL-64 | chr5-117535168-DEL-56 |
| chr5-117536082-DEL-165 | chr5-117536083-DEL-165 |
| chr5-117538187-DEL-541 | chr5-117538188-DEL-541 |
| chr5-117548781-DEL-77 | chr5-117548782-DEL-75 |
| chr5-117577771-DEL-402 | chr5-117577772-DEL-402 |
| chr5-117597221-DEL-56 | chr5-117597222-DEL-56 |
| chr5-117617157-DEL-130 | chr5-117617135-DEL-155 |
| chr5-117618489-DEL-52 | chr5-117618490-DEL-52 |
| chr5-117634345-DEL-70 | chr5-117634346-DEL-70 |
| chr5-117645510-DEL-69 | chr5-117645522-DEL-57 |
| chr5-117651898-DEL-206 | chr5-117651899-DEL-206 |
| chr5-117658484-DEL-52 | chr5-117658485-DEL-52 |
| chr5-117697804-DEL-181 | chr5-117697805-DEL-151 |
| chr5-117698582-DEL-121 | chr5-117698583-DEL-121 |
| chr5-117722363-DEL-53 | chr5-117722362-DEL-69 |
| chr5-117818747-DEL-112 | chr5-117818776-DEL-84 |
| chr5-117823161-DEL-64 | chr5-117823162-DEL-64 |
| chr5-117824288-DEL-172 | chr5-117824289-DEL-172 |
| chr5-117915773-DEL-596 | chr5-117915774-DEL-596 |
| chr5-117946794-DEL-103 | chr5-117946795-DEL-103 |
| chr5-117968008-DEL-126 | chr5-117968009-DEL-94 |
| chr5-117982999-DEL-56 | chr5-117983000-DEL-56 |
| chr5-117994006-DEL-123 | chr5-117993959-DEL-164 |
| chr5-118020973-DEL-58 | chr5-118020974-DEL-58 |
| chr5-118053068-DEL-118 | chr5-118053069-DEL-118 |
| chr5-118088202-DEL-302 | chr5-118088203-DEL-295 |
| chr5-118095045-DEL-62 | chr5-118095048-DEL-50 |
| chr5-118161640-DEL-211 | chr5-118161641-DEL-211 |
| chr5-118162799-DEL-219 | chr5-118162800-DEL-219 |
| chr5-118177976-DEL-425 | chr5-118177977-DEL-422 |
| chr5-118190666-DEL-195 | chr5-118190667-DEL-195 |
| chr5-118193324-DEL-250 | chr5-118193325-DEL-250 |
| chr5-118200802-DEL-69 | chr5-118200803-DEL-69 |
| chr5-118226617-DEL-208 | chr5-118226618-DEL-208 |
| chr5-118250629-DEL-221 | chr5-118250630-DEL-221 |
| chr5-118283117-DEL-56 | chr5-118283118-DEL-56 |
| chr5-118304084-DEL-87 | chr5-118304083-DEL-88 |
| chr5-118334036-DEL-80 | chr5-118334037-DEL-80 |
| chr5-118368028-DEL-123 | chr5-118368031-DEL-124 |
| chr5-118399985-DEL-141 | chr5-118399986-DEL-141 |
| chr5-118408208-DEL-61 | chr5-118408209-DEL-61 |
| chr5-118419411-DEL-60 | chr5-118419412-DEL-58 |
| chr5-118430590-DEL-80 | chr5-118430621-DEL-50 |

| | |
|---|---|
| chr5-118449651-DEL-65 | chr5-118449652-DEL-65 |
| chr5-118461891-DEL-169 | chr5-118461883-DEL-136 |
| chr5-118474465-DEL-318 | chr5-118474470-DEL-314 |
| chr5-118475107-DEL-613 | chr5-118475108-DEL-613 |
| chr5-118475893-DEL-83 | chr5-118475925-DEL-88 |
| chr5-118530445-DEL-86 | chr5-118530439-DEL-96 |
| chr5-118546496-DEL-55 | chr5-118546497-DEL-55 |
| chr5-118564384-DEL-212 | chr5-118564385-DEL-212 |
| chr5-118577162-DEL-2806 | chr5-118577163-DEL-2806 |
| chr5-118578169-DEL-1175 | chr5-118578170-DEL-1175 |
| chr5-118581109-DEL-200 | chr5-118581096-DEL-200 |
| chr5-118617496-DEL-177 | chr5-118617497-DEL-177 |
| chr5-118619813-DEL-104 | chr5-118619814-DEL-104 |
| chr5-118625787-DEL-54 | chr5-118625788-DEL-54 |
| chr5-118628170-DEL-209 | chr5-118628171-DEL-209 |
| chr5-118638685-DEL-223 | chr5-118638686-DEL-223 |
| chr5-118712942-DEL-60 | chr5-118712945-DEL-58 |
| chr5-118780093-DEL-491 | chr5-118780094-DEL-491 |
| chr5-118789577-DEL-87 | chr5-118789578-DEL-87 |
| chr5-118800776-DEL-183 | chr5-118800777-DEL-183 |
| chr5-118852179-DEL-95 | chr5-118852180-DEL-95 |
| chr5-118910613-DEL-51 | chr5-118910614-DEL-51 |
| chr5-118914242-DEL-362 | chr5-118914243-DEL-362 |
| chr5-118925843-DEL-147 | chr5-118925844-DEL-147 |
| chr5-118937768-DEL-76 | chr5-118937773-DEL-72 |
| chr5-118944704-DEL-54 | chr5-118944715-DEL-62 |
| chr5-118951582-DEL-54 | chr5-118951583-DEL-54 |
| chr5-118969058-DEL-59 | chr5-118969041-DEL-92 |
| chr5-118970204-DEL-210 | chr5-118970205-DEL-210 |
| chr5-118974034-DEL-51 | chr5-118974035-DEL-51 |
| chr5-118991318-DEL-306 | chr5-118991295-DEL-285 |
| chr5-119021638-DEL-434 | chr5-119021639-DEL-434 |
| chr5-119025486-DEL-489 | chr5-119025486-DEL-486 |
| chr5-119036001-DEL-58 | chr5-119036000-DEL-60 |
| chr5-119064808-DEL-57 | chr5-119064803-DEL-63 |
| chr5-119124743-DEL-78 | chr5-119124743-DEL-78 |
| chr5-119130775-DEL-903 | chr5-119130776-DEL-887 |
| chr5-119132034-DEL-103 | chr5-119132033-DEL-101 |
| chr5-119179423-DEL-62 | chr5-119179419-DEL-83 |
| chr5-119183872-DEL-332 | chr5-119183873-DEL-332 |
| chr5-119202228-DEL-1106 | chr5-119202229-DEL-1106 |
| chr5-119223060-DEL-72 | chr5-119223041-DEL-100 |
| chr5-119265667-DEL-60 | chr5-119265664-DEL-64 |
| chr5-119266277-DEL-93 | chr5-119266241-DEL-138 |
| chr5-119282826-DEL-56 | chr5-119282827-DEL-56 |
| chr5-119288926-DEL-93 | chr5-119288927-DEL-93 |
| chr5-119320447-DEL-59 | chr5-119320447-DEL-59 |
| chr5-119336349-DEL-103 | chr5-119336350-DEL-100 |
| chr5-119341577-DEL-54 | chr5-119341582-DEL-50 |
| chr5-119367662-DEL-343 | chr5-119367663-DEL-343 |
| chr5-119383513-DEL-717 | chr5-119383514-DEL-733 |
| chr5-119390163-DEL-170 | chr5-119390164-DEL-170 |
| chr5-119394398-DEL-54 | chr5-119394373-DEL-58 |
| chr5-119416183-DEL-1209 | chr5-119416184-DEL-1209 |
| chr5-119439836-DEL-241 | chr5-119439837-DEL-241 |
| chr5-119461399-DEL-77 | chr5-119461400-DEL-77 |
| chr5-119488117-DEL-69 | chr5-119488118-DEL-69 |
| chr5-119495879-DEL-177 | chr5-119495880-DEL-177 |
| chr5-119504186-DEL-72 | chr5-119504187-DEL-76 |
| chr5-119506471-DEL-228 | chr5-119506472-DEL-266 |
| chr5-119513349-DEL-206 | chr5-119513350-DEL-206 |
| chr5-119566954-DEL-63 | chr5-119566965-DEL-51 |
| chr5-119580793-DEL-1317 | chr5-119580794-DEL-1317 |
| chr5-119677338-DEL-136 | chr5-119677339-DEL-136 |
| chr5-119688199-DEL-180 | chr5-119688200-DEL-180 |
| chr5-119707571-DEL-194 | chr5-119707572-DEL-194 |
| chr5-119729196-DEL-78 | chr5-119729194-DEL-81 |
| chr5-119783509-DEL-225 | chr5-119783514-DEL-195 |
| chr5-119792518-DEL-1548 | chr5-119792519-DEL-1547 |
| chr5-119837420-DEL-400 | chr5-119837421-DEL-400 |
| chr5-119842299-DEL-58 | chr5-119842281-DEL-100 |
| chr5-119858265-DEL-72 | chr5-119858266-DEL-55 |
| chr5-119863273-DEL-1895 | chr5-119863274-DEL-1895 |
| chr5-119891573-DEL-227 | chr5-119891574-DEL-227 |
| chr5-119954838-DEL-63 | chr5-119954839-DEL-66 |
| chr5-119957189-DEL-350 | chr5-119957190-DEL-350 |
| chr5-120003750-DEL-62 | chr5-120003751-DEL-66 |
| chr5-120036523-DEL-7180 | chr5-120036524-DEL-7180 |
| chr5-120055562-DEL-88 | chr5-120055563-DEL-86 |
| chr5-120078373-DEL-1163 | chr5-120078395-DEL-1155 |

| | |
|---|---|
| chr5-120099427-DEL-65 | chr5-120099434-DEL-59 |
| chr5-120099962-DEL-60 | chr5-120099963-DEL-50 |
| chr5-120104951-DEL-55 | chr5-120104952-DEL-55 |
| chr5-120114057-DEL-98 | chr5-120114094-DEL-82 |
| chr5-120117340-DEL-282 | chr5-120117341-DEL-282 |
| chr5-120128397-DEL-69 | chr5-120128397-DEL-69 |
| chr5-120153189-DEL-163 | chr5-120153189-DEL-163 |
| chr5-120164462-DEL-1003 | chr5-120164461-DEL-1004 |
| chr5-120171956-DEL-285 | chr5-120171957-DEL-285 |
| chr5-120264764-DEL-224 | chr5-120264763-DEL-224 |
| chr5-120268465-DEL-60 | chr5-120268466-DEL-60 |
| chr5-120354059-DEL-227 | chr5-120354056-DEL-219 |
| chr5-120357895-DEL-85 | chr5-120357918-DEL-63 |
| chr5-120379130-DEL-243 | chr5-120379131-DEL-243 |
| chr5-120436272-DEL-206 | chr5-120436266-DEL-202 |
| chr5-120466513-DEL-148 | chr5-120466497-DEL-132 |
| chr5-120474158-DEL-60 | chr5-120474145-DEL-98 |
| chr5-120482718-DEL-76 | chr5-120482719-DEL-76 |
| chr5-120501412-DEL-64 | chr5-120501408-DEL-63 |
| chr5-120554980-DEL-578 | chr5-120554981-DEL-578 |
| chr5-120564525-DEL-80 | chr5-120564526-DEL-80 |
| chr5-120567406-DEL-81 | chr5-120567419-DEL-69 |
| chr5-120588521-DEL-69 | chr5-120588526-DEL-65 |
| chr5-120601606-DEL-208 | chr5-120601607-DEL-208 |
| chr5-120613833-DEL-188 | chr5-120613830-DEL-181 |
| chr5-120616334-DEL-93 | chr5-120616320-DEL-83 |
| chr5-120657208-DEL-197 | chr5-120657209-DEL-197 |
| chr5-120661798-DEL-3567 | chr5-120661799-DEL-3567 |
| chr5-120687828-DEL-118 | chr5-120687828-DEL-118 |
| chr5-120692628-DEL-193 | chr5-120692629-DEL-193 |
| chr5-120710156-DEL-67 | chr5-120710155-DEL-57 |
| chr5-120713816-DEL-191 | chr5-120713803-DEL-190 |
| chr5-120741206-DEL-99 | chr5-120741207-DEL-99 |
| chr5-120805340-DEL-88 | chr5-120805341-DEL-88 |
| chr5-120822639-DEL-65 | chr5-120822641-DEL-63 |
| chr5-120837437-DEL-2280 | chr5-120837440-DEL-2281 |
| chr5-120853379-DEL-87 | chr5-120853380-DEL-87 |
| chr5-120963703-DEL-4525 | chr5-120963704-DEL-4525 |
| chr5-120987566-DEL-66 | chr5-120987565-DEL-66 |
| chr5-120988053-DEL-238 | chr5-120988054-DEL-238 |
| chr5-120998433-DEL-79 | chr5-120998432-DEL-79 |
| chr5-121030987-DEL-514 | chr5-121030988-DEL-514 |
| chr5-121048162-DEL-433 | chr5-121048161-DEL-431 |
| chr5-121062861-DEL-55 | chr5-121062862-DEL-55 |
| chr5-121069292-DEL-215 | chr5-121069293-DEL-215 |
| chr5-121131179-DEL-402 | chr5-121131180-DEL-402 |
| chr5-121132365-DEL-66 | chr5-121132366-DEL-66 |
| chr5-121135972-DEL-56 | chr5-121135959-DEL-78 |
| chr5-121163179-DEL-54 | chr5-121163180-DEL-54 |
| chr5-121171280-DEL-412 | chr5-121171265-DEL-446 |
| chr5-121181095-DEL-222 | chr5-121181096-DEL-222 |
| chr5-121199268-DEL-52 | chr5-121199265-DEL-56 |
| chr5-121201616-DEL-68 | chr5-121201617-DEL-68 |
| chr5-121209931-DEL-217 | chr5-121209932-DEL-217 |
| chr5-121251115-DEL-79 | chr5-121251116-DEL-61 |
| chr5-121252728-DEL-63 | chr5-121252729-DEL-63 |
| chr5-121262907-DEL-166 | chr5-121262955-DEL-119 |
| chr5-121291484-DEL-226 | chr5-121291485-DEL-226 |
| chr5-121306825-DEL-204 | chr5-121306824-DEL-205 |
| chr5-121309342-DEL-203 | chr5-121309343-DEL-203 |
| chr5-121339345-DEL-143 | chr5-121339350-DEL-139 |
| chr5-121341162-DEL-131 | chr5-121341160-DEL-132 |
| chr5-121348179-DEL-202 | chr5-121348180-DEL-202 |
| chr5-121349061-DEL-59 | chr5-121349062-DEL-59 |
| chr5-121353857-DEL-204 | chr5-121353853-DEL-205 |
| chr5-121365595-DEL-52 | chr5-121365595-DEL-58 |
| chr5-121372470-DEL-625 | chr5-121372471-DEL-625 |
| chr5-121375804-DEL-197 | chr5-121375805-DEL-197 |
| chr5-121426402-DEL-179 | chr5-121426396-DEL-186 |
| chr5-121436353-DEL-358 | chr5-121436354-DEL-358 |
| chr5-121450519-DEL-166 | chr5-121450520-DEL-166 |
| chr5-121464326-DEL-86 | chr5-121464316-DEL-88 |
| chr5-121468177-DEL-426 | chr5-121468174-DEL-426 |
| chr5-121485482-DEL-234 | chr5-121485483-DEL-234 |
| chr5-121512293-DEL-65 | chr5-121512294-DEL-50 |
| chr5-121529444-DEL-201 | chr5-121529445-DEL-201 |
| chr5-121545733-DEL-153 | chr5-121545734-DEL-153 |
| chr5-121595037-DEL-229 | chr5-121595038-DEL-229 |
| chr5-121618668-DEL-678 | chr5-121618669-DEL-678 |
| chr5-121672893-DEL-220 | chr5-121672894-DEL-220 |

| | |
|---|---|
| chr5-121700621-DEL-153 | chr5-121700622-DEL-153 |
| chr5-121704695-DEL-140 | chr5-121704696-DEL-140 |
| chr5-121712852-DEL-60 | chr5-121712853-DEL-60 |
| chr5-121721083-DEL-503 | chr5-121721084-DEL-503 |
| chr5-121726927-DEL-90 | chr5-121726928-DEL-90 |
| chr5-121736566-DEL-135 | chr5-121736567-DEL-135 |
| chr5-121775864-DEL-219 | chr5-121775865-DEL-219 |
| chr5-121803488-DEL-206 | chr5-121803489-DEL-206 |
| chr5-121826706-DEL-203 | chr5-121826707-DEL-203 |
| chr5-121830799-DEL-204 | chr5-121830800-DEL-204 |
| chr5-121840654-DEL-222 | chr5-121840655-DEL-222 |
| chr5-121876429-DEL-279 | chr5-121876430-DEL-279 |
| chr5-121899750-DEL-101 | chr5-121899751-DEL-101 |
| chr5-121904920-DEL-58 | chr5-121904917-DEL-62 |
| chr5-121918551-DEL-111 | chr5-121918552-DEL-111 |
| chr5-121937510-DEL-75 | chr5-121937506-DEL-76 |
| chr5-122035692-DEL-64 | chr5-122035693-DEL-64 |
| chr5-122078837-DEL-501 | chr5-122078838-DEL-501 |
| chr5-122088905-DEL-68 | chr5-122088906-DEL-68 |
| chr5-122127044-DEL-98 | chr5-122127042-DEL-94 |
| chr5-122144013-DEL-364 | chr5-122144014-DEL-364 |
| chr5-122166574-DEL-56 | chr5-122166561-DEL-56 |
| chr5-122199117-DEL-1421 | chr5-122199119-DEL-1420 |
| chr5-122215012-DEL-655 | chr5-122215011-DEL-656 |
| chr5-122257758-DEL-143 | chr5-122257752-DEL-138 |
| chr5-122262387-DEL-58 | chr5-122262388-DEL-58 |
| chr5-122278557-DEL-110 | chr5-122278560-DEL-108 |
| chr5-122281329-DEL-165 | chr5-122281337-DEL-168 |
| chr5-122284938-DEL-56 | chr5-122284939-DEL-56 |
| chr5-122285802-DEL-108 | chr5-122285796-DEL-107 |
| chr5-122290546-DEL-189 | chr5-122290547-DEL-189 |
| chr5-122318512-DEL-1523 | chr5-122318513-DEL-1523 |
| chr5-122326483-DEL-993 | chr5-122326482-DEL-1002 |
| chr5-122330904-DEL-7366 | chr5-122330905-DEL-7366 |
| chr5-122353054-DEL-209 | chr5-122353055-DEL-209 |
| chr5-122364045-DEL-52 | chr5-122364085-DEL-52 |
| chr5-122364703-DEL-1229 | chr5-122364704-DEL-1229 |
| chr5-122365292-DEL-106 | chr5-122365293-DEL-102 |
| chr5-122409798-DEL-52 | chr5-122409798-DEL-52 |
| chr5-122412300-DEL-199 | chr5-122412301-DEL-199 |
| chr5-122415017-DEL-354 | chr5-122415018-DEL-354 |
| chr5-122420987-DEL-203 | chr5-122421000-DEL-206 |
| chr5-122428222-DEL-133 | chr5-122428223-DEL-133 |
| chr5-122437322-DEL-208 | chr5-122437323-DEL-208 |
| chr5-122447303-DEL-57 | chr5-122447304-DEL-57 |
| chr5-122472606-DEL-217 | chr5-122472606-DEL-217 |
| chr5-122512833-DEL-54 | chr5-122512834-DEL-54 |
| chr5-122517521-DEL-93 | chr5-122517530-DEL-85 |
| chr5-122521335-DEL-6479 | chr5-122521336-DEL-6479 |
| chr5-122547053-DEL-65 | chr5-122547044-DEL-52 |
| chr5-122562007-DEL-223 | chr5-122562008-DEL-223 |
| chr5-122585766-DEL-105 | chr5-122585767-DEL-105 |
| chr5-122627588-DEL-1077 | chr5-122627589-DEL-1077 |
| chr5-122650425-DEL-215 | chr5-122650426-DEL-215 |
| chr5-122677323-DEL-60 | chr5-122677324-DEL-60 |
| chr5-122700788-DEL-216 | chr5-122700837-DEL-168 |
| chr5-122714359-DEL-85 | chr5-122714388-DEL-86 |
| chr5-122765721-DEL-72 | chr5-122765727-DEL-58 |
| chr5-122774808-DEL-55 | chr5-122774809-DEL-55 |
| chr5-122777427-DEL-77 | chr5-122777428-DEL-77 |
| chr5-122795488-DEL-447 | chr5-122795489-DEL-447 |
| chr5-122809264-DEL-60 | chr5-122809269-DEL-56 |
| chr5-122812754-DEL-58 | chr5-122812755-DEL-62 |
| chr5-122814493-DEL-201 | chr5-122814494-DEL-201 |
| chr5-122821403-DEL-70 | chr5-122821404-DEL-70 |
| chr5-122832490-DEL-1000 | chr5-122832506-DEL-1015 |
| chr5-122841282-DEL-58 | chr5-122841286-DEL-55 |
| chr5-122858473-DEL-56 | chr5-122858476-DEL-55 |
| chr5-122859592-DEL-169 | chr5-122859593-DEL-169 |
| chr5-122860387-DEL-169 | chr5-122860388-DEL-169 |
| chr5-122862776-DEL-209 | chr5-122862777-DEL-209 |
| chr5-122873248-DEL-393 | chr5-122873249-DEL-393 |
| chr5-122876467-DEL-408 | chr5-122876468-DEL-408 |
| chr5-122889198-DEL-69 | chr5-122889198-DEL-70 |
| chr5-122890156-DEL-1264 | chr5-122890157-DEL-1264 |
| chr5-122928727-DEL-91 | chr5-122928769-DEL-52 |
| chr5-122965594-DEL-326 | chr5-122965604-DEL-321 |
| chr5-122975834-DEL-1039 | chr5-122975834-DEL-1039 |
| chr5-122982950-DEL-75 | chr5-122982951-DEL-75 |
| chr5-122983444-DEL-183 | chr5-122983445-DEL-183 |

| | |
|---|---|
| chr5-122993487-DEL-1207 | chr5-122993488-DEL-1207 |
| chr5-123003992-DEL-167 | chr5-123003993-DEL-167 |
| chr5-123021783-DEL-423 | chr5-123021782-DEL-423 |
| chr5-123024776-DEL-1287 | chr5-123024777-DEL-1287 |
| chr5-123044584-DEL-123 | chr5-123044623-DEL-85 |
| chr5-123066947-DEL-61 | chr5-123066948-DEL-61 |
| chr5-123067590-DEL-197 | chr5-123067591-DEL-197 |
| chr5-123098300-DEL-154 | chr5-123098302-DEL-128 |
| chr5-123100104-DEL-52 | chr5-123100105-DEL-52 |
| chr5-123102283-DEL-714 | chr5-123102284-DEL-714 |
| chr5-123102997-DEL-146 | chr5-123102998-DEL-146 |
| chr5-123136650-DEL-290 | chr5-123136651-DEL-290 |
| chr5-123137885-DEL-1094 | chr5-123137886-DEL-1094 |
| chr5-123178105-DEL-213 | chr5-123178106-DEL-213 |
| chr5-123193762-DEL-202 | chr5-123193763-DEL-202 |
| chr5-123194508-DEL-66 | chr5-123194477-DEL-50 |
| chr5-123199497-DEL-463 | chr5-123199498-DEL-463 |
| chr5-123201983-DEL-145 | chr5-123201975-DEL-151 |
| chr5-123235202-DEL-80 | chr5-123235203-DEL-80 |
| chr5-123303282-DEL-185 | chr5-123303283-DEL-185 |
| chr5-123323268-DEL-74 | chr5-123323271-DEL-72 |
| chr5-123327391-DEL-726 | chr5-123327392-DEL-726 |
| chr5-123328120-DEL-898 | chr5-123328122-DEL-899 |
| chr5-123337058-DEL-657 | chr5-123337059-DEL-652 |
| chr5-123345782-DEL-6452 | chr5-123345783-DEL-6452 |
| chr5-123352531-DEL-211 | chr5-123352532-DEL-211 |
| chr5-123358445-DEL-53 | chr5-123358440-DEL-53 |
| chr5-123388848-DEL-209 | chr5-123388849-DEL-209 |
| chr5-123393507-DEL-431 | chr5-123393508-DEL-431 |
| chr5-123404716-DEL-1654 | chr5-123404717-DEL-1654 |
| chr5-123413606-DEL-62 | chr5-123413607-DEL-62 |
| chr5-123449748-DEL-234 | chr5-123449749-DEL-234 |
| chr5-123460062-DEL-216 | chr5-123460063-DEL-216 |
| chr5-123521125-DEL-83 | chr5-123521122-DEL-79 |
| chr5-123525607-DEL-84 | chr5-123525578-DEL-112 |
| chr5-123538335-DEL-305 | chr5-123538336-DEL-305 |
| chr5-123540057-DEL-1118 | chr5-123540058-DEL-1118 |
| chr5-123561755-DEL-70 | chr5-123561752-DEL-61 |
| chr5-123572958-DEL-872 | chr5-123572959-DEL-872 |
| chr5-123644967-DEL-267 | chr5-123644968-DEL-267 |
| chr5-123659630-DEL-52 | chr5-123659631-DEL-52 |
| chr5-123676793-DEL-6811 | chr5-123676791-DEL-6811 |
| chr5-123685682-DEL-6406 | chr5-123685682-DEL-6406 |
| chr5-123695046-DEL-532 | chr5-123695047-DEL-532 |
| chr5-123698586-DEL-861 | chr5-123698587-DEL-862 |
| chr5-123706137-DEL-110 | chr5-123706138-DEL-110 |
| chr5-123733010-DEL-178 | chr5-123733014-DEL-177 |
| chr5-123734637-DEL-1043 | chr5-123734640-DEL-1041 |
| chr5-123741397-DEL-65 | chr5-123741400-DEL-63 |
| chr5-123763826-DEL-174 | chr5-123763827-DEL-174 |
| chr5-123781506-DEL-55 | chr5-123781516-DEL-50 |
| chr5-123785127-DEL-90 | chr5-123785128-DEL-89 |
| chr5-123791347-DEL-76 | chr5-123791346-DEL-110 |
| chr5-123803080-DEL-212 | chr5-123803081-DEL-212 |
| chr5-123806833-DEL-84 | chr5-123806801-DEL-85 |
| chr5-123815787-DEL-213 | chr5-123815788-DEL-213 |
| chr5-123827300-DEL-255 | chr5-123827301-DEL-254 |
| chr5-123831849-DEL-52 | chr5-123831850-DEL-52 |
| chr5-123833304-DEL-182 | chr5-123833305-DEL-182 |
| chr5-123841527-DEL-56 | chr5-123841534-DEL-57 |
| chr5-123863872-DEL-57 | chr5-123863876-DEL-54 |
| chr5-123894443-DEL-123 | chr5-123894444-DEL-123 |
| chr5-123899886-DEL-82 | chr5-123899887-DEL-112 |
| chr5-123909886-DEL-68 | chr5-123909889-DEL-66 |
| chr5-123971983-DEL-190 | chr5-123971980-DEL-196 |
| chr5-123976572-DEL-162 | chr5-123976573-DEL-162 |
| chr5-123981276-DEL-78 | chr5-123981294-DEL-69 |
| chr5-123987246-DEL-509 | chr5-123987247-DEL-508 |
| chr5-123988784-DEL-196 | chr5-123988785-DEL-195 |
| chr5-124010627-DEL-72 | chr5-124010628-DEL-72 |
| chr5-124011712-DEL-52 | chr5-124011712-DEL-52 |
| chr5-124025652-DEL-58 | chr5-124025653-DEL-58 |
| chr5-124033203-DEL-216 | chr5-124033199-DEL-209 |
| chr5-124057435-DEL-538 | chr5-124057431-DEL-538 |
| chr5-124073100-DEL-58 | chr5-124073103-DEL-55 |
| chr5-124101331-DEL-209 | chr5-124101325-DEL-210 |
| chr5-124148996-DEL-51 | chr5-124148997-DEL-51 |
| chr5-124189464-DEL-53 | chr5-124189462-DEL-52 |
| chr5-124191754-DEL-140 | chr5-124191755-DEL-105 |
| chr5-124203184-DEL-1118 | chr5-124203185-DEL-1118 |

| | |
|---|---|
| chr5-124225805-DEL-222 | chr5-124225806-DEL-222 |
| chr5-124242550-DEL-6460 | chr5-124242551-DEL-6460 |
| chr5-124313917-DEL-223 | chr5-124313918-DEL-223 |
| chr5-124319931-DEL-60 | chr5-124319932-DEL-60 |
| chr5-124342921-DEL-60 | chr5-124342922-DEL-60 |
| chr5-124347139-DEL-260 | chr5-124347140-DEL-260 |
| chr5-124350636-DEL-386 | chr5-124350637-DEL-386 |
| chr5-124409400-DEL-3597 | chr5-124409401-DEL-3597 |
| chr5-124438975-DEL-144 | chr5-124438976-DEL-144 |
| chr5-124442437-DEL-398 | chr5-124442437-DEL-398 |
| chr5-124462052-DEL-202 | chr5-124462053-DEL-202 |
| chr5-124515773-DEL-130 | chr5-124515774-DEL-130 |
| chr5-124525962-DEL-50 | chr5-124525957-DEL-64 |
| chr5-124553927-DEL-168 | chr5-124553928-DEL-168 |
| chr5-124571851-DEL-151 | chr5-124571863-DEL-167 |
| chr5-124603810-DEL-288 | chr5-124603811-DEL-288 |
| chr5-124610060-DEL-53 | chr5-124610061-DEL-53 |
| chr5-124632601-DEL-356 | chr5-124632601-DEL-358 |
| chr5-124638339-DEL-227 | chr5-124638340-DEL-227 |
| chr5-124643724-DEL-198 | chr5-124643725-DEL-198 |
| chr5-124652845-DEL-496 | chr5-124652846-DEL-496 |
| chr5-124661458-DEL-166 | chr5-124661459-DEL-166 |
| chr5-124677671-DEL-205 | chr5-124677672-DEL-205 |
| chr5-124689077-DEL-197 | chr5-124689074-DEL-201 |
| chr5-124701787-DEL-52 | chr5-124701788-DEL-52 |
| chr5-124784878-DEL-162 | chr5-124784879-DEL-160 |
| chr5-124860120-DEL-134 | chr5-124860162-DEL-107 |
| chr5-124861166-DEL-51 | chr5-124861167-DEL-51 |
| chr5-124863850-DEL-399 | chr5-124863851-DEL-399 |
| chr5-124870639-DEL-102 | chr5-124870640-DEL-102 |
| chr5-124892477-DEL-55 | chr5-124892470-DEL-52 |
| chr5-124973552-DEL-56 | chr5-124973552-DEL-57 |
| chr5-124976046-DEL-1003 | chr5-124976047-DEL-1003 |
| chr5-124984285-DEL-4465 | chr5-124984286-DEL-4465 |
| chr5-124994837-DEL-1571 | chr5-124994838-DEL-1571 |
| chr5-125033022-DEL-239 | chr5-125033023-DEL-240 |
| chr5-125068753-DEL-291 | chr5-125068754-DEL-291 |
| chr5-125121883-DEL-253 | chr5-125121884-DEL-253 |
| chr5-125156195-DEL-126 | chr5-125156196-DEL-82 |
| chr5-125168649-DEL-60 | chr5-125168650-DEL-60 |
| chr5-125243534-DEL-124 | chr5-125243535-DEL-124 |
| chr5-125251357-DEL-207 | chr5-125251358-DEL-207 |
| chr5-125285451-DEL-53 | chr5-125285414-DEL-73 |
| chr5-125295698-DEL-219 | chr5-125295699-DEL-219 |
| chr5-125298030-DEL-199 | chr5-125298031-DEL-199 |
| chr5-125308129-DEL-892 | chr5-125308130-DEL-892 |
| chr5-125327125-DEL-174 | chr5-125327126-DEL-174 |
| chr5-125332948-DEL-201 | chr5-125332949-DEL-201 |
| chr5-125357742-DEL-209 | chr5-125357743-DEL-209 |
| chr5-125359594-DEL-243 | chr5-125359583-DEL-232 |
| chr5-125382219-DEL-134 | chr5-125382220-DEL-134 |
| chr5-125424388-DEL-100 | chr5-125424389-DEL-100 |
| chr5-125429722-DEL-1810 | chr5-125429723-DEL-1810 |
| chr5-125431603-DEL-1144 | chr5-125431604-DEL-1144 |
| chr5-125434243-DEL-68 | chr5-125434244-DEL-68 |
| chr5-125438554-DEL-215 | chr5-125438555-DEL-215 |
| chr5-125443391-DEL-198 | chr5-125443392-DEL-198 |
| chr5-125446776-DEL-219 | chr5-125446777-DEL-219 |
| chr5-125449305-DEL-62 | chr5-125449306-DEL-63 |
| chr5-125484331-DEL-92 | chr5-125484332-DEL-92 |
| chr5-125509357-DEL-206 | chr5-125509353-DEL-205 |
| chr5-125513907-DEL-52 | chr5-125513908-DEL-52 |
| chr5-125541316-DEL-167 | chr5-125541316-DEL-167 |
| chr5-125542676-DEL-93 | chr5-125542677-DEL-92 |
| chr5-125543932-DEL-188 | chr5-125543960-DEL-174 |
| chr5-125555614-DEL-56 | chr5-125555615-DEL-56 |
| chr5-125561210-DEL-216 | chr5-125561211-DEL-216 |
| chr5-125563614-DEL-89 | chr5-125563614-DEL-89 |
| chr5-125566249-DEL-212 | chr5-125566274-DEL-182 |
| chr5-125567324-DEL-197 | chr5-125567325-DEL-197 |
| chr5-125580966-DEL-69 | chr5-125580967-DEL-67 |
| chr5-125584595-DEL-201 | chr5-125584596-DEL-201 |
| chr5-125607537-DEL-103 | chr5-125607538-DEL-103 |
| chr5-125630031-DEL-215 | chr5-125630032-DEL-215 |
| chr5-125640076-DEL-287 | chr5-125640077-DEL-287 |
| chr5-125665394-DEL-64 | chr5-125665394-DEL-63 |
| chr5-125675723-DEL-86 | chr5-125675731-DEL-70 |
| chr5-125680662-DEL-53 | chr5-125680631-DEL-53 |
| chr5-125685921-DEL-73 | chr5-125685922-DEL-73 |
| chr5-125774029-DEL-1498 | chr5-125774042-DEL-1484 |

| | |
|---|---|
| chr5-125790150-DEL-109 | chr5-125790151-DEL-109 |
| chr5-125792439-DEL-7099 | chr5-125792440-DEL-7099 |
| chr5-125800901-DEL-104 | chr5-125800902-DEL-104 |
| chr5-125802375-DEL-394 | chr5-125802376-DEL-394 |
| chr5-125824814-DEL-65 | chr5-125824817-DEL-62 |
| chr5-125831139-DEL-81 | chr5-125831140-DEL-81 |
| chr5-125860833-DEL-179 | chr5-125860834-DEL-179 |
| chr5-125885522-DEL-82 | chr5-125885494-DEL-108 |
| chr5-125931249-DEL-52 | chr5-125931250-DEL-52 |
| chr5-125942020-DEL-145 | chr5-125942021-DEL-145 |
| chr5-125977264-DEL-164 | chr5-125977265-DEL-164 |
| chr5-125983171-DEL-52 | chr5-125983171-DEL-52 |
| chr5-126020845-DEL-415 | chr5-126020845-DEL-415 |
| chr5-126102187-DEL-1305 | chr5-126102188-DEL-1305 |
| chr5-126124720-DEL-3146 | chr5-126124721-DEL-3146 |
| chr5-126129801-DEL-2748 | chr5-126129802-DEL-2748 |
| chr5-126164097-DEL-195 | chr5-126164098-DEL-195 |
| chr5-126166195-DEL-57 | chr5-126166196-DEL-57 |
| chr5-126182257-DEL-833 | chr5-126182258-DEL-833 |
| chr5-126212987-DEL-192 | chr5-126212992-DEL-190 |
| chr5-126223031-DEL-52 | chr5-126223015-DEL-60 |
| chr5-126235619-DEL-136 | chr5-126235620-DEL-136 |
| chr5-126242697-DEL-6455 | chr5-126242698-DEL-6455 |
| chr5-126248859-DEL-202 | chr5-126248860-DEL-202 |
| chr5-126271243-DEL-83 | chr5-126271201-DEL-78 |
| chr5-126287523-DEL-63 | chr5-126287487-DEL-62 |
| chr5-126291092-DEL-66 | chr5-126291084-DEL-60 |
| chr5-126315919-DEL-1137 | chr5-126315920-DEL-1137 |
| chr5-126326636-DEL-1263 | chr5-126326637-DEL-1263 |
| chr5-126338556-DEL-64 | chr5-126338570-DEL-53 |
| chr5-126356861-DEL-64 | chr5-126356845-DEL-60 |
| chr5-126364341-DEL-4816 | chr5-126364342-DEL-4816 |
| chr5-126368276-DEL-170 | chr5-126368288-DEL-185 |
| chr5-126393622-DEL-82 | chr5-126393642-DEL-62 |
| chr5-126396274-DEL-117 | chr5-126396272-DEL-118 |
| chr5-126402937-DEL-74 | chr5-126402926-DEL-76 |
| chr5-126426620-DEL-61 | chr5-126426621-DEL-61 |
| chr5-126439502-DEL-6979 | chr5-126439503-DEL-6979 |
| chr5-126452734-DEL-2100 | chr5-126452735-DEL-2100 |
| chr5-126456638-DEL-54 | chr5-126456664-DEL-60 |
| chr5-126504234-DEL-145 | chr5-126504262-DEL-148 |
| chr5-126511613-DEL-60 | chr5-126511614-DEL-60 |
| chr5-126530597-DEL-53 | chr5-126530598-DEL-53 |
| chr5-126575716-DEL-1200 | chr5-126575717-DEL-1200 |
| chr5-126576983-DEL-204 | chr5-126576984-DEL-200 |
| chr5-126606728-DEL-64 | chr5-126606729-DEL-60 |
| chr5-126616700-DEL-63 | chr5-126616700-DEL-63 |
| chr5-126631194-DEL-401 | chr5-126631195-DEL-401 |
| chr5-126640385-DEL-315 | chr5-126640386-DEL-315 |
| chr5-126665886-DEL-350 | chr5-126665887-DEL-350 |
| chr5-126666726-DEL-6557 | chr5-126666724-DEL-6560 |
| chr5-126690270-DEL-85 | chr5-126690271-DEL-85 |
| chr5-126707210-DEL-61 | chr5-126707216-DEL-61 |
| chr5-126721458-DEL-175 | chr5-126721459-DEL-175 |
| chr5-126731268-DEL-93 | chr5-126731266-DEL-94 |
| chr5-126736536-DEL-200 | chr5-126736537-DEL-200 |
| chr5-126747449-DEL-204 | chr5-126747450-DEL-204 |
| chr5-126791530-DEL-129 | chr5-126791531-DEL-129 |
| chr5-126794731-DEL-79 | chr5-126794731-DEL-79 |
| chr5-126822854-DEL-52 | chr5-126822855-DEL-52 |
| chr5-126823504-DEL-52 | chr5-126823505-DEL-52 |
| chr5-126833920-DEL-113 | chr5-126833921-DEL-113 |
| chr5-126844018-DEL-52 | chr5-126844019-DEL-52 |
| chr5-126857127-DEL-2608 | chr5-126857128-DEL-2608 |
| chr5-126866376-DEL-59 | chr5-126866328-DEL-104 |
| chr5-126867480-DEL-52 | chr5-126867487-DEL-50 |
| chr5-126871151-DEL-182 | chr5-126871152-DEL-182 |
| chr5-126886788-DEL-932 | chr5-126886789-DEL-932 |
| chr5-126886991-DEL-178 | chr5-126886992-DEL-178 |
| chr5-126896499-DEL-6415 | chr5-126896500-DEL-6415 |
| chr5-126904115-DEL-128 | chr5-126904116-DEL-128 |
| chr5-126909554-DEL-1472 | chr5-126909555-DEL-1472 |
| chr5-126925630-DEL-1897 | chr5-126925631-DEL-1897 |
| chr5-126929807-DEL-1291 | chr5-126929808-DEL-1291 |
| chr5-126943641-DEL-521 | chr5-126943642-DEL-521 |
| chr5-126947235-DEL-80 | chr5-126947236-DEL-80 |
| chr5-126970897-DEL-60 | chr5-126970894-DEL-61 |
| chr5-126976936-DEL-745 | chr5-126976937-DEL-745 |
| chr5-126981721-DEL-108 | chr5-126981723-DEL-107 |
| chr5-127002511-DEL-121 | chr5-127002512-DEL-121 |

| | |
|---|---|
| chr5-127004380-DEL-84 | chr5-127004381-DEL-84 |
| chr5-127020686-DEL-666 | chr5-127020687-DEL-666 |
| chr5-127025930-DEL-152 | chr5-127025931-DEL-152 |
| chr5-127029878-DEL-67 | chr5-127029879-DEL-67 |
| chr5-127037688-DEL-3911 | chr5-127037689-DEL-3911 |
| chr5-127051215-DEL-58 | chr5-127051216-DEL-58 |
| chr5-127057793-DEL-60 | chr5-127057794-DEL-60 |
| chr5-127060781-DEL-228 | chr5-127060782-DEL-228 |
| chr5-127064659-DEL-78 | chr5-127064660-DEL-78 |
| chr5-127097942-DEL-530 | chr5-127097943-DEL-522 |
| chr5-127101249-DEL-62 | chr5-127101250-DEL-62 |
| chr5-127117957-DEL-378 | chr5-127117955-DEL-373 |
| chr5-127123233-DEL-204 | chr5-127123234-DEL-204 |
| chr5-127144554-DEL-84 | chr5-127144569-DEL-70 |
| chr5-127173190-DEL-625 | chr5-127173191-DEL-625 |
| chr5-127191093-DEL-204 | chr5-127191094-DEL-204 |
| chr5-127194405-DEL-102 | chr5-127194406-DEL-102 |
| chr5-127194949-DEL-1703 | chr5-127194949-DEL-1703 |
| chr5-127204452-DEL-60 | chr5-127204453-DEL-66 |
| chr5-127230205-DEL-1887 | chr5-127230205-DEL-1888 |
| chr5-127245736-DEL-352 | chr5-127245737-DEL-352 |
| chr5-127279957-DEL-69 | chr5-127279958-DEL-69 |
| chr5-127284008-DEL-1544 | chr5-127284009-DEL-1544 |
| chr5-127327929-DEL-146 | chr5-127327930-DEL-146 |
| chr5-127348607-DEL-6611 | chr5-127348608-DEL-6611 |
| chr5-127354677-DEL-201 | chr5-127354678-DEL-201 |
| chr5-127369241-DEL-137 | chr5-127369245-DEL-136 |
| chr5-127400057-DEL-75 | chr5-127400058-DEL-75 |
| chr5-127420543-DEL-502 | chr5-127420544-DEL-502 |
| chr5-127471047-DEL-1399 | chr5-127471048-DEL-1399 |
| chr5-127474640-DEL-70 | chr5-127474645-DEL-66 |
| chr5-127484006-DEL-91 | chr5-127484007-DEL-91 |
| chr5-127486197-DEL-103 | chr5-127486198-DEL-103 |
| chr5-127515230-DEL-70 | chr5-127515231-DEL-70 |
| chr5-127533979-DEL-545 | chr5-127533980-DEL-545 |
| chr5-127535472-DEL-1659 | chr5-127535473-DEL-1659 |
| chr5-127551079-DEL-108 | chr5-127551080-DEL-108 |
| chr5-127567807-DEL-75 | chr5-127567808-DEL-75 |
| chr5-127615017-DEL-3870 | chr5-127615018-DEL-3870 |
| chr5-127658685-DEL-136 | chr5-127658684-DEL-136 |
| chr5-127668404-DEL-224 | chr5-127668405-DEL-224 |
| chr5-127705675-DEL-155 | chr5-127705676-DEL-155 |
| chr5-127711306-DEL-199 | chr5-127711307-DEL-199 |
| chr5-127722902-DEL-58 | chr5-127722905-DEL-51 |
| chr5-127741301-DEL-69 | chr5-127741302-DEL-66 |
| chr5-127751412-DEL-55 | chr5-127751413-DEL-55 |
| chr5-127753002-DEL-72 | chr5-127753011-DEL-65 |
| chr5-127791407-DEL-140 | chr5-127791408-DEL-140 |
| chr5-127799462-DEL-327 | chr5-127799455-DEL-329 |
| chr5-127807096-DEL-51 | chr5-127807096-DEL-51 |
| chr5-127819044-DEL-57 | chr5-127819045-DEL-57 |
| chr5-127819505-DEL-6851 | chr5-127819506-DEL-6851 |
| chr5-127829914-DEL-80 | chr5-127829943-DEL-60 |
| chr5-127833623-DEL-227 | chr5-127833624-DEL-228 |
| chr5-127846366-DEL-59 | chr5-127846363-DEL-56 |
| chr5-127847220-DEL-56 | chr5-127847216-DEL-59 |
| chr5-127873556-DEL-87 | chr5-127873557-DEL-87 |
| chr5-127893717-DEL-132 | chr5-127893718-DEL-132 |
| chr5-127896592-DEL-1749 | chr5-127896593-DEL-1749 |
| chr5-127927062-DEL-54 | chr5-127927063-DEL-54 |
| chr5-127938495-DEL-476 | chr5-127938496-DEL-476 |
| chr5-127944877-DEL-1663 | chr5-127944878-DEL-1663 |
| chr5-127952005-DEL-8158 | chr5-127951993-DEL-8156 |
| chr5-127992430-DEL-78 | chr5-127992431-DEL-78 |
| chr5-128019907-DEL-52 | chr5-128019903-DEL-62 |
| chr5-128032036-DEL-408 | chr5-128032037-DEL-408 |
| chr5-128037873-DEL-62 | chr5-128037874-DEL-62 |
| chr5-128041300-DEL-545 | chr5-128041297-DEL-549 |
| chr5-128057268-DEL-106 | chr5-128057269-DEL-106 |
| chr5-128071567-DEL-905 | chr5-128071568-DEL-905 |
| chr5-128095213-DEL-718 | chr5-128095214-DEL-718 |
| chr5-128098724-DEL-130 | chr5-128098725-DEL-130 |
| chr5-128101322-DEL-1193 | chr5-128101323-DEL-1193 |
| chr5-128200464-DEL-186 | chr5-128200465-DEL-186 |
| chr5-128265189-DEL-168 | chr5-128265190-DEL-167 |
| chr5-128272085-DEL-1158 | chr5-128272095-DEL-1168 |
| chr5-128302440-DEL-52 | chr5-128302436-DEL-70 |
| chr5-128378825-DEL-1066 | chr5-128378828-DEL-1063 |
| chr5-128379933-DEL-106 | chr5-128379931-DEL-109 |
| chr5-128414657-DEL-212 | chr5-128414656-DEL-213 |

| | |
|---|---|
| chr5-128416633-DEL-1856 | chr5-128416634-DEL-1856 |
| chr5-128429838-DEL-56 | chr5-128429839-DEL-56 |
| chr5-128432399-DEL-56 | chr5-128432400-DEL-56 |
| chr5-128444801-DEL-100 | chr5-128444802-DEL-100 |
| chr5-128490416-DEL-119 | chr5-128490417-DEL-119 |
| chr5-128512361-DEL-68 | chr5-128512362-DEL-68 |
| chr5-128520783-DEL-56 | chr5-128520784-DEL-56 |
| chr5-128564581-DEL-104 | chr5-128564582-DEL-104 |
| chr5-128582960-DEL-63 | chr5-128582961-DEL-63 |
| chr5-128594918-DEL-62 | chr5-128594919-DEL-50 |
| chr5-128598347-DEL-57 | chr5-128598347-DEL-57 |
| chr5-128599542-DEL-134 | chr5-128599542-DEL-128 |
| chr5-128627820-DEL-70 | chr5-128627825-DEL-66 |
| chr5-128632541-DEL-202 | chr5-128632542-DEL-202 |
| chr5-128642812-DEL-74 | chr5-128642813-DEL-74 |
| chr5-128690711-DEL-113 | chr5-128690712-DEL-113 |
| chr5-128711654-DEL-52 | chr5-128711651-DEL-56 |
| chr5-128744074-DEL-73 | chr5-128744073-DEL-74 |
| chr5-128751800-DEL-988 | chr5-128751801-DEL-988 |
| chr5-128756770-DEL-53 | chr5-128756771-DEL-53 |
| chr5-128760464-DEL-64 | chr5-128760465-DEL-65 |
| chr5-128815560-DEL-60 | chr5-128815561-DEL-60 |
| chr5-128844665-DEL-1790 | chr5-128844668-DEL-1788 |
| chr5-128862995-DEL-84 | chr5-128862996-DEL-84 |
| chr5-128867644-DEL-488 | chr5-128867661-DEL-465 |
| chr5-128930532-DEL-385 | chr5-128930533-DEL-385 |
| chr5-128932053-DEL-202 | chr5-128932047-DEL-203 |
| chr5-128937023-DEL-175 | chr5-128937024-DEL-159 |
| chr5-128951644-DEL-59 | chr5-128951628-DEL-60 |
| chr5-128959126-DEL-473 | chr5-128959123-DEL-435 |
| chr5-128961782-DEL-63 | chr5-128961783-DEL-63 |
| chr5-128985481-DEL-208 | chr5-128985483-DEL-241 |
| chr5-128999636-DEL-274 | chr5-128999637-DEL-274 |
| chr5-129000688-DEL-58 | chr5-129000686-DEL-55 |
| chr5-129020995-DEL-1876 | chr5-129020996-DEL-1876 |
| chr5-129025222-DEL-53 | chr5-129025223-DEL-53 |
| chr5-129035660-DEL-79 | chr5-129035661-DEL-79 |
| chr5-129036857-DEL-1092 | chr5-129036858-DEL-1092 |
| chr5-129039898-DEL-320 | chr5-129039898-DEL-320 |
| chr5-129052199-DEL-330 | chr5-129052205-DEL-324 |
| chr5-129055156-DEL-782 | chr5-129055157-DEL-782 |
| chr5-129108589-DEL-54 | chr5-129108590-DEL-54 |
| chr5-129156441-DEL-293 | chr5-129156442-DEL-293 |
| chr5-129164584-DEL-71 | chr5-129164565-DEL-72 |
| chr5-129198370-DEL-167 | chr5-129198371-DEL-167 |
| chr5-129202408-DEL-198 | chr5-129202409-DEL-198 |
| chr5-129205050-DEL-586 | chr5-129205051-DEL-586 |
| chr5-129210105-DEL-51 | chr5-129210106-DEL-51 |
| chr5-129231888-DEL-183 | chr5-129231879-DEL-194 |
| chr5-129248014-DEL-60 | chr5-129248015-DEL-60 |
| chr5-129275068-DEL-114 | chr5-129275069-DEL-114 |
| chr5-129315473-DEL-261 | chr5-129315474-DEL-261 |
| chr5-129328292-DEL-415 | chr5-129328293-DEL-415 |
| chr5-129331855-DEL-57 | chr5-129331856-DEL-57 |
| chr5-129351975-DEL-62 | chr5-129351976-DEL-62 |
| chr5-129365381-DEL-246 | chr5-129365382-DEL-246 |
| chr5-129392771-DEL-210 | chr5-129392772-DEL-168 |
| chr5-129394615-DEL-357 | chr5-129394616-DEL-357 |
| chr5-129395175-DEL-55 | chr5-129395176-DEL-55 |
| chr5-129412831-DEL-52 | chr5-129412831-DEL-52 |
| chr5-129424776-DEL-296 | chr5-129424777-DEL-296 |
| chr5-129460629-DEL-1202 | chr5-129460651-DEL-1196 |
| chr5-129460749-DEL-231 | chr5-129460750-DEL-231 |
| chr5-129491536-DEL-124 | chr5-129491537-DEL-124 |
| chr5-129507930-DEL-167 | chr5-129507931-DEL-167 |
| chr5-129514498-DEL-5439 | chr5-129514499-DEL-5439 |
| chr5-129533831-DEL-110 | chr5-129533832-DEL-110 |
| chr5-129608973-DEL-1066 | chr5-129608974-DEL-1066 |
| chr5-129619938-DEL-90 | chr5-129619941-DEL-88 |
| chr5-129643167-DEL-67 | chr5-129643168-DEL-67 |
| chr5-129648705-DEL-1520 | chr5-129648706-DEL-1520 |
| chr5-129650480-DEL-211 | chr5-129650481-DEL-211 |
| chr5-129652169-DEL-111 | chr5-129652162-DEL-112 |
| chr5-129673083-DEL-269 | chr5-129673082-DEL-269 |
| chr5-129692897-DEL-963 | chr5-129692898-DEL-963 |
| chr5-129692902-DEL-121 | chr5-129692901-DEL-117 |
| chr5-129698784-DEL-205 | chr5-129698785-DEL-205 |
| chr5-129699987-DEL-978 | chr5-129699988-DEL-978 |
| chr5-129710299-DEL-161 | chr5-129710300-DEL-161 |
| chr5-129711672-DEL-217 | chr5-129711673-DEL-217 |

| | |
|---|---|
| chr5-129715742-DEL-1254 | chr5-129715743-DEL-1254 |
| chr5-129721320-DEL-5505 | chr5-129721321-DEL-5505 |
| chr5-129729569-DEL-54 | chr5-129729570-DEL-54 |
| chr5-129752485-DEL-69 | chr5-129752486-DEL-69 |
| chr5-129757924-DEL-73 | chr5-129757925-DEL-73 |
| chr5-129813450-DEL-75 | chr5-129813451-DEL-75 |
| chr5-129824518-DEL-174 | chr5-129824519-DEL-174 |
| chr5-129827961-DEL-200 | chr5-129827962-DEL-200 |
| chr5-129833733-DEL-198 | chr5-129833734-DEL-198 |
| chr5-129850634-DEL-653 | chr5-129850635-DEL-653 |
| chr5-129863744-DEL-110 | chr5-129863745-DEL-110 |
| chr5-129867909-DEL-221 | chr5-129867910-DEL-221 |
| chr5-129873224-DEL-370 | chr5-129873225-DEL-370 |
| chr5-129880706-DEL-254 | chr5-129880707-DEL-254 |
| chr5-129882130-DEL-51 | chr5-129882131-DEL-51 |
| chr5-129887517-DEL-217 | chr5-129887518-DEL-217 |
| chr5-129897025-DEL-336 | chr5-129897026-DEL-336 |
| chr5-129903219-DEL-54 | chr5-129903220-DEL-54 |
| chr5-129919204-DEL-202 | chr5-129919205-DEL-202 |
| chr5-129927096-DEL-200 | chr5-129927097-DEL-200 |
| chr5-129947256-DEL-74 | chr5-129947257-DEL-80 |
| chr5-129951473-DEL-194 | chr5-129951474-DEL-195 |
| chr5-129964740-DEL-186 | chr5-129964742-DEL-185 |
| chr5-129967418-DEL-258 | chr5-129967412-DEL-259 |
| chr5-129981805-DEL-206 | chr5-129981806-DEL-206 |
| chr5-130067853-DEL-336 | chr5-130067854-DEL-336 |
| chr5-130072008-DEL-90 | chr5-130072009-DEL-90 |
| chr5-130076945-DEL-202 | chr5-130076939-DEL-196 |
| chr5-130082013-DEL-89 | chr5-130082012-DEL-89 |
| chr5-130122515-DEL-207 | chr5-130122525-DEL-199 |
| chr5-130125782-DEL-61 | chr5-130125783-DEL-61 |
| chr5-130132035-DEL-633 | chr5-130132036-DEL-633 |
| chr5-130150457-DEL-1495 | chr5-130150458-DEL-1495 |
| chr5-130154423-DEL-163 | chr5-130154424-DEL-163 |
| chr5-130161134-DEL-194 | chr5-130161135-DEL-196 |
| chr5-130196717-DEL-56 | chr5-130196718-DEL-56 |
| chr5-130198516-DEL-189 | chr5-130198517-DEL-188 |
| chr5-130217236-DEL-463 | chr5-130217237-DEL-463 |
| chr5-130280241-DEL-1610 | chr5-130280242-DEL-1610 |
| chr5-130297368-DEL-53 | chr5-130297368-DEL-53 |
| chr5-130297853-DEL-74 | chr5-130297854-DEL-74 |
| chr5-130316752-DEL-334 | chr5-130316752-DEL-334 |
| chr5-130324402-DEL-270 | chr5-130324403-DEL-270 |
| chr5-130373641-DEL-74 | chr5-130373619-DEL-75 |
| chr5-130374290-DEL-106 | chr5-130374291-DEL-106 |
| chr5-130383093-DEL-108 | chr5-130383094-DEL-108 |
| chr5-130384656-DEL-1119 | chr5-130384657-DEL-1119 |
| chr5-130426276-DEL-83 | chr5-130426280-DEL-79 |
| chr5-130466701-DEL-2358 | chr5-130466702-DEL-2358 |
| chr5-130488595-DEL-99 | chr5-130488578-DEL-80 |
| chr5-130502677-DEL-58 | chr5-130502665-DEL-50 |
| chr5-130509743-DEL-52 | chr5-130509744-DEL-52 |
| chr5-130511285-DEL-443 | chr5-130511282-DEL-443 |
| chr5-130521676-DEL-64 | chr5-130521676-DEL-64 |
| chr5-130523104-DEL-2085 | chr5-130523105-DEL-2085 |
| chr5-130547762-DEL-62 | chr5-130547762-DEL-62 |
| chr5-130597185-DEL-176 | chr5-130597175-DEL-177 |
| chr5-130622010-DEL-106 | chr5-130622011-DEL-106 |
| chr5-130667699-DEL-57 | chr5-130667698-DEL-52 |
| chr5-130670175-DEL-202 | chr5-130670176-DEL-202 |
| chr5-130677385-DEL-624 | chr5-130677386-DEL-624 |
| chr5-130684050-DEL-53 | chr5-130684051-DEL-53 |
| chr5-130698030-DEL-105 | chr5-130698029-DEL-126 |
| chr5-130711035-DEL-369 | chr5-130711036-DEL-369 |
| chr5-130744966-DEL-118 | chr5-130744969-DEL-74 |
| chr5-130754055-DEL-210 | chr5-130754056-DEL-210 |
| chr5-130775481-DEL-84 | chr5-130775482-DEL-78 |
| chr5-130776596-DEL-537 | chr5-130776597-DEL-537 |
| chr5-130780210-DEL-52 | chr5-130780211-DEL-52 |
| chr5-130781441-DEL-500 | chr5-130781442-DEL-500 |
| chr5-130785204-DEL-65 | chr5-130785205-DEL-65 |
| chr5-130805109-DEL-334 | chr5-130805110-DEL-334 |
| chr5-130816003-DEL-52 | chr5-130815981-DEL-53 |
| chr5-130835764-DEL-92 | chr5-130835765-DEL-92 |
| chr5-130836602-DEL-54 | chr5-130836603-DEL-60 |
| chr5-130837966-DEL-168 | chr5-130837967-DEL-168 |
| chr5-130842187-DEL-60 | chr5-130842188-DEL-60 |
| chr5-130847437-DEL-79 | chr5-130847438-DEL-79 |
| chr5-130865287-DEL-99 | chr5-130865288-DEL-99 |
| chr5-130913165-DEL-236 | chr5-130913166-DEL-236 |

| | |
|---|---|
| chr5-130944242-DEL-92 | chr5-130944255-DEL-80 |
| chr5-130953896-DEL-618 | chr5-130953897-DEL-618 |
| chr5-130957010-DEL-80 | chr5-130957007-DEL-72 |
| chr5-130964043-DEL-80 | chr5-130964044-DEL-80 |
| chr5-130970248-DEL-53 | chr5-130970202-DEL-53 |
| chr5-130981970-DEL-149 | chr5-130981971-DEL-149 |
| chr5-130987962-DEL-154 | chr5-130987964-DEL-152 |
| chr5-131021964-DEL-60 | chr5-131021965-DEL-57 |
| chr5-131024812-DEL-158 | chr5-131024766-DEL-159 |
| chr5-131063270-DEL-198 | chr5-131063268-DEL-201 |
| chr5-131073898-DEL-216 | chr5-131073899-DEL-216 |
| chr5-131079117-DEL-6585 | chr5-131079118-DEL-6585 |
| chr5-131079258-DEL-208 | chr5-131079259-DEL-208 |
| chr5-131114082-DEL-51 | chr5-131114083-DEL-51 |
| chr5-131118549-DEL-66 | chr5-131118550-DEL-66 |
| chr5-131122179-DEL-77 | chr5-131122177-DEL-78 |
| chr5-131128634-DEL-212 | chr5-131128635-DEL-212 |
| chr5-131132313-DEL-58 | chr5-131132324-DEL-50 |
| chr5-131142644-DEL-66 | chr5-131142645-DEL-66 |
| chr5-131162579-DEL-116 | chr5-131162580-DEL-116 |
| chr5-131212455-DEL-70 | chr5-131212456-DEL-70 |
| chr5-131224257-DEL-397 | chr5-131224258-DEL-397 |
| chr5-131278359-DEL-52 | chr5-131278325-DEL-76 |
| chr5-131288957-DEL-184 | chr5-131288978-DEL-160 |
| chr5-131302981-DEL-57 | chr5-131302982-DEL-57 |
| chr5-131307686-DEL-200 | chr5-131307687-DEL-200 |
| chr5-131325409-DEL-722 | chr5-131325410-DEL-722 |
| chr5-131353174-DEL-372 | chr5-131353175-DEL-372 |
| chr5-131369486-DEL-6366 | chr5-131369487-DEL-6366 |
| chr5-131407452-DEL-173 | chr5-131407453-DEL-173 |
| chr5-131412890-DEL-157 | chr5-131412891-DEL-157 |
| chr5-131443772-DEL-119 | chr5-131443773-DEL-119 |
| chr5-131447729-DEL-69 | chr5-131447750-DEL-62 |
| chr5-131455209-DEL-54 | chr5-131455210-DEL-60 |
| chr5-131481797-DEL-401 | chr5-131481798-DEL-401 |
| chr5-131488432-DEL-78 | chr5-131488424-DEL-64 |
| chr5-131493334-DEL-53 | chr5-131493335-DEL-53 |
| chr5-131526511-DEL-54 | chr5-131526512-DEL-54 |
| chr5-131529690-DEL-6445 | chr5-131529691-DEL-6445 |
| chr5-131574759-DEL-53 | chr5-131574759-DEL-53 |
| chr5-131582324-DEL-322 | chr5-131582325-DEL-322 |
| chr5-131588605-DEL-182 | chr5-131588606-DEL-182 |
| chr5-131634340-DEL-88 | chr5-131634345-DEL-84 |
| chr5-131656433-DEL-286 | chr5-131656434-DEL-286 |
| chr5-131659093-DEL-56 | chr5-131659088-DEL-54 |
| chr5-131664869-DEL-2803 | chr5-131664870-DEL-2803 |
| chr5-131669461-DEL-77 | chr5-131669462-DEL-77 |
| chr5-131694735-DEL-56 | chr5-131694720-DEL-72 |
| chr5-131713309-DEL-75 | chr5-131713325-DEL-69 |
| chr5-131717169-DEL-61 | chr5-131717170-DEL-61 |
| chr5-131767230-DEL-448 | chr5-131767231-DEL-448 |
| chr5-131856920-DEL-99 | chr5-131856921-DEL-99 |
| chr5-131910584-DEL-93 | chr5-131910585-DEL-93 |
| chr5-131928021-DEL-65 | chr5-131928049-DEL-54 |
| chr5-131934768-DEL-786 | chr5-131934769-DEL-786 |
| chr5-131948178-DEL-53 | chr5-131948181-DEL-51 |
| chr5-131949791-DEL-53 | chr5-131949792-DEL-53 |
| chr5-131965013-DEL-775 | chr5-131965014-DEL-775 |
| chr5-131974854-DEL-220 | chr5-131974855-DEL-220 |
| chr5-131976551-DEL-223 | chr5-131976552-DEL-223 |
| chr5-131998363-DEL-116 | chr5-131998364-DEL-116 |
| chr5-132007193-DEL-332 | chr5-132007194-DEL-332 |
| chr5-132011006-DEL-205 | chr5-132011004-DEL-208 |
| chr5-132038301-DEL-29865 | chr5-132038301-DEL-29870 |
| chr5-132043590-DEL-201 | chr5-132043591-DEL-201 |
| chr5-132063866-DEL-283 | chr5-132063866-DEL-284 |
| chr5-132066367-DEL-203 | chr5-132066368-DEL-203 |
| chr5-132093141-DEL-82 | chr5-132093144-DEL-80 |
| chr5-132121364-DEL-64 | chr5-132121333-DEL-62 |
| chr5-132146252-DEL-51 | chr5-132146253-DEL-51 |
| chr5-132147535-DEL-7164 | chr5-132147535-DEL-7164 |
| chr5-132175203-DEL-61 | chr5-132175204-DEL-61 |
| chr5-132178182-DEL-178 | chr5-132178177-DEL-175 |
| chr5-132203978-DEL-6457 | chr5-132203979-DEL-6457 |
| chr5-132217816-DEL-86 | chr5-132217854-DEL-50 |
| chr5-132271669-DEL-2926 | chr5-132271670-DEL-2926 |
| chr5-132330736-DEL-51 | chr5-132330733-DEL-53 |
| chr5-132339161-DEL-54 | chr5-132339189-DEL-64 |
| chr5-132342625-DEL-3472 | chr5-132342626-DEL-3472 |
| chr5-132350089-DEL-341 | chr5-132350090-DEL-332 |

| | |
|---|---|
| chr5-132354417-DEL-73 | chr5-132354403-DEL-85 |
| chr5-132408386-DEL-85 | chr5-132408387-DEL-85 |
| chr5-132412225-DEL-96 | chr5-132412226-DEL-96 |
| chr5-132427369-DEL-195 | chr5-132427370-DEL-195 |
| chr5-132489488-DEL-103 | chr5-132489489-DEL-111 |
| chr5-132499073-DEL-63 | chr5-132499074-DEL-63 |
| chr5-132517096-DEL-183 | chr5-132517097-DEL-183 |
| chr5-132534044-DEL-53 | chr5-132534053-DEL-56 |
| chr5-132562835-DEL-52 | chr5-132562836-DEL-52 |
| chr5-132576019-DEL-56 | chr5-132576019-DEL-56 |
| chr5-132593059-DEL-54 | chr5-132593060-DEL-54 |
| chr5-132613076-DEL-55 | chr5-132613066-DEL-62 |
| chr5-132661270-DEL-53 | chr5-132661302-DEL-63 |
| chr5-132704374-DEL-72 | chr5-132704392-DEL-52 |
| chr5-132709634-DEL-6815 | chr5-132709648-DEL-6820 |
| chr5-132720403-DEL-120 | chr5-132720404-DEL-120 |
| chr5-132744793-DEL-153 | chr5-132744794-DEL-153 |
| chr5-132758253-DEL-6818 | chr5-132758254-DEL-6818 |
| chr5-132771327-DEL-68 | chr5-132771330-DEL-66 |
| chr5-132773389-DEL-84 | chr5-132773387-DEL-87 |
| chr5-132783305-DEL-85 | chr5-132783296-DEL-83 |
| chr5-132786061-DEL-71 | chr5-132786062-DEL-71 |
| chr5-132789806-DEL-66 | chr5-132789806-DEL-66 |
| chr5-132797392-DEL-202 | chr5-132797393-DEL-202 |
| chr5-132805908-DEL-69 | chr5-132805900-DEL-67 |
| chr5-132837205-DEL-117 | chr5-132837206-DEL-116 |
| chr5-132861318-DEL-114 | chr5-132861319-DEL-114 |
| chr5-132869214-DEL-399 | chr5-132869215-DEL-399 |
| chr5-132880545-DEL-100 | chr5-132880540-DEL-106 |
| chr5-132888571-DEL-60 | chr5-132888572-DEL-60 |
| chr5-132917604-DEL-56 | chr5-132917605-DEL-56 |
| chr5-132926165-DEL-422 | chr5-132926169-DEL-423 |
| chr5-132942235-DEL-54 | chr5-132942212-DEL-72 |
| chr5-132958653-DEL-134 | chr5-132958651-DEL-135 |
| chr5-132971908-DEL-191 | chr5-132971909-DEL-191 |
| chr5-133005626-DEL-66 | chr5-133005603-DEL-93 |
| chr5-133025971-DEL-55 | chr5-133025972-DEL-55 |
| chr5-133031751-DEL-6461 | chr5-133031752-DEL-6461 |
| chr5-133061469-DEL-134 | chr5-133061472-DEL-132 |
| chr5-133082751-DEL-56 | chr5-133082714-DEL-94 |
| chr5-133094070-DEL-178 | chr5-133094059-DEL-190 |
| chr5-133126360-DEL-52 | chr5-133126361-DEL-52 |
| chr5-133132823-DEL-93 | chr5-133132824-DEL-93 |
| chr5-133138305-DEL-52 | chr5-133138307-DEL-50 |
| chr5-133140258-DEL-57 | chr5-133140257-DEL-60 |
| chr5-133144328-DEL-59 | chr5-133144329-DEL-59 |
| chr5-133152065-DEL-57 | chr5-133152065-DEL-57 |
| chr5-133158715-DEL-53 | chr5-133158716-DEL-53 |
| chr5-133179284-DEL-333 | chr5-133179285-DEL-333 |
| chr5-133205933-DEL-176 | chr5-133205934-DEL-176 |
| chr5-133211104-DEL-57 | chr5-133211092-DEL-72 |
| chr5-133245139-DEL-67 | chr5-133245141-DEL-65 |
| chr5-133246001-DEL-133 | chr5-133245998-DEL-145 |
| chr5-133258756-DEL-84 | chr5-133258755-DEL-84 |
| chr5-133285289-DEL-6756 | chr5-133285291-DEL-6754 |
| chr5-133295115-DEL-86 | chr5-133295120-DEL-82 |
| chr5-133314438-DEL-173 | chr5-133314439-DEL-173 |
| chr5-133315086-DEL-402 | chr5-133315087-DEL-401 |
| chr5-133320557-DEL-94 | chr5-133320534-DEL-96 |
| chr5-133325277-DEL-60 | chr5-133325267-DEL-50 |
| chr5-133350636-DEL-6997 | chr5-133350636-DEL-6997 |
| chr5-133351210-DEL-208 | chr5-133351210-DEL-208 |
| chr5-133363897-DEL-52 | chr5-133363898-DEL-52 |
| chr5-133439981-DEL-344 | chr5-133439982-DEL-344 |
| chr5-133442793-DEL-70 | chr5-133442770-DEL-102 |
| chr5-133497500-DEL-6982 | chr5-133497501-DEL-6982 |
| chr5-133503912-DEL-558 | chr5-133503913-DEL-556 |
| chr5-133535816-DEL-140 | chr5-133535863-DEL-94 |
| chr5-133555244-DEL-190 | chr5-133555239-DEL-191 |
| chr5-133565691-DEL-1463 | chr5-133565692-DEL-1463 |
| chr5-133588203-DEL-53 | chr5-133588246-DEL-55 |
| chr5-133588671-DEL-6674 | chr5-133588672-DEL-6674 |
| chr5-133612357-DEL-53 | chr5-133612358-DEL-53 |
| chr5-133655334-DEL-204 | chr5-133655330-DEL-205 |
| chr5-133667461-DEL-56 | chr5-133667459-DEL-56 |
| chr5-133685110-DEL-210 | chr5-133685111-DEL-208 |
| chr5-133728176-DEL-3792 | chr5-133728177-DEL-3792 |
| chr5-133734201-DEL-87 | chr5-133734202-DEL-87 |
| chr5-133758766-DEL-241 | chr5-133758767-DEL-241 |
| chr5-133770356-DEL-53 | chr5-133770356-DEL-53 |

| | |
|---|---|
| chr5-133814856-DEL-741 | chr5-133814857-DEL-742 |
| chr5-133818726-DEL-54 | chr5-133818726-DEL-54 |
| chr5-133868048-DEL-3455 | chr5-133868049-DEL-3455 |
| chr5-133899764-DEL-2084 | chr5-133899765-DEL-2084 |
| chr5-133914797-DEL-148 | chr5-133914798-DEL-148 |
| chr5-133946439-DEL-1470 | chr5-133946440-DEL-1470 |
| chr5-133978273-DEL-5373 | chr5-133978274-DEL-5373 |
| chr5-133989141-DEL-605 | chr5-133989142-DEL-605 |
| chr5-134045445-DEL-53 | chr5-134045446-DEL-53 |
| chr5-134055118-DEL-53 | chr5-134055119-DEL-53 |
| chr5-134057058-DEL-7048 | chr5-134057059-DEL-7048 |
| chr5-134064661-DEL-65 | chr5-134064620-DEL-103 |
| chr5-134100020-DEL-75 | chr5-134100034-DEL-59 |
| chr5-134108163-DEL-53 | chr5-134108164-DEL-53 |
| chr5-134131093-DEL-156 | chr5-134131086-DEL-154 |
| chr5-134137210-DEL-106 | chr5-134137211-DEL-106 |
| chr5-134150653-DEL-106 | chr5-134150654-DEL-106 |
| chr5-134159912-DEL-56 | chr5-134159900-DEL-53 |
| chr5-134175398-DEL-470 | chr5-134175399-DEL-470 |
| chr5-134183571-DEL-105 | chr5-134183568-DEL-106 |
| chr5-134188628-DEL-208 | chr5-134188629-DEL-209 |
| chr5-134190081-DEL-252 | chr5-134190082-DEL-252 |
| chr5-134230699-DEL-168 | chr5-134230700-DEL-168 |
| chr5-134252643-DEL-53 | chr5-134252644-DEL-53 |
| chr5-134277162-DEL-122 | chr5-134277163-DEL-121 |
| chr5-134317633-DEL-218 | chr5-134317636-DEL-216 |
| chr5-134322886-DEL-89 | chr5-134322887-DEL-89 |
| chr5-134328611-DEL-225 | chr5-134328600-DEL-207 |
| chr5-134349883-DEL-205 | chr5-134349871-DEL-206 |
| chr5-134356562-DEL-252 | chr5-134356563-DEL-252 |
| chr5-134363894-DEL-52 | chr5-134363894-DEL-52 |
| chr5-134368611-DEL-221 | chr5-134368612-DEL-221 |
| chr5-134373410-DEL-166 | chr5-134373410-DEL-166 |
| chr5-134404112-DEL-51 | chr5-134404113-DEL-51 |
| chr5-134409465-DEL-52 | chr5-134409466-DEL-52 |
| chr5-134413391-DEL-52 | chr5-134413392-DEL-52 |
| chr5-134451169-DEL-89 | chr5-134451163-DEL-88 |
| chr5-134453514-DEL-52 | chr5-134453515-DEL-52 |
| chr5-134455680-DEL-58 | chr5-134455681-DEL-58 |
| chr5-134463980-DEL-190 | chr5-134463981-DEL-190 |
| chr5-134475498-DEL-159 | chr5-134475499-DEL-159 |
| chr5-134477459-DEL-474 | chr5-134477460-DEL-474 |
| chr5-134488776-DEL-62 | chr5-134488791-DEL-52 |
| chr5-134539126-DEL-106 | chr5-134539127-DEL-106 |
| chr5-134550948-DEL-61 | chr5-134550946-DEL-58 |
| chr5-134573146-DEL-99 | chr5-134573147-DEL-99 |
| chr5-134578163-DEL-166 | chr5-134578162-DEL-167 |
| chr5-134578882-DEL-106 | chr5-134578883-DEL-106 |
| chr5-134599118-DEL-178 | chr5-134599119-DEL-179 |
| chr5-134607015-DEL-475 | chr5-134607016-DEL-475 |
| chr5-134608163-DEL-53 | chr5-134608178-DEL-52 |
| chr5-134662665-DEL-1882 | chr5-134662665-DEL-1882 |
| chr5-134663855-DEL-200 | chr5-134663856-DEL-200 |
| chr5-134665787-DEL-208 | chr5-134665788-DEL-208 |
| chr5-134680087-DEL-203 | chr5-134680088-DEL-203 |
| chr5-134682194-DEL-231 | chr5-134682195-DEL-231 |
| chr5-134691429-DEL-214 | chr5-134691430-DEL-214 |
| chr5-134719468-DEL-108 | chr5-134719469-DEL-108 |
| chr5-134747455-DEL-106 | chr5-134747456-DEL-106 |
| chr5-134788310-DEL-56 | chr5-134788311-DEL-56 |
| chr5-134801034-DEL-53 | chr5-134801034-DEL-53 |
| chr5-134804187-DEL-60 | chr5-134804188-DEL-60 |
| chr5-134807122-DEL-64 | chr5-134807125-DEL-62 |
| chr5-134825488-DEL-53 | chr5-134825489-DEL-53 |
| chr5-134830376-DEL-69 | chr5-134830389-DEL-57 |
| chr5-134848951-DEL-83 | chr5-134848966-DEL-86 |
| chr5-134886952-DEL-144 | chr5-134886953-DEL-142 |
| chr5-134892625-DEL-76 | chr5-134892626-DEL-76 |
| chr5-134921060-DEL-51 | chr5-134921061-DEL-51 |
| chr5-134939835-DEL-250 | chr5-134939821-DEL-267 |
| chr5-134960733-DEL-80 | chr5-134960734-DEL-80 |
| chr5-134966946-DEL-60 | chr5-134966946-DEL-60 |
| chr5-135001016-DEL-76 | chr5-135001009-DEL-87 |
| chr5-135011654-DEL-125 | chr5-135011696-DEL-80 |
| chr5-135042316-DEL-206 | chr5-135042317-DEL-206 |
| chr5-135045516-DEL-231 | chr5-135045517-DEL-231 |
| chr5-135048581-DEL-53 | chr5-135048539-DEL-53 |
| chr5-135056505-DEL-159 | chr5-135056506-DEL-159 |
| chr5-135057204-DEL-52 | chr5-135057201-DEL-56 |
| chr5-135060584-DEL-220 | chr5-135060585-DEL-220 |

| | |
|---|---|
| chr5-135072312-DEL-415 | chr5-135072335-DEL-377 |
| chr5-135076907-DEL-209 | chr5-135076908-DEL-209 |
| chr5-135078325-DEL-276 | chr5-135078313-DEL-289 |
| chr5-135115139-DEL-60 | chr5-135115140-DEL-60 |
| chr5-135143648-DEL-194 | chr5-135143649-DEL-194 |
| chr5-135149935-DEL-75 | chr5-135149937-DEL-90 |
| chr5-135186652-DEL-205 | chr5-135186653-DEL-205 |
| chr5-135207246-DEL-60 | chr5-135207247-DEL-56 |
| chr5-135236861-DEL-129 | chr5-135236856-DEL-97 |
| chr5-135258335-DEL-209 | chr5-135258336-DEL-209 |
| chr5-135261431-DEL-180 | chr5-135261432-DEL-180 |
| chr5-135267296-DEL-95 | chr5-135267297-DEL-95 |
| chr5-135287983-DEL-470 | chr5-135287984-DEL-470 |
| chr5-135288990-DEL-88 | chr5-135288991-DEL-104 |
| chr5-135311165-DEL-196 | chr5-135311158-DEL-197 |
| chr5-135334422-DEL-52 | chr5-135334423-DEL-52 |
| chr5-135360887-DEL-441 | chr5-135360888-DEL-441 |
| chr5-135386500-DEL-183 | chr5-135386501-DEL-183 |
| chr5-135392175-DEL-212 | chr5-135392177-DEL-211 |
| chr5-135397102-DEL-663 | chr5-135397103-DEL-663 |
| chr5-135401425-DEL-210 | chr5-135401426-DEL-210 |
| chr5-135416194-DEL-475 | chr5-135416195-DEL-475 |
| chr5-135417324-DEL-95 | chr5-135417325-DEL-95 |
| chr5-135429755-DEL-845 | chr5-135429756-DEL-845 |
| chr5-135430895-DEL-219 | chr5-135430896-DEL-219 |
| chr5-135434152-DEL-206 | chr5-135434153-DEL-206 |
| chr5-135440576-DEL-197 | chr5-135440577-DEL-197 |
| chr5-135444041-DEL-188 | chr5-135444042-DEL-188 |
| chr5-135448124-DEL-53 | chr5-135448118-DEL-54 |
| chr5-135460673-DEL-64 | chr5-135460673-DEL-112 |
| chr5-135461107-DEL-493 | chr5-135461108-DEL-493 |
| chr5-135468567-DEL-114 | chr5-135468568-DEL-102 |
| chr5-135477875-DEL-106 | chr5-135477876-DEL-106 |
| chr5-135482287-DEL-95 | chr5-135482288-DEL-95 |
| chr5-135495319-DEL-159 | chr5-135495320-DEL-159 |
| chr5-135522531-DEL-209 | chr5-135522532-DEL-209 |
| chr5-135541375-DEL-74 | chr5-135541348-DEL-86 |
| chr5-135550414-DEL-152 | chr5-135550401-DEL-169 |
| chr5-135556085-DEL-63 | chr5-135556086-DEL-63 |
| chr5-135579293-DEL-208 | chr5-135579294-DEL-208 |
| chr5-135595964-DEL-310 | chr5-135595965-DEL-310 |
| chr5-135627577-DEL-120 | chr5-135627578-DEL-120 |
| chr5-135645947-DEL-193 | chr5-135645949-DEL-192 |
| chr5-135651096-DEL-793 | chr5-135651097-DEL-793 |
| chr5-135652475-DEL-5362 | chr5-135652476-DEL-5362 |
| chr5-135671164-DEL-193 | chr5-135671166-DEL-187 |
| chr5-135676618-DEL-201 | chr5-135676619-DEL-201 |
| chr5-135690525-DEL-57 | chr5-135690525-DEL-57 |
| chr5-135693552-DEL-351 | chr5-135693559-DEL-347 |
| chr5-135696076-DEL-205 | chr5-135696102-DEL-204 |
| chr5-135710659-DEL-53 | chr5-135710660-DEL-53 |
| chr5-135721644-DEL-220 | chr5-135721645-DEL-220 |
| chr5-135787413-DEL-205 | chr5-135787414-DEL-205 |
| chr5-135842730-DEL-193 | chr5-135842731-DEL-193 |
| chr5-135843486-DEL-53 | chr5-135843487-DEL-53 |
| chr5-135868080-DEL-117 | chr5-135868072-DEL-110 |
| chr5-135920690-DEL-89 | chr5-135920697-DEL-83 |
| chr5-135925509-DEL-65 | chr5-135925514-DEL-66 |
| chr5-135954012-DEL-136 | chr5-135954013-DEL-137 |
| chr5-135972825-DEL-58 | chr5-135972819-DEL-62 |
| chr5-135976673-DEL-205 | chr5-135976674-DEL-205 |
| chr5-135977142-DEL-789 | chr5-135977130-DEL-786 |
| chr5-135983565-DEL-179 | chr5-135983566-DEL-179 |
| chr5-136003766-DEL-201 | chr5-136003767-DEL-201 |
| chr5-136015227-DEL-186 | chr5-136015228-DEL-186 |
| chr5-136031320-DEL-53 | chr5-136031321-DEL-53 |
| chr5-136052225-DEL-161 | chr5-136052248-DEL-137 |
| chr5-136076749-DEL-183 | chr5-136076750-DEL-183 |
| chr5-136079496-DEL-212 | chr5-136079497-DEL-212 |
| chr5-136092358-DEL-56 | chr5-136092350-DEL-62 |
| chr5-136098370-DEL-52 | chr5-136098371-DEL-52 |
| chr5-136104469-DEL-497 | chr5-136104470-DEL-497 |
| chr5-136118962-DEL-72 | chr5-136118966-DEL-96 |
| chr5-136124819-DEL-212 | chr5-136124820-DEL-212 |
| chr5-136127955-DEL-61 | chr5-136127928-DEL-52 |
| chr5-136129474-DEL-77 | chr5-136129475-DEL-75 |
| chr5-136137033-DEL-731 | chr5-136137034-DEL-731 |
| chr5-136142520-DEL-53 | chr5-136142521-DEL-53 |
| chr5-136158663-DEL-54 | chr5-136158668-DEL-58 |
| chr5-136172656-DEL-53 | chr5-136172656-DEL-53 |

| | |
|---|---|
| chr5-136184526-DEL-4538 | chr5-136184527-DEL-4538 |
| chr5-136189422-DEL-60 | chr5-136189423-DEL-60 |
| chr5-136195829-DEL-53 | chr5-136195829-DEL-53 |
| chr5-136204699-DEL-240 | chr5-136204698-DEL-237 |
| chr5-136216501-DEL-204 | chr5-136216502-DEL-204 |
| chr5-136233065-DEL-4378 | chr5-136233066-DEL-4378 |
| chr5-136240932-DEL-105 | chr5-136240933-DEL-105 |
| chr5-136241863-DEL-809 | chr5-136241864-DEL-809 |
| chr5-136243041-DEL-420 | chr5-136243042-DEL-420 |
| chr5-136268002-DEL-166 | chr5-136268050-DEL-151 |
| chr5-136278790-DEL-53 | chr5-136278790-DEL-53 |
| chr5-136365231-DEL-3484 | chr5-136365232-DEL-3484 |
| chr5-136379448-DEL-202 | chr5-136379449-DEL-202 |
| chr5-136406927-DEL-53 | chr5-136406928-DEL-53 |
| chr5-136428751-DEL-157 | chr5-136428752-DEL-157 |
| chr5-136458673-DEL-460 | chr5-136458674-DEL-460 |
| chr5-136472382-DEL-65 | chr5-136472383-DEL-50 |
| chr5-136473716-DEL-53 | chr5-136473716-DEL-53 |
| chr5-136486032-DEL-120 | chr5-136486068-DEL-114 |
| chr5-136500066-DEL-217 | chr5-136500062-DEL-217 |
| chr5-136509306-DEL-52 | chr5-136509307-DEL-50 |
| chr5-136550484-DEL-54 | chr5-136550529-DEL-53 |
| chr5-136551265-DEL-60 | chr5-136551229-DEL-51 |
| chr5-136569088-DEL-576 | chr5-136569089-DEL-576 |
| chr5-136618691-DEL-103 | chr5-136618670-DEL-106 |
| chr5-136619611-DEL-941 | chr5-136619612-DEL-941 |
| chr5-136626939-DEL-107 | chr5-136626940-DEL-107 |
| chr5-136636627-DEL-204 | chr5-136636628-DEL-204 |
| chr5-136670363-DEL-54 | chr5-136670364-DEL-54 |
| chr5-136672771-DEL-75 | chr5-136672774-DEL-73 |
| chr5-136677578-DEL-218 | chr5-136677579-DEL-218 |
| chr5-136690647-DEL-560 | chr5-136690654-DEL-554 |
| chr5-136706556-DEL-195 | chr5-136706550-DEL-196 |
| chr5-136715783-DEL-53 | chr5-136715784-DEL-53 |
| chr5-136716971-DEL-66 | chr5-136716972-DEL-54 |
| chr5-136723721-DEL-140 | chr5-136723722-DEL-140 |
| chr5-136728974-DEL-159 | chr5-136728975-DEL-159 |
| chr5-136741010-DEL-82 | chr5-136741011-DEL-82 |
| chr5-136746795-DEL-73 | chr5-136746795-DEL-73 |
| chr5-136755351-DEL-199 | chr5-136755351-DEL-199 |
| chr5-136756314-DEL-439 | chr5-136756277-DEL-463 |
| chr5-136759198-DEL-6413 | chr5-136759199-DEL-6413 |
| chr5-136782065-DEL-159 | chr5-136782092-DEL-159 |
| chr5-136796055-DEL-262 | chr5-136796056-DEL-262 |
| chr5-136805890-DEL-96 | chr5-136805891-DEL-96 |
| chr5-136814502-DEL-64 | chr5-136814505-DEL-53 |
| chr5-136823754-DEL-108 | chr5-136823755-DEL-108 |
| chr5-136846178-DEL-283 | chr5-136846179-DEL-283 |
| chr5-136861696-DEL-2696 | chr5-136861697-DEL-2696 |
| chr5-136867132-DEL-62 | chr5-136867118-DEL-75 |
| chr5-136879310-DEL-53 | chr5-136879311-DEL-53 |
| chr5-136888271-DEL-53 | chr5-136888272-DEL-53 |
| chr5-136906635-DEL-70 | chr5-136906634-DEL-54 |
| chr5-136926558-DEL-216 | chr5-136926545-DEL-203 |
| chr5-136927492-DEL-824 | chr5-136927477-DEL-825 |
| chr5-136944060-DEL-68 | chr5-136944073-DEL-59 |
| chr5-136972849-DEL-94 | chr5-136972850-DEL-94 |
| chr5-136976181-DEL-134 | chr5-136976182-DEL-134 |
| chr5-136978133-DEL-203 | chr5-136978134-DEL-203 |
| chr5-136987090-DEL-59 | chr5-136987084-DEL-107 |
| chr5-137000889-DEL-51 | chr5-137000890-DEL-51 |
| chr5-137007319-DEL-347 | chr5-137007320-DEL-347 |
| chr5-137028851-DEL-203 | chr5-137028880-DEL-188 |
| chr5-137030063-DEL-2279 | chr5-137030064-DEL-2279 |
| chr5-137044056-DEL-55 | chr5-137044052-DEL-59 |
| chr5-137047508-DEL-86 | chr5-137047515-DEL-72 |
| chr5-137052548-DEL-222 | chr5-137052549-DEL-222 |
| chr5-137056413-DEL-89 | chr5-137056388-DEL-90 |
| chr5-137069725-DEL-77 | chr5-137069757-DEL-56 |
| chr5-137093351-DEL-108 | chr5-137093351-DEL-108 |
| chr5-137110686-DEL-515 | chr5-137110687-DEL-515 |
| chr5-137118654-DEL-51 | chr5-137118611-DEL-72 |
| chr5-137122407-DEL-97 | chr5-137122408-DEL-97 |
| chr5-137140643-DEL-53 | chr5-137140644-DEL-53 |
| chr5-137155335-DEL-295 | chr5-137155318-DEL-313 |
| chr5-137156245-DEL-60 | chr5-137156245-DEL-60 |
| chr5-137161885-DEL-54 | chr5-137161885-DEL-54 |
| chr5-137169059-DEL-56 | chr5-137169060-DEL-56 |
| chr5-137178166-DEL-144 | chr5-137178167-DEL-144 |
| chr5-137181478-DEL-54 | chr5-137181478-DEL-54 |

| | |
|---|---|
| chr5-137189346-DEL-54 | chr5-137189347-DEL-54 |
| chr5-137194680-DEL-211 | chr5-137194682-DEL-210 |
| chr5-137205585-DEL-109 | chr5-137205586-DEL-109 |
| chr5-137215614-DEL-2772 | chr5-137215640-DEL-2772 |
| chr5-137221651-DEL-248 | chr5-137221652-DEL-248 |
| chr5-137224397-DEL-53 | chr5-137224398-DEL-53 |
| chr5-137233734-DEL-54 | chr5-137233735-DEL-54 |
| chr5-137234378-DEL-146 | chr5-137234370-DEL-145 |
| chr5-137252778-DEL-588 | chr5-137252779-DEL-588 |
| chr5-137259233-DEL-146 | chr5-137259227-DEL-144 |
| chr5-137266254-DEL-72 | chr5-137266254-DEL-72 |
| chr5-137269363-DEL-102 | chr5-137269393-DEL-94 |
| chr5-137279492-DEL-51 | chr5-137279493-DEL-51 |
| chr5-137280359-DEL-236 | chr5-137280360-DEL-236 |
| chr5-137322829-DEL-60 | chr5-137322862-DEL-50 |
| chr5-137327423-DEL-227 | chr5-137327424-DEL-227 |
| chr5-137345686-DEL-206 | chr5-137345687-DEL-206 |
| chr5-137351739-DEL-57 | chr5-137351739-DEL-57 |
| chr5-137365661-DEL-52 | chr5-137365615-DEL-51 |
| chr5-137368212-DEL-165 | chr5-137368213-DEL-165 |
| chr5-137373627-DEL-222 | chr5-137373628-DEL-222 |
| chr5-137381576-DEL-124 | chr5-137381557-DEL-144 |
| chr5-137387802-DEL-53 | chr5-137387803-DEL-53 |
| chr5-137425242-DEL-74 | chr5-137425243-DEL-74 |
| chr5-137428983-DEL-189 | chr5-137428984-DEL-189 |
| chr5-137429823-DEL-150 | chr5-137429814-DEL-156 |
| chr5-137430732-DEL-96 | chr5-137430736-DEL-53 |
| chr5-137436297-DEL-67 | chr5-137436295-DEL-68 |
| chr5-137444086-DEL-57 | chr5-137444087-DEL-57 |
| chr5-137459529-DEL-107 | chr5-137459530-DEL-107 |
| chr5-137471675-DEL-53 | chr5-137471689-DEL-50 |
| chr5-137477260-DEL-196 | chr5-137477262-DEL-196 |
| chr5-137478578-DEL-98 | chr5-137478570-DEL-107 |
| chr5-137496295-DEL-208 | chr5-137496296-DEL-208 |
| chr5-137503414-DEL-224 | chr5-137503415-DEL-224 |
| chr5-137587186-DEL-143 | chr5-137587190-DEL-96 |
| chr5-137588561-DEL-212 | chr5-137588562-DEL-212 |
| chr5-137595034-DEL-154 | chr5-137594987-DEL-186 |
| chr5-137664337-DEL-197 | chr5-137664338-DEL-197 |
| chr5-137689715-DEL-371 | chr5-137689716-DEL-371 |
| chr5-137699754-DEL-51 | chr5-137699755-DEL-51 |
| chr5-137705995-DEL-106 | chr5-137705996-DEL-106 |
| chr5-137708586-DEL-213 | chr5-137708587-DEL-213 |
| chr5-137713669-DEL-1720 | chr5-137713669-DEL-1720 |
| chr5-137731704-DEL-104 | chr5-137731706-DEL-105 |
| chr5-137734783-DEL-158 | chr5-137734811-DEL-159 |
| chr5-137748193-DEL-420 | chr5-137748194-DEL-420 |
| chr5-137761695-DEL-348 | chr5-137761696-DEL-348 |
| chr5-137767803-DEL-102 | chr5-137767804-DEL-102 |
| chr5-137769613-DEL-212 | chr5-137769612-DEL-212 |
| chr5-137798450-DEL-301 | chr5-137798432-DEL-308 |
| chr5-137824172-DEL-66 | chr5-137824173-DEL-66 |
| chr5-137846798-DEL-1889 | chr5-137846800-DEL-1903 |
| chr5-137871623-DEL-95 | chr5-137871624-DEL-95 |
| chr5-137871801-DEL-831 | chr5-137871796-DEL-829 |
| chr5-137878738-DEL-221 | chr5-137878739-DEL-221 |
| chr5-137879517-DEL-3618 | chr5-137879518-DEL-3618 |
| chr5-137882800-DEL-143 | chr5-137882804-DEL-140 |
| chr5-137940428-DEL-60 | chr5-137940429-DEL-60 |
| chr5-137942366-DEL-70 | chr5-137942371-DEL-74 |
| chr5-137943488-DEL-2125 | chr5-137943487-DEL-2123 |
| chr5-137958742-DEL-99 | chr5-137958743-DEL-88 |
| chr5-137972386-DEL-118 | chr5-137972364-DEL-118 |
| chr5-137980939-DEL-338 | chr5-137980940-DEL-338 |
| chr5-137991069-DEL-58 | chr5-137991070-DEL-58 |
| chr5-137991239-DEL-500 | chr5-137991240-DEL-500 |
| chr5-137997656-DEL-1890 | chr5-137997657-DEL-1890 |
| chr5-138000621-DEL-71 | chr5-138000622-DEL-71 |
| chr5-138004488-DEL-440 | chr5-138004489-DEL-440 |
| chr5-138011823-DEL-122 | chr5-138011852-DEL-104 |
| chr5-138065768-DEL-6369 | chr5-138065769-DEL-6369 |
| chr5-138090517-DEL-174 | chr5-138090518-DEL-174 |
| chr5-138114897-DEL-54 | chr5-138114898-DEL-54 |
| chr5-138129870-DEL-209 | chr5-138129871-DEL-209 |
| chr5-138216817-DEL-62 | chr5-138216817-DEL-62 |
| chr5-138221488-DEL-96 | chr5-138221487-DEL-96 |
| chr5-138229394-DEL-56 | chr5-138229384-DEL-100 |
| chr5-138241202-DEL-52 | chr5-138241183-DEL-82 |
| chr5-138242753-DEL-91 | chr5-138242750-DEL-91 |
| chr5-138267899-DEL-90 | chr5-138267904-DEL-52 |

| | |
|---|---|
| chr5-138347839-DEL-61 | chr5-138347840-DEL-61 |
| chr5-138361501-DEL-2369 | chr5-138361502-DEL-2369 |
| chr5-138443991-DEL-145 | chr5-138443989-DEL-146 |
| chr5-138459041-DEL-1931 | chr5-138459042-DEL-1931 |
| chr5-138460141-DEL-585 | chr5-138460142-DEL-585 |
| chr5-138463239-DEL-232 | chr5-138463240-DEL-232 |
| chr5-138468008-DEL-75 | chr5-138468006-DEL-84 |
| chr5-138484011-DEL-471 | chr5-138484012-DEL-471 |
| chr5-138486783-DEL-175 | chr5-138486782-DEL-176 |
| chr5-138492923-DEL-2007 | chr5-138492924-DEL-2007 |
| chr5-138502959-DEL-1752 | chr5-138502960-DEL-1752 |
| chr5-138510056-DEL-98 | chr5-138510056-DEL-98 |
| chr5-138541231-DEL-206 | chr5-138541232-DEL-206 |
| chr5-138551476-DEL-80 | chr5-138551477-DEL-80 |
| chr5-138565962-DEL-1323 | chr5-138565963-DEL-1323 |
| chr5-138571279-DEL-234 | chr5-138571278-DEL-234 |
| chr5-138592153-DEL-66 | chr5-138592154-DEL-66 |
| chr5-138607001-DEL-237 | chr5-138606978-DEL-238 |
| chr5-138627160-DEL-110 | chr5-138627161-DEL-107 |
| chr5-138628922-DEL-340 | chr5-138628923-DEL-340 |
| chr5-138630727-DEL-54 | chr5-138630724-DEL-52 |
| chr5-138646891-DEL-84 | chr5-138646892-DEL-78 |
| chr5-138653335-DEL-657 | chr5-138653336-DEL-657 |
| chr5-138656109-DEL-52 | chr5-138656110-DEL-52 |
| chr5-138657696-DEL-2356 | chr5-138657697-DEL-2356 |
| chr5-138664068-DEL-316 | chr5-138664069-DEL-316 |
| chr5-138678111-DEL-72 | chr5-138678111-DEL-72 |
| chr5-138683435-DEL-463 | chr5-138683436-DEL-463 |
| chr5-138686687-DEL-162 | chr5-138686688-DEL-162 |
| chr5-138688007-DEL-320 | chr5-138688008-DEL-320 |
| chr5-138715267-DEL-58 | chr5-138715268-DEL-58 |
| chr5-138725662-DEL-1209 | chr5-138725663-DEL-1209 |
| chr5-138758220-DEL-77 | chr5-138758221-DEL-50 |
| chr5-138759433-DEL-78 | chr5-138759434-DEL-78 |
| chr5-138785767-DEL-76 | chr5-138785751-DEL-80 |
| chr5-138788069-DEL-107 | chr5-138788070-DEL-107 |
| chr5-138823138-DEL-358 | chr5-138823143-DEL-326 |
| chr5-138855917-DEL-126 | chr5-138855918-DEL-126 |
| chr5-138857277-DEL-53 | chr5-138857315-DEL-52 |
| chr5-138878224-DEL-262 | chr5-138878191-DEL-263 |
| chr5-138885894-DEL-106 | chr5-138885895-DEL-106 |
| chr5-138920882-DEL-52 | chr5-138920882-DEL-52 |
| chr5-138929406-DEL-52 | chr5-138929407-DEL-52 |
| chr5-138940637-DEL-118 | chr5-138940638-DEL-118 |
| chr5-138950045-DEL-433 | chr5-138950046-DEL-433 |
| chr5-138950686-DEL-68 | chr5-138950686-DEL-68 |
| chr5-138989921-DEL-108 | chr5-138989922-DEL-108 |
| chr5-139038301-DEL-52 | chr5-139038302-DEL-52 |
| chr5-139060904-DEL-53 | chr5-139060905-DEL-53 |
| chr5-139110580-DEL-112 | chr5-139110589-DEL-92 |
| chr5-139197061-DEL-467 | chr5-139197062-DEL-467 |
| chr5-139203086-DEL-98 | chr5-139203087-DEL-98 |
| chr5-139203531-DEL-1888 | chr5-139203532-DEL-1888 |
| chr5-139205476-DEL-68 | chr5-139205477-DEL-68 |
| chr5-139224858-DEL-104 | chr5-139224859-DEL-104 |
| chr5-139264350-DEL-181 | chr5-139264351-DEL-181 |
| chr5-139273272-DEL-53 | chr5-139273273-DEL-53 |
| chr5-139274785-DEL-184 | chr5-139274786-DEL-184 |
| chr5-139278014-DEL-691 | chr5-139278015-DEL-691 |
| chr5-139301850-DEL-111 | chr5-139301814-DEL-101 |
| chr5-139326705-DEL-212 | chr5-139326706-DEL-212 |
| chr5-139334066-DEL-202 | chr5-139334067-DEL-202 |
| chr5-139372489-DEL-66 | chr5-139372490-DEL-66 |
| chr5-139393675-DEL-88 | chr5-139393678-DEL-86 |
| chr5-139437933-DEL-172 | chr5-139437934-DEL-172 |
| chr5-139448201-DEL-64 | chr5-139448202-DEL-64 |
| chr5-139456073-DEL-233 | chr5-139456075-DEL-231 |
| chr5-139463554-DEL-134 | chr5-139463555-DEL-134 |
| chr5-139465526-DEL-301 | chr5-139465527-DEL-301 |
| chr5-139473306-DEL-497 | chr5-139473307-DEL-497 |
| chr5-139482984-DEL-201 | chr5-139482985-DEL-201 |
| chr5-139494139-DEL-84 | chr5-139494140-DEL-80 |
| chr5-139554583-DEL-54 | chr5-139554566-DEL-74 |
| chr5-139573469-DEL-185 | chr5-139573470-DEL-185 |
| chr5-139592621-DEL-52 | chr5-139592621-DEL-52 |
| chr5-139618870-DEL-563 | chr5-139618871-DEL-563 |
| chr5-139632126-DEL-59 | chr5-139632127-DEL-59 |
| chr5-139637839-DEL-60 | chr5-139637840-DEL-60 |
| chr5-139643329-DEL-171 | chr5-139643365-DEL-133 |
| chr5-139649263-DEL-209 | chr5-139649264-DEL-209 |

| | |
|---|---|
| chr5-139657144-DEL-109 | chr5-139657145-DEL-109 |
| chr5-139663872-DEL-113 | chr5-139663873-DEL-113 |
| chr5-139666148-DEL-312 | chr5-139666149-DEL-312 |
| chr5-139685538-DEL-53 | chr5-139685539-DEL-53 |
| chr5-139690541-DEL-184 | chr5-139690542-DEL-184 |
| chr5-139699851-DEL-209 | chr5-139699852-DEL-209 |
| chr5-139700718-DEL-51 | chr5-139700671-DEL-53 |
| chr5-139764805-DEL-205 | chr5-139764806-DEL-205 |
| chr5-139771265-DEL-52 | chr5-139771266-DEL-52 |
| chr5-139774021-DEL-201 | chr5-139774022-DEL-201 |
| chr5-139787732-DEL-105 | chr5-139787733-DEL-105 |
| chr5-139794832-DEL-53 | chr5-139794831-DEL-55 |
| chr5-139801125-DEL-52 | chr5-139801126-DEL-52 |
| chr5-139833297-DEL-106 | chr5-139833294-DEL-105 |
| chr5-139838413-DEL-52 | chr5-139838413-DEL-52 |
| chr5-139843207-DEL-69 | chr5-139843250-DEL-72 |
| chr5-139847376-DEL-203 | chr5-139847375-DEL-204 |
| chr5-139851966-DEL-56 | chr5-139851980-DEL-63 |
| chr5-139852884-DEL-150 | chr5-139852859-DEL-146 |
| chr5-139853736-DEL-1894 | chr5-139853737-DEL-1894 |
| chr5-139860001-DEL-140 | chr5-139860002-DEL-140 |
| chr5-139875813-DEL-105 | chr5-139875814-DEL-105 |
| chr5-139883778-DEL-160 | chr5-139883779-DEL-160 |
| chr5-139890449-DEL-53 | chr5-139890450-DEL-53 |
| chr5-139903242-DEL-211 | chr5-139903243-DEL-211 |
| chr5-139906456-DEL-158 | chr5-139906457-DEL-158 |
| chr5-139919838-DEL-60 | chr5-139919839-DEL-60 |
| chr5-139936539-DEL-216 | chr5-139936540-DEL-216 |
| chr5-139952794-DEL-135 | chr5-139952789-DEL-136 |
| chr5-139968624-DEL-79 | chr5-139968625-DEL-79 |
| chr5-139972732-DEL-66 | chr5-139972733-DEL-61 |
| chr5-139974380-DEL-55 | chr5-139974381-DEL-55 |
| chr5-139975737-DEL-475 | chr5-139975738-DEL-475 |
| chr5-139983751-DEL-137 | chr5-139983752-DEL-137 |
| chr5-139988198-DEL-70 | chr5-139988197-DEL-72 |
| chr5-139997579-DEL-316 | chr5-139997583-DEL-317 |
| chr5-140017075-DEL-55 | chr5-140017100-DEL-54 |
| chr5-140030409-DEL-162 | chr5-140030409-DEL-160 |
| chr5-140105829-DEL-211 | chr5-140105830-DEL-211 |
| chr5-140120766-DEL-230 | chr5-140120767-DEL-230 |
| chr5-140128458-DEL-514 | chr5-140128459-DEL-514 |
| chr5-140159232-DEL-230 | chr5-140159233-DEL-230 |
| chr5-140170157-DEL-921 | chr5-140170158-DEL-921 |
| chr5-140190985-DEL-63 | chr5-140190986-DEL-66 |
| chr5-140229712-DEL-156 | chr5-140229713-DEL-156 |
| chr5-140254883-DEL-207 | chr5-140254884-DEL-207 |
| chr5-140257911-DEL-190 | chr5-140257910-DEL-192 |
| chr5-140259196-DEL-345 | chr5-140259232-DEL-310 |
| chr5-140312692-DEL-52 | chr5-140312693-DEL-52 |
| chr5-140314741-DEL-616 | chr5-140314742-DEL-616 |
| chr5-140328179-DEL-177 | chr5-140328180-DEL-177 |
| chr5-140350531-DEL-2419 | chr5-140350531-DEL-2419 |
| chr5-140353326-DEL-93 | chr5-140353327-DEL-93 |
| chr5-140364618-DEL-209 | chr5-140364619-DEL-209 |
| chr5-140370669-DEL-197 | chr5-140370670-DEL-197 |
| chr5-140384518-DEL-1885 | chr5-140384504-DEL-1900 |
| chr5-140387775-DEL-186 | chr5-140387776-DEL-186 |
| chr5-140396499-DEL-181 | chr5-140396500-DEL-181 |
| chr5-140398441-DEL-616 | chr5-140398455-DEL-603 |
| chr5-140401008-DEL-237 | chr5-140401009-DEL-237 |
| chr5-140424475-DEL-66 | chr5-140424476-DEL-66 |
| chr5-140432799-DEL-118 | chr5-140432799-DEL-118 |
| chr5-140434219-DEL-190 | chr5-140434219-DEL-190 |
| chr5-140456673-DEL-117 | chr5-140456674-DEL-116 |
| chr5-140468661-DEL-80 | chr5-140468662-DEL-80 |
| chr5-140476058-DEL-59 | chr5-140476070-DEL-53 |
| chr5-140483583-DEL-185 | chr5-140483584-DEL-185 |
| chr5-140489582-DEL-86 | chr5-140489583-DEL-86 |
| chr5-140520795-DEL-200 | chr5-140520796-DEL-200 |
| chr5-140524346-DEL-128 | chr5-140524347-DEL-128 |
| chr5-140532891-DEL-202 | chr5-140532900-DEL-180 |
| chr5-140540845-DEL-363 | chr5-140540846-DEL-344 |
| chr5-140542076-DEL-205 | chr5-140542077-DEL-205 |
| chr5-140544155-DEL-3609 | chr5-140544154-DEL-3610 |
| chr5-140549200-DEL-67 | chr5-140549200-DEL-67 |
| chr5-140580033-DEL-266 | chr5-140580034-DEL-266 |
| chr5-140583451-DEL-107 | chr5-140583452-DEL-107 |
| chr5-140585327-DEL-149 | chr5-140585328-DEL-149 |
| chr5-140589760-DEL-56 | chr5-140589761-DEL-56 |
| chr5-140616277-DEL-272 | chr5-140616278-DEL-272 |

| | |
|---|---|
| chr5-140629275-DEL-840 | chr5-140629277-DEL-839 |
| chr5-140639492-DEL-65 | chr5-140639495-DEL-63 |
| chr5-140662716-DEL-52 | chr5-140662712-DEL-53 |
| chr5-140676526-DEL-52 | chr5-140676526-DEL-52 |
| chr5-140680965-DEL-208 | chr5-140680966-DEL-208 |
| chr5-140684499-DEL-105 | chr5-140684500-DEL-105 |
| chr5-140731332-DEL-52 | chr5-140731333-DEL-52 |
| chr5-140736597-DEL-180 | chr5-140736598-DEL-180 |
| chr5-140743952-DEL-93 | chr5-140743951-DEL-95 |
| chr5-140826403-DEL-57 | chr5-140826404-DEL-57 |
| chr5-140831149-DEL-55 | chr5-140831149-DEL-53 |
| chr5-140846683-DEL-6358 | chr5-140846644-DEL-6397 |
| chr5-140860908-DEL-576 | chr5-140860909-DEL-576 |
| chr5-140885734-DEL-99 | chr5-140885735-DEL-99 |
| chr5-140887637-DEL-158 | chr5-140887662-DEL-158 |
| chr5-140892083-DEL-69 | chr5-140892082-DEL-70 |
| chr5-140893417-DEL-61 | chr5-140893418-DEL-61 |
| chr5-140897121-DEL-53 | chr5-140897122-DEL-53 |
| chr5-140901930-DEL-51 | chr5-140901931-DEL-51 |
| chr5-140910836-DEL-105 | chr5-140910836-DEL-105 |
| chr5-140939259-DEL-52 | chr5-140939260-DEL-52 |
| chr5-140943553-DEL-307 | chr5-140943514-DEL-311 |
| chr5-140949544-DEL-265 | chr5-140949545-DEL-265 |
| chr5-140956302-DEL-215 | chr5-140956253-DEL-217 |
| chr5-140987776-DEL-157 | chr5-140987801-DEL-156 |
| chr5-141032620-DEL-75 | chr5-141032625-DEL-63 |
| chr5-141058898-DEL-497 | chr5-141058899-DEL-497 |
| chr5-141060810-DEL-51 | chr5-141060810-DEL-51 |
| chr5-141062593-DEL-53 | chr5-141062594-DEL-53 |
| chr5-141065939-DEL-60 | chr5-141065924-DEL-60 |
| chr5-141074142-DEL-212 | chr5-141074143-DEL-212 |
| chr5-141121630-DEL-52 | chr5-141121631-DEL-52 |
| chr5-141149787-DEL-149 | chr5-141149788-DEL-149 |
| chr5-141156476-DEL-71 | chr5-141156475-DEL-75 |
| chr5-141158973-DEL-60 | chr5-141159005-DEL-56 |
| chr5-141165299-DEL-325 | chr5-141165300-DEL-325 |
| chr5-141168670-DEL-107 | chr5-141168684-DEL-87 |
| chr5-141188855-DEL-69 | chr5-141188854-DEL-55 |
| chr5-141190529-DEL-53 | chr5-141190530-DEL-53 |
| chr5-141194963-DEL-3049 | chr5-141194964-DEL-3049 |
| chr5-141197912-DEL-313 | chr5-141197913-DEL-313 |
| chr5-141220987-DEL-121 | chr5-141220988-DEL-121 |
| chr5-141223841-DEL-138 | chr5-141223843-DEL-141 |
| chr5-141253273-DEL-963 | chr5-141253274-DEL-963 |
| chr5-141307191-DEL-1212 | chr5-141307192-DEL-1212 |
| chr5-141313622-DEL-6839 | chr5-141313623-DEL-6839 |
| chr5-141478567-DEL-78 | chr5-141478568-DEL-78 |
| chr5-141480280-DEL-55 | chr5-141480281-DEL-55 |
| chr5-141482887-DEL-54 | chr5-141482886-DEL-60 |
| chr5-141513592-DEL-387 | chr5-141513593-DEL-387 |
| chr5-141514767-DEL-91 | chr5-141514768-DEL-91 |
| chr5-141529715-DEL-69 | chr5-141529715-DEL-69 |
| chr5-141574176-DEL-144 | chr5-141574177-DEL-144 |
| chr5-141589212-DEL-179 | chr5-141589213-DEL-179 |
| chr5-141625479-DEL-2878 | chr5-141625480-DEL-2878 |
| chr5-141632757-DEL-141 | chr5-141632758-DEL-141 |
| chr5-141660790-DEL-297 | chr5-141660819-DEL-264 |
| chr5-141668084-DEL-51 | chr5-141668077-DEL-52 |
| chr5-141679844-DEL-188 | chr5-141679877-DEL-156 |
| chr5-141684171-DEL-53 | chr5-141684172-DEL-53 |
| chr5-141686693-DEL-129 | chr5-141686694-DEL-129 |
| chr5-141697466-DEL-66 | chr5-141697469-DEL-64 |
| chr5-141709225-DEL-365 | chr5-141709255-DEL-365 |
| chr5-141723288-DEL-135 | chr5-141723289-DEL-135 |
| chr5-141726321-DEL-3513 | chr5-141726322-DEL-3513 |
| chr5-141753962-DEL-69 | chr5-141753963-DEL-69 |
| chr5-141768489-DEL-211 | chr5-141768489-DEL-212 |
| chr5-141785035-DEL-72 | chr5-141785039-DEL-64 |
| chr5-141814410-DEL-257 | chr5-141814411-DEL-257 |
| chr5-141820160-DEL-442 | chr5-141820161-DEL-442 |
| chr5-141840925-DEL-53 | chr5-141840926-DEL-53 |
| chr5-141886254-DEL-94 | chr5-141886255-DEL-102 |
| chr5-141909475-DEL-6232 | chr5-141909476-DEL-6232 |
| chr5-141935113-DEL-100 | chr5-141935114-DEL-100 |
| chr5-141960933-DEL-76 | chr5-141960943-DEL-74 |
| chr5-141977801-DEL-64 | chr5-141977802-DEL-66 |
| chr5-141981759-DEL-53 | chr5-141981759-DEL-52 |
| chr5-141982363-DEL-56 | chr5-141982364-DEL-50 |
| chr5-141993876-DEL-77 | chr5-141993835-DEL-76 |
| chr5-142021845-DEL-110 | chr5-142021846-DEL-110 |

| | |
|---|---|
| chr5-142044947-DEL-53 | chr5-142044940-DEL-52 |
| chr5-142051606-DEL-3763 | chr5-142051607-DEL-3763 |
| chr5-142056744-DEL-228 | chr5-142056745-DEL-228 |
| chr5-142059059-DEL-52 | chr5-142059060-DEL-52 |
| chr5-142081074-DEL-108 | chr5-142081075-DEL-108 |
| chr5-142116210-DEL-52 | chr5-142116210-DEL-52 |
| chr5-142141026-DEL-82 | chr5-142141028-DEL-81 |
| chr5-142154666-DEL-126 | chr5-142154650-DEL-142 |
| chr5-142175897-DEL-75 | chr5-142175898-DEL-75 |
| chr5-142179845-DEL-52 | chr5-142179846-DEL-52 |
| chr5-142193161-DEL-52 | chr5-142193161-DEL-52 |
| chr5-142203938-DEL-52 | chr5-142203939-DEL-52 |
| chr5-142217574-DEL-58 | chr5-142217575-DEL-58 |
| chr5-142223096-DEL-51 | chr5-142223097-DEL-51 |
| chr5-142238214-DEL-477 | chr5-142238215-DEL-477 |
| chr5-142270624-DEL-52 | chr5-142270625-DEL-52 |
| chr5-142279199-DEL-52 | chr5-142279200-DEL-52 |
| chr5-142282214-DEL-80 | chr5-142282215-DEL-80 |
| chr5-142282796-DEL-433 | chr5-142282791-DEL-434 |
| chr5-142286810-DEL-74 | chr5-142286811-DEL-74 |
| chr5-142316247-DEL-52 | chr5-142316247-DEL-52 |
| chr5-142327240-DEL-83 | chr5-142327241-DEL-83 |
| chr5-142332831-DEL-62 | chr5-142332828-DEL-70 |
| chr5-142343696-DEL-431 | chr5-142343697-DEL-431 |
| chr5-142344687-DEL-120 | chr5-142344725-DEL-89 |
| chr5-142347119-DEL-103 | chr5-142347120-DEL-103 |
| chr5-142356560-DEL-208 | chr5-142356561-DEL-208 |
| chr5-142370944-DEL-61 | chr5-142370945-DEL-61 |
| chr5-142379573-DEL-207 | chr5-142379615-DEL-206 |
| chr5-142408670-DEL-244 | chr5-142408671-DEL-244 |
| chr5-142438063-DEL-206 | chr5-142438062-DEL-204 |
| chr5-142466209-DEL-112 | chr5-142466210-DEL-112 |
| chr5-142517637-DEL-177 | chr5-142517638-DEL-177 |
| chr5-142519337-DEL-533 | chr5-142519340-DEL-532 |
| chr5-142522904-DEL-153 | chr5-142522905-DEL-152 |
| chr5-142525775-DEL-52 | chr5-142525776-DEL-52 |
| chr5-142533100-DEL-139 | chr5-142533101-DEL-139 |
| chr5-142543222-DEL-72 | chr5-142543221-DEL-74 |
| chr5-142574744-DEL-60 | chr5-142574755-DEL-50 |
| chr5-142577609-DEL-61 | chr5-142577610-DEL-57 |
| chr5-142586488-DEL-135 | chr5-142586489-DEL-135 |
| chr5-142599198-DEL-52 | chr5-142599199-DEL-52 |
| chr5-142612092-DEL-64 | chr5-142612093-DEL-64 |
| chr5-142628119-DEL-164 | chr5-142628120-DEL-164 |
| chr5-142631247-DEL-217 | chr5-142631248-DEL-217 |
| chr5-142634499-DEL-52 | chr5-142634500-DEL-52 |
| chr5-142671287-DEL-84 | chr5-142671292-DEL-80 |
| chr5-142705240-DEL-53 | chr5-142705241-DEL-53 |
| chr5-142763618-DEL-60 | chr5-142763619-DEL-60 |
| chr5-142765342-DEL-99 | chr5-142765351-DEL-91 |
| chr5-142767708-DEL-103 | chr5-142767707-DEL-103 |
| chr5-142771584-DEL-207 | chr5-142771566-DEL-204 |
| chr5-142781135-DEL-52 | chr5-142781147-DEL-52 |
| chr5-142818483-DEL-206 | chr5-142818484-DEL-206 |
| chr5-142826191-DEL-109 | chr5-142826195-DEL-110 |
| chr5-142831047-DEL-104 | chr5-142831082-DEL-104 |
| chr5-142833807-DEL-59 | chr5-142833808-DEL-58 |
| chr5-142835113-DEL-53 | chr5-142835113-DEL-53 |
| chr5-142841726-DEL-204 | chr5-142841743-DEL-205 |
| chr5-142845897-DEL-72 | chr5-142845894-DEL-72 |
| chr5-142858742-DEL-151 | chr5-142858743-DEL-151 |
| chr5-142876002-DEL-212 | chr5-142876003-DEL-212 |
| chr5-142915896-DEL-194 | chr5-142915897-DEL-194 |
| chr5-142920009-DEL-199 | chr5-142920010-DEL-199 |
| chr5-142926953-DEL-56 | chr5-142926957-DEL-52 |
| chr5-142928315-DEL-68 | chr5-142928331-DEL-53 |
| chr5-142930706-DEL-85 | chr5-142930688-DEL-90 |
| chr5-142934477-DEL-195 | chr5-142934478-DEL-195 |
| chr5-142937013-DEL-119 | chr5-142937014-DEL-119 |
| chr5-142942770-DEL-52 | chr5-142942771-DEL-52 |
| chr5-142950784-DEL-55 | chr5-142950785-DEL-50 |
| chr5-142951484-DEL-66 | chr5-142951469-DEL-68 |
| chr5-142965522-DEL-401 | chr5-142965522-DEL-401 |
| chr5-142968712-DEL-483 | chr5-142968723-DEL-470 |
| chr5-142969710-DEL-105 | chr5-142969711-DEL-105 |
| chr5-142980404-DEL-1912 | chr5-142980405-DEL-1912 |
| chr5-142990446-DEL-4781 | chr5-142990447-DEL-4781 |
| chr5-143003940-DEL-129 | chr5-143003941-DEL-129 |
| chr5-143008184-DEL-234 | chr5-143008185-DEL-234 |
| chr5-143010278-DEL-220 | chr5-143010279-DEL-220 |

| | |
|---|---|
| chr5-143084967-DEL-565 | chr5-143084968-DEL-566 |
| chr5-143091141-DEL-213 | chr5-143091141-DEL-214 |
| chr5-143134356-DEL-888 | chr5-143134355-DEL-894 |
| chr5-143138078-DEL-67 | chr5-143138109-DEL-68 |
| chr5-143168222-DEL-156 | chr5-143168230-DEL-114 |
| chr5-143184028-DEL-113 | chr5-143184029-DEL-113 |
| chr5-143192990-DEL-455 | chr5-143192986-DEL-453 |
| chr5-143194703-DEL-77 | chr5-143194663-DEL-77 |
| chr5-143203116-DEL-64 | chr5-143203117-DEL-64 |
| chr5-143215155-DEL-53 | chr5-143215151-DEL-53 |
| chr5-143217219-DEL-218 | chr5-143217220-DEL-218 |
| chr5-143222787-DEL-55 | chr5-143222788-DEL-55 |
| chr5-143224722-DEL-131 | chr5-143224723-DEL-131 |
| chr5-143228731-DEL-78 | chr5-143228750-DEL-52 |
| chr5-143264271-DEL-155 | chr5-143264249-DEL-186 |
| chr5-143283008-DEL-60 | chr5-143283009-DEL-60 |
| chr5-143312585-DEL-203 | chr5-143312615-DEL-193 |
| chr5-143322460-DEL-126 | chr5-143322461-DEL-126 |
| chr5-143334821-DEL-499 | chr5-143334822-DEL-499 |
| chr5-143336290-DEL-413 | chr5-143336291-DEL-413 |
| chr5-143346676-DEL-54 | chr5-143346652-DEL-56 |
| chr5-143347904-DEL-78 | chr5-143347905-DEL-54 |
| chr5-143364709-DEL-78 | chr5-143364733-DEL-52 |
| chr5-143374922-DEL-86 | chr5-143374923-DEL-86 |
| chr5-143399223-DEL-210 | chr5-143399224-DEL-210 |
| chr5-143412989-DEL-5290 | chr5-143412990-DEL-5290 |
| chr5-143423643-DEL-360 | chr5-143423644-DEL-360 |
| chr5-143425598-DEL-201 | chr5-143425599-DEL-201 |
| chr5-143426311-DEL-187 | chr5-143426312-DEL-187 |
| chr5-143441308-DEL-109 | chr5-143441309-DEL-109 |
| chr5-143454708-DEL-149 | chr5-143454709-DEL-149 |
| chr5-143458369-DEL-63 | chr5-143458367-DEL-64 |
| chr5-143463778-DEL-97 | chr5-143463771-DEL-96 |
| chr5-143466575-DEL-216 | chr5-143466571-DEL-214 |
| chr5-143546431-DEL-229 | chr5-143546432-DEL-229 |
| chr5-143552556-DEL-171 | chr5-143552557-DEL-171 |
| chr5-143553999-DEL-59 | chr5-143554000-DEL-59 |
| chr5-143568449-DEL-53 | chr5-143568450-DEL-53 |
| chr5-143586510-DEL-54 | chr5-143586494-DEL-71 |
| chr5-143597561-DEL-53 | chr5-143597562-DEL-53 |
| chr5-143601055-DEL-54 | chr5-143601056-DEL-54 |
| chr5-143633181-DEL-58 | chr5-143633196-DEL-56 |
| chr5-143652287-DEL-75 | chr5-143652288-DEL-75 |
| chr5-143654503-DEL-65 | chr5-143654504-DEL-65 |
| chr5-143661506-DEL-57 | chr5-143661509-DEL-59 |
| chr5-143677748-DEL-82 | chr5-143677749-DEL-82 |
| chr5-143734734-DEL-106 | chr5-143734731-DEL-107 |
| chr5-143754172-DEL-61 | chr5-143754173-DEL-61 |
| chr5-143754814-DEL-61 | chr5-143754813-DEL-62 |
| chr5-143773143-DEL-7103 | chr5-143773144-DEL-7103 |
| chr5-143781466-DEL-3271 | chr5-143781467-DEL-3271 |
| chr5-143798932-DEL-205 | chr5-143798931-DEL-206 |
| chr5-143805250-DEL-59 | chr5-143805251-DEL-59 |
| chr5-143818395-DEL-200 | chr5-143818396-DEL-200 |
| chr5-143904711-DEL-82 | chr5-143904712-DEL-82 |
| chr5-143912692-DEL-75 | chr5-143912693-DEL-75 |
| chr5-143915625-DEL-208 | chr5-143915626-DEL-208 |
| chr5-143941222-DEL-216 | chr5-143941213-DEL-220 |
| chr5-143943596-DEL-154 | chr5-143943601-DEL-149 |
| chr5-143946792-DEL-177 | chr5-143946793-DEL-177 |
| chr5-143950445-DEL-260 | chr5-143950446-DEL-260 |
| chr5-143955102-DEL-135 | chr5-143955103-DEL-135 |
| chr5-143959454-DEL-90 | chr5-143959455-DEL-90 |
| chr5-143968489-DEL-137 | chr5-143968502-DEL-150 |
| chr5-143972745-DEL-56 | chr5-143972753-DEL-53 |
| chr5-143986224-DEL-106 | chr5-143986225-DEL-106 |
| chr5-143995875-DEL-640 | chr5-143995876-DEL-640 |
| chr5-143997883-DEL-146 | chr5-143997884-DEL-146 |
| chr5-144001242-DEL-206 | chr5-144001243-DEL-206 |
| chr5-144004714-DEL-1924 | chr5-144004760-DEL-1895 |
| chr5-144011236-DEL-70 | chr5-144011235-DEL-72 |
| chr5-144018558-DEL-204 | chr5-144018559-DEL-204 |
| chr5-144027510-DEL-3007 | chr5-144027511-DEL-3007 |
| chr5-144030883-DEL-431 | chr5-144030884-DEL-385 |
| chr5-144060273-DEL-87 | chr5-144060274-DEL-87 |
| chr5-144082432-DEL-233 | chr5-144082433-DEL-233 |
| chr5-144103833-DEL-51 | chr5-144103834-DEL-51 |
| chr5-144116825-DEL-99 | chr5-144116826-DEL-99 |
| chr5-144149549-DEL-110 | chr5-144149549-DEL-112 |
| chr5-144167850-DEL-156 | chr5-144167851-DEL-156 |

| | |
|---|---|
| chr5-144169070-DEL-206 | chr5-144169070-DEL-206 |
| chr5-144180301-DEL-102 | chr5-144180304-DEL-97 |
| chr5-144200658-DEL-103 | chr5-144200647-DEL-114 |
| chr5-144205511-DEL-193 | chr5-144205512-DEL-193 |
| chr5-144222713-DEL-100 | chr5-144222720-DEL-97 |
| chr5-144270833-DEL-57 | chr5-144270834-DEL-57 |
| chr5-144283236-DEL-69 | chr5-144283254-DEL-51 |
| chr5-144289558-DEL-95 | chr5-144289559-DEL-95 |
| chr5-144307654-DEL-58 | chr5-144307658-DEL-54 |
| chr5-144312940-DEL-54 | chr5-144312940-DEL-54 |
| chr5-144315755-DEL-105 | chr5-144315759-DEL-132 |
| chr5-144317941-DEL-107 | chr5-144317941-DEL-107 |
| chr5-144327393-DEL-197 | chr5-144327394-DEL-197 |
| chr5-144328490-DEL-98 | chr5-144328491-DEL-98 |
| chr5-144333466-DEL-52 | chr5-144333467-DEL-52 |
| chr5-144334137-DEL-68 | chr5-144334137-DEL-69 |
| chr5-144348067-DEL-58 | chr5-144348068-DEL-58 |
| chr5-144354056-DEL-77 | chr5-144354015-DEL-80 |
| chr5-144359502-DEL-390 | chr5-144359503-DEL-390 |
| chr5-144369990-DEL-56 | chr5-144369991-DEL-56 |
| chr5-144383724-DEL-51 | chr5-144383725-DEL-51 |
| chr5-144396223-DEL-111 | chr5-144396219-DEL-112 |
| chr5-144407636-DEL-77 | chr5-144407639-DEL-119 |
| chr5-144411654-DEL-119 | chr5-144411650-DEL-115 |
| chr5-144414758-DEL-58 | chr5-144414755-DEL-56 |
| chr5-144434389-DEL-53 | chr5-144434390-DEL-53 |
| chr5-144441734-DEL-106 | chr5-144441735-DEL-106 |
| chr5-144448230-DEL-963 | chr5-144448231-DEL-963 |
| chr5-144449644-DEL-57 | chr5-144449645-DEL-57 |
| chr5-144452364-DEL-1741 | chr5-144452356-DEL-1729 |
| chr5-144466879-DEL-56 | chr5-144466880-DEL-56 |
| chr5-144490603-DEL-96 | chr5-144490604-DEL-96 |
| chr5-144491717-DEL-82 | chr5-144491718-DEL-82 |
| chr5-144494816-DEL-120 | chr5-144494815-DEL-121 |
| chr5-144527156-DEL-196 | chr5-144527157-DEL-196 |
| chr5-144555074-DEL-60 | chr5-144555074-DEL-60 |
| chr5-144561834-DEL-51 | chr5-144561834-DEL-51 |
| chr5-144591787-DEL-498 | chr5-144591788-DEL-498 |
| chr5-144600309-DEL-74 | chr5-144600309-DEL-74 |
| chr5-144606644-DEL-8985 | chr5-144606645-DEL-8985 |
| chr5-144621464-DEL-122 | chr5-144621464-DEL-122 |
| chr5-144633958-DEL-51 | chr5-144633957-DEL-52 |
| chr5-144640028-DEL-53 | chr5-144640029-DEL-53 |
| chr5-144649060-DEL-211 | chr5-144649051-DEL-212 |
| chr5-144656026-DEL-441 | chr5-144656027-DEL-441 |
| chr5-144661010-DEL-133 | chr5-144661011-DEL-132 |
| chr5-144689716-DEL-158 | chr5-144689717-DEL-158 |
| chr5-144700975-DEL-108 | chr5-144700974-DEL-84 |
| chr5-144704471-DEL-147 | chr5-144704465-DEL-149 |
| chr5-144717121-DEL-96 | chr5-144717122-DEL-96 |
| chr5-144724843-DEL-201 | chr5-144724802-DEL-198 |
| chr5-144755846-DEL-53 | chr5-144755847-DEL-53 |
| chr5-144770432-DEL-84 | chr5-144770433-DEL-84 |
| chr5-144771450-DEL-100 | chr5-144771451-DEL-100 |
| chr5-144862137-DEL-207 | chr5-144862138-DEL-207 |
| chr5-144904484-DEL-122 | chr5-144904485-DEL-122 |
| chr5-144916386-DEL-201 | chr5-144916387-DEL-201 |
| chr5-144918533-DEL-61 | chr5-144918534-DEL-61 |
| chr5-144924033-DEL-55 | chr5-144924029-DEL-58 |
| chr5-144937086-DEL-51 | chr5-144937054-DEL-51 |
| chr5-144967962-DEL-191 | chr5-144967923-DEL-192 |
| chr5-144970394-DEL-56 | chr5-144970406-DEL-54 |
| chr5-144998943-DEL-70 | chr5-144998922-DEL-94 |
| chr5-145010838-DEL-160 | chr5-145010839-DEL-160 |
| chr5-145011339-DEL-105 | chr5-145011341-DEL-103 |
| chr5-145063984-DEL-78 | chr5-145063987-DEL-79 |
| chr5-145091160-DEL-104 | chr5-145091161-DEL-104 |
| chr5-145099142-DEL-111 | chr5-145099143-DEL-111 |
| chr5-145116703-DEL-55 | chr5-145116704-DEL-55 |
| chr5-145133961-DEL-175 | chr5-145133962-DEL-175 |
| chr5-145139756-DEL-67 | chr5-145139757-DEL-67 |
| chr5-145204161-DEL-211 | chr5-145204162-DEL-211 |
| chr5-145214074-DEL-2080 | chr5-145214075-DEL-2080 |
| chr5-145287162-DEL-61 | chr5-145287163-DEL-61 |
| chr5-145306992-DEL-53 | chr5-145306993-DEL-53 |
| chr5-145335541-DEL-1587 | chr5-145335542-DEL-1587 |
| chr5-145418065-DEL-6810 | chr5-145418066-DEL-6810 |
| chr5-145418216-DEL-415 | chr5-145418169-DEL-416 |
| chr5-145443412-DEL-6391 | chr5-145443413-DEL-6391 |
| chr5-145476224-DEL-6436 | chr5-145476225-DEL-6436 |

| | |
|---|---|
| chr5-145497179-DEL-76 | chr5-145497178-DEL-74 |
| chr5-145527128-DEL-209 | chr5-145527128-DEL-209 |
| chr5-145555506-DEL-913 | chr5-145555535-DEL-913 |
| chr5-145557668-DEL-77 | chr5-145557669-DEL-72 |
| chr5-145623466-DEL-203 | chr5-145623465-DEL-203 |
| chr5-145661042-DEL-7925 | chr5-145661043-DEL-7924 |
| chr5-145661556-DEL-78 | chr5-145661557-DEL-78 |
| chr5-145662542-DEL-214 | chr5-145662541-DEL-214 |
| chr5-145679244-DEL-96 | chr5-145679285-DEL-70 |
| chr5-145688563-DEL-204 | chr5-145688564-DEL-204 |
| chr5-145716794-DEL-201 | chr5-145716795-DEL-201 |
| chr5-145756742-DEL-195 | chr5-145756722-DEL-181 |
| chr5-145822508-DEL-198 | chr5-145822509-DEL-198 |
| chr5-145874642-DEL-255 | chr5-145874643-DEL-255 |
| chr5-145894955-DEL-64 | chr5-145894953-DEL-64 |
| chr5-145962480-DEL-369 | chr5-145962481-DEL-369 |
| chr5-145973997-DEL-6628 | chr5-145973981-DEL-6644 |
| chr5-146007847-DEL-52 | chr5-146007851-DEL-51 |
| chr5-146010708-DEL-58 | chr5-146010709-DEL-56 |
| chr5-146012484-DEL-368 | chr5-146012485-DEL-368 |
| chr5-146065796-DEL-1293 | chr5-146065797-DEL-1293 |
| chr5-146071297-DEL-2721 | chr5-146071298-DEL-2721 |
| chr5-146075824-DEL-69 | chr5-146075825-DEL-69 |
| chr5-146091899-DEL-56 | chr5-146091899-DEL-56 |
| chr5-146123136-DEL-74 | chr5-146123137-DEL-74 |
| chr5-146197797-DEL-76 | chr5-146197798-DEL-76 |
| chr5-146200989-DEL-190 | chr5-146200990-DEL-190 |
| chr5-146206717-DEL-380 | chr5-146206717-DEL-380 |
| chr5-146217697-DEL-59 | chr5-146217698-DEL-59 |
| chr5-146258844-DEL-62 | chr5-146258845-DEL-62 |
| chr5-146268833-DEL-317 | chr5-146268834-DEL-317 |
| chr5-146299018-DEL-108 | chr5-146299019-DEL-108 |
| chr5-146300408-DEL-136 | chr5-146300409-DEL-136 |
| chr5-146308740-DEL-1001 | chr5-146308741-DEL-1001 |
| chr5-146311864-DEL-454 | chr5-146311865-DEL-454 |
| chr5-146313875-DEL-125 | chr5-146313876-DEL-125 |
| chr5-146318767-DEL-125 | chr5-146318768-DEL-125 |
| chr5-146355725-DEL-78 | chr5-146355751-DEL-78 |
| chr5-146378123-DEL-577 | chr5-146378124-DEL-577 |
| chr5-146432513-DEL-921 | chr5-146432514-DEL-921 |
| chr5-146487677-DEL-78 | chr5-146487678-DEL-78 |
| chr5-146504965-DEL-548 | chr5-146504966-DEL-548 |
| chr5-146514609-DEL-75 | chr5-146514610-DEL-75 |
| chr5-146544615-DEL-726 | chr5-146544638-DEL-711 |
| chr5-146546949-DEL-212 | chr5-146546943-DEL-208 |
| chr5-146589644-DEL-88 | chr5-146589645-DEL-88 |
| chr5-146600534-DEL-136 | chr5-146600535-DEL-136 |
| chr5-146601215-DEL-210 | chr5-146601216-DEL-210 |
| chr5-146603425-DEL-54 | chr5-146603425-DEL-72 |
| chr5-146604325-DEL-529 | chr5-146604326-DEL-529 |
| chr5-146642591-DEL-62 | chr5-146642585-DEL-90 |
| chr5-146678538-DEL-6436 | chr5-146678539-DEL-6436 |
| chr5-146698227-DEL-78 | chr5-146698228-DEL-72 |
| chr5-146757995-DEL-60 | chr5-146757996-DEL-60 |
| chr5-146774429-DEL-178 | chr5-146774430-DEL-178 |
| chr5-146789378-DEL-1766 | chr5-146789379-DEL-1766 |
| chr5-146791453-DEL-152 | chr5-146791454-DEL-152 |
| chr5-146798040-DEL-85 | chr5-146798042-DEL-81 |
| chr5-146799556-DEL-153 | chr5-146799557-DEL-156 |
| chr5-146800806-DEL-78 | chr5-146800807-DEL-78 |
| chr5-146825432-DEL-156 | chr5-146825433-DEL-156 |
| chr5-146854382-DEL-208 | chr5-146854383-DEL-207 |
| chr5-146855242-DEL-89 | chr5-146855221-DEL-95 |
| chr5-146867786-DEL-102 | chr5-146867787-DEL-102 |
| chr5-146872579-DEL-72 | chr5-146872580-DEL-72 |
| chr5-146878851-DEL-132 | chr5-146878831-DEL-132 |
| chr5-146900995-DEL-92 | chr5-146900996-DEL-92 |
| chr5-146908849-DEL-121 | chr5-146908849-DEL-121 |
| chr5-146909408-DEL-187 | chr5-146909409-DEL-187 |
| chr5-146912834-DEL-52 | chr5-146912832-DEL-53 |
| chr5-146916138-DEL-599 | chr5-146916138-DEL-599 |
| chr5-146923818-DEL-162 | chr5-146923820-DEL-166 |
| chr5-146929671-DEL-84 | chr5-146929671-DEL-85 |
| chr5-146937361-DEL-184 | chr5-146937394-DEL-168 |
| chr5-146947973-DEL-81 | chr5-146947974-DEL-81 |
| chr5-147019962-DEL-54 | chr5-147019963-DEL-54 |
| chr5-147036964-DEL-81 | chr5-147036965-DEL-81 |
| chr5-147059140-DEL-57 | chr5-147059141-DEL-57 |
| chr5-147060958-DEL-54 | chr5-147060977-DEL-50 |
| chr5-147065043-DEL-430 | chr5-147065044-DEL-430 |

| | |
|---|---|
| chr5-147067826-DEL-855 | chr5-147067833-DEL-845 |
| chr5-147083494-DEL-163 | chr5-147083495-DEL-163 |
| chr5-147099427-DEL-51 | chr5-147099405-DEL-52 |
| chr5-147102998-DEL-66 | chr5-147102998-DEL-72 |
| chr5-147106682-DEL-124 | chr5-147106698-DEL-134 |
| chr5-147110068-DEL-227 | chr5-147110064-DEL-226 |
| chr5-147111663-DEL-2301 | chr5-147111664-DEL-2301 |
| chr5-147138358-DEL-65 | chr5-147138357-DEL-65 |
| chr5-147140586-DEL-133 | chr5-147140587-DEL-131 |
| chr5-147144701-DEL-62 | chr5-147144702-DEL-62 |
| chr5-147148690-DEL-876 | chr5-147148691-DEL-876 |
| chr5-147155698-DEL-396 | chr5-147155699-DEL-396 |
| chr5-147166656-DEL-58 | chr5-147166635-DEL-72 |
| chr5-147191596-DEL-79 | chr5-147191597-DEL-79 |
| chr5-147233346-DEL-210 | chr5-147233347-DEL-210 |
| chr5-147259353-DEL-125 | chr5-147259354-DEL-125 |
| chr5-147261838-DEL-189 | chr5-147261839-DEL-189 |
| chr5-147275342-DEL-62 | chr5-147275299-DEL-59 |
| chr5-147282108-DEL-783 | chr5-147282109-DEL-783 |
| chr5-147297320-DEL-54 | chr5-147297315-DEL-60 |
| chr5-147298946-DEL-245 | chr5-147298947-DEL-245 |
| chr5-147313249-DEL-365 | chr5-147313249-DEL-366 |
| chr5-147338883-DEL-169 | chr5-147338884-DEL-168 |
| chr5-147355164-DEL-104 | chr5-147355166-DEL-100 |
| chr5-147357821-DEL-870 | chr5-147357822-DEL-870 |
| chr5-147365155-DEL-214 | chr5-147365156-DEL-214 |
| chr5-147388531-DEL-217 | chr5-147388532-DEL-217 |
| chr5-147416061-DEL-51 | chr5-147416062-DEL-51 |
| chr5-147480110-DEL-59 | chr5-147480096-DEL-56 |
| chr5-147510217-DEL-64 | chr5-147510257-DEL-60 |
| chr5-147512706-DEL-55 | chr5-147512707-DEL-55 |
| chr5-147539974-DEL-154 | chr5-147539975-DEL-154 |
| chr5-147541523-DEL-226 | chr5-147541524-DEL-226 |
| chr5-147542349-DEL-195 | chr5-147542350-DEL-195 |
| chr5-147545141-DEL-841 | chr5-147545142-DEL-841 |
| chr5-147576882-DEL-116 | chr5-147576883-DEL-116 |
| chr5-147584034-DEL-241 | chr5-147584035-DEL-241 |
| chr5-147590240-DEL-61 | chr5-147590229-DEL-73 |
| chr5-147591515-DEL-98 | chr5-147591519-DEL-86 |
| chr5-147619995-DEL-77 | chr5-147619996-DEL-77 |
| chr5-147627058-DEL-196 | chr5-147627067-DEL-197 |
| chr5-147630621-DEL-109 | chr5-147630622-DEL-109 |
| chr5-147635102-DEL-80 | chr5-147635103-DEL-80 |
| chr5-147650272-DEL-51 | chr5-147650272-DEL-52 |
| chr5-147672983-DEL-54 | chr5-147672984-DEL-54 |
| chr5-147675434-DEL-64 | chr5-147675421-DEL-86 |
| chr5-147713583-DEL-230 | chr5-147713584-DEL-270 |
| chr5-147727198-DEL-78 | chr5-147727198-DEL-78 |
| chr5-147732523-DEL-917 | chr5-147732535-DEL-905 |
| chr5-147791686-DEL-153 | chr5-147791685-DEL-162 |
| chr5-147794877-DEL-193 | chr5-147794878-DEL-195 |
| chr5-147806520-DEL-76 | chr5-147806521-DEL-76 |
| chr5-147810688-DEL-84 | chr5-147810689-DEL-84 |
| chr5-147825864-DEL-219 | chr5-147825865-DEL-218 |
| chr5-147840899-DEL-113 | chr5-147840889-DEL-116 |
| chr5-147842130-DEL-134 | chr5-147842133-DEL-124 |
| chr5-147845637-DEL-208 | chr5-147845638-DEL-207 |
| chr5-147849625-DEL-54 | chr5-147849591-DEL-84 |
| chr5-147853444-DEL-112 | chr5-147853443-DEL-79 |
| chr5-147853913-DEL-112 | chr5-147853914-DEL-112 |
| chr5-147863878-DEL-112 | chr5-147863879-DEL-112 |
| chr5-147869323-DEL-63 | chr5-147869324-DEL-63 |
| chr5-147871279-DEL-413 | chr5-147871278-DEL-411 |
| chr5-147878742-DEL-129 | chr5-147878741-DEL-126 |
| chr5-147890206-DEL-513 | chr5-147890207-DEL-513 |
| chr5-147929615-DEL-53 | chr5-147929616-DEL-53 |
| chr5-147931298-DEL-464 | chr5-147931307-DEL-454 |
| chr5-147932191-DEL-401 | chr5-147932192-DEL-401 |
| chr5-147933247-DEL-65 | chr5-147933248-DEL-65 |
| chr5-147945777-DEL-66 | chr5-147945778-DEL-66 |
| chr5-147946081-DEL-331 | chr5-147946082-DEL-331 |
| chr5-147955265-DEL-1356 | chr5-147955266-DEL-1356 |
| chr5-147961538-DEL-51 | chr5-147961539-DEL-51 |
| chr5-147980227-DEL-100 | chr5-147980228-DEL-100 |
| chr5-147983599-DEL-54 | chr5-147983570-DEL-54 |
| chr5-148042552-DEL-289 | chr5-148042553-DEL-289 |
| chr5-148054882-DEL-3417 | chr5-148054895-DEL-3410 |
| chr5-148057037-DEL-69 | chr5-148057012-DEL-75 |
| chr5-148061774-DEL-559 | chr5-148061775-DEL-559 |
| chr5-148094652-DEL-1062 | chr5-148094653-DEL-1062 |

| | |
|---|---|
| chr5-148101761-DEL-131 | chr5-148101764-DEL-129 |
| chr5-148110425-DEL-531 | chr5-148110426-DEL-531 |
| chr5-148119452-DEL-1042 | chr5-148119472-DEL-1023 |
| chr5-148124943-DEL-94 | chr5-148124937-DEL-95 |
| chr5-148134590-DEL-161 | chr5-148134632-DEL-127 |
| chr5-148157810-DEL-1225 | chr5-148157807-DEL-1219 |
| chr5-148196285-DEL-56 | chr5-148196286-DEL-56 |
| chr5-148257651-DEL-61 | chr5-148257652-DEL-61 |
| chr5-148271755-DEL-230 | chr5-148271764-DEL-223 |
| chr5-148291662-DEL-1019 | chr5-148291663-DEL-1019 |
| chr5-148317382-DEL-148 | chr5-148317383-DEL-146 |
| chr5-148341725-DEL-488 | chr5-148341726-DEL-488 |
| chr5-148357191-DEL-501 | chr5-148357190-DEL-502 |
| chr5-148377877-DEL-268 | chr5-148377878-DEL-268 |
| chr5-148402188-DEL-174 | chr5-148402189-DEL-174 |
| chr5-148408087-DEL-60 | chr5-148408077-DEL-74 |
| chr5-148424738-DEL-189 | chr5-148424739-DEL-189 |
| chr5-148505339-DEL-486 | chr5-148505340-DEL-486 |
| chr5-148514672-DEL-57 | chr5-148514673-DEL-57 |
| chr5-148574361-DEL-82 | chr5-148574362-DEL-82 |
| chr5-148575192-DEL-137 | chr5-148575155-DEL-126 |
| chr5-148577617-DEL-994 | chr5-148577618-DEL-994 |
| chr5-148579528-DEL-386 | chr5-148579529-DEL-386 |
| chr5-148586545-DEL-207 | chr5-148586546-DEL-207 |
| chr5-148603587-DEL-64 | chr5-148603558-DEL-94 |
| chr5-148639243-DEL-313 | chr5-148639244-DEL-313 |
| chr5-148654142-DEL-166 | chr5-148654143-DEL-166 |
| chr5-148662255-DEL-69 | chr5-148662256-DEL-69 |
| chr5-148671665-DEL-333 | chr5-148671666-DEL-333 |
| chr5-148704676-DEL-56 | chr5-148704661-DEL-74 |
| chr5-148718985-DEL-51 | chr5-148718986-DEL-51 |
| chr5-148727671-DEL-51 | chr5-148727676-DEL-54 |
| chr5-148731287-DEL-66 | chr5-148731288-DEL-66 |
| chr5-148762215-DEL-106 | chr5-148762216-DEL-106 |
| chr5-148769548-DEL-51 | chr5-148769530-DEL-70 |
| chr5-148779014-DEL-4514 | chr5-148779015-DEL-4514 |
| chr5-148806484-DEL-60 | chr5-148806485-DEL-60 |
| chr5-148807733-DEL-210 | chr5-148807734-DEL-210 |
| chr5-148837281-DEL-70 | chr5-148837278-DEL-72 |
| chr5-148851943-DEL-70 | chr5-148851944-DEL-70 |
| chr5-148884487-DEL-52 | chr5-148884488-DEL-52 |
| chr5-148922724-DEL-75 | chr5-148922725-DEL-75 |
| chr5-148950514-DEL-86 | chr5-148950513-DEL-78 |
| chr5-148975060-DEL-85 | chr5-148975071-DEL-50 |
| chr5-149025297-DEL-116 | chr5-149025298-DEL-107 |
| chr5-149068424-DEL-198 | chr5-149068418-DEL-199 |
| chr5-149077062-DEL-88 | chr5-149077063-DEL-82 |
| chr5-149077929-DEL-215 | chr5-149077930-DEL-215 |
| chr5-149090152-DEL-178 | chr5-149090153-DEL-177 |
| chr5-149090916-DEL-2377 | chr5-149090917-DEL-2377 |
| chr5-149147681-DEL-144 | chr5-149147682-DEL-120 |
| chr5-149156158-DEL-202 | chr5-149156159-DEL-202 |
| chr5-149167028-DEL-68 | chr5-149167029-DEL-68 |
| chr5-149169468-DEL-123 | chr5-149169469-DEL-123 |
| chr5-149230555-DEL-74 | chr5-149230556-DEL-74 |
| chr5-149250250-DEL-896 | chr5-149250251-DEL-896 |
| chr5-149251918-DEL-297 | chr5-149251919-DEL-297 |
| chr5-149287162-DEL-60 | chr5-149287136-DEL-72 |
| chr5-149289691-DEL-96 | chr5-149289692-DEL-96 |
| chr5-149323339-DEL-192 | chr5-149323340-DEL-192 |
| chr5-149343646-DEL-56 | chr5-149343647-DEL-56 |
| chr5-149370615-DEL-205 | chr5-149370616-DEL-205 |
| chr5-149375592-DEL-143 | chr5-149375594-DEL-145 |
| chr5-149395569-DEL-662 | chr5-149395570-DEL-662 |
| chr5-149410909-DEL-156 | chr5-149410928-DEL-138 |
| chr5-149418680-DEL-120 | chr5-149418681-DEL-108 |
| chr5-149435558-DEL-104 | chr5-149435545-DEL-81 |
| chr5-149440956-DEL-76 | chr5-149440994-DEL-68 |
| chr5-149455063-DEL-86 | chr5-149455062-DEL-94 |
| chr5-149464316-DEL-97 | chr5-149464317-DEL-97 |
| chr5-149467524-DEL-204 | chr5-149467525-DEL-204 |
| chr5-149469954-DEL-133 | chr5-149469955-DEL-133 |
| chr5-149497054-DEL-170 | chr5-149497048-DEL-171 |
| chr5-149510382-DEL-56 | chr5-149510383-DEL-56 |
| chr5-149524303-DEL-99 | chr5-149524315-DEL-96 |
| chr5-149551534-DEL-212 | chr5-149551535-DEL-212 |
| chr5-149563761-DEL-316 | chr5-149563762-DEL-316 |
| chr5-149567947-DEL-74 | chr5-149567932-DEL-94 |
| chr5-149574959-DEL-1938 | chr5-149574960-DEL-1937 |
| chr5-149575553-DEL-176 | chr5-149575554-DEL-176 |

| | |
|---|---|
| chr5-149595791-DEL-51 | chr5-149595792-DEL-51 |
| chr5-149613738-DEL-164 | chr5-149613737-DEL-165 |
| chr5-149621862-DEL-72 | chr5-149621875-DEL-50 |
| chr5-149632397-DEL-54 | chr5-149632401-DEL-50 |
| chr5-149641629-DEL-87 | chr5-149641630-DEL-87 |
| chr5-149642063-DEL-204 | chr5-149642064-DEL-204 |
| chr5-149659139-DEL-78 | chr5-149659141-DEL-77 |
| chr5-149662267-DEL-122 | chr5-149662226-DEL-144 |
| chr5-149680713-DEL-90 | chr5-149680745-DEL-60 |
| chr5-149763101-DEL-216 | chr5-149763102-DEL-216 |
| chr5-149804127-DEL-107 | chr5-149804146-DEL-66 |
| chr5-149819779-DEL-448 | chr5-149819780-DEL-448 |
| chr5-149822573-DEL-849 | chr5-149822576-DEL-851 |
| chr5-149829472-DEL-148 | chr5-149829480-DEL-140 |
| chr5-149835326-DEL-72 | chr5-149835327-DEL-72 |
| chr5-149840264-DEL-1050 | chr5-149840265-DEL-1050 |
| chr5-149845280-DEL-431 | chr5-149845281-DEL-431 |
| chr5-149853557-DEL-54 | chr5-149853538-DEL-70 |
| chr5-149862217-DEL-229 | chr5-149862217-DEL-183 |
| chr5-149866507-DEL-70 | chr5-149866508-DEL-70 |
| chr5-149874591-DEL-75 | chr5-149874592-DEL-75 |
| chr5-149906065-DEL-52 | chr5-149906066-DEL-52 |
| chr5-149928291-DEL-59 | chr5-149928292-DEL-58 |
| chr5-149931273-DEL-96 | chr5-149931274-DEL-96 |
| chr5-149993276-DEL-54 | chr5-149993278-DEL-52 |
| chr5-150008060-DEL-151 | chr5-150008020-DEL-146 |
| chr5-150010275-DEL-54 | chr5-150010272-DEL-50 |
| chr5-150071570-DEL-104 | chr5-150071584-DEL-58 |
| chr5-150129566-DEL-51 | chr5-150129567-DEL-51 |
| chr5-150132532-DEL-252 | chr5-150132533-DEL-210 |
| chr5-150150099-DEL-58 | chr5-150150100-DEL-58 |
| chr5-150178074-DEL-121 | chr5-150178050-DEL-153 |
| chr5-150231828-DEL-353 | chr5-150231829-DEL-353 |
| chr5-150246142-DEL-55 | chr5-150246142-DEL-55 |
| chr5-150263355-DEL-55 | chr5-150263352-DEL-55 |
| chr5-150271631-DEL-220 | chr5-150271632-DEL-220 |
| chr5-150277164-DEL-102 | chr5-150277141-DEL-114 |
| chr5-150291325-DEL-152 | chr5-150291326-DEL-152 |
| chr5-150300911-DEL-90 | chr5-150300912-DEL-90 |
| chr5-150349223-DEL-197 | chr5-150349224-DEL-197 |
| chr5-150367958-DEL-316 | chr5-150367959-DEL-316 |
| chr5-150368985-DEL-79 | chr5-150368986-DEL-79 |
| chr5-150397202-DEL-1884 | chr5-150397202-DEL-1884 |
| chr5-150409622-DEL-87 | chr5-150409623-DEL-87 |
| chr5-150499413-DEL-57 | chr5-150499412-DEL-58 |
| chr5-150506982-DEL-98 | chr5-150506983-DEL-98 |
| chr5-150508387-DEL-1031 | chr5-150508388-DEL-1031 |
| chr5-150526296-DEL-465 | chr5-150526297-DEL-465 |
| chr5-150533661-DEL-192 | chr5-150533662-DEL-192 |
| chr5-150538559-DEL-208 | chr5-150538560-DEL-208 |
| chr5-150555251-DEL-204 | chr5-150555252-DEL-204 |
| chr5-150589789-DEL-200 | chr5-150589790-DEL-200 |
| chr5-150607029-DEL-1372 | chr5-150607030-DEL-1372 |
| chr5-150608934-DEL-188 | chr5-150608935-DEL-188 |
| chr5-150616320-DEL-92 | chr5-150616321-DEL-91 |
| chr5-150623462-DEL-1388 | chr5-150623463-DEL-1388 |
| chr5-150638325-DEL-217 | chr5-150638326-DEL-217 |
| chr5-150651655-DEL-60 | chr5-150651656-DEL-60 |
| chr5-150682856-DEL-94 | chr5-150682857-DEL-94 |
| chr5-150683631-DEL-183 | chr5-150683632-DEL-183 |
| chr5-150805430-DEL-102 | chr5-150805440-DEL-60 |
| chr5-150809689-DEL-58 | chr5-150809690-DEL-58 |
| chr5-150824278-DEL-85 | chr5-150824279-DEL-85 |
| chr5-150836693-DEL-189 | chr5-150836694-DEL-189 |
| chr5-150841441-DEL-66 | chr5-150841418-DEL-66 |
| chr5-150845920-DEL-67 | chr5-150845921-DEL-67 |
| chr5-150848928-DEL-1825 | chr5-150848926-DEL-1814 |
| chr5-150866674-DEL-187 | chr5-150866675-DEL-187 |
| chr5-150868411-DEL-57 | chr5-150868393-DEL-75 |
| chr5-150871660-DEL-197 | chr5-150871660-DEL-197 |
| chr5-150887604-DEL-93 | chr5-150887605-DEL-93 |
| chr5-151018583-DEL-69 | chr5-151018575-DEL-68 |
| chr5-151048680-DEL-582 | chr5-151048681-DEL-582 |
| chr5-151054800-DEL-217 | chr5-151054799-DEL-218 |
| chr5-151074359-DEL-116 | chr5-151074360-DEL-116 |
| chr5-151076996-DEL-92 | chr5-151076997-DEL-92 |
| chr5-151107815-DEL-458 | chr5-151107816-DEL-458 |
| chr5-151144493-DEL-75 | chr5-151144494-DEL-75 |
| chr5-151152503-DEL-87 | chr5-151152477-DEL-84 |
| chr5-151190114-DEL-121 | chr5-151190115-DEL-121 |

| | |
|---|---|
| chr5-151204656-DEL-206 | chr5-151204657-DEL-206 |
| chr5-151205364-DEL-194 | chr5-151205365-DEL-194 |
| chr5-151210875-DEL-147 | chr5-151210876-DEL-147 |
| chr5-151223496-DEL-459 | chr5-151223497-DEL-459 |
| chr5-151250628-DEL-202 | chr5-151250629-DEL-202 |
| chr5-151257785-DEL-1535 | chr5-151257786-DEL-1535 |
| chr5-151270477-DEL-83 | chr5-151270475-DEL-80 |
| chr5-151276486-DEL-7044 | chr5-151276487-DEL-7044 |
| chr5-151313510-DEL-112 | chr5-151313511-DEL-112 |
| chr5-151326817-DEL-64 | chr5-151326818-DEL-64 |
| chr5-151348523-DEL-53 | chr5-151348524-DEL-53 |
| chr5-151414847-DEL-84 | chr5-151414848-DEL-84 |
| chr5-151431048-DEL-401 | chr5-151431049-DEL-401 |
| chr5-151446784-DEL-10156 | chr5-151446784-DEL-10183 |
| chr5-151475271-DEL-174 | chr5-151475261-DEL-175 |
| chr5-151480447-DEL-2373 | chr5-151480448-DEL-2373 |
| chr5-151497315-DEL-982 | chr5-151497316-DEL-982 |
| chr5-151501361-DEL-81 | chr5-151501360-DEL-82 |
| chr5-151513086-DEL-252 | chr5-151513082-DEL-253 |
| chr5-151514156-DEL-203 | chr5-151514150-DEL-204 |
| chr5-151525371-DEL-1034 | chr5-151525372-DEL-1034 |
| chr5-151526593-DEL-72 | chr5-151526594-DEL-72 |
| chr5-151532906-DEL-3691 | chr5-151532907-DEL-3691 |
| chr5-151549504-DEL-4727 | chr5-151549505-DEL-4727 |
| chr5-151578563-DEL-7300 | chr5-151578564-DEL-7300 |
| chr5-151603050-DEL-6401 | chr5-151603051-DEL-6401 |
| chr5-151631720-DEL-370 | chr5-151631721-DEL-370 |
| chr5-151640989-DEL-181 | chr5-151640990-DEL-181 |
| chr6-3161074-DEL-1975 | chr6-3161074-DEL-1975 |
| chr6-3174277-DEL-52 | chr6-3174277-DEL-52 |
| chr6-3184640-DEL-7141 | chr6-3184641-DEL-7141 |
| chr6-3192068-DEL-81 | chr6-3192069-DEL-81 |
| chr6-3249606-DEL-5999 | chr6-3249607-DEL-5999 |
| chr6-3258434-DEL-4995 | chr6-3258435-DEL-4995 |
| chr6-3383420-DEL-98 | chr6-3383422-DEL-58 |
| chr6-3398580-DEL-62 | chr6-3398583-DEL-60 |
| chr6-3409031-DEL-134 | chr6-3409036-DEL-130 |
| chr6-3423654-DEL-60 | chr6-3423649-DEL-71 |
| chr6-3428583-DEL-729 | chr6-3428584-DEL-729 |
| chr6-3444926-DEL-65 | chr6-3444927-DEL-65 |
| chr6-3461406-DEL-100 | chr6-3461394-DEL-68 |
| chr6-3508993-DEL-125 | chr6-3508994-DEL-125 |
| chr6-3670149-DEL-1076 | chr6-3670150-DEL-1076 |
| chr6-3676656-DEL-351 | chr6-3676657-DEL-351 |
| chr6-3680931-DEL-1321 | chr6-3680932-DEL-1321 |
| chr6-3685118-DEL-63 | chr6-3685119-DEL-63 |
| chr6-3695962-DEL-3418 | chr6-3695963-DEL-3418 |
| chr6-3787349-DEL-7441 | chr6-3787347-DEL-7442 |
| chr6-3809419-DEL-90 | chr6-3809419-DEL-91 |
| chr6-3814303-DEL-6596 | chr6-3814304-DEL-6596 |
| chr6-3829384-DEL-116 | chr6-3829384-DEL-116 |
| chr6-3918305-DEL-70 | chr6-3918310-DEL-66 |
| chr6-3950803-DEL-1047 | chr6-3950804-DEL-1047 |
| chr6-4037857-DEL-956 | chr6-4037865-DEL-952 |
| chr6-4092396-DEL-214 | chr6-4092397-DEL-214 |
| chr6-4108470-DEL-238 | chr6-4108471-DEL-238 |
| chr6-4117155-DEL-461 | chr6-4117157-DEL-459 |
| chr6-4120254-DEL-117 | chr6-4120287-DEL-107 |
| chr6-4143919-DEL-372 | chr6-4143920-DEL-372 |
| chr6-4157801-DEL-60 | chr6-4157805-DEL-95 |
| chr6-4168420-DEL-839 | chr6-4168421-DEL-839 |
| chr6-4174072-DEL-6405 | chr6-4174101-DEL-6377 |
| chr6-4187970-DEL-68 | chr6-4187971-DEL-68 |
| chr6-4213873-DEL-56 | chr6-4213874-DEL-56 |
| chr6-4216931-DEL-429 | chr6-4216932-DEL-429 |
| chr6-4238828-DEL-51 | chr6-4238829-DEL-51 |
| chr6-4242765-DEL-60 | chr6-4242719-DEL-72 |
| chr6-4249146-DEL-62 | chr6-4249147-DEL-62 |
| chr6-4255527-DEL-6223 | chr6-4255528-DEL-6223 |
| chr6-4282281-DEL-6254 | chr6-4282282-DEL-6254 |
| chr6-4301101-DEL-92 | chr6-4301102-DEL-92 |
| chr6-4303566-DEL-108 | chr6-4303591-DEL-96 |
| chr6-4315945-DEL-6399 | chr6-4315946-DEL-6399 |
| chr6-4329340-DEL-1389 | chr6-4329341-DEL-1389 |
| chr6-4363015-DEL-5302 | chr6-4363016-DEL-5302 |
| chr6-4387162-DEL-80 | chr6-4387179-DEL-64 |
| chr6-4434558-DEL-189 | chr6-4434559-DEL-189 |
| chr6-4475785-DEL-52 | chr6-4475786-DEL-72 |
| chr6-4511911-DEL-60 | chr6-4511914-DEL-56 |
| chr6-4559792-DEL-68 | chr6-4559793-DEL-68 |

| | |
|---|---|
| chr6-4564407-DEL-201 | chr6-4564408-DEL-201 |
| chr6-4582920-DEL-77 | chr6-4582921-DEL-77 |
| chr6-4585217-DEL-56 | chr6-4585218-DEL-56 |
| chr6-4609286-DEL-90 | chr6-4609291-DEL-85 |
| chr6-4650127-DEL-1003 | chr6-4650122-DEL-1007 |
| chr6-4662083-DEL-494 | chr6-4662084-DEL-494 |
| chr6-4674073-DEL-150 | chr6-4674086-DEL-124 |
| chr6-4679012-DEL-52 | chr6-4678996-DEL-60 |
| chr6-4696631-DEL-64 | chr6-4696640-DEL-50 |
| chr6-4714235-DEL-83 | chr6-4714236-DEL-83 |
| chr6-4735551-DEL-56 | chr6-4735552-DEL-56 |
| chr6-4765817-DEL-581 | chr6-4765818-DEL-581 |
| chr6-4777840-DEL-7943 | chr6-4777841-DEL-7944 |
| chr6-4794340-DEL-255 | chr6-4794335-DEL-257 |
| chr6-4797423-DEL-166 | chr6-4797453-DEL-123 |
| chr6-4809959-DEL-95 | chr6-4809941-DEL-96 |
| chr6-4814381-DEL-408 | chr6-4814382-DEL-408 |
| chr6-4839071-DEL-54 | chr6-4839072-DEL-54 |
| chr6-4859333-DEL-58 | chr6-4859334-DEL-58 |
| chr6-4860729-DEL-84 | chr6-4860730-DEL-84 |
| chr6-4872088-DEL-765 | chr6-4872088-DEL-765 |
| chr6-4893215-DEL-691 | chr6-4893216-DEL-691 |
| chr6-4946509-DEL-107 | chr6-4946522-DEL-95 |
| chr6-4961230-DEL-223 | chr6-4961196-DEL-225 |
| chr6-4998338-DEL-57 | chr6-4998338-DEL-57 |
| chr6-4999536-DEL-66 | chr6-4999535-DEL-66 |
| chr6-5002226-DEL-89 | chr6-5002240-DEL-82 |
| chr6-5015562-DEL-234 | chr6-5015563-DEL-234 |
| chr6-5053068-DEL-492 | chr6-5053069-DEL-492 |
| chr6-5104609-DEL-206 | chr6-5104610-DEL-206 |
| chr6-5125730-DEL-400 | chr6-5125731-DEL-400 |
| chr6-5196291-DEL-170 | chr6-5196308-DEL-160 |
| chr6-5201052-DEL-130 | chr6-5201053-DEL-130 |
| chr6-5222972-DEL-199 | chr6-5222973-DEL-199 |
| chr6-5253912-DEL-88 | chr6-5253913-DEL-88 |
| chr6-5263789-DEL-63 | chr6-5263789-DEL-63 |
| chr6-5310901-DEL-1838 | chr6-5310902-DEL-1838 |
| chr6-5321095-DEL-250 | chr6-5321096-DEL-250 |
| chr6-5327295-DEL-4707 | chr6-5327296-DEL-4707 |
| chr6-5336499-DEL-1181 | chr6-5336502-DEL-1179 |
| chr6-5360882-DEL-76 | chr6-5360879-DEL-76 |
| chr6-5367639-DEL-192 | chr6-5367640-DEL-192 |
| chr6-5367755-DEL-1515 | chr6-5367756-DEL-1515 |
| chr6-5428943-DEL-200 | chr6-5428944-DEL-200 |
| chr6-5476577-DEL-3815 | chr6-5476550-DEL-3816 |
| chr6-5480973-DEL-1922 | chr6-5480974-DEL-1922 |
| chr6-5493601-DEL-330 | chr6-5493602-DEL-330 |
| chr6-5506076-DEL-68 | chr6-5506077-DEL-68 |
| chr6-5518093-DEL-1441 | chr6-5518097-DEL-1434 |
| chr6-5518914-DEL-6970 | chr6-5518915-DEL-6970 |
| chr6-5526880-DEL-56 | chr6-5526881-DEL-56 |
| chr6-5552088-DEL-146 | chr6-5552089-DEL-126 |
| chr6-5556425-DEL-203 | chr6-5556426-DEL-203 |
| chr6-5585284-DEL-135 | chr6-5585285-DEL-135 |
| chr6-5602614-DEL-53 | chr6-5602616-DEL-51 |
| chr6-5608048-DEL-7183 | chr6-5608049-DEL-7183 |
| chr6-5641421-DEL-62 | chr6-5641397-DEL-61 |
| chr6-5653133-DEL-203 | chr6-5653143-DEL-203 |
| chr6-5656106-DEL-1890 | chr6-5656109-DEL-1889 |
| chr6-5661023-DEL-498 | chr6-5661024-DEL-498 |
| chr6-5676596-DEL-325 | chr6-5676597-DEL-325 |
| chr6-5730967-DEL-194 | chr6-5730967-DEL-194 |
| chr6-5744510-DEL-188 | chr6-5744558-DEL-141 |
| chr6-5787027-DEL-127 | chr6-5787028-DEL-127 |
| chr6-5804083-DEL-55 | chr6-5804084-DEL-55 |
| chr6-5841116-DEL-211 | chr6-5841117-DEL-211 |
| chr6-5846630-DEL-4128 | chr6-5846631-DEL-4128 |
| chr6-5851777-DEL-724 | chr6-5851753-DEL-710 |
| chr6-5857770-DEL-78 | chr6-5857756-DEL-70 |
| chr6-5872709-DEL-90 | chr6-5872697-DEL-78 |
| chr6-5874487-DEL-75 | chr6-5874470-DEL-56 |
| chr6-5880412-DEL-61 | chr6-5880432-DEL-73 |
| chr6-5938653-DEL-74 | chr6-5938654-DEL-74 |
| chr6-5950992-DEL-201 | chr6-5950993-DEL-201 |
| chr6-5958469-DEL-100 | chr6-5958470-DEL-100 |
| chr6-5965796-DEL-59 | chr6-5965797-DEL-59 |
| chr6-5978721-DEL-752 | chr6-5978728-DEL-752 |
| chr6-5999582-DEL-87 | chr6-5999588-DEL-82 |
| chr6-6055018-DEL-269 | chr6-6055019-DEL-269 |
| chr6-6122796-DEL-94 | chr6-6122797-DEL-94 |

| | |
|---|---|
| chr6-6132175-DEL-53 | chr6-6132176-DEL-53 |
| chr6-6135645-DEL-187 | chr6-6135640-DEL-190 |
| chr6-6145541-DEL-58 | chr6-6145542-DEL-58 |
| chr6-6151936-DEL-70 | chr6-6151975-DEL-53 |
| chr6-6182131-DEL-286 | chr6-6182132-DEL-286 |
| chr6-6184712-DEL-206 | chr6-6184713-DEL-200 |
| chr6-6217960-DEL-51 | chr6-6217961-DEL-51 |
| chr6-6249712-DEL-81 | chr6-6249714-DEL-65 |
| chr6-6252248-DEL-196 | chr6-6252249-DEL-196 |
| chr6-6267116-DEL-3059 | chr6-6267117-DEL-3059 |
| chr6-6279355-DEL-169 | chr6-6279355-DEL-169 |
| chr6-6287440-DEL-168 | chr6-6287441-DEL-168 |
| chr6-6319087-DEL-465 | chr6-6319088-DEL-465 |
| chr6-6327693-DEL-401 | chr6-6327694-DEL-401 |
| chr6-6358538-DEL-238 | chr6-6358539-DEL-238 |
| chr6-6366202-DEL-52 | chr6-6366203-DEL-52 |
| chr6-6366689-DEL-308 | chr6-6366691-DEL-307 |
| chr6-6381084-DEL-55 | chr6-6381085-DEL-55 |
| chr6-6408760-DEL-501 | chr6-6408761-DEL-501 |
| chr6-6424596-DEL-519 | chr6-6424597-DEL-519 |
| chr6-6427601-DEL-329 | chr6-6427602-DEL-329 |
| chr6-6439686-DEL-3193 | chr6-6439687-DEL-3196 |
| chr6-6448285-DEL-55 | chr6-6448278-DEL-59 |
| chr6-6451180-DEL-1371 | chr6-6451181-DEL-1371 |
| chr6-6469187-DEL-205 | chr6-6469188-DEL-205 |
| chr6-6470243-DEL-61 | chr6-6470244-DEL-61 |
| chr6-6501207-DEL-1065 | chr6-6501208-DEL-1065 |
| chr6-6520170-DEL-1791 | chr6-6520171-DEL-1791 |
| chr6-6545448-DEL-210 | chr6-6545449-DEL-210 |
| chr6-6624800-DEL-3124 | chr6-6624799-DEL-3125 |
| chr6-6628852-DEL-56 | chr6-6628853-DEL-56 |
| chr6-6631684-DEL-73 | chr6-6631685-DEL-73 |
| chr6-6636969-DEL-95 | chr6-6636970-DEL-95 |
| chr6-6645737-DEL-390 | chr6-6645770-DEL-358 |
| chr6-6689095-DEL-60 | chr6-6689106-DEL-56 |
| chr6-6699880-DEL-6213 | chr6-6699881-DEL-6213 |
| chr6-6708808-DEL-77 | chr6-6708809-DEL-77 |
| chr6-6721649-DEL-6298 | chr6-6721650-DEL-6298 |
| chr6-6732029-DEL-68 | chr6-6732030-DEL-68 |
| chr6-6748045-DEL-78 | chr6-6748057-DEL-70 |
| chr6-6748317-DEL-1225 | chr6-6748318-DEL-1225 |
| chr6-6767856-DEL-797 | chr6-6767854-DEL-798 |
| chr6-6791485-DEL-208 | chr6-6791486-DEL-208 |
| chr6-6792799-DEL-53 | chr6-6792800-DEL-53 |
| chr6-6797292-DEL-68 | chr6-6797293-DEL-68 |
| chr6-6806573-DEL-104 | chr6-6806573-DEL-109 |
| chr6-6811963-DEL-64 | chr6-6811965-DEL-63 |
| chr6-6817894-DEL-154 | chr6-6817895-DEL-154 |
| chr6-6837190-DEL-52 | chr6-6837206-DEL-52 |
| chr6-6885896-DEL-60 | chr6-6885887-DEL-74 |
| chr6-6910800-DEL-412 | chr6-6910801-DEL-412 |
| chr6-6931715-DEL-76 | chr6-6931716-DEL-76 |
| chr6-6941001-DEL-70 | chr6-6941002-DEL-70 |
| chr6-6949224-DEL-59 | chr6-6949225-DEL-59 |
| chr6-6951895-DEL-78 | chr6-6951896-DEL-78 |
| chr6-6952153-DEL-467 | chr6-6952154-DEL-467 |
| chr6-6959482-DEL-609 | chr6-6959483-DEL-609 |
| chr6-6960523-DEL-53 | chr6-6960524-DEL-53 |
| chr6-6983255-DEL-6807 | chr6-6983256-DEL-6807 |
| chr6-6995003-DEL-221 | chr6-6995004-DEL-221 |
| chr6-6995722-DEL-64 | chr6-6995747-DEL-65 |
| chr6-7023673-DEL-3929 | chr6-7023674-DEL-3929 |
| chr6-7030427-DEL-611 | chr6-7030428-DEL-611 |
| chr6-7040523-DEL-182 | chr6-7040532-DEL-174 |
| chr6-7044185-DEL-134 | chr6-7044186-DEL-134 |
| chr6-7047428-DEL-201 | chr6-7047429-DEL-201 |
| chr6-7100094-DEL-7160 | chr6-7100095-DEL-7160 |
| chr6-7120689-DEL-190 | chr6-7120680-DEL-186 |
| chr6-7148492-DEL-6816 | chr6-7148493-DEL-6816 |
| chr6-7152595-DEL-1714 | chr6-7152596-DEL-1714 |
| chr6-7193139-DEL-63 | chr6-7193140-DEL-63 |
| chr6-7201209-DEL-251 | chr6-7201210-DEL-251 |
| chr6-7207895-DEL-159 | chr6-7207896-DEL-159 |
| chr6-7218963-DEL-78 | chr6-7218969-DEL-80 |
| chr6-7225412-DEL-1571 | chr6-7225412-DEL-1571 |
| chr6-7232483-DEL-186 | chr6-7232483-DEL-186 |
| chr6-7239137-DEL-882 | chr6-7239138-DEL-882 |
| chr6-7247182-DEL-87 | chr6-7247187-DEL-83 |
| chr6-7280424-DEL-434 | chr6-7280425-DEL-434 |
| chr6-7317443-DEL-869 | chr6-7317444-DEL-869 |

| | |
|---|---|
| chr6-7332993-DEL-6829 | chr6-7332984-DEL-6830 |
| chr6-7352927-DEL-464 | chr6-7352928-DEL-464 |
| chr6-7384387-DEL-1697 | chr6-7384388-DEL-1697 |
| chr6-7394267-DEL-6653 | chr6-7394268-DEL-6653 |
| chr6-7431565-DEL-2720 | chr6-7431566-DEL-2720 |
| chr6-7454088-DEL-729 | chr6-7454089-DEL-729 |
| chr6-7460389-DEL-330 | chr6-7460387-DEL-331 |
| chr6-7480528-DEL-93 | chr6-7480535-DEL-87 |
| chr6-7481328-DEL-6466 | chr6-7481329-DEL-6466 |
| chr6-7548973-DEL-423 | chr6-7548974-DEL-423 |
| chr6-7583771-DEL-494 | chr6-7583772-DEL-494 |
| chr6-7614106-DEL-199 | chr6-7614107-DEL-199 |
| chr6-7626750-DEL-138 | chr6-7626751-DEL-138 |
| chr6-7646698-DEL-1285 | chr6-7646699-DEL-1285 |
| chr6-7657913-DEL-122 | chr6-7657914-DEL-122 |
| chr6-7663006-DEL-2920 | chr6-7663007-DEL-2920 |
| chr6-7666598-DEL-1014 | chr6-7666599-DEL-1014 |
| chr6-7668035-DEL-349 | chr6-7668036-DEL-349 |
| chr6-7681571-DEL-80 | chr6-7681571-DEL-80 |
| chr6-7683649-DEL-202 | chr6-7683650-DEL-202 |
| chr6-7693788-DEL-66 | chr6-7693789-DEL-66 |
| chr6-7695324-DEL-646 | chr6-7695325-DEL-644 |
| chr6-7700102-DEL-6333 | chr6-7700103-DEL-6333 |
| chr6-7708366-DEL-130 | chr6-7708367-DEL-130 |
| chr6-7716612-DEL-318 | chr6-7716613-DEL-318 |
| chr6-7719098-DEL-8391 | chr6-7719099-DEL-8391 |
| chr6-7743617-DEL-106 | chr6-7743618-DEL-106 |
| chr6-7750648-DEL-62 | chr6-7750632-DEL-56 |
| chr6-7757008-DEL-3248 | chr6-7757009-DEL-3248 |
| chr6-7757689-DEL-287 | chr6-7757690-DEL-287 |
| chr6-7780488-DEL-60 | chr6-7780482-DEL-58 |
| chr6-7780983-DEL-497 | chr6-7780984-DEL-497 |
| chr6-7781596-DEL-804 | chr6-7781597-DEL-804 |
| chr6-7783631-DEL-66 | chr6-7783631-DEL-66 |
| chr6-7788027-DEL-561 | chr6-7788044-DEL-549 |
| chr6-7806825-DEL-152 | chr6-7806824-DEL-149 |
| chr6-7825947-DEL-2135 | chr6-7825948-DEL-2135 |
| chr6-7842861-DEL-116 | chr6-7842862-DEL-116 |
| chr6-7851280-DEL-108 | chr6-7851300-DEL-80 |
| chr6-7865379-DEL-84 | chr6-7865380-DEL-84 |
| chr6-7887438-DEL-97 | chr6-7887439-DEL-96 |
| chr6-7918185-DEL-6874 | chr6-7918186-DEL-6874 |
| chr6-7942222-DEL-424 | chr6-7942223-DEL-424 |
| chr6-7950921-DEL-106 | chr6-7950920-DEL-105 |
| chr6-7960575-DEL-320 | chr6-7960576-DEL-320 |
| chr6-7981792-DEL-1647 | chr6-7981793-DEL-1647 |
| chr6-7986463-DEL-259 | chr6-7986464-DEL-259 |
| chr6-8007961-DEL-106 | chr6-8007962-DEL-106 |
| chr6-8020362-DEL-749 | chr6-8020363-DEL-749 |
| chr6-8052991-DEL-3094 | chr6-8052992-DEL-3094 |
| chr6-8056295-DEL-138 | chr6-8056296-DEL-138 |
| chr6-8081656-DEL-435 | chr6-8081658-DEL-434 |
| chr6-8083895-DEL-112 | chr6-8083896-DEL-112 |
| chr6-8084459-DEL-69 | chr6-8084452-DEL-70 |
| chr6-8102452-DEL-353 | chr6-8102454-DEL-352 |
| chr6-8129236-DEL-95 | chr6-8129232-DEL-100 |
| chr6-8150112-DEL-59 | chr6-8150119-DEL-53 |
| chr6-8169078-DEL-307 | chr6-8169079-DEL-307 |
| chr6-8177409-DEL-67 | chr6-8177410-DEL-67 |
| chr6-8181728-DEL-157 | chr6-8181729-DEL-157 |
| chr6-8187444-DEL-214 | chr6-8187442-DEL-215 |
| chr6-8243647-DEL-201 | chr6-8243648-DEL-201 |
| chr6-8254565-DEL-63 | chr6-8254566-DEL-63 |
| chr6-8262264-DEL-58 | chr6-8262253-DEL-62 |
| chr6-8264021-DEL-305 | chr6-8264022-DEL-305 |
| chr6-8264701-DEL-180 | chr6-8264702-DEL-180 |
| chr6-8268662-DEL-58 | chr6-8268663-DEL-58 |
| chr6-8298025-DEL-69 | chr6-8298026-DEL-69 |
| chr6-8322870-DEL-7149 | chr6-8322870-DEL-7149 |
| chr6-8325380-DEL-1330 | chr6-8325380-DEL-1330 |
| chr6-8330226-DEL-67 | chr6-8330227-DEL-67 |
| chr6-8335102-DEL-284 | chr6-8335103-DEL-284 |
| chr6-8340949-DEL-61 | chr6-8340950-DEL-61 |
| chr6-8341684-DEL-544 | chr6-8341685-DEL-544 |
| chr6-8371707-DEL-140 | chr6-8371708-DEL-140 |
| chr6-8401089-DEL-4194 | chr6-8401090-DEL-4194 |
| chr6-8434107-DEL-317 | chr6-8434070-DEL-312 |
| chr6-8459751-DEL-63 | chr6-8459752-DEL-63 |
| chr6-8518266-DEL-556 | chr6-8518267-DEL-556 |
| chr6-8525821-DEL-245 | chr6-8525822-DEL-245 |

| | |
|---|---|
| chr6-8528943-DEL-1779 | chr6-8528944-DEL-1779 |
| chr6-8538233-DEL-6251 | chr6-8538234-DEL-6251 |
| chr6-8554053-DEL-124 | chr6-8554054-DEL-124 |
| chr6-8606909-DEL-10420 | chr6-8606910-DEL-10420 |
| chr6-8618822-DEL-234 | chr6-8618823-DEL-234 |
| chr6-8634798-DEL-1546 | chr6-8634799-DEL-1546 |
| chr6-8646582-DEL-78 | chr6-8646583-DEL-78 |
| chr6-8671946-DEL-52 | chr6-8671947-DEL-52 |
| chr6-8696153-DEL-7101 | chr6-8696149-DEL-7100 |
| chr6-8746130-DEL-995 | chr6-8746151-DEL-990 |
| chr6-8758609-DEL-334 | chr6-8758610-DEL-334 |
| chr6-8761302-DEL-146 | chr6-8761303-DEL-146 |
| chr6-8796760-DEL-2530 | chr6-8796761-DEL-2530 |
| chr6-8799832-DEL-447 | chr6-8799833-DEL-447 |
| chr6-8807336-DEL-855 | chr6-8807337-DEL-855 |
| chr6-8851638-DEL-660 | chr6-8851639-DEL-660 |
| chr6-8860213-DEL-114 | chr6-8860249-DEL-71 |
| chr6-8887372-DEL-96 | chr6-8887371-DEL-97 |
| chr6-8890033-DEL-58 | chr6-8890034-DEL-58 |
| chr6-8904844-DEL-149 | chr6-8904845-DEL-149 |
| chr6-8913067-DEL-66 | chr6-8913068-DEL-66 |
| chr6-8920315-DEL-116 | chr6-8920316-DEL-118 |
| chr6-8927563-DEL-4658 | chr6-8927564-DEL-4658 |
| chr6-8990316-DEL-411 | chr6-8990334-DEL-414 |
| chr6-8992026-DEL-161 | chr6-8992027-DEL-161 |
| chr6-8999940-DEL-56 | chr6-8999941-DEL-56 |
| chr6-9001035-DEL-107 | chr6-9001036-DEL-107 |
| chr6-9054754-DEL-60 | chr6-9054771-DEL-55 |
| chr6-9064976-DEL-79 | chr6-9064977-DEL-79 |
| chr6-9091289-DEL-138 | chr6-9091290-DEL-138 |
| chr6-9113779-DEL-77 | chr6-9113780-DEL-77 |
| chr6-9117724-DEL-1275 | chr6-9117725-DEL-1275 |
| chr6-9120879-DEL-56 | chr6-9120880-DEL-56 |
| chr6-9139091-DEL-6446 | chr6-9139092-DEL-6446 |
| chr6-9160025-DEL-429 | chr6-9160026-DEL-429 |
| chr6-9209553-DEL-99 | chr6-9209554-DEL-99 |
| chr6-9277065-DEL-901 | chr6-9277065-DEL-901 |
| chr6-9296235-DEL-8814 | chr6-9296236-DEL-8814 |
| chr6-9308818-DEL-1827 | chr6-9308818-DEL-1827 |
| chr6-9330104-DEL-420 | chr6-9330105-DEL-420 |
| chr6-9340006-DEL-146 | chr6-9340007-DEL-146 |
| chr6-9350683-DEL-56 | chr6-9350684-DEL-56 |
| chr6-9357534-DEL-68 | chr6-9357535-DEL-68 |
| chr6-9359916-DEL-52 | chr6-9359917-DEL-52 |
| chr6-9375259-DEL-73 | chr6-9375266-DEL-55 |
| chr6-9387545-DEL-55 | chr6-9387546-DEL-55 |
| chr6-9400086-DEL-366 | chr6-9400088-DEL-333 |
| chr6-9436075-DEL-2168 | chr6-9436076-DEL-2168 |
| chr6-9492817-DEL-1446 | chr6-9492815-DEL-1448 |
| chr6-9505496-DEL-298 | chr6-9505497-DEL-298 |
| chr6-9527782-DEL-1367 | chr6-9527783-DEL-1367 |
| chr6-9555468-DEL-52 | chr6-9555471-DEL-50 |
| chr6-9580525-DEL-6209 | chr6-9580526-DEL-6209 |
| chr6-9603557-DEL-6641 | chr6-9603546-DEL-6642 |
| chr6-9627152-DEL-388 | chr6-9627153-DEL-388 |
| chr6-9628489-DEL-54 | chr6-9628487-DEL-68 |
| chr6-9634965-DEL-474 | chr6-9634966-DEL-474 |
| chr6-9645534-DEL-6548 | chr6-9645535-DEL-6548 |
| chr6-9675958-DEL-56 | chr6-9675961-DEL-54 |
| chr6-9709433-DEL-1090 | chr6-9709434-DEL-1090 |
| chr6-9738360-DEL-639 | chr6-9738361-DEL-639 |
| chr6-9768294-DEL-72 | chr6-9768295-DEL-70 |
| chr6-9791563-DEL-1111 | chr6-9791564-DEL-1111 |
| chr6-9796762-DEL-60 | chr6-9796768-DEL-52 |
| chr6-9802296-DEL-558 | chr6-9802297-DEL-558 |
| chr6-9802506-DEL-5600 | chr6-9802507-DEL-5600 |
| chr6-9809124-DEL-140 | chr6-9809125-DEL-140 |
| chr6-9810972-DEL-90 | chr6-9810946-DEL-74 |
| chr6-9844236-DEL-2061 | chr6-9844237-DEL-2061 |
| chr6-9850962-DEL-5552 | chr6-9850964-DEL-5550 |
| chr6-9897634-DEL-197 | chr6-9897636-DEL-196 |
| chr6-9905995-DEL-69 | chr6-9905996-DEL-69 |
| chr6-9912721-DEL-205 | chr6-9912722-DEL-205 |
| chr6-9953202-DEL-60 | chr6-9953209-DEL-54 |
| chr6-10038392-DEL-87 | chr6-10038400-DEL-51 |
| chr6-10060296-DEL-200 | chr6-10060297-DEL-196 |
| chr6-10073784-DEL-202 | chr6-10073783-DEL-204 |
| chr6-10108200-DEL-69 | chr6-10108201-DEL-69 |
| chr6-10127214-DEL-253 | chr6-10127215-DEL-253 |
| chr6-10139762-DEL-92 | chr6-10139721-DEL-90 |

| | |
|---|---|
| chr6-10143909-DEL-171 | chr6-10143908-DEL-169 |
| chr6-10146984-DEL-6477 | chr6-10146985-DEL-6477 |
| chr6-10155370-DEL-122 | chr6-10155371-DEL-122 |
| chr6-10158085-DEL-190 | chr6-10158086-DEL-190 |
| chr6-10190440-DEL-319 | chr6-10190441-DEL-319 |
| chr6-10195728-DEL-53 | chr6-10195729-DEL-53 |
| chr6-10202564-DEL-158 | chr6-10202565-DEL-158 |
| chr6-10213020-DEL-58 | chr6-10213021-DEL-58 |
| chr6-10260905-DEL-1390 | chr6-10260906-DEL-1390 |
| chr6-10264875-DEL-475 | chr6-10264876-DEL-475 |
| chr6-10309849-DEL-148 | chr6-10309834-DEL-151 |
| chr6-10358804-DEL-1028 | chr6-10358805-DEL-1028 |
| chr6-10371991-DEL-60 | chr6-10371992-DEL-60 |
| chr6-10396797-DEL-8838 | chr6-10396798-DEL-8838 |
| chr6-10406396-DEL-290 | chr6-10406369-DEL-290 |
| chr6-10437621-DEL-206 | chr6-10437622-DEL-206 |
| chr6-10449601-DEL-99 | chr6-10449602-DEL-99 |
| chr6-10513066-DEL-560 | chr6-10513063-DEL-561 |
| chr6-10564870-DEL-1264 | chr6-10564871-DEL-1264 |
| chr6-10603970-DEL-201 | chr6-10603971-DEL-201 |
| chr6-10623052-DEL-6754 | chr6-10623053-DEL-6754 |
| chr6-10637909-DEL-694 | chr6-10637910-DEL-694 |
| chr6-10689573-DEL-5233 | chr6-10689574-DEL-5233 |
| chr6-10732774-DEL-6240 | chr6-10732775-DEL-6240 |
| chr6-10786011-DEL-62 | chr6-10786012-DEL-60 |
| chr6-10794522-DEL-229 | chr6-10794519-DEL-217 |
| chr6-10798246-DEL-1495 | chr6-10798247-DEL-1495 |
| chr6-10814920-DEL-160 | chr6-10814913-DEL-160 |
| chr6-10835127-DEL-5566 | chr6-10835128-DEL-5566 |
| chr6-10859006-DEL-1689 | chr6-10859006-DEL-1689 |
| chr6-10867363-DEL-169 | chr6-10867359-DEL-170 |
| chr6-10914995-DEL-53 | chr6-10914976-DEL-84 |
| chr6-10938508-DEL-510 | chr6-10938509-DEL-510 |
| chr6-11010293-DEL-92 | chr6-11010294-DEL-92 |
| chr6-11030829-DEL-6525 | chr6-11030829-DEL-6525 |
| chr6-11042442-DEL-52 | chr6-11042443-DEL-52 |
| chr6-11048854-DEL-90 | chr6-11048855-DEL-90 |
| chr6-11120419-DEL-6336 | chr6-11120420-DEL-6336 |
| chr6-11161459-DEL-6342 | chr6-11161460-DEL-6342 |
| chr6-11205448-DEL-6418 | chr6-11205449-DEL-6418 |
| chr6-11263423-DEL-201 | chr6-11263424-DEL-201 |
| chr6-11286334-DEL-201 | chr6-11286335-DEL-201 |
| chr6-11299835-DEL-1439 | chr6-11299836-DEL-1439 |
| chr6-11326198-DEL-117 | chr6-11326223-DEL-93 |
| chr6-11336274-DEL-634 | chr6-11336275-DEL-634 |
| chr6-11338713-DEL-79 | chr6-11338713-DEL-79 |
| chr6-11349159-DEL-56 | chr6-11349156-DEL-56 |
| chr6-11354629-DEL-98 | chr6-11354630-DEL-98 |
| chr6-11393756-DEL-646 | chr6-11393758-DEL-645 |
| chr6-11406372-DEL-1235 | chr6-11406373-DEL-1235 |
| chr6-11450868-DEL-7037 | chr6-11450869-DEL-7037 |
| chr6-11472295-DEL-62 | chr6-11472296-DEL-62 |
| chr6-11542101-DEL-527 | chr6-11542102-DEL-527 |
| chr6-11599143-DEL-68 | chr6-11599143-DEL-74 |
| chr6-11655973-DEL-3672 | chr6-11655974-DEL-3672 |
| chr6-11670127-DEL-596 | chr6-11670133-DEL-593 |
| chr6-11717902-DEL-99 | chr6-11717915-DEL-87 |
| chr6-11724732-DEL-62 | chr6-11724733-DEL-62 |
| chr6-11726443-DEL-52 | chr6-11726443-DEL-50 |
| chr6-11739728-DEL-101 | chr6-11739731-DEL-99 |
| chr6-11742569-DEL-6878 | chr6-11742570-DEL-6878 |
| chr6-11760899-DEL-504 | chr6-11760900-DEL-504 |
| chr6-11779698-DEL-6021 | chr6-11779699-DEL-6021 |
| chr6-11798884-DEL-70 | chr6-11798885-DEL-70 |
| chr6-11801575-DEL-610 | chr6-11801576-DEL-610 |
| chr6-11803485-DEL-62 | chr6-11803486-DEL-62 |
| chr6-11813697-DEL-54 | chr6-11813701-DEL-50 |
| chr6-11832303-DEL-212 | chr6-11832304-DEL-212 |
| chr6-11880911-DEL-290 | chr6-11880912-DEL-290 |
| chr6-11935689-DEL-113 | chr6-11935704-DEL-84 |
| chr6-11963379-DEL-6639 | chr6-11963390-DEL-6640 |
| chr6-11969638-DEL-212 | chr6-11969639-DEL-212 |
| chr6-11986221-DEL-64 | chr6-11986222-DEL-64 |
| chr6-12010476-DEL-109 | chr6-12010477-DEL-109 |
| chr6-12032214-DEL-94 | chr6-12032202-DEL-84 |
| chr6-12050195-DEL-68 | chr6-12050196-DEL-68 |
| chr6-12056707-DEL-58 | chr6-12056708-DEL-58 |
| chr6-12105963-DEL-72 | chr6-12105968-DEL-68 |
| chr6-12109880-DEL-121 | chr6-12109881-DEL-121 |
| chr6-12116833-DEL-58 | chr6-12116834-DEL-58 |

| | |
|---|---|
| chr6-12154425-DEL-3155 | chr6-12154426-DEL-3155 |
| chr6-12182273-DEL-697 | chr6-12182274-DEL-697 |
| chr6-12185093-DEL-3258 | chr6-12185094-DEL-3258 |
| chr6-12185527-DEL-61 | chr6-12185528-DEL-61 |
| chr6-12186598-DEL-6213 | chr6-12186603-DEL-6214 |
| chr6-12194313-DEL-592 | chr6-12194314-DEL-592 |
| chr6-12253187-DEL-57 | chr6-12253177-DEL-55 |
| chr6-12264004-DEL-564 | chr6-12264005-DEL-564 |
| chr6-12278987-DEL-6232 | chr6-12278988-DEL-6232 |
| chr6-12289366-DEL-378 | chr6-12289367-DEL-378 |
| chr6-12292130-DEL-1801 | chr6-12292131-DEL-1801 |
| chr6-12304047-DEL-1207 | chr6-12304048-DEL-1207 |
| chr6-12363183-DEL-2314 | chr6-12363182-DEL-2314 |
| chr6-12388472-DEL-83 | chr6-12388473-DEL-83 |
| chr6-12439903-DEL-52 | chr6-12439904-DEL-52 |
| chr6-12454795-DEL-2223 | chr6-12454792-DEL-2221 |
| chr6-12476673-DEL-51 | chr6-12476674-DEL-51 |
| chr6-12489803-DEL-208 | chr6-12489804-DEL-208 |
| chr6-12510530-DEL-85 | chr6-12510557-DEL-86 |
| chr6-12514148-DEL-308 | chr6-12514149-DEL-308 |
| chr6-12522566-DEL-6296 | chr6-12522567-DEL-6296 |
| chr6-12613311-DEL-235 | chr6-12613312-DEL-235 |
| chr6-12618581-DEL-87 | chr6-12618582-DEL-87 |
| chr6-12620913-DEL-6645 | chr6-12620914-DEL-6645 |
| chr6-12639648-DEL-155 | chr6-12639649-DEL-155 |
| chr6-12640247-DEL-6642 | chr6-12640248-DEL-6642 |
| chr6-12671561-DEL-8936 | chr6-12671562-DEL-8936 |
| chr6-12688084-DEL-156 | chr6-12688086-DEL-153 |
| chr6-12689878-DEL-118 | chr6-12689879-DEL-118 |
| chr6-12699977-DEL-56 | chr6-12699978-DEL-56 |
| chr6-12711071-DEL-90 | chr6-12711059-DEL-102 |
| chr6-12736703-DEL-75 | chr6-12736703-DEL-75 |
| chr6-12761774-DEL-86 | chr6-12761775-DEL-88 |
| chr6-12775593-DEL-194 | chr6-12775577-DEL-194 |
| chr6-12831354-DEL-218 | chr6-12831355-DEL-218 |
| chr6-12837634-DEL-2821 | chr6-12837635-DEL-2821 |
| chr6-12839278-DEL-112 | chr6-12839313-DEL-113 |
| chr6-12866295-DEL-399 | chr6-12866291-DEL-400 |
| chr6-12868960-DEL-7024 | chr6-12868960-DEL-7024 |
| chr6-12868993-DEL-214 | chr6-12868994-DEL-214 |
| chr6-12887433-DEL-392 | chr6-12887434-DEL-392 |
| chr6-12995705-DEL-2855 | chr6-12995711-DEL-2856 |
| chr6-13014660-DEL-4762 | chr6-13014661-DEL-4762 |
| chr6-13042650-DEL-76 | chr6-13042651-DEL-68 |
| chr6-13057241-DEL-1492 | chr6-13057242-DEL-1492 |
| chr6-13175520-DEL-236 | chr6-13175521-DEL-236 |
| chr6-13194114-DEL-135 | chr6-13194093-DEL-157 |
| chr6-13213507-DEL-1398 | chr6-13213508-DEL-1398 |
| chr6-13255840-DEL-285 | chr6-13255841-DEL-285 |
| chr6-13259897-DEL-171 | chr6-13259898-DEL-171 |
| chr6-13265910-DEL-6387 | chr6-13265911-DEL-6387 |
| chr6-13284312-DEL-110 | chr6-13284313-DEL-82 |
| chr6-13318568-DEL-57 | chr6-13318568-DEL-57 |
| chr6-13331493-DEL-63 | chr6-13331494-DEL-63 |
| chr6-13342260-DEL-1138 | chr6-13342259-DEL-1138 |
| chr6-13351863-DEL-1472 | chr6-13351864-DEL-1472 |
| chr6-13362243-DEL-6220 | chr6-13362244-DEL-6220 |
| chr6-13427972-DEL-2509 | chr6-13427973-DEL-2509 |
| chr6-13452051-DEL-8697 | chr6-13452052-DEL-8697 |
| chr6-13465655-DEL-427 | chr6-13465656-DEL-427 |
| chr6-13470025-DEL-51 | chr6-13470026-DEL-51 |
| chr6-13477307-DEL-880 | chr6-13477308-DEL-880 |
| chr6-13481449-DEL-1890 | chr6-13481450-DEL-1890 |
| chr6-13487104-DEL-17997 | chr6-13487103-DEL-17998 |
| chr6-13490250-DEL-711 | chr6-13490251-DEL-711 |
| chr6-13491890-DEL-1219 | chr6-13491891-DEL-1219 |
| chr6-13496490-DEL-6234 | chr6-13496491-DEL-6234 |
| chr6-13515067-DEL-54 | chr6-13515068-DEL-54 |
| chr6-13527629-DEL-5670 | chr6-13527630-DEL-5670 |
| chr6-13549398-DEL-54 | chr6-13549399-DEL-54 |
| chr6-13554710-DEL-70 | chr6-13554711-DEL-70 |
| chr6-13566066-DEL-6594 | chr6-13566067-DEL-6594 |
| chr6-13571978-DEL-202 | chr6-13571979-DEL-202 |
| chr6-13575505-DEL-58 | chr6-13575508-DEL-56 |
| chr6-13616677-DEL-131 | chr6-13616682-DEL-132 |
| chr6-13637132-DEL-457 | chr6-13637151-DEL-436 |
| chr6-13665972-DEL-90 | chr6-13665973-DEL-90 |
| chr6-13671575-DEL-674 | chr6-13671576-DEL-674 |
| chr6-13695435-DEL-203 | chr6-13695436-DEL-203 |
| chr6-13700123-DEL-12568 | chr6-13700114-DEL-12567 |

| | |
|---|---|
| chr6-13708059-DEL-2491 | chr6-13708060-DEL-2491 |
| chr6-13725022-DEL-2448 | chr6-13725023-DEL-2448 |
| chr6-13740611-DEL-3013 | chr6-13740612-DEL-3013 |
| chr6-13749281-DEL-52 | chr6-13749282-DEL-52 |
| chr6-13755439-DEL-56 | chr6-13755440-DEL-54 |
| chr6-13766832-DEL-7091 | chr6-13766833-DEL-7091 |
| chr6-13778168-DEL-935 | chr6-13778169-DEL-935 |
| chr6-13785535-DEL-334 | chr6-13785536-DEL-334 |
| chr6-13790351-DEL-1199 | chr6-13790337-DEL-1208 |
| chr6-13795071-DEL-66 | chr6-13795077-DEL-92 |
| chr6-13797306-DEL-54 | chr6-13797305-DEL-56 |
| chr6-13805838-DEL-825 | chr6-13805839-DEL-825 |
| chr6-13825559-DEL-52 | chr6-13825560-DEL-52 |
| chr6-13894390-DEL-152 | chr6-13894391-DEL-152 |
| chr6-13895594-DEL-103 | chr6-13895595-DEL-103 |
| chr6-13897336-DEL-187 | chr6-13897337-DEL-187 |
| chr6-13946763-DEL-328 | chr6-13946764-DEL-328 |
| chr6-13965530-DEL-1120 | chr6-13965531-DEL-1120 |
| chr6-14031793-DEL-66 | chr6-14031783-DEL-58 |
| chr6-14085036-DEL-7189 | chr6-14085037-DEL-7189 |
| chr6-14103775-DEL-154 | chr6-14103776-DEL-154 |
| chr6-14152242-DEL-131 | chr6-14152243-DEL-131 |
| chr6-14163656-DEL-197 | chr6-14163643-DEL-198 |
| chr6-14168261-DEL-59 | chr6-14168262-DEL-59 |
| chr6-14169096-DEL-275 | chr6-14169097-DEL-275 |
| chr6-14173281-DEL-564 | chr6-14173282-DEL-564 |
| chr6-14197783-DEL-104 | chr6-14197783-DEL-104 |
| chr6-14250998-DEL-207 | chr6-14250999-DEL-207 |
| chr6-14282514-DEL-78 | chr6-14282488-DEL-117 |
| chr6-14302246-DEL-2905 | chr6-14302245-DEL-2906 |
| chr6-14304298-DEL-324 | chr6-14304299-DEL-324 |
| chr6-14318170-DEL-253 | chr6-14318170-DEL-253 |
| chr6-14342493-DEL-71 | chr6-14342494-DEL-71 |
| chr6-14369010-DEL-1165 | chr6-14369008-DEL-1162 |
| chr6-14371280-DEL-88 | chr6-14371283-DEL-85 |
| chr6-14390393-DEL-114 | chr6-14390394-DEL-114 |
| chr6-14392516-DEL-202 | chr6-14392517-DEL-202 |
| chr6-14405205-DEL-538 | chr6-14405210-DEL-534 |
| chr6-14419492-DEL-205 | chr6-14419493-DEL-205 |
| chr6-14423806-DEL-1126 | chr6-14423807-DEL-1126 |
| chr6-14424935-DEL-254 | chr6-14424935-DEL-254 |
| chr6-14457625-DEL-58 | chr6-14457626-DEL-58 |
| chr6-14459832-DEL-7253 | chr6-14459833-DEL-7253 |
| chr6-14519557-DEL-1722 | chr6-14519558-DEL-1721 |
| chr6-14526472-DEL-51 | chr6-14526473-DEL-51 |
| chr6-14527009-DEL-70 | chr6-14527010-DEL-70 |
| chr6-14534459-DEL-62 | chr6-14534460-DEL-62 |
| chr6-14544488-DEL-68 | chr6-14544495-DEL-62 |
| chr6-14546449-DEL-6425 | chr6-14546450-DEL-6425 |
| chr6-14546678-DEL-199 | chr6-14546678-DEL-199 |
| chr6-14553480-DEL-67 | chr6-14553481-DEL-64 |
| chr6-14609064-DEL-150 | chr6-14609065-DEL-150 |
| chr6-14648283-DEL-6600 | chr6-14648284-DEL-6600 |
| chr6-14682794-DEL-547 | chr6-14682793-DEL-547 |
| chr6-14729598-DEL-515 | chr6-14729599-DEL-515 |
| chr6-14745353-DEL-199 | chr6-14745350-DEL-181 |
| chr6-14749619-DEL-672 | chr6-14749620-DEL-672 |
| chr6-14765873-DEL-1031 | chr6-14765874-DEL-1031 |
| chr6-14766421-DEL-101 | chr6-14766422-DEL-101 |
| chr6-14775903-DEL-573 | chr6-14775902-DEL-573 |
| chr6-14794660-DEL-213 | chr6-14794661-DEL-213 |
| chr6-14836363-DEL-502 | chr6-14836364-DEL-502 |
| chr6-14842044-DEL-183 | chr6-14842044-DEL-183 |
| chr6-14851111-DEL-1346 | chr6-14851111-DEL-1346 |
| chr6-14873474-DEL-6474 | chr6-14873475-DEL-6474 |
| chr6-14887674-DEL-528 | chr6-14887675-DEL-528 |
| chr6-14929216-DEL-6246 | chr6-14929217-DEL-6246 |
| chr6-14957298-DEL-117 | chr6-14957299-DEL-117 |
| chr6-14960361-DEL-6560 | chr6-14960362-DEL-6560 |
| chr6-14994914-DEL-262 | chr6-14994915-DEL-262 |
| chr6-14995708-DEL-122 | chr6-14995709-DEL-122 |
| chr6-15053169-DEL-68 | chr6-15053186-DEL-52 |
| chr6-15055003-DEL-1407 | chr6-15055002-DEL-1408 |
| chr6-15140222-DEL-415 | chr6-15140225-DEL-409 |
| chr6-15145441-DEL-383 | chr6-15145442-DEL-383 |
| chr6-15163106-DEL-357 | chr6-15163107-DEL-357 |
| chr6-15208630-DEL-58 | chr6-15208653-DEL-51 |
| chr6-15228986-DEL-57 | chr6-15228987-DEL-57 |
| chr6-15238213-DEL-2846 | chr6-15238214-DEL-2846 |
| chr6-15356350-DEL-338 | chr6-15356355-DEL-328 |

| | |
|---|---|
| chr6-15376015-DEL-79 | chr6-15376001-DEL-86 |
| chr6-15394006-DEL-66 | chr6-15394001-DEL-69 |
| chr6-15495151-DEL-202 | chr6-15495153-DEL-201 |
| chr6-15496756-DEL-305 | chr6-15496757-DEL-305 |
| chr6-15506607-DEL-723 | chr6-15506608-DEL-723 |
| chr6-15517922-DEL-6840 | chr6-15517923-DEL-6840 |
| chr6-15524833-DEL-57 | chr6-15524834-DEL-57 |
| chr6-15548176-DEL-76 | chr6-15548185-DEL-68 |
| chr6-15550375-DEL-7795 | chr6-15550376-DEL-7795 |
| chr6-15555817-DEL-786 | chr6-15555819-DEL-785 |
| chr6-15573601-DEL-6086 | chr6-15573602-DEL-6086 |
| chr6-15600528-DEL-7012 | chr6-15600519-DEL-7013 |
| chr6-15639267-DEL-56 | chr6-15639272-DEL-52 |
| chr6-15664417-DEL-3003 | chr6-15664418-DEL-3003 |
| chr6-15684666-DEL-6614 | chr6-15684667-DEL-6614 |
| chr6-15707287-DEL-396 | chr6-15707288-DEL-396 |
| chr6-15733589-DEL-221 | chr6-15733590-DEL-221 |
| chr6-15804290-DEL-78 | chr6-15804279-DEL-63 |
| chr6-15831819-DEL-3515 | chr6-15831820-DEL-3508 |
| chr6-15836415-DEL-56 | chr6-15836410-DEL-64 |
| chr6-15874520-DEL-784 | chr6-15874521-DEL-784 |
| chr6-15883679-DEL-214 | chr6-15883680-DEL-214 |
| chr6-15884832-DEL-869 | chr6-15884833-DEL-869 |
| chr6-15887630-DEL-147 | chr6-15887631-DEL-147 |
| chr6-15892982-DEL-84 | chr6-15892983-DEL-84 |
| chr6-15910433-DEL-68 | chr6-15910434-DEL-68 |
| chr6-15916506-DEL-64 | chr6-15916507-DEL-64 |
| chr6-15924974-DEL-344 | chr6-15924975-DEL-344 |
| chr6-15969334-DEL-157 | chr6-15969335-DEL-157 |
| chr6-15997456-DEL-6925 | chr6-15997457-DEL-6925 |
| chr6-16012130-DEL-69 | chr6-16012131-DEL-69 |
| chr6-16014042-DEL-154 | chr6-16014035-DEL-132 |
| chr6-16016425-DEL-124 | chr6-16016426-DEL-116 |
| chr6-16034888-DEL-1083 | chr6-16034874-DEL-1098 |
| chr6-16054186-DEL-54 | chr6-16054182-DEL-86 |
| chr6-16104134-DEL-382 | chr6-16104135-DEL-382 |
| chr6-16114961-DEL-143 | chr6-16114962-DEL-143 |
| chr6-16158418-DEL-95 | chr6-16158387-DEL-96 |
| chr6-16163904-DEL-100 | chr6-16163905-DEL-100 |
| chr6-16180910-DEL-60 | chr6-16180910-DEL-52 |
| chr6-16207756-DEL-102 | chr6-16207757-DEL-102 |
| chr6-16224333-DEL-54 | chr6-16224332-DEL-74 |
| chr6-16287146-DEL-62 | chr6-16287147-DEL-62 |
| chr6-16299651-DEL-11480 | chr6-16299651-DEL-11481 |
| chr6-16313775-DEL-59 | chr6-16313771-DEL-63 |
| chr6-16359737-DEL-504 | chr6-16359740-DEL-502 |
| chr6-16360589-DEL-97 | chr6-16360599-DEL-93 |
| chr6-16365908-DEL-114 | chr6-16365909-DEL-114 |
| chr6-16366236-DEL-732 | chr6-16366235-DEL-733 |
| chr6-16378725-DEL-172 | chr6-16378726-DEL-172 |
| chr6-16391057-DEL-6677 | chr6-16391058-DEL-6677 |
| chr6-16505493-DEL-104 | chr6-16505502-DEL-96 |
| chr6-16535042-DEL-61 | chr6-16535043-DEL-61 |
| chr6-16541258-DEL-665 | chr6-16541259-DEL-664 |
| chr6-16542508-DEL-54 | chr6-16542509-DEL-54 |
| chr6-16557378-DEL-596 | chr6-16557379-DEL-596 |
| chr6-16559079-DEL-1707 | chr6-16559080-DEL-1707 |
| chr6-16562167-DEL-222 | chr6-16562168-DEL-222 |
| chr6-16595111-DEL-6844 | chr6-16595112-DEL-6844 |
| chr6-16602609-DEL-2974 | chr6-16602610-DEL-2974 |
| chr6-16665842-DEL-55 | chr6-16665843-DEL-55 |
| chr6-16672579-DEL-84 | chr6-16672600-DEL-96 |
| chr6-16675941-DEL-137 | chr6-16675942-DEL-137 |
| chr6-16734293-DEL-514 | chr6-16734294-DEL-514 |
| chr6-16755088-DEL-73 | chr6-16755097-DEL-66 |
| chr6-16814452-DEL-57 | chr6-16814452-DEL-58 |
| chr6-16820452-DEL-66 | chr6-16820453-DEL-54 |
| chr6-16822400-DEL-68 | chr6-16822399-DEL-71 |
| chr6-16909538-DEL-525 | chr6-16909539-DEL-525 |
| chr6-16976371-DEL-243 | chr6-16976372-DEL-243 |
| chr6-16989256-DEL-404 | chr6-16989257-DEL-404 |
| chr6-17033222-DEL-416 | chr6-17033223-DEL-416 |
| chr6-17035551-DEL-6401 | chr6-17035552-DEL-6401 |
| chr6-17054391-DEL-203 | chr6-17054392-DEL-203 |
| chr6-17058746-DEL-309 | chr6-17058747-DEL-309 |
| chr6-17078238-DEL-870 | chr6-17078226-DEL-871 |
| chr6-17087392-DEL-67 | chr6-17087406-DEL-67 |
| chr6-17124312-DEL-338 | chr6-17124313-DEL-338 |
| chr6-17132667-DEL-813 | chr6-17132670-DEL-811 |
| chr6-17140733-DEL-208 | chr6-17140732-DEL-183 |

| | |
|---|---|
| chr6-17157795-DEL-764 | chr6-17157796-DEL-764 |
| chr6-17165185-DEL-132 | chr6-17165204-DEL-114 |
| chr6-17181626-DEL-386 | chr6-17181627-DEL-386 |
| chr6-17191456-DEL-506 | chr6-17191460-DEL-502 |
| chr6-17199588-DEL-66 | chr6-17199581-DEL-74 |
| chr6-17237724-DEL-1202 | chr6-17237725-DEL-1202 |
| chr6-17244026-DEL-82 | chr6-17244027-DEL-82 |
| chr6-17257231-DEL-78 | chr6-17257224-DEL-84 |
| chr6-17263229-DEL-779 | chr6-17263230-DEL-779 |
| chr6-17274387-DEL-825 | chr6-17274388-DEL-825 |
| chr6-17298816-DEL-3909 | chr6-17298817-DEL-3909 |
| chr6-17315338-DEL-195 | chr6-17315339-DEL-195 |
| chr6-17329400-DEL-187 | chr6-17329394-DEL-188 |
| chr6-17343459-DEL-2525 | chr6-17343460-DEL-2525 |
| chr6-17366736-DEL-4461 | chr6-17366724-DEL-4461 |
| chr6-17393937-DEL-2856 | chr6-17393938-DEL-2856 |
| chr6-17396321-DEL-205 | chr6-17396319-DEL-206 |
| chr6-17432928-DEL-71 | chr6-17432911-DEL-100 |
| chr6-17435233-DEL-89 | chr6-17435234-DEL-89 |
| chr6-17436288-DEL-4806 | chr6-17436289-DEL-4806 |
| chr6-17487367-DEL-115 | chr6-17487368-DEL-115 |
| chr6-17510379-DEL-398 | chr6-17510381-DEL-396 |
| chr6-17536394-DEL-680 | chr6-17536395-DEL-680 |
| chr6-17564737-DEL-73 | chr6-17564748-DEL-58 |
| chr6-17579870-DEL-140 | chr6-17579871-DEL-140 |
| chr6-17580799-DEL-243 | chr6-17580800-DEL-243 |
| chr6-17583255-DEL-75 | chr6-17583256-DEL-75 |
| chr6-17588772-DEL-53 | chr6-17588773-DEL-53 |
| chr6-17593366-DEL-54 | chr6-17593367-DEL-54 |
| chr6-17674195-DEL-65 | chr6-17674204-DEL-55 |
| chr6-17681778-DEL-61 | chr6-17681779-DEL-61 |
| chr6-17703865-DEL-73 | chr6-17703866-DEL-74 |
| chr6-17705376-DEL-947 | chr6-17705377-DEL-947 |
| chr6-17713779-DEL-54 | chr6-17713780-DEL-54 |
| chr6-17766165-DEL-82 | chr6-17766166-DEL-82 |
| chr6-17773758-DEL-180 | chr6-17773759-DEL-180 |
| chr6-17791717-DEL-89 | chr6-17791718-DEL-89 |
| chr6-17794707-DEL-570 | chr6-17794709-DEL-567 |
| chr6-17797004-DEL-83 | chr6-17796961-DEL-84 |
| chr6-17807638-DEL-1891 | chr6-17807639-DEL-1891 |
| chr6-17814725-DEL-79 | chr6-17814727-DEL-76 |
| chr6-17830364-DEL-67 | chr6-17830365-DEL-67 |
| chr6-17854515-DEL-325 | chr6-17854516-DEL-325 |
| chr6-17870167-DEL-64 | chr6-17870168-DEL-64 |
| chr6-17881657-DEL-442 | chr6-17881658-DEL-442 |
| chr6-17902324-DEL-208 | chr6-17902325-DEL-208 |
| chr6-17908215-DEL-139 | chr6-17908215-DEL-138 |
| chr6-17911320-DEL-60 | chr6-17911321-DEL-60 |
| chr6-17949877-DEL-68 | chr6-17949891-DEL-51 |
| chr6-17950371-DEL-78 | chr6-17950372-DEL-78 |
| chr6-17954505-DEL-154 | chr6-17954506-DEL-154 |
| chr6-17981677-DEL-1803 | chr6-17981678-DEL-1803 |
| chr6-17996500-DEL-555 | chr6-17996501-DEL-555 |
| chr6-18006273-DEL-103 | chr6-18006274-DEL-103 |
| chr6-18018528-DEL-52 | chr6-18018506-DEL-79 |
| chr6-18063213-DEL-269 | chr6-18063214-DEL-269 |
| chr6-18067058-DEL-1276 | chr6-18067059-DEL-1276 |
| chr6-18117874-DEL-97 | chr6-18117875-DEL-97 |
| chr6-18137695-DEL-3292 | chr6-18137697-DEL-3290 |
| chr6-18162276-DEL-6742 | chr6-18162277-DEL-6742 |
| chr6-18169647-DEL-100 | chr6-18169648-DEL-100 |
| chr6-18185395-DEL-379 | chr6-18185396-DEL-379 |
| chr6-18193719-DEL-510 | chr6-18193720-DEL-510 |
| chr6-18238600-DEL-133 | chr6-18238601-DEL-133 |
| chr6-18244349-DEL-112 | chr6-18244350-DEL-112 |
| chr6-18254776-DEL-258 | chr6-18254767-DEL-258 |
| chr6-18261879-DEL-2439 | chr6-18261879-DEL-2439 |
| chr6-18264848-DEL-792 | chr6-18264848-DEL-792 |
| chr6-18288014-DEL-63 | chr6-18288015-DEL-63 |
| chr6-18325069-DEL-1583 | chr6-18325070-DEL-1583 |
| chr6-18327842-DEL-2151 | chr6-18327843-DEL-2151 |
| chr6-18391799-DEL-1732 | chr6-18391800-DEL-1732 |
| chr6-18394167-DEL-60 | chr6-18394168-DEL-60 |
| chr6-18409074-DEL-117 | chr6-18409075-DEL-117 |
| chr6-18416658-DEL-330 | chr6-18416659-DEL-330 |
| chr6-18429367-DEL-109 | chr6-18429368-DEL-109 |
| chr6-18445365-DEL-172 | chr6-18445366-DEL-172 |
| chr6-18452358-DEL-61 | chr6-18452357-DEL-63 |
| chr6-18465442-DEL-154 | chr6-18465443-DEL-154 |
| chr6-18484485-DEL-62 | chr6-18484486-DEL-62 |

| | |
|---|---|
| chr6-18489341-DEL-70 | chr6-18489342-DEL-70 |
| chr6-18536420-DEL-95 | chr6-18536418-DEL-97 |
| chr6-18566943-DEL-66 | chr6-18566955-DEL-52 |
| chr6-18568522-DEL-122 | chr6-18568523-DEL-122 |
| chr6-18579400-DEL-8055 | chr6-18579400-DEL-8055 |
| chr6-18597288-DEL-171 | chr6-18597289-DEL-171 |
| chr6-18603670-DEL-563 | chr6-18603671-DEL-563 |
| chr6-18609400-DEL-705 | chr6-18609401-DEL-705 |
| chr6-18623825-DEL-52 | chr6-18623826-DEL-54 |
| chr6-18650098-DEL-63 | chr6-18650099-DEL-63 |
| chr6-18669578-DEL-154 | chr6-18669579-DEL-154 |
| chr6-18716143-DEL-1055 | chr6-18716144-DEL-1055 |
| chr6-18717963-DEL-84 | chr6-18717964-DEL-84 |
| chr6-18749730-DEL-52 | chr6-18749731-DEL-52 |
| chr6-18754948-DEL-59 | chr6-18754949-DEL-59 |
| chr6-18756259-DEL-1114 | chr6-18756260-DEL-1114 |
| chr6-18757478-DEL-5330 | chr6-18757483-DEL-5326 |
| chr6-18780662-DEL-6441 | chr6-18780663-DEL-6441 |
| chr6-18799762-DEL-54 | chr6-18799763-DEL-54 |
| chr6-18811017-DEL-587 | chr6-18811018-DEL-587 |
| chr6-18818477-DEL-60 | chr6-18818482-DEL-56 |
| chr6-18859716-DEL-414 | chr6-18859717-DEL-414 |
| chr6-18871101-DEL-677 | chr6-18871102-DEL-677 |
| chr6-18875875-DEL-1214 | chr6-18875876-DEL-1214 |
| chr6-18881117-DEL-6118 | chr6-18881118-DEL-6118 |
| chr6-18899869-DEL-6608 | chr6-18899870-DEL-6608 |
| chr6-18982220-DEL-7037 | chr6-18982221-DEL-7037 |
| chr6-18992174-DEL-63 | chr6-18992187-DEL-54 |
| chr6-19068866-DEL-6621 | chr6-19068858-DEL-6630 |
| chr6-19078240-DEL-120 | chr6-19078241-DEL-120 |
| chr6-19117735-DEL-63 | chr6-19117729-DEL-52 |
| chr6-19143738-DEL-256 | chr6-19143739-DEL-256 |
| chr6-19145839-DEL-636 | chr6-19145840-DEL-636 |
| chr6-19190514-DEL-342 | chr6-19190515-DEL-342 |
| chr6-19213970-DEL-117 | chr6-19213969-DEL-143 |
| chr6-19216339-DEL-6336 | chr6-19216340-DEL-6336 |
| chr6-19226468-DEL-133 | chr6-19226449-DEL-129 |
| chr6-19227012-DEL-84 | chr6-19227005-DEL-85 |
| chr6-19236323-DEL-917 | chr6-19236324-DEL-917 |
| chr6-19236469-DEL-87 | chr6-19236470-DEL-87 |
| chr6-19239465-DEL-526 | chr6-19239466-DEL-526 |
| chr6-19278516-DEL-1408 | chr6-19278517-DEL-1408 |
| chr6-19305353-DEL-352 | chr6-19305354-DEL-352 |
| chr6-19381610-DEL-2121 | chr6-19381611-DEL-2121 |
| chr6-19385969-DEL-6339 | chr6-19385970-DEL-6339 |
| chr6-19407496-DEL-399 | chr6-19407497-DEL-399 |
| chr6-19413801-DEL-56 | chr6-19413802-DEL-56 |
| chr6-19421920-DEL-3266 | chr6-19421921-DEL-3266 |
| chr6-19458129-DEL-6696 | chr6-19458130-DEL-6696 |
| chr6-19476583-DEL-1154 | chr6-19476584-DEL-1154 |
| chr6-19488257-DEL-94 | chr6-19488258-DEL-94 |
| chr6-19492390-DEL-312 | chr6-19492391-DEL-312 |
| chr6-19526000-DEL-197 | chr6-19526001-DEL-197 |
| chr6-19559036-DEL-57 | chr6-19559037-DEL-57 |
| chr6-19582968-DEL-4081 | chr6-19582977-DEL-4076 |
| chr6-19586759-DEL-53 | chr6-19586739-DEL-50 |
| chr6-19598136-DEL-83 | chr6-19598131-DEL-84 |
| chr6-19644109-DEL-8860 | chr6-19644110-DEL-8860 |
| chr6-19645021-DEL-76 | chr6-19645021-DEL-76 |
| chr6-19681296-DEL-777 | chr6-19681297-DEL-776 |
| chr6-19701015-DEL-839 | chr6-19701016-DEL-839 |
| chr6-19740889-DEL-183 | chr6-19740890-DEL-183 |
| chr6-19746535-DEL-162 | chr6-19746536-DEL-161 |
| chr6-19748165-DEL-61 | chr6-19748164-DEL-61 |
| chr6-19774232-DEL-2825 | chr6-19774233-DEL-2825 |
| chr6-19780012-DEL-51 | chr6-19780013-DEL-51 |
| chr6-19781330-DEL-1650 | chr6-19781331-DEL-1650 |
| chr6-19793459-DEL-160 | chr6-19793460-DEL-160 |
| chr6-19810036-DEL-56 | chr6-19810037-DEL-56 |
| chr6-19811837-DEL-685 | chr6-19811838-DEL-685 |
| chr6-19825788-DEL-182 | chr6-19825789-DEL-182 |
| chr6-19839444-DEL-3167 | chr6-19839445-DEL-3167 |
| chr6-19845509-DEL-150 | chr6-19845497-DEL-129 |
| chr6-19867615-DEL-56 | chr6-19867616-DEL-56 |
| chr6-19872744-DEL-74 | chr6-19872744-DEL-72 |
| chr6-19875064-DEL-521 | chr6-19875060-DEL-520 |
| chr6-19893492-DEL-358 | chr6-19893493-DEL-357 |
| chr6-19903102-DEL-207 | chr6-19903103-DEL-207 |
| chr6-19913227-DEL-52 | chr6-19913224-DEL-78 |
| chr6-19915925-DEL-128 | chr6-19915925-DEL-122 |

| | |
|---|---|
| chr6-19923197-DEL-63 | chr6-19923213-DEL-95 |
| chr6-20017552-DEL-6723 | chr6-20017553-DEL-6723 |
| chr6-20043660-DEL-1316 | chr6-20043661-DEL-1316 |
| chr6-20048878-DEL-72 | chr6-20048879-DEL-72 |
| chr6-20083569-DEL-6920 | chr6-20083569-DEL-6920 |
| chr6-20105027-DEL-80 | chr6-20105028-DEL-75 |
| chr6-20119054-DEL-201 | chr6-20119054-DEL-201 |
| chr6-20119875-DEL-400 | chr6-20119875-DEL-400 |
| chr6-20134107-DEL-64 | chr6-20134108-DEL-64 |
| chr6-20146812-DEL-56 | chr6-20146813-DEL-56 |
| chr6-20148330-DEL-62 | chr6-20148355-DEL-54 |
| chr6-20182103-DEL-7014 | chr6-20182104-DEL-7014 |
| chr6-20232909-DEL-56 | chr6-20232910-DEL-56 |
| chr6-20241218-DEL-57 | chr6-20241219-DEL-57 |
| chr6-20270156-DEL-95 | chr6-20270162-DEL-92 |
| chr6-20306383-DEL-66 | chr6-20306384-DEL-66 |
| chr6-20338659-DEL-72 | chr6-20338660-DEL-72 |
| chr6-20357613-DEL-172 | chr6-20357615-DEL-174 |
| chr6-20359429-DEL-744 | chr6-20359430-DEL-744 |
| chr6-20373801-DEL-1547 | chr6-20373807-DEL-1539 |
| chr6-20377323-DEL-211 | chr6-20377355-DEL-213 |
| chr6-20385214-DEL-1056 | chr6-20385215-DEL-1056 |
| chr6-20395405-DEL-1208 | chr6-20395406-DEL-1208 |
| chr6-20417099-DEL-2123 | chr6-20417100-DEL-2109 |
| chr6-20436315-DEL-55 | chr6-20436316-DEL-55 |
| chr6-20458197-DEL-851 | chr6-20458198-DEL-851 |
| chr6-20485923-DEL-115 | chr6-20485924-DEL-115 |
| chr6-20498010-DEL-113 | chr6-20498011-DEL-113 |
| chr6-20508705-DEL-88 | chr6-20508714-DEL-74 |
| chr6-20512996-DEL-142 | chr6-20512997-DEL-142 |
| chr6-20528950-DEL-3427 | chr6-20528951-DEL-3427 |
| chr6-20534690-DEL-1490 | chr6-20534691-DEL-1490 |
| chr6-20544847-DEL-6241 | chr6-20544848-DEL-6241 |
| chr6-20560452-DEL-62 | chr6-20560453-DEL-62 |
| chr6-20561239-DEL-73 | chr6-20561241-DEL-72 |
| chr6-20565695-DEL-64 | chr6-20565696-DEL-64 |
| chr6-20567422-DEL-182 | chr6-20567423-DEL-182 |
| chr6-20579579-DEL-130 | chr6-20579578-DEL-131 |
| chr6-20588355-DEL-160 | chr6-20588356-DEL-160 |
| chr6-20590937-DEL-106 | chr6-20590938-DEL-106 |
| chr6-20594831-DEL-2146 | chr6-20594832-DEL-2146 |
| chr6-20614990-DEL-265 | chr6-20614991-DEL-265 |
| chr6-20648448-DEL-387 | chr6-20648449-DEL-385 |
| chr6-20697438-DEL-5405 | chr6-20697439-DEL-5405 |
| chr6-20697464-DEL-2459 | chr6-20697463-DEL-2459 |
| chr6-20716027-DEL-1287 | chr6-20716028-DEL-1287 |
| chr6-20731104-DEL-3578 | chr6-20731105-DEL-3578 |
| chr6-20782045-DEL-881 | chr6-20782046-DEL-881 |
| chr6-20791617-DEL-68 | chr6-20791618-DEL-68 |
| chr6-20793425-DEL-6466 | chr6-20793412-DEL-6436 |
| chr6-20803173-DEL-783 | chr6-20803174-DEL-783 |
| chr6-20806483-DEL-1892 | chr6-20806484-DEL-1892 |
| chr6-20820079-DEL-386 | chr6-20820080-DEL-386 |
| chr6-20830479-DEL-213 | chr6-20830481-DEL-212 |
| chr6-20869574-DEL-86 | chr6-20869575-DEL-62 |
| chr6-20889267-DEL-86 | chr6-20889274-DEL-80 |
| chr6-20893771-DEL-251 | chr6-20893772-DEL-251 |
| chr6-20897744-DEL-1789 | chr6-20897744-DEL-1789 |
| chr6-20907651-DEL-2177 | chr6-20907651-DEL-2149 |
| chr6-20913856-DEL-72 | chr6-20913850-DEL-62 |
| chr6-20935598-DEL-232 | chr6-20935599-DEL-232 |
| chr6-20969273-DEL-54 | chr6-20969292-DEL-54 |
| chr6-20995853-DEL-95 | chr6-20995854-DEL-96 |
| chr6-21026508-DEL-428 | chr6-21026509-DEL-428 |
| chr6-21044494-DEL-75 | chr6-21044505-DEL-65 |
| chr6-21126843-DEL-78 | chr6-21126834-DEL-60 |
| chr6-21233036-DEL-430 | chr6-21233037-DEL-430 |
| chr6-21236299-DEL-363 | chr6-21236300-DEL-363 |
| chr6-21247982-DEL-302 | chr6-21247983-DEL-301 |
| chr6-21312338-DEL-112 | chr6-21312338-DEL-74 |
| chr6-21344833-DEL-64 | chr6-21344842-DEL-56 |
| chr6-21393942-DEL-6949 | chr6-21393941-DEL-6950 |
| chr6-21394433-DEL-212 | chr6-21394434-DEL-212 |
| chr6-21451884-DEL-6222 | chr6-21451885-DEL-6222 |
| chr6-21466136-DEL-222 | chr6-21466137-DEL-222 |
| chr6-21468624-DEL-472 | chr6-21468625-DEL-472 |
| chr6-21476598-DEL-1530 | chr6-21476591-DEL-1530 |
| chr6-21507134-DEL-181 | chr6-21507135-DEL-181 |
| chr6-21529671-DEL-474 | chr6-21529672-DEL-474 |
| chr6-21639919-DEL-88 | chr6-21639920-DEL-88 |

| | |
|---|---|
| chr6-21666940-DEL-272 | chr6-21666941-DEL-272 |
| chr6-21730287-DEL-84 | chr6-21730272-DEL-71 |
| chr6-21731918-DEL-690 | chr6-21731919-DEL-690 |
| chr6-21794421-DEL-107 | chr6-21794422-DEL-107 |
| chr6-21801228-DEL-2070 | chr6-21801229-DEL-2070 |
| chr6-21841474-DEL-4110 | chr6-21841475-DEL-4110 |
| chr6-21873270-DEL-3531 | chr6-21873271-DEL-3531 |
| chr6-21891408-DEL-59 | chr6-21891404-DEL-60 |
| chr6-21901414-DEL-434 | chr6-21901415-DEL-434 |
| chr6-21922488-DEL-72 | chr6-21922489-DEL-72 |
| chr6-21927683-DEL-88 | chr6-21927636-DEL-89 |
| chr6-21953092-DEL-307 | chr6-21953093-DEL-307 |
| chr6-22023741-DEL-267 | chr6-22023742-DEL-267 |
| chr6-22028533-DEL-139 | chr6-22028534-DEL-139 |
| chr6-22036696-DEL-72 | chr6-22036689-DEL-82 |
| chr6-22055075-DEL-2079 | chr6-22055075-DEL-2079 |
| chr6-22083469-DEL-174 | chr6-22083470-DEL-174 |
| chr6-22096405-DEL-63 | chr6-22096406-DEL-63 |
| chr6-22100318-DEL-65 | chr6-22100329-DEL-55 |
| chr6-22105769-DEL-3073 | chr6-22105781-DEL-3067 |
| chr6-22141918-DEL-60 | chr6-22141919-DEL-60 |
| chr6-22151465-DEL-448 | chr6-22151466-DEL-448 |
| chr6-22171239-DEL-3136 | chr6-22171240-DEL-3136 |
| chr6-22238058-DEL-162 | chr6-22238056-DEL-163 |
| chr6-22260321-DEL-214 | chr6-22260317-DEL-174 |
| chr6-22285359-DEL-55 | chr6-22285363-DEL-55 |
| chr6-22337713-DEL-233 | chr6-22337714-DEL-233 |
| chr6-22340827-DEL-51 | chr6-22340828-DEL-51 |
| chr6-22378141-DEL-61 | chr6-22378153-DEL-63 |
| chr6-22381748-DEL-203 | chr6-22381744-DEL-197 |
| chr6-22397954-DEL-212 | chr6-22397955-DEL-212 |
| chr6-22401580-DEL-86 | chr6-22401621-DEL-87 |
| chr6-22414921-DEL-1274 | chr6-22414922-DEL-1274 |
| chr6-22434071-DEL-234 | chr6-22434072-DEL-234 |
| chr6-22445824-DEL-935 | chr6-22445825-DEL-935 |
| chr6-22457332-DEL-107 | chr6-22457328-DEL-108 |
| chr6-22462760-DEL-199 | chr6-22462761-DEL-199 |
| chr6-22463841-DEL-111 | chr6-22463835-DEL-125 |
| chr6-22472608-DEL-204 | chr6-22472609-DEL-204 |
| chr6-22482694-DEL-1345 | chr6-22482695-DEL-1345 |
| chr6-22484581-DEL-74 | chr6-22484582-DEL-74 |
| chr6-22503244-DEL-66 | chr6-22503245-DEL-66 |
| chr6-22516137-DEL-270 | chr6-22516156-DEL-252 |
| chr6-22524323-DEL-54 | chr6-22524324-DEL-54 |
| chr6-22528095-DEL-6220 | chr6-22528096-DEL-6220 |
| chr6-22547819-DEL-71 | chr6-22547841-DEL-50 |
| chr6-22592816-DEL-685 | chr6-22592817-DEL-685 |
| chr6-22621547-DEL-72 | chr6-22621548-DEL-72 |
| chr6-22624431-DEL-57 | chr6-22624424-DEL-63 |
| chr6-22624689-DEL-1645 | chr6-22624690-DEL-1645 |
| chr6-22627504-DEL-866 | chr6-22627505-DEL-866 |
| chr6-22629068-DEL-137 | chr6-22629069-DEL-137 |
| chr6-22644749-DEL-7192 | chr6-22644750-DEL-7192 |
| chr6-22681883-DEL-3843 | chr6-22681884-DEL-3843 |
| chr6-22722048-DEL-2468 | chr6-22722049-DEL-2468 |
| chr6-22738745-DEL-2379 | chr6-22738746-DEL-2379 |
| chr6-22743833-DEL-136 | chr6-22743834-DEL-136 |
| chr6-22750418-DEL-2453 | chr6-22750420-DEL-2452 |
| chr6-22775979-DEL-91 | chr6-22775980-DEL-91 |
| chr6-22804864-DEL-279 | chr6-22804865-DEL-279 |
| chr6-22807476-DEL-72 | chr6-22807477-DEL-72 |
| chr6-22830863-DEL-200 | chr6-22830864-DEL-199 |
| chr6-22848575-DEL-400 | chr6-22848576-DEL-400 |
| chr6-22853952-DEL-90 | chr6-22853953-DEL-90 |
| chr6-22914817-DEL-307 | chr6-22914818-DEL-307 |
| chr6-22936960-DEL-54 | chr6-22936961-DEL-54 |
| chr6-23003607-DEL-105 | chr6-23003608-DEL-105 |
| chr6-23025540-DEL-96 | chr6-23025541-DEL-84 |
| chr6-23032525-DEL-84 | chr6-23032526-DEL-80 |
| chr6-23038772-DEL-92 | chr6-23038775-DEL-90 |
| chr6-23051513-DEL-88 | chr6-23051515-DEL-87 |
| chr6-23055440-DEL-58 | chr6-23055441-DEL-58 |
| chr6-23060389-DEL-6369 | chr6-23060390-DEL-6369 |
| chr6-23101434-DEL-254 | chr6-23101435-DEL-254 |
| chr6-23115997-DEL-973 | chr6-23115998-DEL-971 |
| chr6-23125377-DEL-198 | chr6-23125379-DEL-187 |
| chr6-23148064-DEL-1110 | chr6-23148064-DEL-1110 |
| chr6-23148941-DEL-126 | chr6-23148942-DEL-126 |
| chr6-23156840-DEL-120 | chr6-23156850-DEL-110 |
| chr6-23178339-DEL-926 | chr6-23178368-DEL-898 |

| | |
|---|---|
| chr6-23183743-DEL-57 | chr6-23183750-DEL-56 |
| chr6-23185609-DEL-67 | chr6-23185608-DEL-69 |
| chr6-23198622-DEL-185 | chr6-23198623-DEL-185 |
| chr6-23201306-DEL-668 | chr6-23201307-DEL-668 |
| chr6-23220106-DEL-106 | chr6-23220107-DEL-106 |
| chr6-23223635-DEL-667 | chr6-23223630-DEL-673 |
| chr6-23238716-DEL-290 | chr6-23238716-DEL-291 |
| chr6-23266511-DEL-52 | chr6-23266512-DEL-52 |
| chr6-23285415-DEL-111 | chr6-23285416-DEL-111 |
| chr6-23295663-DEL-283 | chr6-23295664-DEL-283 |
| chr6-23344468-DEL-643 | chr6-23344469-DEL-643 |
| chr6-23364874-DEL-6425 | chr6-23364875-DEL-6425 |
| chr6-23391194-DEL-87 | chr6-23391196-DEL-85 |
| chr6-23406654-DEL-322 | chr6-23406655-DEL-322 |
| chr6-23415876-DEL-163 | chr6-23415877-DEL-163 |
| chr6-23436997-DEL-66 | chr6-23436998-DEL-66 |
| chr6-23458808-DEL-932 | chr6-23458794-DEL-900 |
| chr6-23460369-DEL-252 | chr6-23460405-DEL-216 |
| chr6-23472632-DEL-51 | chr6-23472630-DEL-85 |
| chr6-23474277-DEL-7043 | chr6-23474278-DEL-7043 |
| chr6-23480416-DEL-416 | chr6-23480416-DEL-416 |
| chr6-23482047-DEL-430 | chr6-23482048-DEL-430 |
| chr6-23491504-DEL-279 | chr6-23491505-DEL-279 |
| chr6-23505392-DEL-1058 | chr6-23505385-DEL-1061 |
| chr6-23511265-DEL-6655 | chr6-23511266-DEL-6655 |
| chr6-23519744-DEL-61 | chr6-23519730-DEL-62 |
| chr6-23522224-DEL-178 | chr6-23522225-DEL-178 |
| chr6-23534249-DEL-800 | chr6-23534250-DEL-800 |
| chr6-23538758-DEL-2218 | chr6-23538759-DEL-2218 |
| chr6-23557660-DEL-6280 | chr6-23557661-DEL-6280 |
| chr6-23576120-DEL-873 | chr6-23576121-DEL-873 |
| chr6-23576785-DEL-239 | chr6-23576786-DEL-239 |
| chr6-23584187-DEL-950 | chr6-23584186-DEL-951 |
| chr6-23614775-DEL-54 | chr6-23614776-DEL-54 |
| chr6-23642827-DEL-97 | chr6-23642836-DEL-87 |
| chr6-23668542-DEL-6560 | chr6-23668543-DEL-6560 |
| chr6-23727428-DEL-202 | chr6-23727429-DEL-202 |
| chr6-23729993-DEL-202 | chr6-23729997-DEL-201 |
| chr6-23754532-DEL-163 | chr6-23754533-DEL-163 |
| chr6-23767643-DEL-6230 | chr6-23767644-DEL-6230 |
| chr6-23776396-DEL-396 | chr6-23776397-DEL-396 |
| chr6-23778252-DEL-1559 | chr6-23778253-DEL-1559 |
| chr6-23785957-DEL-2264 | chr6-23785958-DEL-2263 |
| chr6-23795385-DEL-1233 | chr6-23795386-DEL-1233 |
| chr6-23815629-DEL-52 | chr6-23815632-DEL-50 |
| chr6-23849536-DEL-190 | chr6-23849537-DEL-190 |
| chr6-23855815-DEL-6371 | chr6-23855816-DEL-6371 |
| chr6-23867818-DEL-68 | chr6-23867831-DEL-61 |
| chr6-23889133-DEL-5132 | chr6-23889134-DEL-5132 |
| chr6-23919865-DEL-72 | chr6-23919883-DEL-54 |
| chr6-23958107-DEL-1281 | chr6-23958107-DEL-1281 |
| chr6-23985515-DEL-68 | chr6-23985516-DEL-68 |
| chr6-24037574-DEL-462 | chr6-24037575-DEL-462 |
| chr6-24049981-DEL-642 | chr6-24049970-DEL-641 |
| chr6-24057911-DEL-3290 | chr6-24057913-DEL-3289 |
| chr6-24062027-DEL-486 | chr6-24062028-DEL-486 |
| chr6-24063109-DEL-6467 | chr6-24063110-DEL-6467 |
| chr6-24069240-DEL-214 | chr6-24069240-DEL-214 |
| chr6-24082967-DEL-57 | chr6-24082968-DEL-57 |
| chr6-24103838-DEL-97 | chr6-24103869-DEL-68 |
| chr6-24129159-DEL-254 | chr6-24129160-DEL-254 |
| chr6-24146432-DEL-56 | chr6-24146433-DEL-56 |
| chr6-24155352-DEL-1430 | chr6-24155353-DEL-1430 |
| chr6-24159665-DEL-58 | chr6-24159664-DEL-59 |
| chr6-24173900-DEL-99 | chr6-24173917-DEL-69 |
| chr6-24202608-DEL-51 | chr6-24202592-DEL-68 |
| chr6-24226743-DEL-1643 | chr6-24226742-DEL-1643 |
| chr6-24234718-DEL-522 | chr6-24234719-DEL-522 |
| chr6-24257965-DEL-2373 | chr6-24257966-DEL-2373 |
| chr6-24264730-DEL-63 | chr6-24264710-DEL-63 |
| chr6-24276058-DEL-429 | chr6-24276059-DEL-429 |
| chr6-24279337-DEL-6430 | chr6-24279338-DEL-6430 |
| chr6-24290960-DEL-124 | chr6-24290960-DEL-124 |
| chr6-24302554-DEL-69 | chr6-24302594-DEL-69 |
| chr6-24303029-DEL-126 | chr6-24303030-DEL-126 |
| chr6-24314905-DEL-1179 | chr6-24314905-DEL-1179 |
| chr6-24330709-DEL-130 | chr6-24330717-DEL-113 |
| chr6-24338384-DEL-64 | chr6-24338393-DEL-58 |
| chr6-24341631-DEL-157 | chr6-24341602-DEL-182 |
| chr6-24360368-DEL-64 | chr6-24360367-DEL-57 |

| | |
|---|---|
| chr6-24390309-DEL-3049 | chr6-24390310-DEL-3049 |
| chr6-24390593-DEL-63 | chr6-24390594-DEL-63 |
| chr6-24409713-DEL-2675 | chr6-24409714-DEL-2675 |
| chr6-24417474-DEL-79 | chr6-24417472-DEL-79 |
| chr6-24429208-DEL-6189 | chr6-24429209-DEL-6189 |
| chr6-24472495-DEL-84 | chr6-24472498-DEL-82 |
| chr6-24473883-DEL-277 | chr6-24473882-DEL-278 |
| chr6-24478479-DEL-80 | chr6-24478488-DEL-72 |
| chr6-24483161-DEL-662 | chr6-24483162-DEL-662 |
| chr6-24485305-DEL-155 | chr6-24485306-DEL-155 |
| chr6-24491728-DEL-6675 | chr6-24491729-DEL-6675 |
| chr6-24517565-DEL-80 | chr6-24517574-DEL-72 |
| chr6-24545650-DEL-363 | chr6-24545652-DEL-360 |
| chr6-24581792-DEL-127 | chr6-24581782-DEL-106 |
| chr6-24588560-DEL-1249 | chr6-24588571-DEL-1243 |
| chr6-24592233-DEL-57 | chr6-24592234-DEL-57 |
| chr6-24649627-DEL-70 | chr6-24649628-DEL-70 |
| chr6-24673335-DEL-6473 | chr6-24673336-DEL-6473 |
| chr6-24712920-DEL-273 | chr6-24712921-DEL-273 |
| chr6-24774086-DEL-56 | chr6-24774087-DEL-55 |
| chr6-24787217-DEL-328 | chr6-24787217-DEL-332 |
| chr6-24835206-DEL-69 | chr6-24835207-DEL-69 |
| chr6-24844847-DEL-1235 | chr6-24844848-DEL-1235 |
| chr6-24848264-DEL-176 | chr6-24848265-DEL-176 |
| chr6-24868178-DEL-1885 | chr6-24868179-DEL-1885 |
| chr6-24892914-DEL-1477 | chr6-24892915-DEL-1477 |
| chr6-24911448-DEL-191 | chr6-24911449-DEL-191 |
| chr6-24927231-DEL-4392 | chr6-24927232-DEL-4392 |
| chr6-24933563-DEL-945 | chr6-24933564-DEL-945 |
| chr6-24941721-DEL-6207 | chr6-24941722-DEL-6207 |
| chr6-24971280-DEL-58 | chr6-24971284-DEL-50 |
| chr6-25016070-DEL-58 | chr6-25016073-DEL-58 |
| chr6-25036377-DEL-94 | chr6-25036378-DEL-94 |
| chr6-25039925-DEL-69 | chr6-25039919-DEL-70 |
| chr6-25050389-DEL-243 | chr6-25050390-DEL-243 |
| chr6-25070489-DEL-6427 | chr6-25070490-DEL-6427 |
| chr6-25076664-DEL-209 | chr6-25076665-DEL-209 |
| chr6-25082891-DEL-112 | chr6-25082892-DEL-112 |
| chr6-25084584-DEL-393 | chr6-25084585-DEL-393 |
| chr6-25085433-DEL-442 | chr6-25085439-DEL-437 |
| chr6-25090006-DEL-234 | chr6-25090007-DEL-233 |
| chr6-25105112-DEL-6685 | chr6-25105113-DEL-6685 |
| chr6-25111429-DEL-212 | chr6-25111430-DEL-212 |
| chr6-25114918-DEL-484 | chr6-25114919-DEL-484 |
| chr6-25118374-DEL-239 | chr6-25118375-DEL-239 |
| chr6-25119584-DEL-60 | chr6-25119585-DEL-60 |
| chr6-25123686-DEL-446 | chr6-25123691-DEL-446 |
| chr6-25127933-DEL-1587 | chr6-25127934-DEL-1587 |
| chr6-25129996-DEL-157 | chr6-25129997-DEL-157 |
| chr6-25135078-DEL-100 | chr6-25135079-DEL-100 |
| chr6-25153126-DEL-1558 | chr6-25153127-DEL-1558 |
| chr6-25156884-DEL-2229 | chr6-25156885-DEL-2229 |
| chr6-25158187-DEL-1152 | chr6-25158188-DEL-1155 |
| chr6-25159254-DEL-126 | chr6-25159255-DEL-126 |
| chr6-25251459-DEL-3978 | chr6-25251460-DEL-3978 |
| chr6-25338462-DEL-3662 | chr6-25338461-DEL-3663 |
| chr6-25380527-DEL-88 | chr6-25380529-DEL-83 |
| chr6-25387381-DEL-224 | chr6-25387383-DEL-224 |
| chr6-25392957-DEL-1434 | chr6-25392958-DEL-1434 |
| chr6-25425441-DEL-56 | chr6-25425448-DEL-50 |
| chr6-25438948-DEL-1026 | chr6-25438949-DEL-1026 |
| chr6-25473922-DEL-80 | chr6-25473923-DEL-80 |
| chr6-25526806-DEL-2702 | chr6-25526812-DEL-2706 |
| chr6-25540348-DEL-3117 | chr6-25540349-DEL-3117 |
| chr6-25550907-DEL-6907 | chr6-25550907-DEL-6908 |
| chr6-25571775-DEL-3899 | chr6-25571776-DEL-3899 |
| chr6-25577516-DEL-3439 | chr6-25577517-DEL-3439 |
| chr6-25600049-DEL-1630 | chr6-25600098-DEL-1594 |
| chr6-25607375-DEL-54 | chr6-25607410-DEL-50 |
| chr6-25617515-DEL-7468 | chr6-25617505-DEL-7478 |
| chr6-25697591-DEL-73 | chr6-25697593-DEL-72 |
| chr6-25713143-DEL-1460 | chr6-25713144-DEL-1460 |
| chr6-25714608-DEL-166 | chr6-25714614-DEL-161 |
| chr6-25745702-DEL-128 | chr6-25745693-DEL-130 |
| chr6-25746569-DEL-748 | chr6-25746570-DEL-748 |
| chr6-25767993-DEL-2975 | chr6-25767994-DEL-2975 |
| chr6-25780811-DEL-1788 | chr6-25780812-DEL-1788 |
| chr6-25805647-DEL-120 | chr6-25805616-DEL-132 |
| chr6-25822338-DEL-65 | chr6-25822338-DEL-60 |
| chr6-25828792-DEL-63 | chr6-25828792-DEL-64 |

| | |
|---|---|
| chr6-25842260-DEL-95 | chr6-25842260-DEL-93 |
| chr6-25852093-DEL-1415 | chr6-25852094-DEL-1415 |
| chr6-25866617-DEL-51 | chr6-25866610-DEL-51 |
| chr6-25878977-DEL-617 | chr6-25878978-DEL-617 |
| chr6-25883328-DEL-4950 | chr6-25883328-DEL-4950 |
| chr6-25902211-DEL-260 | chr6-25902212-DEL-260 |
| chr6-25906558-DEL-55 | chr6-25906557-DEL-57 |
| chr6-25940612-DEL-206 | chr6-25940631-DEL-188 |
| chr6-25947841-DEL-978 | chr6-25947842-DEL-978 |
| chr6-25949614-DEL-326 | chr6-25949615-DEL-326 |
| chr6-25976346-DEL-164 | chr6-25976347-DEL-164 |
| chr6-25982787-DEL-6082 | chr6-25982788-DEL-6082 |
| chr6-26001079-DEL-333 | chr6-26001080-DEL-333 |
| chr6-26014220-DEL-437 | chr6-26014221-DEL-437 |
| chr6-26022534-DEL-70 | chr6-26022535-DEL-70 |
| chr6-26028991-DEL-78 | chr6-26028992-DEL-78 |
| chr6-26035907-DEL-432 | chr6-26035908-DEL-432 |
| chr6-26037725-DEL-287 | chr6-26037726-DEL-287 |
| chr6-26064301-DEL-7724 | chr6-26064302-DEL-7724 |
| chr6-26087957-DEL-383 | chr6-26087958-DEL-383 |
| chr6-26114792-DEL-725 | chr6-26114785-DEL-730 |
| chr6-26135129-DEL-1798 | chr6-26135130-DEL-1798 |
| chr6-26151190-DEL-2668 | chr6-26151191-DEL-2668 |
| chr6-26156333-DEL-6136 | chr6-26156334-DEL-6136 |
| chr6-26165525-DEL-3162 | chr6-26165526-DEL-3162 |
| chr6-26189410-DEL-66 | chr6-26189452-DEL-50 |
| chr6-26191917-DEL-2572 | chr6-26191918-DEL-2572 |
| chr6-26208497-DEL-164 | chr6-26208462-DEL-165 |
| chr6-26210263-DEL-116 | chr6-26210253-DEL-117 |
| chr6-26211917-DEL-85 | chr6-26211893-DEL-109 |
| chr6-26269857-DEL-10508 | chr6-26269858-DEL-10508 |
| chr6-26282119-DEL-73 | chr6-26282120-DEL-73 |
| chr6-26294317-DEL-115 | chr6-26294318-DEL-114 |
| chr6-26302614-DEL-1945 | chr6-26302615-DEL-1945 |
| chr6-26307100-DEL-519 | chr6-26307101-DEL-519 |
| chr6-26402643-DEL-789 | chr6-26402644-DEL-789 |
| chr6-26418220-DEL-101 | chr6-26418221-DEL-101 |
| chr6-26422277-DEL-1842 | chr6-26422278-DEL-1842 |
| chr6-26425919-DEL-1571 | chr6-26425920-DEL-1571 |
| chr6-26436634-DEL-1896 | chr6-26436634-DEL-1896 |
| chr6-26454948-DEL-52 | chr6-26454949-DEL-52 |
| chr6-26463811-DEL-55 | chr6-26463812-DEL-73 |
| chr6-26480744-DEL-59 | chr6-26480745-DEL-59 |
| chr6-26491984-DEL-160 | chr6-26491985-DEL-160 |
| chr6-26529965-DEL-1103 | chr6-26529966-DEL-1102 |
| chr6-26629771-DEL-54 | chr6-26629772-DEL-54 |
| chr6-26646126-DEL-70 | chr6-26646127-DEL-70 |
| chr6-26677137-DEL-55 | chr6-26677138-DEL-55 |
| chr6-26691743-DEL-6236 | chr6-26691744-DEL-6236 |
| chr6-26735841-DEL-373 | chr6-26735840-DEL-372 |
| chr6-26747543-DEL-94 | chr6-26747544-DEL-94 |
| chr6-26751827-DEL-3208 | chr6-26751828-DEL-3208 |
| chr6-26759152-DEL-79 | chr6-26759153-DEL-79 |
| chr6-26800110-DEL-300 | chr6-26800109-DEL-301 |
| chr6-26809723-DEL-73 | chr6-26809708-DEL-89 |
| chr6-26818548-DEL-511 | chr6-26818549-DEL-511 |
| chr6-26834521-DEL-6418 | chr6-26834522-DEL-6418 |
| chr6-26844637-DEL-54 | chr6-26844638-DEL-54 |
| chr6-26861926-DEL-108 | chr6-26861927-DEL-108 |
| chr6-26884436-DEL-53 | chr6-26884435-DEL-54 |
| chr6-26913609-DEL-106 | chr6-26913632-DEL-62 |
| chr6-26950447-DEL-528 | chr6-26950450-DEL-526 |
| chr6-26953650-DEL-445 | chr6-26953651-DEL-445 |
| chr6-26959613-DEL-283 | chr6-26959613-DEL-283 |
| chr6-26960416-DEL-6820 | chr6-26960417-DEL-6820 |
| chr6-26970992-DEL-74 | chr6-26971013-DEL-54 |
| chr6-26986698-DEL-62 | chr6-26986699-DEL-62 |
| chr6-26987516-DEL-51 | chr6-26987517-DEL-51 |
| chr6-26988409-DEL-4506 | chr6-26988410-DEL-4506 |
| chr6-27001893-DEL-86 | chr6-27001894-DEL-86 |
| chr6-27010498-DEL-63 | chr6-27010491-DEL-58 |
| chr6-27044666-DEL-1489 | chr6-27044667-DEL-1489 |
| chr6-27058058-DEL-116 | chr6-27058057-DEL-91 |
| chr6-27059641-DEL-58 | chr6-27059642-DEL-56 |
| chr6-27088298-DEL-1411 | chr6-27088299-DEL-1411 |
| chr6-27198883-DEL-6241 | chr6-27198882-DEL-6241 |
| chr6-27222468-DEL-6175 | chr6-27222469-DEL-6175 |
| chr6-27235209-DEL-2578 | chr6-27235210-DEL-2578 |
| chr6-27259853-DEL-96 | chr6-27259854-DEL-120 |
| chr6-27281810-DEL-261 | chr6-27281811-DEL-261 |

| | |
|---|---|
| chr6-27369132-DEL-124 | chr6-27369133-DEL-124 |
| chr6-27405529-DEL-146 | chr6-27405530-DEL-146 |
| chr6-27409046-DEL-2015 | chr6-27409047-DEL-2015 |
| chr6-27434981-DEL-581 | chr6-27434982-DEL-581 |
| chr6-27438452-DEL-343 | chr6-27438453-DEL-343 |
| chr6-27456463-DEL-122 | chr6-27456464-DEL-122 |
| chr6-27468450-DEL-59 | chr6-27468450-DEL-59 |
| chr6-27498905-DEL-327 | chr6-27498906-DEL-327 |
| chr6-27519292-DEL-7599 | chr6-27519293-DEL-7599 |
| chr6-27540906-DEL-230 | chr6-27540905-DEL-231 |
| chr6-27551635-DEL-584 | chr6-27551636-DEL-584 |
| chr6-27554234-DEL-6688 | chr6-27554235-DEL-6688 |
| chr6-27624227-DEL-1751 | chr6-27624228-DEL-1751 |
| chr6-27637629-DEL-6338 | chr6-27637630-DEL-6338 |
| chr6-27713444-DEL-56 | chr6-27713437-DEL-64 |
| chr6-27722348-DEL-57 | chr6-27722349-DEL-57 |
| chr6-27750931-DEL-1884 | chr6-27750932-DEL-1884 |
| chr6-27819489-DEL-64 | chr6-27819494-DEL-56 |
| chr6-27847821-DEL-465 | chr6-27847822-DEL-465 |
| chr6-27924165-DEL-125 | chr6-27924173-DEL-92 |
| chr6-27939451-DEL-179 | chr6-27939452-DEL-179 |
| chr6-27944888-DEL-86 | chr6-27944923-DEL-57 |
| chr6-27947737-DEL-80 | chr6-27947738-DEL-80 |
| chr6-27949484-DEL-100 | chr6-27949477-DEL-108 |
| chr6-27994250-DEL-101 | chr6-27994246-DEL-102 |
| chr6-27997407-DEL-66 | chr6-27997408-DEL-66 |
| chr6-27998239-DEL-58 | chr6-27998244-DEL-54 |
| chr6-28002643-DEL-67 | chr6-28002648-DEL-72 |
| chr6-28057436-DEL-6354 | chr6-28057437-DEL-6354 |
| chr6-28065708-DEL-160 | chr6-28065709-DEL-159 |
| chr6-28140109-DEL-96 | chr6-28140110-DEL-96 |
| chr6-28147550-DEL-735 | chr6-28147551-DEL-735 |
| chr6-28149483-DEL-493 | chr6-28149483-DEL-493 |
| chr6-28155113-DEL-63 | chr6-28155113-DEL-63 |
| chr6-28167413-DEL-976 | chr6-28167414-DEL-976 |
| chr6-28169104-DEL-78 | chr6-28169080-DEL-65 |
| chr6-28178144-DEL-6371 | chr6-28178145-DEL-6371 |
| chr6-28189847-DEL-175 | chr6-28189842-DEL-218 |
| chr6-28199557-DEL-105 | chr6-28199557-DEL-104 |
| chr6-28199650-DEL-630 | chr6-28199651-DEL-630 |
| chr6-28217440-DEL-55 | chr6-28217441-DEL-55 |
| chr6-28308970-DEL-61 | chr6-28308971-DEL-61 |
| chr6-28322581-DEL-55 | chr6-28322582-DEL-55 |
| chr6-28327207-DEL-207 | chr6-28327208-DEL-207 |
| chr6-28356684-DEL-117 | chr6-28356676-DEL-117 |
| chr6-28393720-DEL-317 | chr6-28393721-DEL-317 |
| chr6-28403302-DEL-447 | chr6-28403303-DEL-447 |
| chr6-28407574-DEL-195 | chr6-28407572-DEL-200 |
| chr6-28412800-DEL-200 | chr6-28412793-DEL-201 |
| chr6-28425922-DEL-103 | chr6-28425923-DEL-103 |
| chr6-28443144-DEL-210 | chr6-28443145-DEL-210 |
| chr6-28466447-DEL-433 | chr6-28466448-DEL-433 |
| chr6-28500385-DEL-610 | chr6-28500386-DEL-610 |
| chr6-28502309-DEL-76 | chr6-28502325-DEL-76 |
| chr6-28503717-DEL-80 | chr6-28503718-DEL-80 |
| chr6-28516848-DEL-72 | chr6-28516848-DEL-71 |
| chr6-28531653-DEL-390 | chr6-28531652-DEL-392 |
| chr6-28570852-DEL-926 | chr6-28570853-DEL-926 |
| chr6-28645543-DEL-6591 | chr6-28645544-DEL-6591 |
| chr6-28651791-DEL-208 | chr6-28651792-DEL-208 |
| chr6-28652677-DEL-196 | chr6-28652678-DEL-196 |
| chr6-28673564-DEL-208 | chr6-28673565-DEL-208 |
| chr6-28678139-DEL-496 | chr6-28678140-DEL-496 |
| chr6-28786134-DEL-220 | chr6-28786135-DEL-220 |
| chr6-28858869-DEL-1653 | chr6-28858870-DEL-1653 |
| chr6-28924166-DEL-5185 | chr6-28924167-DEL-5185 |
| chr6-28978333-DEL-141 | chr6-28978345-DEL-96 |
| chr6-29041612-DEL-60 | chr6-29041616-DEL-59 |
| chr6-29047939-DEL-74 | chr6-29047940-DEL-74 |
| chr6-29063103-DEL-86 | chr6-29063104-DEL-86 |
| chr6-29086797-DEL-152 | chr6-29086794-DEL-202 |
| chr6-29104215-DEL-203 | chr6-29104216-DEL-203 |
| chr6-29135178-DEL-132 | chr6-29135179-DEL-132 |
| chr6-29156979-DEL-74 | chr6-29156965-DEL-75 |
| chr6-29186606-DEL-175 | chr6-29186607-DEL-175 |
| chr6-29271820-DEL-230 | chr6-29271821-DEL-185 |
| chr6-29283615-DEL-160 | chr6-29283614-DEL-162 |
| chr6-29311076-DEL-73 | chr6-29311077-DEL-73 |
| chr6-29340233-DEL-986 | chr6-29340234-DEL-986 |
| chr6-29459920-DEL-185 | chr6-29459916-DEL-190 |

| | |
|---|---|
| chr6-29468419-DEL-126 | chr6-29468442-DEL-103 |
| chr6-29511439-DEL-212 | chr6-29511440-DEL-212 |
| chr6-29519048-DEL-54 | chr6-29519049-DEL-54 |
| chr6-29551356-DEL-61 | chr6-29551357-DEL-61 |
| chr6-29581698-DEL-55 | chr6-29581684-DEL-56 |
| chr6-29598952-DEL-58 | chr6-29598953-DEL-58 |
| chr6-29629477-DEL-216 | chr6-29629478-DEL-216 |
| chr6-29658443-DEL-217 | chr6-29658444-DEL-217 |
| chr6-29661340-DEL-206 | chr6-29661341-DEL-206 |
| chr6-29664922-DEL-117 | chr6-29664923-DEL-117 |
| chr6-29668231-DEL-1085 | chr6-29668232-DEL-1085 |
| chr6-29699728-DEL-602 | chr6-29699729-DEL-602 |
| chr6-29746852-DEL-53 | chr6-29746861-DEL-74 |
| chr6-29766239-DEL-226 | chr6-29766240-DEL-226 |
| chr6-29788286-DEL-194 | chr6-29788287-DEL-194 |
| chr6-29874392-DEL-440 | chr6-29874393-DEL-440 |
| chr6-29876637-DEL-55 | chr6-29876638-DEL-56 |
| chr6-29885658-DEL-215 | chr6-29885659-DEL-215 |
| chr6-29956156-DEL-213 | chr6-29956157-DEL-213 |
| chr6-29973918-DEL-58 | chr6-29973945-DEL-58 |
| chr6-29975155-DEL-170 | chr6-29975155-DEL-170 |
| chr6-29975359-DEL-2883 | chr6-29975360-DEL-2883 |
| chr6-29994397-DEL-287 | chr6-29994398-DEL-287 |
| chr6-30003575-DEL-52 | chr6-30003576-DEL-52 |
| chr6-30004851-DEL-201 | chr6-30004854-DEL-196 |
| chr6-30012918-DEL-397 | chr6-30012919-DEL-397 |
| chr6-30024594-DEL-498 | chr6-30024557-DEL-487 |
| chr6-30026204-DEL-52 | chr6-30026207-DEL-52 |
| chr6-30031963-DEL-118 | chr6-30031963-DEL-119 |
| chr6-30042070-DEL-218 | chr6-30042071-DEL-218 |
| chr6-30055866-DEL-189 | chr6-30055867-DEL-189 |
| chr6-30068896-DEL-2132 | chr6-30068897-DEL-2132 |
| chr6-30075794-DEL-59 | chr6-30075789-DEL-60 |
| chr6-30078349-DEL-265 | chr6-30078350-DEL-265 |
| chr6-30080754-DEL-202 | chr6-30080755-DEL-202 |
| chr6-30081886-DEL-617 | chr6-30081887-DEL-617 |
| chr6-30126689-DEL-175 | chr6-30126690-DEL-175 |
| chr6-30164847-DEL-230 | chr6-30164848-DEL-229 |
| chr6-30187875-DEL-214 | chr6-30187876-DEL-214 |
| chr6-30192224-DEL-1081 | chr6-30192226-DEL-1078 |
| chr6-30201839-DEL-333 | chr6-30201840-DEL-333 |
| chr6-30229822-DEL-83 | chr6-30229807-DEL-50 |
| chr6-30233430-DEL-234 | chr6-30233431-DEL-234 |
| chr6-30234600-DEL-122 | chr6-30234601-DEL-122 |
| chr6-30235894-DEL-200 | chr6-30235895-DEL-200 |
| chr6-30237467-DEL-556 | chr6-30237468-DEL-556 |
| chr6-30283953-DEL-313 | chr6-30283954-DEL-313 |
| chr6-30287472-DEL-185 | chr6-30287473-DEL-185 |
| chr6-30334865-DEL-52 | chr6-30334875-DEL-57 |
| chr6-30337467-DEL-171 | chr6-30337468-DEL-171 |
| chr6-30339440-DEL-91 | chr6-30339441-DEL-91 |
| chr6-30345488-DEL-488 | chr6-30345489-DEL-488 |
| chr6-30362334-DEL-239 | chr6-30362335-DEL-239 |
| chr6-30364604-DEL-174 | chr6-30364605-DEL-174 |
| chr6-30385286-DEL-206 | chr6-30385287-DEL-206 |
| chr6-30390434-DEL-190 | chr6-30390436-DEL-188 |
| chr6-30437109-DEL-138 | chr6-30437110-DEL-138 |
| chr6-30465998-DEL-92 | chr6-30465999-DEL-92 |
| chr6-30476917-DEL-61 | chr6-30476916-DEL-58 |
| chr6-30502868-DEL-203 | chr6-30502869-DEL-203 |
| chr6-30540799-DEL-55 | chr6-30540800-DEL-55 |
| chr6-30567216-DEL-73 | chr6-30567210-DEL-58 |
| chr6-30610025-DEL-94 | chr6-30610026-DEL-94 |
| chr6-30629260-DEL-67 | chr6-30629223-DEL-79 |
| chr6-30727115-DEL-76 | chr6-30727114-DEL-75 |
| chr6-30797975-DEL-613 | chr6-30797976-DEL-613 |
| chr6-30800168-DEL-624 | chr6-30800169-DEL-624 |
| chr6-30840891-DEL-492 | chr6-30840891-DEL-497 |
| chr6-30842140-DEL-119 | chr6-30842141-DEL-119 |
| chr6-30844363-DEL-203 | chr6-30844364-DEL-203 |
| chr6-30862601-DEL-395 | chr6-30862602-DEL-395 |
| chr6-30886644-DEL-56 | chr6-30886645-DEL-54 |
| chr6-30905763-DEL-7104 | chr6-30905764-DEL-7104 |
| chr6-30926658-DEL-62 | chr6-30926651-DEL-72 |
| chr6-30979010-DEL-412 | chr6-30979011-DEL-412 |
| chr6-30991105-DEL-64 | chr6-30991123-DEL-52 |
| chr6-30995410-DEL-2547 | chr6-30995411-DEL-2547 |
| chr6-30999677-DEL-1340 | chr6-30999678-DEL-1337 |
| chr6-31019648-DEL-927 | chr6-31019649-DEL-927 |
| chr6-31025361-DEL-327 | chr6-31025386-DEL-301 |

| | |
|---|---|
| chr6-31031410-DEL-58 | chr6-31031419-DEL-50 |
| chr6-31037199-DEL-52 | chr6-31037200-DEL-52 |
| chr6-31072798-DEL-138 | chr6-31072801-DEL-132 |
| chr6-31073815-DEL-64 | chr6-31073816-DEL-64 |
| chr6-31118772-DEL-890 | chr6-31118773-DEL-890 |
| chr6-31181436-DEL-52 | chr6-31181437-DEL-52 |
| chr6-31218172-DEL-181 | chr6-31218173-DEL-181 |
| chr6-31235330-DEL-52 | chr6-31235326-DEL-50 |
| chr6-31247212-DEL-81 | chr6-31247204-DEL-86 |
| chr6-31256907-DEL-121 | chr6-31256908-DEL-121 |
| chr6-31290238-DEL-60 | chr6-31290236-DEL-51 |
| chr6-31309093-DEL-57 | chr6-31309094-DEL-57 |
| chr6-31390462-DEL-92 | chr6-31390463-DEL-92 |
| chr6-31436999-DEL-192 | chr6-31436998-DEL-194 |
| chr6-31444376-DEL-131 | chr6-31444377-DEL-131 |
| chr6-31467555-DEL-229 | chr6-31467556-DEL-229 |
| chr6-31516921-DEL-160 | chr6-31516922-DEL-160 |
| chr6-31529182-DEL-102 | chr6-31529200-DEL-56 |
| chr6-31543544-DEL-90 | chr6-31543545-DEL-90 |
| chr6-31558509-DEL-60 | chr6-31558515-DEL-54 |
| chr6-31571444-DEL-84 | chr6-31571445-DEL-84 |
| chr6-31580713-DEL-62 | chr6-31580725-DEL-51 |
| chr6-31586043-DEL-434 | chr6-31586045-DEL-435 |
| chr6-31588310-DEL-56 | chr6-31588311-DEL-56 |
| chr6-31607675-DEL-228 | chr6-31607676-DEL-228 |
| chr6-31620791-DEL-392 | chr6-31620792-DEL-392 |
| chr6-31675855-DEL-351 | chr6-31675856-DEL-351 |
| chr6-31683211-DEL-223 | chr6-31683170-DEL-224 |
| chr6-31765100-DEL-682 | chr6-31765096-DEL-682 |
| chr6-31826977-DEL-4311 | chr6-31826978-DEL-4311 |
| chr6-31846434-DEL-1723 | chr6-31846435-DEL-1723 |
| chr6-31873665-DEL-56 | chr6-31873666-DEL-56 |
| chr6-31884327-DEL-4710 | chr6-31884328-DEL-4710 |
| chr6-31908344-DEL-653 | chr6-31908345-DEL-653 |
| chr6-31926809-DEL-64 | chr6-31926812-DEL-62 |
| chr6-31950521-DEL-258 | chr6-31950523-DEL-257 |
| chr6-32070510-DEL-7089 | chr6-32070511-DEL-7089 |
| chr6-32070649-DEL-212 | chr6-32070648-DEL-212 |
| chr6-32086987-DEL-508 | chr6-32086992-DEL-504 |
| chr6-32095746-DEL-460 | chr6-32095747-DEL-460 |
| chr6-32155963-DEL-132 | chr6-32155974-DEL-130 |
| chr6-32199104-DEL-455 | chr6-32199105-DEL-455 |
| chr6-32221440-DEL-64 | chr6-32221407-DEL-76 |
| chr6-32243554-DEL-72 | chr6-32243555-DEL-59 |
| chr6-32269301-DEL-56 | chr6-32269252-DEL-57 |
| chr6-32284672-DEL-194 | chr6-32284673-DEL-194 |
| chr6-32288758-DEL-300 | chr6-32288759-DEL-300 |
| chr6-32411617-DEL-96 | chr6-32411596-DEL-123 |
| chr6-32442680-DEL-670 | chr6-32442681-DEL-670 |
| chr6-32507420-DEL-194 | chr6-32507421-DEL-194 |
| chr6-32538562-DEL-176 | chr6-32538553-DEL-167 |
| chr6-32663511-DEL-3943 | chr6-32663512-DEL-3943 |
| chr6-32697606-DEL-54 | chr6-32697601-DEL-60 |
| chr6-32703299-DEL-62 | chr6-32703303-DEL-55 |
| chr6-32717492-DEL-6312 | chr6-32717493-DEL-6312 |
| chr6-32724097-DEL-220 | chr6-32724085-DEL-232 |
| chr6-32752240-DEL-541 | chr6-32752241-DEL-541 |
| chr6-32765383-DEL-146 | chr6-32765384-DEL-146 |
| chr6-32779834-DEL-396 | chr6-32779835-DEL-396 |
| chr6-32784186-DEL-172 | chr6-32784187-DEL-129 |
| chr6-32839876-DEL-60 | chr6-32839877-DEL-60 |
| chr6-32844528-DEL-1467 | chr6-32844529-DEL-1467 |
| chr6-32872603-DEL-124 | chr6-32872604-DEL-124 |
| chr6-32938482-DEL-75 | chr6-32938483-DEL-75 |
| chr6-32978121-DEL-162 | chr6-32978122-DEL-162 |
| chr6-32989268-DEL-51 | chr6-32989269-DEL-51 |
| chr6-32990241-DEL-1487 | chr6-32990269-DEL-1467 |
| chr6-33024190-DEL-330 | chr6-33024187-DEL-330 |
| chr6-33041597-DEL-71 | chr6-33041589-DEL-71 |
| chr6-33047724-DEL-118 | chr6-33047725-DEL-118 |
| chr6-33059005-DEL-108 | chr6-33059046-DEL-64 |
| chr6-33116374-DEL-53 | chr6-33116366-DEL-50 |
| chr6-33129558-DEL-500 | chr6-33129559-DEL-500 |
| chr6-33182881-DEL-71 | chr6-33182882-DEL-71 |
| chr6-33258719-DEL-779 | chr6-33258720-DEL-780 |
| chr6-33291501-DEL-75 | chr6-33291494-DEL-76 |
| chr6-33304812-DEL-343 | chr6-33304813-DEL-343 |
| chr6-33380934-DEL-5428 | chr6-33380936-DEL-5427 |
| chr6-33392643-DEL-336 | chr6-33392644-DEL-336 |
| chr6-33448517-DEL-122 | chr6-33448517-DEL-120 |

| | |
|---|---|
| chr6-33449372-DEL-64 | chr6-33449373-DEL-64 |
| chr6-33462706-DEL-441 | chr6-33462707-DEL-441 |
| chr6-33463366-DEL-59 | chr6-33463331-DEL-52 |
| chr6-33467216-DEL-3367 | chr6-33467217-DEL-3367 |
| chr6-33471525-DEL-625 | chr6-33471530-DEL-621 |
| chr6-33506286-DEL-592 | chr6-33506288-DEL-591 |
| chr6-33527931-DEL-96 | chr6-33527951-DEL-89 |
| chr6-33535619-DEL-920 | chr6-33535620-DEL-920 |
| chr6-33627358-DEL-1578 | chr6-33627359-DEL-1578 |
| chr6-33635920-DEL-6447 | chr6-33635921-DEL-6447 |
| chr6-33677871-DEL-52 | chr6-33677872-DEL-52 |
| chr6-33720579-DEL-5541 | chr6-33720580-DEL-5541 |
| chr6-33759109-DEL-1499 | chr6-33759109-DEL-1499 |
| chr6-33778472-DEL-69 | chr6-33778473-DEL-69 |
| chr6-33781319-DEL-104 | chr6-33781320-DEL-102 |
| chr6-33827681-DEL-75 | chr6-33827681-DEL-75 |
| chr6-33832280-DEL-220 | chr6-33832281-DEL-220 |
| chr6-33911104-DEL-58 | chr6-33911105-DEL-58 |
| chr6-33980698-DEL-170 | chr6-33980699-DEL-170 |
| chr6-34037560-DEL-1214 | chr6-34037561-DEL-1214 |
| chr6-34079026-DEL-60 | chr6-34079026-DEL-60 |
| chr6-34085772-DEL-691 | chr6-34085773-DEL-691 |
| chr6-34097370-DEL-399 | chr6-34097371-DEL-399 |
| chr6-34099789-DEL-449 | chr6-34099787-DEL-448 |
| chr6-34116909-DEL-56 | chr6-34116910-DEL-56 |
| chr6-34118088-DEL-52 | chr6-34118089-DEL-52 |
| chr6-34136097-DEL-53 | chr6-34136098-DEL-53 |
| chr6-34138903-DEL-78 | chr6-34138904-DEL-78 |
| chr6-34168971-DEL-390 | chr6-34168971-DEL-390 |
| chr6-34203514-DEL-302 | chr6-34203515-DEL-302 |
| chr6-34204917-DEL-250 | chr6-34204918-DEL-250 |
| chr6-34231382-DEL-381 | chr6-34231383-DEL-381 |
| chr6-34296810-DEL-228 | chr6-34296797-DEL-229 |
| chr6-34302443-DEL-288 | chr6-34302436-DEL-296 |
| chr6-34308796-DEL-333 | chr6-34308797-DEL-333 |
| chr6-34336780-DEL-420 | chr6-34336781-DEL-420 |
| chr6-34349694-DEL-926 | chr6-34349695-DEL-926 |
| chr6-34356970-DEL-216 | chr6-34356971-DEL-216 |
| chr6-34391397-DEL-97 | chr6-34391401-DEL-100 |
| chr6-34441240-DEL-918 | chr6-34441241-DEL-918 |
| chr6-34477854-DEL-2569 | chr6-34477855-DEL-2569 |
| chr6-34527340-DEL-56 | chr6-34527340-DEL-56 |
| chr6-34540125-DEL-155 | chr6-34540135-DEL-156 |
| chr6-34550513-DEL-62 | chr6-34550514-DEL-62 |
| chr6-34562430-DEL-1888 | chr6-34562431-DEL-1888 |
| chr6-34601871-DEL-91 | chr6-34601872-DEL-91 |
| chr6-34608392-DEL-85 | chr6-34608401-DEL-62 |
| chr6-34616187-DEL-123 | chr6-34616187-DEL-123 |
| chr6-34659940-DEL-168 | chr6-34659941-DEL-168 |
| chr6-34665361-DEL-193 | chr6-34665362-DEL-193 |
| chr6-34667080-DEL-230 | chr6-34667078-DEL-233 |
| chr6-34670014-DEL-450 | chr6-34670015-DEL-450 |
| chr6-34821234-DEL-69 | chr6-34821235-DEL-69 |
| chr6-34865748-DEL-87 | chr6-34865748-DEL-87 |
| chr6-34875688-DEL-224 | chr6-34875689-DEL-224 |
| chr6-34898390-DEL-217 | chr6-34898391-DEL-217 |
| chr6-34900582-DEL-56 | chr6-34900583-DEL-56 |
| chr6-34946135-DEL-341 | chr6-34946136-DEL-341 |
| chr6-34950421-DEL-1373 | chr6-34950422-DEL-1373 |
| chr6-34961733-DEL-2883 | chr6-34961734-DEL-2883 |
| chr6-34983357-DEL-1892 | chr6-34983357-DEL-1892 |
| chr6-34990329-DEL-84 | chr6-34990371-DEL-69 |
| chr6-35008035-DEL-188 | chr6-35008036-DEL-188 |
| chr6-35069950-DEL-70 | chr6-35069951-DEL-70 |
| chr6-35090919-DEL-208 | chr6-35090920-DEL-208 |
| chr6-35124393-DEL-211 | chr6-35124395-DEL-210 |
| chr6-35156209-DEL-203 | chr6-35156210-DEL-203 |
| chr6-35157007-DEL-82 | chr6-35157008-DEL-82 |
| chr6-35185961-DEL-221 | chr6-35185973-DEL-219 |
| chr6-35251581-DEL-202 | chr6-35251582-DEL-202 |
| chr6-35271623-DEL-131 | chr6-35271624-DEL-131 |
| chr6-35299260-DEL-791 | chr6-35299261-DEL-791 |
| chr6-35306408-DEL-223 | chr6-35306409-DEL-223 |
| chr6-35316393-DEL-7961 | chr6-35316394-DEL-7961 |
| chr6-35334453-DEL-60 | chr6-35334434-DEL-80 |
| chr6-35348041-DEL-208 | chr6-35348042-DEL-208 |
| chr6-35354865-DEL-97 | chr6-35354865-DEL-97 |
| chr6-35366252-DEL-6496 | chr6-35366253-DEL-6496 |
| chr6-35401079-DEL-180 | chr6-35401080-DEL-180 |
| chr6-35423177-DEL-6675 | chr6-35423178-DEL-6675 |

| | |
|---|---|
| chr6-35438471-DEL-445 | chr6-35438472-DEL-445 |
| chr6-35444461-DEL-1168 | chr6-35444462-DEL-1169 |
| chr6-35462345-DEL-6396 | chr6-35462346-DEL-6396 |
| chr6-35511517-DEL-202 | chr6-35511518-DEL-192 |
| chr6-35520756-DEL-200 | chr6-35520757-DEL-200 |
| chr6-35587637-DEL-373 | chr6-35587638-DEL-371 |
| chr6-35599712-DEL-387 | chr6-35599713-DEL-387 |
| chr6-35672616-DEL-134 | chr6-35672617-DEL-134 |
| chr6-35677714-DEL-2784 | chr6-35677715-DEL-2784 |
| chr6-35690789-DEL-345 | chr6-35690790-DEL-345 |
| chr6-35748752-DEL-6630 | chr6-35748753-DEL-6630 |
| chr6-35769690-DEL-1211 | chr6-35769691-DEL-1211 |
| chr6-35780632-DEL-56 | chr6-35780633-DEL-60 |
| chr6-35789983-DEL-148 | chr6-35789937-DEL-195 |
| chr6-35817599-DEL-68 | chr6-35817594-DEL-58 |
| chr6-35840845-DEL-72 | chr6-35840846-DEL-72 |
| chr6-35849743-DEL-468 | chr6-35849743-DEL-466 |
| chr6-35863362-DEL-177 | chr6-35863363-DEL-177 |
| chr6-35872356-DEL-294 | chr6-35872357-DEL-294 |
| chr6-35892683-DEL-775 | chr6-35892684-DEL-775 |
| chr6-35895942-DEL-318 | chr6-35895943-DEL-318 |
| chr6-35903062-DEL-1004 | chr6-35903061-DEL-1004 |
| chr6-35908410-DEL-53 | chr6-35908408-DEL-51 |
| chr6-35910105-DEL-2818 | chr6-35910106-DEL-2818 |
| chr6-35917387-DEL-304 | chr6-35917386-DEL-305 |
| chr6-35923626-DEL-3345 | chr6-35923627-DEL-3345 |
| chr6-35932043-DEL-66 | chr6-35932059-DEL-54 |
| chr6-35935805-DEL-6213 | chr6-35935806-DEL-6213 |
| chr6-35957602-DEL-179 | chr6-35957603-DEL-179 |
| chr6-35959775-DEL-1056 | chr6-35959776-DEL-1056 |
| chr6-35971577-DEL-61 | chr6-35971578-DEL-61 |
| chr6-36020456-DEL-193 | chr6-36020457-DEL-193 |
| chr6-36030656-DEL-138 | chr6-36030649-DEL-119 |
| chr6-36038127-DEL-2770 | chr6-36038128-DEL-2770 |
| chr6-36050394-DEL-1002 | chr6-36050404-DEL-1001 |
| chr6-36059674-DEL-6431 | chr6-36059675-DEL-6431 |
| chr6-36117548-DEL-58 | chr6-36117549-DEL-58 |
| chr6-36149345-DEL-2083 | chr6-36149346-DEL-2083 |
| chr6-36167823-DEL-68 | chr6-36167826-DEL-66 |
| chr6-36201204-DEL-6413 | chr6-36201205-DEL-6413 |
| chr6-36212321-DEL-244 | chr6-36212322-DEL-244 |
| chr6-36216359-DEL-163 | chr6-36216360-DEL-163 |
| chr6-36221060-DEL-145 | chr6-36221061-DEL-145 |
| chr6-36280676-DEL-154 | chr6-36280677-DEL-154 |
| chr6-36284778-DEL-1368 | chr6-36284779-DEL-1368 |
| chr6-36291608-DEL-70 | chr6-36291592-DEL-72 |
| chr6-36296852-DEL-272 | chr6-36296853-DEL-272 |
| chr6-36323887-DEL-88 | chr6-36323884-DEL-86 |
| chr6-36333732-DEL-6416 | chr6-36333733-DEL-6416 |
| chr6-36351859-DEL-163 | chr6-36351860-DEL-163 |
| chr6-36377660-DEL-1590 | chr6-36377661-DEL-1590 |
| chr6-36393385-DEL-126 | chr6-36393386-DEL-126 |
| chr6-36397362-DEL-78 | chr6-36397380-DEL-56 |
| chr6-36410044-DEL-649 | chr6-36410034-DEL-650 |
| chr6-36469128-DEL-82 | chr6-36469113-DEL-72 |
| chr6-36473790-DEL-6671 | chr6-36473791-DEL-6671 |
| chr6-36474063-DEL-212 | chr6-36474064-DEL-212 |
| chr6-36489449-DEL-173 | chr6-36489453-DEL-168 |
| chr6-36548420-DEL-4562 | chr6-36548421-DEL-4562 |
| chr6-36580220-DEL-73 | chr6-36580221-DEL-83 |
| chr6-36632107-DEL-54 | chr6-36632114-DEL-62 |
| chr6-36637351-DEL-52 | chr6-36637352-DEL-52 |
| chr6-36680171-DEL-92 | chr6-36680166-DEL-99 |
| chr6-36682095-DEL-6398 | chr6-36682096-DEL-6398 |
| chr6-36836381-DEL-383 | chr6-36836372-DEL-384 |
| chr6-36899353-DEL-115 | chr6-36899334-DEL-122 |
| chr6-36905678-DEL-72 | chr6-36905679-DEL-66 |
| chr6-36919862-DEL-157 | chr6-36919910-DEL-157 |
| chr6-36933572-DEL-3800 | chr6-36933573-DEL-3800 |
| chr6-36960890-DEL-204 | chr6-36960891-DEL-204 |
| chr6-36974796-DEL-60 | chr6-36974803-DEL-53 |
| chr6-36988364-DEL-851 | chr6-36988365-DEL-851 |
| chr6-36992771-DEL-246 | chr6-36992772-DEL-246 |
| chr6-36995299-DEL-202 | chr6-36995297-DEL-203 |
| chr6-37065760-DEL-51 | chr6-37065793-DEL-58 |
| chr6-37153579-DEL-71 | chr6-37153602-DEL-78 |
| chr6-37183581-DEL-968 | chr6-37183587-DEL-962 |
| chr6-37184325-DEL-195 | chr6-37184341-DEL-180 |
| chr6-37245091-DEL-56 | chr6-37245092-DEL-51 |
| chr6-37249136-DEL-327 | chr6-37249137-DEL-327 |

| | |
|---|---|
| chr6-37293697-DEL-59 | chr6-37293700-DEL-52 |
| chr6-37319464-DEL-62 | chr6-37319462-DEL-60 |
| chr6-37320792-DEL-52 | chr6-37320793-DEL-52 |
| chr6-37394132-DEL-212 | chr6-37394133-DEL-212 |
| chr6-37410718-DEL-62 | chr6-37410716-DEL-50 |
| chr6-37420611-DEL-161 | chr6-37420612-DEL-161 |
| chr6-37420827-DEL-3570 | chr6-37420828-DEL-3570 |
| chr6-37432325-DEL-213 | chr6-37432326-DEL-206 |
| chr6-37436452-DEL-341 | chr6-37436453-DEL-341 |
| chr6-37437222-DEL-55 | chr6-37437223-DEL-55 |
| chr6-37441074-DEL-476 | chr6-37441075-DEL-476 |
| chr6-37471047-DEL-60 | chr6-37471048-DEL-60 |
| chr6-37522602-DEL-4390 | chr6-37522603-DEL-4390 |
| chr6-37554605-DEL-622 | chr6-37554606-DEL-622 |
| chr6-37555607-DEL-134 | chr6-37555607-DEL-134 |
| chr6-37563487-DEL-84 | chr6-37563488-DEL-84 |
| chr6-37575957-DEL-209 | chr6-37575958-DEL-209 |
| chr6-37577330-DEL-106 | chr6-37577330-DEL-152 |
| chr6-37625440-DEL-202 | chr6-37625441-DEL-202 |
| chr6-37641586-DEL-63 | chr6-37641595-DEL-55 |
| chr6-37649412-DEL-908 | chr6-37649413-DEL-908 |
| chr6-37650819-DEL-196 | chr6-37650787-DEL-211 |
| chr6-37656457-DEL-99 | chr6-37656458-DEL-99 |
| chr6-37668192-DEL-826 | chr6-37668193-DEL-826 |
| chr6-37677235-DEL-83 | chr6-37677235-DEL-84 |
| chr6-37686544-DEL-468 | chr6-37686545-DEL-468 |
| chr6-37712872-DEL-51 | chr6-37712873-DEL-51 |
| chr6-37737887-DEL-195 | chr6-37737888-DEL-195 |
| chr6-37751535-DEL-1501 | chr6-37751536-DEL-1501 |
| chr6-37752499-DEL-114 | chr6-37752506-DEL-108 |
| chr6-37761233-DEL-566 | chr6-37761234-DEL-566 |
| chr6-37765013-DEL-322 | chr6-37765014-DEL-322 |
| chr6-37769862-DEL-77 | chr6-37769847-DEL-58 |
| chr6-37770610-DEL-220 | chr6-37770610-DEL-221 |
| chr6-37773976-DEL-349 | chr6-37773977-DEL-305 |
| chr6-37782911-DEL-1144 | chr6-37782912-DEL-1144 |
| chr6-37790194-DEL-194 | chr6-37790195-DEL-194 |
| chr6-37826077-DEL-203 | chr6-37826078-DEL-202 |
| chr6-37859998-DEL-235 | chr6-37860000-DEL-229 |
| chr6-37934917-DEL-64 | chr6-37934915-DEL-67 |
| chr6-37939677-DEL-90 | chr6-37939678-DEL-66 |
| chr6-37950684-DEL-6413 | chr6-37950685-DEL-6413 |
| chr6-37973532-DEL-227 | chr6-37973533-DEL-227 |
| chr6-37974351-DEL-457 | chr6-37974352-DEL-457 |
| chr6-37975821-DEL-69 | chr6-37975822-DEL-69 |
| chr6-37996198-DEL-53 | chr6-37996195-DEL-56 |
| chr6-38004505-DEL-1354 | chr6-38004506-DEL-1350 |
| chr6-38015831-DEL-98 | chr6-38015837-DEL-91 |
| chr6-38019776-DEL-219 | chr6-38019777-DEL-219 |
| chr6-38034690-DEL-200 | chr6-38034689-DEL-201 |
| chr6-38089401-DEL-225 | chr6-38089402-DEL-225 |
| chr6-38127330-DEL-68 | chr6-38127329-DEL-56 |
| chr6-38158988-DEL-81 | chr6-38158991-DEL-79 |
| chr6-38177601-DEL-159 | chr6-38177602-DEL-159 |
| chr6-38186305-DEL-2181 | chr6-38186306-DEL-2181 |
| chr6-38193518-DEL-256 | chr6-38193519-DEL-256 |
| chr6-38212808-DEL-198 | chr6-38212809-DEL-198 |
| chr6-38237535-DEL-212 | chr6-38237536-DEL-212 |
| chr6-38244501-DEL-90 | chr6-38244502-DEL-90 |
| chr6-38251936-DEL-372 | chr6-38251937-DEL-372 |
| chr6-38253367-DEL-6592 | chr6-38253368-DEL-6592 |
| chr6-38274424-DEL-220 | chr6-38274425-DEL-220 |
| chr6-38278423-DEL-113 | chr6-38278412-DEL-114 |
| chr6-38284813-DEL-184 | chr6-38284814-DEL-184 |
| chr6-38291833-DEL-173 | chr6-38291834-DEL-173 |
| chr6-38339908-DEL-212 | chr6-38339919-DEL-213 |
| chr6-38340872-DEL-117 | chr6-38340875-DEL-118 |
| chr6-38353654-DEL-513 | chr6-38353655-DEL-513 |
| chr6-38355973-DEL-239 | chr6-38355974-DEL-239 |
| chr6-38366458-DEL-95 | chr6-38366468-DEL-92 |
| chr6-38377502-DEL-186 | chr6-38377503-DEL-186 |
| chr6-38387502-DEL-78 | chr6-38387503-DEL-78 |
| chr6-38420275-DEL-184 | chr6-38420276-DEL-184 |
| chr6-38431331-DEL-211 | chr6-38431326-DEL-212 |
| chr6-38433429-DEL-107 | chr6-38433461-DEL-69 |
| chr6-38445024-DEL-221 | chr6-38445025-DEL-221 |
| chr6-38446121-DEL-55 | chr6-38446122-DEL-55 |
| chr6-38446690-DEL-296 | chr6-38446691-DEL-296 |
| chr6-38481155-DEL-497 | chr6-38481156-DEL-497 |
| chr6-38496106-DEL-178 | chr6-38496107-DEL-178 |

| | |
|---|---|
| chr6-38519346-DEL-213 | chr6-38519347-DEL-213 |
| chr6-38523366-DEL-177 | chr6-38523367-DEL-177 |
| chr6-38559668-DEL-199 | chr6-38559669-DEL-199 |
| chr6-38562564-DEL-224 | chr6-38562567-DEL-248 |
| chr6-38599749-DEL-1566 | chr6-38599750-DEL-1566 |
| chr6-38630512-DEL-52 | chr6-38630513-DEL-52 |
| chr6-38631728-DEL-173 | chr6-38631729-DEL-173 |
| chr6-38647707-DEL-195 | chr6-38647708-DEL-195 |
| chr6-38655396-DEL-218 | chr6-38655408-DEL-194 |
| chr6-38662525-DEL-225 | chr6-38662526-DEL-214 |
| chr6-38710414-DEL-81 | chr6-38710415-DEL-81 |
| chr6-38746282-DEL-165 | chr6-38746283-DEL-165 |
| chr6-38751589-DEL-56 | chr6-38751594-DEL-86 |
| chr6-38760484-DEL-499 | chr6-38760485-DEL-499 |
| chr6-38778727-DEL-58 | chr6-38778728-DEL-58 |
| chr6-38898805-DEL-73 | chr6-38898806-DEL-73 |
| chr6-38936219-DEL-68 | chr6-38936220-DEL-68 |
| chr6-38958859-DEL-58 | chr6-38958860-DEL-58 |
| chr6-38991889-DEL-195 | chr6-38991878-DEL-193 |
| chr6-39009712-DEL-124 | chr6-39009716-DEL-121 |
| chr6-39033966-DEL-70 | chr6-39033967-DEL-70 |
| chr6-39108735-DEL-1915 | chr6-39108736-DEL-1915 |
| chr6-39113277-DEL-221 | chr6-39113278-DEL-221 |
| chr6-39128000-DEL-140 | chr6-39128001-DEL-140 |
| chr6-39197338-DEL-56 | chr6-39197319-DEL-52 |
| chr6-39198743-DEL-54 | chr6-39198726-DEL-80 |
| chr6-39204460-DEL-180 | chr6-39204468-DEL-178 |
| chr6-39214106-DEL-668 | chr6-39214107-DEL-668 |
| chr6-39229644-DEL-226 | chr6-39229645-DEL-226 |
| chr6-39258358-DEL-201 | chr6-39258359-DEL-201 |
| chr6-39271565-DEL-203 | chr6-39271566-DEL-203 |
| chr6-39274400-DEL-98 | chr6-39274400-DEL-98 |
| chr6-39296633-DEL-211 | chr6-39296634-DEL-211 |
| chr6-39301190-DEL-855 | chr6-39301191-DEL-855 |
| chr6-39342759-DEL-197 | chr6-39342746-DEL-215 |
| chr6-39344000-DEL-194 | chr6-39344001-DEL-194 |
| chr6-39355039-DEL-201 | chr6-39355040-DEL-201 |
| chr6-39371658-DEL-60 | chr6-39371703-DEL-63 |
| chr6-39389146-DEL-147 | chr6-39389147-DEL-147 |
| chr6-39392376-DEL-426 | chr6-39392377-DEL-426 |
| chr6-39404147-DEL-90 | chr6-39404152-DEL-86 |
| chr6-39405448-DEL-91 | chr6-39405449-DEL-91 |
| chr6-39423738-DEL-69 | chr6-39423737-DEL-85 |
| chr6-39430086-DEL-210 | chr6-39430087-DEL-210 |
| chr6-39431031-DEL-575 | chr6-39431029-DEL-574 |
| chr6-39434165-DEL-2891 | chr6-39434166-DEL-2891 |
| chr6-39446132-DEL-212 | chr6-39446133-DEL-212 |
| chr6-39467399-DEL-116 | chr6-39467410-DEL-118 |
| chr6-39477379-DEL-68 | chr6-39477376-DEL-76 |
| chr6-39477847-DEL-215 | chr6-39477838-DEL-216 |
| chr6-39486476-DEL-58 | chr6-39486477-DEL-58 |
| chr6-39502777-DEL-209 | chr6-39502778-DEL-209 |
| chr6-39514650-DEL-210 | chr6-39514651-DEL-210 |
| chr6-39541227-DEL-330 | chr6-39541230-DEL-328 |
| chr6-39547311-DEL-102 | chr6-39547316-DEL-98 |
| chr6-39548627-DEL-400 | chr6-39548628-DEL-400 |
| chr6-39557451-DEL-60 | chr6-39557452-DEL-54 |
| chr6-39559480-DEL-156 | chr6-39559482-DEL-155 |
| chr6-39570769-DEL-209 | chr6-39570770-DEL-209 |
| chr6-39604140-DEL-66 | chr6-39604141-DEL-66 |
| chr6-39613562-DEL-199 | chr6-39613563-DEL-199 |
| chr6-39635514-DEL-322 | chr6-39635515-DEL-322 |
| chr6-39650697-DEL-106 | chr6-39650698-DEL-106 |
| chr6-39664734-DEL-199 | chr6-39664735-DEL-199 |
| chr6-39675513-DEL-355 | chr6-39675514-DEL-355 |
| chr6-39704301-DEL-98 | chr6-39704302-DEL-98 |
| chr6-39710334-DEL-98 | chr6-39710359-DEL-50 |
| chr6-39712991-DEL-374 | chr6-39712992-DEL-374 |
| chr6-39733708-DEL-197 | chr6-39733709-DEL-197 |
| chr6-39735116-DEL-225 | chr6-39735117-DEL-220 |
| chr6-39737983-DEL-515 | chr6-39737984-DEL-515 |
| chr6-39739740-DEL-189 | chr6-39739749-DEL-185 |
| chr6-39750738-DEL-205 | chr6-39750739-DEL-205 |
| chr6-39765718-DEL-60 | chr6-39765744-DEL-59 |
| chr6-39784554-DEL-206 | chr6-39784555-DEL-206 |
| chr6-39800491-DEL-313 | chr6-39800492-DEL-313 |
| chr6-39801580-DEL-56 | chr6-39801581-DEL-56 |
| chr6-39814026-DEL-107 | chr6-39814027-DEL-107 |
| chr6-39818423-DEL-117 | chr6-39818424-DEL-117 |
| chr6-39819421-DEL-209 | chr6-39819422-DEL-208 |

| | |
|---|---|
| chr6-39820619-DEL-107 | chr6-39820616-DEL-108 |
| chr6-39872202-DEL-213 | chr6-39872203-DEL-213 |
| chr6-39896577-DEL-6906 | chr6-39896578-DEL-6906 |
| chr6-39916252-DEL-1321 | chr6-39916253-DEL-1321 |
| chr6-39970645-DEL-55 | chr6-39970639-DEL-56 |
| chr6-40007991-DEL-290 | chr6-40007992-DEL-290 |
| chr6-40024090-DEL-1383 | chr6-40024091-DEL-1383 |
| chr6-40048429-DEL-60 | chr6-40048407-DEL-94 |
| chr6-40062871-DEL-135 | chr6-40062872-DEL-135 |
| chr6-40258971-DEL-404 | chr6-40258961-DEL-413 |
| chr6-40269280-DEL-100 | chr6-40269299-DEL-78 |
| chr6-40271892-DEL-1813 | chr6-40271893-DEL-1813 |
| chr6-40309231-DEL-109 | chr6-40309224-DEL-110 |
| chr6-40401140-DEL-84 | chr6-40401137-DEL-92 |
| chr6-40409041-DEL-87 | chr6-40409044-DEL-85 |
| chr6-40427267-DEL-126 | chr6-40427265-DEL-129 |
| chr6-40475226-DEL-211 | chr6-40475227-DEL-211 |
| chr6-40512378-DEL-6455 | chr6-40512379-DEL-6455 |
| chr6-40560735-DEL-99 | chr6-40560766-DEL-99 |
| chr6-40576240-DEL-6251 | chr6-40576241-DEL-6251 |
| chr6-40582204-DEL-199 | chr6-40582205-DEL-199 |
| chr6-40606352-DEL-54 | chr6-40606366-DEL-54 |
| chr6-40610893-DEL-223 | chr6-40610894-DEL-223 |
| chr6-40626813-DEL-366 | chr6-40626814-DEL-366 |
| chr6-40628977-DEL-90 | chr6-40628981-DEL-83 |
| chr6-40641952-DEL-475 | chr6-40641953-DEL-475 |
| chr6-40654909-DEL-60 | chr6-40654913-DEL-80 |
| chr6-40749062-DEL-890 | chr6-40749063-DEL-890 |
| chr6-40756198-DEL-99 | chr6-40756198-DEL-96 |
| chr6-40770651-DEL-142 | chr6-40770651-DEL-140 |
| chr6-40794473-DEL-751 | chr6-40794474-DEL-751 |
| chr6-40805690-DEL-61 | chr6-40805691-DEL-61 |
| chr6-40825394-DEL-2629 | chr6-40825395-DEL-2629 |
| chr6-40854525-DEL-58 | chr6-40854526-DEL-58 |
| chr6-40883855-DEL-6436 | chr6-40883856-DEL-6435 |
| chr6-40884004-DEL-208 | chr6-40884005-DEL-208 |
| chr6-40933353-DEL-218 | chr6-40933354-DEL-218 |
| chr6-40948535-DEL-208 | chr6-40948536-DEL-208 |
| chr6-40969367-DEL-208 | chr6-40969368-DEL-208 |
| chr6-40982544-DEL-117 | chr6-40982544-DEL-115 |
| chr6-41062620-DEL-68 | chr6-41062621-DEL-68 |
| chr6-41077896-DEL-143 | chr6-41077897-DEL-143 |
| chr6-41084793-DEL-92 | chr6-41084807-DEL-88 |
| chr6-41113092-DEL-79 | chr6-41113078-DEL-50 |
| chr6-41129331-DEL-2615 | chr6-41129332-DEL-2621 |
| chr6-41142388-DEL-75 | chr6-41142389-DEL-75 |
| chr6-41145717-DEL-55 | chr6-41145721-DEL-50 |
| chr6-41174074-DEL-4968 | chr6-41174075-DEL-4968 |
| chr6-41193767-DEL-155 | chr6-41193768-DEL-155 |
| chr6-41197596-DEL-86 | chr6-41197607-DEL-76 |
| chr6-41219773-DEL-4412 | chr6-41219774-DEL-4412 |
| chr6-41228211-DEL-2972 | chr6-41228212-DEL-2972 |
| chr6-41242629-DEL-70 | chr6-41242581-DEL-80 |
| chr6-41245699-DEL-214 | chr6-41245700-DEL-214 |
| chr6-41252262-DEL-801 | chr6-41252263-DEL-801 |
| chr6-41275700-DEL-139 | chr6-41275701-DEL-139 |
| chr6-41292974-DEL-198 | chr6-41292971-DEL-225 |
| chr6-41303644-DEL-460 | chr6-41303645-DEL-460 |
| chr6-41316422-DEL-91 | chr6-41316423-DEL-91 |
| chr6-41322874-DEL-57 | chr6-41322875-DEL-57 |
| chr6-41379788-DEL-236 | chr6-41379788-DEL-236 |
| chr6-41453461-DEL-76 | chr6-41453462-DEL-76 |
| chr6-41459807-DEL-161 | chr6-41459808-DEL-161 |
| chr6-41477023-DEL-69 | chr6-41477021-DEL-68 |
| chr6-41483549-DEL-366 | chr6-41483550-DEL-366 |
| chr6-41483925-DEL-711 | chr6-41483926-DEL-711 |
| chr6-41488356-DEL-1718 | chr6-41488357-DEL-1718 |
| chr6-41491519-DEL-53 | chr6-41491520-DEL-53 |
| chr6-41513744-DEL-70 | chr6-41513744-DEL-71 |
| chr6-41518218-DEL-55 | chr6-41518210-DEL-50 |
| chr6-41530146-DEL-70 | chr6-41530102-DEL-64 |
| chr6-41539922-DEL-477 | chr6-41539923-DEL-477 |
| chr6-41645374-DEL-1073 | chr6-41645375-DEL-1073 |
| chr6-41654780-DEL-57 | chr6-41654776-DEL-54 |
| chr6-41687911-DEL-84 | chr6-41687928-DEL-85 |
| chr6-41698781-DEL-710 | chr6-41698782-DEL-710 |
| chr6-41740776-DEL-873 | chr6-41740778-DEL-871 |
| chr6-41741617-DEL-64 | chr6-41741609-DEL-73 |
| chr6-41759830-DEL-825 | chr6-41759831-DEL-825 |
| chr6-41762771-DEL-65 | chr6-41762751-DEL-78 |

| | |
|---|---|
| chr6-41834758-DEL-59 | chr6-41834759-DEL-59 |
| chr6-41842837-DEL-1016 | chr6-41842838-DEL-1016 |
| chr6-41869327-DEL-198 | chr6-41869328-DEL-198 |
| chr6-41875373-DEL-537 | chr6-41875374-DEL-537 |
| chr6-41889917-DEL-260 | chr6-41889920-DEL-258 |
| chr6-41893614-DEL-486 | chr6-41893615-DEL-486 |
| chr6-41902575-DEL-633 | chr6-41902575-DEL-633 |
| chr6-41903511-DEL-2760 | chr6-41903512-DEL-2760 |
| chr6-41922350-DEL-400 | chr6-41922349-DEL-400 |
| chr6-41929145-DEL-56 | chr6-41929146-DEL-56 |
| chr6-41939965-DEL-94 | chr6-41939967-DEL-78 |
| chr6-41942151-DEL-654 | chr6-41942152-DEL-654 |
| chr6-41953850-DEL-1257 | chr6-41953851-DEL-1257 |
| chr6-41986471-DEL-6431 | chr6-41986464-DEL-6436 |
| chr6-42099691-DEL-7700 | chr6-42099692-DEL-7700 |
| chr6-42127972-DEL-117 | chr6-42127973-DEL-117 |
| chr6-42132977-DEL-6793 | chr6-42132982-DEL-6795 |
| chr6-42151381-DEL-3550 | chr6-42151382-DEL-3550 |
| chr6-42161412-DEL-76 | chr6-42161412-DEL-95 |
| chr6-42180474-DEL-657 | chr6-42180475-DEL-657 |
| chr6-42199858-DEL-64 | chr6-42199859-DEL-64 |
| chr6-42219287-DEL-536 | chr6-42219288-DEL-536 |
| chr6-42278085-DEL-206 | chr6-42278086-DEL-206 |
| chr6-42290544-DEL-61 | chr6-42290547-DEL-59 |
| chr6-42308017-DEL-65 | chr6-42308036-DEL-65 |
| chr6-42315131-DEL-387 | chr6-42315132-DEL-387 |
| chr6-42323470-DEL-1230 | chr6-42323472-DEL-1228 |
| chr6-42329773-DEL-55 | chr6-42329776-DEL-53 |
| chr6-42369612-DEL-224 | chr6-42369573-DEL-264 |
| chr6-42378629-DEL-81 | chr6-42378633-DEL-77 |
| chr6-42391437-DEL-184 | chr6-42391438-DEL-184 |
| chr6-42393892-DEL-2953 | chr6-42393893-DEL-2953 |
| chr6-42469474-DEL-6442 | chr6-42469475-DEL-6442 |
| chr6-42490457-DEL-639 | chr6-42490458-DEL-639 |
| chr6-42492162-DEL-1161 | chr6-42492163-DEL-1161 |
| chr6-42558313-DEL-1084 | chr6-42558314-DEL-1084 |
| chr6-42581421-DEL-71 | chr6-42581438-DEL-51 |
| chr6-42583994-DEL-108 | chr6-42583995-DEL-108 |
| chr6-42602748-DEL-195 | chr6-42602722-DEL-194 |
| chr6-42615065-DEL-51 | chr6-42615066-DEL-51 |
| chr6-42641194-DEL-433 | chr6-42641195-DEL-433 |
| chr6-42705607-DEL-343 | chr6-42705608-DEL-343 |
| chr6-42710613-DEL-3830 | chr6-42710614-DEL-3830 |
| chr6-42713130-DEL-228 | chr6-42713131-DEL-228 |
| chr6-42723964-DEL-204 | chr6-42723965-DEL-204 |
| chr6-42729787-DEL-420 | chr6-42729788-DEL-420 |
| chr6-42740506-DEL-6661 | chr6-42740507-DEL-6661 |
| chr6-42751562-DEL-133 | chr6-42751563-DEL-133 |
| chr6-42760261-DEL-232 | chr6-42760260-DEL-233 |
| chr6-42778909-DEL-58 | chr6-42778909-DEL-51 |
| chr6-42804737-DEL-1457 | chr6-42804735-DEL-1458 |
| chr6-42821197-DEL-498 | chr6-42821198-DEL-498 |
| chr6-42830567-DEL-6682 | chr6-42830568-DEL-6682 |
| chr6-42856287-DEL-576 | chr6-42856288-DEL-574 |
| chr6-42856843-DEL-130 | chr6-42856846-DEL-128 |
| chr6-42875806-DEL-1483 | chr6-42875807-DEL-1483 |
| chr6-42909500-DEL-55537 | chr6-42909498-DEL-55539 |
| chr6-42910266-DEL-561 | chr6-42910229-DEL-577 |
| chr6-42950444-DEL-3085 | chr6-42950445-DEL-3085 |
| chr6-42962271-DEL-360 | chr6-42962278-DEL-356 |
| chr6-42963022-DEL-2341 | chr6-42963021-DEL-2342 |
| chr6-42974901-DEL-705 | chr6-42974902-DEL-704 |
| chr6-42975760-DEL-58 | chr6-42975769-DEL-58 |
| chr6-42976759-DEL-58 | chr6-42976760-DEL-58 |
| chr6-43030724-DEL-434 | chr6-43030725-DEL-434 |
| chr6-43032839-DEL-930 | chr6-43032839-DEL-930 |
| chr6-43059096-DEL-422 | chr6-43059097-DEL-422 |
| chr6-43065609-DEL-84 | chr6-43065603-DEL-72 |
| chr6-43079901-DEL-4076 | chr6-43079902-DEL-4076 |
| chr6-43085015-DEL-193 | chr6-43085016-DEL-193 |
| chr6-43100657-DEL-298 | chr6-43100658-DEL-298 |
| chr6-43168530-DEL-3070 | chr6-43168531-DEL-3070 |
| chr6-43240399-DEL-191 | chr6-43240400-DEL-191 |
| chr6-43275108-DEL-1895 | chr6-43275109-DEL-1895 |
| chr6-43293585-DEL-124 | chr6-43293585-DEL-124 |
| chr6-43297492-DEL-6363 | chr6-43297495-DEL-6364 |
| chr6-43313174-DEL-292 | chr6-43313175-DEL-292 |
| chr6-43325136-DEL-2225 | chr6-43325137-DEL-2225 |
| chr6-43361238-DEL-57 | chr6-43361236-DEL-56 |
| chr6-43399839-DEL-65 | chr6-43399840-DEL-50 |

| | |
|---|---|
| chr6-43401349-DEL-67 | chr6-43401350-DEL-67 |
| chr6-43419893-DEL-71 | chr6-43419912-DEL-71 |
| chr6-43424360-DEL-297 | chr6-43424361-DEL-297 |
| chr6-43427279-DEL-51 | chr6-43427280-DEL-51 |
| chr6-43468628-DEL-421 | chr6-43468629-DEL-421 |
| chr6-43504095-DEL-1247 | chr6-43504096-DEL-1246 |
| chr6-43529230-DEL-475 | chr6-43529231-DEL-475 |
| chr6-43557462-DEL-260 | chr6-43557463-DEL-260 |
| chr6-43578172-DEL-79 | chr6-43578173-DEL-79 |
| chr6-43595674-DEL-1098 | chr6-43595675-DEL-1098 |
| chr6-43600798-DEL-155 | chr6-43600799-DEL-155 |
| chr6-43602641-DEL-6491 | chr6-43602641-DEL-6491 |
| chr6-43612714-DEL-3976 | chr6-43612715-DEL-3976 |
| chr6-43666978-DEL-188 | chr6-43666979-DEL-188 |
| chr6-43676594-DEL-68 | chr6-43676599-DEL-106 |
| chr6-43714556-DEL-6829 | chr6-43714557-DEL-6829 |
| chr6-43727166-DEL-485 | chr6-43727167-DEL-485 |
| chr6-43736893-DEL-2900 | chr6-43736894-DEL-2900 |
| chr6-43742062-DEL-2363 | chr6-43742062-DEL-2363 |
| chr6-43757480-DEL-221 | chr6-43757481-DEL-221 |
| chr6-43837218-DEL-56 | chr6-43837219-DEL-56 |
| chr6-43925256-DEL-1206 | chr6-43925257-DEL-1206 |
| chr6-43927122-DEL-2612 | chr6-43927123-DEL-2612 |
| chr6-43948849-DEL-117 | chr6-43948854-DEL-125 |
| chr6-43960076-DEL-2680 | chr6-43960077-DEL-2680 |
| chr6-43961468-DEL-354 | chr6-43961469-DEL-354 |
| chr6-43963741-DEL-115 | chr6-43963746-DEL-111 |
| chr6-43965172-DEL-6430 | chr6-43965173-DEL-6430 |
| chr6-43976053-DEL-489 | chr6-43976054-DEL-489 |
| chr6-43982104-DEL-1459 | chr6-43982105-DEL-1459 |
| chr6-43986596-DEL-82 | chr6-43986579-DEL-85 |
| chr6-44011638-DEL-750 | chr6-44011639-DEL-750 |
| chr6-44017490-DEL-97 | chr6-44017491-DEL-97 |
| chr6-44021158-DEL-566 | chr6-44021159-DEL-566 |
| chr6-44022801-DEL-1111 | chr6-44022802-DEL-1111 |
| chr6-44027284-DEL-130 | chr6-44027285-DEL-130 |
| chr6-44083597-DEL-6475 | chr6-44083598-DEL-6475 |
| chr6-44090902-DEL-344 | chr6-44090909-DEL-345 |
| chr6-44116984-DEL-2353 | chr6-44116985-DEL-2353 |
| chr6-44128487-DEL-2726 | chr6-44128488-DEL-2726 |
| chr6-44137895-DEL-290 | chr6-44137896-DEL-290 |
| chr6-44145628-DEL-577 | chr6-44145629-DEL-577 |
| chr6-44145875-DEL-936 | chr6-44145877-DEL-933 |
| chr6-44189811-DEL-717 | chr6-44189812-DEL-711 |
| chr6-44256838-DEL-67 | chr6-44256846-DEL-73 |
| chr6-44261318-DEL-615 | chr6-44261319-DEL-615 |
| chr6-44274905-DEL-64 | chr6-44274906-DEL-54 |
| chr6-44283032-DEL-3151 | chr6-44283033-DEL-3151 |
| chr6-44296661-DEL-96 | chr6-44296662-DEL-96 |
| chr6-44343807-DEL-144 | chr6-44343808-DEL-144 |
| chr6-44390826-DEL-6657 | chr6-44390827-DEL-6657 |
| chr6-44429503-DEL-1580 | chr6-44429504-DEL-1580 |
| chr6-44471368-DEL-230 | chr6-44471369-DEL-230 |
| chr6-44520444-DEL-675 | chr6-44520445-DEL-675 |
| chr6-44560169-DEL-2435 | chr6-44560170-DEL-2435 |
| chr6-44577325-DEL-481 | chr6-44577323-DEL-482 |
| chr6-44620164-DEL-1222 | chr6-44620165-DEL-1219 |
| chr6-44631950-DEL-401 | chr6-44631951-DEL-401 |
| chr6-44634758-DEL-1364 | chr6-44634759-DEL-1364 |
| chr6-44643259-DEL-517 | chr6-44643260-DEL-517 |
| chr6-44662540-DEL-197 | chr6-44662541-DEL-196 |
| chr6-44664085-DEL-988 | chr6-44664082-DEL-990 |
| chr6-44668220-DEL-7081 | chr6-44668221-DEL-7081 |
| chr6-44679593-DEL-497 | chr6-44679592-DEL-498 |
| chr6-44683003-DEL-999 | chr6-44683004-DEL-999 |
| chr6-44701999-DEL-6433 | chr6-44702000-DEL-6433 |
| chr6-44741757-DEL-56 | chr6-44741760-DEL-54 |
| chr6-44747805-DEL-99 | chr6-44747806-DEL-99 |
| chr6-44752807-DEL-1757 | chr6-44752808-DEL-1757 |
| chr6-44755831-DEL-634 | chr6-44755831-DEL-635 |
| chr6-44757274-DEL-121 | chr6-44757275-DEL-120 |
| chr6-44762056-DEL-7005 | chr6-44762056-DEL-7005 |
| chr6-44769861-DEL-373 | chr6-44769862-DEL-373 |
| chr6-44783104-DEL-130 | chr6-44783105-DEL-130 |
| chr6-44789739-DEL-55 | chr6-44789740-DEL-55 |
| chr6-44839918-DEL-2180 | chr6-44839919-DEL-2180 |
| chr6-44843747-DEL-534 | chr6-44843748-DEL-532 |
| chr6-44865133-DEL-214 | chr6-44865134-DEL-214 |
| chr6-44895958-DEL-55 | chr6-44895959-DEL-55 |
| chr6-44897127-DEL-61 | chr6-44897130-DEL-54 |

| | |
|---|---|
| chr6-44901541-DEL-69 | chr6-44901540-DEL-67 |
| chr6-44907652-DEL-58 | chr6-44907613-DEL-58 |
| chr6-44916428-DEL-58 | chr6-44916438-DEL-59 |
| chr6-44921130-DEL-237 | chr6-44921131-DEL-237 |
| chr6-44924474-DEL-52 | chr6-44924462-DEL-68 |
| chr6-44939739-DEL-2731 | chr6-44939740-DEL-2731 |
| chr6-44945858-DEL-381 | chr6-44945859-DEL-381 |
| chr6-44951184-DEL-400 | chr6-44951185-DEL-400 |
| chr6-44962342-DEL-92 | chr6-44962343-DEL-92 |
| chr6-44969578-DEL-82 | chr6-44969579-DEL-82 |
| chr6-44970334-DEL-626 | chr6-44970335-DEL-626 |
| chr6-44979144-DEL-200 | chr6-44979147-DEL-193 |
| chr6-44985467-DEL-147 | chr6-44985468-DEL-147 |
| chr6-45025327-DEL-68 | chr6-45025328-DEL-68 |
| chr6-45028658-DEL-128 | chr6-45028648-DEL-152 |
| chr6-45080039-DEL-98 | chr6-45080043-DEL-97 |
| chr6-45081573-DEL-91 | chr6-45081574-DEL-91 |
| chr6-45096132-DEL-606 | chr6-45096133-DEL-606 |
| chr6-45133615-DEL-829 | chr6-45133616-DEL-828 |
| chr6-45145134-DEL-6591 | chr6-45145135-DEL-6591 |
| chr6-45210365-DEL-130 | chr6-45210366-DEL-130 |
| chr6-45213396-DEL-170 | chr6-45213397-DEL-170 |
| chr6-45227181-DEL-3160 | chr6-45227182-DEL-3160 |
| chr6-45231606-DEL-56 | chr6-45231607-DEL-56 |
| chr6-45251490-DEL-183 | chr6-45251491-DEL-183 |
| chr6-45318471-DEL-80 | chr6-45318474-DEL-81 |
| chr6-45392129-DEL-215 | chr6-45392130-DEL-215 |
| chr6-45395998-DEL-6419 | chr6-45395999-DEL-6419 |
| chr6-45421729-DEL-210 | chr6-45421730-DEL-210 |
| chr6-45424068-DEL-528 | chr6-45424069-DEL-528 |
| chr6-45456600-DEL-6700 | chr6-45456601-DEL-6700 |
| chr6-45502255-DEL-77 | chr6-45502253-DEL-86 |
| chr6-45511586-DEL-374 | chr6-45511587-DEL-374 |
| chr6-45534794-DEL-1335 | chr6-45534795-DEL-1335 |
| chr6-45540328-DEL-315 | chr6-45540329-DEL-315 |
| chr6-45550535-DEL-67 | chr6-45550536-DEL-65 |
| chr6-45579316-DEL-87 | chr6-45579327-DEL-76 |
| chr6-45612121-DEL-6408 | chr6-45612122-DEL-6408 |
| chr6-45645135-DEL-61 | chr6-45645136-DEL-54 |
| chr6-45658618-DEL-2295 | chr6-45658619-DEL-2295 |
| chr6-45676855-DEL-113 | chr6-45676848-DEL-110 |
| chr6-45681130-DEL-1079 | chr6-45681131-DEL-1079 |
| chr6-45707806-DEL-196 | chr6-45707807-DEL-196 |
| chr6-45740468-DEL-56 | chr6-45740477-DEL-54 |
| chr6-45749360-DEL-62 | chr6-45749361-DEL-58 |
| chr6-45752431-DEL-1759 | chr6-45752422-DEL-1760 |
| chr6-45758072-DEL-1893 | chr6-45758073-DEL-1893 |
| chr6-45794694-DEL-3732 | chr6-45794695-DEL-3732 |
| chr6-45810461-DEL-6295 | chr6-45810462-DEL-6295 |
| chr6-45834369-DEL-953 | chr6-45834369-DEL-953 |
| chr6-45840256-DEL-56 | chr6-45840257-DEL-51 |
| chr6-45876687-DEL-54 | chr6-45876688-DEL-54 |
| chr6-45878804-DEL-75 | chr6-45878805-DEL-75 |
| chr6-45907146-DEL-623 | chr6-45907141-DEL-624 |
| chr6-45927297-DEL-69 | chr6-45927296-DEL-70 |
| chr6-45927973-DEL-60 | chr6-45927972-DEL-62 |
| chr6-45930355-DEL-1043 | chr6-45930356-DEL-1043 |
| chr6-45936224-DEL-235 | chr6-45936225-DEL-235 |
| chr6-45964266-DEL-66 | chr6-45964267-DEL-66 |
| chr6-45971958-DEL-192 | chr6-45971959-DEL-192 |
| chr6-46026918-DEL-1348 | chr6-46026919-DEL-1348 |
| chr6-46121980-DEL-84 | chr6-46121981-DEL-84 |
| chr6-46128943-DEL-1014 | chr6-46128944-DEL-1014 |
| chr6-46130428-DEL-6260 | chr6-46130429-DEL-6260 |
| chr6-46159004-DEL-999 | chr6-46159005-DEL-999 |
| chr6-46163156-DEL-68 | chr6-46163157-DEL-68 |
| chr6-46197684-DEL-6031 | chr6-46197685-DEL-6031 |
| chr6-46220713-DEL-85 | chr6-46220716-DEL-82 |
| chr6-46229656-DEL-103 | chr6-46229668-DEL-91 |
| chr6-46231087-DEL-60 | chr6-46231088-DEL-60 |
| chr6-46246934-DEL-2112 | chr6-46246935-DEL-2112 |
| chr6-46255911-DEL-148 | chr6-46255912-DEL-148 |
| chr6-46264311-DEL-6685 | chr6-46264312-DEL-6685 |
| chr6-46285239-DEL-72 | chr6-46285240-DEL-72 |
| chr6-46300266-DEL-91 | chr6-46300267-DEL-90 |
| chr6-46303449-DEL-146 | chr6-46303449-DEL-147 |
| chr6-46326032-DEL-661 | chr6-46326033-DEL-661 |
| chr6-46329520-DEL-185 | chr6-46329521-DEL-185 |
| chr6-46377394-DEL-1545 | chr6-46377395-DEL-1545 |
| chr6-46450932-DEL-73 | chr6-46450933-DEL-69 |

| | |
|---|---|
| chr6-46457423-DEL-847 | chr6-46457424-DEL-847 |
| chr6-46472276-DEL-933 | chr6-46472277-DEL-933 |
| chr6-46503628-DEL-8156 | chr6-46503629-DEL-8156 |
| chr6-46503844-DEL-3975 | chr6-46503845-DEL-3975 |
| chr6-46549410-DEL-149 | chr6-46549410-DEL-149 |
| chr6-46608228-DEL-184 | chr6-46608229-DEL-184 |
| chr6-46619782-DEL-1451 | chr6-46619783-DEL-1451 |
| chr6-46646779-DEL-599 | chr6-46646780-DEL-599 |
| chr6-46649597-DEL-203 | chr6-46649598-DEL-203 |
| chr6-46655451-DEL-7141 | chr6-46655452-DEL-7141 |
| chr6-46711649-DEL-66 | chr6-46711650-DEL-64 |
| chr6-46729643-DEL-205 | chr6-46729644-DEL-205 |
| chr6-46730719-DEL-69 | chr6-46730719-DEL-67 |
| chr6-46743038-DEL-144 | chr6-46743039-DEL-144 |
| chr6-46745645-DEL-2002 | chr6-46745646-DEL-2002 |
| chr6-46761883-DEL-130 | chr6-46761886-DEL-132 |
| chr6-46772034-DEL-98 | chr6-46772031-DEL-101 |
| chr6-46789842-DEL-2115 | chr6-46789830-DEL-2116 |
| chr6-46795975-DEL-257 | chr6-46795976-DEL-257 |
| chr6-46808196-DEL-3118 | chr6-46808196-DEL-3103 |
| chr6-46827791-DEL-444 | chr6-46827792-DEL-439 |
| chr6-46857651-DEL-1846 | chr6-46857652-DEL-1846 |
| chr6-46867487-DEL-1580 | chr6-46867488-DEL-1580 |
| chr6-46871047-DEL-292 | chr6-46871048-DEL-292 |
| chr6-46880961-DEL-737 | chr6-46880962-DEL-737 |
| chr6-46907381-DEL-4119 | chr6-46907382-DEL-4119 |
| chr6-46908185-DEL-82 | chr6-46908186-DEL-82 |
| chr6-46976177-DEL-661 | chr6-46976147-DEL-693 |
| chr6-47034378-DEL-447 | chr6-47034379-DEL-468 |
| chr6-47072929-DEL-532 | chr6-47072930-DEL-532 |
| chr6-47078936-DEL-64 | chr6-47078937-DEL-56 |
| chr6-47082959-DEL-223 | chr6-47082960-DEL-223 |
| chr6-47098638-DEL-405 | chr6-47098639-DEL-405 |
| chr6-47102896-DEL-184 | chr6-47102897-DEL-184 |
| chr6-47105181-DEL-585 | chr6-47105172-DEL-586 |
| chr6-47156090-DEL-531 | chr6-47156091-DEL-531 |
| chr6-47178576-DEL-122 | chr6-47178577-DEL-122 |
| chr6-47203726-DEL-84 | chr6-47203727-DEL-84 |
| chr6-47253171-DEL-92 | chr6-47253172-DEL-93 |
| chr6-47270971-DEL-507 | chr6-47270972-DEL-507 |
| chr6-47288230-DEL-472 | chr6-47288238-DEL-474 |
| chr6-47300025-DEL-53 | chr6-47300023-DEL-51 |
| chr6-47304937-DEL-69 | chr6-47304929-DEL-83 |
| chr6-47307735-DEL-1550 | chr6-47307736-DEL-1550 |
| chr6-47311920-DEL-656 | chr6-47311920-DEL-641 |
| chr6-47346402-DEL-2803 | chr6-47346403-DEL-2803 |
| chr6-47362955-DEL-192 | chr6-47362956-DEL-192 |
| chr6-47369619-DEL-193 | chr6-47369620-DEL-193 |
| chr6-47387403-DEL-195 | chr6-47387404-DEL-195 |
| chr6-47408690-DEL-1391 | chr6-47408691-DEL-1391 |
| chr6-47427628-DEL-527 | chr6-47427628-DEL-527 |
| chr6-47452240-DEL-2739 | chr6-47452241-DEL-2739 |
| chr6-47539900-DEL-64 | chr6-47539905-DEL-60 |
| chr6-47561084-DEL-200 | chr6-47561085-DEL-200 |
| chr6-47564201-DEL-222 | chr6-47564202-DEL-222 |
| chr6-47629026-DEL-60 | chr6-47629025-DEL-60 |
| chr6-47750189-DEL-1894 | chr6-47750190-DEL-1894 |
| chr6-47756283-DEL-207 | chr6-47756284-DEL-207 |
| chr6-47775034-DEL-59 | chr6-47775035-DEL-59 |
| chr6-47804822-DEL-78 | chr6-47804859-DEL-54 |
| chr6-47825079-DEL-7024 | chr6-47825080-DEL-7024 |
| chr6-47834635-DEL-208 | chr6-47834636-DEL-208 |
| chr6-47837325-DEL-449 | chr6-47837326-DEL-449 |
| chr6-47872186-DEL-340 | chr6-47872187-DEL-340 |
| chr6-47889126-DEL-209 | chr6-47889127-DEL-209 |
| chr6-47890544-DEL-205 | chr6-47890545-DEL-205 |
| chr6-47910128-DEL-445 | chr6-47910129-DEL-445 |
| chr6-47921858-DEL-127 | chr6-47921857-DEL-129 |
| chr6-47938094-DEL-856 | chr6-47938095-DEL-856 |
| chr6-47945718-DEL-199 | chr6-47945713-DEL-200 |
| chr6-47951263-DEL-120 | chr6-47951263-DEL-120 |
| chr6-47976789-DEL-7195 | chr6-47976790-DEL-7195 |
| chr6-47990735-DEL-102 | chr6-47990736-DEL-102 |
| chr6-48018309-DEL-152 | chr6-48018310-DEL-152 |
| chr6-48077512-DEL-301 | chr6-48077514-DEL-299 |
| chr6-48086163-DEL-881 | chr6-48086164-DEL-881 |
| chr6-48097842-DEL-79 | chr6-48097838-DEL-84 |
| chr6-48159422-DEL-67 | chr6-48159433-DEL-59 |
| chr6-48160986-DEL-848 | chr6-48160987-DEL-848 |
| chr6-48168264-DEL-69 | chr6-48168265-DEL-69 |

| | |
|---|---|
| chr6-48177389-DEL-120 | chr6-48177390-DEL-120 |
| chr6-48188839-DEL-58 | chr6-48188840-DEL-80 |
| chr6-48205689-DEL-6495 | chr6-48205690-DEL-6495 |
| chr6-48266147-DEL-77 | chr6-48266147-DEL-77 |
| chr6-48297737-DEL-7131 | chr6-48297738-DEL-7131 |
| chr6-48349988-DEL-524 | chr6-48349989-DEL-524 |
| chr6-48381597-DEL-277 | chr6-48381595-DEL-273 |
| chr6-48442983-DEL-55 | chr6-48442984-DEL-55 |
| chr6-48478197-DEL-613 | chr6-48478165-DEL-614 |
| chr6-48482379-DEL-95 | chr6-48482380-DEL-95 |
| chr6-48483716-DEL-70 | chr6-48483713-DEL-70 |
| chr6-48489073-DEL-2904 | chr6-48489074-DEL-2904 |
| chr6-48497100-DEL-1679 | chr6-48497101-DEL-1679 |
| chr6-48499312-DEL-2753 | chr6-48499313-DEL-2753 |
| chr6-48500377-DEL-478 | chr6-48500378-DEL-478 |
| chr6-48524705-DEL-63 | chr6-48524706-DEL-63 |
| chr6-48526229-DEL-52 | chr6-48526229-DEL-52 |
| chr6-48530683-DEL-269 | chr6-48530684-DEL-269 |
| chr6-48583952-DEL-189 | chr6-48583953-DEL-189 |
| chr6-48591100-DEL-2423 | chr6-48591101-DEL-2423 |
| chr6-48599336-DEL-92 | chr6-48599337-DEL-92 |
| chr6-48628973-DEL-181 | chr6-48628974-DEL-181 |
| chr6-48637441-DEL-365 | chr6-48637433-DEL-366 |
| chr6-48641491-DEL-491 | chr6-48641492-DEL-491 |
| chr6-48643601-DEL-60 | chr6-48643602-DEL-60 |
| chr6-48645244-DEL-268 | chr6-48645245-DEL-268 |
| chr6-48660610-DEL-201 | chr6-48660611-DEL-201 |
| chr6-48685937-DEL-68 | chr6-48685936-DEL-82 |
| chr6-48686007-DEL-1295 | chr6-48686052-DEL-1251 |
| chr6-48727821-DEL-627 | chr6-48727822-DEL-627 |
| chr6-48735048-DEL-139 | chr6-48735049-DEL-135 |
| chr6-48751661-DEL-5753 | chr6-48751662-DEL-5753 |
| chr6-48771017-DEL-92 | chr6-48771018-DEL-92 |
| chr6-48776237-DEL-1390 | chr6-48776238-DEL-1390 |
| chr6-48796991-DEL-56 | chr6-48797000-DEL-59 |
| chr6-48804458-DEL-111 | chr6-48804459-DEL-111 |
| chr6-48849151-DEL-55 | chr6-48849152-DEL-55 |
| chr6-48851910-DEL-157 | chr6-48851911-DEL-157 |
| chr6-48859778-DEL-1124 | chr6-48859779-DEL-1124 |
| chr6-48897457-DEL-66 | chr6-48897502-DEL-58 |
| chr6-48929172-DEL-58 | chr6-48929177-DEL-59 |
| chr6-48971811-DEL-52 | chr6-48971812-DEL-52 |
| chr6-48972522-DEL-957 | chr6-48972523-DEL-957 |
| chr6-48977371-DEL-213 | chr6-48977372-DEL-213 |
| chr6-48990827-DEL-3150 | chr6-48990828-DEL-3150 |
| chr6-49042689-DEL-196 | chr6-49042690-DEL-195 |
| chr6-49117241-DEL-133 | chr6-49117242-DEL-133 |
| chr6-49135183-DEL-85 | chr6-49135184-DEL-85 |
| chr6-49164784-DEL-244 | chr6-49164785-DEL-243 |
| chr6-49166041-DEL-180 | chr6-49166042-DEL-180 |
| chr6-49168598-DEL-1225 | chr6-49168598-DEL-1225 |
| chr6-49233215-DEL-205 | chr6-49233216-DEL-205 |
| chr6-49242936-DEL-66 | chr6-49242926-DEL-50 |
| chr6-49264999-DEL-232 | chr6-49264999-DEL-233 |
| chr6-49277397-DEL-219 | chr6-49277398-DEL-219 |
| chr6-49285496-DEL-172 | chr6-49285497-DEL-172 |
| chr6-49293238-DEL-401 | chr6-49293239-DEL-401 |
| chr6-49295573-DEL-177 | chr6-49295574-DEL-177 |
| chr6-49326238-DEL-81 | chr6-49326239-DEL-81 |
| chr6-49327452-DEL-204 | chr6-49327453-DEL-204 |
| chr6-49328033-DEL-51 | chr6-49328017-DEL-52 |
| chr6-49328684-DEL-54 | chr6-49328685-DEL-54 |
| chr6-49334884-DEL-199 | chr6-49334885-DEL-199 |
| chr6-49342406-DEL-139 | chr6-49342407-DEL-139 |
| chr6-49346769-DEL-1603 | chr6-49346770-DEL-1603 |
| chr6-49354332-DEL-100 | chr6-49354333-DEL-100 |
| chr6-49360612-DEL-70 | chr6-49360613-DEL-70 |
| chr6-49361023-DEL-232 | chr6-49361010-DEL-258 |
| chr6-49472866-DEL-82 | chr6-49472867-DEL-82 |
| chr6-49484890-DEL-179 | chr6-49484893-DEL-180 |
| chr6-49492594-DEL-1500 | chr6-49492595-DEL-1500 |
| chr6-49496598-DEL-175 | chr6-49496599-DEL-174 |
| chr6-49504670-DEL-6331 | chr6-49504671-DEL-6331 |
| chr6-49513511-DEL-703 | chr6-49513512-DEL-703 |
| chr6-49521853-DEL-68 | chr6-49521854-DEL-68 |
| chr6-49545374-DEL-337 | chr6-49545375-DEL-337 |
| chr6-49573678-DEL-139 | chr6-49573673-DEL-144 |
| chr6-49584289-DEL-241 | chr6-49584290-DEL-241 |
| chr6-49584657-DEL-703 | chr6-49584658-DEL-703 |
| chr6-49593228-DEL-7153 | chr6-49593229-DEL-7153 |

| | |
|---|---|
| chr6-49613637-DEL-91 | chr6-49613638-DEL-92 |
| chr6-49664619-DEL-54 | chr6-49664620-DEL-54 |
| chr6-49679075-DEL-1386 | chr6-49679076-DEL-1385 |
| chr6-49704407-DEL-1063 | chr6-49704408-DEL-1063 |
| chr6-49717696-DEL-357 | chr6-49717697-DEL-357 |
| chr6-49723895-DEL-144 | chr6-49723896-DEL-144 |
| chr6-49732389-DEL-409 | chr6-49732390-DEL-409 |
| chr6-49744536-DEL-203 | chr6-49744537-DEL-203 |
| chr6-49771014-DEL-214 | chr6-49771002-DEL-220 |
| chr6-49820444-DEL-176 | chr6-49820446-DEL-175 |
| chr6-49831357-DEL-202 | chr6-49831355-DEL-203 |
| chr6-49839869-DEL-72 | chr6-49839882-DEL-52 |
| chr6-49847484-DEL-132 | chr6-49847451-DEL-182 |
| chr6-49856159-DEL-89 | chr6-49856159-DEL-89 |
| chr6-49862130-DEL-7703 | chr6-49862131-DEL-7703 |
| chr6-49863678-DEL-1595 | chr6-49863678-DEL-1595 |
| chr6-49892361-DEL-80 | chr6-49892354-DEL-88 |
| chr6-49916492-DEL-1161 | chr6-49916493-DEL-1161 |
| chr6-49928906-DEL-293 | chr6-49928859-DEL-325 |
| chr6-49934986-DEL-112 | chr6-49934987-DEL-112 |
| chr6-49946691-DEL-580 | chr6-49946695-DEL-574 |
| chr6-49948138-DEL-357 | chr6-49948139-DEL-357 |
| chr6-49953921-DEL-97 | chr6-49953922-DEL-97 |
| chr6-49971277-DEL-1888 | chr6-49971278-DEL-1888 |
| chr6-49994327-DEL-85 | chr6-49994310-DEL-78 |
| chr6-49995270-DEL-214 | chr6-49995271-DEL-214 |
| chr6-49995593-DEL-578 | chr6-49995594-DEL-578 |
| chr6-50005554-DEL-74 | chr6-50005555-DEL-74 |
| chr6-50013219-DEL-195 | chr6-50013220-DEL-195 |
| chr6-50039109-DEL-67 | chr6-50039110-DEL-67 |
| chr6-50042824-DEL-68 | chr6-50042825-DEL-68 |
| chr6-50092637-DEL-75 | chr6-50092638-DEL-75 |
| chr6-50116163-DEL-1140 | chr6-50116164-DEL-1140 |
| chr6-50149868-DEL-803 | chr6-50149869-DEL-803 |
| chr6-50201745-DEL-61 | chr6-50201746-DEL-61 |
| chr6-50203356-DEL-123 | chr6-50203357-DEL-123 |
| chr6-50204877-DEL-72 | chr6-50204873-DEL-76 |
| chr6-50248145-DEL-744 | chr6-50248146-DEL-744 |
| chr6-50259244-DEL-234 | chr6-50259248-DEL-188 |
| chr6-50268508-DEL-92 | chr6-50268509-DEL-92 |
| chr6-50284451-DEL-56 | chr6-50284452-DEL-56 |
| chr6-50290035-DEL-152 | chr6-50290028-DEL-153 |
| chr6-50336114-DEL-112 | chr6-50336115-DEL-112 |
| chr6-50337445-DEL-1710 | chr6-50337446-DEL-1710 |
| chr6-50381234-DEL-70 | chr6-50381235-DEL-70 |
| chr6-50392823-DEL-76 | chr6-50392824-DEL-76 |
| chr6-50399163-DEL-429 | chr6-50399164-DEL-429 |
| chr6-50435462-DEL-98 | chr6-50435432-DEL-84 |
| chr6-50439792-DEL-64 | chr6-50439793-DEL-64 |
| chr6-50460963-DEL-190 | chr6-50460964-DEL-190 |
| chr6-50464885-DEL-68 | chr6-50464886-DEL-66 |
| chr6-50503816-DEL-122 | chr6-50503817-DEL-122 |
| chr6-50508975-DEL-2437 | chr6-50508976-DEL-2437 |
| chr6-50518893-DEL-123 | chr6-50518894-DEL-119 |
| chr6-50527418-DEL-70 | chr6-50527419-DEL-70 |
| chr6-50615929-DEL-102 | chr6-50615930-DEL-102 |
| chr6-50625808-DEL-575 | chr6-50625809-DEL-575 |
| chr6-50626799-DEL-198 | chr6-50626798-DEL-199 |
| chr6-50668482-DEL-52 | chr6-50668514-DEL-62 |
| chr6-50706895-DEL-51 | chr6-50706894-DEL-52 |
| chr6-50708998-DEL-92 | chr6-50708999-DEL-92 |
| chr6-50732866-DEL-72 | chr6-50732856-DEL-76 |
| chr6-50747169-DEL-4008 | chr6-50747170-DEL-4008 |
| chr6-50769006-DEL-72 | chr6-50769007-DEL-72 |
| chr6-50803544-DEL-80 | chr6-50803545-DEL-80 |
| chr6-50806642-DEL-455 | chr6-50806643-DEL-455 |
| chr6-50815677-DEL-118 | chr6-50815678-DEL-118 |
| chr6-50822483-DEL-240 | chr6-50822484-DEL-240 |
| chr6-50897036-DEL-1674 | chr6-50897059-DEL-1637 |
| chr6-50899978-DEL-63 | chr6-50899978-DEL-63 |
| chr6-50908295-DEL-191 | chr6-50908296-DEL-191 |
| chr6-50975360-DEL-52 | chr6-50975360-DEL-52 |
| chr6-51001040-DEL-53 | chr6-51001041-DEL-53 |
| chr6-51011515-DEL-64 | chr6-51011516-DEL-64 |
| chr6-51051241-DEL-235 | chr6-51051242-DEL-238 |
| chr6-51062401-DEL-96 | chr6-51062401-DEL-96 |
| chr6-51063950-DEL-883 | chr6-51063951-DEL-883 |
| chr6-51086919-DEL-392 | chr6-51086920-DEL-392 |
| chr6-51094418-DEL-981 | chr6-51094419-DEL-981 |
| chr6-51116115-DEL-68 | chr6-51116108-DEL-76 |

| | |
|---|---|
| chr6-51140826-DEL-53 | chr6-51140827-DEL-53 |
| chr6-51198142-DEL-72 | chr6-51198140-DEL-63 |
| chr6-51204068-DEL-306 | chr6-51204069-DEL-305 |
| chr6-51218025-DEL-87 | chr6-51218026-DEL-87 |
| chr6-51234238-DEL-192 | chr6-51234237-DEL-192 |
| chr6-51263963-DEL-308 | chr6-51263964-DEL-308 |
| chr6-51276467-DEL-88 | chr6-51276466-DEL-84 |
| chr6-51287535-DEL-79 | chr6-51287533-DEL-80 |
| chr6-51346242-DEL-80 | chr6-51346243-DEL-80 |
| chr6-51358048-DEL-52 | chr6-51358049-DEL-51 |
| chr6-51364604-DEL-77 | chr6-51364605-DEL-77 |
| chr6-51373352-DEL-1833 | chr6-51373353-DEL-1833 |
| chr6-51386604-DEL-72 | chr6-51386589-DEL-60 |
| chr6-51387994-DEL-1067 | chr6-51387995-DEL-1067 |
| chr6-51406683-DEL-175 | chr6-51406673-DEL-176 |
| chr6-51407728-DEL-87 | chr6-51407728-DEL-87 |
| chr6-51425900-DEL-250 | chr6-51425901-DEL-250 |
| chr6-51464865-DEL-107 | chr6-51464863-DEL-107 |
| chr6-51471235-DEL-174 | chr6-51471235-DEL-175 |
| chr6-51481038-DEL-196 | chr6-51481039-DEL-197 |
| chr6-51488260-DEL-175 | chr6-51488261-DEL-175 |
| chr6-51496283-DEL-213 | chr6-51496284-DEL-213 |
| chr6-51526769-DEL-212 | chr6-51526770-DEL-212 |
| chr6-51578060-DEL-245 | chr6-51578061-DEL-245 |
| chr6-51600654-DEL-2667 | chr6-51600655-DEL-2667 |
| chr6-51614098-DEL-118 | chr6-51614101-DEL-116 |
| chr6-51622316-DEL-287 | chr6-51622317-DEL-287 |
| chr6-51623587-DEL-368 | chr6-51623591-DEL-365 |
| chr6-51629573-DEL-275 | chr6-51629574-DEL-282 |
| chr6-51654939-DEL-338 | chr6-51654940-DEL-338 |
| chr6-51669885-DEL-183 | chr6-51669886-DEL-183 |
| chr6-51678946-DEL-58 | chr6-51678960-DEL-87 |
| chr6-51681780-DEL-58 | chr6-51681781-DEL-58 |
| chr6-51683481-DEL-70 | chr6-51683490-DEL-62 |
| chr6-51725803-DEL-139 | chr6-51725804-DEL-139 |
| chr6-51740201-DEL-80 | chr6-51740202-DEL-80 |
| chr6-51742054-DEL-116 | chr6-51742055-DEL-116 |
| chr6-51750134-DEL-931 | chr6-51750135-DEL-931 |
| chr6-51771959-DEL-54 | chr6-51771960-DEL-54 |
| chr6-51788920-DEL-127 | chr6-51788911-DEL-128 |
| chr6-51801184-DEL-91 | chr6-51801184-DEL-89 |
| chr6-51922659-DEL-101 | chr6-51922666-DEL-101 |
| chr6-51945058-DEL-963 | chr6-51945054-DEL-964 |
| chr6-51956483-DEL-294 | chr6-51956486-DEL-292 |
| chr6-51963214-DEL-351 | chr6-51963215-DEL-351 |
| chr6-52005369-DEL-187 | chr6-52005353-DEL-153 |
| chr6-52016511-DEL-1586 | chr6-52016512-DEL-1586 |
| chr6-52046998-DEL-476 | chr6-52047027-DEL-448 |
| chr6-52060255-DEL-79 | chr6-52060256-DEL-79 |
| chr6-52089531-DEL-106 | chr6-52089556-DEL-63 |
| chr6-52094993-DEL-138 | chr6-52094994-DEL-138 |
| chr6-52099696-DEL-215 | chr6-52099697-DEL-215 |
| chr6-52108483-DEL-84 | chr6-52108460-DEL-108 |
| chr6-52110746-DEL-53 | chr6-52110747-DEL-53 |
| chr6-52121311-DEL-60 | chr6-52121292-DEL-75 |
| chr6-52122901-DEL-91 | chr6-52122902-DEL-91 |
| chr6-52264366-DEL-874 | chr6-52264367-DEL-874 |
| chr6-52281702-DEL-108 | chr6-52281703-DEL-90 |
| chr6-52307787-DEL-215 | chr6-52307788-DEL-215 |
| chr6-52310643-DEL-2127 | chr6-52310642-DEL-2127 |
| chr6-52348271-DEL-627 | chr6-52348272-DEL-627 |
| chr6-52353653-DEL-945 | chr6-52353654-DEL-945 |
| chr6-52358434-DEL-218 | chr6-52358434-DEL-219 |
| chr6-52363396-DEL-165 | chr6-52363400-DEL-147 |
| chr6-52368427-DEL-66 | chr6-52368451-DEL-92 |
| chr6-52422556-DEL-205 | chr6-52422557-DEL-205 |
| chr6-52520258-DEL-2517 | chr6-52520259-DEL-2517 |
| chr6-52529214-DEL-76 | chr6-52529212-DEL-80 |
| chr6-52571640-DEL-117 | chr6-52571641-DEL-117 |
| chr6-52588603-DEL-229 | chr6-52588604-DEL-228 |
| chr6-52624042-DEL-522 | chr6-52624043-DEL-522 |
| chr6-52642203-DEL-65 | chr6-52642218-DEL-65 |
| chr6-52655702-DEL-55 | chr6-52655703-DEL-55 |
| chr6-52660765-DEL-59 | chr6-52660766-DEL-63 |
| chr6-52677298-DEL-68 | chr6-52677312-DEL-56 |
| chr6-52712603-DEL-444 | chr6-52712604-DEL-444 |
| chr6-52740167-DEL-101 | chr6-52740171-DEL-90 |
| chr6-52762483-DEL-684 | chr6-52762484-DEL-684 |
| chr6-52777618-DEL-409 | chr6-52777619-DEL-409 |
| chr6-52792710-DEL-184 | chr6-52792711-DEL-184 |

| | |
|---|---|
| chr6-52833628-DEL-74 | chr6-52833630-DEL-73 |
| chr6-52844613-DEL-538 | chr6-52844614-DEL-538 |
| chr6-52852266-DEL-479 | chr6-52852266-DEL-479 |
| chr6-52947967-DEL-93 | chr6-52947948-DEL-94 |
| chr6-52958566-DEL-253 | chr6-52958561-DEL-251 |
| chr6-52961694-DEL-58 | chr6-52961695-DEL-58 |
| chr6-53036439-DEL-63 | chr6-53036451-DEL-50 |
| chr6-53061961-DEL-1698 | chr6-53061953-DEL-1705 |
| chr6-53077481-DEL-6432 | chr6-53077482-DEL-6432 |
| chr6-53083388-DEL-201 | chr6-53083389-DEL-201 |
| chr6-53090884-DEL-134 | chr6-53090885-DEL-134 |
| chr6-53098458-DEL-1489 | chr6-53098459-DEL-1489 |
| chr6-53104080-DEL-2390 | chr6-53104081-DEL-2389 |
| chr6-53139265-DEL-116 | chr6-53139266-DEL-140 |
| chr6-53143524-DEL-313 | chr6-53143525-DEL-313 |
| chr6-53144025-DEL-70 | chr6-53144026-DEL-70 |
| chr6-53144916-DEL-52 | chr6-53144918-DEL-52 |
| chr6-53150361-DEL-157 | chr6-53150362-DEL-157 |
| chr6-53151727-DEL-204 | chr6-53151728-DEL-204 |
| chr6-53173110-DEL-328 | chr6-53173111-DEL-328 |
| chr6-53178045-DEL-65 | chr6-53178046-DEL-65 |
| chr6-53196815-DEL-214 | chr6-53196802-DEL-228 |
| chr6-53223635-DEL-56 | chr6-53223636-DEL-56 |
| chr6-53292038-DEL-67 | chr6-53292053-DEL-52 |
| chr6-53293584-DEL-120 | chr6-53293585-DEL-120 |
| chr6-53300086-DEL-285 | chr6-53300087-DEL-285 |
| chr6-53310802-DEL-69 | chr6-53310803-DEL-69 |
| chr6-53322674-DEL-103 | chr6-53322675-DEL-103 |
| chr6-53364038-DEL-60 | chr6-53364041-DEL-52 |
| chr6-53379831-DEL-6339 | chr6-53379832-DEL-6339 |
| chr6-53410501-DEL-66 | chr6-53410502-DEL-66 |
| chr6-53441196-DEL-1868 | chr6-53441197-DEL-1868 |
| chr6-53456554-DEL-142 | chr6-53456585-DEL-122 |
| chr6-53457877-DEL-6677 | chr6-53457878-DEL-6677 |
| chr6-53469845-DEL-139 | chr6-53469846-DEL-151 |
| chr6-53559708-DEL-52 | chr6-53559711-DEL-50 |
| chr6-53584248-DEL-77 | chr6-53584249-DEL-77 |
| chr6-53642540-DEL-80 | chr6-53642541-DEL-80 |
| chr6-53659830-DEL-220 | chr6-53659831-DEL-220 |
| chr6-53712035-DEL-67 | chr6-53712052-DEL-50 |
| chr6-53750000-DEL-90 | chr6-53749996-DEL-95 |
| chr6-53752161-DEL-897 | chr6-53752162-DEL-897 |
| chr6-53763493-DEL-6401 | chr6-53763494-DEL-6401 |
| chr6-53823790-DEL-182 | chr6-53823790-DEL-182 |
| chr6-53843239-DEL-60 | chr6-53843240-DEL-60 |
| chr6-53846478-DEL-107 | chr6-53846481-DEL-108 |
| chr6-53869359-DEL-465 | chr6-53869353-DEL-470 |
| chr6-53913890-DEL-53 | chr6-53913882-DEL-54 |
| chr6-53931743-DEL-1298 | chr6-53931744-DEL-1298 |
| chr6-53960510-DEL-68 | chr6-53960528-DEL-52 |
| chr6-53967835-DEL-172 | chr6-53967836-DEL-172 |
| chr6-53990328-DEL-52 | chr6-53990329-DEL-52 |
| chr6-53992457-DEL-674 | chr6-53992458-DEL-674 |
| chr6-54041707-DEL-56 | chr6-54041679-DEL-56 |
| chr6-54046460-DEL-164 | chr6-54046461-DEL-164 |
| chr6-54106055-DEL-52 | chr6-54106056-DEL-52 |
| chr6-54143293-DEL-382 | chr6-54143293-DEL-382 |
| chr6-54154027-DEL-672 | chr6-54154028-DEL-672 |
| chr6-54163923-DEL-1086 | chr6-54163924-DEL-1086 |
| chr6-54183123-DEL-52 | chr6-54183124-DEL-52 |
| chr6-54184592-DEL-58 | chr6-54184593-DEL-58 |
| chr6-54273437-DEL-399 | chr6-54273438-DEL-399 |
| chr6-54279685-DEL-119 | chr6-54279685-DEL-119 |
| chr6-54319049-DEL-505 | chr6-54319053-DEL-501 |
| chr6-54324386-DEL-79 | chr6-54324387-DEL-79 |
| chr6-54333230-DEL-3324 | chr6-54333232-DEL-3323 |
| chr6-54344752-DEL-247 | chr6-54344754-DEL-207 |
| chr6-54355105-DEL-765 | chr6-54355106-DEL-765 |
| chr6-54377324-DEL-1368 | chr6-54377325-DEL-1368 |
| chr6-54394280-DEL-383 | chr6-54394281-DEL-383 |
| chr6-54406122-DEL-59 | chr6-54406123-DEL-59 |
| chr6-54412453-DEL-51 | chr6-54412454-DEL-51 |
| chr6-54453861-DEL-71 | chr6-54453862-DEL-71 |
| chr6-54492207-DEL-365 | chr6-54492208-DEL-365 |
| chr6-54493641-DEL-138 | chr6-54493615-DEL-164 |
| chr6-54494973-DEL-63 | chr6-54494965-DEL-64 |
| chr6-54510150-DEL-56 | chr6-54510151-DEL-56 |
| chr6-54533200-DEL-333 | chr6-54533201-DEL-333 |
| chr6-54536167-DEL-96 | chr6-54536168-DEL-96 |
| chr6-54593265-DEL-56 | chr6-54593266-DEL-56 |

| | |
|---|---|
| chr6-54627462-DEL-71 | chr6-54627463-DEL-71 |
| chr6-54628244-DEL-216 | chr6-54628245-DEL-216 |
| chr6-54637302-DEL-259 | chr6-54637303-DEL-259 |
| chr6-54638455-DEL-54 | chr6-54638456-DEL-54 |
| chr6-54638877-DEL-1792 | chr6-54638877-DEL-1791 |
| chr6-54651868-DEL-3111 | chr6-54651868-DEL-3112 |
| chr6-54672358-DEL-79 | chr6-54672359-DEL-79 |
| chr6-54686702-DEL-243 | chr6-54686703-DEL-243 |
| chr6-54692011-DEL-55 | chr6-54691990-DEL-74 |
| chr6-54693533-DEL-54 | chr6-54693532-DEL-50 |
| chr6-54696180-DEL-230 | chr6-54696181-DEL-230 |
| chr6-54700647-DEL-134 | chr6-54700648-DEL-134 |
| chr6-54701914-DEL-52 | chr6-54701915-DEL-52 |
| chr6-54742768-DEL-55 | chr6-54742772-DEL-52 |
| chr6-54769923-DEL-52 | chr6-54769924-DEL-52 |
| chr6-54822789-DEL-1037 | chr6-54822790-DEL-1037 |
| chr6-54871586-DEL-213 | chr6-54871587-DEL-213 |
| chr6-54877486-DEL-220 | chr6-54877487-DEL-220 |
| chr6-54896761-DEL-187 | chr6-54896760-DEL-188 |
| chr6-54906296-DEL-200 | chr6-54906297-DEL-200 |
| chr6-54935945-DEL-183 | chr6-54935946-DEL-183 |
| chr6-54947801-DEL-181 | chr6-54947802-DEL-181 |
| chr6-54952229-DEL-3392 | chr6-54952230-DEL-3392 |
| chr6-54952391-DEL-84 | chr6-54952392-DEL-84 |
| chr6-54955289-DEL-338 | chr6-54955290-DEL-338 |
| chr6-54964042-DEL-241 | chr6-54964043-DEL-241 |
| chr6-54985981-DEL-69 | chr6-54985982-DEL-69 |
| chr6-54995639-DEL-77 | chr6-54995640-DEL-77 |
| chr6-54997203-DEL-230 | chr6-54997203-DEL-229 |
| chr6-54999331-DEL-4826 | chr6-54999332-DEL-4826 |
| chr6-55034718-DEL-172 | chr6-55034719-DEL-172 |
| chr6-55047226-DEL-305 | chr6-55047210-DEL-306 |
| chr6-55055872-DEL-70 | chr6-55055895-DEL-70 |
| chr6-55071749-DEL-395 | chr6-55071750-DEL-395 |
| chr6-55135203-DEL-163 | chr6-55135204-DEL-163 |
| chr6-55152847-DEL-52 | chr6-55152878-DEL-55 |
| chr6-55202570-DEL-3070 | chr6-55202571-DEL-3070 |
| chr6-55239182-DEL-54 | chr6-55239183-DEL-54 |
| chr6-55244738-DEL-5039 | chr6-55244739-DEL-5039 |
| chr6-55276237-DEL-78 | chr6-55276244-DEL-72 |
| chr6-55309604-DEL-1193 | chr6-55309605-DEL-1193 |
| chr6-55359857-DEL-57 | chr6-55359841-DEL-54 |
| chr6-55367151-DEL-191 | chr6-55367152-DEL-191 |
| chr6-55382724-DEL-52 | chr6-55382725-DEL-52 |
| chr6-55383973-DEL-80 | chr6-55383962-DEL-92 |
| chr6-55397283-DEL-240 | chr6-55397284-DEL-240 |
| chr6-55404783-DEL-2128 | chr6-55404741-DEL-2128 |
| chr6-55494042-DEL-602 | chr6-55494043-DEL-600 |
| chr6-55497175-DEL-80 | chr6-55497176-DEL-72 |
| chr6-55503682-DEL-152 | chr6-55503683-DEL-152 |
| chr6-55519278-DEL-80 | chr6-55519279-DEL-80 |
| chr6-55528873-DEL-257 | chr6-55528874-DEL-257 |
| chr6-55540516-DEL-61 | chr6-55540516-DEL-52 |
| chr6-55555509-DEL-58 | chr6-55555510-DEL-58 |
| chr6-55591301-DEL-768 | chr6-55591302-DEL-768 |
| chr6-55592126-DEL-121 | chr6-55592127-DEL-121 |
| chr6-55619094-DEL-211 | chr6-55619095-DEL-211 |
| chr6-55630512-DEL-6443 | chr6-55630513-DEL-6443 |
| chr6-55643330-DEL-77 | chr6-55643331-DEL-77 |
| chr6-55672648-DEL-423 | chr6-55672649-DEL-423 |
| chr6-55683437-DEL-59 | chr6-55683437-DEL-59 |
| chr6-55685166-DEL-177 | chr6-55685167-DEL-177 |
| chr6-55689380-DEL-68 | chr6-55689350-DEL-50 |
| chr6-55725272-DEL-60 | chr6-55725269-DEL-60 |
| chr6-55749927-DEL-57 | chr6-55749928-DEL-57 |
| chr6-55754399-DEL-444 | chr6-55754400-DEL-444 |
| chr6-55779363-DEL-62 | chr6-55779364-DEL-52 |
| chr6-55792488-DEL-131 | chr6-55792484-DEL-134 |
| chr6-55827132-DEL-63 | chr6-55827123-DEL-73 |
| chr6-55840842-DEL-81 | chr6-55840843-DEL-90 |
| chr6-55930865-DEL-1602 | chr6-55930865-DEL-1602 |
| chr6-55935328-DEL-6185 | chr6-55935323-DEL-6186 |
| chr6-55974246-DEL-252 | chr6-55974247-DEL-252 |
| chr6-56012173-DEL-96 | chr6-56012174-DEL-96 |
| chr6-56013669-DEL-70 | chr6-56013686-DEL-66 |
| chr6-56024071-DEL-60 | chr6-56024072-DEL-60 |
| chr6-56030770-DEL-68 | chr6-56030766-DEL-69 |
| chr6-56036628-DEL-768 | chr6-56036629-DEL-768 |
| chr6-56041851-DEL-540 | chr6-56041852-DEL-540 |
| chr6-56054606-DEL-60 | chr6-56054616-DEL-51 |

| | |
|---|---|
| chr6-56057448-DEL-58 | chr6-56057449-DEL-58 |
| chr6-56062523-DEL-742 | chr6-56062524-DEL-742 |
| chr6-56065272-DEL-416 | chr6-56065273-DEL-416 |
| chr6-56120264-DEL-2304 | chr6-56120265-DEL-2304 |
| chr6-56148843-DEL-103 | chr6-56148844-DEL-103 |
| chr6-56160542-DEL-6420 | chr6-56160543-DEL-6420 |
| chr6-56160573-DEL-201 | chr6-56160574-DEL-201 |
| chr6-56216363-DEL-174 | chr6-56216364-DEL-174 |
| chr6-56228447-DEL-95 | chr6-56228448-DEL-95 |
| chr6-56251290-DEL-1317 | chr6-56251291-DEL-1317 |
| chr6-56256906-DEL-92 | chr6-56256907-DEL-94 |
| chr6-56257714-DEL-192 | chr6-56257715-DEL-192 |
| chr6-56259155-DEL-1136 | chr6-56259156-DEL-1136 |
| chr6-56270281-DEL-6730 | chr6-56270290-DEL-6731 |
| chr6-56296866-DEL-2003 | chr6-56296866-DEL-2003 |
| chr6-56303254-DEL-235 | chr6-56303255-DEL-235 |
| chr6-56328652-DEL-72 | chr6-56328653-DEL-66 |
| chr6-56379717-DEL-165 | chr6-56379718-DEL-165 |
| chr6-56388801-DEL-56 | chr6-56388802-DEL-56 |
| chr6-56394372-DEL-2968 | chr6-56394373-DEL-2968 |
| chr6-56403544-DEL-55 | chr6-56403545-DEL-55 |
| chr6-56419531-DEL-911 | chr6-56419532-DEL-911 |
| chr6-56432790-DEL-1700 | chr6-56432791-DEL-1700 |
| chr6-56437610-DEL-6005 | chr6-56437611-DEL-6005 |
| chr6-56448959-DEL-509 | chr6-56448960-DEL-509 |
| chr6-56457005-DEL-90 | chr6-56457036-DEL-64 |
| chr6-56458877-DEL-257 | chr6-56458878-DEL-257 |
| chr6-56459927-DEL-523 | chr6-56459928-DEL-523 |
| chr6-56484696-DEL-6958 | chr6-56484697-DEL-6958 |
| chr6-56491658-DEL-333 | chr6-56491657-DEL-334 |
| chr6-56494107-DEL-361 | chr6-56494108-DEL-361 |
| chr6-56502855-DEL-612 | chr6-56502856-DEL-612 |
| chr6-56519348-DEL-400 | chr6-56519349-DEL-400 |
| chr6-56523140-DEL-102 | chr6-56523141-DEL-102 |
| chr6-56575005-DEL-56 | chr6-56575006-DEL-56 |
| chr6-56586674-DEL-51 | chr6-56586674-DEL-51 |
| chr6-56597598-DEL-87 | chr6-56597599-DEL-87 |
| chr6-56612590-DEL-60 | chr6-56612591-DEL-60 |
| chr6-56632711-DEL-540 | chr6-56632712-DEL-540 |
| chr6-56633662-DEL-51 | chr6-56633663-DEL-51 |
| chr6-56637381-DEL-318 | chr6-56637381-DEL-319 |
| chr6-56649740-DEL-70 | chr6-56649741-DEL-70 |
| chr6-56660466-DEL-195 | chr6-56660467-DEL-195 |
| chr6-56664967-DEL-90 | chr6-56664971-DEL-91 |
| chr6-56670397-DEL-6633 | chr6-56670398-DEL-6633 |
| chr6-56671372-DEL-413 | chr6-56671373-DEL-413 |
| chr6-56673150-DEL-1335 | chr6-56673151-DEL-1335 |
| chr6-56683940-DEL-440 | chr6-56683941-DEL-440 |
| chr6-56686634-DEL-677 | chr6-56686635-DEL-677 |
| chr6-56723761-DEL-4360 | chr6-56723762-DEL-4360 |
| chr6-56781831-DEL-133 | chr6-56781832-DEL-133 |
| chr6-56784215-DEL-221 | chr6-56784216-DEL-221 |
| chr6-56800494-DEL-982 | chr6-56800494-DEL-983 |
| chr6-56818445-DEL-107 | chr6-56818445-DEL-107 |
| chr6-56831867-DEL-52 | chr6-56831849-DEL-54 |
| chr6-56845482-DEL-69 | chr6-56845483-DEL-69 |
| chr6-56880358-DEL-156 | chr6-56880359-DEL-156 |
| chr6-56909914-DEL-315 | chr6-56909954-DEL-311 |
| chr6-56929492-DEL-1043 | chr6-56929493-DEL-1043 |
| chr6-56932093-DEL-966 | chr6-56932094-DEL-966 |
| chr6-56971169-DEL-1647 | chr6-56971175-DEL-1648 |
| chr6-56987954-DEL-60 | chr6-56987955-DEL-60 |
| chr6-56988921-DEL-375 | chr6-56988922-DEL-375 |
| chr6-57021152-DEL-136 | chr6-57021153-DEL-148 |
| chr6-57034713-DEL-1533 | chr6-57034714-DEL-1533 |
| chr6-57037075-DEL-2722 | chr6-57037076-DEL-2722 |
| chr6-57176203-DEL-978 | chr6-57176204-DEL-978 |
| chr6-57196828-DEL-6261 | chr6-57196829-DEL-6261 |
| chr6-57206542-DEL-465 | chr6-57206542-DEL-465 |
| chr6-57217815-DEL-57 | chr6-57217815-DEL-57 |
| chr6-57224868-DEL-181 | chr6-57224869-DEL-181 |
| chr6-57250300-DEL-88 | chr6-57250301-DEL-88 |
| chr6-57264600-DEL-6621 | chr6-57264601-DEL-6621 |
| chr6-57307522-DEL-124 | chr6-57307524-DEL-123 |
| chr6-57345583-DEL-6440 | chr6-57345584-DEL-6440 |
| chr6-57352179-DEL-1378 | chr6-57352180-DEL-1378 |
| chr6-57379519-DEL-75 | chr6-57379520-DEL-63 |
| chr6-57416992-DEL-6392 | chr6-57416993-DEL-6392 |
| chr6-57431517-DEL-306 | chr6-57431511-DEL-313 |
| chr6-57444056-DEL-368 | chr6-57444057-DEL-368 |

| | |
|---|---|
| chr6-57499441-DEL-140 | chr6-57499442-DEL-140 |
| chr6-57536010-DEL-60 | chr6-57535974-DEL-97 |
| chr6-57542961-DEL-1105 | chr6-57542962-DEL-1105 |
| chr6-57553877-DEL-88 | chr6-57553910-DEL-56 |
| chr6-57565785-DEL-6150 | chr6-57565785-DEL-6150 |
| chr6-57599161-DEL-990 | chr6-57599162-DEL-990 |
| chr6-57605330-DEL-6208 | chr6-57605331-DEL-6208 |
| chr6-57611237-DEL-201 | chr6-57611238-DEL-201 |
| chr6-57621577-DEL-525 | chr6-57621578-DEL-522 |
| chr6-57658573-DEL-1283 | chr6-57658574-DEL-1283 |
| chr6-57701844-DEL-208 | chr6-57701850-DEL-203 |
| chr6-57712401-DEL-265 | chr6-57712403-DEL-264 |
| chr6-57718173-DEL-740 | chr6-57718176-DEL-741 |
| chr6-57726649-DEL-222 | chr6-57726650-DEL-222 |
| chr6-57769931-DEL-341 | chr6-57769932-DEL-341 |
| chr6-57778130-DEL-401 | chr6-57778131-DEL-401 |
| chr6-57800900-DEL-77 | chr6-57800884-DEL-75 |
| chr6-57826778-DEL-363 | chr6-57826780-DEL-362 |
| chr6-57874408-DEL-772 | chr6-57874405-DEL-772 |
| chr6-57875775-DEL-293 | chr6-57875776-DEL-293 |
| chr6-57896488-DEL-6338 | chr6-57896479-DEL-6343 |
| chr6-57952276-DEL-403 | chr6-57952277-DEL-403 |
| chr6-57968507-DEL-102 | chr6-57968508-DEL-96 |
| chr6-58000657-DEL-58 | chr6-58000660-DEL-56 |
| chr6-58022778-DEL-80 | chr6-58022779-DEL-80 |
| chr6-58045501-DEL-4902 | chr6-58045502-DEL-4902 |
| chr6-58150512-DEL-6777 | chr6-58150513-DEL-6777 |
| chr6-58179485-DEL-77 | chr6-58179484-DEL-93 |
| chr6-58182763-DEL-371 | chr6-58182764-DEL-371 |
| chr6-58272001-DEL-500 | chr6-58272002-DEL-500 |
| chr6-58280413-DEL-59 | chr6-58280414-DEL-63 |
| chr6-58316907-DEL-153 | chr6-58316908-DEL-153 |
| chr6-58320317-DEL-52 | chr6-58320317-DEL-52 |
| chr6-58322751-DEL-6596 | chr6-58322752-DEL-6596 |
| chr6-58414980-DEL-589 | chr6-58414980-DEL-589 |
| chr6-58415937-DEL-947 | chr6-58415937-DEL-947 |
| chr6-58431499-DEL-1853 | chr6-58431500-DEL-1853 |
| chr6-58445835-DEL-1447 | chr6-58445836-DEL-1447 |
| chr6-58448230-DEL-742 | chr6-58448231-DEL-742 |
| chr6-58451345-DEL-68 | chr6-58451346-DEL-54 |
| chr6-58462095-DEL-968 | chr6-58462096-DEL-968 |
| chr6-58485358-DEL-3302 | chr6-58485359-DEL-3302 |
| chr6-58565546-DEL-1030 | chr6-58565548-DEL-1029 |
| chr6-58567827-DEL-807 | chr6-58567828-DEL-807 |
| chr6-58570536-DEL-57 | chr6-58570537-DEL-57 |
| chr6-58616592-DEL-239 | chr6-58616593-DEL-239 |
| chr6-58643437-DEL-264 | chr6-58643438-DEL-220 |
| chr6-58689611-DEL-6195 | chr6-58689612-DEL-6195 |
| chr6-58704753-DEL-155 | chr6-58704753-DEL-155 |
| chr6-58747989-DEL-454 | chr6-58747990-DEL-454 |
| chr6-58764844-DEL-199 | chr6-58764845-DEL-199 |
| chr6-58765775-DEL-6745 | chr6-58765776-DEL-6745 |
| chr6-58794226-DEL-8415 | chr6-58794227-DEL-8415 |
| chr6-58806347-DEL-51 | chr6-58806348-DEL-51 |
| chr6-58843025-DEL-84 | chr6-58843026-DEL-84 |
| chr6-58925525-DEL-239 | chr6-58925524-DEL-240 |
| chr6-59100521-DEL-6452 | chr6-59100522-DEL-6452 |
| chr6-59144456-DEL-68 | chr6-59144457-DEL-68 |
| chr6-59147983-DEL-511 | chr6-59147986-DEL-513 |
| chr6-59159793-DEL-57 | chr6-59159794-DEL-57 |
| chr6-59168587-DEL-64 | chr6-59168579-DEL-58 |
| chr6-59176612-DEL-7169 | chr6-59176613-DEL-7169 |
| chr6-59192021-DEL-59 | chr6-59192022-DEL-57 |
| chr6-59217851-DEL-1067 | chr6-59217852-DEL-1067 |
| chr6-59260486-DEL-9270 | chr6-59260487-DEL-9270 |
| chr6-59298475-DEL-2492 | chr6-59298476-DEL-2492 |
| chr6-59308051-DEL-735 | chr6-59308052-DEL-735 |
| chr6-59311549-DEL-443 | chr6-59311550-DEL-440 |
| chr6-59338425-DEL-200 | chr6-59338426-DEL-200 |
| chr6-59376233-DEL-752 | chr6-59376234-DEL-752 |
| chr6-59378961-DEL-55 | chr6-59378962-DEL-52 |
| chr6-59434486-DEL-60 | chr6-59434482-DEL-62 |
| chr6-59463449-DEL-1902 | chr6-59463447-DEL-1900 |
| chr6-59469169-DEL-58 | chr6-59469170-DEL-58 |
| chr6-59475504-DEL-78 | chr6-59475512-DEL-70 |
| chr6-59495978-DEL-400 | chr6-59495979-DEL-400 |
| chr6-59498270-DEL-66 | chr6-59498271-DEL-66 |
| chr6-59514250-DEL-1977 | chr6-59514251-DEL-1977 |
| chr6-59514740-DEL-68 | chr6-59514741-DEL-64 |
| chr6-59518652-DEL-6174 | chr6-59518653-DEL-6174 |

| | |
|---|---|
| chr6-59588144-DEL-51 | chr6-59588145-DEL-51 |
| chr6-59607031-DEL-8029 | chr6-59607032-DEL-8029 |
| chr6-59669894-DEL-5287 | chr6-59669895-DEL-5287 |
| chr6-59759315-DEL-6399 | chr6-59759316-DEL-6399 |
| chr6-59813441-DEL-1259 | chr6-59813442-DEL-1259 |
| chr6-59840875-DEL-53 | chr6-59840874-DEL-55 |
| chr6-59848650-DEL-2298 | chr6-59848650-DEL-2298 |
| chr6-59852568-DEL-498 | chr6-59852568-DEL-498 |
| chr6-59856377-DEL-199 | chr6-59856376-DEL-199 |
| chr6-59860150-DEL-8807 | chr6-59860153-DEL-8809 |
| chr6-59959048-DEL-1541 | chr6-59959031-DEL-1541 |
| chr6-59961781-DEL-70 | chr6-59961781-DEL-111 |
| chr6-59976916-DEL-86 | chr6-59976950-DEL-50 |
| chr6-59980059-DEL-144 | chr6-59980059-DEL-144 |
| chr6-60004345-DEL-214 | chr6-60004344-DEL-214 |
| chr6-60008615-DEL-62 | chr6-60008616-DEL-52 |
| chr6-60036743-DEL-498 | chr6-60036744-DEL-498 |
| chr6-60067665-DEL-2316 | chr6-60067665-DEL-2316 |
| chr6-60078525-DEL-96 | chr6-60078537-DEL-88 |
| chr6-60081271-DEL-185 | chr6-60081279-DEL-185 |
| chr6-60090395-DEL-564 | chr6-60090396-DEL-564 |
| chr6-60123647-DEL-126 | chr6-60123647-DEL-126 |
| chr6-60133504-DEL-676 | chr6-60133505-DEL-676 |
| chr6-60150278-DEL-58 | chr6-60150280-DEL-64 |
| chr6-60205591-DEL-221 | chr6-60205590-DEL-221 |
| chr6-60207849-DEL-239 | chr6-60207849-DEL-239 |
| chr6-60225685-DEL-128 | chr6-60225685-DEL-128 |
| chr6-60227137-DEL-95 | chr6-60227137-DEL-95 |
| chr6-60230478-DEL-1198 | chr6-60230478-DEL-1188 |
| chr6-60302334-DEL-400 | chr6-60302335-DEL-400 |
| chr6-60329837-DEL-410 | chr6-60329838-DEL-410 |
| chr6-60333616-DEL-76 | chr6-60333624-DEL-50 |
| chr6-60382367-DEL-64 | chr6-60382367-DEL-64 |
| chr6-60392792-DEL-4688 | chr6-60392793-DEL-4688 |
| chr6-60401906-DEL-577 | chr6-60401907-DEL-577 |
| chr6-60403375-DEL-360 | chr6-60403382-DEL-361 |
| chr6-60458928-DEL-70 | chr6-60458929-DEL-58 |
| chr6-60475374-DEL-150 | chr6-60475375-DEL-150 |
| chr6-60476346-DEL-51 | chr6-60476345-DEL-51 |
| chr6-60476715-DEL-966 | chr6-60476716-DEL-966 |
| chr6-60476727-DEL-2382 | chr6-60476728-DEL-2382 |
| chr6-60483801-DEL-6103 | chr6-60483802-DEL-6103 |
| chr6-60498095-DEL-202 | chr6-60498128-DEL-209 |
| chr6-60508255-DEL-142 | chr6-60508256-DEL-142 |
| chr6-60511188-DEL-1235 | chr6-60511188-DEL-1235 |
| chr6-60513016-DEL-2571 | chr6-60513016-DEL-2571 |
| chr6-60518028-DEL-6886 | chr6-60518030-DEL-6886 |
| chr6-60528372-DEL-2305 | chr6-60528373-DEL-2305 |
| chr6-60533840-DEL-88 | chr6-60533841-DEL-88 |
| chr6-60568491-DEL-652 | chr6-60568487-DEL-649 |
| chr6-60570951-DEL-5263 | chr6-60570952-DEL-5263 |
| chr6-60618804-DEL-61 | chr6-60618801-DEL-75 |
| chr6-60635467-DEL-542 | chr6-60635468-DEL-541 |
| chr6-60644096-DEL-61 | chr6-60644096-DEL-61 |
| chr6-60644939-DEL-2132 | chr6-60644939-DEL-2132 |
| chr6-60655897-DEL-81 | chr6-60655898-DEL-59 |
| chr6-60682568-DEL-294 | chr6-60682569-DEL-294 |
| chr6-60689283-DEL-650 | chr6-60689284-DEL-650 |
| chr6-60690586-DEL-927 | chr6-60690587-DEL-927 |
| chr6-60703476-DEL-217 | chr6-60703477-DEL-217 |
| chr6-60711503-DEL-3699 | chr6-60711504-DEL-3699 |
| chr6-60721183-DEL-465 | chr6-60721211-DEL-459 |
| chr6-60733018-DEL-1410 | chr6-60733019-DEL-1410 |
| chr6-60736346-DEL-195 | chr6-60736347-DEL-195 |
| chr6-60746477-DEL-1688 | chr6-60746478-DEL-1688 |
| chr6-60767022-DEL-312 | chr6-60767023-DEL-312 |
| chr6-60830950-DEL-72 | chr6-60830969-DEL-54 |
| chr6-60886601-DEL-54 | chr6-60886602-DEL-54 |
| chr6-60900114-DEL-71 | chr6-60900114-DEL-71 |
| chr6-60901526-DEL-2314 | chr6-60901527-DEL-2314 |
| chr6-60906741-DEL-109 | chr6-60906746-DEL-141 |
| chr6-60916392-DEL-65 | chr6-60916393-DEL-50 |
| chr6-60927924-DEL-2568 | chr6-60927925-DEL-2568 |
| chr6-60966467-DEL-68 | chr6-60966467-DEL-68 |
| chr6-60986593-DEL-99 | chr6-60986594-DEL-99 |
| chr6-60988077-DEL-52 | chr6-60988081-DEL-51 |
| chr6-60999050-DEL-174 | chr6-60999044-DEL-167 |
| chr6-61000080-DEL-6218 | chr6-61000080-DEL-6218 |
| chr6-61007345-DEL-330 | chr6-61007346-DEL-331 |
| chr6-61009824-DEL-540 | chr6-61009825-DEL-540 |

| | |
|---|---|
| chr6-61024888-DEL-114 | chr6-61024890-DEL-113 |
| chr6-61027305-DEL-252 | chr6-61027304-DEL-253 |
| chr6-61047060-DEL-898 | chr6-61047061-DEL-898 |
| chr6-61073703-DEL-99 | chr6-61073704-DEL-99 |
| chr6-61108705-DEL-178 | chr6-61108706-DEL-178 |
| chr6-61110108-DEL-2502 | chr6-61110109-DEL-2502 |
| chr6-61126066-DEL-214 | chr6-61126067-DEL-214 |
| chr6-61141220-DEL-750 | chr6-61141221-DEL-750 |
| chr6-61148756-DEL-105 | chr6-61148754-DEL-108 |
| chr6-61183578-DEL-58 | chr6-61183579-DEL-58 |
| chr6-61195460-DEL-992 | chr6-61195460-DEL-990 |
| chr6-61201856-DEL-535 | chr6-61201857-DEL-535 |
| chr6-61207998-DEL-67 | chr6-61208000-DEL-65 |
| chr6-61213783-DEL-1228 | chr6-61213784-DEL-1228 |
| chr6-61231455-DEL-177 | chr6-61231456-DEL-177 |
| chr6-61231905-DEL-3195 | chr6-61231906-DEL-3195 |
| chr6-61261171-DEL-445 | chr6-61261171-DEL-445 |
| chr6-61294374-DEL-1112 | chr6-61294375-DEL-1112 |
| chr6-61314213-DEL-2858 | chr6-61314216-DEL-2853 |
| chr6-61317207-DEL-1052 | chr6-61317208-DEL-1052 |
| chr6-61322116-DEL-107 | chr6-61322117-DEL-107 |
| chr6-61322448-DEL-552 | chr6-61322439-DEL-538 |
| chr6-61425760-DEL-6149 | chr6-61425761-DEL-6149 |
| chr6-61485112-DEL-4407 | chr6-61485113-DEL-4407 |
| chr6-61502783-DEL-212 | chr6-61502784-DEL-212 |
| chr6-61553432-DEL-75 | chr6-61553435-DEL-76 |
| chr6-61553830-DEL-897 | chr6-61553840-DEL-898 |
| chr6-61567717-DEL-1863 | chr6-61567718-DEL-1863 |
| chr6-61568513-DEL-498 | chr6-61568514-DEL-498 |
| chr6-61616707-DEL-209 | chr6-61616708-DEL-209 |
| chr6-61647137-DEL-220 | chr6-61647138-DEL-220 |
| chr6-61662433-DEL-1677 | chr6-61662436-DEL-1674 |
| chr6-61676386-DEL-119 | chr6-61676387-DEL-116 |
| chr6-61699043-DEL-702 | chr6-61699044-DEL-702 |
| chr6-61737933-DEL-58 | chr6-61737933-DEL-58 |
| chr6-61754279-DEL-64 | chr6-61754279-DEL-64 |
| chr6-61755726-DEL-6543 | chr6-61755727-DEL-6543 |
| chr6-61815074-DEL-63 | chr6-61815075-DEL-63 |
| chr6-61849494-DEL-1146 | chr6-61849500-DEL-1141 |
| chr6-61861540-DEL-432 | chr6-61861541-DEL-432 |
| chr6-61868505-DEL-136 | chr6-61868506-DEL-132 |
| chr6-61873475-DEL-1160 | chr6-61873476-DEL-1160 |
| chr6-61888504-DEL-925 | chr6-61888505-DEL-925 |
| chr6-61889984-DEL-214 | chr6-61889985-DEL-214 |
| chr6-61892216-DEL-688 | chr6-61892217-DEL-688 |
| chr6-61893992-DEL-6482 | chr6-61893993-DEL-6482 |
| chr6-61914059-DEL-64 | chr6-61914060-DEL-64 |
| chr6-61925953-DEL-146 | chr6-61925948-DEL-158 |
| chr6-61950608-DEL-208 | chr6-61950609-DEL-208 |
| chr6-61955888-DEL-6384 | chr6-61955889-DEL-6384 |
| chr6-61977351-DEL-86 | chr6-61977352-DEL-86 |
| chr6-61980678-DEL-84 | chr6-61980679-DEL-84 |
| chr6-62012152-DEL-7016 | chr6-62012153-DEL-7016 |
| chr6-62036907-DEL-824 | chr6-62036946-DEL-802 |
| chr6-62115393-DEL-2257 | chr6-62115394-DEL-2257 |
| chr6-62149028-DEL-1503 | chr6-62149029-DEL-1503 |
| chr6-62159154-DEL-6186 | chr6-62159155-DEL-6186 |
| chr6-62171282-DEL-61 | chr6-62171283-DEL-56 |
| chr6-62208941-DEL-182 | chr6-62208944-DEL-180 |
| chr6-62245876-DEL-2084 | chr6-62245878-DEL-2084 |
| chr6-62287442-DEL-78 | chr6-62287420-DEL-54 |
| chr6-62309591-DEL-259 | chr6-62309592-DEL-259 |
| chr6-62367130-DEL-140 | chr6-62367131-DEL-140 |
| chr6-62368077-DEL-6250 | chr6-62368078-DEL-6250 |
| chr6-62395948-DEL-593 | chr6-62395949-DEL-593 |
| chr6-62414680-DEL-1566 | chr6-62414680-DEL-1566 |
| chr6-62423524-DEL-81 | chr6-62423524-DEL-81 |
| chr6-62434474-DEL-277 | chr6-62434470-DEL-276 |
| chr6-62452746-DEL-1786 | chr6-62452746-DEL-1786 |
| chr6-62490154-DEL-52 | chr6-62490150-DEL-99 |
| chr6-62497043-DEL-157 | chr6-62497053-DEL-150 |
| chr6-62525456-DEL-381 | chr6-62525457-DEL-381 |
| chr6-62586750-DEL-72 | chr6-62586724-DEL-54 |
| chr6-62599123-DEL-60 | chr6-62599124-DEL-60 |
| chr6-62602102-DEL-505 | chr6-62602102-DEL-505 |
| chr6-62653860-DEL-155 | chr6-62653861-DEL-155 |
| chr6-62680551-DEL-6562 | chr6-62680552-DEL-6562 |
| chr6-62780646-DEL-54 | chr6-62780647-DEL-54 |
| chr6-62811259-DEL-1061 | chr6-62811260-DEL-1061 |
| chr6-62822778-DEL-69 | chr6-62822781-DEL-66 |

| | |
|---|---|
| chr6-62855080-DEL-6217 | chr6-62855081-DEL-6217 |
| chr6-62866638-DEL-488 | chr6-62866639-DEL-488 |
| chr6-62894199-DEL-265 | chr6-62894200-DEL-262 |
| chr6-62927438-DEL-553 | chr6-62927438-DEL-549 |
| chr6-62928480-DEL-280 | chr6-62928481-DEL-278 |
| chr6-62939224-DEL-6274 | chr6-62939225-DEL-6274 |
| chr6-62948382-DEL-509 | chr6-62948383-DEL-509 |
| chr6-62965822-DEL-55 | chr6-62965823-DEL-52 |
| chr6-62968564-DEL-1049 | chr6-62968565-DEL-1092 |
| chr6-62991949-DEL-7948 | chr6-62991950-DEL-7948 |
| chr6-63001505-DEL-1284 | chr6-63001506-DEL-1284 |
| chr6-63028223-DEL-206 | chr6-63028217-DEL-207 |
| chr6-63050311-DEL-71 | chr6-63050312-DEL-71 |
| chr6-63053144-DEL-2681 | chr6-63053145-DEL-2681 |
| chr6-63059792-DEL-831 | chr6-63059795-DEL-828 |
| chr6-63071121-DEL-51 | chr6-63071122-DEL-51 |
| chr6-63095839-DEL-915 | chr6-63095839-DEL-915 |
| chr6-63110030-DEL-385 | chr6-63110024-DEL-371 |
| chr6-63115499-DEL-67 | chr6-63115500-DEL-67 |
| chr6-63129872-DEL-147 | chr6-63129866-DEL-153 |
| chr6-63186759-DEL-3082 | chr6-63186760-DEL-3082 |
| chr6-63200520-DEL-68 | chr6-63200521-DEL-68 |
| chr6-63271781-DEL-87 | chr6-63271809-DEL-67 |
| chr6-63274286-DEL-278 | chr6-63274287-DEL-278 |
| chr6-63397449-DEL-583 | chr6-63397450-DEL-583 |
| chr6-63407581-DEL-181 | chr6-63407582-DEL-181 |
| chr6-63421109-DEL-51 | chr6-63421110-DEL-51 |
| chr6-63427054-DEL-66 | chr6-63427059-DEL-68 |
| chr6-63430329-DEL-500 | chr6-63430329-DEL-495 |
| chr6-63440713-DEL-130 | chr6-63440714-DEL-130 |
| chr6-63445687-DEL-136 | chr6-63445688-DEL-136 |
| chr6-63461504-DEL-240 | chr6-63461504-DEL-240 |
| chr6-63462643-DEL-235 | chr6-63462644-DEL-235 |
| chr6-63463506-DEL-72 | chr6-63463523-DEL-54 |
| chr6-63497577-DEL-368 | chr6-63497578-DEL-368 |
| chr6-63501608-DEL-583 | chr6-63501610-DEL-582 |
| chr6-63530195-DEL-84 | chr6-63530173-DEL-107 |
| chr6-63556790-DEL-6504 | chr6-63556791-DEL-6504 |
| chr6-63572423-DEL-604 | chr6-63572424-DEL-604 |
| chr6-63594894-DEL-584 | chr6-63594889-DEL-573 |
| chr6-63606446-DEL-128 | chr6-63606429-DEL-129 |
| chr6-63643722-DEL-366 | chr6-63643723-DEL-366 |
| chr6-63644564-DEL-7700 | chr6-63644565-DEL-7700 |
| chr6-63658821-DEL-695 | chr6-63658822-DEL-695 |
| chr6-63663396-DEL-686 | chr6-63663376-DEL-704 |
| chr6-63669629-DEL-241 | chr6-63669630-DEL-241 |
| chr6-63683959-DEL-2767 | chr6-63683960-DEL-2767 |
| chr6-63736725-DEL-222 | chr6-63736721-DEL-223 |
| chr6-63738349-DEL-60 | chr6-63738350-DEL-60 |
| chr6-63742986-DEL-54 | chr6-63742987-DEL-54 |
| chr6-63751056-DEL-58 | chr6-63751052-DEL-78 |
| chr6-63758289-DEL-1279 | chr6-63758290-DEL-1279 |
| chr6-63760594-DEL-484 | chr6-63760595-DEL-484 |
| chr6-63768574-DEL-137 | chr6-63768575-DEL-137 |
| chr6-63807121-DEL-297 | chr6-63807122-DEL-297 |
| chr6-63856989-DEL-736 | chr6-63856990-DEL-736 |
| chr6-63860677-DEL-92 | chr6-63860636-DEL-86 |
| chr6-63869369-DEL-618 | chr6-63869370-DEL-618 |
| chr6-63886867-DEL-63 | chr6-63886863-DEL-88 |
| chr6-63899402-DEL-3101 | chr6-63899402-DEL-3101 |
| chr6-63921911-DEL-62 | chr6-63921912-DEL-62 |
| chr6-63970303-DEL-400 | chr6-63970304-DEL-400 |
| chr6-63980072-DEL-74 | chr6-63980073-DEL-74 |
| chr6-64005758-DEL-1523 | chr6-64005759-DEL-1523 |
| chr6-64042584-DEL-5202 | chr6-64042585-DEL-5202 |
| chr6-64051811-DEL-161 | chr6-64051812-DEL-161 |
| chr6-64116885-DEL-978 | chr6-64116886-DEL-978 |
| chr6-64118362-DEL-1358 | chr6-64118363-DEL-1358 |
| chr6-64131472-DEL-7130 | chr6-64131473-DEL-7130 |
| chr6-64145950-DEL-266 | chr6-64145930-DEL-285 |
| chr6-64148140-DEL-3421 | chr6-64148141-DEL-3421 |
| chr6-64154962-DEL-1772 | chr6-64154964-DEL-1771 |
| chr6-64173580-DEL-1349 | chr6-64173581-DEL-1349 |
| chr6-64200040-DEL-80 | chr6-64200041-DEL-68 |
| chr6-64214596-DEL-98 | chr6-64214636-DEL-91 |
| chr6-64222895-DEL-75 | chr6-64222920-DEL-78 |
| chr6-64258846-DEL-211 | chr6-64258847-DEL-211 |
| chr6-64276393-DEL-132 | chr6-64276394-DEL-132 |
| chr6-64355011-DEL-6241 | chr6-64355012-DEL-6241 |
| chr6-64369241-DEL-757 | chr6-64369231-DEL-758 |

| | |
|---|---|
| chr6-64396006-DEL-56 | chr6-64396005-DEL-58 |
| chr6-64401869-DEL-96 | chr6-64401882-DEL-78 |
| chr6-64418833-DEL-400 | chr6-64418834-DEL-400 |
| chr6-64453273-DEL-696 | chr6-64453274-DEL-696 |
| chr6-64471470-DEL-1158 | chr6-64471470-DEL-1158 |
| chr6-64524957-DEL-221 | chr6-64524958-DEL-221 |
| chr6-64553103-DEL-80 | chr6-64553150-DEL-76 |
| chr6-64563694-DEL-1149 | chr6-64563695-DEL-1149 |
| chr6-64588426-DEL-818 | chr6-64588425-DEL-819 |
| chr6-64623044-DEL-1678 | chr6-64623045-DEL-1674 |
| chr6-64628734-DEL-198 | chr6-64628735-DEL-198 |
| chr6-64630360-DEL-8266 | chr6-64630361-DEL-8266 |
| chr6-64652528-DEL-116 | chr6-64652529-DEL-116 |
| chr6-64727186-DEL-576 | chr6-64727187-DEL-576 |
| chr6-64743207-DEL-193 | chr6-64743206-DEL-193 |
| chr6-64769390-DEL-872 | chr6-64769391-DEL-872 |
| chr6-64783287-DEL-6378 | chr6-64783301-DEL-6377 |
| chr6-64805613-DEL-327 | chr6-64805614-DEL-327 |
| chr6-64808542-DEL-809 | chr6-64808543-DEL-809 |
| chr6-64834351-DEL-200 | chr6-64834352-DEL-200 |
| chr6-64836128-DEL-422 | chr6-64836134-DEL-418 |
| chr6-64836776-DEL-51 | chr6-64836777-DEL-51 |
| chr6-64842892-DEL-65 | chr6-64842888-DEL-68 |
| chr6-64858291-DEL-7038 | chr6-64858297-DEL-7039 |
| chr6-64858657-DEL-617 | chr6-64858657-DEL-617 |
| chr6-64873922-DEL-2525 | chr6-64873923-DEL-2525 |
| chr6-64892689-DEL-161 | chr6-64892690-DEL-161 |
| chr6-64898698-DEL-9356 | chr6-64898682-DEL-9372 |
| chr6-64941654-DEL-96 | chr6-64941663-DEL-88 |
| chr6-64980021-DEL-70 | chr6-64980035-DEL-64 |
| chr6-64994075-DEL-199 | chr6-64994076-DEL-200 |
| chr6-65004826-DEL-59 | chr6-65004827-DEL-59 |
| chr6-65012831-DEL-324 | chr6-65012832-DEL-324 |
| chr6-65024228-DEL-228 | chr6-65024229-DEL-228 |
| chr6-65066378-DEL-218 | chr6-65066379-DEL-218 |
| chr6-65104257-DEL-61 | chr6-65104258-DEL-61 |
| chr6-65122618-DEL-438 | chr6-65122619-DEL-438 |
| chr6-65141528-DEL-369 | chr6-65141535-DEL-342 |
| chr6-65204968-DEL-4724 | chr6-65204969-DEL-4724 |
| chr6-65231971-DEL-6593 | chr6-65231972-DEL-6593 |
| chr6-65264865-DEL-225 | chr6-65264865-DEL-225 |
| chr6-65270254-DEL-13201 | chr6-65270243-DEL-13206 |
| chr6-65295338-DEL-6266 | chr6-65295339-DEL-6266 |
| chr6-65312480-DEL-51 | chr6-65312481-DEL-51 |
| chr6-65338794-DEL-426 | chr6-65338795-DEL-426 |
| chr6-65384185-DEL-429 | chr6-65384186-DEL-429 |
| chr6-65394720-DEL-2162 | chr6-65394721-DEL-2162 |
| chr6-65408104-DEL-102 | chr6-65408109-DEL-82 |
| chr6-65417100-DEL-967 | chr6-65417101-DEL-967 |
| chr6-65443062-DEL-96 | chr6-65443062-DEL-96 |
| chr6-65467720-DEL-62 | chr6-65467718-DEL-63 |
| chr6-65477752-DEL-6202 | chr6-65477753-DEL-6202 |
| chr6-65557836-DEL-1075 | chr6-65557836-DEL-1075 |
| chr6-65566547-DEL-210 | chr6-65566548-DEL-210 |
| chr6-65615217-DEL-133 | chr6-65615225-DEL-115 |
| chr6-65633481-DEL-69 | chr6-65633481-DEL-66 |
| chr6-65639378-DEL-60 | chr6-65639379-DEL-60 |
| chr6-65670917-DEL-83 | chr6-65670890-DEL-94 |
| chr6-65675956-DEL-134 | chr6-65675957-DEL-134 |
| chr6-65691063-DEL-66 | chr6-65691063-DEL-66 |
| chr6-65771552-DEL-56 | chr6-65771511-DEL-70 |
| chr6-65791614-DEL-59 | chr6-65791615-DEL-59 |
| chr6-65809340-DEL-206 | chr6-65809341-DEL-206 |
| chr6-65843468-DEL-1485 | chr6-65843468-DEL-1485 |
| chr6-65866967-DEL-178 | chr6-65866968-DEL-178 |
| chr6-65939295-DEL-1888 | chr6-65939296-DEL-1888 |
| chr6-65942037-DEL-122 | chr6-65942038-DEL-122 |
| chr6-65964324-DEL-464 | chr6-65964325-DEL-464 |
| chr6-65969505-DEL-1646 | chr6-65969506-DEL-1646 |
| chr6-65979740-DEL-68 | chr6-65979741-DEL-68 |
| chr6-66006487-DEL-198 | chr6-66006488-DEL-198 |
| chr6-66015077-DEL-2817 | chr6-66015078-DEL-2817 |
| chr6-66028728-DEL-2363 | chr6-66028729-DEL-2363 |
| chr6-66046928-DEL-6406 | chr6-66046929-DEL-6406 |
| chr6-66060242-DEL-2288 | chr6-66060243-DEL-2288 |
| chr6-66072913-DEL-104 | chr6-66072914-DEL-104 |
| chr6-66076320-DEL-100 | chr6-66076321-DEL-100 |
| chr6-66095197-DEL-103 | chr6-66095197-DEL-119 |
| chr6-66096774-DEL-96 | chr6-66096775-DEL-96 |
| chr6-66112747-DEL-217 | chr6-66112748-DEL-217 |

| | |
|---|---|
| chr6-66128498-DEL-69 | chr6-66128502-DEL-97 |
| chr6-66141133-DEL-1002 | chr6-66141109-DEL-1026 |
| chr6-66154343-DEL-526 | chr6-66154344-DEL-526 |
| chr6-66156095-DEL-120 | chr6-66156096-DEL-116 |
| chr6-66215682-DEL-2774 | chr6-66215683-DEL-2774 |
| chr6-66264553-DEL-51 | chr6-66264554-DEL-51 |
| chr6-66270580-DEL-872 | chr6-66270583-DEL-861 |
| chr6-66271866-DEL-54 | chr6-66271867-DEL-54 |
| chr6-66281496-DEL-88 | chr6-66281496-DEL-88 |
| chr6-66341683-DEL-1256 | chr6-66341684-DEL-1256 |
| chr6-66343688-DEL-88 | chr6-66343702-DEL-106 |
| chr6-66350272-DEL-62 | chr6-66350273-DEL-62 |
| chr6-66351492-DEL-3386 | chr6-66351490-DEL-3387 |
| chr6-66402164-DEL-174 | chr6-66402165-DEL-174 |
| chr6-66403529-DEL-1023 | chr6-66403530-DEL-1023 |
| chr6-66417074-DEL-240 | chr6-66417075-DEL-240 |
| chr6-66435574-DEL-5967 | chr6-66435575-DEL-5967 |
| chr6-66451320-DEL-1133 | chr6-66451321-DEL-1133 |
| chr6-66463395-DEL-3547 | chr6-66463396-DEL-3547 |
| chr6-66484950-DEL-53 | chr6-66484947-DEL-53 |
| chr6-66596443-DEL-400 | chr6-66596444-DEL-400 |
| chr6-66603425-DEL-197 | chr6-66603426-DEL-197 |
| chr6-66625464-DEL-7686 | chr6-66625465-DEL-7686 |
| chr6-66631613-DEL-208 | chr6-66631614-DEL-208 |
| chr6-66644345-DEL-195 | chr6-66644346-DEL-195 |
| chr6-66654029-DEL-92 | chr6-66654029-DEL-52 |
| chr6-66665644-DEL-801 | chr6-66665638-DEL-805 |
| chr6-66667021-DEL-1443 | chr6-66667051-DEL-1414 |
| chr6-66672837-DEL-836 | chr6-66672846-DEL-833 |
| chr6-66708088-DEL-219 | chr6-66708089-DEL-219 |
| chr6-66718186-DEL-74 | chr6-66718187-DEL-74 |
| chr6-66718595-DEL-1933 | chr6-66718596-DEL-1933 |
| chr6-66722237-DEL-292 | chr6-66722237-DEL-291 |
| chr6-66728192-DEL-73 | chr6-66728194-DEL-71 |
| chr6-66733139-DEL-547 | chr6-66733140-DEL-547 |
| chr6-66743087-DEL-6442 | chr6-66743088-DEL-6442 |
| chr6-66767295-DEL-260 | chr6-66767296-DEL-260 |
| chr6-66792332-DEL-17109 | chr6-66792333-DEL-17109 |
| chr6-66847285-DEL-6939 | chr6-66847268-DEL-6951 |
| chr6-66866290-DEL-977 | chr6-66866275-DEL-992 |
| chr6-66872551-DEL-52 | chr6-66872540-DEL-56 |
| chr6-66885236-DEL-82 | chr6-66885237-DEL-82 |
| chr6-66910097-DEL-241 | chr6-66910098-DEL-241 |
| chr6-66910980-DEL-1326 | chr6-66910981-DEL-1326 |
| chr6-66935146-DEL-1057 | chr6-66935147-DEL-1057 |
| chr6-66953576-DEL-56 | chr6-66953577-DEL-56 |
| chr6-66960228-DEL-350 | chr6-66960229-DEL-350 |
| chr6-66963353-DEL-177 | chr6-66963354-DEL-177 |
| chr6-66977548-DEL-9079 | chr6-66977549-DEL-9079 |
| chr6-66999765-DEL-54 | chr6-66999766-DEL-54 |
| chr6-67010171-DEL-205 | chr6-67010172-DEL-205 |
| chr6-67080946-DEL-61 | chr6-67080981-DEL-55 |
| chr6-67093239-DEL-90 | chr6-67093240-DEL-90 |
| chr6-67131952-DEL-207 | chr6-67131947-DEL-206 |
| chr6-67149333-DEL-219 | chr6-67149327-DEL-220 |
| chr6-67157894-DEL-213 | chr6-67157892-DEL-214 |
| chr6-67159845-DEL-197 | chr6-67159846-DEL-197 |
| chr6-67182457-DEL-53 | chr6-67182444-DEL-63 |
| chr6-67184490-DEL-59 | chr6-67184498-DEL-52 |
| chr6-67198988-DEL-218 | chr6-67198989-DEL-218 |
| chr6-67204528-DEL-225 | chr6-67204528-DEL-225 |
| chr6-67216534-DEL-498 | chr6-67216534-DEL-498 |
| chr6-67224862-DEL-118 | chr6-67224863-DEL-118 |
| chr6-67229373-DEL-131 | chr6-67229374-DEL-137 |
| chr6-67229920-DEL-204 | chr6-67229921-DEL-204 |
| chr6-67237874-DEL-212 | chr6-67237875-DEL-212 |
| chr6-67268913-DEL-56 | chr6-67268914-DEL-50 |
| chr6-67271530-DEL-53 | chr6-67271531-DEL-53 |
| chr6-67297443-DEL-374 | chr6-67297443-DEL-373 |
| chr6-67302059-DEL-58 | chr6-67302060-DEL-58 |
| chr6-67306957-DEL-119 | chr6-67306958-DEL-119 |
| chr6-67326825-DEL-111 | chr6-67326826-DEL-111 |
| chr6-67335052-DEL-228 | chr6-67335053-DEL-228 |
| chr6-67366406-DEL-87 | chr6-67366442-DEL-63 |
| chr6-67397658-DEL-91 | chr6-67397659-DEL-91 |
| chr6-67409981-DEL-55 | chr6-67409982-DEL-55 |
| chr6-67434954-DEL-697 | chr6-67434955-DEL-697 |
| chr6-67440671-DEL-112 | chr6-67440672-DEL-112 |
| chr6-67447528-DEL-85 | chr6-67447518-DEL-94 |
| chr6-67471597-DEL-6484 | chr6-67471598-DEL-6484 |

| | |
|---|---|
| chr6-67479007-DEL-6683 | chr6-67479008-DEL-6683 |
| chr6-67498703-DEL-304 | chr6-67498701-DEL-305 |
| chr6-67503085-DEL-186 | chr6-67503086-DEL-186 |
| chr6-67523873-DEL-408 | chr6-67523874-DEL-408 |
| chr6-67661126-DEL-84 | chr6-67661119-DEL-80 |
| chr6-67669506-DEL-84 | chr6-67669498-DEL-82 |
| chr6-67676116-DEL-332 | chr6-67676116-DEL-332 |
| chr6-67677343-DEL-882 | chr6-67677343-DEL-882 |
| chr6-67684802-DEL-3182 | chr6-67684803-DEL-3182 |
| chr6-67709413-DEL-59 | chr6-67709414-DEL-59 |
| chr6-67751377-DEL-425 | chr6-67751378-DEL-425 |
| chr6-67778816-DEL-1008 | chr6-67778816-DEL-1008 |
| chr6-67855126-DEL-66 | chr6-67855127-DEL-66 |
| chr6-67858084-DEL-387 | chr6-67858085-DEL-387 |
| chr6-67859376-DEL-2659 | chr6-67859377-DEL-2659 |
| chr6-67862516-DEL-54 | chr6-67862517-DEL-54 |
| chr6-67866200-DEL-938 | chr6-67866201-DEL-938 |
| chr6-67893509-DEL-60 | chr6-67893510-DEL-60 |
| chr6-67957595-DEL-119 | chr6-67957596-DEL-119 |
| chr6-67963085-DEL-3274 | chr6-67963086-DEL-3274 |
| chr6-67970947-DEL-17539 | chr6-67970948-DEL-17539 |
| chr6-67995731-DEL-79 | chr6-67995732-DEL-79 |
| chr6-68034943-DEL-52 | chr6-68034943-DEL-52 |
| chr6-68038391-DEL-1086 | chr6-68038391-DEL-1086 |
| chr6-68125143-DEL-68 | chr6-68125143-DEL-76 |
| chr6-68232057-DEL-319 | chr6-68232047-DEL-315 |
| chr6-68259581-DEL-58 | chr6-68259570-DEL-68 |
| chr6-68294130-DEL-249 | chr6-68294130-DEL-249 |
| chr6-68305028-DEL-135 | chr6-68305040-DEL-123 |
| chr6-68306537-DEL-1389 | chr6-68306540-DEL-1386 |
| chr6-68321458-DEL-1892 | chr6-68321458-DEL-1892 |
| chr6-68324528-DEL-75 | chr6-68324539-DEL-79 |
| chr6-68329130-DEL-179 | chr6-68329130-DEL-179 |
| chr6-68346627-DEL-197 | chr6-68346613-DEL-197 |
| chr6-68358671-DEL-2596 | chr6-68358671-DEL-2597 |
| chr6-68366670-DEL-60 | chr6-68366671-DEL-60 |
| chr6-68375548-DEL-6459 | chr6-68375549-DEL-6459 |
| chr6-68400316-DEL-176 | chr6-68400317-DEL-175 |
| chr6-68403821-DEL-284 | chr6-68403822-DEL-284 |
| chr6-68405575-DEL-726 | chr6-68405564-DEL-729 |
| chr6-68414163-DEL-59 | chr6-68414161-DEL-61 |
| chr6-68419298-DEL-346 | chr6-68419299-DEL-346 |
| chr6-68428155-DEL-6619 | chr6-68428156-DEL-6619 |
| chr6-68449559-DEL-1030 | chr6-68449560-DEL-1030 |
| chr6-68465061-DEL-170 | chr6-68465058-DEL-168 |
| chr6-68472235-DEL-193 | chr6-68472236-DEL-193 |
| chr6-68488171-DEL-2493 | chr6-68488167-DEL-2485 |
| chr6-68518253-DEL-91 | chr6-68518254-DEL-91 |
| chr6-68529031-DEL-435 | chr6-68529032-DEL-435 |
| chr6-68537916-DEL-917 | chr6-68537917-DEL-917 |
| chr6-68539052-DEL-474 | chr6-68539053-DEL-473 |
| chr6-68542570-DEL-64 | chr6-68542570-DEL-64 |
| chr6-68550691-DEL-59 | chr6-68550687-DEL-64 |
| chr6-68566012-DEL-128 | chr6-68566022-DEL-121 |
| chr6-68575513-DEL-1135 | chr6-68575514-DEL-1135 |
| chr6-68577200-DEL-816 | chr6-68577198-DEL-817 |
| chr6-68587993-DEL-6445 | chr6-68587994-DEL-6445 |
| chr6-68595693-DEL-58 | chr6-68595691-DEL-58 |
| chr6-68615903-DEL-545 | chr6-68615899-DEL-546 |
| chr6-68618065-DEL-6328 | chr6-68618066-DEL-6327 |
| chr6-68711152-DEL-519 | chr6-68711153-DEL-519 |
| chr6-68714019-DEL-58 | chr6-68714020-DEL-58 |
| chr6-68718191-DEL-75 | chr6-68718206-DEL-62 |
| chr6-68730440-DEL-847 | chr6-68730450-DEL-838 |
| chr6-68744021-DEL-142 | chr6-68744021-DEL-142 |
| chr6-68747520-DEL-422 | chr6-68747512-DEL-430 |
| chr6-68750962-DEL-6000 | chr6-68750950-DEL-6001 |
| chr6-68769816-DEL-1044 | chr6-68769817-DEL-1044 |
| chr6-68789959-DEL-185 | chr6-68789973-DEL-177 |
| chr6-68821905-DEL-6260 | chr6-68821906-DEL-6260 |
| chr6-68829161-DEL-620 | chr6-68829162-DEL-620 |
| chr6-68836165-DEL-1941 | chr6-68836166-DEL-1941 |
| chr6-68842781-DEL-2408 | chr6-68842782-DEL-2408 |
| chr6-68848685-DEL-331 | chr6-68848686-DEL-331 |
| chr6-68858760-DEL-2189 | chr6-68858759-DEL-2190 |
| chr6-68876294-DEL-5732 | chr6-68876295-DEL-5732 |
| chr6-68882295-DEL-52 | chr6-68882296-DEL-52 |
| chr6-68893875-DEL-1557 | chr6-68893876-DEL-1557 |
| chr6-68896505-DEL-126 | chr6-68896506-DEL-126 |
| chr6-68910927-DEL-3778 | chr6-68910928-DEL-3778 |

| | |
|---|---|
| chr6-68918024-DEL-725 | chr6-68918024-DEL-725 |
| chr6-68922114-DEL-516 | chr6-68922115-DEL-516 |
| chr6-68937354-DEL-91 | chr6-68937355-DEL-91 |
| chr6-68961730-DEL-59 | chr6-68961730-DEL-56 |
| chr6-68970132-DEL-3453 | chr6-68970133-DEL-3453 |
| chr6-68984442-DEL-68 | chr6-68984443-DEL-68 |
| chr6-68988612-DEL-1722 | chr6-68988613-DEL-1722 |
| chr6-69026786-DEL-52 | chr6-69026787-DEL-52 |
| chr6-69047336-DEL-3976 | chr6-69047344-DEL-3998 |
| chr6-69057226-DEL-2471 | chr6-69057220-DEL-2474 |
| chr6-69063806-DEL-12667 | chr6-69063807-DEL-12667 |
| chr6-69077124-DEL-387 | chr6-69077125-DEL-387 |
| chr6-69102010-DEL-429 | chr6-69102012-DEL-428 |
| chr6-69106461-DEL-191 | chr6-69106462-DEL-191 |
| chr6-69121311-DEL-167 | chr6-69121312-DEL-167 |
| chr6-69269979-DEL-164 | chr6-69269979-DEL-164 |
| chr6-69270745-DEL-70 | chr6-69270746-DEL-70 |
| chr6-69273984-DEL-747 | chr6-69273984-DEL-747 |
| chr6-69304576-DEL-506 | chr6-69304577-DEL-506 |
| chr6-69327893-DEL-54 | chr6-69327893-DEL-54 |
| chr6-69338389-DEL-281 | chr6-69338390-DEL-281 |
| chr6-69344723-DEL-941 | chr6-69344686-DEL-942 |
| chr6-69363649-DEL-54 | chr6-69363650-DEL-54 |
| chr6-69428780-DEL-61 | chr6-69428781-DEL-61 |
| chr6-69429388-DEL-7310 | chr6-69429389-DEL-7310 |
| chr6-69506085-DEL-868 | chr6-69506086-DEL-868 |
| chr6-69592303-DEL-144 | chr6-69592303-DEL-144 |
| chr6-69635078-DEL-8668 | chr6-69635078-DEL-8668 |
| chr6-69651018-DEL-3788 | chr6-69651018-DEL-3788 |
| chr6-69676997-DEL-82 | chr6-69676992-DEL-86 |
| chr6-69681844-DEL-3981 | chr6-69681845-DEL-3981 |
| chr6-69702334-DEL-56 | chr6-69702334-DEL-54 |
| chr6-69706148-DEL-3200 | chr6-69706142-DEL-3198 |
| chr6-69709563-DEL-431 | chr6-69709560-DEL-431 |
| chr6-69729660-DEL-4153 | chr6-69729661-DEL-4153 |
| chr6-69748702-DEL-208 | chr6-69748702-DEL-208 |
| chr6-69779941-DEL-214 | chr6-69779942-DEL-214 |
| chr6-69806728-DEL-74 | chr6-69806729-DEL-74 |
| chr6-69815875-DEL-1390 | chr6-69815876-DEL-1390 |
| chr6-69845016-DEL-206 | chr6-69845017-DEL-206 |
| chr6-69870235-DEL-953 | chr6-69870236-DEL-953 |
| chr6-69879399-DEL-842 | chr6-69879400-DEL-842 |
| chr6-69882556-DEL-378 | chr6-69882557-DEL-378 |
| chr6-69890665-DEL-59 | chr6-69890666-DEL-59 |
| chr6-69892527-DEL-2892 | chr6-69892527-DEL-2892 |
| chr6-69906799-DEL-638 | chr6-69906797-DEL-640 |
| chr6-69910980-DEL-1210 | chr6-69910981-DEL-1210 |
| chr6-69911203-DEL-366 | chr6-69911204-DEL-366 |
| chr6-69918093-DEL-102 | chr6-69918130-DEL-52 |
| chr6-69929261-DEL-6268 | chr6-69929262-DEL-6268 |
| chr6-69951575-DEL-218 | chr6-69951576-DEL-218 |
| chr6-69961566-DEL-2287 | chr6-69961567-DEL-2287 |
| chr6-69970146-DEL-327 | chr6-69970147-DEL-327 |
| chr6-69977205-DEL-1893 | chr6-69977206-DEL-1893 |
| chr6-69979686-DEL-414 | chr6-69979687-DEL-414 |
| chr6-69985564-DEL-340 | chr6-69985565-DEL-340 |
| chr6-69997606-DEL-136 | chr6-69997568-DEL-128 |
| chr6-70006584-DEL-2171 | chr6-70006585-DEL-2171 |
| chr6-70019721-DEL-1196 | chr6-70019721-DEL-1197 |
| chr6-70024697-DEL-6107 | chr6-70024707-DEL-6137 |
| chr6-70036727-DEL-63 | chr6-70036718-DEL-72 |
| chr6-70046997-DEL-350 | chr6-70046997-DEL-350 |
| chr6-70079518-DEL-409 | chr6-70079519-DEL-409 |
| chr6-70100079-DEL-1409 | chr6-70100080-DEL-1409 |
| chr6-70106121-DEL-3395 | chr6-70106121-DEL-3395 |
| chr6-70116737-DEL-103 | chr6-70116765-DEL-60 |
| chr6-70121997-DEL-261 | chr6-70121998-DEL-261 |
| chr6-70139564-DEL-194 | chr6-70139565-DEL-194 |
| chr6-70187547-DEL-2935 | chr6-70187548-DEL-2935 |
| chr6-70202810-DEL-856 | chr6-70202811-DEL-856 |
| chr6-70207708-DEL-84 | chr6-70207709-DEL-84 |
| chr6-70221495-DEL-199 | chr6-70221496-DEL-199 |
| chr6-70360086-DEL-585 | chr6-70360082-DEL-588 |
| chr6-70423101-DEL-51 | chr6-70423102-DEL-51 |
| chr6-70491730-DEL-356 | chr6-70491731-DEL-356 |
| chr6-70562061-DEL-1826 | chr6-70562062-DEL-1826 |
| chr6-70567613-DEL-875 | chr6-70567613-DEL-875 |
| chr6-70627242-DEL-1465 | chr6-70627246-DEL-1462 |
| chr6-70686919-DEL-94 | chr6-70686920-DEL-94 |
| chr6-70731083-DEL-55 | chr6-70731084-DEL-55 |

| | |
|---|---|
| chr6-70732140-DEL-133 | chr6-70732141-DEL-133 |
| chr6-70732528-DEL-359 | chr6-70732527-DEL-361 |
| chr6-70734251-DEL-2782 | chr6-70734252-DEL-2782 |
| chr6-70772048-DEL-208 | chr6-70772049-DEL-208 |
| chr6-70777302-DEL-434 | chr6-70777303-DEL-434 |
| chr6-70795337-DEL-197 | chr6-70795338-DEL-197 |
| chr6-70801264-DEL-111 | chr6-70801265-DEL-111 |
| chr6-70813639-DEL-85 | chr6-70813640-DEL-65 |
| chr6-70815976-DEL-325 | chr6-70815976-DEL-324 |
| chr6-70839525-DEL-75 | chr6-70839537-DEL-91 |
| chr6-70950592-DEL-150 | chr6-70950593-DEL-150 |
| chr6-70952127-DEL-325 | chr6-70952128-DEL-325 |
| chr6-70955492-DEL-216 | chr6-70955493-DEL-216 |
| chr6-70956068-DEL-50 | chr6-70956063-DEL-56 |
| chr6-70961331-DEL-555 | chr6-70961332-DEL-555 |
| chr6-70979485-DEL-3650 | chr6-70979486-DEL-3650 |
| chr6-70982617-DEL-201 | chr6-70982618-DEL-201 |
| chr6-70983858-DEL-51 | chr6-70983859-DEL-51 |
| chr6-71079032-DEL-215 | chr6-71079033-DEL-215 |
| chr6-71079746-DEL-1978 | chr6-71079736-DEL-1979 |
| chr6-71087543-DEL-76 | chr6-71087544-DEL-76 |
| chr6-71101558-DEL-487 | chr6-71101547-DEL-488 |
| chr6-71135167-DEL-399 | chr6-71135168-DEL-399 |
| chr6-71142960-DEL-486 | chr6-71142961-DEL-486 |
| chr6-71146528-DEL-270 | chr6-71146529-DEL-270 |
| chr6-71194596-DEL-837 | chr6-71194597-DEL-837 |
| chr6-71199994-DEL-57 | chr6-71199989-DEL-58 |
| chr6-71230825-DEL-87 | chr6-71230802-DEL-62 |
| chr6-71238304-DEL-516 | chr6-71238305-DEL-516 |
| chr6-71272355-DEL-1668 | chr6-71272356-DEL-1668 |
| chr6-71314751-DEL-111 | chr6-71314752-DEL-111 |
| chr6-71315297-DEL-68 | chr6-71315295-DEL-71 |
| chr6-71362534-DEL-540 | chr6-71362535-DEL-540 |
| chr6-71397015-DEL-267 | chr6-71397016-DEL-267 |
| chr6-71455924-DEL-1203 | chr6-71455925-DEL-1204 |
| chr6-71456263-DEL-132 | chr6-71456264-DEL-132 |
| chr6-71466849-DEL-114 | chr6-71466859-DEL-99 |
| chr6-71470281-DEL-179 | chr6-71470282-DEL-179 |
| chr6-71485392-DEL-60 | chr6-71485393-DEL-60 |
| chr6-71498529-DEL-52 | chr6-71498530-DEL-52 |
| chr6-71502179-DEL-72 | chr6-71502141-DEL-74 |
| chr6-71524981-DEL-1356 | chr6-71524982-DEL-1356 |
| chr6-71540875-DEL-229 | chr6-71540876-DEL-229 |
| chr6-71634684-DEL-75 | chr6-71634688-DEL-72 |
| chr6-71668655-DEL-830 | chr6-71668656-DEL-830 |
| chr6-71677647-DEL-157 | chr6-71677648-DEL-157 |
| chr6-71703154-DEL-64 | chr6-71703161-DEL-64 |
| chr6-71706549-DEL-51 | chr6-71706538-DEL-52 |
| chr6-71710211-DEL-127 | chr6-71710212-DEL-127 |
| chr6-71716889-DEL-881 | chr6-71716890-DEL-881 |
| chr6-71745384-DEL-283 | chr6-71745385-DEL-283 |
| chr6-71757349-DEL-118 | chr6-71757350-DEL-118 |
| chr6-71759203-DEL-71 | chr6-71759199-DEL-75 |
| chr6-71764866-DEL-61 | chr6-71764864-DEL-59 |
| chr6-71769580-DEL-117 | chr6-71769581-DEL-117 |
| chr6-71790240-DEL-599 | chr6-71790241-DEL-598 |
| chr6-71838781-DEL-72 | chr6-71838782-DEL-68 |
| chr6-71854819-DEL-206 | chr6-71854820-DEL-206 |
| chr6-71866817-DEL-52 | chr6-71866818-DEL-52 |
| chr6-71869900-DEL-269 | chr6-71869901-DEL-269 |
| chr6-71872693-DEL-168 | chr6-71872694-DEL-168 |
| chr6-71887235-DEL-585 | chr6-71887236-DEL-585 |
| chr6-71893113-DEL-201 | chr6-71893114-DEL-201 |
| chr6-71902321-DEL-68 | chr6-71902322-DEL-68 |
| chr6-71916241-DEL-198 | chr6-71916231-DEL-200 |
| chr6-71919942-DEL-154 | chr6-71919943-DEL-154 |
| chr6-71932412-DEL-5330 | chr6-71932403-DEL-5340 |
| chr6-71945469-DEL-334 | chr6-71945489-DEL-336 |
| chr6-71962618-DEL-58 | chr6-71962614-DEL-60 |
| chr6-71969678-DEL-365 | chr6-71969679-DEL-365 |
| chr6-71983909-DEL-117 | chr6-71983919-DEL-98 |
| chr6-71989830-DEL-1947 | chr6-71989831-DEL-1947 |
| chr6-71995583-DEL-56 | chr6-71995602-DEL-78 |
| chr6-71999730-DEL-208 | chr6-71999729-DEL-206 |
| chr6-72013624-DEL-270 | chr6-72013625-DEL-270 |
| chr6-72023271-DEL-96 | chr6-72023272-DEL-90 |
| chr6-72024238-DEL-68 | chr6-72024234-DEL-73 |
| chr6-72025152-DEL-68 | chr6-72025153-DEL-68 |
| chr6-72061617-DEL-3369 | chr6-72061617-DEL-3369 |
| chr6-72085395-DEL-528 | chr6-72085396-DEL-528 |

| | |
|---|---|
| chr6-72090364-DEL-306 | chr6-72090365-DEL-306 |
| chr6-72114669-DEL-1892 | chr6-72114670-DEL-1892 |
| chr6-72127333-DEL-385 | chr6-72127334-DEL-385 |
| chr6-72134522-DEL-69 | chr6-72134523-DEL-68 |
| chr6-72150249-DEL-128 | chr6-72150250-DEL-126 |
| chr6-72155455-DEL-56 | chr6-72155456-DEL-56 |
| chr6-72160276-DEL-169 | chr6-72160277-DEL-169 |
| chr6-72162381-DEL-64 | chr6-72162382-DEL-64 |
| chr6-72165129-DEL-73 | chr6-72165130-DEL-73 |
| chr6-72171720-DEL-202 | chr6-72171721-DEL-202 |
| chr6-72200449-DEL-116 | chr6-72200449-DEL-117 |
| chr6-72234731-DEL-200 | chr6-72234732-DEL-200 |
| chr6-72274658-DEL-213 | chr6-72274659-DEL-213 |
| chr6-72283033-DEL-344 | chr6-72283071-DEL-299 |
| chr6-72291033-DEL-51 | chr6-72291034-DEL-51 |
| chr6-72293923-DEL-208 | chr6-72293924-DEL-209 |
| chr6-72315820-DEL-55 | chr6-72315821-DEL-55 |
| chr6-72357943-DEL-117 | chr6-72357944-DEL-117 |
| chr6-72364271-DEL-223 | chr6-72364260-DEL-208 |
| chr6-72373495-DEL-195 | chr6-72373496-DEL-195 |
| chr6-72381867-DEL-65 | chr6-72381870-DEL-63 |
| chr6-72385423-DEL-341 | chr6-72385424-DEL-341 |
| chr6-72403687-DEL-96 | chr6-72403702-DEL-69 |
| chr6-72422631-DEL-63 | chr6-72422632-DEL-51 |
| chr6-72436883-DEL-134 | chr6-72436889-DEL-131 |
| chr6-72438576-DEL-64 | chr6-72438583-DEL-58 |
| chr6-72443752-DEL-52 | chr6-72443745-DEL-52 |
| chr6-72448733-DEL-198 | chr6-72448734-DEL-198 |
| chr6-72472665-DEL-196 | chr6-72472666-DEL-196 |
| chr6-72482353-DEL-165 | chr6-72482351-DEL-160 |
| chr6-72506503-DEL-567 | chr6-72506504-DEL-567 |
| chr6-72508292-DEL-243 | chr6-72508294-DEL-239 |
| chr6-72515553-DEL-2979 | chr6-72515554-DEL-2979 |
| chr6-72531051-DEL-281 | chr6-72531052-DEL-281 |
| chr6-72540995-DEL-270 | chr6-72540996-DEL-270 |
| chr6-72549797-DEL-123 | chr6-72549798-DEL-123 |
| chr6-72566225-DEL-60 | chr6-72566232-DEL-54 |
| chr6-72589221-DEL-217 | chr6-72589222-DEL-217 |
| chr6-72597795-DEL-90 | chr6-72597796-DEL-90 |
| chr6-72599542-DEL-170 | chr6-72599543-DEL-170 |
| chr6-72616964-DEL-163 | chr6-72616965-DEL-163 |
| chr6-72643274-DEL-187 | chr6-72643279-DEL-182 |
| chr6-72652364-DEL-204 | chr6-72652365-DEL-204 |
| chr6-72653238-DEL-64 | chr6-72653241-DEL-65 |
| chr6-72736555-DEL-611 | chr6-72736554-DEL-611 |
| chr6-72763040-DEL-206 | chr6-72763041-DEL-206 |
| chr6-72768765-DEL-407 | chr6-72768767-DEL-406 |
| chr6-72783935-DEL-281 | chr6-72783937-DEL-280 |
| chr6-72796627-DEL-1832 | chr6-72796627-DEL-1832 |
| chr6-72797423-DEL-170 | chr6-72797424-DEL-170 |
| chr6-72804150-DEL-207 | chr6-72804151-DEL-207 |
| chr6-72806853-DEL-620 | chr6-72806854-DEL-620 |
| chr6-72809343-DEL-1226 | chr6-72809344-DEL-1226 |
| chr6-72812796-DEL-63 | chr6-72812797-DEL-63 |
| chr6-72817046-DEL-90 | chr6-72817044-DEL-91 |
| chr6-72821888-DEL-61 | chr6-72821889-DEL-61 |
| chr6-72850969-DEL-54 | chr6-72850970-DEL-54 |
| chr6-72880379-DEL-75 | chr6-72880380-DEL-75 |
| chr6-72880388-DEL-1095 | chr6-72880389-DEL-1095 |
| chr6-72883194-DEL-205 | chr6-72883195-DEL-205 |
| chr6-72898390-DEL-80 | chr6-72898391-DEL-60 |
| chr6-72913890-DEL-97 | chr6-72913891-DEL-97 |
| chr6-72920127-DEL-51 | chr6-72920111-DEL-55 |
| chr6-72940722-DEL-213 | chr6-72940717-DEL-213 |
| chr6-72943008-DEL-113 | chr6-72943007-DEL-114 |
| chr6-72943204-DEL-1647 | chr6-72943205-DEL-1647 |
| chr6-72960299-DEL-66 | chr6-72960300-DEL-66 |
| chr6-72962170-DEL-1146 | chr6-72962171-DEL-1146 |
| chr6-72971420-DEL-69 | chr6-72971431-DEL-59 |
| chr6-72979567-DEL-135 | chr6-72979557-DEL-136 |
| chr6-72996900-DEL-66 | chr6-72996895-DEL-60 |
| chr6-73005319-DEL-53 | chr6-73005298-DEL-73 |
| chr6-73041132-DEL-183 | chr6-73041133-DEL-183 |
| chr6-73082606-DEL-7826 | chr6-73082607-DEL-7826 |
| chr6-73099187-DEL-58 | chr6-73099188-DEL-58 |
| chr6-73124918-DEL-173 | chr6-73124920-DEL-172 |
| chr6-73180018-DEL-1376 | chr6-73180019-DEL-1376 |
| chr6-73184519-DEL-77 | chr6-73184520-DEL-78 |
| chr6-73189686-DEL-7100 | chr6-73189687-DEL-7100 |
| chr6-73199608-DEL-112 | chr6-73199609-DEL-110 |

| | |
|---|---|
| chr6-73205139-DEL-218 | chr6-73205140-DEL-218 |
| chr6-73206844-DEL-499 | chr6-73206845-DEL-499 |
| chr6-73212545-DEL-166 | chr6-73212546-DEL-166 |
| chr6-73218783-DEL-1647 | chr6-73218784-DEL-1647 |
| chr6-73232891-DEL-199 | chr6-73232892-DEL-199 |
| chr6-73263771-DEL-7363 | chr6-73263772-DEL-7363 |
| chr6-73279370-DEL-352 | chr6-73279369-DEL-354 |
| chr6-73290115-DEL-68 | chr6-73290116-DEL-68 |
| chr6-73321879-DEL-424 | chr6-73321879-DEL-424 |
| chr6-73348654-DEL-309 | chr6-73348655-DEL-309 |
| chr6-73355864-DEL-585 | chr6-73355865-DEL-585 |
| chr6-73368597-DEL-526 | chr6-73368598-DEL-526 |
| chr6-73371642-DEL-616 | chr6-73371643-DEL-616 |
| chr6-73379492-DEL-58 | chr6-73379493-DEL-58 |
| chr6-73382377-DEL-1353 | chr6-73382377-DEL-1353 |
| chr6-73388283-DEL-197 | chr6-73388284-DEL-197 |
| chr6-73401698-DEL-108 | chr6-73401699-DEL-108 |
| chr6-73403274-DEL-268 | chr6-73403275-DEL-268 |
| chr6-73406650-DEL-110 | chr6-73406618-DEL-146 |
| chr6-73411160-DEL-839 | chr6-73411161-DEL-839 |
| chr6-73417893-DEL-13534 | chr6-73417894-DEL-13534 |
| chr6-73466043-DEL-166 | chr6-73466044-DEL-166 |
| chr6-73470359-DEL-3123 | chr6-73470359-DEL-3124 |
| chr6-73478606-DEL-6444 | chr6-73478607-DEL-6444 |
| chr6-73508501-DEL-54 | chr6-73508482-DEL-100 |
| chr6-73522699-DEL-8941 | chr6-73522700-DEL-8941 |
| chr6-73527091-DEL-675 | chr6-73527092-DEL-675 |
| chr6-73549665-DEL-951 | chr6-73549666-DEL-949 |
| chr6-73567427-DEL-489 | chr6-73567427-DEL-489 |
| chr6-73611497-DEL-99 | chr6-73611498-DEL-99 |
| chr6-73660211-DEL-60 | chr6-73660214-DEL-57 |
| chr6-73672884-DEL-133 | chr6-73672885-DEL-133 |
| chr6-73679113-DEL-6579 | chr6-73679114-DEL-6579 |
| chr6-73702514-DEL-958 | chr6-73702515-DEL-958 |
| chr6-73720721-DEL-621 | chr6-73720722-DEL-621 |
| chr6-73736283-DEL-195 | chr6-73736290-DEL-188 |
| chr6-73750943-DEL-1030 | chr6-73750944-DEL-1030 |
| chr6-73782026-DEL-52 | chr6-73781984-DEL-50 |
| chr6-73800388-DEL-77 | chr6-73800386-DEL-80 |
| chr6-73839363-DEL-1895 | chr6-73839364-DEL-1895 |
| chr6-73847876-DEL-1107 | chr6-73847878-DEL-1106 |
| chr6-73848513-DEL-352 | chr6-73848514-DEL-352 |
| chr6-73851243-DEL-55 | chr6-73851244-DEL-56 |
| chr6-73852583-DEL-1193 | chr6-73852584-DEL-1193 |
| chr6-73859537-DEL-366 | chr6-73859538-DEL-366 |
| chr6-73869512-DEL-53 | chr6-73869557-DEL-50 |
| chr6-73883502-DEL-83 | chr6-73883511-DEL-75 |
| chr6-73888778-DEL-60 | chr6-73888779-DEL-60 |
| chr6-73915251-DEL-2880 | chr6-73915252-DEL-2880 |
| chr6-73936309-DEL-75 | chr6-73936310-DEL-75 |
| chr6-73947196-DEL-1522 | chr6-73947195-DEL-1520 |
| chr6-73962472-DEL-76 | chr6-73962491-DEL-55 |
| chr6-73965085-DEL-232 | chr6-73965086-DEL-232 |
| chr6-73976342-DEL-81 | chr6-73976343-DEL-81 |
| chr6-73982576-DEL-56 | chr6-73982577-DEL-56 |
| chr6-73999333-DEL-66 | chr6-73999346-DEL-60 |
| chr6-74022174-DEL-74 | chr6-74022175-DEL-98 |
| chr6-74049721-DEL-910 | chr6-74049718-DEL-905 |
| chr6-74060279-DEL-59 | chr6-74060283-DEL-57 |
| chr6-74062935-DEL-243 | chr6-74062957-DEL-239 |
| chr6-74064340-DEL-3598 | chr6-74064341-DEL-3598 |
| chr6-74071138-DEL-381 | chr6-74071139-DEL-381 |
| chr6-74100442-DEL-90 | chr6-74100443-DEL-65 |
| chr6-74110465-DEL-68 | chr6-74110466-DEL-68 |
| chr6-74112442-DEL-72 | chr6-74112443-DEL-72 |
| chr6-74113601-DEL-54 | chr6-74113602-DEL-54 |
| chr6-74118625-DEL-1721 | chr6-74118625-DEL-1721 |
| chr6-74144691-DEL-141 | chr6-74144692-DEL-141 |
| chr6-74213925-DEL-108 | chr6-74213926-DEL-109 |
| chr6-74215058-DEL-361 | chr6-74215059-DEL-361 |
| chr6-74215667-DEL-1211 | chr6-74215668-DEL-1211 |
| chr6-74219583-DEL-1172 | chr6-74219584-DEL-1172 |
| chr6-74220347-DEL-292 | chr6-74220348-DEL-292 |
| chr6-74254459-DEL-58 | chr6-74254456-DEL-90 |
| chr6-74278605-DEL-1824 | chr6-74278606-DEL-1824 |
| chr6-74287702-DEL-78 | chr6-74287687-DEL-94 |
| chr6-74310064-DEL-778 | chr6-74310065-DEL-773 |
| chr6-74340383-DEL-6921 | chr6-74340384-DEL-6921 |
| chr6-74340852-DEL-243 | chr6-74340831-DEL-225 |
| chr6-74355630-DEL-56 | chr6-74355631-DEL-56 |

| | |
|---|---|
| chr6-74360638-DEL-1873 | chr6-74360639-DEL-1873 |
| chr6-74424021-DEL-4865 | chr6-74424022-DEL-4865 |
| chr6-74441202-DEL-53 | chr6-74441169-DEL-87 |
| chr6-74441296-DEL-2316 | chr6-74441297-DEL-2316 |
| chr6-74451037-DEL-9856 | chr6-74451038-DEL-9856 |
| chr6-74487695-DEL-66 | chr6-74487696-DEL-66 |
| chr6-74508052-DEL-119 | chr6-74508052-DEL-118 |
| chr6-74512720-DEL-125 | chr6-74512690-DEL-119 |
| chr6-74519362-DEL-907 | chr6-74519363-DEL-907 |
| chr6-74556087-DEL-743 | chr6-74556088-DEL-743 |
| chr6-74556919-DEL-1205 | chr6-74556920-DEL-1205 |
| chr6-74562870-DEL-1414 | chr6-74562871-DEL-1414 |
| chr6-74562883-DEL-106 | chr6-74562884-DEL-106 |
| chr6-74574533-DEL-4578 | chr6-74574534-DEL-4578 |
| chr6-74607456-DEL-98 | chr6-74607475-DEL-82 |
| chr6-74635816-DEL-536 | chr6-74635817-DEL-536 |
| chr6-74639368-DEL-1384 | chr6-74639360-DEL-1399 |
| chr6-74677065-DEL-6271 | chr6-74677066-DEL-6271 |
| chr6-74709244-DEL-1161 | chr6-74709245-DEL-1161 |
| chr6-74714032-DEL-2582 | chr6-74714033-DEL-2582 |
| chr6-74714834-DEL-68 | chr6-74714835-DEL-68 |
| chr6-74743333-DEL-58 | chr6-74743334-DEL-58 |
| chr6-74763631-DEL-82 | chr6-74763632-DEL-82 |
| chr6-74764588-DEL-135 | chr6-74764589-DEL-135 |
| chr6-74798647-DEL-817 | chr6-74798658-DEL-812 |
| chr6-74859330-DEL-6164 | chr6-74859331-DEL-6164 |
| chr6-74872693-DEL-52 | chr6-74872693-DEL-52 |
| chr6-74890841-DEL-399 | chr6-74890841-DEL-400 |
| chr6-74899461-DEL-58 | chr6-74899462-DEL-60 |
| chr6-74910295-DEL-440 | chr6-74910296-DEL-440 |
| chr6-74935085-DEL-62 | chr6-74935086-DEL-62 |
| chr6-74952120-DEL-2662 | chr6-74952121-DEL-2662 |
| chr6-74964922-DEL-120 | chr6-74964923-DEL-120 |
| chr6-74979785-DEL-51 | chr6-74979750-DEL-52 |
| chr6-74983394-DEL-275 | chr6-74983417-DEL-262 |
| chr6-74988005-DEL-6217 | chr6-74988006-DEL-6217 |
| chr6-74998453-DEL-3474 | chr6-74998454-DEL-3474 |
| chr6-75004687-DEL-143 | chr6-75004688-DEL-143 |
| chr6-75007893-DEL-200 | chr6-75007894-DEL-200 |
| chr6-75012859-DEL-67 | chr6-75012859-DEL-68 |
| chr6-75035973-DEL-6421 | chr6-75035974-DEL-6421 |
| chr6-75045308-DEL-88 | chr6-75045309-DEL-88 |
| chr6-75077353-DEL-72 | chr6-75077398-DEL-50 |
| chr6-75080154-DEL-567 | chr6-75080155-DEL-567 |
| chr6-75086538-DEL-66 | chr6-75086539-DEL-66 |
| chr6-75089053-DEL-490 | chr6-75089054-DEL-489 |
| chr6-75119288-DEL-59 | chr6-75119297-DEL-55 |
| chr6-75122327-DEL-691 | chr6-75122327-DEL-691 |
| chr6-75138935-DEL-321 | chr6-75138936-DEL-321 |
| chr6-75157390-DEL-222 | chr6-75157401-DEL-212 |
| chr6-75209856-DEL-2120 | chr6-75209857-DEL-2120 |
| chr6-75210845-DEL-81 | chr6-75210864-DEL-57 |
| chr6-75250531-DEL-69 | chr6-75250532-DEL-69 |
| chr6-75251197-DEL-142 | chr6-75251198-DEL-142 |
| chr6-75262293-DEL-485 | chr6-75262294-DEL-485 |
| chr6-75277861-DEL-386 | chr6-75277866-DEL-382 |
| chr6-75302482-DEL-63 | chr6-75302483-DEL-63 |
| chr6-75304190-DEL-165 | chr6-75304221-DEL-166 |
| chr6-75306972-DEL-2333 | chr6-75306972-DEL-2333 |
| chr6-75308365-DEL-78 | chr6-75308365-DEL-78 |
| chr6-75310708-DEL-3682 | chr6-75310708-DEL-3682 |
| chr6-75346687-DEL-79 | chr6-75346688-DEL-79 |
| chr6-75353595-DEL-6036 | chr6-75353577-DEL-6028 |
| chr6-75368487-DEL-84 | chr6-75368488-DEL-84 |
| chr6-75378579-DEL-73 | chr6-75378582-DEL-72 |
| chr6-75400125-DEL-64 | chr6-75400126-DEL-64 |
| chr6-75406933-DEL-194 | chr6-75406903-DEL-195 |
| chr6-75407733-DEL-57 | chr6-75407740-DEL-51 |
| chr6-75413686-DEL-54 | chr6-75413687-DEL-54 |
| chr6-75432731-DEL-6221 | chr6-75432732-DEL-6221 |
| chr6-75466551-DEL-3307 | chr6-75466552-DEL-3307 |
| chr6-75483390-DEL-157 | chr6-75483391-DEL-157 |
| chr6-75521673-DEL-102 | chr6-75521674-DEL-102 |
| chr6-75522308-DEL-63 | chr6-75522309-DEL-63 |
| chr6-75536543-DEL-139 | chr6-75536544-DEL-139 |
| chr6-75538639-DEL-208 | chr6-75538640-DEL-208 |
| chr6-75540044-DEL-382 | chr6-75540045-DEL-383 |
| chr6-75542707-DEL-885 | chr6-75542708-DEL-885 |
| chr6-75553593-DEL-56 | chr6-75553594-DEL-56 |
| chr6-75572864-DEL-957 | chr6-75572865-DEL-957 |

| | |
|---|---|
| chr6-75575583-DEL-244 | chr6-75575584-DEL-244 |
| chr6-75581602-DEL-52 | chr6-75581594-DEL-53 |
| chr6-75599565-DEL-454 | chr6-75599566-DEL-454 |
| chr6-75624866-DEL-230 | chr6-75624866-DEL-230 |
| chr6-75663009-DEL-178 | chr6-75663010-DEL-178 |
| chr6-75673316-DEL-176 | chr6-75673317-DEL-175 |
| chr6-75681870-DEL-550 | chr6-75681871-DEL-550 |
| chr6-75693272-DEL-61 | chr6-75693276-DEL-68 |
| chr6-75702871-DEL-53 | chr6-75702872-DEL-53 |
| chr6-75722892-DEL-246 | chr6-75722893-DEL-246 |
| chr6-75744763-DEL-406 | chr6-75744764-DEL-406 |
| chr6-75745761-DEL-283 | chr6-75745760-DEL-284 |
| chr6-75756678-DEL-84 | chr6-75756679-DEL-84 |
| chr6-75759855-DEL-59 | chr6-75759856-DEL-59 |
| chr6-75808170-DEL-120 | chr6-75808171-DEL-120 |
| chr6-75810400-DEL-5653 | chr6-75810400-DEL-5649 |
| chr6-75827073-DEL-63 | chr6-75827074-DEL-63 |
| chr6-75842648-DEL-1014 | chr6-75842648-DEL-1014 |
| chr6-75846932-DEL-479 | chr6-75846933-DEL-479 |
| chr6-75883478-DEL-62 | chr6-75883479-DEL-62 |
| chr6-75888319-DEL-68 | chr6-75888319-DEL-68 |
| chr6-75916134-DEL-301 | chr6-75916135-DEL-301 |
| chr6-75933309-DEL-2451 | chr6-75933306-DEL-2452 |
| chr6-75937081-DEL-1365 | chr6-75937082-DEL-1365 |
| chr6-75942950-DEL-979 | chr6-75942951-DEL-979 |
| chr6-75949125-DEL-358 | chr6-75949126-DEL-358 |
| chr6-75954630-DEL-446 | chr6-75954631-DEL-446 |
| chr6-75958422-DEL-55 | chr6-75958421-DEL-54 |
| chr6-75977127-DEL-348 | chr6-75977128-DEL-348 |
| chr6-75999776-DEL-1291 | chr6-75999777-DEL-1291 |
| chr6-76049739-DEL-6392 | chr6-76049740-DEL-6392 |
| chr6-76049799-DEL-199 | chr6-76049800-DEL-199 |
| chr6-76059389-DEL-220 | chr6-76059389-DEL-220 |
| chr6-76059894-DEL-607 | chr6-76059895-DEL-607 |
| chr6-76067100-DEL-62 | chr6-76067101-DEL-62 |
| chr6-76068517-DEL-140 | chr6-76068518-DEL-140 |
| chr6-76075436-DEL-349 | chr6-76075437-DEL-349 |
| chr6-76075815-DEL-2169 | chr6-76075815-DEL-2169 |
| chr6-76082604-DEL-351 | chr6-76082605-DEL-351 |
| chr6-76086011-DEL-1499 | chr6-76086012-DEL-1499 |
| chr6-76087987-DEL-859 | chr6-76087988-DEL-859 |
| chr6-76100406-DEL-469 | chr6-76100406-DEL-469 |
| chr6-76115197-DEL-438 | chr6-76115198-DEL-438 |
| chr6-76133416-DEL-692 | chr6-76133417-DEL-692 |
| chr6-76139032-DEL-90 | chr6-76139033-DEL-90 |
| chr6-76147042-DEL-859 | chr6-76147054-DEL-852 |
| chr6-76150492-DEL-516 | chr6-76150493-DEL-516 |
| chr6-76158731-DEL-57 | chr6-76158732-DEL-57 |
| chr6-76176289-DEL-59 | chr6-76176266-DEL-89 |
| chr6-76178613-DEL-250 | chr6-76178614-DEL-250 |
| chr6-76182283-DEL-68 | chr6-76182284-DEL-68 |
| chr6-76186009-DEL-199 | chr6-76186010-DEL-199 |
| chr6-76189551-DEL-604 | chr6-76189552-DEL-604 |
| chr6-76195518-DEL-855 | chr6-76195475-DEL-854 |
| chr6-76208611-DEL-4853 | chr6-76208612-DEL-4853 |
| chr6-76222504-DEL-1828 | chr6-76222505-DEL-1828 |
| chr6-76268175-DEL-178 | chr6-76268176-DEL-162 |
| chr6-76299614-DEL-52 | chr6-76299615-DEL-52 |
| chr6-76309511-DEL-875 | chr6-76309512-DEL-875 |
| chr6-76342276-DEL-384 | chr6-76342277-DEL-384 |
| chr6-76356573-DEL-433 | chr6-76356574-DEL-433 |
| chr6-76388379-DEL-3436 | chr6-76388380-DEL-3436 |
| chr6-76404123-DEL-709 | chr6-76404123-DEL-709 |
| chr6-76405110-DEL-6713 | chr6-76405111-DEL-6713 |
| chr6-76408811-DEL-714 | chr6-76408812-DEL-714 |
| chr6-76412611-DEL-385 | chr6-76412612-DEL-385 |
| chr6-76420383-DEL-1268 | chr6-76420384-DEL-1268 |
| chr6-76426241-DEL-54 | chr6-76426242-DEL-54 |
| chr6-76444718-DEL-105 | chr6-76444719-DEL-105 |
| chr6-76448178-DEL-6371 | chr6-76448179-DEL-6371 |
| chr6-76456805-DEL-1033 | chr6-76456806-DEL-1033 |
| chr6-76469329-DEL-7326 | chr6-76469330-DEL-7326 |
| chr6-76470115-DEL-548 | chr6-76470116-DEL-548 |
| chr6-76471598-DEL-191 | chr6-76471599-DEL-192 |
| chr6-76472402-DEL-2688 | chr6-76472403-DEL-2688 |
| chr6-76492448-DEL-56 | chr6-76492449-DEL-56 |
| chr6-76519887-DEL-60 | chr6-76519890-DEL-58 |
| chr6-76536610-DEL-67 | chr6-76536630-DEL-63 |
| chr6-76542669-DEL-1367 | chr6-76542670-DEL-1358 |
| chr6-76542818-DEL-638 | chr6-76542819-DEL-638 |

| | |
|---|---|
| chr6-76561929-DEL-94 | chr6-76561930-DEL-94 |
| chr6-76568270-DEL-70 | chr6-76568271-DEL-70 |
| chr6-76595416-DEL-194 | chr6-76595417-DEL-194 |
| chr6-76598549-DEL-75 | chr6-76598534-DEL-65 |
| chr6-76608810-DEL-705 | chr6-76608811-DEL-705 |
| chr6-76628244-DEL-81 | chr6-76628240-DEL-80 |
| chr6-76643349-DEL-60 | chr6-76643350-DEL-60 |
| chr6-76654941-DEL-6385 | chr6-76654942-DEL-6385 |
| chr6-76675738-DEL-645 | chr6-76675739-DEL-645 |
| chr6-76697520-DEL-54 | chr6-76697521-DEL-54 |
| chr6-76737333-DEL-113 | chr6-76737338-DEL-110 |
| chr6-76769206-DEL-3388 | chr6-76769207-DEL-3388 |
| chr6-76771715-DEL-195 | chr6-76771714-DEL-196 |
| chr6-76772666-DEL-400 | chr6-76772667-DEL-400 |
| chr6-76774729-DEL-2821 | chr6-76774770-DEL-2805 |
| chr6-76780032-DEL-54 | chr6-76780033-DEL-54 |
| chr6-76793855-DEL-399 | chr6-76793856-DEL-399 |
| chr6-76824116-DEL-208 | chr6-76824117-DEL-208 |
| chr6-76824454-DEL-416 | chr6-76824455-DEL-416 |
| chr6-76888252-DEL-52 | chr6-76888253-DEL-52 |
| chr6-76921988-DEL-102 | chr6-76921999-DEL-98 |
| chr6-76927984-DEL-329 | chr6-76927986-DEL-320 |
| chr6-76956698-DEL-77 | chr6-76956699-DEL-77 |
| chr6-76995794-DEL-313 | chr6-76995795-DEL-312 |
| chr6-77138094-DEL-88 | chr6-77138095-DEL-76 |
| chr6-77183803-DEL-1123 | chr6-77183804-DEL-1123 |
| chr6-77186871-DEL-131 | chr6-77186876-DEL-127 |
| chr6-77223013-DEL-140 | chr6-77223060-DEL-94 |
| chr6-77243707-DEL-1359 | chr6-77243708-DEL-1359 |
| chr6-77343693-DEL-1120 | chr6-77343694-DEL-1118 |
| chr6-77349716-DEL-175 | chr6-77349717-DEL-191 |
| chr6-77352612-DEL-178 | chr6-77352596-DEL-206 |
| chr6-77354210-DEL-762 | chr6-77354211-DEL-762 |
| chr6-77387584-DEL-1411 | chr6-77387583-DEL-1411 |
| chr6-77397744-DEL-63 | chr6-77397739-DEL-59 |
| chr6-77413991-DEL-1206 | chr6-77413992-DEL-1206 |
| chr6-77451555-DEL-54 | chr6-77451556-DEL-54 |
| chr6-77486285-DEL-515 | chr6-77486286-DEL-515 |
| chr6-77499138-DEL-148 | chr6-77499138-DEL-149 |
| chr6-77508794-DEL-2626 | chr6-77508795-DEL-2626 |
| chr6-77526952-DEL-187 | chr6-77526953-DEL-186 |
| chr6-77593601-DEL-6840 | chr6-77593602-DEL-6840 |
| chr6-77594126-DEL-208 | chr6-77594127-DEL-208 |
| chr6-77614083-DEL-482 | chr6-77614084-DEL-478 |
| chr6-77651503-DEL-204 | chr6-77651504-DEL-204 |
| chr6-77664779-DEL-897 | chr6-77664780-DEL-896 |
| chr6-77677470-DEL-6476 | chr6-77677429-DEL-6476 |
| chr6-77684243-DEL-77 | chr6-77684244-DEL-77 |
| chr6-77708793-DEL-287 | chr6-77708794-DEL-287 |
| chr6-77727230-DEL-54 | chr6-77727230-DEL-51 |
| chr6-77750655-DEL-5796 | chr6-77750623-DEL-5797 |
| chr6-77796991-DEL-399 | chr6-77796992-DEL-399 |
| chr6-77826997-DEL-1201 | chr6-77826998-DEL-1196 |
| chr6-77874185-DEL-54 | chr6-77874187-DEL-52 |
| chr6-77877166-DEL-1544 | chr6-77877167-DEL-1544 |
| chr6-77903946-DEL-66 | chr6-77903947-DEL-62 |
| chr6-77938911-DEL-76 | chr6-77938906-DEL-92 |
| chr6-77948258-DEL-386 | chr6-77948259-DEL-386 |
| chr6-77968009-DEL-60 | chr6-77968010-DEL-60 |
| chr6-77970353-DEL-197 | chr6-77970357-DEL-194 |
| chr6-77991436-DEL-1695 | chr6-77991437-DEL-1695 |
| chr6-78003942-DEL-6507 | chr6-78003943-DEL-6507 |
| chr6-78021119-DEL-200 | chr6-78021120-DEL-200 |
| chr6-78022597-DEL-63 | chr6-78022598-DEL-55 |
| chr6-78039439-DEL-52 | chr6-78039440-DEL-52 |
| chr6-78053066-DEL-55 | chr6-78053067-DEL-55 |
| chr6-78101767-DEL-2429 | chr6-78101768-DEL-2429 |
| chr6-78103946-DEL-1463 | chr6-78103947-DEL-1463 |
| chr6-78106352-DEL-74 | chr6-78106353-DEL-74 |
| chr6-78131907-DEL-6610 | chr6-78131908-DEL-6610 |
| chr6-78209900-DEL-573 | chr6-78209901-DEL-573 |
| chr6-78235392-DEL-65 | chr6-78235405-DEL-53 |
| chr6-78240507-DEL-6213 | chr6-78240508-DEL-6213 |
| chr6-78251562-DEL-60 | chr6-78251563-DEL-60 |
| chr6-78256716-DEL-130 | chr6-78256716-DEL-131 |
| chr6-78267663-DEL-1794 | chr6-78267664-DEL-1794 |
| chr6-78270031-DEL-107 | chr6-78270032-DEL-107 |
| chr6-78273899-DEL-65 | chr6-78273922-DEL-68 |
| chr6-78298298-DEL-68 | chr6-78298299-DEL-68 |
| chr6-78305365-DEL-65 | chr6-78305361-DEL-72 |

| | |
|---|---|
| chr6-78316348-DEL-130 | chr6-78316349-DEL-130 |
| chr6-78329743-DEL-6888 | chr6-78329751-DEL-6882 |
| chr6-78330092-DEL-201 | chr6-78330093-DEL-201 |
| chr6-78365382-DEL-659 | chr6-78365387-DEL-664 |
| chr6-78370591-DEL-429 | chr6-78370592-DEL-429 |
| chr6-78372375-DEL-8311 | chr6-78372376-DEL-8311 |
| chr6-78433121-DEL-1562 | chr6-78433122-DEL-1562 |
| chr6-78449884-DEL-1882 | chr6-78449885-DEL-1882 |
| chr6-78461679-DEL-6990 | chr6-78461685-DEL-6990 |
| chr6-78469068-DEL-106 | chr6-78469069-DEL-106 |
| chr6-78475651-DEL-2643 | chr6-78475652-DEL-2643 |
| chr6-78477775-DEL-59 | chr6-78477783-DEL-50 |
| chr6-78493892-DEL-50 | chr6-78493867-DEL-50 |
| chr6-78501998-DEL-199 | chr6-78501999-DEL-199 |
| chr6-78510702-DEL-63 | chr6-78510703-DEL-63 |
| chr6-78516629-DEL-116 | chr6-78516630-DEL-116 |
| chr6-78537916-DEL-4308 | chr6-78537917-DEL-4308 |
| chr6-78548808-DEL-76 | chr6-78548801-DEL-80 |
| chr6-78555913-DEL-572 | chr6-78555956-DEL-561 |
| chr6-78561310-DEL-132 | chr6-78561311-DEL-132 |
| chr6-78572047-DEL-55 | chr6-78572052-DEL-58 |
| chr6-78572589-DEL-348 | chr6-78572586-DEL-349 |
| chr6-78579496-DEL-2597 | chr6-78579497-DEL-2599 |
| chr6-78590874-DEL-86 | chr6-78590877-DEL-84 |
| chr6-78606570-DEL-58 | chr6-78606571-DEL-58 |
| chr6-78613509-DEL-72 | chr6-78613510-DEL-72 |
| chr6-78614307-DEL-6024 | chr6-78614308-DEL-6024 |
| chr6-78619345-DEL-367 | chr6-78619346-DEL-367 |
| chr6-78634759-DEL-3590 | chr6-78634760-DEL-3590 |
| chr6-78658062-DEL-64 | chr6-78658061-DEL-76 |
| chr6-78672484-DEL-124 | chr6-78672477-DEL-132 |
| chr6-78693274-DEL-6228 | chr6-78693275-DEL-6228 |
| chr6-78705799-DEL-120 | chr6-78705827-DEL-93 |
| chr6-78711540-DEL-289 | chr6-78711541-DEL-288 |
| chr6-78797653-DEL-67 | chr6-78797662-DEL-65 |
| chr6-78825465-DEL-1731 | chr6-78825466-DEL-1731 |
| chr6-78849722-DEL-332 | chr6-78849723-DEL-332 |
| chr6-78853343-DEL-6405 | chr6-78853344-DEL-6405 |
| chr6-78867826-DEL-423 | chr6-78867827-DEL-423 |
| chr6-78868641-DEL-610 | chr6-78868642-DEL-609 |
| chr6-78882029-DEL-52 | chr6-78882028-DEL-54 |
| chr6-78896920-DEL-6606 | chr6-78896921-DEL-6606 |
| chr6-78915080-DEL-432 | chr6-78915081-DEL-432 |
| chr6-78920868-DEL-1891 | chr6-78920869-DEL-1891 |
| chr6-78953605-DEL-76 | chr6-78953606-DEL-76 |
| chr6-78983823-DEL-477 | chr6-78983823-DEL-472 |
| chr6-79008098-DEL-80 | chr6-79008099-DEL-79 |
| chr6-79033134-DEL-165 | chr6-79033135-DEL-165 |
| chr6-79058574-DEL-165 | chr6-79058575-DEL-165 |
| chr6-79104368-DEL-6449 | chr6-79104368-DEL-6451 |
| chr6-79133611-DEL-234 | chr6-79133612-DEL-234 |
| chr6-79143311-DEL-203 | chr6-79143312-DEL-203 |
| chr6-79151114-DEL-659 | chr6-79151115-DEL-659 |
| chr6-79152773-DEL-226 | chr6-79152774-DEL-226 |
| chr6-79158637-DEL-2011 | chr6-79158638-DEL-2011 |
| chr6-79182059-DEL-2596 | chr6-79182060-DEL-2596 |
| chr6-79201688-DEL-2034 | chr6-79201689-DEL-2034 |
| chr6-79206034-DEL-1270 | chr6-79206035-DEL-1270 |
| chr6-79226529-DEL-52 | chr6-79226527-DEL-71 |
| chr6-79246180-DEL-4549 | chr6-79246181-DEL-4549 |
| chr6-79251194-DEL-92 | chr6-79251194-DEL-93 |
| chr6-79254596-DEL-7071 | chr6-79254593-DEL-7075 |
| chr6-79273234-DEL-390 | chr6-79273235-DEL-390 |
| chr6-79273801-DEL-79 | chr6-79273804-DEL-81 |
| chr6-79275232-DEL-377 | chr6-79275237-DEL-373 |
| chr6-79278904-DEL-196 | chr6-79278905-DEL-196 |
| chr6-79284235-DEL-201 | chr6-79284236-DEL-201 |
| chr6-79287864-DEL-3092 | chr6-79287865-DEL-3092 |
| chr6-79314290-DEL-8196 | chr6-79314283-DEL-8196 |
| chr6-79328353-DEL-53 | chr6-79328353-DEL-54 |
| chr6-79348256-DEL-494 | chr6-79348257-DEL-494 |
| chr6-79349719-DEL-6155 | chr6-79349720-DEL-6155 |
| chr6-79395634-DEL-7116 | chr6-79395635-DEL-7116 |
| chr6-79402425-DEL-218 | chr6-79402426-DEL-218 |
| chr6-79404659-DEL-139 | chr6-79404660-DEL-139 |
| chr6-79452755-DEL-60 | chr6-79452756-DEL-60 |
| chr6-79493764-DEL-6598 | chr6-79493765-DEL-6598 |
| chr6-79501068-DEL-62 | chr6-79501075-DEL-56 |
| chr6-79519247-DEL-1657 | chr6-79519248-DEL-1657 |
| chr6-79540518-DEL-182 | chr6-79540519-DEL-182 |

| | |
|---|---|
| chr6-79549239-DEL-66 | chr6-79549246-DEL-50 |
| chr6-79553763-DEL-1786 | chr6-79553807-DEL-1767 |
| chr6-79604540-DEL-8575 | chr6-79604541-DEL-8575 |
| chr6-79621573-DEL-1726 | chr6-79621574-DEL-1726 |
| chr6-79634881-DEL-7003 | chr6-79634882-DEL-7003 |
| chr6-79646489-DEL-88 | chr6-79646490-DEL-87 |
| chr6-79675749-DEL-2849 | chr6-79675749-DEL-2847 |
| chr6-79754507-DEL-861 | chr6-79754508-DEL-861 |
| chr6-79805753-DEL-177 | chr6-79805754-DEL-177 |
| chr6-79829510-DEL-54 | chr6-79829511-DEL-54 |
| chr6-79847124-DEL-54 | chr6-79847125-DEL-54 |
| chr6-79876026-DEL-7070 | chr6-79876027-DEL-7070 |
| chr6-79941583-DEL-1889 | chr6-79941584-DEL-1889 |
| chr6-79945523-DEL-64 | chr6-79945524-DEL-64 |
| chr6-79986213-DEL-6603 | chr6-79986211-DEL-6604 |
| chr6-80019220-DEL-82 | chr6-80019221-DEL-82 |
| chr6-80076150-DEL-481 | chr6-80076151-DEL-481 |
| chr6-80123382-DEL-108 | chr6-80123383-DEL-112 |
| chr6-80132527-DEL-97 | chr6-80132528-DEL-97 |
| chr6-80161100-DEL-60 | chr6-80161105-DEL-56 |
| chr6-80188214-DEL-704 | chr6-80188215-DEL-704 |
| chr6-80201530-DEL-93 | chr6-80201531-DEL-93 |
| chr6-80210272-DEL-1457 | chr6-80210273-DEL-1457 |
| chr6-80214508-DEL-1083 | chr6-80214509-DEL-1083 |
| chr6-80219772-DEL-118 | chr6-80219773-DEL-118 |
| chr6-80223041-DEL-101 | chr6-80223042-DEL-102 |
| chr6-80244964-DEL-418 | chr6-80244977-DEL-396 |
| chr6-80269139-DEL-139 | chr6-80269140-DEL-139 |
| chr6-80276848-DEL-52 | chr6-80276849-DEL-52 |
| chr6-80309652-DEL-821 | chr6-80309653-DEL-821 |
| chr6-80335089-DEL-596 | chr6-80335090-DEL-594 |
| chr6-80355945-DEL-6472 | chr6-80355936-DEL-6474 |
| chr6-80366022-DEL-501 | chr6-80366023-DEL-501 |
| chr6-80389338-DEL-1370 | chr6-80389339-DEL-1370 |
| chr6-80391419-DEL-82 | chr6-80391420-DEL-82 |
| chr6-80400416-DEL-51 | chr6-80400416-DEL-52 |
| chr6-80407051-DEL-170 | chr6-80407050-DEL-171 |
| chr6-80425121-DEL-56 | chr6-80425120-DEL-62 |
| chr6-80426549-DEL-186 | chr6-80426550-DEL-186 |
| chr6-80475733-DEL-83 | chr6-80475734-DEL-83 |
| chr6-80485818-DEL-1139 | chr6-80485818-DEL-1140 |
| chr6-80501377-DEL-64 | chr6-80501380-DEL-62 |
| chr6-80527866-DEL-117 | chr6-80527867-DEL-119 |
| chr6-80570131-DEL-57 | chr6-80570131-DEL-55 |
| chr6-80594378-DEL-1644 | chr6-80594379-DEL-1624 |
| chr6-80611764-DEL-1206 | chr6-80611765-DEL-1206 |
| chr6-80616298-DEL-1434 | chr6-80616299-DEL-1434 |
| chr6-80627541-DEL-7180 | chr6-80627542-DEL-7180 |
| chr6-80668295-DEL-1742 | chr6-80668296-DEL-1742 |
| chr6-80692054-DEL-95 | chr6-80692019-DEL-84 |
| chr6-80708517-DEL-189 | chr6-80708516-DEL-190 |
| chr6-80751253-DEL-77 | chr6-80751254-DEL-77 |
| chr6-80825839-DEL-1392 | chr6-80825840-DEL-1392 |
| chr6-80857992-DEL-57 | chr6-80857993-DEL-57 |
| chr6-80987457-DEL-485 | chr6-80987455-DEL-485 |
| chr6-80997750-DEL-208 | chr6-80997751-DEL-208 |
| chr6-81011848-DEL-1372 | chr6-81011849-DEL-1372 |
| chr6-81012184-DEL-484 | chr6-81012185-DEL-484 |
| chr6-81018614-DEL-99 | chr6-81018623-DEL-99 |
| chr6-81057976-DEL-53 | chr6-81057977-DEL-53 |
| chr6-81083862-DEL-55 | chr6-81083843-DEL-85 |
| chr6-81099237-DEL-817 | chr6-81099238-DEL-817 |
| chr6-81105971-DEL-3557 | chr6-81105954-DEL-3567 |
| chr6-81112917-DEL-72 | chr6-81112918-DEL-72 |
| chr6-81171311-DEL-106 | chr6-81171318-DEL-100 |
| chr6-81177756-DEL-116 | chr6-81177753-DEL-104 |
| chr6-81212074-DEL-3326 | chr6-81212075-DEL-3326 |
| chr6-81217191-DEL-439 | chr6-81217192-DEL-439 |
| chr6-81231691-DEL-79 | chr6-81231692-DEL-79 |
| chr6-81248946-DEL-2369 | chr6-81248947-DEL-2369 |
| chr6-81252104-DEL-7134 | chr6-81252105-DEL-7134 |
| chr6-81260396-DEL-193 | chr6-81260397-DEL-193 |
| chr6-81268474-DEL-106 | chr6-81268475-DEL-106 |
| chr6-81277794-DEL-1433 | chr6-81277794-DEL-1433 |
| chr6-81292949-DEL-1536 | chr6-81292952-DEL-1534 |
| chr6-81313215-DEL-2730 | chr6-81313216-DEL-2730 |
| chr6-81360231-DEL-174 | chr6-81360232-DEL-174 |
| chr6-81364173-DEL-626 | chr6-81364176-DEL-624 |
| chr6-81373848-DEL-697 | chr6-81373849-DEL-697 |
| chr6-81391869-DEL-52 | chr6-81391870-DEL-52 |

| | |
|---|---|
| chr6-81401869-DEL-64 | chr6-81401870-DEL-64 |
| chr6-81467601-DEL-1992 | chr6-81467601-DEL-1992 |
| chr6-81480074-DEL-75 | chr6-81480076-DEL-80 |
| chr6-81481165-DEL-348 | chr6-81481166-DEL-348 |
| chr6-81524761-DEL-422 | chr6-81524762-DEL-422 |
| chr6-81536150-DEL-93 | chr6-81536148-DEL-88 |
| chr6-81552755-DEL-171 | chr6-81552756-DEL-171 |
| chr6-81565732-DEL-64 | chr6-81565741-DEL-56 |
| chr6-81579597-DEL-145 | chr6-81579598-DEL-145 |
| chr6-81600892-DEL-75 | chr6-81600893-DEL-75 |
| chr6-81609010-DEL-58 | chr6-81609011-DEL-58 |
| chr6-81613131-DEL-842 | chr6-81613132-DEL-842 |
| chr6-81660626-DEL-580 | chr6-81660627-DEL-580 |
| chr6-81662854-DEL-80 | chr6-81662855-DEL-80 |
| chr6-81669611-DEL-70 | chr6-81669612-DEL-76 |
| chr6-81676590-DEL-64 | chr6-81676591-DEL-68 |
| chr6-81699343-DEL-1084 | chr6-81699344-DEL-1084 |
| chr6-81705947-DEL-64 | chr6-81705950-DEL-62 |
| chr6-81749856-DEL-90 | chr6-81749857-DEL-90 |
| chr6-81775179-DEL-77 | chr6-81775180-DEL-77 |
| chr6-81806359-DEL-6232 | chr6-81806360-DEL-6232 |
| chr6-81814482-DEL-344 | chr6-81814483-DEL-344 |
| chr6-81820975-DEL-188 | chr6-81820976-DEL-188 |
| chr6-81823394-DEL-627 | chr6-81823395-DEL-627 |
| chr6-81828300-DEL-22013 | chr6-81828295-DEL-22018 |
| chr6-81831404-DEL-6591 | chr6-81831405-DEL-6591 |
| chr6-81861905-DEL-2076 | chr6-81861906-DEL-2076 |
| chr6-81887945-DEL-52 | chr6-81887946-DEL-52 |
| chr6-81889547-DEL-137 | chr6-81889548-DEL-137 |
| chr6-81901197-DEL-61 | chr6-81901198-DEL-61 |
| chr6-81924607-DEL-56 | chr6-81924608-DEL-56 |
| chr6-81930097-DEL-72 | chr6-81930098-DEL-72 |
| chr6-81931004-DEL-52 | chr6-81931005-DEL-52 |
| chr6-81938556-DEL-180 | chr6-81938557-DEL-180 |
| chr6-81947423-DEL-117 | chr6-81947424-DEL-117 |
| chr6-81954354-DEL-140 | chr6-81954329-DEL-154 |
| chr6-81959117-DEL-51 | chr6-81959106-DEL-61 |
| chr6-81969714-DEL-78 | chr6-81969718-DEL-74 |
| chr6-81970410-DEL-52 | chr6-81970411-DEL-52 |
| chr6-81972635-DEL-8232 | chr6-81972636-DEL-8232 |
| chr6-81982165-DEL-6444 | chr6-81982166-DEL-6444 |
| chr6-81989234-DEL-52 | chr6-81989234-DEL-50 |
| chr6-81996915-DEL-66 | chr6-81996905-DEL-64 |
| chr6-82000163-DEL-315 | chr6-82000164-DEL-315 |
| chr6-82011457-DEL-109 | chr6-82011458-DEL-108 |
| chr6-82014576-DEL-55 | chr6-82014577-DEL-55 |
| chr6-82022793-DEL-63 | chr6-82022841-DEL-62 |
| chr6-82064895-DEL-85 | chr6-82064896-DEL-85 |
| chr6-82077295-DEL-4643 | chr6-82077296-DEL-4643 |
| chr6-82083870-DEL-68 | chr6-82083871-DEL-64 |
| chr6-82085740-DEL-90 | chr6-82085741-DEL-74 |
| chr6-82089015-DEL-56 | chr6-82089015-DEL-57 |
| chr6-82093588-DEL-52 | chr6-82093589-DEL-52 |
| chr6-82104134-DEL-64 | chr6-82104134-DEL-64 |
| chr6-82114601-DEL-213 | chr6-82114602-DEL-213 |
| chr6-82115992-DEL-67 | chr6-82115993-DEL-67 |
| chr6-82126316-DEL-211 | chr6-82126317-DEL-211 |
| chr6-82138740-DEL-204 | chr6-82138741-DEL-204 |
| chr6-82174340-DEL-1426 | chr6-82174341-DEL-1426 |
| chr6-82186966-DEL-375 | chr6-82186967-DEL-375 |
| chr6-82189589-DEL-6243 | chr6-82189580-DEL-6244 |
| chr6-82197477-DEL-6019 | chr6-82197478-DEL-6019 |
| chr6-82205225-DEL-328 | chr6-82205226-DEL-328 |
| chr6-82224274-DEL-349 | chr6-82224278-DEL-353 |
| chr6-82229743-DEL-74 | chr6-82229764-DEL-54 |
| chr6-82232909-DEL-6771 | chr6-82232912-DEL-6772 |
| chr6-82246494-DEL-192 | chr6-82246495-DEL-192 |
| chr6-82281765-DEL-56 | chr6-82281766-DEL-56 |
| chr6-82293146-DEL-53 | chr6-82293133-DEL-54 |
| chr6-82295068-DEL-588 | chr6-82295069-DEL-588 |
| chr6-82339935-DEL-85 | chr6-82339915-DEL-78 |
| chr6-82343151-DEL-124 | chr6-82343151-DEL-124 |
| chr6-82344559-DEL-60 | chr6-82344560-DEL-80 |
| chr6-82372433-DEL-2890 | chr6-82372434-DEL-2890 |
| chr6-82377100-DEL-82 | chr6-82377101-DEL-82 |
| chr6-82385272-DEL-679 | chr6-82385295-DEL-667 |
| chr6-82398365-DEL-382 | chr6-82398366-DEL-382 |
| chr6-82408033-DEL-79 | chr6-82408018-DEL-79 |
| chr6-82412040-DEL-244 | chr6-82412039-DEL-245 |
| chr6-82412653-DEL-69 | chr6-82412654-DEL-69 |

| | |
|---|---|
| chr6-82452518-DEL-167 | chr6-82452519-DEL-167 |
| chr6-82489412-DEL-56 | chr6-82489413-DEL-56 |
| chr6-82493735-DEL-54 | chr6-82493736-DEL-54 |
| chr6-82506016-DEL-51 | chr6-82506017-DEL-51 |
| chr6-82508506-DEL-107 | chr6-82508507-DEL-107 |
| chr6-82518247-DEL-84 | chr6-82518248-DEL-84 |
| chr6-82530617-DEL-54 | chr6-82530618-DEL-54 |
| chr6-82535043-DEL-101 | chr6-82535041-DEL-102 |
| chr6-82541620-DEL-659 | chr6-82541621-DEL-659 |
| chr6-82549419-DEL-62 | chr6-82549420-DEL-62 |
| chr6-82554378-DEL-50 | chr6-82554370-DEL-62 |
| chr6-82562650-DEL-3791 | chr6-82562645-DEL-3791 |
| chr6-82583707-DEL-71 | chr6-82583707-DEL-71 |
| chr6-82599409-DEL-479 | chr6-82599410-DEL-476 |
| chr6-82602579-DEL-6679 | chr6-82602580-DEL-6679 |
| chr6-82613592-DEL-203 | chr6-82613591-DEL-204 |
| chr6-82616796-DEL-63 | chr6-82616784-DEL-63 |
| chr6-82619659-DEL-194 | chr6-82619660-DEL-194 |
| chr6-82627719-DEL-179 | chr6-82627738-DEL-178 |
| chr6-82645494-DEL-1024 | chr6-82645494-DEL-1025 |
| chr6-82655406-DEL-74 | chr6-82655407-DEL-74 |
| chr6-82661717-DEL-959 | chr6-82661717-DEL-959 |
| chr6-82664114-DEL-58 | chr6-82664115-DEL-58 |
| chr6-82685732-DEL-442 | chr6-82685733-DEL-442 |
| chr6-82695348-DEL-682 | chr6-82695349-DEL-682 |
| chr6-82709571-DEL-671 | chr6-82709572-DEL-671 |
| chr6-82721271-DEL-88 | chr6-82721278-DEL-76 |
| chr6-82725987-DEL-65 | chr6-82725988-DEL-65 |
| chr6-82755468-DEL-56 | chr6-82755469-DEL-56 |
| chr6-82781454-DEL-51 | chr6-82781438-DEL-52 |
| chr6-82782480-DEL-82 | chr6-82782500-DEL-84 |
| chr6-82821613-DEL-74 | chr6-82821614-DEL-74 |
| chr6-82825333-DEL-58 | chr6-82825334-DEL-58 |
| chr6-82831649-DEL-74 | chr6-82831650-DEL-74 |
| chr6-82848828-DEL-58 | chr6-82848821-DEL-56 |
| chr6-82881301-DEL-50 | chr6-82881286-DEL-66 |
| chr6-82910909-DEL-76 | chr6-82910910-DEL-76 |
| chr6-82913816-DEL-73 | chr6-82913815-DEL-72 |
| chr6-82919191-DEL-210 | chr6-82919192-DEL-210 |
| chr6-82937574-DEL-66 | chr6-82937575-DEL-66 |
| chr6-82951180-DEL-86 | chr6-82951181-DEL-86 |
| chr6-82962173-DEL-912 | chr6-82962177-DEL-903 |
| chr6-82965817-DEL-204 | chr6-82965818-DEL-204 |
| chr6-82979738-DEL-399 | chr6-82979739-DEL-399 |
| chr6-83002574-DEL-214 | chr6-83002565-DEL-224 |
| chr6-83029359-DEL-67 | chr6-83029357-DEL-68 |
| chr6-83089882-DEL-316 | chr6-83089883-DEL-316 |
| chr6-83103697-DEL-221 | chr6-83103713-DEL-206 |
| chr6-83107120-DEL-218 | chr6-83107121-DEL-218 |
| chr6-83158997-DEL-199 | chr6-83158998-DEL-199 |
| chr6-83178491-DEL-68 | chr6-83178492-DEL-68 |
| chr6-83193699-DEL-485 | chr6-83193700-DEL-485 |
| chr6-83203627-DEL-57 | chr6-83203628-DEL-57 |
| chr6-83206155-DEL-144 | chr6-83206156-DEL-144 |
| chr6-83218147-DEL-112 | chr6-83218148-DEL-112 |
| chr6-83223690-DEL-534 | chr6-83223690-DEL-534 |
| chr6-83231093-DEL-66 | chr6-83231094-DEL-66 |
| chr6-83268946-DEL-53 | chr6-83268947-DEL-53 |
| chr6-83281553-DEL-79 | chr6-83281569-DEL-79 |
| chr6-83301634-DEL-64 | chr6-83301649-DEL-62 |
| chr6-83366626-DEL-85 | chr6-83366627-DEL-85 |
| chr6-83400191-DEL-79 | chr6-83400192-DEL-79 |
| chr6-83414620-DEL-76 | chr6-83414621-DEL-76 |
| chr6-83441618-DEL-56 | chr6-83441594-DEL-68 |
| chr6-83496252-DEL-144 | chr6-83496253-DEL-144 |
| chr6-83498387-DEL-217 | chr6-83498388-DEL-217 |
| chr6-83508498-DEL-154 | chr6-83508498-DEL-154 |
| chr6-83522676-DEL-108 | chr6-83522677-DEL-108 |
| chr6-83524948-DEL-182 | chr6-83524949-DEL-182 |
| chr6-83525745-DEL-2727 | chr6-83525746-DEL-2727 |
| chr6-83529239-DEL-264 | chr6-83529235-DEL-260 |
| chr6-83539668-DEL-863 | chr6-83539669-DEL-863 |
| chr6-83576349-DEL-84 | chr6-83576315-DEL-85 |
| chr6-83577619-DEL-110 | chr6-83577617-DEL-72 |
| chr6-83579657-DEL-3554 | chr6-83579654-DEL-3555 |
| chr6-83596997-DEL-1456 | chr6-83596998-DEL-1456 |
| chr6-83615656-DEL-853 | chr6-83615657-DEL-853 |
| chr6-83618987-DEL-902 | chr6-83618986-DEL-904 |
| chr6-83636590-DEL-140 | chr6-83636591-DEL-140 |
| chr6-83649336-DEL-103 | chr6-83649336-DEL-103 |

| | |
|---|---|
| chr6-83650661-DEL-173 | chr6-83650662-DEL-173 |
| chr6-83655985-DEL-56 | chr6-83655986-DEL-56 |
| chr6-83657016-DEL-1213 | chr6-83657017-DEL-1213 |
| chr6-83678909-DEL-110 | chr6-83678930-DEL-87 |
| chr6-83682039-DEL-232 | chr6-83682040-DEL-232 |
| chr6-83781786-DEL-180 | chr6-83781787-DEL-180 |
| chr6-83807062-DEL-291 | chr6-83807063-DEL-291 |
| chr6-83811827-DEL-182 | chr6-83811828-DEL-182 |
| chr6-83818087-DEL-2772 | chr6-83818097-DEL-2773 |
| chr6-83822329-DEL-52 | chr6-83822332-DEL-50 |
| chr6-83829912-DEL-193 | chr6-83829913-DEL-193 |
| chr6-83851161-DEL-1274 | chr6-83851162-DEL-1274 |
| chr6-83852732-DEL-119 | chr6-83852725-DEL-119 |
| chr6-83856431-DEL-114 | chr6-83856430-DEL-115 |
| chr6-83860210-DEL-140 | chr6-83860211-DEL-140 |
| chr6-83893071-DEL-218 | chr6-83893072-DEL-218 |
| chr6-83913527-DEL-222 | chr6-83913528-DEL-222 |
| chr6-83970517-DEL-191 | chr6-83970518-DEL-191 |
| chr6-83974951-DEL-426 | chr6-83974952-DEL-426 |
| chr6-83978913-DEL-54 | chr6-83978914-DEL-54 |
| chr6-83980704-DEL-602 | chr6-83980705-DEL-602 |
| chr6-83981901-DEL-2939 | chr6-83981901-DEL-2939 |
| chr6-83990866-DEL-169 | chr6-83990867-DEL-169 |
| chr6-83992118-DEL-53 | chr6-83992119-DEL-53 |
| chr6-83994158-DEL-197 | chr6-83994159-DEL-197 |
| chr6-84005615-DEL-51 | chr6-84005616-DEL-51 |
| chr6-84016275-DEL-56 | chr6-84016262-DEL-88 |
| chr6-84025313-DEL-55 | chr6-84025314-DEL-63 |
| chr6-84036160-DEL-72 | chr6-84036161-DEL-72 |
| chr6-84042602-DEL-194 | chr6-84042603-DEL-194 |
| chr6-84046101-DEL-53 | chr6-84046101-DEL-53 |
| chr6-84050195-DEL-75 | chr6-84050196-DEL-75 |
| chr6-84053398-DEL-516 | chr6-84053399-DEL-516 |
| chr6-84095322-DEL-557 | chr6-84095323-DEL-557 |
| chr6-84136428-DEL-442 | chr6-84136429-DEL-442 |
| chr6-84157371-DEL-214 | chr6-84157372-DEL-214 |
| chr6-84179489-DEL-129 | chr6-84179488-DEL-130 |
| chr6-84256052-DEL-401 | chr6-84256053-DEL-401 |
| chr6-84258522-DEL-94 | chr6-84258523-DEL-94 |
| chr6-84260365-DEL-2817 | chr6-84260366-DEL-2817 |
| chr6-84264112-DEL-92 | chr6-84264080-DEL-92 |
| chr6-84278367-DEL-3803 | chr6-84278368-DEL-3803 |
| chr6-84336026-DEL-57 | chr6-84336027-DEL-57 |
| chr6-84345228-DEL-51 | chr6-84345229-DEL-51 |
| chr6-84365260-DEL-197 | chr6-84365259-DEL-198 |
| chr6-84369224-DEL-60 | chr6-84369225-DEL-60 |
| chr6-84374205-DEL-131 | chr6-84374206-DEL-131 |
| chr6-84422842-DEL-51 | chr6-84422843-DEL-51 |
| chr6-84429924-DEL-62 | chr6-84429925-DEL-62 |
| chr6-84433623-DEL-72 | chr6-84433624-DEL-72 |
| chr6-84435252-DEL-86 | chr6-84435297-DEL-57 |
| chr6-84487647-DEL-204 | chr6-84487648-DEL-204 |
| chr6-84498941-DEL-240 | chr6-84498942-DEL-240 |
| chr6-84630310-DEL-83 | chr6-84630298-DEL-60 |
| chr6-84653358-DEL-184 | chr6-84653359-DEL-182 |
| chr6-84658530-DEL-6677 | chr6-84658531-DEL-6677 |
| chr6-84690460-DEL-6802 | chr6-84690461-DEL-6802 |
| chr6-84736832-DEL-2860 | chr6-84736833-DEL-2860 |
| chr6-84741293-DEL-6924 | chr6-84741294-DEL-6924 |
| chr6-84741309-DEL-214 | chr6-84741310-DEL-214 |
| chr6-84892402-DEL-73 | chr6-84892403-DEL-73 |
| chr6-84903885-DEL-685 | chr6-84903877-DEL-687 |
| chr6-85009988-DEL-66 | chr6-85009989-DEL-66 |
| chr6-85053152-DEL-916 | chr6-85053153-DEL-916 |
| chr6-85079419-DEL-92 | chr6-85079420-DEL-92 |
| chr6-85105028-DEL-84 | chr6-85105014-DEL-82 |
| chr6-85129851-DEL-424 | chr6-85129852-DEL-424 |
| chr6-85139242-DEL-58 | chr6-85139243-DEL-56 |
| chr6-85215935-DEL-61 | chr6-85215934-DEL-63 |
| chr6-85216942-DEL-87 | chr6-85216943-DEL-87 |
| chr6-85273720-DEL-147 | chr6-85273705-DEL-147 |
| chr6-85275581-DEL-60 | chr6-85275582-DEL-60 |
| chr6-85359010-DEL-54 | chr6-85359011-DEL-54 |
| chr6-85361491-DEL-578 | chr6-85361492-DEL-578 |
| chr6-85365886-DEL-185 | chr6-85365889-DEL-183 |
| chr6-85376278-DEL-51 | chr6-85376279-DEL-51 |
| chr6-85388374-DEL-182 | chr6-85388385-DEL-175 |
| chr6-85396475-DEL-241 | chr6-85396476-DEL-241 |
| chr6-85454156-DEL-105 | chr6-85454176-DEL-69 |
| chr6-85472914-DEL-66 | chr6-85472915-DEL-66 |

| | |
|---|---|
| chr6-85503754-DEL-199 | chr6-85503765-DEL-188 |
| chr6-85505004-DEL-55 | chr6-85504999-DEL-53 |
| chr6-85511263-DEL-211 | chr6-85511264-DEL-211 |
| chr6-85538041-DEL-107 | chr6-85538038-DEL-111 |
| chr6-85571350-DEL-3499 | chr6-85571341-DEL-3509 |
| chr6-85573526-DEL-80 | chr6-85573529-DEL-78 |
| chr6-85603038-DEL-202 | chr6-85603039-DEL-202 |
| chr6-85614323-DEL-1964 | chr6-85614324-DEL-1964 |
| chr6-85647546-DEL-5100 | chr6-85647546-DEL-5100 |
| chr6-85809294-DEL-52 | chr6-85809294-DEL-52 |
| chr6-85820868-DEL-1519 | chr6-85820830-DEL-1547 |
| chr6-85847920-DEL-6452 | chr6-85847921-DEL-6452 |
| chr6-85857491-DEL-202 | chr6-85857492-DEL-202 |
| chr6-85867802-DEL-2612 | chr6-85867803-DEL-2612 |
| chr6-85999480-DEL-271 | chr6-85999481-DEL-271 |
| chr6-86026940-DEL-97 | chr6-86026941-DEL-96 |
| chr6-86082024-DEL-172 | chr6-86082025-DEL-172 |
| chr6-86116627-DEL-453 | chr6-86116628-DEL-453 |
| chr6-86142133-DEL-61 | chr6-86142134-DEL-61 |
| chr6-86143999-DEL-58 | chr6-86143994-DEL-56 |
| chr6-86184702-DEL-57 | chr6-86184703-DEL-57 |
| chr6-86194817-DEL-84 | chr6-86194818-DEL-84 |
| chr6-86268836-DEL-60 | chr6-86268836-DEL-60 |
| chr6-86271495-DEL-176 | chr6-86271493-DEL-176 |
| chr6-86275330-DEL-66 | chr6-86275331-DEL-66 |
| chr6-86298844-DEL-93 | chr6-86298836-DEL-94 |
| chr6-86308742-DEL-183 | chr6-86308743-DEL-183 |
| chr6-86310025-DEL-97 | chr6-86310027-DEL-96 |
| chr6-86310987-DEL-330 | chr6-86310988-DEL-330 |
| chr6-86324613-DEL-1484 | chr6-86324614-DEL-1484 |
| chr6-86326999-DEL-78 | chr6-86327020-DEL-70 |
| chr6-86357342-DEL-227 | chr6-86357343-DEL-227 |
| chr6-86386014-DEL-499 | chr6-86386015-DEL-499 |
| chr6-86397596-DEL-56 | chr6-86397595-DEL-56 |
| chr6-86422054-DEL-221 | chr6-86422013-DEL-241 |
| chr6-86468588-DEL-446 | chr6-86468588-DEL-442 |
| chr6-86482090-DEL-150 | chr6-86482091-DEL-150 |
| chr6-86484841-DEL-75 | chr6-86484842-DEL-71 |
| chr6-86490600-DEL-221 | chr6-86490601-DEL-195 |
| chr6-86496798-DEL-70 | chr6-86496796-DEL-50 |
| chr6-86505767-DEL-619 | chr6-86505771-DEL-616 |
| chr6-86536268-DEL-440 | chr6-86536269-DEL-440 |
| chr6-86551497-DEL-72 | chr6-86551498-DEL-80 |
| chr6-86553276-DEL-53 | chr6-86553254-DEL-54 |
| chr6-86566051-DEL-391 | chr6-86566052-DEL-391 |
| chr6-86569561-DEL-55 | chr6-86569562-DEL-55 |
| chr6-86579872-DEL-66 | chr6-86579879-DEL-60 |
| chr6-86620148-DEL-96 | chr6-86620154-DEL-95 |
| chr6-86629691-DEL-66 | chr6-86629692-DEL-52 |
| chr6-86634205-DEL-222 | chr6-86634206-DEL-222 |
| chr6-86663339-DEL-77 | chr6-86663335-DEL-78 |
| chr6-86693926-DEL-178 | chr6-86693926-DEL-170 |
| chr6-86725584-DEL-72 | chr6-86725585-DEL-72 |
| chr6-86743323-DEL-199 | chr6-86743324-DEL-199 |
| chr6-86760489-DEL-216 | chr6-86760490-DEL-216 |
| chr6-86762261-DEL-62 | chr6-86762262-DEL-62 |
| chr6-86772221-DEL-188 | chr6-86772222-DEL-188 |
| chr6-86782552-DEL-411 | chr6-86782553-DEL-411 |
| chr6-86785780-DEL-53 | chr6-86785779-DEL-53 |
| chr6-86787309-DEL-52 | chr6-86787280-DEL-52 |
| chr6-86813697-DEL-1149 | chr6-86813674-DEL-1182 |
| chr6-86829995-DEL-70 | chr6-86829995-DEL-70 |
| chr6-86834337-DEL-53 | chr6-86834338-DEL-53 |
| chr6-86860238-DEL-168 | chr6-86860239-DEL-168 |
| chr6-86889304-DEL-2560 | chr6-86889305-DEL-2560 |
| chr6-86899185-DEL-209 | chr6-86899186-DEL-209 |
| chr6-86919389-DEL-205 | chr6-86919390-DEL-205 |
| chr6-86994172-DEL-254 | chr6-86994173-DEL-254 |
| chr6-86994689-DEL-228 | chr6-86994690-DEL-228 |
| chr6-87010483-DEL-214 | chr6-87010457-DEL-193 |
| chr6-87028293-DEL-577 | chr6-87028294-DEL-577 |
| chr6-87043314-DEL-214 | chr6-87043315-DEL-214 |
| chr6-87052009-DEL-50 | chr6-87052010-DEL-55 |
| chr6-87063734-DEL-203 | chr6-87063734-DEL-201 |
| chr6-87080489-DEL-68 | chr6-87080489-DEL-68 |
| chr6-87096420-DEL-57 | chr6-87096421-DEL-57 |
| chr6-87099837-DEL-122 | chr6-87099838-DEL-122 |
| chr6-87146239-DEL-54 | chr6-87146240-DEL-56 |
| chr6-87159296-DEL-145 | chr6-87159300-DEL-144 |
| chr6-87171969-DEL-6415 | chr6-87171970-DEL-6415 |

| | |
|---|---|
| chr6-87184983-DEL-97 | chr6-87184994-DEL-66 |
| chr6-87187825-DEL-2280 | chr6-87187826-DEL-2280 |
| chr6-87243029-DEL-311 | chr6-87243030-DEL-311 |
| chr6-87278122-DEL-76 | chr6-87278141-DEL-76 |
| chr6-87279270-DEL-62 | chr6-87279271-DEL-62 |
| chr6-87334092-DEL-52 | chr6-87334093-DEL-52 |
| chr6-87346935-DEL-110 | chr6-87346936-DEL-110 |
| chr6-87357523-DEL-376 | chr6-87357524-DEL-376 |
| chr6-87379074-DEL-7163 | chr6-87379075-DEL-7163 |
| chr6-87398998-DEL-62 | chr6-87398998-DEL-63 |
| chr6-87422804-DEL-64 | chr6-87422805-DEL-64 |
| chr6-87433711-DEL-68 | chr6-87433714-DEL-66 |
| chr6-87450980-DEL-90 | chr6-87451005-DEL-50 |
| chr6-87458008-DEL-70 | chr6-87458009-DEL-70 |
| chr6-87467821-DEL-58 | chr6-87467822-DEL-58 |
| chr6-87498790-DEL-140 | chr6-87498790-DEL-141 |
| chr6-87525145-DEL-1444 | chr6-87525146-DEL-1444 |
| chr6-87550091-DEL-95 | chr6-87550092-DEL-95 |
| chr6-87576521-DEL-202 | chr6-87576523-DEL-202 |
| chr6-87603500-DEL-556 | chr6-87603502-DEL-557 |
| chr6-87604134-DEL-323 | chr6-87604134-DEL-323 |
| chr6-87605572-DEL-409 | chr6-87605573-DEL-409 |
| chr6-87606198-DEL-80 | chr6-87606199-DEL-112 |
| chr6-87634604-DEL-94 | chr6-87634605-DEL-94 |
| chr6-87646554-DEL-189 | chr6-87646563-DEL-181 |
| chr6-87654495-DEL-643 | chr6-87654506-DEL-645 |
| chr6-87667209-DEL-210 | chr6-87667210-DEL-210 |
| chr6-87682469-DEL-74 | chr6-87682469-DEL-74 |
| chr6-87741860-DEL-1353 | chr6-87741861-DEL-1353 |
| chr6-87768640-DEL-99 | chr6-87768648-DEL-91 |
| chr6-87772166-DEL-69 | chr6-87772167-DEL-70 |
| chr6-87798019-DEL-226 | chr6-87798020-DEL-226 |
| chr6-87809802-DEL-222 | chr6-87809805-DEL-222 |
| chr6-87811020-DEL-163 | chr6-87811019-DEL-165 |
| chr6-87846925-DEL-203 | chr6-87846926-DEL-203 |
| chr6-87870260-DEL-319 | chr6-87870261-DEL-319 |
| chr6-87882705-DEL-212 | chr6-87882706-DEL-212 |
| chr6-87905088-DEL-73 | chr6-87905098-DEL-64 |
| chr6-87925856-DEL-519 | chr6-87925857-DEL-519 |
| chr6-87934063-DEL-251 | chr6-87934064-DEL-251 |
| chr6-87938443-DEL-59 | chr6-87938444-DEL-59 |
| chr6-87948055-DEL-6440 | chr6-87948056-DEL-6440 |
| chr6-87988464-DEL-76 | chr6-87988460-DEL-93 |
| chr6-88010878-DEL-472 | chr6-88010849-DEL-486 |
| chr6-88032163-DEL-61 | chr6-88032164-DEL-57 |
| chr6-88050989-DEL-203 | chr6-88050990-DEL-203 |
| chr6-88060587-DEL-74 | chr6-88060585-DEL-52 |
| chr6-88074127-DEL-54 | chr6-88074128-DEL-54 |
| chr6-88141184-DEL-90 | chr6-88141184-DEL-90 |
| chr6-88161757-DEL-108 | chr6-88161736-DEL-107 |
| chr6-88213869-DEL-211 | chr6-88213870-DEL-211 |
| chr6-88224251-DEL-53 | chr6-88224220-DEL-53 |
| chr6-88224783-DEL-190 | chr6-88224784-DEL-190 |
| chr6-88226760-DEL-75 | chr6-88226717-DEL-105 |
| chr6-88227664-DEL-58 | chr6-88227633-DEL-50 |
| chr6-88285440-DEL-52 | chr6-88285441-DEL-52 |
| chr6-88302173-DEL-74 | chr6-88302174-DEL-74 |
| chr6-88431903-DEL-687 | chr6-88431904-DEL-687 |
| chr6-88444850-DEL-86 | chr6-88444851-DEL-86 |
| chr6-88471479-DEL-1004 | chr6-88471480-DEL-1004 |
| chr6-88498301-DEL-173 | chr6-88498302-DEL-173 |
| chr6-88505133-DEL-56 | chr6-88505134-DEL-54 |
| chr6-88529076-DEL-1680 | chr6-88529077-DEL-1680 |
| chr6-88544124-DEL-6308 | chr6-88544125-DEL-6308 |
| chr6-88554864-DEL-353 | chr6-88554863-DEL-351 |
| chr6-88558422-DEL-425 | chr6-88558423-DEL-425 |
| chr6-88562897-DEL-87 | chr6-88562890-DEL-95 |
| chr6-88578040-DEL-54 | chr6-88578041-DEL-54 |
| chr6-88590335-DEL-76 | chr6-88590336-DEL-76 |
| chr6-88720672-DEL-60 | chr6-88720673-DEL-60 |
| chr6-88722665-DEL-62 | chr6-88722666-DEL-77 |
| chr6-88767456-DEL-57 | chr6-88767463-DEL-57 |
| chr6-88794084-DEL-51 | chr6-88794085-DEL-51 |
| chr6-88797683-DEL-137 | chr6-88797684-DEL-137 |
| chr6-88808760-DEL-640 | chr6-88808763-DEL-638 |
| chr6-88857712-DEL-392 | chr6-88857713-DEL-392 |
| chr6-88972663-DEL-80 | chr6-88972659-DEL-84 |
| chr6-88974274-DEL-84 | chr6-88974275-DEL-84 |
| chr6-88982688-DEL-70 | chr6-88982688-DEL-70 |
| chr6-89034083-DEL-65 | chr6-89034084-DEL-65 |

| | |
|---|---|
| chr6-89037461-DEL-87 | chr6-89037462-DEL-87 |
| chr6-89042553-DEL-57 | chr6-89042554-DEL-57 |
| chr6-89052157-DEL-119 | chr6-89052188-DEL-93 |
| chr6-89055836-DEL-113 | chr6-89055796-DEL-153 |
| chr6-89068053-DEL-125 | chr6-89068054-DEL-125 |
| chr6-89082680-DEL-51 | chr6-89082679-DEL-52 |
| chr6-89085553-DEL-94 | chr6-89085554-DEL-94 |
| chr6-89091408-DEL-2932 | chr6-89091406-DEL-2933 |
| chr6-89092434-DEL-54 | chr6-89092435-DEL-54 |
| chr6-89102766-DEL-104 | chr6-89102767-DEL-104 |
| chr6-89109304-DEL-153 | chr6-89109305-DEL-153 |
| chr6-89110798-DEL-56 | chr6-89110799-DEL-56 |
| chr6-89129290-DEL-583 | chr6-89129291-DEL-583 |
| chr6-89177685-DEL-2547 | chr6-89177686-DEL-2547 |
| chr6-89205765-DEL-322 | chr6-89205771-DEL-325 |
| chr6-89208000-DEL-576 | chr6-89208003-DEL-572 |
| chr6-89232755-DEL-58 | chr6-89232754-DEL-60 |
| chr6-89250464-DEL-84 | chr6-89250465-DEL-84 |
| chr6-89273617-DEL-128 | chr6-89273618-DEL-128 |
| chr6-89347167-DEL-64 | chr6-89347168-DEL-64 |
| chr6-89350461-DEL-206 | chr6-89350462-DEL-206 |
| chr6-89388695-DEL-396 | chr6-89388699-DEL-372 |
| chr6-89479758-DEL-154 | chr6-89479759-DEL-158 |
| chr6-89504354-DEL-6891 | chr6-89504355-DEL-6891 |
| chr6-89554720-DEL-580 | chr6-89554721-DEL-580 |
| chr6-89568500-DEL-68 | chr6-89568501-DEL-68 |
| chr6-89644656-DEL-1051 | chr6-89644659-DEL-1052 |
| chr6-89659703-DEL-1820 | chr6-89659704-DEL-1820 |
| chr6-89662269-DEL-490 | chr6-89662270-DEL-490 |
| chr6-89666431-DEL-6992 | chr6-89666429-DEL-6994 |
| chr6-89672773-DEL-601 | chr6-89672773-DEL-601 |
| chr6-89700133-DEL-6195 | chr6-89700127-DEL-6196 |
| chr6-89711393-DEL-619 | chr6-89711394-DEL-619 |
| chr6-89774922-DEL-253 | chr6-89774921-DEL-230 |
| chr6-89805833-DEL-6202 | chr6-89805813-DEL-6216 |
| chr6-89810375-DEL-572 | chr6-89810376-DEL-572 |
| chr6-89944133-DEL-54 | chr6-89944133-DEL-86 |
| chr6-89982139-DEL-3376 | chr6-89982140-DEL-3376 |
| chr6-90004466-DEL-8395 | chr6-90004467-DEL-8395 |
| chr6-90010757-DEL-212 | chr6-90010758-DEL-212 |
| chr6-90011864-DEL-212 | chr6-90011865-DEL-212 |
| chr6-90048413-DEL-293 | chr6-90048414-DEL-293 |
| chr6-90064234-DEL-76 | chr6-90064251-DEL-56 |
| chr6-90128415-DEL-57 | chr6-90128402-DEL-71 |
| chr6-90168694-DEL-414 | chr6-90168695-DEL-414 |
| chr6-90186587-DEL-124 | chr6-90186588-DEL-124 |
| chr6-90210637-DEL-2030 | chr6-90210638-DEL-2030 |
| chr6-90222733-DEL-51 | chr6-90222734-DEL-51 |
| chr6-90231800-DEL-492 | chr6-90231797-DEL-490 |
| chr6-90244023-DEL-892 | chr6-90244024-DEL-892 |
| chr6-90250311-DEL-1981 | chr6-90250312-DEL-1981 |
| chr6-90284111-DEL-560 | chr6-90284112-DEL-559 |
| chr6-90289759-DEL-2861 | chr6-90289760-DEL-2861 |
| chr6-90391079-DEL-60 | chr6-90391080-DEL-60 |
| chr6-90414953-DEL-7109 | chr6-90414954-DEL-7109 |
| chr6-90509531-DEL-178 | chr6-90509532-DEL-178 |
| chr6-90519647-DEL-51 | chr6-90519648-DEL-51 |
| chr6-90551563-DEL-218 | chr6-90551564-DEL-218 |
| chr6-90562047-DEL-180 | chr6-90562048-DEL-180 |
| chr6-90564575-DEL-69 | chr6-90564576-DEL-69 |
| chr6-90612436-DEL-54 | chr6-90612439-DEL-52 |
| chr6-90671319-DEL-67 | chr6-90671320-DEL-67 |
| chr6-90673538-DEL-124 | chr6-90673539-DEL-124 |
| chr6-90691348-DEL-51 | chr6-90691326-DEL-73 |
| chr6-90749027-DEL-51 | chr6-90749028-DEL-51 |
| chr6-90770970-DEL-272 | chr6-90770971-DEL-272 |
| chr6-90776708-DEL-66 | chr6-90776709-DEL-66 |
| chr6-90782620-DEL-56 | chr6-90782621-DEL-56 |
| chr6-90869206-DEL-206 | chr6-90869207-DEL-206 |
| chr6-90884769-DEL-192 | chr6-90884768-DEL-192 |
| chr6-90917409-DEL-95 | chr6-90917410-DEL-95 |
| chr6-90958489-DEL-1022 | chr6-90958490-DEL-1022 |
| chr6-90992117-DEL-90 | chr6-90992118-DEL-90 |
| chr6-90995089-DEL-91 | chr6-90995092-DEL-92 |
| chr6-90998561-DEL-327 | chr6-90998562-DEL-327 |
| chr6-91004081-DEL-82 | chr6-91004082-DEL-82 |
| chr6-91026076-DEL-90 | chr6-91026077-DEL-90 |
| chr6-91038658-DEL-213 | chr6-91038659-DEL-213 |
| chr6-91095474-DEL-78 | chr6-91095477-DEL-76 |
| chr6-91097962-DEL-66 | chr6-91097963-DEL-66 |

| | |
|---|---|
| chr6-91100516-DEL-196 | chr6-91100517-DEL-196 |
| chr6-91121755-DEL-59 | chr6-91121754-DEL-60 |
| chr6-91154366-DEL-92 | chr6-91154337-DEL-71 |
| chr6-91169220-DEL-79 | chr6-91169195-DEL-55 |
| chr6-91172316-DEL-616 | chr6-91172317-DEL-616 |
| chr6-91180553-DEL-88 | chr6-91180554-DEL-88 |
| chr6-91316330-DEL-91 | chr6-91316336-DEL-60 |
| chr6-91318938-DEL-445 | chr6-91318939-DEL-445 |
| chr6-91348293-DEL-100 | chr6-91348324-DEL-82 |
| chr6-91359060-DEL-71 | chr6-91359048-DEL-72 |
| chr6-91402819-DEL-208 | chr6-91402820-DEL-208 |
| chr6-91405776-DEL-1288 | chr6-91405777-DEL-1288 |
| chr6-91412043-DEL-2946 | chr6-91412044-DEL-2946 |
| chr6-91418307-DEL-1040 | chr6-91418308-DEL-1040 |
| chr6-91422636-DEL-2158 | chr6-91422637-DEL-2158 |
| chr6-91425509-DEL-1394 | chr6-91425510-DEL-1394 |
| chr6-91472542-DEL-179 | chr6-91472543-DEL-179 |
| chr6-91493597-DEL-132 | chr6-91493637-DEL-83 |
| chr6-91565210-DEL-77 | chr6-91565211-DEL-77 |
| chr6-91616952-DEL-498 | chr6-91616953-DEL-498 |
| chr6-91618414-DEL-208 | chr6-91618415-DEL-208 |
| chr6-91622324-DEL-125 | chr6-91622325-DEL-125 |
| chr6-91634569-DEL-197 | chr6-91634570-DEL-197 |
| chr6-91641134-DEL-226 | chr6-91641135-DEL-226 |
| chr6-91648110-DEL-171 | chr6-91648111-DEL-171 |
| chr6-91664231-DEL-215 | chr6-91664232-DEL-215 |
| chr6-91777979-DEL-98 | chr6-91777999-DEL-60 |
| chr6-91802531-DEL-89 | chr6-91802527-DEL-58 |
| chr6-91812551-DEL-160 | chr6-91812560-DEL-136 |
| chr6-91844608-DEL-172 | chr6-91844610-DEL-173 |
| chr6-91851817-DEL-200 | chr6-91851818-DEL-200 |
| chr6-91868071-DEL-776 | chr6-91868072-DEL-776 |
| chr6-91891291-DEL-123 | chr6-91891290-DEL-124 |
| chr6-91928840-DEL-93 | chr6-91928841-DEL-96 |
| chr6-91934834-DEL-324 | chr6-91934835-DEL-324 |
| chr6-91973969-DEL-78 | chr6-91973976-DEL-72 |
| chr6-91996054-DEL-720 | chr6-91996055-DEL-720 |
| chr6-92002535-DEL-196 | chr6-92002536-DEL-174 |
| chr6-92009554-DEL-72 | chr6-92009555-DEL-72 |
| chr6-92033835-DEL-85 | chr6-92033836-DEL-85 |
| chr6-92074413-DEL-184 | chr6-92074414-DEL-184 |
| chr6-92092579-DEL-1293 | chr6-92092579-DEL-1293 |
| chr6-92189770-DEL-168 | chr6-92189771-DEL-168 |
| chr6-92226956-DEL-182 | chr6-92226957-DEL-181 |
| chr6-92227786-DEL-213 | chr6-92227783-DEL-214 |
| chr6-92242528-DEL-252 | chr6-92242529-DEL-252 |
| chr6-92251341-DEL-146 | chr6-92251350-DEL-156 |
| chr6-92255001-DEL-1943 | chr6-92254993-DEL-1944 |
| chr6-92292920-DEL-67 | chr6-92292929-DEL-59 |
| chr6-92294171-DEL-194 | chr6-92294172-DEL-194 |
| chr6-92298552-DEL-6796 | chr6-92298553-DEL-6796 |
| chr6-92319315-DEL-175 | chr6-92319316-DEL-175 |
| chr6-92321164-DEL-96 | chr6-92321179-DEL-82 |
| chr6-92362837-DEL-100 | chr6-92362838-DEL-100 |
| chr6-92367985-DEL-389 | chr6-92367986-DEL-389 |
| chr6-92379533-DEL-63 | chr6-92379534-DEL-87 |
| chr6-92383112-DEL-104 | chr6-92383113-DEL-104 |
| chr6-92460253-DEL-66 | chr6-92460254-DEL-66 |
| chr6-92492220-DEL-756 | chr6-92492221-DEL-756 |
| chr6-92536522-DEL-487 | chr6-92536523-DEL-487 |
| chr6-92571245-DEL-152 | chr6-92571246-DEL-152 |
| chr6-92575362-DEL-52 | chr6-92575375-DEL-58 |
| chr6-92581163-DEL-59 | chr6-92581164-DEL-59 |
| chr6-92602397-DEL-56 | chr6-92602354-DEL-100 |
| chr6-92626769-DEL-111 | chr6-92626771-DEL-109 |
| chr6-92635688-DEL-219 | chr6-92635689-DEL-219 |
| chr6-92655951-DEL-56 | chr6-92655945-DEL-53 |
| chr6-92667011-DEL-52 | chr6-92667012-DEL-52 |
| chr6-92684288-DEL-1120 | chr6-92684289-DEL-1120 |
| chr6-92700697-DEL-83 | chr6-92700698-DEL-83 |
| chr6-92722463-DEL-56 | chr6-92722475-DEL-65 |
| chr6-92731138-DEL-1105 | chr6-92731139-DEL-1105 |
| chr6-92760740-DEL-84 | chr6-92760740-DEL-84 |
| chr6-92765141-DEL-79 | chr6-92765142-DEL-79 |
| chr6-92766286-DEL-112 | chr6-92766299-DEL-112 |
| chr6-92773608-DEL-186 | chr6-92773609-DEL-187 |
| chr6-92820369-DEL-78 | chr6-92820370-DEL-78 |
| chr6-92883578-DEL-1173 | chr6-92883579-DEL-1173 |
| chr6-92916222-DEL-107 | chr6-92916223-DEL-107 |
| chr6-92922744-DEL-57 | chr6-92922745-DEL-57 |

| | |
|---|---|
| chr6-92960035-DEL-275 | chr6-92960020-DEL-322 |
| chr6-92990104-DEL-1531 | chr6-92990105-DEL-1531 |
| chr6-93031824-DEL-75 | chr6-93031825-DEL-56 |
| chr6-93033803-DEL-447 | chr6-93033804-DEL-447 |
| chr6-93045381-DEL-454 | chr6-93045380-DEL-452 |
| chr6-93094660-DEL-163 | chr6-93094668-DEL-157 |
| chr6-93128338-DEL-294 | chr6-93128339-DEL-294 |
| chr6-93128977-DEL-104 | chr6-93128991-DEL-90 |
| chr6-93151010-DEL-53 | chr6-93151013-DEL-59 |
| chr6-93155943-DEL-146 | chr6-93155944-DEL-146 |
| chr6-93158092-DEL-88 | chr6-93158099-DEL-78 |
| chr6-93181240-DEL-89 | chr6-93181236-DEL-90 |
| chr6-93189324-DEL-57 | chr6-93189324-DEL-57 |
| chr6-93197604-DEL-83 | chr6-93197594-DEL-55 |
| chr6-93207797-DEL-58 | chr6-93207796-DEL-54 |
| chr6-93227732-DEL-1610 | chr6-93227733-DEL-1610 |
| chr6-93246549-DEL-220 | chr6-93246550-DEL-220 |
| chr6-93273890-DEL-51 | chr6-93273891-DEL-54 |
| chr6-93327798-DEL-135 | chr6-93327799-DEL-135 |
| chr6-93353112-DEL-7641 | chr6-93353113-DEL-7641 |
| chr6-93353664-DEL-256 | chr6-93353665-DEL-256 |
| chr6-93361692-DEL-440 | chr6-93361693-DEL-440 |
| chr6-93376895-DEL-189 | chr6-93376896-DEL-189 |
| chr6-93388750-DEL-92 | chr6-93388746-DEL-90 |
| chr6-93408948-DEL-60 | chr6-93408937-DEL-72 |
| chr6-93420547-DEL-75 | chr6-93420548-DEL-75 |
| chr6-93442830-DEL-953 | chr6-93442831-DEL-953 |
| chr6-93458598-DEL-1479 | chr6-93458599-DEL-1479 |
| chr6-93468946-DEL-54 | chr6-93468929-DEL-66 |
| chr6-93484985-DEL-53 | chr6-93484986-DEL-53 |
| chr6-93502662-DEL-72 | chr6-93502632-DEL-118 |
| chr6-93522445-DEL-96 | chr6-93522446-DEL-96 |
| chr6-93529354-DEL-232 | chr6-93529355-DEL-232 |
| chr6-93552848-DEL-68 | chr6-93552849-DEL-68 |
| chr6-93587836-DEL-389 | chr6-93587848-DEL-407 |
| chr6-93596102-DEL-97 | chr6-93596101-DEL-90 |
| chr6-93606777-DEL-59 | chr6-93606778-DEL-59 |
| chr6-93702175-DEL-601 | chr6-93702176-DEL-600 |
| chr6-93731371-DEL-131 | chr6-93731372-DEL-131 |
| chr6-93744148-DEL-60 | chr6-93744149-DEL-52 |
| chr6-93811738-DEL-152 | chr6-93811738-DEL-152 |
| chr6-93829355-DEL-94 | chr6-93829355-DEL-78 |
| chr6-93912503-DEL-54 | chr6-93912504-DEL-55 |
| chr6-93915841-DEL-60 | chr6-93915840-DEL-60 |
| chr6-93938796-DEL-52 | chr6-93938786-DEL-62 |
| chr6-93957525-DEL-70 | chr6-93957526-DEL-70 |
| chr6-93989827-DEL-175 | chr6-93989828-DEL-174 |
| chr6-94038970-DEL-323 | chr6-94038971-DEL-323 |
| chr6-94065736-DEL-57 | chr6-94065758-DEL-52 |
| chr6-94085307-DEL-87 | chr6-94085293-DEL-116 |
| chr6-94143665-DEL-77 | chr6-94143666-DEL-77 |
| chr6-94173284-DEL-107 | chr6-94173285-DEL-107 |
| chr6-94210608-DEL-425 | chr6-94210609-DEL-425 |
| chr6-94240032-DEL-205 | chr6-94240033-DEL-205 |
| chr6-94264258-DEL-107 | chr6-94264259-DEL-107 |
| chr6-94281703-DEL-203 | chr6-94281704-DEL-203 |
| chr6-94316582-DEL-63 | chr6-94316589-DEL-57 |
| chr6-94332088-DEL-115 | chr6-94332091-DEL-120 |
| chr6-94335092-DEL-67 | chr6-94335093-DEL-67 |
| chr6-94351173-DEL-104 | chr6-94351174-DEL-104 |
| chr6-94358909-DEL-369 | chr6-94358910-DEL-369 |
| chr6-94376144-DEL-748 | chr6-94376145-DEL-748 |
| chr6-94411259-DEL-59 | chr6-94411260-DEL-59 |
| chr6-94504632-DEL-76 | chr6-94504630-DEL-77 |
| chr6-94561266-DEL-132 | chr6-94561267-DEL-132 |
| chr6-94567959-DEL-323 | chr6-94567960-DEL-323 |
| chr6-94583582-DEL-64 | chr6-94583583-DEL-64 |
| chr6-94592683-DEL-512 | chr6-94592684-DEL-512 |
| chr6-94632656-DEL-68 | chr6-94632657-DEL-68 |
| chr6-94639338-DEL-1190 | chr6-94639339-DEL-1190 |
| chr6-94685642-DEL-218 | chr6-94685643-DEL-218 |
| chr6-94719746-DEL-80 | chr6-94719747-DEL-70 |
| chr6-94721049-DEL-104 | chr6-94721042-DEL-88 |
| chr6-94721477-DEL-254 | chr6-94721478-DEL-254 |
| chr6-94759414-DEL-605 | chr6-94759415-DEL-605 |
| chr6-94779197-DEL-60 | chr6-94779197-DEL-54 |
| chr6-94794739-DEL-95 | chr6-94794734-DEL-101 |
| chr6-94804562-DEL-55 | chr6-94804562-DEL-55 |
| chr6-94808991-DEL-52 | chr6-94808992-DEL-54 |
| chr6-94827100-DEL-70 | chr6-94827101-DEL-70 |

| | |
|---|---|
| chr6-94830360-DEL-65 | chr6-94830346-DEL-69 |
| chr6-94844529-DEL-55 | chr6-94844530-DEL-55 |
| chr6-94858821-DEL-58 | chr6-94858822-DEL-58 |
| chr6-94903975-DEL-51 | chr6-94903976-DEL-51 |
| chr6-94906760-DEL-169 | chr6-94906761-DEL-169 |
| chr6-94959932-DEL-211 | chr6-94959923-DEL-212 |
| chr6-94974982-DEL-956 | chr6-94974983-DEL-956 |
| chr6-94985061-DEL-235 | chr6-94985058-DEL-236 |
| chr6-94987051-DEL-936 | chr6-94987052-DEL-936 |
| chr6-95015327-DEL-5409 | chr6-95015328-DEL-5409 |
| chr6-95023074-DEL-498 | chr6-95023075-DEL-498 |
| chr6-95024565-DEL-82 | chr6-95024566-DEL-82 |
| chr6-95037936-DEL-110 | chr6-95037931-DEL-110 |
| chr6-95072652-DEL-198 | chr6-95072653-DEL-198 |
| chr6-95074047-DEL-77 | chr6-95074043-DEL-78 |
| chr6-95117460-DEL-4686 | chr6-95117461-DEL-4686 |
| chr6-95157933-DEL-370 | chr6-95157934-DEL-370 |
| chr6-95187088-DEL-6499 | chr6-95187089-DEL-6499 |
| chr6-95207646-DEL-1201 | chr6-95207647-DEL-1201 |
| chr6-95289275-DEL-52 | chr6-95289276-DEL-52 |
| chr6-95296181-DEL-306 | chr6-95296182-DEL-306 |
| chr6-95300192-DEL-388 | chr6-95300194-DEL-387 |
| chr6-95306402-DEL-841 | chr6-95306402-DEL-841 |
| chr6-95322706-DEL-71 | chr6-95322707-DEL-64 |
| chr6-95332406-DEL-124 | chr6-95332415-DEL-134 |
| chr6-95387708-DEL-216 | chr6-95387709-DEL-216 |
| chr6-95429552-DEL-52 | chr6-95429521-DEL-74 |
| chr6-95452527-DEL-4210 | chr6-95452528-DEL-4210 |
| chr6-95519788-DEL-385 | chr6-95519829-DEL-336 |
| chr6-95565760-DEL-124 | chr6-95565761-DEL-123 |
| chr6-95580880-DEL-1615 | chr6-95580881-DEL-1615 |
| chr6-95626224-DEL-297 | chr6-95626224-DEL-298 |
| chr6-95631499-DEL-99 | chr6-95631500-DEL-99 |
| chr6-95635032-DEL-6710 | chr6-95635033-DEL-6710 |
| chr6-95673165-DEL-895 | chr6-95673166-DEL-895 |
| chr6-95698740-DEL-468 | chr6-95698772-DEL-425 |
| chr6-95736234-DEL-58 | chr6-95736234-DEL-58 |
| chr6-95750823-DEL-93 | chr6-95750824-DEL-93 |
| chr6-95756250-DEL-359 | chr6-95756251-DEL-359 |
| chr6-95771330-DEL-6441 | chr6-95771331-DEL-6441 |
| chr6-95786208-DEL-78 | chr6-95786209-DEL-78 |
| chr6-95811673-DEL-6625 | chr6-95811674-DEL-6625 |
| chr6-95822380-DEL-923 | chr6-95822381-DEL-923 |
| chr6-95845969-DEL-264 | chr6-95845970-DEL-264 |
| chr6-95865325-DEL-60 | chr6-95865326-DEL-60 |
| chr6-95873820-DEL-66 | chr6-95873804-DEL-64 |
| chr6-95899991-DEL-629 | chr6-95899992-DEL-629 |
| chr6-95908554-DEL-753 | chr6-95908555-DEL-753 |
| chr6-95925847-DEL-507 | chr6-95925848-DEL-507 |
| chr6-95928397-DEL-227 | chr6-95928398-DEL-227 |
| chr6-95928882-DEL-6271 | chr6-95928883-DEL-6271 |
| chr6-95957674-DEL-6432 | chr6-95957701-DEL-6446 |
| chr6-95968491-DEL-116 | chr6-95968492-DEL-116 |
| chr6-95979949-DEL-1675 | chr6-95979950-DEL-1675 |
| chr6-95981098-DEL-817 | chr6-95981099-DEL-817 |
| chr6-95996979-DEL-2702 | chr6-95996980-DEL-2702 |
| chr6-96040966-DEL-119 | chr6-96040967-DEL-119 |
| chr6-96060379-DEL-113 | chr6-96060380-DEL-108 |
| chr6-96070652-DEL-60 | chr6-96070653-DEL-60 |
| chr6-96098841-DEL-199 | chr6-96098842-DEL-199 |
| chr6-96104384-DEL-70 | chr6-96104385-DEL-70 |
| chr6-96117933-DEL-71 | chr6-96117934-DEL-71 |
| chr6-96128267-DEL-1837 | chr6-96128261-DEL-1842 |
| chr6-96128267-DEL-338 | chr6-96128267-DEL-338 |
| chr6-96156917-DEL-151 | chr6-96156918-DEL-151 |
| chr6-96158374-DEL-360 | chr6-96158376-DEL-342 |
| chr6-96200473-DEL-73 | chr6-96200485-DEL-72 |
| chr6-96227236-DEL-94 | chr6-96227237-DEL-94 |
| chr6-96267632-DEL-6464 | chr6-96267627-DEL-6465 |
| chr6-96274285-DEL-397 | chr6-96274286-DEL-398 |
| chr6-96279579-DEL-333 | chr6-96279581-DEL-332 |
| chr6-96293445-DEL-198 | chr6-96293446-DEL-198 |
| chr6-96327369-DEL-96 | chr6-96327408-DEL-58 |
| chr6-96340838-DEL-480 | chr6-96340839-DEL-480 |
| chr6-96346345-DEL-76 | chr6-96346346-DEL-76 |
| chr6-96364321-DEL-64 | chr6-96364292-DEL-94 |
| chr6-96425217-DEL-6627 | chr6-96425218-DEL-6627 |
| chr6-96473090-DEL-128 | chr6-96473094-DEL-146 |
| chr6-96492838-DEL-1464 | chr6-96492839-DEL-1464 |
| chr6-96529214-DEL-164 | chr6-96529215-DEL-134 |

| | |
|---|---|
| chr6-96545898-DEL-192 | chr6-96545899-DEL-192 |
| chr6-96560516-DEL-851 | chr6-96560517-DEL-851 |
| chr6-96572410-DEL-7439 | chr6-96572404-DEL-7440 |
| chr6-96572573-DEL-212 | chr6-96572574-DEL-212 |
| chr6-96607468-DEL-259 | chr6-96607469-DEL-259 |
| chr6-96651781-DEL-76 | chr6-96651786-DEL-72 |
| chr6-96655814-DEL-54 | chr6-96655813-DEL-56 |
| chr6-96674930-DEL-399 | chr6-96674931-DEL-399 |
| chr6-96680728-DEL-2917 | chr6-96680729-DEL-2917 |
| chr6-96714264-DEL-62 | chr6-96714245-DEL-54 |
| chr6-96736844-DEL-140 | chr6-96736845-DEL-140 |
| chr6-96739035-DEL-6356 | chr6-96739032-DEL-6357 |
| chr6-96751533-DEL-938 | chr6-96751534-DEL-938 |
| chr6-96756897-DEL-3015 | chr6-96756898-DEL-3015 |
| chr6-96764202-DEL-6429 | chr6-96764203-DEL-6429 |
| chr6-96779843-DEL-69 | chr6-96779840-DEL-66 |
| chr6-96815744-DEL-476 | chr6-96815745-DEL-476 |
| chr6-96837487-DEL-61 | chr6-96837488-DEL-61 |
| chr6-96853677-DEL-6648 | chr6-96853678-DEL-6648 |
| chr6-96938989-DEL-6420 | chr6-96938989-DEL-6420 |
| chr6-96946571-DEL-6484 | chr6-96946572-DEL-6484 |
| chr6-96946601-DEL-212 | chr6-96946602-DEL-212 |
| chr6-96959549-DEL-57 | chr6-96959549-DEL-57 |
| chr6-96961287-DEL-78 | chr6-96961274-DEL-51 |
| chr6-96992718-DEL-218 | chr6-96992719-DEL-218 |
| chr6-97113097-DEL-84 | chr6-97113097-DEL-84 |
| chr6-97113840-DEL-1950 | chr6-97113841-DEL-1950 |
| chr6-97136280-DEL-637 | chr6-97136281-DEL-637 |
| chr6-97197526-DEL-63 | chr6-97197523-DEL-103 |
| chr6-97214611-DEL-56 | chr6-97214612-DEL-56 |
| chr6-97236997-DEL-81 | chr6-97236998-DEL-90 |
| chr6-97240759-DEL-257 | chr6-97240760-DEL-257 |
| chr6-97291953-DEL-52 | chr6-97291954-DEL-52 |
| chr6-97349212-DEL-7199 | chr6-97349212-DEL-7199 |
| chr6-97363950-DEL-125 | chr6-97363950-DEL-125 |
| chr6-97367710-DEL-512 | chr6-97367714-DEL-507 |
| chr6-97376953-DEL-53 | chr6-97376957-DEL-50 |
| chr6-97380050-DEL-87 | chr6-97380051-DEL-88 |
| chr6-97431070-DEL-182 | chr6-97431046-DEL-176 |
| chr6-97443700-DEL-119 | chr6-97443701-DEL-119 |
| chr6-97465134-DEL-65 | chr6-97465126-DEL-70 |
| chr6-97558811-DEL-68 | chr6-97558814-DEL-64 |
| chr6-97567103-DEL-56 | chr6-97567104-DEL-56 |
| chr6-97576947-DEL-60 | chr6-97576948-DEL-60 |
| chr6-97624729-DEL-7088 | chr6-97624730-DEL-7088 |
| chr6-97639769-DEL-2704 | chr6-97639763-DEL-2701 |
| chr6-97656923-DEL-118 | chr6-97656924-DEL-151 |
| chr6-97666332-DEL-61 | chr6-97666332-DEL-59 |
| chr6-97681180-DEL-157 | chr6-97681193-DEL-145 |
| chr6-97699752-DEL-94 | chr6-97699777-DEL-68 |
| chr6-97706066-DEL-104 | chr6-97706084-DEL-102 |
| chr6-97707383-DEL-406 | chr6-97707386-DEL-403 |
| chr6-97712180-DEL-103 | chr6-97712181-DEL-103 |
| chr6-97716332-DEL-54 | chr6-97716333-DEL-54 |
| chr6-97723015-DEL-62 | chr6-97723016-DEL-62 |
| chr6-97739322-DEL-58 | chr6-97739324-DEL-56 |
| chr6-97745849-DEL-65 | chr6-97745845-DEL-70 |
| chr6-97752990-DEL-128 | chr6-97753003-DEL-118 |
| chr6-97757414-DEL-164 | chr6-97757415-DEL-164 |
| chr6-97760232-DEL-2465 | chr6-97760225-DEL-2468 |
| chr6-97787865-DEL-52 | chr6-97787865-DEL-52 |
| chr6-97789096-DEL-1888 | chr6-97789097-DEL-1888 |
| chr6-97828031-DEL-96 | chr6-97828032-DEL-93 |
| chr6-97851825-DEL-108 | chr6-97851826-DEL-114 |
| chr6-97871217-DEL-124 | chr6-97871218-DEL-164 |
| chr6-97881770-DEL-4527 | chr6-97881771-DEL-4527 |
| chr6-97897220-DEL-56 | chr6-97897221-DEL-56 |
| chr6-97959359-DEL-126 | chr6-97959358-DEL-126 |
| chr6-97960779-DEL-739 | chr6-97960780-DEL-739 |
| chr6-98016617-DEL-1149 | chr6-98016618-DEL-1149 |
| chr6-98103301-DEL-60 | chr6-98103302-DEL-60 |
| chr6-98106992-DEL-73 | chr6-98106993-DEL-73 |
| chr6-98138310-DEL-1029 | chr6-98138311-DEL-1029 |
| chr6-98142582-DEL-519 | chr6-98142583-DEL-519 |
| chr6-98144762-DEL-174 | chr6-98144763-DEL-174 |
| chr6-98202861-DEL-1893 | chr6-98202862-DEL-1893 |
| chr6-98276150-DEL-2456 | chr6-98276151-DEL-2456 |
| chr6-98350311-DEL-66 | chr6-98350312-DEL-66 |
| chr6-98356123-DEL-75 | chr6-98356124-DEL-77 |
| chr6-98437388-DEL-52 | chr6-98437387-DEL-53 |

| | |
|---|---|
| chr6-98490911-DEL-51 | chr6-98490908-DEL-54 |
| chr6-98512573-DEL-80 | chr6-98512576-DEL-78 |
| chr6-98520394-DEL-185 | chr6-98520395-DEL-185 |
| chr6-98564356-DEL-179 | chr6-98564348-DEL-176 |
| chr6-98605542-DEL-522 | chr6-98605543-DEL-522 |
| chr6-98627687-DEL-84 | chr6-98627688-DEL-84 |
| chr6-98647562-DEL-858 | chr6-98647564-DEL-857 |
| chr6-98651619-DEL-85 | chr6-98651620-DEL-85 |
| chr6-98673447-DEL-57 | chr6-98673448-DEL-57 |
| chr6-98681203-DEL-55 | chr6-98681204-DEL-55 |
| chr6-98684433-DEL-72 | chr6-98684434-DEL-72 |
| chr6-98722148-DEL-85 | chr6-98722149-DEL-85 |
| chr6-98733753-DEL-188 | chr6-98733756-DEL-176 |
| chr6-98738231-DEL-1921 | chr6-98738232-DEL-1921 |
| chr6-98794367-DEL-7177 | chr6-98794368-DEL-7177 |
| chr6-98856858-DEL-645 | chr6-98856859-DEL-595 |
| chr6-98872787-DEL-383 | chr6-98872788-DEL-383 |
| chr6-98873828-DEL-183 | chr6-98873829-DEL-183 |
| chr6-98882315-DEL-1290 | chr6-98882316-DEL-1291 |
| chr6-98883861-DEL-69 | chr6-98883862-DEL-69 |
| chr6-98890492-DEL-229 | chr6-98890493-DEL-229 |
| chr6-98919974-DEL-143 | chr6-98919975-DEL-143 |
| chr6-98944498-DEL-65 | chr6-98944510-DEL-58 |
| chr6-99063305-DEL-62 | chr6-99063306-DEL-62 |
| chr6-99075677-DEL-57 | chr6-99075678-DEL-57 |
| chr6-99102580-DEL-99 | chr6-99102581-DEL-99 |
| chr6-99140034-DEL-67 | chr6-99140035-DEL-62 |
| chr6-99339891-DEL-96 | chr6-99339892-DEL-88 |
| chr6-99348733-DEL-1259 | chr6-99348734-DEL-1259 |
| chr6-99394261-DEL-248 | chr6-99394262-DEL-248 |
| chr6-99419774-DEL-109 | chr6-99419780-DEL-84 |
| chr6-99456932-DEL-59 | chr6-99456902-DEL-60 |
| chr6-99542843-DEL-572 | chr6-99542841-DEL-566 |
| chr6-99546126-DEL-51 | chr6-99546126-DEL-51 |
| chr6-99562224-DEL-61 | chr6-99562225-DEL-61 |
| chr6-99581028-DEL-86 | chr6-99581029-DEL-86 |
| chr6-99595663-DEL-87 | chr6-99595654-DEL-99 |
| chr6-99624603-DEL-137 | chr6-99624604-DEL-137 |
| chr6-99738719-DEL-60 | chr6-99738720-DEL-60 |
| chr6-99780030-DEL-287 | chr6-99780031-DEL-287 |
| chr6-99814929-DEL-54 | chr6-99814930-DEL-54 |
| chr6-99819055-DEL-132 | chr6-99819054-DEL-133 |
| chr6-99822101-DEL-58 | chr6-99822102-DEL-56 |
| chr6-99831749-DEL-852 | chr6-99831750-DEL-852 |
| chr6-99837432-DEL-96 | chr6-99837433-DEL-96 |
| chr6-99842714-DEL-57 | chr6-99842715-DEL-56 |
| chr6-99847488-DEL-2996 | chr6-99847489-DEL-2996 |
| chr6-99854556-DEL-838 | chr6-99854557-DEL-838 |
| chr6-99885904-DEL-58 | chr6-99885905-DEL-58 |
| chr6-99892466-DEL-86 | chr6-99892467-DEL-118 |
| chr6-99912740-DEL-580 | chr6-99912741-DEL-580 |
| chr6-99940315-DEL-61 | chr6-99940316-DEL-61 |
| chr6-99942733-DEL-208 | chr6-99942734-DEL-208 |
| chr6-99945040-DEL-173 | chr6-99945041-DEL-173 |
| chr6-99958566-DEL-56 | chr6-99958566-DEL-54 |
| chr6-100005403-DEL-57 | chr6-100005403-DEL-57 |
| chr6-100011965-DEL-272 | chr6-100011966-DEL-272 |
| chr6-100043606-DEL-54 | chr6-100043607-DEL-54 |
| chr6-100083027-DEL-114 | chr6-100083028-DEL-114 |
| chr6-100102382-DEL-73 | chr6-100102377-DEL-79 |
| chr6-100113132-DEL-298 | chr6-100113133-DEL-298 |
| chr6-100134773-DEL-278 | chr6-100134774-DEL-278 |
| chr6-100141807-DEL-214 | chr6-100141808-DEL-214 |
| chr6-100162108-DEL-111 | chr6-100162109-DEL-111 |
| chr6-100178210-DEL-53 | chr6-100178211-DEL-53 |
| chr6-100201568-DEL-201 | chr6-100201569-DEL-201 |
| chr6-100239944-DEL-211 | chr6-100239945-DEL-211 |
| chr6-100248705-DEL-105 | chr6-100248750-DEL-63 |
| chr6-100250279-DEL-198 | chr6-100250278-DEL-199 |
| chr6-100289387-DEL-225 | chr6-100289388-DEL-225 |
| chr6-100292285-DEL-58 | chr6-100292248-DEL-88 |
| chr6-100308212-DEL-131 | chr6-100308213-DEL-131 |
| chr6-100310416-DEL-114 | chr6-100310417-DEL-114 |
| chr6-100326492-DEL-217 | chr6-100326492-DEL-218 |
| chr6-100328221-DEL-433 | chr6-100328222-DEL-433 |
| chr6-100347008-DEL-209 | chr6-100347009-DEL-209 |
| chr6-100348140-DEL-523 | chr6-100348139-DEL-523 |
| chr6-100352231-DEL-89 | chr6-100352238-DEL-83 |
| chr6-100386530-DEL-435 | chr6-100386531-DEL-435 |
| chr6-100398130-DEL-96 | chr6-100398143-DEL-84 |

| | |
|---|---|
| chr6-100413275-DEL-126 | chr6-100413264-DEL-149 |
| chr6-100430113-DEL-104 | chr6-100430117-DEL-101 |
| chr6-100434383-DEL-147 | chr6-100434381-DEL-145 |
| chr6-100435979-DEL-332 | chr6-100435987-DEL-349 |
| chr6-100439113-DEL-63 | chr6-100439114-DEL-63 |
| chr6-100440402-DEL-64 | chr6-100440403-DEL-63 |
| chr6-100445404-DEL-187 | chr6-100445405-DEL-187 |
| chr6-100448144-DEL-422 | chr6-100448145-DEL-422 |
| chr6-100463211-DEL-69 | chr6-100463212-DEL-69 |
| chr6-100474103-DEL-92 | chr6-100474101-DEL-93 |
| chr6-100497971-DEL-68 | chr6-100497972-DEL-68 |
| chr6-100500508-DEL-87 | chr6-100500509-DEL-87 |
| chr6-100531632-DEL-159 | chr6-100531627-DEL-165 |
| chr6-100547922-DEL-658 | chr6-100547913-DEL-648 |
| chr6-100581590-DEL-1700 | chr6-100581591-DEL-1700 |
| chr6-100596449-DEL-288 | chr6-100596450-DEL-288 |
| chr6-100613067-DEL-305 | chr6-100613068-DEL-305 |
| chr6-100614960-DEL-976 | chr6-100614960-DEL-976 |
| chr6-100621463-DEL-251 | chr6-100621464-DEL-251 |
| chr6-100631535-DEL-178 | chr6-100631536-DEL-178 |
| chr6-100633057-DEL-396 | chr6-100633058-DEL-396 |
| chr6-100638397-DEL-76 | chr6-100638398-DEL-76 |
| chr6-100690239-DEL-416 | chr6-100690240-DEL-416 |
| chr6-100701711-DEL-62 | chr6-100701680-DEL-104 |
| chr6-100713428-DEL-4637 | chr6-100713429-DEL-4637 |
| chr6-100728090-DEL-7068 | chr6-100728091-DEL-7068 |
| chr6-100765615-DEL-112 | chr6-100765650-DEL-78 |
| chr6-100781381-DEL-53 | chr6-100781382-DEL-53 |
| chr6-100789085-DEL-6315 | chr6-100789106-DEL-6315 |
| chr6-100816156-DEL-60 | chr6-100816176-DEL-66 |
| chr6-100840150-DEL-64 | chr6-100840151-DEL-56 |
| chr6-100847316-DEL-78 | chr6-100847317-DEL-78 |
| chr6-100853532-DEL-104 | chr6-100853533-DEL-104 |
| chr6-100892392-DEL-658 | chr6-100892393-DEL-658 |
| chr6-100896238-DEL-202 | chr6-100896239-DEL-202 |
| chr6-100908733-DEL-697 | chr6-100908739-DEL-693 |
| chr6-100913607-DEL-64 | chr6-100913608-DEL-52 |
| chr6-100928518-DEL-54 | chr6-100928519-DEL-54 |
| chr6-100939097-DEL-66 | chr6-100939110-DEL-63 |
| chr6-100940014-DEL-51 | chr6-100940006-DEL-54 |
| chr6-100946414-DEL-57 | chr6-100946415-DEL-57 |
| chr6-100984439-DEL-6602 | chr6-100984440-DEL-6602 |
| chr6-100991807-DEL-653 | chr6-100991808-DEL-653 |
| chr6-101000178-DEL-150 | chr6-101000171-DEL-194 |
| chr6-101003387-DEL-64 | chr6-101003382-DEL-70 |
| chr6-101004965-DEL-74 | chr6-101004966-DEL-74 |
| chr6-101017613-DEL-358 | chr6-101017614-DEL-358 |
| chr6-101056769-DEL-60 | chr6-101056770-DEL-60 |
| chr6-101093204-DEL-504 | chr6-101093205-DEL-504 |
| chr6-101105989-DEL-487 | chr6-101105991-DEL-483 |
| chr6-101115310-DEL-5424 | chr6-101115311-DEL-5424 |
| chr6-101140402-DEL-108 | chr6-101140403-DEL-108 |
| chr6-101247951-DEL-598 | chr6-101247952-DEL-598 |
| chr6-101249653-DEL-1115 | chr6-101249654-DEL-1115 |
| chr6-101254812-DEL-76 | chr6-101254813-DEL-76 |
| chr6-101299148-DEL-68 | chr6-101299149-DEL-68 |
| chr6-101299821-DEL-1225 | chr6-101299822-DEL-1225 |
| chr6-101308226-DEL-234 | chr6-101308227-DEL-234 |
| chr6-101310143-DEL-2715 | chr6-101310144-DEL-2715 |
| chr6-101321227-DEL-64 | chr6-101321224-DEL-65 |
| chr6-101325721-DEL-361 | chr6-101325724-DEL-359 |
| chr6-101334226-DEL-976 | chr6-101334227-DEL-976 |
| chr6-101342167-DEL-196 | chr6-101342168-DEL-147 |
| chr6-101355672-DEL-54 | chr6-101355674-DEL-57 |
| chr6-101415441-DEL-6222 | chr6-101415442-DEL-6222 |
| chr6-101423025-DEL-54 | chr6-101423029-DEL-51 |
| chr6-101429702-DEL-1791 | chr6-101429703-DEL-1791 |
| chr6-101432652-DEL-1688 | chr6-101432653-DEL-1688 |
| chr6-101445765-DEL-267 | chr6-101445766-DEL-267 |
| chr6-101446549-DEL-74 | chr6-101446550-DEL-74 |
| chr6-101458724-DEL-152 | chr6-101458737-DEL-132 |
| chr6-101514276-DEL-60 | chr6-101514277-DEL-60 |
| chr6-101516872-DEL-61 | chr6-101516873-DEL-61 |
| chr6-101519875-DEL-54 | chr6-101519876-DEL-52 |
| chr6-101535769-DEL-222 | chr6-101535770-DEL-222 |
| chr6-101570021-DEL-92 | chr6-101570022-DEL-84 |
| chr6-101574462-DEL-157 | chr6-101574463-DEL-157 |
| chr6-101600616-DEL-236 | chr6-101600617-DEL-236 |
| chr6-101604923-DEL-58 | chr6-101604924-DEL-56 |
| chr6-101614339-DEL-54 | chr6-101614294-DEL-85 |

| | |
|---|---|
| chr6-101620141-DEL-7292 | chr6-101620142-DEL-7292 |
| chr6-101620467-DEL-212 | chr6-101620468-DEL-212 |
| chr6-101641563-DEL-842 | chr6-101641564-DEL-842 |
| chr6-101655560-DEL-644 | chr6-101655561-DEL-644 |
| chr6-101656263-DEL-125 | chr6-101656264-DEL-125 |
| chr6-101663227-DEL-62 | chr6-101663228-DEL-68 |
| chr6-101665686-DEL-1910 | chr6-101665687-DEL-1910 |
| chr6-101711129-DEL-583 | chr6-101711130-DEL-583 |
| chr6-101779668-DEL-342 | chr6-101779669-DEL-342 |
| chr6-101796236-DEL-6197 | chr6-101796237-DEL-6197 |
| chr6-101809086-DEL-561 | chr6-101809087-DEL-561 |
| chr6-101836886-DEL-70 | chr6-101836887-DEL-70 |
| chr6-101845548-DEL-729 | chr6-101845549-DEL-729 |
| chr6-101876890-DEL-57 | chr6-101876890-DEL-62 |
| chr6-101895004-DEL-232 | chr6-101895005-DEL-232 |
| chr6-101898943-DEL-178 | chr6-101898944-DEL-178 |
| chr6-101970354-DEL-689 | chr6-101970361-DEL-687 |
| chr6-101981378-DEL-905 | chr6-101981379-DEL-905 |
| chr6-101996307-DEL-458 | chr6-101996308-DEL-458 |
| chr6-102010795-DEL-56 | chr6-102010801-DEL-51 |
| chr6-102011999-DEL-6823 | chr6-102012000-DEL-6823 |
| chr6-102024625-DEL-73 | chr6-102024625-DEL-73 |
| chr6-102028195-DEL-885 | chr6-102028196-DEL-885 |
| chr6-102030429-DEL-121 | chr6-102030439-DEL-112 |
| chr6-102032099-DEL-62 | chr6-102032100-DEL-62 |
| chr6-102032990-DEL-808 | chr6-102032990-DEL-811 |
| chr6-102060138-DEL-70 | chr6-102060138-DEL-90 |
| chr6-102061977-DEL-2089 | chr6-102061978-DEL-2089 |
| chr6-102078275-DEL-216 | chr6-102078276-DEL-216 |
| chr6-102093306-DEL-538 | chr6-102093307-DEL-538 |
| chr6-102103387-DEL-1982 | chr6-102103388-DEL-1982 |
| chr6-102109603-DEL-72 | chr6-102109604-DEL-72 |
| chr6-102111904-DEL-63 | chr6-102111919-DEL-64 |
| chr6-102114138-DEL-1697 | chr6-102114139-DEL-1697 |
| chr6-102116971-DEL-2288 | chr6-102116970-DEL-2289 |
| chr6-102123118-DEL-444 | chr6-102123119-DEL-444 |
| chr6-102135709-DEL-153 | chr6-102135710-DEL-153 |
| chr6-102152136-DEL-6317 | chr6-102152137-DEL-6317 |
| chr6-102174271-DEL-560 | chr6-102174272-DEL-560 |
| chr6-102175968-DEL-54 | chr6-102175969-DEL-54 |
| chr6-102209565-DEL-782 | chr6-102209566-DEL-782 |
| chr6-102214624-DEL-56 | chr6-102214623-DEL-58 |
| chr6-102230775-DEL-284 | chr6-102230776-DEL-284 |
| chr6-102231875-DEL-74 | chr6-102231876-DEL-74 |
| chr6-102242522-DEL-83 | chr6-102242522-DEL-83 |
| chr6-102253252-DEL-141 | chr6-102253253-DEL-141 |
| chr6-102279933-DEL-81 | chr6-102279934-DEL-81 |
| chr6-102288922-DEL-1016 | chr6-102288923-DEL-1016 |
| chr6-102307882-DEL-350 | chr6-102307882-DEL-350 |
| chr6-102324423-DEL-1116 | chr6-102324424-DEL-1116 |
| chr6-102333013-DEL-70 | chr6-102333022-DEL-62 |
| chr6-102356533-DEL-6393 | chr6-102356534-DEL-6393 |
| chr6-102385965-DEL-451 | chr6-102385965-DEL-451 |
| chr6-102392635-DEL-1793 | chr6-102392636-DEL-1793 |
| chr6-102432753-DEL-677 | chr6-102432753-DEL-678 |
| chr6-102442150-DEL-128 | chr6-102442151-DEL-128 |
| chr6-102459790-DEL-2207 | chr6-102459791-DEL-2207 |
| chr6-102474844-DEL-62 | chr6-102474857-DEL-50 |
| chr6-102480483-DEL-2169 | chr6-102480484-DEL-2169 |
| chr6-102515562-DEL-64 | chr6-102515563-DEL-64 |
| chr6-102516385-DEL-2286 | chr6-102516386-DEL-2286 |
| chr6-102526628-DEL-152 | chr6-102526629-DEL-152 |
| chr6-102549922-DEL-7179 | chr6-102549922-DEL-7179 |
| chr6-102562620-DEL-195 | chr6-102562621-DEL-195 |
| chr6-102588949-DEL-56 | chr6-102588956-DEL-50 |
| chr6-102620067-DEL-359 | chr6-102620068-DEL-359 |
| chr6-102634687-DEL-354 | chr6-102634688-DEL-354 |
| chr6-102661201-DEL-214 | chr6-102661202-DEL-214 |
| chr6-102682269-DEL-202 | chr6-102682270-DEL-202 |
| chr6-102706206-DEL-67 | chr6-102706208-DEL-66 |
| chr6-102730657-DEL-1482 | chr6-102730658-DEL-1482 |
| chr6-102741575-DEL-310 | chr6-102741575-DEL-310 |
| chr6-102743245-DEL-315 | chr6-102743246-DEL-315 |
| chr6-102744406-DEL-6497 | chr6-102744407-DEL-6497 |
| chr6-102779216-DEL-1179 | chr6-102779217-DEL-1179 |
| chr6-102795692-DEL-6350 | chr6-102795693-DEL-6350 |
| chr6-102813636-DEL-141 | chr6-102813635-DEL-142 |
| chr6-102849718-DEL-195 | chr6-102849719-DEL-195 |
| chr6-102862087-DEL-123 | chr6-102862088-DEL-123 |
| chr6-102896453-DEL-199 | chr6-102896454-DEL-199 |

| | |
|---|---|
| chr6-102897001-DEL-8596 | chr6-102897002-DEL-8596 |
| chr6-102914836-DEL-528 | chr6-102914837-DEL-528 |
| chr6-102958154-DEL-661 | chr6-102958157-DEL-662 |
| chr6-102960358-DEL-56 | chr6-102960359-DEL-56 |
| chr6-102962696-DEL-358 | chr6-102962697-DEL-358 |
| chr6-102973622-DEL-995 | chr6-102973623-DEL-995 |
| chr6-102990159-DEL-382 | chr6-102990158-DEL-383 |
| chr6-103000645-DEL-259 | chr6-103000645-DEL-257 |
| chr6-103039256-DEL-6474 | chr6-103039257-DEL-6474 |
| chr6-103046150-DEL-392 | chr6-103046151-DEL-392 |
| chr6-103059422-DEL-63 | chr6-103059423-DEL-63 |
| chr6-103090487-DEL-156 | chr6-103090488-DEL-144 |
| chr6-103093971-DEL-57 | chr6-103093978-DEL-60 |
| chr6-103109695-DEL-185 | chr6-103109696-DEL-185 |
| chr6-103119992-DEL-147 | chr6-103119960-DEL-142 |
| chr6-103126239-DEL-1469 | chr6-103126240-DEL-1469 |
| chr6-103133570-DEL-54 | chr6-103133571-DEL-54 |
| chr6-103141144-DEL-135 | chr6-103141145-DEL-135 |
| chr6-103156847-DEL-52 | chr6-103156851-DEL-52 |
| chr6-103192339-DEL-2650 | chr6-103192340-DEL-2650 |
| chr6-103200308-DEL-52 | chr6-103200309-DEL-52 |
| chr6-103224080-DEL-98 | chr6-103224081-DEL-98 |
| chr6-103232625-DEL-66 | chr6-103232626-DEL-66 |
| chr6-103236450-DEL-411 | chr6-103236451-DEL-411 |
| chr6-103255776-DEL-471 | chr6-103255782-DEL-476 |
| chr6-103259168-DEL-85 | chr6-103259169-DEL-85 |
| chr6-103291560-DEL-112 | chr6-103291561-DEL-112 |
| chr6-103301802-DEL-159 | chr6-103301803-DEL-159 |
| chr6-103319189-DEL-116 | chr6-103319142-DEL-154 |
| chr6-103331562-DEL-102 | chr6-103331563-DEL-102 |
| chr6-103333231-DEL-60 | chr6-103333235-DEL-52 |
| chr6-103344501-DEL-69 | chr6-103344502-DEL-63 |
| chr6-103346925-DEL-588 | chr6-103346925-DEL-588 |
| chr6-103350655-DEL-100 | chr6-103350627-DEL-84 |
| chr6-103383784-DEL-6292 | chr6-103383785-DEL-6292 |
| chr6-103423837-DEL-70 | chr6-103423844-DEL-69 |
| chr6-103431399-DEL-522 | chr6-103431400-DEL-521 |
| chr6-103464788-DEL-137 | chr6-103464789-DEL-137 |
| chr6-103483185-DEL-66 | chr6-103483181-DEL-55 |
| chr6-103506831-DEL-164 | chr6-103506853-DEL-135 |
| chr6-103513425-DEL-144 | chr6-103513426-DEL-144 |
| chr6-103516700-DEL-592 | chr6-103516700-DEL-592 |
| chr6-103528266-DEL-606 | chr6-103528267-DEL-606 |
| chr6-103551930-DEL-7265 | chr6-103551931-DEL-7265 |
| chr6-103584341-DEL-85 | chr6-103584342-DEL-85 |
| chr6-103624883-DEL-448 | chr6-103624884-DEL-448 |
| chr6-103625996-DEL-248 | chr6-103625995-DEL-244 |
| chr6-103630988-DEL-56 | chr6-103630971-DEL-62 |
| chr6-103657313-DEL-70 | chr6-103657321-DEL-50 |
| chr6-103676030-DEL-66 | chr6-103676031-DEL-66 |
| chr6-103684252-DEL-217 | chr6-103684253-DEL-218 |
| chr6-103696487-DEL-6951 | chr6-103696488-DEL-6951 |
| chr6-103716345-DEL-51 | chr6-103716346-DEL-51 |
| chr6-103723810-DEL-150 | chr6-103723822-DEL-126 |
| chr6-103731484-DEL-2262 | chr6-103731485-DEL-2262 |
| chr6-103741687-DEL-2105 | chr6-103741688-DEL-2105 |
| chr6-103751878-DEL-1936 | chr6-103751879-DEL-1936 |
| chr6-103758732-DEL-191 | chr6-103758734-DEL-190 |
| chr6-103761704-DEL-211 | chr6-103761705-DEL-211 |
| chr6-103769048-DEL-545 | chr6-103769049-DEL-545 |
| chr6-103808318-DEL-411 | chr6-103808319-DEL-411 |
| chr6-103813643-DEL-73 | chr6-103813644-DEL-73 |
| chr6-103831372-DEL-51 | chr6-103831385-DEL-81 |
| chr6-103875023-DEL-873 | chr6-103875024-DEL-873 |
| chr6-103967270-DEL-378 | chr6-103967271-DEL-378 |
| chr6-104041416-DEL-7120 | chr6-104041417-DEL-7120 |
| chr6-104041598-DEL-214 | chr6-104041598-DEL-214 |
| chr6-104061685-DEL-86 | chr6-104061701-DEL-73 |
| chr6-104076353-DEL-134 | chr6-104076358-DEL-126 |
| chr6-104092612-DEL-79 | chr6-104092613-DEL-79 |
| chr6-104105853-DEL-1365 | chr6-104105859-DEL-1366 |
| chr6-104163160-DEL-168 | chr6-104163161-DEL-168 |
| chr6-104170420-DEL-386 | chr6-104170421-DEL-386 |
| chr6-104182904-DEL-854 | chr6-104182905-DEL-854 |
| chr6-104185390-DEL-57 | chr6-104185391-DEL-57 |
| chr6-104193610-DEL-349 | chr6-104193611-DEL-349 |
| chr6-104195098-DEL-8851 | chr6-104195097-DEL-8852 |
| chr6-104219329-DEL-52 | chr6-104219324-DEL-56 |
| chr6-104232473-DEL-102 | chr6-104232508-DEL-62 |
| chr6-104234030-DEL-7353 | chr6-104234031-DEL-7353 |

| | |
|---|---|
| chr6-104248980-DEL-66 | chr6-104248981-DEL-66 |
| chr6-104255402-DEL-97 | chr6-104255405-DEL-81 |
| chr6-104291337-DEL-1926 | chr6-104291338-DEL-1926 |
| chr6-104300501-DEL-2141 | chr6-104300502-DEL-2141 |
| chr6-104317738-DEL-100 | chr6-104317739-DEL-100 |
| chr6-104332954-DEL-6654 | chr6-104332955-DEL-6654 |
| chr6-104356082-DEL-6478 | chr6-104356083-DEL-6478 |
| chr6-104390918-DEL-67 | chr6-104390919-DEL-66 |
| chr6-104407273-DEL-78 | chr6-104407294-DEL-70 |
| chr6-104416980-DEL-52 | chr6-104416981-DEL-52 |
| chr6-104418157-DEL-407 | chr6-104418158-DEL-407 |
| chr6-104428005-DEL-260 | chr6-104428029-DEL-245 |
| chr6-104454695-DEL-400 | chr6-104454684-DEL-401 |
| chr6-104499083-DEL-58 | chr6-104499084-DEL-58 |
| chr6-104522814-DEL-1010 | chr6-104522815-DEL-1010 |
| chr6-104613680-DEL-111 | chr6-104613690-DEL-99 |
| chr6-104636579-DEL-2725 | chr6-104636580-DEL-2725 |
| chr6-104647644-DEL-167 | chr6-104647645-DEL-166 |
| chr6-104760139-DEL-100 | chr6-104760140-DEL-100 |
| chr6-104775390-DEL-1880 | chr6-104775391-DEL-1880 |
| chr6-104776750-DEL-493 | chr6-104776751-DEL-493 |
| chr6-104795612-DEL-2380 | chr6-104795613-DEL-2380 |
| chr6-104878965-DEL-1336 | chr6-104878966-DEL-1336 |
| chr6-104956075-DEL-67 | chr6-104956073-DEL-68 |
| chr6-104987880-DEL-72 | chr6-104987881-DEL-72 |
| chr6-105007543-DEL-2736 | chr6-105007544-DEL-2736 |
| chr6-105030561-DEL-131 | chr6-105030525-DEL-170 |
| chr6-105049216-DEL-6403 | chr6-105049209-DEL-6404 |
| chr6-105157685-DEL-103 | chr6-105157686-DEL-103 |
| chr6-105171488-DEL-6403 | chr6-105171489-DEL-6403 |
| chr6-105230457-DEL-4292 | chr6-105230458-DEL-4292 |
| chr6-105242996-DEL-972 | chr6-105242997-DEL-972 |
| chr6-105310663-DEL-227 | chr6-105310662-DEL-223 |
| chr6-105315794-DEL-112 | chr6-105315772-DEL-113 |
| chr6-105383561-DEL-7539 | chr6-105383562-DEL-7539 |
| chr6-105387658-DEL-1442 | chr6-105387657-DEL-1444 |
| chr6-105390510-DEL-54 | chr6-105390503-DEL-62 |
| chr6-105403631-DEL-317 | chr6-105403629-DEL-318 |
| chr6-105404138-DEL-200 | chr6-105404128-DEL-201 |
| chr6-105421451-DEL-73 | chr6-105421452-DEL-73 |
| chr6-105463875-DEL-63 | chr6-105463873-DEL-66 |
| chr6-105465685-DEL-4166 | chr6-105465686-DEL-4166 |
| chr6-105481769-DEL-924 | chr6-105481773-DEL-919 |
| chr6-105486459-DEL-417 | chr6-105486458-DEL-418 |
| chr6-105487861-DEL-2608 | chr6-105487864-DEL-2605 |
| chr6-105503826-DEL-304 | chr6-105503829-DEL-302 |
| chr6-105540734-DEL-102 | chr6-105540735-DEL-102 |
| chr6-105552895-DEL-365 | chr6-105552896-DEL-365 |
| chr6-105560833-DEL-9562 | chr6-105560834-DEL-9562 |
| chr6-105566738-DEL-6240 | chr6-105566728-DEL-6241 |
| chr6-105578853-DEL-56 | chr6-105578854-DEL-56 |
| chr6-105580267-DEL-57 | chr6-105580268-DEL-57 |
| chr6-105592141-DEL-143 | chr6-105592140-DEL-108 |
| chr6-105597119-DEL-6366 | chr6-105597120-DEL-6366 |
| chr6-105639911-DEL-6901 | chr6-105639912-DEL-6901 |
| chr6-105675104-DEL-2531 | chr6-105675105-DEL-2531 |
| chr6-105713340-DEL-552 | chr6-105713341-DEL-551 |
| chr6-105731854-DEL-54 | chr6-105731855-DEL-50 |
| chr6-105759694-DEL-56 | chr6-105759697-DEL-54 |
| chr6-105765888-DEL-52 | chr6-105765877-DEL-57 |
| chr6-105786213-DEL-127 | chr6-105786214-DEL-127 |
| chr6-105787027-DEL-533 | chr6-105787028-DEL-533 |
| chr6-105791303-DEL-660 | chr6-105791307-DEL-662 |
| chr6-105799166-DEL-1132 | chr6-105799167-DEL-1132 |
| chr6-105805445-DEL-74 | chr6-105805446-DEL-74 |
| chr6-105881678-DEL-807 | chr6-105881679-DEL-807 |
| chr6-105924807-DEL-70 | chr6-105924805-DEL-68 |
| chr6-105930975-DEL-6364 | chr6-105930976-DEL-6364 |
| chr6-105939501-DEL-52 | chr6-105939502-DEL-52 |
| chr6-105984351-DEL-62 | chr6-105984354-DEL-62 |
| chr6-105985432-DEL-133 | chr6-105985434-DEL-132 |
| chr6-105996686-DEL-870 | chr6-105996687-DEL-869 |
| chr6-105999974-DEL-212 | chr6-105999975-DEL-212 |
| chr6-106043941-DEL-57 | chr6-106043946-DEL-61 |
| chr6-106069726-DEL-8206 | chr6-106069727-DEL-8206 |
| chr6-106180905-DEL-54 | chr6-106180902-DEL-54 |
| chr6-106198727-DEL-498 | chr6-106198728-DEL-496 |
| chr6-106257139-DEL-295 | chr6-106257143-DEL-292 |
| chr6-106273644-DEL-56 | chr6-106273645-DEL-56 |
| chr6-106276836-DEL-63 | chr6-106276816-DEL-76 |

| | |
|---|---|
| chr6-106292257-DEL-6683 | chr6-106292258-DEL-6683 |
| chr6-106315563-DEL-203 | chr6-106315564-DEL-203 |
| chr6-106341895-DEL-357 | chr6-106341896-DEL-357 |
| chr6-106342420-DEL-705 | chr6-106342421-DEL-705 |
| chr6-106374352-DEL-6608 | chr6-106374352-DEL-6609 |
| chr6-106383639-DEL-55 | chr6-106383640-DEL-53 |
| chr6-106443917-DEL-55 | chr6-106443918-DEL-50 |
| chr6-106451443-DEL-176 | chr6-106451444-DEL-176 |
| chr6-106451986-DEL-734 | chr6-106451989-DEL-735 |
| chr6-106454298-DEL-63 | chr6-106454274-DEL-68 |
| chr6-106455229-DEL-2653 | chr6-106455230-DEL-2653 |
| chr6-106475905-DEL-133 | chr6-106475905-DEL-133 |
| chr6-106498574-DEL-353 | chr6-106498575-DEL-353 |
| chr6-106517445-DEL-228 | chr6-106517448-DEL-226 |
| chr6-106525469-DEL-131 | chr6-106525470-DEL-131 |
| chr6-106562860-DEL-115 | chr6-106562864-DEL-113 |
| chr6-106564481-DEL-74 | chr6-106564502-DEL-54 |
| chr6-106580376-DEL-93 | chr6-106580377-DEL-93 |
| chr6-106614807-DEL-54 | chr6-106614782-DEL-60 |
| chr6-106625925-DEL-887 | chr6-106625926-DEL-887 |
| chr6-106632155-DEL-54 | chr6-106632156-DEL-54 |
| chr6-106644901-DEL-55 | chr6-106644902-DEL-60 |
| chr6-106645778-DEL-1093 | chr6-106645779-DEL-1093 |
| chr6-106654257-DEL-125 | chr6-106654258-DEL-125 |
| chr6-106664824-DEL-63 | chr6-106664825-DEL-63 |
| chr6-106679712-DEL-613 | chr6-106679713-DEL-613 |
| chr6-106741396-DEL-940 | chr6-106741397-DEL-940 |
| chr6-106744395-DEL-829 | chr6-106744396-DEL-829 |
| chr6-106746659-DEL-113 | chr6-106746660-DEL-113 |
| chr6-106770176-DEL-63 | chr6-106770177-DEL-63 |
| chr6-106822412-DEL-71 | chr6-106822413-DEL-71 |
| chr6-106842562-DEL-3514 | chr6-106842563-DEL-3514 |
| chr6-106842798-DEL-995 | chr6-106842787-DEL-1003 |
| chr6-106855303-DEL-400 | chr6-106855304-DEL-400 |
| chr6-106859467-DEL-2312 | chr6-106859468-DEL-2312 |
| chr6-106873787-DEL-6221 | chr6-106873788-DEL-6221 |
| chr6-106878153-DEL-95 | chr6-106878158-DEL-86 |
| chr6-106882049-DEL-52 | chr6-106882050-DEL-52 |
| chr6-106905312-DEL-534 | chr6-106905313-DEL-534 |
| chr6-106906958-DEL-68 | chr6-106906959-DEL-66 |
| chr6-106908288-DEL-343 | chr6-106908288-DEL-346 |
| chr6-106957844-DEL-222 | chr6-106957845-DEL-222 |
| chr6-106973737-DEL-657 | chr6-106973738-DEL-657 |
| chr6-106998601-DEL-134 | chr6-106998601-DEL-134 |
| chr6-107000835-DEL-6781 | chr6-107000836-DEL-6782 |
| chr6-107025270-DEL-59 | chr6-107025271-DEL-59 |
| chr6-107047078-DEL-74 | chr6-107047079-DEL-70 |
| chr6-107047475-DEL-6198 | chr6-107047476-DEL-6198 |
| chr6-107055641-DEL-471 | chr6-107055642-DEL-471 |
| chr6-107067809-DEL-561 | chr6-107067799-DEL-562 |
| chr6-107069189-DEL-380 | chr6-107069190-DEL-381 |
| chr6-107093183-DEL-1037 | chr6-107093184-DEL-1037 |
| chr6-107095080-DEL-1120 | chr6-107095081-DEL-1120 |
| chr6-107116728-DEL-6692 | chr6-107116728-DEL-6692 |
| chr6-107116952-DEL-215 | chr6-107116953-DEL-215 |
| chr6-107130891-DEL-84 | chr6-107130892-DEL-84 |
| chr6-107131559-DEL-58 | chr6-107131561-DEL-51 |
| chr6-107151599-DEL-1173 | chr6-107151600-DEL-1173 |
| chr6-107189848-DEL-60 | chr6-107189857-DEL-55 |
| chr6-107208331-DEL-495 | chr6-107208332-DEL-495 |
| chr6-107212203-DEL-358 | chr6-107212204-DEL-358 |
| chr6-107215390-DEL-1169 | chr6-107215391-DEL-1169 |
| chr6-107249091-DEL-821 | chr6-107249092-DEL-821 |
| chr6-107295485-DEL-63 | chr6-107295487-DEL-76 |
| chr6-107306610-DEL-6375 | chr6-107306611-DEL-6375 |
| chr6-107326438-DEL-81 | chr6-107326439-DEL-81 |
| chr6-107389338-DEL-54 | chr6-107389327-DEL-68 |
| chr6-107405575-DEL-440 | chr6-107405576-DEL-440 |
| chr6-107408393-DEL-72 | chr6-107408391-DEL-74 |
| chr6-107447464-DEL-232 | chr6-107447456-DEL-233 |
| chr6-107461940-DEL-3093 | chr6-107461941-DEL-3093 |
| chr6-107479397-DEL-93 | chr6-107479397-DEL-85 |
| chr6-107487517-DEL-64 | chr6-107487495-DEL-65 |
| chr6-107495486-DEL-78 | chr6-107495487-DEL-78 |
| chr6-107499562-DEL-163 | chr6-107499563-DEL-163 |
| chr6-107505261-DEL-529 | chr6-107505262-DEL-529 |
| chr6-107542130-DEL-57 | chr6-107542131-DEL-57 |
| chr6-107552449-DEL-967 | chr6-107552450-DEL-967 |
| chr6-107564464-DEL-4871 | chr6-107564465-DEL-4871 |
| chr6-107583477-DEL-2259 | chr6-107583478-DEL-2259 |

| | |
|---|---|
| chr6-107680347-DEL-491 | chr6-107680348-DEL-491 |
| chr6-107694333-DEL-2385 | chr6-107694334-DEL-2385 |
| chr6-107739345-DEL-234 | chr6-107739346-DEL-234 |
| chr6-107802858-DEL-719 | chr6-107802858-DEL-719 |
| chr6-107900247-DEL-201 | chr6-107900248-DEL-201 |
| chr6-107905880-DEL-66 | chr6-107905881-DEL-66 |
| chr6-107922016-DEL-6627 | chr6-107922017-DEL-6627 |
| chr6-107928292-DEL-201 | chr6-107928293-DEL-201 |
| chr6-107950681-DEL-503 | chr6-107950669-DEL-514 |
| chr6-108018396-DEL-6416 | chr6-108018396-DEL-6416 |
| chr6-108032271-DEL-695 | chr6-108032270-DEL-694 |
| chr6-108059330-DEL-5518 | chr6-108059331-DEL-5518 |
| chr6-108117081-DEL-400 | chr6-108117072-DEL-401 |
| chr6-108242833-DEL-7344 | chr6-108242834-DEL-7344 |
| chr6-108260150-DEL-184 | chr6-108260151-DEL-184 |
| chr6-108262699-DEL-489 | chr6-108262700-DEL-489 |
| chr6-108274276-DEL-98 | chr6-108274277-DEL-103 |
| chr6-108280581-DEL-59 | chr6-108280582-DEL-59 |
| chr6-108283732-DEL-51 | chr6-108283733-DEL-51 |
| chr6-108318564-DEL-60 | chr6-108318562-DEL-57 |
| chr6-108320483-DEL-53 | chr6-108320500-DEL-58 |
| chr6-108336365-DEL-130 | chr6-108336366-DEL-130 |
| chr6-108362495-DEL-453 | chr6-108362496-DEL-453 |
| chr6-108374912-DEL-320 | chr6-108374913-DEL-320 |
| chr6-108386678-DEL-74 | chr6-108386679-DEL-74 |
| chr6-108410523-DEL-96 | chr6-108410524-DEL-80 |
| chr6-108415948-DEL-104 | chr6-108415981-DEL-64 |
| chr6-108436404-DEL-668 | chr6-108436405-DEL-668 |
| chr6-108437601-DEL-51 | chr6-108437602-DEL-51 |
| chr6-108451783-DEL-137 | chr6-108451784-DEL-137 |
| chr6-108455301-DEL-1752 | chr6-108455302-DEL-1752 |
| chr6-108476428-DEL-156 | chr6-108476429-DEL-156 |
| chr6-108530821-DEL-66 | chr6-108530822-DEL-64 |
| chr6-108540388-DEL-2271 | chr6-108540389-DEL-2271 |
| chr6-108566735-DEL-114 | chr6-108566726-DEL-106 |
| chr6-108612066-DEL-432 | chr6-108612067-DEL-432 |
| chr6-108612640-DEL-82 | chr6-108612619-DEL-70 |
| chr6-108676209-DEL-64 | chr6-108676210-DEL-64 |
| chr6-108685162-DEL-323 | chr6-108685163-DEL-323 |
| chr6-108703998-DEL-69 | chr6-108703996-DEL-75 |
| chr6-108723035-DEL-559 | chr6-108723033-DEL-560 |
| chr6-108727895-DEL-162 | chr6-108727903-DEL-158 |
| chr6-108741900-DEL-3627 | chr6-108741901-DEL-3627 |
| chr6-108749527-DEL-208 | chr6-108749560-DEL-192 |
| chr6-108765490-DEL-99 | chr6-108765491-DEL-99 |
| chr6-108781043-DEL-107 | chr6-108781044-DEL-87 |
| chr6-108893171-DEL-565 | chr6-108893172-DEL-565 |
| chr6-108931555-DEL-6712 | chr6-108931556-DEL-6712 |
| chr6-108931863-DEL-205 | chr6-108931864-DEL-205 |
| chr6-108954615-DEL-57 | chr6-108954616-DEL-57 |
| chr6-108954944-DEL-8501 | chr6-108954948-DEL-8502 |
| chr6-108978119-DEL-551 | chr6-108978142-DEL-542 |
| chr6-108990066-DEL-6390 | chr6-108990067-DEL-6390 |
| chr6-109033040-DEL-60 | chr6-109033041-DEL-60 |
| chr6-109105768-DEL-4540 | chr6-109105769-DEL-4540 |
| chr6-109111154-DEL-669 | chr6-109111155-DEL-669 |
| chr6-109165687-DEL-154 | chr6-109165688-DEL-154 |
| chr6-109173136-DEL-6403 | chr6-109173137-DEL-6403 |
| chr6-109179208-DEL-214 | chr6-109179209-DEL-214 |
| chr6-109188668-DEL-141 | chr6-109188669-DEL-141 |
| chr6-109192486-DEL-718 | chr6-109192487-DEL-718 |
| chr6-109227413-DEL-169 | chr6-109227414-DEL-169 |
| chr6-109229622-DEL-525 | chr6-109229623-DEL-526 |
| chr6-109274547-DEL-463 | chr6-109274554-DEL-458 |
| chr6-109281208-DEL-815 | chr6-109281209-DEL-815 |
| chr6-109305043-DEL-2905 | chr6-109305044-DEL-2905 |
| chr6-109309670-DEL-56 | chr6-109309671-DEL-56 |
| chr6-109318834-DEL-320 | chr6-109318835-DEL-320 |
| chr6-109338585-DEL-4325 | chr6-109338585-DEL-4325 |
| chr6-109354639-DEL-72 | chr6-109354639-DEL-66 |
| chr6-109365634-DEL-1293 | chr6-109365635-DEL-1293 |
| chr6-109442409-DEL-73 | chr6-109442410-DEL-69 |
| chr6-109459677-DEL-675 | chr6-109459678-DEL-675 |
| chr6-109465553-DEL-197 | chr6-109465554-DEL-197 |
| chr6-109475187-DEL-1888 | chr6-109475188-DEL-1888 |
| chr6-109488980-DEL-54 | chr6-109488980-DEL-54 |
| chr6-109518671-DEL-368 | chr6-109518672-DEL-368 |
| chr6-109526506-DEL-562 | chr6-109526506-DEL-558 |
| chr6-109536898-DEL-58 | chr6-109536899-DEL-58 |
| chr6-109545213-DEL-74 | chr6-109545226-DEL-62 |

| | |
|---|---|
| chr6-109550967-DEL-173 | chr6-109550932-DEL-209 |
| chr6-109562371-DEL-7164 | chr6-109562372-DEL-7164 |
| chr6-109568459-DEL-214 | chr6-109568460-DEL-214 |
| chr6-109597038-DEL-488 | chr6-109597039-DEL-488 |
| chr6-109618694-DEL-6384 | chr6-109618695-DEL-6384 |
| chr6-109674284-DEL-54 | chr6-109674285-DEL-54 |
| chr6-109692924-DEL-494 | chr6-109692929-DEL-490 |
| chr6-109714151-DEL-66 | chr6-109714154-DEL-88 |
| chr6-109742445-DEL-84 | chr6-109742446-DEL-84 |
| chr6-109755925-DEL-80 | chr6-109755914-DEL-58 |
| chr6-109759751-DEL-84 | chr6-109759752-DEL-84 |
| chr6-109771402-DEL-114 | chr6-109771403-DEL-114 |
| chr6-109774639-DEL-1107 | chr6-109774640-DEL-1107 |
| chr6-109800484-DEL-193 | chr6-109800485-DEL-193 |
| chr6-109817922-DEL-235 | chr6-109817925-DEL-233 |
| chr6-109818760-DEL-1077 | chr6-109818770-DEL-1072 |
| chr6-109820979-DEL-792 | chr6-109820978-DEL-792 |
| chr6-109825704-DEL-84 | chr6-109825705-DEL-84 |
| chr6-109826365-DEL-50 | chr6-109826329-DEL-82 |
| chr6-109828903-DEL-66 | chr6-109828904-DEL-100 |
| chr6-109837307-DEL-200 | chr6-109837308-DEL-200 |
| chr6-109868849-DEL-354 | chr6-109868850-DEL-354 |
| chr6-109881842-DEL-135 | chr6-109881843-DEL-134 |
| chr6-109915364-DEL-556 | chr6-109915365-DEL-556 |
| chr6-109931911-DEL-56 | chr6-109931911-DEL-56 |
| chr6-109952539-DEL-60 | chr6-109952540-DEL-60 |
| chr6-109965811-DEL-575 | chr6-109965811-DEL-575 |
| chr6-109971302-DEL-784 | chr6-109971303-DEL-784 |
| chr6-110001440-DEL-70 | chr6-110001441-DEL-70 |
| chr6-110016608-DEL-65 | chr6-110016612-DEL-52 |
| chr6-110065491-DEL-3878 | chr6-110065492-DEL-3878 |
| chr6-110088029-DEL-166 | chr6-110088030-DEL-166 |
| chr6-110118177-DEL-1020 | chr6-110118178-DEL-1020 |
| chr6-110142006-DEL-76 | chr6-110142007-DEL-76 |
| chr6-110166140-DEL-54 | chr6-110166141-DEL-54 |
| chr6-110167704-DEL-173 | chr6-110167705-DEL-173 |
| chr6-110168920-DEL-72 | chr6-110168925-DEL-68 |
| chr6-110170342-DEL-80 | chr6-110170343-DEL-80 |
| chr6-110206880-DEL-445 | chr6-110206881-DEL-445 |
| chr6-110207492-DEL-1109 | chr6-110207493-DEL-1109 |
| chr6-110229254-DEL-3248 | chr6-110229255-DEL-3248 |
| chr6-110229522-DEL-515 | chr6-110229523-DEL-515 |
| chr6-110234274-DEL-528 | chr6-110234273-DEL-528 |
| chr6-110275367-DEL-56 | chr6-110275340-DEL-52 |
| chr6-110306039-DEL-84 | chr6-110306032-DEL-64 |
| chr6-110311190-DEL-4166 | chr6-110311191-DEL-4166 |
| chr6-110340379-DEL-1191 | chr6-110340380-DEL-1191 |
| chr6-110351947-DEL-55 | chr6-110351948-DEL-55 |
| chr6-110367893-DEL-330 | chr6-110367894-DEL-330 |
| chr6-110370128-DEL-1124 | chr6-110370122-DEL-1125 |
| chr6-110389994-DEL-362 | chr6-110389982-DEL-364 |
| chr6-110400094-DEL-662 | chr6-110400094-DEL-660 |
| chr6-110415779-DEL-491 | chr6-110415780-DEL-491 |
| chr6-110425853-DEL-868 | chr6-110425851-DEL-869 |
| chr6-110461851-DEL-72 | chr6-110461860-DEL-64 |
| chr6-110476435-DEL-56 | chr6-110476437-DEL-95 |
| chr6-110485124-DEL-1731 | chr6-110485125-DEL-1731 |
| chr6-110546462-DEL-67 | chr6-110546412-DEL-98 |
| chr6-110548002-DEL-510 | chr6-110548003-DEL-510 |
| chr6-110562227-DEL-84 | chr6-110562228-DEL-80 |
| chr6-110578489-DEL-186 | chr6-110578490-DEL-186 |
| chr6-110587258-DEL-203 | chr6-110587258-DEL-197 |
| chr6-110614150-DEL-199 | chr6-110614145-DEL-200 |
| chr6-110617755-DEL-592 | chr6-110617756-DEL-592 |
| chr6-110628930-DEL-52 | chr6-110628915-DEL-53 |
| chr6-110673615-DEL-1737 | chr6-110673616-DEL-1737 |
| chr6-110686020-DEL-66 | chr6-110686021-DEL-66 |
| chr6-110707963-DEL-62 | chr6-110707930-DEL-71 |
| chr6-110715668-DEL-66 | chr6-110715706-DEL-50 |
| chr6-110725277-DEL-61 | chr6-110725280-DEL-60 |
| chr6-110735362-DEL-1587 | chr6-110735363-DEL-1587 |
| chr6-110738232-DEL-126 | chr6-110738232-DEL-132 |
| chr6-110779754-DEL-262 | chr6-110779755-DEL-266 |
| chr6-110797789-DEL-212 | chr6-110797790-DEL-212 |
| chr6-110900132-DEL-197 | chr6-110900148-DEL-193 |
| chr6-110913308-DEL-158 | chr6-110913309-DEL-158 |
| chr6-110917791-DEL-280 | chr6-110917792-DEL-280 |
| chr6-110922134-DEL-627 | chr6-110922134-DEL-626 |
| chr6-110924066-DEL-71 | chr6-110924058-DEL-72 |
| chr6-110982040-DEL-376 | chr6-110982041-DEL-376 |

| | |
|---|---|
| chr6-110990490-DEL-6348 | chr6-110990491-DEL-6347 |
| chr6-111006246-DEL-52 | chr6-111006247-DEL-52 |
| chr6-111013966-DEL-695 | chr6-111013967-DEL-695 |
| chr6-111016673-DEL-2114 | chr6-111016674-DEL-2114 |
| chr6-111019342-DEL-774 | chr6-111019343-DEL-774 |
| chr6-111020789-DEL-52 | chr6-111020785-DEL-51 |
| chr6-111024277-DEL-562 | chr6-111024278-DEL-562 |
| chr6-111039702-DEL-53 | chr6-111039706-DEL-60 |
| chr6-111042080-DEL-203 | chr6-111042064-DEL-204 |
| chr6-111056161-DEL-54 | chr6-111056162-DEL-54 |
| chr6-111069665-DEL-866 | chr6-111069666-DEL-865 |
| chr6-111086709-DEL-580 | chr6-111086710-DEL-580 |
| chr6-111090396-DEL-52 | chr6-111090399-DEL-56 |
| chr6-111104872-DEL-635 | chr6-111104874-DEL-636 |
| chr6-111107396-DEL-2657 | chr6-111107397-DEL-2657 |
| chr6-111130944-DEL-6351 | chr6-111130945-DEL-6351 |
| chr6-111136913-DEL-658 | chr6-111136914-DEL-658 |
| chr6-111153374-DEL-813 | chr6-111153375-DEL-813 |
| chr6-111154801-DEL-55 | chr6-111154791-DEL-56 |
| chr6-111156064-DEL-977 | chr6-111156065-DEL-977 |
| chr6-111178505-DEL-2024 | chr6-111178506-DEL-2023 |
| chr6-111189457-DEL-275 | chr6-111189470-DEL-263 |
| chr6-111210428-DEL-56 | chr6-111210385-DEL-106 |
| chr6-111265417-DEL-380 | chr6-111265420-DEL-381 |
| chr6-111275263-DEL-60 | chr6-111275264-DEL-70 |
| chr6-111292081-DEL-50 | chr6-111292077-DEL-55 |
| chr6-111293684-DEL-477 | chr6-111293685-DEL-477 |
| chr6-111296812-DEL-363 | chr6-111296822-DEL-357 |
| chr6-111333415-DEL-189 | chr6-111333416-DEL-189 |
| chr6-111351917-DEL-94 | chr6-111351907-DEL-105 |
| chr6-111354010-DEL-379 | chr6-111354011-DEL-379 |
| chr6-111357316-DEL-994 | chr6-111357314-DEL-994 |
| chr6-111358884-DEL-72 | chr6-111358885-DEL-72 |
| chr6-111420362-DEL-52 | chr6-111420359-DEL-58 |
| chr6-111429066-DEL-89 | chr6-111429079-DEL-77 |
| chr6-111431948-DEL-51 | chr6-111431900-DEL-73 |
| chr6-111435060-DEL-102 | chr6-111435053-DEL-69 |
| chr6-111439642-DEL-6594 | chr6-111439643-DEL-6594 |
| chr6-111452775-DEL-1295 | chr6-111452776-DEL-1293 |
| chr6-111453226-DEL-85 | chr6-111453227-DEL-85 |
| chr6-111462559-DEL-363 | chr6-111462555-DEL-358 |
| chr6-111464236-DEL-94 | chr6-111464237-DEL-94 |
| chr6-111473590-DEL-432 | chr6-111473591-DEL-432 |
| chr6-111501088-DEL-681 | chr6-111501089-DEL-673 |
| chr6-111505407-DEL-120 | chr6-111505408-DEL-120 |
| chr6-111518925-DEL-529 | chr6-111518926-DEL-523 |
| chr6-111522961-DEL-6369 | chr6-111522962-DEL-6369 |
| chr6-111532890-DEL-245 | chr6-111532891-DEL-245 |
| chr6-111540192-DEL-1028 | chr6-111540193-DEL-1028 |
| chr6-111618987-DEL-64 | chr6-111618988-DEL-64 |
| chr6-111641315-DEL-134 | chr6-111641316-DEL-132 |
| chr6-111664280-DEL-298 | chr6-111664282-DEL-299 |
| chr6-111676065-DEL-66 | chr6-111676066-DEL-66 |
| chr6-111678473-DEL-236 | chr6-111678476-DEL-234 |
| chr6-111679696-DEL-55 | chr6-111679697-DEL-55 |
| chr6-111680896-DEL-621 | chr6-111680897-DEL-621 |
| chr6-111699278-DEL-84 | chr6-111699307-DEL-56 |
| chr6-111703744-DEL-497 | chr6-111703745-DEL-496 |
| chr6-111739168-DEL-62 | chr6-111739169-DEL-61 |
| chr6-111752352-DEL-2914 | chr6-111752353-DEL-2914 |
| chr6-111760441-DEL-78 | chr6-111760428-DEL-62 |
| chr6-111812534-DEL-980 | chr6-111812535-DEL-980 |
| chr6-111837134-DEL-365 | chr6-111837131-DEL-366 |
| chr6-111837742-DEL-787 | chr6-111837743-DEL-787 |
| chr6-111848019-DEL-96 | chr6-111848017-DEL-99 |
| chr6-111873922-DEL-2833 | chr6-111873923-DEL-2833 |
| chr6-111882376-DEL-66 | chr6-111882383-DEL-62 |
| chr6-111886027-DEL-53 | chr6-111886028-DEL-53 |
| chr6-111886778-DEL-90 | chr6-111886779-DEL-90 |
| chr6-111892490-DEL-125 | chr6-111892492-DEL-137 |
| chr6-111912434-DEL-4731 | chr6-111912435-DEL-4731 |
| chr6-111981292-DEL-63 | chr6-111981293-DEL-63 |
| chr6-111986074-DEL-599 | chr6-111986075-DEL-599 |
| chr6-111988886-DEL-905 | chr6-111988887-DEL-905 |
| chr6-112020831-DEL-54 | chr6-112020813-DEL-59 |
| chr6-112044342-DEL-6423 | chr6-112044343-DEL-6423 |
| chr6-112060598-DEL-69 | chr6-112060599-DEL-69 |
| chr6-112072535-DEL-54 | chr6-112072536-DEL-54 |
| chr6-112076944-DEL-345 | chr6-112076945-DEL-345 |
| chr6-112095607-DEL-130 | chr6-112095608-DEL-130 |

| | |
|---|---|
| chr6-112101954-DEL-206 | chr6-112101955-DEL-218 |
| chr6-112102912-DEL-1435 | chr6-112102913-DEL-1435 |
| chr6-112124633-DEL-75 | chr6-112124634-DEL-75 |
| chr6-112169409-DEL-322 | chr6-112169409-DEL-322 |
| chr6-112203093-DEL-76 | chr6-112203094-DEL-76 |
| chr6-112211788-DEL-84 | chr6-112211789-DEL-84 |
| chr6-112214531-DEL-4399 | chr6-112214532-DEL-4399 |
| chr6-112232312-DEL-54 | chr6-112232313-DEL-54 |
| chr6-112251763-DEL-85 | chr6-112251764-DEL-85 |
| chr6-112262530-DEL-58 | chr6-112262524-DEL-70 |
| chr6-112263112-DEL-162 | chr6-112263113-DEL-162 |
| chr6-112330967-DEL-56 | chr6-112330972-DEL-62 |
| chr6-112334240-DEL-246 | chr6-112334241-DEL-246 |
| chr6-112335163-DEL-154 | chr6-112335167-DEL-153 |
| chr6-112338850-DEL-316 | chr6-112338851-DEL-316 |
| chr6-112340368-DEL-937 | chr6-112340369-DEL-937 |
| chr6-112349953-DEL-62 | chr6-112349954-DEL-62 |
| chr6-112380493-DEL-478 | chr6-112380494-DEL-478 |
| chr6-112383979-DEL-66 | chr6-112383979-DEL-50 |
| chr6-112403720-DEL-56 | chr6-112403732-DEL-65 |
| chr6-112413394-DEL-3848 | chr6-112413395-DEL-3848 |
| chr6-112432413-DEL-57 | chr6-112432413-DEL-57 |
| chr6-112470096-DEL-141 | chr6-112470097-DEL-141 |
| chr6-112498482-DEL-762 | chr6-112498483-DEL-762 |
| chr6-112508208-DEL-280 | chr6-112508209-DEL-280 |
| chr6-112509411-DEL-400 | chr6-112509412-DEL-400 |
| chr6-112511319-DEL-399 | chr6-112511320-DEL-399 |
| chr6-112511891-DEL-6419 | chr6-112511898-DEL-6419 |
| chr6-112565204-DEL-299 | chr6-112565205-DEL-304 |
| chr6-112582293-DEL-3476 | chr6-112582294-DEL-3476 |
| chr6-112587270-DEL-526 | chr6-112587271-DEL-526 |
| chr6-112629619-DEL-7000 | chr6-112629620-DEL-7000 |
| chr6-112681794-DEL-7195 | chr6-112681797-DEL-7196 |
| chr6-112782871-DEL-442 | chr6-112782872-DEL-442 |
| chr6-112811694-DEL-600 | chr6-112811696-DEL-597 |
| chr6-112821475-DEL-53 | chr6-112821476-DEL-53 |
| chr6-112824258-DEL-340 | chr6-112824265-DEL-337 |
| chr6-112836069-DEL-84 | chr6-112836070-DEL-84 |
| chr6-112932964-DEL-73 | chr6-112932963-DEL-74 |
| chr6-112939636-DEL-2253 | chr6-112939637-DEL-2252 |
| chr6-112942235-DEL-111 | chr6-112942236-DEL-111 |
| chr6-112968367-DEL-95 | chr6-112968368-DEL-95 |
| chr6-112969992-DEL-345 | chr6-112969993-DEL-345 |
| chr6-113000311-DEL-207 | chr6-113000312-DEL-207 |
| chr6-113010639-DEL-52 | chr6-113010638-DEL-54 |
| chr6-113141040-DEL-1867 | chr6-113141041-DEL-1867 |
| chr6-113146196-DEL-88 | chr6-113146197-DEL-89 |
| chr6-113187289-DEL-66 | chr6-113187251-DEL-54 |
| chr6-113190930-DEL-222 | chr6-113190929-DEL-223 |
| chr6-113196774-DEL-1346 | chr6-113196775-DEL-1346 |
| chr6-113198479-DEL-233 | chr6-113198480-DEL-233 |
| chr6-113204227-DEL-90 | chr6-113204228-DEL-60 |
| chr6-113223064-DEL-2822 | chr6-113223065-DEL-2822 |
| chr6-113272842-DEL-60 | chr6-113272843-DEL-60 |
| chr6-113313661-DEL-236 | chr6-113313662-DEL-236 |
| chr6-113342873-DEL-186 | chr6-113342874-DEL-186 |
| chr6-113420952-DEL-376 | chr6-113420953-DEL-376 |
| chr6-113423190-DEL-55 | chr6-113423191-DEL-55 |
| chr6-113520596-DEL-422 | chr6-113520597-DEL-422 |
| chr6-113523878-DEL-242 | chr6-113523878-DEL-243 |
| chr6-113534040-DEL-210 | chr6-113534041-DEL-210 |
| chr6-113549069-DEL-78 | chr6-113549080-DEL-54 |
| chr6-113550312-DEL-84 | chr6-113550313-DEL-84 |
| chr6-113552401-DEL-217 | chr6-113552402-DEL-217 |
| chr6-113571143-DEL-78 | chr6-113571144-DEL-78 |
| chr6-113587014-DEL-201 | chr6-113586996-DEL-180 |
| chr6-113588137-DEL-438 | chr6-113588138-DEL-438 |
| chr6-113592233-DEL-60 | chr6-113592234-DEL-60 |
| chr6-113719629-DEL-59 | chr6-113719630-DEL-59 |
| chr6-113734612-DEL-174 | chr6-113734613-DEL-174 |
| chr6-113809600-DEL-58 | chr6-113809591-DEL-52 |
| chr6-113813655-DEL-60 | chr6-113813656-DEL-60 |
| chr6-113822559-DEL-108 | chr6-113822562-DEL-58 |
| chr6-113843842-DEL-1366 | chr6-113843843-DEL-1366 |
| chr6-113854198-DEL-123 | chr6-113854211-DEL-111 |
| chr6-113913768-DEL-62 | chr6-113913769-DEL-62 |
| chr6-113918731-DEL-410 | chr6-113918736-DEL-416 |
| chr6-113923801-DEL-64 | chr6-113923802-DEL-64 |
| chr6-113948225-DEL-58 | chr6-113948224-DEL-70 |
| chr6-113987235-DEL-97 | chr6-113987231-DEL-98 |

| | |
|---|---|
| chr6-113987794-DEL-322 | chr6-113987795-DEL-322 |
| chr6-113987808-DEL-826 | chr6-113987809-DEL-826 |
| chr6-113995074-DEL-110 | chr6-113995075-DEL-110 |
| chr6-114008433-DEL-1175 | chr6-114008434-DEL-1175 |
| chr6-114025307-DEL-200 | chr6-114025308-DEL-200 |
| chr6-114030845-DEL-70 | chr6-114030846-DEL-70 |
| chr6-114044599-DEL-58 | chr6-114044600-DEL-50 |
| chr6-114046267-DEL-57 | chr6-114046267-DEL-57 |
| chr6-114064571-DEL-263 | chr6-114064572-DEL-263 |
| chr6-114077361-DEL-53 | chr6-114077361-DEL-53 |
| chr6-114079765-DEL-200 | chr6-114079766-DEL-200 |
| chr6-114150377-DEL-114 | chr6-114150378-DEL-114 |
| chr6-114167815-DEL-334 | chr6-114167816-DEL-331 |
| chr6-114252701-DEL-247 | chr6-114252702-DEL-247 |
| chr6-114286590-DEL-432 | chr6-114286591-DEL-432 |
| chr6-114289490-DEL-391 | chr6-114289502-DEL-385 |
| chr6-114318067-DEL-100 | chr6-114318068-DEL-100 |
| chr6-114319114-DEL-255 | chr6-114319113-DEL-250 |
| chr6-114329080-DEL-62 | chr6-114329128-DEL-52 |
| chr6-114339133-DEL-66 | chr6-114339134-DEL-66 |
| chr6-114360082-DEL-306 | chr6-114360083-DEL-306 |
| chr6-114382994-DEL-59 | chr6-114382998-DEL-56 |
| chr6-114423292-DEL-95 | chr6-114423289-DEL-118 |
| chr6-114454095-DEL-70 | chr6-114454092-DEL-68 |
| chr6-114494823-DEL-52 | chr6-114494823-DEL-52 |
| chr6-114562926-DEL-64 | chr6-114562931-DEL-58 |
| chr6-114590029-DEL-54 | chr6-114590035-DEL-54 |
| chr6-114597439-DEL-53 | chr6-114597419-DEL-96 |
| chr6-114607344-DEL-62 | chr6-114607317-DEL-67 |
| chr6-114609860-DEL-74 | chr6-114609815-DEL-108 |
| chr6-114615721-DEL-70 | chr6-114615722-DEL-70 |
| chr6-114623577-DEL-55 | chr6-114623578-DEL-55 |
| chr6-114631205-DEL-63 | chr6-114631194-DEL-64 |
| chr6-114650962-DEL-275 | chr6-114650961-DEL-276 |
| chr6-114652512-DEL-488 | chr6-114652513-DEL-488 |
| chr6-114677368-DEL-90 | chr6-114677369-DEL-90 |
| chr6-114693586-DEL-64 | chr6-114693585-DEL-66 |
| chr6-114698958-DEL-60 | chr6-114698951-DEL-56 |
| chr6-114717793-DEL-320 | chr6-114717794-DEL-319 |
| chr6-114736370-DEL-6185 | chr6-114736371-DEL-6185 |
| chr6-114746537-DEL-128 | chr6-114746510-DEL-114 |
| chr6-114819569-DEL-98 | chr6-114819570-DEL-98 |
| chr6-114841704-DEL-1232 | chr6-114841705-DEL-1232 |
| chr6-114842008-DEL-219 | chr6-114842009-DEL-219 |
| chr6-114861394-DEL-86 | chr6-114861352-DEL-129 |
| chr6-114875617-DEL-123 | chr6-114875618-DEL-124 |
| chr6-114887451-DEL-203 | chr6-114887452-DEL-203 |
| chr6-114899197-DEL-204 | chr6-114899198-DEL-204 |
| chr6-114901048-DEL-371 | chr6-114901049-DEL-371 |
| chr6-114924299-DEL-336 | chr6-114924300-DEL-322 |
| chr6-114927733-DEL-118 | chr6-114927734-DEL-118 |
| chr6-114967866-DEL-268 | chr6-114967867-DEL-268 |
| chr6-114979991-DEL-57 | chr6-114979992-DEL-57 |
| chr6-115018830-DEL-78 | chr6-115018831-DEL-78 |
| chr6-115025676-DEL-58 | chr6-115025665-DEL-50 |
| chr6-115038953-DEL-64 | chr6-115038958-DEL-56 |
| chr6-115043335-DEL-56 | chr6-115043336-DEL-56 |
| chr6-115069073-DEL-51 | chr6-115069074-DEL-51 |
| chr6-115070916-DEL-120 | chr6-115070917-DEL-106 |
| chr6-115072289-DEL-60 | chr6-115072330-DEL-58 |
| chr6-115074776-DEL-103 | chr6-115074769-DEL-94 |
| chr6-115127082-DEL-4031 | chr6-115127083-DEL-4031 |
| chr6-115150264-DEL-745 | chr6-115150265-DEL-745 |
| chr6-115240334-DEL-59 | chr6-115240335-DEL-60 |
| chr6-115244557-DEL-372 | chr6-115244558-DEL-372 |
| chr6-115266499-DEL-94 | chr6-115266500-DEL-94 |
| chr6-115275430-DEL-229 | chr6-115275431-DEL-229 |
| chr6-115311660-DEL-878 | chr6-115311661-DEL-876 |
| chr6-115370767-DEL-128 | chr6-115370768-DEL-88 |
| chr6-115371598-DEL-93 | chr6-115371598-DEL-93 |
| chr6-115387060-DEL-60 | chr6-115387061-DEL-60 |
| chr6-115460083-DEL-64 | chr6-115460096-DEL-52 |
| chr6-115471543-DEL-236 | chr6-115471544-DEL-236 |
| chr6-115491886-DEL-1893 | chr6-115491884-DEL-1893 |
| chr6-115498174-DEL-82 | chr6-115498187-DEL-70 |
| chr6-115518908-DEL-321 | chr6-115518906-DEL-322 |
| chr6-115535782-DEL-78 | chr6-115535791-DEL-70 |
| chr6-115550578-DEL-144 | chr6-115550578-DEL-144 |
| chr6-115565124-DEL-274 | chr6-115565141-DEL-252 |
| chr6-115565778-DEL-56 | chr6-115565769-DEL-50 |

| | |
|---|---|
| chr6-115566687-DEL-630 | chr6-115566688-DEL-630 |
| chr6-115582067-DEL-173 | chr6-115582068-DEL-173 |
| chr6-115589607-DEL-52 | chr6-115589608-DEL-52 |
| chr6-115605555-DEL-98 | chr6-115605564-DEL-90 |
| chr6-115636144-DEL-83 | chr6-115636145-DEL-83 |
| chr6-115647105-DEL-168 | chr6-115647106-DEL-168 |
| chr6-115681335-DEL-184 | chr6-115681336-DEL-184 |
| chr6-115696042-DEL-81 | chr6-115696043-DEL-81 |
| chr6-115708698-DEL-620 | chr6-115708700-DEL-619 |
| chr6-115726331-DEL-483 | chr6-115726332-DEL-483 |
| chr6-115737268-DEL-70 | chr6-115737269-DEL-70 |
| chr6-115745020-DEL-886 | chr6-115745021-DEL-886 |
| chr6-115751306-DEL-86 | chr6-115751307-DEL-88 |
| chr6-115754364-DEL-197 | chr6-115754365-DEL-197 |
| chr6-115764684-DEL-57 | chr6-115764685-DEL-56 |
| chr6-115770246-DEL-78 | chr6-115770280-DEL-94 |
| chr6-115780372-DEL-174 | chr6-115780373-DEL-174 |
| chr6-115782086-DEL-173 | chr6-115782071-DEL-174 |
| chr6-115783254-DEL-92 | chr6-115783255-DEL-92 |
| chr6-115801240-DEL-5512 | chr6-115801241-DEL-5512 |
| chr6-115812798-DEL-213 | chr6-115812799-DEL-214 |
| chr6-115823861-DEL-215 | chr6-115823862-DEL-215 |
| chr6-115845430-DEL-4150 | chr6-115845466-DEL-4104 |
| chr6-115866315-DEL-76 | chr6-115866341-DEL-86 |
| chr6-115894596-DEL-199 | chr6-115894597-DEL-199 |
| chr6-115982008-DEL-53 | chr6-115982008-DEL-53 |
| chr6-115986774-DEL-204 | chr6-115986775-DEL-204 |
| chr6-115989054-DEL-64 | chr6-115989075-DEL-57 |
| chr6-116023487-DEL-2908 | chr6-116023488-DEL-2908 |
| chr6-116024865-DEL-163 | chr6-116024866-DEL-163 |
| chr6-116076678-DEL-80 | chr6-116076671-DEL-76 |
| chr6-116093069-DEL-206 | chr6-116093070-DEL-206 |
| chr6-116150576-DEL-897 | chr6-116150577-DEL-897 |
| chr6-116200418-DEL-187 | chr6-116200419-DEL-187 |
| chr6-116225897-DEL-1927 | chr6-116225898-DEL-1927 |
| chr6-116244851-DEL-3100 | chr6-116244852-DEL-3100 |
| chr6-116252878-DEL-865 | chr6-116252879-DEL-865 |
| chr6-116296894-DEL-1910 | chr6-116296895-DEL-1910 |
| chr6-116354953-DEL-71 | chr6-116354947-DEL-69 |
| chr6-116364864-DEL-3246 | chr6-116364865-DEL-3246 |
| chr6-116405146-DEL-664 | chr6-116405136-DEL-664 |
| chr6-116406281-DEL-104 | chr6-116406282-DEL-104 |
| chr6-116409716-DEL-328 | chr6-116409717-DEL-328 |
| chr6-116443103-DEL-459 | chr6-116443103-DEL-459 |
| chr6-116451106-DEL-6885 | chr6-116451107-DEL-6885 |
| chr6-116486152-DEL-3032 | chr6-116486149-DEL-3032 |
| chr6-116507269-DEL-3983 | chr6-116507270-DEL-3983 |
| chr6-116539286-DEL-801 | chr6-116539287-DEL-801 |
| chr6-116543117-DEL-1123 | chr6-116543118-DEL-1123 |
| chr6-116549484-DEL-6498 | chr6-116549501-DEL-6482 |
| chr6-116567839-DEL-408 | chr6-116567840-DEL-408 |
| chr6-116593179-DEL-53 | chr6-116593180-DEL-53 |
| chr6-116618847-DEL-80 | chr6-116618848-DEL-80 |
| chr6-116636981-DEL-72 | chr6-116636982-DEL-72 |
| chr6-116641596-DEL-80 | chr6-116641597-DEL-80 |
| chr6-116662108-DEL-6422 | chr6-116662109-DEL-6422 |
| chr6-116710890-DEL-126 | chr6-116710891-DEL-126 |
| chr6-116717178-DEL-7263 | chr6-116717179-DEL-7263 |
| chr6-116740335-DEL-120 | chr6-116740332-DEL-81 |
| chr6-116741302-DEL-148 | chr6-116741303-DEL-148 |
| chr6-116749697-DEL-430 | chr6-116749698-DEL-431 |
| chr6-116755892-DEL-52 | chr6-116755893-DEL-52 |
| chr6-116757187-DEL-97 | chr6-116757188-DEL-97 |
| chr6-116816704-DEL-67 | chr6-116816699-DEL-78 |
| chr6-116825920-DEL-54 | chr6-116825921-DEL-54 |
| chr6-116835516-DEL-117 | chr6-116835512-DEL-118 |
| chr6-116848527-DEL-2929 | chr6-116848528-DEL-2929 |
| chr6-116854068-DEL-6908 | chr6-116854068-DEL-6908 |
| chr6-116901601-DEL-320 | chr6-116901602-DEL-320 |
| chr6-116903625-DEL-6453 | chr6-116903626-DEL-6453 |
| chr6-116914111-DEL-1301 | chr6-116914099-DEL-1316 |
| chr6-116927447-DEL-235 | chr6-116927448-DEL-235 |
| chr6-116949159-DEL-181 | chr6-116949160-DEL-181 |
| chr6-116976918-DEL-66 | chr6-116976919-DEL-66 |
| chr6-117038955-DEL-976 | chr6-117038956-DEL-976 |
| chr6-117051433-DEL-198 | chr6-117051434-DEL-198 |
| chr6-117060999-DEL-95 | chr6-117061018-DEL-60 |
| chr6-117064132-DEL-7144 | chr6-117064133-DEL-7144 |
| chr6-117088676-DEL-714 | chr6-117088677-DEL-714 |
| chr6-117097368-DEL-57 | chr6-117097370-DEL-56 |

| | |
|---|---|
| chr6-117100713-DEL-60 | chr6-117100712-DEL-58 |
| chr6-117121979-DEL-105 | chr6-117121980-DEL-105 |
| chr6-117167033-DEL-225 | chr6-117167034-DEL-225 |
| chr6-117181933-DEL-77 | chr6-117181934-DEL-77 |
| chr6-117183411-DEL-60 | chr6-117183400-DEL-72 |
| chr6-117229263-DEL-200 | chr6-117229264-DEL-200 |
| chr6-117253202-DEL-54 | chr6-117253189-DEL-76 |
| chr6-117261174-DEL-98 | chr6-117261175-DEL-98 |
| chr6-117266124-DEL-920 | chr6-117266139-DEL-894 |
| chr6-117287322-DEL-1672 | chr6-117287323-DEL-1672 |
| chr6-117311725-DEL-5223 | chr6-117311726-DEL-5223 |
| chr6-117334787-DEL-65 | chr6-117334765-DEL-113 |
| chr6-117360552-DEL-258 | chr6-117360553-DEL-258 |
| chr6-117368452-DEL-54 | chr6-117368453-DEL-54 |
| chr6-117370970-DEL-6466 | chr6-117370972-DEL-6466 |
| chr6-117377623-DEL-66 | chr6-117377621-DEL-50 |
| chr6-117395827-DEL-1577 | chr6-117395828-DEL-1577 |
| chr6-117405985-DEL-211 | chr6-117405986-DEL-211 |
| chr6-117453796-DEL-1118 | chr6-117453797-DEL-1118 |
| chr6-117497380-DEL-2090 | chr6-117497379-DEL-2090 |
| chr6-117504992-DEL-3401 | chr6-117504993-DEL-3401 |
| chr6-117521971-DEL-293 | chr6-117521972-DEL-293 |
| chr6-117531117-DEL-122 | chr6-117531118-DEL-120 |
| chr6-117562052-DEL-7327 | chr6-117562052-DEL-7327 |
| chr6-117608954-DEL-334 | chr6-117608955-DEL-334 |
| chr6-117611748-DEL-6401 | chr6-117611749-DEL-6401 |
| chr6-117648970-DEL-105 | chr6-117648971-DEL-105 |
| chr6-117660832-DEL-569 | chr6-117660833-DEL-569 |
| chr6-117669024-DEL-7326 | chr6-117669025-DEL-7326 |
| chr6-117683342-DEL-59 | chr6-117683329-DEL-73 |
| chr6-117684959-DEL-98 | chr6-117684960-DEL-98 |
| chr6-117693288-DEL-2559 | chr6-117693289-DEL-2559 |
| chr6-117703999-DEL-116 | chr6-117704016-DEL-116 |
| chr6-117708163-DEL-1485 | chr6-117708164-DEL-1485 |
| chr6-117720445-DEL-9532 | chr6-117720446-DEL-9528 |
| chr6-117773079-DEL-2486 | chr6-117773080-DEL-2486 |
| chr6-117779058-DEL-327 | chr6-117779059-DEL-327 |
| chr6-117779870-DEL-87 | chr6-117779871-DEL-87 |
| chr6-117815520-DEL-202 | chr6-117815519-DEL-176 |
| chr6-117816126-DEL-56 | chr6-117816127-DEL-56 |
| chr6-117826620-DEL-56 | chr6-117826621-DEL-56 |
| chr6-117841915-DEL-475 | chr6-117841916-DEL-475 |
| chr6-117844959-DEL-82 | chr6-117844955-DEL-87 |
| chr6-117846716-DEL-220 | chr6-117846702-DEL-221 |
| chr6-117855187-DEL-200 | chr6-117855188-DEL-200 |
| chr6-117887619-DEL-66 | chr6-117887625-DEL-61 |
| chr6-117892027-DEL-201 | chr6-117892028-DEL-201 |
| chr6-117907686-DEL-210 | chr6-117907687-DEL-210 |
| chr6-117921712-DEL-2505 | chr6-117921713-DEL-2505 |
| chr6-117929155-DEL-171 | chr6-117929156-DEL-171 |
| chr6-117935741-DEL-212 | chr6-117935742-DEL-212 |
| chr6-117938975-DEL-108 | chr6-117938975-DEL-109 |
| chr6-117940810-DEL-293 | chr6-117940811-DEL-293 |
| chr6-117963407-DEL-411 | chr6-117963408-DEL-411 |
| chr6-117978758-DEL-61 | chr6-117978748-DEL-51 |
| chr6-117986488-DEL-244 | chr6-117986487-DEL-244 |
| chr6-118049823-DEL-55 | chr6-118049805-DEL-66 |
| chr6-118079233-DEL-52 | chr6-118079234-DEL-52 |
| chr6-118136940-DEL-60 | chr6-118136941-DEL-60 |
| chr6-118143010-DEL-180 | chr6-118143011-DEL-180 |
| chr6-118165293-DEL-85 | chr6-118165294-DEL-85 |
| chr6-118186089-DEL-6657 | chr6-118186090-DEL-6657 |
| chr6-118217359-DEL-80 | chr6-118217360-DEL-80 |
| chr6-118217997-DEL-204 | chr6-118217998-DEL-204 |
| chr6-118249579-DEL-82 | chr6-118249580-DEL-82 |
| chr6-118258498-DEL-592 | chr6-118258499-DEL-592 |
| chr6-118264663-DEL-2944 | chr6-118264664-DEL-2944 |
| chr6-118274925-DEL-269 | chr6-118274926-DEL-269 |
| chr6-118332536-DEL-56 | chr6-118332537-DEL-56 |
| chr6-118333855-DEL-336 | chr6-118333873-DEL-319 |
| chr6-118339536-DEL-298 | chr6-118339537-DEL-298 |
| chr6-118346380-DEL-2669 | chr6-118346381-DEL-2669 |
| chr6-118376585-DEL-528 | chr6-118376586-DEL-528 |
| chr6-118379818-DEL-78 | chr6-118379822-DEL-75 |
| chr6-118384300-DEL-58 | chr6-118384306-DEL-56 |
| chr6-118399808-DEL-194 | chr6-118399809-DEL-194 |
| chr6-118400622-DEL-215 | chr6-118400623-DEL-215 |
| chr6-118412631-DEL-3448 | chr6-118412632-DEL-3448 |
| chr6-118424023-DEL-1893 | chr6-118424024-DEL-1893 |
| chr6-118443931-DEL-70 | chr6-118443932-DEL-70 |

| | |
|---|---|
| chr6-118462784-DEL-200 | chr6-118462785-DEL-200 |
| chr6-118465880-DEL-6401 | chr6-118465881-DEL-6401 |
| chr6-118475015-DEL-285 | chr6-118475016-DEL-285 |
| chr6-118478003-DEL-112 | chr6-118478043-DEL-75 |
| chr6-118497200-DEL-52 | chr6-118497201-DEL-52 |
| chr6-118521339-DEL-2921 | chr6-118521340-DEL-2921 |
| chr6-118546571-DEL-4157 | chr6-118546572-DEL-4157 |
| chr6-118553521-DEL-81 | chr6-118553508-DEL-82 |
| chr6-118554690-DEL-59 | chr6-118554689-DEL-59 |
| chr6-118556348-DEL-128 | chr6-118556355-DEL-122 |
| chr6-118598184-DEL-63 | chr6-118598222-DEL-56 |
| chr6-118638122-DEL-416 | chr6-118638123-DEL-416 |
| chr6-118739554-DEL-2100 | chr6-118739555-DEL-2100 |
| chr6-118782775-DEL-761 | chr6-118782776-DEL-761 |
| chr6-118812395-DEL-86 | chr6-118812396-DEL-86 |
| chr6-118865725-DEL-474 | chr6-118865725-DEL-472 |
| chr6-118882440-DEL-172 | chr6-118882444-DEL-171 |
| chr6-118945398-DEL-184 | chr6-118945399-DEL-184 |
| chr6-118975144-DEL-187 | chr6-118975145-DEL-187 |
| chr6-118978322-DEL-55 | chr6-118978330-DEL-94 |
| chr6-118991727-DEL-82 | chr6-118991728-DEL-88 |
| chr6-118993754-DEL-94 | chr6-118993755-DEL-94 |
| chr6-118999307-DEL-89 | chr6-118999318-DEL-78 |
| chr6-119053330-DEL-65 | chr6-119053330-DEL-66 |
| chr6-119063486-DEL-52 | chr6-119063487-DEL-52 |
| chr6-119097769-DEL-156 | chr6-119097770-DEL-156 |
| chr6-119118555-DEL-200 | chr6-119118556-DEL-200 |
| chr6-119129117-DEL-214 | chr6-119129118-DEL-214 |
| chr6-119179986-DEL-376 | chr6-119179980-DEL-377 |
| chr6-119245809-DEL-196 | chr6-119245810-DEL-196 |
| chr6-119250386-DEL-207 | chr6-119250387-DEL-207 |
| chr6-119305602-DEL-88 | chr6-119305603-DEL-88 |
| chr6-119373852-DEL-60 | chr6-119373892-DEL-62 |
| chr6-119375518-DEL-233 | chr6-119375519-DEL-233 |
| chr6-119414384-DEL-207 | chr6-119414391-DEL-173 |
| chr6-119420305-DEL-85 | chr6-119420306-DEL-85 |
| chr6-119424872-DEL-132 | chr6-119424873-DEL-88 |
| chr6-119525468-DEL-63 | chr6-119525469-DEL-63 |
| chr6-119530937-DEL-89 | chr6-119530938-DEL-89 |
| chr6-119559709-DEL-199 | chr6-119559710-DEL-199 |
| chr6-119579226-DEL-214 | chr6-119579232-DEL-214 |
| chr6-119583969-DEL-3651 | chr6-119583970-DEL-3651 |
| chr6-119614134-DEL-1449 | chr6-119614134-DEL-1449 |
| chr6-119620153-DEL-54 | chr6-119620148-DEL-50 |
| chr6-119655067-DEL-54 | chr6-119655068-DEL-54 |
| chr6-119672666-DEL-1251 | chr6-119672667-DEL-1251 |
| chr6-119721569-DEL-3360 | chr6-119721562-DEL-3352 |
| chr6-119740050-DEL-70 | chr6-119740051-DEL-70 |
| chr6-119759938-DEL-52 | chr6-119759939-DEL-52 |
| chr6-119815254-DEL-265 | chr6-119815255-DEL-265 |
| chr6-119828962-DEL-235 | chr6-119828963-DEL-235 |
| chr6-119836087-DEL-53 | chr6-119836088-DEL-53 |
| chr6-119850001-DEL-2827 | chr6-119850002-DEL-2827 |
| chr6-119855384-DEL-410 | chr6-119855385-DEL-410 |
| chr6-119880567-DEL-2000 | chr6-119880568-DEL-2000 |
| chr6-119885668-DEL-209 | chr6-119885669-DEL-209 |
| chr6-119894767-DEL-64 | chr6-119894768-DEL-64 |
| chr6-119919216-DEL-160 | chr6-119919218-DEL-154 |
| chr6-119963003-DEL-62 | chr6-119963016-DEL-50 |
| chr6-119982958-DEL-342 | chr6-119982959-DEL-342 |
| chr6-119994307-DEL-78 | chr6-119994304-DEL-82 |
| chr6-120032345-DEL-221 | chr6-120032346-DEL-221 |
| chr6-120052830-DEL-71 | chr6-120052831-DEL-71 |
| chr6-120064220-DEL-209 | chr6-120064221-DEL-209 |
| chr6-120064869-DEL-391 | chr6-120064870-DEL-391 |
| chr6-120099779-DEL-60 | chr6-120099784-DEL-55 |
| chr6-120101318-DEL-219 | chr6-120101319-DEL-219 |
| chr6-120102537-DEL-216 | chr6-120102538-DEL-216 |
| chr6-120104558-DEL-7071 | chr6-120104559-DEL-7071 |
| chr6-120117168-DEL-53 | chr6-120117169-DEL-53 |
| chr6-120129922-DEL-183 | chr6-120129923-DEL-183 |
| chr6-120142569-DEL-272 | chr6-120142570-DEL-272 |
| chr6-120149996-DEL-58 | chr6-120150005-DEL-50 |
| chr6-120159655-DEL-185 | chr6-120159656-DEL-185 |
| chr6-120173600-DEL-78 | chr6-120173601-DEL-78 |
| chr6-120198232-DEL-134 | chr6-120198233-DEL-134 |
| chr6-120204655-DEL-213 | chr6-120204656-DEL-213 |
| chr6-120221019-DEL-60 | chr6-120221009-DEL-58 |
| chr6-120228447-DEL-163 | chr6-120228448-DEL-163 |
| chr6-120244686-DEL-167 | chr6-120244685-DEL-168 |

| | |
|---|---|
| chr6-120278480-DEL-128 | chr6-120278481-DEL-130 |
| chr6-120303084-DEL-2148 | chr6-120303083-DEL-2149 |
| chr6-120308417-DEL-68 | chr6-120308448-DEL-68 |
| chr6-120354665-DEL-204 | chr6-120354666-DEL-204 |
| chr6-120355738-DEL-181 | chr6-120355739-DEL-181 |
| chr6-120360277-DEL-1802 | chr6-120360270-DEL-1803 |
| chr6-120381197-DEL-104 | chr6-120381198-DEL-104 |
| chr6-120382543-DEL-77 | chr6-120382544-DEL-83 |
| chr6-120406728-DEL-204 | chr6-120406729-DEL-204 |
| chr6-120421751-DEL-652 | chr6-120421751-DEL-652 |
| chr6-120423917-DEL-192 | chr6-120423909-DEL-193 |
| chr6-120429658-DEL-136 | chr6-120429659-DEL-136 |
| chr6-120467711-DEL-64 | chr6-120467712-DEL-64 |
| chr6-120477083-DEL-56 | chr6-120477083-DEL-56 |
| chr6-120495830-DEL-161 | chr6-120495829-DEL-161 |
| chr6-120514462-DEL-52 | chr6-120514463-DEL-52 |
| chr6-120522029-DEL-196 | chr6-120522030-DEL-196 |
| chr6-120528154-DEL-56 | chr6-120528155-DEL-56 |
| chr6-120529761-DEL-100 | chr6-120529742-DEL-80 |
| chr6-120534180-DEL-214 | chr6-120534181-DEL-214 |
| chr6-120538416-DEL-137 | chr6-120538415-DEL-138 |
| chr6-120542968-DEL-218 | chr6-120542969-DEL-218 |
| chr6-120545693-DEL-1298 | chr6-120545694-DEL-1298 |
| chr6-120550125-DEL-164 | chr6-120550095-DEL-164 |
| chr6-120554230-DEL-51 | chr6-120554230-DEL-51 |
| chr6-120557509-DEL-54 | chr6-120557510-DEL-58 |
| chr6-120561799-DEL-233 | chr6-120561802-DEL-231 |
| chr6-120571310-DEL-166 | chr6-120571311-DEL-166 |
| chr6-120574172-DEL-172 | chr6-120574166-DEL-173 |
| chr6-120582788-DEL-54 | chr6-120582789-DEL-54 |
| chr6-120586029-DEL-91 | chr6-120586030-DEL-91 |
| chr6-120586955-DEL-313 | chr6-120586956-DEL-313 |
| chr6-120588383-DEL-198 | chr6-120588384-DEL-183 |
| chr6-120603556-DEL-60 | chr6-120603557-DEL-60 |
| chr6-120613859-DEL-96 | chr6-120613872-DEL-84 |
| chr6-120624519-DEL-209 | chr6-120624520-DEL-209 |
| chr6-120627215-DEL-413 | chr6-120627216-DEL-413 |
| chr6-120679830-DEL-3972 | chr6-120679830-DEL-3972 |
| chr6-120704746-DEL-197 | chr6-120704747-DEL-197 |
| chr6-120718372-DEL-205 | chr6-120718373-DEL-205 |
| chr6-120719579-DEL-54 | chr6-120719580-DEL-54 |
| chr6-120724372-DEL-109 | chr6-120724373-DEL-109 |
| chr6-120736008-DEL-203 | chr6-120736009-DEL-203 |
| chr6-120760856-DEL-203 | chr6-120760858-DEL-202 |
| chr6-120764076-DEL-66 | chr6-120764077-DEL-66 |
| chr6-120778298-DEL-182 | chr6-120778299-DEL-182 |
| chr6-120788560-DEL-168 | chr6-120788561-DEL-168 |
| chr6-120789216-DEL-5131 | chr6-120789217-DEL-5131 |
| chr6-120807434-DEL-200 | chr6-120807435-DEL-200 |
| chr6-120810777-DEL-126 | chr6-120810778-DEL-126 |
| chr6-120818186-DEL-218 | chr6-120818187-DEL-218 |
| chr6-120876544-DEL-71 | chr6-120876545-DEL-71 |
| chr6-120943206-DEL-1890 | chr6-120943207-DEL-1890 |
| chr6-120950921-DEL-1894 | chr6-120950922-DEL-1894 |
| chr6-120954393-DEL-56 | chr6-120954394-DEL-56 |
| chr6-120962881-DEL-108 | chr6-120962882-DEL-108 |
| chr6-120968074-DEL-175 | chr6-120968075-DEL-175 |
| chr6-120969284-DEL-886 | chr6-120969285-DEL-886 |
| chr6-120979113-DEL-188 | chr6-120979098-DEL-187 |
| chr6-121048869-DEL-246 | chr6-121048870-DEL-246 |
| chr6-121052272-DEL-257 | chr6-121052273-DEL-257 |
| chr6-121071691-DEL-50 | chr6-121071689-DEL-51 |
| chr6-121090835-DEL-93 | chr6-121090836-DEL-93 |
| chr6-121093375-DEL-74 | chr6-121093376-DEL-74 |
| chr6-121102122-DEL-62 | chr6-121102129-DEL-56 |
| chr6-121117396-DEL-61 | chr6-121117397-DEL-61 |
| chr6-121163141-DEL-636 | chr6-121163142-DEL-636 |
| chr6-121177342-DEL-180 | chr6-121177342-DEL-180 |
| chr6-121201700-DEL-261 | chr6-121201677-DEL-288 |
| chr6-121207721-DEL-210 | chr6-121207707-DEL-202 |
| chr6-121260347-DEL-389 | chr6-121260364-DEL-381 |
| chr6-121278045-DEL-90 | chr6-121278044-DEL-91 |
| chr6-121289153-DEL-112 | chr6-121289154-DEL-112 |
| chr6-121297716-DEL-74 | chr6-121297719-DEL-72 |
| chr6-121341917-DEL-76 | chr6-121341918-DEL-76 |
| chr6-121377124-DEL-70 | chr6-121377125-DEL-70 |
| chr6-121462595-DEL-77 | chr6-121462634-DEL-50 |
| chr6-121482829-DEL-203 | chr6-121482830-DEL-203 |
| chr6-121483553-DEL-74 | chr6-121483556-DEL-52 |
| chr6-121517534-DEL-52 | chr6-121517535-DEL-52 |

| | |
|---|---|
| chr6-121531566-DEL-68 | chr6-121531555-DEL-64 |
| chr6-121542552-DEL-793 | chr6-121542553-DEL-793 |
| chr6-121565573-DEL-194 | chr6-121565574-DEL-194 |
| chr6-121598088-DEL-328 | chr6-121598104-DEL-321 |
| chr6-121603339-DEL-623 | chr6-121603340-DEL-623 |
| chr6-121604510-DEL-78 | chr6-121604546-DEL-78 |
| chr6-121620117-DEL-297 | chr6-121620117-DEL-297 |
| chr6-121627665-DEL-336 | chr6-121627666-DEL-336 |
| chr6-121638839-DEL-216 | chr6-121638840-DEL-216 |
| chr6-121687107-DEL-72 | chr6-121687108-DEL-108 |
| chr6-121795598-DEL-161 | chr6-121795599-DEL-161 |
| chr6-121797230-DEL-1632 | chr6-121797231-DEL-1632 |
| chr6-121799059-DEL-619 | chr6-121799060-DEL-619 |
| chr6-121809065-DEL-311 | chr6-121809066-DEL-311 |
| chr6-121809673-DEL-76 | chr6-121809658-DEL-72 |
| chr6-121811209-DEL-440 | chr6-121811210-DEL-440 |
| chr6-121814358-DEL-61 | chr6-121814359-DEL-61 |
| chr6-121852290-DEL-56 | chr6-121852291-DEL-56 |
| chr6-121870083-DEL-940 | chr6-121870084-DEL-940 |
| chr6-121946817-DEL-69 | chr6-121946818-DEL-69 |
| chr6-121959238-DEL-98 | chr6-121959252-DEL-60 |
| chr6-121997222-DEL-100 | chr6-121997209-DEL-82 |
| chr6-122019888-DEL-214 | chr6-122019888-DEL-214 |
| chr6-122073372-DEL-1481 | chr6-122073372-DEL-1481 |
| chr6-122091018-DEL-75 | chr6-122091019-DEL-75 |
| chr6-122123201-DEL-86 | chr6-122123204-DEL-84 |
| chr6-122226137-DEL-73 | chr6-122226138-DEL-73 |
| chr6-122226999-DEL-55 | chr6-122227000-DEL-83 |
| chr6-122262166-DEL-54 | chr6-122262167-DEL-54 |
| chr6-122264311-DEL-199 | chr6-122264312-DEL-199 |
| chr6-122351665-DEL-140 | chr6-122351666-DEL-140 |
| chr6-122352843-DEL-215 | chr6-122352841-DEL-217 |
| chr6-122397381-DEL-208 | chr6-122397384-DEL-206 |
| chr6-122433807-DEL-571 | chr6-122433808-DEL-571 |
| chr6-122439304-DEL-68 | chr6-122439312-DEL-64 |
| chr6-122452741-DEL-205 | chr6-122452742-DEL-205 |
| chr6-122454631-DEL-226 | chr6-122454632-DEL-226 |
| chr6-122460181-DEL-57 | chr6-122460182-DEL-57 |
| chr6-122498388-DEL-388 | chr6-122498389-DEL-388 |
| chr6-122513660-DEL-1616 | chr6-122513661-DEL-1616 |
| chr6-122520272-DEL-81 | chr6-122520273-DEL-81 |
| chr6-122527413-DEL-82 | chr6-122527414-DEL-82 |
| chr6-122560205-DEL-55 | chr6-122560206-DEL-55 |
| chr6-122563447-DEL-214 | chr6-122563448-DEL-214 |
| chr6-122573300-DEL-54 | chr6-122573300-DEL-65 |
| chr6-122589116-DEL-208 | chr6-122589117-DEL-208 |
| chr6-122608922-DEL-53 | chr6-122608928-DEL-54 |
| chr6-122617039-DEL-124 | chr6-122617040-DEL-123 |
| chr6-122633303-DEL-197 | chr6-122633304-DEL-197 |
| chr6-122644171-DEL-1583 | chr6-122644173-DEL-1580 |
| chr6-122650790-DEL-281 | chr6-122650791-DEL-281 |
| chr6-122660286-DEL-218 | chr6-122660287-DEL-218 |
| chr6-122670584-DEL-88 | chr6-122670560-DEL-128 |
| chr6-122698158-DEL-257 | chr6-122698159-DEL-257 |
| chr6-122708750-DEL-173 | chr6-122708751-DEL-173 |
| chr6-122761052-DEL-102 | chr6-122761053-DEL-102 |
| chr6-122803915-DEL-109 | chr6-122803916-DEL-109 |
| chr6-122819268-DEL-177 | chr6-122819257-DEL-176 |
| chr6-122823179-DEL-58 | chr6-122823180-DEL-58 |
| chr6-122870480-DEL-348 | chr6-122870481-DEL-348 |
| chr6-122886411-DEL-6886 | chr6-122886398-DEL-6888 |
| chr6-122892910-DEL-381 | chr6-122892911-DEL-381 |
| chr6-122903338-DEL-410 | chr6-122903339-DEL-410 |
| chr6-122908335-DEL-201 | chr6-122908336-DEL-201 |
| chr6-122913177-DEL-213 | chr6-122913178-DEL-213 |
| chr6-122925090-DEL-69 | chr6-122925065-DEL-60 |
| chr6-122931474-DEL-289 | chr6-122931475-DEL-289 |
| chr6-122932823-DEL-6586 | chr6-122932824-DEL-6586 |
| chr6-122945653-DEL-418 | chr6-122945654-DEL-418 |
| chr6-122971401-DEL-1319 | chr6-122971402-DEL-1319 |
| chr6-122988250-DEL-119 | chr6-122988250-DEL-119 |
| chr6-122991717-DEL-1502 | chr6-122991718-DEL-1502 |
| chr6-123013070-DEL-6508 | chr6-123013056-DEL-6522 |
| chr6-123020739-DEL-279 | chr6-123020740-DEL-279 |
| chr6-123024958-DEL-54 | chr6-123024914-DEL-55 |
| chr6-123030346-DEL-127 | chr6-123030346-DEL-128 |
| chr6-123034982-DEL-254 | chr6-123034983-DEL-254 |
| chr6-123040372-DEL-182 | chr6-123040373-DEL-182 |
| chr6-123046265-DEL-55 | chr6-123046269-DEL-60 |
| chr6-123070398-DEL-355 | chr6-123070399-DEL-355 |

| | |
|---|---|
| chr6-123124290-DEL-1021 | chr6-123124278-DEL-1020 |
| chr6-123132143-DEL-70 | chr6-123132185-DEL-51 |
| chr6-123135270-DEL-5708 | chr6-123135271-DEL-5708 |
| chr6-123171327-DEL-2943 | chr6-123171328-DEL-2943 |
| chr6-123172620-DEL-53 | chr6-123172621-DEL-53 |
| chr6-123173556-DEL-600 | chr6-123173557-DEL-600 |
| chr6-123196495-DEL-169 | chr6-123196496-DEL-169 |
| chr6-123210798-DEL-150 | chr6-123210799-DEL-150 |
| chr6-123214957-DEL-6208 | chr6-123214957-DEL-6208 |
| chr6-123248690-DEL-288 | chr6-123248691-DEL-288 |
| chr6-123249267-DEL-58 | chr6-123249217-DEL-59 |
| chr6-123252630-DEL-1113 | chr6-123252631-DEL-1113 |
| chr6-123252906-DEL-236 | chr6-123252907-DEL-236 |
| chr6-123261395-DEL-712 | chr6-123261396-DEL-712 |
| chr6-123265384-DEL-51 | chr6-123265385-DEL-51 |
| chr6-123269487-DEL-293 | chr6-123269488-DEL-292 |
| chr6-123275941-DEL-1473 | chr6-123275935-DEL-1490 |
| chr6-123280573-DEL-344 | chr6-123280578-DEL-345 |
| chr6-123289506-DEL-121 | chr6-123289512-DEL-120 |
| chr6-123295308-DEL-401 | chr6-123295309-DEL-401 |
| chr6-123307157-DEL-235 | chr6-123307158-DEL-235 |
| chr6-123363829-DEL-4768 | chr6-123363829-DEL-4768 |
| chr6-123398668-DEL-82 | chr6-123398669-DEL-82 |
| chr6-123452654-DEL-169 | chr6-123452679-DEL-146 |
| chr6-123473169-DEL-244 | chr6-123473170-DEL-244 |
| chr6-123548187-DEL-926 | chr6-123548186-DEL-926 |
| chr6-123567918-DEL-96 | chr6-123567918-DEL-88 |
| chr6-123579029-DEL-1888 | chr6-123579029-DEL-1888 |
| chr6-123590394-DEL-244 | chr6-123590393-DEL-244 |
| chr6-123612417-DEL-69 | chr6-123612418-DEL-69 |
| chr6-123619791-DEL-1804 | chr6-123619792-DEL-1804 |
| chr6-123654325-DEL-7192 | chr6-123654326-DEL-7192 |
| chr6-123687029-DEL-79 | chr6-123687031-DEL-63 |
| chr6-123696230-DEL-65 | chr6-123696213-DEL-61 |
| chr6-123701716-DEL-231 | chr6-123701717-DEL-231 |
| chr6-123703674-DEL-128 | chr6-123703675-DEL-128 |
| chr6-123716902-DEL-54 | chr6-123716874-DEL-53 |
| chr6-123726869-DEL-544 | chr6-123726870-DEL-544 |
| chr6-123731387-DEL-413 | chr6-123731388-DEL-413 |
| chr6-123733187-DEL-1337 | chr6-123733194-DEL-1338 |
| chr6-123781228-DEL-776 | chr6-123781229-DEL-776 |
| chr6-123781801-DEL-85 | chr6-123781802-DEL-85 |
| chr6-123812082-DEL-272 | chr6-123812083-DEL-272 |
| chr6-123877464-DEL-457 | chr6-123877464-DEL-457 |
| chr6-123917754-DEL-204 | chr6-123917755-DEL-204 |
| chr6-123933486-DEL-60 | chr6-123933487-DEL-60 |
| chr6-123936092-DEL-146 | chr6-123936093-DEL-146 |
| chr6-123948838-DEL-178 | chr6-123948839-DEL-178 |
| chr6-123998589-DEL-71 | chr6-123998554-DEL-56 |
| chr6-124011688-DEL-53 | chr6-124011689-DEL-55 |
| chr6-124012691-DEL-520 | chr6-124012692-DEL-520 |
| chr6-124044764-DEL-58 | chr6-124044769-DEL-59 |
| chr6-124082851-DEL-969 | chr6-124082852-DEL-969 |
| chr6-124108702-DEL-6203 | chr6-124108703-DEL-6203 |
| chr6-124139540-DEL-58 | chr6-124139541-DEL-58 |
| chr6-124167290-DEL-433 | chr6-124167291-DEL-433 |
| chr6-124188338-DEL-7161 | chr6-124188339-DEL-7161 |
| chr6-124207602-DEL-444 | chr6-124207603-DEL-438 |
| chr6-124293191-DEL-229 | chr6-124293192-DEL-207 |
| chr6-124308662-DEL-6440 | chr6-124308663-DEL-6440 |
| chr6-124358859-DEL-425 | chr6-124358860-DEL-425 |
| chr6-124419222-DEL-716 | chr6-124419223-DEL-716 |
| chr6-124424063-DEL-218 | chr6-124424064-DEL-218 |
| chr6-124434530-DEL-56 | chr6-124434532-DEL-60 |
| chr6-124445501-DEL-1846 | chr6-124445502-DEL-1846 |
| chr6-124446267-DEL-60 | chr6-124446268-DEL-56 |
| chr6-124461087-DEL-80 | chr6-124461088-DEL-80 |
| chr6-124462480-DEL-131 | chr6-124462481-DEL-131 |
| chr6-124467765-DEL-54 | chr6-124467766-DEL-54 |
| chr6-124474303-DEL-58 | chr6-124474304-DEL-58 |
| chr6-124490118-DEL-54 | chr6-124490118-DEL-55 |
| chr6-124490917-DEL-531 | chr6-124490918-DEL-531 |
| chr6-124500566-DEL-249 | chr6-124500564-DEL-251 |
| chr6-124501087-DEL-129 | chr6-124501092-DEL-112 |
| chr6-124505732-DEL-84 | chr6-124505733-DEL-84 |
| chr6-124552723-DEL-75 | chr6-124552722-DEL-70 |
| chr6-124559556-DEL-371 | chr6-124559557-DEL-371 |
| chr6-124685438-DEL-72 | chr6-124685421-DEL-64 |
| chr6-124702499-DEL-74 | chr6-124702500-DEL-73 |
| chr6-124703694-DEL-204 | chr6-124703695-DEL-204 |

| | |
|---|---|
| chr6-124749212-DEL-160 | chr6-124749213-DEL-160 |
| chr6-124759094-DEL-209 | chr6-124759095-DEL-209 |
| chr6-124762422-DEL-202 | chr6-124762423-DEL-202 |
| chr6-124765132-DEL-886 | chr6-124765118-DEL-918 |
| chr6-124768672-DEL-194 | chr6-124768673-DEL-194 |
| chr6-124773986-DEL-5512 | chr6-124773987-DEL-5512 |
| chr6-124827011-DEL-93 | chr6-124827012-DEL-93 |
| chr6-124851692-DEL-114 | chr6-124851693-DEL-114 |
| chr6-124915907-DEL-217 | chr6-124915908-DEL-217 |
| chr6-124919236-DEL-225 | chr6-124919236-DEL-225 |
| chr6-124926985-DEL-219 | chr6-124926985-DEL-219 |
| chr6-124951026-DEL-7108 | chr6-124951027-DEL-7108 |
| chr6-124966081-DEL-52 | chr6-124966076-DEL-56 |
| chr6-125020089-DEL-228 | chr6-125020090-DEL-228 |
| chr6-125022354-DEL-216 | chr6-125022355-DEL-216 |
| chr6-125042414-DEL-221 | chr6-125042443-DEL-239 |
| chr6-125092655-DEL-86 | chr6-125092660-DEL-86 |
| chr6-125107842-DEL-80 | chr6-125107811-DEL-80 |
| chr6-125115911-DEL-201 | chr6-125115912-DEL-201 |
| chr6-125135259-DEL-207 | chr6-125135260-DEL-207 |
| chr6-125145346-DEL-60 | chr6-125145347-DEL-60 |
| chr6-125160136-DEL-211 | chr6-125160137-DEL-211 |
| chr6-125182975-DEL-156 | chr6-125182976-DEL-156 |
| chr6-125191729-DEL-72 | chr6-125191730-DEL-72 |
| chr6-125245569-DEL-317 | chr6-125245570-DEL-317 |
| chr6-125275304-DEL-387 | chr6-125275305-DEL-387 |
| chr6-125281665-DEL-212 | chr6-125281672-DEL-216 |
| chr6-125311769-DEL-54 | chr6-125311770-DEL-54 |
| chr6-125342305-DEL-91 | chr6-125342306-DEL-91 |
| chr6-125360612-DEL-232 | chr6-125360607-DEL-231 |
| chr6-125385220-DEL-241 | chr6-125385239-DEL-223 |
| chr6-125389653-DEL-194 | chr6-125389654-DEL-194 |
| chr6-125395446-DEL-181 | chr6-125395427-DEL-200 |
| chr6-125410322-DEL-70 | chr6-125410323-DEL-70 |
| chr6-125426230-DEL-422 | chr6-125426226-DEL-420 |
| chr6-125428807-DEL-82 | chr6-125428808-DEL-82 |
| chr6-125484977-DEL-316 | chr6-125484983-DEL-299 |
| chr6-125487091-DEL-78 | chr6-125487105-DEL-52 |
| chr6-125522372-DEL-186 | chr6-125522373-DEL-186 |
| chr6-125548564-DEL-722 | chr6-125548565-DEL-722 |
| chr6-125571522-DEL-59 | chr6-125571565-DEL-59 |
| chr6-125587890-DEL-223 | chr6-125587889-DEL-224 |
| chr6-125604771-DEL-58 | chr6-125604768-DEL-62 |
| chr6-125605752-DEL-144 | chr6-125605753-DEL-144 |
| chr6-125610631-DEL-574 | chr6-125610632-DEL-574 |
| chr6-125615431-DEL-108 | chr6-125615432-DEL-108 |
| chr6-125620990-DEL-91 | chr6-125620991-DEL-91 |
| chr6-125709539-DEL-194 | chr6-125709541-DEL-190 |
| chr6-125712106-DEL-945 | chr6-125712107-DEL-945 |
| chr6-125759625-DEL-52 | chr6-125759626-DEL-52 |
| chr6-125779466-DEL-401 | chr6-125779467-DEL-401 |
| chr6-125796880-DEL-108 | chr6-125796881-DEL-108 |
| chr6-125805124-DEL-1891 | chr6-125805125-DEL-1891 |
| chr6-125813495-DEL-75 | chr6-125813481-DEL-66 |
| chr6-125815826-DEL-504 | chr6-125815827-DEL-504 |
| chr6-125864461-DEL-1008 | chr6-125864462-DEL-1008 |
| chr6-125867940-DEL-2101 | chr6-125867941-DEL-2101 |
| chr6-125934886-DEL-1479 | chr6-125934887-DEL-1479 |
| chr6-125944622-DEL-82 | chr6-125944622-DEL-83 |
| chr6-125986515-DEL-58 | chr6-125986516-DEL-58 |
| chr6-125989051-DEL-52 | chr6-125989054-DEL-50 |
| chr6-125994599-DEL-56 | chr6-125994611-DEL-51 |
| chr6-126001301-DEL-7229 | chr6-126001302-DEL-7229 |
| chr6-126029004-DEL-62 | chr6-126029015-DEL-63 |
| chr6-126035472-DEL-939 | chr6-126035473-DEL-939 |
| chr6-126052059-DEL-85 | chr6-126052084-DEL-61 |
| chr6-126076783-DEL-95 | chr6-126076763-DEL-92 |
| chr6-126086754-DEL-100 | chr6-126086755-DEL-100 |
| chr6-126105174-DEL-77 | chr6-126105148-DEL-56 |
| chr6-126109055-DEL-297 | chr6-126109056-DEL-297 |
| chr6-126112281-DEL-1892 | chr6-126112282-DEL-1892 |
| chr6-126151305-DEL-2097 | chr6-126151306-DEL-2097 |
| chr6-126156066-DEL-218 | chr6-126156067-DEL-218 |
| chr6-126183634-DEL-150 | chr6-126183635-DEL-150 |
| chr6-126189364-DEL-52 | chr6-126189367-DEL-52 |
| chr6-126206068-DEL-72 | chr6-126206069-DEL-72 |
| chr6-126210485-DEL-56 | chr6-126210468-DEL-58 |
| chr6-126225789-DEL-214 | chr6-126225790-DEL-214 |
| chr6-126266389-DEL-234 | chr6-126266390-DEL-234 |
| chr6-126315750-DEL-1131 | chr6-126315751-DEL-1131 |

| | |
|---|---|
| chr6-126336894-DEL-137 | chr6-126336895-DEL-137 |
| chr6-126350029-DEL-56 | chr6-126350030-DEL-56 |
| chr6-126388915-DEL-6279 | chr6-126388916-DEL-6279 |
| chr6-126396148-DEL-55 | chr6-126396149-DEL-55 |
| chr6-126432821-DEL-6894 | chr6-126432822-DEL-6894 |
| chr6-126433034-DEL-424 | chr6-126433034-DEL-424 |
| chr6-126453134-DEL-77 | chr6-126453135-DEL-77 |
| chr6-126454104-DEL-1685 | chr6-126454105-DEL-1685 |
| chr6-126484479-DEL-342 | chr6-126484480-DEL-342 |
| chr6-126489548-DEL-60 | chr6-126489549-DEL-60 |
| chr6-126495309-DEL-80 | chr6-126495310-DEL-80 |
| chr6-126554066-DEL-79 | chr6-126554064-DEL-79 |
| chr6-126565476-DEL-162 | chr6-126565477-DEL-165 |
| chr6-126625652-DEL-67 | chr6-126625627-DEL-92 |
| chr6-126631191-DEL-54 | chr6-126631192-DEL-54 |
| chr6-126635550-DEL-52 | chr6-126635542-DEL-54 |
| chr6-126640855-DEL-86 | chr6-126640813-DEL-129 |
| chr6-126665025-DEL-326 | chr6-126665014-DEL-327 |
| chr6-126679625-DEL-139 | chr6-126679626-DEL-139 |
| chr6-126690642-DEL-260 | chr6-126690648-DEL-255 |
| chr6-126722210-DEL-80 | chr6-126722215-DEL-56 |
| chr6-126776887-DEL-429 | chr6-126776892-DEL-426 |
| chr6-126780734-DEL-212 | chr6-126780733-DEL-207 |
| chr6-126862592-DEL-188 | chr6-126862593-DEL-188 |
| chr6-126865001-DEL-64 | chr6-126865002-DEL-64 |
| chr6-126879304-DEL-3927 | chr6-126879305-DEL-3927 |
| chr6-126919718-DEL-78 | chr6-126919719-DEL-78 |
| chr6-127035480-DEL-54 | chr6-127035481-DEL-54 |
| chr6-127072297-DEL-5156 | chr6-127072298-DEL-5156 |
| chr6-127093297-DEL-116 | chr6-127093299-DEL-112 |
| chr6-127169404-DEL-52 | chr6-127169406-DEL-51 |
| chr6-127204982-DEL-54 | chr6-127204983-DEL-54 |
| chr6-127310461-DEL-70 | chr6-127310461-DEL-72 |
| chr6-127326140-DEL-343 | chr6-127326141-DEL-343 |
| chr6-127354392-DEL-208 | chr6-127354382-DEL-209 |
| chr6-127355945-DEL-471 | chr6-127355946-DEL-471 |
| chr6-127363101-DEL-363 | chr6-127363102-DEL-363 |
| chr6-127371897-DEL-995 | chr6-127371898-DEL-995 |
| chr6-127375173-DEL-80 | chr6-127375173-DEL-80 |
| chr6-127398185-DEL-90 | chr6-127398160-DEL-110 |
| chr6-127412624-DEL-108 | chr6-127412625-DEL-108 |
| chr6-127472574-DEL-126 | chr6-127472574-DEL-136 |
| chr6-127490985-DEL-6947 | chr6-127490986-DEL-6947 |
| chr6-127504080-DEL-120 | chr6-127504081-DEL-120 |
| chr6-127507580-DEL-382 | chr6-127507614-DEL-367 |
| chr6-127519649-DEL-727 | chr6-127519650-DEL-727 |
| chr6-127565563-DEL-497 | chr6-127565564-DEL-497 |
| chr6-127575028-DEL-72 | chr6-127575023-DEL-78 |
| chr6-127603504-DEL-66 | chr6-127603498-DEL-81 |
| chr6-127613962-DEL-53 | chr6-127613950-DEL-54 |
| chr6-127642697-DEL-51 | chr6-127642698-DEL-51 |
| chr6-127682726-DEL-100 | chr6-127682719-DEL-108 |
| chr6-127714126-DEL-443 | chr6-127714133-DEL-442 |
| chr6-127752311-DEL-85 | chr6-127752312-DEL-85 |
| chr6-127836894-DEL-66 | chr6-127836912-DEL-57 |
| chr6-127841875-DEL-196 | chr6-127841876-DEL-196 |
| chr6-127852131-DEL-64 | chr6-127852132-DEL-64 |
| chr6-127857587-DEL-131 | chr6-127857564-DEL-119 |
| chr6-127859013-DEL-62 | chr6-127859020-DEL-59 |
| chr6-127880685-DEL-54 | chr6-127880686-DEL-54 |
| chr6-127890949-DEL-379 | chr6-127890959-DEL-349 |
| chr6-127913438-DEL-269 | chr6-127913439-DEL-269 |
| chr6-127922577-DEL-72 | chr6-127922577-DEL-72 |
| chr6-127947085-DEL-135 | chr6-127947088-DEL-133 |
| chr6-127948575-DEL-190 | chr6-127948577-DEL-189 |
| chr6-127973705-DEL-275 | chr6-127973706-DEL-275 |
| chr6-128017837-DEL-98 | chr6-128017838-DEL-98 |
| chr6-128030527-DEL-92 | chr6-128030528-DEL-92 |
| chr6-128095086-DEL-211 | chr6-128095087-DEL-211 |
| chr6-128101891-DEL-669 | chr6-128101892-DEL-669 |
| chr6-128141282-DEL-598 | chr6-128141283-DEL-598 |
| chr6-128178906-DEL-1362 | chr6-128178907-DEL-1362 |
| chr6-128211432-DEL-84 | chr6-128211433-DEL-84 |
| chr6-128212623-DEL-93 | chr6-128212624-DEL-87 |
| chr6-128299897-DEL-54 | chr6-128299898-DEL-54 |
| chr6-128308587-DEL-242 | chr6-128308598-DEL-243 |
| chr6-128318278-DEL-213 | chr6-128318279-DEL-213 |
| chr6-128330465-DEL-206 | chr6-128330466-DEL-206 |
| chr6-128333446-DEL-284 | chr6-128333447-DEL-283 |
| chr6-128346842-DEL-58 | chr6-128346830-DEL-76 |

| | |
|---|---|
| chr6-128358073-DEL-64 | chr6-128358074-DEL-64 |
| chr6-128368453-DEL-51 | chr6-128368453-DEL-77 |
| chr6-128384602-DEL-280 | chr6-128384603-DEL-280 |
| chr6-128397666-DEL-56 | chr6-128397667-DEL-56 |
| chr6-128428855-DEL-72 | chr6-128428855-DEL-73 |
| chr6-128449603-DEL-766 | chr6-128449604-DEL-766 |
| chr6-128461032-DEL-237 | chr6-128461033-DEL-237 |
| chr6-128481390-DEL-221 | chr6-128481391-DEL-221 |
| chr6-128497446-DEL-93 | chr6-128497421-DEL-94 |
| chr6-128502522-DEL-63 | chr6-128502522-DEL-63 |
| chr6-128508460-DEL-400 | chr6-128508461-DEL-400 |
| chr6-128509057-DEL-115 | chr6-128509028-DEL-130 |
| chr6-128512015-DEL-66 | chr6-128512026-DEL-62 |
| chr6-128515538-DEL-294 | chr6-128515539-DEL-294 |
| chr6-128518839-DEL-119 | chr6-128518840-DEL-119 |
| chr6-128542650-DEL-399 | chr6-128542651-DEL-399 |
| chr6-128586545-DEL-137 | chr6-128586551-DEL-132 |
| chr6-128597417-DEL-274 | chr6-128597418-DEL-274 |
| chr6-128598855-DEL-195 | chr6-128598856-DEL-195 |
| chr6-128606076-DEL-59 | chr6-128606074-DEL-61 |
| chr6-128610516-DEL-196 | chr6-128610517-DEL-196 |
| chr6-128611467-DEL-321 | chr6-128611468-DEL-321 |
| chr6-128621785-DEL-2294 | chr6-128621786-DEL-2294 |
| chr6-128625245-DEL-110 | chr6-128625242-DEL-111 |
| chr6-128628068-DEL-511 | chr6-128628069-DEL-511 |
| chr6-128630255-DEL-1642 | chr6-128630255-DEL-1642 |
| chr6-128636240-DEL-1281 | chr6-128636241-DEL-1281 |
| chr6-128657037-DEL-7596 | chr6-128657038-DEL-7596 |
| chr6-128659773-DEL-823 | chr6-128659773-DEL-823 |
| chr6-128666423-DEL-4373 | chr6-128666425-DEL-4372 |
| chr6-128685608-DEL-1104 | chr6-128685609-DEL-1104 |
| chr6-128694640-DEL-462 | chr6-128694639-DEL-462 |
| chr6-128697195-DEL-208 | chr6-128697195-DEL-208 |
| chr6-128702723-DEL-254 | chr6-128702711-DEL-255 |
| chr6-128710251-DEL-411 | chr6-128710252-DEL-411 |
| chr6-128729683-DEL-324 | chr6-128729684-DEL-324 |
| chr6-128741645-DEL-317 | chr6-128741646-DEL-317 |
| chr6-128746252-DEL-133 | chr6-128746252-DEL-132 |
| chr6-128748608-DEL-803 | chr6-128748609-DEL-803 |
| chr6-128757698-DEL-217 | chr6-128757699-DEL-217 |
| chr6-128759617-DEL-712 | chr6-128759623-DEL-712 |
| chr6-128763089-DEL-246 | chr6-128763091-DEL-232 |
| chr6-128764533-DEL-183 | chr6-128764532-DEL-184 |
| chr6-128798363-DEL-430 | chr6-128798374-DEL-427 |
| chr6-128822903-DEL-202 | chr6-128822904-DEL-202 |
| chr6-128828447-DEL-94 | chr6-128828447-DEL-94 |
| chr6-128829398-DEL-196 | chr6-128829348-DEL-202 |
| chr6-128833676-DEL-360 | chr6-128833677-DEL-360 |
| chr6-128837052-DEL-7037 | chr6-128837051-DEL-7037 |
| chr6-128881507-DEL-7273 | chr6-128881508-DEL-7273 |
| chr6-128898700-DEL-75 | chr6-128898741-DEL-75 |
| chr6-128903823-DEL-145 | chr6-128903824-DEL-145 |
| chr6-128910702-DEL-515 | chr6-128910703-DEL-515 |
| chr6-128913223-DEL-5373 | chr6-128913224-DEL-5373 |
| chr6-128931942-DEL-297 | chr6-128931943-DEL-297 |
| chr6-128947896-DEL-374 | chr6-128947891-DEL-375 |
| chr6-128977524-DEL-65 | chr6-128977524-DEL-65 |
| chr6-128981238-DEL-209 | chr6-128981239-DEL-209 |
| chr6-128988977-DEL-489 | chr6-128988978-DEL-489 |
| chr6-128994821-DEL-453 | chr6-128994822-DEL-453 |
| chr6-129021872-DEL-72 | chr6-129021885-DEL-58 |
| chr6-129025650-DEL-1567 | chr6-129025651-DEL-1567 |
| chr6-129065677-DEL-53 | chr6-129065678-DEL-53 |
| chr6-129073005-DEL-54 | chr6-129073005-DEL-104 |
| chr6-129090668-DEL-115 | chr6-129090669-DEL-115 |
| chr6-129097536-DEL-214 | chr6-129097537-DEL-214 |
| chr6-129098026-DEL-1145 | chr6-129098027-DEL-1145 |
| chr6-129099005-DEL-344 | chr6-129099006-DEL-344 |
| chr6-129141853-DEL-5977 | chr6-129141854-DEL-5977 |
| chr6-129153143-DEL-1323 | chr6-129153144-DEL-1323 |
| chr6-129164678-DEL-66 | chr6-129164676-DEL-60 |
| chr6-129177055-DEL-84 | chr6-129177056-DEL-84 |
| chr6-129178108-DEL-970 | chr6-129178104-DEL-974 |
| chr6-129179818-DEL-465 | chr6-129179819-DEL-465 |
| chr6-129182646-DEL-2340 | chr6-129182647-DEL-2340 |
| chr6-129193860-DEL-4568 | chr6-129193861-DEL-4568 |
| chr6-129200858-DEL-91 | chr6-129200859-DEL-91 |
| chr6-129206421-DEL-64 | chr6-129206427-DEL-57 |
| chr6-129211268-DEL-136 | chr6-129211265-DEL-142 |
| chr6-129228082-DEL-3741 | chr6-129228083-DEL-3741 |

| | |
|---|---|
| chr6-129232733-DEL-60 | chr6-129232735-DEL-59 |
| chr6-129241218-DEL-86 | chr6-129241204-DEL-87 |
| chr6-129263013-DEL-58 | chr6-129263014-DEL-58 |
| chr6-129276647-DEL-56 | chr6-129276647-DEL-52 |
| chr6-129283884-DEL-53 | chr6-129283883-DEL-52 |
| chr6-129286982-DEL-2528 | chr6-129286983-DEL-2528 |
| chr6-129316354-DEL-1443 | chr6-129316355-DEL-1443 |
| chr6-129335997-DEL-71 | chr6-129335998-DEL-71 |
| chr6-129337011-DEL-80 | chr6-129337012-DEL-80 |
| chr6-129338797-DEL-65 | chr6-129338780-DEL-53 |
| chr6-129343711-DEL-187 | chr6-129343712-DEL-187 |
| chr6-129354971-DEL-76 | chr6-129354972-DEL-77 |
| chr6-129368138-DEL-97 | chr6-129368139-DEL-97 |
| chr6-129369755-DEL-68 | chr6-129369760-DEL-64 |
| chr6-129409933-DEL-106 | chr6-129409936-DEL-107 |
| chr6-129436309-DEL-199 | chr6-129436310-DEL-199 |
| chr6-129507545-DEL-824 | chr6-129507545-DEL-825 |
| chr6-129507668-DEL-197 | chr6-129507669-DEL-197 |
| chr6-129546291-DEL-1274 | chr6-129546277-DEL-1276 |
| chr6-129554350-DEL-510 | chr6-129554351-DEL-510 |
| chr6-129566035-DEL-54 | chr6-129566036-DEL-54 |
| chr6-129571429-DEL-202 | chr6-129571430-DEL-202 |
| chr6-129574504-DEL-2385 | chr6-129574505-DEL-2384 |
| chr6-129617483-DEL-52 | chr6-129617484-DEL-52 |
| chr6-129647751-DEL-205 | chr6-129647752-DEL-205 |
| chr6-129647868-DEL-978 | chr6-129647869-DEL-978 |
| chr6-129671164-DEL-154 | chr6-129671165-DEL-154 |
| chr6-129686574-DEL-210 | chr6-129686574-DEL-210 |
| chr6-129703768-DEL-920 | chr6-129703769-DEL-920 |
| chr6-129714013-DEL-1121 | chr6-129714014-DEL-1121 |
| chr6-129716470-DEL-1686 | chr6-129716471-DEL-1686 |
| chr6-129725024-DEL-668 | chr6-129725025-DEL-668 |
| chr6-129727573-DEL-108 | chr6-129727574-DEL-108 |
| chr6-129729202-DEL-57 | chr6-129729239-DEL-58 |
| chr6-129752029-DEL-60 | chr6-129752030-DEL-62 |
| chr6-129754804-DEL-361 | chr6-129754805-DEL-361 |
| chr6-129773931-DEL-3603 | chr6-129773932-DEL-3602 |
| chr6-129780314-DEL-56 | chr6-129780315-DEL-56 |
| chr6-129780853-DEL-8758 | chr6-129780854-DEL-8758 |
| chr6-129815316-DEL-238 | chr6-129815317-DEL-238 |
| chr6-129818499-DEL-344 | chr6-129818500-DEL-344 |
| chr6-129824513-DEL-8989 | chr6-129824514-DEL-8989 |
| chr6-129833754-DEL-7066 | chr6-129833740-DEL-7080 |
| chr6-129859343-DEL-224 | chr6-129859350-DEL-248 |
| chr6-129867396-DEL-50 | chr6-129867397-DEL-50 |
| chr6-129868605-DEL-289 | chr6-129868606-DEL-289 |
| chr6-129893369-DEL-63 | chr6-129893370-DEL-63 |
| chr6-129895495-DEL-118 | chr6-129895495-DEL-118 |
| chr6-129912620-DEL-7081 | chr6-129912620-DEL-7081 |
| chr6-129932532-DEL-4210 | chr6-129932533-DEL-4210 |
| chr6-129954468-DEL-143 | chr6-129954468-DEL-143 |
| chr6-129965071-DEL-939 | chr6-129965072-DEL-939 |
| chr6-129976714-DEL-289 | chr6-129976715-DEL-289 |
| chr6-129983099-DEL-84 | chr6-129983100-DEL-84 |
| chr6-130011028-DEL-229 | chr6-130011027-DEL-229 |
| chr6-130016441-DEL-226 | chr6-130016408-DEL-259 |
| chr6-130075539-DEL-559 | chr6-130075539-DEL-552 |
| chr6-130084113-DEL-921 | chr6-130084114-DEL-921 |
| chr6-130115536-DEL-430 | chr6-130115536-DEL-430 |
| chr6-130174230-DEL-86 | chr6-130174228-DEL-84 |
| chr6-130190896-DEL-64 | chr6-130190910-DEL-50 |
| chr6-130221056-DEL-416 | chr6-130221057-DEL-416 |
| chr6-130232974-DEL-348 | chr6-130232974-DEL-348 |
| chr6-130238688-DEL-112 | chr6-130238708-DEL-80 |
| chr6-130287811-DEL-83 | chr6-130287835-DEL-58 |
| chr6-130290902-DEL-380 | chr6-130290901-DEL-380 |
| chr6-130302324-DEL-63 | chr6-130302324-DEL-63 |
| chr6-130326709-DEL-56 | chr6-130326722-DEL-50 |
| chr6-130335058-DEL-2656 | chr6-130335057-DEL-2656 |
| chr6-130344868-DEL-68 | chr6-130344867-DEL-68 |
| chr6-130357874-DEL-92 | chr6-130357875-DEL-92 |
| chr6-130369922-DEL-76 | chr6-130369923-DEL-76 |
| chr6-130395140-DEL-54 | chr6-130395141-DEL-54 |
| chr6-130415841-DEL-20942 | chr6-130415841-DEL-20942 |
| chr6-130419408-DEL-108 | chr6-130419409-DEL-108 |
| chr6-130449107-DEL-66 | chr6-130449108-DEL-66 |
| chr6-130450873-DEL-52 | chr6-130450882-DEL-50 |
| chr6-130477834-DEL-64 | chr6-130477833-DEL-64 |
| chr6-130497902-DEL-83 | chr6-130497903-DEL-83 |
| chr6-130499540-DEL-2915 | chr6-130499541-DEL-2915 |

| | |
|---|---|
| chr6-130524183-DEL-238 | chr6-130524184-DEL-238 |
| chr6-130575126-DEL-267 | chr6-130575127-DEL-267 |
| chr6-130585138-DEL-522 | chr6-130585139-DEL-522 |
| chr6-130633386-DEL-76 | chr6-130633383-DEL-56 |
| chr6-130651456-DEL-204 | chr6-130651457-DEL-204 |
| chr6-130667052-DEL-1018 | chr6-130667053-DEL-1018 |
| chr6-130673294-DEL-84 | chr6-130673284-DEL-72 |
| chr6-130693966-DEL-810 | chr6-130693967-DEL-810 |
| chr6-130704026-DEL-68 | chr6-130704026-DEL-72 |
| chr6-130742263-DEL-1454 | chr6-130742263-DEL-1454 |
| chr6-130827723-DEL-60 | chr6-130827720-DEL-60 |
| chr6-130845373-DEL-252 | chr6-130845374-DEL-252 |
| chr6-130848304-DEL-70 | chr6-130848282-DEL-60 |
| chr6-130868495-DEL-52 | chr6-130868496-DEL-52 |
| chr6-130883545-DEL-812 | chr6-130883546-DEL-812 |
| chr6-130915989-DEL-84 | chr6-130915988-DEL-84 |
| chr6-130920970-DEL-58 | chr6-130920970-DEL-54 |
| chr6-130931630-DEL-7090 | chr6-130931631-DEL-7090 |
| chr6-130964649-DEL-168 | chr6-130964648-DEL-168 |
| chr6-130979983-DEL-399 | chr6-130979982-DEL-399 |
| chr6-130986209-DEL-1137 | chr6-130986210-DEL-1137 |
| chr6-130993474-DEL-1456 | chr6-130993473-DEL-1456 |
| chr6-131014262-DEL-84 | chr6-131014263-DEL-84 |
| chr6-131033755-DEL-7491 | chr6-131033756-DEL-7491 |
| chr6-131049916-DEL-68 | chr6-131049917-DEL-53 |
| chr6-131074025-DEL-74 | chr6-131074026-DEL-73 |
| chr6-131081946-DEL-2539 | chr6-131081947-DEL-2539 |
| chr6-131114856-DEL-140 | chr6-131114857-DEL-140 |
| chr6-131117693-DEL-2108 | chr6-131117693-DEL-2108 |
| chr6-131146183-DEL-87 | chr6-131146184-DEL-79 |
| chr6-131155391-DEL-1825 | chr6-131155392-DEL-1825 |
| chr6-131193262-DEL-582 | chr6-131193263-DEL-582 |
| chr6-131229213-DEL-2445 | chr6-131229213-DEL-2446 |
| chr6-131241431-DEL-62 | chr6-131241432-DEL-50 |
| chr6-131284367-DEL-56 | chr6-131284368-DEL-56 |
| chr6-131326208-DEL-399 | chr6-131326209-DEL-399 |
| chr6-131327374-DEL-55 | chr6-131327375-DEL-55 |
| chr6-131328199-DEL-216 | chr6-131328203-DEL-217 |
| chr6-131333348-DEL-1237 | chr6-131333349-DEL-1237 |
| chr6-131335250-DEL-445 | chr6-131335251-DEL-445 |
| chr6-131350812-DEL-190 | chr6-131350813-DEL-190 |
| chr6-131375433-DEL-344 | chr6-131375434-DEL-344 |
| chr6-131401476-DEL-942 | chr6-131401476-DEL-942 |
| chr6-131402532-DEL-1776 | chr6-131402532-DEL-1776 |
| chr6-131403631-DEL-199 | chr6-131403632-DEL-199 |
| chr6-131405907-DEL-2247 | chr6-131405908-DEL-2248 |
| chr6-131417368-DEL-4346 | chr6-131417369-DEL-4346 |
| chr6-131425450-DEL-497 | chr6-131425451-DEL-497 |
| chr6-131455861-DEL-60 | chr6-131455869-DEL-56 |
| chr6-131460337-DEL-95 | chr6-131460343-DEL-90 |
| chr6-131474388-DEL-1013 | chr6-131474389-DEL-1013 |
| chr6-131478684-DEL-4248 | chr6-131478685-DEL-4248 |
| chr6-131479162-DEL-282 | chr6-131479163-DEL-282 |
| chr6-131492006-DEL-1924 | chr6-131492011-DEL-1922 |
| chr6-131497740-DEL-360 | chr6-131497740-DEL-360 |
| chr6-131523175-DEL-4937 | chr6-131523175-DEL-4937 |
| chr6-131610764-DEL-641 | chr6-131610765-DEL-641 |
| chr6-131620213-DEL-531 | chr6-131620214-DEL-531 |
| chr6-131632721-DEL-3005 | chr6-131632711-DEL-3006 |
| chr6-131715826-DEL-2998 | chr6-131715827-DEL-2998 |
| chr6-131736589-DEL-6420 | chr6-131736590-DEL-6413 |
| chr6-131774483-DEL-6385 | chr6-131774484-DEL-6385 |
| chr6-131840022-DEL-51 | chr6-131840023-DEL-51 |
| chr6-131860610-DEL-192 | chr6-131860609-DEL-193 |
| chr6-131879106-DEL-1063 | chr6-131879110-DEL-1062 |
| chr6-131894140-DEL-147 | chr6-131894141-DEL-147 |
| chr6-131901797-DEL-7015 | chr6-131901798-DEL-7015 |
| chr6-131928566-DEL-112 | chr6-131928568-DEL-111 |
| chr6-131936715-DEL-53 | chr6-131936716-DEL-53 |
| chr6-131940679-DEL-2811 | chr6-131940679-DEL-2811 |
| chr6-131944644-DEL-7775 | chr6-131944643-DEL-7775 |
| chr6-131956362-DEL-436 | chr6-131956363-DEL-436 |
| chr6-131965430-DEL-91 | chr6-131965450-DEL-83 |
| chr6-131971669-DEL-3463 | chr6-131971670-DEL-3463 |
| chr6-131992655-DEL-62 | chr6-131992656-DEL-62 |
| chr6-131996594-DEL-1519 | chr6-131996595-DEL-1519 |
| chr6-131999658-DEL-458 | chr6-131999659-DEL-458 |
| chr6-132000723-DEL-920 | chr6-132000724-DEL-921 |
| chr6-132028196-DEL-109 | chr6-132028181-DEL-94 |
| chr6-132031474-DEL-2942 | chr6-132031475-DEL-2942 |

| | |
|---|---|
| chr6-132043953-DEL-54 | chr6-132043956-DEL-55 |
| chr6-132118987-DEL-17660 | chr6-132118988-DEL-17660 |
| chr6-132120039-DEL-236 | chr6-132120040-DEL-235 |
| chr6-132129542-DEL-631 | chr6-132129541-DEL-632 |
| chr6-132133371-DEL-307 | chr6-132133372-DEL-307 |
| chr6-132134680-DEL-52 | chr6-132134680-DEL-52 |
| chr6-132148871-DEL-60 | chr6-132148872-DEL-60 |
| chr6-132179153-DEL-1836 | chr6-132179153-DEL-1836 |
| chr6-132188732-DEL-926 | chr6-132188733-DEL-926 |
| chr6-132197721-DEL-69 | chr6-132197718-DEL-66 |
| chr6-132221602-DEL-968 | chr6-132221603-DEL-968 |
| chr6-132222343-DEL-3929 | chr6-132222344-DEL-3929 |
| chr6-132231086-DEL-65 | chr6-132231096-DEL-72 |
| chr6-132232257-DEL-317 | chr6-132232257-DEL-317 |
| chr6-132243543-DEL-73 | chr6-132243544-DEL-73 |
| chr6-132244156-DEL-241 | chr6-132244143-DEL-245 |
| chr6-132263834-DEL-54 | chr6-132263831-DEL-58 |
| chr6-132266859-DEL-18978 | chr6-132266852-DEL-18979 |
| chr6-132337027-DEL-67 | chr6-132337028-DEL-67 |
| chr6-132381843-DEL-204 | chr6-132381842-DEL-205 |
| chr6-132405509-DEL-594 | chr6-132405510-DEL-594 |
| chr6-132415993-DEL-54 | chr6-132415994-DEL-54 |
| chr6-132425359-DEL-81 | chr6-132425360-DEL-81 |
| chr6-132432143-DEL-5192 | chr6-132432144-DEL-5192 |
| chr6-132456771-DEL-143 | chr6-132456772-DEL-135 |
| chr6-132458225-DEL-2572 | chr6-132458226-DEL-2572 |
| chr6-132465381-DEL-254 | chr6-132465382-DEL-254 |
| chr6-132472333-DEL-7298 | chr6-132472334-DEL-7298 |
| chr6-132498279-DEL-6926 | chr6-132498280-DEL-6926 |
| chr6-132531780-DEL-54 | chr6-132531781-DEL-54 |
| chr6-132533085-DEL-4972 | chr6-132533086-DEL-4972 |
| chr6-132547057-DEL-390 | chr6-132547058-DEL-390 |
| chr6-132566911-DEL-52 | chr6-132566912-DEL-52 |
| chr6-132573949-DEL-2005 | chr6-132573949-DEL-2005 |
| chr6-132607499-DEL-13492 | chr6-132607500-DEL-13492 |
| chr6-132632350-DEL-65 | chr6-132632351-DEL-65 |
| chr6-132637159-DEL-126 | chr6-132637109-DEL-126 |
| chr6-132657381-DEL-58 | chr6-132657382-DEL-58 |
| chr6-132683300-DEL-1906 | chr6-132683301-DEL-1906 |
| chr6-132692755-DEL-1179 | chr6-132692756-DEL-1178 |
| chr6-132708651-DEL-57 | chr6-132708636-DEL-58 |
| chr6-132717244-DEL-212 | chr6-132717230-DEL-213 |
| chr6-132740421-DEL-79 | chr6-132740422-DEL-79 |
| chr6-132756109-DEL-1558 | chr6-132756110-DEL-1558 |
| chr6-132762944-DEL-203 | chr6-132762945-DEL-199 |
| chr6-132785920-DEL-1035 | chr6-132785921-DEL-1035 |
| chr6-132804924-DEL-765 | chr6-132804925-DEL-765 |
| chr6-132811811-DEL-6712 | chr6-132811812-DEL-6712 |
| chr6-132852813-DEL-6362 | chr6-132852807-DEL-6363 |
| chr6-132865003-DEL-55 | chr6-132865004-DEL-55 |
| chr6-132894079-DEL-162 | chr6-132894080-DEL-162 |
| chr6-132895904-DEL-453 | chr6-132895905-DEL-453 |
| chr6-132917738-DEL-64 | chr6-132917739-DEL-64 |
| chr6-132930520-DEL-998 | chr6-132930503-DEL-1011 |
| chr6-132931790-DEL-73 | chr6-132931791-DEL-73 |
| chr6-132937188-DEL-1666 | chr6-132937189-DEL-1666 |
| chr6-132940827-DEL-6330 | chr6-132940831-DEL-6326 |
| chr6-132988320-DEL-1035 | chr6-132988321-DEL-1035 |
| chr6-133011232-DEL-343 | chr6-133011233-DEL-343 |
| chr6-133017145-DEL-818 | chr6-133017146-DEL-818 |
| chr6-133023084-DEL-1879 | chr6-133023078-DEL-1880 |
| chr6-133029457-DEL-208 | chr6-133029458-DEL-208 |
| chr6-133073609-DEL-4109 | chr6-133073610-DEL-4109 |
| chr6-133080696-DEL-127 | chr6-133080691-DEL-129 |
| chr6-133111741-DEL-1136 | chr6-133111741-DEL-1136 |
| chr6-133113079-DEL-260 | chr6-133113079-DEL-260 |
| chr6-133118745-DEL-683 | chr6-133118745-DEL-683 |
| chr6-133119902-DEL-1105 | chr6-133119902-DEL-1105 |
| chr6-133140707-DEL-109 | chr6-133140707-DEL-109 |
| chr6-133143982-DEL-212 | chr6-133143982-DEL-212 |
| chr6-133183035-DEL-345 | chr6-133183035-DEL-345 |
| chr6-133244002-DEL-446 | chr6-133244003-DEL-446 |
| chr6-133295679-DEL-480 | chr6-133295680-DEL-480 |
| chr6-133311498-DEL-185 | chr6-133311498-DEL-185 |
| chr6-133362182-DEL-1076 | chr6-133362183-DEL-1076 |
| chr6-133378216-DEL-6079 | chr6-133378208-DEL-6080 |
| chr6-133396643-DEL-186 | chr6-133396644-DEL-186 |
| chr6-133399343-DEL-6233 | chr6-133399344-DEL-6233 |
| chr6-133419728-DEL-403 | chr6-133419729-DEL-403 |
| chr6-133439452-DEL-666 | chr6-133439444-DEL-674 |

| | |
|---|---|
| chr6-133521667-DEL-728 | chr6-133521673-DEL-724 |
| chr6-133522388-DEL-2162 | chr6-133522385-DEL-2166 |
| chr6-133525763-DEL-1031 | chr6-133525764-DEL-1031 |
| chr6-133531956-DEL-214 | chr6-133531957-DEL-214 |
| chr6-133562613-DEL-3694 | chr6-133562614-DEL-3694 |
| chr6-133567385-DEL-309 | chr6-133567386-DEL-308 |
| chr6-133573493-DEL-85 | chr6-133573494-DEL-85 |
| chr6-133576438-DEL-58 | chr6-133576439-DEL-58 |
| chr6-133585615-DEL-70 | chr6-133585609-DEL-71 |
| chr6-133589885-DEL-99 | chr6-133589882-DEL-100 |
| chr6-133591042-DEL-1775 | chr6-133591042-DEL-1775 |
| chr6-133600250-DEL-66 | chr6-133600251-DEL-66 |
| chr6-133683827-DEL-333 | chr6-133683828-DEL-333 |
| chr6-133689154-DEL-664 | chr6-133689155-DEL-664 |
| chr6-133696040-DEL-4496 | chr6-133696041-DEL-4496 |
| chr6-133701362-DEL-1159 | chr6-133701363-DEL-1159 |
| chr6-133702965-DEL-54 | chr6-133702962-DEL-84 |
| chr6-133707086-DEL-74 | chr6-133707068-DEL-75 |
| chr6-133722175-DEL-201 | chr6-133722174-DEL-201 |
| chr6-133739319-DEL-509 | chr6-133739318-DEL-509 |
| chr6-133783352-DEL-2648 | chr6-133783352-DEL-2648 |
| chr6-133792364-DEL-14164 | chr6-133792365-DEL-14164 |
| chr6-133842433-DEL-203 | chr6-133842434-DEL-203 |
| chr6-133856806-DEL-249 | chr6-133856806-DEL-235 |
| chr6-133868294-DEL-8687 | chr6-133868282-DEL-8699 |
| chr6-133870469-DEL-2498 | chr6-133870469-DEL-2498 |
| chr6-133898556-DEL-56 | chr6-133898561-DEL-62 |
| chr6-133898811-DEL-6199 | chr6-133898812-DEL-6199 |
| chr6-133908592-DEL-290 | chr6-133908593-DEL-290 |
| chr6-133909263-DEL-350 | chr6-133909267-DEL-368 |
| chr6-133919869-DEL-6497 | chr6-133919870-DEL-6497 |
| chr6-133941320-DEL-204 | chr6-133941321-DEL-204 |
| chr6-133951581-DEL-125 | chr6-133951557-DEL-168 |
| chr6-133956455-DEL-60 | chr6-133956456-DEL-54 |
| chr6-133967484-DEL-150 | chr6-133967485-DEL-148 |
| chr6-133986161-DEL-5522 | chr6-133986163-DEL-5521 |
| chr6-133994460-DEL-147 | chr6-133994461-DEL-147 |
| chr6-134008037-DEL-151 | chr6-134008038-DEL-151 |
| chr6-134071985-DEL-70 | chr6-134071988-DEL-68 |
| chr6-134103786-DEL-161 | chr6-134103787-DEL-161 |
| chr6-134131065-DEL-107 | chr6-134131066-DEL-107 |
| chr6-134137770-DEL-87 | chr6-134137776-DEL-83 |
| chr6-134143935-DEL-955 | chr6-134143936-DEL-955 |
| chr6-134163877-DEL-426 | chr6-134163878-DEL-426 |
| chr6-134218390-DEL-54 | chr6-134218391-DEL-54 |
| chr6-134229922-DEL-111 | chr6-134229931-DEL-103 |
| chr6-134257767-DEL-54 | chr6-134257768-DEL-54 |
| chr6-134261242-DEL-110 | chr6-134261243-DEL-110 |
| chr6-134265471-DEL-1519 | chr6-134265472-DEL-1519 |
| chr6-134284290-DEL-126 | chr6-134284327-DEL-90 |
| chr6-134295147-DEL-343 | chr6-134295148-DEL-343 |
| chr6-134303667-DEL-1557 | chr6-134303668-DEL-1556 |
| chr6-134305607-DEL-60 | chr6-134305609-DEL-59 |
| chr6-134315149-DEL-159 | chr6-134315111-DEL-160 |
| chr6-134342635-DEL-120 | chr6-134342636-DEL-120 |
| chr6-134367582-DEL-810 | chr6-134367572-DEL-814 |
| chr6-134368454-DEL-364 | chr6-134368457-DEL-360 |
| chr6-134393997-DEL-88 | chr6-134394000-DEL-87 |
| chr6-134419746-DEL-215 | chr6-134419747-DEL-215 |
| chr6-134424795-DEL-63 | chr6-134424796-DEL-63 |
| chr6-134439880-DEL-188 | chr6-134439881-DEL-188 |
| chr6-134456266-DEL-51 | chr6-134456267-DEL-51 |
| chr6-134463144-DEL-62 | chr6-134463145-DEL-62 |
| chr6-134474926-DEL-123 | chr6-134474927-DEL-123 |
| chr6-134493266-DEL-64 | chr6-134493266-DEL-64 |
| chr6-134498901-DEL-114 | chr6-134498902-DEL-114 |
| chr6-134502368-DEL-401 | chr6-134502369-DEL-401 |
| chr6-134523950-DEL-150 | chr6-134523951-DEL-150 |
| chr6-134524688-DEL-196 | chr6-134524689-DEL-196 |
| chr6-134526493-DEL-65 | chr6-134526494-DEL-65 |
| chr6-134535212-DEL-290 | chr6-134535213-DEL-290 |
| chr6-134571170-DEL-205 | chr6-134571171-DEL-205 |
| chr6-134579900-DEL-60 | chr6-134579895-DEL-75 |
| chr6-134602280-DEL-54 | chr6-134602285-DEL-55 |
| chr6-134603615-DEL-3349 | chr6-134603616-DEL-3349 |
| chr6-134622520-DEL-53 | chr6-134622521-DEL-53 |
| chr6-134634966-DEL-182 | chr6-134634967-DEL-182 |
| chr6-134644186-DEL-112 | chr6-134644187-DEL-112 |
| chr6-134653020-DEL-86 | chr6-134653021-DEL-86 |
| chr6-134653615-DEL-227 | chr6-134653616-DEL-227 |

| | |
|---|---|
| chr6-134666374-DEL-3118 | chr6-134666375-DEL-3118 |
| chr6-134675012-DEL-198 | chr6-134675012-DEL-198 |
| chr6-134680910-DEL-82 | chr6-134680912-DEL-83 |
| chr6-134682371-DEL-207 | chr6-134682372-DEL-207 |
| chr6-134688308-DEL-1770 | chr6-134688309-DEL-1770 |
| chr6-134706944-DEL-172 | chr6-134706917-DEL-152 |
| chr6-134736610-DEL-206 | chr6-134736602-DEL-207 |
| chr6-134750117-DEL-68 | chr6-134750118-DEL-68 |
| chr6-134752820-DEL-374 | chr6-134752821-DEL-374 |
| chr6-134773699-DEL-621 | chr6-134773694-DEL-622 |
| chr6-134792943-DEL-62 | chr6-134792944-DEL-62 |
| chr6-134795696-DEL-729 | chr6-134795697-DEL-729 |
| chr6-134799720-DEL-180 | chr6-134799721-DEL-180 |
| chr6-134820798-DEL-52 | chr6-134820799-DEL-52 |
| chr6-134826653-DEL-943 | chr6-134826654-DEL-943 |
| chr6-134856619-DEL-216 | chr6-134856620-DEL-216 |
| chr6-134878035-DEL-142 | chr6-134878035-DEL-140 |
| chr6-134881712-DEL-68 | chr6-134881689-DEL-68 |
| chr6-134895859-DEL-68 | chr6-134895865-DEL-64 |
| chr6-134923486-DEL-62 | chr6-134923485-DEL-62 |
| chr6-134924181-DEL-126 | chr6-134924182-DEL-126 |
| chr6-134931931-DEL-125 | chr6-134931932-DEL-125 |
| chr6-134935699-DEL-206 | chr6-134935690-DEL-199 |
| chr6-134950067-DEL-93 | chr6-134950049-DEL-94 |
| chr6-134980038-DEL-2843 | chr6-134980038-DEL-2839 |
| chr6-134989703-DEL-514 | chr6-134989698-DEL-517 |
| chr6-134999160-DEL-207 | chr6-134999184-DEL-184 |
| chr6-135001692-DEL-108 | chr6-135001693-DEL-108 |
| chr6-135037478-DEL-60 | chr6-135037479-DEL-60 |
| chr6-135039008-DEL-183 | chr6-135038963-DEL-201 |
| chr6-135045549-DEL-147 | chr6-135045549-DEL-147 |
| chr6-135058793-DEL-84 | chr6-135058798-DEL-68 |
| chr6-135062517-DEL-202 | chr6-135062516-DEL-203 |
| chr6-135064184-DEL-219 | chr6-135064185-DEL-219 |
| chr6-135077717-DEL-170 | chr6-135077717-DEL-166 |
| chr6-135079512-DEL-113 | chr6-135079513-DEL-113 |
| chr6-135085689-DEL-234 | chr6-135085685-DEL-226 |
| chr6-135106241-DEL-220 | chr6-135106240-DEL-222 |
| chr6-135108712-DEL-72 | chr6-135108713-DEL-72 |
| chr6-135109802-DEL-55 | chr6-135109803-DEL-55 |
| chr6-135123845-DEL-63 | chr6-135123846-DEL-63 |
| chr6-135129295-DEL-79 | chr6-135129298-DEL-77 |
| chr6-135132720-DEL-72 | chr6-135132720-DEL-68 |
| chr6-135137612-DEL-100 | chr6-135137613-DEL-100 |
| chr6-135143096-DEL-51 | chr6-135143095-DEL-53 |
| chr6-135157055-DEL-128 | chr6-135157056-DEL-124 |
| chr6-135159482-DEL-99 | chr6-135159483-DEL-99 |
| chr6-135162145-DEL-82 | chr6-135162146-DEL-82 |
| chr6-135165982-DEL-69 | chr6-135165983-DEL-67 |
| chr6-135189356-DEL-132 | chr6-135189373-DEL-96 |
| chr6-135190371-DEL-66 | chr6-135190366-DEL-63 |
| chr6-135190452-DEL-924 | chr6-135190452-DEL-924 |
| chr6-135201433-DEL-60 | chr6-135201434-DEL-60 |
| chr6-135250108-DEL-137 | chr6-135250109-DEL-137 |
| chr6-135251047-DEL-148 | chr6-135251048-DEL-148 |
| chr6-135256558-DEL-121 | chr6-135256559-DEL-121 |
| chr6-135258369-DEL-6470 | chr6-135258371-DEL-6460 |
| chr6-135291795-DEL-72 | chr6-135291797-DEL-71 |
| chr6-135308799-DEL-62 | chr6-135308800-DEL-62 |
| chr6-135326064-DEL-723 | chr6-135326068-DEL-715 |
| chr6-135338449-DEL-54 | chr6-135338454-DEL-50 |
| chr6-135338955-DEL-62 | chr6-135338956-DEL-62 |
| chr6-135361598-DEL-78 | chr6-135361603-DEL-70 |
| chr6-135380440-DEL-163 | chr6-135380438-DEL-163 |
| chr6-135392656-DEL-200 | chr6-135392655-DEL-202 |
| chr6-135393689-DEL-402 | chr6-135393690-DEL-402 |
| chr6-135401817-DEL-2802 | chr6-135401830-DEL-2806 |
| chr6-135414002-DEL-54 | chr6-135414003-DEL-52 |
| chr6-135431469-DEL-2419 | chr6-135431470-DEL-2419 |
| chr6-135443092-DEL-63 | chr6-135443093-DEL-63 |
| chr6-135470821-DEL-52 | chr6-135470818-DEL-56 |
| chr6-135494152-DEL-549 | chr6-135494153-DEL-549 |
| chr6-135541046-DEL-68 | chr6-135541033-DEL-69 |
| chr6-135573430-DEL-60 | chr6-135573431-DEL-60 |
| chr6-135576212-DEL-1078 | chr6-135576213-DEL-1078 |
| chr6-135586261-DEL-54 | chr6-135586264-DEL-52 |
| chr6-135590578-DEL-67 | chr6-135590579-DEL-67 |
| chr6-135644189-DEL-2243 | chr6-135644190-DEL-2243 |
| chr6-135654371-DEL-72 | chr6-135654372-DEL-72 |
| chr6-135656245-DEL-70 | chr6-135656246-DEL-58 |

| | |
|---|---|
| chr6-135681896-DEL-53 | chr6-135681897-DEL-53 |
| chr6-135701083-DEL-291 | chr6-135701084-DEL-291 |
| chr6-135738076-DEL-71 | chr6-135738077-DEL-73 |
| chr6-135742981-DEL-56 | chr6-135742982-DEL-52 |
| chr6-135845109-DEL-106 | chr6-135845070-DEL-102 |
| chr6-135861802-DEL-60 | chr6-135861803-DEL-60 |
| chr6-135928399-DEL-72 | chr6-135928400-DEL-72 |
| chr6-135947226-DEL-103 | chr6-135947180-DEL-114 |
| chr6-135968654-DEL-51 | chr6-135968655-DEL-51 |
| chr6-135979950-DEL-174 | chr6-135979951-DEL-174 |
| chr6-135984298-DEL-235 | chr6-135984299-DEL-235 |
| chr6-136011961-DEL-4223 | chr6-136011961-DEL-4222 |
| chr6-136026438-DEL-192 | chr6-136026435-DEL-193 |
| chr6-136040542-DEL-58 | chr6-136040543-DEL-62 |
| chr6-136043609-DEL-90 | chr6-136043610-DEL-90 |
| chr6-136073294-DEL-1760 | chr6-136073295-DEL-1760 |
| chr6-136089084-DEL-52 | chr6-136089079-DEL-66 |
| chr6-136100126-DEL-60 | chr6-136100103-DEL-80 |
| chr6-136114092-DEL-59 | chr6-136114082-DEL-56 |
| chr6-136151044-DEL-112 | chr6-136151047-DEL-110 |
| chr6-136155409-DEL-58 | chr6-136155409-DEL-54 |
| chr6-136176713-DEL-89 | chr6-136176695-DEL-107 |
| chr6-136242771-DEL-314 | chr6-136242772-DEL-314 |
| chr6-136247000-DEL-1361 | chr6-136247001-DEL-1361 |
| chr6-136255107-DEL-68 | chr6-136255108-DEL-68 |
| chr6-136261257-DEL-1295 | chr6-136261258-DEL-1295 |
| chr6-136287535-DEL-56 | chr6-136287538-DEL-54 |
| chr6-136288290-DEL-3597 | chr6-136288291-DEL-3597 |
| chr6-136296731-DEL-70 | chr6-136296732-DEL-70 |
| chr6-136299623-DEL-51 | chr6-136299623-DEL-51 |
| chr6-136301177-DEL-55 | chr6-136301182-DEL-51 |
| chr6-136309105-DEL-164 | chr6-136309104-DEL-164 |
| chr6-136310641-DEL-83 | chr6-136310637-DEL-72 |
| chr6-136323792-DEL-59 | chr6-136323793-DEL-59 |
| chr6-136343631-DEL-218 | chr6-136343632-DEL-218 |
| chr6-136344289-DEL-308 | chr6-136344290-DEL-308 |
| chr6-136363162-DEL-89 | chr6-136363198-DEL-63 |
| chr6-136369914-DEL-74 | chr6-136369915-DEL-74 |
| chr6-136383384-DEL-335 | chr6-136383416-DEL-304 |
| chr6-136385674-DEL-70 | chr6-136385669-DEL-62 |
| chr6-136390189-DEL-70 | chr6-136390190-DEL-70 |
| chr6-136394459-DEL-75 | chr6-136394460-DEL-75 |
| chr6-136395759-DEL-72 | chr6-136395760-DEL-72 |
| chr6-136406074-DEL-52 | chr6-136406068-DEL-62 |
| chr6-136407515-DEL-434 | chr6-136407516-DEL-434 |
| chr6-136434811-DEL-207 | chr6-136434812-DEL-207 |
| chr6-136438457-DEL-181 | chr6-136438457-DEL-179 |
| chr6-136450209-DEL-621 | chr6-136450210-DEL-621 |
| chr6-136457049-DEL-51 | chr6-136457058-DEL-59 |
| chr6-136506502-DEL-135 | chr6-136506503-DEL-135 |
| chr6-136546529-DEL-197 | chr6-136546530-DEL-197 |
| chr6-136560148-DEL-58 | chr6-136560141-DEL-51 |
| chr6-136560508-DEL-836 | chr6-136560499-DEL-858 |
| chr6-136561500-DEL-220 | chr6-136561500-DEL-219 |
| chr6-136584359-DEL-60 | chr6-136584359-DEL-68 |
| chr6-136592327-DEL-68 | chr6-136592328-DEL-68 |
| chr6-136600353-DEL-1634 | chr6-136600346-DEL-1635 |
| chr6-136603024-DEL-59 | chr6-136603015-DEL-60 |
| chr6-136608076-DEL-68 | chr6-136608077-DEL-68 |
| chr6-136626286-DEL-78 | chr6-136626293-DEL-72 |
| chr6-136629378-DEL-69 | chr6-136629393-DEL-74 |
| chr6-136635932-DEL-366 | chr6-136635933-DEL-366 |
| chr6-136641564-DEL-51 | chr6-136641564-DEL-51 |
| chr6-136643584-DEL-150 | chr6-136643539-DEL-148 |
| chr6-136644121-DEL-81 | chr6-136644108-DEL-79 |
| chr6-136683555-DEL-77 | chr6-136683570-DEL-77 |
| chr6-136692040-DEL-72 | chr6-136692041-DEL-71 |
| chr6-136697176-DEL-64 | chr6-136697177-DEL-64 |
| chr6-136702141-DEL-555 | chr6-136702142-DEL-555 |
| chr6-136703970-DEL-67 | chr6-136703971-DEL-67 |
| chr6-136715642-DEL-56 | chr6-136715643-DEL-56 |
| chr6-136723712-DEL-94 | chr6-136723713-DEL-94 |
| chr6-136730784-DEL-93 | chr6-136730785-DEL-93 |
| chr6-136747706-DEL-2104 | chr6-136747707-DEL-2104 |
| chr6-136752185-DEL-76 | chr6-136752186-DEL-76 |
| chr6-136754998-DEL-76 | chr6-136754999-DEL-57 |
| chr6-136767715-DEL-51 | chr6-136767716-DEL-51 |
| chr6-136769471-DEL-52 | chr6-136769472-DEL-52 |
| chr6-136772543-DEL-76 | chr6-136772548-DEL-72 |
| chr6-136778743-DEL-690 | chr6-136778744-DEL-690 |

| | |
|---|---|
| chr6-136791876-DEL-253 | chr6-136791877-DEL-252 |
| chr6-136793683-DEL-175 | chr6-136793683-DEL-210 |
| chr6-136798855-DEL-291 | chr6-136798856-DEL-291 |
| chr6-136803261-DEL-179 | chr6-136803262-DEL-179 |
| chr6-136814623-DEL-68 | chr6-136814622-DEL-68 |
| chr6-136823711-DEL-125 | chr6-136823712-DEL-119 |
| chr6-136843900-DEL-55 | chr6-136843901-DEL-55 |
| chr6-136846794-DEL-999 | chr6-136846795-DEL-999 |
| chr6-136914984-DEL-217 | chr6-136914980-DEL-218 |
| chr6-136931171-DEL-86 | chr6-136931176-DEL-82 |
| chr6-136941032-DEL-92 | chr6-136941033-DEL-92 |
| chr6-136944768-DEL-135 | chr6-136944769-DEL-172 |
| chr6-136946156-DEL-469 | chr6-136946154-DEL-469 |
| chr6-136948349-DEL-6331 | chr6-136948350-DEL-6331 |
| chr6-136959842-DEL-64 | chr6-136959843-DEL-64 |
| chr6-136966287-DEL-182 | chr6-136966286-DEL-178 |
| chr6-136977455-DEL-66 | chr6-136977456-DEL-56 |
| chr6-136987619-DEL-349 | chr6-136987620-DEL-349 |
| chr6-136988148-DEL-67 | chr6-136988144-DEL-68 |
| chr6-136994854-DEL-112 | chr6-136994855-DEL-112 |
| chr6-136999306-DEL-54 | chr6-136999306-DEL-54 |
| chr6-137000590-DEL-54 | chr6-137000591-DEL-54 |
| chr6-137012536-DEL-74 | chr6-137012547-DEL-61 |
| chr6-137020957-DEL-415 | chr6-137020960-DEL-416 |
| chr6-137034175-DEL-74 | chr6-137034209-DEL-51 |
| chr6-137036198-DEL-51 | chr6-137036199-DEL-51 |
| chr6-137037132-DEL-64 | chr6-137037133-DEL-64 |
| chr6-137063428-DEL-683 | chr6-137063429-DEL-683 |
| chr6-137073577-DEL-56 | chr6-137073578-DEL-56 |
| chr6-137095129-DEL-168 | chr6-137095130-DEL-140 |
| chr6-137098778-DEL-60 | chr6-137098787-DEL-61 |
| chr6-137101444-DEL-58 | chr6-137101445-DEL-58 |
| chr6-137105739-DEL-51 | chr6-137105731-DEL-52 |
| chr6-137120480-DEL-52 | chr6-137120483-DEL-52 |
| chr6-137124896-DEL-98 | chr6-137124897-DEL-98 |
| chr6-137169901-DEL-108 | chr6-137169902-DEL-108 |
| chr6-137175867-DEL-2098 | chr6-137175868-DEL-2098 |
| chr6-137204649-DEL-6439 | chr6-137204650-DEL-6439 |
| chr6-137216952-DEL-184 | chr6-137216974-DEL-178 |
| chr6-137270558-DEL-67 | chr6-137270559-DEL-67 |
| chr6-137274355-DEL-415 | chr6-137274356-DEL-415 |
| chr6-137275551-DEL-65 | chr6-137275552-DEL-65 |
| chr6-137279317-DEL-82 | chr6-137279318-DEL-82 |
| chr6-137286012-DEL-299 | chr6-137286013-DEL-299 |
| chr6-137317531-DEL-186 | chr6-137317532-DEL-186 |
| chr6-137324546-DEL-97 | chr6-137324547-DEL-97 |
| chr6-137327816-DEL-98 | chr6-137327817-DEL-98 |
| chr6-137338888-DEL-63 | chr6-137338889-DEL-63 |
| chr6-137352352-DEL-128 | chr6-137352353-DEL-128 |
| chr6-137365054-DEL-243 | chr6-137365055-DEL-243 |
| chr6-137378209-DEL-194 | chr6-137378210-DEL-194 |
| chr6-137403579-DEL-72 | chr6-137403580-DEL-72 |
| chr6-137407409-DEL-120 | chr6-137407410-DEL-120 |
| chr6-137412475-DEL-1193 | chr6-137412476-DEL-1193 |
| chr6-137430380-DEL-138 | chr6-137430381-DEL-138 |
| chr6-137444152-DEL-144 | chr6-137444160-DEL-136 |
| chr6-137458558-DEL-59 | chr6-137458559-DEL-59 |
| chr6-137459427-DEL-96 | chr6-137459437-DEL-80 |
| chr6-137463690-DEL-58 | chr6-137463695-DEL-54 |
| chr6-137464984-DEL-54 | chr6-137464985-DEL-54 |
| chr6-137473690-DEL-55 | chr6-137473691-DEL-55 |
| chr6-137514188-DEL-142 | chr6-137514189-DEL-142 |
| chr6-137521392-DEL-505 | chr6-137521393-DEL-505 |
| chr6-137551771-DEL-115 | chr6-137551772-DEL-115 |
| chr6-137576470-DEL-84 | chr6-137576469-DEL-84 |
| chr6-137595934-DEL-2038 | chr6-137595935-DEL-2038 |
| chr6-137615647-DEL-62 | chr6-137615634-DEL-76 |
| chr6-137616316-DEL-80 | chr6-137616317-DEL-80 |
| chr6-137640390-DEL-191 | chr6-137640391-DEL-191 |
| chr6-137645887-DEL-2526 | chr6-137645888-DEL-2526 |
| chr6-137684643-DEL-112 | chr6-137684644-DEL-112 |
| chr6-137685649-DEL-56 | chr6-137685649-DEL-56 |
| chr6-137685890-DEL-520 | chr6-137685891-DEL-520 |
| chr6-137690736-DEL-90 | chr6-137690737-DEL-90 |
| chr6-137692828-DEL-479 | chr6-137692829-DEL-479 |
| chr6-137706298-DEL-236 | chr6-137706299-DEL-236 |
| chr6-137714673-DEL-192 | chr6-137714674-DEL-192 |
| chr6-137747859-DEL-51 | chr6-137747841-DEL-52 |
| chr6-137760081-DEL-329 | chr6-137760082-DEL-329 |
| chr6-137769134-DEL-142 | chr6-137769143-DEL-140 |

| | |
|---|---|
| chr6-137777448-DEL-135 | chr6-137777449-DEL-135 |
| chr6-137788745-DEL-121 | chr6-137788754-DEL-119 |
| chr6-137794908-DEL-465 | chr6-137794909-DEL-465 |
| chr6-137813221-DEL-174 | chr6-137813222-DEL-145 |
| chr6-137826078-DEL-238 | chr6-137826079-DEL-238 |
| chr6-137830371-DEL-56 | chr6-137830372-DEL-56 |
| chr6-137868238-DEL-60 | chr6-137868238-DEL-60 |
| chr6-137870222-DEL-555 | chr6-137870223-DEL-555 |
| chr6-137870894-DEL-835 | chr6-137870895-DEL-835 |
| chr6-137892251-DEL-345 | chr6-137892252-DEL-345 |
| chr6-137902699-DEL-1480 | chr6-137902700-DEL-1480 |
| chr6-137925246-DEL-155 | chr6-137925247-DEL-155 |
| chr6-137949233-DEL-56 | chr6-137949234-DEL-56 |
| chr6-137955062-DEL-792 | chr6-137955063-DEL-792 |
| chr6-137995571-DEL-56 | chr6-137995572-DEL-56 |
| chr6-138001061-DEL-7977 | chr6-138001062-DEL-7977 |
| chr6-138031436-DEL-58 | chr6-138031437-DEL-58 |
| chr6-138032452-DEL-65 | chr6-138032453-DEL-65 |
| chr6-138039981-DEL-6366 | chr6-138039982-DEL-6366 |
| chr6-138053561-DEL-2512 | chr6-138053562-DEL-2512 |
| chr6-138060782-DEL-363 | chr6-138060782-DEL-363 |
| chr6-138063288-DEL-208 | chr6-138063299-DEL-198 |
| chr6-138086437-DEL-90 | chr6-138086438-DEL-94 |
| chr6-138124934-DEL-187 | chr6-138124962-DEL-159 |
| chr6-138140949-DEL-69 | chr6-138140952-DEL-74 |
| chr6-138146786-DEL-190 | chr6-138146787-DEL-187 |
| chr6-138152526-DEL-100 | chr6-138152532-DEL-91 |
| chr6-138154930-DEL-6534 | chr6-138154930-DEL-6535 |
| chr6-138161858-DEL-63 | chr6-138161860-DEL-59 |
| chr6-138164301-DEL-145 | chr6-138164302-DEL-145 |
| chr6-138176645-DEL-56 | chr6-138176646-DEL-56 |
| chr6-138193055-DEL-6188 | chr6-138193056-DEL-6188 |
| chr6-138199920-DEL-435 | chr6-138199921-DEL-435 |
| chr6-138245313-DEL-205 | chr6-138245314-DEL-205 |
| chr6-138269923-DEL-224 | chr6-138269918-DEL-228 |
| chr6-138279373-DEL-111 | chr6-138279379-DEL-106 |
| chr6-138301924-DEL-54 | chr6-138301893-DEL-50 |
| chr6-138330675-DEL-52 | chr6-138330676-DEL-52 |
| chr6-138375327-DEL-181 | chr6-138375326-DEL-178 |
| chr6-138429595-DEL-412 | chr6-138429596-DEL-412 |
| chr6-138466569-DEL-485 | chr6-138466569-DEL-485 |
| chr6-138508476-DEL-958 | chr6-138508477-DEL-958 |
| chr6-138553418-DEL-101 | chr6-138553418-DEL-102 |
| chr6-138555752-DEL-159 | chr6-138555753-DEL-159 |
| chr6-138614211-DEL-93 | chr6-138614212-DEL-93 |
| chr6-138624232-DEL-1885 | chr6-138624233-DEL-1885 |
| chr6-138640268-DEL-92 | chr6-138640269-DEL-92 |
| chr6-138641124-DEL-176 | chr6-138641128-DEL-169 |
| chr6-138662383-DEL-2628 | chr6-138662371-DEL-2641 |
| chr6-138665753-DEL-70 | chr6-138665754-DEL-70 |
| chr6-138672415-DEL-6185 | chr6-138672416-DEL-6185 |
| chr6-138692291-DEL-58 | chr6-138692291-DEL-58 |
| chr6-138718460-DEL-6659 | chr6-138718453-DEL-6660 |
| chr6-138724702-DEL-214 | chr6-138724702-DEL-214 |
| chr6-138780371-DEL-52 | chr6-138780372-DEL-52 |
| chr6-138793203-DEL-156 | chr6-138793204-DEL-156 |
| chr6-138794262-DEL-3354 | chr6-138794262-DEL-3354 |
| chr6-138809473-DEL-116 | chr6-138809474-DEL-116 |
| chr6-138881386-DEL-65 | chr6-138881387-DEL-65 |
| chr6-138930330-DEL-53 | chr6-138930312-DEL-54 |
| chr6-138942248-DEL-81 | chr6-138942251-DEL-65 |
| chr6-138956483-DEL-1233 | chr6-138956483-DEL-1233 |
| chr6-138961535-DEL-907 | chr6-138961536-DEL-907 |
| chr6-138991542-DEL-60 | chr6-138991542-DEL-60 |
| chr6-139017086-DEL-6176 | chr6-139017086-DEL-6176 |
| chr6-139024416-DEL-66 | chr6-139024417-DEL-66 |
| chr6-139050790-DEL-653 | chr6-139050791-DEL-653 |
| chr6-139078528-DEL-52 | chr6-139078531-DEL-51 |
| chr6-139112723-DEL-62 | chr6-139112724-DEL-60 |
| chr6-139113909-DEL-5130 | chr6-139113910-DEL-5130 |
| chr6-139185837-DEL-84 | chr6-139185838-DEL-84 |
| chr6-139194452-DEL-2373 | chr6-139194453-DEL-2374 |
| chr6-139198472-DEL-1570 | chr6-139198471-DEL-1571 |
| chr6-139219922-DEL-53 | chr6-139219875-DEL-52 |
| chr6-139233199-DEL-51 | chr6-139233199-DEL-51 |
| chr6-139235862-DEL-55 | chr6-139235863-DEL-55 |
| chr6-139261730-DEL-280 | chr6-139261731-DEL-280 |
| chr6-139306924-DEL-66 | chr6-139306925-DEL-66 |
| chr6-139318202-DEL-654 | chr6-139318203-DEL-654 |
| chr6-139372566-DEL-1210 | chr6-139372567-DEL-1210 |

| | |
|---|---|
| chr6-139382883-DEL-223 | chr6-139382878-DEL-223 |
| chr6-139384334-DEL-1238 | chr6-139384335-DEL-1238 |
| chr6-139386814-DEL-2203 | chr6-139386806-DEL-2204 |
| chr6-139451628-DEL-60 | chr6-139451637-DEL-52 |
| chr6-139482869-DEL-56 | chr6-139482863-DEL-56 |
| chr6-139499167-DEL-221 | chr6-139499168-DEL-221 |
| chr6-139508208-DEL-2557 | chr6-139508209-DEL-2557 |
| chr6-139540822-DEL-72 | chr6-139540823-DEL-72 |
| chr6-139561385-DEL-82 | chr6-139561386-DEL-82 |
| chr6-139578672-DEL-80 | chr6-139578673-DEL-80 |
| chr6-139585248-DEL-84 | chr6-139585243-DEL-75 |
| chr6-139619365-DEL-506 | chr6-139619366-DEL-506 |
| chr6-139642174-DEL-948 | chr6-139642174-DEL-947 |
| chr6-139686965-DEL-179 | chr6-139686966-DEL-179 |
| chr6-139694890-DEL-91 | chr6-139694891-DEL-91 |
| chr6-139722166-DEL-211 | chr6-139722167-DEL-211 |
| chr6-139742746-DEL-58 | chr6-139742747-DEL-58 |
| chr6-139762934-DEL-3174 | chr6-139762935-DEL-3174 |
| chr6-139771767-DEL-5334 | chr6-139771768-DEL-5334 |
| chr6-139787686-DEL-181 | chr6-139787656-DEL-189 |
| chr6-139794767-DEL-497 | chr6-139794768-DEL-497 |
| chr6-139799197-DEL-72 | chr6-139799198-DEL-72 |
| chr6-139820915-DEL-67 | chr6-139820915-DEL-68 |
| chr6-139828927-DEL-80 | chr6-139828930-DEL-81 |
| chr6-139864170-DEL-120 | chr6-139864169-DEL-128 |
| chr6-139867216-DEL-852 | chr6-139867217-DEL-852 |
| chr6-139869757-DEL-333 | chr6-139869759-DEL-332 |
| chr6-139876989-DEL-1035 | chr6-139876990-DEL-1035 |
| chr6-139877592-DEL-318 | chr6-139877593-DEL-318 |
| chr6-139915466-DEL-291 | chr6-139915467-DEL-291 |
| chr6-139927666-DEL-83 | chr6-139927667-DEL-77 |
| chr6-139953627-DEL-193 | chr6-139953628-DEL-193 |
| chr6-139971888-DEL-52 | chr6-139971888-DEL-52 |
| chr6-140020527-DEL-128 | chr6-140020528-DEL-128 |
| chr6-140044600-DEL-172 | chr6-140044603-DEL-170 |
| chr6-140048604-DEL-1394 | chr6-140048605-DEL-1394 |
| chr6-140051410-DEL-425 | chr6-140051411-DEL-435 |
| chr6-140056507-DEL-66 | chr6-140056508-DEL-66 |
| chr6-140060207-DEL-114 | chr6-140060208-DEL-114 |
| chr6-140067570-DEL-239 | chr6-140067571-DEL-239 |
| chr6-140071668-DEL-83 | chr6-140071667-DEL-84 |
| chr6-140095537-DEL-79 | chr6-140095538-DEL-79 |
| chr6-140100433-DEL-90 | chr6-140100434-DEL-88 |
| chr6-140104367-DEL-65 | chr6-140104368-DEL-65 |
| chr6-140114811-DEL-399 | chr6-140114812-DEL-399 |
| chr6-140136353-DEL-61 | chr6-140136358-DEL-57 |
| chr6-140154566-DEL-57 | chr6-140154567-DEL-57 |
| chr6-140179955-DEL-191 | chr6-140179939-DEL-192 |
| chr6-140197965-DEL-52 | chr6-140197968-DEL-50 |
| chr6-140206736-DEL-80 | chr6-140206736-DEL-80 |
| chr6-140235880-DEL-76 | chr6-140235846-DEL-77 |
| chr6-140257059-DEL-468 | chr6-140257060-DEL-468 |
| chr6-140267799-DEL-4760 | chr6-140267800-DEL-4760 |
| chr6-140273133-DEL-62 | chr6-140273137-DEL-62 |
| chr6-140279465-DEL-640 | chr6-140279466-DEL-640 |
| chr6-140317313-DEL-100 | chr6-140317314-DEL-100 |
| chr6-140327616-DEL-226 | chr6-140327617-DEL-226 |
| chr6-140335228-DEL-1351 | chr6-140335229-DEL-1351 |
| chr6-140346853-DEL-227 | chr6-140346854-DEL-227 |
| chr6-140379258-DEL-65 | chr6-140379259-DEL-65 |
| chr6-140397049-DEL-390 | chr6-140397050-DEL-390 |
| chr6-140404988-DEL-963 | chr6-140404989-DEL-963 |
| chr6-140420514-DEL-65 | chr6-140420515-DEL-65 |
| chr6-140428783-DEL-65 | chr6-140428784-DEL-65 |
| chr6-140446134-DEL-268 | chr6-140446135-DEL-268 |
| chr6-140475338-DEL-255 | chr6-140475339-DEL-255 |
| chr6-140485940-DEL-56 | chr6-140485941-DEL-56 |
| chr6-140509691-DEL-58 | chr6-140509692-DEL-58 |
| chr6-140530548-DEL-207 | chr6-140530549-DEL-206 |
| chr6-140548508-DEL-347 | chr6-140548509-DEL-347 |
| chr6-140553461-DEL-60 | chr6-140553461-DEL-60 |
| chr6-140556416-DEL-72 | chr6-140556417-DEL-72 |
| chr6-140561977-DEL-184 | chr6-140561978-DEL-184 |
| chr6-140564319-DEL-51 | chr6-140564320-DEL-51 |
| chr6-140573305-DEL-60 | chr6-140573306-DEL-60 |
| chr6-140582439-DEL-76 | chr6-140582439-DEL-72 |
| chr6-140585047-DEL-224 | chr6-140585048-DEL-224 |
| chr6-140592788-DEL-86 | chr6-140592789-DEL-86 |
| chr6-140627276-DEL-173 | chr6-140627277-DEL-173 |
| chr6-140637073-DEL-747 | chr6-140637074-DEL-747 |

| | |
|---|---|
| chr6-140661677-DEL-169 | chr6-140661678-DEL-169 |
| chr6-140665383-DEL-574 | chr6-140665387-DEL-572 |
| chr6-140681990-DEL-76 | chr6-140681988-DEL-76 |
| chr6-140683134-DEL-200 | chr6-140683135-DEL-200 |
| chr6-140687718-DEL-73 | chr6-140687708-DEL-74 |
| chr6-140691843-DEL-144 | chr6-140691852-DEL-137 |
| chr6-140692609-DEL-334 | chr6-140692610-DEL-334 |
| chr6-140702428-DEL-561 | chr6-140702428-DEL-554 |
| chr6-140703922-DEL-91 | chr6-140703915-DEL-94 |
| chr6-140725050-DEL-56 | chr6-140725051-DEL-56 |
| chr6-140732474-DEL-244 | chr6-140732505-DEL-242 |
| chr6-140734794-DEL-65 | chr6-140734750-DEL-75 |
| chr6-140737509-DEL-112 | chr6-140737510-DEL-112 |
| chr6-140742393-DEL-128 | chr6-140742394-DEL-126 |
| chr6-140758987-DEL-672 | chr6-140758988-DEL-672 |
| chr6-140777116-DEL-502 | chr6-140777117-DEL-502 |
| chr6-140780430-DEL-6592 | chr6-140780431-DEL-6592 |
| chr6-140788862-DEL-56 | chr6-140788863-DEL-56 |
| chr6-140793798-DEL-76 | chr6-140793783-DEL-92 |
| chr6-140803787-DEL-292 | chr6-140803792-DEL-290 |
| chr6-140809021-DEL-6483 | chr6-140809022-DEL-6483 |
| chr6-140830093-DEL-104 | chr6-140830109-DEL-60 |
| chr6-140831215-DEL-52 | chr6-140831206-DEL-66 |
| chr6-140847590-DEL-120 | chr6-140847591-DEL-120 |
| chr6-140852838-DEL-58 | chr6-140852841-DEL-56 |
| chr6-140864146-DEL-121 | chr6-140864144-DEL-141 |
| chr6-140866491-DEL-148 | chr6-140866492-DEL-148 |
| chr6-140874286-DEL-98 | chr6-140874289-DEL-101 |
| chr6-140875571-DEL-432 | chr6-140875572-DEL-432 |
| chr6-140877257-DEL-477 | chr6-140877258-DEL-477 |
| chr6-140879588-DEL-193 | chr6-140879584-DEL-196 |
| chr6-140911671-DEL-55 | chr6-140911672-DEL-55 |
| chr6-140931017-DEL-6190 | chr6-140931018-DEL-6190 |
| chr6-140982697-DEL-316 | chr6-140982698-DEL-316 |
| chr6-141003658-DEL-286 | chr6-141003659-DEL-286 |
| chr6-141022407-DEL-126 | chr6-141022410-DEL-124 |
| chr6-141023545-DEL-883 | chr6-141023546-DEL-883 |
| chr6-141035539-DEL-103 | chr6-141035540-DEL-103 |
| chr6-141038733-DEL-77 | chr6-141038734-DEL-77 |
| chr6-141049965-DEL-123 | chr6-141049955-DEL-124 |
| chr6-141054428-DEL-126 | chr6-141054429-DEL-126 |
| chr6-141076386-DEL-777 | chr6-141076387-DEL-777 |
| chr6-141081224-DEL-704 | chr6-141081225-DEL-704 |
| chr6-141082059-DEL-116 | chr6-141082040-DEL-104 |
| chr6-141084756-DEL-180 | chr6-141084757-DEL-140 |
| chr6-141085699-DEL-200 | chr6-141085700-DEL-200 |
| chr6-141086837-DEL-319 | chr6-141086838-DEL-319 |
| chr6-141091540-DEL-1761 | chr6-141091541-DEL-1761 |
| chr6-141105769-DEL-62 | chr6-141105745-DEL-52 |
| chr6-141108864-DEL-390 | chr6-141108865-DEL-390 |
| chr6-141125410-DEL-2004 | chr6-141125411-DEL-2004 |
| chr6-141142272-DEL-52 | chr6-141142273-DEL-52 |
| chr6-141149718-DEL-287 | chr6-141149717-DEL-287 |
| chr6-141156713-DEL-6391 | chr6-141156714-DEL-6391 |
| chr6-141177299-DEL-62 | chr6-141177293-DEL-65 |
| chr6-141185567-DEL-2262 | chr6-141185568-DEL-2262 |
| chr6-141220284-DEL-114 | chr6-141220285-DEL-114 |
| chr6-141278823-DEL-56 | chr6-141278823-DEL-60 |
| chr6-141282831-DEL-915 | chr6-141282832-DEL-915 |
| chr6-141285663-DEL-91 | chr6-141285658-DEL-73 |
| chr6-141301666-DEL-7098 | chr6-141301667-DEL-7098 |
| chr6-141307942-DEL-428 | chr6-141307943-DEL-428 |
| chr6-141313908-DEL-74 | chr6-141313904-DEL-90 |
| chr6-141314528-DEL-212 | chr6-141314529-DEL-212 |
| chr6-141318155-DEL-73 | chr6-141318147-DEL-61 |
| chr6-141347180-DEL-56 | chr6-141347181-DEL-56 |
| chr6-141352579-DEL-57 | chr6-141352580-DEL-57 |
| chr6-141376588-DEL-220 | chr6-141376589-DEL-220 |
| chr6-141382520-DEL-389 | chr6-141382521-DEL-389 |
| chr6-141384887-DEL-71 | chr6-141384888-DEL-71 |
| chr6-141386028-DEL-800 | chr6-141386029-DEL-800 |
| chr6-141393143-DEL-60 | chr6-141393144-DEL-60 |
| chr6-141417948-DEL-78 | chr6-141417949-DEL-76 |
| chr6-141489164-DEL-90 | chr6-141489165-DEL-90 |
| chr6-141498034-DEL-62 | chr6-141498033-DEL-64 |
| chr6-141498259-DEL-500 | chr6-141498260-DEL-500 |
| chr6-141509135-DEL-950 | chr6-141509117-DEL-993 |
| chr6-141523755-DEL-6464 | chr6-141523756-DEL-6464 |
| chr6-141534896-DEL-54 | chr6-141534899-DEL-52 |
| chr6-141543273-DEL-155 | chr6-141543274-DEL-155 |

| | |
|---|---|
| chr6-141545740-DEL-254 | chr6-141545750-DEL-251 |
| chr6-141578426-DEL-60 | chr6-141578427-DEL-60 |
| chr6-141598542-DEL-52 | chr6-141598551-DEL-56 |
| chr6-141674627-DEL-408 | chr6-141674628-DEL-408 |
| chr6-141679570-DEL-316 | chr6-141679571-DEL-316 |
| chr6-141684943-DEL-53 | chr6-141684943-DEL-65 |
| chr6-141693191-DEL-6645 | chr6-141693192-DEL-6645 |
| chr6-141704061-DEL-410 | chr6-141704062-DEL-410 |
| chr6-141746993-DEL-479 | chr6-141746993-DEL-480 |
| chr6-141759865-DEL-137 | chr6-141759854-DEL-135 |
| chr6-141762808-DEL-81 | chr6-141762809-DEL-79 |
| chr6-141777544-DEL-888 | chr6-141777545-DEL-888 |
| chr6-141793748-DEL-72 | chr6-141793757-DEL-64 |
| chr6-141806966-DEL-2729 | chr6-141806967-DEL-2729 |
| chr6-141817146-DEL-1693 | chr6-141817147-DEL-1693 |
| chr6-141842699-DEL-55 | chr6-141842699-DEL-55 |
| chr6-141847952-DEL-119 | chr6-141847953-DEL-119 |
| chr6-141869452-DEL-125 | chr6-141869452-DEL-125 |
| chr6-141920475-DEL-56 | chr6-141920476-DEL-56 |
| chr6-141928629-DEL-1522 | chr6-141928630-DEL-1521 |
| chr6-141946647-DEL-82 | chr6-141946648-DEL-82 |
| chr6-141957768-DEL-54 | chr6-141957764-DEL-51 |
| chr6-141958971-DEL-2218 | chr6-141958971-DEL-2218 |
| chr6-141985207-DEL-606 | chr6-141985208-DEL-606 |
| chr6-142005323-DEL-6410 | chr6-142005325-DEL-6411 |
| chr6-142076633-DEL-210 | chr6-142076634-DEL-210 |
| chr6-142082423-DEL-277 | chr6-142082424-DEL-277 |
| chr6-142094691-DEL-7098 | chr6-142094692-DEL-7098 |
| chr6-142113111-DEL-5439 | chr6-142113112-DEL-5439 |
| chr6-142122969-DEL-86 | chr6-142122970-DEL-80 |
| chr6-142191735-DEL-1057 | chr6-142191736-DEL-1057 |
| chr6-142196631-DEL-144 | chr6-142196632-DEL-144 |
| chr6-142202298-DEL-117 | chr6-142202293-DEL-123 |
| chr6-142203190-DEL-78 | chr6-142203191-DEL-78 |
| chr6-142215639-DEL-2847 | chr6-142215640-DEL-2847 |
| chr6-142240533-DEL-86 | chr6-142240534-DEL-86 |
| chr6-142254371-DEL-52 | chr6-142254368-DEL-52 |
| chr6-142260560-DEL-1402 | chr6-142260547-DEL-1403 |
| chr6-142286829-DEL-63 | chr6-142286830-DEL-63 |
| chr6-142319425-DEL-96 | chr6-142319426-DEL-96 |
| chr6-142360792-DEL-437 | chr6-142360793-DEL-437 |
| chr6-142363051-DEL-75 | chr6-142363052-DEL-75 |
| chr6-142367421-DEL-107 | chr6-142367421-DEL-108 |
| chr6-142369548-DEL-213 | chr6-142369548-DEL-213 |
| chr6-142378979-DEL-6212 | chr6-142378980-DEL-6212 |
| chr6-142387217-DEL-69 | chr6-142387217-DEL-69 |
| chr6-142399442-DEL-60 | chr6-142399443-DEL-60 |
| chr6-142405597-DEL-233 | chr6-142405610-DEL-221 |
| chr6-142412244-DEL-844 | chr6-142412245-DEL-844 |
| chr6-142416690-DEL-170 | chr6-142416691-DEL-170 |
| chr6-142419565-DEL-120 | chr6-142419566-DEL-120 |
| chr6-142422828-DEL-72 | chr6-142422839-DEL-62 |
| chr6-142427214-DEL-496 | chr6-142427215-DEL-496 |
| chr6-142428179-DEL-73 | chr6-142428180-DEL-73 |
| chr6-142431877-DEL-407 | chr6-142431878-DEL-407 |
| chr6-142434445-DEL-557 | chr6-142434446-DEL-557 |
| chr6-142460856-DEL-435 | chr6-142460857-DEL-435 |
| chr6-142474685-DEL-91 | chr6-142474686-DEL-91 |
| chr6-142482788-DEL-93 | chr6-142482789-DEL-93 |
| chr6-142494802-DEL-51 | chr6-142494781-DEL-50 |
| chr6-142501384-DEL-56 | chr6-142501385-DEL-56 |
| chr6-142520068-DEL-572 | chr6-142520069-DEL-572 |
| chr6-142527660-DEL-139 | chr6-142527661-DEL-139 |
| chr6-142529489-DEL-76 | chr6-142529486-DEL-84 |
| chr6-142531660-DEL-60 | chr6-142531661-DEL-60 |
| chr6-142540796-DEL-67 | chr6-142540797-DEL-67 |
| chr6-142545683-DEL-318 | chr6-142545684-DEL-318 |
| chr6-142546242-DEL-184 | chr6-142546243-DEL-184 |
| chr6-142550336-DEL-57 | chr6-142550341-DEL-53 |
| chr6-142562779-DEL-58 | chr6-142562778-DEL-76 |
| chr6-142566644-DEL-368 | chr6-142566645-DEL-368 |
| chr6-142574754-DEL-69 | chr6-142574754-DEL-69 |
| chr6-142607965-DEL-348 | chr6-142607966-DEL-348 |
| chr6-142615334-DEL-245 | chr6-142615335-DEL-245 |
| chr6-142622590-DEL-59 | chr6-142622591-DEL-59 |
| chr6-142627340-DEL-222 | chr6-142627341-DEL-222 |
| chr6-142651900-DEL-1917 | chr6-142651901-DEL-1917 |
| chr6-142664365-DEL-105 | chr6-142664345-DEL-87 |
| chr6-142668599-DEL-80 | chr6-142668600-DEL-80 |
| chr6-142675326-DEL-94 | chr6-142675327-DEL-70 |

| | |
|---|---|
| chr6-142699485-DEL-77 | chr6-142699489-DEL-74 |
| chr6-142701900-DEL-64 | chr6-142701883-DEL-82 |
| chr6-142723932-DEL-219 | chr6-142723933-DEL-219 |
| chr6-142747339-DEL-66 | chr6-142747329-DEL-67 |
| chr6-142807169-DEL-159 | chr6-142807169-DEL-157 |
| chr6-142814571-DEL-803 | chr6-142814572-DEL-803 |
| chr6-142849412-DEL-4414 | chr6-142849413-DEL-4414 |
| chr6-142864057-DEL-66 | chr6-142864033-DEL-107 |
| chr6-142868855-DEL-215 | chr6-142868874-DEL-203 |
| chr6-142891841-DEL-80 | chr6-142891842-DEL-80 |
| chr6-142898846-DEL-182 | chr6-142898847-DEL-182 |
| chr6-142911308-DEL-176 | chr6-142911299-DEL-177 |
| chr6-142915804-DEL-1333 | chr6-142915805-DEL-1333 |
| chr6-142943494-DEL-99 | chr6-142943458-DEL-91 |
| chr6-142947514-DEL-251 | chr6-142947515-DEL-251 |
| chr6-142950884-DEL-210 | chr6-142950885-DEL-210 |
| chr6-142953951-DEL-211 | chr6-142953952-DEL-211 |
| chr6-142968863-DEL-209 | chr6-142968864-DEL-209 |
| chr6-143005764-DEL-56 | chr6-143005765-DEL-56 |
| chr6-143011430-DEL-79 | chr6-143011431-DEL-79 |
| chr6-143054587-DEL-371 | chr6-143054588-DEL-371 |
| chr6-143057464-DEL-179 | chr6-143057465-DEL-179 |
| chr6-143064208-DEL-60 | chr6-143064209-DEL-60 |
| chr6-143074838-DEL-154 | chr6-143074839-DEL-154 |
| chr6-143102107-DEL-229 | chr6-143102103-DEL-214 |
| chr6-143143437-DEL-226 | chr6-143143436-DEL-227 |
| chr6-143159085-DEL-380 | chr6-143159086-DEL-403 |
| chr6-143196800-DEL-98 | chr6-143196793-DEL-110 |
| chr6-143198116-DEL-579 | chr6-143198117-DEL-579 |
| chr6-143207080-DEL-58 | chr6-143207081-DEL-58 |
| chr6-143226182-DEL-103 | chr6-143226183-DEL-103 |
| chr6-143237207-DEL-92 | chr6-143237207-DEL-92 |
| chr6-143245059-DEL-63 | chr6-143245079-DEL-60 |
| chr6-143283650-DEL-70 | chr6-143283655-DEL-66 |
| chr6-143292626-DEL-82 | chr6-143292624-DEL-65 |
| chr6-143293195-DEL-3591 | chr6-143293196-DEL-3591 |
| chr6-143306454-DEL-153 | chr6-143306455-DEL-153 |
| chr6-143328201-DEL-148 | chr6-143328201-DEL-111 |
| chr6-143342716-DEL-50 | chr6-143342681-DEL-74 |
| chr6-143362484-DEL-140 | chr6-143362479-DEL-148 |
| chr6-143372602-DEL-66 | chr6-143372595-DEL-58 |
| chr6-143373442-DEL-143 | chr6-143373439-DEL-144 |
| chr6-143380199-DEL-396 | chr6-143380200-DEL-396 |
| chr6-143387587-DEL-446 | chr6-143387588-DEL-446 |
| chr6-143412200-DEL-6331 | chr6-143412201-DEL-6331 |
| chr6-143426000-DEL-476 | chr6-143426041-DEL-438 |
| chr6-143451446-DEL-7273 | chr6-143451447-DEL-7273 |
| chr6-143473248-DEL-98 | chr6-143473244-DEL-102 |
| chr6-143474477-DEL-68 | chr6-143474478-DEL-68 |
| chr6-143519880-DEL-270 | chr6-143519879-DEL-272 |
| chr6-143521028-DEL-128 | chr6-143521040-DEL-88 |
| chr6-143538030-DEL-51 | chr6-143538031-DEL-51 |
| chr6-143567319-DEL-104 | chr6-143567320-DEL-54 |
| chr6-143576218-DEL-113 | chr6-143576219-DEL-113 |
| chr6-143604980-DEL-90 | chr6-143604981-DEL-75 |
| chr6-143618312-DEL-74 | chr6-143618313-DEL-74 |
| chr6-143622793-DEL-64 | chr6-143622795-DEL-52 |
| chr6-143623784-DEL-53 | chr6-143623785-DEL-53 |
| chr6-143662168-DEL-405 | chr6-143662169-DEL-404 |
| chr6-143720837-DEL-6399 | chr6-143720837-DEL-6399 |
| chr6-143727403-DEL-53 | chr6-143727404-DEL-53 |
| chr6-143746310-DEL-2727 | chr6-143746311-DEL-2727 |
| chr6-143752127-DEL-124 | chr6-143752128-DEL-124 |
| chr6-143779634-DEL-69 | chr6-143779635-DEL-69 |
| chr6-143785464-DEL-64 | chr6-143785465-DEL-64 |
| chr6-143788290-DEL-276 | chr6-143788281-DEL-241 |
| chr6-143829140-DEL-3412 | chr6-143829141-DEL-3412 |
| chr6-143890419-DEL-54 | chr6-143890420-DEL-54 |
| chr6-143896642-DEL-51 | chr6-143896642-DEL-51 |
| chr6-143930574-DEL-98 | chr6-143930575-DEL-98 |
| chr6-143984192-DEL-56 | chr6-143984193-DEL-56 |
| chr6-144014944-DEL-80 | chr6-144014945-DEL-80 |
| chr6-144016712-DEL-51 | chr6-144016713-DEL-51 |
| chr6-144037904-DEL-69 | chr6-144037900-DEL-73 |
| chr6-144040259-DEL-83 | chr6-144040260-DEL-83 |
| chr6-144076930-DEL-63 | chr6-144076933-DEL-61 |
| chr6-144107972-DEL-151 | chr6-144107997-DEL-127 |
| chr6-144113143-DEL-67 | chr6-144113097-DEL-68 |
| chr6-144121385-DEL-75 | chr6-144121386-DEL-75 |
| chr6-144139894-DEL-590 | chr6-144139895-DEL-590 |

| | |
|---|---|
| chr6-144162074-DEL-75 | chr6-144162073-DEL-76 |
| chr6-144188642-DEL-71 | chr6-144188616-DEL-84 |
| chr6-144199902-DEL-54 | chr6-144199905-DEL-52 |
| chr6-144207720-DEL-84 | chr6-144207721-DEL-84 |
| chr6-144269804-DEL-74 | chr6-144269805-DEL-74 |
| chr6-144275186-DEL-123 | chr6-144275157-DEL-124 |
| chr6-144292857-DEL-96 | chr6-144292858-DEL-96 |
| chr6-144323012-DEL-2496 | chr6-144323013-DEL-2496 |
| chr6-144338512-DEL-178 | chr6-144338513-DEL-178 |
| chr6-144357319-DEL-249 | chr6-144357320-DEL-246 |
| chr6-144358600-DEL-407 | chr6-144358601-DEL-407 |
| chr6-144373981-DEL-148 | chr6-144373982-DEL-148 |
| chr6-144381250-DEL-93 | chr6-144381251-DEL-93 |
| chr6-144381876-DEL-6272 | chr6-144381877-DEL-6272 |
| chr6-144392353-DEL-63 | chr6-144392354-DEL-63 |
| chr6-144397262-DEL-100 | chr6-144397263-DEL-100 |
| chr6-144417446-DEL-68 | chr6-144417447-DEL-68 |
| chr6-144421029-DEL-54 | chr6-144421026-DEL-58 |
| chr6-144438495-DEL-82 | chr6-144438496-DEL-82 |
| chr6-144455430-DEL-104 | chr6-144455431-DEL-104 |
| chr6-144471045-DEL-51 | chr6-144471046-DEL-51 |
| chr6-144498293-DEL-79 | chr6-144498294-DEL-79 |
| chr6-144516815-DEL-96 | chr6-144516816-DEL-96 |
| chr6-144534357-DEL-2292 | chr6-144534358-DEL-2292 |
| chr6-144577559-DEL-56 | chr6-144577560-DEL-56 |
| chr6-144598450-DEL-194 | chr6-144598451-DEL-194 |
| chr6-144600818-DEL-1640 | chr6-144600818-DEL-1640 |
| chr6-144603073-DEL-65 | chr6-144603072-DEL-64 |
| chr6-144613663-DEL-2016 | chr6-144613664-DEL-2016 |
| chr6-144646358-DEL-145 | chr6-144646359-DEL-145 |
| chr6-144704120-DEL-154 | chr6-144704127-DEL-148 |
| chr6-144712186-DEL-849 | chr6-144712187-DEL-849 |
| chr6-144751883-DEL-156 | chr6-144751887-DEL-155 |
| chr6-144761902-DEL-60 | chr6-144761899-DEL-50 |
| chr6-144765455-DEL-83 | chr6-144765450-DEL-84 |
| chr6-144767710-DEL-206 | chr6-144767695-DEL-208 |
| chr6-144771720-DEL-7028 | chr6-144771721-DEL-7028 |
| chr6-144796780-DEL-168 | chr6-144796781-DEL-168 |
| chr6-144803144-DEL-54 | chr6-144803143-DEL-55 |
| chr6-144848744-DEL-76 | chr6-144848745-DEL-76 |
| chr6-144864565-DEL-95 | chr6-144864566-DEL-95 |
| chr6-144889304-DEL-120 | chr6-144889284-DEL-120 |
| chr6-144897628-DEL-1994 | chr6-144897629-DEL-1994 |
| chr6-144900074-DEL-554 | chr6-144900074-DEL-555 |
| chr6-144912219-DEL-64 | chr6-144912220-DEL-64 |
| chr6-144920754-DEL-6572 | chr6-144920755-DEL-6572 |
| chr6-144930290-DEL-51 | chr6-144930291-DEL-51 |
| chr6-144932055-DEL-209 | chr6-144932056-DEL-209 |
| chr6-144955320-DEL-61 | chr6-144955321-DEL-61 |
| chr6-144958799-DEL-60 | chr6-144958800-DEL-60 |
| chr6-144963512-DEL-198 | chr6-144963513-DEL-198 |
| chr6-144968597-DEL-91 | chr6-144968570-DEL-92 |
| chr6-144974711-DEL-763 | chr6-144974712-DEL-763 |
| chr6-144987139-DEL-84 | chr6-144987140-DEL-84 |
| chr6-145009629-DEL-223 | chr6-145009630-DEL-223 |
| chr6-145013061-DEL-62 | chr6-145013062-DEL-62 |
| chr6-145016750-DEL-168 | chr6-145016751-DEL-168 |
| chr6-145019753-DEL-98 | chr6-145019754-DEL-98 |
| chr6-145033197-DEL-626 | chr6-145033183-DEL-620 |
| chr6-145039006-DEL-147 | chr6-145039007-DEL-147 |
| chr6-145039662-DEL-160 | chr6-145039663-DEL-160 |
| chr6-145042957-DEL-51 | chr6-145042957-DEL-53 |
| chr6-145045771-DEL-187 | chr6-145045769-DEL-190 |
| chr6-145057388-DEL-484 | chr6-145057389-DEL-484 |
| chr6-145072599-DEL-182 | chr6-145072600-DEL-182 |
| chr6-145129291-DEL-51 | chr6-145129256-DEL-52 |
| chr6-145140421-DEL-348 | chr6-145140422-DEL-348 |
| chr6-145212034-DEL-521 | chr6-145212035-DEL-521 |
| chr6-145216357-DEL-272 | chr6-145216370-DEL-273 |
| chr6-145221852-DEL-246 | chr6-145221853-DEL-246 |
| chr6-145229439-DEL-66 | chr6-145229435-DEL-70 |
| chr6-145246640-DEL-182 | chr6-145246641-DEL-182 |
| chr6-145251113-DEL-202 | chr6-145251114-DEL-202 |
| chr6-145278401-DEL-88 | chr6-145278402-DEL-88 |
| chr6-145298469-DEL-51 | chr6-145298470-DEL-51 |
| chr6-145304307-DEL-83 | chr6-145304302-DEL-82 |
| chr6-145316941-DEL-62 | chr6-145316942-DEL-62 |
| chr6-145317858-DEL-155 | chr6-145317871-DEL-152 |
| chr6-145328111-DEL-72 | chr6-145328112-DEL-72 |
| chr6-145338285-DEL-68 | chr6-145338286-DEL-68 |

| | |
|---|---|
| chr6-145371489-DEL-54 | chr6-145371490-DEL-54 |
| chr6-145385308-DEL-60 | chr6-145385309-DEL-60 |
| chr6-145387171-DEL-62 | chr6-145387172-DEL-62 |
| chr6-145396407-DEL-51 | chr6-145396408-DEL-51 |
| chr6-145400673-DEL-113 | chr6-145400715-DEL-73 |
| chr6-145406316-DEL-1210 | chr6-145406317-DEL-1210 |
| chr6-145411702-DEL-73 | chr6-145411703-DEL-73 |
| chr6-145434770-DEL-4327 | chr6-145434771-DEL-4327 |
| chr6-145446484-DEL-157 | chr6-145446483-DEL-158 |
| chr6-145455361-DEL-232 | chr6-145455362-DEL-232 |
| chr6-145465639-DEL-639 | chr6-145465640-DEL-639 |
| chr6-145471190-DEL-432 | chr6-145471191-DEL-432 |
| chr6-145475437-DEL-72 | chr6-145475430-DEL-70 |
| chr6-145477775-DEL-336 | chr6-145477776-DEL-336 |
| chr6-145486865-DEL-87 | chr6-145486865-DEL-85 |
| chr6-145487257-DEL-450 | chr6-145487258-DEL-450 |
| chr6-145525476-DEL-65 | chr6-145525468-DEL-66 |
| chr6-145538717-DEL-525 | chr6-145538718-DEL-525 |
| chr6-145555962-DEL-207 | chr6-145555963-DEL-207 |
| chr6-145570541-DEL-129 | chr6-145570542-DEL-129 |
| chr6-145587717-DEL-7446 | chr6-145587717-DEL-7446 |
| chr6-145593120-DEL-1029 | chr6-145593121-DEL-1029 |
| chr6-145599165-DEL-165 | chr6-145599166-DEL-165 |
| chr6-145614041-DEL-174 | chr6-145614042-DEL-174 |
| chr6-145620171-DEL-76 | chr6-145620152-DEL-60 |
| chr6-145624869-DEL-175 | chr6-145624870-DEL-175 |
| chr6-145639973-DEL-252 | chr6-145639974-DEL-252 |
| chr6-145662627-DEL-1414 | chr6-145662628-DEL-1414 |
| chr6-145703482-DEL-54 | chr6-145703483-DEL-54 |
| chr6-145710096-DEL-81 | chr6-145710096-DEL-54 |
| chr6-145715170-DEL-420 | chr6-145715171-DEL-378 |
| chr6-145721476-DEL-196 | chr6-145721477-DEL-196 |
| chr6-145729854-DEL-149 | chr6-145729855-DEL-149 |
| chr6-145753292-DEL-200 | chr6-145753293-DEL-200 |
| chr6-145779613-DEL-706 | chr6-145779614-DEL-706 |
| chr6-145794753-DEL-782 | chr6-145794754-DEL-782 |
| chr6-145794861-DEL-53 | chr6-145794862-DEL-53 |
| chr6-145832915-DEL-199 | chr6-145832916-DEL-199 |
| chr6-145841544-DEL-7112 | chr6-145841579-DEL-7085 |
| chr6-145849237-DEL-240 | chr6-145849238-DEL-216 |
| chr6-145855231-DEL-76 | chr6-145855232-DEL-68 |
| chr6-145875817-DEL-53 | chr6-145875814-DEL-56 |
| chr6-145878479-DEL-275 | chr6-145878474-DEL-276 |
| chr6-145887631-DEL-84 | chr6-145887628-DEL-76 |
| chr6-145910551-DEL-72 | chr6-145910552-DEL-72 |
| chr6-145940637-DEL-3114 | chr6-145940638-DEL-3114 |
| chr6-145948891-DEL-68 | chr6-145948877-DEL-68 |
| chr6-145962225-DEL-169 | chr6-145962226-DEL-169 |
| chr6-145965598-DEL-738 | chr6-145965599-DEL-738 |
| chr6-145989356-DEL-63 | chr6-145989357-DEL-63 |
| chr6-145992227-DEL-106 | chr6-145992228-DEL-106 |
| chr6-146003980-DEL-327 | chr6-146003981-DEL-327 |
| chr6-146017622-DEL-74 | chr6-146017616-DEL-86 |
| chr6-146064977-DEL-70 | chr6-146064978-DEL-70 |
| chr6-146099472-DEL-126 | chr6-146099473-DEL-126 |
| chr6-146101231-DEL-118 | chr6-146101233-DEL-111 |
| chr6-146103971-DEL-847 | chr6-146103972-DEL-847 |
| chr6-146115013-DEL-740 | chr6-146115014-DEL-740 |
| chr6-146139204-DEL-70 | chr6-146139217-DEL-58 |
| chr6-146157778-DEL-64 | chr6-146157779-DEL-64 |
| chr6-146158464-DEL-472 | chr6-146158464-DEL-472 |
| chr6-146168626-DEL-797 | chr6-146168627-DEL-797 |
| chr6-146177044-DEL-200 | chr6-146177037-DEL-201 |
| chr6-146189355-DEL-245 | chr6-146189356-DEL-245 |
| chr6-146190738-DEL-80 | chr6-146190739-DEL-80 |
| chr6-146198404-DEL-98 | chr6-146198400-DEL-94 |
| chr6-146205517-DEL-1464 | chr6-146205512-DEL-1434 |
| chr6-146223726-DEL-180 | chr6-146223727-DEL-180 |
| chr6-146236646-DEL-63 | chr6-146236638-DEL-60 |
| chr6-146277706-DEL-1897 | chr6-146277707-DEL-1897 |
| chr6-146285694-DEL-108 | chr6-146285697-DEL-100 |
| chr6-146303900-DEL-152 | chr6-146303901-DEL-152 |
| chr6-146307677-DEL-3262 | chr6-146307677-DEL-3262 |
| chr6-146325588-DEL-331 | chr6-146325589-DEL-331 |
| chr6-146330538-DEL-102 | chr6-146330557-DEL-56 |
| chr6-146352839-DEL-57 | chr6-146352840-DEL-57 |
| chr6-146354045-DEL-68 | chr6-146354046-DEL-68 |
| chr6-146383859-DEL-120 | chr6-146383860-DEL-120 |
| chr6-146409114-DEL-66 | chr6-146409117-DEL-68 |
| chr6-146425842-DEL-194 | chr6-146425843-DEL-194 |

| | |
|---|---|
| chr6-146437438-DEL-433 | chr6-146437439-DEL-433 |
| chr6-146455737-DEL-53 | chr6-146455738-DEL-53 |
| chr6-146458303-DEL-223 | chr6-146458304-DEL-223 |
| chr6-146470308-DEL-2001 | chr6-146470309-DEL-2001 |
| chr6-146512472-DEL-486 | chr6-146512473-DEL-486 |
| chr6-146527140-DEL-383 | chr6-146527141-DEL-383 |
| chr6-146532540-DEL-262 | chr6-146532541-DEL-262 |
| chr6-146536571-DEL-91 | chr6-146536582-DEL-80 |
| chr6-146538407-DEL-198 | chr6-146538408-DEL-198 |
| chr6-146542732-DEL-371 | chr6-146542733-DEL-371 |
| chr6-146550199-DEL-208 | chr6-146550200-DEL-208 |
| chr6-146553937-DEL-474 | chr6-146553938-DEL-474 |
| chr6-146558059-DEL-168 | chr6-146558060-DEL-168 |
| chr6-146574061-DEL-52 | chr6-146574062-DEL-52 |
| chr6-146591196-DEL-5021 | chr6-146591196-DEL-5021 |
| chr6-146605665-DEL-207 | chr6-146605666-DEL-207 |
| chr6-146619714-DEL-174 | chr6-146619715-DEL-174 |
| chr6-146623020-DEL-2682 | chr6-146623021-DEL-2682 |
| chr6-146624915-DEL-200 | chr6-146624914-DEL-201 |
| chr6-146641911-DEL-230 | chr6-146641912-DEL-230 |
| chr6-146654142-DEL-72 | chr6-146654143-DEL-72 |
| chr6-146660327-DEL-255 | chr6-146660328-DEL-255 |
| chr6-146661926-DEL-54 | chr6-146661911-DEL-55 |
| chr6-146665774-DEL-52 | chr6-146665775-DEL-52 |
| chr6-146687658-DEL-5699 | chr6-146687658-DEL-5699 |
| chr6-146710857-DEL-54 | chr6-146710859-DEL-53 |
| chr6-146718506-DEL-449 | chr6-146718507-DEL-449 |
| chr6-146720718-DEL-52 | chr6-146720719-DEL-52 |
| chr6-146736252-DEL-79 | chr6-146736253-DEL-79 |
| chr6-146763319-DEL-155 | chr6-146763320-DEL-155 |
| chr6-146765828-DEL-104 | chr6-146765829-DEL-80 |
| chr6-146772088-DEL-148 | chr6-146772089-DEL-148 |
| chr6-146782858-DEL-54 | chr6-146782858-DEL-54 |
| chr6-146796647-DEL-502 | chr6-146796648-DEL-502 |
| chr6-146820279-DEL-6278 | chr6-146820280-DEL-6278 |
| chr6-146850187-DEL-117 | chr6-146850193-DEL-113 |
| chr6-146892560-DEL-208 | chr6-146892561-DEL-208 |
| chr6-146895292-DEL-110 | chr6-146895293-DEL-110 |
| chr6-146900073-DEL-60 | chr6-146900072-DEL-62 |
| chr6-146963295-DEL-92 | chr6-146963296-DEL-92 |
| chr6-146977754-DEL-268 | chr6-146977757-DEL-269 |
| chr6-146978683-DEL-2757 | chr6-146978684-DEL-2757 |
| chr6-146990634-DEL-57 | chr6-146990611-DEL-52 |
| chr6-147015120-DEL-88 | chr6-147015113-DEL-100 |
| chr6-147025904-DEL-111 | chr6-147025905-DEL-111 |
| chr6-147030599-DEL-866 | chr6-147030600-DEL-866 |
| chr6-147060203-DEL-142 | chr6-147060192-DEL-154 |
| chr6-147067490-DEL-140 | chr6-147067491-DEL-140 |
| chr6-147068129-DEL-58 | chr6-147068130-DEL-58 |
| chr6-147069932-DEL-69 | chr6-147069933-DEL-69 |
| chr6-147076679-DEL-506 | chr6-147076680-DEL-506 |
| chr6-147084772-DEL-381 | chr6-147084773-DEL-381 |
| chr6-147091967-DEL-168 | chr6-147091968-DEL-174 |
| chr6-147093957-DEL-82 | chr6-147093958-DEL-82 |
| chr6-147095595-DEL-141 | chr6-147095596-DEL-135 |
| chr6-147099076-DEL-70 | chr6-147099077-DEL-70 |
| chr6-147112780-DEL-53 | chr6-147112781-DEL-53 |
| chr6-147140739-DEL-56 | chr6-147140740-DEL-56 |
| chr6-147204928-DEL-2972 | chr6-147204927-DEL-2973 |
| chr6-147223231-DEL-91 | chr6-147223232-DEL-87 |
| chr6-147240810-DEL-68 | chr6-147240785-DEL-51 |
| chr6-147259409-DEL-1014 | chr6-147259409-DEL-1014 |
| chr6-147267534-DEL-172 | chr6-147267535-DEL-172 |
| chr6-147320918-DEL-57 | chr6-147320894-DEL-66 |
| chr6-147361449-DEL-4919 | chr6-147361450-DEL-4919 |
| chr6-147371259-DEL-184 | chr6-147371214-DEL-184 |
| chr6-147382625-DEL-61 | chr6-147382626-DEL-61 |
| chr6-147396037-DEL-255 | chr6-147396038-DEL-255 |
| chr6-147423668-DEL-56 | chr6-147423669-DEL-56 |
| chr6-147430599-DEL-205 | chr6-147430599-DEL-203 |
| chr6-147451244-DEL-5152 | chr6-147451245-DEL-5152 |
| chr6-147523414-DEL-408 | chr6-147523415-DEL-407 |
| chr6-147531136-DEL-952 | chr6-147531137-DEL-952 |
| chr6-147536567-DEL-95 | chr6-147536529-DEL-130 |
| chr6-147556457-DEL-1872 | chr6-147556458-DEL-1872 |
| chr6-147569246-DEL-207 | chr6-147569247-DEL-207 |
| chr6-147578717-DEL-252 | chr6-147578717-DEL-249 |
| chr6-147603204-DEL-374 | chr6-147603205-DEL-374 |
| chr6-147612530-DEL-6440 | chr6-147612531-DEL-6440 |
| chr6-147637834-DEL-218 | chr6-147637835-DEL-218 |

| | |
|---|---|
| chr6-147638208-DEL-988 | chr6-147638209-DEL-988 |
| chr6-147651801-DEL-236 | chr6-147651800-DEL-240 |
| chr6-147654566-DEL-785 | chr6-147654567-DEL-785 |
| chr6-147669754-DEL-188 | chr6-147669787-DEL-153 |
| chr6-147698291-DEL-52 | chr6-147698292-DEL-52 |
| chr6-147734515-DEL-498 | chr6-147734516-DEL-516 |
| chr6-147744613-DEL-342 | chr6-147744614-DEL-342 |
| chr6-147772084-DEL-64 | chr6-147772085-DEL-64 |
| chr6-147795121-DEL-14725 | chr6-147795122-DEL-14725 |
| chr6-147816975-DEL-76 | chr6-147816976-DEL-76 |
| chr6-147856074-DEL-52 | chr6-147856075-DEL-52 |
| chr6-147870198-DEL-687 | chr6-147870199-DEL-685 |
| chr6-147905713-DEL-54 | chr6-147905711-DEL-56 |
| chr6-147915101-DEL-704 | chr6-147915102-DEL-704 |
| chr6-147916927-DEL-64 | chr6-147916926-DEL-64 |
| chr6-147921476-DEL-4370 | chr6-147921477-DEL-4370 |
| chr6-147973908-DEL-177 | chr6-147973912-DEL-169 |
| chr6-148001471-DEL-806 | chr6-148001472-DEL-806 |
| chr6-148025368-DEL-1692 | chr6-148025369-DEL-1692 |
| chr6-148027775-DEL-53 | chr6-148027776-DEL-53 |
| chr6-148056342-DEL-459 | chr6-148056343-DEL-459 |
| chr6-148062377-DEL-192 | chr6-148062378-DEL-192 |
| chr6-148067846-DEL-5127 | chr6-148067841-DEL-5128 |
| chr6-148085942-DEL-88 | chr6-148085947-DEL-84 |
| chr6-148129124-DEL-144 | chr6-148129116-DEL-145 |
| chr6-148190666-DEL-54 | chr6-148190667-DEL-54 |
| chr6-148199812-DEL-76 | chr6-148199813-DEL-76 |
| chr6-148217815-DEL-902 | chr6-148217816-DEL-902 |
| chr6-148259719-DEL-161 | chr6-148259720-DEL-161 |
| chr6-148269359-DEL-1737 | chr6-148269360-DEL-1737 |
| chr6-148278001-DEL-101 | chr6-148277999-DEL-82 |
| chr6-148291461-DEL-74 | chr6-148291462-DEL-74 |
| chr6-148329164-DEL-194 | chr6-148329165-DEL-192 |
| chr6-148333514-DEL-88 | chr6-148333514-DEL-88 |
| chr6-148335495-DEL-442 | chr6-148335496-DEL-442 |
| chr6-148337004-DEL-198 | chr6-148337005-DEL-198 |
| chr6-148338675-DEL-76 | chr6-148338676-DEL-76 |
| chr6-148366643-DEL-185 | chr6-148366644-DEL-185 |
| chr6-148388620-DEL-59 | chr6-148388602-DEL-60 |
| chr6-148390308-DEL-530 | chr6-148390309-DEL-530 |
| chr6-148408246-DEL-277 | chr6-148408247-DEL-277 |
| chr6-148445916-DEL-1016 | chr6-148445917-DEL-1016 |
| chr6-148454527-DEL-60 | chr6-148454528-DEL-60 |
| chr6-148547328-DEL-53 | chr6-148547329-DEL-53 |
| chr6-148558581-DEL-65 | chr6-148558582-DEL-63 |
| chr6-148564479-DEL-1868 | chr6-148564480-DEL-1868 |
| chr6-148578513-DEL-186 | chr6-148578514-DEL-186 |
| chr6-148582040-DEL-926 | chr6-148582041-DEL-926 |
| chr6-148587140-DEL-1712 | chr6-148587141-DEL-1712 |
| chr6-148597842-DEL-118 | chr6-148597799-DEL-123 |
| chr6-148602654-DEL-62 | chr6-148602655-DEL-62 |
| chr6-148631268-DEL-1896 | chr6-148631253-DEL-1897 |
| chr6-148649568-DEL-146 | chr6-148649569-DEL-135 |
| chr6-148661157-DEL-78 | chr6-148661158-DEL-78 |
| chr6-148669641-DEL-84 | chr6-148669646-DEL-80 |
| chr6-148670673-DEL-75 | chr6-148670674-DEL-75 |
| chr6-148685220-DEL-59 | chr6-148685200-DEL-80 |
| chr6-148698987-DEL-61 | chr6-148698986-DEL-58 |
| chr6-148715891-DEL-151 | chr6-148715892-DEL-151 |
| chr6-148724446-DEL-384 | chr6-148724448-DEL-382 |
| chr6-148725282-DEL-148 | chr6-148725278-DEL-149 |
| chr6-148748713-DEL-213 | chr6-148748714-DEL-213 |
| chr6-148793785-DEL-105 | chr6-148793796-DEL-60 |
| chr6-148810853-DEL-196 | chr6-148810854-DEL-196 |
| chr6-148813106-DEL-198 | chr6-148813108-DEL-197 |
| chr6-148824932-DEL-55 | chr6-148824922-DEL-56 |
| chr6-148834026-DEL-170 | chr6-148834027-DEL-170 |
| chr6-148849641-DEL-247 | chr6-148849642-DEL-247 |
| chr6-148854066-DEL-83 | chr6-148854064-DEL-83 |
| chr6-148854719-DEL-315 | chr6-148854720-DEL-313 |
| chr6-148856216-DEL-246 | chr6-148856217-DEL-242 |
| chr6-148863671-DEL-223 | chr6-148863671-DEL-224 |
| chr6-148870100-DEL-207 | chr6-148870101-DEL-207 |
| chr6-148882213-DEL-258 | chr6-148882214-DEL-258 |
| chr6-148882755-DEL-204 | chr6-148882756-DEL-199 |
| chr6-148891543-DEL-220 | chr6-148891544-DEL-220 |
| chr6-148909846-DEL-201 | chr6-148909847-DEL-201 |
| chr6-148926406-DEL-1415 | chr6-148926407-DEL-1415 |
| chr6-148997391-DEL-195 | chr6-148997392-DEL-195 |
| chr6-149023843-DEL-82 | chr6-149023844-DEL-82 |

| | |
|---|---|
| chr6-149026766-DEL-1505 | chr6-149026768-DEL-1504 |
| chr6-149029007-DEL-69 | chr6-149028998-DEL-59 |
| chr6-149031730-DEL-3908 | chr6-149031749-DEL-3907 |
| chr6-149049193-DEL-80 | chr6-149049205-DEL-66 |
| chr6-149079096-DEL-1959 | chr6-149079097-DEL-1959 |
| chr6-149124623-DEL-219 | chr6-149124624-DEL-219 |
| chr6-149125768-DEL-572 | chr6-149125769-DEL-572 |
| chr6-149126287-DEL-3037 | chr6-149126288-DEL-3037 |
| chr6-149126909-DEL-6469 | chr6-149126910-DEL-6469 |
| chr6-149148652-DEL-211 | chr6-149148653-DEL-211 |
| chr6-149170794-DEL-5246 | chr6-149170795-DEL-5246 |
| chr6-149177039-DEL-341 | chr6-149177040-DEL-341 |
| chr6-149177724-DEL-293 | chr6-149177725-DEL-293 |
| chr6-149255461-DEL-176 | chr6-149255462-DEL-176 |
| chr6-149256212-DEL-148 | chr6-149256213-DEL-148 |
| chr6-149261189-DEL-560 | chr6-149261190-DEL-560 |
| chr6-149263363-DEL-55 | chr6-149263364-DEL-55 |
| chr6-149279481-DEL-229 | chr6-149279482-DEL-229 |
| chr6-149293012-DEL-629 | chr6-149293013-DEL-629 |
| chr6-149384360-DEL-723 | chr6-149384361-DEL-723 |
| chr6-149465252-DEL-56 | chr6-149465252-DEL-58 |
| chr6-149485338-DEL-84 | chr6-149485339-DEL-84 |
| chr7-3131958-DEL-104 | chr7-3131957-DEL-107 |
| chr7-3142372-DEL-53 | chr7-3142373-DEL-53 |
| chr7-3156845-DEL-419 | chr7-3156846-DEL-419 |
| chr7-3182686-DEL-52 | chr7-3182687-DEL-52 |
| chr7-3201147-DEL-194 | chr7-3201148-DEL-194 |
| chr7-3210202-DEL-211 | chr7-3210202-DEL-211 |
| chr7-3220671-DEL-179 | chr7-3220672-DEL-179 |
| chr7-3222689-DEL-224 | chr7-3222690-DEL-224 |
| chr7-3223061-DEL-802 | chr7-3223062-DEL-802 |
| chr7-3230773-DEL-944 | chr7-3230792-DEL-927 |
| chr7-3236958-DEL-192 | chr7-3236959-DEL-192 |
| chr7-3239483-DEL-173 | chr7-3239484-DEL-173 |
| chr7-3268624-DEL-84 | chr7-3268641-DEL-80 |
| chr7-3291159-DEL-102 | chr7-3291160-DEL-102 |
| chr7-3301738-DEL-198 | chr7-3301739-DEL-198 |
| chr7-3303371-DEL-139 | chr7-3303373-DEL-138 |
| chr7-3304115-DEL-1232 | chr7-3304116-DEL-1232 |
| chr7-3304582-DEL-205 | chr7-3304583-DEL-205 |
| chr7-3314032-DEL-479 | chr7-3314033-DEL-479 |
| chr7-3317751-DEL-67 | chr7-3317738-DEL-64 |
| chr7-3322348-DEL-102 | chr7-3322349-DEL-102 |
| chr7-3327526-DEL-338 | chr7-3327527-DEL-338 |
| chr7-3399291-DEL-113 | chr7-3399292-DEL-113 |
| chr7-3407527-DEL-1215 | chr7-3407528-DEL-1215 |
| chr7-3407951-DEL-465 | chr7-3407952-DEL-455 |
| chr7-3420410-DEL-206 | chr7-3420411-DEL-206 |
| chr7-3424513-DEL-206 | chr7-3424505-DEL-207 |
| chr7-3436653-DEL-88 | chr7-3436654-DEL-88 |
| chr7-3454075-DEL-214 | chr7-3454079-DEL-211 |
| chr7-3455756-DEL-173 | chr7-3455752-DEL-171 |
| chr7-3467843-DEL-240 | chr7-3467844-DEL-240 |
| chr7-3514954-DEL-76 | chr7-3514948-DEL-83 |
| chr7-3531437-DEL-486 | chr7-3531438-DEL-486 |
| chr7-3536956-DEL-57 | chr7-3536960-DEL-54 |
| chr7-3588065-DEL-175 | chr7-3588066-DEL-175 |
| chr7-3595651-DEL-202 | chr7-3595651-DEL-202 |
| chr7-3598449-DEL-150 | chr7-3598416-DEL-151 |
| chr7-3605220-DEL-51 | chr7-3605221-DEL-51 |
| chr7-3632178-DEL-72 | chr7-3632179-DEL-72 |
| chr7-3662621-DEL-53 | chr7-3662622-DEL-53 |
| chr7-3684533-DEL-80 | chr7-3684534-DEL-80 |
| chr7-3691197-DEL-93 | chr7-3691198-DEL-93 |
| chr7-3701286-DEL-80 | chr7-3701287-DEL-80 |
| chr7-3710368-DEL-52 | chr7-3710369-DEL-52 |
| chr7-3718098-DEL-52 | chr7-3718085-DEL-60 |
| chr7-3721397-DEL-205 | chr7-3721410-DEL-214 |
| chr7-3726348-DEL-180 | chr7-3726334-DEL-156 |
| chr7-3746180-DEL-72 | chr7-3746181-DEL-72 |
| chr7-3807598-DEL-1981 | chr7-3807599-DEL-1981 |
| chr7-3987552-DEL-90 | chr7-3987553-DEL-90 |
| chr7-4020904-DEL-6639 | chr7-4020905-DEL-6639 |
| chr7-4020927-DEL-208 | chr7-4020928-DEL-208 |
| chr7-4037874-DEL-823 | chr7-4037875-DEL-823 |
| chr7-4159798-DEL-183 | chr7-4159799-DEL-183 |
| chr7-4188375-DEL-14859 | chr7-4188376-DEL-14859 |
| chr7-4215038-DEL-90 | chr7-4215039-DEL-90 |
| chr7-4223566-DEL-213 | chr7-4223567-DEL-213 |
| chr7-4228867-DEL-1830 | chr7-4228868-DEL-1830 |

| | |
|---|---|
| chr7-4231822-DEL-6045 | chr7-4231823-DEL-6045 |
| chr7-4253583-DEL-511 | chr7-4253583-DEL-511 |
| chr7-4279845-DEL-4427 | chr7-4279846-DEL-4427 |
| chr7-4288279-DEL-57 | chr7-4288280-DEL-57 |
| chr7-4290798-DEL-2805 | chr7-4290798-DEL-2805 |
| chr7-4303971-DEL-201 | chr7-4303972-DEL-201 |
| chr7-4305354-DEL-93 | chr7-4305328-DEL-94 |
| chr7-4306955-DEL-7418 | chr7-4306956-DEL-7418 |
| chr7-4332866-DEL-1995 | chr7-4332867-DEL-1995 |
| chr7-4346195-DEL-173 | chr7-4346196-DEL-172 |
| chr7-4415243-DEL-66 | chr7-4415244-DEL-66 |
| chr7-4415852-DEL-52 | chr7-4415853-DEL-52 |
| chr7-4420959-DEL-328 | chr7-4420960-DEL-328 |
| chr7-4425760-DEL-129 | chr7-4425761-DEL-129 |
| chr7-4440412-DEL-252 | chr7-4440413-DEL-252 |
| chr7-4458434-DEL-5267 | chr7-4458435-DEL-5267 |
| chr7-4555967-DEL-175 | chr7-4555968-DEL-140 |
| chr7-4561215-DEL-1893 | chr7-4561218-DEL-1890 |
| chr7-4594043-DEL-4809 | chr7-4594043-DEL-4809 |
| chr7-4605095-DEL-69 | chr7-4605096-DEL-69 |
| chr7-4609156-DEL-123 | chr7-4609164-DEL-93 |
| chr7-4666072-DEL-59 | chr7-4666072-DEL-87 |
| chr7-4672474-DEL-276 | chr7-4672475-DEL-276 |
| chr7-4681267-DEL-178 | chr7-4681268-DEL-178 |
| chr7-4682563-DEL-210 | chr7-4682557-DEL-211 |
| chr7-4712541-DEL-58 | chr7-4712544-DEL-56 |
| chr7-4721067-DEL-133 | chr7-4721072-DEL-131 |
| chr7-4729259-DEL-63 | chr7-4729260-DEL-63 |
| chr7-4740800-DEL-56 | chr7-4740809-DEL-50 |
| chr7-4759659-DEL-111 | chr7-4759660-DEL-111 |
| chr7-4788707-DEL-145 | chr7-4788708-DEL-145 |
| chr7-4793488-DEL-216 | chr7-4793489-DEL-216 |
| chr7-4798598-DEL-214 | chr7-4798600-DEL-213 |
| chr7-4802906-DEL-192 | chr7-4802907-DEL-192 |
| chr7-4803471-DEL-192 | chr7-4803463-DEL-193 |
| chr7-4819065-DEL-216 | chr7-4819066-DEL-216 |
| chr7-4828350-DEL-96 | chr7-4828351-DEL-96 |
| chr7-4855506-DEL-360 | chr7-4855507-DEL-360 |
| chr7-4871898-DEL-194 | chr7-4871908-DEL-189 |
| chr7-4882818-DEL-697 | chr7-4882819-DEL-697 |
| chr7-4887560-DEL-243 | chr7-4887561-DEL-220 |
| chr7-4906973-DEL-211 | chr7-4906974-DEL-211 |
| chr7-4911481-DEL-248 | chr7-4911482-DEL-248 |
| chr7-4931751-DEL-58 | chr7-4931751-DEL-58 |
| chr7-4934546-DEL-59 | chr7-4934546-DEL-59 |
| chr7-4938722-DEL-219 | chr7-4938723-DEL-219 |
| chr7-4963055-DEL-80 | chr7-4963056-DEL-80 |
| chr7-4971833-DEL-66 | chr7-4971834-DEL-66 |
| chr7-4977214-DEL-64 | chr7-4977205-DEL-58 |
| chr7-5000207-DEL-212 | chr7-5000208-DEL-212 |
| chr7-5104205-DEL-98 | chr7-5104206-DEL-98 |
| chr7-5156133-DEL-6190 | chr7-5156137-DEL-6183 |
| chr7-5224665-DEL-5585 | chr7-5224666-DEL-5585 |
| chr7-5293147-DEL-3094 | chr7-5293148-DEL-3094 |
| chr7-5319692-DEL-401 | chr7-5319672-DEL-415 |
| chr7-5323146-DEL-55 | chr7-5323132-DEL-52 |
| chr7-5367364-DEL-148 | chr7-5367369-DEL-150 |
| chr7-5371302-DEL-200 | chr7-5371268-DEL-201 |
| chr7-5378356-DEL-65 | chr7-5378332-DEL-70 |
| chr7-5420540-DEL-1893 | chr7-5420541-DEL-1893 |
| chr7-5444279-DEL-6680 | chr7-5444280-DEL-6680 |
| chr7-5482921-DEL-179 | chr7-5482922-DEL-179 |
| chr7-5612758-DEL-81 | chr7-5612759-DEL-81 |
| chr7-5834860-DEL-4749 | chr7-5834861-DEL-4749 |
| chr7-5862762-DEL-7069 | chr7-5862757-DEL-7070 |
| chr7-5909853-DEL-111 | chr7-5909854-DEL-111 |
| chr7-5920192-DEL-504 | chr7-5920192-DEL-504 |
| chr7-5946175-DEL-68 | chr7-5946176-DEL-68 |
| chr7-5955213-DEL-216 | chr7-5955214-DEL-216 |
| chr7-5958166-DEL-152 | chr7-5958167-DEL-152 |
| chr7-5994617-DEL-6682 | chr7-5994618-DEL-6682 |
| chr7-6018343-DEL-63 | chr7-6018344-DEL-63 |
| chr7-6041240-DEL-70 | chr7-6041241-DEL-61 |
| chr7-6058397-DEL-66 | chr7-6058398-DEL-66 |
| chr7-6089117-DEL-451 | chr7-6089118-DEL-451 |
| chr7-6097721-DEL-141 | chr7-6097722-DEL-141 |
| chr7-6131580-DEL-200 | chr7-6131581-DEL-200 |
| chr7-6169370-DEL-78 | chr7-6169371-DEL-78 |
| chr7-6173670-DEL-179 | chr7-6173679-DEL-179 |
| chr7-6179036-DEL-152 | chr7-6179037-DEL-152 |

| | |
|---|---|
| chr7-6182468-DEL-174 | chr7-6182469-DEL-174 |
| chr7-6198484-DEL-327 | chr7-6198441-DEL-328 |
| chr7-6202182-DEL-50 | chr7-6202160-DEL-71 |
| chr7-6212289-DEL-170 | chr7-6212289-DEL-170 |
| chr7-6217774-DEL-60 | chr7-6217774-DEL-60 |
| chr7-6240152-DEL-163 | chr7-6240135-DEL-163 |
| chr7-6252360-DEL-3099 | chr7-6252361-DEL-3099 |
| chr7-6260268-DEL-353 | chr7-6260269-DEL-353 |
| chr7-6265612-DEL-142 | chr7-6265613-DEL-142 |
| chr7-6279990-DEL-255 | chr7-6279991-DEL-255 |
| chr7-6282925-DEL-794 | chr7-6282926-DEL-794 |
| chr7-6295806-DEL-261 | chr7-6295807-DEL-261 |
| chr7-6311534-DEL-177 | chr7-6311535-DEL-177 |
| chr7-6312149-DEL-91 | chr7-6312150-DEL-58 |
| chr7-6313917-DEL-211 | chr7-6313918-DEL-211 |
| chr7-6350359-DEL-71 | chr7-6350360-DEL-71 |
| chr7-6350811-DEL-377 | chr7-6350812-DEL-377 |
| chr7-6353734-DEL-124 | chr7-6353783-DEL-104 |
| chr7-6358338-DEL-86 | chr7-6358332-DEL-85 |
| chr7-6358779-DEL-1173 | chr7-6358780-DEL-1173 |
| chr7-6364669-DEL-122 | chr7-6364670-DEL-122 |
| chr7-6390300-DEL-128 | chr7-6390301-DEL-128 |
| chr7-6423595-DEL-94 | chr7-6423599-DEL-91 |
| chr7-6436362-DEL-52 | chr7-6436365-DEL-50 |
| chr7-6469396-DEL-171 | chr7-6469397-DEL-169 |
| chr7-6474666-DEL-74 | chr7-6474664-DEL-59 |
| chr7-6514124-DEL-59 | chr7-6514144-DEL-65 |
| chr7-6537562-DEL-3751 | chr7-6537564-DEL-3750 |
| chr7-6553789-DEL-6602 | chr7-6553790-DEL-6602 |
| chr7-6633809-DEL-84 | chr7-6633822-DEL-73 |
| chr7-6640341-DEL-181 | chr7-6640342-DEL-181 |
| chr7-6644681-DEL-681 | chr7-6644682-DEL-681 |
| chr7-6657059-DEL-3022 | chr7-6657064-DEL-3011 |
| chr7-6668007-DEL-597 | chr7-6668008-DEL-597 |
| chr7-6676603-DEL-74 | chr7-6676604-DEL-74 |
| chr7-6759096-DEL-187 | chr7-6759097-DEL-187 |
| chr7-6773558-DEL-98 | chr7-6773559-DEL-98 |
| chr7-6801394-DEL-76 | chr7-6801389-DEL-73 |
| chr7-6828586-DEL-153 | chr7-6828587-DEL-153 |
| chr7-6867118-DEL-55 | chr7-6867117-DEL-52 |
| chr7-6870686-DEL-66 | chr7-6870687-DEL-66 |
| chr7-6884420-DEL-88 | chr7-6884421-DEL-88 |
| chr7-6886467-DEL-54 | chr7-6886456-DEL-55 |
| chr7-6978904-DEL-230 | chr7-6978905-DEL-230 |
| chr7-6992740-DEL-85 | chr7-6992741-DEL-85 |
| chr7-7009857-DEL-387 | chr7-7009858-DEL-387 |
| chr7-7021069-DEL-75 | chr7-7021070-DEL-76 |
| chr7-7029126-DEL-375 | chr7-7029127-DEL-375 |
| chr7-7031456-DEL-1544 | chr7-7031457-DEL-1544 |
| chr7-7042742-DEL-511 | chr7-7042743-DEL-511 |
| chr7-7051923-DEL-1257 | chr7-7051902-DEL-1263 |
| chr7-7056417-DEL-81 | chr7-7056418-DEL-81 |
| chr7-7063205-DEL-85 | chr7-7063206-DEL-70 |
| chr7-7064514-DEL-352 | chr7-7064515-DEL-352 |
| chr7-7065681-DEL-433 | chr7-7065682-DEL-433 |
| chr7-7071939-DEL-140 | chr7-7071940-DEL-140 |
| chr7-7072904-DEL-8986 | chr7-7072905-DEL-8986 |
| chr7-7083465-DEL-7446 | chr7-7083466-DEL-7446 |
| chr7-7095396-DEL-55 | chr7-7095397-DEL-55 |
| chr7-7098654-DEL-258 | chr7-7098657-DEL-259 |
| chr7-7102588-DEL-258 | chr7-7102589-DEL-258 |
| chr7-7141327-DEL-104 | chr7-7141332-DEL-100 |
| chr7-7149494-DEL-6296 | chr7-7149495-DEL-6296 |
| chr7-7158084-DEL-72 | chr7-7158081-DEL-52 |
| chr7-7188430-DEL-70 | chr7-7188431-DEL-71 |
| chr7-7252148-DEL-1842 | chr7-7252147-DEL-1836 |
| chr7-7291416-DEL-210 | chr7-7291417-DEL-210 |
| chr7-7300687-DEL-248 | chr7-7300652-DEL-242 |
| chr7-7333692-DEL-471 | chr7-7333693-DEL-471 |
| chr7-8499354-DEL-52 | chr7-8499355-DEL-52 |
| chr7-9810973-DEL-981 | chr7-9810974-DEL-981 |
| chr7-9842001-DEL-922 | chr7-9842001-DEL-922 |
| chr7-9872445-DEL-212 | chr7-9872446-DEL-212 |
| chr7-9884417-DEL-6720 | chr7-9884418-DEL-6720 |
| chr7-9894025-DEL-106 | chr7-9894019-DEL-94 |
| chr7-9915889-DEL-1045 | chr7-9915890-DEL-1045 |
| chr7-9984053-DEL-1570 | chr7-9984042-DEL-1581 |
| chr7-10023876-DEL-154 | chr7-10023880-DEL-155 |
| chr7-10125981-DEL-56 | chr7-10125969-DEL-68 |
| chr7-10135685-DEL-6282 | chr7-10135686-DEL-6282 |

| | |
|---|---|
| chr7-10155698-DEL-57 | chr7-10155695-DEL-58 |
| chr7-10160372-DEL-84 | chr7-10160373-DEL-84 |
| chr7-10184207-DEL-69 | chr7-10184208-DEL-68 |
| chr7-10205242-DEL-7466 | chr7-10205243-DEL-7466 |
| chr7-10251638-DEL-1136 | chr7-10251639-DEL-1136 |
| chr7-10252693-DEL-65 | chr7-10252694-DEL-65 |
| chr7-10272931-DEL-1689 | chr7-10272932-DEL-1689 |
| chr7-10412049-DEL-421 | chr7-10412048-DEL-421 |
| chr7-10419401-DEL-430 | chr7-10419402-DEL-424 |
| chr7-10486166-DEL-219 | chr7-10486167-DEL-219 |
| chr7-10496191-DEL-190 | chr7-10496192-DEL-190 |
| chr7-10509444-DEL-467 | chr7-10509445-DEL-467 |
| chr7-10919258-DEL-100 | chr7-10919259-DEL-100 |
| chr7-10940208-DEL-697 | chr7-10940210-DEL-693 |
| chr7-10958374-DEL-222 | chr7-10958375-DEL-222 |
| chr7-10998661-DEL-84 | chr7-10998662-DEL-83 |
| chr7-11039166-DEL-556 | chr7-11039166-DEL-556 |
| chr7-11041703-DEL-1231 | chr7-11041704-DEL-1231 |
| chr7-11056768-DEL-117 | chr7-11056769-DEL-117 |
| chr7-11128813-DEL-210 | chr7-11128812-DEL-201 |
| chr7-11290433-DEL-194 | chr7-11290434-DEL-194 |
| chr7-11378364-DEL-6794 | chr7-11378365-DEL-6794 |
| chr7-11417296-DEL-170 | chr7-11417297-DEL-168 |
| chr7-11421362-DEL-84 | chr7-11421363-DEL-84 |
| chr7-11427708-DEL-1419 | chr7-11427709-DEL-1419 |
| chr7-11453433-DEL-1020 | chr7-11453434-DEL-1020 |
| chr7-11557123-DEL-442 | chr7-11557124-DEL-442 |
| chr7-11560686-DEL-3960 | chr7-11560687-DEL-3960 |
| chr7-11565503-DEL-338 | chr7-11565504-DEL-338 |
| chr7-11566725-DEL-72 | chr7-11566705-DEL-54 |
| chr7-11568336-DEL-139 | chr7-11568337-DEL-139 |
| chr7-11586286-DEL-88 | chr7-11586237-DEL-89 |
| chr7-11601211-DEL-207 | chr7-11601212-DEL-207 |
| chr7-11603316-DEL-4045 | chr7-11603317-DEL-4045 |
| chr7-11611458-DEL-1232 | chr7-11611459-DEL-1232 |
| chr7-11615786-DEL-185 | chr7-11615774-DEL-186 |
| chr7-11619191-DEL-610 | chr7-11619192-DEL-610 |
| chr7-11676411-DEL-742 | chr7-11676411-DEL-743 |
| chr7-11683257-DEL-486 | chr7-11683258-DEL-486 |
| chr7-11700294-DEL-886 | chr7-11700295-DEL-886 |
| chr7-11719340-DEL-1396 | chr7-11719342-DEL-1400 |
| chr7-11757651-DEL-86 | chr7-11757659-DEL-81 |
| chr7-11769730-DEL-125 | chr7-11769775-DEL-109 |
| chr7-11833200-DEL-56 | chr7-11833163-DEL-88 |
| chr7-11871950-DEL-225 | chr7-11871951-DEL-225 |
| chr7-11872963-DEL-75 | chr7-11872964-DEL-75 |
| chr7-11875713-DEL-62 | chr7-11875706-DEL-70 |
| chr7-11889404-DEL-102 | chr7-11889405-DEL-102 |
| chr7-11919198-DEL-2220 | chr7-11919199-DEL-2220 |
| chr7-11922582-DEL-52 | chr7-11922583-DEL-52 |
| chr7-11941567-DEL-1545 | chr7-11941567-DEL-1545 |
| chr7-11952229-DEL-901 | chr7-11952229-DEL-902 |
| chr7-11954379-DEL-52 | chr7-11954380-DEL-52 |
| chr7-11968095-DEL-952 | chr7-11968096-DEL-952 |
| chr7-11978395-DEL-64 | chr7-11978404-DEL-56 |
| chr7-11979371-DEL-1376 | chr7-11979372-DEL-1376 |
| chr7-11994871-DEL-1520 | chr7-11994872-DEL-1520 |
| chr7-12006712-DEL-400 | chr7-12006713-DEL-400 |
| chr7-12017080-DEL-7123 | chr7-12017081-DEL-7123 |
| chr7-12023468-DEL-212 | chr7-12023469-DEL-212 |
| chr7-12035168-DEL-71 | chr7-12035168-DEL-71 |
| chr7-12042566-DEL-64 | chr7-12042568-DEL-60 |
| chr7-12046896-DEL-327 | chr7-12046897-DEL-327 |
| chr7-12053094-DEL-831 | chr7-12053095-DEL-831 |
| chr7-12058010-DEL-1010 | chr7-12058011-DEL-1010 |
| chr7-12063957-DEL-832 | chr7-12063958-DEL-832 |
| chr7-12070163-DEL-3609 | chr7-12070145-DEL-3625 |
| chr7-12084913-DEL-6998 | chr7-12084914-DEL-6998 |
| chr7-12113456-DEL-4987 | chr7-12113457-DEL-4987 |
| chr7-12121631-DEL-203 | chr7-12121632-DEL-203 |
| chr7-12123226-DEL-52 | chr7-12123227-DEL-52 |
| chr7-12177107-DEL-219 | chr7-12177108-DEL-219 |
| chr7-12196419-DEL-90 | chr7-12196424-DEL-86 |
| chr7-12233831-DEL-87 | chr7-12233842-DEL-89 |
| chr7-12238168-DEL-75 | chr7-12238173-DEL-71 |
| chr7-12260356-DEL-51 | chr7-12260352-DEL-52 |
| chr7-12276529-DEL-5324 | chr7-12276530-DEL-5324 |
| chr7-12322292-DEL-64 | chr7-12322293-DEL-64 |
| chr7-12406010-DEL-10281 | chr7-12405993-DEL-10296 |
| chr7-12418061-DEL-61 | chr7-12418062-DEL-61 |

| | |
|---|---|
| chr7-12420388-DEL-54 | chr7-12420389-DEL-54 |
| chr7-12431442-DEL-203 | chr7-12431431-DEL-204 |
| chr7-12438816-DEL-332 | chr7-12438817-DEL-332 |
| chr7-12452047-DEL-165 | chr7-12452048-DEL-165 |
| chr7-12453514-DEL-1740 | chr7-12453514-DEL-1740 |
| chr7-12455744-DEL-6397 | chr7-12455745-DEL-6397 |
| chr7-12475851-DEL-84 | chr7-12475835-DEL-66 |
| chr7-12487003-DEL-6934 | chr7-12487005-DEL-6933 |
| chr7-12487084-DEL-204 | chr7-12487085-DEL-204 |
| chr7-12631245-DEL-114 | chr7-12631284-DEL-64 |
| chr7-12651774-DEL-994 | chr7-12651775-DEL-994 |
| chr7-12674986-DEL-388 | chr7-12674987-DEL-388 |
| chr7-12681501-DEL-201 | chr7-12681495-DEL-201 |
| chr7-12689846-DEL-218 | chr7-12689847-DEL-218 |
| chr7-12690517-DEL-66 | chr7-12690536-DEL-50 |
| chr7-12691063-DEL-171 | chr7-12691064-DEL-171 |
| chr7-12727441-DEL-72 | chr7-12727442-DEL-72 |
| chr7-12733866-DEL-212 | chr7-12733867-DEL-212 |
| chr7-12735989-DEL-1792 | chr7-12735990-DEL-1792 |
| chr7-12739244-DEL-216 | chr7-12739245-DEL-216 |
| chr7-12752492-DEL-56 | chr7-12752495-DEL-54 |
| chr7-12754029-DEL-52 | chr7-12754066-DEL-50 |
| chr7-12755272-DEL-536 | chr7-12755247-DEL-550 |
| chr7-12780798-DEL-128 | chr7-12780792-DEL-129 |
| chr7-12800374-DEL-275 | chr7-12800375-DEL-275 |
| chr7-12802837-DEL-180 | chr7-12802838-DEL-180 |
| chr7-12805470-DEL-75 | chr7-12805471-DEL-75 |
| chr7-12837098-DEL-1478 | chr7-12837099-DEL-1478 |
| chr7-12872220-DEL-729 | chr7-12872221-DEL-729 |
| chr7-12875899-DEL-6417 | chr7-12875901-DEL-6418 |
| chr7-12882700-DEL-197 | chr7-12882701-DEL-197 |
| chr7-12933550-DEL-72 | chr7-12933551-DEL-72 |
| chr7-12957105-DEL-1715 | chr7-12957106-DEL-1715 |
| chr7-12958699-DEL-58 | chr7-12958688-DEL-70 |
| chr7-12959277-DEL-332 | chr7-12959274-DEL-344 |
| chr7-12966002-DEL-204 | chr7-12966003-DEL-204 |
| chr7-12970001-DEL-202 | chr7-12970002-DEL-202 |
| chr7-12978864-DEL-810 | chr7-12978898-DEL-781 |
| chr7-12983643-DEL-99 | chr7-12983646-DEL-91 |
| chr7-12984715-DEL-141 | chr7-12984710-DEL-147 |
| chr7-12985903-DEL-6547 | chr7-12985904-DEL-6547 |
| chr7-13053624-DEL-107 | chr7-13053629-DEL-104 |
| chr7-13066480-DEL-726 | chr7-13066481-DEL-726 |
| chr7-13079395-DEL-146 | chr7-13079387-DEL-158 |
| chr7-13091169-DEL-120 | chr7-13091170-DEL-120 |
| chr7-13100164-DEL-558 | chr7-13100165-DEL-558 |
| chr7-13109514-DEL-66 | chr7-13109515-DEL-64 |
| chr7-13110189-DEL-211 | chr7-13110190-DEL-211 |
| chr7-13122527-DEL-265 | chr7-13122528-DEL-265 |
| chr7-13143340-DEL-14011 | chr7-13143341-DEL-14011 |
| chr7-13158797-DEL-205 | chr7-13158798-DEL-205 |
| chr7-13160347-DEL-136 | chr7-13160348-DEL-136 |
| chr7-13161199-DEL-336 | chr7-13161200-DEL-336 |
| chr7-13161899-DEL-212 | chr7-13161900-DEL-210 |
| chr7-13170497-DEL-104 | chr7-13170500-DEL-102 |
| chr7-13187857-DEL-560 | chr7-13187858-DEL-560 |
| chr7-13224212-DEL-370 | chr7-13224213-DEL-370 |
| chr7-13237249-DEL-199 | chr7-13237250-DEL-199 |
| chr7-13247328-DEL-421 | chr7-13247328-DEL-421 |
| chr7-13269167-DEL-93 | chr7-13269153-DEL-107 |
| chr7-13270880-DEL-3288 | chr7-13270881-DEL-3288 |
| chr7-13283896-DEL-112 | chr7-13283897-DEL-112 |
| chr7-13291817-DEL-81 | chr7-13291790-DEL-109 |
| chr7-13325076-DEL-6570 | chr7-13325060-DEL-6586 |
| chr7-13331313-DEL-208 | chr7-13331314-DEL-208 |
| chr7-13403512-DEL-128 | chr7-13403515-DEL-131 |
| chr7-13437515-DEL-66 | chr7-13437516-DEL-66 |
| chr7-13571728-DEL-160 | chr7-13571725-DEL-164 |
| chr7-13584598-DEL-2414 | chr7-13584598-DEL-2413 |
| chr7-13592630-DEL-1349 | chr7-13592631-DEL-1349 |
| chr7-13638512-DEL-829 | chr7-13638513-DEL-829 |
| chr7-13641824-DEL-311 | chr7-13641825-DEL-311 |
| chr7-13644441-DEL-95 | chr7-13644442-DEL-95 |
| chr7-13667153-DEL-506 | chr7-13667154-DEL-506 |
| chr7-13680159-DEL-6235 | chr7-13680159-DEL-6235 |
| chr7-13688867-DEL-1583 | chr7-13688867-DEL-1583 |
| chr7-13756206-DEL-974 | chr7-13756207-DEL-974 |
| chr7-13763408-DEL-55 | chr7-13763409-DEL-50 |
| chr7-13767177-DEL-1026 | chr7-13767178-DEL-1026 |
| chr7-13785039-DEL-6198 | chr7-13785040-DEL-6198 |

| | |
|---|---|
| chr7-13796533-DEL-859 | chr7-13796534-DEL-859 |
| chr7-13820236-DEL-8148 | chr7-13820237-DEL-8148 |
| chr7-13821130-DEL-212 | chr7-13821131-DEL-212 |
| chr7-13859452-DEL-4276 | chr7-13859453-DEL-4276 |
| chr7-13875938-DEL-62 | chr7-13875930-DEL-66 |
| chr7-13885898-DEL-6224 | chr7-13885899-DEL-6224 |
| chr7-13904023-DEL-935 | chr7-13904024-DEL-935 |
| chr7-13910211-DEL-2870 | chr7-13910211-DEL-2870 |
| chr7-13954274-DEL-493 | chr7-13954275-DEL-493 |
| chr7-13967537-DEL-57 | chr7-13967539-DEL-56 |
| chr7-13969098-DEL-919 | chr7-13969099-DEL-919 |
| chr7-14012865-DEL-539 | chr7-14012866-DEL-539 |
| chr7-14020731-DEL-70 | chr7-14020732-DEL-70 |
| chr7-14025453-DEL-2571 | chr7-14025454-DEL-2571 |
| chr7-14027080-DEL-3856 | chr7-14027080-DEL-3856 |
| chr7-14033165-DEL-4627 | chr7-14033166-DEL-4627 |
| chr7-14054766-DEL-112 | chr7-14054750-DEL-86 |
| chr7-14073923-DEL-666 | chr7-14073924-DEL-666 |
| chr7-14077892-DEL-435 | chr7-14077893-DEL-435 |
| chr7-14097585-DEL-64 | chr7-14097585-DEL-64 |
| chr7-14137633-DEL-602 | chr7-14137592-DEL-603 |
| chr7-14149262-DEL-175 | chr7-14149263-DEL-175 |
| chr7-14166581-DEL-846 | chr7-14166582-DEL-845 |
| chr7-14210537-DEL-111 | chr7-14210522-DEL-124 |
| chr7-14215726-DEL-4307 | chr7-14215725-DEL-4308 |
| chr7-14240480-DEL-56 | chr7-14240481-DEL-56 |
| chr7-14278558-DEL-317 | chr7-14278559-DEL-317 |
| chr7-14286622-DEL-85 | chr7-14286661-DEL-74 |
| chr7-14294210-DEL-187 | chr7-14294211-DEL-187 |
| chr7-14316427-DEL-54 | chr7-14316428-DEL-54 |
| chr7-14369875-DEL-1892 | chr7-14369876-DEL-1892 |
| chr7-14377782-DEL-773 | chr7-14377783-DEL-773 |
| chr7-14476092-DEL-154 | chr7-14476092-DEL-154 |
| chr7-14815373-DEL-3216 | chr7-14815374-DEL-3216 |
| chr7-15022183-DEL-936 | chr7-15022182-DEL-936 |
| chr7-15327157-DEL-1795 | chr7-15327158-DEL-1795 |
| chr7-15348437-DEL-1268 | chr7-15348438-DEL-1268 |
| chr7-15362800-DEL-2317 | chr7-15362800-DEL-2317 |
| chr7-15367192-DEL-235 | chr7-15367188-DEL-236 |
| chr7-15437064-DEL-5306 | chr7-15437065-DEL-5306 |
| chr7-15464943-DEL-9431 | chr7-15464942-DEL-9431 |
| chr7-15594800-DEL-245 | chr7-15594801-DEL-245 |
| chr7-15610222-DEL-383 | chr7-15610223-DEL-383 |
| chr7-15618062-DEL-1310 | chr7-15618063-DEL-1310 |
| chr7-15644256-DEL-551 | chr7-15644257-DEL-552 |
| chr7-15647788-DEL-209 | chr7-15647789-DEL-209 |
| chr7-15657785-DEL-1330 | chr7-15657786-DEL-1330 |
| chr7-15666556-DEL-56 | chr7-15666556-DEL-56 |
| chr7-15696850-DEL-189 | chr7-15696851-DEL-189 |
| chr7-15700937-DEL-93 | chr7-15700938-DEL-93 |
| chr7-15705070-DEL-58 | chr7-15705046-DEL-84 |
| chr7-15707939-DEL-51 | chr7-15707937-DEL-52 |
| chr7-15712270-DEL-107 | chr7-15712271-DEL-107 |
| chr7-15723484-DEL-296 | chr7-15723485-DEL-295 |
| chr7-15732476-DEL-52 | chr7-15732477-DEL-52 |
| chr7-15739662-DEL-55 | chr7-15739663-DEL-55 |
| chr7-15747036-DEL-51 | chr7-15747036-DEL-51 |
| chr7-15748290-DEL-53 | chr7-15748291-DEL-53 |
| chr7-15888517-DEL-136 | chr7-15888518-DEL-136 |
| chr7-15897932-DEL-53 | chr7-15897932-DEL-53 |
| chr7-15924124-DEL-212 | chr7-15924125-DEL-212 |
| chr7-15926666-DEL-55 | chr7-15926663-DEL-55 |
| chr7-15928493-DEL-158 | chr7-15928494-DEL-158 |
| chr7-15954985-DEL-111 | chr7-15954989-DEL-116 |
| chr7-15991091-DEL-53 | chr7-15991134-DEL-53 |
| chr7-16001470-DEL-183 | chr7-16001471-DEL-183 |
| chr7-16003420-DEL-68 | chr7-16003414-DEL-69 |
| chr7-16022742-DEL-108 | chr7-16022735-DEL-100 |
| chr7-16025876-DEL-244 | chr7-16025877-DEL-244 |
| chr7-16036332-DEL-57 | chr7-16036333-DEL-57 |
| chr7-16037677-DEL-209 | chr7-16037678-DEL-209 |
| chr7-16065962-DEL-1640 | chr7-16065962-DEL-1641 |
| chr7-16076825-DEL-53 | chr7-16076826-DEL-53 |
| chr7-16084593-DEL-205 | chr7-16084594-DEL-205 |
| chr7-16085013-DEL-1804 | chr7-16085014-DEL-1804 |
| chr7-16106051-DEL-72 | chr7-16106052-DEL-72 |
| chr7-16122829-DEL-209 | chr7-16122830-DEL-209 |
| chr7-16123919-DEL-306 | chr7-16123920-DEL-306 |
| chr7-16126137-DEL-630 | chr7-16126138-DEL-630 |
| chr7-16127044-DEL-126 | chr7-16127045-DEL-126 |

| | |
|---|---|
| chr7-16144989-DEL-84 | chr7-16144990-DEL-84 |
| chr7-16193478-DEL-265 | chr7-16193479-DEL-265 |
| chr7-16217714-DEL-234 | chr7-16217715-DEL-234 |
| chr7-16224834-DEL-199 | chr7-16224828-DEL-198 |
| chr7-16275057-DEL-212 | chr7-16275058-DEL-212 |
| chr7-16295074-DEL-212 | chr7-16295075-DEL-212 |
| chr7-16311291-DEL-132 | chr7-16311290-DEL-125 |
| chr7-16367817-DEL-253 | chr7-16367818-DEL-252 |
| chr7-16376134-DEL-92 | chr7-16376135-DEL-92 |
| chr7-16381827-DEL-213 | chr7-16381821-DEL-213 |
| chr7-16392796-DEL-57 | chr7-16392797-DEL-53 |
| chr7-16395280-DEL-6483 | chr7-16395279-DEL-6483 |
| chr7-16402556-DEL-86 | chr7-16402557-DEL-86 |
| chr7-16410061-DEL-155 | chr7-16410062-DEL-155 |
| chr7-16419433-DEL-194 | chr7-16419434-DEL-194 |
| chr7-16421601-DEL-197 | chr7-16421602-DEL-198 |
| chr7-16440782-DEL-56 | chr7-16440757-DEL-104 |
| chr7-16447667-DEL-159 | chr7-16447668-DEL-159 |
| chr7-16454058-DEL-205 | chr7-16454059-DEL-205 |
| chr7-16460247-DEL-73 | chr7-16460250-DEL-69 |
| chr7-16468882-DEL-66 | chr7-16468883-DEL-66 |
| chr7-16484355-DEL-186 | chr7-16484356-DEL-186 |
| chr7-16488404-DEL-295 | chr7-16488405-DEL-295 |
| chr7-16493537-DEL-200 | chr7-16493538-DEL-199 |
| chr7-16501559-DEL-85 | chr7-16501560-DEL-85 |
| chr7-16506356-DEL-305 | chr7-16506357-DEL-305 |
| chr7-16510645-DEL-196 | chr7-16510641-DEL-201 |
| chr7-16533710-DEL-53 | chr7-16533711-DEL-53 |
| chr7-16535966-DEL-73 | chr7-16535967-DEL-73 |
| chr7-16536532-DEL-62 | chr7-16536564-DEL-58 |
| chr7-16537256-DEL-146 | chr7-16537259-DEL-128 |
| chr7-16570556-DEL-185 | chr7-16570557-DEL-185 |
| chr7-16588921-DEL-53 | chr7-16588921-DEL-53 |
| chr7-16591275-DEL-157 | chr7-16591290-DEL-158 |
| chr7-16598367-DEL-53 | chr7-16598368-DEL-53 |
| chr7-16629814-DEL-290 | chr7-16629814-DEL-290 |
| chr7-16630915-DEL-159 | chr7-16630892-DEL-160 |
| chr7-16632319-DEL-351 | chr7-16632320-DEL-351 |
| chr7-16687429-DEL-196 | chr7-16687430-DEL-196 |
| chr7-16700709-DEL-214 | chr7-16700710-DEL-214 |
| chr7-16713723-DEL-53 | chr7-16713724-DEL-53 |
| chr7-16718847-DEL-204 | chr7-16718848-DEL-204 |
| chr7-16741463-DEL-62 | chr7-16741464-DEL-62 |
| chr7-16775971-DEL-121 | chr7-16775976-DEL-117 |
| chr7-16798206-DEL-714 | chr7-16798207-DEL-714 |
| chr7-16819772-DEL-120 | chr7-16819773-DEL-120 |
| chr7-16836016-DEL-194 | chr7-16836017-DEL-194 |
| chr7-16868874-DEL-626 | chr7-16868875-DEL-626 |
| chr7-16870432-DEL-665 | chr7-16870433-DEL-665 |
| chr7-16943825-DEL-54 | chr7-16943806-DEL-55 |
| chr7-16949887-DEL-1506 | chr7-16949888-DEL-1506 |
| chr7-16999210-DEL-9382 | chr7-16999209-DEL-9382 |
| chr7-17023965-DEL-66 | chr7-17023966-DEL-66 |
| chr7-17039968-DEL-128 | chr7-17039969-DEL-128 |
| chr7-17049315-DEL-293 | chr7-17049316-DEL-293 |
| chr7-17105312-DEL-479 | chr7-17105311-DEL-479 |
| chr7-17149150-DEL-128 | chr7-17149151-DEL-128 |
| chr7-17153409-DEL-527 | chr7-17153410-DEL-527 |
| chr7-17160762-DEL-135 | chr7-17160762-DEL-129 |
| chr7-17180991-DEL-62 | chr7-17180992-DEL-66 |
| chr7-17195034-DEL-530 | chr7-17195030-DEL-535 |
| chr7-17197562-DEL-135 | chr7-17197561-DEL-116 |
| chr7-17198919-DEL-1664 | chr7-17198920-DEL-1664 |
| chr7-17285147-DEL-473 | chr7-17285146-DEL-473 |
| chr7-17344921-DEL-7404 | chr7-17344920-DEL-7404 |
| chr7-17366644-DEL-473 | chr7-17366645-DEL-473 |
| chr7-17374525-DEL-6442 | chr7-17374526-DEL-6442 |
| chr7-17393080-DEL-8830 | chr7-17393081-DEL-8830 |
| chr7-17488649-DEL-64 | chr7-17488650-DEL-64 |
| chr7-17517802-DEL-68 | chr7-17517803-DEL-68 |
| chr7-17533898-DEL-6818 | chr7-17533899-DEL-6818 |
| chr7-17534135-DEL-208 | chr7-17534136-DEL-208 |
| chr7-17606564-DEL-95 | chr7-17606545-DEL-119 |
| chr7-17627908-DEL-1566 | chr7-17627910-DEL-1566 |
| chr7-17632825-DEL-60 | chr7-17632826-DEL-50 |
| chr7-17641197-DEL-672 | chr7-17641196-DEL-670 |
| chr7-17665879-DEL-58 | chr7-17665880-DEL-58 |
| chr7-17667659-DEL-1712 | chr7-17667660-DEL-1708 |
| chr7-17689362-DEL-1452 | chr7-17689361-DEL-1452 |
| chr7-17722025-DEL-550 | chr7-17722026-DEL-550 |

| | |
|---|---|
| chr7-17737652-DEL-434 | chr7-17737648-DEL-435 |
| chr7-17805950-DEL-3472 | chr7-17805951-DEL-3472 |
| chr7-17826347-DEL-57 | chr7-17826348-DEL-57 |
| chr7-17851115-DEL-5221 | chr7-17851116-DEL-5221 |
| chr7-17856825-DEL-58 | chr7-17856826-DEL-58 |
| chr7-17870998-DEL-124 | chr7-17871037-DEL-84 |
| chr7-17878395-DEL-4261 | chr7-17878396-DEL-4261 |
| chr7-17902054-DEL-70 | chr7-17902019-DEL-81 |
| chr7-17928964-DEL-2227 | chr7-17928965-DEL-2226 |
| chr7-17935971-DEL-115 | chr7-17935972-DEL-115 |
| chr7-17966603-DEL-58 | chr7-17966604-DEL-58 |
| chr7-17978219-DEL-104 | chr7-17978208-DEL-78 |
| chr7-18026110-DEL-65 | chr7-18026111-DEL-65 |
| chr7-18045054-DEL-7554 | chr7-18045055-DEL-7554 |
| chr7-18046372-DEL-1340 | chr7-18046373-DEL-1340 |
| chr7-18099511-DEL-142 | chr7-18099509-DEL-140 |
| chr7-18101220-DEL-689 | chr7-18101221-DEL-689 |
| chr7-18105426-DEL-60 | chr7-18105441-DEL-60 |
| chr7-18122861-DEL-212 | chr7-18122861-DEL-212 |
| chr7-18135181-DEL-106 | chr7-18135182-DEL-106 |
| chr7-18136797-DEL-147 | chr7-18136798-DEL-147 |
| chr7-18214990-DEL-70 | chr7-18214990-DEL-70 |
| chr7-18220952-DEL-6222 | chr7-18220953-DEL-6222 |
| chr7-18227546-DEL-56 | chr7-18227547-DEL-56 |
| chr7-18231178-DEL-78 | chr7-18231171-DEL-67 |
| chr7-18234164-DEL-126 | chr7-18234165-DEL-96 |
| chr7-18279548-DEL-67 | chr7-18279548-DEL-71 |
| chr7-18299909-DEL-68 | chr7-18299910-DEL-68 |
| chr7-18320449-DEL-364 | chr7-18320450-DEL-364 |
| chr7-18322639-DEL-438 | chr7-18322640-DEL-438 |
| chr7-18340204-DEL-486 | chr7-18340201-DEL-487 |
| chr7-18352909-DEL-1851 | chr7-18352909-DEL-1851 |
| chr7-18407203-DEL-94 | chr7-18407204-DEL-94 |
| chr7-18409440-DEL-2992 | chr7-18409441-DEL-2992 |
| chr7-18424186-DEL-471 | chr7-18424187-DEL-471 |
| chr7-18426705-DEL-142 | chr7-18426706-DEL-142 |
| chr7-18428447-DEL-10878 | chr7-18428448-DEL-10878 |
| chr7-18439277-DEL-740 | chr7-18439278-DEL-740 |
| chr7-18445772-DEL-58 | chr7-18445772-DEL-56 |
| chr7-18448767-DEL-858 | chr7-18448760-DEL-872 |
| chr7-18476411-DEL-983 | chr7-18476412-DEL-983 |
| chr7-18488320-DEL-358 | chr7-18488321-DEL-358 |
| chr7-18497580-DEL-6401 | chr7-18497581-DEL-6401 |
| chr7-18572653-DEL-169 | chr7-18572654-DEL-169 |
| chr7-18579170-DEL-187 | chr7-18579157-DEL-188 |
| chr7-18587676-DEL-92 | chr7-18587664-DEL-57 |
| chr7-18600111-DEL-1904 | chr7-18600112-DEL-1904 |
| chr7-18603386-DEL-2043 | chr7-18603387-DEL-2043 |
| chr7-18613150-DEL-100 | chr7-18613151-DEL-100 |
| chr7-18617259-DEL-764 | chr7-18617260-DEL-764 |
| chr7-18638709-DEL-133 | chr7-18638710-DEL-152 |
| chr7-18641424-DEL-75 | chr7-18641425-DEL-75 |
| chr7-18649429-DEL-204 | chr7-18649430-DEL-204 |
| chr7-18654219-DEL-198 | chr7-18654220-DEL-198 |
| chr7-18663362-DEL-53 | chr7-18663363-DEL-53 |
| chr7-18715055-DEL-205 | chr7-18715056-DEL-205 |
| chr7-18719242-DEL-168 | chr7-18719243-DEL-168 |
| chr7-18730982-DEL-265 | chr7-18730983-DEL-265 |
| chr7-18737463-DEL-236 | chr7-18737464-DEL-236 |
| chr7-18749826-DEL-106 | chr7-18749826-DEL-106 |
| chr7-18752977-DEL-159 | chr7-18752978-DEL-159 |
| chr7-18767489-DEL-108 | chr7-18767491-DEL-102 |
| chr7-18795627-DEL-158 | chr7-18795630-DEL-156 |
| chr7-18805912-DEL-218 | chr7-18805899-DEL-219 |
| chr7-18895197-DEL-202 | chr7-18895208-DEL-205 |
| chr7-18905027-DEL-159 | chr7-18905028-DEL-159 |
| chr7-18909263-DEL-62 | chr7-18909227-DEL-60 |
| chr7-18923396-DEL-210 | chr7-18923409-DEL-198 |
| chr7-18926865-DEL-212 | chr7-18926866-DEL-212 |
| chr7-18939148-DEL-52 | chr7-18939103-DEL-53 |
| chr7-18941623-DEL-73 | chr7-18941624-DEL-73 |
| chr7-18951344-DEL-54 | chr7-18951295-DEL-60 |
| chr7-19007481-DEL-306 | chr7-19007482-DEL-311 |
| chr7-19019664-DEL-88 | chr7-19019673-DEL-80 |
| chr7-19032913-DEL-730 | chr7-19032917-DEL-727 |
| chr7-19058845-DEL-53 | chr7-19058846-DEL-53 |
| chr7-19070213-DEL-180 | chr7-19070254-DEL-140 |
| chr7-19074403-DEL-430 | chr7-19074403-DEL-426 |
| chr7-19085145-DEL-212 | chr7-19085145-DEL-212 |
| chr7-19087849-DEL-159 | chr7-19087850-DEL-159 |

| | |
|---|---|
| chr7-19089520-DEL-247 | chr7-19089521-DEL-247 |
| chr7-19121515-DEL-88 | chr7-19121516-DEL-88 |
| chr7-19132490-DEL-369 | chr7-19132491-DEL-369 |
| chr7-19161208-DEL-54 | chr7-19161196-DEL-52 |
| chr7-19172758-DEL-221 | chr7-19172759-DEL-221 |
| chr7-19176457-DEL-53 | chr7-19176458-DEL-53 |
| chr7-19179929-DEL-321 | chr7-19179930-DEL-321 |
| chr7-19186968-DEL-171 | chr7-19186969-DEL-171 |
| chr7-19189527-DEL-108 | chr7-19189528-DEL-108 |
| chr7-19214278-DEL-212 | chr7-19214279-DEL-212 |
| chr7-19228282-DEL-52 | chr7-19228283-DEL-52 |
| chr7-19246087-DEL-222 | chr7-19246088-DEL-222 |
| chr7-19259784-DEL-56 | chr7-19259798-DEL-57 |
| chr7-19279097-DEL-613 | chr7-19279098-DEL-612 |
| chr7-19300466-DEL-179 | chr7-19300465-DEL-180 |
| chr7-19319886-DEL-145 | chr7-19319887-DEL-145 |
| chr7-19330292-DEL-71 | chr7-19330293-DEL-71 |
| chr7-19338699-DEL-212 | chr7-19338700-DEL-212 |
| chr7-19402009-DEL-229 | chr7-19402026-DEL-230 |
| chr7-19407586-DEL-55 | chr7-19407583-DEL-55 |
| chr7-19440820-DEL-109 | chr7-19440822-DEL-108 |
| chr7-19459224-DEL-53 | chr7-19459225-DEL-53 |
| chr7-19462865-DEL-53 | chr7-19462827-DEL-53 |
| chr7-19469507-DEL-106 | chr7-19469508-DEL-106 |
| chr7-19472487-DEL-218 | chr7-19472488-DEL-218 |
| chr7-19475555-DEL-53 | chr7-19475555-DEL-53 |
| chr7-19480957-DEL-472 | chr7-19480958-DEL-472 |
| chr7-19491363-DEL-319 | chr7-19491364-DEL-319 |
| chr7-19497017-DEL-53 | chr7-19497017-DEL-53 |
| chr7-19520595-DEL-401 | chr7-19520596-DEL-401 |
| chr7-19525397-DEL-203 | chr7-19525398-DEL-203 |
| chr7-19560311-DEL-60 | chr7-19560312-DEL-60 |
| chr7-19562132-DEL-254 | chr7-19562133-DEL-254 |
| chr7-19562687-DEL-213 | chr7-19562688-DEL-213 |
| chr7-19571294-DEL-53 | chr7-19571295-DEL-53 |
| chr7-19593111-DEL-53 | chr7-19593112-DEL-53 |
| chr7-19601428-DEL-212 | chr7-19601429-DEL-212 |
| chr7-19608580-DEL-84 | chr7-19608581-DEL-84 |
| chr7-19614011-DEL-123 | chr7-19614012-DEL-123 |
| chr7-19616984-DEL-108 | chr7-19616985-DEL-108 |
| chr7-19622164-DEL-199 | chr7-19622165-DEL-199 |
| chr7-19631011-DEL-6358 | chr7-19631013-DEL-6359 |
| chr7-19640076-DEL-93 | chr7-19640077-DEL-93 |
| chr7-19640853-DEL-132 | chr7-19640894-DEL-133 |
| chr7-19642092-DEL-531 | chr7-19642093-DEL-531 |
| chr7-19642362-DEL-478 | chr7-19642363-DEL-478 |
| chr7-19649243-DEL-66 | chr7-19649244-DEL-66 |
| chr7-19663179-DEL-94 | chr7-19663180-DEL-94 |
| chr7-19665026-DEL-230 | chr7-19665012-DEL-229 |
| chr7-19666858-DEL-102 | chr7-19666857-DEL-104 |
| chr7-19674028-DEL-150 | chr7-19674031-DEL-148 |
| chr7-19710899-DEL-102 | chr7-19710900-DEL-102 |
| chr7-19720297-DEL-105 | chr7-19720298-DEL-105 |
| chr7-19727801-DEL-201 | chr7-19727801-DEL-201 |
| chr7-19735881-DEL-511 | chr7-19735880-DEL-512 |
| chr7-19741574-DEL-815 | chr7-19741569-DEL-818 |
| chr7-19742425-DEL-1360 | chr7-19742425-DEL-1360 |
| chr7-19771638-DEL-1262 | chr7-19771639-DEL-1262 |
| chr7-19772261-DEL-204 | chr7-19772262-DEL-204 |
| chr7-19778518-DEL-96 | chr7-19778519-DEL-96 |
| chr7-19786516-DEL-6804 | chr7-19786517-DEL-6804 |
| chr7-19792679-DEL-208 | chr7-19792679-DEL-208 |
| chr7-19807290-DEL-60 | chr7-19807297-DEL-60 |
| chr7-19809467-DEL-8941 | chr7-19809467-DEL-8941 |
| chr7-20225712-DEL-57 | chr7-20225713-DEL-57 |
| chr7-20229413-DEL-57 | chr7-20229412-DEL-57 |
| chr7-20635393-DEL-82 | chr7-20635402-DEL-72 |
| chr7-20710309-DEL-175 | chr7-20710330-DEL-155 |
| chr7-20723468-DEL-134 | chr7-20723469-DEL-134 |
| chr7-22490688-DEL-329 | chr7-22490688-DEL-329 |
| chr7-22925316-DEL-341 | chr7-22925317-DEL-341 |
| chr7-22979527-DEL-196 | chr7-22979527-DEL-196 |
| chr7-22981943-DEL-89 | chr7-22981943-DEL-89 |
| chr7-22995059-DEL-60 | chr7-22995051-DEL-68 |
| chr7-22996625-DEL-1912 | chr7-22996625-DEL-1912 |
| chr7-23065684-DEL-2633 | chr7-23065685-DEL-2633 |
| chr7-23107732-DEL-721 | chr7-23107733-DEL-721 |
| chr7-23114396-DEL-638 | chr7-23114397-DEL-638 |
| chr7-23126692-DEL-90 | chr7-23126693-DEL-90 |
| chr7-23129797-DEL-2093 | chr7-23129798-DEL-2093 |

| | |
|---|---|
| chr7-23162098-DEL-170 | chr7-23162099-DEL-170 |
| chr7-23176272-DEL-372 | chr7-23176273-DEL-372 |
| chr7-23194110-DEL-8964 | chr7-23194111-DEL-8964 |
| chr7-23246175-DEL-112 | chr7-23246223-DEL-69 |
| chr7-23251401-DEL-7077 | chr7-23251402-DEL-7077 |
| chr7-23288583-DEL-864 | chr7-23288584-DEL-864 |
| chr7-23320789-DEL-85 | chr7-23320790-DEL-85 |
| chr7-23326775-DEL-1172 | chr7-23326776-DEL-1172 |
| chr7-23337657-DEL-491 | chr7-23337658-DEL-491 |
| chr7-23340451-DEL-62 | chr7-23340452-DEL-62 |
| chr7-23392071-DEL-67 | chr7-23392062-DEL-53 |
| chr7-23420652-DEL-1172 | chr7-23420652-DEL-1172 |
| chr7-23443636-DEL-61 | chr7-23443637-DEL-61 |
| chr7-23495201-DEL-244 | chr7-23495201-DEL-244 |
| chr7-23510613-DEL-9706 | chr7-23510614-DEL-9706 |
| chr7-23590142-DEL-52 | chr7-23590143-DEL-52 |
| chr7-23603979-DEL-2467 | chr7-23603980-DEL-2467 |
| chr7-23606642-DEL-572 | chr7-23606643-DEL-572 |
| chr7-23646045-DEL-671 | chr7-23646062-DEL-671 |
| chr7-23674230-DEL-119 | chr7-23674231-DEL-119 |
| chr7-23690605-DEL-101 | chr7-23690605-DEL-98 |
| chr7-23691968-DEL-56 | chr7-23691969-DEL-56 |
| chr7-23738312-DEL-142 | chr7-23738313-DEL-142 |
| chr7-23743425-DEL-248 | chr7-23743378-DEL-248 |
| chr7-23818638-DEL-224 | chr7-23818639-DEL-224 |
| chr7-23819425-DEL-304 | chr7-23819426-DEL-339 |
| chr7-23820010-DEL-105 | chr7-23819995-DEL-76 |
| chr7-23847137-DEL-821 | chr7-23847138-DEL-821 |
| chr7-23851618-DEL-89 | chr7-23851603-DEL-120 |
| chr7-23859848-DEL-3630 | chr7-23859849-DEL-3630 |
| chr7-23863866-DEL-6293 | chr7-23863867-DEL-6293 |
| chr7-23871934-DEL-139 | chr7-23871925-DEL-141 |
| chr7-23873208-DEL-52 | chr7-23873201-DEL-53 |
| chr7-23886214-DEL-194 | chr7-23886215-DEL-194 |
| chr7-23913415-DEL-139 | chr7-23913416-DEL-139 |
| chr7-23918295-DEL-3045 | chr7-23918296-DEL-3044 |
| chr7-23921081-DEL-56 | chr7-23921082-DEL-56 |
| chr7-23942722-DEL-65 | chr7-23942735-DEL-63 |
| chr7-23952104-DEL-322 | chr7-23952105-DEL-322 |
| chr7-23958317-DEL-75 | chr7-23958318-DEL-75 |
| chr7-23965797-DEL-214 | chr7-23965798-DEL-214 |
| chr7-23977868-DEL-108 | chr7-23977869-DEL-108 |
| chr7-23984365-DEL-243 | chr7-23984414-DEL-204 |
| chr7-23987759-DEL-529 | chr7-23987771-DEL-536 |
| chr7-23990353-DEL-59 | chr7-23990355-DEL-57 |
| chr7-24004412-DEL-54 | chr7-24004413-DEL-66 |
| chr7-24031668-DEL-438 | chr7-24031669-DEL-438 |
| chr7-24034857-DEL-80 | chr7-24034867-DEL-70 |
| chr7-24037636-DEL-70 | chr7-24037639-DEL-67 |
| chr7-24062150-DEL-300 | chr7-24062151-DEL-300 |
| chr7-24070359-DEL-105 | chr7-24070360-DEL-105 |
| chr7-24074498-DEL-53 | chr7-24074498-DEL-53 |
| chr7-24074570-DEL-214 | chr7-24074570-DEL-214 |
| chr7-24091776-DEL-52 | chr7-24091768-DEL-53 |
| chr7-24103472-DEL-62 | chr7-24103479-DEL-56 |
| chr7-24103879-DEL-232 | chr7-24103880-DEL-232 |
| chr7-24145502-DEL-106 | chr7-24145467-DEL-107 |
| chr7-24177784-DEL-97 | chr7-24177789-DEL-98 |
| chr7-24216955-DEL-62 | chr7-24216956-DEL-62 |
| chr7-24227644-DEL-52 | chr7-24227645-DEL-52 |
| chr7-24236240-DEL-53 | chr7-24236240-DEL-53 |
| chr7-24238516-DEL-111 | chr7-24238516-DEL-111 |
| chr7-24239627-DEL-192 | chr7-24239628-DEL-192 |
| chr7-24251554-DEL-57 | chr7-24251555-DEL-56 |
| chr7-24252960-DEL-106 | chr7-24252961-DEL-106 |
| chr7-24262080-DEL-54 | chr7-24262080-DEL-54 |
| chr7-24277584-DEL-5971 | chr7-24277585-DEL-5971 |
| chr7-24296917-DEL-52 | chr7-24296918-DEL-52 |
| chr7-24311528-DEL-147 | chr7-24311529-DEL-147 |
| chr7-24313304-DEL-192 | chr7-24313305-DEL-192 |
| chr7-24315255-DEL-52 | chr7-24315256-DEL-52 |
| chr7-24319931-DEL-58 | chr7-24319948-DEL-57 |
| chr7-24368380-DEL-56 | chr7-24368381-DEL-56 |
| chr7-24385253-DEL-458 | chr7-24385254-DEL-458 |
| chr7-24399333-DEL-54 | chr7-24399334-DEL-54 |
| chr7-24455191-DEL-53 | chr7-24455192-DEL-53 |
| chr7-24479573-DEL-205 | chr7-24479574-DEL-205 |
| chr7-24494219-DEL-203 | chr7-24494220-DEL-203 |
| chr7-24503980-DEL-54 | chr7-24503981-DEL-54 |
| chr7-24507209-DEL-55 | chr7-24507210-DEL-55 |

| | |
|---|---|
| chr7-24512990-DEL-64 | chr7-24512991-DEL-62 |
| chr7-24538692-DEL-408 | chr7-24538679-DEL-414 |
| chr7-24544831-DEL-69 | chr7-24544832-DEL-69 |
| chr7-24561613-DEL-507 | chr7-24561614-DEL-507 |
| chr7-24580603-DEL-72 | chr7-24580604-DEL-72 |
| chr7-24613194-DEL-2225 | chr7-24613195-DEL-2225 |
| chr7-24615661-DEL-104 | chr7-24615662-DEL-104 |
| chr7-24616824-DEL-71 | chr7-24616825-DEL-71 |
| chr7-24619945-DEL-220 | chr7-24619946-DEL-220 |
| chr7-24658650-DEL-236 | chr7-24658655-DEL-231 |
| chr7-24667052-DEL-208 | chr7-24667053-DEL-208 |
| chr7-24675384-DEL-206 | chr7-24675385-DEL-206 |
| chr7-24676376-DEL-225 | chr7-24676376-DEL-223 |
| chr7-24683555-DEL-204 | chr7-24683556-DEL-204 |
| chr7-24693365-DEL-150 | chr7-24693364-DEL-151 |
| chr7-24738723-DEL-185 | chr7-24738712-DEL-197 |
| chr7-24754684-DEL-53 | chr7-24754719-DEL-54 |
| chr7-24758772-DEL-197 | chr7-24758758-DEL-202 |
| chr7-24764072-DEL-122 | chr7-24764073-DEL-122 |
| chr7-24798189-DEL-195 | chr7-24798190-DEL-195 |
| chr7-24802811-DEL-55 | chr7-24802812-DEL-55 |
| chr7-24843923-DEL-53 | chr7-24843924-DEL-53 |
| chr7-24859598-DEL-212 | chr7-24859599-DEL-212 |
| chr7-24884710-DEL-69 | chr7-24884708-DEL-78 |
| chr7-24885225-DEL-50 | chr7-24885226-DEL-58 |
| chr7-24906931-DEL-462 | chr7-24906932-DEL-462 |
| chr7-24962037-DEL-196 | chr7-24962038-DEL-196 |
| chr7-24962835-DEL-184 | chr7-24962836-DEL-184 |
| chr7-25018887-DEL-206 | chr7-25018888-DEL-206 |
| chr7-25021969-DEL-112 | chr7-25021962-DEL-109 |
| chr7-25024876-DEL-53 | chr7-25024876-DEL-53 |
| chr7-25038347-DEL-53 | chr7-25038348-DEL-53 |
| chr7-25082307-DEL-59 | chr7-25082307-DEL-60 |
| chr7-25098674-DEL-221 | chr7-25098675-DEL-221 |
| chr7-25109366-DEL-725 | chr7-25109341-DEL-751 |
| chr7-25178493-DEL-240 | chr7-25178484-DEL-246 |
| chr7-25179100-DEL-124 | chr7-25179101-DEL-124 |
| chr7-25179442-DEL-550 | chr7-25179443-DEL-550 |
| chr7-25183681-DEL-60 | chr7-25183682-DEL-60 |
| chr7-25185897-DEL-164 | chr7-25185898-DEL-164 |
| chr7-25195126-DEL-6441 | chr7-25195127-DEL-6441 |
| chr7-25195365-DEL-53 | chr7-25195366-DEL-53 |
| chr7-25209303-DEL-1577 | chr7-25209301-DEL-1577 |
| chr7-25241757-DEL-6403 | chr7-25241758-DEL-6403 |
| chr7-25295533-DEL-62 | chr7-25295534-DEL-62 |
| chr7-25298586-DEL-74 | chr7-25298591-DEL-70 |
| chr7-25350192-DEL-181 | chr7-25350193-DEL-181 |
| chr7-25376788-DEL-199 | chr7-25376789-DEL-199 |
| chr7-25401277-DEL-206 | chr7-25401280-DEL-203 |
| chr7-25402607-DEL-57 | chr7-25402609-DEL-56 |
| chr7-25426570-DEL-217 | chr7-25426571-DEL-217 |
| chr7-25456072-DEL-162 | chr7-25456073-DEL-162 |
| chr7-25471477-DEL-60 | chr7-25471488-DEL-54 |
| chr7-25491583-DEL-1571 | chr7-25491584-DEL-1571 |
| chr7-25509050-DEL-211 | chr7-25509050-DEL-215 |
| chr7-25533657-DEL-158 | chr7-25533634-DEL-160 |
| chr7-25552976-DEL-69 | chr7-25552971-DEL-60 |
| chr7-25553410-DEL-1078 | chr7-25553411-DEL-1078 |
| chr7-25578696-DEL-137 | chr7-25578697-DEL-137 |
| chr7-25595550-DEL-52 | chr7-25595553-DEL-54 |
| chr7-25617140-DEL-700 | chr7-25617141-DEL-700 |
| chr7-25631843-DEL-641 | chr7-25631844-DEL-641 |
| chr7-25636074-DEL-203 | chr7-25636075-DEL-203 |
| chr7-25645083-DEL-282 | chr7-25645084-DEL-282 |
| chr7-25646730-DEL-55 | chr7-25646731-DEL-54 |
| chr7-25652592-DEL-138 | chr7-25652593-DEL-130 |
| chr7-25667037-DEL-6794 | chr7-25667038-DEL-6794 |
| chr7-25673365-DEL-409 | chr7-25673366-DEL-409 |
| chr7-25698411-DEL-1269 | chr7-25698412-DEL-1269 |
| chr7-25757588-DEL-2009 | chr7-25757589-DEL-2009 |
| chr7-25771212-DEL-3971 | chr7-25771213-DEL-3971 |
| chr7-25791685-DEL-1453 | chr7-25791686-DEL-1453 |
| chr7-25796849-DEL-144 | chr7-25796849-DEL-144 |
| chr7-25811025-DEL-420 | chr7-25811025-DEL-420 |
| chr7-25823748-DEL-490 | chr7-25823749-DEL-490 |
| chr7-25945715-DEL-964 | chr7-25945716-DEL-964 |
| chr7-25983282-DEL-212 | chr7-25983234-DEL-212 |
| chr7-26016411-DEL-1934 | chr7-26016412-DEL-1934 |
| chr7-26048795-DEL-52 | chr7-26048794-DEL-50 |
| chr7-26066280-DEL-3918 | chr7-26066281-DEL-3918 |

| | |
|---|---|
| chr7-26115715-DEL-78 | chr7-26115715-DEL-78 |
| chr7-26201895-DEL-2937 | chr7-26201896-DEL-2937 |
| chr7-26240216-DEL-200 | chr7-26240217-DEL-200 |
| chr7-26253687-DEL-380 | chr7-26253688-DEL-380 |
| chr7-26278023-DEL-191 | chr7-26278005-DEL-192 |
| chr7-26279628-DEL-91 | chr7-26279629-DEL-91 |
| chr7-26366681-DEL-87 | chr7-26366730-DEL-63 |
| chr7-26416716-DEL-200 | chr7-26416717-DEL-200 |
| chr7-26454488-DEL-60 | chr7-26454493-DEL-56 |
| chr7-26462590-DEL-70 | chr7-26462584-DEL-76 |
| chr7-26475451-DEL-62 | chr7-26475452-DEL-62 |
| chr7-26497646-DEL-156 | chr7-26497660-DEL-143 |
| chr7-26516647-DEL-942 | chr7-26516648-DEL-942 |
| chr7-26524086-DEL-113 | chr7-26524088-DEL-106 |
| chr7-26528230-DEL-93 | chr7-26528231-DEL-105 |
| chr7-26577125-DEL-60 | chr7-26577126-DEL-60 |
| chr7-26603886-DEL-7335 | chr7-26603887-DEL-7335 |
| chr7-26648639-DEL-199 | chr7-26648640-DEL-199 |
| chr7-26654894-DEL-72 | chr7-26654892-DEL-72 |
| chr7-26657292-DEL-2989 | chr7-26657293-DEL-2989 |
| chr7-26668833-DEL-180 | chr7-26668834-DEL-180 |
| chr7-26674516-DEL-381 | chr7-26674515-DEL-382 |
| chr7-26685871-DEL-2110 | chr7-26685872-DEL-2110 |
| chr7-26714197-DEL-1891 | chr7-26714198-DEL-1891 |
| chr7-26738881-DEL-1280 | chr7-26738882-DEL-1280 |
| chr7-26741629-DEL-736 | chr7-26741617-DEL-737 |
| chr7-26757253-DEL-420 | chr7-26757254-DEL-420 |
| chr7-26780993-DEL-64 | chr7-26780994-DEL-64 |
| chr7-26781530-DEL-1793 | chr7-26781531-DEL-1793 |
| chr7-26824972-DEL-175 | chr7-26825010-DEL-168 |
| chr7-26825517-DEL-52 | chr7-26825507-DEL-53 |
| chr7-26836826-DEL-1002 | chr7-26836827-DEL-1002 |
| chr7-26846009-DEL-890 | chr7-26846010-DEL-890 |
| chr7-26853849-DEL-215 | chr7-26853850-DEL-215 |
| chr7-26876443-DEL-96 | chr7-26876444-DEL-92 |
| chr7-26901404-DEL-54 | chr7-26901405-DEL-54 |
| chr7-26947852-DEL-202 | chr7-26947853-DEL-202 |
| chr7-26956215-DEL-51 | chr7-26956216-DEL-51 |
| chr7-26963477-DEL-60 | chr7-26963478-DEL-60 |
| chr7-27002181-DEL-54 | chr7-27002182-DEL-54 |
| chr7-27011797-DEL-52 | chr7-27011798-DEL-52 |
| chr7-27017843-DEL-184 | chr7-27017841-DEL-185 |
| chr7-27077282-DEL-70 | chr7-27077293-DEL-60 |
| chr7-27091697-DEL-380 | chr7-27091695-DEL-379 |
| chr7-27104728-DEL-82 | chr7-27104744-DEL-70 |
| chr7-27113857-DEL-71 | chr7-27113858-DEL-71 |
| chr7-27129483-DEL-1905 | chr7-27129484-DEL-1905 |
| chr7-27131082-DEL-68 | chr7-27131083-DEL-68 |
| chr7-27135581-DEL-473 | chr7-27135582-DEL-473 |
| chr7-27137381-DEL-405 | chr7-27137382-DEL-405 |
| chr7-27142109-DEL-209 | chr7-27142110-DEL-208 |
| chr7-27164737-DEL-188 | chr7-27164738-DEL-188 |
| chr7-27168263-DEL-1782 | chr7-27168295-DEL-1751 |
| chr7-27187312-DEL-181 | chr7-27187313-DEL-181 |
| chr7-27197907-DEL-51 | chr7-27197899-DEL-54 |
| chr7-27201913-DEL-51 | chr7-27201954-DEL-57 |
| chr7-27202882-DEL-207 | chr7-27202883-DEL-207 |
| chr7-27213687-DEL-225 | chr7-27213688-DEL-225 |
| chr7-27234258-DEL-106 | chr7-27234259-DEL-106 |
| chr7-27248864-DEL-102 | chr7-27248865-DEL-102 |
| chr7-27330000-DEL-198 | chr7-27330001-DEL-198 |
| chr7-27347472-DEL-62 | chr7-27347472-DEL-62 |
| chr7-27363140-DEL-66 | chr7-27363140-DEL-67 |
| chr7-27367191-DEL-106 | chr7-27367192-DEL-106 |
| chr7-27379616-DEL-222 | chr7-27379617-DEL-222 |
| chr7-27381784-DEL-76 | chr7-27381785-DEL-76 |
| chr7-27383179-DEL-53 | chr7-27383202-DEL-53 |
| chr7-27384459-DEL-178 | chr7-27384460-DEL-178 |
| chr7-27387652-DEL-1542 | chr7-27387653-DEL-1542 |
| chr7-27393103-DEL-200 | chr7-27393104-DEL-200 |
| chr7-27395355-DEL-220 | chr7-27395355-DEL-220 |
| chr7-27406098-DEL-144 | chr7-27406095-DEL-142 |
| chr7-27415836-DEL-70 | chr7-27415837-DEL-71 |
| chr7-27420474-DEL-524 | chr7-27420475-DEL-524 |
| chr7-27421065-DEL-264 | chr7-27421066-DEL-264 |
| chr7-27432024-DEL-53 | chr7-27432025-DEL-53 |
| chr7-27444471-DEL-153 | chr7-27444472-DEL-153 |
| chr7-27449593-DEL-82 | chr7-27449594-DEL-82 |
| chr7-27455322-DEL-494 | chr7-27455323-DEL-494 |
| chr7-27465218-DEL-472 | chr7-27465218-DEL-472 |

| | |
|---|---|
| chr7-27465423-DEL-2226 | chr7-27465424-DEL-2226 |
| chr7-27466785-DEL-214 | chr7-27466786-DEL-214 |
| chr7-27479734-DEL-327 | chr7-27479735-DEL-327 |
| chr7-27496952-DEL-194 | chr7-27496952-DEL-194 |
| chr7-27521268-DEL-222 | chr7-27521262-DEL-204 |
| chr7-27561165-DEL-24890 | chr7-27561165-DEL-24890 |
| chr7-27563621-DEL-171 | chr7-27563621-DEL-171 |
| chr7-27576817-DEL-186 | chr7-27576818-DEL-186 |
| chr7-27584542-DEL-195 | chr7-27584543-DEL-195 |
| chr7-27585180-DEL-198 | chr7-27585181-DEL-198 |
| chr7-27587125-DEL-420 | chr7-27587130-DEL-407 |
| chr7-27606069-DEL-871 | chr7-27606070-DEL-871 |
| chr7-27618197-DEL-54 | chr7-27618195-DEL-54 |
| chr7-27623146-DEL-100 | chr7-27623147-DEL-120 |
| chr7-27630728-DEL-836 | chr7-27630698-DEL-831 |
| chr7-27633164-DEL-8326 | chr7-27633165-DEL-8326 |
| chr7-27648348-DEL-872 | chr7-27648349-DEL-872 |
| chr7-27654192-DEL-4016 | chr7-27654193-DEL-4016 |
| chr7-27696096-DEL-8497 | chr7-27696097-DEL-8497 |
| chr7-27716683-DEL-189 | chr7-27716684-DEL-189 |
| chr7-27718631-DEL-8812 | chr7-27718632-DEL-8812 |
| chr7-27743670-DEL-2496 | chr7-27743671-DEL-2496 |
| chr7-27746294-DEL-79 | chr7-27746295-DEL-79 |
| chr7-27756456-DEL-59 | chr7-27756456-DEL-59 |
| chr7-27764397-DEL-65 | chr7-27764403-DEL-60 |
| chr7-27767315-DEL-763 | chr7-27767316-DEL-763 |
| chr7-27769561-DEL-111 | chr7-27769562-DEL-111 |
| chr7-27788072-DEL-84 | chr7-27788073-DEL-84 |
| chr7-27809023-DEL-3219 | chr7-27809024-DEL-3219 |
| chr7-27818815-DEL-216 | chr7-27818852-DEL-189 |
| chr7-27820448-DEL-204 | chr7-27820482-DEL-188 |
| chr7-27834354-DEL-67 | chr7-27834355-DEL-67 |
| chr7-27838314-DEL-110 | chr7-27838341-DEL-124 |
| chr7-27900571-DEL-1925 | chr7-27900572-DEL-1924 |
| chr7-27904286-DEL-135 | chr7-27904288-DEL-90 |
| chr7-27952037-DEL-58 | chr7-27952035-DEL-51 |
| chr7-27953389-DEL-371 | chr7-27953390-DEL-371 |
| chr7-27955693-DEL-5496 | chr7-27955694-DEL-5496 |
| chr7-27970065-DEL-85 | chr7-27970066-DEL-85 |
| chr7-27981707-DEL-211 | chr7-27981708-DEL-210 |
| chr7-27983091-DEL-78 | chr7-27983092-DEL-73 |
| chr7-28003679-DEL-52 | chr7-28003680-DEL-52 |
| chr7-28046295-DEL-192 | chr7-28046296-DEL-192 |
| chr7-28055160-DEL-167 | chr7-28055124-DEL-191 |
| chr7-28081749-DEL-54 | chr7-28081752-DEL-52 |
| chr7-28157077-DEL-210 | chr7-28157078-DEL-210 |
| chr7-28177412-DEL-586 | chr7-28177413-DEL-586 |
| chr7-28189071-DEL-242 | chr7-28189072-DEL-242 |
| chr7-28214018-DEL-53 | chr7-28214019-DEL-53 |
| chr7-28220355-DEL-60 | chr7-28220335-DEL-58 |
| chr7-28224075-DEL-55 | chr7-28224076-DEL-52 |
| chr7-28231703-DEL-239 | chr7-28231704-DEL-239 |
| chr7-28242025-DEL-52 | chr7-28241986-DEL-53 |
| chr7-28253172-DEL-53 | chr7-28253173-DEL-53 |
| chr7-28300715-DEL-56 | chr7-28300716-DEL-56 |
| chr7-28312089-DEL-114 | chr7-28312099-DEL-74 |
| chr7-28315499-DEL-64 | chr7-28315499-DEL-64 |
| chr7-28325452-DEL-98 | chr7-28325433-DEL-120 |
| chr7-28346985-DEL-206 | chr7-28346986-DEL-206 |
| chr7-28356979-DEL-1866 | chr7-28356980-DEL-1866 |
| chr7-28367673-DEL-212 | chr7-28367674-DEL-212 |
| chr7-28380620-DEL-518 | chr7-28380619-DEL-519 |
| chr7-28390453-DEL-62 | chr7-28390454-DEL-62 |
| chr7-28401051-DEL-150 | chr7-28401052-DEL-150 |
| chr7-28407373-DEL-193 | chr7-28407383-DEL-182 |
| chr7-28411268-DEL-69 | chr7-28411269-DEL-69 |
| chr7-28448568-DEL-383 | chr7-28448569-DEL-383 |
| chr7-28450810-DEL-65 | chr7-28450830-DEL-66 |
| chr7-28452188-DEL-167 | chr7-28452189-DEL-167 |
| chr7-28496744-DEL-177 | chr7-28496745-DEL-177 |
| chr7-28509798-DEL-61 | chr7-28509799-DEL-61 |
| chr7-28518738-DEL-206 | chr7-28518739-DEL-206 |
| chr7-28526349-DEL-207 | chr7-28526350-DEL-207 |
| chr7-28535513-DEL-89 | chr7-28535513-DEL-89 |
| chr7-28540800-DEL-58 | chr7-28540797-DEL-53 |
| chr7-28545183-DEL-207 | chr7-28545181-DEL-208 |
| chr7-28551451-DEL-181 | chr7-28551452-DEL-181 |
| chr7-28552798-DEL-220 | chr7-28552799-DEL-220 |
| chr7-28579416-DEL-183 | chr7-28579417-DEL-183 |
| chr7-28581507-DEL-70 | chr7-28581507-DEL-70 |

| | |
|---|---|
| chr7-28610308-DEL-75 | chr7-28610294-DEL-76 |
| chr7-28632781-DEL-198 | chr7-28632782-DEL-198 |
| chr7-28633512-DEL-57 | chr7-28633513-DEL-57 |
| chr7-28679057-DEL-243 | chr7-28679058-DEL-243 |
| chr7-28715720-DEL-87 | chr7-28715721-DEL-95 |
| chr7-28763300-DEL-52 | chr7-28763300-DEL-52 |
| chr7-28765388-DEL-54 | chr7-28765389-DEL-54 |
| chr7-28777764-DEL-60 | chr7-28777763-DEL-61 |
| chr7-28779011-DEL-58 | chr7-28779012-DEL-58 |
| chr7-28783967-DEL-127 | chr7-28783968-DEL-127 |
| chr7-28794285-DEL-53 | chr7-28794286-DEL-53 |
| chr7-28796103-DEL-111 | chr7-28796104-DEL-111 |
| chr7-28798572-DEL-263 | chr7-28798573-DEL-263 |
| chr7-28800569-DEL-159 | chr7-28800587-DEL-160 |
| chr7-28806309-DEL-57 | chr7-28806310-DEL-58 |
| chr7-28827478-DEL-59 | chr7-28827465-DEL-70 |
| chr7-28837449-DEL-175 | chr7-28837449-DEL-175 |
| chr7-28838902-DEL-218 | chr7-28838904-DEL-206 |
| chr7-28891209-DEL-212 | chr7-28891210-DEL-212 |
| chr7-28908992-DEL-181 | chr7-28908993-DEL-181 |
| chr7-28911759-DEL-144 | chr7-28911760-DEL-144 |
| chr7-28912427-DEL-72 | chr7-28912428-DEL-72 |
| chr7-28920431-DEL-68 | chr7-28920432-DEL-68 |
| chr7-28966108-DEL-221 | chr7-28966109-DEL-221 |
| chr7-28996773-DEL-816 | chr7-28996774-DEL-816 |
| chr7-29022822-DEL-229 | chr7-29022823-DEL-229 |
| chr7-29026399-DEL-105 | chr7-29026400-DEL-105 |
| chr7-29027622-DEL-78 | chr7-29027623-DEL-78 |
| chr7-29043645-DEL-76 | chr7-29043662-DEL-74 |
| chr7-29060632-DEL-53 | chr7-29060633-DEL-53 |
| chr7-29088092-DEL-60 | chr7-29088079-DEL-74 |
| chr7-29114408-DEL-155 | chr7-29114409-DEL-155 |
| chr7-29117308-DEL-100 | chr7-29117309-DEL-100 |
| chr7-29163886-DEL-62 | chr7-29163887-DEL-62 |
| chr7-29176916-DEL-97 | chr7-29176915-DEL-97 |
| chr7-29199175-DEL-219 | chr7-29199176-DEL-219 |
| chr7-29208663-DEL-1565 | chr7-29208664-DEL-1565 |
| chr7-29225674-DEL-222 | chr7-29225675-DEL-222 |
| chr7-29243566-DEL-225 | chr7-29243567-DEL-225 |
| chr7-29272428-DEL-80 | chr7-29272427-DEL-71 |
| chr7-29314626-DEL-8703 | chr7-29314627-DEL-8703 |
| chr7-29330085-DEL-429 | chr7-29330086-DEL-424 |
| chr7-29342629-DEL-5537 | chr7-29342630-DEL-5537 |
| chr7-29361190-DEL-6433 | chr7-29361191-DEL-6433 |
| chr7-29425545-DEL-70 | chr7-29425546-DEL-70 |
| chr7-29431585-DEL-370 | chr7-29431586-DEL-370 |
| chr7-29497647-DEL-1647 | chr7-29497648-DEL-1647 |
| chr7-29501725-DEL-436 | chr7-29501726-DEL-436 |
| chr7-29511327-DEL-100 | chr7-29511328-DEL-100 |
| chr7-29523766-DEL-66 | chr7-29523767-DEL-62 |
| chr7-29536724-DEL-414 | chr7-29536720-DEL-408 |
| chr7-29540545-DEL-109 | chr7-29540546-DEL-109 |
| chr7-29549255-DEL-231 | chr7-29549256-DEL-231 |
| chr7-29551984-DEL-51 | chr7-29551983-DEL-52 |
| chr7-29555267-DEL-683 | chr7-29555268-DEL-683 |
| chr7-29561577-DEL-1339 | chr7-29561578-DEL-1339 |
| chr7-29564830-DEL-103 | chr7-29564830-DEL-124 |
| chr7-29571384-DEL-56 | chr7-29571388-DEL-59 |
| chr7-29577108-DEL-294 | chr7-29577109-DEL-294 |
| chr7-29579698-DEL-285 | chr7-29579699-DEL-285 |
| chr7-29589433-DEL-1460 | chr7-29589434-DEL-1460 |
| chr7-29590033-DEL-60 | chr7-29590034-DEL-60 |
| chr7-29603538-DEL-883 | chr7-29603539-DEL-883 |
| chr7-29604663-DEL-85 | chr7-29604663-DEL-50 |
| chr7-29612187-DEL-220 | chr7-29612188-DEL-220 |
| chr7-29622357-DEL-80 | chr7-29622358-DEL-80 |
| chr7-29623480-DEL-196 | chr7-29623481-DEL-196 |
| chr7-29626139-DEL-3735 | chr7-29626140-DEL-3735 |
| chr7-29646920-DEL-1162 | chr7-29646903-DEL-1198 |
| chr7-29646999-DEL-96 | chr7-29647020-DEL-68 |
| chr7-29652383-DEL-199 | chr7-29652384-DEL-199 |
| chr7-29657120-DEL-1158 | chr7-29657121-DEL-1158 |
| chr7-29658729-DEL-226 | chr7-29658730-DEL-226 |
| chr7-29660890-DEL-1608 | chr7-29660891-DEL-1605 |
| chr7-29664483-DEL-594 | chr7-29664484-DEL-594 |
| chr7-29672789-DEL-1453 | chr7-29672790-DEL-1453 |
| chr7-29678818-DEL-512 | chr7-29678819-DEL-512 |
| chr7-29687919-DEL-273 | chr7-29687920-DEL-273 |
| chr7-29693195-DEL-439 | chr7-29693196-DEL-439 |
| chr7-29703677-DEL-1266 | chr7-29703678-DEL-1266 |

| | |
|---|---|
| chr7-29718719-DEL-179 | chr7-29718720-DEL-179 |
| chr7-29732856-DEL-178 | chr7-29732857-DEL-178 |
| chr7-29735433-DEL-80 | chr7-29735430-DEL-80 |
| chr7-29739530-DEL-64 | chr7-29739519-DEL-62 |
| chr7-29740318-DEL-139 | chr7-29740319-DEL-139 |
| chr7-29741345-DEL-225 | chr7-29741346-DEL-225 |
| chr7-29743714-DEL-55 | chr7-29743713-DEL-56 |
| chr7-29746172-DEL-215 | chr7-29746173-DEL-215 |
| chr7-29765400-DEL-177 | chr7-29765400-DEL-178 |
| chr7-29766638-DEL-53 | chr7-29766624-DEL-56 |
| chr7-29773159-DEL-234 | chr7-29773160-DEL-233 |
| chr7-29778276-DEL-202 | chr7-29778277-DEL-200 |
| chr7-29779168-DEL-133 | chr7-29779123-DEL-151 |
| chr7-29781117-DEL-173 | chr7-29781102-DEL-148 |
| chr7-29796218-DEL-216 | chr7-29796219-DEL-216 |
| chr7-29796893-DEL-61 | chr7-29796894-DEL-61 |
| chr7-29814594-DEL-104 | chr7-29814577-DEL-100 |
| chr7-29823496-DEL-941 | chr7-29823497-DEL-941 |
| chr7-29843788-DEL-202 | chr7-29843789-DEL-202 |
| chr7-29853542-DEL-996 | chr7-29853543-DEL-996 |
| chr7-29855467-DEL-1780 | chr7-29855468-DEL-1780 |
| chr7-29865614-DEL-112 | chr7-29865616-DEL-111 |
| chr7-29882620-DEL-93 | chr7-29882621-DEL-93 |
| chr7-29889120-DEL-215 | chr7-29889121-DEL-215 |
| chr7-29889659-DEL-166 | chr7-29889660-DEL-166 |
| chr7-29902245-DEL-58 | chr7-29902246-DEL-58 |
| chr7-29907594-DEL-211 | chr7-29907595-DEL-211 |
| chr7-29908664-DEL-101 | chr7-29908665-DEL-101 |
| chr7-29914917-DEL-205 | chr7-29914918-DEL-205 |
| chr7-29927971-DEL-211 | chr7-29927972-DEL-211 |
| chr7-29929540-DEL-276 | chr7-29929541-DEL-276 |
| chr7-29936971-DEL-96 | chr7-29936972-DEL-96 |
| chr7-29937916-DEL-59 | chr7-29937917-DEL-59 |
| chr7-29957871-DEL-60 | chr7-29957872-DEL-58 |
| chr7-29965237-DEL-62 | chr7-29965238-DEL-62 |
| chr7-29975385-DEL-824 | chr7-29975386-DEL-824 |
| chr7-29977017-DEL-237 | chr7-29977018-DEL-237 |
| chr7-30025782-DEL-86 | chr7-30025792-DEL-74 |
| chr7-30050845-DEL-2076 | chr7-30050846-DEL-2076 |
| chr7-30052831-DEL-60 | chr7-30052832-DEL-60 |
| chr7-30084719-DEL-7139 | chr7-30084720-DEL-7139 |
| chr7-30095131-DEL-1420 | chr7-30095132-DEL-1420 |
| chr7-30097299-DEL-124 | chr7-30097289-DEL-156 |
| chr7-30101722-DEL-102 | chr7-30101697-DEL-100 |
| chr7-30109970-DEL-113 | chr7-30109937-DEL-114 |
| chr7-30111354-DEL-72 | chr7-30111355-DEL-72 |
| chr7-30115482-DEL-52 | chr7-30115483-DEL-52 |
| chr7-30127303-DEL-202 | chr7-30127300-DEL-201 |
| chr7-30142395-DEL-61 | chr7-30142396-DEL-61 |
| chr7-30175426-DEL-158 | chr7-30175411-DEL-159 |
| chr7-30176618-DEL-208 | chr7-30176619-DEL-208 |
| chr7-30177381-DEL-87 | chr7-30177382-DEL-87 |
| chr7-30179986-DEL-184 | chr7-30179941-DEL-203 |
| chr7-30184076-DEL-533 | chr7-30184077-DEL-533 |
| chr7-30203782-DEL-53 | chr7-30203783-DEL-53 |
| chr7-30207215-DEL-52 | chr7-30207207-DEL-53 |
| chr7-30216768-DEL-106 | chr7-30216769-DEL-106 |
| chr7-30224454-DEL-81 | chr7-30224455-DEL-81 |
| chr7-30224966-DEL-75 | chr7-30224967-DEL-75 |
| chr7-30238339-DEL-176 | chr7-30238340-DEL-176 |
| chr7-30243526-DEL-112 | chr7-30243528-DEL-116 |
| chr7-30244497-DEL-272 | chr7-30244498-DEL-272 |
| chr7-30267414-DEL-215 | chr7-30267415-DEL-215 |
| chr7-30299731-DEL-52 | chr7-30299732-DEL-52 |
| chr7-30319292-DEL-204 | chr7-30319293-DEL-204 |
| chr7-30340101-DEL-265 | chr7-30340102-DEL-265 |
| chr7-30374203-DEL-273 | chr7-30374204-DEL-273 |
| chr7-30375963-DEL-1294 | chr7-30375961-DEL-1294 |
| chr7-30378358-DEL-55 | chr7-30378326-DEL-52 |
| chr7-30384359-DEL-768 | chr7-30384350-DEL-769 |
| chr7-30388060-DEL-8985 | chr7-30388061-DEL-8985 |
| chr7-30405673-DEL-89 | chr7-30405674-DEL-89 |
| chr7-30429424-DEL-105 | chr7-30429411-DEL-108 |
| chr7-30434724-DEL-56 | chr7-30434725-DEL-56 |
| chr7-30458160-DEL-64 | chr7-30458161-DEL-64 |
| chr7-30467085-DEL-105 | chr7-30467086-DEL-86 |
| chr7-30472662-DEL-159 | chr7-30472663-DEL-159 |
| chr7-30477386-DEL-59 | chr7-30477387-DEL-59 |
| chr7-30483155-DEL-130 | chr7-30483166-DEL-126 |
| chr7-30493676-DEL-200 | chr7-30493677-DEL-200 |

| | |
|---|---|
| chr7-30502392-DEL-194 | chr7-30502393-DEL-194 |
| chr7-30505054-DEL-525 | chr7-30505055-DEL-525 |
| chr7-30508962-DEL-188 | chr7-30508963-DEL-188 |
| chr7-30525360-DEL-401 | chr7-30525361-DEL-401 |
| chr7-30534336-DEL-262 | chr7-30534337-DEL-262 |
| chr7-30545481-DEL-104 | chr7-30545482-DEL-104 |
| chr7-30578324-DEL-68 | chr7-30578325-DEL-68 |
| chr7-30597313-DEL-54 | chr7-30597313-DEL-54 |
| chr7-30627193-DEL-6547 | chr7-30627193-DEL-6547 |
| chr7-30638111-DEL-421 | chr7-30638138-DEL-392 |
| chr7-30641152-DEL-183 | chr7-30641104-DEL-199 |
| chr7-30710413-DEL-652 | chr7-30710414-DEL-652 |
| chr7-30719183-DEL-660 | chr7-30719184-DEL-660 |
| chr7-30720844-DEL-177 | chr7-30720844-DEL-178 |
| chr7-30739052-DEL-431 | chr7-30739053-DEL-431 |
| chr7-30765599-DEL-106 | chr7-30765600-DEL-106 |
| chr7-30774714-DEL-53 | chr7-30774679-DEL-53 |
| chr7-30806555-DEL-106 | chr7-30806556-DEL-106 |
| chr7-30812180-DEL-53 | chr7-30812181-DEL-53 |
| chr7-30822895-DEL-52 | chr7-30822895-DEL-52 |
| chr7-30824002-DEL-116 | chr7-30824003-DEL-116 |
| chr7-30835920-DEL-80 | chr7-30835934-DEL-80 |
| chr7-30852856-DEL-117 | chr7-30852857-DEL-117 |
| chr7-30858707-DEL-68 | chr7-30858708-DEL-68 |
| chr7-30887489-DEL-2118 | chr7-30887490-DEL-2118 |
| chr7-30898402-DEL-944 | chr7-30898403-DEL-944 |
| chr7-30921775-DEL-61 | chr7-30921776-DEL-61 |
| chr7-30946625-DEL-1564 | chr7-30946626-DEL-1564 |
| chr7-30970903-DEL-8789 | chr7-30970904-DEL-8789 |
| chr7-30974982-DEL-2925 | chr7-30975015-DEL-2925 |
| chr7-30992758-DEL-1166 | chr7-30992758-DEL-1166 |
| chr7-31006503-DEL-58 | chr7-31006552-DEL-50 |
| chr7-31016754-DEL-877 | chr7-31016755-DEL-877 |
| chr7-31035596-DEL-57 | chr7-31035597-DEL-57 |
| chr7-31053509-DEL-76 | chr7-31053510-DEL-76 |
| chr7-31060841-DEL-121 | chr7-31060862-DEL-86 |
| chr7-31064832-DEL-6757 | chr7-31064833-DEL-6757 |
| chr7-31111175-DEL-96 | chr7-31111176-DEL-74 |
| chr7-31130663-DEL-212 | chr7-31130664-DEL-212 |
| chr7-31161758-DEL-1145 | chr7-31161759-DEL-1145 |
| chr7-31171256-DEL-615 | chr7-31171257-DEL-615 |
| chr7-31212926-DEL-1205 | chr7-31212927-DEL-1204 |
| chr7-31214217-DEL-65 | chr7-31214218-DEL-65 |
| chr7-31227010-DEL-75 | chr7-31227011-DEL-75 |
| chr7-31252373-DEL-63 | chr7-31252374-DEL-63 |
| chr7-31300767-DEL-95 | chr7-31300768-DEL-95 |
| chr7-31492535-DEL-159 | chr7-31492543-DEL-145 |
| chr7-31504625-DEL-70 | chr7-31504610-DEL-90 |
| chr7-31601034-DEL-1648 | chr7-31601035-DEL-1648 |
| chr7-31625956-DEL-61 | chr7-31625947-DEL-70 |
| chr7-31671249-DEL-52 | chr7-31671247-DEL-52 |
| chr7-31700563-DEL-58 | chr7-31700564-DEL-56 |
| chr7-31778059-DEL-4268 | chr7-31778060-DEL-4268 |
| chr7-31797299-DEL-190 | chr7-31797300-DEL-190 |
| chr7-31802559-DEL-76 | chr7-31802560-DEL-76 |
| chr7-31833549-DEL-196 | chr7-31833550-DEL-196 |
| chr7-31880452-DEL-434 | chr7-31880453-DEL-434 |
| chr7-31997181-DEL-428 | chr7-31997182-DEL-428 |
| chr7-32068378-DEL-124 | chr7-32068378-DEL-124 |
| chr7-32821011-DEL-131 | chr7-32821012-DEL-131 |
| chr7-32845866-DEL-434 | chr7-32845867-DEL-434 |
| chr7-32856400-DEL-2880 | chr7-32856401-DEL-2880 |
| chr7-32879511-DEL-505 | chr7-32879512-DEL-505 |
| chr7-32889343-DEL-76 | chr7-32889329-DEL-99 |
| chr7-32899440-DEL-943 | chr7-32899440-DEL-943 |
| chr7-32906208-DEL-76 | chr7-32906197-DEL-73 |
| chr7-32910620-DEL-231 | chr7-32910621-DEL-231 |
| chr7-32950359-DEL-214 | chr7-32950360-DEL-214 |
| chr7-32950453-DEL-630 | chr7-32950453-DEL-630 |
| chr7-32976458-DEL-57 | chr7-32976459-DEL-57 |
| chr7-32991722-DEL-62 | chr7-32991685-DEL-90 |
| chr7-33025139-DEL-241 | chr7-33025140-DEL-241 |
| chr7-33035292-DEL-109 | chr7-33035293-DEL-109 |
| chr7-33046988-DEL-67 | chr7-33046989-DEL-67 |
| chr7-33104119-DEL-4541 | chr7-33104120-DEL-4541 |
| chr7-33126078-DEL-90 | chr7-33126089-DEL-102 |
| chr7-33158992-DEL-169 | chr7-33158989-DEL-170 |
| chr7-33181056-DEL-401 | chr7-33181057-DEL-401 |
| chr7-33192599-DEL-64 | chr7-33192596-DEL-68 |
| chr7-33206994-DEL-268 | chr7-33206995-DEL-268 |

| | |
|---|---|
| chr7-33208628-DEL-7079 | chr7-33208629-DEL-7079 |
| chr7-33234566-DEL-3401 | chr7-33234567-DEL-3401 |
| chr7-33251093-DEL-1815 | chr7-33251094-DEL-1815 |
| chr7-33259826-DEL-52 | chr7-33259827-DEL-52 |
| chr7-33273633-DEL-90 | chr7-33273613-DEL-91 |
| chr7-33329127-DEL-154 | chr7-33329128-DEL-154 |
| chr7-33330683-DEL-169 | chr7-33330681-DEL-170 |
| chr7-33367017-DEL-60 | chr7-33367018-DEL-60 |
| chr7-33375702-DEL-585 | chr7-33375661-DEL-623 |
| chr7-33384459-DEL-70 | chr7-33384460-DEL-70 |
| chr7-33406601-DEL-52 | chr7-33406602-DEL-52 |
| chr7-33415944-DEL-54 | chr7-33415945-DEL-54 |
| chr7-33446955-DEL-467 | chr7-33446956-DEL-467 |
| chr7-33451573-DEL-6584 | chr7-33451574-DEL-6584 |
| chr7-33496549-DEL-75 | chr7-33496586-DEL-77 |
| chr7-33516198-DEL-203 | chr7-33516199-DEL-203 |
| chr7-33547091-DEL-1268 | chr7-33547092-DEL-1268 |
| chr7-33560750-DEL-6395 | chr7-33560751-DEL-6395 |
| chr7-33560766-DEL-201 | chr7-33560767-DEL-201 |
| chr7-33604028-DEL-117 | chr7-33604029-DEL-117 |
| chr7-33648879-DEL-6918 | chr7-33648880-DEL-6918 |
| chr7-33698891-DEL-214 | chr7-33698891-DEL-214 |
| chr7-33715927-DEL-386 | chr7-33715928-DEL-386 |
| chr7-33740242-DEL-6680 | chr7-33740243-DEL-6680 |
| chr7-33762844-DEL-2605 | chr7-33762845-DEL-2605 |
| chr7-33783073-DEL-164 | chr7-33783074-DEL-164 |
| chr7-33791253-DEL-2954 | chr7-33791249-DEL-2962 |
| chr7-33805995-DEL-122 | chr7-33805996-DEL-122 |
| chr7-33810196-DEL-54 | chr7-33810197-DEL-54 |
| chr7-33810870-DEL-582 | chr7-33810871-DEL-582 |
| chr7-33812219-DEL-214 | chr7-33812220-DEL-214 |
| chr7-33827962-DEL-70 | chr7-33827963-DEL-72 |
| chr7-33837493-DEL-82 | chr7-33837503-DEL-72 |
| chr7-33859245-DEL-211 | chr7-33859246-DEL-211 |
| chr7-33863535-DEL-230 | chr7-33863536-DEL-230 |
| chr7-33882332-DEL-208 | chr7-33882333-DEL-208 |
| chr7-33898262-DEL-1607 | chr7-33898263-DEL-1607 |
| chr7-33910315-DEL-59 | chr7-33910316-DEL-59 |
| chr7-33949089-DEL-232 | chr7-33949090-DEL-232 |
| chr7-33949387-DEL-542 | chr7-33949388-DEL-542 |
| chr7-33973331-DEL-211 | chr7-33973332-DEL-211 |
| chr7-33989593-DEL-181 | chr7-33989594-DEL-181 |
| chr7-34014279-DEL-207 | chr7-34014280-DEL-207 |
| chr7-34019806-DEL-248 | chr7-34019827-DEL-228 |
| chr7-34028148-DEL-668 | chr7-34028149-DEL-653 |
| chr7-34029181-DEL-1567 | chr7-34029182-DEL-1567 |
| chr7-34031335-DEL-188 | chr7-34031336-DEL-188 |
| chr7-34055945-DEL-207 | chr7-34055960-DEL-193 |
| chr7-34115697-DEL-195 | chr7-34115698-DEL-156 |
| chr7-34132260-DEL-64 | chr7-34132261-DEL-64 |
| chr7-34136831-DEL-63 | chr7-34136832-DEL-63 |
| chr7-34158754-DEL-59 | chr7-34158769-DEL-57 |
| chr7-34166540-DEL-203 | chr7-34166541-DEL-203 |
| chr7-34169189-DEL-440 | chr7-34169190-DEL-440 |
| chr7-34212182-DEL-173 | chr7-34212183-DEL-170 |
| chr7-34223942-DEL-6439 | chr7-34223940-DEL-6440 |
| chr7-34238520-DEL-222 | chr7-34238560-DEL-178 |
| chr7-34239636-DEL-71 | chr7-34239637-DEL-71 |
| chr7-34268642-DEL-210 | chr7-34268643-DEL-210 |
| chr7-34280792-DEL-207 | chr7-34280793-DEL-207 |
| chr7-34281278-DEL-396 | chr7-34281279-DEL-396 |
| chr7-34381554-DEL-211 | chr7-34381555-DEL-211 |
| chr7-34393667-DEL-77 | chr7-34393669-DEL-75 |
| chr7-34400211-DEL-215 | chr7-34400212-DEL-215 |
| chr7-34405644-DEL-79 | chr7-34405641-DEL-83 |
| chr7-34429662-DEL-370 | chr7-34429632-DEL-371 |
| chr7-34431354-DEL-130 | chr7-34431345-DEL-139 |
| chr7-34431712-DEL-519 | chr7-34431713-DEL-519 |
| chr7-34462551-DEL-68 | chr7-34462552-DEL-68 |
| chr7-34472257-DEL-94 | chr7-34472258-DEL-94 |
| chr7-34508335-DEL-146 | chr7-34508334-DEL-147 |
| chr7-34550908-DEL-55 | chr7-34550909-DEL-55 |
| chr7-34551911-DEL-734 | chr7-34551912-DEL-734 |
| chr7-34571068-DEL-200 | chr7-34571069-DEL-200 |
| chr7-34622108-DEL-583 | chr7-34622109-DEL-583 |
| chr7-34631476-DEL-383 | chr7-34631478-DEL-375 |
| chr7-34641193-DEL-139 | chr7-34641194-DEL-139 |
| chr7-34662342-DEL-53 | chr7-34662343-DEL-53 |
| chr7-34682636-DEL-195 | chr7-34682637-DEL-195 |
| chr7-34705347-DEL-203 | chr7-34705348-DEL-203 |

| | |
|---|---|
| chr7-34764849-DEL-140 | chr7-34764850-DEL-140 |
| chr7-34768706-DEL-58 | chr7-34768705-DEL-60 |
| chr7-34800270-DEL-76 | chr7-34800271-DEL-76 |
| chr7-34801890-DEL-56 | chr7-34801891-DEL-52 |
| chr7-34842570-DEL-58 | chr7-34842570-DEL-59 |
| chr7-34859213-DEL-1084 | chr7-34859214-DEL-1084 |
| chr7-34892631-DEL-66 | chr7-34892602-DEL-58 |
| chr7-34921334-DEL-56 | chr7-34921336-DEL-55 |
| chr7-34928447-DEL-52 | chr7-34928444-DEL-56 |
| chr7-34928856-DEL-2029 | chr7-34928857-DEL-2029 |
| chr7-34934743-DEL-56 | chr7-34934737-DEL-73 |
| chr7-34937581-DEL-216 | chr7-34937582-DEL-216 |
| chr7-34941546-DEL-63 | chr7-34941547-DEL-63 |
| chr7-34952066-DEL-263 | chr7-34952081-DEL-246 |
| chr7-34973302-DEL-57 | chr7-34973303-DEL-57 |
| chr7-34990002-DEL-119 | chr7-34990003-DEL-119 |
| chr7-34997692-DEL-192 | chr7-34997693-DEL-192 |
| chr7-35013827-DEL-92 | chr7-35013828-DEL-92 |
| chr7-35016424-DEL-59 | chr7-35016425-DEL-60 |
| chr7-35020991-DEL-60 | chr7-35020995-DEL-57 |
| chr7-35025715-DEL-218 | chr7-35025731-DEL-218 |
| chr7-35027935-DEL-206 | chr7-35027935-DEL-206 |
| chr7-35028249-DEL-599 | chr7-35028250-DEL-599 |
| chr7-35043722-DEL-119 | chr7-35043704-DEL-101 |
| chr7-35060917-DEL-197 | chr7-35060921-DEL-197 |
| chr7-35069537-DEL-361 | chr7-35069531-DEL-362 |
| chr7-35084335-DEL-193 | chr7-35084336-DEL-193 |
| chr7-35089320-DEL-67 | chr7-35089321-DEL-67 |
| chr7-35121429-DEL-185 | chr7-35121430-DEL-185 |
| chr7-35132430-DEL-378 | chr7-35132431-DEL-378 |
| chr7-35195645-DEL-200 | chr7-35195646-DEL-200 |
| chr7-35224583-DEL-87 | chr7-35224597-DEL-73 |
| chr7-35242649-DEL-106 | chr7-35242650-DEL-106 |
| chr7-35249923-DEL-89 | chr7-35249924-DEL-89 |
| chr7-35265648-DEL-117 | chr7-35265649-DEL-117 |
| chr7-35272799-DEL-64 | chr7-35272800-DEL-64 |
| chr7-35279909-DEL-56 | chr7-35279910-DEL-56 |
| chr7-35304471-DEL-132 | chr7-35304472-DEL-132 |
| chr7-35309560-DEL-82 | chr7-35309561-DEL-82 |
| chr7-35320770-DEL-61 | chr7-35320744-DEL-62 |
| chr7-35329744-DEL-130 | chr7-35329745-DEL-145 |
| chr7-35335342-DEL-76 | chr7-35335310-DEL-78 |
| chr7-35341077-DEL-73 | chr7-35341078-DEL-73 |
| chr7-35342891-DEL-104 | chr7-35342892-DEL-104 |
| chr7-35364604-DEL-73 | chr7-35364605-DEL-73 |
| chr7-35378748-DEL-205 | chr7-35378749-DEL-205 |
| chr7-35382853-DEL-53 | chr7-35382854-DEL-53 |
| chr7-35384031-DEL-309 | chr7-35384032-DEL-309 |
| chr7-35404478-DEL-87 | chr7-35404479-DEL-87 |
| chr7-35425445-DEL-106 | chr7-35425464-DEL-79 |
| chr7-35449434-DEL-107 | chr7-35449435-DEL-107 |
| chr7-35462494-DEL-66 | chr7-35462498-DEL-63 |
| chr7-35463627-DEL-72 | chr7-35463628-DEL-72 |
| chr7-35478324-DEL-185 | chr7-35478325-DEL-185 |
| chr7-35508956-DEL-559 | chr7-35508957-DEL-559 |
| chr7-35544714-DEL-106 | chr7-35544720-DEL-101 |
| chr7-35589731-DEL-73 | chr7-35589732-DEL-73 |
| chr7-35600942-DEL-66 | chr7-35600941-DEL-68 |
| chr7-35612426-DEL-355 | chr7-35612430-DEL-351 |
| chr7-35632444-DEL-1865 | chr7-35632445-DEL-1865 |
| chr7-35655824-DEL-53 | chr7-35655825-DEL-53 |
| chr7-35667679-DEL-64 | chr7-35667681-DEL-60 |
| chr7-35670141-DEL-6553 | chr7-35670142-DEL-6553 |
| chr7-35699160-DEL-843 | chr7-35699155-DEL-844 |
| chr7-35708388-DEL-112 | chr7-35708389-DEL-111 |
| chr7-35724363-DEL-106 | chr7-35724364-DEL-106 |
| chr7-35742755-DEL-200 | chr7-35742755-DEL-197 |
| chr7-35788863-DEL-227 | chr7-35788864-DEL-227 |
| chr7-35797547-DEL-80 | chr7-35797548-DEL-80 |
| chr7-35817649-DEL-655 | chr7-35817650-DEL-655 |
| chr7-35848316-DEL-75 | chr7-35848317-DEL-68 |
| chr7-35852936-DEL-113 | chr7-35852937-DEL-113 |
| chr7-35860506-DEL-149 | chr7-35860503-DEL-150 |
| chr7-35873758-DEL-53 | chr7-35873759-DEL-53 |
| chr7-35874482-DEL-82 | chr7-35874483-DEL-82 |
| chr7-35886087-DEL-178 | chr7-35886088-DEL-178 |
| chr7-35919517-DEL-79 | chr7-35919518-DEL-79 |
| chr7-35927805-DEL-165 | chr7-35927806-DEL-165 |
| chr7-35953956-DEL-53 | chr7-35953937-DEL-76 |
| chr7-36000372-DEL-1894 | chr7-36000373-DEL-1894 |

| | |
|---|---|
| chr7-36004651-DEL-90 | chr7-36004652-DEL-90 |
| chr7-36013961-DEL-728 | chr7-36013962-DEL-728 |
| chr7-36018503-DEL-71 | chr7-36018483-DEL-60 |
| chr7-36078658-DEL-58 | chr7-36078659-DEL-64 |
| chr7-36083211-DEL-60 | chr7-36083212-DEL-60 |
| chr7-36090618-DEL-193 | chr7-36090619-DEL-193 |
| chr7-36097133-DEL-230 | chr7-36097139-DEL-225 |
| chr7-36101752-DEL-59 | chr7-36101753-DEL-59 |
| chr7-36118888-DEL-240 | chr7-36118889-DEL-240 |
| chr7-36122497-DEL-72 | chr7-36122503-DEL-70 |
| chr7-36130045-DEL-60 | chr7-36130046-DEL-60 |
| chr7-36154958-DEL-53 | chr7-36154959-DEL-53 |
| chr7-36236613-DEL-178 | chr7-36236614-DEL-178 |
| chr7-36243047-DEL-69 | chr7-36243048-DEL-70 |
| chr7-36249129-DEL-84 | chr7-36249130-DEL-84 |
| chr7-36258453-DEL-77 | chr7-36258453-DEL-78 |
| chr7-36259663-DEL-105 | chr7-36259664-DEL-105 |
| chr7-36271993-DEL-119 | chr7-36271990-DEL-119 |
| chr7-36273611-DEL-79 | chr7-36273612-DEL-79 |
| chr7-36274765-DEL-282 | chr7-36274760-DEL-288 |
| chr7-36275414-DEL-176 | chr7-36275413-DEL-136 |
| chr7-36289689-DEL-77 | chr7-36289690-DEL-77 |
| chr7-36290777-DEL-208 | chr7-36290778-DEL-208 |
| chr7-36302899-DEL-66 | chr7-36302904-DEL-72 |
| chr7-36307976-DEL-56 | chr7-36307977-DEL-56 |
| chr7-36315777-DEL-142 | chr7-36315763-DEL-144 |
| chr7-36345463-DEL-400 | chr7-36345464-DEL-400 |
| chr7-36369082-DEL-105 | chr7-36369083-DEL-105 |
| chr7-36380793-DEL-60 | chr7-36380796-DEL-55 |
| chr7-36381600-DEL-100 | chr7-36381601-DEL-130 |
| chr7-36385012-DEL-79 | chr7-36385008-DEL-79 |
| chr7-36429000-DEL-252 | chr7-36429001-DEL-264 |
| chr7-36447650-DEL-104 | chr7-36447627-DEL-105 |
| chr7-36459336-DEL-54 | chr7-36459337-DEL-52 |
| chr7-36492649-DEL-68 | chr7-36492629-DEL-103 |
| chr7-36503797-DEL-56 | chr7-36503798-DEL-56 |
| chr7-36525180-DEL-58 | chr7-36525181-DEL-58 |
| chr7-36526275-DEL-51 | chr7-36526227-DEL-56 |
| chr7-36538254-DEL-111 | chr7-36538255-DEL-111 |
| chr7-36541841-DEL-299 | chr7-36541842-DEL-299 |
| chr7-36559314-DEL-52 | chr7-36559315-DEL-52 |
| chr7-36568497-DEL-74 | chr7-36568498-DEL-73 |
| chr7-36570299-DEL-59 | chr7-36570288-DEL-60 |
| chr7-36572012-DEL-52 | chr7-36572012-DEL-52 |
| chr7-36608456-DEL-212 | chr7-36608457-DEL-212 |
| chr7-36636570-DEL-80 | chr7-36636569-DEL-82 |
| chr7-36654772-DEL-62 | chr7-36654773-DEL-62 |
| chr7-36655918-DEL-84 | chr7-36655919-DEL-84 |
| chr7-36690533-DEL-647 | chr7-36690534-DEL-647 |
| chr7-36693791-DEL-168 | chr7-36693794-DEL-166 |
| chr7-36726238-DEL-72 | chr7-36726239-DEL-72 |
| chr7-36732763-DEL-126 | chr7-36732764-DEL-126 |
| chr7-36736366-DEL-6698 | chr7-36736367-DEL-6698 |
| chr7-36762520-DEL-155 | chr7-36762521-DEL-155 |
| chr7-36768812-DEL-1432 | chr7-36768813-DEL-1432 |
| chr7-36799065-DEL-62 | chr7-36799066-DEL-62 |
| chr7-36827484-DEL-107 | chr7-36827484-DEL-107 |
| chr7-36847351-DEL-131 | chr7-36847351-DEL-132 |
| chr7-36856552-DEL-62 | chr7-36856553-DEL-62 |
| chr7-36862461-DEL-3392 | chr7-36862462-DEL-3392 |
| chr7-36879004-DEL-65 | chr7-36879004-DEL-58 |
| chr7-36895212-DEL-77 | chr7-36895213-DEL-77 |
| chr7-36906950-DEL-183 | chr7-36906957-DEL-177 |
| chr7-36910100-DEL-324 | chr7-36910105-DEL-316 |
| chr7-36911383-DEL-60 | chr7-36911384-DEL-60 |
| chr7-36932695-DEL-86 | chr7-36932723-DEL-71 |
| chr7-36936488-DEL-92 | chr7-36936473-DEL-134 |
| chr7-36979504-DEL-93 | chr7-36979470-DEL-94 |
| chr7-36982485-DEL-57 | chr7-36982448-DEL-58 |
| chr7-36994279-DEL-112 | chr7-36994280-DEL-112 |
| chr7-37004601-DEL-107 | chr7-37004595-DEL-99 |
| chr7-37025585-DEL-577 | chr7-37025587-DEL-574 |
| chr7-37044729-DEL-60 | chr7-37044730-DEL-60 |
| chr7-37047368-DEL-58 | chr7-37047368-DEL-52 |
| chr7-37052910-DEL-51 | chr7-37052911-DEL-51 |
| chr7-37076606-DEL-165 | chr7-37076619-DEL-153 |
| chr7-37079671-DEL-416 | chr7-37079672-DEL-416 |
| chr7-37084040-DEL-80 | chr7-37084043-DEL-78 |
| chr7-37104096-DEL-75 | chr7-37104097-DEL-76 |
| chr7-37106817-DEL-58 | chr7-37106818-DEL-58 |

| | |
|---|---|
| chr7-37117678-DEL-292 | chr7-37117679-DEL-288 |
| chr7-37221393-DEL-60 | chr7-37221401-DEL-53 |
| chr7-37222266-DEL-56 | chr7-37222269-DEL-52 |
| chr7-37256114-DEL-51 | chr7-37256106-DEL-60 |
| chr7-37279536-DEL-99 | chr7-37279513-DEL-100 |
| chr7-37312417-DEL-728 | chr7-37312418-DEL-728 |
| chr7-37320752-DEL-126 | chr7-37320753-DEL-120 |
| chr7-37343534-DEL-503 | chr7-37343535-DEL-503 |
| chr7-37364950-DEL-92 | chr7-37364951-DEL-92 |
| chr7-37387068-DEL-158 | chr7-37387069-DEL-158 |
| chr7-37397145-DEL-52 | chr7-37397117-DEL-53 |
| chr7-37408090-DEL-64 | chr7-37408104-DEL-66 |
| chr7-37463771-DEL-72 | chr7-37463798-DEL-95 |
| chr7-37481700-DEL-213 | chr7-37481701-DEL-213 |
| chr7-37483632-DEL-64 | chr7-37483641-DEL-56 |
| chr7-37505172-DEL-126 | chr7-37505173-DEL-126 |
| chr7-37513609-DEL-168 | chr7-37513610-DEL-168 |
| chr7-37544919-DEL-51 | chr7-37544911-DEL-52 |
| chr7-37547060-DEL-52 | chr7-37547059-DEL-54 |
| chr7-37600898-DEL-106 | chr7-37600898-DEL-103 |
| chr7-37647306-DEL-70 | chr7-37647307-DEL-70 |
| chr7-37651016-DEL-109 | chr7-37651017-DEL-109 |
| chr7-37679684-DEL-74 | chr7-37679674-DEL-95 |
| chr7-37689294-DEL-401 | chr7-37689295-DEL-401 |
| chr7-37690313-DEL-138 | chr7-37690308-DEL-132 |
| chr7-37698842-DEL-54 | chr7-37698843-DEL-54 |
| chr7-37706371-DEL-63 | chr7-37706372-DEL-63 |
| chr7-37707430-DEL-62 | chr7-37707441-DEL-57 |
| chr7-37711988-DEL-189 | chr7-37711989-DEL-189 |
| chr7-37727660-DEL-109 | chr7-37727661-DEL-109 |
| chr7-37734682-DEL-183 | chr7-37734683-DEL-183 |
| chr7-37745959-DEL-132 | chr7-37745960-DEL-132 |
| chr7-37766435-DEL-211 | chr7-37766428-DEL-209 |
| chr7-37778608-DEL-213 | chr7-37778609-DEL-213 |
| chr7-37779708-DEL-66 | chr7-37779709-DEL-66 |
| chr7-37819887-DEL-66 | chr7-37819888-DEL-66 |
| chr7-37823666-DEL-66 | chr7-37823667-DEL-66 |
| chr7-37831809-DEL-56 | chr7-37831810-DEL-50 |
| chr7-37834620-DEL-197 | chr7-37834621-DEL-197 |
| chr7-37837493-DEL-394 | chr7-37837494-DEL-394 |
| chr7-37847147-DEL-716 | chr7-37847148-DEL-716 |
| chr7-37855634-DEL-476 | chr7-37855635-DEL-476 |
| chr7-37890277-DEL-213 | chr7-37890263-DEL-214 |
| chr7-37916139-DEL-70 | chr7-37916140-DEL-70 |
| chr7-37921785-DEL-54 | chr7-37921786-DEL-54 |
| chr7-37922955-DEL-601 | chr7-37922956-DEL-601 |
| chr7-37943275-DEL-52 | chr7-37943277-DEL-51 |
| chr7-37949944-DEL-96 | chr7-37949945-DEL-96 |
| chr7-37952514-DEL-176 | chr7-37952515-DEL-176 |
| chr7-37971663-DEL-134 | chr7-37971713-DEL-105 |
| chr7-38047173-DEL-476 | chr7-38047172-DEL-476 |
| chr7-38052182-DEL-4817 | chr7-38052181-DEL-4817 |
| chr7-38081206-DEL-463 | chr7-38081207-DEL-463 |
| chr7-38110044-DEL-185 | chr7-38110044-DEL-185 |
| chr7-38196791-DEL-395 | chr7-38196792-DEL-395 |
| chr7-38203819-DEL-59 | chr7-38203819-DEL-66 |
| chr7-38222737-DEL-63 | chr7-38222736-DEL-63 |
| chr7-38495600-DEL-118 | chr7-38495558-DEL-115 |
| chr7-38520892-DEL-82 | chr7-38520903-DEL-56 |
| chr7-38634438-DEL-694 | chr7-38634436-DEL-695 |
| chr7-38657141-DEL-134 | chr7-38657161-DEL-127 |
| chr7-38824355-DEL-53 | chr7-38824356-DEL-87 |
| chr7-38850587-DEL-222 | chr7-38850588-DEL-222 |
| chr7-38877465-DEL-127 | chr7-38877465-DEL-127 |
| chr7-38886475-DEL-495 | chr7-38886475-DEL-495 |
| chr7-38933389-DEL-106 | chr7-38933388-DEL-106 |
| chr7-38937040-DEL-80 | chr7-38937026-DEL-80 |
| chr7-38983126-DEL-451 | chr7-38983125-DEL-451 |
| chr7-39002911-DEL-62 | chr7-39002911-DEL-62 |
| chr7-39005329-DEL-54 | chr7-39005329-DEL-54 |
| chr7-39010715-DEL-84 | chr7-39010735-DEL-94 |
| chr7-39020666-DEL-127 | chr7-39020665-DEL-127 |
| chr7-39087353-DEL-165 | chr7-39087354-DEL-165 |
| chr7-39089067-DEL-684 | chr7-39089066-DEL-686 |
| chr7-39093844-DEL-532 | chr7-39093845-DEL-532 |
| chr7-39115233-DEL-216 | chr7-39115201-DEL-217 |
| chr7-39118892-DEL-106 | chr7-39118893-DEL-106 |
| chr7-39124560-DEL-51 | chr7-39124559-DEL-51 |
| chr7-39124583-DEL-732 | chr7-39124586-DEL-730 |
| chr7-39127347-DEL-58 | chr7-39127354-DEL-52 |

| | |
|---|---|
| chr7-39144053-DEL-755 | chr7-39144056-DEL-753 |
| chr7-39150524-DEL-59 | chr7-39150526-DEL-60 |
| chr7-39154714-DEL-10317 | chr7-39154715-DEL-10317 |
| chr7-39154938-DEL-156 | chr7-39154939-DEL-156 |
| chr7-39171626-DEL-2279 | chr7-39171627-DEL-2279 |
| chr7-39173804-DEL-108 | chr7-39173805-DEL-108 |
| chr7-39176294-DEL-65 | chr7-39176302-DEL-66 |
| chr7-39187835-DEL-387 | chr7-39187834-DEL-388 |
| chr7-39207727-DEL-52 | chr7-39207728-DEL-52 |
| chr7-39237201-DEL-772 | chr7-39237202-DEL-772 |
| chr7-39237913-DEL-3216 | chr7-39237914-DEL-3216 |
| chr7-39238777-DEL-235 | chr7-39238778-DEL-235 |
| chr7-39285914-DEL-166 | chr7-39285916-DEL-147 |
| chr7-39304658-DEL-497 | chr7-39304659-DEL-497 |
| chr7-39316582-DEL-56 | chr7-39316583-DEL-56 |
| chr7-39316738-DEL-420 | chr7-39316739-DEL-420 |
| chr7-39333214-DEL-6409 | chr7-39333215-DEL-6409 |
| chr7-39339273-DEL-207 | chr7-39339274-DEL-207 |
| chr7-39349674-DEL-1334 | chr7-39349673-DEL-1334 |
| chr7-39369077-DEL-354 | chr7-39369078-DEL-354 |
| chr7-39369644-DEL-296 | chr7-39369645-DEL-296 |
| chr7-39389175-DEL-52 | chr7-39389167-DEL-60 |
| chr7-39399613-DEL-69 | chr7-39399614-DEL-69 |
| chr7-39413089-DEL-63 | chr7-39413068-DEL-105 |
| chr7-39427036-DEL-98 | chr7-39427037-DEL-98 |
| chr7-39427490-DEL-4520 | chr7-39427491-DEL-4520 |
| chr7-39463095-DEL-310 | chr7-39463096-DEL-310 |
| chr7-39483143-DEL-285 | chr7-39483144-DEL-285 |
| chr7-39486729-DEL-440 | chr7-39486730-DEL-440 |
| chr7-39488802-DEL-59 | chr7-39488803-DEL-59 |
| chr7-39502731-DEL-111 | chr7-39502728-DEL-109 |
| chr7-39523875-DEL-3026 | chr7-39523875-DEL-3026 |
| chr7-39576835-DEL-112 | chr7-39576837-DEL-104 |
| chr7-39600798-DEL-52 | chr7-39600792-DEL-60 |
| chr7-39614246-DEL-199 | chr7-39614247-DEL-199 |
| chr7-39646258-DEL-144 | chr7-39646262-DEL-141 |
| chr7-39709864-DEL-493 | chr7-39709865-DEL-493 |
| chr7-39737544-DEL-71 | chr7-39737544-DEL-72 |
| chr7-39739072-DEL-124 | chr7-39739073-DEL-124 |
| chr7-39742180-DEL-350 | chr7-39742181-DEL-350 |
| chr7-39744155-DEL-76 | chr7-39744156-DEL-76 |
| chr7-39751421-DEL-291 | chr7-39751422-DEL-285 |
| chr7-39757433-DEL-174 | chr7-39757434-DEL-174 |
| chr7-39758145-DEL-6567 | chr7-39758146-DEL-6567 |
| chr7-39758556-DEL-208 | chr7-39758557-DEL-208 |
| chr7-39790976-DEL-1296 | chr7-39790977-DEL-1296 |
| chr7-39802879-DEL-679 | chr7-39802880-DEL-679 |
| chr7-39809374-DEL-58 | chr7-39809373-DEL-58 |
| chr7-39833827-DEL-57 | chr7-39833828-DEL-57 |
| chr7-39840288-DEL-490 | chr7-39840289-DEL-490 |
| chr7-39845592-DEL-76 | chr7-39845587-DEL-82 |
| chr7-39847549-DEL-983 | chr7-39847550-DEL-983 |
| chr7-39881017-DEL-6391 | chr7-39881018-DEL-6391 |
| chr7-39950371-DEL-247 | chr7-39950372-DEL-247 |
| chr7-39956645-DEL-56 | chr7-39956646-DEL-56 |
| chr7-39972787-DEL-207 | chr7-39972788-DEL-207 |
| chr7-39986421-DEL-6370 | chr7-39986422-DEL-6370 |
| chr7-40011026-DEL-905 | chr7-40011027-DEL-905 |
| chr7-40032323-DEL-576 | chr7-40032324-DEL-576 |
| chr7-40035706-DEL-60 | chr7-40035707-DEL-60 |
| chr7-40037212-DEL-71 | chr7-40037197-DEL-83 |
| chr7-40077118-DEL-54 | chr7-40077119-DEL-54 |
| chr7-40079592-DEL-111 | chr7-40079611-DEL-91 |
| chr7-40095615-DEL-437 | chr7-40095616-DEL-437 |
| chr7-40100467-DEL-68 | chr7-40100468-DEL-98 |
| chr7-40109144-DEL-6192 | chr7-40109145-DEL-6192 |
| chr7-40118574-DEL-390 | chr7-40118575-DEL-390 |
| chr7-40221514-DEL-1116 | chr7-40221515-DEL-1116 |
| chr7-40224884-DEL-52 | chr7-40224878-DEL-59 |
| chr7-40239527-DEL-2060 | chr7-40239528-DEL-2060 |
| chr7-40248358-DEL-579 | chr7-40248359-DEL-579 |
| chr7-40297911-DEL-128 | chr7-40297912-DEL-128 |
| chr7-40335356-DEL-86 | chr7-40335361-DEL-82 |
| chr7-40344538-DEL-2330 | chr7-40344538-DEL-2330 |
| chr7-40349711-DEL-184 | chr7-40349711-DEL-184 |
| chr7-40397171-DEL-111 | chr7-40397172-DEL-111 |
| chr7-40405861-DEL-191 | chr7-40405862-DEL-191 |
| chr7-40413826-DEL-115 | chr7-40413827-DEL-115 |
| chr7-40425482-DEL-60 | chr7-40425483-DEL-60 |
| chr7-40432001-DEL-67 | chr7-40431982-DEL-73 |

| | |
|---|---|
| chr7-40442922-DEL-75 | chr7-40442928-DEL-55 |
| chr7-40564179-DEL-56 | chr7-40564179-DEL-55 |
| chr7-40568614-DEL-88 | chr7-40568615-DEL-88 |
| chr7-40593572-DEL-408 | chr7-40593572-DEL-408 |
| chr7-40601665-DEL-1382 | chr7-40601666-DEL-1382 |
| chr7-40614079-DEL-579 | chr7-40614080-DEL-579 |
| chr7-40656748-DEL-573 | chr7-40656751-DEL-570 |
| chr7-40657605-DEL-665 | chr7-40657606-DEL-665 |
| chr7-40669119-DEL-2318 | chr7-40669119-DEL-2318 |
| chr7-40683341-DEL-6213 | chr7-40683342-DEL-6213 |
| chr7-40693730-DEL-87 | chr7-40693731-DEL-87 |
| chr7-40698965-DEL-70 | chr7-40698982-DEL-54 |
| chr7-40713933-DEL-1284 | chr7-40713934-DEL-1284 |
| chr7-40776111-DEL-403 | chr7-40776110-DEL-403 |
| chr7-40875159-DEL-4767 | chr7-40875158-DEL-4767 |
| chr7-40914095-DEL-1042 | chr7-40914086-DEL-1043 |
| chr7-40925957-DEL-81 | chr7-40925953-DEL-85 |
| chr7-41062459-DEL-206 | chr7-41062460-DEL-206 |
| chr7-41069143-DEL-2204 | chr7-41069144-DEL-2204 |
| chr7-41073643-DEL-80 | chr7-41073644-DEL-80 |
| chr7-41078641-DEL-212 | chr7-41078642-DEL-212 |
| chr7-41078641-DEL-6495 | chr7-41078629-DEL-6470 |
| chr7-41089880-DEL-553 | chr7-41089881-DEL-553 |
| chr7-41103565-DEL-212 | chr7-41103566-DEL-212 |
| chr7-41109207-DEL-86 | chr7-41109208-DEL-86 |
| chr7-41125361-DEL-106 | chr7-41125362-DEL-106 |
| chr7-41134948-DEL-60 | chr7-41134938-DEL-55 |
| chr7-41150686-DEL-233 | chr7-41150687-DEL-233 |
| chr7-41154580-DEL-149 | chr7-41154581-DEL-149 |
| chr7-41173928-DEL-214 | chr7-41173929-DEL-214 |
| chr7-41184964-DEL-2450 | chr7-41184965-DEL-2450 |
| chr7-41185999-DEL-51 | chr7-41185998-DEL-53 |
| chr7-41188396-DEL-769 | chr7-41188396-DEL-769 |
| chr7-41189799-DEL-432 | chr7-41189800-DEL-432 |
| chr7-41202764-DEL-454 | chr7-41202765-DEL-454 |
| chr7-41208247-DEL-138 | chr7-41208251-DEL-132 |
| chr7-41211799-DEL-193 | chr7-41211800-DEL-193 |
| chr7-41213792-DEL-199 | chr7-41213791-DEL-200 |
| chr7-41225017-DEL-563 | chr7-41225018-DEL-563 |
| chr7-41227997-DEL-56 | chr7-41227954-DEL-82 |
| chr7-41239054-DEL-196 | chr7-41239082-DEL-176 |
| chr7-41290221-DEL-85 | chr7-41290218-DEL-89 |
| chr7-41294260-DEL-164 | chr7-41294260-DEL-165 |
| chr7-41300718-DEL-370 | chr7-41300718-DEL-371 |
| chr7-41365294-DEL-52 | chr7-41365294-DEL-52 |
| chr7-41413150-DEL-1571 | chr7-41413151-DEL-1571 |
| chr7-41441274-DEL-6448 | chr7-41441277-DEL-6448 |
| chr7-41447341-DEL-214 | chr7-41447342-DEL-214 |
| chr7-41448730-DEL-2573 | chr7-41448730-DEL-2573 |
| chr7-41459285-DEL-6461 | chr7-41459286-DEL-6461 |
| chr7-41494570-DEL-6821 | chr7-41494571-DEL-6821 |
| chr7-41524528-DEL-70 | chr7-41524564-DEL-78 |
| chr7-41564225-DEL-293 | chr7-41564219-DEL-295 |
| chr7-41588028-DEL-66 | chr7-41588025-DEL-68 |
| chr7-41597610-DEL-316 | chr7-41597611-DEL-315 |
| chr7-41646203-DEL-473 | chr7-41646204-DEL-473 |
| chr7-41689615-DEL-53 | chr7-41689616-DEL-51 |
| chr7-41713245-DEL-57 | chr7-41713246-DEL-57 |
| chr7-41727701-DEL-658 | chr7-41727702-DEL-658 |
| chr7-41771779-DEL-318 | chr7-41771780-DEL-318 |
| chr7-41785477-DEL-81 | chr7-41785501-DEL-86 |
| chr7-41817192-DEL-153 | chr7-41817193-DEL-153 |
| chr7-41819757-DEL-5362 | chr7-41819758-DEL-5362 |
| chr7-41829713-DEL-239 | chr7-41829714-DEL-239 |
| chr7-41851518-DEL-6635 | chr7-41851519-DEL-6635 |
| chr7-41861386-DEL-2667 | chr7-41861386-DEL-2667 |
| chr7-41897318-DEL-413 | chr7-41897319-DEL-413 |
| chr7-41912134-DEL-249 | chr7-41912135-DEL-249 |
| chr7-41921809-DEL-807 | chr7-41921810-DEL-807 |
| chr7-41942446-DEL-880 | chr7-41942448-DEL-879 |
| chr7-41945297-DEL-274 | chr7-41945298-DEL-274 |
| chr7-41954816-DEL-223 | chr7-41954817-DEL-223 |
| chr7-41991993-DEL-67 | chr7-41991994-DEL-67 |
| chr7-41993606-DEL-203 | chr7-41993607-DEL-203 |
| chr7-42007568-DEL-1725 | chr7-42007568-DEL-1725 |
| chr7-42074327-DEL-105 | chr7-42074328-DEL-105 |
| chr7-42097803-DEL-211 | chr7-42097804-DEL-211 |
| chr7-42098217-DEL-463 | chr7-42098218-DEL-463 |
| chr7-42105304-DEL-213 | chr7-42105305-DEL-213 |
| chr7-42120854-DEL-415 | chr7-42120855-DEL-415 |

| | |
|---|---|
| chr7-42154115-DEL-203 | chr7-42154116-DEL-203 |
| chr7-42164426-DEL-8813 | chr7-42164427-DEL-8813 |
| chr7-42194703-DEL-1084 | chr7-42194696-DEL-1084 |
| chr7-42196243-DEL-4497 | chr7-42196245-DEL-4496 |
| chr7-42201611-DEL-213 | chr7-42201612-DEL-213 |
| chr7-42270175-DEL-67 | chr7-42270179-DEL-52 |
| chr7-42290458-DEL-96 | chr7-42290459-DEL-72 |
| chr7-42306651-DEL-102 | chr7-42306614-DEL-102 |
| chr7-42316120-DEL-2326 | chr7-42316121-DEL-2326 |
| chr7-42325269-DEL-237 | chr7-42325270-DEL-237 |
| chr7-42333136-DEL-1214 | chr7-42333137-DEL-1214 |
| chr7-42384613-DEL-62 | chr7-42384614-DEL-62 |
| chr7-42392351-DEL-163 | chr7-42392350-DEL-163 |
| chr7-42476631-DEL-103 | chr7-42476629-DEL-104 |
| chr7-42488642-DEL-68 | chr7-42488612-DEL-104 |
| chr7-42538154-DEL-2887 | chr7-42538155-DEL-2887 |
| chr7-42612382-DEL-94 | chr7-42612383-DEL-94 |
| chr7-42613487-DEL-1058 | chr7-42613488-DEL-1059 |
| chr7-42714463-DEL-1058 | chr7-42714464-DEL-1058 |
| chr7-42728910-DEL-751 | chr7-42728908-DEL-752 |
| chr7-42764601-DEL-63 | chr7-42764600-DEL-64 |
| chr7-42803773-DEL-84 | chr7-42803774-DEL-84 |
| chr7-42815890-DEL-132 | chr7-42815922-DEL-129 |
| chr7-42838928-DEL-54 | chr7-42838929-DEL-54 |
| chr7-42937797-DEL-85 | chr7-42937794-DEL-100 |
| chr7-42972035-DEL-1134 | chr7-42972036-DEL-1134 |
| chr7-42972275-DEL-214 | chr7-42972276-DEL-214 |
| chr7-42973314-DEL-2295 | chr7-42973316-DEL-2293 |
| chr7-42982356-DEL-184 | chr7-42982347-DEL-185 |
| chr7-42983082-DEL-289 | chr7-42983074-DEL-298 |
| chr7-42986379-DEL-1518 | chr7-42986380-DEL-1518 |
| chr7-42986774-DEL-76 | chr7-42986775-DEL-76 |
| chr7-42988119-DEL-173 | chr7-42988115-DEL-174 |
| chr7-43009345-DEL-89 | chr7-43009346-DEL-89 |
| chr7-43014191-DEL-142 | chr7-43014192-DEL-142 |
| chr7-43031413-DEL-103 | chr7-43031415-DEL-102 |
| chr7-43039583-DEL-64 | chr7-43039561-DEL-87 |
| chr7-43065262-DEL-1615 | chr7-43065263-DEL-1615 |
| chr7-43065262-DEL-177 | chr7-43065263-DEL-177 |
| chr7-43099813-DEL-148 | chr7-43099814-DEL-148 |
| chr7-43102043-DEL-60 | chr7-43102044-DEL-60 |
| chr7-43117209-DEL-84 | chr7-43117186-DEL-69 |
| chr7-43120552-DEL-72 | chr7-43120553-DEL-72 |
| chr7-43135341-DEL-52 | chr7-43135342-DEL-52 |
| chr7-43154218-DEL-78 | chr7-43154216-DEL-86 |
| chr7-43157913-DEL-101 | chr7-43157910-DEL-105 |
| chr7-43159334-DEL-153 | chr7-43159335-DEL-153 |
| chr7-43197199-DEL-4566 | chr7-43197200-DEL-4566 |
| chr7-43201305-DEL-201 | chr7-43201306-DEL-201 |
| chr7-43203772-DEL-150 | chr7-43203774-DEL-149 |
| chr7-43204714-DEL-171 | chr7-43204715-DEL-171 |
| chr7-43205618-DEL-458 | chr7-43205619-DEL-458 |
| chr7-43219387-DEL-631 | chr7-43219388-DEL-631 |
| chr7-43231519-DEL-62 | chr7-43231520-DEL-62 |
| chr7-43244908-DEL-58 | chr7-43244908-DEL-58 |
| chr7-43248348-DEL-1287 | chr7-43248349-DEL-1287 |
| chr7-43249641-DEL-87 | chr7-43249640-DEL-82 |
| chr7-43262459-DEL-52 | chr7-43262460-DEL-52 |
| chr7-43318827-DEL-62 | chr7-43318828-DEL-62 |
| chr7-43335980-DEL-837 | chr7-43335981-DEL-837 |
| chr7-43408294-DEL-88 | chr7-43408295-DEL-88 |
| chr7-43464009-DEL-73 | chr7-43464007-DEL-84 |
| chr7-43474217-DEL-295 | chr7-43474218-DEL-295 |
| chr7-43538652-DEL-68 | chr7-43538670-DEL-55 |
| chr7-43543002-DEL-86 | chr7-43543003-DEL-86 |
| chr7-43545540-DEL-586 | chr7-43545540-DEL-586 |
| chr7-43567478-DEL-334 | chr7-43567479-DEL-333 |
| chr7-43568133-DEL-66 | chr7-43568134-DEL-66 |
| chr7-43572488-DEL-265 | chr7-43572489-DEL-265 |
| chr7-43574477-DEL-56 | chr7-43574497-DEL-57 |
| chr7-43582595-DEL-311 | chr7-43582594-DEL-312 |
| chr7-43613372-DEL-64 | chr7-43613373-DEL-64 |
| chr7-43627274-DEL-68 | chr7-43627320-DEL-51 |
| chr7-43633266-DEL-76 | chr7-43633263-DEL-95 |
| chr7-43635992-DEL-223 | chr7-43635993-DEL-223 |
| chr7-43655816-DEL-59 | chr7-43655821-DEL-55 |
| chr7-43670996-DEL-6369 | chr7-43670997-DEL-6369 |
| chr7-43683451-DEL-53 | chr7-43683452-DEL-53 |
| chr7-43694928-DEL-914 | chr7-43694937-DEL-913 |
| chr7-43702312-DEL-88 | chr7-43702313-DEL-88 |

| | |
|---|---|
| chr7-43735250-DEL-69 | chr7-43735251-DEL-69 |
| chr7-43768327-DEL-58 | chr7-43768328-DEL-58 |
| chr7-43805438-DEL-6440 | chr7-43805439-DEL-6440 |
| chr7-43859897-DEL-58 | chr7-43859897-DEL-58 |
| chr7-43928962-DEL-58 | chr7-43928963-DEL-58 |
| chr7-43938800-DEL-214 | chr7-43938801-DEL-214 |
| chr7-43939348-DEL-228 | chr7-43939349-DEL-228 |
| chr7-43996631-DEL-52 | chr7-43996631-DEL-52 |
| chr7-44008185-DEL-85 | chr7-44008155-DEL-105 |
| chr7-44025903-DEL-205 | chr7-44025903-DEL-206 |
| chr7-44029717-DEL-87 | chr7-44029718-DEL-88 |
| chr7-44030811-DEL-86 | chr7-44030818-DEL-80 |
| chr7-44038723-DEL-168 | chr7-44038730-DEL-167 |
| chr7-44048707-DEL-120 | chr7-44048708-DEL-120 |
| chr7-44063101-DEL-138 | chr7-44063102-DEL-138 |
| chr7-44068797-DEL-258 | chr7-44068798-DEL-258 |
| chr7-44133444-DEL-141 | chr7-44133445-DEL-141 |
| chr7-44142121-DEL-76 | chr7-44142122-DEL-76 |
| chr7-44159616-DEL-436 | chr7-44159595-DEL-430 |
| chr7-44195241-DEL-203 | chr7-44195242-DEL-203 |
| chr7-44198633-DEL-215 | chr7-44198623-DEL-216 |
| chr7-44214277-DEL-742 | chr7-44214278-DEL-742 |
| chr7-44216589-DEL-198 | chr7-44216590-DEL-198 |
| chr7-44229082-DEL-212 | chr7-44229083-DEL-183 |
| chr7-44347809-DEL-1407 | chr7-44347810-DEL-1407 |
| chr7-44349581-DEL-159 | chr7-44349578-DEL-160 |
| chr7-44353274-DEL-247 | chr7-44353275-DEL-247 |
| chr7-44374392-DEL-357 | chr7-44374382-DEL-358 |
| chr7-44427319-DEL-96 | chr7-44427313-DEL-97 |
| chr7-44428674-DEL-360 | chr7-44428665-DEL-364 |
| chr7-44434251-DEL-281 | chr7-44434252-DEL-281 |
| chr7-44443907-DEL-207 | chr7-44443908-DEL-207 |
| chr7-44471960-DEL-210 | chr7-44471961-DEL-210 |
| chr7-44524426-DEL-58 | chr7-44524429-DEL-57 |
| chr7-44567941-DEL-70 | chr7-44567942-DEL-70 |
| chr7-44588516-DEL-203 | chr7-44588510-DEL-204 |
| chr7-44591273-DEL-1012 | chr7-44591274-DEL-1012 |
| chr7-44631924-DEL-217 | chr7-44631925-DEL-217 |
| chr7-44638339-DEL-191 | chr7-44638340-DEL-191 |
| chr7-44648424-DEL-219 | chr7-44648425-DEL-219 |
| chr7-44673252-DEL-212 | chr7-44673253-DEL-212 |
| chr7-44678955-DEL-58 | chr7-44678954-DEL-58 |
| chr7-44743049-DEL-56 | chr7-44743050-DEL-56 |
| chr7-44747377-DEL-121 | chr7-44747378-DEL-121 |
| chr7-44754172-DEL-194 | chr7-44754173-DEL-194 |
| chr7-44764858-DEL-74 | chr7-44764860-DEL-64 |
| chr7-44802026-DEL-206 | chr7-44802027-DEL-206 |
| chr7-44861675-DEL-104 | chr7-44861676-DEL-104 |
| chr7-44880198-DEL-201 | chr7-44880199-DEL-201 |
| chr7-44891971-DEL-199 | chr7-44891972-DEL-197 |
| chr7-44899996-DEL-203 | chr7-44899997-DEL-203 |
| chr7-44903706-DEL-1015 | chr7-44903707-DEL-1015 |
| chr7-44919182-DEL-116 | chr7-44919211-DEL-88 |
| chr7-44944325-DEL-223 | chr7-44944326-DEL-223 |
| chr7-44974365-DEL-760 | chr7-44974366-DEL-763 |
| chr7-45008392-DEL-234 | chr7-45008393-DEL-234 |
| chr7-45021630-DEL-55 | chr7-45021631-DEL-55 |
| chr7-45054118-DEL-181 | chr7-45054119-DEL-181 |
| chr7-45061575-DEL-189 | chr7-45061576-DEL-189 |
| chr7-45107107-DEL-52 | chr7-45107108-DEL-52 |
| chr7-45119131-DEL-202 | chr7-45119132-DEL-202 |
| chr7-45122616-DEL-210 | chr7-45122617-DEL-210 |
| chr7-45127703-DEL-206 | chr7-45127704-DEL-206 |
| chr7-45137283-DEL-59 | chr7-45137284-DEL-59 |
| chr7-45154879-DEL-207 | chr7-45154877-DEL-208 |
| chr7-45169656-DEL-207 | chr7-45169657-DEL-207 |
| chr7-45176837-DEL-221 | chr7-45176838-DEL-221 |
| chr7-45195931-DEL-62 | chr7-45195932-DEL-62 |
| chr7-45202358-DEL-69 | chr7-45202359-DEL-69 |
| chr7-45207204-DEL-52 | chr7-45207205-DEL-52 |
| chr7-45213155-DEL-86 | chr7-45213156-DEL-86 |
| chr7-45227428-DEL-203 | chr7-45227429-DEL-203 |
| chr7-45236199-DEL-86 | chr7-45236200-DEL-86 |
| chr7-45267283-DEL-204 | chr7-45267284-DEL-204 |
| chr7-45281141-DEL-138 | chr7-45281147-DEL-131 |
| chr7-45308276-DEL-174 | chr7-45308277-DEL-174 |
| chr7-45344968-DEL-69 | chr7-45344969-DEL-69 |
| chr7-45389848-DEL-410 | chr7-45389873-DEL-386 |
| chr7-45397269-DEL-101 | chr7-45397301-DEL-117 |
| chr7-45400970-DEL-104 | chr7-45400971-DEL-104 |

| | |
|---|---|
| chr7-45405794-DEL-66 | chr7-45405832-DEL-61 |
| chr7-45410944-DEL-211 | chr7-45410945-DEL-211 |
| chr7-45423743-DEL-63 | chr7-45423735-DEL-57 |
| chr7-45445827-DEL-57 | chr7-45445828-DEL-61 |
| chr7-45473449-DEL-133 | chr7-45473450-DEL-133 |
| chr7-45496398-DEL-111 | chr7-45496406-DEL-108 |
| chr7-45497021-DEL-391 | chr7-45497022-DEL-391 |
| chr7-45498691-DEL-92 | chr7-45498692-DEL-92 |
| chr7-45532027-DEL-57 | chr7-45532027-DEL-55 |
| chr7-45550608-DEL-52 | chr7-45550609-DEL-52 |
| chr7-45552697-DEL-120 | chr7-45552696-DEL-100 |
| chr7-45553540-DEL-95 | chr7-45553511-DEL-103 |
| chr7-45578127-DEL-58 | chr7-45578089-DEL-56 |
| chr7-45581137-DEL-305 | chr7-45581141-DEL-304 |
| chr7-45592605-DEL-217 | chr7-45592606-DEL-217 |
| chr7-45600099-DEL-422 | chr7-45600100-DEL-422 |
| chr7-45601530-DEL-60 | chr7-45601531-DEL-60 |
| chr7-45602442-DEL-655 | chr7-45602443-DEL-655 |
| chr7-45603603-DEL-12900 | chr7-45603604-DEL-12900 |
| chr7-45604330-DEL-72 | chr7-45604331-DEL-72 |
| chr7-45607072-DEL-52 | chr7-45607073-DEL-52 |
| chr7-45623175-DEL-52 | chr7-45623176-DEL-52 |
| chr7-45654002-DEL-161 | chr7-45654003-DEL-161 |
| chr7-45656745-DEL-128 | chr7-45656767-DEL-114 |
| chr7-45659893-DEL-107 | chr7-45659894-DEL-106 |
| chr7-45660688-DEL-126 | chr7-45660689-DEL-126 |
| chr7-45661728-DEL-78 | chr7-45661729-DEL-78 |
| chr7-45667063-DEL-74 | chr7-45667017-DEL-72 |
| chr7-45671823-DEL-88 | chr7-45671824-DEL-88 |
| chr7-45673634-DEL-69 | chr7-45673633-DEL-70 |
| chr7-45678943-DEL-173 | chr7-45678944-DEL-173 |
| chr7-45696601-DEL-401 | chr7-45696602-DEL-401 |
| chr7-45698387-DEL-59 | chr7-45698386-DEL-60 |
| chr7-45700313-DEL-604 | chr7-45700314-DEL-604 |
| chr7-45706355-DEL-75 | chr7-45706356-DEL-75 |
| chr7-45707998-DEL-123 | chr7-45707998-DEL-123 |
| chr7-45750938-DEL-57 | chr7-45750937-DEL-59 |
| chr7-45791245-DEL-342 | chr7-45791246-DEL-342 |
| chr7-45794906-DEL-58 | chr7-45794907-DEL-58 |
| chr7-45798487-DEL-80 | chr7-45798489-DEL-79 |
| chr7-45800542-DEL-200 | chr7-45800543-DEL-200 |
| chr7-45818635-DEL-449 | chr7-45818627-DEL-453 |
| chr7-45830496-DEL-132 | chr7-45830541-DEL-110 |
| chr7-45877562-DEL-872 | chr7-45877565-DEL-870 |
| chr7-45893424-DEL-92 | chr7-45893458-DEL-56 |
| chr7-45895137-DEL-63 | chr7-45895137-DEL-63 |
| chr7-45940122-DEL-66 | chr7-45940123-DEL-66 |
| chr7-45953871-DEL-209 | chr7-45953872-DEL-209 |
| chr7-45956760-DEL-418 | chr7-45956761-DEL-418 |
| chr7-46015825-DEL-82 | chr7-46015826-DEL-82 |
| chr7-46021206-DEL-106 | chr7-46021207-DEL-106 |
| chr7-46102836-DEL-75 | chr7-46102837-DEL-81 |
| chr7-46127039-DEL-51 | chr7-46127040-DEL-51 |
| chr7-46129632-DEL-170 | chr7-46129631-DEL-170 |
| chr7-46161578-DEL-428 | chr7-46161579-DEL-428 |
| chr7-46171298-DEL-72 | chr7-46171295-DEL-72 |
| chr7-46247957-DEL-61 | chr7-46247975-DEL-61 |
| chr7-46281474-DEL-63 | chr7-46281477-DEL-57 |
| chr7-46313739-DEL-157 | chr7-46313745-DEL-146 |
| chr7-46340463-DEL-634 | chr7-46340464-DEL-634 |
| chr7-46371547-DEL-60 | chr7-46371557-DEL-51 |
| chr7-46371798-DEL-831 | chr7-46371799-DEL-831 |
| chr7-46377966-DEL-73 | chr7-46377942-DEL-56 |
| chr7-46406244-DEL-856 | chr7-46406245-DEL-856 |
| chr7-46419540-DEL-119 | chr7-46419541-DEL-119 |
| chr7-46426660-DEL-199 | chr7-46426661-DEL-199 |
| chr7-46436272-DEL-190 | chr7-46436267-DEL-194 |
| chr7-46450700-DEL-109 | chr7-46450700-DEL-104 |
| chr7-46457691-DEL-202 | chr7-46457692-DEL-202 |
| chr7-46460829-DEL-145 | chr7-46460830-DEL-145 |
| chr7-46473299-DEL-56 | chr7-46473300-DEL-56 |
| chr7-46474406-DEL-202 | chr7-46474407-DEL-202 |
| chr7-46476031-DEL-110 | chr7-46476032-DEL-110 |
| chr7-46499239-DEL-197 | chr7-46499240-DEL-197 |
| chr7-46506424-DEL-209 | chr7-46506425-DEL-209 |
| chr7-46507705-DEL-86 | chr7-46507722-DEL-70 |
| chr7-46512747-DEL-1394 | chr7-46512748-DEL-1394 |
| chr7-46531947-DEL-202 | chr7-46531948-DEL-202 |
| chr7-46532059-DEL-926 | chr7-46532060-DEL-926 |
| chr7-46552375-DEL-201 | chr7-46552376-DEL-201 |

| | |
|---|---|
| chr7-46573082-DEL-148 | chr7-46573083-DEL-148 |
| chr7-46579181-DEL-209 | chr7-46579182-DEL-209 |
| chr7-46582306-DEL-220 | chr7-46582308-DEL-214 |
| chr7-46591704-DEL-224 | chr7-46591705-DEL-224 |
| chr7-46599375-DEL-63 | chr7-46599360-DEL-96 |
| chr7-46623153-DEL-382 | chr7-46623154-DEL-382 |
| chr7-46650375-DEL-52 | chr7-46650376-DEL-52 |
| chr7-46668690-DEL-56 | chr7-46668699-DEL-54 |
| chr7-46727078-DEL-208 | chr7-46727079-DEL-208 |
| chr7-46824473-DEL-86 | chr7-46824474-DEL-86 |
| chr7-46835719-DEL-2684 | chr7-46835720-DEL-2684 |
| chr7-46971783-DEL-439 | chr7-46971783-DEL-439 |
| chr7-46981048-DEL-1189 | chr7-46981048-DEL-1189 |
| chr7-46991085-DEL-281 | chr7-46991086-DEL-281 |
| chr7-47009516-DEL-105 | chr7-47009517-DEL-105 |
| chr7-47009918-DEL-458 | chr7-47009918-DEL-458 |
| chr7-47035927-DEL-105 | chr7-47035928-DEL-105 |
| chr7-47053324-DEL-2827 | chr7-47053324-DEL-2827 |
| chr7-47060159-DEL-52 | chr7-47060160-DEL-52 |
| chr7-47068379-DEL-131 | chr7-47068380-DEL-131 |
| chr7-47140026-DEL-60 | chr7-47140027-DEL-60 |
| chr7-47146929-DEL-889 | chr7-47146930-DEL-889 |
| chr7-47148928-DEL-127 | chr7-47148929-DEL-127 |
| chr7-47155280-DEL-7326 | chr7-47155281-DEL-7326 |
| chr7-47172543-DEL-415 | chr7-47172544-DEL-415 |
| chr7-47192858-DEL-100 | chr7-47192879-DEL-100 |
| chr7-47224412-DEL-2327 | chr7-47224413-DEL-2327 |
| chr7-47293752-DEL-756 | chr7-47293753-DEL-756 |
| chr7-47331507-DEL-249 | chr7-47331508-DEL-249 |
| chr7-47341803-DEL-82 | chr7-47341812-DEL-74 |
| chr7-47375643-DEL-108 | chr7-47375658-DEL-92 |
| chr7-47378962-DEL-3285 | chr7-47378960-DEL-3270 |
| chr7-47385129-DEL-297 | chr7-47385129-DEL-297 |
| chr7-47387040-DEL-71 | chr7-47387040-DEL-71 |
| chr7-47403057-DEL-270 | chr7-47403056-DEL-270 |
| chr7-47410994-DEL-118 | chr7-47410997-DEL-110 |
| chr7-47415029-DEL-53 | chr7-47415029-DEL-53 |
| chr7-47419729-DEL-73 | chr7-47419728-DEL-73 |
| chr7-47424344-DEL-377 | chr7-47424344-DEL-378 |
| chr7-47441518-DEL-4131 | chr7-47441518-DEL-4131 |
| chr7-47483386-DEL-66 | chr7-47483338-DEL-65 |
| chr7-47523241-DEL-583 | chr7-47523242-DEL-583 |
| chr7-47574810-DEL-989 | chr7-47574810-DEL-989 |
| chr7-47593827-DEL-405 | chr7-47593827-DEL-405 |
| chr7-47597124-DEL-505 | chr7-47597124-DEL-505 |
| chr7-47613272-DEL-65 | chr7-47613247-DEL-91 |
| chr7-47613339-DEL-1589 | chr7-47613340-DEL-1589 |
| chr7-47615457-DEL-183 | chr7-47615464-DEL-186 |
| chr7-47616571-DEL-546 | chr7-47616572-DEL-546 |
| chr7-47629651-DEL-123 | chr7-47629652-DEL-122 |
| chr7-47646940-DEL-6163 | chr7-47646929-DEL-6164 |
| chr7-47664998-DEL-473 | chr7-47664999-DEL-472 |
| chr7-47675130-DEL-1752 | chr7-47675131-DEL-1752 |
| chr7-47675328-DEL-545 | chr7-47675326-DEL-547 |
| chr7-47689679-DEL-1457 | chr7-47689684-DEL-1454 |
| chr7-47690590-DEL-1134 | chr7-47690591-DEL-1134 |
| chr7-47692493-DEL-436 | chr7-47692494-DEL-436 |
| chr7-47696028-DEL-64 | chr7-47696029-DEL-64 |
| chr7-47698604-DEL-1095 | chr7-47698617-DEL-1081 |
| chr7-47711758-DEL-256 | chr7-47711761-DEL-254 |
| chr7-47720397-DEL-914 | chr7-47720437-DEL-918 |
| chr7-47739829-DEL-140 | chr7-47739828-DEL-138 |
| chr7-47741926-DEL-120 | chr7-47741965-DEL-82 |
| chr7-47752258-DEL-2869 | chr7-47752259-DEL-2869 |
| chr7-47757248-DEL-1383 | chr7-47757248-DEL-1383 |
| chr7-47801523-DEL-399 | chr7-47801524-DEL-399 |
| chr7-47807101-DEL-332 | chr7-47807111-DEL-331 |
| chr7-47810369-DEL-160 | chr7-47810373-DEL-155 |
| chr7-47813440-DEL-670 | chr7-47813441-DEL-670 |
| chr7-47814975-DEL-81 | chr7-47814976-DEL-81 |
| chr7-47822994-DEL-1553 | chr7-47822985-DEL-1563 |
| chr7-47845643-DEL-924 | chr7-47845639-DEL-928 |
| chr7-47863194-DEL-171 | chr7-47863194-DEL-172 |
| chr7-47870057-DEL-135 | chr7-47870061-DEL-133 |
| chr7-47943196-DEL-1005 | chr7-47943198-DEL-1004 |
| chr7-47947608-DEL-7332 | chr7-47947609-DEL-7332 |
| chr7-47999370-DEL-61 | chr7-47999370-DEL-61 |
| chr7-48011484-DEL-66 | chr7-48011485-DEL-66 |
| chr7-48016054-DEL-6588 | chr7-48016055-DEL-6588 |
| chr7-48041192-DEL-703 | chr7-48041195-DEL-701 |

| | |
|---|---|
| chr7-48081379-DEL-399 | chr7-48081380-DEL-399 |
| chr7-48095304-DEL-67 | chr7-48095305-DEL-67 |
| chr7-48098156-DEL-101 | chr7-48098155-DEL-102 |
| chr7-48107683-DEL-3077 | chr7-48107684-DEL-3077 |
| chr7-48132974-DEL-52 | chr7-48132975-DEL-52 |
| chr7-48136951-DEL-5516 | chr7-48136952-DEL-5522 |
| chr7-48156300-DEL-6984 | chr7-48156301-DEL-6984 |
| chr7-48167854-DEL-344 | chr7-48167855-DEL-344 |
| chr7-48191494-DEL-6382 | chr7-48191495-DEL-6382 |
| chr7-48209336-DEL-200 | chr7-48209337-DEL-200 |
| chr7-48247171-DEL-195 | chr7-48247172-DEL-195 |
| chr7-48270401-DEL-242 | chr7-48270402-DEL-242 |
| chr7-48282421-DEL-7361 | chr7-48282422-DEL-7361 |
| chr7-48338362-DEL-57 | chr7-48338363-DEL-57 |
| chr7-48353765-DEL-53 | chr7-48353766-DEL-53 |
| chr7-48354460-DEL-56 | chr7-48354458-DEL-64 |
| chr7-48365915-DEL-129 | chr7-48365911-DEL-127 |
| chr7-48372347-DEL-55 | chr7-48372348-DEL-55 |
| chr7-48388524-DEL-4967 | chr7-48388525-DEL-4967 |
| chr7-48400305-DEL-626 | chr7-48400306-DEL-626 |
| chr7-48401405-DEL-200 | chr7-48401406-DEL-199 |
| chr7-48418015-DEL-327 | chr7-48418012-DEL-328 |
| chr7-48425313-DEL-492 | chr7-48425314-DEL-492 |
| chr7-48502012-DEL-166 | chr7-48502013-DEL-166 |
| chr7-48534908-DEL-68 | chr7-48534928-DEL-50 |
| chr7-48588522-DEL-116 | chr7-48588517-DEL-117 |
| chr7-48589040-DEL-56 | chr7-48589063-DEL-68 |
| chr7-48625342-DEL-620 | chr7-48625343-DEL-620 |
| chr7-48629320-DEL-251 | chr7-48629321-DEL-251 |
| chr7-48648565-DEL-61 | chr7-48648538-DEL-62 |
| chr7-48723580-DEL-124 | chr7-48723581-DEL-124 |
| chr7-48730594-DEL-1119 | chr7-48730595-DEL-1119 |
| chr7-48740950-DEL-123 | chr7-48740951-DEL-123 |
| chr7-48742791-DEL-221 | chr7-48742792-DEL-221 |
| chr7-48752633-DEL-62 | chr7-48752634-DEL-62 |
| chr7-48780930-DEL-57 | chr7-48780926-DEL-58 |
| chr7-48784576-DEL-219 | chr7-48784577-DEL-219 |
| chr7-48788984-DEL-92 | chr7-48788973-DEL-104 |
| chr7-48798935-DEL-119 | chr7-48798939-DEL-89 |
| chr7-48834607-DEL-66 | chr7-48834608-DEL-66 |
| chr7-48867265-DEL-2252 | chr7-48867266-DEL-2252 |
| chr7-48926572-DEL-68 | chr7-48926569-DEL-84 |
| chr7-48934712-DEL-366 | chr7-48934713-DEL-366 |
| chr7-48957021-DEL-54 | chr7-48957022-DEL-54 |
| chr7-48973484-DEL-77 | chr7-48973486-DEL-76 |
| chr7-48995806-DEL-206 | chr7-48995807-DEL-206 |
| chr7-48998921-DEL-206 | chr7-48998922-DEL-206 |
| chr7-48999880-DEL-76 | chr7-48999877-DEL-53 |
| chr7-49009500-DEL-223 | chr7-49009509-DEL-232 |
| chr7-49035258-DEL-145 | chr7-49035259-DEL-145 |
| chr7-49055151-DEL-1314 | chr7-49055152-DEL-1314 |
| chr7-49067290-DEL-297 | chr7-49067291-DEL-297 |
| chr7-49082137-DEL-110 | chr7-49082138-DEL-110 |
| chr7-49095734-DEL-64 | chr7-49095733-DEL-50 |
| chr7-49105097-DEL-228 | chr7-49105098-DEL-228 |
| chr7-49112834-DEL-193 | chr7-49112832-DEL-195 |
| chr7-49113554-DEL-214 | chr7-49113555-DEL-214 |
| chr7-49126228-DEL-178 | chr7-49126213-DEL-179 |
| chr7-49136970-DEL-68 | chr7-49136970-DEL-66 |
| chr7-49187440-DEL-54 | chr7-49187440-DEL-52 |
| chr7-49200040-DEL-78 | chr7-49200041-DEL-78 |
| chr7-49210806-DEL-212 | chr7-49210807-DEL-212 |
| chr7-49211428-DEL-192 | chr7-49211429-DEL-192 |
| chr7-49219950-DEL-210 | chr7-49219937-DEL-211 |
| chr7-49236891-DEL-54 | chr7-49236883-DEL-53 |
| chr7-49271514-DEL-64 | chr7-49271507-DEL-72 |
| chr7-49275740-DEL-204 | chr7-49275741-DEL-204 |
| chr7-49279885-DEL-217 | chr7-49279886-DEL-217 |
| chr7-49294986-DEL-605 | chr7-49294987-DEL-605 |
| chr7-49297763-DEL-52 | chr7-49297764-DEL-52 |
| chr7-49311508-DEL-152 | chr7-49311509-DEL-152 |
| chr7-49314853-DEL-59 | chr7-49314844-DEL-61 |
| chr7-49341516-DEL-64 | chr7-49341517-DEL-64 |
| chr7-49362119-DEL-711 | chr7-49362120-DEL-711 |
| chr7-49364918-DEL-1527 | chr7-49364918-DEL-1527 |
| chr7-49372917-DEL-209 | chr7-49372910-DEL-210 |
| chr7-49380402-DEL-82 | chr7-49380403-DEL-52 |
| chr7-49383009-DEL-58 | chr7-49383010-DEL-58 |
| chr7-49385021-DEL-319 | chr7-49385022-DEL-319 |
| chr7-49385468-DEL-1618 | chr7-49385469-DEL-1618 |

| | |
|---|---|
| chr7-49394683-DEL-900 | chr7-49394684-DEL-900 |
| chr7-49477126-DEL-225 | chr7-49477127-DEL-225 |
| chr7-49485803-DEL-337 | chr7-49485804-DEL-337 |
| chr7-49503051-DEL-86 | chr7-49503024-DEL-87 |
| chr7-49575783-DEL-52 | chr7-49575782-DEL-53 |
| chr7-49600985-DEL-59 | chr7-49600986-DEL-59 |
| chr7-49607637-DEL-160 | chr7-49607638-DEL-160 |
| chr7-49629254-DEL-85 | chr7-49629255-DEL-85 |
| chr7-49634685-DEL-132 | chr7-49634686-DEL-132 |
| chr7-49644551-DEL-515 | chr7-49644552-DEL-515 |
| chr7-49658548-DEL-152 | chr7-49658549-DEL-152 |
| chr7-49679437-DEL-72 | chr7-49679438-DEL-72 |
| chr7-49683617-DEL-356 | chr7-49683618-DEL-356 |
| chr7-49687793-DEL-152 | chr7-49687798-DEL-148 |
| chr7-49691938-DEL-83 | chr7-49691946-DEL-60 |
| chr7-49703753-DEL-6958 | chr7-49703754-DEL-6958 |
| chr7-49720561-DEL-56 | chr7-49720562-DEL-56 |
| chr7-49724002-DEL-473 | chr7-49724002-DEL-473 |
| chr7-49740110-DEL-80 | chr7-49740123-DEL-97 |
| chr7-49758474-DEL-6896 | chr7-49758475-DEL-6896 |
| chr7-49791687-DEL-7435 | chr7-49791688-DEL-7435 |
| chr7-49797678-DEL-832 | chr7-49797678-DEL-832 |
| chr7-49821047-DEL-803 | chr7-49821048-DEL-803 |
| chr7-49845724-DEL-6318 | chr7-49845725-DEL-6318 |
| chr7-49878822-DEL-66 | chr7-49878834-DEL-54 |
| chr7-49882093-DEL-7213 | chr7-49882094-DEL-7213 |
| chr7-49948611-DEL-6449 | chr7-49948612-DEL-6449 |
| chr7-49975068-DEL-1898 | chr7-49975068-DEL-1898 |
| chr7-50009315-DEL-1917 | chr7-50009315-DEL-1917 |
| chr7-50014755-DEL-7234 | chr7-50014755-DEL-7234 |
| chr7-50026718-DEL-528 | chr7-50026715-DEL-532 |
| chr7-50033928-DEL-186 | chr7-50033930-DEL-183 |
| chr7-50041344-DEL-200 | chr7-50041345-DEL-200 |
| chr7-50045233-DEL-238 | chr7-50045234-DEL-238 |
| chr7-50049955-DEL-164 | chr7-50049956-DEL-164 |
| chr7-50055085-DEL-71 | chr7-50055086-DEL-67 |
| chr7-50101345-DEL-6248 | chr7-50101346-DEL-6248 |
| chr7-50125348-DEL-89 | chr7-50125349-DEL-89 |
| chr7-50173458-DEL-6870 | chr7-50173459-DEL-6870 |
| chr7-50179808-DEL-212 | chr7-50179809-DEL-212 |
| chr7-50200587-DEL-67 | chr7-50200588-DEL-67 |
| chr7-50216874-DEL-1548 | chr7-50216875-DEL-1548 |
| chr7-50220671-DEL-305 | chr7-50220672-DEL-305 |
| chr7-50240392-DEL-399 | chr7-50240393-DEL-399 |
| chr7-50244336-DEL-52 | chr7-50244335-DEL-54 |
| chr7-50259806-DEL-265 | chr7-50259807-DEL-265 |
| chr7-50271898-DEL-2949 | chr7-50271899-DEL-2949 |
| chr7-50285494-DEL-53 | chr7-50285497-DEL-51 |
| chr7-50291207-DEL-409 | chr7-50291200-DEL-414 |
| chr7-50318676-DEL-2065 | chr7-50318677-DEL-2065 |
| chr7-50402197-DEL-1967 | chr7-50402198-DEL-1967 |
| chr7-50434800-DEL-79 | chr7-50434809-DEL-55 |
| chr7-50475466-DEL-54 | chr7-50475476-DEL-71 |
| chr7-50491031-DEL-204 | chr7-50491032-DEL-204 |
| chr7-50518067-DEL-63 | chr7-50518067-DEL-63 |
| chr7-50523234-DEL-6312 | chr7-50523234-DEL-6312 |
| chr7-50544671-DEL-4447 | chr7-50544672-DEL-4447 |
| chr7-50556053-DEL-80 | chr7-50556054-DEL-80 |
| chr7-50583322-DEL-51 | chr7-50583315-DEL-51 |
| chr7-50638821-DEL-150 | chr7-50638822-DEL-150 |
| chr7-50640186-DEL-80 | chr7-50640187-DEL-80 |
| chr7-50676719-DEL-259 | chr7-50676720-DEL-259 |
| chr7-50718919-DEL-101 | chr7-50718903-DEL-81 |
| chr7-50721519-DEL-72 | chr7-50721520-DEL-72 |
| chr7-50725215-DEL-6047 | chr7-50725215-DEL-6043 |
| chr7-50767126-DEL-7522 | chr7-50767127-DEL-7522 |
| chr7-50776308-DEL-6618 | chr7-50776309-DEL-6618 |
| chr7-50776441-DEL-402 | chr7-50776442-DEL-402 |
| chr7-50794574-DEL-83 | chr7-50794575-DEL-83 |
| chr7-50810208-DEL-1111 | chr7-50810209-DEL-1111 |
| chr7-50829266-DEL-85 | chr7-50829257-DEL-81 |
| chr7-50829682-DEL-287 | chr7-50829683-DEL-287 |
| chr7-50830439-DEL-1474 | chr7-50830439-DEL-1474 |
| chr7-50831971-DEL-4424 | chr7-50831972-DEL-4424 |
| chr7-50833053-DEL-992 | chr7-50833054-DEL-991 |
| chr7-50865651-DEL-1126 | chr7-50865648-DEL-1127 |
| chr7-50912024-DEL-60 | chr7-50912025-DEL-60 |
| chr7-50920915-DEL-8880 | chr7-50920916-DEL-8880 |
| chr7-50930001-DEL-88 | chr7-50929993-DEL-104 |
| chr7-50951284-DEL-121 | chr7-50951285-DEL-123 |

| | |
|---|---|
| chr7-50956333-DEL-60 | chr7-50956334-DEL-60 |
| chr7-50977122-DEL-957 | chr7-50977123-DEL-986 |
| chr7-50979251-DEL-54 | chr7-50979252-DEL-54 |
| chr7-50979416-DEL-6220 | chr7-50979417-DEL-6220 |
| chr7-50990393-DEL-74 | chr7-50990400-DEL-75 |
| chr7-50993412-DEL-72 | chr7-50993413-DEL-72 |
| chr7-51015230-DEL-86 | chr7-51015224-DEL-93 |
| chr7-51016484-DEL-116 | chr7-51016485-DEL-116 |
| chr7-51021030-DEL-152 | chr7-51021042-DEL-104 |
| chr7-51042271-DEL-68 | chr7-51042272-DEL-68 |
| chr7-51053058-DEL-56 | chr7-51053065-DEL-50 |
| chr7-51068828-DEL-5404 | chr7-51068829-DEL-5404 |
| chr7-51085508-DEL-759 | chr7-51085508-DEL-759 |
| chr7-51118634-DEL-7216 | chr7-51118635-DEL-7216 |
| chr7-51140077-DEL-53 | chr7-51140071-DEL-54 |
| chr7-51140929-DEL-379 | chr7-51140930-DEL-379 |
| chr7-51184457-DEL-236 | chr7-51184458-DEL-236 |
| chr7-51193066-DEL-860 | chr7-51193067-DEL-860 |
| chr7-51217555-DEL-705 | chr7-51217556-DEL-705 |
| chr7-51221638-DEL-70 | chr7-51221639-DEL-70 |
| chr7-51252890-DEL-84 | chr7-51252891-DEL-84 |
| chr7-51259945-DEL-935 | chr7-51259946-DEL-935 |
| chr7-51264937-DEL-253 | chr7-51264938-DEL-253 |
| chr7-51323221-DEL-76 | chr7-51323222-DEL-76 |
| chr7-51325757-DEL-553 | chr7-51325758-DEL-550 |
| chr7-51336626-DEL-90 | chr7-51336642-DEL-75 |
| chr7-51360518-DEL-177 | chr7-51360519-DEL-177 |
| chr7-51379232-DEL-97 | chr7-51379214-DEL-80 |
| chr7-51385358-DEL-502 | chr7-51385359-DEL-502 |
| chr7-51396361-DEL-1273 | chr7-51396362-DEL-1271 |
| chr7-51453216-DEL-4611 | chr7-51453230-DEL-4613 |
| chr7-51468855-DEL-52 | chr7-51468855-DEL-52 |
| chr7-51480945-DEL-240 | chr7-51480946-DEL-240 |
| chr7-51482835-DEL-618 | chr7-51482836-DEL-618 |
| chr7-51505588-DEL-105 | chr7-51505589-DEL-105 |
| chr7-51508555-DEL-597 | chr7-51508556-DEL-597 |
| chr7-51547255-DEL-196 | chr7-51547256-DEL-196 |
| chr7-51553634-DEL-482 | chr7-51553634-DEL-483 |
| chr7-51584962-DEL-52 | chr7-51584963-DEL-52 |
| chr7-51651588-DEL-77 | chr7-51651581-DEL-85 |
| chr7-51673122-DEL-1684 | chr7-51673123-DEL-1684 |
| chr7-51679718-DEL-4976 | chr7-51679719-DEL-4976 |
| chr7-51683437-DEL-1792 | chr7-51683438-DEL-1792 |
| chr7-51700181-DEL-166 | chr7-51700182-DEL-166 |
| chr7-51705941-DEL-58 | chr7-51705942-DEL-58 |
| chr7-51716633-DEL-204 | chr7-51716634-DEL-204 |
| chr7-51717961-DEL-1665 | chr7-51717962-DEL-1665 |
| chr7-51727159-DEL-483 | chr7-51727160-DEL-483 |
| chr7-51731455-DEL-1676 | chr7-51731456-DEL-1676 |
| chr7-51734752-DEL-117 | chr7-51734753-DEL-117 |
| chr7-51739854-DEL-58 | chr7-51739855-DEL-50 |
| chr7-51740740-DEL-6191 | chr7-51740741-DEL-6191 |
| chr7-51750978-DEL-248 | chr7-51750979-DEL-248 |
| chr7-51751725-DEL-2421 | chr7-51751726-DEL-2421 |
| chr7-51774293-DEL-63 | chr7-51774283-DEL-74 |
| chr7-51779887-DEL-343 | chr7-51779888-DEL-343 |
| chr7-51788303-DEL-6403 | chr7-51788304-DEL-6403 |
| chr7-51799734-DEL-59 | chr7-51799734-DEL-59 |
| chr7-51800579-DEL-72 | chr7-51800580-DEL-72 |
| chr7-51804722-DEL-122 | chr7-51804724-DEL-129 |
| chr7-51819193-DEL-55 | chr7-51819194-DEL-55 |
| chr7-51837049-DEL-906 | chr7-51837051-DEL-905 |
| chr7-51866861-DEL-399 | chr7-51866862-DEL-399 |
| chr7-51867528-DEL-52 | chr7-51867514-DEL-76 |
| chr7-51869525-DEL-1173 | chr7-51869526-DEL-1173 |
| chr7-51870080-DEL-88 | chr7-51870086-DEL-77 |
| chr7-51876155-DEL-190 | chr7-51876158-DEL-188 |
| chr7-51878098-DEL-2030 | chr7-51878099-DEL-2030 |
| chr7-51881839-DEL-5260 | chr7-51881840-DEL-5260 |
| chr7-51888563-DEL-597 | chr7-51888564-DEL-597 |
| chr7-51891668-DEL-56 | chr7-51891668-DEL-56 |
| chr7-51907086-DEL-1058 | chr7-51907087-DEL-1058 |
| chr7-51908922-DEL-57 | chr7-51908923-DEL-57 |
| chr7-51911701-DEL-394 | chr7-51911702-DEL-394 |
| chr7-51931551-DEL-765 | chr7-51931552-DEL-763 |
| chr7-51952837-DEL-620 | chr7-51952838-DEL-620 |
| chr7-51963755-DEL-165 | chr7-51963756-DEL-165 |
| chr7-51975499-DEL-8419 | chr7-51975500-DEL-8419 |
| chr7-52022249-DEL-64 | chr7-52022236-DEL-60 |
| chr7-52031527-DEL-898 | chr7-52031527-DEL-898 |

| | |
|---|---|
| chr7-52037842-DEL-160 | chr7-52037843-DEL-160 |
| chr7-52060745-DEL-302 | chr7-52060746-DEL-302 |
| chr7-52071769-DEL-78 | chr7-52071766-DEL-78 |
| chr7-52072941-DEL-199 | chr7-52072942-DEL-199 |
| chr7-52094610-DEL-1618 | chr7-52094606-DEL-1622 |
| chr7-52146420-DEL-88 | chr7-52146421-DEL-102 |
| chr7-52161974-DEL-1783 | chr7-52161975-DEL-1783 |
| chr7-52164289-DEL-2881 | chr7-52164289-DEL-2880 |
| chr7-52172315-DEL-2977 | chr7-52172316-DEL-2977 |
| chr7-52187806-DEL-5966 | chr7-52187807-DEL-5966 |
| chr7-52200156-DEL-90 | chr7-52200157-DEL-90 |
| chr7-52205659-DEL-2436 | chr7-52205660-DEL-2436 |
| chr7-52208977-DEL-1652 | chr7-52208978-DEL-1652 |
| chr7-52220521-DEL-189 | chr7-52220522-DEL-189 |
| chr7-52287570-DEL-222 | chr7-52287571-DEL-222 |
| chr7-52289610-DEL-184 | chr7-52289611-DEL-184 |
| chr7-52292560-DEL-1884 | chr7-52292561-DEL-1884 |
| chr7-52322811-DEL-113 | chr7-52322804-DEL-109 |
| chr7-52358944-DEL-971 | chr7-52358945-DEL-971 |
| chr7-52372671-DEL-90 | chr7-52372640-DEL-123 |
| chr7-52380604-DEL-1215 | chr7-52380605-DEL-1215 |
| chr7-52408606-DEL-68 | chr7-52408607-DEL-62 |
| chr7-52419182-DEL-6461 | chr7-52419164-DEL-6479 |
| chr7-52444472-DEL-214 | chr7-52444473-DEL-214 |
| chr7-52465369-DEL-6271 | chr7-52465370-DEL-6271 |
| chr7-52499523-DEL-6411 | chr7-52499524-DEL-6411 |
| chr7-52513161-DEL-2226 | chr7-52513162-DEL-2226 |
| chr7-52523501-DEL-617 | chr7-52523502-DEL-617 |
| chr7-52583863-DEL-95 | chr7-52583857-DEL-95 |
| chr7-52594644-DEL-1419 | chr7-52594645-DEL-1419 |
| chr7-52603413-DEL-3411 | chr7-52603414-DEL-3411 |
| chr7-52616179-DEL-59 | chr7-52616179-DEL-60 |
| chr7-52628178-DEL-77 | chr7-52628144-DEL-67 |
| chr7-52629738-DEL-70 | chr7-52629739-DEL-70 |
| chr7-52675888-DEL-753 | chr7-52675888-DEL-753 |
| chr7-52680851-DEL-265 | chr7-52680852-DEL-265 |
| chr7-52691356-DEL-497 | chr7-52691357-DEL-497 |
| chr7-52691915-DEL-130 | chr7-52691916-DEL-130 |
| chr7-52704424-DEL-548 | chr7-52704425-DEL-548 |
| chr7-52708295-DEL-126 | chr7-52708296-DEL-126 |
| chr7-52729303-DEL-1459 | chr7-52729304-DEL-1459 |
| chr7-52747353-DEL-486 | chr7-52747354-DEL-486 |
| chr7-52756722-DEL-1873 | chr7-52756723-DEL-1873 |
| chr7-52813843-DEL-1893 | chr7-52813844-DEL-1893 |
| chr7-52840970-DEL-52 | chr7-52840970-DEL-58 |
| chr7-52848999-DEL-372 | chr7-52849000-DEL-371 |
| chr7-52851874-DEL-128 | chr7-52851874-DEL-128 |
| chr7-52863915-DEL-3517 | chr7-52863916-DEL-3517 |
| chr7-52868437-DEL-139 | chr7-52868438-DEL-139 |
| chr7-52886766-DEL-432 | chr7-52886767-DEL-432 |
| chr7-52895448-DEL-1484 | chr7-52895449-DEL-1484 |
| chr7-52903166-DEL-558 | chr7-52903167-DEL-558 |
| chr7-52914913-DEL-64 | chr7-52914918-DEL-60 |
| chr7-52922657-DEL-71 | chr7-52922658-DEL-67 |
| chr7-52981177-DEL-4064 | chr7-52981178-DEL-4064 |
| chr7-52992390-DEL-998 | chr7-52992391-DEL-998 |
| chr7-52998796-DEL-429 | chr7-52998797-DEL-429 |
| chr7-53008541-DEL-407 | chr7-53008542-DEL-407 |
| chr7-53021182-DEL-80 | chr7-53021163-DEL-100 |
| chr7-53023762-DEL-306 | chr7-53023763-DEL-304 |
| chr7-53040938-DEL-2943 | chr7-53040939-DEL-2944 |
| chr7-53061371-DEL-55 | chr7-53061362-DEL-56 |
| chr7-53112097-DEL-2695 | chr7-53112096-DEL-2695 |
| chr7-53123887-DEL-233 | chr7-53123889-DEL-232 |
| chr7-53134104-DEL-454 | chr7-53134105-DEL-454 |
| chr7-53139631-DEL-2559 | chr7-53139632-DEL-2559 |
| chr7-53147683-DEL-2628 | chr7-53147684-DEL-2628 |
| chr7-53174362-DEL-100 | chr7-53174373-DEL-74 |
| chr7-53175328-DEL-6175 | chr7-53175329-DEL-6175 |
| chr7-53197059-DEL-314 | chr7-53197060-DEL-314 |
| chr7-53207285-DEL-6651 | chr7-53207286-DEL-6651 |
| chr7-53240272-DEL-56 | chr7-53240272-DEL-56 |
| chr7-53244180-DEL-580 | chr7-53244180-DEL-580 |
| chr7-53245896-DEL-214 | chr7-53245897-DEL-214 |
| chr7-53322371-DEL-109 | chr7-53322372-DEL-109 |
| chr7-53325725-DEL-77 | chr7-53325738-DEL-65 |
| chr7-53327101-DEL-5813 | chr7-53327095-DEL-5819 |
| chr7-53345942-DEL-690 | chr7-53345943-DEL-690 |
| chr7-53391464-DEL-417 | chr7-53391462-DEL-418 |
| chr7-53418983-DEL-59 | chr7-53418983-DEL-59 |

| | |
|---|---|
| chr7-53447115-DEL-1588 | chr7-53447116-DEL-1588 |
| chr7-53455492-DEL-370 | chr7-53455492-DEL-370 |
| chr7-53463970-DEL-244 | chr7-53463971-DEL-244 |
| chr7-53470353-DEL-451 | chr7-53470354-DEL-451 |
| chr7-53471416-DEL-4550 | chr7-53471417-DEL-4550 |
| chr7-53493503-DEL-1894 | chr7-53493504-DEL-1894 |
| chr7-53499845-DEL-699 | chr7-53499846-DEL-699 |
| chr7-53499870-DEL-101 | chr7-53499871-DEL-101 |
| chr7-53505294-DEL-6367 | chr7-53505295-DEL-6367 |
| chr7-53512121-DEL-6160 | chr7-53512130-DEL-6160 |
| chr7-53521519-DEL-185 | chr7-53521520-DEL-185 |
| chr7-53529520-DEL-750 | chr7-53529521-DEL-750 |
| chr7-53532981-DEL-70 | chr7-53532984-DEL-66 |
| chr7-53534605-DEL-5510 | chr7-53534606-DEL-5510 |
| chr7-53550808-DEL-151 | chr7-53550808-DEL-151 |
| chr7-53573699-DEL-573 | chr7-53573700-DEL-573 |
| chr7-53583353-DEL-220 | chr7-53583354-DEL-220 |
| chr7-53592522-DEL-75 | chr7-53592524-DEL-74 |
| chr7-53595369-DEL-205 | chr7-53595370-DEL-205 |
| chr7-53601062-DEL-819 | chr7-53601067-DEL-820 |
| chr7-53606599-DEL-102 | chr7-53606600-DEL-102 |
| chr7-53622297-DEL-399 | chr7-53622298-DEL-399 |
| chr7-53635914-DEL-177 | chr7-53635915-DEL-177 |
| chr7-53650155-DEL-1191 | chr7-53650156-DEL-1191 |
| chr7-53669314-DEL-8222 | chr7-53669315-DEL-8222 |
| chr7-53673165-DEL-1857 | chr7-53673163-DEL-1858 |
| chr7-53708564-DEL-1668 | chr7-53708565-DEL-1668 |
| chr7-53710302-DEL-2078 | chr7-53710303-DEL-2078 |
| chr7-53734276-DEL-486 | chr7-53734277-DEL-486 |
| chr7-53736214-DEL-6157 | chr7-53736214-DEL-6157 |
| chr7-53758602-DEL-6284 | chr7-53758602-DEL-6284 |
| chr7-53797597-DEL-310 | chr7-53797598-DEL-310 |
| chr7-53813089-DEL-1323 | chr7-53813090-DEL-1323 |
| chr7-53854516-DEL-61 | chr7-53854517-DEL-60 |
| chr7-53874923-DEL-413 | chr7-53874924-DEL-413 |
| chr7-53904654-DEL-94 | chr7-53904655-DEL-94 |
| chr7-53960869-DEL-659 | chr7-53960870-DEL-659 |
| chr7-53970601-DEL-208 | chr7-53970602-DEL-208 |
| chr7-53977136-DEL-70 | chr7-53977141-DEL-52 |
| chr7-53985642-DEL-2382 | chr7-53985642-DEL-2382 |
| chr7-54016553-DEL-54 | chr7-54016547-DEL-51 |
| chr7-54025593-DEL-204 | chr7-54025594-DEL-204 |
| chr7-54031279-DEL-7008 | chr7-54031278-DEL-7008 |
| chr7-54042928-DEL-2155 | chr7-54042929-DEL-2155 |
| chr7-54061620-DEL-652 | chr7-54061621-DEL-652 |
| chr7-54069814-DEL-596 | chr7-54069815-DEL-596 |
| chr7-54091487-DEL-51 | chr7-54091473-DEL-65 |
| chr7-54091886-DEL-2012 | chr7-54091887-DEL-2012 |
| chr7-54127659-DEL-163 | chr7-54127681-DEL-154 |
| chr7-54143794-DEL-863 | chr7-54143795-DEL-863 |
| chr7-54149598-DEL-339 | chr7-54149593-DEL-340 |
| chr7-54175830-DEL-6592 | chr7-54175831-DEL-6592 |
| chr7-54209235-DEL-56 | chr7-54209236-DEL-54 |
| chr7-54231967-DEL-1278 | chr7-54231956-DEL-1290 |
| chr7-54236908-DEL-153 | chr7-54236910-DEL-152 |
| chr7-54237167-DEL-522 | chr7-54237169-DEL-522 |
| chr7-54250659-DEL-775 | chr7-54250660-DEL-775 |
| chr7-54279687-DEL-1059 | chr7-54279688-DEL-1059 |
| chr7-54296265-DEL-192 | chr7-54296244-DEL-194 |
| chr7-54313822-DEL-87 | chr7-54313795-DEL-115 |
| chr7-54332047-DEL-1036 | chr7-54332048-DEL-1036 |
| chr7-54374914-DEL-1047 | chr7-54374910-DEL-1043 |
| chr7-54377728-DEL-1608 | chr7-54377717-DEL-1612 |
| chr7-54389013-DEL-80 | chr7-54389030-DEL-51 |
| chr7-54400785-DEL-2815 | chr7-54400786-DEL-2815 |
| chr7-54415648-DEL-1502 | chr7-54415649-DEL-1502 |
| chr7-54427752-DEL-245 | chr7-54427753-DEL-245 |
| chr7-54457736-DEL-122 | chr7-54457737-DEL-122 |
| chr7-54458616-DEL-1788 | chr7-54458628-DEL-1786 |
| chr7-54517283-DEL-450 | chr7-54517284-DEL-450 |
| chr7-54518821-DEL-749 | chr7-54518821-DEL-742 |
| chr7-54548691-DEL-86 | chr7-54548704-DEL-72 |
| chr7-54580803-DEL-74 | chr7-54580803-DEL-62 |
| chr7-54604716-DEL-593 | chr7-54604718-DEL-592 |
| chr7-54614984-DEL-6366 | chr7-54614986-DEL-6364 |
| chr7-54621019-DEL-220 | chr7-54621020-DEL-220 |
| chr7-54661200-DEL-6303 | chr7-54661201-DEL-6303 |
| chr7-54699820-DEL-298 | chr7-54699815-DEL-297 |
| chr7-54703553-DEL-569 | chr7-54703554-DEL-569 |
| chr7-54729644-DEL-6519 | chr7-54729645-DEL-6519 |

| | |
|---|---|
| chr7-54744591-DEL-52 | chr7-54744591-DEL-52 |
| chr7-54751995-DEL-1331 | chr7-54751999-DEL-1327 |
| chr7-54764517-DEL-6605 | chr7-54764518-DEL-6605 |
| chr7-54783858-DEL-6904 | chr7-54783859-DEL-6904 |
| chr7-54837429-DEL-720 | chr7-54837430-DEL-720 |
| chr7-54845277-DEL-2326 | chr7-54845278-DEL-2326 |
| chr7-54857107-DEL-344 | chr7-54857108-DEL-344 |
| chr7-54868450-DEL-65 | chr7-54868451-DEL-65 |
| chr7-54869062-DEL-1751 | chr7-54869063-DEL-1751 |
| chr7-54893831-DEL-51 | chr7-54893832-DEL-54 |
| chr7-54901400-DEL-99 | chr7-54901403-DEL-95 |
| chr7-54906425-DEL-99 | chr7-54906443-DEL-82 |
| chr7-54945364-DEL-76 | chr7-54945365-DEL-76 |
| chr7-54946240-DEL-2448 | chr7-54946241-DEL-2448 |
| chr7-54959643-DEL-229 | chr7-54959644-DEL-229 |
| chr7-54959970-DEL-1508 | chr7-54959971-DEL-1508 |
| chr7-55047759-DEL-797 | chr7-55047760-DEL-796 |
| chr7-55058969-DEL-3035 | chr7-55058970-DEL-3035 |
| chr7-55059849-DEL-75 | chr7-55059850-DEL-75 |
| chr7-55067576-DEL-462 | chr7-55067577-DEL-456 |
| chr7-55095932-DEL-84 | chr7-55095929-DEL-88 |
| chr7-55103873-DEL-6440 | chr7-55103874-DEL-6440 |
| chr7-55118596-DEL-56 | chr7-55118597-DEL-56 |
| chr7-55160642-DEL-102 | chr7-55160643-DEL-97 |
| chr7-55185303-DEL-173 | chr7-55185304-DEL-173 |
| chr7-55200953-DEL-513 | chr7-55200954-DEL-513 |
| chr7-55231677-DEL-330 | chr7-55231678-DEL-330 |
| chr7-55253375-DEL-1596 | chr7-55253376-DEL-1596 |
| chr7-55307147-DEL-1233 | chr7-55307148-DEL-1233 |
| chr7-55327519-DEL-206 | chr7-55327520-DEL-206 |
| chr7-55330101-DEL-2354 | chr7-55330102-DEL-2354 |
| chr7-55336577-DEL-199 | chr7-55336578-DEL-199 |
| chr7-55338401-DEL-2181 | chr7-55338401-DEL-2181 |
| chr7-55363686-DEL-395 | chr7-55363687-DEL-395 |
| chr7-55378361-DEL-347 | chr7-55378356-DEL-348 |
| chr7-55385285-DEL-581 | chr7-55385285-DEL-581 |
| chr7-55391945-DEL-85 | chr7-55391970-DEL-63 |
| chr7-55392659-DEL-139 | chr7-55392661-DEL-138 |
| chr7-55402621-DEL-1744 | chr7-55402622-DEL-1745 |
| chr7-55415209-DEL-68 | chr7-55415210-DEL-68 |
| chr7-55434987-DEL-266 | chr7-55434988-DEL-266 |
| chr7-55440116-DEL-523 | chr7-55440117-DEL-523 |
| chr7-55444153-DEL-55 | chr7-55444154-DEL-55 |
| chr7-55463976-DEL-350 | chr7-55463977-DEL-350 |
| chr7-55479353-DEL-603 | chr7-55479354-DEL-604 |
| chr7-55503326-DEL-192 | chr7-55503327-DEL-192 |
| chr7-55516438-DEL-69 | chr7-55516442-DEL-68 |
| chr7-55524311-DEL-68 | chr7-55524312-DEL-68 |
| chr7-55542230-DEL-89 | chr7-55542229-DEL-83 |
| chr7-55573355-DEL-56 | chr7-55573328-DEL-54 |
| chr7-55596509-DEL-7192 | chr7-55596510-DEL-7192 |
| chr7-55631372-DEL-51 | chr7-55631373-DEL-51 |
| chr7-55640272-DEL-198 | chr7-55640273-DEL-197 |
| chr7-55663665-DEL-517 | chr7-55663666-DEL-511 |
| chr7-55673288-DEL-67 | chr7-55673289-DEL-67 |
| chr7-55681192-DEL-6555 | chr7-55681193-DEL-6555 |
| chr7-55692866-DEL-109 | chr7-55692867-DEL-109 |
| chr7-55724807-DEL-6702 | chr7-55724808-DEL-6702 |
| chr7-55748386-DEL-55 | chr7-55748387-DEL-55 |
| chr7-55768928-DEL-377 | chr7-55768918-DEL-378 |
| chr7-55779309-DEL-105 | chr7-55779306-DEL-66 |
| chr7-55789782-DEL-189 | chr7-55789783-DEL-189 |
| chr7-55797263-DEL-255 | chr7-55797264-DEL-255 |
| chr7-55866756-DEL-689 | chr7-55866757-DEL-689 |
| chr7-55895593-DEL-86 | chr7-55895598-DEL-98 |
| chr7-55908799-DEL-54 | chr7-55908800-DEL-54 |
| chr7-55909319-DEL-218 | chr7-55909320-DEL-218 |
| chr7-55909992-DEL-211 | chr7-55909993-DEL-211 |
| chr7-55913227-DEL-60 | chr7-55913229-DEL-53 |
| chr7-55920143-DEL-5580 | chr7-55920144-DEL-5580 |
| chr7-55928800-DEL-86 | chr7-55928799-DEL-71 |
| chr7-55942279-DEL-390 | chr7-55942280-DEL-390 |
| chr7-55980246-DEL-211 | chr7-55980247-DEL-211 |
| chr7-55982762-DEL-519 | chr7-55982763-DEL-519 |
| chr7-55986158-DEL-4361 | chr7-55986159-DEL-4361 |
| chr7-55991751-DEL-6060 | chr7-55991752-DEL-6060 |
| chr7-56001657-DEL-98 | chr7-56001658-DEL-98 |
| chr7-56013087-DEL-339 | chr7-56013088-DEL-339 |
| chr7-56016116-DEL-58 | chr7-56016111-DEL-59 |
| chr7-56021603-DEL-614 | chr7-56021604-DEL-614 |

| | |
|---|---|
| chr7-56053113-DEL-66 | chr7-56053115-DEL-65 |
| chr7-56061106-DEL-57 | chr7-56061094-DEL-57 |
| chr7-56071483-DEL-136 | chr7-56071488-DEL-132 |
| chr7-56080169-DEL-119 | chr7-56080170-DEL-119 |
| chr7-56102528-DEL-53 | chr7-56102523-DEL-58 |
| chr7-56112621-DEL-657 | chr7-56112622-DEL-657 |
| chr7-56114233-DEL-514 | chr7-56114234-DEL-514 |
| chr7-56115441-DEL-4057 | chr7-56115442-DEL-4057 |
| chr7-56127307-DEL-894 | chr7-56127308-DEL-894 |
| chr7-56154391-DEL-126 | chr7-56154389-DEL-127 |
| chr7-56161442-DEL-6275 | chr7-56161443-DEL-6275 |
| chr7-56169987-DEL-92 | chr7-56169991-DEL-91 |
| chr7-56182303-DEL-162 | chr7-56182304-DEL-162 |
| chr7-56199659-DEL-462 | chr7-56199660-DEL-458 |
| chr7-56220926-DEL-6742 | chr7-56220927-DEL-6742 |
| chr7-56227112-DEL-424 | chr7-56227112-DEL-424 |
| chr7-56234104-DEL-874 | chr7-56234097-DEL-875 |
| chr7-56264019-DEL-1001 | chr7-56264019-DEL-1001 |
| chr7-56273747-DEL-609 | chr7-56273748-DEL-609 |
| chr7-56310041-DEL-5377 | chr7-56310042-DEL-5377 |
| chr7-56336341-DEL-52 | chr7-56336346-DEL-50 |
| chr7-56336799-DEL-6507 | chr7-56336794-DEL-6512 |
| chr7-56394639-DEL-5445 | chr7-56394640-DEL-5445 |
| chr7-56401097-DEL-1027 | chr7-56401098-DEL-1027 |
| chr7-56418931-DEL-2649 | chr7-56418932-DEL-2649 |
| chr7-56426957-DEL-1460 | chr7-56426958-DEL-1460 |
| chr7-56432840-DEL-181 | chr7-56432841-DEL-181 |
| chr7-56443900-DEL-64 | chr7-56443893-DEL-76 |
| chr7-56451215-DEL-134 | chr7-56451216-DEL-134 |
| chr7-56453932-DEL-1003 | chr7-56453933-DEL-1003 |
| chr7-56464399-DEL-65 | chr7-56464400-DEL-65 |
| chr7-56483210-DEL-52 | chr7-56483211-DEL-52 |
| chr7-56515094-DEL-1495 | chr7-56515095-DEL-1495 |
| chr7-56544296-DEL-190 | chr7-56544297-DEL-190 |
| chr7-56549203-DEL-119 | chr7-56549178-DEL-117 |
| chr7-56562894-DEL-1428 | chr7-56562895-DEL-1428 |
| chr7-56567449-DEL-100 | chr7-56567479-DEL-104 |
| chr7-56590885-DEL-167 | chr7-56590889-DEL-166 |
| chr7-56596426-DEL-494 | chr7-56596426-DEL-494 |
| chr7-56601279-DEL-6574 | chr7-56601279-DEL-6574 |
| chr7-56633215-DEL-91 | chr7-56633216-DEL-91 |
| chr7-56648662-DEL-343 | chr7-56648663-DEL-343 |
| chr7-56656561-DEL-1530 | chr7-56656562-DEL-1530 |
| chr7-56664512-DEL-1526 | chr7-56664513-DEL-1526 |
| chr7-56688948-DEL-572 | chr7-56688948-DEL-573 |
| chr7-56705613-DEL-69 | chr7-56705614-DEL-69 |
| chr7-56711609-DEL-1849 | chr7-56711610-DEL-1849 |
| chr7-56740451-DEL-344 | chr7-56740452-DEL-344 |
| chr7-56751126-DEL-199 | chr7-56751127-DEL-199 |
| chr7-56764731-DEL-4205 | chr7-56764732-DEL-4205 |
| chr7-56777956-DEL-4203 | chr7-56777941-DEL-4219 |
| chr7-56789200-DEL-4124 | chr7-56789201-DEL-4124 |
| chr7-56822903-DEL-92 | chr7-56822904-DEL-92 |
| chr7-56832180-DEL-2667 | chr7-56832181-DEL-2667 |
| chr7-56855158-DEL-721 | chr7-56855159-DEL-718 |
| chr7-56870443-DEL-2857 | chr7-56870444-DEL-2857 |
| chr7-56875660-DEL-1538 | chr7-56875658-DEL-1538 |
| chr7-56900199-DEL-113 | chr7-56900199-DEL-113 |
| chr7-56901371-DEL-4432 | chr7-56901372-DEL-4432 |
| chr7-56910150-DEL-347 | chr7-56910152-DEL-346 |
| chr7-56913805-DEL-1699 | chr7-56913806-DEL-1699 |
| chr7-56952299-DEL-71 | chr7-56952303-DEL-68 |
| chr7-56960748-DEL-94 | chr7-56960749-DEL-94 |
| chr7-56992258-DEL-136 | chr7-56992259-DEL-135 |
| chr7-57005264-DEL-498 | chr7-57005265-DEL-498 |
| chr7-57008834-DEL-919 | chr7-57008835-DEL-919 |
| chr7-57016920-DEL-7542 | chr7-57016921-DEL-7542 |
| chr7-57066433-DEL-115 | chr7-57066436-DEL-116 |
| chr7-57076003-DEL-273 | chr7-57076004-DEL-273 |
| chr7-57089143-DEL-71 | chr7-57089144-DEL-71 |
| chr7-57105454-DEL-496 | chr7-57105455-DEL-496 |
| chr7-57110655-DEL-785 | chr7-57110656-DEL-785 |
| chr7-57115825-DEL-1391 | chr7-57115826-DEL-1392 |
| chr7-57161799-DEL-458 | chr7-57161800-DEL-458 |
| chr7-57168102-DEL-80 | chr7-57168107-DEL-76 |
| chr7-57173409-DEL-141 | chr7-57173415-DEL-136 |
| chr7-57187073-DEL-112 | chr7-57187072-DEL-113 |
| chr7-57193123-DEL-225 | chr7-57193124-DEL-225 |
| chr7-57194580-DEL-416 | chr7-57194581-DEL-416 |
| chr7-57208929-DEL-120 | chr7-57208930-DEL-120 |

| | |
|---|---|
| chr7-57298849-DEL-89 | chr7-57298847-DEL-86 |
| chr7-57299732-DEL-100 | chr7-57299733-DEL-100 |
| chr7-57303239-DEL-142 | chr7-57303239-DEL-142 |
| chr7-57309897-DEL-173 | chr7-57309898-DEL-173 |
| chr7-57345673-DEL-484 | chr7-57345674-DEL-484 |
| chr7-57377660-DEL-138 | chr7-57377658-DEL-148 |
| chr7-57387552-DEL-94 | chr7-57387553-DEL-94 |
| chr7-57400244-DEL-157 | chr7-57400245-DEL-157 |
| chr7-57446088-DEL-338 | chr7-57446089-DEL-338 |
| chr7-57463133-DEL-155 | chr7-57463134-DEL-155 |
| chr7-57504497-DEL-1890 | chr7-57504479-DEL-1911 |
| chr7-57527709-DEL-3350 | chr7-57527708-DEL-3351 |
| chr7-57549287-DEL-265 | chr7-57549288-DEL-265 |
| chr7-57551475-DEL-2222 | chr7-57551476-DEL-2221 |
| chr7-57553967-DEL-136 | chr7-57553968-DEL-136 |
| chr7-57590210-DEL-500 | chr7-57590211-DEL-500 |
| chr7-57590993-DEL-1081 | chr7-57590994-DEL-1081 |
| chr7-57596425-DEL-148 | chr7-57596431-DEL-142 |
| chr7-57637927-DEL-68 | chr7-57637947-DEL-79 |
| chr7-57666009-DEL-951 | chr7-57666010-DEL-951 |
| chr7-57668839-DEL-65 | chr7-57668832-DEL-66 |
| chr7-57688122-DEL-214 | chr7-57688123-DEL-214 |
| chr7-57704179-DEL-640 | chr7-57704180-DEL-633 |
| chr7-57706935-DEL-7022 | chr7-57706927-DEL-7023 |
| chr7-57721787-DEL-2796 | chr7-57721787-DEL-2796 |
| chr7-57731825-DEL-575 | chr7-57731826-DEL-575 |
| chr7-57787126-DEL-229 | chr7-57787126-DEL-229 |
| chr7-57803234-DEL-76 | chr7-57803235-DEL-76 |
| chr7-57806302-DEL-113 | chr7-57806303-DEL-113 |
| chr7-57815320-DEL-1456 | chr7-57815320-DEL-1456 |
| chr7-57877440-DEL-60 | chr7-57877442-DEL-51 |
| chr7-57895074-DEL-74 | chr7-57895097-DEL-72 |
| chr7-57938389-DEL-6173 | chr7-57938390-DEL-6173 |
| chr7-57954979-DEL-378 | chr7-57954980-DEL-378 |
| chr7-57981058-DEL-2042 | chr7-57981059-DEL-2041 |
| chr7-57984777-DEL-66 | chr7-57984778-DEL-68 |
| chr7-58029620-DEL-73 | chr7-58029615-DEL-74 |
| chr7-58037703-DEL-213 | chr7-58037704-DEL-213 |
| chr7-58064339-DEL-264 | chr7-58064340-DEL-264 |
| chr7-58071464-DEL-6637 | chr7-58071465-DEL-6637 |
| chr7-58099313-DEL-56 | chr7-58099315-DEL-52 |
| chr7-58099365-DEL-944 | chr7-58099366-DEL-944 |
| chr7-58110186-DEL-199 | chr7-58110187-DEL-198 |
| chr7-58151632-DEL-74 | chr7-58151633-DEL-74 |
| chr7-58161908-DEL-161 | chr7-58161907-DEL-154 |
| chr7-58185427-DEL-6215 | chr7-58185428-DEL-6215 |
| chr7-58238416-DEL-6689 | chr7-58238417-DEL-6689 |
| chr7-58318448-DEL-95 | chr7-58318446-DEL-96 |
| chr7-58324107-DEL-93 | chr7-58324104-DEL-94 |
| chr7-58370416-DEL-60 | chr7-58370417-DEL-60 |
| chr7-58374339-DEL-445 | chr7-58374340-DEL-445 |
| chr7-58387179-DEL-400 | chr7-58387180-DEL-400 |
| chr7-58407729-DEL-65 | chr7-58407728-DEL-65 |
| chr7-58430921-DEL-858 | chr7-58430920-DEL-859 |
| chr7-58455322-DEL-793 | chr7-58455323-DEL-793 |
| chr7-58458273-DEL-142 | chr7-58458266-DEL-137 |
| chr7-58475844-DEL-190 | chr7-58475853-DEL-182 |
| chr7-58489482-DEL-56 | chr7-58489469-DEL-67 |
| chr7-58497362-DEL-6213 | chr7-58497363-DEL-6213 |
| chr7-58532482-DEL-7213 | chr7-58532483-DEL-7213 |
| chr7-58532831-DEL-208 | chr7-58532832-DEL-208 |
| chr7-58576947-DEL-85 | chr7-58576948-DEL-85 |
| chr7-58603570-DEL-58 | chr7-58603571-DEL-58 |
| chr7-58697272-DEL-56 | chr7-58697273-DEL-66 |
| chr7-58728642-DEL-808 | chr7-58728643-DEL-808 |
| chr7-58741950-DEL-6463 | chr7-58741957-DEL-6459 |
| chr7-58749408-DEL-12421 | chr7-58749409-DEL-12421 |
| chr7-58767912-DEL-152 | chr7-58767913-DEL-152 |
| chr7-58774921-DEL-6404 | chr7-58774922-DEL-6404 |
| chr7-58799875-DEL-409 | chr7-58799876-DEL-409 |
| chr7-58817525-DEL-106 | chr7-58817547-DEL-85 |
| chr7-58853527-DEL-7087 | chr7-58853528-DEL-7087 |
| chr7-58861177-DEL-1189 | chr7-58861178-DEL-1189 |
| chr7-58914216-DEL-125 | chr7-58914217-DEL-125 |
| chr7-58937246-DEL-171 | chr7-58937247-DEL-171 |
| chr7-58944263-DEL-5752 | chr7-58944264-DEL-5752 |
| chr7-59046514-DEL-3539 | chr7-59046485-DEL-3540 |
| chr7-59272602-DEL-385 | chr7-59272603-DEL-385 |
| chr7-59610581-DEL-180 | chr7-59610582-DEL-180 |
| chr7-59644955-DEL-66 | chr7-59644991-DEL-62 |

| | |
|---|---|
| chr7-59666894-DEL-240 | chr7-59666895-DEL-240 |
| chr7-59670260-DEL-384 | chr7-59670261-DEL-379 |
| chr7-59708090-DEL-2500 | chr7-59708091-DEL-2500 |
| chr7-59712594-DEL-110 | chr7-59712591-DEL-105 |
| chr7-59715145-DEL-204 | chr7-59715146-DEL-204 |
| chr7-59724710-DEL-1174 | chr7-59724711-DEL-1174 |
| chr7-59775640-DEL-52 | chr7-59775637-DEL-56 |
| chr7-59788046-DEL-112 | chr7-59788047-DEL-112 |
| chr7-59790657-DEL-68 | chr7-59790658-DEL-68 |
| chr7-59793510-DEL-636 | chr7-59793511-DEL-635 |
| chr7-59797535-DEL-950 | chr7-59797536-DEL-950 |
| chr7-59805272-DEL-8455 | chr7-59805262-DEL-8456 |
| chr7-59828517-DEL-1501 | chr7-59828518-DEL-1500 |
| chr7-59838130-DEL-187 | chr7-59838131-DEL-187 |
| chr7-59851323-DEL-1295 | chr7-59851324-DEL-1295 |
| chr7-59859106-DEL-6230 | chr7-59859093-DEL-6243 |
| chr7-59882812-DEL-751 | chr7-59882813-DEL-751 |
| chr7-59883939-DEL-2542 | chr7-59883940-DEL-2542 |
| chr7-59972111-DEL-6300 | chr7-59972112-DEL-6300 |
| chr7-60053058-DEL-51 | chr7-60053059-DEL-51 |
| chr7-60057102-DEL-84 | chr7-60057103-DEL-86 |
| chr7-60095206-DEL-852 | chr7-60095207-DEL-852 |
| chr7-60109353-DEL-4804 | chr7-60109323-DEL-4804 |
| chr7-60112724-DEL-535 | chr7-60112725-DEL-535 |
| chr7-60149070-DEL-53 | chr7-60149038-DEL-54 |
| chr7-60158888-DEL-201 | chr7-60158887-DEL-201 |
| chr7-60180771-DEL-7630 | chr7-60180772-DEL-7630 |
| chr7-60191572-DEL-56 | chr7-60191573-DEL-56 |
| chr7-60193270-DEL-66 | chr7-60193271-DEL-66 |
| chr7-60231213-DEL-1170 | chr7-60231212-DEL-1170 |
| chr7-60251565-DEL-510 | chr7-60251566-DEL-510 |
| chr7-60259070-DEL-58 | chr7-60259071-DEL-58 |
| chr7-60263597-DEL-98 | chr7-60263598-DEL-98 |
| chr7-60278973-DEL-63 | chr7-60278974-DEL-63 |
| chr7-60304525-DEL-92 | chr7-60304525-DEL-92 |
| chr7-60309630-DEL-61 | chr7-60309637-DEL-62 |
| chr7-60310407-DEL-5165 | chr7-60310408-DEL-5165 |
| chr7-60313101-DEL-61 | chr7-60313101-DEL-61 |
| chr7-60326005-DEL-6601 | chr7-60326008-DEL-6597 |
| chr7-60332141-DEL-208 | chr7-60332142-DEL-208 |
| chr7-60346126-DEL-540 | chr7-60346127-DEL-540 |
| chr7-60386727-DEL-56 | chr7-60386728-DEL-56 |
| chr7-60391982-DEL-875 | chr7-60391983-DEL-875 |
| chr7-60425285-DEL-6815 | chr7-60425286-DEL-6815 |
| chr7-60425330-DEL-636 | chr7-60425331-DEL-636 |
| chr7-60473134-DEL-346 | chr7-60473134-DEL-342 |
| chr7-60514713-DEL-238 | chr7-60514714-DEL-238 |
| chr7-60541729-DEL-195 | chr7-60541730-DEL-195 |
| chr7-60562105-DEL-1622 | chr7-60562106-DEL-1622 |
| chr7-60570188-DEL-214 | chr7-60570189-DEL-214 |
| chr7-60578524-DEL-54 | chr7-60578525-DEL-54 |
| chr7-60606053-DEL-1637 | chr7-60606053-DEL-1637 |
| chr7-60623117-DEL-90 | chr7-60623118-DEL-90 |
| chr7-60638558-DEL-501 | chr7-60638559-DEL-501 |
| chr7-60686173-DEL-2024 | chr7-60686174-DEL-2024 |
| chr7-60691033-DEL-235 | chr7-60691034-DEL-235 |
| chr7-60698503-DEL-55 | chr7-60698500-DEL-69 |
| chr7-60719552-DEL-154 | chr7-60719553-DEL-154 |
| chr7-60725155-DEL-95 | chr7-60725183-DEL-65 |
| chr7-60751104-DEL-86 | chr7-60751105-DEL-86 |
| chr7-60798745-DEL-4029 | chr7-60798746-DEL-4029 |
| chr7-60811515-DEL-94 | chr7-60811505-DEL-67 |
| chr7-60832040-DEL-7361 | chr7-60832040-DEL-7361 |
| chr7-60832608-DEL-214 | chr7-60832607-DEL-214 |
| chr7-60834658-DEL-250 | chr7-60834658-DEL-250 |
| chr7-60845634-DEL-124 | chr7-60845613-DEL-96 |
| chr7-60853592-DEL-219 | chr7-60853593-DEL-219 |
| chr7-60882437-DEL-338 | chr7-60882438-DEL-338 |
| chr7-60896826-DEL-312 | chr7-60896818-DEL-308 |
| chr7-60899072-DEL-933 | chr7-60899073-DEL-933 |
| chr7-60933356-DEL-58 | chr7-60933355-DEL-64 |
| chr7-60988165-DEL-507 | chr7-60988166-DEL-507 |
| chr7-61042183-DEL-2977 | chr7-61042182-DEL-2978 |
| chr7-61047593-DEL-1060 | chr7-61047594-DEL-1060 |
| chr7-61108769-DEL-8079 | chr7-61108769-DEL-8079 |
| chr7-61131236-DEL-72 | chr7-61131278-DEL-50 |
| chr7-61157094-DEL-6257 | chr7-61157095-DEL-6257 |
| chr7-61180625-DEL-51 | chr7-61180626-DEL-51 |
| chr7-61207297-DEL-82 | chr7-61207280-DEL-100 |
| chr7-61228842-DEL-51 | chr7-61228861-DEL-51 |

| | |
|---|---|
| chr7-61246956-DEL-82 | chr7-61246945-DEL-90 |
| chr7-61251406-DEL-68 | chr7-61251385-DEL-68 |
| chr7-61257421-DEL-307 | chr7-61257419-DEL-300 |
| chr7-61288756-DEL-4911 | chr7-61288754-DEL-4909 |
| chr7-61295530-DEL-80 | chr7-61295513-DEL-90 |
| chr7-61378565-DEL-75 | chr7-61378561-DEL-80 |
| chr7-61402905-DEL-55 | chr7-61402906-DEL-55 |
| chr7-61417135-DEL-698 | chr7-61417136-DEL-696 |
| chr7-61448862-DEL-53 | chr7-61448863-DEL-53 |
| chr7-61450134-DEL-2185 | chr7-61450135-DEL-2185 |
| chr7-61494289-DEL-2675 | chr7-61494289-DEL-2675 |
| chr7-61503556-DEL-245 | chr7-61503556-DEL-245 |
| chr7-61508043-DEL-85 | chr7-61508043-DEL-85 |
| chr7-61509760-DEL-73 | chr7-61509760-DEL-85 |
| chr7-61512579-DEL-10567 | chr7-61512579-DEL-10567 |
| chr7-61516864-DEL-498 | chr7-61516865-DEL-498 |
| chr7-61525082-DEL-677 | chr7-61525080-DEL-677 |
| chr7-61526837-DEL-205 | chr7-61526838-DEL-205 |
| chr7-61533851-DEL-112 | chr7-61533894-DEL-91 |
| chr7-61538287-DEL-56 | chr7-61538288-DEL-56 |
| chr7-61546796-DEL-824 | chr7-61546797-DEL-824 |
| chr7-61600607-DEL-415 | chr7-61600608-DEL-415 |
| chr7-61615283-DEL-858 | chr7-61615284-DEL-858 |
| chr7-61627200-DEL-1517 | chr7-61627200-DEL-1517 |
| chr7-61648478-DEL-65 | chr7-61648474-DEL-70 |
| chr7-61683696-DEL-62 | chr7-61683673-DEL-70 |
| chr7-61691452-DEL-72 | chr7-61691453-DEL-71 |
| chr7-61695071-DEL-153 | chr7-61695066-DEL-153 |
| chr7-61696646-DEL-64 | chr7-61696647-DEL-66 |
| chr7-61732086-DEL-52 | chr7-61732087-DEL-52 |
| chr7-61737548-DEL-64 | chr7-61737549-DEL-64 |
| chr7-61741128-DEL-56 | chr7-61741129-DEL-88 |
| chr7-61743563-DEL-155 | chr7-61743564-DEL-145 |
| chr7-61747756-DEL-55 | chr7-61747757-DEL-55 |
| chr7-61748045-DEL-543 | chr7-61748046-DEL-543 |
| chr7-61784907-DEL-2893 | chr7-61784909-DEL-2893 |
| chr7-61794297-DEL-62 | chr7-61794298-DEL-62 |
| chr7-61802949-DEL-64 | chr7-61802950-DEL-64 |
| chr7-61816924-DEL-619 | chr7-61816933-DEL-619 |
| chr7-61825006-DEL-505 | chr7-61825003-DEL-500 |
| chr7-61829794-DEL-60 | chr7-61829795-DEL-60 |
| chr7-61834048-DEL-617 | chr7-61834050-DEL-616 |
| chr7-61835998-DEL-242 | chr7-61835999-DEL-242 |
| chr7-61839298-DEL-165 | chr7-61839299-DEL-165 |
| chr7-61911246-DEL-66 | chr7-61911249-DEL-65 |
| chr7-61978708-DEL-6766 | chr7-61978708-DEL-6766 |
| chr7-62140505-DEL-470 | chr7-62140497-DEL-471 |
| chr7-62175763-DEL-80 | chr7-62175764-DEL-80 |
| chr7-62224300-DEL-89 | chr7-62224301-DEL-92 |
| chr7-62306792-DEL-14397 | chr7-62306792-DEL-14397 |
| chr7-62318254-DEL-1617 | chr7-62318255-DEL-1617 |
| chr7-62336046-DEL-7050 | chr7-62336047-DEL-7050 |
| chr7-62336059-DEL-636 | chr7-62336060-DEL-636 |
| chr7-62360796-DEL-155 | chr7-62360797-DEL-155 |
| chr7-62377445-DEL-187 | chr7-62377446-DEL-187 |
| chr7-62384246-DEL-103 | chr7-62384247-DEL-103 |
| chr7-62459771-DEL-248 | chr7-62459772-DEL-248 |
| chr7-62483814-DEL-189 | chr7-62483815-DEL-189 |
| chr7-62489833-DEL-5113 | chr7-62489834-DEL-5113 |
| chr7-62503524-DEL-198 | chr7-62503523-DEL-199 |
| chr7-62531017-DEL-4997 | chr7-62531008-DEL-4998 |
| chr7-62591063-DEL-54 | chr7-62591066-DEL-50 |
| chr7-62616031-DEL-3577 | chr7-62616032-DEL-3577 |
| chr7-62622722-DEL-1352 | chr7-62622723-DEL-1352 |
| chr7-62673558-DEL-7231 | chr7-62673559-DEL-7231 |
| chr7-62682784-DEL-146 | chr7-62682781-DEL-150 |
| chr7-62686675-DEL-488 | chr7-62686676-DEL-488 |
| chr7-62705902-DEL-168 | chr7-62705887-DEL-184 |
| chr7-62727669-DEL-85 | chr7-62727674-DEL-81 |
| chr7-62747991-DEL-64 | chr7-62747992-DEL-60 |
| chr7-62754221-DEL-111 | chr7-62754222-DEL-111 |
| chr7-62794784-DEL-132 | chr7-62794785-DEL-132 |
| chr7-62808568-DEL-166 | chr7-62808569-DEL-166 |
| chr7-62810680-DEL-462 | chr7-62810681-DEL-462 |
| chr7-62813392-DEL-216 | chr7-62813401-DEL-208 |
| chr7-62820804-DEL-848 | chr7-62820804-DEL-849 |
| chr7-62821125-DEL-1162 | chr7-62821126-DEL-1162 |
| chr7-62882599-DEL-69 | chr7-62882600-DEL-69 |
| chr7-62893705-DEL-652 | chr7-62893706-DEL-652 |
| chr7-62903460-DEL-868 | chr7-62903461-DEL-868 |

| | |
|---|---|
| chr7-62952714-DEL-287 | chr7-62952715-DEL-287 |
| chr7-62954105-DEL-78 | chr7-62954146-DEL-56 |
| chr7-62954694-DEL-170 | chr7-62954695-DEL-170 |
| chr7-62955241-DEL-82 | chr7-62955242-DEL-82 |
| chr7-62957656-DEL-428 | chr7-62957655-DEL-431 |
| chr7-62984602-DEL-52 | chr7-62984603-DEL-52 |
| chr7-63012504-DEL-62 | chr7-63012505-DEL-62 |
| chr7-63016657-DEL-52 | chr7-63016658-DEL-52 |
| chr7-63020875-DEL-8794 | chr7-63020876-DEL-8794 |
| chr7-63022716-DEL-497 | chr7-63022717-DEL-497 |
| chr7-63084216-DEL-95 | chr7-63084209-DEL-96 |
| chr7-63126475-DEL-200 | chr7-63126476-DEL-200 |
| chr7-63173817-DEL-67 | chr7-63173813-DEL-68 |
| chr7-63178955-DEL-362 | chr7-63178979-DEL-339 |
| chr7-63211638-DEL-75 | chr7-63211638-DEL-75 |
| chr7-63250541-DEL-555 | chr7-63250542-DEL-555 |
| chr7-63260892-DEL-451 | chr7-63260896-DEL-417 |
| chr7-63261968-DEL-378 | chr7-63261974-DEL-378 |
| chr7-63274428-DEL-424 | chr7-63274429-DEL-424 |
| chr7-63277254-DEL-355 | chr7-63277252-DEL-356 |
| chr7-63330809-DEL-1169 | chr7-63330810-DEL-1169 |
| chr7-63350807-DEL-284 | chr7-63350816-DEL-276 |
| chr7-63354181-DEL-6704 | chr7-63354182-DEL-6704 |
| chr7-63375837-DEL-272 | chr7-63375838-DEL-272 |
| chr7-63390843-DEL-279 | chr7-63390841-DEL-285 |
| chr7-63424893-DEL-97 | chr7-63424900-DEL-91 |
| chr7-63429526-DEL-186 | chr7-63429527-DEL-186 |
| chr7-63438729-DEL-57 | chr7-63438730-DEL-57 |
| chr7-63453411-DEL-7101 | chr7-63453413-DEL-7102 |
| chr7-63464546-DEL-59 | chr7-63464566-DEL-55 |
| chr7-63474760-DEL-344 | chr7-63474761-DEL-344 |
| chr7-63478021-DEL-2832 | chr7-63478022-DEL-2832 |
| chr7-63480969-DEL-1888 | chr7-63480970-DEL-1888 |
| chr7-63521819-DEL-71 | chr7-63521820-DEL-71 |
| chr7-63526306-DEL-105 | chr7-63526307-DEL-105 |
| chr7-63541838-DEL-103 | chr7-63541840-DEL-102 |
| chr7-63544022-DEL-72 | chr7-63544071-DEL-50 |
| chr7-63616638-DEL-146 | chr7-63616676-DEL-107 |
| chr7-63625675-DEL-63 | chr7-63625670-DEL-69 |
| chr7-63628360-DEL-2090 | chr7-63628361-DEL-2090 |
| chr7-63633768-DEL-90 | chr7-63633769-DEL-90 |
| chr7-63650975-DEL-68 | chr7-63650956-DEL-72 |
| chr7-63653191-DEL-66 | chr7-63653192-DEL-66 |
| chr7-63662139-DEL-347 | chr7-63662140-DEL-347 |
| chr7-63665217-DEL-64 | chr7-63665170-DEL-111 |
| chr7-63666423-DEL-52 | chr7-63666424-DEL-52 |
| chr7-63688050-DEL-84 | chr7-63688051-DEL-84 |
| chr7-63718797-DEL-93 | chr7-63718807-DEL-81 |
| chr7-63729548-DEL-165 | chr7-63729549-DEL-165 |
| chr7-63730333-DEL-499 | chr7-63730333-DEL-499 |
| chr7-63732852-DEL-63 | chr7-63732853-DEL-63 |
| chr7-63735212-DEL-335 | chr7-63735213-DEL-335 |
| chr7-63737630-DEL-84 | chr7-63737631-DEL-84 |
| chr7-63738371-DEL-1044 | chr7-63738372-DEL-1044 |
| chr7-63739589-DEL-84 | chr7-63739590-DEL-84 |
| chr7-63739682-DEL-627 | chr7-63739682-DEL-627 |
| chr7-63776305-DEL-9045 | chr7-63776306-DEL-9045 |
| chr7-63821603-DEL-58 | chr7-63821563-DEL-58 |
| chr7-63843834-DEL-732 | chr7-63843835-DEL-732 |
| chr7-63850792-DEL-186 | chr7-63850793-DEL-186 |
| chr7-63885156-DEL-210 | chr7-63885122-DEL-252 |
| chr7-63913491-DEL-541 | chr7-63913492-DEL-541 |
| chr7-63949899-DEL-72 | chr7-63949900-DEL-72 |
| chr7-63992674-DEL-196 | chr7-63992675-DEL-196 |
| chr7-63994975-DEL-90 | chr7-63994975-DEL-76 |
| chr7-63996229-DEL-58 | chr7-63996230-DEL-58 |
| chr7-64008951-DEL-184 | chr7-64008924-DEL-166 |
| chr7-64040843-DEL-173 | chr7-64040844-DEL-173 |
| chr7-64053894-DEL-748 | chr7-64053895-DEL-748 |
| chr7-64070017-DEL-240 | chr7-64070018-DEL-240 |
| chr7-64072630-DEL-134 | chr7-64072635-DEL-130 |
| chr7-64074179-DEL-202 | chr7-64074180-DEL-202 |
| chr7-64097318-DEL-396 | chr7-64097319-DEL-396 |
| chr7-64147646-DEL-53 | chr7-64147648-DEL-51 |
| chr7-64148689-DEL-67 | chr7-64148691-DEL-67 |
| chr7-64177235-DEL-89 | chr7-64177231-DEL-90 |
| chr7-64220128-DEL-51 | chr7-64220129-DEL-51 |
| chr7-64249824-DEL-416 | chr7-64249824-DEL-416 |
| chr7-64271313-DEL-2702 | chr7-64271312-DEL-2703 |
| chr7-64308247-DEL-437 | chr7-64308248-DEL-437 |

| | |
|---|---|
| chr7-64322854-DEL-6500 | chr7-64322855-DEL-6500 |
| chr7-64332815-DEL-568 | chr7-64332816-DEL-568 |
| chr7-64385136-DEL-62 | chr7-64385137-DEL-62 |
| chr7-64442282-DEL-329 | chr7-64442284-DEL-324 |
| chr7-64459100-DEL-328 | chr7-64459101-DEL-328 |
| chr7-64466369-DEL-226 | chr7-64466370-DEL-226 |
| chr7-64470241-DEL-87 | chr7-64470241-DEL-87 |
| chr7-64473060-DEL-3455 | chr7-64473061-DEL-3455 |
| chr7-64484144-DEL-109 | chr7-64484139-DEL-113 |
| chr7-64485595-DEL-692 | chr7-64485596-DEL-692 |
| chr7-64510540-DEL-840 | chr7-64510541-DEL-840 |
| chr7-64615605-DEL-234 | chr7-64615606-DEL-234 |
| chr7-64651277-DEL-123 | chr7-64651278-DEL-123 |
| chr7-64652648-DEL-58 | chr7-64652649-DEL-56 |
| chr7-64656642-DEL-970 | chr7-64656635-DEL-968 |
| chr7-64684683-DEL-82 | chr7-64684683-DEL-60 |
| chr7-64727354-DEL-84 | chr7-64727355-DEL-84 |
| chr7-64733834-DEL-53 | chr7-64733835-DEL-53 |
| chr7-64777223-DEL-86 | chr7-64777260-DEL-50 |
| chr7-64783454-DEL-216 | chr7-64783455-DEL-216 |
| chr7-64792906-DEL-992 | chr7-64792907-DEL-991 |
| chr7-64804172-DEL-58 | chr7-64804170-DEL-60 |
| chr7-64829437-DEL-72 | chr7-64829438-DEL-72 |
| chr7-64836920-DEL-6731 | chr7-64836921-DEL-6731 |
| chr7-64887216-DEL-193 | chr7-64887217-DEL-193 |
| chr7-64909829-DEL-6475 | chr7-64909830-DEL-6475 |
| chr7-64922535-DEL-1633 | chr7-64922536-DEL-1633 |
| chr7-64923743-DEL-131 | chr7-64923744-DEL-132 |
| chr7-64924422-DEL-135 | chr7-64924423-DEL-135 |
| chr7-64931362-DEL-203 | chr7-64931363-DEL-203 |
| chr7-64935936-DEL-64 | chr7-64935937-DEL-64 |
| chr7-64953218-DEL-202 | chr7-64953219-DEL-202 |
| chr7-65003210-DEL-201 | chr7-65003211-DEL-201 |
| chr7-65046776-DEL-778 | chr7-65046773-DEL-775 |
| chr7-65052400-DEL-185 | chr7-65052401-DEL-185 |
| chr7-65081896-DEL-71 | chr7-65081897-DEL-71 |
| chr7-65108842-DEL-58 | chr7-65108847-DEL-54 |
| chr7-65123907-DEL-354 | chr7-65123908-DEL-354 |
| chr7-65128596-DEL-207 | chr7-65128597-DEL-207 |
| chr7-65129135-DEL-1888 | chr7-65129136-DEL-1888 |
| chr7-65137877-DEL-54 | chr7-65137878-DEL-54 |
| chr7-65142118-DEL-59 | chr7-65142117-DEL-60 |
| chr7-65144887-DEL-1904 | chr7-65144888-DEL-1904 |
| chr7-65156711-DEL-92 | chr7-65156676-DEL-138 |
| chr7-65170284-DEL-2126 | chr7-65170285-DEL-2126 |
| chr7-65186620-DEL-163 | chr7-65186616-DEL-137 |
| chr7-65188915-DEL-106 | chr7-65188950-DEL-105 |
| chr7-65199746-DEL-78 | chr7-65199747-DEL-75 |
| chr7-65216350-DEL-342 | chr7-65216351-DEL-342 |
| chr7-65252364-DEL-3086 | chr7-65252365-DEL-3086 |
| chr7-65262770-DEL-2678 | chr7-65262771-DEL-2678 |
| chr7-65281600-DEL-103 | chr7-65281646-DEL-98 |
| chr7-65287373-DEL-646 | chr7-65287374-DEL-646 |
| chr7-65304546-DEL-60 | chr7-65304550-DEL-56 |
| chr7-65305796-DEL-1706 | chr7-65305797-DEL-1706 |
| chr7-65320018-DEL-713 | chr7-65320019-DEL-713 |
| chr7-65330332-DEL-56 | chr7-65330333-DEL-56 |
| chr7-65340027-DEL-115 | chr7-65340028-DEL-115 |
| chr7-65349552-DEL-686 | chr7-65349551-DEL-685 |
| chr7-65364437-DEL-294 | chr7-65364438-DEL-294 |
| chr7-65368907-DEL-206 | chr7-65368907-DEL-206 |
| chr7-65379023-DEL-51 | chr7-65379024-DEL-51 |
| chr7-65396399-DEL-60 | chr7-65396400-DEL-60 |
| chr7-65428208-DEL-72 | chr7-65428201-DEL-88 |
| chr7-65434560-DEL-1502 | chr7-65434540-DEL-1522 |
| chr7-65457862-DEL-537 | chr7-65457863-DEL-537 |
| chr7-65466085-DEL-1636 | chr7-65466086-DEL-1636 |
| chr7-65473151-DEL-211 | chr7-65473153-DEL-211 |
| chr7-65497336-DEL-102 | chr7-65497337-DEL-102 |
| chr7-65502918-DEL-1320 | chr7-65502919-DEL-1320 |
| chr7-65527787-DEL-138 | chr7-65527788-DEL-138 |
| chr7-65542742-DEL-431 | chr7-65542743-DEL-431 |
| chr7-65544095-DEL-56 | chr7-65544096-DEL-56 |
| chr7-65615344-DEL-151 | chr7-65615311-DEL-151 |
| chr7-65617305-DEL-353 | chr7-65617306-DEL-353 |
| chr7-65646327-DEL-280 | chr7-65646328-DEL-277 |
| chr7-65704448-DEL-184 | chr7-65704449-DEL-184 |
| chr7-65709177-DEL-165 | chr7-65709178-DEL-165 |
| chr7-65731869-DEL-52 | chr7-65731870-DEL-52 |
| chr7-65784200-DEL-144 | chr7-65784201-DEL-144 |

| | |
|---|---|
| chr7-65798808-DEL-212 | chr7-65798809-DEL-212 |
| chr7-65817732-DEL-76 | chr7-65817733-DEL-75 |
| chr7-65840585-DEL-169 | chr7-65840586-DEL-169 |
| chr7-65843959-DEL-56 | chr7-65843960-DEL-56 |
| chr7-65844767-DEL-60 | chr7-65844768-DEL-60 |
| chr7-65855206-DEL-925 | chr7-65855207-DEL-925 |
| chr7-65862904-DEL-890 | chr7-65862897-DEL-895 |
| chr7-65882703-DEL-63 | chr7-65882689-DEL-112 |
| chr7-65885066-DEL-233 | chr7-65885067-DEL-233 |
| chr7-65914690-DEL-70 | chr7-65914734-DEL-73 |
| chr7-65930474-DEL-68 | chr7-65930474-DEL-68 |
| chr7-65960378-DEL-145 | chr7-65960379-DEL-145 |
| chr7-65961978-DEL-66 | chr7-65961979-DEL-66 |
| chr7-65963358-DEL-7714 | chr7-65963359-DEL-7714 |
| chr7-65968990-DEL-216 | chr7-65968991-DEL-216 |
| chr7-65969306-DEL-752 | chr7-65969307-DEL-752 |
| chr7-65983844-DEL-100 | chr7-65983845-DEL-100 |
| chr7-65994812-DEL-90 | chr7-65994828-DEL-50 |
| chr7-65996467-DEL-53 | chr7-65996468-DEL-53 |
| chr7-66014569-DEL-128 | chr7-66014570-DEL-126 |
| chr7-66016431-DEL-52 | chr7-66016440-DEL-60 |
| chr7-66027370-DEL-636 | chr7-66027371-DEL-636 |
| chr7-66058247-DEL-199 | chr7-66058248-DEL-199 |
| chr7-66068092-DEL-51 | chr7-66068093-DEL-51 |
| chr7-66080871-DEL-179 | chr7-66080872-DEL-179 |
| chr7-66140459-DEL-63 | chr7-66140460-DEL-63 |
| chr7-66149330-DEL-80 | chr7-66149331-DEL-80 |
| chr7-66150462-DEL-63 | chr7-66150463-DEL-63 |
| chr7-66162982-DEL-64 | chr7-66162983-DEL-64 |
| chr7-66185090-DEL-6840 | chr7-66185091-DEL-6840 |
| chr7-66192992-DEL-950 | chr7-66192993-DEL-950 |
| chr7-66202312-DEL-96 | chr7-66202327-DEL-94 |
| chr7-66209980-DEL-51 | chr7-66209981-DEL-51 |
| chr7-66256979-DEL-153 | chr7-66256980-DEL-153 |
| chr7-66265621-DEL-338 | chr7-66265622-DEL-338 |
| chr7-66277658-DEL-506 | chr7-66277659-DEL-506 |
| chr7-66304196-DEL-53 | chr7-66304193-DEL-54 |
| chr7-66340749-DEL-181 | chr7-66340750-DEL-181 |
| chr7-66348904-DEL-60 | chr7-66348905-DEL-60 |
| chr7-66371582-DEL-302 | chr7-66371589-DEL-292 |
| chr7-66372239-DEL-114 | chr7-66372240-DEL-114 |
| chr7-66378272-DEL-56 | chr7-66378265-DEL-64 |
| chr7-66390985-DEL-311 | chr7-66390986-DEL-311 |
| chr7-66391542-DEL-171 | chr7-66391543-DEL-171 |
| chr7-66415716-DEL-2849 | chr7-66415713-DEL-2852 |
| chr7-66426582-DEL-1091 | chr7-66426583-DEL-1091 |
| chr7-66432253-DEL-54 | chr7-66432224-DEL-55 |
| chr7-66483379-DEL-90 | chr7-66483380-DEL-90 |
| chr7-66496329-DEL-89 | chr7-66496341-DEL-70 |
| chr7-66529516-DEL-751 | chr7-66529517-DEL-751 |
| chr7-66578673-DEL-52 | chr7-66578674-DEL-52 |
| chr7-66640706-DEL-3031 | chr7-66640707-DEL-3031 |
| chr7-66646834-DEL-402 | chr7-66646835-DEL-402 |
| chr7-66654461-DEL-54 | chr7-66654460-DEL-56 |
| chr7-66659367-DEL-54 | chr7-66659368-DEL-54 |
| chr7-66678410-DEL-93 | chr7-66678414-DEL-81 |
| chr7-66682842-DEL-63 | chr7-66682846-DEL-59 |
| chr7-66715904-DEL-81 | chr7-66715905-DEL-81 |
| chr7-66717274-DEL-102 | chr7-66717273-DEL-102 |
| chr7-66724193-DEL-70 | chr7-66724173-DEL-59 |
| chr7-66728897-DEL-440 | chr7-66728898-DEL-440 |
| chr7-66745863-DEL-153 | chr7-66745864-DEL-153 |
| chr7-66773054-DEL-115 | chr7-66773055-DEL-115 |
| chr7-66794793-DEL-199 | chr7-66794794-DEL-199 |
| chr7-66804705-DEL-7145 | chr7-66804706-DEL-7145 |
| chr7-66865435-DEL-70 | chr7-66865436-DEL-70 |
| chr7-66923664-DEL-6535 | chr7-66923665-DEL-6535 |
| chr7-66942659-DEL-68 | chr7-66942660-DEL-68 |
| chr7-66947860-DEL-168 | chr7-66947861-DEL-164 |
| chr7-66987598-DEL-89 | chr7-66987599-DEL-89 |
| chr7-66991015-DEL-2322 | chr7-66991016-DEL-2322 |
| chr7-67027269-DEL-64 | chr7-67027270-DEL-64 |
| chr7-67029372-DEL-250 | chr7-67029371-DEL-251 |
| chr7-67031584-DEL-257 | chr7-67031585-DEL-257 |
| chr7-67033762-DEL-70 | chr7-67033763-DEL-70 |
| chr7-67052093-DEL-61 | chr7-67052094-DEL-61 |
| chr7-67056596-DEL-569 | chr7-67056595-DEL-569 |
| chr7-67065000-DEL-2778 | chr7-67065001-DEL-2778 |
| chr7-67080206-DEL-122 | chr7-67080213-DEL-116 |
| chr7-67106257-DEL-68 | chr7-67106251-DEL-76 |

| | |
|---|---|
| chr7-67147312-DEL-120 | chr7-67147326-DEL-96 |
| chr7-67191796-DEL-341 | chr7-67191797-DEL-341 |
| chr7-67193792-DEL-203 | chr7-67193793-DEL-203 |
| chr7-67242593-DEL-171 | chr7-67242594-DEL-171 |
| chr7-67248523-DEL-54 | chr7-67248519-DEL-54 |
| chr7-67281443-DEL-198 | chr7-67281463-DEL-179 |
| chr7-67313173-DEL-84 | chr7-67313162-DEL-96 |
| chr7-67317229-DEL-68 | chr7-67317230-DEL-68 |
| chr7-67324893-DEL-56 | chr7-67324897-DEL-52 |
| chr7-67346331-DEL-70 | chr7-67346332-DEL-70 |
| chr7-67366186-DEL-84 | chr7-67366208-DEL-62 |
| chr7-67404953-DEL-165 | chr7-67404952-DEL-165 |
| chr7-67428037-DEL-492 | chr7-67428038-DEL-492 |
| chr7-67436104-DEL-1588 | chr7-67436105-DEL-1588 |
| chr7-67460378-DEL-1958 | chr7-67460379-DEL-1958 |
| chr7-67463235-DEL-205 | chr7-67463236-DEL-205 |
| chr7-67539874-DEL-66 | chr7-67539879-DEL-102 |
| chr7-67568260-DEL-67 | chr7-67568257-DEL-64 |
| chr7-67619737-DEL-408 | chr7-67619738-DEL-412 |
| chr7-67628779-DEL-120 | chr7-67628759-DEL-93 |
| chr7-67660603-DEL-52 | chr7-67660603-DEL-52 |
| chr7-67681928-DEL-54 | chr7-67681929-DEL-54 |
| chr7-67693166-DEL-103 | chr7-67693167-DEL-103 |
| chr7-67727922-DEL-206 | chr7-67727923-DEL-206 |
| chr7-67744105-DEL-267 | chr7-67744106-DEL-267 |
| chr7-67768045-DEL-70 | chr7-67768046-DEL-70 |
| chr7-67808380-DEL-351 | chr7-67808379-DEL-352 |
| chr7-67810886-DEL-54 | chr7-67810887-DEL-54 |
| chr7-67820022-DEL-155 | chr7-67820023-DEL-153 |
| chr7-67858182-DEL-60 | chr7-67858183-DEL-60 |
| chr7-67905032-DEL-226 | chr7-67905033-DEL-226 |
| chr7-67908840-DEL-138 | chr7-67908841-DEL-138 |
| chr7-68019409-DEL-76 | chr7-68019410-DEL-80 |
| chr7-68024842-DEL-65 | chr7-68024859-DEL-66 |
| chr7-68031565-DEL-52 | chr7-68031566-DEL-52 |
| chr7-68037726-DEL-94 | chr7-68037773-DEL-65 |
| chr7-68045338-DEL-91 | chr7-68045339-DEL-91 |
| chr7-68047967-DEL-90 | chr7-68047961-DEL-95 |
| chr7-68095575-DEL-145 | chr7-68095576-DEL-145 |
| chr7-68099179-DEL-99 | chr7-68099159-DEL-121 |
| chr7-68114884-DEL-66 | chr7-68114887-DEL-64 |
| chr7-68165325-DEL-486 | chr7-68165326-DEL-486 |
| chr7-68169030-DEL-62 | chr7-68169039-DEL-63 |
| chr7-68173160-DEL-166 | chr7-68173161-DEL-166 |
| chr7-68184485-DEL-3051 | chr7-68184486-DEL-3051 |
| chr7-68263126-DEL-92 | chr7-68263127-DEL-92 |
| chr7-68274777-DEL-204 | chr7-68274778-DEL-204 |
| chr7-68297814-DEL-1276 | chr7-68297861-DEL-1237 |
| chr7-68319578-DEL-152 | chr7-68319579-DEL-152 |
| chr7-68324233-DEL-80 | chr7-68324223-DEL-87 |
| chr7-68351119-DEL-763 | chr7-68351120-DEL-763 |
| chr7-68374151-DEL-80 | chr7-68374152-DEL-74 |
| chr7-68402907-DEL-497 | chr7-68402908-DEL-497 |
| chr7-68423598-DEL-6179 | chr7-68423606-DEL-6161 |
| chr7-68432104-DEL-367 | chr7-68432105-DEL-367 |
| chr7-68436397-DEL-899 | chr7-68436398-DEL-899 |
| chr7-68456996-DEL-591 | chr7-68456997-DEL-591 |
| chr7-68469137-DEL-53 | chr7-68469121-DEL-64 |
| chr7-68470539-DEL-62 | chr7-68470536-DEL-50 |
| chr7-68504248-DEL-300 | chr7-68504249-DEL-300 |
| chr7-68538061-DEL-58 | chr7-68538016-DEL-104 |
| chr7-68539229-DEL-126 | chr7-68539230-DEL-126 |
| chr7-68597136-DEL-1768 | chr7-68597137-DEL-1768 |
| chr7-68642298-DEL-2143 | chr7-68642299-DEL-2143 |
| chr7-68647695-DEL-6582 | chr7-68647696-DEL-6582 |
| chr7-68653829-DEL-208 | chr7-68653830-DEL-208 |
| chr7-68668442-DEL-997 | chr7-68668443-DEL-997 |
| chr7-68673127-DEL-80 | chr7-68673136-DEL-66 |
| chr7-68680231-DEL-211 | chr7-68680232-DEL-211 |
| chr7-68730623-DEL-96 | chr7-68730629-DEL-91 |
| chr7-68735285-DEL-119 | chr7-68735286-DEL-117 |
| chr7-68736155-DEL-1538 | chr7-68736156-DEL-1538 |
| chr7-68750826-DEL-962 | chr7-68750827-DEL-962 |
| chr7-68793349-DEL-181 | chr7-68793350-DEL-181 |
| chr7-68794156-DEL-60 | chr7-68794157-DEL-60 |
| chr7-68812727-DEL-76 | chr7-68812728-DEL-76 |
| chr7-68866698-DEL-68 | chr7-68866727-DEL-84 |
| chr7-68905493-DEL-53 | chr7-68905496-DEL-51 |
| chr7-68966208-DEL-153 | chr7-68966194-DEL-160 |
| chr7-68985417-DEL-90 | chr7-68985418-DEL-90 |

| | |
|---|---|
| chr7-68988524-DEL-426 | chr7-68988524-DEL-426 |
| chr7-68996043-DEL-299 | chr7-68996044-DEL-257 |
| chr7-69015205-DEL-9309 | chr7-69015206-DEL-9309 |
| chr7-69048813-DEL-372 | chr7-69048814-DEL-372 |
| chr7-69078656-DEL-6436 | chr7-69078657-DEL-6436 |
| chr7-69090187-DEL-68 | chr7-69090184-DEL-70 |
| chr7-69113343-DEL-65 | chr7-69113348-DEL-65 |
| chr7-69115416-DEL-722 | chr7-69115417-DEL-722 |
| chr7-69122550-DEL-106 | chr7-69122551-DEL-106 |
| chr7-69141362-DEL-54 | chr7-69141348-DEL-78 |
| chr7-69145583-DEL-312 | chr7-69145584-DEL-312 |
| chr7-69147973-DEL-472 | chr7-69147963-DEL-471 |
| chr7-69174348-DEL-63 | chr7-69174349-DEL-63 |
| chr7-69245184-DEL-2262 | chr7-69245185-DEL-2262 |
| chr7-69256963-DEL-504 | chr7-69256964-DEL-504 |
| chr7-69265086-DEL-779 | chr7-69265087-DEL-779 |
| chr7-69288878-DEL-68 | chr7-69288890-DEL-84 |
| chr7-69307577-DEL-1037 | chr7-69307578-DEL-1037 |
| chr7-69340398-DEL-62 | chr7-69340357-DEL-79 |
| chr7-69357602-DEL-77 | chr7-69357603-DEL-76 |
| chr7-69369116-DEL-69 | chr7-69369117-DEL-69 |
| chr7-69373911-DEL-2949 | chr7-69373952-DEL-2917 |
| chr7-69381568-DEL-176 | chr7-69381568-DEL-176 |
| chr7-69434814-DEL-1621 | chr7-69434815-DEL-1621 |
| chr7-69440826-DEL-1164 | chr7-69440827-DEL-1164 |
| chr7-69466177-DEL-99 | chr7-69466184-DEL-93 |
| chr7-69500318-DEL-2545 | chr7-69500319-DEL-2545 |
| chr7-69721868-DEL-353 | chr7-69721869-DEL-353 |
| chr7-69722708-DEL-1228 | chr7-69722709-DEL-1228 |
| chr7-69762033-DEL-319 | chr7-69762034-DEL-319 |
| chr7-69790557-DEL-53 | chr7-69790550-DEL-54 |
| chr7-69842976-DEL-6785 | chr7-69842977-DEL-6785 |
| chr7-69890133-DEL-1862 | chr7-69890134-DEL-1862 |
| chr7-69924840-DEL-337 | chr7-69924841-DEL-337 |
| chr7-69962782-DEL-197 | chr7-69962783-DEL-197 |
| chr7-70062124-DEL-51 | chr7-70062121-DEL-52 |
| chr7-70086601-DEL-975 | chr7-70086602-DEL-975 |
| chr7-70147876-DEL-57 | chr7-70147877-DEL-57 |
| chr7-70284936-DEL-69 | chr7-70284937-DEL-69 |
| chr7-70292516-DEL-54 | chr7-70292547-DEL-66 |
| chr7-70314349-DEL-77 | chr7-70314350-DEL-77 |
| chr7-70327720-DEL-6668 | chr7-70327721-DEL-6671 |
| chr7-70370237-DEL-214 | chr7-70370234-DEL-214 |
| chr7-70383919-DEL-65 | chr7-70383919-DEL-65 |
| chr7-70407663-DEL-7132 | chr7-70407664-DEL-7132 |
| chr7-70433722-DEL-99 | chr7-70433723-DEL-99 |
| chr7-70501168-DEL-150 | chr7-70501169-DEL-149 |
| chr7-70515488-DEL-341 | chr7-70515489-DEL-341 |
| chr7-70630504-DEL-4169 | chr7-70630505-DEL-4169 |
| chr7-70666843-DEL-1241 | chr7-70666844-DEL-1241 |
| chr7-70804872-DEL-225 | chr7-70804873-DEL-225 |
| chr7-70807408-DEL-52 | chr7-70807409-DEL-57 |
| chr7-70818870-DEL-55 | chr7-70818871-DEL-52 |
| chr7-70847173-DEL-58 | chr7-70847174-DEL-55 |
| chr7-70912909-DEL-307 | chr7-70912910-DEL-307 |
| chr7-70917106-DEL-6575 | chr7-70917098-DEL-6576 |
| chr7-70941661-DEL-50 | chr7-70941630-DEL-52 |
| chr7-70944244-DEL-407 | chr7-70944239-DEL-408 |
| chr7-70970033-DEL-574 | chr7-70970034-DEL-574 |
| chr7-70991325-DEL-98 | chr7-70991365-DEL-78 |
| chr7-71006233-DEL-319 | chr7-71006234-DEL-319 |
| chr7-71066828-DEL-58 | chr7-71066815-DEL-84 |
| chr7-71102242-DEL-1942 | chr7-71102243-DEL-1942 |
| chr7-71154649-DEL-194 | chr7-71154650-DEL-194 |
| chr7-71160652-DEL-220 | chr7-71160653-DEL-220 |
| chr7-71183418-DEL-109 | chr7-71183419-DEL-109 |
| chr7-71200357-DEL-4127 | chr7-71200358-DEL-4127 |
| chr7-71220601-DEL-165 | chr7-71220602-DEL-165 |
| chr7-71248969-DEL-105 | chr7-71248972-DEL-103 |
| chr7-71282535-DEL-261 | chr7-71282536-DEL-261 |
| chr7-71283352-DEL-53 | chr7-71283353-DEL-52 |
| chr7-71344094-DEL-140 | chr7-71344095-DEL-140 |
| chr7-71344777-DEL-1208 | chr7-71344778-DEL-1208 |
| chr7-71359378-DEL-483 | chr7-71359356-DEL-507 |
| chr7-71374092-DEL-84 | chr7-71374093-DEL-84 |
| chr7-71383753-DEL-54 | chr7-71383758-DEL-50 |
| chr7-71399184-DEL-213 | chr7-71399185-DEL-213 |
| chr7-71415413-DEL-212 | chr7-71415414-DEL-212 |
| chr7-71450311-DEL-83 | chr7-71450312-DEL-83 |
| chr7-71451820-DEL-84 | chr7-71451821-DEL-84 |

| | |
|---|---|
| chr7-71468080-DEL-6708 | chr7-71468081-DEL-6708 |
| chr7-71476162-DEL-60 | chr7-71476163-DEL-60 |
| chr7-71520155-DEL-53 | chr7-71520156-DEL-53 |
| chr7-71565322-DEL-892 | chr7-71565322-DEL-892 |
| chr7-71566530-DEL-306 | chr7-71566532-DEL-298 |
| chr7-71568177-DEL-293 | chr7-71568177-DEL-292 |
| chr7-71620830-DEL-4055 | chr7-71620831-DEL-4055 |
| chr7-71627513-DEL-6754 | chr7-71627514-DEL-6754 |
| chr7-71627557-DEL-155 | chr7-71627555-DEL-161 |
| chr7-71633709-DEL-208 | chr7-71633710-DEL-208 |
| chr7-71636405-DEL-505 | chr7-71636406-DEL-505 |
| chr7-71642725-DEL-60 | chr7-71642726-DEL-60 |
| chr7-71647276-DEL-260 | chr7-71647254-DEL-277 |
| chr7-71738185-DEL-72 | chr7-71738186-DEL-72 |
| chr7-71756326-DEL-998 | chr7-71756327-DEL-998 |
| chr7-71757388-DEL-674 | chr7-71757389-DEL-674 |
| chr7-71772674-DEL-2757 | chr7-71772675-DEL-2757 |
| chr7-71786728-DEL-90 | chr7-71786747-DEL-50 |
| chr7-71789439-DEL-63 | chr7-71789446-DEL-63 |
| chr7-71791667-DEL-74 | chr7-71791649-DEL-93 |
| chr7-71805704-DEL-99 | chr7-71805705-DEL-99 |
| chr7-71824729-DEL-159 | chr7-71824730-DEL-159 |
| chr7-71892084-DEL-189 | chr7-71892085-DEL-189 |
| chr7-71903166-DEL-2263 | chr7-71903167-DEL-2263 |
| chr7-71914664-DEL-1017 | chr7-71914665-DEL-1017 |
| chr7-71931840-DEL-224 | chr7-71931829-DEL-236 |
| chr7-71939647-DEL-1249 | chr7-71939614-DEL-1282 |
| chr7-71950589-DEL-1315 | chr7-71950590-DEL-1316 |
| chr7-71960298-DEL-56 | chr7-71960299-DEL-56 |
| chr7-71961485-DEL-70 | chr7-71961466-DEL-72 |
| chr7-71967152-DEL-1246 | chr7-71967153-DEL-1246 |
| chr7-71982663-DEL-6155 | chr7-71982664-DEL-6155 |
| chr7-71998695-DEL-57 | chr7-71998696-DEL-57 |
| chr7-72001623-DEL-380 | chr7-72001624-DEL-380 |
| chr7-72023987-DEL-56 | chr7-72023988-DEL-56 |
| chr7-72028816-DEL-628 | chr7-72028817-DEL-628 |
| chr7-72030754-DEL-1002 | chr7-72030755-DEL-1002 |
| chr7-72031956-DEL-440 | chr7-72031957-DEL-440 |
| chr7-72043006-DEL-638 | chr7-72043007-DEL-638 |
| chr7-72046563-DEL-246 | chr7-72046564-DEL-246 |
| chr7-72058783-DEL-1415 | chr7-72058786-DEL-1413 |
| chr7-72132676-DEL-6420 | chr7-72132677-DEL-6420 |
| chr7-72171007-DEL-6616 | chr7-72171008-DEL-6616 |
| chr7-72205045-DEL-1045 | chr7-72205046-DEL-1045 |
| chr7-72206665-DEL-229 | chr7-72206666-DEL-229 |
| chr7-72214717-DEL-225 | chr7-72214718-DEL-225 |
| chr7-72222946-DEL-59 | chr7-72222947-DEL-59 |
| chr7-72225271-DEL-199 | chr7-72225272-DEL-199 |
| chr7-72257113-DEL-127 | chr7-72257114-DEL-127 |
| chr7-72258530-DEL-52 | chr7-72258531-DEL-52 |
| chr7-72279726-DEL-1889 | chr7-72279727-DEL-1889 |
| chr7-72290808-DEL-7088 | chr7-72290809-DEL-7088 |
| chr7-72298446-DEL-651 | chr7-72298447-DEL-651 |
| chr7-72299978-DEL-10657 | chr7-72299979-DEL-10657 |
| chr7-72320725-DEL-1272 | chr7-72320725-DEL-1272 |
| chr7-72339317-DEL-60 | chr7-72339308-DEL-78 |
| chr7-72356317-DEL-961 | chr7-72356318-DEL-961 |
| chr7-72359435-DEL-119 | chr7-72359436-DEL-119 |
| chr7-72368339-DEL-6172 | chr7-72368340-DEL-6172 |
| chr7-72394805-DEL-2648 | chr7-72394806-DEL-2648 |
| chr7-72398621-DEL-257 | chr7-72398582-DEL-255 |
| chr7-72399127-DEL-433 | chr7-72399127-DEL-434 |
| chr7-72404836-DEL-70 | chr7-72404843-DEL-64 |
| chr7-72428384-DEL-66 | chr7-72428384-DEL-60 |
| chr7-72430107-DEL-1407 | chr7-72430108-DEL-1407 |
| chr7-72470590-DEL-1123 | chr7-72470590-DEL-1123 |
| chr7-72474498-DEL-64 | chr7-72474496-DEL-65 |
| chr7-72493682-DEL-729 | chr7-72493683-DEL-729 |
| chr7-72511428-DEL-2265 | chr7-72511429-DEL-2265 |
| chr7-72534612-DEL-1897 | chr7-72534619-DEL-1890 |
| chr7-72577454-DEL-6900 | chr7-72577455-DEL-6900 |
| chr7-72596373-DEL-472 | chr7-72596374-DEL-472 |
| chr7-72600708-DEL-245 | chr7-72600709-DEL-245 |
| chr7-72610338-DEL-66 | chr7-72610338-DEL-66 |
| chr7-72611587-DEL-192 | chr7-72611588-DEL-192 |
| chr7-72631781-DEL-84 | chr7-72631769-DEL-84 |
| chr7-72633570-DEL-54 | chr7-72633571-DEL-54 |
| chr7-72649704-DEL-498 | chr7-72649705-DEL-498 |
| chr7-72657285-DEL-6496 | chr7-72657286-DEL-6496 |
| chr7-72657296-DEL-212 | chr7-72657297-DEL-212 |

| | |
|---|---|
| chr7-72676401-DEL-102 | chr7-72676402-DEL-102 |
| chr7-72698363-DEL-6624 | chr7-72698364-DEL-6624 |
| chr7-72796876-DEL-63 | chr7-72796877-DEL-63 |
| chr7-72802380-DEL-188 | chr7-72802363-DEL-206 |
| chr7-72810301-DEL-54 | chr7-72810300-DEL-56 |
| chr7-72820280-DEL-53 | chr7-72820279-DEL-53 |
| chr7-72824805-DEL-7070 | chr7-72824805-DEL-7070 |
| chr7-72848414-DEL-53 | chr7-72848415-DEL-53 |
| chr7-72849517-DEL-56 | chr7-72849515-DEL-54 |
| chr7-72861652-DEL-7106 | chr7-72861653-DEL-7106 |
| chr7-72864954-DEL-1039 | chr7-72864955-DEL-1039 |
| chr7-72869475-DEL-144 | chr7-72869475-DEL-137 |
| chr7-72878485-DEL-323 | chr7-72878491-DEL-318 |
| chr7-72886669-DEL-249 | chr7-72886647-DEL-272 |
| chr7-72913332-DEL-295 | chr7-72913327-DEL-267 |
| chr7-72966930-DEL-59 | chr7-72966931-DEL-59 |
| chr7-73002136-DEL-69 | chr7-73002137-DEL-70 |
| chr7-73062165-DEL-63 | chr7-73062166-DEL-63 |
| chr7-73070938-DEL-192 | chr7-73070939-DEL-192 |
| chr7-73127501-DEL-102 | chr7-73127502-DEL-102 |
| chr7-73128686-DEL-271 | chr7-73128687-DEL-271 |
| chr7-73142287-DEL-84 | chr7-73142286-DEL-83 |
| chr7-73182044-DEL-72 | chr7-73182045-DEL-72 |
| chr7-73212568-DEL-52 | chr7-73212530-DEL-76 |
| chr7-73256039-DEL-168 | chr7-73256040-DEL-168 |
| chr7-73269399-DEL-140 | chr7-73269417-DEL-140 |
| chr7-73272738-DEL-86 | chr7-73272739-DEL-85 |
| chr7-73295863-DEL-234 | chr7-73295855-DEL-201 |
| chr7-73302376-DEL-186 | chr7-73302377-DEL-186 |
| chr7-73305816-DEL-118 | chr7-73305817-DEL-131 |
| chr7-73338325-DEL-273 | chr7-73338326-DEL-273 |
| chr7-73372513-DEL-54 | chr7-73372514-DEL-54 |
| chr7-73397279-DEL-67 | chr7-73397280-DEL-67 |
| chr7-73417485-DEL-882 | chr7-73417486-DEL-882 |
| chr7-73495469-DEL-52 | chr7-73495470-DEL-52 |
| chr7-73528348-DEL-52 | chr7-73528353-DEL-52 |
| chr7-73550232-DEL-55 | chr7-73550195-DEL-59 |
| chr7-73552436-DEL-2924 | chr7-73552437-DEL-2924 |
| chr7-73562950-DEL-63 | chr7-73562937-DEL-61 |
| chr7-73581825-DEL-66 | chr7-73581840-DEL-52 |
| chr7-73622043-DEL-91 | chr7-73621993-DEL-92 |
| chr7-73687576-DEL-76 | chr7-73687577-DEL-76 |
| chr7-73725296-DEL-93 | chr7-73725297-DEL-62 |
| chr7-73729155-DEL-145 | chr7-73729154-DEL-147 |
| chr7-73774266-DEL-54 | chr7-73774267-DEL-54 |
| chr7-73784811-DEL-102 | chr7-73784814-DEL-109 |
| chr7-73785546-DEL-690 | chr7-73785547-DEL-690 |
| chr7-73794835-DEL-64 | chr7-73794799-DEL-96 |
| chr7-73809554-DEL-288 | chr7-73809555-DEL-282 |
| chr7-73815920-DEL-749 | chr7-73815924-DEL-748 |
| chr7-73824954-DEL-708 | chr7-73824955-DEL-708 |
| chr7-73832197-DEL-80 | chr7-73832198-DEL-80 |
| chr7-73885186-DEL-201 | chr7-73885187-DEL-201 |
| chr7-73969426-DEL-217 | chr7-73969427-DEL-217 |
| chr7-73974543-DEL-60 | chr7-73974544-DEL-60 |
| chr7-73988652-DEL-210 | chr7-73988653-DEL-210 |
| chr7-73991764-DEL-74 | chr7-73991781-DEL-58 |
| chr7-74040413-DEL-106 | chr7-74040414-DEL-106 |
| chr7-74047828-DEL-656 | chr7-74047829-DEL-656 |
| chr7-74064579-DEL-65 | chr7-74064582-DEL-61 |
| chr7-74080628-DEL-135 | chr7-74080629-DEL-135 |
| chr7-74088567-DEL-52 | chr7-74088568-DEL-52 |
| chr7-74152577-DEL-62 | chr7-74152578-DEL-62 |
| chr7-74173381-DEL-203 | chr7-74173376-DEL-226 |
| chr7-74189690-DEL-114 | chr7-74189691-DEL-114 |
| chr7-74191998-DEL-99 | chr7-74191999-DEL-99 |
| chr7-74225394-DEL-711 | chr7-74225395-DEL-711 |
| chr7-74228959-DEL-60 | chr7-74228940-DEL-76 |
| chr7-74237491-DEL-1199 | chr7-74237492-DEL-1199 |
| chr7-74251024-DEL-6292 | chr7-74251025-DEL-6292 |
| chr7-74263753-DEL-282 | chr7-74263754-DEL-282 |
| chr7-74331015-DEL-767 | chr7-74331013-DEL-765 |
| chr7-74374520-DEL-484 | chr7-74374521-DEL-484 |
| chr7-74407176-DEL-136 | chr7-74407177-DEL-136 |
| chr7-74414929-DEL-131 | chr7-74414914-DEL-147 |
| chr7-74449050-DEL-6935 | chr7-74449051-DEL-6935 |
| chr7-74457700-DEL-227 | chr7-74457701-DEL-227 |
| chr7-74503864-DEL-139 | chr7-74503865-DEL-139 |
| chr7-74542708-DEL-6407 | chr7-74542709-DEL-6407 |
| chr7-74550615-DEL-57 | chr7-74550616-DEL-57 |

| | |
|---|---|
| chr7-74554207-DEL-2702 | chr7-74554211-DEL-2699 |
| chr7-74587176-DEL-107 | chr7-74587177-DEL-107 |
| chr7-74603732-DEL-99 | chr7-74603733-DEL-99 |
| chr7-74605083-DEL-6581 | chr7-74605073-DEL-6577 |
| chr7-74637725-DEL-67 | chr7-74637730-DEL-63 |
| chr7-74652861-DEL-4631 | chr7-74652862-DEL-4631 |
| chr7-74660370-DEL-270 | chr7-74660371-DEL-270 |
| chr7-74661727-DEL-52 | chr7-74661728-DEL-52 |
| chr7-74673228-DEL-355 | chr7-74673229-DEL-355 |
| chr7-74706185-DEL-1174 | chr7-74706186-DEL-1174 |
| chr7-74737451-DEL-372 | chr7-74737452-DEL-372 |
| chr7-74743619-DEL-124 | chr7-74743620-DEL-124 |
| chr7-74754916-DEL-77 | chr7-74754917-DEL-77 |
| chr7-74759887-DEL-269 | chr7-74759888-DEL-269 |
| chr7-74793822-DEL-480 | chr7-74793823-DEL-492 |
| chr7-74830511-DEL-6440 | chr7-74830512-DEL-6440 |
| chr7-74879062-DEL-85 | chr7-74879045-DEL-87 |
| chr7-74900714-DEL-976 | chr7-74900715-DEL-976 |
| chr7-74905182-DEL-70 | chr7-74905183-DEL-71 |
| chr7-74922574-DEL-215 | chr7-74922599-DEL-191 |
| chr7-74938194-DEL-1727 | chr7-74938194-DEL-1727 |
| chr7-74954066-DEL-62 | chr7-74954059-DEL-52 |
| chr7-74962872-DEL-165 | chr7-74962857-DEL-181 |
| chr7-74966748-DEL-123 | chr7-74966749-DEL-123 |
| chr7-74984419-DEL-125 | chr7-74984421-DEL-126 |
| chr7-75002345-DEL-126 | chr7-75002346-DEL-126 |
| chr7-75008593-DEL-192 | chr7-75008590-DEL-199 |
| chr7-75023830-DEL-51 | chr7-75023831-DEL-51 |
| chr7-75053756-DEL-693 | chr7-75053765-DEL-682 |
| chr7-75097898-DEL-1204 | chr7-75097899-DEL-1204 |
| chr7-75099400-DEL-220 | chr7-75099401-DEL-220 |
| chr7-75139376-DEL-191 | chr7-75139364-DEL-193 |
| chr7-75141607-DEL-56 | chr7-75141581-DEL-88 |
| chr7-75144883-DEL-51 | chr7-75144880-DEL-61 |
| chr7-75169513-DEL-53 | chr7-75169514-DEL-53 |
| chr7-75171940-DEL-596 | chr7-75171941-DEL-596 |
| chr7-75191309-DEL-6329 | chr7-75191310-DEL-6329 |
| chr7-75206312-DEL-132 | chr7-75206311-DEL-132 |
| chr7-75207119-DEL-215 | chr7-75207120-DEL-215 |
| chr7-75211863-DEL-214 | chr7-75211864-DEL-214 |
| chr7-75224048-DEL-172 | chr7-75224042-DEL-173 |
| chr7-75242913-DEL-321 | chr7-75242914-DEL-321 |
| chr7-75255914-DEL-669 | chr7-75255915-DEL-669 |
| chr7-75285666-DEL-1799 | chr7-75285667-DEL-1799 |
| chr7-75373808-DEL-58 | chr7-75373809-DEL-58 |
| chr7-75383618-DEL-129 | chr7-75383634-DEL-114 |
| chr7-75406344-DEL-68 | chr7-75406359-DEL-79 |
| chr7-75406804-DEL-530 | chr7-75406805-DEL-530 |
| chr7-75407916-DEL-397 | chr7-75407917-DEL-397 |
| chr7-75453407-DEL-1843 | chr7-75453405-DEL-1846 |
| chr7-75457053-DEL-74 | chr7-75457054-DEL-74 |
| chr7-75459508-DEL-638 | chr7-75459540-DEL-614 |
| chr7-75460227-DEL-7133 | chr7-75460228-DEL-7133 |
| chr7-75467493-DEL-107 | chr7-75467492-DEL-108 |
| chr7-75480952-DEL-136 | chr7-75480952-DEL-136 |
| chr7-75481972-DEL-401 | chr7-75481973-DEL-401 |
| chr7-75487379-DEL-65 | chr7-75487380-DEL-65 |
| chr7-75497072-DEL-126 | chr7-75497073-DEL-126 |
| chr7-75518478-DEL-82 | chr7-75518479-DEL-82 |
| chr7-75536120-DEL-301 | chr7-75536121-DEL-301 |
| chr7-75583389-DEL-1523 | chr7-75583390-DEL-1523 |
| chr7-75589824-DEL-80 | chr7-75589825-DEL-80 |
| chr7-75595540-DEL-5716 | chr7-75595541-DEL-5716 |
| chr7-75600673-DEL-214 | chr7-75600674-DEL-214 |
| chr7-75623085-DEL-6510 | chr7-75623072-DEL-6511 |
| chr7-75666345-DEL-7255 | chr7-75666346-DEL-7255 |
| chr7-75690917-DEL-2845 | chr7-75690918-DEL-2845 |
| chr7-75694767-DEL-69 | chr7-75694766-DEL-67 |
| chr7-75701245-DEL-62 | chr7-75701246-DEL-62 |
| chr7-75717519-DEL-6216 | chr7-75717519-DEL-6216 |
| chr7-75730000-DEL-72 | chr7-75729998-DEL-75 |
| chr7-75731589-DEL-91 | chr7-75731587-DEL-92 |
| chr7-75755937-DEL-773 | chr7-75755937-DEL-773 |
| chr7-75760834-DEL-6285 | chr7-75760835-DEL-6285 |
| chr7-75783171-DEL-457 | chr7-75783170-DEL-413 |
| chr7-75810396-DEL-2183 | chr7-75810397-DEL-2183 |
| chr7-75811878-DEL-78 | chr7-75811866-DEL-90 |
| chr7-75817820-DEL-90 | chr7-75817821-DEL-90 |
| chr7-75846122-DEL-6175 | chr7-75846123-DEL-6175 |
| chr7-75866008-DEL-63 | chr7-75866009-DEL-63 |

| | |
|---|---|
| chr7-75892898-DEL-2970 | chr7-75892899-DEL-2970 |
| chr7-75912476-DEL-3357 | chr7-75912477-DEL-3357 |
| chr7-75918160-DEL-273 | chr7-75918161-DEL-273 |
| chr7-75929061-DEL-89 | chr7-75929062-DEL-89 |
| chr7-75977780-DEL-4298 | chr7-75977781-DEL-4298 |
| chr7-75986158-DEL-890 | chr7-75986159-DEL-890 |
| chr7-76115287-DEL-127 | chr7-76115284-DEL-128 |
| chr7-76124949-DEL-181 | chr7-76124950-DEL-181 |
| chr7-76154505-DEL-406 | chr7-76154506-DEL-406 |
| chr7-76159835-DEL-52 | chr7-76159836-DEL-52 |
| chr7-76175075-DEL-128 | chr7-76175075-DEL-129 |
| chr7-76201406-DEL-65 | chr7-76201418-DEL-62 |
| chr7-76213821-DEL-1093 | chr7-76213823-DEL-1092 |
| chr7-76215963-DEL-192 | chr7-76215964-DEL-192 |
| chr7-76216495-DEL-6252 | chr7-76216511-DEL-6218 |
| chr7-76256158-DEL-87 | chr7-76256171-DEL-96 |
| chr7-76281830-DEL-56 | chr7-76281830-DEL-56 |
| chr7-76285393-DEL-68 | chr7-76285392-DEL-80 |
| chr7-76335066-DEL-58 | chr7-76335068-DEL-52 |
| chr7-76353021-DEL-6825 | chr7-76353022-DEL-6825 |
| chr7-76409145-DEL-52 | chr7-76409146-DEL-52 |
| chr7-76452980-DEL-269 | chr7-76452981-DEL-269 |
| chr7-76458428-DEL-97 | chr7-76458420-DEL-98 |
| chr7-76478784-DEL-71 | chr7-76478785-DEL-70 |
| chr7-76482236-DEL-6993 | chr7-76482237-DEL-6993 |
| chr7-76502955-DEL-231 | chr7-76502956-DEL-231 |
| chr7-76513596-DEL-628 | chr7-76513600-DEL-623 |
| chr7-76517796-DEL-70 | chr7-76517797-DEL-68 |
| chr7-76548935-DEL-50 | chr7-76548925-DEL-53 |
| chr7-76555099-DEL-75 | chr7-76555100-DEL-77 |
| chr7-76560609-DEL-1104 | chr7-76560610-DEL-1104 |
| chr7-76570306-DEL-56 | chr7-76570307-DEL-56 |
| chr7-76605805-DEL-159 | chr7-76605832-DEL-132 |
| chr7-76617907-DEL-62 | chr7-76617912-DEL-61 |
| chr7-76618584-DEL-218 | chr7-76618585-DEL-218 |
| chr7-76621694-DEL-83 | chr7-76621684-DEL-88 |
| chr7-76623808-DEL-76 | chr7-76623803-DEL-83 |
| chr7-76650594-DEL-212 | chr7-76650599-DEL-208 |
| chr7-76682323-DEL-57 | chr7-76682303-DEL-68 |
| chr7-76744116-DEL-69 | chr7-76744117-DEL-69 |
| chr7-76760339-DEL-197 | chr7-76760340-DEL-197 |
| chr7-76765446-DEL-76 | chr7-76765470-DEL-91 |
| chr7-76767689-DEL-78 | chr7-76767702-DEL-75 |
| chr7-76788029-DEL-907 | chr7-76788030-DEL-907 |
| chr7-76804197-DEL-94 | chr7-76804198-DEL-86 |
| chr7-76862405-DEL-7095 | chr7-76862404-DEL-7096 |
| chr7-76916294-DEL-72 | chr7-76916295-DEL-72 |
| chr7-76926413-DEL-915 | chr7-76926414-DEL-914 |
| chr7-76953580-DEL-78 | chr7-76953571-DEL-84 |
| chr7-76955934-DEL-457 | chr7-76955935-DEL-456 |
| chr7-76961128-DEL-127 | chr7-76961129-DEL-127 |
| chr7-76978276-DEL-6643 | chr7-76978271-DEL-6649 |
| chr7-76992896-DEL-406 | chr7-76992897-DEL-406 |
| chr7-76997348-DEL-631 | chr7-76997349-DEL-631 |
| chr7-77019767-DEL-212 | chr7-77019768-DEL-212 |
| chr7-77045453-DEL-52 | chr7-77045454-DEL-52 |
| chr7-77055469-DEL-1018 | chr7-77055470-DEL-1018 |
| chr7-77134009-DEL-111 | chr7-77134010-DEL-111 |
| chr7-77170169-DEL-6554 | chr7-77170170-DEL-6555 |
| chr7-77213720-DEL-121 | chr7-77213733-DEL-102 |
| chr7-77287530-DEL-64 | chr7-77287527-DEL-54 |
| chr7-77297461-DEL-60 | chr7-77297463-DEL-68 |
| chr7-77297848-DEL-7076 | chr7-77297861-DEL-7072 |
| chr7-77304609-DEL-212 | chr7-77304610-DEL-212 |
| chr7-77324450-DEL-153 | chr7-77324459-DEL-154 |
| chr7-77348338-DEL-62 | chr7-77348338-DEL-62 |
| chr7-77351697-DEL-63 | chr7-77351682-DEL-60 |
| chr7-77388160-DEL-68 | chr7-77388160-DEL-71 |
| chr7-77411394-DEL-413 | chr7-77411395-DEL-413 |
| chr7-77413845-DEL-54 | chr7-77413846-DEL-54 |
| chr7-77440634-DEL-345 | chr7-77440644-DEL-336 |
| chr7-77447137-DEL-67 | chr7-77447138-DEL-67 |
| chr7-77466747-DEL-63 | chr7-77466746-DEL-64 |
| chr7-77510541-DEL-80 | chr7-77510542-DEL-80 |
| chr7-77524774-DEL-6476 | chr7-77524775-DEL-6476 |
| chr7-77551431-DEL-55 | chr7-77551406-DEL-52 |
| chr7-77569105-DEL-84 | chr7-77569106-DEL-84 |
| chr7-77571828-DEL-366 | chr7-77571829-DEL-366 |
| chr7-77580369-DEL-53 | chr7-77580362-DEL-52 |
| chr7-77583620-DEL-72 | chr7-77583662-DEL-72 |

| | |
|---|---|
| chr7-77584963-DEL-51 | chr7-77584964-DEL-51 |
| chr7-77603651-DEL-6948 | chr7-77603651-DEL-6948 |
| chr7-77621377-DEL-90 | chr7-77621380-DEL-80 |
| chr7-77641007-DEL-185 | chr7-77641008-DEL-185 |
| chr7-77669552-DEL-2014 | chr7-77669553-DEL-2014 |
| chr7-77763847-DEL-1283 | chr7-77763848-DEL-1283 |
| chr7-77795582-DEL-59 | chr7-77795584-DEL-60 |
| chr7-77797042-DEL-86 | chr7-77797055-DEL-92 |
| chr7-77912625-DEL-104 | chr7-77912640-DEL-100 |
| chr7-78065772-DEL-6783 | chr7-78065773-DEL-6783 |
| chr7-78131990-DEL-267 | chr7-78131991-DEL-267 |
| chr7-78142358-DEL-194 | chr7-78142359-DEL-194 |
| chr7-78166161-DEL-114 | chr7-78166157-DEL-109 |
| chr7-78236230-DEL-164 | chr7-78236231-DEL-162 |
| chr7-78264594-DEL-201 | chr7-78264595-DEL-201 |
| chr7-78271469-DEL-74 | chr7-78271465-DEL-108 |
| chr7-78274063-DEL-77 | chr7-78274058-DEL-82 |
| chr7-78285179-DEL-1126 | chr7-78285177-DEL-1123 |
| chr7-78298468-DEL-71 | chr7-78298468-DEL-67 |
| chr7-78313655-DEL-143 | chr7-78313656-DEL-139 |
| chr7-78323954-DEL-866 | chr7-78323955-DEL-867 |
| chr7-78334371-DEL-62 | chr7-78334372-DEL-62 |
| chr7-78353118-DEL-57 | chr7-78353119-DEL-57 |
| chr7-78377281-DEL-79 | chr7-78377277-DEL-82 |
| chr7-78389974-DEL-207 | chr7-78389975-DEL-207 |
| chr7-78400416-DEL-616 | chr7-78400417-DEL-616 |
| chr7-78501977-DEL-52 | chr7-78501978-DEL-52 |
| chr7-78517957-DEL-66 | chr7-78517971-DEL-50 |
| chr7-78548703-DEL-2876 | chr7-78548704-DEL-2876 |
| chr7-78570687-DEL-211 | chr7-78570688-DEL-211 |
| chr7-78600041-DEL-54 | chr7-78600044-DEL-54 |
| chr7-78618494-DEL-108 | chr7-78618491-DEL-111 |
| chr7-78623666-DEL-66 | chr7-78623667-DEL-66 |
| chr7-78633965-DEL-148 | chr7-78633974-DEL-154 |
| chr7-78654694-DEL-157 | chr7-78654693-DEL-153 |
| chr7-78664782-DEL-1226 | chr7-78664767-DEL-1204 |
| chr7-78665391-DEL-2354 | chr7-78665392-DEL-2354 |
| chr7-78666547-DEL-56 | chr7-78666562-DEL-64 |
| chr7-78700357-DEL-62 | chr7-78700358-DEL-62 |
| chr7-78719833-DEL-185 | chr7-78719834-DEL-185 |
| chr7-78815672-DEL-68 | chr7-78815673-DEL-68 |
| chr7-78849147-DEL-56 | chr7-78849148-DEL-56 |
| chr7-78849809-DEL-194 | chr7-78849810-DEL-193 |
| chr7-78876402-DEL-342 | chr7-78876403-DEL-342 |
| chr7-78884272-DEL-217 | chr7-78884273-DEL-217 |
| chr7-78885056-DEL-175 | chr7-78885057-DEL-174 |
| chr7-78888269-DEL-119 | chr7-78888279-DEL-113 |
| chr7-78890136-DEL-143 | chr7-78890128-DEL-110 |
| chr7-78898442-DEL-230 | chr7-78898443-DEL-230 |
| chr7-78916393-DEL-87 | chr7-78916434-DEL-50 |
| chr7-78928675-DEL-67 | chr7-78928676-DEL-67 |
| chr7-78940086-DEL-219 | chr7-78940087-DEL-209 |
| chr7-78945694-DEL-301 | chr7-78945695-DEL-301 |
| chr7-78979537-DEL-1198 | chr7-78979538-DEL-1198 |
| chr7-78986806-DEL-5537 | chr7-78986807-DEL-5537 |
| chr7-78995801-DEL-827 | chr7-78995801-DEL-827 |
| chr7-79018017-DEL-115 | chr7-79018018-DEL-115 |
| chr7-79030934-DEL-137 | chr7-79030935-DEL-137 |
| chr7-79039798-DEL-52 | chr7-79039799-DEL-52 |
| chr7-79059319-DEL-243 | chr7-79059323-DEL-244 |
| chr7-79086519-DEL-195 | chr7-79086520-DEL-195 |
| chr7-79112522-DEL-221 | chr7-79112523-DEL-221 |
| chr7-79127736-DEL-1892 | chr7-79127737-DEL-1892 |
| chr7-79129237-DEL-231 | chr7-79129237-DEL-231 |
| chr7-79130864-DEL-54 | chr7-79130870-DEL-50 |
| chr7-79174428-DEL-73 | chr7-79174429-DEL-73 |
| chr7-79237058-DEL-377 | chr7-79237059-DEL-377 |
| chr7-79273964-DEL-199 | chr7-79273965-DEL-199 |
| chr7-79282346-DEL-217 | chr7-79282347-DEL-217 |
| chr7-79286331-DEL-200 | chr7-79286317-DEL-201 |
| chr7-79302944-DEL-248 | chr7-79302945-DEL-248 |
| chr7-79305136-DEL-174 | chr7-79305137-DEL-174 |
| chr7-79308056-DEL-97 | chr7-79308057-DEL-97 |
| chr7-79321669-DEL-69 | chr7-79321670-DEL-71 |
| chr7-79364285-DEL-99 | chr7-79364286-DEL-99 |
| chr7-79383402-DEL-56 | chr7-79383379-DEL-59 |
| chr7-79390388-DEL-186 | chr7-79390389-DEL-186 |
| chr7-79399519-DEL-81 | chr7-79399520-DEL-81 |
| chr7-79410057-DEL-52 | chr7-79410058-DEL-52 |
| chr7-79412952-DEL-801 | chr7-79412953-DEL-801 |

| | |
|---|---|
| chr7-79466053-DEL-1041 | chr7-79466054-DEL-1041 |
| chr7-79527736-DEL-721 | chr7-79527735-DEL-722 |
| chr7-79548538-DEL-204 | chr7-79548539-DEL-204 |
| chr7-79580513-DEL-161 | chr7-79580514-DEL-161 |
| chr7-79598459-DEL-225 | chr7-79598460-DEL-225 |
| chr7-79604790-DEL-133 | chr7-79604782-DEL-144 |
| chr7-79608575-DEL-51 | chr7-79608577-DEL-52 |
| chr7-79612817-DEL-354 | chr7-79612818-DEL-354 |
| chr7-79630288-DEL-201 | chr7-79630289-DEL-201 |
| chr7-79645763-DEL-984 | chr7-79645764-DEL-984 |
| chr7-79654016-DEL-208 | chr7-79654017-DEL-208 |
| chr7-79656120-DEL-52 | chr7-79656121-DEL-52 |
| chr7-79658781-DEL-341 | chr7-79658782-DEL-341 |
| chr7-79661740-DEL-52 | chr7-79661729-DEL-64 |
| chr7-79676533-DEL-73 | chr7-79676524-DEL-90 |
| chr7-79693605-DEL-182 | chr7-79693606-DEL-182 |
| chr7-79754773-DEL-196 | chr7-79754774-DEL-196 |
| chr7-79781074-DEL-73 | chr7-79781051-DEL-101 |
| chr7-79797502-DEL-110 | chr7-79797490-DEL-126 |
| chr7-79799968-DEL-53 | chr7-79799995-DEL-60 |
| chr7-79812993-DEL-210 | chr7-79812994-DEL-210 |
| chr7-79834158-DEL-193 | chr7-79834159-DEL-193 |
| chr7-79876198-DEL-120 | chr7-79876198-DEL-114 |
| chr7-79883294-DEL-328 | chr7-79883295-DEL-328 |
| chr7-79912532-DEL-1364 | chr7-79912533-DEL-1364 |
| chr7-79919090-DEL-118 | chr7-79919091-DEL-118 |
| chr7-79921751-DEL-54 | chr7-79921752-DEL-54 |
| chr7-79941059-DEL-72 | chr7-79941030-DEL-74 |
| chr7-79950380-DEL-120 | chr7-79950381-DEL-120 |
| chr7-79966282-DEL-157 | chr7-79966287-DEL-151 |
| chr7-79967715-DEL-322 | chr7-79967716-DEL-322 |
| chr7-79982216-DEL-94 | chr7-79982217-DEL-94 |
| chr7-79989462-DEL-195 | chr7-79989464-DEL-196 |
| chr7-79993835-DEL-1384 | chr7-79993836-DEL-1384 |
| chr7-80044148-DEL-247 | chr7-80044149-DEL-247 |
| chr7-80078640-DEL-210 | chr7-80078654-DEL-209 |
| chr7-80081703-DEL-61 | chr7-80081704-DEL-61 |
| chr7-80085678-DEL-58 | chr7-80085678-DEL-58 |
| chr7-80157688-DEL-56 | chr7-80157689-DEL-56 |
| chr7-80164807-DEL-359 | chr7-80164803-DEL-360 |
| chr7-80198004-DEL-70 | chr7-80198005-DEL-70 |
| chr7-80202847-DEL-369 | chr7-80202841-DEL-356 |
| chr7-80213242-DEL-1545 | chr7-80213245-DEL-1539 |
| chr7-80213411-DEL-217 | chr7-80213412-DEL-217 |
| chr7-80217359-DEL-203 | chr7-80217360-DEL-203 |
| chr7-80255946-DEL-80 | chr7-80255927-DEL-90 |
| chr7-80257812-DEL-85 | chr7-80257809-DEL-85 |
| chr7-80301896-DEL-64 | chr7-80301897-DEL-64 |
| chr7-80330555-DEL-62 | chr7-80330570-DEL-72 |
| chr7-80365772-DEL-205 | chr7-80365773-DEL-205 |
| chr7-80370427-DEL-52 | chr7-80370428-DEL-52 |
| chr7-80374206-DEL-140 | chr7-80374194-DEL-140 |
| chr7-80378339-DEL-87 | chr7-80378336-DEL-102 |
| chr7-80404527-DEL-56 | chr7-80404511-DEL-64 |
| chr7-80407782-DEL-77 | chr7-80407781-DEL-78 |
| chr7-80422117-DEL-201 | chr7-80422118-DEL-201 |
| chr7-80470132-DEL-72 | chr7-80470133-DEL-72 |
| chr7-80479908-DEL-209 | chr7-80479905-DEL-207 |
| chr7-80480830-DEL-170 | chr7-80480849-DEL-170 |
| chr7-80483114-DEL-221 | chr7-80483115-DEL-221 |
| chr7-80486791-DEL-165 | chr7-80486792-DEL-165 |
| chr7-80501124-DEL-203 | chr7-80501125-DEL-203 |
| chr7-80528639-DEL-188 | chr7-80528648-DEL-185 |
| chr7-80544725-DEL-87 | chr7-80544726-DEL-87 |
| chr7-80560454-DEL-126 | chr7-80560455-DEL-126 |
| chr7-80580030-DEL-80 | chr7-80580030-DEL-80 |
| chr7-80585079-DEL-357 | chr7-80585080-DEL-357 |
| chr7-80601721-DEL-107 | chr7-80601752-DEL-74 |
| chr7-80610424-DEL-1893 | chr7-80610425-DEL-1893 |
| chr7-80623022-DEL-65 | chr7-80623023-DEL-65 |
| chr7-80624863-DEL-107 | chr7-80624864-DEL-101 |
| chr7-80626729-DEL-362 | chr7-80626730-DEL-362 |
| chr7-80627968-DEL-88 | chr7-80627969-DEL-88 |
| chr7-80631167-DEL-69 | chr7-80631171-DEL-60 |
| chr7-80763358-DEL-90 | chr7-80763359-DEL-90 |
| chr7-80797289-DEL-1888 | chr7-80797290-DEL-1888 |
| chr7-80809957-DEL-59 | chr7-80809958-DEL-59 |
| chr7-80836376-DEL-210 | chr7-80836377-DEL-210 |
| chr7-80887702-DEL-104 | chr7-80887703-DEL-104 |
| chr7-80904882-DEL-51 | chr7-80904883-DEL-51 |

| | |
|---|---|
| chr7-80956852-DEL-56 | chr7-80956853-DEL-56 |
| chr7-80961956-DEL-196 | chr7-80961957-DEL-196 |
| chr7-81024933-DEL-131 | chr7-81024934-DEL-131 |
| chr7-81045338-DEL-72 | chr7-81045347-DEL-80 |
| chr7-81055934-DEL-144 | chr7-81055933-DEL-145 |
| chr7-81063064-DEL-179 | chr7-81063065-DEL-179 |
| chr7-81070189-DEL-904 | chr7-81070190-DEL-905 |
| chr7-81132035-DEL-198 | chr7-81132028-DEL-199 |
| chr7-81146389-DEL-52 | chr7-81146390-DEL-52 |
| chr7-81152238-DEL-54 | chr7-81152239-DEL-54 |
| chr7-81182969-DEL-52 | chr7-81182968-DEL-52 |
| chr7-81204530-DEL-125 | chr7-81204531-DEL-125 |
| chr7-81232752-DEL-209 | chr7-81232753-DEL-209 |
| chr7-81238693-DEL-60 | chr7-81238689-DEL-70 |
| chr7-81239688-DEL-112 | chr7-81239720-DEL-81 |
| chr7-81253272-DEL-51 | chr7-81253273-DEL-51 |
| chr7-81266349-DEL-51 | chr7-81266350-DEL-51 |
| chr7-81275013-DEL-218 | chr7-81275014-DEL-218 |
| chr7-81284014-DEL-1865 | chr7-81284015-DEL-1865 |
| chr7-81286446-DEL-305 | chr7-81286448-DEL-306 |
| chr7-81287308-DEL-95 | chr7-81287310-DEL-64 |
| chr7-81288060-DEL-63 | chr7-81288061-DEL-63 |
| chr7-81297490-DEL-116 | chr7-81297491-DEL-118 |
| chr7-81311215-DEL-60 | chr7-81311216-DEL-60 |
| chr7-81351901-DEL-70 | chr7-81351902-DEL-70 |
| chr7-81367106-DEL-532 | chr7-81367107-DEL-532 |
| chr7-81390533-DEL-881 | chr7-81390534-DEL-881 |
| chr7-81492369-DEL-62 | chr7-81492370-DEL-62 |
| chr7-81502166-DEL-202 | chr7-81502168-DEL-201 |
| chr7-81526445-DEL-457 | chr7-81526446-DEL-457 |
| chr7-81542163-DEL-484 | chr7-81542164-DEL-484 |
| chr7-81592021-DEL-518 | chr7-81592022-DEL-518 |
| chr7-81607266-DEL-55 | chr7-81607269-DEL-72 |
| chr7-81665743-DEL-203 | chr7-81665744-DEL-203 |
| chr7-81687705-DEL-86 | chr7-81687706-DEL-86 |
| chr7-81740078-DEL-390 | chr7-81740079-DEL-390 |
| chr7-81804739-DEL-6244 | chr7-81804740-DEL-6244 |
| chr7-81811542-DEL-9715 | chr7-81811543-DEL-9715 |
| chr7-81838852-DEL-156 | chr7-81838848-DEL-135 |
| chr7-81863087-DEL-6902 | chr7-81863088-DEL-6902 |
| chr7-81882297-DEL-191 | chr7-81882298-DEL-191 |
| chr7-81885550-DEL-434 | chr7-81885551-DEL-434 |
| chr7-81887953-DEL-512 | chr7-81887954-DEL-512 |
| chr7-81892305-DEL-102 | chr7-81892306-DEL-90 |
| chr7-81900362-DEL-52 | chr7-81900362-DEL-52 |
| chr7-81902461-DEL-85 | chr7-81902462-DEL-85 |
| chr7-81903434-DEL-182 | chr7-81903435-DEL-182 |
| chr7-81910265-DEL-185 | chr7-81910266-DEL-185 |
| chr7-81922684-DEL-66 | chr7-81922685-DEL-66 |
| chr7-81936142-DEL-88 | chr7-81936109-DEL-78 |
| chr7-81946329-DEL-2649 | chr7-81946330-DEL-2649 |
| chr7-81951922-DEL-86 | chr7-81951923-DEL-86 |
| chr7-81954864-DEL-65 | chr7-81954865-DEL-65 |
| chr7-81968224-DEL-189 | chr7-81968212-DEL-190 |
| chr7-81970398-DEL-628 | chr7-81970399-DEL-628 |
| chr7-81971412-DEL-67 | chr7-81971419-DEL-62 |
| chr7-81977645-DEL-1012 | chr7-81977635-DEL-1012 |
| chr7-81978433-DEL-324 | chr7-81978434-DEL-324 |
| chr7-82001529-DEL-170 | chr7-82001516-DEL-147 |
| chr7-82011220-DEL-68 | chr7-82011221-DEL-68 |
| chr7-82017849-DEL-54 | chr7-82017849-DEL-54 |
| chr7-82037013-DEL-70 | chr7-82036985-DEL-84 |
| chr7-82081490-DEL-271 | chr7-82081491-DEL-271 |
| chr7-82082176-DEL-120 | chr7-82082221-DEL-94 |
| chr7-82092538-DEL-112 | chr7-82092539-DEL-112 |
| chr7-82103268-DEL-157 | chr7-82103269-DEL-157 |
| chr7-82117257-DEL-6192 | chr7-82117258-DEL-6192 |
| chr7-82147689-DEL-92 | chr7-82147690-DEL-92 |
| chr7-82158447-DEL-2370 | chr7-82158448-DEL-2370 |
| chr7-82188335-DEL-395 | chr7-82188336-DEL-395 |
| chr7-82198059-DEL-1574 | chr7-82198060-DEL-1574 |
| chr7-82218962-DEL-1789 | chr7-82218963-DEL-1789 |
| chr7-82245289-DEL-64 | chr7-82245290-DEL-64 |
| chr7-82268646-DEL-74 | chr7-82268647-DEL-74 |
| chr7-82271980-DEL-252 | chr7-82271981-DEL-252 |
| chr7-82273839-DEL-201 | chr7-82273840-DEL-201 |
| chr7-82298223-DEL-214 | chr7-82298224-DEL-214 |
| chr7-82301986-DEL-92 | chr7-82301987-DEL-92 |
| chr7-82303162-DEL-256 | chr7-82303163-DEL-256 |
| chr7-82331141-DEL-508 | chr7-82331142-DEL-508 |

| | |
|---|---|
| chr7-82353592-DEL-1248 | chr7-82353593-DEL-1248 |
| chr7-82363477-DEL-150 | chr7-82363477-DEL-151 |
| chr7-82386347-DEL-4757 | chr7-82386339-DEL-4769 |
| chr7-82407440-DEL-214 | chr7-82407441-DEL-214 |
| chr7-82420211-DEL-1074 | chr7-82420212-DEL-1074 |
| chr7-82433388-DEL-74 | chr7-82433389-DEL-74 |
| chr7-82448030-DEL-599 | chr7-82448031-DEL-599 |
| chr7-82449473-DEL-3083 | chr7-82449474-DEL-3084 |
| chr7-82475161-DEL-1430 | chr7-82475162-DEL-1430 |
| chr7-82478047-DEL-227 | chr7-82478048-DEL-227 |
| chr7-82489213-DEL-74 | chr7-82489214-DEL-74 |
| chr7-82522829-DEL-74 | chr7-82522830-DEL-74 |
| chr7-82531148-DEL-1172 | chr7-82531149-DEL-1172 |
| chr7-82542287-DEL-254 | chr7-82542288-DEL-254 |
| chr7-82587569-DEL-82 | chr7-82587570-DEL-82 |
| chr7-82645934-DEL-6512 | chr7-82645934-DEL-6512 |
| chr7-82663915-DEL-122 | chr7-82663890-DEL-132 |
| chr7-82685494-DEL-61 | chr7-82685495-DEL-71 |
| chr7-82727847-DEL-56 | chr7-82727848-DEL-56 |
| chr7-82763781-DEL-211 | chr7-82763734-DEL-218 |
| chr7-82828838-DEL-6748 | chr7-82828839-DEL-6748 |
| chr7-82835075-DEL-212 | chr7-82835076-DEL-212 |
| chr7-82883594-DEL-599 | chr7-82883595-DEL-599 |
| chr7-82915263-DEL-7180 | chr7-82915264-DEL-7180 |
| chr7-82923123-DEL-98 | chr7-82923124-DEL-97 |
| chr7-82947035-DEL-70 | chr7-82947036-DEL-70 |
| chr7-82999560-DEL-408 | chr7-82999561-DEL-408 |
| chr7-83003415-DEL-520 | chr7-83003416-DEL-520 |
| chr7-83115370-DEL-70 | chr7-83115338-DEL-87 |
| chr7-83125676-DEL-346 | chr7-83125677-DEL-346 |
| chr7-83130015-DEL-206 | chr7-83130016-DEL-206 |
| chr7-83142217-DEL-2315 | chr7-83142218-DEL-2315 |
| chr7-83163084-DEL-88 | chr7-83163084-DEL-88 |
| chr7-83165425-DEL-68 | chr7-83165383-DEL-70 |
| chr7-83177710-DEL-192 | chr7-83177715-DEL-188 |
| chr7-83182855-DEL-538 | chr7-83182855-DEL-539 |
| chr7-83204742-DEL-215 | chr7-83204743-DEL-215 |
| chr7-83321034-DEL-291 | chr7-83321042-DEL-287 |
| chr7-83338828-DEL-5553 | chr7-83338829-DEL-5553 |
| chr7-83356816-DEL-60 | chr7-83356817-DEL-66 |
| chr7-83391133-DEL-55 | chr7-83391134-DEL-55 |
| chr7-83405015-DEL-1995 | chr7-83405014-DEL-1995 |
| chr7-83406024-DEL-239 | chr7-83406025-DEL-239 |
| chr7-83409295-DEL-92 | chr7-83409292-DEL-51 |
| chr7-83416334-DEL-52 | chr7-83416327-DEL-53 |
| chr7-83493047-DEL-55 | chr7-83493048-DEL-55 |
| chr7-83524244-DEL-345 | chr7-83524245-DEL-345 |
| chr7-83539413-DEL-74 | chr7-83539413-DEL-75 |
| chr7-83564835-DEL-102 | chr7-83564835-DEL-102 |
| chr7-83566081-DEL-64 | chr7-83566082-DEL-64 |
| chr7-83572858-DEL-5333 | chr7-83572859-DEL-5333 |
| chr7-83587225-DEL-58 | chr7-83587226-DEL-58 |
| chr7-83620168-DEL-129 | chr7-83620169-DEL-129 |
| chr7-83642067-DEL-1395 | chr7-83642061-DEL-1396 |
| chr7-83648743-DEL-85 | chr7-83648743-DEL-86 |
| chr7-83663607-DEL-58 | chr7-83663608-DEL-58 |
| chr7-83680741-DEL-54 | chr7-83680742-DEL-52 |
| chr7-83720053-DEL-167 | chr7-83720061-DEL-166 |
| chr7-83738519-DEL-219 | chr7-83738520-DEL-219 |
| chr7-83765189-DEL-83 | chr7-83765172-DEL-82 |
| chr7-83810374-DEL-84 | chr7-83810375-DEL-84 |
| chr7-83833958-DEL-310 | chr7-83833959-DEL-310 |
| chr7-83843531-DEL-53 | chr7-83843532-DEL-53 |
| chr7-83846944-DEL-508 | chr7-83846945-DEL-508 |
| chr7-83856564-DEL-532 | chr7-83856563-DEL-534 |
| chr7-83864820-DEL-302 | chr7-83864820-DEL-303 |
| chr7-83872667-DEL-2356 | chr7-83872668-DEL-2356 |
| chr7-83879107-DEL-362 | chr7-83879110-DEL-326 |
| chr7-83885556-DEL-341 | chr7-83885557-DEL-341 |
| chr7-83886244-DEL-87 | chr7-83886244-DEL-50 |
| chr7-83927179-DEL-209 | chr7-83927180-DEL-209 |
| chr7-83941747-DEL-182 | chr7-83941748-DEL-181 |
| chr7-83971686-DEL-937 | chr7-83971687-DEL-937 |
| chr7-84017531-DEL-1890 | chr7-84017532-DEL-1890 |
| chr7-84023240-DEL-183 | chr7-84023241-DEL-183 |
| chr7-84116861-DEL-7611 | chr7-84116862-DEL-7611 |
| chr7-84138996-DEL-1950 | chr7-84138995-DEL-1951 |
| chr7-84169811-DEL-189 | chr7-84169812-DEL-189 |
| chr7-84216761-DEL-96 | chr7-84216762-DEL-96 |
| chr7-84217472-DEL-6074 | chr7-84217473-DEL-6074 |

| | |
|---|---|
| chr7-84231174-DEL-294 | chr7-84231175-DEL-294 |
| chr7-84251863-DEL-57 | chr7-84251874-DEL-83 |
| chr7-84290697-DEL-164 | chr7-84290698-DEL-164 |
| chr7-84359042-DEL-55 | chr7-84359030-DEL-55 |
| chr7-84362393-DEL-7330 | chr7-84362394-DEL-7330 |
| chr7-84368991-DEL-424 | chr7-84368991-DEL-424 |
| chr7-84373628-DEL-1892 | chr7-84373629-DEL-1892 |
| chr7-84399610-DEL-82 | chr7-84399568-DEL-84 |
| chr7-84400249-DEL-498 | chr7-84400250-DEL-498 |
| chr7-84422389-DEL-3968 | chr7-84422390-DEL-3967 |
| chr7-84437492-DEL-58 | chr7-84437493-DEL-58 |
| chr7-84438732-DEL-65 | chr7-84438729-DEL-67 |
| chr7-84439807-DEL-4107 | chr7-84439808-DEL-4107 |
| chr7-84444294-DEL-233 | chr7-84444295-DEL-233 |
| chr7-84461376-DEL-65 | chr7-84461377-DEL-65 |
| chr7-84495782-DEL-80 | chr7-84495771-DEL-55 |
| chr7-84510820-DEL-58 | chr7-84510821-DEL-58 |
| chr7-84544905-DEL-5854 | chr7-84544906-DEL-5854 |
| chr7-84577333-DEL-1337 | chr7-84577334-DEL-1337 |
| chr7-84611377-DEL-54 | chr7-84611378-DEL-54 |
| chr7-84624487-DEL-6836 | chr7-84624488-DEL-6836 |
| chr7-84624597-DEL-364 | chr7-84624598-DEL-364 |
| chr7-84666887-DEL-6883 | chr7-84666888-DEL-6883 |
| chr7-84674324-DEL-201 | chr7-84674325-DEL-201 |
| chr7-84688969-DEL-246 | chr7-84688970-DEL-245 |
| chr7-84736911-DEL-52 | chr7-84736912-DEL-52 |
| chr7-84739087-DEL-293 | chr7-84739088-DEL-293 |
| chr7-84746062-DEL-4863 | chr7-84746059-DEL-4860 |
| chr7-84803670-DEL-57 | chr7-84803670-DEL-57 |
| chr7-84807271-DEL-324 | chr7-84807271-DEL-324 |
| chr7-84827082-DEL-6399 | chr7-84827083-DEL-6399 |
| chr7-84865318-DEL-253 | chr7-84865318-DEL-253 |
| chr7-84921080-DEL-596 | chr7-84921080-DEL-597 |
| chr7-84934659-DEL-74 | chr7-84934651-DEL-75 |
| chr7-84958706-DEL-60 | chr7-84958707-DEL-60 |
| chr7-85009888-DEL-5639 | chr7-85009889-DEL-5639 |
| chr7-85012747-DEL-3900 | chr7-85012747-DEL-3900 |
| chr7-85036648-DEL-6435 | chr7-85036649-DEL-6435 |
| chr7-85064599-DEL-105 | chr7-85064600-DEL-105 |
| chr7-85082704-DEL-332 | chr7-85082705-DEL-332 |
| chr7-85150238-DEL-2350 | chr7-85150239-DEL-2350 |
| chr7-85158101-DEL-899 | chr7-85158102-DEL-899 |
| chr7-85171352-DEL-107 | chr7-85171353-DEL-107 |
| chr7-85174834-DEL-3359 | chr7-85174835-DEL-3359 |
| chr7-85190560-DEL-321 | chr7-85190562-DEL-320 |
| chr7-85210713-DEL-209 | chr7-85210714-DEL-209 |
| chr7-85216174-DEL-66 | chr7-85216152-DEL-78 |
| chr7-85221637-DEL-74 | chr7-85221634-DEL-78 |
| chr7-85249031-DEL-483 | chr7-85249032-DEL-483 |
| chr7-85272752-DEL-102 | chr7-85272753-DEL-102 |
| chr7-85315134-DEL-81 | chr7-85315135-DEL-81 |
| chr7-85366357-DEL-6389 | chr7-85366358-DEL-6389 |
| chr7-85375607-DEL-82 | chr7-85375616-DEL-74 |
| chr7-85409122-DEL-6585 | chr7-85409119-DEL-6586 |
| chr7-85428194-DEL-697 | chr7-85428194-DEL-698 |
| chr7-85485808-DEL-375 | chr7-85485803-DEL-376 |
| chr7-85515111-DEL-142 | chr7-85515113-DEL-143 |
| chr7-85553160-DEL-1219 | chr7-85553160-DEL-1219 |
| chr7-85557363-DEL-54 | chr7-85557364-DEL-54 |
| chr7-85558895-DEL-1093 | chr7-85558896-DEL-1093 |
| chr7-85575615-DEL-7088 | chr7-85575616-DEL-7088 |
| chr7-85657370-DEL-135 | chr7-85657371-DEL-135 |
| chr7-85659818-DEL-88 | chr7-85659819-DEL-88 |
| chr7-85666632-DEL-90 | chr7-85666609-DEL-132 |
| chr7-85680998-DEL-7097 | chr7-85680999-DEL-7097 |
| chr7-85699672-DEL-96 | chr7-85699673-DEL-96 |
| chr7-85714614-DEL-1418 | chr7-85714615-DEL-1418 |
| chr7-85734685-DEL-2773 | chr7-85734686-DEL-2773 |
| chr7-85753794-DEL-111 | chr7-85753793-DEL-112 |
| chr7-85759494-DEL-1697 | chr7-85759495-DEL-1697 |
| chr7-85767870-DEL-7217 | chr7-85767871-DEL-7217 |
| chr7-85783830-DEL-4654 | chr7-85783831-DEL-4654 |
| chr7-85835932-DEL-631 | chr7-85835933-DEL-629 |
| chr7-85837895-DEL-928 | chr7-85837888-DEL-929 |
| chr7-85840480-DEL-112 | chr7-85840481-DEL-112 |
| chr7-85852600-DEL-56 | chr7-85852601-DEL-56 |
| chr7-85889797-DEL-235 | chr7-85889798-DEL-235 |
| chr7-85908338-DEL-723 | chr7-85908339-DEL-723 |
| chr7-85993843-DEL-1800 | chr7-85993844-DEL-1800 |
| chr7-85996573-DEL-241 | chr7-85996574-DEL-241 |

| | |
|---|---|
| chr7-86024951-DEL-75 | chr7-86024980-DEL-82 |
| chr7-86026595-DEL-131 | chr7-86026596-DEL-130 |
| chr7-86030422-DEL-173 | chr7-86030423-DEL-173 |
| chr7-86036114-DEL-1143 | chr7-86036115-DEL-1143 |
| chr7-86041402-DEL-1627 | chr7-86041401-DEL-1628 |
| chr7-86061956-DEL-90 | chr7-86061957-DEL-90 |
| chr7-86063978-DEL-3304 | chr7-86063979-DEL-3304 |
| chr7-86070699-DEL-54 | chr7-86070700-DEL-54 |
| chr7-86071881-DEL-4022 | chr7-86071877-DEL-4023 |
| chr7-86077914-DEL-86 | chr7-86077939-DEL-76 |
| chr7-86081226-DEL-126 | chr7-86081227-DEL-126 |
| chr7-86083208-DEL-184 | chr7-86083209-DEL-184 |
| chr7-86105650-DEL-231 | chr7-86105651-DEL-231 |
| chr7-86114245-DEL-95 | chr7-86114249-DEL-91 |
| chr7-86146635-DEL-6484 | chr7-86146636-DEL-6484 |
| chr7-86163292-DEL-581 | chr7-86163293-DEL-581 |
| chr7-86207767-DEL-228 | chr7-86207768-DEL-228 |
| chr7-86220783-DEL-636 | chr7-86220784-DEL-636 |
| chr7-86254724-DEL-630 | chr7-86254725-DEL-630 |
| chr7-86266970-DEL-6438 | chr7-86266971-DEL-6438 |
| chr7-86278271-DEL-2167 | chr7-86278272-DEL-2167 |
| chr7-86281637-DEL-159 | chr7-86281638-DEL-159 |
| chr7-86314267-DEL-480 | chr7-86314268-DEL-480 |
| chr7-86348347-DEL-54 | chr7-86348349-DEL-50 |
| chr7-86404750-DEL-162 | chr7-86404748-DEL-162 |
| chr7-86421329-DEL-70 | chr7-86421330-DEL-70 |
| chr7-86480297-DEL-3353 | chr7-86480304-DEL-3347 |
| chr7-86632919-DEL-63 | chr7-86632919-DEL-63 |
| chr7-86633972-DEL-6596 | chr7-86633973-DEL-6596 |
| chr7-86634065-DEL-208 | chr7-86634066-DEL-208 |
| chr7-86654845-DEL-98 | chr7-86654846-DEL-98 |
| chr7-86668424-DEL-6499 | chr7-86668424-DEL-6499 |
| chr7-86735346-DEL-516 | chr7-86735349-DEL-514 |
| chr7-86742201-DEL-55 | chr7-86742195-DEL-56 |
| chr7-86742606-DEL-402 | chr7-86742607-DEL-401 |
| chr7-86744085-DEL-689 | chr7-86744086-DEL-689 |
| chr7-86765640-DEL-489 | chr7-86765641-DEL-489 |
| chr7-86776485-DEL-80 | chr7-86776486-DEL-80 |
| chr7-86801488-DEL-293 | chr7-86801489-DEL-293 |
| chr7-86859771-DEL-81 | chr7-86859771-DEL-81 |
| chr7-86868832-DEL-1326 | chr7-86868833-DEL-1326 |
| chr7-86888915-DEL-357 | chr7-86888916-DEL-358 |
| chr7-86893298-DEL-137 | chr7-86893299-DEL-135 |
| chr7-86916409-DEL-62 | chr7-86916409-DEL-50 |
| chr7-86919656-DEL-310 | chr7-86919657-DEL-310 |
| chr7-86930390-DEL-197 | chr7-86930391-DEL-197 |
| chr7-86951169-DEL-1172 | chr7-86951170-DEL-1172 |
| chr7-86960846-DEL-55 | chr7-86960847-DEL-55 |
| chr7-86964049-DEL-159 | chr7-86964050-DEL-159 |
| chr7-87025884-DEL-7974 | chr7-87025885-DEL-7974 |
| chr7-87052659-DEL-69 | chr7-87052660-DEL-68 |
| chr7-87062443-DEL-11072 | chr7-87062444-DEL-11072 |
| chr7-87064523-DEL-214 | chr7-87064523-DEL-214 |
| chr7-87072125-DEL-358 | chr7-87072126-DEL-358 |
| chr7-87091932-DEL-76 | chr7-87091932-DEL-50 |
| chr7-87126180-DEL-617 | chr7-87126181-DEL-617 |
| chr7-87143181-DEL-78 | chr7-87143196-DEL-50 |
| chr7-87161106-DEL-6283 | chr7-87161092-DEL-6297 |
| chr7-87173051-DEL-1032 | chr7-87173051-DEL-1032 |
| chr7-87174312-DEL-399 | chr7-87174312-DEL-399 |
| chr7-87198199-DEL-6201 | chr7-87198200-DEL-6201 |
| chr7-87208004-DEL-114 | chr7-87208028-DEL-138 |
| chr7-87239624-DEL-107 | chr7-87239609-DEL-88 |
| chr7-87263924-DEL-6440 | chr7-87263925-DEL-6440 |
| chr7-87296333-DEL-391 | chr7-87296334-DEL-391 |
| chr7-87298466-DEL-73 | chr7-87298466-DEL-74 |
| chr7-87316065-DEL-838 | chr7-87316066-DEL-838 |
| chr7-87318668-DEL-252 | chr7-87318674-DEL-246 |
| chr7-87404689-DEL-256 | chr7-87404690-DEL-256 |
| chr7-87440546-DEL-768 | chr7-87440546-DEL-767 |
| chr7-87477427-DEL-162 | chr7-87477428-DEL-162 |
| chr7-87551769-DEL-7296 | chr7-87551770-DEL-7296 |
| chr7-87638249-DEL-12413 | chr7-87638250-DEL-12413 |
| chr7-87806140-DEL-1322 | chr7-87806140-DEL-1320 |
| chr7-87838338-DEL-2577 | chr7-87838339-DEL-2577 |
| chr7-87845663-DEL-58 | chr7-87845664-DEL-56 |
| chr7-87850106-DEL-265 | chr7-87850107-DEL-265 |
| chr7-87857641-DEL-168 | chr7-87857642-DEL-169 |
| chr7-87877781-DEL-143 | chr7-87877758-DEL-163 |
| chr7-87882812-DEL-53 | chr7-87882814-DEL-52 |

| | |
|---|---|
| chr7-87886350-DEL-99 | chr7-87886351-DEL-99 |
| chr7-87889341-DEL-126 | chr7-87889366-DEL-90 |
| chr7-87904648-DEL-684 | chr7-87904649-DEL-684 |
| chr7-87912676-DEL-99 | chr7-87912677-DEL-99 |
| chr7-87925132-DEL-78 | chr7-87925132-DEL-78 |
| chr7-87973403-DEL-150 | chr7-87973404-DEL-148 |
| chr7-87975424-DEL-1076 | chr7-87975425-DEL-1076 |
| chr7-88014366-DEL-237 | chr7-88014397-DEL-232 |
| chr7-88017626-DEL-66 | chr7-88017627-DEL-66 |
| chr7-88020265-DEL-6172 | chr7-88020264-DEL-6173 |
| chr7-88027799-DEL-185 | chr7-88027800-DEL-137 |
| chr7-88091719-DEL-72 | chr7-88091718-DEL-78 |
| chr7-88104730-DEL-298 | chr7-88104702-DEL-331 |
| chr7-88120045-DEL-52 | chr7-88120019-DEL-58 |
| chr7-88179573-DEL-560 | chr7-88179574-DEL-560 |
| chr7-88190061-DEL-1670 | chr7-88190078-DEL-1651 |
| chr7-88225943-DEL-2060 | chr7-88225944-DEL-2060 |
| chr7-88240781-DEL-300 | chr7-88240782-DEL-300 |
| chr7-88262236-DEL-170 | chr7-88262237-DEL-170 |
| chr7-88281896-DEL-4858 | chr7-88281897-DEL-4857 |
| chr7-88291089-DEL-55 | chr7-88291090-DEL-55 |
| chr7-88349383-DEL-115 | chr7-88349380-DEL-116 |
| chr7-88355146-DEL-447 | chr7-88355155-DEL-429 |
| chr7-88362507-DEL-6451 | chr7-88362508-DEL-6451 |
| chr7-88385510-DEL-474 | chr7-88385511-DEL-474 |
| chr7-88432629-DEL-382 | chr7-88432630-DEL-382 |
| chr7-88442121-DEL-133 | chr7-88442122-DEL-130 |
| chr7-88482163-DEL-6331 | chr7-88482175-DEL-6332 |
| chr7-88490791-DEL-1517 | chr7-88490792-DEL-1517 |
| chr7-88505785-DEL-203 | chr7-88505786-DEL-203 |
| chr7-88526429-DEL-56 | chr7-88526430-DEL-58 |
| chr7-88547747-DEL-1739 | chr7-88547748-DEL-1739 |
| chr7-88550212-DEL-181 | chr7-88550214-DEL-182 |
| chr7-88571559-DEL-427 | chr7-88571560-DEL-427 |
| chr7-88574824-DEL-900 | chr7-88574825-DEL-900 |
| chr7-88655999-DEL-66 | chr7-88656003-DEL-60 |
| chr7-88660743-DEL-52 | chr7-88660745-DEL-50 |
| chr7-88662755-DEL-1015 | chr7-88662756-DEL-1015 |
| chr7-88677149-DEL-1743 | chr7-88677150-DEL-1743 |
| chr7-88682646-DEL-60 | chr7-88682647-DEL-60 |
| chr7-88708314-DEL-381 | chr7-88708315-DEL-377 |
| chr7-88731707-DEL-462 | chr7-88731705-DEL-463 |
| chr7-88781774-DEL-651 | chr7-88781775-DEL-651 |
| chr7-88784032-DEL-521 | chr7-88784033-DEL-521 |
| chr7-88808707-DEL-377 | chr7-88808708-DEL-377 |
| chr7-88843922-DEL-52 | chr7-88843922-DEL-52 |
| chr7-88877905-DEL-221 | chr7-88877906-DEL-221 |
| chr7-88922342-DEL-3864 | chr7-88922343-DEL-3864 |
| chr7-88940189-DEL-5092 | chr7-88940190-DEL-5092 |
| chr7-89009927-DEL-76 | chr7-89009928-DEL-84 |
| chr7-89023227-DEL-6440 | chr7-89023228-DEL-6440 |
| chr7-89036341-DEL-164 | chr7-89036340-DEL-166 |
| chr7-89137743-DEL-3544 | chr7-89137744-DEL-3544 |
| chr7-89141432-DEL-2839 | chr7-89141432-DEL-2839 |
| chr7-89185403-DEL-92 | chr7-89185360-DEL-96 |
| chr7-89196112-DEL-65 | chr7-89196113-DEL-65 |
| chr7-89217799-DEL-5176 | chr7-89217800-DEL-5176 |
| chr7-89247329-DEL-97 | chr7-89247330-DEL-97 |
| chr7-89256825-DEL-1275 | chr7-89256826-DEL-1275 |
| chr7-89292261-DEL-486 | chr7-89292262-DEL-486 |
| chr7-89298082-DEL-81 | chr7-89298082-DEL-81 |
| chr7-89299394-DEL-4423 | chr7-89299395-DEL-4423 |
| chr7-89332491-DEL-5706 | chr7-89332476-DEL-5719 |
| chr7-89338681-DEL-7862 | chr7-89338682-DEL-7862 |
| chr7-89350793-DEL-177 | chr7-89350794-DEL-177 |
| chr7-89352018-DEL-2809 | chr7-89352019-DEL-2809 |
| chr7-89368069-DEL-62 | chr7-89368070-DEL-62 |
| chr7-89382279-DEL-141 | chr7-89382280-DEL-141 |
| chr7-89391818-DEL-118 | chr7-89391819-DEL-118 |
| chr7-89400127-DEL-8486 | chr7-89400128-DEL-8486 |
| chr7-89410569-DEL-53 | chr7-89410549-DEL-54 |
| chr7-89454065-DEL-122 | chr7-89454073-DEL-99 |
| chr7-89467996-DEL-7206 | chr7-89467996-DEL-7206 |
| chr7-89507084-DEL-6440 | chr7-89507085-DEL-6440 |
| chr7-89517106-DEL-131 | chr7-89517075-DEL-157 |
| chr7-89520070-DEL-62 | chr7-89520071-DEL-62 |
| chr7-89553547-DEL-74 | chr7-89553524-DEL-102 |
| chr7-89560935-DEL-146 | chr7-89560937-DEL-144 |
| chr7-89570026-DEL-56 | chr7-89570027-DEL-56 |
| chr7-89584431-DEL-69 | chr7-89584430-DEL-60 |

| | |
|---|---|
| chr7-89599739-DEL-212 | chr7-89599734-DEL-207 |
| chr7-89615826-DEL-210 | chr7-89615827-DEL-210 |
| chr7-89637289-DEL-62 | chr7-89637290-DEL-62 |
| chr7-89648454-DEL-99 | chr7-89648455-DEL-99 |
| chr7-89663350-DEL-204 | chr7-89663351-DEL-204 |
| chr7-89728251-DEL-104 | chr7-89728252-DEL-104 |
| chr7-89729871-DEL-1432 | chr7-89729872-DEL-1432 |
| chr7-89736495-DEL-1929 | chr7-89736496-DEL-1929 |
| chr7-89742656-DEL-60 | chr7-89742657-DEL-60 |
| chr7-89752784-DEL-934 | chr7-89752785-DEL-934 |
| chr7-89757650-DEL-56 | chr7-89757651-DEL-52 |
| chr7-89802857-DEL-657 | chr7-89802858-DEL-657 |
| chr7-89807380-DEL-1979 | chr7-89807381-DEL-1979 |
| chr7-89911717-DEL-60 | chr7-89911718-DEL-60 |
| chr7-89923529-DEL-86 | chr7-89923530-DEL-86 |
| chr7-89932407-DEL-148 | chr7-89932376-DEL-152 |
| chr7-89953098-DEL-71 | chr7-89953121-DEL-60 |
| chr7-89955722-DEL-284 | chr7-89955723-DEL-284 |
| chr7-89977942-DEL-106 | chr7-89977943-DEL-106 |
| chr7-90060015-DEL-9057 | chr7-90060016-DEL-9057 |
| chr7-90080986-DEL-191 | chr7-90080998-DEL-174 |
| chr7-90085102-DEL-70 | chr7-90085095-DEL-52 |
| chr7-90090841-DEL-226 | chr7-90090842-DEL-226 |
| chr7-90111836-DEL-121 | chr7-90111837-DEL-121 |
| chr7-90134435-DEL-62 | chr7-90134437-DEL-68 |
| chr7-90159703-DEL-1497 | chr7-90159704-DEL-1497 |
| chr7-90163118-DEL-3921 | chr7-90163119-DEL-3921 |
| chr7-90172109-DEL-1514 | chr7-90172110-DEL-1514 |
| chr7-90176207-DEL-6871 | chr7-90176208-DEL-6871 |
| chr7-90197407-DEL-52 | chr7-90197413-DEL-54 |
| chr7-90206662-DEL-209 | chr7-90206662-DEL-206 |
| chr7-90216970-DEL-54 | chr7-90216975-DEL-50 |
| chr7-90222532-DEL-728 | chr7-90222533-DEL-728 |
| chr7-90239583-DEL-97 | chr7-90239585-DEL-135 |
| chr7-90244574-DEL-162 | chr7-90244575-DEL-162 |
| chr7-90267741-DEL-8688 | chr7-90267742-DEL-8688 |
| chr7-90285128-DEL-255 | chr7-90285129-DEL-255 |
| chr7-90311603-DEL-92 | chr7-90311604-DEL-92 |
| chr7-90313898-DEL-62 | chr7-90313899-DEL-62 |
| chr7-90314454-DEL-79 | chr7-90314471-DEL-82 |
| chr7-90319147-DEL-186 | chr7-90319148-DEL-186 |
| chr7-90377922-DEL-2717 | chr7-90377923-DEL-2717 |
| chr7-90395124-DEL-52 | chr7-90395127-DEL-52 |
| chr7-90406381-DEL-74 | chr7-90406381-DEL-74 |
| chr7-90417772-DEL-63 | chr7-90417773-DEL-61 |
| chr7-90426367-DEL-276 | chr7-90426368-DEL-276 |
| chr7-90459928-DEL-60 | chr7-90459927-DEL-58 |
| chr7-90471802-DEL-100 | chr7-90471803-DEL-100 |
| chr7-90488875-DEL-62 | chr7-90488851-DEL-77 |
| chr7-90507246-DEL-7129 | chr7-90507249-DEL-7130 |
| chr7-90517782-DEL-3118 | chr7-90517783-DEL-3118 |
| chr7-90548396-DEL-2008 | chr7-90548397-DEL-2008 |
| chr7-90563132-DEL-137 | chr7-90563133-DEL-140 |
| chr7-90597746-DEL-599 | chr7-90597757-DEL-589 |
| chr7-90607027-DEL-739 | chr7-90607028-DEL-739 |
| chr7-90624167-DEL-172 | chr7-90624168-DEL-172 |
| chr7-90652040-DEL-400 | chr7-90652041-DEL-400 |
| chr7-90654805-DEL-202 | chr7-90654806-DEL-202 |
| chr7-90665336-DEL-163 | chr7-90665337-DEL-163 |
| chr7-90668769-DEL-819 | chr7-90668770-DEL-819 |
| chr7-90715787-DEL-92 | chr7-90715789-DEL-84 |
| chr7-90764245-DEL-191 | chr7-90764246-DEL-191 |
| chr7-90765537-DEL-678 | chr7-90765538-DEL-678 |
| chr7-90796288-DEL-52 | chr7-90796289-DEL-52 |
| chr7-90812074-DEL-120 | chr7-90812077-DEL-118 |
| chr7-90812432-DEL-286 | chr7-90812433-DEL-286 |
| chr7-90826251-DEL-64 | chr7-90826264-DEL-85 |
| chr7-90830306-DEL-1362 | chr7-90830307-DEL-1362 |
| chr7-90833636-DEL-57 | chr7-90833637-DEL-56 |
| chr7-90834881-DEL-400 | chr7-90834882-DEL-400 |
| chr7-90852337-DEL-122 | chr7-90852336-DEL-117 |
| chr7-90854016-DEL-85 | chr7-90854017-DEL-85 |
| chr7-90871094-DEL-148 | chr7-90871095-DEL-148 |
| chr7-90929850-DEL-3249 | chr7-90929851-DEL-3249 |
| chr7-90939025-DEL-1599 | chr7-90939026-DEL-1599 |
| chr7-90953861-DEL-65 | chr7-90953845-DEL-69 |
| chr7-90957228-DEL-761 | chr7-90957229-DEL-761 |
| chr7-91016220-DEL-54 | chr7-91016223-DEL-50 |
| chr7-91053890-DEL-68 | chr7-91053891-DEL-68 |
| chr7-91077936-DEL-537 | chr7-91077937-DEL-537 |

| | |
|---|---|
| chr7-91081486-DEL-210 | chr7-91081487-DEL-210 |
| chr7-91103212-DEL-133 | chr7-91103213-DEL-133 |
| chr7-91152353-DEL-347 | chr7-91152359-DEL-341 |
| chr7-91159472-DEL-77 | chr7-91159463-DEL-87 |
| chr7-91161842-DEL-58 | chr7-91161843-DEL-58 |
| chr7-91164183-DEL-52 | chr7-91164184-DEL-52 |
| chr7-91174169-DEL-87 | chr7-91174171-DEL-84 |
| chr7-91197628-DEL-84 | chr7-91197629-DEL-84 |
| chr7-91251774-DEL-55 | chr7-91251775-DEL-55 |
| chr7-91254915-DEL-104 | chr7-91254916-DEL-103 |
| chr7-91255773-DEL-967 | chr7-91255774-DEL-967 |
| chr7-91287886-DEL-2025 | chr7-91287887-DEL-2025 |
| chr7-91292831-DEL-586 | chr7-91292832-DEL-586 |
| chr7-91327780-DEL-120 | chr7-91327793-DEL-105 |
| chr7-91369096-DEL-98 | chr7-91369097-DEL-114 |
| chr7-91390786-DEL-6848 | chr7-91390787-DEL-6848 |
| chr7-91404138-DEL-78 | chr7-91404139-DEL-78 |
| chr7-91437354-DEL-94 | chr7-91437355-DEL-95 |
| chr7-91442716-DEL-57 | chr7-91442721-DEL-53 |
| chr7-91444916-DEL-1610 | chr7-91444917-DEL-1610 |
| chr7-91448667-DEL-179 | chr7-91448668-DEL-179 |
| chr7-91478470-DEL-657 | chr7-91478469-DEL-658 |
| chr7-91497604-DEL-60 | chr7-91497605-DEL-60 |
| chr7-91502454-DEL-78 | chr7-91502455-DEL-78 |
| chr7-91516260-DEL-149 | chr7-91516261-DEL-149 |
| chr7-91541800-DEL-79 | chr7-91541817-DEL-63 |
| chr7-91577220-DEL-208 | chr7-91577228-DEL-211 |
| chr7-91615443-DEL-387 | chr7-91615444-DEL-387 |
| chr7-91619129-DEL-699 | chr7-91619130-DEL-699 |
| chr7-91651298-DEL-54 | chr7-91651297-DEL-56 |
| chr7-91660212-DEL-52 | chr7-91660213-DEL-52 |
| chr7-91671188-DEL-400 | chr7-91671189-DEL-400 |
| chr7-91697851-DEL-93 | chr7-91697851-DEL-93 |
| chr7-91721108-DEL-6359 | chr7-91721109-DEL-6359 |
| chr7-91728117-DEL-76 | chr7-91728122-DEL-72 |
| chr7-91733855-DEL-77 | chr7-91733858-DEL-66 |
| chr7-91758312-DEL-378 | chr7-91758313-DEL-393 |
| chr7-91785826-DEL-60 | chr7-91785826-DEL-66 |
| chr7-91820521-DEL-171 | chr7-91820510-DEL-151 |
| chr7-91841685-DEL-3586 | chr7-91841686-DEL-3586 |
| chr7-91874509-DEL-7096 | chr7-91874509-DEL-7096 |
| chr7-91889357-DEL-905 | chr7-91889358-DEL-905 |
| chr7-91911201-DEL-64 | chr7-91911202-DEL-64 |
| chr7-91912208-DEL-93 | chr7-91912209-DEL-93 |
| chr7-91927132-DEL-2073 | chr7-91927133-DEL-2073 |
| chr7-91945401-DEL-177 | chr7-91945402-DEL-177 |
| chr7-91948053-DEL-1439 | chr7-91948054-DEL-1439 |
| chr7-91985874-DEL-338 | chr7-91985875-DEL-338 |
| chr7-91991149-DEL-82 | chr7-91991149-DEL-82 |
| chr7-91993790-DEL-70 | chr7-91993749-DEL-80 |
| chr7-91999508-DEL-6194 | chr7-91999509-DEL-6194 |
| chr7-92086057-DEL-129 | chr7-92086060-DEL-128 |
| chr7-92135889-DEL-84 | chr7-92135922-DEL-52 |
| chr7-92136670-DEL-58 | chr7-92136642-DEL-87 |
| chr7-92184467-DEL-734 | chr7-92184468-DEL-734 |
| chr7-92211502-DEL-994 | chr7-92211503-DEL-994 |
| chr7-92220716-DEL-515 | chr7-92220717-DEL-515 |
| chr7-92224915-DEL-128 | chr7-92224942-DEL-84 |
| chr7-92236089-DEL-186 | chr7-92236090-DEL-186 |
| chr7-92237674-DEL-1243 | chr7-92237675-DEL-1243 |
| chr7-92247632-DEL-6195 | chr7-92247633-DEL-6195 |
| chr7-92257832-DEL-252 | chr7-92257833-DEL-252 |
| chr7-92279056-DEL-407 | chr7-92279057-DEL-407 |
| chr7-92281777-DEL-163 | chr7-92281796-DEL-145 |
| chr7-92284506-DEL-110 | chr7-92284555-DEL-70 |
| chr7-92289655-DEL-255 | chr7-92289656-DEL-255 |
| chr7-92304356-DEL-798 | chr7-92304357-DEL-798 |
| chr7-92360537-DEL-668 | chr7-92360538-DEL-668 |
| chr7-92361520-DEL-83 | chr7-92361521-DEL-83 |
| chr7-92362925-DEL-403 | chr7-92362926-DEL-403 |
| chr7-92363625-DEL-93 | chr7-92363669-DEL-70 |
| chr7-92367752-DEL-50 | chr7-92367739-DEL-64 |
| chr7-92376861-DEL-103 | chr7-92376862-DEL-103 |
| chr7-92401147-DEL-208 | chr7-92401148-DEL-208 |
| chr7-92431808-DEL-85 | chr7-92431809-DEL-85 |
| chr7-92436141-DEL-7427 | chr7-92436142-DEL-7427 |
| chr7-92467827-DEL-215 | chr7-92467828-DEL-215 |
| chr7-92482908-DEL-171 | chr7-92482909-DEL-171 |
| chr7-92499506-DEL-2851 | chr7-92499507-DEL-2851 |
| chr7-92515338-DEL-200 | chr7-92515339-DEL-200 |

| | |
|---|---|
| chr7-92536017-DEL-100 | chr7-92536018-DEL-100 |
| chr7-92550130-DEL-81 | chr7-92550120-DEL-72 |
| chr7-92559851-DEL-6460 | chr7-92559852-DEL-6460 |
| chr7-92574807-DEL-2030 | chr7-92574808-DEL-2030 |
| chr7-92584458-DEL-128 | chr7-92584458-DEL-128 |
| chr7-92591277-DEL-215 | chr7-92591279-DEL-214 |
| chr7-92602006-DEL-2532 | chr7-92602007-DEL-2532 |
| chr7-92605862-DEL-192 | chr7-92605865-DEL-193 |
| chr7-92607200-DEL-54 | chr7-92607201-DEL-54 |
| chr7-92610805-DEL-2336 | chr7-92610805-DEL-2335 |
| chr7-92617450-DEL-6442 | chr7-92617451-DEL-6442 |
| chr7-92638336-DEL-52 | chr7-92638337-DEL-52 |
| chr7-92660584-DEL-64 | chr7-92660585-DEL-64 |
| chr7-92665216-DEL-826 | chr7-92665217-DEL-826 |
| chr7-92671355-DEL-1036 | chr7-92671356-DEL-1036 |
| chr7-92689361-DEL-58 | chr7-92689360-DEL-59 |
| chr7-92704085-DEL-548 | chr7-92704086-DEL-548 |
| chr7-92736930-DEL-387 | chr7-92736931-DEL-387 |
| chr7-92762401-DEL-3353 | chr7-92762402-DEL-3353 |
| chr7-92766779-DEL-68 | chr7-92766782-DEL-68 |
| chr7-92772282-DEL-70 | chr7-92772287-DEL-66 |
| chr7-92789060-DEL-6383 | chr7-92789061-DEL-6383 |
| chr7-92862088-DEL-6334 | chr7-92862089-DEL-6334 |
| chr7-92870537-DEL-566 | chr7-92870534-DEL-567 |
| chr7-92876213-DEL-1887 | chr7-92876214-DEL-1887 |
| chr7-92909044-DEL-2287 | chr7-92909045-DEL-2287 |
| chr7-92912316-DEL-195 | chr7-92912317-DEL-195 |
| chr7-92921132-DEL-6186 | chr7-92921133-DEL-6186 |
| chr7-92929456-DEL-352 | chr7-92929457-DEL-352 |
| chr7-92979099-DEL-477 | chr7-92979100-DEL-477 |
| chr7-92994754-DEL-56 | chr7-92994740-DEL-80 |
| chr7-93061289-DEL-6841 | chr7-93061290-DEL-6841 |
| chr7-93067465-DEL-424 | chr7-93067466-DEL-424 |
| chr7-93077114-DEL-79 | chr7-93077145-DEL-54 |
| chr7-93098436-DEL-4253 | chr7-93098437-DEL-4253 |
| chr7-93112413-DEL-63 | chr7-93112415-DEL-106 |
| chr7-93123388-DEL-1478 | chr7-93123389-DEL-1478 |
| chr7-93123804-DEL-674 | chr7-93123805-DEL-674 |
| chr7-93131564-DEL-871 | chr7-93131565-DEL-871 |
| chr7-93136427-DEL-506 | chr7-93136428-DEL-506 |
| chr7-93166354-DEL-2108 | chr7-93166355-DEL-2108 |
| chr7-93216360-DEL-51 | chr7-93216361-DEL-51 |
| chr7-93247027-DEL-72 | chr7-93247006-DEL-73 |
| chr7-93253013-DEL-113 | chr7-93253005-DEL-97 |
| chr7-93280052-DEL-1355 | chr7-93280053-DEL-1355 |
| chr7-93294338-DEL-1236 | chr7-93294339-DEL-1236 |
| chr7-93300129-DEL-54 | chr7-93300130-DEL-54 |
| chr7-93310460-DEL-1141 | chr7-93310461-DEL-1141 |
| chr7-93341588-DEL-66 | chr7-93341589-DEL-66 |
| chr7-93349647-DEL-1487 | chr7-93349647-DEL-1487 |
| chr7-93377347-DEL-720 | chr7-93377348-DEL-720 |
| chr7-93384655-DEL-266 | chr7-93384656-DEL-265 |
| chr7-93389541-DEL-309 | chr7-93389541-DEL-309 |
| chr7-93393171-DEL-6609 | chr7-93393172-DEL-6609 |
| chr7-93413850-DEL-1564 | chr7-93413856-DEL-1559 |
| chr7-93529961-DEL-1380 | chr7-93529961-DEL-1380 |
| chr7-93541682-DEL-195 | chr7-93541683-DEL-195 |
| chr7-93564686-DEL-429 | chr7-93564687-DEL-429 |
| chr7-93585913-DEL-59 | chr7-93585914-DEL-66 |
| chr7-93587358-DEL-57 | chr7-93587359-DEL-57 |
| chr7-93588510-DEL-98 | chr7-93588513-DEL-60 |
| chr7-93607006-DEL-54 | chr7-93607010-DEL-53 |
| chr7-93608081-DEL-249 | chr7-93608082-DEL-249 |
| chr7-93659219-DEL-245 | chr7-93659220-DEL-245 |
| chr7-93677771-DEL-62 | chr7-93677751-DEL-62 |
| chr7-93697337-DEL-177 | chr7-93697343-DEL-170 |
| chr7-93719955-DEL-58 | chr7-93719956-DEL-56 |
| chr7-93746879-DEL-135 | chr7-93746880-DEL-135 |
| chr7-93758291-DEL-66 | chr7-93758285-DEL-72 |
| chr7-93785719-DEL-672 | chr7-93785720-DEL-672 |
| chr7-93789471-DEL-1202 | chr7-93789471-DEL-1200 |
| chr7-93793835-DEL-60 | chr7-93793836-DEL-59 |
| chr7-93794203-DEL-1416 | chr7-93794204-DEL-1416 |
| chr7-93808789-DEL-83 | chr7-93808790-DEL-81 |
| chr7-93824542-DEL-1857 | chr7-93824543-DEL-1857 |
| chr7-93829746-DEL-2797 | chr7-93829747-DEL-2796 |
| chr7-93850946-DEL-59 | chr7-93850938-DEL-63 |
| chr7-93851857-DEL-1713 | chr7-93851858-DEL-1713 |
| chr7-93891792-DEL-202 | chr7-93891793-DEL-202 |
| chr7-93897620-DEL-725 | chr7-93897621-DEL-725 |

| | |
|---|---|
| chr7-93923190-DEL-1696 | chr7-93923191-DEL-1696 |
| chr7-93928317-DEL-72 | chr7-93928318-DEL-72 |
| chr7-93942140-DEL-6434 | chr7-93942141-DEL-6434 |
| chr7-93960937-DEL-513 | chr7-93960938-DEL-513 |
| chr7-93969603-DEL-1634 | chr7-93969604-DEL-1634 |
| chr7-93969835-DEL-398 | chr7-93969836-DEL-398 |
| chr7-93973742-DEL-61 | chr7-93973743-DEL-61 |
| chr7-93980656-DEL-188 | chr7-93980657-DEL-188 |
| chr7-94020863-DEL-2660 | chr7-94020864-DEL-2660 |
| chr7-94031647-DEL-613 | chr7-94031648-DEL-613 |
| chr7-94034459-DEL-625 | chr7-94034461-DEL-624 |
| chr7-94038209-DEL-6262 | chr7-94038210-DEL-6257 |
| chr7-94102027-DEL-93 | chr7-94102038-DEL-88 |
| chr7-94115566-DEL-60 | chr7-94115567-DEL-60 |
| chr7-94130081-DEL-56 | chr7-94130066-DEL-72 |
| chr7-94137703-DEL-377 | chr7-94137704-DEL-377 |
| chr7-94150715-DEL-456 | chr7-94150716-DEL-456 |
| chr7-94159561-DEL-1029 | chr7-94159562-DEL-1029 |
| chr7-94165480-DEL-297 | chr7-94165481-DEL-297 |
| chr7-94169731-DEL-144 | chr7-94169733-DEL-138 |
| chr7-94226269-DEL-67 | chr7-94226270-DEL-65 |
| chr7-94238996-DEL-60 | chr7-94238997-DEL-60 |
| chr7-94242949-DEL-92 | chr7-94242950-DEL-92 |
| chr7-94244917-DEL-57 | chr7-94244918-DEL-57 |
| chr7-94261919-DEL-215 | chr7-94261920-DEL-215 |
| chr7-94275920-DEL-1579 | chr7-94275922-DEL-1579 |
| chr7-94281003-DEL-1317 | chr7-94281004-DEL-1317 |
| chr7-94303078-DEL-234 | chr7-94303078-DEL-234 |
| chr7-94319614-DEL-52 | chr7-94319591-DEL-76 |
| chr7-94348576-DEL-304 | chr7-94348585-DEL-295 |
| chr7-94427056-DEL-88 | chr7-94427057-DEL-88 |
| chr7-94427694-DEL-623 | chr7-94427695-DEL-623 |
| chr7-94429545-DEL-64 | chr7-94429546-DEL-68 |
| chr7-94431188-DEL-100 | chr7-94431189-DEL-100 |
| chr7-94436861-DEL-100 | chr7-94436862-DEL-100 |
| chr7-94458379-DEL-81 | chr7-94458387-DEL-60 |
| chr7-94466123-DEL-234 | chr7-94466124-DEL-234 |
| chr7-94479304-DEL-1221 | chr7-94479305-DEL-1221 |
| chr7-94491486-DEL-6574 | chr7-94491487-DEL-6574 |
| chr7-94523607-DEL-62 | chr7-94523608-DEL-62 |
| chr7-94526308-DEL-906 | chr7-94526309-DEL-906 |
| chr7-94531430-DEL-502 | chr7-94531431-DEL-502 |
| chr7-94567046-DEL-2767 | chr7-94567047-DEL-2767 |
| chr7-94593924-DEL-1506 | chr7-94593925-DEL-1506 |
| chr7-94595768-DEL-1630 | chr7-94595769-DEL-1630 |
| chr7-94598604-DEL-74 | chr7-94598608-DEL-71 |
| chr7-94632640-DEL-5344 | chr7-94632640-DEL-5344 |
| chr7-94654338-DEL-126 | chr7-94654339-DEL-126 |
| chr7-94681297-DEL-6490 | chr7-94681293-DEL-6491 |
| chr7-94714368-DEL-6015 | chr7-94714369-DEL-6015 |
| chr7-94721228-DEL-6804 | chr7-94721229-DEL-6804 |
| chr7-94739781-DEL-177 | chr7-94739782-DEL-177 |
| chr7-94742379-DEL-107 | chr7-94742417-DEL-105 |
| chr7-94754426-DEL-418 | chr7-94754427-DEL-418 |
| chr7-94781849-DEL-60 | chr7-94781849-DEL-60 |
| chr7-94826404-DEL-208 | chr7-94826405-DEL-208 |
| chr7-94838377-DEL-2069 | chr7-94838378-DEL-2069 |
| chr7-94851497-DEL-108 | chr7-94851504-DEL-78 |
| chr7-94867115-DEL-140 | chr7-94867116-DEL-140 |
| chr7-94886581-DEL-80 | chr7-94886582-DEL-74 |
| chr7-94902620-DEL-60 | chr7-94902619-DEL-62 |
| chr7-94908409-DEL-290 | chr7-94908410-DEL-289 |
| chr7-94915017-DEL-391 | chr7-94915017-DEL-390 |
| chr7-94934892-DEL-6319 | chr7-94934893-DEL-6319 |
| chr7-94934981-DEL-201 | chr7-94934982-DEL-201 |
| chr7-94960950-DEL-453 | chr7-94960951-DEL-453 |
| chr7-94967325-DEL-61 | chr7-94967323-DEL-72 |
| chr7-94994887-DEL-193 | chr7-94994889-DEL-192 |
| chr7-94999995-DEL-371 | chr7-94999996-DEL-371 |
| chr7-95015301-DEL-62 | chr7-95015302-DEL-62 |
| chr7-95042692-DEL-132 | chr7-95042692-DEL-132 |
| chr7-95051830-DEL-6861 | chr7-95051831-DEL-6861 |
| chr7-95098717-DEL-52 | chr7-95098702-DEL-52 |
| chr7-95104078-DEL-74 | chr7-95104079-DEL-74 |
| chr7-95120846-DEL-436 | chr7-95120847-DEL-436 |
| chr7-95122762-DEL-104 | chr7-95122763-DEL-104 |
| chr7-95128572-DEL-74 | chr7-95128572-DEL-74 |
| chr7-95142434-DEL-607 | chr7-95142435-DEL-607 |
| chr7-95173091-DEL-58 | chr7-95173092-DEL-57 |
| chr7-95176342-DEL-6354 | chr7-95176343-DEL-6354 |

| | |
|---|---|
| chr7-95182441-DEL-200 | chr7-95182442-DEL-200 |
| chr7-95185072-DEL-93 | chr7-95185039-DEL-94 |
| chr7-95186694-DEL-987 | chr7-95186695-DEL-987 |
| chr7-95201644-DEL-81 | chr7-95201645-DEL-81 |
| chr7-95209792-DEL-98 | chr7-95209793-DEL-98 |
| chr7-95220520-DEL-2838 | chr7-95220521-DEL-2838 |
| chr7-95253244-DEL-1107 | chr7-95253245-DEL-1107 |
| chr7-95299976-DEL-51 | chr7-95299977-DEL-51 |
| chr7-95311560-DEL-507 | chr7-95311561-DEL-507 |
| chr7-95321207-DEL-53 | chr7-95321208-DEL-53 |
| chr7-95330678-DEL-485 | chr7-95330674-DEL-483 |
| chr7-95362825-DEL-1611 | chr7-95362826-DEL-1610 |
| chr7-95373162-DEL-1079 | chr7-95373163-DEL-1079 |
| chr7-95396972-DEL-118 | chr7-95396973-DEL-102 |
| chr7-95428901-DEL-54 | chr7-95428902-DEL-54 |
| chr7-95441332-DEL-64 | chr7-95441333-DEL-58 |
| chr7-95443108-DEL-80 | chr7-95443109-DEL-80 |
| chr7-95449970-DEL-210 | chr7-95449971-DEL-210 |
| chr7-95456252-DEL-214 | chr7-95456253-DEL-214 |
| chr7-95485303-DEL-174 | chr7-95485312-DEL-170 |
| chr7-95518556-DEL-54 | chr7-95518557-DEL-54 |
| chr7-95562522-DEL-75 | chr7-95562523-DEL-75 |
| chr7-95590586-DEL-399 | chr7-95590587-DEL-399 |
| chr7-95608904-DEL-296 | chr7-95608905-DEL-296 |
| chr7-95616268-DEL-64 | chr7-95616269-DEL-60 |
| chr7-95627130-DEL-190 | chr7-95627130-DEL-190 |
| chr7-95629746-DEL-1403 | chr7-95629747-DEL-1403 |
| chr7-95657983-DEL-3471 | chr7-95657983-DEL-3471 |
| chr7-95681792-DEL-57 | chr7-95681793-DEL-57 |
| chr7-95705867-DEL-355 | chr7-95705867-DEL-355 |
| chr7-95716459-DEL-3223 | chr7-95716460-DEL-3223 |
| chr7-95721571-DEL-57 | chr7-95721587-DEL-52 |
| chr7-95732189-DEL-324 | chr7-95732190-DEL-324 |
| chr7-95741077-DEL-828 | chr7-95741078-DEL-828 |
| chr7-95776033-DEL-140 | chr7-95776034-DEL-140 |
| chr7-95790296-DEL-431 | chr7-95790297-DEL-431 |
| chr7-95791060-DEL-55 | chr7-95791060-DEL-50 |
| chr7-95799656-DEL-82 | chr7-95799650-DEL-66 |
| chr7-95803667-DEL-64 | chr7-95803668-DEL-64 |
| chr7-95840702-DEL-7023 | chr7-95840703-DEL-7023 |
| chr7-95848900-DEL-52 | chr7-95848901-DEL-52 |
| chr7-95851569-DEL-1280 | chr7-95851570-DEL-1280 |
| chr7-95871620-DEL-71 | chr7-95871603-DEL-63 |
| chr7-95936369-DEL-56 | chr7-95936370-DEL-56 |
| chr7-95964309-DEL-59 | chr7-95964315-DEL-54 |
| chr7-95966808-DEL-7107 | chr7-95966809-DEL-7107 |
| chr7-95974444-DEL-361 | chr7-95974445-DEL-361 |
| chr7-95993480-DEL-74 | chr7-95993481-DEL-74 |
| chr7-96024291-DEL-358 | chr7-96024292-DEL-358 |
| chr7-96038565-DEL-68 | chr7-96038570-DEL-64 |
| chr7-96040834-DEL-124 | chr7-96040839-DEL-120 |
| chr7-96076757-DEL-699 | chr7-96076747-DEL-700 |
| chr7-96083368-DEL-72 | chr7-96083352-DEL-88 |
| chr7-96110358-DEL-58 | chr7-96110359-DEL-74 |
| chr7-96210576-DEL-74 | chr7-96210577-DEL-74 |
| chr7-96273482-DEL-88 | chr7-96273477-DEL-89 |
| chr7-96281164-DEL-504 | chr7-96281165-DEL-504 |
| chr7-96289529-DEL-82 | chr7-96289530-DEL-82 |
| chr7-96290395-DEL-482 | chr7-96290396-DEL-482 |
| chr7-96313256-DEL-54 | chr7-96313276-DEL-55 |
| chr7-96335988-DEL-207 | chr7-96335989-DEL-207 |
| chr7-96363784-DEL-60 | chr7-96363780-DEL-59 |
| chr7-96390857-DEL-211 | chr7-96390858-DEL-211 |
| chr7-96411039-DEL-753 | chr7-96411040-DEL-753 |
| chr7-96433594-DEL-409 | chr7-96433595-DEL-409 |
| chr7-96434520-DEL-62 | chr7-96434516-DEL-67 |
| chr7-96438941-DEL-70 | chr7-96438942-DEL-70 |
| chr7-96527133-DEL-117 | chr7-96527134-DEL-117 |
| chr7-96542919-DEL-109 | chr7-96542920-DEL-109 |
| chr7-96573126-DEL-200 | chr7-96573127-DEL-200 |
| chr7-96585044-DEL-2684 | chr7-96585032-DEL-2683 |
| chr7-96677068-DEL-70 | chr7-96677081-DEL-99 |
| chr7-96687083-DEL-1247 | chr7-96687084-DEL-1247 |
| chr7-96719654-DEL-495 | chr7-96719655-DEL-495 |
| chr7-96751427-DEL-66 | chr7-96751428-DEL-66 |
| chr7-96763613-DEL-6474 | chr7-96763614-DEL-6474 |
| chr7-96790146-DEL-6443 | chr7-96790147-DEL-6443 |
| chr7-96816402-DEL-108 | chr7-96816407-DEL-104 |
| chr7-96816542-DEL-2936 | chr7-96816543-DEL-2936 |
| chr7-96863990-DEL-1471 | chr7-96863990-DEL-1471 |

| | |
|---|---|
| chr7-96876514-DEL-8808 | chr7-96876515-DEL-8808 |
| chr7-96913150-DEL-402 | chr7-96913151-DEL-402 |
| chr7-96923202-DEL-4357 | chr7-96923203-DEL-4357 |
| chr7-96956207-DEL-79 | chr7-96956208-DEL-79 |
| chr7-96972187-DEL-136 | chr7-96972188-DEL-136 |
| chr7-97008854-DEL-70 | chr7-97008855-DEL-70 |
| chr7-97011362-DEL-51 | chr7-97011358-DEL-52 |
| chr7-97011553-DEL-2898 | chr7-97011554-DEL-2898 |
| chr7-97064993-DEL-70 | chr7-97064994-DEL-70 |
| chr7-97108054-DEL-572 | chr7-97108055-DEL-572 |
| chr7-97116366-DEL-116 | chr7-97116367-DEL-116 |
| chr7-97135240-DEL-77 | chr7-97135235-DEL-77 |
| chr7-97136955-DEL-198 | chr7-97136956-DEL-198 |
| chr7-97137957-DEL-59 | chr7-97137960-DEL-57 |
| chr7-97142815-DEL-83 | chr7-97142818-DEL-81 |
| chr7-97147522-DEL-104 | chr7-97147523-DEL-104 |
| chr7-97153901-DEL-213 | chr7-97153902-DEL-213 |
| chr7-97157318-DEL-587 | chr7-97157321-DEL-559 |
| chr7-97165987-DEL-344 | chr7-97165988-DEL-345 |
| chr7-97180729-DEL-132 | chr7-97180758-DEL-132 |
| chr7-97202505-DEL-207 | chr7-97202506-DEL-207 |
| chr7-97245332-DEL-64 | chr7-97245333-DEL-64 |
| chr7-97296286-DEL-82 | chr7-97296286-DEL-80 |
| chr7-97322799-DEL-56 | chr7-97322800-DEL-56 |
| chr7-97329514-DEL-166 | chr7-97329515-DEL-166 |
| chr7-97364309-DEL-776 | chr7-97364309-DEL-773 |
| chr7-97403222-DEL-63 | chr7-97403223-DEL-63 |
| chr7-97417530-DEL-64 | chr7-97417517-DEL-86 |
| chr7-97423189-DEL-133 | chr7-97423188-DEL-137 |
| chr7-97445303-DEL-213 | chr7-97445300-DEL-202 |
| chr7-97446741-DEL-67 | chr7-97446720-DEL-58 |
| chr7-97450311-DEL-200 | chr7-97450312-DEL-200 |
| chr7-97459427-DEL-233 | chr7-97459428-DEL-233 |
| chr7-97469571-DEL-157 | chr7-97469572-DEL-157 |
| chr7-97477643-DEL-213 | chr7-97477644-DEL-213 |
| chr7-97489978-DEL-72 | chr7-97489979-DEL-70 |
| chr7-97497387-DEL-70 | chr7-97497387-DEL-70 |
| chr7-97554513-DEL-105 | chr7-97554514-DEL-106 |
| chr7-97572856-DEL-246 | chr7-97572856-DEL-247 |
| chr7-97576884-DEL-1108 | chr7-97576885-DEL-1108 |
| chr7-97579000-DEL-400 | chr7-97579001-DEL-400 |
| chr7-97611451-DEL-56 | chr7-97611452-DEL-56 |
| chr7-97643225-DEL-69 | chr7-97643226-DEL-69 |
| chr7-97656050-DEL-7077 | chr7-97656042-DEL-7078 |
| chr7-97676462-DEL-93 | chr7-97676483-DEL-93 |
| chr7-97681179-DEL-246 | chr7-97681180-DEL-246 |
| chr7-97750873-DEL-801 | chr7-97750874-DEL-801 |
| chr7-97788554-DEL-63 | chr7-97788555-DEL-63 |
| chr7-97790300-DEL-108 | chr7-97790293-DEL-118 |
| chr7-97798033-DEL-166 | chr7-97798034-DEL-166 |
| chr7-97813693-DEL-70 | chr7-97813692-DEL-71 |
| chr7-97818065-DEL-135 | chr7-97818062-DEL-139 |
| chr7-97835147-DEL-373 | chr7-97835148-DEL-373 |
| chr7-97885390-DEL-215 | chr7-97885352-DEL-223 |
| chr7-97893741-DEL-59 | chr7-97893742-DEL-59 |
| chr7-97908492-DEL-70 | chr7-97908493-DEL-105 |
| chr7-97919867-DEL-869 | chr7-97919868-DEL-869 |
| chr7-97939100-DEL-2323 | chr7-97939100-DEL-2320 |
| chr7-97943865-DEL-245 | chr7-97943856-DEL-246 |
| chr7-97945228-DEL-6445 | chr7-97945229-DEL-6445 |
| chr7-97967213-DEL-206 | chr7-97967214-DEL-206 |
| chr7-97968088-DEL-306 | chr7-97968093-DEL-302 |
| chr7-97996378-DEL-56 | chr7-97996379-DEL-56 |
| chr7-98002478-DEL-87 | chr7-98002479-DEL-58 |
| chr7-98012895-DEL-518 | chr7-98012896-DEL-518 |
| chr7-98024160-DEL-490 | chr7-98024161-DEL-490 |
| chr7-98045695-DEL-531 | chr7-98045696-DEL-531 |
| chr7-98051029-DEL-79 | chr7-98051055-DEL-70 |
| chr7-98077159-DEL-175 | chr7-98077160-DEL-175 |
| chr7-98112158-DEL-57 | chr7-98112159-DEL-57 |
| chr7-98159525-DEL-164 | chr7-98159525-DEL-164 |
| chr7-98173560-DEL-112 | chr7-98173561-DEL-105 |
| chr7-98175635-DEL-316 | chr7-98175636-DEL-316 |
| chr7-98179271-DEL-87 | chr7-98179272-DEL-87 |
| chr7-98200685-DEL-53 | chr7-98200685-DEL-54 |
| chr7-98218352-DEL-52 | chr7-98218353-DEL-52 |
| chr7-98254995-DEL-59 | chr7-98254996-DEL-59 |
| chr7-98272235-DEL-374 | chr7-98272236-DEL-374 |
| chr7-98277721-DEL-72 | chr7-98277722-DEL-72 |
| chr7-98311766-DEL-75 | chr7-98311767-DEL-75 |

| | |
|---|---|
| chr7-98326354-DEL-54 | chr7-98326355-DEL-54 |
| chr7-98379660-DEL-450 | chr7-98379661-DEL-450 |
| chr7-98394657-DEL-481 | chr7-98394658-DEL-481 |
| chr7-98409643-DEL-208 | chr7-98409646-DEL-208 |
| chr7-98412465-DEL-305 | chr7-98412466-DEL-305 |
| chr7-98415418-DEL-70 | chr7-98415419-DEL-70 |
| chr7-98426070-DEL-393 | chr7-98426071-DEL-391 |
| chr7-98509688-DEL-470 | chr7-98509689-DEL-470 |
| chr7-98547584-DEL-934 | chr7-98547585-DEL-934 |
| chr7-98550045-DEL-64 | chr7-98550046-DEL-64 |
| chr7-98590750-DEL-184 | chr7-98590751-DEL-184 |
| chr7-98611701-DEL-54 | chr7-98611702-DEL-54 |
| chr7-98612778-DEL-53 | chr7-98612779-DEL-53 |
| chr7-98620480-DEL-62 | chr7-98620481-DEL-62 |
| chr7-98667704-DEL-281 | chr7-98667705-DEL-281 |
| chr7-98714273-DEL-223 | chr7-98714274-DEL-223 |
| chr7-98745611-DEL-241 | chr7-98745612-DEL-241 |
| chr7-98754975-DEL-196 | chr7-98754976-DEL-196 |
| chr7-98800622-DEL-64 | chr7-98800623-DEL-64 |
| chr7-98846836-DEL-224 | chr7-98846837-DEL-224 |
| chr7-98858509-DEL-140 | chr7-98858522-DEL-128 |
| chr7-98880346-DEL-54 | chr7-98880347-DEL-66 |
| chr7-98889066-DEL-53 | chr7-98889065-DEL-58 |
| chr7-98901843-DEL-64 | chr7-98901844-DEL-64 |
| chr7-98907579-DEL-179 | chr7-98907580-DEL-179 |
| chr7-98910489-DEL-108 | chr7-98910487-DEL-107 |
| chr7-98911538-DEL-211 | chr7-98911539-DEL-211 |
| chr7-98914136-DEL-74 | chr7-98914137-DEL-74 |
| chr7-98953806-DEL-52 | chr7-98953807-DEL-52 |
| chr7-98955522-DEL-282 | chr7-98955523-DEL-282 |
| chr7-98959979-DEL-209 | chr7-98959992-DEL-208 |
| chr7-98980307-DEL-132 | chr7-98980303-DEL-123 |
| chr7-99012101-DEL-171 | chr7-99012102-DEL-171 |
| chr7-99054310-DEL-1398 | chr7-99054311-DEL-1398 |
| chr7-99054910-DEL-149 | chr7-99054892-DEL-125 |
| chr7-99061149-DEL-138 | chr7-99061150-DEL-186 |
| chr7-99067074-DEL-859 | chr7-99067075-DEL-859 |
| chr7-99080821-DEL-243 | chr7-99080822-DEL-243 |
| chr7-99089349-DEL-187 | chr7-99089350-DEL-187 |
| chr7-99113426-DEL-123 | chr7-99113427-DEL-123 |
| chr7-99134671-DEL-107 | chr7-99134672-DEL-107 |
| chr7-99146933-DEL-76 | chr7-99146935-DEL-75 |
| chr7-99147762-DEL-185 | chr7-99147777-DEL-205 |
| chr7-99154311-DEL-208 | chr7-99154324-DEL-208 |
| chr7-99167306-DEL-87 | chr7-99167307-DEL-87 |
| chr7-99223540-DEL-93 | chr7-99223541-DEL-93 |
| chr7-99225419-DEL-108 | chr7-99225399-DEL-109 |
| chr7-99234232-DEL-204 | chr7-99234233-DEL-204 |
| chr7-99247920-DEL-56 | chr7-99247918-DEL-57 |
| chr7-99269190-DEL-58 | chr7-99269203-DEL-64 |
| chr7-99284496-DEL-99 | chr7-99284522-DEL-73 |
| chr7-99355249-DEL-52 | chr7-99355249-DEL-50 |
| chr7-99358444-DEL-53 | chr7-99358427-DEL-71 |
| chr7-99386495-DEL-219 | chr7-99386496-DEL-219 |
| chr7-99424587-DEL-198 | chr7-99424588-DEL-198 |
| chr7-99428717-DEL-53 | chr7-99428718-DEL-53 |
| chr7-99446569-DEL-101 | chr7-99446570-DEL-101 |
| chr7-99447439-DEL-110 | chr7-99447439-DEL-111 |
| chr7-99450684-DEL-1355 | chr7-99450685-DEL-1355 |
| chr7-99456112-DEL-576 | chr7-99456113-DEL-576 |
| chr7-99465037-DEL-390 | chr7-99465038-DEL-390 |
| chr7-99466950-DEL-1622 | chr7-99466951-DEL-1622 |
| chr7-99490334-DEL-770 | chr7-99490335-DEL-770 |
| chr7-99531814-DEL-189 | chr7-99531816-DEL-188 |
| chr7-99547152-DEL-53 | chr7-99547153-DEL-53 |
| chr7-99595337-DEL-54 | chr7-99595338-DEL-54 |
| chr7-99647325-DEL-70 | chr7-99647326-DEL-70 |
| chr7-99648351-DEL-3547 | chr7-99648352-DEL-3547 |
| chr7-99702731-DEL-68 | chr7-99702732-DEL-67 |
| chr7-99729995-DEL-112 | chr7-99729996-DEL-112 |
| chr7-99739175-DEL-396 | chr7-99739176-DEL-396 |
| chr7-99760232-DEL-336 | chr7-99760233-DEL-336 |
| chr7-99809662-DEL-63 | chr7-99809663-DEL-63 |
| chr7-99823492-DEL-108 | chr7-99823492-DEL-106 |
| chr7-99844350-DEL-65 | chr7-99844347-DEL-74 |
| chr7-99846747-DEL-166 | chr7-99846749-DEL-166 |
| chr7-99858603-DEL-484 | chr7-99858604-DEL-484 |
| chr7-99908718-DEL-189 | chr7-99908719-DEL-189 |
| chr7-99913973-DEL-54 | chr7-99913974-DEL-54 |
| chr7-99916460-DEL-90 | chr7-99916461-DEL-90 |

| | |
|---|---|
| chr7-99917990-DEL-234 | chr7-99917991-DEL-234 |
| chr7-99945573-DEL-97 | chr7-99945574-DEL-97 |
| chr7-99945972-DEL-240 | chr7-99945973-DEL-240 |
| chr7-99966407-DEL-163 | chr7-99966409-DEL-162 |
| chr7-99979314-DEL-645 | chr7-99979315-DEL-645 |
| chr7-99986971-DEL-205 | chr7-99986972-DEL-205 |
| chr7-100033169-DEL-85 | chr7-100033170-DEL-128 |
| chr7-100036203-DEL-406 | chr7-100036215-DEL-404 |
| chr7-100050810-DEL-338 | chr7-100050811-DEL-338 |
| chr7-100099124-DEL-835 | chr7-100099125-DEL-835 |
| chr7-100105100-DEL-67 | chr7-100105101-DEL-67 |
| chr7-100117736-DEL-431 | chr7-100117737-DEL-431 |
| chr7-100168270-DEL-1152 | chr7-100168271-DEL-1152 |
| chr7-100168613-DEL-451 | chr7-100168630-DEL-435 |
| chr7-100178746-DEL-193 | chr7-100178727-DEL-213 |
| chr7-100187291-DEL-203 | chr7-100187292-DEL-203 |
| chr7-100203512-DEL-219 | chr7-100203513-DEL-219 |
| chr7-100262967-DEL-63 | chr7-100262989-DEL-57 |
| chr7-100270829-DEL-209 | chr7-100270830-DEL-209 |
| chr7-100301994-DEL-68 | chr7-100301995-DEL-68 |
| chr7-100330807-DEL-1184 | chr7-100330801-DEL-1185 |
| chr7-100332492-DEL-779 | chr7-100332493-DEL-779 |
| chr7-100338642-DEL-276 | chr7-100338643-DEL-276 |
| chr7-100344084-DEL-88 | chr7-100344056-DEL-84 |
| chr7-100362434-DEL-262 | chr7-100362435-DEL-262 |
| chr7-100376610-DEL-156 | chr7-100376611-DEL-156 |
| chr7-100388358-DEL-185 | chr7-100388359-DEL-185 |
| chr7-100391492-DEL-70 | chr7-100391493-DEL-70 |
| chr7-100394902-DEL-177 | chr7-100394898-DEL-178 |
| chr7-100396540-DEL-59 | chr7-100396541-DEL-59 |
| chr7-100397104-DEL-59 | chr7-100397106-DEL-54 |
| chr7-100398407-DEL-2994 | chr7-100398408-DEL-2994 |
| chr7-100403480-DEL-210 | chr7-100403481-DEL-210 |
| chr7-100409802-DEL-1282 | chr7-100409803-DEL-1282 |
| chr7-100411045-DEL-109 | chr7-100411046-DEL-109 |
| chr7-100416816-DEL-74 | chr7-100416817-DEL-74 |
| chr7-100426088-DEL-207 | chr7-100426086-DEL-208 |
| chr7-100430042-DEL-924 | chr7-100430042-DEL-924 |
| chr7-100431341-DEL-52 | chr7-100431358-DEL-52 |
| chr7-100447302-DEL-203 | chr7-100447303-DEL-217 |
| chr7-100463310-DEL-180 | chr7-100463311-DEL-180 |
| chr7-100481462-DEL-212 | chr7-100481463-DEL-212 |
| chr7-100482050-DEL-196 | chr7-100482051-DEL-196 |
| chr7-100492155-DEL-106 | chr7-100492156-DEL-106 |
| chr7-100512924-DEL-162 | chr7-100512927-DEL-160 |
| chr7-100514125-DEL-54 | chr7-100514114-DEL-55 |
| chr7-100540600-DEL-98 | chr7-100540601-DEL-98 |
| chr7-100598205-DEL-55 | chr7-100598201-DEL-56 |
| chr7-100601885-DEL-75 | chr7-100601886-DEL-75 |
| chr7-100603662-DEL-83 | chr7-100603666-DEL-84 |
| chr7-100611106-DEL-196 | chr7-100611107-DEL-196 |
| chr7-100625428-DEL-195 | chr7-100625429-DEL-156 |
| chr7-100626746-DEL-165 | chr7-100626747-DEL-165 |
| chr7-100649294-DEL-83 | chr7-100649295-DEL-83 |
| chr7-100678099-DEL-90 | chr7-100678102-DEL-72 |
| chr7-100682319-DEL-64 | chr7-100682320-DEL-64 |
| chr7-100699421-DEL-79 | chr7-100699416-DEL-60 |
| chr7-100726030-DEL-88 | chr7-100726031-DEL-88 |
| chr7-100798606-DEL-51 | chr7-100798607-DEL-51 |
| chr7-100851261-DEL-217 | chr7-100851262-DEL-217 |
| chr7-100858296-DEL-51 | chr7-100858297-DEL-51 |
| chr7-100876217-DEL-118 | chr7-100876218-DEL-118 |
| chr7-100920121-DEL-214 | chr7-100920122-DEL-214 |
| chr7-100952219-DEL-206 | chr7-100952220-DEL-206 |
| chr7-100964473-DEL-174 | chr7-100964474-DEL-174 |
| chr7-100995964-DEL-95 | chr7-100995965-DEL-95 |
| chr7-101004602-DEL-110 | chr7-101004603-DEL-111 |
| chr7-101086947-DEL-268 | chr7-101086948-DEL-268 |
| chr7-101098439-DEL-72 | chr7-101098449-DEL-75 |
| chr7-101159746-DEL-57 | chr7-101159763-DEL-57 |
| chr7-101194623-DEL-144 | chr7-101194624-DEL-144 |
| chr7-101202336-DEL-72 | chr7-101202337-DEL-72 |
| chr7-101228407-DEL-60 | chr7-101228408-DEL-60 |
| chr7-101246658-DEL-152 | chr7-101246657-DEL-153 |
| chr7-101287019-DEL-60 | chr7-101287000-DEL-84 |
| chr7-101297142-DEL-82 | chr7-101297110-DEL-86 |
| chr7-101299404-DEL-441 | chr7-101299405-DEL-441 |
| chr7-101300576-DEL-185 | chr7-101300577-DEL-185 |
| chr7-101377070-DEL-123 | chr7-101377071-DEL-123 |
| chr7-101409459-DEL-227 | chr7-101409460-DEL-227 |

| | |
|---|---|
| chr7-101433364-DEL-447 | chr7-101433365-DEL-447 |
| chr7-101443711-DEL-397 | chr7-101443712-DEL-397 |
| chr7-101446545-DEL-215 | chr7-101446536-DEL-211 |
| chr7-101455056-DEL-80 | chr7-101455057-DEL-80 |
| chr7-101491848-DEL-54 | chr7-101491849-DEL-54 |
| chr7-101515443-DEL-68 | chr7-101515435-DEL-52 |
| chr7-101526192-DEL-54 | chr7-101526193-DEL-54 |
| chr7-101540202-DEL-83 | chr7-101540204-DEL-81 |
| chr7-101574774-DEL-219 | chr7-101574775-DEL-219 |
| chr7-101589476-DEL-249 | chr7-101589477-DEL-249 |
| chr7-101601273-DEL-51 | chr7-101601249-DEL-52 |
| chr7-101602542-DEL-202 | chr7-101602543-DEL-200 |
| chr7-101606951-DEL-53 | chr7-101606947-DEL-67 |
| chr7-101610168-DEL-208 | chr7-101610169-DEL-208 |
| chr7-101646593-DEL-55 | chr7-101646594-DEL-55 |
| chr7-101675770-DEL-290 | chr7-101675771-DEL-290 |
| chr7-101701411-DEL-906 | chr7-101701412-DEL-906 |
| chr7-101717407-DEL-63 | chr7-101717415-DEL-52 |
| chr7-101798096-DEL-212 | chr7-101798097-DEL-212 |
| chr7-101813809-DEL-201 | chr7-101813810-DEL-201 |
| chr7-101831874-DEL-222 | chr7-101831875-DEL-222 |
| chr7-101877993-DEL-189 | chr7-101877989-DEL-195 |
| chr7-101906884-DEL-64 | chr7-101906885-DEL-64 |
| chr7-101907554-DEL-119 | chr7-101907555-DEL-119 |
| chr7-101908452-DEL-692 | chr7-101908456-DEL-688 |
| chr7-101932497-DEL-60 | chr7-101932498-DEL-58 |
| chr7-101936456-DEL-74 | chr7-101936457-DEL-74 |
| chr7-101943810-DEL-207 | chr7-101943811-DEL-207 |
| chr7-101953582-DEL-59 | chr7-101953577-DEL-60 |
| chr7-101956454-DEL-2307 | chr7-101956455-DEL-2307 |
| chr7-101985400-DEL-93 | chr7-101985401-DEL-93 |
| chr7-102011158-DEL-424 | chr7-102011159-DEL-424 |
| chr7-102047316-DEL-62 | chr7-102047317-DEL-62 |
| chr7-102050705-DEL-56 | chr7-102050706-DEL-56 |
| chr7-102055448-DEL-563 | chr7-102055450-DEL-561 |
| chr7-102066183-DEL-242 | chr7-102066184-DEL-242 |
| chr7-102068685-DEL-52 | chr7-102068681-DEL-57 |
| chr7-102107272-DEL-868 | chr7-102107273-DEL-868 |
| chr7-102113701-DEL-207 | chr7-102113690-DEL-208 |
| chr7-102119282-DEL-5762 | chr7-102119283-DEL-5762 |
| chr7-102128083-DEL-51 | chr7-102128084-DEL-51 |
| chr7-102145432-DEL-136 | chr7-102145433-DEL-136 |
| chr7-102168386-DEL-2668 | chr7-102168388-DEL-2667 |
| chr7-102173101-DEL-1244 | chr7-102173101-DEL-1244 |
| chr7-102174762-DEL-496 | chr7-102174758-DEL-500 |
| chr7-102189309-DEL-177 | chr7-102189310-DEL-177 |
| chr7-102205255-DEL-1614 | chr7-102205256-DEL-1614 |
| chr7-102240232-DEL-94 | chr7-102240233-DEL-94 |
| chr7-102248711-DEL-240 | chr7-102248712-DEL-240 |
| chr7-102278508-DEL-91 | chr7-102278508-DEL-91 |
| chr7-102281896-DEL-236 | chr7-102281897-DEL-236 |
| chr7-102302996-DEL-194 | chr7-102302997-DEL-194 |
| chr7-102310698-DEL-398 | chr7-102310697-DEL-399 |
| chr7-102337114-DEL-156 | chr7-102337115-DEL-156 |
| chr7-102339449-DEL-914 | chr7-102339450-DEL-912 |
| chr7-102344040-DEL-88 | chr7-102344041-DEL-87 |
| chr7-102362239-DEL-63 | chr7-102362249-DEL-61 |
| chr7-102379702-DEL-794 | chr7-102379703-DEL-794 |
| chr7-102383904-DEL-317 | chr7-102383905-DEL-317 |
| chr7-102411221-DEL-109 | chr7-102411222-DEL-109 |
| chr7-102427905-DEL-391 | chr7-102427906-DEL-391 |
| chr7-102443884-DEL-6495 | chr7-102443885-DEL-6495 |
| chr7-102452456-DEL-703 | chr7-102452456-DEL-703 |
| chr7-102475729-DEL-790 | chr7-102475730-DEL-790 |
| chr7-102479682-DEL-12889 | chr7-102479684-DEL-12889 |
| chr7-102482502-DEL-6276 | chr7-102482503-DEL-6276 |
| chr7-102492130-DEL-200 | chr7-102492131-DEL-200 |
| chr7-102496069-DEL-228 | chr7-102496070-DEL-228 |
| chr7-102498332-DEL-708 | chr7-102498333-DEL-708 |
| chr7-102499908-DEL-3209 | chr7-102499909-DEL-3209 |
| chr7-102514577-DEL-1062 | chr7-102514578-DEL-1062 |
| chr7-102516112-DEL-1695 | chr7-102516100-DEL-1696 |
| chr7-102517862-DEL-2607 | chr7-102517863-DEL-2607 |
| chr7-102519245-DEL-294 | chr7-102519246-DEL-294 |
| chr7-102544054-DEL-1555 | chr7-102544055-DEL-1555 |
| chr7-102545104-DEL-119 | chr7-102545145-DEL-79 |
| chr7-102567673-DEL-190 | chr7-102567674-DEL-190 |
| chr7-102569185-DEL-837 | chr7-102569186-DEL-837 |
| chr7-102599584-DEL-2732 | chr7-102599586-DEL-2731 |
| chr7-102635065-DEL-99 | chr7-102635066-DEL-99 |

| | |
|---|---|
| chr7-102646096-DEL-82 | chr7-102646097-DEL-82 |
| chr7-102647884-DEL-7043 | chr7-102647885-DEL-7043 |
| chr7-102662595-DEL-418 | chr7-102662596-DEL-418 |
| chr7-102688027-DEL-83 | chr7-102688028-DEL-83 |
| chr7-102692548-DEL-58 | chr7-102692549-DEL-58 |
| chr7-102705719-DEL-99 | chr7-102705719-DEL-141 |
| chr7-102709777-DEL-2169 | chr7-102709777-DEL-2169 |
| chr7-102717620-DEL-1360 | chr7-102717621-DEL-1360 |
| chr7-102722440-DEL-143 | chr7-102722441-DEL-143 |
| chr7-102723536-DEL-117 | chr7-102723537-DEL-117 |
| chr7-102729889-DEL-57 | chr7-102729890-DEL-57 |
| chr7-102750112-DEL-14125 | chr7-102750113-DEL-14125 |
| chr7-102754469-DEL-212 | chr7-102754470-DEL-212 |
| chr7-102767699-DEL-554 | chr7-102767700-DEL-554 |
| chr7-102769771-DEL-513 | chr7-102769772-DEL-513 |
| chr7-102807593-DEL-6197 | chr7-102807594-DEL-6197 |
| chr7-102829785-DEL-225 | chr7-102829786-DEL-225 |
| chr7-102835639-DEL-80 | chr7-102835642-DEL-78 |
| chr7-102844680-DEL-6674 | chr7-102844667-DEL-6674 |
| chr7-102863198-DEL-58 | chr7-102863197-DEL-57 |
| chr7-102880272-DEL-114 | chr7-102880273-DEL-114 |
| chr7-102915168-DEL-306 | chr7-102915169-DEL-306 |
| chr7-102920697-DEL-297 | chr7-102920698-DEL-297 |
| chr7-102931857-DEL-6782 | chr7-102931857-DEL-6782 |
| chr7-102953250-DEL-1178 | chr7-102953250-DEL-1178 |
| chr7-102970766-DEL-107 | chr7-102970764-DEL-107 |
| chr7-102974326-DEL-3289 | chr7-102974327-DEL-3289 |
| chr7-102978366-DEL-72 | chr7-102978367-DEL-72 |
| chr7-102993875-DEL-60 | chr7-102993876-DEL-60 |
| chr7-103002842-DEL-774 | chr7-103002844-DEL-773 |
| chr7-103013806-DEL-204 | chr7-103013800-DEL-205 |
| chr7-103053378-DEL-161 | chr7-103053379-DEL-161 |
| chr7-103064754-DEL-492 | chr7-103064754-DEL-492 |
| chr7-103084770-DEL-198 | chr7-103084770-DEL-198 |
| chr7-103085712-DEL-2178 | chr7-103085713-DEL-2178 |
| chr7-103103984-DEL-545 | chr7-103103985-DEL-545 |
| chr7-103107758-DEL-652 | chr7-103107759-DEL-652 |
| chr7-103115177-DEL-55 | chr7-103115172-DEL-60 |
| chr7-103126446-DEL-307 | chr7-103126447-DEL-307 |
| chr7-103131849-DEL-6371 | chr7-103131850-DEL-6371 |
| chr7-103198911-DEL-1190 | chr7-103198912-DEL-1190 |
| chr7-103255277-DEL-52 | chr7-103255280-DEL-50 |
| chr7-103257685-DEL-52 | chr7-103257677-DEL-50 |
| chr7-103265044-DEL-3221 | chr7-103265026-DEL-3222 |
| chr7-103270659-DEL-2604 | chr7-103270660-DEL-2604 |
| chr7-103274846-DEL-446 | chr7-103274847-DEL-446 |
| chr7-103280310-DEL-782 | chr7-103280311-DEL-782 |
| chr7-103293494-DEL-829 | chr7-103293495-DEL-829 |
| chr7-103297010-DEL-6223 | chr7-103297011-DEL-6223 |
| chr7-103314785-DEL-1884 | chr7-103314786-DEL-1884 |
| chr7-103329226-DEL-74 | chr7-103329227-DEL-74 |
| chr7-103368502-DEL-58 | chr7-103368495-DEL-66 |
| chr7-103382174-DEL-60 | chr7-103382177-DEL-58 |
| chr7-103388130-DEL-265 | chr7-103388129-DEL-260 |
| chr7-103398551-DEL-100 | chr7-103398541-DEL-101 |
| chr7-103407609-DEL-54 | chr7-103407610-DEL-58 |
| chr7-103411448-DEL-1425 | chr7-103411434-DEL-1406 |
| chr7-103413522-DEL-398 | chr7-103413523-DEL-398 |
| chr7-103446095-DEL-2669 | chr7-103446083-DEL-2670 |
| chr7-103454911-DEL-122 | chr7-103454912-DEL-122 |
| chr7-103463997-DEL-161 | chr7-103463999-DEL-160 |
| chr7-103479968-DEL-69 | chr7-103479968-DEL-59 |
| chr7-103493507-DEL-213 | chr7-103493511-DEL-217 |
| chr7-103523093-DEL-143 | chr7-103523094-DEL-143 |
| chr7-103524495-DEL-486 | chr7-103524503-DEL-481 |
| chr7-103531749-DEL-585 | chr7-103531750-DEL-585 |
| chr7-103536184-DEL-357 | chr7-103536189-DEL-353 |
| chr7-103539199-DEL-73 | chr7-103539225-DEL-77 |
| chr7-103560255-DEL-1118 | chr7-103560256-DEL-1118 |
| chr7-103567730-DEL-61 | chr7-103567731-DEL-61 |
| chr7-103569473-DEL-136 | chr7-103569474-DEL-136 |
| chr7-103570584-DEL-6920 | chr7-103570585-DEL-6921 |
| chr7-103580040-DEL-1382 | chr7-103580041-DEL-1382 |
| chr7-103591197-DEL-56 | chr7-103591198-DEL-56 |
| chr7-103599488-DEL-401 | chr7-103599489-DEL-401 |
| chr7-103609996-DEL-123 | chr7-103609996-DEL-124 |
| chr7-103616296-DEL-115 | chr7-103616297-DEL-115 |
| chr7-103617692-DEL-585 | chr7-103617693-DEL-583 |
| chr7-103640810-DEL-343 | chr7-103640811-DEL-343 |
| chr7-103655517-DEL-676 | chr7-103655508-DEL-685 |

| | |
|---|---|
| chr7-103657751-DEL-3008 | chr7-103657752-DEL-3008 |
| chr7-103676784-DEL-7096 | chr7-103676785-DEL-7096 |
| chr7-103683227-DEL-424 | chr7-103683228-DEL-424 |
| chr7-103713068-DEL-232 | chr7-103713069-DEL-232 |
| chr7-103738712-DEL-86 | chr7-103738712-DEL-86 |
| chr7-103740141-DEL-2001 | chr7-103740141-DEL-2001 |
| chr7-103741076-DEL-192 | chr7-103741077-DEL-192 |
| chr7-103752895-DEL-66 | chr7-103752850-DEL-66 |
| chr7-103758006-DEL-541 | chr7-103758007-DEL-541 |
| chr7-103769259-DEL-6710 | chr7-103769260-DEL-6710 |
| chr7-103789282-DEL-62 | chr7-103789283-DEL-62 |
| chr7-103805448-DEL-3508 | chr7-103805449-DEL-3508 |
| chr7-103810339-DEL-283 | chr7-103810340-DEL-283 |
| chr7-103826703-DEL-938 | chr7-103826704-DEL-938 |
| chr7-103852240-DEL-543 | chr7-103852241-DEL-543 |
| chr7-103857455-DEL-170 | chr7-103857456-DEL-161 |
| chr7-103860300-DEL-77 | chr7-103860301-DEL-77 |
| chr7-103862416-DEL-138 | chr7-103862417-DEL-126 |
| chr7-103872256-DEL-67 | chr7-103872257-DEL-67 |
| chr7-103876372-DEL-202 | chr7-103876373-DEL-202 |
| chr7-103877908-DEL-1020 | chr7-103877909-DEL-1020 |
| chr7-103881106-DEL-59 | chr7-103881107-DEL-59 |
| chr7-103882657-DEL-90 | chr7-103882658-DEL-90 |
| chr7-103892795-DEL-54 | chr7-103892810-DEL-77 |
| chr7-103895336-DEL-171 | chr7-103895337-DEL-171 |
| chr7-103908299-DEL-101 | chr7-103908272-DEL-151 |
| chr7-103951532-DEL-238 | chr7-103951527-DEL-238 |
| chr7-103960211-DEL-62 | chr7-103960212-DEL-62 |
| chr7-103981945-DEL-56 | chr7-103981925-DEL-61 |
| chr7-103983495-DEL-111 | chr7-103983513-DEL-81 |
| chr7-103986293-DEL-204 | chr7-103986294-DEL-204 |
| chr7-103996664-DEL-383 | chr7-103996664-DEL-383 |
| chr7-104005700-DEL-186 | chr7-104005700-DEL-184 |
| chr7-104020834-DEL-64 | chr7-104020870-DEL-65 |
| chr7-104053858-DEL-72 | chr7-104053870-DEL-54 |
| chr7-104072465-DEL-4829 | chr7-104072466-DEL-4829 |
| chr7-104081212-DEL-1497 | chr7-104081212-DEL-1497 |
| chr7-104108785-DEL-91 | chr7-104108786-DEL-91 |
| chr7-104131766-DEL-54 | chr7-104131767-DEL-54 |
| chr7-104197800-DEL-183 | chr7-104197802-DEL-172 |
| chr7-104210627-DEL-102 | chr7-104210628-DEL-102 |
| chr7-104232790-DEL-183 | chr7-104232791-DEL-183 |
| chr7-104248376-DEL-71 | chr7-104248362-DEL-90 |
| chr7-104304874-DEL-456 | chr7-104304883-DEL-456 |
| chr7-104364106-DEL-6817 | chr7-104364107-DEL-6817 |
| chr7-104372529-DEL-4496 | chr7-104372530-DEL-4496 |
| chr7-104384830-DEL-222 | chr7-104384831-DEL-222 |
| chr7-104464564-DEL-69 | chr7-104464570-DEL-53 |
| chr7-104472961-DEL-567 | chr7-104472962-DEL-567 |
| chr7-104481286-DEL-212 | chr7-104481287-DEL-212 |
| chr7-104482867-DEL-260 | chr7-104482868-DEL-260 |
| chr7-104505750-DEL-64 | chr7-104505751-DEL-64 |
| chr7-104548122-DEL-68 | chr7-104548123-DEL-68 |
| chr7-104551808-DEL-54 | chr7-104551795-DEL-58 |
| chr7-104598362-DEL-1330 | chr7-104598363-DEL-1330 |
| chr7-104610710-DEL-110 | chr7-104610711-DEL-110 |
| chr7-104621708-DEL-102 | chr7-104621719-DEL-92 |
| chr7-104675404-DEL-298 | chr7-104675405-DEL-298 |
| chr7-104676030-DEL-321 | chr7-104676031-DEL-321 |
| chr7-104687331-DEL-2252 | chr7-104687332-DEL-2252 |
| chr7-104720314-DEL-54 | chr7-104720319-DEL-50 |
| chr7-104736639-DEL-250 | chr7-104736640-DEL-250 |
| chr7-104749030-DEL-373 | chr7-104749031-DEL-373 |
| chr7-104757271-DEL-53 | chr7-104757267-DEL-57 |
| chr7-104860755-DEL-89 | chr7-104860756-DEL-89 |
| chr7-104864907-DEL-3464 | chr7-104864912-DEL-3461 |
| chr7-104871786-DEL-56 | chr7-104871787-DEL-56 |
| chr7-104901076-DEL-2417 | chr7-104901077-DEL-2417 |
| chr7-104912550-DEL-198 | chr7-104912551-DEL-198 |
| chr7-104926449-DEL-80 | chr7-104926419-DEL-89 |
| chr7-104981307-DEL-55 | chr7-104981308-DEL-50 |
| chr7-104999398-DEL-6464 | chr7-104999397-DEL-6464 |
| chr7-105005734-DEL-131 | chr7-105005729-DEL-132 |
| chr7-105008871-DEL-1232 | chr7-105008872-DEL-1232 |
| chr7-105064029-DEL-99 | chr7-105064036-DEL-69 |
| chr7-105080434-DEL-283 | chr7-105080435-DEL-283 |
| chr7-105109976-DEL-275 | chr7-105109977-DEL-275 |
| chr7-105155199-DEL-65 | chr7-105155201-DEL-63 |
| chr7-105167790-DEL-209 | chr7-105167791-DEL-209 |
| chr7-105175107-DEL-52 | chr7-105175106-DEL-52 |

| | |
|---|---|
| chr7-105181180-DEL-459 | chr7-105181181-DEL-459 |
| chr7-105181771-DEL-337 | chr7-105181772-DEL-337 |
| chr7-105192521-DEL-64 | chr7-105192518-DEL-95 |
| chr7-105196834-DEL-60 | chr7-105196835-DEL-60 |
| chr7-105212162-DEL-172 | chr7-105212163-DEL-172 |
| chr7-105238105-DEL-219 | chr7-105238090-DEL-220 |
| chr7-105244983-DEL-61 | chr7-105244984-DEL-61 |
| chr7-105320145-DEL-78 | chr7-105320184-DEL-66 |
| chr7-105321157-DEL-981 | chr7-105321158-DEL-981 |
| chr7-105329283-DEL-204 | chr7-105329284-DEL-203 |
| chr7-105339211-DEL-574 | chr7-105339212-DEL-574 |
| chr7-105349600-DEL-66 | chr7-105349625-DEL-81 |
| chr7-105396385-DEL-94 | chr7-105396386-DEL-94 |
| chr7-105440208-DEL-76 | chr7-105440209-DEL-76 |
| chr7-105446967-DEL-97 | chr7-105446968-DEL-97 |
| chr7-105491958-DEL-1318 | chr7-105491959-DEL-1318 |
| chr7-105497292-DEL-84 | chr7-105497291-DEL-68 |
| chr7-105510570-DEL-78 | chr7-105510571-DEL-78 |
| chr7-105511289-DEL-216 | chr7-105511290-DEL-216 |
| chr7-105661775-DEL-87 | chr7-105661775-DEL-87 |
| chr7-105664161-DEL-302 | chr7-105664161-DEL-302 |
| chr7-105824994-DEL-52 | chr7-105824995-DEL-52 |
| chr7-105957670-DEL-5411 | chr7-105957671-DEL-5411 |
| chr7-105968059-DEL-8815 | chr7-105968060-DEL-8815 |
| chr7-105983316-DEL-211 | chr7-105983317-DEL-211 |
| chr7-105991727-DEL-95 | chr7-105991728-DEL-103 |
| chr7-106025281-DEL-68 | chr7-106025280-DEL-66 |
| chr7-106045895-DEL-480 | chr7-106045882-DEL-504 |
| chr7-106061099-DEL-6192 | chr7-106061100-DEL-6192 |
| chr7-106068760-DEL-1896 | chr7-106068761-DEL-1896 |
| chr7-106084784-DEL-3369 | chr7-106084785-DEL-3369 |
| chr7-106096859-DEL-2109 | chr7-106096860-DEL-2109 |
| chr7-106116704-DEL-62 | chr7-106116705-DEL-62 |
| chr7-106122637-DEL-1733 | chr7-106122638-DEL-1733 |
| chr7-106122916-DEL-54 | chr7-106122917-DEL-54 |
| chr7-106133014-DEL-481 | chr7-106133015-DEL-481 |
| chr7-106166143-DEL-1367 | chr7-106166144-DEL-1367 |
| chr7-106174695-DEL-2171 | chr7-106174696-DEL-2171 |
| chr7-106179759-DEL-7750 | chr7-106179759-DEL-7750 |
| chr7-106195488-DEL-103 | chr7-106195492-DEL-104 |
| chr7-106203398-DEL-150 | chr7-106203399-DEL-150 |
| chr7-106215577-DEL-4163 | chr7-106215564-DEL-4171 |
| chr7-106220111-DEL-208 | chr7-106220112-DEL-208 |
| chr7-106233562-DEL-2821 | chr7-106233565-DEL-2819 |
| chr7-106236542-DEL-91 | chr7-106236543-DEL-91 |
| chr7-106291706-DEL-550 | chr7-106291707-DEL-548 |
| chr7-106359550-DEL-69 | chr7-106359541-DEL-68 |
| chr7-106425171-DEL-71 | chr7-106425176-DEL-67 |
| chr7-106429198-DEL-79 | chr7-106429217-DEL-73 |
| chr7-106436637-DEL-1382 | chr7-106436638-DEL-1382 |
| chr7-106492473-DEL-1204 | chr7-106492466-DEL-1205 |
| chr7-106519055-DEL-3175 | chr7-106519056-DEL-3175 |
| chr7-106552375-DEL-80 | chr7-106552375-DEL-78 |
| chr7-106589708-DEL-84 | chr7-106589709-DEL-84 |
| chr7-106599348-DEL-78 | chr7-106599343-DEL-88 |
| chr7-106616110-DEL-71 | chr7-106616111-DEL-71 |
| chr7-106659492-DEL-82 | chr7-106659493-DEL-82 |
| chr7-106664831-DEL-2016 | chr7-106664832-DEL-2016 |
| chr7-106734255-DEL-260 | chr7-106734256-DEL-260 |
| chr7-106749424-DEL-420 | chr7-106749425-DEL-420 |
| chr7-106762662-DEL-141 | chr7-106762663-DEL-141 |
| chr7-106778670-DEL-80 | chr7-106778670-DEL-80 |
| chr7-106797618-DEL-52 | chr7-106797615-DEL-64 |
| chr7-106813891-DEL-114 | chr7-106813892-DEL-90 |
| chr7-106853600-DEL-106 | chr7-106853601-DEL-106 |
| chr7-106863610-DEL-7086 | chr7-106863611-DEL-7086 |
| chr7-106878513-DEL-98 | chr7-106878514-DEL-98 |
| chr7-106888942-DEL-58 | chr7-106888943-DEL-58 |
| chr7-106896303-DEL-1681 | chr7-106896327-DEL-1654 |
| chr7-106901000-DEL-6275 | chr7-106901001-DEL-6275 |
| chr7-106922340-DEL-134 | chr7-106922341-DEL-134 |
| chr7-106930429-DEL-189 | chr7-106930430-DEL-189 |
| chr7-106974012-DEL-165 | chr7-106974011-DEL-166 |
| chr7-107013576-DEL-184 | chr7-107013577-DEL-184 |
| chr7-107030827-DEL-188 | chr7-107030828-DEL-188 |
| chr7-107039564-DEL-63 | chr7-107039565-DEL-63 |
| chr7-107067099-DEL-101 | chr7-107067078-DEL-102 |
| chr7-107070402-DEL-6742 | chr7-107070403-DEL-6742 |
| chr7-107082799-DEL-228 | chr7-107082758-DEL-224 |
| chr7-107095397-DEL-75 | chr7-107095398-DEL-75 |

| | |
|---|---|
| chr7-107106901-DEL-267 | chr7-107106902-DEL-267 |
| chr7-107142059-DEL-64 | chr7-107142061-DEL-59 |
| chr7-107155769-DEL-57 | chr7-107155770-DEL-57 |
| chr7-107157334-DEL-1353 | chr7-107157335-DEL-1353 |
| chr7-107172553-DEL-58 | chr7-107172554-DEL-58 |
| chr7-107174758-DEL-135 | chr7-107174765-DEL-129 |
| chr7-107177341-DEL-110 | chr7-107177342-DEL-110 |
| chr7-107184898-DEL-2088 | chr7-107184899-DEL-2088 |
| chr7-107200779-DEL-208 | chr7-107200780-DEL-208 |
| chr7-107228396-DEL-1932 | chr7-107228397-DEL-1932 |
| chr7-107273251-DEL-242 | chr7-107273252-DEL-242 |
| chr7-107278062-DEL-62 | chr7-107278063-DEL-62 |
| chr7-107310485-DEL-113 | chr7-107310486-DEL-113 |
| chr7-107330728-DEL-360 | chr7-107330729-DEL-360 |
| chr7-107333329-DEL-160 | chr7-107333330-DEL-160 |
| chr7-107340928-DEL-1163 | chr7-107340929-DEL-1163 |
| chr7-107357230-DEL-1600 | chr7-107357231-DEL-1600 |
| chr7-107357813-DEL-155 | chr7-107357814-DEL-155 |
| chr7-107379272-DEL-51 | chr7-107379255-DEL-52 |
| chr7-107379898-DEL-405 | chr7-107379899-DEL-405 |
| chr7-107383365-DEL-52 | chr7-107383368-DEL-50 |
| chr7-107404173-DEL-390 | chr7-107404174-DEL-390 |
| chr7-107431556-DEL-1010 | chr7-107431557-DEL-1010 |
| chr7-107453173-DEL-7095 | chr7-107453174-DEL-7095 |
| chr7-107462164-DEL-1186 | chr7-107462163-DEL-1186 |
| chr7-107464473-DEL-66 | chr7-107464470-DEL-62 |
| chr7-107483049-DEL-777 | chr7-107483051-DEL-776 |
| chr7-107486065-DEL-6149 | chr7-107486054-DEL-6160 |
| chr7-107497734-DEL-235 | chr7-107497735-DEL-235 |
| chr7-107540570-DEL-1889 | chr7-107540571-DEL-1889 |
| chr7-107556547-DEL-204 | chr7-107556548-DEL-204 |
| chr7-107562613-DEL-60 | chr7-107562614-DEL-60 |
| chr7-107612293-DEL-3781 | chr7-107612292-DEL-3781 |
| chr7-107612397-DEL-74 | chr7-107612398-DEL-74 |
| chr7-107619652-DEL-7651 | chr7-107619653-DEL-7651 |
| chr7-107629727-DEL-51 | chr7-107629728-DEL-51 |
| chr7-107655792-DEL-70 | chr7-107655793-DEL-70 |
| chr7-107681813-DEL-87 | chr7-107681812-DEL-89 |
| chr7-107691718-DEL-66 | chr7-107691712-DEL-72 |
| chr7-107692380-DEL-1408 | chr7-107692380-DEL-1408 |
| chr7-107712678-DEL-119 | chr7-107712679-DEL-119 |
| chr7-107729508-DEL-76 | chr7-107729505-DEL-80 |
| chr7-107870978-DEL-219 | chr7-107870973-DEL-190 |
| chr7-107889465-DEL-134 | chr7-107889466-DEL-134 |
| chr7-107893230-DEL-3466 | chr7-107893230-DEL-3464 |
| chr7-107925216-DEL-182 | chr7-107925217-DEL-181 |
| chr7-107963252-DEL-98 | chr7-107963220-DEL-89 |
| chr7-107971711-DEL-268 | chr7-107971712-DEL-268 |
| chr7-107975210-DEL-343 | chr7-107975211-DEL-343 |
| chr7-107976843-DEL-203 | chr7-107976844-DEL-203 |
| chr7-107980338-DEL-99 | chr7-107980339-DEL-99 |
| chr7-107986695-DEL-74 | chr7-107986716-DEL-52 |
| chr7-108025365-DEL-437 | chr7-108025366-DEL-437 |
| chr7-108025924-DEL-1363 | chr7-108025925-DEL-1363 |
| chr7-108028416-DEL-5258 | chr7-108028417-DEL-5258 |
| chr7-108117828-DEL-351 | chr7-108117829-DEL-350 |
| chr7-108126852-DEL-57 | chr7-108126853-DEL-57 |
| chr7-108144668-DEL-491 | chr7-108144669-DEL-491 |
| chr7-108158663-DEL-198 | chr7-108158664-DEL-198 |
| chr7-108170982-DEL-109 | chr7-108170975-DEL-117 |
| chr7-108173414-DEL-385 | chr7-108173415-DEL-385 |
| chr7-108183794-DEL-1356 | chr7-108183795-DEL-1356 |
| chr7-108186022-DEL-1860 | chr7-108186013-DEL-1861 |
| chr7-108194207-DEL-61 | chr7-108194208-DEL-61 |
| chr7-108214190-DEL-750 | chr7-108214189-DEL-751 |
| chr7-108262747-DEL-132 | chr7-108262748-DEL-131 |
| chr7-108269413-DEL-134 | chr7-108269415-DEL-135 |
| chr7-108278879-DEL-1434 | chr7-108278880-DEL-1434 |
| chr7-108278881-DEL-60 | chr7-108278888-DEL-58 |
| chr7-108281775-DEL-56 | chr7-108281794-DEL-71 |
| chr7-108290788-DEL-58 | chr7-108290800-DEL-51 |
| chr7-108352442-DEL-567 | chr7-108352443-DEL-567 |
| chr7-108361132-DEL-72 | chr7-108361133-DEL-72 |
| chr7-108367439-DEL-147 | chr7-108367440-DEL-147 |
| chr7-108370696-DEL-169 | chr7-108370697-DEL-163 |
| chr7-108381854-DEL-7232 | chr7-108381855-DEL-7232 |
| chr7-108399855-DEL-166 | chr7-108399856-DEL-166 |
| chr7-108427013-DEL-2901 | chr7-108427014-DEL-2901 |
| chr7-108457660-DEL-150 | chr7-108457661-DEL-151 |
| chr7-108471176-DEL-65 | chr7-108471177-DEL-69 |

| | |
|---|---|
| chr7-108492379-DEL-148 | chr7-108492388-DEL-112 |
| chr7-108544319-DEL-240 | chr7-108544320-DEL-240 |
| chr7-108569686-DEL-225 | chr7-108569687-DEL-225 |
| chr7-108580262-DEL-548 | chr7-108580263-DEL-544 |
| chr7-108585528-DEL-56 | chr7-108585560-DEL-65 |
| chr7-108671377-DEL-1680 | chr7-108671374-DEL-1675 |
| chr7-108673912-DEL-398 | chr7-108673913-DEL-398 |
| chr7-108690642-DEL-105 | chr7-108690671-DEL-91 |
| chr7-108717041-DEL-6473 | chr7-108717042-DEL-6473 |
| chr7-108734053-DEL-92 | chr7-108734064-DEL-82 |
| chr7-108806009-DEL-433 | chr7-108806010-DEL-433 |
| chr7-108807079-DEL-1104 | chr7-108807080-DEL-1104 |
| chr7-108828154-DEL-73 | chr7-108828155-DEL-73 |
| chr7-108857588-DEL-209 | chr7-108857589-DEL-209 |
| chr7-108889853-DEL-103 | chr7-108889846-DEL-109 |
| chr7-108901842-DEL-71 | chr7-108901839-DEL-75 |
| chr7-108965200-DEL-6218 | chr7-108965201-DEL-6218 |
| chr7-109019084-DEL-1226 | chr7-109019085-DEL-1226 |
| chr7-109055543-DEL-192 | chr7-109055577-DEL-150 |
| chr7-109057722-DEL-57 | chr7-109057746-DEL-56 |
| chr7-109087368-DEL-369 | chr7-109087369-DEL-369 |
| chr7-109137842-DEL-53 | chr7-109137843-DEL-53 |
| chr7-109192230-DEL-197 | chr7-109192231-DEL-197 |
| chr7-109212560-DEL-129 | chr7-109212561-DEL-129 |
| chr7-109222819-DEL-318 | chr7-109222820-DEL-318 |
| chr7-109245998-DEL-74 | chr7-109245998-DEL-65 |
| chr7-109249931-DEL-223 | chr7-109249932-DEL-223 |
| chr7-109267278-DEL-219 | chr7-109267279-DEL-210 |
| chr7-109282919-DEL-804 | chr7-109282920-DEL-804 |
| chr7-109296381-DEL-398 | chr7-109296383-DEL-397 |
| chr7-109331595-DEL-74 | chr7-109331612-DEL-58 |
| chr7-109334586-DEL-84 | chr7-109334585-DEL-121 |
| chr7-109370428-DEL-200 | chr7-109370429-DEL-200 |
| chr7-109396563-DEL-207 | chr7-109396564-DEL-207 |
| chr7-109405633-DEL-309 | chr7-109405634-DEL-309 |
| chr7-109427832-DEL-212 | chr7-109427833-DEL-212 |
| chr7-109439657-DEL-240 | chr7-109439695-DEL-243 |
| chr7-109479303-DEL-124 | chr7-109479319-DEL-94 |
| chr7-109490055-DEL-643 | chr7-109490056-DEL-643 |
| chr7-109504010-DEL-99 | chr7-109504011-DEL-99 |
| chr7-109508128-DEL-430 | chr7-109508127-DEL-431 |
| chr7-109520816-DEL-62 | chr7-109520817-DEL-62 |
| chr7-109535893-DEL-1500 | chr7-109535894-DEL-1500 |
| chr7-109537073-DEL-1395 | chr7-109537074-DEL-1395 |
| chr7-109551918-DEL-231 | chr7-109551919-DEL-231 |
| chr7-109589926-DEL-184 | chr7-109589927-DEL-184 |
| chr7-109609780-DEL-75 | chr7-109609781-DEL-75 |
| chr7-109632028-DEL-142 | chr7-109632029-DEL-142 |
| chr7-109632977-DEL-210 | chr7-109632987-DEL-211 |
| chr7-109634337-DEL-65 | chr7-109634338-DEL-65 |
| chr7-109636479-DEL-412 | chr7-109636480-DEL-412 |
| chr7-109664028-DEL-56 | chr7-109664001-DEL-96 |
| chr7-109707477-DEL-137 | chr7-109707485-DEL-143 |
| chr7-109710609-DEL-53 | chr7-109710610-DEL-53 |
| chr7-109710735-DEL-2047 | chr7-109710736-DEL-2047 |
| chr7-109745328-DEL-59 | chr7-109745328-DEL-59 |
| chr7-109753879-DEL-365 | chr7-109753871-DEL-367 |
| chr7-109759537-DEL-194 | chr7-109759538-DEL-194 |
| chr7-109760393-DEL-206 | chr7-109760394-DEL-207 |
| chr7-109766184-DEL-500 | chr7-109766185-DEL-500 |
| chr7-109766837-DEL-195 | chr7-109766838-DEL-195 |
| chr7-109769366-DEL-3696 | chr7-109769367-DEL-3696 |
| chr7-109788648-DEL-118 | chr7-109788620-DEL-114 |
| chr7-109798774-DEL-205 | chr7-109798775-DEL-205 |
| chr7-109805736-DEL-203 | chr7-109805737-DEL-203 |
| chr7-109824248-DEL-66 | chr7-109824249-DEL-66 |
| chr7-109830284-DEL-176 | chr7-109830285-DEL-176 |
| chr7-109840463-DEL-98 | chr7-109840483-DEL-89 |
| chr7-109856180-DEL-58 | chr7-109856181-DEL-58 |
| chr7-109869292-DEL-76 | chr7-109869293-DEL-76 |
| chr7-109899414-DEL-63 | chr7-109899410-DEL-64 |
| chr7-109955912-DEL-848 | chr7-109955913-DEL-847 |
| chr7-110011563-DEL-62 | chr7-110011564-DEL-62 |
| chr7-110013262-DEL-194 | chr7-110013263-DEL-194 |
| chr7-110016720-DEL-406 | chr7-110016719-DEL-407 |
| chr7-110032706-DEL-87 | chr7-110032707-DEL-87 |
| chr7-110045138-DEL-165 | chr7-110045138-DEL-165 |
| chr7-110066283-DEL-86 | chr7-110066284-DEL-82 |
| chr7-110098787-DEL-8491 | chr7-110098788-DEL-8491 |
| chr7-110110749-DEL-199 | chr7-110110750-DEL-199 |

| | |
|---|---|
| chr7-110117554-DEL-326 | chr7-110117555-DEL-326 |
| chr7-110177770-DEL-59 | chr7-110177771-DEL-59 |
| chr7-110239434-DEL-53 | chr7-110239441-DEL-54 |
| chr7-110243345-DEL-875 | chr7-110243346-DEL-875 |
| chr7-110283734-DEL-75 | chr7-110283743-DEL-94 |
| chr7-110311421-DEL-120 | chr7-110311422-DEL-120 |
| chr7-110333338-DEL-110 | chr7-110333339-DEL-110 |
| chr7-110334526-DEL-54 | chr7-110334524-DEL-53 |
| chr7-110334681-DEL-465 | chr7-110334684-DEL-463 |
| chr7-110358261-DEL-2567 | chr7-110358262-DEL-2567 |
| chr7-110371359-DEL-56 | chr7-110371359-DEL-56 |
| chr7-110404244-DEL-124 | chr7-110404245-DEL-124 |
| chr7-110428403-DEL-1963 | chr7-110428404-DEL-1964 |
| chr7-110429040-DEL-138 | chr7-110429041-DEL-138 |
| chr7-110431107-DEL-566 | chr7-110431108-DEL-558 |
| chr7-110438164-DEL-51 | chr7-110438165-DEL-51 |
| chr7-110440501-DEL-511 | chr7-110440502-DEL-511 |
| chr7-110484615-DEL-88 | chr7-110484609-DEL-53 |
| chr7-110488832-DEL-933 | chr7-110488833-DEL-930 |
| chr7-110530518-DEL-1891 | chr7-110530519-DEL-1891 |
| chr7-110543871-DEL-340 | chr7-110543872-DEL-340 |
| chr7-110563220-DEL-65 | chr7-110563223-DEL-66 |
| chr7-110571099-DEL-172 | chr7-110571100-DEL-172 |
| chr7-110574521-DEL-1775 | chr7-110574522-DEL-1775 |
| chr7-110605230-DEL-3897 | chr7-110605231-DEL-3897 |
| chr7-110704259-DEL-177 | chr7-110704258-DEL-178 |
| chr7-110718911-DEL-175 | chr7-110718912-DEL-175 |
| chr7-110777597-DEL-51 | chr7-110777598-DEL-51 |
| chr7-110782362-DEL-217 | chr7-110782363-DEL-217 |
| chr7-110793211-DEL-106 | chr7-110793212-DEL-106 |
| chr7-110795192-DEL-1869 | chr7-110795199-DEL-1863 |
| chr7-110807533-DEL-486 | chr7-110807534-DEL-486 |
| chr7-110825091-DEL-68 | chr7-110825064-DEL-84 |
| chr7-110828072-DEL-87 | chr7-110828040-DEL-130 |
| chr7-110840308-DEL-148 | chr7-110840309-DEL-148 |
| chr7-110852979-DEL-831 | chr7-110852980-DEL-831 |
| chr7-110871094-DEL-128 | chr7-110871095-DEL-128 |
| chr7-110873857-DEL-316 | chr7-110873858-DEL-316 |
| chr7-110886306-DEL-637 | chr7-110886307-DEL-637 |
| chr7-110913308-DEL-88 | chr7-110913277-DEL-89 |
| chr7-110943887-DEL-152 | chr7-110943888-DEL-152 |
| chr7-110968871-DEL-52 | chr7-110968872-DEL-52 |
| chr7-110998250-DEL-335 | chr7-110998251-DEL-335 |
| chr7-111013715-DEL-54 | chr7-111013716-DEL-54 |
| chr7-111039880-DEL-613 | chr7-111039881-DEL-613 |
| chr7-111054958-DEL-269 | chr7-111054959-DEL-269 |
| chr7-111065067-DEL-60 | chr7-111065068-DEL-60 |
| chr7-111122246-DEL-1832 | chr7-111122247-DEL-1832 |
| chr7-111157839-DEL-637 | chr7-111157837-DEL-638 |
| chr7-111213879-DEL-121 | chr7-111213880-DEL-121 |
| chr7-111243882-DEL-2187 | chr7-111243883-DEL-2187 |
| chr7-111250434-DEL-121 | chr7-111250435-DEL-120 |
| chr7-111280436-DEL-63 | chr7-111280437-DEL-63 |
| chr7-111336509-DEL-58 | chr7-111336510-DEL-58 |
| chr7-111357791-DEL-84 | chr7-111357819-DEL-54 |
| chr7-111441499-DEL-923 | chr7-111441500-DEL-923 |
| chr7-111447721-DEL-332 | chr7-111447722-DEL-332 |
| chr7-111449745-DEL-216 | chr7-111449746-DEL-216 |
| chr7-111483948-DEL-95 | chr7-111483925-DEL-73 |
| chr7-111510345-DEL-6380 | chr7-111510346-DEL-6380 |
| chr7-111525329-DEL-72 | chr7-111525330-DEL-54 |
| chr7-111529551-DEL-63 | chr7-111529554-DEL-55 |
| chr7-111554972-DEL-131 | chr7-111554973-DEL-131 |
| chr7-111566998-DEL-146 | chr7-111566976-DEL-122 |
| chr7-111591571-DEL-3266 | chr7-111591572-DEL-3266 |
| chr7-111637462-DEL-93 | chr7-111637463-DEL-93 |
| chr7-111733368-DEL-751 | chr7-111733371-DEL-749 |
| chr7-111734046-DEL-75 | chr7-111734045-DEL-75 |
| chr7-111740829-DEL-185 | chr7-111740823-DEL-186 |
| chr7-111742954-DEL-66 | chr7-111742955-DEL-66 |
| chr7-111755953-DEL-121 | chr7-111755954-DEL-121 |
| chr7-111764660-DEL-198 | chr7-111764660-DEL-187 |
| chr7-111799028-DEL-83 | chr7-111799028-DEL-84 |
| chr7-111820707-DEL-775 | chr7-111820709-DEL-774 |
| chr7-111820730-DEL-70 | chr7-111820721-DEL-92 |
| chr7-111852343-DEL-133 | chr7-111852346-DEL-130 |
| chr7-111876495-DEL-78 | chr7-111876495-DEL-78 |
| chr7-111891624-DEL-251 | chr7-111891623-DEL-252 |
| chr7-111895148-DEL-1280 | chr7-111895149-DEL-1280 |
| chr7-111903150-DEL-218 | chr7-111903151-DEL-218 |

| | |
|---|---|
| chr7-111940994-DEL-57 | chr7-111940991-DEL-58 |
| chr7-111984655-DEL-107 | chr7-111984656-DEL-107 |
| chr7-112024972-DEL-515 | chr7-112024973-DEL-515 |
| chr7-112029405-DEL-4576 | chr7-112029406-DEL-4576 |
| chr7-112048367-DEL-167 | chr7-112048368-DEL-167 |
| chr7-112092052-DEL-201 | chr7-112092053-DEL-201 |
| chr7-112102222-DEL-101 | chr7-112102223-DEL-101 |
| chr7-112136422-DEL-3123 | chr7-112136423-DEL-3123 |
| chr7-112150190-DEL-452 | chr7-112150191-DEL-452 |
| chr7-112162083-DEL-54 | chr7-112162084-DEL-54 |
| chr7-112169575-DEL-52 | chr7-112169575-DEL-52 |
| chr7-112179607-DEL-91 | chr7-112179605-DEL-68 |
| chr7-112180597-DEL-116 | chr7-112180598-DEL-116 |
| chr7-112193667-DEL-144 | chr7-112193668-DEL-144 |
| chr7-112204441-DEL-132 | chr7-112204438-DEL-132 |
| chr7-112205147-DEL-6394 | chr7-112205148-DEL-6394 |
| chr7-112219762-DEL-75 | chr7-112219763-DEL-75 |
| chr7-112224561-DEL-58 | chr7-112224561-DEL-59 |
| chr7-112236315-DEL-2010 | chr7-112236318-DEL-1980 |
| chr7-112245581-DEL-55 | chr7-112245583-DEL-54 |
| chr7-112355243-DEL-84 | chr7-112355244-DEL-82 |
| chr7-112359141-DEL-274 | chr7-112359142-DEL-274 |
| chr7-112360468-DEL-94 | chr7-112360469-DEL-94 |
| chr7-112423441-DEL-233 | chr7-112423442-DEL-233 |
| chr7-112434232-DEL-82 | chr7-112434233-DEL-82 |
| chr7-112442123-DEL-222 | chr7-112442122-DEL-221 |
| chr7-112444444-DEL-408 | chr7-112444445-DEL-408 |
| chr7-112458598-DEL-202 | chr7-112458599-DEL-202 |
| chr7-112468513-DEL-55 | chr7-112468514-DEL-55 |
| chr7-112489139-DEL-210 | chr7-112489140-DEL-210 |
| chr7-112530679-DEL-68 | chr7-112530680-DEL-68 |
| chr7-112535704-DEL-56 | chr7-112535706-DEL-58 |
| chr7-112551043-DEL-1122 | chr7-112551044-DEL-1122 |
| chr7-112585904-DEL-742 | chr7-112585905-DEL-742 |
| chr7-112596263-DEL-203 | chr7-112596264-DEL-203 |
| chr7-112624448-DEL-77 | chr7-112624449-DEL-77 |
| chr7-112637860-DEL-70 | chr7-112637861-DEL-70 |
| chr7-112640792-DEL-101 | chr7-112640793-DEL-101 |
| chr7-112641489-DEL-731 | chr7-112641490-DEL-731 |
| chr7-112678990-DEL-235 | chr7-112678991-DEL-235 |
| chr7-112679388-DEL-342 | chr7-112679389-DEL-342 |
| chr7-112684139-DEL-52 | chr7-112684140-DEL-52 |
| chr7-112700926-DEL-96 | chr7-112700923-DEL-98 |
| chr7-112714327-DEL-54 | chr7-112714327-DEL-90 |
| chr7-112722203-DEL-77 | chr7-112722195-DEL-78 |
| chr7-112749093-DEL-322 | chr7-112749094-DEL-322 |
| chr7-112752981-DEL-71 | chr7-112752977-DEL-64 |
| chr7-112761853-DEL-213 | chr7-112761854-DEL-213 |
| chr7-112772230-DEL-194 | chr7-112772230-DEL-194 |
| chr7-112786685-DEL-54 | chr7-112786690-DEL-50 |
| chr7-112803477-DEL-196 | chr7-112803478-DEL-196 |
| chr7-112811741-DEL-564 | chr7-112811742-DEL-564 |
| chr7-112836397-DEL-78 | chr7-112836398-DEL-78 |
| chr7-112858222-DEL-51 | chr7-112858223-DEL-51 |
| chr7-112866154-DEL-3977 | chr7-112866155-DEL-3977 |
| chr7-112871025-DEL-197 | chr7-112871026-DEL-197 |
| chr7-112878078-DEL-136 | chr7-112878042-DEL-127 |
| chr7-112881133-DEL-191 | chr7-112881134-DEL-191 |
| chr7-112891770-DEL-361 | chr7-112891770-DEL-360 |
| chr7-112897428-DEL-308 | chr7-112897429-DEL-308 |
| chr7-112899989-DEL-199 | chr7-112899990-DEL-199 |
| chr7-112914585-DEL-182 | chr7-112914586-DEL-182 |
| chr7-112956486-DEL-105 | chr7-112956479-DEL-131 |
| chr7-112980704-DEL-1584 | chr7-112980705-DEL-1583 |
| chr7-113001799-DEL-76 | chr7-113001800-DEL-76 |
| chr7-113023289-DEL-6808 | chr7-113023290-DEL-6808 |
| chr7-113083290-DEL-76 | chr7-113083283-DEL-76 |
| chr7-113083802-DEL-6887 | chr7-113083798-DEL-6892 |
| chr7-113089810-DEL-424 | chr7-113089811-DEL-424 |
| chr7-113132369-DEL-156 | chr7-113132370-DEL-117 |
| chr7-113172831-DEL-765 | chr7-113172832-DEL-765 |
| chr7-113197234-DEL-92 | chr7-113197235-DEL-92 |
| chr7-113228910-DEL-100 | chr7-113228910-DEL-100 |
| chr7-113249956-DEL-415 | chr7-113249957-DEL-415 |
| chr7-113257008-DEL-163 | chr7-113257009-DEL-163 |
| chr7-113302817-DEL-170 | chr7-113302822-DEL-158 |
| chr7-113309511-DEL-79 | chr7-113309531-DEL-68 |
| chr7-113318686-DEL-56 | chr7-113318674-DEL-66 |
| chr7-113336639-DEL-52 | chr7-113336640-DEL-52 |
| chr7-113437889-DEL-126 | chr7-113437890-DEL-126 |

| | |
|---|---|
| chr7-113467211-DEL-56 | chr7-113467214-DEL-54 |
| chr7-113471774-DEL-98 | chr7-113471775-DEL-98 |
| chr7-113474840-DEL-62 | chr7-113474822-DEL-68 |
| chr7-113494033-DEL-91 | chr7-113494034-DEL-91 |
| chr7-113496046-DEL-426 | chr7-113496047-DEL-426 |
| chr7-113501960-DEL-54 | chr7-113501957-DEL-58 |
| chr7-113517850-DEL-102 | chr7-113517851-DEL-100 |
| chr7-113541516-DEL-4335 | chr7-113541511-DEL-4336 |
| chr7-113598539-DEL-1351 | chr7-113598540-DEL-1351 |
| chr7-113619154-DEL-63 | chr7-113619144-DEL-60 |
| chr7-113682201-DEL-349 | chr7-113682202-DEL-349 |
| chr7-113719188-DEL-6249 | chr7-113719189-DEL-6249 |
| chr7-113725773-DEL-163 | chr7-113725774-DEL-163 |
| chr7-113733548-DEL-68 | chr7-113733549-DEL-68 |
| chr7-113746325-DEL-97 | chr7-113746326-DEL-97 |
| chr7-113779333-DEL-356 | chr7-113779334-DEL-356 |
| chr7-113785469-DEL-200 | chr7-113785470-DEL-200 |
| chr7-113800666-DEL-200 | chr7-113800667-DEL-200 |
| chr7-113827692-DEL-83 | chr7-113827683-DEL-93 |
| chr7-113837032-DEL-111 | chr7-113837033-DEL-111 |
| chr7-113855289-DEL-53 | chr7-113855290-DEL-53 |
| chr7-113875722-DEL-76 | chr7-113875725-DEL-60 |
| chr7-113887632-DEL-55 | chr7-113887672-DEL-52 |
| chr7-113902475-DEL-212 | chr7-113902476-DEL-212 |
| chr7-113903381-DEL-1118 | chr7-113903382-DEL-1118 |
| chr7-113928716-DEL-165 | chr7-113928717-DEL-165 |
| chr7-113978890-DEL-54 | chr7-113978891-DEL-54 |
| chr7-113986421-DEL-679 | chr7-113986422-DEL-679 |
| chr7-114003718-DEL-7135 | chr7-114003719-DEL-7135 |
| chr7-114040580-DEL-499 | chr7-114040584-DEL-496 |
| chr7-114042314-DEL-75 | chr7-114042311-DEL-75 |
| chr7-114052208-DEL-54 | chr7-114052212-DEL-54 |
| chr7-114056725-DEL-239 | chr7-114056728-DEL-238 |
| chr7-114067017-DEL-70 | chr7-114067018-DEL-70 |
| chr7-114091469-DEL-88 | chr7-114091470-DEL-88 |
| chr7-114164790-DEL-1850 | chr7-114164791-DEL-1850 |
| chr7-114168208-DEL-452 | chr7-114168209-DEL-452 |
| chr7-114193285-DEL-241 | chr7-114193286-DEL-241 |
| chr7-114194517-DEL-1964 | chr7-114194518-DEL-1964 |
| chr7-114261072-DEL-374 | chr7-114261050-DEL-382 |
| chr7-114262163-DEL-600 | chr7-114262164-DEL-600 |
| chr7-114282331-DEL-894 | chr7-114282332-DEL-895 |
| chr7-114289155-DEL-124 | chr7-114289156-DEL-124 |
| chr7-114304423-DEL-1054 | chr7-114304424-DEL-1054 |
| chr7-114434550-DEL-223 | chr7-114434551-DEL-223 |
| chr7-114475379-DEL-68 | chr7-114475380-DEL-68 |
| chr7-114515224-DEL-106 | chr7-114515261-DEL-63 |
| chr7-114539924-DEL-1048 | chr7-114539925-DEL-1048 |
| chr7-114546777-DEL-57 | chr7-114546779-DEL-51 |
| chr7-114548661-DEL-471 | chr7-114548662-DEL-471 |
| chr7-114557756-DEL-82 | chr7-114557738-DEL-50 |
| chr7-114576162-DEL-75 | chr7-114576154-DEL-60 |
| chr7-114583295-DEL-65 | chr7-114583277-DEL-71 |
| chr7-114592529-DEL-108 | chr7-114592530-DEL-108 |
| chr7-114596201-DEL-86 | chr7-114596205-DEL-80 |
| chr7-114602076-DEL-478 | chr7-114602077-DEL-478 |
| chr7-114611955-DEL-143 | chr7-114611956-DEL-143 |
| chr7-114615167-DEL-1223 | chr7-114615168-DEL-1223 |
| chr7-114620777-DEL-81 | chr7-114620778-DEL-81 |
| chr7-114660123-DEL-408 | chr7-114660124-DEL-408 |
| chr7-114664973-DEL-90 | chr7-114664974-DEL-90 |
| chr7-114670592-DEL-149 | chr7-114670593-DEL-149 |
| chr7-114676536-DEL-6610 | chr7-114676537-DEL-6610 |
| chr7-114711487-DEL-88 | chr7-114711439-DEL-125 |
| chr7-114714300-DEL-110 | chr7-114714301-DEL-110 |
| chr7-114724993-DEL-130 | chr7-114724994-DEL-130 |
| chr7-114749154-DEL-400 | chr7-114749155-DEL-400 |
| chr7-114762364-DEL-111 | chr7-114762365-DEL-111 |
| chr7-114821960-DEL-77 | chr7-114821961-DEL-77 |
| chr7-114853836-DEL-273 | chr7-114853834-DEL-274 |
| chr7-114881721-DEL-135 | chr7-114881714-DEL-142 |
| chr7-114882916-DEL-55 | chr7-114882917-DEL-60 |
| chr7-114933737-DEL-55 | chr7-114933733-DEL-61 |
| chr7-114936204-DEL-131 | chr7-114936205-DEL-132 |
| chr7-114961250-DEL-54 | chr7-114961251-DEL-54 |
| chr7-114976751-DEL-82 | chr7-114976752-DEL-82 |
| chr7-115057433-DEL-64 | chr7-115057450-DEL-76 |
| chr7-115060267-DEL-64 | chr7-115060268-DEL-64 |
| chr7-115064430-DEL-170 | chr7-115064431-DEL-170 |
| chr7-115084072-DEL-752 | chr7-115084073-DEL-752 |

| | |
|---|---|
| chr7-115119232-DEL-1026 | chr7-115119233-DEL-1026 |
| chr7-115131600-DEL-143 | chr7-115131601-DEL-143 |
| chr7-115169302-DEL-53 | chr7-115169303-DEL-89 |
| chr7-115188120-DEL-6908 | chr7-115188121-DEL-6908 |
| chr7-115188175-DEL-206 | chr7-115188176-DEL-206 |
| chr7-115199936-DEL-1640 | chr7-115199937-DEL-1640 |
| chr7-115243017-DEL-179 | chr7-115243018-DEL-179 |
| chr7-115269957-DEL-88 | chr7-115269958-DEL-88 |
| chr7-115282171-DEL-73 | chr7-115282179-DEL-66 |
| chr7-115327775-DEL-150 | chr7-115327772-DEL-151 |
| chr7-115367711-DEL-6279 | chr7-115367712-DEL-6279 |
| chr7-115409545-DEL-195 | chr7-115409546-DEL-195 |
| chr7-115410868-DEL-7322 | chr7-115410869-DEL-7322 |
| chr7-115422009-DEL-296 | chr7-115422010-DEL-296 |
| chr7-115425977-DEL-81 | chr7-115425944-DEL-83 |
| chr7-115437640-DEL-282 | chr7-115437641-DEL-270 |
| chr7-115508401-DEL-97 | chr7-115508414-DEL-85 |
| chr7-115526761-DEL-80 | chr7-115526762-DEL-80 |
| chr7-115533981-DEL-306 | chr7-115533982-DEL-306 |
| chr7-115548677-DEL-52 | chr7-115548678-DEL-52 |
| chr7-115578203-DEL-79 | chr7-115578210-DEL-80 |
| chr7-115593176-DEL-53 | chr7-115593177-DEL-53 |
| chr7-115594948-DEL-184 | chr7-115594949-DEL-184 |
| chr7-115665496-DEL-207 | chr7-115665497-DEL-207 |
| chr7-115681831-DEL-2027 | chr7-115681832-DEL-2027 |
| chr7-115685272-DEL-204 | chr7-115685273-DEL-204 |
| chr7-115706520-DEL-567 | chr7-115706521-DEL-567 |
| chr7-115713214-DEL-204 | chr7-115713215-DEL-204 |
| chr7-115738314-DEL-56 | chr7-115738315-DEL-56 |
| chr7-115773284-DEL-123 | chr7-115773285-DEL-123 |
| chr7-115804116-DEL-1008 | chr7-115804117-DEL-1008 |
| chr7-115817869-DEL-230 | chr7-115817870-DEL-230 |
| chr7-115828806-DEL-203 | chr7-115828806-DEL-203 |
| chr7-115864034-DEL-157 | chr7-115864035-DEL-157 |
| chr7-115876007-DEL-526 | chr7-115876008-DEL-526 |
| chr7-115892992-DEL-115 | chr7-115892993-DEL-115 |
| chr7-115902492-DEL-181 | chr7-115902493-DEL-181 |
| chr7-115903848-DEL-221 | chr7-115903849-DEL-221 |
| chr7-115905083-DEL-421 | chr7-115905074-DEL-415 |
| chr7-115909178-DEL-77 | chr7-115909176-DEL-76 |
| chr7-115911390-DEL-1561 | chr7-115911391-DEL-1561 |
| chr7-115919290-DEL-156 | chr7-115919252-DEL-181 |
| chr7-115919379-DEL-670 | chr7-115919379-DEL-670 |
| chr7-115926063-DEL-3622 | chr7-115926064-DEL-3622 |
| chr7-115926956-DEL-209 | chr7-115926957-DEL-207 |
| chr7-115928532-DEL-211 | chr7-115928533-DEL-211 |
| chr7-115930229-DEL-134 | chr7-115930230-DEL-133 |
| chr7-115945548-DEL-177 | chr7-115945549-DEL-177 |
| chr7-115950664-DEL-74 | chr7-115950665-DEL-74 |
| chr7-115962272-DEL-211 | chr7-115962273-DEL-211 |
| chr7-115996341-DEL-1469 | chr7-115996342-DEL-1469 |
| chr7-115999862-DEL-966 | chr7-115999863-DEL-966 |
| chr7-116021843-DEL-208 | chr7-116021844-DEL-208 |
| chr7-116028078-DEL-423 | chr7-116028079-DEL-423 |
| chr7-116030746-DEL-79 | chr7-116030744-DEL-80 |
| chr7-116031339-DEL-5678 | chr7-116031340-DEL-5678 |
| chr7-116080300-DEL-63 | chr7-116080289-DEL-63 |
| chr7-116082659-DEL-82 | chr7-116082657-DEL-79 |
| chr7-116088055-DEL-105 | chr7-116088055-DEL-105 |
| chr7-116107392-DEL-59 | chr7-116107382-DEL-58 |
| chr7-116119065-DEL-465 | chr7-116119066-DEL-465 |
| chr7-116121320-DEL-67 | chr7-116121321-DEL-67 |
| chr7-116122245-DEL-162 | chr7-116122259-DEL-142 |
| chr7-116136200-DEL-1054 | chr7-116136201-DEL-1054 |
| chr7-116141852-DEL-217 | chr7-116141853-DEL-217 |
| chr7-116162980-DEL-372 | chr7-116162981-DEL-364 |
| chr7-116183937-DEL-212 | chr7-116183938-DEL-212 |
| chr7-116202456-DEL-63 | chr7-116202457-DEL-63 |
| chr7-116238808-DEL-257 | chr7-116238809-DEL-257 |
| chr7-116263823-DEL-191 | chr7-116263820-DEL-193 |
| chr7-116311452-DEL-113 | chr7-116311453-DEL-113 |
| chr7-116315807-DEL-2407 | chr7-116315808-DEL-2410 |
| chr7-116319513-DEL-136 | chr7-116319514-DEL-136 |
| chr7-116328692-DEL-60 | chr7-116328691-DEL-62 |
| chr7-116340063-DEL-62 | chr7-116340064-DEL-58 |
| chr7-116356343-DEL-6423 | chr7-116356344-DEL-6423 |
| chr7-116380113-DEL-111 | chr7-116380134-DEL-90 |
| chr7-116385285-DEL-6713 | chr7-116385290-DEL-6709 |
| chr7-116386944-DEL-57 | chr7-116386945-DEL-57 |
| chr7-116398733-DEL-3552 | chr7-116398732-DEL-3556 |

| | |
|---|---|
| chr7-116401741-DEL-82 | chr7-116401741-DEL-82 |
| chr7-116404395-DEL-62 | chr7-116404396-DEL-62 |
| chr7-116408655-DEL-57 | chr7-116408644-DEL-52 |
| chr7-116418196-DEL-70 | chr7-116418197-DEL-70 |
| chr7-116422783-DEL-93 | chr7-116422784-DEL-93 |
| chr7-116423628-DEL-1459 | chr7-116423629-DEL-1459 |
| chr7-116435296-DEL-52 | chr7-116435297-DEL-52 |
| chr7-116436751-DEL-121 | chr7-116436751-DEL-114 |
| chr7-116441681-DEL-69 | chr7-116441676-DEL-65 |
| chr7-116505726-DEL-194 | chr7-116505727-DEL-194 |
| chr7-116507123-DEL-317 | chr7-116507124-DEL-316 |
| chr7-116525938-DEL-203 | chr7-116525939-DEL-203 |
| chr7-116549465-DEL-503 | chr7-116549466-DEL-503 |
| chr7-116559809-DEL-54 | chr7-116559810-DEL-54 |
| chr7-116602214-DEL-116 | chr7-116602214-DEL-116 |
| chr7-116609668-DEL-6228 | chr7-116609669-DEL-6228 |
| chr7-116615629-DEL-101 | chr7-116615630-DEL-101 |
| chr7-116660876-DEL-59 | chr7-116660877-DEL-59 |
| chr7-116675723-DEL-143 | chr7-116675722-DEL-148 |
| chr7-116698065-DEL-58 | chr7-116698066-DEL-58 |
| chr7-116723484-DEL-1056 | chr7-116723485-DEL-1056 |
| chr7-116724717-DEL-136 | chr7-116724710-DEL-144 |
| chr7-116732908-DEL-56 | chr7-116732909-DEL-56 |
| chr7-116737676-DEL-60 | chr7-116737677-DEL-60 |
| chr7-116739529-DEL-82 | chr7-116739528-DEL-84 |
| chr7-116748726-DEL-56 | chr7-116748727-DEL-56 |
| chr7-116750771-DEL-230 | chr7-116750776-DEL-229 |
| chr7-116755573-DEL-78 | chr7-116755574-DEL-78 |
| chr7-116773202-DEL-74 | chr7-116773203-DEL-74 |
| chr7-116806510-DEL-211 | chr7-116806473-DEL-210 |
| chr7-116900003-DEL-656 | chr7-116900004-DEL-656 |
| chr7-116910423-DEL-55 | chr7-116910419-DEL-51 |
| chr7-116924738-DEL-63 | chr7-116924745-DEL-56 |
| chr7-116931448-DEL-268 | chr7-116931449-DEL-268 |
| chr7-116935972-DEL-4326 | chr7-116935973-DEL-4326 |
| chr7-116954156-DEL-56 | chr7-116954159-DEL-54 |
| chr7-117015926-DEL-1184 | chr7-117015927-DEL-1184 |
| chr7-117033545-DEL-96 | chr7-117033558-DEL-93 |
| chr7-117049980-DEL-77 | chr7-117049981-DEL-77 |
| chr7-117175087-DEL-189 | chr7-117175088-DEL-189 |
| chr7-117191425-DEL-72 | chr7-117191426-DEL-72 |
| chr7-117192234-DEL-114 | chr7-117192235-DEL-114 |
| chr7-117214555-DEL-56 | chr7-117214556-DEL-57 |
| chr7-117221488-DEL-82 | chr7-117221503-DEL-68 |
| chr7-117225233-DEL-100 | chr7-117225242-DEL-96 |
| chr7-117232078-DEL-972 | chr7-117232079-DEL-972 |
| chr7-117240644-DEL-89 | chr7-117240645-DEL-89 |
| chr7-117252840-DEL-57 | chr7-117252837-DEL-60 |
| chr7-117260245-DEL-5233 | chr7-117260246-DEL-5233 |
| chr7-117276146-DEL-94 | chr7-117276162-DEL-91 |
| chr7-117284870-DEL-111 | chr7-117284901-DEL-89 |
| chr7-117302927-DEL-53 | chr7-117302926-DEL-52 |
| chr7-117318835-DEL-136 | chr7-117318836-DEL-136 |
| chr7-117340287-DEL-2406 | chr7-117340288-DEL-2406 |
| chr7-117363391-DEL-53 | chr7-117363392-DEL-53 |
| chr7-117430996-DEL-178 | chr7-117430997-DEL-178 |
| chr7-117484109-DEL-190 | chr7-117484110-DEL-190 |
| chr7-117564806-DEL-476 | chr7-117564807-DEL-476 |
| chr7-117573858-DEL-98 | chr7-117573859-DEL-98 |
| chr7-117577897-DEL-273 | chr7-117577898-DEL-268 |
| chr7-117610958-DEL-77 | chr7-117610957-DEL-78 |
| chr7-117664595-DEL-80 | chr7-117664596-DEL-80 |
| chr7-117665795-DEL-102 | chr7-117665796-DEL-102 |
| chr7-117668428-DEL-6123 | chr7-117668429-DEL-6123 |
| chr7-117680858-DEL-57 | chr7-117680874-DEL-71 |
| chr7-117682999-DEL-77 | chr7-117683000-DEL-77 |
| chr7-117683207-DEL-1138 | chr7-117683208-DEL-1138 |
| chr7-117685635-DEL-64 | chr7-117685636-DEL-64 |
| chr7-117709448-DEL-63 | chr7-117709449-DEL-63 |
| chr7-117716099-DEL-109 | chr7-117716100-DEL-109 |
| chr7-117724100-DEL-837 | chr7-117724101-DEL-837 |
| chr7-117763859-DEL-74 | chr7-117763852-DEL-82 |
| chr7-117781217-DEL-206 | chr7-117781218-DEL-206 |
| chr7-117787068-DEL-160 | chr7-117787043-DEL-136 |
| chr7-117787720-DEL-63 | chr7-117787722-DEL-67 |
| chr7-117804262-DEL-64 | chr7-117804263-DEL-64 |
| chr7-117808162-DEL-1295 | chr7-117808163-DEL-1295 |
| chr7-117839975-DEL-209 | chr7-117839976-DEL-209 |
| chr7-117856722-DEL-99 | chr7-117856723-DEL-99 |
| chr7-117867675-DEL-5106 | chr7-117867676-DEL-5106 |

| | |
|---|---|
| chr7-117885037-DEL-7084 | chr7-117885038-DEL-7084 |
| chr7-117899524-DEL-52 | chr7-117899524-DEL-52 |
| chr7-117902279-DEL-57 | chr7-117902280-DEL-57 |
| chr7-117906037-DEL-970 | chr7-117906038-DEL-970 |
| chr7-117925350-DEL-161 | chr7-117925333-DEL-147 |
| chr7-117939137-DEL-496 | chr7-117939138-DEL-496 |
| chr7-117956557-DEL-1840 | chr7-117956558-DEL-1840 |
| chr7-117994748-DEL-377 | chr7-117994749-DEL-387 |
| chr7-117998602-DEL-165 | chr7-117998603-DEL-161 |
| chr7-118002481-DEL-308 | chr7-118002477-DEL-308 |
| chr7-118032092-DEL-934 | chr7-118032093-DEL-934 |
| chr7-118034091-DEL-480 | chr7-118034092-DEL-480 |
| chr7-118055875-DEL-120 | chr7-118055876-DEL-120 |
| chr7-118068865-DEL-136 | chr7-118068856-DEL-137 |
| chr7-118070212-DEL-81 | chr7-118070213-DEL-81 |
| chr7-118073638-DEL-347 | chr7-118073636-DEL-342 |
| chr7-118116269-DEL-74 | chr7-118116238-DEL-84 |
| chr7-118117181-DEL-71 | chr7-118117182-DEL-50 |
| chr7-118175217-DEL-329 | chr7-118175214-DEL-326 |
| chr7-118188488-DEL-150 | chr7-118188497-DEL-143 |
| chr7-118189908-DEL-56 | chr7-118189901-DEL-57 |
| chr7-118191795-DEL-227 | chr7-118191796-DEL-227 |
| chr7-118215716-DEL-120 | chr7-118215717-DEL-120 |
| chr7-118216356-DEL-66 | chr7-118216357-DEL-68 |
| chr7-118221482-DEL-511 | chr7-118221483-DEL-509 |
| chr7-118237175-DEL-134 | chr7-118237176-DEL-134 |
| chr7-118242794-DEL-124 | chr7-118242827-DEL-134 |
| chr7-118273718-DEL-56 | chr7-118273719-DEL-56 |
| chr7-118277222-DEL-610 | chr7-118277223-DEL-610 |
| chr7-118283738-DEL-233 | chr7-118283739-DEL-233 |
| chr7-118285969-DEL-1624 | chr7-118285970-DEL-1624 |
| chr7-118286366-DEL-172 | chr7-118286367-DEL-172 |
| chr7-118343501-DEL-1559 | chr7-118343502-DEL-1559 |
| chr7-118343922-DEL-178 | chr7-118343921-DEL-177 |
| chr7-118349367-DEL-200 | chr7-118349368-DEL-200 |
| chr7-118353400-DEL-285 | chr7-118353401-DEL-285 |
| chr7-118364677-DEL-150 | chr7-118364678-DEL-150 |
| chr7-118377112-DEL-52 | chr7-118377113-DEL-52 |
| chr7-118382689-DEL-54 | chr7-118382688-DEL-51 |
| chr7-118394884-DEL-192 | chr7-118394885-DEL-192 |
| chr7-118402368-DEL-126 | chr7-118402369-DEL-127 |
| chr7-118404744-DEL-754 | chr7-118404745-DEL-754 |
| chr7-118412528-DEL-68 | chr7-118412530-DEL-66 |
| chr7-118418730-DEL-151 | chr7-118418731-DEL-151 |
| chr7-118440303-DEL-54 | chr7-118440304-DEL-54 |
| chr7-118451433-DEL-62 | chr7-118451434-DEL-62 |
| chr7-118465801-DEL-1904 | chr7-118465802-DEL-1904 |
| chr7-118481634-DEL-62 | chr7-118481611-DEL-60 |
| chr7-118485171-DEL-890 | chr7-118485172-DEL-890 |
| chr7-118493109-DEL-66 | chr7-118493110-DEL-64 |
| chr7-118504559-DEL-215 | chr7-118504560-DEL-215 |
| chr7-118512926-DEL-115 | chr7-118512962-DEL-80 |
| chr7-118515261-DEL-190 | chr7-118515285-DEL-189 |
| chr7-118527089-DEL-76 | chr7-118527066-DEL-76 |
| chr7-118535564-DEL-80 | chr7-118535565-DEL-80 |
| chr7-118537888-DEL-193 | chr7-118537889-DEL-193 |
| chr7-118547174-DEL-62 | chr7-118547175-DEL-62 |
| chr7-118550508-DEL-95 | chr7-118550496-DEL-97 |
| chr7-118556730-DEL-141 | chr7-118556731-DEL-141 |
| chr7-118559987-DEL-51 | chr7-118559988-DEL-51 |
| chr7-118598274-DEL-351 | chr7-118598275-DEL-351 |
| chr7-118639076-DEL-88 | chr7-118639077-DEL-88 |
| chr7-118648958-DEL-6782 | chr7-118648959-DEL-6782 |
| chr7-118663562-DEL-107 | chr7-118663563-DEL-107 |
| chr7-118699262-DEL-85 | chr7-118699263-DEL-85 |
| chr7-118711350-DEL-4369 | chr7-118711351-DEL-4369 |
| chr7-118797818-DEL-105 | chr7-118797829-DEL-80 |
| chr7-118798599-DEL-54 | chr7-118798600-DEL-54 |
| chr7-118813797-DEL-70 | chr7-118813802-DEL-66 |
| chr7-118822669-DEL-68 | chr7-118822670-DEL-68 |
| chr7-118834714-DEL-125 | chr7-118834715-DEL-125 |
| chr7-118859703-DEL-54 | chr7-118859704-DEL-54 |
| chr7-118887632-DEL-5207 | chr7-118887633-DEL-5207 |
| chr7-118906126-DEL-206 | chr7-118906127-DEL-206 |
| chr7-118917053-DEL-8238 | chr7-118917049-DEL-8238 |
| chr7-118934483-DEL-1288 | chr7-118934484-DEL-1288 |
| chr7-118966991-DEL-90 | chr7-118966992-DEL-90 |
| chr7-118986202-DEL-314 | chr7-118986203-DEL-314 |
| chr7-118997154-DEL-498 | chr7-118997155-DEL-498 |
| chr7-119001494-DEL-1011 | chr7-119001495-DEL-1011 |

| | |
|---|---|
| chr7-119011947-DEL-5195 | chr7-119011948-DEL-5195 |
| chr7-119029474-DEL-498 | chr7-119029475-DEL-498 |
| chr7-119032011-DEL-90 | chr7-119032000-DEL-94 |
| chr7-119033098-DEL-6250 | chr7-119033099-DEL-6250 |
| chr7-119047928-DEL-222 | chr7-119047929-DEL-221 |
| chr7-119079765-DEL-51 | chr7-119079766-DEL-51 |
| chr7-119090427-DEL-62 | chr7-119090440-DEL-50 |
| chr7-119094981-DEL-67 | chr7-119094984-DEL-51 |
| chr7-119104025-DEL-6273 | chr7-119104026-DEL-6273 |
| chr7-119130757-DEL-121 | chr7-119130804-DEL-72 |
| chr7-119132905-DEL-401 | chr7-119132906-DEL-401 |
| chr7-119198338-DEL-223 | chr7-119198339-DEL-223 |
| chr7-119213847-DEL-66 | chr7-119213839-DEL-66 |
| chr7-119234768-DEL-56 | chr7-119234769-DEL-56 |
| chr7-119259447-DEL-109 | chr7-119259448-DEL-109 |
| chr7-119265118-DEL-344 | chr7-119265114-DEL-342 |
| chr7-119294621-DEL-64 | chr7-119294630-DEL-56 |
| chr7-119314573-DEL-349 | chr7-119314574-DEL-349 |
| chr7-119320183-DEL-2214 | chr7-119320184-DEL-2214 |
| chr7-119320864-DEL-4591 | chr7-119320865-DEL-4591 |
| chr7-119323936-DEL-765 | chr7-119323953-DEL-770 |
| chr7-119325750-DEL-177 | chr7-119325751-DEL-177 |
| chr7-119339117-DEL-2345 | chr7-119339118-DEL-2345 |
| chr7-119365105-DEL-393 | chr7-119365106-DEL-393 |
| chr7-119378389-DEL-83 | chr7-119378390-DEL-83 |
| chr7-119387225-DEL-174 | chr7-119387226-DEL-174 |
| chr7-119424219-DEL-211 | chr7-119424220-DEL-211 |
| chr7-119436222-DEL-401 | chr7-119436223-DEL-401 |
| chr7-119523574-DEL-68 | chr7-119523575-DEL-73 |
| chr7-119526663-DEL-73 | chr7-119526664-DEL-73 |
| chr7-119562574-DEL-289 | chr7-119562575-DEL-289 |
| chr7-119563497-DEL-109 | chr7-119563498-DEL-109 |
| chr7-119572691-DEL-78 | chr7-119572696-DEL-78 |
| chr7-119578958-DEL-105 | chr7-119578959-DEL-105 |
| chr7-119584119-DEL-1917 | chr7-119584120-DEL-1917 |
| chr7-119593099-DEL-120 | chr7-119593110-DEL-120 |
| chr7-119596729-DEL-427 | chr7-119596722-DEL-427 |
| chr7-119597987-DEL-6319 | chr7-119597991-DEL-6321 |
| chr7-119604049-DEL-188 | chr7-119604050-DEL-188 |
| chr7-119608080-DEL-62 | chr7-119608088-DEL-54 |
| chr7-119611407-DEL-3590 | chr7-119611408-DEL-3590 |
| chr7-119624182-DEL-78 | chr7-119624185-DEL-75 |
| chr7-119645206-DEL-70 | chr7-119645209-DEL-72 |
| chr7-119664810-DEL-172 | chr7-119664811-DEL-172 |
| chr7-119712962-DEL-694 | chr7-119712963-DEL-694 |
| chr7-119742945-DEL-67 | chr7-119742943-DEL-67 |
| chr7-119787593-DEL-51 | chr7-119787594-DEL-51 |
| chr7-119808056-DEL-93 | chr7-119808057-DEL-93 |
| chr7-119811053-DEL-57 | chr7-119811054-DEL-57 |
| chr7-119813315-DEL-83 | chr7-119813305-DEL-84 |
| chr7-119821687-DEL-810 | chr7-119821688-DEL-810 |
| chr7-119832692-DEL-733 | chr7-119832691-DEL-735 |
| chr7-119834980-DEL-839 | chr7-119834980-DEL-839 |
| chr7-119847889-DEL-578 | chr7-119847890-DEL-578 |
| chr7-119854941-DEL-100 | chr7-119854936-DEL-106 |
| chr7-119857408-DEL-2049 | chr7-119857409-DEL-2049 |
| chr7-119876287-DEL-56 | chr7-119876288-DEL-56 |
| chr7-119892192-DEL-52 | chr7-119892192-DEL-59 |
| chr7-119893606-DEL-6485 | chr7-119893607-DEL-6485 |
| chr7-119899778-DEL-214 | chr7-119899779-DEL-214 |
| chr7-119925150-DEL-1455 | chr7-119925151-DEL-1455 |
| chr7-119940930-DEL-94 | chr7-119940931-DEL-94 |
| chr7-119969530-DEL-2868 | chr7-119969542-DEL-2872 |
| chr7-119979834-DEL-981 | chr7-119979835-DEL-981 |
| chr7-120011346-DEL-54 | chr7-120011347-DEL-54 |
| chr7-120040779-DEL-1334 | chr7-120040780-DEL-1334 |
| chr7-120047618-DEL-199 | chr7-120047619-DEL-199 |
| chr7-120059598-DEL-2095 | chr7-120059614-DEL-2083 |
| chr7-120105225-DEL-7070 | chr7-120105226-DEL-7070 |
| chr7-120134702-DEL-2030 | chr7-120134703-DEL-2030 |
| chr7-120156270-DEL-3206 | chr7-120156271-DEL-3206 |
| chr7-120156483-DEL-66 | chr7-120156484-DEL-66 |
| chr7-120161481-DEL-1133 | chr7-120161482-DEL-1133 |
| chr7-120185239-DEL-1692 | chr7-120185240-DEL-1692 |
| chr7-120187631-DEL-314 | chr7-120187632-DEL-314 |
| chr7-120191778-DEL-64 | chr7-120191779-DEL-64 |
| chr7-120192562-DEL-398 | chr7-120192563-DEL-398 |
| chr7-120193980-DEL-51 | chr7-120193981-DEL-51 |
| chr7-120194158-DEL-608 | chr7-120194159-DEL-608 |
| chr7-120194845-DEL-187 | chr7-120194846-DEL-187 |

| | |
|---|---|
| chr7-120199261-DEL-515 | chr7-120199262-DEL-515 |
| chr7-120204269-DEL-1307 | chr7-120204270-DEL-1307 |
| chr7-120211988-DEL-137 | chr7-120211989-DEL-137 |
| chr7-120265490-DEL-54 | chr7-120265491-DEL-54 |
| chr7-120267214-DEL-85 | chr7-120267213-DEL-86 |
| chr7-120270984-DEL-208 | chr7-120270985-DEL-208 |
| chr7-120280288-DEL-552 | chr7-120280290-DEL-550 |
| chr7-120354204-DEL-204 | chr7-120354211-DEL-178 |
| chr7-120364899-DEL-376 | chr7-120364900-DEL-376 |
| chr7-120367530-DEL-57 | chr7-120367531-DEL-59 |
| chr7-120369207-DEL-54 | chr7-120369208-DEL-54 |
| chr7-120380522-DEL-187 | chr7-120380523-DEL-187 |
| chr7-120402070-DEL-298 | chr7-120402071-DEL-298 |
| chr7-120417151-DEL-57 | chr7-120417152-DEL-57 |
| chr7-120420246-DEL-107 | chr7-120420246-DEL-107 |
| chr7-120453024-DEL-110 | chr7-120453025-DEL-110 |
| chr7-120467941-DEL-71 | chr7-120467942-DEL-71 |
| chr7-120473753-DEL-207 | chr7-120473743-DEL-208 |
| chr7-120493032-DEL-52 | chr7-120493033-DEL-52 |
| chr7-120500474-DEL-60 | chr7-120500475-DEL-60 |
| chr7-120509745-DEL-182 | chr7-120509746-DEL-182 |
| chr7-120511796-DEL-51 | chr7-120511797-DEL-51 |
| chr7-120560156-DEL-79 | chr7-120560157-DEL-78 |
| chr7-120600252-DEL-188 | chr7-120600253-DEL-188 |
| chr7-120600971-DEL-86 | chr7-120600980-DEL-50 |
| chr7-120605987-DEL-892 | chr7-120605988-DEL-892 |
| chr7-120620468-DEL-1379 | chr7-120620472-DEL-1381 |
| chr7-120679158-DEL-191 | chr7-120679159-DEL-191 |
| chr7-120727510-DEL-76 | chr7-120727511-DEL-76 |
| chr7-120765846-DEL-60 | chr7-120765847-DEL-60 |
| chr7-120784600-DEL-645 | chr7-120784601-DEL-645 |
| chr7-120803871-DEL-353 | chr7-120803872-DEL-353 |
| chr7-120896711-DEL-70 | chr7-120896665-DEL-71 |
| chr7-120909916-DEL-490 | chr7-120909916-DEL-491 |
| chr7-120911195-DEL-276 | chr7-120911196-DEL-276 |
| chr7-120911700-DEL-483 | chr7-120911701-DEL-483 |
| chr7-120915439-DEL-143 | chr7-120915440-DEL-117 |
| chr7-120944690-DEL-55 | chr7-120944691-DEL-55 |
| chr7-120952582-DEL-211 | chr7-120952583-DEL-211 |
| chr7-121021068-DEL-6268 | chr7-121021069-DEL-6268 |
| chr7-121036088-DEL-67 | chr7-121036089-DEL-67 |
| chr7-121045282-DEL-168 | chr7-121045283-DEL-168 |
| chr7-121073534-DEL-80 | chr7-121073538-DEL-78 |
| chr7-121104686-DEL-324 | chr7-121104696-DEL-317 |
| chr7-121118834-DEL-73 | chr7-121118835-DEL-73 |
| chr7-121164081-DEL-1849 | chr7-121164082-DEL-1849 |
| chr7-121183508-DEL-60 | chr7-121183499-DEL-54 |
| chr7-121196500-DEL-3079 | chr7-121196501-DEL-3079 |
| chr7-121217345-DEL-111 | chr7-121217346-DEL-111 |
| chr7-121316157-DEL-131 | chr7-121316158-DEL-131 |
| chr7-121320299-DEL-813 | chr7-121320300-DEL-813 |
| chr7-121375292-DEL-64 | chr7-121375285-DEL-72 |
| chr7-121399359-DEL-1143 | chr7-121399360-DEL-1143 |
| chr7-121410589-DEL-111 | chr7-121410590-DEL-111 |
| chr7-121412852-DEL-76 | chr7-121412853-DEL-76 |
| chr7-121416864-DEL-1100 | chr7-121416865-DEL-1100 |
| chr7-121450077-DEL-5143 | chr7-121450078-DEL-5143 |
| chr7-121455399-DEL-119 | chr7-121455399-DEL-119 |
| chr7-121457145-DEL-55 | chr7-121457142-DEL-59 |
| chr7-121544189-DEL-408 | chr7-121544202-DEL-408 |
| chr7-121559271-DEL-52 | chr7-121559272-DEL-50 |
| chr7-121566593-DEL-104 | chr7-121566612-DEL-56 |
| chr7-121598964-DEL-53 | chr7-121598955-DEL-61 |
| chr7-121634362-DEL-106 | chr7-121634363-DEL-106 |
| chr7-121651109-DEL-599 | chr7-121651097-DEL-600 |
| chr7-121683418-DEL-209 | chr7-121683419-DEL-209 |
| chr7-121692517-DEL-215 | chr7-121692518-DEL-215 |
| chr7-121696495-DEL-154 | chr7-121696496-DEL-154 |
| chr7-121699672-DEL-199 | chr7-121699673-DEL-199 |
| chr7-121704884-DEL-209 | chr7-121704885-DEL-209 |
| chr7-121714775-DEL-212 | chr7-121714776-DEL-212 |
| chr7-121736117-DEL-74 | chr7-121736116-DEL-74 |
| chr7-121746554-DEL-201 | chr7-121746555-DEL-201 |
| chr7-121747061-DEL-68 | chr7-121747059-DEL-68 |
| chr7-121764393-DEL-173 | chr7-121764394-DEL-173 |
| chr7-121788205-DEL-1095 | chr7-121788206-DEL-1095 |
| chr7-121799884-DEL-60 | chr7-121799885-DEL-60 |
| chr7-121817928-DEL-75 | chr7-121817929-DEL-75 |
| chr7-121824454-DEL-60 | chr7-121824455-DEL-60 |
| chr7-121826071-DEL-68 | chr7-121826072-DEL-68 |

| | |
|---|---|
| chr7-121832288-DEL-497 | chr7-121832289-DEL-497 |
| chr7-121860005-DEL-125 | chr7-121860006-DEL-125 |
| chr7-121870093-DEL-76 | chr7-121870098-DEL-68 |
| chr7-121882163-DEL-1079 | chr7-121882164-DEL-1079 |
| chr7-121885549-DEL-70 | chr7-121885579-DEL-69 |
| chr7-121904264-DEL-60 | chr7-121904265-DEL-60 |
| chr7-121947080-DEL-68 | chr7-121947081-DEL-68 |
| chr7-121961721-DEL-623 | chr7-121961722-DEL-623 |
| chr7-121965648-DEL-82 | chr7-121965669-DEL-90 |
| chr7-121967051-DEL-62 | chr7-121967052-DEL-62 |
| chr7-121989252-DEL-134 | chr7-121989229-DEL-123 |
| chr7-122041213-DEL-1430 | chr7-122041214-DEL-1430 |
| chr7-122094118-DEL-355 | chr7-122094119-DEL-355 |
| chr7-122095588-DEL-50 | chr7-122095594-DEL-58 |
| chr7-122117930-DEL-69 | chr7-122117904-DEL-69 |
| chr7-122119446-DEL-72 | chr7-122119449-DEL-70 |
| chr7-122127449-DEL-62 | chr7-122127450-DEL-62 |
| chr7-122134532-DEL-7583 | chr7-122134533-DEL-7583 |
| chr7-122140587-DEL-624 | chr7-122140588-DEL-624 |
| chr7-122141391-DEL-624 | chr7-122141392-DEL-624 |
| chr7-122162638-DEL-89 | chr7-122162628-DEL-90 |
| chr7-122164492-DEL-118 | chr7-122164493-DEL-118 |
| chr7-122198953-DEL-57 | chr7-122198954-DEL-57 |
| chr7-122265286-DEL-948 | chr7-122265287-DEL-948 |
| chr7-122290468-DEL-94 | chr7-122290469-DEL-98 |
| chr7-122326077-DEL-386 | chr7-122326084-DEL-380 |
| chr7-122388626-DEL-2224 | chr7-122388627-DEL-2224 |
| chr7-122405433-DEL-97 | chr7-122405434-DEL-97 |
| chr7-122407657-DEL-1056 | chr7-122407658-DEL-1056 |
| chr7-122408225-DEL-129 | chr7-122408226-DEL-129 |
| chr7-122466055-DEL-62 | chr7-122466056-DEL-62 |
| chr7-122479097-DEL-59 | chr7-122479098-DEL-59 |
| chr7-122492289-DEL-2734 | chr7-122492290-DEL-2734 |
| chr7-122510131-DEL-6487 | chr7-122510132-DEL-6487 |
| chr7-122510375-DEL-53 | chr7-122510376-DEL-53 |
| chr7-122539014-DEL-1019 | chr7-122539015-DEL-1019 |
| chr7-122561621-DEL-498 | chr7-122561622-DEL-498 |
| chr7-122571342-DEL-68 | chr7-122571343-DEL-68 |
| chr7-122594945-DEL-208 | chr7-122594946-DEL-208 |
| chr7-122618922-DEL-58 | chr7-122618923-DEL-58 |
| chr7-122631093-DEL-83 | chr7-122631099-DEL-68 |
| chr7-122635908-DEL-166 | chr7-122635907-DEL-167 |
| chr7-122645264-DEL-214 | chr7-122645265-DEL-214 |
| chr7-122653239-DEL-196 | chr7-122653240-DEL-196 |
| chr7-122663538-DEL-144 | chr7-122663539-DEL-144 |
| chr7-122673081-DEL-111 | chr7-122673068-DEL-125 |
| chr7-122676992-DEL-287 | chr7-122676993-DEL-287 |
| chr7-122689101-DEL-159 | chr7-122689102-DEL-159 |
| chr7-122690002-DEL-71 | chr7-122689983-DEL-54 |
| chr7-122702585-DEL-90 | chr7-122702595-DEL-51 |
| chr7-122703420-DEL-192 | chr7-122703419-DEL-191 |
| chr7-122712232-DEL-94 | chr7-122712244-DEL-81 |
| chr7-122786128-DEL-139 | chr7-122786129-DEL-101 |
| chr7-122812030-DEL-213 | chr7-122812014-DEL-245 |
| chr7-122826057-DEL-69 | chr7-122826058-DEL-69 |
| chr7-122826665-DEL-471 | chr7-122826666-DEL-471 |
| chr7-122869532-DEL-484 | chr7-122869522-DEL-488 |
| chr7-122924093-DEL-64 | chr7-122924121-DEL-64 |
| chr7-122939015-DEL-341 | chr7-122939016-DEL-341 |
| chr7-122980979-DEL-155 | chr7-122980980-DEL-155 |
| chr7-123018879-DEL-63 | chr7-123018880-DEL-63 |
| chr7-123038754-DEL-206 | chr7-123038755-DEL-206 |
| chr7-123047652-DEL-66 | chr7-123047652-DEL-84 |
| chr7-123060060-DEL-80 | chr7-123060061-DEL-80 |
| chr7-123092538-DEL-63 | chr7-123092539-DEL-63 |
| chr7-123116366-DEL-270 | chr7-123116367-DEL-270 |
| chr7-123127584-DEL-213 | chr7-123127541-DEL-234 |
| chr7-123133884-DEL-1101 | chr7-123133885-DEL-1101 |
| chr7-123202905-DEL-1920 | chr7-123202906-DEL-1920 |
| chr7-123204924-DEL-6391 | chr7-123204925-DEL-6391 |
| chr7-123211608-DEL-5710 | chr7-123211609-DEL-5710 |
| chr7-123225096-DEL-1162 | chr7-123225097-DEL-1162 |
| chr7-123264632-DEL-1761 | chr7-123264633-DEL-1761 |
| chr7-123276813-DEL-2611 | chr7-123276814-DEL-2611 |
| chr7-123292768-DEL-100 | chr7-123292769-DEL-100 |
| chr7-123295998-DEL-564 | chr7-123295997-DEL-563 |
| chr7-123320209-DEL-760 | chr7-123320210-DEL-760 |
| chr7-123343937-DEL-759 | chr7-123343938-DEL-759 |
| chr7-123345518-DEL-1043 | chr7-123345519-DEL-1043 |
| chr7-123360140-DEL-74 | chr7-123360141-DEL-74 |

| | |
|---|---|
| chr7-123362877-DEL-51 | chr7-123362878-DEL-51 |
| chr7-123363487-DEL-7070 | chr7-123363488-DEL-7070 |
| chr7-123385232-DEL-735 | chr7-123385233-DEL-735 |
| chr7-123388699-DEL-74 | chr7-123388700-DEL-74 |
| chr7-123452703-DEL-136 | chr7-123452689-DEL-115 |
| chr7-123470834-DEL-6331 | chr7-123470835-DEL-6331 |
| chr7-123498816-DEL-967 | chr7-123498817-DEL-967 |
| chr7-123516131-DEL-532 | chr7-123516133-DEL-533 |
| chr7-123546523-DEL-94 | chr7-123546517-DEL-90 |
| chr7-123670945-DEL-2071 | chr7-123670946-DEL-2071 |
| chr7-123679344-DEL-368 | chr7-123679345-DEL-368 |
| chr7-123698788-DEL-350 | chr7-123698789-DEL-350 |
| chr7-123701821-DEL-58 | chr7-123701850-DEL-56 |
| chr7-123706680-DEL-2907 | chr7-123706681-DEL-2907 |
| chr7-123765897-DEL-54 | chr7-123765883-DEL-52 |
| chr7-123789403-DEL-120 | chr7-123789404-DEL-120 |
| chr7-123804462-DEL-119 | chr7-123804463-DEL-119 |
| chr7-123835801-DEL-436 | chr7-123835802-DEL-436 |
| chr7-123852771-DEL-1321 | chr7-123852772-DEL-1321 |
| chr7-124015907-DEL-550 | chr7-124015908-DEL-550 |
| chr7-124026133-DEL-70 | chr7-124026134-DEL-70 |
| chr7-124052159-DEL-797 | chr7-124052159-DEL-797 |
| chr7-124062870-DEL-3922 | chr7-124062878-DEL-3923 |
| chr7-124071345-DEL-1944 | chr7-124071346-DEL-1944 |
| chr7-124112124-DEL-61 | chr7-124112125-DEL-61 |
| chr7-124112550-DEL-143 | chr7-124112548-DEL-138 |
| chr7-124120700-DEL-6924 | chr7-124120701-DEL-6924 |
| chr7-124139595-DEL-414 | chr7-124139596-DEL-414 |
| chr7-124153476-DEL-401 | chr7-124153477-DEL-401 |
| chr7-124189767-DEL-72 | chr7-124189768-DEL-72 |
| chr7-124197317-DEL-1011 | chr7-124197318-DEL-1011 |
| chr7-124219668-DEL-184 | chr7-124219669-DEL-184 |
| chr7-124259157-DEL-83 | chr7-124259158-DEL-83 |
| chr7-124263443-DEL-306 | chr7-124263444-DEL-306 |
| chr7-124271944-DEL-99 | chr7-124271945-DEL-99 |
| chr7-124287652-DEL-501 | chr7-124287653-DEL-501 |
| chr7-124299174-DEL-552 | chr7-124299175-DEL-552 |
| chr7-124302503-DEL-118 | chr7-124302492-DEL-103 |
| chr7-124303343-DEL-90 | chr7-124303344-DEL-90 |
| chr7-124310216-DEL-27274 | chr7-124310216-DEL-27274 |
| chr7-124325265-DEL-58 | chr7-124325250-DEL-70 |
| chr7-124337562-DEL-74 | chr7-124337563-DEL-74 |
| chr7-124356586-DEL-273 | chr7-124356587-DEL-273 |
| chr7-124380757-DEL-59 | chr7-124380751-DEL-60 |
| chr7-124392243-DEL-104 | chr7-124392244-DEL-104 |
| chr7-124405413-DEL-11536 | chr7-124405414-DEL-11536 |
| chr7-124405914-DEL-5131 | chr7-124405915-DEL-5131 |
| chr7-124416544-DEL-204 | chr7-124416545-DEL-204 |
| chr7-124431730-DEL-420 | chr7-124431731-DEL-420 |
| chr7-124465168-DEL-67 | chr7-124465169-DEL-67 |
| chr7-124482293-DEL-280 | chr7-124482294-DEL-280 |
| chr7-124494345-DEL-7111 | chr7-124494346-DEL-7111 |
| chr7-124511672-DEL-119 | chr7-124511666-DEL-166 |
| chr7-124516607-DEL-70 | chr7-124516608-DEL-70 |
| chr7-124523928-DEL-74 | chr7-124523905-DEL-75 |
| chr7-124536425-DEL-328 | chr7-124536426-DEL-336 |
| chr7-124542849-DEL-59 | chr7-124542849-DEL-59 |
| chr7-124559305-DEL-274 | chr7-124559306-DEL-274 |
| chr7-124568255-DEL-1517 | chr7-124568256-DEL-1517 |
| chr7-124577401-DEL-1182 | chr7-124577402-DEL-1182 |
| chr7-124582943-DEL-2814 | chr7-124582944-DEL-2814 |
| chr7-124589640-DEL-92 | chr7-124589649-DEL-80 |
| chr7-124597065-DEL-60 | chr7-124597066-DEL-60 |
| chr7-124608957-DEL-6906 | chr7-124608958-DEL-6906 |
| chr7-124615462-DEL-212 | chr7-124615463-DEL-212 |
| chr7-124639851-DEL-392 | chr7-124639807-DEL-391 |
| chr7-124656332-DEL-142 | chr7-124656333-DEL-142 |
| chr7-124663433-DEL-146 | chr7-124663434-DEL-146 |
| chr7-124677509-DEL-2687 | chr7-124677510-DEL-2687 |
| chr7-124684569-DEL-327 | chr7-124684570-DEL-327 |
| chr7-124689701-DEL-195 | chr7-124689702-DEL-195 |
| chr7-124693003-DEL-455 | chr7-124693004-DEL-455 |
| chr7-124703781-DEL-1034 | chr7-124703782-DEL-1034 |
| chr7-124713591-DEL-456 | chr7-124713592-DEL-456 |
| chr7-124748936-DEL-1485 | chr7-124748937-DEL-1485 |
| chr7-124770199-DEL-90 | chr7-124770200-DEL-90 |
| chr7-124774478-DEL-68 | chr7-124774459-DEL-88 |
| chr7-124775054-DEL-153 | chr7-124775099-DEL-144 |
| chr7-124778635-DEL-62 | chr7-124778636-DEL-62 |
| chr7-124779858-DEL-321 | chr7-124779859-DEL-321 |

| | |
|---|---|
| chr7-124787550-DEL-703 | chr7-124787551-DEL-703 |
| chr7-124812646-DEL-56 | chr7-124812647-DEL-56 |
| chr7-124813580-DEL-56 | chr7-124813581-DEL-56 |
| chr7-124821031-DEL-152 | chr7-124821032-DEL-152 |
| chr7-124837454-DEL-70 | chr7-124837455-DEL-70 |
| chr7-124852173-DEL-63 | chr7-124852175-DEL-61 |
| chr7-124866298-DEL-78 | chr7-124866277-DEL-66 |
| chr7-124877415-DEL-956 | chr7-124877416-DEL-956 |
| chr7-124880221-DEL-66 | chr7-124880218-DEL-69 |
| chr7-124897536-DEL-6413 | chr7-124897537-DEL-6413 |
| chr7-124940753-DEL-83 | chr7-124940754-DEL-83 |
| chr7-124947878-DEL-148 | chr7-124947879-DEL-148 |
| chr7-124965781-DEL-596 | chr7-124965782-DEL-593 |
| chr7-124966820-DEL-406 | chr7-124966821-DEL-406 |
| chr7-124970821-DEL-68 | chr7-124970822-DEL-68 |
| chr7-124992171-DEL-169 | chr7-124992172-DEL-169 |
| chr7-125018875-DEL-52 | chr7-125018843-DEL-85 |
| chr7-125033264-DEL-65 | chr7-125033265-DEL-65 |
| chr7-125052522-DEL-133 | chr7-125052523-DEL-131 |
| chr7-125056968-DEL-55 | chr7-125056949-DEL-92 |
| chr7-125063273-DEL-655 | chr7-125063274-DEL-655 |
| chr7-125105492-DEL-386 | chr7-125105486-DEL-393 |
| chr7-125107761-DEL-51 | chr7-125107762-DEL-52 |
| chr7-125108843-DEL-167 | chr7-125108844-DEL-167 |
| chr7-125113038-DEL-69 | chr7-125113039-DEL-69 |
| chr7-125115024-DEL-651 | chr7-125115025-DEL-651 |
| chr7-125120964-DEL-142 | chr7-125120965-DEL-142 |
| chr7-125128869-DEL-4486 | chr7-125128870-DEL-4486 |
| chr7-125145563-DEL-74 | chr7-125145573-DEL-50 |
| chr7-125185544-DEL-235 | chr7-125185540-DEL-245 |
| chr7-125186852-DEL-142 | chr7-125186887-DEL-108 |
| chr7-125193955-DEL-835 | chr7-125193956-DEL-835 |
| chr7-125233837-DEL-66 | chr7-125233820-DEL-69 |
| chr7-125236241-DEL-66 | chr7-125236241-DEL-66 |
| chr7-125243781-DEL-60 | chr7-125243782-DEL-60 |
| chr7-125254058-DEL-51 | chr7-125254059-DEL-51 |
| chr7-125270975-DEL-63 | chr7-125270960-DEL-96 |
| chr7-125278338-DEL-463 | chr7-125278339-DEL-463 |
| chr7-125356485-DEL-116 | chr7-125356486-DEL-116 |
| chr7-125445214-DEL-3041 | chr7-125445215-DEL-3041 |
| chr7-125501497-DEL-6399 | chr7-125501498-DEL-6399 |
| chr7-125512353-DEL-339 | chr7-125512374-DEL-319 |
| chr7-125559378-DEL-88 | chr7-125559376-DEL-88 |
| chr7-125631302-DEL-75 | chr7-125631289-DEL-75 |
| chr7-125672623-DEL-229 | chr7-125672624-DEL-229 |
| chr7-125696221-DEL-224 | chr7-125696222-DEL-224 |
| chr7-125719991-DEL-94 | chr7-125720012-DEL-62 |
| chr7-125736308-DEL-64 | chr7-125736309-DEL-64 |
| chr7-125737683-DEL-203 | chr7-125737684-DEL-203 |
| chr7-125745166-DEL-205 | chr7-125745157-DEL-207 |
| chr7-125760594-DEL-224 | chr7-125760595-DEL-224 |
| chr7-125791177-DEL-203 | chr7-125791178-DEL-203 |
| chr7-125791652-DEL-729 | chr7-125791653-DEL-729 |
| chr7-125793199-DEL-199 | chr7-125793200-DEL-199 |
| chr7-125807703-DEL-203 | chr7-125807704-DEL-203 |
| chr7-125809682-DEL-94 | chr7-125809683-DEL-94 |
| chr7-125820197-DEL-58 | chr7-125820198-DEL-58 |
| chr7-125820772-DEL-165 | chr7-125820773-DEL-165 |
| chr7-125829363-DEL-219 | chr7-125829364-DEL-219 |
| chr7-125831479-DEL-206 | chr7-125831480-DEL-206 |
| chr7-125839025-DEL-188 | chr7-125839037-DEL-177 |
| chr7-125850100-DEL-128 | chr7-125850101-DEL-128 |
| chr7-125857381-DEL-168 | chr7-125857380-DEL-168 |
| chr7-125882034-DEL-204 | chr7-125882035-DEL-204 |
| chr7-125886563-DEL-191 | chr7-125886564-DEL-191 |
| chr7-125909778-DEL-364 | chr7-125909779-DEL-364 |
| chr7-125915134-DEL-232 | chr7-125915138-DEL-228 |
| chr7-125918393-DEL-56 | chr7-125918394-DEL-56 |
| chr7-125927137-DEL-1599 | chr7-125927138-DEL-1599 |
| chr7-125927963-DEL-204 | chr7-125927964-DEL-204 |
| chr7-125929793-DEL-78 | chr7-125929791-DEL-79 |
| chr7-125944569-DEL-211 | chr7-125944570-DEL-211 |
| chr7-125946977-DEL-182 | chr7-125946978-DEL-182 |
| chr7-125948364-DEL-242 | chr7-125948365-DEL-242 |
| chr7-125953373-DEL-56 | chr7-125953376-DEL-54 |
| chr7-125986157-DEL-171 | chr7-125986158-DEL-171 |
| chr7-125993510-DEL-317 | chr7-125993511-DEL-317 |
| chr7-126001918-DEL-201 | chr7-126001919-DEL-201 |
| chr7-126017893-DEL-629 | chr7-126017894-DEL-629 |
| chr7-126021667-DEL-216 | chr7-126021668-DEL-216 |

| | |
|---|---|
| chr7-126022736-DEL-51 | chr7-126022737-DEL-51 |
| chr7-126030152-DEL-60 | chr7-126030145-DEL-54 |
| chr7-126082884-DEL-236 | chr7-126082885-DEL-236 |
| chr7-126108395-DEL-197 | chr7-126108396-DEL-197 |
| chr7-126114915-DEL-70 | chr7-126114916-DEL-70 |
| chr7-126133300-DEL-202 | chr7-126133301-DEL-202 |
| chr7-126135973-DEL-811 | chr7-126135975-DEL-807 |
| chr7-126138736-DEL-283 | chr7-126138714-DEL-283 |
| chr7-126152808-DEL-71 | chr7-126152807-DEL-72 |
| chr7-126158555-DEL-227 | chr7-126158556-DEL-227 |
| chr7-126164531-DEL-178 | chr7-126164532-DEL-178 |
| chr7-126168399-DEL-362 | chr7-126168396-DEL-363 |
| chr7-126173193-DEL-200 | chr7-126173194-DEL-200 |
| chr7-126179234-DEL-66 | chr7-126179237-DEL-65 |
| chr7-126191098-DEL-61 | chr7-126191099-DEL-61 |
| chr7-126215273-DEL-206 | chr7-126215274-DEL-206 |
| chr7-126223587-DEL-85 | chr7-126223588-DEL-85 |
| chr7-126229511-DEL-225 | chr7-126229512-DEL-225 |
| chr7-126240721-DEL-200 | chr7-126240722-DEL-200 |
| chr7-126242353-DEL-202 | chr7-126242347-DEL-203 |
| chr7-126252561-DEL-168 | chr7-126252562-DEL-169 |
| chr7-126257557-DEL-2002 | chr7-126257558-DEL-2002 |
| chr7-126261377-DEL-202 | chr7-126261378-DEL-202 |
| chr7-126281621-DEL-2683 | chr7-126281622-DEL-2683 |
| chr7-126289623-DEL-211 | chr7-126289624-DEL-211 |
| chr7-126325451-DEL-205 | chr7-126325452-DEL-205 |
| chr7-126336004-DEL-793 | chr7-126336005-DEL-793 |
| chr7-126401130-DEL-53 | chr7-126401134-DEL-52 |
| chr7-126432811-DEL-55 | chr7-126432813-DEL-55 |
| chr7-126473046-DEL-132 | chr7-126473049-DEL-104 |
| chr7-126481557-DEL-203 | chr7-126481591-DEL-153 |
| chr7-126502923-DEL-114 | chr7-126502942-DEL-96 |
| chr7-126590683-DEL-238 | chr7-126590684-DEL-236 |
| chr7-126635582-DEL-74 | chr7-126635610-DEL-54 |
| chr7-126636560-DEL-57 | chr7-126636546-DEL-63 |
| chr7-126682650-DEL-210 | chr7-126682651-DEL-210 |
| chr7-126727082-DEL-88 | chr7-126727083-DEL-88 |
| chr7-126747172-DEL-316 | chr7-126747173-DEL-316 |
| chr7-126765315-DEL-120 | chr7-126765316-DEL-120 |
| chr7-126767418-DEL-177 | chr7-126767419-DEL-177 |
| chr7-126804023-DEL-82 | chr7-126804024-DEL-82 |
| chr7-126805046-DEL-145 | chr7-126805047-DEL-145 |
| chr7-126833409-DEL-97 | chr7-126833410-DEL-97 |
| chr7-126842188-DEL-181 | chr7-126842188-DEL-182 |
| chr7-126865020-DEL-63 | chr7-126865021-DEL-63 |
| chr7-126866080-DEL-53 | chr7-126866067-DEL-50 |
| chr7-126880923-DEL-223 | chr7-126880924-DEL-223 |
| chr7-126882344-DEL-1369 | chr7-126882343-DEL-1371 |
| chr7-126884323-DEL-72 | chr7-126884318-DEL-82 |
| chr7-126894413-DEL-209 | chr7-126894411-DEL-212 |
| chr7-126896662-DEL-84 | chr7-126896663-DEL-81 |
| chr7-126902400-DEL-421 | chr7-126902401-DEL-421 |
| chr7-126928765-DEL-120 | chr7-126928766-DEL-120 |
| chr7-126968345-DEL-1776 | chr7-126968368-DEL-1761 |
| chr7-127006759-DEL-1151 | chr7-127006760-DEL-1151 |
| chr7-127028572-DEL-182 | chr7-127028573-DEL-182 |
| chr7-127035449-DEL-71 | chr7-127035432-DEL-72 |
| chr7-127038375-DEL-230 | chr7-127038376-DEL-230 |
| chr7-127057828-DEL-220 | chr7-127057829-DEL-220 |
| chr7-127058392-DEL-208 | chr7-127058393-DEL-208 |
| chr7-127062469-DEL-214 | chr7-127062470-DEL-214 |
| chr7-127087491-DEL-209 | chr7-127087492-DEL-209 |
| chr7-127107767-DEL-246 | chr7-127107768-DEL-246 |
| chr7-127149359-DEL-77 | chr7-127149360-DEL-76 |
| chr7-127217839-DEL-7462 | chr7-127217839-DEL-7462 |
| chr7-127231002-DEL-64 | chr7-127231002-DEL-64 |
| chr7-127233514-DEL-1700 | chr7-127233515-DEL-1700 |
| chr7-127244075-DEL-60 | chr7-127244076-DEL-58 |
| chr7-127253190-DEL-195 | chr7-127253184-DEL-208 |
| chr7-127269535-DEL-201 | chr7-127269531-DEL-184 |
| chr7-127270712-DEL-76 | chr7-127270663-DEL-89 |
| chr7-127291279-DEL-207 | chr7-127291270-DEL-211 |
| chr7-127295355-DEL-203 | chr7-127295356-DEL-203 |
| chr7-127299542-DEL-2288 | chr7-127299543-DEL-2288 |
| chr7-127313246-DEL-135 | chr7-127313247-DEL-135 |
| chr7-127319433-DEL-214 | chr7-127319434-DEL-214 |
| chr7-127327376-DEL-185 | chr7-127327377-DEL-185 |
| chr7-127329735-DEL-171 | chr7-127329736-DEL-171 |
| chr7-127331136-DEL-6451 | chr7-127331135-DEL-6452 |
| chr7-127362179-DEL-210 | chr7-127362183-DEL-212 |

| | |
|---|---|
| chr7-127416073-DEL-180 | chr7-127416074-DEL-180 |
| chr7-127427901-DEL-210 | chr7-127427902-DEL-210 |
| chr7-127429019-DEL-210 | chr7-127429020-DEL-210 |
| chr7-127438477-DEL-217 | chr7-127438478-DEL-217 |
| chr7-127452016-DEL-4306 | chr7-127452017-DEL-4306 |
| chr7-127477029-DEL-57 | chr7-127477030-DEL-57 |
| chr7-127502856-DEL-209 | chr7-127502857-DEL-209 |
| chr7-127538987-DEL-197 | chr7-127539012-DEL-175 |
| chr7-127549777-DEL-126 | chr7-127549778-DEL-126 |
| chr7-127554739-DEL-210 | chr7-127554740-DEL-210 |
| chr7-127560392-DEL-52 | chr7-127560393-DEL-52 |
| chr7-127562133-DEL-212 | chr7-127562134-DEL-212 |
| chr7-127617088-DEL-65 | chr7-127617085-DEL-69 |
| chr7-127619704-DEL-176 | chr7-127619704-DEL-177 |
| chr7-127620516-DEL-177 | chr7-127620517-DEL-177 |
| chr7-127633821-DEL-635 | chr7-127633824-DEL-633 |
| chr7-127635856-DEL-100 | chr7-127635857-DEL-100 |
| chr7-127636805-DEL-5241 | chr7-127636805-DEL-5242 |
| chr7-127659494-DEL-60 | chr7-127659495-DEL-60 |
| chr7-127690731-DEL-1221 | chr7-127690731-DEL-1221 |
| chr7-127716717-DEL-56 | chr7-127716687-DEL-50 |
| chr7-127722536-DEL-3076 | chr7-127722537-DEL-3076 |
| chr7-127750333-DEL-312 | chr7-127750334-DEL-312 |
| chr7-127759095-DEL-139 | chr7-127759096-DEL-139 |
| chr7-127784444-DEL-68 | chr7-127784445-DEL-60 |
| chr7-127786047-DEL-1254 | chr7-127786031-DEL-1289 |
| chr7-127795184-DEL-82 | chr7-127795185-DEL-82 |
| chr7-127799098-DEL-64 | chr7-127799091-DEL-72 |
| chr7-127803486-DEL-133 | chr7-127803529-DEL-107 |
| chr7-127825101-DEL-73 | chr7-127825100-DEL-58 |
| chr7-127842676-DEL-274 | chr7-127842677-DEL-274 |
| chr7-127868637-DEL-185 | chr7-127868638-DEL-185 |
| chr7-127882434-DEL-531 | chr7-127882435-DEL-531 |
| chr7-127917244-DEL-449 | chr7-127917245-DEL-449 |
| chr7-127926587-DEL-82 | chr7-127926588-DEL-80 |
| chr7-127944119-DEL-439 | chr7-127944120-DEL-439 |
| chr7-127954214-DEL-135 | chr7-127954215-DEL-135 |
| chr7-127996176-DEL-52 | chr7-127996177-DEL-52 |
| chr7-128001094-DEL-227 | chr7-128001095-DEL-227 |
| chr7-128009860-DEL-54 | chr7-128009861-DEL-54 |
| chr7-128066372-DEL-55 | chr7-128066373-DEL-55 |
| chr7-128091287-DEL-219 | chr7-128091277-DEL-220 |
| chr7-128096629-DEL-473 | chr7-128096630-DEL-474 |
| chr7-128172277-DEL-260 | chr7-128172278-DEL-260 |
| chr7-128184417-DEL-56 | chr7-128184416-DEL-56 |
| chr7-128191426-DEL-204 | chr7-128191427-DEL-204 |
| chr7-128206323-DEL-5468 | chr7-128206326-DEL-5459 |
| chr7-128267666-DEL-66 | chr7-128267667-DEL-66 |
| chr7-128281713-DEL-613 | chr7-128281712-DEL-615 |
| chr7-128285722-DEL-197 | chr7-128285723-DEL-197 |
| chr7-128300746-DEL-64 | chr7-128300735-DEL-50 |
| chr7-128301597-DEL-216 | chr7-128301598-DEL-216 |
| chr7-128306477-DEL-198 | chr7-128306478-DEL-197 |
| chr7-128322695-DEL-60 | chr7-128322696-DEL-60 |
| chr7-128329844-DEL-218 | chr7-128329845-DEL-218 |
| chr7-128361558-DEL-82 | chr7-128361559-DEL-82 |
| chr7-128364275-DEL-213 | chr7-128364269-DEL-214 |
| chr7-128371818-DEL-112 | chr7-128371818-DEL-112 |
| chr7-128385884-DEL-211 | chr7-128385885-DEL-211 |
| chr7-128387212-DEL-62 | chr7-128387213-DEL-62 |
| chr7-128427828-DEL-473 | chr7-128427829-DEL-473 |
| chr7-128447380-DEL-401 | chr7-128447379-DEL-402 |
| chr7-128454549-DEL-62 | chr7-128454558-DEL-54 |
| chr7-128462971-DEL-53 | chr7-128462972-DEL-53 |
| chr7-128479709-DEL-54 | chr7-128479688-DEL-66 |
| chr7-128500543-DEL-176 | chr7-128500544-DEL-176 |
| chr7-128504270-DEL-106 | chr7-128504299-DEL-56 |
| chr7-128511112-DEL-310 | chr7-128511079-DEL-344 |
| chr7-128526466-DEL-244 | chr7-128526465-DEL-246 |
| chr7-128543258-DEL-60 | chr7-128543259-DEL-60 |
| chr7-128586181-DEL-7121 | chr7-128586182-DEL-7121 |
| chr7-128616047-DEL-1830 | chr7-128616046-DEL-1831 |
| chr7-128638506-DEL-6190 | chr7-128638507-DEL-6190 |
| chr7-128654778-DEL-74 | chr7-128654779-DEL-74 |
| chr7-128664479-DEL-65 | chr7-128664467-DEL-66 |
| chr7-128667433-DEL-67 | chr7-128667434-DEL-67 |
| chr7-128688347-DEL-213 | chr7-128688348-DEL-213 |
| chr7-128699246-DEL-145 | chr7-128699247-DEL-145 |
| chr7-128766721-DEL-90 | chr7-128766722-DEL-90 |
| chr7-128794321-DEL-60 | chr7-128794306-DEL-50 |

| | |
|---|---|
| chr7-128826248-DEL-1788 | chr7-128826249-DEL-1787 |
| chr7-128834375-DEL-96 | chr7-128834376-DEL-96 |
| chr7-128867899-DEL-52 | chr7-128867900-DEL-52 |
| chr7-128904430-DEL-301 | chr7-128904431-DEL-301 |
| chr7-128935464-DEL-53 | chr7-128935448-DEL-81 |
| chr7-128944301-DEL-219 | chr7-128944336-DEL-201 |
| chr7-128949492-DEL-2737 | chr7-128949493-DEL-2737 |
| chr7-129018175-DEL-2810 | chr7-129018174-DEL-2811 |
| chr7-129034260-DEL-73 | chr7-129034246-DEL-73 |
| chr7-129054125-DEL-402 | chr7-129054122-DEL-406 |
| chr7-129058133-DEL-979 | chr7-129058134-DEL-979 |
| chr7-129072418-DEL-257 | chr7-129072422-DEL-258 |
| chr7-129078835-DEL-73 | chr7-129078836-DEL-73 |
| chr7-129129112-DEL-60 | chr7-129129113-DEL-60 |
| chr7-129152077-DEL-1115 | chr7-129152078-DEL-1115 |
| chr7-129161285-DEL-7142 | chr7-129161286-DEL-7142 |
| chr7-129172846-DEL-100 | chr7-129172849-DEL-98 |
| chr7-129174468-DEL-123 | chr7-129174460-DEL-131 |
| chr7-129183124-DEL-1907 | chr7-129183123-DEL-1906 |
| chr7-129199467-DEL-727 | chr7-129199468-DEL-727 |
| chr7-129238570-DEL-1642 | chr7-129238564-DEL-1643 |
| chr7-129246470-DEL-162 | chr7-129246471-DEL-162 |
| chr7-129251545-DEL-120 | chr7-129251546-DEL-120 |
| chr7-129257260-DEL-134 | chr7-129257261-DEL-135 |
| chr7-129259397-DEL-107 | chr7-129259398-DEL-105 |
| chr7-129268831-DEL-505 | chr7-129268832-DEL-505 |
| chr7-129307493-DEL-680 | chr7-129307494-DEL-680 |
| chr7-129311661-DEL-69 | chr7-129311662-DEL-53 |
| chr7-129335268-DEL-7129 | chr7-129335269-DEL-7129 |
| chr7-129351941-DEL-64 | chr7-129351942-DEL-64 |
| chr7-129355522-DEL-56 | chr7-129355527-DEL-52 |
| chr7-129371693-DEL-180 | chr7-129371679-DEL-180 |
| chr7-129409405-DEL-53 | chr7-129409406-DEL-53 |
| chr7-129415326-DEL-594 | chr7-129415327-DEL-594 |
| chr7-129438284-DEL-114 | chr7-129438285-DEL-96 |
| chr7-129438368-DEL-1252 | chr7-129438364-DEL-1257 |
| chr7-129441277-DEL-99 | chr7-129441278-DEL-99 |
| chr7-129443569-DEL-60 | chr7-129443566-DEL-63 |
| chr7-129463632-DEL-99 | chr7-129463645-DEL-59 |
| chr7-129493159-DEL-60 | chr7-129493132-DEL-93 |
| chr7-129497564-DEL-401 | chr7-129497565-DEL-401 |
| chr7-129501819-DEL-201 | chr7-129501833-DEL-185 |
| chr7-129519568-DEL-61 | chr7-129519569-DEL-61 |
| chr7-129572095-DEL-212 | chr7-129572091-DEL-213 |
| chr7-129623044-DEL-57 | chr7-129623032-DEL-58 |
| chr7-129633223-DEL-136 | chr7-129633224-DEL-136 |
| chr7-129634587-DEL-62 | chr7-129634590-DEL-60 |
| chr7-129637503-DEL-78 | chr7-129637504-DEL-78 |
| chr7-129642398-DEL-77 | chr7-129642399-DEL-77 |
| chr7-129676814-DEL-82 | chr7-129676800-DEL-129 |
| chr7-129722218-DEL-87 | chr7-129722216-DEL-78 |
| chr7-129722927-DEL-99 | chr7-129722925-DEL-99 |
| chr7-129732503-DEL-465 | chr7-129732504-DEL-465 |
| chr7-129735074-DEL-59 | chr7-129735065-DEL-61 |
| chr7-129748390-DEL-142 | chr7-129748389-DEL-134 |
| chr7-129781201-DEL-174 | chr7-129781204-DEL-185 |
| chr7-129781741-DEL-234 | chr7-129781742-DEL-234 |
| chr7-129833895-DEL-185 | chr7-129833896-DEL-185 |
| chr7-129886077-DEL-57 | chr7-129886078-DEL-57 |
| chr7-129896279-DEL-228 | chr7-129896286-DEL-222 |
| chr7-129921141-DEL-63 | chr7-129921127-DEL-64 |
| chr7-129946942-DEL-6444 | chr7-129946943-DEL-6444 |
| chr7-129962159-DEL-63 | chr7-129962156-DEL-66 |
| chr7-129979000-DEL-195 | chr7-129979001-DEL-195 |
| chr7-129991653-DEL-269 | chr7-129991641-DEL-266 |
| chr7-130001072-DEL-373 | chr7-130001073-DEL-373 |
| chr7-130006934-DEL-53 | chr7-130006935-DEL-53 |
| chr7-130042639-DEL-6403 | chr7-130042640-DEL-6403 |
| chr7-130080794-DEL-195 | chr7-130080795-DEL-195 |
| chr7-130081663-DEL-384 | chr7-130081664-DEL-384 |
| chr7-130160781-DEL-2555 | chr7-130160782-DEL-2555 |
| chr7-130190052-DEL-191 | chr7-130190053-DEL-191 |
| chr7-130202981-DEL-118 | chr7-130202981-DEL-85 |
| chr7-130205751-DEL-97 | chr7-130205747-DEL-89 |
| chr7-130211437-DEL-168 | chr7-130211438-DEL-168 |
| chr7-130216600-DEL-254 | chr7-130216601-DEL-240 |
| chr7-130294023-DEL-56 | chr7-130294018-DEL-64 |
| chr7-130303515-DEL-146 | chr7-130303516-DEL-146 |
| chr7-130307937-DEL-58 | chr7-130307938-DEL-58 |
| chr7-130338432-DEL-201 | chr7-130338433-DEL-201 |

| | |
|---|---|
| chr7-130353428-DEL-64 | chr7-130353429-DEL-64 |
| chr7-130377819-DEL-130 | chr7-130377794-DEL-105 |
| chr7-130404411-DEL-197 | chr7-130404412-DEL-196 |
| chr7-130411077-DEL-319 | chr7-130411078-DEL-319 |
| chr7-130428083-DEL-258 | chr7-130428084-DEL-258 |
| chr7-130437674-DEL-216 | chr7-130437675-DEL-216 |
| chr7-130450676-DEL-221 | chr7-130450677-DEL-218 |
| chr7-130456186-DEL-77 | chr7-130456164-DEL-66 |
| chr7-130475026-DEL-209 | chr7-130475028-DEL-208 |
| chr7-130477702-DEL-99 | chr7-130477703-DEL-100 |
| chr7-130534494-DEL-91 | chr7-130534495-DEL-90 |
| chr7-130551556-DEL-68 | chr7-130551557-DEL-68 |
| chr7-130557009-DEL-119 | chr7-130557001-DEL-91 |
| chr7-130611755-DEL-78 | chr7-130611756-DEL-78 |
| chr7-130656464-DEL-358 | chr7-130656465-DEL-358 |
| chr7-130657193-DEL-91 | chr7-130657196-DEL-89 |
| chr7-130661959-DEL-964 | chr7-130661960-DEL-964 |
| chr7-130679792-DEL-236 | chr7-130679793-DEL-236 |
| chr7-130708903-DEL-102 | chr7-130708904-DEL-102 |
| chr7-130736355-DEL-418 | chr7-130736356-DEL-418 |
| chr7-130750017-DEL-3578 | chr7-130750018-DEL-3578 |
| chr7-130777905-DEL-246 | chr7-130777906-DEL-246 |
| chr7-130794157-DEL-1140 | chr7-130794157-DEL-1141 |
| chr7-130803522-DEL-1044 | chr7-130803523-DEL-1044 |
| chr7-130806040-DEL-2622 | chr7-130806041-DEL-2622 |
| chr7-130813861-DEL-251 | chr7-130813844-DEL-289 |
| chr7-130840365-DEL-687 | chr7-130840366-DEL-687 |
| chr7-130878350-DEL-594 | chr7-130878381-DEL-587 |
| chr7-130891742-DEL-65 | chr7-130891750-DEL-69 |
| chr7-130927373-DEL-56 | chr7-130927364-DEL-84 |
| chr7-130943662-DEL-66 | chr7-130943663-DEL-66 |
| chr7-130976830-DEL-6263 | chr7-130976831-DEL-6263 |
| chr7-130983643-DEL-1903 | chr7-130983644-DEL-1903 |
| chr7-131089148-DEL-7387 | chr7-131089149-DEL-7387 |
| chr7-131103213-DEL-64 | chr7-131103185-DEL-60 |
| chr7-131119038-DEL-104 | chr7-131119018-DEL-96 |
| chr7-131126249-DEL-292 | chr7-131126250-DEL-292 |
| chr7-131134101-DEL-66 | chr7-131134102-DEL-66 |
| chr7-131140345-DEL-242 | chr7-131140346-DEL-242 |
| chr7-131205616-DEL-64 | chr7-131205612-DEL-71 |
| chr7-131215852-DEL-883 | chr7-131215853-DEL-883 |
| chr7-131260346-DEL-56 | chr7-131260347-DEL-56 |
| chr7-131285974-DEL-60 | chr7-131285975-DEL-60 |
| chr7-131296618-DEL-56 | chr7-131296620-DEL-54 |
| chr7-131300118-DEL-58 | chr7-131300119-DEL-58 |
| chr7-131308618-DEL-83 | chr7-131308610-DEL-101 |
| chr7-131308699-DEL-891 | chr7-131308663-DEL-915 |
| chr7-131343651-DEL-104 | chr7-131343662-DEL-94 |
| chr7-131355969-DEL-236 | chr7-131355970-DEL-236 |
| chr7-131359996-DEL-64 | chr7-131359997-DEL-64 |
| chr7-131372930-DEL-171 | chr7-131372931-DEL-169 |
| chr7-131375991-DEL-366 | chr7-131375992-DEL-366 |
| chr7-131458077-DEL-54 | chr7-131458078-DEL-54 |
| chr7-131468005-DEL-190 | chr7-131468005-DEL-190 |
| chr7-131472865-DEL-82 | chr7-131472858-DEL-59 |
| chr7-131473392-DEL-85 | chr7-131473393-DEL-85 |
| chr7-131519445-DEL-394 | chr7-131519446-DEL-394 |
| chr7-131542439-DEL-58 | chr7-131542440-DEL-50 |
| chr7-131556886-DEL-197 | chr7-131556903-DEL-181 |
| chr7-131592137-DEL-211 | chr7-131592138-DEL-211 |
| chr7-131597310-DEL-4159 | chr7-131597311-DEL-4159 |
| chr7-131625154-DEL-497 | chr7-131625155-DEL-497 |
| chr7-131640725-DEL-90 | chr7-131640726-DEL-90 |
| chr7-131659804-DEL-130 | chr7-131659793-DEL-136 |
| chr7-131661810-DEL-134 | chr7-131661811-DEL-134 |
| chr7-131673471-DEL-52 | chr7-131673472-DEL-52 |
| chr7-131674872-DEL-144 | chr7-131674872-DEL-120 |
| chr7-131691891-DEL-54 | chr7-131691892-DEL-50 |
| chr7-131703147-DEL-666 | chr7-131703148-DEL-666 |
| chr7-131715708-DEL-273 | chr7-131715711-DEL-275 |
| chr7-131716135-DEL-1887 | chr7-131716136-DEL-1887 |
| chr7-131724007-DEL-54 | chr7-131724007-DEL-54 |
| chr7-131744197-DEL-514 | chr7-131744196-DEL-514 |
| chr7-131752988-DEL-60 | chr7-131752989-DEL-60 |
| chr7-131777355-DEL-80 | chr7-131777360-DEL-64 |
| chr7-131789831-DEL-53 | chr7-131789832-DEL-53 |
| chr7-131794238-DEL-326 | chr7-131794239-DEL-326 |
| chr7-131807075-DEL-82 | chr7-131807064-DEL-82 |
| chr7-131812745-DEL-474 | chr7-131812746-DEL-474 |
| chr7-131821780-DEL-54 | chr7-131821781-DEL-54 |

| | |
|---|---|
| chr7-131841693-DEL-864 | chr7-131841693-DEL-865 |
| chr7-131896852-DEL-76 | chr7-131896853-DEL-76 |
| chr7-131939514-DEL-65 | chr7-131939515-DEL-65 |
| chr7-131949149-DEL-100 | chr7-131949150-DEL-100 |
| chr7-131956610-DEL-205 | chr7-131956611-DEL-205 |
| chr7-131983205-DEL-138 | chr7-131983206-DEL-138 |
| chr7-132032690-DEL-608 | chr7-132032691-DEL-606 |
| chr7-132080610-DEL-268 | chr7-132080610-DEL-276 |
| chr7-132088742-DEL-216 | chr7-132088727-DEL-222 |
| chr7-132198217-DEL-74 | chr7-132198218-DEL-74 |
| chr7-132308278-DEL-225 | chr7-132308279-DEL-225 |
| chr7-132345270-DEL-1893 | chr7-132345271-DEL-1893 |
| chr7-132421533-DEL-493 | chr7-132421534-DEL-493 |
| chr7-132435046-DEL-140 | chr7-132435047-DEL-140 |
| chr7-132450038-DEL-163 | chr7-132450039-DEL-163 |
| chr7-132475286-DEL-130 | chr7-132475282-DEL-138 |
| chr7-132507622-DEL-98 | chr7-132507623-DEL-98 |
| chr7-132509447-DEL-2512 | chr7-132509442-DEL-2517 |
| chr7-132560134-DEL-61 | chr7-132560135-DEL-61 |
| chr7-132606896-DEL-169 | chr7-132606897-DEL-169 |
| chr7-132677347-DEL-215 | chr7-132677348-DEL-215 |
| chr7-132692607-DEL-57 | chr7-132692616-DEL-59 |
| chr7-132697350-DEL-146 | chr7-132697351-DEL-146 |
| chr7-132698004-DEL-233 | chr7-132698005-DEL-233 |
| chr7-132735688-DEL-121 | chr7-132735690-DEL-119 |
| chr7-132770095-DEL-53 | chr7-132770094-DEL-77 |
| chr7-132796865-DEL-84 | chr7-132796834-DEL-132 |
| chr7-132808825-DEL-66 | chr7-132808834-DEL-58 |
| chr7-132838269-DEL-71 | chr7-132838270-DEL-71 |
| chr7-132870579-DEL-6604 | chr7-132870580-DEL-6604 |
| chr7-132881221-DEL-51 | chr7-132881222-DEL-51 |
| chr7-132908936-DEL-291 | chr7-132908955-DEL-272 |
| chr7-132915689-DEL-137 | chr7-132915690-DEL-137 |
| chr7-132923059-DEL-160 | chr7-132923060-DEL-160 |
| chr7-132943833-DEL-54 | chr7-132943834-DEL-54 |
| chr7-132956483-DEL-91 | chr7-132956484-DEL-91 |
| chr7-132958825-DEL-109 | chr7-132958825-DEL-109 |
| chr7-132967877-DEL-786 | chr7-132967878-DEL-786 |
| chr7-133011645-DEL-789 | chr7-133011636-DEL-790 |
| chr7-133029637-DEL-201 | chr7-133029638-DEL-201 |
| chr7-133033445-DEL-141 | chr7-133033446-DEL-141 |
| chr7-133066136-DEL-3165 | chr7-133066137-DEL-3165 |
| chr7-133108652-DEL-77 | chr7-133108665-DEL-81 |
| chr7-133114817-DEL-75 | chr7-133114818-DEL-75 |
| chr7-133121158-DEL-641 | chr7-133121152-DEL-638 |
| chr7-133123552-DEL-220 | chr7-133123549-DEL-224 |
| chr7-133130568-DEL-7377 | chr7-133130569-DEL-7377 |
| chr7-133179051-DEL-53 | chr7-133179052-DEL-53 |
| chr7-133332482-DEL-74 | chr7-133332483-DEL-74 |
| chr7-133335410-DEL-70 | chr7-133335403-DEL-54 |
| chr7-133338532-DEL-115 | chr7-133338533-DEL-115 |
| chr7-133358186-DEL-1287 | chr7-133358187-DEL-1287 |
| chr7-133371589-DEL-216 | chr7-133371590-DEL-216 |
| chr7-133381354-DEL-61 | chr7-133381355-DEL-61 |
| chr7-133420277-DEL-76 | chr7-133420282-DEL-72 |
| chr7-133427364-DEL-230 | chr7-133427377-DEL-218 |
| chr7-133428669-DEL-192 | chr7-133428667-DEL-193 |
| chr7-133429436-DEL-352 | chr7-133429437-DEL-352 |
| chr7-133439919-DEL-152 | chr7-133439920-DEL-152 |
| chr7-133462815-DEL-381 | chr7-133462816-DEL-381 |
| chr7-133499288-DEL-165 | chr7-133499289-DEL-165 |
| chr7-133500321-DEL-208 | chr7-133500322-DEL-208 |
| chr7-133529572-DEL-217 | chr7-133529573-DEL-217 |
| chr7-133557675-DEL-72 | chr7-133557684-DEL-72 |
| chr7-133567862-DEL-52 | chr7-133567863-DEL-52 |
| chr7-133603692-DEL-2944 | chr7-133603693-DEL-2944 |
| chr7-133632012-DEL-62 | chr7-133632014-DEL-63 |
| chr7-133634472-DEL-253 | chr7-133634473-DEL-253 |
| chr7-133637976-DEL-1400 | chr7-133637977-DEL-1400 |
| chr7-133663927-DEL-67 | chr7-133663928-DEL-67 |
| chr7-133702161-DEL-7037 | chr7-133702162-DEL-7037 |
| chr7-133788916-DEL-628 | chr7-133788917-DEL-628 |
| chr7-133821205-DEL-91 | chr7-133821195-DEL-52 |
| chr7-133822095-DEL-56 | chr7-133822096-DEL-56 |
| chr7-133840225-DEL-12856 | chr7-133840226-DEL-12856 |
| chr7-133862652-DEL-84 | chr7-133862653-DEL-84 |
| chr7-133894377-DEL-60 | chr7-133894377-DEL-68 |
| chr7-133929468-DEL-17070 | chr7-133929470-DEL-17069 |
| chr7-133960086-DEL-400 | chr7-133960087-DEL-400 |
| chr7-133974465-DEL-57 | chr7-133974466-DEL-57 |

| | |
|---|---|
| chr7-134025729-DEL-96 | chr7-134025710-DEL-100 |
| chr7-134081612-DEL-1888 | chr7-134081613-DEL-1888 |
| chr7-134087225-DEL-66 | chr7-134087233-DEL-63 |
| chr7-134095830-DEL-70 | chr7-134095831-DEL-70 |
| chr7-134102695-DEL-60 | chr7-134102692-DEL-64 |
| chr7-134129515-DEL-56 | chr7-134129516-DEL-56 |
| chr7-134130481-DEL-1449 | chr7-134130482-DEL-1449 |
| chr7-134144604-DEL-174 | chr7-134144583-DEL-174 |
| chr7-134154427-DEL-836 | chr7-134154428-DEL-833 |
| chr7-134176652-DEL-88 | chr7-134176653-DEL-88 |
| chr7-134193242-DEL-54 | chr7-134193243-DEL-53 |
| chr7-134216334-DEL-386 | chr7-134216335-DEL-386 |
| chr7-134239436-DEL-249 | chr7-134239437-DEL-249 |
| chr7-134261205-DEL-118 | chr7-134261206-DEL-118 |
| chr7-134261910-DEL-125 | chr7-134261911-DEL-123 |
| chr7-134298745-DEL-112 | chr7-134298734-DEL-132 |
| chr7-134303274-DEL-204 | chr7-134303275-DEL-204 |
| chr7-134314306-DEL-1172 | chr7-134314307-DEL-1172 |
| chr7-134355680-DEL-56 | chr7-134355671-DEL-54 |
| chr7-134385630-DEL-77 | chr7-134385625-DEL-78 |
| chr7-134416596-DEL-116 | chr7-134416593-DEL-136 |
| chr7-134427570-DEL-64 | chr7-134427569-DEL-52 |
| chr7-134442429-DEL-527 | chr7-134442430-DEL-513 |
| chr7-134452321-DEL-70 | chr7-134452322-DEL-70 |
| chr7-134501669-DEL-115 | chr7-134501668-DEL-116 |
| chr7-134507089-DEL-1517 | chr7-134507090-DEL-1517 |
| chr7-134550307-DEL-1890 | chr7-134550308-DEL-1890 |
| chr7-134569800-DEL-748 | chr7-134569801-DEL-748 |
| chr7-134583872-DEL-2765 | chr7-134583873-DEL-2765 |
| chr7-134611226-DEL-61 | chr7-134611228-DEL-60 |
| chr7-134648444-DEL-104 | chr7-134648445-DEL-70 |
| chr7-134748564-DEL-148 | chr7-134748605-DEL-108 |
| chr7-134750522-DEL-116 | chr7-134750521-DEL-90 |
| chr7-134809901-DEL-86 | chr7-134809902-DEL-86 |
| chr7-134811271-DEL-635 | chr7-134811272-DEL-635 |
| chr7-134858002-DEL-54 | chr7-134857997-DEL-60 |
| chr7-134871816-DEL-67 | chr7-134871817-DEL-67 |
| chr7-134910864-DEL-67 | chr7-134910864-DEL-67 |
| chr7-134939431-DEL-3143 | chr7-134939423-DEL-3144 |
| chr7-134957852-DEL-6801 | chr7-134957853-DEL-6801 |
| chr7-134971160-DEL-51 | chr7-134971159-DEL-54 |
| chr7-134976902-DEL-200 | chr7-134976938-DEL-208 |
| chr7-134986964-DEL-95 | chr7-134986965-DEL-74 |
| chr7-134997812-DEL-143 | chr7-134997809-DEL-144 |
| chr7-135031437-DEL-60 | chr7-135031438-DEL-60 |
| chr7-135092844-DEL-522 | chr7-135092845-DEL-522 |
| chr7-135098535-DEL-1493 | chr7-135098536-DEL-1493 |
| chr7-135151652-DEL-266 | chr7-135151653-DEL-266 |
| chr7-135162610-DEL-237 | chr7-135162611-DEL-237 |
| chr7-135163174-DEL-104 | chr7-135163174-DEL-104 |
| chr7-135193861-DEL-2617 | chr7-135193862-DEL-2617 |
| chr7-135203895-DEL-80 | chr7-135203896-DEL-80 |
| chr7-135206456-DEL-64 | chr7-135206457-DEL-64 |
| chr7-135214985-DEL-76 | chr7-135214996-DEL-66 |
| chr7-135218464-DEL-114 | chr7-135218465-DEL-114 |
| chr7-135224612-DEL-822 | chr7-135224613-DEL-822 |
| chr7-135265712-DEL-51 | chr7-135265713-DEL-51 |
| chr7-135275643-DEL-111 | chr7-135275643-DEL-111 |
| chr7-135286043-DEL-79 | chr7-135286048-DEL-77 |
| chr7-135311783-DEL-231 | chr7-135311780-DEL-227 |
| chr7-135312746-DEL-53 | chr7-135312747-DEL-53 |
| chr7-135330096-DEL-5606 | chr7-135330097-DEL-5606 |
| chr7-135353940-DEL-167 | chr7-135353941-DEL-167 |
| chr7-135384515-DEL-58 | chr7-135384516-DEL-66 |
| chr7-135399196-DEL-92 | chr7-135399197-DEL-92 |
| chr7-135404046-DEL-220 | chr7-135404047-DEL-220 |
| chr7-135408317-DEL-61 | chr7-135408318-DEL-63 |
| chr7-135424592-DEL-104 | chr7-135424593-DEL-104 |
| chr7-135441510-DEL-86 | chr7-135441511-DEL-86 |
| chr7-135461382-DEL-161 | chr7-135461382-DEL-162 |
| chr7-135467334-DEL-86 | chr7-135467335-DEL-86 |
| chr7-135486071-DEL-60 | chr7-135486072-DEL-60 |
| chr7-135509613-DEL-156 | chr7-135509614-DEL-156 |
| chr7-135514327-DEL-197 | chr7-135514369-DEL-156 |
| chr7-135522728-DEL-2876 | chr7-135522729-DEL-2876 |
| chr7-135538016-DEL-60 | chr7-135538016-DEL-60 |
| chr7-135612109-DEL-59 | chr7-135612110-DEL-59 |
| chr7-135613866-DEL-103 | chr7-135613861-DEL-119 |
| chr7-135707233-DEL-6169 | chr7-135707274-DEL-6133 |
| chr7-135718960-DEL-672 | chr7-135718961-DEL-672 |

| | |
|---|---|
| chr7-135754675-DEL-234 | chr7-135754676-DEL-234 |
| chr7-135786973-DEL-52 | chr7-135786974-DEL-52 |
| chr7-135802196-DEL-76 | chr7-135802196-DEL-76 |
| chr7-135812507-DEL-54 | chr7-135812492-DEL-55 |
| chr7-135822941-DEL-94 | chr7-135822942-DEL-94 |
| chr7-135841632-DEL-5544 | chr7-135841633-DEL-5544 |
| chr7-135856504-DEL-57 | chr7-135856506-DEL-56 |
| chr7-135861295-DEL-52 | chr7-135861296-DEL-52 |
| chr7-135887809-DEL-596 | chr7-135887810-DEL-596 |
| chr7-135901873-DEL-88 | chr7-135901900-DEL-62 |
| chr7-135927165-DEL-564 | chr7-135927164-DEL-565 |
| chr7-135969138-DEL-503 | chr7-135969139-DEL-503 |
| chr7-135973670-DEL-1071 | chr7-135973671-DEL-1071 |
| chr7-136019730-DEL-53 | chr7-136019716-DEL-52 |
| chr7-136064790-DEL-212 | chr7-136064791-DEL-212 |
| chr7-136135722-DEL-287 | chr7-136135723-DEL-287 |
| chr7-136138672-DEL-117 | chr7-136138673-DEL-117 |
| chr7-136218994-DEL-95 | chr7-136218995-DEL-95 |
| chr7-136224430-DEL-97 | chr7-136224431-DEL-97 |
| chr7-136229285-DEL-444 | chr7-136229286-DEL-444 |
| chr7-136230073-DEL-667 | chr7-136230066-DEL-674 |
| chr7-136239826-DEL-70 | chr7-136239825-DEL-70 |
| chr7-136297133-DEL-283 | chr7-136297134-DEL-283 |
| chr7-136300911-DEL-61 | chr7-136300912-DEL-61 |
| chr7-136325172-DEL-122 | chr7-136325198-DEL-81 |
| chr7-136326979-DEL-178 | chr7-136326980-DEL-178 |
| chr7-136417059-DEL-104 | chr7-136417074-DEL-90 |
| chr7-136474235-DEL-66 | chr7-136474235-DEL-75 |
| chr7-136483107-DEL-103 | chr7-136483108-DEL-103 |
| chr7-136535434-DEL-58 | chr7-136535435-DEL-58 |
| chr7-136543378-DEL-74 | chr7-136543377-DEL-76 |
| chr7-136544998-DEL-354 | chr7-136544999-DEL-354 |
| chr7-136746947-DEL-199 | chr7-136746948-DEL-199 |
| chr7-136884058-DEL-77 | chr7-136884063-DEL-78 |
| chr7-136900792-DEL-73 | chr7-136900793-DEL-73 |
| chr7-136909383-DEL-59 | chr7-136909384-DEL-59 |
| chr7-136952869-DEL-198 | chr7-136952870-DEL-198 |
| chr7-136957993-DEL-150 | chr7-136957994-DEL-150 |
| chr7-136975131-DEL-657 | chr7-136975132-DEL-656 |
| chr7-136984255-DEL-51 | chr7-136984258-DEL-52 |
| chr7-136990374-DEL-180 | chr7-136990382-DEL-177 |
| chr7-137007177-DEL-87 | chr7-137007178-DEL-87 |
| chr7-137013608-DEL-369 | chr7-137013603-DEL-370 |
| chr7-137016741-DEL-5269 | chr7-137016742-DEL-5269 |
| chr7-137022671-DEL-101 | chr7-137022661-DEL-84 |
| chr7-137024267-DEL-167 | chr7-137024268-DEL-167 |
| chr7-137032021-DEL-160 | chr7-137031986-DEL-151 |
| chr7-137040936-DEL-207 | chr7-137040937-DEL-207 |
| chr7-137063406-DEL-80 | chr7-137063404-DEL-80 |
| chr7-137077159-DEL-70 | chr7-137077155-DEL-67 |
| chr7-137097671-DEL-218 | chr7-137097672-DEL-218 |
| chr7-137132035-DEL-63 | chr7-137132036-DEL-63 |
| chr7-137144490-DEL-123 | chr7-137144491-DEL-123 |
| chr7-137180855-DEL-108 | chr7-137180856-DEL-108 |
| chr7-137241722-DEL-205 | chr7-137241723-DEL-205 |
| chr7-137243386-DEL-5330 | chr7-137243387-DEL-5330 |
| chr7-137259065-DEL-51 | chr7-137259035-DEL-70 |
| chr7-137261352-DEL-6415 | chr7-137261353-DEL-6415 |
| chr7-137351397-DEL-51 | chr7-137351398-DEL-51 |
| chr7-137383161-DEL-809 | chr7-137383162-DEL-809 |
| chr7-137394197-DEL-1324 | chr7-137394198-DEL-1324 |
| chr7-137400915-DEL-733 | chr7-137400928-DEL-721 |
| chr7-137425593-DEL-1439 | chr7-137425594-DEL-1439 |
| chr7-137427681-DEL-70 | chr7-137427728-DEL-68 |
| chr7-137448939-DEL-110 | chr7-137448940-DEL-110 |
| chr7-137458920-DEL-644 | chr7-137458921-DEL-644 |
| chr7-137496092-DEL-5448 | chr7-137496093-DEL-5448 |
| chr7-137524587-DEL-60 | chr7-137524588-DEL-60 |
| chr7-137534994-DEL-158 | chr7-137534993-DEL-157 |
| chr7-137538759-DEL-564 | chr7-137538760-DEL-564 |
| chr7-137562751-DEL-1459 | chr7-137562752-DEL-1459 |
| chr7-137565131-DEL-61 | chr7-137565132-DEL-61 |
| chr7-137585732-DEL-116 | chr7-137585733-DEL-116 |
| chr7-137625970-DEL-159 | chr7-137625971-DEL-160 |
| chr7-137626984-DEL-52 | chr7-137627000-DEL-56 |
| chr7-137646486-DEL-60 | chr7-137646487-DEL-60 |
| chr7-137649942-DEL-169 | chr7-137649945-DEL-167 |
| chr7-137712223-DEL-86 | chr7-137712224-DEL-86 |
| chr7-137716266-DEL-80 | chr7-137716266-DEL-80 |
| chr7-137787606-DEL-1605 | chr7-137787607-DEL-1604 |

| | |
|---|---|
| chr7-137792527-DEL-66 | chr7-137792528-DEL-66 |
| chr7-137793929-DEL-287 | chr7-137793930-DEL-287 |
| chr7-137798266-DEL-56 | chr7-137798267-DEL-56 |
| chr7-137830031-DEL-59 | chr7-137830032-DEL-59 |
| chr7-137853060-DEL-66 | chr7-137853061-DEL-60 |
| chr7-137863470-DEL-5309 | chr7-137863471-DEL-5309 |
| chr7-137883045-DEL-104 | chr7-137883046-DEL-64 |
| chr7-137884663-DEL-55 | chr7-137884663-DEL-55 |
| chr7-137954128-DEL-108 | chr7-137954083-DEL-113 |
| chr7-137955152-DEL-372 | chr7-137955153-DEL-372 |
| chr7-137977705-DEL-892 | chr7-137977706-DEL-892 |
| chr7-137982592-DEL-102 | chr7-137982593-DEL-102 |
| chr7-138012116-DEL-66 | chr7-138012117-DEL-66 |
| chr7-138025782-DEL-179 | chr7-138025783-DEL-179 |
| chr7-138030597-DEL-60 | chr7-138030598-DEL-60 |
| chr7-138065222-DEL-387 | chr7-138065223-DEL-387 |
| chr7-138124679-DEL-88 | chr7-138124695-DEL-87 |
| chr7-138128362-DEL-59 | chr7-138128363-DEL-59 |
| chr7-138134725-DEL-557 | chr7-138134723-DEL-558 |
| chr7-138138050-DEL-63 | chr7-138138051-DEL-63 |
| chr7-138154927-DEL-59 | chr7-138154928-DEL-59 |
| chr7-138166220-DEL-153 | chr7-138166221-DEL-153 |
| chr7-138174813-DEL-1166 | chr7-138174814-DEL-1166 |
| chr7-138175760-DEL-3191 | chr7-138175761-DEL-3191 |
| chr7-138192082-DEL-1592 | chr7-138192083-DEL-1592 |
| chr7-138215658-DEL-977 | chr7-138215654-DEL-978 |
| chr7-138229724-DEL-162 | chr7-138229725-DEL-162 |
| chr7-138231659-DEL-175 | chr7-138231660-DEL-175 |
| chr7-138259259-DEL-66 | chr7-138259260-DEL-66 |
| chr7-138267244-DEL-341 | chr7-138267245-DEL-341 |
| chr7-138279285-DEL-84 | chr7-138279282-DEL-78 |
| chr7-138290667-DEL-7297 | chr7-138290668-DEL-7297 |
| chr7-138301620-DEL-117 | chr7-138301621-DEL-117 |
| chr7-138315069-DEL-160 | chr7-138315070-DEL-141 |
| chr7-138315962-DEL-78 | chr7-138315963-DEL-78 |
| chr7-138320029-DEL-698 | chr7-138320030-DEL-697 |
| chr7-138323000-DEL-152 | chr7-138323001-DEL-152 |
| chr7-138355702-DEL-203 | chr7-138355702-DEL-203 |
| chr7-138384277-DEL-63 | chr7-138384278-DEL-63 |
| chr7-138396240-DEL-56 | chr7-138396241-DEL-56 |
| chr7-138400694-DEL-1894 | chr7-138400695-DEL-1894 |
| chr7-138402954-DEL-617 | chr7-138402955-DEL-617 |
| chr7-138423627-DEL-207 | chr7-138423628-DEL-207 |
| chr7-138439053-DEL-79 | chr7-138439054-DEL-79 |
| chr7-138442752-DEL-94 | chr7-138442753-DEL-92 |
| chr7-138495082-DEL-60 | chr7-138495063-DEL-80 |
| chr7-138512294-DEL-262 | chr7-138512295-DEL-262 |
| chr7-138518658-DEL-498 | chr7-138518658-DEL-498 |
| chr7-138520449-DEL-199 | chr7-138520446-DEL-198 |
| chr7-138529788-DEL-56 | chr7-138529792-DEL-53 |
| chr7-138531970-DEL-102 | chr7-138532011-DEL-62 |
| chr7-138534090-DEL-65 | chr7-138534094-DEL-51 |
| chr7-138553069-DEL-62 | chr7-138553047-DEL-61 |
| chr7-138554259-DEL-82 | chr7-138554257-DEL-83 |
| chr7-138556305-DEL-416 | chr7-138556306-DEL-416 |
| chr7-138557025-DEL-647 | chr7-138557026-DEL-647 |
| chr7-138575570-DEL-137 | chr7-138575571-DEL-132 |
| chr7-138594256-DEL-227 | chr7-138594257-DEL-227 |
| chr7-138604956-DEL-1886 | chr7-138604957-DEL-1886 |
| chr7-138622828-DEL-96 | chr7-138622829-DEL-96 |
| chr7-138627329-DEL-233 | chr7-138627329-DEL-233 |
| chr7-138664564-DEL-84 | chr7-138664565-DEL-84 |
| chr7-138691846-DEL-6401 | chr7-138691847-DEL-6401 |
| chr7-138761956-DEL-211 | chr7-138761957-DEL-211 |
| chr7-138763358-DEL-999 | chr7-138763359-DEL-999 |
| chr7-138775274-DEL-81 | chr7-138775248-DEL-108 |
| chr7-138797483-DEL-86 | chr7-138797484-DEL-86 |
| chr7-138800797-DEL-54 | chr7-138800798-DEL-54 |
| chr7-138817730-DEL-140 | chr7-138817732-DEL-139 |
| chr7-138870851-DEL-198 | chr7-138870852-DEL-198 |
| chr7-138939277-DEL-209 | chr7-138939278-DEL-209 |
| chr7-138953647-DEL-203 | chr7-138953647-DEL-203 |
| chr7-138959088-DEL-200 | chr7-138959089-DEL-200 |
| chr7-139017970-DEL-96 | chr7-139017971-DEL-96 |
| chr7-139051161-DEL-86 | chr7-139051162-DEL-86 |
| chr7-139078982-DEL-237 | chr7-139078983-DEL-237 |
| chr7-139082539-DEL-212 | chr7-139082536-DEL-231 |
| chr7-139104377-DEL-363 | chr7-139104378-DEL-363 |
| chr7-139105085-DEL-198 | chr7-139105090-DEL-194 |
| chr7-139126098-DEL-69 | chr7-139126095-DEL-68 |

| | |
|---|---|
| chr7-139185250-DEL-100 | chr7-139185258-DEL-97 |
| chr7-139214078-DEL-160 | chr7-139214079-DEL-160 |
| chr7-139239873-DEL-52 | chr7-139239874-DEL-52 |
| chr7-139252900-DEL-93 | chr7-139252909-DEL-84 |
| chr7-139344278-DEL-148 | chr7-139344279-DEL-148 |
| chr7-139372180-DEL-53 | chr7-139372175-DEL-54 |
| chr7-139378977-DEL-52 | chr7-139378985-DEL-51 |
| chr7-139394782-DEL-88 | chr7-139394799-DEL-64 |
| chr7-139447236-DEL-89 | chr7-139447272-DEL-69 |
| chr7-139454449-DEL-73 | chr7-139454450-DEL-73 |
| chr7-139473695-DEL-59 | chr7-139473696-DEL-59 |
| chr7-139480232-DEL-52 | chr7-139480233-DEL-52 |
| chr7-139487130-DEL-130 | chr7-139487097-DEL-131 |
| chr7-139507942-DEL-200 | chr7-139507943-DEL-200 |
| chr7-139532791-DEL-114 | chr7-139532780-DEL-112 |
| chr7-139579260-DEL-176 | chr7-139579261-DEL-176 |
| chr7-139582009-DEL-68 | chr7-139581982-DEL-112 |
| chr7-139594628-DEL-13020 | chr7-139594629-DEL-13020 |
| chr7-139632534-DEL-200 | chr7-139632535-DEL-200 |
| chr7-139662687-DEL-87 | chr7-139662704-DEL-89 |
| chr7-139665913-DEL-172 | chr7-139665902-DEL-167 |
| chr7-139668203-DEL-136 | chr7-139668204-DEL-136 |
| chr7-139687460-DEL-300 | chr7-139687461-DEL-300 |
| chr7-139700857-DEL-166 | chr7-139700858-DEL-166 |
| chr7-139713351-DEL-281 | chr7-139713352-DEL-281 |
| chr7-139726291-DEL-91 | chr7-139726289-DEL-90 |
| chr7-139728195-DEL-60 | chr7-139728195-DEL-60 |
| chr7-139789439-DEL-66 | chr7-139789442-DEL-64 |
| chr7-139792709-DEL-82 | chr7-139792710-DEL-82 |
| chr7-139795778-DEL-97 | chr7-139795779-DEL-97 |
| chr7-139799075-DEL-78 | chr7-139799070-DEL-78 |
| chr7-139801220-DEL-109 | chr7-139801215-DEL-127 |
| chr7-139815176-DEL-117 | chr7-139815177-DEL-117 |
| chr7-139893271-DEL-530 | chr7-139893272-DEL-530 |
| chr7-139917746-DEL-81 | chr7-139917747-DEL-81 |
| chr7-139929079-DEL-498 | chr7-139929080-DEL-498 |
| chr7-139940548-DEL-72 | chr7-139940549-DEL-72 |
| chr7-139970475-DEL-433 | chr7-139970476-DEL-433 |
| chr7-139973856-DEL-2260 | chr7-139973857-DEL-2260 |
| chr7-140001230-DEL-106 | chr7-140001199-DEL-129 |
| chr7-140034799-DEL-51 | chr7-140034787-DEL-52 |
| chr7-140036838-DEL-242 | chr7-140036832-DEL-249 |
| chr7-140051565-DEL-90 | chr7-140051566-DEL-90 |
| chr7-140086452-DEL-144 | chr7-140086448-DEL-145 |
| chr7-140088981-DEL-88 | chr7-140088941-DEL-104 |
| chr7-140089501-DEL-696 | chr7-140089502-DEL-696 |
| chr7-140097201-DEL-2376 | chr7-140097201-DEL-2376 |
| chr7-140102746-DEL-6193 | chr7-140102731-DEL-6208 |
| chr7-140110863-DEL-2400 | chr7-140110863-DEL-2400 |
| chr7-140116035-DEL-66 | chr7-140116028-DEL-60 |
| chr7-140125652-DEL-804 | chr7-140125653-DEL-804 |
| chr7-140139877-DEL-125 | chr7-140139848-DEL-146 |
| chr7-140194138-DEL-6593 | chr7-140194139-DEL-6593 |
| chr7-140194382-DEL-213 | chr7-140194381-DEL-214 |
| chr7-140215156-DEL-438 | chr7-140215157-DEL-438 |
| chr7-140250444-DEL-2901 | chr7-140250445-DEL-2901 |
| chr7-140264509-DEL-60 | chr7-140264510-DEL-60 |
| chr7-140273043-DEL-3068 | chr7-140273044-DEL-3068 |
| chr7-140275199-DEL-219 | chr7-140275200-DEL-218 |
| chr7-140283114-DEL-96 | chr7-140283119-DEL-92 |
| chr7-140285698-DEL-51 | chr7-140285699-DEL-51 |
| chr7-140303766-DEL-807 | chr7-140303767-DEL-807 |
| chr7-140318109-DEL-3189 | chr7-140318110-DEL-3189 |
| chr7-140369963-DEL-912 | chr7-140369964-DEL-912 |
| chr7-140376613-DEL-209 | chr7-140376614-DEL-209 |
| chr7-140389320-DEL-112 | chr7-140389321-DEL-112 |
| chr7-140405139-DEL-200 | chr7-140405140-DEL-200 |
| chr7-140443988-DEL-53 | chr7-140443989-DEL-53 |
| chr7-140470661-DEL-1514 | chr7-140470662-DEL-1514 |
| chr7-140485228-DEL-170 | chr7-140485229-DEL-170 |
| chr7-140505932-DEL-166 | chr7-140505933-DEL-166 |
| chr7-140541653-DEL-217 | chr7-140541654-DEL-217 |
| chr7-140545459-DEL-60 | chr7-140545460-DEL-90 |
| chr7-140558254-DEL-676 | chr7-140558247-DEL-672 |
| chr7-140562742-DEL-211 | chr7-140562743-DEL-211 |
| chr7-140573977-DEL-119 | chr7-140573978-DEL-113 |
| chr7-140575909-DEL-78 | chr7-140575910-DEL-78 |
| chr7-140607588-DEL-371 | chr7-140607612-DEL-362 |
| chr7-140630036-DEL-1531 | chr7-140630037-DEL-1531 |
| chr7-140631404-DEL-52 | chr7-140631405-DEL-52 |

| | |
|---|---|
| chr7-140664725-DEL-237 | chr7-140664733-DEL-236 |
| chr7-140701796-DEL-232 | chr7-140701797-DEL-232 |
| chr7-140713567-DEL-93 | chr7-140713578-DEL-83 |
| chr7-140720666-DEL-233 | chr7-140720667-DEL-233 |
| chr7-140737537-DEL-221 | chr7-140737538-DEL-221 |
| chr7-140804553-DEL-215 | chr7-140804554-DEL-215 |
| chr7-140815748-DEL-211 | chr7-140815749-DEL-211 |
| chr7-140818266-DEL-1159 | chr7-140818267-DEL-1159 |
| chr7-140826038-DEL-207 | chr7-140826039-DEL-207 |
| chr7-140832638-DEL-244 | chr7-140832639-DEL-244 |
| chr7-140835049-DEL-190 | chr7-140835050-DEL-190 |
| chr7-140846455-DEL-66 | chr7-140846456-DEL-66 |
| chr7-140859812-DEL-204 | chr7-140859813-DEL-204 |
| chr7-140892094-DEL-117 | chr7-140892095-DEL-117 |
| chr7-140928346-DEL-224 | chr7-140928348-DEL-223 |
| chr7-140936620-DEL-225 | chr7-140936621-DEL-225 |
| chr7-140955210-DEL-189 | chr7-140955211-DEL-189 |
| chr7-140977263-DEL-190 | chr7-140977264-DEL-190 |
| chr7-140985177-DEL-168 | chr7-140985178-DEL-168 |
| chr7-140986410-DEL-848 | chr7-140986411-DEL-848 |
| chr7-140995356-DEL-178 | chr7-140995352-DEL-179 |
| chr7-141051746-DEL-195 | chr7-141051747-DEL-195 |
| chr7-141080357-DEL-285 | chr7-141080358-DEL-285 |
| chr7-141091772-DEL-387 | chr7-141091772-DEL-387 |
| chr7-141094282-DEL-191 | chr7-141094281-DEL-193 |
| chr7-141101264-DEL-161 | chr7-141101263-DEL-163 |
| chr7-141122298-DEL-209 | chr7-141122299-DEL-209 |
| chr7-141151940-DEL-67 | chr7-141151891-DEL-99 |
| chr7-141218118-DEL-519 | chr7-141218119-DEL-519 |
| chr7-141220519-DEL-300 | chr7-141220519-DEL-300 |
| chr7-141222246-DEL-208 | chr7-141222247-DEL-207 |
| chr7-141245456-DEL-229 | chr7-141245457-DEL-229 |
| chr7-141256980-DEL-206 | chr7-141256981-DEL-206 |
| chr7-141271176-DEL-73 | chr7-141271169-DEL-60 |
| chr7-141283016-DEL-59 | chr7-141283017-DEL-59 |
| chr7-141296177-DEL-272 | chr7-141296161-DEL-273 |
| chr7-141297283-DEL-144 | chr7-141297283-DEL-144 |
| chr7-141303798-DEL-207 | chr7-141303823-DEL-200 |
| chr7-141313069-DEL-83 | chr7-141313062-DEL-60 |
| chr7-141313755-DEL-4044 | chr7-141313757-DEL-4043 |
| chr7-141363243-DEL-96 | chr7-141363243-DEL-96 |
| chr7-141413518-DEL-54 | chr7-141413519-DEL-54 |
| chr7-141417106-DEL-54 | chr7-141417107-DEL-54 |
| chr7-141490316-DEL-70 | chr7-141490317-DEL-70 |
| chr7-141520503-DEL-1589 | chr7-141520504-DEL-1589 |
| chr7-141525117-DEL-426 | chr7-141525118-DEL-426 |
| chr7-141582183-DEL-92 | chr7-141582208-DEL-62 |
| chr7-141625733-DEL-1894 | chr7-141625734-DEL-1894 |
| chr7-141666832-DEL-71 | chr7-141666836-DEL-50 |
| chr7-141672028-DEL-62 | chr7-141672018-DEL-86 |
| chr7-141674180-DEL-60 | chr7-141674180-DEL-60 |
| chr7-141697536-DEL-78 | chr7-141697537-DEL-60 |
| chr7-141708373-DEL-60 | chr7-141708373-DEL-60 |
| chr7-141711668-DEL-217 | chr7-141711691-DEL-195 |
| chr7-141749479-DEL-168 | chr7-141749480-DEL-168 |
| chr7-141749867-DEL-4672 | chr7-141749868-DEL-4672 |
| chr7-141755268-DEL-66 | chr7-141755267-DEL-66 |
| chr7-141768626-DEL-5854 | chr7-141768627-DEL-5854 |
| chr7-141780849-DEL-215 | chr7-141780850-DEL-215 |
| chr7-141782952-DEL-90 | chr7-141782953-DEL-90 |
| chr7-141794524-DEL-60 | chr7-141794525-DEL-60 |
| chr7-141805029-DEL-69 | chr7-141805030-DEL-69 |
| chr7-141813480-DEL-418 | chr7-141813481-DEL-418 |
| chr7-141821403-DEL-323 | chr7-141821404-DEL-323 |
| chr7-141832017-DEL-58 | chr7-141832018-DEL-58 |
| chr7-141859874-DEL-146 | chr7-141859875-DEL-146 |
| chr7-141867041-DEL-61 | chr7-141867038-DEL-79 |
| chr7-141897390-DEL-52 | chr7-141897391-DEL-52 |
| chr7-141902321-DEL-93 | chr7-141902363-DEL-57 |
| chr7-141958906-DEL-112 | chr7-141958907-DEL-112 |
| chr7-141960052-DEL-288 | chr7-141960094-DEL-247 |
| chr7-141961952-DEL-72 | chr7-141961961-DEL-62 |
| chr7-142008244-DEL-65 | chr7-142008208-DEL-63 |
| chr7-142012423-DEL-60 | chr7-142012406-DEL-78 |
| chr7-142065331-DEL-245 | chr7-142065333-DEL-244 |
| chr7-142096090-DEL-58 | chr7-142096091-DEL-58 |
| chr7-142264008-DEL-192 | chr7-142264009-DEL-192 |
| chr7-142280435-DEL-174 | chr7-142280436-DEL-174 |
| chr7-142285559-DEL-414 | chr7-142285560-DEL-413 |
| chr7-142290334-DEL-267 | chr7-142290335-DEL-267 |

| | |
|---|---|
| chr7-142310897-DEL-724 | chr7-142310907-DEL-727 |
| chr7-142321201-DEL-350 | chr7-142321202-DEL-350 |
| chr7-142328212-DEL-7090 | chr7-142328213-DEL-7090 |
| chr7-142377162-DEL-58 | chr7-142377167-DEL-54 |
| chr7-142377829-DEL-456 | chr7-142377830-DEL-456 |
| chr7-142386113-DEL-222 | chr7-142386114-DEL-222 |
| chr7-142404423-DEL-123 | chr7-142404425-DEL-120 |
| chr7-142422576-DEL-371 | chr7-142422577-DEL-371 |
| chr7-142428320-DEL-1691 | chr7-142428321-DEL-1691 |
| chr7-142477724-DEL-72 | chr7-142477741-DEL-52 |
| chr7-142478517-DEL-556 | chr7-142478518-DEL-556 |
| chr7-142481519-DEL-65 | chr7-142481520-DEL-65 |
| chr7-142509174-DEL-57 | chr7-142509168-DEL-67 |
| chr7-142651332-DEL-182 | chr7-142651333-DEL-183 |
| chr7-142714615-DEL-195 | chr7-142714611-DEL-192 |
| chr7-142725871-DEL-244 | chr7-142725868-DEL-248 |
| chr7-142743635-DEL-77 | chr7-142743594-DEL-78 |
| chr7-142964850-DEL-56 | chr7-142964850-DEL-57 |
| chr7-142986063-DEL-89 | chr7-142986064-DEL-89 |
| chr7-143059437-DEL-210 | chr7-143059432-DEL-216 |
| chr7-143080490-DEL-214 | chr7-143080491-DEL-214 |
| chr7-143118303-DEL-69 | chr7-143118304-DEL-69 |
| chr7-143121697-DEL-184 | chr7-143121698-DEL-184 |
| chr7-143201162-DEL-138 | chr7-143201174-DEL-127 |
| chr7-143220412-DEL-520 | chr7-143220413-DEL-520 |
| chr7-143225391-DEL-6586 | chr7-143225386-DEL-6587 |
| chr7-143241101-DEL-51 | chr7-143241101-DEL-51 |
| chr7-143251766-DEL-102 | chr7-143251767-DEL-84 |
| chr7-143298779-DEL-56 | chr7-143298780-DEL-56 |
| chr7-143324294-DEL-88 | chr7-143324272-DEL-66 |
| chr7-143333716-DEL-68 | chr7-143333681-DEL-72 |
| chr7-143370647-DEL-79 | chr7-143370640-DEL-63 |
| chr7-143371413-DEL-166 | chr7-143371415-DEL-167 |
| chr7-143377447-DEL-1400 | chr7-143377448-DEL-1400 |
| chr7-143377895-DEL-497 | chr7-143377896-DEL-497 |
| chr7-143381836-DEL-807 | chr7-143381837-DEL-805 |
| chr7-143403083-DEL-6401 | chr7-143403084-DEL-6401 |
| chr7-143409620-DEL-7660 | chr7-143409621-DEL-7660 |
| chr7-143429076-DEL-504 | chr7-143429077-DEL-504 |
| chr7-143463201-DEL-52 | chr7-143463181-DEL-70 |
| chr7-143505906-DEL-58 | chr7-143505907-DEL-58 |
| chr7-143554156-DEL-59 | chr7-143554155-DEL-60 |
| chr7-143611057-DEL-70 | chr7-143611073-DEL-55 |
| chr7-143633614-DEL-54 | chr7-143633595-DEL-50 |
| chr7-143635288-DEL-72 | chr7-143635288-DEL-72 |
| chr7-143642147-DEL-80 | chr7-143642142-DEL-100 |
| chr7-143652496-DEL-60 | chr7-143652497-DEL-60 |
| chr7-143707293-DEL-114 | chr7-143707294-DEL-114 |
| chr7-143708589-DEL-1301 | chr7-143708590-DEL-1301 |
| chr7-143710068-DEL-1353 | chr7-143710069-DEL-1353 |
| chr7-143737922-DEL-60 | chr7-143737923-DEL-60 |
| chr7-143758813-DEL-85 | chr7-143758811-DEL-50 |
| chr7-143779246-DEL-189 | chr7-143779247-DEL-185 |
| chr7-143824218-DEL-308 | chr7-143824197-DEL-296 |
| chr7-143864289-DEL-68 | chr7-143864250-DEL-81 |
| chr7-143869130-DEL-236 | chr7-143869131-DEL-236 |
| chr7-143880813-DEL-157 | chr7-143880821-DEL-150 |
| chr7-143903292-DEL-5041 | chr7-143903293-DEL-5041 |
| chr7-143913544-DEL-1548 | chr7-143913545-DEL-1548 |
| chr7-143930731-DEL-180 | chr7-143930732-DEL-180 |
| chr7-143972925-DEL-894 | chr7-143972925-DEL-894 |
| chr7-143981542-DEL-590 | chr7-143981543-DEL-586 |
| chr7-143988349-DEL-763 | chr7-143988350-DEL-763 |
| chr7-144053378-DEL-128 | chr7-144053379-DEL-144 |
| chr7-144064144-DEL-314 | chr7-144064145-DEL-314 |
| chr7-144124744-DEL-112 | chr7-144124745-DEL-127 |
| chr7-144196794-DEL-108 | chr7-144196794-DEL-108 |
| chr7-144234194-DEL-215 | chr7-144234195-DEL-215 |
| chr7-144241444-DEL-63 | chr7-144241445-DEL-63 |
| chr7-144296061-DEL-76 | chr7-144296062-DEL-76 |
| chr7-144330836-DEL-92 | chr7-144330830-DEL-93 |
| chr7-144399303-DEL-204 | chr7-144399304-DEL-204 |
| chr7-144421831-DEL-127 | chr7-144421832-DEL-127 |
| chr7-144424017-DEL-235 | chr7-144424018-DEL-235 |
| chr7-144448242-DEL-51 | chr7-144448234-DEL-59 |
| chr7-144532578-DEL-146 | chr7-144532579-DEL-195 |
| chr7-144574394-DEL-52 | chr7-144574395-DEL-52 |
| chr7-144607613-DEL-68 | chr7-144607587-DEL-108 |
| chr7-144678305-DEL-201 | chr7-144678306-DEL-201 |
| chr7-144689219-DEL-71 | chr7-144689232-DEL-64 |

| | |
|---|---|
| chr7-144691261-DEL-111 | chr7-144691262-DEL-111 |
| chr7-144765118-DEL-58 | chr7-144765119-DEL-54 |
| chr7-144867153-DEL-87 | chr7-144867153-DEL-88 |
| chr7-144869905-DEL-52 | chr7-144869906-DEL-52 |
| chr7-144874063-DEL-80 | chr7-144874082-DEL-50 |
| chr7-144884200-DEL-76 | chr7-144884201-DEL-56 |
| chr8-3135397-DEL-76 | chr8-3135398-DEL-76 |
| chr8-3246571-DEL-72 | chr8-3246547-DEL-96 |
| chr8-3290669-DEL-121 | chr8-3290670-DEL-121 |
| chr8-3307483-DEL-56 | chr8-3307484-DEL-60 |
| chr8-3311663-DEL-54 | chr8-3311664-DEL-54 |
| chr8-3323161-DEL-202 | chr8-3323162-DEL-202 |
| chr8-3340553-DEL-64 | chr8-3340539-DEL-74 |
| chr8-3363446-DEL-62 | chr8-3363447-DEL-62 |
| chr8-3388280-DEL-7140 | chr8-3388281-DEL-7140 |
| chr8-3401352-DEL-314 | chr8-3401353-DEL-314 |
| chr8-3423386-DEL-1149 | chr8-3423387-DEL-1149 |
| chr8-3423889-DEL-1664 | chr8-3423890-DEL-1664 |
| chr8-3440009-DEL-343 | chr8-3440010-DEL-343 |
| chr8-3468141-DEL-65 | chr8-3468142-DEL-65 |
| chr8-3485918-DEL-57 | chr8-3485919-DEL-57 |
| chr8-3534964-DEL-225 | chr8-3534965-DEL-225 |
| chr8-3535906-DEL-55 | chr8-3535909-DEL-53 |
| chr8-3538756-DEL-59 | chr8-3538739-DEL-57 |
| chr8-3602020-DEL-1581 | chr8-3602021-DEL-1581 |
| chr8-3603095-DEL-203 | chr8-3603096-DEL-203 |
| chr8-3605793-DEL-193 | chr8-3605794-DEL-193 |
| chr8-3619156-DEL-66 | chr8-3619157-DEL-66 |
| chr8-3631852-DEL-225 | chr8-3631853-DEL-225 |
| chr8-3633735-DEL-515 | chr8-3633714-DEL-537 |
| chr8-3667130-DEL-210 | chr8-3667131-DEL-210 |
| chr8-3690512-DEL-216 | chr8-3690513-DEL-216 |
| chr8-3712650-DEL-54 | chr8-3712637-DEL-55 |
| chr8-3727601-DEL-87 | chr8-3727602-DEL-87 |
| chr8-3754664-DEL-87 | chr8-3754665-DEL-87 |
| chr8-3775020-DEL-183 | chr8-3775021-DEL-183 |
| chr8-3776246-DEL-56 | chr8-3776246-DEL-57 |
| chr8-3804967-DEL-7105 | chr8-3804968-DEL-7105 |
| chr8-3828796-DEL-53 | chr8-3828797-DEL-53 |
| chr8-3841474-DEL-221 | chr8-3841475-DEL-221 |
| chr8-3873606-DEL-5651 | chr8-3873607-DEL-5651 |
| chr8-3884986-DEL-1009 | chr8-3884986-DEL-1009 |
| chr8-3911548-DEL-182 | chr8-3911549-DEL-182 |
| chr8-3941690-DEL-51 | chr8-3941691-DEL-51 |
| chr8-3957140-DEL-7507 | chr8-3957141-DEL-7507 |
| chr8-3992610-DEL-224 | chr8-3992611-DEL-224 |
| chr8-3995957-DEL-6974 | chr8-3995958-DEL-6974 |
| chr8-4010232-DEL-208 | chr8-4010233-DEL-208 |
| chr8-4116519-DEL-82 | chr8-4116518-DEL-83 |
| chr8-4119592-DEL-400 | chr8-4119593-DEL-400 |
| chr8-4132859-DEL-71 | chr8-4132860-DEL-71 |
| chr8-4162083-DEL-56 | chr8-4162084-DEL-56 |
| chr8-4162702-DEL-111 | chr8-4162694-DEL-112 |
| chr8-4163711-DEL-109 | chr8-4163704-DEL-105 |
| chr8-4203849-DEL-210 | chr8-4203850-DEL-210 |
| chr8-4207649-DEL-196 | chr8-4207650-DEL-196 |
| chr8-4239708-DEL-91 | chr8-4239735-DEL-64 |
| chr8-4262034-DEL-86 | chr8-4261996-DEL-132 |
| chr8-4279523-DEL-196 | chr8-4279526-DEL-198 |
| chr8-4322215-DEL-220 | chr8-4322216-DEL-220 |
| chr8-4369750-DEL-192 | chr8-4369751-DEL-192 |
| chr8-4385206-DEL-242 | chr8-4385207-DEL-242 |
| chr8-4393300-DEL-52 | chr8-4393300-DEL-52 |
| chr8-4405003-DEL-204 | chr8-4405016-DEL-202 |
| chr8-4423780-DEL-6618 | chr8-4423781-DEL-6618 |
| chr8-4432324-DEL-212 | chr8-4432325-DEL-212 |
| chr8-4458165-DEL-123 | chr8-4458166-DEL-123 |
| chr8-4474225-DEL-86 | chr8-4474226-DEL-86 |
| chr8-4496837-DEL-203 | chr8-4496838-DEL-203 |
| chr8-4505814-DEL-1891 | chr8-4505810-DEL-1892 |
| chr8-4516009-DEL-411 | chr8-4516010-DEL-411 |
| chr8-4536394-DEL-127 | chr8-4536391-DEL-128 |
| chr8-4541330-DEL-160 | chr8-4541331-DEL-160 |
| chr8-4581941-DEL-1688 | chr8-4581901-DEL-1689 |
| chr8-4600938-DEL-207 | chr8-4600939-DEL-207 |
| chr8-4614941-DEL-58 | chr8-4614946-DEL-54 |
| chr8-4621170-DEL-7332 | chr8-4621171-DEL-7332 |
| chr8-4633843-DEL-84 | chr8-4633844-DEL-84 |
| chr8-4644316-DEL-485 | chr8-4644317-DEL-485 |
| chr8-4648391-DEL-205 | chr8-4648392-DEL-205 |

| | |
|---|---|
| chr8-4648986-DEL-208 | chr8-4648987-DEL-208 |
| chr8-4658896-DEL-76 | chr8-4658897-DEL-76 |
| chr8-4669989-DEL-224 | chr8-4669990-DEL-224 |
| chr8-4687685-DEL-203 | chr8-4687686-DEL-203 |
| chr8-4711670-DEL-405 | chr8-4711671-DEL-405 |
| chr8-4717820-DEL-177 | chr8-4717821-DEL-165 |
| chr8-4766203-DEL-52 | chr8-4766204-DEL-52 |
| chr8-4768114-DEL-78 | chr8-4768115-DEL-78 |
| chr8-4780113-DEL-87 | chr8-4780119-DEL-84 |
| chr8-4803966-DEL-80 | chr8-4803955-DEL-78 |
| chr8-4804719-DEL-1031 | chr8-4804720-DEL-1031 |
| chr8-4804878-DEL-191 | chr8-4804879-DEL-191 |
| chr8-4805445-DEL-77 | chr8-4805451-DEL-75 |
| chr8-4807195-DEL-77 | chr8-4807195-DEL-77 |
| chr8-4831188-DEL-1646 | chr8-4831188-DEL-1645 |
| chr8-4852405-DEL-6269 | chr8-4852406-DEL-6269 |
| chr8-4863152-DEL-153 | chr8-4863153-DEL-141 |
| chr8-4878388-DEL-85 | chr8-4878431-DEL-54 |
| chr8-4885173-DEL-330 | chr8-4885174-DEL-330 |
| chr8-4968660-DEL-1358 | chr8-4968661-DEL-1358 |
| chr8-5003969-DEL-587 | chr8-5003970-DEL-587 |
| chr8-5009274-DEL-8738 | chr8-5009275-DEL-8738 |
| chr8-5026622-DEL-52 | chr8-5026623-DEL-52 |
| chr8-5073206-DEL-178 | chr8-5073207-DEL-178 |
| chr8-5077752-DEL-542 | chr8-5077753-DEL-542 |
| chr8-5130153-DEL-546 | chr8-5130152-DEL-546 |
| chr8-5135460-DEL-61 | chr8-5135461-DEL-61 |
| chr8-5139459-DEL-1399 | chr8-5139460-DEL-1399 |
| chr8-5150168-DEL-64 | chr8-5150169-DEL-57 |
| chr8-5161558-DEL-753 | chr8-5161559-DEL-753 |
| chr8-5180494-DEL-1091 | chr8-5180461-DEL-1089 |
| chr8-5194500-DEL-53 | chr8-5194497-DEL-51 |
| chr8-5228263-DEL-2193 | chr8-5228264-DEL-2193 |
| chr8-5263134-DEL-92 | chr8-5263137-DEL-90 |
| chr8-5267509-DEL-6720 | chr8-5267510-DEL-6720 |
| chr8-5291888-DEL-6519 | chr8-5291889-DEL-6519 |
| chr8-5311585-DEL-112 | chr8-5311586-DEL-112 |
| chr8-5314963-DEL-6286 | chr8-5314959-DEL-6287 |
| chr8-5337936-DEL-2607 | chr8-5337937-DEL-2607 |
| chr8-5352422-DEL-6643 | chr8-5352423-DEL-6650 |
| chr8-5384594-DEL-70 | chr8-5384595-DEL-70 |
| chr8-5393613-DEL-170 | chr8-5393614-DEL-170 |
| chr8-5423263-DEL-286 | chr8-5423264-DEL-286 |
| chr8-5434200-DEL-51 | chr8-5434201-DEL-51 |
| chr8-5484266-DEL-1464 | chr8-5484267-DEL-1463 |
| chr8-5490840-DEL-498 | chr8-5490841-DEL-498 |
| chr8-5500889-DEL-6580 | chr8-5500890-DEL-6580 |
| chr8-5502781-DEL-339 | chr8-5502777-DEL-340 |
| chr8-5533818-DEL-275 | chr8-5533819-DEL-275 |
| chr8-5558850-DEL-883 | chr8-5558851-DEL-883 |
| chr8-5579484-DEL-2821 | chr8-5579485-DEL-2821 |
| chr8-5604448-DEL-398 | chr8-5604449-DEL-398 |
| chr8-5619177-DEL-7146 | chr8-5619178-DEL-7146 |
| chr8-5652421-DEL-68 | chr8-5652424-DEL-66 |
| chr8-5659784-DEL-6288 | chr8-5659785-DEL-6288 |
| chr8-5689492-DEL-1813 | chr8-5689493-DEL-1813 |
| chr8-5695864-DEL-95 | chr8-5695858-DEL-108 |
| chr8-5770911-DEL-670 | chr8-5770912-DEL-670 |
| chr8-5778899-DEL-55 | chr8-5778900-DEL-55 |
| chr8-5818199-DEL-202 | chr8-5818200-DEL-202 |
| chr8-5834651-DEL-383 | chr8-5834652-DEL-383 |
| chr8-5838738-DEL-146 | chr8-5838739-DEL-146 |
| chr8-5842512-DEL-53 | chr8-5842513-DEL-53 |
| chr8-5845418-DEL-169 | chr8-5845421-DEL-167 |
| chr8-5846684-DEL-2760 | chr8-5846685-DEL-2760 |
| chr8-5864214-DEL-90 | chr8-5864215-DEL-90 |
| chr8-5867880-DEL-82 | chr8-5867881-DEL-82 |
| chr8-5871038-DEL-1132 | chr8-5871039-DEL-1132 |
| chr8-5891450-DEL-55 | chr8-5891451-DEL-55 |
| chr8-5905691-DEL-1899 | chr8-5905693-DEL-1899 |
| chr8-5909573-DEL-292 | chr8-5909574-DEL-292 |
| chr8-5910736-DEL-879 | chr8-5910737-DEL-879 |
| chr8-5912549-DEL-77 | chr8-5912542-DEL-78 |
| chr8-5921746-DEL-87 | chr8-5921746-DEL-88 |
| chr8-5921956-DEL-1327 | chr8-5921957-DEL-1327 |
| chr8-5927563-DEL-2364 | chr8-5927564-DEL-2364 |
| chr8-5934847-DEL-244 | chr8-5934848-DEL-244 |
| chr8-5935470-DEL-209 | chr8-5935471-DEL-209 |
| chr8-5939273-DEL-371 | chr8-5939274-DEL-371 |
| chr8-5966872-DEL-51 | chr8-5966861-DEL-63 |

| | |
|---|---|
| chr8-5970610-DEL-474 | chr8-5970611-DEL-474 |
| chr8-5976432-DEL-63 | chr8-5976431-DEL-64 |
| chr8-5983629-DEL-974 | chr8-5983630-DEL-974 |
| chr8-5994380-DEL-172 | chr8-5994381-DEL-172 |
| chr8-5998551-DEL-736 | chr8-5998552-DEL-734 |
| chr8-6044335-DEL-118 | chr8-6044336-DEL-118 |
| chr8-6045201-DEL-1572 | chr8-6045194-DEL-1573 |
| chr8-6052287-DEL-233 | chr8-6052288-DEL-225 |
| chr8-6057705-DEL-99 | chr8-6057706-DEL-99 |
| chr8-6064165-DEL-6597 | chr8-6064155-DEL-6607 |
| chr8-6110761-DEL-1421 | chr8-6110762-DEL-1421 |
| chr8-6127985-DEL-350 | chr8-6127986-DEL-350 |
| chr8-6143221-DEL-83 | chr8-6143245-DEL-50 |
| chr8-6203090-DEL-6257 | chr8-6203091-DEL-6257 |
| chr8-6221029-DEL-52 | chr8-6221011-DEL-71 |
| chr8-6230093-DEL-52 | chr8-6230094-DEL-52 |
| chr8-6240694-DEL-50 | chr8-6240682-DEL-62 |
| chr8-6248787-DEL-76 | chr8-6248788-DEL-76 |
| chr8-6324021-DEL-459 | chr8-6324022-DEL-459 |
| chr8-6325957-DEL-1253 | chr8-6325958-DEL-1253 |
| chr8-6329707-DEL-164 | chr8-6329708-DEL-164 |
| chr8-6352261-DEL-6189 | chr8-6352262-DEL-6189 |
| chr8-6372848-DEL-2295 | chr8-6372845-DEL-2299 |
| chr8-6381299-DEL-55 | chr8-6381320-DEL-67 |
| chr8-6389106-DEL-385 | chr8-6389109-DEL-384 |
| chr8-6402650-DEL-791 | chr8-6402651-DEL-791 |
| chr8-6404500-DEL-75 | chr8-6404499-DEL-76 |
| chr8-6406654-DEL-135 | chr8-6406655-DEL-135 |
| chr8-6420674-DEL-66 | chr8-6420675-DEL-66 |
| chr8-6501626-DEL-639 | chr8-6501627-DEL-639 |
| chr8-6509184-DEL-153 | chr8-6509185-DEL-153 |
| chr8-6523669-DEL-358 | chr8-6523670-DEL-358 |
| chr8-6525596-DEL-6326 | chr8-6525597-DEL-6326 |
| chr8-6543428-DEL-7083 | chr8-6543429-DEL-7083 |
| chr8-6558694-DEL-180 | chr8-6558695-DEL-180 |
| chr8-6567096-DEL-66 | chr8-6567097-DEL-66 |
| chr8-6595905-DEL-1305 | chr8-6595906-DEL-1304 |
| chr8-6609187-DEL-53 | chr8-6609188-DEL-53 |
| chr8-6645042-DEL-296 | chr8-6645043-DEL-296 |
| chr8-6659511-DEL-60 | chr8-6659512-DEL-60 |
| chr8-6681120-DEL-1966 | chr8-6681118-DEL-1965 |
| chr8-6687571-DEL-6893 | chr8-6687559-DEL-6894 |
| chr8-6705892-DEL-69 | chr8-6705882-DEL-60 |
| chr8-6730007-DEL-6129 | chr8-6730008-DEL-6129 |
| chr8-6773654-DEL-272 | chr8-6773655-DEL-272 |
| chr8-6800498-DEL-53 | chr8-6800499-DEL-53 |
| chr8-6801464-DEL-6246 | chr8-6801465-DEL-6246 |
| chr8-6866228-DEL-53 | chr8-6866229-DEL-53 |
| chr8-6873281-DEL-374 | chr8-6873282-DEL-374 |
| chr8-6888512-DEL-565 | chr8-6888513-DEL-565 |
| chr8-6921403-DEL-65 | chr8-6921373-DEL-100 |
| chr8-6935283-DEL-56 | chr8-6935276-DEL-54 |
| chr8-6937426-DEL-605 | chr8-6937415-DEL-606 |
| chr8-6942140-DEL-56 | chr8-6942118-DEL-57 |
| chr8-6942706-DEL-202 | chr8-6942707-DEL-202 |
| chr8-6956719-DEL-110 | chr8-6956698-DEL-114 |
| chr8-7032431-DEL-5424 | chr8-7032432-DEL-5424 |
| chr8-7042968-DEL-2393 | chr8-7042969-DEL-2390 |
| chr8-7058751-DEL-1032 | chr8-7058752-DEL-1032 |
| chr8-7063109-DEL-1218 | chr8-7063107-DEL-1216 |
| chr8-7070952-DEL-1475 | chr8-7070963-DEL-1474 |
| chr8-7079611-DEL-6195 | chr8-7079612-DEL-6195 |
| chr8-7089759-DEL-3675 | chr8-7089760-DEL-3675 |
| chr8-7150444-DEL-62 | chr8-7150445-DEL-62 |
| chr8-7151045-DEL-151 | chr8-7151046-DEL-151 |
| chr8-7153945-DEL-601 | chr8-7153945-DEL-601 |
| chr8-7155274-DEL-263 | chr8-7155275-DEL-263 |
| chr8-7190556-DEL-50 | chr8-7190555-DEL-52 |
| chr8-7191643-DEL-255 | chr8-7191643-DEL-253 |
| chr8-7201099-DEL-1073 | chr8-7201101-DEL-1072 |
| chr8-7203219-DEL-1490 | chr8-7203219-DEL-1490 |
| chr8-7205798-DEL-106 | chr8-7205799-DEL-106 |
| chr8-7216585-DEL-1958 | chr8-7216586-DEL-1958 |
| chr8-7223880-DEL-199 | chr8-7223881-DEL-199 |
| chr8-7229210-DEL-180 | chr8-7229211-DEL-180 |
| chr8-7295522-DEL-101 | chr8-7295523-DEL-99 |
| chr8-7309963-DEL-6401 | chr8-7309964-DEL-6401 |
| chr8-7322406-DEL-196 | chr8-7322408-DEL-195 |
| chr8-7374363-DEL-6389 | chr8-7374364-DEL-6389 |
| chr8-7400861-DEL-6566 | chr8-7400862-DEL-6566 |

| | |
|---|---|
| chr8-7412264-DEL-264 | chr8-7412265-DEL-264 |
| chr8-7425050-DEL-60 | chr8-7425048-DEL-59 |
| chr8-7433224-DEL-72 | chr8-7433267-DEL-74 |
| chr8-7465734-DEL-262 | chr8-7465745-DEL-258 |
| chr8-7477482-DEL-845 | chr8-7477483-DEL-845 |
| chr8-7500532-DEL-154 | chr8-7500533-DEL-154 |
| chr8-7586069-DEL-12638 | chr8-7586070-DEL-12635 |
| chr8-7658598-DEL-493 | chr8-7658604-DEL-490 |
| chr8-7665139-DEL-289 | chr8-7665141-DEL-282 |
| chr8-7700619-DEL-117 | chr8-7700620-DEL-117 |
| chr8-7701061-DEL-1406 | chr8-7701062-DEL-1406 |
| chr8-7731844-DEL-4326 | chr8-7731845-DEL-4326 |
| chr8-7749791-DEL-113 | chr8-7749792-DEL-113 |
| chr8-7802867-DEL-1612 | chr8-7802868-DEL-1612 |
| chr8-7837892-DEL-6066 | chr8-7837893-DEL-6066 |
| chr8-7859712-DEL-62 | chr8-7859713-DEL-58 |
| chr8-7874780-DEL-145 | chr8-7874791-DEL-143 |
| chr8-7897974-DEL-938 | chr8-7897969-DEL-939 |
| chr8-7901831-DEL-430 | chr8-7901832-DEL-430 |
| chr8-7917477-DEL-787 | chr8-7917478-DEL-787 |
| chr8-7928335-DEL-187 | chr8-7928336-DEL-187 |
| chr8-7936326-DEL-414 | chr8-7936327-DEL-414 |
| chr8-7942138-DEL-150 | chr8-7942143-DEL-146 |
| chr8-7945175-DEL-6338 | chr8-7945176-DEL-6338 |
| chr8-7989153-DEL-74 | chr8-7989154-DEL-74 |
| chr8-8041332-DEL-54 | chr8-8041333-DEL-54 |
| chr8-8076265-DEL-55 | chr8-8076262-DEL-55 |
| chr8-8138233-DEL-2101 | chr8-8138234-DEL-2101 |
| chr8-8280141-DEL-396 | chr8-8280142-DEL-396 |
| chr8-8316928-DEL-630 | chr8-8316929-DEL-630 |
| chr8-8360026-DEL-69 | chr8-8360027-DEL-117 |
| chr8-8365018-DEL-157 | chr8-8365018-DEL-157 |
| chr8-8365261-DEL-5408 | chr8-8365261-DEL-5408 |
| chr8-8382800-DEL-75 | chr8-8382801-DEL-74 |
| chr8-8430507-DEL-90 | chr8-8430470-DEL-80 |
| chr8-8476952-DEL-57 | chr8-8476953-DEL-57 |
| chr8-8482901-DEL-62 | chr8-8482892-DEL-60 |
| chr8-8483892-DEL-69 | chr8-8483893-DEL-69 |
| chr8-8494202-DEL-89 | chr8-8494199-DEL-90 |
| chr8-8496628-DEL-149 | chr8-8496629-DEL-149 |
| chr8-8516329-DEL-56 | chr8-8516330-DEL-55 |
| chr8-8534978-DEL-64 | chr8-8534979-DEL-64 |
| chr8-8555877-DEL-82 | chr8-8555878-DEL-123 |
| chr8-8568308-DEL-56 | chr8-8568309-DEL-56 |
| chr8-8570618-DEL-81 | chr8-8570619-DEL-81 |
| chr8-8634680-DEL-58 | chr8-8634681-DEL-58 |
| chr8-8639372-DEL-66 | chr8-8639373-DEL-66 |
| chr8-8648709-DEL-72 | chr8-8648701-DEL-68 |
| chr8-8676824-DEL-388 | chr8-8676825-DEL-388 |
| chr8-8751021-DEL-87 | chr8-8751020-DEL-88 |
| chr8-8751692-DEL-74 | chr8-8751693-DEL-74 |
| chr8-8768102-DEL-67 | chr8-8768120-DEL-70 |
| chr8-8791687-DEL-61 | chr8-8791688-DEL-61 |
| chr8-8794575-DEL-792 | chr8-8794576-DEL-792 |
| chr8-8797418-DEL-219 | chr8-8797418-DEL-219 |
| chr8-8814271-DEL-210 | chr8-8814272-DEL-210 |
| chr8-8837215-DEL-194 | chr8-8837230-DEL-180 |
| chr8-8857431-DEL-208 | chr8-8857430-DEL-209 |
| chr8-8863145-DEL-449 | chr8-8863145-DEL-449 |
| chr8-8865754-DEL-168 | chr8-8865753-DEL-170 |
| chr8-8886147-DEL-264 | chr8-8886148-DEL-264 |
| chr8-8909558-DEL-750 | chr8-8909544-DEL-729 |
| chr8-8913949-DEL-82 | chr8-8913952-DEL-57 |
| chr8-9013586-DEL-66 | chr8-9013587-DEL-66 |
| chr8-9039824-DEL-400 | chr8-9039825-DEL-400 |
| chr8-9088028-DEL-369 | chr8-9088029-DEL-369 |
| chr8-9096707-DEL-765 | chr8-9096708-DEL-765 |
| chr8-9104202-DEL-232 | chr8-9104202-DEL-231 |
| chr8-9129449-DEL-4102 | chr8-9129450-DEL-4102 |
| chr8-9129539-DEL-96 | chr8-9129540-DEL-96 |
| chr8-9136030-DEL-78 | chr8-9136031-DEL-78 |
| chr8-9169001-DEL-155 | chr8-9169002-DEL-155 |
| chr8-9188596-DEL-165 | chr8-9188597-DEL-165 |
| chr8-9195120-DEL-519 | chr8-9195121-DEL-519 |
| chr8-9271421-DEL-136 | chr8-9271422-DEL-136 |
| chr8-9274503-DEL-120 | chr8-9274505-DEL-130 |
| chr8-9286544-DEL-86 | chr8-9286531-DEL-78 |
| chr8-9300093-DEL-1891 | chr8-9300094-DEL-1891 |
| chr8-9312304-DEL-58 | chr8-9312292-DEL-59 |
| chr8-9320363-DEL-63 | chr8-9320364-DEL-63 |

| | |
|---|---|
| chr8-9345499-DEL-242 | chr8-9345500-DEL-242 |
| chr8-9346766-DEL-836 | chr8-9346795-DEL-810 |
| chr8-9350957-DEL-56 | chr8-9350917-DEL-78 |
| chr8-9401767-DEL-6830 | chr8-9401768-DEL-6830 |
| chr8-9419631-DEL-120 | chr8-9419634-DEL-103 |
| chr8-9439952-DEL-2151 | chr8-9439953-DEL-2151 |
| chr8-9440462-DEL-316 | chr8-9440463-DEL-316 |
| chr8-9480806-DEL-128 | chr8-9480788-DEL-105 |
| chr8-9483035-DEL-65 | chr8-9483036-DEL-65 |
| chr8-9497827-DEL-73 | chr8-9497828-DEL-73 |
| chr8-9545719-DEL-185 | chr8-9545720-DEL-185 |
| chr8-9558681-DEL-6384 | chr8-9558682-DEL-6384 |
| chr8-9573777-DEL-1497 | chr8-9573778-DEL-1497 |
| chr8-9601173-DEL-7004 | chr8-9601174-DEL-7004 |
| chr8-9611370-DEL-723 | chr8-9611371-DEL-723 |
| chr8-9612414-DEL-228 | chr8-9612415-DEL-228 |
| chr8-9632892-DEL-99 | chr8-9632893-DEL-99 |
| chr8-9682255-DEL-464 | chr8-9682247-DEL-463 |
| chr8-9710245-DEL-126 | chr8-9710246-DEL-126 |
| chr8-9726557-DEL-155 | chr8-9726604-DEL-110 |
| chr8-9732457-DEL-90 | chr8-9732487-DEL-60 |
| chr8-9738704-DEL-85 | chr8-9738705-DEL-85 |
| chr8-9739365-DEL-84 | chr8-9739366-DEL-84 |
| chr8-9761705-DEL-176 | chr8-9761704-DEL-177 |
| chr8-9766361-DEL-167 | chr8-9766362-DEL-167 |
| chr8-9786937-DEL-88 | chr8-9786938-DEL-88 |
| chr8-9788809-DEL-2356 | chr8-9788810-DEL-2356 |
| chr8-9828577-DEL-134 | chr8-9828578-DEL-134 |
| chr8-9848224-DEL-141 | chr8-9848225-DEL-141 |
| chr8-9859715-DEL-240 | chr8-9859730-DEL-233 |
| chr8-9880275-DEL-265 | chr8-9880275-DEL-265 |
| chr8-9891237-DEL-6187 | chr8-9891238-DEL-6187 |
| chr8-9921491-DEL-450 | chr8-9921492-DEL-450 |
| chr8-9922191-DEL-56 | chr8-9922192-DEL-52 |
| chr8-9952527-DEL-2352 | chr8-9952520-DEL-2358 |
| chr8-9966027-DEL-178 | chr8-9966028-DEL-178 |
| chr8-9973418-DEL-252 | chr8-9973419-DEL-252 |
| chr8-9987478-DEL-51 | chr8-9987479-DEL-51 |
| chr8-9988587-DEL-116 | chr8-9988588-DEL-114 |
| chr8-9996384-DEL-634 | chr8-9996385-DEL-634 |
| chr8-9997231-DEL-329 | chr8-9997232-DEL-329 |
| chr8-10048800-DEL-167 | chr8-10048801-DEL-167 |
| chr8-10052645-DEL-401 | chr8-10052646-DEL-401 |
| chr8-10079713-DEL-209 | chr8-10079714-DEL-209 |
| chr8-10092459-DEL-96 | chr8-10092440-DEL-145 |
| chr8-10094218-DEL-63 | chr8-10094219-DEL-62 |
| chr8-10130302-DEL-6296 | chr8-10130303-DEL-6296 |
| chr8-10146498-DEL-993 | chr8-10146499-DEL-993 |
| chr8-10150773-DEL-190 | chr8-10150764-DEL-187 |
| chr8-10184065-DEL-72 | chr8-10184066-DEL-72 |
| chr8-10191692-DEL-544 | chr8-10191691-DEL-545 |
| chr8-10240174-DEL-58 | chr8-10240167-DEL-54 |
| chr8-10269594-DEL-54 | chr8-10269595-DEL-54 |
| chr8-10270804-DEL-1115 | chr8-10270805-DEL-1115 |
| chr8-10304018-DEL-781 | chr8-10304019-DEL-781 |
| chr8-10315476-DEL-202 | chr8-10315477-DEL-202 |
| chr8-10343530-DEL-52 | chr8-10343531-DEL-52 |
| chr8-10353475-DEL-527 | chr8-10353476-DEL-527 |
| chr8-10384547-DEL-501 | chr8-10384547-DEL-501 |
| chr8-10406716-DEL-71 | chr8-10406696-DEL-60 |
| chr8-10423223-DEL-95 | chr8-10423256-DEL-63 |
| chr8-10433625-DEL-69 | chr8-10433602-DEL-52 |
| chr8-10475671-DEL-114 | chr8-10475629-DEL-115 |
| chr8-10483013-DEL-122 | chr8-10482987-DEL-168 |
| chr8-10487023-DEL-67 | chr8-10487024-DEL-66 |
| chr8-10505172-DEL-452 | chr8-10505173-DEL-452 |
| chr8-10522123-DEL-56 | chr8-10522124-DEL-56 |
| chr8-10545498-DEL-52 | chr8-10545519-DEL-52 |
| chr8-10565557-DEL-92 | chr8-10565558-DEL-92 |
| chr8-10570229-DEL-82 | chr8-10570230-DEL-82 |
| chr8-10573300-DEL-105 | chr8-10573301-DEL-70 |
| chr8-10599760-DEL-104 | chr8-10599781-DEL-101 |
| chr8-10602972-DEL-498 | chr8-10602973-DEL-498 |
| chr8-10741211-DEL-62 | chr8-10741209-DEL-60 |
| chr8-10742227-DEL-76 | chr8-10742240-DEL-98 |
| chr8-10763204-DEL-210 | chr8-10763197-DEL-218 |
| chr8-10776639-DEL-133 | chr8-10776631-DEL-141 |
| chr8-10779054-DEL-2526 | chr8-10779103-DEL-2477 |
| chr8-10791526-DEL-447 | chr8-10791527-DEL-447 |
| chr8-10812655-DEL-74 | chr8-10812638-DEL-70 |

| | |
|---|---|
| chr8-10817923-DEL-167 | chr8-10817924-DEL-167 |
| chr8-10832896-DEL-100 | chr8-10832897-DEL-100 |
| chr8-10878245-DEL-258 | chr8-10878246-DEL-258 |
| chr8-10918892-DEL-61 | chr8-10918889-DEL-61 |
| chr8-10926497-DEL-94 | chr8-10926502-DEL-93 |
| chr8-10940224-DEL-61 | chr8-10940224-DEL-69 |
| chr8-10968213-DEL-90 | chr8-10968214-DEL-82 |
| chr8-10972681-DEL-144 | chr8-10972682-DEL-144 |
| chr8-11003389-DEL-77 | chr8-11003390-DEL-77 |
| chr8-11027375-DEL-100 | chr8-11027362-DEL-98 |
| chr8-11062630-DEL-51 | chr8-11062602-DEL-64 |
| chr8-11065585-DEL-96 | chr8-11065585-DEL-96 |
| chr8-11069679-DEL-113 | chr8-11069680-DEL-113 |
| chr8-11087481-DEL-151 | chr8-11087483-DEL-147 |
| chr8-11120983-DEL-52 | chr8-11120984-DEL-52 |
| chr8-11123415-DEL-408 | chr8-11123416-DEL-408 |
| chr8-11126251-DEL-108 | chr8-11126296-DEL-80 |
| chr8-11145782-DEL-1565 | chr8-11145783-DEL-1565 |
| chr8-11183727-DEL-209 | chr8-11183728-DEL-209 |
| chr8-11188401-DEL-604 | chr8-11188402-DEL-604 |
| chr8-11201227-DEL-68 | chr8-11201220-DEL-76 |
| chr8-11217016-DEL-137 | chr8-11217017-DEL-137 |
| chr8-11367222-DEL-112 | chr8-11367223-DEL-112 |
| chr8-11378724-DEL-75 | chr8-11378725-DEL-81 |
| chr8-11388431-DEL-59 | chr8-11388432-DEL-59 |
| chr8-11415663-DEL-64 | chr8-11415623-DEL-60 |
| chr8-11425496-DEL-52 | chr8-11425457-DEL-58 |
| chr8-11516797-DEL-194 | chr8-11516786-DEL-193 |
| chr8-11531288-DEL-241 | chr8-11531289-DEL-241 |
| chr8-11543348-DEL-238 | chr8-11543349-DEL-238 |
| chr8-11544508-DEL-79 | chr8-11544509-DEL-79 |
| chr8-11550862-DEL-130 | chr8-11550860-DEL-131 |
| chr8-11551366-DEL-86 | chr8-11551367-DEL-86 |
| chr8-11594289-DEL-212 | chr8-11594291-DEL-212 |
| chr8-11660130-DEL-654 | chr8-11660130-DEL-652 |
| chr8-11692006-DEL-194 | chr8-11692007-DEL-194 |
| chr8-11699747-DEL-54 | chr8-11699750-DEL-50 |
| chr8-11704555-DEL-197 | chr8-11704556-DEL-197 |
| chr8-11724660-DEL-566 | chr8-11724661-DEL-566 |
| chr8-11733380-DEL-880 | chr8-11733383-DEL-879 |
| chr8-11738878-DEL-183 | chr8-11738869-DEL-190 |
| chr8-11760111-DEL-78 | chr8-11760112-DEL-78 |
| chr8-11764284-DEL-268 | chr8-11764285-DEL-268 |
| chr8-11781299-DEL-634 | chr8-11781300-DEL-634 |
| chr8-11800489-DEL-88 | chr8-11800490-DEL-88 |
| chr8-11862227-DEL-60 | chr8-11862228-DEL-60 |
| chr8-11910811-DEL-597 | chr8-11910812-DEL-597 |
| chr8-11966283-DEL-86 | chr8-11966284-DEL-86 |
| chr8-11970313-DEL-63 | chr8-11970314-DEL-63 |
| chr8-11994509-DEL-188 | chr8-11994510-DEL-188 |
| chr8-11996465-DEL-121 | chr8-11996466-DEL-121 |
| chr8-11998461-DEL-96 | chr8-11998459-DEL-96 |
| chr8-12003969-DEL-696 | chr8-12003970-DEL-696 |
| chr8-12017199-DEL-389 | chr8-12017187-DEL-389 |
| chr8-12024543-DEL-84 | chr8-12024544-DEL-84 |
| chr8-12058848-DEL-65 | chr8-12058841-DEL-75 |
| chr8-12069573-DEL-406 | chr8-12069574-DEL-406 |
| chr8-12086869-DEL-79 | chr8-12086915-DEL-78 |
| chr8-12095756-DEL-55 | chr8-12095757-DEL-55 |
| chr8-12106671-DEL-319 | chr8-12106672-DEL-319 |
| chr8-12115166-DEL-106 | chr8-12115167-DEL-106 |
| chr8-12121291-DEL-95 | chr8-12121324-DEL-63 |
| chr8-12140401-DEL-170 | chr8-12140402-DEL-170 |
| chr8-12165749-DEL-93 | chr8-12165750-DEL-93 |
| chr8-12218100-DEL-98 | chr8-12218101-DEL-98 |
| chr8-12276185-DEL-773 | chr8-12276186-DEL-773 |
| chr8-12298643-DEL-54 | chr8-12298652-DEL-52 |
| chr8-12339126-DEL-76 | chr8-12339137-DEL-78 |
| chr8-12390640-DEL-60 | chr8-12390641-DEL-60 |
| chr8-12397955-DEL-67 | chr8-12397960-DEL-60 |
| chr8-12406960-DEL-415 | chr8-12406961-DEL-415 |
| chr8-12429836-DEL-64 | chr8-12429837-DEL-64 |
| chr8-12462059-DEL-52 | chr8-12462060-DEL-52 |
| chr8-12471288-DEL-64 | chr8-12471263-DEL-56 |
| chr8-12491600-DEL-176 | chr8-12491601-DEL-176 |
| chr8-12492276-DEL-302 | chr8-12492277-DEL-302 |
| chr8-12516459-DEL-628 | chr8-12516460-DEL-628 |
| chr8-12526336-DEL-114 | chr8-12526358-DEL-101 |
| chr8-12546207-DEL-59 | chr8-12546196-DEL-60 |
| chr8-12573041-DEL-72 | chr8-12573025-DEL-92 |

| | |
|---|---|
| chr8-12578121-DEL-113 | chr8-12578122-DEL-113 |
| chr8-12612995-DEL-113 | chr8-12612996-DEL-113 |
| chr8-12619996-DEL-1314 | chr8-12619997-DEL-1314 |
| chr8-12643837-DEL-60 | chr8-12643838-DEL-60 |
| chr8-12683361-DEL-74 | chr8-12683370-DEL-66 |
| chr8-12695543-DEL-546 | chr8-12695544-DEL-546 |
| chr8-12704003-DEL-202 | chr8-12703978-DEL-203 |
| chr8-12737447-DEL-70 | chr8-12737448-DEL-70 |
| chr8-12746909-DEL-54 | chr8-12746914-DEL-65 |
| chr8-12762589-DEL-213 | chr8-12762590-DEL-213 |
| chr8-12768401-DEL-200 | chr8-12768402-DEL-200 |
| chr8-12769773-DEL-245 | chr8-12769800-DEL-196 |
| chr8-12773276-DEL-54 | chr8-12773277-DEL-54 |
| chr8-12783696-DEL-167 | chr8-12783697-DEL-167 |
| chr8-12787545-DEL-114 | chr8-12787543-DEL-81 |
| chr8-12795722-DEL-373 | chr8-12795723-DEL-373 |
| chr8-12860987-DEL-81 | chr8-12860993-DEL-59 |
| chr8-12922881-DEL-181 | chr8-12922882-DEL-181 |
| chr8-12947719-DEL-55 | chr8-12947720-DEL-55 |
| chr8-13026080-DEL-56 | chr8-13026081-DEL-56 |
| chr8-13048274-DEL-75 | chr8-13048277-DEL-71 |
| chr8-13068377-DEL-54 | chr8-13068378-DEL-54 |
| chr8-13084561-DEL-122 | chr8-13084538-DEL-146 |
| chr8-13121042-DEL-112 | chr8-13121043-DEL-112 |
| chr8-13130087-DEL-1020 | chr8-13130088-DEL-1020 |
| chr8-13133105-DEL-301 | chr8-13133111-DEL-294 |
| chr8-13133629-DEL-515 | chr8-13133661-DEL-475 |
| chr8-13134313-DEL-87 | chr8-13134314-DEL-87 |
| chr8-13144772-DEL-682 | chr8-13144773-DEL-683 |
| chr8-13147028-DEL-195 | chr8-13147030-DEL-196 |
| chr8-13157957-DEL-75 | chr8-13157963-DEL-70 |
| chr8-13181855-DEL-108 | chr8-13181856-DEL-108 |
| chr8-13194611-DEL-84 | chr8-13194612-DEL-84 |
| chr8-13201984-DEL-217 | chr8-13201985-DEL-217 |
| chr8-13203122-DEL-248 | chr8-13203123-DEL-248 |
| chr8-13253893-DEL-66 | chr8-13253894-DEL-51 |
| chr8-13262605-DEL-70 | chr8-13262606-DEL-70 |
| chr8-13270946-DEL-210 | chr8-13270947-DEL-210 |
| chr8-13286878-DEL-1881 | chr8-13286879-DEL-1881 |
| chr8-13356767-DEL-74 | chr8-13356736-DEL-75 |
| chr8-13366523-DEL-108 | chr8-13366524-DEL-108 |
| chr8-13387262-DEL-78 | chr8-13387263-DEL-78 |
| chr8-13434317-DEL-54 | chr8-13434318-DEL-60 |
| chr8-13440907-DEL-1894 | chr8-13440908-DEL-1894 |
| chr8-13459931-DEL-65 | chr8-13459932-DEL-65 |
| chr8-13466906-DEL-367 | chr8-13466907-DEL-367 |
| chr8-13481986-DEL-434 | chr8-13481987-DEL-434 |
| chr8-13500766-DEL-68 | chr8-13500767-DEL-67 |
| chr8-13501338-DEL-60 | chr8-13501339-DEL-60 |
| chr8-13515177-DEL-69 | chr8-13515177-DEL-68 |
| chr8-13560736-DEL-53 | chr8-13560737-DEL-53 |
| chr8-13561906-DEL-82 | chr8-13561917-DEL-80 |
| chr8-13592370-DEL-2031 | chr8-13592370-DEL-2028 |
| chr8-13606261-DEL-54 | chr8-13606259-DEL-54 |
| chr8-13624538-DEL-89 | chr8-13624553-DEL-51 |
| chr8-13648857-DEL-106 | chr8-13648856-DEL-105 |
| chr8-13693331-DEL-112 | chr8-13693332-DEL-112 |
| chr8-13708449-DEL-72 | chr8-13708450-DEL-52 |
| chr8-13724663-DEL-84 | chr8-13724664-DEL-84 |
| chr8-13758177-DEL-268 | chr8-13758178-DEL-268 |
| chr8-13777207-DEL-192 | chr8-13777208-DEL-192 |
| chr8-13795901-DEL-94 | chr8-13795902-DEL-94 |
| chr8-13801968-DEL-201 | chr8-13801956-DEL-202 |
| chr8-13826554-DEL-187 | chr8-13826555-DEL-187 |
| chr8-13841434-DEL-148 | chr8-13841435-DEL-148 |
| chr8-13861807-DEL-234 | chr8-13861808-DEL-234 |
| chr8-13910717-DEL-6440 | chr8-13910718-DEL-6440 |
| chr8-13923980-DEL-198 | chr8-13923981-DEL-198 |
| chr8-13966560-DEL-146 | chr8-13966548-DEL-147 |
| chr8-13969028-DEL-184 | chr8-13969029-DEL-184 |
| chr8-13984048-DEL-385 | chr8-13984047-DEL-386 |
| chr8-14013911-DEL-54 | chr8-14013888-DEL-54 |
| chr8-14030551-DEL-3338 | chr8-14030552-DEL-3338 |
| chr8-14043476-DEL-61 | chr8-14043477-DEL-61 |
| chr8-14045170-DEL-252 | chr8-14045172-DEL-250 |
| chr8-14048961-DEL-66 | chr8-14048940-DEL-88 |
| chr8-14065664-DEL-78 | chr8-14065650-DEL-100 |
| chr8-14068228-DEL-225 | chr8-14068226-DEL-226 |
| chr8-14069875-DEL-1945 | chr8-14069876-DEL-1945 |
| chr8-14086593-DEL-84 | chr8-14086614-DEL-64 |

| | |
|---|---|
| chr8-14092377-DEL-205 | chr8-14092378-DEL-205 |
| chr8-14099335-DEL-201 | chr8-14099336-DEL-201 |
| chr8-14119389-DEL-1892 | chr8-14119389-DEL-1892 |
| chr8-14126303-DEL-58 | chr8-14126304-DEL-58 |
| chr8-14127876-DEL-57 | chr8-14127883-DEL-51 |
| chr8-14212287-DEL-199 | chr8-14212288-DEL-199 |
| chr8-14224608-DEL-118 | chr8-14224609-DEL-118 |
| chr8-14225551-DEL-90 | chr8-14225550-DEL-87 |
| chr8-14246240-DEL-6417 | chr8-14246241-DEL-6417 |
| chr8-14256696-DEL-183 | chr8-14256697-DEL-183 |
| chr8-14282184-DEL-90 | chr8-14282192-DEL-86 |
| chr8-14290980-DEL-1812 | chr8-14290981-DEL-1812 |
| chr8-14319580-DEL-165 | chr8-14319578-DEL-168 |
| chr8-14332898-DEL-194 | chr8-14332899-DEL-194 |
| chr8-14339389-DEL-363 | chr8-14339393-DEL-362 |
| chr8-14344889-DEL-2444 | chr8-14344890-DEL-2444 |
| chr8-14346479-DEL-825 | chr8-14346480-DEL-825 |
| chr8-14407388-DEL-60 | chr8-14407387-DEL-54 |
| chr8-14418746-DEL-89 | chr8-14418737-DEL-50 |
| chr8-14459152-DEL-84 | chr8-14459153-DEL-84 |
| chr8-14498474-DEL-51 | chr8-14498467-DEL-59 |
| chr8-14502903-DEL-200 | chr8-14502909-DEL-197 |
| chr8-14521256-DEL-81 | chr8-14521291-DEL-53 |
| chr8-14574869-DEL-177 | chr8-14574870-DEL-177 |
| chr8-14584536-DEL-120 | chr8-14584522-DEL-96 |
| chr8-14635542-DEL-124 | chr8-14635537-DEL-170 |
| chr8-14737977-DEL-60 | chr8-14737970-DEL-60 |
| chr8-14786827-DEL-215 | chr8-14786828-DEL-215 |
| chr8-14856444-DEL-78 | chr8-14856445-DEL-78 |
| chr8-14885313-DEL-97 | chr8-14885314-DEL-97 |
| chr8-14893710-DEL-128 | chr8-14893711-DEL-128 |
| chr8-14911475-DEL-109 | chr8-14911476-DEL-109 |
| chr8-14953384-DEL-94 | chr8-14953380-DEL-94 |
| chr8-14981607-DEL-88 | chr8-14981608-DEL-88 |
| chr8-15003499-DEL-682 | chr8-15003504-DEL-676 |
| chr8-15006158-DEL-74 | chr8-15006156-DEL-77 |
| chr8-15014930-DEL-78 | chr8-15014911-DEL-78 |
| chr8-15017269-DEL-163 | chr8-15017273-DEL-159 |
| chr8-15061944-DEL-221 | chr8-15061945-DEL-221 |
| chr8-15099052-DEL-65 | chr8-15099052-DEL-65 |
| chr8-15104035-DEL-625 | chr8-15104022-DEL-623 |
| chr8-15107953-DEL-167 | chr8-15107954-DEL-167 |
| chr8-15129549-DEL-72 | chr8-15129558-DEL-64 |
| chr8-15139417-DEL-6216 | chr8-15139418-DEL-6216 |
| chr8-15155120-DEL-269 | chr8-15155121-DEL-269 |
| chr8-15167197-DEL-161 | chr8-15167166-DEL-153 |
| chr8-15175743-DEL-52 | chr8-15175744-DEL-52 |
| chr8-15179814-DEL-74 | chr8-15179815-DEL-74 |
| chr8-15187804-DEL-58 | chr8-15187805-DEL-58 |
| chr8-15194291-DEL-196 | chr8-15194291-DEL-196 |
| chr8-15195884-DEL-119 | chr8-15195885-DEL-119 |
| chr8-15200523-DEL-68 | chr8-15200484-DEL-66 |
| chr8-15204056-DEL-60 | chr8-15204057-DEL-60 |
| chr8-15204712-DEL-354 | chr8-15204713-DEL-354 |
| chr8-15235137-DEL-54 | chr8-15235138-DEL-54 |
| chr8-15275346-DEL-2227 | chr8-15275347-DEL-2227 |
| chr8-15310725-DEL-179 | chr8-15310726-DEL-179 |
| chr8-15318915-DEL-57 | chr8-15318916-DEL-63 |
| chr8-15326982-DEL-57 | chr8-15326983-DEL-57 |
| chr8-15333086-DEL-68 | chr8-15333087-DEL-68 |
| chr8-15359339-DEL-72 | chr8-15359332-DEL-80 |
| chr8-15360353-DEL-217 | chr8-15360354-DEL-217 |
| chr8-15364482-DEL-156 | chr8-15364483-DEL-156 |
| chr8-15366039-DEL-75 | chr8-15366043-DEL-72 |
| chr8-15463763-DEL-5083 | chr8-15463764-DEL-5083 |
| chr8-15489380-DEL-78 | chr8-15489381-DEL-78 |
| chr8-15492194-DEL-7483 | chr8-15492195-DEL-7483 |
| chr8-15504533-DEL-63 | chr8-15504526-DEL-64 |
| chr8-15510304-DEL-869 | chr8-15510305-DEL-826 |
| chr8-15514900-DEL-60 | chr8-15514894-DEL-66 |
| chr8-15530666-DEL-485 | chr8-15530667-DEL-485 |
| chr8-15531310-DEL-75 | chr8-15531311-DEL-75 |
| chr8-15557701-DEL-6461 | chr8-15557702-DEL-6461 |
| chr8-15569778-DEL-341 | chr8-15569779-DEL-340 |
| chr8-15593769-DEL-101 | chr8-15593770-DEL-99 |
| chr8-15610049-DEL-59 | chr8-15610050-DEL-59 |
| chr8-15616178-DEL-51 | chr8-15616179-DEL-51 |
| chr8-15623169-DEL-2963 | chr8-15623170-DEL-2963 |
| chr8-15636025-DEL-55 | chr8-15636026-DEL-55 |
| chr8-15637960-DEL-247 | chr8-15637961-DEL-247 |

| | |
|---|---|
| chr8-15639671-DEL-320 | chr8-15639648-DEL-326 |
| chr8-15644055-DEL-1894 | chr8-15644056-DEL-1894 |
| chr8-15702253-DEL-6450 | chr8-15702244-DEL-6451 |
| chr8-15717855-DEL-801 | chr8-15717856-DEL-801 |
| chr8-15731724-DEL-231 | chr8-15731725-DEL-231 |
| chr8-15739479-DEL-376 | chr8-15739480-DEL-376 |
| chr8-15742018-DEL-6403 | chr8-15742019-DEL-6403 |
| chr8-15763610-DEL-6668 | chr8-15763611-DEL-6669 |
| chr8-15788301-DEL-3001 | chr8-15788302-DEL-3001 |
| chr8-15804064-DEL-7221 | chr8-15804065-DEL-7221 |
| chr8-15860995-DEL-196 | chr8-15860996-DEL-196 |
| chr8-15877456-DEL-260 | chr8-15877457-DEL-260 |
| chr8-15929293-DEL-83 | chr8-15929294-DEL-83 |
| chr8-15947158-DEL-174 | chr8-15947159-DEL-174 |
| chr8-15965449-DEL-187 | chr8-15965450-DEL-187 |
| chr8-16017712-DEL-6965 | chr8-16017713-DEL-6965 |
| chr8-16040195-DEL-62 | chr8-16040196-DEL-62 |
| chr8-16068004-DEL-98 | chr8-16068047-DEL-56 |
| chr8-16091762-DEL-6191 | chr8-16091763-DEL-6191 |
| chr8-16099488-DEL-936 | chr8-16099489-DEL-936 |
| chr8-16109859-DEL-52 | chr8-16109850-DEL-62 |
| chr8-16127724-DEL-135 | chr8-16127725-DEL-135 |
| chr8-16135901-DEL-52 | chr8-16135915-DEL-53 |
| chr8-16188104-DEL-6125 | chr8-16188095-DEL-6124 |
| chr8-16228446-DEL-58 | chr8-16228447-DEL-58 |
| chr8-16229823-DEL-1221 | chr8-16229824-DEL-1221 |
| chr8-16254750-DEL-102 | chr8-16254751-DEL-102 |
| chr8-16266386-DEL-6501 | chr8-16266387-DEL-6501 |
| chr8-16307501-DEL-72 | chr8-16307502-DEL-72 |
| chr8-16315644-DEL-773 | chr8-16315645-DEL-773 |
| chr8-16329019-DEL-87 | chr8-16329027-DEL-80 |
| chr8-16329265-DEL-526 | chr8-16329266-DEL-526 |
| chr8-16342878-DEL-144 | chr8-16342882-DEL-141 |
| chr8-16347431-DEL-1939 | chr8-16347431-DEL-1939 |
| chr8-16359837-DEL-50 | chr8-16359826-DEL-62 |
| chr8-16360915-DEL-770 | chr8-16360916-DEL-770 |
| chr8-16363910-DEL-111 | chr8-16363910-DEL-111 |
| chr8-16374896-DEL-79 | chr8-16374876-DEL-89 |
| chr8-16377288-DEL-64 | chr8-16377289-DEL-64 |
| chr8-16411137-DEL-6012 | chr8-16411138-DEL-6012 |
| chr8-16467764-DEL-58 | chr8-16467765-DEL-58 |
| chr8-16496351-DEL-643 | chr8-16496352-DEL-643 |
| chr8-16517047-DEL-82 | chr8-16517048-DEL-82 |
| chr8-16542287-DEL-81 | chr8-16542288-DEL-81 |
| chr8-16573072-DEL-5057 | chr8-16573072-DEL-5057 |
| chr8-16584453-DEL-60 | chr8-16584454-DEL-60 |
| chr8-16659687-DEL-793 | chr8-16659688-DEL-793 |
| chr8-16703258-DEL-63 | chr8-16703272-DEL-52 |
| chr8-16764121-DEL-622 | chr8-16764120-DEL-617 |
| chr8-16765343-DEL-72 | chr8-16765344-DEL-72 |
| chr8-16769131-DEL-395 | chr8-16769132-DEL-395 |
| chr8-16785212-DEL-70 | chr8-16785213-DEL-70 |
| chr8-16803271-DEL-55 | chr8-16803272-DEL-55 |
| chr8-16836379-DEL-130 | chr8-16836380-DEL-130 |
| chr8-16840186-DEL-5403 | chr8-16840187-DEL-5403 |
| chr8-16860276-DEL-2379 | chr8-16860277-DEL-2379 |
| chr8-16867381-DEL-57 | chr8-16867382-DEL-57 |
| chr8-16946369-DEL-7111 | chr8-16946370-DEL-7111 |
| chr8-16970280-DEL-7782 | chr8-16970281-DEL-7782 |
| chr8-16996315-DEL-829 | chr8-16996316-DEL-829 |
| chr8-17003633-DEL-214 | chr8-17003634-DEL-213 |
| chr8-17029633-DEL-122 | chr8-17029634-DEL-122 |
| chr8-17051258-DEL-1030 | chr8-17051265-DEL-1024 |
| chr8-17091885-DEL-345 | chr8-17091885-DEL-345 |
| chr8-17100493-DEL-2569 | chr8-17100492-DEL-2570 |
| chr8-17108115-DEL-100 | chr8-17108116-DEL-100 |
| chr8-17110136-DEL-172 | chr8-17110139-DEL-168 |
| chr8-17112861-DEL-134 | chr8-17112862-DEL-134 |
| chr8-17116825-DEL-6487 | chr8-17116826-DEL-6487 |
| chr8-17146805-DEL-437 | chr8-17146806-DEL-437 |
| chr8-17200435-DEL-490 | chr8-17200436-DEL-490 |
| chr8-17204014-DEL-351 | chr8-17204015-DEL-351 |
| chr8-17234458-DEL-77 | chr8-17234459-DEL-73 |
| chr8-17236343-DEL-142 | chr8-17236362-DEL-124 |
| chr8-17240281-DEL-6412 | chr8-17240282-DEL-6412 |
| chr8-17250994-DEL-55 | chr8-17250995-DEL-57 |
| chr8-17257777-DEL-109 | chr8-17257788-DEL-99 |
| chr8-17264649-DEL-66 | chr8-17264680-DEL-69 |
| chr8-17320153-DEL-404 | chr8-17320154-DEL-404 |
| chr8-17320836-DEL-215 | chr8-17320837-DEL-215 |

| | |
|---|---|
| chr8-17352211-DEL-830 | chr8-17352212-DEL-830 |
| chr8-17387231-DEL-63 | chr8-17387232-DEL-63 |
| chr8-17388502-DEL-3052 | chr8-17388503-DEL-3052 |
| chr8-17432428-DEL-3072 | chr8-17432429-DEL-3072 |
| chr8-17453255-DEL-377 | chr8-17453260-DEL-375 |
| chr8-17464387-DEL-66 | chr8-17464382-DEL-60 |
| chr8-17466272-DEL-60 | chr8-17466273-DEL-60 |
| chr8-17491639-DEL-69 | chr8-17491639-DEL-69 |
| chr8-17502014-DEL-60 | chr8-17502015-DEL-60 |
| chr8-17542339-DEL-6852 | chr8-17542340-DEL-6852 |
| chr8-17554308-DEL-56 | chr8-17554309-DEL-56 |
| chr8-17630538-DEL-133 | chr8-17630539-DEL-133 |
| chr8-17633178-DEL-1360 | chr8-17633179-DEL-1360 |
| chr8-17647901-DEL-142 | chr8-17647902-DEL-148 |
| chr8-17649256-DEL-72 | chr8-17649227-DEL-73 |
| chr8-17661595-DEL-693 | chr8-17661596-DEL-692 |
| chr8-17666315-DEL-433 | chr8-17666315-DEL-434 |
| chr8-17679028-DEL-1300 | chr8-17679029-DEL-1300 |
| chr8-17687531-DEL-6400 | chr8-17687531-DEL-6400 |
| chr8-17699951-DEL-212 | chr8-17699952-DEL-212 |
| chr8-17702180-DEL-51 | chr8-17702181-DEL-51 |
| chr8-17702715-DEL-132 | chr8-17702716-DEL-132 |
| chr8-17727343-DEL-110 | chr8-17727297-DEL-117 |
| chr8-17793520-DEL-59 | chr8-17793521-DEL-59 |
| chr8-17797324-DEL-401 | chr8-17797325-DEL-401 |
| chr8-17808831-DEL-56 | chr8-17808832-DEL-56 |
| chr8-17812721-DEL-167 | chr8-17812722-DEL-167 |
| chr8-17813808-DEL-5098 | chr8-17813809-DEL-5098 |
| chr8-17822715-DEL-427 | chr8-17822716-DEL-427 |
| chr8-17845647-DEL-78 | chr8-17845648-DEL-78 |
| chr8-17858899-DEL-53 | chr8-17858887-DEL-52 |
| chr8-17877088-DEL-3143 | chr8-17877089-DEL-3143 |
| chr8-17895442-DEL-55 | chr8-17895443-DEL-55 |
| chr8-17913003-DEL-183 | chr8-17913004-DEL-183 |
| chr8-17917777-DEL-5104 | chr8-17917778-DEL-5104 |
| chr8-17938785-DEL-532 | chr8-17938786-DEL-532 |
| chr8-17941677-DEL-62 | chr8-17941688-DEL-52 |
| chr8-17985633-DEL-88 | chr8-17985655-DEL-80 |
| chr8-17989199-DEL-113 | chr8-17989200-DEL-113 |
| chr8-18009308-DEL-103 | chr8-18009309-DEL-103 |
| chr8-18014208-DEL-6136 | chr8-18014195-DEL-6137 |
| chr8-18033578-DEL-1335 | chr8-18033579-DEL-1335 |
| chr8-18048552-DEL-257 | chr8-18048553-DEL-257 |
| chr8-18071688-DEL-1390 | chr8-18071689-DEL-1390 |
| chr8-18084140-DEL-6482 | chr8-18084141-DEL-6482 |
| chr8-18091405-DEL-1646 | chr8-18091406-DEL-1646 |
| chr8-18101965-DEL-202 | chr8-18101966-DEL-202 |
| chr8-18118453-DEL-124 | chr8-18118454-DEL-124 |
| chr8-18122366-DEL-66 | chr8-18122367-DEL-66 |
| chr8-18151085-DEL-60 | chr8-18151086-DEL-60 |
| chr8-18151573-DEL-345 | chr8-18151574-DEL-345 |
| chr8-18155926-DEL-110 | chr8-18155927-DEL-110 |
| chr8-18182178-DEL-1629 | chr8-18182179-DEL-1629 |
| chr8-18188219-DEL-4201 | chr8-18188220-DEL-4201 |
| chr8-18213126-DEL-90 | chr8-18213127-DEL-90 |
| chr8-18228561-DEL-614 | chr8-18228562-DEL-614 |
| chr8-18272411-DEL-127 | chr8-18272412-DEL-126 |
| chr8-18299607-DEL-1664 | chr8-18299608-DEL-1664 |
| chr8-18299908-DEL-303 | chr8-18299909-DEL-303 |
| chr8-18303455-DEL-105 | chr8-18303456-DEL-105 |
| chr8-18305533-DEL-226 | chr8-18305534-DEL-226 |
| chr8-18309128-DEL-6185 | chr8-18309129-DEL-6185 |
| chr8-18344743-DEL-119 | chr8-18344744-DEL-119 |
| chr8-18355620-DEL-1333 | chr8-18355621-DEL-1333 |
| chr8-18356332-DEL-958 | chr8-18356333-DEL-958 |
| chr8-18357256-DEL-1234 | chr8-18357257-DEL-1234 |
| chr8-18417464-DEL-188 | chr8-18417465-DEL-188 |
| chr8-18428671-DEL-60 | chr8-18428661-DEL-71 |
| chr8-18432684-DEL-85 | chr8-18432671-DEL-117 |
| chr8-18442294-DEL-355 | chr8-18442295-DEL-355 |
| chr8-18468363-DEL-62 | chr8-18468364-DEL-62 |
| chr8-18471901-DEL-66 | chr8-18471902-DEL-66 |
| chr8-18477082-DEL-1231 | chr8-18477083-DEL-1231 |
| chr8-18482267-DEL-398 | chr8-18482256-DEL-399 |
| chr8-18493940-DEL-67 | chr8-18493941-DEL-67 |
| chr8-18521173-DEL-799 | chr8-18521174-DEL-799 |
| chr8-18532219-DEL-221 | chr8-18532220-DEL-221 |
| chr8-18545379-DEL-52 | chr8-18545376-DEL-70 |
| chr8-18546841-DEL-151 | chr8-18546856-DEL-147 |
| chr8-18557819-DEL-534 | chr8-18557823-DEL-531 |

| | |
|---|---|
| chr8-18582607-DEL-2242 | chr8-18582608-DEL-2242 |
| chr8-18590392-DEL-141 | chr8-18590392-DEL-141 |
| chr8-18598128-DEL-1177 | chr8-18598129-DEL-1177 |
| chr8-18616803-DEL-1980 | chr8-18616803-DEL-1981 |
| chr8-18625264-DEL-4858 | chr8-18625249-DEL-4826 |
| chr8-18636080-DEL-72 | chr8-18636081-DEL-54 |
| chr8-18660806-DEL-60 | chr8-18660807-DEL-60 |
| chr8-18666379-DEL-758 | chr8-18666380-DEL-758 |
| chr8-18698269-DEL-202 | chr8-18698270-DEL-202 |
| chr8-18876353-DEL-626 | chr8-18876354-DEL-626 |
| chr8-18887486-DEL-312 | chr8-18887497-DEL-263 |
| chr8-18912792-DEL-200 | chr8-18912793-DEL-200 |
| chr8-18946411-DEL-337 | chr8-18946412-DEL-337 |
| chr8-18963628-DEL-109 | chr8-18963629-DEL-109 |
| chr8-19011796-DEL-148 | chr8-19011790-DEL-150 |
| chr8-19020592-DEL-537 | chr8-19020593-DEL-537 |
| chr8-19051998-DEL-2182 | chr8-19051998-DEL-2183 |
| chr8-19057128-DEL-766 | chr8-19057128-DEL-766 |
| chr8-19064583-DEL-163 | chr8-19064584-DEL-163 |
| chr8-19070464-DEL-206 | chr8-19070465-DEL-206 |
| chr8-19088448-DEL-6899 | chr8-19088449-DEL-6899 |
| chr8-19094755-DEL-428 | chr8-19094756-DEL-428 |
| chr8-19104694-DEL-5107 | chr8-19104694-DEL-5107 |
| chr8-19111195-DEL-118 | chr8-19111196-DEL-116 |
| chr8-19125799-DEL-62 | chr8-19125800-DEL-62 |
| chr8-19134675-DEL-6615 | chr8-19134660-DEL-6630 |
| chr8-19140889-DEL-201 | chr8-19140889-DEL-201 |
| chr8-19166320-DEL-3733 | chr8-19166321-DEL-3733 |
| chr8-19210153-DEL-69 | chr8-19210154-DEL-69 |
| chr8-19223923-DEL-7085 | chr8-19223924-DEL-7085 |
| chr8-19249325-DEL-333 | chr8-19249373-DEL-286 |
| chr8-19321796-DEL-108 | chr8-19321797-DEL-108 |
| chr8-19323545-DEL-141 | chr8-19323542-DEL-141 |
| chr8-19327737-DEL-206 | chr8-19327738-DEL-206 |
| chr8-19341829-DEL-124 | chr8-19341830-DEL-124 |
| chr8-19461360-DEL-157 | chr8-19461361-DEL-157 |
| chr8-19475761-DEL-327 | chr8-19475762-DEL-327 |
| chr8-19484209-DEL-549 | chr8-19484210-DEL-549 |
| chr8-19497509-DEL-3082 | chr8-19497510-DEL-3067 |
| chr8-19516003-DEL-5912 | chr8-19516004-DEL-5912 |
| chr8-19516156-DEL-471 | chr8-19516157-DEL-471 |
| chr8-19516682-DEL-432 | chr8-19516683-DEL-432 |
| chr8-19520752-DEL-399 | chr8-19520753-DEL-399 |
| chr8-19536697-DEL-113 | chr8-19536697-DEL-113 |
| chr8-19537608-DEL-4510 | chr8-19537609-DEL-4510 |
| chr8-19551747-DEL-90 | chr8-19551729-DEL-93 |
| chr8-19571970-DEL-202 | chr8-19571971-DEL-202 |
| chr8-19589391-DEL-401 | chr8-19589392-DEL-401 |
| chr8-19609751-DEL-53 | chr8-19609752-DEL-59 |
| chr8-19651568-DEL-60 | chr8-19651569-DEL-60 |
| chr8-19657693-DEL-548 | chr8-19657694-DEL-548 |
| chr8-19684591-DEL-138 | chr8-19684592-DEL-138 |
| chr8-19695628-DEL-123 | chr8-19695622-DEL-88 |
| chr8-19721290-DEL-3812 | chr8-19721290-DEL-3812 |
| chr8-20011645-DEL-200 | chr8-20011646-DEL-200 |
| chr8-20015526-DEL-634 | chr8-20015527-DEL-639 |
| chr8-20052379-DEL-210 | chr8-20052380-DEL-210 |
| chr8-20539256-DEL-122 | chr8-20539227-DEL-135 |
| chr8-20612223-DEL-198 | chr8-20612223-DEL-198 |
| chr8-20681803-DEL-72 | chr8-20681804-DEL-50 |
| chr8-21368748-DEL-400 | chr8-21368749-DEL-400 |
| chr8-21372861-DEL-343 | chr8-21372862-DEL-343 |
| chr8-21396286-DEL-750 | chr8-21396287-DEL-751 |
| chr8-21435646-DEL-3361 | chr8-21435647-DEL-3361 |
| chr8-21447636-DEL-6224 | chr8-21447637-DEL-6224 |
| chr8-21492038-DEL-2692 | chr8-21492039-DEL-2692 |
| chr8-21499758-DEL-98 | chr8-21499757-DEL-84 |
| chr8-21506854-DEL-413 | chr8-21506855-DEL-413 |
| chr8-21525226-DEL-244 | chr8-21525222-DEL-245 |
| chr8-21589341-DEL-159 | chr8-21589340-DEL-159 |
| chr8-21735990-DEL-736 | chr8-21735991-DEL-736 |
| chr8-21738367-DEL-841 | chr8-21738367-DEL-841 |
| chr8-21745341-DEL-831 | chr8-21745342-DEL-831 |
| chr8-22265160-DEL-52 | chr8-22265161-DEL-52 |
| chr8-22276691-DEL-3032 | chr8-22276692-DEL-3032 |
| chr8-22290617-DEL-401 | chr8-22290618-DEL-401 |
| chr8-22315205-DEL-1278 | chr8-22315206-DEL-1278 |
| chr8-22336045-DEL-95 | chr8-22336035-DEL-90 |
| chr8-22365496-DEL-167 | chr8-22365509-DEL-164 |
| chr8-22375071-DEL-61 | chr8-22375072-DEL-68 |

| | |
|---|---|
| chr8-22398124-DEL-374 | chr8-22398125-DEL-374 |
| chr8-22404720-DEL-72 | chr8-22404715-DEL-66 |
| chr8-22441406-DEL-498 | chr8-22441407-DEL-498 |
| chr8-22473508-DEL-6053 | chr8-22473509-DEL-6053 |
| chr8-22523864-DEL-55 | chr8-22523868-DEL-75 |
| chr8-22543065-DEL-60 | chr8-22543066-DEL-60 |
| chr8-22568812-DEL-56 | chr8-22568802-DEL-60 |
| chr8-22578263-DEL-407 | chr8-22578264-DEL-407 |
| chr8-22586188-DEL-77 | chr8-22586188-DEL-77 |
| chr8-22588627-DEL-5243 | chr8-22588628-DEL-5243 |
| chr8-22601292-DEL-113 | chr8-22601293-DEL-113 |
| chr8-22611883-DEL-132 | chr8-22611884-DEL-132 |
| chr8-22619555-DEL-792 | chr8-22619556-DEL-792 |
| chr8-22626980-DEL-2759 | chr8-22626981-DEL-2759 |
| chr8-22642707-DEL-249 | chr8-22642708-DEL-249 |
| chr8-22653716-DEL-287 | chr8-22653733-DEL-250 |
| chr8-22663877-DEL-105 | chr8-22663878-DEL-105 |
| chr8-22693021-DEL-143 | chr8-22693022-DEL-143 |
| chr8-22694640-DEL-51 | chr8-22694641-DEL-51 |
| chr8-22703577-DEL-989 | chr8-22703578-DEL-989 |
| chr8-22738602-DEL-193 | chr8-22738603-DEL-193 |
| chr8-22739772-DEL-189 | chr8-22739773-DEL-189 |
| chr8-22740422-DEL-51 | chr8-22740423-DEL-51 |
| chr8-22743157-DEL-66 | chr8-22743158-DEL-66 |
| chr8-22751957-DEL-102 | chr8-22751958-DEL-102 |
| chr8-22759762-DEL-55 | chr8-22759762-DEL-83 |
| chr8-22761142-DEL-638 | chr8-22761143-DEL-638 |
| chr8-22769271-DEL-1066 | chr8-22769272-DEL-1066 |
| chr8-22779326-DEL-6625 | chr8-22779327-DEL-6625 |
| chr8-22836132-DEL-62 | chr8-22836133-DEL-62 |
| chr8-22838328-DEL-784 | chr8-22838332-DEL-777 |
| chr8-22929751-DEL-210 | chr8-22929756-DEL-209 |
| chr8-22948916-DEL-181 | chr8-22948917-DEL-181 |
| chr8-22986674-DEL-219 | chr8-22986675-DEL-219 |
| chr8-23019283-DEL-196 | chr8-23019284-DEL-196 |
| chr8-23028135-DEL-54 | chr8-23028136-DEL-54 |
| chr8-23037359-DEL-162 | chr8-23037360-DEL-162 |
| chr8-23041177-DEL-7132 | chr8-23041178-DEL-7132 |
| chr8-23076474-DEL-212 | chr8-23076475-DEL-212 |
| chr8-23101727-DEL-147 | chr8-23101727-DEL-148 |
| chr8-23123052-DEL-82 | chr8-23123053-DEL-82 |
| chr8-23136026-DEL-226 | chr8-23136027-DEL-226 |
| chr8-23187802-DEL-123 | chr8-23187798-DEL-124 |
| chr8-23190932-DEL-136 | chr8-23190968-DEL-108 |
| chr8-23194413-DEL-157 | chr8-23194414-DEL-157 |
| chr8-23200977-DEL-106 | chr8-23200986-DEL-98 |
| chr8-23207497-DEL-852 | chr8-23207498-DEL-852 |
| chr8-23239836-DEL-55 | chr8-23239837-DEL-55 |
| chr8-23246387-DEL-74 | chr8-23246391-DEL-72 |
| chr8-23320099-DEL-196 | chr8-23320092-DEL-197 |
| chr8-23440107-DEL-69 | chr8-23440108-DEL-69 |
| chr8-23469238-DEL-58 | chr8-23469239-DEL-58 |
| chr8-23512147-DEL-164 | chr8-23512172-DEL-120 |
| chr8-23547616-DEL-416 | chr8-23547620-DEL-414 |
| chr8-23575907-DEL-629 | chr8-23575886-DEL-629 |
| chr8-23585491-DEL-328 | chr8-23585492-DEL-328 |
| chr8-23602061-DEL-174 | chr8-23602062-DEL-174 |
| chr8-23607355-DEL-191 | chr8-23607356-DEL-191 |
| chr8-23631465-DEL-54 | chr8-23631466-DEL-54 |
| chr8-23633317-DEL-127 | chr8-23633318-DEL-127 |
| chr8-23658304-DEL-51 | chr8-23658305-DEL-51 |
| chr8-23666144-DEL-217 | chr8-23666145-DEL-217 |
| chr8-23726497-DEL-82 | chr8-23726484-DEL-94 |
| chr8-23743609-DEL-206 | chr8-23743610-DEL-206 |
| chr8-23752362-DEL-54 | chr8-23752367-DEL-50 |
| chr8-23756420-DEL-3204 | chr8-23756417-DEL-3203 |
| chr8-23762983-DEL-386 | chr8-23762984-DEL-386 |
| chr8-23763435-DEL-2885 | chr8-23763436-DEL-2885 |
| chr8-23792846-DEL-66 | chr8-23792847-DEL-60 |
| chr8-23815954-DEL-75 | chr8-23815955-DEL-75 |
| chr8-23836251-DEL-268 | chr8-23836252-DEL-268 |
| chr8-23859344-DEL-3741 | chr8-23859345-DEL-3741 |
| chr8-23930052-DEL-176 | chr8-23930053-DEL-176 |
| chr8-23944570-DEL-63 | chr8-23944576-DEL-58 |
| chr8-24003610-DEL-1492 | chr8-24003611-DEL-1492 |
| chr8-24018696-DEL-53 | chr8-24018697-DEL-53 |
| chr8-24019271-DEL-585 | chr8-24019272-DEL-585 |
| chr8-24032177-DEL-207 | chr8-24032178-DEL-207 |
| chr8-24033550-DEL-293 | chr8-24033551-DEL-293 |
| chr8-24035582-DEL-5405 | chr8-24035583-DEL-5405 |

| | |
|---|---|
| chr8-24049711-DEL-58 | chr8-24049718-DEL-58 |
| chr8-24054073-DEL-158 | chr8-24054074-DEL-158 |
| chr8-24060238-DEL-535 | chr8-24060239-DEL-535 |
| chr8-24061078-DEL-409 | chr8-24061079-DEL-409 |
| chr8-24074184-DEL-1389 | chr8-24074185-DEL-1389 |
| chr8-24090379-DEL-329 | chr8-24090380-DEL-329 |
| chr8-24114866-DEL-60 | chr8-24114867-DEL-80 |
| chr8-24137086-DEL-64 | chr8-24137099-DEL-52 |
| chr8-24143426-DEL-206 | chr8-24143422-DEL-207 |
| chr8-24153230-DEL-96 | chr8-24153259-DEL-105 |
| chr8-24172078-DEL-90 | chr8-24172079-DEL-90 |
| chr8-24195226-DEL-119 | chr8-24195224-DEL-122 |
| chr8-24216662-DEL-62 | chr8-24216663-DEL-62 |
| chr8-24244245-DEL-408 | chr8-24244246-DEL-408 |
| chr8-24245344-DEL-56 | chr8-24245345-DEL-56 |
| chr8-24274482-DEL-495 | chr8-24274483-DEL-495 |
| chr8-24276219-DEL-56 | chr8-24276220-DEL-56 |
| chr8-24321019-DEL-6451 | chr8-24321020-DEL-6451 |
| chr8-24338440-DEL-181 | chr8-24338441-DEL-181 |
| chr8-24360097-DEL-6202 | chr8-24360098-DEL-6202 |
| chr8-24421753-DEL-210 | chr8-24421756-DEL-208 |
| chr8-24470037-DEL-136 | chr8-24470046-DEL-128 |
| chr8-24543498-DEL-535 | chr8-24543485-DEL-522 |
| chr8-24621753-DEL-128 | chr8-24621787-DEL-112 |
| chr8-24648458-DEL-112 | chr8-24648464-DEL-100 |
| chr8-24667595-DEL-504 | chr8-24667596-DEL-504 |
| chr8-24697911-DEL-99 | chr8-24697912-DEL-99 |
| chr8-24739678-DEL-1709 | chr8-24739679-DEL-1709 |
| chr8-24775514-DEL-1165 | chr8-24775515-DEL-1165 |
| chr8-24867176-DEL-53 | chr8-24867177-DEL-53 |
| chr8-24883380-DEL-86 | chr8-24883381-DEL-86 |
| chr8-24915198-DEL-92 | chr8-24915215-DEL-76 |
| chr8-24926308-DEL-104 | chr8-24926309-DEL-104 |
| chr8-24949593-DEL-63 | chr8-24949594-DEL-62 |
| chr8-24959099-DEL-179 | chr8-24959100-DEL-179 |
| chr8-24959956-DEL-70 | chr8-24959957-DEL-70 |
| chr8-24961649-DEL-1893 | chr8-24961650-DEL-1893 |
| chr8-24980299-DEL-63 | chr8-24980287-DEL-75 |
| chr8-24989095-DEL-111 | chr8-24989087-DEL-112 |
| chr8-24996988-DEL-380 | chr8-24996989-DEL-380 |
| chr8-25007949-DEL-322 | chr8-25007950-DEL-322 |
| chr8-25020860-DEL-701 | chr8-25020835-DEL-693 |
| chr8-25024323-DEL-100 | chr8-25024324-DEL-97 |
| chr8-25037945-DEL-86 | chr8-25037918-DEL-87 |
| chr8-25041504-DEL-609 | chr8-25041504-DEL-600 |
| chr8-25051581-DEL-550 | chr8-25051582-DEL-549 |
| chr8-25052567-DEL-124 | chr8-25052566-DEL-125 |
| chr8-25059840-DEL-227 | chr8-25059841-DEL-227 |
| chr8-25062193-DEL-144 | chr8-25062194-DEL-144 |
| chr8-25088557-DEL-79 | chr8-25088558-DEL-79 |
| chr8-25094875-DEL-543 | chr8-25094876-DEL-543 |
| chr8-25110558-DEL-3610 | chr8-25110548-DEL-3620 |
| chr8-25126072-DEL-140 | chr8-25126038-DEL-175 |
| chr8-25150438-DEL-3627 | chr8-25150439-DEL-3627 |
| chr8-25162676-DEL-52 | chr8-25162680-DEL-54 |
| chr8-25165478-DEL-401 | chr8-25165480-DEL-402 |
| chr8-25169568-DEL-121 | chr8-25169572-DEL-126 |
| chr8-25208431-DEL-607 | chr8-25208432-DEL-607 |
| chr8-25250117-DEL-3237 | chr8-25250118-DEL-3237 |
| chr8-25256534-DEL-65 | chr8-25256515-DEL-54 |
| chr8-25257422-DEL-7190 | chr8-25257423-DEL-7190 |
| chr8-25333178-DEL-6546 | chr8-25333179-DEL-6546 |
| chr8-25356330-DEL-410 | chr8-25356331-DEL-410 |
| chr8-25392309-DEL-205 | chr8-25392310-DEL-205 |
| chr8-25401676-DEL-206 | chr8-25401677-DEL-206 |
| chr8-25450334-DEL-1038 | chr8-25450335-DEL-1038 |
| chr8-25452879-DEL-54 | chr8-25452880-DEL-54 |
| chr8-25491763-DEL-1286 | chr8-25491764-DEL-1286 |
| chr8-25538646-DEL-59 | chr8-25538647-DEL-59 |
| chr8-25559660-DEL-74 | chr8-25559662-DEL-73 |
| chr8-25606633-DEL-1245 | chr8-25606634-DEL-1245 |
| chr8-25642125-DEL-968 | chr8-25642126-DEL-968 |
| chr8-25651464-DEL-65 | chr8-25651465-DEL-65 |
| chr8-25666067-DEL-143 | chr8-25666068-DEL-143 |
| chr8-25675503-DEL-144 | chr8-25675505-DEL-141 |
| chr8-25678788-DEL-977 | chr8-25678789-DEL-977 |
| chr8-25697189-DEL-67 | chr8-25697189-DEL-67 |
| chr8-25697908-DEL-3569 | chr8-25697909-DEL-3569 |
| chr8-25704371-DEL-132 | chr8-25704372-DEL-132 |
| chr8-25708821-DEL-74 | chr8-25708804-DEL-84 |

| | |
|---|---|
| chr8-25723722-DEL-79 | chr8-25723723-DEL-76 |
| chr8-25748699-DEL-62 | chr8-25748700-DEL-62 |
| chr8-25750416-DEL-440 | chr8-25750417-DEL-440 |
| chr8-25751068-DEL-615 | chr8-25751071-DEL-651 |
| chr8-25754683-DEL-63 | chr8-25754696-DEL-51 |
| chr8-25756672-DEL-414 | chr8-25756678-DEL-398 |
| chr8-25758170-DEL-158 | chr8-25758171-DEL-158 |
| chr8-25804173-DEL-100 | chr8-25804176-DEL-101 |
| chr8-25805802-DEL-91 | chr8-25805803-DEL-91 |
| chr8-25831657-DEL-164 | chr8-25831658-DEL-164 |
| chr8-25854593-DEL-93 | chr8-25854594-DEL-93 |
| chr8-25876973-DEL-82 | chr8-25876974-DEL-82 |
| chr8-25883620-DEL-64 | chr8-25883621-DEL-64 |
| chr8-25905422-DEL-71 | chr8-25905423-DEL-70 |
| chr8-25914735-DEL-348 | chr8-25914736-DEL-348 |
| chr8-25929016-DEL-86 | chr8-25928995-DEL-88 |
| chr8-25956648-DEL-205 | chr8-25956649-DEL-205 |
| chr8-25960229-DEL-199 | chr8-25960260-DEL-167 |
| chr8-25962368-DEL-105 | chr8-25962369-DEL-101 |
| chr8-25967116-DEL-180 | chr8-25967115-DEL-181 |
| chr8-25976322-DEL-82 | chr8-25976322-DEL-82 |
| chr8-25979581-DEL-70 | chr8-25979582-DEL-68 |
| chr8-25990210-DEL-64 | chr8-25990224-DEL-50 |
| chr8-25992947-DEL-185 | chr8-25992948-DEL-185 |
| chr8-26009860-DEL-74 | chr8-26009833-DEL-72 |
| chr8-26020130-DEL-68 | chr8-26020141-DEL-58 |
| chr8-26038408-DEL-95 | chr8-26038409-DEL-95 |
| chr8-26041116-DEL-90 | chr8-26041095-DEL-61 |
| chr8-26073497-DEL-217 | chr8-26073498-DEL-217 |
| chr8-26084441-DEL-189 | chr8-26084442-DEL-189 |
| chr8-26118843-DEL-228 | chr8-26118844-DEL-228 |
| chr8-26147615-DEL-115 | chr8-26147652-DEL-100 |
| chr8-26183968-DEL-88 | chr8-26184013-DEL-63 |
| chr8-26241144-DEL-98 | chr8-26241144-DEL-98 |
| chr8-26252724-DEL-216 | chr8-26252725-DEL-216 |
| chr8-26267995-DEL-332 | chr8-26267996-DEL-328 |
| chr8-26308871-DEL-323 | chr8-26308865-DEL-320 |
| chr8-26310845-DEL-1859 | chr8-26310846-DEL-1859 |
| chr8-26326393-DEL-1890 | chr8-26326393-DEL-1890 |
| chr8-26352676-DEL-77 | chr8-26352677-DEL-50 |
| chr8-26361476-DEL-7091 | chr8-26361477-DEL-7091 |
| chr8-26369173-DEL-198 | chr8-26369174-DEL-198 |
| chr8-26409035-DEL-67 | chr8-26409018-DEL-83 |
| chr8-26439369-DEL-184 | chr8-26439370-DEL-184 |
| chr8-26455878-DEL-711 | chr8-26455879-DEL-711 |
| chr8-26462291-DEL-192 | chr8-26462292-DEL-192 |
| chr8-26478639-DEL-265 | chr8-26478642-DEL-264 |
| chr8-26491152-DEL-109 | chr8-26491153-DEL-109 |
| chr8-26493542-DEL-146 | chr8-26493553-DEL-105 |
| chr8-26516855-DEL-54 | chr8-26516856-DEL-54 |
| chr8-26528405-DEL-213 | chr8-26528406-DEL-213 |
| chr8-26529993-DEL-213 | chr8-26529994-DEL-213 |
| chr8-26540136-DEL-88 | chr8-26540169-DEL-86 |
| chr8-26554170-DEL-207 | chr8-26554171-DEL-207 |
| chr8-26566556-DEL-118 | chr8-26566557-DEL-118 |
| chr8-26568220-DEL-77 | chr8-26568221-DEL-77 |
| chr8-26583268-DEL-140 | chr8-26583269-DEL-140 |
| chr8-26589712-DEL-181 | chr8-26589713-DEL-181 |
| chr8-26612106-DEL-207 | chr8-26612128-DEL-177 |
| chr8-26621596-DEL-214 | chr8-26621597-DEL-214 |
| chr8-26656773-DEL-86 | chr8-26656774-DEL-86 |
| chr8-26657880-DEL-61 | chr8-26657877-DEL-65 |
| chr8-26660846-DEL-251 | chr8-26660831-DEL-247 |
| chr8-26671709-DEL-193 | chr8-26671708-DEL-195 |
| chr8-26690568-DEL-236 | chr8-26690569-DEL-236 |
| chr8-26726158-DEL-52 | chr8-26726159-DEL-52 |
| chr8-26740518-DEL-91 | chr8-26740517-DEL-64 |
| chr8-26746528-DEL-72 | chr8-26746531-DEL-70 |
| chr8-26774528-DEL-113 | chr8-26774529-DEL-113 |
| chr8-26796432-DEL-179 | chr8-26796415-DEL-186 |
| chr8-26816571-DEL-409 | chr8-26816572-DEL-409 |
| chr8-26818216-DEL-432 | chr8-26818217-DEL-432 |
| chr8-26864290-DEL-56 | chr8-26864291-DEL-56 |
| chr8-26886422-DEL-171 | chr8-26886423-DEL-163 |
| chr8-26887775-DEL-137 | chr8-26887776-DEL-137 |
| chr8-26918141-DEL-82 | chr8-26918139-DEL-82 |
| chr8-26928266-DEL-76 | chr8-26928267-DEL-76 |
| chr8-26945164-DEL-572 | chr8-26945165-DEL-572 |
| chr8-26945809-DEL-1405 | chr8-26945810-DEL-1404 |
| chr8-26988032-DEL-319 | chr8-26988033-DEL-319 |

| | |
|---|---|
| chr8-27000145-DEL-123 | chr8-27000146-DEL-123 |
| chr8-27005301-DEL-465 | chr8-27005302-DEL-465 |
| chr8-27007073-DEL-99 | chr8-27007074-DEL-99 |
| chr8-27012815-DEL-413 | chr8-27012816-DEL-413 |
| chr8-27065986-DEL-74 | chr8-27065994-DEL-71 |
| chr8-27070612-DEL-7121 | chr8-27070613-DEL-7121 |
| chr8-27101035-DEL-729 | chr8-27101036-DEL-729 |
| chr8-27111721-DEL-109 | chr8-27111722-DEL-109 |
| chr8-27132355-DEL-163 | chr8-27132356-DEL-163 |
| chr8-27157091-DEL-822 | chr8-27157092-DEL-822 |
| chr8-27187072-DEL-291 | chr8-27187065-DEL-292 |
| chr8-27211334-DEL-86 | chr8-27211341-DEL-100 |
| chr8-27217404-DEL-661 | chr8-27217405-DEL-661 |
| chr8-27224750-DEL-314 | chr8-27224751-DEL-314 |
| chr8-27230413-DEL-91 | chr8-27230401-DEL-93 |
| chr8-27270342-DEL-105 | chr8-27270349-DEL-99 |
| chr8-27272588-DEL-157 | chr8-27272589-DEL-157 |
| chr8-27277667-DEL-400 | chr8-27277668-DEL-400 |
| chr8-27296222-DEL-56 | chr8-27296221-DEL-58 |
| chr8-27308517-DEL-92 | chr8-27308540-DEL-70 |
| chr8-27358097-DEL-75 | chr8-27358098-DEL-75 |
| chr8-27407544-DEL-1206 | chr8-27407544-DEL-1206 |
| chr8-27412155-DEL-540 | chr8-27412156-DEL-540 |
| chr8-27426032-DEL-81 | chr8-27426033-DEL-81 |
| chr8-27436631-DEL-91 | chr8-27436632-DEL-91 |
| chr8-27441538-DEL-6534 | chr8-27441539-DEL-6534 |
| chr8-27493566-DEL-163 | chr8-27493612-DEL-153 |
| chr8-27505007-DEL-2554 | chr8-27505008-DEL-2554 |
| chr8-27517410-DEL-153 | chr8-27517417-DEL-135 |
| chr8-27544054-DEL-1031 | chr8-27544055-DEL-1031 |
| chr8-27555783-DEL-72 | chr8-27555796-DEL-60 |
| chr8-27561490-DEL-225 | chr8-27561491-DEL-225 |
| chr8-27563875-DEL-418 | chr8-27563876-DEL-418 |
| chr8-27564930-DEL-192 | chr8-27564961-DEL-152 |
| chr8-27619824-DEL-89 | chr8-27619825-DEL-89 |
| chr8-27620491-DEL-209 | chr8-27620492-DEL-209 |
| chr8-27625457-DEL-163 | chr8-27625458-DEL-163 |
| chr8-27635762-DEL-70 | chr8-27635763-DEL-70 |
| chr8-27648730-DEL-210 | chr8-27648731-DEL-210 |
| chr8-27653558-DEL-87 | chr8-27653560-DEL-84 |
| chr8-27655971-DEL-178 | chr8-27656017-DEL-133 |
| chr8-27668146-DEL-175 | chr8-27668147-DEL-175 |
| chr8-27668956-DEL-181 | chr8-27668957-DEL-181 |
| chr8-27671269-DEL-55 | chr8-27671270-DEL-50 |
| chr8-27702956-DEL-73 | chr8-27702957-DEL-73 |
| chr8-27738633-DEL-184 | chr8-27738634-DEL-184 |
| chr8-27758368-DEL-52 | chr8-27758369-DEL-52 |
| chr8-27772217-DEL-54 | chr8-27772218-DEL-54 |
| chr8-27780420-DEL-1178 | chr8-27780412-DEL-1179 |
| chr8-27807846-DEL-1403 | chr8-27807847-DEL-1403 |
| chr8-27820210-DEL-201 | chr8-27820211-DEL-201 |
| chr8-27821499-DEL-209 | chr8-27821500-DEL-209 |
| chr8-27842059-DEL-64 | chr8-27842047-DEL-62 |
| chr8-27880878-DEL-249 | chr8-27880879-DEL-249 |
| chr8-27885224-DEL-51 | chr8-27885221-DEL-54 |
| chr8-27890699-DEL-325 | chr8-27890700-DEL-325 |
| chr8-27891327-DEL-336 | chr8-27891328-DEL-336 |
| chr8-27928904-DEL-256 | chr8-27928905-DEL-256 |
| chr8-27952416-DEL-72 | chr8-27952418-DEL-72 |
| chr8-27952929-DEL-73 | chr8-27952930-DEL-73 |
| chr8-27954655-DEL-7062 | chr8-27954656-DEL-7062 |
| chr8-27965310-DEL-358 | chr8-27965311-DEL-358 |
| chr8-27995742-DEL-204 | chr8-27995743-DEL-203 |
| chr8-27997712-DEL-6914 | chr8-27997713-DEL-6914 |
| chr8-28005490-DEL-567 | chr8-28005491-DEL-567 |
| chr8-28014337-DEL-289 | chr8-28014333-DEL-288 |
| chr8-28016477-DEL-66 | chr8-28016478-DEL-66 |
| chr8-28039074-DEL-400 | chr8-28039075-DEL-400 |
| chr8-28040503-DEL-415 | chr8-28040504-DEL-415 |
| chr8-28062210-DEL-62 | chr8-28062207-DEL-63 |
| chr8-28076826-DEL-6247 | chr8-28076827-DEL-6247 |
| chr8-28112810-DEL-78 | chr8-28112811-DEL-78 |
| chr8-28127818-DEL-1926 | chr8-28127819-DEL-1926 |
| chr8-28133282-DEL-196 | chr8-28133283-DEL-196 |
| chr8-28146074-DEL-486 | chr8-28146074-DEL-486 |
| chr8-28185962-DEL-6371 | chr8-28185963-DEL-6371 |
| chr8-28197406-DEL-58 | chr8-28197385-DEL-94 |
| chr8-28219224-DEL-6806 | chr8-28219225-DEL-6806 |
| chr8-28219595-DEL-208 | chr8-28219596-DEL-208 |
| chr8-28232053-DEL-2246 | chr8-28232054-DEL-2246 |

| | |
|---|---|
| chr8-28243388-DEL-1240 | chr8-28243389-DEL-1240 |
| chr8-28246592-DEL-134 | chr8-28246595-DEL-130 |
| chr8-28288900-DEL-62 | chr8-28288899-DEL-62 |
| chr8-28292217-DEL-60 | chr8-28292218-DEL-60 |
| chr8-28292984-DEL-7181 | chr8-28292985-DEL-7181 |
| chr8-28298417-DEL-623 | chr8-28298418-DEL-623 |
| chr8-28303514-DEL-267 | chr8-28303510-DEL-262 |
| chr8-28367495-DEL-224 | chr8-28367496-DEL-224 |
| chr8-28424096-DEL-1401 | chr8-28424097-DEL-1401 |
| chr8-28426229-DEL-52 | chr8-28426272-DEL-52 |
| chr8-28427043-DEL-194 | chr8-28427044-DEL-194 |
| chr8-28445311-DEL-58 | chr8-28445312-DEL-58 |
| chr8-28446525-DEL-3436 | chr8-28446526-DEL-3436 |
| chr8-28500305-DEL-96 | chr8-28500306-DEL-92 |
| chr8-28512321-DEL-211 | chr8-28512322-DEL-211 |
| chr8-28530221-DEL-220 | chr8-28530222-DEL-220 |
| chr8-28541727-DEL-54 | chr8-28541717-DEL-54 |
| chr8-28553346-DEL-1792 | chr8-28553347-DEL-1792 |
| chr8-28560743-DEL-120 | chr8-28560748-DEL-121 |
| chr8-28589675-DEL-9312 | chr8-28589667-DEL-9311 |
| chr8-28697536-DEL-81 | chr8-28697537-DEL-81 |
| chr8-28698634-DEL-2738 | chr8-28698641-DEL-2737 |
| chr8-28734993-DEL-83 | chr8-28734999-DEL-60 |
| chr8-28755488-DEL-393 | chr8-28755489-DEL-393 |
| chr8-28772801-DEL-1161 | chr8-28772802-DEL-1161 |
| chr8-28791103-DEL-1822 | chr8-28791104-DEL-1822 |
| chr8-28829358-DEL-373 | chr8-28829359-DEL-373 |
| chr8-28836460-DEL-52 | chr8-28836460-DEL-52 |
| chr8-28838168-DEL-57 | chr8-28838169-DEL-57 |
| chr8-28876651-DEL-6643 | chr8-28876652-DEL-6643 |
| chr8-28883018-DEL-212 | chr8-28883019-DEL-212 |
| chr8-28924230-DEL-6425 | chr8-28924231-DEL-6425 |
| chr8-28941882-DEL-1880 | chr8-28941883-DEL-1880 |
| chr8-28982038-DEL-1054 | chr8-28982039-DEL-1056 |
| chr8-29020135-DEL-6649 | chr8-29020136-DEL-6649 |
| chr8-29028729-DEL-290 | chr8-29028730-DEL-290 |
| chr8-29031148-DEL-320 | chr8-29031149-DEL-320 |
| chr8-29032939-DEL-918 | chr8-29032940-DEL-917 |
| chr8-29035368-DEL-965 | chr8-29035369-DEL-965 |
| chr8-29036389-DEL-442 | chr8-29036390-DEL-442 |
| chr8-29049257-DEL-76 | chr8-29049258-DEL-75 |
| chr8-29094898-DEL-232 | chr8-29094899-DEL-232 |
| chr8-29103841-DEL-6173 | chr8-29103842-DEL-6173 |
| chr8-29130393-DEL-343 | chr8-29130394-DEL-343 |
| chr8-29211524-DEL-52 | chr8-29211554-DEL-60 |
| chr8-29216712-DEL-294 | chr8-29216713-DEL-294 |
| chr8-29221834-DEL-2649 | chr8-29221835-DEL-2649 |
| chr8-29265317-DEL-80 | chr8-29265318-DEL-80 |
| chr8-29266318-DEL-382 | chr8-29266312-DEL-383 |
| chr8-29288884-DEL-60 | chr8-29288883-DEL-62 |
| chr8-29315718-DEL-543 | chr8-29315716-DEL-544 |
| chr8-29331828-DEL-60 | chr8-29331829-DEL-60 |
| chr8-29351068-DEL-6438 | chr8-29351069-DEL-6438 |
| chr8-29357644-DEL-82 | chr8-29357643-DEL-84 |
| chr8-29374146-DEL-518 | chr8-29374147-DEL-518 |
| chr8-29377404-DEL-1116 | chr8-29377405-DEL-1116 |
| chr8-29394271-DEL-6208 | chr8-29394272-DEL-6208 |
| chr8-29403446-DEL-97 | chr8-29403447-DEL-97 |
| chr8-29412913-DEL-78 | chr8-29412882-DEL-106 |
| chr8-29428305-DEL-500 | chr8-29428310-DEL-496 |
| chr8-29447099-DEL-72 | chr8-29447118-DEL-62 |
| chr8-29452703-DEL-1466 | chr8-29452704-DEL-1466 |
| chr8-29454344-DEL-6887 | chr8-29454345-DEL-6887 |
| chr8-29460767-DEL-424 | chr8-29460767-DEL-424 |
| chr8-29482183-DEL-72 | chr8-29482179-DEL-77 |
| chr8-29489488-DEL-2092 | chr8-29489488-DEL-2090 |
| chr8-29496541-DEL-51 | chr8-29496542-DEL-51 |
| chr8-29501692-DEL-185 | chr8-29501693-DEL-185 |
| chr8-29509115-DEL-350 | chr8-29509116-DEL-350 |
| chr8-29541057-DEL-60 | chr8-29541058-DEL-60 |
| chr8-29571238-DEL-54 | chr8-29571239-DEL-54 |
| chr8-29583427-DEL-85 | chr8-29583458-DEL-55 |
| chr8-29586055-DEL-965 | chr8-29586056-DEL-965 |
| chr8-29615074-DEL-84 | chr8-29615075-DEL-84 |
| chr8-29638982-DEL-2031 | chr8-29638983-DEL-2031 |
| chr8-29667335-DEL-96 | chr8-29667336-DEL-96 |
| chr8-29684049-DEL-6938 | chr8-29684050-DEL-6938 |
| chr8-29692924-DEL-86 | chr8-29692941-DEL-66 |
| chr8-29701509-DEL-90 | chr8-29701510-DEL-90 |
| chr8-29747206-DEL-6413 | chr8-29747207-DEL-6413 |

| | |
|---|---|
| chr8-29754206-DEL-55 | chr8-29754207-DEL-55 |
| chr8-29754818-DEL-250 | chr8-29754819-DEL-250 |
| chr8-29776198-DEL-1101 | chr8-29776199-DEL-1101 |
| chr8-29787995-DEL-2531 | chr8-29788006-DEL-2523 |
| chr8-29795828-DEL-52 | chr8-29795829-DEL-52 |
| chr8-29800485-DEL-53 | chr8-29800486-DEL-53 |
| chr8-29818971-DEL-121 | chr8-29818970-DEL-128 |
| chr8-29828745-DEL-71 | chr8-29828754-DEL-63 |
| chr8-29844029-DEL-6746 | chr8-29844030-DEL-6746 |
| chr8-29871678-DEL-327 | chr8-29871679-DEL-327 |
| chr8-29899881-DEL-6495 | chr8-29899882-DEL-6495 |
| chr8-29905923-DEL-213 | chr8-29905923-DEL-213 |
| chr8-29943054-DEL-845 | chr8-29943055-DEL-845 |
| chr8-29948882-DEL-1025 | chr8-29948883-DEL-1025 |
| chr8-29977657-DEL-168 | chr8-29977644-DEL-154 |
| chr8-29993246-DEL-92 | chr8-29993263-DEL-62 |
| chr8-29995875-DEL-8175 | chr8-29995859-DEL-8191 |
| chr8-30005508-DEL-5159 | chr8-30005509-DEL-5159 |
| chr8-30016131-DEL-921 | chr8-30016132-DEL-921 |
| chr8-30038442-DEL-6392 | chr8-30038443-DEL-6392 |
| chr8-30049536-DEL-539 | chr8-30049537-DEL-539 |
| chr8-30082128-DEL-101 | chr8-30082142-DEL-87 |
| chr8-30093981-DEL-68 | chr8-30093982-DEL-68 |
| chr8-30100441-DEL-56 | chr8-30100442-DEL-56 |
| chr8-30113636-DEL-35387 | chr8-30113636-DEL-35387 |
| chr8-30164335-DEL-326 | chr8-30164342-DEL-326 |
| chr8-30206181-DEL-1624 | chr8-30206182-DEL-1624 |
| chr8-30228495-DEL-77 | chr8-30228493-DEL-67 |
| chr8-30234140-DEL-364 | chr8-30234141-DEL-364 |
| chr8-30245998-DEL-53 | chr8-30245958-DEL-57 |
| chr8-30253176-DEL-258 | chr8-30253177-DEL-258 |
| chr8-30254161-DEL-72 | chr8-30254126-DEL-73 |
| chr8-30276394-DEL-462 | chr8-30276395-DEL-462 |
| chr8-30280475-DEL-424 | chr8-30280484-DEL-398 |
| chr8-30286770-DEL-854 | chr8-30286771-DEL-854 |
| chr8-30288082-DEL-6216 | chr8-30288083-DEL-6217 |
| chr8-30335803-DEL-4868 | chr8-30335804-DEL-4868 |
| chr8-30351378-DEL-4034 | chr8-30351379-DEL-4034 |
| chr8-30361444-DEL-152 | chr8-30361445-DEL-152 |
| chr8-30424756-DEL-6645 | chr8-30424757-DEL-6645 |
| chr8-30440836-DEL-1381 | chr8-30440838-DEL-1370 |
| chr8-30465192-DEL-61 | chr8-30465189-DEL-61 |
| chr8-30470585-DEL-6475 | chr8-30470586-DEL-6475 |
| chr8-30487902-DEL-57 | chr8-30487900-DEL-60 |
| chr8-30504715-DEL-559 | chr8-30504716-DEL-559 |
| chr8-30506783-DEL-7199 | chr8-30506784-DEL-7199 |
| chr8-30546408-DEL-96 | chr8-30546408-DEL-96 |
| chr8-30590471-DEL-214 | chr8-30590472-DEL-214 |
| chr8-30617947-DEL-379 | chr8-30617946-DEL-379 |
| chr8-30635303-DEL-114 | chr8-30635304-DEL-114 |
| chr8-30645423-DEL-13556 | chr8-30645424-DEL-13556 |
| chr8-30661030-DEL-280 | chr8-30661031-DEL-280 |
| chr8-30667919-DEL-95 | chr8-30667920-DEL-95 |
| chr8-30672692-DEL-185 | chr8-30672693-DEL-185 |
| chr8-30677504-DEL-304 | chr8-30677466-DEL-304 |
| chr8-30689376-DEL-2982 | chr8-30689377-DEL-2982 |
| chr8-30756922-DEL-128 | chr8-30756923-DEL-125 |
| chr8-30772441-DEL-96 | chr8-30772445-DEL-92 |
| chr8-30805202-DEL-558 | chr8-30805203-DEL-558 |
| chr8-30889042-DEL-6194 | chr8-30889043-DEL-6194 |
| chr8-30900888-DEL-142 | chr8-30900889-DEL-142 |
| chr8-30915841-DEL-1204 | chr8-30915842-DEL-1204 |
| chr8-30931467-DEL-194 | chr8-30931468-DEL-194 |
| chr8-30937812-DEL-2505 | chr8-30937813-DEL-2505 |
| chr8-30958462-DEL-83 | chr8-30958463-DEL-83 |
| chr8-30974988-DEL-343 | chr8-30974989-DEL-343 |
| chr8-30979794-DEL-241 | chr8-30979795-DEL-241 |
| chr8-31016817-DEL-61 | chr8-31016817-DEL-59 |
| chr8-31026906-DEL-3178 | chr8-31026907-DEL-3178 |
| chr8-31058931-DEL-127 | chr8-31058932-DEL-127 |
| chr8-31084677-DEL-73 | chr8-31084670-DEL-77 |
| chr8-31094756-DEL-62 | chr8-31094759-DEL-58 |
| chr8-31115147-DEL-619 | chr8-31115148-DEL-619 |
| chr8-31135435-DEL-96 | chr8-31135437-DEL-95 |
| chr8-31141585-DEL-59 | chr8-31141586-DEL-59 |
| chr8-31156839-DEL-211 | chr8-31156840-DEL-211 |
| chr8-31167811-DEL-1083 | chr8-31167812-DEL-1083 |
| chr8-31195170-DEL-6792 | chr8-31195170-DEL-6793 |
| chr8-31202192-DEL-70 | chr8-31202193-DEL-70 |
| chr8-31203086-DEL-1115 | chr8-31203087-DEL-1115 |

| | |
|---|---|
| chr8-31214124-DEL-86 | chr8-31214125-DEL-86 |
| chr8-31215329-DEL-1232 | chr8-31215330-DEL-1228 |
| chr8-31237825-DEL-88 | chr8-31237832-DEL-64 |
| chr8-31244498-DEL-184 | chr8-31244499-DEL-184 |
| chr8-31250670-DEL-79 | chr8-31250671-DEL-87 |
| chr8-31263203-DEL-87 | chr8-31263204-DEL-87 |
| chr8-31286152-DEL-60 | chr8-31286152-DEL-60 |
| chr8-31289052-DEL-209 | chr8-31289053-DEL-209 |
| chr8-31301538-DEL-166 | chr8-31301539-DEL-166 |
| chr8-31308596-DEL-69 | chr8-31308596-DEL-70 |
| chr8-31319066-DEL-98 | chr8-31319066-DEL-93 |
| chr8-31335310-DEL-167 | chr8-31335310-DEL-165 |
| chr8-31345197-DEL-68 | chr8-31345198-DEL-68 |
| chr8-31370395-DEL-66 | chr8-31370395-DEL-64 |
| chr8-31371100-DEL-136 | chr8-31371093-DEL-133 |
| chr8-31374912-DEL-4937 | chr8-31374913-DEL-4937 |
| chr8-31377175-DEL-2195 | chr8-31377176-DEL-2195 |
| chr8-31380416-DEL-92 | chr8-31380417-DEL-92 |
| chr8-31382182-DEL-204 | chr8-31382179-DEL-208 |
| chr8-31406936-DEL-74 | chr8-31406937-DEL-72 |
| chr8-31415392-DEL-85 | chr8-31415423-DEL-50 |
| chr8-31428659-DEL-56 | chr8-31428660-DEL-56 |
| chr8-31432979-DEL-285 | chr8-31432977-DEL-282 |
| chr8-31434683-DEL-268 | chr8-31434684-DEL-267 |
| chr8-31438728-DEL-375 | chr8-31438729-DEL-375 |
| chr8-31451069-DEL-196 | chr8-31451070-DEL-196 |
| chr8-31452717-DEL-160 | chr8-31452718-DEL-160 |
| chr8-31460268-DEL-51 | chr8-31460266-DEL-61 |
| chr8-31465579-DEL-57 | chr8-31465586-DEL-76 |
| chr8-31466492-DEL-1780 | chr8-31466495-DEL-1781 |
| chr8-31468935-DEL-233 | chr8-31468936-DEL-233 |
| chr8-31504843-DEL-567 | chr8-31504844-DEL-567 |
| chr8-31511037-DEL-2770 | chr8-31511038-DEL-2770 |
| chr8-31530421-DEL-68 | chr8-31530376-DEL-78 |
| chr8-31536099-DEL-323 | chr8-31536100-DEL-321 |
| chr8-31538615-DEL-131 | chr8-31538597-DEL-132 |
| chr8-31544227-DEL-78 | chr8-31544228-DEL-82 |
| chr8-31558517-DEL-83 | chr8-31558511-DEL-70 |
| chr8-31590946-DEL-2333 | chr8-31590947-DEL-2333 |
| chr8-31647650-DEL-430 | chr8-31647651-DEL-430 |
| chr8-31659092-DEL-2345 | chr8-31659093-DEL-2345 |
| chr8-31674070-DEL-1796 | chr8-31674071-DEL-1796 |
| chr8-31679053-DEL-75 | chr8-31679054-DEL-75 |
| chr8-31686634-DEL-83 | chr8-31686635-DEL-83 |
| chr8-31692710-DEL-102 | chr8-31692711-DEL-102 |
| chr8-31706535-DEL-72 | chr8-31706529-DEL-76 |
| chr8-31727822-DEL-799 | chr8-31727822-DEL-799 |
| chr8-31728975-DEL-1751 | chr8-31728967-DEL-1749 |
| chr8-31735251-DEL-66 | chr8-31735242-DEL-68 |
| chr8-31741024-DEL-89 | chr8-31741023-DEL-89 |
| chr8-31757106-DEL-54 | chr8-31757107-DEL-54 |
| chr8-31820346-DEL-650 | chr8-31820349-DEL-649 |
| chr8-31829264-DEL-67 | chr8-31829278-DEL-54 |
| chr8-31837512-DEL-732 | chr8-31837513-DEL-732 |
| chr8-31841542-DEL-1963 | chr8-31841526-DEL-1979 |
| chr8-31845219-DEL-54 | chr8-31845216-DEL-62 |
| chr8-31851763-DEL-232 | chr8-31851764-DEL-232 |
| chr8-31860928-DEL-90 | chr8-31860929-DEL-90 |
| chr8-31873455-DEL-58 | chr8-31873454-DEL-76 |
| chr8-31880938-DEL-6327 | chr8-31880939-DEL-6327 |
| chr8-31905085-DEL-322 | chr8-31905086-DEL-322 |
| chr8-31905887-DEL-114 | chr8-31905888-DEL-114 |
| chr8-31936274-DEL-130 | chr8-31936275-DEL-130 |
| chr8-31948077-DEL-713 | chr8-31948078-DEL-713 |
| chr8-31958607-DEL-81 | chr8-31958608-DEL-81 |
| chr8-31967431-DEL-59 | chr8-31967431-DEL-59 |
| chr8-31971172-DEL-56 | chr8-31971172-DEL-56 |
| chr8-31981709-DEL-57 | chr8-31981677-DEL-57 |
| chr8-31995656-DEL-6390 | chr8-31995657-DEL-6390 |
| chr8-32002897-DEL-712 | chr8-32002898-DEL-712 |
| chr8-32004099-DEL-256 | chr8-32004100-DEL-256 |
| chr8-32032778-DEL-565 | chr8-32032779-DEL-565 |
| chr8-32035230-DEL-291 | chr8-32035231-DEL-291 |
| chr8-32049199-DEL-402 | chr8-32049200-DEL-402 |
| chr8-32052925-DEL-67 | chr8-32052926-DEL-67 |
| chr8-32079092-DEL-60 | chr8-32079093-DEL-60 |
| chr8-32085206-DEL-76 | chr8-32085215-DEL-60 |
| chr8-32103719-DEL-58 | chr8-32103688-DEL-90 |
| chr8-32121378-DEL-66 | chr8-32121391-DEL-60 |
| chr8-32123827-DEL-3173 | chr8-32123828-DEL-3173 |

| | |
|---|---|
| chr8-32140765-DEL-341 | chr8-32140766-DEL-341 |
| chr8-32151308-DEL-1774 | chr8-32151309-DEL-1774 |
| chr8-32159845-DEL-2234 | chr8-32159846-DEL-2234 |
| chr8-32160893-DEL-224 | chr8-32160894-DEL-216 |
| chr8-32165219-DEL-6584 | chr8-32165220-DEL-6584 |
| chr8-32173299-DEL-59 | chr8-32173300-DEL-59 |
| chr8-32198127-DEL-59 | chr8-32198128-DEL-59 |
| chr8-32202067-DEL-692 | chr8-32202068-DEL-692 |
| chr8-32207756-DEL-66 | chr8-32207757-DEL-66 |
| chr8-32270633-DEL-294 | chr8-32270630-DEL-295 |
| chr8-32278152-DEL-3613 | chr8-32278153-DEL-3597 |
| chr8-32288721-DEL-831 | chr8-32288722-DEL-831 |
| chr8-32320663-DEL-56 | chr8-32320648-DEL-87 |
| chr8-32339864-DEL-91 | chr8-32339865-DEL-91 |
| chr8-32362927-DEL-369 | chr8-32362928-DEL-366 |
| chr8-32379321-DEL-74 | chr8-32379322-DEL-74 |
| chr8-32393749-DEL-52 | chr8-32393750-DEL-60 |
| chr8-32404635-DEL-210 | chr8-32404636-DEL-210 |
| chr8-32409116-DEL-60 | chr8-32409117-DEL-60 |
| chr8-32417442-DEL-97 | chr8-32417443-DEL-97 |
| chr8-32434072-DEL-201 | chr8-32434073-DEL-201 |
| chr8-32459138-DEL-1833 | chr8-32459139-DEL-1833 |
| chr8-32480255-DEL-74 | chr8-32480256-DEL-74 |
| chr8-32504225-DEL-77 | chr8-32504210-DEL-60 |
| chr8-32514787-DEL-111 | chr8-32514788-DEL-111 |
| chr8-32545826-DEL-99 | chr8-32545825-DEL-100 |
| chr8-32550405-DEL-69 | chr8-32550404-DEL-70 |
| chr8-32583760-DEL-136 | chr8-32583759-DEL-134 |
| chr8-32610087-DEL-50 | chr8-32610087-DEL-50 |
| chr8-32620381-DEL-67 | chr8-32620382-DEL-67 |
| chr8-32641658-DEL-53 | chr8-32641654-DEL-51 |
| chr8-32651034-DEL-117 | chr8-32651035-DEL-117 |
| chr8-32675045-DEL-194 | chr8-32675046-DEL-195 |
| chr8-32682137-DEL-6329 | chr8-32682138-DEL-6329 |
| chr8-32716964-DEL-65 | chr8-32716973-DEL-105 |
| chr8-32718983-DEL-64 | chr8-32718984-DEL-64 |
| chr8-32725191-DEL-68 | chr8-32725192-DEL-68 |
| chr8-32747675-DEL-135 | chr8-32747676-DEL-135 |
| chr8-32749515-DEL-56 | chr8-32749514-DEL-58 |
| chr8-32802079-DEL-610 | chr8-32802080-DEL-610 |
| chr8-32844659-DEL-350 | chr8-32844660-DEL-350 |
| chr8-32856957-DEL-200 | chr8-32856958-DEL-200 |
| chr8-32876820-DEL-58 | chr8-32876820-DEL-56 |
| chr8-32880498-DEL-136 | chr8-32880498-DEL-136 |
| chr8-32899265-DEL-604 | chr8-32899266-DEL-604 |
| chr8-32925372-DEL-106 | chr8-32925373-DEL-106 |
| chr8-32930035-DEL-52 | chr8-32930036-DEL-50 |
| chr8-32943173-DEL-180 | chr8-32943174-DEL-180 |
| chr8-32958156-DEL-76 | chr8-32958157-DEL-76 |
| chr8-32966839-DEL-541 | chr8-32966840-DEL-541 |
| chr8-33047049-DEL-773 | chr8-33047050-DEL-773 |
| chr8-33051739-DEL-400 | chr8-33051740-DEL-400 |
| chr8-33062120-DEL-395 | chr8-33062121-DEL-395 |
| chr8-33065491-DEL-6475 | chr8-33065492-DEL-6475 |
| chr8-33104135-DEL-53 | chr8-33104135-DEL-53 |
| chr8-33120825-DEL-2217 | chr8-33120826-DEL-2217 |
| chr8-33125499-DEL-68 | chr8-33125500-DEL-68 |
| chr8-33141982-DEL-439 | chr8-33141979-DEL-439 |
| chr8-33175802-DEL-132 | chr8-33175805-DEL-102 |
| chr8-33197808-DEL-52 | chr8-33197809-DEL-52 |
| chr8-33201070-DEL-485 | chr8-33201071-DEL-485 |
| chr8-33204267-DEL-6530 | chr8-33204268-DEL-6530 |
| chr8-33226058-DEL-91 | chr8-33226059-DEL-87 |
| chr8-33230502-DEL-6387 | chr8-33230502-DEL-6387 |
| chr8-33255785-DEL-62 | chr8-33255769-DEL-66 |
| chr8-33264777-DEL-210 | chr8-33264778-DEL-210 |
| chr8-33286583-DEL-202 | chr8-33286578-DEL-207 |
| chr8-33287777-DEL-2698 | chr8-33287778-DEL-2698 |
| chr8-33288190-DEL-262 | chr8-33288191-DEL-262 |
| chr8-33290358-DEL-69 | chr8-33290359-DEL-69 |
| chr8-33291896-DEL-2987 | chr8-33291897-DEL-2987 |
| chr8-33321571-DEL-80 | chr8-33321572-DEL-80 |
| chr8-33341216-DEL-60 | chr8-33341243-DEL-55 |
| chr8-33343294-DEL-6596 | chr8-33343295-DEL-6596 |
| chr8-33358738-DEL-935 | chr8-33358739-DEL-935 |
| chr8-33362331-DEL-224 | chr8-33362332-DEL-224 |
| chr8-33381928-DEL-57 | chr8-33381930-DEL-50 |
| chr8-33382560-DEL-3434 | chr8-33382561-DEL-3434 |
| chr8-33411792-DEL-1611 | chr8-33411793-DEL-1611 |
| chr8-33419145-DEL-237 | chr8-33419146-DEL-237 |

| | |
|---|---|
| chr8-33455793-DEL-121 | chr8-33455794-DEL-121 |
| chr8-33471274-DEL-315 | chr8-33471275-DEL-315 |
| chr8-33497213-DEL-73 | chr8-33497212-DEL-74 |
| chr8-33503758-DEL-659 | chr8-33503759-DEL-659 |
| chr8-33504863-DEL-129 | chr8-33504864-DEL-129 |
| chr8-33528937-DEL-144 | chr8-33528938-DEL-144 |
| chr8-33541726-DEL-75 | chr8-33541734-DEL-67 |
| chr8-33545625-DEL-89 | chr8-33545626-DEL-89 |
| chr8-33547149-DEL-184 | chr8-33547150-DEL-184 |
| chr8-33619498-DEL-1580 | chr8-33619499-DEL-1580 |
| chr8-33630513-DEL-1402 | chr8-33630514-DEL-1402 |
| chr8-33632327-DEL-199 | chr8-33632328-DEL-199 |
| chr8-33641184-DEL-6787 | chr8-33641185-DEL-6787 |
| chr8-33670615-DEL-175 | chr8-33670616-DEL-175 |
| chr8-33679371-DEL-2990 | chr8-33679372-DEL-2990 |
| chr8-33690209-DEL-138 | chr8-33690169-DEL-159 |
| chr8-33704775-DEL-264 | chr8-33704776-DEL-264 |
| chr8-33712400-DEL-139 | chr8-33712407-DEL-131 |
| chr8-33726950-DEL-1887 | chr8-33726951-DEL-1887 |
| chr8-33729076-DEL-68 | chr8-33729076-DEL-68 |
| chr8-33744252-DEL-1050 | chr8-33744253-DEL-1050 |
| chr8-33751494-DEL-68 | chr8-33751497-DEL-66 |
| chr8-33756419-DEL-139 | chr8-33756420-DEL-137 |
| chr8-33772667-DEL-127 | chr8-33772668-DEL-127 |
| chr8-33773551-DEL-56 | chr8-33773552-DEL-56 |
| chr8-33775422-DEL-206 | chr8-33775423-DEL-206 |
| chr8-33780326-DEL-83 | chr8-33780286-DEL-84 |
| chr8-33796833-DEL-58 | chr8-33796840-DEL-52 |
| chr8-33819268-DEL-65 | chr8-33819297-DEL-52 |
| chr8-33831197-DEL-205 | chr8-33831201-DEL-204 |
| chr8-33835947-DEL-559 | chr8-33835948-DEL-559 |
| chr8-33841766-DEL-196 | chr8-33841767-DEL-196 |
| chr8-33847122-DEL-164 | chr8-33847123-DEL-164 |
| chr8-33859850-DEL-3392 | chr8-33859850-DEL-3392 |
| chr8-33863263-DEL-1344 | chr8-33863265-DEL-1343 |
| chr8-33887326-DEL-263 | chr8-33887311-DEL-263 |
| chr8-33892611-DEL-69 | chr8-33892611-DEL-70 |
| chr8-33935735-DEL-550 | chr8-33935736-DEL-550 |
| chr8-33938403-DEL-154 | chr8-33938404-DEL-154 |
| chr8-33940370-DEL-204 | chr8-33940371-DEL-204 |
| chr8-33962417-DEL-107 | chr8-33962437-DEL-80 |
| chr8-33978569-DEL-2756 | chr8-33978570-DEL-2756 |
| chr8-33982826-DEL-488 | chr8-33982827-DEL-488 |
| chr8-33986909-DEL-62 | chr8-33986910-DEL-62 |
| chr8-33993248-DEL-97 | chr8-33993249-DEL-97 |
| chr8-33997384-DEL-59 | chr8-33997385-DEL-59 |
| chr8-34017323-DEL-589 | chr8-34017324-DEL-589 |
| chr8-34035573-DEL-369 | chr8-34035574-DEL-369 |
| chr8-34094423-DEL-181 | chr8-34094424-DEL-181 |
| chr8-34123355-DEL-66 | chr8-34123356-DEL-64 |
| chr8-34141122-DEL-177 | chr8-34141123-DEL-177 |
| chr8-34154211-DEL-56 | chr8-34154212-DEL-56 |
| chr8-34154485-DEL-573 | chr8-34154486-DEL-573 |
| chr8-34155278-DEL-71 | chr8-34155279-DEL-71 |
| chr8-34158836-DEL-52 | chr8-34158837-DEL-52 |
| chr8-34160977-DEL-217 | chr8-34160978-DEL-217 |
| chr8-34169841-DEL-210 | chr8-34169842-DEL-210 |
| chr8-34188640-DEL-195 | chr8-34188641-DEL-170 |
| chr8-34190489-DEL-52 | chr8-34190490-DEL-52 |
| chr8-34197370-DEL-62 | chr8-34197361-DEL-92 |
| chr8-34200177-DEL-67 | chr8-34200178-DEL-67 |
| chr8-34203767-DEL-5001 | chr8-34203768-DEL-5001 |
| chr8-34210776-DEL-74 | chr8-34210789-DEL-62 |
| chr8-34215059-DEL-213 | chr8-34215060-DEL-213 |
| chr8-34229005-DEL-223 | chr8-34229006-DEL-223 |
| chr8-34249329-DEL-65 | chr8-34249330-DEL-66 |
| chr8-34254944-DEL-85 | chr8-34254945-DEL-85 |
| chr8-34256951-DEL-222 | chr8-34256952-DEL-222 |
| chr8-34287895-DEL-58 | chr8-34287898-DEL-56 |
| chr8-34320970-DEL-59 | chr8-34320971-DEL-59 |
| chr8-34324922-DEL-161 | chr8-34324923-DEL-161 |
| chr8-34336930-DEL-72 | chr8-34336922-DEL-56 |
| chr8-34343328-DEL-90 | chr8-34343341-DEL-78 |
| chr8-34346414-DEL-186 | chr8-34346418-DEL-183 |
| chr8-34350396-DEL-51 | chr8-34350396-DEL-52 |
| chr8-34382129-DEL-112 | chr8-34382130-DEL-112 |
| chr8-34383243-DEL-198 | chr8-34383244-DEL-198 |
| chr8-34422190-DEL-98 | chr8-34422191-DEL-98 |
| chr8-34422635-DEL-328 | chr8-34422637-DEL-327 |
| chr8-34425634-DEL-221 | chr8-34425635-DEL-221 |

| | |
|---|---|
| chr8-34432152-DEL-201 | chr8-34432153-DEL-201 |
| chr8-34437729-DEL-851 | chr8-34437730-DEL-851 |
| chr8-34446390-DEL-210 | chr8-34446391-DEL-210 |
| chr8-34451068-DEL-103 | chr8-34451069-DEL-103 |
| chr8-34459169-DEL-884 | chr8-34459170-DEL-884 |
| chr8-34459241-DEL-209 | chr8-34459242-DEL-209 |
| chr8-34460565-DEL-215 | chr8-34460569-DEL-212 |
| chr8-34466911-DEL-56 | chr8-34466911-DEL-54 |
| chr8-34473155-DEL-55 | chr8-34473105-DEL-66 |
| chr8-34488277-DEL-106 | chr8-34488280-DEL-110 |
| chr8-34492474-DEL-197 | chr8-34492474-DEL-197 |
| chr8-34499117-DEL-73 | chr8-34499126-DEL-71 |
| chr8-34499832-DEL-411 | chr8-34499833-DEL-411 |
| chr8-34513481-DEL-189 | chr8-34513482-DEL-189 |
| chr8-34529462-DEL-616 | chr8-34529462-DEL-616 |
| chr8-34531813-DEL-225 | chr8-34531814-DEL-225 |
| chr8-34533831-DEL-204 | chr8-34533832-DEL-204 |
| chr8-34554309-DEL-1904 | chr8-34554310-DEL-1904 |
| chr8-34562440-DEL-993 | chr8-34562441-DEL-993 |
| chr8-34570509-DEL-1158 | chr8-34570510-DEL-1158 |
| chr8-34578102-DEL-789 | chr8-34578103-DEL-789 |
| chr8-34585587-DEL-365 | chr8-34585588-DEL-365 |
| chr8-34597000-DEL-95 | chr8-34596997-DEL-96 |
| chr8-34619549-DEL-292 | chr8-34619552-DEL-291 |
| chr8-34628577-DEL-299 | chr8-34628578-DEL-299 |
| chr8-34631127-DEL-95 | chr8-34631129-DEL-94 |
| chr8-34642381-DEL-579 | chr8-34642382-DEL-579 |
| chr8-34644570-DEL-126 | chr8-34644581-DEL-129 |
| chr8-34650932-DEL-75 | chr8-34650934-DEL-73 |
| chr8-34656780-DEL-174 | chr8-34656781-DEL-174 |
| chr8-34658037-DEL-60 | chr8-34658038-DEL-60 |
| chr8-34672370-DEL-332 | chr8-34672363-DEL-333 |
| chr8-34676765-DEL-66 | chr8-34676766-DEL-66 |
| chr8-34708333-DEL-393 | chr8-34708338-DEL-389 |
| chr8-34711027-DEL-2845 | chr8-34711028-DEL-2845 |
| chr8-34715199-DEL-1204 | chr8-34715200-DEL-1204 |
| chr8-34727538-DEL-88 | chr8-34727555-DEL-104 |
| chr8-34729589-DEL-53 | chr8-34729557-DEL-54 |
| chr8-34740428-DEL-52 | chr8-34740428-DEL-52 |
| chr8-34745495-DEL-1459 | chr8-34745496-DEL-1459 |
| chr8-34750116-DEL-202 | chr8-34750117-DEL-202 |
| chr8-34752308-DEL-325 | chr8-34752309-DEL-325 |
| chr8-34759124-DEL-146 | chr8-34759125-DEL-146 |
| chr8-34774025-DEL-194 | chr8-34774024-DEL-192 |
| chr8-34785588-DEL-458 | chr8-34785593-DEL-454 |
| chr8-34790117-DEL-116 | chr8-34790122-DEL-116 |
| chr8-34796439-DEL-72 | chr8-34796430-DEL-72 |
| chr8-34802600-DEL-452 | chr8-34802601-DEL-452 |
| chr8-34808068-DEL-51 | chr8-34808053-DEL-96 |
| chr8-34809858-DEL-74 | chr8-34809878-DEL-54 |
| chr8-34811973-DEL-67 | chr8-34811974-DEL-67 |
| chr8-34812259-DEL-358 | chr8-34812260-DEL-358 |
| chr8-34846286-DEL-102 | chr8-34846287-DEL-102 |
| chr8-34861699-DEL-58 | chr8-34861702-DEL-59 |
| chr8-34862643-DEL-56 | chr8-34862644-DEL-56 |
| chr8-34884590-DEL-56 | chr8-34884591-DEL-56 |
| chr8-34903313-DEL-2977 | chr8-34903314-DEL-2977 |
| chr8-34908530-DEL-176 | chr8-34908531-DEL-186 |
| chr8-34943387-DEL-105 | chr8-34943402-DEL-84 |
| chr8-34950032-DEL-1191 | chr8-34950033-DEL-1190 |
| chr8-34952306-DEL-2696 | chr8-34952307-DEL-2696 |
| chr8-34957274-DEL-57 | chr8-34957275-DEL-57 |
| chr8-35008182-DEL-84 | chr8-35008191-DEL-75 |
| chr8-35013176-DEL-64 | chr8-35013185-DEL-56 |
| chr8-35022853-DEL-209 | chr8-35022854-DEL-209 |
| chr8-35066818-DEL-432 | chr8-35066849-DEL-402 |
| chr8-35075603-DEL-1347 | chr8-35075604-DEL-1347 |
| chr8-35098959-DEL-52 | chr8-35098960-DEL-52 |
| chr8-35114372-DEL-55 | chr8-35114368-DEL-56 |
| chr8-35124319-DEL-3924 | chr8-35124320-DEL-3924 |
| chr8-35132952-DEL-731 | chr8-35132953-DEL-731 |
| chr8-35136493-DEL-654 | chr8-35136494-DEL-654 |
| chr8-35140291-DEL-90 | chr8-35140292-DEL-90 |
| chr8-35141030-DEL-66 | chr8-35141031-DEL-66 |
| chr8-35141665-DEL-491 | chr8-35141666-DEL-491 |
| chr8-35146540-DEL-74 | chr8-35146541-DEL-74 |
| chr8-35166060-DEL-58 | chr8-35166022-DEL-77 |
| chr8-35184817-DEL-1469 | chr8-35184818-DEL-1469 |
| chr8-35189777-DEL-116 | chr8-35189778-DEL-124 |
| chr8-35209884-DEL-56 | chr8-35209884-DEL-56 |

| | |
|---|---|
| chr8-35212299-DEL-180 | chr8-35212329-DEL-190 |
| chr8-35222090-DEL-108 | chr8-35222088-DEL-111 |
| chr8-35225985-DEL-171 | chr8-35225986-DEL-171 |
| chr8-35242643-DEL-59 | chr8-35242642-DEL-60 |
| chr8-35254377-DEL-132 | chr8-35254405-DEL-94 |
| chr8-35259369-DEL-83 | chr8-35259370-DEL-83 |
| chr8-35315067-DEL-89 | chr8-35315098-DEL-59 |
| chr8-35335447-DEL-168 | chr8-35335448-DEL-168 |
| chr8-35349942-DEL-2252 | chr8-35349943-DEL-2252 |
| chr8-35353113-DEL-249 | chr8-35353116-DEL-247 |
| chr8-35387246-DEL-54 | chr8-35387247-DEL-54 |
| chr8-35428289-DEL-609 | chr8-35428291-DEL-597 |
| chr8-35456841-DEL-56 | chr8-35456841-DEL-56 |
| chr8-35459445-DEL-56 | chr8-35459444-DEL-62 |
| chr8-35469124-DEL-54 | chr8-35469097-DEL-72 |
| chr8-35503474-DEL-82 | chr8-35503475-DEL-82 |
| chr8-35512904-DEL-123 | chr8-35512889-DEL-124 |
| chr8-35519090-DEL-64 | chr8-35519095-DEL-64 |
| chr8-35533186-DEL-753 | chr8-35533187-DEL-753 |
| chr8-35540835-DEL-57 | chr8-35540835-DEL-57 |
| chr8-35550923-DEL-54 | chr8-35550924-DEL-54 |
| chr8-35561550-DEL-52 | chr8-35561551-DEL-52 |
| chr8-35602053-DEL-66 | chr8-35602054-DEL-60 |
| chr8-35621919-DEL-664 | chr8-35621920-DEL-664 |
| chr8-35623439-DEL-50 | chr8-35623440-DEL-50 |
| chr8-35626366-DEL-58 | chr8-35626367-DEL-54 |
| chr8-35639658-DEL-136 | chr8-35639659-DEL-136 |
| chr8-35651979-DEL-80 | chr8-35651980-DEL-80 |
| chr8-35653089-DEL-76 | chr8-35653090-DEL-76 |
| chr8-35654745-DEL-64 | chr8-35654746-DEL-64 |
| chr8-35677012-DEL-146 | chr8-35676985-DEL-148 |
| chr8-35702471-DEL-68 | chr8-35702432-DEL-66 |
| chr8-35715378-DEL-51 | chr8-35715379-DEL-51 |
| chr8-35733855-DEL-55 | chr8-35733857-DEL-101 |
| chr8-35736879-DEL-73 | chr8-35736882-DEL-74 |
| chr8-35745016-DEL-4169 | chr8-35745017-DEL-4169 |
| chr8-35756518-DEL-60 | chr8-35756519-DEL-60 |
| chr8-35776673-DEL-52 | chr8-35776674-DEL-52 |
| chr8-35777633-DEL-685 | chr8-35777634-DEL-685 |
| chr8-35779881-DEL-429 | chr8-35779882-DEL-429 |
| chr8-35822369-DEL-59 | chr8-35822370-DEL-59 |
| chr8-35829077-DEL-191 | chr8-35829078-DEL-191 |
| chr8-35869531-DEL-464 | chr8-35869532-DEL-464 |
| chr8-35911663-DEL-1104 | chr8-35911663-DEL-1104 |
| chr8-35951573-DEL-6170 | chr8-35951574-DEL-6170 |
| chr8-35965278-DEL-60 | chr8-35965279-DEL-60 |
| chr8-35966607-DEL-6394 | chr8-35966608-DEL-6394 |
| chr8-35975158-DEL-58 | chr8-35975159-DEL-58 |
| chr8-35991635-DEL-55 | chr8-35991636-DEL-55 |
| chr8-35998606-DEL-74 | chr8-35998607-DEL-74 |
| chr8-36010417-DEL-114 | chr8-36010418-DEL-114 |
| chr8-36029083-DEL-55 | chr8-36029080-DEL-57 |
| chr8-36033058-DEL-70 | chr8-36033068-DEL-68 |
| chr8-36091220-DEL-393 | chr8-36091221-DEL-393 |
| chr8-36104685-DEL-1946 | chr8-36104686-DEL-1946 |
| chr8-36110797-DEL-90 | chr8-36110798-DEL-90 |
| chr8-36124517-DEL-389 | chr8-36124518-DEL-389 |
| chr8-36153766-DEL-70 | chr8-36153767-DEL-70 |
| chr8-36157409-DEL-177 | chr8-36157416-DEL-186 |
| chr8-36160610-DEL-69 | chr8-36160602-DEL-69 |
| chr8-36163956-DEL-68 | chr8-36163957-DEL-68 |
| chr8-36189157-DEL-108 | chr8-36189158-DEL-108 |
| chr8-36194086-DEL-68 | chr8-36194087-DEL-68 |
| chr8-36199139-DEL-216 | chr8-36199140-DEL-216 |
| chr8-36222208-DEL-55 | chr8-36222218-DEL-56 |
| chr8-36231756-DEL-59 | chr8-36231756-DEL-59 |
| chr8-36247690-DEL-155 | chr8-36247690-DEL-147 |
| chr8-36250826-DEL-178 | chr8-36250827-DEL-178 |
| chr8-36253927-DEL-80 | chr8-36253928-DEL-76 |
| chr8-36263606-DEL-60 | chr8-36263607-DEL-60 |
| chr8-36277152-DEL-60 | chr8-36277167-DEL-50 |
| chr8-36280396-DEL-53 | chr8-36280390-DEL-56 |
| chr8-36305312-DEL-70 | chr8-36305310-DEL-74 |
| chr8-36362834-DEL-512 | chr8-36362835-DEL-512 |
| chr8-36363580-DEL-121 | chr8-36363579-DEL-122 |
| chr8-36370106-DEL-857 | chr8-36370110-DEL-858 |
| chr8-36373318-DEL-992 | chr8-36373319-DEL-992 |
| chr8-36380467-DEL-59 | chr8-36380468-DEL-59 |
| chr8-36387818-DEL-51 | chr8-36387815-DEL-51 |
| chr8-36389338-DEL-100 | chr8-36389368-DEL-69 |

| | |
|---|---|
| chr8-36393352-DEL-900 | chr8-36393353-DEL-900 |
| chr8-36395639-DEL-330 | chr8-36395640-DEL-330 |
| chr8-36440418-DEL-286 | chr8-36440419-DEL-286 |
| chr8-36443497-DEL-829 | chr8-36443498-DEL-829 |
| chr8-36490739-DEL-83 | chr8-36490691-DEL-87 |
| chr8-36495097-DEL-156 | chr8-36495141-DEL-129 |
| chr8-36527624-DEL-92 | chr8-36527620-DEL-93 |
| chr8-36565440-DEL-87 | chr8-36565441-DEL-87 |
| chr8-36598415-DEL-55 | chr8-36598416-DEL-55 |
| chr8-36606281-DEL-119 | chr8-36606277-DEL-123 |
| chr8-36607726-DEL-57 | chr8-36607727-DEL-57 |
| chr8-36613161-DEL-75 | chr8-36613169-DEL-58 |
| chr8-36671225-DEL-139 | chr8-36671226-DEL-130 |
| chr8-36710230-DEL-409 | chr8-36710231-DEL-409 |
| chr8-36732224-DEL-84 | chr8-36732225-DEL-84 |
| chr8-36738884-DEL-215 | chr8-36738880-DEL-190 |
| chr8-36740448-DEL-509 | chr8-36740449-DEL-509 |
| chr8-36742183-DEL-469 | chr8-36742160-DEL-470 |
| chr8-36764326-DEL-7693 | chr8-36764327-DEL-7693 |
| chr8-36782343-DEL-88 | chr8-36782344-DEL-88 |
| chr8-36790211-DEL-2006 | chr8-36790212-DEL-2006 |
| chr8-36825086-DEL-87 | chr8-36825087-DEL-87 |
| chr8-36847601-DEL-83 | chr8-36847602-DEL-83 |
| chr8-36854328-DEL-6211 | chr8-36854329-DEL-6211 |
| chr8-36865869-DEL-612 | chr8-36865870-DEL-612 |
| chr8-36876112-DEL-222 | chr8-36876103-DEL-221 |
| chr8-36900844-DEL-144 | chr8-36900834-DEL-146 |
| chr8-36919353-DEL-68 | chr8-36919354-DEL-68 |
| chr8-36933038-DEL-399 | chr8-36933028-DEL-400 |
| chr8-36960651-DEL-115 | chr8-36960654-DEL-113 |
| chr8-36967210-DEL-216 | chr8-36967211-DEL-216 |
| chr8-36973927-DEL-138 | chr8-36973921-DEL-134 |
| chr8-36985008-DEL-815 | chr8-36985009-DEL-815 |
| chr8-36990621-DEL-1372 | chr8-36990622-DEL-1372 |
| chr8-36991526-DEL-105 | chr8-36991531-DEL-104 |
| chr8-36998120-DEL-173 | chr8-36998114-DEL-174 |
| chr8-37001920-DEL-169 | chr8-37001921-DEL-169 |
| chr8-37005147-DEL-132 | chr8-37005154-DEL-122 |
| chr8-37025432-DEL-62 | chr8-37025433-DEL-62 |
| chr8-37043259-DEL-144 | chr8-37043260-DEL-144 |
| chr8-37046164-DEL-52 | chr8-37046165-DEL-52 |
| chr8-37059450-DEL-117 | chr8-37059468-DEL-114 |
| chr8-37077448-DEL-79 | chr8-37077449-DEL-79 |
| chr8-37087843-DEL-54 | chr8-37087844-DEL-58 |
| chr8-37092002-DEL-57 | chr8-37092003-DEL-57 |
| chr8-37104686-DEL-98 | chr8-37104687-DEL-98 |
| chr8-37115974-DEL-123 | chr8-37115978-DEL-128 |
| chr8-37120928-DEL-66 | chr8-37120900-DEL-50 |
| chr8-37139558-DEL-200 | chr8-37139559-DEL-200 |
| chr8-37158982-DEL-60 | chr8-37158987-DEL-56 |
| chr8-37166990-DEL-341 | chr8-37166989-DEL-341 |
| chr8-37204280-DEL-1304 | chr8-37204281-DEL-1304 |
| chr8-37209972-DEL-488 | chr8-37209973-DEL-488 |
| chr8-37224620-DEL-398 | chr8-37224621-DEL-398 |
| chr8-37232212-DEL-122 | chr8-37232213-DEL-122 |
| chr8-37236815-DEL-72 | chr8-37236820-DEL-68 |
| chr8-37246788-DEL-50 | chr8-37246789-DEL-50 |
| chr8-37259249-DEL-559 | chr8-37259250-DEL-559 |
| chr8-37274535-DEL-324 | chr8-37274536-DEL-324 |
| chr8-37325611-DEL-58 | chr8-37325612-DEL-50 |
| chr8-37348031-DEL-129 | chr8-37348030-DEL-126 |
| chr8-37359866-DEL-72 | chr8-37359876-DEL-57 |
| chr8-37372831-DEL-203 | chr8-37372834-DEL-204 |
| chr8-37396837-DEL-60 | chr8-37396819-DEL-71 |
| chr8-37466729-DEL-53 | chr8-37466730-DEL-53 |
| chr8-37530319-DEL-80 | chr8-37530315-DEL-73 |
| chr8-37532583-DEL-6370 | chr8-37532584-DEL-6370 |
| chr8-37552869-DEL-162 | chr8-37552870-DEL-162 |
| chr8-37611358-DEL-524 | chr8-37611359-DEL-524 |
| chr8-37612526-DEL-67 | chr8-37612510-DEL-68 |
| chr8-37614005-DEL-65 | chr8-37614008-DEL-63 |
| chr8-37621277-DEL-136 | chr8-37621278-DEL-136 |
| chr8-37647421-DEL-79 | chr8-37647422-DEL-79 |
| chr8-37650544-DEL-160 | chr8-37650545-DEL-160 |
| chr8-37671679-DEL-97 | chr8-37671680-DEL-97 |
| chr8-37686463-DEL-923 | chr8-37686464-DEL-923 |
| chr8-37688861-DEL-316 | chr8-37688859-DEL-317 |
| chr8-37712412-DEL-703 | chr8-37712409-DEL-659 |
| chr8-37726452-DEL-583 | chr8-37726453-DEL-583 |
| chr8-37734303-DEL-1842 | chr8-37734310-DEL-1828 |

| | |
|---|---|
| chr8-37777292-DEL-3278 | chr8-37777293-DEL-3274 |
| chr8-37781719-DEL-1248 | chr8-37781720-DEL-1248 |
| chr8-37791508-DEL-2472 | chr8-37791507-DEL-2472 |
| chr8-37804297-DEL-2300 | chr8-37804298-DEL-2300 |
| chr8-37809083-DEL-3069 | chr8-37809084-DEL-3069 |
| chr8-37826394-DEL-111 | chr8-37826395-DEL-111 |
| chr8-37832890-DEL-51 | chr8-37832891-DEL-51 |
| chr8-37881324-DEL-80 | chr8-37881325-DEL-80 |
| chr8-37887812-DEL-96 | chr8-37887811-DEL-96 |
| chr8-37906355-DEL-110 | chr8-37906356-DEL-110 |
| chr8-37918891-DEL-468 | chr8-37918892-DEL-468 |
| chr8-37934704-DEL-1672 | chr8-37934705-DEL-1672 |
| chr8-37943964-DEL-983 | chr8-37943956-DEL-988 |
| chr8-37948143-DEL-200 | chr8-37948144-DEL-200 |
| chr8-37948531-DEL-1046 | chr8-37948532-DEL-1046 |
| chr8-37998510-DEL-60 | chr8-37998511-DEL-60 |
| chr8-38021090-DEL-6497 | chr8-38021091-DEL-6497 |
| chr8-38039439-DEL-6360 | chr8-38039445-DEL-6355 |
| chr8-38049178-DEL-960 | chr8-38049176-DEL-962 |
| chr8-38075835-DEL-55 | chr8-38075836-DEL-55 |
| chr8-38131846-DEL-6220 | chr8-38131847-DEL-6220 |
| chr8-38144498-DEL-507 | chr8-38144499-DEL-507 |
| chr8-38148668-DEL-139 | chr8-38148688-DEL-150 |
| chr8-38221775-DEL-2267 | chr8-38221776-DEL-2267 |
| chr8-38244783-DEL-275 | chr8-38244775-DEL-275 |
| chr8-38248301-DEL-6190 | chr8-38248302-DEL-6190 |
| chr8-38259355-DEL-170 | chr8-38259356-DEL-170 |
| chr8-38272441-DEL-78 | chr8-38272438-DEL-82 |
| chr8-38286685-DEL-3694 | chr8-38286686-DEL-3694 |
| chr8-38294313-DEL-3573 | chr8-38294314-DEL-3573 |
| chr8-38299205-DEL-56 | chr8-38299206-DEL-56 |
| chr8-38319979-DEL-136 | chr8-38320001-DEL-95 |
| chr8-38320964-DEL-53 | chr8-38320965-DEL-53 |
| chr8-38322999-DEL-198 | chr8-38322992-DEL-196 |
| chr8-38327543-DEL-75 | chr8-38327549-DEL-76 |
| chr8-38335990-DEL-84 | chr8-38335991-DEL-56 |
| chr8-38364512-DEL-259 | chr8-38364513-DEL-259 |
| chr8-38394676-DEL-53 | chr8-38394683-DEL-51 |
| chr8-38396041-DEL-51 | chr8-38396037-DEL-56 |
| chr8-38434494-DEL-1037 | chr8-38434495-DEL-1037 |
| chr8-38434615-DEL-212 | chr8-38434616-DEL-212 |
| chr8-38440567-DEL-205 | chr8-38440568-DEL-205 |
| chr8-38464594-DEL-4436 | chr8-38464583-DEL-4446 |
| chr8-38475339-DEL-4265 | chr8-38475338-DEL-4266 |
| chr8-38500682-DEL-132 | chr8-38500685-DEL-130 |
| chr8-38504334-DEL-56 | chr8-38504334-DEL-56 |
| chr8-38510167-DEL-54 | chr8-38510168-DEL-54 |
| chr8-38516583-DEL-1004 | chr8-38516584-DEL-1004 |
| chr8-38518490-DEL-889 | chr8-38518490-DEL-889 |
| chr8-38527029-DEL-1079 | chr8-38527030-DEL-1079 |
| chr8-38539379-DEL-67 | chr8-38539380-DEL-67 |
| chr8-38541244-DEL-65 | chr8-38541245-DEL-65 |
| chr8-38544721-DEL-159 | chr8-38544718-DEL-160 |
| chr8-38550435-DEL-389 | chr8-38550445-DEL-385 |
| chr8-38567757-DEL-108 | chr8-38567758-DEL-108 |
| chr8-38574324-DEL-1490 | chr8-38574324-DEL-1490 |
| chr8-38584166-DEL-633 | chr8-38584166-DEL-633 |
| chr8-38608047-DEL-185 | chr8-38608051-DEL-187 |
| chr8-38609333-DEL-210 | chr8-38609334-DEL-210 |
| chr8-38629706-DEL-238 | chr8-38629700-DEL-239 |
| chr8-38642626-DEL-69 | chr8-38642641-DEL-50 |
| chr8-38670221-DEL-366 | chr8-38670222-DEL-366 |
| chr8-38674511-DEL-5983 | chr8-38674497-DEL-5992 |
| chr8-38686771-DEL-64 | chr8-38686772-DEL-64 |
| chr8-38693916-DEL-711 | chr8-38693917-DEL-711 |
| chr8-38711489-DEL-87 | chr8-38711490-DEL-87 |
| chr8-38720707-DEL-2780 | chr8-38720708-DEL-2780 |
| chr8-38733163-DEL-75 | chr8-38733163-DEL-75 |
| chr8-38737500-DEL-59 | chr8-38737501-DEL-59 |
| chr8-38738452-DEL-1552 | chr8-38738453-DEL-1552 |
| chr8-38762763-DEL-193 | chr8-38762764-DEL-193 |
| chr8-38798141-DEL-241 | chr8-38798142-DEL-241 |
| chr8-38848205-DEL-1312 | chr8-38848209-DEL-1309 |
| chr8-38851041-DEL-83 | chr8-38851041-DEL-84 |
| chr8-38853013-DEL-228 | chr8-38853014-DEL-227 |
| chr8-38875366-DEL-203 | chr8-38875367-DEL-203 |
| chr8-38884665-DEL-1182 | chr8-38884665-DEL-1182 |
| chr8-38895475-DEL-6360 | chr8-38895476-DEL-6360 |
| chr8-38913212-DEL-57 | chr8-38913210-DEL-60 |
| chr8-38921810-DEL-334 | chr8-38921811-DEL-334 |

| | |
|---|---|
| chr8-38925119-DEL-56 | chr8-38925122-DEL-54 |
| chr8-38926026-DEL-64 | chr8-38926027-DEL-64 |
| chr8-38929275-DEL-688 | chr8-38929276-DEL-688 |
| chr8-38952125-DEL-160 | chr8-38952126-DEL-160 |
| chr8-38985506-DEL-101 | chr8-38985507-DEL-101 |
| chr8-38997564-DEL-52 | chr8-38997565-DEL-52 |
| chr8-39019175-DEL-2583 | chr8-39019176-DEL-2583 |
| chr8-39039696-DEL-72 | chr8-39039697-DEL-72 |
| chr8-39062584-DEL-172 | chr8-39062585-DEL-172 |
| chr8-39114273-DEL-104 | chr8-39114283-DEL-102 |
| chr8-39138921-DEL-182 | chr8-39138922-DEL-182 |
| chr8-39159775-DEL-765 | chr8-39159776-DEL-765 |
| chr8-39169912-DEL-1286 | chr8-39169913-DEL-1286 |
| chr8-39225579-DEL-80 | chr8-39225580-DEL-80 |
| chr8-39245020-DEL-6619 | chr8-39245021-DEL-6619 |
| chr8-39286197-DEL-6584 | chr8-39286198-DEL-6584 |
| chr8-39318316-DEL-86 | chr8-39318316-DEL-87 |
| chr8-39327131-DEL-223 | chr8-39327132-DEL-223 |
| chr8-39339602-DEL-2333 | chr8-39339603-DEL-2333 |
| chr8-39342171-DEL-6467 | chr8-39342172-DEL-6467 |
| chr8-39364425-DEL-177 | chr8-39364426-DEL-177 |
| chr8-39397851-DEL-65 | chr8-39397831-DEL-57 |
| chr8-39407928-DEL-590 | chr8-39407929-DEL-590 |
| chr8-39417242-DEL-66 | chr8-39417243-DEL-66 |
| chr8-39438792-DEL-387 | chr8-39438793-DEL-387 |
| chr8-39493398-DEL-1308 | chr8-39493399-DEL-1308 |
| chr8-39494887-DEL-96 | chr8-39494888-DEL-96 |
| chr8-39509463-DEL-104 | chr8-39509460-DEL-132 |
| chr8-39512298-DEL-288 | chr8-39512299-DEL-288 |
| chr8-39542810-DEL-314 | chr8-39542811-DEL-314 |
| chr8-39544102-DEL-64 | chr8-39544103-DEL-64 |
| chr8-39577298-DEL-510 | chr8-39577299-DEL-510 |
| chr8-39626857-DEL-62 | chr8-39626858-DEL-62 |
| chr8-39646328-DEL-451 | chr8-39646329-DEL-451 |
| chr8-39647154-DEL-6299 | chr8-39647155-DEL-6299 |
| chr8-39686659-DEL-108 | chr8-39686660-DEL-108 |
| chr8-39695295-DEL-1154 | chr8-39695296-DEL-1154 |
| chr8-39699750-DEL-6640 | chr8-39699751-DEL-6640 |
| chr8-39737122-DEL-282 | chr8-39737123-DEL-279 |
| chr8-39740428-DEL-63 | chr8-39740408-DEL-83 |
| chr8-39757610-DEL-54 | chr8-39757611-DEL-54 |
| chr8-39782867-DEL-3393 | chr8-39782867-DEL-3397 |
| chr8-39794779-DEL-76 | chr8-39794780-DEL-76 |
| chr8-39806263-DEL-79 | chr8-39806264-DEL-79 |
| chr8-39826925-DEL-222 | chr8-39826955-DEL-214 |
| chr8-39829370-DEL-72 | chr8-39829371-DEL-72 |
| chr8-39832103-DEL-7079 | chr8-39832104-DEL-7079 |
| chr8-39853126-DEL-55 | chr8-39853114-DEL-56 |
| chr8-39887699-DEL-81 | chr8-39887700-DEL-81 |
| chr8-39901273-DEL-422 | chr8-39901275-DEL-420 |
| chr8-39916214-DEL-1127 | chr8-39916215-DEL-1127 |
| chr8-39929740-DEL-711 | chr8-39929741-DEL-711 |
| chr8-39933850-DEL-60 | chr8-39933851-DEL-60 |
| chr8-39936306-DEL-56 | chr8-39936293-DEL-55 |
| chr8-39958504-DEL-58 | chr8-39958508-DEL-59 |
| chr8-39994381-DEL-82 | chr8-39994411-DEL-80 |
| chr8-39997176-DEL-6584 | chr8-39997177-DEL-6584 |
| chr8-40048208-DEL-163 | chr8-40048221-DEL-151 |
| chr8-40050035-DEL-73 | chr8-40050036-DEL-73 |
| chr8-40083318-DEL-248 | chr8-40083321-DEL-263 |
| chr8-40089784-DEL-166 | chr8-40089803-DEL-148 |
| chr8-40108791-DEL-810 | chr8-40108792-DEL-810 |
| chr8-40115466-DEL-2857 | chr8-40115467-DEL-2857 |
| chr8-40141378-DEL-55 | chr8-40141379-DEL-55 |
| chr8-40143413-DEL-56 | chr8-40143425-DEL-57 |
| chr8-40149430-DEL-2803 | chr8-40149430-DEL-2803 |
| chr8-40176923-DEL-6367 | chr8-40176924-DEL-6367 |
| chr8-40185229-DEL-60 | chr8-40185230-DEL-60 |
| chr8-40193537-DEL-57 | chr8-40193538-DEL-61 |
| chr8-40201631-DEL-64 | chr8-40201632-DEL-60 |
| chr8-40209341-DEL-274 | chr8-40209342-DEL-273 |
| chr8-40229505-DEL-102 | chr8-40229506-DEL-102 |
| chr8-40251426-DEL-58 | chr8-40251427-DEL-58 |
| chr8-40252245-DEL-231 | chr8-40252238-DEL-229 |
| chr8-40258847-DEL-1747 | chr8-40258847-DEL-1747 |
| chr8-40277128-DEL-6361 | chr8-40277129-DEL-6362 |
| chr8-40299527-DEL-52 | chr8-40299529-DEL-51 |
| chr8-40318737-DEL-396 | chr8-40318737-DEL-397 |
| chr8-40336200-DEL-378 | chr8-40336201-DEL-378 |
| chr8-40338842-DEL-8921 | chr8-40338843-DEL-8921 |

| | |
|---|---|
| chr8-40341213-DEL-2538 | chr8-40341202-DEL-2534 |
| chr8-40350833-DEL-59 | chr8-40350831-DEL-60 |
| chr8-40398885-DEL-660 | chr8-40398886-DEL-660 |
| chr8-40407363-DEL-63 | chr8-40407364-DEL-69 |
| chr8-40419972-DEL-56 | chr8-40419974-DEL-52 |
| chr8-40429078-DEL-175 | chr8-40429079-DEL-175 |
| chr8-40441190-DEL-2247 | chr8-40441191-DEL-2247 |
| chr8-40447269-DEL-365 | chr8-40447270-DEL-365 |
| chr8-40447791-DEL-143 | chr8-40447809-DEL-125 |
| chr8-40486914-DEL-328 | chr8-40486915-DEL-328 |
| chr8-40490310-DEL-71 | chr8-40490311-DEL-71 |
| chr8-40501254-DEL-3446 | chr8-40501255-DEL-3444 |
| chr8-40501888-DEL-411 | chr8-40501889-DEL-411 |
| chr8-40506398-DEL-60 | chr8-40506399-DEL-60 |
| chr8-40509197-DEL-6377 | chr8-40509198-DEL-6377 |
| chr8-40521562-DEL-268 | chr8-40521563-DEL-268 |
| chr8-40582446-DEL-1564 | chr8-40582447-DEL-1564 |
| chr8-40599979-DEL-76 | chr8-40599980-DEL-76 |
| chr8-40617507-DEL-125 | chr8-40617509-DEL-126 |
| chr8-40628793-DEL-1254 | chr8-40628794-DEL-1254 |
| chr8-40642593-DEL-175 | chr8-40642594-DEL-175 |
| chr8-40650157-DEL-6432 | chr8-40650158-DEL-6432 |
| chr8-40659403-DEL-2858 | chr8-40659404-DEL-2858 |
| chr8-40668816-DEL-192 | chr8-40668817-DEL-192 |
| chr8-40670572-DEL-55 | chr8-40670573-DEL-55 |
| chr8-40695581-DEL-57 | chr8-40695582-DEL-57 |
| chr8-40698830-DEL-6207 | chr8-40698822-DEL-6208 |
| chr8-40712041-DEL-289 | chr8-40712040-DEL-290 |
| chr8-40714437-DEL-58 | chr8-40714438-DEL-58 |
| chr8-40733595-DEL-54 | chr8-40733596-DEL-54 |
| chr8-40735427-DEL-56 | chr8-40735428-DEL-56 |
| chr8-40757095-DEL-63 | chr8-40757096-DEL-63 |
| chr8-40815680-DEL-202 | chr8-40815681-DEL-202 |
| chr8-40821318-DEL-6426 | chr8-40821319-DEL-6426 |
| chr8-40869151-DEL-570 | chr8-40869152-DEL-570 |
| chr8-40883181-DEL-236 | chr8-40883182-DEL-236 |
| chr8-40889547-DEL-178 | chr8-40889547-DEL-177 |
| chr8-40894048-DEL-210 | chr8-40894049-DEL-210 |
| chr8-40904111-DEL-230 | chr8-40904110-DEL-231 |
| chr8-40921999-DEL-180 | chr8-40922000-DEL-180 |
| chr8-40930810-DEL-4066 | chr8-40930810-DEL-4066 |
| chr8-40950823-DEL-51 | chr8-40950823-DEL-51 |
| chr8-40953708-DEL-61 | chr8-40953710-DEL-60 |
| chr8-40983655-DEL-397 | chr8-40983656-DEL-397 |
| chr8-41019733-DEL-5509 | chr8-41019731-DEL-5510 |
| chr8-41054436-DEL-239 | chr8-41054437-DEL-239 |
| chr8-41062979-DEL-403 | chr8-41062980-DEL-403 |
| chr8-41102715-DEL-176 | chr8-41102716-DEL-176 |
| chr8-41115666-DEL-76 | chr8-41115667-DEL-76 |
| chr8-41128612-DEL-51 | chr8-41128613-DEL-58 |
| chr8-41150061-DEL-123 | chr8-41150062-DEL-123 |
| chr8-41186742-DEL-160 | chr8-41186743-DEL-160 |
| chr8-41228913-DEL-6581 | chr8-41228913-DEL-6581 |
| chr8-41273338-DEL-66 | chr8-41273339-DEL-66 |
| chr8-41291867-DEL-1129 | chr8-41291867-DEL-1129 |
| chr8-41297284-DEL-309 | chr8-41297285-DEL-309 |
| chr8-41300939-DEL-135 | chr8-41300947-DEL-136 |
| chr8-41313318-DEL-944 | chr8-41313319-DEL-944 |
| chr8-41322810-DEL-873 | chr8-41322803-DEL-874 |
| chr8-41387730-DEL-436 | chr8-41387731-DEL-436 |
| chr8-41396092-DEL-195 | chr8-41396093-DEL-195 |
| chr8-41400412-DEL-273 | chr8-41400423-DEL-272 |
| chr8-41405045-DEL-366 | chr8-41405046-DEL-366 |
| chr8-41411038-DEL-127 | chr8-41411039-DEL-127 |
| chr8-41413382-DEL-98 | chr8-41413383-DEL-98 |
| chr8-41415828-DEL-61 | chr8-41415829-DEL-61 |
| chr8-41423046-DEL-77 | chr8-41423031-DEL-89 |
| chr8-41431498-DEL-225 | chr8-41431499-DEL-225 |
| chr8-41432299-DEL-1602 | chr8-41432300-DEL-1602 |
| chr8-41441402-DEL-155 | chr8-41441403-DEL-155 |
| chr8-41477077-DEL-201 | chr8-41477078-DEL-201 |
| chr8-41515857-DEL-135 | chr8-41515848-DEL-136 |
| chr8-41523562-DEL-56 | chr8-41523563-DEL-56 |
| chr8-41526640-DEL-52 | chr8-41526641-DEL-52 |
| chr8-41543201-DEL-751 | chr8-41543202-DEL-751 |
| chr8-41555720-DEL-202 | chr8-41555721-DEL-202 |
| chr8-41560230-DEL-149 | chr8-41560231-DEL-149 |
| chr8-41561880-DEL-172 | chr8-41561881-DEL-172 |
| chr8-41617241-DEL-6298 | chr8-41617242-DEL-6298 |
| chr8-41627138-DEL-1384 | chr8-41627139-DEL-1384 |

| | |
|---|---|
| chr8-41628793-DEL-220 | chr8-41628793-DEL-221 |
| chr8-41629848-DEL-439 | chr8-41629844-DEL-442 |
| chr8-41632719-DEL-195 | chr8-41632720-DEL-195 |
| chr8-41638522-DEL-67 | chr8-41638523-DEL-67 |
| chr8-41651577-DEL-372 | chr8-41651578-DEL-373 |
| chr8-41657275-DEL-210 | chr8-41657276-DEL-210 |
| chr8-41660313-DEL-60 | chr8-41660314-DEL-60 |
| chr8-41665740-DEL-566 | chr8-41665741-DEL-566 |
| chr8-41679184-DEL-6599 | chr8-41679181-DEL-6600 |
| chr8-41700746-DEL-70 | chr8-41700747-DEL-70 |
| chr8-41708650-DEL-68 | chr8-41708651-DEL-68 |
| chr8-41737792-DEL-1142 | chr8-41737793-DEL-1142 |
| chr8-41761428-DEL-235 | chr8-41761429-DEL-235 |
| chr8-41770609-DEL-163 | chr8-41770610-DEL-163 |
| chr8-41771183-DEL-143 | chr8-41771184-DEL-143 |
| chr8-41809874-DEL-183 | chr8-41809875-DEL-183 |
| chr8-41811876-DEL-54 | chr8-41811881-DEL-50 |
| chr8-41817036-DEL-83 | chr8-41817031-DEL-84 |
| chr8-41819597-DEL-117 | chr8-41819598-DEL-117 |
| chr8-41822593-DEL-129 | chr8-41822594-DEL-129 |
| chr8-41824911-DEL-173 | chr8-41824912-DEL-173 |
| chr8-41831137-DEL-83 | chr8-41831138-DEL-83 |
| chr8-41851569-DEL-78 | chr8-41851570-DEL-78 |
| chr8-41858837-DEL-52 | chr8-41858838-DEL-52 |
| chr8-41868488-DEL-54 | chr8-41868489-DEL-54 |
| chr8-41872825-DEL-6839 | chr8-41872826-DEL-6839 |
| chr8-41888517-DEL-172 | chr8-41888518-DEL-172 |
| chr8-41889772-DEL-58 | chr8-41889773-DEL-58 |
| chr8-41899353-DEL-3360 | chr8-41899354-DEL-3360 |
| chr8-41908852-DEL-103 | chr8-41908877-DEL-79 |
| chr8-41910776-DEL-2249 | chr8-41910777-DEL-2249 |
| chr8-41939764-DEL-66 | chr8-41939765-DEL-66 |
| chr8-41954736-DEL-1334 | chr8-41954735-DEL-1332 |
| chr8-41959960-DEL-6872 | chr8-41959961-DEL-6872 |
| chr8-41975866-DEL-2780 | chr8-41975867-DEL-2780 |
| chr8-41978928-DEL-52 | chr8-41978929-DEL-52 |
| chr8-42005666-DEL-74 | chr8-42005666-DEL-75 |
| chr8-42032379-DEL-1244 | chr8-42032380-DEL-1244 |
| chr8-42043574-DEL-92 | chr8-42043581-DEL-86 |
| chr8-42052042-DEL-53 | chr8-42052041-DEL-53 |
| chr8-42059074-DEL-79 | chr8-42059059-DEL-80 |
| chr8-42068171-DEL-438 | chr8-42068164-DEL-429 |
| chr8-42070264-DEL-6213 | chr8-42070265-DEL-6213 |
| chr8-42109250-DEL-7346 | chr8-42109251-DEL-7346 |
| chr8-42142737-DEL-366 | chr8-42142738-DEL-366 |
| chr8-42158150-DEL-100 | chr8-42158151-DEL-100 |
| chr8-42184242-DEL-347 | chr8-42184243-DEL-347 |
| chr8-42239650-DEL-194 | chr8-42239651-DEL-194 |
| chr8-42241698-DEL-215 | chr8-42241699-DEL-215 |
| chr8-42250788-DEL-58 | chr8-42250788-DEL-56 |
| chr8-42256982-DEL-140 | chr8-42256983-DEL-140 |
| chr8-42286442-DEL-70 | chr8-42286440-DEL-71 |
| chr8-42295771-DEL-322 | chr8-42295772-DEL-322 |
| chr8-42304519-DEL-150 | chr8-42304519-DEL-151 |
| chr8-42314353-DEL-76 | chr8-42314354-DEL-76 |
| chr8-42319079-DEL-140 | chr8-42319083-DEL-136 |
| chr8-42348960-DEL-4023 | chr8-42348961-DEL-4023 |
| chr8-42361760-DEL-51 | chr8-42361761-DEL-51 |
| chr8-42362049-DEL-6326 | chr8-42362050-DEL-6326 |
| chr8-42381215-DEL-372 | chr8-42381216-DEL-372 |
| chr8-42390368-DEL-262 | chr8-42390369-DEL-262 |
| chr8-42416385-DEL-203 | chr8-42416386-DEL-203 |
| chr8-42418793-DEL-326 | chr8-42418794-DEL-326 |
| chr8-42426546-DEL-100 | chr8-42426546-DEL-100 |
| chr8-42442738-DEL-56 | chr8-42442739-DEL-56 |
| chr8-42445569-DEL-779 | chr8-42445541-DEL-778 |
| chr8-42455597-DEL-261 | chr8-42455604-DEL-254 |
| chr8-42484267-DEL-6587 | chr8-42484278-DEL-6585 |
| chr8-42502037-DEL-1697 | chr8-42502035-DEL-1698 |
| chr8-42529085-DEL-3167 | chr8-42529086-DEL-3167 |
| chr8-42611835-DEL-1056 | chr8-42611836-DEL-1056 |
| chr8-42638606-DEL-331 | chr8-42638607-DEL-331 |
| chr8-42654178-DEL-2790 | chr8-42654179-DEL-2790 |
| chr8-42682644-DEL-51 | chr8-42682629-DEL-66 |
| chr8-42691382-DEL-331 | chr8-42691383-DEL-331 |
| chr8-42697564-DEL-5587 | chr8-42697565-DEL-5587 |
| chr8-42707853-DEL-57 | chr8-42707853-DEL-57 |
| chr8-42714698-DEL-127 | chr8-42714702-DEL-124 |
| chr8-42719252-DEL-6385 | chr8-42719253-DEL-6385 |
| chr8-42725197-DEL-401 | chr8-42725198-DEL-402 |

| | |
|---|---|
| chr8-42738664-DEL-94 | chr8-42738667-DEL-92 |
| chr8-42767084-DEL-54 | chr8-42767085-DEL-54 |
| chr8-42768908-DEL-58 | chr8-42768907-DEL-58 |
| chr8-42791409-DEL-124 | chr8-42791409-DEL-124 |
| chr8-42805385-DEL-107 | chr8-42805385-DEL-107 |
| chr8-42817044-DEL-963 | chr8-42817034-DEL-964 |
| chr8-42818367-DEL-1669 | chr8-42818370-DEL-1666 |
| chr8-42860864-DEL-203 | chr8-42860865-DEL-203 |
| chr8-42883677-DEL-66 | chr8-42883690-DEL-54 |
| chr8-42887917-DEL-699 | chr8-42887918-DEL-699 |
| chr8-42922972-DEL-101 | chr8-42922965-DEL-100 |
| chr8-42923403-DEL-2627 | chr8-42923401-DEL-2625 |
| chr8-42939965-DEL-6640 | chr8-42939966-DEL-6639 |
| chr8-42971045-DEL-552 | chr8-42971046-DEL-552 |
| chr8-43011193-DEL-52 | chr8-43011194-DEL-52 |
| chr8-43043035-DEL-66 | chr8-43043035-DEL-66 |
| chr8-43061536-DEL-100 | chr8-43061537-DEL-100 |
| chr8-43072363-DEL-6416 | chr8-43072365-DEL-6415 |
| chr8-43084633-DEL-1872 | chr8-43084634-DEL-1872 |
| chr8-43091086-DEL-426 | chr8-43091087-DEL-426 |
| chr8-43117534-DEL-473 | chr8-43117535-DEL-473 |
| chr8-43156692-DEL-59 | chr8-43156693-DEL-59 |
| chr8-43207558-DEL-54 | chr8-43207559-DEL-54 |
| chr8-43269745-DEL-6572 | chr8-43269745-DEL-6572 |
| chr8-43287840-DEL-61 | chr8-43287841-DEL-61 |
| chr8-43297233-DEL-556 | chr8-43297234-DEL-555 |
| chr8-43349368-DEL-52 | chr8-43349369-DEL-52 |
| chr8-43356236-DEL-196 | chr8-43356240-DEL-194 |
| chr8-43370737-DEL-322 | chr8-43370738-DEL-322 |
| chr8-43384137-DEL-880 | chr8-43384138-DEL-880 |
| chr8-43402259-DEL-130 | chr8-43402260-DEL-130 |
| chr8-43403671-DEL-942 | chr8-43403672-DEL-942 |
| chr8-43418718-DEL-100 | chr8-43418719-DEL-96 |
| chr8-43427974-DEL-52 | chr8-43427975-DEL-53 |
| chr8-43428451-DEL-300 | chr8-43428452-DEL-300 |
| chr8-43449834-DEL-457 | chr8-43449835-DEL-457 |
| chr8-43450480-DEL-1437 | chr8-43450481-DEL-1437 |
| chr8-43458836-DEL-6620 | chr8-43458837-DEL-6620 |
| chr8-43467047-DEL-2059 | chr8-43467048-DEL-2059 |
| chr8-43474626-DEL-73 | chr8-43474627-DEL-73 |
| chr8-43508718-DEL-136 | chr8-43508719-DEL-136 |
| chr8-43518780-DEL-449 | chr8-43518781-DEL-449 |
| chr8-43521109-DEL-1742 | chr8-43521110-DEL-1742 |
| chr8-43527348-DEL-2036 | chr8-43527349-DEL-2036 |
| chr8-43533661-DEL-2673 | chr8-43533662-DEL-2673 |
| chr8-43541728-DEL-274 | chr8-43541729-DEL-274 |
| chr8-43545337-DEL-211 | chr8-43545338-DEL-211 |
| chr8-43553971-DEL-60 | chr8-43553972-DEL-66 |
| chr8-43559198-DEL-54 | chr8-43559191-DEL-50 |
| chr8-43572813-DEL-92 | chr8-43572817-DEL-88 |
| chr8-43588713-DEL-409 | chr8-43588714-DEL-409 |
| chr8-43598561-DEL-577 | chr8-43598562-DEL-577 |
| chr8-43603071-DEL-6848 | chr8-43603066-DEL-6849 |
| chr8-43615774-DEL-79 | chr8-43615775-DEL-79 |
| chr8-43627325-DEL-68 | chr8-43627326-DEL-72 |
| chr8-43632920-DEL-64 | chr8-43632909-DEL-51 |
| chr8-43636415-DEL-72 | chr8-43636416-DEL-72 |
| chr8-43665839-DEL-88 | chr8-43665840-DEL-88 |
| chr8-43695247-DEL-672 | chr8-43695248-DEL-672 |
| chr8-43698899-DEL-223 | chr8-43698875-DEL-203 |
| chr8-43702865-DEL-1156 | chr8-43702866-DEL-1156 |
| chr8-43719563-DEL-254 | chr8-43719564-DEL-254 |
| chr8-43729789-DEL-51 | chr8-43729790-DEL-51 |
| chr8-43731182-DEL-52 | chr8-43731183-DEL-52 |
| chr8-43751873-DEL-194 | chr8-43751874-DEL-194 |
| chr8-43764762-DEL-8737 | chr8-43764763-DEL-8737 |
| chr8-43775891-DEL-133 | chr8-43775892-DEL-137 |
| chr8-43777149-DEL-118 | chr8-43777150-DEL-118 |
| chr8-43779615-DEL-283 | chr8-43779616-DEL-283 |
| chr8-43784434-DEL-94 | chr8-43784462-DEL-69 |
| chr8-43786914-DEL-123 | chr8-43786907-DEL-131 |
| chr8-43794353-DEL-188 | chr8-43794354-DEL-188 |
| chr8-43797953-DEL-66 | chr8-43797953-DEL-66 |
| chr8-43804939-DEL-199 | chr8-43804940-DEL-199 |
| chr8-43816549-DEL-384 | chr8-43816550-DEL-384 |
| chr8-43821084-DEL-52 | chr8-43821085-DEL-58 |
| chr8-43839450-DEL-94 | chr8-43839451-DEL-94 |
| chr8-43846469-DEL-226 | chr8-43846470-DEL-226 |
| chr8-43850501-DEL-99 | chr8-43850502-DEL-99 |
| chr8-43892843-DEL-101 | chr8-43892840-DEL-102 |

| | |
|---|---|
| chr8-43896961-DEL-55 | chr8-43896981-DEL-67 |
| chr8-43915713-DEL-232 | chr8-43915714-DEL-232 |
| chr8-43937077-DEL-100 | chr8-43937078-DEL-98 |
| chr8-43970551-DEL-2317 | chr8-43970551-DEL-2310 |
| chr8-43995893-DEL-393 | chr8-43995894-DEL-393 |
| chr8-44026478-DEL-107 | chr8-44026479-DEL-105 |
| chr8-44033478-DEL-2839 | chr8-44033479-DEL-2839 |
| chr8-44041941-DEL-6490 | chr8-44041942-DEL-6490 |
| chr8-44056741-DEL-194 | chr8-44056702-DEL-195 |
| chr8-44086740-DEL-1412 | chr8-44086741-DEL-1412 |
| chr8-44176543-DEL-2908 | chr8-44176544-DEL-2908 |
| chr8-44191398-DEL-729 | chr8-44191366-DEL-730 |
| chr8-44202557-DEL-73 | chr8-44202565-DEL-67 |
| chr8-44203562-DEL-788 | chr8-44203563-DEL-788 |
| chr8-44225535-DEL-170 | chr8-44225525-DEL-171 |
| chr8-44244115-DEL-290 | chr8-44244116-DEL-290 |
| chr8-44244739-DEL-6618 | chr8-44244729-DEL-6628 |
| chr8-44289228-DEL-6427 | chr8-44289208-DEL-6401 |
| chr8-44327518-DEL-89 | chr8-44327519-DEL-89 |
| chr8-44367464-DEL-6401 | chr8-44367465-DEL-6401 |
| chr8-44379636-DEL-608 | chr8-44379637-DEL-608 |
| chr8-44380536-DEL-315 | chr8-44380537-DEL-315 |
| chr8-44390379-DEL-132 | chr8-44390380-DEL-132 |
| chr8-44413005-DEL-146 | chr8-44413006-DEL-146 |
| chr8-44414266-DEL-6463 | chr8-44414267-DEL-6463 |
| chr8-44440306-DEL-3285 | chr8-44440289-DEL-3286 |
| chr8-44457284-DEL-175 | chr8-44457285-DEL-175 |
| chr8-44461597-DEL-75 | chr8-44461561-DEL-120 |
| chr8-44490999-DEL-289 | chr8-44490993-DEL-290 |
| chr8-44493510-DEL-528 | chr8-44493511-DEL-528 |
| chr8-44517944-DEL-60 | chr8-44517945-DEL-52 |
| chr8-44526645-DEL-275 | chr8-44526647-DEL-286 |
| chr8-44601819-DEL-271 | chr8-44601820-DEL-271 |
| chr8-44622145-DEL-72 | chr8-44622146-DEL-70 |
| chr8-44653393-DEL-139 | chr8-44653394-DEL-139 |
| chr8-44663043-DEL-1508 | chr8-44663044-DEL-1507 |
| chr8-44664989-DEL-52 | chr8-44664990-DEL-52 |
| chr8-44702820-DEL-226 | chr8-44702821-DEL-226 |
| chr8-44713942-DEL-58 | chr8-44713943-DEL-58 |
| chr8-44726016-DEL-56 | chr8-44726017-DEL-56 |
| chr8-44742988-DEL-67 | chr8-44742987-DEL-68 |
| chr8-44804093-DEL-455 | chr8-44804094-DEL-455 |
| chr8-44816472-DEL-645 | chr8-44816473-DEL-645 |
| chr8-44866160-DEL-158 | chr8-44866161-DEL-158 |
| chr8-44874483-DEL-86 | chr8-44874484-DEL-78 |
| chr8-44875171-DEL-57 | chr8-44875172-DEL-57 |
| chr8-44879108-DEL-2665 | chr8-44879109-DEL-2665 |
| chr8-44894898-DEL-354 | chr8-44894899-DEL-354 |
| chr8-44922683-DEL-68 | chr8-44922684-DEL-68 |
| chr8-44960513-DEL-574 | chr8-44960514-DEL-574 |
| chr8-44994191-DEL-701 | chr8-44994192-DEL-701 |
| chr8-45006504-DEL-103 | chr8-45006501-DEL-71 |
| chr8-45020226-DEL-63 | chr8-45020227-DEL-63 |
| chr8-45036144-DEL-6238 | chr8-45036143-DEL-6260 |
| chr8-45047067-DEL-2382 | chr8-45047068-DEL-2382 |
| chr8-45059244-DEL-497 | chr8-45059245-DEL-497 |
| chr8-45119551-DEL-67 | chr8-45119552-DEL-67 |
| chr8-45169978-DEL-1413 | chr8-45169979-DEL-1413 |
| chr8-45175549-DEL-561 | chr8-45175546-DEL-577 |
| chr8-45187974-DEL-160 | chr8-45187970-DEL-164 |
| chr8-45199878-DEL-2235 | chr8-45199879-DEL-2235 |
| chr8-45236249-DEL-1864 | chr8-45236250-DEL-1864 |
| chr8-45376147-DEL-206 | chr8-45376148-DEL-206 |
| chr8-45397503-DEL-148 | chr8-45397504-DEL-148 |
| chr8-45421522-DEL-190 | chr8-45421523-DEL-190 |
| chr8-45440604-DEL-89 | chr8-45440599-DEL-78 |
| chr8-45462361-DEL-357 | chr8-45462362-DEL-357 |
| chr8-45509341-DEL-597 | chr8-45509342-DEL-597 |
| chr8-45512817-DEL-76 | chr8-45512801-DEL-92 |
| chr8-45514985-DEL-488 | chr8-45514972-DEL-478 |
| chr8-45518538-DEL-93 | chr8-45518538-DEL-93 |
| chr8-45521551-DEL-54 | chr8-45521552-DEL-54 |
| chr8-45537824-DEL-85 | chr8-45537870-DEL-51 |
| chr8-45548616-DEL-734 | chr8-45548613-DEL-722 |
| chr8-45563845-DEL-83 | chr8-45563846-DEL-83 |
| chr8-45579502-DEL-402 | chr8-45579503-DEL-402 |
| chr8-45610868-DEL-414 | chr8-45610869-DEL-414 |
| chr8-45617395-DEL-58 | chr8-45617396-DEL-58 |
| chr8-45629356-DEL-319 | chr8-45629360-DEL-316 |
| chr8-45632001-DEL-6466 | chr8-45632002-DEL-6466 |

| | |
|---|---|
| chr8-45636622-DEL-56 | chr8-45636602-DEL-57 |
| chr8-45643428-DEL-320 | chr8-45643429-DEL-320 |
| chr8-45647800-DEL-320 | chr8-45647801-DEL-320 |
| chr8-45663626-DEL-116 | chr8-45663636-DEL-106 |
| chr8-45676875-DEL-62 | chr8-45676886-DEL-92 |
| chr8-45696821-DEL-1679 | chr8-45696822-DEL-1679 |
| chr8-45707559-DEL-61 | chr8-45707560-DEL-61 |
| chr8-45716260-DEL-175 | chr8-45716268-DEL-146 |
| chr8-45767912-DEL-106 | chr8-45767913-DEL-106 |
| chr8-45787801-DEL-203 | chr8-45787802-DEL-203 |
| chr8-45829750-DEL-83 | chr8-45829751-DEL-83 |
| chr8-45836675-DEL-184 | chr8-45836672-DEL-163 |
| chr8-45843012-DEL-202 | chr8-45843013-DEL-202 |
| chr8-45844902-DEL-63 | chr8-45844903-DEL-64 |
| chr8-45852470-DEL-55 | chr8-45852471-DEL-55 |
| chr8-45858141-DEL-81 | chr8-45858142-DEL-82 |
| chr8-45872380-DEL-449 | chr8-45872381-DEL-449 |
| chr8-45873910-DEL-8178 | chr8-45873911-DEL-8178 |
| chr8-45900006-DEL-56 | chr8-45900007-DEL-56 |
| chr8-45910794-DEL-87 | chr8-45910780-DEL-57 |
| chr8-45911310-DEL-90 | chr8-45911311-DEL-90 |
| chr8-45916449-DEL-816 | chr8-45916449-DEL-816 |
| chr8-45918851-DEL-86 | chr8-45918838-DEL-60 |
| chr8-45929662-DEL-235 | chr8-45929663-DEL-235 |
| chr8-45942300-DEL-2621 | chr8-45942300-DEL-2622 |
| chr8-45952843-DEL-781 | chr8-45952844-DEL-781 |
| chr8-45967238-DEL-60 | chr8-45967239-DEL-60 |
| chr8-45972256-DEL-163 | chr8-45972257-DEL-163 |
| chr8-45982544-DEL-917 | chr8-45982545-DEL-917 |
| chr8-45985823-DEL-2253 | chr8-45985816-DEL-2247 |
| chr8-45986039-DEL-914 | chr8-45986043-DEL-911 |
| chr8-45988073-DEL-396 | chr8-45988079-DEL-397 |
| chr8-46009309-DEL-194 | chr8-46009302-DEL-202 |
| chr8-46030234-DEL-360 | chr8-46030237-DEL-363 |
| chr8-46031866-DEL-75 | chr8-46031829-DEL-66 |
| chr8-46031915-DEL-839 | chr8-46031914-DEL-837 |
| chr8-46032559-DEL-183 | chr8-46032558-DEL-183 |
| chr8-46034541-DEL-51 | chr8-46034526-DEL-51 |
| chr8-46069321-DEL-72 | chr8-46069339-DEL-55 |
| chr8-46210847-DEL-52 | chr8-46210848-DEL-51 |
| chr8-46215222-DEL-126 | chr8-46215223-DEL-126 |
| chr8-46230855-DEL-125 | chr8-46230871-DEL-110 |
| chr8-46254102-DEL-913 | chr8-46254103-DEL-913 |
| chr8-46277755-DEL-86 | chr8-46277756-DEL-86 |
| chr8-46283708-DEL-52 | chr8-46283709-DEL-52 |
| chr8-46303178-DEL-52 | chr8-46303171-DEL-68 |
| chr8-46306944-DEL-67 | chr8-46306941-DEL-79 |
| chr8-46308613-DEL-1690 | chr8-46308614-DEL-1690 |
| chr8-46309455-DEL-498 | chr8-46309449-DEL-494 |
| chr8-46326882-DEL-585 | chr8-46326891-DEL-578 |
| chr8-46329517-DEL-583 | chr8-46329518-DEL-583 |
| chr8-46337443-DEL-88 | chr8-46337444-DEL-87 |
| chr8-46350902-DEL-1895 | chr8-46350903-DEL-1895 |
| chr8-46369777-DEL-207 | chr8-46369778-DEL-207 |
| chr8-46375733-DEL-7112 | chr8-46375734-DEL-7112 |
| chr8-46426041-DEL-54 | chr8-46426042-DEL-54 |
| chr8-46433706-DEL-51 | chr8-46433707-DEL-51 |
| chr8-46438797-DEL-80 | chr8-46438798-DEL-75 |
| chr8-46460617-DEL-371 | chr8-46460613-DEL-372 |
| chr8-46484312-DEL-121 | chr8-46484313-DEL-121 |
| chr8-46503818-DEL-252 | chr8-46503811-DEL-263 |
| chr8-46513660-DEL-266 | chr8-46513659-DEL-266 |
| chr8-46547885-DEL-61 | chr8-46547886-DEL-61 |
| chr8-46578256-DEL-2593 | chr8-46578257-DEL-2593 |
| chr8-46657029-DEL-208 | chr8-46657030-DEL-208 |
| chr8-46658610-DEL-130 | chr8-46658611-DEL-130 |
| chr8-46667593-DEL-90 | chr8-46667594-DEL-90 |
| chr8-46675770-DEL-213 | chr8-46675768-DEL-213 |
| chr8-46677218-DEL-53 | chr8-46677213-DEL-53 |
| chr8-46680286-DEL-70 | chr8-46680287-DEL-70 |
| chr8-46706187-DEL-410 | chr8-46706196-DEL-386 |
| chr8-46706449-DEL-1768 | chr8-46706450-DEL-1768 |
| chr8-46726061-DEL-56 | chr8-46726061-DEL-56 |
| chr8-46737837-DEL-400 | chr8-46737838-DEL-400 |
| chr8-46738976-DEL-695 | chr8-46738977-DEL-695 |
| chr8-46757710-DEL-306 | chr8-46757711-DEL-306 |
| chr8-46759297-DEL-61 | chr8-46759298-DEL-61 |
| chr8-46796453-DEL-80 | chr8-46796454-DEL-80 |
| chr8-46798416-DEL-5426 | chr8-46798417-DEL-5426 |
| chr8-46822438-DEL-962 | chr8-46822439-DEL-962 |

| | |
|---|---|
| chr8-46824641-DEL-164 | chr8-46824642-DEL-162 |
| chr8-46831675-DEL-6235 | chr8-46831676-DEL-6235 |
| chr8-46858183-DEL-112 | chr8-46858182-DEL-109 |
| chr8-46859227-DEL-2090 | chr8-46859228-DEL-2090 |
| chr8-46863780-DEL-52 | chr8-46863781-DEL-52 |
| chr8-46883636-DEL-91 | chr8-46883638-DEL-87 |
| chr8-46935365-DEL-440 | chr8-46935366-DEL-418 |
| chr8-46949250-DEL-186 | chr8-46949251-DEL-186 |
| chr8-46989209-DEL-102 | chr8-46989210-DEL-101 |
| chr8-47016818-DEL-1066 | chr8-47016819-DEL-1066 |
| chr8-47057182-DEL-216 | chr8-47057183-DEL-216 |
| chr8-47074746-DEL-200 | chr8-47074749-DEL-199 |
| chr8-47084256-DEL-172 | chr8-47084257-DEL-172 |
| chr8-47100790-DEL-376 | chr8-47100791-DEL-376 |
| chr8-47115375-DEL-389 | chr8-47115376-DEL-389 |
| chr8-47127143-DEL-78 | chr8-47127144-DEL-78 |
| chr8-47134625-DEL-56 | chr8-47134628-DEL-60 |
| chr8-47139211-DEL-69 | chr8-47139252-DEL-78 |
| chr8-47178839-DEL-395 | chr8-47178861-DEL-371 |
| chr8-47197143-DEL-84 | chr8-47197144-DEL-84 |
| chr8-47198006-DEL-195 | chr8-47198007-DEL-195 |
| chr8-47211716-DEL-65 | chr8-47211710-DEL-66 |
| chr8-47263448-DEL-104 | chr8-47263449-DEL-84 |
| chr8-47331590-DEL-135 | chr8-47331591-DEL-135 |
| chr8-47354784-DEL-2931 | chr8-47354785-DEL-2931 |
| chr8-47365706-DEL-117 | chr8-47365707-DEL-117 |
| chr8-47409220-DEL-6396 | chr8-47409221-DEL-6396 |
| chr8-47422222-DEL-221 | chr8-47422223-DEL-221 |
| chr8-47444467-DEL-69 | chr8-47444468-DEL-69 |
| chr8-47466449-DEL-70 | chr8-47466450-DEL-70 |
| chr8-47473274-DEL-143 | chr8-47473256-DEL-152 |
| chr8-47484928-DEL-76 | chr8-47484929-DEL-76 |
| chr8-47512654-DEL-56 | chr8-47512655-DEL-56 |
| chr8-47524718-DEL-3153 | chr8-47524719-DEL-3153 |
| chr8-47573669-DEL-490 | chr8-47573669-DEL-490 |
| chr8-47582541-DEL-182 | chr8-47582542-DEL-182 |
| chr8-47583942-DEL-701 | chr8-47583943-DEL-701 |
| chr8-47598576-DEL-6420 | chr8-47598577-DEL-6420 |
| chr8-47608134-DEL-85 | chr8-47608140-DEL-59 |
| chr8-47611118-DEL-401 | chr8-47611119-DEL-401 |
| chr8-47614162-DEL-75 | chr8-47614163-DEL-75 |
| chr8-47620604-DEL-76 | chr8-47620605-DEL-76 |
| chr8-47670855-DEL-61 | chr8-47670843-DEL-73 |
| chr8-47754356-DEL-98 | chr8-47754357-DEL-98 |
| chr8-47801411-DEL-53 | chr8-47801412-DEL-53 |
| chr8-47810927-DEL-87 | chr8-47810928-DEL-87 |
| chr8-47823313-DEL-88 | chr8-47823314-DEL-88 |
| chr8-47862458-DEL-123 | chr8-47862459-DEL-123 |
| chr8-47893263-DEL-869 | chr8-47893264-DEL-869 |
| chr8-47904965-DEL-2963 | chr8-47904966-DEL-2963 |
| chr8-47918659-DEL-13791 | chr8-47918660-DEL-13791 |
| chr8-47941608-DEL-56 | chr8-47941610-DEL-54 |
| chr8-47964091-DEL-60 | chr8-47964107-DEL-60 |
| chr8-47987365-DEL-186 | chr8-47987366-DEL-186 |
| chr8-47994125-DEL-441 | chr8-47994127-DEL-428 |
| chr8-48001519-DEL-1249 | chr8-48001520-DEL-1249 |
| chr8-48002723-DEL-700 | chr8-48002723-DEL-700 |
| chr8-48018340-DEL-95 | chr8-48018344-DEL-91 |
| chr8-48038559-DEL-58 | chr8-48038562-DEL-59 |
| chr8-48038899-DEL-520 | chr8-48038900-DEL-520 |
| chr8-48039127-DEL-1689 | chr8-48039128-DEL-1689 |
| chr8-48040094-DEL-2375 | chr8-48040095-DEL-2375 |
| chr8-48048834-DEL-82 | chr8-48048835-DEL-82 |
| chr8-48050905-DEL-199 | chr8-48050906-DEL-199 |
| chr8-48097553-DEL-845 | chr8-48097554-DEL-845 |
| chr8-48107085-DEL-306 | chr8-48107086-DEL-306 |
| chr8-48138764-DEL-242 | chr8-48138765-DEL-242 |
| chr8-48152395-DEL-99 | chr8-48152396-DEL-99 |
| chr8-48178184-DEL-52 | chr8-48178185-DEL-52 |
| chr8-48209711-DEL-92 | chr8-48209722-DEL-70 |
| chr8-48219122-DEL-51 | chr8-48219123-DEL-51 |
| chr8-48220575-DEL-64 | chr8-48220575-DEL-80 |
| chr8-48226094-DEL-117 | chr8-48226095-DEL-117 |
| chr8-48228009-DEL-58 | chr8-48228010-DEL-55 |
| chr8-48303206-DEL-852 | chr8-48303207-DEL-852 |
| chr8-48356322-DEL-75 | chr8-48356333-DEL-93 |
| chr8-48363251-DEL-3355 | chr8-48363252-DEL-3355 |
| chr8-48377355-DEL-196 | chr8-48377356-DEL-196 |
| chr8-48378367-DEL-203 | chr8-48378368-DEL-203 |
| chr8-48385279-DEL-4158 | chr8-48385280-DEL-4158 |

| | |
|---|---|
| chr8-48413634-DEL-151 | chr8-48413635-DEL-151 |
| chr8-48454037-DEL-72 | chr8-48454038-DEL-72 |
| chr8-48484148-DEL-79 | chr8-48484134-DEL-65 |
| chr8-48515940-DEL-85 | chr8-48515941-DEL-85 |
| chr8-48534538-DEL-332 | chr8-48534539-DEL-331 |
| chr8-48589033-DEL-68 | chr8-48589034-DEL-68 |
| chr8-48620642-DEL-577 | chr8-48620643-DEL-577 |
| chr8-48641943-DEL-7105 | chr8-48641947-DEL-7104 |
| chr8-48650075-DEL-197 | chr8-48650076-DEL-197 |
| chr8-48651940-DEL-62 | chr8-48651940-DEL-62 |
| chr8-48662566-DEL-515 | chr8-48662567-DEL-515 |
| chr8-48683681-DEL-351 | chr8-48683693-DEL-347 |
| chr8-48687354-DEL-54 | chr8-48687355-DEL-56 |
| chr8-48706561-DEL-118 | chr8-48706562-DEL-118 |
| chr8-48715213-DEL-68 | chr8-48715214-DEL-68 |
| chr8-48723676-DEL-84 | chr8-48723677-DEL-84 |
| chr8-48772062-DEL-676 | chr8-48772063-DEL-676 |
| chr8-48795920-DEL-66 | chr8-48795921-DEL-66 |
| chr8-48812135-DEL-355 | chr8-48812136-DEL-355 |
| chr8-48836012-DEL-54 | chr8-48836013-DEL-54 |
| chr8-48843715-DEL-78 | chr8-48843716-DEL-78 |
| chr8-48849165-DEL-56 | chr8-48849166-DEL-96 |
| chr8-48886554-DEL-451 | chr8-48886555-DEL-451 |
| chr8-48889851-DEL-68 | chr8-48889852-DEL-68 |
| chr8-48945193-DEL-68 | chr8-48945201-DEL-65 |
| chr8-48945898-DEL-54 | chr8-48945899-DEL-54 |
| chr8-48970005-DEL-72 | chr8-48970002-DEL-70 |
| chr8-48976473-DEL-69 | chr8-48976474-DEL-69 |
| chr8-48986845-DEL-69 | chr8-48986846-DEL-69 |
| chr8-48997382-DEL-72 | chr8-48997379-DEL-76 |
| chr8-49050342-DEL-78 | chr8-49050343-DEL-78 |
| chr8-49083834-DEL-1641 | chr8-49083835-DEL-1641 |
| chr8-49103411-DEL-833 | chr8-49103412-DEL-833 |
| chr8-49128222-DEL-233 | chr8-49128223-DEL-232 |
| chr8-49147266-DEL-101 | chr8-49147270-DEL-100 |
| chr8-49180496-DEL-120 | chr8-49180530-DEL-84 |
| chr8-49220961-DEL-570 | chr8-49220962-DEL-570 |
| chr8-49314676-DEL-87 | chr8-49314676-DEL-92 |
| chr8-49318916-DEL-54 | chr8-49318920-DEL-67 |
| chr8-49352602-DEL-65 | chr8-49352603-DEL-65 |
| chr8-49371355-DEL-60 | chr8-49371355-DEL-60 |
| chr8-49378151-DEL-211 | chr8-49378152-DEL-211 |
| chr8-49389843-DEL-77 | chr8-49389843-DEL-67 |
| chr8-49437210-DEL-7078 | chr8-49437211-DEL-7078 |
| chr8-49468649-DEL-121 | chr8-49468650-DEL-121 |
| chr8-49469711-DEL-299 | chr8-49469712-DEL-299 |
| chr8-49525789-DEL-7008 | chr8-49525790-DEL-7008 |
| chr8-49540389-DEL-168 | chr8-49540389-DEL-167 |
| chr8-49553156-DEL-58 | chr8-49553157-DEL-58 |
| chr8-49622315-DEL-59 | chr8-49622316-DEL-59 |
| chr8-49654428-DEL-484 | chr8-49654428-DEL-481 |
| chr8-49725765-DEL-53 | chr8-49725766-DEL-53 |
| chr8-49758793-DEL-196 | chr8-49758794-DEL-196 |
| chr8-49859437-DEL-458 | chr8-49859438-DEL-458 |
| chr8-49890158-DEL-89 | chr8-49890159-DEL-89 |
| chr8-49922463-DEL-579 | chr8-49922464-DEL-579 |
| chr8-49925179-DEL-124 | chr8-49925184-DEL-120 |
| chr8-49950805-DEL-72 | chr8-49950803-DEL-72 |
| chr8-49984011-DEL-98 | chr8-49984012-DEL-98 |
| chr8-49984543-DEL-63 | chr8-49984544-DEL-63 |
| chr8-50026724-DEL-7022 | chr8-50026725-DEL-7022 |
| chr8-50026739-DEL-636 | chr8-50026740-DEL-636 |
| chr8-50043013-DEL-60 | chr8-50043022-DEL-52 |
| chr8-50045101-DEL-372 | chr8-50045102-DEL-372 |
| chr8-50053514-DEL-6272 | chr8-50053515-DEL-6272 |
| chr8-50092294-DEL-104 | chr8-50092285-DEL-102 |
| chr8-50112437-DEL-632 | chr8-50112438-DEL-632 |
| chr8-50167697-DEL-68 | chr8-50167690-DEL-76 |
| chr8-50168379-DEL-55 | chr8-50168387-DEL-56 |
| chr8-50230864-DEL-72 | chr8-50230872-DEL-56 |
| chr8-50287711-DEL-5895 | chr8-50287712-DEL-5895 |
| chr8-50313382-DEL-52 | chr8-50313383-DEL-52 |
| chr8-50327173-DEL-455 | chr8-50327174-DEL-455 |
| chr8-50348578-DEL-151 | chr8-50348580-DEL-148 |
| chr8-50371676-DEL-89 | chr8-50371677-DEL-89 |
| chr8-50386251-DEL-354 | chr8-50386254-DEL-355 |
| chr8-50400033-DEL-111 | chr8-50400034-DEL-111 |
| chr8-50426399-DEL-77 | chr8-50426397-DEL-76 |
| chr8-50430863-DEL-202 | chr8-50430864-DEL-202 |
| chr8-50432549-DEL-262 | chr8-50432579-DEL-260 |

| | |
|---|---|
| chr8-50434877-DEL-6900 | chr8-50434878-DEL-6900 |
| chr8-50435115-DEL-636 | chr8-50435116-DEL-636 |
| chr8-50472062-DEL-473 | chr8-50472051-DEL-479 |
| chr8-50490522-DEL-232 | chr8-50490505-DEL-250 |
| chr8-50494887-DEL-498 | chr8-50494896-DEL-499 |
| chr8-50514511-DEL-3540 | chr8-50514511-DEL-3540 |
| chr8-50549997-DEL-6288 | chr8-50549998-DEL-6288 |
| chr8-50574473-DEL-536 | chr8-50574474-DEL-527 |
| chr8-50596886-DEL-685 | chr8-50596887-DEL-685 |
| chr8-50625674-DEL-1365 | chr8-50625675-DEL-1365 |
| chr8-50640683-DEL-208 | chr8-50640684-DEL-208 |
| chr8-50660750-DEL-229 | chr8-50660744-DEL-248 |
| chr8-50671430-DEL-625 | chr8-50671431-DEL-625 |
| chr8-50674109-DEL-103 | chr8-50674110-DEL-103 |
| chr8-50696914-DEL-6165 | chr8-50696915-DEL-6165 |
| chr8-50703642-DEL-162 | chr8-50703643-DEL-162 |
| chr8-50706683-DEL-67 | chr8-50706684-DEL-67 |
| chr8-50713984-DEL-154 | chr8-50713985-DEL-154 |
| chr8-50718598-DEL-62 | chr8-50718606-DEL-62 |
| chr8-50730496-DEL-209 | chr8-50730497-DEL-209 |
| chr8-50761339-DEL-98 | chr8-50761340-DEL-98 |
| chr8-50788418-DEL-245 | chr8-50788419-DEL-245 |
| chr8-50795488-DEL-1624 | chr8-50795494-DEL-1619 |
| chr8-50806618-DEL-117 | chr8-50806619-DEL-117 |
| chr8-50854261-DEL-150 | chr8-50854259-DEL-151 |
| chr8-50880937-DEL-91 | chr8-50880938-DEL-91 |
| chr8-50956779-DEL-1373 | chr8-50956780-DEL-1373 |
| chr8-50976639-DEL-77 | chr8-50976640-DEL-77 |
| chr8-51033664-DEL-157 | chr8-51033665-DEL-157 |
| chr8-51050214-DEL-7210 | chr8-51050215-DEL-7210 |
| chr8-51070469-DEL-67 | chr8-51070471-DEL-70 |
| chr8-51074627-DEL-655 | chr8-51074628-DEL-655 |
| chr8-51091850-DEL-57 | chr8-51091846-DEL-58 |
| chr8-51095025-DEL-119 | chr8-51095070-DEL-114 |
| chr8-51141135-DEL-61 | chr8-51141142-DEL-56 |
| chr8-51213337-DEL-401 | chr8-51213338-DEL-402 |
| chr8-51216459-DEL-1172 | chr8-51216460-DEL-1172 |
| chr8-51281705-DEL-389 | chr8-51281706-DEL-389 |
| chr8-51284169-DEL-4153 | chr8-51284170-DEL-4153 |
| chr8-51323124-DEL-211 | chr8-51323118-DEL-205 |
| chr8-51343261-DEL-140 | chr8-51343262-DEL-139 |
| chr8-51346938-DEL-56 | chr8-51346943-DEL-62 |
| chr8-51359848-DEL-61 | chr8-51359823-DEL-60 |
| chr8-51412052-DEL-934 | chr8-51412053-DEL-934 |
| chr8-51449143-DEL-4640 | chr8-51449144-DEL-4640 |
| chr8-51464757-DEL-197 | chr8-51464758-DEL-198 |
| chr8-51494615-DEL-412 | chr8-51494616-DEL-412 |
| chr8-51503703-DEL-2350 | chr8-51503704-DEL-2350 |
| chr8-51520027-DEL-127 | chr8-51520028-DEL-121 |
| chr8-51559938-DEL-105 | chr8-51559944-DEL-100 |
| chr8-51566187-DEL-78 | chr8-51566186-DEL-77 |
| chr8-51579183-DEL-1037 | chr8-51579187-DEL-1032 |
| chr8-51580789-DEL-779 | chr8-51580790-DEL-779 |
| chr8-51582750-DEL-2993 | chr8-51582796-DEL-2949 |
| chr8-51594906-DEL-53 | chr8-51594906-DEL-52 |
| chr8-51607329-DEL-340 | chr8-51607330-DEL-340 |
| chr8-51620405-DEL-725 | chr8-51620406-DEL-725 |
| chr8-51632071-DEL-69 | chr8-51632059-DEL-68 |
| chr8-51646468-DEL-312 | chr8-51646480-DEL-293 |
| chr8-51652360-DEL-57 | chr8-51652361-DEL-57 |
| chr8-51692063-DEL-77 | chr8-51692064-DEL-79 |
| chr8-51719964-DEL-6399 | chr8-51719964-DEL-6399 |
| chr8-51721759-DEL-440 | chr8-51721759-DEL-438 |
| chr8-51732236-DEL-61 | chr8-51732248-DEL-50 |
| chr8-51737583-DEL-60 | chr8-51737583-DEL-60 |
| chr8-51744981-DEL-522 | chr8-51744982-DEL-522 |
| chr8-51757537-DEL-68 | chr8-51757530-DEL-69 |
| chr8-51758552-DEL-178 | chr8-51758513-DEL-179 |
| chr8-51799660-DEL-6922 | chr8-51799661-DEL-6922 |
| chr8-51839843-DEL-79 | chr8-51839844-DEL-79 |
| chr8-51841252-DEL-609 | chr8-51841243-DEL-610 |
| chr8-51842701-DEL-178 | chr8-51842702-DEL-178 |
| chr8-51847882-DEL-2842 | chr8-51847882-DEL-2843 |
| chr8-51897287-DEL-68 | chr8-51897288-DEL-68 |
| chr8-51922496-DEL-55 | chr8-51922497-DEL-55 |
| chr8-51969798-DEL-331 | chr8-51969799-DEL-331 |
| chr8-51987576-DEL-562 | chr8-51987577-DEL-562 |
| chr8-51996441-DEL-6111 | chr8-51996442-DEL-6111 |
| chr8-52006116-DEL-51 | chr8-52006116-DEL-52 |
| chr8-52006442-DEL-498 | chr8-52006443-DEL-498 |

| | |
|---|---|
| chr8-52016785-DEL-6515 | chr8-52016786-DEL-6515 |
| chr8-52023027-DEL-204 | chr8-52023028-DEL-204 |
| chr8-52065867-DEL-5228 | chr8-52065868-DEL-5228 |
| chr8-52084967-DEL-54 | chr8-52084972-DEL-54 |
| chr8-52099581-DEL-82 | chr8-52099582-DEL-82 |
| chr8-52138820-DEL-60 | chr8-52138821-DEL-68 |
| chr8-52157850-DEL-95 | chr8-52157851-DEL-95 |
| chr8-52164700-DEL-6419 | chr8-52164701-DEL-6419 |
| chr8-52177198-DEL-72 | chr8-52177199-DEL-72 |
| chr8-52184613-DEL-4156 | chr8-52184614-DEL-4156 |
| chr8-52189672-DEL-163 | chr8-52189673-DEL-163 |
| chr8-52205292-DEL-6583 | chr8-52205293-DEL-6583 |
| chr8-52220848-DEL-1253 | chr8-52220849-DEL-1253 |
| chr8-52220911-DEL-372 | chr8-52220912-DEL-372 |
| chr8-52235304-DEL-3228 | chr8-52235305-DEL-3228 |
| chr8-52241077-DEL-2242 | chr8-52241083-DEL-2237 |
| chr8-52256859-DEL-412 | chr8-52256860-DEL-412 |
| chr8-52289650-DEL-685 | chr8-52289651-DEL-685 |
| chr8-52291131-DEL-2119 | chr8-52291132-DEL-2119 |
| chr8-52291587-DEL-677 | chr8-52291592-DEL-674 |
| chr8-52296552-DEL-2836 | chr8-52296553-DEL-2836 |
| chr8-52317083-DEL-281 | chr8-52317084-DEL-281 |
| chr8-52328017-DEL-62 | chr8-52328018-DEL-62 |
| chr8-52330839-DEL-83 | chr8-52330838-DEL-83 |
| chr8-52337664-DEL-123 | chr8-52337665-DEL-123 |
| chr8-52352665-DEL-2779 | chr8-52352663-DEL-2779 |
| chr8-52363796-DEL-77 | chr8-52363797-DEL-77 |
| chr8-52370083-DEL-196 | chr8-52370084-DEL-196 |
| chr8-52372823-DEL-64 | chr8-52372823-DEL-68 |
| chr8-52376172-DEL-7225 | chr8-52376173-DEL-7225 |
| chr8-52397885-DEL-194 | chr8-52397886-DEL-194 |
| chr8-52407874-DEL-72 | chr8-52407875-DEL-72 |
| chr8-52411648-DEL-72 | chr8-52411649-DEL-72 |
| chr8-52422015-DEL-1498 | chr8-52422016-DEL-1498 |
| chr8-52427846-DEL-492 | chr8-52427847-DEL-492 |
| chr8-52474402-DEL-63 | chr8-52474442-DEL-59 |
| chr8-52514456-DEL-6190 | chr8-52514456-DEL-6190 |
| chr8-52560038-DEL-135 | chr8-52560039-DEL-135 |
| chr8-52564806-DEL-56 | chr8-52564807-DEL-56 |
| chr8-52570638-DEL-132 | chr8-52570639-DEL-132 |
| chr8-52590039-DEL-101 | chr8-52590089-DEL-51 |
| chr8-52599576-DEL-336 | chr8-52599577-DEL-336 |
| chr8-52602584-DEL-57 | chr8-52602585-DEL-57 |
| chr8-52642629-DEL-69 | chr8-52642630-DEL-69 |
| chr8-52645321-DEL-413 | chr8-52645322-DEL-413 |
| chr8-52648792-DEL-1426 | chr8-52648793-DEL-1426 |
| chr8-52682453-DEL-52 | chr8-52682454-DEL-52 |
| chr8-52684580-DEL-54 | chr8-52684576-DEL-51 |
| chr8-52687146-DEL-2867 | chr8-52687147-DEL-2867 |
| chr8-52692099-DEL-57 | chr8-52692100-DEL-57 |
| chr8-52709105-DEL-3622 | chr8-52709106-DEL-3622 |
| chr8-52712967-DEL-76 | chr8-52712967-DEL-74 |
| chr8-52715991-DEL-883 | chr8-52715993-DEL-881 |
| chr8-52724679-DEL-1590 | chr8-52724679-DEL-1590 |
| chr8-52745732-DEL-56 | chr8-52745769-DEL-59 |
| chr8-52756591-DEL-11110 | chr8-52756585-DEL-11111 |
| chr8-52760084-DEL-424 | chr8-52760083-DEL-424 |
| chr8-52789725-DEL-416 | chr8-52789724-DEL-417 |
| chr8-52792876-DEL-66 | chr8-52792872-DEL-65 |
| chr8-52818217-DEL-385 | chr8-52818218-DEL-385 |
| chr8-52833155-DEL-1057 | chr8-52833156-DEL-1057 |
| chr8-52874121-DEL-136 | chr8-52874122-DEL-136 |
| chr8-52879840-DEL-6444 | chr8-52879841-DEL-6444 |
| chr8-52891482-DEL-57 | chr8-52891483-DEL-57 |
| chr8-52910820-DEL-80 | chr8-52910821-DEL-78 |
| chr8-52939802-DEL-62 | chr8-52939803-DEL-62 |
| chr8-53032546-DEL-70 | chr8-53032546-DEL-70 |
| chr8-53039304-DEL-345 | chr8-53039305-DEL-345 |
| chr8-53047861-DEL-6390 | chr8-53047862-DEL-6390 |
| chr8-53064590-DEL-332 | chr8-53064591-DEL-332 |
| chr8-53065950-DEL-69 | chr8-53065946-DEL-70 |
| chr8-53091098-DEL-756 | chr8-53091099-DEL-756 |
| chr8-53164220-DEL-304 | chr8-53164221-DEL-304 |
| chr8-53178216-DEL-78 | chr8-53178216-DEL-73 |
| chr8-53183706-DEL-79 | chr8-53183707-DEL-79 |
| chr8-53188898-DEL-2076 | chr8-53188899-DEL-2076 |
| chr8-53197168-DEL-76 | chr8-53197169-DEL-76 |
| chr8-53201975-DEL-122 | chr8-53201975-DEL-123 |
| chr8-53208238-DEL-1891 | chr8-53208239-DEL-1891 |
| chr8-53212743-DEL-73 | chr8-53212706-DEL-64 |

| | |
|---|---|
| chr8-53223440-DEL-96 | chr8-53223441-DEL-96 |
| chr8-53230699-DEL-4286 | chr8-53230700-DEL-4286 |
| chr8-53240891-DEL-456 | chr8-53240891-DEL-456 |
| chr8-53244690-DEL-52 | chr8-53244678-DEL-64 |
| chr8-53268483-DEL-229 | chr8-53268484-DEL-229 |
| chr8-53289180-DEL-7748 | chr8-53289181-DEL-7748 |
| chr8-53319332-DEL-365 | chr8-53319332-DEL-365 |
| chr8-53329430-DEL-75 | chr8-53329431-DEL-75 |
| chr8-53356107-DEL-549 | chr8-53356114-DEL-536 |
| chr8-53356810-DEL-6368 | chr8-53356811-DEL-6368 |
| chr8-53356862-DEL-214 | chr8-53356863-DEL-214 |
| chr8-53415950-DEL-7198 | chr8-53415950-DEL-7198 |
| chr8-53433270-DEL-65 | chr8-53433246-DEL-60 |
| chr8-53452629-DEL-60 | chr8-53452630-DEL-60 |
| chr8-53532908-DEL-90 | chr8-53532908-DEL-86 |
| chr8-53536210-DEL-1447 | chr8-53536211-DEL-1447 |
| chr8-53537852-DEL-93 | chr8-53537853-DEL-93 |
| chr8-53547088-DEL-412 | chr8-53547088-DEL-412 |
| chr8-53555252-DEL-3008 | chr8-53555253-DEL-3008 |
| chr8-53566805-DEL-6231 | chr8-53566806-DEL-6231 |
| chr8-53609316-DEL-2736 | chr8-53609317-DEL-2736 |
| chr8-53636318-DEL-136 | chr8-53636317-DEL-138 |
| chr8-53661763-DEL-822 | chr8-53661758-DEL-789 |
| chr8-53673629-DEL-575 | chr8-53673630-DEL-575 |
| chr8-53684774-DEL-105 | chr8-53684773-DEL-106 |
| chr8-53691121-DEL-70 | chr8-53691122-DEL-70 |
| chr8-53717586-DEL-3176 | chr8-53717587-DEL-3176 |
| chr8-53736879-DEL-63 | chr8-53736842-DEL-64 |
| chr8-53749353-DEL-320 | chr8-53749354-DEL-320 |
| chr8-53758211-DEL-100 | chr8-53758215-DEL-85 |
| chr8-53759345-DEL-56 | chr8-53759317-DEL-63 |
| chr8-53780053-DEL-289 | chr8-53780054-DEL-289 |
| chr8-53787103-DEL-2647 | chr8-53787104-DEL-2647 |
| chr8-53793685-DEL-868 | chr8-53793686-DEL-868 |
| chr8-53830312-DEL-2376 | chr8-53830310-DEL-2366 |
| chr8-53854677-DEL-367 | chr8-53854691-DEL-366 |
| chr8-53873519-DEL-90 | chr8-53873520-DEL-90 |
| chr8-53879982-DEL-168 | chr8-53879983-DEL-168 |
| chr8-53901916-DEL-57 | chr8-53901919-DEL-55 |
| chr8-53904017-DEL-6251 | chr8-53904017-DEL-6251 |
| chr8-53913558-DEL-71 | chr8-53913559-DEL-71 |
| chr8-53917544-DEL-345 | chr8-53917545-DEL-345 |
| chr8-53928790-DEL-52 | chr8-53928787-DEL-56 |
| chr8-54004256-DEL-79 | chr8-54004245-DEL-80 |
| chr8-54007364-DEL-1946 | chr8-54007365-DEL-1946 |
| chr8-54066666-DEL-64 | chr8-54066667-DEL-64 |
| chr8-54068437-DEL-1886 | chr8-54068419-DEL-1906 |
| chr8-54071393-DEL-91 | chr8-54071394-DEL-91 |
| chr8-54098313-DEL-388 | chr8-54098314-DEL-388 |
| chr8-54155747-DEL-1439 | chr8-54155748-DEL-1439 |
| chr8-54180862-DEL-165 | chr8-54180863-DEL-165 |
| chr8-54184846-DEL-475 | chr8-54184847-DEL-475 |
| chr8-54292161-DEL-80 | chr8-54292152-DEL-85 |
| chr8-54298059-DEL-157 | chr8-54298075-DEL-142 |
| chr8-54316298-DEL-3460 | chr8-54316298-DEL-3460 |
| chr8-54355065-DEL-1656 | chr8-54355066-DEL-1656 |
| chr8-54377933-DEL-1533 | chr8-54377932-DEL-1533 |
| chr8-54395261-DEL-1350 | chr8-54395262-DEL-1350 |
| chr8-54452105-DEL-52 | chr8-54452083-DEL-55 |
| chr8-54463808-DEL-735 | chr8-54463809-DEL-735 |
| chr8-54492600-DEL-284 | chr8-54492601-DEL-284 |
| chr8-54493513-DEL-80 | chr8-54493538-DEL-78 |
| chr8-54526357-DEL-92 | chr8-54526356-DEL-92 |
| chr8-54544751-DEL-2561 | chr8-54544752-DEL-2561 |
| chr8-54560066-DEL-707 | chr8-54560067-DEL-707 |
| chr8-54562577-DEL-680 | chr8-54562578-DEL-680 |
| chr8-54595915-DEL-7154 | chr8-54595916-DEL-7154 |
| chr8-54608387-DEL-54 | chr8-54608387-DEL-54 |
| chr8-54608910-DEL-167 | chr8-54608910-DEL-161 |
| chr8-54632717-DEL-86 | chr8-54632706-DEL-60 |
| chr8-54640416-DEL-62 | chr8-54640417-DEL-62 |
| chr8-54650975-DEL-512 | chr8-54650975-DEL-512 |
| chr8-54692208-DEL-770 | chr8-54692209-DEL-770 |
| chr8-54717134-DEL-152 | chr8-54717135-DEL-152 |
| chr8-54754228-DEL-200 | chr8-54754229-DEL-200 |
| chr8-54774968-DEL-514 | chr8-54774969-DEL-514 |
| chr8-54802066-DEL-62 | chr8-54802058-DEL-72 |
| chr8-54838668-DEL-1662 | chr8-54838669-DEL-1662 |
| chr8-54843436-DEL-60 | chr8-54843432-DEL-61 |
| chr8-54856669-DEL-78 | chr8-54856667-DEL-79 |

| | |
|---|---|
| chr8-54861825-DEL-238 | chr8-54861826-DEL-238 |
| chr8-54868187-DEL-183 | chr8-54868188-DEL-183 |
| chr8-54876628-DEL-333 | chr8-54876629-DEL-333 |
| chr8-54878018-DEL-56 | chr8-54877995-DEL-60 |
| chr8-54878378-DEL-1508 | chr8-54878380-DEL-1506 |
| chr8-54900908-DEL-209 | chr8-54900909-DEL-209 |
| chr8-54924387-DEL-168 | chr8-54924388-DEL-168 |
| chr8-54934444-DEL-55 | chr8-54934445-DEL-55 |
| chr8-54937369-DEL-918 | chr8-54937364-DEL-916 |
| chr8-54953555-DEL-2204 | chr8-54953556-DEL-2201 |
| chr8-54965492-DEL-1893 | chr8-54965493-DEL-1893 |
| chr8-54992984-DEL-426 | chr8-54992992-DEL-426 |
| chr8-55007283-DEL-58 | chr8-55007282-DEL-60 |
| chr8-55011974-DEL-397 | chr8-55011975-DEL-397 |
| chr8-55028985-DEL-72 | chr8-55029002-DEL-62 |
| chr8-55048465-DEL-514 | chr8-55048466-DEL-514 |
| chr8-55076129-DEL-400 | chr8-55076130-DEL-400 |
| chr8-55090871-DEL-53 | chr8-55090872-DEL-53 |
| chr8-55108707-DEL-56 | chr8-55108708-DEL-56 |
| chr8-55122914-DEL-75 | chr8-55122913-DEL-76 |
| chr8-55133912-DEL-62 | chr8-55133913-DEL-62 |
| chr8-55154248-DEL-325 | chr8-55154249-DEL-325 |
| chr8-55161691-DEL-3329 | chr8-55161692-DEL-3329 |
| chr8-55170321-DEL-351 | chr8-55170328-DEL-345 |
| chr8-55186314-DEL-864 | chr8-55186316-DEL-837 |
| chr8-55187028-DEL-285 | chr8-55186993-DEL-317 |
| chr8-55194664-DEL-406 | chr8-55194665-DEL-406 |
| chr8-55228677-DEL-315 | chr8-55228678-DEL-315 |
| chr8-55245993-DEL-3722 | chr8-55245994-DEL-3722 |
| chr8-55262994-DEL-362 | chr8-55262995-DEL-362 |
| chr8-55296001-DEL-6402 | chr8-55296002-DEL-6402 |
| chr8-55321171-DEL-6701 | chr8-55321172-DEL-6701 |
| chr8-55359072-DEL-79 | chr8-55359073-DEL-79 |
| chr8-55361911-DEL-52 | chr8-55361912-DEL-52 |
| chr8-55397498-DEL-126 | chr8-55397505-DEL-142 |
| chr8-55430592-DEL-450 | chr8-55430593-DEL-450 |
| chr8-55434608-DEL-263 | chr8-55434608-DEL-263 |
| chr8-55435400-DEL-58 | chr8-55435401-DEL-58 |
| chr8-55436750-DEL-497 | chr8-55436751-DEL-497 |
| chr8-55456920-DEL-400 | chr8-55456921-DEL-400 |
| chr8-55516961-DEL-73 | chr8-55516962-DEL-73 |
| chr8-55519830-DEL-60 | chr8-55519831-DEL-60 |
| chr8-55862259-DEL-80 | chr8-55862296-DEL-70 |
| chr8-55895541-DEL-66 | chr8-55895542-DEL-66 |
| chr8-55898790-DEL-194 | chr8-55898791-DEL-194 |
| chr8-55979317-DEL-3017 | chr8-55979318-DEL-3017 |
| chr8-55999809-DEL-5259 | chr8-55999810-DEL-5259 |
| chr8-56077977-DEL-69 | chr8-56078006-DEL-74 |
| chr8-56111448-DEL-135 | chr8-56111449-DEL-135 |
| chr8-56121975-DEL-95 | chr8-56121976-DEL-102 |
| chr8-56161652-DEL-138 | chr8-56161653-DEL-138 |
| chr8-56163728-DEL-1310 | chr8-56163733-DEL-1313 |
| chr8-56169411-DEL-637 | chr8-56169412-DEL-637 |
| chr8-56188511-DEL-644 | chr8-56188505-DEL-645 |
| chr8-56194799-DEL-225 | chr8-56194800-DEL-225 |
| chr8-56196652-DEL-201 | chr8-56196653-DEL-201 |
| chr8-56211290-DEL-70 | chr8-56211312-DEL-68 |
| chr8-56241903-DEL-52 | chr8-56241894-DEL-62 |
| chr8-56245196-DEL-2108 | chr8-56245197-DEL-2110 |
| chr8-56261690-DEL-60 | chr8-56261691-DEL-60 |
| chr8-56292603-DEL-6262 | chr8-56292604-DEL-6262 |
| chr8-56337548-DEL-623 | chr8-56337549-DEL-623 |
| chr8-56366528-DEL-201 | chr8-56366529-DEL-201 |
| chr8-56374772-DEL-224 | chr8-56374773-DEL-224 |
| chr8-56386877-DEL-610 | chr8-56386878-DEL-610 |
| chr8-56404256-DEL-67 | chr8-56404262-DEL-56 |
| chr8-56410511-DEL-114 | chr8-56410512-DEL-114 |
| chr8-56424806-DEL-1661 | chr8-56424806-DEL-1662 |
| chr8-56433407-DEL-101 | chr8-56433408-DEL-101 |
| chr8-56446601-DEL-746 | chr8-56446602-DEL-746 |
| chr8-56499945-DEL-116 | chr8-56499946-DEL-116 |
| chr8-56501275-DEL-544 | chr8-56501276-DEL-544 |
| chr8-56519031-DEL-51 | chr8-56519032-DEL-51 |
| chr8-56587397-DEL-6232 | chr8-56587383-DEL-6232 |
| chr8-56627898-DEL-7095 | chr8-56627899-DEL-7095 |
| chr8-56643022-DEL-1616 | chr8-56643022-DEL-1621 |
| chr8-56787872-DEL-99 | chr8-56787873-DEL-87 |
| chr8-56803467-DEL-6392 | chr8-56803468-DEL-6392 |
| chr8-56824429-DEL-6212 | chr8-56824430-DEL-6212 |
| chr8-56868410-DEL-134 | chr8-56868410-DEL-132 |

| | |
|---|---|
| chr8-56876678-DEL-60 | chr8-56876679-DEL-60 |
| chr8-56882527-DEL-59 | chr8-56882528-DEL-59 |
| chr8-56885999-DEL-773 | chr8-56886000-DEL-775 |
| chr8-56892913-DEL-52 | chr8-56892913-DEL-52 |
| chr8-56896660-DEL-58 | chr8-56896661-DEL-58 |
| chr8-56916526-DEL-59 | chr8-56916528-DEL-58 |
| chr8-56974246-DEL-56 | chr8-56974219-DEL-50 |
| chr8-56975115-DEL-65 | chr8-56975143-DEL-54 |
| chr8-57015987-DEL-72 | chr8-57015979-DEL-52 |
| chr8-57081248-DEL-182 | chr8-57081249-DEL-182 |
| chr8-57094411-DEL-57 | chr8-57094384-DEL-85 |
| chr8-57105454-DEL-2214 | chr8-57105455-DEL-2214 |
| chr8-57139434-DEL-164 | chr8-57139435-DEL-164 |
| chr8-57140103-DEL-84 | chr8-57140084-DEL-104 |
| chr8-57144932-DEL-66 | chr8-57144933-DEL-66 |
| chr8-57145786-DEL-219 | chr8-57145787-DEL-219 |
| chr8-57173877-DEL-567 | chr8-57173872-DEL-561 |
| chr8-57196802-DEL-6470 | chr8-57196797-DEL-6470 |
| chr8-57203686-DEL-127 | chr8-57203687-DEL-127 |
| chr8-57206883-DEL-558 | chr8-57206884-DEL-558 |
| chr8-57215282-DEL-74 | chr8-57215282-DEL-72 |
| chr8-57217219-DEL-60 | chr8-57217220-DEL-60 |
| chr8-57220744-DEL-73 | chr8-57220777-DEL-74 |
| chr8-57271031-DEL-1024 | chr8-57271032-DEL-1024 |
| chr8-57274281-DEL-6846 | chr8-57274282-DEL-6846 |
| chr8-57319794-DEL-5171 | chr8-57319795-DEL-5171 |
| chr8-57333708-DEL-439 | chr8-57333709-DEL-439 |
| chr8-57337654-DEL-177 | chr8-57337655-DEL-177 |
| chr8-57339403-DEL-74 | chr8-57339404-DEL-74 |
| chr8-57381608-DEL-56 | chr8-57381609-DEL-56 |
| chr8-57396138-DEL-428 | chr8-57396139-DEL-428 |
| chr8-57487301-DEL-6653 | chr8-57487302-DEL-6653 |
| chr8-57493433-DEL-424 | chr8-57493434-DEL-424 |
| chr8-57497445-DEL-114 | chr8-57497446-DEL-115 |
| chr8-57508141-DEL-75 | chr8-57508147-DEL-70 |
| chr8-57519535-DEL-5063 | chr8-57519536-DEL-5063 |
| chr8-57545776-DEL-72 | chr8-57545777-DEL-72 |
| chr8-57551412-DEL-6671 | chr8-57551412-DEL-6671 |
| chr8-57589185-DEL-309 | chr8-57589186-DEL-309 |
| chr8-57668573-DEL-6217 | chr8-57668574-DEL-6217 |
| chr8-57675683-DEL-1437 | chr8-57675684-DEL-1437 |
| chr8-57683446-DEL-91 | chr8-57683429-DEL-85 |
| chr8-57696433-DEL-75 | chr8-57696432-DEL-69 |
| chr8-57716469-DEL-1225 | chr8-57716470-DEL-1225 |
| chr8-57737268-DEL-84 | chr8-57737277-DEL-68 |
| chr8-57739624-DEL-1126 | chr8-57739625-DEL-1126 |
| chr8-57743002-DEL-206 | chr8-57743003-DEL-206 |
| chr8-57758101-DEL-136 | chr8-57758102-DEL-109 |
| chr8-57824498-DEL-1750 | chr8-57824499-DEL-1750 |
| chr8-57836609-DEL-233 | chr8-57836610-DEL-233 |
| chr8-57841060-DEL-82 | chr8-57841061-DEL-82 |
| chr8-57843870-DEL-6382 | chr8-57843871-DEL-6382 |
| chr8-57883402-DEL-72 | chr8-57883406-DEL-74 |
| chr8-57898984-DEL-60 | chr8-57898987-DEL-58 |
| chr8-57903522-DEL-210 | chr8-57903523-DEL-210 |
| chr8-57943927-DEL-198 | chr8-57943913-DEL-201 |
| chr8-58003244-DEL-1295 | chr8-58003256-DEL-1284 |
| chr8-58014846-DEL-199 | chr8-58014847-DEL-199 |
| chr8-58024682-DEL-318 | chr8-58024683-DEL-318 |
| chr8-58025404-DEL-110 | chr8-58025405-DEL-110 |
| chr8-58034282-DEL-67 | chr8-58034266-DEL-68 |
| chr8-58056262-DEL-179 | chr8-58056263-DEL-179 |
| chr8-58065749-DEL-185 | chr8-58065750-DEL-185 |
| chr8-58092087-DEL-135 | chr8-58092088-DEL-135 |
| chr8-58099364-DEL-57 | chr8-58099365-DEL-57 |
| chr8-58122694-DEL-51 | chr8-58122695-DEL-51 |
| chr8-58123182-DEL-6483 | chr8-58123183-DEL-6483 |
| chr8-58151338-DEL-425 | chr8-58151339-DEL-425 |
| chr8-58155397-DEL-154 | chr8-58155398-DEL-154 |
| chr8-58159875-DEL-58 | chr8-58159876-DEL-58 |
| chr8-58171761-DEL-856 | chr8-58171762-DEL-825 |
| chr8-58174244-DEL-1453 | chr8-58174245-DEL-1453 |
| chr8-58177675-DEL-335 | chr8-58177669-DEL-333 |
| chr8-58185129-DEL-181 | chr8-58185130-DEL-181 |
| chr8-58187357-DEL-210 | chr8-58187358-DEL-210 |
| chr8-58203301-DEL-512 | chr8-58203302-DEL-512 |
| chr8-58210836-DEL-154 | chr8-58210837-DEL-154 |
| chr8-58212147-DEL-55 | chr8-58212151-DEL-54 |
| chr8-58242178-DEL-52 | chr8-58242179-DEL-50 |
| chr8-58272467-DEL-53 | chr8-58272465-DEL-51 |

| | |
|---|---|
| chr8-58296416-DEL-62 | chr8-58296418-DEL-58 |
| chr8-58314941-DEL-2139 | chr8-58314942-DEL-2139 |
| chr8-58318296-DEL-74 | chr8-58318297-DEL-74 |
| chr8-58321335-DEL-54 | chr8-58321324-DEL-54 |
| chr8-58323602-DEL-71 | chr8-58323603-DEL-71 |
| chr8-58367158-DEL-558 | chr8-58367168-DEL-555 |
| chr8-58378228-DEL-76 | chr8-58378229-DEL-80 |
| chr8-58392047-DEL-76 | chr8-58392048-DEL-76 |
| chr8-58423302-DEL-94 | chr8-58423303-DEL-94 |
| chr8-58432805-DEL-6397 | chr8-58432806-DEL-6397 |
| chr8-58439933-DEL-139 | chr8-58439944-DEL-128 |
| chr8-58444189-DEL-73 | chr8-58444168-DEL-72 |
| chr8-58454272-DEL-513 | chr8-58454273-DEL-513 |
| chr8-58469720-DEL-66 | chr8-58469727-DEL-60 |
| chr8-58474568-DEL-125 | chr8-58474569-DEL-125 |
| chr8-58497935-DEL-79 | chr8-58497938-DEL-71 |
| chr8-58546082-DEL-56 | chr8-58546083-DEL-55 |
| chr8-58548294-DEL-7445 | chr8-58548295-DEL-7445 |
| chr8-58548673-DEL-1040 | chr8-58548673-DEL-1040 |
| chr8-58567894-DEL-63 | chr8-58567895-DEL-63 |
| chr8-58568910-DEL-6332 | chr8-58568911-DEL-6332 |
| chr8-58589974-DEL-4845 | chr8-58589975-DEL-4845 |
| chr8-58627216-DEL-182 | chr8-58627217-DEL-182 |
| chr8-58643703-DEL-166 | chr8-58643704-DEL-166 |
| chr8-58651583-DEL-118 | chr8-58651584-DEL-118 |
| chr8-58652390-DEL-376 | chr8-58652392-DEL-377 |
| chr8-58655073-DEL-489 | chr8-58655074-DEL-489 |
| chr8-58681636-DEL-2829 | chr8-58681637-DEL-2829 |
| chr8-58698482-DEL-51 | chr8-58698483-DEL-51 |
| chr8-58700707-DEL-167 | chr8-58700708-DEL-167 |
| chr8-58702863-DEL-138 | chr8-58702864-DEL-138 |
| chr8-58721215-DEL-114 | chr8-58721212-DEL-78 |
| chr8-58745101-DEL-6798 | chr8-58745102-DEL-6798 |
| chr8-58780420-DEL-69 | chr8-58780399-DEL-70 |
| chr8-58789276-DEL-512 | chr8-58789277-DEL-512 |
| chr8-58802545-DEL-416 | chr8-58802546-DEL-416 |
| chr8-58808667-DEL-111 | chr8-58808668-DEL-111 |
| chr8-58810127-DEL-70 | chr8-58810140-DEL-58 |
| chr8-58825088-DEL-52 | chr8-58825091-DEL-50 |
| chr8-58841354-DEL-7312 | chr8-58841355-DEL-7312 |
| chr8-58841412-DEL-424 | chr8-58841413-DEL-424 |
| chr8-58854453-DEL-59 | chr8-58854456-DEL-52 |
| chr8-58868967-DEL-587 | chr8-58868968-DEL-587 |
| chr8-58892788-DEL-202 | chr8-58892789-DEL-202 |
| chr8-58900699-DEL-361 | chr8-58900700-DEL-361 |
| chr8-58915823-DEL-8820 | chr8-58915824-DEL-8820 |
| chr8-58928587-DEL-74 | chr8-58928588-DEL-74 |
| chr8-58937114-DEL-69 | chr8-58937115-DEL-69 |
| chr8-58938044-DEL-1879 | chr8-58938044-DEL-1879 |
| chr8-58942227-DEL-1834 | chr8-58942230-DEL-1831 |
| chr8-58944428-DEL-878 | chr8-58944428-DEL-878 |
| chr8-58946219-DEL-80 | chr8-58946218-DEL-58 |
| chr8-58947089-DEL-102 | chr8-58947088-DEL-102 |
| chr8-58954913-DEL-57 | chr8-58954914-DEL-57 |
| chr8-58970875-DEL-56 | chr8-58970875-DEL-88 |
| chr8-58984269-DEL-666 | chr8-58984270-DEL-666 |
| chr8-59036879-DEL-97 | chr8-59036880-DEL-97 |
| chr8-59044650-DEL-104 | chr8-59044650-DEL-102 |
| chr8-59064896-DEL-282 | chr8-59064901-DEL-270 |
| chr8-59065575-DEL-56 | chr8-59065576-DEL-56 |
| chr8-59068471-DEL-459 | chr8-59068472-DEL-460 |
| chr8-59077441-DEL-3469 | chr8-59077441-DEL-3469 |
| chr8-59082109-DEL-664 | chr8-59082110-DEL-664 |
| chr8-59087382-DEL-51 | chr8-59087383-DEL-51 |
| chr8-59094191-DEL-54 | chr8-59094192-DEL-54 |
| chr8-59101449-DEL-229 | chr8-59101455-DEL-228 |
| chr8-59109395-DEL-54 | chr8-59109395-DEL-54 |
| chr8-59124965-DEL-254 | chr8-59124966-DEL-254 |
| chr8-59179395-DEL-5948 | chr8-59179396-DEL-5948 |
| chr8-59186781-DEL-78 | chr8-59186782-DEL-78 |
| chr8-59194479-DEL-54 | chr8-59194480-DEL-55 |
| chr8-59218378-DEL-113 | chr8-59218379-DEL-113 |
| chr8-59224583-DEL-94 | chr8-59224585-DEL-91 |
| chr8-59255573-DEL-1892 | chr8-59255574-DEL-1892 |
| chr8-59261168-DEL-66 | chr8-59261169-DEL-66 |
| chr8-59263609-DEL-408 | chr8-59263610-DEL-408 |
| chr8-59264403-DEL-53 | chr8-59264404-DEL-53 |
| chr8-59278761-DEL-561 | chr8-59278762-DEL-561 |
| chr8-59312394-DEL-634 | chr8-59312395-DEL-634 |
| chr8-59373844-DEL-71 | chr8-59373845-DEL-71 |

| | |
|---|---|
| chr8-59374941-DEL-84 | chr8-59374935-DEL-92 |
| chr8-59375987-DEL-111 | chr8-59375986-DEL-80 |
| chr8-59383562-DEL-94 | chr8-59383563-DEL-94 |
| chr8-59426001-DEL-6214 | chr8-59426002-DEL-6214 |
| chr8-59435663-DEL-60 | chr8-59435660-DEL-85 |
| chr8-59448077-DEL-56 | chr8-59448078-DEL-56 |
| chr8-59467754-DEL-102 | chr8-59467779-DEL-60 |
| chr8-59470664-DEL-232 | chr8-59470664-DEL-232 |
| chr8-59499938-DEL-6307 | chr8-59499937-DEL-6309 |
| chr8-59557744-DEL-670 | chr8-59557753-DEL-662 |
| chr8-59574874-DEL-56 | chr8-59574873-DEL-72 |
| chr8-59576592-DEL-513 | chr8-59576592-DEL-513 |
| chr8-59584832-DEL-7090 | chr8-59584833-DEL-7090 |
| chr8-59592709-DEL-838 | chr8-59592710-DEL-838 |
| chr8-59643523-DEL-111 | chr8-59643524-DEL-111 |
| chr8-59701202-DEL-441 | chr8-59701206-DEL-439 |
| chr8-59709227-DEL-5347 | chr8-59709227-DEL-5347 |
| chr8-59723414-DEL-399 | chr8-59723406-DEL-400 |
| chr8-59742912-DEL-1158 | chr8-59742913-DEL-1158 |
| chr8-59752353-DEL-51 | chr8-59752354-DEL-51 |
| chr8-59787456-DEL-1656 | chr8-59787463-DEL-1656 |
| chr8-59790081-DEL-6424 | chr8-59790082-DEL-6424 |
| chr8-59818422-DEL-205 | chr8-59818413-DEL-196 |
| chr8-59841144-DEL-98 | chr8-59841145-DEL-98 |
| chr8-59870031-DEL-283 | chr8-59870032-DEL-283 |
| chr8-59871794-DEL-1100 | chr8-59871795-DEL-1100 |
| chr8-59878620-DEL-52 | chr8-59878611-DEL-66 |
| chr8-59880767-DEL-50 | chr8-59880782-DEL-67 |
| chr8-59908029-DEL-58 | chr8-59908030-DEL-58 |
| chr8-59949262-DEL-1548 | chr8-59949263-DEL-1548 |
| chr8-60003851-DEL-1434 | chr8-60003852-DEL-1434 |
| chr8-60028838-DEL-112 | chr8-60028839-DEL-112 |
| chr8-60039012-DEL-2688 | chr8-60039013-DEL-2688 |
| chr8-60044251-DEL-64 | chr8-60044271-DEL-66 |
| chr8-60048940-DEL-76 | chr8-60048941-DEL-76 |
| chr8-60068770-DEL-6467 | chr8-60068771-DEL-6467 |
| chr8-60074916-DEL-200 | chr8-60074917-DEL-200 |
| chr8-60080624-DEL-1418 | chr8-60080625-DEL-1418 |
| chr8-60088012-DEL-58 | chr8-60088013-DEL-58 |
| chr8-60102706-DEL-6879 | chr8-60102707-DEL-6879 |
| chr8-60103117-DEL-212 | chr8-60103118-DEL-212 |
| chr8-60116252-DEL-142 | chr8-60116254-DEL-141 |
| chr8-60118800-DEL-64 | chr8-60118805-DEL-60 |
| chr8-60130553-DEL-7364 | chr8-60130554-DEL-7364 |
| chr8-60159645-DEL-1524 | chr8-60159631-DEL-1538 |
| chr8-60188718-DEL-6025 | chr8-60188718-DEL-6025 |
| chr8-60205973-DEL-2154 | chr8-60205974-DEL-2154 |
| chr8-60209998-DEL-51 | chr8-60209999-DEL-51 |
| chr8-60212752-DEL-267 | chr8-60212753-DEL-267 |
| chr8-60218139-DEL-126 | chr8-60218140-DEL-126 |
| chr8-60228721-DEL-800 | chr8-60228722-DEL-800 |
| chr8-60286823-DEL-6403 | chr8-60286824-DEL-6403 |
| chr8-60303378-DEL-159 | chr8-60303379-DEL-159 |
| chr8-60306688-DEL-51 | chr8-60306677-DEL-63 |
| chr8-60308156-DEL-1214 | chr8-60308157-DEL-1214 |
| chr8-60329249-DEL-450 | chr8-60329250-DEL-450 |
| chr8-60332453-DEL-4071 | chr8-60332454-DEL-4073 |
| chr8-60386057-DEL-3728 | chr8-60386058-DEL-3728 |
| chr8-60413901-DEL-55 | chr8-60413901-DEL-55 |
| chr8-60414290-DEL-468 | chr8-60414291-DEL-468 |
| chr8-60431442-DEL-253 | chr8-60431443-DEL-253 |
| chr8-60438717-DEL-2590 | chr8-60438718-DEL-2590 |
| chr8-60440088-DEL-698 | chr8-60440087-DEL-699 |
| chr8-60446147-DEL-108 | chr8-60446148-DEL-108 |
| chr8-60456789-DEL-201 | chr8-60456790-DEL-201 |
| chr8-60466174-DEL-730 | chr8-60466175-DEL-730 |
| chr8-60497135-DEL-62 | chr8-60497136-DEL-62 |
| chr8-60498627-DEL-92 | chr8-60498614-DEL-86 |
| chr8-60501856-DEL-58 | chr8-60501842-DEL-58 |
| chr8-60509328-DEL-6424 | chr8-60509329-DEL-6424 |
| chr8-60528138-DEL-3394 | chr8-60528125-DEL-3395 |
| chr8-60533478-DEL-207 | chr8-60533479-DEL-207 |
| chr8-60555774-DEL-5589 | chr8-60555775-DEL-5589 |
| chr8-60559081-DEL-543 | chr8-60559082-DEL-543 |
| chr8-60571167-DEL-69 | chr8-60571150-DEL-72 |
| chr8-60580965-DEL-1367 | chr8-60580966-DEL-1367 |
| chr8-60641324-DEL-68 | chr8-60641329-DEL-64 |
| chr8-60656666-DEL-1796 | chr8-60656667-DEL-1796 |
| chr8-60686343-DEL-119 | chr8-60686344-DEL-119 |
| chr8-60695354-DEL-68 | chr8-60695353-DEL-70 |

| | |
|---|---|
| chr8-60711844-DEL-979 | chr8-60711845-DEL-979 |
| chr8-60712828-DEL-63 | chr8-60712828-DEL-63 |
| chr8-60713634-DEL-1457 | chr8-60713635-DEL-1457 |
| chr8-60719420-DEL-233 | chr8-60719421-DEL-236 |
| chr8-60723542-DEL-85 | chr8-60723543-DEL-85 |
| chr8-60725091-DEL-1504 | chr8-60725093-DEL-1503 |
| chr8-60734388-DEL-7020 | chr8-60734389-DEL-7020 |
| chr8-60771960-DEL-7015 | chr8-60771961-DEL-7015 |
| chr8-60793571-DEL-498 | chr8-60793572-DEL-498 |
| chr8-60802183-DEL-1732 | chr8-60802184-DEL-1732 |
| chr8-60806157-DEL-86 | chr8-60806155-DEL-86 |
| chr8-60821415-DEL-874 | chr8-60821416-DEL-874 |
| chr8-60850986-DEL-204 | chr8-60850987-DEL-204 |
| chr8-60860346-DEL-470 | chr8-60860347-DEL-470 |
| chr8-60870683-DEL-4174 | chr8-60870684-DEL-4174 |
| chr8-60872507-DEL-724 | chr8-60872507-DEL-724 |
| chr8-60884305-DEL-68 | chr8-60884306-DEL-68 |
| chr8-60887889-DEL-310 | chr8-60887899-DEL-309 |
| chr8-60903240-DEL-1938 | chr8-60903241-DEL-1938 |
| chr8-60923074-DEL-105 | chr8-60923079-DEL-101 |
| chr8-60923464-DEL-200 | chr8-60923454-DEL-199 |
| chr8-60925889-DEL-6405 | chr8-60925890-DEL-6405 |
| chr8-60932482-DEL-512 | chr8-60932483-DEL-512 |
| chr8-60938151-DEL-548 | chr8-60938152-DEL-548 |
| chr8-60970959-DEL-224 | chr8-60970960-DEL-224 |
| chr8-60996623-DEL-139 | chr8-60996624-DEL-139 |
| chr8-61012314-DEL-138 | chr8-61012315-DEL-138 |
| chr8-61013345-DEL-402 | chr8-61013345-DEL-400 |
| chr8-61050648-DEL-8572 | chr8-61050649-DEL-8572 |
| chr8-61069109-DEL-77 | chr8-61069101-DEL-78 |
| chr8-61094008-DEL-123 | chr8-61094005-DEL-120 |
| chr8-61099805-DEL-186 | chr8-61099806-DEL-186 |
| chr8-61104682-DEL-323 | chr8-61104683-DEL-323 |
| chr8-61137681-DEL-216 | chr8-61137682-DEL-216 |
| chr8-61138466-DEL-7097 | chr8-61138467-DEL-7097 |
| chr8-61149480-DEL-80 | chr8-61149437-DEL-96 |
| chr8-61160260-DEL-403 | chr8-61160249-DEL-401 |
| chr8-61163437-DEL-60 | chr8-61163438-DEL-60 |
| chr8-61181092-DEL-162 | chr8-61181085-DEL-163 |
| chr8-61191121-DEL-6934 | chr8-61191122-DEL-6934 |
| chr8-61201021-DEL-1004 | chr8-61201022-DEL-1004 |
| chr8-61213586-DEL-127 | chr8-61213587-DEL-127 |
| chr8-61215545-DEL-1041 | chr8-61215546-DEL-1040 |
| chr8-61227642-DEL-322 | chr8-61227643-DEL-322 |
| chr8-61229275-DEL-67 | chr8-61229271-DEL-71 |
| chr8-61232733-DEL-2211 | chr8-61232734-DEL-2211 |
| chr8-61242173-DEL-2893 | chr8-61242174-DEL-2893 |
| chr8-61250554-DEL-136 | chr8-61250553-DEL-137 |
| chr8-61259040-DEL-180 | chr8-61259041-DEL-180 |
| chr8-61269206-DEL-292 | chr8-61269207-DEL-292 |
| chr8-61270442-DEL-7125 | chr8-61270443-DEL-7125 |
| chr8-61281112-DEL-379 | chr8-61281113-DEL-379 |
| chr8-61282454-DEL-58 | chr8-61282455-DEL-58 |
| chr8-61303806-DEL-6568 | chr8-61303807-DEL-6568 |
| chr8-61304049-DEL-214 | chr8-61304050-DEL-214 |
| chr8-61307219-DEL-115 | chr8-61307220-DEL-115 |
| chr8-61331442-DEL-1361 | chr8-61331457-DEL-1347 |
| chr8-61333681-DEL-186 | chr8-61333682-DEL-186 |
| chr8-61354830-DEL-354 | chr8-61354831-DEL-354 |
| chr8-61389071-DEL-170 | chr8-61389072-DEL-170 |
| chr8-61393350-DEL-100 | chr8-61393350-DEL-101 |
| chr8-61412523-DEL-124 | chr8-61412524-DEL-124 |
| chr8-61443680-DEL-701 | chr8-61443681-DEL-701 |
| chr8-61451998-DEL-7099 | chr8-61451999-DEL-7099 |
| chr8-61548225-DEL-166 | chr8-61548226-DEL-166 |
| chr8-61553137-DEL-80 | chr8-61553138-DEL-79 |
| chr8-61566355-DEL-206 | chr8-61566356-DEL-206 |
| chr8-61604031-DEL-81 | chr8-61604026-DEL-81 |
| chr8-61606793-DEL-174 | chr8-61606780-DEL-166 |
| chr8-61640916-DEL-126 | chr8-61640917-DEL-121 |
| chr8-61641675-DEL-52 | chr8-61641676-DEL-52 |
| chr8-61652941-DEL-223 | chr8-61652942-DEL-223 |
| chr8-61653186-DEL-3899 | chr8-61653186-DEL-3899 |
| chr8-61672250-DEL-54 | chr8-61672251-DEL-54 |
| chr8-61673324-DEL-84 | chr8-61673325-DEL-84 |
| chr8-61732048-DEL-591 | chr8-61732049-DEL-591 |
| chr8-61786413-DEL-96 | chr8-61786414-DEL-96 |
| chr8-61791736-DEL-979 | chr8-61791737-DEL-980 |
| chr8-61799638-DEL-302 | chr8-61799629-DEL-306 |
| chr8-61820049-DEL-4511 | chr8-61820050-DEL-4511 |

| | |
|---|---|
| chr8-61833477-DEL-979 | chr8-61833478-DEL-979 |
| chr8-61862659-DEL-190 | chr8-61862669-DEL-181 |
| chr8-61871289-DEL-128 | chr8-61871290-DEL-127 |
| chr8-61873172-DEL-59 | chr8-61873162-DEL-60 |
| chr8-61902958-DEL-56 | chr8-61902965-DEL-50 |
| chr8-61906049-DEL-60 | chr8-61906049-DEL-65 |
| chr8-61908072-DEL-90 | chr8-61908047-DEL-117 |
| chr8-61931438-DEL-72 | chr8-61931439-DEL-72 |
| chr8-61948786-DEL-193 | chr8-61948786-DEL-193 |
| chr8-61963459-DEL-80 | chr8-61963460-DEL-80 |
| chr8-62022448-DEL-137 | chr8-62022449-DEL-137 |
| chr8-62030075-DEL-68 | chr8-62030075-DEL-68 |
| chr8-62040304-DEL-236 | chr8-62040305-DEL-236 |
| chr8-62043553-DEL-52 | chr8-62043536-DEL-70 |
| chr8-62127373-DEL-168 | chr8-62127374-DEL-168 |
| chr8-62181202-DEL-83 | chr8-62181203-DEL-83 |
| chr8-62191523-DEL-3997 | chr8-62191524-DEL-3997 |
| chr8-62288364-DEL-60 | chr8-62288365-DEL-58 |
| chr8-62301264-DEL-277 | chr8-62301265-DEL-277 |
| chr8-62314586-DEL-64 | chr8-62314587-DEL-64 |
| chr8-62327998-DEL-69 | chr8-62327999-DEL-69 |
| chr8-62328718-DEL-199 | chr8-62328719-DEL-199 |
| chr8-62337147-DEL-284 | chr8-62337148-DEL-285 |
| chr8-62344433-DEL-77 | chr8-62344433-DEL-77 |
| chr8-62346019-DEL-343 | chr8-62346019-DEL-343 |
| chr8-62370643-DEL-397 | chr8-62370644-DEL-397 |
| chr8-62386218-DEL-57 | chr8-62386219-DEL-57 |
| chr8-62396828-DEL-1048 | chr8-62396829-DEL-1048 |
| chr8-62398004-DEL-299 | chr8-62398004-DEL-300 |
| chr8-62405087-DEL-751 | chr8-62405085-DEL-752 |
| chr8-62483068-DEL-937 | chr8-62483069-DEL-937 |
| chr8-62555322-DEL-6382 | chr8-62555323-DEL-6382 |
| chr8-62609381-DEL-6553 | chr8-62609382-DEL-6553 |
| chr8-62697246-DEL-6439 | chr8-62697247-DEL-6439 |
| chr8-62783927-DEL-81 | chr8-62783928-DEL-81 |
| chr8-62803463-DEL-204 | chr8-62803464-DEL-204 |
| chr8-62851792-DEL-60 | chr8-62851793-DEL-60 |
| chr8-62853239-DEL-137 | chr8-62853240-DEL-137 |
| chr8-62869461-DEL-402 | chr8-62869462-DEL-402 |
| chr8-62873592-DEL-186 | chr8-62873593-DEL-186 |
| chr8-62895414-DEL-4710 | chr8-62895415-DEL-4710 |
| chr8-62900754-DEL-428 | chr8-62900754-DEL-426 |
| chr8-62904636-DEL-200 | chr8-62904637-DEL-200 |
| chr8-62949125-DEL-1165 | chr8-62949126-DEL-1165 |
| chr8-62952545-DEL-174 | chr8-62952546-DEL-174 |
| chr8-62972118-DEL-103 | chr8-62972119-DEL-103 |
| chr8-62995549-DEL-2932 | chr8-62995543-DEL-2899 |
| chr8-63015320-DEL-4419 | chr8-63015320-DEL-4419 |
| chr8-63049770-DEL-600 | chr8-63049771-DEL-600 |
| chr8-63050553-DEL-211 | chr8-63050554-DEL-211 |
| chr8-63068293-DEL-407 | chr8-63068294-DEL-407 |
| chr8-63071825-DEL-60 | chr8-63071824-DEL-107 |
| chr8-63097579-DEL-209 | chr8-63097586-DEL-203 |
| chr8-63101957-DEL-6591 | chr8-63101958-DEL-6591 |
| chr8-63112149-DEL-69 | chr8-63112139-DEL-72 |
| chr8-63113443-DEL-304 | chr8-63113444-DEL-304 |
| chr8-63138851-DEL-183 | chr8-63138843-DEL-183 |
| chr8-63204905-DEL-513 | chr8-63204906-DEL-513 |
| chr8-63212156-DEL-1246 | chr8-63212157-DEL-1246 |
| chr8-63217259-DEL-101 | chr8-63217265-DEL-96 |
| chr8-63238490-DEL-516 | chr8-63238488-DEL-514 |
| chr8-63258247-DEL-7105 | chr8-63258248-DEL-7105 |
| chr8-63282879-DEL-404 | chr8-63282880-DEL-404 |
| chr8-63317008-DEL-4358 | chr8-63317009-DEL-4358 |
| chr8-63368607-DEL-3982 | chr8-63368608-DEL-3982 |
| chr8-63393367-DEL-2211 | chr8-63393368-DEL-2211 |
| chr8-63408134-DEL-52 | chr8-63408135-DEL-52 |
| chr8-63509679-DEL-628 | chr8-63509680-DEL-628 |
| chr8-63517256-DEL-829 | chr8-63517257-DEL-829 |
| chr8-63521816-DEL-7076 | chr8-63521817-DEL-7076 |
| chr8-63528250-DEL-212 | chr8-63528251-DEL-212 |
| chr8-63569297-DEL-56 | chr8-63569298-DEL-56 |
| chr8-63579922-DEL-1833 | chr8-63579923-DEL-1833 |
| chr8-63597858-DEL-110 | chr8-63597884-DEL-98 |
| chr8-63607372-DEL-396 | chr8-63607373-DEL-396 |
| chr8-63642847-DEL-556 | chr8-63642848-DEL-556 |
| chr8-63657874-DEL-824 | chr8-63657875-DEL-824 |
| chr8-63673265-DEL-74 | chr8-63673266-DEL-68 |
| chr8-63676771-DEL-214 | chr8-63676772-DEL-214 |
| chr8-63678295-DEL-1921 | chr8-63678296-DEL-1921 |

| | |
|---|---|
| chr8-63682869-DEL-515 | chr8-63682871-DEL-514 |
| chr8-63686648-DEL-54 | chr8-63686646-DEL-60 |
| chr8-63693857-DEL-1281 | chr8-63693858-DEL-1281 |
| chr8-63696276-DEL-66 | chr8-63696277-DEL-66 |
| chr8-63744671-DEL-997 | chr8-63744672-DEL-997 |
| chr8-63747621-DEL-119 | chr8-63747622-DEL-119 |
| chr8-63760819-DEL-1253 | chr8-63760819-DEL-1253 |
| chr8-63762075-DEL-8809 | chr8-63762075-DEL-8809 |
| chr8-63785270-DEL-65 | chr8-63785271-DEL-65 |
| chr8-63786370-DEL-1537 | chr8-63786371-DEL-1537 |
| chr8-63796142-DEL-539 | chr8-63796146-DEL-535 |
| chr8-63799783-DEL-757 | chr8-63799784-DEL-757 |
| chr8-63812493-DEL-485 | chr8-63812494-DEL-485 |
| chr8-63816327-DEL-106 | chr8-63816328-DEL-106 |
| chr8-63825150-DEL-3765 | chr8-63825150-DEL-3765 |
| chr8-63837064-DEL-231 | chr8-63837065-DEL-235 |
| chr8-63838680-DEL-208 | chr8-63838679-DEL-209 |
| chr8-63857146-DEL-758 | chr8-63857142-DEL-759 |
| chr8-63917236-DEL-399 | chr8-63917207-DEL-432 |
| chr8-63920669-DEL-56 | chr8-63920669-DEL-56 |
| chr8-63984617-DEL-91 | chr8-63984610-DEL-92 |
| chr8-63985694-DEL-119 | chr8-63985691-DEL-123 |
| chr8-64006956-DEL-395 | chr8-64006957-DEL-395 |
| chr8-64013759-DEL-253 | chr8-64013760-DEL-253 |
| chr8-64018144-DEL-56 | chr8-64018141-DEL-60 |
| chr8-64072452-DEL-3416 | chr8-64072453-DEL-3416 |
| chr8-64083888-DEL-3673 | chr8-64083894-DEL-3667 |
| chr8-64093601-DEL-106 | chr8-64093602-DEL-106 |
| chr8-64098873-DEL-843 | chr8-64098871-DEL-847 |
| chr8-64114057-DEL-1107 | chr8-64114059-DEL-1104 |
| chr8-64118818-DEL-84 | chr8-64118851-DEL-52 |
| chr8-64119624-DEL-325 | chr8-64119625-DEL-325 |
| chr8-64126361-DEL-645 | chr8-64126361-DEL-645 |
| chr8-64127064-DEL-4701 | chr8-64127065-DEL-4701 |
| chr8-64136252-DEL-6191 | chr8-64136253-DEL-6191 |
| chr8-64163656-DEL-420 | chr8-64163657-DEL-420 |
| chr8-64182504-DEL-847 | chr8-64182505-DEL-847 |
| chr8-64192478-DEL-7108 | chr8-64192478-DEL-7108 |
| chr8-64231745-DEL-4982 | chr8-64231746-DEL-4982 |
| chr8-64232387-DEL-76 | chr8-64232382-DEL-58 |
| chr8-64258102-DEL-2105 | chr8-64258103-DEL-2105 |
| chr8-64285125-DEL-182 | chr8-64285126-DEL-180 |
| chr8-64315931-DEL-618 | chr8-64315920-DEL-619 |
| chr8-64329189-DEL-250 | chr8-64329190-DEL-250 |
| chr8-64343141-DEL-100 | chr8-64343141-DEL-96 |
| chr8-64356265-DEL-1273 | chr8-64356265-DEL-1273 |
| chr8-64359665-DEL-101 | chr8-64359666-DEL-99 |
| chr8-64362083-DEL-119 | chr8-64362084-DEL-119 |
| chr8-64362852-DEL-902 | chr8-64362845-DEL-901 |
| chr8-64394945-DEL-130 | chr8-64394946-DEL-130 |
| chr8-64399134-DEL-164 | chr8-64399135-DEL-163 |
| chr8-64399803-DEL-470 | chr8-64399804-DEL-470 |
| chr8-64402735-DEL-1405 | chr8-64402736-DEL-1405 |
| chr8-64403344-DEL-271 | chr8-64403345-DEL-271 |
| chr8-64436348-DEL-100 | chr8-64436349-DEL-100 |
| chr8-64484842-DEL-252 | chr8-64484842-DEL-252 |
| chr8-64485240-DEL-1127 | chr8-64485241-DEL-1127 |
| chr8-64492696-DEL-72 | chr8-64492705-DEL-60 |
| chr8-64511525-DEL-60 | chr8-64511532-DEL-54 |
| chr8-64533932-DEL-94 | chr8-64533933-DEL-94 |
| chr8-64546425-DEL-74 | chr8-64546426-DEL-74 |
| chr8-64632471-DEL-57 | chr8-64632466-DEL-69 |
| chr8-64668985-DEL-59 | chr8-64668986-DEL-57 |
| chr8-64687697-DEL-82 | chr8-64687698-DEL-82 |
| chr8-64688964-DEL-52 | chr8-64688964-DEL-52 |
| chr8-64690538-DEL-357 | chr8-64690539-DEL-357 |
| chr8-64698279-DEL-5667 | chr8-64698279-DEL-5667 |
| chr8-64714107-DEL-2756 | chr8-64714108-DEL-2756 |
| chr8-64720102-DEL-60 | chr8-64720097-DEL-72 |
| chr8-64738762-DEL-52 | chr8-64738763-DEL-52 |
| chr8-64740765-DEL-6258 | chr8-64740765-DEL-6258 |
| chr8-64761701-DEL-2830 | chr8-64761702-DEL-2830 |
| chr8-64769402-DEL-333 | chr8-64769403-DEL-347 |
| chr8-64789151-DEL-65 | chr8-64789152-DEL-65 |
| chr8-64872325-DEL-51 | chr8-64872326-DEL-52 |
| chr8-64883513-DEL-68 | chr8-64883513-DEL-68 |
| chr8-64884565-DEL-231 | chr8-64884577-DEL-222 |
| chr8-64893224-DEL-2124 | chr8-64893225-DEL-2124 |
| chr8-64898855-DEL-7292 | chr8-64898856-DEL-7292 |
| chr8-64911305-DEL-122 | chr8-64911310-DEL-118 |

| | |
|---|---|
| chr8-64916752-DEL-2042 | chr8-64916753-DEL-2042 |
| chr8-64946185-DEL-2140 | chr8-64946186-DEL-2140 |
| chr8-64974310-DEL-80 | chr8-64974331-DEL-76 |
| chr8-64998637-DEL-64 | chr8-64998638-DEL-64 |
| chr8-65009859-DEL-56 | chr8-65009860-DEL-60 |
| chr8-65010224-DEL-6235 | chr8-65010225-DEL-6235 |
| chr8-65025631-DEL-1895 | chr8-65025632-DEL-1895 |
| chr8-65042298-DEL-266 | chr8-65042299-DEL-266 |
| chr8-65057543-DEL-66 | chr8-65057543-DEL-66 |
| chr8-65062342-DEL-59 | chr8-65062343-DEL-59 |
| chr8-65086389-DEL-509 | chr8-65086387-DEL-510 |
| chr8-65137306-DEL-61 | chr8-65137307-DEL-61 |
| chr8-65179803-DEL-207 | chr8-65179804-DEL-207 |
| chr8-65208467-DEL-215 | chr8-65208468-DEL-215 |
| chr8-65209330-DEL-144 | chr8-65209331-DEL-144 |
| chr8-65221630-DEL-72 | chr8-65221626-DEL-70 |
| chr8-65224387-DEL-112 | chr8-65224390-DEL-110 |
| chr8-65231228-DEL-286 | chr8-65231229-DEL-286 |
| chr8-65235486-DEL-3953 | chr8-65235479-DEL-3961 |
| chr8-65249235-DEL-197 | chr8-65249236-DEL-197 |
| chr8-65261252-DEL-59 | chr8-65261246-DEL-77 |
| chr8-65269968-DEL-104 | chr8-65269969-DEL-104 |
| chr8-65297006-DEL-107 | chr8-65297007-DEL-107 |
| chr8-65306394-DEL-235 | chr8-65306395-DEL-235 |
| chr8-65318302-DEL-118 | chr8-65318315-DEL-86 |
| chr8-65343198-DEL-6246 | chr8-65343199-DEL-6246 |
| chr8-65410094-DEL-56 | chr8-65410095-DEL-56 |
| chr8-65414656-DEL-439 | chr8-65414657-DEL-439 |
| chr8-65425057-DEL-6684 | chr8-65425058-DEL-6684 |
| chr8-65441493-DEL-2124 | chr8-65441494-DEL-2124 |
| chr8-65463613-DEL-188 | chr8-65463614-DEL-188 |
| chr8-65472756-DEL-1749 | chr8-65472757-DEL-1748 |
| chr8-65499281-DEL-82 | chr8-65499282-DEL-82 |
| chr8-65502819-DEL-70 | chr8-65502831-DEL-69 |
| chr8-65509063-DEL-52 | chr8-65509075-DEL-53 |
| chr8-65522187-DEL-55 | chr8-65522178-DEL-53 |
| chr8-65549026-DEL-1382 | chr8-65549027-DEL-1379 |
| chr8-65558782-DEL-52 | chr8-65558783-DEL-52 |
| chr8-65562573-DEL-116 | chr8-65562590-DEL-100 |
| chr8-65565029-DEL-1350 | chr8-65565030-DEL-1350 |
| chr8-65567173-DEL-1160 | chr8-65567174-DEL-1160 |
| chr8-65568533-DEL-196 | chr8-65568549-DEL-169 |
| chr8-65585600-DEL-52 | chr8-65585601-DEL-52 |
| chr8-65594108-DEL-186 | chr8-65594109-DEL-186 |
| chr8-65616095-DEL-66 | chr8-65616114-DEL-52 |
| chr8-65617953-DEL-194 | chr8-65617953-DEL-187 |
| chr8-65637508-DEL-6324 | chr8-65637509-DEL-6324 |
| chr8-65661298-DEL-1619 | chr8-65661299-DEL-1619 |
| chr8-65667416-DEL-192 | chr8-65667425-DEL-184 |
| chr8-65674935-DEL-313 | chr8-65674934-DEL-313 |
| chr8-65675988-DEL-315 | chr8-65675989-DEL-315 |
| chr8-65685228-DEL-785 | chr8-65685229-DEL-785 |
| chr8-65687530-DEL-4297 | chr8-65687531-DEL-4297 |
| chr8-65694733-DEL-6242 | chr8-65694734-DEL-6242 |
| chr8-65710303-DEL-210 | chr8-65710304-DEL-210 |
| chr8-65718585-DEL-115 | chr8-65718586-DEL-115 |
| chr8-65727446-DEL-69 | chr8-65727447-DEL-70 |
| chr8-65729896-DEL-68 | chr8-65729901-DEL-72 |
| chr8-65775181-DEL-214 | chr8-65775182-DEL-214 |
| chr8-65812577-DEL-480 | chr8-65812578-DEL-480 |
| chr8-65836476-DEL-109 | chr8-65836477-DEL-109 |
| chr8-65836995-DEL-52 | chr8-65836988-DEL-58 |
| chr8-65856094-DEL-424 | chr8-65856095-DEL-424 |
| chr8-65906554-DEL-9482 | chr8-65906555-DEL-9481 |
| chr8-65906574-DEL-214 | chr8-65906575-DEL-214 |
| chr8-65916569-DEL-62 | chr8-65916570-DEL-62 |
| chr8-65932407-DEL-237 | chr8-65932408-DEL-237 |
| chr8-65952800-DEL-410 | chr8-65952801-DEL-408 |
| chr8-65970964-DEL-89 | chr8-65970958-DEL-81 |
| chr8-65995242-DEL-6421 | chr8-65995243-DEL-6424 |
| chr8-65995355-DEL-199 | chr8-65995356-DEL-199 |
| chr8-66010942-DEL-226 | chr8-66010971-DEL-204 |
| chr8-66021511-DEL-2207 | chr8-66021512-DEL-2207 |
| chr8-66036260-DEL-1319 | chr8-66036261-DEL-1319 |
| chr8-66039003-DEL-175 | chr8-66039004-DEL-175 |
| chr8-66043517-DEL-6606 | chr8-66043518-DEL-6606 |
| chr8-66094380-DEL-67 | chr8-66094381-DEL-67 |
| chr8-66104493-DEL-72 | chr8-66104500-DEL-66 |
| chr8-66107579-DEL-420 | chr8-66107580-DEL-420 |
| chr8-66114144-DEL-184 | chr8-66114145-DEL-183 |

| | |
|---|---|
| chr8-66115811-DEL-1664 | chr8-66115812-DEL-1664 |
| chr8-66154667-DEL-207 | chr8-66154668-DEL-207 |
| chr8-66156742-DEL-7920 | chr8-66156742-DEL-7921 |
| chr8-66176992-DEL-117 | chr8-66176993-DEL-117 |
| chr8-66177630-DEL-1468 | chr8-66177631-DEL-1468 |
| chr8-66209610-DEL-300 | chr8-66209611-DEL-300 |
| chr8-66227012-DEL-206 | chr8-66227013-DEL-206 |
| chr8-66231321-DEL-2681 | chr8-66231322-DEL-2681 |
| chr8-66234275-DEL-54 | chr8-66234276-DEL-54 |
| chr8-66237354-DEL-54 | chr8-66237355-DEL-54 |
| chr8-66260358-DEL-505 | chr8-66260350-DEL-502 |
| chr8-66264181-DEL-60 | chr8-66264170-DEL-76 |
| chr8-66266195-DEL-543 | chr8-66266196-DEL-544 |
| chr8-66283130-DEL-6589 | chr8-66283131-DEL-6589 |
| chr8-66306797-DEL-598 | chr8-66306798-DEL-598 |
| chr8-66364421-DEL-64 | chr8-66364422-DEL-64 |
| chr8-66371950-DEL-1558 | chr8-66371951-DEL-1558 |
| chr8-66377346-DEL-5927 | chr8-66377347-DEL-5927 |
| chr8-66426138-DEL-2326 | chr8-66426139-DEL-2326 |
| chr8-66446827-DEL-288 | chr8-66446828-DEL-288 |
| chr8-66458951-DEL-516 | chr8-66458952-DEL-516 |
| chr8-66471388-DEL-223 | chr8-66471389-DEL-223 |
| chr8-66478209-DEL-76 | chr8-66478210-DEL-76 |
| chr8-66485627-DEL-498 | chr8-66485628-DEL-498 |
| chr8-66501275-DEL-96 | chr8-66501276-DEL-96 |
| chr8-66509596-DEL-127 | chr8-66509596-DEL-127 |
| chr8-66511565-DEL-84 | chr8-66511609-DEL-61 |
| chr8-66512453-DEL-736 | chr8-66512454-DEL-736 |
| chr8-66535437-DEL-73 | chr8-66535485-DEL-72 |
| chr8-66554462-DEL-1165 | chr8-66554463-DEL-1165 |
| chr8-66558280-DEL-3338 | chr8-66558281-DEL-3338 |
| chr8-66589763-DEL-52 | chr8-66589764-DEL-52 |
| chr8-66606480-DEL-73 | chr8-66606483-DEL-73 |
| chr8-66619476-DEL-74 | chr8-66619513-DEL-69 |
| chr8-66640733-DEL-349 | chr8-66640734-DEL-349 |
| chr8-66787861-DEL-2136 | chr8-66787862-DEL-2136 |
| chr8-66805698-DEL-170 | chr8-66805699-DEL-170 |
| chr8-66811287-DEL-363 | chr8-66811288-DEL-363 |
| chr8-66827714-DEL-3426 | chr8-66827715-DEL-3426 |
| chr8-66834799-DEL-477 | chr8-66834800-DEL-477 |
| chr8-66907209-DEL-188 | chr8-66907210-DEL-188 |
| chr8-66918622-DEL-52 | chr8-66918623-DEL-52 |
| chr8-66924666-DEL-92 | chr8-66924684-DEL-52 |
| chr8-66951698-DEL-109 | chr8-66951699-DEL-109 |
| chr8-66956896-DEL-2428 | chr8-66956897-DEL-2428 |
| chr8-66984413-DEL-54 | chr8-66984414-DEL-54 |
| chr8-67011616-DEL-163 | chr8-67011617-DEL-163 |
| chr8-67030659-DEL-1011 | chr8-67030660-DEL-1011 |
| chr8-67049064-DEL-7747 | chr8-67049065-DEL-7747 |
| chr8-67092993-DEL-2025 | chr8-67092994-DEL-2025 |
| chr8-67105023-DEL-6284 | chr8-67105024-DEL-6284 |
| chr8-67119400-DEL-134 | chr8-67119401-DEL-134 |
| chr8-67136301-DEL-80 | chr8-67136294-DEL-84 |
| chr8-67166304-DEL-214 | chr8-67166305-DEL-214 |
| chr8-67181870-DEL-258 | chr8-67181871-DEL-258 |
| chr8-67196003-DEL-400 | chr8-67196004-DEL-400 |
| chr8-67261552-DEL-84 | chr8-67261583-DEL-54 |
| chr8-67270767-DEL-1020 | chr8-67270765-DEL-1021 |
| chr8-67301964-DEL-206 | chr8-67301965-DEL-206 |
| chr8-67303544-DEL-62 | chr8-67303554-DEL-52 |
| chr8-67306824-DEL-102 | chr8-67306825-DEL-102 |
| chr8-67324093-DEL-176 | chr8-67324094-DEL-176 |
| chr8-67356497-DEL-625 | chr8-67356501-DEL-622 |
| chr8-67364766-DEL-1329 | chr8-67364767-DEL-1329 |
| chr8-67393570-DEL-991 | chr8-67393571-DEL-991 |
| chr8-67406123-DEL-6806 | chr8-67406112-DEL-6807 |
| chr8-67406661-DEL-207 | chr8-67406691-DEL-208 |
| chr8-67442101-DEL-56 | chr8-67442106-DEL-50 |
| chr8-67459607-DEL-850 | chr8-67459607-DEL-850 |
| chr8-67487998-DEL-7194 | chr8-67487999-DEL-7194 |
| chr8-67502578-DEL-390 | chr8-67502579-DEL-390 |
| chr8-67547654-DEL-981 | chr8-67547656-DEL-980 |
| chr8-67566854-DEL-441 | chr8-67566855-DEL-441 |
| chr8-67608224-DEL-750 | chr8-67608225-DEL-750 |
| chr8-67628816-DEL-3235 | chr8-67628817-DEL-3235 |
| chr8-67635802-DEL-64 | chr8-67635821-DEL-50 |
| chr8-67638883-DEL-6241 | chr8-67638876-DEL-6242 |
| chr8-67638975-DEL-201 | chr8-67638976-DEL-201 |
| chr8-67673405-DEL-894 | chr8-67673406-DEL-894 |
| chr8-67675619-DEL-96 | chr8-67675626-DEL-90 |

| | |
|---|---|
| chr8-67681130-DEL-61 | chr8-67681083-DEL-62 |
| chr8-67706621-DEL-1816 | chr8-67706622-DEL-1816 |
| chr8-67727845-DEL-665 | chr8-67727846-DEL-665 |
| chr8-67750936-DEL-2620 | chr8-67750937-DEL-2620 |
| chr8-67782957-DEL-2404 | chr8-67782958-DEL-2404 |
| chr8-67816635-DEL-20926 | chr8-67816635-DEL-20926 |
| chr8-67831504-DEL-178 | chr8-67831505-DEL-178 |
| chr8-67836179-DEL-69 | chr8-67836180-DEL-69 |
| chr8-67853240-DEL-195 | chr8-67853241-DEL-195 |
| chr8-67873006-DEL-423 | chr8-67873007-DEL-423 |
| chr8-67878817-DEL-63 | chr8-67878818-DEL-63 |
| chr8-67927253-DEL-288 | chr8-67927254-DEL-288 |
| chr8-67962613-DEL-70 | chr8-67962614-DEL-70 |
| chr8-67966612-DEL-1698 | chr8-67966613-DEL-1698 |
| chr8-67977638-DEL-1209 | chr8-67977639-DEL-1209 |
| chr8-68056310-DEL-3389 | chr8-68056311-DEL-3390 |
| chr8-68060424-DEL-747 | chr8-68060425-DEL-747 |
| chr8-68062451-DEL-138 | chr8-68062476-DEL-114 |
| chr8-68063593-DEL-6391 | chr8-68063594-DEL-6391 |
| chr8-68102836-DEL-1865 | chr8-68102834-DEL-1866 |
| chr8-68106640-DEL-6450 | chr8-68106641-DEL-6450 |
| chr8-68139693-DEL-54 | chr8-68139694-DEL-54 |
| chr8-68216032-DEL-79 | chr8-68216034-DEL-77 |
| chr8-68222847-DEL-377 | chr8-68222848-DEL-377 |
| chr8-68224834-DEL-52 | chr8-68224835-DEL-52 |
| chr8-68259840-DEL-198 | chr8-68259841-DEL-198 |
| chr8-68279342-DEL-55 | chr8-68279346-DEL-56 |
| chr8-68291460-DEL-66 | chr8-68291461-DEL-66 |
| chr8-68371640-DEL-270 | chr8-68371659-DEL-231 |
| chr8-68397636-DEL-207 | chr8-68397637-DEL-207 |
| chr8-68466556-DEL-2001 | chr8-68466557-DEL-2001 |
| chr8-68499513-DEL-51 | chr8-68499482-DEL-51 |
| chr8-68510797-DEL-224 | chr8-68510798-DEL-224 |
| chr8-68523875-DEL-1260 | chr8-68523876-DEL-1260 |
| chr8-68553547-DEL-442 | chr8-68553547-DEL-439 |
| chr8-68561382-DEL-453 | chr8-68561383-DEL-453 |
| chr8-68576644-DEL-401 | chr8-68576645-DEL-401 |
| chr8-68612811-DEL-411 | chr8-68612812-DEL-411 |
| chr8-68617685-DEL-977 | chr8-68617686-DEL-977 |
| chr8-68630840-DEL-550 | chr8-68630841-DEL-550 |
| chr8-68639465-DEL-366 | chr8-68639466-DEL-366 |
| chr8-68668432-DEL-62 | chr8-68668433-DEL-62 |
| chr8-68670668-DEL-938 | chr8-68670669-DEL-938 |
| chr8-68673507-DEL-6586 | chr8-68673508-DEL-6586 |
| chr8-68735669-DEL-67 | chr8-68735670-DEL-67 |
| chr8-68736698-DEL-57 | chr8-68736698-DEL-57 |
| chr8-68754510-DEL-1892 | chr8-68754511-DEL-1892 |
| chr8-68764667-DEL-98 | chr8-68764671-DEL-96 |
| chr8-68803871-DEL-1685 | chr8-68803872-DEL-1685 |
| chr8-68835962-DEL-1490 | chr8-68835962-DEL-1490 |
| chr8-68837871-DEL-1881 | chr8-68837882-DEL-1871 |
| chr8-68857750-DEL-6419 | chr8-68857742-DEL-6419 |
| chr8-68867672-DEL-227 | chr8-68867673-DEL-227 |
| chr8-68868881-DEL-291 | chr8-68868882-DEL-291 |
| chr8-68870537-DEL-52 | chr8-68870538-DEL-52 |
| chr8-68952061-DEL-6575 | chr8-68952062-DEL-6575 |
| chr8-68959175-DEL-202 | chr8-68959176-DEL-202 |
| chr8-68968365-DEL-176 | chr8-68968366-DEL-176 |
| chr8-68984661-DEL-205 | chr8-68984662-DEL-204 |
| chr8-68992719-DEL-67 | chr8-68992720-DEL-67 |
| chr8-68999069-DEL-291 | chr8-68999070-DEL-291 |
| chr8-69012977-DEL-195 | chr8-69012978-DEL-195 |
| chr8-69022576-DEL-6207 | chr8-69022577-DEL-6207 |
| chr8-69062390-DEL-62 | chr8-69062397-DEL-68 |
| chr8-69090591-DEL-225 | chr8-69090592-DEL-225 |
| chr8-69096503-DEL-63 | chr8-69096504-DEL-72 |
| chr8-69125487-DEL-338 | chr8-69125483-DEL-333 |
| chr8-69127753-DEL-605 | chr8-69127762-DEL-597 |
| chr8-69129321-DEL-7078 | chr8-69129318-DEL-7076 |
| chr8-69163675-DEL-1653 | chr8-69163676-DEL-1653 |
| chr8-69168216-DEL-67 | chr8-69168217-DEL-67 |
| chr8-69173918-DEL-96 | chr8-69173944-DEL-64 |
| chr8-69175328-DEL-122 | chr8-69175319-DEL-128 |
| chr8-69191546-DEL-338 | chr8-69191547-DEL-338 |
| chr8-69206974-DEL-854 | chr8-69206975-DEL-854 |
| chr8-69216154-DEL-53 | chr8-69216157-DEL-51 |
| chr8-69220055-DEL-1307 | chr8-69220056-DEL-1306 |
| chr8-69248899-DEL-68 | chr8-69248899-DEL-68 |
| chr8-69258123-DEL-186 | chr8-69258127-DEL-182 |
| chr8-69274069-DEL-196 | chr8-69274070-DEL-196 |

| | |
|---|---|
| chr8-69280807-DEL-202 | chr8-69280808-DEL-202 |
| chr8-69298717-DEL-55 | chr8-69298716-DEL-51 |
| chr8-69304646-DEL-58 | chr8-69304656-DEL-81 |
| chr8-69328094-DEL-328 | chr8-69328090-DEL-333 |
| chr8-69340608-DEL-52 | chr8-69340609-DEL-52 |
| chr8-69352804-DEL-76 | chr8-69352797-DEL-82 |
| chr8-69360545-DEL-56 | chr8-69360542-DEL-57 |
| chr8-69375236-DEL-52 | chr8-69375237-DEL-68 |
| chr8-69397407-DEL-298 | chr8-69397408-DEL-295 |
| chr8-69418552-DEL-4621 | chr8-69418553-DEL-4621 |
| chr8-69444367-DEL-207 | chr8-69444368-DEL-207 |
| chr8-69461872-DEL-202 | chr8-69461873-DEL-202 |
| chr8-69485973-DEL-66 | chr8-69485974-DEL-66 |
| chr8-69502969-DEL-80 | chr8-69502985-DEL-88 |
| chr8-69515478-DEL-96 | chr8-69515479-DEL-114 |
| chr8-69521613-DEL-56 | chr8-69521614-DEL-56 |
| chr8-69548548-DEL-195 | chr8-69548549-DEL-195 |
| chr8-69551676-DEL-77 | chr8-69551681-DEL-77 |
| chr8-69599217-DEL-80 | chr8-69599218-DEL-80 |
| chr8-69619087-DEL-87 | chr8-69619088-DEL-87 |
| chr8-69625282-DEL-57 | chr8-69625280-DEL-55 |
| chr8-69631247-DEL-70 | chr8-69631248-DEL-70 |
| chr8-69641134-DEL-52 | chr8-69641134-DEL-54 |
| chr8-69671498-DEL-199 | chr8-69671499-DEL-199 |
| chr8-69677434-DEL-108 | chr8-69677415-DEL-128 |
| chr8-69692865-DEL-2299 | chr8-69692872-DEL-2286 |
| chr8-69697488-DEL-185 | chr8-69697489-DEL-185 |
| chr8-69698628-DEL-2441 | chr8-69698629-DEL-2441 |
| chr8-69704051-DEL-213 | chr8-69704052-DEL-213 |
| chr8-69718689-DEL-59 | chr8-69718690-DEL-59 |
| chr8-69719761-DEL-1050 | chr8-69719767-DEL-1045 |
| chr8-69731774-DEL-542 | chr8-69731775-DEL-542 |
| chr8-69733125-DEL-365 | chr8-69733126-DEL-365 |
| chr8-69737057-DEL-110 | chr8-69737059-DEL-61 |
| chr8-69742896-DEL-172 | chr8-69742897-DEL-172 |
| chr8-69744816-DEL-826 | chr8-69744817-DEL-826 |
| chr8-69749471-DEL-182 | chr8-69749472-DEL-182 |
| chr8-69770514-DEL-59 | chr8-69770515-DEL-59 |
| chr8-69780231-DEL-189 | chr8-69780237-DEL-189 |
| chr8-69782415-DEL-113 | chr8-69782418-DEL-111 |
| chr8-69786614-DEL-87 | chr8-69786615-DEL-84 |
| chr8-69811986-DEL-2330 | chr8-69811987-DEL-2330 |
| chr8-69834263-DEL-2788 | chr8-69834264-DEL-2788 |
| chr8-69853389-DEL-250 | chr8-69853390-DEL-250 |
| chr8-69854265-DEL-59 | chr8-69854265-DEL-59 |
| chr8-69909392-DEL-58 | chr8-69909393-DEL-58 |
| chr8-69979164-DEL-188 | chr8-69979161-DEL-188 |
| chr8-69982260-DEL-2103 | chr8-69982261-DEL-2103 |
| chr8-70022467-DEL-336 | chr8-70022468-DEL-336 |
| chr8-70075386-DEL-409 | chr8-70075390-DEL-406 |
| chr8-70090241-DEL-54 | chr8-70090242-DEL-54 |
| chr8-70094207-DEL-7980 | chr8-70094206-DEL-7980 |
| chr8-70104952-DEL-1303 | chr8-70104953-DEL-1303 |
| chr8-70106555-DEL-252 | chr8-70106556-DEL-251 |
| chr8-70131219-DEL-1049 | chr8-70131220-DEL-1049 |
| chr8-70137660-DEL-197 | chr8-70137661-DEL-197 |
| chr8-70142523-DEL-66 | chr8-70142536-DEL-58 |
| chr8-70144215-DEL-9034 | chr8-70144216-DEL-9034 |
| chr8-70151241-DEL-1578 | chr8-70151240-DEL-1579 |
| chr8-70180949-DEL-595 | chr8-70180950-DEL-595 |
| chr8-70187430-DEL-203 | chr8-70187431-DEL-203 |
| chr8-70207697-DEL-447 | chr8-70207698-DEL-447 |
| chr8-70219615-DEL-169 | chr8-70219616-DEL-168 |
| chr8-70232070-DEL-212 | chr8-70232085-DEL-196 |
| chr8-70239403-DEL-207 | chr8-70239404-DEL-207 |
| chr8-70248446-DEL-54 | chr8-70248449-DEL-52 |
| chr8-70262188-DEL-55 | chr8-70262161-DEL-56 |
| chr8-70267587-DEL-81 | chr8-70267588-DEL-78 |
| chr8-70283848-DEL-203 | chr8-70283849-DEL-203 |
| chr8-70302862-DEL-60 | chr8-70302863-DEL-60 |
| chr8-70306107-DEL-185 | chr8-70306108-DEL-185 |
| chr8-70313034-DEL-54 | chr8-70313023-DEL-53 |
| chr8-70326560-DEL-115 | chr8-70326561-DEL-113 |
| chr8-70351037-DEL-201 | chr8-70351036-DEL-202 |
| chr8-70389707-DEL-99 | chr8-70389708-DEL-99 |
| chr8-70414940-DEL-3426 | chr8-70414941-DEL-3426 |
| chr8-70434971-DEL-796 | chr8-70434963-DEL-784 |
| chr8-70452840-DEL-51 | chr8-70452791-DEL-51 |
| chr8-70507037-DEL-76 | chr8-70507040-DEL-68 |
| chr8-70554071-DEL-171 | chr8-70554074-DEL-170 |

| | |
|---|---|
| chr8-70570802-DEL-210 | chr8-70570803-DEL-210 |
| chr8-70594004-DEL-220 | chr8-70594005-DEL-220 |
| chr8-70614435-DEL-71 | chr8-70614436-DEL-71 |
| chr8-70617504-DEL-75 | chr8-70617505-DEL-75 |
| chr8-70619129-DEL-61 | chr8-70619106-DEL-60 |
| chr8-70637884-DEL-58 | chr8-70637878-DEL-64 |
| chr8-70657970-DEL-102 | chr8-70657978-DEL-84 |
| chr8-70658102-DEL-649 | chr8-70658135-DEL-617 |
| chr8-70670796-DEL-87 | chr8-70670778-DEL-88 |
| chr8-70679556-DEL-228 | chr8-70679557-DEL-228 |
| chr8-70685012-DEL-1101 | chr8-70685013-DEL-1101 |
| chr8-70692728-DEL-261 | chr8-70692729-DEL-261 |
| chr8-70697199-DEL-113 | chr8-70697200-DEL-113 |
| chr8-70722613-DEL-322 | chr8-70722614-DEL-319 |
| chr8-70730043-DEL-104 | chr8-70730044-DEL-104 |
| chr8-70885867-DEL-52 | chr8-70885868-DEL-52 |
| chr8-70897091-DEL-314 | chr8-70897088-DEL-318 |
| chr8-70901757-DEL-62 | chr8-70901760-DEL-62 |
| chr8-70926707-DEL-99 | chr8-70926708-DEL-99 |
| chr8-70932692-DEL-115 | chr8-70932693-DEL-115 |
| chr8-70976738-DEL-200 | chr8-70976739-DEL-200 |
| chr8-71030290-DEL-105 | chr8-71030291-DEL-105 |
| chr8-71031092-DEL-204 | chr8-71031093-DEL-204 |
| chr8-71094968-DEL-208 | chr8-71094969-DEL-208 |
| chr8-71116956-DEL-130 | chr8-71116943-DEL-111 |
| chr8-71120273-DEL-61 | chr8-71120281-DEL-56 |
| chr8-71147014-DEL-199 | chr8-71147032-DEL-180 |
| chr8-71175656-DEL-92 | chr8-71175657-DEL-92 |
| chr8-71187108-DEL-195 | chr8-71187109-DEL-195 |
| chr8-71280164-DEL-205 | chr8-71280165-DEL-205 |
| chr8-71283995-DEL-76 | chr8-71283998-DEL-56 |
| chr8-71305778-DEL-66 | chr8-71305779-DEL-66 |
| chr8-71325205-DEL-104 | chr8-71325210-DEL-100 |
| chr8-71353125-DEL-199 | chr8-71353126-DEL-199 |
| chr8-71356175-DEL-56 | chr8-71356164-DEL-56 |
| chr8-71379604-DEL-422 | chr8-71379605-DEL-422 |
| chr8-71381084-DEL-270 | chr8-71381085-DEL-270 |
| chr8-71382229-DEL-56 | chr8-71382230-DEL-56 |
| chr8-71391405-DEL-1592 | chr8-71391406-DEL-1592 |
| chr8-71391867-DEL-207 | chr8-71391868-DEL-207 |
| chr8-71401317-DEL-77 | chr8-71401318-DEL-77 |
| chr8-71404418-DEL-2356 | chr8-71404405-DEL-2356 |
| chr8-71409936-DEL-85 | chr8-71409937-DEL-85 |
| chr8-71418133-DEL-189 | chr8-71418134-DEL-189 |
| chr8-71418572-DEL-1520 | chr8-71418573-DEL-1520 |
| chr8-71421972-DEL-1292 | chr8-71421972-DEL-1292 |
| chr8-71424000-DEL-1292 | chr8-71424001-DEL-1292 |
| chr8-71425260-DEL-381 | chr8-71425231-DEL-382 |
| chr8-71426442-DEL-187 | chr8-71426443-DEL-187 |
| chr8-71443088-DEL-57 | chr8-71443089-DEL-55 |
| chr8-71450530-DEL-645 | chr8-71450514-DEL-645 |
| chr8-71451310-DEL-304 | chr8-71451311-DEL-304 |
| chr8-71452267-DEL-75 | chr8-71452268-DEL-75 |
| chr8-71452437-DEL-418 | chr8-71452438-DEL-418 |
| chr8-71457557-DEL-564 | chr8-71457556-DEL-564 |
| chr8-71470697-DEL-2060 | chr8-71470697-DEL-2060 |
| chr8-71511677-DEL-213 | chr8-71511678-DEL-213 |
| chr8-71529881-DEL-370 | chr8-71529882-DEL-370 |
| chr8-71598319-DEL-563 | chr8-71598320-DEL-563 |
| chr8-71657784-DEL-404 | chr8-71657785-DEL-404 |
| chr8-71666520-DEL-51 | chr8-71666521-DEL-51 |
| chr8-71681626-DEL-5389 | chr8-71681627-DEL-5389 |
| chr8-71698163-DEL-206 | chr8-71698164-DEL-206 |
| chr8-71714255-DEL-444 | chr8-71714256-DEL-444 |
| chr8-71736000-DEL-56 | chr8-71736005-DEL-52 |
| chr8-71852884-DEL-199 | chr8-71852885-DEL-199 |
| chr8-71863726-DEL-173 | chr8-71863718-DEL-174 |
| chr8-71881435-DEL-202 | chr8-71881436-DEL-202 |
| chr8-71893897-DEL-176 | chr8-71893898-DEL-176 |
| chr8-71896408-DEL-305 | chr8-71896409-DEL-305 |
| chr8-71946719-DEL-777 | chr8-71946720-DEL-777 |
| chr8-71954204-DEL-202 | chr8-71954205-DEL-197 |
| chr8-71972156-DEL-51 | chr8-71972139-DEL-52 |
| chr8-71981951-DEL-181 | chr8-71981953-DEL-157 |
| chr8-71991633-DEL-66 | chr8-71991634-DEL-66 |
| chr8-71998919-DEL-203 | chr8-71998920-DEL-203 |
| chr8-72002559-DEL-201 | chr8-72002560-DEL-201 |
| chr8-72017889-DEL-202 | chr8-72017890-DEL-202 |
| chr8-72042590-DEL-63 | chr8-72042584-DEL-88 |
| chr8-72062927-DEL-199 | chr8-72062928-DEL-199 |

| | |
|---|---|
| chr8-72126257-DEL-209 | chr8-72126258-DEL-209 |
| chr8-72127162-DEL-74 | chr8-72127163-DEL-74 |
| chr8-72150757-DEL-66 | chr8-72150758-DEL-66 |
| chr8-72171458-DEL-80 | chr8-72171459-DEL-80 |
| chr8-72232849-DEL-537 | chr8-72232850-DEL-537 |
| chr8-72247153-DEL-498 | chr8-72247154-DEL-498 |
| chr8-72269064-DEL-209 | chr8-72269065-DEL-209 |
| chr8-72680841-DEL-61 | chr8-72680842-DEL-61 |
| chr8-72708892-DEL-98 | chr8-72708893-DEL-98 |
| chr8-72757457-DEL-68 | chr8-72757501-DEL-58 |
| chr8-72758279-DEL-280 | chr8-72758280-DEL-277 |
| chr8-72772732-DEL-371 | chr8-72772733-DEL-371 |
| chr8-72773322-DEL-264 | chr8-72773322-DEL-262 |
| chr8-72775705-DEL-806 | chr8-72775705-DEL-805 |
| chr8-72778628-DEL-8300 | chr8-72778620-DEL-8306 |
| chr8-72788991-DEL-821 | chr8-72788992-DEL-821 |
| chr8-72790056-DEL-202 | chr8-72790057-DEL-202 |
| chr8-72801949-DEL-60 | chr8-72801980-DEL-60 |
| chr8-72805524-DEL-78 | chr8-72805525-DEL-78 |
| chr8-72817585-DEL-206 | chr8-72817585-DEL-201 |
| chr8-72828903-DEL-138 | chr8-72828903-DEL-138 |
| chr8-72839789-DEL-68 | chr8-72839788-DEL-68 |
| chr8-72849186-DEL-643 | chr8-72849187-DEL-643 |
| chr8-72855850-DEL-226 | chr8-72855851-DEL-226 |
| chr8-72865266-DEL-612 | chr8-72865267-DEL-612 |
| chr8-72867488-DEL-70 | chr8-72867481-DEL-64 |
| chr8-72873618-DEL-3344 | chr8-72873618-DEL-3344 |
| chr8-72882345-DEL-102 | chr8-72882342-DEL-106 |
| chr8-72898912-DEL-81 | chr8-72898901-DEL-83 |
| chr8-72902373-DEL-211 | chr8-72902374-DEL-211 |
| chr8-72921521-DEL-85 | chr8-72921522-DEL-85 |
| chr8-72973002-DEL-202 | chr8-72973003-DEL-202 |
| chr8-72978404-DEL-14513 | chr8-72978405-DEL-14513 |
| chr8-72993247-DEL-90 | chr8-72993248-DEL-90 |
| chr8-73031870-DEL-249 | chr8-73031871-DEL-243 |
| chr8-73035919-DEL-80 | chr8-73035921-DEL-65 |
| chr8-73045424-DEL-172 | chr8-73045417-DEL-137 |
| chr8-73047859-DEL-56 | chr8-73047859-DEL-56 |
| chr8-73052033-DEL-205 | chr8-73052034-DEL-205 |
| chr8-73053139-DEL-76 | chr8-73053127-DEL-84 |
| chr8-73057910-DEL-1248 | chr8-73057910-DEL-1249 |
| chr8-73065417-DEL-200 | chr8-73065418-DEL-200 |
| chr8-73071147-DEL-60 | chr8-73071148-DEL-60 |
| chr8-73075546-DEL-60 | chr8-73075581-DEL-54 |
| chr8-73147697-DEL-55 | chr8-73147698-DEL-55 |
| chr8-73176258-DEL-210 | chr8-73176259-DEL-210 |
| chr8-73186320-DEL-116 | chr8-73186306-DEL-128 |
| chr8-73187455-DEL-907 | chr8-73187456-DEL-907 |
| chr8-73187515-DEL-174 | chr8-73187516-DEL-174 |
| chr8-73198110-DEL-211 | chr8-73198111-DEL-211 |
| chr8-73199906-DEL-205 | chr8-73199907-DEL-205 |
| chr8-73209980-DEL-53 | chr8-73209981-DEL-53 |
| chr8-73262262-DEL-569 | chr8-73262263-DEL-569 |
| chr8-73282622-DEL-64 | chr8-73282623-DEL-64 |
| chr8-73292895-DEL-187 | chr8-73292896-DEL-187 |
| chr8-73324197-DEL-182 | chr8-73324190-DEL-183 |
| chr8-73343091-DEL-51 | chr8-73343090-DEL-52 |
| chr8-73349495-DEL-80 | chr8-73349519-DEL-90 |
| chr8-73362972-DEL-178 | chr8-73362973-DEL-178 |
| chr8-73379913-DEL-65 | chr8-73379907-DEL-69 |
| chr8-73402950-DEL-2957 | chr8-73402951-DEL-2957 |
| chr8-73410658-DEL-192 | chr8-73410659-DEL-192 |
| chr8-73425104-DEL-156 | chr8-73425127-DEL-130 |
| chr8-73428681-DEL-127 | chr8-73428682-DEL-127 |
| chr8-73529233-DEL-208 | chr8-73529245-DEL-208 |
| chr8-73565089-DEL-6718 | chr8-73565090-DEL-6718 |
| chr8-73575286-DEL-55 | chr8-73575287-DEL-55 |
| chr8-73576745-DEL-947 | chr8-73576746-DEL-947 |
| chr8-73588944-DEL-9672 | chr8-73588945-DEL-9672 |
| chr8-73612990-DEL-123 | chr8-73612996-DEL-118 |
| chr8-73616056-DEL-6356 | chr8-73616057-DEL-6356 |
| chr8-73640122-DEL-61 | chr8-73640127-DEL-63 |
| chr8-73648897-DEL-10491 | chr8-73648898-DEL-10491 |
| chr8-73694569-DEL-6544 | chr8-73694570-DEL-6544 |
| chr8-73714533-DEL-595 | chr8-73714534-DEL-595 |
| chr8-73717579-DEL-84 | chr8-73717565-DEL-69 |
| chr8-73718991-DEL-85 | chr8-73719013-DEL-56 |
| chr8-73725405-DEL-81 | chr8-73725404-DEL-82 |
| chr8-73745531-DEL-195 | chr8-73745526-DEL-199 |
| chr8-73769240-DEL-192 | chr8-73769241-DEL-192 |

| | |
|---|---|
| chr8-73790595-DEL-120 | chr8-73790596-DEL-120 |
| chr8-73799104-DEL-1376 | chr8-73799105-DEL-1376 |
| chr8-73878071-DEL-6403 | chr8-73878072-DEL-6403 |
| chr8-73901363-DEL-61 | chr8-73901341-DEL-62 |
| chr8-73903137-DEL-361 | chr8-73903138-DEL-361 |
| chr8-73904924-DEL-4214 | chr8-73904925-DEL-4214 |
| chr8-73909172-DEL-78 | chr8-73909173-DEL-78 |
| chr8-73934547-DEL-546 | chr8-73934548-DEL-546 |
| chr8-73963307-DEL-132 | chr8-73963308-DEL-132 |
| chr8-73972290-DEL-8690 | chr8-73972291-DEL-8690 |
| chr8-74002188-DEL-120 | chr8-74002189-DEL-120 |
| chr8-74003031-DEL-195 | chr8-74003032-DEL-195 |
| chr8-74005121-DEL-224 | chr8-74005122-DEL-224 |
| chr8-74010655-DEL-1469 | chr8-74010656-DEL-1469 |
| chr8-74032044-DEL-54 | chr8-74031998-DEL-62 |
| chr8-74045880-DEL-521 | chr8-74045881-DEL-521 |
| chr8-74074093-DEL-57 | chr8-74074090-DEL-51 |
| chr8-74108022-DEL-585 | chr8-74108023-DEL-585 |
| chr8-74189613-DEL-65 | chr8-74189584-DEL-82 |
| chr8-74203847-DEL-67 | chr8-74203848-DEL-67 |
| chr8-74224683-DEL-199 | chr8-74224684-DEL-199 |
| chr8-74247318-DEL-462 | chr8-74247319-DEL-462 |
| chr8-74250772-DEL-7230 | chr8-74250773-DEL-7230 |
| chr8-74300620-DEL-62 | chr8-74300621-DEL-62 |
| chr8-74319262-DEL-498 | chr8-74319263-DEL-498 |
| chr8-74339045-DEL-207 | chr8-74339046-DEL-207 |
| chr8-74383320-DEL-607 | chr8-74383321-DEL-607 |
| chr8-74393484-DEL-66 | chr8-74393485-DEL-66 |
| chr8-74394155-DEL-204 | chr8-74394148-DEL-204 |
| chr8-74403707-DEL-108 | chr8-74403709-DEL-109 |
| chr8-74405454-DEL-357 | chr8-74405455-DEL-357 |
| chr8-74410013-DEL-2423 | chr8-74410014-DEL-2423 |
| chr8-74423303-DEL-66 | chr8-74423304-DEL-66 |
| chr8-74424297-DEL-694 | chr8-74424298-DEL-694 |
| chr8-74451326-DEL-64 | chr8-74451329-DEL-65 |
| chr8-74456739-DEL-145 | chr8-74456745-DEL-140 |
| chr8-74481308-DEL-1928 | chr8-74481313-DEL-1924 |
| chr8-74488557-DEL-332 | chr8-74488558-DEL-332 |
| chr8-74490031-DEL-2253 | chr8-74490032-DEL-2253 |
| chr8-74496286-DEL-1133 | chr8-74496287-DEL-1133 |
| chr8-74502641-DEL-264 | chr8-74502642-DEL-264 |
| chr8-74518845-DEL-87 | chr8-74518846-DEL-87 |
| chr8-74521930-DEL-530 | chr8-74521931-DEL-530 |
| chr8-74569616-DEL-55 | chr8-74569584-DEL-56 |
| chr8-74589041-DEL-1668 | chr8-74589042-DEL-1668 |
| chr8-74591291-DEL-58 | chr8-74591292-DEL-58 |
| chr8-74641470-DEL-522 | chr8-74641471-DEL-522 |
| chr8-74665615-DEL-311 | chr8-74665616-DEL-311 |
| chr8-74722840-DEL-115 | chr8-74722841-DEL-115 |
| chr8-74770284-DEL-6440 | chr8-74770285-DEL-6440 |
| chr8-74793276-DEL-75 | chr8-74793274-DEL-58 |
| chr8-74805802-DEL-104 | chr8-74805803-DEL-103 |
| chr8-74831445-DEL-268 | chr8-74831450-DEL-272 |
| chr8-74832339-DEL-85 | chr8-74832353-DEL-70 |
| chr8-74841194-DEL-939 | chr8-74841194-DEL-939 |
| chr8-74846441-DEL-490 | chr8-74846444-DEL-491 |
| chr8-74875573-DEL-692 | chr8-74875574-DEL-692 |
| chr8-74895841-DEL-74 | chr8-74895842-DEL-74 |
| chr8-74917682-DEL-6431 | chr8-74917683-DEL-6431 |
| chr8-74936092-DEL-105 | chr8-74936091-DEL-105 |
| chr8-74938343-DEL-203 | chr8-74938344-DEL-203 |
| chr8-74944454-DEL-176 | chr8-74944455-DEL-176 |
| chr8-74970071-DEL-6213 | chr8-74970072-DEL-6213 |
| chr8-74988036-DEL-2119 | chr8-74988037-DEL-2119 |
| chr8-74991238-DEL-761 | chr8-74991239-DEL-761 |
| chr8-74997314-DEL-2933 | chr8-74997315-DEL-2933 |
| chr8-75026393-DEL-56 | chr8-75026394-DEL-56 |
| chr8-75030137-DEL-1432 | chr8-75030138-DEL-1432 |
| chr8-75060208-DEL-197 | chr8-75060209-DEL-195 |
| chr8-75106316-DEL-6345 | chr8-75106317-DEL-6345 |
| chr8-75153956-DEL-4865 | chr8-75153957-DEL-4865 |
| chr8-75175400-DEL-60 | chr8-75175401-DEL-50 |
| chr8-75211500-DEL-6275 | chr8-75211501-DEL-6275 |
| chr8-75228727-DEL-88 | chr8-75228728-DEL-88 |
| chr8-75230327-DEL-116 | chr8-75230327-DEL-116 |
| chr8-75241977-DEL-4194 | chr8-75241978-DEL-4194 |
| chr8-75246654-DEL-195 | chr8-75246655-DEL-195 |
| chr8-75273664-DEL-64 | chr8-75273664-DEL-64 |
| chr8-75310097-DEL-81 | chr8-75310098-DEL-81 |
| chr8-75312466-DEL-2790 | chr8-75312467-DEL-2790 |

| | |
|---|---|
| chr8-75330205-DEL-2785 | chr8-75330206-DEL-2785 |
| chr8-75358049-DEL-504 | chr8-75358050-DEL-504 |
| chr8-75375013-DEL-80 | chr8-75375012-DEL-82 |
| chr8-75407428-DEL-159 | chr8-75407429-DEL-159 |
| chr8-75416000-DEL-1146 | chr8-75416001-DEL-1146 |
| chr8-75437790-DEL-91 | chr8-75437791-DEL-91 |
| chr8-75438503-DEL-96 | chr8-75438504-DEL-95 |
| chr8-75439516-DEL-120 | chr8-75439517-DEL-120 |
| chr8-75443754-DEL-13594 | chr8-75443755-DEL-13594 |
| chr8-75472505-DEL-6830 | chr8-75472506-DEL-6830 |
| chr8-75481078-DEL-140 | chr8-75481079-DEL-140 |
| chr8-75541464-DEL-511 | chr8-75541468-DEL-512 |
| chr8-75551679-DEL-66 | chr8-75551680-DEL-65 |
| chr8-75555987-DEL-209 | chr8-75555985-DEL-209 |
| chr8-75594372-DEL-187 | chr8-75594373-DEL-187 |
| chr8-75630607-DEL-952 | chr8-75630608-DEL-952 |
| chr8-75634524-DEL-341 | chr8-75634525-DEL-341 |
| chr8-75655425-DEL-540 | chr8-75655426-DEL-540 |
| chr8-75664247-DEL-1531 | chr8-75664246-DEL-1532 |
| chr8-75665112-DEL-96 | chr8-75665113-DEL-96 |
| chr8-75666843-DEL-171 | chr8-75666828-DEL-170 |
| chr8-75680890-DEL-201 | chr8-75680891-DEL-201 |
| chr8-75681549-DEL-746 | chr8-75681550-DEL-746 |
| chr8-75717118-DEL-57 | chr8-75717117-DEL-59 |
| chr8-75753133-DEL-248 | chr8-75753133-DEL-247 |
| chr8-75768227-DEL-86 | chr8-75768232-DEL-78 |
| chr8-75774820-DEL-161 | chr8-75774821-DEL-161 |
| chr8-75793558-DEL-64 | chr8-75793562-DEL-65 |
| chr8-75798501-DEL-80 | chr8-75798502-DEL-80 |
| chr8-75810313-DEL-105 | chr8-75810314-DEL-105 |
| chr8-75833965-DEL-231 | chr8-75833964-DEL-228 |
| chr8-75854970-DEL-1389 | chr8-75854970-DEL-1389 |
| chr8-75856509-DEL-916 | chr8-75856509-DEL-916 |
| chr8-75866626-DEL-6724 | chr8-75866627-DEL-6724 |
| chr8-75899913-DEL-198 | chr8-75899914-DEL-198 |
| chr8-75900923-DEL-374 | chr8-75900926-DEL-372 |
| chr8-75902343-DEL-6401 | chr8-75902344-DEL-6401 |
| chr8-75909997-DEL-84 | chr8-75910015-DEL-56 |
| chr8-75932090-DEL-52 | chr8-75932119-DEL-54 |
| chr8-75933576-DEL-182 | chr8-75933577-DEL-182 |
| chr8-75949384-DEL-58 | chr8-75949363-DEL-59 |
| chr8-75971924-DEL-506 | chr8-75971925-DEL-506 |
| chr8-75991973-DEL-75 | chr8-75991974-DEL-75 |
| chr8-76034476-DEL-1488 | chr8-76034476-DEL-1488 |
| chr8-76059219-DEL-7260 | chr8-76059220-DEL-7260 |
| chr8-76070945-DEL-1429 | chr8-76070946-DEL-1429 |
| chr8-76131919-DEL-1297 | chr8-76131920-DEL-1297 |
| chr8-76148048-DEL-64 | chr8-76148049-DEL-64 |
| chr8-76150192-DEL-1572 | chr8-76150193-DEL-1576 |
| chr8-76176033-DEL-246 | chr8-76176048-DEL-233 |
| chr8-76182514-DEL-79 | chr8-76182509-DEL-56 |
| chr8-76254202-DEL-224 | chr8-76254203-DEL-224 |
| chr8-76264616-DEL-208 | chr8-76264616-DEL-208 |
| chr8-76265085-DEL-3013 | chr8-76265085-DEL-3013 |
| chr8-76294356-DEL-51 | chr8-76294357-DEL-51 |
| chr8-76308144-DEL-6670 | chr8-76308145-DEL-6670 |
| chr8-76314305-DEL-214 | chr8-76314306-DEL-214 |
| chr8-76323632-DEL-354 | chr8-76323628-DEL-354 |
| chr8-76379911-DEL-1018 | chr8-76379912-DEL-1018 |
| chr8-76380780-DEL-96 | chr8-76380744-DEL-125 |
| chr8-76387100-DEL-61 | chr8-76387101-DEL-61 |
| chr8-76416093-DEL-7151 | chr8-76416094-DEL-7151 |
| chr8-76435212-DEL-1416 | chr8-76435213-DEL-1416 |
| chr8-76438068-DEL-1847 | chr8-76438083-DEL-1847 |
| chr8-76459386-DEL-1729 | chr8-76459384-DEL-1728 |
| chr8-76469754-DEL-489 | chr8-76469755-DEL-489 |
| chr8-76480689-DEL-58 | chr8-76480687-DEL-60 |
| chr8-76494832-DEL-55 | chr8-76494833-DEL-55 |
| chr8-76511116-DEL-409 | chr8-76511123-DEL-412 |
| chr8-76519743-DEL-1592 | chr8-76519744-DEL-1592 |
| chr8-76528703-DEL-3115 | chr8-76528704-DEL-3115 |
| chr8-76537332-DEL-3888 | chr8-76537333-DEL-3888 |
| chr8-76548381-DEL-497 | chr8-76548382-DEL-497 |
| chr8-76578194-DEL-5965 | chr8-76578195-DEL-5965 |
| chr8-76598143-DEL-3407 | chr8-76598144-DEL-3407 |
| chr8-76638793-DEL-451 | chr8-76638794-DEL-451 |
| chr8-76646131-DEL-979 | chr8-76646132-DEL-979 |
| chr8-76671132-DEL-722 | chr8-76671132-DEL-722 |
| chr8-76697294-DEL-129 | chr8-76697295-DEL-129 |
| chr8-76752221-DEL-2225 | chr8-76752216-DEL-2225 |

| | |
|---|---|
| chr8-76781237-DEL-127 | chr8-76781276-DEL-85 |
| chr8-76824404-DEL-6867 | chr8-76824405-DEL-6867 |
| chr8-76926726-DEL-6174 | chr8-76926727-DEL-6174 |
| chr8-76998027-DEL-52 | chr8-76998028-DEL-53 |
| chr8-77037779-DEL-3218 | chr8-77037780-DEL-3218 |
| chr8-77079699-DEL-75 | chr8-77079700-DEL-75 |
| chr8-77097639-DEL-716 | chr8-77097640-DEL-716 |
| chr8-77117151-DEL-66 | chr8-77117146-DEL-72 |
| chr8-77125366-DEL-67 | chr8-77125367-DEL-67 |
| chr8-77138461-DEL-8436 | chr8-77138462-DEL-8436 |
| chr8-77161054-DEL-865 | chr8-77161064-DEL-856 |
| chr8-77225192-DEL-360 | chr8-77225193-DEL-360 |
| chr8-77226229-DEL-923 | chr8-77226230-DEL-923 |
| chr8-77279194-DEL-64 | chr8-77279195-DEL-56 |
| chr8-77325615-DEL-127 | chr8-77325616-DEL-127 |
| chr8-77335843-DEL-62 | chr8-77335850-DEL-56 |
| chr8-77356384-DEL-54 | chr8-77356381-DEL-54 |
| chr8-77372080-DEL-60 | chr8-77372064-DEL-63 |
| chr8-77388639-DEL-346 | chr8-77388639-DEL-346 |
| chr8-77390021-DEL-52 | chr8-77390036-DEL-51 |
| chr8-77468263-DEL-58 | chr8-77468272-DEL-94 |
| chr8-77477386-DEL-80 | chr8-77477384-DEL-74 |
| chr8-77490380-DEL-87 | chr8-77490381-DEL-87 |
| chr8-77492399-DEL-72 | chr8-77492399-DEL-72 |
| chr8-77504790-DEL-2566 | chr8-77504791-DEL-2566 |
| chr8-77531859-DEL-712 | chr8-77531860-DEL-712 |
| chr8-77544880-DEL-63 | chr8-77544881-DEL-63 |
| chr8-77569684-DEL-1028 | chr8-77569685-DEL-1028 |
| chr8-77586887-DEL-322 | chr8-77586888-DEL-322 |
| chr8-77594097-DEL-64 | chr8-77594077-DEL-51 |
| chr8-77647296-DEL-107 | chr8-77647297-DEL-107 |
| chr8-77659452-DEL-52 | chr8-77659421-DEL-76 |
| chr8-77672410-DEL-2566 | chr8-77672411-DEL-2566 |
| chr8-77688119-DEL-574 | chr8-77688120-DEL-585 |
| chr8-77700341-DEL-103 | chr8-77700342-DEL-103 |
| chr8-77728293-DEL-54 | chr8-77728294-DEL-54 |
| chr8-77785292-DEL-378 | chr8-77785293-DEL-378 |
| chr8-77791373-DEL-62 | chr8-77791374-DEL-62 |
| chr8-77815334-DEL-51 | chr8-77815335-DEL-51 |
| chr8-77828855-DEL-57 | chr8-77828856-DEL-57 |
| chr8-77829428-DEL-97 | chr8-77829421-DEL-98 |
| chr8-77840041-DEL-594 | chr8-77840042-DEL-594 |
| chr8-77863524-DEL-6343 | chr8-77863522-DEL-6344 |
| chr8-77880743-DEL-302 | chr8-77880744-DEL-302 |
| chr8-77884368-DEL-99 | chr8-77884369-DEL-99 |
| chr8-77885197-DEL-138 | chr8-77885190-DEL-137 |
| chr8-77897826-DEL-62 | chr8-77897829-DEL-60 |
| chr8-77908901-DEL-1691 | chr8-77908902-DEL-1691 |
| chr8-77924614-DEL-90 | chr8-77924632-DEL-60 |
| chr8-77964485-DEL-440 | chr8-77964476-DEL-441 |
| chr8-77979905-DEL-99 | chr8-77979914-DEL-99 |
| chr8-77998061-DEL-97 | chr8-77998062-DEL-97 |
| chr8-78027404-DEL-3909 | chr8-78027405-DEL-3909 |
| chr8-78049122-DEL-359 | chr8-78049123-DEL-359 |
| chr8-78075366-DEL-74 | chr8-78075367-DEL-74 |
| chr8-78084719-DEL-62 | chr8-78084719-DEL-63 |
| chr8-78089957-DEL-94 | chr8-78089958-DEL-94 |
| chr8-78100116-DEL-702 | chr8-78100113-DEL-669 |
| chr8-78106074-DEL-545 | chr8-78106075-DEL-545 |
| chr8-78108833-DEL-82 | chr8-78108834-DEL-82 |
| chr8-78113566-DEL-152 | chr8-78113616-DEL-136 |
| chr8-78115825-DEL-6438 | chr8-78115826-DEL-6438 |
| chr8-78125621-DEL-903 | chr8-78125622-DEL-903 |
| chr8-78138060-DEL-78 | chr8-78138058-DEL-78 |
| chr8-78150486-DEL-69 | chr8-78150487-DEL-69 |
| chr8-78171867-DEL-56 | chr8-78171872-DEL-72 |
| chr8-78179168-DEL-56 | chr8-78179169-DEL-56 |
| chr8-78184629-DEL-55 | chr8-78184589-DEL-80 |
| chr8-78207689-DEL-117 | chr8-78207690-DEL-117 |
| chr8-78219825-DEL-61 | chr8-78219826-DEL-61 |
| chr8-78221858-DEL-76 | chr8-78221853-DEL-82 |
| chr8-78231655-DEL-199 | chr8-78231656-DEL-199 |
| chr8-78258341-DEL-215 | chr8-78258342-DEL-215 |
| chr8-78259975-DEL-161 | chr8-78259976-DEL-161 |
| chr8-78264216-DEL-104 | chr8-78264216-DEL-104 |
| chr8-78264733-DEL-2276 | chr8-78264734-DEL-2276 |
| chr8-78266899-DEL-169 | chr8-78266899-DEL-171 |
| chr8-78277162-DEL-512 | chr8-78277163-DEL-512 |
| chr8-78292819-DEL-109 | chr8-78292820-DEL-109 |
| chr8-78294251-DEL-442 | chr8-78294252-DEL-442 |

| | |
|---|---|
| chr8-78298400-DEL-85 | chr8-78298401-DEL-85 |
| chr8-78333983-DEL-823 | chr8-78333984-DEL-823 |
| chr8-78352632-DEL-201 | chr8-78352601-DEL-191 |
| chr8-78369932-DEL-7119 | chr8-78369933-DEL-7119 |
| chr8-78378468-DEL-71 | chr8-78378469-DEL-71 |
| chr8-78382715-DEL-558 | chr8-78382716-DEL-558 |
| chr8-78425244-DEL-52 | chr8-78425245-DEL-52 |
| chr8-78439134-DEL-58 | chr8-78439135-DEL-58 |
| chr8-78449363-DEL-76 | chr8-78449369-DEL-75 |
| chr8-78453260-DEL-66 | chr8-78453260-DEL-66 |
| chr8-78477630-DEL-77 | chr8-78477626-DEL-125 |
| chr8-78489425-DEL-161 | chr8-78489426-DEL-161 |
| chr8-78499700-DEL-143 | chr8-78499701-DEL-143 |
| chr8-78506838-DEL-372 | chr8-78506839-DEL-373 |
| chr8-78514644-DEL-177 | chr8-78514645-DEL-177 |
| chr8-78525476-DEL-3663 | chr8-78525477-DEL-3663 |
| chr8-78538877-DEL-334 | chr8-78538878-DEL-334 |
| chr8-78547785-DEL-457 | chr8-78547786-DEL-457 |
| chr8-78550831-DEL-81 | chr8-78550832-DEL-81 |
| chr8-78564324-DEL-158 | chr8-78564325-DEL-158 |
| chr8-78589153-DEL-186 | chr8-78589154-DEL-185 |
| chr8-78590624-DEL-4960 | chr8-78590624-DEL-4950 |
| chr8-78607609-DEL-93 | chr8-78607612-DEL-91 |
| chr8-78609470-DEL-264 | chr8-78609471-DEL-263 |
| chr8-78611663-DEL-6956 | chr8-78611664-DEL-6956 |
| chr8-78624503-DEL-83 | chr8-78624504-DEL-93 |
| chr8-78668570-DEL-513 | chr8-78668571-DEL-513 |
| chr8-78701795-DEL-1451 | chr8-78701796-DEL-1451 |
| chr8-78718537-DEL-2349 | chr8-78718538-DEL-2349 |
| chr8-78724305-DEL-1465 | chr8-78724306-DEL-1465 |
| chr8-78747245-DEL-67 | chr8-78747238-DEL-68 |
| chr8-78769323-DEL-204 | chr8-78769324-DEL-204 |
| chr8-78775760-DEL-52 | chr8-78775761-DEL-52 |
| chr8-78779061-DEL-51 | chr8-78779067-DEL-58 |
| chr8-78781272-DEL-69 | chr8-78781273-DEL-69 |
| chr8-78783433-DEL-72 | chr8-78783434-DEL-72 |
| chr8-78798568-DEL-95 | chr8-78798569-DEL-95 |
| chr8-78831610-DEL-149 | chr8-78831599-DEL-142 |
| chr8-78834815-DEL-503 | chr8-78834816-DEL-503 |
| chr8-78852681-DEL-68 | chr8-78852674-DEL-66 |
| chr8-78866377-DEL-7483 | chr8-78866378-DEL-7483 |
| chr8-78896029-DEL-379 | chr8-78896030-DEL-379 |
| chr8-78911288-DEL-3603 | chr8-78911289-DEL-3602 |
| chr8-78949722-DEL-122 | chr8-78949749-DEL-100 |
| chr8-78992241-DEL-72 | chr8-78992242-DEL-72 |
| chr8-79003891-DEL-54 | chr8-79003866-DEL-94 |
| chr8-79008432-DEL-341 | chr8-79008433-DEL-341 |
| chr8-79065992-DEL-1759 | chr8-79065992-DEL-1759 |
| chr8-79069914-DEL-61 | chr8-79069914-DEL-61 |
| chr8-79079689-DEL-90 | chr8-79079642-DEL-136 |
| chr8-79083508-DEL-120 | chr8-79083517-DEL-96 |
| chr8-79089742-DEL-401 | chr8-79089743-DEL-401 |
| chr8-79142925-DEL-204 | chr8-79142926-DEL-204 |
| chr8-79195768-DEL-52 | chr8-79195742-DEL-78 |
| chr8-79196334-DEL-1235 | chr8-79196336-DEL-1234 |
| chr8-79197695-DEL-562 | chr8-79197696-DEL-562 |
| chr8-79209378-DEL-291 | chr8-79209379-DEL-279 |
| chr8-79212515-DEL-584 | chr8-79212516-DEL-584 |
| chr8-79254231-DEL-1520 | chr8-79254232-DEL-1520 |
| chr8-79260321-DEL-78 | chr8-79260322-DEL-78 |
| chr8-79269406-DEL-132 | chr8-79269426-DEL-99 |
| chr8-79271753-DEL-327 | chr8-79271754-DEL-327 |
| chr8-79282008-DEL-261 | chr8-79282009-DEL-261 |
| chr8-79292211-DEL-744 | chr8-79292210-DEL-740 |
| chr8-79306435-DEL-78 | chr8-79306410-DEL-104 |
| chr8-79309430-DEL-104 | chr8-79309458-DEL-78 |
| chr8-79315054-DEL-105 | chr8-79315055-DEL-70 |
| chr8-79344702-DEL-174 | chr8-79344704-DEL-173 |
| chr8-79358419-DEL-114 | chr8-79358418-DEL-115 |
| chr8-79359246-DEL-65 | chr8-79359227-DEL-99 |
| chr8-79361689-DEL-81 | chr8-79361681-DEL-82 |
| chr8-79363354-DEL-234 | chr8-79363355-DEL-234 |
| chr8-79375897-DEL-432 | chr8-79375898-DEL-432 |
| chr8-79395118-DEL-1910 | chr8-79395119-DEL-1910 |
| chr8-79397466-DEL-1520 | chr8-79397467-DEL-1520 |
| chr8-79404955-DEL-167 | chr8-79404913-DEL-169 |
| chr8-79421095-DEL-1388 | chr8-79421096-DEL-1388 |
| chr8-79435302-DEL-456 | chr8-79435303-DEL-456 |
| chr8-79440489-DEL-259 | chr8-79440490-DEL-259 |
| chr8-79443898-DEL-127 | chr8-79443899-DEL-127 |

| | |
|---|---|
| chr8-79486366-DEL-75 | chr8-79486372-DEL-72 |
| chr8-79509870-DEL-1023 | chr8-79509871-DEL-1023 |
| chr8-79527164-DEL-264 | chr8-79527165-DEL-263 |
| chr8-79528776-DEL-189 | chr8-79528777-DEL-162 |
| chr8-79554515-DEL-82 | chr8-79554516-DEL-82 |
| chr8-79578891-DEL-650 | chr8-79578892-DEL-650 |
| chr8-79588685-DEL-89 | chr8-79588686-DEL-89 |
| chr8-79591732-DEL-90 | chr8-79591733-DEL-90 |
| chr8-79597474-DEL-2211 | chr8-79597459-DEL-2226 |
| chr8-79601465-DEL-63 | chr8-79601433-DEL-87 |
| chr8-79610853-DEL-90 | chr8-79610820-DEL-127 |
| chr8-79622915-DEL-485 | chr8-79622916-DEL-485 |
| chr8-79635048-DEL-70 | chr8-79635055-DEL-64 |
| chr8-79665049-DEL-52 | chr8-79665017-DEL-55 |
| chr8-79671927-DEL-53 | chr8-79671922-DEL-58 |
| chr8-79744637-DEL-167 | chr8-79744635-DEL-158 |
| chr8-79772064-DEL-783 | chr8-79772065-DEL-767 |
| chr8-79825941-DEL-73 | chr8-79825922-DEL-74 |
| chr8-79833565-DEL-272 | chr8-79833566-DEL-272 |
| chr8-79836166-DEL-1366 | chr8-79836166-DEL-1366 |
| chr8-79856269-DEL-106 | chr8-79856270-DEL-106 |
| chr8-79860800-DEL-110 | chr8-79860802-DEL-104 |
| chr8-79893770-DEL-97 | chr8-79893771-DEL-97 |
| chr8-79923607-DEL-62 | chr8-79923606-DEL-59 |
| chr8-79934334-DEL-1451 | chr8-79934335-DEL-1451 |
| chr8-79982374-DEL-75 | chr8-79982376-DEL-74 |
| chr8-80012976-DEL-63 | chr8-80012967-DEL-81 |
| chr8-80019709-DEL-154 | chr8-80019710-DEL-154 |
| chr8-80036157-DEL-59 | chr8-80036158-DEL-59 |
| chr8-80062965-DEL-323 | chr8-80062966-DEL-323 |
| chr8-80104253-DEL-103 | chr8-80104254-DEL-103 |
| chr8-80106016-DEL-286 | chr8-80106003-DEL-320 |
| chr8-80156356-DEL-180 | chr8-80156371-DEL-176 |
| chr8-80185471-DEL-78 | chr8-80185475-DEL-89 |
| chr8-80189435-DEL-324 | chr8-80189436-DEL-325 |
| chr8-80205375-DEL-184 | chr8-80205376-DEL-184 |
| chr8-80222625-DEL-170 | chr8-80222626-DEL-170 |
| chr8-80250419-DEL-227 | chr8-80250420-DEL-227 |
| chr8-80254451-DEL-78 | chr8-80254452-DEL-78 |
| chr8-80280875-DEL-130 | chr8-80280884-DEL-110 |
| chr8-80296986-DEL-98 | chr8-80296972-DEL-142 |
| chr8-80297457-DEL-790 | chr8-80297458-DEL-790 |
| chr8-80297535-DEL-172 | chr8-80297536-DEL-172 |
| chr8-80298512-DEL-449 | chr8-80298510-DEL-450 |
| chr8-80305724-DEL-416 | chr8-80305725-DEL-416 |
| chr8-80329510-DEL-150 | chr8-80329511-DEL-150 |
| chr8-80330332-DEL-285 | chr8-80330333-DEL-273 |
| chr8-80334421-DEL-238 | chr8-80334422-DEL-238 |
| chr8-80335971-DEL-396 | chr8-80335972-DEL-396 |
| chr8-80385296-DEL-185 | chr8-80385297-DEL-185 |
| chr8-80404633-DEL-138 | chr8-80404634-DEL-136 |
| chr8-80409597-DEL-148 | chr8-80409598-DEL-148 |
| chr8-80412790-DEL-216 | chr8-80412791-DEL-216 |
| chr8-80441908-DEL-399 | chr8-80441909-DEL-399 |
| chr8-80455395-DEL-114 | chr8-80455396-DEL-114 |
| chr8-80475913-DEL-52 | chr8-80475914-DEL-52 |
| chr8-80486372-DEL-6446 | chr8-80486373-DEL-6446 |
| chr8-80509152-DEL-1350 | chr8-80509153-DEL-1350 |
| chr8-80529380-DEL-56 | chr8-80529380-DEL-56 |
| chr8-80539326-DEL-226 | chr8-80539327-DEL-226 |
| chr8-80542767-DEL-256 | chr8-80542768-DEL-256 |
| chr8-80545489-DEL-6154 | chr8-80545490-DEL-6154 |
| chr8-80580563-DEL-224 | chr8-80580564-DEL-224 |
| chr8-80597070-DEL-68 | chr8-80597071-DEL-68 |
| chr8-80628734-DEL-101 | chr8-80628735-DEL-101 |
| chr8-80629990-DEL-81 | chr8-80629991-DEL-81 |
| chr8-80639209-DEL-2563 | chr8-80639210-DEL-2562 |
| chr8-80668928-DEL-86 | chr8-80668929-DEL-86 |
| chr8-80671425-DEL-69 | chr8-80671426-DEL-69 |
| chr8-80686230-DEL-1794 | chr8-80686231-DEL-1794 |
| chr8-80704077-DEL-65 | chr8-80704078-DEL-65 |
| chr8-80766546-DEL-52 | chr8-80766547-DEL-52 |
| chr8-80771551-DEL-69 | chr8-80771551-DEL-69 |
| chr8-80794880-DEL-1908 | chr8-80794881-DEL-1908 |
| chr8-80828189-DEL-331 | chr8-80828185-DEL-332 |
| chr8-80849135-DEL-52 | chr8-80849136-DEL-52 |
| chr8-80851883-DEL-1844 | chr8-80851884-DEL-1844 |
| chr8-80853338-DEL-231 | chr8-80853339-DEL-231 |
| chr8-80885282-DEL-1038 | chr8-80885283-DEL-1038 |
| chr8-80929493-DEL-102 | chr8-80929497-DEL-98 |

| | |
|---|---|
| chr8-80947957-DEL-1284 | chr8-80947958-DEL-1284 |
| chr8-80954686-DEL-52 | chr8-80954687-DEL-52 |
| chr8-80961504-DEL-59 | chr8-80961465-DEL-92 |
| chr8-80972242-DEL-971 | chr8-80972243-DEL-971 |
| chr8-81001790-DEL-766 | chr8-81001812-DEL-764 |
| chr8-81072582-DEL-6499 | chr8-81072583-DEL-6499 |
| chr8-81091467-DEL-825 | chr8-81091468-DEL-825 |
| chr8-81097222-DEL-506 | chr8-81097223-DEL-507 |
| chr8-81100061-DEL-3201 | chr8-81100065-DEL-3200 |
| chr8-81133986-DEL-687 | chr8-81133984-DEL-688 |
| chr8-81135026-DEL-6478 | chr8-81135027-DEL-6478 |
| chr8-81169090-DEL-204 | chr8-81169091-DEL-204 |
| chr8-81180773-DEL-5174 | chr8-81180774-DEL-5174 |
| chr8-81205803-DEL-1766 | chr8-81205804-DEL-1766 |
| chr8-81221707-DEL-424 | chr8-81221704-DEL-425 |
| chr8-81241339-DEL-373 | chr8-81241340-DEL-373 |
| chr8-81259221-DEL-87 | chr8-81259222-DEL-87 |
| chr8-81281972-DEL-168 | chr8-81281959-DEL-169 |
| chr8-81288393-DEL-81 | chr8-81288390-DEL-81 |
| chr8-81314369-DEL-464 | chr8-81314370-DEL-464 |
| chr8-81334791-DEL-1455 | chr8-81334790-DEL-1456 |
| chr8-81347454-DEL-649 | chr8-81347443-DEL-662 |
| chr8-81353318-DEL-1494 | chr8-81353314-DEL-1495 |
| chr8-81355300-DEL-366 | chr8-81355301-DEL-366 |
| chr8-81355923-DEL-3267 | chr8-81355933-DEL-3268 |
| chr8-81366699-DEL-2054 | chr8-81366700-DEL-2054 |
| chr8-81379204-DEL-305 | chr8-81379205-DEL-305 |
| chr8-81403785-DEL-4641 | chr8-81403786-DEL-4641 |
| chr8-81412897-DEL-557 | chr8-81412898-DEL-557 |
| chr8-81445167-DEL-60 | chr8-81445168-DEL-56 |
| chr8-81481755-DEL-194 | chr8-81481756-DEL-194 |
| chr8-81484493-DEL-180 | chr8-81484494-DEL-180 |
| chr8-81529746-DEL-63 | chr8-81529747-DEL-63 |
| chr8-81536669-DEL-200 | chr8-81536670-DEL-200 |
| chr8-81590775-DEL-209 | chr8-81590776-DEL-209 |
| chr8-81593688-DEL-649 | chr8-81593689-DEL-649 |
| chr8-81598675-DEL-80 | chr8-81598676-DEL-80 |
| chr8-81622750-DEL-196 | chr8-81622751-DEL-196 |
| chr8-81632228-DEL-129 | chr8-81632213-DEL-132 |
| chr8-81634104-DEL-54 | chr8-81634105-DEL-54 |
| chr8-81642472-DEL-93 | chr8-81642473-DEL-93 |
| chr8-81651577-DEL-80 | chr8-81651564-DEL-76 |
| chr8-81654284-DEL-56 | chr8-81654285-DEL-56 |
| chr8-81677903-DEL-206 | chr8-81677942-DEL-179 |
| chr8-81691929-DEL-82 | chr8-81691934-DEL-78 |
| chr8-81722890-DEL-63 | chr8-81722891-DEL-63 |
| chr8-81752828-DEL-52 | chr8-81752829-DEL-52 |
| chr8-81766962-DEL-108 | chr8-81766963-DEL-108 |
| chr8-81767881-DEL-186 | chr8-81767882-DEL-186 |
| chr8-81799524-DEL-6486 | chr8-81799523-DEL-6487 |
| chr8-81822241-DEL-186 | chr8-81822276-DEL-150 |
| chr8-81839245-DEL-2557 | chr8-81839246-DEL-2557 |
| chr8-81848812-DEL-3228 | chr8-81848813-DEL-3228 |
| chr8-81852924-DEL-120 | chr8-81852925-DEL-116 |
| chr8-81879535-DEL-434 | chr8-81879536-DEL-434 |
| chr8-81889632-DEL-88 | chr8-81889651-DEL-91 |
| chr8-81916249-DEL-7119 | chr8-81916250-DEL-7119 |
| chr8-81927145-DEL-4592 | chr8-81927146-DEL-4592 |
| chr8-81943928-DEL-1174 | chr8-81943929-DEL-1174 |
| chr8-81957217-DEL-68 | chr8-81957184-DEL-74 |
| chr8-81990833-DEL-228 | chr8-81990828-DEL-234 |
| chr8-82002078-DEL-61 | chr8-82002052-DEL-63 |
| chr8-82003675-DEL-556 | chr8-82003676-DEL-556 |
| chr8-82019895-DEL-117 | chr8-82019896-DEL-117 |
| chr8-82028256-DEL-51 | chr8-82028257-DEL-51 |
| chr8-82044854-DEL-53 | chr8-82044842-DEL-54 |
| chr8-82076960-DEL-70 | chr8-82076954-DEL-60 |
| chr8-82103559-DEL-306 | chr8-82103560-DEL-306 |
| chr8-82108442-DEL-96 | chr8-82108443-DEL-96 |
| chr8-82110559-DEL-1655 | chr8-82110560-DEL-1655 |
| chr8-82122237-DEL-198 | chr8-82122238-DEL-198 |
| chr8-82124817-DEL-82 | chr8-82124824-DEL-80 |
| chr8-82180794-DEL-157 | chr8-82180795-DEL-157 |
| chr8-82184321-DEL-937 | chr8-82184322-DEL-937 |
| chr8-82195547-DEL-63 | chr8-82195548-DEL-63 |
| chr8-82215396-DEL-784 | chr8-82215397-DEL-784 |
| chr8-82220343-DEL-2790 | chr8-82220344-DEL-2790 |
| chr8-82236389-DEL-72 | chr8-82236394-DEL-75 |
| chr8-82236988-DEL-198 | chr8-82236989-DEL-198 |
| chr8-82306571-DEL-207 | chr8-82306572-DEL-207 |

| | |
|---|---|
| chr8-82323219-DEL-58 | chr8-82323227-DEL-59 |
| chr8-82345311-DEL-239 | chr8-82345312-DEL-239 |
| chr8-82396056-DEL-455 | chr8-82396057-DEL-455 |
| chr8-82399930-DEL-1580 | chr8-82399931-DEL-1580 |
| chr8-82432266-DEL-75 | chr8-82432264-DEL-54 |
| chr8-82435087-DEL-114 | chr8-82435088-DEL-112 |
| chr8-82447937-DEL-4203 | chr8-82447937-DEL-4203 |
| chr8-82462865-DEL-124 | chr8-82462866-DEL-124 |
| chr8-82472517-DEL-171 | chr8-82472518-DEL-171 |
| chr8-82498984-DEL-97 | chr8-82498951-DEL-84 |
| chr8-82502728-DEL-202 | chr8-82502729-DEL-202 |
| chr8-82507332-DEL-166 | chr8-82507333-DEL-166 |
| chr8-82508488-DEL-222 | chr8-82508485-DEL-223 |
| chr8-82513072-DEL-211 | chr8-82513073-DEL-211 |
| chr8-82553410-DEL-181 | chr8-82553411-DEL-181 |
| chr8-82555925-DEL-54 | chr8-82555926-DEL-57 |
| chr8-82557824-DEL-58 | chr8-82557825-DEL-58 |
| chr8-82563824-DEL-147 | chr8-82563825-DEL-147 |
| chr8-82572536-DEL-349 | chr8-82572537-DEL-349 |
| chr8-82600009-DEL-123 | chr8-82599959-DEL-133 |
| chr8-82618656-DEL-595 | chr8-82618657-DEL-595 |
| chr8-82637604-DEL-6365 | chr8-82637605-DEL-6365 |
| chr8-82650861-DEL-79 | chr8-82650860-DEL-88 |
| chr8-82651409-DEL-72 | chr8-82651410-DEL-72 |
| chr8-82655421-DEL-64 | chr8-82655411-DEL-74 |
| chr8-82669674-DEL-1247 | chr8-82669675-DEL-1247 |
| chr8-82672242-DEL-70 | chr8-82672243-DEL-70 |
| chr8-82681198-DEL-6895 | chr8-82681199-DEL-6895 |
| chr8-82697011-DEL-53 | chr8-82697010-DEL-54 |
| chr8-82724181-DEL-52 | chr8-82724182-DEL-56 |
| chr8-82732353-DEL-1084 | chr8-82732354-DEL-1084 |
| chr8-82757503-DEL-64 | chr8-82757504-DEL-64 |
| chr8-82761145-DEL-6321 | chr8-82761146-DEL-6321 |
| chr8-82777179-DEL-1499 | chr8-82777180-DEL-1499 |
| chr8-82793555-DEL-83 | chr8-82793555-DEL-83 |
| chr8-82830970-DEL-54 | chr8-82830968-DEL-56 |
| chr8-82838396-DEL-249 | chr8-82838402-DEL-250 |
| chr8-82861770-DEL-86 | chr8-82861771-DEL-86 |
| chr8-82866259-DEL-70 | chr8-82866260-DEL-71 |
| chr8-82898189-DEL-52 | chr8-82898186-DEL-56 |
| chr8-82908188-DEL-118 | chr8-82908188-DEL-116 |
| chr8-82911194-DEL-207 | chr8-82911195-DEL-207 |
| chr8-82934013-DEL-60 | chr8-82934014-DEL-60 |
| chr8-82935366-DEL-4092 | chr8-82935367-DEL-4093 |
| chr8-82973754-DEL-57 | chr8-82973755-DEL-57 |
| chr8-82994745-DEL-6332 | chr8-82994746-DEL-6332 |
| chr8-83015419-DEL-1084 | chr8-83015420-DEL-1084 |
| chr8-83016950-DEL-1166 | chr8-83016951-DEL-1166 |
| chr8-83030872-DEL-481 | chr8-83030873-DEL-481 |
| chr8-83043682-DEL-64 | chr8-83043683-DEL-62 |
| chr8-83107516-DEL-1142 | chr8-83107517-DEL-1142 |
| chr8-83161924-DEL-3821 | chr8-83161957-DEL-3789 |
| chr8-83177699-DEL-1430 | chr8-83177700-DEL-1430 |
| chr8-83184287-DEL-1932 | chr8-83184288-DEL-1932 |
| chr8-83213906-DEL-801 | chr8-83213906-DEL-801 |
| chr8-83315689-DEL-95 | chr8-83315681-DEL-96 |
| chr8-83318265-DEL-84 | chr8-83318266-DEL-84 |
| chr8-83323747-DEL-386 | chr8-83323748-DEL-386 |
| chr8-83344593-DEL-54 | chr8-83344594-DEL-54 |
| chr8-83350659-DEL-357 | chr8-83350660-DEL-357 |
| chr8-83379691-DEL-983 | chr8-83379692-DEL-983 |
| chr8-83384235-DEL-892 | chr8-83384236-DEL-892 |
| chr8-83429750-DEL-219 | chr8-83429751-DEL-219 |
| chr8-83439424-DEL-7304 | chr8-83439422-DEL-7305 |
| chr8-83482342-DEL-2210 | chr8-83482343-DEL-2210 |
| chr8-83507675-DEL-61 | chr8-83507675-DEL-61 |
| chr8-83518836-DEL-54 | chr8-83518815-DEL-55 |
| chr8-83527639-DEL-78 | chr8-83527624-DEL-64 |
| chr8-83557692-DEL-66 | chr8-83557693-DEL-66 |
| chr8-83559015-DEL-58 | chr8-83559005-DEL-60 |
| chr8-83576541-DEL-875 | chr8-83576542-DEL-874 |
| chr8-83589009-DEL-104 | chr8-83589010-DEL-104 |
| chr8-83659183-DEL-68 | chr8-83659158-DEL-64 |
| chr8-83662480-DEL-177 | chr8-83662481-DEL-177 |
| chr8-83697636-DEL-162 | chr8-83697637-DEL-162 |
| chr8-83731189-DEL-232 | chr8-83731190-DEL-220 |
| chr8-83732329-DEL-72 | chr8-83732330-DEL-72 |
| chr8-83737223-DEL-184 | chr8-83737224-DEL-184 |
| chr8-83755355-DEL-83 | chr8-83755347-DEL-91 |
| chr8-83760125-DEL-677 | chr8-83760126-DEL-677 |

| | |
|---|---|
| chr8-83788780-DEL-2061 | chr8-83788781-DEL-2061 |
| chr8-83836950-DEL-125 | chr8-83836981-DEL-84 |
| chr8-83864197-DEL-59 | chr8-83864198-DEL-53 |
| chr8-83869842-DEL-524 | chr8-83869846-DEL-521 |
| chr8-83871699-DEL-452 | chr8-83871698-DEL-453 |
| chr8-83872515-DEL-175 | chr8-83872516-DEL-175 |
| chr8-83880241-DEL-72 | chr8-83880242-DEL-72 |
| chr8-83911189-DEL-198 | chr8-83911188-DEL-193 |
| chr8-83912318-DEL-614 | chr8-83912348-DEL-602 |
| chr8-83916927-DEL-792 | chr8-83916928-DEL-792 |
| chr8-83924001-DEL-63 | chr8-83924006-DEL-61 |
| chr8-83932973-DEL-186 | chr8-83932982-DEL-178 |
| chr8-83945745-DEL-95 | chr8-83945744-DEL-71 |
| chr8-83955172-DEL-2972 | chr8-83955168-DEL-2979 |
| chr8-83963420-DEL-1344 | chr8-83963421-DEL-1344 |
| chr8-83965800-DEL-59 | chr8-83965801-DEL-59 |
| chr8-83973731-DEL-194 | chr8-83973732-DEL-194 |
| chr8-83974294-DEL-81 | chr8-83974295-DEL-81 |
| chr8-83999723-DEL-165 | chr8-83999724-DEL-165 |
| chr8-84001730-DEL-209 | chr8-84001731-DEL-209 |
| chr8-84013150-DEL-164 | chr8-84013151-DEL-164 |
| chr8-84030003-DEL-433 | chr8-84030004-DEL-433 |
| chr8-84032801-DEL-580 | chr8-84032802-DEL-580 |
| chr8-84033943-DEL-59 | chr8-84033944-DEL-59 |
| chr8-84034717-DEL-60 | chr8-84034720-DEL-58 |
| chr8-84064545-DEL-458 | chr8-84064546-DEL-458 |
| chr8-84075580-DEL-114 | chr8-84075581-DEL-114 |
| chr8-84093604-DEL-114 | chr8-84093605-DEL-114 |
| chr8-84119708-DEL-67 | chr8-84119694-DEL-68 |
| chr8-84128276-DEL-51 | chr8-84128266-DEL-51 |
| chr8-84132901-DEL-64 | chr8-84132901-DEL-64 |
| chr8-84166121-DEL-86 | chr8-84166102-DEL-101 |
| chr8-84178212-DEL-2100 | chr8-84178213-DEL-2100 |
| chr8-84222945-DEL-166 | chr8-84222946-DEL-165 |
| chr8-84228728-DEL-1774 | chr8-84228729-DEL-1774 |
| chr8-84236547-DEL-372 | chr8-84236547-DEL-372 |
| chr8-84239492-DEL-62 | chr8-84239493-DEL-62 |
| chr8-84259798-DEL-123 | chr8-84259800-DEL-121 |
| chr8-84283988-DEL-218 | chr8-84283989-DEL-218 |
| chr8-84286556-DEL-208 | chr8-84286557-DEL-208 |
| chr8-84297670-DEL-60 | chr8-84297671-DEL-60 |
| chr8-84309629-DEL-439 | chr8-84309630-DEL-439 |
| chr8-84317694-DEL-138 | chr8-84317695-DEL-138 |
| chr8-84329705-DEL-534 | chr8-84329706-DEL-534 |
| chr8-84344642-DEL-67 | chr8-84344643-DEL-67 |
| chr8-84354764-DEL-536 | chr8-84354765-DEL-536 |
| chr8-84360146-DEL-53 | chr8-84360148-DEL-52 |
| chr8-84362527-DEL-52 | chr8-84362528-DEL-51 |
| chr8-84362745-DEL-998 | chr8-84362746-DEL-998 |
| chr8-84370283-DEL-70 | chr8-84370284-DEL-50 |
| chr8-84381010-DEL-55 | chr8-84380999-DEL-51 |
| chr8-84429190-DEL-80 | chr8-84429191-DEL-80 |
| chr8-84434846-DEL-1600 | chr8-84434847-DEL-1600 |
| chr8-84481138-DEL-58 | chr8-84481137-DEL-59 |
| chr8-84485281-DEL-60 | chr8-84485282-DEL-60 |
| chr8-84494922-DEL-112 | chr8-84494904-DEL-106 |
| chr8-84499037-DEL-56 | chr8-84499042-DEL-52 |
| chr8-84508409-DEL-204 | chr8-84508422-DEL-203 |
| chr8-84521593-DEL-98 | chr8-84521621-DEL-96 |
| chr8-84523146-DEL-66 | chr8-84523147-DEL-66 |
| chr8-84524328-DEL-62 | chr8-84524314-DEL-74 |
| chr8-84554619-DEL-68 | chr8-84554619-DEL-68 |
| chr8-84588636-DEL-56 | chr8-84588637-DEL-60 |
| chr8-84592843-DEL-5809 | chr8-84592844-DEL-5809 |
| chr8-84604052-DEL-65 | chr8-84604053-DEL-65 |
| chr8-84676177-DEL-1203 | chr8-84676178-DEL-1203 |
| chr8-84685134-DEL-223 | chr8-84685135-DEL-223 |
| chr8-84709493-DEL-390 | chr8-84709485-DEL-394 |
| chr8-84734707-DEL-193 | chr8-84734708-DEL-193 |
| chr8-84735992-DEL-205 | chr8-84735979-DEL-206 |
| chr8-84743213-DEL-233 | chr8-84743214-DEL-237 |
| chr8-84754588-DEL-64 | chr8-84754589-DEL-64 |
| chr8-84772358-DEL-76 | chr8-84772375-DEL-56 |
| chr8-84773928-DEL-67 | chr8-84773929-DEL-67 |
| chr8-84785527-DEL-211 | chr8-84785528-DEL-211 |
| chr8-84787753-DEL-54 | chr8-84787749-DEL-54 |
| chr8-84827424-DEL-302 | chr8-84827437-DEL-302 |
| chr8-84833623-DEL-54 | chr8-84833624-DEL-54 |
| chr8-84834882-DEL-108 | chr8-84834883-DEL-102 |
| chr8-84838082-DEL-395 | chr8-84838083-DEL-395 |

| | |
|---|---|
| chr8-84843823-DEL-175 | chr8-84843824-DEL-175 |
| chr8-84850218-DEL-1420 | chr8-84850219-DEL-1438 |
| chr8-84858407-DEL-55 | chr8-84858408-DEL-55 |
| chr8-84872326-DEL-154 | chr8-84872327-DEL-154 |
| chr8-84880451-DEL-79 | chr8-84880450-DEL-83 |
| chr8-84882254-DEL-1365 | chr8-84882258-DEL-1367 |
| chr8-84921368-DEL-207 | chr8-84921369-DEL-207 |
| chr8-84923481-DEL-94 | chr8-84923482-DEL-94 |
| chr8-84933104-DEL-204 | chr8-84933105-DEL-204 |
| chr8-84936737-DEL-137 | chr8-84936738-DEL-139 |
| chr8-84947881-DEL-198 | chr8-84947882-DEL-198 |
| chr8-84975031-DEL-52 | chr8-84975032-DEL-52 |
| chr8-84979743-DEL-60 | chr8-84979744-DEL-60 |
| chr8-85004969-DEL-702 | chr8-85004975-DEL-697 |
| chr8-85013198-DEL-1861 | chr8-85013191-DEL-1892 |
| chr8-85016698-DEL-288 | chr8-85016707-DEL-291 |
| chr8-85017217-DEL-203 | chr8-85017218-DEL-203 |
| chr8-85035734-DEL-50 | chr8-85035735-DEL-50 |
| chr8-85035756-DEL-3077 | chr8-85035759-DEL-3075 |
| chr8-85036392-DEL-348 | chr8-85036411-DEL-330 |
| chr8-85042719-DEL-257 | chr8-85042722-DEL-248 |
| chr8-85049068-DEL-89 | chr8-85049066-DEL-95 |
| chr8-85053422-DEL-871 | chr8-85053423-DEL-871 |
| chr8-85068964-DEL-205 | chr8-85068965-DEL-205 |
| chr8-85084466-DEL-57 | chr8-85084467-DEL-57 |
| chr8-85096990-DEL-64 | chr8-85096946-DEL-68 |
| chr8-85098424-DEL-83 | chr8-85098431-DEL-58 |
| chr8-85100841-DEL-82 | chr8-85100826-DEL-54 |
| chr8-85119478-DEL-246 | chr8-85119479-DEL-240 |
| chr8-85123800-DEL-555 | chr8-85123801-DEL-555 |
| chr8-85125893-DEL-280 | chr8-85125924-DEL-278 |
| chr8-85160999-DEL-54 | chr8-85161000-DEL-54 |
| chr8-85161396-DEL-297 | chr8-85161397-DEL-297 |
| chr8-85183577-DEL-68 | chr8-85183576-DEL-69 |
| chr8-85202275-DEL-97 | chr8-85202276-DEL-97 |
| chr8-85227298-DEL-96 | chr8-85227299-DEL-96 |
| chr8-85236516-DEL-68 | chr8-85236517-DEL-68 |
| chr8-85240431-DEL-51 | chr8-85240432-DEL-51 |
| chr8-85243193-DEL-65 | chr8-85243199-DEL-64 |
| chr8-85350454-DEL-62 | chr8-85350444-DEL-72 |
| chr8-85352379-DEL-193 | chr8-85352380-DEL-193 |
| chr8-85354383-DEL-52 | chr8-85354384-DEL-52 |
| chr8-85355539-DEL-73 | chr8-85355539-DEL-73 |
| chr8-85377445-DEL-102 | chr8-85377446-DEL-102 |
| chr8-85390416-DEL-56 | chr8-85390417-DEL-56 |
| chr8-85400700-DEL-66 | chr8-85400700-DEL-68 |
| chr8-85441921-DEL-75 | chr8-85441953-DEL-79 |
| chr8-85528069-DEL-66 | chr8-85528064-DEL-86 |
| chr8-85538501-DEL-60 | chr8-85538501-DEL-60 |
| chr8-85546972-DEL-205 | chr8-85546973-DEL-205 |
| chr8-85580523-DEL-133 | chr8-85580524-DEL-133 |
| chr8-85589140-DEL-195 | chr8-85589142-DEL-191 |
| chr8-85601217-DEL-173 | chr8-85601218-DEL-173 |
| chr8-85615573-DEL-211 | chr8-85615574-DEL-211 |
| chr8-85633933-DEL-124 | chr8-85633928-DEL-127 |
| chr8-85640489-DEL-52 | chr8-85640490-DEL-74 |
| chr8-85660146-DEL-79 | chr8-85660146-DEL-67 |
| chr8-85676217-DEL-72 | chr8-85676207-DEL-90 |
| chr8-85687191-DEL-218 | chr8-85687192-DEL-218 |
| chr8-85691351-DEL-194 | chr8-85691352-DEL-194 |
| chr8-85700092-DEL-69 | chr8-85700093-DEL-69 |
| chr8-85725956-DEL-199 | chr8-85725958-DEL-200 |
| chr8-85731083-DEL-196 | chr8-85731084-DEL-196 |
| chr8-85738827-DEL-56 | chr8-85738826-DEL-56 |
| chr8-85749638-DEL-146 | chr8-85749639-DEL-146 |
| chr8-85790062-DEL-121 | chr8-85790063-DEL-121 |
| chr8-85799439-DEL-140 | chr8-85799437-DEL-141 |
| chr8-85816714-DEL-717 | chr8-85816715-DEL-717 |
| chr8-85829655-DEL-71 | chr8-85829655-DEL-80 |
| chr8-85834396-DEL-201 | chr8-85834397-DEL-201 |
| chr8-85839195-DEL-225 | chr8-85839196-DEL-225 |
| chr8-85845132-DEL-82 | chr8-85845133-DEL-80 |
| chr8-85848710-DEL-59 | chr8-85848696-DEL-74 |
| chr8-85873128-DEL-81 | chr8-85873129-DEL-81 |
| chr8-85874582-DEL-661 | chr8-85874583-DEL-661 |
| chr8-85875459-DEL-210 | chr8-85875460-DEL-210 |
| chr8-85880403-DEL-138 | chr8-85880404-DEL-138 |
| chr8-85883275-DEL-363 | chr8-85883276-DEL-363 |
| chr8-85888408-DEL-6286 | chr8-85888409-DEL-6286 |
| chr8-85903417-DEL-3581 | chr8-85903416-DEL-3581 |

| | |
|---|---|
| chr8-85913028-DEL-57 | chr8-85913012-DEL-59 |
| chr8-85925890-DEL-1448 | chr8-85925891-DEL-1448 |
| chr8-85942591-DEL-206 | chr8-85942592-DEL-206 |
| chr8-85946256-DEL-59 | chr8-85946257-DEL-59 |
| chr8-85958847-DEL-76 | chr8-85958848-DEL-76 |
| chr8-85960059-DEL-6188 | chr8-85960060-DEL-6188 |
| chr8-85970019-DEL-4633 | chr8-85970020-DEL-4633 |
| chr8-85978042-DEL-205 | chr8-85978037-DEL-206 |
| chr8-85988910-DEL-82 | chr8-85988911-DEL-82 |
| chr8-85992147-DEL-67 | chr8-85992148-DEL-67 |
| chr8-86002983-DEL-222 | chr8-86003000-DEL-211 |
| chr8-86037795-DEL-56 | chr8-86037796-DEL-56 |
| chr8-86055788-DEL-189 | chr8-86055789-DEL-189 |
| chr8-86070237-DEL-6197 | chr8-86070231-DEL-6198 |
| chr8-86081052-DEL-213 | chr8-86081053-DEL-213 |
| chr8-86083334-DEL-55 | chr8-86083335-DEL-55 |
| chr8-86105321-DEL-60 | chr8-86105322-DEL-60 |
| chr8-86111742-DEL-202 | chr8-86111743-DEL-202 |
| chr8-86122317-DEL-1014 | chr8-86122318-DEL-1014 |
| chr8-86142729-DEL-178 | chr8-86142730-DEL-178 |
| chr8-86162165-DEL-212 | chr8-86162165-DEL-213 |
| chr8-86175319-DEL-214 | chr8-86175320-DEL-214 |
| chr8-86178962-DEL-52 | chr8-86178963-DEL-52 |
| chr8-86194490-DEL-94 | chr8-86194491-DEL-94 |
| chr8-86210637-DEL-63 | chr8-86210638-DEL-63 |
| chr8-86224307-DEL-195 | chr8-86224308-DEL-195 |
| chr8-86230608-DEL-194 | chr8-86230609-DEL-194 |
| chr8-86342124-DEL-57 | chr8-86342124-DEL-67 |
| chr8-86362842-DEL-63 | chr8-86362842-DEL-58 |
| chr8-86379818-DEL-491 | chr8-86379820-DEL-490 |
| chr8-86429016-DEL-498 | chr8-86429017-DEL-498 |
| chr8-86433343-DEL-198 | chr8-86433344-DEL-198 |
| chr8-86506473-DEL-346 | chr8-86506474-DEL-346 |
| chr8-86515343-DEL-53 | chr8-86515371-DEL-50 |
| chr8-86546143-DEL-286 | chr8-86546144-DEL-286 |
| chr8-86548683-DEL-54 | chr8-86548684-DEL-53 |
| chr8-86552407-DEL-346 | chr8-86552408-DEL-346 |
| chr8-86583765-DEL-332 | chr8-86583766-DEL-332 |
| chr8-86584652-DEL-157 | chr8-86584653-DEL-154 |
| chr8-86585392-DEL-1210 | chr8-86585393-DEL-1204 |
| chr8-86609859-DEL-66 | chr8-86609860-DEL-66 |
| chr8-86616668-DEL-4623 | chr8-86616669-DEL-4623 |
| chr8-86624066-DEL-54 | chr8-86624067-DEL-54 |
| chr8-86624896-DEL-476 | chr8-86624901-DEL-459 |
| chr8-86630600-DEL-95 | chr8-86630612-DEL-72 |
| chr8-86633402-DEL-850 | chr8-86633403-DEL-850 |
| chr8-86640508-DEL-6639 | chr8-86640511-DEL-6640 |
| chr8-86670601-DEL-85 | chr8-86670600-DEL-55 |
| chr8-86674976-DEL-6670 | chr8-86674982-DEL-6671 |
| chr8-86674992-DEL-428 | chr8-86674992-DEL-428 |
| chr8-86718886-DEL-628 | chr8-86718887-DEL-628 |
| chr8-86721271-DEL-118 | chr8-86721272-DEL-118 |
| chr8-86725728-DEL-102 | chr8-86725729-DEL-102 |
| chr8-86726567-DEL-490 | chr8-86726568-DEL-490 |
| chr8-86738973-DEL-57 | chr8-86738973-DEL-57 |
| chr8-86763429-DEL-81 | chr8-86763432-DEL-82 |
| chr8-86793654-DEL-1408 | chr8-86793655-DEL-1408 |
| chr8-86808113-DEL-2471 | chr8-86808114-DEL-2471 |
| chr8-86834357-DEL-6210 | chr8-86834350-DEL-6211 |
| chr8-86841303-DEL-55 | chr8-86841304-DEL-55 |
| chr8-86853420-DEL-210 | chr8-86853421-DEL-210 |
| chr8-86864399-DEL-647 | chr8-86864400-DEL-647 |
| chr8-86870888-DEL-75 | chr8-86870890-DEL-72 |
| chr8-86901570-DEL-5358 | chr8-86901571-DEL-5358 |
| chr8-86908288-DEL-95 | chr8-86908264-DEL-96 |
| chr8-86914765-DEL-2058 | chr8-86914766-DEL-2058 |
| chr8-86921429-DEL-174 | chr8-86921430-DEL-174 |
| chr8-86932232-DEL-181 | chr8-86932242-DEL-182 |
| chr8-86942607-DEL-6231 | chr8-86942608-DEL-6231 |
| chr8-86957834-DEL-3037 | chr8-86957835-DEL-3037 |
| chr8-86966539-DEL-1252 | chr8-86966540-DEL-1252 |
| chr8-86971885-DEL-6209 | chr8-86971886-DEL-6209 |
| chr8-87003909-DEL-521 | chr8-87003910-DEL-521 |
| chr8-87013285-DEL-1862 | chr8-87013286-DEL-1862 |
| chr8-87018811-DEL-54 | chr8-87018812-DEL-54 |
| chr8-87024489-DEL-55 | chr8-87024495-DEL-50 |
| chr8-87062533-DEL-109 | chr8-87062538-DEL-105 |
| chr8-87099116-DEL-6644 | chr8-87099117-DEL-6644 |
| chr8-87135506-DEL-196 | chr8-87135507-DEL-196 |
| chr8-87183381-DEL-386 | chr8-87183383-DEL-383 |

| | |
|---|---|
| chr8-87247895-DEL-55 | chr8-87247896-DEL-55 |
| chr8-87294037-DEL-6338 | chr8-87294038-DEL-6338 |
| chr8-87317738-DEL-285 | chr8-87317739-DEL-285 |
| chr8-87353954-DEL-188 | chr8-87353957-DEL-184 |
| chr8-87368527-DEL-159 | chr8-87368528-DEL-159 |
| chr8-87387014-DEL-203 | chr8-87387015-DEL-203 |
| chr8-87390095-DEL-207 | chr8-87390096-DEL-207 |
| chr8-87393160-DEL-239 | chr8-87393160-DEL-239 |
| chr8-87405579-DEL-6441 | chr8-87405580-DEL-6441 |
| chr8-87423815-DEL-337 | chr8-87423816-DEL-337 |
| chr8-87488909-DEL-59 | chr8-87488906-DEL-86 |
| chr8-87500719-DEL-122 | chr8-87500720-DEL-118 |
| chr8-87504086-DEL-400 | chr8-87504094-DEL-352 |
| chr8-87522910-DEL-88 | chr8-87522901-DEL-83 |
| chr8-87538607-DEL-506 | chr8-87538608-DEL-506 |
| chr8-87576246-DEL-208 | chr8-87576247-DEL-198 |
| chr8-87596004-DEL-181 | chr8-87596005-DEL-181 |
| chr8-87629709-DEL-62 | chr8-87629710-DEL-62 |
| chr8-87649745-DEL-63 | chr8-87649731-DEL-68 |
| chr8-87663623-DEL-203 | chr8-87663655-DEL-217 |
| chr8-87671821-DEL-84 | chr8-87671822-DEL-84 |
| chr8-87707152-DEL-75 | chr8-87707153-DEL-73 |
| chr8-87714667-DEL-6402 | chr8-87714668-DEL-6402 |
| chr8-87754419-DEL-300 | chr8-87754420-DEL-300 |
| chr8-87757812-DEL-65 | chr8-87757813-DEL-64 |
| chr8-87770231-DEL-82 | chr8-87770234-DEL-80 |
| chr8-87774321-DEL-549 | chr8-87774322-DEL-549 |
| chr8-87788860-DEL-1019 | chr8-87788861-DEL-1019 |
| chr8-87789426-DEL-187 | chr8-87789427-DEL-187 |
| chr8-87818611-DEL-114 | chr8-87818588-DEL-138 |
| chr8-87844350-DEL-406 | chr8-87844351-DEL-404 |
| chr8-87890434-DEL-97 | chr8-87890435-DEL-97 |
| chr8-87903490-DEL-66 | chr8-87903491-DEL-66 |
| chr8-87909097-DEL-74 | chr8-87909098-DEL-74 |
| chr8-87924476-DEL-57 | chr8-87924477-DEL-57 |
| chr8-87938590-DEL-58 | chr8-87938593-DEL-86 |
| chr8-87963547-DEL-343 | chr8-87963548-DEL-343 |
| chr8-87987664-DEL-203 | chr8-87987658-DEL-204 |
| chr8-88017766-DEL-177 | chr8-88017767-DEL-177 |
| chr8-88041076-DEL-112 | chr8-88041077-DEL-112 |
| chr8-88052524-DEL-286 | chr8-88052525-DEL-286 |
| chr8-88055055-DEL-159 | chr8-88055076-DEL-159 |
| chr8-88064124-DEL-64 | chr8-88064125-DEL-64 |
| chr8-88087470-DEL-88 | chr8-88087487-DEL-53 |
| chr8-88090251-DEL-60 | chr8-88090245-DEL-102 |
| chr8-88130039-DEL-67 | chr8-88130040-DEL-67 |
| chr8-88136867-DEL-54 | chr8-88136868-DEL-54 |
| chr8-88140499-DEL-1241 | chr8-88140500-DEL-1241 |
| chr8-88155827-DEL-248 | chr8-88155828-DEL-248 |
| chr8-88173189-DEL-781 | chr8-88173190-DEL-781 |
| chr8-88208525-DEL-336 | chr8-88208526-DEL-336 |
| chr8-88223078-DEL-87 | chr8-88223076-DEL-58 |
| chr8-88228239-DEL-95 | chr8-88228240-DEL-95 |
| chr8-88234796-DEL-198 | chr8-88234799-DEL-196 |
| chr8-88258571-DEL-171 | chr8-88258579-DEL-164 |
| chr8-88262829-DEL-84 | chr8-88262831-DEL-83 |
| chr8-88297376-DEL-53 | chr8-88297347-DEL-54 |
| chr8-88304638-DEL-392 | chr8-88304639-DEL-392 |
| chr8-88320869-DEL-79 | chr8-88320871-DEL-74 |
| chr8-88361015-DEL-59 | chr8-88361019-DEL-67 |
| chr8-88379254-DEL-57 | chr8-88379255-DEL-57 |
| chr8-88397278-DEL-214 | chr8-88397279-DEL-214 |
| chr8-88410567-DEL-60 | chr8-88410567-DEL-56 |
| chr8-88452466-DEL-254 | chr8-88452467-DEL-254 |
| chr8-88490279-DEL-72 | chr8-88490279-DEL-72 |
| chr8-88519150-DEL-201 | chr8-88519151-DEL-201 |
| chr8-88522029-DEL-56 | chr8-88522004-DEL-57 |
| chr8-88540302-DEL-91 | chr8-88540301-DEL-92 |
| chr8-88541904-DEL-391 | chr8-88541905-DEL-391 |
| chr8-88605598-DEL-54 | chr8-88605599-DEL-50 |
| chr8-88625183-DEL-409 | chr8-88625184-DEL-409 |
| chr8-88626406-DEL-73 | chr8-88626411-DEL-69 |
| chr8-88633549-DEL-211 | chr8-88633550-DEL-211 |
| chr8-88649484-DEL-132 | chr8-88649480-DEL-135 |
| chr8-88669804-DEL-55 | chr8-88669804-DEL-55 |
| chr8-88704945-DEL-199 | chr8-88704946-DEL-199 |
| chr8-88705894-DEL-189 | chr8-88705895-DEL-189 |
| chr8-88729997-DEL-79 | chr8-88729998-DEL-79 |
| chr8-88744081-DEL-342 | chr8-88744082-DEL-342 |
| chr8-88753287-DEL-326 | chr8-88753288-DEL-326 |

| | |
|---|---|
| chr8-88756567-DEL-64 | chr8-88756568-DEL-64 |
| chr8-88761326-DEL-213 | chr8-88761327-DEL-213 |
| chr8-88764897-DEL-110 | chr8-88764898-DEL-110 |
| chr8-88774272-DEL-89 | chr8-88774272-DEL-58 |
| chr8-88789722-DEL-188 | chr8-88789723-DEL-188 |
| chr8-88794472-DEL-569 | chr8-88794475-DEL-567 |
| chr8-88809945-DEL-528 | chr8-88809946-DEL-528 |
| chr8-88810832-DEL-206 | chr8-88810833-DEL-206 |
| chr8-88811898-DEL-188 | chr8-88811898-DEL-188 |
| chr8-88835742-DEL-495 | chr8-88835743-DEL-495 |
| chr8-88839487-DEL-186 | chr8-88839488-DEL-186 |
| chr8-88854728-DEL-60 | chr8-88854729-DEL-60 |
| chr8-88869116-DEL-1140 | chr8-88869117-DEL-1140 |
| chr8-88873537-DEL-209 | chr8-88873538-DEL-209 |
| chr8-88887496-DEL-215 | chr8-88887497-DEL-215 |
| chr8-88892208-DEL-215 | chr8-88892209-DEL-215 |
| chr8-88903622-DEL-144 | chr8-88903623-DEL-144 |
| chr8-88904252-DEL-194 | chr8-88904253-DEL-194 |
| chr8-88906601-DEL-507 | chr8-88906602-DEL-507 |
| chr8-88909374-DEL-93 | chr8-88909379-DEL-89 |
| chr8-88913054-DEL-173 | chr8-88913055-DEL-173 |
| chr8-88915465-DEL-146 | chr8-88915465-DEL-146 |
| chr8-88917719-DEL-52 | chr8-88917720-DEL-52 |
| chr8-88922237-DEL-56 | chr8-88922238-DEL-56 |
| chr8-88936860-DEL-219 | chr8-88936861-DEL-219 |
| chr8-88970614-DEL-218 | chr8-88970615-DEL-218 |
| chr8-88972760-DEL-67 | chr8-88972761-DEL-67 |
| chr8-88976330-DEL-58 | chr8-88976307-DEL-103 |
| chr8-88996472-DEL-58 | chr8-88996473-DEL-58 |
| chr8-89008458-DEL-102 | chr8-89008459-DEL-102 |
| chr8-89013541-DEL-213 | chr8-89013542-DEL-213 |
| chr8-89033884-DEL-201 | chr8-89033885-DEL-201 |
| chr8-89064176-DEL-364 | chr8-89064177-DEL-364 |
| chr8-89091178-DEL-100 | chr8-89091179-DEL-100 |
| chr8-89099526-DEL-197 | chr8-89099522-DEL-198 |
| chr8-89101346-DEL-363 | chr8-89101346-DEL-363 |
| chr8-89130672-DEL-1879 | chr8-89130673-DEL-1879 |
| chr8-89198601-DEL-74 | chr8-89198602-DEL-74 |
| chr8-89226006-DEL-93 | chr8-89226007-DEL-93 |
| chr8-89230412-DEL-2153 | chr8-89230427-DEL-2144 |
| chr8-89242960-DEL-847 | chr8-89242961-DEL-847 |
| chr8-89303967-DEL-163 | chr8-89303947-DEL-183 |
| chr8-89317740-DEL-84 | chr8-89317741-DEL-84 |
| chr8-89343643-DEL-150 | chr8-89343644-DEL-150 |
| chr8-89357246-DEL-95 | chr8-89357247-DEL-95 |
| chr8-89370769-DEL-59 | chr8-89370770-DEL-59 |
| chr8-89409430-DEL-426 | chr8-89409437-DEL-420 |
| chr8-89411578-DEL-56 | chr8-89411573-DEL-62 |
| chr8-89457589-DEL-73 | chr8-89457593-DEL-72 |
| chr8-89486628-DEL-892 | chr8-89486629-DEL-892 |
| chr8-89488330-DEL-179 | chr8-89488331-DEL-179 |
| chr8-89490222-DEL-1879 | chr8-89490223-DEL-1879 |
| chr8-89496683-DEL-109 | chr8-89496684-DEL-109 |
| chr8-89500482-DEL-97 | chr8-89500483-DEL-97 |
| chr8-89528102-DEL-399 | chr8-89528101-DEL-399 |
| chr8-89529316-DEL-54 | chr8-89529317-DEL-54 |
| chr8-89532980-DEL-834 | chr8-89532981-DEL-834 |
| chr8-89547637-DEL-84 | chr8-89547632-DEL-83 |
| chr8-89548109-DEL-878 | chr8-89548108-DEL-879 |
| chr8-89551671-DEL-281 | chr8-89551672-DEL-281 |
| chr8-89566344-DEL-55 | chr8-89566340-DEL-56 |
| chr8-89567947-DEL-55 | chr8-89567953-DEL-50 |
| chr8-89568507-DEL-88 | chr8-89568508-DEL-88 |
| chr8-89580758-DEL-87 | chr8-89580759-DEL-87 |
| chr8-89593544-DEL-124 | chr8-89593541-DEL-136 |
| chr8-89605943-DEL-89 | chr8-89605941-DEL-90 |
| chr8-89613121-DEL-181 | chr8-89613122-DEL-181 |
| chr8-89646339-DEL-120 | chr8-89646345-DEL-115 |
| chr8-89659608-DEL-607 | chr8-89659609-DEL-607 |
| chr8-89661678-DEL-238 | chr8-89661679-DEL-238 |
| chr8-89685440-DEL-54 | chr8-89685434-DEL-55 |
| chr8-89698503-DEL-57 | chr8-89698498-DEL-54 |
| chr8-89739371-DEL-54 | chr8-89739368-DEL-53 |
| chr8-89742814-DEL-237 | chr8-89742815-DEL-237 |
| chr8-89788876-DEL-768 | chr8-89788877-DEL-768 |
| chr8-89822067-DEL-1564 | chr8-89822068-DEL-1564 |
| chr8-89829641-DEL-201 | chr8-89829642-DEL-201 |
| chr8-89845600-DEL-3173 | chr8-89845599-DEL-3171 |
| chr8-89846973-DEL-58 | chr8-89846992-DEL-52 |
| chr8-89866796-DEL-157 | chr8-89866797-DEL-157 |

| | |
|---|---|
| chr8-89872465-DEL-131 | chr8-89872432-DEL-121 |
| chr8-89889571-DEL-68 | chr8-89889572-DEL-66 |
| chr8-89890712-DEL-102 | chr8-89890707-DEL-108 |
| chr8-89915885-DEL-1083 | chr8-89915886-DEL-1083 |
| chr8-89946406-DEL-69 | chr8-89946407-DEL-69 |
| chr8-89972676-DEL-105 | chr8-89972677-DEL-56 |
| chr8-89974073-DEL-56 | chr8-89974074-DEL-56 |
| chr8-89984880-DEL-680 | chr8-89984880-DEL-680 |
| chr8-90054276-DEL-122 | chr8-90054277-DEL-122 |
| chr8-90092551-DEL-2600 | chr8-90092552-DEL-2600 |
| chr8-90120154-DEL-82 | chr8-90120155-DEL-82 |
| chr8-90125327-DEL-56 | chr8-90125328-DEL-56 |
| chr8-90128049-DEL-53 | chr8-90128011-DEL-90 |
| chr8-90145689-DEL-165 | chr8-90145690-DEL-165 |
| chr8-90158152-DEL-305 | chr8-90158154-DEL-306 |
| chr8-90163611-DEL-74 | chr8-90163611-DEL-76 |
| chr8-90168533-DEL-52 | chr8-90168517-DEL-56 |
| chr8-90210991-DEL-181 | chr8-90211000-DEL-173 |
| chr8-90348646-DEL-122 | chr8-90348647-DEL-122 |
| chr8-90387696-DEL-82 | chr8-90387702-DEL-54 |
| chr8-90402759-DEL-90 | chr8-90402760-DEL-90 |
| chr8-90431834-DEL-54 | chr8-90431835-DEL-66 |
| chr8-90445818-DEL-89 | chr8-90445808-DEL-62 |
| chr8-90458010-DEL-61 | chr8-90458011-DEL-61 |
| chr8-90461225-DEL-59 | chr8-90461234-DEL-50 |
| chr8-90479249-DEL-92 | chr8-90479234-DEL-96 |
| chr8-90487284-DEL-76 | chr8-90487284-DEL-76 |
| chr8-90532602-DEL-350 | chr8-90532602-DEL-350 |
| chr8-90569639-DEL-152 | chr8-90569639-DEL-152 |
| chr8-90582779-DEL-201 | chr8-90582780-DEL-201 |
| chr8-90644648-DEL-7257 | chr8-90644649-DEL-7257 |
| chr8-90672801-DEL-2030 | chr8-90672802-DEL-2030 |
| chr8-90675999-DEL-1494 | chr8-90676000-DEL-1494 |
| chr8-90682494-DEL-125 | chr8-90682495-DEL-125 |
| chr8-90697741-DEL-1070 | chr8-90697742-DEL-1070 |
| chr8-90717160-DEL-52 | chr8-90717149-DEL-69 |
| chr8-90717777-DEL-116 | chr8-90717778-DEL-114 |
| chr8-90721575-DEL-262 | chr8-90721576-DEL-262 |
| chr8-90754296-DEL-55 | chr8-90754297-DEL-55 |
| chr8-90759947-DEL-1611 | chr8-90759948-DEL-1611 |
| chr8-90800076-DEL-67 | chr8-90800066-DEL-69 |
| chr8-90809715-DEL-777 | chr8-90809716-DEL-777 |
| chr8-90810854-DEL-2124 | chr8-90810855-DEL-2124 |
| chr8-90814317-DEL-7263 | chr8-90814318-DEL-7263 |
| chr8-90825762-DEL-462 | chr8-90825763-DEL-462 |
| chr8-90889677-DEL-3698 | chr8-90889678-DEL-3698 |
| chr8-90942159-DEL-6174 | chr8-90942160-DEL-6174 |
| chr8-90983511-DEL-267 | chr8-90983495-DEL-288 |
| chr8-90987484-DEL-69 | chr8-90987485-DEL-69 |
| chr8-90989197-DEL-104 | chr8-90989198-DEL-104 |
| chr8-91010199-DEL-639 | chr8-91010200-DEL-639 |
| chr8-91047298-DEL-400 | chr8-91047299-DEL-400 |
| chr8-91100931-DEL-80 | chr8-91100932-DEL-80 |
| chr8-91108725-DEL-7118 | chr8-91108726-DEL-7118 |
| chr8-91172497-DEL-65 | chr8-91172503-DEL-65 |
| chr8-91174989-DEL-56 | chr8-91174978-DEL-68 |
| chr8-91225621-DEL-51 | chr8-91225620-DEL-58 |
| chr8-91233736-DEL-105 | chr8-91233736-DEL-105 |
| chr8-91290876-DEL-229 | chr8-91290877-DEL-229 |
| chr8-91293464-DEL-76 | chr8-91293465-DEL-76 |
| chr8-91299625-DEL-130 | chr8-91299626-DEL-130 |
| chr8-91340039-DEL-85 | chr8-91340040-DEL-85 |
| chr8-91341127-DEL-87 | chr8-91341128-DEL-87 |
| chr8-91343728-DEL-1031 | chr8-91343729-DEL-1031 |
| chr8-91354220-DEL-220 | chr8-91354221-DEL-220 |
| chr8-91360164-DEL-65 | chr8-91360148-DEL-81 |
| chr8-91379591-DEL-146 | chr8-91379592-DEL-146 |
| chr8-91381198-DEL-64 | chr8-91381199-DEL-64 |
| chr8-91399406-DEL-58 | chr8-91399407-DEL-58 |
| chr8-91405108-DEL-115 | chr8-91405109-DEL-115 |
| chr8-91408459-DEL-82 | chr8-91408462-DEL-68 |
| chr8-91416761-DEL-126 | chr8-91416766-DEL-122 |
| chr8-91418345-DEL-53 | chr8-91418352-DEL-53 |
| chr8-91436094-DEL-123 | chr8-91436095-DEL-124 |
| chr8-91437926-DEL-95 | chr8-91437931-DEL-91 |
| chr8-91440704-DEL-63 | chr8-91440705-DEL-63 |
| chr8-91455211-DEL-66 | chr8-91455215-DEL-63 |
| chr8-91461021-DEL-348 | chr8-91461023-DEL-346 |
| chr8-91471565-DEL-84 | chr8-91471566-DEL-84 |
| chr8-91494991-DEL-90 | chr8-91494992-DEL-90 |

| | |
|---|---|
| chr8-91517399-DEL-144 | chr8-91517400-DEL-144 |
| chr8-91518202-DEL-88 | chr8-91518225-DEL-66 |
| chr8-91519150-DEL-443 | chr8-91519151-DEL-443 |
| chr8-91520980-DEL-54 | chr8-91520981-DEL-54 |
| chr8-91532272-DEL-255 | chr8-91532271-DEL-256 |
| chr8-91533812-DEL-202 | chr8-91533806-DEL-203 |
| chr8-91553973-DEL-57 | chr8-91553974-DEL-54 |
| chr8-91561682-DEL-54 | chr8-91561683-DEL-54 |
| chr8-91570703-DEL-114 | chr8-91570703-DEL-114 |
| chr8-91582740-DEL-131 | chr8-91582747-DEL-115 |
| chr8-91611023-DEL-195 | chr8-91611024-DEL-195 |
| chr8-91612914-DEL-5672 | chr8-91612915-DEL-5672 |
| chr8-91642395-DEL-66 | chr8-91642396-DEL-52 |
| chr8-91672623-DEL-204 | chr8-91672617-DEL-205 |
| chr8-91673715-DEL-191 | chr8-91673711-DEL-189 |
| chr8-91708168-DEL-81 | chr8-91708218-DEL-51 |
| chr8-91713037-DEL-1211 | chr8-91713038-DEL-1211 |
| chr8-91719409-DEL-58 | chr8-91719411-DEL-57 |
| chr8-91729404-DEL-213 | chr8-91729405-DEL-213 |
| chr8-91740961-DEL-83 | chr8-91740962-DEL-82 |
| chr8-91744212-DEL-51 | chr8-91744201-DEL-52 |
| chr8-91768403-DEL-198 | chr8-91768404-DEL-198 |
| chr8-91772342-DEL-125 | chr8-91772359-DEL-124 |
| chr8-91815366-DEL-931 | chr8-91815367-DEL-931 |
| chr8-91824377-DEL-199 | chr8-91824378-DEL-199 |
| chr8-91858789-DEL-208 | chr8-91858790-DEL-208 |
| chr8-91863960-DEL-134 | chr8-91863956-DEL-135 |
| chr8-91866710-DEL-115 | chr8-91866727-DEL-103 |
| chr8-91878850-DEL-75 | chr8-91878831-DEL-78 |
| chr8-91880835-DEL-68 | chr8-91880836-DEL-68 |
| chr8-91882533-DEL-109 | chr8-91882534-DEL-109 |
| chr8-91899380-DEL-74 | chr8-91899379-DEL-76 |
| chr8-91913154-DEL-62 | chr8-91913155-DEL-62 |
| chr8-91970565-DEL-51 | chr8-91970565-DEL-51 |
| chr8-92021626-DEL-89 | chr8-92021626-DEL-87 |
| chr8-92052567-DEL-196 | chr8-92052568-DEL-196 |
| chr8-92060156-DEL-585 | chr8-92060169-DEL-588 |
| chr8-92061740-DEL-189 | chr8-92061741-DEL-189 |
| chr8-92063387-DEL-172 | chr8-92063388-DEL-129 |
| chr8-92076132-DEL-72 | chr8-92076133-DEL-72 |
| chr8-92083435-DEL-58 | chr8-92083439-DEL-69 |
| chr8-92088817-DEL-283 | chr8-92088818-DEL-283 |
| chr8-92124481-DEL-57 | chr8-92124482-DEL-57 |
| chr8-92140964-DEL-513 | chr8-92140965-DEL-513 |
| chr8-92142252-DEL-118 | chr8-92142253-DEL-118 |
| chr8-92143578-DEL-6915 | chr8-92143579-DEL-6915 |
| chr8-92155769-DEL-64 | chr8-92155770-DEL-64 |
| chr8-92157701-DEL-3426 | chr8-92157702-DEL-3426 |
| chr8-92210409-DEL-105 | chr8-92210426-DEL-65 |
| chr8-92218978-DEL-495 | chr8-92218979-DEL-495 |
| chr8-92239782-DEL-213 | chr8-92239783-DEL-213 |
| chr8-92264780-DEL-61 | chr8-92264780-DEL-62 |
| chr8-92278005-DEL-114 | chr8-92278006-DEL-114 |
| chr8-92313906-DEL-54 | chr8-92313887-DEL-50 |
| chr8-92315996-DEL-64 | chr8-92315999-DEL-62 |
| chr8-92330011-DEL-1863 | chr8-92330012-DEL-1863 |
| chr8-92345942-DEL-51 | chr8-92345943-DEL-51 |
| chr8-92347879-DEL-90 | chr8-92347880-DEL-55 |
| chr8-92376282-DEL-63 | chr8-92376283-DEL-63 |
| chr8-92407350-DEL-87 | chr8-92407351-DEL-87 |
| chr8-92428629-DEL-208 | chr8-92428630-DEL-208 |
| chr8-92433644-DEL-58 | chr8-92433616-DEL-76 |
| chr8-92451935-DEL-108 | chr8-92451941-DEL-90 |
| chr8-92452566-DEL-62 | chr8-92452522-DEL-75 |
| chr8-92456372-DEL-686 | chr8-92456373-DEL-686 |
| chr8-92465553-DEL-113 | chr8-92465566-DEL-89 |
| chr8-92467467-DEL-2829 | chr8-92467468-DEL-2829 |
| chr8-92470982-DEL-93 | chr8-92470976-DEL-55 |
| chr8-92475368-DEL-115 | chr8-92475369-DEL-115 |
| chr8-92487042-DEL-1608 | chr8-92487043-DEL-1608 |
| chr8-92501155-DEL-53 | chr8-92501156-DEL-52 |
| chr8-92548140-DEL-119 | chr8-92548141-DEL-119 |
| chr8-92551124-DEL-66 | chr8-92551125-DEL-66 |
| chr8-92583584-DEL-195 | chr8-92583600-DEL-180 |
| chr8-92622760-DEL-56 | chr8-92622761-DEL-56 |
| chr8-92647355-DEL-69 | chr8-92647343-DEL-81 |
| chr8-92656260-DEL-181 | chr8-92656307-DEL-163 |
| chr8-92660745-DEL-54 | chr8-92660746-DEL-54 |
| chr8-92661296-DEL-1349 | chr8-92661296-DEL-1349 |
| chr8-92740170-DEL-311 | chr8-92740170-DEL-312 |

| | |
|---|---|
| chr8-92779153-DEL-56 | chr8-92779154-DEL-61 |
| chr8-92797958-DEL-56 | chr8-92797959-DEL-56 |
| chr8-92811121-DEL-103 | chr8-92811122-DEL-103 |
| chr8-92819823-DEL-58 | chr8-92819830-DEL-52 |
| chr8-92820642-DEL-89 | chr8-92820643-DEL-94 |
| chr8-92833693-DEL-6006 | chr8-92833694-DEL-6006 |
| chr8-92869421-DEL-399 | chr8-92869422-DEL-399 |
| chr8-92901331-DEL-52 | chr8-92901319-DEL-65 |
| chr8-92905398-DEL-74 | chr8-92905399-DEL-74 |
| chr8-92905896-DEL-99 | chr8-92905897-DEL-99 |
| chr8-92911272-DEL-91 | chr8-92911273-DEL-91 |
| chr8-92935803-DEL-1286 | chr8-92935801-DEL-1286 |
| chr8-92937178-DEL-315 | chr8-92937178-DEL-315 |
| chr8-92977688-DEL-304 | chr8-92977689-DEL-304 |
| chr8-92980599-DEL-299 | chr8-92980600-DEL-299 |
| chr8-92987965-DEL-138 | chr8-92987953-DEL-139 |
| chr8-93009946-DEL-70 | chr8-93009928-DEL-67 |
| chr8-93040826-DEL-211 | chr8-93040827-DEL-211 |
| chr8-93055930-DEL-69 | chr8-93055931-DEL-55 |
| chr8-93065739-DEL-401 | chr8-93065740-DEL-401 |
| chr8-93097512-DEL-322 | chr8-93097513-DEL-322 |
| chr8-93133938-DEL-258 | chr8-93133939-DEL-258 |
| chr8-93138338-DEL-327 | chr8-93138339-DEL-327 |
| chr8-93153360-DEL-96 | chr8-93153361-DEL-114 |
| chr8-93160671-DEL-52 | chr8-93160672-DEL-60 |
| chr8-93174710-DEL-51 | chr8-93174711-DEL-51 |
| chr8-93192653-DEL-61 | chr8-93192654-DEL-61 |
| chr8-93202848-DEL-434 | chr8-93202849-DEL-434 |
| chr8-93204480-DEL-54 | chr8-93204492-DEL-51 |
| chr8-93211738-DEL-58 | chr8-93211735-DEL-62 |
| chr8-93215961-DEL-94 | chr8-93215962-DEL-70 |
| chr8-93243699-DEL-506 | chr8-93243700-DEL-506 |
| chr8-93286214-DEL-497 | chr8-93286189-DEL-492 |
| chr8-93307104-DEL-1693 | chr8-93307105-DEL-1693 |
| chr8-93312111-DEL-6545 | chr8-93312112-DEL-6545 |
| chr8-93339358-DEL-102 | chr8-93339343-DEL-105 |
| chr8-93368772-DEL-133 | chr8-93368768-DEL-93 |
| chr8-93386865-DEL-217 | chr8-93386866-DEL-217 |
| chr8-93394828-DEL-59 | chr8-93394824-DEL-56 |
| chr8-93415219-DEL-66 | chr8-93415218-DEL-56 |
| chr8-93438043-DEL-2551 | chr8-93438044-DEL-2551 |
| chr8-93442342-DEL-1287 | chr8-93442343-DEL-1287 |
| chr8-93475319-DEL-145 | chr8-93475320-DEL-145 |
| chr8-93490683-DEL-132 | chr8-93490692-DEL-130 |
| chr8-93496130-DEL-201 | chr8-93496129-DEL-202 |
| chr8-93542981-DEL-60 | chr8-93542982-DEL-60 |
| chr8-93555586-DEL-1525 | chr8-93555587-DEL-1525 |
| chr8-93569566-DEL-54 | chr8-93569567-DEL-72 |
| chr8-93585433-DEL-389 | chr8-93585434-DEL-389 |
| chr8-93586366-DEL-217 | chr8-93586367-DEL-217 |
| chr8-93590272-DEL-217 | chr8-93590273-DEL-217 |
| chr8-93607682-DEL-175 | chr8-93607684-DEL-161 |
| chr8-93615295-DEL-92 | chr8-93615296-DEL-92 |
| chr8-93624749-DEL-99 | chr8-93624747-DEL-101 |
| chr8-93639906-DEL-85 | chr8-93639863-DEL-112 |
| chr8-93641562-DEL-202 | chr8-93641562-DEL-202 |
| chr8-93648898-DEL-81 | chr8-93648870-DEL-72 |
| chr8-93653205-DEL-122 | chr8-93653206-DEL-122 |
| chr8-93666694-DEL-64 | chr8-93666695-DEL-68 |
| chr8-93703736-DEL-64 | chr8-93703737-DEL-64 |
| chr8-93708236-DEL-53 | chr8-93708237-DEL-53 |
| chr8-93714983-DEL-79 | chr8-93714984-DEL-76 |
| chr8-93723174-DEL-184 | chr8-93723181-DEL-178 |
| chr8-93728640-DEL-218 | chr8-93728636-DEL-214 |
| chr8-93754219-DEL-57 | chr8-93754220-DEL-61 |
| chr8-93769759-DEL-1053 | chr8-93769760-DEL-1053 |
| chr8-93776874-DEL-959 | chr8-93776875-DEL-959 |
| chr8-93789461-DEL-53 | chr8-93789449-DEL-54 |
| chr8-93809800-DEL-2830 | chr8-93809800-DEL-2830 |
| chr8-93815379-DEL-139 | chr8-93815380-DEL-139 |
| chr8-93825996-DEL-1020 | chr8-93825997-DEL-1020 |
| chr8-93829936-DEL-230 | chr8-93829937-DEL-231 |
| chr8-93837823-DEL-6584 | chr8-93837824-DEL-6584 |
| chr8-93848212-DEL-57 | chr8-93848213-DEL-63 |
| chr8-93853818-DEL-178 | chr8-93853819-DEL-178 |
| chr8-93883724-DEL-1588 | chr8-93883725-DEL-1588 |
| chr8-93926532-DEL-164 | chr8-93926494-DEL-206 |
| chr8-93930184-DEL-133 | chr8-93930185-DEL-133 |
| chr8-93964111-DEL-1454 | chr8-93964120-DEL-1452 |
| chr8-93968491-DEL-478 | chr8-93968492-DEL-478 |

| | |
|---|---|
| chr8-93977806-DEL-486 | chr8-93977808-DEL-486 |
| chr8-94019458-DEL-6361 | chr8-94019459-DEL-6361 |
| chr8-94045214-DEL-55 | chr8-94045208-DEL-75 |
| chr8-94049099-DEL-65 | chr8-94049094-DEL-63 |
| chr8-94055880-DEL-105 | chr8-94055880-DEL-105 |
| chr8-94068266-DEL-6612 | chr8-94068260-DEL-6613 |
| chr8-94090570-DEL-84 | chr8-94090549-DEL-55 |
| chr8-94117531-DEL-419 | chr8-94117532-DEL-419 |
| chr8-94176558-DEL-198 | chr8-94176559-DEL-198 |
| chr8-94184758-DEL-3126 | chr8-94184759-DEL-3126 |
| chr8-94214321-DEL-81 | chr8-94214325-DEL-78 |
| chr8-94217964-DEL-89 | chr8-94217921-DEL-90 |
| chr8-94232351-DEL-6305 | chr8-94232352-DEL-6305 |
| chr8-94304003-DEL-7063 | chr8-94304004-DEL-7063 |
| chr8-94315738-DEL-126 | chr8-94315723-DEL-126 |
| chr8-94329894-DEL-475 | chr8-94329895-DEL-475 |
| chr8-94338027-DEL-103 | chr8-94338028-DEL-103 |
| chr8-94367620-DEL-56 | chr8-94367606-DEL-56 |
| chr8-94399285-DEL-59 | chr8-94399292-DEL-53 |
| chr8-94405259-DEL-929 | chr8-94405260-DEL-929 |
| chr8-94406272-DEL-62 | chr8-94406273-DEL-62 |
| chr8-94412234-DEL-63 | chr8-94412235-DEL-63 |
| chr8-94442752-DEL-71 | chr8-94442755-DEL-70 |
| chr8-94446071-DEL-880 | chr8-94446073-DEL-878 |
| chr8-94446572-DEL-56 | chr8-94446573-DEL-58 |
| chr8-94447913-DEL-440 | chr8-94447914-DEL-440 |
| chr8-94463062-DEL-60 | chr8-94463063-DEL-54 |
| chr8-94601068-DEL-746 | chr8-94601059-DEL-747 |
| chr8-94636482-DEL-53 | chr8-94636483-DEL-53 |
| chr8-94653516-DEL-207 | chr8-94653516-DEL-207 |
| chr8-94685584-DEL-127 | chr8-94685579-DEL-128 |
| chr8-94714923-DEL-83 | chr8-94714924-DEL-83 |
| chr8-94735391-DEL-69 | chr8-94735354-DEL-57 |
| chr8-94767930-DEL-63 | chr8-94767931-DEL-63 |
| chr8-94802673-DEL-231 | chr8-94802675-DEL-230 |
| chr8-94804222-DEL-207 | chr8-94804223-DEL-207 |
| chr8-94822464-DEL-149 | chr8-94822461-DEL-157 |
| chr8-94838353-DEL-52 | chr8-94838354-DEL-52 |
| chr8-94840370-DEL-56 | chr8-94840371-DEL-56 |
| chr8-94842514-DEL-182 | chr8-94842514-DEL-180 |
| chr8-94844591-DEL-62 | chr8-94844609-DEL-62 |
| chr8-94872054-DEL-86 | chr8-94872042-DEL-60 |
| chr8-94873336-DEL-193 | chr8-94873337-DEL-193 |
| chr8-94908052-DEL-205 | chr8-94908066-DEL-207 |
| chr8-94910977-DEL-67 | chr8-94910978-DEL-67 |
| chr8-94911607-DEL-219 | chr8-94911606-DEL-219 |
| chr8-94922437-DEL-149 | chr8-94922438-DEL-149 |
| chr8-94925310-DEL-544 | chr8-94925337-DEL-522 |
| chr8-94933844-DEL-192 | chr8-94933845-DEL-192 |
| chr8-94949564-DEL-794 | chr8-94949565-DEL-794 |
| chr8-94985742-DEL-85 | chr8-94985740-DEL-88 |
| chr8-95095788-DEL-86 | chr8-95095797-DEL-78 |
| chr8-95130200-DEL-2519 | chr8-95130201-DEL-2519 |
| chr8-95145943-DEL-414 | chr8-95145944-DEL-414 |
| chr8-95152009-DEL-944 | chr8-95152010-DEL-944 |
| chr8-95173017-DEL-510 | chr8-95173018-DEL-510 |
| chr8-95176436-DEL-85 | chr8-95176437-DEL-60 |
| chr8-95279230-DEL-75 | chr8-95279232-DEL-74 |
| chr8-95343133-DEL-5367 | chr8-95343134-DEL-5367 |
| chr8-95377146-DEL-193 | chr8-95377147-DEL-193 |
| chr8-95378815-DEL-203 | chr8-95378816-DEL-203 |
| chr8-95426762-DEL-52 | chr8-95426763-DEL-52 |
| chr8-95428268-DEL-975 | chr8-95428274-DEL-969 |
| chr8-95447698-DEL-355 | chr8-95447699-DEL-355 |
| chr8-95474824-DEL-62 | chr8-95474825-DEL-62 |
| chr8-95521887-DEL-4854 | chr8-95521888-DEL-4854 |
| chr8-95553354-DEL-235 | chr8-95553355-DEL-235 |
| chr8-95560856-DEL-195 | chr8-95560856-DEL-196 |
| chr8-95561549-DEL-151 | chr8-95561550-DEL-151 |
| chr8-95566171-DEL-244 | chr8-95566172-DEL-244 |
| chr8-95578132-DEL-204 | chr8-95578131-DEL-205 |
| chr8-95608914-DEL-76 | chr8-95608915-DEL-76 |
| chr8-95658506-DEL-171 | chr8-95658507-DEL-171 |
| chr8-95680502-DEL-117 | chr8-95680503-DEL-117 |
| chr8-95683278-DEL-68 | chr8-95683278-DEL-72 |
| chr8-95699124-DEL-70 | chr8-95699125-DEL-70 |
| chr8-95748151-DEL-1020 | chr8-95748152-DEL-1020 |
| chr8-95754650-DEL-71 | chr8-95754651-DEL-71 |
| chr8-95765022-DEL-546 | chr8-95765023-DEL-546 |
| chr8-95775967-DEL-326 | chr8-95775968-DEL-326 |

| | |
|---|---|
| chr8-95779120-DEL-129 | chr8-95779119-DEL-130 |
| chr8-95841300-DEL-6439 | chr8-95841301-DEL-6439 |
| chr8-95870666-DEL-176 | chr8-95870667-DEL-176 |
| chr8-95887767-DEL-192 | chr8-95887768-DEL-192 |
| chr8-95894864-DEL-201 | chr8-95894865-DEL-201 |
| chr8-95907348-DEL-52 | chr8-95907349-DEL-52 |
| chr8-95916955-DEL-64 | chr8-95916959-DEL-61 |
| chr8-95925142-DEL-110 | chr8-95925123-DEL-130 |
| chr8-95954621-DEL-4110 | chr8-95954601-DEL-4131 |
| chr8-95962015-DEL-365 | chr8-95962012-DEL-366 |
| chr8-95985057-DEL-66 | chr8-95985062-DEL-62 |
| chr8-96002532-DEL-76 | chr8-96002511-DEL-76 |
| chr8-96010114-DEL-194 | chr8-96010115-DEL-194 |
| chr8-96091490-DEL-65 | chr8-96091490-DEL-52 |
| chr8-96126604-DEL-72 | chr8-96126605-DEL-72 |
| chr8-96138743-DEL-6446 | chr8-96138752-DEL-6438 |
| chr8-96150325-DEL-157 | chr8-96150326-DEL-157 |
| chr8-96170432-DEL-196 | chr8-96170433-DEL-196 |
| chr8-96180896-DEL-158 | chr8-96180897-DEL-158 |
| chr8-96238420-DEL-95 | chr8-96238421-DEL-95 |
| chr8-96241990-DEL-464 | chr8-96241967-DEL-484 |
| chr8-96244281-DEL-574 | chr8-96244282-DEL-562 |
| chr8-96245521-DEL-3160 | chr8-96245522-DEL-3160 |
| chr8-96276356-DEL-63 | chr8-96276357-DEL-63 |
| chr8-96290162-DEL-58 | chr8-96290163-DEL-58 |
| chr8-96308763-DEL-1489 | chr8-96308763-DEL-1489 |
| chr8-96321851-DEL-51 | chr8-96321852-DEL-51 |
| chr8-96325489-DEL-211 | chr8-96325490-DEL-211 |
| chr8-96361532-DEL-84 | chr8-96361535-DEL-94 |
| chr8-96363493-DEL-220 | chr8-96363494-DEL-220 |
| chr8-96378581-DEL-58 | chr8-96378582-DEL-58 |
| chr8-96381679-DEL-203 | chr8-96381680-DEL-203 |
| chr8-96384454-DEL-68 | chr8-96384458-DEL-64 |
| chr8-96409110-DEL-218 | chr8-96409111-DEL-218 |
| chr8-96413178-DEL-502 | chr8-96413178-DEL-502 |
| chr8-96420035-DEL-54 | chr8-96420036-DEL-60 |
| chr8-96436768-DEL-56 | chr8-96436769-DEL-56 |
| chr8-96476447-DEL-314 | chr8-96476448-DEL-314 |
| chr8-96520663-DEL-221 | chr8-96520664-DEL-221 |
| chr8-96524061-DEL-168 | chr8-96524064-DEL-170 |
| chr8-96539384-DEL-219 | chr8-96539416-DEL-194 |
| chr8-96551968-DEL-817 | chr8-96551968-DEL-817 |
| chr8-96573005-DEL-340 | chr8-96573006-DEL-340 |
| chr8-96581609-DEL-745 | chr8-96581610-DEL-745 |
| chr8-96582903-DEL-191 | chr8-96582906-DEL-189 |
| chr8-96603144-DEL-204 | chr8-96603145-DEL-203 |
| chr8-96620672-DEL-198 | chr8-96620673-DEL-198 |
| chr8-96627022-DEL-167 | chr8-96627014-DEL-168 |
| chr8-96638439-DEL-1865 | chr8-96638440-DEL-1865 |
| chr8-96643548-DEL-211 | chr8-96643561-DEL-199 |
| chr8-96647674-DEL-204 | chr8-96647675-DEL-204 |
| chr8-96650888-DEL-185 | chr8-96650889-DEL-185 |
| chr8-96654133-DEL-116 | chr8-96654134-DEL-112 |
| chr8-96654884-DEL-565 | chr8-96654876-DEL-568 |
| chr8-96657746-DEL-83 | chr8-96657747-DEL-83 |
| chr8-96678798-DEL-133 | chr8-96678814-DEL-122 |
| chr8-96681570-DEL-574 | chr8-96681571-DEL-574 |
| chr8-96686996-DEL-197 | chr8-96686997-DEL-197 |
| chr8-96712702-DEL-88 | chr8-96712699-DEL-90 |
| chr8-96723808-DEL-2447 | chr8-96723809-DEL-2447 |
| chr8-96734966-DEL-441 | chr8-96734967-DEL-441 |
| chr8-96745436-DEL-104 | chr8-96745437-DEL-104 |
| chr8-96769653-DEL-403 | chr8-96769654-DEL-400 |
| chr8-96779266-DEL-4084 | chr8-96779267-DEL-4084 |
| chr8-96786954-DEL-6591 | chr8-96786955-DEL-6591 |
| chr8-96815343-DEL-168 | chr8-96815344-DEL-168 |
| chr8-96838602-DEL-2124 | chr8-96838603-DEL-2124 |
| chr8-96841160-DEL-540 | chr8-96841161-DEL-540 |
| chr8-96875585-DEL-144 | chr8-96875628-DEL-102 |
| chr8-96919929-DEL-6284 | chr8-96919930-DEL-6284 |
| chr8-96959900-DEL-436 | chr8-96959901-DEL-436 |
| chr8-96995973-DEL-112 | chr8-96995984-DEL-109 |
| chr8-97005665-DEL-190 | chr8-97005666-DEL-190 |
| chr8-97065956-DEL-64 | chr8-97065957-DEL-64 |
| chr8-97083769-DEL-67 | chr8-97083770-DEL-67 |
| chr8-97137323-DEL-805 | chr8-97137324-DEL-805 |
| chr8-97151569-DEL-360 | chr8-97151570-DEL-360 |
| chr8-97190821-DEL-92 | chr8-97190822-DEL-92 |
| chr8-97228104-DEL-6217 | chr8-97228105-DEL-6217 |
| chr8-97238970-DEL-427 | chr8-97238971-DEL-427 |

| | |
|---|---|
| chr8-97258325-DEL-896 | chr8-97258326-DEL-896 |
| chr8-97306325-DEL-267 | chr8-97306326-DEL-267 |
| chr8-97330706-DEL-53 | chr8-97330709-DEL-51 |
| chr8-97339379-DEL-70 | chr8-97339384-DEL-66 |
| chr8-97367446-DEL-212 | chr8-97367450-DEL-211 |
| chr8-97382639-DEL-56 | chr8-97382641-DEL-54 |
| chr8-97385781-DEL-665 | chr8-97385782-DEL-665 |
| chr8-97397297-DEL-825 | chr8-97397297-DEL-825 |
| chr8-97399436-DEL-909 | chr8-97399436-DEL-909 |
| chr8-97416555-DEL-122 | chr8-97416576-DEL-130 |
| chr8-97472310-DEL-52 | chr8-97472311-DEL-52 |
| chr8-97479796-DEL-90 | chr8-97479797-DEL-90 |
| chr8-97492228-DEL-8415 | chr8-97492229-DEL-8415 |
| chr8-97516110-DEL-133 | chr8-97516111-DEL-133 |
| chr8-97516681-DEL-58 | chr8-97516682-DEL-58 |
| chr8-97525807-DEL-161 | chr8-97525808-DEL-161 |
| chr8-97538127-DEL-91 | chr8-97538127-DEL-91 |
| chr8-97549550-DEL-1421 | chr8-97549551-DEL-1421 |
| chr8-97563610-DEL-201 | chr8-97563611-DEL-201 |
| chr8-97571784-DEL-56 | chr8-97571784-DEL-52 |
| chr8-97577792-DEL-235 | chr8-97577760-DEL-268 |
| chr8-97582504-DEL-114 | chr8-97582505-DEL-114 |
| chr8-97632986-DEL-1947 | chr8-97632987-DEL-1947 |
| chr8-97649867-DEL-105 | chr8-97649868-DEL-105 |
| chr8-97655891-DEL-92 | chr8-97655904-DEL-86 |
| chr8-97660019-DEL-6224 | chr8-97660006-DEL-6225 |
| chr8-97675430-DEL-419 | chr8-97675431-DEL-419 |
| chr8-97703002-DEL-62 | chr8-97703003-DEL-62 |
| chr8-97712578-DEL-282 | chr8-97712579-DEL-282 |
| chr8-97713907-DEL-242 | chr8-97713908-DEL-242 |
| chr8-97726149-DEL-60 | chr8-97726150-DEL-60 |
| chr8-97772290-DEL-1780 | chr8-97772291-DEL-1780 |
| chr8-97772854-DEL-89 | chr8-97772855-DEL-89 |
| chr8-97774454-DEL-475 | chr8-97774455-DEL-475 |
| chr8-97775176-DEL-562 | chr8-97775177-DEL-562 |
| chr8-97804509-DEL-1083 | chr8-97804510-DEL-1083 |
| chr8-97807808-DEL-68 | chr8-97807809-DEL-68 |
| chr8-97830740-DEL-788 | chr8-97830741-DEL-788 |
| chr8-97836691-DEL-6583 | chr8-97836692-DEL-6583 |
| chr8-97848754-DEL-106 | chr8-97848755-DEL-103 |
| chr8-97896177-DEL-52 | chr8-97896178-DEL-52 |
| chr8-97896569-DEL-247 | chr8-97896570-DEL-247 |
| chr8-97900107-DEL-287 | chr8-97900126-DEL-287 |
| chr8-97939360-DEL-1134 | chr8-97939350-DEL-1152 |
| chr8-97941786-DEL-191 | chr8-97941833-DEL-143 |
| chr8-97965915-DEL-6400 | chr8-97965916-DEL-6400 |
| chr8-97999448-DEL-219 | chr8-97999457-DEL-203 |
| chr8-98005221-DEL-173 | chr8-98005222-DEL-172 |
| chr8-98010663-DEL-325 | chr8-98010664-DEL-323 |
| chr8-98038754-DEL-55 | chr8-98038747-DEL-70 |
| chr8-98043999-DEL-99 | chr8-98044000-DEL-99 |
| chr8-98069078-DEL-173 | chr8-98069040-DEL-169 |
| chr8-98116972-DEL-140 | chr8-98117009-DEL-104 |
| chr8-98128012-DEL-51 | chr8-98128013-DEL-51 |
| chr8-98170075-DEL-64 | chr8-98170076-DEL-64 |
| chr8-98229800-DEL-2607 | chr8-98229800-DEL-2608 |
| chr8-98232152-DEL-255 | chr8-98232153-DEL-255 |
| chr8-98235477-DEL-1604 | chr8-98235478-DEL-1604 |
| chr8-98269263-DEL-98 | chr8-98269303-DEL-84 |
| chr8-98271908-DEL-6704 | chr8-98271909-DEL-6704 |
| chr8-98299611-DEL-98 | chr8-98299612-DEL-98 |
| chr8-98313432-DEL-56 | chr8-98313419-DEL-96 |
| chr8-98317611-DEL-993 | chr8-98317614-DEL-991 |
| chr8-98320075-DEL-51 | chr8-98320076-DEL-51 |
| chr8-98349555-DEL-2541 | chr8-98349556-DEL-2541 |
| chr8-98358097-DEL-54 | chr8-98358080-DEL-76 |
| chr8-98392175-DEL-79 | chr8-98392176-DEL-79 |
| chr8-98392916-DEL-309 | chr8-98392918-DEL-308 |
| chr8-98395985-DEL-55 | chr8-98395951-DEL-52 |
| chr8-98402393-DEL-68 | chr8-98402418-DEL-57 |
| chr8-98407126-DEL-272 | chr8-98407125-DEL-259 |
| chr8-98423189-DEL-3801 | chr8-98423190-DEL-3801 |
| chr8-98444621-DEL-4057 | chr8-98444622-DEL-4057 |
| chr8-98472171-DEL-504 | chr8-98472172-DEL-504 |
| chr8-98501388-DEL-369 | chr8-98501422-DEL-330 |
| chr8-98532585-DEL-417 | chr8-98532586-DEL-417 |
| chr8-98535138-DEL-826 | chr8-98535133-DEL-826 |
| chr8-98537220-DEL-171 | chr8-98537221-DEL-171 |
| chr8-98561790-DEL-446 | chr8-98561791-DEL-446 |
| chr8-98580937-DEL-69 | chr8-98580938-DEL-69 |

| | |
|---|---|
| chr8-98582128-DEL-52 | chr8-98582113-DEL-68 |
| chr8-98648417-DEL-194 | chr8-98648418-DEL-194 |
| chr8-98649466-DEL-370 | chr8-98649467-DEL-370 |
| chr8-98704127-DEL-1095 | chr8-98704128-DEL-1095 |
| chr8-98713396-DEL-325 | chr8-98713397-DEL-325 |
| chr8-98734315-DEL-2216 | chr8-98734316-DEL-2216 |
| chr8-98750443-DEL-170 | chr8-98750444-DEL-170 |
| chr8-98777386-DEL-58 | chr8-98777384-DEL-88 |
| chr8-98814355-DEL-91 | chr8-98814355-DEL-90 |
| chr8-98830664-DEL-144 | chr8-98830660-DEL-149 |
| chr8-98831635-DEL-52 | chr8-98831637-DEL-62 |
| chr8-98852297-DEL-3712 | chr8-98852298-DEL-3712 |
| chr8-98865938-DEL-56 | chr8-98865927-DEL-68 |
| chr8-98866427-DEL-980 | chr8-98866428-DEL-980 |
| chr8-98867812-DEL-152 | chr8-98867792-DEL-173 |
| chr8-98874051-DEL-646 | chr8-98874093-DEL-602 |
| chr8-98881276-DEL-58 | chr8-98881252-DEL-74 |
| chr8-98892884-DEL-747 | chr8-98892885-DEL-747 |
| chr8-98896640-DEL-60 | chr8-98896650-DEL-51 |
| chr8-98900962-DEL-7073 | chr8-98900963-DEL-7073 |
| chr8-98931568-DEL-109 | chr8-98931569-DEL-110 |
| chr8-98966832-DEL-65 | chr8-98966823-DEL-66 |
| chr8-98975315-DEL-70 | chr8-98975333-DEL-52 |
| chr8-98976217-DEL-6437 | chr8-98976218-DEL-6437 |
| chr8-98990778-DEL-72 | chr8-98990779-DEL-72 |
| chr8-99009477-DEL-7240 | chr8-99009477-DEL-7240 |
| chr8-99037085-DEL-1041 | chr8-99037086-DEL-1041 |
| chr8-99049875-DEL-66 | chr8-99049872-DEL-62 |
| chr8-99058413-DEL-361 | chr8-99058424-DEL-351 |
| chr8-99086477-DEL-2801 | chr8-99086478-DEL-2801 |
| chr8-99089656-DEL-188 | chr8-99089657-DEL-188 |
| chr8-99102159-DEL-1925 | chr8-99102160-DEL-1925 |
| chr8-99155743-DEL-6360 | chr8-99155744-DEL-6360 |
| chr8-99175254-DEL-85 | chr8-99175255-DEL-86 |
| chr8-99178842-DEL-778 | chr8-99178843-DEL-778 |
| chr8-99208386-DEL-98 | chr8-99208386-DEL-98 |
| chr8-99237445-DEL-1250 | chr8-99237446-DEL-1250 |
| chr8-99280406-DEL-2847 | chr8-99280407-DEL-2847 |
| chr8-99280518-DEL-1136 | chr8-99280519-DEL-1136 |
| chr8-99368631-DEL-154 | chr8-99368632-DEL-154 |
| chr8-99410197-DEL-771 | chr8-99410198-DEL-771 |
| chr8-99419396-DEL-6848 | chr8-99419397-DEL-6848 |
| chr8-99425746-DEL-424 | chr8-99425747-DEL-424 |
| chr8-99448718-DEL-130 | chr8-99448719-DEL-84 |
| chr8-99456486-DEL-225 | chr8-99456487-DEL-225 |
| chr8-99467220-DEL-66 | chr8-99467221-DEL-76 |
| chr8-99469010-DEL-2186 | chr8-99469011-DEL-2185 |
| chr8-99498532-DEL-490 | chr8-99498532-DEL-490 |
| chr8-99499228-DEL-87 | chr8-99499236-DEL-101 |
| chr8-99565102-DEL-7411 | chr8-99565090-DEL-7412 |
| chr8-99621825-DEL-167 | chr8-99621826-DEL-167 |
| chr8-99631105-DEL-205 | chr8-99631106-DEL-205 |
| chr8-99666165-DEL-1865 | chr8-99666167-DEL-1864 |
| chr8-99679069-DEL-62 | chr8-99679070-DEL-62 |
| chr8-99679614-DEL-626 | chr8-99679615-DEL-626 |
| chr8-99688453-DEL-101 | chr8-99688497-DEL-91 |
| chr8-99690941-DEL-702 | chr8-99690942-DEL-702 |
| chr8-99710282-DEL-446 | chr8-99710283-DEL-446 |
| chr8-99732210-DEL-88 | chr8-99732211-DEL-58 |
| chr8-99734681-DEL-321 | chr8-99734682-DEL-321 |
| chr8-99736428-DEL-320 | chr8-99736429-DEL-320 |
| chr8-99749286-DEL-58 | chr8-99749287-DEL-58 |
| chr8-99774461-DEL-265 | chr8-99774462-DEL-265 |
| chr8-99799078-DEL-4827 | chr8-99799079-DEL-4827 |
| chr8-99805712-DEL-1184 | chr8-99805714-DEL-1185 |
| chr8-99856581-DEL-386 | chr8-99856582-DEL-386 |
| chr8-99861238-DEL-471 | chr8-99861239-DEL-471 |
| chr8-99947982-DEL-374 | chr8-99947983-DEL-374 |
| chr8-100007891-DEL-230 | chr8-100007892-DEL-230 |
| chr8-100014837-DEL-96 | chr8-100014838-DEL-96 |
| chr8-100015512-DEL-60 | chr8-100015513-DEL-60 |
| chr8-100024628-DEL-1510 | chr8-100024630-DEL-1482 |
| chr8-100034263-DEL-164 | chr8-100034264-DEL-164 |
| chr8-100079756-DEL-58 | chr8-100079755-DEL-56 |
| chr8-100099121-DEL-495 | chr8-100099122-DEL-495 |
| chr8-100106273-DEL-61 | chr8-100106275-DEL-60 |
| chr8-100173475-DEL-60 | chr8-100173486-DEL-50 |
| chr8-100192892-DEL-7135 | chr8-100192893-DEL-7135 |
| chr8-100231600-DEL-61 | chr8-100231595-DEL-61 |
| chr8-100241062-DEL-188 | chr8-100241063-DEL-188 |

| | |
|---|---|
| chr8-100298669-DEL-70 | chr8-100298688-DEL-50 |
| chr8-100346379-DEL-166 | chr8-100346380-DEL-166 |
| chr8-100383983-DEL-1875 | chr8-100383983-DEL-1875 |
| chr8-100396067-DEL-6341 | chr8-100396071-DEL-6339 |
| chr8-100402074-DEL-201 | chr8-100402075-DEL-201 |
| chr8-100422427-DEL-1232 | chr8-100422404-DEL-1231 |
| chr8-100439740-DEL-451 | chr8-100439743-DEL-443 |
| chr8-100456961-DEL-90 | chr8-100457009-DEL-52 |
| chr8-100497390-DEL-62 | chr8-100497379-DEL-51 |
| chr8-100563100-DEL-81 | chr8-100563101-DEL-81 |
| chr8-100566927-DEL-58 | chr8-100566928-DEL-58 |
| chr8-100574471-DEL-94 | chr8-100574472-DEL-92 |
| chr8-100596330-DEL-51 | chr8-100596330-DEL-54 |
| chr8-100601599-DEL-63 | chr8-100601600-DEL-61 |
| chr8-100608680-DEL-61 | chr8-100608680-DEL-62 |
| chr8-100614451-DEL-155 | chr8-100614452-DEL-155 |
| chr8-100665156-DEL-74 | chr8-100665157-DEL-74 |
| chr8-100668911-DEL-59 | chr8-100668912-DEL-59 |
| chr8-100680601-DEL-2228 | chr8-100680602-DEL-2228 |
| chr8-100685872-DEL-324 | chr8-100685873-DEL-324 |
| chr8-100687018-DEL-52 | chr8-100687016-DEL-50 |
| chr8-100722493-DEL-86 | chr8-100722494-DEL-86 |
| chr8-100757103-DEL-1696 | chr8-100757104-DEL-1696 |
| chr8-100775770-DEL-528 | chr8-100775757-DEL-526 |
| chr8-100816991-DEL-54 | chr8-100816992-DEL-54 |
| chr8-100836659-DEL-59 | chr8-100836660-DEL-59 |
| chr8-100837547-DEL-6741 | chr8-100837548-DEL-6741 |
| chr8-100857958-DEL-3084 | chr8-100857959-DEL-3084 |
| chr8-100861707-DEL-67 | chr8-100861713-DEL-66 |
| chr8-100874726-DEL-421 | chr8-100874727-DEL-421 |
| chr8-100876088-DEL-62 | chr8-100876073-DEL-75 |
| chr8-100890170-DEL-4383 | chr8-100890171-DEL-4383 |
| chr8-100943394-DEL-505 | chr8-100943395-DEL-505 |
| chr8-100944222-DEL-900 | chr8-100944222-DEL-898 |
| chr8-100946323-DEL-1492 | chr8-100946324-DEL-1492 |
| chr8-100963143-DEL-71 | chr8-100963144-DEL-53 |
| chr8-100975374-DEL-6625 | chr8-100975375-DEL-6625 |
| chr8-101001491-DEL-6442 | chr8-101001492-DEL-6442 |
| chr8-101019782-DEL-189 | chr8-101019789-DEL-184 |
| chr8-101021728-DEL-66 | chr8-101021729-DEL-66 |
| chr8-101034050-DEL-2602 | chr8-101034051-DEL-2602 |
| chr8-101072503-DEL-615 | chr8-101072504-DEL-615 |
| chr8-101075557-DEL-73 | chr8-101075558-DEL-73 |
| chr8-101080995-DEL-6508 | chr8-101080996-DEL-6508 |
| chr8-101111493-DEL-54 | chr8-101111494-DEL-50 |
| chr8-101113643-DEL-73 | chr8-101113644-DEL-73 |
| chr8-101147300-DEL-56 | chr8-101147301-DEL-56 |
| chr8-101152972-DEL-152 | chr8-101152972-DEL-152 |
| chr8-101154350-DEL-57 | chr8-101154350-DEL-52 |
| chr8-101155234-DEL-212 | chr8-101155235-DEL-212 |
| chr8-101165240-DEL-735 | chr8-101165241-DEL-735 |
| chr8-101171287-DEL-52 | chr8-101171288-DEL-52 |
| chr8-101180970-DEL-68 | chr8-101180971-DEL-74 |
| chr8-101195463-DEL-6213 | chr8-101195464-DEL-6213 |
| chr8-101197457-DEL-476 | chr8-101197458-DEL-476 |
| chr8-101245258-DEL-4363 | chr8-101245259-DEL-4363 |
| chr8-101287954-DEL-52 | chr8-101287955-DEL-52 |
| chr8-101289635-DEL-181 | chr8-101289636-DEL-181 |
| chr8-101301157-DEL-304 | chr8-101301157-DEL-304 |
| chr8-101306969-DEL-62 | chr8-101306962-DEL-60 |
| chr8-101313245-DEL-91 | chr8-101313246-DEL-91 |
| chr8-101319085-DEL-56 | chr8-101319082-DEL-56 |
| chr8-101324344-DEL-88 | chr8-101324344-DEL-88 |
| chr8-101324752-DEL-4518 | chr8-101324753-DEL-4518 |
| chr8-101344378-DEL-4054 | chr8-101344379-DEL-4054 |
| chr8-101381929-DEL-1565 | chr8-101381930-DEL-1565 |
| chr8-101387024-DEL-579 | chr8-101387022-DEL-580 |
| chr8-101400420-DEL-210 | chr8-101400421-DEL-210 |
| chr8-101403107-DEL-58 | chr8-101403106-DEL-60 |
| chr8-101407303-DEL-100 | chr8-101407304-DEL-100 |
| chr8-101441469-DEL-56 | chr8-101441470-DEL-56 |
| chr8-101491375-DEL-411 | chr8-101491376-DEL-411 |
| chr8-101506843-DEL-379 | chr8-101506843-DEL-367 |
| chr8-101537407-DEL-454 | chr8-101537408-DEL-454 |
| chr8-101565724-DEL-612 | chr8-101565727-DEL-610 |
| chr8-101586773-DEL-68 | chr8-101586774-DEL-58 |
| chr8-101590435-DEL-76 | chr8-101590438-DEL-74 |
| chr8-101591337-DEL-627 | chr8-101591332-DEL-628 |
| chr8-101691313-DEL-1953 | chr8-101691314-DEL-1954 |
| chr8-101700756-DEL-1266 | chr8-101700757-DEL-1267 |

| | |
|---|---|
| chr8-101701957-DEL-67 | chr8-101701962-DEL-63 |
| chr8-101712582-DEL-60 | chr8-101712582-DEL-58 |
| chr8-101715224-DEL-3982 | chr8-101715214-DEL-3960 |
| chr8-101732007-DEL-1466 | chr8-101732008-DEL-1466 |
| chr8-101741242-DEL-72 | chr8-101741244-DEL-70 |
| chr8-101758829-DEL-132 | chr8-101758828-DEL-131 |
| chr8-101768683-DEL-512 | chr8-101768684-DEL-512 |
| chr8-101831520-DEL-1200 | chr8-101831521-DEL-1200 |
| chr8-101852311-DEL-107 | chr8-101852312-DEL-107 |
| chr8-101880642-DEL-1862 | chr8-101880643-DEL-1862 |
| chr8-101882623-DEL-74 | chr8-101882622-DEL-74 |
| chr8-101883948-DEL-68 | chr8-101883951-DEL-54 |
| chr8-101927392-DEL-121 | chr8-101927393-DEL-121 |
| chr8-101934963-DEL-6780 | chr8-101934964-DEL-6780 |
| chr8-101966918-DEL-8191 | chr8-101966919-DEL-8191 |
| chr8-102030835-DEL-19036 | chr8-102030836-DEL-19036 |
| chr8-102062327-DEL-729 | chr8-102062328-DEL-729 |
| chr8-102070763-DEL-1434 | chr8-102070764-DEL-1434 |
| chr8-102087231-DEL-5285 | chr8-102087232-DEL-5285 |
| chr8-102110088-DEL-238 | chr8-102110089-DEL-238 |
| chr8-102137676-DEL-4541 | chr8-102137677-DEL-4540 |
| chr8-102145196-DEL-627 | chr8-102145197-DEL-627 |
| chr8-102202883-DEL-780 | chr8-102202884-DEL-780 |
| chr8-102218749-DEL-78 | chr8-102218749-DEL-77 |
| chr8-102225193-DEL-555 | chr8-102225194-DEL-555 |
| chr8-102265958-DEL-180 | chr8-102265980-DEL-165 |
| chr8-102358220-DEL-1090 | chr8-102358221-DEL-1090 |
| chr8-102361506-DEL-72 | chr8-102361507-DEL-72 |
| chr8-102392236-DEL-319 | chr8-102392237-DEL-319 |
| chr8-102398625-DEL-4392 | chr8-102398628-DEL-4389 |
| chr8-102400692-DEL-512 | chr8-102400693-DEL-512 |
| chr8-102427815-DEL-1099 | chr8-102427816-DEL-1099 |
| chr8-102433661-DEL-56 | chr8-102433662-DEL-56 |
| chr8-102434601-DEL-58 | chr8-102434601-DEL-54 |
| chr8-102437378-DEL-6372 | chr8-102437379-DEL-6372 |
| chr8-102448527-DEL-300 | chr8-102448528-DEL-300 |
| chr8-102453210-DEL-56 | chr8-102453211-DEL-54 |
| chr8-102474515-DEL-270 | chr8-102474516-DEL-270 |
| chr8-102475119-DEL-60 | chr8-102475120-DEL-60 |
| chr8-102487540-DEL-78 | chr8-102487545-DEL-74 |
| chr8-102500657-DEL-56 | chr8-102500658-DEL-56 |
| chr8-102526095-DEL-61 | chr8-102526093-DEL-64 |
| chr8-102526315-DEL-807 | chr8-102526316-DEL-807 |
| chr8-102530876-DEL-55 | chr8-102530879-DEL-53 |
| chr8-102534926-DEL-77 | chr8-102534944-DEL-50 |
| chr8-102550587-DEL-607 | chr8-102550587-DEL-607 |
| chr8-102551956-DEL-563 | chr8-102551957-DEL-563 |
| chr8-102558880-DEL-434 | chr8-102558881-DEL-434 |
| chr8-102589216-DEL-55 | chr8-102589209-DEL-56 |
| chr8-102590446-DEL-1540 | chr8-102590447-DEL-1540 |
| chr8-102598163-DEL-455 | chr8-102598163-DEL-455 |
| chr8-102603385-DEL-130 | chr8-102603386-DEL-130 |
| chr8-102604719-DEL-77 | chr8-102604720-DEL-77 |
| chr8-102620043-DEL-60 | chr8-102620044-DEL-60 |
| chr8-102622271-DEL-1938 | chr8-102622278-DEL-1937 |
| chr8-102627883-DEL-221 | chr8-102627884-DEL-221 |
| chr8-102654017-DEL-400 | chr8-102654057-DEL-361 |
| chr8-102670286-DEL-92 | chr8-102670264-DEL-111 |
| chr8-102671162-DEL-96 | chr8-102671163-DEL-96 |
| chr8-102690410-DEL-3129 | chr8-102690411-DEL-3129 |
| chr8-102698235-DEL-742 | chr8-102698230-DEL-712 |
| chr8-102698346-DEL-54 | chr8-102698347-DEL-54 |
| chr8-102706473-DEL-2775 | chr8-102706474-DEL-2775 |
| chr8-102718980-DEL-56 | chr8-102718966-DEL-56 |
| chr8-102724879-DEL-219 | chr8-102724882-DEL-217 |
| chr8-102731542-DEL-5368 | chr8-102731543-DEL-5368 |
| chr8-102739593-DEL-420 | chr8-102739594-DEL-420 |
| chr8-102761425-DEL-404 | chr8-102761426-DEL-404 |
| chr8-102782239-DEL-80 | chr8-102782255-DEL-65 |
| chr8-102799479-DEL-130 | chr8-102799471-DEL-139 |
| chr8-102816436-DEL-156 | chr8-102816433-DEL-156 |
| chr8-102834680-DEL-754 | chr8-102834681-DEL-754 |
| chr8-102839300-DEL-1018 | chr8-102839301-DEL-1017 |
| chr8-102841246-DEL-1893 | chr8-102841247-DEL-1893 |
| chr8-102849331-DEL-82 | chr8-102849338-DEL-76 |
| chr8-102868077-DEL-7135 | chr8-102868079-DEL-7135 |
| chr8-102897728-DEL-76 | chr8-102897729-DEL-76 |
| chr8-102898856-DEL-55 | chr8-102898857-DEL-50 |
| chr8-102901009-DEL-63 | chr8-102901010-DEL-63 |
| chr8-102919840-DEL-594 | chr8-102919841-DEL-594 |

| | |
|---|---|
| chr8-102938175-DEL-967 | chr8-102938176-DEL-967 |
| chr8-102938225-DEL-371 | chr8-102938226-DEL-371 |
| chr8-102972959-DEL-6502 | chr8-102972960-DEL-6502 |
| chr8-102982179-DEL-90 | chr8-102982180-DEL-90 |
| chr8-102983764-DEL-545 | chr8-102983765-DEL-545 |
| chr8-102996803-DEL-61 | chr8-102996814-DEL-72 |
| chr8-103008985-DEL-101 | chr8-103008986-DEL-101 |
| chr8-103018182-DEL-1452 | chr8-103018183-DEL-1452 |
| chr8-103028989-DEL-52 | chr8-103028990-DEL-52 |
| chr8-103038173-DEL-75 | chr8-103038174-DEL-75 |
| chr8-103046704-DEL-506 | chr8-103046705-DEL-505 |
| chr8-103061590-DEL-124 | chr8-103061591-DEL-124 |
| chr8-103064297-DEL-3496 | chr8-103064298-DEL-3496 |
| chr8-103076960-DEL-57 | chr8-103076961-DEL-57 |
| chr8-103092082-DEL-6305 | chr8-103092083-DEL-6305 |
| chr8-103120846-DEL-1780 | chr8-103120832-DEL-1794 |
| chr8-103134792-DEL-883 | chr8-103134793-DEL-882 |
| chr8-103190352-DEL-75 | chr8-103190369-DEL-50 |
| chr8-103192650-DEL-665 | chr8-103192643-DEL-667 |
| chr8-103195555-DEL-1337 | chr8-103195556-DEL-1337 |
| chr8-103208705-DEL-659 | chr8-103208705-DEL-659 |
| chr8-103210487-DEL-106 | chr8-103210488-DEL-108 |
| chr8-103211628-DEL-730 | chr8-103211629-DEL-730 |
| chr8-103212602-DEL-537 | chr8-103212585-DEL-575 |
| chr8-103216413-DEL-106 | chr8-103216414-DEL-106 |
| chr8-103227275-DEL-191 | chr8-103227276-DEL-191 |
| chr8-103263684-DEL-81 | chr8-103263695-DEL-56 |
| chr8-103281419-DEL-72 | chr8-103281410-DEL-78 |
| chr8-103349178-DEL-160 | chr8-103349179-DEL-160 |
| chr8-103387085-DEL-2873 | chr8-103387086-DEL-2873 |
| chr8-103388648-DEL-68 | chr8-103388649-DEL-68 |
| chr8-103403051-DEL-127 | chr8-103403052-DEL-127 |
| chr8-103408147-DEL-657 | chr8-103408146-DEL-659 |
| chr8-103471498-DEL-68 | chr8-103471499-DEL-68 |
| chr8-103500348-DEL-114 | chr8-103500355-DEL-120 |
| chr8-103543366-DEL-482 | chr8-103543367-DEL-478 |
| chr8-103591371-DEL-1144 | chr8-103591363-DEL-1145 |
| chr8-103598803-DEL-2376 | chr8-103598804-DEL-2376 |
| chr8-103614044-DEL-5968 | chr8-103614077-DEL-5979 |
| chr8-103620343-DEL-256 | chr8-103620344-DEL-256 |
| chr8-103624252-DEL-96 | chr8-103624260-DEL-105 |
| chr8-103625055-DEL-297 | chr8-103625050-DEL-303 |
| chr8-103630571-DEL-54 | chr8-103630574-DEL-52 |
| chr8-103638585-DEL-69 | chr8-103638594-DEL-60 |
| chr8-103667240-DEL-571 | chr8-103667241-DEL-571 |
| chr8-103674383-DEL-6165 | chr8-103674384-DEL-6165 |
| chr8-103686468-DEL-1381 | chr8-103686469-DEL-1381 |
| chr8-103695842-DEL-922 | chr8-103695847-DEL-922 |
| chr8-103708249-DEL-87 | chr8-103708274-DEL-63 |
| chr8-103744279-DEL-124 | chr8-103744280-DEL-124 |
| chr8-103774196-DEL-58 | chr8-103774213-DEL-80 |
| chr8-103785564-DEL-173 | chr8-103785569-DEL-167 |
| chr8-103791201-DEL-90 | chr8-103791186-DEL-106 |
| chr8-103805938-DEL-4268 | chr8-103805939-DEL-4268 |
| chr8-103841889-DEL-159 | chr8-103841887-DEL-157 |
| chr8-103858386-DEL-121 | chr8-103858338-DEL-127 |
| chr8-103986835-DEL-80 | chr8-103986835-DEL-81 |
| chr8-104087145-DEL-170 | chr8-104087146-DEL-170 |
| chr8-104089792-DEL-78 | chr8-104089774-DEL-64 |
| chr8-104108615-DEL-122 | chr8-104108616-DEL-122 |
| chr8-104117324-DEL-163 | chr8-104117348-DEL-159 |
| chr8-104120550-DEL-630 | chr8-104120549-DEL-631 |
| chr8-104123035-DEL-509 | chr8-104123036-DEL-509 |
| chr8-104154192-DEL-778 | chr8-104154193-DEL-778 |
| chr8-104212085-DEL-851 | chr8-104212086-DEL-851 |
| chr8-104229532-DEL-54 | chr8-104229511-DEL-76 |
| chr8-104276869-DEL-56 | chr8-104276870-DEL-56 |
| chr8-104290140-DEL-377 | chr8-104290140-DEL-378 |
| chr8-104292609-DEL-414 | chr8-104292610-DEL-414 |
| chr8-104296240-DEL-125 | chr8-104296241-DEL-125 |
| chr8-104358193-DEL-555 | chr8-104358194-DEL-555 |
| chr8-104373780-DEL-1226 | chr8-104373781-DEL-1226 |
| chr8-104379234-DEL-253 | chr8-104379235-DEL-254 |
| chr8-104399208-DEL-6132 | chr8-104399216-DEL-6143 |
| chr8-104413190-DEL-77 | chr8-104413191-DEL-77 |
| chr8-104421505-DEL-240 | chr8-104421506-DEL-240 |
| chr8-104443030-DEL-242 | chr8-104443031-DEL-242 |
| chr8-104446524-DEL-662 | chr8-104446525-DEL-662 |
| chr8-104447348-DEL-52 | chr8-104447349-DEL-52 |
| chr8-104471508-DEL-87 | chr8-104471509-DEL-87 |

| | |
|---|---|
| chr8-104481492-DEL-6495 | chr8-104481478-DEL-6496 |
| chr8-104501220-DEL-761 | chr8-104501221-DEL-761 |
| chr8-104502366-DEL-131 | chr8-104502367-DEL-131 |
| chr8-104552622-DEL-124 | chr8-104552623-DEL-124 |
| chr8-104589889-DEL-180 | chr8-104589890-DEL-178 |
| chr8-104676955-DEL-76 | chr8-104676956-DEL-76 |
| chr8-104719509-DEL-56 | chr8-104719510-DEL-60 |
| chr8-104730584-DEL-245 | chr8-104730585-DEL-245 |
| chr8-104744294-DEL-64 | chr8-104744342-DEL-64 |
| chr8-104775102-DEL-68 | chr8-104775103-DEL-68 |
| chr8-104836087-DEL-238 | chr8-104836088-DEL-238 |
| chr8-104858571-DEL-1295 | chr8-104858572-DEL-1295 |
| chr8-104860413-DEL-244 | chr8-104860418-DEL-240 |
| chr8-104879133-DEL-1892 | chr8-104879134-DEL-1892 |
| chr8-104882048-DEL-3007 | chr8-104882049-DEL-3007 |
| chr8-104890317-DEL-166 | chr8-104890339-DEL-123 |
| chr8-104910040-DEL-92 | chr8-104910041-DEL-68 |
| chr8-104913005-DEL-1787 | chr8-104913006-DEL-1787 |
| chr8-104915769-DEL-94 | chr8-104915770-DEL-94 |
| chr8-104918658-DEL-313 | chr8-104918659-DEL-313 |
| chr8-104950109-DEL-1377 | chr8-104950110-DEL-1377 |
| chr8-104992273-DEL-1903 | chr8-104992274-DEL-1903 |
| chr8-104996290-DEL-3261 | chr8-104996278-DEL-3267 |
| chr8-105001689-DEL-216 | chr8-105001690-DEL-216 |
| chr8-105002800-DEL-78 | chr8-105002801-DEL-78 |
| chr8-105025898-DEL-211 | chr8-105025899-DEL-211 |
| chr8-105048377-DEL-57 | chr8-105048378-DEL-57 |
| chr8-105050267-DEL-69 | chr8-105050265-DEL-72 |
| chr8-105054094-DEL-120 | chr8-105054075-DEL-92 |
| chr8-105076806-DEL-59 | chr8-105076831-DEL-58 |
| chr8-105098846-DEL-220 | chr8-105098847-DEL-220 |
| chr8-105103958-DEL-56 | chr8-105103959-DEL-56 |
| chr8-105119696-DEL-183 | chr8-105119697-DEL-183 |
| chr8-105130863-DEL-194 | chr8-105130859-DEL-195 |
| chr8-105151233-DEL-186 | chr8-105151234-DEL-186 |
| chr8-105153274-DEL-245 | chr8-105153273-DEL-246 |
| chr8-105186325-DEL-5698 | chr8-105186326-DEL-5698 |
| chr8-105210270-DEL-230 | chr8-105210271-DEL-230 |
| chr8-105223693-DEL-84 | chr8-105223694-DEL-84 |
| chr8-105227167-DEL-400 | chr8-105227168-DEL-400 |
| chr8-105282105-DEL-334 | chr8-105282106-DEL-334 |
| chr8-105287991-DEL-199 | chr8-105287992-DEL-199 |
| chr8-105296900-DEL-96 | chr8-105296883-DEL-120 |
| chr8-105303664-DEL-98 | chr8-105303665-DEL-98 |
| chr8-105326708-DEL-192 | chr8-105326707-DEL-194 |
| chr8-105373126-DEL-6232 | chr8-105373127-DEL-6232 |
| chr8-105380186-DEL-276 | chr8-105380187-DEL-276 |
| chr8-105391219-DEL-633 | chr8-105391246-DEL-589 |
| chr8-105412883-DEL-851 | chr8-105412884-DEL-851 |
| chr8-105415858-DEL-6475 | chr8-105415859-DEL-6475 |
| chr8-105436354-DEL-123 | chr8-105436355-DEL-107 |
| chr8-105440188-DEL-368 | chr8-105440189-DEL-368 |
| chr8-105473182-DEL-335 | chr8-105473183-DEL-335 |
| chr8-105505144-DEL-4930 | chr8-105505145-DEL-4930 |
| chr8-105514876-DEL-400 | chr8-105514876-DEL-400 |
| chr8-105545762-DEL-6439 | chr8-105545763-DEL-6439 |
| chr8-105552908-DEL-52 | chr8-105552909-DEL-52 |
| chr8-105581512-DEL-970 | chr8-105581513-DEL-970 |
| chr8-105582408-DEL-74 | chr8-105582424-DEL-58 |
| chr8-105727856-DEL-7287 | chr8-105727857-DEL-7287 |
| chr8-105752364-DEL-1836 | chr8-105752365-DEL-1836 |
| chr8-105860898-DEL-212 | chr8-105860899-DEL-212 |
| chr8-105862123-DEL-217 | chr8-105862124-DEL-217 |
| chr8-105865655-DEL-77 | chr8-105865641-DEL-91 |
| chr8-105892416-DEL-454 | chr8-105892416-DEL-453 |
| chr8-105899112-DEL-280 | chr8-105899113-DEL-280 |
| chr8-105902296-DEL-202 | chr8-105902301-DEL-198 |
| chr8-105905502-DEL-822 | chr8-105905503-DEL-822 |
| chr8-105923951-DEL-66 | chr8-105923952-DEL-66 |
| chr8-105931451-DEL-125 | chr8-105931452-DEL-125 |
| chr8-105937978-DEL-203 | chr8-105937979-DEL-203 |
| chr8-105967180-DEL-211 | chr8-105967181-DEL-211 |
| chr8-106058198-DEL-66 | chr8-106058207-DEL-57 |
| chr8-106091491-DEL-113 | chr8-106091492-DEL-113 |
| chr8-106131508-DEL-241 | chr8-106131509-DEL-241 |
| chr8-106138183-DEL-212 | chr8-106138184-DEL-212 |
| chr8-106141707-DEL-2030 | chr8-106141708-DEL-2030 |
| chr8-106145042-DEL-102 | chr8-106145043-DEL-102 |
| chr8-106164915-DEL-426 | chr8-106164916-DEL-426 |
| chr8-106224927-DEL-199 | chr8-106224945-DEL-199 |

| | |
|---|---|
| chr8-106267009-DEL-193 | chr8-106267010-DEL-193 |
| chr8-106275860-DEL-105 | chr8-106275853-DEL-112 |
| chr8-106282348-DEL-96 | chr8-106282349-DEL-92 |
| chr8-106297695-DEL-171 | chr8-106297696-DEL-171 |
| chr8-106302891-DEL-104 | chr8-106302892-DEL-104 |
| chr8-106304206-DEL-104 | chr8-106304175-DEL-105 |
| chr8-106316880-DEL-59 | chr8-106316866-DEL-56 |
| chr8-106327031-DEL-70 | chr8-106327032-DEL-70 |
| chr8-106327795-DEL-139 | chr8-106327788-DEL-164 |
| chr8-106340005-DEL-178 | chr8-106340006-DEL-178 |
| chr8-106347353-DEL-54 | chr8-106347354-DEL-54 |
| chr8-106356623-DEL-78 | chr8-106356624-DEL-78 |
| chr8-106367392-DEL-58 | chr8-106367393-DEL-58 |
| chr8-106406375-DEL-105 | chr8-106406376-DEL-105 |
| chr8-106456613-DEL-382 | chr8-106456614-DEL-382 |
| chr8-106463683-DEL-72 | chr8-106463665-DEL-93 |
| chr8-106475066-DEL-56 | chr8-106475060-DEL-90 |
| chr8-106523611-DEL-210 | chr8-106523612-DEL-210 |
| chr8-106550241-DEL-70 | chr8-106550257-DEL-91 |
| chr8-106555816-DEL-83 | chr8-106555817-DEL-83 |
| chr8-106560464-DEL-176 | chr8-106560469-DEL-175 |
| chr8-106589527-DEL-205 | chr8-106589528-DEL-205 |
| chr8-106596418-DEL-362 | chr8-106596419-DEL-362 |
| chr8-106597588-DEL-200 | chr8-106597589-DEL-200 |
| chr8-106626627-DEL-69 | chr8-106626628-DEL-69 |
| chr8-106630942-DEL-107 | chr8-106630943-DEL-107 |
| chr8-106631691-DEL-191 | chr8-106631692-DEL-191 |
| chr8-106637325-DEL-204 | chr8-106637320-DEL-210 |
| chr8-106672863-DEL-179 | chr8-106672864-DEL-179 |
| chr8-106681261-DEL-220 | chr8-106681262-DEL-219 |
| chr8-106685319-DEL-54 | chr8-106685320-DEL-54 |
| chr8-106693067-DEL-92 | chr8-106693068-DEL-92 |
| chr8-106726734-DEL-221 | chr8-106726735-DEL-221 |
| chr8-106739077-DEL-205 | chr8-106739078-DEL-205 |
| chr8-106779414-DEL-64 | chr8-106779415-DEL-64 |
| chr8-106827830-DEL-154 | chr8-106827831-DEL-154 |
| chr8-106838788-DEL-70 | chr8-106838785-DEL-72 |
| chr8-106840941-DEL-176 | chr8-106840942-DEL-176 |
| chr8-106842034-DEL-224 | chr8-106842035-DEL-224 |
| chr8-106876345-DEL-56 | chr8-106876353-DEL-53 |
| chr8-106887062-DEL-76 | chr8-106887063-DEL-76 |
| chr8-106898011-DEL-213 | chr8-106898012-DEL-213 |
| chr8-106932894-DEL-217 | chr8-106932895-DEL-217 |
| chr8-106934863-DEL-213 | chr8-106934864-DEL-213 |
| chr8-106935645-DEL-106 | chr8-106935646-DEL-118 |
| chr8-106949499-DEL-209 | chr8-106949500-DEL-209 |
| chr8-107082621-DEL-199 | chr8-107082622-DEL-199 |
| chr8-107158927-DEL-359 | chr8-107158928-DEL-359 |
| chr8-107167933-DEL-158 | chr8-107167934-DEL-158 |
| chr8-107170525-DEL-867 | chr8-107170526-DEL-867 |
| chr8-107206126-DEL-13937 | chr8-107206127-DEL-13937 |
| chr8-107224662-DEL-191 | chr8-107224663-DEL-191 |
| chr8-107227463-DEL-56 | chr8-107227477-DEL-60 |
| chr8-107229678-DEL-347 | chr8-107229658-DEL-370 |
| chr8-107269260-DEL-140 | chr8-107269261-DEL-140 |
| chr8-107280368-DEL-211 | chr8-107280369-DEL-211 |
| chr8-107291713-DEL-167 | chr8-107291709-DEL-166 |
| chr8-107305891-DEL-227 | chr8-107305892-DEL-227 |
| chr8-107314153-DEL-53 | chr8-107314129-DEL-52 |
| chr8-107320517-DEL-68 | chr8-107320544-DEL-75 |
| chr8-107334654-DEL-199 | chr8-107334655-DEL-199 |
| chr8-107340479-DEL-101 | chr8-107340480-DEL-101 |
| chr8-107379043-DEL-210 | chr8-107379044-DEL-210 |
| chr8-107383846-DEL-143 | chr8-107383847-DEL-143 |
| chr8-107393859-DEL-82 | chr8-107393860-DEL-82 |
| chr8-107401286-DEL-180 | chr8-107401286-DEL-180 |
| chr8-107402902-DEL-5445 | chr8-107402903-DEL-5445 |
| chr8-107409254-DEL-210 | chr8-107409255-DEL-210 |
| chr8-107414462-DEL-60 | chr8-107414465-DEL-58 |
| chr8-107416209-DEL-180 | chr8-107416210-DEL-180 |
| chr8-107428649-DEL-93 | chr8-107428681-DEL-71 |
| chr8-107509999-DEL-103 | chr8-107510000-DEL-103 |
| chr8-107510260-DEL-469 | chr8-107510261-DEL-469 |
| chr8-107514166-DEL-82 | chr8-107514167-DEL-82 |
| chr8-107529268-DEL-232 | chr8-107529269-DEL-232 |
| chr8-107551242-DEL-210 | chr8-107551243-DEL-210 |
| chr8-107554516-DEL-52 | chr8-107554518-DEL-52 |
| chr8-107559354-DEL-229 | chr8-107559374-DEL-219 |
| chr8-107579017-DEL-185 | chr8-107579018-DEL-185 |
| chr8-107588731-DEL-198 | chr8-107588732-DEL-198 |

| | |
|---|---|
| chr8-107648681-DEL-62 | chr8-107648674-DEL-66 |
| chr8-107650508-DEL-207 | chr8-107650509-DEL-207 |
| chr8-107653075-DEL-231 | chr8-107653076-DEL-231 |
| chr8-107661460-DEL-209 | chr8-107661461-DEL-209 |
| chr8-107686861-DEL-211 | chr8-107686862-DEL-211 |
| chr8-107693530-DEL-1205 | chr8-107693533-DEL-1203 |
| chr8-107707031-DEL-202 | chr8-107707032-DEL-202 |
| chr8-107717378-DEL-191 | chr8-107717379-DEL-191 |
| chr8-107721610-DEL-837 | chr8-107721608-DEL-837 |
| chr8-107732649-DEL-389 | chr8-107732601-DEL-388 |
| chr8-107748076-DEL-227 | chr8-107748077-DEL-227 |
| chr8-107754044-DEL-203 | chr8-107754045-DEL-203 |
| chr8-107785996-DEL-1092 | chr8-107785997-DEL-1092 |
| chr8-107789112-DEL-52 | chr8-107789113-DEL-52 |
| chr8-107816605-DEL-189 | chr8-107816606-DEL-189 |
| chr8-107820768-DEL-242 | chr8-107820788-DEL-220 |
| chr8-107838239-DEL-174 | chr8-107838236-DEL-175 |
| chr8-107861719-DEL-245 | chr8-107861720-DEL-245 |
| chr8-107864006-DEL-205 | chr8-107864007-DEL-205 |
| chr8-107874978-DEL-322 | chr8-107874982-DEL-322 |
| chr8-107878575-DEL-410 | chr8-107878577-DEL-409 |
| chr8-107897884-DEL-193 | chr8-107897885-DEL-193 |
| chr8-107944130-DEL-329 | chr8-107944131-DEL-329 |
| chr8-107951374-DEL-224 | chr8-107951375-DEL-223 |
| chr8-107953366-DEL-194 | chr8-107953367-DEL-194 |
| chr8-107954088-DEL-581 | chr8-107954089-DEL-581 |
| chr8-107970262-DEL-140 | chr8-107970260-DEL-145 |
| chr8-107974300-DEL-70 | chr8-107974302-DEL-67 |
| chr8-107989360-DEL-570 | chr8-107989361-DEL-570 |
| chr8-108061337-DEL-210 | chr8-108061338-DEL-209 |
| chr8-108084172-DEL-163 | chr8-108084179-DEL-163 |
| chr8-108125001-DEL-70 | chr8-108125002-DEL-70 |
| chr8-108143959-DEL-55 | chr8-108143962-DEL-50 |
| chr8-108158114-DEL-180 | chr8-108158115-DEL-180 |
| chr8-108165748-DEL-471 | chr8-108165749-DEL-471 |
| chr8-108170656-DEL-243 | chr8-108170688-DEL-214 |
| chr8-108177349-DEL-55 | chr8-108177350-DEL-55 |
| chr8-108212402-DEL-68 | chr8-108212403-DEL-68 |
| chr8-108223694-DEL-80 | chr8-108223695-DEL-80 |
| chr8-108236318-DEL-159 | chr8-108236319-DEL-159 |
| chr8-108239498-DEL-75 | chr8-108239498-DEL-75 |
| chr8-108280206-DEL-226 | chr8-108280184-DEL-243 |
| chr8-108293330-DEL-920 | chr8-108293329-DEL-916 |
| chr8-108312156-DEL-124 | chr8-108312157-DEL-124 |
| chr8-108315396-DEL-200 | chr8-108315397-DEL-200 |
| chr8-108317888-DEL-224 | chr8-108317889-DEL-224 |
| chr8-108331551-DEL-123 | chr8-108331552-DEL-123 |
| chr8-108341033-DEL-3070 | chr8-108341034-DEL-3070 |
| chr8-108368629-DEL-581 | chr8-108368624-DEL-582 |
| chr8-108373069-DEL-345 | chr8-108373070-DEL-344 |
| chr8-108380958-DEL-612 | chr8-108380959-DEL-612 |
| chr8-108395476-DEL-119 | chr8-108395477-DEL-119 |
| chr8-108427458-DEL-579 | chr8-108427459-DEL-579 |
| chr8-108442349-DEL-90 | chr8-108442360-DEL-72 |
| chr8-108458744-DEL-6475 | chr8-108458745-DEL-6475 |
| chr8-108478684-DEL-1578 | chr8-108478685-DEL-1578 |
| chr8-108484795-DEL-68 | chr8-108484796-DEL-68 |
| chr8-108492850-DEL-79 | chr8-108492851-DEL-79 |
| chr8-108500496-DEL-54 | chr8-108500501-DEL-50 |
| chr8-108505079-DEL-66 | chr8-108505080-DEL-66 |
| chr8-108505931-DEL-52 | chr8-108505932-DEL-52 |
| chr8-108512903-DEL-142 | chr8-108512904-DEL-142 |
| chr8-108528932-DEL-2107 | chr8-108528933-DEL-2107 |
| chr8-108528940-DEL-159 | chr8-108528941-DEL-159 |
| chr8-108530805-DEL-403 | chr8-108530806-DEL-403 |
| chr8-108548831-DEL-346 | chr8-108548832-DEL-346 |
| chr8-108550282-DEL-54 | chr8-108550282-DEL-54 |
| chr8-108567514-DEL-1894 | chr8-108567515-DEL-1894 |
| chr8-108574220-DEL-75 | chr8-108574203-DEL-77 |
| chr8-108607930-DEL-60 | chr8-108607931-DEL-60 |
| chr8-108610907-DEL-90 | chr8-108610917-DEL-56 |
| chr8-108620229-DEL-72 | chr8-108620243-DEL-60 |
| chr8-108629539-DEL-204 | chr8-108629540-DEL-204 |
| chr8-108691377-DEL-93 | chr8-108691378-DEL-95 |
| chr8-108710295-DEL-58 | chr8-108710295-DEL-60 |
| chr8-108713981-DEL-114 | chr8-108713981-DEL-152 |
| chr8-108730677-DEL-53 | chr8-108730678-DEL-53 |
| chr8-108739162-DEL-56 | chr8-108739162-DEL-56 |
| chr8-108765662-DEL-68 | chr8-108765674-DEL-89 |
| chr8-108822144-DEL-62 | chr8-108822147-DEL-61 |

| | |
|---|---|
| chr8-108830774-DEL-76 | chr8-108830779-DEL-68 |
| chr8-108833739-DEL-206 | chr8-108833740-DEL-206 |
| chr8-108843699-DEL-6403 | chr8-108843700-DEL-6403 |
| chr8-108858736-DEL-195 | chr8-108858737-DEL-195 |
| chr8-108869413-DEL-700 | chr8-108869414-DEL-700 |
| chr8-108871210-DEL-262 | chr8-108871211-DEL-262 |
| chr8-108953698-DEL-254 | chr8-108953699-DEL-254 |
| chr8-108977514-DEL-51 | chr8-108977515-DEL-51 |
| chr8-109019129-DEL-558 | chr8-109019130-DEL-558 |
| chr8-109064412-DEL-7001 | chr8-109064413-DEL-7001 |
| chr8-109070645-DEL-208 | chr8-109070646-DEL-208 |
| chr8-109085226-DEL-1889 | chr8-109085227-DEL-1889 |
| chr8-109092689-DEL-156 | chr8-109092690-DEL-156 |
| chr8-109094761-DEL-124 | chr8-109094762-DEL-124 |
| chr8-109108118-DEL-94 | chr8-109108119-DEL-94 |
| chr8-109169402-DEL-546 | chr8-109169403-DEL-546 |
| chr8-109213766-DEL-375 | chr8-109213767-DEL-375 |
| chr8-109215570-DEL-76 | chr8-109215563-DEL-81 |
| chr8-109227823-DEL-65 | chr8-109227824-DEL-65 |
| chr8-109242688-DEL-208 | chr8-109242689-DEL-208 |
| chr8-109304742-DEL-52 | chr8-109304743-DEL-52 |
| chr8-109362750-DEL-414 | chr8-109362750-DEL-414 |
| chr8-109390405-DEL-89 | chr8-109390406-DEL-89 |
| chr8-109414810-DEL-54 | chr8-109414811-DEL-54 |
| chr8-109495984-DEL-51 | chr8-109495985-DEL-51 |
| chr8-109561389-DEL-70 | chr8-109561390-DEL-70 |
| chr8-109568721-DEL-68 | chr8-109568721-DEL-68 |
| chr8-109610936-DEL-94 | chr8-109610937-DEL-94 |
| chr8-109641573-DEL-64 | chr8-109641574-DEL-56 |
| chr8-109651788-DEL-119 | chr8-109651789-DEL-119 |
| chr8-109665815-DEL-135 | chr8-109665775-DEL-184 |
| chr8-109717899-DEL-105 | chr8-109717913-DEL-109 |
| chr8-109732626-DEL-64 | chr8-109732635-DEL-50 |
| chr8-109735587-DEL-190 | chr8-109735588-DEL-190 |
| chr8-109744426-DEL-162 | chr8-109744430-DEL-163 |
| chr8-109755702-DEL-109 | chr8-109755703-DEL-109 |
| chr8-109756270-DEL-57 | chr8-109756271-DEL-57 |
| chr8-109760302-DEL-103 | chr8-109760302-DEL-99 |
| chr8-109773421-DEL-510 | chr8-109773422-DEL-510 |
| chr8-109792473-DEL-496 | chr8-109792474-DEL-496 |
| chr8-109806317-DEL-73 | chr8-109806318-DEL-73 |
| chr8-109852718-DEL-6325 | chr8-109852715-DEL-6321 |
| chr8-109863773-DEL-283 | chr8-109863774-DEL-283 |
| chr8-109883619-DEL-2915 | chr8-109883620-DEL-2915 |
| chr8-109887160-DEL-77 | chr8-109887148-DEL-76 |
| chr8-109930222-DEL-400 | chr8-109930223-DEL-400 |
| chr8-109967598-DEL-220 | chr8-109967599-DEL-220 |
| chr8-109998355-DEL-560 | chr8-109998356-DEL-560 |
| chr8-110004052-DEL-78 | chr8-110004058-DEL-72 |
| chr8-110022614-DEL-591 | chr8-110022613-DEL-592 |
| chr8-110053716-DEL-1888 | chr8-110053717-DEL-1888 |
| chr8-110066405-DEL-1136 | chr8-110066406-DEL-1136 |
| chr8-110073648-DEL-737 | chr8-110073649-DEL-737 |
| chr8-110082470-DEL-224 | chr8-110082471-DEL-224 |
| chr8-110121106-DEL-72 | chr8-110121107-DEL-86 |
| chr8-110140907-DEL-82 | chr8-110140908-DEL-82 |
| chr8-110152309-DEL-64 | chr8-110152316-DEL-50 |
| chr8-110157198-DEL-363 | chr8-110157200-DEL-361 |
| chr8-110160718-DEL-63 | chr8-110160719-DEL-63 |
| chr8-110171937-DEL-80 | chr8-110171940-DEL-58 |
| chr8-110173511-DEL-144 | chr8-110173512-DEL-108 |
| chr8-110196897-DEL-82 | chr8-110196898-DEL-50 |
| chr8-110237050-DEL-62 | chr8-110237051-DEL-62 |
| chr8-110271375-DEL-474 | chr8-110271376-DEL-474 |
| chr8-110315887-DEL-179 | chr8-110315905-DEL-171 |
| chr8-110322931-DEL-84 | chr8-110322932-DEL-63 |
| chr8-110327072-DEL-54 | chr8-110327073-DEL-54 |
| chr8-110331114-DEL-390 | chr8-110331115-DEL-390 |
| chr8-110350426-DEL-468 | chr8-110350427-DEL-468 |
| chr8-110393134-DEL-94 | chr8-110393135-DEL-94 |
| chr8-110396767-DEL-57 | chr8-110396773-DEL-52 |
| chr8-110410592-DEL-465 | chr8-110410593-DEL-465 |
| chr8-110430687-DEL-415 | chr8-110430718-DEL-383 |
| chr8-110449404-DEL-101 | chr8-110449405-DEL-101 |
| chr8-110465676-DEL-7155 | chr8-110465680-DEL-7154 |
| chr8-110491048-DEL-6003 | chr8-110491049-DEL-6003 |
| chr8-110496296-DEL-197 | chr8-110496271-DEL-215 |
| chr8-110498230-DEL-106 | chr8-110498212-DEL-128 |
| chr8-110503549-DEL-137 | chr8-110503544-DEL-137 |
| chr8-110528275-DEL-207 | chr8-110528276-DEL-207 |

| | |
|---|---|
| chr8-110537167-DEL-3462 | chr8-110537168-DEL-3462 |
| chr8-110574991-DEL-193 | chr8-110574992-DEL-193 |
| chr8-110604102-DEL-209 | chr8-110604103-DEL-209 |
| chr8-110624596-DEL-81 | chr8-110624597-DEL-81 |
| chr8-110632898-DEL-81 | chr8-110632906-DEL-89 |
| chr8-110753170-DEL-169 | chr8-110753158-DEL-170 |
| chr8-110770695-DEL-54 | chr8-110770695-DEL-54 |
| chr8-110781705-DEL-57 | chr8-110781658-DEL-75 |
| chr8-110791235-DEL-60 | chr8-110791247-DEL-58 |
| chr8-110794225-DEL-441 | chr8-110794226-DEL-441 |
| chr8-110802026-DEL-116 | chr8-110802027-DEL-116 |
| chr8-110803937-DEL-261 | chr8-110803938-DEL-261 |
| chr8-110814619-DEL-56 | chr8-110814645-DEL-52 |
| chr8-110898185-DEL-55 | chr8-110898186-DEL-55 |
| chr8-110900183-DEL-76 | chr8-110900180-DEL-80 |
| chr8-110962421-DEL-110 | chr8-110962421-DEL-65 |
| chr8-110976489-DEL-446 | chr8-110976490-DEL-446 |
| chr8-110997681-DEL-90 | chr8-110997711-DEL-83 |
| chr8-111046774-DEL-115 | chr8-111046772-DEL-115 |
| chr8-111054007-DEL-68 | chr8-111054008-DEL-52 |
| chr8-111088080-DEL-75 | chr8-111088081-DEL-75 |
| chr8-111088570-DEL-72 | chr8-111088580-DEL-63 |
| chr8-111156948-DEL-144 | chr8-111156923-DEL-123 |
| chr8-111211229-DEL-204 | chr8-111211230-DEL-204 |
| chr8-111218526-DEL-171 | chr8-111218527-DEL-169 |
| chr8-111236119-DEL-178 | chr8-111236120-DEL-178 |
| chr8-111241672-DEL-128 | chr8-111241709-DEL-92 |
| chr8-111243268-DEL-86 | chr8-111243269-DEL-86 |
| chr8-111275086-DEL-206 | chr8-111275087-DEL-206 |
| chr8-111489859-DEL-216 | chr8-111489860-DEL-216 |
| chr8-111507817-DEL-184 | chr8-111507818-DEL-184 |
| chr8-111511277-DEL-62 | chr8-111511278-DEL-62 |
| chr8-111528785-DEL-193 | chr8-111528781-DEL-195 |
| chr8-111601404-DEL-225 | chr8-111601405-DEL-225 |
| chr8-111617681-DEL-217 | chr8-111617682-DEL-217 |
| chr8-111618476-DEL-193 | chr8-111618477-DEL-193 |
| chr8-111643560-DEL-172 | chr8-111643561-DEL-172 |
| chr8-111651634-DEL-162 | chr8-111651630-DEL-162 |
| chr8-111653268-DEL-135 | chr8-111653269-DEL-135 |
| chr8-111674696-DEL-350 | chr8-111674710-DEL-354 |
| chr8-111676454-DEL-219 | chr8-111676454-DEL-220 |
| chr8-111678544-DEL-318 | chr8-111678545-DEL-318 |
| chr8-111699326-DEL-1551 | chr8-111699324-DEL-1552 |
| chr8-111717481-DEL-186 | chr8-111717482-DEL-186 |
| chr8-111721480-DEL-213 | chr8-111721481-DEL-213 |
| chr8-111722198-DEL-97 | chr8-111722179-DEL-94 |
| chr8-111722999-DEL-367 | chr8-111723000-DEL-367 |
| chr8-111761411-DEL-77 | chr8-111761412-DEL-77 |
| chr8-111765594-DEL-62 | chr8-111765595-DEL-62 |
| chr8-111803072-DEL-144 | chr8-111803074-DEL-134 |
| chr8-111815558-DEL-1277 | chr8-111815559-DEL-1277 |
| chr8-111826473-DEL-185 | chr8-111826464-DEL-195 |
| chr8-111832274-DEL-53 | chr8-111832312-DEL-61 |
| chr8-111839846-DEL-1108 | chr8-111839858-DEL-1103 |
| chr8-111872324-DEL-170 | chr8-111872325-DEL-170 |
| chr8-111917233-DEL-217 | chr8-111917234-DEL-217 |
| chr8-111929020-DEL-117 | chr8-111929021-DEL-117 |
| chr8-111942441-DEL-190 | chr8-111942442-DEL-190 |
| chr8-111946544-DEL-77 | chr8-111946551-DEL-82 |
| chr8-111949472-DEL-208 | chr8-111949473-DEL-208 |
| chr8-111979720-DEL-628 | chr8-111979721-DEL-623 |
| chr8-111980747-DEL-118 | chr8-111980748-DEL-118 |
| chr8-111990364-DEL-214 | chr8-111990365-DEL-214 |
| chr8-111991245-DEL-209 | chr8-111991246-DEL-209 |
| chr8-112022563-DEL-216 | chr8-112022568-DEL-214 |
| chr8-112056176-DEL-63 | chr8-112056177-DEL-61 |
| chr8-112079773-DEL-183 | chr8-112079773-DEL-183 |
| chr8-112106571-DEL-92 | chr8-112106571-DEL-96 |
| chr8-112137517-DEL-73 | chr8-112137518-DEL-73 |
| chr8-112162856-DEL-358 | chr8-112162857-DEL-358 |
| chr8-112193069-DEL-203 | chr8-112193060-DEL-204 |
| chr8-112203061-DEL-420 | chr8-112203062-DEL-420 |
| chr8-112215038-DEL-90 | chr8-112215039-DEL-90 |
| chr8-112221220-DEL-93 | chr8-112221221-DEL-93 |
| chr8-112245913-DEL-207 | chr8-112245914-DEL-207 |
| chr8-112250413-DEL-213 | chr8-112250414-DEL-213 |
| chr8-112257187-DEL-204 | chr8-112257188-DEL-204 |
| chr8-112270410-DEL-222 | chr8-112270411-DEL-222 |
| chr8-112275665-DEL-185 | chr8-112275666-DEL-185 |
| chr8-112277880-DEL-51 | chr8-112277881-DEL-51 |

| | |
|---|---|
| chr8-112279908-DEL-89 | chr8-112279915-DEL-57 |
| chr8-112282368-DEL-222 | chr8-112282369-DEL-222 |
| chr8-112282926-DEL-229 | chr8-112282927-DEL-229 |
| chr8-112330206-DEL-108 | chr8-112330207-DEL-108 |
| chr8-112342449-DEL-2496 | chr8-112342449-DEL-2496 |
| chr8-112395208-DEL-90 | chr8-112395209-DEL-90 |
| chr8-112410433-DEL-238 | chr8-112410434-DEL-238 |
| chr8-112430574-DEL-568 | chr8-112430575-DEL-545 |
| chr8-112431162-DEL-164 | chr8-112431163-DEL-164 |
| chr8-112433790-DEL-283 | chr8-112433797-DEL-277 |
| chr8-112479304-DEL-481 | chr8-112479305-DEL-481 |
| chr8-112485922-DEL-622 | chr8-112485923-DEL-622 |
| chr8-112492624-DEL-65 | chr8-112492625-DEL-65 |
| chr8-112495986-DEL-78 | chr8-112495987-DEL-78 |
| chr8-112518877-DEL-197 | chr8-112518869-DEL-198 |
| chr8-112521518-DEL-91 | chr8-112521511-DEL-68 |
| chr8-112536829-DEL-215 | chr8-112536830-DEL-215 |
| chr8-112538021-DEL-53 | chr8-112538022-DEL-53 |
| chr8-112547205-DEL-147 | chr8-112547206-DEL-147 |
| chr8-112562842-DEL-57 | chr8-112562842-DEL-51 |
| chr8-112563689-DEL-1857 | chr8-112563690-DEL-1857 |
| chr8-112576268-DEL-76 | chr8-112576269-DEL-76 |
| chr8-112595769-DEL-207 | chr8-112595770-DEL-207 |
| chr8-112596931-DEL-213 | chr8-112596932-DEL-213 |
| chr8-112610703-DEL-197 | chr8-112610690-DEL-199 |
| chr8-112615628-DEL-58 | chr8-112615629-DEL-58 |
| chr8-112619686-DEL-71 | chr8-112619687-DEL-71 |
| chr8-112620652-DEL-214 | chr8-112620653-DEL-212 |
| chr8-112624425-DEL-766 | chr8-112624404-DEL-771 |
| chr8-112626448-DEL-215 | chr8-112626450-DEL-211 |
| chr8-112630814-DEL-111 | chr8-112630815-DEL-105 |
| chr8-112635508-DEL-216 | chr8-112635533-DEL-192 |
| chr8-112644003-DEL-180 | chr8-112644016-DEL-181 |
| chr8-112649405-DEL-95 | chr8-112649402-DEL-96 |
| chr8-112677225-DEL-115 | chr8-112677225-DEL-163 |
| chr8-112684752-DEL-359 | chr8-112684753-DEL-360 |
| chr8-112685464-DEL-541 | chr8-112685465-DEL-541 |
| chr8-112688157-DEL-204 | chr8-112688157-DEL-204 |
| chr8-112696943-DEL-208 | chr8-112696944-DEL-208 |
| chr8-112699979-DEL-54 | chr8-112699980-DEL-54 |
| chr8-112713296-DEL-201 | chr8-112713297-DEL-201 |
| chr8-112722868-DEL-70 | chr8-112722869-DEL-58 |
| chr8-112731038-DEL-188 | chr8-112731045-DEL-182 |
| chr8-112731760-DEL-149 | chr8-112731761-DEL-149 |
| chr8-112740184-DEL-1509 | chr8-112740185-DEL-1509 |
| chr8-112743802-DEL-64 | chr8-112743803-DEL-64 |
| chr8-112777729-DEL-586 | chr8-112777727-DEL-585 |
| chr8-112779166-DEL-132 | chr8-112779175-DEL-123 |
| chr8-112780763-DEL-162 | chr8-112780763-DEL-170 |
| chr8-112791388-DEL-190 | chr8-112791387-DEL-190 |
| chr8-112808689-DEL-52 | chr8-112808688-DEL-54 |
| chr8-112821928-DEL-155 | chr8-112821913-DEL-171 |
| chr8-112855589-DEL-90 | chr8-112855590-DEL-90 |
| chr8-112867755-DEL-67 | chr8-112867756-DEL-67 |
| chr8-112871672-DEL-66 | chr8-112871673-DEL-66 |
| chr8-112892416-DEL-135 | chr8-112892441-DEL-105 |
| chr8-112953259-DEL-399 | chr8-112953260-DEL-399 |
| chr8-112956045-DEL-242 | chr8-112956046-DEL-242 |
| chr8-112964103-DEL-57 | chr8-112964079-DEL-58 |
| chr8-112978524-DEL-52 | chr8-112978513-DEL-66 |
| chr8-113016860-DEL-98 | chr8-113016861-DEL-98 |
| chr8-113067541-DEL-56 | chr8-113067529-DEL-95 |
| chr8-113068564-DEL-129 | chr8-113068561-DEL-120 |
| chr8-113094469-DEL-294 | chr8-113094470-DEL-294 |
| chr8-113113052-DEL-84 | chr8-113113085-DEL-52 |
| chr8-113165516-DEL-70 | chr8-113165517-DEL-70 |
| chr8-113169739-DEL-69 | chr8-113169740-DEL-69 |
| chr8-113179048-DEL-2014 | chr8-113179049-DEL-2014 |
| chr8-113206272-DEL-401 | chr8-113206273-DEL-401 |
| chr8-113209031-DEL-63 | chr8-113209032-DEL-63 |
| chr8-113210204-DEL-6710 | chr8-113210205-DEL-6710 |
| chr8-113225610-DEL-53 | chr8-113225609-DEL-55 |
| chr8-113246626-DEL-55 | chr8-113246627-DEL-55 |
| chr8-113258950-DEL-60 | chr8-113258951-DEL-60 |
| chr8-113275178-DEL-99 | chr8-113275179-DEL-99 |
| chr8-113284847-DEL-172 | chr8-113284848-DEL-172 |
| chr8-113310455-DEL-118 | chr8-113310465-DEL-108 |
| chr8-113333131-DEL-277 | chr8-113333132-DEL-277 |
| chr8-113371958-DEL-6618 | chr8-113371959-DEL-6618 |
| chr8-113402772-DEL-70 | chr8-113402751-DEL-86 |

| | |
|---|---|
| chr8-113420423-DEL-65 | chr8-113420424-DEL-65 |
| chr8-113438480-DEL-52 | chr8-113438481-DEL-52 |
| chr8-113446964-DEL-919 | chr8-113446965-DEL-919 |
| chr8-113463315-DEL-52 | chr8-113463316-DEL-56 |
| chr8-113481054-DEL-66 | chr8-113481054-DEL-66 |
| chr8-113507108-DEL-88 | chr8-113507145-DEL-56 |
| chr8-113516325-DEL-76 | chr8-113516330-DEL-50 |
| chr8-113517702-DEL-6765 | chr8-113517703-DEL-6765 |
| chr8-113553289-DEL-6448 | chr8-113553290-DEL-6448 |
| chr8-113559350-DEL-214 | chr8-113559351-DEL-214 |
| chr8-113565610-DEL-54 | chr8-113565611-DEL-54 |
| chr8-113576109-DEL-370 | chr8-113576110-DEL-366 |
| chr8-113580988-DEL-88 | chr8-113580998-DEL-52 |
| chr8-113582164-DEL-58 | chr8-113582177-DEL-54 |
| chr8-113636620-DEL-1486 | chr8-113636621-DEL-1486 |
| chr8-113664779-DEL-360 | chr8-113664780-DEL-360 |
| chr8-113687979-DEL-60 | chr8-113687987-DEL-56 |
| chr8-113714063-DEL-104 | chr8-113714061-DEL-97 |
| chr8-113730046-DEL-61 | chr8-113730047-DEL-61 |
| chr8-113783023-DEL-67 | chr8-113783024-DEL-67 |
| chr8-113811627-DEL-205 | chr8-113811626-DEL-206 |
| chr8-113826142-DEL-1763 | chr8-113826143-DEL-1763 |
| chr8-113841437-DEL-1174 | chr8-113841438-DEL-1174 |
| chr8-113860121-DEL-67 | chr8-113860122-DEL-67 |
| chr8-113867053-DEL-345 | chr8-113867054-DEL-345 |
| chr8-113881700-DEL-570 | chr8-113881701-DEL-570 |
| chr8-113959255-DEL-279 | chr8-113959256-DEL-279 |
| chr8-113982834-DEL-383 | chr8-113982835-DEL-383 |
| chr8-113995229-DEL-85 | chr8-113995234-DEL-81 |
| chr8-113996734-DEL-307 | chr8-113996735-DEL-307 |
| chr8-114006982-DEL-7182 | chr8-114006980-DEL-7184 |
| chr8-114015920-DEL-67 | chr8-114015919-DEL-116 |
| chr8-114022036-DEL-299 | chr8-114022037-DEL-299 |
| chr8-114023771-DEL-78 | chr8-114023784-DEL-96 |
| chr8-114050809-DEL-113 | chr8-114050838-DEL-78 |
| chr8-114054640-DEL-152 | chr8-114054641-DEL-152 |
| chr8-114065467-DEL-790 | chr8-114065468-DEL-790 |
| chr8-114106976-DEL-51 | chr8-114106977-DEL-51 |
| chr8-114112359-DEL-58 | chr8-114112364-DEL-54 |
| chr8-114113513-DEL-52 | chr8-114113514-DEL-52 |
| chr8-114154957-DEL-53 | chr8-114154958-DEL-53 |
| chr8-114184604-DEL-74 | chr8-114184605-DEL-74 |
| chr8-114185876-DEL-727 | chr8-114185874-DEL-726 |
| chr8-114195187-DEL-199 | chr8-114195187-DEL-200 |
| chr8-114248475-DEL-68 | chr8-114248476-DEL-68 |
| chr8-114256247-DEL-216 | chr8-114256208-DEL-264 |
| chr8-114260733-DEL-204 | chr8-114260733-DEL-204 |
| chr8-114287381-DEL-15719 | chr8-114287382-DEL-15719 |
| chr8-114291292-DEL-6881 | chr8-114291293-DEL-6881 |
| chr8-114303558-DEL-232 | chr8-114303559-DEL-232 |
| chr8-114315796-DEL-3180 | chr8-114315797-DEL-3180 |
| chr8-114327870-DEL-119 | chr8-114327871-DEL-119 |
| chr8-114328380-DEL-6200 | chr8-114328381-DEL-6200 |
| chr8-114356324-DEL-305 | chr8-114356325-DEL-304 |
| chr8-114377571-DEL-187 | chr8-114377560-DEL-168 |
| chr8-114390584-DEL-74 | chr8-114390583-DEL-73 |
| chr8-114398769-DEL-197 | chr8-114398770-DEL-197 |
| chr8-114415817-DEL-74 | chr8-114415818-DEL-74 |
| chr8-114438923-DEL-86 | chr8-114438924-DEL-86 |
| chr8-114445300-DEL-251 | chr8-114445301-DEL-251 |
| chr8-114492864-DEL-67 | chr8-114492866-DEL-66 |
| chr8-114513776-DEL-62 | chr8-114513809-DEL-52 |
| chr8-114524851-DEL-54 | chr8-114524806-DEL-58 |
| chr8-114527323-DEL-132 | chr8-114527324-DEL-132 |
| chr8-114544840-DEL-55 | chr8-114544860-DEL-58 |
| chr8-114547069-DEL-59 | chr8-114547071-DEL-50 |
| chr8-114555916-DEL-110 | chr8-114555939-DEL-88 |
| chr8-114627921-DEL-1055 | chr8-114627922-DEL-1055 |
| chr8-114646388-DEL-84 | chr8-114646389-DEL-84 |
| chr8-114649759-DEL-70 | chr8-114649754-DEL-71 |
| chr8-114651373-DEL-68 | chr8-114651374-DEL-68 |
| chr8-114672253-DEL-742 | chr8-114672254-DEL-742 |
| chr8-114688883-DEL-87 | chr8-114688884-DEL-87 |
| chr8-114693568-DEL-64 | chr8-114693559-DEL-86 |
| chr8-114697203-DEL-288 | chr8-114697204-DEL-288 |
| chr8-114700347-DEL-286 | chr8-114700348-DEL-286 |
| chr8-114707595-DEL-51 | chr8-114707596-DEL-51 |
| chr8-114713181-DEL-237 | chr8-114713182-DEL-237 |
| chr8-114719083-DEL-52 | chr8-114719084-DEL-52 |
| chr8-114746730-DEL-95 | chr8-114746714-DEL-96 |

| | |
|---|---|
| chr8-114768001-DEL-410 | chr8-114768002-DEL-410 |
| chr8-114772726-DEL-124 | chr8-114772738-DEL-112 |
| chr8-114782557-DEL-224 | chr8-114782558-DEL-224 |
| chr8-114790671-DEL-400 | chr8-114790672-DEL-400 |
| chr8-114795105-DEL-60 | chr8-114795110-DEL-56 |
| chr8-114814533-DEL-59 | chr8-114814529-DEL-86 |
| chr8-114839267-DEL-204 | chr8-114839268-DEL-204 |
| chr8-114842858-DEL-1189 | chr8-114842859-DEL-1189 |
| chr8-114859748-DEL-84 | chr8-114859722-DEL-110 |
| chr8-114875548-DEL-68 | chr8-114875549-DEL-68 |
| chr8-114881659-DEL-59 | chr8-114881659-DEL-57 |
| chr8-114882876-DEL-65 | chr8-114882837-DEL-66 |
| chr8-114887580-DEL-102 | chr8-114887581-DEL-102 |
| chr8-114896933-DEL-52 | chr8-114896934-DEL-52 |
| chr8-114913962-DEL-323 | chr8-114913963-DEL-323 |
| chr8-114917033-DEL-78 | chr8-114917020-DEL-68 |
| chr8-114928593-DEL-76 | chr8-114928594-DEL-76 |
| chr8-114939367-DEL-105 | chr8-114939367-DEL-117 |
| chr8-114988635-DEL-90 | chr8-114988631-DEL-95 |
| chr8-115028635-DEL-393 | chr8-115028636-DEL-393 |
| chr8-115029209-DEL-890 | chr8-115029210-DEL-890 |
| chr8-115040593-DEL-55 | chr8-115040625-DEL-56 |
| chr8-115043899-DEL-73 | chr8-115043862-DEL-110 |
| chr8-115050896-DEL-5006 | chr8-115050897-DEL-5006 |
| chr8-115056558-DEL-581 | chr8-115056559-DEL-581 |
| chr8-115064599-DEL-241 | chr8-115064600-DEL-241 |
| chr8-115065386-DEL-58 | chr8-115065387-DEL-58 |
| chr8-115068801-DEL-218 | chr8-115068799-DEL-219 |
| chr8-115074223-DEL-52 | chr8-115074224-DEL-52 |
| chr8-115077635-DEL-100 | chr8-115077590-DEL-116 |
| chr8-115117838-DEL-7160 | chr8-115117839-DEL-7160 |
| chr8-115127031-DEL-170 | chr8-115127032-DEL-170 |
| chr8-115131337-DEL-93 | chr8-115131342-DEL-94 |
| chr8-115135767-DEL-52 | chr8-115135778-DEL-54 |
| chr8-115155948-DEL-182 | chr8-115155946-DEL-186 |
| chr8-115190549-DEL-1240 | chr8-115190550-DEL-1240 |
| chr8-115197807-DEL-168 | chr8-115197808-DEL-168 |
| chr8-115204136-DEL-54 | chr8-115204137-DEL-54 |
| chr8-115214477-DEL-88 | chr8-115214478-DEL-88 |
| chr8-115255824-DEL-52 | chr8-115255825-DEL-52 |
| chr8-115278355-DEL-137 | chr8-115278356-DEL-137 |
| chr8-115447355-DEL-118 | chr8-115447355-DEL-124 |
| chr8-115462609-DEL-752 | chr8-115462610-DEL-752 |
| chr8-115462690-DEL-55 | chr8-115462691-DEL-55 |
| chr8-115467030-DEL-1288 | chr8-115467031-DEL-1288 |
| chr8-115467109-DEL-82 | chr8-115467109-DEL-82 |
| chr8-115487210-DEL-51 | chr8-115487216-DEL-51 |
| chr8-115488079-DEL-201 | chr8-115488080-DEL-201 |
| chr8-115507578-DEL-333 | chr8-115507596-DEL-323 |
| chr8-115557967-DEL-58 | chr8-115557968-DEL-58 |
| chr8-115590614-DEL-230 | chr8-115590615-DEL-230 |
| chr8-115629826-DEL-399 | chr8-115629827-DEL-399 |
| chr8-115630405-DEL-356 | chr8-115630406-DEL-356 |
| chr8-115640778-DEL-59 | chr8-115640779-DEL-59 |
| chr8-115681063-DEL-313 | chr8-115681064-DEL-313 |
| chr8-115700539-DEL-168 | chr8-115700540-DEL-168 |
| chr8-115734376-DEL-68 | chr8-115734377-DEL-64 |
| chr8-115737794-DEL-53 | chr8-115737795-DEL-53 |
| chr8-115744808-DEL-100 | chr8-115744829-DEL-60 |
| chr8-115757586-DEL-57 | chr8-115757587-DEL-56 |
| chr8-115771699-DEL-58 | chr8-115771689-DEL-80 |
| chr8-115789586-DEL-74 | chr8-115789587-DEL-74 |
| chr8-115792972-DEL-66 | chr8-115792973-DEL-66 |
| chr8-115827619-DEL-54 | chr8-115827620-DEL-54 |
| chr8-115844965-DEL-1677 | chr8-115844966-DEL-1677 |
| chr8-115855712-DEL-99 | chr8-115855709-DEL-100 |
| chr8-115863983-DEL-81 | chr8-115863984-DEL-81 |
| chr8-115867317-DEL-360 | chr8-115867319-DEL-358 |
| chr8-115874930-DEL-1293 | chr8-115874933-DEL-1291 |
| chr8-115891164-DEL-57 | chr8-115891165-DEL-57 |
| chr8-115901543-DEL-1708 | chr8-115901544-DEL-1708 |
| chr8-115920873-DEL-98 | chr8-115920874-DEL-98 |
| chr8-115931432-DEL-102 | chr8-115931433-DEL-102 |
| chr8-115944417-DEL-70 | chr8-115944406-DEL-82 |
| chr8-115981052-DEL-77 | chr8-115981050-DEL-78 |
| chr8-116003398-DEL-52 | chr8-116003383-DEL-68 |
| chr8-116007472-DEL-498 | chr8-116007472-DEL-498 |
| chr8-116008655-DEL-5838 | chr8-116008656-DEL-5838 |
| chr8-116052861-DEL-106 | chr8-116052862-DEL-106 |
| chr8-116117081-DEL-60 | chr8-116117082-DEL-60 |

| | |
|---|---|
| chr8-116118328-DEL-485 | chr8-116118329-DEL-485 |
| chr8-116161674-DEL-818 | chr8-116161675-DEL-818 |
| chr8-116182035-DEL-72 | chr8-116182056-DEL-60 |
| chr8-116186388-DEL-775 | chr8-116186389-DEL-775 |
| chr8-116194856-DEL-564 | chr8-116194857-DEL-564 |
| chr8-116208384-DEL-239 | chr8-116208385-DEL-239 |
| chr8-116249651-DEL-442 | chr8-116249652-DEL-442 |
| chr8-116282711-DEL-568 | chr8-116282712-DEL-568 |
| chr8-116284151-DEL-66 | chr8-116284152-DEL-66 |
| chr8-116294061-DEL-871 | chr8-116294062-DEL-871 |
| chr8-116299250-DEL-139 | chr8-116299257-DEL-99 |
| chr8-116328215-DEL-374 | chr8-116328201-DEL-375 |
| chr8-116350844-DEL-400 | chr8-116350845-DEL-400 |
| chr8-116373065-DEL-62 | chr8-116373066-DEL-62 |
| chr8-116442527-DEL-74 | chr8-116442528-DEL-68 |
| chr8-116457907-DEL-59 | chr8-116457908-DEL-59 |
| chr8-116461574-DEL-2719 | chr8-116461575-DEL-2719 |
| chr8-116476792-DEL-51 | chr8-116476821-DEL-68 |
| chr8-116489458-DEL-91 | chr8-116489411-DEL-135 |
| chr8-116523703-DEL-6450 | chr8-116523703-DEL-6450 |
| chr8-116538110-DEL-82 | chr8-116538111-DEL-82 |
| chr8-116595886-DEL-71 | chr8-116595887-DEL-71 |
| chr8-116654463-DEL-173 | chr8-116654464-DEL-173 |
| chr8-116657978-DEL-55 | chr8-116657979-DEL-55 |
| chr8-116660123-DEL-52 | chr8-116660119-DEL-56 |
| chr8-116698379-DEL-423 | chr8-116698380-DEL-407 |
| chr8-116699501-DEL-58 | chr8-116699503-DEL-56 |
| chr8-116715647-DEL-248 | chr8-116715650-DEL-246 |
| chr8-116737965-DEL-52 | chr8-116737966-DEL-64 |
| chr8-116745935-DEL-141 | chr8-116745936-DEL-141 |
| chr8-116748875-DEL-92 | chr8-116748884-DEL-56 |
| chr8-116757316-DEL-66 | chr8-116757317-DEL-66 |
| chr8-116778660-DEL-77 | chr8-116778661-DEL-77 |
| chr8-116833516-DEL-335 | chr8-116833517-DEL-335 |
| chr8-116833822-DEL-1566 | chr8-116833823-DEL-1566 |
| chr8-116840198-DEL-70 | chr8-116840241-DEL-62 |
| chr8-116842264-DEL-116 | chr8-116842280-DEL-100 |
| chr8-116868984-DEL-118 | chr8-116869019-DEL-70 |
| chr8-116946250-DEL-342 | chr8-116946251-DEL-342 |
| chr8-116948799-DEL-401 | chr8-116948800-DEL-401 |
| chr8-116954829-DEL-302 | chr8-116954830-DEL-294 |
| chr8-117014322-DEL-85 | chr8-117014323-DEL-85 |
| chr8-117069869-DEL-63 | chr8-117069847-DEL-56 |
| chr8-117077775-DEL-392 | chr8-117077776-DEL-392 |
| chr8-117093905-DEL-185 | chr8-117093953-DEL-138 |
| chr8-117128001-DEL-209 | chr8-117128002-DEL-209 |
| chr8-117150104-DEL-70 | chr8-117150105-DEL-70 |
| chr8-117159075-DEL-249 | chr8-117159076-DEL-249 |
| chr8-117209042-DEL-63 | chr8-117209043-DEL-63 |
| chr8-117234563-DEL-210 | chr8-117234602-DEL-183 |
| chr8-117257022-DEL-1721 | chr8-117257023-DEL-1721 |
| chr8-117257710-DEL-177 | chr8-117257702-DEL-186 |
| chr8-117268426-DEL-122 | chr8-117268430-DEL-119 |
| chr8-117274333-DEL-92 | chr8-117274340-DEL-86 |
| chr8-117305181-DEL-124 | chr8-117305182-DEL-124 |
| chr8-117327227-DEL-61 | chr8-117327227-DEL-93 |
| chr8-117331926-DEL-54 | chr8-117331927-DEL-54 |
| chr8-117342655-DEL-56 | chr8-117342649-DEL-64 |
| chr8-117373276-DEL-439 | chr8-117373277-DEL-439 |
| chr8-117379079-DEL-202 | chr8-117379080-DEL-202 |
| chr8-117411852-DEL-68 | chr8-117411853-DEL-68 |
| chr8-117492929-DEL-315 | chr8-117492930-DEL-315 |
| chr8-117499370-DEL-88 | chr8-117499371-DEL-88 |
| chr8-117509822-DEL-96 | chr8-117509823-DEL-96 |
| chr8-117510502-DEL-180 | chr8-117510503-DEL-180 |
| chr8-117518639-DEL-400 | chr8-117518640-DEL-400 |
| chr8-117521169-DEL-147 | chr8-117521170-DEL-147 |
| chr8-117522832-DEL-397 | chr8-117522833-DEL-397 |
| chr8-117525363-DEL-72 | chr8-117525384-DEL-62 |
| chr8-117531619-DEL-448 | chr8-117531620-DEL-448 |
| chr8-117534186-DEL-154 | chr8-117534187-DEL-154 |
| chr8-117541881-DEL-63 | chr8-117541886-DEL-64 |
| chr8-117542008-DEL-1837 | chr8-117541999-DEL-1838 |
| chr8-117567703-DEL-76 | chr8-117567697-DEL-52 |
| chr8-117576638-DEL-90 | chr8-117576641-DEL-68 |
| chr8-117595790-DEL-51 | chr8-117595791-DEL-51 |
| chr8-117596762-DEL-56 | chr8-117596752-DEL-65 |
| chr8-117605493-DEL-178 | chr8-117605499-DEL-169 |
| chr8-117608321-DEL-68 | chr8-117608322-DEL-53 |
| chr8-117608826-DEL-268 | chr8-117608827-DEL-268 |

| | |
|---|---|
| chr8-117622998-DEL-7167 | chr8-117622999-DEL-7167 |
| chr8-117673911-DEL-3725 | chr8-117673912-DEL-3725 |
| chr8-117678466-DEL-77 | chr8-117678487-DEL-73 |
| chr8-117687744-DEL-208 | chr8-117687745-DEL-208 |
| chr8-117701425-DEL-86 | chr8-117701426-DEL-86 |
| chr8-117729298-DEL-5153 | chr8-117729299-DEL-5153 |
| chr8-117744823-DEL-398 | chr8-117744824-DEL-398 |
| chr8-117748079-DEL-54 | chr8-117748080-DEL-52 |
| chr8-117752624-DEL-190 | chr8-117752625-DEL-190 |
| chr8-117768161-DEL-98 | chr8-117768190-DEL-70 |
| chr8-117784910-DEL-62 | chr8-117784911-DEL-62 |
| chr8-117798797-DEL-154 | chr8-117798798-DEL-154 |
| chr8-117870472-DEL-54 | chr8-117870473-DEL-54 |
| chr8-117872219-DEL-52 | chr8-117872220-DEL-52 |
| chr8-117913200-DEL-62 | chr8-117913219-DEL-55 |
| chr8-117949045-DEL-270 | chr8-117949046-DEL-270 |
| chr8-117975533-DEL-221 | chr8-117975534-DEL-221 |
| chr8-117982350-DEL-70 | chr8-117982350-DEL-70 |
| chr8-118008127-DEL-203 | chr8-118008128-DEL-203 |
| chr8-118040423-DEL-199 | chr8-118040424-DEL-199 |
| chr8-118054643-DEL-86 | chr8-118054648-DEL-70 |
| chr8-118080854-DEL-75 | chr8-118080855-DEL-75 |
| chr8-118154801-DEL-205 | chr8-118154814-DEL-209 |
| chr8-118189542-DEL-258 | chr8-118189576-DEL-225 |
| chr8-118191597-DEL-195 | chr8-118191585-DEL-193 |
| chr8-118194876-DEL-79 | chr8-118194877-DEL-79 |
| chr8-118200521-DEL-248 | chr8-118200502-DEL-246 |
| chr8-118202009-DEL-172 | chr8-118202010-DEL-172 |
| chr8-118240245-DEL-64 | chr8-118240246-DEL-64 |
| chr8-118247190-DEL-178 | chr8-118247191-DEL-179 |
| chr8-118287474-DEL-160 | chr8-118287453-DEL-161 |
| chr8-118296529-DEL-54 | chr8-118296529-DEL-54 |
| chr8-118301080-DEL-64 | chr8-118301078-DEL-68 |
| chr8-118306187-DEL-60 | chr8-118306188-DEL-60 |
| chr8-118314958-DEL-94 | chr8-118314959-DEL-94 |
| chr8-118319541-DEL-2903 | chr8-118319542-DEL-2903 |
| chr8-118334173-DEL-379 | chr8-118334174-DEL-379 |
| chr8-118335735-DEL-68 | chr8-118335736-DEL-68 |
| chr8-118356290-DEL-112 | chr8-118356295-DEL-107 |
| chr8-118379857-DEL-76 | chr8-118379858-DEL-76 |
| chr8-118417325-DEL-66 | chr8-118417326-DEL-66 |
| chr8-118421673-DEL-52 | chr8-118421674-DEL-52 |
| chr8-118457392-DEL-55 | chr8-118457393-DEL-55 |
| chr8-118458816-DEL-144 | chr8-118458817-DEL-144 |
| chr8-118472738-DEL-84 | chr8-118472741-DEL-81 |
| chr8-118479315-DEL-488 | chr8-118479313-DEL-489 |
| chr8-118494000-DEL-404 | chr8-118494001-DEL-404 |
| chr8-118500102-DEL-96 | chr8-118500103-DEL-96 |
| chr8-118523418-DEL-51 | chr8-118523419-DEL-51 |
| chr8-118584687-DEL-207 | chr8-118584688-DEL-207 |
| chr8-118585895-DEL-75 | chr8-118585896-DEL-77 |
| chr8-118603082-DEL-68 | chr8-118603083-DEL-68 |
| chr8-118623623-DEL-53 | chr8-118623624-DEL-53 |
| chr8-118640844-DEL-58 | chr8-118640845-DEL-58 |
| chr8-118642513-DEL-58 | chr8-118642514-DEL-58 |
| chr8-118670091-DEL-674 | chr8-118670092-DEL-674 |
| chr8-118676080-DEL-124 | chr8-118676081-DEL-124 |
| chr8-118704808-DEL-6675 | chr8-118704809-DEL-6675 |
| chr8-118704824-DEL-214 | chr8-118704825-DEL-214 |
| chr8-118711871-DEL-712 | chr8-118711872-DEL-712 |
| chr8-118712957-DEL-303 | chr8-118712958-DEL-303 |
| chr8-118717450-DEL-55 | chr8-118717451-DEL-55 |
| chr8-118720194-DEL-80 | chr8-118720197-DEL-87 |
| chr8-118746172-DEL-168 | chr8-118746173-DEL-168 |
| chr8-118761794-DEL-112 | chr8-118761798-DEL-100 |
| chr8-118770692-DEL-6189 | chr8-118770686-DEL-6190 |
| chr8-118791481-DEL-746 | chr8-118791482-DEL-746 |
| chr8-118830329-DEL-125 | chr8-118830355-DEL-100 |
| chr8-118884676-DEL-377 | chr8-118884677-DEL-375 |
| chr8-118887614-DEL-55 | chr8-118887607-DEL-53 |
| chr8-118902793-DEL-75 | chr8-118902794-DEL-75 |
| chr8-118903808-DEL-68 | chr8-118903812-DEL-53 |
| chr8-118922645-DEL-401 | chr8-118922646-DEL-401 |
| chr8-118923181-DEL-468 | chr8-118923182-DEL-468 |
| chr8-118950486-DEL-399 | chr8-118950487-DEL-399 |
| chr8-118993537-DEL-112 | chr8-118993538-DEL-112 |
| chr8-119104107-DEL-74 | chr8-119104150-DEL-62 |
| chr8-119109844-DEL-156 | chr8-119109845-DEL-153 |
| chr8-119160127-DEL-164 | chr8-119160128-DEL-164 |
| chr8-119231032-DEL-81 | chr8-119231036-DEL-53 |

| | |
|---|---|
| chr8-119238030-DEL-140 | chr8-119238031-DEL-140 |
| chr8-119294135-DEL-54 | chr8-119294136-DEL-54 |
| chr8-119350588-DEL-59 | chr8-119350589-DEL-59 |
| chr8-119356397-DEL-64 | chr8-119356403-DEL-60 |
| chr8-119370914-DEL-803 | chr8-119370915-DEL-803 |
| chr8-119384900-DEL-1264 | chr8-119384901-DEL-1264 |
| chr8-119397204-DEL-685 | chr8-119397195-DEL-663 |
| chr8-119485193-DEL-6223 | chr8-119485194-DEL-6223 |
| chr8-119493697-DEL-86 | chr8-119493698-DEL-86 |
| chr8-119513461-DEL-71 | chr8-119513461-DEL-71 |
| chr8-119561030-DEL-187 | chr8-119561031-DEL-187 |
| chr8-119626093-DEL-635 | chr8-119626137-DEL-600 |
| chr8-119631939-DEL-349 | chr8-119631940-DEL-349 |
| chr8-119633532-DEL-58 | chr8-119633520-DEL-59 |
| chr8-119637975-DEL-88 | chr8-119637976-DEL-88 |
| chr8-119647240-DEL-176 | chr8-119647233-DEL-140 |
| chr8-119686405-DEL-60 | chr8-119686406-DEL-60 |
| chr8-119717875-DEL-333 | chr8-119717876-DEL-333 |
| chr8-119719020-DEL-169 | chr8-119719020-DEL-135 |
| chr8-119771929-DEL-7166 | chr8-119771930-DEL-7166 |
| chr8-119794457-DEL-64 | chr8-119794466-DEL-56 |
| chr8-119795135-DEL-71 | chr8-119795135-DEL-71 |
| chr8-119841029-DEL-60 | chr8-119841019-DEL-64 |
| chr8-119841276-DEL-477 | chr8-119841277-DEL-477 |
| chr8-119848903-DEL-62 | chr8-119848904-DEL-62 |
| chr8-119855433-DEL-968 | chr8-119855434-DEL-968 |
| chr8-119886058-DEL-82 | chr8-119886059-DEL-108 |
| chr8-119906923-DEL-208 | chr8-119906924-DEL-208 |
| chr8-119921671-DEL-52 | chr8-119921676-DEL-53 |
| chr8-119931364-DEL-77 | chr8-119931375-DEL-66 |
| chr8-119968789-DEL-71 | chr8-119968790-DEL-71 |
| chr8-119973763-DEL-76 | chr8-119973792-DEL-57 |
| chr8-119988367-DEL-79 | chr8-119988338-DEL-75 |
| chr8-119998097-DEL-288 | chr8-119998098-DEL-288 |
| chr8-120057266-DEL-56 | chr8-120057267-DEL-56 |
| chr8-120057729-DEL-7091 | chr8-120057730-DEL-7091 |
| chr8-120076685-DEL-344 | chr8-120076686-DEL-344 |
| chr8-120083105-DEL-440 | chr8-120083106-DEL-440 |
| chr8-120088637-DEL-75 | chr8-120088638-DEL-75 |
| chr8-120097150-DEL-114 | chr8-120097175-DEL-114 |
| chr8-120098692-DEL-130 | chr8-120098693-DEL-130 |
| chr8-120118811-DEL-218 | chr8-120118811-DEL-218 |
| chr8-120201078-DEL-1530 | chr8-120201079-DEL-1530 |
| chr8-120209919-DEL-126 | chr8-120209919-DEL-126 |
| chr8-120229609-DEL-79 | chr8-120229610-DEL-79 |
| chr8-120288064-DEL-81 | chr8-120288066-DEL-80 |
| chr8-120338250-DEL-168 | chr8-120338258-DEL-163 |
| chr8-120368514-DEL-96 | chr8-120368515-DEL-96 |
| chr8-120399230-DEL-1858 | chr8-120399231-DEL-1858 |
| chr8-120403420-DEL-257 | chr8-120403421-DEL-257 |
| chr8-120416967-DEL-200 | chr8-120416968-DEL-200 |
| chr8-120464048-DEL-212 | chr8-120464049-DEL-212 |
| chr8-120478917-DEL-72 | chr8-120478918-DEL-72 |
| chr8-120496166-DEL-204 | chr8-120496167-DEL-204 |
| chr8-120563184-DEL-80 | chr8-120563185-DEL-80 |
| chr8-120581912-DEL-205 | chr8-120581913-DEL-205 |
| chr8-120599460-DEL-233 | chr8-120599461-DEL-233 |
| chr8-120610455-DEL-110 | chr8-120610439-DEL-111 |
| chr8-120616300-DEL-7480 | chr8-120616300-DEL-7480 |
| chr8-120636098-DEL-78 | chr8-120636104-DEL-58 |
| chr8-120709570-DEL-55 | chr8-120709571-DEL-55 |
| chr8-120712900-DEL-57 | chr8-120712906-DEL-54 |
| chr8-120716152-DEL-114 | chr8-120716180-DEL-114 |
| chr8-120726276-DEL-159 | chr8-120726277-DEL-159 |
| chr8-120735396-DEL-98 | chr8-120735394-DEL-99 |
| chr8-120787401-DEL-56 | chr8-120787393-DEL-56 |
| chr8-120809632-DEL-129 | chr8-120809633-DEL-129 |
| chr8-120838688-DEL-54 | chr8-120838689-DEL-54 |
| chr8-120855037-DEL-231 | chr8-120855038-DEL-231 |
| chr8-120868987-DEL-98 | chr8-120868988-DEL-98 |
| chr8-120874055-DEL-76 | chr8-120874075-DEL-56 |
| chr8-120881970-DEL-208 | chr8-120881971-DEL-208 |
| chr8-120896887-DEL-263 | chr8-120896888-DEL-263 |
| chr8-120919323-DEL-7132 | chr8-120919324-DEL-7132 |
| chr8-120927379-DEL-184 | chr8-120927398-DEL-204 |
| chr8-120935622-DEL-60 | chr8-120935629-DEL-50 |
| chr8-120971802-DEL-220 | chr8-120971803-DEL-220 |
| chr8-120975455-DEL-70 | chr8-120975454-DEL-71 |
| chr8-120989661-DEL-101 | chr8-120989682-DEL-81 |
| chr8-120998705-DEL-208 | chr8-120998706-DEL-208 |

| | |
|---|---|
| chr8-121006468-DEL-170 | chr8-121006456-DEL-171 |
| chr8-121042992-DEL-74 | chr8-121042993-DEL-76 |
| chr8-121052733-DEL-196 | chr8-121052735-DEL-195 |
| chr8-121085094-DEL-54 | chr8-121085098-DEL-52 |
| chr8-121158649-DEL-78 | chr8-121158604-DEL-116 |
| chr8-121182651-DEL-220 | chr8-121182650-DEL-221 |
| chr8-121183517-DEL-106 | chr8-121183518-DEL-106 |
| chr8-121243690-DEL-60 | chr8-121243691-DEL-50 |
| chr8-121272936-DEL-162 | chr8-121272937-DEL-162 |
| chr8-121310452-DEL-198 | chr8-121310453-DEL-198 |
| chr8-121317713-DEL-210 | chr8-121317714-DEL-210 |
| chr8-121327821-DEL-427 | chr8-121327822-DEL-427 |
| chr8-121339284-DEL-180 | chr8-121339285-DEL-180 |
| chr8-121350294-DEL-115 | chr8-121350269-DEL-125 |
| chr8-121370117-DEL-57 | chr8-121370118-DEL-57 |
| chr8-121403367-DEL-189 | chr8-121403368-DEL-189 |
| chr8-121448950-DEL-53 | chr8-121448951-DEL-53 |
| chr8-121459085-DEL-241 | chr8-121459086-DEL-241 |
| chr8-121503489-DEL-68 | chr8-121503490-DEL-68 |
| chr8-121513071-DEL-75 | chr8-121513065-DEL-75 |
| chr8-121571271-DEL-56 | chr8-121571272-DEL-56 |
| chr8-121655985-DEL-59 | chr8-121655983-DEL-60 |
| chr8-121667932-DEL-105 | chr8-121667933-DEL-105 |
| chr8-121678895-DEL-58 | chr8-121678896-DEL-58 |
| chr8-121713317-DEL-83 | chr8-121713312-DEL-80 |
| chr8-121766577-DEL-100 | chr8-121766561-DEL-81 |
| chr8-121774886-DEL-153 | chr8-121774907-DEL-141 |
| chr8-121821645-DEL-74 | chr8-121821648-DEL-65 |
| chr8-121872435-DEL-216 | chr8-121872436-DEL-216 |
| chr8-121878629-DEL-100 | chr8-121878626-DEL-97 |
| chr8-121919836-DEL-107 | chr8-121919837-DEL-107 |
| chr8-121972521-DEL-349 | chr8-121972521-DEL-350 |
| chr8-121974350-DEL-75 | chr8-121974351-DEL-75 |
| chr8-121974884-DEL-151 | chr8-121974860-DEL-176 |
| chr8-121977075-DEL-113 | chr8-121977092-DEL-103 |
| chr8-122048292-DEL-56 | chr8-122048293-DEL-56 |
| chr8-122056505-DEL-126 | chr8-122056506-DEL-126 |
| chr8-122060611-DEL-128 | chr8-122060593-DEL-129 |
| chr8-122066482-DEL-63 | chr8-122066473-DEL-62 |
| chr8-122074829-DEL-80 | chr8-122074830-DEL-80 |
| chr8-122125464-DEL-54 | chr8-122125446-DEL-92 |
| chr8-122132024-DEL-7252 | chr8-122132025-DEL-7252 |
| chr8-122142748-DEL-73 | chr8-122142757-DEL-62 |
| chr8-122159199-DEL-60 | chr8-122159240-DEL-55 |
| chr8-122167935-DEL-59 | chr8-122167906-DEL-85 |
| chr8-122171183-DEL-60 | chr8-122171184-DEL-60 |
| chr8-122180743-DEL-75 | chr8-122180744-DEL-50 |
| chr8-122186984-DEL-63 | chr8-122186985-DEL-63 |
| chr8-122189808-DEL-393 | chr8-122189827-DEL-377 |
| chr8-122193791-DEL-70 | chr8-122193791-DEL-70 |
| chr8-122220468-DEL-84 | chr8-122220469-DEL-96 |
| chr8-122247312-DEL-64 | chr8-122247313-DEL-63 |
| chr8-122251263-DEL-66 | chr8-122251248-DEL-67 |
| chr8-122261047-DEL-244 | chr8-122261067-DEL-205 |
| chr8-122284119-DEL-224 | chr8-122284111-DEL-226 |
| chr8-122288364-DEL-52 | chr8-122288365-DEL-52 |
| chr8-122303347-DEL-102 | chr8-122303348-DEL-102 |
| chr8-122330217-DEL-206 | chr8-122330218-DEL-206 |
| chr8-122351747-DEL-68 | chr8-122351748-DEL-55 |
| chr8-122394349-DEL-68 | chr8-122394350-DEL-68 |
| chr8-122417345-DEL-72 | chr8-122417346-DEL-72 |
| chr8-122425781-DEL-226 | chr8-122425780-DEL-242 |
| chr8-122439042-DEL-200 | chr8-122439043-DEL-200 |
| chr8-122475748-DEL-55 | chr8-122475743-DEL-52 |
| chr8-122528704-DEL-204 | chr8-122528705-DEL-204 |
| chr8-122534540-DEL-851 | chr8-122534543-DEL-850 |
| chr8-122540967-DEL-102 | chr8-122540968-DEL-102 |
| chr8-122543129-DEL-64 | chr8-122543130-DEL-64 |
| chr8-122544625-DEL-1008 | chr8-122544626-DEL-1008 |
| chr8-122559579-DEL-8424 | chr8-122559580-DEL-8424 |
| chr8-122575981-DEL-179 | chr8-122575982-DEL-179 |
| chr8-122580633-DEL-72 | chr8-122580634-DEL-72 |
| chr8-122588308-DEL-254 | chr8-122588309-DEL-254 |
| chr8-122595546-DEL-135 | chr8-122595547-DEL-135 |
| chr8-122601365-DEL-88 | chr8-122601366-DEL-88 |
| chr8-122605322-DEL-200 | chr8-122605324-DEL-199 |
| chr8-122611379-DEL-95 | chr8-122611380-DEL-95 |
| chr8-122629072-DEL-68 | chr8-122629073-DEL-68 |
| chr8-122638082-DEL-117 | chr8-122638083-DEL-117 |
| chr8-122669574-DEL-88 | chr8-122669575-DEL-85 |

| | |
|---|---|
| chr8-122697750-DEL-126 | chr8-122697751-DEL-126 |
| chr8-122732841-DEL-229 | chr8-122732842-DEL-230 |
| chr8-122736716-DEL-58 | chr8-122736717-DEL-58 |
| chr8-122740354-DEL-156 | chr8-122740355-DEL-156 |
| chr8-122774898-DEL-70 | chr8-122774913-DEL-58 |
| chr8-122787720-DEL-73 | chr8-122787721-DEL-73 |
| chr8-122808355-DEL-877 | chr8-122808356-DEL-877 |
| chr8-122834496-DEL-425 | chr8-122834492-DEL-426 |
| chr8-122858679-DEL-72 | chr8-122858680-DEL-72 |
| chr8-122858769-DEL-682 | chr8-122858770-DEL-682 |
| chr8-122862058-DEL-2808 | chr8-122862059-DEL-2808 |
| chr8-122882739-DEL-87 | chr8-122882733-DEL-88 |
| chr8-122884987-DEL-1892 | chr8-122884988-DEL-1892 |
| chr8-122906543-DEL-80 | chr8-122906544-DEL-80 |
| chr8-122934282-DEL-58 | chr8-122934302-DEL-56 |
| chr8-122945942-DEL-154 | chr8-122945942-DEL-154 |
| chr8-122953718-DEL-94 | chr8-122953743-DEL-70 |
| chr8-123005501-DEL-145 | chr8-123005502-DEL-145 |
| chr8-123050859-DEL-82 | chr8-123050860-DEL-82 |
| chr8-123069075-DEL-367 | chr8-123069076-DEL-367 |
| chr8-123083501-DEL-77 | chr8-123083502-DEL-77 |
| chr8-123089742-DEL-63 | chr8-123089743-DEL-63 |
| chr8-123128888-DEL-66 | chr8-123128889-DEL-66 |
| chr8-123174053-DEL-78 | chr8-123174054-DEL-78 |
| chr8-123176225-DEL-62 | chr8-123176226-DEL-62 |
| chr8-123178693-DEL-187 | chr8-123178694-DEL-187 |
| chr8-123205159-DEL-64 | chr8-123205160-DEL-64 |
| chr8-123209486-DEL-67 | chr8-123209487-DEL-67 |
| chr8-123223868-DEL-63 | chr8-123223853-DEL-85 |
| chr8-123256474-DEL-764 | chr8-123256475-DEL-764 |
| chr8-123279462-DEL-77 | chr8-123279461-DEL-78 |
| chr8-123282089-DEL-52 | chr8-123282090-DEL-52 |
| chr8-123297807-DEL-61 | chr8-123297808-DEL-61 |
| chr8-123316337-DEL-60 | chr8-123316338-DEL-60 |
| chr8-123321168-DEL-1001 | chr8-123321169-DEL-1001 |
| chr8-123333841-DEL-198 | chr8-123333842-DEL-198 |
| chr8-123343692-DEL-209 | chr8-123343693-DEL-209 |
| chr8-123354870-DEL-63 | chr8-123354870-DEL-61 |
| chr8-123357947-DEL-608 | chr8-123357948-DEL-608 |
| chr8-123410341-DEL-726 | chr8-123410342-DEL-726 |
| chr8-123416887-DEL-77 | chr8-123416876-DEL-78 |
| chr8-123430222-DEL-78 | chr8-123430223-DEL-78 |
| chr8-123465306-DEL-55 | chr8-123465307-DEL-55 |
| chr8-123468054-DEL-60 | chr8-123468055-DEL-60 |
| chr8-123475680-DEL-98 | chr8-123475681-DEL-99 |
| chr8-123484093-DEL-111 | chr8-123484094-DEL-111 |
| chr8-123555937-DEL-53 | chr8-123555938-DEL-53 |
| chr8-123568313-DEL-703 | chr8-123568314-DEL-703 |
| chr8-123569599-DEL-203 | chr8-123569600-DEL-203 |
| chr8-123589221-DEL-52 | chr8-123589221-DEL-52 |
| chr8-123609906-DEL-82 | chr8-123609907-DEL-82 |
| chr8-123613447-DEL-134 | chr8-123613448-DEL-134 |
| chr8-123742633-DEL-2484 | chr8-123742634-DEL-2484 |
| chr8-123750208-DEL-184 | chr8-123750209-DEL-184 |
| chr8-123760022-DEL-51 | chr8-123759991-DEL-68 |
| chr8-123771467-DEL-100 | chr8-123771468-DEL-100 |
| chr8-123778044-DEL-71 | chr8-123778015-DEL-90 |
| chr8-123779667-DEL-205 | chr8-123779668-DEL-205 |
| chr8-123785905-DEL-217 | chr8-123785906-DEL-217 |
| chr8-123796121-DEL-147 | chr8-123796122-DEL-147 |
| chr8-123797666-DEL-232 | chr8-123797666-DEL-231 |
| chr8-123817741-DEL-348 | chr8-123817742-DEL-348 |
| chr8-123819361-DEL-520 | chr8-123819362-DEL-520 |
| chr8-123835313-DEL-70 | chr8-123835314-DEL-70 |
| chr8-123858611-DEL-96 | chr8-123858611-DEL-94 |
| chr8-123862841-DEL-200 | chr8-123862842-DEL-200 |
| chr8-123870020-DEL-172 | chr8-123870021-DEL-172 |
| chr8-123876887-DEL-71 | chr8-123876888-DEL-71 |
| chr8-123879010-DEL-131 | chr8-123879009-DEL-82 |
| chr8-123887618-DEL-267 | chr8-123887619-DEL-267 |
| chr8-123889355-DEL-54 | chr8-123889355-DEL-54 |
| chr8-123891450-DEL-7366 | chr8-123891451-DEL-7366 |
| chr8-123900775-DEL-1931 | chr8-123900776-DEL-1931 |
| chr8-123906124-DEL-131 | chr8-123906132-DEL-88 |
| chr8-123916727-DEL-55 | chr8-123916728-DEL-55 |
| chr8-123918387-DEL-328 | chr8-123918388-DEL-328 |
| chr8-123919592-DEL-102 | chr8-123919596-DEL-107 |
| chr8-123923899-DEL-60 | chr8-123923899-DEL-60 |
| chr8-123930369-DEL-108 | chr8-123930370-DEL-108 |
| chr8-123933245-DEL-559 | chr8-123933246-DEL-559 |

| | |
|---|---|
| chr8-123944869-DEL-52 | chr8-123944870-DEL-50 |
| chr8-123966412-DEL-1126 | chr8-123966413-DEL-1126 |
| chr8-123989463-DEL-111 | chr8-123989464-DEL-111 |
| chr8-123991289-DEL-168 | chr8-123991290-DEL-168 |
| chr8-124004653-DEL-66 | chr8-124004652-DEL-64 |
| chr8-124035438-DEL-1590 | chr8-124035439-DEL-1590 |
| chr8-124038026-DEL-196 | chr8-124038027-DEL-196 |
| chr8-124042117-DEL-172 | chr8-124042118-DEL-172 |
| chr8-124048136-DEL-216 | chr8-124048137-DEL-216 |
| chr8-124050413-DEL-569 | chr8-124050420-DEL-560 |
| chr8-124051332-DEL-505 | chr8-124051325-DEL-509 |
| chr8-124055468-DEL-81 | chr8-124055469-DEL-81 |
| chr8-124062798-DEL-217 | chr8-124062799-DEL-217 |
| chr8-124093932-DEL-52 | chr8-124093932-DEL-52 |
| chr8-124102113-DEL-53 | chr8-124102113-DEL-53 |
| chr8-124104958-DEL-53 | chr8-124104959-DEL-53 |
| chr8-124114669-DEL-132 | chr8-124114670-DEL-133 |
| chr8-124121349-DEL-104 | chr8-124121350-DEL-104 |
| chr8-124151802-DEL-56 | chr8-124151796-DEL-57 |
| chr8-124152538-DEL-8732 | chr8-124152539-DEL-8732 |
| chr8-124167062-DEL-1322 | chr8-124167062-DEL-1322 |
| chr8-124180747-DEL-64 | chr8-124180748-DEL-64 |
| chr8-124185552-DEL-61 | chr8-124185553-DEL-65 |
| chr8-124191380-DEL-74 | chr8-124191381-DEL-74 |
| chr8-124194462-DEL-9176 | chr8-124194463-DEL-9176 |
| chr8-124362707-DEL-2864 | chr8-124362708-DEL-2864 |
| chr8-124371241-DEL-60 | chr8-124371242-DEL-60 |
| chr8-124371691-DEL-226 | chr8-124371714-DEL-205 |
| chr8-124372662-DEL-208 | chr8-124372663-DEL-208 |
| chr8-124384926-DEL-96 | chr8-124384927-DEL-96 |
| chr8-124403484-DEL-58 | chr8-124403485-DEL-58 |
| chr8-124407244-DEL-438 | chr8-124407245-DEL-438 |
| chr8-124434460-DEL-52 | chr8-124434452-DEL-60 |
| chr8-124449213-DEL-117 | chr8-124449214-DEL-117 |
| chr8-124458711-DEL-79 | chr8-124458712-DEL-79 |
| chr8-124487243-DEL-71 | chr8-124487282-DEL-66 |
| chr8-124514982-DEL-183 | chr8-124514983-DEL-182 |
| chr8-124537901-DEL-116 | chr8-124537902-DEL-116 |
| chr8-124541880-DEL-201 | chr8-124541881-DEL-201 |
| chr8-124548428-DEL-419 | chr8-124548429-DEL-419 |
| chr8-124564016-DEL-757 | chr8-124564017-DEL-757 |
| chr8-124570973-DEL-162 | chr8-124570967-DEL-136 |
| chr8-124572210-DEL-501 | chr8-124572211-DEL-501 |
| chr8-124597528-DEL-199 | chr8-124597529-DEL-198 |
| chr8-124630521-DEL-559 | chr8-124630522-DEL-559 |
| chr8-124655758-DEL-79 | chr8-124655759-DEL-79 |
| chr8-124682032-DEL-95 | chr8-124682033-DEL-95 |
| chr8-124688247-DEL-75 | chr8-124688227-DEL-70 |
| chr8-124690440-DEL-70 | chr8-124690448-DEL-67 |
| chr8-124692703-DEL-56 | chr8-124692704-DEL-56 |
| chr8-124694969-DEL-57 | chr8-124694970-DEL-57 |
| chr8-124699051-DEL-168 | chr8-124699052-DEL-168 |
| chr8-124700668-DEL-205 | chr8-124700669-DEL-205 |
| chr8-124703458-DEL-210 | chr8-124703459-DEL-210 |
| chr8-124717919-DEL-216 | chr8-124717909-DEL-220 |
| chr8-124739415-DEL-206 | chr8-124739416-DEL-206 |
| chr8-124759354-DEL-534 | chr8-124759355-DEL-536 |
| chr8-124770820-DEL-56 | chr8-124770821-DEL-56 |
| chr8-124771435-DEL-62 | chr8-124771436-DEL-62 |
| chr8-124796065-DEL-63 | chr8-124796051-DEL-63 |
| chr8-124799546-DEL-156 | chr8-124799583-DEL-148 |
| chr8-124801214-DEL-198 | chr8-124801214-DEL-199 |
| chr8-124863052-DEL-88 | chr8-124863053-DEL-66 |
| chr8-124880376-DEL-58 | chr8-124880375-DEL-59 |
| chr8-124885868-DEL-297 | chr8-124885869-DEL-297 |
| chr8-124887950-DEL-256 | chr8-124887951-DEL-256 |
| chr8-124889022-DEL-694 | chr8-124889022-DEL-695 |
| chr8-124894808-DEL-96 | chr8-124894844-DEL-64 |
| chr8-124897026-DEL-343 | chr8-124897027-DEL-343 |
| chr8-124900813-DEL-78 | chr8-124900814-DEL-78 |
| chr8-124905657-DEL-64 | chr8-124905658-DEL-64 |
| chr8-124911879-DEL-52 | chr8-124911879-DEL-52 |
| chr8-124932968-DEL-440 | chr8-124932969-DEL-440 |
| chr8-124940559-DEL-60 | chr8-124940560-DEL-60 |
| chr8-124981589-DEL-333 | chr8-124981590-DEL-333 |
| chr8-125025320-DEL-60 | chr8-125025304-DEL-52 |
| chr8-125027498-DEL-60 | chr8-125027499-DEL-60 |
| chr8-125063742-DEL-139 | chr8-125063743-DEL-139 |
| chr8-125078223-DEL-58 | chr8-125078224-DEL-58 |
| chr8-125093265-DEL-217 | chr8-125093266-DEL-217 |

| | |
|---|---|
| chr8-125105173-DEL-83 | chr8-125105174-DEL-83 |
| chr8-125132894-DEL-61 | chr8-125132895-DEL-61 |
| chr8-125136624-DEL-280 | chr8-125136625-DEL-280 |
| chr8-125137968-DEL-277 | chr8-125137969-DEL-277 |
| chr8-125143774-DEL-58 | chr8-125143771-DEL-62 |
| chr8-125181047-DEL-66 | chr8-125181048-DEL-66 |
| chr8-125182827-DEL-228 | chr8-125182828-DEL-184 |
| chr8-125193638-DEL-2808 | chr8-125193639-DEL-2808 |
| chr8-125204816-DEL-62 | chr8-125204814-DEL-62 |
| chr8-125206508-DEL-116 | chr8-125206509-DEL-116 |
| chr8-125209041-DEL-190 | chr8-125209042-DEL-190 |
| chr8-125210796-DEL-1042 | chr8-125210796-DEL-1042 |
| chr8-125211877-DEL-1075 | chr8-125211877-DEL-1075 |
| chr8-125219333-DEL-124 | chr8-125219334-DEL-124 |
| chr8-125282842-DEL-54 | chr8-125282843-DEL-54 |
| chr8-125299509-DEL-149 | chr8-125299510-DEL-149 |
| chr8-125305108-DEL-4766 | chr8-125305109-DEL-4766 |
| chr8-125311104-DEL-53 | chr8-125311105-DEL-53 |
| chr8-125327562-DEL-116 | chr8-125327543-DEL-100 |
| chr8-125329672-DEL-54 | chr8-125329673-DEL-54 |
| chr8-125345873-DEL-135 | chr8-125345874-DEL-135 |
| chr8-125346578-DEL-136 | chr8-125346580-DEL-137 |
| chr8-125367235-DEL-102 | chr8-125367236-DEL-102 |
| chr8-125375123-DEL-216 | chr8-125375124-DEL-216 |
| chr8-125414264-DEL-76 | chr8-125414265-DEL-76 |
| chr8-125415079-DEL-86 | chr8-125415080-DEL-86 |
| chr8-125418118-DEL-89 | chr8-125418119-DEL-89 |
| chr8-125433527-DEL-100 | chr8-125433559-DEL-100 |
| chr8-125450721-DEL-348 | chr8-125450722-DEL-348 |
| chr8-125451361-DEL-206 | chr8-125451362-DEL-206 |
| chr8-125466643-DEL-294 | chr8-125466644-DEL-294 |
| chr8-125473842-DEL-74 | chr8-125473825-DEL-88 |
| chr8-125490770-DEL-2343 | chr8-125490753-DEL-2344 |
| chr8-125495316-DEL-427 | chr8-125495317-DEL-427 |
| chr8-125506772-DEL-216 | chr8-125506773-DEL-216 |
| chr8-125511641-DEL-414 | chr8-125511642-DEL-414 |
| chr8-125533216-DEL-4597 | chr8-125533218-DEL-4596 |
| chr8-125553121-DEL-66 | chr8-125553146-DEL-54 |
| chr8-125565613-DEL-91 | chr8-125565614-DEL-91 |
| chr8-125572760-DEL-6442 | chr8-125572761-DEL-6442 |
| chr8-125609192-DEL-73 | chr8-125609193-DEL-73 |
| chr8-125628029-DEL-52 | chr8-125628030-DEL-52 |
| chr8-125673018-DEL-279 | chr8-125673008-DEL-274 |
| chr8-125685686-DEL-100 | chr8-125685687-DEL-100 |
| chr8-125699222-DEL-208 | chr8-125699225-DEL-206 |
| chr8-125706036-DEL-177 | chr8-125706037-DEL-177 |
| chr8-125710882-DEL-180 | chr8-125710922-DEL-138 |
| chr8-125720397-DEL-1662 | chr8-125720402-DEL-1647 |
| chr8-125721898-DEL-254 | chr8-125721911-DEL-249 |
| chr8-125731324-DEL-281 | chr8-125731325-DEL-281 |
| chr8-125742575-DEL-130 | chr8-125742575-DEL-130 |
| chr8-125746439-DEL-480 | chr8-125746440-DEL-448 |
| chr8-125747121-DEL-3382 | chr8-125747122-DEL-3382 |
| chr8-125756559-DEL-112 | chr8-125756560-DEL-112 |
| chr8-125762269-DEL-52 | chr8-125762270-DEL-52 |
| chr8-125774419-DEL-89 | chr8-125774392-DEL-95 |
| chr8-125775046-DEL-179 | chr8-125775047-DEL-179 |
| chr8-125782965-DEL-470 | chr8-125782990-DEL-492 |
| chr8-125783834-DEL-193 | chr8-125783836-DEL-231 |
| chr8-125790693-DEL-62 | chr8-125790693-DEL-62 |
| chr8-125792949-DEL-79 | chr8-125792950-DEL-79 |
| chr8-125799802-DEL-80 | chr8-125799805-DEL-78 |
| chr8-125839568-DEL-57 | chr8-125839568-DEL-57 |
| chr8-125859107-DEL-84 | chr8-125859108-DEL-56 |
| chr8-125873264-DEL-1263 | chr8-125873265-DEL-1263 |
| chr8-125880823-DEL-235 | chr8-125880832-DEL-236 |
| chr8-125897649-DEL-56 | chr8-125897650-DEL-56 |
| chr8-125902305-DEL-59 | chr8-125902305-DEL-60 |
| chr8-125910128-DEL-132 | chr8-125910129-DEL-132 |
| chr8-125916154-DEL-162 | chr8-125916155-DEL-162 |
| chr8-125918124-DEL-437 | chr8-125918143-DEL-418 |
| chr8-125919635-DEL-77 | chr8-125919636-DEL-77 |
| chr8-125971670-DEL-87 | chr8-125971670-DEL-87 |
| chr8-125975408-DEL-83 | chr8-125975409-DEL-83 |
| chr8-126003103-DEL-76 | chr8-126003104-DEL-60 |
| chr8-126028297-DEL-103 | chr8-126028298-DEL-103 |
| chr8-126043457-DEL-687 | chr8-126043456-DEL-688 |
| chr8-126058917-DEL-539 | chr8-126058918-DEL-539 |
| chr8-126090778-DEL-60 | chr8-126090779-DEL-52 |
| chr8-126092129-DEL-208 | chr8-126092130-DEL-208 |

| | |
|---|---|
| chr8-126096035-DEL-187 | chr8-126096036-DEL-187 |
| chr8-126110971-DEL-52 | chr8-126110970-DEL-52 |
| chr8-126128256-DEL-100 | chr8-126128257-DEL-100 |
| chr8-126139343-DEL-391 | chr8-126139344-DEL-391 |
| chr8-126158013-DEL-54 | chr8-126158014-DEL-54 |
| chr8-126265759-DEL-73 | chr8-126265760-DEL-73 |
| chr8-126266589-DEL-236 | chr8-126266589-DEL-235 |
| chr8-126319538-DEL-400 | chr8-126319539-DEL-400 |
| chr8-126335956-DEL-98 | chr8-126335946-DEL-70 |
| chr8-126356056-DEL-7101 | chr8-126356057-DEL-7101 |
| chr8-126363869-DEL-60 | chr8-126363870-DEL-60 |
| chr8-126369343-DEL-2887 | chr8-126369344-DEL-2887 |
| chr8-126371096-DEL-445 | chr8-126371097-DEL-445 |
| chr8-126388604-DEL-211 | chr8-126388605-DEL-211 |
| chr8-126411758-DEL-115 | chr8-126411790-DEL-87 |
| chr8-126418872-DEL-175 | chr8-126418873-DEL-175 |
| chr8-126501165-DEL-499 | chr8-126501168-DEL-498 |
| chr8-126508160-DEL-60 | chr8-126508166-DEL-60 |
| chr8-126515623-DEL-62 | chr8-126515624-DEL-62 |
| chr8-126537460-DEL-162 | chr8-126537454-DEL-168 |
| chr8-126554370-DEL-60 | chr8-126554362-DEL-82 |
| chr8-126558756-DEL-80 | chr8-126558757-DEL-80 |
| chr8-126560536-DEL-174 | chr8-126560537-DEL-172 |
| chr8-126598242-DEL-75 | chr8-126598253-DEL-60 |
| chr8-126626512-DEL-400 | chr8-126626513-DEL-400 |
| chr8-126632431-DEL-104 | chr8-126632432-DEL-104 |
| chr8-126641950-DEL-74 | chr8-126641965-DEL-76 |
| chr8-126649570-DEL-102 | chr8-126649571-DEL-102 |
| chr8-126656710-DEL-160 | chr8-126656711-DEL-160 |
| chr8-126662664-DEL-73 | chr8-126662665-DEL-73 |
| chr8-126669411-DEL-582 | chr8-126669413-DEL-581 |
| chr8-126703365-DEL-51 | chr8-126703376-DEL-51 |
| chr8-126732592-DEL-85 | chr8-126732593-DEL-85 |
| chr8-126738257-DEL-80 | chr8-126738258-DEL-80 |
| chr8-126749926-DEL-253 | chr8-126749927-DEL-253 |
| chr8-126798136-DEL-212 | chr8-126798139-DEL-209 |
| chr8-126802442-DEL-82 | chr8-126802408-DEL-119 |
| chr8-126832683-DEL-5214 | chr8-126832684-DEL-5214 |
| chr8-126847472-DEL-381 | chr8-126847473-DEL-375 |
| chr8-126854569-DEL-136 | chr8-126854558-DEL-124 |
| chr8-126873398-DEL-94 | chr8-126873398-DEL-95 |
| chr8-126910370-DEL-54 | chr8-126910383-DEL-54 |
| chr8-126931153-DEL-204 | chr8-126931154-DEL-204 |
| chr8-126956490-DEL-200 | chr8-126956480-DEL-201 |
| chr8-126995037-DEL-351 | chr8-126995038-DEL-351 |
| chr8-127017952-DEL-110 | chr8-127017975-DEL-76 |
| chr8-127036375-DEL-253 | chr8-127036353-DEL-253 |
| chr8-127063099-DEL-112 | chr8-127063100-DEL-112 |
| chr8-127064925-DEL-130 | chr8-127064920-DEL-136 |
| chr8-127067060-DEL-57 | chr8-127067061-DEL-57 |
| chr8-127090850-DEL-304 | chr8-127090857-DEL-302 |
| chr8-127093809-DEL-120 | chr8-127093810-DEL-120 |
| chr8-127126021-DEL-1888 | chr8-127126022-DEL-1888 |
| chr8-127140388-DEL-5160 | chr8-127140389-DEL-5160 |
| chr8-127170414-DEL-155 | chr8-127170415-DEL-155 |
| chr8-127185167-DEL-56 | chr8-127185167-DEL-56 |
| chr8-127190207-DEL-571 | chr8-127190208-DEL-571 |
| chr8-127211751-DEL-117 | chr8-127211752-DEL-117 |
| chr8-127212751-DEL-440 | chr8-127212752-DEL-440 |
| chr8-127214123-DEL-58 | chr8-127214113-DEL-59 |
| chr8-127218952-DEL-121 | chr8-127218953-DEL-114 |
| chr8-127230074-DEL-111 | chr8-127230067-DEL-112 |
| chr8-127232477-DEL-72 | chr8-127232466-DEL-73 |
| chr8-127238821-DEL-60 | chr8-127238821-DEL-60 |
| chr8-127245698-DEL-74 | chr8-127245699-DEL-74 |
| chr8-127250670-DEL-105 | chr8-127250692-DEL-84 |
| chr8-127259930-DEL-3069 | chr8-127259931-DEL-3069 |
| chr8-127265212-DEL-68 | chr8-127265213-DEL-68 |
| chr8-127277088-DEL-102 | chr8-127277095-DEL-96 |
| chr8-127331002-DEL-98 | chr8-127331003-DEL-98 |
| chr8-127335846-DEL-106 | chr8-127335853-DEL-94 |
| chr8-127351079-DEL-114 | chr8-127351074-DEL-111 |
| chr8-127416057-DEL-67 | chr8-127416054-DEL-63 |
| chr8-127434985-DEL-60 | chr8-127434986-DEL-60 |
| chr8-127443870-DEL-56 | chr8-127443871-DEL-56 |
| chr8-127449791-DEL-57 | chr8-127449792-DEL-57 |
| chr8-127459636-DEL-101 | chr8-127459600-DEL-93 |
| chr8-127484353-DEL-144 | chr8-127484354-DEL-144 |
| chr8-127510056-DEL-52 | chr8-127510057-DEL-52 |
| chr8-127536864-DEL-202 | chr8-127536856-DEL-187 |

| | |
|---|---|
| chr8-127580397-DEL-253 | chr8-127580398-DEL-253 |
| chr8-127607297-DEL-51 | chr8-127607298-DEL-51 |
| chr8-127642444-DEL-64 | chr8-127642445-DEL-64 |
| chr8-127646986-DEL-52 | chr8-127646986-DEL-52 |
| chr8-127651656-DEL-56 | chr8-127651657-DEL-56 |
| chr8-127659876-DEL-95 | chr8-127659877-DEL-95 |
| chr8-127665600-DEL-202 | chr8-127665601-DEL-202 |
| chr8-127669935-DEL-201 | chr8-127669936-DEL-201 |
| chr8-127690551-DEL-105 | chr8-127690552-DEL-105 |
| chr8-127731337-DEL-140 | chr8-127731338-DEL-140 |
| chr8-127740951-DEL-246 | chr8-127740952-DEL-246 |
| chr8-127759320-DEL-85 | chr8-127759321-DEL-85 |
| chr8-127814588-DEL-51 | chr8-127814589-DEL-51 |
| chr8-127828745-DEL-121 | chr8-127828747-DEL-120 |
| chr8-127878370-DEL-56 | chr8-127878371-DEL-56 |
| chr8-127890758-DEL-9645 | chr8-127890759-DEL-9645 |
| chr8-127896520-DEL-73 | chr8-127896505-DEL-76 |
| chr8-127910018-DEL-369 | chr8-127910019-DEL-369 |
| chr8-127930924-DEL-361 | chr8-127930925-DEL-361 |
| chr8-127935018-DEL-241 | chr8-127935019-DEL-241 |
| chr8-127943764-DEL-141 | chr8-127943765-DEL-141 |
| chr8-127950911-DEL-121 | chr8-127950912-DEL-121 |
| chr8-127954526-DEL-68 | chr8-127954519-DEL-54 |
| chr8-127956575-DEL-83 | chr8-127956576-DEL-83 |
| chr8-127959008-DEL-560 | chr8-127959009-DEL-560 |
| chr8-127963426-DEL-4414 | chr8-127963426-DEL-4415 |
| chr8-128008689-DEL-188 | chr8-128008684-DEL-189 |
| chr8-128040691-DEL-163 | chr8-128040691-DEL-163 |
| chr8-128056062-DEL-619 | chr8-128056063-DEL-619 |
| chr8-128111277-DEL-357 | chr8-128111273-DEL-361 |
| chr8-128119719-DEL-192 | chr8-128119724-DEL-188 |
| chr8-128131497-DEL-96 | chr8-128131498-DEL-96 |
| chr8-128195456-DEL-646 | chr8-128195457-DEL-673 |
| chr8-128203286-DEL-799 | chr8-128203287-DEL-799 |
| chr8-128211173-DEL-467 | chr8-128211174-DEL-467 |
| chr8-128218200-DEL-52 | chr8-128218201-DEL-52 |
| chr8-128234466-DEL-146 | chr8-128234457-DEL-158 |
| chr8-128262842-DEL-104 | chr8-128262841-DEL-102 |
| chr8-128263819-DEL-1877 | chr8-128263820-DEL-1877 |
| chr8-128310862-DEL-124 | chr8-128310863-DEL-124 |
| chr8-128352117-DEL-1784 | chr8-128352118-DEL-1784 |
| chr8-128352791-DEL-118 | chr8-128352792-DEL-116 |
| chr8-128357857-DEL-367 | chr8-128357858-DEL-367 |
| chr8-128449177-DEL-60 | chr8-128449180-DEL-58 |
| chr8-128459011-DEL-155 | chr8-128458985-DEL-156 |
| chr8-128471186-DEL-57 | chr8-128471187-DEL-57 |
| chr8-128474299-DEL-56 | chr8-128474300-DEL-56 |
| chr8-128495883-DEL-59 | chr8-128495887-DEL-60 |
| chr8-128555237-DEL-112 | chr8-128555238-DEL-112 |
| chr8-128557742-DEL-163 | chr8-128557739-DEL-167 |
| chr8-128568199-DEL-1203 | chr8-128568200-DEL-1203 |
| chr8-128572656-DEL-57 | chr8-128572658-DEL-56 |
| chr8-128579095-DEL-1282 | chr8-128579096-DEL-1282 |
| chr8-128579706-DEL-59 | chr8-128579708-DEL-58 |
| chr8-128594328-DEL-4356 | chr8-128594329-DEL-4356 |
| chr8-128604244-DEL-116 | chr8-128604286-DEL-97 |
| chr8-128625056-DEL-6336 | chr8-128625043-DEL-6337 |
| chr8-128650729-DEL-54 | chr8-128650730-DEL-54 |
| chr8-128675603-DEL-88 | chr8-128675624-DEL-66 |
| chr8-128681833-DEL-72 | chr8-128681834-DEL-72 |
| chr8-128767253-DEL-327 | chr8-128767254-DEL-327 |
| chr8-128801719-DEL-478 | chr8-128801720-DEL-478 |
| chr8-128816011-DEL-1962 | chr8-128816012-DEL-1961 |
| chr8-128822481-DEL-287 | chr8-128822482-DEL-287 |
| chr8-128832653-DEL-72 | chr8-128832656-DEL-70 |
| chr8-128852782-DEL-60 | chr8-128852783-DEL-60 |
| chr8-128916461-DEL-159 | chr8-128916462-DEL-159 |
| chr8-128968416-DEL-510 | chr8-128968417-DEL-510 |
| chr8-129004246-DEL-193 | chr8-129004241-DEL-196 |
| chr8-129037013-DEL-200 | chr8-129037014-DEL-199 |
| chr8-129039248-DEL-67 | chr8-129039233-DEL-78 |
| chr8-129056507-DEL-426 | chr8-129056508-DEL-426 |
| chr8-129113323-DEL-76 | chr8-129113328-DEL-73 |
| chr8-129158992-DEL-83 | chr8-129158990-DEL-92 |
| chr8-129179079-DEL-86 | chr8-129179085-DEL-62 |
| chr8-129275527-DEL-168 | chr8-129275528-DEL-168 |
| chr8-129279154-DEL-5321 | chr8-129279155-DEL-5321 |
| chr8-129288067-DEL-54 | chr8-129288068-DEL-54 |
| chr8-129346238-DEL-616 | chr8-129346239-DEL-616 |
| chr8-129353925-DEL-4113 | chr8-129353926-DEL-4113 |

| | |
|---|---|
| chr8-129364223-DEL-1582 | chr8-129364224-DEL-1583 |
| chr8-129373894-DEL-74 | chr8-129373897-DEL-68 |
| chr8-129405166-DEL-440 | chr8-129405167-DEL-440 |
| chr8-129406048-DEL-84 | chr8-129406068-DEL-52 |
| chr8-129430662-DEL-125 | chr8-129430643-DEL-145 |
| chr8-129434376-DEL-73 | chr8-129434377-DEL-73 |
| chr8-129447227-DEL-354 | chr8-129447228-DEL-354 |
| chr8-129472929-DEL-7483 | chr8-129472930-DEL-7483 |
| chr8-129497079-DEL-847 | chr8-129497082-DEL-848 |
| chr8-129502942-DEL-2260 | chr8-129502941-DEL-2260 |
| chr8-129510863-DEL-138 | chr8-129510864-DEL-138 |
| chr8-129539783-DEL-6428 | chr8-129539784-DEL-6428 |
| chr8-129572508-DEL-298 | chr8-129572509-DEL-296 |
| chr8-129583325-DEL-66 | chr8-129583330-DEL-62 |
| chr8-129637473-DEL-429 | chr8-129637474-DEL-429 |
| chr8-129644725-DEL-63 | chr8-129644726-DEL-63 |
| chr8-129663289-DEL-4076 | chr8-129663290-DEL-4076 |
| chr8-129680161-DEL-260 | chr8-129680161-DEL-260 |
| chr8-129705722-DEL-191 | chr8-129705723-DEL-191 |
| chr8-129716975-DEL-120 | chr8-129717001-DEL-90 |
| chr8-129757421-DEL-81 | chr8-129757422-DEL-66 |
| chr8-129763241-DEL-2798 | chr8-129763242-DEL-2798 |
| chr8-129777538-DEL-229 | chr8-129777543-DEL-228 |
| chr8-129800142-DEL-60 | chr8-129800143-DEL-61 |
| chr8-129865104-DEL-6679 | chr8-129865105-DEL-6679 |
| chr8-129871230-DEL-214 | chr8-129871231-DEL-214 |
| chr8-129898012-DEL-236 | chr8-129898013-DEL-236 |
| chr8-129970894-DEL-11466 | chr8-129970896-DEL-11467 |
| chr8-129999360-DEL-6382 | chr8-129999361-DEL-6382 |
| chr8-130015797-DEL-66 | chr8-130015798-DEL-66 |
| chr9-3046518-DEL-100 | chr9-3046519-DEL-100 |
| chr9-3047204-DEL-100 | chr9-3047231-DEL-120 |
| chr9-3053553-DEL-62 | chr9-3053554-DEL-62 |
| chr9-3055846-DEL-391 | chr9-3055847-DEL-391 |
| chr9-3058579-DEL-64 | chr9-3058575-DEL-62 |
| chr9-3079388-DEL-4565 | chr9-3079389-DEL-4565 |
| chr9-3122502-DEL-872 | chr9-3122503-DEL-872 |
| chr9-3153012-DEL-108 | chr9-3153025-DEL-84 |
| chr9-3163180-DEL-13158 | chr9-3163181-DEL-13158 |
| chr9-3197569-DEL-80 | chr9-3197574-DEL-76 |
| chr9-3246638-DEL-105 | chr9-3246671-DEL-72 |
| chr9-3302144-DEL-205 | chr9-3302145-DEL-205 |
| chr9-3315559-DEL-499 | chr9-3315560-DEL-499 |
| chr9-3324146-DEL-65 | chr9-3324153-DEL-54 |
| chr9-3332502-DEL-204 | chr9-3332503-DEL-204 |
| chr9-3364589-DEL-61 | chr9-3364612-DEL-76 |
| chr9-3416224-DEL-73 | chr9-3416226-DEL-72 |
| chr9-3461286-DEL-5185 | chr9-3461287-DEL-5185 |
| chr9-3647858-DEL-1324 | chr9-3647858-DEL-1324 |
| chr9-3649787-DEL-659 | chr9-3649788-DEL-659 |
| chr9-3658249-DEL-1830 | chr9-3658250-DEL-1830 |
| chr9-3681038-DEL-818 | chr9-3681039-DEL-818 |
| chr9-3698096-DEL-2867 | chr9-3698097-DEL-2867 |
| chr9-3764946-DEL-551 | chr9-3764961-DEL-554 |
| chr9-3783263-DEL-2740 | chr9-3783264-DEL-2740 |
| chr9-3800052-DEL-520 | chr9-3800053-DEL-520 |
| chr9-3885753-DEL-718 | chr9-3885755-DEL-717 |
| chr9-3887755-DEL-600 | chr9-3887755-DEL-600 |
| chr9-3931560-DEL-145 | chr9-3931561-DEL-145 |
| chr9-4002175-DEL-1159 | chr9-4002176-DEL-1159 |
| chr9-4024090-DEL-1068 | chr9-4024090-DEL-1068 |
| chr9-4112777-DEL-6024 | chr9-4112778-DEL-6024 |
| chr9-4155975-DEL-158 | chr9-4155976-DEL-158 |
| chr9-4177150-DEL-416 | chr9-4177151-DEL-416 |
| chr9-4182494-DEL-6566 | chr9-4182495-DEL-6566 |
| chr9-4270667-DEL-519 | chr9-4270668-DEL-505 |
| chr9-4359854-DEL-501 | chr9-4359855-DEL-501 |
| chr9-4377248-DEL-52 | chr9-4377249-DEL-52 |
| chr9-4377391-DEL-902 | chr9-4377392-DEL-902 |
| chr9-4394177-DEL-65 | chr9-4394185-DEL-52 |
| chr9-4408491-DEL-72 | chr9-4408492-DEL-72 |
| chr9-4445206-DEL-1559 | chr9-4445190-DEL-1560 |
| chr9-4491279-DEL-7176 | chr9-4491280-DEL-7176 |
| chr9-4491841-DEL-75 | chr9-4491842-DEL-75 |
| chr9-4574599-DEL-989 | chr9-4574600-DEL-989 |
| chr9-4580453-DEL-95 | chr9-4580441-DEL-87 |
| chr9-4607338-DEL-1557 | chr9-4607338-DEL-1557 |
| chr9-4678545-DEL-54 | chr9-4678546-DEL-54 |
| chr9-4693868-DEL-2030 | chr9-4693869-DEL-2030 |
| chr9-4726716-DEL-113 | chr9-4726717-DEL-113 |

| | |
|---|---|
| chr9-4731094-DEL-52 | chr9-4731095-DEL-52 |
| chr9-4765323-DEL-6826 | chr9-4765324-DEL-6826 |
| chr9-4782063-DEL-100 | chr9-4782064-DEL-100 |
| chr9-4807668-DEL-66 | chr9-4807668-DEL-66 |
| chr9-4808896-DEL-400 | chr9-4808897-DEL-400 |
| chr9-4813167-DEL-1571 | chr9-4813135-DEL-1565 |
| chr9-4814873-DEL-971 | chr9-4814874-DEL-972 |
| chr9-4815038-DEL-209 | chr9-4815039-DEL-209 |
| chr9-4831073-DEL-111 | chr9-4831074-DEL-111 |
| chr9-4854587-DEL-76 | chr9-4854571-DEL-76 |
| chr9-4858662-DEL-197 | chr9-4858660-DEL-197 |
| chr9-4871505-DEL-452 | chr9-4871506-DEL-452 |
| chr9-4874462-DEL-997 | chr9-4874463-DEL-997 |
| chr9-4889868-DEL-1360 | chr9-4889869-DEL-1360 |
| chr9-4896690-DEL-400 | chr9-4896690-DEL-400 |
| chr9-4907742-DEL-251 | chr9-4907743-DEL-251 |
| chr9-4914902-DEL-325 | chr9-4914902-DEL-325 |
| chr9-4916573-DEL-5510 | chr9-4916574-DEL-5510 |
| chr9-4979092-DEL-122 | chr9-4979093-DEL-122 |
| chr9-5027656-DEL-2364 | chr9-5027657-DEL-2364 |
| chr9-5051252-DEL-62 | chr9-5051267-DEL-50 |
| chr9-5079266-DEL-1925 | chr9-5079267-DEL-1925 |
| chr9-5101957-DEL-67 | chr9-5101958-DEL-66 |
| chr9-5117161-DEL-99 | chr9-5117159-DEL-148 |
| chr9-5168841-DEL-134 | chr9-5168842-DEL-132 |
| chr9-5179254-DEL-649 | chr9-5179244-DEL-650 |
| chr9-5198445-DEL-186 | chr9-5198446-DEL-186 |
| chr9-5211837-DEL-99 | chr9-5211838-DEL-98 |
| chr9-5219301-DEL-133 | chr9-5219302-DEL-133 |
| chr9-5247897-DEL-6131 | chr9-5247898-DEL-6131 |
| chr9-5280105-DEL-4455 | chr9-5280106-DEL-4455 |
| chr9-5292786-DEL-1664 | chr9-5292787-DEL-1664 |
| chr9-5319164-DEL-78 | chr9-5319165-DEL-78 |
| chr9-5335443-DEL-431 | chr9-5335444-DEL-430 |
| chr9-5343136-DEL-56 | chr9-5343111-DEL-58 |
| chr9-5359120-DEL-6187 | chr9-5359121-DEL-6187 |
| chr9-5440652-DEL-257 | chr9-5440649-DEL-255 |
| chr9-5460171-DEL-473 | chr9-5460172-DEL-473 |
| chr9-5478192-DEL-725 | chr9-5478193-DEL-725 |
| chr9-5494095-DEL-779 | chr9-5494096-DEL-779 |
| chr9-5502342-DEL-1663 | chr9-5502343-DEL-1663 |
| chr9-5503771-DEL-197 | chr9-5503772-DEL-197 |
| chr9-5548464-DEL-7523 | chr9-5548465-DEL-7523 |
| chr9-5562328-DEL-313 | chr9-5562329-DEL-313 |
| chr9-5563339-DEL-80 | chr9-5563352-DEL-68 |
| chr9-5577139-DEL-1129 | chr9-5577140-DEL-1129 |
| chr9-5577199-DEL-190 | chr9-5577200-DEL-190 |
| chr9-5596684-DEL-233 | chr9-5596685-DEL-233 |
| chr9-5610245-DEL-2594 | chr9-5610246-DEL-2594 |
| chr9-5614293-DEL-214 | chr9-5614283-DEL-215 |
| chr9-5639445-DEL-58 | chr9-5639446-DEL-60 |
| chr9-5695129-DEL-60 | chr9-5695130-DEL-60 |
| chr9-5697691-DEL-59 | chr9-5697690-DEL-60 |
| chr9-5698788-DEL-81 | chr9-5698789-DEL-81 |
| chr9-5701980-DEL-267 | chr9-5701980-DEL-267 |
| chr9-5703722-DEL-800 | chr9-5703723-DEL-800 |
| chr9-5704117-DEL-1397 | chr9-5704118-DEL-1397 |
| chr9-5708046-DEL-52 | chr9-5708017-DEL-58 |
| chr9-5732412-DEL-75 | chr9-5732378-DEL-75 |
| chr9-5780968-DEL-6575 | chr9-5780959-DEL-6576 |
| chr9-5809882-DEL-52 | chr9-5809874-DEL-60 |
| chr9-5855934-DEL-1313 | chr9-5855935-DEL-1313 |
| chr9-5857920-DEL-225 | chr9-5857921-DEL-225 |
| chr9-5860721-DEL-82 | chr9-5860719-DEL-68 |
| chr9-5862959-DEL-56 | chr9-5862959-DEL-56 |
| chr9-5871898-DEL-217 | chr9-5871898-DEL-217 |
| chr9-5877362-DEL-2158 | chr9-5877363-DEL-2158 |
| chr9-5880230-DEL-400 | chr9-5880231-DEL-400 |
| chr9-5923439-DEL-64 | chr9-5923440-DEL-64 |
| chr9-5934500-DEL-7025 | chr9-5934501-DEL-7025 |
| chr9-6024771-DEL-78 | chr9-6024772-DEL-90 |
| chr9-6039699-DEL-90 | chr9-6039700-DEL-62 |
| chr9-6040499-DEL-78 | chr9-6040493-DEL-56 |
| chr9-6043224-DEL-86 | chr9-6043225-DEL-86 |
| chr9-6047609-DEL-2156 | chr9-6047608-DEL-2157 |
| chr9-6057428-DEL-206 | chr9-6057429-DEL-206 |
| chr9-6063569-DEL-133 | chr9-6063570-DEL-133 |
| chr9-6069421-DEL-209 | chr9-6069422-DEL-209 |
| chr9-6106258-DEL-62 | chr9-6106259-DEL-62 |
| chr9-6120495-DEL-115 | chr9-6120496-DEL-115 |

| | |
|---|---|
| chr9-6143340-DEL-462 | chr9-6143341-DEL-462 |
| chr9-6146022-DEL-72 | chr9-6146023-DEL-72 |
| chr9-6154656-DEL-173 | chr9-6154657-DEL-173 |
| chr9-6229990-DEL-736 | chr9-6229992-DEL-736 |
| chr9-6237797-DEL-233 | chr9-6237798-DEL-233 |
| chr9-6258388-DEL-65 | chr9-6258389-DEL-65 |
| chr9-6278223-DEL-254 | chr9-6278224-DEL-254 |
| chr9-6307875-DEL-420 | chr9-6307876-DEL-420 |
| chr9-6316183-DEL-2338 | chr9-6316181-DEL-2336 |
| chr9-6323641-DEL-576 | chr9-6323610-DEL-572 |
| chr9-6339296-DEL-128 | chr9-6339297-DEL-128 |
| chr9-6428180-DEL-358 | chr9-6428181-DEL-358 |
| chr9-6431480-DEL-1294 | chr9-6431481-DEL-1294 |
| chr9-6442500-DEL-6413 | chr9-6442500-DEL-6413 |
| chr9-6442615-DEL-201 | chr9-6442615-DEL-201 |
| chr9-6458149-DEL-81 | chr9-6458136-DEL-61 |
| chr9-6462135-DEL-6328 | chr9-6462136-DEL-6328 |
| chr9-6500627-DEL-388 | chr9-6500628-DEL-388 |
| chr9-6525540-DEL-79 | chr9-6525541-DEL-79 |
| chr9-6594427-DEL-1089 | chr9-6594428-DEL-1089 |
| chr9-6597291-DEL-177 | chr9-6597292-DEL-177 |
| chr9-6609607-DEL-142 | chr9-6609620-DEL-136 |
| chr9-6624416-DEL-207 | chr9-6624417-DEL-207 |
| chr9-6627343-DEL-60 | chr9-6627344-DEL-69 |
| chr9-6631841-DEL-4224 | chr9-6631842-DEL-4224 |
| chr9-6640615-DEL-517 | chr9-6640616-DEL-517 |
| chr9-6647548-DEL-2525 | chr9-6647549-DEL-2525 |
| chr9-6657611-DEL-6508 | chr9-6657612-DEL-6508 |
| chr9-6694755-DEL-8657 | chr9-6694756-DEL-8656 |
| chr9-6708154-DEL-96 | chr9-6708153-DEL-119 |
| chr9-6757522-DEL-1783 | chr9-6757516-DEL-1783 |
| chr9-6780311-DEL-5341 | chr9-6780309-DEL-5342 |
| chr9-6804764-DEL-8778 | chr9-6804765-DEL-8777 |
| chr9-6819745-DEL-203 | chr9-6819748-DEL-203 |
| chr9-6826067-DEL-65 | chr9-6826065-DEL-66 |
| chr9-6837489-DEL-205 | chr9-6837490-DEL-205 |
| chr9-6846358-DEL-978 | chr9-6846359-DEL-978 |
| chr9-6849713-DEL-375 | chr9-6849714-DEL-375 |
| chr9-6860878-DEL-224 | chr9-6860879-DEL-224 |
| chr9-6886793-DEL-363 | chr9-6886794-DEL-363 |
| chr9-6888018-DEL-130 | chr9-6888019-DEL-130 |
| chr9-6889194-DEL-102 | chr9-6889195-DEL-102 |
| chr9-6896067-DEL-673 | chr9-6896068-DEL-673 |
| chr9-6916779-DEL-73 | chr9-6916814-DEL-78 |
| chr9-6937456-DEL-64 | chr9-6937457-DEL-64 |
| chr9-6945420-DEL-6357 | chr9-6945420-DEL-6357 |
| chr9-6954746-DEL-741 | chr9-6954747-DEL-741 |
| chr9-6964063-DEL-69 | chr9-6964064-DEL-69 |
| chr9-6983871-DEL-849 | chr9-6983872-DEL-849 |
| chr9-7003274-DEL-276 | chr9-7003275-DEL-276 |
| chr9-7024810-DEL-52 | chr9-7024811-DEL-52 |
| chr9-7034072-DEL-75 | chr9-7034092-DEL-64 |
| chr9-7064289-DEL-6395 | chr9-7064290-DEL-6395 |
| chr9-7151313-DEL-79 | chr9-7151314-DEL-79 |
| chr9-7152124-DEL-2218 | chr9-7152125-DEL-2218 |
| chr9-7171129-DEL-339 | chr9-7171130-DEL-339 |
| chr9-7209137-DEL-98 | chr9-7209138-DEL-98 |
| chr9-7211758-DEL-54 | chr9-7211747-DEL-50 |
| chr9-7245658-DEL-2045 | chr9-7245659-DEL-2045 |
| chr9-7267402-DEL-1893 | chr9-7267403-DEL-1893 |
| chr9-7294534-DEL-6575 | chr9-7294535-DEL-6575 |
| chr9-7313148-DEL-132 | chr9-7313149-DEL-136 |
| chr9-7320541-DEL-91 | chr9-7320544-DEL-65 |
| chr9-7371132-DEL-4395 | chr9-7371133-DEL-4395 |
| chr9-7406320-DEL-3146 | chr9-7406321-DEL-3146 |
| chr9-7407035-DEL-8261 | chr9-7407035-DEL-8261 |
| chr9-7461587-DEL-375 | chr9-7461612-DEL-351 |
| chr9-7462604-DEL-734 | chr9-7462605-DEL-734 |
| chr9-7485244-DEL-3496 | chr9-7485245-DEL-3496 |
| chr9-7515032-DEL-2338 | chr9-7515039-DEL-2333 |
| chr9-7520830-DEL-5119 | chr9-7520831-DEL-5119 |
| chr9-7543047-DEL-121 | chr9-7543048-DEL-121 |
| chr9-7602118-DEL-217 | chr9-7602121-DEL-218 |
| chr9-7605129-DEL-56 | chr9-7605129-DEL-60 |
| chr9-7609894-DEL-5080 | chr9-7609895-DEL-5080 |
| chr9-7623737-DEL-196 | chr9-7623738-DEL-196 |
| chr9-7646843-DEL-535 | chr9-7646844-DEL-535 |
| chr9-7671861-DEL-82 | chr9-7671855-DEL-83 |
| chr9-7699927-DEL-210 | chr9-7699934-DEL-202 |
| chr9-7714100-DEL-51 | chr9-7714089-DEL-56 |

| | |
|---|---|
| chr9-7721054-DEL-2304 | chr9-7721055-DEL-2304 |
| chr9-7722617-DEL-116 | chr9-7722624-DEL-112 |
| chr9-7723027-DEL-1734 | chr9-7723028-DEL-1734 |
| chr9-7728093-DEL-194 | chr9-7728094-DEL-194 |
| chr9-7744781-DEL-120 | chr9-7744812-DEL-104 |
| chr9-7755279-DEL-158 | chr9-7755262-DEL-140 |
| chr9-7756838-DEL-919 | chr9-7756839-DEL-919 |
| chr9-7779590-DEL-554 | chr9-7779591-DEL-561 |
| chr9-7787064-DEL-60 | chr9-7787065-DEL-60 |
| chr9-7797302-DEL-52 | chr9-7797298-DEL-56 |
| chr9-7798962-DEL-6226 | chr9-7798962-DEL-6226 |
| chr9-7813774-DEL-567 | chr9-7813775-DEL-567 |
| chr9-7824430-DEL-400 | chr9-7824431-DEL-400 |
| chr9-7839934-DEL-233 | chr9-7839918-DEL-218 |
| chr9-7859315-DEL-211 | chr9-7859316-DEL-211 |
| chr9-7892737-DEL-220 | chr9-7892738-DEL-220 |
| chr9-7894259-DEL-51 | chr9-7894224-DEL-51 |
| chr9-7899205-DEL-54 | chr9-7899195-DEL-50 |
| chr9-7908140-DEL-74 | chr9-7908141-DEL-70 |
| chr9-7909166-DEL-2240 | chr9-7909167-DEL-2240 |
| chr9-7911590-DEL-155 | chr9-7911590-DEL-155 |
| chr9-7915801-DEL-8179 | chr9-7915801-DEL-8179 |
| chr9-7934972-DEL-270 | chr9-7934973-DEL-270 |
| chr9-7942374-DEL-100 | chr9-7942375-DEL-90 |
| chr9-7977778-DEL-58 | chr9-7977779-DEL-58 |
| chr9-8009009-DEL-75 | chr9-8009010-DEL-75 |
| chr9-8029394-DEL-281 | chr9-8029395-DEL-281 |
| chr9-8037758-DEL-104 | chr9-8037759-DEL-104 |
| chr9-8046528-DEL-1333 | chr9-8046529-DEL-1333 |
| chr9-8048213-DEL-6450 | chr9-8048214-DEL-6450 |
| chr9-8063513-DEL-4582 | chr9-8063514-DEL-4582 |
| chr9-8073925-DEL-132 | chr9-8073926-DEL-132 |
| chr9-8080937-DEL-65 | chr9-8080938-DEL-65 |
| chr9-8132870-DEL-208 | chr9-8132871-DEL-208 |
| chr9-8141533-DEL-187 | chr9-8141534-DEL-187 |
| chr9-8189541-DEL-95 | chr9-8189544-DEL-95 |
| chr9-8231322-DEL-6201 | chr9-8231321-DEL-6199 |
| chr9-8251776-DEL-2179 | chr9-8251777-DEL-2179 |
| chr9-8276086-DEL-7125 | chr9-8276087-DEL-7125 |
| chr9-8290526-DEL-1797 | chr9-8290527-DEL-1797 |
| chr9-8326334-DEL-110 | chr9-8326335-DEL-110 |
| chr9-8342828-DEL-65 | chr9-8342828-DEL-70 |
| chr9-8344396-DEL-399 | chr9-8344397-DEL-399 |
| chr9-8346428-DEL-950 | chr9-8346429-DEL-950 |
| chr9-8390594-DEL-1430 | chr9-8390595-DEL-1430 |
| chr9-8396775-DEL-2920 | chr9-8396776-DEL-2920 |
| chr9-8405209-DEL-6015 | chr9-8405210-DEL-6015 |
| chr9-8445241-DEL-170 | chr9-8445242-DEL-170 |
| chr9-8447696-DEL-154 | chr9-8447697-DEL-154 |
| chr9-8452668-DEL-1093 | chr9-8452669-DEL-1093 |
| chr9-8458474-DEL-292 | chr9-8458476-DEL-280 |
| chr9-8462148-DEL-203 | chr9-8462149-DEL-203 |
| chr9-8466894-DEL-7596 | chr9-8466895-DEL-7596 |
| chr9-8486221-DEL-86 | chr9-8486201-DEL-107 |
| chr9-8540270-DEL-59 | chr9-8540271-DEL-59 |
| chr9-8597740-DEL-66 | chr9-8597743-DEL-64 |
| chr9-8606688-DEL-58 | chr9-8606689-DEL-54 |
| chr9-8628261-DEL-5343 | chr9-8628262-DEL-5343 |
| chr9-8663548-DEL-6085 | chr9-8663549-DEL-6085 |
| chr9-8681342-DEL-649 | chr9-8681325-DEL-665 |
| chr9-8708665-DEL-109 | chr9-8708666-DEL-109 |
| chr9-8767870-DEL-140 | chr9-8767871-DEL-140 |
| chr9-8770829-DEL-119 | chr9-8770830-DEL-119 |
| chr9-8771224-DEL-1932 | chr9-8771225-DEL-1932 |
| chr9-8782885-DEL-7209 | chr9-8782886-DEL-7209 |
| chr9-8822092-DEL-92 | chr9-8822093-DEL-92 |
| chr9-8859659-DEL-6090 | chr9-8859660-DEL-6089 |
| chr9-8981073-DEL-217 | chr9-8981074-DEL-217 |
| chr9-9017626-DEL-521 | chr9-9017627-DEL-521 |
| chr9-9038992-DEL-135 | chr9-9038993-DEL-135 |
| chr9-9040751-DEL-83 | chr9-9040752-DEL-83 |
| chr9-9041662-DEL-790 | chr9-9041686-DEL-775 |
| chr9-9053520-DEL-660 | chr9-9053523-DEL-659 |
| chr9-9059349-DEL-100 | chr9-9059350-DEL-100 |
| chr9-9069503-DEL-56 | chr9-9069504-DEL-56 |
| chr9-9083034-DEL-507 | chr9-9083035-DEL-507 |
| chr9-9089783-DEL-229 | chr9-9089784-DEL-229 |
| chr9-9119392-DEL-839 | chr9-9119393-DEL-839 |
| chr9-9128868-DEL-197 | chr9-9128869-DEL-197 |
| chr9-9135847-DEL-62 | chr9-9135850-DEL-60 |

| | |
|---|---|
| chr9-9155640-DEL-59 | chr9-9155638-DEL-60 |
| chr9-9185952-DEL-152 | chr9-9185971-DEL-106 |
| chr9-9191295-DEL-221 | chr9-9191296-DEL-221 |
| chr9-9234955-DEL-173 | chr9-9234955-DEL-179 |
| chr9-9244166-DEL-231 | chr9-9244197-DEL-191 |
| chr9-9246775-DEL-107 | chr9-9246727-DEL-105 |
| chr9-9262937-DEL-147 | chr9-9262916-DEL-164 |
| chr9-9277906-DEL-183 | chr9-9277907-DEL-183 |
| chr9-9292713-DEL-7228 | chr9-9292714-DEL-7228 |
| chr9-9302670-DEL-203 | chr9-9302671-DEL-203 |
| chr9-9306198-DEL-639 | chr9-9306199-DEL-639 |
| chr9-9312797-DEL-530 | chr9-9312798-DEL-530 |
| chr9-9318381-DEL-82 | chr9-9318415-DEL-82 |
| chr9-9324070-DEL-355 | chr9-9324071-DEL-354 |
| chr9-9371582-DEL-81 | chr9-9371583-DEL-81 |
| chr9-9388448-DEL-560 | chr9-9388449-DEL-560 |
| chr9-9389609-DEL-573 | chr9-9389610-DEL-573 |
| chr9-9396972-DEL-51 | chr9-9396959-DEL-65 |
| chr9-9417749-DEL-1213 | chr9-9417750-DEL-1213 |
| chr9-9443816-DEL-200 | chr9-9443819-DEL-200 |
| chr9-9495670-DEL-2739 | chr9-9495671-DEL-2739 |
| chr9-9508972-DEL-509 | chr9-9508985-DEL-507 |
| chr9-9511942-DEL-83 | chr9-9511943-DEL-83 |
| chr9-9513519-DEL-200 | chr9-9513520-DEL-200 |
| chr9-9516354-DEL-87 | chr9-9516381-DEL-55 |
| chr9-9516951-DEL-56 | chr9-9516929-DEL-60 |
| chr9-9527640-DEL-324 | chr9-9527641-DEL-324 |
| chr9-9538668-DEL-6691 | chr9-9538667-DEL-6692 |
| chr9-9545027-DEL-212 | chr9-9545028-DEL-212 |
| chr9-9550469-DEL-379 | chr9-9550470-DEL-379 |
| chr9-9598744-DEL-2666 | chr9-9598745-DEL-2666 |
| chr9-9606606-DEL-563 | chr9-9606606-DEL-565 |
| chr9-9619800-DEL-6990 | chr9-9619790-DEL-6991 |
| chr9-9655120-DEL-86 | chr9-9655129-DEL-78 |
| chr9-9661968-DEL-361 | chr9-9661968-DEL-359 |
| chr9-9667833-DEL-100 | chr9-9667833-DEL-101 |
| chr9-9702673-DEL-638 | chr9-9702674-DEL-638 |
| chr9-9739965-DEL-118 | chr9-9739966-DEL-118 |
| chr9-9777920-DEL-109 | chr9-9777921-DEL-109 |
| chr9-9788584-DEL-393 | chr9-9788577-DEL-394 |
| chr9-9812051-DEL-165 | chr9-9812052-DEL-165 |
| chr9-9843544-DEL-158 | chr9-9843545-DEL-158 |
| chr9-9846765-DEL-6217 | chr9-9846766-DEL-6217 |
| chr9-9865337-DEL-2703 | chr9-9865338-DEL-2702 |
| chr9-9887470-DEL-4428 | chr9-9887471-DEL-4428 |
| chr9-9921895-DEL-5965 | chr9-9921895-DEL-5965 |
| chr9-9973085-DEL-887 | chr9-9973088-DEL-885 |
| chr9-10021250-DEL-1034 | chr9-10021251-DEL-1034 |
| chr9-10052221-DEL-730 | chr9-10052222-DEL-730 |
| chr9-10109471-DEL-213 | chr9-10109472-DEL-213 |
| chr9-10135387-DEL-1524 | chr9-10135388-DEL-1524 |
| chr9-10158509-DEL-5327 | chr9-10158495-DEL-5341 |
| chr9-10193307-DEL-5401 | chr9-10193308-DEL-5401 |
| chr9-10234787-DEL-52 | chr9-10234788-DEL-52 |
| chr9-10290275-DEL-5094 | chr9-10290276-DEL-5094 |
| chr9-10320191-DEL-2473 | chr9-10320192-DEL-2473 |
| chr9-10329578-DEL-431 | chr9-10329579-DEL-431 |
| chr9-10349038-DEL-233 | chr9-10349039-DEL-233 |
| chr9-10390700-DEL-54 | chr9-10390705-DEL-52 |
| chr9-10452467-DEL-1465 | chr9-10452468-DEL-1465 |
| chr9-10492623-DEL-61 | chr9-10492628-DEL-71 |
| chr9-10541196-DEL-233 | chr9-10541197-DEL-233 |
| chr9-10549318-DEL-55 | chr9-10549319-DEL-55 |
| chr9-10552134-DEL-6258 | chr9-10552135-DEL-6258 |
| chr9-10635779-DEL-863 | chr9-10635780-DEL-863 |
| chr9-10671638-DEL-60 | chr9-10671649-DEL-50 |
| chr9-10672229-DEL-161 | chr9-10672225-DEL-187 |
| chr9-10687436-DEL-736 | chr9-10687428-DEL-736 |
| chr9-10701200-DEL-6243 | chr9-10701201-DEL-6243 |
| chr9-10732552-DEL-2388 | chr9-10732553-DEL-2388 |
| chr9-10739947-DEL-330 | chr9-10739941-DEL-330 |
| chr9-10742149-DEL-4128 | chr9-10742148-DEL-4129 |
| chr9-10758958-DEL-1810 | chr9-10758958-DEL-1808 |
| chr9-10763841-DEL-455 | chr9-10763842-DEL-455 |
| chr9-10787437-DEL-70 | chr9-10787441-DEL-68 |
| chr9-10812569-DEL-86 | chr9-10812570-DEL-86 |
| chr9-10827012-DEL-371 | chr9-10827013-DEL-371 |
| chr9-10868846-DEL-731 | chr9-10868847-DEL-731 |
| chr9-10924458-DEL-402 | chr9-10924459-DEL-402 |
| chr9-10964167-DEL-7290 | chr9-10964167-DEL-7290 |

| | |
|---|---|
| chr9-10964306-DEL-214 | chr9-10964262-DEL-214 |
| chr9-11012188-DEL-365 | chr9-11012189-DEL-365 |
| chr9-11066884-DEL-353 | chr9-11066885-DEL-353 |
| chr9-11070079-DEL-533 | chr9-11070072-DEL-530 |
| chr9-11076706-DEL-5660 | chr9-11076707-DEL-5660 |
| chr9-11087403-DEL-3198 | chr9-11087400-DEL-3199 |
| chr9-11102243-DEL-215 | chr9-11102244-DEL-215 |
| chr9-11108833-DEL-6202 | chr9-11108834-DEL-6202 |
| chr9-11117048-DEL-1163 | chr9-11117049-DEL-1163 |
| chr9-11118440-DEL-107 | chr9-11118441-DEL-107 |
| chr9-11134302-DEL-6665 | chr9-11134303-DEL-6665 |
| chr9-11180198-DEL-432 | chr9-11180199-DEL-432 |
| chr9-11205219-DEL-375 | chr9-11205215-DEL-376 |
| chr9-11207985-DEL-843 | chr9-11207985-DEL-843 |
| chr9-11211125-DEL-94 | chr9-11211125-DEL-102 |
| chr9-11214038-DEL-4878 | chr9-11214039-DEL-4878 |
| chr9-11224305-DEL-285 | chr9-11224298-DEL-302 |
| chr9-11268400-DEL-86 | chr9-11268422-DEL-85 |
| chr9-11270379-DEL-61 | chr9-11270382-DEL-54 |
| chr9-11305290-DEL-4367 | chr9-11305291-DEL-4367 |
| chr9-11319124-DEL-1098 | chr9-11319125-DEL-1098 |
| chr9-11319577-DEL-425 | chr9-11319578-DEL-425 |
| chr9-11341984-DEL-3276 | chr9-11341985-DEL-3276 |
| chr9-11352374-DEL-445 | chr9-11352375-DEL-445 |
| chr9-11358239-DEL-96 | chr9-11358272-DEL-60 |
| chr9-11373912-DEL-222 | chr9-11373913-DEL-222 |
| chr9-11400580-DEL-1880 | chr9-11400581-DEL-1880 |
| chr9-11410621-DEL-6670 | chr9-11410622-DEL-6670 |
| chr9-11498671-DEL-6589 | chr9-11498672-DEL-6589 |
| chr9-11508731-DEL-623 | chr9-11508732-DEL-623 |
| chr9-11521746-DEL-1655 | chr9-11521747-DEL-1655 |
| chr9-11554043-DEL-3619 | chr9-11554044-DEL-3619 |
| chr9-11612263-DEL-63 | chr9-11612264-DEL-63 |
| chr9-11633584-DEL-873 | chr9-11633588-DEL-869 |
| chr9-11642167-DEL-24147 | chr9-11642166-DEL-24148 |
| chr9-11652026-DEL-6316 | chr9-11652027-DEL-6316 |
| chr9-11652039-DEL-214 | chr9-11652040-DEL-214 |
| chr9-11716361-DEL-86 | chr9-11716362-DEL-86 |
| chr9-11719906-DEL-55 | chr9-11719907-DEL-55 |
| chr9-11722523-DEL-6259 | chr9-11722524-DEL-6259 |
| chr9-11722536-DEL-212 | chr9-11722537-DEL-212 |
| chr9-11729193-DEL-59 | chr9-11729193-DEL-59 |
| chr9-11736906-DEL-107 | chr9-11736877-DEL-97 |
| chr9-11792939-DEL-5316 | chr9-11792940-DEL-5316 |
| chr9-11806383-DEL-202 | chr9-11806384-DEL-202 |
| chr9-11876452-DEL-427 | chr9-11876453-DEL-427 |
| chr9-11890160-DEL-778 | chr9-11890154-DEL-778 |
| chr9-11919445-DEL-115 | chr9-11919444-DEL-115 |
| chr9-11920879-DEL-786 | chr9-11920872-DEL-787 |
| chr9-11945590-DEL-6642 | chr9-11945591-DEL-6642 |
| chr9-12019500-DEL-103 | chr9-12019497-DEL-81 |
| chr9-12066268-DEL-4082 | chr9-12066269-DEL-4082 |
| chr9-12086781-DEL-793 | chr9-12086782-DEL-793 |
| chr9-12129514-DEL-9962 | chr9-12129514-DEL-9962 |
| chr9-12132487-DEL-1889 | chr9-12132488-DEL-1889 |
| chr9-12332200-DEL-54 | chr9-12332202-DEL-54 |
| chr9-12373120-DEL-6605 | chr9-12373121-DEL-6605 |
| chr9-12444033-DEL-3567 | chr9-12444034-DEL-3567 |
| chr9-12469140-DEL-4237 | chr9-12469141-DEL-4237 |
| chr9-12481788-DEL-64 | chr9-12481793-DEL-60 |
| chr9-12524905-DEL-587 | chr9-12524906-DEL-587 |
| chr9-12557558-DEL-103 | chr9-12557559-DEL-103 |
| chr9-12596524-DEL-790 | chr9-12596525-DEL-790 |
| chr9-12644669-DEL-96 | chr9-12644670-DEL-96 |
| chr9-12665906-DEL-328 | chr9-12665907-DEL-328 |
| chr9-12682395-DEL-74 | chr9-12682396-DEL-74 |
| chr9-12685707-DEL-81 | chr9-12685690-DEL-82 |
| chr9-12720069-DEL-75 | chr9-12720059-DEL-60 |
| chr9-12739962-DEL-6872 | chr9-12739963-DEL-6872 |
| chr9-12839788-DEL-12700 | chr9-12839789-DEL-12700 |
| chr9-12884731-DEL-199 | chr9-12884732-DEL-199 |
| chr9-12903523-DEL-826 | chr9-12903524-DEL-826 |
| chr9-12916122-DEL-6205 | chr9-12916119-DEL-6206 |
| chr9-12963717-DEL-662 | chr9-12963718-DEL-662 |
| chr9-13069367-DEL-370 | chr9-13069368-DEL-370 |
| chr9-13124481-DEL-1090 | chr9-13124482-DEL-1090 |
| chr9-13138401-DEL-2345 | chr9-13138401-DEL-2345 |
| chr9-13220502-DEL-2533 | chr9-13220500-DEL-2532 |
| chr9-13334560-DEL-61 | chr9-13334566-DEL-50 |
| chr9-13421735-DEL-89 | chr9-13421742-DEL-83 |

| | |
|---|---|
| chr9-13438294-DEL-176 | chr9-13438295-DEL-176 |
| chr9-13453102-DEL-610 | chr9-13453105-DEL-608 |
| chr9-13478811-DEL-95 | chr9-13478824-DEL-82 |
| chr9-13527064-DEL-149 | chr9-13527053-DEL-161 |
| chr9-13555668-DEL-335 | chr9-13555669-DEL-335 |
| chr9-13574784-DEL-53 | chr9-13574789-DEL-51 |
| chr9-13586571-DEL-177 | chr9-13586572-DEL-177 |
| chr9-13609668-DEL-188 | chr9-13609661-DEL-224 |
| chr9-13635820-DEL-1641 | chr9-13635821-DEL-1641 |
| chr9-13665525-DEL-4172 | chr9-13665526-DEL-4172 |
| chr9-13690533-DEL-723 | chr9-13690534-DEL-723 |
| chr9-13702385-DEL-215 | chr9-13702386-DEL-215 |
| chr9-13775069-DEL-51 | chr9-13775069-DEL-51 |
| chr9-13822134-DEL-66 | chr9-13822137-DEL-66 |
| chr9-13849246-DEL-6210 | chr9-13849247-DEL-6210 |
| chr9-13906589-DEL-77 | chr9-13906590-DEL-77 |
| chr9-13940002-DEL-927 | chr9-13939997-DEL-924 |
| chr9-13971980-DEL-72 | chr9-13971973-DEL-67 |
| chr9-13985062-DEL-245 | chr9-13985063-DEL-244 |
| chr9-14006275-DEL-7017 | chr9-14006276-DEL-7017 |
| chr9-14015500-DEL-57 | chr9-14015484-DEL-58 |
| chr9-14053960-DEL-57 | chr9-14053958-DEL-57 |
| chr9-14120015-DEL-54 | chr9-14120017-DEL-51 |
| chr9-14135409-DEL-2373 | chr9-14135410-DEL-2373 |
| chr9-14145532-DEL-59 | chr9-14145531-DEL-56 |
| chr9-14154824-DEL-75 | chr9-14154825-DEL-75 |
| chr9-14170083-DEL-59 | chr9-14170084-DEL-59 |
| chr9-14204761-DEL-63 | chr9-14204762-DEL-63 |
| chr9-14233032-DEL-57 | chr9-14233036-DEL-58 |
| chr9-14253931-DEL-239 | chr9-14253924-DEL-240 |
| chr9-14265118-DEL-160 | chr9-14265119-DEL-120 |
| chr9-14277892-DEL-132 | chr9-14277893-DEL-132 |
| chr9-14288991-DEL-299 | chr9-14288992-DEL-299 |
| chr9-14296104-DEL-411 | chr9-14296105-DEL-411 |
| chr9-14300850-DEL-450 | chr9-14300851-DEL-450 |
| chr9-14310846-DEL-201 | chr9-14310847-DEL-201 |
| chr9-14329139-DEL-222 | chr9-14329140-DEL-222 |
| chr9-14330606-DEL-6452 | chr9-14330607-DEL-6452 |
| chr9-14356545-DEL-114 | chr9-14356542-DEL-118 |
| chr9-14422018-DEL-144 | chr9-14422019-DEL-144 |
| chr9-14427822-DEL-169 | chr9-14427823-DEL-169 |
| chr9-14431989-DEL-147 | chr9-14431990-DEL-147 |
| chr9-14439803-DEL-494 | chr9-14439804-DEL-494 |
| chr9-14450477-DEL-58 | chr9-14450490-DEL-58 |
| chr9-14473785-DEL-884 | chr9-14473786-DEL-884 |
| chr9-14487895-DEL-73 | chr9-14487912-DEL-58 |
| chr9-14488485-DEL-2107 | chr9-14488486-DEL-2107 |
| chr9-14494443-DEL-1267 | chr9-14494444-DEL-1267 |
| chr9-14499023-DEL-88 | chr9-14499024-DEL-88 |
| chr9-14509173-DEL-704 | chr9-14509174-DEL-704 |
| chr9-14518483-DEL-111 | chr9-14518484-DEL-111 |
| chr9-14521212-DEL-232 | chr9-14521188-DEL-233 |
| chr9-14540232-DEL-19074 | chr9-14540233-DEL-19074 |
| chr9-14552575-DEL-93 | chr9-14552576-DEL-93 |
| chr9-14573275-DEL-98 | chr9-14573276-DEL-98 |
| chr9-14598699-DEL-394 | chr9-14598700-DEL-394 |
| chr9-14607305-DEL-57 | chr9-14607306-DEL-57 |
| chr9-14657972-DEL-266 | chr9-14657979-DEL-260 |
| chr9-14664016-DEL-196 | chr9-14664017-DEL-196 |
| chr9-14687881-DEL-204 | chr9-14687882-DEL-203 |
| chr9-14690832-DEL-134 | chr9-14690833-DEL-134 |
| chr9-14718833-DEL-80 | chr9-14718834-DEL-80 |
| chr9-14721904-DEL-65 | chr9-14721905-DEL-65 |
| chr9-14738857-DEL-199 | chr9-14738859-DEL-198 |
| chr9-14752910-DEL-207 | chr9-14752911-DEL-207 |
| chr9-14758591-DEL-80 | chr9-14758592-DEL-80 |
| chr9-14790362-DEL-88 | chr9-14790363-DEL-73 |
| chr9-14820012-DEL-1196 | chr9-14820014-DEL-1195 |
| chr9-14840136-DEL-488 | chr9-14840137-DEL-488 |
| chr9-14872463-DEL-580 | chr9-14872461-DEL-576 |
| chr9-14880657-DEL-97 | chr9-14880658-DEL-116 |
| chr9-14896752-DEL-1891 | chr9-14896753-DEL-1891 |
| chr9-14922757-DEL-901 | chr9-14922758-DEL-901 |
| chr9-15000602-DEL-329 | chr9-15000603-DEL-329 |
| chr9-15008820-DEL-184 | chr9-15008821-DEL-184 |
| chr9-15041328-DEL-633 | chr9-15041328-DEL-631 |
| chr9-15050747-DEL-53 | chr9-15050743-DEL-54 |
| chr9-15059637-DEL-585 | chr9-15059638-DEL-585 |
| chr9-15063423-DEL-119 | chr9-15063424-DEL-119 |
| chr9-15092390-DEL-269 | chr9-15092391-DEL-269 |

| | |
|---|---|
| chr9-15106953-DEL-214 | chr9-15106953-DEL-214 |
| chr9-15110320-DEL-837 | chr9-15110318-DEL-839 |
| chr9-15111798-DEL-233 | chr9-15111799-DEL-233 |
| chr9-15122578-DEL-195 | chr9-15122584-DEL-198 |
| chr9-15136872-DEL-251 | chr9-15136873-DEL-250 |
| chr9-15144141-DEL-51 | chr9-15144142-DEL-51 |
| chr9-15145367-DEL-303 | chr9-15145368-DEL-303 |
| chr9-15146473-DEL-637 | chr9-15146474-DEL-637 |
| chr9-15182768-DEL-101 | chr9-15182780-DEL-79 |
| chr9-15213414-DEL-195 | chr9-15213415-DEL-195 |
| chr9-15252408-DEL-200 | chr9-15252409-DEL-200 |
| chr9-15274521-DEL-314 | chr9-15274522-DEL-314 |
| chr9-15302423-DEL-200 | chr9-15302424-DEL-200 |
| chr9-15321764-DEL-342 | chr9-15321765-DEL-342 |
| chr9-15322439-DEL-204 | chr9-15322441-DEL-203 |
| chr9-15335909-DEL-65 | chr9-15335910-DEL-65 |
| chr9-15345645-DEL-347 | chr9-15345646-DEL-347 |
| chr9-15363462-DEL-336 | chr9-15363463-DEL-336 |
| chr9-15366193-DEL-236 | chr9-15366194-DEL-236 |
| chr9-15369086-DEL-873 | chr9-15369087-DEL-873 |
| chr9-15376617-DEL-116 | chr9-15376618-DEL-116 |
| chr9-15385133-DEL-1733 | chr9-15385134-DEL-1733 |
| chr9-15393037-DEL-491 | chr9-15393038-DEL-491 |
| chr9-15403706-DEL-66 | chr9-15403707-DEL-66 |
| chr9-15418031-DEL-52 | chr9-15417996-DEL-57 |
| chr9-15423140-DEL-113 | chr9-15423104-DEL-150 |
| chr9-15437131-DEL-441 | chr9-15437132-DEL-441 |
| chr9-15454033-DEL-84 | chr9-15454034-DEL-85 |
| chr9-15455395-DEL-119 | chr9-15455396-DEL-119 |
| chr9-15457109-DEL-76 | chr9-15457110-DEL-76 |
| chr9-15484362-DEL-110 | chr9-15484363-DEL-82 |
| chr9-15503506-DEL-78 | chr9-15503507-DEL-74 |
| chr9-15519612-DEL-272 | chr9-15519613-DEL-272 |
| chr9-15539816-DEL-366 | chr9-15539817-DEL-366 |
| chr9-15565931-DEL-94 | chr9-15565932-DEL-94 |
| chr9-15611001-DEL-1142 | chr9-15611002-DEL-1142 |
| chr9-15638502-DEL-111 | chr9-15638503-DEL-111 |
| chr9-15692423-DEL-159 | chr9-15692424-DEL-159 |
| chr9-15699150-DEL-269 | chr9-15699153-DEL-267 |
| chr9-15726101-DEL-96 | chr9-15726102-DEL-96 |
| chr9-15741601-DEL-102 | chr9-15741603-DEL-106 |
| chr9-15760255-DEL-643 | chr9-15760256-DEL-643 |
| chr9-15765454-DEL-3446 | chr9-15765455-DEL-3446 |
| chr9-15780449-DEL-73 | chr9-15780450-DEL-73 |
| chr9-15787032-DEL-52 | chr9-15787033-DEL-52 |
| chr9-15799798-DEL-987 | chr9-15799799-DEL-987 |
| chr9-15801616-DEL-4107 | chr9-15801615-DEL-4107 |
| chr9-15810035-DEL-161 | chr9-15810036-DEL-161 |
| chr9-15813593-DEL-597 | chr9-15813594-DEL-597 |
| chr9-15840376-DEL-1888 | chr9-15840377-DEL-1888 |
| chr9-15857149-DEL-122 | chr9-15857117-DEL-166 |
| chr9-15889811-DEL-69 | chr9-15889812-DEL-69 |
| chr9-15911672-DEL-130 | chr9-15911673-DEL-118 |
| chr9-15922129-DEL-154 | chr9-15922130-DEL-154 |
| chr9-15923623-DEL-55 | chr9-15923643-DEL-50 |
| chr9-15961690-DEL-399 | chr9-15961684-DEL-400 |
| chr9-15971577-DEL-80 | chr9-15971612-DEL-72 |
| chr9-15991692-DEL-2202 | chr9-15991693-DEL-2202 |
| chr9-16027625-DEL-84 | chr9-16027625-DEL-98 |
| chr9-16029131-DEL-57 | chr9-16029132-DEL-57 |
| chr9-16089110-DEL-474 | chr9-16089111-DEL-474 |
| chr9-16125588-DEL-167 | chr9-16125589-DEL-167 |
| chr9-16130451-DEL-128 | chr9-16130452-DEL-128 |
| chr9-16140430-DEL-1303 | chr9-16140431-DEL-1304 |
| chr9-16190808-DEL-109 | chr9-16190809-DEL-107 |
| chr9-16199899-DEL-63 | chr9-16199908-DEL-53 |
| chr9-16228449-DEL-58 | chr9-16228450-DEL-52 |
| chr9-16230085-DEL-87 | chr9-16230086-DEL-87 |
| chr9-16245663-DEL-107 | chr9-16245659-DEL-107 |
| chr9-16249300-DEL-194 | chr9-16249301-DEL-194 |
| chr9-16251483-DEL-84 | chr9-16251484-DEL-84 |
| chr9-16269944-DEL-101 | chr9-16269945-DEL-101 |
| chr9-16281319-DEL-119 | chr9-16281308-DEL-103 |
| chr9-16315840-DEL-211 | chr9-16315841-DEL-212 |
| chr9-16351669-DEL-70 | chr9-16351670-DEL-70 |
| chr9-16359030-DEL-224 | chr9-16359029-DEL-225 |
| chr9-16363996-DEL-6392 | chr9-16363997-DEL-6392 |
| chr9-16380458-DEL-57 | chr9-16380453-DEL-64 |
| chr9-16395155-DEL-665 | chr9-16395156-DEL-665 |
| chr9-16424663-DEL-133 | chr9-16424664-DEL-133 |

| | |
|---|---|
| chr9-16475896-DEL-583 | chr9-16475897-DEL-583 |
| chr9-16505824-DEL-464 | chr9-16505825-DEL-463 |
| chr9-16515811-DEL-196 | chr9-16515812-DEL-196 |
| chr9-16516208-DEL-1505 | chr9-16516209-DEL-1505 |
| chr9-16517941-DEL-6842 | chr9-16517942-DEL-6842 |
| chr9-16527145-DEL-203 | chr9-16527146-DEL-203 |
| chr9-16530071-DEL-58 | chr9-16530072-DEL-58 |
| chr9-16536747-DEL-1615 | chr9-16536758-DEL-1615 |
| chr9-16555883-DEL-74 | chr9-16555884-DEL-74 |
| chr9-16560709-DEL-4758 | chr9-16560710-DEL-4758 |
| chr9-16594813-DEL-58 | chr9-16594768-DEL-62 |
| chr9-16597084-DEL-6206 | chr9-16597085-DEL-6206 |
| chr9-16607375-DEL-786 | chr9-16607376-DEL-786 |
| chr9-16633525-DEL-889 | chr9-16633525-DEL-888 |
| chr9-16660110-DEL-55 | chr9-16660111-DEL-50 |
| chr9-16665319-DEL-154 | chr9-16665299-DEL-155 |
| chr9-16695810-DEL-97 | chr9-16695843-DEL-62 |
| chr9-16725267-DEL-480 | chr9-16725268-DEL-480 |
| chr9-16744933-DEL-6737 | chr9-16744934-DEL-6737 |
| chr9-16777741-DEL-4506 | chr9-16777742-DEL-4506 |
| chr9-16798705-DEL-85 | chr9-16798706-DEL-85 |
| chr9-16827060-DEL-54 | chr9-16827061-DEL-54 |
| chr9-16895564-DEL-1099 | chr9-16895565-DEL-1099 |
| chr9-16897004-DEL-371 | chr9-16897005-DEL-371 |
| chr9-16904590-DEL-489 | chr9-16904591-DEL-489 |
| chr9-16944344-DEL-1551 | chr9-16944342-DEL-1554 |
| chr9-16948132-DEL-6140 | chr9-16948120-DEL-6152 |
| chr9-16956434-DEL-58 | chr9-16956435-DEL-58 |
| chr9-16964562-DEL-467 | chr9-16964563-DEL-467 |
| chr9-16970280-DEL-6576 | chr9-16970281-DEL-6576 |
| chr9-16987514-DEL-95 | chr9-16987498-DEL-72 |
| chr9-16994765-DEL-4359 | chr9-16994766-DEL-4359 |
| chr9-17014819-DEL-60 | chr9-17014824-DEL-54 |
| chr9-17015991-DEL-494 | chr9-17015992-DEL-494 |
| chr9-17037833-DEL-276 | chr9-17037834-DEL-276 |
| chr9-17051948-DEL-146 | chr9-17051949-DEL-146 |
| chr9-17065915-DEL-270 | chr9-17065916-DEL-270 |
| chr9-17091376-DEL-67 | chr9-17091369-DEL-75 |
| chr9-17106772-DEL-204 | chr9-17106773-DEL-204 |
| chr9-17107899-DEL-11568 | chr9-17107900-DEL-11568 |
| chr9-17125968-DEL-55 | chr9-17125950-DEL-59 |
| chr9-17131803-DEL-483 | chr9-17131804-DEL-483 |
| chr9-17135590-DEL-455 | chr9-17135590-DEL-479 |
| chr9-17147932-DEL-98 | chr9-17147933-DEL-98 |
| chr9-17151016-DEL-5187 | chr9-17151017-DEL-5187 |
| chr9-17172719-DEL-1864 | chr9-17172720-DEL-1864 |
| chr9-17182664-DEL-68 | chr9-17182665-DEL-68 |
| chr9-17198116-DEL-134 | chr9-17198152-DEL-128 |
| chr9-17198566-DEL-363 | chr9-17198567-DEL-363 |
| chr9-17201274-DEL-163 | chr9-17201263-DEL-166 |
| chr9-17205493-DEL-113 | chr9-17205494-DEL-113 |
| chr9-17207934-DEL-62 | chr9-17207929-DEL-68 |
| chr9-17208869-DEL-2028 | chr9-17208870-DEL-2028 |
| chr9-17250870-DEL-371 | chr9-17250871-DEL-371 |
| chr9-17271622-DEL-1043 | chr9-17271623-DEL-1043 |
| chr9-17278278-DEL-52 | chr9-17278279-DEL-51 |
| chr9-17298598-DEL-4338 | chr9-17298591-DEL-4345 |
| chr9-17310838-DEL-820 | chr9-17310839-DEL-820 |
| chr9-17312529-DEL-122 | chr9-17312530-DEL-122 |
| chr9-17313331-DEL-6233 | chr9-17313332-DEL-6233 |
| chr9-17334542-DEL-52 | chr9-17334543-DEL-52 |
| chr9-17346369-DEL-447 | chr9-17346370-DEL-447 |
| chr9-17351207-DEL-357 | chr9-17351208-DEL-357 |
| chr9-17362843-DEL-343 | chr9-17362844-DEL-343 |
| chr9-17372292-DEL-133 | chr9-17372292-DEL-133 |
| chr9-17373017-DEL-358 | chr9-17373018-DEL-358 |
| chr9-17375098-DEL-2339 | chr9-17375099-DEL-2339 |
| chr9-17384330-DEL-118 | chr9-17384319-DEL-119 |
| chr9-17387372-DEL-259 | chr9-17387373-DEL-260 |
| chr9-17413351-DEL-248 | chr9-17413355-DEL-241 |
| chr9-17413913-DEL-90 | chr9-17413914-DEL-54 |
| chr9-17414865-DEL-63 | chr9-17414877-DEL-52 |
| chr9-17418486-DEL-650 | chr9-17418486-DEL-650 |
| chr9-17420789-DEL-62 | chr9-17420765-DEL-86 |
| chr9-17443566-DEL-131 | chr9-17443575-DEL-126 |
| chr9-17505629-DEL-53 | chr9-17505629-DEL-53 |
| chr9-17513037-DEL-520 | chr9-17513038-DEL-520 |
| chr9-17538270-DEL-408 | chr9-17538271-DEL-408 |
| chr9-17539374-DEL-888 | chr9-17539375-DEL-888 |
| chr9-17540589-DEL-793 | chr9-17540590-DEL-793 |

| | |
|---|---|
| chr9-17543389-DEL-3759 | chr9-17543389-DEL-3759 |
| chr9-17559993-DEL-183 | chr9-17559994-DEL-177 |
| chr9-17562253-DEL-176 | chr9-17562254-DEL-186 |
| chr9-17599868-DEL-605 | chr9-17599869-DEL-605 |
| chr9-17625383-DEL-6203 | chr9-17625384-DEL-6203 |
| chr9-17665880-DEL-1876 | chr9-17665881-DEL-1876 |
| chr9-17671972-DEL-96 | chr9-17671973-DEL-96 |
| chr9-17673523-DEL-1862 | chr9-17673524-DEL-1862 |
| chr9-17676192-DEL-212 | chr9-17676184-DEL-214 |
| chr9-17677627-DEL-1405 | chr9-17677628-DEL-1405 |
| chr9-17687055-DEL-61 | chr9-17687056-DEL-61 |
| chr9-17696560-DEL-580 | chr9-17696560-DEL-581 |
| chr9-17701617-DEL-1978 | chr9-17701612-DEL-1976 |
| chr9-17722149-DEL-101 | chr9-17722150-DEL-101 |
| chr9-17726202-DEL-74 | chr9-17726218-DEL-69 |
| chr9-17761349-DEL-6631 | chr9-17761350-DEL-6631 |
| chr9-17775201-DEL-397 | chr9-17775202-DEL-395 |
| chr9-17826624-DEL-6085 | chr9-17826628-DEL-6082 |
| chr9-17857332-DEL-87 | chr9-17857333-DEL-87 |
| chr9-17859653-DEL-465 | chr9-17859654-DEL-465 |
| chr9-17861760-DEL-274 | chr9-17861761-DEL-274 |
| chr9-17864489-DEL-1829 | chr9-17864492-DEL-1828 |
| chr9-17867561-DEL-1788 | chr9-17867562-DEL-1788 |
| chr9-17872833-DEL-138 | chr9-17872834-DEL-138 |
| chr9-17878826-DEL-6344 | chr9-17878827-DEL-6344 |
| chr9-17886416-DEL-195 | chr9-17886417-DEL-195 |
| chr9-17908423-DEL-86 | chr9-17908424-DEL-86 |
| chr9-17913123-DEL-140 | chr9-17913124-DEL-140 |
| chr9-17918616-DEL-658 | chr9-17918617-DEL-658 |
| chr9-17920286-DEL-7033 | chr9-17920287-DEL-7033 |
| chr9-17934944-DEL-70 | chr9-17934943-DEL-72 |
| chr9-17940469-DEL-242 | chr9-17940470-DEL-242 |
| chr9-17946239-DEL-64 | chr9-17946240-DEL-64 |
| chr9-17967868-DEL-209 | chr9-17967918-DEL-159 |
| chr9-17971650-DEL-8863 | chr9-17971654-DEL-8864 |
| chr9-17987109-DEL-57 | chr9-17987114-DEL-85 |
| chr9-17993958-DEL-6588 | chr9-17993959-DEL-6588 |
| chr9-18017410-DEL-76 | chr9-18017413-DEL-74 |
| chr9-18028157-DEL-135 | chr9-18028159-DEL-134 |
| chr9-18051752-DEL-4460 | chr9-18051753-DEL-4460 |
| chr9-18067132-DEL-261 | chr9-18067133-DEL-261 |
| chr9-18067274-DEL-6252 | chr9-18067275-DEL-6252 |
| chr9-18073504-DEL-208 | chr9-18073504-DEL-210 |
| chr9-18074136-DEL-96 | chr9-18074142-DEL-90 |
| chr9-18077275-DEL-730 | chr9-18077276-DEL-730 |
| chr9-18096970-DEL-6412 | chr9-18096971-DEL-6412 |
| chr9-18106661-DEL-81 | chr9-18106665-DEL-63 |
| chr9-18139198-DEL-330 | chr9-18139199-DEL-330 |
| chr9-18173263-DEL-197 | chr9-18173244-DEL-198 |
| chr9-18246608-DEL-2221 | chr9-18246608-DEL-2217 |
| chr9-18278747-DEL-176 | chr9-18278748-DEL-177 |
| chr9-18287163-DEL-195 | chr9-18287164-DEL-194 |
| chr9-18292763-DEL-64 | chr9-18292776-DEL-58 |
| chr9-18320099-DEL-215 | chr9-18320100-DEL-215 |
| chr9-18346830-DEL-76 | chr9-18346831-DEL-76 |
| chr9-18347975-DEL-273 | chr9-18347976-DEL-273 |
| chr9-18371561-DEL-116 | chr9-18371562-DEL-114 |
| chr9-18468620-DEL-59 | chr9-18468625-DEL-55 |
| chr9-18502195-DEL-643 | chr9-18502189-DEL-642 |
| chr9-18505137-DEL-66 | chr9-18505138-DEL-66 |
| chr9-18526003-DEL-169 | chr9-18526004-DEL-169 |
| chr9-18539672-DEL-7840 | chr9-18539673-DEL-7840 |
| chr9-18588363-DEL-623 | chr9-18588364-DEL-623 |
| chr9-18592517-DEL-651 | chr9-18592520-DEL-648 |
| chr9-18606215-DEL-371 | chr9-18606215-DEL-372 |
| chr9-18616593-DEL-93 | chr9-18616574-DEL-85 |
| chr9-18627307-DEL-116 | chr9-18627302-DEL-76 |
| chr9-18630568-DEL-52 | chr9-18630571-DEL-50 |
| chr9-18659143-DEL-247 | chr9-18659144-DEL-247 |
| chr9-18687766-DEL-70 | chr9-18687763-DEL-66 |
| chr9-18709771-DEL-129 | chr9-18709736-DEL-172 |
| chr9-18747790-DEL-196 | chr9-18747785-DEL-197 |
| chr9-18751379-DEL-984 | chr9-18751375-DEL-988 |
| chr9-18761788-DEL-437 | chr9-18761789-DEL-437 |
| chr9-18771411-DEL-505 | chr9-18771412-DEL-505 |
| chr9-18815086-DEL-109 | chr9-18815087-DEL-109 |
| chr9-18856994-DEL-8523 | chr9-18856994-DEL-8521 |
| chr9-18891682-DEL-7079 | chr9-18891683-DEL-7079 |
| chr9-18902487-DEL-3507 | chr9-18902486-DEL-3508 |
| chr9-18926886-DEL-66 | chr9-18926887-DEL-66 |

| | |
|---|---|
| chr9-18939489-DEL-54 | chr9-18939490-DEL-54 |
| chr9-18949016-DEL-56 | chr9-18949034-DEL-62 |
| chr9-18957386-DEL-2890 | chr9-18957388-DEL-2890 |
| chr9-18983070-DEL-201 | chr9-18983071-DEL-201 |
| chr9-18992598-DEL-654 | chr9-18992593-DEL-654 |
| chr9-18993548-DEL-69 | chr9-18993548-DEL-69 |
| chr9-19024909-DEL-1830 | chr9-19024910-DEL-1830 |
| chr9-19126161-DEL-490 | chr9-19126162-DEL-490 |
| chr9-19170032-DEL-126 | chr9-19170048-DEL-131 |
| chr9-19170894-DEL-7254 | chr9-19170892-DEL-7255 |
| chr9-19178062-DEL-474 | chr9-19178063-DEL-474 |
| chr9-19191386-DEL-76 | chr9-19191387-DEL-76 |
| chr9-19196269-DEL-13899 | chr9-19196271-DEL-13901 |
| chr9-19199731-DEL-6369 | chr9-19199732-DEL-6369 |
| chr9-19232814-DEL-310 | chr9-19232815-DEL-310 |
| chr9-19238271-DEL-52 | chr9-19238256-DEL-52 |
| chr9-19254381-DEL-262 | chr9-19254382-DEL-262 |
| chr9-19339822-DEL-385 | chr9-19339823-DEL-385 |
| chr9-19346793-DEL-267 | chr9-19346793-DEL-267 |
| chr9-19378160-DEL-1568 | chr9-19378161-DEL-1568 |
| chr9-19397119-DEL-125 | chr9-19397120-DEL-125 |
| chr9-19414934-DEL-6298 | chr9-19414927-DEL-6299 |
| chr9-19443888-DEL-54 | chr9-19443880-DEL-51 |
| chr9-19498502-DEL-100 | chr9-19498503-DEL-72 |
| chr9-19532934-DEL-64 | chr9-19532935-DEL-64 |
| chr9-19542489-DEL-88 | chr9-19542490-DEL-88 |
| chr9-19543330-DEL-483 | chr9-19543329-DEL-484 |
| chr9-19544154-DEL-62 | chr9-19544155-DEL-62 |
| chr9-19568811-DEL-646 | chr9-19568812-DEL-646 |
| chr9-19631857-DEL-4831 | chr9-19631858-DEL-4831 |
| chr9-19707425-DEL-54 | chr9-19707426-DEL-54 |
| chr9-19745414-DEL-60 | chr9-19745415-DEL-60 |
| chr9-19812221-DEL-5087 | chr9-19812221-DEL-5088 |
| chr9-19822424-DEL-63 | chr9-19822425-DEL-63 |
| chr9-19845232-DEL-664 | chr9-19845233-DEL-664 |
| chr9-19854179-DEL-85 | chr9-19854132-DEL-95 |
| chr9-19961000-DEL-149 | chr9-19961001-DEL-149 |
| chr9-19964702-DEL-101 | chr9-19964703-DEL-99 |
| chr9-19970911-DEL-511 | chr9-19970911-DEL-511 |
| chr9-19973447-DEL-6371 | chr9-19973448-DEL-6371 |
| chr9-19984035-DEL-138 | chr9-19984042-DEL-132 |
| chr9-20043338-DEL-3139 | chr9-20043338-DEL-3139 |
| chr9-20057860-DEL-60 | chr9-20057873-DEL-97 |
| chr9-20070941-DEL-58 | chr9-20070944-DEL-50 |
| chr9-20075805-DEL-161 | chr9-20075798-DEL-169 |
| chr9-20095012-DEL-7095 | chr9-20095013-DEL-7095 |
| chr9-20103746-DEL-5520 | chr9-20103730-DEL-5536 |
| chr9-20121297-DEL-74 | chr9-20121298-DEL-72 |
| chr9-20142431-DEL-234 | chr9-20142432-DEL-234 |
| chr9-20198628-DEL-2351 | chr9-20198629-DEL-2351 |
| chr9-20232966-DEL-6581 | chr9-20232966-DEL-6581 |
| chr9-20242995-DEL-52 | chr9-20242996-DEL-52 |
| chr9-20244709-DEL-84 | chr9-20244726-DEL-68 |
| chr9-20291331-DEL-1374 | chr9-20291331-DEL-1374 |
| chr9-20299683-DEL-1824 | chr9-20299685-DEL-1817 |
| chr9-20304244-DEL-207 | chr9-20304245-DEL-207 |
| chr9-20312250-DEL-218 | chr9-20312251-DEL-218 |
| chr9-20322717-DEL-328 | chr9-20322718-DEL-328 |
| chr9-20323139-DEL-1959 | chr9-20323138-DEL-1943 |
| chr9-20328415-DEL-504 | chr9-20328416-DEL-504 |
| chr9-20339972-DEL-309 | chr9-20339992-DEL-290 |
| chr9-20342385-DEL-66 | chr9-20342386-DEL-66 |
| chr9-20368245-DEL-85 | chr9-20368247-DEL-86 |
| chr9-20376113-DEL-84 | chr9-20376120-DEL-70 |
| chr9-20420694-DEL-61 | chr9-20420695-DEL-61 |
| chr9-20437381-DEL-193 | chr9-20437374-DEL-194 |
| chr9-20439832-DEL-152 | chr9-20439833-DEL-152 |
| chr9-20440254-DEL-324 | chr9-20440255-DEL-360 |
| chr9-20444122-DEL-57 | chr9-20444123-DEL-73 |
| chr9-20473012-DEL-64 | chr9-20473028-DEL-77 |
| chr9-20485453-DEL-363 | chr9-20485454-DEL-363 |
| chr9-20494422-DEL-87 | chr9-20494418-DEL-87 |
| chr9-20509868-DEL-91 | chr9-20509869-DEL-91 |
| chr9-20530478-DEL-6401 | chr9-20530479-DEL-6401 |
| chr9-20551872-DEL-73 | chr9-20551851-DEL-89 |
| chr9-20595209-DEL-68 | chr9-20595210-DEL-68 |
| chr9-20600529-DEL-254 | chr9-20600530-DEL-254 |
| chr9-20661045-DEL-82 | chr9-20661051-DEL-56 |
| chr9-20664891-DEL-63 | chr9-20664896-DEL-63 |
| chr9-20732130-DEL-52 | chr9-20732131-DEL-52 |

| | |
|---|---|
| chr9-20749107-DEL-58 | chr9-20749119-DEL-56 |
| chr9-20772187-DEL-216 | chr9-20772188-DEL-213 |
| chr9-20796915-DEL-201 | chr9-20796916-DEL-201 |
| chr9-20800398-DEL-206 | chr9-20800399-DEL-206 |
| chr9-20898562-DEL-143 | chr9-20898563-DEL-143 |
| chr9-20905334-DEL-106 | chr9-20905335-DEL-106 |
| chr9-20912777-DEL-169 | chr9-20912778-DEL-169 |
| chr9-20915331-DEL-212 | chr9-20915332-DEL-212 |
| chr9-20920655-DEL-176 | chr9-20920656-DEL-176 |
| chr9-20922445-DEL-217 | chr9-20922446-DEL-217 |
| chr9-20934651-DEL-241 | chr9-20934652-DEL-241 |
| chr9-20937417-DEL-77 | chr9-20937418-DEL-77 |
| chr9-20945306-DEL-217 | chr9-20945307-DEL-217 |
| chr9-20953702-DEL-695 | chr9-20953703-DEL-695 |
| chr9-20966256-DEL-710 | chr9-20966257-DEL-710 |
| chr9-20971342-DEL-203 | chr9-20971343-DEL-203 |
| chr9-20989344-DEL-118 | chr9-20989316-DEL-98 |
| chr9-21015044-DEL-55 | chr9-21015006-DEL-52 |
| chr9-21017678-DEL-62 | chr9-21017679-DEL-62 |
| chr9-21023976-DEL-211 | chr9-21023977-DEL-211 |
| chr9-21028780-DEL-211 | chr9-21028772-DEL-212 |
| chr9-21030287-DEL-174 | chr9-21030288-DEL-174 |
| chr9-21047046-DEL-982 | chr9-21047047-DEL-982 |
| chr9-21048530-DEL-59 | chr9-21048531-DEL-59 |
| chr9-21064801-DEL-89 | chr9-21064801-DEL-89 |
| chr9-21081311-DEL-95 | chr9-21081312-DEL-83 |
| chr9-21091305-DEL-1099 | chr9-21091306-DEL-1099 |
| chr9-21099426-DEL-83 | chr9-21099427-DEL-84 |
| chr9-21134724-DEL-201 | chr9-21134724-DEL-201 |
| chr9-21139803-DEL-71 | chr9-21139804-DEL-71 |
| chr9-21145635-DEL-67 | chr9-21145636-DEL-67 |
| chr9-21153813-DEL-178 | chr9-21153814-DEL-178 |
| chr9-21166499-DEL-2404 | chr9-21166500-DEL-2404 |
| chr9-21192030-DEL-201 | chr9-21192031-DEL-201 |
| chr9-21220570-DEL-53 | chr9-21220571-DEL-53 |
| chr9-21243662-DEL-398 | chr9-21243663-DEL-398 |
| chr9-21262380-DEL-117 | chr9-21262382-DEL-115 |
| chr9-21267624-DEL-352 | chr9-21267634-DEL-343 |
| chr9-21282536-DEL-211 | chr9-21282537-DEL-211 |
| chr9-21321397-DEL-100 | chr9-21321398-DEL-100 |
| chr9-21321908-DEL-170 | chr9-21321909-DEL-170 |
| chr9-21331056-DEL-64 | chr9-21331057-DEL-64 |
| chr9-21346290-DEL-206 | chr9-21346291-DEL-206 |
| chr9-21399770-DEL-214 | chr9-21399787-DEL-219 |
| chr9-21402747-DEL-87 | chr9-21402750-DEL-84 |
| chr9-21411715-DEL-1240 | chr9-21411671-DEL-1285 |
| chr9-21423051-DEL-200 | chr9-21423052-DEL-200 |
| chr9-21429782-DEL-54 | chr9-21429783-DEL-50 |
| chr9-21430392-DEL-2976 | chr9-21430393-DEL-2976 |
| chr9-21438604-DEL-198 | chr9-21438605-DEL-198 |
| chr9-21444138-DEL-80 | chr9-21444147-DEL-72 |
| chr9-21460879-DEL-220 | chr9-21460880-DEL-220 |
| chr9-21466665-DEL-65 | chr9-21466637-DEL-66 |
| chr9-21508531-DEL-196 | chr9-21508532-DEL-196 |
| chr9-21509190-DEL-240 | chr9-21509191-DEL-240 |
| chr9-21513833-DEL-73 | chr9-21513834-DEL-73 |
| chr9-21525614-DEL-124 | chr9-21525615-DEL-124 |
| chr9-21533353-DEL-137 | chr9-21533345-DEL-135 |
| chr9-21540447-DEL-207 | chr9-21540448-DEL-207 |
| chr9-21555429-DEL-215 | chr9-21555430-DEL-215 |
| chr9-21569468-DEL-134 | chr9-21569466-DEL-132 |
| chr9-21621771-DEL-2909 | chr9-21621772-DEL-2909 |
| chr9-21625641-DEL-178 | chr9-21625642-DEL-178 |
| chr9-21638603-DEL-79 | chr9-21638603-DEL-79 |
| chr9-21651635-DEL-56 | chr9-21651637-DEL-57 |
| chr9-21654516-DEL-406 | chr9-21654517-DEL-406 |
| chr9-21662133-DEL-64 | chr9-21662134-DEL-64 |
| chr9-21666897-DEL-196 | chr9-21666898-DEL-196 |
| chr9-21676355-DEL-222 | chr9-21676356-DEL-222 |
| chr9-21682402-DEL-579 | chr9-21682403-DEL-579 |
| chr9-21682985-DEL-181 | chr9-21682986-DEL-181 |
| chr9-21687177-DEL-218 | chr9-21687187-DEL-213 |
| chr9-21688511-DEL-150 | chr9-21688518-DEL-149 |
| chr9-21702193-DEL-203 | chr9-21702185-DEL-204 |
| chr9-21733902-DEL-92 | chr9-21733903-DEL-92 |
| chr9-21734586-DEL-71 | chr9-21734587-DEL-69 |
| chr9-21739604-DEL-81 | chr9-21739605-DEL-81 |
| chr9-21777780-DEL-104 | chr9-21777781-DEL-104 |
| chr9-21790487-DEL-7077 | chr9-21790480-DEL-7077 |
| chr9-21801170-DEL-205 | chr9-21801171-DEL-176 |

| | |
|---|---|
| chr9-21801818-DEL-201 | chr9-21801819-DEL-201 |
| chr9-21816228-DEL-74 | chr9-21816229-DEL-74 |
| chr9-21897138-DEL-172 | chr9-21897158-DEL-157 |
| chr9-21931811-DEL-61 | chr9-21931812-DEL-61 |
| chr9-21932592-DEL-54 | chr9-21932592-DEL-54 |
| chr9-21933255-DEL-138 | chr9-21933238-DEL-132 |
| chr9-21960337-DEL-140 | chr9-21960338-DEL-138 |
| chr9-21979919-DEL-219 | chr9-21979920-DEL-219 |
| chr9-21987018-DEL-239 | chr9-21987019-DEL-239 |
| chr9-22002093-DEL-576 | chr9-22002094-DEL-576 |
| chr9-22026500-DEL-203 | chr9-22026501-DEL-203 |
| chr9-22038637-DEL-131 | chr9-22038638-DEL-131 |
| chr9-22047468-DEL-53 | chr9-22047470-DEL-52 |
| chr9-22052443-DEL-210 | chr9-22052444-DEL-209 |
| chr9-22053175-DEL-208 | chr9-22053176-DEL-208 |
| chr9-22066744-DEL-71 | chr9-22066745-DEL-71 |
| chr9-22070770-DEL-104 | chr9-22070771-DEL-104 |
| chr9-22072198-DEL-4799 | chr9-22072199-DEL-4799 |
| chr9-22088781-DEL-7102 | chr9-22088782-DEL-7102 |
| chr9-22127102-DEL-160 | chr9-22127103-DEL-160 |
| chr9-22131529-DEL-108 | chr9-22131530-DEL-100 |
| chr9-22144335-DEL-594 | chr9-22144336-DEL-594 |
| chr9-22181403-DEL-76 | chr9-22181404-DEL-76 |
| chr9-22182347-DEL-187 | chr9-22182365-DEL-188 |
| chr9-22183991-DEL-189 | chr9-22183992-DEL-189 |
| chr9-22199676-DEL-60 | chr9-22199675-DEL-59 |
| chr9-22213491-DEL-406 | chr9-22213497-DEL-396 |
| chr9-22216913-DEL-192 | chr9-22216914-DEL-192 |
| chr9-22219528-DEL-205 | chr9-22219523-DEL-215 |
| chr9-22237885-DEL-154 | chr9-22237886-DEL-154 |
| chr9-22251979-DEL-724 | chr9-22251980-DEL-724 |
| chr9-22257196-DEL-184 | chr9-22257197-DEL-184 |
| chr9-22284709-DEL-209 | chr9-22284710-DEL-208 |
| chr9-22302293-DEL-234 | chr9-22302286-DEL-235 |
| chr9-22319096-DEL-213 | chr9-22319097-DEL-213 |
| chr9-22326670-DEL-76 | chr9-22326699-DEL-63 |
| chr9-22352723-DEL-70 | chr9-22352724-DEL-70 |
| chr9-22431978-DEL-72 | chr9-22431979-DEL-54 |
| chr9-22435724-DEL-1456 | chr9-22435725-DEL-1459 |
| chr9-22465476-DEL-140 | chr9-22465500-DEL-113 |
| chr9-22472593-DEL-1074 | chr9-22472591-DEL-1075 |
| chr9-22475681-DEL-210 | chr9-22475682-DEL-210 |
| chr9-22477659-DEL-66 | chr9-22477660-DEL-66 |
| chr9-22479627-DEL-66 | chr9-22479622-DEL-67 |
| chr9-22532908-DEL-60 | chr9-22532909-DEL-60 |
| chr9-22537645-DEL-217 | chr9-22537646-DEL-217 |
| chr9-22564611-DEL-210 | chr9-22564612-DEL-210 |
| chr9-22571479-DEL-527 | chr9-22571480-DEL-527 |
| chr9-22602633-DEL-62 | chr9-22602642-DEL-54 |
| chr9-22616366-DEL-100 | chr9-22616385-DEL-94 |
| chr9-22630161-DEL-6440 | chr9-22630162-DEL-6440 |
| chr9-22636677-DEL-6455 | chr9-22636678-DEL-6455 |
| chr9-22712478-DEL-782 | chr9-22712479-DEL-782 |
| chr9-22791436-DEL-522 | chr9-22791429-DEL-530 |
| chr9-22792740-DEL-560 | chr9-22792735-DEL-561 |
| chr9-22801977-DEL-3581 | chr9-22801972-DEL-3582 |
| chr9-22808524-DEL-6962 | chr9-22808521-DEL-6966 |
| chr9-22831307-DEL-6401 | chr9-22831308-DEL-6401 |
| chr9-22853358-DEL-622 | chr9-22853359-DEL-622 |
| chr9-22859866-DEL-126 | chr9-22859867-DEL-126 |
| chr9-22905991-DEL-64 | chr9-22905992-DEL-64 |
| chr9-22931530-DEL-186 | chr9-22931531-DEL-186 |
| chr9-22932111-DEL-215 | chr9-22932112-DEL-215 |
| chr9-22958256-DEL-94 | chr9-22958266-DEL-85 |
| chr9-22996450-DEL-95 | chr9-22996451-DEL-95 |
| chr9-23012677-DEL-76 | chr9-23012675-DEL-68 |
| chr9-23021620-DEL-163 | chr9-23021621-DEL-163 |
| chr9-23030802-DEL-63 | chr9-23030796-DEL-70 |
| chr9-23039527-DEL-68 | chr9-23039528-DEL-68 |
| chr9-23039758-DEL-999 | chr9-23039759-DEL-999 |
| chr9-23073673-DEL-68 | chr9-23073676-DEL-64 |
| chr9-23093588-DEL-101 | chr9-23093589-DEL-97 |
| chr9-23143302-DEL-78 | chr9-23143314-DEL-78 |
| chr9-23160572-DEL-54 | chr9-23160563-DEL-55 |
| chr9-23188993-DEL-6678 | chr9-23188994-DEL-6678 |
| chr9-23232909-DEL-72 | chr9-23232917-DEL-71 |
| chr9-23241611-DEL-391 | chr9-23241612-DEL-391 |
| chr9-23269960-DEL-178 | chr9-23269961-DEL-178 |
| chr9-23273592-DEL-54 | chr9-23273593-DEL-54 |
| chr9-23346195-DEL-206 | chr9-23346196-DEL-206 |

| | |
|---|---|
| chr9-23354661-DEL-1717 | chr9-23354662-DEL-1717 |
| chr9-23357202-DEL-945 | chr9-23357203-DEL-945 |
| chr9-23379098-DEL-256 | chr9-23379099-DEL-256 |
| chr9-23391840-DEL-57 | chr9-23391822-DEL-62 |
| chr9-23419444-DEL-2790 | chr9-23419445-DEL-2790 |
| chr9-23423506-DEL-430 | chr9-23423507-DEL-430 |
| chr9-23426133-DEL-269 | chr9-23426134-DEL-269 |
| chr9-23427505-DEL-445 | chr9-23427506-DEL-445 |
| chr9-23457307-DEL-325 | chr9-23457307-DEL-325 |
| chr9-23459787-DEL-72 | chr9-23459787-DEL-72 |
| chr9-23462400-DEL-56 | chr9-23462400-DEL-56 |
| chr9-23468791-DEL-58 | chr9-23468792-DEL-58 |
| chr9-23515583-DEL-73 | chr9-23515584-DEL-73 |
| chr9-23519555-DEL-4038 | chr9-23519547-DEL-4041 |
| chr9-23541157-DEL-598 | chr9-23541158-DEL-598 |
| chr9-23548252-DEL-1025 | chr9-23548253-DEL-1025 |
| chr9-23552977-DEL-76 | chr9-23552968-DEL-76 |
| chr9-23553637-DEL-1490 | chr9-23553601-DEL-1489 |
| chr9-23570182-DEL-5209 | chr9-23570183-DEL-5209 |
| chr9-23571415-DEL-1996 | chr9-23571416-DEL-1996 |
| chr9-23579510-DEL-54 | chr9-23579511-DEL-54 |
| chr9-23585808-DEL-436 | chr9-23585809-DEL-436 |
| chr9-23625028-DEL-622 | chr9-23625029-DEL-622 |
| chr9-23627910-DEL-5442 | chr9-23627912-DEL-5441 |
| chr9-23632604-DEL-168 | chr9-23632605-DEL-168 |
| chr9-23634655-DEL-447 | chr9-23634656-DEL-447 |
| chr9-23652228-DEL-3915 | chr9-23652229-DEL-3915 |
| chr9-23660599-DEL-96 | chr9-23660635-DEL-52 |
| chr9-23679662-DEL-421 | chr9-23679663-DEL-421 |
| chr9-23720654-DEL-249 | chr9-23720656-DEL-250 |
| chr9-23724933-DEL-52 | chr9-23724934-DEL-52 |
| chr9-23728763-DEL-288 | chr9-23728741-DEL-311 |
| chr9-23758011-DEL-131 | chr9-23758012-DEL-131 |
| chr9-23758458-DEL-893 | chr9-23758459-DEL-893 |
| chr9-23764386-DEL-1173 | chr9-23764387-DEL-1173 |
| chr9-23771268-DEL-487 | chr9-23771265-DEL-488 |
| chr9-23804269-DEL-4926 | chr9-23804270-DEL-4926 |
| chr9-23816987-DEL-54 | chr9-23816970-DEL-68 |
| chr9-23847615-DEL-520 | chr9-23847616-DEL-520 |
| chr9-23858323-DEL-66 | chr9-23858327-DEL-104 |
| chr9-23866883-DEL-51 | chr9-23866865-DEL-51 |
| chr9-23876395-DEL-354 | chr9-23876372-DEL-376 |
| chr9-23882150-DEL-55 | chr9-23882151-DEL-55 |
| chr9-23883030-DEL-53 | chr9-23883031-DEL-53 |
| chr9-23890435-DEL-426 | chr9-23890436-DEL-426 |
| chr9-23892792-DEL-6614 | chr9-23892793-DEL-6614 |
| chr9-23892895-DEL-214 | chr9-23892895-DEL-214 |
| chr9-23913268-DEL-54 | chr9-23913282-DEL-82 |
| chr9-23951709-DEL-6237 | chr9-23951708-DEL-6238 |
| chr9-23961862-DEL-76 | chr9-23961856-DEL-100 |
| chr9-23970930-DEL-55 | chr9-23970931-DEL-57 |
| chr9-23974303-DEL-1114 | chr9-23974304-DEL-1114 |
| chr9-23974476-DEL-84 | chr9-23974477-DEL-84 |
| chr9-23975735-DEL-54 | chr9-23975736-DEL-54 |
| chr9-23980348-DEL-588 | chr9-23980346-DEL-589 |
| chr9-23996639-DEL-368 | chr9-23996640-DEL-368 |
| chr9-24023754-DEL-1689 | chr9-24023755-DEL-1689 |
| chr9-24032750-DEL-502 | chr9-24032751-DEL-502 |
| chr9-24058621-DEL-7703 | chr9-24058622-DEL-7703 |
| chr9-24067487-DEL-80 | chr9-24067489-DEL-93 |
| chr9-24178085-DEL-6761 | chr9-24178086-DEL-6761 |
| chr9-24210811-DEL-327 | chr9-24210812-DEL-327 |
| chr9-24213474-DEL-115 | chr9-24213507-DEL-121 |
| chr9-24216795-DEL-872 | chr9-24216801-DEL-870 |
| chr9-24220573-DEL-123 | chr9-24220574-DEL-123 |
| chr9-24222135-DEL-239 | chr9-24222136-DEL-239 |
| chr9-24225936-DEL-52 | chr9-24225892-DEL-51 |
| chr9-24239630-DEL-214 | chr9-24239631-DEL-214 |
| chr9-24244873-DEL-981 | chr9-24244874-DEL-981 |
| chr9-24265645-DEL-457 | chr9-24265646-DEL-457 |
| chr9-24281635-DEL-388 | chr9-24281636-DEL-388 |
| chr9-24284380-DEL-112 | chr9-24284393-DEL-92 |
| chr9-24287689-DEL-72 | chr9-24287688-DEL-72 |
| chr9-24300671-DEL-1111 | chr9-24300672-DEL-1111 |
| chr9-24387220-DEL-120 | chr9-24387221-DEL-120 |
| chr9-24419298-DEL-284 | chr9-24419299-DEL-284 |
| chr9-24434528-DEL-1469 | chr9-24434529-DEL-1469 |
| chr9-24436849-DEL-6259 | chr9-24436845-DEL-6260 |
| chr9-24453304-DEL-161 | chr9-24453304-DEL-161 |
| chr9-24455487-DEL-1539 | chr9-24455488-DEL-1539 |

| | |
|---|---|
| chr9-24461008-DEL-3324 | chr9-24461009-DEL-3324 |
| chr9-24479204-DEL-348 | chr9-24479205-DEL-348 |
| chr9-24482726-DEL-4492 | chr9-24482727-DEL-4492 |
| chr9-24509741-DEL-1014 | chr9-24509741-DEL-1015 |
| chr9-24556418-DEL-162 | chr9-24556440-DEL-162 |
| chr9-24564688-DEL-2386 | chr9-24564689-DEL-2386 |
| chr9-24613661-DEL-2905 | chr9-24613662-DEL-2905 |
| chr9-24620735-DEL-416 | chr9-24620736-DEL-416 |
| chr9-24640545-DEL-81 | chr9-24640556-DEL-82 |
| chr9-24644447-DEL-52 | chr9-24644448-DEL-52 |
| chr9-24654107-DEL-135 | chr9-24654108-DEL-133 |
| chr9-24657311-DEL-462 | chr9-24657312-DEL-462 |
| chr9-24671601-DEL-289 | chr9-24671602-DEL-289 |
| chr9-24673239-DEL-55 | chr9-24673225-DEL-76 |
| chr9-24733419-DEL-205 | chr9-24733420-DEL-205 |
| chr9-24739356-DEL-191 | chr9-24739357-DEL-191 |
| chr9-24751223-DEL-122 | chr9-24751224-DEL-122 |
| chr9-24777166-DEL-238 | chr9-24777167-DEL-238 |
| chr9-24786830-DEL-610 | chr9-24786831-DEL-610 |
| chr9-24814088-DEL-116 | chr9-24814134-DEL-93 |
| chr9-24840145-DEL-1952 | chr9-24840146-DEL-1952 |
| chr9-24843493-DEL-51 | chr9-24843494-DEL-51 |
| chr9-24853178-DEL-708 | chr9-24853179-DEL-708 |
| chr9-24928408-DEL-310 | chr9-24928409-DEL-310 |
| chr9-24946445-DEL-218 | chr9-24946446-DEL-218 |
| chr9-24955964-DEL-195 | chr9-24955964-DEL-195 |
| chr9-24958188-DEL-1319 | chr9-24958209-DEL-1319 |
| chr9-24965466-DEL-397 | chr9-24965467-DEL-398 |
| chr9-24977890-DEL-188 | chr9-24977891-DEL-188 |
| chr9-24988518-DEL-1264 | chr9-24988519-DEL-1264 |
| chr9-25076312-DEL-2548 | chr9-25076303-DEL-2549 |
| chr9-25092147-DEL-844 | chr9-25092148-DEL-844 |
| chr9-25144783-DEL-6404 | chr9-25144784-DEL-6404 |
| chr9-25152881-DEL-77 | chr9-25152882-DEL-77 |
| chr9-25208558-DEL-145 | chr9-25208546-DEL-148 |
| chr9-25215338-DEL-86 | chr9-25215339-DEL-86 |
| chr9-25228980-DEL-83 | chr9-25228985-DEL-79 |
| chr9-25229692-DEL-2230 | chr9-25229688-DEL-2231 |
| chr9-25247043-DEL-291 | chr9-25247044-DEL-291 |
| chr9-25266251-DEL-207 | chr9-25266250-DEL-183 |
| chr9-25267899-DEL-74 | chr9-25267900-DEL-74 |
| chr9-25273737-DEL-109 | chr9-25273739-DEL-106 |
| chr9-25275867-DEL-181 | chr9-25275861-DEL-165 |
| chr9-25278031-DEL-52 | chr9-25278032-DEL-52 |
| chr9-25312310-DEL-60 | chr9-25312311-DEL-60 |
| chr9-25319075-DEL-389 | chr9-25319076-DEL-389 |
| chr9-25329420-DEL-1102 | chr9-25329421-DEL-1102 |
| chr9-25331397-DEL-6401 | chr9-25331398-DEL-6401 |
| chr9-25377117-DEL-655 | chr9-25377117-DEL-655 |
| chr9-25387563-DEL-291 | chr9-25387564-DEL-291 |
| chr9-25388931-DEL-96 | chr9-25388932-DEL-96 |
| chr9-25409047-DEL-688 | chr9-25409036-DEL-664 |
| chr9-25458995-DEL-58 | chr9-25458996-DEL-58 |
| chr9-25499488-DEL-211 | chr9-25499489-DEL-211 |
| chr9-25507708-DEL-6353 | chr9-25507709-DEL-6353 |
| chr9-25574760-DEL-223 | chr9-25574761-DEL-223 |
| chr9-25577269-DEL-51 | chr9-25577274-DEL-53 |
| chr9-25603153-DEL-99 | chr9-25603150-DEL-103 |
| chr9-25624431-DEL-384 | chr9-25624432-DEL-384 |
| chr9-25663483-DEL-6673 | chr9-25663484-DEL-6673 |
| chr9-25669753-DEL-214 | chr9-25669754-DEL-214 |
| chr9-25671828-DEL-1411 | chr9-25671818-DEL-1412 |
| chr9-25677182-DEL-110 | chr9-25677187-DEL-111 |
| chr9-25683756-DEL-116 | chr9-25683757-DEL-116 |
| chr9-25687305-DEL-4603 | chr9-25687306-DEL-4603 |
| chr9-25693399-DEL-166 | chr9-25693379-DEL-160 |
| chr9-25719871-DEL-173 | chr9-25719872-DEL-173 |
| chr9-25720147-DEL-1490 | chr9-25720148-DEL-1489 |
| chr9-25766521-DEL-8419 | chr9-25766522-DEL-8419 |
| chr9-25778251-DEL-220 | chr9-25778252-DEL-220 |
| chr9-25782940-DEL-1204 | chr9-25782941-DEL-1204 |
| chr9-25791220-DEL-1533 | chr9-25791221-DEL-1533 |
| chr9-25793258-DEL-1566 | chr9-25793259-DEL-1563 |
| chr9-25797804-DEL-218 | chr9-25797805-DEL-218 |
| chr9-25800489-DEL-204 | chr9-25800490-DEL-204 |
| chr9-25811648-DEL-57 | chr9-25811645-DEL-91 |
| chr9-25831658-DEL-52 | chr9-25831617-DEL-94 |
| chr9-25857500-DEL-66 | chr9-25857459-DEL-80 |
| chr9-25863432-DEL-58 | chr9-25863431-DEL-59 |
| chr9-25877773-DEL-3590 | chr9-25877774-DEL-3590 |

| | |
|---|---|
| chr9-25881425-DEL-2544 | chr9-25881426-DEL-2544 |
| chr9-25902640-DEL-4835 | chr9-25902641-DEL-4836 |
| chr9-25923999-DEL-402 | chr9-25924000-DEL-402 |
| chr9-25981142-DEL-1421 | chr9-25981143-DEL-1421 |
| chr9-25998310-DEL-76 | chr9-25998331-DEL-58 |
| chr9-26007532-DEL-7129 | chr9-26007533-DEL-7129 |
| chr9-26028965-DEL-190 | chr9-26028966-DEL-190 |
| chr9-26032688-DEL-324 | chr9-26032685-DEL-325 |
| chr9-26040428-DEL-67 | chr9-26040398-DEL-66 |
| chr9-26052123-DEL-201 | chr9-26052124-DEL-201 |
| chr9-26058666-DEL-60 | chr9-26058667-DEL-60 |
| chr9-26066439-DEL-692 | chr9-26066441-DEL-691 |
| chr9-26067470-DEL-3326 | chr9-26067471-DEL-3326 |
| chr9-26068515-DEL-7665 | chr9-26068515-DEL-7665 |
| chr9-26115135-DEL-3610 | chr9-26115136-DEL-3610 |
| chr9-26147701-DEL-440 | chr9-26147700-DEL-441 |
| chr9-26168613-DEL-58 | chr9-26168614-DEL-58 |
| chr9-26205791-DEL-122 | chr9-26205792-DEL-122 |
| chr9-26303117-DEL-790 | chr9-26303117-DEL-790 |
| chr9-26311774-DEL-7936 | chr9-26311775-DEL-7936 |
| chr9-26318705-DEL-847 | chr9-26318702-DEL-848 |
| chr9-26322735-DEL-88 | chr9-26322755-DEL-63 |
| chr9-26347448-DEL-197 | chr9-26347438-DEL-193 |
| chr9-26358197-DEL-134 | chr9-26358196-DEL-135 |
| chr9-26416178-DEL-742 | chr9-26416177-DEL-743 |
| chr9-26427539-DEL-51 | chr9-26427528-DEL-54 |
| chr9-26470047-DEL-1893 | chr9-26470048-DEL-1893 |
| chr9-26474634-DEL-411 | chr9-26474635-DEL-411 |
| chr9-26478403-DEL-1449 | chr9-26478404-DEL-1450 |
| chr9-26504022-DEL-2118 | chr9-26504023-DEL-2119 |
| chr9-26521873-DEL-61 | chr9-26521875-DEL-59 |
| chr9-26579637-DEL-190 | chr9-26579638-DEL-190 |
| chr9-26591371-DEL-95 | chr9-26591372-DEL-95 |
| chr9-26612178-DEL-7107 | chr9-26612179-DEL-7107 |
| chr9-26680736-DEL-51 | chr9-26680737-DEL-51 |
| chr9-26684451-DEL-106 | chr9-26684452-DEL-106 |
| chr9-26700296-DEL-59 | chr9-26700314-DEL-51 |
| chr9-26763324-DEL-217 | chr9-26763325-DEL-217 |
| chr9-26797420-DEL-868 | chr9-26797421-DEL-868 |
| chr9-26822787-DEL-71 | chr9-26822788-DEL-71 |
| chr9-26859595-DEL-82 | chr9-26859597-DEL-81 |
| chr9-26876491-DEL-224 | chr9-26876492-DEL-224 |
| chr9-26882014-DEL-1118 | chr9-26882015-DEL-1118 |
| chr9-26931953-DEL-51 | chr9-26931926-DEL-58 |
| chr9-26936699-DEL-108 | chr9-26936700-DEL-108 |
| chr9-26964038-DEL-2592 | chr9-26964039-DEL-2592 |
| chr9-26971390-DEL-210 | chr9-26971391-DEL-210 |
| chr9-26974429-DEL-100 | chr9-26974430-DEL-100 |
| chr9-26977409-DEL-76 | chr9-26977409-DEL-60 |
| chr9-27030275-DEL-184 | chr9-27030260-DEL-174 |
| chr9-27056172-DEL-129 | chr9-27056173-DEL-129 |
| chr9-27056657-DEL-204 | chr9-27056658-DEL-204 |
| chr9-27146844-DEL-70 | chr9-27146845-DEL-60 |
| chr9-27148403-DEL-406 | chr9-27148404-DEL-406 |
| chr9-27176754-DEL-60 | chr9-27176755-DEL-60 |
| chr9-27190219-DEL-110 | chr9-27190220-DEL-110 |
| chr9-27191843-DEL-74 | chr9-27191839-DEL-81 |
| chr9-27236805-DEL-62 | chr9-27236774-DEL-96 |
| chr9-27288709-DEL-600 | chr9-27288710-DEL-600 |
| chr9-27319424-DEL-423 | chr9-27319424-DEL-421 |
| chr9-27320824-DEL-63 | chr9-27320849-DEL-51 |
| chr9-27326967-DEL-80 | chr9-27326967-DEL-81 |
| chr9-27365986-DEL-189 | chr9-27365987-DEL-189 |
| chr9-27376244-DEL-6209 | chr9-27376276-DEL-6190 |
| chr9-27385673-DEL-116 | chr9-27385675-DEL-115 |
| chr9-27389760-DEL-118 | chr9-27389761-DEL-118 |
| chr9-27465975-DEL-93 | chr9-27465997-DEL-60 |
| chr9-27511876-DEL-76 | chr9-27511877-DEL-74 |
| chr9-27513502-DEL-63 | chr9-27513499-DEL-63 |
| chr9-27524176-DEL-974 | chr9-27524177-DEL-974 |
| chr9-27551154-DEL-143 | chr9-27551156-DEL-142 |
| chr9-27571523-DEL-62 | chr9-27571524-DEL-60 |
| chr9-27577251-DEL-80 | chr9-27577250-DEL-81 |
| chr9-27646080-DEL-72 | chr9-27646081-DEL-72 |
| chr9-27656672-DEL-78 | chr9-27656673-DEL-78 |
| chr9-27691169-DEL-60 | chr9-27691170-DEL-60 |
| chr9-27708630-DEL-1543 | chr9-27708631-DEL-1543 |
| chr9-27727412-DEL-55 | chr9-27727413-DEL-55 |
| chr9-27859299-DEL-794 | chr9-27859300-DEL-794 |
| chr9-27866552-DEL-67 | chr9-27866554-DEL-63 |

| | |
|---|---|
| chr9-27881226-DEL-56 | chr9-27881227-DEL-56 |
| chr9-27885779-DEL-7125 | chr9-27885780-DEL-7125 |
| chr9-27906428-DEL-58 | chr9-27906429-DEL-58 |
| chr9-27939292-DEL-63 | chr9-27939289-DEL-67 |
| chr9-27958685-DEL-186 | chr9-27958686-DEL-185 |
| chr9-27991881-DEL-177 | chr9-27991879-DEL-178 |
| chr9-28028326-DEL-59 | chr9-28028329-DEL-57 |
| chr9-28036925-DEL-66 | chr9-28036916-DEL-64 |
| chr9-28171514-DEL-211 | chr9-28171501-DEL-212 |
| chr9-28182257-DEL-661 | chr9-28182258-DEL-661 |
| chr9-28290704-DEL-86 | chr9-28290705-DEL-86 |
| chr9-28356879-DEL-142 | chr9-28356916-DEL-120 |
| chr9-28374252-DEL-181 | chr9-28374253-DEL-181 |
| chr9-28381289-DEL-66 | chr9-28381290-DEL-66 |
| chr9-28408789-DEL-58 | chr9-28408790-DEL-58 |
| chr9-28409539-DEL-58 | chr9-28409540-DEL-58 |
| chr9-28422919-DEL-138 | chr9-28422919-DEL-134 |
| chr9-28432932-DEL-909 | chr9-28432932-DEL-909 |
| chr9-28436452-DEL-1240 | chr9-28436453-DEL-1240 |
| chr9-28470466-DEL-62 | chr9-28470467-DEL-60 |
| chr9-28491577-DEL-76 | chr9-28491576-DEL-78 |
| chr9-28499646-DEL-54 | chr9-28499646-DEL-54 |
| chr9-28515109-DEL-84 | chr9-28515134-DEL-52 |
| chr9-28551441-DEL-131 | chr9-28551442-DEL-127 |
| chr9-28709525-DEL-54 | chr9-28709526-DEL-54 |
| chr9-28714625-DEL-106 | chr9-28714626-DEL-105 |
| chr9-28739030-DEL-66 | chr9-28739023-DEL-78 |
| chr9-28748855-DEL-70 | chr9-28748856-DEL-60 |
| chr9-28751227-DEL-3169 | chr9-28751228-DEL-3169 |
| chr9-28809983-DEL-74 | chr9-28809977-DEL-75 |
| chr9-28825279-DEL-57 | chr9-28825238-DEL-93 |
| chr9-28826898-DEL-72 | chr9-28826906-DEL-70 |
| chr9-28855481-DEL-2281 | chr9-28855482-DEL-2281 |
| chr9-28907225-DEL-65 | chr9-28907226-DEL-65 |
| chr9-28912819-DEL-59 | chr9-28912820-DEL-59 |
| chr9-28950033-DEL-60 | chr9-28950034-DEL-60 |
| chr9-29027091-DEL-157 | chr9-29027092-DEL-157 |
| chr9-29030447-DEL-3738 | chr9-29030448-DEL-3738 |
| chr9-29131381-DEL-54 | chr9-29131382-DEL-54 |
| chr9-29137912-DEL-69 | chr9-29137913-DEL-69 |
| chr9-29139966-DEL-70 | chr9-29139967-DEL-70 |
| chr9-29149217-DEL-75 | chr9-29149207-DEL-73 |
| chr9-29170139-DEL-54 | chr9-29170140-DEL-54 |
| chr9-29188662-DEL-2013 | chr9-29188663-DEL-2013 |
| chr9-29234743-DEL-63 | chr9-29234744-DEL-56 |
| chr9-29241132-DEL-419 | chr9-29241133-DEL-419 |
| chr9-29262913-DEL-92 | chr9-29262903-DEL-74 |
| chr9-29268493-DEL-55 | chr9-29268494-DEL-55 |
| chr9-29294201-DEL-57 | chr9-29294202-DEL-57 |
| chr9-29316127-DEL-288 | chr9-29316128-DEL-288 |
| chr9-29316740-DEL-52 | chr9-29316741-DEL-52 |
| chr9-29338636-DEL-60 | chr9-29338637-DEL-54 |
| chr9-29387129-DEL-93 | chr9-29387130-DEL-91 |
| chr9-29457931-DEL-127 | chr9-29457932-DEL-127 |
| chr9-29516240-DEL-229 | chr9-29516244-DEL-226 |
| chr9-29613118-DEL-76 | chr9-29613119-DEL-76 |
| chr9-29619030-DEL-136 | chr9-29619047-DEL-112 |
| chr9-29648251-DEL-340 | chr9-29648251-DEL-340 |
| chr9-29651122-DEL-1863 | chr9-29651123-DEL-1863 |
| chr9-29655982-DEL-401 | chr9-29655983-DEL-401 |
| chr9-29659120-DEL-220 | chr9-29659121-DEL-220 |
| chr9-29741859-DEL-51 | chr9-29741861-DEL-63 |
| chr9-29756359-DEL-66 | chr9-29756357-DEL-52 |
| chr9-29788142-DEL-65 | chr9-29788142-DEL-63 |
| chr9-29795026-DEL-76 | chr9-29795027-DEL-76 |
| chr9-29801856-DEL-443 | chr9-29801857-DEL-443 |
| chr9-29850497-DEL-60 | chr9-29850508-DEL-50 |
| chr9-29851928-DEL-67 | chr9-29851929-DEL-67 |
| chr9-29877019-DEL-51 | chr9-29877016-DEL-52 |
| chr9-29881382-DEL-1417 | chr9-29881383-DEL-1417 |
| chr9-29898935-DEL-56 | chr9-29898936-DEL-56 |
| chr9-29907060-DEL-263 | chr9-29907061-DEL-263 |
| chr9-29914867-DEL-68 | chr9-29914864-DEL-72 |
| chr9-29939344-DEL-1708 | chr9-29939345-DEL-1708 |
| chr9-29967104-DEL-401 | chr9-29967105-DEL-401 |
| chr9-30021637-DEL-51 | chr9-30021672-DEL-51 |
| chr9-30059595-DEL-439 | chr9-30059596-DEL-439 |
| chr9-30107469-DEL-78 | chr9-30107461-DEL-68 |
| chr9-30137987-DEL-205 | chr9-30137988-DEL-205 |
| chr9-30233519-DEL-224 | chr9-30233520-DEL-224 |

| | |
|---|---|
| chr9-30240431-DEL-63 | chr9-30240432-DEL-63 |
| chr9-30248212-DEL-137 | chr9-30248221-DEL-129 |
| chr9-30285967-DEL-79 | chr9-30285968-DEL-80 |
| chr9-30319895-DEL-2904 | chr9-30319896-DEL-2904 |
| chr9-30329984-DEL-116 | chr9-30329987-DEL-140 |
| chr9-30342961-DEL-332 | chr9-30342962-DEL-332 |
| chr9-30353238-DEL-73 | chr9-30353239-DEL-73 |
| chr9-30394730-DEL-218 | chr9-30394731-DEL-218 |
| chr9-30426576-DEL-71 | chr9-30426589-DEL-59 |
| chr9-30480850-DEL-250 | chr9-30480851-DEL-249 |
| chr9-30485877-DEL-719 | chr9-30485878-DEL-719 |
| chr9-30495217-DEL-1206 | chr9-30495218-DEL-1206 |
| chr9-30503320-DEL-68 | chr9-30503321-DEL-56 |
| chr9-30509995-DEL-317 | chr9-30509996-DEL-317 |
| chr9-30515747-DEL-1894 | chr9-30515748-DEL-1894 |
| chr9-30550041-DEL-676 | chr9-30550040-DEL-676 |
| chr9-30553111-DEL-7134 | chr9-30553112-DEL-7134 |
| chr9-30573480-DEL-382 | chr9-30573481-DEL-379 |
| chr9-30574112-DEL-2532 | chr9-30574113-DEL-2532 |
| chr9-30583839-DEL-53 | chr9-30583838-DEL-64 |
| chr9-30593903-DEL-1171 | chr9-30593904-DEL-1171 |
| chr9-30601697-DEL-6295 | chr9-30601698-DEL-6295 |
| chr9-30608027-DEL-561 | chr9-30608027-DEL-561 |
| chr9-30664587-DEL-62 | chr9-30664584-DEL-60 |
| chr9-30701025-DEL-238 | chr9-30701027-DEL-237 |
| chr9-30736443-DEL-309 | chr9-30736484-DEL-261 |
| chr9-30741987-DEL-139 | chr9-30741983-DEL-109 |
| chr9-30744268-DEL-57 | chr9-30744269-DEL-57 |
| chr9-30749921-DEL-425 | chr9-30749922-DEL-425 |
| chr9-30752090-DEL-104 | chr9-30752088-DEL-105 |
| chr9-30771203-DEL-108 | chr9-30771212-DEL-99 |
| chr9-30777283-DEL-6603 | chr9-30777284-DEL-6603 |
| chr9-30785080-DEL-726 | chr9-30785081-DEL-726 |
| chr9-30801749-DEL-127 | chr9-30801757-DEL-123 |
| chr9-30841131-DEL-1171 | chr9-30841132-DEL-1171 |
| chr9-30849398-DEL-166 | chr9-30849366-DEL-206 |
| chr9-30871570-DEL-75 | chr9-30871571-DEL-66 |
| chr9-30877635-DEL-99 | chr9-30877621-DEL-114 |
| chr9-30886816-DEL-58 | chr9-30886789-DEL-54 |
| chr9-30890914-DEL-421 | chr9-30890962-DEL-372 |
| chr9-30896270-DEL-128 | chr9-30896271-DEL-128 |
| chr9-30915464-DEL-59 | chr9-30915465-DEL-59 |
| chr9-30919564-DEL-72 | chr9-30919583-DEL-54 |
| chr9-30962753-DEL-197 | chr9-30962754-DEL-197 |
| chr9-30977287-DEL-84 | chr9-30977288-DEL-84 |
| chr9-31005325-DEL-62 | chr9-31005326-DEL-62 |
| chr9-31007140-DEL-147 | chr9-31007141-DEL-98 |
| chr9-31046640-DEL-1449 | chr9-31046640-DEL-1449 |
| chr9-31064162-DEL-72 | chr9-31064163-DEL-70 |
| chr9-31066241-DEL-96 | chr9-31066242-DEL-96 |
| chr9-31090670-DEL-213 | chr9-31090671-DEL-213 |
| chr9-31097797-DEL-1230 | chr9-31097798-DEL-1230 |
| chr9-31100017-DEL-117 | chr9-31100018-DEL-117 |
| chr9-31105310-DEL-135 | chr9-31105311-DEL-135 |
| chr9-31108890-DEL-178 | chr9-31108891-DEL-178 |
| chr9-31139318-DEL-185 | chr9-31139319-DEL-185 |
| chr9-31154370-DEL-52 | chr9-31154371-DEL-52 |
| chr9-31157306-DEL-189 | chr9-31157316-DEL-190 |
| chr9-31175611-DEL-85 | chr9-31175615-DEL-82 |
| chr9-31195127-DEL-174 | chr9-31195128-DEL-174 |
| chr9-31196630-DEL-1074 | chr9-31196631-DEL-1074 |
| chr9-31202753-DEL-248 | chr9-31202754-DEL-248 |
| chr9-31229271-DEL-84 | chr9-31229240-DEL-67 |
| chr9-31251890-DEL-187 | chr9-31251891-DEL-187 |
| chr9-31280108-DEL-407 | chr9-31280109-DEL-406 |
| chr9-31311043-DEL-62 | chr9-31311043-DEL-62 |
| chr9-31329254-DEL-207 | chr9-31329255-DEL-207 |
| chr9-31341828-DEL-531 | chr9-31341829-DEL-531 |
| chr9-31349535-DEL-143 | chr9-31349560-DEL-138 |
| chr9-31364887-DEL-383 | chr9-31364888-DEL-383 |
| chr9-31365426-DEL-169 | chr9-31365427-DEL-169 |
| chr9-31385233-DEL-81 | chr9-31385234-DEL-81 |
| chr9-31393222-DEL-329 | chr9-31393223-DEL-329 |
| chr9-31401576-DEL-92 | chr9-31401577-DEL-92 |
| chr9-31404671-DEL-204 | chr9-31404672-DEL-204 |
| chr9-31416651-DEL-69 | chr9-31416652-DEL-69 |
| chr9-31443775-DEL-70 | chr9-31443776-DEL-70 |
| chr9-31444429-DEL-3470 | chr9-31444430-DEL-3470 |
| chr9-31456494-DEL-73 | chr9-31456495-DEL-59 |
| chr9-31467605-DEL-88 | chr9-31467610-DEL-78 |

| | |
|---|---|
| chr9-31472933-DEL-91 | chr9-31472934-DEL-91 |
| chr9-31479551-DEL-52 | chr9-31479552-DEL-52 |
| chr9-31485913-DEL-65 | chr9-31485903-DEL-97 |
| chr9-31532924-DEL-353 | chr9-31532930-DEL-352 |
| chr9-31544840-DEL-131 | chr9-31544844-DEL-127 |
| chr9-31585242-DEL-76 | chr9-31585243-DEL-76 |
| chr9-31595343-DEL-119 | chr9-31595344-DEL-119 |
| chr9-31605152-DEL-65 | chr9-31605153-DEL-65 |
| chr9-31627670-DEL-68 | chr9-31627671-DEL-68 |
| chr9-31646235-DEL-6197 | chr9-31646237-DEL-6211 |
| chr9-31678683-DEL-333 | chr9-31678684-DEL-333 |
| chr9-31690350-DEL-1261 | chr9-31690351-DEL-1261 |
| chr9-31712311-DEL-70 | chr9-31712312-DEL-70 |
| chr9-31716770-DEL-116 | chr9-31716766-DEL-121 |
| chr9-31724276-DEL-722 | chr9-31724276-DEL-721 |
| chr9-31737231-DEL-59 | chr9-31737195-DEL-59 |
| chr9-31803202-DEL-186 | chr9-31803203-DEL-186 |
| chr9-31839205-DEL-454 | chr9-31839206-DEL-454 |
| chr9-31863799-DEL-74 | chr9-31863800-DEL-74 |
| chr9-31865172-DEL-194 | chr9-31865173-DEL-194 |
| chr9-31868290-DEL-180 | chr9-31868291-DEL-180 |
| chr9-31911024-DEL-100 | chr9-31911068-DEL-71 |
| chr9-31985681-DEL-320 | chr9-31985682-DEL-320 |
| chr9-31986186-DEL-115 | chr9-31986187-DEL-115 |
| chr9-31987339-DEL-270 | chr9-31987340-DEL-270 |
| chr9-31990272-DEL-4492 | chr9-31990273-DEL-4492 |
| chr9-32009965-DEL-215 | chr9-32009963-DEL-216 |
| chr9-32013595-DEL-116 | chr9-32013596-DEL-116 |
| chr9-32019413-DEL-112 | chr9-32019406-DEL-110 |
| chr9-32106328-DEL-199 | chr9-32106329-DEL-199 |
| chr9-32110626-DEL-210 | chr9-32110627-DEL-210 |
| chr9-32113004-DEL-636 | chr9-32113005-DEL-636 |
| chr9-32113822-DEL-76 | chr9-32113823-DEL-76 |
| chr9-32124336-DEL-216 | chr9-32124337-DEL-216 |
| chr9-32129681-DEL-52 | chr9-32129682-DEL-52 |
| chr9-32155023-DEL-236 | chr9-32155062-DEL-235 |
| chr9-32161620-DEL-269 | chr9-32161615-DEL-262 |
| chr9-32189178-DEL-92 | chr9-32189211-DEL-85 |
| chr9-32198045-DEL-280 | chr9-32198057-DEL-278 |
| chr9-32214964-DEL-659 | chr9-32214965-DEL-659 |
| chr9-32218901-DEL-645 | chr9-32218883-DEL-645 |
| chr9-32248583-DEL-138 | chr9-32248584-DEL-138 |
| chr9-32261307-DEL-52 | chr9-32261309-DEL-52 |
| chr9-32265594-DEL-2207 | chr9-32265594-DEL-2207 |
| chr9-32322277-DEL-65 | chr9-32322285-DEL-55 |
| chr9-32353840-DEL-80 | chr9-32353845-DEL-72 |
| chr9-32373781-DEL-56 | chr9-32373784-DEL-54 |
| chr9-32475025-DEL-108 | chr9-32475000-DEL-144 |
| chr9-32495562-DEL-1199 | chr9-32495563-DEL-1199 |
| chr9-32497590-DEL-127 | chr9-32497591-DEL-127 |
| chr9-32546747-DEL-161 | chr9-32546748-DEL-161 |
| chr9-32550818-DEL-57 | chr9-32550817-DEL-64 |
| chr9-32576845-DEL-54 | chr9-32576842-DEL-58 |
| chr9-32585029-DEL-71 | chr9-32585012-DEL-72 |
| chr9-32683283-DEL-324 | chr9-32683284-DEL-324 |
| chr9-32695146-DEL-110 | chr9-32695146-DEL-110 |
| chr9-32702141-DEL-59 | chr9-32702142-DEL-59 |
| chr9-32739781-DEL-59 | chr9-32739782-DEL-59 |
| chr9-32750599-DEL-7206 | chr9-32750600-DEL-7206 |
| chr9-32800014-DEL-7180 | chr9-32799996-DEL-7185 |
| chr9-32816888-DEL-68 | chr9-32816889-DEL-68 |
| chr9-32831477-DEL-86 | chr9-32831484-DEL-80 |
| chr9-32832067-DEL-72 | chr9-32832071-DEL-70 |
| chr9-32840601-DEL-125 | chr9-32840602-DEL-125 |
| chr9-32849257-DEL-51 | chr9-32849250-DEL-52 |
| chr9-32862772-DEL-203 | chr9-32862773-DEL-203 |
| chr9-32864380-DEL-4898 | chr9-32864381-DEL-4898 |
| chr9-32872150-DEL-283 | chr9-32872150-DEL-282 |
| chr9-32874585-DEL-223 | chr9-32874586-DEL-223 |
| chr9-32903503-DEL-6794 | chr9-32903504-DEL-6794 |
| chr9-32919595-DEL-1482 | chr9-32919596-DEL-1482 |
| chr9-32921359-DEL-109 | chr9-32921357-DEL-109 |
| chr9-32926094-DEL-787 | chr9-32926095-DEL-787 |
| chr9-32935717-DEL-116 | chr9-32935718-DEL-116 |
| chr9-32951893-DEL-7839 | chr9-32951894-DEL-7839 |
| chr9-32952007-DEL-1248 | chr9-32952008-DEL-1248 |
| chr9-32998015-DEL-56 | chr9-32998021-DEL-50 |
| chr9-33006616-DEL-98 | chr9-33006617-DEL-98 |
| chr9-33007616-DEL-76 | chr9-33007614-DEL-77 |
| chr9-33020608-DEL-421 | chr9-33020609-DEL-421 |

| | |
|---|---|
| chr9-33039428-DEL-2260 | chr9-33039429-DEL-2260 |
| chr9-33065089-DEL-7126 | chr9-33065090-DEL-7126 |
| chr9-33128888-DEL-2588 | chr9-33128889-DEL-2588 |
| chr9-33142204-DEL-132 | chr9-33142205-DEL-132 |
| chr9-33160395-DEL-378 | chr9-33160396-DEL-378 |
| chr9-33161024-DEL-53 | chr9-33161025-DEL-53 |
| chr9-33195484-DEL-178 | chr9-33195485-DEL-177 |
| chr9-33201342-DEL-72 | chr9-33201336-DEL-82 |
| chr9-33207833-DEL-594 | chr9-33207834-DEL-594 |
| chr9-33209287-DEL-1247 | chr9-33209288-DEL-1247 |
| chr9-33222561-DEL-254 | chr9-33222562-DEL-254 |
| chr9-33266525-DEL-66 | chr9-33266525-DEL-67 |
| chr9-33282091-DEL-55 | chr9-33282092-DEL-55 |
| chr9-33339711-DEL-55 | chr9-33339712-DEL-53 |
| chr9-33357137-DEL-65 | chr9-33357147-DEL-52 |
| chr9-33375814-DEL-62 | chr9-33375823-DEL-54 |
| chr9-33387213-DEL-52 | chr9-33387200-DEL-66 |
| chr9-33436576-DEL-51 | chr9-33436606-DEL-51 |
| chr9-33483630-DEL-521 | chr9-33483631-DEL-519 |
| chr9-33501191-DEL-52 | chr9-33501194-DEL-50 |
| chr9-33551000-DEL-495 | chr9-33551001-DEL-495 |
| chr9-33583831-DEL-94 | chr9-33583854-DEL-62 |
| chr9-33600141-DEL-2826 | chr9-33600142-DEL-2826 |
| chr9-33605444-DEL-68 | chr9-33605443-DEL-70 |
| chr9-33637043-DEL-1928 | chr9-33637056-DEL-1926 |
| chr9-33639438-DEL-2030 | chr9-33639439-DEL-2030 |
| chr9-33676407-DEL-731 | chr9-33676408-DEL-731 |
| chr9-33715411-DEL-214 | chr9-33715412-DEL-214 |
| chr9-33729551-DEL-64 | chr9-33729552-DEL-64 |
| chr9-33740173-DEL-1066 | chr9-33740174-DEL-1066 |
| chr9-33797609-DEL-96 | chr9-33797611-DEL-76 |
| chr9-33818665-DEL-6793 | chr9-33818666-DEL-6793 |
| chr9-33853341-DEL-362 | chr9-33853342-DEL-362 |
| chr9-33859389-DEL-326 | chr9-33859390-DEL-326 |
| chr9-33860848-DEL-1886 | chr9-33860849-DEL-1886 |
| chr9-33866993-DEL-6201 | chr9-33866994-DEL-6201 |
| chr9-33904433-DEL-1891 | chr9-33904434-DEL-1891 |
| chr9-33907792-DEL-281 | chr9-33907791-DEL-321 |
| chr9-33921869-DEL-469 | chr9-33921870-DEL-469 |
| chr9-33940054-DEL-102 | chr9-33940055-DEL-102 |
| chr9-33942021-DEL-62 | chr9-33942022-DEL-62 |
| chr9-33947058-DEL-2516 | chr9-33947059-DEL-2516 |
| chr9-33950170-DEL-125 | chr9-33950205-DEL-91 |
| chr9-33982305-DEL-453 | chr9-33982306-DEL-453 |
| chr9-33987071-DEL-65 | chr9-33987082-DEL-62 |
| chr9-33990541-DEL-1791 | chr9-33990542-DEL-1791 |
| chr9-34005381-DEL-130 | chr9-34005382-DEL-130 |
| chr9-34016929-DEL-80 | chr9-34016930-DEL-72 |
| chr9-34019850-DEL-81 | chr9-34019851-DEL-81 |
| chr9-34050324-DEL-85 | chr9-34050325-DEL-85 |
| chr9-34065598-DEL-106 | chr9-34065599-DEL-106 |
| chr9-34073374-DEL-154 | chr9-34073374-DEL-154 |
| chr9-34083428-DEL-91 | chr9-34083429-DEL-91 |
| chr9-34106647-DEL-60 | chr9-34106649-DEL-54 |
| chr9-34107685-DEL-252 | chr9-34107688-DEL-249 |
| chr9-34118004-DEL-292 | chr9-34118005-DEL-292 |
| chr9-34120958-DEL-983 | chr9-34120957-DEL-985 |
| chr9-34140081-DEL-52 | chr9-34140082-DEL-52 |
| chr9-34186130-DEL-613 | chr9-34186131-DEL-612 |
| chr9-34215949-DEL-68 | chr9-34215950-DEL-68 |
| chr9-34227260-DEL-68 | chr9-34227261-DEL-65 |
| chr9-34399003-DEL-107 | chr9-34399004-DEL-107 |
| chr9-34439048-DEL-1893 | chr9-34439049-DEL-1893 |
| chr9-34468203-DEL-81 | chr9-34468204-DEL-78 |
| chr9-34507940-DEL-56 | chr9-34507941-DEL-56 |
| chr9-34513253-DEL-72 | chr9-34513266-DEL-68 |
| chr9-34556579-DEL-62 | chr9-34556580-DEL-62 |
| chr9-34562336-DEL-415 | chr9-34562337-DEL-415 |
| chr9-34565121-DEL-62 | chr9-34565122-DEL-62 |
| chr9-34711495-DEL-75 | chr9-34711496-DEL-75 |
| chr9-34718870-DEL-831 | chr9-34718871-DEL-831 |
| chr9-34719851-DEL-173 | chr9-34719851-DEL-177 |
| chr9-34774598-DEL-90 | chr9-34774599-DEL-90 |
| chr9-34791739-DEL-60 | chr9-34791738-DEL-64 |
| chr9-34815320-DEL-84 | chr9-34815320-DEL-91 |
| chr9-34818584-DEL-61 | chr9-34818585-DEL-61 |
| chr9-34843786-DEL-2018 | chr9-34843787-DEL-2018 |
| chr9-34846341-DEL-54 | chr9-34846342-DEL-54 |
| chr9-34979142-DEL-167 | chr9-34979141-DEL-167 |
| chr9-35044852-DEL-205 | chr9-35044853-DEL-205 |

| | |
|---|---|
| chr9-35112438-DEL-258 | chr9-35112439-DEL-258 |
| chr9-35147820-DEL-146 | chr9-35147821-DEL-146 |
| chr9-35215385-DEL-492 | chr9-35215386-DEL-492 |
| chr9-35216511-DEL-376 | chr9-35216512-DEL-376 |
| chr9-35239500-DEL-218 | chr9-35239501-DEL-218 |
| chr9-35243260-DEL-76 | chr9-35243261-DEL-76 |
| chr9-35267200-DEL-1890 | chr9-35267201-DEL-1890 |
| chr9-35295653-DEL-2740 | chr9-35295653-DEL-2740 |
| chr9-35299654-DEL-6436 | chr9-35299655-DEL-6436 |
| chr9-35320894-DEL-84 | chr9-35320895-DEL-84 |
| chr9-35326139-DEL-132 | chr9-35326140-DEL-132 |
| chr9-35347443-DEL-1349 | chr9-35347444-DEL-1349 |
| chr9-35368304-DEL-51 | chr9-35368305-DEL-51 |
| chr9-35403677-DEL-80 | chr9-35403686-DEL-72 |
| chr9-35453516-DEL-78 | chr9-35453517-DEL-78 |
| chr9-35614635-DEL-1776 | chr9-35614634-DEL-1776 |
| chr9-35619081-DEL-109 | chr9-35619082-DEL-109 |
| chr9-35625974-DEL-72 | chr9-35625967-DEL-108 |
| chr9-35675311-DEL-70 | chr9-35675288-DEL-63 |
| chr9-35678413-DEL-6577 | chr9-35678414-DEL-6577 |
| chr9-35692270-DEL-65 | chr9-35692284-DEL-58 |
| chr9-35725731-DEL-457 | chr9-35725732-DEL-457 |
| chr9-35759897-DEL-9960 | chr9-35759898-DEL-9960 |
| chr9-35767192-DEL-8303 | chr9-35767193-DEL-8303 |
| chr9-35788195-DEL-412 | chr9-35788196-DEL-412 |
| chr9-35791194-DEL-7514 | chr9-35791195-DEL-7514 |
| chr9-35856443-DEL-973 | chr9-35856444-DEL-973 |
| chr9-35857832-DEL-1295 | chr9-35857833-DEL-1295 |
| chr9-35889168-DEL-952 | chr9-35889170-DEL-949 |
| chr9-35891113-DEL-52 | chr9-35891114-DEL-52 |
| chr9-35922137-DEL-204 | chr9-35922138-DEL-204 |
| chr9-35925070-DEL-6395 | chr9-35925071-DEL-6395 |
| chr9-35936585-DEL-1063 | chr9-35936586-DEL-1063 |
| chr9-35956662-DEL-2235 | chr9-35956663-DEL-2234 |
| chr9-35962071-DEL-990 | chr9-35962072-DEL-990 |
| chr9-35979928-DEL-197 | chr9-35979929-DEL-197 |
| chr9-35989502-DEL-11380 | chr9-35989500-DEL-11382 |
| chr9-36014407-DEL-70 | chr9-36014408-DEL-70 |
| chr9-36034535-DEL-2999 | chr9-36034536-DEL-2999 |
| chr9-36051585-DEL-254 | chr9-36051586-DEL-254 |
| chr9-36090265-DEL-51 | chr9-36090266-DEL-51 |
| chr9-36124945-DEL-1005 | chr9-36124946-DEL-1005 |
| chr9-36142121-DEL-96 | chr9-36142122-DEL-96 |
| chr9-36146539-DEL-6678 | chr9-36146540-DEL-6678 |
| chr9-36178633-DEL-85 | chr9-36178634-DEL-85 |
| chr9-36180462-DEL-54 | chr9-36180463-DEL-54 |
| chr9-36204748-DEL-82 | chr9-36204747-DEL-84 |
| chr9-36279481-DEL-453 | chr9-36279482-DEL-453 |
| chr9-36293669-DEL-141 | chr9-36293673-DEL-140 |
| chr9-36305353-DEL-6453 | chr9-36305354-DEL-6453 |
| chr9-36332548-DEL-1336 | chr9-36332549-DEL-1336 |
| chr9-36396827-DEL-6194 | chr9-36396828-DEL-6194 |
| chr9-36451335-DEL-60 | chr9-36451342-DEL-54 |
| chr9-36453881-DEL-56 | chr9-36453866-DEL-72 |
| chr9-36464885-DEL-62 | chr9-36464886-DEL-62 |
| chr9-36467850-DEL-154 | chr9-36467851-DEL-154 |
| chr9-36468981-DEL-74 | chr9-36468975-DEL-74 |
| chr9-36470787-DEL-289 | chr9-36470791-DEL-286 |
| chr9-36480083-DEL-196 | chr9-36480084-DEL-196 |
| chr9-36502303-DEL-3035 | chr9-36502303-DEL-3035 |
| chr9-36514495-DEL-54 | chr9-36514495-DEL-54 |
| chr9-36563871-DEL-7429 | chr9-36563872-DEL-7429 |
| chr9-36574287-DEL-4699 | chr9-36574288-DEL-4699 |
| chr9-36583007-DEL-66 | chr9-36583008-DEL-66 |
| chr9-36627045-DEL-113 | chr9-36627046-DEL-113 |
| chr9-36636170-DEL-171 | chr9-36636171-DEL-171 |
| chr9-36651530-DEL-52 | chr9-36651530-DEL-52 |
| chr9-36666821-DEL-2570 | chr9-36666822-DEL-2570 |
| chr9-36666857-DEL-234 | chr9-36666871-DEL-198 |
| chr9-36667994-DEL-134 | chr9-36667995-DEL-134 |
| chr9-36672910-DEL-288 | chr9-36672911-DEL-288 |
| chr9-36680752-DEL-65 | chr9-36680751-DEL-67 |
| chr9-36683963-DEL-266 | chr9-36683964-DEL-266 |
| chr9-36686953-DEL-1383 | chr9-36686954-DEL-1383 |
| chr9-36695243-DEL-52 | chr9-36695244-DEL-52 |
| chr9-36707330-DEL-249 | chr9-36707331-DEL-249 |
| chr9-36719112-DEL-76 | chr9-36719102-DEL-81 |
| chr9-36769602-DEL-52 | chr9-36769599-DEL-72 |
| chr9-36776215-DEL-69 | chr9-36776226-DEL-58 |
| chr9-36782409-DEL-83 | chr9-36782410-DEL-83 |

| | |
|---|---|
| chr9-36800086-DEL-194 | chr9-36800087-DEL-194 |
| chr9-36814177-DEL-670 | chr9-36814190-DEL-666 |
| chr9-36870851-DEL-122 | chr9-36870876-DEL-108 |
| chr9-36891728-DEL-488 | chr9-36891729-DEL-488 |
| chr9-36905352-DEL-60 | chr9-36905352-DEL-60 |
| chr9-36912340-DEL-54 | chr9-36912341-DEL-51 |
| chr9-36931786-DEL-62 | chr9-36931786-DEL-63 |
| chr9-37049101-DEL-56 | chr9-37049102-DEL-56 |
| chr9-37049733-DEL-68 | chr9-37049721-DEL-80 |
| chr9-37122931-DEL-175 | chr9-37122910-DEL-197 |
| chr9-37164203-DEL-458 | chr9-37164204-DEL-458 |
| chr9-37208960-DEL-2586 | chr9-37208961-DEL-2586 |
| chr9-37211692-DEL-175 | chr9-37211651-DEL-176 |
| chr9-37242659-DEL-7113 | chr9-37242660-DEL-7113 |
| chr9-37258266-DEL-121 | chr9-37258267-DEL-121 |
| chr9-37285440-DEL-217 | chr9-37285441-DEL-217 |
| chr9-37300520-DEL-169 | chr9-37300520-DEL-170 |
| chr9-37305329-DEL-190 | chr9-37305330-DEL-190 |
| chr9-37307010-DEL-59 | chr9-37306980-DEL-50 |
| chr9-37350725-DEL-75 | chr9-37350722-DEL-81 |
| chr9-37353057-DEL-6394 | chr9-37353058-DEL-6394 |
| chr9-37365001-DEL-122 | chr9-37365002-DEL-122 |
| chr9-37378522-DEL-81 | chr9-37378517-DEL-75 |
| chr9-37379446-DEL-206 | chr9-37379447-DEL-206 |
| chr9-37380259-DEL-176 | chr9-37380260-DEL-176 |
| chr9-37396110-DEL-204 | chr9-37396111-DEL-204 |
| chr9-37397997-DEL-179 | chr9-37397998-DEL-179 |
| chr9-37403684-DEL-185 | chr9-37403685-DEL-185 |
| chr9-37421579-DEL-125 | chr9-37421580-DEL-125 |
| chr9-37432984-DEL-55 | chr9-37432980-DEL-73 |
| chr9-37503035-DEL-204 | chr9-37503036-DEL-204 |
| chr9-37540187-DEL-58 | chr9-37540192-DEL-56 |
| chr9-37593422-DEL-98 | chr9-37593423-DEL-98 |
| chr9-37595384-DEL-141 | chr9-37595384-DEL-141 |
| chr9-37597643-DEL-704 | chr9-37597644-DEL-704 |
| chr9-37601089-DEL-54 | chr9-37601089-DEL-54 |
| chr9-37629099-DEL-2477 | chr9-37629099-DEL-2478 |
| chr9-37648557-DEL-400 | chr9-37648557-DEL-400 |
| chr9-37651329-DEL-88 | chr9-37651300-DEL-116 |
| chr9-37686248-DEL-217 | chr9-37686247-DEL-237 |
| chr9-37697144-DEL-527 | chr9-37697131-DEL-528 |
| chr9-37698684-DEL-90 | chr9-37698707-DEL-87 |
| chr9-37727035-DEL-357 | chr9-37727036-DEL-357 |
| chr9-37774294-DEL-58 | chr9-37774294-DEL-58 |
| chr9-37776164-DEL-147 | chr9-37776161-DEL-145 |
| chr9-37784199-DEL-78 | chr9-37784200-DEL-78 |
| chr9-37786929-DEL-56 | chr9-37786930-DEL-56 |
| chr9-37789423-DEL-257 | chr9-37789424-DEL-257 |
| chr9-37793161-DEL-6198 | chr9-37793158-DEL-6199 |
| chr9-37801283-DEL-77 | chr9-37801284-DEL-77 |
| chr9-37841904-DEL-409 | chr9-37841904-DEL-408 |
| chr9-37890000-DEL-499 | chr9-37890001-DEL-499 |
| chr9-37916520-DEL-6205 | chr9-37916521-DEL-6205 |
| chr9-37935985-DEL-54 | chr9-37935986-DEL-54 |
| chr9-37940651-DEL-69 | chr9-37940652-DEL-69 |
| chr9-37992037-DEL-58 | chr9-37992032-DEL-67 |
| chr9-38078479-DEL-1112 | chr9-38078480-DEL-1112 |
| chr9-38082424-DEL-484 | chr9-38082425-DEL-484 |
| chr9-38099724-DEL-54 | chr9-38099750-DEL-52 |
| chr9-38100511-DEL-85 | chr9-38100511-DEL-85 |
| chr9-38106756-DEL-1327 | chr9-38106757-DEL-1327 |
| chr9-38110047-DEL-81 | chr9-38110087-DEL-54 |
| chr9-38152010-DEL-8626 | chr9-38152011-DEL-8626 |
| chr9-38192648-DEL-800 | chr9-38192649-DEL-800 |
| chr9-38204880-DEL-66 | chr9-38204872-DEL-82 |
| chr9-38205755-DEL-541 | chr9-38205751-DEL-542 |
| chr9-38209354-DEL-1724 | chr9-38209355-DEL-1724 |
| chr9-38224801-DEL-3728 | chr9-38224801-DEL-3728 |
| chr9-38230032-DEL-2516 | chr9-38230031-DEL-2517 |
| chr9-38261745-DEL-192 | chr9-38261755-DEL-181 |
| chr9-38262732-DEL-373 | chr9-38262726-DEL-374 |
| chr9-38275956-DEL-1012 | chr9-38275956-DEL-1012 |
| chr9-38283725-DEL-806 | chr9-38283725-DEL-806 |
| chr9-38309213-DEL-304 | chr9-38309212-DEL-304 |
| chr9-38324381-DEL-4819 | chr9-38324382-DEL-4820 |
| chr9-38330880-DEL-184 | chr9-38330875-DEL-185 |
| chr9-38366273-DEL-1740 | chr9-38366274-DEL-1721 |
| chr9-38368142-DEL-56 | chr9-38368092-DEL-105 |
| chr9-38380164-DEL-995 | chr9-38380165-DEL-995 |
| chr9-38388565-DEL-440 | chr9-38388566-DEL-440 |

| | |
|---|---|
| chr9-38400819-DEL-54 | chr9-38400814-DEL-64 |
| chr9-38407592-DEL-497 | chr9-38407591-DEL-497 |
| chr9-38411088-DEL-6488 | chr9-38411089-DEL-6488 |
| chr9-38443196-DEL-56 | chr9-38443197-DEL-56 |
| chr9-38462889-DEL-432 | chr9-38462890-DEL-432 |
| chr9-38489153-DEL-188 | chr9-38489154-DEL-188 |
| chr9-38506297-DEL-909 | chr9-38506298-DEL-909 |
| chr9-38559033-DEL-58 | chr9-38559043-DEL-50 |
| chr9-38559954-DEL-113 | chr9-38559955-DEL-113 |
| chr9-38566704-DEL-575 | chr9-38566705-DEL-575 |
| chr9-38567253-DEL-112 | chr9-38567255-DEL-111 |
| chr9-38572002-DEL-52 | chr9-38572003-DEL-52 |
| chr9-38576891-DEL-858 | chr9-38576891-DEL-858 |
| chr9-38582876-DEL-2780 | chr9-38582877-DEL-2780 |
| chr9-38610550-DEL-229 | chr9-38610551-DEL-229 |
| chr9-38617394-DEL-203 | chr9-38617393-DEL-204 |
| chr9-38624169-DEL-919 | chr9-38624169-DEL-919 |
| chr9-38627699-DEL-142 | chr9-38627700-DEL-142 |
| chr9-38642736-DEL-52 | chr9-38642729-DEL-60 |
| chr9-38654744-DEL-62 | chr9-38654739-DEL-60 |
| chr9-38683575-DEL-230 | chr9-38683576-DEL-230 |
| chr9-38711968-DEL-368 | chr9-38711969-DEL-368 |
| chr9-38719832-DEL-1154 | chr9-38719833-DEL-1154 |
| chr9-38745133-DEL-6851 | chr9-38745116-DEL-6868 |
| chr9-38754248-DEL-76 | chr9-38754249-DEL-76 |
| chr9-38805090-DEL-174 | chr9-38805091-DEL-174 |
| chr9-38807821-DEL-6604 | chr9-38807822-DEL-6604 |
| chr9-38817595-DEL-334 | chr9-38817596-DEL-334 |
| chr9-38842752-DEL-68 | chr9-38842753-DEL-68 |
| chr9-38844801-DEL-56 | chr9-38844798-DEL-52 |
| chr9-38860018-DEL-106 | chr9-38860019-DEL-106 |
| chr9-38869909-DEL-6659 | chr9-38869910-DEL-6659 |
| chr9-38886872-DEL-63 | chr9-38886873-DEL-63 |
| chr9-38915933-DEL-61 | chr9-38915933-DEL-61 |
| chr9-38939778-DEL-73 | chr9-38939761-DEL-90 |
| chr9-38947733-DEL-6408 | chr9-38947734-DEL-6408 |
| chr9-38953817-DEL-268 | chr9-38953818-DEL-268 |
| chr9-38967484-DEL-1647 | chr9-38967489-DEL-1644 |
| chr9-38986191-DEL-508 | chr9-38986192-DEL-508 |
| chr9-39004205-DEL-330 | chr9-39004206-DEL-330 |
| chr9-39036181-DEL-102 | chr9-39036182-DEL-102 |
| chr9-39134190-DEL-65 | chr9-39134191-DEL-65 |
| chr9-39142913-DEL-90 | chr9-39142914-DEL-88 |
| chr9-39221735-DEL-204 | chr9-39221736-DEL-204 |
| chr9-39223654-DEL-259 | chr9-39223655-DEL-259 |
| chr9-39279479-DEL-1654 | chr9-39279479-DEL-1654 |
| chr9-39287320-DEL-987 | chr9-39287321-DEL-987 |
| chr9-39354519-DEL-6411 | chr9-39354520-DEL-6411 |
| chr9-39384638-DEL-439 | chr9-39384639-DEL-439 |
| chr9-39386039-DEL-4426 | chr9-39386039-DEL-4423 |
| chr9-39389448-DEL-168 | chr9-39389449-DEL-168 |
| chr9-39405661-DEL-2376 | chr9-39405662-DEL-2376 |
| chr9-39407724-DEL-6229 | chr9-39407725-DEL-6229 |
| chr9-39409640-DEL-327 | chr9-39409641-DEL-327 |
| chr9-39414672-DEL-2884 | chr9-39414673-DEL-2884 |
| chr9-39419460-DEL-56 | chr9-39419461-DEL-52 |
| chr9-39427891-DEL-197 | chr9-39427892-DEL-197 |
| chr9-39449579-DEL-981 | chr9-39449580-DEL-981 |
| chr9-39451415-DEL-133 | chr9-39451416-DEL-133 |
| chr9-39538254-DEL-1227 | chr9-39538255-DEL-1227 |
| chr9-39552546-DEL-92 | chr9-39552585-DEL-70 |
| chr9-39594973-DEL-230 | chr9-39594974-DEL-230 |
| chr9-39601579-DEL-220 | chr9-39601580-DEL-220 |
| chr9-39602251-DEL-1169 | chr9-39602252-DEL-1169 |
| chr9-39612640-DEL-348 | chr9-39612641-DEL-348 |
| chr9-39650584-DEL-597 | chr9-39650585-DEL-597 |
| chr9-39654393-DEL-88 | chr9-39654390-DEL-92 |
| chr9-39735565-DEL-487 | chr9-39735566-DEL-487 |
| chr9-39746487-DEL-1555 | chr9-39746488-DEL-1555 |
| chr9-39752813-DEL-830 | chr9-39752814-DEL-830 |
| chr9-39757094-DEL-1455 | chr9-39757095-DEL-1455 |
| chr9-39763265-DEL-1056 | chr9-39763266-DEL-1056 |
| chr9-39768539-DEL-4749 | chr9-39768540-DEL-4749 |
| chr9-39776613-DEL-66 | chr9-39776608-DEL-76 |
| chr9-39804889-DEL-5301 | chr9-39804890-DEL-5301 |
| chr9-39831494-DEL-6222 | chr9-39831481-DEL-6235 |
| chr9-39838603-DEL-202 | chr9-39838604-DEL-202 |
| chr9-39842843-DEL-60 | chr9-39842830-DEL-73 |
| chr9-39860336-DEL-631 | chr9-39860337-DEL-631 |
| chr9-39871981-DEL-708 | chr9-39871982-DEL-708 |

| | |
|---|---|
| chr9-39907045-DEL-136 | chr9-39907040-DEL-126 |
| chr9-39931718-DEL-56 | chr9-39931719-DEL-56 |
| chr9-39973072-DEL-241 | chr9-39973071-DEL-242 |
| chr9-39975398-DEL-6591 | chr9-39975404-DEL-6590 |
| chr9-39983492-DEL-52 | chr9-39983493-DEL-52 |
| chr9-40027273-DEL-132 | chr9-40027313-DEL-96 |
| chr9-40046010-DEL-204 | chr9-40046011-DEL-204 |
| chr9-40056953-DEL-649 | chr9-40056954-DEL-649 |
| chr9-40059186-DEL-51 | chr9-40059187-DEL-51 |
| chr9-40075482-DEL-70 | chr9-40075482-DEL-58 |
| chr9-40076347-DEL-51 | chr9-40076345-DEL-60 |
| chr9-40080768-DEL-747 | chr9-40080769-DEL-747 |
| chr9-40097975-DEL-79 | chr9-40097950-DEL-92 |
| chr9-40100911-DEL-698 | chr9-40100912-DEL-698 |
| chr9-40114879-DEL-574 | chr9-40114881-DEL-573 |
| chr9-40133507-DEL-139 | chr9-40133508-DEL-141 |
| chr9-40138136-DEL-4724 | chr9-40138137-DEL-4724 |
| chr9-40154020-DEL-159 | chr9-40154021-DEL-159 |
| chr9-40162032-DEL-66 | chr9-40162033-DEL-80 |
| chr9-40165464-DEL-6507 | chr9-40165465-DEL-6507 |
| chr9-40165563-DEL-204 | chr9-40165564-DEL-204 |
| chr9-40195392-DEL-394 | chr9-40195393-DEL-394 |
| chr9-40198535-DEL-589 | chr9-40198529-DEL-590 |
| chr9-40210039-DEL-172 | chr9-40210040-DEL-172 |
| chr9-40241347-DEL-202 | chr9-40241348-DEL-202 |
| chr9-40294890-DEL-56 | chr9-40294850-DEL-96 |
| chr9-40299487-DEL-204 | chr9-40299490-DEL-203 |
| chr9-40314191-DEL-120 | chr9-40314193-DEL-117 |
| chr9-40314799-DEL-52 | chr9-40314790-DEL-66 |
| chr9-40338806-DEL-52 | chr9-40338807-DEL-52 |
| chr9-40403194-DEL-6473 | chr9-40403195-DEL-6473 |
| chr9-40413377-DEL-89 | chr9-40413364-DEL-85 |
| chr9-40433483-DEL-66 | chr9-40433484-DEL-50 |
| chr9-40448921-DEL-54 | chr9-40448922-DEL-60 |
| chr9-40468098-DEL-103 | chr9-40468099-DEL-107 |
| chr9-40499648-DEL-840 | chr9-40499641-DEL-825 |
| chr9-40515213-DEL-58 | chr9-40515220-DEL-52 |
| chr9-40532369-DEL-370 | chr9-40532335-DEL-415 |
| chr9-40540540-DEL-89 | chr9-40540540-DEL-73 |
| chr9-40543217-DEL-1348 | chr9-40543218-DEL-1349 |
| chr9-40544335-DEL-480 | chr9-40544336-DEL-480 |
| chr9-40552359-DEL-123 | chr9-40552360-DEL-123 |
| chr9-40569052-DEL-115 | chr9-40569053-DEL-125 |
| chr9-40623714-DEL-58 | chr9-40623715-DEL-58 |
| chr9-40644512-DEL-162 | chr9-40644518-DEL-157 |
| chr9-40653881-DEL-62 | chr9-40653882-DEL-62 |
| chr9-40680788-DEL-534 | chr9-40680789-DEL-534 |
| chr9-40683945-DEL-83 | chr9-40683946-DEL-60 |
| chr9-40689028-DEL-233 | chr9-40689029-DEL-233 |
| chr9-40690888-DEL-64 | chr9-40690883-DEL-70 |
| chr9-40692842-DEL-54 | chr9-40692843-DEL-72 |
| chr9-40700357-DEL-292 | chr9-40700358-DEL-292 |
| chr9-40703126-DEL-56 | chr9-40703126-DEL-53 |
| chr9-40704317-DEL-204 | chr9-40704327-DEL-212 |
| chr9-40717079-DEL-478 | chr9-40717080-DEL-478 |
| chr9-40724261-DEL-63 | chr9-40724266-DEL-64 |
| chr9-40725928-DEL-205 | chr9-40725929-DEL-205 |
| chr9-40727887-DEL-396 | chr9-40727887-DEL-390 |
| chr9-40728504-DEL-233 | chr9-40728505-DEL-233 |
| chr9-40730535-DEL-225 | chr9-40730536-DEL-225 |
| chr9-40788362-DEL-62 | chr9-40788363-DEL-62 |
| chr9-40809865-DEL-6385 | chr9-40809867-DEL-6384 |
| chr9-40906302-DEL-70 | chr9-40906307-DEL-72 |
| chr9-40909093-DEL-85 | chr9-40909092-DEL-94 |
| chr9-40910702-DEL-225 | chr9-40910703-DEL-225 |
| chr9-40928820-DEL-128 | chr9-40928821-DEL-128 |
| chr9-40941476-DEL-63 | chr9-40941477-DEL-63 |
| chr9-40954451-DEL-54 | chr9-40954452-DEL-54 |
| chr9-40970684-DEL-53 | chr9-40970685-DEL-53 |
| chr9-40976509-DEL-53 | chr9-40976533-DEL-54 |
| chr9-40982597-DEL-83 | chr9-40982586-DEL-84 |
| chr9-40997038-DEL-117 | chr9-40997039-DEL-117 |
| chr9-41015170-DEL-68 | chr9-41015171-DEL-68 |
| chr9-41018375-DEL-57 | chr9-41018376-DEL-57 |
| chr9-41037277-DEL-82 | chr9-41037278-DEL-82 |
| chr9-41039022-DEL-76 | chr9-41039019-DEL-84 |
| chr9-41045589-DEL-212 | chr9-41045590-DEL-212 |
| chr9-41051595-DEL-116 | chr9-41051594-DEL-127 |
| chr9-41065848-DEL-327 | chr9-41065849-DEL-327 |
| chr9-41078934-DEL-1435 | chr9-41078935-DEL-1435 |

| | |
|---|---|
| chr9-41088890-DEL-222 | chr9-41088891-DEL-222 |
| chr9-41121047-DEL-62 | chr9-41121047-DEL-62 |
| chr9-41164770-DEL-501 | chr9-41164771-DEL-501 |
| chr9-41183554-DEL-51 | chr9-41183565-DEL-51 |
| chr9-41189639-DEL-83 | chr9-41189640-DEL-83 |
| chr9-41204110-DEL-52 | chr9-41204111-DEL-52 |
| chr9-41208855-DEL-68 | chr9-41208814-DEL-69 |
| chr9-41220228-DEL-210 | chr9-41220229-DEL-210 |
| chr9-41227771-DEL-4592 | chr9-41227772-DEL-4592 |
| chr9-41244600-DEL-67 | chr9-41244602-DEL-66 |
| chr9-41245193-DEL-78 | chr9-41245194-DEL-78 |
| chr9-41262840-DEL-76 | chr9-41262840-DEL-75 |
| chr9-41320133-DEL-64 | chr9-41320134-DEL-64 |
| chr9-41324362-DEL-74 | chr9-41324377-DEL-72 |
| chr9-41350535-DEL-70 | chr9-41350555-DEL-75 |
| chr9-41377242-DEL-340 | chr9-41377243-DEL-340 |
| chr9-41379988-DEL-74 | chr9-41379990-DEL-72 |
| chr9-41388185-DEL-100 | chr9-41388186-DEL-100 |
| chr9-41410283-DEL-84 | chr9-41410284-DEL-84 |
| chr9-41459631-DEL-419 | chr9-41459632-DEL-419 |
| chr9-41470470-DEL-11475 | chr9-41470471-DEL-11475 |
| chr9-41486881-DEL-50 | chr9-41486881-DEL-60 |
| chr9-41493371-DEL-66 | chr9-41493372-DEL-66 |
| chr9-41495922-DEL-51 | chr9-41495923-DEL-51 |
| chr9-41529305-DEL-159 | chr9-41529306-DEL-159 |
| chr9-41533932-DEL-60 | chr9-41533933-DEL-60 |
| chr9-41564202-DEL-85 | chr9-41564203-DEL-83 |
| chr9-41569324-DEL-1152 | chr9-41569325-DEL-1152 |
| chr9-41570589-DEL-574 | chr9-41570590-DEL-574 |
| chr9-41621709-DEL-437 | chr9-41621710-DEL-437 |
| chr9-41653859-DEL-51 | chr9-41653853-DEL-53 |
| chr9-41667699-DEL-54 | chr9-41667700-DEL-52 |
| chr9-41712206-DEL-153 | chr9-41712206-DEL-154 |
| chr9-41749504-DEL-478 | chr9-41749505-DEL-478 |
| chr9-41753314-DEL-1234 | chr9-41753315-DEL-1234 |
| chr9-41815596-DEL-827 | chr9-41815597-DEL-827 |
| chr9-41816778-DEL-6788 | chr9-41816779-DEL-6788 |
| chr9-41848222-DEL-261 | chr9-41848215-DEL-269 |
| chr9-41873824-DEL-83 | chr9-41873820-DEL-88 |
| chr9-41900833-DEL-58 | chr9-41900836-DEL-56 |
| chr9-41908644-DEL-66 | chr9-41908645-DEL-66 |
| chr9-41940824-DEL-57 | chr9-41940825-DEL-57 |
| chr9-41962935-DEL-68 | chr9-41962932-DEL-68 |
| chr9-41977500-DEL-150 | chr9-41977493-DEL-138 |
| chr9-41979723-DEL-72 | chr9-41979718-DEL-67 |
| chr9-41988558-DEL-79 | chr9-41988559-DEL-79 |
| chr9-42053565-DEL-52 | chr9-42053566-DEL-52 |
| chr9-42079372-DEL-1422 | chr9-42079373-DEL-1422 |
| chr9-42100866-DEL-71 | chr9-42100864-DEL-70 |
| chr9-42106888-DEL-63 | chr9-42106888-DEL-60 |
| chr9-42110197-DEL-166 | chr9-42110198-DEL-166 |
| chr9-42126278-DEL-52 | chr9-42126261-DEL-70 |
| chr9-42145425-DEL-62 | chr9-42145426-DEL-62 |
| chr9-42194003-DEL-549 | chr9-42194004-DEL-549 |
| chr9-42210769-DEL-1063 | chr9-42210770-DEL-1063 |
| chr9-42236159-DEL-100 | chr9-42236160-DEL-100 |
| chr9-42278626-DEL-96 | chr9-42278626-DEL-96 |
| chr9-42285523-DEL-120 | chr9-42285523-DEL-120 |
| chr9-42337400-DEL-112 | chr9-42337401-DEL-84 |
| chr9-42376361-DEL-62 | chr9-42376362-DEL-62 |
| chr9-42404807-DEL-132 | chr9-42404808-DEL-132 |
| chr9-42415493-DEL-56 | chr9-42415492-DEL-54 |
| chr9-42415684-DEL-928 | chr9-42415685-DEL-928 |
| chr9-42467564-DEL-66 | chr9-42467534-DEL-78 |
| chr9-42520317-DEL-693 | chr9-42520318-DEL-693 |
| chr9-42542418-DEL-67 | chr9-42542419-DEL-67 |
| chr9-42564251-DEL-2303 | chr9-42564252-DEL-2303 |
| chr9-42608810-DEL-56 | chr9-42608811-DEL-56 |
| chr9-42626249-DEL-100 | chr9-42626254-DEL-96 |
| chr9-42636728-DEL-109 | chr9-42636724-DEL-103 |
| chr9-42766895-DEL-1311 | chr9-42766896-DEL-1311 |
| chr9-42773330-DEL-69 | chr9-42773331-DEL-69 |
| chr9-42817235-DEL-55 | chr9-42817224-DEL-60 |
| chr9-42834403-DEL-177 | chr9-42834428-DEL-172 |
| chr9-42902484-DEL-604 | chr9-42902485-DEL-604 |
| chr9-42914118-DEL-209 | chr9-42914119-DEL-209 |
| chr9-42945849-DEL-111 | chr9-42945850-DEL-111 |
| chr9-43008097-DEL-241 | chr9-43008098-DEL-241 |
| chr9-43051851-DEL-102 | chr9-43051852-DEL-102 |
| chr9-43067051-DEL-332 | chr9-43067052-DEL-332 |

| | |
|---|---|
| chr9-43072654-DEL-68 | chr9-43072663-DEL-60 |
| chr9-43077048-DEL-110 | chr9-43077049-DEL-110 |
| chr9-43097389-DEL-51 | chr9-43097389-DEL-53 |
| chr9-43105785-DEL-54 | chr9-43105783-DEL-56 |
| chr9-43108656-DEL-167 | chr9-43108662-DEL-170 |
| chr9-43121100-DEL-60 | chr9-43121102-DEL-58 |
| chr9-43156312-DEL-80 | chr9-43156313-DEL-80 |
| chr9-43172382-DEL-74 | chr9-43172403-DEL-80 |
| chr9-43200529-DEL-2105 | chr9-43200530-DEL-2105 |
| chr9-43255445-DEL-52 | chr9-43255445-DEL-52 |
| chr9-43321695-DEL-66 | chr9-43321676-DEL-52 |
| chr9-43340627-DEL-470 | chr9-43340628-DEL-470 |
| chr9-43342973-DEL-305 | chr9-43342978-DEL-294 |
| chr9-43362901-DEL-150 | chr9-43362902-DEL-150 |
| chr9-43388906-DEL-204 | chr9-43388907-DEL-204 |
| chr9-43395175-DEL-171 | chr9-43395178-DEL-171 |
| chr9-43404666-DEL-77 | chr9-43404619-DEL-87 |
| chr9-43408289-DEL-52 | chr9-43408241-DEL-56 |
| chr9-43464059-DEL-82 | chr9-43464060-DEL-82 |
| chr9-43548723-DEL-209 | chr9-43548724-DEL-209 |
| chr9-43549330-DEL-128 | chr9-43549319-DEL-102 |
| chr9-43554874-DEL-61 | chr9-43554872-DEL-60 |
| chr9-43573867-DEL-81 | chr9-43573867-DEL-81 |
| chr9-43577616-DEL-320 | chr9-43577617-DEL-320 |
| chr9-43578542-DEL-66 | chr9-43578543-DEL-66 |
| chr9-43586676-DEL-87 | chr9-43586674-DEL-88 |
| chr9-43596323-DEL-1653 | chr9-43596324-DEL-1653 |
| chr9-43614850-DEL-117 | chr9-43614851-DEL-117 |
| chr9-43695670-DEL-76 | chr9-43695671-DEL-74 |
| chr9-43697000-DEL-225 | chr9-43697001-DEL-225 |
| chr9-43724142-DEL-97 | chr9-43724144-DEL-95 |
| chr9-43727151-DEL-76 | chr9-43727145-DEL-57 |
| chr9-43743313-DEL-103 | chr9-43743300-DEL-72 |
| chr9-43768888-DEL-53 | chr9-43768851-DEL-66 |
| chr9-43793113-DEL-123 | chr9-43793114-DEL-123 |
| chr9-43794101-DEL-224 | chr9-43794102-DEL-228 |
| chr9-43799262-DEL-64 | chr9-43799263-DEL-62 |
| chr9-43813839-DEL-77 | chr9-43813852-DEL-94 |
| chr9-43832008-DEL-217 | chr9-43832007-DEL-220 |
| chr9-43838541-DEL-58 | chr9-43838542-DEL-58 |
| chr9-43858045-DEL-341 | chr9-43858047-DEL-337 |
| chr9-43975701-DEL-109 | chr9-43975702-DEL-109 |
| chr9-44012556-DEL-80 | chr9-44012561-DEL-80 |
| chr9-44027867-DEL-171 | chr9-44027871-DEL-168 |
| chr9-44028524-DEL-169 | chr9-44028525-DEL-169 |
| chr9-44068437-DEL-237 | chr9-44068438-DEL-237 |
| chr9-44074338-DEL-56 | chr9-44074339-DEL-56 |
| chr9-44125588-DEL-6452 | chr9-44125589-DEL-6452 |
| chr9-44141993-DEL-194 | chr9-44141988-DEL-200 |
| chr9-44146572-DEL-165 | chr9-44146573-DEL-165 |
| chr9-44196717-DEL-199 | chr9-44196718-DEL-199 |
| chr9-44197508-DEL-62 | chr9-44197509-DEL-62 |
| chr9-44258047-DEL-91 | chr9-44258048-DEL-91 |
| chr9-44263768-DEL-78 | chr9-44263769-DEL-78 |
| chr9-44269791-DEL-213 | chr9-44269792-DEL-213 |
| chr9-44276516-DEL-7368 | chr9-44276517-DEL-7368 |
| chr9-44314921-DEL-81 | chr9-44314922-DEL-81 |
| chr9-44343018-DEL-379 | chr9-44343019-DEL-372 |
| chr9-44343693-DEL-155 | chr9-44343681-DEL-170 |
| chr9-44375441-DEL-1393 | chr9-44375442-DEL-1393 |
| chr9-44425931-DEL-209 | chr9-44425932-DEL-209 |
| chr9-44438930-DEL-56 | chr9-44438931-DEL-56 |
| chr9-44440642-DEL-55 | chr9-44440643-DEL-55 |
| chr9-44450507-DEL-72 | chr9-44450508-DEL-72 |
| chr9-44457836-DEL-223 | chr9-44457837-DEL-223 |
| chr9-44458801-DEL-108 | chr9-44458802-DEL-108 |
| chr9-44480578-DEL-104 | chr9-44480579-DEL-104 |
| chr9-44481541-DEL-125 | chr9-44481530-DEL-126 |
| chr9-44507477-DEL-90 | chr9-44507477-DEL-86 |
| chr9-44522655-DEL-213 | chr9-44522655-DEL-213 |
| chr9-44567448-DEL-81 | chr9-44567449-DEL-81 |
| chr9-44591308-DEL-60 | chr9-44591309-DEL-61 |
| chr9-44640767-DEL-202 | chr9-44640768-DEL-202 |
| chr9-44665767-DEL-99 | chr9-44665768-DEL-99 |
| chr9-44679285-DEL-218 | chr9-44679286-DEL-218 |
| chr9-44683137-DEL-507 | chr9-44683138-DEL-507 |
| chr9-44688951-DEL-187 | chr9-44688957-DEL-182 |
| chr9-44737414-DEL-64 | chr9-44737415-DEL-64 |
| chr9-44738929-DEL-828 | chr9-44738924-DEL-829 |
| chr9-44795004-DEL-209 | chr9-44795005-DEL-209 |

| | |
|---|---|
| chr9-44799206-DEL-200 | chr9-44799196-DEL-201 |
| chr9-44808508-DEL-1420 | chr9-44808509-DEL-1420 |
| chr9-44809621-DEL-211 | chr9-44809614-DEL-212 |
| chr9-44814073-DEL-104 | chr9-44814074-DEL-104 |
| chr9-44820084-DEL-81 | chr9-44820085-DEL-81 |
| chr9-44827854-DEL-198 | chr9-44827855-DEL-198 |
| chr9-44849994-DEL-218 | chr9-44849995-DEL-218 |
| chr9-44869419-DEL-188 | chr9-44869420-DEL-188 |
| chr9-44870358-DEL-211 | chr9-44870359-DEL-211 |
| chr9-44879838-DEL-144 | chr9-44879834-DEL-141 |
| chr9-44902810-DEL-204 | chr9-44902811-DEL-204 |
| chr9-44904011-DEL-432 | chr9-44904012-DEL-432 |
| chr9-44962464-DEL-512 | chr9-44962465-DEL-512 |
| chr9-44998742-DEL-82 | chr9-44998739-DEL-85 |
| chr9-45003295-DEL-62 | chr9-45003296-DEL-62 |
| chr9-45008441-DEL-174 | chr9-45008442-DEL-174 |
| chr9-45022601-DEL-56 | chr9-45022602-DEL-56 |
| chr9-45173087-DEL-117 | chr9-45173106-DEL-116 |
| chr9-45190145-DEL-222 | chr9-45190146-DEL-222 |
| chr9-45226990-DEL-84 | chr9-45227037-DEL-52 |
| chr9-45227933-DEL-415 | chr9-45227934-DEL-415 |
| chr9-45249184-DEL-85 | chr9-45249189-DEL-81 |
| chr9-45272454-DEL-100 | chr9-45272455-DEL-100 |
| chr9-45275562-DEL-111 | chr9-45275580-DEL-97 |
| chr9-45278410-DEL-822 | chr9-45278411-DEL-822 |
| chr9-45283254-DEL-58 | chr9-45283255-DEL-58 |
| chr9-45289367-DEL-346 | chr9-45289367-DEL-346 |
| chr9-45373111-DEL-3443 | chr9-45373112-DEL-3443 |
| chr9-45400594-DEL-188 | chr9-45400595-DEL-188 |
| chr9-45507243-DEL-81 | chr9-45507244-DEL-81 |
| chr9-45517396-DEL-52 | chr9-45517387-DEL-60 |
| chr9-45526248-DEL-71 | chr9-45526251-DEL-69 |
| chr9-45570872-DEL-60 | chr9-45570859-DEL-61 |
| chr9-45619483-DEL-84 | chr9-45619484-DEL-84 |
| chr9-45622045-DEL-62 | chr9-45622054-DEL-60 |
| chr9-45637550-DEL-185 | chr9-45637562-DEL-173 |
| chr9-45654780-DEL-312 | chr9-45654780-DEL-313 |
| chr9-45659632-DEL-168 | chr9-45659676-DEL-148 |
| chr9-45662861-DEL-62 | chr9-45662862-DEL-62 |
| chr9-45670607-DEL-101 | chr9-45670608-DEL-101 |
| chr9-45676451-DEL-313 | chr9-45676452-DEL-313 |
| chr9-45705851-DEL-188 | chr9-45705852-DEL-188 |
| chr9-45722122-DEL-55 | chr9-45722123-DEL-55 |
| chr9-45738473-DEL-112 | chr9-45738472-DEL-113 |
| chr9-45741799-DEL-216 | chr9-45741800-DEL-216 |
| chr9-45846516-DEL-60 | chr9-45846543-DEL-62 |
| chr9-45851509-DEL-387 | chr9-45851510-DEL-387 |
| chr9-45870651-DEL-218 | chr9-45870652-DEL-218 |
| chr9-45904354-DEL-1343 | chr9-45904348-DEL-1344 |
| chr9-45906607-DEL-864 | chr9-45906608-DEL-864 |
| chr9-45948703-DEL-224 | chr9-45948704-DEL-224 |
| chr9-45955436-DEL-52 | chr9-45955433-DEL-58 |
| chr9-46002584-DEL-110 | chr9-46002585-DEL-110 |
| chr9-46009108-DEL-56 | chr9-46009109-DEL-56 |
| chr9-46046922-DEL-442 | chr9-46046923-DEL-442 |
| chr9-46158395-DEL-122 | chr9-46158398-DEL-120 |
| chr9-46183690-DEL-224 | chr9-46183689-DEL-212 |
| chr9-46205033-DEL-160 | chr9-46205034-DEL-166 |
| chr9-46214080-DEL-68 | chr9-46214085-DEL-50 |
| chr9-46247843-DEL-178 | chr9-46247844-DEL-178 |
| chr9-46279685-DEL-1026 | chr9-46279686-DEL-1026 |
| chr9-46295760-DEL-96 | chr9-46295734-DEL-139 |
| chr9-46315701-DEL-58 | chr9-46315699-DEL-60 |
| chr9-46367466-DEL-174 | chr9-46367467-DEL-174 |
| chr9-46380325-DEL-126 | chr9-46380326-DEL-126 |
| chr9-46392120-DEL-150 | chr9-46392121-DEL-150 |
| chr9-46400091-DEL-61 | chr9-46400092-DEL-61 |
| chr9-46491690-DEL-72 | chr9-46491691-DEL-72 |
| chr9-46496453-DEL-419 | chr9-46496454-DEL-419 |
| chr9-46507089-DEL-2707 | chr9-46507090-DEL-2707 |
| chr9-46515283-DEL-54 | chr9-46515267-DEL-70 |
| chr9-46521170-DEL-67 | chr9-46521171-DEL-67 |
| chr9-46524257-DEL-118 | chr9-46524239-DEL-117 |
| chr9-46541423-DEL-97 | chr9-46541424-DEL-97 |
| chr9-46605026-DEL-58 | chr9-46605027-DEL-58 |
| chr9-46612914-DEL-104 | chr9-46612919-DEL-100 |
| chr9-46617039-DEL-57 | chr9-46617040-DEL-51 |
| chr9-46680293-DEL-52 | chr9-46680294-DEL-52 |
| chr9-46785447-DEL-56 | chr9-46785448-DEL-50 |
| chr9-46913811-DEL-392 | chr9-46913814-DEL-390 |

| | |
|---|---|
| chr9-46931288-DEL-1155 | chr9-46931289-DEL-1155 |
| chr9-46939802-DEL-715 | chr9-46939803-DEL-715 |
| chr9-46947578-DEL-54 | chr9-46947579-DEL-54 |
| chr9-46959753-DEL-52 | chr9-46959759-DEL-55 |
| chr9-46960672-DEL-54 | chr9-46960663-DEL-64 |
| chr9-46996176-DEL-280 | chr9-46996177-DEL-280 |
| chr9-47017551-DEL-54 | chr9-47017552-DEL-60 |
| chr9-47034062-DEL-54 | chr9-47034059-DEL-58 |
| chr9-47043853-DEL-72 | chr9-47043853-DEL-70 |
| chr9-47061151-DEL-152 | chr9-47061162-DEL-135 |
| chr9-47107031-DEL-2421 | chr9-47107032-DEL-2421 |
| chr9-47118924-DEL-74 | chr9-47118925-DEL-74 |
| chr9-47132429-DEL-66 | chr9-47132412-DEL-64 |
| chr9-47156518-DEL-74 | chr9-47156519-DEL-74 |
| chr9-47183060-DEL-327 | chr9-47183061-DEL-327 |
| chr9-47209802-DEL-479 | chr9-47209846-DEL-465 |
| chr9-47238496-DEL-390 | chr9-47238497-DEL-390 |
| chr9-47246403-DEL-60 | chr9-47246396-DEL-58 |
| chr9-47293422-DEL-217 | chr9-47293423-DEL-217 |
| chr9-47341397-DEL-67 | chr9-47341366-DEL-59 |
| chr9-47365757-DEL-122 | chr9-47365758-DEL-133 |
| chr9-47470797-DEL-565 | chr9-47470798-DEL-565 |
| chr9-47494933-DEL-73 | chr9-47494934-DEL-75 |
| chr9-47577911-DEL-362 | chr9-47577912-DEL-362 |
| chr9-47588228-DEL-107 | chr9-47588220-DEL-118 |
| chr9-47614748-DEL-106 | chr9-47614749-DEL-106 |
| chr9-47631546-DEL-1238 | chr9-47631547-DEL-1238 |
| chr9-47643014-DEL-88 | chr9-47643015-DEL-88 |
| chr9-47677773-DEL-1571 | chr9-47677774-DEL-1571 |
| chr9-47736319-DEL-58 | chr9-47736320-DEL-58 |
| chr9-47774141-DEL-58 | chr9-47774146-DEL-54 |
| chr9-47789953-DEL-101 | chr9-47789954-DEL-101 |
| chr9-47798628-DEL-72 | chr9-47798629-DEL-72 |
| chr9-47814720-DEL-154 | chr9-47814721-DEL-154 |
| chr9-47837592-DEL-52 | chr9-47837592-DEL-52 |
| chr9-47893478-DEL-2451 | chr9-47893479-DEL-2449 |
| chr9-47904279-DEL-110 | chr9-47904280-DEL-108 |
| chr9-47906953-DEL-71 | chr9-47906945-DEL-68 |
| chr9-47915042-DEL-55 | chr9-47915045-DEL-56 |
| chr9-47968536-DEL-69 | chr9-47968537-DEL-69 |
| chr9-47971993-DEL-74 | chr9-47971977-DEL-87 |
| chr9-47973364-DEL-71 | chr9-47973365-DEL-71 |
| chr9-48006426-DEL-62 | chr9-48006427-DEL-62 |
| chr9-48013930-DEL-443 | chr9-48013931-DEL-443 |
| chr9-48051626-DEL-100 | chr9-48051627-DEL-100 |
| chr9-48070000-DEL-378 | chr9-48070001-DEL-378 |
| chr9-48075618-DEL-230 | chr9-48075619-DEL-230 |
| chr9-48110656-DEL-738 | chr9-48110657-DEL-738 |
| chr9-48112405-DEL-381 | chr9-48112406-DEL-381 |
| chr9-48118911-DEL-66 | chr9-48118912-DEL-66 |
| chr9-48128917-DEL-167 | chr9-48128904-DEL-188 |
| chr9-48167980-DEL-140 | chr9-48168015-DEL-136 |
| chr9-48183965-DEL-554 | chr9-48183966-DEL-554 |
| chr9-48203056-DEL-95 | chr9-48203056-DEL-96 |
| chr9-48222845-DEL-358 | chr9-48222846-DEL-358 |
| chr9-48225693-DEL-654 | chr9-48225694-DEL-654 |
| chr9-48264337-DEL-3744 | chr9-48264338-DEL-3744 |
| chr9-48274172-DEL-65 | chr9-48274173-DEL-65 |
| chr9-48275455-DEL-230 | chr9-48275447-DEL-239 |
| chr9-48279392-DEL-266 | chr9-48279393-DEL-259 |
| chr9-48286354-DEL-221 | chr9-48286355-DEL-221 |
| chr9-48292301-DEL-490 | chr9-48292302-DEL-490 |
| chr9-48302589-DEL-72 | chr9-48302590-DEL-72 |
| chr9-48319943-DEL-1691 | chr9-48319944-DEL-1691 |
| chr9-48322387-DEL-62 | chr9-48322388-DEL-62 |
| chr9-48323766-DEL-54 | chr9-48323767-DEL-54 |
| chr9-48329367-DEL-105 | chr9-48329368-DEL-105 |
| chr9-48346756-DEL-90 | chr9-48346757-DEL-90 |
| chr9-48350317-DEL-95 | chr9-48350310-DEL-84 |
| chr9-48361545-DEL-1313 | chr9-48361546-DEL-1313 |
| chr9-48373499-DEL-225 | chr9-48373500-DEL-225 |
| chr9-48376405-DEL-137 | chr9-48376406-DEL-137 |
| chr9-48388869-DEL-331 | chr9-48388870-DEL-331 |
| chr9-48423509-DEL-148 | chr9-48423528-DEL-130 |
| chr9-48429924-DEL-5212 | chr9-48429925-DEL-5212 |
| chr9-48446902-DEL-413 | chr9-48446904-DEL-412 |
| chr9-48448254-DEL-63 | chr9-48448261-DEL-57 |
| chr9-48449211-DEL-86 | chr9-48449192-DEL-82 |
| chr9-48513428-DEL-92 | chr9-48513429-DEL-92 |
| chr9-48534882-DEL-56 | chr9-48534882-DEL-57 |

| | |
|---|---|
| chr9-48550300-DEL-99 | chr9-48550301-DEL-99 |
| chr9-48562683-DEL-311 | chr9-48562684-DEL-311 |
| chr9-48571343-DEL-97 | chr9-48571344-DEL-97 |
| chr9-48623718-DEL-53 | chr9-48623719-DEL-53 |
| chr9-48659116-DEL-139 | chr9-48659117-DEL-139 |
| chr9-48664720-DEL-53 | chr9-48664720-DEL-53 |
| chr9-48681793-DEL-82 | chr9-48681794-DEL-82 |
| chr9-48691891-DEL-171 | chr9-48691892-DEL-171 |
| chr9-48697626-DEL-112 | chr9-48697631-DEL-113 |
| chr9-48713554-DEL-84 | chr9-48713555-DEL-84 |
| chr9-48721793-DEL-51 | chr9-48721791-DEL-51 |
| chr9-48738541-DEL-52 | chr9-48738542-DEL-52 |
| chr9-48754758-DEL-52 | chr9-48754759-DEL-52 |
| chr9-48756284-DEL-1480 | chr9-48756285-DEL-1480 |
| chr9-48759899-DEL-99 | chr9-48759900-DEL-99 |
| chr9-48767007-DEL-59 | chr9-48767004-DEL-63 |
| chr9-48787561-DEL-209 | chr9-48787562-DEL-209 |
| chr9-48792653-DEL-345 | chr9-48792654-DEL-345 |
| chr9-48795806-DEL-55 | chr9-48795813-DEL-54 |
| chr9-48806040-DEL-90 | chr9-48806039-DEL-89 |
| chr9-48814197-DEL-115 | chr9-48814198-DEL-115 |
| chr9-48826298-DEL-72 | chr9-48826281-DEL-104 |
| chr9-48831504-DEL-204 | chr9-48831505-DEL-204 |
| chr9-48865201-DEL-95 | chr9-48865206-DEL-91 |
| chr9-48878887-DEL-52 | chr9-48878887-DEL-52 |
| chr9-48881213-DEL-83 | chr9-48881214-DEL-83 |
| chr9-48885503-DEL-864 | chr9-48885504-DEL-864 |
| chr9-48888924-DEL-399 | chr9-48888925-DEL-399 |
| chr9-48895229-DEL-68 | chr9-48895230-DEL-68 |
| chr9-48900787-DEL-207 | chr9-48900787-DEL-207 |
| chr9-48957632-DEL-135 | chr9-48957633-DEL-135 |
| chr9-48961635-DEL-160 | chr9-48961636-DEL-176 |
| chr9-48969329-DEL-197 | chr9-48969330-DEL-197 |
| chr9-48971015-DEL-56 | chr9-48971016-DEL-56 |
| chr9-49008961-DEL-90 | chr9-49008956-DEL-96 |
| chr9-49062863-DEL-1820 | chr9-49062864-DEL-1820 |
| chr9-49065384-DEL-59 | chr9-49065386-DEL-62 |
| chr9-49084280-DEL-66 | chr9-49084281-DEL-66 |
| chr9-49146712-DEL-89 | chr9-49146713-DEL-89 |
| chr9-49156506-DEL-84 | chr9-49156511-DEL-72 |
| chr9-49188614-DEL-78 | chr9-49188641-DEL-52 |
| chr9-49206109-DEL-86 | chr9-49206100-DEL-78 |
| chr9-49215467-DEL-453 | chr9-49215468-DEL-453 |
| chr9-49220521-DEL-58 | chr9-49220522-DEL-56 |
| chr9-49320516-DEL-54 | chr9-49320517-DEL-52 |
| chr9-49405910-DEL-73 | chr9-49405917-DEL-72 |
| chr9-49428652-DEL-58 | chr9-49428657-DEL-70 |
| chr9-49431849-DEL-52 | chr9-49431855-DEL-53 |
| chr9-49446976-DEL-111 | chr9-49446941-DEL-112 |
| chr9-49485545-DEL-95 | chr9-49485546-DEL-95 |
| chr9-49486899-DEL-72 | chr9-49486902-DEL-68 |
| chr9-49522718-DEL-60 | chr9-49522711-DEL-76 |
| chr9-49566397-DEL-90 | chr9-49566398-DEL-78 |
| chr9-49575857-DEL-173 | chr9-49575858-DEL-173 |
| chr9-49577828-DEL-96 | chr9-49577813-DEL-112 |
| chr9-49589884-DEL-67 | chr9-49589885-DEL-67 |
| chr9-49644490-DEL-60 | chr9-49644491-DEL-60 |
| chr9-49779826-DEL-261 | chr9-49779827-DEL-261 |
| chr9-49814751-DEL-327 | chr9-49814752-DEL-327 |
| chr9-49863557-DEL-480 | chr9-49863558-DEL-480 |
| chr9-49904647-DEL-400 | chr9-49904648-DEL-400 |
| chr9-49935589-DEL-66 | chr9-49935584-DEL-56 |
| chr9-49946817-DEL-67 | chr9-49946813-DEL-68 |
| chr9-49968478-DEL-93 | chr9-49968479-DEL-93 |
| chr9-49970932-DEL-366 | chr9-49970933-DEL-366 |
| chr9-49993280-DEL-6413 | chr9-49993281-DEL-6413 |
| chr9-50028425-DEL-64 | chr9-50028426-DEL-64 |
| chr9-50041400-DEL-341 | chr9-50041396-DEL-342 |
| chr9-50051724-DEL-119 | chr9-50051753-DEL-83 |
| chr9-50058455-DEL-63 | chr9-50058456-DEL-63 |
| chr9-50067293-DEL-2046 | chr9-50067293-DEL-2046 |
| chr9-50074058-DEL-1495 | chr9-50074059-DEL-1496 |
| chr9-50122513-DEL-67 | chr9-50122492-DEL-66 |
| chr9-50166951-DEL-86 | chr9-50166952-DEL-86 |
| chr9-50195242-DEL-6697 | chr9-50195243-DEL-6697 |
| chr9-50210516-DEL-326 | chr9-50210517-DEL-326 |
| chr9-50218871-DEL-602 | chr9-50218872-DEL-600 |
| chr9-50242996-DEL-52 | chr9-50242949-DEL-52 |
| chr9-50255487-DEL-796 | chr9-50255488-DEL-796 |
| chr9-50264850-DEL-1141 | chr9-50264851-DEL-1140 |

| | |
|---|---|
| chr9-50283006-DEL-164 | chr9-50283006-DEL-165 |
| chr9-50303665-DEL-1891 | chr9-50303666-DEL-1891 |
| chr9-50309046-DEL-131 | chr9-50309044-DEL-132 |
| chr9-50311865-DEL-246 | chr9-50311861-DEL-239 |
| chr9-50329627-DEL-1782 | chr9-50329628-DEL-1782 |
| chr9-50342909-DEL-77 | chr9-50342902-DEL-100 |
| chr9-50351206-DEL-433 | chr9-50351207-DEL-433 |
| chr9-50357400-DEL-70 | chr9-50357401-DEL-66 |
| chr9-50373382-DEL-144 | chr9-50373383-DEL-144 |
| chr9-50483366-DEL-56 | chr9-50483367-DEL-56 |
| chr9-50484286-DEL-74 | chr9-50484287-DEL-74 |
| chr9-50488209-DEL-304 | chr9-50488183-DEL-292 |
| chr9-50489541-DEL-202 | chr9-50489542-DEL-202 |
| chr9-50533828-DEL-766 | chr9-50533829-DEL-766 |
| chr9-50537573-DEL-5736 | chr9-50537574-DEL-5736 |
| chr9-50540667-DEL-217 | chr9-50540668-DEL-217 |
| chr9-50544675-DEL-418 | chr9-50544676-DEL-418 |
| chr9-50582404-DEL-422 | chr9-50582405-DEL-422 |
| chr9-50584904-DEL-308 | chr9-50584905-DEL-308 |
| chr9-50603106-DEL-198 | chr9-50603107-DEL-198 |
| chr9-50620104-DEL-254 | chr9-50620114-DEL-242 |
| chr9-50642415-DEL-209 | chr9-50642416-DEL-209 |
| chr9-50673922-DEL-53 | chr9-50673924-DEL-54 |
| chr9-50704366-DEL-199 | chr9-50704367-DEL-199 |
| chr9-50726063-DEL-408 | chr9-50726064-DEL-408 |
| chr9-50740698-DEL-519 | chr9-50740699-DEL-520 |
| chr9-50780951-DEL-66 | chr9-50780957-DEL-60 |
| chr9-50786274-DEL-189 | chr9-50786275-DEL-189 |
| chr9-50871976-DEL-51 | chr9-50871964-DEL-52 |
| chr9-50901587-DEL-942 | chr9-50901588-DEL-942 |
| chr9-50901715-DEL-253 | chr9-50901716-DEL-253 |
| chr9-50959805-DEL-53 | chr9-50959795-DEL-54 |
| chr9-50963961-DEL-2773 | chr9-50963962-DEL-2773 |
| chr9-50976638-DEL-51 | chr9-50976639-DEL-51 |
| chr9-51001434-DEL-137 | chr9-51001434-DEL-137 |
| chr9-51082999-DEL-84 | chr9-51083000-DEL-84 |
| chr9-51099455-DEL-825 | chr9-51099456-DEL-823 |
| chr9-51166111-DEL-1013 | chr9-51166112-DEL-1013 |
| chr9-51167505-DEL-52 | chr9-51167488-DEL-76 |
| chr9-51200313-DEL-63 | chr9-51200318-DEL-64 |
| chr9-51248693-DEL-865 | chr9-51248689-DEL-863 |
| chr9-51273529-DEL-51 | chr9-51273526-DEL-55 |
| chr9-51287910-DEL-1306 | chr9-51287911-DEL-1306 |
| chr9-51290365-DEL-56 | chr9-51290360-DEL-62 |
| chr9-51293403-DEL-90 | chr9-51293404-DEL-90 |
| chr9-51342576-DEL-97 | chr9-51342573-DEL-77 |
| chr9-51350191-DEL-94 | chr9-51350230-DEL-102 |
| chr9-51355796-DEL-465 | chr9-51355800-DEL-466 |
| chr9-51366612-DEL-217 | chr9-51366613-DEL-217 |
| chr9-51369838-DEL-161 | chr9-51369839-DEL-161 |
| chr9-51422837-DEL-1132 | chr9-51422835-DEL-1133 |
| chr9-51464809-DEL-96 | chr9-51464810-DEL-96 |
| chr9-51475872-DEL-6667 | chr9-51475873-DEL-6667 |
| chr9-51523614-DEL-8148 | chr9-51523615-DEL-8148 |
| chr9-51578375-DEL-1856 | chr9-51578376-DEL-1856 |
| chr9-51616544-DEL-5896 | chr9-51616545-DEL-5896 |
| chr9-51649202-DEL-806 | chr9-51649203-DEL-806 |
| chr9-51654058-DEL-206 | chr9-51654052-DEL-200 |
| chr9-51666815-DEL-629 | chr9-51666816-DEL-629 |
| chr9-51672479-DEL-99 | chr9-51672506-DEL-50 |
| chr9-51695051-DEL-966 | chr9-51695052-DEL-966 |
| chr9-51714843-DEL-300 | chr9-51714844-DEL-298 |
| chr9-51719520-DEL-835 | chr9-51719521-DEL-835 |
| chr9-51733881-DEL-64 | chr9-51733894-DEL-52 |
| chr9-51757910-DEL-225 | chr9-51757911-DEL-225 |
| chr9-51762793-DEL-111 | chr9-51762794-DEL-111 |
| chr9-51807864-DEL-333 | chr9-51807878-DEL-310 |
| chr9-51829177-DEL-70 | chr9-51829185-DEL-70 |
| chr9-51842114-DEL-56 | chr9-51842115-DEL-56 |
| chr9-51894516-DEL-1131 | chr9-51894516-DEL-1131 |
| chr9-51926780-DEL-731 | chr9-51926781-DEL-731 |
| chr9-51946308-DEL-165 | chr9-51946268-DEL-160 |
| chr9-51956561-DEL-58 | chr9-51956562-DEL-58 |
| chr9-51982645-DEL-56 | chr9-51982645-DEL-56 |
| chr9-51985657-DEL-374 | chr9-51985658-DEL-374 |
| chr9-52035496-DEL-208 | chr9-52035497-DEL-208 |
| chr9-52088023-DEL-62 | chr9-52088015-DEL-63 |
| chr9-52093871-DEL-1111 | chr9-52093872-DEL-1111 |
| chr9-52107825-DEL-971 | chr9-52107826-DEL-971 |
| chr9-52230133-DEL-97 | chr9-52230131-DEL-98 |

| | |
|---|---|
| chr9-52247397-DEL-62 | chr9-52247398-DEL-61 |
| chr9-52259208-DEL-3999 | chr9-52259209-DEL-3999 |
| chr9-52309434-DEL-84 | chr9-52309438-DEL-80 |
| chr9-52311613-DEL-403 | chr9-52311614-DEL-403 |
| chr9-52312118-DEL-257 | chr9-52312119-DEL-257 |
| chr9-52325308-DEL-88 | chr9-52325309-DEL-92 |
| chr9-52330110-DEL-3514 | chr9-52330111-DEL-3514 |
| chr9-52348149-DEL-1813 | chr9-52348150-DEL-1814 |
| chr9-52356732-DEL-52 | chr9-52356744-DEL-60 |
| chr9-52364636-DEL-196 | chr9-52364637-DEL-196 |
| chr9-52367796-DEL-1668 | chr9-52367797-DEL-1668 |
| chr9-52379543-DEL-1983 | chr9-52379544-DEL-1983 |
| chr9-52381232-DEL-56 | chr9-52381233-DEL-56 |
| chr9-52382411-DEL-952 | chr9-52382412-DEL-952 |
| chr9-52410040-DEL-98 | chr9-52410041-DEL-98 |
| chr9-52445454-DEL-56 | chr9-52445451-DEL-60 |
| chr9-52472589-DEL-200 | chr9-52472590-DEL-201 |
| chr9-52474759-DEL-80 | chr9-52474757-DEL-96 |
| chr9-52478472-DEL-97 | chr9-52478470-DEL-81 |
| chr9-52497614-DEL-653 | chr9-52497614-DEL-653 |
| chr9-52522093-DEL-6382 | chr9-52522093-DEL-6382 |
| chr9-52559243-DEL-666 | chr9-52559244-DEL-666 |
| chr9-52607103-DEL-142 | chr9-52607104-DEL-142 |
| chr9-52642875-DEL-69 | chr9-52642876-DEL-69 |
| chr9-52661015-DEL-6434 | chr9-52661016-DEL-6434 |
| chr9-52670230-DEL-262 | chr9-52670231-DEL-262 |
| chr9-52719841-DEL-216 | chr9-52719841-DEL-217 |
| chr9-52721544-DEL-130 | chr9-52721545-DEL-130 |
| chr9-52760452-DEL-2076 | chr9-52760453-DEL-2076 |
| chr9-52832594-DEL-498 | chr9-52832595-DEL-498 |
| chr9-52857093-DEL-62 | chr9-52857093-DEL-63 |
| chr9-52865844-DEL-98 | chr9-52865845-DEL-98 |
| chr9-52903844-DEL-6384 | chr9-52903845-DEL-6384 |
| chr9-52915901-DEL-129 | chr9-52915902-DEL-129 |
| chr9-52924765-DEL-6228 | chr9-52924766-DEL-6228 |
| chr9-52937507-DEL-91 | chr9-52937508-DEL-91 |
| chr9-52944705-DEL-126 | chr9-52944706-DEL-126 |
| chr9-52949889-DEL-978 | chr9-52949890-DEL-978 |
| chr9-52969690-DEL-647 | chr9-52969691-DEL-647 |
| chr9-52975163-DEL-145 | chr9-52975164-DEL-145 |
| chr9-52986755-DEL-187 | chr9-52986756-DEL-187 |
| chr9-53026935-DEL-145 | chr9-53026936-DEL-145 |
| chr9-53058035-DEL-1303 | chr9-53058036-DEL-1303 |
| chr9-53079078-DEL-135 | chr9-53079079-DEL-135 |
| chr9-53166042-DEL-203 | chr9-53166043-DEL-203 |
| chr9-53184310-DEL-189 | chr9-53184311-DEL-189 |
| chr9-53199381-DEL-285 | chr9-53199388-DEL-279 |
| chr9-53241470-DEL-74 | chr9-53241472-DEL-83 |
| chr9-53265090-DEL-680 | chr9-53265091-DEL-680 |
| chr9-53308354-DEL-176 | chr9-53308355-DEL-176 |
| chr9-53324414-DEL-154 | chr9-53324416-DEL-153 |
| chr9-53340381-DEL-179 | chr9-53340382-DEL-179 |
| chr9-53369460-DEL-233 | chr9-53369475-DEL-235 |
| chr9-53372415-DEL-58 | chr9-53372425-DEL-56 |
| chr9-53380655-DEL-204 | chr9-53380652-DEL-205 |
| chr9-53385103-DEL-1313 | chr9-53385104-DEL-1313 |
| chr9-53388302-DEL-223 | chr9-53388303-DEL-223 |
| chr9-53419783-DEL-119 | chr9-53419824-DEL-79 |
| chr9-53426945-DEL-74 | chr9-53426930-DEL-68 |
| chr9-53430318-DEL-268 | chr9-53430323-DEL-267 |
| chr9-53431785-DEL-189 | chr9-53431786-DEL-189 |
| chr9-53454089-DEL-201 | chr9-53454090-DEL-201 |
| chr9-53464542-DEL-60 | chr9-53464552-DEL-75 |
| chr9-53507569-DEL-1687 | chr9-53507570-DEL-1689 |
| chr9-53507635-DEL-293 | chr9-53507636-DEL-293 |
| chr9-53514021-DEL-202 | chr9-53514022-DEL-202 |
| chr9-53525742-DEL-83 | chr9-53525743-DEL-83 |
| chr9-53531366-DEL-88 | chr9-53531367-DEL-88 |
| chr9-53538657-DEL-435 | chr9-53538657-DEL-435 |
| chr9-53574114-DEL-214 | chr9-53574117-DEL-212 |
| chr9-53600777-DEL-869 | chr9-53600781-DEL-884 |
| chr9-53606243-DEL-62 | chr9-53606243-DEL-62 |
| chr9-53620306-DEL-141 | chr9-53620276-DEL-123 |
| chr9-53627096-DEL-86 | chr9-53627106-DEL-77 |
| chr9-53634375-DEL-173 | chr9-53634370-DEL-155 |
| chr9-53635502-DEL-7197 | chr9-53635503-DEL-7197 |
| chr9-53652706-DEL-53 | chr9-53652706-DEL-53 |
| chr9-53663167-DEL-1229 | chr9-53663168-DEL-1229 |
| chr9-53685412-DEL-206 | chr9-53685413-DEL-206 |
| chr9-53711320-DEL-1293 | chr9-53711321-DEL-1293 |

| | |
|---|---|
| chr9-53718785-DEL-721 | chr9-53718786-DEL-722 |
| chr9-53728419-DEL-75 | chr9-53728424-DEL-71 |
| chr9-53741745-DEL-88 | chr9-53741702-DEL-132 |
| chr9-53749585-DEL-65 | chr9-53749586-DEL-65 |
| chr9-53750549-DEL-54 | chr9-53750577-DEL-76 |
| chr9-53753530-DEL-60 | chr9-53753531-DEL-82 |
| chr9-53772335-DEL-204 | chr9-53772336-DEL-204 |
| chr9-53785089-DEL-1063 | chr9-53785090-DEL-1063 |
| chr9-53804809-DEL-349 | chr9-53804810-DEL-349 |
| chr9-53824663-DEL-95 | chr9-53824664-DEL-95 |
| chr9-53841167-DEL-53 | chr9-53841163-DEL-54 |
| chr9-53866448-DEL-6226 | chr9-53866449-DEL-6226 |
| chr9-53923602-DEL-4456 | chr9-53923603-DEL-4456 |
| chr9-53939690-DEL-1129 | chr9-53939691-DEL-1129 |
| chr9-53946120-DEL-1654 | chr9-53946121-DEL-1654 |
| chr9-54022796-DEL-81 | chr9-54022797-DEL-81 |
| chr9-54032538-DEL-6552 | chr9-54032539-DEL-6552 |
| chr9-54046687-DEL-5670 | chr9-54046688-DEL-5670 |
| chr9-54072474-DEL-328 | chr9-54072475-DEL-328 |
| chr9-54073488-DEL-104 | chr9-54073491-DEL-102 |
| chr9-54096590-DEL-80 | chr9-54096591-DEL-84 |
| chr9-54198398-DEL-4765 | chr9-54198399-DEL-4765 |
| chr9-54223822-DEL-1212 | chr9-54223823-DEL-1212 |
| chr9-54264335-DEL-6904 | chr9-54264336-DEL-6904 |
| chr9-54271860-DEL-189 | chr9-54271861-DEL-189 |
| chr9-54326870-DEL-61 | chr9-54326871-DEL-61 |
| chr9-54337854-DEL-55 | chr9-54337850-DEL-60 |
| chr9-54340847-DEL-442 | chr9-54340802-DEL-476 |
| chr9-54355575-DEL-97 | chr9-54355576-DEL-97 |
| chr9-54386213-DEL-211 | chr9-54386214-DEL-211 |
| chr9-54404723-DEL-55 | chr9-54404728-DEL-54 |
| chr9-54460993-DEL-78 | chr9-54460989-DEL-83 |
| chr9-54491660-DEL-203 | chr9-54491661-DEL-203 |
| chr9-54507425-DEL-66 | chr9-54507444-DEL-76 |
| chr9-54529608-DEL-55 | chr9-54529646-DEL-55 |
| chr9-54533370-DEL-172 | chr9-54533371-DEL-172 |
| chr9-54559280-DEL-74 | chr9-54559281-DEL-74 |
| chr9-54589571-DEL-231 | chr9-54589593-DEL-209 |
| chr9-54648491-DEL-96 | chr9-54648492-DEL-96 |
| chr9-54650637-DEL-70 | chr9-54650638-DEL-70 |
| chr9-54665851-DEL-181 | chr9-54665852-DEL-181 |
| chr9-54671006-DEL-52 | chr9-54671007-DEL-52 |
| chr9-54680645-DEL-69 | chr9-54680644-DEL-67 |
| chr9-54688029-DEL-113 | chr9-54688018-DEL-125 |
| chr9-54737926-DEL-1592 | chr9-54737927-DEL-1592 |
| chr9-54743331-DEL-338 | chr9-54743308-DEL-350 |
| chr9-54748124-DEL-61 | chr9-54748116-DEL-52 |
| chr9-54750248-DEL-60 | chr9-54750249-DEL-60 |
| chr9-54812234-DEL-668 | chr9-54812235-DEL-668 |
| chr9-54822309-DEL-82 | chr9-54822322-DEL-70 |
| chr9-54830183-DEL-6231 | chr9-54830184-DEL-6231 |
| chr9-54878136-DEL-421 | chr9-54878137-DEL-421 |
| chr9-54878842-DEL-381 | chr9-54878843-DEL-381 |
| chr9-54894403-DEL-419 | chr9-54894404-DEL-419 |
| chr9-54905941-DEL-120 | chr9-54905942-DEL-120 |
| chr9-54910201-DEL-88 | chr9-54910202-DEL-88 |
| chr9-54924940-DEL-54 | chr9-54924897-DEL-73 |
| chr9-54963058-DEL-120 | chr9-54963059-DEL-120 |
| chr9-54967262-DEL-186 | chr9-54967267-DEL-186 |
| chr9-54971638-DEL-120 | chr9-54971669-DEL-114 |
| chr9-54978047-DEL-7167 | chr9-54978048-DEL-7167 |
| chr9-55012646-DEL-58 | chr9-55012647-DEL-58 |
| chr9-55016911-DEL-487 | chr9-55016912-DEL-487 |
| chr9-55018337-DEL-207 | chr9-55018338-DEL-207 |
| chr9-55023972-DEL-220 | chr9-55023973-DEL-220 |
| chr9-55026540-DEL-1863 | chr9-55026540-DEL-1863 |
| chr9-55028010-DEL-222 | chr9-55028011-DEL-222 |
| chr9-55039515-DEL-208 | chr9-55039516-DEL-208 |
| chr9-55144482-DEL-155 | chr9-55144483-DEL-155 |
| chr9-55167696-DEL-132 | chr9-55167697-DEL-132 |
| chr9-55170218-DEL-60 | chr9-55170218-DEL-60 |
| chr9-55201212-DEL-117 | chr9-55201211-DEL-118 |
| chr9-55206672-DEL-1465 | chr9-55206673-DEL-1465 |
| chr9-55278504-DEL-64 | chr9-55278505-DEL-65 |
| chr9-55303361-DEL-54 | chr9-55303362-DEL-54 |
| chr9-55326758-DEL-83 | chr9-55326759-DEL-83 |
| chr9-55348169-DEL-78 | chr9-55348170-DEL-78 |
| chr9-55349041-DEL-1049 | chr9-55349042-DEL-1049 |
| chr9-55349319-DEL-223 | chr9-55349320-DEL-223 |
| chr9-55386905-DEL-66 | chr9-55386882-DEL-90 |

| | |
|---|---|
| chr9-55388263-DEL-169 | chr9-55388257-DEL-170 |
| chr9-55393388-DEL-162 | chr9-55393389-DEL-162 |
| chr9-55421746-DEL-55 | chr9-55421747-DEL-55 |
| chr9-55428452-DEL-497 | chr9-55428453-DEL-497 |
| chr9-55480414-DEL-425 | chr9-55480415-DEL-425 |
| chr9-55539023-DEL-6587 | chr9-55539024-DEL-6587 |
| chr9-55586150-DEL-992 | chr9-55586151-DEL-992 |
| chr9-55681399-DEL-146 | chr9-55681400-DEL-146 |
| chr9-55700686-DEL-4985 | chr9-55700687-DEL-4985 |
| chr9-55717523-DEL-66 | chr9-55717524-DEL-66 |
| chr9-55735912-DEL-6552 | chr9-55735912-DEL-6552 |
| chr9-55758823-DEL-6329 | chr9-55758822-DEL-6336 |
| chr9-55763681-DEL-196 | chr9-55763682-DEL-196 |
| chr9-55789549-DEL-90 | chr9-55789550-DEL-90 |
| chr9-55833951-DEL-60 | chr9-55833952-DEL-60 |
| chr9-55835627-DEL-1697 | chr9-55835628-DEL-1697 |
| chr9-55853012-DEL-71 | chr9-55852996-DEL-50 |
| chr9-55873570-DEL-6385 | chr9-55873569-DEL-6385 |
| chr9-55884788-DEL-1654 | chr9-55884789-DEL-1654 |
| chr9-55927408-DEL-53 | chr9-55927379-DEL-65 |
| chr9-55937922-DEL-862 | chr9-55937923-DEL-861 |
| chr9-55941626-DEL-164 | chr9-55941627-DEL-164 |
| chr9-55989286-DEL-399 | chr9-55989287-DEL-399 |
| chr9-55989869-DEL-56 | chr9-55989870-DEL-56 |
| chr9-56033557-DEL-184 | chr9-56033546-DEL-183 |
| chr9-56130775-DEL-466 | chr9-56130775-DEL-464 |
| chr9-56136796-DEL-202 | chr9-56136797-DEL-202 |
| chr9-56141104-DEL-185 | chr9-56141105-DEL-185 |
| chr9-56152822-DEL-226 | chr9-56152823-DEL-226 |
| chr9-56162949-DEL-891 | chr9-56162950-DEL-891 |
| chr9-56259369-DEL-331 | chr9-56259370-DEL-331 |
| chr9-56260825-DEL-81 | chr9-56260826-DEL-81 |
| chr9-56267650-DEL-79 | chr9-56267651-DEL-79 |
| chr9-56336629-DEL-1894 | chr9-56336630-DEL-1894 |
| chr9-56344695-DEL-1039 | chr9-56344696-DEL-1039 |
| chr9-56346755-DEL-56 | chr9-56346756-DEL-56 |
| chr9-56358570-DEL-219 | chr9-56358571-DEL-219 |
| chr9-56359681-DEL-169 | chr9-56359687-DEL-172 |
| chr9-56425847-DEL-174 | chr9-56425848-DEL-174 |
| chr9-56436491-DEL-55 | chr9-56436492-DEL-55 |
| chr9-56474872-DEL-171 | chr9-56474873-DEL-171 |
| chr9-56507621-DEL-76 | chr9-56507622-DEL-76 |
| chr9-56511192-DEL-72 | chr9-56511193-DEL-72 |
| chr9-56532591-DEL-192 | chr9-56532592-DEL-192 |
| chr9-56586066-DEL-200 | chr9-56586082-DEL-199 |
| chr9-56637851-DEL-58 | chr9-56637852-DEL-58 |
| chr9-56643648-DEL-59 | chr9-56643649-DEL-59 |
| chr9-56649724-DEL-217 | chr9-56649725-DEL-217 |
| chr9-56679562-DEL-98 | chr9-56679551-DEL-98 |
| chr9-56724525-DEL-61 | chr9-56724545-DEL-58 |
| chr9-56780867-DEL-79 | chr9-56780868-DEL-79 |
| chr9-56810260-DEL-215 | chr9-56810261-DEL-215 |
| chr9-56855956-DEL-170 | chr9-56855957-DEL-170 |
| chr9-56896252-DEL-216 | chr9-56896253-DEL-216 |
| chr9-56899042-DEL-1886 | chr9-56899039-DEL-1887 |
| chr9-56915026-DEL-101 | chr9-56915024-DEL-57 |
| chr9-56945359-DEL-219 | chr9-56945360-DEL-219 |
| chr9-56949706-DEL-71 | chr9-56949707-DEL-69 |
| chr9-56991548-DEL-1911 | chr9-56991549-DEL-1911 |
| chr9-57016351-DEL-355 | chr9-57016352-DEL-355 |
| chr9-57059051-DEL-186 | chr9-57059052-DEL-186 |
| chr9-57065730-DEL-120 | chr9-57065730-DEL-120 |
| chr9-57073128-DEL-3640 | chr9-57073130-DEL-3638 |
| chr9-57078694-DEL-90 | chr9-57078695-DEL-90 |
| chr9-57081950-DEL-192 | chr9-57081951-DEL-192 |
| chr9-57103813-DEL-1115 | chr9-57103814-DEL-1115 |
| chr9-57111210-DEL-72 | chr9-57111210-DEL-72 |
| chr9-57114671-DEL-146 | chr9-57114688-DEL-140 |
| chr9-57129389-DEL-359 | chr9-57129395-DEL-355 |
| chr9-57132433-DEL-317 | chr9-57132434-DEL-317 |
| chr9-57172230-DEL-162 | chr9-57172231-DEL-162 |
| chr9-57233795-DEL-1109 | chr9-57233796-DEL-1109 |
| chr9-57241347-DEL-759 | chr9-57241348-DEL-759 |
| chr9-57275211-DEL-414 | chr9-57275212-DEL-414 |
| chr9-57291267-DEL-145 | chr9-57291268-DEL-145 |
| chr9-57315269-DEL-1483 | chr9-57315270-DEL-1483 |
| chr9-57356041-DEL-204 | chr9-57356042-DEL-204 |
| chr9-57361606-DEL-63 | chr9-57361607-DEL-63 |
| chr9-57366470-DEL-68 | chr9-57366471-DEL-68 |
| chr9-57379109-DEL-220 | chr9-57379110-DEL-220 |

| | |
|---|---|
| chr9-57382255-DEL-169 | chr9-57382262-DEL-167 |
| chr9-57384379-DEL-4243 | chr9-57384380-DEL-4243 |
| chr9-57387774-DEL-166 | chr9-57387775-DEL-166 |
| chr9-57397188-DEL-60 | chr9-57397191-DEL-53 |
| chr9-57420329-DEL-100 | chr9-57420326-DEL-104 |
| chr9-57435059-DEL-213 | chr9-57435060-DEL-213 |
| chr9-57463862-DEL-174 | chr9-57463863-DEL-182 |
| chr9-57528990-DEL-285 | chr9-57528991-DEL-285 |
| chr9-57556241-DEL-228 | chr9-57556251-DEL-229 |
| chr9-57591885-DEL-143 | chr9-57591891-DEL-143 |
| chr9-57608434-DEL-57 | chr9-57608431-DEL-57 |
| chr9-57614870-DEL-184 | chr9-57614871-DEL-184 |
| chr9-57624760-DEL-212 | chr9-57624761-DEL-212 |
| chr9-57675170-DEL-823 | chr9-57675171-DEL-823 |
| chr9-57721058-DEL-71 | chr9-57721059-DEL-71 |
| chr9-57722526-DEL-149 | chr9-57722575-DEL-101 |
| chr9-57726887-DEL-1453 | chr9-57726888-DEL-1453 |
| chr9-57748374-DEL-307 | chr9-57748375-DEL-307 |
| chr9-57749654-DEL-104 | chr9-57749655-DEL-104 |
| chr9-57766315-DEL-2121 | chr9-57766316-DEL-2121 |
| chr9-57768529-DEL-234 | chr9-57768530-DEL-234 |
| chr9-57777555-DEL-56 | chr9-57777558-DEL-54 |
| chr9-57785056-DEL-191 | chr9-57785057-DEL-191 |
| chr9-57801636-DEL-55 | chr9-57801633-DEL-55 |
| chr9-57803070-DEL-54 | chr9-57803071-DEL-54 |
| chr9-57820678-DEL-431 | chr9-57820679-DEL-431 |
| chr9-57828083-DEL-102 | chr9-57828075-DEL-132 |
| chr9-57832283-DEL-59 | chr9-57832284-DEL-59 |
| chr9-57832743-DEL-233 | chr9-57832744-DEL-233 |
| chr9-57836231-DEL-88 | chr9-57836232-DEL-88 |
| chr9-57877997-DEL-192 | chr9-57877998-DEL-192 |
| chr9-57882203-DEL-401 | chr9-57882204-DEL-401 |
| chr9-57891741-DEL-202 | chr9-57891742-DEL-202 |
| chr9-57941932-DEL-107 | chr9-57941975-DEL-110 |
| chr9-58015336-DEL-320 | chr9-58015337-DEL-310 |
| chr9-58035359-DEL-202 | chr9-58035360-DEL-202 |
| chr9-58077153-DEL-197 | chr9-58077154-DEL-197 |
| chr9-58099877-DEL-213 | chr9-58099878-DEL-213 |
| chr9-58166715-DEL-200 | chr9-58166716-DEL-200 |
| chr9-58168365-DEL-57 | chr9-58168365-DEL-57 |
| chr9-58170482-DEL-243 | chr9-58170483-DEL-243 |
| chr9-58201258-DEL-203 | chr9-58201259-DEL-203 |
| chr9-58233125-DEL-199 | chr9-58233126-DEL-199 |
| chr9-58239674-DEL-456 | chr9-58239675-DEL-456 |
| chr9-58270929-DEL-104 | chr9-58270907-DEL-109 |
| chr9-58323088-DEL-56 | chr9-58323088-DEL-57 |
| chr9-58337554-DEL-56 | chr9-58337555-DEL-50 |
| chr9-58361605-DEL-144 | chr9-58361606-DEL-144 |
| chr9-58388130-DEL-77 | chr9-58388127-DEL-74 |
| chr9-58468193-DEL-885 | chr9-58468194-DEL-885 |
| chr9-58480568-DEL-131 | chr9-58480585-DEL-129 |
| chr9-58482463-DEL-260 | chr9-58482468-DEL-261 |
| chr9-58483636-DEL-235 | chr9-58483637-DEL-235 |
| chr9-58487873-DEL-56 | chr9-58487874-DEL-56 |
| chr9-58495118-DEL-758 | chr9-58495118-DEL-758 |
| chr9-58503407-DEL-166 | chr9-58503408-DEL-166 |
| chr9-58513464-DEL-157 | chr9-58513465-DEL-152 |
| chr9-58515508-DEL-94 | chr9-58515545-DEL-53 |
| chr9-58531416-DEL-200 | chr9-58531417-DEL-200 |
| chr9-58532063-DEL-60 | chr9-58532052-DEL-72 |
| chr9-58535564-DEL-202 | chr9-58535565-DEL-203 |
| chr9-58573936-DEL-209 | chr9-58573937-DEL-209 |
| chr9-58586002-DEL-84 | chr9-58586003-DEL-84 |
| chr9-58593042-DEL-1252 | chr9-58593043-DEL-1252 |
| chr9-58599072-DEL-199 | chr9-58599073-DEL-199 |
| chr9-58603886-DEL-1889 | chr9-58603887-DEL-1889 |
| chr9-58616311-DEL-67 | chr9-58616312-DEL-67 |
| chr9-58621106-DEL-226 | chr9-58621107-DEL-226 |
| chr9-58636141-DEL-54 | chr9-58636142-DEL-54 |
| chr9-58664793-DEL-152 | chr9-58664794-DEL-152 |
| chr9-58668624-DEL-66 | chr9-58668618-DEL-64 |
| chr9-58674772-DEL-101 | chr9-58674773-DEL-101 |
| chr9-58678918-DEL-197 | chr9-58678919-DEL-197 |
| chr9-58685651-DEL-7316 | chr9-58685653-DEL-7315 |
| chr9-58720184-DEL-54 | chr9-58720185-DEL-56 |
| chr9-58744274-DEL-156 | chr9-58744275-DEL-156 |
| chr9-58835279-DEL-592 | chr9-58835280-DEL-592 |
| chr9-58847382-DEL-1302 | chr9-58847383-DEL-1302 |
| chr9-58856917-DEL-1576 | chr9-58856918-DEL-1576 |
| chr9-58886467-DEL-60 | chr9-58886468-DEL-60 |

| | |
|---|---|
| chr9-58926917-DEL-518 | chr9-58926918-DEL-518 |
| chr9-58953547-DEL-6452 | chr9-58953548-DEL-6452 |
| chr9-59000351-DEL-270 | chr9-59000386-DEL-259 |
| chr9-59025687-DEL-70 | chr9-59025688-DEL-56 |
| chr9-59026430-DEL-6176 | chr9-59026431-DEL-6176 |
| chr9-59065154-DEL-63 | chr9-59065156-DEL-61 |
| chr9-59083866-DEL-319 | chr9-59083867-DEL-319 |
| chr9-59094618-DEL-86 | chr9-59094608-DEL-92 |
| chr9-59118352-DEL-205 | chr9-59118353-DEL-205 |
| chr9-59119148-DEL-2433 | chr9-59119157-DEL-2433 |
| chr9-59131631-DEL-234 | chr9-59131632-DEL-234 |
| chr9-59149003-DEL-218 | chr9-59149004-DEL-218 |
| chr9-59150035-DEL-62 | chr9-59150044-DEL-54 |
| chr9-59160900-DEL-76 | chr9-59160890-DEL-86 |
| chr9-59165763-DEL-160 | chr9-59165764-DEL-160 |
| chr9-59172340-DEL-64 | chr9-59172341-DEL-58 |
| chr9-59202005-DEL-55 | chr9-59201974-DEL-55 |
| chr9-59242839-DEL-61 | chr9-59242840-DEL-61 |
| chr9-59251605-DEL-76 | chr9-59251606-DEL-76 |
| chr9-59283419-DEL-217 | chr9-59283420-DEL-217 |
| chr9-59347805-DEL-136 | chr9-59347806-DEL-136 |
| chr9-59409521-DEL-79 | chr9-59409522-DEL-79 |
| chr9-59490339-DEL-285 | chr9-59490344-DEL-284 |
| chr9-59576812-DEL-135 | chr9-59576813-DEL-147 |
| chr9-59584443-DEL-201 | chr9-59584444-DEL-201 |
| chr9-59620993-DEL-57 | chr9-59620994-DEL-57 |
| chr9-59625755-DEL-60 | chr9-59625756-DEL-60 |
| chr9-59634388-DEL-80 | chr9-59634389-DEL-80 |
| chr9-59664240-DEL-211 | chr9-59664241-DEL-211 |
| chr9-59674577-DEL-198 | chr9-59674578-DEL-198 |
| chr9-59681938-DEL-58 | chr9-59681916-DEL-64 |
| chr9-59699293-DEL-207 | chr9-59699294-DEL-207 |
| chr9-59715487-DEL-155 | chr9-59715488-DEL-155 |
| chr9-59738305-DEL-216 | chr9-59738306-DEL-216 |
| chr9-59781481-DEL-57 | chr9-59781482-DEL-57 |
| chr9-59784608-DEL-355 | chr9-59784609-DEL-355 |
| chr9-59863590-DEL-130 | chr9-59863616-DEL-94 |
| chr9-59865117-DEL-1145 | chr9-59865118-DEL-1145 |
| chr9-59920082-DEL-327 | chr9-59920083-DEL-327 |
| chr9-59952770-DEL-6320 | chr9-59952771-DEL-6320 |
| chr9-59976721-DEL-506 | chr9-59976722-DEL-506 |
| chr9-60077896-DEL-104 | chr9-60077907-DEL-103 |
| chr9-60081352-DEL-241 | chr9-60081353-DEL-241 |
| chr9-60100669-DEL-195 | chr9-60100670-DEL-195 |
| chr9-60121257-DEL-80 | chr9-60121287-DEL-52 |
| chr9-60124241-DEL-630 | chr9-60124242-DEL-630 |
| chr9-60169632-DEL-115 | chr9-60169633-DEL-115 |
| chr9-60225112-DEL-358 | chr9-60225113-DEL-358 |
| chr9-60235154-DEL-70 | chr9-60235161-DEL-62 |
| chr9-60235279-DEL-451 | chr9-60235280-DEL-451 |
| chr9-60278381-DEL-217 | chr9-60278382-DEL-217 |
| chr9-60294623-DEL-633 | chr9-60294624-DEL-633 |
| chr9-60307349-DEL-86 | chr9-60307350-DEL-86 |
| chr9-60316682-DEL-1677 | chr9-60316683-DEL-1677 |
| chr9-60329701-DEL-418 | chr9-60329702-DEL-418 |
| chr9-60333981-DEL-504 | chr9-60333982-DEL-504 |
| chr9-60366355-DEL-62 | chr9-60366338-DEL-58 |
| chr9-60366602-DEL-349 | chr9-60366594-DEL-349 |
| chr9-60418925-DEL-74 | chr9-60418924-DEL-74 |
| chr9-60419575-DEL-60 | chr9-60419572-DEL-56 |
| chr9-60422926-DEL-67 | chr9-60422926-DEL-68 |
| chr9-60431772-DEL-56 | chr9-60431772-DEL-56 |
| chr9-60454446-DEL-967 | chr9-60454447-DEL-967 |
| chr9-60457085-DEL-1427 | chr9-60457079-DEL-1429 |
| chr9-60483579-DEL-493 | chr9-60483580-DEL-493 |
| chr9-60500671-DEL-148 | chr9-60500672-DEL-148 |
| chr9-60504531-DEL-65 | chr9-60504531-DEL-66 |
| chr9-60525262-DEL-229 | chr9-60525266-DEL-232 |
| chr9-60530829-DEL-76 | chr9-60530829-DEL-95 |
| chr9-60538977-DEL-262 | chr9-60538979-DEL-255 |
| chr9-60541654-DEL-169 | chr9-60541655-DEL-169 |
| chr9-60542812-DEL-59 | chr9-60542813-DEL-59 |
| chr9-60544948-DEL-102 | chr9-60544947-DEL-103 |
| chr9-60564267-DEL-450 | chr9-60564290-DEL-415 |
| chr9-60571806-DEL-56 | chr9-60571807-DEL-56 |
| chr9-60588671-DEL-205 | chr9-60588672-DEL-205 |
| chr9-60606445-DEL-56 | chr9-60606445-DEL-56 |
| chr9-60613710-DEL-55 | chr9-60613713-DEL-53 |
| chr9-60614529-DEL-252 | chr9-60614530-DEL-252 |
| chr9-60633492-DEL-7099 | chr9-60633493-DEL-7099 |

| | |
|---|---|
| chr9-60659036-DEL-139 | chr9-60659037-DEL-139 |
| chr9-60688594-DEL-154 | chr9-60688595-DEL-154 |
| chr9-60722001-DEL-119 | chr9-60722002-DEL-119 |
| chr9-60725669-DEL-398 | chr9-60725670-DEL-398 |
| chr9-60754999-DEL-118 | chr9-60755000-DEL-118 |
| chr9-60755629-DEL-88 | chr9-60755639-DEL-70 |
| chr9-60769391-DEL-699 | chr9-60769377-DEL-700 |
| chr9-60792914-DEL-71 | chr9-60792914-DEL-72 |
| chr9-60795115-DEL-235 | chr9-60795116-DEL-235 |
| chr9-60815043-DEL-4336 | chr9-60815044-DEL-4336 |
| chr9-60848934-DEL-206 | chr9-60848935-DEL-206 |
| chr9-60947363-DEL-82 | chr9-60947379-DEL-82 |
| chr9-60971347-DEL-62 | chr9-60971348-DEL-62 |
| chr9-61007679-DEL-93 | chr9-61007680-DEL-93 |
| chr9-61030555-DEL-60 | chr9-61030558-DEL-52 |
| chr9-61030575-DEL-879 | chr9-61030574-DEL-880 |
| chr9-61060555-DEL-221 | chr9-61060556-DEL-221 |
| chr9-61062403-DEL-60 | chr9-61062404-DEL-60 |
| chr9-61106956-DEL-82 | chr9-61106973-DEL-60 |
| chr9-61151088-DEL-51 | chr9-61151089-DEL-51 |
| chr9-61163809-DEL-75 | chr9-61163810-DEL-75 |
| chr9-61238936-DEL-115 | chr9-61238937-DEL-115 |
| chr9-61253340-DEL-105 | chr9-61253341-DEL-105 |
| chr9-61327692-DEL-51 | chr9-61327693-DEL-51 |
| chr9-61361727-DEL-86 | chr9-61361698-DEL-72 |
| chr9-61369577-DEL-51 | chr9-61369574-DEL-54 |
| chr9-61376162-DEL-62 | chr9-61376163-DEL-73 |
| chr9-61420735-DEL-149 | chr9-61420736-DEL-149 |
| chr9-61459401-DEL-157 | chr9-61459402-DEL-157 |
| chr9-61476238-DEL-98 | chr9-61476279-DEL-80 |
| chr9-61487653-DEL-80 | chr9-61487681-DEL-64 |
| chr9-61489509-DEL-130 | chr9-61489510-DEL-130 |
| chr9-61494819-DEL-62 | chr9-61494864-DEL-50 |
| chr9-61517742-DEL-579 | chr9-61517741-DEL-581 |
| chr9-61519584-DEL-101 | chr9-61519585-DEL-101 |
| chr9-61530879-DEL-193 | chr9-61530880-DEL-193 |
| chr9-61549938-DEL-58 | chr9-61549938-DEL-64 |
| chr9-61557810-DEL-99 | chr9-61557811-DEL-99 |
| chr9-61568736-DEL-76 | chr9-61568737-DEL-71 |
| chr9-61588633-DEL-54 | chr9-61588633-DEL-54 |
| chr9-61613291-DEL-248 | chr9-61613293-DEL-246 |
| chr9-61638838-DEL-93 | chr9-61638838-DEL-93 |
| chr9-61639215-DEL-334 | chr9-61639201-DEL-345 |
| chr9-61646195-DEL-162 | chr9-61646196-DEL-162 |
| chr9-61781180-DEL-218 | chr9-61781181-DEL-218 |
| chr9-61825312-DEL-398 | chr9-61825313-DEL-398 |
| chr9-61882244-DEL-53 | chr9-61882245-DEL-53 |
| chr9-61908610-DEL-55 | chr9-61908594-DEL-56 |
| chr9-61913848-DEL-105 | chr9-61913841-DEL-113 |
| chr9-61952041-DEL-213 | chr9-61952042-DEL-213 |
| chr9-61953070-DEL-167 | chr9-61953072-DEL-163 |
| chr9-62014542-DEL-270 | chr9-62014543-DEL-270 |
| chr9-62102024-DEL-187 | chr9-62102025-DEL-187 |
| chr9-62143302-DEL-432 | chr9-62143303-DEL-432 |
| chr9-62150907-DEL-146 | chr9-62150908-DEL-146 |
| chr9-62155849-DEL-336 | chr9-62155850-DEL-336 |
| chr9-62206909-DEL-54 | chr9-62206910-DEL-52 |
| chr9-62224559-DEL-56 | chr9-62224560-DEL-56 |
| chr9-62233698-DEL-209 | chr9-62233699-DEL-209 |
| chr9-62240408-DEL-198 | chr9-62240409-DEL-198 |
| chr9-62259884-DEL-232 | chr9-62259893-DEL-235 |
| chr9-62272138-DEL-95 | chr9-62272139-DEL-95 |
| chr9-62308482-DEL-90 | chr9-62308483-DEL-90 |
| chr9-62315986-DEL-1588 | chr9-62315987-DEL-1588 |
| chr9-62322956-DEL-64 | chr9-62322961-DEL-60 |
| chr9-62330977-DEL-123 | chr9-62330970-DEL-120 |
| chr9-62346452-DEL-8763 | chr9-62346452-DEL-8763 |
| chr9-62356155-DEL-99 | chr9-62356156-DEL-99 |
| chr9-62356365-DEL-434 | chr9-62356366-DEL-434 |
| chr9-62358739-DEL-644 | chr9-62358740-DEL-644 |
| chr9-62365364-DEL-52 | chr9-62365365-DEL-52 |
| chr9-62400213-DEL-70 | chr9-62400216-DEL-68 |
| chr9-62401142-DEL-55 | chr9-62401143-DEL-55 |
| chr9-62419812-DEL-90 | chr9-62419812-DEL-90 |
| chr9-62452602-DEL-2122 | chr9-62452603-DEL-2122 |
| chr9-62455469-DEL-672 | chr9-62455470-DEL-672 |
| chr9-62479600-DEL-323 | chr9-62479601-DEL-323 |
| chr9-62506612-DEL-6006 | chr9-62506613-DEL-6006 |
| chr9-62546632-DEL-3069 | chr9-62546634-DEL-3068 |
| chr9-62583395-DEL-96 | chr9-62583396-DEL-96 |

| | |
|---|---|
| chr9-62597171-DEL-87 | chr9-62597172-DEL-87 |
| chr9-62620913-DEL-226 | chr9-62620914-DEL-226 |
| chr9-62631121-DEL-105 | chr9-62631121-DEL-106 |
| chr9-62633509-DEL-226 | chr9-62633510-DEL-226 |
| chr9-62638668-DEL-144 | chr9-62638660-DEL-153 |
| chr9-62647147-DEL-465 | chr9-62647148-DEL-465 |
| chr9-62650403-DEL-65 | chr9-62650404-DEL-65 |
| chr9-62653299-DEL-66 | chr9-62653305-DEL-66 |
| chr9-62677120-DEL-356 | chr9-62677121-DEL-356 |
| chr9-62679341-DEL-88 | chr9-62679354-DEL-76 |
| chr9-62709420-DEL-201 | chr9-62709421-DEL-201 |
| chr9-62731754-DEL-197 | chr9-62731767-DEL-185 |
| chr9-62734520-DEL-107 | chr9-62734555-DEL-73 |
| chr9-62735100-DEL-79 | chr9-62735107-DEL-73 |
| chr9-62741141-DEL-387 | chr9-62741142-DEL-387 |
| chr9-62769106-DEL-6473 | chr9-62769107-DEL-6473 |
| chr9-62780775-DEL-217 | chr9-62780780-DEL-213 |
| chr9-62808346-DEL-190 | chr9-62808347-DEL-190 |
| chr9-62835852-DEL-209 | chr9-62835853-DEL-209 |
| chr9-62990596-DEL-208 | chr9-62990597-DEL-208 |
| chr9-63118327-DEL-898 | chr9-63118328-DEL-898 |
| chr9-63130016-DEL-92 | chr9-63130053-DEL-103 |
| chr9-63189436-DEL-51 | chr9-63189437-DEL-51 |
| chr9-63198988-DEL-61 | chr9-63198982-DEL-62 |
| chr9-63229904-DEL-72 | chr9-63229911-DEL-52 |
| chr9-63325140-DEL-223 | chr9-63325141-DEL-223 |
| chr9-63356151-DEL-68 | chr9-63356152-DEL-68 |
| chr9-63365493-DEL-220 | chr9-63365494-DEL-220 |
| chr9-63383486-DEL-108 | chr9-63383505-DEL-90 |
| chr9-63429882-DEL-206 | chr9-63429883-DEL-206 |
| chr9-63457949-DEL-4026 | chr9-63457950-DEL-4026 |
| chr9-63502226-DEL-54 | chr9-63502179-DEL-98 |
| chr9-63513036-DEL-249 | chr9-63513034-DEL-250 |
| chr9-63534984-DEL-231 | chr9-63534985-DEL-231 |
| chr9-63543938-DEL-333 | chr9-63543939-DEL-333 |
| chr9-63669439-DEL-139 | chr9-63669440-DEL-139 |
| chr9-63708361-DEL-86 | chr9-63708362-DEL-84 |
| chr9-63720187-DEL-190 | chr9-63720188-DEL-190 |
| chr9-63721326-DEL-65 | chr9-63721327-DEL-65 |
| chr9-63722892-DEL-254 | chr9-63722893-DEL-254 |
| chr9-63737791-DEL-180 | chr9-63737795-DEL-177 |
| chr9-63749033-DEL-327 | chr9-63749034-DEL-327 |
| chr9-63753586-DEL-207 | chr9-63753588-DEL-206 |
| chr9-63762763-DEL-94 | chr9-63762741-DEL-78 |
| chr9-63766857-DEL-729 | chr9-63766862-DEL-725 |
| chr9-63807224-DEL-277 | chr9-63807225-DEL-277 |
| chr9-63841959-DEL-84 | chr9-63841953-DEL-82 |
| chr9-63846372-DEL-266 | chr9-63846373-DEL-266 |
| chr9-63847115-DEL-60 | chr9-63847116-DEL-54 |
| chr9-63864222-DEL-78 | chr9-63864223-DEL-78 |
| chr9-63942519-DEL-200 | chr9-63942513-DEL-198 |
| chr9-63945044-DEL-258 | chr9-63945045-DEL-258 |
| chr9-63959271-DEL-210 | chr9-63959272-DEL-210 |
| chr9-63971491-DEL-56 | chr9-63971498-DEL-50 |
| chr9-64032019-DEL-119 | chr9-64032020-DEL-119 |
| chr9-64034761-DEL-204 | chr9-64034762-DEL-204 |
| chr9-64037648-DEL-201 | chr9-64037649-DEL-201 |
| chr9-64077207-DEL-194 | chr9-64077208-DEL-194 |
| chr9-64088056-DEL-89 | chr9-64088057-DEL-89 |
| chr9-64118135-DEL-177 | chr9-64118136-DEL-177 |
| chr9-64125013-DEL-738 | chr9-64125007-DEL-745 |
| chr9-64143906-DEL-1888 | chr9-64143907-DEL-1888 |
| chr9-64151836-DEL-216 | chr9-64151837-DEL-216 |
| chr9-64165745-DEL-1239 | chr9-64165744-DEL-1240 |
| chr9-64168768-DEL-101 | chr9-64168769-DEL-101 |
| chr9-64182857-DEL-407 | chr9-64182861-DEL-411 |
| chr9-64184428-DEL-168 | chr9-64184429-DEL-168 |
| chr9-64185484-DEL-366 | chr9-64185490-DEL-362 |
| chr9-64223903-DEL-213 | chr9-64223904-DEL-213 |
| chr9-64232458-DEL-69 | chr9-64232455-DEL-69 |
| chr9-64268454-DEL-123 | chr9-64268455-DEL-123 |
| chr9-64273638-DEL-56 | chr9-64273638-DEL-56 |
| chr9-64275500-DEL-57 | chr9-64275501-DEL-67 |
| chr9-64276785-DEL-56 | chr9-64276786-DEL-56 |
| chr9-64280192-DEL-421 | chr9-64280189-DEL-423 |
| chr9-64287585-DEL-307 | chr9-64287586-DEL-307 |
| chr9-64290249-DEL-105 | chr9-64290265-DEL-56 |
| chr9-64301968-DEL-54 | chr9-64301969-DEL-54 |
| chr9-64310181-DEL-55 | chr9-64310182-DEL-55 |
| chr9-64316880-DEL-73 | chr9-64316881-DEL-73 |

| | |
|---|---|
| chr9-64317776-DEL-7115 | chr9-64317777-DEL-7115 |
| chr9-64326621-DEL-81 | chr9-64326622-DEL-81 |
| chr9-64330605-DEL-72 | chr9-64330606-DEL-68 |
| chr9-64345246-DEL-54 | chr9-64345247-DEL-54 |
| chr9-64364109-DEL-399 | chr9-64364110-DEL-399 |
| chr9-64386101-DEL-56 | chr9-64386081-DEL-76 |
| chr9-64507092-DEL-92 | chr9-64507093-DEL-92 |
| chr9-64509206-DEL-183 | chr9-64509207-DEL-183 |
| chr9-64534261-DEL-857 | chr9-64534262-DEL-857 |
| chr9-64544096-DEL-111 | chr9-64544097-DEL-111 |
| chr9-64564159-DEL-390 | chr9-64564160-DEL-390 |
| chr9-64592563-DEL-72 | chr9-64592582-DEL-64 |
| chr9-64650628-DEL-272 | chr9-64650629-DEL-272 |
| chr9-64651620-DEL-631 | chr9-64651622-DEL-632 |
| chr9-64661105-DEL-55 | chr9-64661111-DEL-50 |
| chr9-64667977-DEL-2763 | chr9-64667978-DEL-2763 |
| chr9-64668813-DEL-63 | chr9-64668800-DEL-86 |
| chr9-64670215-DEL-121 | chr9-64670220-DEL-75 |
| chr9-64676448-DEL-53 | chr9-64676449-DEL-53 |
| chr9-64691230-DEL-541 | chr9-64691231-DEL-541 |
| chr9-64694550-DEL-90 | chr9-64694539-DEL-90 |
| chr9-64696108-DEL-165 | chr9-64696109-DEL-165 |
| chr9-64697417-DEL-164 | chr9-64697418-DEL-164 |
| chr9-64706486-DEL-75 | chr9-64706487-DEL-75 |
| chr9-64753905-DEL-201 | chr9-64753906-DEL-201 |
| chr9-64782815-DEL-92 | chr9-64782824-DEL-80 |
| chr9-64808133-DEL-133 | chr9-64808134-DEL-133 |
| chr9-64847313-DEL-208 | chr9-64847314-DEL-208 |
| chr9-64865717-DEL-225 | chr9-64865718-DEL-225 |
| chr9-64904243-DEL-54 | chr9-64904244-DEL-54 |
| chr9-64951185-DEL-68 | chr9-64951186-DEL-68 |
| chr9-64966830-DEL-212 | chr9-64966831-DEL-212 |
| chr9-64968437-DEL-212 | chr9-64968438-DEL-212 |
| chr9-65108171-DEL-63 | chr9-65108171-DEL-63 |
| chr9-65156426-DEL-212 | chr9-65156427-DEL-212 |
| chr9-65164083-DEL-217 | chr9-65164091-DEL-218 |
| chr9-65170618-DEL-168 | chr9-65170619-DEL-168 |
| chr9-65195805-DEL-53 | chr9-65195806-DEL-53 |
| chr9-65211259-DEL-947 | chr9-65211255-DEL-947 |
| chr9-65226900-DEL-204 | chr9-65226901-DEL-204 |
| chr9-65235378-DEL-82 | chr9-65235379-DEL-82 |
| chr9-65236031-DEL-62 | chr9-65236015-DEL-94 |
| chr9-65239373-DEL-125 | chr9-65239374-DEL-125 |
| chr9-65275146-DEL-91 | chr9-65275147-DEL-91 |
| chr9-65276938-DEL-406 | chr9-65276939-DEL-406 |
| chr9-65292325-DEL-1016 | chr9-65292321-DEL-1013 |
| chr9-65313695-DEL-54 | chr9-65313684-DEL-66 |
| chr9-65328771-DEL-473 | chr9-65328772-DEL-473 |
| chr9-65335710-DEL-168 | chr9-65335711-DEL-168 |
| chr9-65410403-DEL-53 | chr9-65410404-DEL-53 |
| chr9-65478726-DEL-198 | chr9-65478727-DEL-198 |
| chr9-65479333-DEL-88 | chr9-65479334-DEL-96 |
| chr9-65481295-DEL-381 | chr9-65481296-DEL-381 |
| chr9-65489368-DEL-196 | chr9-65489369-DEL-196 |
| chr9-65505351-DEL-90 | chr9-65505352-DEL-90 |
| chr9-65511999-DEL-209 | chr9-65512000-DEL-209 |
| chr9-65552655-DEL-71 | chr9-65552656-DEL-71 |
| chr9-65573873-DEL-203 | chr9-65573874-DEL-203 |
| chr9-65612376-DEL-256 | chr9-65612377-DEL-256 |
| chr9-65692135-DEL-155 | chr9-65692136-DEL-155 |
| chr9-65713632-DEL-282 | chr9-65713633-DEL-282 |
| chr9-65716757-DEL-218 | chr9-65716758-DEL-218 |
| chr9-65767681-DEL-176 | chr9-65767682-DEL-176 |
| chr9-65772740-DEL-211 | chr9-65772741-DEL-211 |
| chr9-65775065-DEL-1016 | chr9-65775074-DEL-982 |
| chr9-65781827-DEL-211 | chr9-65781828-DEL-211 |
| chr9-65784230-DEL-195 | chr9-65784232-DEL-174 |
| chr9-65848928-DEL-5373 | chr9-65848929-DEL-5373 |
| chr9-65885797-DEL-219 | chr9-65885798-DEL-219 |
| chr9-65888942-DEL-3345 | chr9-65888943-DEL-3344 |
| chr9-65891479-DEL-185 | chr9-65891476-DEL-178 |
| chr9-65902123-DEL-70 | chr9-65902116-DEL-76 |
| chr9-65932111-DEL-858 | chr9-65932112-DEL-858 |
| chr9-65954266-DEL-224 | chr9-65954267-DEL-224 |
| chr9-65972013-DEL-425 | chr9-65972014-DEL-425 |
| chr9-66012358-DEL-78 | chr9-66012359-DEL-78 |
| chr9-66020862-DEL-172 | chr9-66020863-DEL-172 |
| chr9-66022866-DEL-79 | chr9-66022867-DEL-79 |
| chr9-66038580-DEL-141 | chr9-66038565-DEL-157 |
| chr9-66047483-DEL-1017 | chr9-66047484-DEL-1017 |

| | |
|---|---|
| chr9-66048610-DEL-713 | chr9-66048605-DEL-713 |
| chr9-66057164-DEL-95 | chr9-66057159-DEL-89 |
| chr9-66081554-DEL-61 | chr9-66081555-DEL-61 |
| chr9-66093561-DEL-82 | chr9-66093562-DEL-82 |
| chr9-66140186-DEL-197 | chr9-66140187-DEL-197 |
| chr9-66154524-DEL-147 | chr9-66154525-DEL-147 |
| chr9-66161916-DEL-61 | chr9-66161917-DEL-61 |
| chr9-66196530-DEL-64 | chr9-66196531-DEL-64 |
| chr9-66203387-DEL-738 | chr9-66203388-DEL-738 |
| chr9-66273991-DEL-228 | chr9-66273992-DEL-208 |
| chr9-66314644-DEL-126 | chr9-66314649-DEL-126 |
| chr9-66390102-DEL-216 | chr9-66390103-DEL-216 |
| chr9-66395511-DEL-209 | chr9-66395512-DEL-209 |
| chr9-66398130-DEL-303 | chr9-66398131-DEL-303 |
| chr9-66410502-DEL-54 | chr9-66410503-DEL-54 |
| chr9-66456076-DEL-1253 | chr9-66456077-DEL-1253 |
| chr9-66506742-DEL-142 | chr9-66506743-DEL-142 |
| chr9-66540227-DEL-6602 | chr9-66540228-DEL-6602 |
| chr9-66558156-DEL-96 | chr9-66558177-DEL-72 |
| chr9-66596152-DEL-173 | chr9-66596153-DEL-173 |
| chr9-66613683-DEL-944 | chr9-66613682-DEL-944 |
| chr9-66637237-DEL-213 | chr9-66637238-DEL-213 |
| chr9-66648858-DEL-345 | chr9-66648862-DEL-340 |
| chr9-66749715-DEL-64 | chr9-66749716-DEL-64 |
| chr9-66757412-DEL-205 | chr9-66757413-DEL-205 |
| chr9-66759325-DEL-52 | chr9-66759322-DEL-56 |
| chr9-66760326-DEL-92 | chr9-66760327-DEL-92 |
| chr9-66805903-DEL-70 | chr9-66805904-DEL-70 |
| chr9-66815955-DEL-61 | chr9-66815956-DEL-61 |
| chr9-66827694-DEL-63 | chr9-66827672-DEL-88 |
| chr9-66839263-DEL-214 | chr9-66839264-DEL-214 |
| chr9-66851232-DEL-214 | chr9-66851227-DEL-215 |
| chr9-66857420-DEL-1123 | chr9-66857421-DEL-1123 |
| chr9-66887086-DEL-199 | chr9-66887087-DEL-199 |
| chr9-66906667-DEL-420 | chr9-66906668-DEL-420 |
| chr9-66909849-DEL-53 | chr9-66909821-DEL-54 |
| chr9-66921814-DEL-54 | chr9-66921815-DEL-57 |
| chr9-66924946-DEL-1564 | chr9-66924947-DEL-1564 |
| chr9-66957557-DEL-240 | chr9-66957558-DEL-240 |
| chr9-66962978-DEL-266 | chr9-66962979-DEL-266 |
| chr9-66968762-DEL-54 | chr9-66968763-DEL-54 |
| chr9-66969508-DEL-79 | chr9-66969505-DEL-106 |
| chr9-66970021-DEL-60 | chr9-66970022-DEL-60 |
| chr9-66986301-DEL-794 | chr9-66986302-DEL-794 |
| chr9-67016175-DEL-397 | chr9-67016178-DEL-401 |
| chr9-67033334-DEL-93 | chr9-67033366-DEL-57 |
| chr9-67045532-DEL-60 | chr9-67045529-DEL-56 |
| chr9-67046930-DEL-80 | chr9-67046928-DEL-81 |
| chr9-67048894-DEL-220 | chr9-67048895-DEL-220 |
| chr9-67053030-DEL-224 | chr9-67053031-DEL-224 |
| chr9-67057947-DEL-235 | chr9-67057948-DEL-235 |
| chr9-67064888-DEL-75 | chr9-67064898-DEL-62 |
| chr9-67073229-DEL-90 | chr9-67073230-DEL-90 |
| chr9-67105900-DEL-6387 | chr9-67105901-DEL-6387 |
| chr9-67116964-DEL-60 | chr9-67116965-DEL-60 |
| chr9-67138737-DEL-62 | chr9-67138738-DEL-62 |
| chr9-67144737-DEL-64 | chr9-67144738-DEL-64 |
| chr9-67147148-DEL-57 | chr9-67147149-DEL-57 |
| chr9-67160186-DEL-197 | chr9-67160187-DEL-197 |
| chr9-67170912-DEL-73 | chr9-67170913-DEL-73 |
| chr9-67212800-DEL-108 | chr9-67212804-DEL-92 |
| chr9-67216517-DEL-217 | chr9-67216515-DEL-218 |
| chr9-67282338-DEL-100 | chr9-67282343-DEL-88 |
| chr9-67331274-DEL-1319 | chr9-67331275-DEL-1319 |
| chr9-67355047-DEL-198 | chr9-67355048-DEL-198 |
| chr9-67356692-DEL-1213 | chr9-67356692-DEL-1211 |
| chr9-67357263-DEL-115 | chr9-67357268-DEL-108 |
| chr9-67414155-DEL-66 | chr9-67414156-DEL-66 |
| chr9-67445767-DEL-497 | chr9-67445768-DEL-497 |
| chr9-67505978-DEL-60 | chr9-67505979-DEL-60 |
| chr9-67507926-DEL-88 | chr9-67507927-DEL-88 |
| chr9-67527418-DEL-189 | chr9-67527419-DEL-189 |
| chr9-67532325-DEL-65 | chr9-67532323-DEL-67 |
| chr9-67552031-DEL-328 | chr9-67552032-DEL-328 |
| chr9-67555728-DEL-2723 | chr9-67555729-DEL-2723 |
| chr9-67580950-DEL-340 | chr9-67580951-DEL-340 |
| chr9-67583649-DEL-125 | chr9-67583650-DEL-125 |
| chr9-67587408-DEL-63 | chr9-67587457-DEL-60 |
| chr9-67589470-DEL-94 | chr9-67589471-DEL-94 |
| chr9-67599713-DEL-62 | chr9-67599712-DEL-64 |

| | |
|---|---|
| chr9-67601449-DEL-71 | chr9-67601450-DEL-71 |
| chr9-67605804-DEL-4837 | chr9-67605805-DEL-4837 |
| chr9-67611808-DEL-445 | chr9-67611809-DEL-445 |
| chr9-67622735-DEL-1465 | chr9-67622736-DEL-1465 |
| chr9-67647641-DEL-150 | chr9-67647617-DEL-141 |
| chr9-67672762-DEL-65 | chr9-67672758-DEL-70 |
| chr9-67682829-DEL-1296 | chr9-67682830-DEL-1296 |
| chr9-67720762-DEL-223 | chr9-67720763-DEL-223 |
| chr9-67743511-DEL-112 | chr9-67743512-DEL-112 |
| chr9-67763622-DEL-101 | chr9-67763652-DEL-51 |
| chr9-67769754-DEL-58 | chr9-67769755-DEL-58 |
| chr9-67883400-DEL-6198 | chr9-67883401-DEL-6198 |
| chr9-67918451-DEL-54 | chr9-67918451-DEL-54 |
| chr9-67948178-DEL-1807 | chr9-67948179-DEL-1807 |
| chr9-67977026-DEL-337 | chr9-67977027-DEL-337 |
| chr9-67979413-DEL-76 | chr9-67979374-DEL-78 |
| chr9-67991377-DEL-822 | chr9-67991378-DEL-822 |
| chr9-68019812-DEL-78 | chr9-68019813-DEL-78 |
| chr9-68026116-DEL-1735 | chr9-68026117-DEL-1735 |
| chr9-68043810-DEL-532 | chr9-68043811-DEL-532 |
| chr9-68050093-DEL-68 | chr9-68050124-DEL-60 |
| chr9-68052971-DEL-6449 | chr9-68052972-DEL-6449 |
| chr9-68061586-DEL-52 | chr9-68061587-DEL-52 |
| chr9-68065365-DEL-2257 | chr9-68065366-DEL-2257 |
| chr9-68080804-DEL-214 | chr9-68080805-DEL-214 |
| chr9-68082606-DEL-217 | chr9-68082607-DEL-211 |
| chr9-68085636-DEL-351 | chr9-68085632-DEL-352 |
| chr9-68090415-DEL-3218 | chr9-68090416-DEL-3218 |
| chr9-68093898-DEL-193 | chr9-68093899-DEL-190 |
| chr9-68107876-DEL-103 | chr9-68107889-DEL-103 |
| chr9-68144542-DEL-80 | chr9-68144525-DEL-123 |
| chr9-68145084-DEL-1341 | chr9-68145086-DEL-1343 |
| chr9-68155712-DEL-800 | chr9-68155724-DEL-798 |
| chr9-68164436-DEL-183 | chr9-68164437-DEL-183 |
| chr9-68168094-DEL-108 | chr9-68168095-DEL-108 |
| chr9-68181823-DEL-65 | chr9-68181824-DEL-65 |
| chr9-68199726-DEL-166 | chr9-68199727-DEL-165 |
| chr9-68219646-DEL-1932 | chr9-68219647-DEL-1932 |
| chr9-68283903-DEL-140 | chr9-68283904-DEL-126 |
| chr9-68285910-DEL-101 | chr9-68285911-DEL-101 |
| chr9-68321728-DEL-552 | chr9-68321729-DEL-552 |
| chr9-68326001-DEL-186 | chr9-68326002-DEL-186 |
| chr9-68326511-DEL-67 | chr9-68326512-DEL-67 |
| chr9-68350652-DEL-134 | chr9-68350653-DEL-134 |
| chr9-68355889-DEL-85 | chr9-68355842-DEL-87 |
| chr9-68357582-DEL-2150 | chr9-68357583-DEL-2150 |
| chr9-68377906-DEL-85 | chr9-68377895-DEL-78 |
| chr9-68384142-DEL-237 | chr9-68384143-DEL-237 |
| chr9-68394739-DEL-51 | chr9-68394735-DEL-70 |
| chr9-68429979-DEL-118 | chr9-68429980-DEL-118 |
| chr9-68444062-DEL-74 | chr9-68444063-DEL-74 |
| chr9-68460347-DEL-240 | chr9-68460348-DEL-240 |
| chr9-68463241-DEL-392 | chr9-68463232-DEL-404 |
| chr9-68476170-DEL-56 | chr9-68476171-DEL-50 |
| chr9-68479213-DEL-68 | chr9-68479214-DEL-62 |
| chr9-68484854-DEL-916 | chr9-68484855-DEL-916 |
| chr9-68504799-DEL-151 | chr9-68504796-DEL-155 |
| chr9-68590239-DEL-72 | chr9-68590240-DEL-72 |
| chr9-68594472-DEL-70 | chr9-68594462-DEL-68 |
| chr9-68630872-DEL-57 | chr9-68630872-DEL-57 |
| chr9-68634119-DEL-61 | chr9-68634120-DEL-61 |
| chr9-68661739-DEL-68 | chr9-68661704-DEL-68 |
| chr9-68680929-DEL-207 | chr9-68680930-DEL-207 |
| chr9-68690833-DEL-73 | chr9-68690849-DEL-70 |
| chr9-68708921-DEL-59 | chr9-68708922-DEL-59 |
| chr9-68755999-DEL-236 | chr9-68756000-DEL-236 |
| chr9-68798258-DEL-184 | chr9-68798259-DEL-184 |
| chr9-68826394-DEL-193 | chr9-68826395-DEL-193 |
| chr9-68837478-DEL-112 | chr9-68837487-DEL-80 |
| chr9-68841849-DEL-66 | chr9-68841850-DEL-66 |
| chr9-68861760-DEL-60 | chr9-68861761-DEL-60 |
| chr9-68864172-DEL-84 | chr9-68864176-DEL-63 |
| chr9-68870569-DEL-58 | chr9-68870570-DEL-58 |
| chr9-68968409-DEL-86 | chr9-68968450-DEL-86 |
| chr9-68986859-DEL-1315 | chr9-68986860-DEL-1315 |
| chr9-68993018-DEL-60 | chr9-68993017-DEL-60 |
| chr9-69001293-DEL-54 | chr9-69001294-DEL-54 |
| chr9-69023675-DEL-52 | chr9-69023676-DEL-52 |
| chr9-69027509-DEL-62 | chr9-69027506-DEL-52 |
| chr9-69051928-DEL-54 | chr9-69051929-DEL-54 |

| | |
|---|---|
| chr9-69054992-DEL-83 | chr9-69054993-DEL-83 |
| chr9-69079315-DEL-57 | chr9-69079306-DEL-60 |
| chr9-69116649-DEL-54 | chr9-69116650-DEL-54 |
| chr9-69134923-DEL-57 | chr9-69134922-DEL-61 |
| chr9-69157192-DEL-526 | chr9-69157197-DEL-522 |
| chr9-69168071-DEL-63 | chr9-69168062-DEL-64 |
| chr9-69230314-DEL-252 | chr9-69230315-DEL-252 |
| chr9-69235274-DEL-72 | chr9-69235275-DEL-72 |
| chr9-69267795-DEL-52 | chr9-69267796-DEL-52 |
| chr9-69283065-DEL-102 | chr9-69283062-DEL-56 |
| chr9-69292063-DEL-56 | chr9-69292063-DEL-56 |
| chr9-69297718-DEL-644 | chr9-69297719-DEL-644 |
| chr9-69302186-DEL-68 | chr9-69302199-DEL-64 |
| chr9-69310120-DEL-206 | chr9-69310121-DEL-206 |
| chr9-69310944-DEL-116 | chr9-69310945-DEL-116 |
| chr9-69313919-DEL-184 | chr9-69313920-DEL-184 |
| chr9-69341098-DEL-202 | chr9-69341099-DEL-202 |
| chr9-69384557-DEL-263 | chr9-69384558-DEL-263 |
| chr9-69386076-DEL-87 | chr9-69386063-DEL-78 |
| chr9-69433198-DEL-68 | chr9-69433197-DEL-68 |
| chr9-69434709-DEL-117 | chr9-69434710-DEL-117 |
| chr9-69444826-DEL-1165 | chr9-69444827-DEL-1165 |
| chr9-69473089-DEL-643 | chr9-69473090-DEL-643 |
| chr9-69478966-DEL-829 | chr9-69478967-DEL-828 |
| chr9-69496930-DEL-66 | chr9-69496921-DEL-64 |
| chr9-69507543-DEL-56 | chr9-69507514-DEL-68 |
| chr9-69525647-DEL-205 | chr9-69525648-DEL-205 |
| chr9-69526967-DEL-58 | chr9-69526968-DEL-58 |
| chr9-69566795-DEL-109 | chr9-69566796-DEL-109 |
| chr9-69572328-DEL-52 | chr9-69572329-DEL-52 |
| chr9-69582632-DEL-426 | chr9-69582662-DEL-400 |
| chr9-69584698-DEL-56 | chr9-69584699-DEL-54 |
| chr9-69602146-DEL-607 | chr9-69602147-DEL-607 |
| chr9-69608934-DEL-609 | chr9-69608933-DEL-610 |
| chr9-69646206-DEL-68 | chr9-69646206-DEL-68 |
| chr9-69657740-DEL-158 | chr9-69657741-DEL-152 |
| chr9-69690457-DEL-721 | chr9-69690455-DEL-721 |
| chr9-69694238-DEL-60 | chr9-69694239-DEL-60 |
| chr9-69702334-DEL-55 | chr9-69702335-DEL-55 |
| chr9-69706187-DEL-762 | chr9-69706185-DEL-762 |
| chr9-69728098-DEL-75 | chr9-69728102-DEL-72 |
| chr9-69764261-DEL-406 | chr9-69764262-DEL-406 |
| chr9-69784346-DEL-129 | chr9-69784319-DEL-129 |
| chr9-69803113-DEL-806 | chr9-69803114-DEL-806 |
| chr9-69825686-DEL-496 | chr9-69825687-DEL-496 |
| chr9-69882747-DEL-131 | chr9-69882748-DEL-131 |
| chr9-69891442-DEL-263 | chr9-69891443-DEL-263 |
| chr9-69902373-DEL-70 | chr9-69902374-DEL-70 |
| chr9-69954426-DEL-295 | chr9-69954427-DEL-295 |
| chr9-69990349-DEL-583 | chr9-69990350-DEL-583 |
| chr9-70070611-DEL-130 | chr9-70070612-DEL-130 |
| chr9-70072970-DEL-1005 | chr9-70072971-DEL-1005 |
| chr9-70102835-DEL-210 | chr9-70102836-DEL-210 |
| chr9-70110842-DEL-135 | chr9-70110843-DEL-135 |
| chr9-70133052-DEL-4037 | chr9-70133053-DEL-4037 |
| chr9-70157972-DEL-54 | chr9-70157973-DEL-58 |
| chr9-70181835-DEL-62 | chr9-70181837-DEL-61 |
| chr9-70217982-DEL-209 | chr9-70217983-DEL-209 |
| chr9-70241348-DEL-65 | chr9-70241349-DEL-65 |
| chr9-70255345-DEL-1018 | chr9-70255346-DEL-1018 |
| chr9-70287430-DEL-54 | chr9-70287431-DEL-66 |
| chr9-70305408-DEL-113 | chr9-70305407-DEL-109 |
| chr9-70356089-DEL-235 | chr9-70356090-DEL-235 |
| chr9-70430293-DEL-210 | chr9-70430294-DEL-210 |
| chr9-70448565-DEL-77 | chr9-70448566-DEL-77 |
| chr9-70464661-DEL-174 | chr9-70464662-DEL-172 |
| chr9-70477843-DEL-325 | chr9-70477844-DEL-325 |
| chr9-70486123-DEL-64 | chr9-70486124-DEL-64 |
| chr9-70528700-DEL-826 | chr9-70528703-DEL-823 |
| chr9-70532493-DEL-203 | chr9-70532494-DEL-203 |
| chr9-70535300-DEL-216 | chr9-70535301-DEL-216 |
| chr9-70544622-DEL-203 | chr9-70544623-DEL-203 |
| chr9-70553563-DEL-62 | chr9-70553564-DEL-62 |
| chr9-70566126-DEL-98 | chr9-70566119-DEL-92 |
| chr9-70609529-DEL-116 | chr9-70609531-DEL-104 |
| chr9-70697310-DEL-72 | chr9-70697329-DEL-54 |
| chr9-70751605-DEL-164 | chr9-70751606-DEL-152 |
| chr9-70761671-DEL-110 | chr9-70761672-DEL-110 |
| chr9-70789082-DEL-72 | chr9-70789081-DEL-72 |
| chr9-70791191-DEL-80 | chr9-70791193-DEL-64 |

| | |
|---|---|
| chr9-70810998-DEL-6522 | chr9-70810999-DEL-6522 |
| chr9-70811209-DEL-212 | chr9-70811210-DEL-212 |
| chr9-70856084-DEL-143 | chr9-70856085-DEL-143 |
| chr9-70864473-DEL-99 | chr9-70864472-DEL-99 |
| chr9-70917567-DEL-54 | chr9-70917568-DEL-54 |
| chr9-70986736-DEL-72 | chr9-70986737-DEL-72 |
| chr9-70987672-DEL-10881 | chr9-70987673-DEL-10881 |
| chr9-71006191-DEL-120 | chr9-71006193-DEL-119 |
| chr9-71009706-DEL-247 | chr9-71009707-DEL-247 |
| chr9-71021515-DEL-51 | chr9-71021516-DEL-51 |
| chr9-71021708-DEL-6440 | chr9-71021709-DEL-6440 |
| chr9-71028553-DEL-53 | chr9-71028554-DEL-53 |
| chr9-71030590-DEL-172 | chr9-71030589-DEL-174 |
| chr9-71034258-DEL-52 | chr9-71034259-DEL-52 |
| chr9-71048299-DEL-428 | chr9-71048300-DEL-428 |
| chr9-71050371-DEL-70 | chr9-71050371-DEL-70 |
| chr9-71055029-DEL-54 | chr9-71055036-DEL-58 |
| chr9-71083992-DEL-536 | chr9-71083993-DEL-536 |
| chr9-71084606-DEL-187 | chr9-71084606-DEL-187 |
| chr9-71096914-DEL-7338 | chr9-71096915-DEL-7338 |
| chr9-71130979-DEL-362 | chr9-71130980-DEL-362 |
| chr9-71148273-DEL-84 | chr9-71148262-DEL-96 |
| chr9-71175842-DEL-213 | chr9-71175843-DEL-212 |
| chr9-71238417-DEL-270 | chr9-71238369-DEL-270 |
| chr9-71273567-DEL-55 | chr9-71273558-DEL-55 |
| chr9-71277990-DEL-1613 | chr9-71277995-DEL-1616 |
| chr9-71294518-DEL-82 | chr9-71294520-DEL-62 |
| chr9-71298892-DEL-321 | chr9-71298893-DEL-321 |
| chr9-71327887-DEL-54 | chr9-71327888-DEL-54 |
| chr9-71328351-DEL-13548 | chr9-71328352-DEL-13548 |
| chr9-71382883-DEL-139 | chr9-71382884-DEL-139 |
| chr9-71410493-DEL-64 | chr9-71410493-DEL-72 |
| chr9-71413068-DEL-186 | chr9-71413069-DEL-186 |
| chr9-71433868-DEL-104 | chr9-71433888-DEL-86 |
| chr9-71477187-DEL-50 | chr9-71477201-DEL-57 |
| chr9-71517350-DEL-119 | chr9-71517351-DEL-119 |
| chr9-71527176-DEL-198 | chr9-71527177-DEL-198 |
| chr9-71550787-DEL-198 | chr9-71550788-DEL-198 |
| chr9-71552618-DEL-55 | chr9-71552622-DEL-84 |
| chr9-71569313-DEL-67 | chr9-71569323-DEL-86 |
| chr9-71580226-DEL-69 | chr9-71580227-DEL-69 |
| chr9-71580271-DEL-947 | chr9-71580248-DEL-971 |
| chr9-71584009-DEL-84 | chr9-71584010-DEL-84 |
| chr9-71585119-DEL-183 | chr9-71585120-DEL-183 |
| chr9-71594560-DEL-123 | chr9-71594561-DEL-123 |
| chr9-71599394-DEL-98 | chr9-71599389-DEL-91 |
| chr9-71604943-DEL-438 | chr9-71604944-DEL-438 |
| chr9-71667633-DEL-89 | chr9-71667633-DEL-89 |
| chr9-71682257-DEL-56 | chr9-71682258-DEL-56 |
| chr9-71688793-DEL-2547 | chr9-71688794-DEL-2547 |
| chr9-71702251-DEL-871 | chr9-71702252-DEL-871 |
| chr9-71830906-DEL-202 | chr9-71830907-DEL-202 |
| chr9-71838275-DEL-54 | chr9-71838276-DEL-54 |
| chr9-71872273-DEL-7237 | chr9-71872274-DEL-7237 |
| chr9-71894028-DEL-155 | chr9-71894029-DEL-155 |
| chr9-71904981-DEL-205 | chr9-71904982-DEL-205 |
| chr9-71919574-DEL-87 | chr9-71919575-DEL-87 |
| chr9-71945535-DEL-61 | chr9-71945536-DEL-61 |
| chr9-71989580-DEL-721 | chr9-71989581-DEL-721 |
| chr9-72060339-DEL-53 | chr9-72060339-DEL-53 |
| chr9-72068697-DEL-356 | chr9-72068698-DEL-356 |
| chr9-72080438-DEL-65 | chr9-72080439-DEL-65 |
| chr9-72081766-DEL-521 | chr9-72081767-DEL-521 |
| chr9-72094344-DEL-77 | chr9-72094355-DEL-68 |
| chr9-72121537-DEL-205 | chr9-72121538-DEL-205 |
| chr9-72143795-DEL-220 | chr9-72143796-DEL-220 |
| chr9-72148521-DEL-119 | chr9-72148522-DEL-119 |
| chr9-72155828-DEL-1200 | chr9-72155829-DEL-1200 |
| chr9-72157167-DEL-184 | chr9-72157168-DEL-184 |
| chr9-72184749-DEL-205 | chr9-72184750-DEL-205 |
| chr9-72199539-DEL-1114 | chr9-72199540-DEL-1114 |
| chr9-72209390-DEL-99 | chr9-72209391-DEL-99 |
| chr9-72249151-DEL-96 | chr9-72249152-DEL-96 |
| chr9-72264626-DEL-52 | chr9-72264627-DEL-52 |
| chr9-72271612-DEL-1833 | chr9-72271613-DEL-1833 |
| chr9-72274888-DEL-191 | chr9-72274889-DEL-191 |
| chr9-72339134-DEL-95 | chr9-72339138-DEL-95 |
| chr9-72348588-DEL-242 | chr9-72348589-DEL-242 |
| chr9-72356852-DEL-93 | chr9-72356853-DEL-93 |
| chr9-72373593-DEL-368 | chr9-72373594-DEL-368 |

| | |
|---|---|
| chr9-72379213-DEL-80 | chr9-72379214-DEL-80 |
| chr9-72382500-DEL-206 | chr9-72382501-DEL-206 |
| chr9-72383741-DEL-241 | chr9-72383742-DEL-241 |
| chr9-72426828-DEL-56 | chr9-72426829-DEL-56 |
| chr9-72464820-DEL-212 | chr9-72464816-DEL-213 |
| chr9-72478031-DEL-77 | chr9-72478032-DEL-77 |
| chr9-72503411-DEL-1330 | chr9-72503412-DEL-1330 |
| chr9-72512263-DEL-210 | chr9-72512264-DEL-210 |
| chr9-72537919-DEL-214 | chr9-72537920-DEL-214 |
| chr9-72573872-DEL-221 | chr9-72573873-DEL-221 |
| chr9-72575944-DEL-205 | chr9-72575945-DEL-205 |
| chr9-72578067-DEL-220 | chr9-72578068-DEL-220 |
| chr9-72587595-DEL-200 | chr9-72587596-DEL-200 |
| chr9-72618282-DEL-73 | chr9-72618283-DEL-73 |
| chr9-72636813-DEL-218 | chr9-72636814-DEL-218 |
| chr9-72659768-DEL-160 | chr9-72659766-DEL-161 |
| chr9-72662025-DEL-129 | chr9-72662026-DEL-129 |
| chr9-72672655-DEL-254 | chr9-72672656-DEL-254 |
| chr9-72674677-DEL-72 | chr9-72674722-DEL-70 |
| chr9-72685712-DEL-156 | chr9-72685713-DEL-156 |
| chr9-72686700-DEL-224 | chr9-72686701-DEL-224 |
| chr9-72708961-DEL-214 | chr9-72708962-DEL-214 |
| chr9-72731195-DEL-54 | chr9-72731196-DEL-54 |
| chr9-72754735-DEL-163 | chr9-72754736-DEL-163 |
| chr9-72755859-DEL-68 | chr9-72755860-DEL-68 |
| chr9-72768363-DEL-218 | chr9-72768364-DEL-218 |
| chr9-72793258-DEL-54 | chr9-72793259-DEL-54 |
| chr9-72796707-DEL-195 | chr9-72796708-DEL-195 |
| chr9-72803129-DEL-4991 | chr9-72803130-DEL-4991 |
| chr9-72826465-DEL-81 | chr9-72826466-DEL-84 |
| chr9-72878423-DEL-215 | chr9-72878424-DEL-215 |
| chr9-72883415-DEL-3190 | chr9-72883416-DEL-3190 |
| chr9-72890157-DEL-122 | chr9-72890159-DEL-124 |
| chr9-72939963-DEL-169 | chr9-72939964-DEL-169 |
| chr9-72941997-DEL-957 | chr9-72941998-DEL-957 |
| chr9-72944176-DEL-201 | chr9-72944166-DEL-205 |
| chr9-72960682-DEL-55 | chr9-72960670-DEL-56 |
| chr9-72977372-DEL-198 | chr9-72977373-DEL-198 |
| chr9-72983210-DEL-210 | chr9-72983211-DEL-210 |
| chr9-72986868-DEL-198 | chr9-72986869-DEL-198 |
| chr9-72990610-DEL-107 | chr9-72990611-DEL-107 |
| chr9-73001049-DEL-232 | chr9-73001049-DEL-233 |
| chr9-73016311-DEL-988 | chr9-73016312-DEL-988 |
| chr9-73024213-DEL-197 | chr9-73024214-DEL-197 |
| chr9-73051388-DEL-59 | chr9-73051389-DEL-59 |
| chr9-73060953-DEL-77 | chr9-73060954-DEL-77 |
| chr9-73064130-DEL-98 | chr9-73064131-DEL-98 |
| chr9-73065385-DEL-90 | chr9-73065365-DEL-107 |
| chr9-73065976-DEL-110 | chr9-73065977-DEL-110 |
| chr9-73082645-DEL-588 | chr9-73082660-DEL-598 |
| chr9-73100189-DEL-467 | chr9-73100190-DEL-467 |
| chr9-73110899-DEL-69 | chr9-73110904-DEL-65 |
| chr9-73136686-DEL-51 | chr9-73136687-DEL-51 |
| chr9-73136913-DEL-6900 | chr9-73136914-DEL-6900 |
| chr9-73148944-DEL-54 | chr9-73148945-DEL-54 |
| chr9-73176320-DEL-56 | chr9-73176321-DEL-56 |
| chr9-73190077-DEL-55 | chr9-73190078-DEL-55 |
| chr9-73203615-DEL-5286 | chr9-73203616-DEL-5286 |
| chr9-73211467-DEL-198 | chr9-73211468-DEL-198 |
| chr9-73238323-DEL-105 | chr9-73238324-DEL-105 |
| chr9-73258906-DEL-51 | chr9-73258907-DEL-51 |
| chr9-73260096-DEL-646 | chr9-73260097-DEL-646 |
| chr9-73264656-DEL-571 | chr9-73264684-DEL-543 |
| chr9-73290657-DEL-100 | chr9-73290644-DEL-113 |
| chr9-73296798-DEL-82 | chr9-73296800-DEL-102 |
| chr9-73303980-DEL-6805 | chr9-73303981-DEL-6805 |
| chr9-73304141-DEL-208 | chr9-73304142-DEL-208 |
| chr9-73312593-DEL-7071 | chr9-73312594-DEL-7071 |
| chr9-73329264-DEL-179 | chr9-73329265-DEL-179 |
| chr9-73331520-DEL-1286 | chr9-73331521-DEL-1286 |
| chr9-73358429-DEL-103 | chr9-73358430-DEL-103 |
| chr9-73370758-DEL-774 | chr9-73370759-DEL-774 |
| chr9-73374105-DEL-192 | chr9-73374106-DEL-192 |
| chr9-73391309-DEL-70 | chr9-73391310-DEL-70 |
| chr9-73396658-DEL-67 | chr9-73396659-DEL-67 |
| chr9-73397784-DEL-284 | chr9-73397785-DEL-284 |
| chr9-73423954-DEL-97 | chr9-73423955-DEL-97 |
| chr9-73426001-DEL-56 | chr9-73426002-DEL-56 |
| chr9-73426860-DEL-51 | chr9-73426861-DEL-51 |
| chr9-73442310-DEL-362 | chr9-73442305-DEL-355 |

| | |
|---|---|
| chr9-73453461-DEL-58 | chr9-73453481-DEL-60 |
| chr9-73454949-DEL-71 | chr9-73454950-DEL-67 |
| chr9-73475319-DEL-98 | chr9-73475320-DEL-139 |
| chr9-73489747-DEL-3403 | chr9-73489750-DEL-3401 |
| chr9-73493422-DEL-625 | chr9-73493423-DEL-625 |
| chr9-73500820-DEL-1597 | chr9-73500821-DEL-1597 |
| chr9-73504614-DEL-6250 | chr9-73504605-DEL-6251 |
| chr9-73538111-DEL-1680 | chr9-73538112-DEL-1680 |
| chr9-73553851-DEL-1259 | chr9-73553852-DEL-1259 |
| chr9-73582184-DEL-93 | chr9-73582185-DEL-93 |
| chr9-73595539-DEL-1485 | chr9-73595540-DEL-1485 |
| chr9-73623558-DEL-276 | chr9-73623574-DEL-242 |
| chr9-73631414-DEL-857 | chr9-73631415-DEL-857 |
| chr9-73679541-DEL-167 | chr9-73679542-DEL-167 |
| chr9-73680690-DEL-76 | chr9-73680691-DEL-76 |
| chr9-73701689-DEL-100 | chr9-73701690-DEL-100 |
| chr9-73702706-DEL-195 | chr9-73702707-DEL-195 |
| chr9-73716651-DEL-74 | chr9-73716652-DEL-56 |
| chr9-73725202-DEL-193 | chr9-73725199-DEL-192 |
| chr9-73731942-DEL-6399 | chr9-73731943-DEL-6399 |
| chr9-73742737-DEL-62 | chr9-73742762-DEL-58 |
| chr9-73768677-DEL-1888 | chr9-73768678-DEL-1888 |
| chr9-73786651-DEL-371 | chr9-73786652-DEL-371 |
| chr9-73807431-DEL-4505 | chr9-73807432-DEL-4505 |
| chr9-73816991-DEL-574 | chr9-73816992-DEL-574 |
| chr9-73842826-DEL-6612 | chr9-73842827-DEL-6612 |
| chr9-73850693-DEL-3326 | chr9-73850694-DEL-3326 |
| chr9-73851170-DEL-1110 | chr9-73851171-DEL-1110 |
| chr9-73874644-DEL-51 | chr9-73874636-DEL-52 |
| chr9-73943914-DEL-7067 | chr9-73943915-DEL-7067 |
| chr9-73970306-DEL-456 | chr9-73970303-DEL-422 |
| chr9-73994253-DEL-455 | chr9-73994254-DEL-455 |
| chr9-73995343-DEL-450 | chr9-73995331-DEL-464 |
| chr9-74019452-DEL-199 | chr9-74019453-DEL-199 |
| chr9-74024722-DEL-198 | chr9-74024719-DEL-197 |
| chr9-74040721-DEL-206 | chr9-74040722-DEL-206 |
| chr9-74110550-DEL-1307 | chr9-74110551-DEL-1307 |
| chr9-74169600-DEL-189 | chr9-74169601-DEL-189 |
| chr9-74247063-DEL-324 | chr9-74247064-DEL-324 |
| chr9-74271302-DEL-351 | chr9-74271303-DEL-351 |
| chr9-74286093-DEL-961 | chr9-74286094-DEL-961 |
| chr9-74312619-DEL-409 | chr9-74312620-DEL-409 |
| chr9-74317450-DEL-6221 | chr9-74317447-DEL-6220 |
| chr9-74379899-DEL-63 | chr9-74379899-DEL-61 |
| chr9-74390349-DEL-312 | chr9-74390350-DEL-312 |
| chr9-74402509-DEL-987 | chr9-74402509-DEL-987 |
| chr9-74403823-DEL-64 | chr9-74403823-DEL-64 |
| chr9-74417517-DEL-493 | chr9-74417518-DEL-493 |
| chr9-74434519-DEL-941 | chr9-74434518-DEL-942 |
| chr9-74455278-DEL-60 | chr9-74455283-DEL-61 |
| chr9-74460965-DEL-84 | chr9-74460976-DEL-52 |
| chr9-74467221-DEL-436 | chr9-74467222-DEL-436 |
| chr9-74473393-DEL-109 | chr9-74473407-DEL-95 |
| chr9-74475745-DEL-64 | chr9-74475735-DEL-63 |
| chr9-74535409-DEL-13586 | chr9-74535410-DEL-13586 |
| chr9-74571995-DEL-6127 | chr9-74571996-DEL-6127 |
| chr9-74591847-DEL-6257 | chr9-74591848-DEL-6257 |
| chr9-74591999-DEL-199 | chr9-74592005-DEL-198 |
| chr9-74604148-DEL-75 | chr9-74604128-DEL-75 |
| chr9-74620907-DEL-354 | chr9-74620908-DEL-354 |
| chr9-74622581-DEL-73 | chr9-74622589-DEL-90 |
| chr9-74632514-DEL-75 | chr9-74632512-DEL-66 |
| chr9-74640411-DEL-1738 | chr9-74640412-DEL-1738 |
| chr9-74663592-DEL-749 | chr9-74663593-DEL-749 |
| chr9-74669543-DEL-92 | chr9-74669545-DEL-93 |
| chr9-74675331-DEL-57 | chr9-74675326-DEL-63 |
| chr9-74675862-DEL-6175 | chr9-74675848-DEL-6182 |
| chr9-74695252-DEL-270 | chr9-74695260-DEL-271 |
| chr9-74723782-DEL-112 | chr9-74723784-DEL-100 |
| chr9-74736849-DEL-93 | chr9-74736850-DEL-93 |
| chr9-74807177-DEL-210 | chr9-74807171-DEL-209 |
| chr9-74820850-DEL-90 | chr9-74820871-DEL-91 |
| chr9-74849479-DEL-88 | chr9-74849494-DEL-58 |
| chr9-74890145-DEL-99 | chr9-74890145-DEL-66 |
| chr9-74918987-DEL-64 | chr9-74918990-DEL-62 |
| chr9-74921679-DEL-110 | chr9-74921680-DEL-110 |
| chr9-74925637-DEL-111 | chr9-74925638-DEL-111 |
| chr9-74934709-DEL-213 | chr9-74934710-DEL-213 |
| chr9-74951731-DEL-205 | chr9-74951732-DEL-205 |
| chr9-74968643-DEL-684 | chr9-74968647-DEL-680 |

| | |
|---|---|
| chr9-74989780-DEL-680 | chr9-74989739-DEL-698 |
| chr9-75003352-DEL-709 | chr9-75003353-DEL-709 |
| chr9-75029090-DEL-110 | chr9-75029091-DEL-110 |
| chr9-75032111-DEL-52 | chr9-75032084-DEL-80 |
| chr9-75087611-DEL-53 | chr9-75087612-DEL-53 |
| chr9-75133236-DEL-56 | chr9-75133237-DEL-56 |
| chr9-75140130-DEL-54 | chr9-75140131-DEL-56 |
| chr9-75154810-DEL-294 | chr9-75154811-DEL-294 |
| chr9-75161484-DEL-175 | chr9-75161483-DEL-176 |
| chr9-75206077-DEL-66 | chr9-75206078-DEL-66 |
| chr9-75210108-DEL-169 | chr9-75210109-DEL-169 |
| chr9-75212801-DEL-534 | chr9-75212801-DEL-534 |
| chr9-75230425-DEL-50 | chr9-75230416-DEL-71 |
| chr9-75348065-DEL-208 | chr9-75348076-DEL-195 |
| chr9-75354075-DEL-222 | chr9-75354076-DEL-222 |
| chr9-75358145-DEL-247 | chr9-75358146-DEL-247 |
| chr9-75376814-DEL-52 | chr9-75376815-DEL-52 |
| chr9-75390989-DEL-399 | chr9-75390990-DEL-399 |
| chr9-75400208-DEL-62 | chr9-75400209-DEL-61 |
| chr9-75412601-DEL-1582 | chr9-75412602-DEL-1582 |
| chr9-75427585-DEL-176 | chr9-75427585-DEL-201 |
| chr9-75433659-DEL-639 | chr9-75433660-DEL-639 |
| chr9-75452680-DEL-206 | chr9-75452681-DEL-206 |
| chr9-75463145-DEL-672 | chr9-75463146-DEL-672 |
| chr9-75477845-DEL-101 | chr9-75477846-DEL-101 |
| chr9-75480382-DEL-62 | chr9-75480383-DEL-62 |
| chr9-75500290-DEL-51 | chr9-75500291-DEL-51 |
| chr9-75524918-DEL-139 | chr9-75524919-DEL-139 |
| chr9-75525263-DEL-1762 | chr9-75525264-DEL-1762 |
| chr9-75537860-DEL-836 | chr9-75537861-DEL-836 |
| chr9-75543648-DEL-189 | chr9-75543646-DEL-186 |
| chr9-75549857-DEL-54 | chr9-75549858-DEL-54 |
| chr9-75554059-DEL-139 | chr9-75554060-DEL-139 |
| chr9-75557041-DEL-73 | chr9-75557031-DEL-73 |
| chr9-75558661-DEL-100 | chr9-75558662-DEL-99 |
| chr9-75571197-DEL-185 | chr9-75571198-DEL-185 |
| chr9-75581332-DEL-91 | chr9-75581333-DEL-91 |
| chr9-75598058-DEL-63 | chr9-75598058-DEL-63 |
| chr9-75648856-DEL-771 | chr9-75648857-DEL-771 |
| chr9-75665337-DEL-171 | chr9-75665338-DEL-171 |
| chr9-75712972-DEL-67 | chr9-75712975-DEL-65 |
| chr9-75818710-DEL-99 | chr9-75818711-DEL-99 |
| chr9-75825210-DEL-6857 | chr9-75825209-DEL-6858 |
| chr9-75831587-DEL-424 | chr9-75831588-DEL-424 |
| chr9-75843173-DEL-63 | chr9-75843174-DEL-63 |
| chr9-75877634-DEL-128 | chr9-75877635-DEL-128 |
| chr9-75880371-DEL-6184 | chr9-75880372-DEL-6184 |
| chr9-75918139-DEL-129 | chr9-75918141-DEL-126 |
| chr9-75955630-DEL-1246 | chr9-75955631-DEL-1246 |
| chr9-75962867-DEL-231 | chr9-75962868-DEL-231 |
| chr9-76013730-DEL-497 | chr9-76013731-DEL-497 |
| chr9-76047706-DEL-54 | chr9-76047707-DEL-52 |
| chr9-76061218-DEL-87 | chr9-76061219-DEL-88 |
| chr9-76063645-DEL-96 | chr9-76063646-DEL-96 |
| chr9-76090326-DEL-6469 | chr9-76090327-DEL-6469 |
| chr9-76096450-DEL-214 | chr9-76096451-DEL-214 |
| chr9-76164437-DEL-203 | chr9-76164438-DEL-203 |
| chr9-76201227-DEL-60 | chr9-76201228-DEL-60 |
| chr9-76279138-DEL-258 | chr9-76279139-DEL-258 |
| chr9-76285214-DEL-106 | chr9-76285215-DEL-106 |
| chr9-76319577-DEL-51 | chr9-76319578-DEL-51 |
| chr9-76330473-DEL-114 | chr9-76330482-DEL-99 |
| chr9-76347548-DEL-521 | chr9-76347549-DEL-521 |
| chr9-76353752-DEL-431 | chr9-76353753-DEL-431 |
| chr9-76361524-DEL-1342 | chr9-76361525-DEL-1342 |
| chr9-76364853-DEL-67 | chr9-76364806-DEL-68 |
| chr9-76377941-DEL-54 | chr9-76377942-DEL-54 |
| chr9-76454507-DEL-96 | chr9-76454496-DEL-68 |
| chr9-76484290-DEL-147 | chr9-76484298-DEL-120 |
| chr9-76498381-DEL-281 | chr9-76498383-DEL-284 |
| chr9-76502655-DEL-1079 | chr9-76502656-DEL-1079 |
| chr9-76513980-DEL-4393 | chr9-76513981-DEL-4393 |
| chr9-76519107-DEL-1119 | chr9-76519108-DEL-1119 |
| chr9-76532806-DEL-350 | chr9-76532807-DEL-350 |
| chr9-76551770-DEL-933 | chr9-76551771-DEL-933 |
| chr9-76573419-DEL-72 | chr9-76573420-DEL-72 |
| chr9-76579874-DEL-2769 | chr9-76579875-DEL-2769 |
| chr9-76587854-DEL-208 | chr9-76587864-DEL-199 |
| chr9-76589758-DEL-76 | chr9-76589759-DEL-76 |
| chr9-76608364-DEL-127 | chr9-76608374-DEL-118 |

| | |
|---|---|
| chr9-76642034-DEL-124 | chr9-76642035-DEL-124 |
| chr9-76642652-DEL-3906 | chr9-76642653-DEL-3906 |
| chr9-76662236-DEL-57 | chr9-76662237-DEL-57 |
| chr9-76662405-DEL-2424 | chr9-76662404-DEL-2425 |
| chr9-76716877-DEL-6266 | chr9-76716878-DEL-6266 |
| chr9-76775789-DEL-2155 | chr9-76775790-DEL-2155 |
| chr9-76780508-DEL-64 | chr9-76780509-DEL-64 |
| chr9-76780697-DEL-1677 | chr9-76780698-DEL-1677 |
| chr9-76790028-DEL-249 | chr9-76790026-DEL-250 |
| chr9-76829415-DEL-70 | chr9-76829411-DEL-95 |
| chr9-76856335-DEL-755 | chr9-76856336-DEL-755 |
| chr9-76865898-DEL-386 | chr9-76865900-DEL-385 |
| chr9-76879649-DEL-1415 | chr9-76879655-DEL-1413 |
| chr9-76890862-DEL-125 | chr9-76890863-DEL-125 |
| chr9-76928028-DEL-686 | chr9-76928029-DEL-686 |
| chr9-77008911-DEL-176 | chr9-77008912-DEL-176 |
| chr9-77036635-DEL-4630 | chr9-77036636-DEL-4630 |
| chr9-77100589-DEL-71 | chr9-77100590-DEL-71 |
| chr9-77107673-DEL-141 | chr9-77107674-DEL-141 |
| chr9-77113739-DEL-63 | chr9-77113696-DEL-70 |
| chr9-77126644-DEL-125 | chr9-77126645-DEL-125 |
| chr9-77149221-DEL-814 | chr9-77149222-DEL-814 |
| chr9-77150487-DEL-117 | chr9-77150484-DEL-84 |
| chr9-77158250-DEL-58 | chr9-77158246-DEL-60 |
| chr9-77191097-DEL-1101 | chr9-77191098-DEL-1101 |
| chr9-77201210-DEL-256 | chr9-77201210-DEL-256 |
| chr9-77205282-DEL-124 | chr9-77205283-DEL-124 |
| chr9-77239122-DEL-2575 | chr9-77239123-DEL-2575 |
| chr9-77242653-DEL-687 | chr9-77242654-DEL-687 |
| chr9-77257180-DEL-356 | chr9-77257181-DEL-356 |
| chr9-77282398-DEL-150 | chr9-77282401-DEL-148 |
| chr9-77285749-DEL-1236 | chr9-77285750-DEL-1236 |
| chr9-77309157-DEL-497 | chr9-77309158-DEL-497 |
| chr9-77311652-DEL-66 | chr9-77311653-DEL-66 |
| chr9-77324560-DEL-130 | chr9-77324573-DEL-118 |
| chr9-77347324-DEL-52 | chr9-77347327-DEL-50 |
| chr9-77367407-DEL-111 | chr9-77367408-DEL-111 |
| chr9-77398253-DEL-619 | chr9-77398254-DEL-619 |
| chr9-77407640-DEL-63 | chr9-77407637-DEL-63 |
| chr9-77411965-DEL-2923 | chr9-77411966-DEL-2923 |
| chr9-77459408-DEL-110 | chr9-77459409-DEL-110 |
| chr9-77479700-DEL-2423 | chr9-77479702-DEL-2422 |
| chr9-77479857-DEL-129 | chr9-77479851-DEL-133 |
| chr9-77516080-DEL-195 | chr9-77516082-DEL-194 |
| chr9-77517480-DEL-400 | chr9-77517481-DEL-400 |
| chr9-77588664-DEL-112 | chr9-77588665-DEL-112 |
| chr9-77589901-DEL-60 | chr9-77589902-DEL-60 |
| chr9-77603668-DEL-1865 | chr9-77603658-DEL-1866 |
| chr9-77619527-DEL-203 | chr9-77619528-DEL-203 |
| chr9-77648236-DEL-74 | chr9-77648237-DEL-74 |
| chr9-77664214-DEL-225 | chr9-77664215-DEL-225 |
| chr9-77672270-DEL-71 | chr9-77672271-DEL-71 |
| chr9-77725182-DEL-65 | chr9-77725153-DEL-81 |
| chr9-77738892-DEL-182 | chr9-77738893-DEL-181 |
| chr9-77749911-DEL-213 | chr9-77749912-DEL-213 |
| chr9-77793180-DEL-106 | chr9-77793181-DEL-106 |
| chr9-77813385-DEL-81 | chr9-77813386-DEL-81 |
| chr9-77843818-DEL-173 | chr9-77843819-DEL-174 |
| chr9-77858859-DEL-420 | chr9-77858860-DEL-420 |
| chr9-77865243-DEL-135 | chr9-77865244-DEL-135 |
| chr9-77869087-DEL-1305 | chr9-77869088-DEL-1305 |
| chr9-77892826-DEL-81 | chr9-77892832-DEL-75 |
| chr9-77894651-DEL-100 | chr9-77894664-DEL-92 |
| chr9-77914527-DEL-56 | chr9-77914528-DEL-56 |
| chr9-77945529-DEL-66 | chr9-77945509-DEL-50 |
| chr9-77963524-DEL-205 | chr9-77963525-DEL-205 |
| chr9-77975234-DEL-206 | chr9-77975235-DEL-206 |
| chr9-77993719-DEL-146 | chr9-77993720-DEL-146 |
| chr9-77995575-DEL-56 | chr9-77995576-DEL-56 |
| chr9-78003587-DEL-179 | chr9-78003588-DEL-179 |
| chr9-78046691-DEL-74 | chr9-78046681-DEL-54 |
| chr9-78058380-DEL-69 | chr9-78058392-DEL-62 |
| chr9-78070915-DEL-176 | chr9-78070916-DEL-176 |
| chr9-78086966-DEL-3038 | chr9-78086967-DEL-3038 |
| chr9-78091965-DEL-80 | chr9-78091966-DEL-80 |
| chr9-78100374-DEL-58 | chr9-78100332-DEL-52 |
| chr9-78103548-DEL-1874 | chr9-78103547-DEL-1875 |
| chr9-78129764-DEL-59 | chr9-78129750-DEL-60 |
| chr9-78190041-DEL-6401 | chr9-78190042-DEL-6401 |
| chr9-78230878-DEL-205 | chr9-78230879-DEL-205 |

| | |
|---|---|
| chr9-78232956-DEL-569 | chr9-78232955-DEL-568 |
| chr9-78242074-DEL-211 | chr9-78242084-DEL-212 |
| chr9-78254776-DEL-102 | chr9-78254783-DEL-96 |
| chr9-78257037-DEL-180 | chr9-78257038-DEL-180 |
| chr9-78257757-DEL-233 | chr9-78257758-DEL-233 |
| chr9-78265980-DEL-185 | chr9-78265981-DEL-185 |
| chr9-78305896-DEL-215 | chr9-78305897-DEL-215 |
| chr9-78334764-DEL-60 | chr9-78334765-DEL-60 |
| chr9-78352714-DEL-141 | chr9-78352673-DEL-183 |
| chr9-78354604-DEL-63 | chr9-78354605-DEL-63 |
| chr9-78364024-DEL-123 | chr9-78364025-DEL-123 |
| chr9-78367169-DEL-209 | chr9-78367170-DEL-209 |
| chr9-78373687-DEL-58 | chr9-78373688-DEL-58 |
| chr9-78375580-DEL-198 | chr9-78375581-DEL-198 |
| chr9-78402494-DEL-75 | chr9-78402461-DEL-115 |
| chr9-78412235-DEL-194 | chr9-78412235-DEL-194 |
| chr9-78415178-DEL-63 | chr9-78415188-DEL-53 |
| chr9-78417139-DEL-197 | chr9-78417135-DEL-198 |
| chr9-78417958-DEL-332 | chr9-78417959-DEL-332 |
| chr9-78418642-DEL-216 | chr9-78418643-DEL-216 |
| chr9-78424726-DEL-202 | chr9-78424727-DEL-202 |
| chr9-78429404-DEL-210 | chr9-78429449-DEL-199 |
| chr9-78430207-DEL-104 | chr9-78430209-DEL-102 |
| chr9-78436132-DEL-491 | chr9-78436134-DEL-490 |
| chr9-78442645-DEL-269 | chr9-78442646-DEL-269 |
| chr9-78447274-DEL-188 | chr9-78447285-DEL-178 |
| chr9-78451002-DEL-209 | chr9-78451003-DEL-209 |
| chr9-78455510-DEL-456 | chr9-78455512-DEL-454 |
| chr9-78456134-DEL-168 | chr9-78456135-DEL-168 |
| chr9-78462072-DEL-1895 | chr9-78462074-DEL-1895 |
| chr9-78465683-DEL-320 | chr9-78465684-DEL-320 |
| chr9-78470152-DEL-5133 | chr9-78470153-DEL-5133 |
| chr9-78499575-DEL-59 | chr9-78499576-DEL-56 |
| chr9-78527028-DEL-243 | chr9-78527029-DEL-243 |
| chr9-78546514-DEL-100 | chr9-78546515-DEL-50 |
| chr9-78547911-DEL-66 | chr9-78547912-DEL-80 |
| chr9-78554600-DEL-62 | chr9-78554601-DEL-62 |
| chr9-78619176-DEL-53 | chr9-78619179-DEL-51 |
| chr9-78632408-DEL-138 | chr9-78632409-DEL-92 |
| chr9-78642067-DEL-331 | chr9-78642062-DEL-323 |
| chr9-78650876-DEL-341 | chr9-78650870-DEL-342 |
| chr9-78652822-DEL-120 | chr9-78652823-DEL-120 |
| chr9-78674099-DEL-81 | chr9-78674103-DEL-74 |
| chr9-78719662-DEL-60 | chr9-78719663-DEL-66 |
| chr9-78722406-DEL-60 | chr9-78722417-DEL-50 |
| chr9-78725846-DEL-3351 | chr9-78725847-DEL-3351 |
| chr9-78730578-DEL-7077 | chr9-78730579-DEL-7077 |
| chr9-78761258-DEL-417 | chr9-78761262-DEL-419 |
| chr9-78775541-DEL-64 | chr9-78775564-DEL-57 |
| chr9-78776022-DEL-304 | chr9-78776023-DEL-304 |
| chr9-78790105-DEL-7472 | chr9-78790106-DEL-7472 |
| chr9-78803093-DEL-52 | chr9-78803088-DEL-60 |
| chr9-78809597-DEL-52 | chr9-78809598-DEL-52 |
| chr9-78837396-DEL-1717 | chr9-78837396-DEL-1717 |
| chr9-78838278-DEL-2104 | chr9-78838278-DEL-2104 |
| chr9-78852046-DEL-355 | chr9-78852047-DEL-355 |
| chr9-78854535-DEL-1209 | chr9-78854534-DEL-1209 |
| chr9-78855777-DEL-954 | chr9-78855777-DEL-954 |
| chr9-78868379-DEL-192 | chr9-78868380-DEL-192 |
| chr9-78881254-DEL-1223 | chr9-78881255-DEL-1223 |
| chr9-78907907-DEL-55 | chr9-78907908-DEL-55 |
| chr9-78912682-DEL-1051 | chr9-78912683-DEL-1051 |
| chr9-78972685-DEL-488 | chr9-78972686-DEL-488 |
| chr9-78999915-DEL-8798 | chr9-78999916-DEL-8798 |
| chr9-79020194-DEL-2907 | chr9-79020188-DEL-2913 |
| chr9-79027803-DEL-514 | chr9-79027804-DEL-513 |
| chr9-79058918-DEL-230 | chr9-79058919-DEL-260 |
| chr9-79071617-DEL-813 | chr9-79071618-DEL-813 |
| chr9-79125941-DEL-638 | chr9-79125942-DEL-638 |
| chr9-79134106-DEL-505 | chr9-79134102-DEL-506 |
| chr9-79161736-DEL-7136 | chr9-79161737-DEL-7136 |
| chr9-79181987-DEL-82 | chr9-79181988-DEL-82 |
| chr9-79200352-DEL-623 | chr9-79200353-DEL-623 |
| chr9-79205375-DEL-586 | chr9-79205375-DEL-587 |
| chr9-79226936-DEL-910 | chr9-79226937-DEL-910 |
| chr9-79229666-DEL-173 | chr9-79229667-DEL-173 |
| chr9-79243608-DEL-289 | chr9-79243609-DEL-289 |
| chr9-79274313-DEL-168 | chr9-79274314-DEL-168 |
| chr9-79278055-DEL-1222 | chr9-79278056-DEL-1222 |
| chr9-79293842-DEL-52 | chr9-79293843-DEL-52 |

| | |
|---|---|
| chr9-79310154-DEL-6602 | chr9-79310155-DEL-6602 |
| chr9-79359969-DEL-52 | chr9-79360007-DEL-52 |
| chr9-79371048-DEL-2103 | chr9-79371049-DEL-2102 |
| chr9-79375432-DEL-3674 | chr9-79375433-DEL-3674 |
| chr9-79383223-DEL-55 | chr9-79383224-DEL-55 |
| chr9-79392375-DEL-1318 | chr9-79392375-DEL-1318 |
| chr9-79413548-DEL-779 | chr9-79413549-DEL-779 |
| chr9-79432278-DEL-87 | chr9-79432279-DEL-87 |
| chr9-79452554-DEL-510 | chr9-79452555-DEL-510 |
| chr9-79542971-DEL-1011 | chr9-79542972-DEL-1011 |
| chr9-79566610-DEL-68 | chr9-79566611-DEL-68 |
| chr9-79609801-DEL-309 | chr9-79609802-DEL-309 |
| chr9-79618398-DEL-56 | chr9-79618399-DEL-56 |
| chr9-79719617-DEL-54 | chr9-79719617-DEL-60 |
| chr9-79728074-DEL-60 | chr9-79728075-DEL-60 |
| chr9-79763323-DEL-94 | chr9-79763324-DEL-92 |
| chr9-79807045-DEL-4249 | chr9-79807046-DEL-4249 |
| chr9-79808235-DEL-409 | chr9-79808236-DEL-409 |
| chr9-79822161-DEL-106 | chr9-79822162-DEL-106 |
| chr9-79832574-DEL-72 | chr9-79832575-DEL-108 |
| chr9-79836258-DEL-62 | chr9-79836259-DEL-62 |
| chr9-79853667-DEL-659 | chr9-79853668-DEL-659 |
| chr9-79869702-DEL-106 | chr9-79869699-DEL-110 |
| chr9-79898911-DEL-239 | chr9-79898889-DEL-231 |
| chr9-79903659-DEL-51 | chr9-79903660-DEL-51 |
| chr9-79905174-DEL-66 | chr9-79905190-DEL-51 |
| chr9-79907571-DEL-168 | chr9-79907572-DEL-168 |
| chr9-79931193-DEL-138 | chr9-79931194-DEL-138 |
| chr9-79936376-DEL-59 | chr9-79936377-DEL-59 |
| chr9-79940023-DEL-2683 | chr9-79940024-DEL-2683 |
| chr9-79948727-DEL-1059 | chr9-79948728-DEL-1059 |
| chr9-79991227-DEL-396 | chr9-79991228-DEL-396 |
| chr9-80043287-DEL-56 | chr9-80043288-DEL-56 |
| chr9-80049747-DEL-138 | chr9-80049748-DEL-138 |
| chr9-80077968-DEL-78 | chr9-80077969-DEL-72 |
| chr9-80080481-DEL-92 | chr9-80080472-DEL-86 |
| chr9-80083823-DEL-136 | chr9-80083824-DEL-140 |
| chr9-80105979-DEL-497 | chr9-80105980-DEL-497 |
| chr9-80117291-DEL-62 | chr9-80117292-DEL-62 |
| chr9-80137530-DEL-246 | chr9-80137531-DEL-246 |
| chr9-80143603-DEL-144 | chr9-80143605-DEL-143 |
| chr9-80172467-DEL-54 | chr9-80172450-DEL-72 |
| chr9-80226205-DEL-1571 | chr9-80226206-DEL-1571 |
| chr9-80232307-DEL-234 | chr9-80232309-DEL-228 |
| chr9-80234442-DEL-205 | chr9-80234443-DEL-205 |
| chr9-80235197-DEL-103 | chr9-80235198-DEL-103 |
| chr9-80244221-DEL-697 | chr9-80244222-DEL-697 |
| chr9-80246703-DEL-61 | chr9-80246704-DEL-61 |
| chr9-80269377-DEL-73 | chr9-80269378-DEL-73 |
| chr9-80297871-DEL-91 | chr9-80297876-DEL-87 |
| chr9-80300739-DEL-190 | chr9-80300740-DEL-190 |
| chr9-80302790-DEL-1030 | chr9-80302807-DEL-1039 |
| chr9-80315718-DEL-236 | chr9-80315726-DEL-187 |
| chr9-80335485-DEL-64 | chr9-80335490-DEL-60 |
| chr9-80356596-DEL-63 | chr9-80356597-DEL-63 |
| chr9-80359805-DEL-580 | chr9-80359806-DEL-580 |
| chr9-80382683-DEL-605 | chr9-80382684-DEL-605 |
| chr9-80401936-DEL-6518 | chr9-80401937-DEL-6518 |
| chr9-80409274-DEL-527 | chr9-80409275-DEL-527 |
| chr9-80438657-DEL-275 | chr9-80438658-DEL-275 |
| chr9-80444111-DEL-66 | chr9-80444112-DEL-66 |
| chr9-80449650-DEL-69 | chr9-80449650-DEL-69 |
| chr9-80452091-DEL-448 | chr9-80452092-DEL-448 |
| chr9-80452841-DEL-879 | chr9-80452842-DEL-879 |
| chr9-80460254-DEL-70 | chr9-80460255-DEL-70 |
| chr9-80470660-DEL-6407 | chr9-80470661-DEL-6407 |
| chr9-80485798-DEL-784 | chr9-80485799-DEL-784 |
| chr9-80499850-DEL-586 | chr9-80499851-DEL-586 |
| chr9-80509645-DEL-387 | chr9-80509646-DEL-387 |
| chr9-80524495-DEL-418 | chr9-80524497-DEL-412 |
| chr9-80538225-DEL-339 | chr9-80538243-DEL-329 |
| chr9-80542794-DEL-5976 | chr9-80542795-DEL-5976 |
| chr9-80568663-DEL-228 | chr9-80568664-DEL-228 |
| chr9-80602560-DEL-81 | chr9-80602561-DEL-81 |
| chr9-80603122-DEL-2656 | chr9-80603123-DEL-2656 |
| chr9-80624482-DEL-52 | chr9-80624483-DEL-52 |
| chr9-80626474-DEL-84 | chr9-80626475-DEL-84 |
| chr9-80665989-DEL-174 | chr9-80665990-DEL-174 |
| chr9-80699108-DEL-84 | chr9-80699112-DEL-62 |
| chr9-80722376-DEL-81 | chr9-80722333-DEL-89 |

| | |
|---|---|
| chr9-80787021-DEL-100 | chr9-80787021-DEL-99 |
| chr9-80789793-DEL-223 | chr9-80789794-DEL-221 |
| chr9-80814801-DEL-87 | chr9-80814802-DEL-87 |
| chr9-80831076-DEL-60 | chr9-80831072-DEL-62 |
| chr9-80832023-DEL-8572 | chr9-80832024-DEL-8572 |
| chr9-80865368-DEL-1052 | chr9-80865369-DEL-1052 |
| chr9-80887795-DEL-73 | chr9-80887796-DEL-73 |
| chr9-80896981-DEL-1056 | chr9-80896982-DEL-1056 |
| chr9-80917579-DEL-1661 | chr9-80917580-DEL-1661 |
| chr9-80929351-DEL-130 | chr9-80929359-DEL-123 |
| chr9-80932229-DEL-6177 | chr9-80932230-DEL-6177 |
| chr9-80982999-DEL-1533 | chr9-80983042-DEL-1490 |
| chr9-80999178-DEL-203 | chr9-80999178-DEL-204 |
| chr9-81007972-DEL-152 | chr9-81007971-DEL-154 |
| chr9-81042008-DEL-641 | chr9-81042009-DEL-641 |
| chr9-81058193-DEL-1683 | chr9-81058194-DEL-1683 |
| chr9-81064690-DEL-6514 | chr9-81064691-DEL-6514 |
| chr9-81083405-DEL-59 | chr9-81083402-DEL-55 |
| chr9-81095506-DEL-54 | chr9-81095462-DEL-62 |
| chr9-81124461-DEL-70 | chr9-81124477-DEL-60 |
| chr9-81132408-DEL-9932 | chr9-81132409-DEL-9932 |
| chr9-81151910-DEL-173 | chr9-81151911-DEL-173 |
| chr9-81155874-DEL-699 | chr9-81155875-DEL-699 |
| chr9-81161639-DEL-9073 | chr9-81161640-DEL-9073 |
| chr9-81171228-DEL-119 | chr9-81171229-DEL-119 |
| chr9-81178255-DEL-83 | chr9-81178256-DEL-83 |
| chr9-81180380-DEL-4676 | chr9-81180381-DEL-4676 |
| chr9-81190599-DEL-327 | chr9-81190600-DEL-327 |
| chr9-81201368-DEL-65 | chr9-81201363-DEL-66 |
| chr9-81209978-DEL-1061 | chr9-81209979-DEL-1061 |
| chr9-81220366-DEL-52 | chr9-81220363-DEL-56 |
| chr9-81236814-DEL-394 | chr9-81236815-DEL-394 |
| chr9-81308291-DEL-2468 | chr9-81308292-DEL-2468 |
| chr9-81324133-DEL-957 | chr9-81324133-DEL-958 |
| chr9-81355209-DEL-531 | chr9-81355210-DEL-531 |
| chr9-81366875-DEL-6776 | chr9-81366876-DEL-6776 |
| chr9-81383096-DEL-60 | chr9-81383097-DEL-60 |
| chr9-81387000-DEL-58 | chr9-81387001-DEL-58 |
| chr9-81419782-DEL-642 | chr9-81419783-DEL-642 |
| chr9-81425677-DEL-934 | chr9-81425678-DEL-934 |
| chr9-81430981-DEL-55 | chr9-81430982-DEL-60 |
| chr9-81450340-DEL-102 | chr9-81450344-DEL-99 |
| chr9-81459294-DEL-204 | chr9-81459295-DEL-204 |
| chr9-81463822-DEL-358 | chr9-81463823-DEL-358 |
| chr9-81477865-DEL-102 | chr9-81477820-DEL-129 |
| chr9-81496144-DEL-6592 | chr9-81496145-DEL-6592 |
| chr9-81547199-DEL-400 | chr9-81547200-DEL-400 |
| chr9-81549697-DEL-1691 | chr9-81549698-DEL-1691 |
| chr9-81622307-DEL-606 | chr9-81622308-DEL-606 |
| chr9-81639224-DEL-139 | chr9-81639225-DEL-139 |
| chr9-81645034-DEL-107 | chr9-81645035-DEL-107 |
| chr9-81654901-DEL-1483 | chr9-81654900-DEL-1483 |
| chr9-81687031-DEL-52 | chr9-81687032-DEL-52 |
| chr9-81690837-DEL-137 | chr9-81690838-DEL-137 |
| chr9-81743274-DEL-208 | chr9-81743275-DEL-208 |
| chr9-81780404-DEL-53 | chr9-81780392-DEL-54 |
| chr9-81788494-DEL-6488 | chr9-81788495-DEL-6488 |
| chr9-81808898-DEL-75 | chr9-81808899-DEL-75 |
| chr9-81812432-DEL-6865 | chr9-81812433-DEL-6865 |
| chr9-81855320-DEL-68 | chr9-81855321-DEL-68 |
| chr9-81870953-DEL-439 | chr9-81870954-DEL-439 |
| chr9-81873064-DEL-168 | chr9-81873071-DEL-161 |
| chr9-81882144-DEL-135 | chr9-81882145-DEL-135 |
| chr9-81887027-DEL-387 | chr9-81887028-DEL-387 |
| chr9-81891542-DEL-159 | chr9-81891543-DEL-160 |
| chr9-81897632-DEL-65 | chr9-81897632-DEL-66 |
| chr9-81904341-DEL-113 | chr9-81904341-DEL-110 |
| chr9-81915314-DEL-194 | chr9-81915314-DEL-194 |
| chr9-81915564-DEL-1063 | chr9-81915565-DEL-1063 |
| chr9-81924436-DEL-173 | chr9-81924437-DEL-173 |
| chr9-81925536-DEL-52 | chr9-81925537-DEL-52 |
| chr9-81966333-DEL-973 | chr9-81966332-DEL-974 |
| chr9-81974683-DEL-231 | chr9-81974684-DEL-231 |
| chr9-81987246-DEL-3272 | chr9-81987247-DEL-3272 |
| chr9-82008165-DEL-1036 | chr9-82008166-DEL-1036 |
| chr9-82069246-DEL-205 | chr9-82069247-DEL-205 |
| chr9-82097692-DEL-91 | chr9-82097693-DEL-91 |
| chr9-82103340-DEL-93 | chr9-82103309-DEL-139 |
| chr9-82124050-DEL-82 | chr9-82124047-DEL-86 |
| chr9-82171204-DEL-1973 | chr9-82171205-DEL-1973 |

| | |
|---|---|
| chr9-82192191-DEL-525 | chr9-82192192-DEL-525 |
| chr9-82194371-DEL-197 | chr9-82194371-DEL-197 |
| chr9-82204846-DEL-386 | chr9-82204847-DEL-386 |
| chr9-82209720-DEL-1211 | chr9-82209721-DEL-1211 |
| chr9-82227019-DEL-259 | chr9-82227020-DEL-260 |
| chr9-82232994-DEL-50 | chr9-82233005-DEL-58 |
| chr9-82234278-DEL-6691 | chr9-82234279-DEL-6691 |
| chr9-82247137-DEL-301 | chr9-82247138-DEL-301 |
| chr9-82248894-DEL-3524 | chr9-82248895-DEL-3524 |
| chr9-82253089-DEL-1648 | chr9-82253090-DEL-1648 |
| chr9-82276546-DEL-221 | chr9-82276556-DEL-224 |
| chr9-82277175-DEL-713 | chr9-82277171-DEL-710 |
| chr9-82336263-DEL-270 | chr9-82336267-DEL-269 |
| chr9-82336599-DEL-505 | chr9-82336600-DEL-505 |
| chr9-82349482-DEL-54 | chr9-82349483-DEL-54 |
| chr9-82351913-DEL-6725 | chr9-82351914-DEL-6725 |
| chr9-82394047-DEL-234 | chr9-82394048-DEL-234 |
| chr9-82412867-DEL-52 | chr9-82412868-DEL-52 |
| chr9-82434163-DEL-68 | chr9-82434142-DEL-72 |
| chr9-82446954-DEL-6808 | chr9-82446955-DEL-6808 |
| chr9-82447192-DEL-208 | chr9-82447193-DEL-208 |
| chr9-82495838-DEL-321 | chr9-82495839-DEL-321 |
| chr9-82503338-DEL-53 | chr9-82503339-DEL-53 |
| chr9-82508210-DEL-324 | chr9-82508211-DEL-324 |
| chr9-82511820-DEL-77 | chr9-82511854-DEL-77 |
| chr9-82551852-DEL-269 | chr9-82551853-DEL-269 |
| chr9-82554278-DEL-1143 | chr9-82554278-DEL-1143 |
| chr9-82580878-DEL-862 | chr9-82580879-DEL-862 |
| chr9-82583690-DEL-2272 | chr9-82583691-DEL-2272 |
| chr9-82601384-DEL-54 | chr9-82601385-DEL-54 |
| chr9-82605976-DEL-196 | chr9-82605977-DEL-208 |
| chr9-82607112-DEL-467 | chr9-82607113-DEL-467 |
| chr9-82612339-DEL-589 | chr9-82612340-DEL-589 |
| chr9-82617161-DEL-143 | chr9-82617161-DEL-142 |
| chr9-82626368-DEL-113 | chr9-82626374-DEL-108 |
| chr9-82636089-DEL-68 | chr9-82636089-DEL-71 |
| chr9-82641598-DEL-63 | chr9-82641595-DEL-67 |
| chr9-82646609-DEL-1318 | chr9-82646610-DEL-1318 |
| chr9-82649970-DEL-12368 | chr9-82649971-DEL-12368 |
| chr9-82665423-DEL-139 | chr9-82665424-DEL-139 |
| chr9-82687340-DEL-150 | chr9-82687326-DEL-198 |
| chr9-82688625-DEL-66 | chr9-82688633-DEL-65 |
| chr9-82706041-DEL-90 | chr9-82706042-DEL-90 |
| chr9-82725164-DEL-233 | chr9-82725165-DEL-233 |
| chr9-82726861-DEL-54 | chr9-82726862-DEL-54 |
| chr9-82842242-DEL-504 | chr9-82842243-DEL-504 |
| chr9-82850190-DEL-97 | chr9-82850190-DEL-97 |
| chr9-82882843-DEL-237 | chr9-82882844-DEL-237 |
| chr9-82907080-DEL-308 | chr9-82907081-DEL-308 |
| chr9-82911152-DEL-296 | chr9-82911153-DEL-296 |
| chr9-82990464-DEL-59 | chr9-82990417-DEL-76 |
| chr9-83030006-DEL-2214 | chr9-83030007-DEL-2214 |
| chr9-83055127-DEL-4984 | chr9-83055128-DEL-4984 |
| chr9-83092921-DEL-96 | chr9-83092917-DEL-97 |
| chr9-83093605-DEL-882 | chr9-83093606-DEL-882 |
| chr9-83109791-DEL-102 | chr9-83109769-DEL-102 |
| chr9-83124174-DEL-75 | chr9-83124163-DEL-50 |
| chr9-83128355-DEL-114 | chr9-83128325-DEL-110 |
| chr9-83148117-DEL-92 | chr9-83148118-DEL-92 |
| chr9-83151194-DEL-59 | chr9-83151195-DEL-59 |
| chr9-83160850-DEL-3942 | chr9-83160850-DEL-3942 |
| chr9-83172394-DEL-804 | chr9-83172395-DEL-804 |
| chr9-83173557-DEL-448 | chr9-83173558-DEL-448 |
| chr9-83177123-DEL-52 | chr9-83177123-DEL-54 |
| chr9-83227985-DEL-51 | chr9-83227986-DEL-75 |
| chr9-83234663-DEL-74 | chr9-83234663-DEL-74 |
| chr9-83291040-DEL-1499 | chr9-83291041-DEL-1499 |
| chr9-83291667-DEL-557 | chr9-83291659-DEL-560 |
| chr9-83342671-DEL-6400 | chr9-83342654-DEL-6400 |
| chr9-83354837-DEL-551 | chr9-83354837-DEL-551 |
| chr9-83357868-DEL-1951 | chr9-83357869-DEL-1951 |
| chr9-83364324-DEL-163 | chr9-83364325-DEL-163 |
| chr9-83386508-DEL-504 | chr9-83386509-DEL-504 |
| chr9-83389231-DEL-60 | chr9-83389232-DEL-60 |
| chr9-83394464-DEL-1893 | chr9-83394465-DEL-1893 |
| chr9-83447394-DEL-1050 | chr9-83447395-DEL-1050 |
| chr9-83498449-DEL-501 | chr9-83498448-DEL-502 |
| chr9-83499692-DEL-118 | chr9-83499689-DEL-122 |
| chr9-83525753-DEL-66 | chr9-83525755-DEL-57 |
| chr9-83538492-DEL-627 | chr9-83538493-DEL-627 |

| | |
|---|---|
| chr9-83540033-DEL-121 | chr9-83540066-DEL-72 |
| chr9-83570200-DEL-341 | chr9-83570201-DEL-360 |
| chr9-83576517-DEL-215 | chr9-83576518-DEL-215 |
| chr9-83577899-DEL-96 | chr9-83577900-DEL-96 |
| chr9-83591486-DEL-156 | chr9-83591487-DEL-156 |
| chr9-83615166-DEL-216 | chr9-83615157-DEL-214 |
| chr9-83622161-DEL-3507 | chr9-83622161-DEL-3507 |
| chr9-83626308-DEL-201 | chr9-83626274-DEL-198 |
| chr9-83626445-DEL-2394 | chr9-83626445-DEL-2394 |
| chr9-83682339-DEL-200 | chr9-83682342-DEL-195 |
| chr9-83702122-DEL-175 | chr9-83702123-DEL-175 |
| chr9-83723897-DEL-57 | chr9-83723898-DEL-57 |
| chr9-83734722-DEL-221 | chr9-83734723-DEL-221 |
| chr9-83763554-DEL-112 | chr9-83763555-DEL-112 |
| chr9-83777158-DEL-135 | chr9-83777159-DEL-135 |
| chr9-83808888-DEL-164 | chr9-83808889-DEL-164 |
| chr9-83824594-DEL-7327 | chr9-83824595-DEL-7327 |
| chr9-83834828-DEL-56 | chr9-83834829-DEL-56 |
| chr9-83855639-DEL-85 | chr9-83855640-DEL-85 |
| chr9-83860059-DEL-414 | chr9-83860060-DEL-414 |
| chr9-83860894-DEL-130 | chr9-83860895-DEL-130 |
| chr9-83861807-DEL-70 | chr9-83861814-DEL-66 |
| chr9-83869739-DEL-51 | chr9-83869735-DEL-52 |
| chr9-83874508-DEL-6511 | chr9-83874509-DEL-6511 |
| chr9-83895407-DEL-64 | chr9-83895366-DEL-63 |
| chr9-83904920-DEL-142 | chr9-83904955-DEL-141 |
| chr9-83906717-DEL-195 | chr9-83906718-DEL-195 |
| chr9-83921412-DEL-236 | chr9-83921433-DEL-215 |
| chr9-83936044-DEL-654 | chr9-83936045-DEL-654 |
| chr9-83940611-DEL-68 | chr9-83940612-DEL-51 |
| chr9-83981829-DEL-327 | chr9-83981830-DEL-327 |
| chr9-83990419-DEL-215 | chr9-83990420-DEL-215 |
| chr9-84010884-DEL-143 | chr9-84010921-DEL-142 |
| chr9-84014469-DEL-460 | chr9-84014465-DEL-466 |
| chr9-84046963-DEL-123 | chr9-84046964-DEL-123 |
| chr9-84093946-DEL-1319 | chr9-84093947-DEL-1319 |
| chr9-84102004-DEL-3647 | chr9-84102005-DEL-3647 |
| chr9-84108809-DEL-214 | chr9-84108810-DEL-214 |
| chr9-84129003-DEL-75 | chr9-84129004-DEL-75 |
| chr9-84145305-DEL-174 | chr9-84145306-DEL-174 |
| chr9-84157466-DEL-78 | chr9-84157449-DEL-51 |
| chr9-84186186-DEL-3506 | chr9-84186190-DEL-3503 |
| chr9-84209083-DEL-199 | chr9-84209084-DEL-199 |
| chr9-84243754-DEL-474 | chr9-84243755-DEL-474 |
| chr9-84250720-DEL-1432 | chr9-84250721-DEL-1432 |
| chr9-84254494-DEL-96 | chr9-84254495-DEL-96 |
| chr9-84289046-DEL-58 | chr9-84289049-DEL-56 |
| chr9-84305994-DEL-902 | chr9-84305995-DEL-902 |
| chr9-84312715-DEL-117 | chr9-84312716-DEL-117 |
| chr9-84316245-DEL-59 | chr9-84316246-DEL-59 |
| chr9-84318910-DEL-56 | chr9-84318911-DEL-52 |
| chr9-84337032-DEL-1186 | chr9-84337033-DEL-1186 |
| chr9-84357712-DEL-66 | chr9-84357697-DEL-72 |
| chr9-84387069-DEL-1736 | chr9-84387069-DEL-1737 |
| chr9-84389425-DEL-887 | chr9-84389426-DEL-887 |
| chr9-84395523-DEL-2721 | chr9-84395524-DEL-2721 |
| chr9-84405329-DEL-56 | chr9-84405330-DEL-56 |
| chr9-84406958-DEL-719 | chr9-84406959-DEL-719 |
| chr9-84411840-DEL-1243 | chr9-84411839-DEL-1242 |
| chr9-84416228-DEL-145 | chr9-84416227-DEL-146 |
| chr9-84454139-DEL-628 | chr9-84454140-DEL-628 |
| chr9-84486896-DEL-54 | chr9-84486898-DEL-52 |
| chr9-84490653-DEL-638 | chr9-84490655-DEL-637 |
| chr9-84507814-DEL-84 | chr9-84507815-DEL-84 |
| chr9-84528436-DEL-185 | chr9-84528437-DEL-185 |
| chr9-84535104-DEL-866 | chr9-84535105-DEL-866 |
| chr9-84540988-DEL-204 | chr9-84540967-DEL-204 |
| chr9-84554295-DEL-94 | chr9-84554296-DEL-94 |
| chr9-84589172-DEL-3075 | chr9-84589172-DEL-3075 |
| chr9-84646214-DEL-482 | chr9-84646215-DEL-482 |
| chr9-84649252-DEL-1381 | chr9-84649245-DEL-1382 |
| chr9-84667490-DEL-146 | chr9-84667491-DEL-146 |
| chr9-84678270-DEL-2048 | chr9-84678271-DEL-2048 |
| chr9-84688466-DEL-150 | chr9-84688459-DEL-151 |
| chr9-84700949-DEL-63 | chr9-84700939-DEL-64 |
| chr9-84715228-DEL-816 | chr9-84715229-DEL-816 |
| chr9-84721751-DEL-192 | chr9-84721752-DEL-192 |
| chr9-84821921-DEL-54 | chr9-84821909-DEL-64 |
| chr9-84839060-DEL-114 | chr9-84839061-DEL-114 |
| chr9-84876174-DEL-1866 | chr9-84876175-DEL-1866 |

| | |
|---|---|
| chr9-84878102-DEL-1537 | chr9-84878103-DEL-1537 |
| chr9-84884208-DEL-63 | chr9-84884212-DEL-60 |
| chr9-84890763-DEL-1546 | chr9-84890764-DEL-1546 |
| chr9-84954035-DEL-6223 | chr9-84954037-DEL-6223 |
| chr9-84961026-DEL-81 | chr9-84961047-DEL-63 |
| chr9-84974351-DEL-520 | chr9-84974371-DEL-501 |
| chr9-84975840-DEL-1372 | chr9-84975838-DEL-1371 |
| chr9-84992603-DEL-55 | chr9-84992602-DEL-57 |
| chr9-84993772-DEL-73 | chr9-84993774-DEL-72 |
| chr9-84995843-DEL-148 | chr9-84995844-DEL-148 |
| chr9-85014696-DEL-695 | chr9-85014697-DEL-695 |
| chr9-85018553-DEL-66 | chr9-85018554-DEL-66 |
| chr9-85019769-DEL-54 | chr9-85019770-DEL-54 |
| chr9-85027056-DEL-64 | chr9-85027057-DEL-64 |
| chr9-85053508-DEL-146 | chr9-85053509-DEL-146 |
| chr9-85079026-DEL-463 | chr9-85079027-DEL-463 |
| chr9-85091845-DEL-53 | chr9-85091846-DEL-53 |
| chr9-85093373-DEL-52 | chr9-85093371-DEL-54 |
| chr9-85124402-DEL-197 | chr9-85124403-DEL-173 |
| chr9-85130889-DEL-2703 | chr9-85130890-DEL-2703 |
| chr9-85145172-DEL-343 | chr9-85145173-DEL-343 |
| chr9-85159866-DEL-66 | chr9-85159875-DEL-50 |
| chr9-85164288-DEL-54 | chr9-85164287-DEL-58 |
| chr9-85175427-DEL-87 | chr9-85175449-DEL-66 |
| chr9-85195432-DEL-57 | chr9-85195432-DEL-58 |
| chr9-85262102-DEL-51 | chr9-85262103-DEL-51 |
| chr9-85284524-DEL-108 | chr9-85284552-DEL-81 |
| chr9-85298230-DEL-757 | chr9-85298234-DEL-754 |
| chr9-85313424-DEL-1030 | chr9-85313425-DEL-1030 |
| chr9-85340375-DEL-303 | chr9-85340376-DEL-303 |
| chr9-85357764-DEL-63 | chr9-85357765-DEL-63 |
| chr9-85358717-DEL-123 | chr9-85358747-DEL-112 |
| chr9-85388433-DEL-64 | chr9-85388426-DEL-72 |
| chr9-85390123-DEL-131 | chr9-85390124-DEL-131 |
| chr9-85393033-DEL-83 | chr9-85393052-DEL-64 |
| chr9-85395917-DEL-2406 | chr9-85395918-DEL-2406 |
| chr9-85411236-DEL-335 | chr9-85411237-DEL-335 |
| chr9-85422765-DEL-880 | chr9-85422766-DEL-880 |
| chr9-85460618-DEL-2247 | chr9-85460619-DEL-2247 |
| chr9-85496612-DEL-70 | chr9-85496591-DEL-76 |
| chr9-85530007-DEL-149 | chr9-85530008-DEL-149 |
| chr9-85539429-DEL-160 | chr9-85539431-DEL-159 |
| chr9-85542167-DEL-120 | chr9-85542168-DEL-120 |
| chr9-85586294-DEL-166 | chr9-85586286-DEL-162 |
| chr9-85628374-DEL-236 | chr9-85628375-DEL-236 |
| chr9-85637549-DEL-346 | chr9-85637550-DEL-346 |
| chr9-85642606-DEL-486 | chr9-85642607-DEL-486 |
| chr9-85663832-DEL-902 | chr9-85663833-DEL-902 |
| chr9-85674004-DEL-101 | chr9-85674005-DEL-101 |
| chr9-85693776-DEL-60 | chr9-85693763-DEL-61 |
| chr9-85746392-DEL-179 | chr9-85746393-DEL-179 |
| chr9-85755717-DEL-52 | chr9-85755716-DEL-50 |
| chr9-85780912-DEL-6662 | chr9-85780913-DEL-6662 |
| chr9-85792613-DEL-565 | chr9-85792613-DEL-565 |
| chr9-85796386-DEL-54 | chr9-85796357-DEL-84 |
| chr9-85798649-DEL-2791 | chr9-85798650-DEL-2791 |
| chr9-85801497-DEL-52 | chr9-85801496-DEL-50 |
| chr9-85817438-DEL-58 | chr9-85817439-DEL-58 |
| chr9-85821394-DEL-4168 | chr9-85821395-DEL-4168 |
| chr9-85847124-DEL-77 | chr9-85847124-DEL-78 |
| chr9-85848675-DEL-1357 | chr9-85848676-DEL-1357 |
| chr9-85869816-DEL-64 | chr9-85869817-DEL-64 |
| chr9-85878000-DEL-167 | chr9-85878001-DEL-167 |
| chr9-85882295-DEL-57 | chr9-85882287-DEL-63 |
| chr9-85886346-DEL-180 | chr9-85886347-DEL-180 |
| chr9-85891340-DEL-100 | chr9-85891341-DEL-98 |
| chr9-85939311-DEL-56 | chr9-85939312-DEL-56 |
| chr9-85944803-DEL-7121 | chr9-85944804-DEL-7121 |
| chr9-85985524-DEL-68 | chr9-85985525-DEL-68 |
| chr9-86002127-DEL-6605 | chr9-86002128-DEL-6605 |
| chr9-86016005-DEL-167 | chr9-86016006-DEL-167 |
| chr9-86017905-DEL-1891 | chr9-86017906-DEL-1891 |
| chr9-86063223-DEL-123 | chr9-86063224-DEL-123 |
| chr9-86070455-DEL-78 | chr9-86070456-DEL-78 |
| chr9-86075290-DEL-6240 | chr9-86075291-DEL-6240 |
| chr9-86109574-DEL-115 | chr9-86109575-DEL-115 |
| chr9-86118628-DEL-96 | chr9-86118629-DEL-88 |
| chr9-86159389-DEL-56 | chr9-86159390-DEL-56 |
| chr9-86161288-DEL-173 | chr9-86161290-DEL-172 |
| chr9-86171960-DEL-6480 | chr9-86171961-DEL-6480 |

| | |
|---|---|
| chr9-86199117-DEL-67 | chr9-86199118-DEL-73 |
| chr9-86229866-DEL-516 | chr9-86229867-DEL-516 |
| chr9-86263006-DEL-2507 | chr9-86263007-DEL-2507 |
| chr9-86268200-DEL-497 | chr9-86268201-DEL-497 |
| chr9-86296273-DEL-4750 | chr9-86296274-DEL-4750 |
| chr9-86319208-DEL-88 | chr9-86319209-DEL-96 |
| chr9-86322044-DEL-203 | chr9-86322045-DEL-203 |
| chr9-86340220-DEL-4984 | chr9-86340220-DEL-4984 |
| chr9-86353070-DEL-77 | chr9-86353071-DEL-77 |
| chr9-86406392-DEL-121 | chr9-86406393-DEL-121 |
| chr9-86438067-DEL-57 | chr9-86438068-DEL-57 |
| chr9-86470853-DEL-98 | chr9-86470885-DEL-100 |
| chr9-86473853-DEL-60 | chr9-86473850-DEL-58 |
| chr9-86479899-DEL-144 | chr9-86479900-DEL-144 |
| chr9-86495240-DEL-143 | chr9-86495243-DEL-143 |
| chr9-86544010-DEL-177 | chr9-86544011-DEL-177 |
| chr9-86553801-DEL-60 | chr9-86553802-DEL-60 |
| chr9-86560833-DEL-907 | chr9-86560834-DEL-907 |
| chr9-86576439-DEL-188 | chr9-86576440-DEL-188 |
| chr9-86629411-DEL-74 | chr9-86629412-DEL-74 |
| chr9-86634636-DEL-90 | chr9-86634637-DEL-90 |
| chr9-86635862-DEL-453 | chr9-86635863-DEL-453 |
| chr9-86714750-DEL-521 | chr9-86714751-DEL-521 |
| chr9-86731340-DEL-646 | chr9-86731341-DEL-646 |
| chr9-86745747-DEL-83 | chr9-86745748-DEL-83 |
| chr9-86770125-DEL-78 | chr9-86770126-DEL-117 |
| chr9-86782367-DEL-364 | chr9-86782368-DEL-364 |
| chr9-86792860-DEL-4758 | chr9-86792861-DEL-4758 |
| chr9-86823417-DEL-6323 | chr9-86823418-DEL-6323 |
| chr9-86842664-DEL-63 | chr9-86842665-DEL-57 |
| chr9-86849133-DEL-96 | chr9-86849114-DEL-116 |
| chr9-86869645-DEL-114 | chr9-86869646-DEL-114 |
| chr9-86887281-DEL-92 | chr9-86887282-DEL-92 |
| chr9-86906175-DEL-114 | chr9-86906176-DEL-114 |
| chr9-86929734-DEL-175 | chr9-86929735-DEL-175 |
| chr9-86933225-DEL-115 | chr9-86933228-DEL-113 |
| chr9-86962256-DEL-646 | chr9-86962257-DEL-646 |
| chr9-86972791-DEL-94 | chr9-86972792-DEL-94 |
| chr9-86980854-DEL-7133 | chr9-86980856-DEL-7135 |
| chr9-86988082-DEL-6331 | chr9-86988082-DEL-6331 |
| chr9-87015442-DEL-60 | chr9-87015468-DEL-60 |
| chr9-87037409-DEL-7113 | chr9-87037410-DEL-7113 |
| chr9-87087371-DEL-293 | chr9-87087372-DEL-293 |
| chr9-87146345-DEL-386 | chr9-87146346-DEL-386 |
| chr9-87159147-DEL-6394 | chr9-87159148-DEL-6394 |
| chr9-87166384-DEL-374 | chr9-87166385-DEL-374 |
| chr9-87174138-DEL-318 | chr9-87174138-DEL-312 |
| chr9-87229877-DEL-2654 | chr9-87229878-DEL-2654 |
| chr9-87254154-DEL-74 | chr9-87254157-DEL-72 |
| chr9-87283383-DEL-186 | chr9-87283384-DEL-186 |
| chr9-87294123-DEL-755 | chr9-87294124-DEL-755 |
| chr9-87317830-DEL-1347 | chr9-87317831-DEL-1347 |
| chr9-87328769-DEL-735 | chr9-87328770-DEL-735 |
| chr9-87332423-DEL-83 | chr9-87332424-DEL-83 |
| chr9-87337037-DEL-728 | chr9-87337038-DEL-728 |
| chr9-87356504-DEL-84 | chr9-87356505-DEL-60 |
| chr9-87360006-DEL-309 | chr9-87360007-DEL-314 |
| chr9-87373672-DEL-7085 | chr9-87373673-DEL-7085 |
| chr9-87415697-DEL-101 | chr9-87415698-DEL-99 |
| chr9-87447764-DEL-51 | chr9-87447764-DEL-51 |
| chr9-87464841-DEL-214 | chr9-87464841-DEL-214 |
| chr9-87469691-DEL-5528 | chr9-87469692-DEL-5528 |
| chr9-87535040-DEL-55 | chr9-87535041-DEL-55 |
| chr9-87536196-DEL-66 | chr9-87536197-DEL-66 |
| chr9-87552993-DEL-6866 | chr9-87552994-DEL-6866 |
| chr9-87594331-DEL-52 | chr9-87594332-DEL-52 |
| chr9-87598708-DEL-216 | chr9-87598702-DEL-224 |
| chr9-87627398-DEL-197 | chr9-87627399-DEL-197 |
| chr9-87639186-DEL-1081 | chr9-87639187-DEL-1081 |
| chr9-87640615-DEL-388 | chr9-87640616-DEL-388 |
| chr9-87669939-DEL-6586 | chr9-87669940-DEL-6586 |
| chr9-87671785-DEL-3243 | chr9-87671786-DEL-3243 |
| chr9-87722113-DEL-208 | chr9-87722114-DEL-208 |
| chr9-87812010-DEL-54 | chr9-87812014-DEL-55 |
| chr9-87827561-DEL-6885 | chr9-87827562-DEL-6885 |
| chr9-87841071-DEL-375 | chr9-87841072-DEL-375 |
| chr9-87872308-DEL-765 | chr9-87872309-DEL-765 |
| chr9-87889914-DEL-6195 | chr9-87889899-DEL-6196 |
| chr9-87903137-DEL-58 | chr9-87903138-DEL-58 |
| chr9-87933108-DEL-51 | chr9-87933106-DEL-52 |

| | |
|---|---|
| chr9-87978452-DEL-112 | chr9-87978453-DEL-96 |
| chr9-87987875-DEL-188 | chr9-87987876-DEL-188 |
| chr9-87990929-DEL-1357 | chr9-87990929-DEL-1357 |
| chr9-88011639-DEL-6449 | chr9-88011640-DEL-6449 |
| chr9-88021297-DEL-101 | chr9-88021298-DEL-101 |
| chr9-88026091-DEL-344 | chr9-88026092-DEL-344 |
| chr9-88057882-DEL-115 | chr9-88057883-DEL-115 |
| chr9-88105720-DEL-53 | chr9-88105721-DEL-53 |
| chr9-88117559-DEL-980 | chr9-88117560-DEL-980 |
| chr9-88181521-DEL-1374 | chr9-88181504-DEL-1392 |
| chr9-88196402-DEL-1319 | chr9-88196403-DEL-1319 |
| chr9-88199803-DEL-6891 | chr9-88199797-DEL-6891 |
| chr9-88203165-DEL-2755 | chr9-88203166-DEL-2755 |
| chr9-88272838-DEL-585 | chr9-88272839-DEL-585 |
| chr9-88297157-DEL-250 | chr9-88297161-DEL-240 |
| chr9-88389161-DEL-88 | chr9-88389162-DEL-88 |
| chr9-88389884-DEL-1889 | chr9-88389885-DEL-1889 |
| chr9-88403730-DEL-268 | chr9-88403731-DEL-268 |
| chr9-88421175-DEL-2777 | chr9-88421176-DEL-2777 |
| chr9-88423006-DEL-205 | chr9-88423007-DEL-204 |
| chr9-88437392-DEL-200 | chr9-88437393-DEL-200 |
| chr9-88529708-DEL-1451 | chr9-88529708-DEL-1451 |
| chr9-88554485-DEL-208 | chr9-88554486-DEL-208 |
| chr9-88569507-DEL-977 | chr9-88569503-DEL-978 |
| chr9-88576480-DEL-280 | chr9-88576467-DEL-245 |
| chr9-88619576-DEL-1223 | chr9-88619588-DEL-1240 |
| chr9-88630708-DEL-3005 | chr9-88630709-DEL-3005 |
| chr9-88671142-DEL-206 | chr9-88671143-DEL-206 |
| chr9-88703405-DEL-7124 | chr9-88703406-DEL-7124 |
| chr9-88723082-DEL-555 | chr9-88723083-DEL-555 |
| chr9-88755285-DEL-507 | chr9-88755286-DEL-507 |
| chr9-88775841-DEL-78 | chr9-88775841-DEL-78 |
| chr9-88916314-DEL-418 | chr9-88916315-DEL-418 |
| chr9-88983858-DEL-158 | chr9-88983858-DEL-156 |
| chr9-89106442-DEL-55 | chr9-89106440-DEL-56 |
| chr9-89155079-DEL-84 | chr9-89155078-DEL-86 |
| chr9-89185460-DEL-454 | chr9-89185459-DEL-454 |
| chr9-89203406-DEL-1681 | chr9-89203407-DEL-1681 |
| chr9-89218563-DEL-1495 | chr9-89218564-DEL-1494 |
| chr9-89221781-DEL-6017 | chr9-89221781-DEL-6017 |
| chr9-89248045-DEL-194 | chr9-89248046-DEL-192 |
| chr9-89332306-DEL-54 | chr9-89332316-DEL-54 |
| chr9-89350887-DEL-96 | chr9-89350888-DEL-96 |
| chr9-89354311-DEL-6442 | chr9-89354312-DEL-6442 |
| chr9-89405345-DEL-63 | chr9-89405358-DEL-65 |
| chr9-89438779-DEL-724 | chr9-89438780-DEL-724 |
| chr9-89440626-DEL-428 | chr9-89440627-DEL-425 |
| chr9-89443254-DEL-76 | chr9-89443255-DEL-76 |
| chr9-89448563-DEL-263 | chr9-89448564-DEL-263 |
| chr9-89473768-DEL-78 | chr9-89473743-DEL-107 |
| chr9-89515037-DEL-58 | chr9-89515038-DEL-58 |
| chr9-89527879-DEL-193 | chr9-89527896-DEL-186 |
| chr9-89558099-DEL-195 | chr9-89558076-DEL-219 |
| chr9-89565036-DEL-712 | chr9-89565038-DEL-713 |
| chr9-89608711-DEL-1204 | chr9-89608712-DEL-1204 |
| chr9-89630452-DEL-211 | chr9-89630453-DEL-211 |
| chr9-89692802-DEL-78 | chr9-89692803-DEL-78 |
| chr9-89699717-DEL-777 | chr9-89699718-DEL-777 |
| chr9-89716536-DEL-1858 | chr9-89716537-DEL-1858 |
| chr9-89746941-DEL-116 | chr9-89746942-DEL-116 |
| chr9-89766431-DEL-53 | chr9-89766428-DEL-66 |
| chr9-89776753-DEL-1942 | chr9-89776754-DEL-1942 |
| chr9-89799135-DEL-91 | chr9-89799136-DEL-67 |
| chr9-89843629-DEL-224 | chr9-89843628-DEL-224 |
| chr9-89844066-DEL-373 | chr9-89844067-DEL-373 |
| chr9-89845802-DEL-54 | chr9-89845827-DEL-78 |
| chr9-89855730-DEL-176 | chr9-89855731-DEL-176 |
| chr9-89880265-DEL-64 | chr9-89880261-DEL-68 |
| chr9-89885813-DEL-88 | chr9-89885814-DEL-88 |
| chr9-89931577-DEL-82 | chr9-89931576-DEL-119 |
| chr9-89946798-DEL-151 | chr9-89946799-DEL-151 |
| chr9-89956277-DEL-69 | chr9-89956275-DEL-50 |
| chr9-89960372-DEL-191 | chr9-89960373-DEL-191 |
| chr9-89963839-DEL-196 | chr9-89963840-DEL-196 |
| chr9-89967960-DEL-1835 | chr9-89967953-DEL-1836 |
| chr9-90000918-DEL-2250 | chr9-90000919-DEL-2250 |
| chr9-90028529-DEL-81 | chr9-90028487-DEL-96 |
| chr9-90029797-DEL-1411 | chr9-90029821-DEL-1388 |
| chr9-90032600-DEL-79 | chr9-90032594-DEL-71 |
| chr9-90050101-DEL-194 | chr9-90050107-DEL-195 |

| | |
|---|---|
| chr9-90086445-DEL-57 | chr9-90086446-DEL-57 |
| chr9-90103293-DEL-470 | chr9-90103294-DEL-470 |
| chr9-90135551-DEL-83 | chr9-90135552-DEL-83 |
| chr9-90160769-DEL-51 | chr9-90160770-DEL-51 |
| chr9-90163863-DEL-358 | chr9-90163864-DEL-358 |
| chr9-90179338-DEL-340 | chr9-90179339-DEL-340 |
| chr9-90192976-DEL-422 | chr9-90192977-DEL-422 |
| chr9-90239412-DEL-75 | chr9-90239413-DEL-75 |
| chr9-90268873-DEL-1588 | chr9-90268876-DEL-1586 |
| chr9-90286677-DEL-1584 | chr9-90286678-DEL-1579 |
| chr9-90308631-DEL-106 | chr9-90308632-DEL-106 |
| chr9-90314613-DEL-828 | chr9-90314614-DEL-828 |
| chr9-90371893-DEL-52 | chr9-90371894-DEL-52 |
| chr9-90387005-DEL-117 | chr9-90387006-DEL-117 |
| chr9-90395169-DEL-95 | chr9-90395127-DEL-119 |
| chr9-90418742-DEL-51 | chr9-90418733-DEL-50 |
| chr9-90461991-DEL-2138 | chr9-90461992-DEL-2138 |
| chr9-90465161-DEL-161 | chr9-90465162-DEL-161 |
| chr9-90537358-DEL-74 | chr9-90537383-DEL-60 |
| chr9-90621444-DEL-103 | chr9-90621463-DEL-75 |
| chr9-90630366-DEL-243 | chr9-90630367-DEL-243 |
| chr9-90680894-DEL-84 | chr9-90680895-DEL-86 |
| chr9-90690193-DEL-6436 | chr9-90690194-DEL-6436 |
| chr9-90739379-DEL-7138 | chr9-90739380-DEL-7138 |
| chr9-90748410-DEL-418 | chr9-90748411-DEL-418 |
| chr9-90809291-DEL-14922 | chr9-90809292-DEL-14922 |
| chr9-90857037-DEL-975 | chr9-90857038-DEL-975 |
| chr9-90881635-DEL-2205 | chr9-90881636-DEL-2205 |
| chr9-90888472-DEL-62 | chr9-90888473-DEL-62 |
| chr9-90902898-DEL-116 | chr9-90902899-DEL-116 |
| chr9-90979085-DEL-1628 | chr9-90979086-DEL-1628 |
| chr9-90999464-DEL-969 | chr9-90999465-DEL-969 |
| chr9-91024877-DEL-3872 | chr9-91024878-DEL-3872 |
| chr9-91032580-DEL-339 | chr9-91032581-DEL-339 |
| chr9-91044211-DEL-334 | chr9-91044212-DEL-334 |
| chr9-91047187-DEL-962 | chr9-91047188-DEL-962 |
| chr9-91063905-DEL-156 | chr9-91063906-DEL-156 |
| chr9-91099466-DEL-6353 | chr9-91099467-DEL-6353 |
| chr9-91113135-DEL-98 | chr9-91113136-DEL-98 |
| chr9-91119468-DEL-6590 | chr9-91119469-DEL-6590 |
| chr9-91143296-DEL-7750 | chr9-91143297-DEL-7750 |
| chr9-91160209-DEL-2322 | chr9-91160210-DEL-2322 |
| chr9-91160966-DEL-782 | chr9-91160966-DEL-782 |
| chr9-91170213-DEL-324 | chr9-91170214-DEL-324 |
| chr9-91176095-DEL-259 | chr9-91176096-DEL-258 |
| chr9-91188879-DEL-89 | chr9-91188880-DEL-89 |
| chr9-91193141-DEL-1576 | chr9-91193142-DEL-1576 |
| chr9-91219388-DEL-376 | chr9-91219389-DEL-376 |
| chr9-91221718-DEL-226 | chr9-91221719-DEL-226 |
| chr9-91235479-DEL-86 | chr9-91235480-DEL-86 |
| chr9-91301128-DEL-53 | chr9-91301129-DEL-53 |
| chr9-91336242-DEL-84 | chr9-91336233-DEL-85 |
| chr9-91373876-DEL-59 | chr9-91373864-DEL-61 |
| chr9-91462072-DEL-64 | chr9-91462074-DEL-62 |
| chr9-91501046-DEL-212 | chr9-91501047-DEL-212 |
| chr9-91572118-DEL-1038 | chr9-91572119-DEL-1038 |
| chr9-91594411-DEL-1384 | chr9-91594412-DEL-1384 |
| chr9-91632791-DEL-196 | chr9-91632792-DEL-196 |
| chr9-91670695-DEL-214 | chr9-91670696-DEL-214 |
| chr9-91687094-DEL-4345 | chr9-91687094-DEL-4345 |
| chr9-91691977-DEL-1893 | chr9-91691978-DEL-1893 |
| chr9-91724369-DEL-1020 | chr9-91724370-DEL-1020 |
| chr9-91731553-DEL-1284 | chr9-91731554-DEL-1284 |
| chr9-91742828-DEL-2792 | chr9-91742829-DEL-2792 |
| chr9-91745820-DEL-271 | chr9-91745821-DEL-271 |
| chr9-91779174-DEL-55 | chr9-91779175-DEL-66 |
| chr9-91855144-DEL-73 | chr9-91855103-DEL-89 |
| chr9-91867960-DEL-196 | chr9-91867946-DEL-197 |
| chr9-91876605-DEL-229 | chr9-91876606-DEL-228 |
| chr9-91952492-DEL-62 | chr9-91952493-DEL-62 |
| chr9-91994105-DEL-1158 | chr9-91994106-DEL-1158 |
| chr9-92035543-DEL-86 | chr9-92035543-DEL-90 |
| chr9-92063556-DEL-136 | chr9-92063591-DEL-137 |
| chr9-92087127-DEL-52 | chr9-92087094-DEL-84 |
| chr9-92090585-DEL-59 | chr9-92090591-DEL-57 |
| chr9-92121842-DEL-3050 | chr9-92121843-DEL-3050 |
| chr9-92138285-DEL-100 | chr9-92138296-DEL-70 |
| chr9-92146745-DEL-108 | chr9-92146746-DEL-108 |
| chr9-92200271-DEL-688 | chr9-92200272-DEL-688 |
| chr9-92284252-DEL-99 | chr9-92284257-DEL-104 |

| | |
|---|---|
| chr9-92288069-DEL-3280 | chr9-92288070-DEL-3280 |
| chr9-92313737-DEL-775 | chr9-92313738-DEL-775 |
| chr9-92328443-DEL-7076 | chr9-92328434-DEL-7073 |
| chr9-92344945-DEL-143 | chr9-92344946-DEL-133 |
| chr9-92376538-DEL-626 | chr9-92376539-DEL-626 |
| chr9-92395191-DEL-908 | chr9-92395192-DEL-908 |
| chr9-92399686-DEL-1357 | chr9-92399687-DEL-1357 |
| chr9-92408082-DEL-168 | chr9-92408083-DEL-168 |
| chr9-92429577-DEL-213 | chr9-92429579-DEL-212 |
| chr9-92463849-DEL-1115 | chr9-92463861-DEL-1109 |
| chr9-92501656-DEL-52 | chr9-92501657-DEL-52 |
| chr9-92506941-DEL-6588 | chr9-92506942-DEL-6588 |
| chr9-92523037-DEL-445 | chr9-92523037-DEL-445 |
| chr9-92524652-DEL-2757 | chr9-92524653-DEL-2757 |
| chr9-92529417-DEL-7140 | chr9-92529418-DEL-7140 |
| chr9-92540109-DEL-53 | chr9-92540097-DEL-54 |
| chr9-92574262-DEL-75 | chr9-92574266-DEL-74 |
| chr9-92595545-DEL-1690 | chr9-92595546-DEL-1690 |
| chr9-92611757-DEL-152 | chr9-92611778-DEL-131 |
| chr9-92616712-DEL-7439 | chr9-92616713-DEL-7439 |
| chr9-92630657-DEL-88 | chr9-92630658-DEL-88 |
| chr9-92631323-DEL-75 | chr9-92631324-DEL-75 |
| chr9-92638108-DEL-53 | chr9-92638109-DEL-54 |
| chr9-92690707-DEL-377 | chr9-92690708-DEL-377 |
| chr9-92698198-DEL-2548 | chr9-92698199-DEL-2548 |
| chr9-92704594-DEL-3283 | chr9-92704595-DEL-3283 |
| chr9-92740501-DEL-2477 | chr9-92740502-DEL-2477 |
| chr9-92786477-DEL-75 | chr9-92786478-DEL-75 |
| chr9-92846399-DEL-1393 | chr9-92846400-DEL-1393 |
| chr9-92858275-DEL-329 | chr9-92858276-DEL-328 |
| chr9-92881849-DEL-696 | chr9-92881850-DEL-696 |
| chr9-92895321-DEL-8028 | chr9-92895322-DEL-8028 |
| chr9-92901944-DEL-642 | chr9-92901945-DEL-642 |
| chr9-92925035-DEL-436 | chr9-92925035-DEL-436 |
| chr9-92928927-DEL-420 | chr9-92928928-DEL-420 |
| chr9-92931688-DEL-390 | chr9-92931689-DEL-390 |
| chr9-92937136-DEL-352 | chr9-92937137-DEL-352 |
| chr9-92940394-DEL-54 | chr9-92940404-DEL-60 |
| chr9-92945463-DEL-431 | chr9-92945464-DEL-431 |
| chr9-92951663-DEL-299 | chr9-92951664-DEL-274 |
| chr9-92968448-DEL-80 | chr9-92968449-DEL-80 |
| chr9-92991761-DEL-475 | chr9-92991762-DEL-475 |
| chr9-93038006-DEL-72 | chr9-93038007-DEL-69 |
| chr9-93052956-DEL-425 | chr9-93052956-DEL-425 |
| chr9-93086938-DEL-211 | chr9-93086939-DEL-211 |
| chr9-93164115-DEL-308 | chr9-93164121-DEL-302 |
| chr9-93167331-DEL-6621 | chr9-93167332-DEL-6621 |
| chr9-93202399-DEL-2355 | chr9-93202401-DEL-2356 |
| chr9-93205119-DEL-80 | chr9-93205120-DEL-80 |
| chr9-93205570-DEL-4173 | chr9-93205571-DEL-4173 |
| chr9-93276859-DEL-6475 | chr9-93276860-DEL-6475 |
| chr9-93316831-DEL-7309 | chr9-93316832-DEL-7309 |
| chr9-93346347-DEL-58 | chr9-93346348-DEL-58 |
| chr9-93356566-DEL-1461 | chr9-93356567-DEL-1460 |
| chr9-93360421-DEL-554 | chr9-93360422-DEL-554 |
| chr9-93365857-DEL-104 | chr9-93365858-DEL-104 |
| chr9-93385364-DEL-67 | chr9-93385365-DEL-67 |
| chr9-93395497-DEL-909 | chr9-93395498-DEL-909 |
| chr9-93400670-DEL-56 | chr9-93400671-DEL-56 |
| chr9-93487125-DEL-52 | chr9-93487126-DEL-52 |
| chr9-93506411-DEL-5725 | chr9-93506412-DEL-5725 |
| chr9-93519075-DEL-629 | chr9-93519076-DEL-629 |
| chr9-93526916-DEL-75 | chr9-93526917-DEL-75 |
| chr9-93565536-DEL-60 | chr9-93565537-DEL-60 |
| chr9-93570624-DEL-811 | chr9-93570624-DEL-811 |
| chr9-93583617-DEL-944 | chr9-93583618-DEL-944 |
| chr9-93614866-DEL-657 | chr9-93614867-DEL-657 |
| chr9-93646925-DEL-152 | chr9-93646926-DEL-152 |
| chr9-93652905-DEL-336 | chr9-93652906-DEL-336 |
| chr9-93661025-DEL-561 | chr9-93661026-DEL-561 |
| chr9-93696209-DEL-1580 | chr9-93696209-DEL-1580 |
| chr9-93717713-DEL-129 | chr9-93717714-DEL-119 |
| chr9-93736017-DEL-430 | chr9-93736018-DEL-430 |
| chr9-93776832-DEL-77 | chr9-93776833-DEL-77 |
| chr9-93779747-DEL-87 | chr9-93779748-DEL-87 |
| chr9-93782730-DEL-58 | chr9-93782731-DEL-58 |
| chr9-93784561-DEL-733 | chr9-93784562-DEL-733 |
| chr9-93793041-DEL-7118 | chr9-93793042-DEL-7118 |
| chr9-93799210-DEL-617 | chr9-93799211-DEL-617 |
| chr9-93825765-DEL-62 | chr9-93825766-DEL-62 |

| | |
|---|---|
| chr9-93840783-DEL-4499 | chr9-93840784-DEL-4499 |
| chr9-93844483-DEL-53 | chr9-93844484-DEL-53 |
| chr9-93861624-DEL-72 | chr9-93861625-DEL-72 |
| chr9-93873592-DEL-120 | chr9-93873618-DEL-80 |
| chr9-93881102-DEL-65 | chr9-93881103-DEL-65 |
| chr9-93892951-DEL-1145 | chr9-93892952-DEL-1143 |
| chr9-93902775-DEL-137 | chr9-93902776-DEL-137 |
| chr9-93910086-DEL-465 | chr9-93910087-DEL-465 |
| chr9-93975258-DEL-64 | chr9-93975259-DEL-62 |
| chr9-93990878-DEL-168 | chr9-93990856-DEL-163 |
| chr9-94025859-DEL-72 | chr9-94025860-DEL-72 |
| chr9-94027717-DEL-499 | chr9-94027718-DEL-497 |
| chr9-94058312-DEL-54 | chr9-94058313-DEL-54 |
| chr9-94059467-DEL-454 | chr9-94059468-DEL-454 |
| chr9-94063779-DEL-178 | chr9-94063780-DEL-177 |
| chr9-94068891-DEL-70 | chr9-94068894-DEL-112 |
| chr9-94075157-DEL-74 | chr9-94075178-DEL-52 |
| chr9-94092217-DEL-117 | chr9-94092228-DEL-109 |
| chr9-94104759-DEL-2435 | chr9-94104760-DEL-2435 |
| chr9-94158764-DEL-71 | chr9-94158765-DEL-71 |
| chr9-94164557-DEL-1606 | chr9-94164558-DEL-1606 |
| chr9-94185927-DEL-62 | chr9-94185928-DEL-54 |
| chr9-94186711-DEL-70 | chr9-94186712-DEL-70 |
| chr9-94188128-DEL-78 | chr9-94188129-DEL-78 |
| chr9-94197664-DEL-147 | chr9-94197665-DEL-147 |
| chr9-94245336-DEL-68 | chr9-94245337-DEL-62 |
| chr9-94251708-DEL-107 | chr9-94251708-DEL-107 |
| chr9-94268885-DEL-6469 | chr9-94268886-DEL-6469 |
| chr9-94300608-DEL-79 | chr9-94300599-DEL-80 |
| chr9-94314462-DEL-51 | chr9-94314463-DEL-51 |
| chr9-94331236-DEL-182 | chr9-94331237-DEL-182 |
| chr9-94366796-DEL-56 | chr9-94366797-DEL-56 |
| chr9-94372217-DEL-1129 | chr9-94372218-DEL-1129 |
| chr9-94378759-DEL-348 | chr9-94378760-DEL-348 |
| chr9-94381352-DEL-4644 | chr9-94381353-DEL-4644 |
| chr9-94392548-DEL-678 | chr9-94392549-DEL-678 |
| chr9-94411874-DEL-64 | chr9-94411865-DEL-52 |
| chr9-94470083-DEL-67 | chr9-94470084-DEL-67 |
| chr9-94479319-DEL-4314 | chr9-94479309-DEL-4315 |
| chr9-94491036-DEL-59 | chr9-94491037-DEL-59 |
| chr9-94515646-DEL-117 | chr9-94515647-DEL-117 |
| chr9-94518083-DEL-6590 | chr9-94518098-DEL-6576 |
| chr9-94547193-DEL-1749 | chr9-94547194-DEL-1749 |
| chr9-94554179-DEL-96 | chr9-94554180-DEL-96 |
| chr9-94610396-DEL-72 | chr9-94610399-DEL-84 |
| chr9-94611416-DEL-72 | chr9-94611417-DEL-60 |
| chr9-94657330-DEL-170 | chr9-94657331-DEL-170 |
| chr9-94663219-DEL-6379 | chr9-94663220-DEL-6379 |
| chr9-94703736-DEL-6400 | chr9-94703737-DEL-6400 |
| chr9-94719682-DEL-200 | chr9-94719683-DEL-200 |
| chr9-94752133-DEL-1282 | chr9-94752134-DEL-1282 |
| chr9-94769306-DEL-56 | chr9-94769307-DEL-56 |
| chr9-94797914-DEL-5583 | chr9-94797915-DEL-5583 |
| chr9-94843903-DEL-60 | chr9-94843903-DEL-60 |
| chr9-94861603-DEL-56 | chr9-94861637-DEL-56 |
| chr9-94868791-DEL-74 | chr9-94868796-DEL-66 |
| chr9-94872740-DEL-275 | chr9-94872747-DEL-272 |
| chr9-94893599-DEL-120 | chr9-94893609-DEL-111 |
| chr9-94918475-DEL-241 | chr9-94918476-DEL-241 |
| chr9-94944063-DEL-66 | chr9-94944088-DEL-57 |
| chr9-94997694-DEL-58 | chr9-94997722-DEL-58 |
| chr9-95071862-DEL-400 | chr9-95071863-DEL-400 |
| chr9-95080236-DEL-470 | chr9-95080237-DEL-468 |
| chr9-95123803-DEL-157 | chr9-95123804-DEL-157 |
| chr9-95151241-DEL-6411 | chr9-95151242-DEL-6412 |
| chr9-95164372-DEL-72 | chr9-95164373-DEL-72 |
| chr9-95169638-DEL-90 | chr9-95169639-DEL-90 |
| chr9-95174098-DEL-7562 | chr9-95174099-DEL-7562 |
| chr9-95174117-DEL-212 | chr9-95174118-DEL-212 |
| chr9-95185689-DEL-87 | chr9-95185690-DEL-87 |
| chr9-95226965-DEL-63 | chr9-95226963-DEL-66 |
| chr9-95228001-DEL-786 | chr9-95228002-DEL-786 |
| chr9-95234898-DEL-83 | chr9-95234899-DEL-83 |
| chr9-95237918-DEL-127 | chr9-95237917-DEL-109 |
| chr9-95246569-DEL-92 | chr9-95246561-DEL-96 |
| chr9-95250433-DEL-65 | chr9-95250434-DEL-65 |
| chr9-95262147-DEL-95 | chr9-95262148-DEL-95 |
| chr9-95282591-DEL-112 | chr9-95282592-DEL-112 |
| chr9-95313315-DEL-94 | chr9-95313316-DEL-94 |
| chr9-95314154-DEL-57 | chr9-95314155-DEL-57 |

| | |
|---|---|
| chr9-95330234-DEL-1309 | chr9-95330235-DEL-1309 |
| chr9-95356855-DEL-354 | chr9-95356856-DEL-354 |
| chr9-95399820-DEL-198 | chr9-95399821-DEL-198 |
| chr9-95401101-DEL-74 | chr9-95401101-DEL-74 |
| chr9-95403985-DEL-52 | chr9-95403983-DEL-55 |
| chr9-95405126-DEL-75 | chr9-95405127-DEL-74 |
| chr9-95406210-DEL-65 | chr9-95406257-DEL-54 |
| chr9-95410781-DEL-259 | chr9-95410781-DEL-259 |
| chr9-95485143-DEL-952 | chr9-95485143-DEL-953 |
| chr9-95494544-DEL-57 | chr9-95494544-DEL-57 |
| chr9-95502996-DEL-236 | chr9-95502996-DEL-236 |
| chr9-95516413-DEL-240 | chr9-95516414-DEL-238 |
| chr9-95531588-DEL-155 | chr9-95531589-DEL-155 |
| chr9-95552436-DEL-159 | chr9-95552437-DEL-159 |
| chr9-95559067-DEL-420 | chr9-95559068-DEL-420 |
| chr9-95560615-DEL-139 | chr9-95560616-DEL-139 |
| chr9-95574320-DEL-68 | chr9-95574321-DEL-68 |
| chr9-95575442-DEL-66 | chr9-95575435-DEL-67 |
| chr9-95589998-DEL-129 | chr9-95589995-DEL-123 |
| chr9-95592839-DEL-491 | chr9-95592840-DEL-491 |
| chr9-95645279-DEL-96 | chr9-95645280-DEL-96 |
| chr9-95648149-DEL-467 | chr9-95648150-DEL-467 |
| chr9-95673619-DEL-56 | chr9-95673620-DEL-58 |
| chr9-95675068-DEL-511 | chr9-95675064-DEL-530 |
| chr9-95676757-DEL-196 | chr9-95676778-DEL-176 |
| chr9-95679055-DEL-2478 | chr9-95679056-DEL-2478 |
| chr9-95695417-DEL-200 | chr9-95695418-DEL-200 |
| chr9-95708430-DEL-56 | chr9-95708431-DEL-56 |
| chr9-95711447-DEL-1629 | chr9-95711449-DEL-1644 |
| chr9-95850150-DEL-114 | chr9-95850151-DEL-114 |
| chr9-95857572-DEL-320 | chr9-95857573-DEL-320 |
| chr9-95872683-DEL-54 | chr9-95872684-DEL-54 |
| chr9-95942582-DEL-218 | chr9-95942583-DEL-218 |
| chr9-95961415-DEL-1039 | chr9-95961401-DEL-1021 |
| chr9-95963753-DEL-3071 | chr9-95963754-DEL-3071 |
| chr9-95983237-DEL-54 | chr9-95983238-DEL-54 |
| chr9-95986940-DEL-6473 | chr9-95986941-DEL-6473 |
| chr9-95995712-DEL-118 | chr9-95995713-DEL-118 |
| chr9-96008190-DEL-98 | chr9-96008193-DEL-97 |
| chr9-96008878-DEL-56 | chr9-96008886-DEL-54 |
| chr9-96018352-DEL-212 | chr9-96018353-DEL-212 |
| chr9-96053380-DEL-94 | chr9-96053381-DEL-94 |
| chr9-96054727-DEL-433 | chr9-96054728-DEL-433 |
| chr9-96074926-DEL-66 | chr9-96074927-DEL-66 |
| chr9-96091389-DEL-1675 | chr9-96091403-DEL-1671 |
| chr9-96109978-DEL-55 | chr9-96109956-DEL-58 |
| chr9-96122606-DEL-164 | chr9-96122603-DEL-165 |
| chr9-96137668-DEL-63 | chr9-96137674-DEL-59 |
| chr9-96150896-DEL-86 | chr9-96150895-DEL-88 |
| chr9-96167412-DEL-748 | chr9-96167413-DEL-743 |
| chr9-96168044-DEL-126 | chr9-96168082-DEL-94 |
| chr9-96176433-DEL-115 | chr9-96176434-DEL-115 |
| chr9-96232079-DEL-52 | chr9-96232080-DEL-56 |
| chr9-96237227-DEL-219 | chr9-96237228-DEL-219 |
| chr9-96249664-DEL-219 | chr9-96249665-DEL-219 |
| chr9-96256835-DEL-107 | chr9-96256835-DEL-108 |
| chr9-96261442-DEL-56 | chr9-96261403-DEL-92 |
| chr9-96287475-DEL-52 | chr9-96287476-DEL-57 |
| chr9-96309143-DEL-66 | chr9-96309143-DEL-66 |
| chr9-96325957-DEL-506 | chr9-96325958-DEL-506 |
| chr9-96331043-DEL-211 | chr9-96331044-DEL-211 |
| chr9-96332207-DEL-64 | chr9-96332208-DEL-64 |
| chr9-96335609-DEL-191 | chr9-96335610-DEL-191 |
| chr9-96355006-DEL-73 | chr9-96355007-DEL-73 |
| chr9-96358052-DEL-220 | chr9-96358053-DEL-220 |
| chr9-96371537-DEL-180 | chr9-96371538-DEL-180 |
| chr9-96383950-DEL-206 | chr9-96383950-DEL-206 |
| chr9-96388938-DEL-400 | chr9-96388939-DEL-400 |
| chr9-96392829-DEL-213 | chr9-96392830-DEL-213 |
| chr9-96398862-DEL-51 | chr9-96398863-DEL-51 |
| chr9-96416369-DEL-250 | chr9-96416370-DEL-250 |
| chr9-96416757-DEL-752 | chr9-96416734-DEL-776 |
| chr9-96452044-DEL-127 | chr9-96452045-DEL-127 |
| chr9-96457510-DEL-604 | chr9-96457511-DEL-604 |
| chr9-96471811-DEL-659 | chr9-96471812-DEL-659 |
| chr9-96485719-DEL-488 | chr9-96485720-DEL-488 |
| chr9-96499203-DEL-518 | chr9-96499204-DEL-518 |
| chr9-96507315-DEL-220 | chr9-96507316-DEL-220 |
| chr9-96507823-DEL-94 | chr9-96507824-DEL-94 |
| chr9-96523436-DEL-417 | chr9-96523432-DEL-443 |

| | |
|---|---|
| chr9-96524776-DEL-71 | chr9-96524783-DEL-70 |
| chr9-96541129-DEL-54 | chr9-96541130-DEL-54 |
| chr9-96546522-DEL-3554 | chr9-96546523-DEL-3554 |
| chr9-96600384-DEL-60 | chr9-96600385-DEL-60 |
| chr9-96618647-DEL-89 | chr9-96618638-DEL-86 |
| chr9-96636954-DEL-52 | chr9-96636955-DEL-52 |
| chr9-96638278-DEL-75 | chr9-96638279-DEL-60 |
| chr9-96639537-DEL-90 | chr9-96639538-DEL-90 |
| chr9-96640075-DEL-60 | chr9-96640076-DEL-60 |
| chr9-96643234-DEL-209 | chr9-96643234-DEL-209 |
| chr9-96647760-DEL-54 | chr9-96647761-DEL-54 |
| chr9-96676961-DEL-84 | chr9-96676962-DEL-84 |
| chr9-96681892-DEL-1561 | chr9-96681893-DEL-1561 |
| chr9-96692324-DEL-159 | chr9-96692327-DEL-157 |
| chr9-96699133-DEL-468 | chr9-96699149-DEL-453 |
| chr9-96699755-DEL-569 | chr9-96699756-DEL-569 |
| chr9-96702425-DEL-163 | chr9-96702426-DEL-163 |
| chr9-96715117-DEL-64 | chr9-96715165-DEL-66 |
| chr9-96726551-DEL-54 | chr9-96726552-DEL-54 |
| chr9-96734989-DEL-240 | chr9-96734990-DEL-240 |
| chr9-96756573-DEL-89 | chr9-96756574-DEL-89 |
| chr9-96757248-DEL-53 | chr9-96757249-DEL-53 |
| chr9-96766199-DEL-51 | chr9-96766200-DEL-51 |
| chr9-96777650-DEL-772 | chr9-96777651-DEL-772 |
| chr9-96793290-DEL-72 | chr9-96793291-DEL-72 |
| chr9-96881678-DEL-98 | chr9-96881709-DEL-68 |
| chr9-96888548-DEL-58 | chr9-96888554-DEL-54 |
| chr9-96908518-DEL-191 | chr9-96908519-DEL-191 |
| chr9-96910097-DEL-179 | chr9-96910098-DEL-179 |
| chr9-96914009-DEL-1137 | chr9-96914010-DEL-1137 |
| chr9-96923591-DEL-54 | chr9-96923591-DEL-56 |
| chr9-96929285-DEL-191 | chr9-96929286-DEL-198 |
| chr9-96955486-DEL-197 | chr9-96955487-DEL-197 |
| chr9-96961908-DEL-200 | chr9-96961909-DEL-200 |
| chr9-96994836-DEL-96 | chr9-96994837-DEL-96 |
| chr9-97008550-DEL-236 | chr9-97008551-DEL-236 |
| chr9-97037265-DEL-57 | chr9-97037266-DEL-57 |
| chr9-97047074-DEL-199 | chr9-97047075-DEL-199 |
| chr9-97055414-DEL-56 | chr9-97055414-DEL-56 |
| chr9-97056690-DEL-83 | chr9-97056690-DEL-83 |
| chr9-97061195-DEL-886 | chr9-97061195-DEL-886 |
| chr9-97062137-DEL-216 | chr9-97062137-DEL-216 |
| chr9-97074079-DEL-144 | chr9-97074079-DEL-144 |
| chr9-97102369-DEL-400 | chr9-97102369-DEL-400 |
| chr9-97113108-DEL-61 | chr9-97113108-DEL-61 |
| chr9-97121467-DEL-65 | chr9-97121483-DEL-50 |
| chr9-97150381-DEL-862 | chr9-97150382-DEL-862 |
| chr9-97153658-DEL-408 | chr9-97153660-DEL-406 |
| chr9-97154682-DEL-616 | chr9-97154683-DEL-616 |
| chr9-97158511-DEL-58 | chr9-97158512-DEL-58 |
| chr9-97206714-DEL-665 | chr9-97206715-DEL-665 |
| chr9-97243925-DEL-92 | chr9-97243938-DEL-80 |
| chr9-97269309-DEL-100 | chr9-97269310-DEL-100 |
| chr9-97291927-DEL-259 | chr9-97291928-DEL-259 |
| chr9-97292989-DEL-463 | chr9-97292990-DEL-463 |
| chr9-97320810-DEL-64 | chr9-97320810-DEL-62 |
| chr9-97384127-DEL-62 | chr9-97384109-DEL-52 |
| chr9-97476334-DEL-747 | chr9-97476355-DEL-733 |
| chr9-97487599-DEL-54 | chr9-97487600-DEL-54 |
| chr9-97493959-DEL-2276 | chr9-97493960-DEL-2276 |
| chr9-97553528-DEL-99 | chr9-97553529-DEL-99 |
| chr9-97571042-DEL-618 | chr9-97571044-DEL-619 |
| chr9-97618783-DEL-425 | chr9-97618784-DEL-425 |
| chr9-97713671-DEL-351 | chr9-97713672-DEL-351 |
| chr9-97750332-DEL-1530 | chr9-97750333-DEL-1530 |
| chr9-97763514-DEL-235 | chr9-97763515-DEL-235 |
| chr9-97784566-DEL-78 | chr9-97784559-DEL-85 |
| chr9-97794865-DEL-1171 | chr9-97794866-DEL-1171 |
| chr9-97798837-DEL-84 | chr9-97798838-DEL-84 |
| chr9-97810546-DEL-52 | chr9-97810547-DEL-50 |
| chr9-97814703-DEL-52 | chr9-97814702-DEL-56 |
| chr9-97845388-DEL-52 | chr9-97845389-DEL-50 |
| chr9-97982924-DEL-78 | chr9-97982924-DEL-75 |
| chr9-97990919-DEL-8044 | chr9-97990920-DEL-8044 |
| chr9-98002527-DEL-2338 | chr9-98002528-DEL-2338 |
| chr9-98020248-DEL-92 | chr9-98020249-DEL-92 |
| chr9-98026209-DEL-430 | chr9-98026210-DEL-430 |
| chr9-98037882-DEL-82 | chr9-98037883-DEL-82 |
| chr9-98044984-DEL-437 | chr9-98044984-DEL-437 |
| chr9-98048393-DEL-6594 | chr9-98048394-DEL-6594 |

| | |
|---|---|
| chr9-98132092-DEL-7741 | chr9-98132093-DEL-7741 |
| chr9-98132408-DEL-424 | chr9-98132409-DEL-424 |
| chr9-98224872-DEL-2869 | chr9-98224873-DEL-2869 |
| chr9-98247857-DEL-108 | chr9-98247856-DEL-109 |
| chr9-98296196-DEL-138 | chr9-98296197-DEL-138 |
| chr9-98326035-DEL-202 | chr9-98326036-DEL-202 |
| chr9-98333442-DEL-7179 | chr9-98333443-DEL-7179 |
| chr9-98343692-DEL-60 | chr9-98343693-DEL-60 |
| chr9-98350300-DEL-74 | chr9-98350297-DEL-60 |
| chr9-98354382-DEL-430 | chr9-98354381-DEL-431 |
| chr9-98385424-DEL-71 | chr9-98385425-DEL-71 |
| chr9-98403238-DEL-89 | chr9-98403223-DEL-117 |
| chr9-98438308-DEL-226 | chr9-98438307-DEL-227 |
| chr9-98484261-DEL-175 | chr9-98484262-DEL-176 |
| chr9-98484987-DEL-127 | chr9-98484988-DEL-128 |
| chr9-98495831-DEL-54 | chr9-98495826-DEL-54 |
| chr9-98499551-DEL-52 | chr9-98499552-DEL-58 |
| chr9-98533426-DEL-64 | chr9-98533427-DEL-64 |
| chr9-98554202-DEL-328 | chr9-98554203-DEL-328 |
| chr9-98601491-DEL-61 | chr9-98601492-DEL-60 |
| chr9-98602756-DEL-119 | chr9-98602757-DEL-119 |
| chr9-98604203-DEL-184 | chr9-98604204-DEL-184 |
| chr9-98627855-DEL-3034 | chr9-98627856-DEL-3034 |
| chr9-98657058-DEL-108 | chr9-98657059-DEL-108 |
| chr9-98660267-DEL-1395 | chr9-98660294-DEL-1369 |
| chr9-98666330-DEL-56 | chr9-98666331-DEL-56 |
| chr9-98671019-DEL-94 | chr9-98671019-DEL-93 |
| chr9-98707057-DEL-312 | chr9-98707058-DEL-312 |
| chr9-98731745-DEL-1421 | chr9-98731745-DEL-1421 |
| chr9-98750768-DEL-1454 | chr9-98750737-DEL-1468 |
| chr9-98850293-DEL-180 | chr9-98850276-DEL-174 |
| chr9-98865368-DEL-2731 | chr9-98865369-DEL-2732 |
| chr9-98906163-DEL-62 | chr9-98906192-DEL-69 |
| chr9-98907121-DEL-67 | chr9-98907122-DEL-67 |
| chr9-98927610-DEL-106 | chr9-98927611-DEL-106 |
| chr9-98942686-DEL-92 | chr9-98942687-DEL-92 |
| chr9-98946735-DEL-126 | chr9-98946736-DEL-126 |
| chr9-98956272-DEL-78 | chr9-98956273-DEL-78 |
| chr9-98988177-DEL-206 | chr9-98988178-DEL-206 |
| chr9-99039469-DEL-53 | chr9-99039464-DEL-57 |
| chr9-99050540-DEL-100 | chr9-99050541-DEL-101 |
| chr9-99080604-DEL-126 | chr9-99080605-DEL-126 |
| chr9-99134049-DEL-54 | chr9-99134050-DEL-54 |
| chr9-99138996-DEL-69 | chr9-99138997-DEL-69 |
| chr9-99141539-DEL-154 | chr9-99141540-DEL-154 |
| chr9-99144176-DEL-198 | chr9-99144176-DEL-198 |
| chr9-99148760-DEL-6413 | chr9-99148761-DEL-6413 |
| chr9-99164942-DEL-234 | chr9-99164927-DEL-226 |
| chr9-99194704-DEL-2091 | chr9-99194705-DEL-2091 |
| chr9-99200989-DEL-51 | chr9-99200990-DEL-51 |
| chr9-99208740-DEL-703 | chr9-99208741-DEL-703 |
| chr9-99220436-DEL-140 | chr9-99220437-DEL-140 |
| chr9-99230825-DEL-159 | chr9-99230826-DEL-159 |
| chr9-99232814-DEL-112 | chr9-99232815-DEL-112 |
| chr9-99245964-DEL-186 | chr9-99245965-DEL-186 |
| chr9-99251026-DEL-68 | chr9-99251027-DEL-68 |
| chr9-99295372-DEL-62 | chr9-99295373-DEL-62 |
| chr9-99301384-DEL-96 | chr9-99301415-DEL-66 |
| chr9-99322787-DEL-81 | chr9-99322788-DEL-54 |
| chr9-99324217-DEL-83 | chr9-99324218-DEL-83 |
| chr9-99325042-DEL-114 | chr9-99325043-DEL-114 |
| chr9-99359065-DEL-1190 | chr9-99359050-DEL-1190 |
| chr9-99367978-DEL-96 | chr9-99367979-DEL-96 |
| chr9-99375969-DEL-368 | chr9-99375970-DEL-368 |
| chr9-99422055-DEL-1444 | chr9-99422057-DEL-1443 |
| chr9-99435900-DEL-196 | chr9-99435901-DEL-196 |
| chr9-99437904-DEL-4522 | chr9-99437905-DEL-4522 |
| chr9-99444649-DEL-182 | chr9-99444650-DEL-182 |
| chr9-99454353-DEL-87 | chr9-99454354-DEL-87 |
| chr9-99475555-DEL-107 | chr9-99475543-DEL-152 |
| chr9-99490782-DEL-129 | chr9-99490783-DEL-129 |
| chr9-99492579-DEL-85 | chr9-99492580-DEL-85 |
| chr9-99498573-DEL-74 | chr9-99498574-DEL-74 |
| chr9-99501536-DEL-325 | chr9-99501537-DEL-325 |
| chr9-99504600-DEL-319 | chr9-99504601-DEL-319 |
| chr9-99528660-DEL-78 | chr9-99528643-DEL-52 |
| chr9-99557452-DEL-212 | chr9-99557453-DEL-212 |
| chr9-99575692-DEL-68 | chr9-99575690-DEL-70 |
| chr9-99578690-DEL-125 | chr9-99578689-DEL-124 |
| chr9-99582283-DEL-52 | chr9-99582284-DEL-52 |

| | |
|---|---|
| chr9-99601285-DEL-113 | chr9-99601282-DEL-114 |
| chr9-99617828-DEL-7131 | chr9-99617829-DEL-7131 |
| chr9-99626009-DEL-135 | chr9-99626010-DEL-135 |
| chr9-99631886-DEL-62 | chr9-99631887-DEL-54 |
| chr9-99638938-DEL-6440 | chr9-99638939-DEL-6440 |
| chr9-99646394-DEL-352 | chr9-99646395-DEL-352 |
| chr9-99672732-DEL-69 | chr9-99672740-DEL-81 |
| chr9-99690214-DEL-72 | chr9-99690179-DEL-108 |
| chr9-99724098-DEL-54 | chr9-99724099-DEL-54 |
| chr9-99731143-DEL-143 | chr9-99731193-DEL-94 |
| chr9-99737264-DEL-194 | chr9-99737302-DEL-153 |
| chr9-99765069-DEL-206 | chr9-99765060-DEL-204 |
| chr9-99775930-DEL-200 | chr9-99775931-DEL-200 |
| chr9-99806314-DEL-271 | chr9-99806330-DEL-259 |
| chr9-99815735-DEL-6223 | chr9-99815735-DEL-6224 |
| chr9-99829427-DEL-97 | chr9-99829428-DEL-97 |
| chr9-99840160-DEL-494 | chr9-99840161-DEL-494 |
| chr9-99847354-DEL-166 | chr9-99847355-DEL-166 |
| chr9-99862490-DEL-188 | chr9-99862491-DEL-188 |
| chr9-99863518-DEL-137 | chr9-99863519-DEL-137 |
| chr9-99946249-DEL-88 | chr9-99946250-DEL-88 |
| chr9-99951177-DEL-830 | chr9-99951174-DEL-832 |
| chr9-99955864-DEL-6866 | chr9-99955865-DEL-6866 |
| chr9-99955892-DEL-212 | chr9-99955893-DEL-212 |
| chr9-99965416-DEL-68 | chr9-99965416-DEL-68 |
| chr9-99983695-DEL-727 | chr9-99983696-DEL-727 |
| chr9-99993824-DEL-714 | chr9-99993825-DEL-714 |
| chr9-100028186-DEL-10981 | chr9-100028188-DEL-10979 |
| chr9-100042067-DEL-102 | chr9-100042068-DEL-102 |
| chr9-100101244-DEL-212 | chr9-100101247-DEL-224 |
| chr9-100106645-DEL-980 | chr9-100106646-DEL-978 |
| chr9-100113766-DEL-716 | chr9-100113767-DEL-716 |
| chr9-100121294-DEL-52 | chr9-100121295-DEL-52 |
| chr9-100128917-DEL-205 | chr9-100128918-DEL-205 |
| chr9-100150151-DEL-107 | chr9-100150172-DEL-87 |
| chr9-100158998-DEL-63 | chr9-100158999-DEL-63 |
| chr9-100196371-DEL-1211 | chr9-100196372-DEL-1211 |
| chr9-100222676-DEL-60 | chr9-100222677-DEL-56 |
| chr9-100226515-DEL-903 | chr9-100226516-DEL-903 |
| chr9-100242839-DEL-96 | chr9-100242840-DEL-96 |
| chr9-100244024-DEL-400 | chr9-100244025-DEL-400 |
| chr9-100258281-DEL-323 | chr9-100258282-DEL-323 |
| chr9-100264816-DEL-146 | chr9-100264816-DEL-146 |
| chr9-100274318-DEL-1766 | chr9-100274317-DEL-1757 |
| chr9-100283156-DEL-91 | chr9-100283157-DEL-91 |
| chr9-100289681-DEL-69 | chr9-100289682-DEL-69 |
| chr9-100290993-DEL-2171 | chr9-100290994-DEL-2171 |
| chr9-100300165-DEL-67 | chr9-100300143-DEL-96 |
| chr9-100306383-DEL-52 | chr9-100306384-DEL-52 |
| chr9-100308514-DEL-51 | chr9-100308515-DEL-51 |
| chr9-100312950-DEL-52 | chr9-100312951-DEL-52 |
| chr9-100323537-DEL-540 | chr9-100323539-DEL-537 |
| chr9-100324143-DEL-1274 | chr9-100324144-DEL-1274 |
| chr9-100331815-DEL-343 | chr9-100331816-DEL-343 |
| chr9-100352682-DEL-226 | chr9-100352685-DEL-224 |
| chr9-100362490-DEL-52 | chr9-100362491-DEL-56 |
| chr9-100369304-DEL-69 | chr9-100369305-DEL-69 |
| chr9-100383667-DEL-168 | chr9-100383668-DEL-168 |
| chr9-100385457-DEL-195 | chr9-100385458-DEL-195 |
| chr9-100433354-DEL-65 | chr9-100433345-DEL-66 |
| chr9-100444633-DEL-119 | chr9-100444634-DEL-116 |
| chr9-100448657-DEL-965 | chr9-100448657-DEL-965 |
| chr9-100467427-DEL-160 | chr9-100467428-DEL-160 |
| chr9-100468274-DEL-72 | chr9-100468275-DEL-72 |
| chr9-100495482-DEL-116 | chr9-100495472-DEL-117 |
| chr9-100509222-DEL-221 | chr9-100509223-DEL-221 |
| chr9-100513855-DEL-3396 | chr9-100513856-DEL-3396 |
| chr9-100518308-DEL-1745 | chr9-100518307-DEL-1746 |
| chr9-100518840-DEL-90 | chr9-100518838-DEL-85 |
| chr9-100519163-DEL-503 | chr9-100519164-DEL-503 |
| chr9-100531579-DEL-80 | chr9-100531580-DEL-80 |
| chr9-100558463-DEL-87 | chr9-100558464-DEL-87 |
| chr9-100576680-DEL-221 | chr9-100576681-DEL-221 |
| chr9-100653153-DEL-193 | chr9-100653155-DEL-192 |
| chr9-100682195-DEL-439 | chr9-100682196-DEL-439 |
| chr9-100701983-DEL-88 | chr9-100701984-DEL-88 |
| chr9-100767634-DEL-121 | chr9-100767635-DEL-121 |
| chr9-100788994-DEL-237 | chr9-100788995-DEL-237 |
| chr9-100789435-DEL-184 | chr9-100789437-DEL-183 |
| chr9-100795506-DEL-237 | chr9-100795507-DEL-237 |

| | |
|---|---|
| chr9-100802455-DEL-1920 | chr9-100802456-DEL-1920 |
| chr9-100820746-DEL-762 | chr9-100820756-DEL-757 |
| chr9-100823880-DEL-143 | chr9-100823881-DEL-143 |
| chr9-100850609-DEL-62 | chr9-100850610-DEL-62 |
| chr9-100865549-DEL-61 | chr9-100865550-DEL-61 |
| chr9-100870474-DEL-60 | chr9-100870474-DEL-61 |
| chr9-100890425-DEL-163 | chr9-100890426-DEL-163 |
| chr9-100891582-DEL-147 | chr9-100891583-DEL-147 |
| chr9-100896766-DEL-224 | chr9-100896767-DEL-224 |
| chr9-100902942-DEL-194 | chr9-100902943-DEL-194 |
| chr9-100909848-DEL-186 | chr9-100909849-DEL-185 |
| chr9-100914797-DEL-511 | chr9-100914773-DEL-495 |
| chr9-100927218-DEL-80 | chr9-100927219-DEL-80 |
| chr9-100939553-DEL-1462 | chr9-100939564-DEL-1466 |
| chr9-100944954-DEL-431 | chr9-100944955-DEL-431 |
| chr9-100972144-DEL-212 | chr9-100972145-DEL-212 |
| chr9-100988904-DEL-229 | chr9-100988905-DEL-229 |
| chr9-100994690-DEL-201 | chr9-100994691-DEL-201 |
| chr9-100998845-DEL-278 | chr9-100998875-DEL-258 |
| chr9-101030318-DEL-130 | chr9-101030319-DEL-130 |
| chr9-101040189-DEL-6169 | chr9-101040190-DEL-6169 |
| chr9-101101084-DEL-1200 | chr9-101101085-DEL-1200 |
| chr9-101102985-DEL-250 | chr9-101102986-DEL-250 |
| chr9-101140702-DEL-600 | chr9-101140703-DEL-600 |
| chr9-101169803-DEL-5733 | chr9-101169804-DEL-5733 |
| chr9-101179942-DEL-84 | chr9-101179943-DEL-83 |
| chr9-101187037-DEL-1893 | chr9-101187038-DEL-1893 |
| chr9-101193776-DEL-400 | chr9-101193777-DEL-400 |
| chr9-101196959-DEL-250 | chr9-101196961-DEL-249 |
| chr9-101208019-DEL-117 | chr9-101208020-DEL-117 |
| chr9-101213966-DEL-65 | chr9-101213963-DEL-69 |
| chr9-101286735-DEL-710 | chr9-101286736-DEL-710 |
| chr9-101304672-DEL-7288 | chr9-101304673-DEL-7288 |
| chr9-101304778-DEL-1060 | chr9-101304778-DEL-1060 |
| chr9-101314136-DEL-118 | chr9-101314140-DEL-115 |
| chr9-101316752-DEL-72 | chr9-101316755-DEL-72 |
| chr9-101343303-DEL-59 | chr9-101343304-DEL-59 |
| chr9-101352601-DEL-80 | chr9-101352602-DEL-80 |
| chr9-101366769-DEL-58 | chr9-101366770-DEL-58 |
| chr9-101367158-DEL-854 | chr9-101367158-DEL-860 |
| chr9-101388445-DEL-60 | chr9-101388446-DEL-60 |
| chr9-101426969-DEL-303 | chr9-101426970-DEL-303 |
| chr9-101439401-DEL-649 | chr9-101439403-DEL-647 |
| chr9-101483623-DEL-1634 | chr9-101483633-DEL-1624 |
| chr9-101486167-DEL-230 | chr9-101486168-DEL-230 |
| chr9-101506183-DEL-95 | chr9-101506184-DEL-95 |
| chr9-101547307-DEL-52 | chr9-101547308-DEL-52 |
| chr9-101636518-DEL-197 | chr9-101636556-DEL-198 |
| chr9-101714567-DEL-150 | chr9-101714568-DEL-150 |
| chr9-101730432-DEL-66 | chr9-101730435-DEL-64 |
| chr9-101806856-DEL-56 | chr9-101806857-DEL-56 |
| chr9-101835748-DEL-130 | chr9-101835749-DEL-128 |
| chr9-101977597-DEL-52 | chr9-101977599-DEL-50 |
| chr9-101984378-DEL-54 | chr9-101984377-DEL-54 |
| chr9-102031696-DEL-53 | chr9-102031693-DEL-57 |
| chr9-102079033-DEL-56 | chr9-102079034-DEL-56 |
| chr9-102128089-DEL-85 | chr9-102128081-DEL-91 |
| chr9-102154562-DEL-6432 | chr9-102154567-DEL-6433 |
| chr9-102240779-DEL-118 | chr9-102240780-DEL-118 |
| chr9-102279449-DEL-116 | chr9-102279448-DEL-106 |
| chr9-102305217-DEL-191 | chr9-102305218-DEL-191 |
| chr9-102336218-DEL-56 | chr9-102336223-DEL-52 |
| chr9-102348050-DEL-6598 | chr9-102348051-DEL-6598 |
| chr9-102387093-DEL-81 | chr9-102387094-DEL-81 |
| chr9-102441412-DEL-1847 | chr9-102441413-DEL-1847 |
| chr9-102458941-DEL-56 | chr9-102458938-DEL-60 |
| chr9-102481640-DEL-57 | chr9-102481641-DEL-57 |
| chr9-102499619-DEL-87 | chr9-102499617-DEL-88 |
| chr9-102517680-DEL-211 | chr9-102517682-DEL-211 |
| chr9-102523165-DEL-82 | chr9-102523115-DEL-122 |
| chr9-102551562-DEL-101 | chr9-102551558-DEL-102 |
| chr9-102556841-DEL-80 | chr9-102556846-DEL-75 |
| chr9-102621639-DEL-208 | chr9-102621640-DEL-208 |
| chr9-102641730-DEL-90 | chr9-102641746-DEL-69 |
| chr9-102642017-DEL-1891 | chr9-102642018-DEL-1891 |
| chr9-102669891-DEL-184 | chr9-102669892-DEL-184 |
| chr9-102691080-DEL-215 | chr9-102691060-DEL-189 |
| chr9-102691766-DEL-201 | chr9-102691767-DEL-201 |
| chr9-102694212-DEL-205 | chr9-102694213-DEL-205 |
| chr9-102697043-DEL-1699 | chr9-102697044-DEL-1699 |

| | |
|---|---|
| chr9-102746020-DEL-270 | chr9-102746021-DEL-270 |
| chr9-102759593-DEL-63 | chr9-102759588-DEL-55 |
| chr9-102760672-DEL-68 | chr9-102760673-DEL-64 |
| chr9-102778512-DEL-76 | chr9-102778513-DEL-81 |
| chr9-102786945-DEL-145 | chr9-102786946-DEL-145 |
| chr9-102790520-DEL-399 | chr9-102790521-DEL-399 |
| chr9-102794540-DEL-192 | chr9-102794541-DEL-192 |
| chr9-102820423-DEL-107 | chr9-102820418-DEL-113 |
| chr9-102833606-DEL-1886 | chr9-102833607-DEL-1886 |
| chr9-102840795-DEL-59 | chr9-102840796-DEL-59 |
| chr9-102860759-DEL-206 | chr9-102860760-DEL-206 |
| chr9-102905945-DEL-157 | chr9-102905942-DEL-158 |
| chr9-102922975-DEL-202 | chr9-102922976-DEL-202 |
| chr9-102961299-DEL-60 | chr9-102961300-DEL-60 |
| chr9-102972253-DEL-81 | chr9-102972254-DEL-81 |
| chr9-102985793-DEL-517 | chr9-102985794-DEL-517 |
| chr9-103002250-DEL-209 | chr9-103002251-DEL-209 |
| chr9-103029063-DEL-196 | chr9-103029065-DEL-195 |
| chr9-103032812-DEL-76 | chr9-103032813-DEL-74 |
| chr9-103066683-DEL-98 | chr9-103066666-DEL-147 |
| chr9-103092740-DEL-90 | chr9-103092741-DEL-90 |
| chr9-103121695-DEL-180 | chr9-103121704-DEL-180 |
| chr9-103122463-DEL-54 | chr9-103122453-DEL-95 |
| chr9-103160637-DEL-201 | chr9-103160638-DEL-201 |
| chr9-103163249-DEL-236 | chr9-103163250-DEL-236 |
| chr9-103164708-DEL-83 | chr9-103164709-DEL-83 |
| chr9-103172674-DEL-54 | chr9-103172676-DEL-52 |
| chr9-103196232-DEL-192 | chr9-103196233-DEL-192 |
| chr9-103216479-DEL-110 | chr9-103216480-DEL-110 |
| chr9-103224827-DEL-54 | chr9-103224828-DEL-52 |
| chr9-103259084-DEL-81 | chr9-103259084-DEL-96 |
| chr9-103272723-DEL-1263 | chr9-103272724-DEL-1264 |
| chr9-103277263-DEL-117 | chr9-103277282-DEL-87 |
| chr9-103293037-DEL-52 | chr9-103293037-DEL-52 |
| chr9-103294602-DEL-189 | chr9-103294596-DEL-186 |
| chr9-103296559-DEL-180 | chr9-103296560-DEL-180 |
| chr9-103312093-DEL-182 | chr9-103312094-DEL-182 |
| chr9-103319896-DEL-323 | chr9-103319873-DEL-331 |
| chr9-103328006-DEL-70 | chr9-103328007-DEL-70 |
| chr9-103330316-DEL-6297 | chr9-103330317-DEL-6297 |
| chr9-103334294-DEL-176 | chr9-103334295-DEL-176 |
| chr9-103336314-DEL-132 | chr9-103336313-DEL-132 |
| chr9-103370751-DEL-485 | chr9-103370784-DEL-451 |
| chr9-103374767-DEL-154 | chr9-103374770-DEL-152 |
| chr9-103388253-DEL-56 | chr9-103388254-DEL-56 |
| chr9-103396623-DEL-990 | chr9-103396626-DEL-951 |
| chr9-103413491-DEL-76 | chr9-103413491-DEL-76 |
| chr9-103419796-DEL-91 | chr9-103419797-DEL-91 |
| chr9-103433764-DEL-62 | chr9-103433765-DEL-62 |
| chr9-103456693-DEL-51 | chr9-103456694-DEL-51 |
| chr9-103463291-DEL-1559 | chr9-103463292-DEL-1559 |
| chr9-103466146-DEL-234 | chr9-103466147-DEL-234 |
| chr9-103480094-DEL-102 | chr9-103480120-DEL-84 |
| chr9-103486159-DEL-214 | chr9-103486160-DEL-214 |
| chr9-103491385-DEL-63 | chr9-103491386-DEL-63 |
| chr9-103499020-DEL-51 | chr9-103499021-DEL-51 |
| chr9-103547932-DEL-136 | chr9-103547933-DEL-136 |
| chr9-103577893-DEL-51 | chr9-103577894-DEL-51 |
| chr9-103587376-DEL-173 | chr9-103587363-DEL-145 |
| chr9-103662493-DEL-298 | chr9-103662494-DEL-298 |
| chr9-103685444-DEL-111 | chr9-103685445-DEL-111 |
| chr9-103686470-DEL-2237 | chr9-103686471-DEL-2237 |
| chr9-103724916-DEL-134 | chr9-103724925-DEL-126 |
| chr9-103732021-DEL-109 | chr9-103732022-DEL-109 |
| chr9-103749736-DEL-217 | chr9-103749737-DEL-217 |
| chr9-103755413-DEL-88 | chr9-103755414-DEL-88 |
| chr9-103782614-DEL-67 | chr9-103782615-DEL-68 |
| chr9-103852425-DEL-59 | chr9-103852423-DEL-56 |
| chr9-103877610-DEL-125 | chr9-103877631-DEL-105 |
| chr9-103887746-DEL-60 | chr9-103887765-DEL-61 |
| chr9-103889012-DEL-89 | chr9-103889012-DEL-89 |
| chr9-103907698-DEL-102 | chr9-103907685-DEL-94 |
| chr9-103924353-DEL-64 | chr9-103924354-DEL-64 |
| chr9-103930489-DEL-208 | chr9-103930490-DEL-208 |
| chr9-103947559-DEL-61 | chr9-103947558-DEL-61 |
| chr9-103982420-DEL-60 | chr9-103982396-DEL-60 |
| chr9-103991101-DEL-69 | chr9-103991105-DEL-66 |
| chr9-103995954-DEL-88 | chr9-103995955-DEL-88 |
| chr9-104006789-DEL-376 | chr9-104006790-DEL-376 |
| chr9-104012716-DEL-214 | chr9-104012717-DEL-214 |

| | |
|---|---|
| chr9-104050820-DEL-77 | chr9-104050821-DEL-77 |
| chr9-104078351-DEL-619 | chr9-104078397-DEL-620 |
| chr9-104088760-DEL-114 | chr9-104088754-DEL-87 |
| chr9-104099226-DEL-6248 | chr9-104099227-DEL-6248 |
| chr9-104187019-DEL-187 | chr9-104187020-DEL-187 |
| chr9-104189050-DEL-246 | chr9-104189051-DEL-246 |
| chr9-104190716-DEL-203 | chr9-104190717-DEL-203 |
| chr9-104223552-DEL-196 | chr9-104223553-DEL-196 |
| chr9-104234854-DEL-58 | chr9-104234873-DEL-51 |
| chr9-104239412-DEL-68 | chr9-104239404-DEL-75 |
| chr9-104258981-DEL-159 | chr9-104258982-DEL-159 |
| chr9-104271036-DEL-106 | chr9-104271027-DEL-86 |
| chr9-104273440-DEL-182 | chr9-104273427-DEL-183 |
| chr9-104274614-DEL-195 | chr9-104274615-DEL-195 |
| chr9-104278099-DEL-52 | chr9-104278100-DEL-52 |
| chr9-104281902-DEL-75 | chr9-104281903-DEL-74 |
| chr9-104282969-DEL-272 | chr9-104282956-DEL-250 |
| chr9-104288567-DEL-219 | chr9-104288568-DEL-219 |
| chr9-104298631-DEL-60 | chr9-104298632-DEL-60 |
| chr9-104301875-DEL-78 | chr9-104301873-DEL-67 |
| chr9-104308593-DEL-176 | chr9-104308594-DEL-176 |
| chr9-104339844-DEL-336 | chr9-104339837-DEL-334 |
| chr9-104372965-DEL-205 | chr9-104372966-DEL-205 |
| chr9-104382284-DEL-65 | chr9-104382256-DEL-60 |
| chr9-104394006-DEL-666 | chr9-104394007-DEL-666 |
| chr9-104398192-DEL-875 | chr9-104398193-DEL-875 |
| chr9-104399188-DEL-130 | chr9-104399179-DEL-134 |
| chr9-104401100-DEL-52 | chr9-104401101-DEL-52 |
| chr9-104436940-DEL-1870 | chr9-104436941-DEL-1870 |
| chr9-104438727-DEL-83 | chr9-104438728-DEL-83 |
| chr9-104441353-DEL-248 | chr9-104441322-DEL-242 |
| chr9-104454272-DEL-648 | chr9-104454273-DEL-648 |
| chr9-104473602-DEL-140 | chr9-104473636-DEL-127 |
| chr9-104494721-DEL-224 | chr9-104494722-DEL-224 |
| chr9-104509557-DEL-129 | chr9-104509559-DEL-98 |
| chr9-104511547-DEL-52 | chr9-104511548-DEL-52 |
| chr9-104527361-DEL-52 | chr9-104527362-DEL-52 |
| chr9-104536220-DEL-80 | chr9-104536221-DEL-80 |
| chr9-104543534-DEL-310 | chr9-104543535-DEL-310 |
| chr9-104553546-DEL-67 | chr9-104553546-DEL-68 |
| chr9-104569941-DEL-50 | chr9-104569934-DEL-62 |
| chr9-104589308-DEL-63 | chr9-104589309-DEL-63 |
| chr9-104611909-DEL-74 | chr9-104611910-DEL-74 |
| chr9-104616764-DEL-87 | chr9-104616753-DEL-99 |
| chr9-104675385-DEL-213 | chr9-104675377-DEL-214 |
| chr9-104698822-DEL-240 | chr9-104698823-DEL-240 |
| chr9-104742722-DEL-52 | chr9-104742723-DEL-52 |
| chr9-104754518-DEL-392 | chr9-104754519-DEL-392 |
| chr9-104755624-DEL-91 | chr9-104755625-DEL-91 |
| chr9-104765644-DEL-166 | chr9-104765645-DEL-166 |
| chr9-104779915-DEL-144 | chr9-104779916-DEL-144 |
| chr9-104782076-DEL-168 | chr9-104782077-DEL-168 |
| chr9-104789607-DEL-199 | chr9-104789608-DEL-199 |
| chr9-104794880-DEL-64 | chr9-104794881-DEL-64 |
| chr9-104801088-DEL-62 | chr9-104801089-DEL-66 |
| chr9-104825165-DEL-52 | chr9-104825166-DEL-52 |
| chr9-104833162-DEL-77 | chr9-104833163-DEL-77 |
| chr9-104834715-DEL-110 | chr9-104834715-DEL-119 |
| chr9-104854440-DEL-62 | chr9-104854441-DEL-62 |
| chr9-104864822-DEL-187 | chr9-104864823-DEL-187 |
| chr9-104883708-DEL-113 | chr9-104883709-DEL-113 |
| chr9-104897387-DEL-52 | chr9-104897352-DEL-56 |
| chr9-104934749-DEL-101 | chr9-104934751-DEL-100 |
| chr9-104937916-DEL-186 | chr9-104937917-DEL-186 |
| chr9-104962578-DEL-6217 | chr9-104962579-DEL-6217 |
| chr9-104980147-DEL-74 | chr9-104980148-DEL-74 |
| chr9-104980663-DEL-200 | chr9-104980664-DEL-200 |
| chr9-104984951-DEL-163 | chr9-104984953-DEL-158 |
| chr9-104994620-DEL-1892 | chr9-104994621-DEL-1892 |
| chr9-105005464-DEL-576 | chr9-105005465-DEL-576 |
| chr9-105014978-DEL-52 | chr9-105014979-DEL-52 |
| chr9-105015739-DEL-69 | chr9-105015738-DEL-70 |
| chr9-105017823-DEL-63 | chr9-105017824-DEL-63 |
| chr9-105032580-DEL-942 | chr9-105032581-DEL-942 |
| chr9-105050672-DEL-58 | chr9-105050677-DEL-54 |
| chr9-105076645-DEL-115 | chr9-105076619-DEL-116 |
| chr9-105080718-DEL-180 | chr9-105080757-DEL-152 |
| chr9-105085110-DEL-87 | chr9-105085111-DEL-87 |
| chr9-105086425-DEL-1072 | chr9-105086426-DEL-1072 |
| chr9-105091140-DEL-2098 | chr9-105091141-DEL-2098 |

| | |
|---|---|
| chr9-105096130-DEL-635 | chr9-105096131-DEL-635 |
| chr9-105113513-DEL-507 | chr9-105113514-DEL-507 |
| chr9-105136740-DEL-56 | chr9-105136741-DEL-50 |
| chr9-105147068-DEL-361 | chr9-105147069-DEL-361 |
| chr9-105161298-DEL-56 | chr9-105161299-DEL-56 |
| chr9-105173832-DEL-62 | chr9-105173837-DEL-58 |
| chr9-105174743-DEL-58 | chr9-105174744-DEL-58 |
| chr9-105202837-DEL-2575 | chr9-105202838-DEL-2575 |
| chr9-105210930-DEL-6996 | chr9-105210931-DEL-6996 |
| chr9-105221726-DEL-1766 | chr9-105221727-DEL-1766 |
| chr9-105234680-DEL-70 | chr9-105234675-DEL-70 |
| chr9-105250873-DEL-52 | chr9-105250874-DEL-52 |
| chr9-105260795-DEL-293 | chr9-105260796-DEL-293 |
| chr9-105268216-DEL-88 | chr9-105268212-DEL-88 |
| chr9-105269395-DEL-53 | chr9-105269396-DEL-54 |
| chr9-105286524-DEL-1642 | chr9-105286526-DEL-1639 |
| chr9-105316240-DEL-157 | chr9-105316243-DEL-125 |
| chr9-105317097-DEL-962 | chr9-105317098-DEL-962 |
| chr9-105318960-DEL-170 | chr9-105318961-DEL-170 |
| chr9-105332175-DEL-1869 | chr9-105332176-DEL-1869 |
| chr9-105344142-DEL-61 | chr9-105344127-DEL-59 |
| chr9-105385231-DEL-68 | chr9-105385232-DEL-68 |
| chr9-105390761-DEL-260 | chr9-105390762-DEL-260 |
| chr9-105416516-DEL-60 | chr9-105416517-DEL-60 |
| chr9-105419509-DEL-55 | chr9-105419510-DEL-50 |
| chr9-105423387-DEL-396 | chr9-105423388-DEL-397 |
| chr9-105469900-DEL-225 | chr9-105469901-DEL-225 |
| chr9-105484431-DEL-3876 | chr9-105484469-DEL-3829 |
| chr9-105494815-DEL-60 | chr9-105494816-DEL-80 |
| chr9-105500694-DEL-6395 | chr9-105500695-DEL-6395 |
| chr9-105515152-DEL-72 | chr9-105515153-DEL-66 |
| chr9-105522733-DEL-142 | chr9-105522734-DEL-142 |
| chr9-105552622-DEL-55 | chr9-105552597-DEL-56 |
| chr9-105554666-DEL-60 | chr9-105554667-DEL-60 |
| chr9-105583853-DEL-360 | chr9-105583815-DEL-363 |
| chr9-105587491-DEL-237 | chr9-105587513-DEL-204 |
| chr9-105615589-DEL-62 | chr9-105615562-DEL-90 |
| chr9-105647015-DEL-676 | chr9-105647016-DEL-676 |
| chr9-105681633-DEL-150 | chr9-105681643-DEL-147 |
| chr9-105700149-DEL-191 | chr9-105700150-DEL-191 |
| chr9-105722967-DEL-88 | chr9-105722968-DEL-88 |
| chr9-105724785-DEL-56 | chr9-105724786-DEL-56 |
| chr9-105726028-DEL-61 | chr9-105726029-DEL-61 |
| chr9-105776749-DEL-169 | chr9-105776750-DEL-167 |
| chr9-105796339-DEL-66 | chr9-105796340-DEL-66 |
| chr9-105800076-DEL-201 | chr9-105800077-DEL-201 |
| chr9-105819003-DEL-384 | chr9-105819004-DEL-384 |
| chr9-105819623-DEL-91 | chr9-105819624-DEL-90 |
| chr9-105821174-DEL-151 | chr9-105821175-DEL-151 |
| chr9-105823375-DEL-115 | chr9-105823377-DEL-114 |
| chr9-105825810-DEL-155 | chr9-105825811-DEL-155 |
| chr9-105835599-DEL-588 | chr9-105835600-DEL-588 |
| chr9-105843552-DEL-80 | chr9-105843549-DEL-84 |
| chr9-105860481-DEL-192 | chr9-105860469-DEL-189 |
| chr9-105868814-DEL-73 | chr9-105868815-DEL-73 |
| chr9-105873921-DEL-140 | chr9-105873922-DEL-140 |
| chr9-105916020-DEL-54 | chr9-105916019-DEL-56 |
| chr9-105931421-DEL-752 | chr9-105931420-DEL-753 |
| chr9-105946021-DEL-68 | chr9-105946022-DEL-68 |
| chr9-105974491-DEL-135 | chr9-105974492-DEL-135 |
| chr9-105979861-DEL-196 | chr9-105979862-DEL-196 |
| chr9-105982801-DEL-223 | chr9-105982791-DEL-224 |
| chr9-105983274-DEL-340 | chr9-105983275-DEL-340 |
| chr9-105993652-DEL-62 | chr9-105993653-DEL-62 |
| chr9-106005431-DEL-54 | chr9-106005432-DEL-54 |
| chr9-106012568-DEL-179 | chr9-106012561-DEL-184 |
| chr9-106014221-DEL-192 | chr9-106014209-DEL-193 |
| chr9-106018025-DEL-72 | chr9-106018026-DEL-72 |
| chr9-106029195-DEL-58 | chr9-106029200-DEL-59 |
| chr9-106038232-DEL-54 | chr9-106038232-DEL-54 |
| chr9-106045926-DEL-208 | chr9-106045927-DEL-208 |
| chr9-106062819-DEL-211 | chr9-106062820-DEL-211 |
| chr9-106068858-DEL-55 | chr9-106068859-DEL-55 |
| chr9-106076928-DEL-51 | chr9-106076929-DEL-51 |
| chr9-106077891-DEL-82 | chr9-106077892-DEL-82 |
| chr9-106121779-DEL-103 | chr9-106121815-DEL-72 |
| chr9-106123183-DEL-54 | chr9-106123184-DEL-54 |
| chr9-106130232-DEL-53 | chr9-106130233-DEL-53 |
| chr9-106136588-DEL-387 | chr9-106136589-DEL-387 |
| chr9-106148546-DEL-60 | chr9-106148547-DEL-60 |

| | |
|---|---|
| chr9-106149356-DEL-53 | chr9-106149333-DEL-54 |
| chr9-106164803-DEL-85 | chr9-106164804-DEL-85 |
| chr9-106168919-DEL-68 | chr9-106168916-DEL-63 |
| chr9-106180467-DEL-52 | chr9-106180468-DEL-52 |
| chr9-106198367-DEL-118 | chr9-106198368-DEL-118 |
| chr9-106212824-DEL-70 | chr9-106212825-DEL-70 |
| chr9-106234097-DEL-51 | chr9-106234055-DEL-93 |
| chr9-106253332-DEL-436 | chr9-106253333-DEL-436 |
| chr9-106259698-DEL-493 | chr9-106259699-DEL-494 |
| chr9-106265676-DEL-324 | chr9-106265677-DEL-324 |
| chr9-106275809-DEL-60 | chr9-106275793-DEL-84 |
| chr9-106305632-DEL-240 | chr9-106305633-DEL-240 |
| chr9-106322850-DEL-118 | chr9-106322858-DEL-115 |
| chr9-106338596-DEL-150 | chr9-106338597-DEL-150 |
| chr9-106344682-DEL-476 | chr9-106344683-DEL-476 |
| chr9-106363508-DEL-2175 | chr9-106363509-DEL-2175 |
| chr9-106363918-DEL-196 | chr9-106363919-DEL-196 |
| chr9-106370159-DEL-6422 | chr9-106370160-DEL-6422 |
| chr9-106414840-DEL-115 | chr9-106414833-DEL-116 |
| chr9-106416321-DEL-391 | chr9-106416322-DEL-391 |
| chr9-106423331-DEL-199 | chr9-106423332-DEL-199 |
| chr9-106426409-DEL-377 | chr9-106426410-DEL-373 |
| chr9-106452600-DEL-101 | chr9-106452598-DEL-94 |
| chr9-106456866-DEL-237 | chr9-106456867-DEL-237 |
| chr9-106461597-DEL-6512 | chr9-106461595-DEL-6514 |
| chr9-106480147-DEL-56 | chr9-106480157-DEL-50 |
| chr9-106484847-DEL-73 | chr9-106484853-DEL-68 |
| chr9-106486739-DEL-662 | chr9-106486738-DEL-663 |
| chr9-106490771-DEL-76 | chr9-106490772-DEL-76 |
| chr9-106512381-DEL-94 | chr9-106512382-DEL-94 |
| chr9-106571234-DEL-335 | chr9-106571235-DEL-335 |
| chr9-106609064-DEL-89 | chr9-106609069-DEL-85 |
| chr9-106612043-DEL-202 | chr9-106612042-DEL-202 |
| chr9-106622187-DEL-60 | chr9-106622187-DEL-66 |
| chr9-106647750-DEL-61 | chr9-106647751-DEL-61 |
| chr9-106671931-DEL-209 | chr9-106671932-DEL-209 |
| chr9-106676521-DEL-56 | chr9-106676522-DEL-56 |
| chr9-106688611-DEL-1568 | chr9-106688612-DEL-1567 |
| chr9-106697932-DEL-84 | chr9-106697933-DEL-84 |
| chr9-106722948-DEL-170 | chr9-106722949-DEL-170 |
| chr9-106728378-DEL-548 | chr9-106728379-DEL-548 |
| chr9-106748200-DEL-623 | chr9-106748201-DEL-623 |
| chr9-106750545-DEL-66 | chr9-106750524-DEL-68 |
| chr9-106757414-DEL-52 | chr9-106757415-DEL-52 |
| chr9-106794248-DEL-200 | chr9-106794249-DEL-200 |
| chr9-106795621-DEL-6313 | chr9-106795622-DEL-6313 |
| chr9-106811129-DEL-130 | chr9-106811130-DEL-126 |
| chr9-106855054-DEL-215 | chr9-106855055-DEL-215 |
| chr9-106859621-DEL-112 | chr9-106859622-DEL-112 |
| chr9-106886902-DEL-63 | chr9-106886911-DEL-58 |
| chr9-106894789-DEL-91 | chr9-106894787-DEL-91 |
| chr9-106902304-DEL-88 | chr9-106902307-DEL-86 |
| chr9-106906363-DEL-72 | chr9-106906352-DEL-100 |
| chr9-106927884-DEL-200 | chr9-106927885-DEL-200 |
| chr9-106949476-DEL-241 | chr9-106949477-DEL-241 |
| chr9-106950027-DEL-6252 | chr9-106950028-DEL-6252 |
| chr9-106958998-DEL-68 | chr9-106958999-DEL-66 |
| chr9-106980911-DEL-2125 | chr9-106980912-DEL-2125 |
| chr9-106991826-DEL-67 | chr9-106991823-DEL-68 |
| chr9-106992253-DEL-336 | chr9-106992254-DEL-336 |
| chr9-107017745-DEL-3417 | chr9-107017745-DEL-3417 |
| chr9-107043392-DEL-352 | chr9-107043393-DEL-352 |
| chr9-107087782-DEL-1749 | chr9-107087783-DEL-1749 |
| chr9-107090810-DEL-6062 | chr9-107090811-DEL-6062 |
| chr9-107097988-DEL-222 | chr9-107097989-DEL-222 |
| chr9-107172412-DEL-60 | chr9-107172413-DEL-65 |
| chr9-107184295-DEL-110 | chr9-107184294-DEL-110 |
| chr9-107218851-DEL-140 | chr9-107218852-DEL-140 |
| chr9-107222413-DEL-237 | chr9-107222416-DEL-235 |
| chr9-107228196-DEL-99 | chr9-107228198-DEL-93 |
| chr9-107233919-DEL-198 | chr9-107233920-DEL-198 |
| chr9-107239946-DEL-52 | chr9-107239946-DEL-57 |
| chr9-107243745-DEL-87 | chr9-107243746-DEL-87 |
| chr9-107260682-DEL-60 | chr9-107260674-DEL-59 |
| chr9-107272774-DEL-65 | chr9-107272777-DEL-53 |
| chr9-107292995-DEL-63 | chr9-107293005-DEL-50 |
| chr9-107315302-DEL-245 | chr9-107315303-DEL-245 |
| chr9-107341005-DEL-54 | chr9-107341006-DEL-54 |
| chr9-107409747-DEL-101 | chr9-107409748-DEL-99 |
| chr9-107451528-DEL-51 | chr9-107451529-DEL-51 |

| | |
|---|---|
| chr9-107454491-DEL-94 | chr9-107454492-DEL-94 |
| chr9-107513703-DEL-175 | chr9-107513704-DEL-175 |
| chr9-107530214-DEL-170 | chr9-107530213-DEL-172 |
| chr9-107555258-DEL-203 | chr9-107555259-DEL-203 |
| chr9-107630682-DEL-232 | chr9-107630683-DEL-232 |
| chr9-107632394-DEL-89 | chr9-107632402-DEL-69 |
| chr9-107655873-DEL-51 | chr9-107655871-DEL-52 |
| chr9-107665848-DEL-205 | chr9-107665849-DEL-205 |
| chr9-107668493-DEL-210 | chr9-107668494-DEL-210 |
| chr9-107684620-DEL-3087 | chr9-107684621-DEL-3087 |
| chr9-107696749-DEL-82 | chr9-107696730-DEL-102 |
| chr9-107705681-DEL-117 | chr9-107705682-DEL-117 |
| chr9-107743070-DEL-156 | chr9-107743092-DEL-108 |
| chr9-107753370-DEL-226 | chr9-107753371-DEL-226 |
| chr9-107764353-DEL-92 | chr9-107764340-DEL-94 |
| chr9-107829790-DEL-56 | chr9-107829791-DEL-56 |
| chr9-107903190-DEL-195 | chr9-107903191-DEL-195 |
| chr9-107913956-DEL-120 | chr9-107913958-DEL-114 |
| chr9-107914841-DEL-206 | chr9-107914842-DEL-206 |
| chr9-107952200-DEL-193 | chr9-107952204-DEL-172 |
| chr9-107953866-DEL-72 | chr9-107953863-DEL-76 |
| chr9-107995721-DEL-736 | chr9-107995722-DEL-736 |
| chr9-108000274-DEL-124 | chr9-108000277-DEL-115 |
| chr9-108071446-DEL-103 | chr9-108071447-DEL-103 |
| chr9-108073674-DEL-197 | chr9-108073675-DEL-197 |
| chr9-108125149-DEL-53 | chr9-108125150-DEL-53 |
| chr9-108129578-DEL-206 | chr9-108129579-DEL-206 |
| chr9-108133049-DEL-211 | chr9-108133050-DEL-211 |
| chr9-108163037-DEL-56 | chr9-108163038-DEL-56 |
| chr9-108164987-DEL-205 | chr9-108164986-DEL-206 |
| chr9-108190260-DEL-114 | chr9-108190255-DEL-120 |
| chr9-108197260-DEL-64 | chr9-108197261-DEL-64 |
| chr9-108203715-DEL-181 | chr9-108203716-DEL-181 |
| chr9-108220497-DEL-63 | chr9-108220498-DEL-63 |
| chr9-108222127-DEL-216 | chr9-108222128-DEL-217 |
| chr9-108223900-DEL-71 | chr9-108223912-DEL-92 |
| chr9-108251652-DEL-69 | chr9-108251653-DEL-69 |
| chr9-108252881-DEL-213 | chr9-108252882-DEL-213 |
| chr9-108280802-DEL-114 | chr9-108280803-DEL-114 |
| chr9-108304091-DEL-202 | chr9-108304090-DEL-202 |
| chr9-108314838-DEL-69 | chr9-108314839-DEL-69 |
| chr9-108332791-DEL-54 | chr9-108332792-DEL-54 |
| chr9-108387958-DEL-79 | chr9-108387959-DEL-82 |
| chr9-108412078-DEL-203 | chr9-108412079-DEL-203 |
| chr9-108416802-DEL-231 | chr9-108416800-DEL-232 |
| chr9-108425405-DEL-220 | chr9-108425406-DEL-220 |
| chr9-108462178-DEL-124 | chr9-108462179-DEL-124 |
| chr9-108469247-DEL-87 | chr9-108469248-DEL-87 |
| chr9-108472560-DEL-531 | chr9-108472561-DEL-531 |
| chr9-108495669-DEL-977 | chr9-108495674-DEL-974 |
| chr9-108499344-DEL-60 | chr9-108499345-DEL-60 |
| chr9-108537071-DEL-210 | chr9-108537072-DEL-210 |
| chr9-108541559-DEL-1729 | chr9-108541588-DEL-1713 |
| chr9-108545953-DEL-152 | chr9-108545951-DEL-152 |
| chr9-108552194-DEL-201 | chr9-108552195-DEL-201 |
| chr9-108584912-DEL-1002 | chr9-108584913-DEL-1002 |
| chr9-108606028-DEL-78 | chr9-108606029-DEL-78 |
| chr9-108623466-DEL-52 | chr9-108623467-DEL-52 |
| chr9-108630012-DEL-528 | chr9-108630013-DEL-528 |
| chr9-108634439-DEL-112 | chr9-108634442-DEL-90 |
| chr9-108635272-DEL-178 | chr9-108635273-DEL-178 |
| chr9-108639922-DEL-201 | chr9-108639923-DEL-202 |
| chr9-108648755-DEL-1204 | chr9-108648756-DEL-1204 |
| chr9-108741034-DEL-59 | chr9-108741035-DEL-59 |
| chr9-108756482-DEL-192 | chr9-108756468-DEL-188 |
| chr9-108766825-DEL-205 | chr9-108766826-DEL-205 |
| chr9-108772362-DEL-620 | chr9-108772363-DEL-620 |
| chr9-108809358-DEL-82 | chr9-108809359-DEL-82 |
| chr9-108824937-DEL-91 | chr9-108824938-DEL-91 |
| chr9-108890872-DEL-205 | chr9-108890873-DEL-205 |
| chr9-108914963-DEL-151 | chr9-108914950-DEL-171 |
| chr9-108957613-DEL-106 | chr9-108957614-DEL-106 |
| chr9-109029516-DEL-1684 | chr9-109029517-DEL-1684 |
| chr9-109033862-DEL-114 | chr9-109033863-DEL-114 |
| chr9-109074047-DEL-7087 | chr9-109074048-DEL-7087 |
| chr9-109106746-DEL-112 | chr9-109106747-DEL-112 |
| chr9-109204628-DEL-109 | chr9-109204629-DEL-108 |
| chr9-109242218-DEL-188 | chr9-109242218-DEL-181 |
| chr9-109245624-DEL-1980 | chr9-109245624-DEL-1980 |
| chr9-109284503-DEL-112 | chr9-109284482-DEL-134 |

| | |
|---|---|
| chr9-109300315-DEL-4580 | chr9-109300316-DEL-4580 |
| chr9-109356930-DEL-1144 | chr9-109356918-DEL-1145 |
| chr9-109394968-DEL-86 | chr9-109394990-DEL-84 |
| chr9-109405406-DEL-69 | chr9-109405407-DEL-69 |
| chr9-109414810-DEL-203 | chr9-109414807-DEL-203 |
| chr9-109415676-DEL-2313 | chr9-109415677-DEL-2313 |
| chr9-109519184-DEL-80 | chr9-109519185-DEL-80 |
| chr9-109551710-DEL-83 | chr9-109551711-DEL-83 |
| chr9-109555608-DEL-121 | chr9-109555609-DEL-121 |
| chr9-109597931-DEL-177 | chr9-109597966-DEL-143 |
| chr9-109604137-DEL-1430 | chr9-109604138-DEL-1430 |
| chr9-109607491-DEL-1062 | chr9-109607492-DEL-1062 |
| chr9-109665103-DEL-261 | chr9-109665104-DEL-254 |
| chr9-109680592-DEL-236 | chr9-109680593-DEL-236 |
| chr9-109681957-DEL-206 | chr9-109681958-DEL-206 |
| chr9-109684997-DEL-56 | chr9-109684996-DEL-57 |
| chr9-109686434-DEL-146 | chr9-109686435-DEL-146 |
| chr9-109693626-DEL-93 | chr9-109693615-DEL-59 |
| chr9-109694374-DEL-217 | chr9-109694375-DEL-217 |
| chr9-109702384-DEL-94 | chr9-109702385-DEL-94 |
| chr9-109717470-DEL-345 | chr9-109717471-DEL-345 |
| chr9-109737668-DEL-233 | chr9-109737709-DEL-196 |
| chr9-109745593-DEL-214 | chr9-109745594-DEL-214 |
| chr9-109775671-DEL-337 | chr9-109775672-DEL-337 |
| chr9-109778540-DEL-108 | chr9-109778541-DEL-108 |
| chr9-109796869-DEL-101 | chr9-109796884-DEL-76 |
| chr9-109809277-DEL-170 | chr9-109809277-DEL-170 |
| chr9-109823926-DEL-184 | chr9-109823926-DEL-184 |
| chr9-109840608-DEL-210 | chr9-109840609-DEL-210 |
| chr9-109845407-DEL-80 | chr9-109845408-DEL-80 |
| chr9-109847706-DEL-123 | chr9-109847707-DEL-123 |
| chr9-109854127-DEL-635 | chr9-109854128-DEL-635 |
| chr9-109856979-DEL-208 | chr9-109856980-DEL-208 |
| chr9-109865221-DEL-194 | chr9-109865222-DEL-194 |
| chr9-109887587-DEL-173 | chr9-109887588-DEL-173 |
| chr9-109905482-DEL-218 | chr9-109905483-DEL-218 |
| chr9-109941153-DEL-166 | chr9-109941154-DEL-166 |
| chr9-109949389-DEL-111 | chr9-109949390-DEL-111 |
| chr9-109951714-DEL-56 | chr9-109951715-DEL-56 |
| chr9-109956243-DEL-208 | chr9-109956244-DEL-208 |
| chr9-109977312-DEL-206 | chr9-109977313-DEL-206 |
| chr9-109983147-DEL-211 | chr9-109983148-DEL-211 |
| chr9-109983654-DEL-216 | chr9-109983655-DEL-216 |
| chr9-109984303-DEL-1525 | chr9-109984303-DEL-1525 |
| chr9-109997818-DEL-211 | chr9-109997819-DEL-211 |
| chr9-109998866-DEL-198 | chr9-109998867-DEL-197 |
| chr9-110001089-DEL-50 | chr9-110001055-DEL-66 |
| chr9-110016124-DEL-205 | chr9-110016125-DEL-205 |
| chr9-110023599-DEL-206 | chr9-110023600-DEL-206 |
| chr9-110043431-DEL-131 | chr9-110043432-DEL-131 |
| chr9-110070609-DEL-57 | chr9-110070610-DEL-57 |
| chr9-110081138-DEL-3006 | chr9-110081139-DEL-3006 |
| chr9-110130180-DEL-206 | chr9-110130181-DEL-206 |
| chr9-110132840-DEL-202 | chr9-110132841-DEL-202 |
| chr9-110137585-DEL-197 | chr9-110137586-DEL-197 |
| chr9-110137979-DEL-493 | chr9-110137963-DEL-484 |
| chr9-110141795-DEL-242 | chr9-110141806-DEL-235 |
| chr9-110158751-DEL-204 | chr9-110158752-DEL-204 |
| chr9-110192595-DEL-2564 | chr9-110192596-DEL-2564 |
| chr9-110206842-DEL-123 | chr9-110206847-DEL-121 |
| chr9-110232618-DEL-202 | chr9-110232619-DEL-202 |
| chr9-110244816-DEL-422 | chr9-110244817-DEL-422 |
| chr9-110254639-DEL-224 | chr9-110254640-DEL-224 |
| chr9-110263177-DEL-187 | chr9-110263178-DEL-187 |
| chr9-110281309-DEL-80 | chr9-110281310-DEL-80 |
| chr9-110283155-DEL-1000 | chr9-110283143-DEL-994 |
| chr9-110287728-DEL-74 | chr9-110287740-DEL-52 |
| chr9-110292235-DEL-196 | chr9-110292236-DEL-196 |
| chr9-110301435-DEL-255 | chr9-110301436-DEL-255 |
| chr9-110304189-DEL-189 | chr9-110304190-DEL-189 |
| chr9-110318035-DEL-337 | chr9-110318036-DEL-337 |
| chr9-110328348-DEL-53 | chr9-110328349-DEL-53 |
| chr9-110332124-DEL-55 | chr9-110332125-DEL-55 |
| chr9-110390660-DEL-205 | chr9-110390661-DEL-205 |
| chr9-110394423-DEL-206 | chr9-110394424-DEL-206 |
| chr9-110410625-DEL-134 | chr9-110410626-DEL-134 |
| chr9-110433639-DEL-964 | chr9-110433640-DEL-964 |
| chr9-110445562-DEL-90 | chr9-110445519-DEL-136 |
| chr9-110528194-DEL-237 | chr9-110528195-DEL-237 |
| chr9-110534227-DEL-204 | chr9-110534228-DEL-204 |

| | |
|---|---|
| chr9-110534965-DEL-56 | chr9-110534966-DEL-56 |
| chr9-110540743-DEL-68 | chr9-110540744-DEL-68 |
| chr9-110552211-DEL-118 | chr9-110552212-DEL-118 |
| chr9-110566968-DEL-208 | chr9-110566969-DEL-208 |
| chr9-110614612-DEL-64 | chr9-110614607-DEL-69 |
| chr9-110621024-DEL-174 | chr9-110621031-DEL-134 |
| chr9-110635388-DEL-358 | chr9-110635389-DEL-358 |
| chr9-110636287-DEL-101 | chr9-110636337-DEL-87 |
| chr9-110639708-DEL-209 | chr9-110639701-DEL-209 |
| chr9-110663203-DEL-61 | chr9-110663204-DEL-61 |
| chr9-110694074-DEL-260 | chr9-110694075-DEL-260 |
| chr9-110696782-DEL-214 | chr9-110696783-DEL-214 |
| chr9-110713255-DEL-74 | chr9-110713240-DEL-75 |
| chr9-110730846-DEL-203 | chr9-110730847-DEL-203 |
| chr9-110731830-DEL-159 | chr9-110731830-DEL-159 |
| chr9-110751301-DEL-56 | chr9-110751302-DEL-56 |
| chr9-110765885-DEL-236 | chr9-110765885-DEL-236 |
| chr9-110766576-DEL-210 | chr9-110766577-DEL-212 |
| chr9-110771991-DEL-52 | chr9-110771980-DEL-58 |
| chr9-110781512-DEL-138 | chr9-110781513-DEL-138 |
| chr9-110797604-DEL-211 | chr9-110797605-DEL-210 |
| chr9-110799478-DEL-5456 | chr9-110799467-DEL-5468 |
| chr9-110805457-DEL-119 | chr9-110805457-DEL-119 |
| chr9-110807819-DEL-192 | chr9-110807833-DEL-166 |
| chr9-110809691-DEL-58 | chr9-110809692-DEL-58 |
| chr9-110812423-DEL-215 | chr9-110812424-DEL-215 |
| chr9-110820382-DEL-237 | chr9-110820383-DEL-237 |
| chr9-110823005-DEL-97 | chr9-110823003-DEL-84 |
| chr9-110844208-DEL-133 | chr9-110844209-DEL-133 |
| chr9-110858388-DEL-56 | chr9-110858389-DEL-56 |
| chr9-110901481-DEL-279 | chr9-110901482-DEL-279 |
| chr9-110919282-DEL-2010 | chr9-110919283-DEL-2010 |
| chr9-110929054-DEL-223 | chr9-110929050-DEL-225 |
| chr9-110939823-DEL-456 | chr9-110939824-DEL-456 |
| chr9-110954023-DEL-148 | chr9-110954024-DEL-148 |
| chr9-110954870-DEL-62 | chr9-110954871-DEL-58 |
| chr9-110960317-DEL-78 | chr9-110960318-DEL-78 |
| chr9-110986486-DEL-108 | chr9-110986479-DEL-124 |
| chr9-111022927-DEL-312 | chr9-111022928-DEL-302 |
| chr9-111039855-DEL-69 | chr9-111039856-DEL-64 |
| chr9-111045054-DEL-54 | chr9-111045055-DEL-54 |
| chr9-111079751-DEL-426 | chr9-111079752-DEL-426 |
| chr9-111097492-DEL-87 | chr9-111097493-DEL-87 |
| chr9-111097534-DEL-754 | chr9-111097535-DEL-754 |
| chr9-111112130-DEL-80 | chr9-111112131-DEL-80 |
| chr9-111121753-DEL-504 | chr9-111121754-DEL-504 |
| chr9-111127082-DEL-99 | chr9-111127087-DEL-132 |
| chr9-111159629-DEL-786 | chr9-111159630-DEL-786 |
| chr9-111166494-DEL-67 | chr9-111166495-DEL-67 |
| chr9-111201161-DEL-56 | chr9-111201162-DEL-56 |
| chr9-111246265-DEL-88 | chr9-111246275-DEL-125 |
| chr9-111280626-DEL-156 | chr9-111280627-DEL-156 |
| chr9-111311399-DEL-178 | chr9-111311419-DEL-150 |
| chr9-111317192-DEL-53 | chr9-111317196-DEL-54 |
| chr9-111355602-DEL-702 | chr9-111355603-DEL-702 |
| chr9-111377180-DEL-143 | chr9-111377181-DEL-143 |
| chr9-111397027-DEL-74 | chr9-111397026-DEL-78 |
| chr9-111408651-DEL-89 | chr9-111408653-DEL-87 |
| chr9-111409281-DEL-1236 | chr9-111409282-DEL-1236 |
| chr9-111437579-DEL-76 | chr9-111437576-DEL-80 |
| chr9-111459043-DEL-66 | chr9-111459075-DEL-76 |
| chr9-111472799-DEL-60 | chr9-111472800-DEL-60 |
| chr9-111477371-DEL-628 | chr9-111477372-DEL-628 |
| chr9-111490795-DEL-2567 | chr9-111490796-DEL-2567 |
| chr9-111494929-DEL-103 | chr9-111494930-DEL-84 |
| chr9-111496482-DEL-1328 | chr9-111496483-DEL-1325 |
| chr9-111509147-DEL-87 | chr9-111509148-DEL-87 |
| chr9-111530154-DEL-3100 | chr9-111530155-DEL-3100 |
| chr9-111542223-DEL-90 | chr9-111542224-DEL-90 |
| chr9-111548474-DEL-60 | chr9-111548474-DEL-60 |
| chr9-111581514-DEL-363 | chr9-111581515-DEL-363 |
| chr9-111584158-DEL-94 | chr9-111584155-DEL-90 |
| chr9-111603131-DEL-76 | chr9-111603116-DEL-66 |
| chr9-111609976-DEL-675 | chr9-111609970-DEL-675 |
| chr9-111621719-DEL-152 | chr9-111621720-DEL-154 |
| chr9-111634356-DEL-6596 | chr9-111634356-DEL-6597 |
| chr9-111656989-DEL-195 | chr9-111656989-DEL-195 |
| chr9-111660898-DEL-59 | chr9-111660900-DEL-61 |
| chr9-111675457-DEL-60 | chr9-111675438-DEL-60 |
| chr9-111675637-DEL-6164 | chr9-111675638-DEL-6164 |

| | |
|---|---|
| chr9-111676294-DEL-66 | chr9-111676295-DEL-66 |
| chr9-111705152-DEL-200 | chr9-111705150-DEL-201 |
| chr9-111715332-DEL-72 | chr9-111715333-DEL-72 |
| chr9-111717086-DEL-1644 | chr9-111717086-DEL-1644 |
| chr9-111746819-DEL-66 | chr9-111746827-DEL-59 |
| chr9-111759122-DEL-90 | chr9-111759123-DEL-92 |
| chr9-111782736-DEL-70 | chr9-111782737-DEL-70 |
| chr9-111798322-DEL-1982 | chr9-111798334-DEL-1986 |
| chr9-111802301-DEL-133 | chr9-111802306-DEL-129 |
| chr9-111819518-DEL-107 | chr9-111819519-DEL-107 |
| chr9-111830954-DEL-151 | chr9-111830955-DEL-148 |
| chr9-111863056-DEL-324 | chr9-111863057-DEL-324 |
| chr9-111869979-DEL-108 | chr9-111869989-DEL-101 |
| chr9-111888822-DEL-295 | chr9-111888823-DEL-295 |
| chr9-111923645-DEL-364 | chr9-111923646-DEL-364 |
| chr9-111926392-DEL-209 | chr9-111926395-DEL-228 |
| chr9-111979268-DEL-215 | chr9-111979289-DEL-180 |
| chr9-111988956-DEL-116 | chr9-111988966-DEL-84 |
| chr9-111992291-DEL-56 | chr9-111992293-DEL-55 |
| chr9-112022964-DEL-56 | chr9-112022965-DEL-56 |
| chr9-112035520-DEL-1213 | chr9-112035521-DEL-1213 |
| chr9-112050236-DEL-116 | chr9-112050271-DEL-104 |
| chr9-112096508-DEL-62 | chr9-112096508-DEL-62 |
| chr9-112124011-DEL-98 | chr9-112124012-DEL-95 |
| chr9-112124877-DEL-1243 | chr9-112124878-DEL-1243 |
| chr9-112126651-DEL-70 | chr9-112126656-DEL-66 |
| chr9-112134188-DEL-1875 | chr9-112134187-DEL-1875 |
| chr9-112159108-DEL-6473 | chr9-112159109-DEL-6473 |
| chr9-112197658-DEL-51 | chr9-112197654-DEL-50 |
| chr9-112252192-DEL-56 | chr9-112252193-DEL-56 |
| chr9-112267404-DEL-156 | chr9-112267404-DEL-153 |
| chr9-112285110-DEL-308 | chr9-112285108-DEL-320 |
| chr9-112295371-DEL-87 | chr9-112295372-DEL-87 |
| chr9-112304764-DEL-55 | chr9-112304755-DEL-65 |
| chr9-112306127-DEL-68 | chr9-112306146-DEL-50 |
| chr9-112311059-DEL-80 | chr9-112311060-DEL-80 |
| chr9-112311822-DEL-747 | chr9-112311823-DEL-744 |
| chr9-112312554-DEL-6595 | chr9-112312555-DEL-6595 |
| chr9-112318915-DEL-207 | chr9-112318919-DEL-201 |
| chr9-112323619-DEL-57 | chr9-112323620-DEL-57 |
| chr9-112327381-DEL-70 | chr9-112327382-DEL-70 |
| chr9-112379309-DEL-117 | chr9-112379310-DEL-117 |
| chr9-112443260-DEL-92 | chr9-112443249-DEL-104 |
| chr9-112448024-DEL-6242 | chr9-112448022-DEL-6243 |
| chr9-112459898-DEL-56 | chr9-112459899-DEL-57 |
| chr9-112490266-DEL-6340 | chr9-112490270-DEL-6339 |
| chr9-112501197-DEL-54 | chr9-112501198-DEL-54 |
| chr9-112506524-DEL-178 | chr9-112506525-DEL-178 |
| chr9-112514998-DEL-565 | chr9-112514999-DEL-565 |
| chr9-112551220-DEL-70 | chr9-112551227-DEL-71 |
| chr9-112611507-DEL-64 | chr9-112611508-DEL-64 |
| chr9-112659827-DEL-60 | chr9-112659843-DEL-55 |
| chr9-112675597-DEL-52 | chr9-112675597-DEL-52 |
| chr9-112677696-DEL-107 | chr9-112677697-DEL-107 |
| chr9-112680169-DEL-5127 | chr9-112680170-DEL-5127 |
| chr9-112699225-DEL-139 | chr9-112699226-DEL-139 |
| chr9-112740941-DEL-184 | chr9-112740942-DEL-184 |
| chr9-112741709-DEL-105 | chr9-112741710-DEL-135 |
| chr9-112743639-DEL-60 | chr9-112743640-DEL-60 |
| chr9-112774680-DEL-81 | chr9-112774681-DEL-81 |
| chr9-112777449-DEL-86 | chr9-112777450-DEL-82 |
| chr9-112787897-DEL-6891 | chr9-112787898-DEL-6891 |
| chr9-112788504-DEL-212 | chr9-112788505-DEL-212 |
| chr9-112813490-DEL-201 | chr9-112813491-DEL-201 |
| chr9-112831850-DEL-68 | chr9-112831852-DEL-52 |
| chr9-112838520-DEL-73 | chr9-112838514-DEL-121 |
| chr9-112846337-DEL-120 | chr9-112846338-DEL-120 |
| chr9-112884407-DEL-53 | chr9-112884411-DEL-53 |
| chr9-112928120-DEL-78 | chr9-112928119-DEL-56 |
| chr9-113010764-DEL-1579 | chr9-113010765-DEL-1579 |
| chr9-113027648-DEL-60 | chr9-113027649-DEL-60 |
| chr9-113041137-DEL-253 | chr9-113041138-DEL-253 |
| chr9-113102613-DEL-76 | chr9-113102614-DEL-76 |
| chr9-113129457-DEL-63 | chr9-113129452-DEL-64 |
| chr9-113183795-DEL-75 | chr9-113183800-DEL-71 |
| chr9-113190185-DEL-6235 | chr9-113190186-DEL-6235 |
| chr9-113203302-DEL-58 | chr9-113203303-DEL-58 |
| chr9-113212276-DEL-391 | chr9-113212277-DEL-388 |
| chr9-113221862-DEL-80 | chr9-113221864-DEL-82 |
| chr9-113226355-DEL-100 | chr9-113226356-DEL-78 |

| | |
|---|---|
| chr9-113241922-DEL-90 | chr9-113241945-DEL-68 |
| chr9-113249621-DEL-54 | chr9-113249622-DEL-54 |
| chr9-113254406-DEL-974 | chr9-113254407-DEL-972 |
| chr9-113257417-DEL-64 | chr9-113257434-DEL-58 |
| chr9-113310305-DEL-137 | chr9-113310306-DEL-137 |
| chr9-113311574-DEL-166 | chr9-113311575-DEL-166 |
| chr9-113325295-DEL-135 | chr9-113325296-DEL-120 |
| chr9-113343091-DEL-94 | chr9-113343116-DEL-71 |
| chr9-113345018-DEL-90 | chr9-113345016-DEL-80 |
| chr9-113364319-DEL-6217 | chr9-113364320-DEL-6217 |
| chr9-113395251-DEL-52 | chr9-113395252-DEL-64 |
| chr9-113399450-DEL-220 | chr9-113399447-DEL-214 |
| chr9-113425199-DEL-66 | chr9-113425157-DEL-66 |
| chr9-113442372-DEL-995 | chr9-113442373-DEL-995 |
| chr9-113523829-DEL-212 | chr9-113523828-DEL-212 |
| chr9-113528206-DEL-240 | chr9-113528207-DEL-240 |
| chr9-113541431-DEL-108 | chr9-113541428-DEL-109 |
| chr9-113567458-DEL-165 | chr9-113567459-DEL-152 |
| chr9-113584035-DEL-250 | chr9-113584016-DEL-260 |
| chr9-113586519-DEL-60 | chr9-113586519-DEL-82 |
| chr9-113629135-DEL-97 | chr9-113629108-DEL-84 |
| chr9-113662415-DEL-423 | chr9-113662416-DEL-423 |
| chr9-113711996-DEL-73 | chr9-113711997-DEL-73 |
| chr9-113712936-DEL-151 | chr9-113712937-DEL-151 |
| chr9-113733649-DEL-223 | chr9-113733650-DEL-223 |
| chr9-113816950-DEL-189 | chr9-113816955-DEL-168 |
| chr9-113829697-DEL-199 | chr9-113829698-DEL-199 |
| chr9-113834283-DEL-190 | chr9-113834284-DEL-190 |
| chr9-113839212-DEL-70 | chr9-113839181-DEL-71 |
| chr9-113840564-DEL-1164 | chr9-113840565-DEL-1164 |
| chr9-113843730-DEL-620 | chr9-113843731-DEL-620 |
| chr9-113863058-DEL-347 | chr9-113863059-DEL-347 |
| chr9-113865897-DEL-198 | chr9-113865898-DEL-197 |
| chr9-113930621-DEL-7082 | chr9-113930622-DEL-7082 |
| chr9-113953588-DEL-61 | chr9-113953589-DEL-61 |
| chr9-114025129-DEL-1890 | chr9-114025130-DEL-1890 |
| chr9-114048108-DEL-6475 | chr9-114048109-DEL-6475 |
| chr9-114139489-DEL-61 | chr9-114139490-DEL-61 |
| chr9-114141776-DEL-99 | chr9-114141777-DEL-99 |
| chr9-114147913-DEL-449 | chr9-114147914-DEL-449 |
| chr9-114149742-DEL-1557 | chr9-114149743-DEL-1557 |
| chr9-114173888-DEL-207 | chr9-114173889-DEL-207 |
| chr9-114190471-DEL-212 | chr9-114190472-DEL-212 |
| chr9-114208811-DEL-68 | chr9-114208806-DEL-100 |
| chr9-114215885-DEL-153 | chr9-114215885-DEL-153 |
| chr9-114220527-DEL-97 | chr9-114220528-DEL-97 |
| chr9-114229288-DEL-51 | chr9-114229263-DEL-69 |
| chr9-114236482-DEL-193 | chr9-114236483-DEL-193 |
| chr9-114247650-DEL-82 | chr9-114247646-DEL-52 |
| chr9-114253350-DEL-114 | chr9-114253361-DEL-102 |
| chr9-114254713-DEL-140 | chr9-114254714-DEL-140 |
| chr9-114266256-DEL-163 | chr9-114266252-DEL-164 |
| chr9-114277159-DEL-168 | chr9-114277160-DEL-168 |
| chr9-114278792-DEL-204 | chr9-114278793-DEL-204 |
| chr9-114282831-DEL-59 | chr9-114282827-DEL-60 |
| chr9-114322356-DEL-218 | chr9-114322357-DEL-218 |
| chr9-114323604-DEL-192 | chr9-114323605-DEL-192 |
| chr9-114331530-DEL-195 | chr9-114331528-DEL-198 |
| chr9-114356963-DEL-79 | chr9-114356978-DEL-75 |
| chr9-114369110-DEL-162 | chr9-114369111-DEL-162 |
| chr9-114372557-DEL-106 | chr9-114372558-DEL-106 |
| chr9-114397116-DEL-704 | chr9-114397117-DEL-704 |
| chr9-114400322-DEL-170 | chr9-114400323-DEL-170 |
| chr9-114407203-DEL-76 | chr9-114407195-DEL-84 |
| chr9-114409038-DEL-130 | chr9-114409039-DEL-130 |
| chr9-114409647-DEL-214 | chr9-114409648-DEL-214 |
| chr9-114439356-DEL-56 | chr9-114439357-DEL-56 |
| chr9-114440569-DEL-101 | chr9-114440564-DEL-99 |
| chr9-114445113-DEL-211 | chr9-114445114-DEL-211 |
| chr9-114464852-DEL-199 | chr9-114464853-DEL-199 |
| chr9-114465488-DEL-72 | chr9-114465489-DEL-72 |
| chr9-114465899-DEL-209 | chr9-114465900-DEL-209 |
| chr9-114475040-DEL-211 | chr9-114475040-DEL-211 |
| chr9-114477466-DEL-206 | chr9-114477467-DEL-200 |
| chr9-114495791-DEL-157 | chr9-114495792-DEL-157 |
| chr9-114499546-DEL-62 | chr9-114499547-DEL-62 |
| chr9-114502851-DEL-61 | chr9-114502851-DEL-61 |
| chr9-114507292-DEL-82 | chr9-114507293-DEL-82 |
| chr9-114530291-DEL-1174 | chr9-114530292-DEL-1174 |
| chr9-114532356-DEL-96 | chr9-114532349-DEL-87 |

| | |
|---|---|
| chr9-114543645-DEL-90 | chr9-114543646-DEL-90 |
| chr9-114547865-DEL-171 | chr9-114547866-DEL-171 |
| chr9-114554951-DEL-144 | chr9-114554970-DEL-126 |
| chr9-114568539-DEL-520 | chr9-114568540-DEL-520 |
| chr9-114587185-DEL-422 | chr9-114587192-DEL-415 |
| chr9-114598194-DEL-137 | chr9-114598195-DEL-137 |
| chr9-114600381-DEL-162 | chr9-114600382-DEL-160 |
| chr9-114607700-DEL-73 | chr9-114607700-DEL-70 |
| chr9-114623651-DEL-133 | chr9-114623681-DEL-107 |
| chr9-114666183-DEL-52 | chr9-114666184-DEL-52 |
| chr9-114678688-DEL-53 | chr9-114678678-DEL-72 |
| chr9-114680108-DEL-126 | chr9-114680109-DEL-126 |
| chr9-114694959-DEL-54 | chr9-114694960-DEL-54 |
| chr9-114701441-DEL-99 | chr9-114701442-DEL-97 |
| chr9-114703162-DEL-4953 | chr9-114703163-DEL-4953 |
| chr9-114727380-DEL-347 | chr9-114727381-DEL-347 |
| chr9-114743666-DEL-1025 | chr9-114743647-DEL-1026 |
| chr9-114757406-DEL-80 | chr9-114757401-DEL-56 |
| chr9-114765157-DEL-196 | chr9-114765158-DEL-196 |
| chr9-114770353-DEL-2304 | chr9-114770354-DEL-2304 |
| chr9-114776304-DEL-60 | chr9-114776305-DEL-60 |
| chr9-114778595-DEL-224 | chr9-114778596-DEL-224 |
| chr9-114793337-DEL-136 | chr9-114793338-DEL-136 |
| chr9-114794703-DEL-189 | chr9-114794704-DEL-189 |
| chr9-114795741-DEL-100 | chr9-114795742-DEL-100 |
| chr9-114803756-DEL-54 | chr9-114803757-DEL-54 |
| chr9-114804149-DEL-416 | chr9-114804150-DEL-416 |
| chr9-114811419-DEL-388 | chr9-114811420-DEL-388 |
| chr9-114814995-DEL-62 | chr9-114814961-DEL-62 |
| chr9-114820054-DEL-146 | chr9-114820045-DEL-112 |
| chr9-114820625-DEL-224 | chr9-114820623-DEL-225 |
| chr9-114837555-DEL-884 | chr9-114837556-DEL-884 |
| chr9-114847551-DEL-337 | chr9-114847552-DEL-337 |
| chr9-114860065-DEL-100 | chr9-114860066-DEL-100 |
| chr9-114860484-DEL-303 | chr9-114860485-DEL-303 |
| chr9-114864284-DEL-511 | chr9-114864282-DEL-500 |
| chr9-114877100-DEL-529 | chr9-114877101-DEL-529 |
| chr9-114878119-DEL-84 | chr9-114878124-DEL-56 |
| chr9-114880605-DEL-126 | chr9-114880606-DEL-126 |
| chr9-114888328-DEL-83 | chr9-114888329-DEL-83 |
| chr9-114891429-DEL-74 | chr9-114891429-DEL-74 |
| chr9-114894939-DEL-197 | chr9-114894940-DEL-197 |
| chr9-114901811-DEL-478 | chr9-114901812-DEL-478 |
| chr9-114905036-DEL-144 | chr9-114905011-DEL-126 |
| chr9-114907088-DEL-7187 | chr9-114907089-DEL-7187 |
| chr9-114919902-DEL-78 | chr9-114919903-DEL-77 |
| chr9-114920372-DEL-287 | chr9-114920373-DEL-287 |
| chr9-114941092-DEL-128 | chr9-114941097-DEL-124 |
| chr9-114958538-DEL-57 | chr9-114958524-DEL-60 |
| chr9-114964558-DEL-507 | chr9-114964557-DEL-506 |
| chr9-114970524-DEL-69 | chr9-114970523-DEL-78 |
| chr9-114971484-DEL-252 | chr9-114971483-DEL-253 |
| chr9-114972610-DEL-208 | chr9-114972611-DEL-208 |
| chr9-114980196-DEL-204 | chr9-114980197-DEL-204 |
| chr9-114991642-DEL-110 | chr9-114991643-DEL-110 |
| chr9-115019723-DEL-58 | chr9-115019724-DEL-58 |
| chr9-115023511-DEL-370 | chr9-115023512-DEL-370 |
| chr9-115038533-DEL-478 | chr9-115038534-DEL-478 |
| chr9-115039096-DEL-217 | chr9-115039097-DEL-217 |
| chr9-115047710-DEL-473 | chr9-115047711-DEL-473 |
| chr9-115060088-DEL-55 | chr9-115060089-DEL-55 |
| chr9-115074630-DEL-60 | chr9-115074631-DEL-60 |
| chr9-115075645-DEL-194 | chr9-115075609-DEL-190 |
| chr9-115085763-DEL-86 | chr9-115085764-DEL-86 |
| chr9-115092607-DEL-208 | chr9-115092608-DEL-208 |
| chr9-115106386-DEL-210 | chr9-115106387-DEL-210 |
| chr9-115112396-DEL-144 | chr9-115112445-DEL-96 |
| chr9-115157242-DEL-393 | chr9-115157243-DEL-392 |
| chr9-115206613-DEL-127 | chr9-115206614-DEL-127 |
| chr9-115210986-DEL-137 | chr9-115210987-DEL-137 |
| chr9-115266023-DEL-205 | chr9-115266024-DEL-205 |
| chr9-115301643-DEL-65 | chr9-115301634-DEL-71 |
| chr9-115304600-DEL-57 | chr9-115304601-DEL-57 |
| chr9-115308926-DEL-65 | chr9-115308917-DEL-73 |
| chr9-115314954-DEL-99 | chr9-115314953-DEL-102 |
| chr9-115315875-DEL-102 | chr9-115315876-DEL-102 |
| chr9-115319013-DEL-135 | chr9-115319017-DEL-138 |
| chr9-115326024-DEL-311 | chr9-115326055-DEL-272 |
| chr9-115327730-DEL-156 | chr9-115327731-DEL-156 |
| chr9-115331606-DEL-96 | chr9-115331607-DEL-96 |

| | |
|---|---|
| chr9-115346070-DEL-52 | chr9-115346070-DEL-53 |
| chr9-115381171-DEL-660 | chr9-115381171-DEL-660 |
| chr9-115389005-DEL-119 | chr9-115389026-DEL-113 |
| chr9-115394104-DEL-576 | chr9-115394105-DEL-576 |
| chr9-115401863-DEL-84 | chr9-115401868-DEL-80 |
| chr9-115406976-DEL-579 | chr9-115406977-DEL-577 |
| chr9-115413061-DEL-181 | chr9-115413062-DEL-181 |
| chr9-115458023-DEL-143 | chr9-115458024-DEL-143 |
| chr9-115467978-DEL-96 | chr9-115467979-DEL-96 |
| chr9-115488183-DEL-62 | chr9-115488184-DEL-62 |
| chr9-115524756-DEL-223 | chr9-115524751-DEL-228 |
| chr9-115539893-DEL-112 | chr9-115539894-DEL-112 |
| chr9-115544070-DEL-78 | chr9-115544071-DEL-78 |
| chr9-115550823-DEL-239 | chr9-115550824-DEL-239 |
| chr9-115556231-DEL-58 | chr9-115556232-DEL-58 |
| chr9-115619957-DEL-6272 | chr9-115619958-DEL-6272 |
| chr9-115628227-DEL-134 | chr9-115628225-DEL-129 |
| chr9-115657499-DEL-71 | chr9-115657500-DEL-71 |
| chr9-115665264-DEL-226 | chr9-115665265-DEL-214 |
| chr9-115705483-DEL-1132 | chr9-115705484-DEL-1132 |
| chr9-115706484-DEL-1336 | chr9-115706485-DEL-1336 |
| chr9-115724443-DEL-115 | chr9-115724444-DEL-115 |
| chr9-115782833-DEL-61 | chr9-115782830-DEL-61 |
| chr9-115814048-DEL-86 | chr9-115814059-DEL-76 |
| chr9-115828246-DEL-52 | chr9-115828247-DEL-52 |
| chr9-115857788-DEL-171 | chr9-115857789-DEL-171 |
| chr9-115869598-DEL-609 | chr9-115869588-DEL-610 |
| chr9-115874521-DEL-59 | chr9-115874516-DEL-60 |
| chr9-115875019-DEL-176 | chr9-115875020-DEL-173 |
| chr9-115881573-DEL-59 | chr9-115881574-DEL-60 |
| chr9-115884772-DEL-64 | chr9-115884773-DEL-64 |
| chr9-115893987-DEL-72 | chr9-115893965-DEL-90 |
| chr9-115900963-DEL-88 | chr9-115900989-DEL-69 |
| chr9-115905646-DEL-54 | chr9-115905647-DEL-55 |
| chr9-115926526-DEL-61 | chr9-115926527-DEL-61 |
| chr9-115943497-DEL-61 | chr9-115943498-DEL-61 |
| chr9-115967217-DEL-70 | chr9-115967214-DEL-66 |
| chr9-115970926-DEL-78 | chr9-115970903-DEL-61 |
| chr9-115972033-DEL-599 | chr9-115972034-DEL-599 |
| chr9-115984227-DEL-59 | chr9-115984228-DEL-59 |
| chr9-116031291-DEL-507 | chr9-116031292-DEL-507 |
| chr9-116049244-DEL-3483 | chr9-116049245-DEL-3483 |
| chr9-116052942-DEL-74 | chr9-116052951-DEL-60 |
| chr9-116089524-DEL-4729 | chr9-116089529-DEL-4725 |
| chr9-116093915-DEL-209 | chr9-116093916-DEL-209 |
| chr9-116116630-DEL-158 | chr9-116116630-DEL-158 |
| chr9-116209156-DEL-491 | chr9-116209157-DEL-491 |
| chr9-116232357-DEL-66 | chr9-116232358-DEL-66 |
| chr9-116271048-DEL-4178 | chr9-116271059-DEL-4168 |
| chr9-116305791-DEL-100 | chr9-116305753-DEL-101 |
| chr9-116430941-DEL-857 | chr9-116430931-DEL-858 |
| chr9-116432212-DEL-177 | chr9-116432221-DEL-169 |
| chr9-116447473-DEL-118 | chr9-116447515-DEL-109 |
| chr9-116457878-DEL-64 | chr9-116457879-DEL-64 |
| chr9-116477947-DEL-218 | chr9-116477954-DEL-212 |
| chr9-116502170-DEL-62 | chr9-116502171-DEL-62 |
| chr9-116519144-DEL-291 | chr9-116519145-DEL-291 |
| chr9-116522962-DEL-99 | chr9-116522940-DEL-87 |
| chr9-116533953-DEL-156 | chr9-116533960-DEL-150 |
| chr9-116534428-DEL-357 | chr9-116534427-DEL-353 |
| chr9-116558726-DEL-685 | chr9-116558727-DEL-685 |
| chr9-116584520-DEL-69 | chr9-116584521-DEL-69 |
| chr9-116588775-DEL-84 | chr9-116588776-DEL-84 |
| chr9-116591634-DEL-64 | chr9-116591635-DEL-64 |
| chr9-116605062-DEL-111 | chr9-116605061-DEL-108 |
| chr9-116631311-DEL-52 | chr9-116631307-DEL-56 |
| chr9-116645330-DEL-95 | chr9-116645331-DEL-95 |
| chr9-116668517-DEL-58 | chr9-116668518-DEL-58 |
| chr9-116668872-DEL-580 | chr9-116668873-DEL-580 |
| chr9-116704504-DEL-347 | chr9-116704505-DEL-347 |
| chr9-116732982-DEL-78 | chr9-116732983-DEL-78 |
| chr9-116750238-DEL-52 | chr9-116750239-DEL-52 |
| chr9-116790507-DEL-73 | chr9-116790507-DEL-50 |
| chr9-116810565-DEL-7161 | chr9-116810566-DEL-7161 |
| chr9-116834576-DEL-52 | chr9-116834538-DEL-70 |
| chr9-116839360-DEL-72 | chr9-116839361-DEL-67 |
| chr9-116870559-DEL-78 | chr9-116870530-DEL-78 |
| chr9-116872370-DEL-528 | chr9-116872370-DEL-528 |
| chr9-116937999-DEL-75 | chr9-116937980-DEL-95 |
| chr9-116984228-DEL-83 | chr9-116984229-DEL-83 |

| | |
|---|---|
| chr9-116991602-DEL-122 | chr9-116991578-DEL-109 |
| chr9-117009002-DEL-96 | chr9-117009003-DEL-96 |
| chr9-117020122-DEL-58 | chr9-117020122-DEL-58 |
| chr9-117029970-DEL-275 | chr9-117029971-DEL-275 |
| chr9-117059883-DEL-191 | chr9-117059884-DEL-191 |
| chr9-117091318-DEL-57 | chr9-117091350-DEL-59 |
| chr9-117132184-DEL-178 | chr9-117132183-DEL-179 |
| chr9-117139815-DEL-269 | chr9-117139816-DEL-269 |
| chr9-117210242-DEL-235 | chr9-117210243-DEL-235 |
| chr9-117220340-DEL-58 | chr9-117220341-DEL-58 |
| chr9-117232494-DEL-603 | chr9-117232495-DEL-603 |
| chr9-117269787-DEL-136 | chr9-117269788-DEL-136 |
| chr9-117313302-DEL-58 | chr9-117313303-DEL-58 |
| chr9-117316191-DEL-54 | chr9-117316192-DEL-54 |
| chr9-117336967-DEL-407 | chr9-117336968-DEL-407 |
| chr9-117365363-DEL-78 | chr9-117365363-DEL-84 |
| chr9-117375334-DEL-146 | chr9-117375321-DEL-160 |
| chr9-117388476-DEL-119 | chr9-117388466-DEL-107 |
| chr9-117416922-DEL-715 | chr9-117416923-DEL-715 |
| chr9-117467010-DEL-63 | chr9-117466968-DEL-56 |
| chr9-117491447-DEL-60 | chr9-117491458-DEL-52 |
| chr9-117493056-DEL-67 | chr9-117493055-DEL-69 |
| chr9-117498390-DEL-95 | chr9-117498391-DEL-95 |
| chr9-117499111-DEL-872 | chr9-117499112-DEL-872 |
| chr9-117541031-DEL-84 | chr9-117541032-DEL-84 |
| chr9-117543136-DEL-92 | chr9-117543137-DEL-92 |
| chr9-117546045-DEL-62 | chr9-117546045-DEL-62 |
| chr9-117547063-DEL-72 | chr9-117547064-DEL-72 |
| chr9-117554409-DEL-125 | chr9-117554410-DEL-125 |
| chr9-117585371-DEL-445 | chr9-117585372-DEL-445 |
| chr9-117597832-DEL-1091 | chr9-117597833-DEL-1091 |
| chr9-117622458-DEL-88 | chr9-117622479-DEL-92 |
| chr9-117628186-DEL-56 | chr9-117628187-DEL-56 |
| chr9-117642961-DEL-56 | chr9-117642946-DEL-54 |
| chr9-117646379-DEL-60 | chr9-117646366-DEL-52 |
| chr9-117665363-DEL-54 | chr9-117665362-DEL-52 |
| chr9-117672576-DEL-1571 | chr9-117672577-DEL-1571 |
| chr9-117675674-DEL-51 | chr9-117675676-DEL-50 |
| chr9-117691514-DEL-72 | chr9-117691515-DEL-72 |
| chr9-117713266-DEL-244 | chr9-117713267-DEL-244 |
| chr9-117715331-DEL-54 | chr9-117715332-DEL-54 |
| chr9-117738558-DEL-192 | chr9-117738559-DEL-192 |
| chr9-117748399-DEL-52 | chr9-117748400-DEL-52 |
| chr9-117756125-DEL-54 | chr9-117756126-DEL-54 |
| chr9-117778367-DEL-2341 | chr9-117778367-DEL-2341 |
| chr9-117787040-DEL-65 | chr9-117787041-DEL-65 |
| chr9-117800511-DEL-78 | chr9-117800529-DEL-66 |
| chr9-117833884-DEL-6569 | chr9-117833885-DEL-6569 |
| chr9-117904826-DEL-1810 | chr9-117904827-DEL-1810 |
| chr9-117919845-DEL-80 | chr9-117919846-DEL-80 |
| chr9-117920560-DEL-1146 | chr9-117920560-DEL-1146 |
| chr9-117941784-DEL-568 | chr9-117941761-DEL-569 |
| chr9-117942172-DEL-1494 | chr9-117942173-DEL-1494 |
| chr9-117957231-DEL-6179 | chr9-117957232-DEL-6179 |
| chr9-117965668-DEL-65 | chr9-117965669-DEL-65 |
| chr9-117975713-DEL-452 | chr9-117975698-DEL-494 |
| chr9-117988734-DEL-53 | chr9-117988701-DEL-53 |
| chr9-117995639-DEL-177 | chr9-117995640-DEL-177 |
| chr9-118003479-DEL-101 | chr9-118003480-DEL-96 |
| chr9-118015224-DEL-105 | chr9-118015225-DEL-105 |
| chr9-118021696-DEL-154 | chr9-118021697-DEL-154 |
| chr9-118037504-DEL-2856 | chr9-118037505-DEL-2856 |
| chr9-118040896-DEL-70 | chr9-118040897-DEL-70 |
| chr9-118055966-DEL-69 | chr9-118055979-DEL-56 |
| chr9-118065802-DEL-180 | chr9-118065814-DEL-169 |
| chr9-118067150-DEL-150 | chr9-118067153-DEL-104 |
| chr9-118092699-DEL-79 | chr9-118092700-DEL-79 |
| chr9-118112957-DEL-176 | chr9-118112956-DEL-164 |
| chr9-118124572-DEL-58 | chr9-118124573-DEL-58 |
| chr9-118125438-DEL-63 | chr9-118125434-DEL-53 |
| chr9-118133354-DEL-156 | chr9-118133355-DEL-156 |
| chr9-118134560-DEL-170 | chr9-118134561-DEL-170 |
| chr9-118141283-DEL-84 | chr9-118141285-DEL-83 |
| chr9-118142076-DEL-432 | chr9-118142078-DEL-428 |
| chr9-118144875-DEL-54 | chr9-118144875-DEL-54 |
| chr9-118154209-DEL-61 | chr9-118154210-DEL-61 |
| chr9-118159432-DEL-141 | chr9-118159406-DEL-151 |
| chr9-118186140-DEL-440 | chr9-118186141-DEL-440 |
| chr9-118194315-DEL-55 | chr9-118194316-DEL-54 |
| chr9-118197085-DEL-210 | chr9-118197086-DEL-210 |

| | |
|---|---|
| chr9-118215754-DEL-130 | chr9-118215755-DEL-130 |
| chr9-118230449-DEL-51 | chr9-118230450-DEL-51 |
| chr9-118259546-DEL-401 | chr9-118259547-DEL-401 |
| chr9-118269126-DEL-99 | chr9-118269129-DEL-97 |
| chr9-118301237-DEL-100 | chr9-118301238-DEL-100 |
| chr9-118436978-DEL-84 | chr9-118436979-DEL-84 |
| chr9-118443925-DEL-61 | chr9-118443955-DEL-59 |
| chr9-118447229-DEL-95 | chr9-118447230-DEL-95 |
| chr9-118482316-DEL-109 | chr9-118482316-DEL-105 |
| chr9-118532662-DEL-423 | chr9-118532659-DEL-427 |
| chr9-118533272-DEL-73 | chr9-118533275-DEL-81 |
| chr9-118536967-DEL-176 | chr9-118536968-DEL-176 |
| chr9-118537664-DEL-86 | chr9-118537690-DEL-63 |
| chr9-118538163-DEL-135 | chr9-118538209-DEL-90 |
| chr9-118585244-DEL-114 | chr9-118585242-DEL-115 |
| chr9-118590479-DEL-51 | chr9-118590480-DEL-51 |
| chr9-118622446-DEL-198 | chr9-118622447-DEL-198 |
| chr9-118626838-DEL-934 | chr9-118626839-DEL-934 |
| chr9-118628964-DEL-53 | chr9-118628965-DEL-53 |
| chr9-118633021-DEL-53 | chr9-118633022-DEL-53 |
| chr9-118646540-DEL-140 | chr9-118646541-DEL-140 |
| chr9-118653431-DEL-59 | chr9-118653432-DEL-59 |
| chr9-118678492-DEL-74 | chr9-118678492-DEL-74 |
| chr9-118696593-DEL-87 | chr9-118696594-DEL-87 |
| chr9-118745245-DEL-475 | chr9-118745246-DEL-475 |
| chr9-118747512-DEL-72 | chr9-118747502-DEL-98 |
| chr9-118768119-DEL-185 | chr9-118768120-DEL-185 |
| chr9-118775277-DEL-434 | chr9-118775273-DEL-435 |
| chr9-118776327-DEL-55 | chr9-118776328-DEL-54 |
| chr9-118791422-DEL-6666 | chr9-118791423-DEL-6666 |
| chr9-118884314-DEL-168 | chr9-118884363-DEL-168 |
| chr9-118888031-DEL-62 | chr9-118888031-DEL-62 |
| chr9-118944592-DEL-194 | chr9-118944593-DEL-194 |
| chr9-119015214-DEL-230 | chr9-119015215-DEL-230 |
| chr9-119023043-DEL-84 | chr9-119023044-DEL-84 |
| chr9-119057001-DEL-75 | chr9-119057002-DEL-75 |
| chr9-119063841-DEL-171 | chr9-119063842-DEL-172 |
| chr9-119064343-DEL-77 | chr9-119064344-DEL-77 |
| chr9-119094965-DEL-160 | chr9-119094966-DEL-160 |
| chr9-119100318-DEL-51 | chr9-119100319-DEL-51 |
| chr9-119103743-DEL-201 | chr9-119103744-DEL-201 |
| chr9-119121490-DEL-117 | chr9-119121491-DEL-117 |
| chr9-119139865-DEL-204 | chr9-119139866-DEL-204 |
| chr9-119142138-DEL-114 | chr9-119142139-DEL-114 |
| chr9-119154853-DEL-77 | chr9-119154862-DEL-69 |
| chr9-119206852-DEL-51 | chr9-119206853-DEL-51 |
| chr9-119208709-DEL-134 | chr9-119208710-DEL-134 |
| chr9-119219028-DEL-245 | chr9-119219029-DEL-245 |
| chr9-119235776-DEL-129 | chr9-119235777-DEL-129 |
| chr9-119274518-DEL-68 | chr9-119274519-DEL-68 |
| chr9-119278393-DEL-64 | chr9-119278394-DEL-62 |
| chr9-119288510-DEL-61 | chr9-119288511-DEL-61 |
| chr9-119304825-DEL-187 | chr9-119304826-DEL-187 |
| chr9-119309360-DEL-1355 | chr9-119309361-DEL-1355 |
| chr9-119337044-DEL-52 | chr9-119337069-DEL-50 |
| chr9-119345506-DEL-62 | chr9-119345509-DEL-60 |
| chr9-119367978-DEL-156 | chr9-119367978-DEL-156 |
| chr9-119374869-DEL-406 | chr9-119374871-DEL-405 |
| chr9-119442877-DEL-129 | chr9-119442876-DEL-131 |
| chr9-119448994-DEL-2553 | chr9-119448988-DEL-2554 |
| chr9-119452312-DEL-52 | chr9-119452313-DEL-52 |
| chr9-119478766-DEL-321 | chr9-119478767-DEL-320 |
| chr9-119484462-DEL-270 | chr9-119484463-DEL-270 |
| chr9-119539019-DEL-57 | chr9-119539020-DEL-57 |
| chr9-119581252-DEL-148 | chr9-119581253-DEL-148 |
| chr9-119638995-DEL-82 | chr9-119638997-DEL-94 |
| chr9-119639534-DEL-432 | chr9-119639497-DEL-429 |
| chr9-119645218-DEL-63 | chr9-119645219-DEL-63 |
| chr9-119661993-DEL-114 | chr9-119662022-DEL-92 |
| chr9-119679772-DEL-96 | chr9-119679773-DEL-96 |
| chr9-119827478-DEL-60 | chr9-119827479-DEL-58 |
| chr9-119832707-DEL-207 | chr9-119832708-DEL-207 |
| chr9-119835604-DEL-206 | chr9-119835605-DEL-206 |
| chr9-119862971-DEL-58 | chr9-119862972-DEL-58 |
| chr9-119871250-DEL-82 | chr9-119871293-DEL-64 |
| chr9-119881488-DEL-54 | chr9-119881488-DEL-54 |
| chr9-119885148-DEL-237 | chr9-119885122-DEL-237 |
| chr9-119888914-DEL-60 | chr9-119888915-DEL-60 |
| chr9-119903362-DEL-63 | chr9-119903363-DEL-63 |
| chr9-119912616-DEL-63 | chr9-119912617-DEL-63 |

| | |
|---|---|
| chr9-119925551-DEL-61 | chr9-119925552-DEL-61 |
| chr9-119935560-DEL-205 | chr9-119935560-DEL-206 |
| chr9-119936560-DEL-486 | chr9-119936561-DEL-486 |
| chr9-119937201-DEL-492 | chr9-119937202-DEL-492 |
| chr9-119950808-DEL-227 | chr9-119950809-DEL-227 |
| chr9-119953780-DEL-557 | chr9-119953781-DEL-557 |
| chr9-119955196-DEL-96 | chr9-119955208-DEL-91 |
| chr9-119979455-DEL-163 | chr9-119979455-DEL-164 |
| chr9-119981911-DEL-52 | chr9-119981912-DEL-52 |
| chr9-120015683-DEL-226 | chr9-120015678-DEL-227 |
| chr9-120020310-DEL-88 | chr9-120020311-DEL-88 |
| chr9-120023795-DEL-86 | chr9-120023808-DEL-70 |
| chr9-120024547-DEL-70 | chr9-120024548-DEL-70 |
| chr9-120038600-DEL-66 | chr9-120038601-DEL-66 |
| chr9-120040681-DEL-66 | chr9-120040684-DEL-56 |
| chr9-120045456-DEL-64 | chr9-120045451-DEL-70 |
| chr9-120066920-DEL-484 | chr9-120066925-DEL-480 |
| chr9-120092327-DEL-238 | chr9-120092328-DEL-238 |
| chr9-120094396-DEL-6399 | chr9-120094397-DEL-6399 |
| chr9-120121145-DEL-77 | chr9-120121152-DEL-71 |
| chr9-120137767-DEL-206 | chr9-120137768-DEL-206 |
| chr9-120169581-DEL-52 | chr9-120169582-DEL-52 |
| chr9-120173682-DEL-203 | chr9-120173684-DEL-203 |
| chr9-120186952-DEL-51 | chr9-120186953-DEL-51 |
| chr9-120194812-DEL-340 | chr9-120194813-DEL-340 |
| chr9-120199972-DEL-96 | chr9-120199973-DEL-96 |
| chr9-120216649-DEL-99 | chr9-120216652-DEL-105 |
| chr9-120226482-DEL-432 | chr9-120226483-DEL-432 |
| chr9-120228843-DEL-392 | chr9-120228843-DEL-385 |
| chr9-120234311-DEL-123 | chr9-120234312-DEL-123 |
| chr9-120239823-DEL-327 | chr9-120239853-DEL-297 |
| chr9-120262620-DEL-202 | chr9-120262639-DEL-190 |
| chr9-120280122-DEL-57 | chr9-120280123-DEL-57 |
| chr9-120294413-DEL-76 | chr9-120294430-DEL-82 |
| chr9-120343493-DEL-373 | chr9-120343504-DEL-363 |
| chr9-120369399-DEL-54 | chr9-120369400-DEL-50 |
| chr9-120370246-DEL-69 | chr9-120370282-DEL-50 |
| chr9-120384271-DEL-64 | chr9-120384290-DEL-63 |
| chr9-120390193-DEL-66 | chr9-120390194-DEL-66 |
| chr9-120405396-DEL-289 | chr9-120405423-DEL-269 |
| chr9-120414443-DEL-182 | chr9-120414427-DEL-216 |
| chr9-120425554-DEL-176 | chr9-120425555-DEL-132 |
| chr9-120429362-DEL-203 | chr9-120429363-DEL-203 |
| chr9-120435657-DEL-203 | chr9-120435658-DEL-203 |
| chr9-120443677-DEL-462 | chr9-120443678-DEL-462 |
| chr9-120451045-DEL-214 | chr9-120451046-DEL-214 |
| chr9-120463040-DEL-7270 | chr9-120463041-DEL-7270 |
| chr9-120473971-DEL-54 | chr9-120473972-DEL-54 |
| chr9-120498427-DEL-136 | chr9-120498421-DEL-163 |
| chr9-120509917-DEL-184 | chr9-120509918-DEL-184 |
| chr9-120514026-DEL-80 | chr9-120514012-DEL-94 |
| chr9-120516370-DEL-189 | chr9-120516371-DEL-189 |
| chr9-120517340-DEL-122 | chr9-120517341-DEL-122 |
| chr9-120519083-DEL-90 | chr9-120519084-DEL-90 |
| chr9-120520613-DEL-2293 | chr9-120520614-DEL-2293 |
| chr9-120538721-DEL-1418 | chr9-120538722-DEL-1418 |
| chr9-120543609-DEL-92 | chr9-120543594-DEL-93 |
| chr9-120560867-DEL-61 | chr9-120560869-DEL-60 |
| chr9-120565443-DEL-161 | chr9-120565444-DEL-161 |
| chr9-120571821-DEL-143 | chr9-120571818-DEL-147 |
| chr9-120581211-DEL-102 | chr9-120581212-DEL-56 |
| chr9-120583525-DEL-76 | chr9-120583526-DEL-72 |
| chr9-120584542-DEL-226 | chr9-120584543-DEL-226 |
| chr9-120599699-DEL-85 | chr9-120599700-DEL-85 |
| chr9-120608693-DEL-101 | chr9-120608707-DEL-110 |
| chr9-120624149-DEL-360 | chr9-120624150-DEL-360 |
| chr9-120648266-DEL-64 | chr9-120648251-DEL-62 |
| chr9-120655701-DEL-60 | chr9-120655702-DEL-60 |
| chr9-120671288-DEL-124 | chr9-120671287-DEL-125 |
| chr9-120673869-DEL-199 | chr9-120673870-DEL-199 |
| chr9-120706746-DEL-192 | chr9-120706747-DEL-192 |
| chr9-120720875-DEL-71 | chr9-120720876-DEL-71 |
| chr9-120740549-DEL-301 | chr9-120740553-DEL-291 |
| chr9-120746485-DEL-340 | chr9-120746486-DEL-340 |
| chr9-120766741-DEL-924 | chr9-120766742-DEL-924 |
| chr9-120839082-DEL-280 | chr9-120839084-DEL-270 |
| chr9-120853738-DEL-4801 | chr9-120853739-DEL-4801 |
| chr9-120883061-DEL-64 | chr9-120883062-DEL-64 |
| chr9-120909448-DEL-358 | chr9-120909449-DEL-358 |
| chr9-120913577-DEL-60 | chr9-120913578-DEL-60 |

| | |
|---|---|
| chr9-120935638-DEL-100 | chr9-120935639-DEL-98 |
| chr9-120947408-DEL-62 | chr9-120947409-DEL-62 |
| chr9-120957314-DEL-54 | chr9-120957288-DEL-50 |
| chr9-120960513-DEL-118 | chr9-120960507-DEL-120 |
| chr9-120977199-DEL-52 | chr9-120977201-DEL-50 |
| chr9-120981481-DEL-213 | chr9-120981482-DEL-213 |
| chr9-120982803-DEL-84 | chr9-120982803-DEL-84 |
| chr9-120983070-DEL-865 | chr9-120983071-DEL-865 |
| chr9-120984314-DEL-106 | chr9-120984315-DEL-106 |
| chr9-120997705-DEL-204 | chr9-120997706-DEL-204 |
| chr9-121007440-DEL-6158 | chr9-121007441-DEL-6158 |
| chr9-121017123-DEL-102 | chr9-121017124-DEL-102 |
| chr9-121019156-DEL-57 | chr9-121019157-DEL-61 |
| chr9-121026678-DEL-51 | chr9-121026678-DEL-51 |
| chr9-121040514-DEL-105 | chr9-121040515-DEL-105 |
| chr9-121045978-DEL-74 | chr9-121045979-DEL-74 |
| chr9-121048120-DEL-511 | chr9-121048121-DEL-511 |
| chr9-121072165-DEL-210 | chr9-121072166-DEL-210 |
| chr9-121075711-DEL-119 | chr9-121075712-DEL-119 |
| chr9-121087640-DEL-67 | chr9-121087640-DEL-67 |
| chr9-121089782-DEL-465 | chr9-121089782-DEL-465 |
| chr9-121091158-DEL-55 | chr9-121091121-DEL-56 |
| chr9-121098865-DEL-62 | chr9-121098866-DEL-62 |
| chr9-121108860-DEL-193 | chr9-121108861-DEL-193 |
| chr9-121112412-DEL-203 | chr9-121112413-DEL-203 |
| chr9-121118895-DEL-112 | chr9-121118910-DEL-98 |
| chr9-121120723-DEL-200 | chr9-121120724-DEL-200 |
| chr9-121148229-DEL-57 | chr9-121148230-DEL-57 |
| chr9-121151090-DEL-65 | chr9-121151080-DEL-104 |
| chr9-121151437-DEL-6112 | chr9-121151438-DEL-6112 |
| chr9-121179815-DEL-85 | chr9-121179816-DEL-85 |
| chr9-121189337-DEL-54 | chr9-121189369-DEL-54 |
| chr9-121219406-DEL-59 | chr9-121219406-DEL-59 |
| chr9-121228878-DEL-80 | chr9-121228879-DEL-80 |
| chr9-121235466-DEL-7104 | chr9-121235467-DEL-7104 |
| chr9-121247042-DEL-144 | chr9-121247043-DEL-144 |
| chr9-121247792-DEL-173 | chr9-121247793-DEL-173 |
| chr9-121262081-DEL-160 | chr9-121262082-DEL-160 |
| chr9-121267392-DEL-382 | chr9-121267393-DEL-382 |
| chr9-121268291-DEL-76 | chr9-121268287-DEL-80 |
| chr9-121288227-DEL-337 | chr9-121288228-DEL-337 |
| chr9-121289113-DEL-76 | chr9-121289113-DEL-76 |
| chr9-121298354-DEL-62 | chr9-121298355-DEL-62 |
| chr9-121309174-DEL-205 | chr9-121309175-DEL-205 |
| chr9-121319820-DEL-57 | chr9-121319821-DEL-57 |
| chr9-121334882-DEL-439 | chr9-121334883-DEL-439 |
| chr9-121360319-DEL-114 | chr9-121360320-DEL-114 |
| chr9-121361411-DEL-232 | chr9-121361412-DEL-232 |
| chr9-121370746-DEL-79 | chr9-121370718-DEL-79 |
| chr9-121422807-DEL-142 | chr9-121422808-DEL-142 |
| chr9-121435647-DEL-82 | chr9-121435648-DEL-82 |
| chr9-121440356-DEL-2230 | chr9-121440356-DEL-2230 |
| chr9-121447384-DEL-63 | chr9-121447426-DEL-60 |
| chr9-121451708-DEL-343 | chr9-121451709-DEL-343 |
| chr9-121458809-DEL-56 | chr9-121458804-DEL-56 |
| chr9-121473629-DEL-78 | chr9-121473630-DEL-78 |
| chr9-121500696-DEL-120 | chr9-121500697-DEL-120 |
| chr9-121510106-DEL-51 | chr9-121510106-DEL-51 |
| chr9-121515535-DEL-88 | chr9-121515502-DEL-72 |
| chr9-121525093-DEL-498 | chr9-121525094-DEL-498 |
| chr9-121551407-DEL-171 | chr9-121551408-DEL-171 |
| chr9-121613455-DEL-81 | chr9-121613408-DEL-82 |
| chr9-121626327-DEL-287 | chr9-121626327-DEL-285 |
| chr9-121627490-DEL-134 | chr9-121627503-DEL-137 |
| chr9-121635297-DEL-90 | chr9-121635328-DEL-104 |
| chr9-121653523-DEL-50 | chr9-121653482-DEL-94 |
| chr9-121654824-DEL-218 | chr9-121654825-DEL-218 |
| chr9-121672118-DEL-58 | chr9-121672119-DEL-58 |
| chr9-121690305-DEL-400 | chr9-121690306-DEL-400 |
| chr9-121691955-DEL-59 | chr9-121691956-DEL-59 |
| chr9-121721562-DEL-1768 | chr9-121721563-DEL-1768 |
| chr9-121758581-DEL-2438 | chr9-121758572-DEL-2437 |
| chr9-121761706-DEL-103 | chr9-121761707-DEL-104 |
| chr9-121763500-DEL-126 | chr9-121763495-DEL-128 |
| chr9-121787747-DEL-451 | chr9-121787748-DEL-451 |
| chr9-121802013-DEL-109 | chr9-121802058-DEL-65 |
| chr9-121820870-DEL-68 | chr9-121820871-DEL-68 |
| chr9-121831920-DEL-58 | chr9-121831921-DEL-58 |
| chr9-121847357-DEL-59 | chr9-121847358-DEL-59 |
| chr9-121864053-DEL-86 | chr9-121864063-DEL-98 |

| | |
|---|---|
| chr9-121871808-DEL-175 | chr9-121871809-DEL-169 |
| chr9-121889384-DEL-473 | chr9-121889385-DEL-473 |
| chr9-121892822-DEL-64 | chr9-121892827-DEL-60 |
| chr9-121897269-DEL-412 | chr9-121897229-DEL-428 |
| chr9-121902436-DEL-267 | chr9-121902437-DEL-267 |
| chr9-121905809-DEL-756 | chr9-121905810-DEL-756 |
| chr9-121914576-DEL-62 | chr9-121914579-DEL-64 |
| chr9-121969395-DEL-1146 | chr9-121969396-DEL-1146 |
| chr9-121971118-DEL-882 | chr9-121971119-DEL-882 |
| chr9-121991865-DEL-92 | chr9-121991866-DEL-92 |
| chr9-122029117-DEL-82 | chr9-122029118-DEL-82 |
| chr9-122039622-DEL-428 | chr9-122039587-DEL-464 |
| chr9-122056999-DEL-73 | chr9-122057002-DEL-51 |
| chr9-122065001-DEL-188 | chr9-122065002-DEL-188 |
| chr9-122086595-DEL-60 | chr9-122086596-DEL-60 |
| chr9-122087717-DEL-178 | chr9-122087718-DEL-178 |
| chr9-122106598-DEL-2651 | chr9-122106599-DEL-2651 |
| chr9-122110836-DEL-434 | chr9-122110837-DEL-434 |
| chr9-122112850-DEL-1627 | chr9-122112851-DEL-1627 |
| chr9-122125897-DEL-132 | chr9-122125898-DEL-132 |
| chr9-122126409-DEL-6233 | chr9-122126415-DEL-6244 |
| chr9-122142076-DEL-120 | chr9-122142077-DEL-86 |
| chr9-122146065-DEL-157 | chr9-122146062-DEL-124 |
| chr9-122157653-DEL-6473 | chr9-122157654-DEL-6473 |
| chr9-122175755-DEL-186 | chr9-122175757-DEL-185 |
| chr9-122188128-DEL-3255 | chr9-122188129-DEL-3255 |
| chr9-122190988-DEL-75 | chr9-122191015-DEL-75 |
| chr9-122230735-DEL-7070 | chr9-122230736-DEL-7070 |
| chr9-122244032-DEL-106 | chr9-122244011-DEL-140 |
| chr9-122244899-DEL-377 | chr9-122244900-DEL-377 |
| chr9-122262773-DEL-87 | chr9-122262774-DEL-87 |
| chr9-122267172-DEL-268 | chr9-122267173-DEL-268 |
| chr9-122267846-DEL-3128 | chr9-122267847-DEL-3128 |
| chr9-122272262-DEL-530 | chr9-122272263-DEL-530 |
| chr9-122278048-DEL-78 | chr9-122278049-DEL-75 |
| chr9-122286517-DEL-423 | chr9-122286518-DEL-423 |
| chr9-122326074-DEL-81 | chr9-122326093-DEL-69 |
| chr9-122370528-DEL-77 | chr9-122370528-DEL-77 |
| chr9-122392700-DEL-76 | chr9-122392703-DEL-57 |
| chr9-122407930-DEL-367 | chr9-122407931-DEL-367 |
| chr9-122454412-DEL-56 | chr9-122454413-DEL-56 |
| chr9-122468047-DEL-269 | chr9-122468048-DEL-269 |
| chr9-122484852-DEL-100 | chr9-122484843-DEL-70 |
| chr9-122486882-DEL-399 | chr9-122486882-DEL-399 |
| chr9-122489717-DEL-6444 | chr9-122489718-DEL-6444 |
| chr9-122498097-DEL-65 | chr9-122498109-DEL-64 |
| chr9-122528586-DEL-59 | chr9-122528587-DEL-59 |
| chr9-122532492-DEL-6450 | chr9-122532493-DEL-6450 |
| chr9-122559000-DEL-72 | chr9-122559001-DEL-72 |
| chr9-122660475-DEL-106 | chr9-122660457-DEL-87 |
| chr9-122661906-DEL-61 | chr9-122661907-DEL-61 |
| chr9-122707399-DEL-1045 | chr9-122707400-DEL-1045 |
| chr9-122770165-DEL-386 | chr9-122770166-DEL-386 |
| chr9-122772672-DEL-63 | chr9-122772673-DEL-63 |
| chr9-122867499-DEL-373 | chr9-122867500-DEL-373 |
| chr9-122869221-DEL-146 | chr9-122869222-DEL-146 |
| chr9-122872393-DEL-296 | chr9-122872392-DEL-297 |
| chr9-122876280-DEL-82 | chr9-122876269-DEL-83 |
| chr9-122916851-DEL-74 | chr9-122916852-DEL-74 |
| chr9-122921120-DEL-2306 | chr9-122921121-DEL-2306 |
| chr9-122922335-DEL-183 | chr9-122922336-DEL-183 |
| chr9-122928324-DEL-170 | chr9-122928325-DEL-170 |
| chr9-122933222-DEL-147 | chr9-122933223-DEL-147 |
| chr9-122946769-DEL-66 | chr9-122946770-DEL-66 |
| chr9-122986485-DEL-221 | chr9-122986486-DEL-221 |
| chr9-123008285-DEL-81 | chr9-123008286-DEL-81 |
| chr9-123019890-DEL-7382 | chr9-123019891-DEL-7382 |
| chr9-123025837-DEL-81 | chr9-123025838-DEL-81 |
| chr9-123038207-DEL-68 | chr9-123038208-DEL-68 |
| chr9-123039952-DEL-83 | chr9-123039953-DEL-83 |
| chr9-123057549-DEL-58 | chr9-123057550-DEL-59 |
| chr9-123118810-DEL-53 | chr9-123118811-DEL-53 |
| chr9-123132686-DEL-659 | chr9-123132687-DEL-659 |
| chr9-123133621-DEL-329 | chr9-123133622-DEL-329 |
| chr9-123135623-DEL-1429 | chr9-123135624-DEL-1429 |
| chr9-123147359-DEL-3977 | chr9-123147360-DEL-3977 |
| chr9-123171359-DEL-389 | chr9-123171360-DEL-389 |
| chr9-123184390-DEL-358 | chr9-123184391-DEL-358 |
| chr9-123194148-DEL-68 | chr9-123194149-DEL-68 |
| chr9-123217484-DEL-354 | chr9-123217485-DEL-346 |

| | |
|---|---|
| chr9-123230856-DEL-55 | chr9-123230857-DEL-55 |
| chr9-123237374-DEL-265 | chr9-123237375-DEL-265 |
| chr9-123251320-DEL-58 | chr9-123251321-DEL-58 |
| chr9-123275238-DEL-88 | chr9-123275239-DEL-88 |
| chr9-123275284-DEL-1467 | chr9-123275288-DEL-1465 |
| chr9-123307627-DEL-76 | chr9-123307627-DEL-76 |
| chr9-123310713-DEL-53 | chr9-123310715-DEL-52 |
| chr9-123324124-DEL-201 | chr9-123324125-DEL-200 |
| chr9-123338631-DEL-86 | chr9-123338639-DEL-79 |
| chr9-123343255-DEL-204 | chr9-123343256-DEL-204 |
| chr9-123353990-DEL-90 | chr9-123353984-DEL-88 |
| chr9-123376759-DEL-159 | chr9-123376760-DEL-159 |
| chr9-123388749-DEL-224 | chr9-123388750-DEL-192 |
| chr9-123476057-DEL-56 | chr9-123476058-DEL-72 |
| chr9-123479593-DEL-158 | chr9-123479594-DEL-158 |
| chr9-123489230-DEL-76 | chr9-123489231-DEL-76 |
| chr9-123497104-DEL-52 | chr9-123497093-DEL-52 |
| chr9-123502007-DEL-72 | chr9-123502008-DEL-72 |
| chr9-123517915-DEL-1425 | chr9-123517916-DEL-1425 |
| chr9-123529304-DEL-146 | chr9-123529296-DEL-147 |
| chr9-123545166-DEL-464 | chr9-123545167-DEL-464 |
| chr9-123547265-DEL-901 | chr9-123547266-DEL-901 |
| chr9-123547931-DEL-196 | chr9-123547932-DEL-196 |
| chr9-123557706-DEL-96 | chr9-123557753-DEL-71 |
| chr9-123571775-DEL-7100 | chr9-123571776-DEL-7100 |
| chr9-123579892-DEL-2676 | chr9-123579888-DEL-2676 |
| chr9-123604173-DEL-126 | chr9-123604174-DEL-126 |
| chr9-123664155-DEL-57 | chr9-123664156-DEL-56 |
| chr9-123680960-DEL-120 | chr9-123680961-DEL-90 |
| chr9-123693907-DEL-201 | chr9-123693908-DEL-201 |
| chr9-123699121-DEL-63 | chr9-123699121-DEL-63 |
| chr9-123725884-DEL-1607 | chr9-123725884-DEL-1607 |
| chr9-123734489-DEL-881 | chr9-123734499-DEL-871 |
| chr9-123818281-DEL-128 | chr9-123818282-DEL-128 |
| chr9-123820661-DEL-614 | chr9-123820662-DEL-614 |
| chr9-123836612-DEL-69 | chr9-123836627-DEL-52 |
| chr9-123846055-DEL-1065 | chr9-123846056-DEL-1065 |
| chr9-123847179-DEL-2042 | chr9-123847180-DEL-2042 |
| chr9-123857063-DEL-797 | chr9-123857096-DEL-783 |
| chr9-123896565-DEL-429 | chr9-123896566-DEL-427 |
| chr9-123914059-DEL-81 | chr9-123914060-DEL-81 |
| chr9-123923729-DEL-50 | chr9-123923722-DEL-58 |
| chr9-123934576-DEL-5594 | chr9-123934574-DEL-5592 |
| chr9-123942530-DEL-58 | chr9-123942530-DEL-58 |
| chr9-123947527-DEL-201 | chr9-123947528-DEL-201 |
| chr9-123948962-DEL-111 | chr9-123948964-DEL-109 |
| chr9-124101989-DEL-212 | chr9-124101990-DEL-212 |
| chr9-124110166-DEL-100 | chr9-124110167-DEL-79 |
| chr9-124141452-DEL-147 | chr9-124141450-DEL-148 |
| chr9-124143072-DEL-133 | chr9-124143073-DEL-133 |
| chr9-124182137-DEL-203 | chr9-124182138-DEL-203 |
| chr9-124188211-DEL-187 | chr9-124188212-DEL-187 |
| chr9-124211093-DEL-52 | chr9-124211094-DEL-52 |
| chrX-3687360-DEL-51 | chrX-3687361-DEL-51 |
| chrX-5300589-DEL-1434 | chrX-5300590-DEL-1434 |
| chrX-5315095-DEL-1867 | chrX-5315096-DEL-1867 |
| chrX-5339982-DEL-707 | chrX-5339938-DEL-732 |
| chrX-5388694-DEL-58 | chrX-5388694-DEL-58 |
| chrX-5430634-DEL-9163 | chrX-5430635-DEL-9163 |
| chrX-5498578-DEL-1120 | chrX-5498579-DEL-1120 |
| chrX-5507568-DEL-68 | chrX-5507569-DEL-68 |
| chrX-5518660-DEL-1441 | chrX-5518661-DEL-1441 |
| chrX-5538462-DEL-1149 | chrX-5538463-DEL-1149 |
| chrX-5554125-DEL-6041 | chrX-5554126-DEL-6041 |
| chrX-5588578-DEL-633 | chrX-5588579-DEL-633 |
| chrX-5589706-DEL-1418 | chrX-5589707-DEL-1418 |
| chrX-5600578-DEL-3425 | chrX-5600579-DEL-3425 |
| chrX-5625321-DEL-96 | chrX-5625303-DEL-110 |
| chrX-5732730-DEL-210 | chrX-5732731-DEL-210 |
| chrX-5733767-DEL-888 | chrX-5733768-DEL-888 |
| chrX-5739922-DEL-2571 | chrX-5739923-DEL-2571 |
| chrX-5764602-DEL-295 | chrX-5764603-DEL-295 |
| chrX-5789757-DEL-2003 | chrX-5789758-DEL-2003 |
| chrX-5821294-DEL-213 | chrX-5821295-DEL-213 |
| chrX-5887187-DEL-54 | chrX-5887188-DEL-54 |
| chrX-5904746-DEL-3697 | chrX-5904747-DEL-3697 |
| chrX-5969919-DEL-104 | chrX-5969920-DEL-104 |
| chrX-6029609-DEL-54 | chrX-6029610-DEL-50 |
| chrX-6043383-DEL-54 | chrX-6043387-DEL-50 |
| chrX-6090297-DEL-223 | chrX-6090298-DEL-223 |

| | |
|---|---|
| chrX-6134533-DEL-1511 | chrX-6134549-DEL-1507 |
| chrX-6155243-DEL-867 | chrX-6155244-DEL-867 |
| chrX-6217870-DEL-53 | chrX-6217871-DEL-53 |
| chrX-6340011-DEL-614 | chrX-6340012-DEL-614 |
| chrX-6343306-DEL-186 | chrX-6343307-DEL-186 |
| chrX-6360590-DEL-55 | chrX-6360591-DEL-55 |
| chrX-6388614-DEL-719 | chrX-6388613-DEL-719 |
| chrX-6428809-DEL-89 | chrX-6428796-DEL-115 |
| chrX-6430869-DEL-114 | chrX-6430870-DEL-114 |
| chrX-6462840-DEL-1194 | chrX-6462852-DEL-1148 |
| chrX-6478380-DEL-6599 | chrX-6478381-DEL-6599 |
| chrX-6484407-DEL-212 | chrX-6484408-DEL-212 |
| chrX-6504565-DEL-1176 | chrX-6504566-DEL-1176 |
| chrX-6534699-DEL-56 | chrX-6534676-DEL-60 |
| chrX-6604722-DEL-65 | chrX-6604723-DEL-65 |
| chrX-6611697-DEL-779 | chrX-6611686-DEL-788 |
| chrX-6667852-DEL-526 | chrX-6667853-DEL-526 |
| chrX-6712012-DEL-208 | chrX-6712014-DEL-207 |
| chrX-6736394-DEL-54 | chrX-6736395-DEL-54 |
| chrX-6781387-DEL-1558 | chrX-6781388-DEL-1558 |
| chrX-6843156-DEL-1016 | chrX-6843157-DEL-1016 |
| chrX-6894567-DEL-695 | chrX-6894568-DEL-695 |
| chrX-6937931-DEL-119 | chrX-6937932-DEL-119 |
| chrX-6954696-DEL-71 | chrX-6954696-DEL-73 |
| chrX-7039957-DEL-79 | chrX-7040005-DEL-50 |
| chrX-7075516-DEL-208 | chrX-7075517-DEL-208 |
| chrX-7134098-DEL-154 | chrX-7134099-DEL-154 |
| chrX-7233275-DEL-811 | chrX-7233276-DEL-811 |
| chrX-7305021-DEL-297 | chrX-7304993-DEL-326 |
| chrX-7346934-DEL-154 | chrX-7346934-DEL-155 |
| chrX-7366293-DEL-440 | chrX-7366294-DEL-440 |
| chrX-7372888-DEL-1335 | chrX-7372894-DEL-1328 |
| chrX-7410683-DEL-51 | chrX-7410684-DEL-51 |
| chrX-7448898-DEL-69 | chrX-7448905-DEL-64 |
| chrX-7488994-DEL-160 | chrX-7488995-DEL-160 |
| chrX-7506868-DEL-57 | chrX-7506822-DEL-70 |
| chrX-7516208-DEL-69 | chrX-7516209-DEL-69 |
| chrX-7548353-DEL-207 | chrX-7548354-DEL-207 |
| chrX-7558399-DEL-202 | chrX-7558397-DEL-199 |
| chrX-7561035-DEL-274 | chrX-7561054-DEL-249 |
| chrX-7623580-DEL-191 | chrX-7623581-DEL-191 |
| chrX-7626564-DEL-175 | chrX-7626577-DEL-136 |
| chrX-7633134-DEL-53 | chrX-7633135-DEL-53 |
| chrX-7663280-DEL-212 | chrX-7663281-DEL-212 |
| chrX-7693224-DEL-193 | chrX-7693225-DEL-193 |
| chrX-7774763-DEL-216 | chrX-7774764-DEL-216 |
| chrX-7800793-DEL-58 | chrX-7800794-DEL-54 |
| chrX-7827680-DEL-205 | chrX-7827681-DEL-205 |
| chrX-7845741-DEL-86 | chrX-7845741-DEL-85 |
| chrX-7899731-DEL-194 | chrX-7899741-DEL-211 |
| chrX-7949442-DEL-56 | chrX-7949447-DEL-52 |
| chrX-7961973-DEL-67 | chrX-7961974-DEL-65 |
| chrX-8018498-DEL-264 | chrX-8018499-DEL-264 |
| chrX-8026892-DEL-62 | chrX-8026921-DEL-74 |
| chrX-8033626-DEL-74 | chrX-8033618-DEL-62 |
| chrX-8046775-DEL-209 | chrX-8046776-DEL-209 |
| chrX-8050118-DEL-185 | chrX-8050119-DEL-185 |
| chrX-8079334-DEL-96 | chrX-8079335-DEL-96 |
| chrX-8082300-DEL-200 | chrX-8082301-DEL-200 |
| chrX-8091983-DEL-52 | chrX-8091993-DEL-51 |
| chrX-8102323-DEL-218 | chrX-8102324-DEL-218 |
| chrX-8104601-DEL-238 | chrX-8104602-DEL-238 |
| chrX-8135407-DEL-54 | chrX-8135408-DEL-54 |
| chrX-8153228-DEL-238 | chrX-8153229-DEL-238 |
| chrX-8154384-DEL-224 | chrX-8154386-DEL-220 |
| chrX-8384076-DEL-5222 | chrX-8384077-DEL-5222 |
| chrX-8400685-DEL-474 | chrX-8400685-DEL-475 |
| chrX-8561124-DEL-2008 | chrX-8561125-DEL-2008 |
| chrX-8578425-DEL-3668 | chrX-8578426-DEL-3668 |
| chrX-8610378-DEL-56 | chrX-8610379-DEL-56 |
| chrX-8610665-DEL-1865 | chrX-8610664-DEL-1865 |
| chrX-8622816-DEL-6614 | chrX-8622816-DEL-6614 |
| chrX-8632311-DEL-3839 | chrX-8632312-DEL-3839 |
| chrX-8786200-DEL-55 | chrX-8786201-DEL-55 |
| chrX-8804898-DEL-1947 | chrX-8804899-DEL-1947 |
| chrX-8863687-DEL-6600 | chrX-8863688-DEL-6600 |
| chrX-8869861-DEL-205 | chrX-8869862-DEL-205 |
| chrX-8873284-DEL-481 | chrX-8873285-DEL-480 |
| chrX-8894058-DEL-405 | chrX-8894052-DEL-412 |
| chrX-8928980-DEL-97 | chrX-8928959-DEL-97 |

| | |
|---|---|
| chrX-8954990-DEL-2126 | chrX-8954991-DEL-2126 |
| chrX-8982343-DEL-69 | chrX-8982344-DEL-71 |
| chrX-8989438-DEL-56 | chrX-8989438-DEL-56 |
| chrX-8995516-DEL-52 | chrX-8995517-DEL-52 |
| chrX-9031992-DEL-192 | chrX-9031993-DEL-192 |
| chrX-9081716-DEL-108 | chrX-9081755-DEL-92 |
| chrX-9094172-DEL-301 | chrX-9094173-DEL-300 |
| chrX-9134439-DEL-60 | chrX-9134440-DEL-60 |
| chrX-9138301-DEL-89 | chrX-9138302-DEL-89 |
| chrX-9148946-DEL-56 | chrX-9148947-DEL-56 |
| chrX-9169608-DEL-53 | chrX-9169609-DEL-53 |
| chrX-9172971-DEL-577 | chrX-9172972-DEL-577 |
| chrX-9208915-DEL-108 | chrX-9208916-DEL-108 |
| chrX-9212150-DEL-4154 | chrX-9212151-DEL-4154 |
| chrX-9223251-DEL-64 | chrX-9223251-DEL-64 |
| chrX-9227468-DEL-1108 | chrX-9227467-DEL-1109 |
| chrX-9248177-DEL-1136 | chrX-9248178-DEL-1136 |
| chrX-9265706-DEL-54 | chrX-9265706-DEL-54 |
| chrX-9266700-DEL-1021 | chrX-9266700-DEL-1021 |
| chrX-9267544-DEL-106 | chrX-9267545-DEL-106 |
| chrX-9272170-DEL-267 | chrX-9272171-DEL-267 |
| chrX-9277883-DEL-54 | chrX-9277864-DEL-74 |
| chrX-9280084-DEL-88 | chrX-9280101-DEL-72 |
| chrX-9285787-DEL-76 | chrX-9285790-DEL-77 |
| chrX-9293511-DEL-7000 | chrX-9293512-DEL-7000 |
| chrX-9366355-DEL-9412 | chrX-9366356-DEL-9412 |
| chrX-9375253-DEL-201 | chrX-9375254-DEL-201 |
| chrX-9382118-DEL-191 | chrX-9382119-DEL-191 |
| chrX-9384476-DEL-538 | chrX-9384477-DEL-538 |
| chrX-9408940-DEL-197 | chrX-9408941-DEL-197 |
| chrX-9445374-DEL-53 | chrX-9445373-DEL-53 |
| chrX-9448965-DEL-54 | chrX-9448951-DEL-54 |
| chrX-9452480-DEL-160 | chrX-9452481-DEL-160 |
| chrX-9455454-DEL-60 | chrX-9455455-DEL-60 |
| chrX-9456028-DEL-6589 | chrX-9456029-DEL-6589 |
| chrX-9462289-DEL-214 | chrX-9462290-DEL-214 |
| chrX-9484024-DEL-1823 | chrX-9484022-DEL-1825 |
| chrX-9488617-DEL-54 | chrX-9488617-DEL-54 |
| chrX-9489261-DEL-9001 | chrX-9489245-DEL-9017 |
| chrX-9498902-DEL-162 | chrX-9498902-DEL-162 |
| chrX-9504765-DEL-108 | chrX-9504766-DEL-108 |
| chrX-9508329-DEL-4080 | chrX-9508330-DEL-4080 |
| chrX-9513162-DEL-1337 | chrX-9513162-DEL-1337 |
| chrX-9607128-DEL-54 | chrX-9607129-DEL-53 |
| chrX-9616821-DEL-109 | chrX-9616822-DEL-109 |
| chrX-9627349-DEL-65 | chrX-9627356-DEL-58 |
| chrX-9643068-DEL-152 | chrX-9643069-DEL-152 |
| chrX-9661027-DEL-171 | chrX-9661028-DEL-171 |
| chrX-9725312-DEL-653 | chrX-9725313-DEL-653 |
| chrX-9762010-DEL-208 | chrX-9762011-DEL-208 |
| chrX-9791446-DEL-213 | chrX-9791430-DEL-214 |
| chrX-9826580-DEL-286 | chrX-9826581-DEL-286 |
| chrX-9846445-DEL-86 | chrX-9846485-DEL-75 |
| chrX-9881846-DEL-1277 | chrX-9881846-DEL-1277 |
| chrX-9917964-DEL-1560 | chrX-9917965-DEL-1560 |
| chrX-9961940-DEL-60 | chrX-9961941-DEL-60 |
| chrX-9970132-DEL-130 | chrX-9970171-DEL-114 |
| chrX-9987538-DEL-1545 | chrX-9987539-DEL-1545 |
| chrX-10003776-DEL-385 | chrX-10003777-DEL-385 |
| chrX-10004494-DEL-550 | chrX-10004495-DEL-550 |
| chrX-10202543-DEL-127 | chrX-10202530-DEL-128 |
| chrX-10209914-DEL-973 | chrX-10209915-DEL-973 |
| chrX-10233749-DEL-62 | chrX-10233752-DEL-60 |
| chrX-10238380-DEL-822 | chrX-10238381-DEL-822 |
| chrX-10268701-DEL-165 | chrX-10268702-DEL-153 |
| chrX-10269291-DEL-6562 | chrX-10269292-DEL-6562 |
| chrX-10286844-DEL-860 | chrX-10286827-DEL-837 |
| chrX-10385196-DEL-214 | chrX-10385197-DEL-214 |
| chrX-10385827-DEL-969 | chrX-10385828-DEL-969 |
| chrX-10391014-DEL-6522 | chrX-10391015-DEL-6522 |
| chrX-10408577-DEL-552 | chrX-10408578-DEL-552 |
| chrX-10493557-DEL-228 | chrX-10493558-DEL-228 |
| chrX-10494224-DEL-341 | chrX-10494225-DEL-341 |
| chrX-10496021-DEL-65 | chrX-10496028-DEL-58 |
| chrX-10551208-DEL-71 | chrX-10551209-DEL-71 |
| chrX-10557523-DEL-87 | chrX-10557512-DEL-84 |
| chrX-10572301-DEL-730 | chrX-10572300-DEL-730 |
| chrX-10751904-DEL-1043 | chrX-10751905-DEL-1043 |
| chrX-10766646-DEL-102 | chrX-10766648-DEL-104 |
| chrX-10799052-DEL-155 | chrX-10799053-DEL-155 |

| | |
|---|---|
| chrX-10814044-DEL-1493 | chrX-10814045-DEL-1493 |
| chrX-10831443-DEL-7508 | chrX-10831444-DEL-7508 |
| chrX-10853777-DEL-75 | chrX-10853775-DEL-77 |
| chrX-10871417-DEL-92 | chrX-10871448-DEL-62 |
| chrX-10886390-DEL-224 | chrX-10886391-DEL-224 |
| chrX-10892120-DEL-1409 | chrX-10892121-DEL-1409 |
| chrX-10900937-DEL-1527 | chrX-10900938-DEL-1527 |
| chrX-10926359-DEL-93 | chrX-10926371-DEL-87 |
| chrX-10985675-DEL-720 | chrX-10985676-DEL-720 |
| chrX-11011028-DEL-6802 | chrX-11011043-DEL-6802 |
| chrX-11020981-DEL-62 | chrX-11020982-DEL-62 |
| chrX-11149472-DEL-557 | chrX-11149473-DEL-557 |
| chrX-11197625-DEL-71 | chrX-11197626-DEL-71 |
| chrX-11323621-DEL-429 | chrX-11323622-DEL-429 |
| chrX-11350733-DEL-332 | chrX-11350734-DEL-332 |
| chrX-11381894-DEL-187 | chrX-11381895-DEL-187 |
| chrX-11426096-DEL-52 | chrX-11426099-DEL-62 |
| chrX-11436143-DEL-155 | chrX-11436144-DEL-155 |
| chrX-11469605-DEL-140 | chrX-11469606-DEL-140 |
| chrX-11562581-DEL-99 | chrX-11562582-DEL-99 |
| chrX-11745167-DEL-203 | chrX-11745168-DEL-203 |
| chrX-11816880-DEL-295 | chrX-11816881-DEL-295 |
| chrX-11833881-DEL-55 | chrX-11833875-DEL-65 |
| chrX-11880578-DEL-92 | chrX-11880574-DEL-86 |
| chrX-11898917-DEL-84 | chrX-11898914-DEL-62 |
| chrX-11900006-DEL-61 | chrX-11899993-DEL-62 |
| chrX-11905586-DEL-109 | chrX-11905587-DEL-109 |
| chrX-11968175-DEL-87 | chrX-11968152-DEL-111 |
| chrX-12002389-DEL-66 | chrX-12002385-DEL-64 |
| chrX-12014645-DEL-396 | chrX-12014649-DEL-396 |
| chrX-12066395-DEL-230 | chrX-12066396-DEL-230 |
| chrX-12090400-DEL-220 | chrX-12090393-DEL-220 |
| chrX-12093404-DEL-805 | chrX-12093425-DEL-797 |
| chrX-12097632-DEL-7148 | chrX-12097633-DEL-7148 |
| chrX-12111603-DEL-815 | chrX-12111604-DEL-815 |
| chrX-12296683-DEL-183 | chrX-12296684-DEL-183 |
| chrX-12322582-DEL-51 | chrX-12322583-DEL-51 |
| chrX-12348380-DEL-1037 | chrX-12348381-DEL-1037 |
| chrX-12359705-DEL-92 | chrX-12359706-DEL-92 |
| chrX-12373334-DEL-191 | chrX-12373335-DEL-191 |
| chrX-12390677-DEL-312 | chrX-12390699-DEL-281 |
| chrX-12397975-DEL-2132 | chrX-12397976-DEL-2132 |
| chrX-12408543-DEL-65 | chrX-12408544-DEL-65 |
| chrX-12413913-DEL-174 | chrX-12413914-DEL-174 |
| chrX-12423087-DEL-228 | chrX-12423088-DEL-228 |
| chrX-12432412-DEL-91 | chrX-12432413-DEL-91 |
| chrX-12434986-DEL-1359 | chrX-12434987-DEL-1359 |
| chrX-12447242-DEL-55 | chrX-12447235-DEL-56 |
| chrX-12449965-DEL-1888 | chrX-12449966-DEL-1888 |
| chrX-12513521-DEL-197 | chrX-12513518-DEL-198 |
| chrX-12517872-DEL-220 | chrX-12517873-DEL-220 |
| chrX-12577768-DEL-60 | chrX-12577769-DEL-60 |
| chrX-12593621-DEL-196 | chrX-12593622-DEL-196 |
| chrX-12630607-DEL-260 | chrX-12630608-DEL-260 |
| chrX-12662746-DEL-477 | chrX-12662747-DEL-477 |
| chrX-12664801-DEL-194 | chrX-12664796-DEL-195 |
| chrX-12670935-DEL-126 | chrX-12670936-DEL-126 |
| chrX-12679161-DEL-774 | chrX-12679160-DEL-775 |
| chrX-12729679-DEL-213 | chrX-12729690-DEL-203 |
| chrX-12749452-DEL-221 | chrX-12749453-DEL-221 |
| chrX-12758461-DEL-202 | chrX-12758462-DEL-202 |
| chrX-12801582-DEL-432 | chrX-12801578-DEL-432 |
| chrX-12809203-DEL-320 | chrX-12809204-DEL-320 |
| chrX-12817903-DEL-193 | chrX-12817906-DEL-192 |
| chrX-12856000-DEL-72 | chrX-12855980-DEL-110 |
| chrX-12864924-DEL-93 | chrX-12864897-DEL-82 |
| chrX-12869244-DEL-208 | chrX-12869238-DEL-209 |
| chrX-12887539-DEL-84 | chrX-12887540-DEL-84 |
| chrX-12904786-DEL-215 | chrX-12904787-DEL-215 |
| chrX-12926813-DEL-76 | chrX-12926823-DEL-95 |
| chrX-13067545-DEL-473 | chrX-13067546-DEL-473 |
| chrX-13068779-DEL-1334 | chrX-13068780-DEL-1334 |
| chrX-13071655-DEL-117 | chrX-13071656-DEL-117 |
| chrX-13073774-DEL-83 | chrX-13073775-DEL-83 |
| chrX-13075876-DEL-231 | chrX-13075877-DEL-231 |
| chrX-13084137-DEL-757 | chrX-13084138-DEL-757 |
| chrX-13091599-DEL-173 | chrX-13091600-DEL-173 |
| chrX-13094236-DEL-203 | chrX-13094237-DEL-203 |
| chrX-13114386-DEL-73 | chrX-13114387-DEL-73 |
| chrX-13115183-DEL-945 | chrX-13115184-DEL-945 |

| | |
|---|---|
| chrX-13118271-DEL-208 | chrX-13118272-DEL-208 |
| chrX-13123381-DEL-3105 | chrX-13123382-DEL-3105 |
| chrX-13137990-DEL-250 | chrX-13137991-DEL-250 |
| chrX-13142768-DEL-241 | chrX-13142775-DEL-238 |
| chrX-13183843-DEL-6436 | chrX-13183844-DEL-6436 |
| chrX-13241418-DEL-341 | chrX-13241419-DEL-341 |
| chrX-13263729-DEL-2945 | chrX-13263730-DEL-2945 |
| chrX-13267702-DEL-408 | chrX-13267679-DEL-432 |
| chrX-13271836-DEL-406 | chrX-13271837-DEL-406 |
| chrX-13287180-DEL-56 | chrX-13287181-DEL-56 |
| chrX-13287486-DEL-353 | chrX-13287487-DEL-353 |
| chrX-13304606-DEL-111 | chrX-13304607-DEL-111 |
| chrX-13318969-DEL-57 | chrX-13318970-DEL-57 |
| chrX-13354064-DEL-2527 | chrX-13354065-DEL-2527 |
| chrX-13381537-DEL-51 | chrX-13381538-DEL-51 |
| chrX-13409929-DEL-201 | chrX-13409930-DEL-201 |
| chrX-13412028-DEL-587 | chrX-13412035-DEL-582 |
| chrX-13499435-DEL-61 | chrX-13499436-DEL-61 |
| chrX-13507500-DEL-4762 | chrX-13507501-DEL-4762 |
| chrX-13661800-DEL-90 | chrX-13661801-DEL-66 |
| chrX-13719388-DEL-5713 | chrX-13719389-DEL-5713 |
| chrX-13736438-DEL-397 | chrX-13736439-DEL-397 |
| chrX-13747672-DEL-55 | chrX-13747673-DEL-55 |
| chrX-13762336-DEL-178 | chrX-13762337-DEL-178 |
| chrX-13816533-DEL-6693 | chrX-13816534-DEL-6690 |
| chrX-13841745-DEL-400 | chrX-13841746-DEL-400 |
| chrX-13877012-DEL-180 | chrX-13877014-DEL-176 |
| chrX-13878424-DEL-77 | chrX-13878425-DEL-77 |
| chrX-13895698-DEL-201 | chrX-13895695-DEL-184 |
| chrX-13918408-DEL-1733 | chrX-13918409-DEL-1733 |
| chrX-13936122-DEL-10018 | chrX-13936124-DEL-10019 |
| chrX-13967530-DEL-73 | chrX-13967531-DEL-73 |
| chrX-13981805-DEL-62 | chrX-13981800-DEL-75 |
| chrX-14009847-DEL-6826 | chrX-14009847-DEL-6826 |
| chrX-14034987-DEL-268 | chrX-14034988-DEL-268 |
| chrX-14043536-DEL-204 | chrX-14043537-DEL-204 |
| chrX-14056880-DEL-6961 | chrX-14056881-DEL-6961 |
| chrX-14106903-DEL-2790 | chrX-14106904-DEL-2790 |
| chrX-14115113-DEL-760 | chrX-14115114-DEL-760 |
| chrX-14268091-DEL-198 | chrX-14268092-DEL-198 |
| chrX-14276776-DEL-1634 | chrX-14276776-DEL-1634 |
| chrX-14278494-DEL-5007 | chrX-14278494-DEL-5007 |
| chrX-14287209-DEL-80 | chrX-14287234-DEL-56 |
| chrX-14355418-DEL-2743 | chrX-14355419-DEL-2743 |
| chrX-14355690-DEL-212 | chrX-14355691-DEL-212 |
| chrX-14414890-DEL-68 | chrX-14414891-DEL-62 |
| chrX-14465634-DEL-51 | chrX-14465634-DEL-50 |
| chrX-14474677-DEL-459 | chrX-14474678-DEL-459 |
| chrX-14516593-DEL-70 | chrX-14516627-DEL-68 |
| chrX-14523078-DEL-317 | chrX-14523079-DEL-317 |
| chrX-14553946-DEL-327 | chrX-14553947-DEL-327 |
| chrX-14614052-DEL-244 | chrX-14614053-DEL-244 |
| chrX-14619851-DEL-799 | chrX-14619852-DEL-799 |
| chrX-14645411-DEL-2293 | chrX-14645412-DEL-2291 |
| chrX-14651470-DEL-383 | chrX-14651471-DEL-383 |
| chrX-14652648-DEL-129 | chrX-14652649-DEL-129 |
| chrX-14655491-DEL-554 | chrX-14655493-DEL-555 |
| chrX-14670946-DEL-119 | chrX-14670947-DEL-119 |
| chrX-14674417-DEL-242 | chrX-14674418-DEL-242 |
| chrX-14677070-DEL-7451 | chrX-14677071-DEL-7451 |
| chrX-14686793-DEL-2515 | chrX-14686794-DEL-2515 |
| chrX-14701193-DEL-120 | chrX-14701195-DEL-118 |
| chrX-14820833-DEL-519 | chrX-14820834-DEL-519 |
| chrX-14832309-DEL-2312 | chrX-14832310-DEL-2312 |
| chrX-14879595-DEL-1336 | chrX-14879596-DEL-1336 |
| chrX-14960795-DEL-73 | chrX-14960796-DEL-68 |
| chrX-15039453-DEL-568 | chrX-15039455-DEL-566 |
| chrX-15047306-DEL-804 | chrX-15047307-DEL-804 |
| chrX-15129243-DEL-6235 | chrX-15129235-DEL-6236 |
| chrX-15159042-DEL-1028 | chrX-15159043-DEL-1028 |
| chrX-15165818-DEL-124 | chrX-15165819-DEL-124 |
| chrX-15258830-DEL-8262 | chrX-15258831-DEL-8262 |
| chrX-15265863-DEL-618 | chrX-15265862-DEL-618 |
| chrX-15287467-DEL-506 | chrX-15287468-DEL-506 |
| chrX-15299118-DEL-199 | chrX-15299119-DEL-199 |
| chrX-15340282-DEL-8985 | chrX-15340283-DEL-8985 |
| chrX-15453198-DEL-6749 | chrX-15453199-DEL-6749 |
| chrX-15467269-DEL-185 | chrX-15467270-DEL-184 |
| chrX-15492463-DEL-264 | chrX-15492464-DEL-264 |
| chrX-15522399-DEL-3720 | chrX-15522400-DEL-3720 |

| | |
|---|---|
| chrX-15584192-DEL-246 | chrX-15584193-DEL-246 |
| chrX-15597696-DEL-2442 | chrX-15597697-DEL-2442 |
| chrX-15617637-DEL-2127 | chrX-15617638-DEL-2127 |
| chrX-15625924-DEL-187 | chrX-15625921-DEL-188 |
| chrX-15655099-DEL-84 | chrX-15655055-DEL-126 |
| chrX-15656260-DEL-6548 | chrX-15656261-DEL-6548 |
| chrX-15677126-DEL-254 | chrX-15677127-DEL-254 |
| chrX-15706669-DEL-54 | chrX-15706670-DEL-54 |
| chrX-15717901-DEL-406 | chrX-15717902-DEL-406 |
| chrX-15733650-DEL-2895 | chrX-15733650-DEL-2895 |
| chrX-15760452-DEL-469 | chrX-15760428-DEL-470 |
| chrX-15769049-DEL-6700 | chrX-15769050-DEL-6700 |
| chrX-15775496-DEL-214 | chrX-15775497-DEL-214 |
| chrX-15792401-DEL-73 | chrX-15792407-DEL-52 |
| chrX-15864784-DEL-501 | chrX-15864785-DEL-501 |
| chrX-15886896-DEL-1307 | chrX-15886896-DEL-1307 |
| chrX-15910671-DEL-687 | chrX-15910672-DEL-687 |
| chrX-15978409-DEL-153 | chrX-15978398-DEL-165 |
| chrX-16002312-DEL-70 | chrX-16002353-DEL-52 |
| chrX-16168306-DEL-51 | chrX-16168307-DEL-51 |
| chrX-16180100-DEL-1740 | chrX-16180101-DEL-1739 |
| chrX-16253915-DEL-62 | chrX-16253890-DEL-52 |
| chrX-16306222-DEL-6841 | chrX-16306223-DEL-6841 |
| chrX-16335299-DEL-1102 | chrX-16335300-DEL-1102 |
| chrX-16337398-DEL-127 | chrX-16337381-DEL-128 |
| chrX-16401247-DEL-83 | chrX-16401262-DEL-82 |
| chrX-16405038-DEL-304 | chrX-16405039-DEL-304 |
| chrX-16415909-DEL-119 | chrX-16415910-DEL-119 |
| chrX-16425308-DEL-177 | chrX-16425309-DEL-173 |
| chrX-16456677-DEL-6819 | chrX-16456678-DEL-6819 |
| chrX-16463840-DEL-96 | chrX-16463855-DEL-80 |
| chrX-16473817-DEL-90 | chrX-16473818-DEL-90 |
| chrX-16532351-DEL-6234 | chrX-16532352-DEL-6234 |
| chrX-16540987-DEL-621 | chrX-16540988-DEL-621 |
| chrX-16639826-DEL-869 | chrX-16639827-DEL-869 |
| chrX-16719252-DEL-277 | chrX-16719253-DEL-288 |
| chrX-16735102-DEL-7161 | chrX-16735103-DEL-7161 |
| chrX-16789865-DEL-56 | chrX-16789888-DEL-55 |
| chrX-16806915-DEL-508 | chrX-16806916-DEL-508 |
| chrX-16939360-DEL-70 | chrX-16939311-DEL-77 |
| chrX-16998785-DEL-108 | chrX-16998786-DEL-108 |
| chrX-17036898-DEL-627 | chrX-17036899-DEL-627 |
| chrX-17045088-DEL-400 | chrX-17045089-DEL-400 |
| chrX-17062501-DEL-1251 | chrX-17062502-DEL-1251 |
| chrX-17136522-DEL-453 | chrX-17136523-DEL-453 |
| chrX-17200463-DEL-145 | chrX-17200464-DEL-145 |
| chrX-17285908-DEL-6375 | chrX-17285909-DEL-6375 |
| chrX-17314980-DEL-501 | chrX-17314981-DEL-499 |
| chrX-17330017-DEL-157 | chrX-17330014-DEL-161 |
| chrX-17338649-DEL-498 | chrX-17338650-DEL-498 |
| chrX-17341861-DEL-1865 | chrX-17341862-DEL-1865 |
| chrX-17361950-DEL-9553 | chrX-17361951-DEL-9553 |
| chrX-17376964-DEL-68 | chrX-17376965-DEL-68 |
| chrX-17454975-DEL-53 | chrX-17454976-DEL-53 |
| chrX-17462456-DEL-488 | chrX-17462457-DEL-488 |
| chrX-17474756-DEL-1757 | chrX-17474757-DEL-1757 |
| chrX-17562919-DEL-504 | chrX-17562920-DEL-504 |
| chrX-17620568-DEL-112 | chrX-17620581-DEL-104 |
| chrX-17628384-DEL-162 | chrX-17628385-DEL-162 |
| chrX-17633624-DEL-208 | chrX-17633625-DEL-208 |
| chrX-17641417-DEL-492 | chrX-17641418-DEL-493 |
| chrX-17647492-DEL-7409 | chrX-17647493-DEL-7409 |
| chrX-17816438-DEL-204 | chrX-17816471-DEL-204 |
| chrX-17861679-DEL-724 | chrX-17861680-DEL-724 |
| chrX-17904522-DEL-397 | chrX-17904523-DEL-396 |
| chrX-17930270-DEL-80 | chrX-17930271-DEL-80 |
| chrX-18097478-DEL-55 | chrX-18097475-DEL-58 |
| chrX-18149668-DEL-152 | chrX-18149669-DEL-151 |
| chrX-18197809-DEL-1489 | chrX-18197810-DEL-1489 |
| chrX-18227603-DEL-6427 | chrX-18227604-DEL-6427 |
| chrX-18254628-DEL-62 | chrX-18254629-DEL-62 |
| chrX-18294881-DEL-108 | chrX-18294882-DEL-108 |
| chrX-18312103-DEL-224 | chrX-18312104-DEL-224 |
| chrX-18358390-DEL-2531 | chrX-18358391-DEL-2531 |
| chrX-18371654-DEL-6502 | chrX-18371655-DEL-6502 |
| chrX-18371669-DEL-214 | chrX-18371669-DEL-214 |
| chrX-18390110-DEL-152 | chrX-18390111-DEL-152 |
| chrX-18436802-DEL-63 | chrX-18436809-DEL-58 |
| chrX-18472688-DEL-69 | chrX-18472689-DEL-69 |
| chrX-18491531-DEL-6610 | chrX-18491532-DEL-6610 |

| | |
|---|---|
| chrX-18515116-DEL-1920 | chrX-18515117-DEL-1920 |
| chrX-18522489-DEL-74 | chrX-18522445-DEL-110 |
| chrX-18547408-DEL-1187 | chrX-18547409-DEL-1187 |
| chrX-18556093-DEL-521 | chrX-18556102-DEL-527 |
| chrX-18560963-DEL-2719 | chrX-18560964-DEL-2719 |
| chrX-18564869-DEL-154 | chrX-18564870-DEL-154 |
| chrX-18581795-DEL-7301 | chrX-18581786-DEL-7302 |
| chrX-18625472-DEL-178 | chrX-18625473-DEL-178 |
| chrX-18629156-DEL-91 | chrX-18629157-DEL-91 |
| chrX-18633245-DEL-4098 | chrX-18633246-DEL-4098 |
| chrX-18651504-DEL-58 | chrX-18651505-DEL-58 |
| chrX-18656696-DEL-58 | chrX-18656695-DEL-61 |
| chrX-18662990-DEL-8297 | chrX-18662991-DEL-8297 |
| chrX-18769149-DEL-2516 | chrX-18769150-DEL-2516 |
| chrX-18800794-DEL-4810 | chrX-18800795-DEL-4810 |
| chrX-18861258-DEL-197 | chrX-18861259-DEL-197 |
| chrX-18871864-DEL-117 | chrX-18871865-DEL-115 |
| chrX-18897786-DEL-170 | chrX-18897787-DEL-170 |
| chrX-18899223-DEL-3365 | chrX-18899223-DEL-3365 |
| chrX-18903308-DEL-248 | chrX-18903308-DEL-246 |
| chrX-18905057-DEL-78 | chrX-18905042-DEL-68 |
| chrX-18914150-DEL-201 | chrX-18914151-DEL-201 |
| chrX-18928949-DEL-3441 | chrX-18928950-DEL-3441 |
| chrX-18935679-DEL-935 | chrX-18935680-DEL-935 |
| chrX-18940096-DEL-60 | chrX-18940101-DEL-56 |
| chrX-18949698-DEL-103 | chrX-18949701-DEL-101 |
| chrX-18994103-DEL-3666 | chrX-18994103-DEL-3663 |
| chrX-19025202-DEL-62 | chrX-19025217-DEL-50 |
| chrX-19052791-DEL-6731 | chrX-19052793-DEL-6729 |
| chrX-19059055-DEL-212 | chrX-19059056-DEL-212 |
| chrX-19082965-DEL-201 | chrX-19082966-DEL-201 |
| chrX-19092664-DEL-175 | chrX-19092665-DEL-175 |
| chrX-19111834-DEL-87 | chrX-19111835-DEL-87 |
| chrX-19152412-DEL-138 | chrX-19152425-DEL-126 |
| chrX-19189339-DEL-74 | chrX-19189340-DEL-74 |
| chrX-19222514-DEL-62 | chrX-19222515-DEL-62 |
| chrX-19235604-DEL-1506 | chrX-19235605-DEL-1506 |
| chrX-19302167-DEL-6306 | chrX-19302158-DEL-6305 |
| chrX-19311746-DEL-275 | chrX-19311746-DEL-275 |
| chrX-19336800-DEL-1204 | chrX-19336801-DEL-1204 |
| chrX-19364999-DEL-230 | chrX-19365000-DEL-230 |
| chrX-19367317-DEL-227 | chrX-19367318-DEL-227 |
| chrX-19384315-DEL-6753 | chrX-19384316-DEL-6753 |
| chrX-19400596-DEL-60 | chrX-19400597-DEL-60 |
| chrX-19408001-DEL-60 | chrX-19408002-DEL-60 |
| chrX-19425475-DEL-6173 | chrX-19425480-DEL-6169 |
| chrX-19492454-DEL-3148 | chrX-19492455-DEL-3148 |
| chrX-19496403-DEL-1899 | chrX-19496404-DEL-1899 |
| chrX-19527920-DEL-7134 | chrX-19527921-DEL-7134 |
| chrX-19562563-DEL-400 | chrX-19562566-DEL-401 |
| chrX-19580927-DEL-559 | chrX-19580928-DEL-559 |
| chrX-19613771-DEL-603 | chrX-19613772-DEL-603 |
| chrX-19638638-DEL-1260 | chrX-19638637-DEL-1261 |
| chrX-19651251-DEL-498 | chrX-19651252-DEL-498 |
| chrX-19663191-DEL-2629 | chrX-19663192-DEL-2629 |
| chrX-19673283-DEL-1363 | chrX-19673284-DEL-1363 |
| chrX-19697262-DEL-7086 | chrX-19697263-DEL-7086 |
| chrX-19719749-DEL-709 | chrX-19719750-DEL-709 |
| chrX-19726225-DEL-6091 | chrX-19726226-DEL-6092 |
| chrX-19744121-DEL-615 | chrX-19744122-DEL-615 |
| chrX-19803938-DEL-349 | chrX-19803939-DEL-349 |
| chrX-19805835-DEL-1860 | chrX-19805836-DEL-1860 |
| chrX-19810216-DEL-97 | chrX-19810217-DEL-97 |
| chrX-19818887-DEL-1231 | chrX-19818888-DEL-1231 |
| chrX-19825101-DEL-229 | chrX-19825102-DEL-229 |
| chrX-19830798-DEL-3462 | chrX-19830799-DEL-3462 |
| chrX-19834071-DEL-132 | chrX-19834072-DEL-132 |
| chrX-19834842-DEL-74 | chrX-19834843-DEL-74 |
| chrX-19837460-DEL-4415 | chrX-19837462-DEL-4413 |
| chrX-19841904-DEL-1645 | chrX-19841905-DEL-1645 |
| chrX-19844379-DEL-660 | chrX-19844380-DEL-660 |
| chrX-19853084-DEL-138 | chrX-19853085-DEL-138 |
| chrX-19861296-DEL-6418 | chrX-19861297-DEL-6418 |
| chrX-19890631-DEL-53 | chrX-19890632-DEL-53 |
| chrX-19900176-DEL-6415 | chrX-19900177-DEL-6415 |
| chrX-19940885-DEL-60 | chrX-19940879-DEL-66 |
| chrX-19973553-DEL-688 | chrX-19973554-DEL-688 |
| chrX-20061579-DEL-518 | chrX-20061580-DEL-518 |
| chrX-20073439-DEL-504 | chrX-20073456-DEL-488 |
| chrX-20099746-DEL-5455 | chrX-20099744-DEL-5456 |

| | |
|---|---|
| chrX-20170381-DEL-227 | chrX-20170382-DEL-227 |
| chrX-20178392-DEL-296 | chrX-20178399-DEL-278 |
| chrX-20179430-DEL-2310 | chrX-20179431-DEL-2310 |
| chrX-20183738-DEL-144 | chrX-20183739-DEL-144 |
| chrX-20188688-DEL-531 | chrX-20188689-DEL-531 |
| chrX-20189987-DEL-213 | chrX-20189988-DEL-213 |
| chrX-20196296-DEL-126 | chrX-20196297-DEL-126 |
| chrX-20215880-DEL-123 | chrX-20215881-DEL-126 |
| chrX-20231737-DEL-60 | chrX-20231738-DEL-60 |
| chrX-20239110-DEL-252 | chrX-20239111-DEL-252 |
| chrX-20256530-DEL-81 | chrX-20256531-DEL-81 |
| chrX-20268569-DEL-205 | chrX-20268570-DEL-205 |
| chrX-20289283-DEL-219 | chrX-20289284-DEL-219 |
| chrX-20307616-DEL-106 | chrX-20307617-DEL-106 |
| chrX-20308246-DEL-4849 | chrX-20308248-DEL-4848 |
| chrX-20323823-DEL-197 | chrX-20323824-DEL-197 |
| chrX-20326875-DEL-184 | chrX-20326876-DEL-184 |
| chrX-20340082-DEL-2765 | chrX-20340083-DEL-2765 |
| chrX-20378501-DEL-447 | chrX-20378502-DEL-447 |
| chrX-20401139-DEL-2501 | chrX-20401139-DEL-2501 |
| chrX-20446773-DEL-360 | chrX-20446774-DEL-357 |
| chrX-20454775-DEL-251 | chrX-20454769-DEL-234 |
| chrX-20490719-DEL-1155 | chrX-20490720-DEL-1155 |
| chrX-20491397-DEL-213 | chrX-20491398-DEL-213 |
| chrX-20520583-DEL-65 | chrX-20520584-DEL-65 |
| chrX-20544686-DEL-94 | chrX-20544686-DEL-70 |
| chrX-20566432-DEL-183 | chrX-20566415-DEL-186 |
| chrX-20568591-DEL-141 | chrX-20568588-DEL-131 |
| chrX-20581978-DEL-218 | chrX-20581979-DEL-218 |
| chrX-20591350-DEL-64 | chrX-20591321-DEL-60 |
| chrX-20605942-DEL-342 | chrX-20605950-DEL-314 |
| chrX-20612161-DEL-197 | chrX-20612162-DEL-197 |
| chrX-20612895-DEL-106 | chrX-20612882-DEL-120 |
| chrX-20622376-DEL-89 | chrX-20622377-DEL-89 |
| chrX-20626122-DEL-186 | chrX-20626121-DEL-189 |
| chrX-20638671-DEL-564 | chrX-20638672-DEL-564 |
| chrX-20646143-DEL-200 | chrX-20646144-DEL-200 |
| chrX-20646606-DEL-372 | chrX-20646607-DEL-372 |
| chrX-20647761-DEL-333 | chrX-20647762-DEL-333 |
| chrX-20650516-DEL-165 | chrX-20650517-DEL-165 |
| chrX-20688714-DEL-94 | chrX-20688715-DEL-94 |
| chrX-20699937-DEL-206 | chrX-20699942-DEL-201 |
| chrX-20722045-DEL-51 | chrX-20722046-DEL-51 |
| chrX-20768053-DEL-427 | chrX-20768054-DEL-427 |
| chrX-20835182-DEL-520 | chrX-20835183-DEL-520 |
| chrX-20849064-DEL-353 | chrX-20849065-DEL-353 |
| chrX-20873568-DEL-504 | chrX-20873569-DEL-504 |
| chrX-20916386-DEL-667 | chrX-20916387-DEL-667 |
| chrX-20936141-DEL-4537 | chrX-20936142-DEL-4537 |
| chrX-20958066-DEL-302 | chrX-20958087-DEL-326 |
| chrX-20959841-DEL-589 | chrX-20959842-DEL-589 |
| chrX-20967844-DEL-6397 | chrX-20967845-DEL-6397 |
| chrX-21019437-DEL-4145 | chrX-21019438-DEL-4145 |
| chrX-21049523-DEL-8118 | chrX-21049524-DEL-8118 |
| chrX-21072651-DEL-62 | chrX-21072652-DEL-62 |
| chrX-21153368-DEL-95 | chrX-21153388-DEL-72 |
| chrX-21177932-DEL-52 | chrX-21177933-DEL-52 |
| chrX-21213151-DEL-70 | chrX-21213152-DEL-70 |
| chrX-21213391-DEL-2872 | chrX-21213392-DEL-2872 |
| chrX-21252671-DEL-104 | chrX-21252655-DEL-105 |
| chrX-21266790-DEL-67 | chrX-21266817-DEL-59 |
| chrX-21269430-DEL-6684 | chrX-21269431-DEL-6684 |
| chrX-21332713-DEL-6661 | chrX-21332714-DEL-6661 |
| chrX-21338576-DEL-229 | chrX-21338577-DEL-229 |
| chrX-21378448-DEL-1253 | chrX-21378449-DEL-1253 |
| chrX-21382033-DEL-398 | chrX-21382034-DEL-398 |
| chrX-21400059-DEL-1005 | chrX-21400060-DEL-1005 |
| chrX-21421621-DEL-411 | chrX-21421622-DEL-411 |
| chrX-21471434-DEL-355 | chrX-21471434-DEL-373 |
| chrX-21478049-DEL-449 | chrX-21478050-DEL-449 |
| chrX-21480491-DEL-92 | chrX-21480492-DEL-90 |
| chrX-21485099-DEL-7049 | chrX-21485100-DEL-7049 |
| chrX-21495401-DEL-242 | chrX-21495402-DEL-242 |
| chrX-21523695-DEL-1328 | chrX-21523696-DEL-1328 |
| chrX-21622081-DEL-6668 | chrX-21622082-DEL-6668 |
| chrX-21663718-DEL-3759 | chrX-21663719-DEL-3759 |
| chrX-21670361-DEL-134 | chrX-21670362-DEL-134 |
| chrX-21731332-DEL-6557 | chrX-21731333-DEL-6557 |
| chrX-21731391-DEL-212 | chrX-21731391-DEL-212 |
| chrX-21760792-DEL-1648 | chrX-21760793-DEL-1648 |

| | |
|---|---|
| chrX-21785513-DEL-506 | chrX-21785514-DEL-506 |
| chrX-21843715-DEL-203 | chrX-21843716-DEL-203 |
| chrX-21854642-DEL-360 | chrX-21854642-DEL-360 |
| chrX-21933320-DEL-53 | chrX-21933321-DEL-53 |
| chrX-21939131-DEL-52 | chrX-21939116-DEL-53 |
| chrX-21943832-DEL-66 | chrX-21943833-DEL-66 |
| chrX-21996348-DEL-416 | chrX-21996348-DEL-416 |
| chrX-22029398-DEL-351 | chrX-22029399-DEL-351 |
| chrX-22052981-DEL-71 | chrX-22052982-DEL-71 |
| chrX-22128032-DEL-96 | chrX-22128033-DEL-96 |
| chrX-22149889-DEL-62 | chrX-22149888-DEL-62 |
| chrX-22160516-DEL-439 | chrX-22160517-DEL-439 |
| chrX-22250675-DEL-7492 | chrX-22250676-DEL-7492 |
| chrX-22270311-DEL-152 | chrX-22270312-DEL-152 |
| chrX-22293725-DEL-56 | chrX-22293726-DEL-56 |
| chrX-22345438-DEL-461 | chrX-22345439-DEL-461 |
| chrX-22355238-DEL-60 | chrX-22355243-DEL-52 |
| chrX-22363489-DEL-233 | chrX-22363490-DEL-233 |
| chrX-22405536-DEL-59 | chrX-22405536-DEL-60 |
| chrX-22510608-DEL-6385 | chrX-22510609-DEL-6385 |
| chrX-22518505-DEL-545 | chrX-22518506-DEL-545 |
| chrX-22523008-DEL-217 | chrX-22523009-DEL-217 |
| chrX-22531370-DEL-134 | chrX-22531371-DEL-128 |
| chrX-22537680-DEL-199 | chrX-22537681-DEL-199 |
| chrX-22540933-DEL-52 | chrX-22540933-DEL-52 |
| chrX-22601544-DEL-127 | chrX-22601545-DEL-127 |
| chrX-22616818-DEL-1806 | chrX-22616820-DEL-1803 |
| chrX-22642954-DEL-834 | chrX-22642955-DEL-834 |
| chrX-22708135-DEL-1684 | chrX-22708136-DEL-1684 |
| chrX-22754177-DEL-214 | chrX-22754178-DEL-211 |
| chrX-22762287-DEL-59 | chrX-22762288-DEL-59 |
| chrX-22781419-DEL-1718 | chrX-22781422-DEL-1716 |
| chrX-22781798-DEL-184 | chrX-22781799-DEL-184 |
| chrX-22833786-DEL-1021 | chrX-22833778-DEL-1022 |
| chrX-22846654-DEL-78 | chrX-22846655-DEL-78 |
| chrX-22867627-DEL-227 | chrX-22867628-DEL-227 |
| chrX-22906614-DEL-2838 | chrX-22906615-DEL-2838 |
| chrX-22908539-DEL-310 | chrX-22908540-DEL-310 |
| chrX-22950151-DEL-334 | chrX-22950152-DEL-334 |
| chrX-22970852-DEL-244 | chrX-22970831-DEL-232 |
| chrX-22997187-DEL-108 | chrX-22997188-DEL-108 |
| chrX-23006245-DEL-77 | chrX-23006233-DEL-69 |
| chrX-23078246-DEL-66 | chrX-23078247-DEL-84 |
| chrX-23176450-DEL-89 | chrX-23176451-DEL-89 |
| chrX-23219169-DEL-448 | chrX-23219170-DEL-448 |
| chrX-23331345-DEL-57 | chrX-23331345-DEL-57 |
| chrX-23425312-DEL-177 | chrX-23425312-DEL-178 |
| chrX-23428120-DEL-470 | chrX-23428121-DEL-470 |
| chrX-23430052-DEL-1856 | chrX-23430053-DEL-1856 |
| chrX-23478251-DEL-1877 | chrX-23478252-DEL-1846 |
| chrX-23640656-DEL-2494 | chrX-23640657-DEL-2494 |
| chrX-23667459-DEL-199 | chrX-23667460-DEL-199 |
| chrX-23687334-DEL-83 | chrX-23687335-DEL-83 |
| chrX-23840545-DEL-6962 | chrX-23840546-DEL-6962 |
| chrX-23841026-DEL-208 | chrX-23841026-DEL-208 |
| chrX-23861360-DEL-400 | chrX-23861361-DEL-399 |
| chrX-23867151-DEL-57 | chrX-23867144-DEL-62 |
| chrX-23935673-DEL-99 | chrX-23935673-DEL-99 |
| chrX-23958456-DEL-67 | chrX-23958457-DEL-67 |
| chrX-25360272-DEL-63 | chrX-25360281-DEL-54 |
| chrX-26076882-DEL-607 | chrX-26076883-DEL-607 |
| chrX-30903382-DEL-93 | chrX-30903416-DEL-60 |
| chrX-35045972-DEL-466 | chrX-35045973-DEL-466 |
| chrX-35086173-DEL-6962 | chrX-35086174-DEL-6962 |
| chrX-35094001-DEL-139 | chrX-35094002-DEL-139 |
| chrX-35123753-DEL-7300 | chrX-35123753-DEL-7299 |
| chrX-35148963-DEL-214 | chrX-35148962-DEL-216 |
| chrX-35170941-DEL-212 | chrX-35170942-DEL-212 |
| chrX-35178663-DEL-4524 | chrX-35178664-DEL-4524 |
| chrX-35221928-DEL-179 | chrX-35221929-DEL-179 |
| chrX-35278831-DEL-58 | chrX-35278830-DEL-58 |
| chrX-35293870-DEL-93 | chrX-35293871-DEL-93 |
| chrX-35418082-DEL-107 | chrX-35418083-DEL-107 |
| chrX-35440584-DEL-211 | chrX-35440582-DEL-212 |
| chrX-35444624-DEL-193 | chrX-35444625-DEL-193 |
| chrX-35485349-DEL-193 | chrX-35485349-DEL-194 |
| chrX-35569874-DEL-55 | chrX-35569875-DEL-55 |
| chrX-35739873-DEL-1393 | chrX-35739874-DEL-1393 |
| chrX-35750806-DEL-1396 | chrX-35750807-DEL-1396 |
| chrX-35808718-DEL-99 | chrX-35808719-DEL-99 |

| | |
|---|---|
| chrX-35860124-DEL-205 | chrX-35860125-DEL-204 |
| chrX-35913192-DEL-3029 | chrX-35913193-DEL-3029 |
| chrX-35919919-DEL-54 | chrX-35919917-DEL-54 |
| chrX-35929223-DEL-95 | chrX-35929224-DEL-95 |
| chrX-35942540-DEL-432 | chrX-35942542-DEL-431 |
| chrX-35951070-DEL-2109 | chrX-35951071-DEL-2109 |
| chrX-35954205-DEL-1351 | chrX-35954206-DEL-1351 |
| chrX-35985792-DEL-1171 | chrX-35985793-DEL-1171 |
| chrX-36002545-DEL-82 | chrX-36002546-DEL-82 |
| chrX-36020457-DEL-68 | chrX-36020458-DEL-68 |
| chrX-36034373-DEL-110 | chrX-36034374-DEL-110 |
| chrX-36041063-DEL-227 | chrX-36041064-DEL-227 |
| chrX-36086574-DEL-202 | chrX-36086575-DEL-202 |
| chrX-36139733-DEL-214 | chrX-36139725-DEL-214 |
| chrX-36142436-DEL-150 | chrX-36142436-DEL-150 |
| chrX-36250124-DEL-71 | chrX-36250125-DEL-71 |
| chrX-36268435-DEL-99 | chrX-36268434-DEL-99 |
| chrX-36285714-DEL-604 | chrX-36285715-DEL-604 |
| chrX-36302237-DEL-86 | chrX-36302236-DEL-87 |
| chrX-36303110-DEL-1525 | chrX-36303111-DEL-1525 |
| chrX-36318673-DEL-5847 | chrX-36318674-DEL-5847 |
| chrX-36344209-DEL-202 | chrX-36344210-DEL-201 |
| chrX-36384947-DEL-214 | chrX-36384948-DEL-214 |
| chrX-36387257-DEL-209 | chrX-36387258-DEL-208 |
| chrX-36407603-DEL-989 | chrX-36407604-DEL-989 |
| chrX-36424974-DEL-174 | chrX-36424975-DEL-174 |
| chrX-36439475-DEL-81 | chrX-36439476-DEL-81 |
| chrX-36442108-DEL-56 | chrX-36442079-DEL-56 |
| chrX-36443633-DEL-209 | chrX-36443634-DEL-209 |
| chrX-36456973-DEL-179 | chrX-36456974-DEL-179 |
| chrX-36862059-DEL-120 | chrX-36862060-DEL-120 |
| chrX-36934127-DEL-52 | chrX-36934128-DEL-52 |
| chrX-36934622-DEL-117 | chrX-36934631-DEL-105 |
| chrX-36998584-DEL-6685 | chrX-36998585-DEL-6685 |
| chrX-37057347-DEL-58 | chrX-37057348-DEL-58 |
| chrX-37120916-DEL-1798 | chrX-37120917-DEL-1798 |
| chrX-37183678-DEL-96 | chrX-37183683-DEL-92 |
| chrX-37222965-DEL-229 | chrX-37222966-DEL-229 |
| chrX-37254738-DEL-79 | chrX-37254727-DEL-91 |
| chrX-37264255-DEL-54 | chrX-37264250-DEL-64 |
| chrX-37299358-DEL-3207 | chrX-37299357-DEL-3204 |
| chrX-37364707-DEL-97 | chrX-37364708-DEL-97 |
| chrX-37463774-DEL-56 | chrX-37463775-DEL-56 |
| chrX-37539822-DEL-188 | chrX-37539823-DEL-188 |
| chrX-37546123-DEL-55 | chrX-37546124-DEL-55 |
| chrX-37554454-DEL-78 | chrX-37554455-DEL-72 |
| chrX-37587113-DEL-788 | chrX-37587114-DEL-788 |
| chrX-37600807-DEL-1906 | chrX-37600808-DEL-1906 |
| chrX-37601456-DEL-57 | chrX-37601457-DEL-54 |
| chrX-37604120-DEL-51 | chrX-37604092-DEL-56 |
| chrX-37612012-DEL-210 | chrX-37612013-DEL-210 |
| chrX-37616451-DEL-878 | chrX-37616452-DEL-878 |
| chrX-37623694-DEL-293 | chrX-37623695-DEL-293 |
| chrX-37626519-DEL-710 | chrX-37626520-DEL-710 |
| chrX-37635009-DEL-112 | chrX-37635011-DEL-111 |
| chrX-37640003-DEL-101 | chrX-37640004-DEL-101 |
| chrX-37656517-DEL-220 | chrX-37656516-DEL-220 |
| chrX-37732772-DEL-778 | chrX-37732756-DEL-779 |
| chrX-37738062-DEL-1336 | chrX-37738062-DEL-1342 |
| chrX-37788998-DEL-179 | chrX-37788999-DEL-182 |
| chrX-37856541-DEL-2482 | chrX-37856542-DEL-2482 |
| chrX-37859901-DEL-90 | chrX-37859932-DEL-64 |
| chrX-37873005-DEL-6623 | chrX-37873006-DEL-6623 |
| chrX-37881057-DEL-206 | chrX-37881058-DEL-206 |
| chrX-37924709-DEL-8464 | chrX-37924709-DEL-8464 |
| chrX-37938242-DEL-1866 | chrX-37938243-DEL-1866 |
| chrX-38019620-DEL-200 | chrX-38019621-DEL-200 |
| chrX-38058660-DEL-445 | chrX-38058661-DEL-445 |
| chrX-38135745-DEL-2873 | chrX-38135746-DEL-2873 |
| chrX-38158348-DEL-6353 | chrX-38158349-DEL-6353 |
| chrX-38183945-DEL-340 | chrX-38183946-DEL-340 |
| chrX-38212662-DEL-4004 | chrX-38212663-DEL-4004 |
| chrX-38259053-DEL-2656 | chrX-38259054-DEL-2656 |
| chrX-38294651-DEL-214 | chrX-38294652-DEL-214 |
| chrX-38301258-DEL-9555 | chrX-38301259-DEL-9555 |
| chrX-38373205-DEL-154 | chrX-38373206-DEL-154 |
| chrX-38420731-DEL-6908 | chrX-38420717-DEL-6911 |
| chrX-38576675-DEL-1789 | chrX-38576676-DEL-1789 |
| chrX-38617089-DEL-185 | chrX-38617090-DEL-185 |
| chrX-38631306-DEL-487 | chrX-38631315-DEL-479 |

| | |
|---|---|
| chrX-38632097-DEL-96 | chrX-38632116-DEL-68 |
| chrX-38653347-DEL-213 | chrX-38653348-DEL-213 |
| chrX-38682844-DEL-7043 | chrX-38682845-DEL-7043 |
| chrX-38710070-DEL-51 | chrX-38710061-DEL-67 |
| chrX-38750018-DEL-6469 | chrX-38750019-DEL-6469 |
| chrX-38780774-DEL-245 | chrX-38780763-DEL-243 |
| chrX-38783318-DEL-75 | chrX-38783319-DEL-75 |
| chrX-38843490-DEL-61 | chrX-38843475-DEL-69 |
| chrX-38863254-DEL-1995 | chrX-38863255-DEL-1995 |
| chrX-38884961-DEL-473 | chrX-38884962-DEL-473 |
| chrX-38916383-DEL-608 | chrX-38916384-DEL-608 |
| chrX-38998051-DEL-113 | chrX-38998052-DEL-113 |
| chrX-39022489-DEL-6591 | chrX-39022490-DEL-6591 |
| chrX-39078828-DEL-719 | chrX-39078816-DEL-729 |
| chrX-39085881-DEL-6051 | chrX-39085882-DEL-6051 |
| chrX-39106922-DEL-1700 | chrX-39106923-DEL-1700 |
| chrX-39191906-DEL-56 | chrX-39191907-DEL-56 |
| chrX-39198277-DEL-69 | chrX-39198275-DEL-71 |
| chrX-39200846-DEL-192 | chrX-39200847-DEL-195 |
| chrX-39234641-DEL-503 | chrX-39234642-DEL-503 |
| chrX-39271189-DEL-6420 | chrX-39271190-DEL-6420 |
| chrX-39271219-DEL-208 | chrX-39271220-DEL-208 |
| chrX-39330747-DEL-6380 | chrX-39330748-DEL-6380 |
| chrX-39358190-DEL-558 | chrX-39358191-DEL-558 |
| chrX-39374739-DEL-66 | chrX-39374760-DEL-56 |
| chrX-39393537-DEL-3470 | chrX-39393538-DEL-3469 |
| chrX-39562414-DEL-769 | chrX-39562415-DEL-769 |
| chrX-39728369-DEL-199 | chrX-39728331-DEL-201 |
| chrX-39800521-DEL-970 | chrX-39800522-DEL-970 |
| chrX-39856009-DEL-62 | chrX-39856014-DEL-58 |
| chrX-39864725-DEL-96 | chrX-39864726-DEL-96 |
| chrX-39873420-DEL-485 | chrX-39873421-DEL-491 |
| chrX-39922188-DEL-6922 | chrX-39922189-DEL-6922 |
| chrX-39953781-DEL-199 | chrX-39953782-DEL-199 |
| chrX-40021629-DEL-1894 | chrX-40021630-DEL-1894 |
| chrX-40037030-DEL-7498 | chrX-40037031-DEL-7498 |
| chrX-40101076-DEL-51 | chrX-40101077-DEL-51 |
| chrX-40114146-DEL-57 | chrX-40114147-DEL-57 |
| chrX-40172276-DEL-3785 | chrX-40172277-DEL-3785 |
| chrX-40178034-DEL-900 | chrX-40178035-DEL-900 |
| chrX-40215272-DEL-56 | chrX-40215289-DEL-58 |
| chrX-40262995-DEL-6896 | chrX-40262981-DEL-6892 |
| chrX-40269576-DEL-212 | chrX-40269576-DEL-212 |
| chrX-40312112-DEL-57 | chrX-40312113-DEL-57 |
| chrX-40313582-DEL-869 | chrX-40313583-DEL-869 |
| chrX-40320615-DEL-6761 | chrX-40320616-DEL-6761 |
| chrX-40393943-DEL-6428 | chrX-40393944-DEL-6428 |
| chrX-40550534-DEL-194 | chrX-40550535-DEL-194 |
| chrX-40571324-DEL-7206 | chrX-40571325-DEL-7205 |
| chrX-40609858-DEL-3169 | chrX-40609859-DEL-3169 |
| chrX-40642117-DEL-69 | chrX-40642118-DEL-69 |
| chrX-40713559-DEL-525 | chrX-40713546-DEL-536 |
| chrX-40728075-DEL-4035 | chrX-40728082-DEL-4036 |
| chrX-40740353-DEL-129 | chrX-40740354-DEL-129 |
| chrX-40781562-DEL-55 | chrX-40781612-DEL-50 |
| chrX-40787883-DEL-6903 | chrX-40787884-DEL-6903 |
| chrX-40809993-DEL-7088 | chrX-40809994-DEL-7088 |
| chrX-40821619-DEL-68 | chrX-40821604-DEL-84 |
| chrX-40841329-DEL-56 | chrX-40841330-DEL-56 |
| chrX-40924769-DEL-56 | chrX-40924770-DEL-56 |
| chrX-40949897-DEL-1486 | chrX-40949898-DEL-1486 |
| chrX-40963285-DEL-132 | chrX-40963286-DEL-132 |
| chrX-40975396-DEL-6403 | chrX-40975397-DEL-6403 |
| chrX-40998124-DEL-108 | chrX-40998132-DEL-80 |
| chrX-41028239-DEL-1894 | chrX-41028240-DEL-1894 |
| chrX-41038554-DEL-202 | chrX-41038555-DEL-202 |
| chrX-41077880-DEL-6792 | chrX-41077881-DEL-6792 |
| chrX-41106128-DEL-172 | chrX-41106129-DEL-172 |
| chrX-41123701-DEL-338 | chrX-41123700-DEL-339 |
| chrX-41128269-DEL-171 | chrX-41128270-DEL-171 |
| chrX-41132446-DEL-185 | chrX-41132447-DEL-185 |
| chrX-41133028-DEL-405 | chrX-41133006-DEL-425 |
| chrX-41141390-DEL-331 | chrX-41141395-DEL-327 |
| chrX-41162705-DEL-628 | chrX-41162706-DEL-628 |
| chrX-41166548-DEL-60 | chrX-41166530-DEL-90 |
| chrX-41253777-DEL-51 | chrX-41253778-DEL-51 |
| chrX-41285232-DEL-200 | chrX-41285233-DEL-200 |
| chrX-41655622-DEL-6883 | chrX-41655623-DEL-6883 |
| chrX-41681307-DEL-212 | chrX-41681299-DEL-214 |
| chrX-41721815-DEL-6192 | chrX-41721816-DEL-6192 |

| | |
|---|---|
| chrX-41728186-DEL-715 | chrX-41728186-DEL-715 |
| chrX-41767847-DEL-4453 | chrX-41767848-DEL-4453 |
| chrX-41769185-DEL-454 | chrX-41769186-DEL-454 |
| chrX-41774878-DEL-2064 | chrX-41774879-DEL-2064 |
| chrX-41798288-DEL-910 | chrX-41798289-DEL-910 |
| chrX-41802889-DEL-69 | chrX-41802895-DEL-64 |
| chrX-41850003-DEL-6477 | chrX-41850004-DEL-6477 |
| chrX-41970668-DEL-849 | chrX-41970669-DEL-849 |
| chrX-41972322-DEL-492 | chrX-41972325-DEL-490 |
| chrX-41993977-DEL-603 | chrX-41993979-DEL-602 |
| chrX-42009543-DEL-6466 | chrX-42009544-DEL-6466 |
| chrX-42156830-DEL-1438 | chrX-42156831-DEL-1438 |
| chrX-42208717-DEL-172 | chrX-42208718-DEL-172 |
| chrX-42249950-DEL-903 | chrX-42249951-DEL-903 |
| chrX-42253578-DEL-478 | chrX-42253579-DEL-478 |
| chrX-42259575-DEL-644 | chrX-42259576-DEL-644 |
| chrX-42265958-DEL-356 | chrX-42265959-DEL-356 |
| chrX-42266624-DEL-2611 | chrX-42266625-DEL-2611 |
| chrX-42418775-DEL-208 | chrX-42418776-DEL-208 |
| chrX-42427359-DEL-790 | chrX-42427360-DEL-790 |
| chrX-42446401-DEL-468 | chrX-42446402-DEL-468 |
| chrX-42452032-DEL-72 | chrX-42452033-DEL-72 |
| chrX-42467763-DEL-3245 | chrX-42467771-DEL-3229 |
| chrX-42530090-DEL-387 | chrX-42530091-DEL-397 |
| chrX-42530769-DEL-467 | chrX-42530770-DEL-467 |
| chrX-42542407-DEL-207 | chrX-42542408-DEL-207 |
| chrX-42559865-DEL-4897 | chrX-42559866-DEL-4897 |
| chrX-42634779-DEL-140 | chrX-42634760-DEL-160 |
| chrX-42749545-DEL-214 | chrX-42749546-DEL-214 |
| chrX-42761974-DEL-7554 | chrX-42761975-DEL-7554 |
| chrX-42955964-DEL-1122 | chrX-42955965-DEL-1122 |
| chrX-43036534-DEL-201 | chrX-43036535-DEL-201 |
| chrX-43047345-DEL-2833 | chrX-43047346-DEL-2833 |
| chrX-43101235-DEL-4692 | chrX-43101236-DEL-4692 |
| chrX-43115626-DEL-78 | chrX-43115627-DEL-78 |
| chrX-43121835-DEL-99 | chrX-43121836-DEL-101 |
| chrX-43132306-DEL-6682 | chrX-43132307-DEL-6682 |
| chrX-43141868-DEL-84 | chrX-43141869-DEL-66 |
| chrX-43167649-DEL-535 | chrX-43167650-DEL-535 |
| chrX-43189666-DEL-359 | chrX-43189667-DEL-359 |
| chrX-43206841-DEL-2588 | chrX-43206841-DEL-2588 |
| chrX-43219467-DEL-1059 | chrX-43219468-DEL-1057 |
| chrX-43222129-DEL-2143 | chrX-43222130-DEL-2143 |
| chrX-43286517-DEL-7008 | chrX-43286518-DEL-7008 |
| chrX-43357500-DEL-208 | chrX-43357501-DEL-208 |
| chrX-43358181-DEL-208 | chrX-43358182-DEL-208 |
| chrX-43388530-DEL-390 | chrX-43388531-DEL-390 |
| chrX-43408973-DEL-1226 | chrX-43408974-DEL-1226 |
| chrX-43475830-DEL-6594 | chrX-43475831-DEL-6594 |
| chrX-43475870-DEL-208 | chrX-43475871-DEL-208 |
| chrX-43505291-DEL-6397 | chrX-43505292-DEL-6397 |
| chrX-43525234-DEL-6500 | chrX-43525235-DEL-6500 |
| chrX-43532111-DEL-1705 | chrX-43532111-DEL-1709 |
| chrX-43578084-DEL-6796 | chrX-43578085-DEL-6796 |
| chrX-43593005-DEL-109 | chrX-43593006-DEL-109 |
| chrX-43655822-DEL-1384 | chrX-43655823-DEL-1384 |
| chrX-43680855-DEL-2282 | chrX-43680856-DEL-2282 |
| chrX-43684800-DEL-2091 | chrX-43684795-DEL-2091 |
| chrX-43702201-DEL-1920 | chrX-43702202-DEL-1920 |
| chrX-43733795-DEL-2364 | chrX-43733796-DEL-2364 |
| chrX-43771046-DEL-82 | chrX-43771037-DEL-84 |
| chrX-43774714-DEL-67 | chrX-43774730-DEL-55 |
| chrX-43804114-DEL-66 | chrX-43804114-DEL-66 |
| chrX-43920633-DEL-691 | chrX-43920634-DEL-691 |
| chrX-43975442-DEL-2430 | chrX-43975445-DEL-2431 |
| chrX-43983439-DEL-68 | chrX-43983440-DEL-68 |
| chrX-44060519-DEL-368 | chrX-44060521-DEL-370 |
| chrX-44100031-DEL-6693 | chrX-44100032-DEL-6693 |
| chrX-44126931-DEL-1116 | chrX-44126932-DEL-1116 |
| chrX-44136187-DEL-3143 | chrX-44136188-DEL-3143 |
| chrX-44262420-DEL-158 | chrX-44262420-DEL-159 |
| chrX-44301245-DEL-84 | chrX-44301252-DEL-60 |
| chrX-44341931-DEL-6591 | chrX-44341931-DEL-6591 |
| chrX-44460426-DEL-683 | chrX-44460427-DEL-682 |
| chrX-44557259-DEL-5109 | chrX-44557260-DEL-5109 |
| chrX-44609707-DEL-1111 | chrX-44609708-DEL-1111 |
| chrX-44641304-DEL-69 | chrX-44641316-DEL-60 |
| chrX-44656751-DEL-405 | chrX-44656752-DEL-405 |
| chrX-44703620-DEL-78 | chrX-44703621-DEL-78 |
| chrX-44710346-DEL-82 | chrX-44710347-DEL-82 |

| | |
|---|---|
| chrX-44714812-DEL-282 | chrX-44714813-DEL-282 |
| chrX-44729066-DEL-7080 | chrX-44729067-DEL-7080 |
| chrX-44753202-DEL-9971 | chrX-44753204-DEL-9970 |
| chrX-44768575-DEL-199 | chrX-44768576-DEL-199 |
| chrX-44809688-DEL-2232 | chrX-44809689-DEL-2232 |
| chrX-44904448-DEL-1890 | chrX-44904449-DEL-1890 |
| chrX-44968239-DEL-58 | chrX-44968249-DEL-62 |
| chrX-44996244-DEL-60 | chrX-44996245-DEL-60 |
| chrX-45121328-DEL-129 | chrX-45121329-DEL-129 |
| chrX-45263926-DEL-51 | chrX-45263926-DEL-52 |
| chrX-45305019-DEL-2296 | chrX-45305014-DEL-2299 |
| chrX-45306238-DEL-1378 | chrX-45306239-DEL-1378 |
| chrX-45361576-DEL-6386 | chrX-45361577-DEL-6386 |
| chrX-45398986-DEL-3673 | chrX-45398987-DEL-3673 |
| chrX-45435051-DEL-100 | chrX-45435052-DEL-97 |
| chrX-45448761-DEL-3325 | chrX-45448762-DEL-3325 |
| chrX-45473206-DEL-323 | chrX-45473207-DEL-323 |
| chrX-45507012-DEL-1942 | chrX-45507013-DEL-1942 |
| chrX-45509206-DEL-224 | chrX-45509207-DEL-224 |
| chrX-45537167-DEL-626 | chrX-45537168-DEL-626 |
| chrX-45556782-DEL-526 | chrX-45556783-DEL-526 |
| chrX-45557845-DEL-67 | chrX-45557843-DEL-53 |
| chrX-45612630-DEL-6211 | chrX-45612631-DEL-6211 |
| chrX-45619886-DEL-697 | chrX-45619874-DEL-698 |
| chrX-45631291-DEL-6420 | chrX-45631292-DEL-6420 |
| chrX-45734440-DEL-356 | chrX-45734441-DEL-356 |
| chrX-45749358-DEL-5937 | chrX-45749359-DEL-5937 |
| chrX-45775133-DEL-1071 | chrX-45775134-DEL-1071 |
| chrX-45777945-DEL-722 | chrX-45777946-DEL-722 |
| chrX-45795346-DEL-3614 | chrX-45795347-DEL-3614 |
| chrX-45816532-DEL-451 | chrX-45816533-DEL-451 |
| chrX-45888137-DEL-6750 | chrX-45888138-DEL-6750 |
| chrX-45957505-DEL-58 | chrX-45957506-DEL-58 |
| chrX-46127923-DEL-65 | chrX-46127924-DEL-65 |
| chrX-46202784-DEL-1757 | chrX-46202785-DEL-1757 |
| chrX-46215871-DEL-229 | chrX-46215872-DEL-229 |
| chrX-46218686-DEL-7266 | chrX-46218687-DEL-7266 |
| chrX-46285823-DEL-1101 | chrX-46285819-DEL-1102 |
| chrX-46435999-DEL-3794 | chrX-46436000-DEL-3794 |
| chrX-46441507-DEL-819 | chrX-46441505-DEL-814 |
| chrX-46444675-DEL-6332 | chrX-46444676-DEL-6332 |
| chrX-46513307-DEL-208 | chrX-46513308-DEL-208 |
| chrX-46524763-DEL-7311 | chrX-46524764-DEL-7311 |
| chrX-46537750-DEL-2473 | chrX-46537752-DEL-2472 |
| chrX-46542834-DEL-358 | chrX-46542835-DEL-358 |
| chrX-46588915-DEL-54 | chrX-46588926-DEL-52 |
| chrX-46613749-DEL-1110 | chrX-46613750-DEL-1110 |
| chrX-46663297-DEL-103 | chrX-46663298-DEL-103 |
| chrX-46703693-DEL-400 | chrX-46703694-DEL-400 |
| chrX-46721862-DEL-1888 | chrX-46721863-DEL-1888 |
| chrX-46725764-DEL-197 | chrX-46725765-DEL-197 |
| chrX-46744037-DEL-681 | chrX-46744038-DEL-681 |
| chrX-46759396-DEL-84 | chrX-46759429-DEL-60 |
| chrX-46765244-DEL-891 | chrX-46765245-DEL-891 |
| chrX-46825283-DEL-6445 | chrX-46825284-DEL-6445 |
| chrX-46831735-DEL-63 | chrX-46831734-DEL-62 |
| chrX-46855463-DEL-250 | chrX-46855451-DEL-250 |
| chrX-46856206-DEL-1787 | chrX-46856207-DEL-1787 |
| chrX-46878381-DEL-6414 | chrX-46878382-DEL-6414 |
| chrX-46926824-DEL-2643 | chrX-46926825-DEL-2643 |
| chrX-46955242-DEL-7357 | chrX-46955243-DEL-7357 |
| chrX-47048573-DEL-212 | chrX-47048574-DEL-212 |
| chrX-47058762-DEL-348 | chrX-47058763-DEL-348 |
| chrX-47072146-DEL-55 | chrX-47072149-DEL-51 |
| chrX-47118403-DEL-204 | chrX-47118404-DEL-204 |
| chrX-47170186-DEL-221 | chrX-47170187-DEL-221 |
| chrX-47214475-DEL-108 | chrX-47214476-DEL-108 |
| chrX-47298849-DEL-203 | chrX-47298836-DEL-204 |
| chrX-47313317-DEL-72 | chrX-47313318-DEL-72 |
| chrX-47315981-DEL-156 | chrX-47315982-DEL-156 |
| chrX-47317150-DEL-238 | chrX-47317151-DEL-238 |
| chrX-47320953-DEL-54 | chrX-47320954-DEL-54 |
| chrX-47353982-DEL-354 | chrX-47353983-DEL-354 |
| chrX-47376832-DEL-76 | chrX-47376833-DEL-76 |
| chrX-47379892-DEL-165 | chrX-47379893-DEL-165 |
| chrX-47389525-DEL-1018 | chrX-47389526-DEL-1018 |
| chrX-47402387-DEL-146 | chrX-47402388-DEL-146 |
| chrX-47412281-DEL-1226 | chrX-47412282-DEL-1226 |
| chrX-47423132-DEL-202 | chrX-47423133-DEL-202 |
| chrX-47465559-DEL-252 | chrX-47465560-DEL-252 |

| | |
|---|---|
| chrX-47478671-DEL-93 | chrX-47478698-DEL-80 |
| chrX-47524768-DEL-52 | chrX-47524769-DEL-52 |
| chrX-47549187-DEL-82 | chrX-47549188-DEL-82 |
| chrX-47567055-DEL-1337 | chrX-47567056-DEL-1335 |
| chrX-47567900-DEL-209 | chrX-47567901-DEL-209 |
| chrX-47583734-DEL-51 | chrX-47583735-DEL-51 |
| chrX-47635920-DEL-68 | chrX-47635921-DEL-68 |
| chrX-47642129-DEL-70 | chrX-47642132-DEL-68 |
| chrX-47644065-DEL-205 | chrX-47644066-DEL-205 |
| chrX-47655238-DEL-88 | chrX-47655239-DEL-88 |
| chrX-47660293-DEL-422 | chrX-47660294-DEL-422 |
| chrX-47681863-DEL-74 | chrX-47681864-DEL-72 |
| chrX-47688411-DEL-182 | chrX-47688412-DEL-182 |
| chrX-47699981-DEL-128 | chrX-47699958-DEL-140 |
| chrX-47700663-DEL-80 | chrX-47700649-DEL-95 |
| chrX-47727250-DEL-164 | chrX-47727251-DEL-164 |
| chrX-47742557-DEL-112 | chrX-47742558-DEL-112 |
| chrX-47756012-DEL-209 | chrX-47756013-DEL-209 |
| chrX-47758223-DEL-207 | chrX-47758224-DEL-207 |
| chrX-47787844-DEL-6669 | chrX-47787845-DEL-6669 |
| chrX-47793984-DEL-212 | chrX-47793985-DEL-212 |
| chrX-47818485-DEL-90 | chrX-47818487-DEL-89 |
| chrX-47853191-DEL-120 | chrX-47853187-DEL-124 |
| chrX-47865592-DEL-180 | chrX-47865593-DEL-180 |
| chrX-47913980-DEL-681 | chrX-47913981-DEL-681 |
| chrX-47935400-DEL-199 | chrX-47935401-DEL-199 |
| chrX-47955620-DEL-102 | chrX-47955621-DEL-68 |
| chrX-48005082-DEL-61 | chrX-48005079-DEL-63 |
| chrX-48006856-DEL-2056 | chrX-48006857-DEL-2056 |
| chrX-48018448-DEL-417 | chrX-48018449-DEL-417 |
| chrX-48028191-DEL-123 | chrX-48028189-DEL-109 |
| chrX-48041325-DEL-71 | chrX-48041347-DEL-76 |
| chrX-48051840-DEL-8689 | chrX-48051841-DEL-8689 |
| chrX-48097836-DEL-150 | chrX-48097835-DEL-118 |
| chrX-48155258-DEL-56 | chrX-48155260-DEL-55 |
| chrX-48228188-DEL-2711 | chrX-48228189-DEL-2711 |
| chrX-48265170-DEL-1292 | chrX-48265171-DEL-1292 |
| chrX-48267108-DEL-99 | chrX-48267079-DEL-81 |
| chrX-48282540-DEL-77 | chrX-48282541-DEL-77 |
| chrX-48295437-DEL-2565 | chrX-48295438-DEL-2565 |
| chrX-48332210-DEL-51 | chrX-48332211-DEL-51 |
| chrX-48340495-DEL-145 | chrX-48340496-DEL-145 |
| chrX-48385256-DEL-7144 | chrX-48385257-DEL-7144 |
| chrX-48412568-DEL-341 | chrX-48412569-DEL-341 |
| chrX-48440506-DEL-13549 | chrX-48440507-DEL-13549 |
| chrX-48470073-DEL-65 | chrX-48470074-DEL-65 |
| chrX-48513927-DEL-7681 | chrX-48513941-DEL-7681 |
| chrX-48514321-DEL-1248 | chrX-48514322-DEL-1248 |
| chrX-48562185-DEL-90 | chrX-48562223-DEL-55 |
| chrX-48571527-DEL-137 | chrX-48571528-DEL-137 |
| chrX-48572510-DEL-66 | chrX-48572521-DEL-66 |
| chrX-48649962-DEL-6430 | chrX-48649963-DEL-6430 |
| chrX-48649973-DEL-56 | chrX-48649974-DEL-55 |
| chrX-48658255-DEL-284 | chrX-48658256-DEL-283 |
| chrX-48695336-DEL-7571 | chrX-48695337-DEL-7571 |
| chrX-48728987-DEL-2103 | chrX-48729024-DEL-2101 |
| chrX-48749694-DEL-730 | chrX-48749695-DEL-731 |
| chrX-48754452-DEL-1509 | chrX-48754452-DEL-1509 |
| chrX-48759762-DEL-382 | chrX-48759763-DEL-382 |
| chrX-48782689-DEL-6847 | chrX-48782690-DEL-6847 |
| chrX-48804506-DEL-1602 | chrX-48804507-DEL-1602 |
| chrX-48893966-DEL-323 | chrX-48893967-DEL-323 |
| chrX-48920476-DEL-624 | chrX-48920477-DEL-624 |
| chrX-48966093-DEL-58 | chrX-48966094-DEL-58 |
| chrX-49022163-DEL-54 | chrX-49022164-DEL-54 |
| chrX-49030240-DEL-1215 | chrX-49030241-DEL-1215 |
| chrX-49039242-DEL-6591 | chrX-49039243-DEL-6591 |
| chrX-49054638-DEL-3245 | chrX-49054639-DEL-3245 |
| chrX-49059763-DEL-55 | chrX-49059764-DEL-55 |
| chrX-49172236-DEL-7129 | chrX-49172237-DEL-7129 |
| chrX-49188346-DEL-98 | chrX-49188347-DEL-98 |
| chrX-49198427-DEL-385 | chrX-49198428-DEL-385 |
| chrX-49247370-DEL-4161 | chrX-49247370-DEL-4161 |
| chrX-49259865-DEL-2044 | chrX-49259865-DEL-2044 |
| chrX-49269110-DEL-5387 | chrX-49269111-DEL-5387 |
| chrX-49300081-DEL-29015 | chrX-49300073-DEL-29023 |
| chrX-49300544-DEL-6504 | chrX-49300545-DEL-6504 |
| chrX-49317022-DEL-1895 | chrX-49317023-DEL-1895 |
| chrX-49320215-DEL-171 | chrX-49320215-DEL-169 |
| chrX-49359050-DEL-241 | chrX-49359051-DEL-241 |

| | |
|---|---|
| chrX-49502998-DEL-4442 | chrX-49502999-DEL-4442 |
| chrX-49549102-DEL-6606 | chrX-49549103-DEL-6606 |
| chrX-49555353-DEL-201 | chrX-49555353-DEL-201 |
| chrX-49595544-DEL-3598 | chrX-49595545-DEL-3598 |
| chrX-49601119-DEL-198 | chrX-49601115-DEL-199 |
| chrX-49677525-DEL-203 | chrX-49677526-DEL-203 |
| chrX-49711647-DEL-207 | chrX-49711648-DEL-207 |
| chrX-49713269-DEL-87 | chrX-49713270-DEL-87 |
| chrX-49733847-DEL-415 | chrX-49733848-DEL-415 |
| chrX-49792774-DEL-1294 | chrX-49792775-DEL-1294 |
| chrX-49800020-DEL-505 | chrX-49800021-DEL-505 |
| chrX-49836307-DEL-4984 | chrX-49836308-DEL-4984 |
| chrX-49836541-DEL-193 | chrX-49836541-DEL-193 |
| chrX-49839088-DEL-320 | chrX-49839089-DEL-320 |
| chrX-49861621-DEL-213 | chrX-49861622-DEL-213 |
| chrX-49899272-DEL-7680 | chrX-49899259-DEL-7681 |
| chrX-49996281-DEL-555 | chrX-49996293-DEL-529 |
| chrX-50013829-DEL-107 | chrX-50013830-DEL-107 |
| chrX-50015475-DEL-59 | chrX-50015474-DEL-62 |
| chrX-50017682-DEL-76 | chrX-50017679-DEL-108 |
| chrX-50267994-DEL-85 | chrX-50267995-DEL-85 |
| chrX-50274838-DEL-217 | chrX-50274839-DEL-217 |
| chrX-50280102-DEL-70 | chrX-50280126-DEL-53 |
| chrX-50319780-DEL-110 | chrX-50319801-DEL-90 |
| chrX-50359474-DEL-6582 | chrX-50359474-DEL-6583 |
| chrX-50378194-DEL-1064 | chrX-50378195-DEL-1064 |
| chrX-50395738-DEL-54 | chrX-50395739-DEL-54 |
| chrX-50440179-DEL-66 | chrX-50440180-DEL-66 |
| chrX-50465982-DEL-436 | chrX-50465983-DEL-436 |
| chrX-50496646-DEL-108 | chrX-50496657-DEL-98 |
| chrX-50525568-DEL-6409 | chrX-50525556-DEL-6410 |
| chrX-50592340-DEL-60 | chrX-50592343-DEL-50 |
| chrX-50597273-DEL-69 | chrX-50597274-DEL-66 |
| chrX-50611802-DEL-86 | chrX-50611802-DEL-86 |
| chrX-50634426-DEL-52 | chrX-50634426-DEL-52 |
| chrX-50655534-DEL-6442 | chrX-50655535-DEL-6442 |
| chrX-50715364-DEL-534 | chrX-50715365-DEL-534 |
| chrX-50796113-DEL-105 | chrX-50796114-DEL-105 |
| chrX-50818400-DEL-6221 | chrX-50818385-DEL-6236 |
| chrX-50824934-DEL-113 | chrX-50824935-DEL-113 |
| chrX-50859651-DEL-1217 | chrX-50859651-DEL-1217 |
| chrX-50880680-DEL-59 | chrX-50880655-DEL-60 |
| chrX-50883912-DEL-201 | chrX-50883913-DEL-201 |
| chrX-50893113-DEL-857 | chrX-50893111-DEL-859 |
| chrX-50926065-DEL-96 | chrX-50926066-DEL-96 |
| chrX-51004999-DEL-2736 | chrX-51005000-DEL-2736 |
| chrX-51013340-DEL-9449 | chrX-51013341-DEL-9449 |
| chrX-51016079-DEL-3042 | chrX-51016080-DEL-3042 |
| chrX-51020389-DEL-169 | chrX-51020390-DEL-169 |
| chrX-51025801-DEL-54 | chrX-51025800-DEL-59 |
| chrX-51030764-DEL-386 | chrX-51030765-DEL-386 |
| chrX-51075876-DEL-1890 | chrX-51075877-DEL-1890 |
| chrX-51082964-DEL-4501 | chrX-51082965-DEL-4501 |
| chrX-51126288-DEL-1710 | chrX-51126289-DEL-1710 |
| chrX-51133939-DEL-448 | chrX-51133940-DEL-448 |
| chrX-51214683-DEL-76 | chrX-51214686-DEL-72 |
| chrX-51222422-DEL-197 | chrX-51222423-DEL-197 |
| chrX-51258923-DEL-547 | chrX-51258924-DEL-547 |
| chrX-51262011-DEL-113 | chrX-51262012-DEL-113 |
| chrX-51264530-DEL-197 | chrX-51264531-DEL-197 |
| chrX-51323766-DEL-104 | chrX-51323776-DEL-66 |
| chrX-51379786-DEL-75 | chrX-51379786-DEL-75 |
| chrX-51434977-DEL-194 | chrX-51434978-DEL-194 |
| chrX-51444300-DEL-6402 | chrX-51444301-DEL-6402 |
| chrX-51461946-DEL-210 | chrX-51461947-DEL-210 |
| chrX-51592220-DEL-72 | chrX-51592221-DEL-72 |
| chrX-51659578-DEL-224 | chrX-51659579-DEL-224 |
| chrX-51681004-DEL-121 | chrX-51681005-DEL-121 |
| chrX-51745743-DEL-70 | chrX-51745743-DEL-52 |
| chrX-51761546-DEL-197 | chrX-51761547-DEL-197 |
| chrX-51851072-DEL-157 | chrX-51851073-DEL-157 |
| chrX-51859235-DEL-736 | chrX-51859236-DEL-736 |
| chrX-51861155-DEL-494 | chrX-51861156-DEL-494 |
| chrX-51873689-DEL-293 | chrX-51873690-DEL-293 |
| chrX-51878880-DEL-486 | chrX-51878882-DEL-485 |
| chrX-51922392-DEL-551 | chrX-51922393-DEL-551 |
| chrX-51938376-DEL-106 | chrX-51938387-DEL-74 |
| chrX-51938761-DEL-1076 | chrX-51938762-DEL-1076 |
| chrX-51987422-DEL-986 | chrX-51987423-DEL-986 |
| chrX-52027929-DEL-172 | chrX-52027930-DEL-172 |

| | |
|---|---|
| chrX-52032339-DEL-732 | chrX-52032334-DEL-744 |
| chrX-52054772-DEL-176 | chrX-52054774-DEL-175 |
| chrX-52069822-DEL-229 | chrX-52069823-DEL-229 |
| chrX-52074896-DEL-62 | chrX-52074897-DEL-62 |
| chrX-52157208-DEL-63 | chrX-52157205-DEL-67 |
| chrX-52226313-DEL-52 | chrX-52226314-DEL-50 |
| chrX-52283904-DEL-489 | chrX-52283904-DEL-489 |
| chrX-52368225-DEL-86 | chrX-52368225-DEL-86 |
| chrX-52377486-DEL-51 | chrX-52377509-DEL-56 |
| chrX-52493916-DEL-1467 | chrX-52493917-DEL-1467 |
| chrX-52506349-DEL-252 | chrX-52506350-DEL-252 |
| chrX-52507211-DEL-943 | chrX-52507212-DEL-943 |
| chrX-52567346-DEL-5175 | chrX-52567347-DEL-5175 |
| chrX-52589936-DEL-56 | chrX-52589937-DEL-56 |
| chrX-52591984-DEL-789 | chrX-52591985-DEL-789 |
| chrX-52593587-DEL-60 | chrX-52593588-DEL-60 |
| chrX-52595761-DEL-150 | chrX-52595761-DEL-150 |
| chrX-52627072-DEL-56 | chrX-52627073-DEL-56 |
| chrX-52636229-DEL-99 | chrX-52636230-DEL-99 |
| chrX-52660413-DEL-6527 | chrX-52660411-DEL-6528 |
| chrX-52674663-DEL-60 | chrX-52674665-DEL-58 |
| chrX-52688135-DEL-369 | chrX-52688136-DEL-369 |
| chrX-52706182-DEL-108 | chrX-52706183-DEL-108 |
| chrX-52756629-DEL-1193 | chrX-52756630-DEL-1194 |
| chrX-52840481-DEL-80 | chrX-52840486-DEL-76 |
| chrX-52901392-DEL-3472 | chrX-52901392-DEL-3472 |
| chrX-52912970-DEL-3546 | chrX-52912970-DEL-3546 |
| chrX-53024738-DEL-1971 | chrX-53024739-DEL-1971 |
| chrX-53110340-DEL-3642 | chrX-53110341-DEL-3642 |
| chrX-53143516-DEL-171 | chrX-53143517-DEL-171 |
| chrX-53175088-DEL-89 | chrX-53175086-DEL-70 |
| chrX-53293797-DEL-332 | chrX-53293798-DEL-332 |
| chrX-54412510-DEL-1406 | chrX-54412511-DEL-1406 |
| chrX-54516328-DEL-664 | chrX-54516322-DEL-664 |
| chrX-54727095-DEL-4304 | chrX-54727096-DEL-4304 |
| chrX-54873918-DEL-68 | chrX-54873936-DEL-73 |
| chrX-54909198-DEL-688 | chrX-54909199-DEL-688 |
| chrX-54965893-DEL-10753 | chrX-54965894-DEL-10753 |
| chrX-55043982-DEL-5067 | chrX-55043983-DEL-5067 |
| chrX-55093238-DEL-126 | chrX-55093239-DEL-126 |
| chrX-55105699-DEL-5681 | chrX-55105700-DEL-5682 |
| chrX-55126961-DEL-6590 | chrX-55126962-DEL-6590 |
| chrX-55196276-DEL-398 | chrX-55196277-DEL-398 |
| chrX-55269044-DEL-2020 | chrX-55269054-DEL-2019 |
| chrX-55285856-DEL-214 | chrX-55285857-DEL-214 |
| chrX-55297311-DEL-90 | chrX-55297312-DEL-90 |
| chrX-55310451-DEL-3902 | chrX-55310452-DEL-3902 |
| chrX-55333984-DEL-740 | chrX-55333985-DEL-740 |
| chrX-55351991-DEL-359 | chrX-55351988-DEL-363 |
| chrX-55368775-DEL-89 | chrX-55368776-DEL-89 |
| chrX-55386045-DEL-206 | chrX-55386046-DEL-206 |
| chrX-55417578-DEL-207 | chrX-55417579-DEL-207 |
| chrX-55443124-DEL-216 | chrX-55443125-DEL-216 |
| chrX-55450342-DEL-325 | chrX-55450343-DEL-325 |
| chrX-55453885-DEL-935 | chrX-55453886-DEL-935 |
| chrX-55463476-DEL-204 | chrX-55463476-DEL-205 |
| chrX-55470479-DEL-113 | chrX-55470500-DEL-70 |
| chrX-55473567-DEL-402 | chrX-55473584-DEL-387 |
| chrX-55581653-DEL-207 | chrX-55581654-DEL-207 |
| chrX-55597930-DEL-66 | chrX-55597922-DEL-50 |
| chrX-55600587-DEL-55 | chrX-55600586-DEL-56 |
| chrX-55605703-DEL-288 | chrX-55605704-DEL-288 |
| chrX-55628112-DEL-833 | chrX-55628113-DEL-833 |
| chrX-55689932-DEL-722 | chrX-55689933-DEL-722 |
| chrX-55753693-DEL-62 | chrX-55753694-DEL-62 |
| chrX-55772123-DEL-887 | chrX-55772124-DEL-887 |
| chrX-55883042-DEL-58 | chrX-55883043-DEL-58 |
| chrX-55899475-DEL-507 | chrX-55899476-DEL-507 |
| chrX-55912431-DEL-205 | chrX-55912432-DEL-205 |
| chrX-55926172-DEL-6851 | chrX-55926173-DEL-6851 |
| chrX-55926186-DEL-424 | chrX-55926187-DEL-424 |
| chrX-55940646-DEL-1197 | chrX-55940659-DEL-1185 |
| chrX-55948419-DEL-741 | chrX-55948420-DEL-741 |
| chrX-56036319-DEL-81 | chrX-56036320-DEL-81 |
| chrX-56056120-DEL-1480 | chrX-56056118-DEL-1483 |
| chrX-56121009-DEL-226 | chrX-56121010-DEL-226 |
| chrX-56154533-DEL-52 | chrX-56154534-DEL-52 |
| chrX-56165540-DEL-414 | chrX-56165541-DEL-414 |
| chrX-56169730-DEL-1005 | chrX-56169731-DEL-1005 |
| chrX-56214407-DEL-3095 | chrX-56214408-DEL-3095 |

| | |
|---|---|
| chrX-56216092-DEL-69 | chrX-56216093-DEL-69 |
| chrX-56243001-DEL-66 | chrX-56243002-DEL-66 |
| chrX-56245095-DEL-97 | chrX-56245136-DEL-73 |
| chrX-56344167-DEL-68 | chrX-56344168-DEL-68 |
| chrX-56361783-DEL-1349 | chrX-56361784-DEL-1349 |
| chrX-56364955-DEL-990 | chrX-56364980-DEL-974 |
| chrX-56495555-DEL-1560 | chrX-56495556-DEL-1560 |
| chrX-56510186-DEL-385 | chrX-56510187-DEL-385 |
| chrX-56515478-DEL-114 | chrX-56515479-DEL-114 |
| chrX-56580391-DEL-60 | chrX-56580392-DEL-60 |
| chrX-56588823-DEL-6580 | chrX-56588824-DEL-6580 |
| chrX-56613869-DEL-3464 | chrX-56613870-DEL-3464 |
| chrX-56618885-DEL-7197 | chrX-56618886-DEL-7197 |
| chrX-56643680-DEL-7113 | chrX-56643681-DEL-7113 |
| chrX-56651089-DEL-555 | chrX-56651090-DEL-555 |
| chrX-56688077-DEL-6243 | chrX-56688078-DEL-6243 |
| chrX-56704731-DEL-196 | chrX-56704732-DEL-196 |
| chrX-56744224-DEL-135 | chrX-56744225-DEL-135 |
| chrX-56795243-DEL-233 | chrX-56795244-DEL-233 |
| chrX-56802083-DEL-1747 | chrX-56802084-DEL-1747 |
| chrX-56818746-DEL-233 | chrX-56818747-DEL-233 |
| chrX-56838856-DEL-98 | chrX-56838861-DEL-94 |
| chrX-56864756-DEL-4924 | chrX-56864757-DEL-4924 |
| chrX-56878870-DEL-54 | chrX-56878873-DEL-52 |
| chrX-56898940-DEL-2217 | chrX-56898941-DEL-2213 |
| chrX-56916840-DEL-54 | chrX-56916882-DEL-52 |
| chrX-56928270-DEL-149 | chrX-56928265-DEL-155 |
| chrX-57017764-DEL-218 | chrX-57017765-DEL-218 |
| chrX-57039000-DEL-162 | chrX-57038968-DEL-167 |
| chrX-57044417-DEL-1679 | chrX-57044418-DEL-1679 |
| chrX-57050866-DEL-1959 | chrX-57050866-DEL-1959 |
| chrX-57069292-DEL-80 | chrX-57069288-DEL-50 |
| chrX-57104295-DEL-231 | chrX-57104281-DEL-218 |
| chrX-57123099-DEL-92 | chrX-57123119-DEL-106 |
| chrX-57155284-DEL-52 | chrX-57155284-DEL-56 |
| chrX-57204906-DEL-76 | chrX-57204923-DEL-50 |
| chrX-57252313-DEL-13181 | chrX-57252314-DEL-13181 |
| chrX-57302589-DEL-6436 | chrX-57302590-DEL-6436 |
| chrX-57333787-DEL-6287 | chrX-57333788-DEL-6287 |
| chrX-57347714-DEL-403 | chrX-57347715-DEL-403 |
| chrX-57384325-DEL-1568 | chrX-57384325-DEL-1568 |
| chrX-57403961-DEL-7191 | chrX-57403962-DEL-7191 |
| chrX-57428998-DEL-6378 | chrX-57428999-DEL-6378 |
| chrX-57457308-DEL-5219 | chrX-57457309-DEL-5219 |
| chrX-57498918-DEL-166 | chrX-57498919-DEL-166 |
| chrX-57520309-DEL-3564 | chrX-57520310-DEL-3564 |
| chrX-57639471-DEL-186 | chrX-57639472-DEL-186 |
| chrX-57643191-DEL-193 | chrX-57643189-DEL-196 |
| chrX-57644414-DEL-198 | chrX-57644415-DEL-198 |
| chrX-57674573-DEL-2195 | chrX-57674572-DEL-2196 |
| chrX-57687828-DEL-63 | chrX-57687826-DEL-75 |
| chrX-57725139-DEL-205 | chrX-57725140-DEL-205 |
| chrX-57764949-DEL-98 | chrX-57764950-DEL-98 |
| chrX-57839798-DEL-201 | chrX-57839798-DEL-201 |
| chrX-57840483-DEL-6475 | chrX-57840484-DEL-6475 |
| chrX-57847986-DEL-68 | chrX-57847978-DEL-76 |
| chrX-57850755-DEL-1853 | chrX-57850756-DEL-1853 |
| chrX-57890099-DEL-2231 | chrX-57890100-DEL-2231 |
| chrX-57906977-DEL-6653 | chrX-57906978-DEL-6653 |
| chrX-57926000-DEL-491 | chrX-57926001-DEL-491 |
| chrX-57959088-DEL-4385 | chrX-57959090-DEL-4365 |
| chrX-58034869-DEL-89 | chrX-58034870-DEL-89 |
| chrX-58035299-DEL-829 | chrX-58035300-DEL-829 |
| chrX-58082166-DEL-55 | chrX-58082167-DEL-55 |
| chrX-58118875-DEL-52 | chrX-58118866-DEL-50 |
| chrX-58127488-DEL-58 | chrX-58127489-DEL-58 |
| chrX-58164986-DEL-1000 | chrX-58164987-DEL-1000 |
| chrX-58165796-DEL-217 | chrX-58165797-DEL-217 |
| chrX-58216902-DEL-62 | chrX-58216890-DEL-50 |
| chrX-58253539-DEL-60 | chrX-58253529-DEL-51 |
| chrX-58257295-DEL-350 | chrX-58257296-DEL-350 |
| chrX-58302043-DEL-1021 | chrX-58302044-DEL-1021 |
| chrX-58310855-DEL-106 | chrX-58310856-DEL-106 |
| chrX-58329213-DEL-7024 | chrX-58329214-DEL-7024 |
| chrX-58329914-DEL-204 | chrX-58329915-DEL-204 |
| chrX-58386037-DEL-301 | chrX-58386040-DEL-298 |
| chrX-58400246-DEL-212 | chrX-58400247-DEL-212 |
| chrX-58413979-DEL-951 | chrX-58413980-DEL-951 |
| chrX-58458596-DEL-84 | chrX-58458597-DEL-84 |
| chrX-58505466-DEL-6444 | chrX-58505467-DEL-6444 |

| | |
|---|---|
| chrX-58538172-DEL-96 | chrX-58538173-DEL-96 |
| chrX-58547075-DEL-324 | chrX-58547076-DEL-324 |
| chrX-58549191-DEL-139 | chrX-58549192-DEL-137 |
| chrX-58683627-DEL-6662 | chrX-58683628-DEL-6662 |
| chrX-58694678-DEL-1121 | chrX-58694679-DEL-1120 |
| chrX-58750812-DEL-66 | chrX-58750812-DEL-66 |
| chrX-58768028-DEL-6202 | chrX-58768029-DEL-6202 |
| chrX-58803522-DEL-674 | chrX-58803523-DEL-674 |
| chrX-58832737-DEL-9562 | chrX-58832738-DEL-9562 |
| chrX-58853814-DEL-62 | chrX-58853815-DEL-62 |
| chrX-58873825-DEL-635 | chrX-58873826-DEL-635 |
| chrX-58894977-DEL-6392 | chrX-58894976-DEL-6392 |
| chrX-58900022-DEL-52 | chrX-58900023-DEL-52 |
| chrX-58974576-DEL-226 | chrX-58974577-DEL-226 |
| chrX-59040676-DEL-195 | chrX-59040677-DEL-195 |
| chrX-59081424-DEL-1078 | chrX-59081425-DEL-1078 |
| chrX-59093664-DEL-745 | chrX-59093665-DEL-745 |
| chrX-59098943-DEL-60 | chrX-59098920-DEL-50 |
| chrX-59138415-DEL-144 | chrX-59138407-DEL-153 |
| chrX-59140670-DEL-1751 | chrX-59140670-DEL-1751 |
| chrX-59147733-DEL-196 | chrX-59147734-DEL-196 |
| chrX-59164665-DEL-386 | chrX-59164666-DEL-386 |
| chrX-59205752-DEL-114 | chrX-59205753-DEL-114 |
| chrX-59234110-DEL-99 | chrX-59234099-DEL-100 |
| chrX-59246204-DEL-6498 | chrX-59246205-DEL-6498 |
| chrX-59255281-DEL-72 | chrX-59255282-DEL-72 |
| chrX-59309231-DEL-1005 | chrX-59309232-DEL-1005 |
| chrX-59403789-DEL-116 | chrX-59403790-DEL-116 |
| chrX-59488698-DEL-582 | chrX-59488699-DEL-582 |
| chrX-59506450-DEL-102 | chrX-59506451-DEL-102 |
| chrX-59516688-DEL-52 | chrX-59516689-DEL-52 |
| chrX-59521162-DEL-464 | chrX-59521163-DEL-464 |
| chrX-59529166-DEL-100 | chrX-59529167-DEL-100 |
| chrX-59530178-DEL-58 | chrX-59530149-DEL-67 |
| chrX-59530955-DEL-392 | chrX-59530956-DEL-392 |
| chrX-59603274-DEL-2264 | chrX-59603275-DEL-2264 |
| chrX-59629698-DEL-571 | chrX-59629699-DEL-571 |
| chrX-59635825-DEL-133 | chrX-59635821-DEL-125 |
| chrX-59643247-DEL-1274 | chrX-59643248-DEL-1274 |
| chrX-59661996-DEL-6652 | chrX-59661997-DEL-6652 |
| chrX-59709031-DEL-6172 | chrX-59709032-DEL-6172 |
| chrX-59731718-DEL-56 | chrX-59731725-DEL-50 |
| chrX-59800461-DEL-141 | chrX-59800472-DEL-124 |
| chrX-59831804-DEL-119 | chrX-59831839-DEL-118 |
| chrX-59898655-DEL-54 | chrX-59898644-DEL-67 |
| chrX-59906091-DEL-87 | chrX-59906092-DEL-87 |
| chrX-59913250-DEL-57 | chrX-59913251-DEL-57 |
| chrX-59917177-DEL-56 | chrX-59917175-DEL-56 |
| chrX-59949393-DEL-221 | chrX-59949394-DEL-221 |
| chrX-59969333-DEL-397 | chrX-59969334-DEL-397 |
| chrX-59993603-DEL-91 | chrX-59993604-DEL-91 |
| chrX-60005269-DEL-214 | chrX-60005269-DEL-215 |
| chrX-60051541-DEL-60 | chrX-60051542-DEL-60 |
| chrX-60094074-DEL-73 | chrX-60094074-DEL-78 |
| chrX-60102771-DEL-4166 | chrX-60102772-DEL-4166 |
| chrX-60110314-DEL-2117 | chrX-60110314-DEL-2117 |
| chrX-60258714-DEL-181 | chrX-60258715-DEL-181 |
| chrX-60266506-DEL-112 | chrX-60266546-DEL-72 |
| chrX-60277373-DEL-123 | chrX-60277373-DEL-120 |
| chrX-60295421-DEL-7102 | chrX-60295422-DEL-7102 |
| chrX-60302782-DEL-7302 | chrX-60302782-DEL-7302 |
| chrX-60394671-DEL-1326 | chrX-60394672-DEL-1326 |
| chrX-60425969-DEL-1049 | chrX-60425970-DEL-1049 |
| chrX-60428442-DEL-95 | chrX-60428435-DEL-96 |
| chrX-60483650-DEL-6808 | chrX-60483651-DEL-6808 |
| chrX-60520530-DEL-613 | chrX-60520529-DEL-614 |
| chrX-60735655-DEL-218 | chrX-60735656-DEL-218 |
| chrX-60811338-DEL-4341 | chrX-60811339-DEL-4341 |
| chrX-60846251-DEL-68 | chrX-60846246-DEL-60 |
| chrX-60866291-DEL-764 | chrX-60866292-DEL-764 |
| chrX-60895705-DEL-61 | chrX-60895706-DEL-61 |
| chrX-60908273-DEL-9453 | chrX-60908274-DEL-9453 |
| chrX-60953934-DEL-124 | chrX-60953944-DEL-118 |
| chrX-60982479-DEL-4860 | chrX-60982480-DEL-4860 |
| chrX-61006029-DEL-7201 | chrX-61006030-DEL-7201 |
| chrX-61061260-DEL-2322 | chrX-61061261-DEL-2322 |
| chrX-61064349-DEL-79 | chrX-61064327-DEL-56 |
| chrX-61070167-DEL-936 | chrX-61070168-DEL-941 |
| chrX-61080727-DEL-75 | chrX-61080728-DEL-75 |
| chrX-61090937-DEL-109 | chrX-61090930-DEL-100 |

| | |
|---|---|
| chrX-61121359-DEL-7220 | chrX-61121360-DEL-7220 |
| chrX-61151229-DEL-201 | chrX-61151204-DEL-201 |
| chrX-61240890-DEL-6973 | chrX-61240891-DEL-6973 |
| chrX-61241391-DEL-221 | chrX-61241391-DEL-221 |
| chrX-61254941-DEL-198 | chrX-61254942-DEL-198 |
| chrX-61320641-DEL-95 | chrX-61320642-DEL-95 |
| chrX-61340964-DEL-80 | chrX-61340965-DEL-80 |
| chrX-61395582-DEL-81 | chrX-61395583-DEL-81 |
| chrX-61396641-DEL-174 | chrX-61396642-DEL-174 |
| chrX-61407747-DEL-1851 | chrX-61407748-DEL-1851 |
| chrX-61459208-DEL-58 | chrX-61459214-DEL-53 |
| chrX-61492718-DEL-420 | chrX-61492719-DEL-421 |
| chrX-61534100-DEL-70 | chrX-61534109-DEL-50 |
| chrX-61535198-DEL-545 | chrX-61535199-DEL-545 |
| chrX-61535465-DEL-811 | chrX-61535466-DEL-811 |
| chrX-61536564-DEL-88 | chrX-61536565-DEL-88 |
| chrX-61542306-DEL-54 | chrX-61542307-DEL-54 |
| chrX-61546992-DEL-901 | chrX-61546993-DEL-901 |
| chrX-61548200-DEL-189 | chrX-61548201-DEL-189 |
| chrX-61624012-DEL-63 | chrX-61624013-DEL-63 |
| chrX-61673836-DEL-212 | chrX-61673835-DEL-212 |
| chrX-61691420-DEL-3543 | chrX-61691422-DEL-3527 |
| chrX-61715448-DEL-52 | chrX-61715449-DEL-52 |
| chrX-61733841-DEL-111 | chrX-61733842-DEL-111 |
| chrX-61786939-DEL-70 | chrX-61786934-DEL-91 |
| chrX-61791078-DEL-6938 | chrX-61791079-DEL-6937 |
| chrX-61791173-DEL-208 | chrX-61791174-DEL-208 |
| chrX-61796162-DEL-563 | chrX-61796163-DEL-563 |
| chrX-61814525-DEL-5824 | chrX-61814526-DEL-5824 |
| chrX-61884224-DEL-6620 | chrX-61884225-DEL-6620 |
| chrX-61884392-DEL-212 | chrX-61884393-DEL-212 |
| chrX-61909160-DEL-6175 | chrX-61909161-DEL-6175 |
| chrX-61920767-DEL-107 | chrX-61920767-DEL-98 |
| chrX-61923980-DEL-112 | chrX-61923999-DEL-72 |
| chrX-61925832-DEL-677 | chrX-61925833-DEL-677 |
| chrX-61955146-DEL-1873 | chrX-61955147-DEL-1873 |
| chrX-62018214-DEL-113 | chrX-62018215-DEL-113 |
| chrX-62025524-DEL-78 | chrX-62025525-DEL-78 |
| chrX-62164033-DEL-6518 | chrX-62164034-DEL-6518 |
| chrX-62183016-DEL-1250 | chrX-62183016-DEL-1250 |
| chrX-62184904-DEL-5119 | chrX-62184905-DEL-5119 |
| chrX-62190974-DEL-1422 | chrX-62190975-DEL-1422 |
| chrX-62201736-DEL-106 | chrX-62201737-DEL-106 |
| chrX-62238142-DEL-945 | chrX-62238143-DEL-945 |
| chrX-62260477-DEL-183 | chrX-62260478-DEL-183 |
| chrX-62281145-DEL-3042 | chrX-62281146-DEL-3042 |
| chrX-62318131-DEL-571 | chrX-62318132-DEL-571 |
| chrX-62329129-DEL-235 | chrX-62329130-DEL-235 |
| chrX-62335943-DEL-96 | chrX-62335943-DEL-96 |
| chrX-62357975-DEL-202 | chrX-62357986-DEL-203 |
| chrX-62413180-DEL-53 | chrX-62413181-DEL-53 |
| chrX-62428324-DEL-1849 | chrX-62428325-DEL-1849 |
| chrX-62452352-DEL-140 | chrX-62452353-DEL-140 |
| chrX-62473351-DEL-402 | chrX-62473352-DEL-401 |
| chrX-62490654-DEL-6260 | chrX-62490640-DEL-6261 |
| chrX-62502150-DEL-166 | chrX-62502169-DEL-169 |
| chrX-62515586-DEL-2456 | chrX-62515587-DEL-2455 |
| chrX-62552697-DEL-6799 | chrX-62552716-DEL-6784 |
| chrX-62597304-DEL-1882 | chrX-62597305-DEL-1882 |
| chrX-62619034-DEL-3921 | chrX-62619035-DEL-3921 |
| chrX-62631069-DEL-1602 | chrX-62631070-DEL-1602 |
| chrX-62703233-DEL-85 | chrX-62703234-DEL-85 |
| chrX-62792876-DEL-82 | chrX-62792885-DEL-54 |
| chrX-62803639-DEL-579 | chrX-62803640-DEL-579 |
| chrX-62836519-DEL-964 | chrX-62836503-DEL-980 |
| chrX-62843685-DEL-4141 | chrX-62843686-DEL-4141 |
| chrX-62857147-DEL-70 | chrX-62857150-DEL-54 |
| chrX-62864291-DEL-870 | chrX-62864292-DEL-870 |
| chrX-62898771-DEL-66 | chrX-62898772-DEL-66 |
| chrX-62911372-DEL-72 | chrX-62911370-DEL-67 |
| chrX-62914531-DEL-54 | chrX-62914532-DEL-54 |
| chrX-62936424-DEL-206 | chrX-62936401-DEL-207 |
| chrX-62975475-DEL-198 | chrX-62975476-DEL-198 |
| chrX-62981390-DEL-311 | chrX-62981391-DEL-311 |
| chrX-62989545-DEL-6637 | chrX-62989546-DEL-6637 |
| chrX-62989643-DEL-212 | chrX-62989644-DEL-212 |
| chrX-63074047-DEL-620 | chrX-63074047-DEL-620 |
| chrX-63086580-DEL-291 | chrX-63086581-DEL-291 |
| chrX-63169573-DEL-6267 | chrX-63169574-DEL-6267 |
| chrX-63191320-DEL-129 | chrX-63191321-DEL-129 |

| | |
|---|---|
| chrX-63269805-DEL-72 | chrX-63269802-DEL-56 |
| chrX-63304094-DEL-1219 | chrX-63304095-DEL-1219 |
| chrX-63312435-DEL-63 | chrX-63312436-DEL-63 |
| chrX-63464591-DEL-71 | chrX-63464592-DEL-57 |
| chrX-63514733-DEL-121 | chrX-63514734-DEL-121 |
| chrX-63532764-DEL-6778 | chrX-63532765-DEL-6778 |
| chrX-63598937-DEL-6464 | chrX-63598938-DEL-6464 |
| chrX-63627717-DEL-643 | chrX-63627718-DEL-643 |
| chrX-63669597-DEL-7377 | chrX-63669598-DEL-7377 |
| chrX-63675782-DEL-206 | chrX-63675783-DEL-206 |
| chrX-63791921-DEL-187 | chrX-63791922-DEL-187 |
| chrX-63797297-DEL-2755 | chrX-63797298-DEL-2755 |
| chrX-63927303-DEL-6795 | chrX-63927304-DEL-6795 |
| chrX-63955087-DEL-334 | chrX-63955088-DEL-334 |
| chrX-63965465-DEL-204 | chrX-63965466-DEL-204 |
| chrX-63976703-DEL-5583 | chrX-63976704-DEL-5583 |
| chrX-64039596-DEL-6605 | chrX-64039597-DEL-6605 |
| chrX-64045736-DEL-208 | chrX-64045737-DEL-208 |
| chrX-64125981-DEL-562 | chrX-64125982-DEL-562 |
| chrX-64183837-DEL-52 | chrX-64183803-DEL-50 |
| chrX-64196650-DEL-7432 | chrX-64196651-DEL-7432 |
| chrX-64216435-DEL-1951 | chrX-64216436-DEL-1951 |
| chrX-64256451-DEL-132 | chrX-64256452-DEL-132 |
| chrX-64296543-DEL-63 | chrX-64296544-DEL-63 |
| chrX-64330406-DEL-80 | chrX-64330423-DEL-60 |
| chrX-64367684-DEL-57 | chrX-64367678-DEL-65 |
| chrX-64390557-DEL-852 | chrX-64390558-DEL-852 |
| chrX-64410593-DEL-102 | chrX-64410594-DEL-102 |
| chrX-64435530-DEL-67 | chrX-64435511-DEL-65 |
| chrX-64450192-DEL-6796 | chrX-64450193-DEL-6796 |
| chrX-64450223-DEL-623 | chrX-64450206-DEL-624 |
| chrX-64457225-DEL-123 | chrX-64457226-DEL-123 |
| chrX-64466371-DEL-72 | chrX-64466390-DEL-54 |
| chrX-64513498-DEL-285 | chrX-64513499-DEL-285 |
| chrX-64583549-DEL-6865 | chrX-64583550-DEL-6865 |
| chrX-64710303-DEL-7467 | chrX-64710304-DEL-7467 |
| chrX-64710316-DEL-1357 | chrX-64710317-DEL-1349 |
| chrX-64750524-DEL-7069 | chrX-64750525-DEL-7069 |
| chrX-64758950-DEL-58 | chrX-64758951-DEL-58 |
| chrX-64790017-DEL-1956 | chrX-64790018-DEL-1956 |
| chrX-64818650-DEL-52 | chrX-64818650-DEL-52 |
| chrX-64825211-DEL-1891 | chrX-64825212-DEL-1891 |
| chrX-64862491-DEL-6588 | chrX-64862492-DEL-6588 |
| chrX-64870185-DEL-188 | chrX-64870186-DEL-188 |
| chrX-64894095-DEL-3649 | chrX-64894096-DEL-3649 |
| chrX-64902666-DEL-64 | chrX-64902667-DEL-64 |
| chrX-64913946-DEL-66 | chrX-64913995-DEL-54 |
| chrX-64921583-DEL-266 | chrX-64921584-DEL-266 |
| chrX-64935503-DEL-627 | chrX-64935510-DEL-621 |
| chrX-65018496-DEL-6989 | chrX-65018497-DEL-6989 |
| chrX-65027414-DEL-10482 | chrX-65027417-DEL-10480 |
| chrX-65079355-DEL-139 | chrX-65079356-DEL-139 |
| chrX-65111415-DEL-72 | chrX-65111388-DEL-64 |
| chrX-65112610-DEL-992 | chrX-65112611-DEL-992 |
| chrX-65121074-DEL-6568 | chrX-65121075-DEL-6568 |
| chrX-65142880-DEL-324 | chrX-65142880-DEL-323 |
| chrX-65183687-DEL-5443 | chrX-65183688-DEL-5443 |
| chrX-65218149-DEL-2794 | chrX-65218150-DEL-2794 |
| chrX-65252110-DEL-234 | chrX-65252111-DEL-234 |
| chrX-65300632-DEL-186 | chrX-65300633-DEL-186 |
| chrX-65328204-DEL-415 | chrX-65328205-DEL-415 |
| chrX-65374914-DEL-6453 | chrX-65374915-DEL-6453 |
| chrX-65495522-DEL-6561 | chrX-65495523-DEL-6561 |
| chrX-65530466-DEL-1826 | chrX-65530467-DEL-1826 |
| chrX-65591089-DEL-80 | chrX-65591102-DEL-68 |
| chrX-65591275-DEL-675 | chrX-65591276-DEL-675 |
| chrX-65607786-DEL-300 | chrX-65607787-DEL-300 |
| chrX-65608848-DEL-329 | chrX-65608849-DEL-329 |
| chrX-65612110-DEL-1628 | chrX-65612111-DEL-1628 |
| chrX-65637506-DEL-526 | chrX-65637507-DEL-526 |
| chrX-65652224-DEL-6627 | chrX-65652225-DEL-6627 |
| chrX-65672888-DEL-81 | chrX-65672889-DEL-75 |
| chrX-65762219-DEL-104 | chrX-65762220-DEL-104 |
| chrX-65768065-DEL-193 | chrX-65768066-DEL-193 |
| chrX-65801880-DEL-4604 | chrX-65801881-DEL-4604 |
| chrX-65822019-DEL-463 | chrX-65822024-DEL-457 |
| chrX-65826669-DEL-2995 | chrX-65826670-DEL-2995 |
| chrX-65867475-DEL-13958 | chrX-65867475-DEL-13958 |
| chrX-65894485-DEL-54 | chrX-65894486-DEL-54 |
| chrX-65936297-DEL-54 | chrX-65936296-DEL-68 |

| | |
|---|---|
| chrX-65946048-DEL-161 | chrX-65946049-DEL-161 |
| chrX-65992904-DEL-715 | chrX-65992905-DEL-715 |
| chrX-66010694-DEL-102 | chrX-66010701-DEL-96 |
| chrX-66047858-DEL-54 | chrX-66047859-DEL-72 |
| chrX-66057386-DEL-424 | chrX-66057386-DEL-424 |
| chrX-66091298-DEL-4434 | chrX-66091299-DEL-4434 |
| chrX-66119583-DEL-425 | chrX-66119584-DEL-425 |
| chrX-66145703-DEL-60 | chrX-66145703-DEL-60 |
| chrX-66186929-DEL-6825 | chrX-66186929-DEL-6825 |
| chrX-66223099-DEL-118 | chrX-66223100-DEL-118 |
| chrX-66268101-DEL-5154 | chrX-66268102-DEL-5154 |
| chrX-66268324-DEL-1518 | chrX-66268325-DEL-1518 |
| chrX-66268974-DEL-212 | chrX-66268975-DEL-212 |
| chrX-66305451-DEL-1297 | chrX-66305452-DEL-1297 |
| chrX-66351588-DEL-1295 | chrX-66351589-DEL-1295 |
| chrX-66373734-DEL-120 | chrX-66373735-DEL-120 |
| chrX-66386250-DEL-401 | chrX-66386249-DEL-401 |
| chrX-66417043-DEL-7101 | chrX-66417044-DEL-7101 |
| chrX-66456687-DEL-531 | chrX-66456686-DEL-532 |
| chrX-66475238-DEL-1977 | chrX-66475239-DEL-1977 |
| chrX-66494986-DEL-2620 | chrX-66494987-DEL-2620 |
| chrX-66508640-DEL-6472 | chrX-66508641-DEL-6472 |
| chrX-66583179-DEL-2821 | chrX-66583180-DEL-2821 |
| chrX-66587323-DEL-1892 | chrX-66587324-DEL-1892 |
| chrX-66762260-DEL-201 | chrX-66762261-DEL-201 |
| chrX-66860908-DEL-117 | chrX-66860891-DEL-99 |
| chrX-67063049-DEL-342 | chrX-67063049-DEL-342 |
| chrX-67074029-DEL-4837 | chrX-67074030-DEL-4837 |
| chrX-67100898-DEL-109 | chrX-67100899-DEL-109 |
| chrX-67104448-DEL-66 | chrX-67104433-DEL-102 |
| chrX-67119983-DEL-233 | chrX-67119966-DEL-225 |
| chrX-67132175-DEL-62 | chrX-67132176-DEL-62 |
| chrX-67145337-DEL-199 | chrX-67145328-DEL-201 |
| chrX-67220488-DEL-6428 | chrX-67220489-DEL-6428 |
| chrX-67277142-DEL-61 | chrX-67277142-DEL-61 |
| chrX-67308684-DEL-1998 | chrX-67308685-DEL-1998 |
| chrX-67314678-DEL-91 | chrX-67314728-DEL-55 |
| chrX-67345254-DEL-1520 | chrX-67345255-DEL-1520 |
| chrX-67364291-DEL-55 | chrX-67364292-DEL-55 |
| chrX-67383089-DEL-6452 | chrX-67383090-DEL-6452 |
| chrX-67391244-DEL-243 | chrX-67391245-DEL-243 |
| chrX-67396401-DEL-93 | chrX-67396402-DEL-93 |
| chrX-67397828-DEL-2857 | chrX-67397829-DEL-2857 |
| chrX-67407005-DEL-76 | chrX-67407027-DEL-60 |
| chrX-67409248-DEL-7747 | chrX-67409213-DEL-7748 |
| chrX-67411091-DEL-3588 | chrX-67411092-DEL-3588 |
| chrX-67441871-DEL-2830 | chrX-67441872-DEL-2830 |
| chrX-67483756-DEL-1148 | chrX-67483757-DEL-1148 |
| chrX-67560273-DEL-6395 | chrX-67560274-DEL-6395 |
| chrX-67584224-DEL-1770 | chrX-67584225-DEL-1770 |
| chrX-67622339-DEL-530 | chrX-67622340-DEL-530 |
| chrX-67650941-DEL-6219 | chrX-67650942-DEL-6219 |
| chrX-67767468-DEL-208 | chrX-67767473-DEL-203 |
| chrX-67768232-DEL-2264 | chrX-67768232-DEL-2262 |
| chrX-67770962-DEL-109 | chrX-67770963-DEL-109 |
| chrX-67814379-DEL-205 | chrX-67814380-DEL-205 |
| chrX-67838521-DEL-110 | chrX-67838536-DEL-95 |
| chrX-67843430-DEL-496 | chrX-67843431-DEL-496 |
| chrX-67853283-DEL-96 | chrX-67853284-DEL-96 |
| chrX-67864688-DEL-4249 | chrX-67864689-DEL-4249 |
| chrX-67917357-DEL-6214 | chrX-67917358-DEL-6214 |
| chrX-67921139-DEL-2122 | chrX-67921140-DEL-2122 |
| chrX-67954223-DEL-175 | chrX-67954224-DEL-175 |
| chrX-67992419-DEL-478 | chrX-67992420-DEL-478 |
| chrX-68012660-DEL-3071 | chrX-68012661-DEL-3071 |
| chrX-68035223-DEL-2782 | chrX-68035209-DEL-2797 |
| chrX-68044844-DEL-1446 | chrX-68044845-DEL-1446 |
| chrX-68046631-DEL-235 | chrX-68046632-DEL-235 |
| chrX-68055180-DEL-2724 | chrX-68055181-DEL-2724 |
| chrX-68061479-DEL-6720 | chrX-68061480-DEL-6720 |
| chrX-68090601-DEL-58 | chrX-68090602-DEL-57 |
| chrX-68091572-DEL-80 | chrX-68091573-DEL-80 |
| chrX-68096415-DEL-54 | chrX-68096416-DEL-54 |
| chrX-68107362-DEL-786 | chrX-68107349-DEL-768 |
| chrX-68112213-DEL-2624 | chrX-68112214-DEL-2624 |
| chrX-68139498-DEL-144 | chrX-68139499-DEL-144 |
| chrX-68210543-DEL-3975 | chrX-68210544-DEL-3975 |
| chrX-68223994-DEL-6317 | chrX-68223995-DEL-6317 |
| chrX-68224035-DEL-200 | chrX-68224027-DEL-201 |
| chrX-68251042-DEL-698 | chrX-68251043-DEL-698 |

| | |
|---|---|
| chrX-68380559-DEL-449 | chrX-68380560-DEL-449 |
| chrX-68448507-DEL-2610 | chrX-68448508-DEL-2610 |
| chrX-68467390-DEL-879 | chrX-68467376-DEL-893 |
| chrX-68507996-DEL-6201 | chrX-68507997-DEL-6201 |
| chrX-68522387-DEL-52 | chrX-68522388-DEL-52 |
| chrX-68600427-DEL-975 | chrX-68600427-DEL-975 |
| chrX-68601425-DEL-348 | chrX-68601425-DEL-348 |
| chrX-68616481-DEL-6838 | chrX-68616482-DEL-6838 |
| chrX-68634654-DEL-66 | chrX-68634654-DEL-66 |
| chrX-68644047-DEL-2084 | chrX-68644048-DEL-2084 |
| chrX-68648480-DEL-1413 | chrX-68648481-DEL-1413 |
| chrX-68673152-DEL-6868 | chrX-68673153-DEL-6868 |
| chrX-68679161-DEL-424 | chrX-68679162-DEL-424 |
| chrX-68697724-DEL-282 | chrX-68697725-DEL-282 |
| chrX-68700310-DEL-291 | chrX-68700311-DEL-291 |
| chrX-68760259-DEL-170 | chrX-68760258-DEL-171 |
| chrX-68818771-DEL-93 | chrX-68818772-DEL-93 |
| chrX-68898220-DEL-126 | chrX-68898221-DEL-126 |
| chrX-68969899-DEL-64 | chrX-68969900-DEL-50 |
| chrX-68974482-DEL-7645 | chrX-68974483-DEL-7645 |
| chrX-68980506-DEL-424 | chrX-68980507-DEL-424 |
| chrX-68980507-DEL-1272 | chrX-68980507-DEL-1272 |
| chrX-68990154-DEL-87 | chrX-68990164-DEL-87 |
| chrX-69030484-DEL-621 | chrX-69030485-DEL-621 |
| chrX-69067933-DEL-90 | chrX-69067934-DEL-90 |
| chrX-69084562-DEL-344 | chrX-69084563-DEL-344 |
| chrX-69116312-DEL-1596 | chrX-69116312-DEL-1596 |
| chrX-69152224-DEL-311 | chrX-69152225-DEL-311 |
| chrX-69172772-DEL-421 | chrX-69172773-DEL-421 |
| chrX-69237833-DEL-445 | chrX-69237834-DEL-445 |
| chrX-69267885-DEL-52 | chrX-69267886-DEL-52 |
| chrX-69322087-DEL-52 | chrX-69322079-DEL-60 |
| chrX-69347470-DEL-2770 | chrX-69347471-DEL-2770 |
| chrX-69360741-DEL-56 | chrX-69360742-DEL-56 |
| chrX-69392526-DEL-871 | chrX-69392527-DEL-871 |
| chrX-69521511-DEL-156 | chrX-69521512-DEL-156 |
| chrX-69613523-DEL-91 | chrX-69613524-DEL-91 |
| chrX-69621236-DEL-105 | chrX-69621237-DEL-105 |
| chrX-69654555-DEL-2838 | chrX-69654556-DEL-2838 |
| chrX-69677993-DEL-2116 | chrX-69677994-DEL-2116 |
| chrX-69699261-DEL-1057 | chrX-69699262-DEL-1057 |
| chrX-69707087-DEL-248 | chrX-69707088-DEL-248 |
| chrX-69748534-DEL-57 | chrX-69748535-DEL-57 |
| chrX-69758878-DEL-354 | chrX-69758879-DEL-354 |
| chrX-69764927-DEL-380 | chrX-69764928-DEL-380 |
| chrX-69785761-DEL-6590 | chrX-69785762-DEL-6590 |
| chrX-69809877-DEL-105 | chrX-69809862-DEL-150 |
| chrX-69879246-DEL-1324 | chrX-69879247-DEL-1324 |
| chrX-69962252-DEL-433 | chrX-69962253-DEL-433 |
| chrX-69987643-DEL-405 | chrX-69987644-DEL-405 |
| chrX-70027604-DEL-2698 | chrX-70027605-DEL-2698 |
| chrX-70028012-DEL-1131 | chrX-70028013-DEL-1131 |
| chrX-70052129-DEL-56 | chrX-70052130-DEL-56 |
| chrX-70067195-DEL-189 | chrX-70067196-DEL-189 |
| chrX-70197255-DEL-133 | chrX-70197256-DEL-129 |
| chrX-70204118-DEL-206 | chrX-70204119-DEL-206 |
| chrX-70216873-DEL-97 | chrX-70216891-DEL-80 |
| chrX-70242529-DEL-59 | chrX-70242530-DEL-59 |
| chrX-70247410-DEL-53 | chrX-70247401-DEL-54 |
| chrX-70314083-DEL-400 | chrX-70314084-DEL-400 |
| chrX-70500262-DEL-387 | chrX-70500263-DEL-387 |
| chrX-70500894-DEL-204 | chrX-70500895-DEL-204 |
| chrX-70511920-DEL-274 | chrX-70511921-DEL-274 |
| chrX-70515458-DEL-133 | chrX-70515459-DEL-133 |
| chrX-70539137-DEL-168 | chrX-70539138-DEL-174 |
| chrX-70547521-DEL-62 | chrX-70547549-DEL-66 |
| chrX-70551414-DEL-807 | chrX-70551415-DEL-807 |
| chrX-70557118-DEL-75 | chrX-70557119-DEL-75 |
| chrX-70576432-DEL-598 | chrX-70576433-DEL-598 |
| chrX-70578632-DEL-210 | chrX-70578633-DEL-210 |
| chrX-70589631-DEL-214 | chrX-70589632-DEL-214 |
| chrX-70621711-DEL-199 | chrX-70621708-DEL-198 |
| chrX-70630349-DEL-118 | chrX-70630340-DEL-117 |
| chrX-70634052-DEL-58 | chrX-70634056-DEL-54 |
| chrX-70648208-DEL-67 | chrX-70648209-DEL-67 |
| chrX-70655753-DEL-400 | chrX-70655754-DEL-400 |
| chrX-70659685-DEL-2354 | chrX-70659686-DEL-2354 |
| chrX-70677193-DEL-111 | chrX-70677194-DEL-111 |
| chrX-70726609-DEL-194 | chrX-70726610-DEL-194 |
| chrX-70727671-DEL-4032 | chrX-70727672-DEL-4032 |

| | |
|---|---|
| chrX-70732354-DEL-105 | chrX-70732355-DEL-105 |
| chrX-70732726-DEL-2642 | chrX-70732716-DEL-2654 |
| chrX-70742130-DEL-1100 | chrX-70742131-DEL-1100 |
| chrX-70745939-DEL-3114 | chrX-70745940-DEL-3114 |
| chrX-70750038-DEL-64 | chrX-70749999-DEL-64 |
| chrX-70763117-DEL-64 | chrX-70763118-DEL-56 |
| chrX-70851171-DEL-527 | chrX-70851172-DEL-527 |
| chrX-70853748-DEL-184 | chrX-70853731-DEL-202 |
| chrX-70878571-DEL-697 | chrX-70878572-DEL-697 |
| chrX-70879592-DEL-461 | chrX-70879593-DEL-461 |
| chrX-70914020-DEL-1639 | chrX-70914021-DEL-1639 |
| chrX-70923786-DEL-3979 | chrX-70923787-DEL-3979 |
| chrX-70934648-DEL-315 | chrX-70934648-DEL-316 |
| chrX-70963750-DEL-80 | chrX-70963764-DEL-57 |
| chrX-71047145-DEL-52 | chrX-71047137-DEL-60 |
| chrX-71093507-DEL-214 | chrX-71093508-DEL-213 |
| chrX-71106241-DEL-340 | chrX-71106243-DEL-341 |
| chrX-71122014-DEL-7717 | chrX-71122015-DEL-7717 |
| chrX-71130108-DEL-78 | chrX-71130109-DEL-78 |
| chrX-71140180-DEL-615 | chrX-71140179-DEL-615 |
| chrX-71141351-DEL-786 | chrX-71141352-DEL-786 |
| chrX-71142440-DEL-239 | chrX-71142441-DEL-239 |
| chrX-71144349-DEL-6644 | chrX-71144350-DEL-6644 |
| chrX-71166341-DEL-6136 | chrX-71166337-DEL-6137 |
| chrX-71187406-DEL-6264 | chrX-71187393-DEL-6265 |
| chrX-71221733-DEL-2882 | chrX-71221732-DEL-2882 |
| chrX-71268266-DEL-278 | chrX-71268264-DEL-282 |
| chrX-71313874-DEL-84 | chrX-71313875-DEL-84 |
| chrX-71350025-DEL-569 | chrX-71350026-DEL-569 |
| chrX-71353891-DEL-768 | chrX-71353892-DEL-768 |
| chrX-71373173-DEL-57 | chrX-71373174-DEL-57 |
| chrX-71484467-DEL-83 | chrX-71484468-DEL-83 |
| chrX-71524992-DEL-98 | chrX-71524985-DEL-92 |
| chrX-71557131-DEL-3057 | chrX-71557132-DEL-3057 |
| chrX-71580280-DEL-111 | chrX-71580298-DEL-114 |
| chrX-71582673-DEL-101 | chrX-71582674-DEL-101 |
| chrX-71616444-DEL-13643 | chrX-71616445-DEL-13643 |
| chrX-71631910-DEL-249 | chrX-71631911-DEL-249 |
| chrX-71656460-DEL-6915 | chrX-71656461-DEL-6915 |
| chrX-71714864-DEL-468 | chrX-71714865-DEL-468 |
| chrX-71717421-DEL-68 | chrX-71717379-DEL-94 |
| chrX-71718524-DEL-51 | chrX-71718525-DEL-51 |
| chrX-71722785-DEL-6517 | chrX-71722785-DEL-6517 |
| chrX-71728799-DEL-212 | chrX-71728800-DEL-212 |
| chrX-71738991-DEL-206 | chrX-71738991-DEL-206 |
| chrX-71741536-DEL-578 | chrX-71741537-DEL-578 |
| chrX-71753324-DEL-3902 | chrX-71753325-DEL-3902 |
| chrX-71774362-DEL-431 | chrX-71774363-DEL-431 |
| chrX-71784519-DEL-8776 | chrX-71784520-DEL-8776 |
| chrX-71796156-DEL-1892 | chrX-71796157-DEL-1892 |
| chrX-71860832-DEL-1479 | chrX-71860833-DEL-1479 |
| chrX-71907442-DEL-2715 | chrX-71907443-DEL-2715 |
| chrX-71917728-DEL-4930 | chrX-71917729-DEL-4930 |
| chrX-71936465-DEL-948 | chrX-71936466-DEL-948 |
| chrX-71941137-DEL-111 | chrX-71941153-DEL-82 |
| chrX-71966190-DEL-78 | chrX-71966231-DEL-78 |
| chrX-71969530-DEL-285 | chrX-71969531-DEL-285 |
| chrX-71985552-DEL-7160 | chrX-71985553-DEL-7160 |
| chrX-72014986-DEL-607 | chrX-72014987-DEL-607 |
| chrX-72020952-DEL-492 | chrX-72020953-DEL-492 |
| chrX-72054127-DEL-130 | chrX-72054128-DEL-130 |
| chrX-72072259-DEL-127 | chrX-72072260-DEL-127 |
| chrX-72211895-DEL-6768 | chrX-72211896-DEL-6768 |
| chrX-72231765-DEL-61 | chrX-72231812-DEL-58 |
| chrX-72255595-DEL-1345 | chrX-72255595-DEL-1345 |
| chrX-72279970-DEL-920 | chrX-72279971-DEL-918 |
| chrX-72290841-DEL-53 | chrX-72290841-DEL-53 |
| chrX-72343591-DEL-7728 | chrX-72343592-DEL-7728 |
| chrX-72364191-DEL-203 | chrX-72364192-DEL-203 |
| chrX-72378543-DEL-427 | chrX-72378543-DEL-427 |
| chrX-72422259-DEL-236 | chrX-72422238-DEL-234 |
| chrX-72461158-DEL-165 | chrX-72461159-DEL-165 |
| chrX-72467671-DEL-60 | chrX-72467671-DEL-64 |
| chrX-72481680-DEL-470 | chrX-72481681-DEL-470 |
| chrX-72504889-DEL-2881 | chrX-72504890-DEL-2881 |
| chrX-72531864-DEL-53 | chrX-72531850-DEL-50 |
| chrX-72634341-DEL-182 | chrX-72634342-DEL-182 |
| chrX-72679807-DEL-76 | chrX-72679808-DEL-76 |
| chrX-72698170-DEL-208 | chrX-72698171-DEL-207 |
| chrX-72742560-DEL-161 | chrX-72742560-DEL-161 |

| | |
|---|---|
| chrX-72785659-DEL-94 | chrX-72785660-DEL-94 |
| chrX-72791901-DEL-211 | chrX-72791902-DEL-211 |
| chrX-72876554-DEL-209 | chrX-72876555-DEL-209 |
| chrX-72891866-DEL-56 | chrX-72891867-DEL-56 |
| chrX-72896631-DEL-172 | chrX-72896626-DEL-166 |
| chrX-72917784-DEL-197 | chrX-72917785-DEL-197 |
| chrX-72921867-DEL-145 | chrX-72921862-DEL-185 |
| chrX-72981361-DEL-189 | chrX-72981362-DEL-188 |
| chrX-73038800-DEL-201 | chrX-73038801-DEL-201 |
| chrX-73049697-DEL-179 | chrX-73049698-DEL-179 |
| chrX-73052503-DEL-170 | chrX-73052514-DEL-168 |
| chrX-73054134-DEL-207 | chrX-73054135-DEL-207 |
| chrX-73086677-DEL-951 | chrX-73086678-DEL-951 |
| chrX-73094331-DEL-161 | chrX-73094339-DEL-160 |
| chrX-73113715-DEL-54 | chrX-73113716-DEL-54 |
| chrX-73210402-DEL-1013 | chrX-73210403-DEL-1013 |
| chrX-73212924-DEL-66 | chrX-73212902-DEL-87 |
| chrX-73223012-DEL-203 | chrX-73223013-DEL-203 |
| chrX-73235582-DEL-550 | chrX-73235583-DEL-550 |
| chrX-73252464-DEL-1176 | chrX-73252465-DEL-1176 |
| chrX-73255325-DEL-2292 | chrX-73255326-DEL-2292 |
| chrX-73305633-DEL-389 | chrX-73305634-DEL-389 |
| chrX-73311759-DEL-167 | chrX-73311760-DEL-167 |
| chrX-73333997-DEL-60 | chrX-73333998-DEL-60 |
| chrX-73360560-DEL-208 | chrX-73360561-DEL-208 |
| chrX-73400878-DEL-226 | chrX-73400879-DEL-226 |
| chrX-73418834-DEL-85 | chrX-73418793-DEL-126 |
| chrX-73420953-DEL-1476 | chrX-73420954-DEL-1476 |
| chrX-73439276-DEL-203 | chrX-73439277-DEL-203 |
| chrX-73446809-DEL-51 | chrX-73446810-DEL-51 |
| chrX-73463864-DEL-979 | chrX-73463865-DEL-979 |
| chrX-73482660-DEL-84 | chrX-73482661-DEL-84 |
| chrX-73487759-DEL-2054 | chrX-73487760-DEL-2054 |
| chrX-73496137-DEL-205 | chrX-73496138-DEL-205 |
| chrX-73509201-DEL-218 | chrX-73509202-DEL-218 |
| chrX-73536907-DEL-777 | chrX-73536908-DEL-777 |
| chrX-73563716-DEL-2838 | chrX-73563716-DEL-2838 |
| chrX-73578272-DEL-6442 | chrX-73578273-DEL-6442 |
| chrX-73584935-DEL-112 | chrX-73584933-DEL-113 |
| chrX-73591630-DEL-85 | chrX-73591644-DEL-70 |
| chrX-73618973-DEL-103 | chrX-73618943-DEL-115 |
| chrX-73659417-DEL-6469 | chrX-73659424-DEL-6468 |
| chrX-73685902-DEL-355 | chrX-73685903-DEL-355 |
| chrX-73722491-DEL-58 | chrX-73722491-DEL-58 |
| chrX-73750757-DEL-1489 | chrX-73750758-DEL-1489 |
| chrX-73756617-DEL-863 | chrX-73756617-DEL-863 |
| chrX-73758617-DEL-191 | chrX-73758618-DEL-191 |
| chrX-73786458-DEL-157 | chrX-73786447-DEL-141 |
| chrX-73788454-DEL-78 | chrX-73788455-DEL-78 |
| chrX-73795684-DEL-83 | chrX-73795685-DEL-83 |
| chrX-73830228-DEL-54 | chrX-73830228-DEL-54 |
| chrX-73850912-DEL-66 | chrX-73850913-DEL-66 |
| chrX-73863264-DEL-55 | chrX-73863258-DEL-54 |
| chrX-73908034-DEL-8518 | chrX-73908035-DEL-8518 |
| chrX-73945198-DEL-250 | chrX-73945199-DEL-250 |
| chrX-73946263-DEL-1320 | chrX-73946257-DEL-1320 |
| chrX-73951771-DEL-2711 | chrX-73951772-DEL-2711 |
| chrX-73976591-DEL-6270 | chrX-73976591-DEL-6270 |
| chrX-74004031-DEL-150 | chrX-74004028-DEL-152 |
| chrX-74013286-DEL-521 | chrX-74013287-DEL-521 |
| chrX-74033436-DEL-118 | chrX-74033437-DEL-118 |
| chrX-74108859-DEL-222 | chrX-74108860-DEL-222 |
| chrX-74185756-DEL-178 | chrX-74185757-DEL-178 |
| chrX-74196300-DEL-403 | chrX-74196303-DEL-401 |
| chrX-74237898-DEL-7144 | chrX-74237899-DEL-7144 |
| chrX-74244462-DEL-212 | chrX-74244462-DEL-212 |
| chrX-74346469-DEL-7295 | chrX-74346470-DEL-7295 |
| chrX-74398073-DEL-70 | chrX-74398074-DEL-70 |
| chrX-74401710-DEL-2731 | chrX-74401703-DEL-2732 |
| chrX-74473764-DEL-697 | chrX-74473765-DEL-697 |
| chrX-74482119-DEL-1081 | chrX-74482120-DEL-1081 |
| chrX-74484327-DEL-2881 | chrX-74484327-DEL-2881 |
| chrX-74515634-DEL-2088 | chrX-74515631-DEL-2086 |
| chrX-74602243-DEL-1139 | chrX-74602244-DEL-1139 |
| chrX-74625785-DEL-4692 | chrX-74625786-DEL-4692 |
| chrX-74633172-DEL-2469 | chrX-74633173-DEL-2469 |
| chrX-74644053-DEL-530 | chrX-74644054-DEL-530 |
| chrX-74653355-DEL-3280 | chrX-74653356-DEL-3280 |
| chrX-74700813-DEL-230 | chrX-74700805-DEL-228 |
| chrX-74759440-DEL-1244 | chrX-74759441-DEL-1243 |

| | |
|---|---|
| chrX-74765712-DEL-555 | chrX-74765713-DEL-555 |
| chrX-74767300-DEL-93 | chrX-74767301-DEL-93 |
| chrX-74812110-DEL-850 | chrX-74812139-DEL-843 |
| chrX-74813005-DEL-74 | chrX-74813006-DEL-74 |
| chrX-74865493-DEL-117 | chrX-74865494-DEL-117 |
| chrX-74872184-DEL-195 | chrX-74872185-DEL-195 |
| chrX-74880095-DEL-1973 | chrX-74880096-DEL-1973 |
| chrX-74928028-DEL-231 | chrX-74928029-DEL-231 |
| chrX-74929101-DEL-51 | chrX-74929101-DEL-51 |
| chrX-74935135-DEL-66 | chrX-74935136-DEL-90 |
| chrX-74952511-DEL-52 | chrX-74952512-DEL-52 |
| chrX-75001007-DEL-3015 | chrX-75001008-DEL-3015 |
| chrX-75023645-DEL-481 | chrX-75023646-DEL-481 |
| chrX-75029337-DEL-103 | chrX-75029338-DEL-103 |
| chrX-75035312-DEL-1602 | chrX-75035313-DEL-1602 |
| chrX-75062049-DEL-797 | chrX-75062050-DEL-797 |
| chrX-75076716-DEL-5583 | chrX-75076717-DEL-5583 |
| chrX-75110229-DEL-214 | chrX-75110230-DEL-214 |
| chrX-75130534-DEL-749 | chrX-75130488-DEL-794 |
| chrX-75153122-DEL-447 | chrX-75153123-DEL-447 |
| chrX-75162152-DEL-1351 | chrX-75162153-DEL-1351 |
| chrX-75190261-DEL-6578 | chrX-75190262-DEL-6578 |
| chrX-75211781-DEL-945 | chrX-75211782-DEL-945 |
| chrX-75215156-DEL-1137 | chrX-75215157-DEL-1137 |
| chrX-75267496-DEL-2475 | chrX-75267496-DEL-2475 |
| chrX-75280072-DEL-6405 | chrX-75280073-DEL-6405 |
| chrX-75315395-DEL-8420 | chrX-75315396-DEL-8420 |
| chrX-75341923-DEL-690 | chrX-75341924-DEL-690 |
| chrX-75354951-DEL-185 | chrX-75354944-DEL-186 |
| chrX-75361674-DEL-1037 | chrX-75361675-DEL-1037 |
| chrX-75383578-DEL-206 | chrX-75383579-DEL-206 |
| chrX-75394134-DEL-659 | chrX-75394135-DEL-659 |
| chrX-75405666-DEL-7279 | chrX-75405667-DEL-7279 |
| chrX-75417384-DEL-198 | chrX-75417385-DEL-198 |
| chrX-75438503-DEL-210 | chrX-75438504-DEL-210 |
| chrX-75441657-DEL-199 | chrX-75441641-DEL-199 |
| chrX-75471044-DEL-65 | chrX-75471045-DEL-65 |
| chrX-75592028-DEL-1185 | chrX-75592025-DEL-1185 |
| chrX-75668430-DEL-5206 | chrX-75668430-DEL-5206 |
| chrX-75680091-DEL-4400 | chrX-75680092-DEL-4401 |
| chrX-75690392-DEL-277 | chrX-75690393-DEL-277 |
| chrX-75698732-DEL-305 | chrX-75698733-DEL-305 |
| chrX-75704905-DEL-56 | chrX-75704906-DEL-65 |
| chrX-75718596-DEL-2018 | chrX-75718596-DEL-2018 |
| chrX-75744950-DEL-6232 | chrX-75744951-DEL-6232 |
| chrX-75752583-DEL-391 | chrX-75752584-DEL-391 |
| chrX-75804238-DEL-6420 | chrX-75804239-DEL-6420 |
| chrX-75851208-DEL-1154 | chrX-75851209-DEL-1154 |
| chrX-75911138-DEL-640 | chrX-75911139-DEL-640 |
| chrX-75937523-DEL-8833 | chrX-75937524-DEL-8833 |
| chrX-75956791-DEL-54 | chrX-75956790-DEL-55 |
| chrX-75970063-DEL-799 | chrX-75970062-DEL-800 |
| chrX-75990599-DEL-60 | chrX-75990596-DEL-68 |
| chrX-76000531-DEL-1889 | chrX-76000532-DEL-1889 |
| chrX-76025943-DEL-1000 | chrX-76025944-DEL-1000 |
| chrX-76067639-DEL-1893 | chrX-76067640-DEL-1893 |
| chrX-76086693-DEL-406 | chrX-76086694-DEL-406 |
| chrX-76117223-DEL-4932 | chrX-76117224-DEL-4932 |
| chrX-76146132-DEL-458 | chrX-76146133-DEL-458 |
| chrX-76170478-DEL-55 | chrX-76170466-DEL-54 |
| chrX-76215974-DEL-12956 | chrX-76215969-DEL-12957 |
| chrX-76255449-DEL-1951 | chrX-76255450-DEL-1951 |
| chrX-76294424-DEL-999 | chrX-76294425-DEL-999 |
| chrX-76358856-DEL-6872 | chrX-76358857-DEL-6872 |
| chrX-76438855-DEL-464 | chrX-76438856-DEL-457 |
| chrX-76441351-DEL-6686 | chrX-76441352-DEL-6686 |
| chrX-76508771-DEL-103 | chrX-76508771-DEL-64 |
| chrX-76514907-DEL-579 | chrX-76514908-DEL-579 |
| chrX-76517653-DEL-454 | chrX-76517672-DEL-446 |
| chrX-76526411-DEL-996 | chrX-76526412-DEL-996 |
| chrX-76562244-DEL-201 | chrX-76562245-DEL-201 |
| chrX-76593537-DEL-1182 | chrX-76593538-DEL-1182 |
| chrX-76622197-DEL-138 | chrX-76622198-DEL-138 |
| chrX-76630860-DEL-54 | chrX-76630859-DEL-54 |
| chrX-76635824-DEL-196 | chrX-76635825-DEL-196 |
| chrX-76721250-DEL-726 | chrX-76721251-DEL-720 |
| chrX-76769945-DEL-2274 | chrX-76769945-DEL-2274 |
| chrX-76775251-DEL-58 | chrX-76775252-DEL-58 |
| chrX-76794844-DEL-618 | chrX-76794845-DEL-616 |
| chrX-76798980-DEL-66 | chrX-76798951-DEL-66 |

| | |
|---|---|
| chrX-76815477-DEL-80 | chrX-76815478-DEL-80 |
| chrX-76823304-DEL-139 | chrX-76823307-DEL-137 |
| chrX-76846690-DEL-498 | chrX-76846691-DEL-498 |
| chrX-76922097-DEL-1679 | chrX-76922097-DEL-1679 |
| chrX-76927085-DEL-51 | chrX-76927077-DEL-59 |
| chrX-76928038-DEL-3158 | chrX-76928038-DEL-3158 |
| chrX-76986018-DEL-461 | chrX-76986019-DEL-461 |
| chrX-77012239-DEL-6861 | chrX-77012240-DEL-6861 |
| chrX-77051510-DEL-413 | chrX-77051516-DEL-411 |
| chrX-77072108-DEL-70 | chrX-77072109-DEL-70 |
| chrX-77088503-DEL-382 | chrX-77088504-DEL-382 |
| chrX-77104674-DEL-663 | chrX-77104675-DEL-663 |
| chrX-77161158-DEL-1076 | chrX-77161159-DEL-1076 |
| chrX-77183039-DEL-6235 | chrX-77183043-DEL-6234 |
| chrX-77188625-DEL-176 | chrX-77188626-DEL-176 |
| chrX-77211865-DEL-1238 | chrX-77211866-DEL-1238 |
| chrX-77246877-DEL-221 | chrX-77246878-DEL-221 |
| chrX-77274462-DEL-6713 | chrX-77274443-DEL-6736 |
| chrX-77274464-DEL-424 | chrX-77274465-DEL-424 |
| chrX-77300398-DEL-2977 | chrX-77300399-DEL-2977 |
| chrX-77311519-DEL-1859 | chrX-77311520-DEL-1859 |
| chrX-77356907-DEL-1234 | chrX-77356908-DEL-1234 |
| chrX-77376872-DEL-4872 | chrX-77376873-DEL-4872 |
| chrX-77416560-DEL-78 | chrX-77416561-DEL-78 |
| chrX-77417560-DEL-2969 | chrX-77417561-DEL-2969 |
| chrX-77437203-DEL-2781 | chrX-77437204-DEL-2781 |
| chrX-77444033-DEL-544 | chrX-77444014-DEL-518 |
| chrX-77463653-DEL-487 | chrX-77463654-DEL-487 |
| chrX-77473715-DEL-3686 | chrX-77473716-DEL-3684 |
| chrX-77500579-DEL-105 | chrX-77500580-DEL-105 |
| chrX-77507858-DEL-859 | chrX-77507859-DEL-859 |
| chrX-77581803-DEL-153 | chrX-77581817-DEL-112 |
| chrX-77674565-DEL-203 | chrX-77674566-DEL-203 |
| chrX-77679424-DEL-123 | chrX-77679425-DEL-123 |
| chrX-77687993-DEL-741 | chrX-77687994-DEL-741 |
| chrX-77694266-DEL-482 | chrX-77694267-DEL-482 |
| chrX-77722131-DEL-204 | chrX-77722132-DEL-204 |
| chrX-77727840-DEL-170 | chrX-77727839-DEL-165 |
| chrX-77737050-DEL-3639 | chrX-77737051-DEL-3639 |
| chrX-77800029-DEL-54 | chrX-77800030-DEL-62 |
| chrX-77817539-DEL-396 | chrX-77817540-DEL-396 |
| chrX-77853282-DEL-495 | chrX-77853283-DEL-495 |
| chrX-77931163-DEL-8318 | chrX-77931163-DEL-8318 |
| chrX-77968634-DEL-67 | chrX-77968646-DEL-55 |
| chrX-77981798-DEL-6402 | chrX-77981799-DEL-6402 |
| chrX-78035905-DEL-750 | chrX-78035906-DEL-750 |
| chrX-78042663-DEL-23455 | chrX-78042663-DEL-23455 |
| chrX-78073099-DEL-6012 | chrX-78073100-DEL-6012 |
| chrX-78098141-DEL-2301 | chrX-78098141-DEL-2301 |
| chrX-78100550-DEL-246 | chrX-78100550-DEL-246 |
| chrX-78110742-DEL-8852 | chrX-78110743-DEL-8852 |
| chrX-78136224-DEL-2966 | chrX-78136225-DEL-2966 |
| chrX-78181701-DEL-60 | chrX-78181702-DEL-67 |
| chrX-78186749-DEL-7176 | chrX-78186750-DEL-7176 |
| chrX-78289440-DEL-66 | chrX-78289448-DEL-52 |
| chrX-78319116-DEL-133 | chrX-78319117-DEL-133 |
| chrX-78352441-DEL-73 | chrX-78352442-DEL-73 |
| chrX-78441403-DEL-108 | chrX-78441406-DEL-90 |
| chrX-78475804-DEL-58 | chrX-78475805-DEL-56 |
| chrX-78504260-DEL-94 | chrX-78504227-DEL-135 |
| chrX-78542148-DEL-511 | chrX-78542149-DEL-511 |
| chrX-78574204-DEL-3066 | chrX-78574205-DEL-3066 |
| chrX-78651923-DEL-78 | chrX-78651882-DEL-92 |
| chrX-78717848-DEL-2581 | chrX-78717849-DEL-2581 |
| chrX-78722613-DEL-1692 | chrX-78722614-DEL-1692 |
| chrX-78731966-DEL-5492 | chrX-78731967-DEL-5492 |
| chrX-78755297-DEL-6537 | chrX-78755298-DEL-6537 |
| chrX-78766377-DEL-218 | chrX-78766378-DEL-218 |
| chrX-78781398-DEL-7106 | chrX-78781399-DEL-7106 |
| chrX-78802002-DEL-117 | chrX-78802003-DEL-117 |
| chrX-78825036-DEL-197 | chrX-78825037-DEL-197 |
| chrX-78829677-DEL-8017 | chrX-78829678-DEL-8017 |
| chrX-78840881-DEL-112 | chrX-78840881-DEL-112 |
| chrX-78846543-DEL-1057 | chrX-78846544-DEL-1057 |
| chrX-78851150-DEL-261 | chrX-78851134-DEL-262 |
| chrX-78860584-DEL-56 | chrX-78860585-DEL-56 |
| chrX-78863160-DEL-6692 | chrX-78863161-DEL-6692 |
| chrX-78898664-DEL-7114 | chrX-78898665-DEL-7114 |
| chrX-78904786-DEL-348 | chrX-78904787-DEL-348 |
| chrX-79021453-DEL-1780 | chrX-79021454-DEL-1780 |

| | |
|---|---|
| chrX-79132857-DEL-1510 | chrX-79132858-DEL-1510 |
| chrX-79134895-DEL-897 | chrX-79134895-DEL-897 |
| chrX-79145352-DEL-2716 | chrX-79145352-DEL-2716 |
| chrX-79182818-DEL-6550 | chrX-79182819-DEL-6550 |
| chrX-79225827-DEL-1003 | chrX-79225828-DEL-1003 |
| chrX-79237246-DEL-1290 | chrX-79237247-DEL-1290 |
| chrX-79251709-DEL-6917 | chrX-79251710-DEL-6917 |
| chrX-79280680-DEL-52 | chrX-79280681-DEL-58 |
| chrX-79325241-DEL-529 | chrX-79325242-DEL-529 |
| chrX-79328653-DEL-756 | chrX-79328654-DEL-755 |
| chrX-79336585-DEL-135 | chrX-79336580-DEL-120 |
| chrX-79381339-DEL-52 | chrX-79381329-DEL-53 |
| chrX-79389529-DEL-98 | chrX-79389530-DEL-98 |
| chrX-79415450-DEL-201 | chrX-79415451-DEL-201 |
| chrX-79439191-DEL-4964 | chrX-79439192-DEL-4964 |
| chrX-79440763-DEL-295 | chrX-79440764-DEL-295 |
| chrX-79461786-DEL-253 | chrX-79461775-DEL-256 |
| chrX-79463786-DEL-72 | chrX-79463786-DEL-72 |
| chrX-79500277-DEL-1893 | chrX-79500278-DEL-1893 |
| chrX-79510255-DEL-5502 | chrX-79510256-DEL-5502 |
| chrX-79510397-DEL-209 | chrX-79510397-DEL-209 |
| chrX-79562547-DEL-196 | chrX-79562558-DEL-188 |
| chrX-79651529-DEL-54 | chrX-79651530-DEL-54 |
| chrX-79661724-DEL-69 | chrX-79661722-DEL-72 |
| chrX-79665284-DEL-3475 | chrX-79665285-DEL-3475 |
| chrX-79744018-DEL-203 | chrX-79744019-DEL-203 |
| chrX-79747031-DEL-167 | chrX-79747029-DEL-166 |
| chrX-79775459-DEL-80 | chrX-79775460-DEL-82 |
| chrX-79798578-DEL-214 | chrX-79798578-DEL-214 |
| chrX-79807670-DEL-6220 | chrX-79807671-DEL-6220 |
| chrX-79838527-DEL-525 | chrX-79838528-DEL-525 |
| chrX-79859294-DEL-6552 | chrX-79859295-DEL-6552 |
| chrX-79868140-DEL-625 | chrX-79868141-DEL-625 |
| chrX-79902547-DEL-666 | chrX-79902548-DEL-666 |
| chrX-79907643-DEL-901 | chrX-79907644-DEL-901 |
| chrX-79952099-DEL-1105 | chrX-79952099-DEL-1105 |
| chrX-79964741-DEL-256 | chrX-79964742-DEL-256 |
| chrX-79977136-DEL-100 | chrX-79977124-DEL-64 |
| chrX-80002692-DEL-7740 | chrX-80002693-DEL-7740 |
| chrX-80059821-DEL-100 | chrX-80059822-DEL-100 |
| chrX-80093758-DEL-3282 | chrX-80093759-DEL-3282 |
| chrX-80156274-DEL-282 | chrX-80156275-DEL-282 |
| chrX-80198169-DEL-183 | chrX-80198170-DEL-183 |
| chrX-80214079-DEL-953 | chrX-80214080-DEL-953 |
| chrX-80238417-DEL-16947 | chrX-80238418-DEL-16947 |
| chrX-80286732-DEL-6299 | chrX-80286733-DEL-6299 |
| chrX-80296948-DEL-535 | chrX-80296949-DEL-535 |
| chrX-80307948-DEL-926 | chrX-80307949-DEL-926 |
| chrX-80325053-DEL-79 | chrX-80325054-DEL-79 |
| chrX-80327440-DEL-698 | chrX-80327441-DEL-698 |
| chrX-80332030-DEL-110 | chrX-80332031-DEL-110 |
| chrX-80361580-DEL-1595 | chrX-80361581-DEL-1595 |
| chrX-80419496-DEL-65 | chrX-80419497-DEL-65 |
| chrX-80441697-DEL-87 | chrX-80441698-DEL-87 |
| chrX-80456556-DEL-6583 | chrX-80456557-DEL-6583 |
| chrX-80478212-DEL-228 | chrX-80478213-DEL-228 |
| chrX-80482033-DEL-501 | chrX-80482044-DEL-502 |
| chrX-80487034-DEL-54 | chrX-80487034-DEL-54 |
| chrX-80562168-DEL-496 | chrX-80562168-DEL-496 |
| chrX-80610731-DEL-5881 | chrX-80610732-DEL-5881 |
| chrX-80690249-DEL-5424 | chrX-80690250-DEL-5424 |
| chrX-80719541-DEL-12845 | chrX-80719541-DEL-12845 |
| chrX-80736070-DEL-101 | chrX-80736077-DEL-93 |
| chrX-80804053-DEL-5443 | chrX-80804054-DEL-5443 |
| chrX-80844236-DEL-88 | chrX-80844237-DEL-68 |
| chrX-80883065-DEL-4512 | chrX-80883065-DEL-4512 |
| chrX-80898366-DEL-824 | chrX-80898367-DEL-825 |
| chrX-80906355-DEL-6381 | chrX-80906356-DEL-6381 |
| chrX-80954807-DEL-197 | chrX-80954807-DEL-197 |
| chrX-80998620-DEL-68 | chrX-80998621-DEL-68 |
| chrX-81013659-DEL-1145 | chrX-81013660-DEL-1145 |
| chrX-81025761-DEL-6600 | chrX-81025762-DEL-6600 |
| chrX-81026141-DEL-208 | chrX-81026141-DEL-208 |
| chrX-81076665-DEL-6050 | chrX-81076666-DEL-6050 |
| chrX-81088538-DEL-3801 | chrX-81088539-DEL-3801 |
| chrX-81098590-DEL-1888 | chrX-81098591-DEL-1888 |
| chrX-81158096-DEL-85 | chrX-81158095-DEL-85 |
| chrX-81189783-DEL-439 | chrX-81189784-DEL-439 |
| chrX-81200488-DEL-416 | chrX-81200489-DEL-416 |
| chrX-81203307-DEL-2336 | chrX-81203308-DEL-2336 |

| | |
|---|---|
| chrX-81245271-DEL-7443 | chrX-81245272-DEL-7443 |
| chrX-81254605-DEL-2638 | chrX-81254606-DEL-2638 |
| chrX-81282375-DEL-214 | chrX-81282376-DEL-214 |
| chrX-81286697-DEL-4791 | chrX-81286698-DEL-4791 |
| chrX-81297207-DEL-224 | chrX-81297200-DEL-225 |
| chrX-81306315-DEL-6600 | chrX-81306316-DEL-6600 |
| chrX-81312353-DEL-208 | chrX-81312353-DEL-208 |
| chrX-81336861-DEL-55 | chrX-81336857-DEL-55 |
| chrX-81364943-DEL-56 | chrX-81364936-DEL-64 |
| chrX-81425591-DEL-92 | chrX-81425592-DEL-91 |
| chrX-81432804-DEL-6172 | chrX-81432805-DEL-6172 |
| chrX-81467282-DEL-164 | chrX-81467282-DEL-165 |
| chrX-81519361-DEL-6442 | chrX-81519361-DEL-6442 |
| chrX-81551647-DEL-486 | chrX-81551648-DEL-486 |
| chrX-81585324-DEL-461 | chrX-81585327-DEL-458 |
| chrX-81619730-DEL-404 | chrX-81619731-DEL-404 |
| chrX-81631521-DEL-146 | chrX-81631522-DEL-146 |
| chrX-81632736-DEL-1548 | chrX-81632737-DEL-1548 |
| chrX-81646202-DEL-6387 | chrX-81646203-DEL-6387 |
| chrX-81664175-DEL-51 | chrX-81664165-DEL-69 |
| chrX-81698329-DEL-3722 | chrX-81698330-DEL-3722 |
| chrX-81702728-DEL-1093 | chrX-81702728-DEL-1093 |
| chrX-81894299-DEL-247 | chrX-81894300-DEL-247 |
| chrX-81978254-DEL-6667 | chrX-81978255-DEL-6667 |
| chrX-82014612-DEL-6240 | chrX-82014613-DEL-6240 |
| chrX-82048555-DEL-107 | chrX-82048556-DEL-107 |
| chrX-82079546-DEL-1615 | chrX-82079547-DEL-1615 |
| chrX-82129065-DEL-123 | chrX-82129066-DEL-123 |
| chrX-82129171-DEL-859 | chrX-82129172-DEL-859 |
| chrX-82131375-DEL-671 | chrX-82131341-DEL-666 |
| chrX-82137708-DEL-52 | chrX-82137709-DEL-52 |
| chrX-82143326-DEL-994 | chrX-82143327-DEL-994 |
| chrX-82186548-DEL-594 | chrX-82186549-DEL-594 |
| chrX-82223903-DEL-179 | chrX-82223904-DEL-179 |
| chrX-82260753-DEL-123 | chrX-82260754-DEL-123 |
| chrX-82289131-DEL-1900 | chrX-82289140-DEL-1897 |
| chrX-82309288-DEL-195 | chrX-82309289-DEL-195 |
| chrX-82369918-DEL-72 | chrX-82369918-DEL-72 |
| chrX-82407318-DEL-743 | chrX-82407318-DEL-741 |
| chrX-82436879-DEL-338 | chrX-82436880-DEL-338 |
| chrX-82441839-DEL-180 | chrX-82441840-DEL-180 |
| chrX-82449730-DEL-202 | chrX-82449731-DEL-202 |
| chrX-82541422-DEL-101 | chrX-82541422-DEL-101 |
| chrX-82562853-DEL-526 | chrX-82562854-DEL-526 |
| chrX-82567188-DEL-678 | chrX-82567187-DEL-677 |
| chrX-82579623-DEL-4226 | chrX-82579624-DEL-4226 |
| chrX-82593846-DEL-3728 | chrX-82593846-DEL-3728 |
| chrX-82690138-DEL-472 | chrX-82690141-DEL-468 |
| chrX-82706684-DEL-238 | chrX-82706685-DEL-238 |
| chrX-82719911-DEL-1051 | chrX-82719912-DEL-1051 |
| chrX-82758746-DEL-187 | chrX-82758747-DEL-187 |
| chrX-82795480-DEL-62 | chrX-82795479-DEL-52 |
| chrX-82826770-DEL-1051 | chrX-82826771-DEL-1051 |
| chrX-82834262-DEL-1316 | chrX-82834262-DEL-1316 |
| chrX-82843737-DEL-871 | chrX-82843738-DEL-871 |
| chrX-82867791-DEL-404 | chrX-82867792-DEL-404 |
| chrX-82869647-DEL-310 | chrX-82869648-DEL-310 |
| chrX-82874716-DEL-2318 | chrX-82874714-DEL-2320 |
| chrX-82891629-DEL-69 | chrX-82891621-DEL-78 |
| chrX-82932221-DEL-6656 | chrX-82932222-DEL-6656 |
| chrX-82981753-DEL-72 | chrX-82981754-DEL-72 |
| chrX-83010009-DEL-59 | chrX-83010010-DEL-59 |
| chrX-83026978-DEL-158 | chrX-83026979-DEL-158 |
| chrX-83044515-DEL-54 | chrX-83044538-DEL-54 |
| chrX-83045444-DEL-702 | chrX-83045445-DEL-702 |
| chrX-83048340-DEL-1888 | chrX-83048341-DEL-1888 |
| chrX-83106057-DEL-172 | chrX-83106058-DEL-172 |
| chrX-83157544-DEL-686 | chrX-83157545-DEL-686 |
| chrX-83196931-DEL-7234 | chrX-83196932-DEL-7234 |
| chrX-83202943-DEL-424 | chrX-83202943-DEL-424 |
| chrX-83247921-DEL-62 | chrX-83247921-DEL-62 |
| chrX-83292701-DEL-746 | chrX-83292702-DEL-748 |
| chrX-83319308-DEL-158 | chrX-83319313-DEL-154 |
| chrX-83384909-DEL-2454 | chrX-83384910-DEL-2453 |
| chrX-83389163-DEL-282 | chrX-83389164-DEL-282 |
| chrX-83423912-DEL-1029 | chrX-83423913-DEL-1029 |
| chrX-83432950-DEL-6418 | chrX-83432947-DEL-6419 |
| chrX-83447776-DEL-2300 | chrX-83447777-DEL-2279 |
| chrX-83470334-DEL-64 | chrX-83470335-DEL-64 |
| chrX-83502180-DEL-3263 | chrX-83502181-DEL-3263 |

| | |
|---|---|
| chrX-83506011-DEL-636 | chrX-83506012-DEL-626 |
| chrX-83552217-DEL-4481 | chrX-83552218-DEL-4481 |
| chrX-83557563-DEL-108 | chrX-83557564-DEL-108 |
| chrX-83582741-DEL-6921 | chrX-83582742-DEL-6921 |
| chrX-83651199-DEL-6355 | chrX-83651200-DEL-6355 |
| chrX-83786190-DEL-743 | chrX-83786191-DEL-743 |
| chrX-83851739-DEL-6189 | chrX-83851740-DEL-6189 |
| chrX-83878727-DEL-126 | chrX-83878728-DEL-126 |
| chrX-83893020-DEL-471 | chrX-83893021-DEL-471 |
| chrX-83897866-DEL-697 | chrX-83897867-DEL-697 |
| chrX-83899288-DEL-60 | chrX-83899289-DEL-60 |
| chrX-83913137-DEL-118 | chrX-83913138-DEL-118 |
| chrX-83917223-DEL-1455 | chrX-83917224-DEL-1455 |
| chrX-84077378-DEL-54 | chrX-84077379-DEL-54 |
| chrX-84082244-DEL-54 | chrX-84082245-DEL-54 |
| chrX-84109388-DEL-897 | chrX-84109389-DEL-897 |
| chrX-84141622-DEL-403 | chrX-84141623-DEL-403 |
| chrX-84211581-DEL-189 | chrX-84211582-DEL-189 |
| chrX-84249532-DEL-211 | chrX-84249533-DEL-211 |
| chrX-84261531-DEL-8091 | chrX-84261529-DEL-8091 |
| chrX-84284307-DEL-207 | chrX-84284308-DEL-207 |
| chrX-84293699-DEL-343 | chrX-84293677-DEL-344 |
| chrX-84316686-DEL-2394 | chrX-84316687-DEL-2394 |
| chrX-84340037-DEL-911 | chrX-84340037-DEL-912 |
| chrX-84386256-DEL-313 | chrX-84386257-DEL-313 |
| chrX-84389712-DEL-1614 | chrX-84389712-DEL-1614 |
| chrX-84399278-DEL-92 | chrX-84399277-DEL-94 |
| chrX-84422600-DEL-73 | chrX-84422601-DEL-73 |
| chrX-84444034-DEL-115 | chrX-84444035-DEL-115 |
| chrX-84463834-DEL-56 | chrX-84463831-DEL-52 |
| chrX-84554902-DEL-199 | chrX-84554884-DEL-200 |
| chrX-84561162-DEL-89 | chrX-84561165-DEL-82 |
| chrX-84568203-DEL-429 | chrX-84568204-DEL-429 |
| chrX-84638258-DEL-791 | chrX-84638259-DEL-791 |
| chrX-84705249-DEL-6039 | chrX-84705250-DEL-6039 |
| chrX-84744147-DEL-55 | chrX-84744148-DEL-55 |
| chrX-84766962-DEL-2178 | chrX-84766963-DEL-2178 |
| chrX-84788921-DEL-707 | chrX-84788922-DEL-707 |
| chrX-84801691-DEL-236 | chrX-84801692-DEL-236 |
| chrX-84813156-DEL-82 | chrX-84813157-DEL-74 |
| chrX-84852605-DEL-204 | chrX-84852606-DEL-204 |
| chrX-84907796-DEL-191 | chrX-84907796-DEL-189 |
| chrX-84925896-DEL-400 | chrX-84925897-DEL-400 |
| chrX-84953715-DEL-504 | chrX-84953715-DEL-504 |
| chrX-85009064-DEL-2983 | chrX-85009072-DEL-2978 |
| chrX-85030659-DEL-108 | chrX-85030660-DEL-108 |
| chrX-85032440-DEL-172 | chrX-85032441-DEL-172 |
| chrX-85037632-DEL-211 | chrX-85037633-DEL-211 |
| chrX-85085374-DEL-65 | chrX-85085375-DEL-65 |
| chrX-85163557-DEL-368 | chrX-85163558-DEL-368 |
| chrX-85164783-DEL-135 | chrX-85164784-DEL-135 |
| chrX-85166039-DEL-468 | chrX-85166039-DEL-468 |
| chrX-85177390-DEL-60 | chrX-85177387-DEL-59 |
| chrX-85181578-DEL-318 | chrX-85181579-DEL-318 |
| chrX-85184636-DEL-640 | chrX-85184637-DEL-640 |
| chrX-85221177-DEL-53 | chrX-85221166-DEL-68 |
| chrX-85226506-DEL-51 | chrX-85226495-DEL-52 |
| chrX-85261428-DEL-6408 | chrX-85261429-DEL-6408 |
| chrX-85270804-DEL-1354 | chrX-85270805-DEL-1354 |
| chrX-85281932-DEL-216 | chrX-85281933-DEL-216 |
| chrX-85296346-DEL-68 | chrX-85296346-DEL-68 |
| chrX-85327627-DEL-66 | chrX-85327628-DEL-66 |
| chrX-85350709-DEL-3292 | chrX-85350710-DEL-3292 |
| chrX-85358797-DEL-117 | chrX-85358798-DEL-117 |
| chrX-85390927-DEL-473 | chrX-85390928-DEL-473 |
| chrX-85412713-DEL-74 | chrX-85412714-DEL-74 |
| chrX-85431099-DEL-96 | chrX-85431100-DEL-96 |
| chrX-85454012-DEL-1786 | chrX-85454013-DEL-1786 |
| chrX-85500017-DEL-138 | chrX-85500018-DEL-134 |
| chrX-85505097-DEL-1921 | chrX-85505098-DEL-1921 |
| chrX-85567753-DEL-386 | chrX-85567754-DEL-383 |
| chrX-85569609-DEL-98 | chrX-85569610-DEL-98 |
| chrX-85583509-DEL-2258 | chrX-85583510-DEL-2258 |
| chrX-85584696-DEL-67 | chrX-85584697-DEL-67 |
| chrX-85589214-DEL-98 | chrX-85589217-DEL-98 |
| chrX-85595299-DEL-96 | chrX-85595319-DEL-54 |
| chrX-85606105-DEL-139 | chrX-85606106-DEL-139 |
| chrX-85610599-DEL-62 | chrX-85610604-DEL-66 |
| chrX-85620604-DEL-6376 | chrX-85620605-DEL-6376 |
| chrX-85643564-DEL-433 | chrX-85643565-DEL-434 |

| | |
|---|---|
| chrX-85671201-DEL-114 | chrX-85671202-DEL-114 |
| chrX-85676842-DEL-54 | chrX-85676843-DEL-54 |
| chrX-85687423-DEL-2128 | chrX-85687424-DEL-2128 |
| chrX-85715042-DEL-96 | chrX-85715043-DEL-96 |
| chrX-85753208-DEL-804 | chrX-85753209-DEL-804 |
| chrX-85824643-DEL-54 | chrX-85824630-DEL-52 |
| chrX-85865178-DEL-7027 | chrX-85865179-DEL-7027 |
| chrX-85865199-DEL-208 | chrX-85865199-DEL-208 |
| chrX-85909490-DEL-218 | chrX-85909491-DEL-218 |
| chrX-85935714-DEL-6640 | chrX-85935715-DEL-6640 |
| chrX-85975516-DEL-112 | chrX-85975517-DEL-112 |
| chrX-85981085-DEL-6211 | chrX-85981086-DEL-6211 |
| chrX-86000344-DEL-88 | chrX-86000345-DEL-88 |
| chrX-86006511-DEL-400 | chrX-86006512-DEL-400 |
| chrX-86017029-DEL-792 | chrX-86017030-DEL-792 |
| chrX-86047363-DEL-1394 | chrX-86047364-DEL-1394 |
| chrX-86066954-DEL-231 | chrX-86066955-DEL-231 |
| chrX-86072644-DEL-1311 | chrX-86072645-DEL-1311 |
| chrX-86099490-DEL-2013 | chrX-86099490-DEL-2013 |
| chrX-86121151-DEL-7170 | chrX-86121152-DEL-7170 |
| chrX-86150196-DEL-314 | chrX-86150197-DEL-314 |
| chrX-86168394-DEL-107 | chrX-86168396-DEL-106 |
| chrX-86172056-DEL-739 | chrX-86172057-DEL-739 |
| chrX-86185521-DEL-57 | chrX-86185542-DEL-51 |
| chrX-86194358-DEL-188 | chrX-86194359-DEL-188 |
| chrX-86230139-DEL-811 | chrX-86230140-DEL-811 |
| chrX-86231042-DEL-266 | chrX-86231043-DEL-266 |
| chrX-86242861-DEL-348 | chrX-86242862-DEL-348 |
| chrX-86246658-DEL-85 | chrX-86246656-DEL-85 |
| chrX-86265912-DEL-1740 | chrX-86265913-DEL-1740 |
| chrX-86298739-DEL-7502 | chrX-86298740-DEL-7502 |
| chrX-86303649-DEL-1290 | chrX-86303650-DEL-1290 |
| chrX-86376995-DEL-400 | chrX-86376996-DEL-400 |
| chrX-86399191-DEL-192 | chrX-86399192-DEL-192 |
| chrX-86402203-DEL-919 | chrX-86402206-DEL-917 |
| chrX-86421084-DEL-51 | chrX-86421085-DEL-51 |
| chrX-86437076-DEL-92 | chrX-86437077-DEL-92 |
| chrX-86472403-DEL-3031 | chrX-86472404-DEL-3031 |
| chrX-86482873-DEL-60 | chrX-86482874-DEL-56 |
| chrX-86503624-DEL-6700 | chrX-86503625-DEL-6700 |
| chrX-86513187-DEL-51 | chrX-86513187-DEL-51 |
| chrX-86521236-DEL-2311 | chrX-86521237-DEL-2311 |
| chrX-86538738-DEL-171 | chrX-86538739-DEL-171 |
| chrX-86570796-DEL-270 | chrX-86570792-DEL-277 |
| chrX-86595056-DEL-1200 | chrX-86595045-DEL-1206 |
| chrX-86638272-DEL-943 | chrX-86638273-DEL-943 |
| chrX-86694801-DEL-932 | chrX-86694802-DEL-932 |
| chrX-86711395-DEL-1939 | chrX-86711396-DEL-1939 |
| chrX-86774257-DEL-591 | chrX-86774258-DEL-591 |
| chrX-86788018-DEL-63 | chrX-86788019-DEL-63 |
| chrX-86842304-DEL-54 | chrX-86842305-DEL-54 |
| chrX-86882101-DEL-6739 | chrX-86882102-DEL-6739 |
| chrX-86911534-DEL-64 | chrX-86911543-DEL-83 |
| chrX-86963853-DEL-386 | chrX-86963854-DEL-386 |
| chrX-86969034-DEL-6993 | chrX-86969035-DEL-6993 |
| chrX-86979010-DEL-979 | chrX-86979011-DEL-979 |
| chrX-87069715-DEL-1609 | chrX-87069716-DEL-1608 |
| chrX-87110387-DEL-899 | chrX-87110387-DEL-900 |
| chrX-87112080-DEL-599 | chrX-87112081-DEL-599 |
| chrX-87113227-DEL-195 | chrX-87113228-DEL-195 |
| chrX-87198918-DEL-417 | chrX-87198919-DEL-417 |
| chrX-87200685-DEL-638 | chrX-87200686-DEL-638 |
| chrX-87304092-DEL-80 | chrX-87304118-DEL-54 |
| chrX-87390405-DEL-199 | chrX-87390398-DEL-200 |
| chrX-87421050-DEL-84 | chrX-87421051-DEL-85 |
| chrX-87430590-DEL-56 | chrX-87430601-DEL-58 |
| chrX-87433163-DEL-404 | chrX-87433164-DEL-404 |
| chrX-87437897-DEL-7525 | chrX-87437898-DEL-7523 |
| chrX-87476426-DEL-173 | chrX-87476432-DEL-175 |
| chrX-87531356-DEL-184 | chrX-87531357-DEL-184 |
| chrX-87553074-DEL-827 | chrX-87553075-DEL-827 |
| chrX-87554246-DEL-1625 | chrX-87554247-DEL-1625 |
| chrX-87701089-DEL-826 | chrX-87701090-DEL-826 |
| chrX-87710099-DEL-3882 | chrX-87710100-DEL-3882 |
| chrX-87722619-DEL-1865 | chrX-87722618-DEL-1865 |
| chrX-87741706-DEL-436 | chrX-87741706-DEL-436 |
| chrX-87803042-DEL-884 | chrX-87803042-DEL-884 |
| chrX-87806790-DEL-425 | chrX-87806791-DEL-425 |
| chrX-87862817-DEL-71 | chrX-87862818-DEL-71 |
| chrX-87877879-DEL-213 | chrX-87877880-DEL-213 |

| | |
|---|---|
| chrX-87879633-DEL-60 | chrX-87879638-DEL-56 |
| chrX-87888880-DEL-1561 | chrX-87888881-DEL-1561 |
| chrX-87924077-DEL-3204 | chrX-87924077-DEL-3204 |
| chrX-87929876-DEL-6953 | chrX-87929877-DEL-6953 |
| chrX-87974023-DEL-789 | chrX-87974024-DEL-789 |
| chrX-88035248-DEL-6670 | chrX-88035249-DEL-6670 |
| chrX-88054873-DEL-8818 | chrX-88054874-DEL-8818 |
| chrX-88117624-DEL-859 | chrX-88117625-DEL-859 |
| chrX-88125832-DEL-1112 | chrX-88125833-DEL-1112 |
| chrX-88138181-DEL-361 | chrX-88138190-DEL-384 |
| chrX-88144083-DEL-2587 | chrX-88144084-DEL-2587 |
| chrX-88146140-DEL-217 | chrX-88146141-DEL-217 |
| chrX-88149236-DEL-222 | chrX-88149237-DEL-222 |
| chrX-88177802-DEL-742 | chrX-88177803-DEL-741 |
| chrX-88181998-DEL-874 | chrX-88181999-DEL-874 |
| chrX-88223807-DEL-1077 | chrX-88223808-DEL-1077 |
| chrX-88226408-DEL-2921 | chrX-88226409-DEL-2921 |
| chrX-88321528-DEL-306 | chrX-88321529-DEL-306 |
| chrX-88343860-DEL-7139 | chrX-88343861-DEL-7139 |
| chrX-88395381-DEL-524 | chrX-88395382-DEL-524 |
| chrX-88465875-DEL-6389 | chrX-88465876-DEL-6389 |
| chrX-88500969-DEL-2119 | chrX-88500970-DEL-2119 |
| chrX-88517545-DEL-54 | chrX-88517546-DEL-54 |
| chrX-88564164-DEL-4561 | chrX-88564165-DEL-4561 |
| chrX-88624398-DEL-58 | chrX-88624397-DEL-62 |
| chrX-88721189-DEL-4196 | chrX-88721190-DEL-4196 |
| chrX-88911664-DEL-2210 | chrX-88911665-DEL-2210 |
| chrX-88990541-DEL-2887 | chrX-88990542-DEL-2887 |
| chrX-89019462-DEL-52 | chrX-89019462-DEL-52 |
| chrX-89035506-DEL-1717 | chrX-89035507-DEL-1717 |
| chrX-89044642-DEL-627 | chrX-89044643-DEL-627 |
| chrX-89076345-DEL-1487 | chrX-89076346-DEL-1487 |
| chrX-89098087-DEL-161 | chrX-89098088-DEL-161 |
| chrX-89125063-DEL-66 | chrX-89125064-DEL-64 |
| chrX-89140481-DEL-486 | chrX-89140482-DEL-486 |
| chrX-89169235-DEL-3339 | chrX-89169236-DEL-3339 |
| chrX-89227375-DEL-887 | chrX-89227376-DEL-887 |
| chrX-89232324-DEL-218 | chrX-89232325-DEL-218 |
| chrX-89292562-DEL-4964 | chrX-89292563-DEL-4964 |
| chrX-89298313-DEL-467 | chrX-89298314-DEL-466 |
| chrX-89330908-DEL-1186 | chrX-89330909-DEL-1186 |
| chrX-89373567-DEL-6649 | chrX-89373568-DEL-6649 |
| chrX-89388721-DEL-45481 | chrX-89388721-DEL-45481 |
| chrX-89394454-DEL-160 | chrX-89394455-DEL-145 |
| chrX-89397631-DEL-839 | chrX-89397632-DEL-839 |
| chrX-89422240-DEL-6671 | chrX-89422241-DEL-6671 |
| chrX-89568545-DEL-6213 | chrX-89568546-DEL-6213 |
| chrX-89598096-DEL-436 | chrX-89598097-DEL-436 |
| chrX-89649525-DEL-6228 | chrX-89649526-DEL-6228 |
| chrX-89668241-DEL-3027 | chrX-89668242-DEL-3027 |
| chrX-89683889-DEL-1473 | chrX-89683890-DEL-1473 |
| chrX-89702482-DEL-425 | chrX-89702479-DEL-426 |
| chrX-89708766-DEL-6839 | chrX-89708767-DEL-6839 |
| chrX-89755387-DEL-6345 | chrX-89755388-DEL-6345 |
| chrX-89767730-DEL-1474 | chrX-89767731-DEL-1474 |
| chrX-89777236-DEL-551 | chrX-89777237-DEL-551 |
| chrX-89788011-DEL-1706 | chrX-89788030-DEL-1711 |
| chrX-89818725-DEL-2141 | chrX-89818726-DEL-2141 |
| chrX-89836017-DEL-132 | chrX-89836018-DEL-132 |
| chrX-89882574-DEL-6708 | chrX-89882575-DEL-6708 |
| chrX-89906385-DEL-3254 | chrX-89906386-DEL-3254 |
| chrX-89928074-DEL-60 | chrX-89928071-DEL-64 |
| chrX-89961302-DEL-93 | chrX-89961273-DEL-94 |
| chrX-90030827-DEL-1525 | chrX-90030828-DEL-1525 |
| chrX-90044281-DEL-58 | chrX-90044284-DEL-56 |
| chrX-90056733-DEL-7544 | chrX-90056734-DEL-7544 |
| chrX-90070807-DEL-1775 | chrX-90070808-DEL-1775 |
| chrX-90072525-DEL-62 | chrX-90072526-DEL-50 |
| chrX-90093566-DEL-6057 | chrX-90093550-DEL-6071 |
| chrX-90105998-DEL-53 | chrX-90105999-DEL-53 |
| chrX-90109973-DEL-61 | chrX-90109971-DEL-62 |
| chrX-90131264-DEL-279 | chrX-90131265-DEL-279 |
| chrX-90169444-DEL-2843 | chrX-90169445-DEL-2843 |
| chrX-90180720-DEL-72 | chrX-90180735-DEL-82 |
| chrX-90192926-DEL-60 | chrX-90192931-DEL-56 |
| chrX-90200811-DEL-6279 | chrX-90200812-DEL-6280 |
| chrX-90262616-DEL-80 | chrX-90262617-DEL-80 |
| chrX-90327356-DEL-77 | chrX-90327357-DEL-77 |
| chrX-90335467-DEL-56 | chrX-90335468-DEL-56 |
| chrX-90396785-DEL-148 | chrX-90396786-DEL-148 |

| | |
|---|---|
| chrX-90398358-DEL-489 | chrX-90398389-DEL-459 |
| chrX-90546513-DEL-73 | chrX-90546513-DEL-73 |
| chrX-90575034-DEL-60 | chrX-90575027-DEL-66 |
| chrX-90652315-DEL-6245 | chrX-90652316-DEL-6245 |
| chrX-90666279-DEL-178 | chrX-90666280-DEL-178 |
| chrX-90672758-DEL-64 | chrX-90672761-DEL-74 |
| chrX-90815496-DEL-84 | chrX-90815496-DEL-84 |
| chrX-90839409-DEL-4121 | chrX-90839410-DEL-4121 |
| chrX-90860576-DEL-7140 | chrX-90860577-DEL-7140 |
| chrX-90884989-DEL-2120 | chrX-90884988-DEL-2120 |
| chrX-90912499-DEL-1894 | chrX-90912500-DEL-1894 |
| chrX-90917231-DEL-201 | chrX-90917231-DEL-201 |
| chrX-90923020-DEL-277 | chrX-90923021-DEL-277 |
| chrX-90938447-DEL-7032 | chrX-90938451-DEL-7033 |
| chrX-90949024-DEL-377 | chrX-90949025-DEL-377 |
| chrX-90964300-DEL-52 | chrX-90964301-DEL-50 |
| chrX-91091874-DEL-58 | chrX-91091875-DEL-58 |
| chrX-91167179-DEL-1565 | chrX-91167183-DEL-1562 |
| chrX-91273693-DEL-348 | chrX-91273694-DEL-348 |
| chrX-91280272-DEL-2589 | chrX-91280273-DEL-2589 |
| chrX-91285981-DEL-201 | chrX-91285982-DEL-201 |
| chrX-91291975-DEL-6706 | chrX-91291976-DEL-6706 |
| chrX-91312196-DEL-6587 | chrX-91312200-DEL-6586 |
| chrX-91327114-DEL-64 | chrX-91327111-DEL-68 |
| chrX-91342990-DEL-3641 | chrX-91342990-DEL-3641 |
| chrX-91376138-DEL-125 | chrX-91376101-DEL-123 |
| chrX-91405289-DEL-840 | chrX-91405290-DEL-840 |
| chrX-91414329-DEL-74 | chrX-91414372-DEL-75 |
| chrX-91429850-DEL-213 | chrX-91429851-DEL-213 |
| chrX-91510156-DEL-115 | chrX-91510154-DEL-76 |
| chrX-91537259-DEL-107 | chrX-91537260-DEL-107 |
| chrX-91539369-DEL-242 | chrX-91539370-DEL-242 |
| chrX-91556401-DEL-74 | chrX-91556410-DEL-82 |
| chrX-91575329-DEL-162 | chrX-91575330-DEL-162 |
| chrX-91610908-DEL-630 | chrX-91610909-DEL-630 |
| chrX-91611846-DEL-3563 | chrX-91611847-DEL-3564 |
| chrX-91615878-DEL-6311 | chrX-91615879-DEL-6311 |
| chrX-91658444-DEL-6413 | chrX-91658445-DEL-6413 |
| chrX-91727963-DEL-560 | chrX-91727964-DEL-560 |
| chrX-91748630-DEL-62 | chrX-91748643-DEL-52 |
| chrX-91755832-DEL-6398 | chrX-91755833-DEL-6398 |
| chrX-91808791-DEL-2824 | chrX-91808792-DEL-2824 |
| chrX-91844227-DEL-6649 | chrX-91844228-DEL-6649 |
| chrX-91859028-DEL-1943 | chrX-91859029-DEL-1943 |
| chrX-91915366-DEL-1248 | chrX-91915367-DEL-1248 |
| chrX-91952600-DEL-6606 | chrX-91952601-DEL-6606 |
| chrX-91987011-DEL-6705 | chrX-91987012-DEL-6705 |
| chrX-92043353-DEL-399 | chrX-92043354-DEL-399 |
| chrX-92127860-DEL-1379 | chrX-92127861-DEL-1379 |
| chrX-92170760-DEL-410 | chrX-92170761-DEL-410 |
| chrX-92209474-DEL-51 | chrX-92209475-DEL-51 |
| chrX-92232669-DEL-1554 | chrX-92232670-DEL-1554 |
| chrX-92254351-DEL-99 | chrX-92254352-DEL-99 |
| chrX-92296107-DEL-102 | chrX-92296108-DEL-102 |
| chrX-92377241-DEL-60 | chrX-92377242-DEL-61 |
| chrX-92439352-DEL-538 | chrX-92439353-DEL-538 |
| chrX-92512130-DEL-1381 | chrX-92512126-DEL-1380 |
| chrX-92562290-DEL-181 | chrX-92562291-DEL-181 |
| chrX-92572384-DEL-524 | chrX-92572385-DEL-524 |
| chrX-92594169-DEL-206 | chrX-92594170-DEL-206 |
| chrX-92611457-DEL-3373 | chrX-92611458-DEL-3373 |
| chrX-92650217-DEL-5527 | chrX-92650218-DEL-5527 |
| chrX-92656447-DEL-3564 | chrX-92656448-DEL-3564 |
| chrX-92661342-DEL-2578 | chrX-92661343-DEL-2578 |
| chrX-92682010-DEL-2242 | chrX-92682013-DEL-2243 |
| chrX-92704231-DEL-109 | chrX-92704231-DEL-110 |
| chrX-92721601-DEL-78 | chrX-92721594-DEL-78 |
| chrX-92765477-DEL-6239 | chrX-92765478-DEL-6239 |
| chrX-92817732-DEL-665 | chrX-92817733-DEL-665 |
| chrX-92829462-DEL-205 | chrX-92829462-DEL-205 |
| chrX-92835871-DEL-208 | chrX-92835872-DEL-208 |
| chrX-92846824-DEL-284 | chrX-92846825-DEL-284 |
| chrX-92853715-DEL-2438 | chrX-92853716-DEL-2438 |
| chrX-92869638-DEL-733 | chrX-92869639-DEL-733 |
| chrX-92903589-DEL-67 | chrX-92903573-DEL-68 |
| chrX-92905388-DEL-120 | chrX-92905383-DEL-126 |
| chrX-92985759-DEL-401 | chrX-92985766-DEL-399 |
| chrX-93019396-DEL-246 | chrX-93019397-DEL-246 |
| chrX-93107161-DEL-152 | chrX-93107162-DEL-152 |
| chrX-93107625-DEL-497 | chrX-93107626-DEL-497 |

| | |
|---|---|
| chrX-93129398-DEL-366 | chrX-93129401-DEL-365 |
| chrX-93133715-DEL-80 | chrX-93133698-DEL-88 |
| chrX-93150725-DEL-206 | chrX-93150726-DEL-206 |
| chrX-93207068-DEL-233 | chrX-93207069-DEL-233 |
| chrX-93272357-DEL-84 | chrX-93272358-DEL-84 |
| chrX-93305403-DEL-370 | chrX-93305404-DEL-370 |
| chrX-93329866-DEL-205 | chrX-93329867-DEL-205 |
| chrX-93357911-DEL-145 | chrX-93357912-DEL-145 |
| chrX-93390026-DEL-71 | chrX-93390027-DEL-71 |
| chrX-93423072-DEL-54 | chrX-93423073-DEL-54 |
| chrX-93428716-DEL-184 | chrX-93428717-DEL-184 |
| chrX-93441048-DEL-56 | chrX-93441049-DEL-56 |
| chrX-93605822-DEL-75 | chrX-93605813-DEL-64 |
| chrX-93614641-DEL-1782 | chrX-93614642-DEL-1782 |
| chrX-93618874-DEL-4261 | chrX-93618875-DEL-4261 |
| chrX-93677413-DEL-60 | chrX-93677414-DEL-60 |
| chrX-93721855-DEL-433 | chrX-93721856-DEL-433 |
| chrX-93723122-DEL-202 | chrX-93723122-DEL-202 |
| chrX-93730379-DEL-2870 | chrX-93730380-DEL-2870 |
| chrX-93733587-DEL-808 | chrX-93733588-DEL-808 |
| chrX-93745432-DEL-256 | chrX-93745433-DEL-256 |
| chrX-93895396-DEL-203 | chrX-93895397-DEL-203 |
| chrX-94065023-DEL-93 | chrX-94065016-DEL-81 |
| chrX-94117814-DEL-208 | chrX-94117815-DEL-208 |
| chrX-94163012-DEL-567 | chrX-94163013-DEL-567 |
| chrX-94184495-DEL-1390 | chrX-94184496-DEL-1390 |
| chrX-94227502-DEL-1182 | chrX-94227503-DEL-1182 |
| chrX-94254224-DEL-82 | chrX-94254227-DEL-80 |
| chrX-94282866-DEL-78 | chrX-94282875-DEL-86 |
| chrX-94291803-DEL-64 | chrX-94291806-DEL-62 |
| chrX-94334919-DEL-208 | chrX-94334920-DEL-208 |
| chrX-94354148-DEL-577 | chrX-94354149-DEL-577 |
| chrX-94356375-DEL-1368 | chrX-94356376-DEL-1368 |
| chrX-94372189-DEL-54 | chrX-94372189-DEL-58 |
| chrX-94393474-DEL-4974 | chrX-94393475-DEL-4974 |
| chrX-94427744-DEL-206 | chrX-94427745-DEL-206 |
| chrX-94456034-DEL-6356 | chrX-94456035-DEL-6356 |
| chrX-94477396-DEL-54 | chrX-94477397-DEL-54 |
| chrX-94576593-DEL-113 | chrX-94576594-DEL-113 |
| chrX-94581160-DEL-333 | chrX-94581161-DEL-330 |
| chrX-94646615-DEL-216 | chrX-94646616-DEL-216 |
| chrX-94672421-DEL-493 | chrX-94672422-DEL-493 |
| chrX-94690630-DEL-105 | chrX-94690631-DEL-105 |
| chrX-94707408-DEL-1422 | chrX-94707409-DEL-1422 |
| chrX-94736891-DEL-6429 | chrX-94736892-DEL-6429 |
| chrX-94824142-DEL-74 | chrX-94824143-DEL-74 |
| chrX-94865858-DEL-6427 | chrX-94865859-DEL-6427 |
| chrX-94882235-DEL-306 | chrX-94882236-DEL-306 |
| chrX-94902494-DEL-1392 | chrX-94902495-DEL-1392 |
| chrX-94906513-DEL-735 | chrX-94906514-DEL-735 |
| chrX-94909554-DEL-225 | chrX-94909555-DEL-225 |
| chrX-94914042-DEL-6584 | chrX-94914043-DEL-6584 |
| chrX-94914056-DEL-208 | chrX-94914057-DEL-208 |
| chrX-94928321-DEL-221 | chrX-94928322-DEL-221 |
| chrX-94935645-DEL-90 | chrX-94935670-DEL-66 |
| chrX-95015485-DEL-1454 | chrX-95015458-DEL-1454 |
| chrX-95017572-DEL-102 | chrX-95017573-DEL-54 |
| chrX-95023917-DEL-836 | chrX-95023917-DEL-836 |
| chrX-95026086-DEL-201 | chrX-95026087-DEL-201 |
| chrX-95057099-DEL-66 | chrX-95057100-DEL-65 |
| chrX-95079648-DEL-82 | chrX-95079657-DEL-74 |
| chrX-95088827-DEL-786 | chrX-95088828-DEL-786 |
| chrX-95094558-DEL-1142 | chrX-95094559-DEL-1142 |
| chrX-95104097-DEL-1694 | chrX-95104097-DEL-1694 |
| chrX-95143270-DEL-143 | chrX-95143271-DEL-143 |
| chrX-95221505-DEL-126 | chrX-95221495-DEL-128 |
| chrX-95230519-DEL-1186 | chrX-95230537-DEL-1180 |
| chrX-95325090-DEL-308 | chrX-95325087-DEL-309 |
| chrX-95389531-DEL-50 | chrX-95389528-DEL-54 |
| chrX-95391409-DEL-5209 | chrX-95391410-DEL-5209 |
| chrX-95397146-DEL-67 | chrX-95397125-DEL-102 |
| chrX-95412158-DEL-180 | chrX-95412159-DEL-180 |
| chrX-95528449-DEL-661 | chrX-95528450-DEL-661 |
| chrX-95584544-DEL-2168 | chrX-95584536-DEL-2169 |
| chrX-95614672-DEL-57 | chrX-95614677-DEL-53 |
| chrX-95640562-DEL-2198 | chrX-95640563-DEL-2198 |
| chrX-95673829-DEL-799 | chrX-95673830-DEL-799 |
| chrX-95683545-DEL-53 | chrX-95683510-DEL-54 |
| chrX-95687680-DEL-181 | chrX-95687661-DEL-201 |
| chrX-95696898-DEL-6639 | chrX-95696899-DEL-6639 |

| | |
|---|---|
| chrX-95713416-DEL-740 | chrX-95713417-DEL-740 |
| chrX-95725634-DEL-5062 | chrX-95725635-DEL-5062 |
| chrX-95788873-DEL-400 | chrX-95788874-DEL-400 |
| chrX-95794918-DEL-64 | chrX-95794927-DEL-56 |
| chrX-95869802-DEL-8434 | chrX-95869803-DEL-8434 |
| chrX-95963849-DEL-833 | chrX-95963850-DEL-833 |
| chrX-96001294-DEL-3282 | chrX-96001295-DEL-3282 |
| chrX-96043899-DEL-4362 | chrX-96043909-DEL-4359 |
| chrX-96058387-DEL-1491 | chrX-96058388-DEL-1491 |
| chrX-96093302-DEL-6584 | chrX-96093303-DEL-6584 |
| chrX-96093312-DEL-208 | chrX-96093313-DEL-208 |
| chrX-96111108-DEL-58 | chrX-96111100-DEL-80 |
| chrX-96179475-DEL-460 | chrX-96179476-DEL-460 |
| chrX-96185341-DEL-481 | chrX-96185342-DEL-481 |
| chrX-96212718-DEL-1738 | chrX-96212719-DEL-1738 |
| chrX-96226564-DEL-4369 | chrX-96226565-DEL-4369 |
| chrX-96275319-DEL-82 | chrX-96275316-DEL-86 |
| chrX-96286215-DEL-183 | chrX-96286210-DEL-187 |
| chrX-96310485-DEL-445 | chrX-96310486-DEL-445 |
| chrX-96338290-DEL-642 | chrX-96338273-DEL-679 |
| chrX-96347777-DEL-5348 | chrX-96347778-DEL-5348 |
| chrX-96406236-DEL-316 | chrX-96406243-DEL-305 |
| chrX-96422591-DEL-106 | chrX-96422592-DEL-106 |
| chrX-96501411-DEL-2071 | chrX-96501412-DEL-2071 |
| chrX-96520363-DEL-308 | chrX-96520364-DEL-308 |
| chrX-96529180-DEL-3877 | chrX-96529181-DEL-3877 |
| chrX-96546396-DEL-60 | chrX-96546385-DEL-62 |
| chrX-96595704-DEL-89 | chrX-96595696-DEL-90 |
| chrX-96630385-DEL-3793 | chrX-96630386-DEL-3793 |
| chrX-96645506-DEL-112 | chrX-96645507-DEL-112 |
| chrX-96647819-DEL-7138 | chrX-96647820-DEL-7138 |
| chrX-96731372-DEL-1950 | chrX-96731373-DEL-1950 |
| chrX-96753279-DEL-482 | chrX-96753280-DEL-481 |
| chrX-96759850-DEL-553 | chrX-96759851-DEL-553 |
| chrX-96771495-DEL-6885 | chrX-96771496-DEL-6885 |
| chrX-96801373-DEL-2883 | chrX-96801418-DEL-2846 |
| chrX-96877200-DEL-6519 | chrX-96877201-DEL-6519 |
| chrX-96892436-DEL-369 | chrX-96892437-DEL-369 |
| chrX-97003753-DEL-7661 | chrX-97003754-DEL-7661 |
| chrX-97012662-DEL-7608 | chrX-97012648-DEL-7622 |
| chrX-97051617-DEL-2865 | chrX-97051618-DEL-2865 |
| chrX-97076288-DEL-212 | chrX-97076288-DEL-212 |
| chrX-97100346-DEL-62 | chrX-97100347-DEL-62 |
| chrX-97124619-DEL-225 | chrX-97124620-DEL-225 |
| chrX-97131397-DEL-226 | chrX-97131398-DEL-226 |
| chrX-97133666-DEL-52 | chrX-97133660-DEL-50 |
| chrX-97136094-DEL-788 | chrX-97136095-DEL-788 |
| chrX-97205786-DEL-7171 | chrX-97205787-DEL-7171 |
| chrX-97209759-DEL-83 | chrX-97209803-DEL-52 |
| chrX-97266161-DEL-6643 | chrX-97266162-DEL-6643 |
| chrX-97284047-DEL-255 | chrX-97284051-DEL-251 |
| chrX-97305541-DEL-60 | chrX-97305538-DEL-64 |
| chrX-97459090-DEL-859 | chrX-97459091-DEL-859 |
| chrX-97468171-DEL-6061 | chrX-97468172-DEL-6061 |
| chrX-97492796-DEL-340 | chrX-97492796-DEL-340 |
| chrX-97498395-DEL-109 | chrX-97498395-DEL-109 |
| chrX-97507925-DEL-6209 | chrX-97507926-DEL-6209 |
| chrX-97517083-DEL-411 | chrX-97517084-DEL-411 |
| chrX-97556237-DEL-51 | chrX-97556238-DEL-51 |
| chrX-97626603-DEL-849 | chrX-97626603-DEL-849 |
| chrX-97646777-DEL-1843 | chrX-97646778-DEL-1843 |
| chrX-97732811-DEL-495 | chrX-97732812-DEL-495 |
| chrX-97745146-DEL-58 | chrX-97745147-DEL-58 |
| chrX-97762172-DEL-770 | chrX-97762173-DEL-770 |
| chrX-97778158-DEL-349 | chrX-97778159-DEL-350 |
| chrX-97785368-DEL-204 | chrX-97785369-DEL-204 |
| chrX-97797733-DEL-80 | chrX-97797713-DEL-100 |
| chrX-97811792-DEL-251 | chrX-97811793-DEL-251 |
| chrX-97816517-DEL-497 | chrX-97816518-DEL-497 |
| chrX-97821227-DEL-381 | chrX-97821229-DEL-383 |
| chrX-97844943-DEL-977 | chrX-97844944-DEL-977 |
| chrX-97847116-DEL-1960 | chrX-97847117-DEL-1960 |
| chrX-97883324-DEL-163 | chrX-97883325-DEL-163 |
| chrX-97921573-DEL-693 | chrX-97921574-DEL-693 |
| chrX-97939269-DEL-204 | chrX-97939270-DEL-204 |
| chrX-97946219-DEL-241 | chrX-97946220-DEL-241 |
| chrX-98017562-DEL-468 | chrX-98017563-DEL-468 |
| chrX-98019058-DEL-1822 | chrX-98019059-DEL-1822 |
| chrX-98026967-DEL-90 | chrX-98026968-DEL-72 |
| chrX-98067289-DEL-235 | chrX-98067290-DEL-228 |

| | |
|---|---|
| chrX-98148341-DEL-157 | chrX-98148342-DEL-157 |
| chrX-98151193-DEL-205 | chrX-98151194-DEL-205 |
| chrX-98163412-DEL-56 | chrX-98163413-DEL-56 |
| chrX-98168512-DEL-64 | chrX-98168513-DEL-64 |
| chrX-98349637-DEL-287 | chrX-98349638-DEL-287 |
| chrX-98353362-DEL-205 | chrX-98353363-DEL-205 |
| chrX-98501460-DEL-52 | chrX-98501452-DEL-55 |
| chrX-98582336-DEL-129 | chrX-98582337-DEL-128 |
| chrX-98631519-DEL-183 | chrX-98631520-DEL-183 |
| chrX-98685654-DEL-981 | chrX-98685655-DEL-981 |
| chrX-98752545-DEL-897 | chrX-98752546-DEL-897 |
| chrX-98807936-DEL-3000 | chrX-98807937-DEL-3000 |
| chrX-98821090-DEL-80 | chrX-98821091-DEL-80 |
| chrX-98826153-DEL-1691 | chrX-98826154-DEL-1691 |
| chrX-98837585-DEL-478 | chrX-98837586-DEL-478 |
| chrX-98945174-DEL-7134 | chrX-98945175-DEL-7134 |
| chrX-98975353-DEL-116 | chrX-98975354-DEL-116 |
| chrX-98979858-DEL-747 | chrX-98979861-DEL-745 |
| chrX-99014582-DEL-162 | chrX-99014583-DEL-162 |
| chrX-99044725-DEL-6651 | chrX-99044726-DEL-6651 |
| chrX-99057550-DEL-932 | chrX-99057551-DEL-932 |
| chrX-99126580-DEL-794 | chrX-99126581-DEL-794 |
| chrX-99142719-DEL-78 | chrX-99142720-DEL-75 |
| chrX-99151651-DEL-7031 | chrX-99151652-DEL-7031 |
| chrX-99214440-DEL-6989 | chrX-99214434-DEL-6994 |
| chrX-99230165-DEL-56 | chrX-99230163-DEL-56 |
| chrX-99245617-DEL-56 | chrX-99245610-DEL-64 |
| chrX-99261138-DEL-227 | chrX-99261139-DEL-227 |
| chrX-99338302-DEL-7081 | chrX-99338303-DEL-7081 |
| chrX-99413861-DEL-190 | chrX-99413862-DEL-190 |
| chrX-99470069-DEL-58 | chrX-99470062-DEL-56 |
| chrX-99486055-DEL-143 | chrX-99486056-DEL-143 |
| chrX-99503836-DEL-617 | chrX-99503837-DEL-617 |
| chrX-99522103-DEL-424 | chrX-99522104-DEL-424 |
| chrX-99527299-DEL-580 | chrX-99527300-DEL-580 |
| chrX-99556607-DEL-210 | chrX-99556608-DEL-210 |
| chrX-99567755-DEL-437 | chrX-99567756-DEL-437 |
| chrX-99608110-DEL-658 | chrX-99608099-DEL-672 |
| chrX-99676912-DEL-101 | chrX-99676902-DEL-114 |
| chrX-99689046-DEL-60 | chrX-99689045-DEL-50 |
| chrX-99705316-DEL-876 | chrX-99705319-DEL-877 |
| chrX-99749603-DEL-95 | chrX-99749604-DEL-95 |
| chrX-99758496-DEL-86 | chrX-99758497-DEL-86 |
| chrX-99801400-DEL-52 | chrX-99801389-DEL-50 |
| chrX-99802586-DEL-6474 | chrX-99802587-DEL-6474 |
| chrX-99868131-DEL-679 | chrX-99868153-DEL-679 |
| chrX-99902648-DEL-6876 | chrX-99902649-DEL-6876 |
| chrX-99943006-DEL-231 | chrX-99943007-DEL-231 |
| chrX-99984393-DEL-116 | chrX-99984394-DEL-116 |
| chrX-100024583-DEL-3497 | chrX-100024571-DEL-3498 |
| chrX-100030433-DEL-803 | chrX-100030434-DEL-803 |
| chrX-100043098-DEL-169 | chrX-100043112-DEL-140 |
| chrX-100095710-DEL-147 | chrX-100095711-DEL-147 |
| chrX-100133384-DEL-145 | chrX-100133385-DEL-145 |
| chrX-100134362-DEL-63 | chrX-100134363-DEL-63 |
| chrX-100135519-DEL-527 | chrX-100135521-DEL-524 |
| chrX-100143061-DEL-205 | chrX-100143062-DEL-205 |
| chrX-100163671-DEL-52 | chrX-100163672-DEL-52 |
| chrX-100167519-DEL-203 | chrX-100167520-DEL-203 |
| chrX-100176760-DEL-211 | chrX-100176761-DEL-211 |
| chrX-100232241-DEL-208 | chrX-100232242-DEL-208 |
| chrX-100290817-DEL-72 | chrX-100290818-DEL-72 |
| chrX-100371166-DEL-212 | chrX-100371167-DEL-212 |
| chrX-100424656-DEL-100 | chrX-100424657-DEL-100 |
| chrX-100436163-DEL-247 | chrX-100436164-DEL-250 |
| chrX-100475571-DEL-61 | chrX-100475572-DEL-61 |
| chrX-100476248-DEL-104 | chrX-100476249-DEL-104 |
| chrX-100483564-DEL-195 | chrX-100483565-DEL-195 |
| chrX-100511690-DEL-421 | chrX-100511698-DEL-404 |
| chrX-100530435-DEL-212 | chrX-100530436-DEL-209 |
| chrX-100537059-DEL-56 | chrX-100537060-DEL-56 |
| chrX-100537920-DEL-256 | chrX-100537921-DEL-256 |
| chrX-100538526-DEL-165 | chrX-100538528-DEL-162 |
| chrX-100543794-DEL-6679 | chrX-100543795-DEL-6679 |
| chrX-100567646-DEL-178 | chrX-100567647-DEL-178 |
| chrX-100623637-DEL-201 | chrX-100623638-DEL-201 |
| chrX-100626950-DEL-211 | chrX-100626951-DEL-211 |
| chrX-100633171-DEL-346 | chrX-100633172-DEL-346 |
| chrX-100642232-DEL-148 | chrX-100642233-DEL-148 |
| chrX-100654543-DEL-60 | chrX-100654544-DEL-60 |

| | |
|---|---|
| chrX-100675140-DEL-108 | chrX-100675137-DEL-112 |
| chrX-100675749-DEL-209 | chrX-100675750-DEL-209 |
| chrX-100688239-DEL-193 | chrX-100688232-DEL-196 |
| chrX-100741313-DEL-83 | chrX-100741282-DEL-129 |
| chrX-100749257-DEL-198 | chrX-100749294-DEL-170 |
| chrX-100751927-DEL-182 | chrX-100751928-DEL-182 |
| chrX-100842480-DEL-72 | chrX-100842481-DEL-72 |
| chrX-100879671-DEL-55 | chrX-100879671-DEL-56 |
| chrX-100880223-DEL-154 | chrX-100880224-DEL-154 |
| chrX-100881197-DEL-70 | chrX-100881198-DEL-70 |
| chrX-100881577-DEL-1568 | chrX-100881578-DEL-1568 |
| chrX-100882560-DEL-360 | chrX-100882561-DEL-360 |
| chrX-100953010-DEL-96 | chrX-100953027-DEL-80 |
| chrX-101098131-DEL-496 | chrX-101098132-DEL-496 |
| chrX-101134703-DEL-75 | chrX-101134688-DEL-51 |
| chrX-101148570-DEL-386 | chrX-101148571-DEL-386 |
| chrX-101174703-DEL-199 | chrX-101174704-DEL-199 |
| chrX-101178206-DEL-1204 | chrX-101178207-DEL-1204 |
| chrX-101185263-DEL-386 | chrX-101185264-DEL-386 |
| chrX-101223394-DEL-274 | chrX-101223395-DEL-274 |
| chrX-101234309-DEL-83 | chrX-101234310-DEL-83 |
| chrX-101247956-DEL-227 | chrX-101247957-DEL-227 |
| chrX-101277688-DEL-225 | chrX-101277689-DEL-225 |
| chrX-101289333-DEL-74 | chrX-101289334-DEL-70 |
| chrX-101308468-DEL-59 | chrX-101308451-DEL-60 |
| chrX-101343501-DEL-196 | chrX-101343502-DEL-196 |
| chrX-101354577-DEL-1489 | chrX-101354578-DEL-1489 |
| chrX-101439620-DEL-787 | chrX-101439621-DEL-787 |
| chrX-101484607-DEL-67 | chrX-101484612-DEL-57 |
| chrX-101492921-DEL-341 | chrX-101492918-DEL-341 |
| chrX-101503962-DEL-84 | chrX-101503963-DEL-84 |
| chrX-101508651-DEL-2440 | chrX-101508652-DEL-2440 |
| chrX-101534909-DEL-215 | chrX-101534910-DEL-215 |
| chrX-101569910-DEL-146 | chrX-101569911-DEL-146 |
| chrX-101584055-DEL-54 | chrX-101584046-DEL-57 |
| chrX-101599370-DEL-90 | chrX-101599371-DEL-90 |
| chrX-101604295-DEL-6612 | chrX-101604296-DEL-6612 |
| chrX-101629331-DEL-198 | chrX-101629332-DEL-198 |
| chrX-101631543-DEL-192 | chrX-101631544-DEL-192 |
| chrX-101670608-DEL-78 | chrX-101670609-DEL-78 |
| chrX-101691992-DEL-70 | chrX-101691993-DEL-70 |
| chrX-101737922-DEL-102 | chrX-101737923-DEL-102 |
| chrX-101828669-DEL-1551 | chrX-101828670-DEL-1551 |
| chrX-101833608-DEL-52 | chrX-101833609-DEL-52 |
| chrX-101872944-DEL-58 | chrX-101872945-DEL-58 |
| chrX-101905639-DEL-258 | chrX-101905657-DEL-256 |
| chrX-101935278-DEL-97 | chrX-101935279-DEL-97 |
| chrX-102022537-DEL-688 | chrX-102022538-DEL-688 |
| chrX-102148035-DEL-754 | chrX-102148036-DEL-754 |
| chrX-102150826-DEL-658 | chrX-102150827-DEL-658 |
| chrX-102206424-DEL-2752 | chrX-102206425-DEL-2752 |
| chrX-102244186-DEL-173 | chrX-102244187-DEL-173 |
| chrX-102294911-DEL-56 | chrX-102294912-DEL-56 |
| chrX-102309432-DEL-56 | chrX-102309413-DEL-106 |
| chrX-102315378-DEL-203 | chrX-102315379-DEL-203 |
| chrX-102318837-DEL-190 | chrX-102318841-DEL-187 |
| chrX-102359987-DEL-405 | chrX-102359988-DEL-405 |
| chrX-102385774-DEL-75 | chrX-102385742-DEL-112 |
| chrX-102422483-DEL-206 | chrX-102422484-DEL-206 |
| chrX-102446139-DEL-193 | chrX-102446132-DEL-194 |
| chrX-102461275-DEL-169 | chrX-102461273-DEL-170 |
| chrX-102468419-DEL-177 | chrX-102468420-DEL-177 |
| chrX-102477596-DEL-322 | chrX-102477597-DEL-323 |
| chrX-102532360-DEL-58 | chrX-102532361-DEL-58 |
| chrX-102556867-DEL-228 | chrX-102556868-DEL-228 |
| chrX-102559698-DEL-56 | chrX-102559699-DEL-56 |
| chrX-102592630-DEL-234 | chrX-102592631-DEL-234 |
| chrX-102598682-DEL-107 | chrX-102598683-DEL-107 |
| chrX-102602427-DEL-186 | chrX-102602428-DEL-186 |
| chrX-102608345-DEL-551 | chrX-102608346-DEL-551 |
| chrX-102611346-DEL-64 | chrX-102611346-DEL-64 |
| chrX-102631302-DEL-213 | chrX-102631303-DEL-213 |
| chrX-102632887-DEL-59 | chrX-102632888-DEL-59 |
| chrX-102662875-DEL-160 | chrX-102662874-DEL-160 |
| chrX-102684022-DEL-230 | chrX-102684023-DEL-230 |
| chrX-102702013-DEL-126 | chrX-102702006-DEL-162 |
| chrX-102707716-DEL-142 | chrX-102707733-DEL-126 |
| chrX-102711538-DEL-72 | chrX-102711539-DEL-72 |
| chrX-102801308-DEL-319 | chrX-102801298-DEL-318 |
| chrX-102807155-DEL-2568 | chrX-102807156-DEL-2568 |

| | |
|---|---|
| chrX-102855549-DEL-1853 | chrX-102855550-DEL-1853 |
| chrX-102857957-DEL-3354 | chrX-102857958-DEL-3354 |
| chrX-102948022-DEL-210 | chrX-102948023-DEL-210 |
| chrX-102973822-DEL-204 | chrX-102973823-DEL-204 |
| chrX-103018548-DEL-219 | chrX-103018549-DEL-219 |
| chrX-103034560-DEL-308 | chrX-103034561-DEL-308 |
| chrX-103044326-DEL-138 | chrX-103044355-DEL-110 |
| chrX-103067020-DEL-80 | chrX-103067005-DEL-95 |
| chrX-103069026-DEL-2932 | chrX-103069027-DEL-2932 |
| chrX-103091834-DEL-2830 | chrX-103091835-DEL-2830 |
| chrX-103136318-DEL-53 | chrX-103136320-DEL-77 |
| chrX-103140745-DEL-140 | chrX-103140750-DEL-124 |
| chrX-103210932-DEL-6801 | chrX-103210933-DEL-6801 |
| chrX-103297103-DEL-393 | chrX-103297104-DEL-392 |
| chrX-103298305-DEL-349 | chrX-103298305-DEL-349 |
| chrX-103301908-DEL-674 | chrX-103301909-DEL-674 |
| chrX-103306981-DEL-132 | chrX-103306982-DEL-132 |
| chrX-103310177-DEL-6514 | chrX-103310178-DEL-6514 |
| chrX-103336798-DEL-144 | chrX-103336804-DEL-127 |
| chrX-103388856-DEL-3261 | chrX-103388857-DEL-3261 |
| chrX-103568183-DEL-62 | chrX-103568184-DEL-62 |
| chrX-103666431-DEL-175 | chrX-103666432-DEL-175 |
| chrX-103691342-DEL-64 | chrX-103691341-DEL-58 |
| chrX-103725088-DEL-120 | chrX-103725089-DEL-120 |
| chrX-103749312-DEL-818 | chrX-103749313-DEL-818 |
| chrX-103764953-DEL-6467 | chrX-103764954-DEL-6467 |
| chrX-103790050-DEL-958 | chrX-103790051-DEL-958 |
| chrX-103820922-DEL-3564 | chrX-103820923-DEL-3564 |
| chrX-103841884-DEL-6349 | chrX-103841872-DEL-6361 |
| chrX-103841980-DEL-212 | chrX-103841981-DEL-212 |
| chrX-103905170-DEL-8354 | chrX-103905171-DEL-8354 |
| chrX-103934285-DEL-493 | chrX-103934286-DEL-493 |
| chrX-103954347-DEL-868 | chrX-103954348-DEL-868 |
| chrX-103989841-DEL-76 | chrX-103989842-DEL-76 |
| chrX-104061629-DEL-207 | chrX-104061630-DEL-207 |
| chrX-104087785-DEL-94 | chrX-104087796-DEL-84 |
| chrX-104093171-DEL-66 | chrX-104093172-DEL-66 |
| chrX-104109170-DEL-159 | chrX-104109171-DEL-159 |
| chrX-104117829-DEL-535 | chrX-104117830-DEL-535 |
| chrX-104136732-DEL-3438 | chrX-104136733-DEL-3438 |
| chrX-104147057-DEL-519 | chrX-104147058-DEL-519 |
| chrX-104193901-DEL-2103 | chrX-104193902-DEL-2103 |
| chrX-104207974-DEL-1031 | chrX-104207974-DEL-1034 |
| chrX-104226479-DEL-615 | chrX-104226486-DEL-609 |
| chrX-104241790-DEL-880 | chrX-104241794-DEL-864 |
| chrX-104246237-DEL-196 | chrX-104246238-DEL-197 |
| chrX-104286173-DEL-1203 | chrX-104286174-DEL-1203 |
| chrX-104292881-DEL-6382 | chrX-104292881-DEL-6382 |
| chrX-104296599-DEL-2701 | chrX-104296600-DEL-2701 |
| chrX-104322166-DEL-506 | chrX-104322167-DEL-506 |
| chrX-104328080-DEL-198 | chrX-104328081-DEL-198 |
| chrX-104335344-DEL-7101 | chrX-104335343-DEL-7101 |
| chrX-104356619-DEL-6214 | chrX-104356620-DEL-6214 |
| chrX-104365588-DEL-239 | chrX-104365589-DEL-239 |
| chrX-104469720-DEL-538 | chrX-104469721-DEL-538 |
| chrX-104471376-DEL-83 | chrX-104471381-DEL-78 |
| chrX-104524448-DEL-276 | chrX-104524420-DEL-263 |
| chrX-104551366-DEL-616 | chrX-104551367-DEL-616 |
| chrX-104553748-DEL-65 | chrX-104553749-DEL-64 |
| chrX-104582926-DEL-90 | chrX-104582928-DEL-86 |
| chrX-104596772-DEL-1927 | chrX-104596780-DEL-1934 |
| chrX-104616116-DEL-8950 | chrX-104616117-DEL-8949 |
| chrX-104658500-DEL-6432 | chrX-104658501-DEL-6433 |
| chrX-104687001-DEL-639 | chrX-104687001-DEL-644 |
| chrX-104691627-DEL-5107 | chrX-104691628-DEL-5107 |
| chrX-104709430-DEL-90 | chrX-104709431-DEL-90 |
| chrX-104719811-DEL-111 | chrX-104719812-DEL-111 |
| chrX-104720776-DEL-818 | chrX-104720776-DEL-818 |
| chrX-104732723-DEL-6224 | chrX-104732724-DEL-6224 |
| chrX-104767346-DEL-76 | chrX-104767351-DEL-72 |
| chrX-104769973-DEL-456 | chrX-104769991-DEL-447 |
| chrX-104779912-DEL-13512 | chrX-104779912-DEL-13512 |
| chrX-104805483-DEL-221 | chrX-104805478-DEL-215 |
| chrX-104813564-DEL-607 | chrX-104813565-DEL-607 |
| chrX-104951206-DEL-56 | chrX-104951180-DEL-73 |
| chrX-104963715-DEL-374 | chrX-104963716-DEL-374 |
| chrX-104965354-DEL-55 | chrX-104965354-DEL-55 |
| chrX-104995298-DEL-200 | chrX-104995299-DEL-200 |
| chrX-105001793-DEL-168 | chrX-105001798-DEL-172 |
| chrX-105005107-DEL-204 | chrX-105005108-DEL-204 |

| | |
|---|---|
| chrX-105008436-DEL-214 | chrX-105008437-DEL-214 |
| chrX-105011116-DEL-165 | chrX-105011117-DEL-165 |
| chrX-105022543-DEL-178 | chrX-105022590-DEL-150 |
| chrX-105051163-DEL-216 | chrX-105051178-DEL-215 |
| chrX-105051521-DEL-1224 | chrX-105051522-DEL-1224 |
| chrX-105058547-DEL-207 | chrX-105058548-DEL-207 |
| chrX-105062279-DEL-214 | chrX-105062280-DEL-214 |
| chrX-105097051-DEL-2099 | chrX-105097051-DEL-2097 |
| chrX-105097212-DEL-205 | chrX-105097213-DEL-205 |
| chrX-105107956-DEL-71 | chrX-105107955-DEL-72 |
| chrX-105139898-DEL-281 | chrX-105139899-DEL-281 |
| chrX-105140587-DEL-57 | chrX-105140548-DEL-57 |
| chrX-105141829-DEL-186 | chrX-105141830-DEL-186 |
| chrX-105164505-DEL-205 | chrX-105164506-DEL-205 |
| chrX-105175575-DEL-165 | chrX-105175576-DEL-164 |
| chrX-105180638-DEL-412 | chrX-105180639-DEL-412 |
| chrX-105224686-DEL-1892 | chrX-105224687-DEL-1892 |
| chrX-105311644-DEL-444 | chrX-105311645-DEL-444 |
| chrX-105326437-DEL-2282 | chrX-105326438-DEL-2282 |
| chrX-105351968-DEL-66 | chrX-105351969-DEL-66 |
| chrX-105367341-DEL-3179 | chrX-105367342-DEL-3179 |
| chrX-105368128-DEL-1292 | chrX-105368128-DEL-1292 |
| chrX-105376559-DEL-510 | chrX-105376560-DEL-510 |
| chrX-105390083-DEL-181 | chrX-105390084-DEL-181 |
| chrX-105472454-DEL-1690 | chrX-105472455-DEL-1690 |
| chrX-105544951-DEL-989 | chrX-105544952-DEL-989 |
| chrX-105597181-DEL-52 | chrX-105597182-DEL-52 |
| chrX-105601134-DEL-131 | chrX-105601135-DEL-131 |
| chrX-105643980-DEL-403 | chrX-105643981-DEL-403 |
| chrX-105671744-DEL-613 | chrX-105671745-DEL-613 |
| chrX-105673335-DEL-1751 | chrX-105673327-DEL-1752 |
| chrX-105688413-DEL-236 | chrX-105688414-DEL-236 |
| chrX-105707793-DEL-996 | chrX-105707793-DEL-996 |
| chrX-105709026-DEL-1808 | chrX-105709027-DEL-1808 |
| chrX-105734505-DEL-4087 | chrX-105734506-DEL-4087 |
| chrX-105762750-DEL-2342 | chrX-105762753-DEL-2341 |
| chrX-105772251-DEL-214 | chrX-105772252-DEL-214 |
| chrX-105778888-DEL-1224 | chrX-105778888-DEL-1224 |
| chrX-105784299-DEL-472 | chrX-105784299-DEL-474 |
| chrX-105799788-DEL-52 | chrX-105799789-DEL-80 |
| chrX-105809610-DEL-892 | chrX-105809612-DEL-891 |
| chrX-105828536-DEL-205 | chrX-105828537-DEL-205 |
| chrX-105830016-DEL-658 | chrX-105830017-DEL-658 |
| chrX-105885526-DEL-71 | chrX-105885514-DEL-72 |
| chrX-105945120-DEL-594 | chrX-105945121-DEL-594 |
| chrX-105953828-DEL-122 | chrX-105953829-DEL-122 |
| chrX-105954044-DEL-942 | chrX-105954045-DEL-942 |
| chrX-105959898-DEL-223 | chrX-105959899-DEL-223 |
| chrX-105961706-DEL-436 | chrX-105961707-DEL-436 |
| chrX-105989559-DEL-64 | chrX-105989574-DEL-50 |
| chrX-106038924-DEL-196 | chrX-106038925-DEL-196 |
| chrX-106045870-DEL-1285 | chrX-106045871-DEL-1285 |
| chrX-106089608-DEL-11537 | chrX-106089609-DEL-11537 |
| chrX-106103069-DEL-2995 | chrX-106103070-DEL-2995 |
| chrX-106130854-DEL-53 | chrX-106130844-DEL-54 |
| chrX-106158120-DEL-15967 | chrX-106158121-DEL-15967 |
| chrX-106168255-DEL-268 | chrX-106168256-DEL-268 |
| chrX-106189003-DEL-65 | chrX-106189004-DEL-65 |
| chrX-106237203-DEL-155 | chrX-106237199-DEL-156 |
| chrX-106280382-DEL-1214 | chrX-106280379-DEL-1215 |
| chrX-106287520-DEL-3065 | chrX-106287521-DEL-3065 |
| chrX-106347893-DEL-152 | chrX-106347892-DEL-154 |
| chrX-106358066-DEL-1704 | chrX-106358067-DEL-1704 |
| chrX-106363357-DEL-53 | chrX-106363358-DEL-53 |
| chrX-106364482-DEL-615 | chrX-106364483-DEL-615 |
| chrX-106374006-DEL-6249 | chrX-106374007-DEL-6249 |
| chrX-106443438-DEL-170 | chrX-106443439-DEL-170 |
| chrX-106473358-DEL-98 | chrX-106473359-DEL-98 |
| chrX-106475148-DEL-179 | chrX-106475148-DEL-175 |
| chrX-106522368-DEL-76 | chrX-106522369-DEL-114 |
| chrX-106530888-DEL-66 | chrX-106530903-DEL-62 |
| chrX-106559717-DEL-5982 | chrX-106559718-DEL-5982 |
| chrX-106633124-DEL-71 | chrX-106633121-DEL-74 |
| chrX-106636448-DEL-171 | chrX-106636449-DEL-171 |
| chrX-106645582-DEL-6722 | chrX-106645583-DEL-6722 |
| chrX-106685490-DEL-428 | chrX-106685491-DEL-428 |
| chrX-106774503-DEL-7595 | chrX-106774504-DEL-7595 |
| chrX-106779829-DEL-1397 | chrX-106779830-DEL-1397 |
| chrX-106963373-DEL-62 | chrX-106963377-DEL-56 |
| chrX-107001594-DEL-6003 | chrX-107001594-DEL-6003 |

| | |
|---|---|
| chrX-107022144-DEL-8728 | chrX-107022145-DEL-8728 |
| chrX-107030491-DEL-201 | chrX-107030447-DEL-201 |
| chrX-107061579-DEL-4761 | chrX-107061580-DEL-4761 |
| chrX-107114570-DEL-6791 | chrX-107114570-DEL-6791 |
| chrX-107142320-DEL-185 | chrX-107142321-DEL-185 |
| chrX-107174585-DEL-142 | chrX-107174586-DEL-142 |
| chrX-107197633-DEL-62 | chrX-107197630-DEL-62 |
| chrX-107219297-DEL-2395 | chrX-107219298-DEL-2395 |
| chrX-107246605-DEL-358 | chrX-107246606-DEL-358 |
| chrX-107254839-DEL-96 | chrX-107254840-DEL-96 |
| chrX-107316733-DEL-60 | chrX-107316736-DEL-58 |
| chrX-107429039-DEL-2207 | chrX-107429040-DEL-2207 |
| chrX-107518944-DEL-289 | chrX-107518945-DEL-289 |
| chrX-107539680-DEL-6374 | chrX-107539681-DEL-6374 |
| chrX-107561505-DEL-1744 | chrX-107561506-DEL-1744 |
| chrX-107565463-DEL-152 | chrX-107565464-DEL-152 |
| chrX-107654588-DEL-1615 | chrX-107654588-DEL-1615 |
| chrX-107671362-DEL-1452 | chrX-107671363-DEL-1452 |
| chrX-107697343-DEL-1294 | chrX-107697332-DEL-1293 |
| chrX-107709212-DEL-385 | chrX-107709213-DEL-383 |
| chrX-107791014-DEL-63 | chrX-107791015-DEL-63 |
| chrX-107823908-DEL-105 | chrX-107823946-DEL-94 |
| chrX-107858824-DEL-269 | chrX-107858825-DEL-269 |
| chrX-107861041-DEL-145 | chrX-107861042-DEL-145 |
| chrX-107900355-DEL-168 | chrX-107900350-DEL-166 |
| chrX-107927289-DEL-53 | chrX-107927257-DEL-56 |
| chrX-107931032-DEL-58 | chrX-107931038-DEL-55 |
| chrX-107948520-DEL-17607 | chrX-107948521-DEL-17606 |
| chrX-107967040-DEL-225 | chrX-107967040-DEL-225 |
| chrX-108018116-DEL-193 | chrX-108018116-DEL-193 |
| chrX-108068657-DEL-53 | chrX-108068658-DEL-53 |
| chrX-108096422-DEL-7590 | chrX-108096423-DEL-7590 |
| chrX-108101737-DEL-1235 | chrX-108101736-DEL-1235 |
| chrX-108105549-DEL-113 | chrX-108105550-DEL-113 |
| chrX-108112254-DEL-1850 | chrX-108112242-DEL-1861 |
| chrX-108122880-DEL-209 | chrX-108122881-DEL-209 |
| chrX-108172337-DEL-63 | chrX-108172342-DEL-59 |
| chrX-108253480-DEL-356 | chrX-108253481-DEL-356 |
| chrX-108261400-DEL-1068 | chrX-108261401-DEL-1068 |
| chrX-108282749-DEL-57 | chrX-108282750-DEL-57 |
| chrX-108290233-DEL-1474 | chrX-108290234-DEL-1470 |
| chrX-108328217-DEL-416 | chrX-108328217-DEL-416 |
| chrX-108363710-DEL-57 | chrX-108363711-DEL-57 |
| chrX-108367810-DEL-62 | chrX-108367815-DEL-58 |
| chrX-108382630-DEL-285 | chrX-108382631-DEL-285 |
| chrX-108389308-DEL-282 | chrX-108389309-DEL-280 |
| chrX-108412621-DEL-6204 | chrX-108412622-DEL-6204 |
| chrX-108479044-DEL-5070 | chrX-108479045-DEL-5070 |
| chrX-108491498-DEL-425 | chrX-108491471-DEL-427 |
| chrX-108499145-DEL-3067 | chrX-108499146-DEL-3067 |
| chrX-108536371-DEL-6346 | chrX-108536371-DEL-6346 |
| chrX-108542846-DEL-1379 | chrX-108542847-DEL-1379 |
| chrX-108548397-DEL-1144 | chrX-108548398-DEL-1144 |
| chrX-108580793-DEL-6194 | chrX-108580794-DEL-6194 |
| chrX-108647982-DEL-64 | chrX-108647983-DEL-64 |
| chrX-108747608-DEL-6244 | chrX-108747613-DEL-6242 |
| chrX-108824857-DEL-1492 | chrX-108824858-DEL-1492 |
| chrX-108877846-DEL-5071 | chrX-108877847-DEL-5071 |
| chrX-108919290-DEL-161 | chrX-108919291-DEL-161 |
| chrX-108944192-DEL-605 | chrX-108944193-DEL-605 |
| chrX-108964608-DEL-89 | chrX-108964609-DEL-89 |
| chrX-108992413-DEL-76 | chrX-108992414-DEL-98 |
| chrX-109021139-DEL-7032 | chrX-109021140-DEL-7032 |
| chrX-109027340-DEL-208 | chrX-109027341-DEL-208 |
| chrX-109130198-DEL-6640 | chrX-109130198-DEL-6640 |
| chrX-109157435-DEL-515 | chrX-109157436-DEL-515 |
| chrX-109267448-DEL-494 | chrX-109267449-DEL-494 |
| chrX-109303871-DEL-538 | chrX-109303872-DEL-538 |
| chrX-109358963-DEL-164 | chrX-109358964-DEL-164 |
| chrX-109372509-DEL-2061 | chrX-109372510-DEL-2061 |
| chrX-109424295-DEL-473 | chrX-109424296-DEL-473 |
| chrX-109430082-DEL-101 | chrX-109430083-DEL-102 |
| chrX-109436050-DEL-201 | chrX-109436051-DEL-201 |
| chrX-109440190-DEL-54 | chrX-109440191-DEL-54 |
| chrX-109443619-DEL-82 | chrX-109443620-DEL-82 |
| chrX-109450418-DEL-248 | chrX-109450419-DEL-248 |
| chrX-109460410-DEL-7053 | chrX-109460411-DEL-7053 |
| chrX-109529133-DEL-1551 | chrX-109529134-DEL-1552 |
| chrX-109536608-DEL-855 | chrX-109536609-DEL-855 |
| chrX-109558302-DEL-6402 | chrX-109558303-DEL-6402 |

| | |
|---|---|
| chrX-109566420-DEL-1265 | chrX-109566421-DEL-1265 |
| chrX-109610902-DEL-54 | chrX-109610869-DEL-66 |
| chrX-109627263-DEL-184 | chrX-109627264-DEL-184 |
| chrX-109655511-DEL-6352 | chrX-109655512-DEL-6352 |
| chrX-109662894-DEL-498 | chrX-109662895-DEL-498 |
| chrX-109728358-DEL-54 | chrX-109728359-DEL-53 |
| chrX-109769102-DEL-6413 | chrX-109769103-DEL-6413 |
| chrX-109777403-DEL-3131 | chrX-109777404-DEL-3131 |
| chrX-109875088-DEL-63 | chrX-109875082-DEL-69 |
| chrX-109906706-DEL-4800 | chrX-109906707-DEL-4800 |
| chrX-109923670-DEL-7839 | chrX-109923671-DEL-7839 |
| chrX-109924011-DEL-208 | chrX-109924012-DEL-208 |
| chrX-109988865-DEL-8768 | chrX-109988866-DEL-8768 |
| chrX-110032840-DEL-396 | chrX-110032841-DEL-396 |
| chrX-110038160-DEL-822 | chrX-110038161-DEL-822 |
| chrX-110039477-DEL-258 | chrX-110039478-DEL-258 |
| chrX-110074323-DEL-527 | chrX-110074324-DEL-528 |
| chrX-110142446-DEL-6438 | chrX-110142447-DEL-6438 |
| chrX-110156228-DEL-55 | chrX-110156229-DEL-55 |
| chrX-110159075-DEL-1484 | chrX-110159076-DEL-1484 |
| chrX-110181963-DEL-338 | chrX-110181964-DEL-338 |
| chrX-110200627-DEL-320 | chrX-110200617-DEL-328 |
| chrX-110209027-DEL-1153 | chrX-110209028-DEL-1153 |
| chrX-110212741-DEL-210 | chrX-110212742-DEL-210 |
| chrX-110214907-DEL-1478 | chrX-110214908-DEL-1478 |
| chrX-110248430-DEL-846 | chrX-110248431-DEL-846 |
| chrX-110270343-DEL-6648 | chrX-110270344-DEL-6648 |
| chrX-110293113-DEL-505 | chrX-110293114-DEL-505 |
| chrX-110375814-DEL-54 | chrX-110375815-DEL-54 |
| chrX-110388622-DEL-4575 | chrX-110388623-DEL-4575 |
| chrX-110503812-DEL-288 | chrX-110503807-DEL-291 |
| chrX-110627995-DEL-6217 | chrX-110627996-DEL-6217 |
| chrX-110641321-DEL-100 | chrX-110641335-DEL-68 |
| chrX-110668337-DEL-172 | chrX-110668338-DEL-172 |
| chrX-110688026-DEL-76 | chrX-110688027-DEL-76 |
| chrX-110691309-DEL-6823 | chrX-110691310-DEL-6823 |
| chrX-110697558-DEL-214 | chrX-110697559-DEL-214 |
| chrX-110733876-DEL-58 | chrX-110733877-DEL-58 |
| chrX-110762799-DEL-336 | chrX-110762800-DEL-336 |
| chrX-110815145-DEL-192 | chrX-110815146-DEL-192 |
| chrX-110945203-DEL-311 | chrX-110945204-DEL-311 |
| chrX-110958887-DEL-1325 | chrX-110958888-DEL-1325 |
| chrX-110979635-DEL-105 | chrX-110979636-DEL-105 |
| chrX-110986754-DEL-1950 | chrX-110986753-DEL-1950 |
| chrX-111017047-DEL-78 | chrX-111017048-DEL-78 |
| chrX-111054735-DEL-6777 | chrX-111054736-DEL-6777 |
| chrX-111107099-DEL-1326 | chrX-111107100-DEL-1326 |
| chrX-111110395-DEL-944 | chrX-111110396-DEL-944 |
| chrX-111143912-DEL-79 | chrX-111143899-DEL-80 |
| chrX-111166401-DEL-72 | chrX-111166402-DEL-54 |
| chrX-111280399-DEL-550 | chrX-111280394-DEL-550 |
| chrX-111287043-DEL-1849 | chrX-111287044-DEL-1849 |
| chrX-111295350-DEL-3177 | chrX-111295351-DEL-3177 |
| chrX-111332150-DEL-681 | chrX-111332151-DEL-681 |
| chrX-111336674-DEL-56 | chrX-111336675-DEL-56 |
| chrX-111389774-DEL-214 | chrX-111389775-DEL-214 |
| chrX-111443673-DEL-3085 | chrX-111443674-DEL-3085 |
| chrX-111487179-DEL-6285 | chrX-111487180-DEL-6285 |
| chrX-111497442-DEL-275 | chrX-111497443-DEL-275 |
| chrX-111595344-DEL-425 | chrX-111595345-DEL-425 |
| chrX-111619913-DEL-310 | chrX-111619911-DEL-311 |
| chrX-111707259-DEL-524 | chrX-111707260-DEL-524 |
| chrX-111730612-DEL-64 | chrX-111730617-DEL-60 |
| chrX-111732477-DEL-72 | chrX-111732477-DEL-54 |
| chrX-111749799-DEL-6413 | chrX-111749800-DEL-6413 |
| chrX-111768828-DEL-60 | chrX-111768825-DEL-72 |
| chrX-111782448-DEL-60 | chrX-111782449-DEL-60 |
| chrX-111829008-DEL-1889 | chrX-111829009-DEL-1889 |
| chrX-111847093-DEL-112 | chrX-111847094-DEL-112 |
| chrX-111848451-DEL-4406 | chrX-111848452-DEL-4406 |
| chrX-111868791-DEL-2304 | chrX-111868792-DEL-2304 |
| chrX-111872479-DEL-1123 | chrX-111872468-DEL-1132 |
| chrX-111928295-DEL-128 | chrX-111928296-DEL-128 |
| chrX-112000888-DEL-1932 | chrX-112000889-DEL-1932 |
| chrX-112027057-DEL-74 | chrX-112027020-DEL-75 |
| chrX-112046361-DEL-191 | chrX-112046362-DEL-191 |
| chrX-112081003-DEL-55 | chrX-112081004-DEL-55 |
| chrX-112105295-DEL-6205 | chrX-112105296-DEL-6205 |
| chrX-112122000-DEL-55 | chrX-112122001-DEL-54 |
| chrX-112130987-DEL-1533 | chrX-112130988-DEL-1533 |

| | |
|---|---|
| chrX-112153844-DEL-6463 | chrX-112153845-DEL-6463 |
| chrX-112195750-DEL-2692 | chrX-112195751-DEL-2692 |
| chrX-112229874-DEL-3839 | chrX-112229875-DEL-3839 |
| chrX-112246265-DEL-747 | chrX-112246266-DEL-747 |
| chrX-112251889-DEL-4030 | chrX-112251890-DEL-4030 |
| chrX-112261571-DEL-6698 | chrX-112261572-DEL-6698 |
| chrX-112287731-DEL-506 | chrX-112287732-DEL-506 |
| chrX-112294385-DEL-76 | chrX-112294386-DEL-76 |
| chrX-112321963-DEL-76 | chrX-112321956-DEL-84 |
| chrX-112406829-DEL-3969 | chrX-112406830-DEL-3969 |
| chrX-112414255-DEL-969 | chrX-112414256-DEL-969 |
| chrX-112418626-DEL-540 | chrX-112418627-DEL-540 |
| chrX-112444436-DEL-51 | chrX-112444431-DEL-57 |
| chrX-112445284-DEL-56 | chrX-112445285-DEL-56 |
| chrX-112470883-DEL-1076 | chrX-112470884-DEL-1076 |
| chrX-112519121-DEL-201 | chrX-112519122-DEL-201 |
| chrX-112559735-DEL-3330 | chrX-112559731-DEL-3334 |
| chrX-112615817-DEL-171 | chrX-112615818-DEL-171 |
| chrX-112629489-DEL-106 | chrX-112629490-DEL-106 |
| chrX-112637837-DEL-57 | chrX-112637846-DEL-60 |
| chrX-112653883-DEL-1766 | chrX-112653884-DEL-1766 |
| chrX-112709806-DEL-5217 | chrX-112709807-DEL-5217 |
| chrX-112727537-DEL-64 | chrX-112727538-DEL-58 |
| chrX-112737558-DEL-1189 | chrX-112737559-DEL-1189 |
| chrX-112765774-DEL-820 | chrX-112765775-DEL-820 |
| chrX-112820081-DEL-5199 | chrX-112820082-DEL-5199 |
| chrX-112840968-DEL-95 | chrX-112840969-DEL-95 |
| chrX-112841808-DEL-3187 | chrX-112841809-DEL-3187 |
| chrX-112854131-DEL-135 | chrX-112854132-DEL-133 |
| chrX-112914422-DEL-479 | chrX-112914423-DEL-479 |
| chrX-112985261-DEL-55 | chrX-112985287-DEL-50 |
| chrX-112999855-DEL-132 | chrX-112999856-DEL-132 |
| chrX-113005884-DEL-55 | chrX-113005884-DEL-55 |
| chrX-113011737-DEL-6510 | chrX-113011738-DEL-6510 |
| chrX-113088101-DEL-205 | chrX-113088102-DEL-205 |
| chrX-113093021-DEL-589 | chrX-113093013-DEL-580 |
| chrX-113096918-DEL-2795 | chrX-113096919-DEL-2795 |
| chrX-113105155-DEL-249 | chrX-113105156-DEL-249 |
| chrX-113150075-DEL-674 | chrX-113150076-DEL-674 |
| chrX-113161150-DEL-198 | chrX-113161135-DEL-199 |
| chrX-113176546-DEL-1060 | chrX-113176546-DEL-1060 |
| chrX-113194944-DEL-134 | chrX-113194945-DEL-134 |
| chrX-113205619-DEL-5621 | chrX-113205620-DEL-5621 |
| chrX-113236868-DEL-5568 | chrX-113236869-DEL-5568 |
| chrX-113299551-DEL-931 | chrX-113299552-DEL-931 |
| chrX-113321283-DEL-5023 | chrX-113321284-DEL-5023 |
| chrX-113381676-DEL-95 | chrX-113381676-DEL-95 |
| chrX-113406865-DEL-6401 | chrX-113406866-DEL-6401 |
| chrX-113442786-DEL-6423 | chrX-113442787-DEL-6423 |
| chrX-113494354-DEL-102 | chrX-113494367-DEL-96 |
| chrX-113508155-DEL-76 | chrX-113508156-DEL-76 |
| chrX-113550211-DEL-92 | chrX-113550214-DEL-92 |
| chrX-113550895-DEL-1130 | chrX-113550897-DEL-1130 |
| chrX-113566513-DEL-4726 | chrX-113566514-DEL-4726 |
| chrX-113666196-DEL-214 | chrX-113666197-DEL-214 |
| chrX-113707775-DEL-90 | chrX-113707780-DEL-86 |
| chrX-113782377-DEL-5468 | chrX-113782378-DEL-5468 |
| chrX-113799695-DEL-4540 | chrX-113799696-DEL-4540 |
| chrX-113813841-DEL-205 | chrX-113813842-DEL-205 |
| chrX-113886580-DEL-205 | chrX-113886581-DEL-205 |
| chrX-113888067-DEL-520 | chrX-113888068-DEL-520 |
| chrX-113934220-DEL-279 | chrX-113934221-DEL-279 |
| chrX-113942279-DEL-478 | chrX-113942280-DEL-478 |
| chrX-113953608-DEL-169 | chrX-113953609-DEL-169 |
| chrX-113956400-DEL-1484 | chrX-113956401-DEL-1481 |
| chrX-113958557-DEL-52 | chrX-113958560-DEL-52 |
| chrX-113985399-DEL-6538 | chrX-113985400-DEL-6538 |
| chrX-113991479-DEL-208 | chrX-113991480-DEL-208 |
| chrX-114000721-DEL-2107 | chrX-114000722-DEL-2107 |
| chrX-114020617-DEL-200 | chrX-114020617-DEL-200 |
| chrX-114024605-DEL-1468 | chrX-114024605-DEL-1468 |
| chrX-114030208-DEL-117 | chrX-114030208-DEL-117 |
| chrX-114037055-DEL-198 | chrX-114037056-DEL-198 |
| chrX-114043187-DEL-240 | chrX-114043189-DEL-236 |
| chrX-114052924-DEL-6696 | chrX-114052925-DEL-6696 |
| chrX-114085789-DEL-96 | chrX-114085778-DEL-110 |
| chrX-114093190-DEL-2820 | chrX-114093191-DEL-2820 |
| chrX-114096106-DEL-922 | chrX-114096107-DEL-922 |
| chrX-114098688-DEL-501 | chrX-114098689-DEL-501 |
| chrX-114101241-DEL-52 | chrX-114101244-DEL-50 |

| | |
|---|---|
| chrX-114136523-DEL-60 | chrX-114136524-DEL-60 |
| chrX-114139353-DEL-172 | chrX-114139343-DEL-173 |
| chrX-114190216-DEL-162 | chrX-114190217-DEL-162 |
| chrX-114227652-DEL-224 | chrX-114227653-DEL-224 |
| chrX-114228474-DEL-1170 | chrX-114228475-DEL-1170 |
| chrX-114271358-DEL-64 | chrX-114271359-DEL-64 |
| chrX-114271993-DEL-66 | chrX-114271994-DEL-62 |
| chrX-114294122-DEL-6586 | chrX-114294123-DEL-6586 |
| chrX-114324205-DEL-7559 | chrX-114324206-DEL-7560 |
| chrX-114338925-DEL-66 | chrX-114338926-DEL-66 |
| chrX-114404785-DEL-3601 | chrX-114404786-DEL-3601 |
| chrX-114425006-DEL-2504 | chrX-114425007-DEL-2504 |
| chrX-114435548-DEL-58 | chrX-114435549-DEL-58 |
| chrX-114459504-DEL-7117 | chrX-114459505-DEL-7117 |
| chrX-114475641-DEL-1544 | chrX-114475642-DEL-1544 |
| chrX-114519165-DEL-77 | chrX-114519166-DEL-77 |
| chrX-114528922-DEL-55 | chrX-114528903-DEL-77 |
| chrX-114583959-DEL-574 | chrX-114583958-DEL-574 |
| chrX-114600899-DEL-1659 | chrX-114600900-DEL-1659 |
| chrX-114622526-DEL-414 | chrX-114622527-DEL-414 |
| chrX-114638490-DEL-448 | chrX-114638490-DEL-440 |
| chrX-114773396-DEL-1488 | chrX-114773443-DEL-1488 |
| chrX-114800427-DEL-515 | chrX-114800428-DEL-515 |
| chrX-114902148-DEL-2857 | chrX-114902149-DEL-2857 |
| chrX-114908947-DEL-50 | chrX-114908944-DEL-57 |
| chrX-114910184-DEL-177 | chrX-114910185-DEL-177 |
| chrX-114924622-DEL-374 | chrX-114924623-DEL-374 |
| chrX-114957275-DEL-553 | chrX-114957276-DEL-553 |
| chrX-115001508-DEL-851 | chrX-115001509-DEL-851 |
| chrX-115013006-DEL-97 | chrX-115013007-DEL-97 |
| chrX-115077206-DEL-74 | chrX-115077207-DEL-74 |
| chrX-115078575-DEL-62 | chrX-115078536-DEL-70 |
| chrX-115114259-DEL-608 | chrX-115114260-DEL-609 |
| chrX-115134861-DEL-201 | chrX-115134862-DEL-201 |
| chrX-115179751-DEL-873 | chrX-115179752-DEL-873 |
| chrX-115207630-DEL-1145 | chrX-115207631-DEL-1145 |
| chrX-115246532-DEL-64 | chrX-115246533-DEL-64 |
| chrX-115267843-DEL-226 | chrX-115267844-DEL-226 |
| chrX-115285076-DEL-52 | chrX-115285077-DEL-52 |
| chrX-115285406-DEL-1538 | chrX-115285407-DEL-1538 |
| chrX-115290787-DEL-823 | chrX-115290788-DEL-823 |
| chrX-115297368-DEL-139 | chrX-115297369-DEL-139 |
| chrX-115314543-DEL-8428 | chrX-115314545-DEL-8427 |
| chrX-115371210-DEL-1166 | chrX-115371210-DEL-1166 |
| chrX-115373991-DEL-2002 | chrX-115373991-DEL-2002 |
| chrX-115390984-DEL-55 | chrX-115390983-DEL-56 |
| chrX-115397597-DEL-202 | chrX-115397598-DEL-202 |
| chrX-115437499-DEL-207 | chrX-115437500-DEL-207 |
| chrX-115441751-DEL-108 | chrX-115441752-DEL-104 |
| chrX-115444581-DEL-696 | chrX-115444582-DEL-696 |
| chrX-115499554-DEL-82 | chrX-115499522-DEL-97 |
| chrX-115546705-DEL-85 | chrX-115546706-DEL-85 |
| chrX-115548209-DEL-617 | chrX-115548210-DEL-617 |
| chrX-115561412-DEL-4569 | chrX-115561413-DEL-4569 |
| chrX-115585051-DEL-2886 | chrX-115585052-DEL-2886 |
| chrX-115604942-DEL-52 | chrX-115604943-DEL-52 |
| chrX-115610104-DEL-58 | chrX-115610111-DEL-76 |
| chrX-115623307-DEL-6048 | chrX-115623308-DEL-6048 |
| chrX-115631701-DEL-2703 | chrX-115631702-DEL-2703 |
| chrX-115712433-DEL-222 | chrX-115712434-DEL-222 |
| chrX-115728504-DEL-210 | chrX-115728505-DEL-210 |
| chrX-115765746-DEL-1786 | chrX-115765747-DEL-1786 |
| chrX-115818896-DEL-2010 | chrX-115818897-DEL-2010 |
| chrX-115831089-DEL-203 | chrX-115831090-DEL-203 |
| chrX-115885819-DEL-456 | chrX-115885820-DEL-456 |
| chrX-115887337-DEL-4760 | chrX-115887338-DEL-4760 |
| chrX-115939536-DEL-351 | chrX-115939537-DEL-351 |
| chrX-115996845-DEL-8762 | chrX-115996846-DEL-8762 |
| chrX-116050914-DEL-726 | chrX-116050915-DEL-726 |
| chrX-116061073-DEL-13690 | chrX-116061077-DEL-13688 |
| chrX-116116284-DEL-468 | chrX-116116285-DEL-468 |
| chrX-116119366-DEL-6414 | chrX-116119364-DEL-6415 |
| chrX-116226546-DEL-5186 | chrX-116226547-DEL-5186 |
| chrX-116346401-DEL-56 | chrX-116346401-DEL-56 |
| chrX-116364514-DEL-80 | chrX-116364514-DEL-81 |
| chrX-116368324-DEL-182 | chrX-116368325-DEL-182 |
| chrX-116380891-DEL-340 | chrX-116380889-DEL-329 |
| chrX-116384291-DEL-56 | chrX-116384292-DEL-56 |
| chrX-116386274-DEL-2888 | chrX-116386275-DEL-2888 |
| chrX-116413650-DEL-64 | chrX-116413647-DEL-65 |

| | |
|---|---|
| chrX-116421628-DEL-103 | chrX-116421631-DEL-101 |
| chrX-116496142-DEL-3121 | chrX-116496143-DEL-3121 |
| chrX-116497521-DEL-52 | chrX-116497522-DEL-52 |
| chrX-116500018-DEL-456 | chrX-116500019-DEL-456 |
| chrX-116544675-DEL-1017 | chrX-116544676-DEL-1017 |
| chrX-116590674-DEL-690 | chrX-116590675-DEL-690 |
| chrX-116676570-DEL-726 | chrX-116676571-DEL-726 |
| chrX-116709703-DEL-358 | chrX-116709704-DEL-358 |
| chrX-116750139-DEL-4327 | chrX-116750138-DEL-4329 |
| chrX-116764086-DEL-7641 | chrX-116764087-DEL-7641 |
| chrX-116764102-DEL-416 | chrX-116764103-DEL-416 |
| chrX-116794637-DEL-69 | chrX-116794644-DEL-63 |
| chrX-116831750-DEL-1187 | chrX-116831751-DEL-1187 |
| chrX-116919825-DEL-82 | chrX-116919814-DEL-56 |
| chrX-116942915-DEL-89 | chrX-116942916-DEL-89 |
| chrX-116960766-DEL-562 | chrX-116960767-DEL-562 |
| chrX-117003539-DEL-3344 | chrX-117003540-DEL-3344 |
| chrX-117028427-DEL-495 | chrX-117028428-DEL-495 |
| chrX-117046458-DEL-9487 | chrX-117046459-DEL-9487 |
| chrX-117068273-DEL-650 | chrX-117068274-DEL-650 |
| chrX-117077683-DEL-500 | chrX-117077684-DEL-500 |
| chrX-117107692-DEL-57 | chrX-117107673-DEL-58 |
| chrX-117109696-DEL-6436 | chrX-117109697-DEL-6436 |
| chrX-117154157-DEL-2014 | chrX-117154158-DEL-2014 |
| chrX-117316207-DEL-7145 | chrX-117316208-DEL-7145 |
| chrX-117351462-DEL-200 | chrX-117351463-DEL-198 |
| chrX-117417970-DEL-52 | chrX-117417984-DEL-53 |
| chrX-117460102-DEL-2278 | chrX-117460103-DEL-2278 |
| chrX-117479637-DEL-7455 | chrX-117479638-DEL-7455 |
| chrX-117523416-DEL-2920 | chrX-117523417-DEL-2920 |
| chrX-117582851-DEL-5136 | chrX-117582851-DEL-5136 |
| chrX-117591748-DEL-78 | chrX-117591711-DEL-88 |
| chrX-117617308-DEL-3260 | chrX-117617309-DEL-3260 |
| chrX-117627079-DEL-352 | chrX-117627080-DEL-352 |
| chrX-117628841-DEL-107 | chrX-117628842-DEL-107 |
| chrX-117676032-DEL-88 | chrX-117676033-DEL-88 |
| chrX-117682774-DEL-497 | chrX-117682775-DEL-497 |
| chrX-117713208-DEL-70 | chrX-117713209-DEL-70 |
| chrX-117728905-DEL-65 | chrX-117728906-DEL-65 |
| chrX-117746304-DEL-6014 | chrX-117746305-DEL-6014 |
| chrX-117752621-DEL-55 | chrX-117752621-DEL-55 |
| chrX-117756479-DEL-51 | chrX-117756480-DEL-51 |
| chrX-117758000-DEL-144 | chrX-117758001-DEL-144 |
| chrX-117759184-DEL-124 | chrX-117759185-DEL-124 |
| chrX-117792481-DEL-165 | chrX-117792481-DEL-162 |
| chrX-117808962-DEL-5302 | chrX-117808963-DEL-5302 |
| chrX-117851721-DEL-6343 | chrX-117851722-DEL-6343 |
| chrX-117888254-DEL-1152 | chrX-117888255-DEL-1152 |
| chrX-117907772-DEL-495 | chrX-117907773-DEL-495 |
| chrX-117931087-DEL-2492 | chrX-117931088-DEL-2492 |
| chrX-117969767-DEL-162 | chrX-117969768-DEL-162 |
| chrX-117999387-DEL-595 | chrX-117999388-DEL-595 |
| chrX-118049140-DEL-2347 | chrX-118049141-DEL-2347 |
| chrX-118051602-DEL-541 | chrX-118051603-DEL-541 |
| chrX-118072187-DEL-379 | chrX-118072188-DEL-379 |
| chrX-118076438-DEL-51 | chrX-118076439-DEL-51 |
| chrX-118081182-DEL-2248 | chrX-118081183-DEL-2248 |
| chrX-118090367-DEL-1552 | chrX-118090368-DEL-1552 |
| chrX-118128832-DEL-1438 | chrX-118128833-DEL-1438 |
| chrX-118161133-DEL-6444 | chrX-118161134-DEL-6444 |
| chrX-118163716-DEL-115 | chrX-118163717-DEL-116 |
| chrX-118171929-DEL-113 | chrX-118171930-DEL-113 |
| chrX-118197562-DEL-1765 | chrX-118197563-DEL-1765 |
| chrX-118233722-DEL-434 | chrX-118233723-DEL-434 |
| chrX-118270158-DEL-365 | chrX-118270159-DEL-365 |
| chrX-118271180-DEL-2554 | chrX-118271184-DEL-2555 |
| chrX-118289905-DEL-2973 | chrX-118289906-DEL-2973 |
| chrX-118296892-DEL-79 | chrX-118296903-DEL-76 |
| chrX-118320900-DEL-1264 | chrX-118320901-DEL-1264 |
| chrX-118343799-DEL-6076 | chrX-118343800-DEL-6076 |
| chrX-118373850-DEL-1364 | chrX-118373851-DEL-1364 |
| chrX-118383273-DEL-1593 | chrX-118383274-DEL-1593 |
| chrX-118412250-DEL-7062 | chrX-118412251-DEL-7062 |
| chrX-118467795-DEL-1718 | chrX-118467796-DEL-1718 |
| chrX-118505441-DEL-390 | chrX-118505450-DEL-394 |
| chrX-118585589-DEL-7099 | chrX-118585590-DEL-7099 |
| chrX-118622503-DEL-303 | chrX-118622504-DEL-303 |
| chrX-118630582-DEL-696 | chrX-118630583-DEL-696 |
| chrX-118660770-DEL-395 | chrX-118660760-DEL-401 |
| chrX-118679565-DEL-60 | chrX-118679566-DEL-60 |

| | |
|---|---|
| chrX-118700756-DEL-3273 | chrX-118700757-DEL-3273 |
| chrX-118719594-DEL-1394 | chrX-118719595-DEL-1394 |
| chrX-118764727-DEL-4851 | chrX-118764728-DEL-4851 |
| chrX-118772445-DEL-110 | chrX-118772446-DEL-110 |
| chrX-118858834-DEL-201 | chrX-118858835-DEL-201 |
| chrX-118872250-DEL-56 | chrX-118872241-DEL-57 |
| chrX-118898843-DEL-1855 | chrX-118898844-DEL-1855 |
| chrX-118912285-DEL-1458 | chrX-118912288-DEL-1459 |
| chrX-118989349-DEL-8827 | chrX-118989349-DEL-8827 |
| chrX-118997177-DEL-76 | chrX-118997178-DEL-76 |
| chrX-119036597-DEL-478 | chrX-119036598-DEL-478 |
| chrX-119038560-DEL-85 | chrX-119038563-DEL-77 |
| chrX-119059155-DEL-59 | chrX-119059154-DEL-61 |
| chrX-119130897-DEL-52 | chrX-119130898-DEL-52 |
| chrX-119162940-DEL-62 | chrX-119162940-DEL-60 |
| chrX-119245019-DEL-938 | chrX-119245020-DEL-938 |
| chrX-119295185-DEL-6503 | chrX-119295186-DEL-6503 |
| chrX-119347307-DEL-936 | chrX-119347308-DEL-937 |
| chrX-119444658-DEL-3780 | chrX-119444659-DEL-3780 |
| chrX-119489782-DEL-149 | chrX-119489783-DEL-149 |
| chrX-119492085-DEL-136 | chrX-119492094-DEL-128 |
| chrX-119516200-DEL-55 | chrX-119516201-DEL-55 |
| chrX-119583684-DEL-6654 | chrX-119583685-DEL-6654 |
| chrX-119607841-DEL-1434 | chrX-119607842-DEL-1434 |
| chrX-119697146-DEL-555 | chrX-119697147-DEL-555 |
| chrX-119741734-DEL-5200 | chrX-119741735-DEL-5200 |
| chrX-119786306-DEL-894 | chrX-119786307-DEL-894 |
| chrX-119823400-DEL-704 | chrX-119823401-DEL-704 |
| chrX-119830688-DEL-693 | chrX-119830689-DEL-693 |
| chrX-119868861-DEL-6495 | chrX-119868862-DEL-6495 |
| chrX-119886349-DEL-7355 | chrX-119886350-DEL-7355 |
| chrX-119906823-DEL-65 | chrX-119906824-DEL-65 |
| chrX-119950368-DEL-215 | chrX-119950372-DEL-212 |
| chrX-119959760-DEL-8086 | chrX-119959761-DEL-8086 |
| chrX-119970053-DEL-53 | chrX-119970054-DEL-53 |
| chrX-119972198-DEL-73 | chrX-119972191-DEL-72 |
| chrX-119978002-DEL-97 | chrX-119978003-DEL-97 |
| chrX-119985426-DEL-219 | chrX-119985427-DEL-219 |
| chrX-120060410-DEL-199 | chrX-120060411-DEL-199 |
| chrX-120061794-DEL-1880 | chrX-120061795-DEL-1880 |
| chrX-120085327-DEL-2261 | chrX-120085328-DEL-2261 |
| chrX-120125733-DEL-55 | chrX-120125744-DEL-56 |
| chrX-120129138-DEL-612 | chrX-120129139-DEL-612 |
| chrX-120164558-DEL-14130 | chrX-120164560-DEL-14129 |
| chrX-120219909-DEL-5656 | chrX-120219909-DEL-5656 |
| chrX-120222704-DEL-421 | chrX-120222705-DEL-421 |
| chrX-120230186-DEL-786 | chrX-120230141-DEL-787 |
| chrX-120237112-DEL-2835 | chrX-120237113-DEL-2835 |
| chrX-120267230-DEL-203 | chrX-120267231-DEL-203 |
| chrX-120276940-DEL-1294 | chrX-120276940-DEL-1294 |
| chrX-120308646-DEL-199 | chrX-120308647-DEL-199 |
| chrX-120348777-DEL-7138 | chrX-120348778-DEL-7138 |
| chrX-120364815-DEL-92 | chrX-120364816-DEL-93 |
| chrX-120365881-DEL-431 | chrX-120365881-DEL-429 |
| chrX-120459801-DEL-7259 | chrX-120459802-DEL-7259 |
| chrX-120479263-DEL-1298 | chrX-120479264-DEL-1298 |
| chrX-120494891-DEL-6654 | chrX-120494892-DEL-6654 |
| chrX-120535951-DEL-55 | chrX-120535948-DEL-63 |
| chrX-120584849-DEL-1891 | chrX-120584850-DEL-1891 |
| chrX-120609863-DEL-2773 | chrX-120609864-DEL-2770 |
| chrX-120634194-DEL-52 | chrX-120634191-DEL-53 |
| chrX-120642690-DEL-7547 | chrX-120642691-DEL-7547 |
| chrX-120656633-DEL-115 | chrX-120656632-DEL-115 |
| chrX-120666469-DEL-719 | chrX-120666470-DEL-719 |
| chrX-120695918-DEL-1101 | chrX-120695919-DEL-1101 |
| chrX-120758025-DEL-1683 | chrX-120758026-DEL-1683 |
| chrX-120807091-DEL-198 | chrX-120807092-DEL-199 |
| chrX-120822736-DEL-76 | chrX-120822735-DEL-75 |
| chrX-120822897-DEL-1613 | chrX-120822897-DEL-1630 |
| chrX-120826966-DEL-452 | chrX-120826967-DEL-452 |
| chrX-120848242-DEL-156 | chrX-120848243-DEL-156 |
| chrX-120849892-DEL-336 | chrX-120849893-DEL-336 |
| chrX-120859255-DEL-6673 | chrX-120859256-DEL-6673 |
| chrX-120888969-DEL-1587 | chrX-120888970-DEL-1587 |
| chrX-120917856-DEL-3095 | chrX-120917866-DEL-3092 |
| chrX-120970885-DEL-6818 | chrX-120970886-DEL-6818 |
| chrX-120980169-DEL-65 | chrX-120980170-DEL-65 |
| chrX-120992372-DEL-64 | chrX-120992347-DEL-56 |
| chrX-121036639-DEL-52 | chrX-121036640-DEL-52 |
| chrX-121082987-DEL-198 | chrX-121082987-DEL-198 |

| | |
|---|---|
| chrX-121089052-DEL-6414 | chrX-121089053-DEL-6414 |
| chrX-121089061-DEL-214 | chrX-121089062-DEL-214 |
| chrX-121093664-DEL-116 | chrX-121093665-DEL-116 |
| chrX-121120788-DEL-52 | chrX-121120789-DEL-52 |
| chrX-121183848-DEL-6135 | chrX-121183849-DEL-6135 |
| chrX-121206337-DEL-2728 | chrX-121206338-DEL-2728 |
| chrX-121228011-DEL-273 | chrX-121228012-DEL-273 |
| chrX-121240496-DEL-64 | chrX-121240497-DEL-64 |
| chrX-121253325-DEL-2921 | chrX-121253326-DEL-2921 |
| chrX-121256991-DEL-1714 | chrX-121256992-DEL-1714 |
| chrX-121262657-DEL-70 | chrX-121262658-DEL-70 |
| chrX-121267559-DEL-3333 | chrX-121267560-DEL-3333 |
| chrX-121283386-DEL-345 | chrX-121283387-DEL-345 |
| chrX-121311127-DEL-515 | chrX-121311128-DEL-515 |
| chrX-121354522-DEL-1655 | chrX-121354523-DEL-1655 |
| chrX-121382744-DEL-936 | chrX-121382745-DEL-936 |
| chrX-121389250-DEL-212 | chrX-121389251-DEL-212 |
| chrX-121411657-DEL-2473 | chrX-121411658-DEL-2473 |
| chrX-121575695-DEL-61 | chrX-121575696-DEL-61 |
| chrX-121580678-DEL-6484 | chrX-121580679-DEL-6484 |
| chrX-121634252-DEL-7258 | chrX-121634253-DEL-7258 |
| chrX-121656197-DEL-6202 | chrX-121656198-DEL-6202 |
| chrX-121663528-DEL-2829 | chrX-121663529-DEL-2829 |
| chrX-121741002-DEL-349 | chrX-121741001-DEL-349 |
| chrX-121767543-DEL-66 | chrX-121767523-DEL-82 |
| chrX-121811315-DEL-6381 | chrX-121811316-DEL-6381 |
| chrX-121875032-DEL-68 | chrX-121875029-DEL-72 |
| chrX-121930980-DEL-2935 | chrX-121930981-DEL-2935 |
| chrX-121944171-DEL-202 | chrX-121944183-DEL-191 |
| chrX-121950405-DEL-84 | chrX-121950438-DEL-53 |
| chrX-121974662-DEL-81 | chrX-121974651-DEL-84 |
| chrX-121977042-DEL-54 | chrX-121977047-DEL-50 |
| chrX-121981833-DEL-78 | chrX-121981803-DEL-86 |
| chrX-122035131-DEL-360 | chrX-122035132-DEL-360 |
| chrX-122394530-DEL-89 | chrX-122394531-DEL-88 |
| chrX-122935218-DEL-131 | chrX-122935219-DEL-131 |
| chrX-122967920-DEL-215 | chrX-122967921-DEL-215 |
| chrX-123101051-DEL-826 | chrX-123101052-DEL-826 |
| chrX-123121436-DEL-101 | chrX-123121437-DEL-101 |
| chrX-123124576-DEL-344 | chrX-123124583-DEL-338 |
| chrX-123149695-DEL-196 | chrX-123149696-DEL-196 |
| chrX-123220976-DEL-998 | chrX-123220977-DEL-998 |
| chrX-123248000-DEL-78 | chrX-123248001-DEL-78 |
| chrX-124975728-DEL-290 | chrX-124975729-DEL-290 |
| chrX-125028283-DEL-96 | chrX-125028284-DEL-96 |
| chrX-125051177-DEL-7125 | chrX-125051178-DEL-7125 |
| chrX-125082585-DEL-4906 | chrX-125082586-DEL-4906 |
| chrX-125103462-DEL-6381 | chrX-125103463-DEL-6381 |
| chrX-125161993-DEL-10811 | chrX-125161993-DEL-10811 |
| chrX-125181380-DEL-3503 | chrX-125181381-DEL-3503 |
| chrX-125226312-DEL-88 | chrX-125226313-DEL-88 |
| chrX-125242033-DEL-3483 | chrX-125242034-DEL-3483 |
| chrX-125310953-DEL-9812 | chrX-125310953-DEL-9812 |
| chrX-125347950-DEL-93 | chrX-125347951-DEL-93 |
| chrX-125372774-DEL-52 | chrX-125372775-DEL-54 |
| chrX-125380838-DEL-4352 | chrX-125380839-DEL-4352 |
| chrX-125393666-DEL-6769 | chrX-125393667-DEL-6769 |
| chrX-125442931-DEL-13746 | chrX-125442932-DEL-13746 |
| chrX-125466676-DEL-201 | chrX-125466639-DEL-239 |
| chrX-125470588-DEL-132 | chrX-125470589-DEL-128 |
| chrX-125597995-DEL-345 | chrX-125597996-DEL-345 |
| chrX-125670027-DEL-181 | chrX-125670028-DEL-181 |
| chrX-125689714-DEL-1415 | chrX-125689715-DEL-1415 |
| chrX-125728745-DEL-6780 | chrX-125728746-DEL-6780 |
| chrX-125738439-DEL-2471 | chrX-125738440-DEL-2470 |
| chrX-125762402-DEL-1603 | chrX-125762402-DEL-1603 |
| chrX-125783170-DEL-6440 | chrX-125783171-DEL-6440 |
| chrX-125795530-DEL-73 | chrX-125795531-DEL-83 |
| chrX-125811527-DEL-100 | chrX-125811541-DEL-56 |
| chrX-125853885-DEL-66 | chrX-125853886-DEL-66 |
| chrX-125891580-DEL-63 | chrX-125891587-DEL-57 |
| chrX-125963330-DEL-68 | chrX-125963331-DEL-68 |
| chrX-126027716-DEL-227 | chrX-126027717-DEL-227 |
| chrX-126032278-DEL-6500 | chrX-126032279-DEL-6500 |
| chrX-126063308-DEL-1267 | chrX-126063309-DEL-1267 |
| chrX-126092966-DEL-585 | chrX-126092967-DEL-585 |
| chrX-126099847-DEL-2910 | chrX-126099848-DEL-2910 |
| chrX-126109668-DEL-58 | chrX-126109669-DEL-58 |
| chrX-126125303-DEL-793 | chrX-126125304-DEL-793 |
| chrX-126160069-DEL-467 | chrX-126160070-DEL-467 |

| | |
|---|---|
| chrX-126163034-DEL-82 | chrX-126163035-DEL-82 |
| chrX-126172331-DEL-58 | chrX-126172339-DEL-57 |
| chrX-126176198-DEL-6455 | chrX-126176199-DEL-6455 |
| chrX-126188766-DEL-745 | chrX-126188767-DEL-745 |
| chrX-126193346-DEL-2032 | chrX-126193347-DEL-2032 |
| chrX-126277673-DEL-7117 | chrX-126277674-DEL-7117 |
| chrX-126333536-DEL-9246 | chrX-126333537-DEL-9246 |
| chrX-126354190-DEL-347 | chrX-126354191-DEL-347 |
| chrX-126367639-DEL-1795 | chrX-126367640-DEL-1795 |
| chrX-126389472-DEL-506 | chrX-126389473-DEL-506 |
| chrX-126403512-DEL-579 | chrX-126403513-DEL-579 |
| chrX-126409419-DEL-78 | chrX-126409420-DEL-78 |
| chrX-126449690-DEL-1666 | chrX-126449691-DEL-1666 |
| chrX-126484952-DEL-68 | chrX-126484953-DEL-66 |
| chrX-126486767-DEL-264 | chrX-126486768-DEL-264 |
| chrX-126530285-DEL-59 | chrX-126530286-DEL-59 |
| chrX-126563832-DEL-266 | chrX-126563829-DEL-261 |
| chrX-126607660-DEL-64 | chrX-126607664-DEL-60 |
| chrX-126612500-DEL-2254 | chrX-126612501-DEL-2254 |
| chrX-126632787-DEL-6613 | chrX-126632787-DEL-6613 |
| chrX-126765252-DEL-52 | chrX-126765253-DEL-52 |
| chrX-126800919-DEL-7903 | chrX-126800920-DEL-7903 |
| chrX-126838703-DEL-5291 | chrX-126838704-DEL-5291 |
| chrX-126858114-DEL-6578 | chrX-126858115-DEL-6569 |
| chrX-126896528-DEL-76 | chrX-126896529-DEL-76 |
| chrX-126938327-DEL-1719 | chrX-126938328-DEL-1719 |
| chrX-126979175-DEL-59 | chrX-126979176-DEL-59 |
| chrX-126981966-DEL-64 | chrX-126981967-DEL-64 |
| chrX-126991193-DEL-5226 | chrX-126991193-DEL-5226 |
| chrX-127048406-DEL-62 | chrX-127048409-DEL-60 |
| chrX-127094725-DEL-453 | chrX-127094726-DEL-453 |
| chrX-127097300-DEL-113 | chrX-127097301-DEL-113 |
| chrX-127123196-DEL-92 | chrX-127123197-DEL-92 |
| chrX-127123999-DEL-4540 | chrX-127124000-DEL-4540 |
| chrX-127134126-DEL-6963 | chrX-127134121-DEL-6964 |
| chrX-127158283-DEL-67 | chrX-127158284-DEL-66 |
| chrX-127214006-DEL-127 | chrX-127214007-DEL-127 |
| chrX-127217142-DEL-4037 | chrX-127217143-DEL-4037 |
| chrX-127260448-DEL-84 | chrX-127260449-DEL-78 |
| chrX-127262243-DEL-2495 | chrX-127262244-DEL-2495 |
| chrX-127338646-DEL-572 | chrX-127338647-DEL-572 |
| chrX-127444371-DEL-1479 | chrX-127444372-DEL-1479 |
| chrX-127446753-DEL-8839 | chrX-127446754-DEL-8839 |
| chrX-127496616-DEL-6530 | chrX-127496618-DEL-6529 |
| chrX-127504357-DEL-1902 | chrX-127504358-DEL-1902 |
| chrX-127507566-DEL-71 | chrX-127507567-DEL-71 |
| chrX-127534971-DEL-927 | chrX-127534972-DEL-927 |
| chrX-127626692-DEL-4517 | chrX-127626693-DEL-4517 |
| chrX-127647654-DEL-68 | chrX-127647681-DEL-68 |
| chrX-127695539-DEL-6256 | chrX-127695540-DEL-6256 |
| chrX-127722463-DEL-253 | chrX-127722465-DEL-251 |
| chrX-127859501-DEL-6191 | chrX-127859501-DEL-6191 |
| chrX-127957315-DEL-58 | chrX-127957322-DEL-52 |
| chrX-127968907-DEL-4510 | chrX-127968910-DEL-4510 |
| chrX-128046273-DEL-7073 | chrX-128046262-DEL-7087 |
| chrX-128108776-DEL-54 | chrX-128108808-DEL-56 |
| chrX-128257660-DEL-6367 | chrX-128257661-DEL-6367 |
| chrX-128297473-DEL-6579 | chrX-128297474-DEL-6577 |
| chrX-128329466-DEL-214 | chrX-128329467-DEL-214 |
| chrX-128342177-DEL-6577 | chrX-128342178-DEL-6577 |
| chrX-128522908-DEL-2125 | chrX-128522862-DEL-2123 |
| chrX-128574338-DEL-383 | chrX-128574339-DEL-383 |
| chrX-128625442-DEL-195 | chrX-128625443-DEL-195 |
| chrX-128660334-DEL-189 | chrX-128660335-DEL-189 |
| chrX-128718453-DEL-197 | chrX-128718454-DEL-197 |
| chrX-128744038-DEL-91 | chrX-128744039-DEL-91 |
| chrX-128756348-DEL-209 | chrX-128756349-DEL-209 |
| chrX-128804685-DEL-6495 | chrX-128804686-DEL-6495 |
| chrX-128821078-DEL-58 | chrX-128821080-DEL-60 |
| chrX-128851414-DEL-524 | chrX-128851415-DEL-524 |
| chrX-128876138-DEL-259 | chrX-128876139-DEL-259 |
| chrX-128900132-DEL-408 | chrX-128900139-DEL-408 |
| chrX-128951962-DEL-136 | chrX-128951963-DEL-136 |
| chrX-128952011-DEL-834 | chrX-128952022-DEL-823 |
| chrX-128964323-DEL-636 | chrX-128964324-DEL-636 |
| chrX-128998989-DEL-2784 | chrX-128998990-DEL-2784 |
| chrX-129098404-DEL-6483 | chrX-129098405-DEL-6483 |
| chrX-129113994-DEL-914 | chrX-129113995-DEL-914 |
| chrX-129149815-DEL-603 | chrX-129149816-DEL-603 |
| chrX-129172312-DEL-63 | chrX-129172313-DEL-63 |

| | |
|---|---|
| chrX-129174239-DEL-211 | chrX-129174260-DEL-205 |
| chrX-129225797-DEL-554 | chrX-129225797-DEL-549 |
| chrX-129231850-DEL-62 | chrX-129231851-DEL-62 |
| chrX-129286781-DEL-1529 | chrX-129286782-DEL-1529 |
| chrX-129394959-DEL-158 | chrX-129394960-DEL-158 |
| chrX-129422498-DEL-62 | chrX-129422499-DEL-61 |
| chrX-129439108-DEL-828 | chrX-129439109-DEL-828 |
| chrX-129455007-DEL-1558 | chrX-129455008-DEL-1558 |
| chrX-129487324-DEL-223 | chrX-129487324-DEL-224 |
| chrX-129510069-DEL-3330 | chrX-129510070-DEL-3330 |
| chrX-129527018-DEL-81 | chrX-129527019-DEL-81 |
| chrX-129548333-DEL-1867 | chrX-129548334-DEL-1867 |
| chrX-129612559-DEL-853 | chrX-129612560-DEL-853 |
| chrX-129619309-DEL-200 | chrX-129619310-DEL-200 |
| chrX-129638406-DEL-193 | chrX-129638407-DEL-193 |
| chrX-129649223-DEL-52 | chrX-129649238-DEL-51 |
| chrX-129656654-DEL-6718 | chrX-129656655-DEL-6718 |
| chrX-129662924-DEL-206 | chrX-129662925-DEL-206 |
| chrX-129700486-DEL-6424 | chrX-129700487-DEL-6424 |
| chrX-129746297-DEL-257 | chrX-129746298-DEL-257 |
| chrX-129854355-DEL-2303 | chrX-129854356-DEL-2303 |
| chrX-129954816-DEL-62 | chrX-129954817-DEL-50 |
| chrX-129970644-DEL-226 | chrX-129970645-DEL-226 |
| chrX-129993513-DEL-70 | chrX-129993492-DEL-55 |
| chrX-130224831-DEL-838 | chrX-130224832-DEL-838 |
| chrX-130242349-DEL-4131 | chrX-130242350-DEL-4131 |
| chrX-130288839-DEL-1029 | chrX-130288840-DEL-1029 |
| chrX-130369999-DEL-6450 | chrX-130370000-DEL-6450 |
| chrX-130386894-DEL-68 | chrX-130386895-DEL-68 |
| chrX-130398322-DEL-3909 | chrX-130398323-DEL-3909 |
| chrX-130470702-DEL-55 | chrX-130470664-DEL-64 |
| chrX-130580082-DEL-90 | chrX-130580083-DEL-90 |
| chrX-130627652-DEL-1039 | chrX-130627653-DEL-1039 |
| chrX-130643101-DEL-207 | chrX-130643102-DEL-207 |
| chrX-130654887-DEL-565 | chrX-130654888-DEL-565 |
| chrX-130700346-DEL-718 | chrX-130700347-DEL-718 |
| chrX-130702003-DEL-197 | chrX-130702004-DEL-197 |
| chrX-130708131-DEL-6287 | chrX-130708132-DEL-6287 |
| chrX-130862567-DEL-6672 | chrX-130862568-DEL-6672 |
| chrX-130863105-DEL-64 | chrX-130863106-DEL-64 |
| chrX-130894328-DEL-6227 | chrX-130894329-DEL-6227 |
| chrX-130907794-DEL-90 | chrX-130907822-DEL-53 |
| chrX-130920351-DEL-455 | chrX-130920352-DEL-455 |
| chrX-130924958-DEL-6218 | chrX-130924959-DEL-6218 |
| chrX-130958606-DEL-838 | chrX-130958607-DEL-838 |
| chrX-131047986-DEL-60 | chrX-131047987-DEL-60 |
| chrX-131108210-DEL-78 | chrX-131108211-DEL-78 |
| chrX-131118043-DEL-2277 | chrX-131118054-DEL-2246 |
| chrX-131128229-DEL-511 | chrX-131128230-DEL-511 |
| chrX-131142605-DEL-115 | chrX-131142606-DEL-115 |
| chrX-131173671-DEL-2835 | chrX-131173671-DEL-2835 |
| chrX-131226233-DEL-56 | chrX-131226234-DEL-56 |
| chrX-131330959-DEL-81 | chrX-131330960-DEL-81 |
| chrX-131394164-DEL-592 | chrX-131394165-DEL-592 |
| chrX-131400177-DEL-393 | chrX-131400178-DEL-393 |
| chrX-131424437-DEL-413 | chrX-131424438-DEL-413 |
| chrX-131435768-DEL-1881 | chrX-131435769-DEL-1873 |
| chrX-131458079-DEL-309 | chrX-131458080-DEL-309 |
| chrX-131463251-DEL-6421 | chrX-131463252-DEL-6421 |
| chrX-131539960-DEL-54 | chrX-131539961-DEL-54 |
| chrX-131563595-DEL-526 | chrX-131563596-DEL-526 |
| chrX-131609282-DEL-964 | chrX-131609283-DEL-964 |
| chrX-131645995-DEL-199 | chrX-131645996-DEL-199 |
| chrX-131650770-DEL-503 | chrX-131650767-DEL-504 |
| chrX-131708200-DEL-210 | chrX-131708201-DEL-210 |
| chrX-131748222-DEL-1125 | chrX-131748223-DEL-1125 |
| chrX-131842692-DEL-72 | chrX-131842696-DEL-65 |
| chrX-131857392-DEL-661 | chrX-131857393-DEL-661 |
| chrX-131993455-DEL-1411 | chrX-131993461-DEL-1406 |
| chrX-132014777-DEL-1457 | chrX-132014778-DEL-1457 |
| chrX-132033104-DEL-6946 | chrX-132033105-DEL-6946 |
| chrX-132033397-DEL-408 | chrX-132033398-DEL-408 |
| chrX-132058604-DEL-54 | chrX-132058604-DEL-54 |
| chrX-132123903-DEL-78 | chrX-132123903-DEL-75 |
| chrX-132140463-DEL-215 | chrX-132140464-DEL-214 |
| chrX-132143904-DEL-56 | chrX-132143905-DEL-56 |
| chrX-132153312-DEL-77 | chrX-132153312-DEL-77 |
| chrX-132161952-DEL-8234 | chrX-132161953-DEL-8234 |
| chrX-132190592-DEL-395 | chrX-132190593-DEL-395 |
| chrX-132240071-DEL-6648 | chrX-132240072-DEL-6648 |

| | |
|---|---|
| chrX-132329111-DEL-6599 | chrX-132329112-DEL-6599 |
| chrX-132380750-DEL-580 | chrX-132380751-DEL-580 |
| chrX-132410989-DEL-59 | chrX-132410990-DEL-59 |
| chrX-132472084-DEL-567 | chrX-132472085-DEL-567 |
| chrX-132491892-DEL-453 | chrX-132491893-DEL-453 |
| chrX-132500976-DEL-199 | chrX-132500977-DEL-199 |
| chrX-132597634-DEL-1140 | chrX-132597635-DEL-1140 |
| chrX-132664967-DEL-70 | chrX-132664966-DEL-72 |
| chrX-132699057-DEL-2177 | chrX-132699058-DEL-2177 |
| chrX-132713380-DEL-354 | chrX-132713381-DEL-354 |
| chrX-132775882-DEL-6801 | chrX-132775882-DEL-6801 |
| chrX-132891527-DEL-54 | chrX-132891528-DEL-54 |
| chrX-132925454-DEL-912 | chrX-132925455-DEL-912 |
| chrX-132990687-DEL-213 | chrX-132990688-DEL-213 |
| chrX-133053539-DEL-211 | chrX-133053556-DEL-209 |
| chrX-133064713-DEL-52 | chrX-133064708-DEL-62 |
| chrX-133095830-DEL-827 | chrX-133095832-DEL-828 |
| chrX-133127175-DEL-486 | chrX-133127176-DEL-486 |
| chrX-133133170-DEL-475 | chrX-133133171-DEL-475 |
| chrX-133148756-DEL-57 | chrX-133148741-DEL-107 |
| chrX-133206899-DEL-221 | chrX-133206900-DEL-221 |
| chrX-133221148-DEL-619 | chrX-133221116-DEL-620 |
| chrX-133224265-DEL-215 | chrX-133224266-DEL-215 |
| chrX-133272305-DEL-399 | chrX-133272342-DEL-360 |
| chrX-133288771-DEL-92 | chrX-133288772-DEL-92 |
| chrX-133313316-DEL-660 | chrX-133313317-DEL-660 |
| chrX-133323177-DEL-82 | chrX-133323188-DEL-78 |
| chrX-133384971-DEL-127 | chrX-133384934-DEL-129 |
| chrX-133389565-DEL-1066 | chrX-133389566-DEL-1066 |
| chrX-133394316-DEL-877 | chrX-133394317-DEL-876 |
| chrX-133398100-DEL-195 | chrX-133398101-DEL-194 |
| chrX-133415951-DEL-204 | chrX-133415951-DEL-204 |
| chrX-133446462-DEL-184 | chrX-133446463-DEL-184 |
| chrX-133508494-DEL-244 | chrX-133508495-DEL-244 |
| chrX-133565165-DEL-81 | chrX-133565166-DEL-81 |
| chrX-133565784-DEL-63 | chrX-133565785-DEL-63 |
| chrX-133569876-DEL-203 | chrX-133569877-DEL-203 |
| chrX-133608854-DEL-58 | chrX-133608843-DEL-61 |
| chrX-133628912-DEL-102 | chrX-133628908-DEL-104 |
| chrX-133633117-DEL-65 | chrX-133633118-DEL-65 |
| chrX-133663105-DEL-84 | chrX-133663100-DEL-56 |
| chrX-133691659-DEL-440 | chrX-133691660-DEL-440 |
| chrX-133698527-DEL-98 | chrX-133698546-DEL-80 |
| chrX-133720829-DEL-339 | chrX-133720831-DEL-337 |
| chrX-133728822-DEL-109 | chrX-133728823-DEL-109 |
| chrX-133747159-DEL-157 | chrX-133747160-DEL-157 |
| chrX-133773499-DEL-130 | chrX-133773500-DEL-130 |
| chrX-133793420-DEL-5637 | chrX-133793421-DEL-5637 |
| chrX-133799602-DEL-200 | chrX-133799603-DEL-200 |
| chrX-133826263-DEL-685 | chrX-133826264-DEL-685 |
| chrX-133913993-DEL-254 | chrX-133913994-DEL-254 |
| chrX-133957851-DEL-1324 | chrX-133957852-DEL-1324 |
| chrX-133958299-DEL-54 | chrX-133958300-DEL-54 |
| chrX-133973561-DEL-88 | chrX-133973562-DEL-88 |
| chrX-134003666-DEL-157 | chrX-134003667-DEL-157 |
| chrX-134004228-DEL-64 | chrX-134004228-DEL-64 |
| chrX-134021408-DEL-64 | chrX-134021409-DEL-64 |
| chrX-134129085-DEL-1475 | chrX-134129086-DEL-1475 |
| chrX-134133354-DEL-205 | chrX-134133355-DEL-205 |
| chrX-134146564-DEL-2607 | chrX-134146565-DEL-2607 |
| chrX-134162831-DEL-116 | chrX-134162832-DEL-116 |
| chrX-134164852-DEL-64 | chrX-134164859-DEL-54 |
| chrX-134184920-DEL-2093 | chrX-134184921-DEL-2093 |
| chrX-134197945-DEL-4605 | chrX-134197946-DEL-4605 |
| chrX-134204287-DEL-288 | chrX-134204288-DEL-287 |
| chrX-134221496-DEL-326 | chrX-134221496-DEL-326 |
| chrX-134228738-DEL-324 | chrX-134228738-DEL-324 |
| chrX-134255397-DEL-52 | chrX-134255398-DEL-52 |
| chrX-134492100-DEL-171 | chrX-134492101-DEL-171 |
| chrX-134521007-DEL-263 | chrX-134521005-DEL-264 |
| chrX-134530771-DEL-8197 | chrX-134530772-DEL-8194 |
| chrX-134532334-DEL-1597 | chrX-134532334-DEL-1597 |
| chrX-134537366-DEL-326 | chrX-134537367-DEL-326 |
| chrX-134575217-DEL-60 | chrX-134575223-DEL-54 |
| chrX-134577296-DEL-62 | chrX-134577296-DEL-94 |
| chrX-134609898-DEL-60 | chrX-134609899-DEL-60 |
| chrX-134612618-DEL-530 | chrX-134612619-DEL-530 |
| chrX-134632442-DEL-1595 | chrX-134632443-DEL-1595 |
| chrX-134636315-DEL-199 | chrX-134636302-DEL-195 |
| chrX-134636843-DEL-397 | chrX-134636844-DEL-397 |

| | |
|---|---|
| chrX-134639693-DEL-260 | chrX-134639694-DEL-260 |
| chrX-134708706-DEL-202 | chrX-134708707-DEL-202 |
| chrX-134747995-DEL-430 | chrX-134747990-DEL-431 |
| chrX-134754001-DEL-430 | chrX-134754002-DEL-431 |
| chrX-134766070-DEL-274 | chrX-134766071-DEL-274 |
| chrX-134767144-DEL-514 | chrX-134767145-DEL-514 |
| chrX-134778692-DEL-217 | chrX-134778693-DEL-217 |
| chrX-134809054-DEL-106 | chrX-134809055-DEL-104 |
| chrX-134809926-DEL-168 | chrX-134809933-DEL-192 |
| chrX-134818216-DEL-54 | chrX-134818217-DEL-54 |
| chrX-134820442-DEL-91 | chrX-134820443-DEL-63 |
| chrX-134848133-DEL-90 | chrX-134848132-DEL-70 |
| chrX-134852058-DEL-1763 | chrX-134852059-DEL-1763 |
| chrX-134902402-DEL-51 | chrX-134902403-DEL-51 |
| chrX-134906250-DEL-79 | chrX-134906251-DEL-79 |
| chrX-134913295-DEL-931 | chrX-134913292-DEL-930 |
| chrX-134917455-DEL-79 | chrX-134917455-DEL-77 |
| chrX-134955148-DEL-58 | chrX-134955149-DEL-58 |
| chrX-134960527-DEL-131 | chrX-134960528-DEL-131 |
| chrX-134982678-DEL-216 | chrX-134982679-DEL-216 |
| chrX-135017663-DEL-1923 | chrX-135017664-DEL-1923 |
| chrX-135084442-DEL-229 | chrX-135084443-DEL-229 |
| chrX-135108294-DEL-204 | chrX-135108295-DEL-204 |
| chrX-135124985-DEL-133 | chrX-135124986-DEL-133 |
| chrX-135132491-DEL-592 | chrX-135132492-DEL-592 |
| chrX-135139234-DEL-165 | chrX-135139235-DEL-165 |
| chrX-135143495-DEL-62 | chrX-135143496-DEL-62 |
| chrX-135184439-DEL-2980 | chrX-135184440-DEL-2980 |
| chrX-135190790-DEL-62 | chrX-135190791-DEL-62 |
| chrX-135199517-DEL-84 | chrX-135199518-DEL-84 |
| chrX-135221861-DEL-134 | chrX-135221862-DEL-146 |
| chrX-135268788-DEL-708 | chrX-135268792-DEL-718 |
| chrX-135277013-DEL-890 | chrX-135277012-DEL-890 |
| chrX-135316296-DEL-173 | chrX-135316297-DEL-173 |
| chrX-135321406-DEL-19135 | chrX-135321407-DEL-19135 |
| chrX-135347238-DEL-84 | chrX-135347239-DEL-84 |
| chrX-135360256-DEL-6237 | chrX-135360257-DEL-6237 |
| chrX-135377640-DEL-1308 | chrX-135377641-DEL-1308 |
| chrX-135391585-DEL-4077 | chrX-135391586-DEL-4077 |
| chrX-135398082-DEL-409 | chrX-135398082-DEL-409 |
| chrX-135431256-DEL-424 | chrX-135431257-DEL-424 |
| chrX-135435274-DEL-66 | chrX-135435292-DEL-50 |
| chrX-135466736-DEL-799 | chrX-135466737-DEL-798 |
| chrX-135488941-DEL-193 | chrX-135488942-DEL-193 |
| chrX-135498702-DEL-446 | chrX-135498703-DEL-446 |
| chrX-135523615-DEL-52 | chrX-135523616-DEL-52 |
| chrX-135532663-DEL-603 | chrX-135532664-DEL-603 |
| chrX-135536288-DEL-631 | chrX-135536292-DEL-627 |
| chrX-135585752-DEL-3003 | chrX-135585753-DEL-3003 |
| chrX-135591618-DEL-186 | chrX-135591619-DEL-186 |
| chrX-135595949-DEL-956 | chrX-135595950-DEL-956 |
| chrX-135603895-DEL-131 | chrX-135603896-DEL-131 |
| chrX-135669633-DEL-1860 | chrX-135669634-DEL-1860 |
| chrX-135707803-DEL-83 | chrX-135707804-DEL-83 |
| chrX-135774192-DEL-465 | chrX-135774196-DEL-465 |
| chrX-135774801-DEL-438 | chrX-135774802-DEL-438 |
| chrX-135780217-DEL-1566 | chrX-135780218-DEL-1566 |
| chrX-135799246-DEL-105 | chrX-135799247-DEL-105 |
| chrX-135814748-DEL-2379 | chrX-135814723-DEL-2380 |
| chrX-135815753-DEL-581 | chrX-135815741-DEL-582 |
| chrX-135819689-DEL-1024 | chrX-135819690-DEL-1024 |
| chrX-135822523-DEL-563 | chrX-135822524-DEL-563 |
| chrX-135828925-DEL-95 | chrX-135828926-DEL-95 |
| chrX-135833583-DEL-97 | chrX-135833579-DEL-85 |
| chrX-135865338-DEL-75 | chrX-135865338-DEL-75 |
| chrX-135874178-DEL-911 | chrX-135874178-DEL-909 |
| chrX-135902305-DEL-51 | chrX-135902306-DEL-51 |
| chrX-135909513-DEL-211 | chrX-135909514-DEL-211 |
| chrX-135961717-DEL-56 | chrX-135961728-DEL-66 |
| chrX-135967475-DEL-234 | chrX-135967475-DEL-235 |
| chrX-135969163-DEL-61 | chrX-135969164-DEL-57 |
| chrX-135984075-DEL-70 | chrX-135984079-DEL-50 |
| chrX-135989838-DEL-95 | chrX-135989867-DEL-53 |
| chrX-136014105-DEL-56 | chrX-136014058-DEL-60 |
| chrX-136017482-DEL-360 | chrX-136017483-DEL-360 |
| chrX-136019779-DEL-76 | chrX-136019780-DEL-76 |
| chrX-136022878-DEL-137 | chrX-136022882-DEL-138 |
| chrX-136038049-DEL-208 | chrX-136038050-DEL-208 |
| chrX-136047190-DEL-68 | chrX-136047191-DEL-64 |
| chrX-136070605-DEL-92 | chrX-136070614-DEL-85 |

| | |
|---|---|
| chrX-136100619-DEL-1923 | chrX-136100620-DEL-1923 |
| chrX-136134320-DEL-948 | chrX-136134320-DEL-948 |
| chrX-136192516-DEL-8917 | chrX-136192517-DEL-8917 |
| chrX-136276610-DEL-1136 | chrX-136276611-DEL-1136 |
| chrX-136300868-DEL-4875 | chrX-136300869-DEL-4875 |
| chrX-136322070-DEL-70 | chrX-136322071-DEL-70 |
| chrX-136327471-DEL-6864 | chrX-136327472-DEL-6864 |
| chrX-136338053-DEL-965 | chrX-136338054-DEL-965 |
| chrX-136357018-DEL-1200 | chrX-136357019-DEL-1200 |
| chrX-136366362-DEL-230 | chrX-136366362-DEL-228 |
| chrX-136401922-DEL-5063 | chrX-136401923-DEL-5063 |
| chrX-136457077-DEL-70 | chrX-136457112-DEL-54 |
| chrX-136475609-DEL-66 | chrX-136475610-DEL-64 |
| chrX-136509187-DEL-56 | chrX-136509188-DEL-56 |
| chrX-136510573-DEL-70 | chrX-136510570-DEL-78 |
| chrX-136517863-DEL-196 | chrX-136517864-DEL-196 |
| chrX-136590092-DEL-7123 | chrX-136590093-DEL-7123 |
| chrX-136621890-DEL-729 | chrX-136621891-DEL-729 |
| chrX-136667536-DEL-80 | chrX-136667497-DEL-120 |
| chrX-136743383-DEL-215 | chrX-136743384-DEL-216 |
| chrX-136757208-DEL-171 | chrX-136757207-DEL-168 |
| chrX-136775070-DEL-688 | chrX-136775071-DEL-688 |
| chrX-136779596-DEL-892 | chrX-136779597-DEL-892 |
| chrX-136814704-DEL-55 | chrX-136814705-DEL-55 |
| chrX-136864336-DEL-4764 | chrX-136864337-DEL-4764 |
| chrX-136872255-DEL-2112 | chrX-136872258-DEL-2111 |
| chrX-136873370-DEL-847 | chrX-136873371-DEL-847 |
| chrX-136928218-DEL-1382 | chrX-136928219-DEL-1382 |
| chrX-136937429-DEL-68 | chrX-136937459-DEL-51 |
| chrX-136951585-DEL-166 | chrX-136951586-DEL-166 |
| chrX-136952549-DEL-236 | chrX-136952572-DEL-220 |
| chrX-136988577-DEL-1479 | chrX-136988578-DEL-1479 |
| chrX-137054542-DEL-84 | chrX-137054543-DEL-84 |
| chrX-137084240-DEL-131 | chrX-137084241-DEL-131 |
| chrX-137117046-DEL-139 | chrX-137117047-DEL-139 |
| chrX-137184666-DEL-443 | chrX-137184667-DEL-443 |
| chrX-137193735-DEL-2481 | chrX-137193733-DEL-2482 |
| chrX-137201057-DEL-727 | chrX-137201057-DEL-728 |
| chrX-137219493-DEL-5947 | chrX-137219494-DEL-5946 |
| chrX-137254825-DEL-2721 | chrX-137254826-DEL-2721 |
| chrX-137259111-DEL-90 | chrX-137259116-DEL-86 |
| chrX-137274279-DEL-103 | chrX-137274284-DEL-99 |
| chrX-137284303-DEL-270 | chrX-137284304-DEL-270 |
| chrX-137288154-DEL-68 | chrX-137288129-DEL-62 |
| chrX-137311267-DEL-98 | chrX-137311264-DEL-57 |
| chrX-137331477-DEL-80 | chrX-137331478-DEL-80 |
| chrX-137409568-DEL-6396 | chrX-137409569-DEL-6396 |
| chrX-137531067-DEL-2875 | chrX-137531068-DEL-2875 |
| chrX-137564259-DEL-7428 | chrX-137564258-DEL-7428 |
| chrX-137582756-DEL-199 | chrX-137582757-DEL-199 |
| chrX-137601364-DEL-276 | chrX-137601365-DEL-276 |
| chrX-137617235-DEL-52 | chrX-137617236-DEL-52 |
| chrX-137625518-DEL-86 | chrX-137625519-DEL-86 |
| chrX-137661358-DEL-238 | chrX-137661358-DEL-226 |
| chrX-137667327-DEL-145 | chrX-137667330-DEL-143 |
| chrX-137709731-DEL-53 | chrX-137709730-DEL-54 |
| chrX-137795152-DEL-67 | chrX-137795153-DEL-67 |
| chrX-137918495-DEL-750 | chrX-137918493-DEL-750 |
| chrX-137957473-DEL-431 | chrX-137957472-DEL-432 |
| chrX-137974645-DEL-1029 | chrX-137974646-DEL-1029 |
| chrX-138004352-DEL-816 | chrX-138004356-DEL-812 |
| chrX-138037516-DEL-66 | chrX-138037531-DEL-66 |
| chrX-138055456-DEL-361 | chrX-138055457-DEL-361 |
| chrX-138099143-DEL-61 | chrX-138099143-DEL-61 |
| chrX-138150742-DEL-72 | chrX-138150743-DEL-72 |
| chrX-138271593-DEL-66 | chrX-138271600-DEL-50 |
| chrX-138283689-DEL-6417 | chrX-138283690-DEL-6417 |
| chrX-138311935-DEL-218 | chrX-138311936-DEL-218 |
| chrX-138328477-DEL-66 | chrX-138328479-DEL-65 |
| chrX-138353446-DEL-391 | chrX-138353447-DEL-391 |
| chrX-138358499-DEL-6415 | chrX-138358500-DEL-6415 |
| chrX-138364688-DEL-208 | chrX-138364689-DEL-208 |
| chrX-138376810-DEL-90 | chrX-138376811-DEL-90 |
| chrX-138385433-DEL-86 | chrX-138385434-DEL-86 |
| chrX-138398120-DEL-52 | chrX-138398121-DEL-52 |
| chrX-138420972-DEL-210 | chrX-138420973-DEL-210 |
| chrX-138522654-DEL-72 | chrX-138522655-DEL-75 |
| chrX-138529476-DEL-735 | chrX-138529477-DEL-735 |
| chrX-138603572-DEL-1436 | chrX-138603572-DEL-1436 |
| chrX-138697492-DEL-69 | chrX-138697493-DEL-69 |

| | |
|---|---|
| chrX-138743873-DEL-72 | chrX-138743904-DEL-82 |
| chrX-138805712-DEL-333 | chrX-138805713-DEL-333 |
| chrX-138814487-DEL-1866 | chrX-138814488-DEL-1866 |
| chrX-138849045-DEL-486 | chrX-138849046-DEL-486 |
| chrX-138962559-DEL-112 | chrX-138962559-DEL-112 |
| chrX-139005515-DEL-352 | chrX-139005512-DEL-356 |
| chrX-139016787-DEL-53 | chrX-139016784-DEL-61 |
| chrX-139101035-DEL-117 | chrX-139101036-DEL-117 |
| chrX-139105891-DEL-198 | chrX-139105892-DEL-198 |
| chrX-139130032-DEL-180 | chrX-139130033-DEL-180 |
| chrX-139312612-DEL-182 | chrX-139312598-DEL-182 |
| chrX-139318810-DEL-52 | chrX-139318808-DEL-50 |
| chrX-139366957-DEL-359 | chrX-139366958-DEL-359 |
| chrX-139383375-DEL-474 | chrX-139383376-DEL-474 |
| chrX-139385108-DEL-195 | chrX-139385109-DEL-195 |
| chrX-139399672-DEL-79 | chrX-139399673-DEL-79 |
| chrX-139410451-DEL-224 | chrX-139410450-DEL-198 |
| chrX-139452270-DEL-105 | chrX-139452221-DEL-105 |
| chrX-139464173-DEL-200 | chrX-139464174-DEL-200 |
| chrX-139542063-DEL-53 | chrX-139542064-DEL-53 |
| chrX-139630283-DEL-383 | chrX-139630284-DEL-383 |
| chrX-139645416-DEL-709 | chrX-139645417-DEL-709 |
| chrX-139657896-DEL-207 | chrX-139657897-DEL-207 |
| chrX-139703456-DEL-794 | chrX-139703457-DEL-786 |
| chrX-139752468-DEL-144 | chrX-139752469-DEL-144 |
| chrX-139777170-DEL-183 | chrX-139777171-DEL-183 |
| chrX-139794565-DEL-4105 | chrX-139794566-DEL-4105 |
| chrX-139840372-DEL-1163 | chrX-139840370-DEL-1164 |
| chrX-139862134-DEL-877 | chrX-139862135-DEL-877 |
| chrX-139871212-DEL-202 | chrX-139871213-DEL-202 |
| chrX-139900944-DEL-100 | chrX-139900945-DEL-98 |
| chrX-139926912-DEL-57 | chrX-139926906-DEL-77 |
| chrX-139928186-DEL-65 | chrX-139928187-DEL-65 |
| chrX-139933465-DEL-630 | chrX-139933420-DEL-630 |
| chrX-139993328-DEL-685 | chrX-139993335-DEL-670 |
| chrX-140031080-DEL-384 | chrX-140031081-DEL-384 |
| chrX-140064372-DEL-61 | chrX-140064373-DEL-61 |
| chrX-140075194-DEL-159 | chrX-140075195-DEL-157 |
| chrX-140089552-DEL-64 | chrX-140089542-DEL-66 |
| chrX-140145563-DEL-64 | chrX-140145564-DEL-64 |
| chrX-140147925-DEL-123 | chrX-140147926-DEL-123 |
| chrX-140177048-DEL-1523 | chrX-140177049-DEL-1523 |
| chrX-140189727-DEL-1300 | chrX-140189728-DEL-1300 |
| chrX-140200909-DEL-202 | chrX-140200910-DEL-202 |
| chrX-140290927-DEL-55 | chrX-140290928-DEL-55 |
| chrX-140304554-DEL-358 | chrX-140304555-DEL-358 |
| chrX-140344098-DEL-7589 | chrX-140344099-DEL-7589 |
| chrX-140376677-DEL-214 | chrX-140376684-DEL-208 |
| chrX-140393580-DEL-103 | chrX-140393570-DEL-97 |
| chrX-140408947-DEL-111 | chrX-140408948-DEL-107 |
| chrX-140436152-DEL-177 | chrX-140436153-DEL-177 |
| chrX-140460011-DEL-223 | chrX-140460016-DEL-219 |
| chrX-140468608-DEL-405 | chrX-140468609-DEL-405 |
| chrX-140499833-DEL-150 | chrX-140499834-DEL-150 |
| chrX-140524345-DEL-6517 | chrX-140524346-DEL-6517 |
| chrX-140603748-DEL-6267 | chrX-140603749-DEL-6267 |
| chrX-140631201-DEL-99 | chrX-140631213-DEL-97 |
| chrX-140662001-DEL-1129 | chrX-140662002-DEL-1129 |
| chrX-140675404-DEL-6677 | chrX-140675405-DEL-6677 |
| chrX-140699406-DEL-1211 | chrX-140699407-DEL-1211 |
| chrX-140733094-DEL-57 | chrX-140733079-DEL-54 |
| chrX-140736234-DEL-57 | chrX-140736234-DEL-57 |
| chrX-140752971-DEL-63 | chrX-140752972-DEL-63 |
| chrX-140762489-DEL-56 | chrX-140762500-DEL-54 |
| chrX-140895083-DEL-407 | chrX-140895084-DEL-407 |
| chrX-140952688-DEL-197 | chrX-140952689-DEL-197 |
| chrX-140974106-DEL-79 | chrX-140974107-DEL-79 |
| chrX-141010400-DEL-291 | chrX-141010401-DEL-291 |
| chrX-141100552-DEL-58 | chrX-141100553-DEL-52 |
| chrX-141195031-DEL-644 | chrX-141195032-DEL-644 |
| chrX-141207921-DEL-144 | chrX-141207922-DEL-143 |
| chrX-141225048-DEL-563 | chrX-141225051-DEL-572 |
| chrX-141254474-DEL-195 | chrX-141254480-DEL-194 |
| chrX-141274992-DEL-127 | chrX-141274993-DEL-127 |
| chrX-141321409-DEL-406 | chrX-141321410-DEL-406 |
| chrX-141339126-DEL-192 | chrX-141339114-DEL-225 |
| chrX-141361641-DEL-89 | chrX-141361635-DEL-87 |
| chrX-141378171-DEL-88 | chrX-141378143-DEL-128 |
| chrX-141387005-DEL-441 | chrX-141387006-DEL-441 |
| chrX-141413692-DEL-2612 | chrX-141413684-DEL-2613 |

| | |
|---|---|
| chrX-141432623-DEL-64 | chrX-141432623-DEL-50 |
| chrX-141474663-DEL-64 | chrX-141474644-DEL-65 |
| chrX-141487339-DEL-108 | chrX-141487340-DEL-108 |
| chrX-141491836-DEL-6293 | chrX-141491837-DEL-6293 |
| chrX-141604126-DEL-58 | chrX-141604127-DEL-58 |
| chrX-141645839-DEL-6322 | chrX-141645840-DEL-6322 |
| chrX-141695437-DEL-892 | chrX-141695438-DEL-892 |
| chrX-141713588-DEL-2500 | chrX-141713589-DEL-2500 |
| chrX-141723384-DEL-280 | chrX-141723385-DEL-280 |
| chrX-141788847-DEL-57 | chrX-141788848-DEL-57 |
| chrX-141808348-DEL-2302 | chrX-141808349-DEL-2302 |
| chrX-141824099-DEL-84 | chrX-141824096-DEL-69 |
| chrX-141830694-DEL-61 | chrX-141830695-DEL-61 |
| chrX-141831520-DEL-64 | chrX-141831521-DEL-64 |
| chrX-141928132-DEL-444 | chrX-141928133-DEL-444 |
| chrX-141933394-DEL-407 | chrX-141933395-DEL-407 |
| chrX-141946677-DEL-975 | chrX-141946676-DEL-975 |
| chrX-141955594-DEL-7070 | chrX-141955595-DEL-7070 |
| chrX-141976379-DEL-65 | chrX-141976380-DEL-65 |
| chrX-141998363-DEL-66 | chrX-141998370-DEL-60 |
| chrX-142025858-DEL-126 | chrX-142025877-DEL-108 |
| chrX-142067251-DEL-890 | chrX-142067252-DEL-889 |
| chrX-142102113-DEL-1038 | chrX-142102114-DEL-1038 |
| chrX-142182062-DEL-52 | chrX-142182062-DEL-71 |
| chrX-142202025-DEL-1123 | chrX-142202026-DEL-1123 |
| chrX-142218400-DEL-370 | chrX-142218401-DEL-370 |
| chrX-142243787-DEL-712 | chrX-142243788-DEL-712 |
| chrX-142265993-DEL-85 | chrX-142265993-DEL-84 |
| chrX-142323343-DEL-1021 | chrX-142323344-DEL-1021 |
| chrX-142324611-DEL-1022 | chrX-142324612-DEL-1022 |
| chrX-142337771-DEL-60 | chrX-142337762-DEL-61 |
| chrX-142385092-DEL-387 | chrX-142385093-DEL-387 |
| chrX-142387306-DEL-449 | chrX-142387307-DEL-449 |
| chrX-142402851-DEL-113 | chrX-142402852-DEL-113 |
| chrX-142436422-DEL-564 | chrX-142436423-DEL-564 |
| chrX-142488311-DEL-205 | chrX-142488312-DEL-205 |
| chrX-142505448-DEL-178 | chrX-142505449-DEL-178 |
| chrX-142507075-DEL-283 | chrX-142507076-DEL-283 |
| chrX-142507240-DEL-1146 | chrX-142507241-DEL-1146 |
| chrX-142544621-DEL-102 | chrX-142544635-DEL-92 |
| chrX-142567189-DEL-1731 | chrX-142567190-DEL-1731 |
| chrX-142602949-DEL-80 | chrX-142602950-DEL-80 |
| chrX-142607423-DEL-92 | chrX-142607423-DEL-92 |
| chrX-142623082-DEL-2615 | chrX-142623083-DEL-2615 |
| chrX-142693760-DEL-58 | chrX-142693761-DEL-58 |
| chrX-142802078-DEL-444 | chrX-142802079-DEL-444 |
| chrX-142828802-DEL-2692 | chrX-142828803-DEL-2692 |
| chrX-142846521-DEL-235 | chrX-142846522-DEL-234 |
| chrX-142898824-DEL-7264 | chrX-142898825-DEL-7264 |
| chrX-142910064-DEL-1588 | chrX-142910070-DEL-1587 |
| chrX-142925402-DEL-2414 | chrX-142925402-DEL-2414 |
| chrX-143005217-DEL-6258 | chrX-143005229-DEL-6257 |
| chrX-143021527-DEL-185 | chrX-143021525-DEL-186 |
| chrX-143030322-DEL-203 | chrX-143030320-DEL-204 |
| chrX-143036009-DEL-196 | chrX-143036009-DEL-196 |
| chrX-143052291-DEL-75 | chrX-143052292-DEL-75 |
| chrX-143055578-DEL-6938 | chrX-143055578-DEL-6938 |
| chrX-143064700-DEL-132 | chrX-143064700-DEL-132 |
| chrX-143071882-DEL-196 | chrX-143071882-DEL-196 |
| chrX-143094340-DEL-328 | chrX-143094343-DEL-326 |
| chrX-143102113-DEL-138 | chrX-143102078-DEL-185 |
| chrX-143109650-DEL-212 | chrX-143109651-DEL-212 |
| chrX-143130160-DEL-682 | chrX-143130161-DEL-682 |
| chrX-143133826-DEL-235 | chrX-143133829-DEL-246 |
| chrX-143161286-DEL-77 | chrX-143161287-DEL-77 |
| chrX-143166736-DEL-82 | chrX-143166737-DEL-82 |
| chrX-143183236-DEL-197 | chrX-143183237-DEL-194 |
| chrX-143200649-DEL-52 | chrX-143200648-DEL-54 |
| chrX-143215182-DEL-55 | chrX-143215183-DEL-50 |
| chrX-143232334-DEL-70 | chrX-143232335-DEL-70 |
| chrX-143239805-DEL-60 | chrX-143239806-DEL-60 |
| chrX-143282704-DEL-175 | chrX-143282703-DEL-177 |
| chrX-143360866-DEL-812 | chrX-143360865-DEL-813 |
| chrX-143372181-DEL-6226 | chrX-143372182-DEL-6226 |
| chrX-143384664-DEL-1686 | chrX-143384666-DEL-1684 |
| chrX-143393052-DEL-849 | chrX-143393058-DEL-837 |
| chrX-143396981-DEL-6395 | chrX-143396982-DEL-6395 |
| chrX-143409331-DEL-72 | chrX-143409332-DEL-72 |
| chrX-143446981-DEL-2451 | chrX-143446982-DEL-2451 |
| chrX-143454376-DEL-547 | chrX-143454377-DEL-547 |

| | |
|---|---|
| chrX-143458039-DEL-292 | chrX-143458040-DEL-292 |
| chrX-143482005-DEL-4409 | chrX-143482006-DEL-4409 |
| chrX-143490882-DEL-84 | chrX-143490884-DEL-83 |
| chrX-143500468-DEL-55 | chrX-143500469-DEL-55 |
| chrX-143544866-DEL-1451 | chrX-143544867-DEL-1451 |
| chrX-143573068-DEL-109 | chrX-143573069-DEL-109 |
| chrX-143597830-DEL-394 | chrX-143597831-DEL-394 |
| chrX-143603968-DEL-6843 | chrX-143603969-DEL-6843 |
| chrX-143612507-DEL-69 | chrX-143612535-DEL-80 |
| chrX-143640880-DEL-584 | chrX-143640881-DEL-584 |
| chrX-143656023-DEL-316 | chrX-143656024-DEL-316 |
| chrX-143683047-DEL-52 | chrX-143683048-DEL-52 |
| chrX-143688292-DEL-772 | chrX-143688293-DEL-772 |
| chrX-143699972-DEL-125 | chrX-143699973-DEL-121 |
| chrX-143710407-DEL-1388 | chrX-143710408-DEL-1388 |
| chrX-143715698-DEL-53 | chrX-143715699-DEL-53 |
| chrX-143754500-DEL-140 | chrX-143754519-DEL-122 |
| chrX-143803831-DEL-6244 | chrX-143803832-DEL-6244 |
| chrX-143822361-DEL-952 | chrX-143822362-DEL-952 |
| chrX-143828165-DEL-6706 | chrX-143828166-DEL-6706 |
| chrX-143836806-DEL-3558 | chrX-143836807-DEL-3558 |
| chrX-143860230-DEL-1494 | chrX-143860231-DEL-1494 |
| chrX-143879058-DEL-78 | chrX-143879059-DEL-78 |
| chrX-143897089-DEL-1705 | chrX-143897090-DEL-1705 |
| chrX-143923191-DEL-2103 | chrX-143923192-DEL-2103 |
| chrX-144016982-DEL-6236 | chrX-144016983-DEL-6236 |
| chrX-144064248-DEL-6463 | chrX-144064249-DEL-6463 |
| chrX-144077407-DEL-2074 | chrX-144077408-DEL-2074 |
| chrX-144093297-DEL-477 | chrX-144093298-DEL-477 |
| chrX-144097709-DEL-6507 | chrX-144097710-DEL-6507 |
| chrX-144105408-DEL-74 | chrX-144105409-DEL-74 |
| chrX-144143489-DEL-136 | chrX-144143490-DEL-136 |
| chrX-144166574-DEL-223 | chrX-144166575-DEL-223 |
| chrX-144220949-DEL-1464 | chrX-144220950-DEL-1464 |
| chrX-144278211-DEL-210 | chrX-144278212-DEL-210 |
| chrX-144301498-DEL-61 | chrX-144301499-DEL-61 |
| chrX-144377481-DEL-1563 | chrX-144377482-DEL-1563 |
| chrX-144379826-DEL-90 | chrX-144379824-DEL-86 |
| chrX-144389546-DEL-60 | chrX-144389547-DEL-60 |
| chrX-144422445-DEL-552 | chrX-144422446-DEL-552 |
| chrX-144426095-DEL-511 | chrX-144426096-DEL-511 |
| chrX-144427452-DEL-51 | chrX-144427455-DEL-55 |
| chrX-144433345-DEL-372 | chrX-144433346-DEL-372 |
| chrX-144452827-DEL-107 | chrX-144452828-DEL-107 |
| chrX-144455354-DEL-193 | chrX-144455358-DEL-193 |
| chrX-144469631-DEL-857 | chrX-144469632-DEL-857 |
| chrX-144492714-DEL-8492 | chrX-144492715-DEL-8492 |
| chrX-144513333-DEL-751 | chrX-144513334-DEL-751 |
| chrX-144516728-DEL-6668 | chrX-144516729-DEL-6668 |
| chrX-144540337-DEL-344 | chrX-144540338-DEL-344 |
| chrX-144563323-DEL-6159 | chrX-144563324-DEL-6159 |
| chrX-144609617-DEL-679 | chrX-144609611-DEL-677 |
| chrX-144611322-DEL-836 | chrX-144611323-DEL-836 |
| chrX-144685119-DEL-6739 | chrX-144685124-DEL-6739 |
| chrX-144691497-DEL-204 | chrX-144691497-DEL-204 |
| chrX-144706311-DEL-485 | chrX-144706312-DEL-485 |
| chrX-144713278-DEL-7116 | chrX-144713279-DEL-7116 |
| chrX-144739289-DEL-1276 | chrX-144739290-DEL-1276 |
| chrX-144775882-DEL-69 | chrX-144775860-DEL-102 |
| chrX-144786127-DEL-58 | chrX-144786123-DEL-51 |
| chrX-144830387-DEL-119 | chrX-144830388-DEL-119 |
| chrX-144837049-DEL-73 | chrX-144837050-DEL-67 |
| chrX-144843851-DEL-64 | chrX-144843852-DEL-64 |
| chrX-144844896-DEL-80 | chrX-144844898-DEL-69 |
| chrX-144862159-DEL-779 | chrX-144862160-DEL-779 |
| chrX-144917932-DEL-169 | chrX-144917933-DEL-169 |
| chrX-144923847-DEL-203 | chrX-144923848-DEL-204 |
| chrX-144925702-DEL-70 | chrX-144925703-DEL-70 |
| chrX-144935785-DEL-134 | chrX-144935786-DEL-134 |
| chrX-145014043-DEL-491 | chrX-145014044-DEL-491 |
| chrX-145020217-DEL-6207 | chrX-145020202-DEL-6222 |
| chrX-145034374-DEL-59 | chrX-145034375-DEL-59 |
| chrX-145041816-DEL-52 | chrX-145041813-DEL-55 |
| chrX-145052770-DEL-53 | chrX-145052772-DEL-52 |
| chrX-145062857-DEL-155 | chrX-145062858-DEL-155 |
| chrX-145084086-DEL-76 | chrX-145084067-DEL-76 |
| chrX-145092349-DEL-6519 | chrX-145092350-DEL-6519 |
| chrX-145107691-DEL-58 | chrX-145107699-DEL-56 |
| chrX-145118463-DEL-74 | chrX-145118427-DEL-62 |
| chrX-145127709-DEL-51 | chrX-145127709-DEL-51 |

| | |
|---|---|
| chrX-145136443-DEL-6035 | chrX-145136444-DEL-6035 |
| chrX-145175147-DEL-52 | chrX-145175148-DEL-51 |
| chrX-145179894-DEL-2373 | chrX-145179886-DEL-2382 |
| chrX-145185353-DEL-197 | chrX-145185354-DEL-197 |
| chrX-145211033-DEL-1317 | chrX-145211037-DEL-1315 |
| chrX-145235281-DEL-152 | chrX-145235282-DEL-152 |
| chrX-145282896-DEL-1600 | chrX-145282897-DEL-1600 |
| chrX-145297543-DEL-3116 | chrX-145297544-DEL-3116 |
| chrX-145325301-DEL-421 | chrX-145325302-DEL-421 |
| chrX-145365456-DEL-6276 | chrX-145365457-DEL-6276 |
| chrX-145365561-DEL-201 | chrX-145365562-DEL-201 |
| chrX-145464952-DEL-229 | chrX-145464953-DEL-229 |
| chrX-145473230-DEL-2651 | chrX-145473231-DEL-2651 |
| chrX-145479567-DEL-4338 | chrX-145479568-DEL-4338 |
| chrX-145504424-DEL-633 | chrX-145504425-DEL-633 |
| chrX-145507736-DEL-62 | chrX-145507737-DEL-58 |
| chrX-145541067-DEL-3790 | chrX-145541068-DEL-3790 |
| chrX-145561869-DEL-447 | chrX-145561870-DEL-447 |
| chrX-145610546-DEL-410 | chrX-145610547-DEL-410 |
| chrX-145612910-DEL-1566 | chrX-145612911-DEL-1566 |
| chrX-145632105-DEL-2575 | chrX-145632106-DEL-2575 |
| chrX-145676536-DEL-1371 | chrX-145676537-DEL-1371 |
| chrX-145747783-DEL-63 | chrX-145747784-DEL-63 |
| chrX-145827632-DEL-322 | chrX-145827633-DEL-322 |
| chrX-145830357-DEL-2886 | chrX-145830358-DEL-2886 |
| chrX-145893093-DEL-6217 | chrX-145893094-DEL-6217 |
| chrX-145899853-DEL-194 | chrX-145899854-DEL-194 |
| chrX-145901931-DEL-78 | chrX-145901932-DEL-78 |
| chrX-145950498-DEL-1249 | chrX-145950499-DEL-1249 |
| chrX-146006849-DEL-72 | chrX-146006848-DEL-73 |
| chrX-146058083-DEL-4478 | chrX-146058084-DEL-4478 |
| chrX-146095940-DEL-747 | chrX-146095941-DEL-747 |
| chrX-146096852-DEL-964 | chrX-146096853-DEL-964 |
| chrX-146126375-DEL-1206 | chrX-146126376-DEL-1206 |
| chrX-146128476-DEL-80 | chrX-146128476-DEL-69 |
| chrX-146137393-DEL-62 | chrX-146137394-DEL-50 |
| chrX-146163785-DEL-147 | chrX-146163786-DEL-147 |
| chrX-146173103-DEL-56 | chrX-146173095-DEL-64 |
| chrX-146204215-DEL-54 | chrX-146204216-DEL-54 |
| chrX-146213417-DEL-58 | chrX-146213425-DEL-50 |
| chrX-146223707-DEL-1700 | chrX-146223708-DEL-1700 |
| chrX-146235554-DEL-1309 | chrX-146235555-DEL-1309 |
| chrX-146320127-DEL-471 | chrX-146320131-DEL-470 |
| chrX-146356454-DEL-1666 | chrX-146356455-DEL-1666 |
| chrX-146435577-DEL-1410 | chrX-146435578-DEL-1410 |
| chrX-146746916-DEL-635 | chrX-146746917-DEL-635 |
| chrX-147355847-DEL-579 | chrX-147355848-DEL-579 |
| chrX-147602003-DEL-296 | chrX-147602008-DEL-293 |
| chrX-147612968-DEL-638 | chrX-147612969-DEL-638 |
| chrX-147732302-DEL-254 | chrX-147732303-DEL-254 |
| chrX-148512667-DEL-108 | chrX-148512668-DEL-108 |
| chrX-148598424-DEL-857 | chrX-148598425-DEL-857 |
| chrX-148906827-DEL-174 | chrX-148906828-DEL-174 |
| chrX-148907827-DEL-1123 | chrX-148907827-DEL-1123 |
| chrX-148917069-DEL-2580 | chrX-148917070-DEL-2580 |
| chrX-148941385-DEL-52 | chrX-148941386-DEL-52 |
| chrX-148976710-DEL-6403 | chrX-148976711-DEL-6403 |
| chrX-149072660-DEL-3843 | chrX-149072661-DEL-3843 |
| chrX-149130052-DEL-117 | chrX-149130052-DEL-117 |
| chrX-149164253-DEL-503 | chrX-149164253-DEL-503 |
| chrX-149209361-DEL-150 | chrX-149209362-DEL-150 |
| chrX-149212433-DEL-2219 | chrX-149212434-DEL-2219 |
| chrX-149235825-DEL-57 | chrX-149235814-DEL-58 |
| chrX-149240101-DEL-333 | chrX-149240102-DEL-333 |
| chrX-149251785-DEL-497 | chrX-149251786-DEL-497 |
| chrX-149268013-DEL-4951 | chrX-149268014-DEL-4951 |
| chrX-149274749-DEL-3132 | chrX-149274739-DEL-3133 |
| chrX-149297461-DEL-582 | chrX-149297462-DEL-582 |
| chrX-149308530-DEL-615 | chrX-149308532-DEL-613 |
| chrX-149312965-DEL-6319 | chrX-149312966-DEL-6319 |
| chrX-149321802-DEL-186 | chrX-149321801-DEL-187 |
| chrX-149373626-DEL-2529 | chrX-149373627-DEL-2529 |
| chrX-149394973-DEL-851 | chrX-149394974-DEL-851 |
| chrX-149397351-DEL-6396 | chrX-149397351-DEL-6396 |
| chrX-149411115-DEL-400 | chrX-149411116-DEL-400 |
| chrX-149413404-DEL-3694 | chrX-149413405-DEL-3694 |
| chrX-149431902-DEL-60 | chrX-149431903-DEL-60 |
| chrX-149444453-DEL-134 | chrX-149444454-DEL-144 |
| chrX-149449135-DEL-68 | chrX-149449136-DEL-68 |
| chrX-149462665-DEL-545 | chrX-149462666-DEL-545 |

| | |
|---|---|
| chrX-149481218-DEL-4495 | chrX-149481219-DEL-4495 |
| chrX-149516034-DEL-3554 | chrX-149516034-DEL-3554 |
| chrX-149520513-DEL-2551 | chrX-149520513-DEL-2551 |
| chrX-149522555-DEL-428 | chrX-149522556-DEL-428 |
| chrX-149566001-DEL-198 | chrX-149566002-DEL-198 |
| chrX-149644205-DEL-6659 | chrX-149644206-DEL-6659 |
| chrX-149654670-DEL-309 | chrX-149654671-DEL-309 |
| chrX-149666460-DEL-166 | chrX-149666461-DEL-166 |
| chrX-149673537-DEL-231 | chrX-149673538-DEL-231 |
| chrX-149689237-DEL-729 | chrX-149689238-DEL-729 |
| chrX-149705734-DEL-156 | chrX-149705743-DEL-149 |
| chrX-149722838-DEL-2772 | chrX-149722839-DEL-2772 |
| chrX-149731581-DEL-58 | chrX-149731586-DEL-50 |
| chrX-149733592-DEL-90 | chrX-149733593-DEL-90 |
| chrX-149744289-DEL-64 | chrX-149744290-DEL-64 |
| chrX-149766753-DEL-168 | chrX-149766736-DEL-159 |
| chrX-149793720-DEL-212 | chrX-149793721-DEL-212 |
| chrX-149812872-DEL-186 | chrX-149812879-DEL-185 |
| chrX-149817932-DEL-162 | chrX-149817933-DEL-162 |
| chrX-149854077-DEL-211 | chrX-149854078-DEL-211 |
| chrX-149860338-DEL-70 | chrX-149860339-DEL-70 |
| chrX-149875424-DEL-198 | chrX-149875425-DEL-198 |
| chrX-149881223-DEL-98 | chrX-149881224-DEL-98 |
| chrX-149910220-DEL-204 | chrX-149910221-DEL-204 |
| chrX-149911958-DEL-394 | chrX-149911959-DEL-394 |
| chrX-149974812-DEL-214 | chrX-149974813-DEL-214 |
| chrX-149997687-DEL-199 | chrX-149997688-DEL-199 |
| chrX-149998120-DEL-576 | chrX-149998121-DEL-575 |
| chrX-150006577-DEL-56 | chrX-150006578-DEL-56 |
| chrX-150021287-DEL-7128 | chrX-150021288-DEL-7128 |
| chrX-150061450-DEL-498 | chrX-150061451-DEL-498 |
| chrX-150065584-DEL-867 | chrX-150065585-DEL-867 |
| chrX-150083126-DEL-229 | chrX-150083127-DEL-229 |
| chrX-150091205-DEL-102 | chrX-150091206-DEL-102 |
| chrX-150096825-DEL-224 | chrX-150096826-DEL-224 |
| chrX-150108216-DEL-200 | chrX-150108217-DEL-200 |
| chrX-150118703-DEL-369 | chrX-150118704-DEL-369 |
| chrX-150170084-DEL-100 | chrX-150170085-DEL-100 |
| chrX-150172356-DEL-91 | chrX-150172357-DEL-91 |
| chrX-150173998-DEL-142 | chrX-150173999-DEL-142 |
| chrX-150196488-DEL-1074 | chrX-150196489-DEL-1075 |
| chrX-150284842-DEL-6389 | chrX-150284843-DEL-6389 |
| chrX-150361136-DEL-7091 | chrX-150361137-DEL-7091 |
| chrX-150394346-DEL-64 | chrX-150394347-DEL-64 |
| chrX-150400767-DEL-175 | chrX-150400768-DEL-175 |
| chrX-150407492-DEL-253 | chrX-150407493-DEL-253 |
| chrX-150419291-DEL-242 | chrX-150419292-DEL-242 |
| chrX-150438385-DEL-2169 | chrX-150438386-DEL-2169 |
| chrX-150443718-DEL-207 | chrX-150443719-DEL-207 |
| chrX-150458749-DEL-2704 | chrX-150458750-DEL-2704 |
| chrX-150481214-DEL-200 | chrX-150481215-DEL-200 |
| chrX-150511273-DEL-189 | chrX-150511235-DEL-190 |
| chrX-150525648-DEL-7421 | chrX-150525649-DEL-7421 |
| chrX-150541464-DEL-328 | chrX-150541465-DEL-328 |
| chrX-150559804-DEL-194 | chrX-150559805-DEL-194 |
| chrX-150591073-DEL-62 | chrX-150591074-DEL-62 |
| chrX-150609880-DEL-224 | chrX-150609881-DEL-224 |
| chrX-150646074-DEL-55 | chrX-150646074-DEL-55 |
| chrX-150655535-DEL-52 | chrX-150655536-DEL-60 |
| chrX-150776568-DEL-66 | chrX-150776569-DEL-66 |
| chrX-150776968-DEL-216 | chrX-150776969-DEL-216 |
| chrX-150805052-DEL-211 | chrX-150805053-DEL-211 |
| chrX-150837833-DEL-113 | chrX-150837834-DEL-113 |
| chrX-150854290-DEL-59 | chrX-150854289-DEL-61 |
| chrX-150923584-DEL-212 | chrX-150923585-DEL-212 |
| chrX-150946719-DEL-84 | chrX-150946724-DEL-80 |
| chrX-150958972-DEL-206 | chrX-150958973-DEL-206 |
| chrX-151068295-DEL-219 | chrX-151068296-DEL-219 |
| chrX-151069494-DEL-812 | chrX-151069495-DEL-812 |
| chrX-151070642-DEL-181 | chrX-151070643-DEL-181 |
| chrX-151083477-DEL-216 | chrX-151083478-DEL-216 |
| chrX-151085516-DEL-1400 | chrX-151085516-DEL-1400 |
| chrX-151159067-DEL-633 | chrX-151159068-DEL-633 |
| chrX-151180649-DEL-58 | chrX-151180650-DEL-58 |
| chrX-151182692-DEL-84 | chrX-151182685-DEL-92 |
| chrX-151190901-DEL-192 | chrX-151190900-DEL-191 |
| chrX-151227598-DEL-193 | chrX-151227599-DEL-193 |
| chrX-151256704-DEL-108 | chrX-151256701-DEL-103 |
| chrX-151259751-DEL-8744 | chrX-151259752-DEL-8744 |
| chrX-151306596-DEL-6903 | chrX-151306597-DEL-6903 |

| | |
|---|---|
| chrX-151340779-DEL-223 | chrX-151340780-DEL-223 |
| chrX-151352189-DEL-4452 | chrX-151352190-DEL-4452 |
| chrX-151388506-DEL-189 | chrX-151388507-DEL-189 |
| chrX-151450873-DEL-1206 | chrX-151450874-DEL-1206 |
| chrX-151495640-DEL-1088 | chrX-151495640-DEL-1088 |
| chrX-151511179-DEL-192 | chrX-151511180-DEL-201 |
| chrX-151539622-DEL-205 | chrX-151539623-DEL-205 |
| chrX-151578283-DEL-58 | chrX-151578282-DEL-60 |
| chrX-151591886-DEL-226 | chrX-151591887-DEL-226 |
| chrX-151612022-DEL-182 | chrX-151612023-DEL-182 |
| chrX-151644575-DEL-3733 | chrX-151644576-DEL-3733 |
| chrX-151677680-DEL-7087 | chrX-151677681-DEL-7087 |
| chrX-151706517-DEL-239 | chrX-151706518-DEL-239 |
| chrX-151746067-DEL-4903 | chrX-151746068-DEL-4903 |
| chrX-151761520-DEL-143 | chrX-151761521-DEL-143 |
| chrX-151825796-DEL-92 | chrX-151825796-DEL-92 |
| chrX-151838437-DEL-1420 | chrX-151838438-DEL-1420 |
| chrX-151839948-DEL-73 | chrX-151839949-DEL-73 |
| chrX-151845507-DEL-71 | chrX-151845502-DEL-67 |
| chrX-151885066-DEL-220 | chrX-151885057-DEL-221 |
| chrX-151956896-DEL-60 | chrX-151956897-DEL-60 |
| chrX-151967070-DEL-406 | chrX-151967071-DEL-406 |
| chrX-151976490-DEL-4379 | chrX-151976491-DEL-4379 |
| chrX-151986157-DEL-206 | chrX-151986163-DEL-207 |
| chrX-152008573-DEL-68 | chrX-152008573-DEL-68 |
| chrX-152042647-DEL-165 | chrX-152042648-DEL-165 |
| chrX-152055296-DEL-171 | chrX-152055297-DEL-171 |
| chrX-152107769-DEL-327 | chrX-152107770-DEL-327 |
| chrX-152146811-DEL-233 | chrX-152146812-DEL-233 |
| chrX-152147970-DEL-211 | chrX-152147971-DEL-211 |
| chrX-152152152-DEL-52 | chrX-152152155-DEL-50 |
| chrX-152221952-DEL-67 | chrX-152221952-DEL-68 |
| chrX-152236865-DEL-77 | chrX-152236866-DEL-77 |
| chrX-152304946-DEL-4811 | chrX-152304947-DEL-4811 |
| chrX-152369999-DEL-90 | chrX-152370000-DEL-74 |
| chrX-152373681-DEL-6661 | chrX-152373682-DEL-6661 |
| chrX-152390555-DEL-64 | chrX-152390556-DEL-64 |
| chrX-152424579-DEL-3164 | chrX-152424580-DEL-3164 |
| chrX-152440801-DEL-76 | chrX-152440802-DEL-76 |
| chrX-152443610-DEL-200 | chrX-152443609-DEL-198 |
| chrX-152605783-DEL-6203 | chrX-152605767-DEL-6203 |
| chrX-152680664-DEL-90 | chrX-152680665-DEL-50 |
| chrX-152690803-DEL-60 | chrX-152690805-DEL-58 |
| chrX-152721251-DEL-2800 | chrX-152721252-DEL-2800 |
| chrX-152930445-DEL-10934 | chrX-152930446-DEL-10934 |
| chrX-152944943-DEL-61 | chrX-152944943-DEL-61 |
| chrX-153088092-DEL-3010 | chrX-153088093-DEL-3009 |
| chrX-153117331-DEL-54 | chrX-153117307-DEL-78 |
| chrX-153147340-DEL-1269 | chrX-153147341-DEL-1269 |
| chrX-153166366-DEL-3993 | chrX-153166367-DEL-3993 |
| chrX-153210218-DEL-33380 | chrX-153210215-DEL-33383 |
| chrX-153283794-DEL-6195 | chrX-153283795-DEL-6195 |
| chrX-153497038-DEL-1027 | chrX-153497039-DEL-1027 |
| chrX-153532002-DEL-87 | chrX-153532003-DEL-87 |
| chrX-153562097-DEL-782 | chrX-153562098-DEL-782 |
| chrX-153587783-DEL-2584 | chrX-153587784-DEL-2584 |
| chrX-153596190-DEL-1496 | chrX-153596191-DEL-1496 |
| chrX-153601401-DEL-4162 | chrX-153601402-DEL-4162 |
| chrX-153624140-DEL-91 | chrX-153624141-DEL-77 |
| chrX-153689223-DEL-351 | chrX-153689224-DEL-351 |
| chrX-153716808-DEL-557 | chrX-153716808-DEL-557 |
| chrX-153759902-DEL-145 | chrX-153759903-DEL-145 |
| chrX-153860142-DEL-4008 | chrX-153860144-DEL-4007 |
| chrX-153884222-DEL-88 | chrX-153884229-DEL-66 |
| chrX-153899229-DEL-275 | chrX-153899230-DEL-276 |
| chrX-153921812-DEL-471 | chrX-153921813-DEL-471 |
| chrX-153936959-DEL-4369 | chrX-153936960-DEL-4369 |
| chrX-153943384-DEL-1462 | chrX-153943381-DEL-1466 |
| chrX-153963425-DEL-1449 | chrX-153963426-DEL-1449 |
| chrX-153971770-DEL-154 | chrX-153971807-DEL-119 |
| chrX-154004301-DEL-7069 | chrX-154004302-DEL-7069 |
| chrX-154036557-DEL-219 | chrX-154036558-DEL-219 |
| chrX-154057012-DEL-109 | chrX-154057018-DEL-107 |
| chrX-154099890-DEL-368 | chrX-154099891-DEL-368 |
| chrX-154100578-DEL-394 | chrX-154100579-DEL-394 |
| chrX-154101803-DEL-1231 | chrX-154101804-DEL-1231 |
| chrX-154113896-DEL-373 | chrX-154113897-DEL-373 |
| chrX-154145011-DEL-57 | chrX-154145010-DEL-60 |
| chrX-154157305-DEL-225 | chrX-154157306-DEL-225 |
| chrX-154165448-DEL-411 | chrX-154165449-DEL-411 |

| | |
|---|---|
| chrX-154176159-DEL-73 | chrX-154176155-DEL-69 |
| chrX-154196237-DEL-1000 | chrX-154196238-DEL-1000 |
| chrX-154235465-DEL-1716 | chrX-154235466-DEL-1716 |
| chrX-154242207-DEL-6882 | chrX-154242208-DEL-6882 |
| chrX-154257812-DEL-71 | chrX-154257813-DEL-71 |
| chrX-154387725-DEL-293 | chrX-154387725-DEL-294 |
| chrX-154397952-DEL-98 | chrX-154397953-DEL-98 |
| chrX-154442803-DEL-201 | chrX-154442804-DEL-201 |
| chrX-154454380-DEL-3755 | chrX-154454381-DEL-3755 |
| chrX-154459450-DEL-52 | chrX-154459409-DEL-72 |
| chrX-154464651-DEL-6525 | chrX-154464652-DEL-6525 |
| chrX-154473445-DEL-59 | chrX-154473438-DEL-58 |
| chrX-154512987-DEL-198 | chrX-154513028-DEL-198 |
| chrX-154561700-DEL-4402 | chrX-154561701-DEL-4402 |
| chrX-154575447-DEL-171 | chrX-154575448-DEL-171 |
| chrX-154580410-DEL-982 | chrX-154580411-DEL-982 |
| chrX-154594068-DEL-87 | chrX-154594069-DEL-95 |
| chrX-154632207-DEL-7077 | chrX-154632208-DEL-7077 |
| chrX-154655758-DEL-6312 | chrX-154655759-DEL-6312 |
| chrX-154686879-DEL-1377 | chrX-154686880-DEL-1377 |
| chrX-154700141-DEL-87 | chrX-154700134-DEL-95 |
| chrX-154707326-DEL-1356 | chrX-154707327-DEL-1356 |
| chrX-154773842-DEL-6401 | chrX-154773843-DEL-6401 |
| chrX-154791810-DEL-152 | chrX-154791810-DEL-151 |
| chrX-154822218-DEL-60 | chrX-154822222-DEL-56 |
| chrX-154846996-DEL-7418 | chrX-154846997-DEL-7418 |
| chrX-154933615-DEL-385 | chrX-154933616-DEL-385 |
| chrX-154935381-DEL-508 | chrX-154935382-DEL-508 |
| chrX-154953096-DEL-213 | chrX-154953097-DEL-213 |
| chrX-154980435-DEL-57 | chrX-154980430-DEL-58 |
| chrX-154990047-DEL-266 | chrX-154990048-DEL-266 |
| chrX-155053083-DEL-281 | chrX-155053084-DEL-281 |
| chrX-155064755-DEL-68 | chrX-155064766-DEL-50 |
| chrX-155088220-DEL-64 | chrX-155088197-DEL-88 |
| chrX-155101482-DEL-4815 | chrX-155101483-DEL-4815 |
| chrX-155110344-DEL-53 | chrX-155110344-DEL-57 |
| chrX-155117351-DEL-67 | chrX-155117352-DEL-67 |
| chrX-155153633-DEL-8614 | chrX-155153635-DEL-8613 |
| chrX-155253451-DEL-203 | chrX-155253452-DEL-203 |
| chrX-155284420-DEL-107 | chrX-155284412-DEL-88 |
| chrX-155287106-DEL-2897 | chrX-155287107-DEL-2897 |
| chrX-155304935-DEL-428 | chrX-155304936-DEL-428 |
| chrX-155307102-DEL-54 | chrX-155307096-DEL-65 |
| chrX-155332138-DEL-52 | chrX-155332139-DEL-52 |
| chrX-155338590-DEL-1481 | chrX-155338591-DEL-1481 |
| chrX-155362860-DEL-7086 | chrX-155362861-DEL-7086 |
| chrX-155381874-DEL-53 | chrX-155381850-DEL-66 |
| chrX-155400848-DEL-154 | chrX-155400850-DEL-153 |
| chrX-155422443-DEL-972 | chrX-155422442-DEL-968 |
| chrX-155438497-DEL-6613 | chrX-155438496-DEL-6613 |
| chrX-155514968-DEL-1932 | chrX-155514968-DEL-1932 |
| chrX-155537947-DEL-239 | chrX-155537948-DEL-239 |
| chrX-155573695-DEL-484 | chrX-155573696-DEL-484 |
| chrX-155620970-DEL-214 | chrX-155620971-DEL-214 |
| chrX-155640670-DEL-661 | chrX-155640671-DEL-661 |
| chrX-155651682-DEL-204 | chrX-155651683-DEL-204 |
| chrX-155678986-DEL-233 | chrX-155678987-DEL-233 |
| chrX-155709852-DEL-1705 | chrX-155709853-DEL-1705 |
| chrX-155811295-DEL-208 | chrX-155811296-DEL-208 |
| chrX-155831890-DEL-995 | chrX-155831891-DEL-995 |
| chrX-155880114-DEL-3240 | chrX-155880115-DEL-3240 |
| chrX-155882787-DEL-6654 | chrX-155882788-DEL-6654 |
| chrX-155894104-DEL-205 | chrX-155894105-DEL-205 |
| chrX-155909798-DEL-90 | chrX-155909799-DEL-90 |
| chrX-155910393-DEL-87 | chrX-155910394-DEL-80 |
| chrX-155934780-DEL-1424 | chrX-155934775-DEL-1429 |
| chrX-155965333-DEL-84 | chrX-155965360-DEL-66 |
| chrX-155974343-DEL-1561 | chrX-155974332-DEL-1534 |
| chrX-155995368-DEL-54 | chrX-155995373-DEL-50 |
| chrX-156028571-DEL-120 | chrX-156028572-DEL-128 |
| chrX-156039715-DEL-454 | chrX-156039715-DEL-454 |
| chrX-156046842-DEL-306 | chrX-156046843-DEL-306 |
| chrX-156065956-DEL-6706 | chrX-156065957-DEL-6706 |
| chrX-156163197-DEL-497 | chrX-156163198-DEL-497 |
| chrX-156164555-DEL-64 | chrX-156164566-DEL-61 |
| chrX-156194461-DEL-189 | chrX-156194462-DEL-189 |
| chrX-156217740-DEL-87 | chrX-156217741-DEL-87 |
| chrX-156218748-DEL-199 | chrX-156218749-DEL-199 |
| chrX-156232947-DEL-222 | chrX-156232948-DEL-222 |
| chrX-156306452-DEL-338 | chrX-156306453-DEL-338 |

| | |
|---|---|
| chrX-156390716-DEL-208 | chrX-156390748-DEL-193 |
| chrX-156402190-DEL-6594 | chrX-156402191-DEL-6594 |
| chrX-156443883-DEL-131 | chrX-156443882-DEL-114 |
| chrX-156447418-DEL-86 | chrX-156447433-DEL-70 |
| chrX-156466677-DEL-1531 | chrX-156466678-DEL-1531 |
| chrX-156482717-DEL-53 | chrX-156482718-DEL-53 |
| chrX-156516602-DEL-68 | chrX-156516613-DEL-58 |
| chrX-156525173-DEL-1802 | chrX-156525174-DEL-1802 |
| chrX-156527497-DEL-104 | chrX-156527498-DEL-104 |
| chrX-156546177-DEL-6625 | chrX-156546178-DEL-6625 |
| chrX-156554064-DEL-554 | chrX-156554065-DEL-554 |
| chrX-156581008-DEL-475 | chrX-156581009-DEL-475 |
| chrX-156656928-DEL-1691 | chrX-156656929-DEL-1691 |
| chrX-156662561-DEL-83 | chrX-156662587-DEL-58 |
| chrX-156690104-DEL-400 | chrX-156690105-DEL-400 |
| chrX-156739656-DEL-841 | chrX-156739657-DEL-841 |
| chrX-156770277-DEL-498 | chrX-156770278-DEL-498 |
| chrX-156794353-DEL-876 | chrX-156794354-DEL-876 |
| chrX-156852159-DEL-1487 | chrX-156852164-DEL-1483 |
| chrX-156880880-DEL-124 | chrX-156880881-DEL-123 |
| chrX-156895369-DEL-6704 | chrX-156895370-DEL-6704 |
| chrX-156895477-DEL-431 | chrX-156895476-DEL-431 |
| chrX-156925175-DEL-6656 | chrX-156925176-DEL-6656 |
| chrX-156932707-DEL-459 | chrX-156932708-DEL-459 |
| chrX-156942443-DEL-2878 | chrX-156942444-DEL-2877 |
| chrX-156985004-DEL-1633 | chrX-156985005-DEL-1633 |
| chrX-157003976-DEL-1623 | chrX-157003977-DEL-1621 |
| chrX-157043853-DEL-101 | chrX-157043889-DEL-75 |
| chrX-157096575-DEL-64 | chrX-157096576-DEL-64 |
| chrX-157107346-DEL-3183 | chrX-157107347-DEL-3183 |
| chrX-157170386-DEL-3322 | chrX-157170387-DEL-3322 |
| chrX-157219144-DEL-90 | chrX-157219145-DEL-90 |
| chrX-157226833-DEL-2447 | chrX-157226834-DEL-2447 |
| chrX-157241291-DEL-1288 | chrX-157241292-DEL-1288 |
| chrX-157346989-DEL-6591 | chrX-157346990-DEL-6591 |
| chrX-157372314-DEL-533 | chrX-157372315-DEL-533 |
| chrX-157383786-DEL-6496 | chrX-157383787-DEL-6496 |
| chrX-157390598-DEL-76 | chrX-157390599-DEL-76 |
| chrX-157417895-DEL-183 | chrX-157417928-DEL-142 |
| chrX-157459984-DEL-7075 | chrX-157459985-DEL-7075 |
| chrX-157508603-DEL-270 | chrX-157508604-DEL-280 |
| chrX-157524054-DEL-2660 | chrX-157524055-DEL-2660 |
| chrX-157609570-DEL-316 | chrX-157609571-DEL-316 |
| chrX-157619729-DEL-55 | chrX-157619730-DEL-55 |
| chrX-157652822-DEL-6677 | chrX-157652823-DEL-6677 |
| chrX-157669817-DEL-201 | chrX-157669818-DEL-201 |
| chrX-157708738-DEL-471 | chrX-157708739-DEL-471 |
| chrX-157732857-DEL-4878 | chrX-157732858-DEL-4878 |
| chrX-157741978-DEL-491 | chrX-157741979-DEL-491 |
| chrX-157751324-DEL-395 | chrX-157751325-DEL-394 |
| chrX-157769459-DEL-383 | chrX-157769463-DEL-378 |
| chrX-157788794-DEL-1277 | chrX-157788795-DEL-1277 |
| chrX-157793561-DEL-7159 | chrX-157793561-DEL-7159 |
| chrX-157818899-DEL-442 | chrX-157818900-DEL-442 |
| chrX-157867115-DEL-454 | chrX-157867102-DEL-450 |
| chrX-157892205-DEL-70 | chrX-157892206-DEL-70 |
| chrX-157932994-DEL-4865 | chrX-157932995-DEL-4865 |
| chrX-157964278-DEL-111 | chrX-157964279-DEL-111 |
| chrX-157976927-DEL-135 | chrX-157976928-DEL-135 |
| chrX-157989069-DEL-117 | chrX-157989052-DEL-114 |
| chrX-158007255-DEL-51 | chrX-158007255-DEL-51 |
| chrX-158014530-DEL-56 | chrX-158014531-DEL-56 |
| chrX-158022802-DEL-431 | chrX-158022802-DEL-431 |
| chrX-158026974-DEL-125 | chrX-158026975-DEL-126 |
| chrX-158081998-DEL-2780 | chrX-158081999-DEL-2780 |
| chrX-158141028-DEL-202 | chrX-158141029-DEL-202 |
| chrX-158209882-DEL-199 | chrX-158209883-DEL-199 |
| chrX-158238022-DEL-53 | chrX-158238023-DEL-53 |
| chrX-158258534-DEL-207 | chrX-158258535-DEL-207 |
| chrX-158276465-DEL-230 | chrX-158276466-DEL-230 |
| chrX-158291816-DEL-93 | chrX-158291817-DEL-94 |
| chrX-158366387-DEL-219 | chrX-158366380-DEL-217 |
| chrX-158370893-DEL-180 | chrX-158370894-DEL-181 |
| chrX-158390958-DEL-360 | chrX-158390959-DEL-360 |
| chrX-158444163-DEL-722 | chrX-158444164-DEL-722 |
| chrX-158445798-DEL-6402 | chrX-158445799-DEL-6402 |
| chrX-158477497-DEL-197 | chrX-158477498-DEL-197 |
| chrX-158541940-DEL-2804 | chrX-158541941-DEL-2804 |
| chrX-158573343-DEL-75 | chrX-158573346-DEL-51 |
| chrX-158574257-DEL-472 | chrX-158574253-DEL-472 |

| | |
|---|---|
| chrX-158609358-DEL-57 | chrX-158609359-DEL-57 |
| chrX-158697613-DEL-280 | chrX-158697614-DEL-280 |
| chrX-158721269-DEL-70 | chrX-158721270-DEL-70 |
| chrX-158728122-DEL-208 | chrX-158728123-DEL-208 |
| chrX-158753071-DEL-193 | chrX-158753072-DEL-193 |
| chrX-158754641-DEL-66 | chrX-158754641-DEL-72 |
| chrX-158755388-DEL-63 | chrX-158755389-DEL-63 |
| chrX-158783748-DEL-56 | chrX-158783749-DEL-56 |
| chrX-158786605-DEL-203 | chrX-158786606-DEL-203 |
| chrX-158808188-DEL-862 | chrX-158808189-DEL-862 |
| chrX-158810224-DEL-407 | chrX-158810225-DEL-407 |
| chrX-158829388-DEL-79 | chrX-158829389-DEL-79 |
| chrX-158840731-DEL-399 | chrX-158840732-DEL-399 |
| chrX-158864908-DEL-243 | chrX-158864909-DEL-243 |
| chrX-158928755-DEL-551 | chrX-158928756-DEL-551 |
| chrX-158948733-DEL-577 | chrX-158948734-DEL-575 |
| chrX-158984857-DEL-416 | chrX-158984858-DEL-416 |
| chrX-158997457-DEL-1407 | chrX-158997458-DEL-1407 |
| chrX-159042124-DEL-133 | chrX-159042125-DEL-133 |
| chrX-159045273-DEL-407 | chrX-159045274-DEL-407 |
| chrX-159063029-DEL-3280 | chrX-159063030-DEL-3280 |
| chrX-159068584-DEL-7199 | chrX-159068585-DEL-7199 |
| chrX-159083782-DEL-2876 | chrX-159083783-DEL-2876 |
| chrX-159222040-DEL-108 | chrX-159222030-DEL-101 |
| chrX-159250784-DEL-535 | chrX-159250785-DEL-535 |
| chrX-159258542-DEL-76 | chrX-159258588-DEL-68 |
| chrX-159262364-DEL-249 | chrX-159262365-DEL-249 |
| chrX-159291008-DEL-685 | chrX-159291009-DEL-685 |
| chrX-159296850-DEL-655 | chrX-159296851-DEL-655 |
| chrX-159305376-DEL-72 | chrX-159305377-DEL-72 |
| chrX-159315020-DEL-500 | chrX-159315021-DEL-500 |
| chrX-159317625-DEL-6203 | chrX-159317626-DEL-6203 |
| chrX-159345089-DEL-387 | chrX-159345090-DEL-387 |
| chrX-159349745-DEL-105 | chrX-159349746-DEL-105 |
| chrX-159386171-DEL-65 | chrX-159386177-DEL-64 |
| chrX-159391962-DEL-201 | chrX-159391955-DEL-202 |
| chrX-159406787-DEL-53 | chrX-159406788-DEL-53 |
| chrX-159410849-DEL-1494 | chrX-159410848-DEL-1494 |
| chrX-159421448-DEL-731 | chrX-159421449-DEL-731 |
| chrX-159436341-DEL-6265 | chrX-159436342-DEL-6265 |
| chrX-159470635-DEL-547 | chrX-159470636-DEL-547 |
| chrX-159526506-DEL-68 | chrX-159526507-DEL-68 |
| chrX-159545741-DEL-155 | chrX-159545742-DEL-155 |
| chrX-159546572-DEL-401 | chrX-159546573-DEL-401 |
| chrX-159552582-DEL-226 | chrX-159552579-DEL-225 |
| chrX-159839267-DEL-6477 | chrX-159839268-DEL-6477 |
| chrX-159910339-DEL-214 | chrX-159910378-DEL-210 |
| chrX-159949092-DEL-3674 | chrX-159949093-DEL-3674 |
| chrX-159962170-DEL-6413 | chrX-159962171-DEL-6413 |
| chrX-159977875-DEL-265 | chrX-159977876-DEL-265 |
| chrX-159983640-DEL-400 | chrX-159983641-DEL-400 |
| chrX-159998883-DEL-88 | chrX-159998884-DEL-88 |
| chrX-160000053-DEL-215 | chrX-160000054-DEL-215 |
| chrX-160008011-DEL-1526 | chrX-160008012-DEL-1528 |
| chrX-160093113-DEL-60 | chrX-160093111-DEL-56 |
| chrX-160093772-DEL-4219 | chrX-160093773-DEL-4219 |
| chrX-160100702-DEL-1792 | chrX-160100703-DEL-1792 |
| chrX-160130273-DEL-1031 | chrX-160130274-DEL-1031 |
| chrX-160158907-DEL-128 | chrX-160158908-DEL-128 |
| chrX-160173464-DEL-373 | chrX-160173465-DEL-373 |
| chrX-160176338-DEL-60 | chrX-160176339-DEL-60 |
| chrX-160191700-DEL-208 | chrX-160191701-DEL-208 |
| chrX-160255960-DEL-59 | chrX-160255961-DEL-59 |
| chrX-160266032-DEL-161 | chrX-160266033-DEL-161 |
| chrX-160277580-DEL-99 | chrX-160277582-DEL-95 |
| chrX-160291895-DEL-235 | chrX-160291896-DEL-235 |
| chrX-160323862-DEL-6670 | chrX-160323863-DEL-6670 |
| chrX-160373089-DEL-81 | chrX-160373095-DEL-87 |
| chrX-160389521-DEL-225 | chrX-160389522-DEL-225 |
| chrX-160405733-DEL-6176 | chrX-160405734-DEL-6176 |
| chrX-160441692-DEL-70 | chrX-160441696-DEL-52 |
| chrX-160590679-DEL-603 | chrX-160590680-DEL-603 |
| chrX-160610992-DEL-99 | chrX-160611002-DEL-123 |
| chrX-160661834-DEL-912 | chrX-160661835-DEL-912 |
| chrX-160729777-DEL-64 | chrX-160729768-DEL-80 |
| chrX-160731294-DEL-767 | chrX-160731295-DEL-767 |
| chrX-160852445-DEL-3092 | chrX-160852446-DEL-3092 |
| chrX-160946683-DEL-70 | chrX-160946684-DEL-70 |
| chrX-160960331-DEL-13542 | chrX-160960332-DEL-13542 |
| chrX-161021594-DEL-1186 | chrX-161021595-DEL-1185 |

| | |
|---|---|
| chrX-161103121-DEL-4634 | chrX-161103122-DEL-4634 |
| chrX-161115811-DEL-321 | chrX-161115812-DEL-321 |
| chrX-161203068-DEL-56 | chrX-161203041-DEL-68 |
| chrX-161209632-DEL-52 | chrX-161209628-DEL-62 |
| chrX-161213073-DEL-2118 | chrX-161213074-DEL-2118 |
| chrX-161264538-DEL-232 | chrX-161264539-DEL-232 |
| chrX-161341056-DEL-1525 | chrX-161341058-DEL-1524 |
| chrX-161376302-DEL-2046 | chrX-161376302-DEL-2046 |
| chrX-161382128-DEL-317 | chrX-161382115-DEL-317 |
| chrX-161385709-DEL-1893 | chrX-161385715-DEL-1893 |
| chrX-161395061-DEL-60 | chrX-161395062-DEL-60 |
| chrX-161451728-DEL-6805 | chrX-161451729-DEL-6805 |
| chrX-161478666-DEL-1711 | chrX-161478667-DEL-1711 |
| chrX-161480996-DEL-1486 | chrX-161480997-DEL-1486 |
| chrX-161484138-DEL-68 | chrX-161484139-DEL-52 |
| chrX-161487374-DEL-302 | chrX-161487375-DEL-302 |
| chrX-161489225-DEL-2109 | chrX-161489226-DEL-2109 |
| chrX-161494089-DEL-197 | chrX-161494090-DEL-197 |
| chrX-161499276-DEL-1697 | chrX-161499277-DEL-1697 |
| chrX-161500163-DEL-212 | chrX-161500163-DEL-212 |
| chrX-161508033-DEL-1264 | chrX-161508080-DEL-1215 |
| chrX-161538599-DEL-198 | chrX-161538599-DEL-199 |
| chrX-161559889-DEL-98 | chrX-161559890-DEL-94 |
| chrX-161565344-DEL-109 | chrX-161565345-DEL-109 |
| chrX-161579174-DEL-200 | chrX-161579172-DEL-195 |
| chrX-161593467-DEL-212 | chrX-161593468-DEL-212 |
| chrX-161596357-DEL-80 | chrX-161596358-DEL-80 |
| chrX-161597656-DEL-534 | chrX-161597660-DEL-525 |
| chrX-161598169-DEL-108 | chrX-161598172-DEL-106 |
| chrX-161599579-DEL-242 | chrX-161599580-DEL-242 |
| chrX-161602044-DEL-87 | chrX-161602050-DEL-81 |
| chrX-161604373-DEL-60 | chrX-161604374-DEL-60 |
| chrX-161618338-DEL-63 | chrX-161618339-DEL-51 |
| chrX-161635270-DEL-231 | chrX-161635263-DEL-227 |
| chrX-161635943-DEL-196 | chrX-161635964-DEL-173 |
| chrX-161645057-DEL-209 | chrX-161645058-DEL-209 |
| chrX-161677448-DEL-213 | chrX-161677449-DEL-213 |
| chrX-161692054-DEL-321 | chrX-161692055-DEL-321 |
| chrX-161693815-DEL-323 | chrX-161693816-DEL-323 |
| chrX-161708420-DEL-92 | chrX-161708389-DEL-124 |
| chrX-161711290-DEL-214 | chrX-161711291-DEL-214 |
| chrX-161727538-DEL-114 | chrX-161727550-DEL-113 |
| chrX-161761621-DEL-7131 | chrX-161761622-DEL-7131 |
| chrX-161796614-DEL-70 | chrX-161796615-DEL-70 |
| chrX-161801336-DEL-182 | chrX-161801337-DEL-182 |
| chrX-161809681-DEL-54 | chrX-161809644-DEL-62 |
| chrX-161836748-DEL-206 | chrX-161836749-DEL-206 |
| chrX-161841448-DEL-204 | chrX-161841449-DEL-204 |
| chrX-161845245-DEL-592 | chrX-161845246-DEL-592 |
| chrX-161864065-DEL-2171 | chrX-161864066-DEL-2171 |
| chrX-161903717-DEL-210 | chrX-161903718-DEL-210 |
| chrX-161913145-DEL-107 | chrX-161913159-DEL-108 |
| chrX-161918725-DEL-86 | chrX-161918722-DEL-51 |
| chrX-161979369-DEL-54 | chrX-161979324-DEL-55 |
| chrX-161986395-DEL-51 | chrX-161986363-DEL-65 |
| chrX-162002357-DEL-4547 | chrX-162002357-DEL-4547 |
| chrX-162062552-DEL-2621 | chrX-162062553-DEL-2620 |
| chrX-162103023-DEL-815 | chrX-162103023-DEL-815 |
| chrX-162104281-DEL-189 | chrX-162104282-DEL-189 |
| chrX-162127950-DEL-445 | chrX-162127951-DEL-445 |
| chrX-162155226-DEL-1423 | chrX-162155219-DEL-1424 |
| chrX-162182563-DEL-203 | chrX-162182564-DEL-203 |
| chrX-162277037-DEL-2883 | chrX-162277038-DEL-2882 |
| chrX-162288641-DEL-54 | chrX-162288638-DEL-50 |
| chrX-162321960-DEL-54 | chrX-162321960-DEL-66 |
| chrX-162361903-DEL-62 | chrX-162361905-DEL-58 |
| chrX-162393433-DEL-412 | chrX-162393434-DEL-412 |
| chrX-162402989-DEL-62 | chrX-162402990-DEL-62 |
| chrX-162422746-DEL-73 | chrX-162422747-DEL-73 |
| chrX-162450029-DEL-254 | chrX-162450030-DEL-254 |
| chrX-162472933-DEL-104 | chrX-162472934-DEL-104 |
| chrX-162525360-DEL-114 | chrX-162525361-DEL-114 |
| chrX-162575856-DEL-103 | chrX-162575857-DEL-103 |
| chrX-162593860-DEL-54 | chrX-162593861-DEL-54 |
| chrX-162642379-DEL-3745 | chrX-162642380-DEL-3745 |
| chrX-162676967-DEL-526 | chrX-162676968-DEL-526 |
| chrX-162738615-DEL-53 | chrX-162738616-DEL-53 |
| chrX-162748235-DEL-119 | chrX-162748236-DEL-119 |
| chrX-162751308-DEL-56 | chrX-162751309-DEL-56 |
| chrX-162754527-DEL-120 | chrX-162754528-DEL-120 |

| | |
|---|---|
| chrX-162761881-DEL-68 | chrX-162761872-DEL-74 |
| chrX-162765922-DEL-1190 | chrX-162765922-DEL-1188 |
| chrX-162790796-DEL-70 | chrX-162790805-DEL-69 |
| chrX-162811834-DEL-108 | chrX-162811835-DEL-96 |
| chrX-162831333-DEL-5639 | chrX-162831334-DEL-5639 |
| chrX-162853265-DEL-168 | chrX-162853266-DEL-168 |
| chrX-162877490-DEL-64 | chrX-162877491-DEL-64 |
| chrX-162931162-DEL-64 | chrX-162931162-DEL-65 |
| chrX-162934688-DEL-679 | chrX-162934689-DEL-679 |
| chrX-162987115-DEL-489 | chrX-162987103-DEL-490 |
| chrX-162995301-DEL-181 | chrX-162995302-DEL-181 |
| chrX-162997589-DEL-459 | chrX-162997586-DEL-461 |
| chrX-163006381-DEL-133 | chrX-163006382-DEL-133 |
| chrX-163048501-DEL-555 | chrX-163048502-DEL-555 |
| chrX-163080787-DEL-1411 | chrX-163080788-DEL-1411 |
| chrX-163097636-DEL-182 | chrX-163097639-DEL-183 |
| chrX-163134192-DEL-6650 | chrX-163134193-DEL-6650 |
| chrX-163166816-DEL-1577 | chrX-163166817-DEL-1577 |
| chrX-163203490-DEL-111 | chrX-163203494-DEL-112 |
| chrX-163208529-DEL-101 | chrX-163208530-DEL-101 |
| chrX-163217715-DEL-216 | chrX-163217716-DEL-216 |
| chrX-163244187-DEL-212 | chrX-163244192-DEL-205 |
| chrX-163249102-DEL-1284 | chrX-163249105-DEL-1282 |
| chrX-163275364-DEL-1218 | chrX-163275365-DEL-1218 |
| chrX-163278937-DEL-916 | chrX-163278938-DEL-916 |
| chrX-163282423-DEL-104 | chrX-163282423-DEL-64 |
| chrX-163315247-DEL-57 | chrX-163315234-DEL-71 |
| chrX-163345645-DEL-6680 | chrX-163345646-DEL-6680 |
| chrX-163372070-DEL-198 | chrX-163372071-DEL-198 |
| chrX-163397946-DEL-7285 | chrX-163397947-DEL-7285 |
| chrX-163441278-DEL-1437 | chrX-163441274-DEL-1437 |
| chrX-163448513-DEL-428 | chrX-163448503-DEL-422 |
| chrX-163451820-DEL-64 | chrX-163451835-DEL-54 |
| chrX-163459805-DEL-66 | chrX-163459806-DEL-66 |
| chrX-163467896-DEL-1094 | chrX-163467897-DEL-1094 |
| chrX-163473403-DEL-58 | chrX-163473404-DEL-58 |
| chrX-163489313-DEL-6770 | chrX-163489314-DEL-6770 |
| chrX-163540385-DEL-385 | chrX-163540385-DEL-386 |
| chrX-163551698-DEL-56 | chrX-163551693-DEL-60 |
| chrX-163552302-DEL-328 | chrX-163552303-DEL-328 |
| chrX-163600119-DEL-61 | chrX-163600112-DEL-60 |
| chrX-163615577-DEL-2366 | chrX-163615578-DEL-2366 |
| chrX-163643675-DEL-80 | chrX-163643680-DEL-76 |
| chrX-163673885-DEL-4173 | chrX-163673886-DEL-4173 |
| chrX-163689371-DEL-65 | chrX-163689391-DEL-54 |
| chrX-163724889-DEL-600 | chrX-163724894-DEL-596 |
| chrX-163727050-DEL-124 | chrX-163727057-DEL-90 |
| chrX-163736527-DEL-1198 | chrX-163736528-DEL-1198 |
| chrX-163802030-DEL-91 | chrX-163802050-DEL-52 |
| chrX-163802949-DEL-201 | chrX-163802950-DEL-201 |
| chrX-163813247-DEL-855 | chrX-163813248-DEL-855 |
| chrX-163855677-DEL-848 | chrX-163855678-DEL-848 |
| chrX-163866459-DEL-57 | chrX-163866460-DEL-57 |
| chrX-163871726-DEL-6505 | chrX-163871727-DEL-6505 |
| chrX-163882629-DEL-889 | chrX-163882630-DEL-889 |
| chrX-163891062-DEL-97 | chrX-163891063-DEL-97 |
| chrX-163934762-DEL-866 | chrX-163934763-DEL-866 |
| chrX-163949048-DEL-360 | chrX-163949049-DEL-360 |
| chrX-163964340-DEL-63 | chrX-163964341-DEL-63 |
| chrX-163997844-DEL-1761 | chrX-163997845-DEL-1761 |
| chrX-164083742-DEL-1725 | chrX-164083743-DEL-1725 |
| chrX-164249228-DEL-6855 | chrX-164249229-DEL-6855 |
| chrX-164265063-DEL-457 | chrX-164265064-DEL-457 |
| chrX-164284790-DEL-547 | chrX-164284791-DEL-547 |
| chrX-164391220-DEL-328 | chrX-164391221-DEL-328 |
| chrX-164417319-DEL-6505 | chrX-164417320-DEL-6505 |
| chrX-164417342-DEL-214 | chrX-164417343-DEL-214 |
| chrX-164425940-DEL-1996 | chrX-164425941-DEL-1996 |
| chrX-164431811-DEL-77 | chrX-164431767-DEL-77 |
| chrX-164439786-DEL-54 | chrX-164439788-DEL-53 |
| chrX-164449779-DEL-481 | chrX-164449780-DEL-481 |
| chrX-164455156-DEL-79 | chrX-164455156-DEL-71 |
| chrX-164516123-DEL-6914 | chrX-164516123-DEL-6914 |
| chrX-164614005-DEL-80 | chrX-164614006-DEL-80 |
| chrX-164652642-DEL-4354 | chrX-164652643-DEL-4354 |
| chrX-164695721-DEL-115 | chrX-164695714-DEL-98 |
| chrX-164796141-DEL-204 | chrX-164796142-DEL-204 |
| chrX-164800096-DEL-259 | chrX-164800097-DEL-259 |
| chrX-164893522-DEL-177 | chrX-164893523-DEL-177 |
| chrX-164900366-DEL-3870 | chrX-164900367-DEL-3870 |

| | |
|---|---|
| chrX-164910188-DEL-548 | chrX-164910188-DEL-548 |
| chrX-164912340-DEL-213 | chrX-164912341-DEL-213 |
| chrX-164931681-DEL-237 | chrX-164931682-DEL-237 |
| chrX-164960016-DEL-473 | chrX-164960025-DEL-465 |
| chrX-165005957-DEL-522 | chrX-165005958-DEL-522 |
| chrX-165044622-DEL-62 | chrX-165044627-DEL-56 |
| chrX-165054291-DEL-65 | chrX-165054292-DEL-65 |
| chrX-165057206-DEL-55 | chrX-165057207-DEL-55 |
| chrX-165127549-DEL-79 | chrX-165127550-DEL-79 |
| chrX-165184550-DEL-230 | chrX-165184551-DEL-230 |
| chrX-165199467-DEL-6240 | chrX-165199468-DEL-6240 |
| chrX-165207411-DEL-61 | chrX-165207425-DEL-62 |
| chrX-165211903-DEL-205 | chrX-165211904-DEL-203 |
| chrX-165275440-DEL-100 | chrX-165275460-DEL-50 |
| chrX-165277988-DEL-54 | chrX-165277989-DEL-54 |
| chrX-165315080-DEL-52 | chrX-165315081-DEL-52 |
| chrX-165337252-DEL-64 | chrX-165337253-DEL-61 |
| chrX-165427291-DEL-61 | chrX-165427293-DEL-58 |
| chrX-165440397-DEL-78 | chrX-165440378-DEL-58 |
| chrX-165518503-DEL-6444 | chrX-165518498-DEL-6445 |
| chrX-165530828-DEL-203 | chrX-165530829-DEL-203 |
| chrX-165556984-DEL-174 | chrX-165557017-DEL-149 |
| chrX-165575380-DEL-454 | chrX-165575381-DEL-454 |
| chrX-165607104-DEL-134 | chrX-165607105-DEL-134 |
| chrX-165607519-DEL-6169 | chrX-165607520-DEL-6169 |
| chrX-165614338-DEL-61 | chrX-165614333-DEL-58 |
| chrX-165616423-DEL-53 | chrX-165616424-DEL-53 |
| chrX-165637169-DEL-889 | chrX-165637170-DEL-889 |
| chrX-165695847-DEL-98 | chrX-165695848-DEL-98 |
| chrX-165736887-DEL-56 | chrX-165736888-DEL-56 |
| chrX-165783150-DEL-704 | chrX-165783147-DEL-705 |
| chrX-165790133-DEL-66 | chrX-165790134-DEL-66 |
| chrX-165798595-DEL-71 | chrX-165798562-DEL-60 |
| chrX-165811280-DEL-60 | chrX-165811281-DEL-60 |
| chrX-165911801-DEL-81 | chrX-165911802-DEL-81 |
| chrX-165996343-DEL-6879 | chrX-165996344-DEL-6879 |
| chrX-166065989-DEL-51 | chrX-166065993-DEL-66 |
| chrX-166071747-DEL-67 | chrX-166071741-DEL-68 |
| chrX-166079284-DEL-3087 | chrX-166079285-DEL-3083 |
| chrX-166093197-DEL-147 | chrX-166093156-DEL-156 |
| chrX-166150719-DEL-235 | chrX-166150720-DEL-233 |
| chrX-166167053-DEL-54 | chrX-166167053-DEL-60 |
| chrX-166188210-DEL-407 | chrX-166188211-DEL-407 |
| chrX-166192129-DEL-1229 | chrX-166192130-DEL-1229 |
| chrX-166268673-DEL-191 | chrX-166268674-DEL-192 |
| chrX-166273263-DEL-71 | chrX-166273264-DEL-71 |
| chrX-166298320-DEL-220 | chrX-166298319-DEL-220 |
| chrX-166327203-DEL-409 | chrX-166327204-DEL-409 |
| chrX-166329558-DEL-766 | chrX-166329559-DEL-766 |
| chrX-166360285-DEL-54 | chrX-166360282-DEL-56 |
| chrX-166366556-DEL-491 | chrX-166366563-DEL-484 |
| chrX-166369700-DEL-1728 | chrX-166369701-DEL-1728 |
| chrX-166379159-DEL-823 | chrX-166379160-DEL-823 |
| chrX-166381544-DEL-585 | chrX-166381545-DEL-585 |
| chrX-166412908-DEL-265 | chrX-166412927-DEL-247 |
| chrX-166532976-DEL-1594 | chrX-166532977-DEL-1594 |
| chrX-166541236-DEL-54 | chrX-166541237-DEL-54 |
| chrX-166542328-DEL-57 | chrX-166542329-DEL-57 |
| chrX-166561363-DEL-109 | chrX-166561364-DEL-109 |
| chrX-166606676-DEL-66 | chrX-166606677-DEL-60 |
| chrX-166696173-DEL-1280 | chrX-166696174-DEL-1280 |
| chrX-166706138-DEL-501 | chrX-166706139-DEL-501 |
| chrX-166713915-DEL-142 | chrX-166713916-DEL-142 |
| chrX-166718188-DEL-70 | chrX-166718205-DEL-54 |
| chrX-166788638-DEL-52 | chrX-166788630-DEL-74 |
| chrX-166799073-DEL-763 | chrX-166799074-DEL-763 |
| chrX-166821924-DEL-564 | chrX-166821925-DEL-564 |
| chrX-166862618-DEL-532 | chrX-166862619-DEL-532 |
| chrX-166864099-DEL-2376 | chrX-166864100-DEL-2375 |
| chrX-166873907-DEL-52 | chrX-166873907-DEL-52 |
| chrX-166897344-DEL-569 | chrX-166897345-DEL-569 |
| chrX-166912955-DEL-2797 | chrX-166912956-DEL-2797 |
| chrX-166955073-DEL-203 | chrX-166955074-DEL-203 |
| chrX-166959602-DEL-5010 | chrX-166959603-DEL-5010 |
| chrX-166979954-DEL-222 | chrX-166979955-DEL-222 |
| chrX-166986915-DEL-170 | chrX-166986916-DEL-170 |
| chrX-167016149-DEL-242 | chrX-167016150-DEL-242 |
| chrX-167026745-DEL-1217 | chrX-167026746-DEL-1218 |
| chrX-167043315-DEL-327 | chrX-167043316-DEL-327 |
| chrX-167135523-DEL-2477 | chrX-167135523-DEL-2477 |

| | |
|---|---|
| chrX-167165162-DEL-746 | chrX-167165163-DEL-746 |
| chrX-167167959-DEL-51 | chrX-167167960-DEL-51 |
| chrX-167278200-DEL-6841 | chrX-167278201-DEL-6841 |
| chrX-167279719-DEL-200 | chrX-167279720-DEL-200 |
| chrX-167312051-DEL-1227 | chrX-167312052-DEL-1227 |
| chrX-167336007-DEL-203 | chrX-167336008-DEL-215 |
| chrX-167402295-DEL-75 | chrX-167402296-DEL-75 |
| chrX-167458739-DEL-145 | chrX-167458740-DEL-145 |
| chrX-167513333-DEL-1724 | chrX-167513334-DEL-1724 |
| chrX-167604770-DEL-570 | chrX-167604782-DEL-563 |
| chrX-167622519-DEL-455 | chrX-167622520-DEL-455 |
| chrX-167641994-DEL-60 | chrX-167641994-DEL-60 |
| chrX-167649233-DEL-152 | chrX-167649234-DEL-152 |
| chrX-167694755-DEL-56 | chrX-167694758-DEL-54 |
| chrX-167705066-DEL-6604 | chrX-167705067-DEL-6604 |
| chrX-167735123-DEL-838 | chrX-167735112-DEL-839 |
| chrX-167855298-DEL-55 | chrX-167855283-DEL-100 |
| chrX-167862612-DEL-117 | chrX-167862629-DEL-100 |
| chrX-167917475-DEL-1216 | chrX-167917475-DEL-1216 |
| chrX-167933228-DEL-54 | chrX-167933229-DEL-54 |
| chrX-167942770-DEL-6648 | chrX-167942771-DEL-6648 |
| chrX-168068181-DEL-6727 | chrX-168068182-DEL-6727 |
| chrX-168118756-DEL-132 | chrX-168118766-DEL-123 |
| chrX-168128236-DEL-73 | chrX-168128237-DEL-73 |
| chrX-168129868-DEL-3128 | chrX-168129869-DEL-3128 |
| chrX-168178199-DEL-63 | chrX-168178200-DEL-63 |
| chrX-168194386-DEL-6888 | chrX-168194387-DEL-6888 |
| chrX-168206284-DEL-219 | chrX-168206284-DEL-213 |
| chrX-168214373-DEL-110 | chrX-168214374-DEL-110 |
| chrX-168223798-DEL-2188 | chrX-168223799-DEL-2188 |
| chrX-168245090-DEL-1415 | chrX-168245091-DEL-1415 |
| chrX-168263405-DEL-79 | chrX-168263406-DEL-79 |
| chrX-168266729-DEL-151 | chrX-168266730-DEL-151 |
| chrX-168274839-DEL-999 | chrX-168274839-DEL-999 |
| chrX-168301923-DEL-335 | chrX-168301924-DEL-335 |
| chrX-168377327-DEL-68 | chrX-168377328-DEL-68 |
| chrX-168380455-DEL-625 | chrX-168380457-DEL-624 |
| chrX-168430752-DEL-70 | chrX-168430753-DEL-70 |
| chrX-168523680-DEL-6267 | chrX-168523676-DEL-6269 |
| chrX-168604348-DEL-295 | chrX-168604349-DEL-295 |
| chrX-168609192-DEL-169 | chrX-168609196-DEL-165 |
| chrX-168619102-DEL-1606 | chrX-168619103-DEL-1606 |
| chrX-168637213-DEL-59 | chrX-168637213-DEL-59 |
| chrX-168714643-DEL-53 | chrX-168714646-DEL-50 |
| chrX-168745757-DEL-91 | chrX-168745758-DEL-91 |
| chrX-168757179-DEL-2063 | chrX-168757180-DEL-2063 |
| chrX-168763444-DEL-763 | chrX-168763445-DEL-763 |
| chrX-168765234-DEL-827 | chrX-168765232-DEL-828 |
| chrX-168767144-DEL-62 | chrX-168767096-DEL-62 |
| chrX-168777275-DEL-93 | chrX-168777286-DEL-92 |
| chrX-168779398-DEL-60 | chrX-168779421-DEL-59 |
| chrX-168792909-DEL-81 | chrX-168792949-DEL-75 |
| chrX-168793312-DEL-183 | chrX-168793293-DEL-177 |
| chrX-169074332-DEL-610 | chrX-169074333-DEL-623 |
| chrX-169081022-DEL-185 | chrX-169081023-DEL-185 |
| chrX-169082469-DEL-123 | chrX-169082428-DEL-124 |
| chrX-169090225-DEL-66 | chrX-169090192-DEL-62 |
| chrX-169097208-DEL-219 | chrX-169097254-DEL-219 |
| chrX-169098050-DEL-58 | chrX-169098051-DEL-58 |
| chrX-169098746-DEL-291 | chrX-169098747-DEL-291 |
| chrX-169106047-DEL-150 | chrX-169106019-DEL-178 |
| chrX-169110601-DEL-89 | chrX-169110601-DEL-89 |
| chrX-169119383-DEL-91 | chrX-169119384-DEL-91 |

**Identified_By_Us**
chr1-3097022-DEL-313
chr1-3107938-DEL-6338
chr1-3115104-DEL-580
chr1-3121610-DEL-123
chr1-3146947-DEL-694
chr1-3195763-DEL-4425
chr1-3220905-DEL-5498
chr1-3232585-DEL-595
chr1-3245734-DEL-481
chr1-3246710-DEL-560
chr1-3248112-DEL-2648
chr1-3252268-DEL-54
chr1-3287426-DEL-339
chr1-3305609-DEL-798
chr1-3308086-DEL-5972
chr1-3335749-DEL-176
chr1-3338371-DEL-623
chr1-3344368-DEL-6292
chr1-3357572-DEL-101
chr1-3360940-DEL-1582
chr1-3372075-DEL-199
chr1-3374827-DEL-1604
chr1-3384758-DEL-185
chr1-3414865-DEL-3897
chr1-3420523-DEL-150
chr1-3460424-DEL-975
chr1-3503669-DEL-226
chr1-3564936-DEL-94
chr1-3602984-DEL-1063
chr1-3625308-DEL-579
chr1-3644508-DEL-180
chr1-3659430-DEL-60
chr1-3660442-DEL-54
chr1-3673712-DEL-1188
chr1-3682482-DEL-2276
chr1-3684835-DEL-733
chr1-3721546-DEL-91
chr1-3764098-DEL-55
chr1-3835603-DEL-60
chr1-3836254-DEL-6124
chr1-3844629-DEL-442
chr1-3859638-DEL-54
chr1-3868927-DEL-5545
chr1-3875550-DEL-749
chr1-3925069-DEL-399
chr1-3991720-DEL-1967
chr1-4026303-DEL-154
chr1-4053139-DEL-71
chr1-4053655-DEL-4576
chr1-4065200-DEL-54
chr1-4070563-DEL-67
chr1-4087229-DEL-504
chr1-4089948-DEL-2898
chr1-4117862-DEL-59
chr1-4124698-DEL-88
chr1-4137859-DEL-7206
chr1-4143036-DEL-120
chr1-4223716-DEL-51
chr1-4238486-DEL-117
chr1-4257617-DEL-9136
chr1-4289610-DEL-51
chr1-4299779-DEL-626
chr1-4326681-DEL-399
chr1-4326746-DEL-2284
chr1-4327451-DEL-245
chr1-4328219-DEL-88
chr1-4328734-DEL-60
chr1-4364025-DEL-797
chr1-4368748-DEL-74
chr1-4444138-DEL-4053
chr1-4467421-DEL-72
chr1-4470232-DEL-1101
chr1-4483917-DEL-72
chr1-4509976-DEL-5168
chr1-4519221-DEL-92
chr1-4521384-DEL-52

chr1-4530989-DEL-52
chr1-4550946-DEL-250
chr1-4579644-DEL-163
chr1-4583088-DEL-126
chr1-4593427-DEL-55
chr1-4599239-DEL-834
chr1-4627196-DEL-10670
chr1-4643979-DEL-276
chr1-4645632-DEL-752
chr1-4676618-DEL-1740
chr1-4746065-DEL-54
chr1-4767003-DEL-105
chr1-4825470-DEL-87
chr1-4835827-DEL-207
chr1-4841192-DEL-52
chr1-4853561-DEL-200
chr1-4860630-DEL-84
chr1-4880205-DEL-123
chr1-4895443-DEL-2031
chr1-4905653-DEL-226
chr1-4918114-DEL-169
chr1-4921441-DEL-443
chr1-4952470-DEL-145
chr1-4972340-DEL-83
chr1-4980033-DEL-92
chr1-4981478-DEL-832
chr1-5001536-DEL-72
chr1-5104947-DEL-6211
chr1-5114425-DEL-670
chr1-5116257-DEL-1339
chr1-5120038-DEL-444
chr1-5127720-DEL-1613
chr1-5129370-DEL-1107
chr1-5142441-DEL-499
chr1-5179570-DEL-4891
chr1-5179654-DEL-4148
chr1-5181960-DEL-2789
chr1-5184008-DEL-741
chr1-5200141-DEL-51
chr1-5208543-DEL-198
chr1-5213076-DEL-52
chr1-5232795-DEL-706
chr1-5291998-DEL-3171
chr1-5297892-DEL-94
chr1-5320219-DEL-6479
chr1-5331802-DEL-291
chr1-5336947-DEL-6335
chr1-5363768-DEL-551
chr1-5394225-DEL-463
chr1-5438213-DEL-494
chr1-5450831-DEL-282
chr1-5455006-DEL-90
chr1-5467827-DEL-2523
chr1-5475837-DEL-503
chr1-5479965-DEL-214
chr1-5486063-DEL-77
chr1-5493657-DEL-6110
chr1-5500042-DEL-6819
chr1-5513161-DEL-6972
chr1-5522476-DEL-127
chr1-5540178-DEL-168
chr1-5561459-DEL-761
chr1-5602108-DEL-196
chr1-5625278-DEL-402
chr1-5664594-DEL-144
chr1-5689368-DEL-302
chr1-5701026-DEL-95
chr1-5703117-DEL-135
chr1-5714637-DEL-399
chr1-5746972-DEL-5486
chr1-5758165-DEL-542
chr1-5759096-DEL-6666
chr1-5795103-DEL-54
chr1-5808997-DEL-4015
chr1-5816935-DEL-2478
chr1-5874567-DEL-7100
chr1-5890202-DEL-6614

chr1-5931435-DEL-112
chr1-5945304-DEL-814
chr1-5969153-DEL-92
chr1-5990345-DEL-1513
chr1-6003999-DEL-341
chr1-6008680-DEL-1667
chr1-6014790-DEL-160
chr1-6052402-DEL-3624
chr1-6060676-DEL-71
chr1-6130841-DEL-181
chr1-6133940-DEL-55
chr1-6195152-DEL-474
chr1-6198803-DEL-245
chr1-6200500-DEL-415
chr1-6221197-DEL-199
chr1-6257385-DEL-85
chr1-6305824-DEL-178
chr1-6361053-DEL-999
chr1-6385758-DEL-1152
chr1-6395915-DEL-1494
chr1-6417255-DEL-276
chr1-6431976-DEL-231
chr1-6460355-DEL-564
chr1-6475108-DEL-328
chr1-6475768-DEL-738
chr1-6491328-DEL-72
chr1-6514334-DEL-1358
chr1-6587866-DEL-422
chr1-6681220-DEL-276
chr1-6715149-DEL-141
chr1-6716110-DEL-989
chr1-6732981-DEL-1891
chr1-6756805-DEL-2088
chr1-6759410-DEL-5849
chr1-6785716-DEL-114
chr1-6819078-DEL-1065
chr1-6868264-DEL-64
chr1-6986709-DEL-7152
chr1-6987361-DEL-200
chr1-7016801-DEL-4189
chr1-7059075-DEL-51
chr1-7122846-DEL-399
chr1-7150225-DEL-60
chr1-7162783-DEL-89
chr1-7164421-DEL-954
chr1-7182084-DEL-86
chr1-7189050-DEL-223
chr1-7196479-DEL-461
chr1-7210428-DEL-92
chr1-7258120-DEL-157
chr1-7300836-DEL-11975
chr1-7341852-DEL-6217
chr1-7350605-DEL-8347
chr1-7380877-DEL-67
chr1-7392787-DEL-186
chr1-7393301-DEL-171
chr1-7418760-DEL-66
chr1-7451000-DEL-2452
chr1-7454381-DEL-57
chr1-7462440-DEL-58
chr1-7472137-DEL-83
chr1-7479241-DEL-52
chr1-7509871-DEL-2728
chr1-7518680-DEL-2607
chr1-7520694-DEL-441
chr1-7538281-DEL-71
chr1-7585436-DEL-2084
chr1-7632766-DEL-172
chr1-7633910-DEL-515
chr1-7708097-DEL-66
chr1-7720923-DEL-1176
chr1-7772239-DEL-11204
chr1-7802122-DEL-82
chr1-7806602-DEL-56
chr1-7816530-DEL-381
chr1-7858147-DEL-4592
chr1-7865192-DEL-1422

chr1-7880379-DEL-298
chr1-7881667-DEL-15200
chr1-7935250-DEL-1223
chr1-7937685-DEL-407
chr1-7952030-DEL-7377
chr1-7975881-DEL-260
chr1-7976182-DEL-2222
chr1-7992966-DEL-17122
chr1-7998230-DEL-524
chr1-7999065-DEL-1312
chr1-7999067-DEL-7263
chr1-8000449-DEL-2636
chr1-8003164-DEL-3187
chr1-8010255-DEL-206
chr1-8033974-DEL-414
chr1-8049238-DEL-1068
chr1-8060598-DEL-1291
chr1-8074237-DEL-329
chr1-8074654-DEL-993
chr1-8088011-DEL-313
chr1-8092415-DEL-81
chr1-8104052-DEL-644
chr1-8108543-DEL-2070
chr1-8110919-DEL-6540
chr1-8163203-DEL-123
chr1-8167707-DEL-201
chr1-8218539-DEL-115
chr1-8226423-DEL-7046
chr1-8238721-DEL-61
chr1-8253869-DEL-61
chr1-8260234-DEL-7119
chr1-8271009-DEL-126
chr1-8279704-DEL-12808
chr1-8301980-DEL-1159
chr1-8304854-DEL-6092
chr1-8308366-DEL-665
chr1-8315778-DEL-147
chr1-8335207-DEL-136
chr1-8351559-DEL-1092
chr1-8362788-DEL-76
chr1-8375445-DEL-781
chr1-8387933-DEL-154
chr1-8392827-DEL-65
chr1-8441760-DEL-51
chr1-8454730-DEL-352
chr1-8459787-DEL-331
chr1-8461307-DEL-273
chr1-8463682-DEL-9914
chr1-8475097-DEL-586
chr1-8486664-DEL-353
chr1-8494638-DEL-75
chr1-8499242-DEL-143
chr1-8532008-DEL-775
chr1-8537138-DEL-2614
chr1-8540963-DEL-96
chr1-8580004-DEL-5939
chr1-8588132-DEL-7289
chr1-8594226-DEL-2991
chr1-8595476-DEL-1720
chr1-8620237-DEL-84
chr1-8638301-DEL-2427
chr1-8641949-DEL-4352
chr1-8674395-DEL-103
chr1-8678886-DEL-1129
chr1-8705217-DEL-6520
chr1-8756640-DEL-53
chr1-8758481-DEL-6997
chr1-8758921-DEL-415
chr1-8769913-DEL-2458
chr1-8776551-DEL-1962
chr1-8779674-DEL-2875
chr1-8788857-DEL-266
chr1-8789145-DEL-2123
chr1-8792217-DEL-141
chr1-8800595-DEL-1887
chr1-8803123-DEL-195
chr1-8807171-DEL-252

chr1-8833656-DEL-75
chr1-8842732-DEL-5056
chr1-8855531-DEL-99
chr1-8856902-DEL-3663
chr1-8880331-DEL-1674
chr1-8882031-DEL-1864
chr1-8903431-DEL-204
chr1-8910660-DEL-81
chr1-8918465-DEL-66
chr1-8922696-DEL-328
chr1-8926869-DEL-585
chr1-8950920-DEL-447
chr1-8960513-DEL-3978
chr1-8979736-DEL-65
chr1-9011349-DEL-75
chr1-9019351-DEL-193
chr1-9073758-DEL-100
chr1-9088243-DEL-571
chr1-9109929-DEL-1610
chr1-9111836-DEL-2010
chr1-9128369-DEL-6406
chr1-9143175-DEL-222
chr1-9144484-DEL-6549
chr1-9165709-DEL-1637
chr1-9276925-DEL-82
chr1-9301764-DEL-2961
chr1-9305255-DEL-12129
chr1-9310726-DEL-6657
chr1-9336111-DEL-95
chr1-9398475-DEL-320
chr1-9406708-DEL-52
chr1-9431939-DEL-77
chr1-9435738-DEL-52
chr1-9548809-DEL-51
chr1-9597241-DEL-1432
chr1-9600710-DEL-251
chr1-9602613-DEL-154
chr1-9668149-DEL-399
chr1-9701214-DEL-87
chr1-9707214-DEL-330
chr1-9766960-DEL-51
chr1-9781087-DEL-200
chr1-9784934-DEL-1800
chr1-9802732-DEL-923
chr1-9804603-DEL-2096
chr1-9810995-DEL-190
chr1-9834254-DEL-204
chr1-9852018-DEL-131
chr1-9854595-DEL-360
chr1-9859118-DEL-54
chr1-9861745-DEL-484
chr1-9912344-DEL-198
chr1-9915876-DEL-195
chr1-9937065-DEL-54
chr1-9965704-DEL-71
chr1-9987764-DEL-58
chr1-9990990-DEL-201
chr1-10011297-DEL-183
chr1-10039521-DEL-171
chr1-10040066-DEL-806
chr1-10054574-DEL-674
chr1-10074912-DEL-54
chr1-10119908-DEL-78
chr1-10132080-DEL-237
chr1-10148637-DEL-180
chr1-10162280-DEL-280
chr1-10170988-DEL-837
chr1-10171080-DEL-158
chr1-10232786-DEL-296
chr1-10238100-DEL-202
chr1-10250480-DEL-191
chr1-10270742-DEL-74
chr1-10285336-DEL-70
chr1-10325081-DEL-79
chr1-10399615-DEL-56
chr1-10407791-DEL-1752
chr1-10452805-DEL-6533

chr1-10452999-DEL-201
chr1-10459665-DEL-117
chr1-10486175-DEL-54
chr1-10487891-DEL-370
chr1-10489167-DEL-1562
chr1-10492354-DEL-5802
chr1-10503357-DEL-135
chr1-10504019-DEL-5173
chr1-10526333-DEL-4665
chr1-10548465-DEL-174
chr1-10581756-DEL-3796
chr1-10586515-DEL-3929
chr1-10598034-DEL-64
chr1-10656909-DEL-80
chr1-10660638-DEL-324
chr1-10696961-DEL-175
chr1-10746800-DEL-81
chr1-10756803-DEL-3045
chr1-10763775-DEL-74
chr1-10785124-DEL-1131
chr1-10812456-DEL-950
chr1-10829211-DEL-556
chr1-10832094-DEL-56
chr1-10846356-DEL-1506
chr1-10854171-DEL-747
chr1-10887086-DEL-5520
chr1-10931110-DEL-92
chr1-10950072-DEL-144
chr1-10991234-DEL-379
chr1-11004579-DEL-1556
chr1-11090676-DEL-51
chr1-11139721-DEL-4651
chr1-11164295-DEL-53
chr1-11204490-DEL-624
chr1-11212061-DEL-103
chr1-11259566-DEL-1005
chr1-11262230-DEL-113
chr1-11264624-DEL-252
chr1-11297142-DEL-2768
chr1-11327956-DEL-6439
chr1-11369857-DEL-122
chr1-11449871-DEL-180
chr1-11467972-DEL-62
chr1-11490723-DEL-114
chr1-11529391-DEL-563
chr1-11542414-DEL-259
chr1-11549259-DEL-58
chr1-11550054-DEL-59
chr1-11555634-DEL-652
chr1-11556494-DEL-1245
chr1-11565235-DEL-3849
chr1-11669124-DEL-67
chr1-11675332-DEL-67
chr1-11689571-DEL-5546
chr1-11725857-DEL-108
chr1-11784616-DEL-6320
chr1-11792243-DEL-609
chr1-11799464-DEL-208
chr1-11875564-DEL-6279
chr1-11897658-DEL-685
chr1-11913250-DEL-975
chr1-11953462-DEL-71
chr1-11965175-DEL-66
chr1-11972708-DEL-737
chr1-11975722-DEL-84
chr1-11992772-DEL-180
chr1-12001199-DEL-74
chr1-12010302-DEL-182
chr1-12092950-DEL-106
chr1-12136248-DEL-109
chr1-12145626-DEL-6184
chr1-12189306-DEL-718
chr1-12198759-DEL-1176
chr1-12217141-DEL-102
chr1-12227578-DEL-1945
chr1-12246321-DEL-195
chr1-12246970-DEL-200

chr1-12247178-DEL-1156
chr1-12268816-DEL-86
chr1-12286106-DEL-1143
chr1-12314025-DEL-155
chr1-12317572-DEL-890
chr1-12318515-DEL-1305
chr1-12332644-DEL-66
chr1-12334166-DEL-6162
chr1-12383959-DEL-307
chr1-12414287-DEL-57
chr1-12430361-DEL-103
chr1-12434280-DEL-99
chr1-12521957-DEL-95
chr1-12533727-DEL-695
chr1-12546404-DEL-56
chr1-12556132-DEL-68
chr1-12604783-DEL-207
chr1-12605631-DEL-1585
chr1-12662665-DEL-335
chr1-12715515-DEL-2817
chr1-12727837-DEL-370
chr1-12787975-DEL-82
chr1-12791405-DEL-51
chr1-12813901-DEL-92
chr1-12826593-DEL-177
chr1-12829716-DEL-128
chr1-12831649-DEL-5597
chr1-12844524-DEL-202
chr1-12869135-DEL-402
chr1-12894732-DEL-400
chr1-12899477-DEL-464
chr1-12903171-DEL-66
chr1-12910853-DEL-187
chr1-12911422-DEL-157
chr1-12912327-DEL-94
chr1-12914635-DEL-670
chr1-12926844-DEL-71
chr1-12946401-DEL-83
chr1-12978563-DEL-63
chr1-13014976-DEL-1534
chr1-13017316-DEL-89
chr1-13022247-DEL-309
chr1-13028115-DEL-120
chr1-13077554-DEL-188
chr1-13085537-DEL-333
chr1-13108173-DEL-212
chr1-13116785-DEL-132
chr1-13121750-DEL-151
chr1-13130892-DEL-144
chr1-13140304-DEL-53
chr1-13164064-DEL-72
chr1-13169469-DEL-350
chr1-13187108-DEL-448
chr1-13215392-DEL-102
chr1-13225891-DEL-63
chr1-13228356-DEL-397
chr1-13245968-DEL-56
chr1-13260703-DEL-415
chr1-13293077-DEL-75
chr1-13305257-DEL-51
chr1-13325234-DEL-62
chr1-13328310-DEL-160
chr1-13357843-DEL-151
chr1-13388336-DEL-254
chr1-13388960-DEL-144
chr1-13397960-DEL-55
chr1-13407821-DEL-427
chr1-13407838-DEL-1408
chr1-13408422-DEL-61
chr1-13408925-DEL-121
chr1-13447495-DEL-1886
chr1-13466346-DEL-61
chr1-13478375-DEL-52
chr1-13483228-DEL-1054
chr1-13511866-DEL-111
chr1-13518549-DEL-63
chr1-13519078-DEL-219

chr1-13542729-DEL-6344
chr1-13552370-DEL-177
chr1-13558852-DEL-138
chr1-13563477-DEL-114
chr1-13569295-DEL-171
chr1-13589539-DEL-1416
chr1-13606777-DEL-196
chr1-13621076-DEL-6880
chr1-13645732-DEL-93
chr1-13684089-DEL-570
chr1-13697783-DEL-88
chr1-13704008-DEL-51
chr1-13740264-DEL-177
chr1-13741632-DEL-77
chr1-13760119-DEL-72
chr1-13772415-DEL-403
chr1-13791180-DEL-900
chr1-13800574-DEL-528
chr1-13809037-DEL-100
chr1-13817806-DEL-182
chr1-13824129-DEL-193
chr1-13872759-DEL-307
chr1-13916738-DEL-52
chr1-13948423-DEL-62
chr1-13959412-DEL-180
chr1-13961231-DEL-6446
chr1-13971622-DEL-190
chr1-13977178-DEL-2390
chr1-13983162-DEL-1166
chr1-14008178-DEL-64
chr1-14065021-DEL-553
chr1-14075680-DEL-117
chr1-14082692-DEL-65
chr1-14093762-DEL-402
chr1-14104396-DEL-85
chr1-14106809-DEL-70
chr1-14121783-DEL-224
chr1-14127593-DEL-912
chr1-14153034-DEL-1492
chr1-14193940-DEL-51
chr1-14219655-DEL-93
chr1-14221712-DEL-108
chr1-14246279-DEL-869
chr1-14267206-DEL-576
chr1-14309422-DEL-396
chr1-14386444-DEL-164
chr1-14400297-DEL-116
chr1-14402398-DEL-2850
chr1-14428629-DEL-95
chr1-14433322-DEL-5159
chr1-14434604-DEL-217
chr1-14435111-DEL-184
chr1-14502806-DEL-62
chr1-14509545-DEL-92
chr1-14554885-DEL-97
chr1-14566916-DEL-84
chr1-14663791-DEL-161
chr1-14672291-DEL-74
chr1-14683696-DEL-168
chr1-14686220-DEL-65
chr1-14759353-DEL-514
chr1-14871994-DEL-362
chr1-14875883-DEL-208
chr1-14889782-DEL-175
chr1-14934971-DEL-906
chr1-14968786-DEL-66
chr1-14981892-DEL-96
chr1-14986861-DEL-83
chr1-15004840-DEL-63
chr1-15009749-DEL-159
chr1-15019904-DEL-122
chr1-15039526-DEL-6938
chr1-15045808-DEL-204
chr1-15053835-DEL-2085
chr1-15058774-DEL-6407
chr1-15106216-DEL-623
chr1-15134847-DEL-119

chr1-15139175-DEL-337
chr1-15154263-DEL-214
chr1-15197687-DEL-218
chr1-15206088-DEL-155
chr1-15211958-DEL-78
chr1-15243876-DEL-137
chr1-15297223-DEL-8789
chr1-15298657-DEL-5994
chr1-15298779-DEL-2150
chr1-15300254-DEL-717
chr1-15306090-DEL-1861
chr1-15338818-DEL-1277
chr1-15340612-DEL-2867
chr1-15354354-DEL-175
chr1-15371446-DEL-161
chr1-15372953-DEL-68
chr1-15389874-DEL-831
chr1-15469823-DEL-164
chr1-15485530-DEL-352
chr1-15491827-DEL-738
chr1-15492749-DEL-198
chr1-15571272-DEL-249
chr1-15578744-DEL-116
chr1-15580841-DEL-614
chr1-15610741-DEL-1393
chr1-15644755-DEL-211
chr1-15657943-DEL-55
chr1-15659980-DEL-186
chr1-15660983-DEL-71
chr1-15694004-DEL-53
chr1-15788128-DEL-52
chr1-15796394-DEL-82
chr1-15803213-DEL-245
chr1-15806561-DEL-8032
chr1-15814900-DEL-4041
chr1-15827886-DEL-3085
chr1-15831446-DEL-6247
chr1-15864681-DEL-370
chr1-15871867-DEL-210
chr1-15884739-DEL-598
chr1-15946347-DEL-400
chr1-15960672-DEL-58
chr1-15977928-DEL-52
chr1-16005400-DEL-64
chr1-16008341-DEL-171
chr1-16030196-DEL-212
chr1-16033848-DEL-85
chr1-16037934-DEL-100
chr1-16066451-DEL-72
chr1-16086311-DEL-68
chr1-16091751-DEL-492
chr1-16092541-DEL-60
chr1-16095388-DEL-199
chr1-16119709-DEL-300
chr1-16125211-DEL-83
chr1-16192470-DEL-214
chr1-16214956-DEL-59
chr1-16233898-DEL-6341
chr1-16245202-DEL-674
chr1-16329789-DEL-93
chr1-16353074-DEL-119
chr1-16354556-DEL-1540
chr1-16397891-DEL-548
chr1-16410775-DEL-776
chr1-16412928-DEL-162
chr1-16487466-DEL-645
chr1-16497750-DEL-640
chr1-16500008-DEL-2327
chr1-16506293-DEL-1142
chr1-16514362-DEL-66
chr1-16532970-DEL-60
chr1-16546236-DEL-240
chr1-16572249-DEL-719
chr1-16589782-DEL-68
chr1-16594876-DEL-400
chr1-16596638-DEL-577
chr1-16637083-DEL-92

chr1-16662923-DEL-663
chr1-16663602-DEL-153
chr1-16668752-DEL-231
chr1-16670572-DEL-340
chr1-16734894-DEL-111
chr1-16765949-DEL-57
chr1-16775416-DEL-702
chr1-16780452-DEL-3880
chr1-16780453-DEL-627
chr1-16797352-DEL-53
chr1-16827649-DEL-573
chr1-16855092-DEL-668
chr1-16856068-DEL-1722
chr1-16874094-DEL-507
chr1-16916051-DEL-74
chr1-16952953-DEL-158
chr1-16953411-DEL-1411
chr1-16955570-DEL-1961
chr1-16971072-DEL-60
chr1-17007010-DEL-5511
chr1-17012613-DEL-68
chr1-17015771-DEL-226
chr1-17023815-DEL-206
chr1-17030061-DEL-645
chr1-17035469-DEL-140
chr1-17038408-DEL-231
chr1-17064764-DEL-194
chr1-17077816-DEL-67
chr1-17093340-DEL-299
chr1-17094805-DEL-153
chr1-17096356-DEL-3438
chr1-17097773-DEL-182
chr1-17114992-DEL-1198
chr1-17131441-DEL-924
chr1-17175429-DEL-65
chr1-17188713-DEL-57
chr1-17205479-DEL-389
chr1-17207695-DEL-964
chr1-17222602-DEL-368
chr1-17241530-DEL-138
chr1-17244467-DEL-53
chr1-17247043-DEL-661
chr1-17262599-DEL-728
chr1-17268267-DEL-216
chr1-17292619-DEL-5549
chr1-17300804-DEL-68
chr1-17335393-DEL-778
chr1-17368325-DEL-84
chr1-17411949-DEL-19788
chr1-17437947-DEL-82
chr1-17452959-DEL-468
chr1-17459528-DEL-212
chr1-17460503-DEL-213
chr1-17466361-DEL-91
chr1-17530794-DEL-428
chr1-17535831-DEL-14979
chr1-17537064-DEL-13100
chr1-17537476-DEL-212
chr1-17572117-DEL-65
chr1-17579594-DEL-3030
chr1-17584457-DEL-75
chr1-17594755-DEL-2431
chr1-17609723-DEL-537
chr1-17646812-DEL-2148
chr1-17711535-DEL-1460
chr1-17713009-DEL-59
chr1-17843730-DEL-9079
chr1-17865545-DEL-125
chr1-17865825-DEL-175
chr1-17871647-DEL-401
chr1-17883216-DEL-63
chr1-17888375-DEL-60
chr1-17965724-DEL-582
chr1-17967840-DEL-6492
chr1-17975993-DEL-2437
chr1-17981681-DEL-53
chr1-17992571-DEL-212

chr1-18000243-DEL-1183
chr1-18012946-DEL-1343
chr1-18023067-DEL-13812
chr1-18038327-DEL-265
chr1-18049118-DEL-132
chr1-18086623-DEL-301
chr1-18096401-DEL-331
chr1-18096784-DEL-2877
chr1-18117866-DEL-376
chr1-18131213-DEL-53
chr1-18134388-DEL-1017
chr1-18140538-DEL-342
chr1-18148780-DEL-428
chr1-18148994-DEL-214
chr1-18184869-DEL-187
chr1-18188655-DEL-208
chr1-18200798-DEL-66
chr1-18257496-DEL-76
chr1-18263173-DEL-519
chr1-18316210-DEL-62
chr1-18333031-DEL-178
chr1-18350875-DEL-72
chr1-18362546-DEL-58
chr1-18386425-DEL-1164
chr1-18400142-DEL-89
chr1-18423623-DEL-2511
chr1-18425208-DEL-110
chr1-18446349-DEL-79
chr1-18484680-DEL-61
chr1-18509346-DEL-67
chr1-18512427-DEL-4448
chr1-18549491-DEL-806
chr1-18570264-DEL-329
chr1-18593268-DEL-711
chr1-18604853-DEL-118
chr1-18678855-DEL-704
chr1-18688579-DEL-210
chr1-18698580-DEL-62
chr1-18741095-DEL-2060
chr1-18752107-DEL-205
chr1-18764970-DEL-78
chr1-18766143-DEL-826
chr1-18769661-DEL-1263
chr1-18795424-DEL-685
chr1-18805926-DEL-310
chr1-18808168-DEL-2292
chr1-18820657-DEL-51
chr1-18822207-DEL-171
chr1-18853448-DEL-66
chr1-18855299-DEL-56
chr1-18870512-DEL-1445
chr1-18881806-DEL-413
chr1-18907242-DEL-367
chr1-18915402-DEL-72
chr1-18930390-DEL-2406
chr1-18942646-DEL-6240
chr1-18961799-DEL-180
chr1-18962710-DEL-704
chr1-18964602-DEL-78
chr1-18979245-DEL-356
chr1-18996212-DEL-147
chr1-18997801-DEL-6239
chr1-19004695-DEL-214
chr1-19008106-DEL-840
chr1-19019756-DEL-485
chr1-19097853-DEL-77
chr1-19099568-DEL-240
chr1-19109006-DEL-363
chr1-19120141-DEL-3004
chr1-19134490-DEL-494
chr1-19137041-DEL-56
chr1-19163308-DEL-79
chr1-19217780-DEL-256
chr1-19223023-DEL-1289
chr1-19241726-DEL-233
chr1-19248708-DEL-105
chr1-19259169-DEL-1562

chr1-19283873-DEL-110
chr1-19303785-DEL-52
chr1-19331038-DEL-69
chr1-19336573-DEL-76
chr1-19385331-DEL-1449
chr1-19387037-DEL-72
chr1-19439191-DEL-207
chr1-19471323-DEL-96
chr1-19473418-DEL-2608
chr1-19495652-DEL-63
chr1-19497244-DEL-79
chr1-19501595-DEL-166
chr1-19504936-DEL-6194
chr1-19514024-DEL-6461
chr1-19593132-DEL-281
chr1-19621607-DEL-238
chr1-19622929-DEL-753
chr1-19689605-DEL-60
chr1-19693047-DEL-79
chr1-19721528-DEL-8721
chr1-19722635-DEL-2224
chr1-19756290-DEL-7628
chr1-19817742-DEL-740
chr1-19823141-DEL-3356
chr1-19858283-DEL-2049
chr1-19904102-DEL-310
chr1-19934845-DEL-80
chr1-19936067-DEL-58
chr1-19954954-DEL-1068
chr1-19956278-DEL-767
chr1-19970198-DEL-107
chr1-19991560-DEL-54
chr1-19997042-DEL-68
chr1-20001074-DEL-156
chr1-20019005-DEL-136
chr1-20027733-DEL-856
chr1-20028400-DEL-194
chr1-20055269-DEL-245
chr1-20069454-DEL-407
chr1-20112433-DEL-66
chr1-20131514-DEL-4459
chr1-20153353-DEL-52
chr1-20161354-DEL-172
chr1-20161589-DEL-1412
chr1-20163770-DEL-1711
chr1-20166505-DEL-6218
chr1-20182033-DEL-56
chr1-20216359-DEL-660
chr1-20272680-DEL-201
chr1-20287373-DEL-1209
chr1-20364995-DEL-452
chr1-20383639-DEL-471
chr1-20406627-DEL-96
chr1-20440342-DEL-54
chr1-20483454-DEL-60
chr1-20497277-DEL-207
chr1-20502846-DEL-212
chr1-20539788-DEL-395
chr1-20542097-DEL-182
chr1-20570653-DEL-140
chr1-20594894-DEL-7070
chr1-20622804-DEL-404
chr1-20634929-DEL-52
chr1-20662675-DEL-526
chr1-20674579-DEL-63
chr1-20679410-DEL-68
chr1-20694434-DEL-96
chr1-20748642-DEL-192
chr1-20758224-DEL-53
chr1-20771534-DEL-202
chr1-20772617-DEL-1490
chr1-20789112-DEL-137
chr1-20794047-DEL-58
chr1-20805876-DEL-287
chr1-20831058-DEL-111
chr1-20832730-DEL-1730
chr1-20851001-DEL-186

chr1-20861059-DEL-131
chr1-20871269-DEL-93
chr1-20895207-DEL-849
chr1-20917941-DEL-195
chr1-20938497-DEL-88
chr1-21028002-DEL-2188
chr1-21072413-DEL-120
chr1-21079874-DEL-68
chr1-21094997-DEL-204
chr1-21114894-DEL-132
chr1-21211756-DEL-344
chr1-21221170-DEL-200
chr1-21229335-DEL-73
chr1-21244572-DEL-80
chr1-21280427-DEL-144
chr1-21287301-DEL-72
chr1-21292491-DEL-80
chr1-21301300-DEL-117
chr1-21339533-DEL-151
chr1-21388374-DEL-52
chr1-21398458-DEL-62
chr1-21404653-DEL-171
chr1-21434734-DEL-14756
chr1-21450754-DEL-472
chr1-21465117-DEL-54
chr1-21467002-DEL-226
chr1-21478114-DEL-54
chr1-21479821-DEL-126
chr1-21482502-DEL-64
chr1-21491688-DEL-122
chr1-21565115-DEL-63
chr1-21573886-DEL-52
chr1-21582070-DEL-1294
chr1-21606615-DEL-108
chr1-21669248-DEL-540
chr1-21670727-DEL-79
chr1-21696103-DEL-1381
chr1-21696321-DEL-2606
chr1-21732056-DEL-117
chr1-21733202-DEL-69
chr1-21764884-DEL-501
chr1-21786241-DEL-185
chr1-21787838-DEL-201
chr1-21793087-DEL-529
chr1-21814019-DEL-224
chr1-21904320-DEL-282
chr1-21911027-DEL-69
chr1-21929269-DEL-56
chr1-21937756-DEL-70
chr1-21954414-DEL-74
chr1-21960210-DEL-15272
chr1-21974740-DEL-275
chr1-21988563-DEL-80
chr1-22028861-DEL-79
chr1-22038986-DEL-196
chr1-22043445-DEL-402
chr1-22080426-DEL-193
chr1-22083273-DEL-2685
chr1-22120054-DEL-682
chr1-22127771-DEL-844
chr1-22158356-DEL-538
chr1-22159208-DEL-458
chr1-22196415-DEL-407
chr1-22208675-DEL-923
chr1-22218201-DEL-1511
chr1-22261250-DEL-635
chr1-22270134-DEL-77
chr1-22292086-DEL-369
chr1-22301074-DEL-340
chr1-22301077-DEL-880
chr1-22303545-DEL-1745
chr1-22305388-DEL-1866
chr1-22328226-DEL-55
chr1-22369109-DEL-184
chr1-22369547-DEL-81
chr1-22397811-DEL-208
chr1-22428368-DEL-4572

chr1-22430975-DEL-56
chr1-22443068-DEL-171
chr1-22451416-DEL-93
chr1-22521397-DEL-113
chr1-22524822-DEL-54
chr1-22532417-DEL-125
chr1-22534495-DEL-73
chr1-22543820-DEL-533
chr1-22579690-DEL-191
chr1-22599140-DEL-316
chr1-22605369-DEL-258
chr1-22606413-DEL-2656
chr1-22618855-DEL-134
chr1-22640614-DEL-69
chr1-22643064-DEL-481
chr1-22648174-DEL-160
chr1-22686812-DEL-101
chr1-22706235-DEL-710
chr1-22707141-DEL-434
chr1-22722194-DEL-193
chr1-22746326-DEL-371
chr1-22760995-DEL-89
chr1-22781642-DEL-1379
chr1-22854647-DEL-592
chr1-22863706-DEL-52
chr1-22920195-DEL-1365
chr1-22933443-DEL-58
chr1-22936190-DEL-72
chr1-22936850-DEL-60
chr1-22962741-DEL-84
chr1-22970304-DEL-118
chr1-22980579-DEL-66
chr1-22985335-DEL-1902
chr1-23010788-DEL-51
chr1-23018533-DEL-118
chr1-23043625-DEL-78
chr1-23058717-DEL-399
chr1-23090329-DEL-60
chr1-23099929-DEL-131
chr1-23142938-DEL-304
chr1-23163516-DEL-282
chr1-23186678-DEL-75
chr1-23253450-DEL-85
chr1-23279012-DEL-719
chr1-23283421-DEL-259
chr1-23347919-DEL-53
chr1-23363658-DEL-58
chr1-23432724-DEL-2212
chr1-23438899-DEL-72
chr1-23442358-DEL-2246
chr1-23453578-DEL-266
chr1-23456106-DEL-152
chr1-23458774-DEL-675
chr1-23487332-DEL-125
chr1-23496783-DEL-388
chr1-23498405-DEL-174
chr1-23511060-DEL-123
chr1-23522466-DEL-154
chr1-23530756-DEL-56
chr1-23535116-DEL-201
chr1-23551768-DEL-100
chr1-23564283-DEL-98
chr1-23567143-DEL-2439
chr1-23569880-DEL-192
chr1-23572554-DEL-184
chr1-23595243-DEL-636
chr1-23612218-DEL-59
chr1-23614652-DEL-70
chr1-23620296-DEL-921
chr1-23633693-DEL-111
chr1-23653421-DEL-6244
chr1-23702201-DEL-6240
chr1-23708880-DEL-329
chr1-23721016-DEL-2261
chr1-23721590-DEL-110
chr1-23746341-DEL-84
chr1-23749268-DEL-178

chr1-23760848-DEL-310
chr1-23763307-DEL-6163
chr1-23773381-DEL-59
chr1-23789965-DEL-52
chr1-23791086-DEL-60
chr1-23858944-DEL-352
chr1-23865614-DEL-90
chr1-23922587-DEL-91
chr1-23973264-DEL-171
chr1-23990380-DEL-716
chr1-23994657-DEL-60
chr1-24021019-DEL-3208
chr1-24084310-DEL-78
chr1-24100251-DEL-60
chr1-24108711-DEL-73
chr1-24127463-DEL-999
chr1-24127717-DEL-54
chr1-24135911-DEL-129
chr1-24143534-DEL-102
chr1-24178909-DEL-67
chr1-24220255-DEL-199
chr1-24226638-DEL-225
chr1-24246707-DEL-52
chr1-24247602-DEL-100
chr1-24255614-DEL-154
chr1-24279739-DEL-759
chr1-24295932-DEL-51
chr1-24307512-DEL-169
chr1-24323380-DEL-187
chr1-24347807-DEL-52
chr1-24366049-DEL-60
chr1-24379086-DEL-5058
chr1-24384148-DEL-147
chr1-24387530-DEL-75
chr1-24398894-DEL-74
chr1-24402800-DEL-725
chr1-24405072-DEL-60
chr1-24411995-DEL-52
chr1-24416248-DEL-52
chr1-24443315-DEL-55
chr1-24446552-DEL-153
chr1-24458258-DEL-2071
chr1-24496748-DEL-67
chr1-24515046-DEL-140
chr1-24527097-DEL-56
chr1-24597749-DEL-79
chr1-24602724-DEL-72
chr1-24636014-DEL-52
chr1-24658947-DEL-195
chr1-24692350-DEL-72
chr1-24728348-DEL-61
chr1-24729482-DEL-63
chr1-24738590-DEL-69
chr1-24739322-DEL-98
chr1-24793084-DEL-3960
chr1-24842762-DEL-4188
chr1-24851359-DEL-286
chr1-24869383-DEL-314
chr1-24920128-DEL-2154
chr1-24923531-DEL-2282
chr1-24961747-DEL-71
chr1-24972020-DEL-79
chr1-24975693-DEL-264
chr1-25008269-DEL-107
chr1-25016234-DEL-478
chr1-25040844-DEL-215
chr1-25041264-DEL-528
chr1-25045067-DEL-69
chr1-25049944-DEL-168
chr1-25053904-DEL-4396
chr1-25059082-DEL-211
chr1-25063535-DEL-63
chr1-25079395-DEL-121
chr1-25091382-DEL-127
chr1-25100898-DEL-427
chr1-25112572-DEL-87
chr1-25119976-DEL-1054

chr1-25159088-DEL-242
chr1-25186872-DEL-96
chr1-25226568-DEL-121
chr1-25234808-DEL-112
chr1-25377111-DEL-91
chr1-25380229-DEL-57
chr1-25394999-DEL-661
chr1-25404807-DEL-74
chr1-25415238-DEL-192
chr1-25423541-DEL-5108
chr1-25442296-DEL-6229
chr1-25476617-DEL-71
chr1-25480414-DEL-84
chr1-25508929-DEL-184
chr1-25570373-DEL-53
chr1-25618564-DEL-699
chr1-25642093-DEL-3292
chr1-25651912-DEL-176
chr1-25657836-DEL-80
chr1-25701148-DEL-432
chr1-25726026-DEL-693
chr1-25767553-DEL-54
chr1-25778677-DEL-1048
chr1-25784193-DEL-124
chr1-25837661-DEL-1045
chr1-25901640-DEL-403
chr1-25906125-DEL-118
chr1-25948822-DEL-9620
chr1-25967427-DEL-2550
chr1-25972222-DEL-185
chr1-25975141-DEL-57
chr1-26022854-DEL-93
chr1-26033667-DEL-65
chr1-26034301-DEL-100
chr1-26037154-DEL-52
chr1-26090446-DEL-55
chr1-26090470-DEL-1401
chr1-26092066-DEL-1666
chr1-26139403-DEL-59
chr1-26140240-DEL-1484
chr1-26141972-DEL-162
chr1-26154451-DEL-188
chr1-26176946-DEL-200
chr1-26182441-DEL-1623
chr1-26191740-DEL-60
chr1-26205924-DEL-1409
chr1-26211475-DEL-305
chr1-26219393-DEL-111
chr1-26229266-DEL-111
chr1-26238435-DEL-57
chr1-26245703-DEL-60
chr1-26302082-DEL-423
chr1-26313694-DEL-256
chr1-26327915-DEL-409
chr1-26333263-DEL-497
chr1-26372207-DEL-63
chr1-26397586-DEL-169
chr1-26408020-DEL-590
chr1-26410141-DEL-819
chr1-26412298-DEL-1245
chr1-26414654-DEL-436
chr1-26446531-DEL-76
chr1-26449714-DEL-132
chr1-26452475-DEL-183
chr1-26478334-DEL-66
chr1-26525732-DEL-310
chr1-26544142-DEL-216
chr1-26564606-DEL-377
chr1-26608528-DEL-509
chr1-26621014-DEL-52
chr1-26635033-DEL-52
chr1-26639449-DEL-58
chr1-26654607-DEL-333
chr1-26681204-DEL-60
chr1-26728174-DEL-568
chr1-26737874-DEL-57
chr1-26743744-DEL-67

chr1-26770772-DEL-178
chr1-26797180-DEL-56
chr1-26802101-DEL-963
chr1-26826803-DEL-52
chr1-26853859-DEL-52
chr1-26890515-DEL-564
chr1-26897815-DEL-399
chr1-26904969-DEL-72
chr1-26938328-DEL-119
chr1-26945799-DEL-54
chr1-26972180-DEL-178
chr1-26988090-DEL-431
chr1-26995143-DEL-77
chr1-27002998-DEL-68
chr1-27049149-DEL-3807
chr1-27062506-DEL-1822
chr1-27088727-DEL-2316
chr1-27093845-DEL-2745
chr1-27097645-DEL-70
chr1-27104224-DEL-1227
chr1-27129268-DEL-1122
chr1-27139868-DEL-55
chr1-27209888-DEL-1628
chr1-27210722-DEL-130
chr1-27212914-DEL-971
chr1-27217942-DEL-66
chr1-27218344-DEL-535
chr1-27228567-DEL-683
chr1-27229548-DEL-728
chr1-27230188-DEL-135
chr1-27232521-DEL-84
chr1-27233079-DEL-7012
chr1-27233454-DEL-208
chr1-27236980-DEL-951
chr1-27239192-DEL-67
chr1-27240641-DEL-6363
chr1-27259868-DEL-574
chr1-27299964-DEL-103
chr1-27369575-DEL-993
chr1-27436250-DEL-463
chr1-27467177-DEL-98
chr1-27482035-DEL-386
chr1-27526204-DEL-584
chr1-27542510-DEL-458
chr1-27553724-DEL-4488
chr1-27559878-DEL-87
chr1-27565240-DEL-52
chr1-27596084-DEL-313
chr1-27612365-DEL-334
chr1-27635963-DEL-76
chr1-27639700-DEL-250
chr1-27652939-DEL-521
chr1-27670191-DEL-55
chr1-27684995-DEL-844
chr1-27685953-DEL-197
chr1-27687662-DEL-1605
chr1-27688265-DEL-1263
chr1-27703292-DEL-4438
chr1-27725332-DEL-55
chr1-27748425-DEL-108
chr1-27754030-DEL-317
chr1-27767167-DEL-14840
chr1-27788341-DEL-236
chr1-27824103-DEL-2723
chr1-27847893-DEL-70
chr1-27856378-DEL-63
chr1-27884830-DEL-2671
chr1-27892139-DEL-52
chr1-27900241-DEL-58
chr1-27908928-DEL-1485
chr1-27952049-DEL-52
chr1-27972584-DEL-105
chr1-27981407-DEL-390
chr1-27986848-DEL-122
chr1-27997665-DEL-427
chr1-28000608-DEL-400
chr1-28014522-DEL-1257

chr1-28025398-DEL-58
chr1-28027686-DEL-1293
chr1-28032337-DEL-1085
chr1-28126317-DEL-1145
chr1-28135146-DEL-665
chr1-28151305-DEL-7629
chr1-28169852-DEL-739
chr1-28174961-DEL-142
chr1-28185775-DEL-54
chr1-28208738-DEL-242
chr1-28215142-DEL-62
chr1-28231485-DEL-212
chr1-28240016-DEL-66
chr1-28249835-DEL-198
chr1-28282544-DEL-63
chr1-28292727-DEL-68
chr1-28302845-DEL-62
chr1-28305040-DEL-78
chr1-28314889-DEL-295
chr1-28323164-DEL-1228
chr1-28326757-DEL-424
chr1-28339493-DEL-64
chr1-28341847-DEL-6481
chr1-28400287-DEL-1484
chr1-28406968-DEL-51
chr1-28423970-DEL-329
chr1-28437988-DEL-6258
chr1-28449834-DEL-620
chr1-28463313-DEL-827
chr1-28470724-DEL-145
chr1-28477037-DEL-611
chr1-28497840-DEL-82
chr1-28506929-DEL-1857
chr1-28507242-DEL-83
chr1-28514090-DEL-108
chr1-28525298-DEL-62
chr1-28525388-DEL-1206
chr1-28533217-DEL-2593
chr1-28558882-DEL-205
chr1-28600356-DEL-56
chr1-28618252-DEL-190
chr1-28630562-DEL-1684
chr1-28639874-DEL-1601
chr1-28664836-DEL-2427
chr1-28699475-DEL-66
chr1-28700017-DEL-53
chr1-28714853-DEL-1391
chr1-28730225-DEL-215
chr1-28732287-DEL-1211
chr1-28759980-DEL-113
chr1-28776316-DEL-151
chr1-28784739-DEL-216
chr1-28807009-DEL-2982
chr1-28812469-DEL-203
chr1-28834498-DEL-347
chr1-28840541-DEL-1218
chr1-28865659-DEL-262
chr1-28867292-DEL-52
chr1-28879209-DEL-347
chr1-28894412-DEL-1053
chr1-28904664-DEL-94
chr1-28936024-DEL-215
chr1-28946102-DEL-11596
chr1-29013396-DEL-225
chr1-29014445-DEL-89
chr1-29048301-DEL-969
chr1-29062465-DEL-359
chr1-29080833-DEL-1216
chr1-29091408-DEL-55
chr1-29092881-DEL-65
chr1-29103778-DEL-54
chr1-29104600-DEL-1284
chr1-29110183-DEL-74
chr1-29121767-DEL-387
chr1-29136620-DEL-85
chr1-29143749-DEL-714
chr1-29177437-DEL-4550

chr1-29179208-DEL-275
chr1-29180323-DEL-86
chr1-29190644-DEL-1135
chr1-29202390-DEL-126
chr1-29203691-DEL-893
chr1-29214025-DEL-717
chr1-29230718-DEL-384
chr1-29259700-DEL-85
chr1-29280458-DEL-69
chr1-29283089-DEL-107
chr1-29297805-DEL-967
chr1-29302369-DEL-1872
chr1-29304012-DEL-563
chr1-29344584-DEL-1661
chr1-29351305-DEL-59
chr1-29371925-DEL-6153
chr1-29400730-DEL-64
chr1-29405268-DEL-259
chr1-29513439-DEL-138
chr1-29514829-DEL-792
chr1-29523596-DEL-1451
chr1-29559358-DEL-62
chr1-29565254-DEL-1944
chr1-29592229-DEL-72
chr1-29667573-DEL-58
chr1-29673413-DEL-154
chr1-29678276-DEL-77
chr1-29693771-DEL-4370
chr1-29701944-DEL-586
chr1-29718202-DEL-488
chr1-29752838-DEL-59
chr1-29774208-DEL-54
chr1-29783116-DEL-2429
chr1-29814386-DEL-207
chr1-29825769-DEL-129
chr1-29827706-DEL-431
chr1-29830312-DEL-363
chr1-29835525-DEL-6440
chr1-29843079-DEL-70
chr1-29846599-DEL-1124
chr1-29872787-DEL-6164
chr1-29880720-DEL-327
chr1-29904012-DEL-54
chr1-29908960-DEL-1052
chr1-29911458-DEL-197
chr1-29916038-DEL-68
chr1-29954774-DEL-195
chr1-29955622-DEL-505
chr1-29958878-DEL-237
chr1-29960972-DEL-399
chr1-29973742-DEL-1455
chr1-29980305-DEL-9253
chr1-29981812-DEL-2372
chr1-30007529-DEL-659
chr1-30030982-DEL-2305
chr1-30033868-DEL-2215
chr1-30046046-DEL-200
chr1-30055630-DEL-638
chr1-30082455-DEL-523
chr1-30087828-DEL-58
chr1-30098616-DEL-197
chr1-30102637-DEL-75
chr1-30105861-DEL-4979
chr1-30111399-DEL-1141
chr1-30138001-DEL-64
chr1-30157998-DEL-443
chr1-30159455-DEL-52
chr1-30166649-DEL-311
chr1-30170653-DEL-84
chr1-30187191-DEL-102
chr1-30198504-DEL-191
chr1-30200159-DEL-2839
chr1-30203076-DEL-694
chr1-30203815-DEL-1057
chr1-30213092-DEL-82
chr1-30218904-DEL-57
chr1-30225908-DEL-96

chr1-30232722-DEL-3517
chr1-30246606-DEL-98
chr1-30249128-DEL-2490
chr1-30258642-DEL-9295
chr1-30260737-DEL-222
chr1-30274452-DEL-726
chr1-30276449-DEL-72
chr1-30280754-DEL-100
chr1-30297646-DEL-56
chr1-30305117-DEL-1099
chr1-30313575-DEL-339
chr1-30330494-DEL-6189
chr1-30337114-DEL-4323
chr1-30353363-DEL-70
chr1-30397796-DEL-100
chr1-30409853-DEL-7050
chr1-30410277-DEL-418
chr1-30464903-DEL-117
chr1-30468795-DEL-128
chr1-30479934-DEL-670
chr1-30490682-DEL-149
chr1-30512885-DEL-53
chr1-30515010-DEL-935
chr1-30525769-DEL-75
chr1-30547577-DEL-211
chr1-30573387-DEL-52
chr1-30613207-DEL-2836
chr1-30649818-DEL-2691
chr1-30652683-DEL-3806
chr1-30666124-DEL-1424
chr1-30682580-DEL-2555
chr1-30683213-DEL-1412
chr1-30706911-DEL-422
chr1-30713919-DEL-1890
chr1-30725840-DEL-87
chr1-30726008-DEL-9241
chr1-30749260-DEL-97
chr1-30761613-DEL-6317
chr1-30769410-DEL-6530
chr1-30776885-DEL-96
chr1-30790747-DEL-62
chr1-30799597-DEL-6391
chr1-30836169-DEL-2581
chr1-30836320-DEL-444
chr1-30850404-DEL-66
chr1-30921096-DEL-2567
chr1-30963969-DEL-570
chr1-30970209-DEL-80
chr1-30977775-DEL-276
chr1-30978385-DEL-360
chr1-30998964-DEL-8241
chr1-31003772-DEL-58
chr1-31009801-DEL-1166
chr1-31044172-DEL-2612
chr1-31077663-DEL-1023
chr1-31094167-DEL-132
chr1-31107974-DEL-970
chr1-31113240-DEL-52
chr1-31114201-DEL-212
chr1-31140269-DEL-5729
chr1-31167161-DEL-373
chr1-31168042-DEL-724
chr1-31190014-DEL-106
chr1-31241551-DEL-71
chr1-31254549-DEL-62
chr1-31281585-DEL-480
chr1-31285294-DEL-1539
chr1-31289299-DEL-788
chr1-31300288-DEL-205
chr1-31304332-DEL-22347
chr1-31304998-DEL-210
chr1-31421007-DEL-61
chr1-31422992-DEL-6196
chr1-31434434-DEL-326
chr1-31435535-DEL-79
chr1-31438177-DEL-117
chr1-31493562-DEL-6173

chr1-31502723-DEL-99
chr1-31509115-DEL-3819
chr1-31537980-DEL-94
chr1-31542130-DEL-1023
chr1-31565085-DEL-309
chr1-31610474-DEL-1431
chr1-31618207-DEL-65
chr1-31637448-DEL-155
chr1-31651850-DEL-213
chr1-31652444-DEL-1557
chr1-31661644-DEL-62
chr1-31668360-DEL-80
chr1-31687397-DEL-76
chr1-31689864-DEL-242
chr1-31718520-DEL-374
chr1-31759598-DEL-172
chr1-31768087-DEL-110
chr1-31822988-DEL-404
chr1-31833486-DEL-2653
chr1-31873999-DEL-188
chr1-31938751-DEL-114
chr1-31942821-DEL-897
chr1-31971113-DEL-75
chr1-31976850-DEL-134
chr1-31988244-DEL-156
chr1-31996357-DEL-8503
chr1-32011439-DEL-814
chr1-32016154-DEL-113
chr1-32026288-DEL-1637
chr1-32027372-DEL-94
chr1-32042468-DEL-1274
chr1-32056597-DEL-312
chr1-32058389-DEL-248
chr1-32067679-DEL-91
chr1-32089408-DEL-7862
chr1-32093665-DEL-311
chr1-32097204-DEL-389
chr1-32103093-DEL-10573
chr1-32156724-DEL-66
chr1-32161406-DEL-183
chr1-32182216-DEL-328
chr1-32185854-DEL-52
chr1-32189067-DEL-438
chr1-32220983-DEL-124
chr1-32230718-DEL-631
chr1-32260894-DEL-85
chr1-32271968-DEL-249
chr1-32284558-DEL-52
chr1-32309305-DEL-134
chr1-32310000-DEL-1233
chr1-32324513-DEL-51
chr1-32351526-DEL-1079
chr1-32370713-DEL-52
chr1-32406360-DEL-4637
chr1-32418688-DEL-65
chr1-32475388-DEL-6531
chr1-32482018-DEL-432
chr1-32482546-DEL-75
chr1-32485101-DEL-66
chr1-32491494-DEL-2535
chr1-32536807-DEL-70
chr1-32544093-DEL-1240
chr1-32582483-DEL-157
chr1-32583598-DEL-184
chr1-32617528-DEL-2628
chr1-32620234-DEL-123
chr1-32657801-DEL-91
chr1-32672348-DEL-216
chr1-32687893-DEL-166
chr1-32699815-DEL-52
chr1-32713879-DEL-797
chr1-32717453-DEL-96
chr1-32778901-DEL-52
chr1-32782114-DEL-193
chr1-32789736-DEL-1923
chr1-32800728-DEL-686
chr1-32821331-DEL-108

chr1-32822387-DEL-153
chr1-32839830-DEL-536
chr1-32903402-DEL-65
chr1-32924759-DEL-297
chr1-32938607-DEL-243
chr1-32947921-DEL-7488
chr1-33001422-DEL-238
chr1-33006243-DEL-379
chr1-33015176-DEL-492
chr1-33015826-DEL-1859
chr1-33017712-DEL-1137
chr1-33019212-DEL-410
chr1-33025179-DEL-85
chr1-33029821-DEL-203
chr1-33048744-DEL-89
chr1-33080803-DEL-386
chr1-33094440-DEL-1342
chr1-33096026-DEL-80
chr1-33109583-DEL-8355
chr1-33122308-DEL-156
chr1-33139396-DEL-247
chr1-33173778-DEL-78
chr1-33221622-DEL-531
chr1-33228444-DEL-163
chr1-33255089-DEL-57
chr1-33266513-DEL-6247
chr1-33279448-DEL-195
chr1-33327887-DEL-82
chr1-33329099-DEL-61
chr1-33344708-DEL-102
chr1-33356725-DEL-641
chr1-33363993-DEL-377
chr1-33365872-DEL-242
chr1-33396944-DEL-77
chr1-33423918-DEL-34672
chr1-33446628-DEL-1258
chr1-33461686-DEL-52
chr1-33482555-DEL-244
chr1-33494923-DEL-587
chr1-33503297-DEL-57
chr1-33508830-DEL-65
chr1-33525289-DEL-229
chr1-33531457-DEL-928
chr1-33554365-DEL-4956
chr1-33561214-DEL-274
chr1-33585050-DEL-1683
chr1-33596373-DEL-2264
chr1-33606207-DEL-3003
chr1-33606320-DEL-312
chr1-33607139-DEL-174
chr1-33608349-DEL-195
chr1-33609516-DEL-117
chr1-33609629-DEL-1422
chr1-33609986-DEL-156
chr1-33626463-DEL-59
chr1-33639787-DEL-796
chr1-33651914-DEL-75
chr1-33670238-DEL-172
chr1-33695818-DEL-669
chr1-33700472-DEL-196
chr1-33720673-DEL-205
chr1-33743907-DEL-74
chr1-33770007-DEL-163
chr1-33790888-DEL-1033
chr1-33864826-DEL-152
chr1-33865361-DEL-379
chr1-33890251-DEL-209
chr1-33894082-DEL-53
chr1-33933399-DEL-90
chr1-33933846-DEL-266
chr1-33954440-DEL-71
chr1-33972818-DEL-107
chr1-34023498-DEL-118
chr1-34060071-DEL-64
chr1-34072768-DEL-62
chr1-34105702-DEL-186
chr1-34114884-DEL-67

chr1-34126473-DEL-59
chr1-34131146-DEL-199
chr1-34132736-DEL-58
chr1-34146541-DEL-549
chr1-34154604-DEL-169
chr1-34183086-DEL-445
chr1-34205701-DEL-86
chr1-34250529-DEL-270
chr1-34269641-DEL-272
chr1-34270490-DEL-70
chr1-34291119-DEL-99
chr1-34296345-DEL-71
chr1-34297104-DEL-1395
chr1-34348140-DEL-1187
chr1-34353155-DEL-215
chr1-34353800-DEL-156
chr1-34361405-DEL-222
chr1-34369189-DEL-55
chr1-34373500-DEL-168
chr1-34396388-DEL-52
chr1-34416321-DEL-65
chr1-34428128-DEL-96
chr1-34456831-DEL-204
chr1-34465322-DEL-176
chr1-34472439-DEL-1374
chr1-34472777-DEL-62
chr1-34481506-DEL-178
chr1-34484668-DEL-69
chr1-34494723-DEL-54
chr1-34521029-DEL-159
chr1-34527871-DEL-1187
chr1-34536718-DEL-144
chr1-34565734-DEL-127
chr1-34568085-DEL-201
chr1-34570747-DEL-132
chr1-34573842-DEL-344
chr1-34594800-DEL-198
chr1-34630414-DEL-72
chr1-34632148-DEL-474
chr1-34682549-DEL-72
chr1-34711367-DEL-79
chr1-34750991-DEL-105
chr1-34770388-DEL-148
chr1-34786808-DEL-111
chr1-34827738-DEL-2075
chr1-34831899-DEL-72
chr1-34837012-DEL-188
chr1-34839428-DEL-108
chr1-34877857-DEL-399
chr1-34925331-DEL-293
chr1-34925982-DEL-1630
chr1-34934504-DEL-333
chr1-34951679-DEL-93
chr1-34965094-DEL-105
chr1-34976701-DEL-158
chr1-34983493-DEL-76
chr1-34990511-DEL-85
chr1-34993440-DEL-173
chr1-35005563-DEL-977
chr1-35012120-DEL-62
chr1-35019879-DEL-114
chr1-35034755-DEL-4631
chr1-35065567-DEL-60
chr1-35071325-DEL-4968
chr1-35077730-DEL-57
chr1-35081813-DEL-6358
chr1-35179947-DEL-53
chr1-35202454-DEL-54
chr1-35208054-DEL-241
chr1-35230841-DEL-2652
chr1-35244716-DEL-5200
chr1-35264968-DEL-58
chr1-35273071-DEL-539
chr1-35273999-DEL-91
chr1-35298200-DEL-774
chr1-35309868-DEL-56
chr1-35310626-DEL-77

chr1-35315848-DEL-78
chr1-35349767-DEL-177
chr1-35389577-DEL-60
chr1-35426777-DEL-66
chr1-35433709-DEL-200
chr1-35462019-DEL-572
chr1-35473207-DEL-53
chr1-35474528-DEL-196
chr1-35524031-DEL-198
chr1-35528516-DEL-1324
chr1-35532731-DEL-170
chr1-35560370-DEL-721
chr1-35584735-DEL-620
chr1-35592878-DEL-415
chr1-35595969-DEL-88
chr1-35605110-DEL-59
chr1-35616708-DEL-1410
chr1-35650236-DEL-115
chr1-35653232-DEL-59
chr1-35695352-DEL-52
chr1-35695833-DEL-172
chr1-35699261-DEL-93
chr1-35701204-DEL-105
chr1-35702798-DEL-56
chr1-35727265-DEL-63
chr1-35736047-DEL-250
chr1-35749066-DEL-54
chr1-35777302-DEL-94
chr1-35785792-DEL-60
chr1-35793780-DEL-1263
chr1-35805685-DEL-74
chr1-35810185-DEL-273
chr1-35835701-DEL-139
chr1-35847693-DEL-253
chr1-35852108-DEL-477
chr1-35853581-DEL-227
chr1-35872705-DEL-235
chr1-35881002-DEL-82
chr1-35913757-DEL-71
chr1-35939376-DEL-56
chr1-35969503-DEL-72
chr1-35975198-DEL-71
chr1-36026804-DEL-56
chr1-36030233-DEL-355
chr1-36036584-DEL-60
chr1-36050327-DEL-845
chr1-36052190-DEL-81
chr1-36083641-DEL-53
chr1-36088314-DEL-324
chr1-36095298-DEL-84
chr1-36113925-DEL-135
chr1-36127122-DEL-56
chr1-36131368-DEL-126
chr1-36136604-DEL-111
chr1-36151297-DEL-649
chr1-36163372-DEL-60
chr1-36168709-DEL-128
chr1-36175856-DEL-329
chr1-36176629-DEL-70
chr1-36204780-DEL-818
chr1-36228162-DEL-1024
chr1-36239288-DEL-67
chr1-36265881-DEL-139
chr1-36267683-DEL-83
chr1-36297838-DEL-63
chr1-36298897-DEL-211
chr1-36302903-DEL-65
chr1-36313114-DEL-210
chr1-36316174-DEL-59
chr1-36326538-DEL-110
chr1-36402530-DEL-289
chr1-36424927-DEL-52
chr1-36454969-DEL-223
chr1-36464172-DEL-122
chr1-36470570-DEL-385
chr1-36533334-DEL-133
chr1-36541077-DEL-78

chr1-36558828-DEL-143
chr1-36584104-DEL-1433
chr1-36609800-DEL-61
chr1-36618546-DEL-153
chr1-36646687-DEL-55
chr1-36649858-DEL-294
chr1-36675611-DEL-1328
chr1-36700941-DEL-104
chr1-36707293-DEL-277
chr1-36724752-DEL-116
chr1-36733566-DEL-1081
chr1-36736878-DEL-73
chr1-36751853-DEL-195
chr1-36755768-DEL-132
chr1-36765850-DEL-444
chr1-36795593-DEL-59
chr1-36822011-DEL-1172
chr1-36830349-DEL-194
chr1-36848482-DEL-53
chr1-36869067-DEL-61
chr1-36869469-DEL-240
chr1-36950335-DEL-3683
chr1-36952961-DEL-102
chr1-36963920-DEL-1332
chr1-36975154-DEL-415
chr1-36989905-DEL-84
chr1-36999046-DEL-56
chr1-37003145-DEL-234
chr1-37011330-DEL-223
chr1-37042057-DEL-110
chr1-37045821-DEL-201
chr1-37050250-DEL-228
chr1-37063554-DEL-152
chr1-37105577-DEL-56
chr1-37107397-DEL-2737
chr1-37147138-DEL-316
chr1-37170511-DEL-167
chr1-37213109-DEL-728
chr1-37220247-DEL-99
chr1-37237740-DEL-446
chr1-37240163-DEL-1657
chr1-37267077-DEL-1218
chr1-37271988-DEL-258
chr1-37287771-DEL-190
chr1-37306080-DEL-1666
chr1-37308584-DEL-125
chr1-37335929-DEL-1365
chr1-37391777-DEL-60
chr1-37413673-DEL-59
chr1-37430719-DEL-320
chr1-37435526-DEL-101
chr1-37458652-DEL-94
chr1-37461756-DEL-128
chr1-37474019-DEL-68
chr1-37500943-DEL-152
chr1-37507319-DEL-1166
chr1-37522554-DEL-2548
chr1-37623821-DEL-259
chr1-37635387-DEL-559
chr1-37664420-DEL-84
chr1-37674535-DEL-203
chr1-37693221-DEL-550
chr1-37695502-DEL-812
chr1-37697229-DEL-68
chr1-37798775-DEL-99
chr1-37883216-DEL-84
chr1-37902686-DEL-88
chr1-37953076-DEL-157
chr1-38005973-DEL-55
chr1-38018214-DEL-1477
chr1-38042257-DEL-2234
chr1-38054079-DEL-347
chr1-38066524-DEL-566
chr1-38068703-DEL-90
chr1-38077821-DEL-1732
chr1-38083618-DEL-105
chr1-38111267-DEL-209

chr1-38117356-DEL-82
chr1-38150350-DEL-395
chr1-38194284-DEL-106
chr1-38198935-DEL-229
chr1-38305399-DEL-62
chr1-38308279-DEL-158
chr1-38313379-DEL-168
chr1-38322731-DEL-497
chr1-38328709-DEL-1388
chr1-38351522-DEL-58
chr1-38365074-DEL-258
chr1-38395891-DEL-81
chr1-38413398-DEL-77
chr1-38414223-DEL-79
chr1-38418017-DEL-329
chr1-38425342-DEL-289
chr1-38469541-DEL-126
chr1-38472717-DEL-1560
chr1-38510327-DEL-383
chr1-38519032-DEL-200
chr1-38535263-DEL-78
chr1-38545097-DEL-164
chr1-38579882-DEL-306
chr1-38590639-DEL-179
chr1-38616050-DEL-92
chr1-38625594-DEL-68
chr1-38649743-DEL-226
chr1-38669682-DEL-113
chr1-38713438-DEL-578
chr1-38732833-DEL-96
chr1-38757888-DEL-93
chr1-38848092-DEL-561
chr1-38849323-DEL-134
chr1-38896163-DEL-173
chr1-38920758-DEL-638
chr1-38925587-DEL-66
chr1-38966980-DEL-582
chr1-38990771-DEL-341
chr1-39012646-DEL-551
chr1-39019043-DEL-184
chr1-39022619-DEL-1297
chr1-39028023-DEL-176
chr1-39031890-DEL-1360
chr1-39035924-DEL-1222
chr1-39057505-DEL-60
chr1-39067575-DEL-190
chr1-39101505-DEL-73
chr1-39135321-DEL-319
chr1-39156563-DEL-2226
chr1-39185849-DEL-122
chr1-39262034-DEL-140
chr1-39283374-DEL-119
chr1-39286261-DEL-220
chr1-39288448-DEL-1212
chr1-39333721-DEL-64
chr1-39354569-DEL-51
chr1-39368741-DEL-85
chr1-39393814-DEL-52
chr1-39416234-DEL-142
chr1-39417743-DEL-222
chr1-39462799-DEL-104
chr1-39466799-DEL-487
chr1-39520112-DEL-194
chr1-39534275-DEL-109
chr1-39536320-DEL-52
chr1-39538807-DEL-53
chr1-39568783-DEL-593
chr1-39571007-DEL-194
chr1-39589057-DEL-197
chr1-39591007-DEL-246
chr1-39619768-DEL-280
chr1-39639974-DEL-1281
chr1-39667928-DEL-84
chr1-39701537-DEL-124
chr1-39715092-DEL-109
chr1-39729975-DEL-62
chr1-39772318-DEL-62

chr1-39778971-DEL-138
chr1-39831552-DEL-474
chr1-39835267-DEL-431
chr1-39861789-DEL-204
chr1-39866367-DEL-252
chr1-39873239-DEL-133
chr1-39894891-DEL-57
chr1-39903197-DEL-181
chr1-39975145-DEL-243
chr1-40003741-DEL-119
chr1-40014098-DEL-214
chr1-40032171-DEL-72
chr1-40033548-DEL-1215
chr1-40107446-DEL-113
chr1-40117982-DEL-400
chr1-40160633-DEL-132
chr1-40176234-DEL-578
chr1-40186454-DEL-85
chr1-40188968-DEL-345
chr1-40190111-DEL-205
chr1-40277113-DEL-1705
chr1-40299001-DEL-126
chr1-40322653-DEL-214
chr1-40324965-DEL-605
chr1-40337625-DEL-129
chr1-40355830-DEL-1137
chr1-40383472-DEL-158
chr1-40400406-DEL-84
chr1-40429961-DEL-439
chr1-40463667-DEL-165
chr1-40521539-DEL-51
chr1-40521985-DEL-923
chr1-40558227-DEL-208
chr1-40583409-DEL-312
chr1-40592020-DEL-1897
chr1-40605989-DEL-2206
chr1-40626970-DEL-72
chr1-40628709-DEL-424
chr1-40637587-DEL-400
chr1-40639064-DEL-375
chr1-40643228-DEL-133
chr1-40647014-DEL-66
chr1-40691193-DEL-357
chr1-40698750-DEL-78
chr1-40712873-DEL-106
chr1-40727664-DEL-87
chr1-40764079-DEL-216
chr1-40779468-DEL-96
chr1-40782106-DEL-69
chr1-40833733-DEL-51
chr1-40851381-DEL-183
chr1-40863293-DEL-432
chr1-40883048-DEL-234
chr1-40888039-DEL-164
chr1-40921567-DEL-292
chr1-40924542-DEL-124
chr1-40944519-DEL-54
chr1-40974971-DEL-518
chr1-40991970-DEL-275
chr1-41019184-DEL-55
chr1-41126491-DEL-149
chr1-41146649-DEL-476
chr1-41199621-DEL-192
chr1-41215391-DEL-1366
chr1-41218526-DEL-56
chr1-41233532-DEL-76
chr1-41246325-DEL-1829
chr1-41269923-DEL-1565
chr1-41327389-DEL-147
chr1-41335391-DEL-82
chr1-41353779-DEL-56
chr1-41374190-DEL-53
chr1-41404313-DEL-210
chr1-41410336-DEL-901
chr1-41411077-DEL-164
chr1-41420220-DEL-62
chr1-41492108-DEL-59

chr1-41496377-DEL-101
chr1-41604649-DEL-90
chr1-41606372-DEL-51
chr1-41668059-DEL-116
chr1-41710363-DEL-56
chr1-41734387-DEL-67
chr1-41796331-DEL-108
chr1-41806967-DEL-111
chr1-41845728-DEL-51
chr1-41868535-DEL-155
chr1-41874363-DEL-82
chr1-41893733-DEL-100
chr1-41894058-DEL-498
chr1-41895229-DEL-66
chr1-41919924-DEL-104
chr1-41950270-DEL-125
chr1-41957078-DEL-332
chr1-41988490-DEL-66
chr1-42012660-DEL-52
chr1-42013534-DEL-626
chr1-42107669-DEL-84
chr1-42120575-DEL-1050
chr1-42128779-DEL-647
chr1-42166551-DEL-362
chr1-42176718-DEL-208
chr1-42185718-DEL-364
chr1-42186221-DEL-2621
chr1-42228410-DEL-82
chr1-42278648-DEL-80
chr1-42282526-DEL-58
chr1-42285787-DEL-58
chr1-42313345-DEL-162
chr1-42315089-DEL-117
chr1-42357793-DEL-56
chr1-42370316-DEL-83
chr1-42399788-DEL-256
chr1-42419850-DEL-2540
chr1-42420123-DEL-331
chr1-42422114-DEL-237
chr1-42443335-DEL-96
chr1-42460252-DEL-389
chr1-42470527-DEL-194
chr1-42477098-DEL-881
chr1-42491050-DEL-250
chr1-42545935-DEL-258
chr1-42596035-DEL-55
chr1-42600427-DEL-82
chr1-42625932-DEL-474
chr1-42640829-DEL-1031
chr1-42671881-DEL-60
chr1-42759755-DEL-72
chr1-42774103-DEL-72
chr1-42804331-DEL-252
chr1-42825990-DEL-52
chr1-42830008-DEL-343
chr1-42862212-DEL-126
chr1-42865164-DEL-80
chr1-42904281-DEL-1046
chr1-42970749-DEL-218
chr1-42998982-DEL-1573
chr1-43003964-DEL-57
chr1-43023751-DEL-158
chr1-43032873-DEL-3661
chr1-43060422-DEL-1388
chr1-43127348-DEL-54
chr1-43151512-DEL-99
chr1-43158077-DEL-440
chr1-43215791-DEL-61
chr1-43237003-DEL-96
chr1-43239753-DEL-708
chr1-43249643-DEL-54
chr1-43265328-DEL-6281
chr1-43317023-DEL-68
chr1-43373893-DEL-70
chr1-43378033-DEL-144
chr1-43428646-DEL-52
chr1-43456712-DEL-206

chr1-43460039-DEL-88
chr1-43604724-DEL-751
chr1-43674040-DEL-122
chr1-43679895-DEL-445
chr1-43715169-DEL-54
chr1-43723248-DEL-121
chr1-43724880-DEL-672
chr1-43729571-DEL-120
chr1-43767185-DEL-163
chr1-43816762-DEL-1281
chr1-43816811-DEL-122
chr1-43817405-DEL-122
chr1-43827251-DEL-107
chr1-43864066-DEL-107
chr1-43881224-DEL-2610
chr1-43886783-DEL-321
chr1-43921211-DEL-360
chr1-43930880-DEL-205
chr1-43956536-DEL-79
chr1-43966670-DEL-1672
chr1-43991527-DEL-127
chr1-43998175-DEL-87
chr1-44018667-DEL-402
chr1-44025200-DEL-88
chr1-44035662-DEL-232
chr1-44066342-DEL-273
chr1-44068647-DEL-104
chr1-44093080-DEL-490
chr1-44128210-DEL-664
chr1-44155739-DEL-85
chr1-44193610-DEL-552
chr1-44234935-DEL-1788
chr1-44239800-DEL-3359
chr1-44249493-DEL-56
chr1-44260565-DEL-378
chr1-44288707-DEL-11432
chr1-44292416-DEL-1965
chr1-44299464-DEL-61
chr1-44306318-DEL-8431
chr1-44310292-DEL-2210
chr1-44311079-DEL-2184
chr1-44329865-DEL-5057
chr1-44335613-DEL-528
chr1-44342618-DEL-207
chr1-44358169-DEL-85
chr1-44360085-DEL-350
chr1-44368489-DEL-63
chr1-44376278-DEL-852
chr1-44380265-DEL-58355
chr1-44381433-DEL-55
chr1-44382169-DEL-590
chr1-44396222-DEL-342
chr1-44399346-DEL-2897
chr1-44406269-DEL-14530
chr1-44436566-DEL-70
chr1-44448658-DEL-344
chr1-44451605-DEL-109
chr1-44479357-DEL-231
chr1-44491501-DEL-232
chr1-44515944-DEL-1942
chr1-44519599-DEL-8223
chr1-44533436-DEL-90
chr1-44535741-DEL-7588
chr1-44535756-DEL-79
chr1-44536455-DEL-6455
chr1-44550658-DEL-205
chr1-44551212-DEL-893
chr1-44582109-DEL-517
chr1-44595159-DEL-378
chr1-44668920-DEL-2229
chr1-44676483-DEL-2151
chr1-44713786-DEL-195
chr1-44725761-DEL-1614
chr1-44731829-DEL-208
chr1-44742169-DEL-85
chr1-44756978-DEL-669
chr1-44790946-DEL-333

chr1-44795696-DEL-1883
chr1-44799865-DEL-771
chr1-44840451-DEL-643
chr1-44841107-DEL-1926
chr1-44866700-DEL-217
chr1-44867797-DEL-190
chr1-44878882-DEL-54
chr1-44897910-DEL-507
chr1-44898553-DEL-2120
chr1-44936211-DEL-12295
chr1-44956320-DEL-311
chr1-44973558-DEL-285
chr1-44999030-DEL-212
chr1-45032251-DEL-3900
chr1-45038704-DEL-1017
chr1-45054759-DEL-95
chr1-45070413-DEL-88
chr1-45073426-DEL-1014
chr1-45075748-DEL-160
chr1-45090228-DEL-80
chr1-45102018-DEL-1554
chr1-45144794-DEL-5950
chr1-45168772-DEL-454
chr1-45174000-DEL-67
chr1-45180670-DEL-683
chr1-45187032-DEL-91
chr1-45201154-DEL-12998
chr1-45216983-DEL-63
chr1-45227011-DEL-544
chr1-45232218-DEL-66
chr1-45259061-DEL-746
chr1-45268731-DEL-230
chr1-45279699-DEL-425
chr1-45281973-DEL-618
chr1-45286019-DEL-6213
chr1-45296440-DEL-105
chr1-45300358-DEL-82
chr1-45352623-DEL-62
chr1-45390714-DEL-1243
chr1-45506758-DEL-52
chr1-45524151-DEL-1319
chr1-45524258-DEL-101
chr1-45534191-DEL-109
chr1-45548814-DEL-266
chr1-45564953-DEL-663
chr1-45579155-DEL-1104
chr1-45609888-DEL-9846
chr1-45648029-DEL-1390
chr1-45653754-DEL-241
chr1-45675722-DEL-353
chr1-45708127-DEL-198
chr1-45717150-DEL-815
chr1-45735493-DEL-185
chr1-45737059-DEL-62
chr1-45750297-DEL-653
chr1-45754106-DEL-622
chr1-45790029-DEL-2084
chr1-45799006-DEL-143
chr1-45812744-DEL-52
chr1-45825095-DEL-6283
chr1-45846280-DEL-156
chr1-45855910-DEL-198
chr1-45875306-DEL-53
chr1-45879256-DEL-116
chr1-45902064-DEL-52
chr1-45937668-DEL-95
chr1-45939144-DEL-98
chr1-45940660-DEL-152
chr1-45988474-DEL-3161
chr1-45994846-DEL-9262
chr1-46003683-DEL-518
chr1-46027466-DEL-690
chr1-46046893-DEL-125
chr1-46081034-DEL-6335
chr1-46094147-DEL-970
chr1-46098342-DEL-70
chr1-46104506-DEL-631

chr1-46128167-DEL-182
chr1-46129132-DEL-6601
chr1-46147148-DEL-1082
chr1-46149662-DEL-71
chr1-46185703-DEL-1871
chr1-46211010-DEL-76
chr1-46236480-DEL-58
chr1-46248099-DEL-301
chr1-46321970-DEL-6473
chr1-46352505-DEL-6416
chr1-46353659-DEL-1609
chr1-46365724-DEL-822
chr1-46368023-DEL-5363
chr1-46381664-DEL-506
chr1-46392957-DEL-300
chr1-46408228-DEL-71
chr1-46428497-DEL-56
chr1-46432178-DEL-82
chr1-46442477-DEL-175
chr1-46451343-DEL-64
chr1-46471053-DEL-219
chr1-46480009-DEL-1739
chr1-46488148-DEL-424
chr1-46500267-DEL-57
chr1-46508355-DEL-1027
chr1-46523855-DEL-7011
chr1-46525272-DEL-5136
chr1-46537797-DEL-60
chr1-46561194-DEL-313
chr1-46570462-DEL-74
chr1-46575537-DEL-96
chr1-46584986-DEL-53
chr1-46586469-DEL-106
chr1-46619120-DEL-106
chr1-46650506-DEL-55
chr1-46674047-DEL-400
chr1-46694989-DEL-76
chr1-46739231-DEL-8169
chr1-46746881-DEL-201
chr1-46766875-DEL-57
chr1-46767820-DEL-1059
chr1-46771714-DEL-70
chr1-46778470-DEL-76
chr1-46790589-DEL-6206
chr1-46801765-DEL-66
chr1-46826067-DEL-1268
chr1-46879963-DEL-391
chr1-46887285-DEL-667
chr1-46906136-DEL-59
chr1-46906863-DEL-60
chr1-46954864-DEL-80
chr1-46999087-DEL-1633
chr1-47026375-DEL-209
chr1-47047580-DEL-440
chr1-47061300-DEL-126
chr1-47081561-DEL-63
chr1-47088030-DEL-125
chr1-47115137-DEL-327
chr1-47119841-DEL-6140
chr1-47153892-DEL-1996
chr1-47164099-DEL-1544
chr1-47167441-DEL-231
chr1-47229718-DEL-146
chr1-47247279-DEL-1251
chr1-47273076-DEL-7738
chr1-47274889-DEL-304
chr1-47304871-DEL-721
chr1-47320866-DEL-60
chr1-47342958-DEL-62
chr1-47351079-DEL-55
chr1-47370173-DEL-121
chr1-47382874-DEL-241
chr1-47383418-DEL-554
chr1-47389657-DEL-54
chr1-47431413-DEL-5390
chr1-47449578-DEL-798
chr1-47507797-DEL-52

chr1-47546683-DEL-95
chr1-47619029-DEL-117
chr1-47633446-DEL-54
chr1-47668521-DEL-412
chr1-47686543-DEL-1196
chr1-47687984-DEL-122
chr1-47700903-DEL-278
chr1-47709683-DEL-456
chr1-47711954-DEL-69
chr1-47760288-DEL-214
chr1-47764101-DEL-1365
chr1-47777795-DEL-938
chr1-47788319-DEL-492
chr1-47789835-DEL-494
chr1-47791623-DEL-8264
chr1-47793158-DEL-294
chr1-47797780-DEL-1929
chr1-47825821-DEL-4324
chr1-47832012-DEL-2384
chr1-47845934-DEL-100
chr1-47873751-DEL-698
chr1-47906663-DEL-206
chr1-47912940-DEL-191
chr1-47929336-DEL-63
chr1-47930607-DEL-410
chr1-47951828-DEL-2596
chr1-47960322-DEL-58
chr1-47974287-DEL-115
chr1-47980250-DEL-79
chr1-47990846-DEL-9145
chr1-48006614-DEL-67
chr1-48008283-DEL-2886
chr1-48022505-DEL-413
chr1-48036556-DEL-84
chr1-48063799-DEL-1882
chr1-48066497-DEL-65
chr1-48067208-DEL-54
chr1-48082955-DEL-5995
chr1-48090256-DEL-70
chr1-48091082-DEL-3589
chr1-48117068-DEL-173
chr1-48118565-DEL-2141
chr1-48150887-DEL-3781
chr1-48170088-DEL-537
chr1-48170854-DEL-66
chr1-48173180-DEL-433
chr1-48189679-DEL-562
chr1-48205165-DEL-3327
chr1-48214100-DEL-64
chr1-48218884-DEL-3223
chr1-48288411-DEL-6201
chr1-48307853-DEL-3280
chr1-48345977-DEL-511
chr1-48355830-DEL-110
chr1-48369092-DEL-288
chr1-48372233-DEL-326
chr1-48384155-DEL-249
chr1-48392663-DEL-15401
chr1-48407642-DEL-214
chr1-48477595-DEL-54
chr1-48500053-DEL-10399
chr1-48515105-DEL-6189
chr1-48518038-DEL-124
chr1-48522566-DEL-62
chr1-48523209-DEL-1144
chr1-48533919-DEL-6392
chr1-48568708-DEL-508
chr1-48569217-DEL-257
chr1-48619121-DEL-54
chr1-48645890-DEL-191
chr1-48657797-DEL-1372
chr1-48664513-DEL-63
chr1-48673856-DEL-161
chr1-48689971-DEL-54
chr1-48700842-DEL-116
chr1-48710142-DEL-3259
chr1-48728455-DEL-6324

chr1-48740503-DEL-628
chr1-48740661-DEL-1220
chr1-48747500-DEL-7030
chr1-48772036-DEL-57
chr1-48804662-DEL-918
chr1-48823793-DEL-521
chr1-48840332-DEL-6780
chr1-48860630-DEL-647
chr1-48908925-DEL-2934
chr1-48917086-DEL-71
chr1-48930191-DEL-1131
chr1-48934546-DEL-61
chr1-48952227-DEL-515
chr1-48985588-DEL-1045
chr1-48992420-DEL-1330
chr1-49034905-DEL-125
chr1-49035134-DEL-926
chr1-49049872-DEL-6015
chr1-49058850-DEL-61
chr1-49065101-DEL-1536
chr1-49147270-DEL-130
chr1-49169792-DEL-107
chr1-49175506-DEL-1389
chr1-49199571-DEL-1286
chr1-49204143-DEL-796
chr1-49307181-DEL-6148
chr1-49313492-DEL-4133
chr1-49338108-DEL-2722
chr1-49340905-DEL-581
chr1-49393354-DEL-1191
chr1-49395582-DEL-75
chr1-49415583-DEL-1221
chr1-49427366-DEL-2522
chr1-49445082-DEL-56
chr1-49446913-DEL-218
chr1-49507450-DEL-53
chr1-49516928-DEL-800
chr1-49547000-DEL-1702
chr1-49556251-DEL-66
chr1-49570222-DEL-628
chr1-49578787-DEL-83
chr1-49579502-DEL-2494
chr1-49593163-DEL-1156
chr1-49597447-DEL-312
chr1-49602932-DEL-198
chr1-49604065-DEL-94
chr1-49608251-DEL-337
chr1-49633043-DEL-5681
chr1-49650036-DEL-64
chr1-49654762-DEL-650
chr1-49665037-DEL-284
chr1-49671047-DEL-78
chr1-49672490-DEL-1271
chr1-49674382-DEL-1845
chr1-49676251-DEL-9459
chr1-49686288-DEL-265
chr1-49690761-DEL-228
chr1-49748600-DEL-5098
chr1-49765449-DEL-386
chr1-49785004-DEL-398
chr1-49788539-DEL-812
chr1-49793348-DEL-278
chr1-49811718-DEL-664
chr1-49826870-DEL-97
chr1-49868915-DEL-1007
chr1-49877225-DEL-127
chr1-49893571-DEL-346
chr1-49896499-DEL-1110
chr1-49905331-DEL-52
chr1-49911233-DEL-1889
chr1-49923474-DEL-4177
chr1-49945700-DEL-54
chr1-49993609-DEL-514
chr1-50015937-DEL-1528
chr1-50021843-DEL-1985
chr1-50048579-DEL-163
chr1-50053789-DEL-2430

chr1-50054734-DEL-2414
chr1-50055509-DEL-2429
chr1-50060843-DEL-4497
chr1-50084991-DEL-817
chr1-50090582-DEL-1340
chr1-50112558-DEL-90
chr1-50116829-DEL-564
chr1-50134132-DEL-164
chr1-50145421-DEL-2727
chr1-50172365-DEL-562
chr1-50175393-DEL-84
chr1-50189889-DEL-75
chr1-50252401-DEL-6146
chr1-50283549-DEL-1811
chr1-50312771-DEL-60
chr1-50360297-DEL-2533
chr1-50401000-DEL-62
chr1-50407692-DEL-319
chr1-50433838-DEL-93
chr1-50461602-DEL-169
chr1-50585552-DEL-104
chr1-50619038-DEL-2019
chr1-50648662-DEL-87
chr1-50692560-DEL-56
chr1-50696099-DEL-159
chr1-50722436-DEL-402
chr1-50757823-DEL-749
chr1-50775982-DEL-2299
chr1-50825970-DEL-1606
chr1-50835440-DEL-64
chr1-50877288-DEL-215
chr1-50891011-DEL-178
chr1-50943529-DEL-52
chr1-50947870-DEL-2300
chr1-50961237-DEL-68
chr1-50967540-DEL-69
chr1-50998564-DEL-325
chr1-51006254-DEL-286
chr1-51011388-DEL-51
chr1-51049039-DEL-99
chr1-51050741-DEL-200
chr1-51058217-DEL-64
chr1-51062183-DEL-356
chr1-51073835-DEL-637
chr1-51075718-DEL-216
chr1-51116370-DEL-327
chr1-51151382-DEL-148
chr1-51165971-DEL-130
chr1-51262164-DEL-104
chr1-51325575-DEL-1212
chr1-51337706-DEL-224
chr1-51358604-DEL-77
chr1-51403568-DEL-168
chr1-51447243-DEL-1397
chr1-51447968-DEL-1325
chr1-51463098-DEL-177
chr1-51471202-DEL-12252
chr1-51472859-DEL-214
chr1-51520937-DEL-161
chr1-51556861-DEL-1201
chr1-51649240-DEL-151
chr1-51682367-DEL-138
chr1-51715018-DEL-73
chr1-51865763-DEL-115
chr1-51890966-DEL-522
chr1-51953278-DEL-53
chr1-52040720-DEL-56
chr1-52050329-DEL-93
chr1-52085527-DEL-610
chr1-52097044-DEL-483
chr1-52101869-DEL-54
chr1-52123774-DEL-189
chr1-52129724-DEL-507
chr1-52134196-DEL-128
chr1-52142614-DEL-118
chr1-52156193-DEL-61
chr1-52165390-DEL-148

chr1-52212020-DEL-80
chr1-52275229-DEL-2500
chr1-52300828-DEL-2634
chr1-52301707-DEL-57
chr1-52324255-DEL-945
chr1-52324927-DEL-170
chr1-52341729-DEL-53
chr1-52370457-DEL-137
chr1-52438321-DEL-8434
chr1-52439018-DEL-7026
chr1-52453755-DEL-777
chr1-52519072-DEL-330
chr1-52547521-DEL-2388
chr1-52620063-DEL-139
chr1-52660702-DEL-1495
chr1-52667511-DEL-408
chr1-52671974-DEL-192
chr1-52694729-DEL-550
chr1-52697295-DEL-58
chr1-52721161-DEL-795
chr1-52726651-DEL-88
chr1-52784893-DEL-963
chr1-52794929-DEL-63
chr1-52805993-DEL-1928
chr1-52810038-DEL-870
chr1-52810121-DEL-288
chr1-52810486-DEL-450
chr1-52826885-DEL-876
chr1-52843299-DEL-54
chr1-52861405-DEL-167
chr1-52866270-DEL-166
chr1-52891136-DEL-316
chr1-52909053-DEL-224
chr1-52916459-DEL-880
chr1-52926284-DEL-163
chr1-52945655-DEL-698
chr1-52972063-DEL-61
chr1-52976843-DEL-838
chr1-53001800-DEL-76
chr1-53008379-DEL-272
chr1-53042726-DEL-54
chr1-53117720-DEL-597
chr1-53134286-DEL-3158
chr1-53147966-DEL-53
chr1-53151334-DEL-129
chr1-53153290-DEL-53
chr1-53176032-DEL-491
chr1-53193555-DEL-57
chr1-53194666-DEL-96
chr1-53215045-DEL-1250
chr1-53218466-DEL-84
chr1-53234876-DEL-217
chr1-53238167-DEL-87
chr1-53254342-DEL-168
chr1-53256459-DEL-77
chr1-53265162-DEL-67
chr1-53275139-DEL-2331
chr1-53302623-DEL-66
chr1-53307790-DEL-78
chr1-53398422-DEL-134
chr1-53435706-DEL-245
chr1-53457738-DEL-115
chr1-53470365-DEL-324
chr1-53471346-DEL-152
chr1-53475064-DEL-188
chr1-53520048-DEL-67
chr1-53527316-DEL-6120
chr1-53548936-DEL-52
chr1-53569232-DEL-97
chr1-53580672-DEL-642
chr1-53581688-DEL-1127
chr1-53606567-DEL-161
chr1-53608351-DEL-209
chr1-53609347-DEL-66
chr1-53613722-DEL-115
chr1-53633167-DEL-8336
chr1-53650732-DEL-811

chr1-53658303-DEL-399
chr1-53704447-DEL-2114
chr1-53738021-DEL-259
chr1-53747810-DEL-451
chr1-53757894-DEL-191
chr1-53758657-DEL-327
chr1-53857533-DEL-56
chr1-53880861-DEL-424
chr1-53898507-DEL-758
chr1-53898629-DEL-326
chr1-53946153-DEL-240
chr1-53957108-DEL-4013
chr1-53974517-DEL-74
chr1-53983245-DEL-96
chr1-53991907-DEL-10766
chr1-54031970-DEL-2376
chr1-54045818-DEL-61
chr1-54083074-DEL-54
chr1-54139900-DEL-62
chr1-54199600-DEL-68
chr1-54203809-DEL-318
chr1-54236245-DEL-138
chr1-54247137-DEL-405
chr1-54297817-DEL-1171
chr1-54347796-DEL-199
chr1-54357697-DEL-1227
chr1-54360374-DEL-127
chr1-54438716-DEL-236
chr1-54445438-DEL-7742
chr1-54493319-DEL-569
chr1-54494117-DEL-2049
chr1-54532907-DEL-198
chr1-54537864-DEL-241
chr1-54539683-DEL-503
chr1-54547432-DEL-803
chr1-54621883-DEL-125
chr1-54650077-DEL-99
chr1-54693185-DEL-197
chr1-54732409-DEL-862
chr1-54748144-DEL-296
chr1-54793501-DEL-263
chr1-54819002-DEL-190
chr1-54824903-DEL-84
chr1-54829602-DEL-88
chr1-54843781-DEL-64
chr1-54850405-DEL-188
chr1-54859871-DEL-364
chr1-54860238-DEL-139
chr1-54873654-DEL-271
chr1-54905724-DEL-69
chr1-54921813-DEL-156
chr1-54982193-DEL-199
chr1-54996291-DEL-364
chr1-54996797-DEL-946
chr1-55017130-DEL-1038
chr1-55026976-DEL-52
chr1-55027465-DEL-934
chr1-55104147-DEL-780
chr1-55113061-DEL-465
chr1-55136498-DEL-411
chr1-55139702-DEL-248
chr1-55154209-DEL-65
chr1-55159003-DEL-210
chr1-55186336-DEL-108
chr1-55196819-DEL-62
chr1-55201091-DEL-777
chr1-55205872-DEL-108
chr1-55214061-DEL-813
chr1-55215008-DEL-198
chr1-55216897-DEL-1826
chr1-55226066-DEL-51
chr1-55244675-DEL-293
chr1-55269175-DEL-1089
chr1-55274822-DEL-173
chr1-55304312-DEL-51
chr1-55314183-DEL-7662
chr1-55392536-DEL-1516

chr1-55423423-DEL-1782
chr1-55428200-DEL-2201
chr1-55501830-DEL-52
chr1-55521499-DEL-212
chr1-55531714-DEL-1036
chr1-55551939-DEL-71
chr1-55590603-DEL-388
chr1-55613240-DEL-91
chr1-55614084-DEL-53
chr1-55623352-DEL-67
chr1-55630830-DEL-60
chr1-55637351-DEL-443
chr1-55680751-DEL-146
chr1-55711071-DEL-279
chr1-55739314-DEL-272
chr1-55788484-DEL-131
chr1-55799246-DEL-1494
chr1-55799346-DEL-822
chr1-55800050-DEL-376
chr1-55809370-DEL-76
chr1-55811798-DEL-11698
chr1-55836279-DEL-5469
chr1-55864503-DEL-178
chr1-55879557-DEL-2001
chr1-55882435-DEL-63
chr1-55908600-DEL-160
chr1-55962075-DEL-61
chr1-55981878-DEL-392
chr1-56051727-DEL-64
chr1-56059848-DEL-55
chr1-56077370-DEL-81
chr1-56081642-DEL-1643
chr1-56106809-DEL-460
chr1-56158839-DEL-1145
chr1-56194977-DEL-266
chr1-56195608-DEL-60
chr1-56199345-DEL-210
chr1-56253732-DEL-68
chr1-56267175-DEL-156
chr1-56273668-DEL-213
chr1-56280756-DEL-220
chr1-56304738-DEL-86
chr1-56360167-DEL-77
chr1-56433469-DEL-64
chr1-56439436-DEL-461
chr1-56441476-DEL-62
chr1-56510704-DEL-51
chr1-56516926-DEL-56
chr1-56520670-DEL-52
chr1-56538649-DEL-192
chr1-56560223-DEL-518
chr1-56569269-DEL-226
chr1-56577910-DEL-56
chr1-56581806-DEL-111
chr1-56599181-DEL-93
chr1-56600887-DEL-486
chr1-56608673-DEL-64
chr1-56630874-DEL-524
chr1-56638006-DEL-116
chr1-56684371-DEL-52
chr1-56694378-DEL-150
chr1-56698369-DEL-129
chr1-56748803-DEL-276
chr1-56770162-DEL-120
chr1-56785389-DEL-90
chr1-56799528-DEL-51
chr1-56841205-DEL-272
chr1-56856384-DEL-2955
chr1-56870877-DEL-365
chr1-56931401-DEL-129
chr1-56956126-DEL-221
chr1-56987791-DEL-775
chr1-57038201-DEL-492
chr1-57053569-DEL-104
chr1-57061541-DEL-171
chr1-57085654-DEL-99
chr1-57100479-DEL-58

chr1-57126383-DEL-115
chr1-57154459-DEL-783
chr1-57155453-DEL-56
chr1-57172677-DEL-411
chr1-57249754-DEL-129
chr1-57295440-DEL-637
chr1-57310345-DEL-120
chr1-57355729-DEL-2072
chr1-57386528-DEL-211
chr1-57388287-DEL-56
chr1-57403854-DEL-77
chr1-57413277-DEL-86
chr1-57442894-DEL-214
chr1-57490495-DEL-196
chr1-57495582-DEL-4384
chr1-57571809-DEL-675
chr1-57581509-DEL-2438
chr1-57627305-DEL-81
chr1-57650983-DEL-67
chr1-57702584-DEL-620
chr1-57718054-DEL-76
chr1-57752669-DEL-133
chr1-57890739-DEL-333
chr1-57893305-DEL-56
chr1-57904643-DEL-127
chr1-57909615-DEL-105
chr1-57955058-DEL-51
chr1-57961402-DEL-303
chr1-57969524-DEL-159
chr1-57975648-DEL-133
chr1-58024003-DEL-184
chr1-58057659-DEL-591
chr1-58074081-DEL-62
chr1-58128904-DEL-93
chr1-58143212-DEL-80
chr1-58151341-DEL-52
chr1-58175609-DEL-307
chr1-58213945-DEL-284
chr1-58254662-DEL-83
chr1-58257355-DEL-97
chr1-58387551-DEL-86
chr1-58399427-DEL-54
chr1-58404910-DEL-66
chr1-58408494-DEL-83
chr1-58409478-DEL-2451
chr1-58411065-DEL-126
chr1-58414014-DEL-90
chr1-58442520-DEL-199
chr1-58447182-DEL-56
chr1-58449500-DEL-60
chr1-58458275-DEL-199
chr1-58473082-DEL-69
chr1-58474548-DEL-197
chr1-58484557-DEL-55
chr1-58488379-DEL-99
chr1-58495075-DEL-211
chr1-58521500-DEL-80
chr1-58599390-DEL-203
chr1-58615399-DEL-1005
chr1-58675392-DEL-63
chr1-58679920-DEL-10164
chr1-58693886-DEL-121
chr1-58694827-DEL-81
chr1-58704203-DEL-64
chr1-58715286-DEL-85
chr1-58716964-DEL-202
chr1-58725404-DEL-1458
chr1-58741855-DEL-452
chr1-58743078-DEL-157
chr1-58796295-DEL-199
chr1-58799887-DEL-1951
chr1-58810562-DEL-330
chr1-58828566-DEL-195
chr1-58837402-DEL-1010
chr1-58840967-DEL-145
chr1-58850844-DEL-7173
chr1-58885467-DEL-172

chr1-58907843-DEL-2924
chr1-58917706-DEL-150
chr1-58918766-DEL-205
chr1-58923862-DEL-343
chr1-58934328-DEL-202
chr1-58938177-DEL-94
chr1-58983547-DEL-54
chr1-59045609-DEL-51
chr1-59074489-DEL-667
chr1-59085546-DEL-239
chr1-59093331-DEL-428
chr1-59095322-DEL-399
chr1-59108152-DEL-326
chr1-59116372-DEL-70
chr1-59122660-DEL-101
chr1-59130394-DEL-60
chr1-59141974-DEL-1573
chr1-59177899-DEL-212
chr1-59178737-DEL-103
chr1-59183357-DEL-77
chr1-59249596-DEL-534
chr1-59280598-DEL-1768
chr1-59333203-DEL-58
chr1-59344126-DEL-393
chr1-59359342-DEL-82
chr1-59375428-DEL-144
chr1-59391701-DEL-176
chr1-59425408-DEL-75
chr1-59430151-DEL-55
chr1-59434344-DEL-53
chr1-59439721-DEL-540
chr1-59440245-DEL-139
chr1-59441593-DEL-145
chr1-59460793-DEL-140
chr1-59465350-DEL-176
chr1-59497485-DEL-83
chr1-59519994-DEL-74
chr1-59536136-DEL-58
chr1-59562575-DEL-135
chr1-59569309-DEL-66
chr1-59586478-DEL-72
chr1-59626168-DEL-198
chr1-59639582-DEL-92
chr1-59640205-DEL-198
chr1-59640739-DEL-126
chr1-59643459-DEL-373
chr1-59663579-DEL-194
chr1-59687175-DEL-106
chr1-59689832-DEL-195
chr1-59701327-DEL-63
chr1-59706231-DEL-198
chr1-59718845-DEL-148
chr1-59745397-DEL-77
chr1-59754391-DEL-202
chr1-59763243-DEL-214
chr1-59767988-DEL-194
chr1-59789896-DEL-68
chr1-59794414-DEL-454
chr1-59801179-DEL-54
chr1-59807728-DEL-184
chr1-59831490-DEL-653
chr1-59881252-DEL-202
chr1-59949847-DEL-256
chr1-59960153-DEL-125
chr1-59975539-DEL-195
chr1-59975645-DEL-1060
chr1-59999988-DEL-1726
chr1-60039185-DEL-79
chr1-60042606-DEL-65
chr1-60047215-DEL-2002
chr1-60048205-DEL-145
chr1-60049552-DEL-60
chr1-60064897-DEL-1072
chr1-60068932-DEL-52
chr1-60072260-DEL-914
chr1-60108723-DEL-193
chr1-60141067-DEL-91

chr1-60205337-DEL-53
chr1-60293865-DEL-59
chr1-60295114-DEL-747
chr1-60329846-DEL-471
chr1-60330373-DEL-111
chr1-60340050-DEL-208
chr1-60388767-DEL-1865
chr1-60406681-DEL-58
chr1-60419133-DEL-134
chr1-60471910-DEL-68
chr1-60484598-DEL-59
chr1-60498313-DEL-202
chr1-60507667-DEL-81
chr1-60594458-DEL-916
chr1-60623711-DEL-1088
chr1-60624843-DEL-2524
chr1-60658294-DEL-252
chr1-60687774-DEL-219
chr1-60700623-DEL-142
chr1-60707977-DEL-61
chr1-60713264-DEL-314
chr1-60747342-DEL-59
chr1-60801126-DEL-191
chr1-60857785-DEL-1337
chr1-60874184-DEL-1588
chr1-60910359-DEL-104
chr1-60923519-DEL-156
chr1-60926903-DEL-60
chr1-60960313-DEL-60
chr1-60965921-DEL-493
chr1-60970986-DEL-225
chr1-60973007-DEL-293
chr1-60988367-DEL-354
chr1-61005323-DEL-64
chr1-61015947-DEL-59
chr1-61026910-DEL-1202
chr1-61048977-DEL-54
chr1-61075396-DEL-98
chr1-61086137-DEL-111
chr1-61107955-DEL-1301
chr1-61209082-DEL-394
chr1-61209836-DEL-59
chr1-61211954-DEL-178
chr1-61232481-DEL-134
chr1-61244009-DEL-142
chr1-61278742-DEL-105
chr1-61291081-DEL-196
chr1-61335435-DEL-134
chr1-61387154-DEL-69
chr1-61407562-DEL-201
chr1-61480140-DEL-1368
chr1-61496689-DEL-68
chr1-61525990-DEL-555
chr1-61549244-DEL-226
chr1-61554061-DEL-424
chr1-61591163-DEL-50
chr1-61606007-DEL-113
chr1-61608470-DEL-199
chr1-61650286-DEL-59
chr1-61652898-DEL-118
chr1-61657429-DEL-73
chr1-61658048-DEL-215
chr1-61671718-DEL-54
chr1-61685539-DEL-382
chr1-61694211-DEL-174
chr1-61763083-DEL-495
chr1-61770538-DEL-54
chr1-61794412-DEL-1367
chr1-61820470-DEL-613
chr1-61838064-DEL-60
chr1-61846591-DEL-113
chr1-61863602-DEL-64
chr1-61885641-DEL-100
chr1-61894990-DEL-63
chr1-61910235-DEL-652
chr1-61924551-DEL-261
chr1-61937361-DEL-599

chr1-61949299-DEL-112
chr1-61986043-DEL-1196
chr1-61994904-DEL-262
chr1-62078379-DEL-1945
chr1-62088886-DEL-56
chr1-62116783-DEL-177
chr1-62126110-DEL-71
chr1-62145945-DEL-557
chr1-62150758-DEL-667
chr1-62175609-DEL-54
chr1-62178648-DEL-87
chr1-62199296-DEL-70
chr1-62217822-DEL-1187
chr1-62285036-DEL-74
chr1-62286398-DEL-1289
chr1-62349857-DEL-777
chr1-62359023-DEL-66
chr1-62381893-DEL-133
chr1-62424173-DEL-2706
chr1-62470472-DEL-71
chr1-62511990-DEL-60
chr1-62640731-DEL-64
chr1-62668166-DEL-132
chr1-62719096-DEL-568
chr1-62784855-DEL-145
chr1-62890083-DEL-69
chr1-63010819-DEL-431
chr1-63045158-DEL-135
chr1-63060337-DEL-122
chr1-63066559-DEL-5677
chr1-63123848-DEL-181
chr1-63138631-DEL-52
chr1-63146316-DEL-228
chr1-63167878-DEL-60
chr1-63170515-DEL-168
chr1-63174027-DEL-82
chr1-63181097-DEL-348
chr1-63212779-DEL-122
chr1-63225861-DEL-96
chr1-63229472-DEL-53
chr1-63231404-DEL-56
chr1-63236808-DEL-58
chr1-63246032-DEL-72
chr1-63267669-DEL-134
chr1-63273224-DEL-215
chr1-63279260-DEL-53
chr1-63282022-DEL-205
chr1-63322046-DEL-122
chr1-63324922-DEL-198
chr1-63365704-DEL-162
chr1-63437055-DEL-201
chr1-63468587-DEL-206
chr1-63528675-DEL-52
chr1-63547763-DEL-63
chr1-63615116-DEL-60
chr1-63664875-DEL-926
chr1-63702478-DEL-252
chr1-63729271-DEL-135
chr1-63743024-DEL-66
chr1-63775185-DEL-522
chr1-63814223-DEL-1036
chr1-63836850-DEL-6691
chr1-63843074-DEL-206
chr1-63848302-DEL-468
chr1-63868315-DEL-58
chr1-63878299-DEL-238
chr1-63878836-DEL-3809
chr1-63898415-DEL-201
chr1-63967935-DEL-133
chr1-63998193-DEL-1227
chr1-64049310-DEL-269
chr1-64208941-DEL-62
chr1-64287373-DEL-379
chr1-64298895-DEL-6304
chr1-64308474-DEL-151
chr1-64311561-DEL-108
chr1-64359488-DEL-96

chr1-64495922-DEL-677
chr1-64529274-DEL-78
chr1-64531194-DEL-53
chr1-64539535-DEL-485
chr1-64585206-DEL-72
chr1-64623082-DEL-389
chr1-64626219-DEL-469
chr1-64662167-DEL-421
chr1-64670394-DEL-90
chr1-64696474-DEL-63
chr1-64721679-DEL-59
chr1-64789562-DEL-201
chr1-64799471-DEL-236
chr1-64802440-DEL-68
chr1-64837588-DEL-357
chr1-64837604-DEL-1938
chr1-64838152-DEL-911
chr1-64887574-DEL-125
chr1-64933290-DEL-2378
chr1-64970203-DEL-194
chr1-64974740-DEL-206
chr1-64978399-DEL-2139
chr1-64982397-DEL-63
chr1-65036155-DEL-100
chr1-65049256-DEL-134
chr1-65080487-DEL-631
chr1-65082144-DEL-224
chr1-65093266-DEL-180
chr1-65093620-DEL-470
chr1-65134295-DEL-654
chr1-65137226-DEL-89
chr1-65148219-DEL-156
chr1-65155191-DEL-415
chr1-65170392-DEL-87
chr1-65182478-DEL-88
chr1-65196325-DEL-88
chr1-65203414-DEL-98
chr1-65240005-DEL-141
chr1-65325172-DEL-959
chr1-65334415-DEL-208
chr1-65372936-DEL-56
chr1-65424418-DEL-114
chr1-65451046-DEL-6443
chr1-65462999-DEL-6159
chr1-65472307-DEL-163
chr1-65472861-DEL-202
chr1-65494680-DEL-2046
chr1-65502852-DEL-981
chr1-65517725-DEL-112
chr1-65544373-DEL-1826
chr1-65552458-DEL-58
chr1-65563280-DEL-1177
chr1-65565497-DEL-5341
chr1-65594185-DEL-1328
chr1-65621483-DEL-207
chr1-65626609-DEL-3258
chr1-65643449-DEL-58
chr1-65648679-DEL-70
chr1-65650448-DEL-299
chr1-65653081-DEL-260
chr1-65658338-DEL-7068
chr1-65672697-DEL-218
chr1-65685637-DEL-572
chr1-65689534-DEL-54
chr1-65702191-DEL-212
chr1-65771303-DEL-65
chr1-65809323-DEL-3659
chr1-65818998-DEL-6594
chr1-65832407-DEL-2016
chr1-65834953-DEL-853
chr1-65849178-DEL-1676
chr1-65861767-DEL-368
chr1-65881576-DEL-1161
chr1-65985528-DEL-1193
chr1-66027257-DEL-51
chr1-66076058-DEL-6242
chr1-66087691-DEL-5365

chr1-66099708-DEL-62
chr1-66109128-DEL-3089
chr1-66113529-DEL-509
chr1-66128350-DEL-2436
chr1-66128707-DEL-67
chr1-66137558-DEL-66
chr1-66158054-DEL-176
chr1-66193322-DEL-402
chr1-66195114-DEL-375
chr1-66205580-DEL-1025
chr1-66210282-DEL-109
chr1-66210971-DEL-96
chr1-66281368-DEL-950
chr1-66295592-DEL-98
chr1-66296770-DEL-447
chr1-66298723-DEL-110
chr1-66299771-DEL-436
chr1-66327441-DEL-906
chr1-66356162-DEL-68
chr1-66543771-DEL-56
chr1-66550217-DEL-66
chr1-66573005-DEL-1938
chr1-66577261-DEL-102
chr1-66603437-DEL-921
chr1-66622608-DEL-1915
chr1-66644990-DEL-56
chr1-66652636-DEL-2299
chr1-66676366-DEL-365
chr1-66692754-DEL-127
chr1-66696629-DEL-107
chr1-66698017-DEL-54
chr1-66699065-DEL-761
chr1-66777044-DEL-56
chr1-66882659-DEL-126
chr1-66883169-DEL-734
chr1-66922682-DEL-1685
chr1-66931974-DEL-61
chr1-66966228-DEL-1518
chr1-66967269-DEL-252
chr1-66967742-DEL-126
chr1-66985892-DEL-472
chr1-67011809-DEL-2442
chr1-67021751-DEL-442
chr1-67054522-DEL-57
chr1-67127329-DEL-1843
chr1-67154411-DEL-81
chr1-67161229-DEL-109
chr1-67266456-DEL-96
chr1-67268820-DEL-62
chr1-67279182-DEL-4828
chr1-67348352-DEL-402
chr1-67348867-DEL-54
chr1-67367147-DEL-52
chr1-67367794-DEL-1106
chr1-67369302-DEL-2511
chr1-67387395-DEL-1146
chr1-67407491-DEL-401
chr1-67410482-DEL-7104
chr1-67431524-DEL-111
chr1-67467678-DEL-114
chr1-67488939-DEL-1213
chr1-67495830-DEL-2334
chr1-67501313-DEL-124
chr1-67506738-DEL-176
chr1-67536034-DEL-428
chr1-67552664-DEL-110
chr1-67575814-DEL-6406
chr1-67581884-DEL-201
chr1-67601352-DEL-73
chr1-67611525-DEL-1057
chr1-67619845-DEL-424
chr1-67653372-DEL-82
chr1-67704151-DEL-360
chr1-67731151-DEL-430
chr1-67735619-DEL-3651
chr1-67745890-DEL-436
chr1-67752833-DEL-7753

chr1-67822840-DEL-267
chr1-67836295-DEL-731
chr1-67872755-DEL-468
chr1-67875060-DEL-256
chr1-67920107-DEL-127
chr1-68022583-DEL-290
chr1-68054954-DEL-898
chr1-68060715-DEL-283
chr1-68062375-DEL-486
chr1-68092516-DEL-60
chr1-68126183-DEL-52
chr1-68138757-DEL-71
chr1-68151106-DEL-1903
chr1-68151196-DEL-739
chr1-68164590-DEL-80
chr1-68199128-DEL-66
chr1-68214409-DEL-74
chr1-68261462-DEL-137
chr1-68268200-DEL-423
chr1-68318640-DEL-792
chr1-68361031-DEL-614
chr1-68433437-DEL-149
chr1-68503895-DEL-141
chr1-68520949-DEL-252
chr1-68548205-DEL-59
chr1-68550678-DEL-82
chr1-68581162-DEL-55
chr1-68608346-DEL-1285
chr1-68637415-DEL-6607
chr1-68708941-DEL-58
chr1-68759526-DEL-219
chr1-68773519-DEL-146
chr1-68773780-DEL-2132
chr1-68787492-DEL-54
chr1-68793416-DEL-52
chr1-68857711-DEL-321
chr1-68895330-DEL-506
chr1-68937631-DEL-1357
chr1-68943015-DEL-1655
chr1-68954387-DEL-60
chr1-68966913-DEL-461
chr1-68986882-DEL-60
chr1-69004262-DEL-56
chr1-69005967-DEL-90
chr1-69026204-DEL-56
chr1-69040348-DEL-69
chr1-69050305-DEL-1140
chr1-69052514-DEL-9369
chr1-69070898-DEL-182
chr1-69109630-DEL-56
chr1-69119317-DEL-214
chr1-69143207-DEL-54
chr1-69193266-DEL-215
chr1-69199221-DEL-181
chr1-69200181-DEL-717
chr1-69214551-DEL-2698
chr1-69220841-DEL-58
chr1-69243194-DEL-97
chr1-69308386-DEL-1343
chr1-69310442-DEL-824
chr1-69339534-DEL-160
chr1-69359870-DEL-85
chr1-69365353-DEL-70
chr1-69380868-DEL-92
chr1-69393726-DEL-126
chr1-69397757-DEL-54
chr1-69409953-DEL-1541
chr1-69460520-DEL-380
chr1-69496052-DEL-67
chr1-69506950-DEL-55
chr1-69551346-DEL-55
chr1-69551953-DEL-316
chr1-69561828-DEL-61
chr1-69568527-DEL-61
chr1-69596244-DEL-1385
chr1-69608260-DEL-532
chr1-69615807-DEL-64

chr1-69627315-DEL-62
chr1-69709897-DEL-64
chr1-69754261-DEL-68
chr1-69761697-DEL-55
chr1-69782288-DEL-95
chr1-69791725-DEL-64
chr1-69825310-DEL-57
chr1-69843612-DEL-4696
chr1-69849145-DEL-1801
chr1-69862359-DEL-135
chr1-69870666-DEL-349
chr1-69877922-DEL-1520
chr1-69891174-DEL-245
chr1-69897980-DEL-994
chr1-69962427-DEL-57
chr1-69971490-DEL-54
chr1-69990149-DEL-64
chr1-70001829-DEL-6459
chr1-70003797-DEL-4490
chr1-70005569-DEL-2656
chr1-70006074-DEL-787
chr1-70008350-DEL-2316
chr1-70023776-DEL-60
chr1-70023839-DEL-309
chr1-70025407-DEL-136
chr1-70044131-DEL-416
chr1-70066140-DEL-66
chr1-70089184-DEL-111
chr1-70092428-DEL-208
chr1-70111901-DEL-128
chr1-70136249-DEL-1033
chr1-70164218-DEL-57
chr1-70191981-DEL-312
chr1-70241038-DEL-21233
chr1-70244704-DEL-76
chr1-70259347-DEL-51
chr1-70274576-DEL-178
chr1-70293747-DEL-53
chr1-70294378-DEL-56
chr1-70305962-DEL-179
chr1-70313401-DEL-1109
chr1-70449276-DEL-1941
chr1-70512653-DEL-143
chr1-70538575-DEL-52
chr1-70540572-DEL-390
chr1-70543311-DEL-4394
chr1-70579182-DEL-58
chr1-70579856-DEL-139
chr1-70594467-DEL-1018
chr1-70595940-DEL-64
chr1-70597209-DEL-555
chr1-70644141-DEL-195
chr1-70647245-DEL-283
chr1-70652360-DEL-241
chr1-70684491-DEL-52
chr1-70714479-DEL-67
chr1-70817730-DEL-1042
chr1-70873935-DEL-2324
chr1-70878883-DEL-112
chr1-70879949-DEL-661
chr1-70909853-DEL-75
chr1-70932317-DEL-217
chr1-70941136-DEL-1094
chr1-70954195-DEL-64
chr1-70963867-DEL-337
chr1-70986990-DEL-1486
chr1-70986993-DEL-668
chr1-70988395-DEL-4920
chr1-71018561-DEL-62
chr1-71064933-DEL-178
chr1-71068086-DEL-283
chr1-71072183-DEL-169
chr1-71138462-DEL-56
chr1-71143865-DEL-514
chr1-71172165-DEL-285
chr1-71177363-DEL-58
chr1-71203025-DEL-80

chr1-71214620-DEL-88
chr1-71241748-DEL-365
chr1-71254569-DEL-1812
chr1-71262880-DEL-120
chr1-71264166-DEL-186
chr1-71272493-DEL-1817
chr1-71322975-DEL-51
chr1-71349482-DEL-145
chr1-71363124-DEL-239
chr1-71380412-DEL-120
chr1-71407342-DEL-56
chr1-71427057-DEL-98
chr1-71448804-DEL-579
chr1-71508261-DEL-80
chr1-71509875-DEL-413
chr1-71518396-DEL-3316
chr1-71536708-DEL-556
chr1-71537914-DEL-230
chr1-71544967-DEL-53
chr1-71557302-DEL-64
chr1-71581852-DEL-188
chr1-71587783-DEL-1008
chr1-71598506-DEL-687
chr1-71648377-DEL-76
chr1-71649424-DEL-341
chr1-71672177-DEL-80
chr1-71707317-DEL-417
chr1-71709010-DEL-92
chr1-71724658-DEL-757
chr1-71748335-DEL-950
chr1-71758967-DEL-91
chr1-71764324-DEL-77
chr1-71764899-DEL-190
chr1-71783164-DEL-66
chr1-71784820-DEL-179
chr1-71800009-DEL-68
chr1-71819150-DEL-1750
chr1-71839737-DEL-241
chr1-71848986-DEL-133
chr1-71902208-DEL-359
chr1-71955859-DEL-176
chr1-71960512-DEL-134
chr1-71971808-DEL-165
chr1-71972539-DEL-1259
chr1-71975195-DEL-195
chr1-71983505-DEL-127
chr1-71985419-DEL-91
chr1-72027734-DEL-188
chr1-72031375-DEL-626
chr1-72058999-DEL-54
chr1-72063360-DEL-1350
chr1-72107426-DEL-466
chr1-72218652-DEL-352
chr1-72221164-DEL-72
chr1-72223756-DEL-1393
chr1-72228656-DEL-227
chr1-72254477-DEL-108
chr1-72261646-DEL-2790
chr1-72291834-DEL-189
chr1-72301285-DEL-433
chr1-72343277-DEL-78
chr1-72361540-DEL-595
chr1-72374420-DEL-466
chr1-72420041-DEL-54
chr1-72478961-DEL-121
chr1-72512602-DEL-1335
chr1-72514420-DEL-1990
chr1-72539978-DEL-86
chr1-72560700-DEL-63
chr1-72688984-DEL-141
chr1-72699145-DEL-55
chr1-72706280-DEL-51
chr1-72735588-DEL-60
chr1-72743750-DEL-83
chr1-72752304-DEL-255
chr1-72756606-DEL-1561
chr1-72760678-DEL-56

chr1-72781109-DEL-84
chr1-72781600-DEL-128
chr1-72813233-DEL-1383
chr1-72816338-DEL-121
chr1-72841753-DEL-204
chr1-72851208-DEL-932
chr1-72881932-DEL-75
chr1-72885120-DEL-656
chr1-72886499-DEL-52
chr1-72932816-DEL-801
chr1-72936395-DEL-83
chr1-72945387-DEL-255
chr1-72976187-DEL-2527
chr1-72988566-DEL-932
chr1-72990815-DEL-66
chr1-73021154-DEL-58
chr1-73046695-DEL-440
chr1-73069718-DEL-53
chr1-73072697-DEL-912
chr1-73130279-DEL-183
chr1-73233971-DEL-1319
chr1-73235639-DEL-1921
chr1-73243831-DEL-62
chr1-73268202-DEL-155
chr1-73272134-DEL-52
chr1-73285344-DEL-78
chr1-73343785-DEL-52
chr1-73373792-DEL-60
chr1-73380156-DEL-52
chr1-73442806-DEL-51
chr1-73479621-DEL-358
chr1-73504986-DEL-150
chr1-73524967-DEL-62
chr1-73655828-DEL-1360
chr1-73746857-DEL-6216
chr1-73767380-DEL-79
chr1-73787194-DEL-90
chr1-73805613-DEL-79
chr1-73813725-DEL-66
chr1-73836822-DEL-60
chr1-73852319-DEL-280
chr1-73870561-DEL-112
chr1-73872461-DEL-159
chr1-73892131-DEL-823
chr1-73915627-DEL-292
chr1-73924398-DEL-6648
chr1-73931670-DEL-64
chr1-73941343-DEL-256
chr1-74023439-DEL-92
chr1-74082413-DEL-60
chr1-74116056-DEL-195
chr1-74140219-DEL-201
chr1-74212034-DEL-71
chr1-74216602-DEL-87
chr1-74237772-DEL-106
chr1-74242206-DEL-160
chr1-74253013-DEL-225
chr1-74308117-DEL-79
chr1-74312826-DEL-54
chr1-74351035-DEL-222
chr1-74353161-DEL-334
chr1-74359886-DEL-226
chr1-74376892-DEL-290
chr1-74379965-DEL-444
chr1-74384282-DEL-789
chr1-74416620-DEL-316
chr1-74425514-DEL-59
chr1-74428491-DEL-63
chr1-74468613-DEL-140
chr1-74483938-DEL-71
chr1-74490229-DEL-73
chr1-74497305-DEL-55
chr1-74503451-DEL-196
chr1-74532332-DEL-84
chr1-74536318-DEL-60
chr1-74550425-DEL-71
chr1-74555696-DEL-109

chr1-74588812-DEL-741
chr1-74591583-DEL-58
chr1-74592839-DEL-115
chr1-74596454-DEL-207
chr1-74651214-DEL-57
chr1-74667524-DEL-577
chr1-74679013-DEL-102
chr1-74689576-DEL-74
chr1-74690381-DEL-206
chr1-74712955-DEL-173
chr1-74717615-DEL-55
chr1-74732064-DEL-195
chr1-74750870-DEL-745
chr1-74756604-DEL-198
chr1-74757703-DEL-503
chr1-74796099-DEL-203
chr1-74808179-DEL-75
chr1-74819010-DEL-150
chr1-74821105-DEL-63
chr1-74831933-DEL-78
chr1-74852398-DEL-120
chr1-74877506-DEL-207
chr1-74902326-DEL-2347
chr1-74912586-DEL-217
chr1-74954693-DEL-69
chr1-74969138-DEL-434
chr1-75011807-DEL-125
chr1-75015560-DEL-71
chr1-75031416-DEL-301
chr1-75043832-DEL-121
chr1-75056740-DEL-56
chr1-75058332-DEL-101
chr1-75064637-DEL-204
chr1-75084951-DEL-334
chr1-75089412-DEL-208
chr1-75095127-DEL-1685
chr1-75114576-DEL-432
chr1-75115732-DEL-74
chr1-75138010-DEL-78
chr1-75138849-DEL-480
chr1-75143675-DEL-162
chr1-75226388-DEL-58
chr1-75228075-DEL-103
chr1-75285757-DEL-81
chr1-75296628-DEL-86
chr1-75323537-DEL-61
chr1-75434023-DEL-169
chr1-75437342-DEL-1312
chr1-75437769-DEL-106
chr1-75437978-DEL-78
chr1-75438595-DEL-98
chr1-75453002-DEL-194
chr1-75489031-DEL-182
chr1-75506304-DEL-62
chr1-75511481-DEL-214
chr1-75577606-DEL-78
chr1-75606576-DEL-212
chr1-75611416-DEL-69
chr1-75612927-DEL-1362
chr1-75621963-DEL-113
chr1-75627320-DEL-126
chr1-75632840-DEL-120
chr1-75642961-DEL-101
chr1-75667620-DEL-106
chr1-75678382-DEL-54
chr1-75684044-DEL-56
chr1-75691297-DEL-212
chr1-75793093-DEL-198
chr1-75814238-DEL-109
chr1-75829977-DEL-52
chr1-75831436-DEL-99
chr1-75836769-DEL-676
chr1-75866322-DEL-220
chr1-75874309-DEL-52
chr1-75897928-DEL-75
chr1-75925463-DEL-105
chr1-75985422-DEL-56

chr1-76028170-DEL-52
chr1-76043328-DEL-91
chr1-76044538-DEL-103
chr1-76056332-DEL-64
chr1-76070889-DEL-310
chr1-76083132-DEL-114
chr1-76137983-DEL-68
chr1-76139905-DEL-79
chr1-76150177-DEL-3302
chr1-76178395-DEL-56
chr1-76186723-DEL-411
chr1-76197852-DEL-182
chr1-76197995-DEL-1258
chr1-76206503-DEL-274
chr1-76235663-DEL-111
chr1-76243810-DEL-2051
chr1-76274088-DEL-902
chr1-76382407-DEL-61
chr1-76418993-DEL-60
chr1-76487855-DEL-61
chr1-76494366-DEL-1655
chr1-76505043-DEL-523
chr1-76546952-DEL-557
chr1-76548452-DEL-403
chr1-76611555-DEL-64
chr1-76660744-DEL-106
chr1-76678602-DEL-57
chr1-76721532-DEL-75
chr1-76729221-DEL-474
chr1-76735070-DEL-101
chr1-76754836-DEL-293
chr1-76774304-DEL-52
chr1-76779779-DEL-52
chr1-76780298-DEL-169
chr1-76874440-DEL-130
chr1-76940200-DEL-390
chr1-76947808-DEL-68
chr1-76967150-DEL-153
chr1-77001340-DEL-72
chr1-77008068-DEL-253
chr1-77011374-DEL-129
chr1-77018524-DEL-82
chr1-77029008-DEL-76
chr1-77047085-DEL-243
chr1-77052501-DEL-265
chr1-77063016-DEL-387
chr1-77083486-DEL-83
chr1-77088049-DEL-79
chr1-77094360-DEL-1569
chr1-77111446-DEL-314
chr1-77129908-DEL-73
chr1-77147392-DEL-149
chr1-77202687-DEL-59
chr1-77226223-DEL-383
chr1-77294499-DEL-52
chr1-77334035-DEL-56
chr1-77432759-DEL-219
chr1-77445119-DEL-103
chr1-77500537-DEL-119
chr1-77513734-DEL-64
chr1-77540486-DEL-94
chr1-77551601-DEL-80
chr1-77582503-DEL-60
chr1-77583825-DEL-74
chr1-77588295-DEL-63
chr1-77639172-DEL-60
chr1-77671137-DEL-85
chr1-77681234-DEL-52
chr1-77682696-DEL-657
chr1-77715270-DEL-125
chr1-77733228-DEL-55
chr1-77753944-DEL-1629
chr1-77773300-DEL-96
chr1-77776634-DEL-321
chr1-77831413-DEL-503
chr1-77832003-DEL-389
chr1-77854932-DEL-58

chr1-77901769-DEL-56
chr1-77937304-DEL-82
chr1-77950069-DEL-63
chr1-77952644-DEL-159
chr1-77976513-DEL-56
chr1-77990404-DEL-104
chr1-77991846-DEL-82
chr1-78028301-DEL-55
chr1-78051764-DEL-175
chr1-78083962-DEL-72
chr1-78131166-DEL-58
chr1-78132910-DEL-51
chr1-78154499-DEL-215
chr1-78156850-DEL-83
chr1-78207717-DEL-120
chr1-78237933-DEL-111
chr1-78281264-DEL-66
chr1-78374979-DEL-66
chr1-78541759-DEL-71
chr1-78557178-DEL-54
chr1-78559724-DEL-146
chr1-78597987-DEL-245
chr1-78601047-DEL-130
chr1-78611565-DEL-278
chr1-78616347-DEL-235
chr1-78633167-DEL-60
chr1-78653338-DEL-1227
chr1-78668284-DEL-52
chr1-78673131-DEL-539
chr1-78690141-DEL-73
chr1-78724696-DEL-78
chr1-78727744-DEL-393
chr1-78739618-DEL-2309
chr1-78744345-DEL-504
chr1-78762880-DEL-1112
chr1-78762886-DEL-844
chr1-78765547-DEL-87
chr1-78777755-DEL-4801
chr1-78777759-DEL-5837
chr1-78783581-DEL-1311
chr1-78802095-DEL-427
chr1-78854530-DEL-160
chr1-78876205-DEL-197
chr1-78889052-DEL-90
chr1-78900795-DEL-53
chr1-78944075-DEL-1739
chr1-78955076-DEL-225
chr1-78969467-DEL-52
chr1-78983643-DEL-51
chr1-79001196-DEL-51
chr1-79004448-DEL-1090
chr1-79008716-DEL-334
chr1-79011976-DEL-3493
chr1-79013154-DEL-334
chr1-79016438-DEL-6738
chr1-79034928-DEL-702
chr1-79047388-DEL-556
chr1-79065198-DEL-2182
chr1-79076471-DEL-201
chr1-79113477-DEL-3420
chr1-79114167-DEL-58
chr1-79119146-DEL-2800
chr1-79155007-DEL-86
chr1-79159606-DEL-1694
chr1-79175236-DEL-144
chr1-79186087-DEL-65
chr1-79214324-DEL-214
chr1-79250244-DEL-164
chr1-79253753-DEL-6152
chr1-79255126-DEL-586
chr1-79266853-DEL-57
chr1-79312590-DEL-194
chr1-79347542-DEL-69
chr1-79424183-DEL-52
chr1-79441840-DEL-85
chr1-79445217-DEL-113
chr1-79450728-DEL-923

chr1-79497951-DEL-190
chr1-79519607-DEL-70
chr1-79570228-DEL-71
chr1-79573485-DEL-338
chr1-79587433-DEL-117
chr1-79606728-DEL-60
chr1-79622746-DEL-2453
chr1-79661554-DEL-3327
chr1-79693270-DEL-7427
chr1-79700702-DEL-2896
chr1-79710996-DEL-298
chr1-79715470-DEL-344
chr1-79731956-DEL-152
chr1-79758244-DEL-55
chr1-79772440-DEL-80
chr1-79773951-DEL-95
chr1-79777990-DEL-68
chr1-79813882-DEL-161
chr1-79924195-DEL-114
chr1-79963979-DEL-402
chr1-79979451-DEL-6459
chr1-80000488-DEL-348
chr1-80023082-DEL-6188
chr1-80091120-DEL-2171
chr1-80104050-DEL-112
chr1-80118222-DEL-6367
chr1-80140166-DEL-567
chr1-80163390-DEL-827
chr1-80189341-DEL-99
chr1-80220905-DEL-238
chr1-80255939-DEL-101
chr1-80260868-DEL-86
chr1-80277748-DEL-127
chr1-80310508-DEL-546
chr1-80324851-DEL-1760
chr1-80335122-DEL-748
chr1-80372572-DEL-125
chr1-80378776-DEL-55
chr1-80408550-DEL-1121
chr1-80445362-DEL-2017
chr1-80515717-DEL-413
chr1-80525810-DEL-245
chr1-80556584-DEL-981
chr1-80584568-DEL-660
chr1-80589281-DEL-805
chr1-80616483-DEL-986
chr1-80665450-DEL-6577
chr1-80682979-DEL-121
chr1-80687586-DEL-144
chr1-80692854-DEL-102
chr1-80695829-DEL-181
chr1-80710348-DEL-443
chr1-80744587-DEL-78
chr1-80752027-DEL-59
chr1-80753799-DEL-67
chr1-80756640-DEL-1599
chr1-80802421-DEL-94
chr1-80806348-DEL-395
chr1-80871206-DEL-553
chr1-80904273-DEL-1420
chr1-80950516-DEL-95
chr1-80976010-DEL-52
chr1-80988192-DEL-329
chr1-81100568-DEL-617
chr1-81116952-DEL-326
chr1-81124432-DEL-110
chr1-81149081-DEL-490
chr1-81150743-DEL-141
chr1-81182988-DEL-165
chr1-81182988-DEL-769
chr1-81184353-DEL-381
chr1-81193385-DEL-60
chr1-81214806-DEL-712
chr1-81234404-DEL-183
chr1-81254592-DEL-350
chr1-81282417-DEL-60
chr1-81288143-DEL-84

chr1-81326163-DEL-82
chr1-81330374-DEL-129
chr1-81340713-DEL-77
chr1-81371181-DEL-5239
chr1-81379486-DEL-187
chr1-81398004-DEL-212
chr1-81421577-DEL-8311
chr1-81430565-DEL-74
chr1-81445703-DEL-1188
chr1-81455270-DEL-1904
chr1-81457673-DEL-54
chr1-81476206-DEL-212
chr1-81502180-DEL-119
chr1-81536276-DEL-1753
chr1-81556825-DEL-434
chr1-81573583-DEL-147
chr1-81582339-DEL-184
chr1-81582891-DEL-112
chr1-81591080-DEL-135
chr1-81597807-DEL-54
chr1-81611896-DEL-75
chr1-81618069-DEL-248
chr1-81637165-DEL-80
chr1-81658247-DEL-51
chr1-81679838-DEL-132
chr1-81683164-DEL-61
chr1-81686993-DEL-75
chr1-81697816-DEL-54
chr1-81719284-DEL-1458
chr1-81739403-DEL-76
chr1-81758860-DEL-367
chr1-81799924-DEL-2443
chr1-81807665-DEL-834
chr1-81812377-DEL-600
chr1-81856798-DEL-588
chr1-81885881-DEL-69
chr1-81897707-DEL-68
chr1-82005593-DEL-120
chr1-82011909-DEL-1220
chr1-82025949-DEL-133
chr1-82036909-DEL-1358
chr1-82049349-DEL-568
chr1-82052428-DEL-174
chr1-82078741-DEL-149
chr1-82081437-DEL-342
chr1-82092393-DEL-84
chr1-82109359-DEL-98
chr1-82130190-DEL-72
chr1-82131565-DEL-1008
chr1-82135412-DEL-496
chr1-82189506-DEL-8122
chr1-82351120-DEL-231
chr1-82351743-DEL-90
chr1-82414304-DEL-460
chr1-82436457-DEL-51
chr1-82460868-DEL-78
chr1-82509818-DEL-675
chr1-82520565-DEL-189
chr1-82530992-DEL-1187
chr1-82539705-DEL-769
chr1-82542161-DEL-90
chr1-82542528-DEL-169
chr1-82546430-DEL-245
chr1-82551515-DEL-117
chr1-82553591-DEL-1872
chr1-82561362-DEL-1636
chr1-82572303-DEL-529
chr1-82584820-DEL-58
chr1-82593809-DEL-58
chr1-82594669-DEL-365
chr1-82617184-DEL-113
chr1-82625411-DEL-74
chr1-82637399-DEL-69
chr1-82639132-DEL-56
chr1-82664677-DEL-1014
chr1-82680781-DEL-3842
chr1-82749540-DEL-313

chr1-82780038-DEL-57
chr1-82805107-DEL-52
chr1-82807388-DEL-63
chr1-82835092-DEL-174
chr1-82848593-DEL-351
chr1-82853190-DEL-77
chr1-82856975-DEL-430
chr1-82910296-DEL-145
chr1-82910776-DEL-283
chr1-82924237-DEL-60
chr1-82942773-DEL-72
chr1-82978482-DEL-147
chr1-82981142-DEL-2784
chr1-82997771-DEL-55
chr1-83007470-DEL-2646
chr1-83021047-DEL-594
chr1-83036904-DEL-54
chr1-83038864-DEL-57
chr1-83071028-DEL-275
chr1-83081063-DEL-84
chr1-83110455-DEL-2826
chr1-83115992-DEL-159
chr1-83141169-DEL-53
chr1-83148724-DEL-70
chr1-83155358-DEL-155
chr1-83158704-DEL-202
chr1-83183891-DEL-62
chr1-83207365-DEL-5820
chr1-83214330-DEL-107
chr1-83278812-DEL-1874
chr1-83293277-DEL-508
chr1-83306687-DEL-583
chr1-83331058-DEL-461
chr1-83360238-DEL-225
chr1-83416830-DEL-60
chr1-83424611-DEL-228
chr1-83444345-DEL-700
chr1-83444500-DEL-1272
chr1-83477150-DEL-2350
chr1-83480326-DEL-230
chr1-83502651-DEL-166
chr1-83550037-DEL-3248
chr1-83587482-DEL-64
chr1-83600811-DEL-2063
chr1-83601945-DEL-62
chr1-83606290-DEL-75
chr1-83650845-DEL-6586
chr1-83707978-DEL-1219
chr1-83717347-DEL-88
chr1-83761568-DEL-66
chr1-83773430-DEL-436
chr1-83831975-DEL-2650
chr1-83888431-DEL-3114
chr1-83901837-DEL-193
chr1-83907295-DEL-220
chr1-83908707-DEL-59
chr1-83908965-DEL-1112
chr1-83913583-DEL-1177
chr1-83931425-DEL-80
chr1-83937940-DEL-59
chr1-84006940-DEL-87
chr1-84009239-DEL-71
chr1-84028420-DEL-241
chr1-84034759-DEL-144
chr1-84048761-DEL-707
chr1-84067921-DEL-51
chr1-84108653-DEL-57
chr1-84114281-DEL-86
chr1-84148427-DEL-59
chr1-84210825-DEL-52
chr1-84233906-DEL-454
chr1-84240073-DEL-104
chr1-84262927-DEL-146
chr1-84270164-DEL-88
chr1-84271810-DEL-225
chr1-84282904-DEL-53
chr1-84286206-DEL-596

chr1-84287233-DEL-219
chr1-84313858-DEL-1562
chr1-84343804-DEL-83
chr1-84362008-DEL-245
chr1-84375756-DEL-59
chr1-84408484-DEL-73
chr1-84422900-DEL-67
chr1-84424034-DEL-585
chr1-84429674-DEL-208
chr1-84438729-DEL-64
chr1-84458331-DEL-343
chr1-84459211-DEL-868
chr1-84485792-DEL-86
chr1-84513364-DEL-65
chr1-84538034-DEL-76
chr1-84641497-DEL-180
chr1-84646595-DEL-870
chr1-84663276-DEL-516
chr1-84663277-DEL-1160
chr1-84663311-DEL-2934
chr1-84663851-DEL-1710
chr1-84664476-DEL-1762
chr1-84689841-DEL-63
chr1-84724635-DEL-74
chr1-84737892-DEL-52
chr1-84742731-DEL-195
chr1-84784004-DEL-335
chr1-84795376-DEL-146
chr1-84797927-DEL-213
chr1-84834782-DEL-495
chr1-84836844-DEL-374
chr1-84837690-DEL-166
chr1-84843136-DEL-201
chr1-84851083-DEL-66
chr1-84966578-DEL-89
chr1-85009983-DEL-3871
chr1-85025863-DEL-259
chr1-85136609-DEL-1494
chr1-85139088-DEL-52
chr1-85198281-DEL-9256
chr1-85215516-DEL-5101
chr1-85244197-DEL-1153
chr1-85250408-DEL-63
chr1-85275566-DEL-192
chr1-85277565-DEL-558
chr1-85304700-DEL-92
chr1-85311650-DEL-6365
chr1-85377760-DEL-185
chr1-85392649-DEL-269
chr1-85395864-DEL-8846
chr1-85414011-DEL-1495
chr1-85435692-DEL-294
chr1-85441086-DEL-102
chr1-85443431-DEL-6665
chr1-85447350-DEL-455
chr1-85470358-DEL-299
chr1-85530992-DEL-660
chr1-85581719-DEL-101
chr1-85583532-DEL-1387
chr1-85595794-DEL-143
chr1-85651673-DEL-1723
chr1-85657073-DEL-86
chr1-85716363-DEL-202
chr1-85720549-DEL-52
chr1-85767303-DEL-60
chr1-85779708-DEL-1003
chr1-85783701-DEL-257
chr1-85788861-DEL-125
chr1-85802657-DEL-1249
chr1-85812461-DEL-299
chr1-85881758-DEL-135
chr1-85903517-DEL-86
chr1-85928883-DEL-282
chr1-85939565-DEL-71
chr1-85941032-DEL-2753
chr1-85963435-DEL-890
chr1-86031136-DEL-79

chr1-86059707-DEL-195
chr1-86069393-DEL-210
chr1-86098388-DEL-210
chr1-86104375-DEL-94
chr1-86156721-DEL-163
chr1-86163324-DEL-810
chr1-86167987-DEL-268
chr1-86184016-DEL-56
chr1-86205773-DEL-73
chr1-86211526-DEL-5677
chr1-86222580-DEL-588
chr1-86225722-DEL-60
chr1-86245292-DEL-2825
chr1-86260981-DEL-93
chr1-86264108-DEL-159
chr1-86281956-DEL-70
chr1-86344417-DEL-88
chr1-86369207-DEL-67
chr1-86383291-DEL-170
chr1-86388635-DEL-469
chr1-86401937-DEL-221
chr1-86406120-DEL-4499
chr1-86440559-DEL-174
chr1-86462175-DEL-681
chr1-86494200-DEL-53
chr1-86524954-DEL-149
chr1-86535313-DEL-66
chr1-86586493-DEL-139
chr1-86610485-DEL-165
chr1-86637024-DEL-313
chr1-86641763-DEL-196
chr1-86644287-DEL-1219
chr1-86646333-DEL-551
chr1-86648959-DEL-91
chr1-86651832-DEL-55
chr1-86678274-DEL-104
chr1-86702855-DEL-6011
chr1-86724543-DEL-67
chr1-86729882-DEL-196
chr1-86756403-DEL-197
chr1-86834508-DEL-783
chr1-86836453-DEL-972
chr1-86849447-DEL-1100
chr1-86903495-DEL-97
chr1-86904359-DEL-59
chr1-86916694-DEL-334
chr1-86945428-DEL-78
chr1-86953217-DEL-203
chr1-87008617-DEL-586
chr1-87058138-DEL-248
chr1-87060894-DEL-132
chr1-87072695-DEL-230
chr1-87105261-DEL-74
chr1-87106492-DEL-80
chr1-87124337-DEL-73
chr1-87130013-DEL-249
chr1-87174435-DEL-186
chr1-87187636-DEL-78
chr1-87194884-DEL-121
chr1-87239607-DEL-108
chr1-87259867-DEL-77
chr1-87264910-DEL-179
chr1-87348477-DEL-93
chr1-87349434-DEL-174
chr1-87365696-DEL-229
chr1-87382581-DEL-111
chr1-87397608-DEL-60
chr1-87406320-DEL-75
chr1-87449130-DEL-153
chr1-87477808-DEL-52
chr1-87561512-DEL-200
chr1-87586178-DEL-3184
chr1-87601255-DEL-209
chr1-87602812-DEL-86
chr1-87609956-DEL-314
chr1-87631628-DEL-74
chr1-87637506-DEL-52

chr1-87688339-DEL-768
chr1-87697477-DEL-97
chr1-87700323-DEL-506
chr1-87733166-DEL-65
chr1-87748765-DEL-191
chr1-87750012-DEL-435
chr1-87750522-DEL-172
chr1-87767011-DEL-67
chr1-87772955-DEL-66
chr1-87801451-DEL-68
chr1-87802454-DEL-84
chr1-87817044-DEL-87
chr1-87827008-DEL-951
chr1-87830425-DEL-765
chr1-87855595-DEL-99
chr1-87859928-DEL-118
chr1-87919736-DEL-56
chr1-88016812-DEL-490
chr1-88017338-DEL-53
chr1-88036472-DEL-52
chr1-88115489-DEL-54
chr1-88136507-DEL-61
chr1-88136776-DEL-1196
chr1-88161904-DEL-69
chr1-88163436-DEL-544
chr1-88198507-DEL-506
chr1-88250263-DEL-55
chr1-88272798-DEL-64
chr1-88313420-DEL-302
chr1-88328135-DEL-134
chr1-88331377-DEL-1283
chr1-88359086-DEL-82
chr1-88363487-DEL-172
chr1-88387853-DEL-75
chr1-88410991-DEL-75
chr1-88430869-DEL-258
chr1-88457355-DEL-135
chr1-88474043-DEL-403
chr1-88482185-DEL-132
chr1-88499307-DEL-297
chr1-88499897-DEL-1558
chr1-88543771-DEL-525
chr1-88603133-DEL-194
chr1-88604480-DEL-59
chr1-88640797-DEL-137
chr1-88644155-DEL-196
chr1-88742661-DEL-400
chr1-88747079-DEL-76
chr1-88750239-DEL-186
chr1-88771068-DEL-75
chr1-88795682-DEL-100
chr1-88798074-DEL-56
chr1-88813082-DEL-536
chr1-88867362-DEL-54
chr1-88898593-DEL-728
chr1-88923250-DEL-237
chr1-89080322-DEL-92
chr1-89086747-DEL-88
chr1-89144342-DEL-6995
chr1-89255945-DEL-63
chr1-89267214-DEL-64
chr1-89298202-DEL-156
chr1-89310751-DEL-62
chr1-89323913-DEL-51
chr1-89325713-DEL-107
chr1-89339130-DEL-201
chr1-89339524-DEL-733
chr1-89355767-DEL-80
chr1-89385943-DEL-155
chr1-89390093-DEL-108
chr1-89411929-DEL-79
chr1-89429563-DEL-1033
chr1-89430512-DEL-96
chr1-89430608-DEL-417
chr1-89442773-DEL-1459
chr1-89489363-DEL-255
chr1-89496454-DEL-316

chr1-89553338-DEL-464
chr1-89559096-DEL-312
chr1-89573824-DEL-80
chr1-89579180-DEL-57
chr1-89609075-DEL-2400
chr1-89609579-DEL-719
chr1-89665988-DEL-68
chr1-89695489-DEL-619
chr1-89734295-DEL-55
chr1-89790918-DEL-60
chr1-89809864-DEL-91
chr1-89831397-DEL-203
chr1-89873363-DEL-733
chr1-89874515-DEL-203
chr1-89885836-DEL-62
chr1-89911393-DEL-89
chr1-89911850-DEL-199
chr1-89927947-DEL-214
chr1-89985325-DEL-431
chr1-89992949-DEL-148
chr1-89995251-DEL-177
chr1-90008807-DEL-200
chr1-90009353-DEL-198
chr1-90017645-DEL-460
chr1-90019153-DEL-175
chr1-90033191-DEL-2143
chr1-90038960-DEL-60
chr1-90089917-DEL-94
chr1-90109452-DEL-155
chr1-90116812-DEL-210
chr1-90118246-DEL-6208
chr1-90120216-DEL-88
chr1-90164114-DEL-401
chr1-90197845-DEL-210
chr1-90205351-DEL-65
chr1-90219319-DEL-62
chr1-90229739-DEL-90
chr1-90244096-DEL-463
chr1-90278721-DEL-64
chr1-90289109-DEL-231
chr1-90300308-DEL-96
chr1-90303494-DEL-114
chr1-90309824-DEL-229
chr1-90328078-DEL-568
chr1-90328258-DEL-2433
chr1-90346542-DEL-124
chr1-90373135-DEL-3198
chr1-90373862-DEL-268
chr1-90373990-DEL-1654
chr1-90374384-DEL-160
chr1-90376118-DEL-133
chr1-90490690-DEL-79
chr1-90505197-DEL-324
chr1-90517599-DEL-463
chr1-90542046-DEL-1149
chr1-90544841-DEL-86
chr1-90548456-DEL-54
chr1-90548689-DEL-9825
chr1-90560754-DEL-318
chr1-90622274-DEL-433
chr1-90626655-DEL-133
chr1-90663771-DEL-102
chr1-90684142-DEL-144
chr1-90703600-DEL-62
chr1-90735899-DEL-1325
chr1-90736056-DEL-115
chr1-90748886-DEL-252
chr1-90759975-DEL-125
chr1-90791718-DEL-141
chr1-90797645-DEL-67
chr1-90832778-DEL-84
chr1-90841397-DEL-242
chr1-90859475-DEL-169
chr1-90877870-DEL-144
chr1-90889078-DEL-82
chr1-90892108-DEL-127
chr1-90905066-DEL-309

chr1-90912550-DEL-2133
chr1-90944386-DEL-102
chr1-90951218-DEL-468
chr1-91070661-DEL-56
chr1-91083615-DEL-187
chr1-91129793-DEL-126
chr1-91172565-DEL-60
chr1-91183443-DEL-124
chr1-91214463-DEL-109
chr1-91231388-DEL-202
chr1-91307270-DEL-496
chr1-91322710-DEL-55
chr1-91349905-DEL-495
chr1-91350182-DEL-117
chr1-91390430-DEL-130
chr1-91393356-DEL-68
chr1-91420049-DEL-60
chr1-91421767-DEL-118
chr1-91466177-DEL-61
chr1-91469682-DEL-78
chr1-91517181-DEL-125
chr1-91521352-DEL-67
chr1-91522145-DEL-89
chr1-91559535-DEL-100
chr1-91569289-DEL-235
chr1-91576511-DEL-347
chr1-91577413-DEL-207
chr1-91609804-DEL-527
chr1-91610590-DEL-506
chr1-91617171-DEL-64
chr1-91646704-DEL-214
chr1-91672135-DEL-57
chr1-91709808-DEL-175
chr1-91732608-DEL-78
chr1-91925203-DEL-96
chr1-91952369-DEL-1097
chr1-92020715-DEL-3595
chr1-92038068-DEL-72
chr1-92047857-DEL-884
chr1-92053269-DEL-1120
chr1-92169219-DEL-74
chr1-92172037-DEL-1443
chr1-92202424-DEL-55
chr1-92226650-DEL-84
chr1-92231533-DEL-67
chr1-92233748-DEL-1202
chr1-92251259-DEL-79
chr1-92258687-DEL-52
chr1-92334692-DEL-1679
chr1-92346976-DEL-208
chr1-92396763-DEL-207
chr1-92404190-DEL-78
chr1-92427764-DEL-1800
chr1-92444881-DEL-177
chr1-92612435-DEL-171
chr1-92620366-DEL-54
chr1-92628192-DEL-70
chr1-92629498-DEL-119
chr1-92695779-DEL-216
chr1-92737818-DEL-53
chr1-92815695-DEL-55
chr1-92819539-DEL-191
chr1-92841157-DEL-60
chr1-92859458-DEL-82
chr1-92884514-DEL-331
chr1-92903094-DEL-443
chr1-92905074-DEL-1049
chr1-92972574-DEL-92
chr1-92976925-DEL-68
chr1-93014960-DEL-1067
chr1-93017574-DEL-109
chr1-93031576-DEL-52
chr1-93047142-DEL-51
chr1-93047737-DEL-114
chr1-93057071-DEL-169
chr1-93057642-DEL-92
chr1-93089166-DEL-184

chr1-93185196-DEL-87
chr1-93255880-DEL-60
chr1-93310577-DEL-168
chr1-93349368-DEL-894
chr1-93360306-DEL-590
chr1-93377973-DEL-860
chr1-93386707-DEL-62
chr1-93417976-DEL-801
chr1-93451703-DEL-212
chr1-93466569-DEL-210
chr1-93470145-DEL-1280
chr1-93471495-DEL-93
chr1-93524370-DEL-86
chr1-93538233-DEL-69
chr1-93557803-DEL-1757
chr1-93573775-DEL-169
chr1-93577013-DEL-60
chr1-93587263-DEL-1735
chr1-93653196-DEL-652
chr1-93692323-DEL-75
chr1-93699157-DEL-72
chr1-93717641-DEL-306
chr1-93719448-DEL-68
chr1-93723938-DEL-1487
chr1-93735369-DEL-68
chr1-93761759-DEL-347
chr1-93766863-DEL-6492
chr1-93775400-DEL-56
chr1-93784573-DEL-493
chr1-93791165-DEL-7426
chr1-93821145-DEL-73
chr1-93829800-DEL-2426
chr1-93838027-DEL-69
chr1-93847691-DEL-844
chr1-93871941-DEL-1031
chr1-93871956-DEL-848
chr1-93884298-DEL-4719
chr1-93893418-DEL-495
chr1-93911265-DEL-841
chr1-93912255-DEL-206
chr1-93924965-DEL-161
chr1-93928346-DEL-239
chr1-93958858-DEL-67
chr1-93993849-DEL-189
chr1-93995406-DEL-201
chr1-94005214-DEL-55
chr1-94006433-DEL-171
chr1-94015019-DEL-91
chr1-94028645-DEL-8673
chr1-94028799-DEL-64
chr1-94030934-DEL-4707
chr1-94030936-DEL-2489
chr1-94056207-DEL-216
chr1-94068235-DEL-2563
chr1-94086866-DEL-79
chr1-94105355-DEL-203
chr1-94112462-DEL-209
chr1-94133426-DEL-258
chr1-94140526-DEL-4691
chr1-94143456-DEL-178
chr1-94147879-DEL-54
chr1-94152518-DEL-187
chr1-94156000-DEL-5677
chr1-94163666-DEL-6410
chr1-94169767-DEL-201
chr1-94177059-DEL-81
chr1-94180110-DEL-55
chr1-94182260-DEL-6660
chr1-94201228-DEL-534
chr1-94238026-DEL-51
chr1-94258455-DEL-910
chr1-94268550-DEL-53
chr1-94272698-DEL-52
chr1-94273713-DEL-68
chr1-94278066-DEL-600
chr1-94305024-DEL-164
chr1-94306172-DEL-179

chr1-94307486-DEL-990
chr1-94315798-DEL-6616
chr1-94327132-DEL-6249
chr1-94357113-DEL-224
chr1-94383564-DEL-197
chr1-94410613-DEL-56
chr1-94452189-DEL-78
chr1-94462083-DEL-327
chr1-94467302-DEL-59
chr1-94470650-DEL-130
chr1-94492776-DEL-1105
chr1-94494459-DEL-195
chr1-94511543-DEL-2155
chr1-94511863-DEL-113
chr1-94515791-DEL-2708
chr1-94518287-DEL-181
chr1-94546079-DEL-487
chr1-94557948-DEL-54
chr1-94572864-DEL-60
chr1-94592315-DEL-112
chr1-94610282-DEL-109
chr1-94618803-DEL-2055
chr1-94670351-DEL-204
chr1-94702047-DEL-51
chr1-94714154-DEL-101
chr1-94727752-DEL-79
chr1-94728848-DEL-77
chr1-94730616-DEL-65
chr1-94733025-DEL-371
chr1-94734417-DEL-688
chr1-94739110-DEL-221
chr1-94757444-DEL-128
chr1-94760155-DEL-78
chr1-94763421-DEL-1149
chr1-94764079-DEL-70
chr1-94793165-DEL-82
chr1-94811724-DEL-80
chr1-94828275-DEL-59
chr1-94843485-DEL-349
chr1-94851785-DEL-60
chr1-94868470-DEL-3602
chr1-94882529-DEL-262
chr1-94889688-DEL-615
chr1-94892811-DEL-51
chr1-94895667-DEL-129
chr1-94900440-DEL-93
chr1-95000104-DEL-58
chr1-95011221-DEL-525
chr1-95021285-DEL-359
chr1-95050066-DEL-53
chr1-95055968-DEL-1809
chr1-95075486-DEL-170
chr1-95091830-DEL-132
chr1-95096603-DEL-358
chr1-95108753-DEL-281
chr1-95111008-DEL-855
chr1-95135415-DEL-380
chr1-95149933-DEL-77
chr1-95200641-DEL-2605
chr1-95218653-DEL-105
chr1-95230027-DEL-1863
chr1-95230056-DEL-226
chr1-95282387-DEL-161
chr1-95283574-DEL-1209
chr1-95296085-DEL-60
chr1-95309880-DEL-1522
chr1-95340370-DEL-548
chr1-95383254-DEL-52
chr1-95384383-DEL-789
chr1-95404973-DEL-1073
chr1-95405020-DEL-386
chr1-95411392-DEL-1085
chr1-95429431-DEL-328
chr1-95459787-DEL-57
chr1-95470622-DEL-2491
chr1-95485858-DEL-564
chr1-95492655-DEL-80

chr1-95509938-DEL-1601
chr1-95527917-DEL-571
chr1-95530639-DEL-55
chr1-95532501-DEL-455
chr1-95538873-DEL-3763
chr1-95540101-DEL-91
chr1-95548833-DEL-664
chr1-95573901-DEL-2590
chr1-95577163-DEL-57
chr1-95600394-DEL-450
chr1-95621388-DEL-68
chr1-95640991-DEL-831
chr1-95655047-DEL-318
chr1-95685170-DEL-217
chr1-95711317-DEL-200
chr1-95723947-DEL-1539
chr1-95725197-DEL-55
chr1-95731422-DEL-2586
chr1-95785475-DEL-139
chr1-95810659-DEL-360
chr1-95816923-DEL-56
chr1-95818974-DEL-80
chr1-95822192-DEL-3473
chr1-95823488-DEL-704
chr1-95827087-DEL-943
chr1-95829084-DEL-81
chr1-95834376-DEL-1158
chr1-95849952-DEL-80
chr1-95852354-DEL-5554
chr1-95879874-DEL-1890
chr1-95880399-DEL-2270
chr1-95883886-DEL-60
chr1-95887960-DEL-69
chr1-95944494-DEL-204
chr1-95962904-DEL-261
chr1-95973772-DEL-397
chr1-95992193-DEL-1551
chr1-96010092-DEL-478
chr1-96012096-DEL-510
chr1-96021181-DEL-6218
chr1-96047817-DEL-450
chr1-96066767-DEL-432
chr1-96073284-DEL-63
chr1-96116542-DEL-183
chr1-96134535-DEL-89
chr1-96148300-DEL-64
chr1-96159168-DEL-77
chr1-96163756-DEL-64
chr1-96167637-DEL-67
chr1-96185897-DEL-2793
chr1-96199127-DEL-77
chr1-96206863-DEL-77
chr1-96255830-DEL-6337
chr1-96262296-DEL-52
chr1-96268868-DEL-85
chr1-96271078-DEL-146
chr1-96272819-DEL-88
chr1-96286268-DEL-801
chr1-96291444-DEL-64
chr1-96293689-DEL-405
chr1-96313421-DEL-52
chr1-96319277-DEL-3630
chr1-96345486-DEL-108
chr1-96357485-DEL-55
chr1-96362995-DEL-401
chr1-96366469-DEL-6179
chr1-96396015-DEL-358
chr1-96398594-DEL-4654
chr1-96447545-DEL-64
chr1-96481853-DEL-614
chr1-96489621-DEL-123
chr1-96510649-DEL-1413
chr1-96519722-DEL-97
chr1-96521659-DEL-577
chr1-96533955-DEL-1188
chr1-96542433-DEL-75
chr1-96544808-DEL-1755

chr1-96570102-DEL-52
chr1-96572026-DEL-544
chr1-96586829-DEL-60
chr1-96648671-DEL-1252
chr1-96668177-DEL-5400
chr1-96672588-DEL-996
chr1-96679816-DEL-1384
chr1-96690039-DEL-74
chr1-96708073-DEL-4478
chr1-96732513-DEL-219
chr1-96738410-DEL-729
chr1-96785762-DEL-908
chr1-96794237-DEL-184
chr1-96819120-DEL-849
chr1-96841022-DEL-98
chr1-96851256-DEL-64
chr1-96864828-DEL-65
chr1-96866465-DEL-854
chr1-96866519-DEL-149
chr1-96872160-DEL-324
chr1-96879603-DEL-551
chr1-96886044-DEL-89
chr1-96894091-DEL-4016
chr1-96903946-DEL-4808
chr1-96910372-DEL-4347
chr1-96918294-DEL-68
chr1-96944679-DEL-103
chr1-96969432-DEL-1105
chr1-96975525-DEL-306
chr1-96985161-DEL-67
chr1-96991370-DEL-104
chr1-97050361-DEL-74
chr1-97085447-DEL-2790
chr1-97118006-DEL-92
chr1-97128236-DEL-1090
chr1-97202475-DEL-798
chr1-97206983-DEL-6367
chr1-97226845-DEL-658
chr1-97259753-DEL-53
chr1-97288264-DEL-472
chr1-97302933-DEL-788
chr1-97315110-DEL-71
chr1-97320032-DEL-6195
chr1-97353532-DEL-1149
chr1-97355666-DEL-2602
chr1-97358762-DEL-108
chr1-97402348-DEL-2608
chr1-97434534-DEL-6640
chr1-97442493-DEL-906
chr1-97450974-DEL-390
chr1-97485762-DEL-108
chr1-97560515-DEL-186
chr1-97591637-DEL-60
chr1-97625922-DEL-100
chr1-97632130-DEL-96
chr1-97638894-DEL-84
chr1-97675442-DEL-200
chr1-97699745-DEL-777
chr1-97714817-DEL-122
chr1-97727345-DEL-288
chr1-97733684-DEL-189
chr1-97744982-DEL-54
chr1-97746611-DEL-705
chr1-97758635-DEL-200
chr1-97847172-DEL-370
chr1-97879096-DEL-836
chr1-97937555-DEL-170
chr1-97950779-DEL-577
chr1-97962035-DEL-276
chr1-97965584-DEL-64
chr1-97968686-DEL-816
chr1-98003921-DEL-128
chr1-98029784-DEL-4253
chr1-98046693-DEL-1986
chr1-98066477-DEL-351
chr1-98070501-DEL-146
chr1-98091820-DEL-51

chr1-98094551-DEL-73
chr1-98143722-DEL-1179
chr1-98157072-DEL-154
chr1-98168627-DEL-980
chr1-98174169-DEL-163
chr1-98178629-DEL-359
chr1-98183492-DEL-545
chr1-98194861-DEL-476
chr1-98196438-DEL-83
chr1-98270047-DEL-420
chr1-98270180-DEL-1691
chr1-98271993-DEL-911
chr1-98287057-DEL-1453
chr1-98302877-DEL-165
chr1-98312854-DEL-6786
chr1-98313015-DEL-428
chr1-98326535-DEL-72
chr1-98337558-DEL-54
chr1-98357333-DEL-483
chr1-98365844-DEL-79
chr1-98396113-DEL-99
chr1-98411697-DEL-245
chr1-98420167-DEL-887
chr1-98436544-DEL-53
chr1-98463129-DEL-147
chr1-98468620-DEL-51
chr1-98475817-DEL-83
chr1-98494089-DEL-177
chr1-98496272-DEL-344
chr1-98514600-DEL-5231
chr1-98517698-DEL-56
chr1-98523174-DEL-817
chr1-98524089-DEL-367
chr1-98524737-DEL-233
chr1-98533499-DEL-60
chr1-98552770-DEL-1393
chr1-98559632-DEL-1423
chr1-98578799-DEL-208
chr1-98580258-DEL-95
chr1-98584710-DEL-2585
chr1-98593344-DEL-113
chr1-98616222-DEL-385
chr1-98625999-DEL-54
chr1-98670942-DEL-2206
chr1-98677778-DEL-2697
chr1-98680303-DEL-200
chr1-98681777-DEL-324
chr1-98682492-DEL-372
chr1-98684051-DEL-63
chr1-98693896-DEL-6487
chr1-98707861-DEL-134
chr1-98711871-DEL-336
chr1-98713966-DEL-144
chr1-98723012-DEL-647
chr1-98724126-DEL-4364
chr1-98731330-DEL-201
chr1-98737067-DEL-62
chr1-98758875-DEL-56
chr1-98764889-DEL-300
chr1-98774671-DEL-381
chr1-98801054-DEL-1998
chr1-98803031-DEL-83
chr1-98810817-DEL-88
chr1-98810886-DEL-6482
chr1-98818542-DEL-680
chr1-98823337-DEL-6206
chr1-98869824-DEL-902
chr1-98873666-DEL-675
chr1-98885862-DEL-1762
chr1-98909715-DEL-964
chr1-98915878-DEL-210
chr1-98918339-DEL-62
chr1-98920853-DEL-160
chr1-98922095-DEL-56
chr1-98979681-DEL-76
chr1-99003014-DEL-1194
chr1-99018137-DEL-64

chr1-99062834-DEL-213
chr1-99106430-DEL-2236
chr1-99108035-DEL-127
chr1-99127568-DEL-391
chr1-99129435-DEL-8485
chr1-99142329-DEL-295
chr1-99163983-DEL-310
chr1-99177134-DEL-506
chr1-99178922-DEL-1214
chr1-99181169-DEL-6170
chr1-99190281-DEL-4786
chr1-99201432-DEL-2118
chr1-99216875-DEL-474
chr1-99217850-DEL-2855
chr1-99264394-DEL-63
chr1-99269283-DEL-329
chr1-99269860-DEL-54
chr1-99295150-DEL-767
chr1-99297754-DEL-165
chr1-99312706-DEL-148
chr1-99321608-DEL-56
chr1-99336811-DEL-100
chr1-99339194-DEL-6232
chr1-99365792-DEL-139
chr1-99365924-DEL-1034
chr1-99369629-DEL-1625
chr1-99397284-DEL-1827
chr1-99403790-DEL-2972
chr1-99422935-DEL-791
chr1-99443879-DEL-55
chr1-99448788-DEL-55
chr1-99473873-DEL-396
chr1-99578347-DEL-174
chr1-99578846-DEL-518
chr1-99587300-DEL-61
chr1-99602959-DEL-58
chr1-99611554-DEL-696
chr1-99618602-DEL-107
chr1-99618912-DEL-1408
chr1-99621629-DEL-627
chr1-99678207-DEL-334
chr1-99690742-DEL-908
chr1-99720434-DEL-58
chr1-99751686-DEL-201
chr1-99757749-DEL-870
chr1-99758630-DEL-3488
chr1-99783881-DEL-510
chr1-99785822-DEL-5109
chr1-99787154-DEL-129
chr1-99811393-DEL-148
chr1-99824927-DEL-851
chr1-99827077-DEL-372
chr1-99848975-DEL-5983
chr1-99855591-DEL-58
chr1-99869964-DEL-1010
chr1-99880625-DEL-516
chr1-99890169-DEL-85
chr1-99899419-DEL-51
chr1-99903646-DEL-1260
chr1-99962399-DEL-4577
chr1-99981496-DEL-515
chr1-100006352-DEL-148
chr1-100007735-DEL-58
chr1-100008083-DEL-794
chr1-100025912-DEL-70
chr1-100027097-DEL-229
chr1-100074793-DEL-536
chr1-100080449-DEL-56
chr1-100101704-DEL-1602
chr1-100115274-DEL-170
chr1-100127177-DEL-1295
chr1-100142214-DEL-2272
chr1-100163466-DEL-124
chr1-100188288-DEL-265
chr1-100197112-DEL-328
chr1-100199995-DEL-71
chr1-100203597-DEL-116

chr1-100228431-DEL-5946
chr1-100271573-DEL-222
chr1-100273610-DEL-316
chr1-100284786-DEL-112
chr1-100311817-DEL-197
chr1-100316137-DEL-226
chr1-100325117-DEL-110
chr1-100326404-DEL-293
chr1-100336705-DEL-8534
chr1-100360991-DEL-2452
chr1-100365300-DEL-1207
chr1-100375983-DEL-58
chr1-100377679-DEL-1949
chr1-100433595-DEL-103
chr1-100456672-DEL-2813
chr1-100472802-DEL-351
chr1-100475251-DEL-560
chr1-100480869-DEL-334
chr1-100482286-DEL-84
chr1-100487306-DEL-6187
chr1-100519730-DEL-71
chr1-100538251-DEL-260
chr1-100550972-DEL-86
chr1-100560863-DEL-212
chr1-100563535-DEL-6741
chr1-100609405-DEL-1687
chr1-100614236-DEL-1518
chr1-100616725-DEL-79
chr1-100618832-DEL-57
chr1-100621919-DEL-396
chr1-100622753-DEL-120
chr1-100635056-DEL-186
chr1-100648068-DEL-6796
chr1-100651480-DEL-2551
chr1-100671324-DEL-78
chr1-100673918-DEL-6391
chr1-100682043-DEL-416
chr1-100689377-DEL-8139
chr1-100691694-DEL-776
chr1-100699323-DEL-68
chr1-100703202-DEL-276
chr1-100703603-DEL-7954
chr1-100743863-DEL-781
chr1-100745968-DEL-72
chr1-100821979-DEL-389
chr1-100841318-DEL-487
chr1-100845465-DEL-187
chr1-100845877-DEL-4332
chr1-100863147-DEL-98
chr1-100896174-DEL-52
chr1-100908056-DEL-6598
chr1-100914633-DEL-7636
chr1-100917322-DEL-5166
chr1-100939539-DEL-217
chr1-100963486-DEL-91
chr1-100965394-DEL-51
chr1-100975389-DEL-946
chr1-100986004-DEL-275
chr1-100993098-DEL-388
chr1-101004900-DEL-57
chr1-101014532-DEL-9270
chr1-101017060-DEL-4156
chr1-101031205-DEL-6026
chr1-101031206-DEL-5205
chr1-101031423-DEL-2605
chr1-101046194-DEL-3037
chr1-101070296-DEL-64
chr1-101086245-DEL-61
chr1-101096966-DEL-1360
chr1-101101030-DEL-2073
chr1-101122238-DEL-76
chr1-101124560-DEL-2745
chr1-101149427-DEL-189
chr1-101165239-DEL-1984
chr1-101169324-DEL-9474
chr1-101182774-DEL-358
chr1-101204532-DEL-436

chr1-101212050-DEL-618
chr1-101214742-DEL-5090
chr1-101228758-DEL-197
chr1-101248933-DEL-296
chr1-101270619-DEL-107
chr1-101303234-DEL-104
chr1-101306217-DEL-734
chr1-101312359-DEL-457
chr1-101319295-DEL-80
chr1-101321208-DEL-744
chr1-101322158-DEL-5059
chr1-101340908-DEL-66
chr1-101347234-DEL-63
chr1-101359410-DEL-52
chr1-101362447-DEL-62
chr1-101375940-DEL-764
chr1-101381462-DEL-101
chr1-101399540-DEL-486
chr1-101410531-DEL-1978
chr1-101416606-DEL-213
chr1-101436759-DEL-156
chr1-101445194-DEL-4830
chr1-101449381-DEL-192
chr1-101467913-DEL-159
chr1-101495971-DEL-390
chr1-101498542-DEL-736
chr1-101510689-DEL-600
chr1-101511916-DEL-88
chr1-101554885-DEL-109
chr1-101555616-DEL-63
chr1-101558496-DEL-194
chr1-101563052-DEL-1719
chr1-101571134-DEL-260
chr1-101571278-DEL-1234
chr1-101579620-DEL-734
chr1-101592626-DEL-576
chr1-101598621-DEL-66
chr1-101614345-DEL-116
chr1-101627492-DEL-60
chr1-101630182-DEL-437
chr1-101657605-DEL-1148
chr1-101698550-DEL-333
chr1-101716489-DEL-1660
chr1-101734943-DEL-93
chr1-101739884-DEL-1146
chr1-101743300-DEL-53
chr1-101744981-DEL-136
chr1-101763799-DEL-173
chr1-101772506-DEL-1874
chr1-101782225-DEL-71
chr1-101804382-DEL-51
chr1-101806631-DEL-129
chr1-101807996-DEL-2265
chr1-101811619-DEL-2318
chr1-101812623-DEL-401
chr1-101815226-DEL-3849
chr1-101825593-DEL-5161
chr1-101834612-DEL-109
chr1-101836067-DEL-98
chr1-101841965-DEL-290
chr1-101855167-DEL-12602
chr1-101856083-DEL-1424
chr1-101857646-DEL-5359
chr1-101857892-DEL-483
chr1-101860026-DEL-160
chr1-101860450-DEL-1037
chr1-101861614-DEL-1379
chr1-101862379-DEL-208
chr1-101870952-DEL-174
chr1-101872045-DEL-1161
chr1-101873631-DEL-5067
chr1-101884660-DEL-52
chr1-101902951-DEL-51
chr1-101903518-DEL-129
chr1-101929805-DEL-6130
chr1-101934061-DEL-403
chr1-101937580-DEL-8973

chr1-101949242-DEL-291
chr1-101981673-DEL-2424
chr1-101991158-DEL-529
chr1-101998802-DEL-55
chr1-102007373-DEL-6078
chr1-102013790-DEL-6859
chr1-102025164-DEL-53
chr1-102043230-DEL-71
chr1-102083692-DEL-398
chr1-102106641-DEL-103
chr1-102159424-DEL-95
chr1-102159802-DEL-1598
chr1-102164724-DEL-4080
chr1-102166544-DEL-7154
chr1-102183162-DEL-846
chr1-102183630-DEL-323
chr1-102196379-DEL-2335
chr1-102199257-DEL-1019
chr1-102223198-DEL-180
chr1-102232579-DEL-155
chr1-102236767-DEL-17995
chr1-102263773-DEL-428
chr1-102284073-DEL-3221
chr1-102288551-DEL-4833
chr1-102300449-DEL-426
chr1-102301821-DEL-80
chr1-102311107-DEL-91
chr1-102312347-DEL-1056
chr1-102330658-DEL-99
chr1-102366064-DEL-1237
chr1-102370967-DEL-1874
chr1-102374350-DEL-52
chr1-102474190-DEL-1459
chr1-102526882-DEL-4068
chr1-102564194-DEL-2123
chr1-102585443-DEL-86
chr1-102597784-DEL-56
chr1-102599959-DEL-72
chr1-102627480-DEL-52
chr1-102657647-DEL-1448
chr1-102672881-DEL-2805
chr1-102673123-DEL-147
chr1-102675768-DEL-1981
chr1-102675811-DEL-2528
chr1-102677788-DEL-1721
chr1-102678352-DEL-1168
chr1-102726406-DEL-174
chr1-102744297-DEL-7354
chr1-102752985-DEL-189
chr1-102806849-DEL-311
chr1-102816032-DEL-3981
chr1-102832435-DEL-486
chr1-102860733-DEL-52
chr1-102865355-DEL-525
chr1-102866224-DEL-4447
chr1-102867170-DEL-1428
chr1-102868999-DEL-3067
chr1-102877242-DEL-68
chr1-102885450-DEL-4552
chr1-102890700-DEL-6439
chr1-102917456-DEL-260
chr1-102941379-DEL-2508
chr1-102946012-DEL-124
chr1-102959384-DEL-7134
chr1-102980642-DEL-84
chr1-102991835-DEL-1919
chr1-103022222-DEL-106
chr1-103022324-DEL-739
chr1-103024596-DEL-896
chr1-103038923-DEL-6797
chr1-103045189-DEL-208
chr1-103058138-DEL-4872
chr1-103066681-DEL-112
chr1-103097689-DEL-105
chr1-103105784-DEL-3062
chr1-103132573-DEL-9932
chr1-103144773-DEL-73

chr1-103151945-DEL-519
chr1-103190745-DEL-101
chr1-103218987-DEL-2529
chr1-103225075-DEL-253
chr1-103226923-DEL-138
chr1-103229899-DEL-120
chr1-103245596-DEL-8272
chr1-103278074-DEL-174
chr1-103288559-DEL-57
chr1-103298739-DEL-212
chr1-103299667-DEL-1647
chr1-103317282-DEL-141
chr1-103322446-DEL-15414
chr1-103331334-DEL-6200
chr1-103335065-DEL-8127
chr1-103340939-DEL-6669
chr1-103355605-DEL-55
chr1-103356756-DEL-1584
chr1-103400852-DEL-122
chr1-103410970-DEL-1708
chr1-103411092-DEL-90
chr1-103416507-DEL-1870
chr1-103430320-DEL-117
chr1-103437443-DEL-4216
chr1-103440934-DEL-1948
chr1-103446699-DEL-100
chr1-103455570-DEL-84
chr1-103461519-DEL-347
chr1-103484299-DEL-186
chr1-103494136-DEL-4615
chr1-103497862-DEL-739
chr1-103540066-DEL-91
chr1-103558858-DEL-151
chr1-103561734-DEL-376
chr1-103566074-DEL-1912
chr1-103578985-DEL-72
chr1-103603510-DEL-119
chr1-103620967-DEL-450
chr1-103621712-DEL-112
chr1-103623594-DEL-1382
chr1-103627456-DEL-68
chr1-103640735-DEL-744
chr1-103651610-DEL-207
chr1-103698577-DEL-1185
chr1-103701177-DEL-993
chr1-103702053-DEL-179
chr1-103726894-DEL-75
chr1-103768640-DEL-109
chr1-103772383-DEL-262
chr1-103775706-DEL-124
chr1-103816029-DEL-58
chr1-103817817-DEL-654
chr1-103853911-DEL-208
chr1-103862759-DEL-1890
chr1-103878069-DEL-1660
chr1-103881268-DEL-170
chr1-103917498-DEL-1871
chr1-103920344-DEL-288
chr1-103944509-DEL-683
chr1-103945405-DEL-340
chr1-103952824-DEL-228
chr1-103956072-DEL-1192
chr1-103989144-DEL-108
chr1-103990630-DEL-1123
chr1-103993599-DEL-60
chr1-104013293-DEL-3631
chr1-104016211-DEL-208
chr1-104017945-DEL-220
chr1-104020520-DEL-1871
chr1-104022372-DEL-100
chr1-104022788-DEL-579
chr1-104025737-DEL-6213
chr1-104032247-DEL-290
chr1-104037818-DEL-883
chr1-104043993-DEL-698
chr1-104051338-DEL-67
chr1-104065535-DEL-288

chr1-104073656-DEL-58
chr1-104079344-DEL-760
chr1-104086476-DEL-6474
chr1-104107029-DEL-540
chr1-104108542-DEL-6059
chr1-104119628-DEL-117
chr1-104121882-DEL-170
chr1-104129580-DEL-58
chr1-104165653-DEL-214
chr1-104194535-DEL-6252
chr1-104212016-DEL-1209
chr1-104216935-DEL-6944
chr1-104223911-DEL-1694
chr1-104267635-DEL-103
chr1-104269376-DEL-4959
chr1-104295490-DEL-276
chr1-104303614-DEL-79
chr1-104307160-DEL-810
chr1-104310643-DEL-6214
chr1-104327261-DEL-1183
chr1-104345114-DEL-1707
chr1-104365083-DEL-1626
chr1-104406431-DEL-65
chr1-104410573-DEL-88
chr1-104413868-DEL-938
chr1-104426276-DEL-496
chr1-104445964-DEL-78
chr1-104449572-DEL-254
chr1-104485810-DEL-59
chr1-104524861-DEL-59
chr1-104562671-DEL-744
chr1-104583229-DEL-2865
chr1-104583240-DEL-60
chr1-104634891-DEL-284
chr1-104647846-DEL-126
chr1-104656098-DEL-170
chr1-104664447-DEL-2779
chr1-104686925-DEL-52
chr1-104709493-DEL-134
chr1-104714886-DEL-76
chr1-104748135-DEL-525
chr1-104761702-DEL-1259
chr1-104763202-DEL-620
chr1-104768595-DEL-88
chr1-104790442-DEL-1880
chr1-104798549-DEL-114
chr1-104848515-DEL-119
chr1-104897777-DEL-51
chr1-104941364-DEL-13742
chr1-104941425-DEL-99
chr1-104970490-DEL-162
chr1-104970763-DEL-1592
chr1-104970989-DEL-1514
chr1-104972441-DEL-213
chr1-104972799-DEL-2872
chr1-104974235-DEL-1190
chr1-104984570-DEL-2122
chr1-104988068-DEL-593
chr1-104992921-DEL-96
chr1-105017866-DEL-73
chr1-105048918-DEL-201
chr1-105057766-DEL-69
chr1-105063732-DEL-60
chr1-105067617-DEL-6161
chr1-105134704-DEL-180
chr1-105140497-DEL-134
chr1-105244540-DEL-64
chr1-105259618-DEL-3410
chr1-105259995-DEL-98
chr1-105262065-DEL-87
chr1-105294754-DEL-58
chr1-105298047-DEL-53
chr1-105298165-DEL-885
chr1-105310191-DEL-65
chr1-105341883-DEL-170
chr1-105356121-DEL-62
chr1-105399068-DEL-6401

chr1-105413397-DEL-338
chr1-105482486-DEL-70
chr1-105482810-DEL-1782
chr1-105490742-DEL-252
chr1-105493145-DEL-229
chr1-105533952-DEL-3335
chr1-105545818-DEL-491
chr1-105561182-DEL-411
chr1-105593468-DEL-485
chr1-105625659-DEL-4171
chr1-105626463-DEL-155
chr1-105707504-DEL-124
chr1-105713321-DEL-71
chr1-105726360-DEL-115
chr1-105743127-DEL-695
chr1-105777722-DEL-64
chr1-105783749-DEL-1186
chr1-105801350-DEL-1570
chr1-105815630-DEL-400
chr1-105845272-DEL-107
chr1-105860070-DEL-152
chr1-105884125-DEL-122
chr1-105890552-DEL-842
chr1-105977787-DEL-113
chr1-105990799-DEL-137
chr1-106001383-DEL-113
chr1-106006716-DEL-635
chr1-106023671-DEL-345
chr1-106052332-DEL-76
chr1-106055643-DEL-82
chr1-106057042-DEL-228
chr1-106067820-DEL-357
chr1-106070111-DEL-355
chr1-106086470-DEL-55
chr1-106094981-DEL-230
chr1-106096542-DEL-82
chr1-106146091-DEL-106
chr1-106176769-DEL-1290
chr1-106178721-DEL-324
chr1-106215453-DEL-58
chr1-106229245-DEL-827
chr1-106260377-DEL-52
chr1-106271836-DEL-108
chr1-106310512-DEL-898
chr1-106310706-DEL-273
chr1-106337853-DEL-132
chr1-106357349-DEL-58
chr1-106388632-DEL-115
chr1-106394695-DEL-129
chr1-106400674-DEL-1605
chr1-106400966-DEL-600
chr1-106438317-DEL-1074
chr1-106443342-DEL-61
chr1-106538207-DEL-178
chr1-106541736-DEL-758
chr1-106588752-DEL-350
chr1-106626052-DEL-71
chr1-106643580-DEL-160
chr1-106645633-DEL-145
chr1-106667411-DEL-210
chr1-106670929-DEL-233
chr1-106730247-DEL-3228
chr1-106757591-DEL-133
chr1-106758009-DEL-1901
chr1-106782584-DEL-212
chr1-106790893-DEL-306
chr1-106836691-DEL-57
chr1-106856123-DEL-334
chr1-106865602-DEL-56
chr1-106891534-DEL-138
chr1-106900785-DEL-192
chr1-106931492-DEL-421
chr1-106934255-DEL-108
chr1-106944400-DEL-65
chr1-106963048-DEL-7064
chr1-106986070-DEL-1230
chr1-107017114-DEL-399

chr1-107025630-DEL-187
chr1-107026305-DEL-83
chr1-107048668-DEL-6275
chr1-107144865-DEL-2280
chr1-107156943-DEL-73
chr1-107208468-DEL-60
chr1-107216173-DEL-3487
chr1-107217005-DEL-3898
chr1-107225098-DEL-569
chr1-107246181-DEL-410
chr1-107325063-DEL-405
chr1-107382614-DEL-425
chr1-107402848-DEL-138
chr1-107403395-DEL-3001
chr1-107418469-DEL-79
chr1-107445259-DEL-1668
chr1-107492583-DEL-600
chr1-107572790-DEL-54
chr1-107614866-DEL-1659
chr1-107667920-DEL-333
chr1-107667974-DEL-342
chr1-107689780-DEL-692
chr1-107717554-DEL-88
chr1-107739013-DEL-512
chr1-107843962-DEL-62
chr1-107888178-DEL-231
chr1-107999310-DEL-1433
chr1-108006739-DEL-760
chr1-108020491-DEL-651
chr1-108030708-DEL-94
chr1-108055587-DEL-70
chr1-108083165-DEL-76
chr1-108111710-DEL-589
chr1-108119124-DEL-70
chr1-108181775-DEL-1882
chr1-108241906-DEL-150
chr1-108293996-DEL-61
chr1-108300188-DEL-708
chr1-108313195-DEL-1714
chr1-108324283-DEL-5934
chr1-108326047-DEL-1287
chr1-108327745-DEL-165
chr1-108329410-DEL-609
chr1-108336579-DEL-7190
chr1-108368902-DEL-1136
chr1-108377428-DEL-148
chr1-108421791-DEL-551
chr1-108432553-DEL-2128
chr1-108443240-DEL-57
chr1-108466202-DEL-332
chr1-108479809-DEL-335
chr1-108486964-DEL-58
chr1-108498455-DEL-124
chr1-108518796-DEL-269
chr1-108535101-DEL-60
chr1-108539165-DEL-52
chr1-108552305-DEL-6471
chr1-108560260-DEL-87
chr1-108616247-DEL-1334
chr1-108622917-DEL-807
chr1-108627851-DEL-184
chr1-108638857-DEL-1209
chr1-108641801-DEL-63
chr1-108650639-DEL-1107
chr1-108675136-DEL-1838
chr1-108690122-DEL-1101
chr1-108697568-DEL-74
chr1-108704470-DEL-497
chr1-108705651-DEL-254
chr1-108716033-DEL-242
chr1-108730975-DEL-2336
chr1-108734809-DEL-552
chr1-108749760-DEL-55
chr1-108764938-DEL-208
chr1-108780012-DEL-3453
chr1-108806813-DEL-997
chr1-108808777-DEL-68

chr1-108815376-DEL-126
chr1-108825698-DEL-72
chr1-108833024-DEL-68
chr1-108855402-DEL-715
chr1-108875870-DEL-75
chr1-108884538-DEL-3778
chr1-108934120-DEL-937
chr1-108935305-DEL-212
chr1-108956640-DEL-1060
chr1-108957189-DEL-189
chr1-108970102-DEL-56
chr1-108972978-DEL-272
chr1-109007033-DEL-64
chr1-109010209-DEL-1111
chr1-109022813-DEL-71
chr1-109039253-DEL-519
chr1-109043271-DEL-66
chr1-109043846-DEL-91
chr1-109056099-DEL-8633
chr1-109080166-DEL-932
chr1-109081376-DEL-487
chr1-109101358-DEL-1860
chr1-109109340-DEL-239
chr1-109123675-DEL-1234
chr1-109128592-DEL-357
chr1-109142147-DEL-58
chr1-109155743-DEL-86
chr1-109178691-DEL-1751
chr1-109180699-DEL-2212
chr1-109183626-DEL-1584
chr1-109185893-DEL-129
chr1-109191879-DEL-6254
chr1-109213536-DEL-597
chr1-109217262-DEL-221
chr1-109221090-DEL-135
chr1-109231475-DEL-64
chr1-109267497-DEL-52
chr1-109304368-DEL-380
chr1-109325247-DEL-52
chr1-109328345-DEL-767
chr1-109366379-DEL-108
chr1-109370166-DEL-2141
chr1-109414793-DEL-67
chr1-109429795-DEL-1769
chr1-109431378-DEL-207
chr1-109431623-DEL-2909
chr1-109461187-DEL-76
chr1-109516986-DEL-2990
chr1-109523001-DEL-81
chr1-109553048-DEL-74
chr1-109581394-DEL-8387
chr1-109581546-DEL-330
chr1-109609961-DEL-408
chr1-109610320-DEL-1918
chr1-109641672-DEL-52
chr1-109651792-DEL-6388
chr1-109662987-DEL-58
chr1-109666287-DEL-960
chr1-109696254-DEL-2707
chr1-109718229-DEL-64
chr1-109722802-DEL-264
chr1-109727401-DEL-95
chr1-109774863-DEL-882
chr1-109782199-DEL-63
chr1-109787753-DEL-1051
chr1-109795756-DEL-1757
chr1-109810674-DEL-240
chr1-109851245-DEL-1267
chr1-109872796-DEL-129
chr1-109875575-DEL-771
chr1-110002816-DEL-152
chr1-110004110-DEL-179
chr1-110032918-DEL-68
chr1-110034427-DEL-152
chr1-110042545-DEL-156
chr1-110043062-DEL-1582
chr1-110051941-DEL-4712

chr1-110082963-DEL-51
chr1-110084627-DEL-476
chr1-110087455-DEL-84
chr1-110090417-DEL-679
chr1-110107253-DEL-183
chr1-110112772-DEL-3605
chr1-110128386-DEL-59
chr1-110138649-DEL-1233
chr1-110158711-DEL-3294
chr1-110180364-DEL-1041
chr1-110185073-DEL-73
chr1-110206522-DEL-58
chr1-110208427-DEL-113
chr1-110216180-DEL-6471
chr1-110231278-DEL-500
chr1-110251398-DEL-52
chr1-110252403-DEL-1769
chr1-110272036-DEL-5172
chr1-110283440-DEL-909
chr1-110299618-DEL-144
chr1-110302397-DEL-74
chr1-110304957-DEL-523
chr1-110313098-DEL-91
chr1-110317081-DEL-155
chr1-110318164-DEL-1845
chr1-110340938-DEL-251
chr1-110352065-DEL-181
chr1-110362304-DEL-348
chr1-110363648-DEL-102
chr1-110367471-DEL-388
chr1-110385702-DEL-117
chr1-110395330-DEL-1684
chr1-110401337-DEL-69
chr1-110412569-DEL-76
chr1-110417760-DEL-373
chr1-110418453-DEL-78
chr1-110419708-DEL-171
chr1-110438268-DEL-5986
chr1-110457182-DEL-673
chr1-110463790-DEL-55
chr1-110469593-DEL-6196
chr1-110488749-DEL-323
chr1-110491919-DEL-76
chr1-110537183-DEL-65
chr1-110556723-DEL-1626
chr1-110566418-DEL-1522
chr1-110587551-DEL-12378
chr1-110605737-DEL-6146
chr1-110616209-DEL-6630
chr1-110622569-DEL-201
chr1-110629543-DEL-451
chr1-110633407-DEL-64
chr1-110640916-DEL-74
chr1-110643399-DEL-681
chr1-110663855-DEL-1164
chr1-110663862-DEL-70
chr1-110676586-DEL-523
chr1-110700280-DEL-910
chr1-110729534-DEL-630
chr1-110735331-DEL-5779
chr1-110759476-DEL-818
chr1-110766230-DEL-115
chr1-110776587-DEL-5571
chr1-110787997-DEL-2151
chr1-110788968-DEL-104
chr1-110814106-DEL-213
chr1-110819557-DEL-56
chr1-110886664-DEL-2084
chr1-110901745-DEL-181
chr1-110904803-DEL-124
chr1-110912858-DEL-657
chr1-110921421-DEL-6145
chr1-110938824-DEL-120
chr1-110943286-DEL-1610
chr1-110952562-DEL-494
chr1-110964736-DEL-259
chr1-110968545-DEL-80

chr1-110973275-DEL-1713
chr1-110992523-DEL-243
chr1-111001477-DEL-59
chr1-111004881-DEL-129
chr1-111007898-DEL-250
chr1-111015676-DEL-299
chr1-111040087-DEL-1236
chr1-111051755-DEL-66
chr1-111058527-DEL-1601
chr1-111084439-DEL-867
chr1-111096492-DEL-936
chr1-111103061-DEL-1992
chr1-111129011-DEL-1138
chr1-111148412-DEL-68
chr1-111153786-DEL-5597
chr1-111155068-DEL-3345
chr1-111165717-DEL-2088
chr1-111194538-DEL-68
chr1-111200300-DEL-6438
chr1-111220237-DEL-204
chr1-111221847-DEL-674
chr1-111253056-DEL-62
chr1-111254487-DEL-3709
chr1-111259403-DEL-392
chr1-111314737-DEL-60
chr1-111314838-DEL-5627
chr1-111346999-DEL-58
chr1-111356643-DEL-2408
chr1-111357253-DEL-2408
chr1-111360005-DEL-4029
chr1-111374826-DEL-1131
chr1-111408044-DEL-119
chr1-111416564-DEL-2093
chr1-111428243-DEL-316
chr1-111452855-DEL-318
chr1-111467457-DEL-7537
chr1-111475174-DEL-1101
chr1-111489944-DEL-6228
chr1-111491370-DEL-2187
chr1-111496773-DEL-228
chr1-111509717-DEL-76
chr1-111554056-DEL-6456
chr1-111606367-DEL-5734
chr1-111618962-DEL-5493
chr1-111627443-DEL-1131
chr1-111631019-DEL-7182
chr1-111661590-DEL-113
chr1-111663126-DEL-84
chr1-111676773-DEL-785
chr1-111681398-DEL-1793
chr1-111714408-DEL-2936
chr1-111733598-DEL-584
chr1-111736722-DEL-679
chr1-111852762-DEL-80
chr1-111854094-DEL-974
chr1-111857502-DEL-804
chr1-111899988-DEL-4718
chr1-111946908-DEL-515
chr1-111954805-DEL-724
chr1-111957462-DEL-270
chr1-111969175-DEL-806
chr1-111974757-DEL-510
chr1-111983562-DEL-202
chr1-112004161-DEL-65
chr1-112031215-DEL-108
chr1-112069598-DEL-117
chr1-112082216-DEL-2039
chr1-112101836-DEL-1688
chr1-112105264-DEL-651
chr1-112119562-DEL-78
chr1-112125474-DEL-549
chr1-112127426-DEL-834
chr1-112130099-DEL-1837
chr1-112151986-DEL-1827
chr1-112156701-DEL-1298
chr1-112160818-DEL-6705
chr1-112160823-DEL-2118

chr1-112160824-DEL-4236
chr1-112192096-DEL-7293
chr1-112200167-DEL-123
chr1-112239282-DEL-106
chr1-112254041-DEL-56
chr1-112293474-DEL-79
chr1-112295146-DEL-3644
chr1-112299049-DEL-1976
chr1-112333829-DEL-311
chr1-112345900-DEL-127
chr1-112367859-DEL-4546
chr1-112380686-DEL-70
chr1-112401118-DEL-54
chr1-112403909-DEL-201
chr1-112404606-DEL-816
chr1-112427256-DEL-513
chr1-112428593-DEL-73
chr1-112444498-DEL-230
chr1-112452243-DEL-779
chr1-112466074-DEL-1094
chr1-112469589-DEL-342
chr1-112478204-DEL-6639
chr1-112491913-DEL-5287
chr1-112505140-DEL-6133
chr1-112506035-DEL-5171
chr1-112579899-DEL-6396
chr1-112588131-DEL-106
chr1-112594071-DEL-120
chr1-112644042-DEL-3051
chr1-112659241-DEL-52
chr1-112664720-DEL-107
chr1-112667413-DEL-358
chr1-112690493-DEL-203
chr1-112699402-DEL-2707
chr1-112725601-DEL-5098
chr1-112731897-DEL-2700
chr1-112754166-DEL-1586
chr1-112758297-DEL-440
chr1-112771921-DEL-8561
chr1-112779441-DEL-61
chr1-112781529-DEL-60
chr1-112792503-DEL-6162
chr1-112799342-DEL-146
chr1-112802831-DEL-514
chr1-112807881-DEL-4042
chr1-112811160-DEL-381
chr1-112843840-DEL-70
chr1-112878155-DEL-52
chr1-112879971-DEL-108
chr1-112890579-DEL-615
chr1-112952359-DEL-179
chr1-112960204-DEL-3394
chr1-112967293-DEL-125
chr1-112968804-DEL-975
chr1-112973800-DEL-60
chr1-112985617-DEL-242
chr1-113015458-DEL-276
chr1-113018062-DEL-1697
chr1-113025315-DEL-56
chr1-113027121-DEL-104
chr1-113046490-DEL-5285
chr1-113062843-DEL-1155
chr1-113064950-DEL-258
chr1-113074179-DEL-88
chr1-113098427-DEL-77
chr1-113154848-DEL-88
chr1-113226791-DEL-54
chr1-113280905-DEL-117
chr1-113282671-DEL-106
chr1-113289025-DEL-859
chr1-113293977-DEL-2445
chr1-113328747-DEL-66
chr1-113343082-DEL-83
chr1-113347462-DEL-2338
chr1-113350230-DEL-1353
chr1-113410405-DEL-66
chr1-113437622-DEL-486

chr1-113443882-DEL-1268
chr1-113445843-DEL-458
chr1-113486484-DEL-155
chr1-113488605-DEL-305
chr1-113497170-DEL-4092
chr1-113505156-DEL-85
chr1-113507200-DEL-66
chr1-113510307-DEL-52
chr1-113557780-DEL-56
chr1-113558467-DEL-170
chr1-113596533-DEL-67
chr1-113601547-DEL-2114
chr1-113613868-DEL-1736
chr1-113616217-DEL-690
chr1-113620789-DEL-272
chr1-113639601-DEL-170
chr1-113639983-DEL-6743
chr1-113648865-DEL-1621
chr1-113649295-DEL-1621
chr1-113660996-DEL-880
chr1-113672293-DEL-6385
chr1-113684948-DEL-6272
chr1-113703261-DEL-70
chr1-113707735-DEL-82
chr1-113743544-DEL-209
chr1-113744744-DEL-350
chr1-113748010-DEL-1229
chr1-113784691-DEL-991
chr1-113808445-DEL-110
chr1-113851839-DEL-3007
chr1-113856868-DEL-53
chr1-113860606-DEL-990
chr1-113864770-DEL-1133
chr1-113878614-DEL-106
chr1-113884898-DEL-58
chr1-113896311-DEL-522
chr1-113921410-DEL-99
chr1-113931247-DEL-344
chr1-113931990-DEL-365
chr1-113980990-DEL-366
chr1-113987278-DEL-919
chr1-113991973-DEL-5932
chr1-113999284-DEL-436
chr1-114031512-DEL-2364
chr1-114034946-DEL-242
chr1-114036051-DEL-2055
chr1-114038346-DEL-5755
chr1-114039544-DEL-2871
chr1-114044157-DEL-5036
chr1-114062113-DEL-207
chr1-114080635-DEL-5921
chr1-114097777-DEL-1228
chr1-114121923-DEL-100
chr1-114132268-DEL-2175
chr1-114148827-DEL-72
chr1-114150140-DEL-55
chr1-114151025-DEL-220
chr1-114179717-DEL-349
chr1-114188185-DEL-2044
chr1-114197059-DEL-145
chr1-114227729-DEL-4312
chr1-114237823-DEL-436
chr1-114246615-DEL-969
chr1-114249214-DEL-1273
chr1-114252243-DEL-473
chr1-114253020-DEL-75
chr1-114275442-DEL-54
chr1-114298235-DEL-3595
chr1-114301684-DEL-314
chr1-114303280-DEL-418
chr1-114304099-DEL-109
chr1-114329770-DEL-381
chr1-114331200-DEL-2298
chr1-114346258-DEL-410
chr1-114360241-DEL-284
chr1-114362920-DEL-218
chr1-114427729-DEL-6223

chr1-114436591-DEL-3279
chr1-114453540-DEL-90
chr1-114504091-DEL-328
chr1-114510185-DEL-1317
chr1-114546357-DEL-124
chr1-114555582-DEL-225
chr1-114567517-DEL-84
chr1-114574229-DEL-106
chr1-114582349-DEL-780
chr1-114594407-DEL-95
chr1-114596909-DEL-106
chr1-114597195-DEL-2008
chr1-114612895-DEL-373
chr1-114614433-DEL-62
chr1-114653115-DEL-580
chr1-114676609-DEL-90
chr1-114692215-DEL-3800
chr1-114696039-DEL-2861
chr1-114700872-DEL-131
chr1-114703443-DEL-422
chr1-114706411-DEL-119
chr1-114716808-DEL-509
chr1-114720893-DEL-52
chr1-114727594-DEL-80
chr1-114758442-DEL-1557
chr1-114759952-DEL-147
chr1-114764126-DEL-226
chr1-114766851-DEL-479
chr1-114808929-DEL-53
chr1-114821008-DEL-1108
chr1-114838888-DEL-392
chr1-114873057-DEL-52
chr1-114883003-DEL-102
chr1-114892065-DEL-630
chr1-114936574-DEL-73
chr1-114952080-DEL-983
chr1-114967013-DEL-1182
chr1-114969986-DEL-1264
chr1-114991281-DEL-429
chr1-114991984-DEL-95
chr1-114994325-DEL-72
chr1-114995545-DEL-830
chr1-114998022-DEL-60
chr1-115019459-DEL-2892
chr1-115027981-DEL-6370
chr1-115028077-DEL-201
chr1-115034345-DEL-5229
chr1-115043389-DEL-159
chr1-115048986-DEL-571
chr1-115056603-DEL-951
chr1-115059876-DEL-338
chr1-115065272-DEL-2885
chr1-115083304-DEL-1627
chr1-115102101-DEL-150
chr1-115103377-DEL-1040
chr1-115114644-DEL-134
chr1-115138519-DEL-467
chr1-115146401-DEL-381
chr1-115165665-DEL-439
chr1-115175823-DEL-96
chr1-115202371-DEL-58
chr1-115217198-DEL-1034
chr1-115238500-DEL-509
chr1-115258612-DEL-1444
chr1-115259963-DEL-92
chr1-115267445-DEL-957
chr1-115315994-DEL-112
chr1-115321376-DEL-2779
chr1-115337787-DEL-1396
chr1-115360270-DEL-1535
chr1-115366533-DEL-637
chr1-115368931-DEL-66
chr1-115370325-DEL-1859
chr1-115420591-DEL-130
chr1-115427951-DEL-80
chr1-115429841-DEL-627
chr1-115434749-DEL-743

chr1-115437985-DEL-267
chr1-115448437-DEL-473
chr1-115453276-DEL-246
chr1-115471532-DEL-147
chr1-115480542-DEL-785
chr1-115495938-DEL-4104
chr1-115503312-DEL-445
chr1-115510219-DEL-1246
chr1-115536495-DEL-6159
chr1-115570035-DEL-101
chr1-115581000-DEL-92
chr1-115601935-DEL-597
chr1-115648353-DEL-1301
chr1-115661084-DEL-80
chr1-115745233-DEL-516
chr1-115778595-DEL-60
chr1-115790256-DEL-2772
chr1-115793067-DEL-1149
chr1-115799184-DEL-105
chr1-115818126-DEL-75
chr1-115822073-DEL-335
chr1-115823052-DEL-57
chr1-115824521-DEL-380
chr1-115829323-DEL-1315
chr1-115833278-DEL-1999
chr1-115895887-DEL-1226
chr1-115903097-DEL-58
chr1-115935542-DEL-255
chr1-115970395-DEL-180
chr1-115984434-DEL-128
chr1-116009084-DEL-139
chr1-116039180-DEL-62
chr1-116052551-DEL-103
chr1-116089223-DEL-923
chr1-116134458-DEL-700
chr1-116136648-DEL-81
chr1-116191643-DEL-657
chr1-116243740-DEL-251
chr1-116272877-DEL-56
chr1-116280901-DEL-76
chr1-116333252-DEL-3315
chr1-116390927-DEL-59
chr1-116392918-DEL-5801
chr1-116441677-DEL-1176
chr1-116443211-DEL-307
chr1-116471109-DEL-64
chr1-116472031-DEL-587
chr1-116488315-DEL-781
chr1-116524161-DEL-123
chr1-116531032-DEL-184
chr1-116538501-DEL-64
chr1-116571647-DEL-27260
chr1-116588920-DEL-7147
chr1-116597405-DEL-1275
chr1-116609109-DEL-424
chr1-116626276-DEL-92
chr1-116646047-DEL-146
chr1-116675545-DEL-365
chr1-116676831-DEL-910
chr1-116677928-DEL-808
chr1-116678260-DEL-3456
chr1-116684492-DEL-6873
chr1-116696067-DEL-6749
chr1-116696079-DEL-4096
chr1-116700882-DEL-2343
chr1-116732986-DEL-3553
chr1-116752757-DEL-68
chr1-116754484-DEL-183
chr1-116781424-DEL-210
chr1-116795673-DEL-405
chr1-116824894-DEL-6258
chr1-116864199-DEL-2424
chr1-116894403-DEL-53
chr1-116935763-DEL-140
chr1-116960260-DEL-342
chr1-116978006-DEL-241
chr1-116978917-DEL-536

chr1-116981411-DEL-599
chr1-116983052-DEL-1974
chr1-117024188-DEL-1240
chr1-117027482-DEL-144
chr1-117044535-DEL-71
chr1-117073886-DEL-778
chr1-117095202-DEL-212
chr1-117118189-DEL-5054
chr1-117131680-DEL-6341
chr1-117141866-DEL-2238
chr1-117144183-DEL-4109
chr1-117156027-DEL-1038
chr1-117159524-DEL-6307
chr1-117167781-DEL-59
chr1-117175317-DEL-399
chr1-117181311-DEL-205
chr1-117216170-DEL-626
chr1-117223103-DEL-2546
chr1-117256897-DEL-74
chr1-117290367-DEL-306
chr1-117303698-DEL-645
chr1-117307056-DEL-212
chr1-117307970-DEL-731
chr1-117309414-DEL-323
chr1-117325119-DEL-442
chr1-117354489-DEL-54
chr1-117366025-DEL-3574
chr1-117374929-DEL-202
chr1-117384384-DEL-86
chr1-117384620-DEL-502
chr1-117413760-DEL-211
chr1-117420383-DEL-408
chr1-117429450-DEL-54
chr1-117431269-DEL-1788
chr1-117444058-DEL-205
chr1-117463855-DEL-51
chr1-117469002-DEL-53
chr1-117511459-DEL-1326
chr1-117515040-DEL-6469
chr1-117538423-DEL-124
chr1-117538654-DEL-308
chr1-117552555-DEL-147
chr1-117553645-DEL-1092
chr1-117559990-DEL-2592
chr1-117564459-DEL-6186
chr1-117574704-DEL-54
chr1-117646465-DEL-787
chr1-117649346-DEL-1666
chr1-117669911-DEL-637
chr1-117673473-DEL-640
chr1-117674897-DEL-2675
chr1-117728948-DEL-168
chr1-117735484-DEL-243
chr1-117743721-DEL-79
chr1-117768895-DEL-356
chr1-117773257-DEL-69
chr1-117774050-DEL-644
chr1-117775571-DEL-65
chr1-117788688-DEL-207
chr1-117790102-DEL-1339
chr1-117801794-DEL-10816
chr1-117809397-DEL-201
chr1-117813268-DEL-197
chr1-117838775-DEL-2341
chr1-117842507-DEL-283
chr1-117871730-DEL-589
chr1-117938695-DEL-474
chr1-117956065-DEL-6195
chr1-118010607-DEL-400
chr1-118024819-DEL-207
chr1-118032325-DEL-348
chr1-118034461-DEL-700
chr1-118038341-DEL-177
chr1-118056816-DEL-84
chr1-118070060-DEL-4806
chr1-118080628-DEL-150
chr1-118086568-DEL-204

chr1-118176985-DEL-197
chr1-118205882-DEL-1519
chr1-118219171-DEL-206
chr1-118221427-DEL-144
chr1-118231242-DEL-443
chr1-118241366-DEL-202
chr1-118250961-DEL-133
chr1-118269381-DEL-191
chr1-118277055-DEL-166
chr1-118279539-DEL-116
chr1-118300474-DEL-180
chr1-118303139-DEL-66
chr1-118305123-DEL-87
chr1-118307522-DEL-84
chr1-118379657-DEL-1537
chr1-118394309-DEL-196
chr1-118417829-DEL-1381
chr1-118418674-DEL-1393
chr1-118424974-DEL-116
chr1-118447478-DEL-59
chr1-118486593-DEL-184
chr1-118541044-DEL-272
chr1-118597285-DEL-65
chr1-118617895-DEL-143
chr1-118637543-DEL-86
chr1-118668660-DEL-109
chr1-118694124-DEL-89
chr1-118698914-DEL-67
chr1-118703924-DEL-107
chr1-118751820-DEL-51
chr1-118788258-DEL-51
chr1-118861264-DEL-209
chr1-118883883-DEL-975
chr1-118898977-DEL-729
chr1-118902250-DEL-71
chr1-118917773-DEL-86
chr1-118924483-DEL-52
chr1-118928262-DEL-60
chr1-118936340-DEL-57
chr1-118967458-DEL-108
chr1-118990154-DEL-146
chr1-118991801-DEL-63
chr1-119008426-DEL-117
chr1-119029605-DEL-54
chr1-119038192-DEL-52
chr1-119042328-DEL-54
chr1-119046403-DEL-170
chr1-119056233-DEL-92
chr1-119081105-DEL-61
chr1-119128328-DEL-175
chr1-119167303-DEL-80
chr1-119208461-DEL-118
chr1-119239641-DEL-56
chr1-119262865-DEL-370
chr1-119267976-DEL-57
chr1-119286891-DEL-66
chr1-119296411-DEL-68
chr1-119323035-DEL-106
chr1-119416567-DEL-364
chr1-119500670-DEL-92
chr1-119575020-DEL-70
chr1-119599908-DEL-209
chr1-119645133-DEL-658
chr1-119658052-DEL-6118
chr1-119672040-DEL-6489
chr1-119699466-DEL-190
chr1-119745415-DEL-150
chr1-119776337-DEL-495
chr1-119777911-DEL-86
chr1-119782941-DEL-293
chr1-119783419-DEL-1397
chr1-119784920-DEL-1519
chr1-119808624-DEL-723
chr1-119843302-DEL-175
chr1-119858141-DEL-116
chr1-119867956-DEL-4898
chr1-119879539-DEL-196

chr1-119880221-DEL-80
chr1-119887274-DEL-64
chr1-119911397-DEL-60
chr1-119916551-DEL-149
chr1-119917947-DEL-69
chr1-119961985-DEL-120
chr1-119968123-DEL-98
chr1-119974865-DEL-85
chr1-119979305-DEL-77
chr1-119980421-DEL-58
chr1-119987857-DEL-414
chr1-120064497-DEL-1301
chr1-120085479-DEL-1576
chr1-120087340-DEL-249
chr1-120153188-DEL-90
chr1-120155899-DEL-181
chr1-120161029-DEL-154
chr1-120166589-DEL-191
chr1-120195562-DEL-458
chr1-120211308-DEL-80
chr1-120255217-DEL-56
chr1-120306292-DEL-456
chr1-120335518-DEL-868
chr1-120348978-DEL-55
chr1-120387278-DEL-98
chr1-120413209-DEL-63
chr1-120423531-DEL-90
chr1-120457079-DEL-66
chr1-120473714-DEL-91
chr1-120497812-DEL-8060
chr1-120586063-DEL-760
chr1-120625595-DEL-120
chr1-120627713-DEL-209
chr1-120635623-DEL-1245
chr1-120760656-DEL-63
chr1-120765103-DEL-106
chr1-120830125-DEL-60
chr1-120830589-DEL-4904
chr1-120834380-DEL-1636
chr1-120855026-DEL-120
chr1-120856507-DEL-389
chr1-120879166-DEL-54
chr1-120920450-DEL-54
chr1-120939468-DEL-131
chr1-120944382-DEL-666
chr1-120971728-DEL-63
chr1-120998443-DEL-77
chr1-121005253-DEL-65
chr1-121019731-DEL-924
chr1-121042050-DEL-537
chr1-121092118-DEL-59
chr1-121092663-DEL-142
chr1-121118229-DEL-76
chr1-121163675-DEL-61
chr1-121182816-DEL-65
chr1-121192352-DEL-99
chr1-121286254-DEL-719
chr1-121286929-DEL-63
chr1-121291431-DEL-83
chr1-121297408-DEL-68
chr1-121304525-DEL-452
chr1-121331728-DEL-195
chr1-121366076-DEL-353
chr1-121411159-DEL-780
chr1-121411160-DEL-104
chr1-121411783-DEL-962
chr1-121412343-DEL-182
chr1-121442557-DEL-59
chr1-121470895-DEL-560
chr1-121496611-DEL-92
chr1-121530223-DEL-55
chr1-121572975-DEL-245
chr1-121626628-DEL-64
chr1-121631838-DEL-370
chr1-121681871-DEL-51
chr1-121688581-DEL-510
chr1-121725327-DEL-64

chr1-121727300-DEL-79
chr1-121732617-DEL-105
chr1-121794033-DEL-235
chr1-121800138-DEL-66
chr1-121839515-DEL-73
chr1-121862870-DEL-91
chr1-121867783-DEL-94
chr1-121877864-DEL-114
chr1-121881601-DEL-701
chr1-121882590-DEL-288
chr1-121888517-DEL-204
chr1-121894354-DEL-56
chr1-121899850-DEL-509
chr1-121901788-DEL-99
chr1-121923092-DEL-239
chr1-121928325-DEL-64
chr1-121942216-DEL-80
chr1-121973647-DEL-896
chr1-121977764-DEL-439
chr1-122000684-DEL-401
chr1-122028017-DEL-289
chr1-122076804-DEL-403
chr1-122091093-DEL-838
chr1-122104350-DEL-107
chr1-122121545-DEL-99
chr1-122132658-DEL-457
chr1-122140818-DEL-1188
chr1-122170347-DEL-78
chr1-122186279-DEL-59
chr1-122203811-DEL-71
chr1-122211821-DEL-120
chr1-122219014-DEL-525
chr1-122243147-DEL-430
chr1-122246221-DEL-448
chr1-122259827-DEL-64
chr1-122273630-DEL-67
chr1-122296959-DEL-128
chr1-122300363-DEL-54
chr1-122305442-DEL-3363
chr1-122338465-DEL-248
chr1-122345277-DEL-282
chr1-122348680-DEL-1263
chr1-122371600-DEL-6284
chr1-122411933-DEL-78
chr1-122431013-DEL-100
chr1-122435644-DEL-190
chr1-122438943-DEL-498
chr1-122453042-DEL-467
chr1-122483145-DEL-401
chr1-122485331-DEL-559
chr1-122487737-DEL-98
chr1-122489530-DEL-65
chr1-122502640-DEL-444
chr1-122533335-DEL-346
chr1-122534360-DEL-512
chr1-122543474-DEL-449
chr1-122544170-DEL-861
chr1-122560899-DEL-519
chr1-122566489-DEL-174
chr1-122599461-DEL-3455
chr1-122599475-DEL-6218
chr1-122602945-DEL-2747
chr1-122641291-DEL-4113
chr1-122643228-DEL-5104
chr1-122646091-DEL-985
chr1-122648974-DEL-3333
chr1-122673700-DEL-158
chr1-122749145-DEL-563
chr1-122772695-DEL-54
chr1-122776944-DEL-231
chr1-122835940-DEL-239
chr1-122846592-DEL-66
chr1-122866551-DEL-567
chr1-122870996-DEL-531
chr1-122900508-DEL-381
chr1-122917357-DEL-4972
chr1-122966306-DEL-363

chr1-122973983-DEL-62
chr1-122988969-DEL-96
chr1-123006548-DEL-3399
chr1-123006785-DEL-468
chr1-123007594-DEL-667
chr1-123008606-DEL-1339
chr1-123017154-DEL-64
chr1-123023078-DEL-58
chr1-123031549-DEL-127
chr1-123075949-DEL-571
chr1-123084153-DEL-85
chr1-123123767-DEL-6015
chr1-123132780-DEL-214
chr1-123143567-DEL-1456
chr1-123151722-DEL-1490
chr1-123185338-DEL-8098
chr1-123202561-DEL-130
chr1-123206283-DEL-763
chr1-123211102-DEL-51
chr1-123223841-DEL-2551
chr1-123226791-DEL-176
chr1-123228010-DEL-650
chr1-123230341-DEL-204
chr1-123242814-DEL-1251
chr1-123272212-DEL-2575
chr1-123276524-DEL-101
chr1-123303630-DEL-118
chr1-123307267-DEL-56
chr1-123314087-DEL-60
chr1-123315126-DEL-66
chr1-123344791-DEL-75
chr1-123384367-DEL-195
chr1-123397225-DEL-247
chr1-123404958-DEL-103
chr1-123412436-DEL-336
chr1-123416794-DEL-220
chr1-123442127-DEL-2345
chr1-123478951-DEL-75
chr1-123484993-DEL-365
chr1-123489456-DEL-123
chr1-123495245-DEL-955
chr1-123496216-DEL-247
chr1-123527982-DEL-619
chr1-123550694-DEL-778
chr1-123555908-DEL-58
chr1-123573635-DEL-1372
chr1-123586480-DEL-119
chr1-123600501-DEL-132
chr1-123618139-DEL-74
chr1-123625503-DEL-62
chr1-123634719-DEL-54
chr1-123640242-DEL-83
chr1-123692737-DEL-70
chr1-123747226-DEL-191
chr1-123748069-DEL-54
chr1-123748929-DEL-62
chr1-123768314-DEL-2050
chr1-123799953-DEL-349
chr1-123801504-DEL-6376
chr1-123801749-DEL-198
chr1-123818653-DEL-4156
chr1-123826342-DEL-65
chr1-123829525-DEL-56
chr1-123838948-DEL-1376
chr1-123840567-DEL-1313
chr1-123848607-DEL-160
chr1-123867089-DEL-52
chr1-123870796-DEL-79
chr1-123879772-DEL-246
chr1-123893594-DEL-86
chr1-123905917-DEL-70
chr1-123920313-DEL-110
chr1-123936334-DEL-184
chr1-123951380-DEL-138
chr1-123955357-DEL-56
chr1-123962127-DEL-91
chr1-124004260-DEL-67

chr1-124007359-DEL-55
chr1-124023585-DEL-61
chr1-124026369-DEL-189
chr1-124048035-DEL-113
chr1-124055711-DEL-56
chr1-124056818-DEL-58
chr1-124083246-DEL-168
chr1-124089637-DEL-56
chr1-124119485-DEL-68
chr1-124152928-DEL-52
chr1-124154711-DEL-63
chr1-124170251-DEL-1116
chr1-124184512-DEL-700
chr1-124186186-DEL-62
chr1-124197665-DEL-1552
chr1-124199084-DEL-451
chr1-124261562-DEL-645
chr1-124264659-DEL-210
chr1-124265901-DEL-80
chr1-124288416-DEL-261
chr1-124320222-DEL-87
chr1-124349983-DEL-63
chr1-124352486-DEL-538
chr1-124353072-DEL-388
chr1-124364700-DEL-51
chr1-124392570-DEL-3812
chr1-124396888-DEL-118
chr1-124397736-DEL-254
chr1-124402965-DEL-3254
chr1-124424635-DEL-66
chr1-124425414-DEL-282
chr1-124487001-DEL-294
chr1-124523172-DEL-1111
chr1-124574377-DEL-132
chr1-124585982-DEL-182
chr1-124608273-DEL-1332
chr1-124670323-DEL-63
chr1-124676195-DEL-59
chr1-124695192-DEL-68
chr1-124711293-DEL-57
chr1-124733892-DEL-118
chr1-124740114-DEL-64
chr1-124769628-DEL-168
chr1-124782167-DEL-129
chr1-124834055-DEL-95
chr1-124843283-DEL-969
chr1-124849652-DEL-7023
chr1-124891731-DEL-56
chr1-124941124-DEL-176
chr1-125001901-DEL-206
chr1-125077662-DEL-113
chr1-125091093-DEL-4184
chr1-125099675-DEL-619
chr1-125113518-DEL-57
chr1-125151729-DEL-1470
chr1-125153402-DEL-1431
chr1-125184149-DEL-56
chr1-125185345-DEL-542
chr1-125196410-DEL-1907
chr1-125207074-DEL-116
chr1-125292171-DEL-570
chr1-125300066-DEL-66
chr1-125306288-DEL-54
chr1-125316771-DEL-227
chr1-125342138-DEL-183
chr1-125351698-DEL-191
chr1-125396577-DEL-74
chr1-125434923-DEL-697
chr1-125438423-DEL-97
chr1-125442113-DEL-184
chr1-125443844-DEL-137
chr1-125545022-DEL-3528
chr1-125555694-DEL-1313
chr1-125580608-DEL-101
chr1-125583540-DEL-1423
chr1-125586645-DEL-74
chr1-125623909-DEL-1149

chr1-125669562-DEL-261
chr1-125680648-DEL-65
chr1-125683591-DEL-51
chr1-125688993-DEL-668
chr1-125728322-DEL-1047
chr1-125729280-DEL-272
chr1-125733421-DEL-52
chr1-125742338-DEL-1131
chr1-125760530-DEL-73
chr1-125772293-DEL-81
chr1-125913555-DEL-58
chr1-125923165-DEL-703
chr1-125926941-DEL-103
chr1-126019603-DEL-230
chr1-126085310-DEL-177
chr1-126131756-DEL-64
chr1-126156779-DEL-67
chr1-126169167-DEL-73
chr1-126185668-DEL-150
chr1-126189073-DEL-143
chr1-126203142-DEL-495
chr1-126218443-DEL-259
chr1-126249614-DEL-72
chr1-126250655-DEL-224
chr1-126327935-DEL-329
chr1-126366783-DEL-117
chr1-126394068-DEL-56
chr1-126431066-DEL-70
chr1-126448536-DEL-525
chr1-126470624-DEL-100
chr1-126481878-DEL-52
chr1-126492270-DEL-207
chr1-126494406-DEL-239
chr1-126511688-DEL-75
chr1-126539401-DEL-143
chr1-126569710-DEL-293
chr1-126610283-DEL-836
chr1-126644707-DEL-70
chr1-126658396-DEL-119
chr1-126666886-DEL-131
chr1-126676055-DEL-59
chr1-126688974-DEL-68
chr1-126695221-DEL-6420
chr1-126706503-DEL-284
chr1-126711925-DEL-56
chr1-126739820-DEL-52
chr1-126757506-DEL-66
chr1-126764764-DEL-288
chr1-126790099-DEL-237
chr1-126811918-DEL-1704
chr1-126837683-DEL-52
chr1-126887181-DEL-859
chr1-126893391-DEL-2257
chr1-126907866-DEL-425
chr1-126921586-DEL-70
chr1-126923190-DEL-1049
chr1-126953357-DEL-1285
chr1-126965159-DEL-155
chr1-126972000-DEL-54
chr1-126981438-DEL-430
chr1-126991883-DEL-133
chr1-126995221-DEL-5874
chr1-127019409-DEL-59
chr1-127037644-DEL-1208
chr1-127084089-DEL-152
chr1-127186999-DEL-86
chr1-127223111-DEL-93
chr1-127259967-DEL-2569
chr1-127277772-DEL-1222
chr1-127309622-DEL-73
chr1-127318063-DEL-82
chr1-127320203-DEL-153
chr1-127331544-DEL-57
chr1-127333759-DEL-62
chr1-127340830-DEL-79
chr1-127374294-DEL-574
chr1-127409080-DEL-81

chr1-127415511-DEL-54
chr1-127422214-DEL-359
chr1-127430750-DEL-66
chr1-127478023-DEL-66
chr1-127502231-DEL-67
chr1-127506898-DEL-55
chr1-127525939-DEL-154
chr1-127525969-DEL-642
chr1-127526402-DEL-3358
chr1-127526519-DEL-106
chr1-127529354-DEL-186
chr1-127558651-DEL-196
chr1-127637287-DEL-958
chr1-127717387-DEL-928
chr1-127723785-DEL-58
chr1-127753561-DEL-1267
chr1-127792534-DEL-77
chr1-127800326-DEL-604
chr1-127802811-DEL-7100
chr1-127819714-DEL-5276
chr1-127826091-DEL-193
chr1-127878225-DEL-153
chr1-127892225-DEL-190
chr1-127907566-DEL-587
chr1-127912297-DEL-234
chr1-127940786-DEL-189
chr1-127957402-DEL-71
chr1-127975535-DEL-56
chr1-127981366-DEL-1194
chr1-127990475-DEL-420
chr1-128020454-DEL-195
chr1-128022473-DEL-63
chr1-128038186-DEL-213
chr1-128039624-DEL-74
chr1-128062643-DEL-1520
chr1-128113005-DEL-53
chr1-128152910-DEL-887
chr1-128156526-DEL-64
chr1-128168498-DEL-401
chr1-128192783-DEL-184
chr1-128194637-DEL-329
chr1-128208505-DEL-1047
chr1-128216939-DEL-1957
chr1-128220011-DEL-399
chr1-128241614-DEL-106
chr1-128243510-DEL-352
chr1-128243914-DEL-654
chr1-128244822-DEL-150
chr1-128266781-DEL-89
chr1-128278558-DEL-184
chr1-128285401-DEL-351
chr1-128296217-DEL-62
chr1-128312962-DEL-52
chr1-128373012-DEL-550
chr1-128376769-DEL-121
chr1-128411607-DEL-6070
chr1-128419140-DEL-58
chr1-128435431-DEL-60
chr1-128444561-DEL-132
chr1-128446091-DEL-241
chr1-128450762-DEL-1073
chr1-128488433-DEL-68
chr1-128539145-DEL-57
chr1-128540198-DEL-230
chr1-128558012-DEL-345
chr1-128578802-DEL-60
chr1-128603785-DEL-55
chr1-128614561-DEL-56
chr1-128643253-DEL-196
chr1-128645232-DEL-66
chr1-128674185-DEL-79
chr1-128718417-DEL-78
chr1-128721896-DEL-93
chr1-128735764-DEL-83
chr1-128815804-DEL-3599
chr1-128849319-DEL-176
chr1-128875904-DEL-54

chr1-128900442-DEL-611
chr1-128938116-DEL-66
chr1-128965293-DEL-114
chr1-128994332-DEL-78
chr1-129003772-DEL-298
chr1-129015311-DEL-128
chr1-129018129-DEL-2556
chr1-129025446-DEL-402
chr1-129027914-DEL-86
chr1-129058260-DEL-3130
chr1-129071363-DEL-83
chr1-129072516-DEL-722
chr1-129074797-DEL-51
chr1-129112220-DEL-1888
chr1-129125630-DEL-54
chr1-129174960-DEL-55
chr1-129184017-DEL-349
chr1-129239374-DEL-90
chr1-129242701-DEL-116
chr1-129257476-DEL-1496
chr1-129262442-DEL-2731
chr1-129267570-DEL-58
chr1-129294009-DEL-1105
chr1-129386428-DEL-1584
chr1-129396700-DEL-93
chr1-129424522-DEL-2982
chr1-129436912-DEL-1006
chr1-129465167-DEL-101
chr1-129534787-DEL-52
chr1-129560393-DEL-196
chr1-129569135-DEL-550
chr1-129579718-DEL-96
chr1-129580420-DEL-737
chr1-129581804-DEL-2300
chr1-129598262-DEL-454
chr1-129658466-DEL-58
chr1-129664660-DEL-626
chr1-129729006-DEL-589
chr1-129767194-DEL-1898
chr1-129784351-DEL-312
chr1-129785136-DEL-1872
chr1-129813390-DEL-70
chr1-129847800-DEL-546
chr1-129849738-DEL-467
chr1-129850940-DEL-73
chr1-129882694-DEL-238
chr1-129925021-DEL-93
chr1-129939508-DEL-228
chr1-129947277-DEL-118
chr1-129952455-DEL-1508
chr1-129959538-DEL-8235
chr1-129967659-DEL-1422
chr1-130033473-DEL-801
chr1-130064379-DEL-96
chr1-130087942-DEL-2370
chr1-130126420-DEL-72
chr1-130134456-DEL-88
chr1-130235755-DEL-1507
chr1-130237235-DEL-72
chr1-130246363-DEL-6311
chr1-130255549-DEL-1850
chr1-130275672-DEL-62
chr1-130288325-DEL-621
chr1-130288790-DEL-108
chr1-130288908-DEL-78
chr1-130319097-DEL-624
chr1-130376845-DEL-51
chr1-130408993-DEL-233
chr1-130442680-DEL-65
chr1-130455562-DEL-92
chr1-130475982-DEL-224
chr1-130484817-DEL-6227
chr1-130508070-DEL-67
chr1-130512945-DEL-281
chr1-130516855-DEL-51
chr1-130518083-DEL-166
chr1-130527658-DEL-816

chr1-130535427-DEL-2265
chr1-130575015-DEL-66
chr1-130613635-DEL-336
chr1-130625133-DEL-72
chr1-130672588-DEL-304
chr1-130690695-DEL-64
chr1-130693112-DEL-87
chr1-130701941-DEL-76
chr1-130718150-DEL-229
chr1-130740893-DEL-6575
chr1-130762436-DEL-91
chr1-130791771-DEL-155
chr1-130816410-DEL-214
chr1-130837595-DEL-701
chr1-130841681-DEL-1897
chr1-130878238-DEL-90
chr1-130900924-DEL-5737
chr1-130911551-DEL-1182
chr1-130944463-DEL-269
chr1-130993243-DEL-778
chr1-131060006-DEL-52
chr1-131072016-DEL-81
chr1-131078445-DEL-53
chr1-131113643-DEL-176
chr1-131126343-DEL-68
chr1-131153629-DEL-871
chr1-131191787-DEL-531
chr1-131236126-DEL-205
chr1-131284186-DEL-55
chr1-131292293-DEL-136
chr1-131295661-DEL-187
chr1-131325551-DEL-120
chr1-131470640-DEL-207
chr1-131493241-DEL-68
chr1-131493793-DEL-324
chr1-131536573-DEL-211
chr1-131570310-DEL-2598
chr1-131572917-DEL-6455
chr1-131622017-DEL-84
chr1-131637770-DEL-126
chr1-131649353-DEL-231
chr1-131654328-DEL-165
chr1-131669067-DEL-52
chr1-131685367-DEL-154
chr1-131694194-DEL-535
chr1-131697924-DEL-59
chr1-131749570-DEL-204
chr1-131754758-DEL-84
chr1-131813257-DEL-65
chr1-131815218-DEL-108
chr1-131821632-DEL-65
chr1-131840741-DEL-251
chr1-131841561-DEL-203
chr1-131867537-DEL-195
chr1-131884149-DEL-180
chr1-131917179-DEL-52
chr1-131922327-DEL-198
chr1-131924684-DEL-188
chr1-131929105-DEL-204
chr1-131955145-DEL-140
chr1-131958653-DEL-83
chr1-131968753-DEL-58
chr1-131975575-DEL-3477
chr1-132014650-DEL-186
chr1-132015555-DEL-136
chr1-132028835-DEL-711
chr1-132035312-DEL-1323
chr1-132058876-DEL-356
chr1-132071802-DEL-105
chr1-132109861-DEL-1051
chr1-132112658-DEL-214
chr1-132151481-DEL-88
chr1-132153434-DEL-69
chr1-132169872-DEL-203
chr1-132174991-DEL-201
chr1-132238125-DEL-73
chr1-132337558-DEL-99

chr1-132406818-DEL-53
chr1-132418720-DEL-77
chr1-132420846-DEL-197
chr1-132445656-DEL-175
chr1-132477760-DEL-828
chr1-132530063-DEL-65
chr1-132532178-DEL-218
chr1-132572307-DEL-1538
chr1-132641622-DEL-116
chr1-132649701-DEL-90
chr1-132665308-DEL-1185
chr1-132677458-DEL-829
chr1-132681263-DEL-726
chr1-132690280-DEL-51
chr1-132704729-DEL-153
chr1-132715830-DEL-204
chr1-132723774-DEL-156
chr1-132762991-DEL-192
chr1-132793849-DEL-109
chr1-132843419-DEL-60
chr1-132844612-DEL-66
chr1-132893106-DEL-70
chr1-132908319-DEL-213
chr1-132909618-DEL-162
chr1-132949974-DEL-191
chr1-132953534-DEL-373
chr1-133030318-DEL-136
chr1-133053729-DEL-1571
chr1-133077484-DEL-183
chr1-133099794-DEL-173
chr1-133107480-DEL-98
chr1-133177759-DEL-201
chr1-133255478-DEL-129
chr1-133260686-DEL-447
chr1-133266881-DEL-210
chr1-133339011-DEL-54
chr1-133372631-DEL-1501
chr1-133377847-DEL-197
chr1-133438756-DEL-215
chr1-133482236-DEL-58
chr1-133493021-DEL-125
chr1-133496521-DEL-199
chr1-133505481-DEL-505
chr1-133511276-DEL-98
chr1-133520555-DEL-51
chr1-133522221-DEL-1674
chr1-133539335-DEL-53
chr1-133542780-DEL-51
chr1-133578309-DEL-210
chr1-133579325-DEL-81
chr1-133580175-DEL-204
chr1-133595869-DEL-1453
chr1-133629601-DEL-60
chr1-133637905-DEL-163
chr1-133649825-DEL-213
chr1-133651244-DEL-68
chr1-133676246-DEL-1143
chr1-133701973-DEL-55
chr1-133735730-DEL-63
chr1-133763225-DEL-887
chr1-133764259-DEL-660
chr1-133821710-DEL-528
chr1-133832481-DEL-73
chr1-133852908-DEL-66
chr1-133871822-DEL-884
chr1-133873657-DEL-92
chr1-133882433-DEL-4112
chr1-133919983-DEL-85
chr1-133950157-DEL-236
chr1-134017284-DEL-653
chr1-134027943-DEL-109
chr1-134031947-DEL-368
chr1-134081797-DEL-99
chr1-134098484-DEL-71
chr1-134106341-DEL-61
chr1-134247546-DEL-100
chr1-134248035-DEL-100

chr1-134252516-DEL-203
chr1-134261377-DEL-151
chr1-134300598-DEL-56
chr1-134310219-DEL-59
chr1-134354458-DEL-760
chr1-134356936-DEL-203
chr1-134369710-DEL-506
chr1-134373860-DEL-143
chr1-134386490-DEL-328
chr1-134394514-DEL-549
chr1-134432256-DEL-56
chr1-134443407-DEL-62
chr1-134445749-DEL-688
chr1-134450836-DEL-147
chr1-134461441-DEL-514
chr1-134467422-DEL-254
chr1-134478073-DEL-167
chr1-134504143-DEL-154
chr1-134511805-DEL-55
chr1-134531319-DEL-208
chr1-134538193-DEL-248
chr1-134544671-DEL-219
chr1-134567608-DEL-860
chr1-134572042-DEL-190
chr1-134595623-DEL-64
chr1-134601000-DEL-209
chr1-134645406-DEL-253
chr1-134696988-DEL-134
chr1-134697528-DEL-768
chr1-134698984-DEL-68
chr1-134714543-DEL-423
chr1-134760685-DEL-52
chr1-134776997-DEL-475
chr1-134777654-DEL-91
chr1-134795609-DEL-62
chr1-134802177-DEL-98
chr1-134806547-DEL-220
chr1-134825493-DEL-68
chr1-134871373-DEL-203
chr1-134880918-DEL-101
chr1-134926486-DEL-355
chr1-134927727-DEL-77
chr1-134973219-DEL-56
chr1-134999608-DEL-86
chr1-135027722-DEL-90
chr1-135081793-DEL-104
chr1-135093460-DEL-53
chr1-135100302-DEL-72
chr1-135119353-DEL-149
chr1-135122602-DEL-187
chr1-135141427-DEL-52
chr1-135142750-DEL-159
chr1-135148731-DEL-218
chr1-135196939-DEL-160
chr1-135205355-DEL-187
chr1-135216467-DEL-455
chr1-135217669-DEL-197
chr1-135224343-DEL-114
chr1-135241679-DEL-1400
chr1-135243415-DEL-401
chr1-135253285-DEL-1070
chr1-135272709-DEL-177
chr1-135281002-DEL-409
chr1-135284421-DEL-104
chr1-135288420-DEL-60
chr1-135301542-DEL-61
chr1-135325088-DEL-63
chr1-135335887-DEL-258
chr1-135417264-DEL-180
chr1-135441790-DEL-52
chr1-135493284-DEL-132
chr1-135500201-DEL-59
chr1-135503362-DEL-641
chr1-135570035-DEL-183
chr1-135579321-DEL-59
chr1-135606527-DEL-136
chr1-135631027-DEL-196

chr1-135635505-DEL-147
chr1-135677281-DEL-63
chr1-135681004-DEL-64
chr1-135688387-DEL-182
chr1-135693350-DEL-53
chr1-135701513-DEL-61
chr1-135717034-DEL-131
chr1-135725872-DEL-59
chr1-135752982-DEL-105
chr1-135849210-DEL-100
chr1-135859119-DEL-231
chr1-135877395-DEL-1104
chr1-135885295-DEL-87
chr1-135946839-DEL-142
chr1-135979628-DEL-60
chr1-135981969-DEL-65
chr1-135989875-DEL-146
chr1-135999388-DEL-56
chr1-136018144-DEL-61
chr1-136074491-DEL-75
chr1-136146433-DEL-57
chr1-136169159-DEL-53
chr1-136264441-DEL-2192
chr1-136264660-DEL-901
chr1-136265171-DEL-433
chr1-136266203-DEL-89
chr1-136266514-DEL-556
chr1-136280767-DEL-93
chr1-136285965-DEL-251
chr1-136292626-DEL-297
chr1-136304000-DEL-204
chr1-136310001-DEL-156
chr1-136313815-DEL-833
chr1-136331872-DEL-1229
chr1-136361251-DEL-91
chr1-136364517-DEL-161
chr1-136379590-DEL-108
chr1-136392175-DEL-1491
chr1-136392743-DEL-222
chr1-136401253-DEL-63
chr1-136411776-DEL-94
chr1-136448715-DEL-120
chr1-136459817-DEL-55
chr1-136477323-DEL-401
chr1-136488706-DEL-190
chr1-136499881-DEL-208
chr1-136575307-DEL-343
chr1-136575885-DEL-108
chr1-136576408-DEL-537
chr1-136590412-DEL-259
chr1-136648981-DEL-316
chr1-136651198-DEL-99
chr1-136663020-DEL-1075
chr1-136666452-DEL-62
chr1-136670944-DEL-58
chr1-136688514-DEL-110
chr1-136695722-DEL-286
chr1-136717431-DEL-230
chr1-136783494-DEL-225
chr1-136790886-DEL-569
chr1-136795727-DEL-61
chr1-136799557-DEL-207
chr1-136814131-DEL-199
chr1-136858137-DEL-68
chr1-136860669-DEL-104
chr1-136865272-DEL-83
chr1-136879535-DEL-72
chr1-136902228-DEL-206
chr1-136909697-DEL-178
chr1-136914771-DEL-114
chr1-136974513-DEL-103
chr1-136979582-DEL-372
chr1-136988160-DEL-51
chr1-136990175-DEL-5911
chr1-137021236-DEL-381
chr1-137058764-DEL-59
chr1-137093093-DEL-55

chr1-137108468-DEL-77
chr1-137111734-DEL-51
chr1-137132460-DEL-965
chr1-137174729-DEL-56
chr1-137218419-DEL-103
chr1-137228071-DEL-450
chr1-137232852-DEL-57
chr1-137248785-DEL-1273
chr1-137251547-DEL-91
chr1-137287524-DEL-467
chr1-137302243-DEL-63
chr1-137320560-DEL-303
chr1-137328791-DEL-60
chr1-137339650-DEL-63
chr1-137339895-DEL-3815
chr1-137343727-DEL-114
chr1-137353647-DEL-78
chr1-137383165-DEL-615
chr1-137395638-DEL-54
chr1-137454600-DEL-3918
chr1-137458563-DEL-2327
chr1-137474635-DEL-515
chr1-137484778-DEL-522
chr1-137501114-DEL-87
chr1-137505141-DEL-708
chr1-137538625-DEL-6188
chr1-137573742-DEL-1755
chr1-137672304-DEL-72
chr1-137684639-DEL-3160
chr1-137689297-DEL-75
chr1-137723837-DEL-391
chr1-137780699-DEL-1013
chr1-137796130-DEL-372
chr1-137805991-DEL-122
chr1-137843970-DEL-499
chr1-137852475-DEL-63
chr1-137862929-DEL-176
chr1-137867931-DEL-56
chr1-137871980-DEL-895
chr1-137882912-DEL-436
chr1-137899632-DEL-70
chr1-137913260-DEL-66
chr1-137977857-DEL-6249
chr1-137988398-DEL-60
chr1-138031799-DEL-1014
chr1-138056611-DEL-52
chr1-138061755-DEL-153
chr1-138132006-DEL-163
chr1-138139167-DEL-1664
chr1-138153976-DEL-5788
chr1-138181314-DEL-1674
chr1-138199036-DEL-351
chr1-138254795-DEL-446
chr1-138278424-DEL-231
chr1-138327323-DEL-669
chr1-138340614-DEL-113
chr1-138356403-DEL-56
chr1-138421473-DEL-205
chr1-138513184-DEL-110
chr1-138516450-DEL-68
chr1-138518363-DEL-101
chr1-138521209-DEL-151
chr1-138525734-DEL-187
chr1-138577098-DEL-685
chr1-138578096-DEL-257
chr1-138581527-DEL-74
chr1-138630334-DEL-62
chr1-138638064-DEL-115
chr1-138682114-DEL-97
chr1-138716542-DEL-6862
chr1-138725012-DEL-222
chr1-138730930-DEL-5885
chr1-138745773-DEL-789
chr1-138751316-DEL-201
chr1-138761372-DEL-76
chr1-138793640-DEL-6416
chr1-138885290-DEL-199

chr1-138895593-DEL-364
chr1-138981414-DEL-95
chr1-139072568-DEL-229
chr1-139106684-DEL-59
chr1-139117907-DEL-344
chr1-139184412-DEL-59
chr1-139193302-DEL-186
chr1-139213531-DEL-361
chr1-139226038-DEL-2754
chr1-139235591-DEL-92
chr1-139248470-DEL-596
chr1-139257914-DEL-1532
chr1-139271596-DEL-201
chr1-139273193-DEL-298
chr1-139282728-DEL-162
chr1-139291722-DEL-339
chr1-139300581-DEL-68
chr1-139327530-DEL-160
chr1-139355015-DEL-64
chr1-139411964-DEL-330
chr1-139413720-DEL-64
chr1-139439584-DEL-540
chr1-139497323-DEL-13146
chr1-139504393-DEL-2500
chr1-139507408-DEL-300
chr1-139515562-DEL-10504
chr1-139515564-DEL-5390
chr1-139522249-DEL-1045
chr1-139568438-DEL-125
chr1-139568861-DEL-12958
chr1-139585850-DEL-112
chr1-139591541-DEL-6182
chr1-139605226-DEL-3333
chr1-139609209-DEL-127
chr1-139644058-DEL-6887
chr1-139652500-DEL-5363
chr1-139783969-DEL-69
chr1-139812501-DEL-1289
chr1-139841746-DEL-173
chr1-139868995-DEL-60
chr1-139873548-DEL-879
chr1-139935322-DEL-599
chr1-139965098-DEL-2301
chr1-139967136-DEL-266
chr1-139979968-DEL-7441
chr1-139980615-DEL-6633
chr1-139987583-DEL-849
chr1-139995211-DEL-63
chr1-140068251-DEL-198
chr1-140093864-DEL-332
chr1-140096917-DEL-52
chr1-140108427-DEL-76
chr1-140120809-DEL-148
chr1-140121982-DEL-683
chr1-140135001-DEL-94
chr1-140157098-DEL-550
chr1-140158208-DEL-112
chr1-140170647-DEL-178
chr1-140197980-DEL-202
chr1-140213638-DEL-806
chr1-140226970-DEL-80
chr1-140239633-DEL-6256
chr1-140253037-DEL-307
chr1-140257829-DEL-544
chr1-140288036-DEL-102
chr1-140295921-DEL-137
chr1-140339294-DEL-89
chr1-140375326-DEL-403
chr1-140438349-DEL-693
chr1-140470108-DEL-305
chr1-140478699-DEL-69
chr1-140489599-DEL-187
chr1-140528206-DEL-276
chr1-140547076-DEL-2904
chr1-140553228-DEL-1636
chr1-140582647-DEL-133
chr1-140599765-DEL-165

chr1-140602900-DEL-276
chr1-140621297-DEL-2928
chr1-140625925-DEL-964
chr1-140633535-DEL-1966
chr1-140637579-DEL-6425
chr1-140644869-DEL-1900
chr1-140647988-DEL-6196
chr1-140648594-DEL-445
chr1-140650295-DEL-6533
chr1-140655868-DEL-189
chr1-140657808-DEL-98
chr1-140668892-DEL-2325
chr1-140704466-DEL-98
chr1-140722163-DEL-2792
chr1-140732685-DEL-396
chr1-140752557-DEL-397
chr1-140762628-DEL-232
chr1-140764895-DEL-170
chr1-140831731-DEL-6306
chr1-140839324-DEL-71
chr1-140845797-DEL-812
chr1-140849819-DEL-74
chr1-140851616-DEL-6528
chr1-140855423-DEL-5495
chr1-140868991-DEL-150
chr1-140877494-DEL-86
chr1-140887610-DEL-15816
chr1-140929589-DEL-141
chr1-140946944-DEL-6182
chr1-140994670-DEL-129
chr1-141006542-DEL-679
chr1-141009033-DEL-876
chr1-141076914-DEL-662
chr1-141078954-DEL-266
chr1-141083202-DEL-118
chr1-141086565-DEL-69
chr1-141107017-DEL-97
chr1-141110957-DEL-73
chr1-141114425-DEL-13589
chr1-141117418-DEL-7061
chr1-141130100-DEL-128
chr1-141134774-DEL-1269
chr1-141141213-DEL-1554
chr1-141152668-DEL-453
chr1-141175835-DEL-1862
chr1-141180510-DEL-158
chr1-141201491-DEL-1059
chr1-141232627-DEL-4796
chr1-141246452-DEL-324
chr1-141250290-DEL-60
chr1-141254150-DEL-276
chr1-141274389-DEL-269
chr1-141275361-DEL-338
chr1-141316945-DEL-537
chr1-141349601-DEL-322
chr1-141384718-DEL-188
chr1-141392816-DEL-205
chr1-141403208-DEL-58
chr1-141412382-DEL-69
chr1-141472228-DEL-5039
chr1-141474601-DEL-383
chr1-141511481-DEL-133
chr1-141524533-DEL-60
chr1-141529808-DEL-416
chr1-141575733-DEL-245
chr1-141589844-DEL-521
chr1-141594291-DEL-6720
chr1-141647142-DEL-289
chr1-141674945-DEL-457
chr1-141692342-DEL-341
chr1-141744358-DEL-70
chr1-141765839-DEL-653
chr1-141787004-DEL-272
chr1-141828208-DEL-483
chr1-141838512-DEL-90
chr1-141915885-DEL-302
chr1-141946349-DEL-62

chr1-141953415-DEL-360
chr1-141957449-DEL-6326
chr1-142017109-DEL-2377
chr1-142020444-DEL-192
chr1-142027809-DEL-1047
chr1-142091651-DEL-1121
chr1-142095196-DEL-75
chr1-142104572-DEL-67
chr1-142150453-DEL-555
chr1-142160392-DEL-493
chr1-142172694-DEL-219
chr1-142175895-DEL-346
chr1-142235520-DEL-1916
chr1-142238601-DEL-92
chr1-142263163-DEL-570
chr1-142284019-DEL-5101
chr1-142298445-DEL-5115
chr1-142308503-DEL-774
chr1-142351798-DEL-68
chr1-142362142-DEL-3184
chr1-142387223-DEL-2519
chr1-142403553-DEL-84
chr1-142404342-DEL-579
chr1-142413878-DEL-1696
chr1-142459164-DEL-652
chr1-142469232-DEL-1992
chr1-142479285-DEL-1492
chr1-142485145-DEL-279
chr1-142511782-DEL-7568
chr1-142532849-DEL-222
chr1-142539774-DEL-3717
chr1-142592321-DEL-104
chr1-142594929-DEL-6189
chr1-142617985-DEL-911
chr1-142621650-DEL-207
chr1-142630120-DEL-636
chr1-142674273-DEL-64
chr1-142690386-DEL-706
chr1-142736610-DEL-66
chr1-142740947-DEL-1462
chr1-142745496-DEL-167
chr1-142756905-DEL-267
chr1-142775439-DEL-197
chr1-142789120-DEL-52
chr1-142789938-DEL-743
chr1-142848446-DEL-1343
chr1-142873083-DEL-1497
chr1-142918142-DEL-3844
chr1-142919136-DEL-206
chr1-142919978-DEL-74
chr1-142941165-DEL-1621
chr1-142972487-DEL-54
chr1-142980240-DEL-636
chr1-143001356-DEL-51
chr1-143033781-DEL-5085
chr1-143039106-DEL-81
chr1-143084426-DEL-115
chr1-143143723-DEL-285
chr1-143172073-DEL-123
chr1-143172870-DEL-71
chr1-143176131-DEL-4768
chr1-143177446-DEL-2494
chr1-143209332-DEL-396
chr1-143211217-DEL-2729
chr1-143220944-DEL-51
chr1-143234710-DEL-324
chr1-143238657-DEL-1014
chr1-143259788-DEL-208
chr1-143270332-DEL-399
chr1-143308243-DEL-6387
chr1-143320561-DEL-8407
chr1-143374709-DEL-444
chr1-143381080-DEL-60
chr1-143381505-DEL-6456
chr1-143415073-DEL-756
chr1-143421003-DEL-3335
chr1-143440723-DEL-105

chr1-143458152-DEL-202
chr1-143466250-DEL-89
chr1-143469375-DEL-66
chr1-143493614-DEL-84
chr1-143558901-DEL-1550
chr1-143637811-DEL-859
chr1-143660993-DEL-95
chr1-143718631-DEL-158
chr1-143722885-DEL-1688
chr1-143723134-DEL-2481
chr1-143737823-DEL-3886
chr1-143741523-DEL-186
chr1-143741772-DEL-2720
chr1-143747486-DEL-108
chr1-143773074-DEL-67
chr1-143776115-DEL-71
chr1-143798114-DEL-56
chr1-143800224-DEL-98
chr1-143835001-DEL-66
chr1-143840704-DEL-8503
chr1-143840745-DEL-62
chr1-143860916-DEL-90
chr1-143926072-DEL-536
chr1-143955551-DEL-123
chr1-143966838-DEL-1761
chr1-143976503-DEL-76
chr1-143984471-DEL-1724
chr1-144000638-DEL-59
chr1-144058617-DEL-132
chr1-144066636-DEL-106
chr1-144072090-DEL-446
chr1-144128153-DEL-3287
chr1-144133362-DEL-79
chr1-144145092-DEL-1874
chr1-144166757-DEL-60
chr1-144189360-DEL-220
chr1-144247026-DEL-507
chr1-144275814-DEL-67
chr1-144286702-DEL-295
chr1-144287737-DEL-1575
chr1-144292621-DEL-521
chr1-144354897-DEL-974
chr1-144400224-DEL-242
chr1-144407548-DEL-633
chr1-144421187-DEL-104
chr1-144437828-DEL-198
chr1-144444996-DEL-75
chr1-144462084-DEL-60
chr1-144474695-DEL-1210
chr1-144490884-DEL-639
chr1-144525051-DEL-1380
chr1-144530373-DEL-1572
chr1-144592557-DEL-95
chr1-144598519-DEL-592
chr1-144599275-DEL-99
chr1-144640786-DEL-51
chr1-144659390-DEL-3352
chr1-144676892-DEL-233
chr1-144720982-DEL-1603
chr1-144741686-DEL-1560
chr1-144775008-DEL-77
chr1-144783073-DEL-78
chr1-144788547-DEL-53
chr1-144814023-DEL-120
chr1-144822122-DEL-2410
chr1-144824584-DEL-674
chr1-144825341-DEL-2243
chr1-144852655-DEL-110
chr1-144859564-DEL-80
chr1-144873258-DEL-95
chr1-144883652-DEL-769
chr1-144901178-DEL-6045
chr1-144925148-DEL-1143
chr1-144938520-DEL-390
chr1-144963316-DEL-1687
chr1-144970223-DEL-97
chr1-144996009-DEL-417

chr1-145044797-DEL-145
chr1-145050000-DEL-4755
chr1-145058794-DEL-358
chr1-145085091-DEL-79
chr1-145093659-DEL-1446
chr1-145105848-DEL-6154
chr1-145118762-DEL-437
chr1-145138423-DEL-53
chr1-145139717-DEL-147
chr1-145143489-DEL-95
chr1-145178072-DEL-123
chr1-145181258-DEL-804
chr1-145189494-DEL-71
chr1-145195785-DEL-58
chr1-145198788-DEL-346
chr1-145205541-DEL-66
chr1-145206687-DEL-419
chr1-145243241-DEL-1093
chr1-145290894-DEL-2992
chr1-145294355-DEL-307
chr1-145329833-DEL-81
chr1-145413757-DEL-74
chr1-145434724-DEL-523
chr1-145438515-DEL-947
chr1-145446693-DEL-2520
chr1-145461906-DEL-142
chr1-145486540-DEL-53
chr1-145492183-DEL-223
chr1-145522569-DEL-309
chr1-145539336-DEL-10413
chr1-145552706-DEL-92
chr1-145553011-DEL-329
chr1-145554259-DEL-57
chr1-145564197-DEL-1444
chr1-145591920-DEL-60
chr1-145594176-DEL-71
chr1-145623095-DEL-538
chr1-145668115-DEL-80
chr1-145683016-DEL-413
chr1-145720748-DEL-302
chr1-145749664-DEL-397
chr1-145760161-DEL-4700
chr1-145783006-DEL-241
chr1-145783844-DEL-963
chr1-145811572-DEL-55
chr1-145827994-DEL-56
chr1-145839031-DEL-55
chr1-145856598-DEL-90
chr1-145869470-DEL-1611
chr1-145893771-DEL-6754
chr1-145900200-DEL-199
chr1-145915651-DEL-202
chr1-145916302-DEL-83
chr1-145965121-DEL-824
chr1-145981247-DEL-492
chr1-145981639-DEL-1694
chr1-145982046-DEL-3448
chr1-145995857-DEL-170
chr1-145996336-DEL-8750
chr1-146030298-DEL-108
chr1-146038702-DEL-60
chr1-146054318-DEL-69
chr1-146071223-DEL-9088
chr1-146137922-DEL-651
chr1-146145077-DEL-352
chr1-146181078-DEL-3186
chr1-146191099-DEL-450
chr1-146220306-DEL-396
chr1-146227869-DEL-61
chr1-146237080-DEL-1398
chr1-146253032-DEL-29119
chr1-146264772-DEL-176
chr1-146281923-DEL-385
chr1-146295558-DEL-625
chr1-146297707-DEL-943
chr1-146315577-DEL-445
chr1-146316230-DEL-522

chr1-146328764-DEL-68
chr1-146351011-DEL-93
chr1-146356789-DEL-1281
chr1-146364683-DEL-227
chr1-146404216-DEL-976
chr1-146491014-DEL-77
chr1-146497912-DEL-1982
chr1-146502449-DEL-90
chr1-146521487-DEL-65
chr1-146539792-DEL-6975
chr1-146547101-DEL-2843
chr1-146547116-DEL-867
chr1-146548076-DEL-1866
chr1-146555132-DEL-73
chr1-146567508-DEL-2026
chr1-146582251-DEL-2381
chr1-146617038-DEL-2703
chr1-146622572-DEL-100
chr1-146626138-DEL-303
chr1-146648404-DEL-951
chr1-146680094-DEL-2078
chr1-146701553-DEL-60
chr1-146734326-DEL-592
chr1-146762917-DEL-55
chr1-146772781-DEL-2051
chr1-146792688-DEL-150
chr1-146801175-DEL-94
chr1-146833298-DEL-88
chr1-146834219-DEL-80
chr1-146854530-DEL-1786
chr1-146892807-DEL-92
chr1-146900940-DEL-82
chr1-146916257-DEL-5230
chr1-146917808-DEL-3049
chr1-146918573-DEL-1524
chr1-146930328-DEL-239
chr1-146986004-DEL-3690
chr1-147001043-DEL-533
chr1-147018579-DEL-1087
chr1-147029273-DEL-430
chr1-147082230-DEL-66
chr1-147105364-DEL-183
chr1-147114509-DEL-58
chr1-147131379-DEL-1956
chr1-147142100-DEL-588
chr1-147200440-DEL-1111
chr1-147201318-DEL-1077
chr1-147201715-DEL-894
chr1-147206041-DEL-99
chr1-147209122-DEL-6188
chr1-147242576-DEL-1896
chr1-147281648-DEL-287
chr1-147283371-DEL-91
chr1-147288454-DEL-796
chr1-147299947-DEL-204
chr1-147315319-DEL-143
chr1-147342178-DEL-302
chr1-147350624-DEL-122
chr1-147367735-DEL-150
chr1-147409736-DEL-2333
chr1-147412627-DEL-66
chr1-147422842-DEL-51
chr1-147430612-DEL-778
chr1-147446073-DEL-1734
chr1-147453024-DEL-617
chr1-147465846-DEL-997
chr1-147475616-DEL-712
chr1-147477087-DEL-182
chr1-147477794-DEL-159
chr1-147478562-DEL-281
chr1-147499413-DEL-2538
chr1-147513072-DEL-562
chr1-147544331-DEL-1499
chr1-147547004-DEL-5398
chr1-147552410-DEL-1077
chr1-147557733-DEL-1619
chr1-147564930-DEL-616

chr1-147603826-DEL-2019
chr1-147609069-DEL-4593
chr1-147622390-DEL-916
chr1-147641149-DEL-55
chr1-147650867-DEL-208
chr1-147652756-DEL-114
chr1-147670503-DEL-144
chr1-147696144-DEL-1476
chr1-147702407-DEL-3284
chr1-147731051-DEL-63
chr1-147736315-DEL-92
chr1-147740584-DEL-402
chr1-147745248-DEL-100
chr1-147789389-DEL-70
chr1-147815502-DEL-62
chr1-147819534-DEL-739
chr1-147826855-DEL-55
chr1-147828637-DEL-283
chr1-147832349-DEL-53
chr1-147868901-DEL-201
chr1-147880929-DEL-871
chr1-147892898-DEL-66
chr1-147915924-DEL-59
chr1-147939821-DEL-189
chr1-147949746-DEL-62
chr1-147959366-DEL-994
chr1-147966248-DEL-745
chr1-148001344-DEL-6261
chr1-148065807-DEL-389
chr1-148114594-DEL-91
chr1-148138270-DEL-285
chr1-148146455-DEL-353
chr1-148184263-DEL-2405
chr1-148194683-DEL-51
chr1-148196327-DEL-58
chr1-148208130-DEL-6935
chr1-148238136-DEL-224
chr1-148254407-DEL-7001
chr1-148270125-DEL-64
chr1-148270799-DEL-1172
chr1-148277452-DEL-147
chr1-148280722-DEL-3265
chr1-148287157-DEL-57
chr1-148296317-DEL-280
chr1-148324213-DEL-1213
chr1-148429002-DEL-6214
chr1-148449521-DEL-70
chr1-148460922-DEL-293
chr1-148462267-DEL-2241
chr1-148480513-DEL-1094
chr1-148530647-DEL-50
chr1-148589188-DEL-92
chr1-148598556-DEL-792
chr1-148599074-DEL-276
chr1-148623447-DEL-283
chr1-148666102-DEL-270
chr1-148672377-DEL-84
chr1-148736615-DEL-1605
chr1-148753677-DEL-6539
chr1-148769112-DEL-273
chr1-148783087-DEL-148
chr1-148787485-DEL-151
chr1-148790134-DEL-800
chr1-148812324-DEL-57
chr1-148843963-DEL-138
chr1-148846990-DEL-555
chr1-148864659-DEL-1414
chr1-148866599-DEL-709
chr1-148873680-DEL-96
chr1-148879281-DEL-111
chr1-148909364-DEL-905
chr1-148920424-DEL-192
chr1-148930052-DEL-230
chr1-148999517-DEL-1428
chr1-149050336-DEL-344
chr1-149052594-DEL-4449
chr1-149085576-DEL-1656

chr1-149160744-DEL-1453
chr1-149170159-DEL-70
chr1-149172195-DEL-208
chr1-149179603-DEL-67
chr1-149190101-DEL-98
chr1-149211281-DEL-69
chr1-149220286-DEL-185
chr1-149249516-DEL-937
chr1-149256450-DEL-177
chr1-149291981-DEL-99
chr1-149294776-DEL-1706
chr1-149305931-DEL-110
chr1-149313005-DEL-157
chr1-149337865-DEL-52
chr1-149341889-DEL-3216
chr1-149368156-DEL-6073
chr1-149378612-DEL-125
chr1-149390023-DEL-7124
chr1-149410037-DEL-135
chr1-149433409-DEL-89
chr1-149457261-DEL-7146
chr1-149475332-DEL-6351
chr1-149487968-DEL-254
chr1-149526060-DEL-392
chr1-149531095-DEL-67
chr1-149578281-DEL-51
chr1-149591683-DEL-681
chr1-149604496-DEL-51
chr1-149665178-DEL-63
chr1-149686267-DEL-1479
chr1-149719448-DEL-123
chr1-149798716-DEL-85
chr1-149846754-DEL-289
chr1-149853163-DEL-112
chr1-149861874-DEL-62
chr1-149908113-DEL-92
chr1-149952080-DEL-540
chr1-149964350-DEL-90
chr1-150002584-DEL-107
chr1-150016333-DEL-231
chr1-150025243-DEL-1805
chr1-150115847-DEL-54
chr1-150119817-DEL-68
chr1-150124888-DEL-58
chr1-150159264-DEL-290
chr1-150179074-DEL-165
chr1-150230099-DEL-6148
chr1-150262847-DEL-52
chr1-150294960-DEL-76
chr1-150331320-DEL-188
chr1-150369417-DEL-61
chr1-150396442-DEL-52
chr1-150404371-DEL-201
chr1-150419317-DEL-125
chr1-150441347-DEL-52
chr1-150467988-DEL-54
chr1-150526555-DEL-1154
chr1-150528737-DEL-127
chr1-150558844-DEL-64
chr1-150568129-DEL-440
chr1-150575488-DEL-121
chr1-150586330-DEL-80
chr1-150611795-DEL-53
chr1-150621915-DEL-678
chr1-150621996-DEL-170
chr1-150622564-DEL-66
chr1-150651426-DEL-52
chr1-150677766-DEL-80
chr1-150703712-DEL-60
chr1-150743360-DEL-66
chr1-150752494-DEL-101
chr1-150762728-DEL-256
chr1-150764386-DEL-1286
chr1-150786460-DEL-149
chr1-150802353-DEL-145
chr1-150812154-DEL-196
chr1-150813333-DEL-55

chr1-150864096-DEL-61
chr1-150893614-DEL-239
chr1-150931059-DEL-70
chr1-150933977-DEL-127
chr1-151002221-DEL-54
chr1-151007780-DEL-71
chr1-151049830-DEL-443
chr1-151122013-DEL-928
chr1-151142250-DEL-7209
chr1-151152174-DEL-52
chr1-151158823-DEL-96
chr1-151166474-DEL-77
chr1-151189545-DEL-98
chr1-151252149-DEL-2651
chr1-151309680-DEL-78
chr1-151312845-DEL-163
chr1-151321539-DEL-57
chr1-151323540-DEL-54
chr1-151328572-DEL-210
chr1-151329396-DEL-222
chr1-151331168-DEL-52
chr1-151337280-DEL-936
chr1-151344648-DEL-113
chr1-151361602-DEL-195
chr1-151381462-DEL-62
chr1-151407914-DEL-70
chr1-151432782-DEL-150
chr1-151434585-DEL-1032
chr1-151450639-DEL-174
chr1-151455192-DEL-492
chr1-151462081-DEL-700
chr1-151463275-DEL-131
chr1-151483626-DEL-53
chr1-151487884-DEL-127
chr1-151498045-DEL-206
chr1-151505053-DEL-51
chr1-151508379-DEL-1026
chr1-151526955-DEL-139
chr1-151550712-DEL-59
chr1-151557813-DEL-176
chr1-151599445-DEL-64
chr1-151618071-DEL-2363
chr1-151625567-DEL-99
chr1-151676060-DEL-2433
chr1-151676191-DEL-138
chr1-151676717-DEL-81
chr1-151677331-DEL-268
chr1-151677864-DEL-254
chr1-151678563-DEL-126
chr1-151716030-DEL-163
chr1-151752473-DEL-704
chr1-151772875-DEL-1886
chr1-151777500-DEL-87
chr1-151780252-DEL-88
chr1-151783168-DEL-104
chr1-151803692-DEL-60
chr1-151830581-DEL-2492
chr1-151897873-DEL-192
chr1-151918584-DEL-137
chr1-151995290-DEL-430
chr1-152054906-DEL-2765
chr1-152066560-DEL-779
chr1-152090042-DEL-64
chr1-152108816-DEL-163
chr1-152143567-DEL-75
chr1-152150178-DEL-1866
chr1-152152964-DEL-138
chr1-152185388-DEL-105
chr1-152216927-DEL-1762
chr1-152219604-DEL-63
chr1-152251758-DEL-70
chr1-152301715-DEL-79
chr1-152342717-DEL-64
chr1-152349784-DEL-63
chr1-152368997-DEL-1599
chr1-152374995-DEL-61
chr1-152398262-DEL-82

chr1-152443392-DEL-68
chr1-152480967-DEL-176
chr1-152484370-DEL-124
chr1-152494372-DEL-67
chr1-152518081-DEL-136
chr1-152526079-DEL-52
chr1-152538620-DEL-5281
chr1-152549354-DEL-304
chr1-152582337-DEL-265
chr1-152598360-DEL-1420
chr1-152602700-DEL-179
chr1-152618311-DEL-249
chr1-152637974-DEL-184
chr1-152639048-DEL-101
chr1-152680203-DEL-124
chr1-152688409-DEL-56
chr1-152701096-DEL-52
chr1-152722366-DEL-198
chr1-152726098-DEL-62
chr1-152749284-DEL-55
chr1-152769642-DEL-83
chr1-152822594-DEL-222
chr1-152856798-DEL-69
chr1-152865312-DEL-280
chr1-152867844-DEL-62
chr1-152897799-DEL-169
chr1-152903332-DEL-3927
chr1-152928870-DEL-151
chr1-152932281-DEL-91
chr1-152943702-DEL-267
chr1-152945101-DEL-59
chr1-152953671-DEL-5454
chr1-152995189-DEL-77
chr1-153038460-DEL-182
chr1-153052098-DEL-139
chr1-153083967-DEL-91
chr1-153091954-DEL-314
chr1-153092970-DEL-187
chr1-153146671-DEL-123
chr1-153189839-DEL-319
chr1-153211395-DEL-120
chr1-153228685-DEL-54
chr1-153246443-DEL-186
chr1-153263089-DEL-7674
chr1-153277599-DEL-221
chr1-153289612-DEL-102
chr1-153304140-DEL-85
chr1-153326066-DEL-197
chr1-153339749-DEL-94
chr1-153343989-DEL-192
chr1-153389498-DEL-255
chr1-153399879-DEL-116
chr1-153408216-DEL-120
chr1-153416729-DEL-1851
chr1-153426001-DEL-224
chr1-153449746-DEL-415
chr1-153457825-DEL-63
chr1-153469277-DEL-122
chr1-153471151-DEL-1351
chr1-153560382-DEL-56
chr1-153605952-DEL-84
chr1-153632694-DEL-941
chr1-153793965-DEL-82
chr1-153794996-DEL-70
chr1-153818461-DEL-161
chr1-153843525-DEL-197
chr1-153876859-DEL-1092
chr1-153893161-DEL-96
chr1-153911340-DEL-66
chr1-153918270-DEL-57
chr1-153929118-DEL-142
chr1-153971451-DEL-219
chr1-154031330-DEL-499
chr1-154039143-DEL-1249
chr1-154062347-DEL-53
chr1-154066942-DEL-4908
chr1-154082750-DEL-1451

chr1-154110809-DEL-1572
chr1-154144541-DEL-58
chr1-154147644-DEL-856
chr1-154208920-DEL-156
chr1-154214636-DEL-99
chr1-154236114-DEL-56
chr1-154243603-DEL-168
chr1-154322890-DEL-188
chr1-154365233-DEL-90
chr1-154381064-DEL-981
chr1-154389221-DEL-951
chr1-154431122-DEL-81
chr1-154469792-DEL-938
chr1-154477031-DEL-401
chr1-154491407-DEL-88
chr1-154530371-DEL-1047
chr1-154545789-DEL-220
chr1-154586044-DEL-73
chr1-154636936-DEL-52
chr1-154668638-DEL-1518
chr1-154674034-DEL-111
chr1-154728365-DEL-284
chr1-154734599-DEL-172
chr1-154739353-DEL-968
chr1-154743869-DEL-59
chr1-154756994-DEL-259
chr1-154757497-DEL-97
chr1-154795160-DEL-150
chr1-154811752-DEL-440
chr1-154845265-DEL-710
chr1-154853686-DEL-93
chr1-154882207-DEL-152
chr1-154895835-DEL-1290
chr1-154897162-DEL-2218
chr1-154933397-DEL-84
chr1-154978235-DEL-78
chr1-154978320-DEL-804
chr1-154988753-DEL-151
chr1-154990864-DEL-144
chr1-155028813-DEL-107
chr1-155036565-DEL-51
chr1-155038110-DEL-93
chr1-155052700-DEL-137
chr1-155093514-DEL-351
chr1-155107964-DEL-100
chr1-155126919-DEL-336
chr1-155132498-DEL-545
chr1-155139161-DEL-91
chr1-155148710-DEL-52
chr1-155164715-DEL-193
chr1-155167335-DEL-722
chr1-155180014-DEL-108
chr1-155191453-DEL-72
chr1-155193266-DEL-2108
chr1-155201351-DEL-2346
chr1-155210951-DEL-1004
chr1-155215046-DEL-232
chr1-155249699-DEL-111
chr1-155267273-DEL-1073
chr1-155270199-DEL-58
chr1-155276668-DEL-183
chr1-155331706-DEL-54
chr1-155333611-DEL-191
chr1-155354033-DEL-822
chr1-155356083-DEL-122
chr1-155372472-DEL-243
chr1-155392668-DEL-73
chr1-155444486-DEL-185
chr1-155516482-DEL-89
chr1-155519235-DEL-52
chr1-155528146-DEL-1366
chr1-155536598-DEL-56
chr1-155553968-DEL-60
chr1-155602819-DEL-223
chr1-155630171-DEL-222
chr1-155673218-DEL-535
chr1-155680499-DEL-100

chr1-155816883-DEL-270
chr1-155855510-DEL-1329
chr1-155858544-DEL-131
chr1-155885097-DEL-271
chr1-155897229-DEL-206
chr1-155957998-DEL-196
chr1-155958727-DEL-155
chr1-155978135-DEL-96
chr1-156022609-DEL-52
chr1-156035835-DEL-565
chr1-156038223-DEL-203
chr1-156051263-DEL-77
chr1-156078019-DEL-183
chr1-156078356-DEL-426
chr1-156080087-DEL-99
chr1-156080647-DEL-399
chr1-156085566-DEL-51
chr1-156094560-DEL-73
chr1-156110598-DEL-200
chr1-156147127-DEL-196
chr1-156157096-DEL-443
chr1-156159913-DEL-55
chr1-156201836-DEL-51
chr1-156204757-DEL-68
chr1-156207296-DEL-62
chr1-156218028-DEL-115
chr1-156222157-DEL-201
chr1-156228099-DEL-322
chr1-156230682-DEL-9122
chr1-156244531-DEL-394
chr1-156278754-DEL-248
chr1-156279589-DEL-2122
chr1-156292059-DEL-6816
chr1-156299243-DEL-1108
chr1-156335780-DEL-3474
chr1-156339565-DEL-64
chr1-156355124-DEL-2000
chr1-156366280-DEL-90
chr1-156383744-DEL-52
chr1-156393752-DEL-100
chr1-156409001-DEL-62
chr1-156423861-DEL-219
chr1-156425023-DEL-226
chr1-156432079-DEL-108
chr1-156433842-DEL-139
chr1-156439259-DEL-55
chr1-156454171-DEL-53
chr1-156476171-DEL-59
chr1-156477607-DEL-88
chr1-156478191-DEL-481
chr1-156478854-DEL-188
chr1-156504227-DEL-210
chr1-156521425-DEL-295
chr1-156534852-DEL-213
chr1-156550561-DEL-376
chr1-156552926-DEL-491
chr1-156614515-DEL-103
chr1-156626543-DEL-344
chr1-156637571-DEL-269
chr1-156719468-DEL-325
chr1-156747271-DEL-350
chr1-156758513-DEL-121
chr1-156789644-DEL-61
chr1-156830157-DEL-68
chr1-156833068-DEL-78
chr1-156869804-DEL-358
chr1-156905356-DEL-241
chr1-156923574-DEL-1433
chr1-156954948-DEL-668
chr1-156962319-DEL-247
chr1-156991016-DEL-138
chr1-157029983-DEL-1587
chr1-157034421-DEL-3605
chr1-157042502-DEL-80
chr1-157044282-DEL-53
chr1-157060944-DEL-79
chr1-157085983-DEL-338

chr1-157088166-DEL-1128
chr1-157096566-DEL-255
chr1-157128328-DEL-654
chr1-157130358-DEL-74
chr1-157180412-DEL-847
chr1-157194393-DEL-371
chr1-157200027-DEL-191
chr1-157223553-DEL-321
chr1-157228533-DEL-1764
chr1-157254192-DEL-207
chr1-157290993-DEL-281
chr1-157343698-DEL-198
chr1-157366454-DEL-64
chr1-157381206-DEL-343
chr1-157386475-DEL-191
chr1-157390817-DEL-220
chr1-157406747-DEL-840
chr1-157440344-DEL-96
chr1-157524325-DEL-66
chr1-157615226-DEL-73
chr1-157635033-DEL-162
chr1-157656825-DEL-99
chr1-157710929-DEL-208
chr1-157720969-DEL-197
chr1-157804798-DEL-438
chr1-157805168-DEL-525
chr1-157807058-DEL-267
chr1-157884857-DEL-88
chr1-157894118-DEL-558
chr1-157914720-DEL-206
chr1-157942693-DEL-53
chr1-157955476-DEL-68
chr1-157970722-DEL-124
chr1-157983826-DEL-64
chr1-158032989-DEL-399
chr1-158050448-DEL-67
chr1-158075002-DEL-174
chr1-158085335-DEL-184
chr1-158089395-DEL-223
chr1-158098237-DEL-382
chr1-158102947-DEL-90
chr1-158132685-DEL-3681
chr1-158200153-DEL-504
chr1-158259549-DEL-1412
chr1-158270662-DEL-1771
chr1-158297954-DEL-61
chr1-158313337-DEL-98
chr1-158316732-DEL-156
chr1-158362126-DEL-54
chr1-158365236-DEL-81
chr1-158386043-DEL-112
chr1-158410078-DEL-6221
chr1-158452148-DEL-136
chr1-158482781-DEL-274
chr1-158514558-DEL-114
chr1-158522674-DEL-242
chr1-158574208-DEL-103
chr1-158642835-DEL-946
chr1-158715038-DEL-66
chr1-158723950-DEL-116
chr1-158726987-DEL-156
chr1-158742452-DEL-400
chr1-158793766-DEL-941
chr1-158810101-DEL-186
chr1-158811266-DEL-160
chr1-158827861-DEL-167
chr1-158849410-DEL-6839
chr1-158910282-DEL-72
chr1-158914670-DEL-1258
chr1-158942731-DEL-90
chr1-158953998-DEL-92
chr1-158957706-DEL-571
chr1-158981820-DEL-62
chr1-158987722-DEL-53
chr1-159007024-DEL-127
chr1-159017475-DEL-502
chr1-159042912-DEL-1671

chr1-159058671-DEL-554
chr1-159130542-DEL-1403
chr1-159161429-DEL-1882
chr1-159193289-DEL-2195
chr1-159225561-DEL-98
chr1-159260562-DEL-79
chr1-159357067-DEL-80
chr1-159380151-DEL-76
chr1-159453436-DEL-400
chr1-159481003-DEL-54
chr1-159536023-DEL-6189
chr1-159578981-DEL-1628
chr1-159608868-DEL-624
chr1-159612315-DEL-94
chr1-159620831-DEL-227
chr1-159678654-DEL-58
chr1-159723956-DEL-236
chr1-159761606-DEL-62
chr1-159767920-DEL-205
chr1-159786212-DEL-3162
chr1-159824170-DEL-2003
chr1-159840144-DEL-300
chr1-159862753-DEL-68
chr1-159984488-DEL-470
chr1-159997597-DEL-227
chr1-160190004-DEL-6427
chr1-160193436-DEL-107
chr1-160217218-DEL-319
chr1-160239336-DEL-1313
chr1-160246605-DEL-1506
chr1-160268562-DEL-72
chr1-160285009-DEL-52
chr1-160287163-DEL-63
chr1-160300324-DEL-70
chr1-160414575-DEL-93
chr1-160430884-DEL-61
chr1-160493667-DEL-269
chr1-160498194-DEL-326
chr1-160519223-DEL-327
chr1-160547360-DEL-135
chr1-160557050-DEL-95
chr1-160584510-DEL-98
chr1-160595019-DEL-455
chr1-160596000-DEL-733
chr1-160609796-DEL-197
chr1-160634564-DEL-329
chr1-160725848-DEL-190
chr1-160733448-DEL-76
chr1-160749482-DEL-421
chr1-160803274-DEL-810
chr1-160818424-DEL-114
chr1-160914559-DEL-309
chr1-160924703-DEL-230
chr1-160956353-DEL-220
chr1-160981148-DEL-329
chr1-161004065-DEL-76
chr1-161019795-DEL-70
chr1-161039992-DEL-331
chr1-161045632-DEL-102
chr1-161048656-DEL-586
chr1-161051186-DEL-112
chr1-161055853-DEL-176
chr1-161060631-DEL-120
chr1-161103154-DEL-739
chr1-161116141-DEL-227
chr1-161119200-DEL-165
chr1-161120050-DEL-263
chr1-161150007-DEL-2424
chr1-161176079-DEL-1311
chr1-161185106-DEL-2803
chr1-161190705-DEL-73
chr1-161198685-DEL-1084
chr1-161238959-DEL-210
chr1-161245882-DEL-329
chr1-161261113-DEL-161
chr1-161269118-DEL-254
chr1-161283568-DEL-68

chr1-161305709-DEL-160
chr1-161306748-DEL-561
chr1-161336458-DEL-1319
chr1-161359741-DEL-4002
chr1-161369732-DEL-279
chr1-161371119-DEL-956
chr1-161391709-DEL-54
chr1-161412540-DEL-295
chr1-161424741-DEL-76
chr1-161441119-DEL-126
chr1-161447183-DEL-401
chr1-161464876-DEL-82
chr1-161467385-DEL-107
chr1-161474668-DEL-2612
chr1-161477319-DEL-1116
chr1-161505866-DEL-1914
chr1-161509629-DEL-189
chr1-161529424-DEL-124
chr1-161579635-DEL-54
chr1-161641235-DEL-295
chr1-161658887-DEL-608
chr1-161694396-DEL-267
chr1-161710141-DEL-124
chr1-161710658-DEL-53
chr1-161826078-DEL-178
chr1-161828166-DEL-83
chr1-161952981-DEL-84
chr1-161957532-DEL-198
chr1-161974372-DEL-274
chr1-162016908-DEL-1411
chr1-162066090-DEL-74
chr1-162078949-DEL-81
chr1-162084146-DEL-134
chr1-162207429-DEL-484
chr1-162242250-DEL-286
chr1-162257563-DEL-91
chr1-162281621-DEL-1711
chr1-162327263-DEL-348
chr1-162356494-DEL-110
chr1-162365552-DEL-199
chr1-162379469-DEL-734
chr1-162381189-DEL-63
chr1-162397151-DEL-236
chr1-162434015-DEL-75
chr1-162450913-DEL-84
chr1-162461144-DEL-96
chr1-162475176-DEL-59
chr1-162484324-DEL-141
chr1-162491288-DEL-266
chr1-162496278-DEL-211
chr1-162518257-DEL-58
chr1-162574833-DEL-1501
chr1-162613259-DEL-378
chr1-162658981-DEL-54
chr1-162662374-DEL-1252
chr1-162664423-DEL-99
chr1-162688097-DEL-150
chr1-162694874-DEL-80
chr1-162740236-DEL-230
chr1-162771686-DEL-95
chr1-162793187-DEL-66
chr1-162797887-DEL-663
chr1-162840296-DEL-955
chr1-162841394-DEL-1214
chr1-162864152-DEL-110
chr1-162909174-DEL-485
chr1-162910335-DEL-2253
chr1-162946106-DEL-398
chr1-162948962-DEL-494
chr1-162951230-DEL-116
chr1-162958086-DEL-179
chr1-163033001-DEL-781
chr1-163061215-DEL-731
chr1-163071680-DEL-70
chr1-163087554-DEL-54
chr1-163093744-DEL-321
chr1-163131031-DEL-121

chr1-163133021-DEL-59
chr1-163213459-DEL-90
chr1-163234642-DEL-82
chr1-163257169-DEL-52
chr1-163264317-DEL-544
chr1-163277138-DEL-962
chr1-163290042-DEL-197
chr1-163323930-DEL-97
chr1-163348428-DEL-1985
chr1-163404487-DEL-129
chr1-163427019-DEL-112
chr1-163464206-DEL-122
chr1-163482744-DEL-61
chr1-163551185-DEL-66
chr1-163577305-DEL-764
chr1-163581944-DEL-886
chr1-163599291-DEL-6196
chr1-163611632-DEL-212
chr1-163661064-DEL-487
chr1-163706010-DEL-2705
chr1-163764802-DEL-193
chr1-163766453-DEL-299
chr1-163771086-DEL-84
chr1-163787845-DEL-54
chr1-163820107-DEL-66
chr1-163840637-DEL-335
chr1-163871856-DEL-220
chr1-163940524-DEL-510
chr1-163951443-DEL-62
chr1-164000591-DEL-56
chr1-164014599-DEL-60
chr1-164068497-DEL-108
chr1-164079883-DEL-60
chr1-164093940-DEL-439
chr1-164098368-DEL-258
chr1-164130752-DEL-74
chr1-164153345-DEL-53
chr1-164157802-DEL-1630
chr1-164158346-DEL-1087
chr1-164190918-DEL-2067
chr1-164203865-DEL-692
chr1-164214558-DEL-197
chr1-164291794-DEL-897
chr1-164353508-DEL-66
chr1-164359065-DEL-198
chr1-164362831-DEL-380
chr1-164363076-DEL-760
chr1-164367019-DEL-792
chr1-164378566-DEL-1532
chr1-164405428-DEL-2522
chr1-164425044-DEL-669
chr1-164427048-DEL-424
chr1-164464912-DEL-120
chr1-164535539-DEL-126
chr1-164536133-DEL-500
chr1-164596440-DEL-50
chr1-164617745-DEL-507
chr1-164678980-DEL-1427
chr1-164769321-DEL-54
chr1-164817149-DEL-2730
chr1-164824418-DEL-169
chr1-164841806-DEL-97
chr1-164861671-DEL-114
chr1-164862743-DEL-70
chr1-164872108-DEL-95
chr1-164894396-DEL-145
chr1-164895106-DEL-127
chr1-164909980-DEL-363
chr1-164914796-DEL-498
chr1-164926004-DEL-697
chr1-164930495-DEL-382
chr1-164943800-DEL-110
chr1-164965772-DEL-67
chr1-165032332-DEL-237
chr1-165036698-DEL-2086
chr1-165067930-DEL-106
chr1-165080112-DEL-2002

chr1-165156377-DEL-304
chr1-165174449-DEL-602
chr1-165180954-DEL-207
chr1-165204125-DEL-56
chr1-165204707-DEL-552
chr1-165207831-DEL-290
chr1-165224622-DEL-1683
chr1-165232673-DEL-212
chr1-165241702-DEL-158
chr1-165271833-DEL-8338
chr1-165318963-DEL-77
chr1-165330936-DEL-132
chr1-165354412-DEL-212
chr1-165365543-DEL-106
chr1-165385412-DEL-81
chr1-165389617-DEL-67
chr1-165396526-DEL-74
chr1-165406203-DEL-241
chr1-165416099-DEL-102
chr1-165427390-DEL-1035
chr1-165448141-DEL-97
chr1-165484367-DEL-512
chr1-165506216-DEL-115
chr1-165507795-DEL-224
chr1-165544488-DEL-102
chr1-165558346-DEL-67
chr1-165594910-DEL-366
chr1-165612846-DEL-200
chr1-165635584-DEL-85
chr1-165667627-DEL-81
chr1-165669482-DEL-54
chr1-165739256-DEL-228
chr1-165743653-DEL-62
chr1-165791176-DEL-208
chr1-165865932-DEL-56
chr1-165886470-DEL-60
chr1-165896931-DEL-220
chr1-165897807-DEL-202
chr1-165899081-DEL-135
chr1-165905666-DEL-211
chr1-165915481-DEL-115
chr1-165933098-DEL-79
chr1-165962708-DEL-144
chr1-165971020-DEL-1186
chr1-166010319-DEL-74
chr1-166030352-DEL-324
chr1-166039525-DEL-7089
chr1-166040216-DEL-80
chr1-166079373-DEL-63
chr1-166179445-DEL-7956
chr1-166206774-DEL-80
chr1-166209173-DEL-421
chr1-166211877-DEL-123
chr1-166223124-DEL-65
chr1-166244534-DEL-56
chr1-166254915-DEL-79
chr1-166260752-DEL-4621
chr1-166263323-DEL-52
chr1-166273813-DEL-66
chr1-166287668-DEL-4771
chr1-166294703-DEL-57
chr1-166320969-DEL-322
chr1-166325464-DEL-591
chr1-166368460-DEL-6167
chr1-166382986-DEL-66
chr1-166395456-DEL-1266
chr1-166396833-DEL-186
chr1-166427909-DEL-12973
chr1-166441813-DEL-2999
chr1-166446842-DEL-116
chr1-166447691-DEL-6388
chr1-166523485-DEL-188
chr1-166532800-DEL-373
chr1-166577156-DEL-268
chr1-166579580-DEL-101
chr1-166626612-DEL-62
chr1-166638194-DEL-100

chr1-166645962-DEL-68
chr1-166649773-DEL-56
chr1-166650299-DEL-57
chr1-166664703-DEL-56
chr1-166664795-DEL-1105
chr1-166711814-DEL-654
chr1-166759296-DEL-70
chr1-166795244-DEL-177
chr1-166796290-DEL-143
chr1-166862940-DEL-318
chr1-166878416-DEL-125
chr1-166927475-DEL-404
chr1-167019856-DEL-129
chr1-167039423-DEL-155
chr1-167042962-DEL-645
chr1-167054758-DEL-52
chr1-167066754-DEL-773
chr1-167070534-DEL-58
chr1-167100708-DEL-56
chr1-167107358-DEL-221
chr1-167107793-DEL-945
chr1-167128118-DEL-197
chr1-167135354-DEL-64
chr1-167147652-DEL-336
chr1-167159183-DEL-100
chr1-167166043-DEL-375
chr1-167224834-DEL-548
chr1-167235726-DEL-113
chr1-167245037-DEL-599
chr1-167246112-DEL-186
chr1-167329395-DEL-55
chr1-167331294-DEL-109
chr1-167364413-DEL-861
chr1-167426125-DEL-174
chr1-167441102-DEL-110
chr1-167513058-DEL-58
chr1-167530729-DEL-56
chr1-167539044-DEL-959
chr1-167593138-DEL-478
chr1-167604296-DEL-113
chr1-167622786-DEL-192
chr1-167651570-DEL-5552
chr1-167661401-DEL-6440
chr1-167681158-DEL-56
chr1-167691828-DEL-208
chr1-167696816-DEL-53
chr1-167711113-DEL-895
chr1-167711349-DEL-72
chr1-167722041-DEL-138
chr1-167745086-DEL-97
chr1-167790016-DEL-61
chr1-167810912-DEL-1605
chr1-167813560-DEL-99
chr1-167819996-DEL-2580
chr1-167824501-DEL-52
chr1-167832211-DEL-1547
chr1-167865549-DEL-206
chr1-167929893-DEL-133
chr1-167957812-DEL-70
chr1-168037980-DEL-607
chr1-168062618-DEL-68
chr1-168088634-DEL-182
chr1-168099738-DEL-1125
chr1-168111380-DEL-52
chr1-168143622-DEL-4482
chr1-168189445-DEL-88
chr1-168231486-DEL-151
chr1-168243401-DEL-71
chr1-168293472-DEL-947
chr1-168329269-DEL-99
chr1-168355944-DEL-109
chr1-168390375-DEL-94
chr1-168437659-DEL-3560
chr1-168439018-DEL-6437
chr1-168444191-DEL-1985
chr1-168457066-DEL-205
chr1-168468578-DEL-3764

chr1-168477335-DEL-1043
chr1-168496394-DEL-272
chr1-168517642-DEL-591
chr1-168537750-DEL-105
chr1-168547459-DEL-225
chr1-168567590-DEL-51
chr1-168574374-DEL-241
chr1-168584477-DEL-941
chr1-168600772-DEL-138
chr1-168623942-DEL-89
chr1-168638541-DEL-218
chr1-168655667-DEL-78
chr1-168681027-DEL-6033
chr1-168687798-DEL-875
chr1-168712388-DEL-328
chr1-168733555-DEL-260
chr1-168787483-DEL-363
chr1-168793455-DEL-80
chr1-168824523-DEL-61
chr1-168825991-DEL-70
chr1-168826738-DEL-395
chr1-168852937-DEL-74
chr1-168952290-DEL-366
chr1-169023633-DEL-907
chr1-169025257-DEL-71
chr1-169027066-DEL-171
chr1-169089470-DEL-53
chr1-169097436-DEL-157
chr1-169101462-DEL-52
chr1-169133931-DEL-52
chr1-169207353-DEL-60
chr1-169233352-DEL-819
chr1-169237354-DEL-52
chr1-169292957-DEL-61
chr1-169322524-DEL-59
chr1-169333957-DEL-64
chr1-169345502-DEL-113
chr1-169347106-DEL-69
chr1-169350136-DEL-57
chr1-169362591-DEL-1357
chr1-169406441-DEL-77
chr1-169415752-DEL-433
chr1-169443103-DEL-62
chr1-169536995-DEL-66
chr1-169545679-DEL-74
chr1-169583403-DEL-261
chr1-169589791-DEL-65
chr1-169598115-DEL-365
chr1-169635416-DEL-760
chr1-169671833-DEL-56
chr1-169685155-DEL-324
chr1-169695677-DEL-70
chr1-169708387-DEL-135
chr1-169709328-DEL-74
chr1-169758304-DEL-88
chr1-169816803-DEL-166
chr1-169824105-DEL-129
chr1-169845682-DEL-129
chr1-169847534-DEL-1098
chr1-169855674-DEL-997
chr1-169872955-DEL-51
chr1-169903538-DEL-84
chr1-169930170-DEL-68
chr1-169952619-DEL-51
chr1-169961171-DEL-82
chr1-169967579-DEL-573
chr1-169979012-DEL-125
chr1-169980430-DEL-2533
chr1-169999274-DEL-79
chr1-170004568-DEL-56
chr1-170041505-DEL-125
chr1-170062212-DEL-162
chr1-170142102-DEL-111
chr1-170179871-DEL-328
chr1-170216043-DEL-221
chr1-170217654-DEL-375
chr1-170251040-DEL-63

chr1-170265367-DEL-68
chr1-170290712-DEL-116
chr1-170298389-DEL-397
chr1-170327034-DEL-209
chr1-170356578-DEL-132
chr1-170409412-DEL-61
chr1-170463842-DEL-52
chr1-170464762-DEL-80
chr1-170498827-DEL-1087
chr1-170508186-DEL-267
chr1-170510267-DEL-6047
chr1-170521901-DEL-66
chr1-170528086-DEL-51
chr1-170597829-DEL-831
chr1-170605528-DEL-56
chr1-170606067-DEL-1500
chr1-170606778-DEL-1552
chr1-170607689-DEL-77
chr1-170611561-DEL-95
chr1-170628064-DEL-387
chr1-170631857-DEL-68
chr1-170652838-DEL-127
chr1-170663354-DEL-366
chr1-170681345-DEL-1245
chr1-170692295-DEL-847
chr1-170718669-DEL-199
chr1-170729115-DEL-164
chr1-170815474-DEL-216
chr1-170821599-DEL-149
chr1-170852528-DEL-138
chr1-170861712-DEL-534
chr1-170862937-DEL-66
chr1-170873570-DEL-455
chr1-170874636-DEL-290
chr1-170882416-DEL-360
chr1-170887881-DEL-284
chr1-170917744-DEL-56
chr1-170927416-DEL-96
chr1-170928034-DEL-532
chr1-170933583-DEL-83
chr1-171004246-DEL-3379
chr1-171027948-DEL-122
chr1-171032973-DEL-69
chr1-171036616-DEL-110
chr1-171043631-DEL-64
chr1-171144076-DEL-165
chr1-171174115-DEL-106
chr1-171191203-DEL-204
chr1-171204904-DEL-181
chr1-171223474-DEL-660
chr1-171233033-DEL-204
chr1-171260603-DEL-199
chr1-171261337-DEL-196
chr1-171274331-DEL-82
chr1-171283476-DEL-2320
chr1-171292240-DEL-201
chr1-171294718-DEL-53
chr1-171299475-DEL-163
chr1-171302807-DEL-167
chr1-171317491-DEL-66
chr1-171346216-DEL-6199
chr1-171359705-DEL-139
chr1-171363157-DEL-63
chr1-171373829-DEL-56
chr1-171374206-DEL-184
chr1-171378200-DEL-5646
chr1-171400697-DEL-116
chr1-171407540-DEL-55
chr1-171413143-DEL-51
chr1-171414888-DEL-107
chr1-171418862-DEL-81
chr1-171427657-DEL-53
chr1-171450280-DEL-52
chr1-171519088-DEL-1686
chr1-171520840-DEL-1510
chr1-171527819-DEL-195
chr1-171580523-DEL-1229

chr1-171586487-DEL-54
chr1-171587537-DEL-264
chr1-171593492-DEL-64
chr1-171607175-DEL-177
chr1-171610856-DEL-52
chr1-171620774-DEL-203
chr1-171642376-DEL-529
chr1-171715644-DEL-7201
chr1-171738834-DEL-1783
chr1-171748091-DEL-55
chr1-171752893-DEL-2140
chr1-171776993-DEL-65
chr1-171805508-DEL-150
chr1-171825167-DEL-209
chr1-171865546-DEL-197
chr1-171879011-DEL-196
chr1-171890617-DEL-59
chr1-171902062-DEL-63
chr1-171928401-DEL-138
chr1-171964309-DEL-178
chr1-171969545-DEL-92
chr1-171985406-DEL-206
chr1-172032696-DEL-56
chr1-172066807-DEL-71
chr1-172077765-DEL-132
chr1-172088802-DEL-82
chr1-172093234-DEL-4061
chr1-172101599-DEL-54
chr1-172103987-DEL-95
chr1-172126394-DEL-54
chr1-172130825-DEL-84
chr1-172136342-DEL-103
chr1-172166499-DEL-79
chr1-172179116-DEL-884
chr1-172206792-DEL-341
chr1-172207320-DEL-469
chr1-172224430-DEL-52
chr1-172225458-DEL-1998
chr1-172237411-DEL-52
chr1-172253510-DEL-277
chr1-172277740-DEL-168
chr1-172278975-DEL-89
chr1-172298151-DEL-103
chr1-172299694-DEL-138
chr1-172378999-DEL-130
chr1-172386594-DEL-67
chr1-172422961-DEL-100
chr1-172436265-DEL-195
chr1-172438544-DEL-288
chr1-172446305-DEL-602
chr1-172453339-DEL-69
chr1-172481043-DEL-148
chr1-172489664-DEL-60
chr1-172492342-DEL-86
chr1-172502167-DEL-1014
chr1-172516156-DEL-1398
chr1-172544252-DEL-6487
chr1-172554143-DEL-1073
chr1-172559378-DEL-6378
chr1-172598788-DEL-52
chr1-172612692-DEL-312
chr1-172623023-DEL-55
chr1-172636517-DEL-155
chr1-172639392-DEL-80
chr1-172642931-DEL-306
chr1-172643772-DEL-1177
chr1-172685213-DEL-338
chr1-172716128-DEL-2122
chr1-172733363-DEL-481
chr1-172734477-DEL-573
chr1-172760409-DEL-560
chr1-172788708-DEL-593
chr1-172793205-DEL-82
chr1-172798283-DEL-6414
chr1-172821540-DEL-52
chr1-172822433-DEL-60
chr1-172831610-DEL-60

chr1-172848150-DEL-461
chr1-172850537-DEL-52
chr1-172882436-DEL-245
chr1-172920642-DEL-277
chr1-172937299-DEL-6224
chr1-172975367-DEL-1223
chr1-172992735-DEL-248
chr1-172995125-DEL-137
chr1-173022000-DEL-1735
chr1-173087911-DEL-201
chr1-173172337-DEL-54
chr1-173186032-DEL-78
chr1-173233543-DEL-1131
chr1-173269446-DEL-271
chr1-173296757-DEL-427
chr1-173308091-DEL-426
chr1-173326648-DEL-73
chr1-173367894-DEL-6073
chr1-173421973-DEL-2576
chr1-173426933-DEL-206
chr1-173436492-DEL-645
chr1-173545730-DEL-772
chr1-173599775-DEL-139
chr1-173614831-DEL-6359
chr1-173660449-DEL-517
chr1-173660451-DEL-7080
chr1-173661016-DEL-6246
chr1-173667457-DEL-60
chr1-173668578-DEL-51
chr1-173691685-DEL-375
chr1-173720190-DEL-2113
chr1-173722170-DEL-216
chr1-173815795-DEL-4833
chr1-173815800-DEL-3399
chr1-173819599-DEL-1022
chr1-173820655-DEL-1309
chr1-173837824-DEL-249
chr1-173892785-DEL-52
chr1-173940962-DEL-241
chr1-173954915-DEL-334
chr1-173954931-DEL-3024
chr1-173955152-DEL-1300
chr1-173955278-DEL-270
chr1-173956579-DEL-458
chr1-173961266-DEL-1677
chr1-174015388-DEL-1450
chr1-174015451-DEL-73
chr1-174031072-DEL-80
chr1-174044322-DEL-194
chr1-174068296-DEL-68
chr1-174081586-DEL-6928
chr1-174103121-DEL-6282
chr1-174110313-DEL-1246
chr1-174115761-DEL-6742
chr1-174153163-DEL-107
chr1-174153184-DEL-3594
chr1-174187336-DEL-2220
chr1-174204045-DEL-7245
chr1-174204337-DEL-66
chr1-174211076-DEL-66
chr1-174221831-DEL-64
chr1-174222173-DEL-6228
chr1-174222174-DEL-11734
chr1-174228605-DEL-6054
chr1-174228723-DEL-5182
chr1-174234922-DEL-193
chr1-174240924-DEL-196
chr1-174292396-DEL-125
chr1-174294929-DEL-269
chr1-174302388-DEL-159
chr1-174303287-DEL-1025
chr1-174309534-DEL-457
chr1-174326627-DEL-52
chr1-174347651-DEL-74
chr1-174353980-DEL-244
chr1-174370376-DEL-54
chr1-174381167-DEL-68

chr1-174382547-DEL-68
chr1-174408256-DEL-586
chr1-174417876-DEL-237
chr1-174459227-DEL-54
chr1-174551245-DEL-479
chr1-174557075-DEL-54
chr1-174656348-DEL-62
chr1-174677153-DEL-60
chr1-174689105-DEL-1062
chr1-174705459-DEL-118
chr1-174715964-DEL-54
chr1-174738109-DEL-303
chr1-174743664-DEL-130
chr1-174749660-DEL-134
chr1-174752262-DEL-666
chr1-174767426-DEL-195
chr1-174786890-DEL-54
chr1-174792362-DEL-162
chr1-174800879-DEL-82
chr1-174835766-DEL-73
chr1-174883741-DEL-174
chr1-174899292-DEL-1197
chr1-174940203-DEL-2960
chr1-174947220-DEL-298
chr1-174948298-DEL-66
chr1-174981937-DEL-164
chr1-175005356-DEL-62
chr1-175022918-DEL-197
chr1-175029466-DEL-85
chr1-175055931-DEL-50
chr1-175056460-DEL-296
chr1-175059649-DEL-6388
chr1-175074474-DEL-67
chr1-175105442-DEL-97
chr1-175126171-DEL-60
chr1-175174382-DEL-203
chr1-175205544-DEL-86
chr1-175220537-DEL-146
chr1-175224235-DEL-6235
chr1-175308841-DEL-298
chr1-175353046-DEL-107
chr1-175367782-DEL-95
chr1-175401129-DEL-73
chr1-175420371-DEL-65
chr1-175461007-DEL-904
chr1-175463391-DEL-55
chr1-175465645-DEL-207
chr1-175478309-DEL-60
chr1-175558876-DEL-121
chr1-175572972-DEL-1193
chr1-175575307-DEL-2735
chr1-175597834-DEL-130
chr1-175608597-DEL-56
chr1-175613665-DEL-475
chr1-175623337-DEL-77
chr1-175629017-DEL-104
chr1-175635386-DEL-459
chr1-175638691-DEL-125
chr1-175681463-DEL-1578
chr1-175718867-DEL-270
chr1-175729955-DEL-55
chr1-175739008-DEL-581
chr1-175781691-DEL-233
chr1-175782364-DEL-183
chr1-175814955-DEL-62
chr1-175819860-DEL-1192
chr1-175831540-DEL-63
chr1-175843135-DEL-60
chr1-175847582-DEL-156
chr1-175859204-DEL-56
chr1-175875239-DEL-1256
chr1-175905830-DEL-1668
chr1-175960840-DEL-60
chr1-175964497-DEL-78
chr1-175969720-DEL-180
chr1-176000532-DEL-575
chr1-176057596-DEL-149

chr1-176077017-DEL-57
chr1-176085499-DEL-59
chr1-176108409-DEL-68
chr1-176110104-DEL-204
chr1-176125783-DEL-58
chr1-176171474-DEL-1356
chr1-176171489-DEL-2035
chr1-176172962-DEL-441
chr1-176205909-DEL-548
chr1-176221027-DEL-1485
chr1-176266785-DEL-73
chr1-176286307-DEL-520
chr1-176301418-DEL-126
chr1-176303647-DEL-122
chr1-176335774-DEL-1145
chr1-176346309-DEL-352
chr1-176390152-DEL-641
chr1-176406358-DEL-1024
chr1-176406384-DEL-1637
chr1-176407452-DEL-308
chr1-176408084-DEL-991
chr1-176410903-DEL-60
chr1-176413045-DEL-1852
chr1-176414514-DEL-262
chr1-176420105-DEL-52
chr1-176465835-DEL-53
chr1-176480728-DEL-270
chr1-176484892-DEL-56
chr1-176549689-DEL-202
chr1-176572303-DEL-65
chr1-176622202-DEL-68
chr1-176630128-DEL-105
chr1-176647186-DEL-108
chr1-176718690-DEL-699
chr1-176729999-DEL-83
chr1-176737492-DEL-571
chr1-176802632-DEL-92
chr1-176812131-DEL-213
chr1-176868045-DEL-289
chr1-176945419-DEL-3184
chr1-176953186-DEL-456
chr1-176955724-DEL-863
chr1-176966715-DEL-57
chr1-176974565-DEL-61
chr1-176987824-DEL-882
chr1-176987983-DEL-219
chr1-177074255-DEL-128
chr1-177116508-DEL-112
chr1-177119474-DEL-58
chr1-177143044-DEL-746
chr1-177164466-DEL-177
chr1-177178945-DEL-513
chr1-177180522-DEL-342
chr1-177289561-DEL-59
chr1-177306352-DEL-56
chr1-177361578-DEL-67
chr1-177398411-DEL-60
chr1-177413994-DEL-1988
chr1-177444827-DEL-418
chr1-177466042-DEL-57
chr1-177495573-DEL-107
chr1-177498267-DEL-148
chr1-177502434-DEL-331
chr1-177505394-DEL-197
chr1-177517355-DEL-52
chr1-177541923-DEL-1575
chr1-177548056-DEL-84
chr1-177573476-DEL-2807
chr1-177581579-DEL-326
chr1-177614791-DEL-81
chr1-177636427-DEL-54
chr1-177660861-DEL-1927
chr1-177660863-DEL-271
chr1-177672791-DEL-749
chr1-177685509-DEL-225
chr1-177698797-DEL-199
chr1-177720525-DEL-1807

chr1-177738567-DEL-90
chr1-177766039-DEL-100
chr1-177777162-DEL-56
chr1-177780120-DEL-348
chr1-177797191-DEL-288
chr1-177805018-DEL-2873
chr1-177838196-DEL-214
chr1-177846307-DEL-182
chr1-177885587-DEL-7105
chr1-177906520-DEL-338
chr1-177915589-DEL-5169
chr1-177937771-DEL-90
chr1-177940266-DEL-133
chr1-177954637-DEL-2384
chr1-177960163-DEL-201
chr1-177961994-DEL-5743
chr1-177969803-DEL-100
chr1-177981486-DEL-77
chr1-177989444-DEL-91
chr1-178035993-DEL-251
chr1-178038759-DEL-3934
chr1-178058290-DEL-83
chr1-178081237-DEL-737
chr1-178100060-DEL-189
chr1-178118292-DEL-144
chr1-178123815-DEL-192
chr1-178194317-DEL-142
chr1-178208021-DEL-1149
chr1-178208260-DEL-581
chr1-178214947-DEL-71
chr1-178221932-DEL-755
chr1-178261870-DEL-1026
chr1-178315180-DEL-148
chr1-178336550-DEL-1489
chr1-178346497-DEL-56
chr1-178354616-DEL-103
chr1-178376700-DEL-66
chr1-178377640-DEL-58
chr1-178440134-DEL-359
chr1-178461692-DEL-64
chr1-178477247-DEL-655
chr1-178596221-DEL-62
chr1-178598541-DEL-74
chr1-178625751-DEL-56
chr1-178637416-DEL-96
chr1-178646162-DEL-121
chr1-178669656-DEL-85
chr1-178685147-DEL-54
chr1-178686639-DEL-338
chr1-178707037-DEL-109
chr1-178732484-DEL-3459
chr1-178804979-DEL-89
chr1-178815883-DEL-345
chr1-178843260-DEL-62
chr1-178857516-DEL-206
chr1-178893422-DEL-2757
chr1-178896657-DEL-861
chr1-178900000-DEL-68
chr1-178931608-DEL-64
chr1-178939099-DEL-165
chr1-178990660-DEL-223
chr1-178990995-DEL-909
chr1-178991598-DEL-143
chr1-179009745-DEL-110
chr1-179014302-DEL-212
chr1-179029103-DEL-100
chr1-179069267-DEL-58
chr1-179070113-DEL-181
chr1-179125395-DEL-642
chr1-179147550-DEL-98
chr1-179149250-DEL-66
chr1-179188414-DEL-144
chr1-179200888-DEL-59
chr1-179225199-DEL-309
chr1-179248748-DEL-1192
chr1-179274638-DEL-346
chr1-179275114-DEL-875

chr1-179283712-DEL-64
chr1-179324594-DEL-132
chr1-179346289-DEL-838
chr1-179347797-DEL-676
chr1-179353455-DEL-363
chr1-179380091-DEL-84
chr1-179392222-DEL-184
chr1-179404629-DEL-51
chr1-179417001-DEL-51
chr1-179420471-DEL-194
chr1-179421442-DEL-374
chr1-179423496-DEL-3136
chr1-179425221-DEL-56
chr1-179425941-DEL-106
chr1-179426819-DEL-207
chr1-179430856-DEL-179
chr1-179434715-DEL-141
chr1-179455757-DEL-716
chr1-179463598-DEL-6342
chr1-179470218-DEL-80
chr1-179492450-DEL-53
chr1-179499181-DEL-67
chr1-179530319-DEL-908
chr1-179534231-DEL-486
chr1-179542537-DEL-652
chr1-179562598-DEL-105
chr1-179618911-DEL-172
chr1-179625072-DEL-218
chr1-179639860-DEL-1139
chr1-179655038-DEL-1120
chr1-179665413-DEL-96
chr1-179667075-DEL-434
chr1-179678425-DEL-96
chr1-179684078-DEL-60
chr1-179695887-DEL-682
chr1-179730057-DEL-2155
chr1-179730142-DEL-310
chr1-179730168-DEL-230
chr1-179730932-DEL-786
chr1-179731129-DEL-1093
chr1-179731907-DEL-367
chr1-179731979-DEL-1012
chr1-179732317-DEL-570
chr1-179733078-DEL-115
chr1-179752444-DEL-567
chr1-179756428-DEL-120
chr1-179769730-DEL-383
chr1-179794356-DEL-117
chr1-179843843-DEL-61
chr1-179844820-DEL-92
chr1-179861231-DEL-481
chr1-179862274-DEL-183
chr1-179891722-DEL-192
chr1-179893300-DEL-55
chr1-179943420-DEL-62
chr1-179948591-DEL-148
chr1-179963072-DEL-410
chr1-180023239-DEL-149
chr1-180063774-DEL-73
chr1-180069547-DEL-825
chr1-180069598-DEL-349
chr1-180070043-DEL-399
chr1-180070102-DEL-1874
chr1-180070994-DEL-324
chr1-180071894-DEL-98
chr1-180090698-DEL-70
chr1-180100782-DEL-58
chr1-180109973-DEL-84
chr1-180124780-DEL-151
chr1-180173660-DEL-133
chr1-180175026-DEL-59
chr1-180180198-DEL-60
chr1-180184437-DEL-53
chr1-180211133-DEL-90
chr1-180239590-DEL-112
chr1-180243584-DEL-402
chr1-180287955-DEL-60

chr1-180302600-DEL-178
chr1-180315303-DEL-152
chr1-180342943-DEL-218
chr1-180349801-DEL-77
chr1-180367688-DEL-66
chr1-180402321-DEL-132
chr1-180413625-DEL-85
chr1-180415030-DEL-374
chr1-180423117-DEL-413
chr1-180425110-DEL-59
chr1-180438275-DEL-70
chr1-180439647-DEL-3970
chr1-180439648-DEL-2402
chr1-180442065-DEL-2017
chr1-180443947-DEL-155
chr1-180478633-DEL-190
chr1-180480777-DEL-327
chr1-180483579-DEL-2086
chr1-180499706-DEL-61
chr1-180500738-DEL-181
chr1-180503720-DEL-93
chr1-180506430-DEL-203
chr1-180522946-DEL-177
chr1-180576705-DEL-84
chr1-180577759-DEL-64
chr1-180578125-DEL-1375
chr1-180593916-DEL-80
chr1-180628153-DEL-93
chr1-180632589-DEL-94
chr1-180638830-DEL-177
chr1-180645795-DEL-1093
chr1-180647206-DEL-182
chr1-180650722-DEL-131
chr1-180651249-DEL-708
chr1-180656747-DEL-146
chr1-180668582-DEL-563
chr1-180673358-DEL-1602
chr1-180679165-DEL-52
chr1-180681466-DEL-686
chr1-180699970-DEL-355
chr1-180717457-DEL-129
chr1-180745819-DEL-794
chr1-180755535-DEL-152
chr1-180757996-DEL-219
chr1-180760935-DEL-83
chr1-180829737-DEL-225
chr1-180833413-DEL-489
chr1-180886108-DEL-178
chr1-180895323-DEL-119
chr1-180907610-DEL-123
chr1-180946543-DEL-198
chr1-180956470-DEL-64
chr1-180978253-DEL-53
chr1-180987137-DEL-56
chr1-181017378-DEL-197
chr1-181021173-DEL-76
chr1-181032646-DEL-62
chr1-181037594-DEL-65
chr1-181114239-DEL-567
chr1-181151678-DEL-125
chr1-181185466-DEL-353
chr1-181186440-DEL-85
chr1-181194346-DEL-60
chr1-181242934-DEL-160
chr1-181337209-DEL-102
chr1-181360202-DEL-70
chr1-181385477-DEL-656
chr1-181386620-DEL-59
chr1-181391375-DEL-444
chr1-181395816-DEL-141
chr1-181418518-DEL-114
chr1-181435236-DEL-82
chr1-181457690-DEL-463
chr1-181458466-DEL-473
chr1-181499152-DEL-1422
chr1-181504354-DEL-112
chr1-181505925-DEL-358

chr1-181540515-DEL-2644
chr1-181560859-DEL-298
chr1-181571026-DEL-480
chr1-181579081-DEL-435
chr1-181588518-DEL-2000
chr1-181614406-DEL-670
chr1-181662107-DEL-214
chr1-181687210-DEL-59
chr1-181693919-DEL-65
chr1-181712689-DEL-295
chr1-181733708-DEL-173
chr1-181745440-DEL-181
chr1-181747403-DEL-84
chr1-181751753-DEL-228
chr1-181754259-DEL-397
chr1-181810585-DEL-340
chr1-181822771-DEL-75
chr1-181827623-DEL-196
chr1-181831897-DEL-100
chr1-181834956-DEL-197
chr1-181890742-DEL-171
chr1-181940238-DEL-98
chr1-181970752-DEL-152
chr1-181975324-DEL-8946
chr1-181978846-DEL-285
chr1-181994674-DEL-211
chr1-182013910-DEL-120
chr1-182062815-DEL-67
chr1-182069167-DEL-1549
chr1-182105127-DEL-428
chr1-182113370-DEL-65
chr1-182169807-DEL-52
chr1-182201497-DEL-51
chr1-182238081-DEL-754
chr1-182250254-DEL-96
chr1-182310950-DEL-135
chr1-182320585-DEL-906
chr1-182322819-DEL-62
chr1-182332709-DEL-154
chr1-182345425-DEL-501
chr1-182369734-DEL-492
chr1-182375051-DEL-67
chr1-182386650-DEL-83
chr1-182393622-DEL-62
chr1-182457116-DEL-72
chr1-182469060-DEL-1335
chr1-182471366-DEL-801
chr1-182489117-DEL-62
chr1-182512256-DEL-56
chr1-182528139-DEL-533
chr1-182605025-DEL-58
chr1-182618618-DEL-103
chr1-182637420-DEL-2408
chr1-182640629-DEL-89
chr1-182663550-DEL-510
chr1-182678891-DEL-51
chr1-182687866-DEL-123
chr1-182691290-DEL-1089
chr1-182721272-DEL-839
chr1-182728239-DEL-59
chr1-182752369-DEL-119
chr1-182756788-DEL-136
chr1-182763351-DEL-65
chr1-182789796-DEL-132
chr1-182793247-DEL-373
chr1-182822942-DEL-91
chr1-182831121-DEL-90
chr1-182834959-DEL-486
chr1-182843479-DEL-833
chr1-182844619-DEL-182
chr1-182887480-DEL-486
chr1-182888860-DEL-60
chr1-182898425-DEL-101
chr1-182906866-DEL-3954
chr1-182924037-DEL-135
chr1-182943302-DEL-179
chr1-182955835-DEL-452

chr1-182962269-DEL-175
chr1-182968015-DEL-56
chr1-182973301-DEL-60
chr1-183038776-DEL-145
chr1-183040045-DEL-922
chr1-183124576-DEL-147
chr1-183172414-DEL-324
chr1-183173254-DEL-152
chr1-183176095-DEL-236
chr1-183180114-DEL-747
chr1-183180887-DEL-391
chr1-183202924-DEL-212
chr1-183210769-DEL-59
chr1-183223543-DEL-296
chr1-183227674-DEL-784
chr1-183257459-DEL-90
chr1-183267863-DEL-75
chr1-183288948-DEL-377
chr1-183307389-DEL-114
chr1-183310144-DEL-182
chr1-183330428-DEL-755
chr1-183337298-DEL-59
chr1-183360865-DEL-80
chr1-183373881-DEL-395
chr1-183391804-DEL-312
chr1-183406404-DEL-64
chr1-183410405-DEL-126
chr1-183414231-DEL-976
chr1-183418804-DEL-69
chr1-183455137-DEL-84
chr1-183470364-DEL-803
chr1-183484875-DEL-264
chr1-183510375-DEL-54
chr1-183521484-DEL-147
chr1-183541940-DEL-6792
chr1-183555803-DEL-338
chr1-183556450-DEL-1657
chr1-183591552-DEL-51
chr1-183602154-DEL-747
chr1-183611829-DEL-51
chr1-183612519-DEL-51
chr1-183651868-DEL-287
chr1-183694548-DEL-106
chr1-183703255-DEL-399
chr1-183752486-DEL-68
chr1-183757633-DEL-400
chr1-183778951-DEL-121
chr1-183792931-DEL-54
chr1-183813564-DEL-201
chr1-183824666-DEL-380
chr1-183836534-DEL-620
chr1-183852770-DEL-510
chr1-183855613-DEL-64
chr1-183871414-DEL-53
chr1-183888501-DEL-135
chr1-183892033-DEL-171
chr1-183896197-DEL-52
chr1-183906903-DEL-67
chr1-183919149-DEL-54
chr1-183921750-DEL-68
chr1-183922812-DEL-703
chr1-183923424-DEL-93
chr1-183976060-DEL-64
chr1-183980794-DEL-208
chr1-183981795-DEL-1270
chr1-183988000-DEL-915
chr1-184001551-DEL-89
chr1-184015763-DEL-75
chr1-184051762-DEL-3166
chr1-184062240-DEL-270
chr1-184077380-DEL-110
chr1-184101989-DEL-893
chr1-184103526-DEL-4074
chr1-184129907-DEL-95
chr1-184140574-DEL-145
chr1-184147857-DEL-122
chr1-184160538-DEL-66

chr1-184164281-DEL-243
chr1-184178755-DEL-55
chr1-184182121-DEL-56
chr1-184201516-DEL-200
chr1-184206292-DEL-288
chr1-184225056-DEL-54
chr1-184238088-DEL-115
chr1-184258765-DEL-111
chr1-184273839-DEL-80
chr1-184305226-DEL-193
chr1-184312160-DEL-96
chr1-184328756-DEL-52
chr1-184331577-DEL-538
chr1-184348578-DEL-53
chr1-184366256-DEL-6283
chr1-184427635-DEL-86
chr1-184434923-DEL-689
chr1-184435520-DEL-189
chr1-184437080-DEL-1153
chr1-184437758-DEL-68
chr1-184461312-DEL-86
chr1-184480630-DEL-70
chr1-184485672-DEL-182
chr1-184495594-DEL-60
chr1-184496802-DEL-742
chr1-184518391-DEL-188
chr1-184523668-DEL-51
chr1-184526608-DEL-158
chr1-184546664-DEL-437
chr1-184547280-DEL-500
chr1-184555529-DEL-962
chr1-184559369-DEL-55
chr1-184594851-DEL-57
chr1-184621064-DEL-86
chr1-184623492-DEL-141
chr1-184649645-DEL-65
chr1-184660696-DEL-105
chr1-184688613-DEL-237
chr1-184693006-DEL-3840
chr1-184703193-DEL-149
chr1-184708075-DEL-104
chr1-184744970-DEL-189
chr1-184755761-DEL-116
chr1-184775449-DEL-428
chr1-184803259-DEL-152
chr1-184808702-DEL-514
chr1-184812918-DEL-1501
chr1-184814745-DEL-334
chr1-184821221-DEL-69
chr1-184840970-DEL-60
chr1-184846529-DEL-131
chr1-184855338-DEL-66
chr1-184874014-DEL-74
chr1-184877712-DEL-2014
chr1-184912513-DEL-1131
chr1-184969603-DEL-871
chr1-184979299-DEL-117
chr1-185001804-DEL-661
chr1-185019469-DEL-82
chr1-185031542-DEL-204
chr1-185051268-DEL-700
chr1-185051922-DEL-149
chr1-185089441-DEL-152
chr1-185124889-DEL-62
chr1-185134286-DEL-65
chr1-185180887-DEL-109
chr1-185185273-DEL-83
chr1-185189378-DEL-87
chr1-185190420-DEL-354
chr1-185217211-DEL-164
chr1-185222887-DEL-147
chr1-185235368-DEL-60
chr1-185243405-DEL-197
chr1-185245840-DEL-126
chr1-185258854-DEL-1439
chr1-185272970-DEL-204
chr1-185285443-DEL-71

chr1-185290840-DEL-60
chr1-185405998-DEL-264
chr1-185407016-DEL-80
chr1-185425380-DEL-254
chr1-185452898-DEL-186
chr1-185482531-DEL-387
chr1-185488661-DEL-90
chr1-185556824-DEL-1458
chr1-185584912-DEL-56
chr1-185590470-DEL-60
chr1-185610097-DEL-1066
chr1-185651252-DEL-51
chr1-185660986-DEL-233
chr1-185664486-DEL-148
chr1-185668011-DEL-54
chr1-185685017-DEL-388
chr1-185691837-DEL-129
chr1-185703520-DEL-1124
chr1-185707664-DEL-122
chr1-185717216-DEL-74
chr1-185720677-DEL-105
chr1-185736845-DEL-2198
chr1-185790981-DEL-116
chr1-185803528-DEL-5760
chr1-185817151-DEL-73
chr1-185826251-DEL-58
chr1-185827103-DEL-399
chr1-185835486-DEL-76
chr1-185836041-DEL-75
chr1-185872323-DEL-91
chr1-185900544-DEL-66
chr1-185916318-DEL-605
chr1-185922293-DEL-178
chr1-185931235-DEL-63
chr1-185945949-DEL-331
chr1-185955334-DEL-83
chr1-185956513-DEL-1123
chr1-185996542-DEL-56
chr1-186000659-DEL-63
chr1-186002794-DEL-130
chr1-186022873-DEL-568
chr1-186065407-DEL-70
chr1-186070283-DEL-60
chr1-186092126-DEL-185
chr1-186117489-DEL-256
chr1-186128439-DEL-147
chr1-186130092-DEL-53
chr1-186149949-DEL-1505
chr1-186180554-DEL-247
chr1-186186005-DEL-117
chr1-186191096-DEL-51
chr1-186227509-DEL-60
chr1-186228547-DEL-82
chr1-186238894-DEL-56
chr1-186257970-DEL-945
chr1-186269391-DEL-280
chr1-186290492-DEL-100
chr1-186293919-DEL-54
chr1-186304768-DEL-113
chr1-186320206-DEL-1481
chr1-186339755-DEL-56
chr1-186347598-DEL-112
chr1-186353931-DEL-72
chr1-186371276-DEL-71
chr1-186375561-DEL-648
chr1-186393766-DEL-789
chr1-186404472-DEL-94
chr1-186434035-DEL-112
chr1-186438518-DEL-78
chr1-186442034-DEL-834
chr1-186442602-DEL-175
chr1-186462167-DEL-3137
chr1-186462581-DEL-93
chr1-186463634-DEL-1701
chr1-186481897-DEL-51
chr1-186534856-DEL-224
chr1-186546156-DEL-152

chr1-186553427-DEL-262
chr1-186557750-DEL-58
chr1-186578040-DEL-517
chr1-186654719-DEL-67
chr1-186725663-DEL-836
chr1-186726856-DEL-84
chr1-186737339-DEL-632
chr1-186771462-DEL-2092
chr1-186793618-DEL-68
chr1-186794509-DEL-99
chr1-186831310-DEL-81
chr1-186835804-DEL-2124
chr1-186841944-DEL-478
chr1-186849851-DEL-167
chr1-186860659-DEL-2299
chr1-186898082-DEL-51
chr1-186903571-DEL-768
chr1-186917493-DEL-62
chr1-186919554-DEL-90
chr1-186924106-DEL-441
chr1-186931420-DEL-62
chr1-186933178-DEL-760
chr1-186936210-DEL-113
chr1-186962521-DEL-52
chr1-187015951-DEL-294
chr1-187041837-DEL-57
chr1-187062904-DEL-290
chr1-187071251-DEL-140
chr1-187071284-DEL-710
chr1-187085332-DEL-1392
chr1-187085683-DEL-1798
chr1-187087267-DEL-243
chr1-187125106-DEL-848
chr1-187135940-DEL-75
chr1-187138455-DEL-1245
chr1-187154336-DEL-151
chr1-187155866-DEL-243
chr1-187197426-DEL-55
chr1-187230153-DEL-737
chr1-187234789-DEL-339
chr1-187267092-DEL-83
chr1-187267688-DEL-52
chr1-187276890-DEL-1183
chr1-187292114-DEL-159
chr1-187323245-DEL-300
chr1-187348709-DEL-75
chr1-187358187-DEL-57
chr1-187475372-DEL-58
chr1-187481870-DEL-56
chr1-187484087-DEL-236
chr1-187494167-DEL-54
chr1-187507313-DEL-172
chr1-187545223-DEL-95
chr1-187546340-DEL-64
chr1-187547904-DEL-164
chr1-187574279-DEL-95
chr1-187575111-DEL-54
chr1-187588829-DEL-53
chr1-187601201-DEL-79
chr1-187606146-DEL-95
chr1-187625666-DEL-136
chr1-187629611-DEL-66
chr1-187639096-DEL-96
chr1-187673413-DEL-89
chr1-187693428-DEL-202
chr1-187693929-DEL-55
chr1-187710374-DEL-67
chr1-187722014-DEL-79
chr1-187723520-DEL-64
chr1-187728485-DEL-52
chr1-187731004-DEL-72
chr1-187744910-DEL-544
chr1-187783375-DEL-55
chr1-187805690-DEL-65
chr1-187817026-DEL-128
chr1-187820569-DEL-94
chr1-187846976-DEL-70

chr1-187852027-DEL-434
chr1-187862812-DEL-264
chr1-187867785-DEL-86
chr1-187872326-DEL-82
chr1-187912416-DEL-58
chr1-187920542-DEL-140
chr1-187934088-DEL-251
chr1-187952061-DEL-112
chr1-187972336-DEL-214
chr1-187980488-DEL-342
chr1-188031674-DEL-142
chr1-188038894-DEL-130
chr1-188040528-DEL-1458
chr1-188044686-DEL-81
chr1-188056495-DEL-800
chr1-188057661-DEL-99
chr1-188057683-DEL-376
chr1-188061870-DEL-206
chr1-188075117-DEL-215
chr1-188082113-DEL-135
chr1-188085272-DEL-71
chr1-188109620-DEL-65
chr1-188156297-DEL-92
chr1-188192344-DEL-338
chr1-188194544-DEL-56
chr1-188218272-DEL-313
chr1-188242433-DEL-139
chr1-188279597-DEL-63
chr1-188300130-DEL-58
chr1-188305180-DEL-2120
chr1-188320711-DEL-64
chr1-188331025-DEL-110
chr1-188376586-DEL-1692
chr1-188379403-DEL-689
chr1-188386064-DEL-171
chr1-188402952-DEL-271
chr1-188414524-DEL-154
chr1-188520378-DEL-337
chr1-188522198-DEL-189
chr1-188525608-DEL-228
chr1-188526362-DEL-176
chr1-188549323-DEL-547
chr1-188570562-DEL-59
chr1-188601491-DEL-150
chr1-188607439-DEL-90
chr1-188613838-DEL-73
chr1-188625665-DEL-195
chr1-188628317-DEL-240
chr1-188663091-DEL-67
chr1-188673627-DEL-484
chr1-188705531-DEL-61
chr1-188721203-DEL-141
chr1-188723910-DEL-80
chr1-188730876-DEL-97
chr1-188742006-DEL-130
chr1-188762744-DEL-126
chr1-188765110-DEL-153
chr1-188799015-DEL-55
chr1-188801093-DEL-173
chr1-188846062-DEL-132
chr1-188859839-DEL-208
chr1-188861620-DEL-61
chr1-188862912-DEL-79
chr1-188880911-DEL-59
chr1-188899405-DEL-3427
chr1-188903719-DEL-344
chr1-188906848-DEL-61
chr1-188926492-DEL-627
chr1-188944583-DEL-97
chr1-188967159-DEL-86
chr1-188975771-DEL-103
chr1-188982252-DEL-143
chr1-189017604-DEL-175
chr1-189030972-DEL-55
chr1-189032885-DEL-256
chr1-189058979-DEL-88
chr1-189088918-DEL-62

chr1-189151852-DEL-133
chr1-189173543-DEL-59
chr1-189211826-DEL-77
chr1-189233281-DEL-145
chr1-189238555-DEL-1496
chr1-189240501-DEL-94
chr1-189244098-DEL-56
chr1-189244675-DEL-68
chr1-189267154-DEL-612
chr1-189269886-DEL-173
chr1-189270452-DEL-1910
chr1-189292134-DEL-159
chr1-189325717-DEL-188
chr1-189327119-DEL-675
chr1-189342303-DEL-534
chr1-189349310-DEL-338
chr1-189372044-DEL-1379
chr1-189425186-DEL-74
chr1-189431185-DEL-53
chr1-189466569-DEL-143
chr1-189469094-DEL-168
chr1-189483385-DEL-314
chr1-189495781-DEL-75
chr1-189498344-DEL-1254
chr1-189521592-DEL-57
chr1-189551164-DEL-131
chr1-189557691-DEL-103
chr1-189588530-DEL-133
chr1-189599597-DEL-67
chr1-189606609-DEL-73
chr1-189637956-DEL-490
chr1-189638636-DEL-127
chr1-189659215-DEL-62
chr1-189659784-DEL-65
chr1-189693472-DEL-146
chr1-189695768-DEL-2638
chr1-189712699-DEL-236
chr1-189716815-DEL-84
chr1-189738390-DEL-69
chr1-189744878-DEL-554
chr1-189748714-DEL-57
chr1-189751650-DEL-938
chr1-189772322-DEL-54
chr1-189773963-DEL-200
chr1-189776735-DEL-124
chr1-189787243-DEL-66
chr1-189790707-DEL-146
chr1-189802405-DEL-112
chr1-189833712-DEL-298
chr1-189837983-DEL-56
chr1-189857675-DEL-75
chr1-189884615-DEL-192
chr1-189915485-DEL-53
chr1-189935194-DEL-111
chr1-189944049-DEL-592
chr1-189964102-DEL-62
chr1-189967999-DEL-270
chr1-190000398-DEL-397
chr1-190004180-DEL-79
chr1-190009830-DEL-407
chr1-190024917-DEL-82
chr1-190025333-DEL-229
chr1-190025814-DEL-771
chr1-190032317-DEL-3924
chr1-190033589-DEL-213
chr1-190042985-DEL-66
chr1-190058362-DEL-55
chr1-190063059-DEL-76
chr1-190063956-DEL-62
chr1-190089206-DEL-104
chr1-190150836-DEL-183
chr1-190162403-DEL-51
chr1-190177783-DEL-75
chr1-190179147-DEL-72
chr1-190242531-DEL-110
chr1-190259293-DEL-97
chr1-190263740-DEL-224

chr1-190268700-DEL-100
chr1-190275628-DEL-111
chr1-190297878-DEL-58
chr1-190339705-DEL-59
chr1-190362409-DEL-124
chr1-190375607-DEL-53
chr1-190394459-DEL-86
chr1-190406942-DEL-249
chr1-190407566-DEL-73
chr1-190446912-DEL-67
chr1-190475207-DEL-54
chr1-190479500-DEL-51
chr1-190482565-DEL-105
chr1-190488221-DEL-620
chr1-190488808-DEL-104
chr1-190501323-DEL-262
chr1-190512661-DEL-147
chr1-190512808-DEL-988
chr1-190513130-DEL-298
chr1-190514037-DEL-177
chr1-190522585-DEL-123
chr1-190534085-DEL-322
chr1-190564686-DEL-68
chr1-190583278-DEL-827
chr1-190583486-DEL-508
chr1-190613331-DEL-82
chr1-190616518-DEL-100
chr1-190618307-DEL-75
chr1-190650183-DEL-71
chr1-190671908-DEL-111
chr1-190679356-DEL-76
chr1-190682508-DEL-90
chr1-190690416-DEL-162
chr1-190707775-DEL-478
chr1-190721016-DEL-65
chr1-190732924-DEL-63
chr1-190745846-DEL-295
chr1-190776035-DEL-409
chr1-190782944-DEL-1388
chr1-190783615-DEL-101
chr1-190784688-DEL-95
chr1-190792206-DEL-77
chr1-190843984-DEL-161
chr1-190867329-DEL-69
chr1-190869965-DEL-298
chr1-190882733-DEL-884
chr1-190902059-DEL-169
chr1-190908665-DEL-54
chr1-190947058-DEL-60
chr1-190985145-DEL-67
chr1-190992523-DEL-266
chr1-191001137-DEL-52
chr1-191006657-DEL-71
chr1-191024186-DEL-83
chr1-191035541-DEL-114
chr1-191040957-DEL-404
chr1-191045819-DEL-234
chr1-191047976-DEL-51
chr1-191050529-DEL-137
chr1-191061969-DEL-444
chr1-191066343-DEL-160
chr1-191071351-DEL-112
chr1-191083477-DEL-207
chr1-191102205-DEL-505
chr1-191111664-DEL-203
chr1-191116354-DEL-83
chr1-191132263-DEL-77
chr1-191150997-DEL-297
chr1-191155228-DEL-118
chr1-191174570-DEL-54
chr1-191184792-DEL-347
chr1-191185468-DEL-191
chr1-191199740-DEL-60
chr1-191205310-DEL-342
chr1-191220901-DEL-98
chr1-191224383-DEL-100
chr1-191257515-DEL-161

chr1-191266603-DEL-183
chr1-191284453-DEL-78
chr1-191287438-DEL-683
chr1-191298097-DEL-196
chr1-191310179-DEL-1349
chr1-191359666-DEL-68
chr1-191368067-DEL-116
chr1-191378868-DEL-276
chr1-191384462-DEL-802
chr1-191401426-DEL-51
chr1-191406133-DEL-1294
chr1-191412598-DEL-328
chr1-191422370-DEL-342
chr1-191448957-DEL-78
chr1-191455731-DEL-823
chr1-191472071-DEL-72
chr1-191491032-DEL-53
chr1-191530031-DEL-871
chr1-191531300-DEL-358
chr1-191598836-DEL-108
chr1-191600502-DEL-288
chr1-191619016-DEL-267
chr1-191620718-DEL-176
chr1-191628372-DEL-56
chr1-191646084-DEL-141
chr1-191683454-DEL-55
chr1-191732481-DEL-689
chr1-191735497-DEL-70
chr1-191741786-DEL-94
chr1-191796025-DEL-144
chr1-191803145-DEL-190
chr1-191811647-DEL-352
chr1-191832226-DEL-634
chr1-191856873-DEL-55
chr1-191864918-DEL-1810
chr1-191868567-DEL-177
chr1-191871909-DEL-77
chr1-191884507-DEL-156
chr1-191896761-DEL-98
chr1-191901083-DEL-72
chr1-191912197-DEL-1151
chr1-191912267-DEL-864
chr1-191912309-DEL-96
chr1-191913854-DEL-143
chr1-191952652-DEL-64
chr1-191983459-DEL-67
chr1-192000038-DEL-76
chr1-192012721-DEL-66
chr1-192034647-DEL-913
chr1-192044640-DEL-56
chr1-192047553-DEL-192
chr1-192048495-DEL-348
chr1-192065280-DEL-108
chr1-192089209-DEL-243
chr1-192092699-DEL-116
chr1-192097302-DEL-288
chr1-192102549-DEL-1099
chr1-192108351-DEL-52
chr1-192115501-DEL-1492
chr1-192130551-DEL-152
chr1-192142677-DEL-115
chr1-192147613-DEL-6650
chr1-192153783-DEL-208
chr1-192173533-DEL-137
chr1-192230146-DEL-108
chr1-192265800-DEL-64
chr1-192280463-DEL-64
chr1-192299146-DEL-945
chr1-192310634-DEL-102
chr1-192311828-DEL-2944
chr1-192341928-DEL-98
chr1-192350790-DEL-926
chr1-192352116-DEL-703
chr1-192352123-DEL-1414
chr1-192352478-DEL-402
chr1-192363180-DEL-52
chr1-192399937-DEL-105

chr1-192404297-DEL-276
chr1-192424398-DEL-58
chr1-192434827-DEL-67
chr1-192457405-DEL-301
chr1-192464248-DEL-1287
chr1-192475737-DEL-89
chr1-192485868-DEL-648
chr1-192560858-DEL-76
chr1-192600852-DEL-292
chr1-192714743-DEL-257
chr1-192729793-DEL-58
chr1-192734160-DEL-770
chr1-192737875-DEL-198
chr1-192742609-DEL-86
chr1-192773806-DEL-54
chr1-192780598-DEL-54
chr1-192804019-DEL-71
chr1-192837254-DEL-177
chr1-192885030-DEL-276
chr1-192902971-DEL-225
chr1-192906479-DEL-104
chr1-192927742-DEL-869
chr1-192934400-DEL-64
chr1-192936425-DEL-227
chr1-192938380-DEL-193
chr1-192983138-DEL-210
chr1-193028031-DEL-68
chr1-193057260-DEL-65
chr1-193058369-DEL-120
chr1-193087531-DEL-116
chr1-193091386-DEL-65
chr1-193099344-DEL-198
chr1-193099973-DEL-271
chr1-193105243-DEL-81
chr1-193109848-DEL-59
chr1-193154629-DEL-286
chr1-193180409-DEL-109
chr1-193206279-DEL-80
chr1-193214719-DEL-408
chr1-193307236-DEL-161
chr1-193325814-DEL-1172
chr1-193341299-DEL-1014
chr1-193424568-DEL-78
chr1-193440414-DEL-75
chr1-193441514-DEL-1178
chr1-193447145-DEL-1121
chr1-193545807-DEL-810
chr1-193561697-DEL-185
chr1-193602686-DEL-57
chr1-193687830-DEL-56
chr1-193688568-DEL-919
chr1-193690344-DEL-175
chr1-193691036-DEL-3802
chr1-193691040-DEL-1399
chr1-193708825-DEL-70
chr1-193715563-DEL-116
chr1-193722954-DEL-114
chr1-193741914-DEL-654
chr1-193746178-DEL-80
chr1-193757801-DEL-640
chr1-193769793-DEL-539
chr1-193789296-DEL-90
chr1-193815022-DEL-140
chr1-193816020-DEL-1977
chr1-193825943-DEL-1785
chr1-193827728-DEL-488
chr1-193837187-DEL-396
chr1-193863640-DEL-74
chr1-193873952-DEL-325
chr1-193885152-DEL-463
chr1-193895662-DEL-53
chr1-193926189-DEL-108
chr1-193966712-DEL-86
chr1-193976209-DEL-243
chr1-194057901-DEL-1108
chr1-194068482-DEL-1292
chr1-194115134-DEL-61

chr1-194144557-DEL-259
chr1-194155205-DEL-502
chr1-194211377-DEL-6817
chr1-194211865-DEL-51
chr1-194259149-DEL-75
chr1-194276941-DEL-1321
chr1-194290548-DEL-400
chr1-194331601-DEL-66
chr1-194372805-DEL-104
chr1-194381924-DEL-576
chr1-194408847-DEL-111
chr1-194417935-DEL-52
chr1-194431857-DEL-488
chr1-194555752-DEL-313
chr1-194575123-DEL-503
chr1-194580274-DEL-75
chr1-194608879-DEL-360
chr1-194632416-DEL-59
chr1-194654722-DEL-190
chr1-194659376-DEL-232
chr1-194707142-DEL-146
chr1-194738562-DEL-304
chr1-194772333-DEL-128
chr1-194827975-DEL-1081
chr1-194843845-DEL-185
chr1-194886031-DEL-2191
chr1-194910380-DEL-439
chr1-194916386-DEL-6166
chr1-194926002-DEL-6143
chr1-194936915-DEL-7465
chr1-194937501-DEL-72
chr1-194959646-DEL-3427
chr1-195001503-DEL-57
chr1-195013257-DEL-655
chr1-195018454-DEL-52
chr1-195020568-DEL-66
chr1-195043190-DEL-173
chr10-3059408-DEL-2190
chr10-3060473-DEL-770
chr10-3062093-DEL-503
chr10-3062792-DEL-2057
chr10-3062810-DEL-148
chr10-3064934-DEL-1295
chr10-3066533-DEL-446
chr10-3071100-DEL-397
chr10-3078021-DEL-127
chr10-3118378-DEL-2176
chr10-3125202-DEL-368
chr10-3126254-DEL-178
chr10-3140442-DEL-171
chr10-3168235-DEL-201
chr10-3216432-DEL-3030
chr10-3259727-DEL-52
chr10-3266237-DEL-5159
chr10-3299540-DEL-1074
chr10-3322680-DEL-490
chr10-3362755-DEL-69
chr10-3375800-DEL-52
chr10-3396644-DEL-84
chr10-3433457-DEL-2041
chr10-3470649-DEL-111
chr10-3482291-DEL-121
chr10-3516435-DEL-2963
chr10-3517533-DEL-620
chr10-3535665-DEL-116
chr10-3559399-DEL-66
chr10-3569373-DEL-203
chr10-3570833-DEL-6911
chr10-3570888-DEL-204
chr10-3604366-DEL-921
chr10-3616815-DEL-69
chr10-3642935-DEL-223
chr10-3671482-DEL-457
chr10-3685752-DEL-702
chr10-3720132-DEL-113
chr10-3731280-DEL-53
chr10-3746318-DEL-72

chr10-3778927-DEL-104
chr10-3787060-DEL-53
chr10-3816732-DEL-491
chr10-3818809-DEL-198
chr10-3824457-DEL-118
chr10-3832638-DEL-52
chr10-3841285-DEL-75
chr10-3878926-DEL-153
chr10-3886924-DEL-132
chr10-3927045-DEL-56
chr10-3938671-DEL-79
chr10-3940599-DEL-196
chr10-3948946-DEL-843
chr10-3950816-DEL-678
chr10-3968767-DEL-94
chr10-4012737-DEL-869
chr10-4051921-DEL-320
chr10-4073253-DEL-305
chr10-4087215-DEL-226
chr10-4108426-DEL-119
chr10-4132977-DEL-184
chr10-4151519-DEL-660
chr10-4167257-DEL-2326
chr10-4187162-DEL-207
chr10-4188410-DEL-203
chr10-4191250-DEL-522
chr10-4194391-DEL-126
chr10-4195285-DEL-548
chr10-4212650-DEL-58
chr10-4219651-DEL-270
chr10-4222471-DEL-196
chr10-4223336-DEL-65
chr10-4226608-DEL-191
chr10-4236844-DEL-75
chr10-4248467-DEL-1444
chr10-4268596-DEL-214
chr10-4292340-DEL-1278
chr10-4325076-DEL-57
chr10-4376834-DEL-90
chr10-4399973-DEL-454
chr10-4419336-DEL-281
chr10-4423012-DEL-63
chr10-4429307-DEL-65
chr10-4448724-DEL-486
chr10-4452768-DEL-2054
chr10-4455128-DEL-56
chr10-4458434-DEL-51
chr10-4488214-DEL-77
chr10-4489094-DEL-622
chr10-4492566-DEL-58
chr10-4500996-DEL-281
chr10-4546136-DEL-505
chr10-4551294-DEL-782
chr10-4569784-DEL-102
chr10-4642779-DEL-60
chr10-4665204-DEL-131
chr10-4674564-DEL-391
chr10-4687591-DEL-72
chr10-4704327-DEL-121
chr10-4711348-DEL-461
chr10-4737771-DEL-6978
chr10-4753205-DEL-317
chr10-4810444-DEL-562
chr10-4814165-DEL-1319
chr10-4822035-DEL-729
chr10-4861715-DEL-212
chr10-4891596-DEL-54
chr10-4896161-DEL-76
chr10-4898291-DEL-1374
chr10-4914475-DEL-132
chr10-4941174-DEL-2236
chr10-4956529-DEL-53
chr10-4966821-DEL-68
chr10-4999964-DEL-895
chr10-5019313-DEL-76
chr10-5030907-DEL-1044
chr10-5031005-DEL-946

chr10-5032000-DEL-1304
chr10-5087483-DEL-428
chr10-5098698-DEL-161
chr10-5107956-DEL-421
chr10-5160029-DEL-642
chr10-5207509-DEL-90
chr10-5269358-DEL-164
chr10-5304302-DEL-91
chr10-5305288-DEL-142
chr10-5310250-DEL-58
chr10-5326220-DEL-183
chr10-5333821-DEL-118
chr10-5338664-DEL-1349
chr10-5339106-DEL-208
chr10-5343038-DEL-536
chr10-5384303-DEL-57
chr10-5406124-DEL-178
chr10-5424333-DEL-204
chr10-5433444-DEL-1928
chr10-5436072-DEL-213
chr10-5450112-DEL-150
chr10-5451687-DEL-3146
chr10-5482805-DEL-567
chr10-5515446-DEL-3176
chr10-5524815-DEL-1554
chr10-5536874-DEL-61
chr10-5558900-DEL-76
chr10-5625233-DEL-789
chr10-5645144-DEL-1328
chr10-5650608-DEL-67
chr10-5732747-DEL-188
chr10-5759589-DEL-184
chr10-5765017-DEL-194
chr10-5770738-DEL-237
chr10-5825150-DEL-178
chr10-5825662-DEL-395
chr10-5827762-DEL-255
chr10-5833313-DEL-9924
chr10-5847807-DEL-266
chr10-5848983-DEL-66
chr10-5858306-DEL-89
chr10-5865447-DEL-304
chr10-5884763-DEL-291
chr10-5887132-DEL-89
chr10-5899314-DEL-515
chr10-5907349-DEL-6229
chr10-5923894-DEL-170
chr10-5929833-DEL-125
chr10-5932594-DEL-475
chr10-5938125-DEL-1488
chr10-5941963-DEL-618
chr10-5959843-DEL-1258
chr10-5991125-DEL-74
chr10-6015986-DEL-488
chr10-6018994-DEL-2819
chr10-6095476-DEL-268
chr10-6125097-DEL-204
chr10-6137953-DEL-124
chr10-6159327-DEL-4938
chr10-6187675-DEL-199
chr10-6213737-DEL-559
chr10-6250135-DEL-2450
chr10-6263556-DEL-95
chr10-6284620-DEL-222
chr10-6295518-DEL-5130
chr10-6312753-DEL-52
chr10-6335560-DEL-572
chr10-6348727-DEL-8814
chr10-6374411-DEL-184
chr10-6390752-DEL-245
chr10-6395973-DEL-673
chr10-6396616-DEL-90
chr10-6397769-DEL-3543
chr10-6398827-DEL-128
chr10-6401356-DEL-245
chr10-6404899-DEL-1258
chr10-6413822-DEL-55

chr10-6461429-DEL-514
chr10-6475823-DEL-57
chr10-6515284-DEL-110
chr10-6550735-DEL-502
chr10-6573647-DEL-6215
chr10-6580501-DEL-308
chr10-6584616-DEL-8536
chr10-6587945-DEL-5929
chr10-6619246-DEL-163
chr10-6643456-DEL-388
chr10-6688435-DEL-56
chr10-6699118-DEL-144
chr10-6717739-DEL-54
chr10-6725844-DEL-2440
chr10-6752952-DEL-137
chr10-6780710-DEL-328
chr10-6799428-DEL-191
chr10-6816039-DEL-2688
chr10-6838587-DEL-54
chr10-6877856-DEL-54
chr10-6921006-DEL-700
chr10-6931478-DEL-54
chr10-6934874-DEL-153
chr10-6937098-DEL-493
chr10-6940015-DEL-54
chr10-6943953-DEL-1230
chr10-6967720-DEL-321
chr10-6999027-DEL-133
chr10-7000325-DEL-53
chr10-7005543-DEL-58
chr10-7018848-DEL-204
chr10-7031017-DEL-4219
chr10-7034826-DEL-198
chr10-7037423-DEL-5594
chr10-7038072-DEL-10739
chr10-7042573-DEL-869
chr10-7055301-DEL-600
chr10-7099254-DEL-191
chr10-7099533-DEL-1391
chr10-7117723-DEL-1398
chr10-7118418-DEL-221
chr10-7119911-DEL-465
chr10-7123464-DEL-504
chr10-7140211-DEL-130
chr10-7147650-DEL-702
chr10-7173007-DEL-1230
chr10-7186498-DEL-69
chr10-7191766-DEL-81
chr10-7224234-DEL-142
chr10-7319826-DEL-81
chr10-7345180-DEL-145
chr10-7345942-DEL-6372
chr10-7354546-DEL-583
chr10-7356089-DEL-527
chr10-7358539-DEL-71
chr10-7361478-DEL-5117
chr10-7361722-DEL-75
chr10-7366355-DEL-70
chr10-7371060-DEL-15006
chr10-7439797-DEL-219
chr10-7449444-DEL-139
chr10-7468514-DEL-94
chr10-7485162-DEL-642
chr10-7491528-DEL-2110
chr10-7509483-DEL-2579
chr10-7509583-DEL-1920
chr10-7520301-DEL-219
chr10-7532251-DEL-364
chr10-7582511-DEL-194
chr10-7602959-DEL-125
chr10-7604469-DEL-101
chr10-7613698-DEL-280
chr10-7649590-DEL-61
chr10-7675887-DEL-83
chr10-7699201-DEL-392
chr10-7708988-DEL-443
chr10-7713075-DEL-202

chr10-7722342-DEL-392
chr10-7741784-DEL-169
chr10-7778417-DEL-347
chr10-7803219-DEL-89
chr10-7805701-DEL-636
chr10-7856475-DEL-1061
chr10-7873969-DEL-54
chr10-7897509-DEL-122
chr10-7905659-DEL-714
chr10-7938986-DEL-85
chr10-7989412-DEL-72
chr10-8015394-DEL-2753
chr10-8077629-DEL-63
chr10-8168778-DEL-550
chr10-8180365-DEL-109
chr10-8195455-DEL-201
chr10-8201360-DEL-57
chr10-8224466-DEL-148
chr10-8241722-DEL-1200
chr10-8254153-DEL-7639
chr10-8289008-DEL-149
chr10-8290569-DEL-164
chr10-8312012-DEL-1395
chr10-8317595-DEL-80
chr10-8322202-DEL-56
chr10-8325663-DEL-130
chr10-8329832-DEL-169
chr10-8358331-DEL-196
chr10-8378329-DEL-52
chr10-8415353-DEL-129
chr10-8436064-DEL-74
chr10-8467480-DEL-63
chr10-8548388-DEL-1110
chr10-8571824-DEL-50
chr10-8594131-DEL-62
chr10-8609944-DEL-2873
chr10-8617804-DEL-74
chr10-8639645-DEL-498
chr10-8688019-DEL-209
chr10-8693153-DEL-139
chr10-8715304-DEL-88
chr10-8725332-DEL-182
chr10-8768970-DEL-3193
chr10-8790741-DEL-81
chr10-8791206-DEL-2126
chr10-8800198-DEL-329
chr10-8801690-DEL-67
chr10-8821998-DEL-196
chr10-8837903-DEL-134
chr10-8893722-DEL-254
chr10-8938605-DEL-144
chr10-8940758-DEL-102
chr10-8949958-DEL-137
chr10-8960580-DEL-606
chr10-8990408-DEL-2853
chr10-9023913-DEL-78
chr10-9037463-DEL-60
chr10-9053932-DEL-168
chr10-9056941-DEL-1387
chr10-9091302-DEL-54
chr10-9179903-DEL-721
chr10-9225803-DEL-170
chr10-9241993-DEL-1732
chr10-9248605-DEL-1851
chr10-9258136-DEL-272
chr10-9278309-DEL-435
chr10-9358429-DEL-80
chr10-9369364-DEL-68
chr10-9406583-DEL-80
chr10-9454466-DEL-250
chr10-9464482-DEL-51
chr10-9493948-DEL-204
chr10-9511795-DEL-68
chr10-9583815-DEL-159
chr10-9587261-DEL-122
chr10-9613662-DEL-94
chr10-9669765-DEL-183

chr10-9715978-DEL-64
chr10-9729361-DEL-1656
chr10-9730802-DEL-252
chr10-9732821-DEL-350
chr10-9758176-DEL-52
chr10-9758984-DEL-83
chr10-9853886-DEL-277
chr10-9861696-DEL-6227
chr10-9935512-DEL-178
chr10-9969559-DEL-81
chr10-10006657-DEL-1276
chr10-10008706-DEL-113
chr10-10008976-DEL-578
chr10-10014697-DEL-699
chr10-10016829-DEL-6198
chr10-10036763-DEL-559
chr10-10046226-DEL-211
chr10-10047502-DEL-861
chr10-10048720-DEL-4022
chr10-10192572-DEL-1055
chr10-10210980-DEL-146
chr10-10260130-DEL-67
chr10-10275506-DEL-59
chr10-10279675-DEL-896
chr10-10308944-DEL-117
chr10-10375358-DEL-9039
chr10-10377054-DEL-1786
chr10-10379196-DEL-769
chr10-10382703-DEL-1684
chr10-10417770-DEL-1476
chr10-10423616-DEL-144
chr10-10440000-DEL-1484
chr10-10488366-DEL-637
chr10-10496434-DEL-6185
chr10-10498858-DEL-212
chr10-10500244-DEL-140
chr10-10503536-DEL-116
chr10-10542903-DEL-150
chr10-10568763-DEL-58
chr10-10604551-DEL-191
chr10-10659150-DEL-799
chr10-10662079-DEL-177
chr10-10696203-DEL-60
chr10-10742462-DEL-65
chr10-10767988-DEL-100
chr10-10796947-DEL-205
chr10-10823818-DEL-72
chr10-10842631-DEL-90
chr10-10847495-DEL-86
chr10-10868822-DEL-428
chr10-10889336-DEL-66
chr10-10899304-DEL-434
chr10-10909035-DEL-63
chr10-10914399-DEL-225
chr10-10960485-DEL-51
chr10-10999475-DEL-244
chr10-11002655-DEL-806
chr10-11055918-DEL-107
chr10-11084704-DEL-163
chr10-11086540-DEL-204
chr10-11100650-DEL-59
chr10-11116914-DEL-1261
chr10-11135540-DEL-229
chr10-11142366-DEL-477
chr10-11143989-DEL-669
chr10-11150718-DEL-195
chr10-11152276-DEL-116
chr10-11183297-DEL-228
chr10-11193914-DEL-111
chr10-11235443-DEL-1897
chr10-11254095-DEL-62
chr10-11275832-DEL-199
chr10-11285432-DEL-534
chr10-11286543-DEL-84
chr10-11323373-DEL-710
chr10-11333298-DEL-56
chr10-11364460-DEL-5203

chr10-11399353-DEL-7620
chr10-11428200-DEL-638
chr10-11431208-DEL-115
chr10-11441139-DEL-192
chr10-11449690-DEL-161
chr10-11450175-DEL-276
chr10-11466463-DEL-78
chr10-11478487-DEL-7734
chr10-11483097-DEL-1005
chr10-11486672-DEL-2234
chr10-11488024-DEL-1103
chr10-11488678-DEL-474
chr10-11497111-DEL-7961
chr10-11508086-DEL-909
chr10-11509773-DEL-53
chr10-11510314-DEL-63
chr10-11511514-DEL-6904
chr10-11511652-DEL-211
chr10-11511734-DEL-213
chr10-11522225-DEL-68
chr10-11524479-DEL-71
chr10-11545466-DEL-219
chr10-11567425-DEL-59
chr10-11568096-DEL-10309
chr10-11569113-DEL-8642
chr10-11584829-DEL-52
chr10-11588026-DEL-72
chr10-11590455-DEL-197
chr10-11617762-DEL-167
chr10-11644757-DEL-63
chr10-11651159-DEL-624
chr10-11664047-DEL-7975
chr10-11668487-DEL-1781
chr10-11676855-DEL-399
chr10-11678046-DEL-164
chr10-11683425-DEL-272
chr10-11727186-DEL-3126
chr10-11731969-DEL-6241
chr10-11751916-DEL-382
chr10-11776152-DEL-1289
chr10-11778525-DEL-75
chr10-11791575-DEL-128
chr10-11825263-DEL-565
chr10-11891169-DEL-88
chr10-11891897-DEL-184
chr10-11891965-DEL-1234
chr10-11894710-DEL-59
chr10-11946619-DEL-6014
chr10-12013053-DEL-72
chr10-12029221-DEL-175
chr10-12041062-DEL-742
chr10-12110824-DEL-81
chr10-12116471-DEL-1195
chr10-12149748-DEL-293
chr10-12214836-DEL-125
chr10-12237831-DEL-110
chr10-12241203-DEL-214
chr10-12245186-DEL-204
chr10-12256483-DEL-901
chr10-12272697-DEL-84
chr10-12330655-DEL-57
chr10-12355207-DEL-1487
chr10-12456624-DEL-128
chr10-12456714-DEL-998
chr10-12483876-DEL-254
chr10-12511897-DEL-174
chr10-12523745-DEL-88
chr10-12529333-DEL-131
chr10-12564657-DEL-152
chr10-12590290-DEL-110
chr10-12599184-DEL-400
chr10-12635892-DEL-293
chr10-12648756-DEL-53
chr10-12653751-DEL-96
chr10-12673664-DEL-197
chr10-12675744-DEL-54
chr10-12691373-DEL-60

chr10-12709643-DEL-60
chr10-12742148-DEL-55
chr10-12754015-DEL-228
chr10-12794326-DEL-226
chr10-12801233-DEL-97
chr10-12815032-DEL-76
chr10-12840656-DEL-83
chr10-12841836-DEL-728
chr10-12874948-DEL-564
chr10-12878257-DEL-1399
chr10-12913341-DEL-519
chr10-12913930-DEL-125
chr10-12940413-DEL-109
chr10-12950859-DEL-58
chr10-13006623-DEL-57
chr10-13007253-DEL-58
chr10-13023019-DEL-62
chr10-13060104-DEL-199
chr10-13105383-DEL-2715
chr10-13107838-DEL-162
chr10-13143940-DEL-57
chr10-13157414-DEL-77
chr10-13174684-DEL-65
chr10-13205171-DEL-90
chr10-13242151-DEL-51
chr10-13277895-DEL-678
chr10-13306047-DEL-96
chr10-13412312-DEL-288
chr10-13414231-DEL-662
chr10-13426508-DEL-61
chr10-13461646-DEL-62
chr10-13467837-DEL-429
chr10-13481434-DEL-214
chr10-13498797-DEL-64
chr10-13510853-DEL-56
chr10-13530383-DEL-189
chr10-13623053-DEL-109
chr10-13633393-DEL-56
chr10-13649996-DEL-717
chr10-13660727-DEL-320
chr10-13747563-DEL-51
chr10-13768956-DEL-212
chr10-13856470-DEL-58
chr10-13858360-DEL-147
chr10-13876401-DEL-466
chr10-13878184-DEL-74
chr10-13937128-DEL-179
chr10-13981646-DEL-80
chr10-14028616-DEL-850
chr10-14035288-DEL-136
chr10-14096181-DEL-76
chr10-14099851-DEL-55
chr10-14102065-DEL-102
chr10-14116313-DEL-3417
chr10-14121809-DEL-1161
chr10-14147239-DEL-195
chr10-14152150-DEL-93
chr10-14215234-DEL-102
chr10-14245559-DEL-255
chr10-14270711-DEL-64
chr10-14271535-DEL-93
chr10-14284330-DEL-632
chr10-14377888-DEL-58
chr10-14378481-DEL-400
chr10-14383095-DEL-60
chr10-14412951-DEL-79
chr10-14413426-DEL-234
chr10-14450403-DEL-6281
chr10-14484521-DEL-5156
chr10-14490344-DEL-160
chr10-14492501-DEL-57
chr10-14494407-DEL-58
chr10-14521257-DEL-2659
chr10-14537444-DEL-429
chr10-14538477-DEL-1483
chr10-14569816-DEL-59
chr10-14573330-DEL-149

chr10-14621017-DEL-2121
chr10-14650972-DEL-577
chr10-14679488-DEL-52
chr10-14686456-DEL-51
chr10-14696657-DEL-134
chr10-14727142-DEL-297
chr10-14731607-DEL-202
chr10-14749676-DEL-96
chr10-14752623-DEL-138
chr10-14757643-DEL-318
chr10-14763509-DEL-121
chr10-14764723-DEL-2510
chr10-14790927-DEL-68
chr10-14791369-DEL-3351
chr10-14839488-DEL-79
chr10-14866857-DEL-223
chr10-14874282-DEL-6210
chr10-14883357-DEL-92
chr10-14923183-DEL-2083
chr10-14934762-DEL-59
chr10-14936905-DEL-52
chr10-14938408-DEL-138
chr10-14947332-DEL-75
chr10-14955866-DEL-5533
chr10-14976374-DEL-94
chr10-14984745-DEL-1457
chr10-14988252-DEL-7977
chr10-15000222-DEL-3347
chr10-15012543-DEL-63
chr10-15028356-DEL-61
chr10-15036080-DEL-6453
chr10-15064157-DEL-181
chr10-15066316-DEL-17539
chr10-15090076-DEL-60
chr10-15096196-DEL-146
chr10-15154943-DEL-1023
chr10-15183896-DEL-101
chr10-15211919-DEL-6641
chr10-15222052-DEL-6391
chr10-15235921-DEL-325
chr10-15248867-DEL-1083
chr10-15248914-DEL-821
chr10-15249863-DEL-304
chr10-15256893-DEL-2970
chr10-15259870-DEL-3354
chr10-15299103-DEL-1560
chr10-15309825-DEL-66
chr10-15355782-DEL-343
chr10-15369186-DEL-313
chr10-15404281-DEL-6076
chr10-15424731-DEL-264
chr10-15426970-DEL-91
chr10-15442497-DEL-6992
chr10-15448633-DEL-409
chr10-15461536-DEL-82
chr10-15469191-DEL-118
chr10-15473400-DEL-216
chr10-15475059-DEL-3205
chr10-15495294-DEL-7398
chr10-15507311-DEL-60
chr10-15507993-DEL-1316
chr10-15508003-DEL-61
chr10-15510545-DEL-7018
chr10-15523533-DEL-71
chr10-15553926-DEL-1641
chr10-15570855-DEL-1354
chr10-15607243-DEL-621
chr10-15659522-DEL-1208
chr10-15662598-DEL-79
chr10-15766190-DEL-63
chr10-15776975-DEL-1359
chr10-15784486-DEL-356
chr10-15813924-DEL-358
chr10-15821501-DEL-195
chr10-15839877-DEL-3382
chr10-15853334-DEL-71
chr10-15879534-DEL-98

chr10-15940043-DEL-115
chr10-15967918-DEL-336
chr10-15978963-DEL-6905
chr10-16010586-DEL-176
chr10-16035259-DEL-304
chr10-16042349-DEL-89
chr10-16044091-DEL-673
chr10-16064251-DEL-366
chr10-16085118-DEL-55
chr10-16111953-DEL-70
chr10-16113480-DEL-1580
chr10-16118619-DEL-251
chr10-16124352-DEL-88
chr10-16156737-DEL-1167
chr10-16158248-DEL-258
chr10-16176865-DEL-198
chr10-16179016-DEL-1533
chr10-16253907-DEL-144
chr10-16272740-DEL-64
chr10-16287361-DEL-617
chr10-16326477-DEL-1920
chr10-16391765-DEL-5723
chr10-16398177-DEL-530
chr10-16405742-DEL-933
chr10-16417573-DEL-402
chr10-16418612-DEL-975
chr10-16433711-DEL-323
chr10-16436750-DEL-1090
chr10-16477759-DEL-993
chr10-16481209-DEL-57
chr10-16498676-DEL-132
chr10-16520521-DEL-386
chr10-16525086-DEL-66
chr10-16553374-DEL-58
chr10-16583269-DEL-155
chr10-16612564-DEL-57
chr10-16642325-DEL-62
chr10-16655963-DEL-159
chr10-16659716-DEL-1226
chr10-16670264-DEL-1575
chr10-16671669-DEL-175
chr10-16677084-DEL-80
chr10-16737542-DEL-112
chr10-16742096-DEL-86
chr10-16750357-DEL-55
chr10-16814832-DEL-97
chr10-16829211-DEL-86
chr10-16843121-DEL-6077
chr10-16852289-DEL-84
chr10-16854296-DEL-58
chr10-16889170-DEL-128
chr10-16948222-DEL-209
chr10-16959731-DEL-53
chr10-16965353-DEL-417
chr10-17013021-DEL-6201
chr10-17013057-DEL-12484
chr10-17028548-DEL-174
chr10-17069948-DEL-134
chr10-17092825-DEL-149
chr10-17133674-DEL-70
chr10-17155181-DEL-66
chr10-17187272-DEL-343
chr10-17215666-DEL-471
chr10-17222684-DEL-66
chr10-17249001-DEL-154
chr10-17286439-DEL-318
chr10-17291521-DEL-71
chr10-17306278-DEL-69
chr10-17326474-DEL-293
chr10-17337642-DEL-2156
chr10-17337670-DEL-857
chr10-17337671-DEL-259
chr10-17338783-DEL-533
chr10-17431529-DEL-53
chr10-17462424-DEL-225
chr10-17482364-DEL-110
chr10-17546734-DEL-80

chr10-17619454-DEL-1592
chr10-17656471-DEL-81
chr10-17665792-DEL-60
chr10-17694536-DEL-61
chr10-17696578-DEL-314
chr10-17734001-DEL-587
chr10-17738303-DEL-59
chr10-17738909-DEL-325
chr10-17744183-DEL-252
chr10-17812576-DEL-2107
chr10-17817819-DEL-651
chr10-17877053-DEL-61
chr10-17915047-DEL-70
chr10-17916142-DEL-390
chr10-17959422-DEL-196
chr10-17969373-DEL-87
chr10-17976581-DEL-202
chr10-18020073-DEL-184
chr10-18058816-DEL-225
chr10-18061973-DEL-58
chr10-18111660-DEL-207
chr10-18114167-DEL-465
chr10-18124919-DEL-846
chr10-18127504-DEL-147
chr10-18154880-DEL-101
chr10-18161120-DEL-66
chr10-18187023-DEL-310
chr10-18242053-DEL-214
chr10-18243403-DEL-54
chr10-18249645-DEL-927
chr10-18254251-DEL-1113
chr10-18257357-DEL-54
chr10-18263824-DEL-65
chr10-18265653-DEL-65
chr10-18290256-DEL-60
chr10-18293824-DEL-200
chr10-18314880-DEL-52
chr10-18317572-DEL-52
chr10-18336387-DEL-428
chr10-18370656-DEL-312
chr10-18396946-DEL-599
chr10-18405189-DEL-73
chr10-18412657-DEL-300
chr10-18425137-DEL-66
chr10-18451245-DEL-2650
chr10-18453424-DEL-86
chr10-18543158-DEL-366
chr10-18558238-DEL-62
chr10-18604184-DEL-641
chr10-18612062-DEL-404
chr10-18613356-DEL-78
chr10-18633203-DEL-1162
chr10-18688850-DEL-115
chr10-18699411-DEL-2013
chr10-18714118-DEL-148
chr10-18788403-DEL-306
chr10-18820810-DEL-75
chr10-18837266-DEL-84
chr10-18928417-DEL-351
chr10-18954474-DEL-1448
chr10-18965015-DEL-378
chr10-18968521-DEL-168
chr10-18998763-DEL-213
chr10-19005936-DEL-83
chr10-19074646-DEL-93
chr10-19079592-DEL-291
chr10-19082852-DEL-139
chr10-19100062-DEL-398
chr10-19151531-DEL-197
chr10-19255616-DEL-1159
chr10-19262651-DEL-422
chr10-19269124-DEL-378
chr10-19281388-DEL-399
chr10-19303115-DEL-774
chr10-19303400-DEL-325
chr10-19307357-DEL-83
chr10-19393475-DEL-201

chr10-19395902-DEL-89
chr10-19401111-DEL-102
chr10-19405208-DEL-84
chr10-19439061-DEL-115
chr10-19456285-DEL-126
chr10-19473519-DEL-184
chr10-19489631-DEL-330
chr10-19497531-DEL-62
chr10-19507709-DEL-86
chr10-19520061-DEL-120
chr10-19577645-DEL-498
chr10-19580825-DEL-1121
chr10-19583722-DEL-62
chr10-19590430-DEL-65
chr10-19599501-DEL-1481
chr10-19613077-DEL-381
chr10-19613629-DEL-231
chr10-19622215-DEL-632
chr10-19629114-DEL-56
chr10-19645778-DEL-79
chr10-19706633-DEL-129
chr10-19790667-DEL-341
chr10-19820645-DEL-154
chr10-19864210-DEL-98
chr10-19938853-DEL-57
chr10-19939867-DEL-316
chr10-19967820-DEL-67
chr10-19976683-DEL-138
chr10-19982480-DEL-204
chr10-20013052-DEL-220
chr10-20022077-DEL-51
chr10-20065565-DEL-310
chr10-20068243-DEL-117
chr10-20098058-DEL-875
chr10-20116299-DEL-188
chr10-20160685-DEL-267
chr10-20163939-DEL-200
chr10-20166740-DEL-91
chr10-20169044-DEL-621
chr10-20171332-DEL-499
chr10-20173093-DEL-518
chr10-20177119-DEL-1975
chr10-20228887-DEL-56
chr10-20233626-DEL-171
chr10-20235146-DEL-845
chr10-20247050-DEL-2191
chr10-20266765-DEL-69
chr10-20269424-DEL-63
chr10-20324944-DEL-87
chr10-20357490-DEL-77
chr10-20363156-DEL-62
chr10-20396474-DEL-64
chr10-20411712-DEL-106
chr10-20415302-DEL-711
chr10-20439012-DEL-73
chr10-20464892-DEL-120
chr10-20467013-DEL-100
chr10-20468532-DEL-70
chr10-20480338-DEL-202
chr10-20499273-DEL-172
chr10-20502624-DEL-281
chr10-20511670-DEL-53
chr10-20573076-DEL-123
chr10-20580757-DEL-1295
chr10-20594142-DEL-56
chr10-20596705-DEL-188
chr10-20605881-DEL-78
chr10-20655328-DEL-2121
chr10-20660622-DEL-122
chr10-20699139-DEL-98
chr10-20708557-DEL-152
chr10-20709251-DEL-131
chr10-20711645-DEL-743
chr10-20716685-DEL-74
chr10-20751269-DEL-342
chr10-20790289-DEL-194
chr10-20802785-DEL-200

chr10-20854687-DEL-75
chr10-20865057-DEL-207
chr10-20865884-DEL-376
chr10-20912455-DEL-53
chr10-20922240-DEL-151
chr10-20927432-DEL-172
chr10-20928846-DEL-68
chr10-20952333-DEL-254
chr10-21079914-DEL-506
chr10-21117275-DEL-52
chr10-21139124-DEL-224
chr10-21167078-DEL-94
chr10-21223690-DEL-58
chr10-21242458-DEL-66
chr10-21255612-DEL-284
chr10-21269127-DEL-101
chr10-21274800-DEL-66
chr10-21294666-DEL-591
chr10-21295279-DEL-62
chr10-21303504-DEL-120
chr10-21318436-DEL-78
chr10-21330812-DEL-88
chr10-21335079-DEL-208
chr10-21342893-DEL-101
chr10-21348049-DEL-112
chr10-21393766-DEL-245
chr10-21413569-DEL-810
chr10-21428336-DEL-106
chr10-21518175-DEL-3393
chr10-21521580-DEL-3840
chr10-21532777-DEL-1883
chr10-21548492-DEL-56
chr10-21556308-DEL-72
chr10-21583022-DEL-52
chr10-21597909-DEL-54
chr10-21703061-DEL-113
chr10-21716055-DEL-59
chr10-21732324-DEL-112
chr10-21738625-DEL-1290
chr10-21739655-DEL-90
chr10-21746287-DEL-63
chr10-21755193-DEL-264
chr10-21762634-DEL-196
chr10-21780047-DEL-63
chr10-21801227-DEL-73
chr10-21821815-DEL-88
chr10-21836224-DEL-158
chr10-21847685-DEL-228
chr10-21849796-DEL-3241
chr10-21886971-DEL-280
chr10-21888463-DEL-103
chr10-21891121-DEL-95
chr10-21899240-DEL-195
chr10-21925549-DEL-102
chr10-21936457-DEL-206
chr10-21938609-DEL-70
chr10-21941364-DEL-180
chr10-21953243-DEL-152
chr10-21955283-DEL-2960
chr10-21958428-DEL-56
chr10-21960049-DEL-3078
chr10-21965430-DEL-6450
chr10-21971969-DEL-807
chr10-21978643-DEL-52
chr10-21979313-DEL-396
chr10-21989330-DEL-372
chr10-21989786-DEL-442
chr10-21993630-DEL-682
chr10-22001346-DEL-589
chr10-22007795-DEL-7483
chr10-22010892-DEL-79
chr10-22015711-DEL-80
chr10-22017772-DEL-76
chr10-22028766-DEL-1727
chr10-22039501-DEL-332
chr10-22048663-DEL-84
chr10-22058273-DEL-1028

chr10-22062007-DEL-56
chr10-22071742-DEL-180
chr10-22075431-DEL-3316
chr10-22079413-DEL-572
chr10-22105279-DEL-2949
chr10-22118968-DEL-348
chr10-22121500-DEL-6333
chr10-22147752-DEL-398
chr10-22154266-DEL-88
chr10-22164856-DEL-3348
chr10-22172243-DEL-415
chr10-22173829-DEL-668
chr10-22178034-DEL-59
chr10-22203866-DEL-17540
chr10-22217907-DEL-851
chr10-22237206-DEL-61
chr10-22261899-DEL-179
chr10-22263990-DEL-87
chr10-22286844-DEL-438
chr10-22290123-DEL-56
chr10-22324103-DEL-141
chr10-22340289-DEL-5577
chr10-22346494-DEL-759
chr10-22372690-DEL-61
chr10-22381714-DEL-11850
chr10-22394322-DEL-2687
chr10-22395747-DEL-23995
chr10-22399153-DEL-8026
chr10-22406416-DEL-783
chr10-22409166-DEL-6490
chr10-22416993-DEL-1817
chr10-22420073-DEL-11478
chr10-22440540-DEL-51
chr10-22465489-DEL-924
chr10-22479686-DEL-92
chr10-22485960-DEL-159
chr10-22491531-DEL-69
chr10-22552089-DEL-76
chr10-22573923-DEL-135
chr10-22586950-DEL-179
chr10-22620553-DEL-474
chr10-22620584-DEL-991
chr10-22621062-DEL-217
chr10-22634808-DEL-6392
chr10-22634993-DEL-198
chr10-22658856-DEL-111
chr10-22684521-DEL-1045
chr10-22732511-DEL-434
chr10-22758978-DEL-68
chr10-22760038-DEL-307
chr10-22772554-DEL-844
chr10-22790671-DEL-58
chr10-22811642-DEL-6448
chr10-22833205-DEL-59
chr10-22855181-DEL-528
chr10-22860920-DEL-62
chr10-22904367-DEL-53
chr10-22914256-DEL-259
chr10-22946822-DEL-168
chr10-22967427-DEL-69
chr10-23019313-DEL-97
chr10-23044992-DEL-71
chr10-23056648-DEL-97
chr10-23135316-DEL-123
chr10-23154341-DEL-103
chr10-23173876-DEL-365
chr10-23204141-DEL-633
chr10-23232710-DEL-64
chr10-23250974-DEL-53
chr10-23254170-DEL-303
chr10-23303199-DEL-72
chr10-23315732-DEL-210
chr10-23342004-DEL-59
chr10-23345470-DEL-6411
chr10-23356854-DEL-6393
chr10-23384190-DEL-631
chr10-23397610-DEL-245

chr10-23422937-DEL-274
chr10-23441301-DEL-132
chr10-23444301-DEL-289
chr10-23469414-DEL-103
chr10-23507430-DEL-4756
chr10-23512352-DEL-1679
chr10-23522242-DEL-138
chr10-23523540-DEL-490
chr10-23528976-DEL-200
chr10-23573845-DEL-877
chr10-23596786-DEL-119
chr10-23645925-DEL-190
chr10-23648122-DEL-298
chr10-23648563-DEL-654
chr10-23658349-DEL-158
chr10-23665799-DEL-142
chr10-23667572-DEL-61
chr10-23676549-DEL-155
chr10-23684321-DEL-502
chr10-23687692-DEL-535
chr10-23690967-DEL-68
chr10-23692906-DEL-292
chr10-23713988-DEL-413
chr10-23714675-DEL-201
chr10-23726628-DEL-54
chr10-23732591-DEL-5702
chr10-23743972-DEL-359
chr10-23747216-DEL-7142
chr10-23761305-DEL-2324
chr10-23769841-DEL-72
chr10-23772233-DEL-157
chr10-23779473-DEL-131
chr10-23792808-DEL-1355
chr10-23794805-DEL-207
chr10-23803862-DEL-370
chr10-23804270-DEL-468
chr10-23811151-DEL-61
chr10-23813082-DEL-207
chr10-23824421-DEL-59
chr10-23840952-DEL-80
chr10-23849026-DEL-327
chr10-23850967-DEL-455
chr10-23893594-DEL-161
chr10-23899021-DEL-548
chr10-23936920-DEL-469
chr10-23944576-DEL-1037
chr10-23956319-DEL-1099
chr10-23960133-DEL-76
chr10-23971686-DEL-1438
chr10-23973694-DEL-392
chr10-24000858-DEL-3249
chr10-24045969-DEL-1533
chr10-24065206-DEL-102
chr10-24069382-DEL-78
chr10-24077536-DEL-106
chr10-24090080-DEL-1347
chr10-24116419-DEL-76
chr10-24121144-DEL-72
chr10-24141343-DEL-101
chr10-24165877-DEL-91
chr10-24181116-DEL-1532
chr10-24200743-DEL-59
chr10-24230141-DEL-1067
chr10-24246198-DEL-124
chr10-24299181-DEL-146
chr10-24302070-DEL-394
chr10-24386866-DEL-6305
chr10-24476513-DEL-184
chr10-24539311-DEL-122
chr10-24541333-DEL-51
chr10-24559525-DEL-92
chr10-24581904-DEL-99
chr10-24585945-DEL-76
chr10-24604433-DEL-7502
chr10-24614671-DEL-75
chr10-24620942-DEL-435
chr10-24621678-DEL-1392

chr10-24623627-DEL-1578
chr10-24632849-DEL-100
chr10-24637584-DEL-68
chr10-24639054-DEL-69
chr10-24676022-DEL-70
chr10-24681981-DEL-53
chr10-24684318-DEL-66
chr10-24687774-DEL-261
chr10-24705125-DEL-122
chr10-24714173-DEL-201
chr10-24721966-DEL-59
chr10-24724192-DEL-6368
chr10-24753556-DEL-82
chr10-24794318-DEL-51
chr10-24795269-DEL-131
chr10-24796051-DEL-79
chr10-24815690-DEL-56
chr10-24831110-DEL-322
chr10-24833697-DEL-150
chr10-24868832-DEL-1766
chr10-24874957-DEL-432
chr10-24881173-DEL-66
chr10-24884733-DEL-436
chr10-24929293-DEL-1417
chr10-24937698-DEL-879
chr10-24961573-DEL-75
chr10-24967515-DEL-985
chr10-24968276-DEL-206
chr10-24973774-DEL-137
chr10-24977457-DEL-4954
chr10-24985107-DEL-393
chr10-25007452-DEL-78
chr10-25019174-DEL-317
chr10-25057047-DEL-112
chr10-25063248-DEL-292
chr10-25134228-DEL-63
chr10-25150729-DEL-59
chr10-25155746-DEL-61
chr10-25160736-DEL-201
chr10-25172795-DEL-77
chr10-25175072-DEL-234
chr10-25192796-DEL-231
chr10-25213181-DEL-233
chr10-25250974-DEL-57
chr10-25275884-DEL-347
chr10-25308695-DEL-63
chr10-25311005-DEL-623
chr10-25318515-DEL-799
chr10-25334359-DEL-847
chr10-25350376-DEL-68
chr10-25351088-DEL-551
chr10-25357989-DEL-174
chr10-25372323-DEL-952
chr10-25427145-DEL-1317
chr10-25436885-DEL-117
chr10-25460741-DEL-63
chr10-25466435-DEL-1563
chr10-25478579-DEL-474
chr10-25479657-DEL-189
chr10-25520053-DEL-1320
chr10-25577792-DEL-93
chr10-25611012-DEL-257
chr10-25628879-DEL-75
chr10-25644371-DEL-609
chr10-25650571-DEL-167
chr10-25658912-DEL-538
chr10-25671104-DEL-2017
chr10-25713311-DEL-245
chr10-25724438-DEL-198
chr10-25725992-DEL-1276
chr10-25728118-DEL-824
chr10-25729537-DEL-1563
chr10-25760960-DEL-1000
chr10-25764925-DEL-192
chr10-25788509-DEL-54
chr10-25827345-DEL-135
chr10-25856538-DEL-3521

chr10-25923893-DEL-53
chr10-26043828-DEL-3032
chr10-26081154-DEL-103
chr10-26145008-DEL-59
chr10-26176209-DEL-56
chr10-26188726-DEL-160
chr10-26237080-DEL-88
chr10-26257519-DEL-406
chr10-26267729-DEL-454
chr10-26297050-DEL-60
chr10-26300289-DEL-706
chr10-26349891-DEL-168
chr10-26381201-DEL-504
chr10-26390531-DEL-2378
chr10-26393009-DEL-514
chr10-26408136-DEL-1923
chr10-26447566-DEL-67
chr10-26486508-DEL-136
chr10-26498660-DEL-57
chr10-26565359-DEL-51
chr10-26572187-DEL-3109
chr10-26603622-DEL-139
chr10-26614308-DEL-57
chr10-26620663-DEL-982
chr10-26622408-DEL-765
chr10-26642732-DEL-160
chr10-26683845-DEL-204
chr10-26713671-DEL-137
chr10-26766102-DEL-208
chr10-26783251-DEL-850
chr10-26796159-DEL-52
chr10-26799713-DEL-317
chr10-26824036-DEL-108
chr10-26868527-DEL-321
chr10-26935171-DEL-633
chr10-26939590-DEL-55
chr10-26955141-DEL-838
chr10-26965762-DEL-68
chr10-26979015-DEL-642
chr10-27008211-DEL-61
chr10-27038796-DEL-1592
chr10-27054247-DEL-170
chr10-27074801-DEL-1797
chr10-27113239-DEL-1086
chr10-27116819-DEL-95
chr10-27180609-DEL-95
chr10-27187872-DEL-2680
chr10-27200837-DEL-153
chr10-27208143-DEL-128
chr10-27211831-DEL-221
chr10-27214067-DEL-337
chr10-27228021-DEL-65
chr10-27253271-DEL-442
chr10-27327976-DEL-1088
chr10-27338027-DEL-2145
chr10-27341026-DEL-101
chr10-27346378-DEL-400
chr10-27394004-DEL-357
chr10-27432260-DEL-2303
chr10-27451328-DEL-145
chr10-27475084-DEL-550
chr10-27485902-DEL-52
chr10-27594920-DEL-5928
chr10-27600448-DEL-290
chr10-27624104-DEL-2678
chr10-27624111-DEL-433
chr10-27653059-DEL-74
chr10-27665357-DEL-6735
chr10-27725384-DEL-708
chr10-27739214-DEL-64
chr10-27741694-DEL-89
chr10-27755690-DEL-493
chr10-27785326-DEL-66
chr10-27805318-DEL-69
chr10-27834926-DEL-376
chr10-27840879-DEL-285
chr10-27867251-DEL-52

chr10-27877910-DEL-63
chr10-27933349-DEL-64
chr10-27946502-DEL-231
chr10-27968066-DEL-65
chr10-27970830-DEL-64
chr10-28011711-DEL-1068
chr10-28013319-DEL-55
chr10-28023502-DEL-878
chr10-28047659-DEL-162
chr10-28114929-DEL-115
chr10-28129217-DEL-650
chr10-28130690-DEL-813
chr10-28147339-DEL-2034
chr10-28168925-DEL-351
chr10-28171288-DEL-70
chr10-28179436-DEL-56
chr10-28181501-DEL-214
chr10-28192939-DEL-317
chr10-28199952-DEL-5912
chr10-28201757-DEL-57
chr10-28228071-DEL-97
chr10-28292204-DEL-79
chr10-28297501-DEL-60
chr10-28335070-DEL-75
chr10-28370676-DEL-166
chr10-28379123-DEL-3478
chr10-28382077-DEL-228
chr10-28412972-DEL-4207
chr10-28483354-DEL-76
chr10-28496420-DEL-134
chr10-28514049-DEL-72
chr10-28518918-DEL-51
chr10-28536291-DEL-76
chr10-28543064-DEL-68
chr10-28547209-DEL-1092
chr10-28548365-DEL-270
chr10-28555102-DEL-1334
chr10-28610536-DEL-70
chr10-28612416-DEL-56
chr10-28619773-DEL-1873
chr10-28640783-DEL-250
chr10-28644649-DEL-6144
chr10-28679415-DEL-105
chr10-28721628-DEL-301
chr10-28727442-DEL-84
chr10-28732839-DEL-382
chr10-28802696-DEL-259
chr10-28809758-DEL-196
chr10-28850911-DEL-63
chr10-28867478-DEL-80
chr10-28874714-DEL-1408
chr10-28883541-DEL-511
chr10-28891900-DEL-72
chr10-28923269-DEL-1046
chr10-28949569-DEL-609
chr10-28953004-DEL-173
chr10-28953031-DEL-1824
chr10-28953224-DEL-1026
chr10-28954948-DEL-90
chr10-28955136-DEL-1038
chr10-28956207-DEL-1836
chr10-28958075-DEL-995
chr10-28965943-DEL-1153
chr10-28991825-DEL-5193
chr10-29002131-DEL-587
chr10-29002927-DEL-58
chr10-29009444-DEL-1483
chr10-29014001-DEL-126
chr10-29015073-DEL-128
chr10-29108707-DEL-1014
chr10-29137326-DEL-213
chr10-29160719-DEL-75
chr10-29202661-DEL-100
chr10-29203108-DEL-621
chr10-29303416-DEL-72
chr10-29313542-DEL-2948
chr10-29316730-DEL-200

chr10-29318303-DEL-486
chr10-29320599-DEL-169
chr10-29424639-DEL-61
chr10-29493558-DEL-1352
chr10-29600280-DEL-82
chr10-29622351-DEL-161
chr10-29641156-DEL-261
chr10-29681198-DEL-74
chr10-29681973-DEL-93
chr10-29744931-DEL-367
chr10-29759505-DEL-1901
chr10-29794160-DEL-583
chr10-29810364-DEL-386
chr10-29829551-DEL-13459
chr10-29834425-DEL-5752
chr10-29857218-DEL-61
chr10-29863945-DEL-57
chr10-29887007-DEL-280
chr10-29893321-DEL-435
chr10-29898933-DEL-140
chr10-29934410-DEL-203
chr10-29945060-DEL-414
chr10-29947829-DEL-736
chr10-30010546-DEL-690
chr10-30013242-DEL-324
chr10-30045003-DEL-63
chr10-30046091-DEL-2152
chr10-30050130-DEL-55
chr10-30061144-DEL-546
chr10-30062007-DEL-195
chr10-30064685-DEL-57
chr10-30066846-DEL-280
chr10-30075572-DEL-106
chr10-30092435-DEL-423
chr10-30119673-DEL-380
chr10-30145262-DEL-195
chr10-30171933-DEL-9816
chr10-30229097-DEL-4005
chr10-30240168-DEL-73
chr10-30287518-DEL-286
chr10-30313351-DEL-214
chr10-30320465-DEL-68
chr10-30326510-DEL-65
chr10-30348883-DEL-6207
chr10-30369923-DEL-268
chr10-30376142-DEL-146
chr10-30382510-DEL-220
chr10-30421476-DEL-2066
chr10-30455727-DEL-3267
chr10-30458502-DEL-666
chr10-30462247-DEL-60
chr10-30466499-DEL-248
chr10-30477156-DEL-263
chr10-30484741-DEL-1533
chr10-30498700-DEL-63
chr10-30554727-DEL-335
chr10-30661863-DEL-80
chr10-30685834-DEL-84
chr10-30750009-DEL-3072
chr10-30764189-DEL-734
chr10-30765231-DEL-333
chr10-30770775-DEL-723
chr10-30776869-DEL-6849
chr10-30805865-DEL-422
chr10-30814771-DEL-58
chr10-30888664-DEL-88
chr10-30900551-DEL-555
chr10-30903472-DEL-408
chr10-30951459-DEL-55
chr10-30980961-DEL-55
chr10-31016927-DEL-334
chr10-31019192-DEL-257
chr10-31030073-DEL-772
chr10-31063333-DEL-63
chr10-31078783-DEL-51
chr10-31132085-DEL-138
chr10-31188224-DEL-112

chr10-31242381-DEL-190
chr10-31275777-DEL-233
chr10-31279888-DEL-83
chr10-31346563-DEL-51
chr10-31392689-DEL-204
chr10-31394983-DEL-764
chr10-31421188-DEL-52
chr10-31454903-DEL-167
chr10-31537221-DEL-151
chr10-31542475-DEL-74
chr10-31551046-DEL-109
chr10-31555393-DEL-5067
chr10-31589584-DEL-101
chr10-31597463-DEL-5921
chr10-31609521-DEL-254
chr10-31676475-DEL-3231
chr10-31681976-DEL-381
chr10-31683999-DEL-596
chr10-31691757-DEL-6306
chr10-31726434-DEL-1306
chr10-31733024-DEL-410
chr10-31755736-DEL-6367
chr10-31814918-DEL-222
chr10-31851513-DEL-60
chr10-31855994-DEL-5989
chr10-31888865-DEL-207
chr10-31913350-DEL-173
chr10-31981548-DEL-2679
chr10-31986575-DEL-6167
chr10-32001673-DEL-215
chr10-32020838-DEL-1573
chr10-32032540-DEL-577
chr10-32051972-DEL-6464
chr10-32066772-DEL-500
chr10-32069542-DEL-746
chr10-32071002-DEL-80
chr10-32072634-DEL-4630
chr10-32102499-DEL-458
chr10-32109639-DEL-54
chr10-32128211-DEL-702
chr10-32132471-DEL-5420
chr10-32195434-DEL-58
chr10-32202408-DEL-53
chr10-32241088-DEL-258
chr10-32270099-DEL-149
chr10-32293881-DEL-840
chr10-32294547-DEL-203
chr10-32299575-DEL-56
chr10-32303133-DEL-73
chr10-32305645-DEL-4450
chr10-32328752-DEL-65
chr10-32343392-DEL-118
chr10-32351057-DEL-101
chr10-32352590-DEL-5674
chr10-32363865-DEL-94
chr10-32373403-DEL-8344
chr10-32375161-DEL-212
chr10-32381772-DEL-1193
chr10-32384922-DEL-51
chr10-32390705-DEL-6060
chr10-32397686-DEL-55
chr10-32398598-DEL-10335
chr10-32401898-DEL-1002
chr10-32403709-DEL-5564
chr10-32426366-DEL-859
chr10-32428514-DEL-69
chr10-32433432-DEL-653
chr10-32443658-DEL-1281
chr10-32449323-DEL-60
chr10-32461493-DEL-114
chr10-32463511-DEL-100
chr10-32470037-DEL-73
chr10-32508742-DEL-2704
chr10-32523881-DEL-141
chr10-32542914-DEL-419
chr10-32565859-DEL-418
chr10-32577192-DEL-665

chr10-32585366-DEL-84
chr10-32597259-DEL-60
chr10-32606524-DEL-228
chr10-32609967-DEL-985
chr10-32619893-DEL-299
chr10-32661268-DEL-987
chr10-32673450-DEL-133
chr10-32713630-DEL-63
chr10-32722602-DEL-234
chr10-32727182-DEL-6270
chr10-32733681-DEL-56
chr10-32767843-DEL-1311
chr10-32777397-DEL-96
chr10-32794533-DEL-6300
chr10-32820257-DEL-4436
chr10-32831675-DEL-53
chr10-32833398-DEL-69
chr10-32839977-DEL-102
chr10-32848321-DEL-212
chr10-32873850-DEL-173
chr10-32877228-DEL-359
chr10-32882709-DEL-6236
chr10-32901794-DEL-53
chr10-32928791-DEL-79
chr10-32942876-DEL-946
chr10-32943856-DEL-365
chr10-32954103-DEL-82
chr10-32977397-DEL-1765
chr10-32984638-DEL-151
chr10-32992494-DEL-4468
chr10-33035430-DEL-64
chr10-33057139-DEL-1268
chr10-33083541-DEL-64
chr10-33134630-DEL-693
chr10-33154109-DEL-109
chr10-33161469-DEL-110
chr10-33233306-DEL-145
chr10-33244572-DEL-121
chr10-33255144-DEL-867
chr10-33288070-DEL-9470
chr10-33310055-DEL-86
chr10-33312607-DEL-390
chr10-33321733-DEL-442
chr10-33333322-DEL-142
chr10-33341428-DEL-710
chr10-33367725-DEL-6275
chr10-33377982-DEL-106
chr10-33394017-DEL-2192
chr10-33424061-DEL-56
chr10-33435776-DEL-6703
chr10-33445727-DEL-56
chr10-33446890-DEL-548
chr10-33461607-DEL-1107
chr10-33468767-DEL-311
chr10-33506913-DEL-153
chr10-33509307-DEL-1333
chr10-33531428-DEL-6204
chr10-33539519-DEL-450
chr10-33551633-DEL-6595
chr10-33552769-DEL-343
chr10-33556659-DEL-410
chr10-33559656-DEL-271
chr10-33584415-DEL-1498
chr10-33614618-DEL-99
chr10-33630725-DEL-812
chr10-33636483-DEL-58
chr10-33641777-DEL-337
chr10-33648377-DEL-4729
chr10-33655255-DEL-386
chr10-33658360-DEL-648
chr10-33668426-DEL-755
chr10-33669732-DEL-249
chr10-33684285-DEL-1280
chr10-33697640-DEL-212
chr10-33707266-DEL-438
chr10-33711913-DEL-3591
chr10-33716221-DEL-369

chr10-33722510-DEL-65
chr10-33731828-DEL-223
chr10-33780420-DEL-150
chr10-33839177-DEL-117
chr10-33847673-DEL-182
chr10-33864474-DEL-2119
chr10-33864989-DEL-407
chr10-33878754-DEL-1234
chr10-33926925-DEL-2035
chr10-33995924-DEL-151
chr10-34062384-DEL-108
chr10-34076419-DEL-69
chr10-34088563-DEL-90
chr10-34107750-DEL-138
chr10-34140472-DEL-70
chr10-34146980-DEL-84
chr10-34164451-DEL-410
chr10-34182724-DEL-183
chr10-34224411-DEL-153
chr10-34225117-DEL-182
chr10-34230062-DEL-362
chr10-34260486-DEL-379
chr10-34317905-DEL-199
chr10-34329535-DEL-359
chr10-34338754-DEL-392
chr10-34348549-DEL-51
chr10-34361470-DEL-58
chr10-34362233-DEL-4752
chr10-34466518-DEL-66
chr10-34473674-DEL-252
chr10-34530309-DEL-507
chr10-34563820-DEL-120
chr10-34571190-DEL-400
chr10-34571191-DEL-1596
chr10-34575363-DEL-555
chr10-34599801-DEL-71
chr10-34612777-DEL-121
chr10-34617814-DEL-7108
chr10-34634529-DEL-180
chr10-34670928-DEL-115
chr10-34705381-DEL-690
chr10-34722637-DEL-172
chr10-34726341-DEL-68
chr10-34749926-DEL-25110
chr10-34767789-DEL-7799
chr10-34777883-DEL-1394
chr10-34793523-DEL-117
chr10-34796400-DEL-6529
chr10-34819718-DEL-4744
chr10-34852820-DEL-336
chr10-34872986-DEL-93
chr10-34892195-DEL-593
chr10-34895041-DEL-513
chr10-34916391-DEL-1583
chr10-34966241-DEL-311
chr10-34982295-DEL-182
chr10-35001804-DEL-62
chr10-35011799-DEL-108
chr10-35058214-DEL-259
chr10-35076660-DEL-188
chr10-35077283-DEL-496
chr10-35079824-DEL-163
chr10-35097342-DEL-135
chr10-35128407-DEL-324
chr10-35156270-DEL-996
chr10-35161565-DEL-447
chr10-35162815-DEL-1078
chr10-35199879-DEL-3503
chr10-35203333-DEL-299
chr10-35216981-DEL-169
chr10-35221566-DEL-87
chr10-35224485-DEL-51
chr10-35260522-DEL-196
chr10-35269037-DEL-72
chr10-35309733-DEL-371
chr10-35314043-DEL-6234
chr10-35339888-DEL-165

chr10-35344401-DEL-1244
chr10-35360461-DEL-159
chr10-35364361-DEL-117
chr10-35366529-DEL-441
chr10-35384812-DEL-14369
chr10-35429024-DEL-10026
chr10-35457459-DEL-64
chr10-35463328-DEL-82
chr10-35468455-DEL-213
chr10-35479456-DEL-62
chr10-35507266-DEL-52
chr10-35519994-DEL-718
chr10-35532095-DEL-191
chr10-35553096-DEL-624
chr10-35567127-DEL-742
chr10-35578973-DEL-206
chr10-35583919-DEL-212
chr10-35584630-DEL-3258
chr10-35647984-DEL-1355
chr10-35710375-DEL-234
chr10-35711536-DEL-452
chr10-35724287-DEL-6367
chr10-35741068-DEL-203
chr10-35748377-DEL-58
chr10-35753944-DEL-2904
chr10-35754910-DEL-78
chr10-35784846-DEL-119
chr10-35808285-DEL-350
chr10-35817205-DEL-1748
chr10-35828619-DEL-486
chr10-35833139-DEL-55
chr10-35842067-DEL-301
chr10-35842833-DEL-1581
chr10-35844460-DEL-1996
chr10-35858477-DEL-542
chr10-35863469-DEL-3934
chr10-35884204-DEL-103
chr10-35887185-DEL-840
chr10-35900007-DEL-62
chr10-35907726-DEL-1636
chr10-35918576-DEL-3983
chr10-35926540-DEL-99
chr10-36006132-DEL-6434
chr10-36014939-DEL-506
chr10-36024091-DEL-360
chr10-36032652-DEL-74
chr10-36035679-DEL-59
chr10-36048404-DEL-74
chr10-36060147-DEL-413
chr10-36076053-DEL-50
chr10-36080783-DEL-334
chr10-36116219-DEL-60
chr10-36152020-DEL-6467
chr10-36178308-DEL-227
chr10-36179670-DEL-1540
chr10-36203771-DEL-169
chr10-36321774-DEL-2580
chr10-36355560-DEL-100
chr10-36357461-DEL-56
chr10-36381695-DEL-118
chr10-36424291-DEL-98
chr10-36429167-DEL-65
chr10-36451214-DEL-51
chr10-36465274-DEL-270
chr10-36468963-DEL-5903
chr10-36476100-DEL-1240
chr10-36476166-DEL-401
chr10-36478472-DEL-997
chr10-36510129-DEL-104
chr10-36619795-DEL-686
chr10-36639720-DEL-3201
chr10-36653310-DEL-2968
chr10-36689144-DEL-2090
chr10-36689265-DEL-659
chr10-36689964-DEL-1270
chr10-36693403-DEL-196
chr10-36696344-DEL-125

chr10-36697442-DEL-2713
chr10-36700936-DEL-59
chr10-36717342-DEL-1085
chr10-36733855-DEL-445
chr10-36744750-DEL-68
chr10-36777505-DEL-528
chr10-36786473-DEL-265
chr10-36794285-DEL-151
chr10-36814890-DEL-77
chr10-36829573-DEL-62
chr10-36844358-DEL-1589
chr10-36913832-DEL-6021
chr10-36951730-DEL-56
chr10-36968717-DEL-235
chr10-36970070-DEL-73
chr10-36976344-DEL-60
chr10-36995661-DEL-128
chr10-37038694-DEL-72
chr10-37061577-DEL-3793
chr10-37076666-DEL-196
chr10-37099779-DEL-73
chr10-37139697-DEL-65
chr10-37163580-DEL-83
chr10-37179376-DEL-978
chr10-37259674-DEL-347
chr10-37279514-DEL-463
chr10-37312415-DEL-58
chr10-37323135-DEL-109
chr10-37326646-DEL-696
chr10-37354974-DEL-298
chr10-37387469-DEL-104
chr10-37422331-DEL-1761
chr10-37452392-DEL-3458
chr10-37483367-DEL-53
chr10-37486296-DEL-152
chr10-37519194-DEL-750
chr10-37575919-DEL-53
chr10-37576997-DEL-60
chr10-37583939-DEL-212
chr10-37587969-DEL-59
chr10-37627282-DEL-2024
chr10-37646883-DEL-4300
chr10-37658992-DEL-75
chr10-37686286-DEL-60
chr10-37694133-DEL-68
chr10-37700012-DEL-290
chr10-37700560-DEL-279
chr10-37731182-DEL-73
chr10-37741456-DEL-382
chr10-37759025-DEL-2188
chr10-37761223-DEL-1492
chr10-37769408-DEL-114
chr10-37788652-DEL-615
chr10-37794968-DEL-371
chr10-37803595-DEL-2737
chr10-37806397-DEL-1444
chr10-37822901-DEL-577
chr10-37834177-DEL-63
chr10-37865659-DEL-109
chr10-37883044-DEL-379
chr10-37890890-DEL-61
chr10-37918479-DEL-105
chr10-37922186-DEL-1625
chr10-37930815-DEL-942
chr10-37935214-DEL-4184
chr10-37940263-DEL-56
chr10-37955097-DEL-256
chr10-37960234-DEL-397
chr10-38009647-DEL-79
chr10-38083434-DEL-514
chr10-38084015-DEL-1152
chr10-38121479-DEL-381
chr10-38130474-DEL-1741
chr10-38156081-DEL-205
chr10-38158889-DEL-280
chr10-38192316-DEL-90
chr10-38236983-DEL-81

chr10-38266817-DEL-162
chr10-38310159-DEL-2124
chr10-38363115-DEL-83
chr10-38465892-DEL-6067
chr10-38490512-DEL-370
chr10-38508860-DEL-68
chr10-38510732-DEL-178
chr10-38521131-DEL-2107
chr10-38533486-DEL-881
chr10-38536244-DEL-1572
chr10-38536296-DEL-773
chr10-38537658-DEL-145
chr10-38546800-DEL-60
chr10-38565783-DEL-56
chr10-38579238-DEL-193
chr10-38582184-DEL-2129
chr10-38591096-DEL-775
chr10-38612802-DEL-6378
chr10-38667639-DEL-325
chr10-38677535-DEL-1192
chr10-38685826-DEL-671
chr10-38688806-DEL-1768
chr10-38701036-DEL-274
chr10-38703335-DEL-193
chr10-38716697-DEL-6318
chr10-38748564-DEL-61
chr10-38752131-DEL-85
chr10-38806045-DEL-52
chr10-38817959-DEL-261
chr10-38824177-DEL-177
chr10-38826463-DEL-97
chr10-38848320-DEL-79
chr10-38852477-DEL-70
chr10-38865669-DEL-53
chr10-38868671-DEL-286
chr10-38884648-DEL-552
chr10-38897136-DEL-328
chr10-38908713-DEL-264
chr10-38926418-DEL-56
chr10-38947131-DEL-75
chr10-38968896-DEL-406
chr10-38976596-DEL-270
chr10-38992875-DEL-64
chr10-39028569-DEL-74
chr10-39036597-DEL-168
chr10-39054924-DEL-81
chr10-39069680-DEL-488
chr10-39099082-DEL-1357
chr10-39105180-DEL-57
chr10-39147837-DEL-69
chr10-39173516-DEL-1200
chr10-39224628-DEL-79
chr10-39225729-DEL-101
chr10-39271944-DEL-244
chr10-39280358-DEL-67
chr10-39321600-DEL-2288
chr10-39327551-DEL-261
chr10-39449790-DEL-177
chr10-39456849-DEL-802
chr10-39467752-DEL-196
chr10-39470592-DEL-64
chr10-39474650-DEL-140
chr10-39483559-DEL-294
chr10-39495112-DEL-192
chr10-39512915-DEL-68
chr10-39541557-DEL-96
chr10-39593686-DEL-51
chr10-39624465-DEL-207
chr10-39651736-DEL-1502
chr10-39675177-DEL-58
chr10-39715841-DEL-74
chr10-39739971-DEL-68
chr10-39749351-DEL-51
chr10-39778665-DEL-402
chr10-39785684-DEL-107
chr10-39787973-DEL-122
chr10-39790723-DEL-212

chr10-39793845-DEL-51
chr10-39795882-DEL-185
chr10-39810438-DEL-146
chr10-39829130-DEL-129
chr10-39847598-DEL-160
chr10-39849655-DEL-3589
chr10-39867160-DEL-79
chr10-39867600-DEL-170
chr10-39884630-DEL-2823
chr10-39892780-DEL-503
chr10-39914336-DEL-1787
chr10-39918448-DEL-195
chr10-39931187-DEL-241
chr10-39960782-DEL-150
chr10-39962934-DEL-136
chr10-39983054-DEL-66
chr10-39991144-DEL-52
chr10-40002640-DEL-162
chr10-40003504-DEL-280
chr10-40024777-DEL-2595
chr10-40031006-DEL-702
chr10-40033428-DEL-932
chr10-40053246-DEL-79
chr10-40056532-DEL-133
chr10-40056846-DEL-2120
chr10-40078457-DEL-1685
chr10-40092380-DEL-209
chr10-40093602-DEL-58
chr10-40099964-DEL-54
chr10-40130694-DEL-1788
chr10-40151156-DEL-88
chr10-40154794-DEL-767
chr10-40157328-DEL-64
chr10-40165930-DEL-130
chr10-40180619-DEL-74
chr10-40190979-DEL-197
chr10-40195003-DEL-56
chr10-40202567-DEL-92
chr10-40208380-DEL-152
chr10-40220256-DEL-66
chr10-40236435-DEL-353
chr10-40252748-DEL-61
chr10-40325070-DEL-216
chr10-40328771-DEL-65
chr10-40336853-DEL-480
chr10-40341341-DEL-210
chr10-40349987-DEL-250
chr10-40374126-DEL-95
chr10-40426308-DEL-56
chr10-40438301-DEL-410
chr10-40470893-DEL-175
chr10-40485053-DEL-1730
chr10-40521173-DEL-92
chr10-40521964-DEL-1112
chr10-40547785-DEL-164
chr10-40552850-DEL-60
chr10-40588967-DEL-78
chr10-40624240-DEL-199
chr10-40708506-DEL-219
chr10-40786672-DEL-107
chr10-40787573-DEL-197
chr10-40794425-DEL-287
chr10-40810917-DEL-266
chr10-40815108-DEL-53
chr10-40821794-DEL-90
chr10-40849180-DEL-6165
chr10-40859608-DEL-53
chr10-40864491-DEL-6182
chr10-40894926-DEL-79
chr10-40899786-DEL-79
chr10-40924623-DEL-401
chr10-40937746-DEL-56
chr10-40944197-DEL-69
chr10-40971179-DEL-74
chr10-40993734-DEL-86
chr10-40997642-DEL-1778
chr10-41000149-DEL-59

chr10-41003855-DEL-101
chr10-41039755-DEL-196
chr10-41044665-DEL-1545
chr10-41052726-DEL-62
chr10-41067904-DEL-54
chr10-41068348-DEL-777
chr10-41068474-DEL-79
chr10-41069111-DEL-542
chr10-41070094-DEL-793
chr10-41086433-DEL-75
chr10-41097575-DEL-1117
chr10-41100093-DEL-2321
chr10-41108531-DEL-145
chr10-41135241-DEL-965
chr10-41184805-DEL-6881
chr10-41184872-DEL-57
chr10-41184980-DEL-1371
chr10-41186395-DEL-5259
chr10-41235991-DEL-53
chr10-41269914-DEL-1057
chr10-41274992-DEL-336
chr10-41280812-DEL-194
chr10-41321779-DEL-104
chr10-41347542-DEL-182
chr10-41414402-DEL-58
chr10-41492434-DEL-243
chr10-41500289-DEL-226
chr10-41505630-DEL-74
chr10-41514633-DEL-185
chr10-41516278-DEL-1199
chr10-41523316-DEL-98
chr10-41535738-DEL-1863
chr10-41550799-DEL-216
chr10-41559679-DEL-86
chr10-41611837-DEL-1195
chr10-41612706-DEL-245
chr10-41618686-DEL-239
chr10-41625308-DEL-289
chr10-41666821-DEL-176
chr10-41738426-DEL-178
chr10-41743212-DEL-64
chr10-41758038-DEL-64
chr10-41775777-DEL-94
chr10-41810312-DEL-52
chr10-41836263-DEL-140
chr10-41974169-DEL-54
chr10-41995979-DEL-70
chr10-42005723-DEL-702
chr10-42008747-DEL-880
chr10-42013073-DEL-304
chr10-42040908-DEL-1657
chr10-42050926-DEL-146
chr10-42070243-DEL-310
chr10-42211667-DEL-395
chr10-42220457-DEL-639
chr10-42233726-DEL-238
chr10-42246999-DEL-184
chr10-42269215-DEL-257
chr10-42277598-DEL-150
chr10-42278416-DEL-300
chr10-42290841-DEL-54
chr10-42316894-DEL-55
chr10-42322739-DEL-132
chr10-42326082-DEL-51
chr10-42333138-DEL-351
chr10-42367526-DEL-519
chr10-42379182-DEL-114
chr10-42393453-DEL-294
chr10-42412793-DEL-54
chr10-42413001-DEL-1356
chr10-42414488-DEL-590
chr10-42415413-DEL-78
chr10-42424431-DEL-72
chr10-42477108-DEL-52
chr10-42488522-DEL-59
chr10-42502093-DEL-139
chr10-42502531-DEL-970

chr10-42531929-DEL-58
chr10-42539361-DEL-1000
chr10-42541285-DEL-97
chr10-42553721-DEL-95
chr10-42560773-DEL-343
chr10-42580048-DEL-109
chr10-42626153-DEL-337
chr10-42649253-DEL-211
chr10-42658604-DEL-182
chr10-42667295-DEL-876
chr10-42728907-DEL-51
chr10-42758538-DEL-87
chr10-42766507-DEL-61
chr10-42767701-DEL-55
chr10-42791680-DEL-150
chr10-42800626-DEL-528
chr10-42826519-DEL-69
chr10-42847664-DEL-173
chr10-42860606-DEL-556
chr10-42867979-DEL-387
chr10-42931375-DEL-66
chr10-42933527-DEL-88
chr10-42939455-DEL-96
chr10-42964611-DEL-71
chr10-43071125-DEL-2537
chr10-43091112-DEL-69
chr10-43104593-DEL-58
chr10-43119675-DEL-715
chr10-43130162-DEL-1864
chr10-43156782-DEL-172
chr10-43165325-DEL-399
chr10-43176041-DEL-60
chr10-43223855-DEL-764
chr10-43273865-DEL-432
chr10-43297858-DEL-54
chr10-43332083-DEL-454
chr10-43347586-DEL-103
chr10-43373405-DEL-197
chr10-43384213-DEL-111
chr10-43423474-DEL-198
chr10-43429097-DEL-286
chr10-43445432-DEL-1266
chr10-43486122-DEL-112
chr10-43494064-DEL-87
chr10-43509586-DEL-56
chr10-43550542-DEL-81
chr10-43554572-DEL-138
chr10-43559966-DEL-136
chr10-43576156-DEL-119
chr10-43604540-DEL-128
chr10-43621520-DEL-51
chr10-43625510-DEL-51
chr10-43632701-DEL-51
chr10-43637749-DEL-53
chr10-43654761-DEL-301
chr10-43732836-DEL-1566
chr10-43780679-DEL-1030
chr10-43785466-DEL-131
chr10-43847284-DEL-247
chr10-43852121-DEL-116
chr10-43868180-DEL-140
chr10-43869473-DEL-77
chr10-43890581-DEL-184
chr10-43893129-DEL-197
chr10-43895812-DEL-59
chr10-43918098-DEL-81
chr10-43931170-DEL-63
chr10-43958883-DEL-1580
chr10-43979725-DEL-508
chr10-43993283-DEL-538
chr10-44007369-DEL-280
chr10-44092474-DEL-185
chr10-44138427-DEL-358
chr10-44140605-DEL-2019
chr10-44143020-DEL-63
chr10-44147142-DEL-58
chr10-44177973-DEL-579

chr10-44184038-DEL-2316
chr10-44243788-DEL-134
chr10-44265880-DEL-55
chr10-44274218-DEL-195
chr10-44283672-DEL-65
chr10-44300344-DEL-157
chr10-44346917-DEL-66
chr10-44348891-DEL-202
chr10-44360528-DEL-107
chr10-44374395-DEL-76
chr10-44437217-DEL-61
chr10-44473387-DEL-51
chr10-44608824-DEL-178
chr10-44610515-DEL-214
chr10-44653408-DEL-651
chr10-44658701-DEL-144
chr10-44665717-DEL-60
chr10-44674889-DEL-51
chr10-44710652-DEL-75
chr10-44715945-DEL-75
chr10-44724566-DEL-687
chr10-44727145-DEL-52
chr10-44743758-DEL-1912
chr10-44778283-DEL-718
chr10-44793520-DEL-178
chr10-44794949-DEL-84
chr10-44798181-DEL-148
chr10-44873266-DEL-54
chr10-44876030-DEL-56
chr10-44887476-DEL-76
chr10-44899131-DEL-243
chr10-44944357-DEL-558
chr10-44973832-DEL-137
chr10-45003529-DEL-63
chr10-45012012-DEL-51
chr10-45044756-DEL-104
chr10-45046342-DEL-178
chr10-45055533-DEL-101
chr10-45078115-DEL-2179
chr10-45101403-DEL-104
chr10-45119916-DEL-72
chr10-45120449-DEL-162
chr10-45121930-DEL-61
chr10-45122988-DEL-76
chr10-45124339-DEL-413
chr10-45156254-DEL-1454
chr10-45164121-DEL-70
chr10-45208456-DEL-62
chr10-45225379-DEL-66
chr10-45236251-DEL-198
chr10-45251788-DEL-63
chr10-45269644-DEL-3395
chr10-45278713-DEL-246
chr10-45282559-DEL-162
chr10-45299683-DEL-2636
chr10-45301953-DEL-277
chr10-45339266-DEL-1030
chr10-45390936-DEL-11327
chr10-45393032-DEL-7895
chr10-45406804-DEL-424
chr10-45407207-DEL-6353
chr10-45407741-DEL-6647
chr10-45414948-DEL-63
chr10-45415844-DEL-1379
chr10-45416243-DEL-84
chr10-45429031-DEL-111
chr10-45454692-DEL-69
chr10-45500993-DEL-1250
chr10-45502312-DEL-120
chr10-45502333-DEL-977
chr10-45503442-DEL-888
chr10-45503451-DEL-147
chr10-45507166-DEL-170
chr10-45520813-DEL-220
chr10-45530932-DEL-58
chr10-45532696-DEL-375
chr10-45545329-DEL-719

chr10-45551227-DEL-62
chr10-45577402-DEL-349
chr10-45579568-DEL-69
chr10-45591198-DEL-101
chr10-45594251-DEL-335
chr10-45631097-DEL-1887
chr10-45635212-DEL-66
chr10-45644995-DEL-310
chr10-45690661-DEL-717
chr10-45722593-DEL-200
chr10-45795465-DEL-72
chr10-45809585-DEL-60
chr10-45822854-DEL-204
chr10-45825364-DEL-944
chr10-45854459-DEL-55
chr10-45920841-DEL-678
chr10-45938380-DEL-100
chr10-45966318-DEL-609
chr10-45985529-DEL-1185
chr10-45998538-DEL-82
chr10-46030994-DEL-717
chr10-46041120-DEL-52
chr10-46052385-DEL-60
chr10-46068201-DEL-55
chr10-46080757-DEL-3995
chr10-46087772-DEL-64
chr10-46094236-DEL-108
chr10-46106440-DEL-498
chr10-46118147-DEL-52
chr10-46183610-DEL-165
chr10-46187228-DEL-411
chr10-46190258-DEL-62
chr10-46214804-DEL-3826
chr10-46234393-DEL-544
chr10-46248125-DEL-178
chr10-46256281-DEL-6478
chr10-46263186-DEL-1505
chr10-46276627-DEL-572
chr10-46290994-DEL-54
chr10-46307962-DEL-896
chr10-46308718-DEL-933
chr10-46318503-DEL-661
chr10-46344361-DEL-3885
chr10-46348057-DEL-445
chr10-46376558-DEL-1567
chr10-46397195-DEL-179
chr10-46402183-DEL-91
chr10-46448760-DEL-59
chr10-46463122-DEL-207
chr10-46465829-DEL-424
chr10-46475928-DEL-443
chr10-46493860-DEL-151
chr10-46525120-DEL-64
chr10-46536106-DEL-393
chr10-46547217-DEL-1816
chr10-46578912-DEL-4067
chr10-46601520-DEL-429
chr10-46607112-DEL-3423
chr10-46614164-DEL-244
chr10-46615453-DEL-1188
chr10-46640377-DEL-3006
chr10-46644940-DEL-66
chr10-46652948-DEL-354
chr10-46655858-DEL-1088
chr10-46661839-DEL-629
chr10-46680676-DEL-515
chr10-46716217-DEL-439
chr10-46735907-DEL-2671
chr10-46771162-DEL-58
chr10-46788592-DEL-60
chr10-46795147-DEL-212
chr10-46808720-DEL-1048
chr10-46836402-DEL-5041
chr10-46855019-DEL-1008
chr10-46860286-DEL-1892
chr10-46862085-DEL-437
chr10-46863902-DEL-148

chr10-46906188-DEL-725
chr10-46938955-DEL-160
chr10-46966978-DEL-93
chr10-46988363-DEL-259
chr10-46989687-DEL-269
chr10-46999710-DEL-5931
chr10-47043966-DEL-165
chr10-47072395-DEL-635
chr10-47119168-DEL-223
chr10-47146132-DEL-86
chr10-47151986-DEL-1871
chr10-47172063-DEL-4092
chr10-47179961-DEL-241
chr10-47194392-DEL-700
chr10-47200813-DEL-75
chr10-47201961-DEL-2789
chr10-47213038-DEL-71
chr10-47231446-DEL-1019
chr10-47232963-DEL-295
chr10-47242186-DEL-139
chr10-47245496-DEL-52
chr10-47251593-DEL-1337
chr10-47255526-DEL-245
chr10-47274436-DEL-217
chr10-47314459-DEL-211
chr10-47324899-DEL-8863
chr10-47359172-DEL-3454
chr10-47363183-DEL-652
chr10-47398977-DEL-919
chr10-47400521-DEL-6129
chr10-47412082-DEL-393
chr10-47433090-DEL-875
chr10-47459926-DEL-1091
chr10-47479055-DEL-231
chr10-47560889-DEL-2997
chr10-47572513-DEL-203
chr10-47650701-DEL-73
chr10-47651646-DEL-429
chr10-47654671-DEL-387
chr10-47692835-DEL-2470
chr10-47711952-DEL-108
chr10-47749933-DEL-8528
chr10-47751291-DEL-6428
chr10-47751429-DEL-201
chr10-47759179-DEL-894
chr10-47761984-DEL-32989
chr10-47766077-DEL-86
chr10-47767188-DEL-3665
chr10-47768830-DEL-63
chr10-47773052-DEL-6369
chr10-47789147-DEL-1816
chr10-47792636-DEL-282
chr10-47801495-DEL-60
chr10-47803465-DEL-1308
chr10-47811433-DEL-187
chr10-47816356-DEL-5847
chr10-47835332-DEL-135
chr10-47853858-DEL-262
chr10-47861521-DEL-3864
chr10-47880745-DEL-242
chr10-47905265-DEL-2879
chr10-47908341-DEL-2305
chr10-47947206-DEL-363
chr10-47959676-DEL-645
chr10-47962757-DEL-894
chr10-47992825-DEL-931
chr10-48056685-DEL-346
chr10-48059046-DEL-1361
chr10-48077117-DEL-115
chr10-48090293-DEL-126
chr10-48132174-DEL-2829
chr10-48149915-DEL-1841
chr10-48166724-DEL-77
chr10-48167112-DEL-4674
chr10-48184360-DEL-98
chr10-48234926-DEL-290
chr10-48245918-DEL-1921

chr10-48272639-DEL-6931
chr10-48299340-DEL-237
chr10-48323049-DEL-1448
chr10-48324532-DEL-52
chr10-48351280-DEL-376
chr10-48356171-DEL-12273
chr10-48406596-DEL-149
chr10-48427846-DEL-491
chr10-48430029-DEL-52
chr10-48449522-DEL-53
chr10-48452785-DEL-4328
chr10-48462447-DEL-867
chr10-48466597-DEL-1148
chr10-48472346-DEL-2639
chr10-48475389-DEL-80
chr10-48480522-DEL-60
chr10-48531361-DEL-54
chr10-48590101-DEL-1986
chr10-48598119-DEL-205
chr10-48607816-DEL-131
chr10-48614579-DEL-1373
chr10-48620579-DEL-1163
chr10-48627002-DEL-3552
chr10-48645374-DEL-92
chr10-48650872-DEL-451
chr10-48662373-DEL-559
chr10-48713725-DEL-2170
chr10-48731247-DEL-400
chr10-48740054-DEL-246
chr10-48740224-DEL-666
chr10-48741710-DEL-1131
chr10-48745052-DEL-69
chr10-48772048-DEL-62
chr10-48808557-DEL-111
chr10-48811187-DEL-7960
chr10-48820558-DEL-3541
chr10-48826388-DEL-90
chr10-48901573-DEL-6103
chr10-48903247-DEL-187
chr10-48921378-DEL-1293
chr10-48964615-DEL-733
chr10-48968779-DEL-256
chr10-48996181-DEL-197
chr10-49004767-DEL-226
chr10-49018527-DEL-534
chr10-49045576-DEL-701
chr10-49054633-DEL-1773
chr10-49056268-DEL-239
chr10-49063521-DEL-58
chr10-49080742-DEL-238
chr10-49109542-DEL-964
chr10-49112829-DEL-879
chr10-49150338-DEL-1684
chr10-49158725-DEL-1102
chr10-49177632-DEL-1643
chr10-49211950-DEL-1019
chr10-49216032-DEL-55
chr10-49245967-DEL-112
chr10-49260927-DEL-179
chr10-49282969-DEL-423
chr10-49290912-DEL-67
chr10-49300379-DEL-257
chr10-49306635-DEL-121
chr10-49326950-DEL-173
chr10-49336833-DEL-175
chr10-49348456-DEL-831
chr10-49352882-DEL-930
chr10-49403820-DEL-52
chr10-49416659-DEL-1155
chr10-49441460-DEL-56
chr10-49456396-DEL-454
chr10-49461584-DEL-552
chr10-49468517-DEL-779
chr10-49486170-DEL-835
chr10-49500877-DEL-418
chr10-49594247-DEL-113
chr10-49629252-DEL-229

chr10-49669175-DEL-52
chr10-49674727-DEL-62
chr10-49712002-DEL-6270
chr10-49729611-DEL-1868
chr10-49763167-DEL-6156
chr10-49770974-DEL-62
chr10-49813541-DEL-2986
chr10-49865406-DEL-55
chr10-49888299-DEL-72
chr10-49898745-DEL-334
chr10-49913004-DEL-122
chr10-49944701-DEL-66
chr10-49982031-DEL-277
chr10-50003785-DEL-125
chr10-50006741-DEL-1513
chr10-50042923-DEL-579
chr10-50051797-DEL-139
chr10-50086032-DEL-442
chr10-50105670-DEL-187
chr10-50115923-DEL-114
chr10-50128885-DEL-9627
chr10-50143127-DEL-2467
chr10-50146006-DEL-6164
chr10-50191975-DEL-1800
chr10-50197134-DEL-237
chr10-50218593-DEL-106
chr10-50247935-DEL-220
chr10-50250781-DEL-660
chr10-50305260-DEL-315
chr10-50308020-DEL-480
chr10-50309191-DEL-144
chr10-50317199-DEL-73
chr10-50318784-DEL-61
chr10-50322964-DEL-2402
chr10-50342634-DEL-170
chr10-50367398-DEL-3817
chr10-50375826-DEL-107
chr10-50424906-DEL-6191
chr10-50440087-DEL-260
chr10-50472810-DEL-76
chr10-50494544-DEL-59
chr10-50495998-DEL-816
chr10-50531057-DEL-102
chr10-50538737-DEL-5286
chr10-50668494-DEL-53
chr10-50690519-DEL-124
chr10-50696025-DEL-60
chr10-50705478-DEL-52
chr10-50730791-DEL-83
chr10-50740124-DEL-998
chr10-50744158-DEL-61
chr10-50745949-DEL-253
chr10-50749979-DEL-1152
chr10-50812305-DEL-64
chr10-50829498-DEL-1620
chr10-50848653-DEL-355
chr10-50849415-DEL-145
chr10-50865404-DEL-149
chr10-50910485-DEL-208
chr10-50934288-DEL-51
chr10-51002126-DEL-14331
chr10-51009250-DEL-7206
chr10-51014807-DEL-2294
chr10-51037631-DEL-58
chr10-51072644-DEL-2915
chr10-51077485-DEL-306
chr10-51078566-DEL-3813
chr10-51082050-DEL-53
chr10-51144463-DEL-89
chr10-51177394-DEL-96
chr10-51178089-DEL-939
chr10-51179050-DEL-391
chr10-51183333-DEL-124
chr10-51185470-DEL-512
chr10-51187695-DEL-68
chr10-51193256-DEL-268
chr10-51213048-DEL-428

chr10-51219066-DEL-5659
chr10-51229255-DEL-75
chr10-51232042-DEL-6011
chr10-51254138-DEL-349
chr10-51279001-DEL-376
chr10-51344680-DEL-7816
chr10-51397525-DEL-157
chr10-51402276-DEL-103
chr10-51408338-DEL-1413
chr10-51413505-DEL-779
chr10-51422833-DEL-976
chr10-51424720-DEL-181
chr10-51430728-DEL-52
chr10-51434675-DEL-2339
chr10-51439172-DEL-618
chr10-51440237-DEL-1270
chr10-51477277-DEL-6332
chr10-51496129-DEL-372
chr10-51614127-DEL-478
chr10-51618283-DEL-8352
chr10-51676514-DEL-316
chr10-51677181-DEL-86
chr10-51681092-DEL-824
chr10-51699169-DEL-1000
chr10-51713057-DEL-174
chr10-51728407-DEL-141
chr10-51770084-DEL-1069
chr10-51830806-DEL-70
chr10-51835600-DEL-569
chr10-51839316-DEL-441
chr10-51851996-DEL-238
chr10-51867368-DEL-218
chr10-51934702-DEL-53
chr10-51935591-DEL-1484
chr10-51935636-DEL-64
chr10-51992756-DEL-200
chr10-51995336-DEL-68
chr10-52064693-DEL-57
chr10-52072276-DEL-211
chr10-52074162-DEL-71
chr10-52081650-DEL-8369
chr10-52081842-DEL-6464
chr10-52088838-DEL-74
chr10-52097476-DEL-60
chr10-52171120-DEL-221
chr10-52181589-DEL-86
chr10-52188628-DEL-174
chr10-52194227-DEL-149
chr10-52199823-DEL-109
chr10-52225777-DEL-51
chr10-52237661-DEL-53
chr10-52242440-DEL-61
chr10-52278383-DEL-60
chr10-52285014-DEL-56
chr10-52291002-DEL-66
chr10-52301082-DEL-82
chr10-52319512-DEL-300
chr10-52321218-DEL-6274
chr10-52367423-DEL-12593
chr10-52397235-DEL-9182
chr10-52397249-DEL-201
chr10-52430898-DEL-202
chr10-52444229-DEL-3479
chr10-52451728-DEL-670
chr10-52457285-DEL-6025
chr10-52457779-DEL-6033
chr10-52468411-DEL-216
chr10-52473090-DEL-2812
chr10-52484780-DEL-2536
chr10-52496282-DEL-132
chr10-52498403-DEL-325
chr10-52512330-DEL-1887
chr10-52524759-DEL-6496
chr10-52581522-DEL-63
chr10-52594900-DEL-399
chr10-52618405-DEL-90
chr10-52627129-DEL-189

chr10-52644736-DEL-396
chr10-52658199-DEL-2271
chr10-52658724-DEL-56
chr10-52670704-DEL-2849
chr10-52672090-DEL-375
chr10-52680681-DEL-176
chr10-52688765-DEL-179
chr10-52689801-DEL-574
chr10-52701663-DEL-2301
chr10-52705158-DEL-70
chr10-52733475-DEL-55
chr10-52751413-DEL-92
chr10-52772886-DEL-82
chr10-52789596-DEL-60
chr10-52812113-DEL-115
chr10-52836027-DEL-201
chr10-52837642-DEL-456
chr10-52852473-DEL-360
chr10-52856128-DEL-555
chr10-52856948-DEL-1514
chr10-52871786-DEL-487
chr10-52907386-DEL-79
chr10-52927636-DEL-2431
chr10-52966832-DEL-64
chr10-52969658-DEL-64
chr10-52973772-DEL-767
chr10-53013834-DEL-4000
chr10-53020839-DEL-2064
chr10-53069588-DEL-270
chr10-53069842-DEL-1375
chr10-53072348-DEL-407
chr10-53077719-DEL-210
chr10-53118329-DEL-51
chr10-53121007-DEL-63
chr10-53138754-DEL-85
chr10-53146192-DEL-493
chr10-53147264-DEL-451
chr10-53152662-DEL-87
chr10-53201704-DEL-63
chr10-53310297-DEL-83
chr10-53327042-DEL-79
chr10-53335413-DEL-93
chr10-53359842-DEL-65
chr10-53390264-DEL-143
chr10-53439840-DEL-65
chr10-53464623-DEL-101
chr10-53467259-DEL-60
chr10-53529838-DEL-195
chr10-53564458-DEL-117
chr10-53570100-DEL-66
chr10-53570433-DEL-282
chr10-53571193-DEL-136
chr10-53597244-DEL-166
chr10-53651277-DEL-68
chr10-53659279-DEL-618
chr10-53660930-DEL-1016
chr10-53678971-DEL-683
chr10-53681487-DEL-217
chr10-53701601-DEL-340
chr10-53708353-DEL-3540
chr10-53713804-DEL-651
chr10-53715097-DEL-204
chr10-53719632-DEL-66
chr10-53735799-DEL-450
chr10-53803187-DEL-133
chr10-53871011-DEL-52
chr10-53874174-DEL-282
chr10-53880307-DEL-89
chr10-53945306-DEL-198
chr10-53959146-DEL-84
chr10-54000660-DEL-77
chr10-54004693-DEL-74
chr10-54027804-DEL-663
chr10-54040086-DEL-191
chr10-54047442-DEL-56
chr10-54079119-DEL-120
chr10-54094739-DEL-3860

chr10-54100701-DEL-58
chr10-54150364-DEL-3662
chr10-54192647-DEL-475
chr10-54218977-DEL-285
chr10-54276495-DEL-437
chr10-54277290-DEL-63
chr10-54286375-DEL-441
chr10-54306789-DEL-195
chr10-54320074-DEL-120
chr10-54380101-DEL-627
chr10-54383714-DEL-862
chr10-54394036-DEL-1121
chr10-54445625-DEL-371
chr10-54455189-DEL-76
chr10-54458360-DEL-96
chr10-54472315-DEL-69
chr10-54511450-DEL-6033
chr10-54530399-DEL-368
chr10-54556794-DEL-674
chr10-54565850-DEL-12411
chr10-54568540-DEL-6371
chr10-54607207-DEL-118
chr10-54609968-DEL-120
chr10-54622176-DEL-81
chr10-54632919-DEL-64
chr10-54684373-DEL-960
chr10-54700036-DEL-393
chr10-54704626-DEL-94
chr10-54704990-DEL-704
chr10-54708945-DEL-72
chr10-54721608-DEL-64
chr10-54750871-DEL-112
chr10-54825192-DEL-197
chr10-54881181-DEL-1134
chr10-54883752-DEL-5974
chr10-54898559-DEL-90
chr10-54910287-DEL-932
chr10-54933090-DEL-953
chr10-54945295-DEL-166
chr10-54952032-DEL-237
chr10-54955418-DEL-4776
chr10-54964818-DEL-1753
chr10-54975654-DEL-716
chr10-54994906-DEL-120
chr10-55003342-DEL-205
chr10-55016040-DEL-1015
chr10-55022176-DEL-64
chr10-55044214-DEL-910
chr10-55044753-DEL-209
chr10-55056393-DEL-198
chr10-55065629-DEL-2123
chr10-55086508-DEL-74
chr10-55121884-DEL-210
chr10-55124115-DEL-726
chr10-55129725-DEL-177
chr10-55138662-DEL-705
chr10-55146645-DEL-368
chr10-55159958-DEL-1059
chr10-55173106-DEL-318
chr10-55180777-DEL-89
chr10-55190975-DEL-658
chr10-55194429-DEL-337
chr10-55227691-DEL-286
chr10-55234646-DEL-76
chr10-55268222-DEL-536
chr10-55279429-DEL-336
chr10-55285715-DEL-173
chr10-55287341-DEL-146
chr10-55317132-DEL-578
chr10-55371760-DEL-201
chr10-55374098-DEL-6820
chr10-55374099-DEL-2082
chr10-55377798-DEL-3123
chr10-55380318-DEL-424
chr10-55390394-DEL-191
chr10-55395353-DEL-81
chr10-55423833-DEL-59

chr10-55426586-DEL-555
chr10-55428672-DEL-554
chr10-55446593-DEL-214
chr10-55502976-DEL-359
chr10-55513939-DEL-159
chr10-55522615-DEL-60
chr10-55529248-DEL-6434
chr10-55571989-DEL-471
chr10-55609918-DEL-6184
chr10-55620850-DEL-6367
chr10-55652403-DEL-683
chr10-55660237-DEL-65
chr10-55704610-DEL-55
chr10-55717771-DEL-6272
chr10-55763457-DEL-1345
chr10-55770615-DEL-6271
chr10-55794481-DEL-705
chr10-55875688-DEL-89
chr10-55897988-DEL-1041
chr10-55902131-DEL-106
chr10-55916694-DEL-66
chr10-55930311-DEL-424
chr10-55934384-DEL-138
chr10-55977136-DEL-1259
chr10-55978094-DEL-747
chr10-56014001-DEL-6466
chr10-56044883-DEL-1014
chr10-56103914-DEL-51
chr10-56136951-DEL-76
chr10-56142482-DEL-9892
chr10-56143953-DEL-1198
chr10-56145241-DEL-305
chr10-56145788-DEL-2115
chr10-56152926-DEL-399
chr10-56158145-DEL-69
chr10-56160050-DEL-180
chr10-56172899-DEL-134
chr10-56206754-DEL-113
chr10-56208786-DEL-96
chr10-56226712-DEL-98
chr10-56279486-DEL-76
chr10-56280524-DEL-59
chr10-56290179-DEL-82
chr10-56331398-DEL-356
chr10-56353061-DEL-57
chr10-56361648-DEL-80
chr10-56364805-DEL-143
chr10-56368178-DEL-53
chr10-56428781-DEL-72
chr10-56431680-DEL-126
chr10-56472176-DEL-318
chr10-56483399-DEL-54
chr10-56490744-DEL-901
chr10-56494310-DEL-6209
chr10-56500734-DEL-75
chr10-56571568-DEL-151
chr10-56583237-DEL-415
chr10-56611908-DEL-864
chr10-56612968-DEL-150
chr10-56617608-DEL-122
chr10-56618038-DEL-540
chr10-56647932-DEL-55
chr10-56678755-DEL-71
chr10-56701890-DEL-2645
chr10-56708968-DEL-223
chr10-56713737-DEL-64
chr10-56770142-DEL-1377
chr10-56771559-DEL-1803
chr10-56806320-DEL-351
chr10-56810311-DEL-835
chr10-56817597-DEL-73
chr10-56828373-DEL-1513
chr10-56910957-DEL-54
chr10-56924656-DEL-66
chr10-56940641-DEL-1150
chr10-56955456-DEL-54
chr10-56959716-DEL-202

chr10-56962614-DEL-454
chr10-56975364-DEL-6417
chr10-57025182-DEL-192
chr10-57027824-DEL-2027
chr10-57065770-DEL-618
chr10-57072325-DEL-381
chr10-57074307-DEL-4264
chr10-57083721-DEL-652
chr10-57089097-DEL-201
chr10-57100258-DEL-167
chr10-57101479-DEL-2197
chr10-57134195-DEL-79
chr10-57137749-DEL-94
chr10-57176891-DEL-62
chr10-57209158-DEL-212
chr10-57212646-DEL-528
chr10-57214443-DEL-80
chr10-57239576-DEL-1257
chr10-57300194-DEL-252
chr10-57310381-DEL-214
chr10-57339641-DEL-275
chr10-57344476-DEL-52
chr10-57444373-DEL-54
chr10-57455151-DEL-3402
chr10-57465375-DEL-224
chr10-57482793-DEL-74
chr10-57490633-DEL-101
chr10-57500430-DEL-901
chr10-57529763-DEL-804
chr10-57530664-DEL-3645
chr10-57532717-DEL-591
chr10-57550437-DEL-614
chr10-57554252-DEL-196
chr10-57572773-DEL-80
chr10-57574915-DEL-656
chr10-57577222-DEL-430
chr10-57589299-DEL-210
chr10-57589757-DEL-325
chr10-57642775-DEL-347
chr10-57649371-DEL-63
chr10-57706402-DEL-60
chr10-57728427-DEL-4846
chr10-57738022-DEL-4336
chr10-58008259-DEL-4218
chr10-58008483-DEL-2875
chr10-58012838-DEL-2655
chr10-58032100-DEL-71
chr10-58072791-DEL-63
chr10-58077085-DEL-86
chr10-58101663-DEL-129
chr10-58109350-DEL-60
chr10-58117534-DEL-68
chr10-58129649-DEL-196
chr10-58138270-DEL-1417
chr10-58145126-DEL-2574
chr10-58145273-DEL-431
chr10-58147841-DEL-989
chr10-58180443-DEL-2049
chr10-58182184-DEL-2611
chr10-58211022-DEL-245
chr10-58212273-DEL-84
chr10-58287309-DEL-70
chr10-58288195-DEL-642
chr10-58306986-DEL-170
chr10-58310215-DEL-109
chr10-58333421-DEL-98
chr10-58336175-DEL-77
chr10-58356021-DEL-3738
chr10-58399671-DEL-78
chr10-58420281-DEL-257
chr10-58433400-DEL-568
chr10-58434714-DEL-867
chr10-58448581-DEL-64
chr10-58461740-DEL-80
chr10-58466158-DEL-60
chr10-58480934-DEL-51
chr10-58500146-DEL-116

chr10-58525452-DEL-68
chr10-58531069-DEL-1418
chr10-58534606-DEL-1121
chr10-58547648-DEL-1935
chr10-58555403-DEL-6217
chr10-58585307-DEL-2345
chr10-58596752-DEL-312
chr10-58604145-DEL-4178
chr10-58667147-DEL-151
chr10-58717531-DEL-117
chr10-58729687-DEL-120
chr10-58761110-DEL-384
chr10-58761549-DEL-112
chr10-58766559-DEL-150
chr10-58798683-DEL-74
chr10-58799214-DEL-1879
chr10-58813015-DEL-58
chr10-58841678-DEL-440
chr10-58859368-DEL-94
chr10-58869106-DEL-804
chr10-58917309-DEL-116
chr10-58962768-DEL-153
chr10-58970839-DEL-51
chr10-58989930-DEL-400
chr10-58994056-DEL-1556
chr10-59042048-DEL-522
chr10-59043687-DEL-351
chr10-59050543-DEL-1072
chr10-59063975-DEL-56
chr10-59094019-DEL-130
chr10-59108557-DEL-97
chr10-59119911-DEL-196
chr10-59123981-DEL-60
chr10-59137784-DEL-4178
chr10-59141979-DEL-1557
chr10-59144378-DEL-193
chr10-59152281-DEL-359
chr10-59163973-DEL-854
chr10-59180431-DEL-384
chr10-59214691-DEL-1472
chr10-59250961-DEL-80
chr10-59255824-DEL-51
chr10-59314202-DEL-63
chr10-59320909-DEL-61
chr10-59322475-DEL-77
chr10-59332546-DEL-117
chr10-59334996-DEL-128
chr10-59358494-DEL-345
chr10-59488131-DEL-747
chr10-59511527-DEL-68
chr10-59521500-DEL-338
chr10-59554325-DEL-188
chr10-59581625-DEL-57
chr10-59584923-DEL-183
chr10-59672769-DEL-127
chr10-59703913-DEL-130
chr10-59740368-DEL-230
chr10-59758949-DEL-166
chr10-59762523-DEL-91
chr10-59763319-DEL-491
chr10-59765493-DEL-221
chr10-59766487-DEL-1701
chr10-59767397-DEL-651
chr10-59773756-DEL-274
chr10-59775386-DEL-51
chr10-59777365-DEL-106
chr10-59781504-DEL-58
chr10-59782877-DEL-188
chr10-59804846-DEL-189
chr10-59901685-DEL-58
chr10-59906078-DEL-216
chr10-59906612-DEL-126
chr10-59907375-DEL-3272
chr10-59917586-DEL-86
chr10-59930984-DEL-60
chr10-59981403-DEL-2580
chr10-59986488-DEL-51

chr10-60057483-DEL-166
chr10-60073016-DEL-106
chr10-60083433-DEL-426
chr10-60092229-DEL-73
chr10-60110013-DEL-174
chr10-60142800-DEL-298
chr10-60176076-DEL-240
chr10-60194401-DEL-56
chr10-60196285-DEL-56
chr10-60237722-DEL-87
chr10-60254344-DEL-185
chr10-60256532-DEL-197
chr10-60259322-DEL-121
chr10-60281122-DEL-134
chr10-60296020-DEL-77
chr10-60304463-DEL-338
chr10-60312522-DEL-65
chr10-60330766-DEL-62
chr10-60419871-DEL-312
chr10-60446895-DEL-113
chr10-60543282-DEL-57
chr10-60548866-DEL-58
chr10-60561776-DEL-1519
chr10-60567261-DEL-83
chr10-60568725-DEL-131
chr10-60595548-DEL-177
chr10-60652243-DEL-406
chr10-60681087-DEL-60
chr10-60686607-DEL-113
chr10-60689754-DEL-504
chr10-60704531-DEL-116
chr10-60811330-DEL-621
chr10-60820460-DEL-64
chr10-60831008-DEL-153
chr10-60845223-DEL-75
chr10-60867626-DEL-115
chr10-60878271-DEL-591
chr10-60917862-DEL-714
chr10-60919472-DEL-398
chr10-60937214-DEL-230
chr10-60940727-DEL-62
chr10-60990744-DEL-166
chr10-61014055-DEL-102
chr10-61021183-DEL-64
chr10-61052904-DEL-106
chr10-61082489-DEL-136
chr10-61095078-DEL-76
chr10-61104965-DEL-85
chr10-61113423-DEL-56
chr10-61162046-DEL-224
chr10-61169852-DEL-54
chr10-61194985-DEL-211
chr10-61209161-DEL-161
chr10-61212196-DEL-514
chr10-61231825-DEL-642
chr10-61239281-DEL-202
chr10-61246496-DEL-79
chr10-61273514-DEL-316
chr10-61295333-DEL-283
chr10-61334827-DEL-259
chr10-61357908-DEL-211
chr10-61358511-DEL-718
chr10-61403528-DEL-166
chr10-61415647-DEL-56
chr10-61428845-DEL-204
chr10-61469299-DEL-61
chr10-61546628-DEL-306
chr10-61554167-DEL-205
chr10-61557842-DEL-52
chr10-61558526-DEL-93
chr10-61569020-DEL-80
chr10-61572152-DEL-62
chr10-61578371-DEL-1910
chr10-61589922-DEL-53
chr10-61596409-DEL-101
chr10-61603944-DEL-178
chr10-61611902-DEL-115

chr10-61637261-DEL-65
chr10-61640461-DEL-133
chr10-61642565-DEL-2704
chr10-61647219-DEL-110
chr10-61653767-DEL-325
chr10-61658248-DEL-142
chr10-61690253-DEL-74
chr10-61727537-DEL-92
chr10-61773954-DEL-146
chr10-61804397-DEL-119
chr10-61837289-DEL-52
chr10-61847336-DEL-65
chr10-61890240-DEL-92
chr10-61906778-DEL-1622
chr10-61951590-DEL-80
chr10-61960020-DEL-1118
chr10-61974185-DEL-60
chr10-61984404-DEL-137
chr10-61987902-DEL-69
chr10-62007320-DEL-278
chr10-62009644-DEL-196
chr10-62019663-DEL-72
chr10-62042497-DEL-532
chr10-62047950-DEL-377
chr10-62063563-DEL-268
chr10-62078299-DEL-132
chr10-62092208-DEL-142
chr10-62112675-DEL-58
chr10-62137228-DEL-156
chr10-62148084-DEL-104
chr10-62167401-DEL-343
chr10-62181968-DEL-410
chr10-62184866-DEL-95
chr10-62192723-DEL-55
chr10-62195345-DEL-71
chr10-62220325-DEL-349
chr10-62234244-DEL-67
chr10-62236941-DEL-88
chr10-62238120-DEL-175
chr10-62239120-DEL-58
chr10-62275132-DEL-945
chr10-62278317-DEL-139
chr10-62287711-DEL-345
chr10-62340447-DEL-1606
chr10-62346363-DEL-102
chr10-62352287-DEL-169
chr10-62359406-DEL-93
chr10-62374199-DEL-104
chr10-62380934-DEL-644
chr10-62390164-DEL-834
chr10-62394029-DEL-51
chr10-62409477-DEL-224
chr10-62414958-DEL-177
chr10-62431981-DEL-417
chr10-62435377-DEL-77
chr10-62436240-DEL-217
chr10-62465290-DEL-81
chr10-62472703-DEL-1493
chr10-62488348-DEL-380
chr10-62494255-DEL-52
chr10-62497919-DEL-81
chr10-62505263-DEL-56
chr10-62512580-DEL-239
chr10-62528927-DEL-216
chr10-62548193-DEL-692
chr10-62567167-DEL-1481
chr10-62584057-DEL-56
chr10-62590800-DEL-97
chr10-62608028-DEL-211
chr10-62657537-DEL-113
chr10-62658617-DEL-59
chr10-62669702-DEL-52
chr10-62689591-DEL-81
chr10-62728521-DEL-53
chr10-62739461-DEL-117
chr10-62741122-DEL-2871
chr10-62741819-DEL-80

chr10-62744028-DEL-2416
chr10-62747365-DEL-572
chr10-62755776-DEL-127
chr10-62759420-DEL-975
chr10-62770577-DEL-98
chr10-62788074-DEL-182
chr10-62803935-DEL-535
chr10-62826918-DEL-857
chr10-62865892-DEL-705
chr10-62873256-DEL-980
chr10-62905464-DEL-925
chr10-62923378-DEL-728
chr10-62939108-DEL-8052
chr10-62953649-DEL-161
chr10-62969943-DEL-179
chr10-62981043-DEL-86
chr10-63000005-DEL-56
chr10-63022188-DEL-57
chr10-63047382-DEL-208
chr10-63050180-DEL-6209
chr10-63101129-DEL-437
chr10-63102725-DEL-213
chr10-63112310-DEL-206
chr10-63116919-DEL-58
chr10-63119672-DEL-1185
chr10-63128550-DEL-203
chr10-63138865-DEL-557
chr10-63140922-DEL-95
chr10-63182030-DEL-56
chr10-63215758-DEL-175
chr10-63221449-DEL-203
chr10-63226033-DEL-551
chr10-63226606-DEL-4473
chr10-63240290-DEL-2397
chr10-63248230-DEL-70
chr10-63250389-DEL-256
chr10-63251486-DEL-288
chr10-63271763-DEL-114
chr10-63342550-DEL-62
chr10-63343192-DEL-114
chr10-63363550-DEL-56
chr10-63372248-DEL-499
chr10-63389451-DEL-223
chr10-63403985-DEL-239
chr10-63414691-DEL-195
chr10-63461263-DEL-65
chr10-63462003-DEL-72
chr10-63481642-DEL-148
chr10-63486648-DEL-75
chr10-63547548-DEL-509
chr10-63577413-DEL-6339
chr10-63586496-DEL-2037
chr10-63638733-DEL-1376
chr10-63649007-DEL-471
chr10-63669556-DEL-686
chr10-63703434-DEL-422
chr10-63744274-DEL-54
chr10-63751635-DEL-152
chr10-63790628-DEL-97
chr10-63823916-DEL-73
chr10-63832377-DEL-1243
chr10-63873271-DEL-68
chr10-63882163-DEL-535
chr10-63886620-DEL-1559
chr10-63891284-DEL-162
chr10-63901606-DEL-2292
chr10-63912641-DEL-143
chr10-63989699-DEL-70
chr10-63999233-DEL-59
chr10-64006832-DEL-62
chr10-64120270-DEL-168
chr10-64171889-DEL-94
chr10-64209992-DEL-91
chr10-64243304-DEL-59
chr10-64247352-DEL-53
chr10-64254438-DEL-1314
chr10-64264593-DEL-405

chr10-64290804-DEL-389
chr10-64335734-DEL-1454
chr10-64474613-DEL-112
chr10-64497573-DEL-78
chr10-64542764-DEL-52
chr10-64563931-DEL-1534
chr10-64573381-DEL-76
chr10-64574432-DEL-467
chr10-64609231-DEL-12314
chr10-64620852-DEL-237
chr10-64635545-DEL-79
chr10-64650153-DEL-52
chr10-64667298-DEL-871
chr10-64675616-DEL-1508
chr10-64760387-DEL-259
chr10-64788628-DEL-749
chr10-64799327-DEL-100
chr10-64813703-DEL-170
chr10-64831819-DEL-56
chr10-64863966-DEL-59
chr10-64945737-DEL-481
chr10-64959842-DEL-130
chr10-64974679-DEL-53
chr10-64975910-DEL-64
chr10-64980718-DEL-94
chr10-64998871-DEL-249
chr10-65001090-DEL-68
chr10-65034980-DEL-118
chr10-65035861-DEL-1637
chr10-65048498-DEL-56
chr10-65062564-DEL-85
chr10-65078356-DEL-262
chr10-65089747-DEL-66
chr10-65107599-DEL-217
chr10-65153045-DEL-238
chr10-65188676-DEL-230
chr10-65205469-DEL-69
chr10-65206389-DEL-571
chr10-65242815-DEL-92
chr10-65267108-DEL-54
chr10-65288134-DEL-4188
chr10-65288253-DEL-5629
chr10-65298771-DEL-66
chr10-65307719-DEL-2113
chr10-65325465-DEL-2716
chr10-65338852-DEL-52
chr10-65344544-DEL-610
chr10-65353984-DEL-78
chr10-65362318-DEL-877
chr10-65366852-DEL-96
chr10-65372483-DEL-147
chr10-65404965-DEL-6228
chr10-65441238-DEL-1827
chr10-65450979-DEL-139
chr10-65459536-DEL-102
chr10-65470238-DEL-3135
chr10-65486969-DEL-97
chr10-65501528-DEL-6200
chr10-65508590-DEL-77
chr10-65540881-DEL-4192
chr10-65556599-DEL-2102
chr10-65560930-DEL-432
chr10-65571895-DEL-196
chr10-65601894-DEL-78
chr10-65624620-DEL-1189
chr10-65640662-DEL-280
chr10-65658194-DEL-438
chr10-65659415-DEL-6371
chr10-65715811-DEL-237
chr10-65719654-DEL-52
chr10-65723517-DEL-659
chr10-65727463-DEL-58
chr10-65773810-DEL-6771
chr10-65773957-DEL-208
chr10-65780979-DEL-284
chr10-65786539-DEL-5428
chr10-65801195-DEL-65

chr10-65815227-DEL-90
chr10-65894404-DEL-441
chr10-65918316-DEL-838
chr10-66045378-DEL-72
chr10-66086800-DEL-1057
chr10-66092509-DEL-500
chr10-66097446-DEL-5511
chr10-66108273-DEL-89
chr10-66132327-DEL-380
chr10-66147312-DEL-526
chr10-66148810-DEL-688
chr10-66150697-DEL-2060
chr10-66202038-DEL-55
chr10-66216223-DEL-530
chr10-66245904-DEL-338
chr10-66253777-DEL-2867
chr10-66329739-DEL-236
chr10-66347370-DEL-162
chr10-66348155-DEL-760
chr10-66348600-DEL-928
chr10-66356318-DEL-227
chr10-66404937-DEL-126
chr10-66405507-DEL-1449
chr10-66407705-DEL-94
chr10-66411616-DEL-56
chr10-66489174-DEL-547
chr10-66510826-DEL-2944
chr10-66514547-DEL-618
chr10-66515233-DEL-1640
chr10-66529430-DEL-63
chr10-66542095-DEL-615
chr10-66550266-DEL-213
chr10-66571844-DEL-59
chr10-66607267-DEL-104
chr10-66609271-DEL-60
chr10-66633876-DEL-261
chr10-66636060-DEL-100
chr10-66637900-DEL-63
chr10-66643627-DEL-3116
chr10-66647899-DEL-64
chr10-66665783-DEL-62
chr10-66676921-DEL-1755
chr10-66692347-DEL-66
chr10-66693428-DEL-824
chr10-66699151-DEL-130
chr10-66718991-DEL-132
chr10-66721775-DEL-55
chr10-66723842-DEL-112
chr10-66749732-DEL-59
chr10-66752091-DEL-76
chr10-66761851-DEL-170
chr10-66765297-DEL-67
chr10-66775126-DEL-1064
chr10-66784704-DEL-386
chr10-66787959-DEL-52
chr10-66792193-DEL-53
chr10-66824113-DEL-59
chr10-66827585-DEL-76
chr10-66831711-DEL-87
chr10-66848253-DEL-55
chr10-66851688-DEL-215
chr10-66864037-DEL-169
chr10-66921592-DEL-130
chr10-66937657-DEL-473
chr10-67036562-DEL-68
chr10-67099159-DEL-354
chr10-67109262-DEL-80
chr10-67138449-DEL-970
chr10-67139436-DEL-302
chr10-67154083-DEL-62
chr10-67163007-DEL-67
chr10-67169626-DEL-136
chr10-67183906-DEL-341
chr10-67194453-DEL-58
chr10-67206668-DEL-127
chr10-67222189-DEL-709
chr10-67230431-DEL-52

chr10-67234643-DEL-51
chr10-67264243-DEL-65
chr10-67278165-DEL-58
chr10-67284662-DEL-122
chr10-67284769-DEL-3218
chr10-67285299-DEL-1333
chr10-67299317-DEL-104
chr10-67315321-DEL-101
chr10-67433051-DEL-121
chr10-67475836-DEL-157
chr10-67480058-DEL-80
chr10-67498039-DEL-509
chr10-67515762-DEL-80
chr10-67541849-DEL-103
chr10-67575614-DEL-69
chr10-67599860-DEL-85
chr10-67611447-DEL-102
chr10-67627243-DEL-67
chr10-67644129-DEL-694
chr10-67644206-DEL-616
chr10-67644954-DEL-499
chr10-67665905-DEL-122
chr10-67688031-DEL-120
chr10-67715393-DEL-64
chr10-67756851-DEL-74
chr10-67778857-DEL-168
chr10-67792057-DEL-55
chr10-67817512-DEL-450
chr10-67834123-DEL-78
chr10-67849562-DEL-214
chr10-67883307-DEL-66
chr10-67890941-DEL-1824
chr10-67926344-DEL-64
chr10-67935695-DEL-175
chr10-67953235-DEL-93
chr10-67976252-DEL-167
chr10-68041650-DEL-87
chr10-68043138-DEL-70
chr10-68050990-DEL-835
chr10-68059817-DEL-78
chr10-68068778-DEL-380
chr10-68101171-DEL-84
chr10-68143081-DEL-396
chr10-68157992-DEL-289
chr10-68173452-DEL-192
chr10-68181558-DEL-221
chr10-68333731-DEL-1038
chr10-68353580-DEL-3615
chr10-68392798-DEL-62
chr10-68395302-DEL-96
chr10-68439509-DEL-991
chr10-68440553-DEL-489
chr10-68454333-DEL-165
chr10-68530780-DEL-1155
chr10-68579126-DEL-1912
chr10-68582791-DEL-123
chr10-68585170-DEL-80
chr10-68597476-DEL-2140
chr10-68598721-DEL-3726
chr10-68606215-DEL-390
chr10-68626536-DEL-74
chr10-68644166-DEL-53
chr10-68645676-DEL-53
chr10-68650229-DEL-757
chr10-68690687-DEL-54
chr10-68697635-DEL-167
chr10-68771139-DEL-150
chr10-68772186-DEL-75
chr10-68775334-DEL-82
chr10-68777723-DEL-98
chr10-68782808-DEL-62
chr10-68789923-DEL-224
chr10-68810134-DEL-52
chr10-68813576-DEL-89
chr10-68825946-DEL-62
chr10-68855600-DEL-147
chr10-68859505-DEL-1067

chr10-68859640-DEL-58
chr10-68882328-DEL-1391
chr10-68886149-DEL-6258
chr10-68903025-DEL-76
chr10-68903947-DEL-76
chr10-68905322-DEL-80
chr10-68915584-DEL-1244
chr10-68938541-DEL-187
chr10-68956380-DEL-2009
chr10-68958290-DEL-234
chr10-68965100-DEL-106
chr10-68975749-DEL-227
chr10-68989331-DEL-76
chr10-69010356-DEL-1365
chr10-69052039-DEL-112
chr10-69053569-DEL-262
chr10-69096598-DEL-53
chr10-69109525-DEL-54
chr10-69150755-DEL-56
chr10-69161859-DEL-123
chr10-69202658-DEL-64
chr10-69232391-DEL-100
chr10-69238523-DEL-111
chr10-69249319-DEL-52
chr10-69263341-DEL-2386
chr10-69298935-DEL-76
chr10-69323372-DEL-363
chr10-69392946-DEL-61
chr10-69395802-DEL-84
chr10-69400853-DEL-59
chr10-69450325-DEL-66
chr10-69497244-DEL-202
chr10-69505990-DEL-63
chr10-69508903-DEL-290
chr10-69512509-DEL-409
chr10-69520130-DEL-62
chr10-69523312-DEL-1482
chr10-69530048-DEL-64
chr10-69571029-DEL-210
chr10-69572474-DEL-1338
chr10-69590669-DEL-147
chr10-69592068-DEL-52
chr10-69601980-DEL-129
chr10-69610588-DEL-574
chr10-69683696-DEL-83
chr10-69713140-DEL-107
chr10-69738566-DEL-92
chr10-69775583-DEL-105
chr10-69799524-DEL-108
chr10-69802527-DEL-907
chr10-69803448-DEL-149
chr10-69833050-DEL-77
chr10-69858115-DEL-60
chr10-69907839-DEL-59
chr10-69996271-DEL-127
chr10-70005973-DEL-60
chr10-70016034-DEL-310
chr10-70020520-DEL-103
chr10-70022911-DEL-56
chr10-70024574-DEL-179
chr10-70030824-DEL-62
chr10-70034666-DEL-121
chr10-70039696-DEL-120
chr10-70067027-DEL-960
chr10-70076698-DEL-212
chr10-70082769-DEL-57
chr10-70101889-DEL-145
chr10-70103566-DEL-708
chr10-70107492-DEL-57
chr10-70135229-DEL-66
chr10-70136226-DEL-362
chr10-70147674-DEL-375
chr10-70181191-DEL-106
chr10-70187416-DEL-246
chr10-70220724-DEL-765
chr10-70249311-DEL-73
chr10-70261085-DEL-116

chr10-70265483-DEL-81
chr10-70305828-DEL-227
chr10-70307447-DEL-78
chr10-70344202-DEL-55
chr10-70537023-DEL-293
chr10-70557873-DEL-238
chr10-70605153-DEL-407
chr10-70666169-DEL-3049
chr10-70687647-DEL-138
chr10-70733079-DEL-98
chr10-70784537-DEL-651
chr10-70832440-DEL-564
chr10-70845272-DEL-195
chr10-70858571-DEL-96
chr10-70874662-DEL-485
chr10-70905451-DEL-65
chr10-70942803-DEL-69
chr10-70997158-DEL-186
chr10-71014720-DEL-150
chr10-71015397-DEL-59
chr10-71024575-DEL-65
chr10-71024973-DEL-966
chr10-71037557-DEL-73
chr10-71076789-DEL-62
chr10-71077599-DEL-92
chr10-71097359-DEL-66
chr10-71107555-DEL-113
chr10-71121506-DEL-73
chr10-71133566-DEL-85
chr10-71140925-DEL-67
chr10-71164758-DEL-437
chr10-71181730-DEL-174
chr10-71203655-DEL-64
chr10-71206616-DEL-181
chr10-71225740-DEL-1252
chr10-71231971-DEL-214
chr10-71233910-DEL-165
chr10-71259542-DEL-132
chr10-71281276-DEL-1733
chr10-71295530-DEL-1191
chr10-71323810-DEL-61
chr10-71324952-DEL-135
chr10-71341149-DEL-61
chr10-71345349-DEL-6381
chr10-71359032-DEL-74
chr10-71361500-DEL-55
chr10-71392243-DEL-68
chr10-71395319-DEL-6346
chr10-71402153-DEL-2769
chr10-71406634-DEL-81
chr10-71406988-DEL-3223
chr10-71406997-DEL-6840
chr10-71413240-DEL-5583
chr10-71413705-DEL-4535
chr10-71419907-DEL-124
chr10-71420999-DEL-6763
chr10-71424726-DEL-2948
chr10-71461406-DEL-12394
chr10-71478153-DEL-242
chr10-71518302-DEL-104
chr10-71544287-DEL-387
chr10-71554086-DEL-126
chr10-71566049-DEL-175
chr10-71579172-DEL-84
chr10-71622940-DEL-57
chr10-71637923-DEL-2243
chr10-71654215-DEL-1503
chr10-71654215-DEL-904
chr10-71655154-DEL-1155
chr10-71656348-DEL-1549
chr10-71659081-DEL-1736
chr10-71676982-DEL-6175
chr10-71698972-DEL-373
chr10-71706565-DEL-68
chr10-71762679-DEL-271
chr10-71765464-DEL-97
chr10-71812806-DEL-120

chr10-71835621-DEL-106
chr10-71855071-DEL-625
chr10-71855737-DEL-410
chr10-71873908-DEL-312
chr10-71914284-DEL-72
chr10-71924694-DEL-383
chr10-71933719-DEL-107
chr10-71953633-DEL-147
chr10-71982259-DEL-107
chr10-71989189-DEL-2128
chr10-72010097-DEL-569
chr10-72010749-DEL-193
chr10-72021230-DEL-1181
chr10-72039124-DEL-267
chr10-72046186-DEL-1277
chr10-72050799-DEL-2266
chr10-72066551-DEL-78
chr10-72071631-DEL-101
chr10-72091785-DEL-199
chr10-72104454-DEL-480
chr10-72138808-DEL-67
chr10-72144797-DEL-6381
chr10-72170567-DEL-100
chr10-72186804-DEL-54
chr10-72193726-DEL-4713
chr10-72215877-DEL-54
chr10-72225582-DEL-104
chr10-72249276-DEL-140
chr10-72292893-DEL-6389
chr10-72302885-DEL-454
chr10-72305798-DEL-76
chr10-72311199-DEL-54
chr10-72339372-DEL-554
chr10-72351816-DEL-1461
chr10-72364929-DEL-71
chr10-72372696-DEL-94
chr10-72387745-DEL-270
chr10-72390049-DEL-145
chr10-72395624-DEL-2119
chr10-72422669-DEL-754
chr10-72436631-DEL-172
chr10-72466577-DEL-52
chr10-72467880-DEL-59
chr10-72472246-DEL-56
chr10-72474520-DEL-83
chr10-72486556-DEL-2158
chr10-72522093-DEL-211
chr10-72526673-DEL-6242
chr10-72533630-DEL-446
chr10-72538061-DEL-7294
chr10-72550979-DEL-56
chr10-72568386-DEL-2258
chr10-72656726-DEL-83
chr10-72685693-DEL-112
chr10-72692141-DEL-1265
chr10-72727649-DEL-146
chr10-72734459-DEL-56
chr10-72765672-DEL-725
chr10-72797204-DEL-6342
chr10-72825229-DEL-81
chr10-72839214-DEL-62
chr10-72883365-DEL-58
chr10-72890122-DEL-59
chr10-72891246-DEL-55
chr10-72914031-DEL-1139
chr10-72928488-DEL-52
chr10-72937320-DEL-92
chr10-72941474-DEL-182
chr10-72961058-DEL-59
chr10-73035499-DEL-78
chr10-73041145-DEL-1235
chr10-73056068-DEL-1918
chr10-73064687-DEL-774
chr10-73066138-DEL-422
chr10-73066388-DEL-1081
chr10-73070401-DEL-70
chr10-73072676-DEL-60

chr10-73084218-DEL-1449
chr10-73104977-DEL-3003
chr10-73178472-DEL-67
chr10-73202427-DEL-71
chr10-73206561-DEL-196
chr10-73214819-DEL-254
chr10-73255693-DEL-66
chr10-73368228-DEL-286
chr10-73383872-DEL-669
chr10-73384562-DEL-1433
chr10-73386052-DEL-1003
chr10-73420241-DEL-126
chr10-73424413-DEL-132
chr10-73425896-DEL-1700
chr10-73427714-DEL-232
chr10-73431628-DEL-6408
chr10-73455628-DEL-753
chr10-73475092-DEL-3493
chr10-73486508-DEL-59
chr10-73486738-DEL-457
chr10-73543039-DEL-146
chr10-73559788-DEL-546
chr10-73607930-DEL-122
chr10-73614382-DEL-1258
chr10-73615843-DEL-1247
chr10-73648048-DEL-87
chr10-73653340-DEL-85
chr10-73660983-DEL-60
chr10-73665740-DEL-60
chr10-73710571-DEL-119
chr10-73730079-DEL-490
chr10-73733885-DEL-58
chr10-73750325-DEL-122
chr10-73763206-DEL-476
chr10-73782477-DEL-70
chr10-73813245-DEL-92
chr10-73831647-DEL-81
chr10-73834446-DEL-135
chr10-73835143-DEL-182
chr10-73842495-DEL-128
chr10-73861260-DEL-69
chr10-73873859-DEL-2730
chr10-73880087-DEL-60
chr10-73901096-DEL-91
chr10-73930239-DEL-77
chr10-73930982-DEL-77
chr10-73972043-DEL-6291
chr10-73980863-DEL-92
chr10-73988598-DEL-6402
chr10-74002294-DEL-576
chr10-74003166-DEL-76
chr10-74008174-DEL-88
chr10-74035604-DEL-78
chr10-74079155-DEL-6417
chr10-74079464-DEL-199
chr10-74097468-DEL-1238
chr10-74130970-DEL-341
chr10-74137740-DEL-76
chr10-74156082-DEL-475
chr10-74176732-DEL-145
chr10-74190883-DEL-61
chr10-74198789-DEL-939
chr10-74235414-DEL-88
chr10-74239728-DEL-400
chr10-74264541-DEL-59
chr10-74268839-DEL-7860
chr10-74269125-DEL-6248
chr10-74306939-DEL-124
chr10-74313862-DEL-183
chr10-74331814-DEL-147
chr10-74364972-DEL-94
chr10-74372934-DEL-1040
chr10-74425109-DEL-373
chr10-74450553-DEL-464
chr10-74470305-DEL-63
chr10-74504798-DEL-6759
chr10-74542263-DEL-103

chr10-74550524-DEL-5648
chr10-74610259-DEL-218
chr10-74611147-DEL-494
chr10-74613773-DEL-56
chr10-74631062-DEL-63
chr10-74639587-DEL-83
chr10-74657770-DEL-55
chr10-74685965-DEL-68
chr10-74704076-DEL-51
chr10-74710887-DEL-223
chr10-74728315-DEL-150
chr10-74734392-DEL-923
chr10-74755449-DEL-849
chr10-74756595-DEL-73
chr10-74786232-DEL-1015
chr10-74789383-DEL-57
chr10-74840638-DEL-211
chr10-74875471-DEL-81
chr10-74947949-DEL-210
chr10-74963347-DEL-85
chr10-74989308-DEL-57
chr10-75038431-DEL-70
chr10-75043276-DEL-253
chr10-75043814-DEL-127
chr10-75056155-DEL-146
chr10-75068924-DEL-469
chr10-75075416-DEL-143
chr10-75114473-DEL-284
chr10-75119671-DEL-57
chr10-75123569-DEL-136
chr10-75141546-DEL-132
chr10-75190775-DEL-80
chr10-75204843-DEL-104
chr10-75214068-DEL-56
chr10-75240594-DEL-84
chr10-75255132-DEL-70
chr10-75261746-DEL-94
chr10-75271114-DEL-82
chr10-75304944-DEL-789
chr10-75341732-DEL-191
chr10-75379360-DEL-1919
chr10-75386584-DEL-97
chr10-75406032-DEL-68
chr10-75406671-DEL-61
chr10-75430123-DEL-127
chr10-75436896-DEL-367
chr10-75497901-DEL-119
chr10-75540959-DEL-538
chr10-75559759-DEL-220
chr10-75581755-DEL-179
chr10-75624813-DEL-72
chr10-75627252-DEL-230
chr10-75667475-DEL-187
chr10-75704318-DEL-103
chr10-75715778-DEL-867
chr10-75742364-DEL-62
chr10-75770273-DEL-228
chr10-75774764-DEL-127
chr10-75787417-DEL-111
chr10-75800526-DEL-170
chr10-75806164-DEL-202
chr10-75813792-DEL-251
chr10-75820462-DEL-148
chr10-75832630-DEL-147
chr10-75836464-DEL-1552
chr10-75855179-DEL-1244
chr10-75857464-DEL-743
chr10-75859016-DEL-198
chr10-75863373-DEL-72
chr10-75882560-DEL-67
chr10-75883645-DEL-120
chr10-75900116-DEL-198
chr10-75908722-DEL-208
chr10-75912995-DEL-1802
chr10-75916145-DEL-144
chr10-75918217-DEL-127
chr10-75926162-DEL-84

chr10-75949415-DEL-431
chr10-75950054-DEL-74
chr10-75971215-DEL-1081
chr10-75991286-DEL-597
chr10-76003183-DEL-71
chr10-76023094-DEL-80
chr10-76030781-DEL-1914
chr10-76061651-DEL-253
chr10-76065594-DEL-56109
chr10-76080303-DEL-61
chr10-76108193-DEL-64
chr10-76146212-DEL-1157
chr10-76146248-DEL-314
chr10-76146645-DEL-136
chr10-76147154-DEL-971
chr10-76147936-DEL-258
chr10-76166455-DEL-570
chr10-76246353-DEL-76
chr10-76272043-DEL-68
chr10-76289608-DEL-200
chr10-76290959-DEL-69
chr10-76295601-DEL-226
chr10-76309222-DEL-78
chr10-76369550-DEL-54
chr10-76401555-DEL-191
chr10-76413616-DEL-62
chr10-76456225-DEL-160
chr10-76461732-DEL-69
chr10-76467484-DEL-70
chr10-76470461-DEL-233
chr10-76474694-DEL-356
chr10-76512316-DEL-449
chr10-76523789-DEL-70
chr10-76539874-DEL-180
chr10-76562381-DEL-264
chr10-76588090-DEL-98
chr10-76647667-DEL-215
chr10-76662462-DEL-470
chr10-76666535-DEL-60
chr10-76690664-DEL-97
chr10-76714777-DEL-60
chr10-76785158-DEL-383
chr10-76822454-DEL-154
chr10-76834861-DEL-75
chr10-76846621-DEL-113
chr10-76876104-DEL-93
chr10-76881786-DEL-366
chr10-76912618-DEL-93
chr10-76920854-DEL-94
chr10-76922503-DEL-228
chr10-76945018-DEL-71
chr10-76951026-DEL-1628
chr10-76961961-DEL-974
chr10-76985593-DEL-92
chr10-77009623-DEL-76
chr10-77010829-DEL-53
chr10-77011880-DEL-183
chr10-77026206-DEL-60
chr10-77063503-DEL-517
chr10-77066709-DEL-152
chr10-77089053-DEL-56
chr10-77089997-DEL-97
chr10-77111537-DEL-60
chr10-77121775-DEL-399
chr10-77139631-DEL-304
chr10-77141172-DEL-128
chr10-77163973-DEL-63
chr10-77213080-DEL-330
chr10-77329018-DEL-1747
chr10-77340308-DEL-201
chr10-77354774-DEL-103
chr10-77389895-DEL-338
chr10-77422014-DEL-966
chr10-77436768-DEL-376
chr10-77464760-DEL-322
chr10-77502600-DEL-75
chr10-77517562-DEL-210

chr10-77519501-DEL-162
chr10-77529353-DEL-80
chr10-77585330-DEL-69
chr10-77591480-DEL-200
chr10-77640709-DEL-2101
chr10-77650652-DEL-59
chr10-77707448-DEL-178
chr10-77770094-DEL-100
chr10-77797076-DEL-224
chr10-77799197-DEL-194
chr10-77806900-DEL-66
chr10-77849728-DEL-151
chr10-77872713-DEL-78
chr10-77952734-DEL-87
chr10-77968075-DEL-195
chr10-77974335-DEL-52
chr10-77993118-DEL-128
chr10-78089482-DEL-129
chr10-78124750-DEL-63
chr10-78181211-DEL-64
chr10-78206763-DEL-204
chr10-78214628-DEL-166
chr10-78216219-DEL-226
chr10-78222870-DEL-58
chr10-78231863-DEL-120
chr10-78241561-DEL-212
chr10-78270747-DEL-1212
chr10-78271808-DEL-367
chr10-78367957-DEL-58
chr10-78378886-DEL-4230
chr10-78385892-DEL-2434
chr10-78395737-DEL-150
chr10-78430004-DEL-94
chr10-78465559-DEL-2408
chr10-78470336-DEL-3599
chr10-78543721-DEL-276
chr10-78548574-DEL-55
chr10-78553843-DEL-54
chr10-78561616-DEL-156
chr10-78574261-DEL-54
chr10-78575318-DEL-70
chr10-78578536-DEL-73
chr10-78610513-DEL-90
chr10-78639288-DEL-6918
chr10-78652298-DEL-6588
chr10-78688704-DEL-5828
chr10-78695263-DEL-8696
chr10-78706046-DEL-492
chr10-78708182-DEL-240
chr10-78716399-DEL-478
chr10-78727239-DEL-9840
chr10-78730592-DEL-214
chr10-78776357-DEL-6496
chr10-78807880-DEL-897
chr10-78855226-DEL-56
chr10-78866044-DEL-59
chr10-78884957-DEL-127
chr10-78939797-DEL-2677
chr10-78973272-DEL-210
chr10-78992450-DEL-206
chr10-79103747-DEL-600
chr10-79133827-DEL-200
chr10-79150089-DEL-8319
chr10-79166832-DEL-6621
chr10-79202750-DEL-1665
chr10-79233685-DEL-2163
chr10-79237228-DEL-414
chr10-79244439-DEL-136
chr10-79255642-DEL-53
chr10-79360973-DEL-156
chr10-79417201-DEL-2101
chr10-79424383-DEL-1117
chr10-79425440-DEL-67
chr10-79426751-DEL-95
chr10-79442257-DEL-128
chr10-79453492-DEL-164
chr10-79477314-DEL-689

chr10-79492585-DEL-198
chr10-79514459-DEL-507
chr10-79521923-DEL-53
chr10-79560671-DEL-60
chr10-79562662-DEL-200
chr10-79627987-DEL-197
chr10-79650634-DEL-95
chr10-79660118-DEL-62
chr10-79720642-DEL-252
chr10-79732084-DEL-78
chr10-79748660-DEL-83
chr10-79760329-DEL-199
chr10-79795076-DEL-131
chr10-79799136-DEL-54
chr10-79805917-DEL-283
chr10-79904904-DEL-196
chr10-79909934-DEL-61
chr10-79921830-DEL-207
chr10-79924575-DEL-86
chr10-79948108-DEL-154
chr10-79976764-DEL-54
chr10-80038937-DEL-178
chr10-80047876-DEL-1634
chr10-80049087-DEL-174
chr10-80067040-DEL-196
chr10-80112373-DEL-178
chr10-80127455-DEL-275
chr10-80179685-DEL-122
chr10-80189675-DEL-176
chr10-80194955-DEL-52
chr10-80197617-DEL-413
chr10-80203204-DEL-99
chr10-80203958-DEL-126
chr10-80204833-DEL-204
chr10-80305889-DEL-83
chr10-80307029-DEL-80
chr10-80319042-DEL-76
chr10-80320617-DEL-1059
chr10-80323826-DEL-182
chr10-80324456-DEL-151
chr10-80337029-DEL-66
chr10-80337854-DEL-79
chr10-80341569-DEL-69
chr10-80344511-DEL-76
chr10-80347208-DEL-56
chr10-80369276-DEL-180
chr10-80373605-DEL-59
chr10-80380349-DEL-146
chr10-80394619-DEL-185
chr10-80395695-DEL-73
chr10-80429435-DEL-1820
chr10-80441527-DEL-188
chr10-80477283-DEL-215
chr10-80567434-DEL-168
chr10-80578933-DEL-205
chr10-80615870-DEL-110
chr10-80625698-DEL-207
chr10-80661247-DEL-65
chr10-80696089-DEL-414
chr10-80717615-DEL-809
chr10-80728445-DEL-203
chr10-80737013-DEL-146
chr10-80747851-DEL-92
chr10-80789923-DEL-210
chr10-80819389-DEL-70
chr10-80844675-DEL-655
chr10-80846508-DEL-208
chr10-80850475-DEL-73
chr10-80881055-DEL-178
chr10-80892681-DEL-369
chr10-80893877-DEL-167
chr10-80896949-DEL-162
chr10-80930847-DEL-67
chr10-80935170-DEL-217
chr10-80936128-DEL-68
chr10-80962803-DEL-86
chr10-81020938-DEL-70

chr10-81024151-DEL-181
chr10-81039709-DEL-96
chr10-81074560-DEL-60
chr10-81092708-DEL-283
chr10-81117424-DEL-576
chr10-81163361-DEL-112
chr10-81171660-DEL-84
chr10-81175083-DEL-233
chr10-81187435-DEL-117
chr10-81191849-DEL-69
chr10-81210813-DEL-206
chr10-81343166-DEL-51
chr10-81350907-DEL-196
chr10-81351818-DEL-196
chr10-81383711-DEL-232
chr10-81415305-DEL-89
chr10-81449439-DEL-188
chr10-81455637-DEL-189
chr10-81464820-DEL-144
chr10-81475473-DEL-159
chr10-81503988-DEL-85
chr10-81505682-DEL-240
chr10-81506406-DEL-7198
chr10-81518078-DEL-62
chr10-81520034-DEL-473
chr10-81556706-DEL-84
chr10-81560795-DEL-76
chr10-81580676-DEL-84
chr10-81634337-DEL-4078
chr10-81641854-DEL-88
chr10-81685536-DEL-598
chr10-81688575-DEL-552
chr10-81746466-DEL-1064
chr10-81793775-DEL-3904
chr10-81801256-DEL-146
chr10-81803734-DEL-7093
chr10-81816374-DEL-64
chr10-81817710-DEL-598
chr10-81821699-DEL-551
chr10-81840215-DEL-293
chr10-81846813-DEL-433
chr10-81853414-DEL-77
chr10-81859886-DEL-5999
chr10-81898259-DEL-139
chr10-81909750-DEL-72
chr10-81911471-DEL-1363
chr10-81925473-DEL-64
chr10-81951273-DEL-51
chr10-81957836-DEL-1086
chr10-82001840-DEL-76
chr10-82017229-DEL-851
chr10-82022142-DEL-867
chr10-82045306-DEL-8382
chr10-82085434-DEL-979
chr10-82087057-DEL-6076
chr10-82106861-DEL-81
chr10-82124463-DEL-126
chr10-82136686-DEL-123
chr10-82164550-DEL-2964
chr10-82168441-DEL-3983
chr10-82221761-DEL-680
chr10-82246428-DEL-167
chr10-82247150-DEL-6500
chr10-82275391-DEL-93
chr10-82280467-DEL-2595
chr10-82293669-DEL-372
chr10-82329808-DEL-451
chr10-82348358-DEL-2311
chr10-82353238-DEL-1506
chr10-82407201-DEL-3820
chr10-82421206-DEL-2176
chr10-82425711-DEL-318
chr10-82435346-DEL-2529
chr10-82442786-DEL-949
chr10-82446470-DEL-3907
chr10-82530806-DEL-142
chr10-82559725-DEL-56

chr10-82609163-DEL-72
chr10-82618870-DEL-2237
chr10-82633370-DEL-88
chr10-82639035-DEL-373
chr10-82652634-DEL-107
chr10-82670547-DEL-53
chr10-82673522-DEL-83
chr10-82688048-DEL-334
chr10-82734688-DEL-202
chr10-82743178-DEL-1002
chr10-82751510-DEL-59
chr10-82758506-DEL-753
chr10-82767021-DEL-105
chr10-82768800-DEL-56
chr10-82811412-DEL-61
chr10-82844694-DEL-2223
chr10-82849470-DEL-60
chr10-82866297-DEL-235
chr10-82884665-DEL-220
chr10-82888087-DEL-60
chr10-82909432-DEL-696
chr10-82922600-DEL-181
chr10-82947589-DEL-92
chr10-82956442-DEL-88
chr10-82957078-DEL-66
chr10-82977895-DEL-56
chr10-82983660-DEL-55
chr10-83050468-DEL-55
chr10-83060803-DEL-88
chr10-83074922-DEL-234
chr10-83084804-DEL-64
chr10-83116796-DEL-105
chr10-83121499-DEL-54
chr10-83132283-DEL-118
chr10-83145285-DEL-110
chr10-83181184-DEL-173
chr10-83191623-DEL-112
chr10-83219304-DEL-66074
chr10-83227178-DEL-77
chr10-83254238-DEL-75
chr10-83290736-DEL-3513
chr10-83295350-DEL-102
chr10-83295812-DEL-14550
chr10-83342008-DEL-169
chr10-83360348-DEL-143
chr10-83363911-DEL-432
chr10-83372779-DEL-56
chr10-83381304-DEL-59
chr10-83382747-DEL-208
chr10-83384649-DEL-57
chr10-83447150-DEL-61
chr10-83451139-DEL-87
chr10-83462754-DEL-106
chr10-83506781-DEL-7082
chr10-83538315-DEL-57
chr10-83545043-DEL-68
chr10-83550016-DEL-199
chr10-83551870-DEL-1623
chr10-83568442-DEL-56
chr10-83589533-DEL-64
chr10-83630032-DEL-183
chr10-83644382-DEL-57
chr10-83680208-DEL-611
chr10-83683967-DEL-317
chr10-83713437-DEL-109
chr10-83739765-DEL-71
chr10-83742123-DEL-59
chr10-83756758-DEL-411
chr10-83772383-DEL-61
chr10-83790323-DEL-65
chr10-83915289-DEL-68
chr10-83958939-DEL-114
chr10-83978422-DEL-56
chr10-84000777-DEL-612
chr10-84020336-DEL-91
chr10-84031354-DEL-969
chr10-84064389-DEL-864

chr10-84065356-DEL-1409
chr10-84067631-DEL-64
chr10-84094056-DEL-174
chr10-84098157-DEL-615
chr10-84126836-DEL-356
chr10-84146483-DEL-1128
chr10-84165766-DEL-92
chr10-84181443-DEL-56
chr10-84191576-DEL-56
chr10-84213155-DEL-62
chr10-84263393-DEL-58
chr10-84284770-DEL-198
chr10-84310598-DEL-198
chr10-84323397-DEL-73
chr10-84351808-DEL-137
chr10-84358268-DEL-66
chr10-84384518-DEL-89
chr10-84429180-DEL-136
chr10-84434813-DEL-243
chr10-84435370-DEL-641
chr10-84462822-DEL-330
chr10-84470287-DEL-82
chr10-84483734-DEL-186
chr10-84491138-DEL-296
chr10-84496775-DEL-159
chr10-84499951-DEL-245
chr10-84538649-DEL-1122
chr10-84553251-DEL-306
chr10-84567607-DEL-60
chr10-84569040-DEL-66
chr10-84573168-DEL-187
chr10-84586742-DEL-231
chr10-84597023-DEL-187
chr10-84597757-DEL-217
chr10-84633653-DEL-201
chr10-84672959-DEL-60
chr10-84677143-DEL-996
chr10-84680395-DEL-57
chr10-84714355-DEL-184
chr10-84717618-DEL-1048
chr10-84779849-DEL-94
chr10-84870320-DEL-53
chr10-84907881-DEL-157
chr10-84917070-DEL-198
chr10-84918234-DEL-109
chr10-84924015-DEL-358
chr10-84977270-DEL-137
chr10-85027339-DEL-462
chr10-85028099-DEL-112
chr10-85101591-DEL-1431
chr10-85117014-DEL-350
chr10-85147008-DEL-63
chr10-85149324-DEL-1000
chr10-85163944-DEL-142
chr10-85198535-DEL-191
chr10-85201053-DEL-175
chr10-85203785-DEL-72
chr10-85221595-DEL-442
chr10-85229032-DEL-56
chr10-85240221-DEL-277
chr10-85249265-DEL-6067
chr10-85249487-DEL-370
chr10-85250227-DEL-5105
chr10-85251411-DEL-3736
chr10-85264539-DEL-55
chr10-85288223-DEL-91
chr10-85308264-DEL-67
chr10-85321537-DEL-746
chr10-85323013-DEL-83
chr10-85351596-DEL-107
chr10-85357251-DEL-145
chr10-85372263-DEL-71
chr10-85403998-DEL-64
chr10-85406078-DEL-58
chr10-85409072-DEL-770
chr10-85437544-DEL-1259
chr10-85446423-DEL-54

chr10-85510208-DEL-545
chr10-85519680-DEL-390
chr10-85532397-DEL-1364
chr10-85567014-DEL-99
chr10-85615794-DEL-543
chr10-85620437-DEL-198
chr10-85690805-DEL-4388
chr10-85695418-DEL-3080
chr10-85702996-DEL-131
chr10-85705162-DEL-902
chr10-85722634-DEL-59
chr10-85732573-DEL-156
chr10-85756472-DEL-69
chr10-85766482-DEL-1276
chr10-85767726-DEL-185
chr10-85794924-DEL-502
chr10-85839142-DEL-170
chr10-85887206-DEL-84
chr10-85894194-DEL-51
chr10-85932911-DEL-381
chr10-85969135-DEL-86
chr10-85981150-DEL-56
chr10-85987885-DEL-215
chr10-86004556-DEL-66
chr10-86041118-DEL-85
chr10-86042058-DEL-73
chr10-86070602-DEL-212
chr10-86080523-DEL-963
chr10-86082054-DEL-191
chr10-86083025-DEL-189
chr10-86092474-DEL-86
chr10-86165778-DEL-52
chr10-86250637-DEL-210
chr10-86308599-DEL-1020
chr10-86314829-DEL-426
chr10-86330114-DEL-1168
chr10-86394767-DEL-139
chr10-86407456-DEL-1284
chr10-86408117-DEL-1281
chr10-86429692-DEL-6172
chr10-86438430-DEL-321
chr10-86447256-DEL-1686
chr10-86464179-DEL-3210
chr10-86493219-DEL-181
chr10-86499297-DEL-176
chr10-86510220-DEL-58
chr10-86511128-DEL-132
chr10-86512048-DEL-8403
chr10-86525079-DEL-54
chr10-86558820-DEL-95
chr10-86567562-DEL-808
chr10-86575786-DEL-2707
chr10-86577720-DEL-2034
chr10-86578521-DEL-244
chr10-86579320-DEL-874
chr10-86585595-DEL-175
chr10-86595244-DEL-60
chr10-86623916-DEL-140
chr10-86640422-DEL-63
chr10-86658651-DEL-112
chr10-86676098-DEL-114
chr10-86696082-DEL-60
chr10-86728572-DEL-123
chr10-86736987-DEL-185
chr10-86740235-DEL-290
chr10-86743794-DEL-66
chr10-86764743-DEL-353
chr10-86774772-DEL-424
chr10-86788854-DEL-58
chr10-86796987-DEL-100
chr10-86800268-DEL-1051
chr10-86802622-DEL-111
chr10-86804014-DEL-276
chr10-86806130-DEL-258
chr10-86825560-DEL-73
chr10-86838241-DEL-380
chr10-86860668-DEL-50

chr10-86890481-DEL-144
chr10-86926555-DEL-80
chr10-86978117-DEL-3516
chr10-86992619-DEL-54
chr10-86999497-DEL-768
chr10-87013026-DEL-122
chr10-87020683-DEL-1174
chr10-87022079-DEL-86
chr10-87022304-DEL-1673
chr10-87037938-DEL-107
chr10-87043210-DEL-528
chr10-87053004-DEL-397
chr10-87084770-DEL-1012
chr10-87088509-DEL-612
chr10-87117581-DEL-125
chr10-87121668-DEL-945
chr10-87127793-DEL-666
chr10-87132940-DEL-99
chr10-87156341-DEL-2271
chr10-87201682-DEL-62
chr10-87209496-DEL-57
chr10-87211061-DEL-3637
chr10-87250223-DEL-379
chr10-87263223-DEL-58
chr10-87272743-DEL-4754
chr10-87272842-DEL-6135
chr10-87284568-DEL-80
chr10-87295656-DEL-406
chr10-87303499-DEL-216
chr10-87306569-DEL-2436
chr10-87306735-DEL-79
chr10-87319242-DEL-912
chr10-87348233-DEL-227
chr10-87361078-DEL-174
chr10-87362405-DEL-60
chr10-87368617-DEL-908
chr10-87379157-DEL-204
chr10-87418241-DEL-358
chr10-87434888-DEL-1322
chr10-87471825-DEL-259
chr10-87474175-DEL-502
chr10-87486367-DEL-1521
chr10-87503993-DEL-60
chr10-87523912-DEL-1050
chr10-87537135-DEL-101
chr10-87541308-DEL-62
chr10-87575209-DEL-298
chr10-87578263-DEL-68
chr10-87616631-DEL-53
chr10-87623070-DEL-163
chr10-87628028-DEL-63
chr10-87635581-DEL-940
chr10-87752412-DEL-168
chr10-87780287-DEL-631
chr10-87802622-DEL-3222
chr10-87825224-DEL-1533
chr10-87833304-DEL-328
chr10-87846207-DEL-626
chr10-87868624-DEL-531
chr10-87886831-DEL-490
chr10-87901837-DEL-983
chr10-87917646-DEL-61
chr10-87973379-DEL-1288
chr10-87973928-DEL-1095
chr10-87992848-DEL-116
chr10-88000487-DEL-76
chr10-88008940-DEL-540
chr10-88028052-DEL-57
chr10-88031063-DEL-56
chr10-88046385-DEL-110
chr10-88090912-DEL-84
chr10-88106352-DEL-546
chr10-88121212-DEL-69
chr10-88141434-DEL-51
chr10-88153236-DEL-1361
chr10-88167781-DEL-90
chr10-88186083-DEL-92

chr10-88195542-DEL-62
chr10-88213370-DEL-153
chr10-88237553-DEL-310
chr10-88276961-DEL-75
chr10-88287761-DEL-143
chr10-88322890-DEL-217
chr10-88347240-DEL-745
chr10-88353424-DEL-147
chr10-88358362-DEL-54
chr10-88365485-DEL-174
chr10-88365965-DEL-228
chr10-88377504-DEL-194
chr10-88398019-DEL-460
chr10-88405282-DEL-81
chr10-88417722-DEL-76
chr10-88423377-DEL-73
chr10-88430136-DEL-87
chr10-88482015-DEL-246
chr10-88494192-DEL-157
chr10-88496813-DEL-180
chr10-88507352-DEL-1388
chr10-88520154-DEL-84
chr10-88532243-DEL-63
chr10-88580657-DEL-61
chr10-88581271-DEL-410
chr10-88591330-DEL-87
chr10-88604044-DEL-386
chr10-88610298-DEL-72
chr10-88617031-DEL-665
chr10-88641249-DEL-185
chr10-88657167-DEL-144
chr10-88686729-DEL-94
chr10-88686970-DEL-402
chr10-88702251-DEL-72
chr10-88708593-DEL-160
chr10-88715783-DEL-328
chr10-88748549-DEL-164
chr10-88783100-DEL-168
chr10-88783714-DEL-299
chr10-88786772-DEL-400
chr10-88816524-DEL-82
chr10-88843567-DEL-239
chr10-88880812-DEL-7133
chr10-88895872-DEL-1391
chr10-88909690-DEL-72
chr10-88917819-DEL-53
chr10-88933193-DEL-1874
chr10-89008196-DEL-906
chr10-89015461-DEL-279
chr10-89027500-DEL-122
chr10-89048009-DEL-147
chr10-89067428-DEL-80
chr10-89076263-DEL-1246
chr10-89132627-DEL-97
chr10-89134093-DEL-63
chr10-89175130-DEL-133
chr10-89175918-DEL-417
chr10-89183941-DEL-60
chr10-89221282-DEL-7084
chr10-89222976-DEL-118
chr10-89242966-DEL-55
chr10-89266472-DEL-185
chr10-89311313-DEL-102
chr10-89353499-DEL-56
chr10-89379554-DEL-67
chr10-89438845-DEL-1605
chr10-89448117-DEL-59
chr10-89448125-DEL-2209
chr10-89450639-DEL-100
chr10-89500250-DEL-149
chr10-89522301-DEL-8363
chr10-89523237-DEL-1670
chr10-89541181-DEL-52
chr10-89545398-DEL-472
chr10-89549704-DEL-227
chr10-89553398-DEL-179
chr10-89566490-DEL-215

chr10-89621503-DEL-448
chr10-89649382-DEL-479
chr10-89652366-DEL-95
chr10-89652652-DEL-461
chr10-89660264-DEL-921
chr10-89661070-DEL-96
chr10-89668609-DEL-229
chr10-89684415-DEL-767
chr10-89685224-DEL-1314
chr10-89688414-DEL-165
chr10-89703246-DEL-94
chr10-89730490-DEL-97
chr10-89734043-DEL-221
chr10-89751178-DEL-225
chr10-89775215-DEL-93
chr10-89785434-DEL-15361
chr10-89802851-DEL-453
chr10-89819688-DEL-293
chr10-89825045-DEL-2495
chr10-89829010-DEL-79
chr10-89911350-DEL-132
chr10-89924337-DEL-312
chr10-89961561-DEL-163
chr10-90007287-DEL-213
chr10-90023176-DEL-219
chr10-90028148-DEL-167
chr10-90052391-DEL-2689
chr10-90092036-DEL-100
chr10-90093633-DEL-1044
chr10-90095274-DEL-182
chr10-90102214-DEL-643
chr10-90140787-DEL-128
chr10-90151791-DEL-651
chr10-90159085-DEL-55
chr10-90182565-DEL-134
chr10-90201210-DEL-80
chr10-90238669-DEL-579
chr10-90262355-DEL-232
chr10-90286255-DEL-1748
chr10-90295520-DEL-52
chr10-90334244-DEL-803
chr10-90341751-DEL-104
chr10-90368884-DEL-89
chr10-90405819-DEL-88
chr10-90421683-DEL-156
chr10-90438532-DEL-60
chr10-90528916-DEL-169
chr10-90530379-DEL-160
chr10-90600253-DEL-357
chr10-90610519-DEL-51
chr10-90610966-DEL-176
chr10-90638912-DEL-157
chr10-90685300-DEL-91
chr10-90690989-DEL-357
chr10-90730229-DEL-175
chr10-90737637-DEL-81
chr10-90740115-DEL-62
chr10-90781374-DEL-51
chr10-90823518-DEL-59
chr10-90846167-DEL-98
chr10-90847666-DEL-1149
chr10-90892233-DEL-95
chr10-90924462-DEL-929
chr10-90953311-DEL-87
chr10-90975659-DEL-692
chr10-90976473-DEL-220
chr10-90980818-DEL-191
chr10-91053523-DEL-84
chr10-91088080-DEL-6295
chr10-91108935-DEL-56
chr10-91110501-DEL-201
chr10-91111834-DEL-158
chr10-91138239-DEL-61
chr10-91200959-DEL-89
chr10-91212213-DEL-123
chr10-91318464-DEL-68
chr10-91339611-DEL-63

chr10-91355349-DEL-176
chr10-91398535-DEL-62
chr10-91454474-DEL-178
chr10-91469495-DEL-56
chr10-91475532-DEL-1004
chr10-91481707-DEL-70
chr10-91503371-DEL-64
chr10-91522710-DEL-494
chr10-91572947-DEL-68
chr10-91590962-DEL-453
chr10-91601484-DEL-588
chr10-91604456-DEL-171
chr10-91607208-DEL-56
chr10-91611809-DEL-6306
chr10-91650510-DEL-2296
chr10-91653704-DEL-907
chr10-91683095-DEL-64
chr10-91714154-DEL-141
chr10-91716571-DEL-64
chr10-91721766-DEL-64
chr10-91741176-DEL-96
chr10-91779119-DEL-1325
chr10-91809596-DEL-114
chr10-91835267-DEL-84
chr10-91887326-DEL-768
chr10-91959933-DEL-89
chr10-92079501-DEL-65
chr10-92143353-DEL-120
chr10-92193306-DEL-84
chr10-92212430-DEL-75
chr10-92242229-DEL-86
chr10-92245880-DEL-670
chr10-92338686-DEL-121
chr10-92361325-DEL-57
chr10-92383266-DEL-59
chr10-92396036-DEL-54
chr10-92430330-DEL-70
chr10-92457093-DEL-52
chr10-92480885-DEL-54
chr10-92481467-DEL-152
chr10-92524898-DEL-273
chr10-92568639-DEL-116
chr10-92579600-DEL-62
chr10-92616818-DEL-1024
chr10-92639066-DEL-6256
chr10-92646032-DEL-101
chr10-92654214-DEL-87
chr10-92675111-DEL-51
chr10-92675754-DEL-101
chr10-92678525-DEL-298
chr10-92686546-DEL-2778
chr10-92693854-DEL-707
chr10-92706206-DEL-122
chr10-92738007-DEL-156
chr10-92738965-DEL-66
chr10-92762059-DEL-62
chr10-92794866-DEL-141
chr10-92814360-DEL-110
chr10-92819686-DEL-324
chr10-92838646-DEL-580
chr10-92861063-DEL-174
chr10-92876521-DEL-52
chr10-92879406-DEL-195
chr10-92886294-DEL-53
chr10-92907583-DEL-62
chr10-92935211-DEL-51
chr10-92943965-DEL-168
chr10-92986505-DEL-275
chr10-92993210-DEL-52
chr10-92999932-DEL-98
chr10-93014157-DEL-74
chr10-93031188-DEL-133
chr10-93034250-DEL-245
chr10-93058197-DEL-140
chr10-93058219-DEL-1243
chr10-93058267-DEL-353
chr10-93058816-DEL-308

chr10-93058964-DEL-752
chr10-93070202-DEL-212
chr10-93082785-DEL-274
chr10-93135358-DEL-54
chr10-93150588-DEL-110
chr10-93170289-DEL-52
chr10-93171158-DEL-89
chr10-93182016-DEL-56
chr10-93185952-DEL-111
chr10-93245458-DEL-6405
chr10-93253944-DEL-63
chr10-93263934-DEL-184
chr10-93269380-DEL-102
chr10-93281183-DEL-449
chr10-93332897-DEL-54
chr10-93353198-DEL-169
chr10-93394009-DEL-395
chr10-93402233-DEL-97
chr10-93435868-DEL-770
chr10-93436671-DEL-1283
chr10-93461634-DEL-124
chr10-93462882-DEL-676
chr10-93463586-DEL-102
chr10-93469228-DEL-167
chr10-93507886-DEL-84
chr10-93529694-DEL-56
chr10-93550060-DEL-75
chr10-93562843-DEL-744
chr10-93562860-DEL-190
chr10-93587592-DEL-208
chr10-93594711-DEL-52
chr10-93595072-DEL-312
chr10-93595460-DEL-483
chr10-93608416-DEL-56
chr10-93611221-DEL-57
chr10-93637873-DEL-434
chr10-93641492-DEL-63
chr10-93674034-DEL-152
chr10-93685664-DEL-64
chr10-93739712-DEL-68
chr10-93757393-DEL-51
chr10-93761641-DEL-594
chr10-93765730-DEL-186
chr10-93775237-DEL-52
chr10-93796992-DEL-51
chr10-93811431-DEL-80
chr10-93821894-DEL-840
chr10-93845287-DEL-131
chr10-93898439-DEL-62
chr10-93904302-DEL-79
chr10-94020223-DEL-211
chr10-94048333-DEL-1413
chr10-94054053-DEL-50
chr10-94079363-DEL-244
chr10-94091986-DEL-53
chr10-94097769-DEL-100
chr10-94120497-DEL-172
chr10-94132437-DEL-60
chr10-94173846-DEL-68
chr10-94215917-DEL-80
chr10-94227652-DEL-204
chr10-94231555-DEL-76
chr10-94249157-DEL-52
chr10-94295381-DEL-97
chr10-94335222-DEL-60
chr10-94358334-DEL-60
chr10-94389306-DEL-4266
chr10-94390791-DEL-204
chr10-94399664-DEL-150
chr10-94433924-DEL-167
chr10-94452698-DEL-54
chr10-94460674-DEL-131
chr10-94461186-DEL-75
chr10-94469737-DEL-457
chr10-94542280-DEL-1484
chr10-94544053-DEL-89
chr10-94569228-DEL-75

chr10-94571288-DEL-275
chr10-94588773-DEL-281
chr10-94698348-DEL-62
chr10-94705236-DEL-70
chr10-94711728-DEL-56
chr10-94750959-DEL-128
chr10-94784646-DEL-432
chr10-94792537-DEL-51
chr10-94822197-DEL-266
chr10-94831982-DEL-475
chr10-94835485-DEL-217
chr10-94857158-DEL-1374
chr10-94859105-DEL-2258
chr10-94926967-DEL-108
chr10-94949998-DEL-768
chr10-94950820-DEL-926
chr10-95015175-DEL-116
chr10-95019651-DEL-154
chr10-95037065-DEL-73
chr10-95044223-DEL-1370
chr10-95071751-DEL-244
chr10-95097510-DEL-1268
chr10-95098588-DEL-164
chr10-95116370-DEL-365
chr10-95121972-DEL-160
chr10-95141517-DEL-297
chr10-95168909-DEL-76
chr10-95179142-DEL-56
chr10-95215997-DEL-327
chr10-95254237-DEL-854
chr10-95259806-DEL-117
chr10-95264705-DEL-294
chr10-95281644-DEL-1314
chr10-95283889-DEL-52
chr10-95291813-DEL-942
chr10-95295974-DEL-399
chr10-95300711-DEL-75
chr10-95334127-DEL-943
chr10-95347568-DEL-99
chr10-95348560-DEL-53
chr10-95371766-DEL-208
chr10-95381748-DEL-81
chr10-95392814-DEL-70
chr10-95396516-DEL-1306
chr10-95396530-DEL-133
chr10-95404881-DEL-66
chr10-95407810-DEL-235
chr10-95414325-DEL-1206
chr10-95422173-DEL-156
chr10-95430243-DEL-110
chr10-95435440-DEL-67
chr10-95455022-DEL-178
chr10-95458034-DEL-159
chr10-95474674-DEL-60
chr10-95487395-DEL-197
chr10-95531035-DEL-292
chr10-95536382-DEL-52
chr10-95546223-DEL-65
chr10-95554885-DEL-85
chr10-95579191-DEL-195
chr10-95634951-DEL-97
chr10-95648424-DEL-58
chr10-95648622-DEL-443
chr10-95670388-DEL-388
chr10-95689591-DEL-204
chr10-95696546-DEL-121
chr10-95702701-DEL-247
chr10-95720760-DEL-55
chr10-95721473-DEL-400
chr10-95746669-DEL-173
chr10-95747462-DEL-96
chr10-95762346-DEL-143
chr10-95762920-DEL-179
chr10-95769432-DEL-325
chr10-95810495-DEL-60
chr10-95890284-DEL-4355
chr10-95904287-DEL-6010

chr10-95921105-DEL-78
chr10-95922791-DEL-139
chr10-96007767-DEL-86
chr10-96010137-DEL-66
chr10-96072215-DEL-60
chr10-96082803-DEL-54
chr10-96113178-DEL-79
chr10-96191094-DEL-54
chr10-96201976-DEL-1372
chr10-96256496-DEL-2926
chr10-96292378-DEL-555
chr10-96297676-DEL-415
chr10-96307063-DEL-95
chr10-96333848-DEL-1347
chr10-96376445-DEL-352
chr10-96389443-DEL-1341
chr10-96411871-DEL-172
chr10-96439822-DEL-67
chr10-96474007-DEL-79
chr10-96476166-DEL-214
chr10-96485973-DEL-354
chr10-96492635-DEL-52
chr10-96511361-DEL-93
chr10-96517917-DEL-56
chr10-96544030-DEL-106
chr10-96544922-DEL-156
chr10-96554819-DEL-167
chr10-96582249-DEL-62
chr10-96610784-DEL-2228
chr10-96615644-DEL-445
chr10-96631754-DEL-216
chr10-96693307-DEL-123
chr10-96795714-DEL-54
chr10-96832389-DEL-172
chr10-96853316-DEL-51
chr10-96871575-DEL-1094
chr10-96916742-DEL-1052
chr10-96946673-DEL-317
chr10-96953834-DEL-55
chr10-96954040-DEL-377
chr10-96956713-DEL-1465
chr10-96960961-DEL-68
chr10-96961139-DEL-1033
chr10-97099901-DEL-54
chr10-97112107-DEL-1578
chr10-97118777-DEL-2240
chr10-97167967-DEL-72
chr10-97179725-DEL-103
chr10-97184643-DEL-907
chr10-97185899-DEL-56
chr10-97196052-DEL-309
chr10-97229907-DEL-85
chr10-97242238-DEL-78
chr10-97243270-DEL-254
chr10-97251039-DEL-123
chr10-97290570-DEL-267
chr10-97299448-DEL-70
chr10-97303456-DEL-63
chr10-97326932-DEL-102
chr10-97332228-DEL-454
chr10-97352090-DEL-54
chr10-97355413-DEL-76
chr10-97381232-DEL-92
chr10-97393442-DEL-111
chr10-97421579-DEL-121
chr10-97435872-DEL-274
chr10-97436901-DEL-174
chr10-97438683-DEL-390
chr10-97439766-DEL-570
chr10-97453691-DEL-5070
chr10-97456145-DEL-2029
chr10-97468418-DEL-92
chr10-97470265-DEL-60
chr10-97519188-DEL-797
chr10-97552573-DEL-198
chr10-97598039-DEL-269
chr10-97608229-DEL-386

chr10-97650621-DEL-1169
chr10-97652690-DEL-1766
chr10-97656771-DEL-871
chr10-97663389-DEL-715
chr10-97664470-DEL-1616
chr10-97675701-DEL-884
chr10-97691157-DEL-6253
chr10-97717077-DEL-63
chr10-97766025-DEL-134
chr10-97880176-DEL-471
chr10-97881166-DEL-6872
chr10-97889949-DEL-78
chr10-97961011-DEL-222
chr10-97989675-DEL-345
chr10-97996638-DEL-6272
chr10-98001579-DEL-68
chr10-98020072-DEL-80
chr10-98059754-DEL-898
chr10-98071539-DEL-443
chr10-98093051-DEL-60
chr10-98133887-DEL-161
chr10-98144260-DEL-58
chr10-98175550-DEL-2750
chr10-98195713-DEL-825
chr10-98202460-DEL-2095
chr10-98214282-DEL-54
chr10-98247415-DEL-84
chr10-98282466-DEL-129
chr10-98286647-DEL-541
chr10-98291684-DEL-238
chr10-98323994-DEL-93
chr10-98326328-DEL-56
chr10-98332968-DEL-1129
chr10-98342113-DEL-1097
chr10-98355240-DEL-857
chr10-98359965-DEL-237
chr10-98365139-DEL-92
chr10-98375051-DEL-84
chr10-98386648-DEL-212
chr10-98412325-DEL-64
chr10-98436206-DEL-7349
chr10-98440970-DEL-1290
chr10-98456307-DEL-1861
chr10-98463049-DEL-82
chr10-98463556-DEL-175
chr10-98489229-DEL-61
chr10-98571146-DEL-99
chr10-98578033-DEL-59
chr10-98588005-DEL-89
chr10-98630334-DEL-414
chr10-98631624-DEL-171
chr10-98641617-DEL-87
chr10-98667061-DEL-1421
chr10-98711697-DEL-73
chr10-98732661-DEL-343
chr10-98736003-DEL-474
chr10-98744828-DEL-212
chr10-98847407-DEL-244
chr10-98856748-DEL-769
chr10-98877584-DEL-129
chr10-98890822-DEL-152
chr10-98924591-DEL-176
chr10-98967936-DEL-73
chr10-99006460-DEL-286
chr10-99022038-DEL-93
chr10-99044099-DEL-157
chr10-99057021-DEL-80
chr10-99142410-DEL-53
chr10-99164271-DEL-86
chr10-99254049-DEL-52
chr10-99269015-DEL-774
chr10-99282201-DEL-282
chr10-99288828-DEL-121
chr10-99302233-DEL-116
chr10-99355004-DEL-111
chr10-99378767-DEL-325
chr10-99381738-DEL-270

chr10-99382054-DEL-1738
chr10-99386931-DEL-354
chr10-99423835-DEL-1322
chr10-99428689-DEL-75
chr10-99482462-DEL-132
chr10-99490776-DEL-63
chr10-99530671-DEL-1055
chr10-99554525-DEL-64
chr10-99554803-DEL-6445
chr10-99562770-DEL-72
chr10-99563221-DEL-229
chr10-99569202-DEL-7106
chr10-99578986-DEL-2931
chr10-99590331-DEL-279
chr10-99590846-DEL-150
chr10-99597678-DEL-179
chr10-99644262-DEL-53
chr10-99678946-DEL-62
chr10-99703267-DEL-608
chr10-99790458-DEL-83
chr10-99810162-DEL-51
chr10-99823866-DEL-460
chr10-99832189-DEL-77
chr10-99835183-DEL-1639
chr10-99872901-DEL-89
chr10-99928486-DEL-287
chr10-99972521-DEL-158
chr10-99984136-DEL-97
chr10-100005976-DEL-71
chr10-100012137-DEL-4848
chr10-100032098-DEL-67
chr10-100088935-DEL-5932
chr10-100108924-DEL-1659
chr10-100121761-DEL-58
chr10-100133677-DEL-81
chr10-100140390-DEL-98
chr10-100141085-DEL-431
chr10-100149791-DEL-533
chr10-100151622-DEL-6054
chr10-100161964-DEL-87
chr10-100213758-DEL-102
chr10-100238347-DEL-59
chr10-100308878-DEL-371
chr10-100342410-DEL-981
chr10-100347240-DEL-55
chr10-100355755-DEL-291
chr10-100355836-DEL-706
chr10-100375332-DEL-62
chr10-100400672-DEL-1005
chr10-100470965-DEL-75
chr10-100497156-DEL-68
chr10-100510263-DEL-7183
chr10-100527424-DEL-56
chr10-100528182-DEL-171
chr10-100568030-DEL-59
chr10-100569382-DEL-173
chr10-100574934-DEL-132
chr10-100580162-DEL-52
chr10-100614892-DEL-77
chr10-100662764-DEL-1591
chr10-100667459-DEL-386
chr10-100671862-DEL-70
chr10-100677395-DEL-92
chr10-100740668-DEL-480
chr10-100744189-DEL-6122
chr10-100806400-DEL-204
chr10-100826318-DEL-83
chr10-100837940-DEL-83
chr10-100857861-DEL-75
chr10-100891853-DEL-57
chr10-100921368-DEL-541
chr10-100935680-DEL-314
chr10-100938312-DEL-220
chr10-100941395-DEL-2135
chr10-100969272-DEL-74553
chr10-100983897-DEL-4368
chr10-101121104-DEL-772

chr10-101153380-DEL-63
chr10-101176758-DEL-168
chr10-101196026-DEL-264
chr10-101223730-DEL-606
chr10-101232720-DEL-808
chr10-101252757-DEL-215
chr10-101254737-DEL-107
chr10-101266304-DEL-74
chr10-101295240-DEL-276
chr10-101311597-DEL-183
chr10-101319466-DEL-54
chr10-101324250-DEL-261
chr10-101331821-DEL-65
chr10-101337824-DEL-75
chr10-101339942-DEL-228
chr10-101356074-DEL-59
chr10-101382166-DEL-89
chr10-101418897-DEL-58
chr10-101438282-DEL-84
chr10-101515967-DEL-52
chr10-101529105-DEL-100
chr10-101531952-DEL-1379
chr10-101557080-DEL-265
chr10-101558085-DEL-157
chr10-101597182-DEL-168
chr10-101626526-DEL-681
chr10-101677538-DEL-209
chr10-101689253-DEL-244
chr10-101701094-DEL-104
chr10-101709536-DEL-1849
chr10-101712601-DEL-53
chr10-101716343-DEL-498
chr10-101731725-DEL-84
chr10-101733878-DEL-73
chr10-101760013-DEL-70
chr10-101763241-DEL-150
chr10-101772225-DEL-636
chr10-101792480-DEL-3996
chr10-101800887-DEL-150
chr10-101808583-DEL-164
chr10-101881215-DEL-105
chr10-101886132-DEL-106
chr10-101887571-DEL-54
chr10-101888114-DEL-115
chr10-101898386-DEL-490
chr10-101899955-DEL-75
chr10-101918041-DEL-127
chr10-101955276-DEL-103
chr10-101973089-DEL-86
chr10-102007171-DEL-145
chr10-102019485-DEL-365
chr10-102061523-DEL-838
chr10-102134966-DEL-615
chr10-102175961-DEL-720
chr10-102215908-DEL-84
chr10-102217285-DEL-91
chr10-102238343-DEL-140
chr10-102279852-DEL-78
chr10-102286116-DEL-52
chr10-102289151-DEL-72
chr10-102306230-DEL-815
chr10-102321803-DEL-142
chr10-102362911-DEL-401
chr10-102379962-DEL-357
chr10-102432915-DEL-370
chr10-102454426-DEL-76
chr10-102457775-DEL-128
chr10-102464878-DEL-66
chr10-102472777-DEL-125
chr10-102496266-DEL-396
chr10-102497367-DEL-1866
chr10-102523689-DEL-72
chr10-102539194-DEL-832
chr10-102557801-DEL-6840
chr10-102567790-DEL-64
chr10-102575416-DEL-186
chr10-102585945-DEL-285

chr10-102599072-DEL-91
chr10-102625840-DEL-755
chr10-102668575-DEL-1033
chr10-102673665-DEL-234
chr10-102676340-DEL-336
chr10-102721417-DEL-414
chr10-102729555-DEL-78
chr10-102745557-DEL-76
chr10-102753497-DEL-80
chr10-102770206-DEL-108
chr10-102780975-DEL-113
chr10-102789369-DEL-776
chr10-102796188-DEL-63
chr10-102831169-DEL-65
chr10-102874859-DEL-595
chr10-102879961-DEL-50
chr10-102910567-DEL-1146
chr10-102941707-DEL-211
chr10-102944584-DEL-153
chr10-102962784-DEL-99
chr10-103028389-DEL-81
chr10-103040631-DEL-273
chr10-103049542-DEL-212
chr10-103098730-DEL-3199
chr10-103124847-DEL-2586
chr10-103140919-DEL-63
chr10-103148193-DEL-73
chr10-103163499-DEL-1134
chr10-103186263-DEL-3737
chr10-103211409-DEL-1642
chr10-103251487-DEL-479
chr10-103283690-DEL-74
chr10-103296464-DEL-119
chr10-103309637-DEL-329
chr10-103325656-DEL-735
chr10-103332332-DEL-147
chr10-103335951-DEL-3204
chr10-103348780-DEL-6214
chr10-103358678-DEL-935
chr10-103362345-DEL-91
chr10-103395381-DEL-96
chr10-103418021-DEL-156
chr10-103445633-DEL-6616
chr10-103453811-DEL-345
chr10-103458292-DEL-66
chr10-103465253-DEL-650
chr10-103466864-DEL-10213
chr10-103484619-DEL-177
chr10-103489456-DEL-271
chr10-103498159-DEL-432
chr10-103500357-DEL-2160
chr10-103506591-DEL-4213
chr10-103507095-DEL-2776
chr10-103514471-DEL-6448
chr10-103546329-DEL-194
chr10-103547094-DEL-862
chr10-103568397-DEL-63
chr10-103618993-DEL-51
chr10-103625614-DEL-788
chr10-103678320-DEL-50
chr10-103689197-DEL-337
chr10-103727983-DEL-52
chr10-103734865-DEL-51
chr10-103772527-DEL-6136
chr10-103809224-DEL-386
chr10-103846759-DEL-75
chr10-103857454-DEL-613
chr10-103858576-DEL-132
chr10-103871797-DEL-68
chr10-103884388-DEL-95
chr10-103906805-DEL-100
chr10-103909176-DEL-186
chr10-103922841-DEL-51
chr10-103925915-DEL-250
chr10-103936803-DEL-88
chr10-103976218-DEL-3680
chr10-104051105-DEL-83

chr10-104052710-DEL-412
chr10-104053257-DEL-397
chr10-104064951-DEL-5828
chr10-104072367-DEL-118
chr10-104075090-DEL-6041
chr10-104082312-DEL-51
chr10-104107033-DEL-63
chr10-104136305-DEL-196
chr10-104159058-DEL-712
chr10-104202028-DEL-69
chr10-104215754-DEL-4380
chr10-104215799-DEL-68
chr10-104247227-DEL-505
chr10-104276549-DEL-89
chr10-104285381-DEL-1611
chr10-104325158-DEL-175
chr10-104353637-DEL-1275
chr10-104357495-DEL-71
chr10-104370185-DEL-100
chr10-104375481-DEL-6502
chr10-104382477-DEL-1790
chr10-104385163-DEL-573
chr10-104391128-DEL-7172
chr10-104438234-DEL-93
chr10-104440225-DEL-158
chr10-104447169-DEL-114
chr10-104450248-DEL-992
chr10-104484634-DEL-641
chr10-104502215-DEL-202
chr10-104543378-DEL-1081
chr10-104599479-DEL-726
chr10-104600970-DEL-6474
chr10-104619863-DEL-68
chr10-104655304-DEL-5398
chr10-104670184-DEL-8982
chr10-104694847-DEL-429
chr10-104695768-DEL-1007
chr10-104716245-DEL-54
chr10-104723554-DEL-76
chr10-104735343-DEL-58
chr10-104743030-DEL-1162
chr10-104760810-DEL-1160
chr10-104790296-DEL-589
chr10-104802515-DEL-1509
chr10-104822069-DEL-66
chr10-104844377-DEL-88
chr10-104858508-DEL-70
chr10-104892238-DEL-752
chr10-104902212-DEL-1708
chr10-104915778-DEL-69
chr10-104979037-DEL-127
chr10-104989731-DEL-181
chr10-105007262-DEL-165
chr10-105051156-DEL-714
chr10-105084769-DEL-55
chr10-105111074-DEL-893
chr10-105142894-DEL-182
chr10-105184056-DEL-106
chr10-105241280-DEL-61
chr10-105254562-DEL-122
chr10-105267629-DEL-54
chr10-105295736-DEL-58
chr10-105340067-DEL-596
chr10-105357381-DEL-79
chr10-105394865-DEL-198
chr10-105419761-DEL-878
chr10-105425111-DEL-170
chr10-105449049-DEL-73
chr10-105493143-DEL-54
chr10-105496291-DEL-1000
chr10-105506198-DEL-64
chr10-105535937-DEL-1022
chr10-105545766-DEL-88
chr10-105589461-DEL-933
chr10-105591461-DEL-5177
chr10-105608148-DEL-52
chr10-105620002-DEL-126

chr10-105627147-DEL-57
chr10-105637715-DEL-81
chr10-105698302-DEL-58
chr10-105726432-DEL-172
chr10-105737536-DEL-741
chr10-105739663-DEL-56
chr10-105740561-DEL-1987
chr10-105751387-DEL-98
chr10-105782110-DEL-3315
chr10-105784719-DEL-2930
chr10-105798476-DEL-92
chr10-105803227-DEL-990
chr10-105806652-DEL-1616
chr10-105811854-DEL-141
chr10-105818172-DEL-595
chr10-105850529-DEL-2256
chr10-105878003-DEL-55
chr10-105895866-DEL-56
chr10-105908974-DEL-54
chr10-105915423-DEL-244
chr10-105916136-DEL-208
chr10-105925135-DEL-4981
chr10-105954016-DEL-1892
chr10-105963111-DEL-465
chr10-105968622-DEL-199
chr10-106018139-DEL-62
chr10-106042780-DEL-636
chr10-106053672-DEL-5348
chr10-106098095-DEL-59
chr10-106103791-DEL-59
chr10-106117634-DEL-194
chr10-106126061-DEL-54
chr10-106138976-DEL-2907
chr10-106181883-DEL-1043
chr10-106185432-DEL-68
chr10-106190993-DEL-103
chr10-106194648-DEL-458
chr10-106226266-DEL-13367
chr10-106232453-DEL-5117
chr10-106257753-DEL-1199
chr10-106263154-DEL-70
chr10-106280759-DEL-59
chr10-106285321-DEL-6177
chr10-106322732-DEL-725
chr10-106329518-DEL-134
chr10-106338479-DEL-822
chr10-106343548-DEL-89
chr10-106365444-DEL-6710
chr10-106381300-DEL-68
chr10-106392915-DEL-52
chr10-106412533-DEL-55
chr10-106418715-DEL-106
chr10-106430144-DEL-180
chr10-106442216-DEL-343
chr10-106450642-DEL-913
chr10-106479196-DEL-713
chr10-106480312-DEL-249
chr10-106481857-DEL-490
chr10-106502838-DEL-865
chr10-106547001-DEL-368
chr10-106561883-DEL-171
chr10-106577177-DEL-56
chr10-106593557-DEL-802
chr10-106596376-DEL-332
chr10-106628417-DEL-174
chr10-106637042-DEL-449
chr10-106755530-DEL-3204
chr10-106783437-DEL-509
chr10-106791005-DEL-208
chr10-106808192-DEL-369
chr10-106817920-DEL-1788
chr10-106826814-DEL-616
chr10-106856889-DEL-59
chr10-106866114-DEL-2037
chr10-106885915-DEL-52
chr10-106895929-DEL-57
chr10-106898040-DEL-62

chr10-106923503-DEL-249
chr10-106945309-DEL-461
chr10-106947764-DEL-54
chr10-106977422-DEL-400
chr10-106990576-DEL-424
chr10-106993199-DEL-52
chr10-107024412-DEL-6982
chr10-107048825-DEL-205
chr10-107050686-DEL-794
chr10-107055506-DEL-77
chr10-107062102-DEL-1679
chr10-107089501-DEL-1008
chr10-107154179-DEL-311
chr10-107168664-DEL-208
chr10-107178539-DEL-92
chr10-107211105-DEL-1110
chr10-107252839-DEL-195
chr10-107273817-DEL-226
chr10-107297236-DEL-839
chr10-107298285-DEL-2654
chr10-107301144-DEL-715
chr10-107308573-DEL-243
chr10-107314477-DEL-63
chr10-107387558-DEL-1748
chr10-107410550-DEL-80
chr10-107434698-DEL-143
chr10-107455359-DEL-527
chr10-107467809-DEL-136
chr10-107468900-DEL-552
chr10-107497029-DEL-384
chr10-107514352-DEL-198
chr10-107549612-DEL-303
chr10-107568053-DEL-84
chr10-107572978-DEL-80
chr10-107574567-DEL-117
chr10-107585227-DEL-191
chr10-107587118-DEL-60
chr10-107596065-DEL-85
chr10-107626889-DEL-177
chr10-107698740-DEL-1094
chr10-107715909-DEL-84
chr10-107729780-DEL-801
chr10-107731934-DEL-170
chr10-107763088-DEL-121
chr10-107790184-DEL-132
chr10-107816590-DEL-442
chr10-107823110-DEL-60
chr10-107864529-DEL-359
chr10-107878938-DEL-61
chr10-107900852-DEL-102
chr10-107901481-DEL-59
chr10-107923281-DEL-60
chr10-107923318-DEL-427
chr10-107950185-DEL-53
chr10-107965396-DEL-118
chr10-107980930-DEL-438
chr10-107990512-DEL-168
chr10-107996119-DEL-312
chr10-108010899-DEL-93
chr10-108016363-DEL-56
chr10-108036430-DEL-144
chr10-108066795-DEL-397
chr10-108091765-DEL-262
chr10-108099548-DEL-209
chr10-108120588-DEL-62
chr10-108124123-DEL-100
chr10-108223483-DEL-267
chr10-108236527-DEL-86
chr10-108265886-DEL-72
chr10-108266773-DEL-213
chr10-108274185-DEL-1890
chr10-108279454-DEL-69
chr10-108291756-DEL-51
chr10-108376528-DEL-69
chr10-108423360-DEL-70
chr10-108465708-DEL-90
chr10-108469191-DEL-637

chr10-108480290-DEL-1543
chr10-108533616-DEL-67
chr10-108549949-DEL-172
chr10-108550878-DEL-475
chr10-108554778-DEL-366
chr10-108561669-DEL-222
chr10-108601176-DEL-121
chr10-108645515-DEL-62
chr10-108672913-DEL-57
chr10-108685908-DEL-69
chr10-108701969-DEL-54
chr10-108712221-DEL-680
chr10-108726327-DEL-66
chr10-108727856-DEL-625
chr10-108745252-DEL-350
chr10-108768222-DEL-96
chr10-108803571-DEL-54
chr10-108831956-DEL-1893
chr10-108841339-DEL-72
chr10-108881457-DEL-66
chr10-108884026-DEL-297
chr10-108889814-DEL-226
chr10-108917832-DEL-155
chr10-108935408-DEL-105
chr10-108967060-DEL-52
chr10-108994456-DEL-60
chr10-109005530-DEL-93
chr10-109007824-DEL-438
chr10-109013499-DEL-67
chr10-109141601-DEL-61
chr10-109177794-DEL-1080
chr10-109183537-DEL-175
chr10-109203141-DEL-2342
chr10-109236847-DEL-72
chr10-109243070-DEL-2562
chr10-109252723-DEL-58
chr10-109340170-DEL-55
chr10-109340799-DEL-228
chr10-109346630-DEL-368
chr10-109401827-DEL-55
chr10-109420908-DEL-51
chr10-109432342-DEL-644
chr10-109433705-DEL-133
chr10-109450517-DEL-126
chr10-109453531-DEL-3044
chr10-109458749-DEL-1393
chr10-109498004-DEL-124
chr10-109529643-DEL-192
chr10-109545718-DEL-1431
chr10-109608662-DEL-162
chr10-109623607-DEL-98
chr10-109624568-DEL-4308
chr10-109632638-DEL-116
chr10-109652489-DEL-118
chr10-109659764-DEL-108
chr10-109686473-DEL-65
chr10-109766265-DEL-86
chr10-109768872-DEL-68
chr10-109777175-DEL-64
chr10-109777447-DEL-6454
chr10-109784133-DEL-89
chr10-109800726-DEL-150
chr10-109807695-DEL-55
chr10-109824403-DEL-148
chr10-109825552-DEL-366
chr10-109853981-DEL-164
chr10-109862675-DEL-65
chr10-109879351-DEL-126
chr10-109896069-DEL-58
chr10-109919949-DEL-90
chr10-109935525-DEL-211
chr10-109959271-DEL-124
chr10-109967854-DEL-619
chr10-109985758-DEL-1293
chr10-110008594-DEL-312
chr10-110024829-DEL-52
chr10-110071858-DEL-115

chr10-110104266-DEL-55
chr10-110104895-DEL-81
chr10-110109719-DEL-210
chr10-110122231-DEL-1986
chr10-110160789-DEL-508
chr10-110163158-DEL-98
chr10-110186656-DEL-674
chr10-110224205-DEL-125
chr10-110241887-DEL-51
chr10-110267167-DEL-173
chr10-110339846-DEL-333
chr10-110339846-DEL-953
chr10-110339867-DEL-1193
chr10-110340269-DEL-6473
chr10-110347615-DEL-2507
chr10-110350204-DEL-965
chr10-110351218-DEL-1284
chr10-110351978-DEL-510
chr10-110353707-DEL-59
chr10-110392715-DEL-879
chr10-110394037-DEL-60
chr10-110454886-DEL-77
chr10-110463298-DEL-6629
chr10-110483315-DEL-1602
chr10-110493619-DEL-80
chr10-110503244-DEL-1493
chr10-110527017-DEL-1986
chr10-110531139-DEL-545
chr10-110550736-DEL-518
chr10-110565830-DEL-78
chr10-110573505-DEL-749
chr10-110611016-DEL-138
chr10-110619688-DEL-100
chr10-110654127-DEL-128
chr10-110661782-DEL-30246
chr10-110665868-DEL-199
chr10-110680085-DEL-669
chr10-110697539-DEL-242
chr10-110701088-DEL-52
chr10-110725721-DEL-53
chr10-110730652-DEL-9344
chr10-110730674-DEL-2758
chr10-110733458-DEL-6538
chr10-110738994-DEL-1304
chr10-110743048-DEL-181
chr10-110750693-DEL-454
chr10-110751675-DEL-192
chr10-110754005-DEL-199
chr10-110773807-DEL-83
chr10-110787547-DEL-75
chr10-110790038-DEL-684
chr10-110818984-DEL-111
chr10-110821470-DEL-57
chr10-110828988-DEL-55
chr10-110858721-DEL-339
chr10-110871567-DEL-162
chr10-110922060-DEL-125
chr10-110934237-DEL-96
chr10-110940508-DEL-100
chr10-110957342-DEL-422
chr10-110974634-DEL-120
chr10-111005811-DEL-1213
chr10-111026602-DEL-480
chr10-111028382-DEL-186
chr10-111036043-DEL-60
chr10-111048546-DEL-861
chr10-111053722-DEL-920
chr10-111061012-DEL-52
chr10-111064901-DEL-2299
chr10-111114836-DEL-195
chr10-111160943-DEL-2386
chr10-111161903-DEL-212
chr10-111170936-DEL-512
chr10-111177430-DEL-77
chr10-111178963-DEL-249
chr10-111185936-DEL-121
chr10-111189981-DEL-483

chr10-111215754-DEL-518
chr10-111253246-DEL-72
chr10-111278257-DEL-56
chr10-111284988-DEL-73
chr10-111287382-DEL-256
chr10-111287493-DEL-833
chr10-111304542-DEL-138
chr10-111368670-DEL-57
chr10-111371894-DEL-156
chr10-111375089-DEL-169
chr10-111377900-DEL-91
chr10-111383278-DEL-108
chr10-111393256-DEL-377
chr10-111399754-DEL-2496
chr10-111407401-DEL-114
chr10-111421404-DEL-822
chr10-111437442-DEL-113
chr10-111448714-DEL-225
chr10-111449371-DEL-151
chr10-111456649-DEL-1552
chr10-111458400-DEL-408
chr10-111466039-DEL-63
chr10-111499827-DEL-3833
chr10-111558619-DEL-605
chr10-111562425-DEL-95
chr10-111598836-DEL-58
chr10-111612634-DEL-281
chr10-111613467-DEL-96
chr10-111662823-DEL-127
chr10-111676740-DEL-171
chr10-111688282-DEL-250
chr10-111722343-DEL-71
chr10-111763227-DEL-98
chr10-111798150-DEL-253
chr10-111849872-DEL-1485
chr10-111851621-DEL-6155
chr10-111887739-DEL-177
chr10-111921728-DEL-69
chr10-111941599-DEL-331
chr10-111948065-DEL-907
chr10-111949640-DEL-1276
chr10-111951683-DEL-166
chr10-111953340-DEL-67
chr10-111953719-DEL-2455
chr10-112003156-DEL-7610
chr10-112009315-DEL-428
chr10-112009957-DEL-214
chr10-112095070-DEL-70
chr10-112119882-DEL-138
chr10-112120526-DEL-763
chr10-112127749-DEL-54
chr10-112131561-DEL-473
chr10-112137273-DEL-6196
chr10-112159092-DEL-6436
chr10-112165562-DEL-1420
chr10-112192756-DEL-378
chr10-112201067-DEL-401
chr10-112205077-DEL-55
chr10-112246511-DEL-102
chr10-112254693-DEL-1152
chr10-112319369-DEL-52
chr10-112369367-DEL-63
chr10-112434456-DEL-6309
chr10-112456015-DEL-839
chr10-112478903-DEL-2089
chr10-112486196-DEL-74
chr10-112490031-DEL-54
chr10-112500129-DEL-6393
chr10-112510174-DEL-694
chr10-112518248-DEL-3183
chr10-112524569-DEL-95
chr10-112527636-DEL-51
chr10-112534998-DEL-427
chr10-112547761-DEL-199
chr10-112558864-DEL-846
chr10-112563768-DEL-1083
chr10-112583736-DEL-329

chr10-112612462-DEL-400
chr10-112620329-DEL-866
chr10-112643494-DEL-58
chr10-112648410-DEL-337
chr10-112663260-DEL-4114
chr10-112693212-DEL-871
chr10-112707989-DEL-212
chr10-112711844-DEL-55
chr10-112750954-DEL-95
chr10-112770186-DEL-66
chr10-112794113-DEL-465
chr10-112810746-DEL-2499
chr10-112838389-DEL-1867
chr10-112934470-DEL-521
chr10-112957530-DEL-54
chr10-112959350-DEL-440
chr10-112966724-DEL-68
chr10-112972480-DEL-111
chr10-112974462-DEL-95
chr10-112980325-DEL-832
chr10-113009742-DEL-579
chr10-113016259-DEL-369
chr10-113022121-DEL-124
chr10-113074912-DEL-195
chr10-113075087-DEL-516
chr10-113084404-DEL-201
chr10-113090688-DEL-618
chr10-113101642-DEL-58
chr10-113114147-DEL-434
chr10-113168013-DEL-167
chr10-113191317-DEL-57
chr10-113210258-DEL-57
chr10-113238865-DEL-66
chr10-113239434-DEL-64
chr10-113343579-DEL-155
chr10-113354965-DEL-186
chr10-113357589-DEL-142
chr10-113360433-DEL-100
chr10-113364984-DEL-1513
chr10-113395258-DEL-367
chr10-113395932-DEL-570
chr10-113405045-DEL-70
chr10-113443310-DEL-983
chr10-113476516-DEL-1053
chr10-113476644-DEL-900
chr10-113476649-DEL-401
chr10-113490606-DEL-72
chr10-113518608-DEL-93
chr10-113528568-DEL-54
chr10-113535934-DEL-83
chr10-113558621-DEL-7079
chr10-113571243-DEL-2245
chr10-113637860-DEL-68
chr10-113640423-DEL-86
chr10-113651793-DEL-70
chr10-113663645-DEL-357
chr10-113672795-DEL-1085
chr10-113711287-DEL-56
chr10-113721418-DEL-63
chr10-113723814-DEL-57
chr10-113740355-DEL-403
chr10-113742352-DEL-297
chr10-113750185-DEL-105
chr10-113752016-DEL-140
chr10-113752218-DEL-126
chr10-113763174-DEL-66
chr10-113799998-DEL-112
chr10-113824773-DEL-1167
chr10-113830925-DEL-250
chr10-113846802-DEL-1390
chr10-113868709-DEL-492
chr10-113874885-DEL-154
chr10-113923853-DEL-60
chr10-113949793-DEL-112
chr10-113956912-DEL-146
chr10-113969415-DEL-70
chr10-113974367-DEL-163

chr10-114002794-DEL-8164
chr10-114008026-DEL-1491
chr10-114012851-DEL-122
chr10-114045725-DEL-1463
chr10-114061069-DEL-348
chr10-114072684-DEL-53
chr10-114103486-DEL-55
chr10-114118102-DEL-56
chr10-114214999-DEL-52
chr10-114224547-DEL-135
chr10-114260341-DEL-121
chr10-114263323-DEL-110
chr10-114265401-DEL-562
chr10-114270030-DEL-55
chr10-114296734-DEL-110
chr10-114330800-DEL-127
chr10-114362048-DEL-6386
chr10-114444125-DEL-834
chr10-114460329-DEL-52
chr10-114484270-DEL-99
chr10-114501060-DEL-75
chr10-114505633-DEL-104
chr10-114512047-DEL-65
chr10-114515042-DEL-54
chr10-114539966-DEL-80
chr10-114544268-DEL-1964
chr10-114558311-DEL-97
chr10-114589052-DEL-277
chr10-114605590-DEL-65
chr10-114651441-DEL-575
chr10-114652284-DEL-1055
chr10-114664378-DEL-87
chr10-114720222-DEL-162
chr10-114722590-DEL-61
chr10-114731593-DEL-55
chr10-114774961-DEL-139
chr10-114784219-DEL-74
chr10-114790276-DEL-78
chr10-114802876-DEL-425
chr10-114840320-DEL-121
chr10-114856433-DEL-66
chr10-114866983-DEL-1277
chr10-114884477-DEL-90
chr10-114913065-DEL-101
chr10-114928612-DEL-55
chr10-114976769-DEL-101
chr10-114982590-DEL-97
chr10-114993615-DEL-57
chr10-115014125-DEL-64
chr10-115022081-DEL-52
chr10-115024783-DEL-57
chr10-115027646-DEL-547
chr10-115027903-DEL-1796
chr10-115029722-DEL-1107
chr10-115036751-DEL-288
chr10-115076123-DEL-151
chr10-115096282-DEL-56
chr10-115119204-DEL-54
chr10-115138189-DEL-190
chr10-115145869-DEL-148
chr10-115165104-DEL-66
chr10-115192076-DEL-1183
chr10-115210118-DEL-55
chr10-115241751-DEL-266
chr10-115275407-DEL-810
chr10-115276511-DEL-196
chr10-115316978-DEL-52
chr10-115318977-DEL-94
chr10-115330432-DEL-84
chr10-115358009-DEL-54
chr10-115380611-DEL-1375
chr10-115402071-DEL-77
chr10-115407978-DEL-56
chr10-115408862-DEL-550
chr10-115428944-DEL-155
chr10-115434850-DEL-1409
chr10-115438256-DEL-58

chr10-115464229-DEL-1094
chr10-115478933-DEL-147
chr10-115480090-DEL-166
chr10-115487759-DEL-400
chr10-115538220-DEL-367
chr10-115541347-DEL-1910
chr10-115541812-DEL-145
chr10-115559153-DEL-1539
chr10-115566023-DEL-70
chr10-115572021-DEL-194
chr10-115582337-DEL-66
chr10-115593718-DEL-93
chr10-115621244-DEL-1294
chr10-115624319-DEL-2398
chr10-115696770-DEL-54
chr10-115711646-DEL-180
chr10-115715886-DEL-195
chr10-115730515-DEL-151
chr10-115734334-DEL-86
chr10-115756517-DEL-195
chr10-115770117-DEL-132
chr10-115824691-DEL-676
chr10-115827937-DEL-1260
chr10-115829744-DEL-124
chr10-115834920-DEL-236
chr10-115852944-DEL-63
chr10-115893636-DEL-134
chr10-115908517-DEL-53
chr10-115945670-DEL-107
chr10-115955019-DEL-141
chr10-115973653-DEL-343
chr10-115988628-DEL-526
chr10-116031714-DEL-60
chr10-116049626-DEL-51
chr10-116058099-DEL-76
chr10-116063939-DEL-108
chr10-116071543-DEL-60
chr10-116075840-DEL-162
chr10-116102012-DEL-54
chr10-116112793-DEL-51
chr10-116118434-DEL-79
chr10-116119511-DEL-105
chr10-116125322-DEL-83
chr10-116147923-DEL-220
chr10-116151443-DEL-53
chr10-116165904-DEL-73
chr10-116170619-DEL-54
chr10-116180934-DEL-203
chr10-116185561-DEL-61
chr10-116192051-DEL-57
chr10-116197528-DEL-55
chr10-116205741-DEL-188
chr10-116243471-DEL-150
chr10-116268055-DEL-93
chr10-116274520-DEL-71
chr10-116293069-DEL-71
chr10-116337479-DEL-179
chr10-116357938-DEL-151
chr10-116370564-DEL-56
chr10-116388167-DEL-68
chr10-116395011-DEL-89
chr10-116399998-DEL-72
chr10-116402368-DEL-198
chr10-116426391-DEL-481
chr10-116432232-DEL-54
chr10-116432454-DEL-500
chr10-116433037-DEL-52
chr10-116454851-DEL-93
chr10-116476188-DEL-153
chr10-116477597-DEL-468
chr10-116485412-DEL-65
chr10-116500392-DEL-131
chr10-116503264-DEL-144
chr10-116518337-DEL-86
chr10-116538838-DEL-108
chr10-116550413-DEL-154
chr10-116560351-DEL-431

chr10-116561595-DEL-188
chr10-116566662-DEL-170
chr10-116570425-DEL-121
chr10-116572016-DEL-51
chr10-116623600-DEL-259
chr10-116625406-DEL-124
chr10-116627974-DEL-356
chr10-116633467-DEL-70
chr10-116641580-DEL-56
chr10-116661810-DEL-102
chr10-116662996-DEL-188
chr10-116669727-DEL-53
chr10-116673769-DEL-142
chr10-116689892-DEL-177
chr10-116714114-DEL-152
chr10-116715121-DEL-542
chr10-116744883-DEL-76
chr10-116759899-DEL-176
chr10-116788358-DEL-182
chr10-116803381-DEL-449
chr10-116846158-DEL-118
chr10-116853220-DEL-480
chr10-116871800-DEL-62
chr10-116872417-DEL-152
chr10-116876837-DEL-192
chr10-116894366-DEL-156
chr10-116896057-DEL-446
chr10-116917700-DEL-199
chr10-116929282-DEL-193
chr10-116946551-DEL-52
chr10-116960158-DEL-181
chr10-116964985-DEL-12682
chr10-116967320-DEL-520
chr10-116977769-DEL-58
chr10-117008591-DEL-74
chr10-117026372-DEL-54
chr10-117028662-DEL-56
chr10-117112426-DEL-147
chr10-117145829-DEL-509
chr10-117195875-DEL-70
chr10-117201384-DEL-201
chr10-117219665-DEL-58
chr10-117257969-DEL-200
chr10-117331628-DEL-106
chr10-117370292-DEL-348
chr10-117409441-DEL-1631
chr10-117412668-DEL-1770
chr10-117432117-DEL-402
chr10-117444252-DEL-86
chr10-117462921-DEL-56
chr10-117513367-DEL-60
chr10-117527839-DEL-195
chr10-117529222-DEL-351
chr10-117539625-DEL-54
chr10-117554780-DEL-72
chr10-117556434-DEL-193
chr10-117583042-DEL-2701
chr10-117605193-DEL-207
chr10-117609673-DEL-60
chr10-117625206-DEL-495
chr10-117632735-DEL-142
chr10-117633389-DEL-213
chr10-117664158-DEL-3315
chr10-117699385-DEL-218
chr10-117704091-DEL-205
chr10-117704498-DEL-1435
chr10-117706322-DEL-72
chr10-117727402-DEL-60
chr10-117732128-DEL-566
chr10-117740263-DEL-66
chr10-117755303-DEL-403
chr10-117764298-DEL-198
chr10-117793987-DEL-1424
chr10-117799814-DEL-57
chr10-117809916-DEL-141
chr10-117850205-DEL-412
chr10-117850408-DEL-747

chr10-117850408-DEL-879
chr10-117851352-DEL-321
chr10-117851557-DEL-932
chr10-117853576-DEL-6473
chr10-117894177-DEL-678
chr10-117905114-DEL-492
chr10-117915953-DEL-102
chr10-117924383-DEL-871
chr10-117931471-DEL-65
chr10-117952554-DEL-89
chr10-118039607-DEL-51
chr10-118074184-DEL-8591
chr10-118093856-DEL-16592
chr10-118110514-DEL-870
chr10-118124459-DEL-63
chr10-118126787-DEL-74
chr10-118209884-DEL-195
chr10-118214003-DEL-380
chr10-118216339-DEL-99
chr10-118218392-DEL-665
chr10-118241460-DEL-696
chr10-118242112-DEL-111
chr10-118254546-DEL-98
chr10-118258303-DEL-4091
chr10-118262416-DEL-1721
chr10-118269550-DEL-124
chr10-118291192-DEL-583
chr10-118324901-DEL-149
chr10-118326173-DEL-178
chr10-118340437-DEL-234
chr10-118356555-DEL-139
chr10-118362934-DEL-77
chr10-118381350-DEL-68
chr10-118385352-DEL-230
chr10-118387991-DEL-64
chr10-118438105-DEL-576
chr10-118461414-DEL-65
chr10-118463113-DEL-395
chr10-118478577-DEL-85
chr10-118485647-DEL-64
chr10-118486923-DEL-178
chr10-118525292-DEL-78
chr10-118526156-DEL-6462
chr10-118526170-DEL-5375
chr10-118526198-DEL-3262
chr10-118531605-DEL-1015
chr10-118531605-DEL-609
chr10-118604557-DEL-76
chr10-118637269-DEL-618
chr10-118650946-DEL-62
chr10-118655495-DEL-233
chr10-118663633-DEL-75
chr10-118744585-DEL-57
chr10-118766157-DEL-62
chr10-118787703-DEL-118
chr10-118788078-DEL-320
chr10-118810458-DEL-56
chr10-118845786-DEL-73
chr10-118846580-DEL-60
chr10-118887956-DEL-125
chr10-118903596-DEL-62
chr10-118914116-DEL-204
chr10-118930673-DEL-70
chr10-118959129-DEL-201
chr10-118965152-DEL-4466
chr10-118980740-DEL-514
chr10-119031851-DEL-673
chr10-119093383-DEL-60
chr10-119094494-DEL-1405
chr10-119125775-DEL-77
chr10-119142498-DEL-100
chr10-119168374-DEL-55
chr10-119170809-DEL-137
chr10-119187068-DEL-61
chr10-119193130-DEL-61
chr10-119233111-DEL-1925
chr10-119244698-DEL-158

chr10-119262856-DEL-288
chr10-119280350-DEL-66
chr10-119320147-DEL-97
chr10-119372836-DEL-68
chr10-119374504-DEL-73
chr10-119386617-DEL-617
chr10-119490956-DEL-1176
chr10-119501160-DEL-71
chr10-119505782-DEL-52
chr10-119508934-DEL-54
chr10-119577576-DEL-52
chr10-119593972-DEL-288
chr10-119607420-DEL-75
chr10-119620534-DEL-304
chr10-119662128-DEL-251
chr10-119672943-DEL-59
chr10-119675884-DEL-58
chr10-119695802-DEL-492
chr10-119713674-DEL-93
chr10-119729400-DEL-672
chr10-119729452-DEL-1494
chr10-119729867-DEL-867
chr10-119730502-DEL-139
chr10-119776268-DEL-206
chr10-119784566-DEL-60
chr10-119789285-DEL-87
chr10-119807083-DEL-232
chr10-119810953-DEL-192
chr10-119874076-DEL-64
chr10-119884686-DEL-151
chr10-119907274-DEL-62
chr10-119929008-DEL-97
chr10-119942589-DEL-74
chr10-119947716-DEL-181
chr10-119949431-DEL-118
chr10-119955785-DEL-71
chr10-119959475-DEL-127
chr10-119965352-DEL-63
chr10-119984794-DEL-3700
chr10-119991517-DEL-612
chr10-119998193-DEL-685
chr10-119999614-DEL-122
chr10-120002997-DEL-54
chr10-120018943-DEL-239
chr10-120036605-DEL-78
chr10-120041350-DEL-68
chr10-120055246-DEL-330
chr10-120066696-DEL-51
chr10-120097934-DEL-57
chr10-120104122-DEL-342
chr10-120114627-DEL-270
chr10-120115922-DEL-53
chr10-120129376-DEL-189
chr10-120131327-DEL-82
chr10-120146163-DEL-82
chr10-120173731-DEL-91
chr10-120176798-DEL-79
chr10-120188949-DEL-2368
chr10-120202238-DEL-119
chr10-120225441-DEL-80
chr10-120246104-DEL-177
chr10-120255573-DEL-115
chr10-120257581-DEL-80
chr10-120286263-DEL-205
chr10-120294643-DEL-284
chr10-120321100-DEL-86
chr10-120336229-DEL-200
chr10-120347562-DEL-52
chr10-120371360-DEL-142
chr10-120385798-DEL-59
chr10-120386656-DEL-55
chr10-120390334-DEL-200
chr10-120424761-DEL-233
chr10-120437798-DEL-191
chr10-120438162-DEL-741
chr10-120444084-DEL-1705
chr10-120451570-DEL-475

chr10-120454497-DEL-53
chr10-120458572-DEL-66
chr10-120466254-DEL-139
chr10-120470691-DEL-60
chr10-120520446-DEL-71
chr10-120568304-DEL-64
chr10-120574706-DEL-62
chr10-120606500-DEL-73
chr10-120654244-DEL-63
chr10-120672189-DEL-1586
chr10-120687729-DEL-72
chr10-120730378-DEL-63
chr10-120753321-DEL-63
chr10-120780376-DEL-247
chr10-120790082-DEL-369
chr10-120793506-DEL-125
chr10-120800189-DEL-117
chr10-120803784-DEL-83
chr10-120804891-DEL-182
chr10-120811022-DEL-1172
chr10-120812484-DEL-145
chr10-120830691-DEL-502
chr10-120832383-DEL-427
chr10-120856075-DEL-90
chr10-120860189-DEL-84
chr10-120862720-DEL-225
chr10-120887788-DEL-65
chr10-120887959-DEL-1077
chr10-120901398-DEL-80
chr10-120906010-DEL-237
chr10-120918463-DEL-1290
chr10-120971732-DEL-54
chr10-120982038-DEL-182
chr10-120982601-DEL-383
chr10-120983527-DEL-132
chr10-120984102-DEL-65
chr10-121020758-DEL-65
chr10-121077225-DEL-89
chr10-121081252-DEL-130
chr10-121113514-DEL-73
chr10-121128357-DEL-150
chr10-121129231-DEL-91
chr10-121129539-DEL-369
chr10-121143022-DEL-72
chr10-121178329-DEL-120
chr10-121183537-DEL-148
chr10-121194263-DEL-52
chr10-121218238-DEL-66
chr10-121226944-DEL-108
chr10-121234590-DEL-2575
chr10-121235387-DEL-1430
chr10-121239867-DEL-53
chr10-121274959-DEL-2464
chr10-121301451-DEL-290
chr10-121320321-DEL-565
chr10-121321240-DEL-155
chr10-121323209-DEL-70
chr10-121327731-DEL-256
chr10-121331521-DEL-103
chr10-121334888-DEL-71
chr10-121335391-DEL-88
chr10-121351408-DEL-129
chr10-121379655-DEL-58
chr10-121396656-DEL-60
chr10-121402009-DEL-1888
chr10-121408487-DEL-860
chr10-121409725-DEL-328
chr10-121412965-DEL-390
chr10-121415421-DEL-498
chr10-121436045-DEL-177
chr10-121510420-DEL-522
chr10-121518996-DEL-726
chr10-121525067-DEL-83
chr10-121553224-DEL-124
chr10-121560973-DEL-203
chr10-121593632-DEL-92
chr10-121632015-DEL-96

chr10-121696724-DEL-176
chr10-121697441-DEL-746
chr10-121712773-DEL-67
chr10-121724917-DEL-536
chr10-121725870-DEL-54
chr10-121789490-DEL-280
chr10-121818784-DEL-292
chr10-121825283-DEL-127
chr10-121858187-DEL-204
chr10-121891569-DEL-398
chr10-121909035-DEL-1455
chr10-121916393-DEL-196
chr10-121923736-DEL-192
chr10-121940317-DEL-873
chr10-121955245-DEL-56
chr10-121966365-DEL-72
chr10-121968911-DEL-53
chr10-121986644-DEL-51
chr10-121994909-DEL-201
chr10-121998218-DEL-278
chr10-122001401-DEL-326
chr10-122002892-DEL-56
chr10-122016301-DEL-60
chr10-122049035-DEL-2851
chr10-122083758-DEL-329
chr10-122093932-DEL-199
chr10-122140484-DEL-3981
chr10-122146438-DEL-201
chr10-122176548-DEL-103
chr10-122181864-DEL-382
chr10-122215123-DEL-137
chr10-122249827-DEL-72
chr10-122252263-DEL-98
chr10-122261826-DEL-125
chr10-122272002-DEL-69
chr10-122283542-DEL-105
chr10-122285744-DEL-62
chr10-122295255-DEL-325
chr10-122300902-DEL-227
chr10-122301427-DEL-1114
chr10-122301587-DEL-954
chr10-122324789-DEL-501
chr10-122329533-DEL-73
chr10-122340530-DEL-321
chr10-122361988-DEL-63
chr10-122363574-DEL-176
chr10-122392485-DEL-648
chr10-122416468-DEL-202
chr10-122431590-DEL-2483
chr10-122467467-DEL-6603
chr10-122486876-DEL-55
chr10-122533451-DEL-83
chr10-122545870-DEL-101
chr10-122550202-DEL-67
chr10-122553140-DEL-124
chr10-122567355-DEL-78
chr10-122576016-DEL-57
chr10-122591433-DEL-331
chr10-122595366-DEL-85
chr10-122616627-DEL-254
chr10-122621913-DEL-331
chr10-122623977-DEL-119
chr10-122635127-DEL-618
chr10-122689328-DEL-56
chr10-122696360-DEL-208
chr10-122746538-DEL-67
chr10-122750759-DEL-172
chr10-122772376-DEL-1971
chr10-122772385-DEL-1961
chr10-122782678-DEL-212
chr10-122802455-DEL-1154
chr10-122857598-DEL-73
chr10-122882991-DEL-1182
chr10-122883221-DEL-648
chr10-122883295-DEL-540
chr10-122918378-DEL-92
chr10-122923538-DEL-56

chr10-122994108-DEL-65
chr10-123043467-DEL-4035
chr10-123050498-DEL-956
chr10-123074827-DEL-56
chr10-123076398-DEL-52
chr10-123143740-DEL-51
chr10-123183442-DEL-141
chr10-123214790-DEL-186
chr10-123266275-DEL-98
chr10-123274670-DEL-63
chr10-123288925-DEL-144
chr10-123319931-DEL-382
chr10-123320422-DEL-361
chr10-123342757-DEL-3153
chr10-123385024-DEL-50
chr10-123392595-DEL-66
chr10-123468960-DEL-153
chr10-123494469-DEL-66
chr10-123497384-DEL-397
chr10-123532876-DEL-342
chr10-123561235-DEL-64
chr10-123606694-DEL-114
chr10-123614175-DEL-201
chr10-123618897-DEL-172
chr10-123654525-DEL-112
chr10-123695259-DEL-7409
chr10-123737148-DEL-55
chr10-123738419-DEL-59
chr10-123747053-DEL-289
chr10-123814931-DEL-1444
chr10-123823662-DEL-73
chr10-123845711-DEL-55
chr10-123898001-DEL-303
chr10-123901752-DEL-72
chr10-123906535-DEL-66
chr10-123907918-DEL-68
chr10-123913701-DEL-55
chr10-123923189-DEL-87
chr10-123941248-DEL-144
chr10-123943465-DEL-64
chr10-123944893-DEL-143
chr10-123949245-DEL-1392
chr10-123961823-DEL-65
chr10-123962666-DEL-1023
chr10-123963657-DEL-5092
chr10-123978976-DEL-164
chr10-124007024-DEL-107
chr10-124017823-DEL-90
chr10-124067142-DEL-2565
chr10-124070438-DEL-175
chr10-124070864-DEL-1046
chr10-124128666-DEL-83
chr10-124149965-DEL-87
chr10-124153915-DEL-438
chr10-124166137-DEL-90
chr10-124170402-DEL-76
chr10-124204332-DEL-68
chr10-124214232-DEL-52
chr10-124271572-DEL-75
chr10-124273838-DEL-213
chr10-124300724-DEL-340
chr10-124304228-DEL-54
chr10-124306018-DEL-404
chr10-124347637-DEL-361
chr10-124374343-DEL-51
chr10-124384679-DEL-3868
chr10-124452682-DEL-327
chr10-124464645-DEL-98
chr10-124505915-DEL-530
chr10-124581971-DEL-69
chr10-124585096-DEL-82
chr10-124586201-DEL-118
chr10-124626288-DEL-75
chr10-124636793-DEL-155
chr10-124644585-DEL-110
chr10-124672910-DEL-1805
chr10-124674455-DEL-273

chr10-124685336-DEL-1075
chr10-124688781-DEL-484
chr10-124698957-DEL-301
chr10-124712592-DEL-700
chr10-124753840-DEL-80
chr10-124800665-DEL-217
chr10-124825244-DEL-207
chr10-124846005-DEL-66
chr10-124861153-DEL-237
chr10-124915630-DEL-856
chr10-124964853-DEL-54
chr10-124966044-DEL-102
chr10-124968528-DEL-56
chr10-124971103-DEL-255
chr10-125006411-DEL-74
chr10-125047129-DEL-434
chr10-125075674-DEL-159
chr10-125076823-DEL-1131
chr10-125077530-DEL-172
chr10-125085488-DEL-1689
chr10-125116060-DEL-112
chr10-125126232-DEL-255
chr10-125160910-DEL-51
chr10-125202888-DEL-134
chr10-125232087-DEL-237
chr10-125236384-DEL-249
chr10-125260422-DEL-141
chr10-125289383-DEL-132
chr10-125301109-DEL-87
chr10-125306286-DEL-52
chr10-125310846-DEL-57
chr10-125315718-DEL-52
chr10-125323431-DEL-243
chr10-125351355-DEL-103
chr10-125370231-DEL-63
chr10-125383779-DEL-96
chr10-125386236-DEL-88
chr10-125388863-DEL-105
chr10-125393399-DEL-82
chr10-125415897-DEL-121
chr10-125420852-DEL-533
chr10-125421828-DEL-490
chr10-125450907-DEL-63
chr10-125452008-DEL-158
chr10-125472130-DEL-60
chr10-125476985-DEL-57
chr10-125489253-DEL-90
chr10-125498891-DEL-52
chr10-125512593-DEL-60
chr10-125513446-DEL-120
chr10-125528025-DEL-122
chr10-125528931-DEL-51
chr10-125537137-DEL-253
chr10-125550465-DEL-6588
chr10-125562125-DEL-199
chr10-125567898-DEL-56
chr10-125665682-DEL-961
chr10-125666440-DEL-251
chr10-125691019-DEL-88
chr10-125691620-DEL-78
chr10-125703509-DEL-3874
chr10-125714268-DEL-201
chr10-125717657-DEL-62
chr10-125720129-DEL-100
chr10-125726659-DEL-1043
chr10-125739422-DEL-848
chr10-125740188-DEL-504
chr10-125755991-DEL-265
chr10-125812098-DEL-90
chr10-125816442-DEL-90
chr10-125822034-DEL-147
chr10-125844636-DEL-96
chr10-125870184-DEL-53
chr10-125872237-DEL-611
chr10-125913977-DEL-249
chr10-125914264-DEL-342
chr10-125928279-DEL-106

chr10-125940251-DEL-184
chr10-125975776-DEL-93
chr10-125984248-DEL-158
chr10-125989226-DEL-492
chr10-126020903-DEL-191
chr10-126036811-DEL-365
chr10-126041040-DEL-502
chr10-126047600-DEL-89
chr10-126061569-DEL-4284
chr10-126073709-DEL-167
chr10-126078748-DEL-239
chr10-126081253-DEL-237
chr10-126082086-DEL-164
chr10-126134084-DEL-57
chr10-126138855-DEL-6155
chr10-126149552-DEL-62
chr10-126164060-DEL-105
chr10-126180214-DEL-4096
chr10-126198213-DEL-699
chr10-126203121-DEL-141
chr10-126231599-DEL-139
chr10-126301606-DEL-161
chr10-126318314-DEL-456
chr10-126363997-DEL-126
chr10-126373011-DEL-379
chr10-126378752-DEL-78
chr10-126401597-DEL-64
chr10-126420537-DEL-3087
chr10-126428682-DEL-100
chr10-126430759-DEL-104
chr10-126431627-DEL-54
chr10-126444753-DEL-1491
chr10-126453510-DEL-240
chr10-126460518-DEL-52
chr10-126462858-DEL-787
chr10-126484750-DEL-81
chr10-126497296-DEL-125
chr10-126498171-DEL-156
chr10-126499567-DEL-194
chr10-126511493-DEL-808
chr10-126518385-DEL-158
chr10-126518704-DEL-3301
chr10-126522125-DEL-133
chr10-126523490-DEL-526
chr10-126524501-DEL-219
chr10-126528099-DEL-495
chr10-126581380-DEL-345
chr10-126595985-DEL-2922
chr10-126606861-DEL-80
chr10-126614118-DEL-60
chr10-126659865-DEL-184
chr10-126679261-DEL-332
chr10-126695281-DEL-2604
chr10-126707186-DEL-54
chr10-126718848-DEL-399
chr10-126730632-DEL-154
chr10-126745641-DEL-152
chr10-126789754-DEL-64
chr10-126794103-DEL-195
chr10-126819419-DEL-81
chr10-126841337-DEL-90
chr10-126906452-DEL-369
chr10-126908093-DEL-195
chr10-126920319-DEL-90
chr10-126942393-DEL-120
chr10-126995043-DEL-2522
chr10-127048195-DEL-98
chr10-127059903-DEL-56
chr10-127085391-DEL-1051
chr10-127135734-DEL-52
chr10-127145165-DEL-147
chr10-127207243-DEL-69
chr10-127209000-DEL-186
chr10-127228311-DEL-421
chr10-127240428-DEL-57
chr10-127245489-DEL-522
chr10-127375900-DEL-53

chr10-127390619-DEL-56
chr10-127394249-DEL-76
chr10-127398327-DEL-373
chr10-127465533-DEL-205
chr10-127473574-DEL-326
chr10-127493694-DEL-173
chr10-127494355-DEL-69
chr10-127508118-DEL-750
chr10-127510713-DEL-125
chr10-127519139-DEL-59
chr10-127522412-DEL-184
chr10-127552301-DEL-308
chr10-127621928-DEL-836
chr10-127651696-DEL-60
chr10-127677116-DEL-174
chr10-127680768-DEL-116
chr10-127683536-DEL-883
chr10-127688685-DEL-1150
chr10-127704258-DEL-1468
chr10-127708430-DEL-62
chr10-127719976-DEL-384
chr10-127738721-DEL-84
chr10-127768119-DEL-234
chr10-127786074-DEL-107
chr10-127790570-DEL-1316
chr10-127792174-DEL-94
chr10-127794980-DEL-55
chr10-127797800-DEL-509
chr10-127804959-DEL-65
chr10-127806932-DEL-1182
chr10-127815877-DEL-280
chr10-127833138-DEL-83
chr10-127847610-DEL-490
chr10-127848686-DEL-183
chr10-127854563-DEL-219
chr10-127890884-DEL-145
chr10-127896888-DEL-2818
chr10-127915877-DEL-194
chr10-127939807-DEL-143
chr10-128010464-DEL-58
chr10-128018514-DEL-590
chr10-128024767-DEL-71
chr10-128069018-DEL-707
chr10-128096187-DEL-203
chr10-128108930-DEL-204
chr10-128111166-DEL-218
chr10-128126582-DEL-216
chr10-128168450-DEL-205
chr10-128171519-DEL-54
chr10-128208582-DEL-227
chr10-128210528-DEL-220
chr10-128242958-DEL-268
chr10-128294928-DEL-79
chr10-128303286-DEL-56
chr10-128313959-DEL-126
chr10-128322666-DEL-181
chr10-128362753-DEL-213
chr10-128373507-DEL-494
chr10-128375582-DEL-59
chr10-128385403-DEL-1547
chr10-128388141-DEL-184
chr10-128390547-DEL-130
chr10-128391716-DEL-195
chr10-128392692-DEL-55
chr10-128398421-DEL-61
chr10-128412145-DEL-101
chr10-128449871-DEL-169
chr10-128453935-DEL-439
chr10-128465237-DEL-105
chr10-128491791-DEL-162
chr10-128520314-DEL-52
chr10-128545019-DEL-159
chr10-128590718-DEL-96
chr10-128611307-DEL-65
chr10-128612154-DEL-1046
chr10-128615805-DEL-52
chr10-128618845-DEL-87

chr10-128623687-DEL-92
chr10-128624978-DEL-158
chr10-128635078-DEL-139
chr10-128645650-DEL-122
chr10-128669700-DEL-84
chr10-128736663-DEL-136
chr10-128798343-DEL-60
chr10-128803526-DEL-512
chr10-128811381-DEL-536
chr10-128838596-DEL-131
chr10-128843610-DEL-267
chr10-128862640-DEL-84
chr10-128877907-DEL-1414
chr10-128886857-DEL-4524
chr10-128891116-DEL-4355
chr10-128892517-DEL-66
chr10-128908641-DEL-66
chr10-128919413-DEL-676
chr10-128932466-DEL-874
chr10-128942192-DEL-1441
chr10-128949513-DEL-537
chr10-128949931-DEL-788
chr10-128950423-DEL-73
chr10-128957071-DEL-54
chr10-128977828-DEL-6503
chr10-129039711-DEL-71
chr10-129044553-DEL-12170
chr10-129044578-DEL-3161
chr10-129093233-DEL-597
chr10-129116375-DEL-2602
chr10-129176208-DEL-5484
chr10-129177628-DEL-160
chr10-129183010-DEL-91
chr10-129220071-DEL-160
chr10-129235107-DEL-404
chr10-129235687-DEL-517
chr10-129260674-DEL-408
chr10-129266795-DEL-63
chr10-129295025-DEL-92
chr10-129319521-DEL-786
chr10-129319564-DEL-58
chr10-129327696-DEL-540
chr10-129349519-DEL-350
chr10-129408775-DEL-148
chr10-129410410-DEL-54
chr10-129412400-DEL-56
chr10-129435068-DEL-156
chr10-129436669-DEL-2524
chr10-129479770-DEL-305
chr10-129489502-DEL-57
chr10-129500722-DEL-1037
chr10-129510624-DEL-99
chr10-129535421-DEL-52
chr10-129536073-DEL-293
chr10-129559438-DEL-4772
chr10-129571346-DEL-60
chr10-129586982-DEL-933
chr10-129599775-DEL-52
chr10-129632801-DEL-64
chr10-129642734-DEL-3268
chr10-129646775-DEL-12971
chr10-129650045-DEL-4974
chr10-129726274-DEL-11033
chr10-129742751-DEL-498
chr10-129752299-DEL-1396
chr10-129804349-DEL-2454
chr10-129804350-DEL-11328
chr10-129828820-DEL-67
chr10-129874654-DEL-6644
chr10-129882650-DEL-6177
chr10-129883048-DEL-116
chr10-129883758-DEL-11453
chr10-129923683-DEL-64
chr10-129949625-DEL-414
chr10-129963725-DEL-161
chr10-129970151-DEL-3627
chr10-130005836-DEL-1957

chr10-130031044-DEL-6150
chr10-130053737-DEL-6541
chr10-130073497-DEL-616
chr10-130084382-DEL-1393
chr10-130107732-DEL-94
chr10-130156301-DEL-743
chr10-130194551-DEL-1180
chr10-130217035-DEL-54
chr10-130242698-DEL-58
chr10-130248102-DEL-1910
chr10-130275647-DEL-383
chr10-130298964-DEL-3378
chr10-130300653-DEL-1489
chr10-130313033-DEL-80
chr10-130321243-DEL-2981
chr10-130345834-DEL-60
chr10-130346527-DEL-691
chr10-130386520-DEL-5499
chr10-130395840-DEL-184
chr10-130404690-DEL-131
chr10-130416989-DEL-5490
chr10-130425665-DEL-395
chr10-130427004-DEL-240
chr11-3052841-DEL-118
chr11-3062947-DEL-2252
chr11-3070627-DEL-498
chr11-3098902-DEL-574
chr11-3101572-DEL-468
chr11-3206114-DEL-334
chr11-3220736-DEL-1992
chr11-3264623-DEL-176
chr11-3281703-DEL-64
chr11-3299782-DEL-182
chr11-3321891-DEL-300
chr11-3355238-DEL-214
chr11-3380781-DEL-60
chr11-3392822-DEL-77
chr11-3405267-DEL-62
chr11-3435414-DEL-62
chr11-3483046-DEL-102
chr11-3510631-DEL-57
chr11-3518437-DEL-81
chr11-3526737-DEL-125
chr11-3526766-DEL-511
chr11-3533757-DEL-53
chr11-3544851-DEL-213
chr11-3549622-DEL-211
chr11-3552137-DEL-367
chr11-3563708-DEL-349
chr11-3567470-DEL-2491
chr11-3585783-DEL-243
chr11-3651826-DEL-93
chr11-3682462-DEL-68
chr11-3695727-DEL-69
chr11-3737357-DEL-105
chr11-3737881-DEL-100
chr11-3763644-DEL-78
chr11-3774661-DEL-599
chr11-3781933-DEL-272
chr11-3816021-DEL-2452
chr11-3820434-DEL-260
chr11-3827304-DEL-83
chr11-3828797-DEL-108
chr11-3890797-DEL-413
chr11-3893063-DEL-192
chr11-3899883-DEL-2330
chr11-3929664-DEL-342
chr11-3935229-DEL-59
chr11-3935815-DEL-56
chr11-3954907-DEL-266
chr11-3963437-DEL-69
chr11-3965883-DEL-6343
chr11-3976041-DEL-2258
chr11-3994763-DEL-368
chr11-3995355-DEL-663
chr11-4004256-DEL-663
chr11-4012102-DEL-74

chr11-4015475-DEL-78
chr11-4021865-DEL-676
chr11-4028323-DEL-112
chr11-4050758-DEL-402
chr11-4054147-DEL-201
chr11-4059885-DEL-58
chr11-4063836-DEL-205
chr11-4134033-DEL-218
chr11-4136093-DEL-62
chr11-4191174-DEL-70
chr11-4204943-DEL-183
chr11-4225713-DEL-84
chr11-4268227-DEL-366
chr11-4277920-DEL-200
chr11-4287121-DEL-66
chr11-4288771-DEL-207
chr11-4298732-DEL-1095
chr11-4306211-DEL-240
chr11-4314730-DEL-197
chr11-4317561-DEL-58
chr11-4320518-DEL-79
chr11-4325194-DEL-200
chr11-4334376-DEL-60
chr11-4344785-DEL-1818
chr11-4373066-DEL-59
chr11-4378586-DEL-62
chr11-4420074-DEL-209
chr11-4441345-DEL-64
chr11-4444575-DEL-68
chr11-4492484-DEL-60
chr11-4496862-DEL-65
chr11-4510027-DEL-178
chr11-4514846-DEL-192
chr11-4522871-DEL-1192
chr11-4523094-DEL-596
chr11-4556157-DEL-84
chr11-4580770-DEL-64
chr11-4580936-DEL-2043
chr11-4585954-DEL-73
chr11-4602798-DEL-434
chr11-4619086-DEL-172
chr11-4621641-DEL-58
chr11-4626016-DEL-737
chr11-4627762-DEL-199
chr11-4640211-DEL-156
chr11-4648395-DEL-135
chr11-4660359-DEL-197
chr11-4677367-DEL-84
chr11-4771952-DEL-209
chr11-4774282-DEL-448
chr11-4788326-DEL-859
chr11-4792688-DEL-92
chr11-4795360-DEL-191
chr11-4803349-DEL-386
chr11-4819141-DEL-85
chr11-4820861-DEL-1058
chr11-4831805-DEL-192
chr11-4848807-DEL-94
chr11-4852213-DEL-55
chr11-4859511-DEL-58
chr11-4859915-DEL-206
chr11-4865636-DEL-215
chr11-4909216-DEL-199
chr11-4911290-DEL-127
chr11-4915497-DEL-70
chr11-4916396-DEL-68
chr11-4938184-DEL-307
chr11-4971456-DEL-214
chr11-4971776-DEL-256
chr11-5008691-DEL-56
chr11-5025998-DEL-1025
chr11-5075000-DEL-450
chr11-5096661-DEL-315
chr11-5109022-DEL-404
chr11-5118251-DEL-343
chr11-5118443-DEL-847
chr11-5119418-DEL-866

chr11-5119816-DEL-235
chr11-5134666-DEL-116
chr11-5136349-DEL-119
chr11-5147069-DEL-98
chr11-5153267-DEL-90
chr11-5168346-DEL-290
chr11-5179002-DEL-73
chr11-5186882-DEL-60
chr11-5207822-DEL-62
chr11-5248754-DEL-950
chr11-5308781-DEL-732
chr11-5325644-DEL-51
chr11-5339975-DEL-67
chr11-5358371-DEL-2885
chr11-5383745-DEL-74
chr11-5390490-DEL-64
chr11-5396483-DEL-206
chr11-5407286-DEL-6790
chr11-5414543-DEL-108
chr11-5416718-DEL-288
chr11-5434520-DEL-137
chr11-5444758-DEL-832
chr11-5459378-DEL-58
chr11-5487391-DEL-231
chr11-5502093-DEL-76
chr11-5558803-DEL-90
chr11-5566540-DEL-82
chr11-5624409-DEL-55
chr11-5631692-DEL-2180
chr11-5634378-DEL-60
chr11-5635046-DEL-577
chr11-5643331-DEL-132
chr11-5646688-DEL-214
chr11-5648972-DEL-159
chr11-5651318-DEL-100
chr11-5675347-DEL-53
chr11-5683358-DEL-66
chr11-5687789-DEL-191
chr11-5706238-DEL-563
chr11-5727594-DEL-70
chr11-5734784-DEL-204
chr11-5764734-DEL-1678
chr11-5765465-DEL-74
chr11-5771286-DEL-88
chr11-5774463-DEL-201
chr11-5777071-DEL-80
chr11-5803155-DEL-193
chr11-5814430-DEL-60
chr11-5841629-DEL-80
chr11-5855933-DEL-104
chr11-5877399-DEL-58
chr11-5890141-DEL-98
chr11-5901163-DEL-195
chr11-5978063-DEL-57
chr11-5986304-DEL-149
chr11-5997061-DEL-78
chr11-6003862-DEL-212
chr11-6033905-DEL-167
chr11-6040986-DEL-356
chr11-6112706-DEL-403
chr11-6117535-DEL-63
chr11-6122572-DEL-496
chr11-6135146-DEL-54
chr11-6146668-DEL-205
chr11-6157813-DEL-204
chr11-6185635-DEL-9439
chr11-6205990-DEL-272
chr11-6221314-DEL-67
chr11-6222496-DEL-82
chr11-6234285-DEL-598
chr11-6278095-DEL-208
chr11-6284707-DEL-228
chr11-6291724-DEL-82
chr11-6300921-DEL-165
chr11-6325922-DEL-119
chr11-6327453-DEL-59
chr11-6329687-DEL-193

chr11-6344968-DEL-62
chr11-6363174-DEL-68
chr11-6372038-DEL-1362
chr11-6385702-DEL-63
chr11-6388364-DEL-104
chr11-6389838-DEL-212
chr11-6393038-DEL-86
chr11-6397273-DEL-55
chr11-6398899-DEL-78
chr11-6404262-DEL-375
chr11-6408849-DEL-1009
chr11-6413066-DEL-183
chr11-6443878-DEL-123
chr11-6481472-DEL-3967
chr11-6481481-DEL-3319
chr11-6481482-DEL-2167
chr11-6484699-DEL-1173
chr11-6484972-DEL-897
chr11-6486361-DEL-570
chr11-6486855-DEL-102
chr11-6524309-DEL-61
chr11-6557932-DEL-212
chr11-6580130-DEL-303
chr11-6592404-DEL-98
chr11-6598809-DEL-939
chr11-6615195-DEL-284
chr11-6639445-DEL-52
chr11-6647400-DEL-204
chr11-6660647-DEL-81
chr11-6694896-DEL-14452
chr11-6700314-DEL-1707
chr11-6733767-DEL-81
chr11-6738156-DEL-105
chr11-6774019-DEL-61
chr11-6779748-DEL-76
chr11-6787998-DEL-175
chr11-6795406-DEL-519
chr11-6834770-DEL-59
chr11-6836234-DEL-572
chr11-6853143-DEL-98
chr11-6888437-DEL-92
chr11-6890219-DEL-95
chr11-6903593-DEL-75
chr11-6930299-DEL-53
chr11-6947776-DEL-78
chr11-6954585-DEL-3125
chr11-6962523-DEL-51
chr11-6972852-DEL-54
chr11-6984803-DEL-121
chr11-6985258-DEL-170
chr11-6987722-DEL-86
chr11-7000558-DEL-327
chr11-7082179-DEL-75
chr11-7112243-DEL-55
chr11-7115743-DEL-52
chr11-7120087-DEL-66
chr11-7134088-DEL-466
chr11-7139181-DEL-150
chr11-7154744-DEL-79
chr11-7179127-DEL-86
chr11-7202697-DEL-304
chr11-7206241-DEL-314
chr11-7209688-DEL-54
chr11-7236711-DEL-87
chr11-7241142-DEL-830
chr11-7248263-DEL-75
chr11-7281370-DEL-346
chr11-7292434-DEL-1999
chr11-7329690-DEL-57
chr11-7392960-DEL-586
chr11-7394518-DEL-302
chr11-7427262-DEL-82
chr11-7553732-DEL-1724
chr11-7571578-DEL-1326
chr11-7583638-DEL-237
chr11-7587005-DEL-214
chr11-7588262-DEL-522

chr11-7603859-DEL-6614
chr11-7680748-DEL-275
chr11-7725579-DEL-58
chr11-7740613-DEL-191
chr11-7741159-DEL-1908
chr11-7784170-DEL-63
chr11-7791614-DEL-3096
chr11-7817233-DEL-64
chr11-7818317-DEL-93
chr11-7819373-DEL-75
chr11-7819757-DEL-6277
chr11-7826880-DEL-63
chr11-7834997-DEL-1733
chr11-7905087-DEL-441
chr11-7934030-DEL-2040
chr11-7981399-DEL-667
chr11-7982383-DEL-5286
chr11-7983940-DEL-3443
chr11-7998122-DEL-328
chr11-7998925-DEL-332
chr11-8043588-DEL-215
chr11-8064993-DEL-55
chr11-8147870-DEL-229
chr11-8174141-DEL-263
chr11-8186576-DEL-196
chr11-8210488-DEL-67
chr11-8234461-DEL-90
chr11-8270658-DEL-254
chr11-8274768-DEL-1920
chr11-8302653-DEL-77
chr11-8315388-DEL-67
chr11-8334535-DEL-757
chr11-8359678-DEL-79
chr11-8371886-DEL-299
chr11-8391147-DEL-73
chr11-8490453-DEL-91
chr11-8526201-DEL-666
chr11-8531167-DEL-82
chr11-8538547-DEL-94
chr11-8592836-DEL-51
chr11-8684169-DEL-5778
chr11-8697817-DEL-70
chr11-8699783-DEL-70
chr11-8701547-DEL-587
chr11-8705364-DEL-74
chr11-8718298-DEL-84
chr11-8723365-DEL-908
chr11-8826492-DEL-552
chr11-8840790-DEL-56
chr11-8859217-DEL-53
chr11-8882598-DEL-87
chr11-8884435-DEL-833
chr11-8898381-DEL-62
chr11-8950930-DEL-400
chr11-9039678-DEL-499
chr11-9087958-DEL-2485
chr11-9109810-DEL-58
chr11-9139483-DEL-316
chr11-9145328-DEL-62
chr11-9159075-DEL-60
chr11-9191380-DEL-56
chr11-9207978-DEL-425
chr11-9236872-DEL-490
chr11-9239509-DEL-120
chr11-9255615-DEL-164
chr11-9315084-DEL-200
chr11-9321854-DEL-1176
chr11-9374924-DEL-112
chr11-9408246-DEL-1453
chr11-9424729-DEL-1277
chr11-9447063-DEL-2096
chr11-9449200-DEL-97
chr11-9485339-DEL-78
chr11-9502833-DEL-255
chr11-9504320-DEL-63
chr11-9530070-DEL-1023
chr11-9550180-DEL-52

chr11-9553910-DEL-3261
chr11-9584006-DEL-52
chr11-9587389-DEL-133
chr11-9598649-DEL-187
chr11-9615842-DEL-1257
chr11-9622244-DEL-54
chr11-9642642-DEL-533
chr11-9694072-DEL-6403
chr11-9710400-DEL-544
chr11-9736797-DEL-73
chr11-9757699-DEL-6440
chr11-9764272-DEL-63
chr11-9766146-DEL-155
chr11-9813259-DEL-317
chr11-9824996-DEL-213
chr11-9826629-DEL-57
chr11-9832286-DEL-8296
chr11-9853247-DEL-1523
chr11-9886891-DEL-3263
chr11-9888420-DEL-276
chr11-9929822-DEL-184
chr11-9931963-DEL-561
chr11-9980674-DEL-440
chr11-9982998-DEL-56
chr11-10000798-DEL-71
chr11-10002315-DEL-184
chr11-10038585-DEL-6284
chr11-10111757-DEL-173
chr11-10117065-DEL-6654
chr11-10141455-DEL-59
chr11-10146305-DEL-195
chr11-10147336-DEL-351
chr11-10185437-DEL-357
chr11-10196559-DEL-84
chr11-10200737-DEL-6164
chr11-10212307-DEL-322
chr11-10213614-DEL-6141
chr11-10221578-DEL-1941
chr11-10242950-DEL-520
chr11-10262970-DEL-2197
chr11-10327976-DEL-215
chr11-10336323-DEL-3239
chr11-10340003-DEL-1497
chr11-10368932-DEL-2557
chr11-10385896-DEL-8834
chr11-10396921-DEL-55
chr11-10398727-DEL-6420
chr11-10410655-DEL-1798
chr11-10459507-DEL-687
chr11-10506247-DEL-71
chr11-10507978-DEL-235
chr11-10538424-DEL-986
chr11-10574261-DEL-4943
chr11-10644337-DEL-59
chr11-10653674-DEL-992
chr11-10656799-DEL-84
chr11-10678835-DEL-227
chr11-10683115-DEL-402
chr11-10687047-DEL-87
chr11-10706509-DEL-58
chr11-10710086-DEL-177
chr11-10710306-DEL-7109
chr11-10719797-DEL-421
chr11-10726336-DEL-2177
chr11-10727131-DEL-398
chr11-10728601-DEL-4907
chr11-10734724-DEL-55
chr11-10820304-DEL-75
chr11-10828445-DEL-246
chr11-10839493-DEL-1021
chr11-10842465-DEL-722
chr11-10862945-DEL-69
chr11-10902551-DEL-124
chr11-10903511-DEL-158
chr11-10939755-DEL-164
chr11-10970373-DEL-809
chr11-10974172-DEL-694

chr11-11011446-DEL-475
chr11-11018268-DEL-435
chr11-11037203-DEL-70
chr11-11049193-DEL-587
chr11-11097803-DEL-214
chr11-11134321-DEL-372
chr11-11149122-DEL-219
chr11-11179606-DEL-105
chr11-11237313-DEL-212
chr11-11237887-DEL-425
chr11-11262022-DEL-2791
chr11-11279947-DEL-1018
chr11-11292720-DEL-6417
chr11-11320282-DEL-214
chr11-11327854-DEL-6249
chr11-11337535-DEL-166
chr11-11338922-DEL-7722
chr11-11349054-DEL-1142
chr11-11373854-DEL-58
chr11-11408442-DEL-53
chr11-11451250-DEL-145
chr11-11471589-DEL-188
chr11-11513111-DEL-177
chr11-11542324-DEL-2311
chr11-11553988-DEL-4898
chr11-11561525-DEL-6627
chr11-11576687-DEL-6902
chr11-11581486-DEL-522
chr11-11583972-DEL-342
chr11-11608444-DEL-89
chr11-11664549-DEL-506
chr11-11688245-DEL-81
chr11-11715947-DEL-68
chr11-11745086-DEL-193
chr11-11749387-DEL-68
chr11-11770006-DEL-58
chr11-11777683-DEL-282
chr11-11868373-DEL-69
chr11-11914542-DEL-72
chr11-12072395-DEL-15375
chr11-12079178-DEL-6805
chr11-12087049-DEL-313
chr11-12089493-DEL-7641
chr11-12193614-DEL-311
chr11-12205867-DEL-134
chr11-12208132-DEL-123
chr11-12228551-DEL-602
chr11-12263477-DEL-58
chr11-12280222-DEL-60
chr11-12283584-DEL-74
chr11-12310855-DEL-77
chr11-12322598-DEL-73
chr11-12381188-DEL-78
chr11-12386769-DEL-180
chr11-12391242-DEL-100
chr11-12434840-DEL-71
chr11-12444558-DEL-150
chr11-12467192-DEL-126
chr11-12483466-DEL-343
chr11-12499721-DEL-558
chr11-12507490-DEL-107
chr11-12510476-DEL-8579
chr11-12546558-DEL-61
chr11-12558977-DEL-91
chr11-12572087-DEL-3586
chr11-12598061-DEL-397
chr11-12640388-DEL-80
chr11-12650203-DEL-67
chr11-12678324-DEL-876
chr11-12683897-DEL-120
chr11-12686360-DEL-743
chr11-12695364-DEL-239
chr11-12699154-DEL-182
chr11-12700424-DEL-6450
chr11-12714929-DEL-206
chr11-12722437-DEL-6169
chr11-12737778-DEL-72

chr11-12777596-DEL-806
chr11-12804847-DEL-119
chr11-12807472-DEL-587
chr11-12811788-DEL-1928
chr11-12812542-DEL-332
chr11-12843742-DEL-94
chr11-12847454-DEL-8938
chr11-12879431-DEL-110
chr11-12888158-DEL-152
chr11-12909477-DEL-1291
chr11-12911826-DEL-18248
chr11-12979442-DEL-1743
chr11-12993251-DEL-1959
chr11-13021535-DEL-903
chr11-13022548-DEL-103
chr11-13040983-DEL-1777
chr11-13043253-DEL-1700
chr11-13083429-DEL-182
chr11-13105248-DEL-189
chr11-13106450-DEL-1666
chr11-13110568-DEL-1618
chr11-13126043-DEL-6285
chr11-13132967-DEL-6180
chr11-13171547-DEL-697
chr11-13184404-DEL-2056
chr11-13195507-DEL-210
chr11-13232155-DEL-218
chr11-13268199-DEL-439
chr11-13270206-DEL-64
chr11-13273634-DEL-168
chr11-13285231-DEL-163
chr11-13294230-DEL-12572
chr11-13328998-DEL-6746
chr11-13363725-DEL-63
chr11-13379941-DEL-64
chr11-13453154-DEL-360
chr11-13488254-DEL-2963
chr11-13489431-DEL-448
chr11-13496474-DEL-409
chr11-13518381-DEL-97
chr11-13518748-DEL-1361
chr11-13523651-DEL-2300
chr11-13565429-DEL-212
chr11-13566176-DEL-212
chr11-13584811-DEL-2046
chr11-13614362-DEL-4507
chr11-13634281-DEL-59
chr11-13656663-DEL-79
chr11-13687835-DEL-58
chr11-13699238-DEL-1368
chr11-13706340-DEL-6179
chr11-13761130-DEL-13472
chr11-13799117-DEL-555
chr11-13813646-DEL-2293
chr11-13815318-DEL-686
chr11-13818463-DEL-68
chr11-13844972-DEL-1478
chr11-13846657-DEL-105
chr11-13906020-DEL-505
chr11-13950935-DEL-3193
chr11-13968804-DEL-211
chr11-13970177-DEL-3437
chr11-14006542-DEL-946
chr11-14030871-DEL-163
chr11-14032542-DEL-2184
chr11-14071833-DEL-6470
chr11-14093135-DEL-502
chr11-14108910-DEL-52
chr11-14135390-DEL-99
chr11-14140568-DEL-884
chr11-14161611-DEL-755
chr11-14162406-DEL-4677
chr11-14169512-DEL-1676
chr11-14205963-DEL-74
chr11-14214568-DEL-2878
chr11-14257078-DEL-5128
chr11-14270672-DEL-54

chr11-14277298-DEL-1210
chr11-14292326-DEL-66
chr11-14302560-DEL-127
chr11-14304816-DEL-174
chr11-14362050-DEL-364
chr11-14363289-DEL-59
chr11-14385744-DEL-65
chr11-14391593-DEL-2682
chr11-14404117-DEL-978
chr11-14445328-DEL-77
chr11-14491409-DEL-93
chr11-14516834-DEL-1033
chr11-14518964-DEL-2588
chr11-14544578-DEL-51
chr11-14567616-DEL-90
chr11-14572679-DEL-309
chr11-14591706-DEL-2981
chr11-14595152-DEL-140
chr11-14597646-DEL-1478
chr11-14600384-DEL-3394
chr11-14601257-DEL-2545
chr11-14625443-DEL-249
chr11-14636673-DEL-139
chr11-14646680-DEL-52
chr11-14725778-DEL-51
chr11-14746766-DEL-399
chr11-14760122-DEL-1345
chr11-14787630-DEL-488
chr11-14817148-DEL-55
chr11-14860331-DEL-78
chr11-14860742-DEL-1922
chr11-14876081-DEL-766
chr11-14888740-DEL-1012
chr11-14895857-DEL-52
chr11-14913119-DEL-74
chr11-14936331-DEL-54
chr11-14996344-DEL-1031
chr11-15012720-DEL-58
chr11-15015986-DEL-92
chr11-15025538-DEL-57
chr11-15043375-DEL-3301
chr11-15046550-DEL-864
chr11-15048422-DEL-261
chr11-15053829-DEL-66
chr11-15071597-DEL-75
chr11-15085442-DEL-180
chr11-15137041-DEL-6215
chr11-15144164-DEL-228
chr11-15168440-DEL-1448
chr11-15213976-DEL-1235
chr11-15214872-DEL-2736
chr11-15216112-DEL-1140
chr11-15229225-DEL-11986
chr11-15241971-DEL-112
chr11-15255581-DEL-70
chr11-15256557-DEL-1610
chr11-15277526-DEL-794
chr11-15280774-DEL-164
chr11-15324617-DEL-82
chr11-15344519-DEL-88
chr11-15359189-DEL-676
chr11-15379358-DEL-560
chr11-15452914-DEL-1329
chr11-15473920-DEL-79
chr11-15519097-DEL-280
chr11-15532609-DEL-56
chr11-15582329-DEL-194
chr11-15582666-DEL-2415
chr11-15615107-DEL-202
chr11-15616453-DEL-692
chr11-15617274-DEL-5345
chr11-15628462-DEL-128
chr11-15695449-DEL-1090
chr11-15715364-DEL-53
chr11-15779905-DEL-437
chr11-15782700-DEL-58
chr11-15790385-DEL-66

chr11-15801418-DEL-115
chr11-15867802-DEL-71
chr11-15877028-DEL-1560
chr11-15887922-DEL-7539
chr11-15888945-DEL-1343
chr11-15900396-DEL-515
chr11-15917681-DEL-80
chr11-15946516-DEL-145
chr11-15962004-DEL-214
chr11-15973383-DEL-612
chr11-15975083-DEL-422
chr11-15990335-DEL-212
chr11-16006074-DEL-54
chr11-16010791-DEL-185
chr11-16013595-DEL-195
chr11-16021191-DEL-6289
chr11-16052672-DEL-107
chr11-16064437-DEL-2043
chr11-16182958-DEL-4671
chr11-16191526-DEL-84
chr11-16192820-DEL-4623
chr11-16204925-DEL-672
chr11-16253623-DEL-485
chr11-16261143-DEL-95
chr11-16274866-DEL-170
chr11-16286460-DEL-1052
chr11-16327184-DEL-84
chr11-16330650-DEL-15616
chr11-16368170-DEL-1449
chr11-16374256-DEL-74
chr11-16402856-DEL-2056
chr11-16402858-DEL-5026
chr11-16404965-DEL-2004
chr11-16407255-DEL-1119
chr11-16407427-DEL-460
chr11-16407967-DEL-328
chr11-16423034-DEL-72
chr11-16459869-DEL-176
chr11-16492174-DEL-83
chr11-16512785-DEL-3528
chr11-16513486-DEL-458
chr11-16542347-DEL-124
chr11-16556307-DEL-104
chr11-16563275-DEL-787
chr11-16613038-DEL-790
chr11-16614088-DEL-224
chr11-16678937-DEL-3772
chr11-16707558-DEL-2439
chr11-16720496-DEL-88
chr11-16761016-DEL-105
chr11-16787067-DEL-575
chr11-16822049-DEL-576
chr11-16826590-DEL-1060
chr11-16829025-DEL-1011
chr11-16829157-DEL-476
chr11-16838719-DEL-202
chr11-16878314-DEL-3309
chr11-16905048-DEL-203
chr11-16907742-DEL-127
chr11-16936229-DEL-53
chr11-16940942-DEL-102
chr11-16945542-DEL-82
chr11-16980858-DEL-122
chr11-16991514-DEL-150
chr11-16993289-DEL-200
chr11-17012934-DEL-94
chr11-17044701-DEL-684
chr11-17045465-DEL-167
chr11-17045683-DEL-1087
chr11-17047032-DEL-586
chr11-17054888-DEL-1112
chr11-17069792-DEL-548
chr11-17070437-DEL-1398
chr11-17102602-DEL-556
chr11-17128560-DEL-529
chr11-17158364-DEL-215
chr11-17164568-DEL-140

chr11-17166928-DEL-786
chr11-17194019-DEL-203
chr11-17219691-DEL-102
chr11-17248043-DEL-1166
chr11-17269773-DEL-108
chr11-17291580-DEL-54
chr11-17306165-DEL-746
chr11-17330164-DEL-62
chr11-17346226-DEL-96
chr11-17368315-DEL-144
chr11-17387354-DEL-612
chr11-17415496-DEL-987
chr11-17416250-DEL-186
chr11-17433858-DEL-67
chr11-17436305-DEL-406
chr11-17446806-DEL-574
chr11-17453123-DEL-659
chr11-17460266-DEL-71
chr11-17461800-DEL-6592
chr11-17467952-DEL-212
chr11-17484456-DEL-4019
chr11-17517632-DEL-185
chr11-17531216-DEL-3211
chr11-17535619-DEL-2501
chr11-17579316-DEL-53
chr11-17583539-DEL-55
chr11-17598376-DEL-217
chr11-17601549-DEL-164
chr11-17691139-DEL-68
chr11-17701183-DEL-202
chr11-17701726-DEL-164
chr11-17806092-DEL-88
chr11-17815240-DEL-54
chr11-17837790-DEL-1361
chr11-17840516-DEL-384
chr11-17858260-DEL-3113
chr11-17880390-DEL-204
chr11-17881820-DEL-77
chr11-17887418-DEL-2414
chr11-17904434-DEL-1365
chr11-17909242-DEL-11835
chr11-17909412-DEL-5565
chr11-17950183-DEL-143
chr11-17951479-DEL-86
chr11-18002671-DEL-706
chr11-18007005-DEL-4460
chr11-18019239-DEL-58
chr11-18045387-DEL-116
chr11-18048956-DEL-117
chr11-18131041-DEL-70
chr11-18138933-DEL-3484
chr11-18139484-DEL-738
chr11-18146830-DEL-1233
chr11-18148354-DEL-212
chr11-18172000-DEL-128
chr11-18181953-DEL-6310
chr11-18183638-DEL-1514
chr11-18202104-DEL-499
chr11-18203508-DEL-723
chr11-18247201-DEL-6233
chr11-18251877-DEL-128
chr11-18281665-DEL-648
chr11-18309301-DEL-65
chr11-18405134-DEL-2358
chr11-18409449-DEL-538
chr11-18410096-DEL-2591
chr11-18458310-DEL-85
chr11-18461671-DEL-6193
chr11-18468722-DEL-187
chr11-18470609-DEL-127
chr11-18470823-DEL-421
chr11-18523611-DEL-808
chr11-18527088-DEL-952
chr11-18560634-DEL-874
chr11-18573162-DEL-6569
chr11-18586153-DEL-6484
chr11-18593215-DEL-809

chr11-18610318-DEL-1323
chr11-18653873-DEL-220
chr11-18657839-DEL-168
chr11-18660412-DEL-59
chr11-18667658-DEL-84
chr11-18695109-DEL-98
chr11-18718566-DEL-938
chr11-18747752-DEL-139
chr11-18769503-DEL-59
chr11-18774961-DEL-478
chr11-18844064-DEL-115
chr11-18930323-DEL-57
chr11-18991413-DEL-69
chr11-19025748-DEL-548
chr11-19029265-DEL-283
chr11-19031888-DEL-7383
chr11-19085474-DEL-185
chr11-19129583-DEL-62
chr11-19134890-DEL-78
chr11-19185723-DEL-260
chr11-19224462-DEL-83
chr11-19235735-DEL-1679
chr11-19272471-DEL-66
chr11-19302851-DEL-58
chr11-19310473-DEL-422
chr11-19333603-DEL-1327
chr11-19346559-DEL-88
chr11-19348367-DEL-52
chr11-19396486-DEL-53
chr11-19407845-DEL-61
chr11-19433162-DEL-66
chr11-19434295-DEL-93
chr11-19442412-DEL-98
chr11-19458206-DEL-344
chr11-19472867-DEL-532
chr11-19475614-DEL-195
chr11-19497833-DEL-52
chr11-19507130-DEL-1196
chr11-19530200-DEL-73
chr11-19571706-DEL-882
chr11-19579340-DEL-154
chr11-19608085-DEL-158
chr11-19643676-DEL-55
chr11-19647192-DEL-199
chr11-19667954-DEL-119
chr11-19670211-DEL-1375
chr11-19683792-DEL-1570
chr11-19692832-DEL-156
chr11-19693571-DEL-2089
chr11-19724605-DEL-82
chr11-19745894-DEL-226
chr11-19755515-DEL-496
chr11-19756548-DEL-1360
chr11-19801366-DEL-78
chr11-19806148-DEL-736
chr11-19824738-DEL-119
chr11-19830261-DEL-351
chr11-19865327-DEL-74
chr11-19883568-DEL-219
chr11-19887062-DEL-51
chr11-19900874-DEL-195
chr11-19905964-DEL-76
chr11-19918228-DEL-420
chr11-19956665-DEL-1687
chr11-19960274-DEL-194
chr11-19964414-DEL-152
chr11-19977670-DEL-100
chr11-19982944-DEL-323
chr11-19991005-DEL-51
chr11-19992692-DEL-178
chr11-19996162-DEL-108
chr11-20020333-DEL-71
chr11-20025803-DEL-197
chr11-20041018-DEL-211
chr11-20044752-DEL-59
chr11-20051083-DEL-291
chr11-20057684-DEL-65

chr11-20083262-DEL-154
chr11-20090623-DEL-1105
chr11-20092502-DEL-90
chr11-20098706-DEL-198
chr11-20099638-DEL-180
chr11-20109845-DEL-115
chr11-20131577-DEL-425
chr11-20133726-DEL-62
chr11-20138979-DEL-65
chr11-20148728-DEL-204
chr11-20192264-DEL-188
chr11-20206201-DEL-1486
chr11-20255182-DEL-71
chr11-20260450-DEL-920
chr11-20260961-DEL-160
chr11-20329481-DEL-199
chr11-20354039-DEL-8337
chr11-20361824-DEL-155
chr11-20363656-DEL-200
chr11-20396394-DEL-462
chr11-20424090-DEL-121
chr11-20435101-DEL-61
chr11-20458943-DEL-4650
chr11-20484394-DEL-172
chr11-20515858-DEL-59
chr11-20532474-DEL-64
chr11-20537529-DEL-4251
chr11-20619072-DEL-80
chr11-20624728-DEL-125
chr11-20667161-DEL-212
chr11-20675249-DEL-106
chr11-20695766-DEL-65
chr11-20712257-DEL-5436
chr11-20721605-DEL-650
chr11-20735943-DEL-379
chr11-20741395-DEL-2076
chr11-20804964-DEL-166
chr11-20827306-DEL-95
chr11-20882334-DEL-467
chr11-20882856-DEL-270
chr11-20886669-DEL-93
chr11-20891060-DEL-139
chr11-20919317-DEL-1348
chr11-20990173-DEL-61
chr11-21024621-DEL-209
chr11-21063983-DEL-337
chr11-21074223-DEL-6443
chr11-21086661-DEL-709
chr11-21101105-DEL-1121
chr11-21116503-DEL-58
chr11-21126484-DEL-76
chr11-21148534-DEL-109
chr11-21167770-DEL-1645
chr11-21179330-DEL-1794
chr11-21183942-DEL-954
chr11-21198270-DEL-168
chr11-21243913-DEL-53
chr11-21282988-DEL-62
chr11-21289752-DEL-2121
chr11-21291925-DEL-891
chr11-21313875-DEL-291
chr11-21327552-DEL-156
chr11-21334623-DEL-214
chr11-21368728-DEL-84
chr11-21447066-DEL-529
chr11-21541054-DEL-1485
chr11-21542692-DEL-2306
chr11-21543825-DEL-1092
chr11-21573150-DEL-1901
chr11-21671868-DEL-106
chr11-21676783-DEL-52
chr11-21679426-DEL-75
chr11-21721571-DEL-126
chr11-21736985-DEL-984
chr11-21756529-DEL-1970
chr11-21793280-DEL-358
chr11-21795552-DEL-64

chr11-21847257-DEL-196
chr11-21888515-DEL-114
chr11-21890147-DEL-951
chr11-21917937-DEL-129
chr11-21930685-DEL-162
chr11-22014199-DEL-628
chr11-22049045-DEL-869
chr11-22059255-DEL-940
chr11-22093662-DEL-66
chr11-22130616-DEL-1331
chr11-22162128-DEL-2503
chr11-22177834-DEL-93
chr11-22184733-DEL-77
chr11-22202991-DEL-1303
chr11-22214458-DEL-495
chr11-22220589-DEL-96
chr11-22231362-DEL-118
chr11-22252609-DEL-2571
chr11-22255742-DEL-445
chr11-22261936-DEL-417
chr11-22298366-DEL-56
chr11-22308428-DEL-170
chr11-22322815-DEL-119
chr11-22336033-DEL-5977
chr11-22368080-DEL-61
chr11-22377134-DEL-54
chr11-22377275-DEL-5993
chr11-22395896-DEL-331
chr11-22405978-DEL-54
chr11-22407764-DEL-468
chr11-22409933-DEL-80
chr11-22410873-DEL-989
chr11-22434433-DEL-1303
chr11-22457106-DEL-187
chr11-22470603-DEL-1605
chr11-22470663-DEL-167
chr11-22470986-DEL-764
chr11-22481070-DEL-775
chr11-22494018-DEL-181
chr11-22505066-DEL-95
chr11-22506663-DEL-121
chr11-22544191-DEL-1885
chr11-22546326-DEL-542
chr11-22589435-DEL-51
chr11-22597883-DEL-5451
chr11-22603570-DEL-2824
chr11-22610087-DEL-406
chr11-22612782-DEL-80
chr11-22688206-DEL-128
chr11-22731272-DEL-2887
chr11-22753250-DEL-410
chr11-22764881-DEL-55
chr11-22779288-DEL-109
chr11-22781190-DEL-80
chr11-22791834-DEL-146
chr11-22793879-DEL-71
chr11-22828656-DEL-60
chr11-22831345-DEL-645
chr11-22858568-DEL-85
chr11-22870602-DEL-53
chr11-22880674-DEL-67
chr11-22894323-DEL-190
chr11-22913637-DEL-142
chr11-22938445-DEL-72
chr11-22976319-DEL-88
chr11-22981471-DEL-52
chr11-23009504-DEL-783
chr11-23021888-DEL-2620
chr11-23023575-DEL-74
chr11-23048181-DEL-273
chr11-23062955-DEL-200
chr11-23073964-DEL-427
chr11-23094819-DEL-95
chr11-23095956-DEL-336
chr11-23103061-DEL-942
chr11-23105884-DEL-78
chr11-23130192-DEL-63

chr11-23135695-DEL-518
chr11-23177374-DEL-12867
chr11-23227226-DEL-551
chr11-23238836-DEL-184
chr11-23242174-DEL-218
chr11-23251471-DEL-491
chr11-23255382-DEL-216
chr11-23299633-DEL-783
chr11-23365752-DEL-209
chr11-23390296-DEL-354
chr11-23401303-DEL-625
chr11-23488926-DEL-644
chr11-23491204-DEL-1144
chr11-23491500-DEL-368
chr11-23506198-DEL-180
chr11-23509347-DEL-1737
chr11-23522252-DEL-168
chr11-23536866-DEL-76
chr11-23569508-DEL-75
chr11-23578765-DEL-60
chr11-23586467-DEL-526
chr11-23620161-DEL-70
chr11-23642700-DEL-665
chr11-23643629-DEL-1256
chr11-23664060-DEL-3321
chr11-23671178-DEL-249
chr11-23741885-DEL-218
chr11-23759539-DEL-56
chr11-23784706-DEL-8811
chr11-23838081-DEL-197
chr11-23854312-DEL-749
chr11-23859451-DEL-2166
chr11-23868975-DEL-1920
chr11-23898727-DEL-400
chr11-23963848-DEL-159
chr11-23968777-DEL-104
chr11-24118889-DEL-117
chr11-24124893-DEL-94
chr11-24298535-DEL-53
chr11-24307770-DEL-140
chr11-24330561-DEL-56
chr11-24350225-DEL-58
chr11-24379599-DEL-798
chr11-24395057-DEL-62
chr11-24415908-DEL-95
chr11-24471720-DEL-796
chr11-24474735-DEL-442
chr11-24539394-DEL-8758
chr11-24616103-DEL-76
chr11-24620225-DEL-61
chr11-24649232-DEL-76
chr11-24707678-DEL-54
chr11-24724315-DEL-185
chr11-24730388-DEL-274
chr11-24777280-DEL-51
chr11-24809072-DEL-1654
chr11-24842866-DEL-84
chr11-24904431-DEL-77
chr11-24930850-DEL-88
chr11-24950771-DEL-66
chr11-25068402-DEL-1309
chr11-25180767-DEL-496
chr11-25203510-DEL-345
chr11-25208473-DEL-860
chr11-25213711-DEL-68
chr11-25242399-DEL-75
chr11-25291168-DEL-67
chr11-25320432-DEL-6210
chr11-25329714-DEL-171
chr11-25357708-DEL-208
chr11-25380881-DEL-1296
chr11-25503269-DEL-637
chr11-25570560-DEL-183
chr11-25575582-DEL-103
chr11-25598900-DEL-320
chr11-25602162-DEL-77
chr11-25606594-DEL-59

chr11-25804182-DEL-191
chr11-25880907-DEL-388
chr11-25903331-DEL-2843
chr11-25917055-DEL-204
chr11-25925943-DEL-541
chr11-25940429-DEL-59
chr11-25988677-DEL-116
chr11-26016134-DEL-561
chr11-26063636-DEL-115
chr11-26078116-DEL-1350
chr11-26084660-DEL-260
chr11-26163698-DEL-182
chr11-26190417-DEL-60
chr11-26195301-DEL-3271
chr11-26206283-DEL-74
chr11-26301865-DEL-70
chr11-26307815-DEL-229
chr11-26313308-DEL-4474
chr11-26319159-DEL-204
chr11-26319969-DEL-517
chr11-26350536-DEL-144
chr11-26354267-DEL-384
chr11-26401013-DEL-525
chr11-26408659-DEL-199
chr11-26409897-DEL-224
chr11-26452193-DEL-660
chr11-26493188-DEL-128
chr11-26494066-DEL-2157
chr11-26544435-DEL-66
chr11-26569075-DEL-1528
chr11-26589107-DEL-313
chr11-26619319-DEL-6539
chr11-26669387-DEL-91
chr11-26679068-DEL-62
chr11-26697543-DEL-1062
chr11-26780032-DEL-62
chr11-26786249-DEL-66
chr11-26805977-DEL-194
chr11-26823764-DEL-199
chr11-26847214-DEL-131
chr11-26848393-DEL-2182
chr11-26867447-DEL-59
chr11-26906556-DEL-90
chr11-26924019-DEL-165
chr11-26937496-DEL-510
chr11-26959092-DEL-3508
chr11-26968151-DEL-634
chr11-26972694-DEL-99
chr11-26987041-DEL-600
chr11-27002085-DEL-177
chr11-27012395-DEL-201
chr11-27034768-DEL-454
chr11-27035048-DEL-1362
chr11-27067427-DEL-576
chr11-27070102-DEL-262
chr11-27073576-DEL-204
chr11-27075336-DEL-1092
chr11-27091664-DEL-473
chr11-27108102-DEL-77
chr11-27119521-DEL-292
chr11-27139149-DEL-1888
chr11-27158028-DEL-910
chr11-27167596-DEL-328
chr11-27209657-DEL-86
chr11-27221129-DEL-98
chr11-27233802-DEL-3563
chr11-27257579-DEL-2484
chr11-27262921-DEL-5474
chr11-27269633-DEL-141
chr11-27273603-DEL-193
chr11-27318824-DEL-63
chr11-27343463-DEL-279
chr11-27395871-DEL-185
chr11-27440091-DEL-4511
chr11-27447169-DEL-901
chr11-27448260-DEL-98
chr11-27455935-DEL-149

chr11-27474459-DEL-958
chr11-27483337-DEL-8151
chr11-27506924-DEL-85
chr11-27509012-DEL-58
chr11-27523029-DEL-2708
chr11-27570122-DEL-165
chr11-27618941-DEL-72
chr11-27634410-DEL-6388
chr11-27670360-DEL-87
chr11-27680930-DEL-102
chr11-27682213-DEL-62
chr11-27704953-DEL-146
chr11-27712127-DEL-64
chr11-27712192-DEL-2507
chr11-27714559-DEL-301
chr11-27723531-DEL-6212
chr11-27730260-DEL-161
chr11-27746301-DEL-3299
chr11-27785024-DEL-645
chr11-27802579-DEL-2656
chr11-27849936-DEL-61
chr11-27872364-DEL-73
chr11-27879562-DEL-213
chr11-27885376-DEL-1088
chr11-27889586-DEL-62
chr11-27914858-DEL-102
chr11-27961111-DEL-1213
chr11-27968033-DEL-1668
chr11-27979949-DEL-96
chr11-28033824-DEL-153
chr11-28040348-DEL-145
chr11-28046731-DEL-118
chr11-28085877-DEL-107
chr11-28088330-DEL-1348
chr11-28090472-DEL-5540
chr11-28112147-DEL-722
chr11-28124087-DEL-689
chr11-28125580-DEL-73
chr11-28159488-DEL-279
chr11-28159732-DEL-389
chr11-28165521-DEL-658
chr11-28179109-DEL-100
chr11-28180518-DEL-2015
chr11-28183675-DEL-6638
chr11-28183720-DEL-337
chr11-28184826-DEL-2826
chr11-28190565-DEL-227
chr11-28203199-DEL-693
chr11-28211245-DEL-149
chr11-28215171-DEL-316
chr11-28242154-DEL-7346
chr11-28249330-DEL-369
chr11-28259684-DEL-340
chr11-28264087-DEL-1563
chr11-28286388-DEL-3788
chr11-28312547-DEL-64
chr11-28375579-DEL-59004
chr11-28386302-DEL-1632
chr11-28437887-DEL-51
chr11-28453737-DEL-199
chr11-28455651-DEL-723
chr11-28461168-DEL-70
chr11-28462177-DEL-77
chr11-28530398-DEL-213
chr11-28536528-DEL-54
chr11-28557232-DEL-74
chr11-28598944-DEL-73
chr11-28634046-DEL-179
chr11-28648963-DEL-93
chr11-28667762-DEL-113
chr11-28677422-DEL-2104
chr11-28730028-DEL-168
chr11-28752269-DEL-501
chr11-28772384-DEL-1153
chr11-28833640-DEL-675
chr11-28841028-DEL-144
chr11-28913088-DEL-119

chr11-28913372-DEL-460
chr11-28914371-DEL-592
chr11-28918065-DEL-216
chr11-28919042-DEL-6331
chr11-28930478-DEL-5671
chr11-28962183-DEL-398
chr11-28965568-DEL-134
chr11-28980525-DEL-214
chr11-28985424-DEL-630
chr11-28987092-DEL-93
chr11-29012775-DEL-75
chr11-29013915-DEL-6212
chr11-29030707-DEL-5412
chr11-29031931-DEL-6153
chr11-29035428-DEL-51
chr11-29072235-DEL-244
chr11-29118756-DEL-73
chr11-29133765-DEL-154
chr11-29149636-DEL-188
chr11-29187194-DEL-101
chr11-29201581-DEL-875
chr11-29216771-DEL-182
chr11-29229967-DEL-1421
chr11-29237901-DEL-188
chr11-29240670-DEL-106
chr11-29245366-DEL-59
chr11-29245774-DEL-61
chr11-29273975-DEL-57
chr11-29275860-DEL-6763
chr11-29280209-DEL-119
chr11-29281866-DEL-190
chr11-29317302-DEL-82
chr11-29348065-DEL-178
chr11-29359400-DEL-61
chr11-29364152-DEL-3630
chr11-29370711-DEL-617
chr11-29431348-DEL-128
chr11-29473386-DEL-548
chr11-29495654-DEL-51
chr11-29560543-DEL-223
chr11-29579323-DEL-51
chr11-29593902-DEL-1310
chr11-29611025-DEL-675
chr11-29618886-DEL-70
chr11-29622869-DEL-53
chr11-29625576-DEL-199
chr11-29636843-DEL-53
chr11-29641658-DEL-216
chr11-29651775-DEL-55
chr11-29654918-DEL-263
chr11-29692938-DEL-78
chr11-29706206-DEL-167
chr11-29712249-DEL-283
chr11-29732284-DEL-91
chr11-29756590-DEL-217
chr11-29762357-DEL-75
chr11-29785761-DEL-69
chr11-29826351-DEL-70
chr11-29860609-DEL-919
chr11-29901124-DEL-69
chr11-29911106-DEL-6399
chr11-29911301-DEL-200
chr11-29923035-DEL-107
chr11-29925914-DEL-866
chr11-29948751-DEL-139
chr11-29997169-DEL-59
chr11-29999373-DEL-60
chr11-30007808-DEL-117
chr11-30011309-DEL-12783
chr11-30014557-DEL-1009
chr11-30015552-DEL-6414
chr11-30019810-DEL-2156
chr11-30026293-DEL-1071
chr11-30053750-DEL-57
chr11-30070018-DEL-78
chr11-30076892-DEL-80
chr11-30139434-DEL-288

chr11-30243193-DEL-56
chr11-30268423-DEL-429
chr11-30314962-DEL-86
chr11-30357992-DEL-6375
chr11-30372244-DEL-611
chr11-30416904-DEL-1967
chr11-30417029-DEL-350
chr11-30417392-DEL-917
chr11-30418487-DEL-244
chr11-30426266-DEL-3317
chr11-30444638-DEL-188
chr11-30455820-DEL-116
chr11-30460952-DEL-3495
chr11-30464194-DEL-252
chr11-30476591-DEL-175
chr11-30484084-DEL-81
chr11-30508507-DEL-129
chr11-30512979-DEL-51
chr11-30543986-DEL-563
chr11-30545552-DEL-1172
chr11-30550742-DEL-68
chr11-30552080-DEL-2858
chr11-30554996-DEL-2343
chr11-30557379-DEL-115
chr11-30600123-DEL-121
chr11-30612884-DEL-3807
chr11-30616716-DEL-5103
chr11-30629867-DEL-58
chr11-30638863-DEL-443
chr11-30651972-DEL-384
chr11-30653267-DEL-648
chr11-30762753-DEL-181
chr11-30804561-DEL-256
chr11-30818283-DEL-2269
chr11-30831690-DEL-96
chr11-30854327-DEL-62
chr11-30854714-DEL-254
chr11-30882357-DEL-54
chr11-30883421-DEL-102
chr11-30915237-DEL-57
chr11-30931040-DEL-84
chr11-30945974-DEL-1886
chr11-30961556-DEL-6267
chr11-30977388-DEL-79
chr11-31009962-DEL-612
chr11-31026577-DEL-68
chr11-31053581-DEL-385
chr11-31057349-DEL-2130
chr11-31073955-DEL-366
chr11-31079606-DEL-497
chr11-31115718-DEL-2229
chr11-31147450-DEL-807
chr11-31153156-DEL-52
chr11-31157399-DEL-78
chr11-31162779-DEL-2833
chr11-31185856-DEL-344
chr11-31189014-DEL-801
chr11-31193449-DEL-400
chr11-31198914-DEL-2940
chr11-31201881-DEL-380
chr11-31202386-DEL-2235
chr11-31217588-DEL-54
chr11-31236184-DEL-201
chr11-31325054-DEL-82
chr11-31358722-DEL-309
chr11-31416336-DEL-131
chr11-31431868-DEL-365
chr11-31437468-DEL-1719
chr11-31445520-DEL-88
chr11-31460836-DEL-96
chr11-31470663-DEL-157
chr11-31471247-DEL-264
chr11-31567564-DEL-60
chr11-31572157-DEL-308
chr11-31574668-DEL-90
chr11-31610492-DEL-170
chr11-31626019-DEL-51

chr11-31678046-DEL-366
chr11-31700282-DEL-889
chr11-31724124-DEL-304
chr11-31724751-DEL-737
chr11-31725515-DEL-1610
chr11-31736651-DEL-343
chr11-31795392-DEL-60
chr11-31901515-DEL-151
chr11-31904173-DEL-57
chr11-31950888-DEL-192
chr11-32003135-DEL-99
chr11-32042363-DEL-1103
chr11-32045055-DEL-58
chr11-32059912-DEL-135
chr11-32074702-DEL-182
chr11-32089192-DEL-262
chr11-32100205-DEL-66
chr11-32112822-DEL-184
chr11-32116316-DEL-60
chr11-32122666-DEL-210
chr11-32127388-DEL-326
chr11-32129041-DEL-1352
chr11-32135794-DEL-88
chr11-32143004-DEL-83
chr11-32145244-DEL-96
chr11-32150944-DEL-56
chr11-32211859-DEL-290
chr11-32231002-DEL-175
chr11-32232502-DEL-270
chr11-32243449-DEL-54
chr11-32247811-DEL-64
chr11-32265121-DEL-932
chr11-32272923-DEL-252
chr11-32293284-DEL-73
chr11-32308762-DEL-88
chr11-32322359-DEL-152
chr11-32332770-DEL-53
chr11-32359649-DEL-109
chr11-32366177-DEL-93
chr11-32389250-DEL-56
chr11-32401698-DEL-53
chr11-32439542-DEL-204
chr11-32442873-DEL-192
chr11-32480878-DEL-590
chr11-32494969-DEL-53
chr11-32498790-DEL-132
chr11-32530240-DEL-123
chr11-32552533-DEL-292
chr11-32553035-DEL-188
chr11-32556467-DEL-60
chr11-32559167-DEL-53
chr11-32613159-DEL-213
chr11-32624566-DEL-498
chr11-32625178-DEL-194
chr11-32633099-DEL-442
chr11-32641683-DEL-375
chr11-32646824-DEL-76
chr11-32675842-DEL-51
chr11-32676452-DEL-153
chr11-32698362-DEL-199
chr11-32702843-DEL-267
chr11-32716319-DEL-615
chr11-32729662-DEL-82
chr11-32746389-DEL-533
chr11-32757124-DEL-749
chr11-32758097-DEL-58
chr11-32770774-DEL-50
chr11-32774267-DEL-98
chr11-32779109-DEL-1808
chr11-32781158-DEL-325
chr11-32792831-DEL-82
chr11-32941697-DEL-61
chr11-32958571-DEL-64
chr11-33010832-DEL-1007
chr11-33055780-DEL-61
chr11-33122866-DEL-606
chr11-33165870-DEL-51

chr11-33219634-DEL-202
chr11-33223518-DEL-342
chr11-33288697-DEL-103
chr11-33357860-DEL-1818
chr11-33368675-DEL-1512
chr11-33420448-DEL-193
chr11-33434846-DEL-183
chr11-33457213-DEL-1156
chr11-33461163-DEL-203
chr11-33488755-DEL-64
chr11-33499812-DEL-96
chr11-33511633-DEL-379
chr11-33552514-DEL-390
chr11-33553152-DEL-60
chr11-33573508-DEL-136
chr11-33582161-DEL-711
chr11-33645371-DEL-92
chr11-33652033-DEL-208
chr11-33660488-DEL-325
chr11-33681283-DEL-125
chr11-33738910-DEL-114
chr11-33761142-DEL-169
chr11-33865107-DEL-117
chr11-33880437-DEL-56
chr11-33896169-DEL-399
chr11-33903991-DEL-64
chr11-33979933-DEL-622
chr11-34025376-DEL-54
chr11-34036837-DEL-120
chr11-34069342-DEL-133
chr11-34071747-DEL-126
chr11-34099079-DEL-66
chr11-34107936-DEL-612
chr11-34117547-DEL-54
chr11-34170429-DEL-160
chr11-34173432-DEL-132
chr11-34176802-DEL-238
chr11-34197063-DEL-79
chr11-34234131-DEL-149
chr11-34270776-DEL-185
chr11-34285976-DEL-96
chr11-34349057-DEL-343
chr11-34402826-DEL-56
chr11-34443487-DEL-62
chr11-34452582-DEL-135
chr11-34490916-DEL-1907
chr11-34495835-DEL-2750
chr11-34497132-DEL-1454
chr11-34497563-DEL-900
chr11-34499374-DEL-122
chr11-34506204-DEL-120
chr11-34513695-DEL-168
chr11-34516060-DEL-193
chr11-34517753-DEL-1126
chr11-34518883-DEL-2787
chr11-34520255-DEL-931
chr11-34520257-DEL-237
chr11-34521514-DEL-277
chr11-34527546-DEL-70
chr11-34531006-DEL-217
chr11-34531166-DEL-386
chr11-34534142-DEL-71
chr11-34535610-DEL-71
chr11-34538359-DEL-1864
chr11-34540452-DEL-97
chr11-34541321-DEL-891
chr11-34543354-DEL-116
chr11-34543665-DEL-1104
chr11-34544785-DEL-2223
chr11-34544905-DEL-85
chr11-34547701-DEL-1096
chr11-34551466-DEL-115
chr11-34551775-DEL-1099
chr11-34553223-DEL-73
chr11-34554723-DEL-217
chr11-34554883-DEL-625
chr11-34555869-DEL-210

chr11-34556052-DEL-239
chr11-34556964-DEL-854
chr11-34557417-DEL-120
chr11-34558462-DEL-71
chr11-34559418-DEL-3468
chr11-34559644-DEL-118
chr11-34561147-DEL-71
chr11-34562349-DEL-122
chr11-34563125-DEL-605
chr11-34563808-DEL-96
chr11-34566711-DEL-187
chr11-34568759-DEL-217
chr11-34637810-DEL-185
chr11-34657171-DEL-64
chr11-34658199-DEL-318
chr11-34698871-DEL-549
chr11-34701601-DEL-74
chr11-34703886-DEL-155
chr11-34705801-DEL-85
chr11-34713684-DEL-55
chr11-34734702-DEL-62
chr11-34763923-DEL-98
chr11-34765576-DEL-916
chr11-34768499-DEL-1388
chr11-34790094-DEL-69
chr11-34804261-DEL-144
chr11-34807381-DEL-93
chr11-34809953-DEL-1426
chr11-34850827-DEL-1352
chr11-34861864-DEL-108
chr11-34877773-DEL-1079
chr11-34880259-DEL-166
chr11-34880443-DEL-1109
chr11-34881877-DEL-400
chr11-34882937-DEL-152
chr11-34887402-DEL-158
chr11-34901096-DEL-158
chr11-34945383-DEL-72
chr11-34954977-DEL-90
chr11-34955681-DEL-54
chr11-35036263-DEL-400
chr11-35036868-DEL-183
chr11-35039454-DEL-153
chr11-35078482-DEL-84
chr11-35096717-DEL-54
chr11-35156311-DEL-56
chr11-35183790-DEL-98
chr11-35189768-DEL-122
chr11-35202472-DEL-174
chr11-35205850-DEL-147
chr11-35216087-DEL-125
chr11-35270263-DEL-53
chr11-35291054-DEL-205
chr11-35353095-DEL-78
chr11-35408672-DEL-232
chr11-35450682-DEL-74
chr11-35457821-DEL-1604
chr11-35504969-DEL-51
chr11-35508852-DEL-145
chr11-35586167-DEL-86
chr11-35588584-DEL-190
chr11-35654253-DEL-980
chr11-35654724-DEL-256
chr11-35654919-DEL-760
chr11-35655227-DEL-1015
chr11-35656054-DEL-158
chr11-35692955-DEL-1504
chr11-35745067-DEL-58
chr11-35834651-DEL-240
chr11-35873434-DEL-143
chr11-35874551-DEL-428
chr11-35906526-DEL-2085
chr11-35907883-DEL-77
chr11-35919861-DEL-312
chr11-35930036-DEL-59
chr11-35951617-DEL-152
chr11-35993335-DEL-258

chr11-36037372-DEL-56
chr11-36038836-DEL-192
chr11-36088699-DEL-281
chr11-36123043-DEL-70
chr11-36126469-DEL-94
chr11-36174744-DEL-215
chr11-36179925-DEL-52
chr11-36187160-DEL-94
chr11-36391734-DEL-85
chr11-36413329-DEL-105
chr11-36423717-DEL-72
chr11-36436298-DEL-112
chr11-36483193-DEL-62
chr11-36525251-DEL-51
chr11-36546278-DEL-115
chr11-36583297-DEL-61
chr11-36602479-DEL-126
chr11-36689439-DEL-111
chr11-36696929-DEL-100
chr11-36771922-DEL-74
chr11-36788068-DEL-464
chr11-36844510-DEL-61
chr11-36868241-DEL-860
chr11-36923836-DEL-79
chr11-37003618-DEL-119
chr11-37016888-DEL-569
chr11-37058029-DEL-608
chr11-37080119-DEL-1358
chr11-37082860-DEL-764
chr11-37096127-DEL-194
chr11-37098769-DEL-58
chr11-37129921-DEL-62
chr11-37157871-DEL-110
chr11-37196008-DEL-222
chr11-37402399-DEL-1205
chr11-37407036-DEL-69
chr11-37441091-DEL-350
chr11-37445705-DEL-174
chr11-37544318-DEL-2101
chr11-37587745-DEL-54
chr11-37588558-DEL-149
chr11-37610469-DEL-93
chr11-37621432-DEL-1030
chr11-37650943-DEL-192
chr11-37699208-DEL-66
chr11-37731431-DEL-64
chr11-37749042-DEL-58
chr11-37760542-DEL-159
chr11-37762588-DEL-56
chr11-37770675-DEL-61
chr11-37782369-DEL-52
chr11-37878820-DEL-903
chr11-37889353-DEL-327
chr11-37891015-DEL-83
chr11-37892726-DEL-667
chr11-37896977-DEL-126
chr11-37897214-DEL-771
chr11-37930485-DEL-52
chr11-38003884-DEL-216
chr11-38005905-DEL-80
chr11-38032356-DEL-50
chr11-38040115-DEL-138
chr11-38066889-DEL-133
chr11-38072180-DEL-445
chr11-38082649-DEL-54
chr11-38087207-DEL-2163
chr11-38124420-DEL-102
chr11-38135050-DEL-102
chr11-38147924-DEL-115
chr11-38151268-DEL-54
chr11-38158635-DEL-1649
chr11-38190049-DEL-200
chr11-38212878-DEL-138
chr11-38213444-DEL-319
chr11-38216337-DEL-69
chr11-38232796-DEL-218
chr11-38241058-DEL-75

chr11-38253524-DEL-338
chr11-38267313-DEL-1495
chr11-38327712-DEL-56
chr11-38350116-DEL-69
chr11-38358443-DEL-53
chr11-38363631-DEL-379
chr11-38371199-DEL-56
chr11-38407881-DEL-120
chr11-38488853-DEL-151
chr11-38496230-DEL-1360
chr11-38498346-DEL-86
chr11-38556177-DEL-3013
chr11-38569480-DEL-749
chr11-38582593-DEL-78
chr11-38620496-DEL-3371
chr11-38623540-DEL-118
chr11-38688446-DEL-658
chr11-38726183-DEL-96
chr11-38733147-DEL-52
chr11-38735907-DEL-57
chr11-38786395-DEL-1088
chr11-38880683-DEL-3324
chr11-38889852-DEL-1163
chr11-38899277-DEL-133
chr11-38920859-DEL-61
chr11-38925909-DEL-148
chr11-38951119-DEL-351
chr11-38966899-DEL-388
chr11-39008829-DEL-63
chr11-39012027-DEL-456
chr11-39024419-DEL-506
chr11-39033872-DEL-2190
chr11-39079223-DEL-80
chr11-39081557-DEL-95
chr11-39096527-DEL-54
chr11-39110274-DEL-249
chr11-39132035-DEL-54
chr11-39147400-DEL-74
chr11-39176886-DEL-177
chr11-39183197-DEL-52
chr11-39220838-DEL-428
chr11-39248026-DEL-233
chr11-39258425-DEL-10952
chr11-39268597-DEL-689
chr11-39269978-DEL-25031
chr11-39274259-DEL-815
chr11-39276110-DEL-51
chr11-39280040-DEL-409
chr11-39289184-DEL-1019
chr11-39309917-DEL-253
chr11-39323392-DEL-118
chr11-39327774-DEL-1371
chr11-39346900-DEL-758
chr11-39352942-DEL-56
chr11-39354403-DEL-51
chr11-39371834-DEL-1530
chr11-39378383-DEL-79
chr11-39413828-DEL-80
chr11-39457697-DEL-109
chr11-39469559-DEL-84
chr11-39477269-DEL-62
chr11-39478149-DEL-66
chr11-39497957-DEL-72
chr11-39513826-DEL-299
chr11-39549581-DEL-198
chr11-39551559-DEL-55
chr11-39573294-DEL-2541
chr11-39577531-DEL-66
chr11-39710373-DEL-612
chr11-39720599-DEL-56
chr11-39737652-DEL-444
chr11-39787672-DEL-390
chr11-39840072-DEL-64
chr11-39853376-DEL-1184
chr11-39873889-DEL-96
chr11-39876892-DEL-1068
chr11-39895936-DEL-244

chr11-39916857-DEL-204
chr11-39950527-DEL-891
chr11-39976320-DEL-1478
chr11-39978571-DEL-1463
chr11-40013269-DEL-1320
chr11-40031622-DEL-52
chr11-40065516-DEL-626
chr11-40073289-DEL-62
chr11-40077488-DEL-400
chr11-40101259-DEL-468
chr11-40121211-DEL-326
chr11-40133646-DEL-6676
chr11-40190793-DEL-591
chr11-40213420-DEL-1050
chr11-40223353-DEL-1826
chr11-40239140-DEL-416
chr11-40248629-DEL-1876
chr11-40263177-DEL-66
chr11-40281141-DEL-708
chr11-40310296-DEL-212
chr11-40332695-DEL-52
chr11-40425832-DEL-1045
chr11-40484375-DEL-1892
chr11-40533532-DEL-401
chr11-40551810-DEL-6236
chr11-40588443-DEL-56
chr11-40590163-DEL-102
chr11-40642672-DEL-68
chr11-40652078-DEL-686
chr11-40658017-DEL-5847
chr11-40682624-DEL-9056
chr11-40688397-DEL-1287
chr11-40694935-DEL-6292
chr11-40707394-DEL-461
chr11-40711526-DEL-52
chr11-40714446-DEL-59
chr11-40745302-DEL-77
chr11-40828123-DEL-180
chr11-40831459-DEL-185
chr11-40876315-DEL-1734
chr11-40876336-DEL-3852
chr11-40880216-DEL-1503
chr11-40882023-DEL-620
chr11-40916423-DEL-876
chr11-40921714-DEL-63
chr11-40923402-DEL-52
chr11-40939903-DEL-99
chr11-40971388-DEL-196
chr11-40983596-DEL-263
chr11-41015346-DEL-250
chr11-41022407-DEL-1307
chr11-41023838-DEL-1005
chr11-41025134-DEL-161
chr11-41060210-DEL-163
chr11-41082624-DEL-112
chr11-41166661-DEL-67
chr11-41180786-DEL-398
chr11-41200329-DEL-196
chr11-41220378-DEL-73
chr11-41225274-DEL-550
chr11-41226793-DEL-1130
chr11-41236395-DEL-782
chr11-41244994-DEL-439
chr11-41246676-DEL-725
chr11-41338591-DEL-77
chr11-41341345-DEL-343
chr11-41350756-DEL-601
chr11-41358691-DEL-642
chr11-41428929-DEL-803
chr11-41457360-DEL-310
chr11-41457974-DEL-772
chr11-41468131-DEL-2708
chr11-41492478-DEL-52
chr11-41505685-DEL-485
chr11-41509725-DEL-857
chr11-41510906-DEL-1758
chr11-41523446-DEL-118

chr11-41573332-DEL-54
chr11-41628686-DEL-1127
chr11-41637800-DEL-958
chr11-41639784-DEL-180
chr11-41644837-DEL-217
chr11-41649195-DEL-311
chr11-41650842-DEL-119
chr11-41653066-DEL-389
chr11-41657092-DEL-1809
chr11-41683560-DEL-117
chr11-41684561-DEL-2119
chr11-41703241-DEL-80
chr11-41713842-DEL-60
chr11-41726375-DEL-117
chr11-41728531-DEL-897
chr11-41738871-DEL-59
chr11-41744970-DEL-125
chr11-41776855-DEL-227
chr11-41780310-DEL-6579
chr11-41832062-DEL-64
chr11-41860281-DEL-314
chr11-41898649-DEL-574
chr11-41911426-DEL-2152
chr11-41950467-DEL-1510
chr11-41993520-DEL-550
chr11-42106560-DEL-62
chr11-42147742-DEL-467
chr11-42156447-DEL-603
chr11-42157073-DEL-70
chr11-42162813-DEL-511
chr11-42188219-DEL-54
chr11-42191639-DEL-1073
chr11-42217619-DEL-74
chr11-42261682-DEL-54
chr11-42264986-DEL-495
chr11-42309461-DEL-62
chr11-42373989-DEL-1033
chr11-42401229-DEL-72
chr11-42407571-DEL-240
chr11-42443033-DEL-338
chr11-42488112-DEL-191
chr11-42574810-DEL-6110
chr11-42574882-DEL-212
chr11-42610035-DEL-4977
chr11-42647820-DEL-1815
chr11-42664412-DEL-73
chr11-42717821-DEL-238
chr11-42723858-DEL-54
chr11-42767948-DEL-527
chr11-42799457-DEL-197
chr11-42805768-DEL-1168
chr11-42806493-DEL-54
chr11-42936236-DEL-187
chr11-42969719-DEL-6167
chr11-42993845-DEL-1231
chr11-43001762-DEL-627
chr11-43045154-DEL-57
chr11-43058902-DEL-648
chr11-43104451-DEL-199
chr11-43110023-DEL-78
chr11-43148521-DEL-62
chr11-43153869-DEL-78
chr11-43170070-DEL-436
chr11-43184946-DEL-9596
chr11-43195409-DEL-8262
chr11-43223699-DEL-1061
chr11-43269664-DEL-92
chr11-43312523-DEL-86
chr11-43340607-DEL-4850
chr11-43342088-DEL-120
chr11-43346214-DEL-6773
chr11-43356598-DEL-1913
chr11-43360358-DEL-64
chr11-43385480-DEL-1395
chr11-43393091-DEL-1515
chr11-43393135-DEL-75
chr11-43398345-DEL-197

chr11-43432568-DEL-68
chr11-43445788-DEL-140
chr11-43467488-DEL-171
chr11-43488737-DEL-227
chr11-43490070-DEL-167
chr11-43535527-DEL-257
chr11-43536096-DEL-896
chr11-43559673-DEL-61
chr11-43601846-DEL-199
chr11-43613993-DEL-58
chr11-43649933-DEL-78
chr11-43707671-DEL-56
chr11-43710883-DEL-217
chr11-43728439-DEL-351
chr11-43728686-DEL-1125
chr11-43799660-DEL-304
chr11-43809230-DEL-54
chr11-43853012-DEL-160
chr11-43919315-DEL-52
chr11-44077174-DEL-54
chr11-44092153-DEL-221
chr11-44115438-DEL-66
chr11-44171175-DEL-71
chr11-44183417-DEL-1016
chr11-44209671-DEL-75
chr11-44211317-DEL-56
chr11-44218469-DEL-66
chr11-44236538-DEL-53
chr11-44250346-DEL-2061
chr11-44275128-DEL-2733
chr11-44288226-DEL-109
chr11-44295803-DEL-781
chr11-44317271-DEL-1162
chr11-44349864-DEL-51
chr11-44353361-DEL-361
chr11-44386433-DEL-325
chr11-44390333-DEL-159
chr11-44391116-DEL-108
chr11-44444646-DEL-56
chr11-44456597-DEL-88
chr11-44471098-DEL-90
chr11-44493906-DEL-54
chr11-44571963-DEL-85
chr11-44586854-DEL-122
chr11-44603105-DEL-66
chr11-44690935-DEL-51
chr11-44699645-DEL-172
chr11-44743005-DEL-136
chr11-44770792-DEL-325
chr11-44775045-DEL-55
chr11-44849022-DEL-56
chr11-44945522-DEL-578
chr11-44950697-DEL-375
chr11-44982935-DEL-82
chr11-45040278-DEL-78
chr11-45063634-DEL-396
chr11-45070603-DEL-68
chr11-45141037-DEL-103
chr11-45167006-DEL-184
chr11-45207472-DEL-61
chr11-45214249-DEL-84
chr11-45218687-DEL-549
chr11-45229055-DEL-260
chr11-45247448-DEL-95
chr11-45258798-DEL-636
chr11-45313431-DEL-2142
chr11-45333856-DEL-264
chr11-45342591-DEL-1014
chr11-45361526-DEL-520
chr11-45401581-DEL-706
chr11-45403591-DEL-242
chr11-45408098-DEL-332
chr11-45473114-DEL-3900
chr11-45477041-DEL-3479
chr11-45480269-DEL-124
chr11-45481045-DEL-66
chr11-45525035-DEL-73

chr11-45547257-DEL-640
chr11-45565218-DEL-101
chr11-45591414-DEL-55
chr11-45602344-DEL-97
chr11-45627852-DEL-72
chr11-45629477-DEL-538
chr11-45637900-DEL-3033
chr11-45640990-DEL-518
chr11-45645580-DEL-52
chr11-45647173-DEL-6749
chr11-45670822-DEL-3451
chr11-45687189-DEL-690
chr11-45739059-DEL-72
chr11-45746657-DEL-68
chr11-45774036-DEL-68
chr11-45795027-DEL-1031
chr11-45802378-DEL-471
chr11-45814219-DEL-194
chr11-45855854-DEL-57
chr11-45859892-DEL-428
chr11-45869572-DEL-168
chr11-45885535-DEL-682
chr11-45893684-DEL-53
chr11-45898932-DEL-6465
chr11-45905134-DEL-201
chr11-45911660-DEL-225
chr11-45935413-DEL-97
chr11-45936495-DEL-474
chr11-45946462-DEL-120
chr11-45958925-DEL-147
chr11-45966019-DEL-107
chr11-46005946-DEL-112
chr11-46030314-DEL-143
chr11-46069700-DEL-848
chr11-46081538-DEL-67
chr11-46115116-DEL-75
chr11-46174898-DEL-62
chr11-46187198-DEL-362
chr11-46207315-DEL-101
chr11-46214872-DEL-86
chr11-46234284-DEL-72
chr11-46272466-DEL-123
chr11-46276627-DEL-183
chr11-46299300-DEL-307
chr11-46308132-DEL-55
chr11-46308542-DEL-176
chr11-46322444-DEL-130
chr11-46372250-DEL-1907
chr11-46380806-DEL-177
chr11-46398726-DEL-230
chr11-46401863-DEL-722
chr11-46407861-DEL-131
chr11-46422738-DEL-66
chr11-46437974-DEL-58
chr11-46448100-DEL-196
chr11-46464163-DEL-189
chr11-46465074-DEL-80
chr11-46473029-DEL-114
chr11-46480354-DEL-184
chr11-46518918-DEL-66
chr11-46524338-DEL-345
chr11-46550967-DEL-84
chr11-46586937-DEL-172
chr11-46592857-DEL-134
chr11-46614146-DEL-5847
chr11-46620853-DEL-85
chr11-46624754-DEL-107
chr11-46685902-DEL-71
chr11-46692461-DEL-57
chr11-46695724-DEL-373
chr11-46703987-DEL-54
chr11-46706973-DEL-92
chr11-46735856-DEL-58
chr11-46741929-DEL-56
chr11-46744742-DEL-71
chr11-46746417-DEL-4168
chr11-46781260-DEL-190

chr11-46794264-DEL-940
chr11-46805195-DEL-90
chr11-46827422-DEL-87
chr11-46838613-DEL-72
chr11-46846584-DEL-113
chr11-46848294-DEL-55
chr11-46901292-DEL-200
chr11-46907659-DEL-349
chr11-46908223-DEL-69
chr11-46962249-DEL-264
chr11-46971423-DEL-327
chr11-46977929-DEL-445
chr11-47006025-DEL-51
chr11-47033074-DEL-176
chr11-47036198-DEL-912
chr11-47078577-DEL-3098
chr11-47094340-DEL-194
chr11-47153932-DEL-204
chr11-47170309-DEL-56
chr11-47193250-DEL-946
chr11-47213232-DEL-76
chr11-47235070-DEL-1276
chr11-47257496-DEL-66
chr11-47294804-DEL-903
chr11-47310063-DEL-15919
chr11-47318683-DEL-7093
chr11-47318815-DEL-208
chr11-47319536-DEL-208
chr11-47335348-DEL-1528
chr11-47338955-DEL-261
chr11-47344549-DEL-150
chr11-47361276-DEL-83
chr11-47362294-DEL-342
chr11-47375334-DEL-66
chr11-47381341-DEL-4708
chr11-47381828-DEL-199
chr11-47407850-DEL-557
chr11-47413242-DEL-648
chr11-47426492-DEL-551
chr11-47427654-DEL-62
chr11-47433198-DEL-86
chr11-47465082-DEL-145
chr11-47497660-DEL-68
chr11-47552685-DEL-377
chr11-47572556-DEL-860
chr11-47579027-DEL-82
chr11-47588641-DEL-87
chr11-47594124-DEL-318
chr11-47622980-DEL-687
chr11-47695717-DEL-85
chr11-47737772-DEL-14196
chr11-47755572-DEL-774
chr11-47779010-DEL-393
chr11-47781668-DEL-134
chr11-47870431-DEL-167
chr11-47896248-DEL-60
chr11-47919204-DEL-100
chr11-48014902-DEL-4221
chr11-48031120-DEL-170
chr11-48087922-DEL-70
chr11-48088601-DEL-57
chr11-48096367-DEL-119
chr11-48128230-DEL-138
chr11-48167739-DEL-59
chr11-48203021-DEL-108
chr11-48206898-DEL-6228
chr11-48214576-DEL-517
chr11-48215083-DEL-6242
chr11-48235676-DEL-143
chr11-48236528-DEL-1734
chr11-48301737-DEL-109
chr11-48327285-DEL-56
chr11-48342666-DEL-6502
chr11-48342827-DEL-213
chr11-48358660-DEL-6511
chr11-48360646-DEL-3457
chr11-48403025-DEL-443

chr11-48496772-DEL-68
chr11-48542572-DEL-93
chr11-48545430-DEL-693
chr11-48546358-DEL-459
chr11-48560177-DEL-1474
chr11-48593783-DEL-64
chr11-48599557-DEL-481
chr11-48602785-DEL-351
chr11-48603558-DEL-798
chr11-48623630-DEL-259
chr11-48624157-DEL-200
chr11-48645505-DEL-54
chr11-48664773-DEL-285
chr11-48675263-DEL-60
chr11-48676447-DEL-2633
chr11-48676666-DEL-432
chr11-48683493-DEL-2231
chr11-48722599-DEL-359
chr11-48725414-DEL-132
chr11-48751486-DEL-80
chr11-48811812-DEL-96
chr11-48812511-DEL-689
chr11-48818570-DEL-71
chr11-48825263-DEL-612
chr11-48836155-DEL-821
chr11-48842081-DEL-196
chr11-48852569-DEL-2780
chr11-48853378-DEL-77
chr11-48866990-DEL-979
chr11-48881639-DEL-664
chr11-48883570-DEL-88
chr11-48884161-DEL-543
chr11-48889692-DEL-58
chr11-48892274-DEL-69
chr11-48894331-DEL-546
chr11-48938831-DEL-978
chr11-48953532-DEL-749
chr11-48966145-DEL-133
chr11-48989473-DEL-62
chr11-49011495-DEL-220
chr11-49028910-DEL-639
chr11-49031890-DEL-74
chr11-49041458-DEL-83
chr11-49077480-DEL-131
chr11-49082266-DEL-71
chr11-49095098-DEL-56
chr11-49118220-DEL-145
chr11-49118755-DEL-992
chr11-49118969-DEL-117
chr11-49164539-DEL-120
chr11-49167331-DEL-327
chr11-49222907-DEL-1125
chr11-49228198-DEL-63
chr11-49251868-DEL-57
chr11-49272178-DEL-1228
chr11-49331998-DEL-78
chr11-49333294-DEL-249
chr11-49387222-DEL-1158
chr11-49411496-DEL-183
chr11-49441279-DEL-98
chr11-49441875-DEL-163
chr11-49454322-DEL-67
chr11-49456577-DEL-464
chr11-49494658-DEL-880
chr11-49496325-DEL-66
chr11-49591233-DEL-195
chr11-49593644-DEL-196
chr11-49622651-DEL-226
chr11-49631717-DEL-128
chr11-49650520-DEL-68
chr11-49701490-DEL-215
chr11-49750712-DEL-158
chr11-49796237-DEL-6054
chr11-49806825-DEL-746
chr11-49808087-DEL-174
chr11-49855732-DEL-200
chr11-49880377-DEL-84

chr11-49886808-DEL-282
chr11-49921416-DEL-78
chr11-49921697-DEL-63
chr11-49945297-DEL-728
chr11-49955451-DEL-123
chr11-49969139-DEL-254
chr11-49973414-DEL-198
chr11-49992092-DEL-56
chr11-50001593-DEL-90
chr11-50020677-DEL-84
chr11-50023854-DEL-290
chr11-50026204-DEL-76
chr11-50056676-DEL-170
chr11-50058225-DEL-358
chr11-50081046-DEL-201
chr11-50092335-DEL-235
chr11-50104230-DEL-123
chr11-50123663-DEL-70
chr11-50126239-DEL-69
chr11-50151224-DEL-327
chr11-50162298-DEL-1287
chr11-50200441-DEL-293
chr11-50238031-DEL-68
chr11-50290750-DEL-52
chr11-50299662-DEL-68
chr11-50303515-DEL-540
chr11-50310178-DEL-54
chr11-50339959-DEL-482
chr11-50364734-DEL-1204
chr11-50381288-DEL-96
chr11-50579868-DEL-122
chr11-50582117-DEL-57
chr11-50588671-DEL-51
chr11-50618276-DEL-183
chr11-50669745-DEL-105
chr11-50709544-DEL-191
chr11-50725315-DEL-364
chr11-50757119-DEL-2863
chr11-50767061-DEL-89
chr11-50783800-DEL-372
chr11-50787907-DEL-229
chr11-50827716-DEL-510
chr11-50836385-DEL-51
chr11-50848274-DEL-221
chr11-50851077-DEL-160
chr11-50903717-DEL-345
chr11-50907537-DEL-1039
chr11-50908959-DEL-6460
chr11-50909041-DEL-542
chr11-50909616-DEL-5824
chr11-50915627-DEL-984
chr11-50918795-DEL-455
chr11-50919433-DEL-80
chr11-50920102-DEL-648
chr11-50924401-DEL-52
chr11-50929278-DEL-3640
chr11-50929964-DEL-894
chr11-50931262-DEL-52
chr11-50931528-DEL-3601
chr11-50952747-DEL-1282
chr11-50975103-DEL-151
chr11-50987677-DEL-57
chr11-51002276-DEL-3139
chr11-51005040-DEL-166
chr11-51005968-DEL-1632
chr11-51009082-DEL-7134
chr11-51046757-DEL-937
chr11-51093337-DEL-2778
chr11-51117773-DEL-127
chr11-51167965-DEL-64
chr11-51212755-DEL-63
chr11-51240766-DEL-491
chr11-51252071-DEL-240
chr11-51261957-DEL-151
chr11-51271379-DEL-255
chr11-51284008-DEL-253
chr11-51297012-DEL-171

chr11-51311433-DEL-80
chr11-51326195-DEL-106
chr11-51328518-DEL-1964
chr11-51330497-DEL-2389
chr11-51332628-DEL-254
chr11-51380528-DEL-939
chr11-51422208-DEL-384
chr11-51423584-DEL-119
chr11-51428406-DEL-109
chr11-51452521-DEL-54
chr11-51456330-DEL-198
chr11-51465768-DEL-256
chr11-51469746-DEL-83
chr11-51503409-DEL-168
chr11-51535723-DEL-66
chr11-51555030-DEL-6882
chr11-51555353-DEL-254
chr11-51582559-DEL-69
chr11-51588304-DEL-341
chr11-51609643-DEL-498
chr11-51628212-DEL-66
chr11-51635929-DEL-322
chr11-51638728-DEL-53
chr11-51641400-DEL-1313
chr11-51649780-DEL-68
chr11-51684175-DEL-144
chr11-51686006-DEL-977
chr11-51694017-DEL-51
chr11-51710096-DEL-60
chr11-51719484-DEL-202
chr11-51736960-DEL-63
chr11-51761203-DEL-53
chr11-51782882-DEL-75
chr11-51832741-DEL-270
chr11-51839265-DEL-61
chr11-51841243-DEL-64
chr11-51842489-DEL-58
chr11-51907525-DEL-929
chr11-51914546-DEL-138
chr11-51935234-DEL-54
chr11-51941039-DEL-78
chr11-51948150-DEL-950
chr11-51996164-DEL-240
chr11-52001344-DEL-292
chr11-52008010-DEL-60
chr11-52023454-DEL-84
chr11-52063136-DEL-54
chr11-52064300-DEL-63
chr11-52071782-DEL-203
chr11-52074735-DEL-69
chr11-52088899-DEL-760
chr11-52094179-DEL-250
chr11-52095095-DEL-1259
chr11-52097320-DEL-128
chr11-52107462-DEL-234
chr11-52112491-DEL-57
chr11-52129493-DEL-207
chr11-52186187-DEL-1488
chr11-52213368-DEL-64
chr11-52262649-DEL-70
chr11-52275078-DEL-181
chr11-52281748-DEL-388
chr11-52296539-DEL-236
chr11-52302982-DEL-4114
chr11-52320076-DEL-73
chr11-52389662-DEL-96
chr11-52398010-DEL-256
chr11-52403675-DEL-68
chr11-52421575-DEL-196
chr11-52444346-DEL-1478
chr11-52474129-DEL-73
chr11-52502697-DEL-389
chr11-52503891-DEL-71
chr11-52548055-DEL-183
chr11-52569837-DEL-178
chr11-52572382-DEL-2654
chr11-52692021-DEL-443

chr11-52715567-DEL-1482
chr11-52735779-DEL-75
chr11-52742804-DEL-68
chr11-52745401-DEL-147
chr11-52784476-DEL-66
chr11-52793031-DEL-118
chr11-52816193-DEL-53
chr11-52831487-DEL-76
chr11-52863807-DEL-110
chr11-52872018-DEL-551
chr11-52881381-DEL-974
chr11-52886538-DEL-65
chr11-52891373-DEL-122
chr11-52898899-DEL-129
chr11-52918124-DEL-63
chr11-52920177-DEL-259
chr11-52953568-DEL-55
chr11-52982222-DEL-66
chr11-53003119-DEL-222
chr11-53022541-DEL-162
chr11-53034014-DEL-629
chr11-53066109-DEL-898
chr11-53066339-DEL-140
chr11-53066762-DEL-308
chr11-53095697-DEL-88
chr11-53102189-DEL-70
chr11-53112552-DEL-606
chr11-53116368-DEL-79
chr11-53116977-DEL-84
chr11-53140265-DEL-216
chr11-53149820-DEL-70
chr11-53167427-DEL-52
chr11-53196956-DEL-216
chr11-53204792-DEL-117
chr11-53226631-DEL-58
chr11-53228923-DEL-208
chr11-53260530-DEL-200
chr11-53275608-DEL-199
chr11-53289359-DEL-461
chr11-53329737-DEL-192
chr11-53373297-DEL-85
chr11-53415283-DEL-104
chr11-53429099-DEL-576
chr11-53549719-DEL-204
chr11-53572604-DEL-1090
chr11-53602859-DEL-203
chr11-53648817-DEL-180
chr11-53671497-DEL-71
chr11-53682600-DEL-56
chr11-53707443-DEL-177
chr11-53710162-DEL-132
chr11-53714276-DEL-58
chr11-53737507-DEL-197
chr11-53772504-DEL-68
chr11-53817634-DEL-1491
chr11-53825081-DEL-199
chr11-53828562-DEL-67
chr11-53871273-DEL-91
chr11-53890252-DEL-109
chr11-53925591-DEL-69
chr11-53933472-DEL-1186
chr11-53947749-DEL-182
chr11-54125497-DEL-51
chr11-54246493-DEL-100
chr11-54265301-DEL-119
chr11-54268510-DEL-3018
chr11-54279812-DEL-214
chr11-54290875-DEL-290
chr11-54297497-DEL-763
chr11-54303570-DEL-51
chr11-54309028-DEL-55
chr11-54333374-DEL-166
chr11-54361145-DEL-141
chr11-54435170-DEL-1561
chr11-54478693-DEL-385
chr11-54492287-DEL-215
chr11-54494575-DEL-60

chr11-54573406-DEL-810
chr11-54577571-DEL-69
chr11-54584073-DEL-189
chr11-54585750-DEL-89
chr11-54635964-DEL-819
chr11-54636879-DEL-95
chr11-54638101-DEL-56
chr11-54657104-DEL-175
chr11-54671746-DEL-208
chr11-54682047-DEL-86
chr11-54706851-DEL-2414
chr11-54728738-DEL-66
chr11-54742777-DEL-173
chr11-54765083-DEL-133
chr11-54767439-DEL-145
chr11-54799203-DEL-129
chr11-54815749-DEL-164
chr11-54856888-DEL-180
chr11-54889631-DEL-80
chr11-54914709-DEL-492
chr11-54929907-DEL-149
chr11-54944037-DEL-91
chr11-54976913-DEL-174
chr11-54981081-DEL-54
chr11-55004899-DEL-57
chr11-55008609-DEL-1770
chr11-55014398-DEL-576
chr11-55028685-DEL-64
chr11-55029632-DEL-253
chr11-55064462-DEL-393
chr11-55067659-DEL-79
chr11-55081783-DEL-51
chr11-55084468-DEL-94
chr11-55231356-DEL-318
chr11-55253894-DEL-310
chr11-55306512-DEL-52
chr11-55330522-DEL-61
chr11-55342793-DEL-728
chr11-55363263-DEL-217
chr11-55366235-DEL-203
chr11-55369227-DEL-230
chr11-55374198-DEL-203
chr11-55432089-DEL-58
chr11-55452383-DEL-52
chr11-55469849-DEL-115
chr11-55476451-DEL-524
chr11-55591558-DEL-1101
chr11-55617174-DEL-1050
chr11-55710630-DEL-54
chr11-55731738-DEL-4177
chr11-55758173-DEL-496
chr11-55789135-DEL-445
chr11-55790314-DEL-53
chr11-55793937-DEL-89
chr11-55802648-DEL-64
chr11-55886862-DEL-54
chr11-55905821-DEL-74
chr11-55906914-DEL-85
chr11-55921883-DEL-86
chr11-55925128-DEL-60
chr11-55928082-DEL-58
chr11-55939367-DEL-148
chr11-55982414-DEL-583
chr11-55987624-DEL-99
chr11-56010088-DEL-51
chr11-56012366-DEL-85
chr11-56047910-DEL-467
chr11-56079585-DEL-317
chr11-56085754-DEL-233
chr11-56095462-DEL-423
chr11-56113448-DEL-68
chr11-56147401-DEL-814
chr11-56177448-DEL-178
chr11-56184750-DEL-3250
chr11-56188591-DEL-54
chr11-56228823-DEL-104
chr11-56236797-DEL-51

chr11-56270298-DEL-585
chr11-56310469-DEL-168
chr11-56331059-DEL-140
chr11-56420759-DEL-582
chr11-56429965-DEL-209
chr11-56442330-DEL-120
chr11-56527954-DEL-70
chr11-56538674-DEL-327
chr11-56570735-DEL-1250
chr11-56574261-DEL-69
chr11-56600807-DEL-142
chr11-56645631-DEL-62
chr11-56679921-DEL-3683
chr11-56701798-DEL-53
chr11-56734788-DEL-53
chr11-56753283-DEL-62
chr11-56761897-DEL-60
chr11-56959828-DEL-56
chr11-56999776-DEL-52
chr11-57028513-DEL-87
chr11-57033644-DEL-111
chr11-57103586-DEL-51
chr11-57193480-DEL-54
chr11-57206003-DEL-70
chr11-57222713-DEL-1497
chr11-57227595-DEL-1074
chr11-57294769-DEL-136
chr11-57296133-DEL-396
chr11-57344187-DEL-62
chr11-57375529-DEL-108
chr11-57392688-DEL-153
chr11-57397308-DEL-89
chr11-57436891-DEL-113
chr11-57456137-DEL-197
chr11-57462872-DEL-81
chr11-57479150-DEL-137
chr11-57564592-DEL-184
chr11-57590599-DEL-112
chr11-57652776-DEL-210
chr11-57658608-DEL-160
chr11-57680472-DEL-81
chr11-57681863-DEL-79
chr11-57687475-DEL-209
chr11-57712543-DEL-150
chr11-57729480-DEL-73
chr11-57734596-DEL-182
chr11-57737231-DEL-177
chr11-57737780-DEL-69
chr11-57740582-DEL-56
chr11-57744992-DEL-56
chr11-57753757-DEL-124
chr11-57766073-DEL-624
chr11-57767643-DEL-213
chr11-57811251-DEL-248
chr11-57818166-DEL-184
chr11-57837695-DEL-207
chr11-57878532-DEL-451
chr11-57888510-DEL-236
chr11-57890576-DEL-283
chr11-57892744-DEL-54
chr11-57961610-DEL-268
chr11-57966602-DEL-153
chr11-57974694-DEL-961
chr11-57975741-DEL-52
chr11-57979275-DEL-115
chr11-57996503-DEL-609
chr11-57997567-DEL-180
chr11-58003199-DEL-114
chr11-58057843-DEL-178
chr11-58082297-DEL-659
chr11-58144618-DEL-71
chr11-58163185-DEL-352
chr11-58196043-DEL-243
chr11-58251406-DEL-183
chr11-58253563-DEL-3636
chr11-58272304-DEL-226
chr11-58273773-DEL-99

chr11-58278712-DEL-195
chr11-58282139-DEL-98
chr11-58284560-DEL-380
chr11-58297342-DEL-362
chr11-58303122-DEL-139
chr11-58324190-DEL-835
chr11-58333602-DEL-198
chr11-58334806-DEL-140
chr11-58352964-DEL-81
chr11-58359530-DEL-51
chr11-58395083-DEL-68
chr11-58396033-DEL-2178
chr11-58399548-DEL-653
chr11-58415477-DEL-227
chr11-58416407-DEL-656
chr11-58443518-DEL-499
chr11-58460954-DEL-79
chr11-58477653-DEL-1438
chr11-58580417-DEL-6730
chr11-58603361-DEL-3043
chr11-58633303-DEL-319
chr11-58642218-DEL-130
chr11-58691796-DEL-58
chr11-58702643-DEL-80
chr11-58704587-DEL-124
chr11-58718641-DEL-159
chr11-58745346-DEL-489
chr11-58757610-DEL-51
chr11-58763939-DEL-66
chr11-58784708-DEL-129
chr11-58815345-DEL-91
chr11-58826083-DEL-334
chr11-58875638-DEL-249
chr11-58895397-DEL-210
chr11-58895846-DEL-1149
chr11-58896116-DEL-750
chr11-58983703-DEL-154
chr11-58987048-DEL-81
chr11-58990668-DEL-3585
chr11-59032525-DEL-2860
chr11-59040559-DEL-61
chr11-59094102-DEL-180
chr11-59133130-DEL-53
chr11-59160242-DEL-69
chr11-59254115-DEL-62
chr11-59283058-DEL-59
chr11-59289892-DEL-230
chr11-59323658-DEL-2108
chr11-59343467-DEL-156
chr11-59372434-DEL-163
chr11-59380775-DEL-1509
chr11-59407108-DEL-62
chr11-59411112-DEL-57
chr11-59453784-DEL-115
chr11-59504104-DEL-199
chr11-59507172-DEL-162
chr11-59539103-DEL-109
chr11-59577862-DEL-66
chr11-59579232-DEL-239
chr11-59625714-DEL-67
chr11-59628592-DEL-96
chr11-59693732-DEL-144
chr11-59694986-DEL-68
chr11-59722776-DEL-171
chr11-59798293-DEL-154
chr11-59802834-DEL-65
chr11-59814392-DEL-460
chr11-59850915-DEL-176
chr11-59871786-DEL-628
chr11-59886133-DEL-616
chr11-59897548-DEL-96
chr11-59948337-DEL-293
chr11-59951033-DEL-130
chr11-60042247-DEL-208
chr11-60098632-DEL-56
chr11-60130236-DEL-78
chr11-60138026-DEL-191

chr11-60175587-DEL-893
chr11-60197106-DEL-87
chr11-60228906-DEL-179
chr11-60247333-DEL-96
chr11-60270258-DEL-157
chr11-60276964-DEL-890
chr11-60341975-DEL-489
chr11-60406741-DEL-106
chr11-60437812-DEL-233
chr11-60455602-DEL-518
chr11-60462746-DEL-206
chr11-60463151-DEL-304
chr11-60466469-DEL-443
chr11-60469470-DEL-7364
chr11-60479141-DEL-81
chr11-60480822-DEL-602
chr11-60496520-DEL-73
chr11-60505963-DEL-2195
chr11-60513075-DEL-445
chr11-60517941-DEL-68
chr11-60518595-DEL-1569
chr11-60519855-DEL-500
chr11-60523949-DEL-443
chr11-60530962-DEL-601
chr11-60532890-DEL-15615
chr11-60542100-DEL-6406
chr11-60556961-DEL-1539
chr11-60562061-DEL-423
chr11-60565768-DEL-75
chr11-60570944-DEL-745
chr11-60627728-DEL-1598
chr11-60686681-DEL-80
chr11-60691669-DEL-215
chr11-60700931-DEL-62
chr11-60702489-DEL-71
chr11-60709655-DEL-1873
chr11-60715262-DEL-218
chr11-60720850-DEL-100
chr11-60734102-DEL-137
chr11-60741765-DEL-68
chr11-60785299-DEL-108
chr11-60814920-DEL-57
chr11-60832565-DEL-52
chr11-60855263-DEL-75
chr11-60862920-DEL-85
chr11-60865098-DEL-63
chr11-60886664-DEL-140
chr11-60931454-DEL-56
chr11-60942104-DEL-59
chr11-60990441-DEL-568
chr11-61035960-DEL-51
chr11-61075210-DEL-981
chr11-61076512-DEL-111
chr11-61101484-DEL-88
chr11-61111650-DEL-52
chr11-61112104-DEL-212
chr11-61130025-DEL-74
chr11-61143242-DEL-1084
chr11-61151691-DEL-328
chr11-61189689-DEL-625
chr11-61209265-DEL-142
chr11-61232498-DEL-92
chr11-61262143-DEL-106
chr11-61287729-DEL-51
chr11-61288291-DEL-51
chr11-61299393-DEL-54
chr11-61358061-DEL-203
chr11-61446978-DEL-199
chr11-61459754-DEL-136
chr11-61473854-DEL-179
chr11-61480586-DEL-181
chr11-61484451-DEL-1033
chr11-61486113-DEL-1144
chr11-61531627-DEL-184
chr11-61653690-DEL-237
chr11-61678633-DEL-379
chr11-61713763-DEL-73

chr11-61787373-DEL-80
chr11-61826651-DEL-727
chr11-61838261-DEL-1526
chr11-61843840-DEL-98
chr11-61922004-DEL-60
chr11-62033725-DEL-63
chr11-62057941-DEL-81
chr11-62066475-DEL-439
chr11-62072347-DEL-2827
chr11-62085592-DEL-96
chr11-62087741-DEL-591
chr11-62090402-DEL-78
chr11-62101290-DEL-1048
chr11-62125483-DEL-60
chr11-62195257-DEL-63
chr11-62252619-DEL-81
chr11-62268231-DEL-214
chr11-62301055-DEL-158
chr11-62325533-DEL-390
chr11-62372989-DEL-183
chr11-62399416-DEL-511
chr11-62405713-DEL-290
chr11-62413619-DEL-175
chr11-62418837-DEL-117
chr11-62434172-DEL-192
chr11-62435405-DEL-68
chr11-62435940-DEL-393
chr11-62437214-DEL-88
chr11-62448379-DEL-68
chr11-62462131-DEL-172
chr11-62523379-DEL-173
chr11-62547940-DEL-114
chr11-62557901-DEL-1321
chr11-62655571-DEL-65
chr11-62722498-DEL-158
chr11-62757037-DEL-63
chr11-62760892-DEL-377
chr11-62790075-DEL-59
chr11-62797955-DEL-1541
chr11-62803954-DEL-192
chr11-62806087-DEL-74
chr11-62851440-DEL-190
chr11-62901647-DEL-1249
chr11-62912247-DEL-115
chr11-62934043-DEL-531
chr11-62947469-DEL-63
chr11-62979912-DEL-64
chr11-63035762-DEL-1831
chr11-63051743-DEL-80
chr11-63078337-DEL-812
chr11-63079411-DEL-55
chr11-63111955-DEL-1607
chr11-63224886-DEL-1891
chr11-63262353-DEL-82
chr11-63297583-DEL-90
chr11-63309969-DEL-79
chr11-63310683-DEL-109
chr11-63342020-DEL-401
chr11-63352224-DEL-182
chr11-63353241-DEL-56
chr11-63375443-DEL-221
chr11-63407714-DEL-373
chr11-63429654-DEL-255
chr11-63453327-DEL-52
chr11-63491459-DEL-51
chr11-63515654-DEL-2169
chr11-63521640-DEL-470
chr11-63522201-DEL-282
chr11-63527414-DEL-1628
chr11-63560413-DEL-170
chr11-63566584-DEL-594
chr11-63615021-DEL-135
chr11-63616127-DEL-989
chr11-63616563-DEL-1023
chr11-63619553-DEL-938
chr11-63632072-DEL-83
chr11-63640881-DEL-60

chr11-63668558-DEL-52
chr11-63690288-DEL-487
chr11-63691274-DEL-682
chr11-63694643-DEL-1865
chr11-63717355-DEL-67
chr11-63742683-DEL-663
chr11-63757296-DEL-123
chr11-63782822-DEL-722
chr11-63795534-DEL-95
chr11-63833604-DEL-57
chr11-63841349-DEL-69
chr11-63918360-DEL-2097
chr11-64008993-DEL-60
chr11-64036364-DEL-67
chr11-64043546-DEL-53
chr11-64071633-DEL-948
chr11-64098642-DEL-76
chr11-64105344-DEL-66
chr11-64127926-DEL-130
chr11-64138969-DEL-600
chr11-64144876-DEL-51
chr11-64151813-DEL-52
chr11-64155729-DEL-2496
chr11-64183915-DEL-92
chr11-64243465-DEL-984
chr11-64279321-DEL-54
chr11-64304984-DEL-102
chr11-64305217-DEL-868
chr11-64307365-DEL-1972
chr11-64314532-DEL-51
chr11-64367313-DEL-82
chr11-64374636-DEL-360
chr11-64388712-DEL-89
chr11-64414309-DEL-99
chr11-64429484-DEL-52
chr11-64442688-DEL-108
chr11-64446900-DEL-66
chr11-64448410-DEL-596
chr11-64458893-DEL-164
chr11-64466789-DEL-75
chr11-64493888-DEL-4030
chr11-64511493-DEL-53
chr11-64546497-DEL-255
chr11-64568668-DEL-147
chr11-64577428-DEL-101
chr11-64595907-DEL-52
chr11-64597383-DEL-118
chr11-64598123-DEL-52
chr11-64604475-DEL-92
chr11-64633818-DEL-64
chr11-64680460-DEL-60
chr11-64689895-DEL-55
chr11-64732766-DEL-236
chr11-64738209-DEL-542
chr11-64767832-DEL-148
chr11-64775796-DEL-56
chr11-64779658-DEL-186
chr11-64782667-DEL-176
chr11-64784046-DEL-192
chr11-64799857-DEL-70
chr11-64811707-DEL-65
chr11-64812487-DEL-54
chr11-64860748-DEL-3771
chr11-64861726-DEL-1917
chr11-64879243-DEL-75
chr11-64911194-DEL-53
chr11-65129954-DEL-70
chr11-65156019-DEL-57
chr11-65187145-DEL-85
chr11-65201919-DEL-63
chr11-65225071-DEL-51
chr11-65241698-DEL-493
chr11-65249265-DEL-106
chr11-65295582-DEL-196
chr11-65300105-DEL-1064
chr11-65319828-DEL-1053
chr11-65324520-DEL-210

chr11-65380440-DEL-63
chr11-65396987-DEL-670
chr11-65405119-DEL-92
chr11-65425274-DEL-115
chr11-65443510-DEL-126
chr11-65549123-DEL-2405
chr11-65560914-DEL-117
chr11-65593751-DEL-65
chr11-65609608-DEL-80
chr11-65613262-DEL-1117
chr11-65650171-DEL-105
chr11-65682187-DEL-171
chr11-65745187-DEL-55
chr11-65755919-DEL-57
chr11-65758342-DEL-186
chr11-65760820-DEL-59
chr11-65781819-DEL-55
chr11-65787976-DEL-6481
chr11-65795488-DEL-57
chr11-65796782-DEL-52
chr11-65799366-DEL-1686
chr11-65864503-DEL-73
chr11-65882435-DEL-680
chr11-65934133-DEL-80
chr11-65957821-DEL-58
chr11-65958509-DEL-141
chr11-65989521-DEL-198
chr11-66060356-DEL-575
chr11-66129180-DEL-96
chr11-66162672-DEL-1343
chr11-66174769-DEL-183
chr11-66216563-DEL-73
chr11-66311796-DEL-162
chr11-66326890-DEL-52
chr11-66328980-DEL-300
chr11-66351542-DEL-6466
chr11-66361973-DEL-56
chr11-66363682-DEL-6194
chr11-66374820-DEL-674
chr11-66380989-DEL-1193
chr11-66388873-DEL-103
chr11-66421446-DEL-91
chr11-66432064-DEL-127
chr11-66444907-DEL-52
chr11-66499231-DEL-476
chr11-66508350-DEL-511
chr11-66555010-DEL-95
chr11-66619492-DEL-52
chr11-66633200-DEL-358
chr11-66656965-DEL-169
chr11-66668259-DEL-66
chr11-66670901-DEL-186
chr11-66699176-DEL-111
chr11-66704029-DEL-152
chr11-66711333-DEL-72
chr11-66734693-DEL-55
chr11-66735820-DEL-66
chr11-66786587-DEL-61
chr11-66797899-DEL-1045
chr11-66818847-DEL-576
chr11-66848932-DEL-191
chr11-66897551-DEL-440
chr11-66902556-DEL-167
chr11-66905124-DEL-756
chr11-66910519-DEL-134
chr11-66914204-DEL-87
chr11-66915554-DEL-161
chr11-66927645-DEL-187
chr11-66983509-DEL-146
chr11-67012221-DEL-171
chr11-67012524-DEL-233
chr11-67051030-DEL-1154
chr11-67051133-DEL-362
chr11-67052444-DEL-799
chr11-67067225-DEL-68
chr11-67116613-DEL-1901
chr11-67191478-DEL-199

chr11-67200308-DEL-838
chr11-67208078-DEL-97
chr11-67256587-DEL-138
chr11-67268972-DEL-153
chr11-67336747-DEL-66
chr11-67358902-DEL-53
chr11-67384481-DEL-274
chr11-67414351-DEL-81
chr11-67463308-DEL-105
chr11-67480622-DEL-126
chr11-67496676-DEL-343
chr11-67506919-DEL-79
chr11-67508715-DEL-88
chr11-67512123-DEL-142
chr11-67546940-DEL-64
chr11-67554063-DEL-96
chr11-67571821-DEL-98
chr11-67578758-DEL-624
chr11-67701013-DEL-222
chr11-67743621-DEL-179
chr11-67791020-DEL-116
chr11-67806932-DEL-54
chr11-67873561-DEL-102
chr11-67938714-DEL-423
chr11-67978457-DEL-194
chr11-68044113-DEL-80
chr11-68097232-DEL-391
chr11-68112760-DEL-72
chr11-68145204-DEL-738
chr11-68176186-DEL-64
chr11-68218747-DEL-62
chr11-68220968-DEL-298
chr11-68253852-DEL-84
chr11-68350755-DEL-54
chr11-68353221-DEL-220
chr11-68357372-DEL-63
chr11-68391617-DEL-54
chr11-68421180-DEL-208
chr11-68435229-DEL-65
chr11-68442504-DEL-77
chr11-68454954-DEL-62
chr11-68459407-DEL-225
chr11-68516807-DEL-1852
chr11-68529999-DEL-175
chr11-68534575-DEL-831
chr11-68537456-DEL-669
chr11-68540029-DEL-196
chr11-68559831-DEL-52
chr11-68614323-DEL-164
chr11-68620089-DEL-393
chr11-68671244-DEL-126
chr11-68687718-DEL-224
chr11-68691268-DEL-58
chr11-68701354-DEL-487
chr11-68701935-DEL-196
chr11-68714225-DEL-56
chr11-68716205-DEL-51
chr11-68728724-DEL-320
chr11-68791249-DEL-104
chr11-68800922-DEL-580
chr11-68805143-DEL-207
chr11-68816523-DEL-188
chr11-68846878-DEL-88
chr11-68860417-DEL-85
chr11-68902607-DEL-68
chr11-68903657-DEL-652
chr11-68984373-DEL-59
chr11-69004602-DEL-164
chr11-69014497-DEL-96
chr11-69023397-DEL-77
chr11-69052788-DEL-58
chr11-69072993-DEL-70
chr11-69083735-DEL-620
chr11-69104077-DEL-219
chr11-69136052-DEL-97
chr11-69141628-DEL-419
chr11-69174515-DEL-242

chr11-69178082-DEL-2318
chr11-69187223-DEL-147
chr11-69188379-DEL-86
chr11-69226463-DEL-191
chr11-69265440-DEL-102
chr11-69270077-DEL-156
chr11-69271929-DEL-956
chr11-69317908-DEL-109
chr11-69340437-DEL-128
chr11-69352306-DEL-94
chr11-69357940-DEL-171
chr11-69369339-DEL-83
chr11-69371475-DEL-1245
chr11-69441580-DEL-1690
chr11-69441598-DEL-590
chr11-69489303-DEL-177
chr11-69510423-DEL-217
chr11-69517589-DEL-80
chr11-69538031-DEL-213
chr11-69560668-DEL-77
chr11-69592829-DEL-1560
chr11-69596428-DEL-408
chr11-69605056-DEL-63
chr11-69615403-DEL-372
chr11-69617163-DEL-75
chr11-69638351-DEL-219
chr11-69683231-DEL-55
chr11-69747114-DEL-51
chr11-69748615-DEL-55
chr11-69752483-DEL-647
chr11-69766047-DEL-611
chr11-69845033-DEL-501
chr11-69849014-DEL-107
chr11-69868766-DEL-116
chr11-69894560-DEL-957
chr11-69924999-DEL-183
chr11-69943404-DEL-334
chr11-69962313-DEL-209
chr11-69974324-DEL-115
chr11-69977271-DEL-3262
chr11-70004435-DEL-1398
chr11-70012311-DEL-2217
chr11-70015909-DEL-372
chr11-70028144-DEL-341
chr11-70046226-DEL-196
chr11-70138907-DEL-81
chr11-70148544-DEL-54
chr11-70194103-DEL-400
chr11-70195678-DEL-838
chr11-70229290-DEL-236
chr11-70233587-DEL-53
chr11-70256460-DEL-268
chr11-70261602-DEL-200
chr11-70264554-DEL-151
chr11-70268743-DEL-57
chr11-70280874-DEL-477
chr11-70284314-DEL-101
chr11-70291334-DEL-588
chr11-70345503-DEL-80
chr11-70349136-DEL-216
chr11-70361348-DEL-135
chr11-70382962-DEL-534
chr11-70394178-DEL-55
chr11-70413506-DEL-59
chr11-70443955-DEL-53
chr11-70449193-DEL-208
chr11-70461597-DEL-66
chr11-70465889-DEL-77
chr11-70483931-DEL-81
chr11-70519636-DEL-374
chr11-70526891-DEL-316
chr11-70554291-DEL-92
chr11-70581697-DEL-210
chr11-70600211-DEL-84
chr11-70627662-DEL-126
chr11-70641762-DEL-89
chr11-70659476-DEL-500

chr11-70665271-DEL-292
chr11-70668847-DEL-169
chr11-70670880-DEL-98
chr11-70697164-DEL-197
chr11-70717561-DEL-1941
chr11-70730139-DEL-55
chr11-70758509-DEL-111
chr11-70798027-DEL-55
chr11-70824235-DEL-240
chr11-70891623-DEL-445
chr11-70911394-DEL-1362
chr11-70948379-DEL-359
chr11-70956545-DEL-1179
chr11-70957603-DEL-269
chr11-70957906-DEL-771
chr11-70959204-DEL-849
chr11-70988187-DEL-84
chr11-70992060-DEL-1586
chr11-71056726-DEL-66
chr11-71057126-DEL-1904
chr11-71061959-DEL-235
chr11-71068364-DEL-2667
chr11-71073767-DEL-149
chr11-71099171-DEL-5146
chr11-71104440-DEL-105
chr11-71164324-DEL-124
chr11-71176497-DEL-61
chr11-71187808-DEL-294
chr11-71190970-DEL-7448
chr11-71210897-DEL-873
chr11-71215657-DEL-1118
chr11-71216819-DEL-2836
chr11-71219411-DEL-1916
chr11-71252701-DEL-57
chr11-71263642-DEL-90
chr11-71264260-DEL-184
chr11-71351573-DEL-2402
chr11-71352442-DEL-99
chr11-71353596-DEL-94
chr11-71355928-DEL-278
chr11-71429056-DEL-2280
chr11-71492244-DEL-191
chr11-71498320-DEL-249
chr11-71557708-DEL-468
chr11-71558465-DEL-72
chr11-71582286-DEL-789
chr11-71623973-DEL-96
chr11-71689208-DEL-93
chr11-71760346-DEL-1391
chr11-71826356-DEL-99
chr11-71880535-DEL-63
chr11-71896407-DEL-52
chr11-71958844-DEL-1011
chr11-71993578-DEL-309
chr11-72033688-DEL-88
chr11-72073328-DEL-152
chr11-72075175-DEL-70
chr11-72078629-DEL-165
chr11-72103918-DEL-233
chr11-72139708-DEL-815
chr11-72140610-DEL-433
chr11-72165017-DEL-87
chr11-72183020-DEL-408
chr11-72184499-DEL-195
chr11-72187317-DEL-1837
chr11-72188613-DEL-2237
chr11-72202854-DEL-287
chr11-72256383-DEL-140
chr11-72264771-DEL-804
chr11-72278558-DEL-53
chr11-72283799-DEL-55
chr11-72289654-DEL-62
chr11-72296364-DEL-52
chr11-72319877-DEL-69
chr11-72339787-DEL-79
chr11-72374820-DEL-417
chr11-72402793-DEL-618

chr11-72404968-DEL-83
chr11-72409033-DEL-56
chr11-72430549-DEL-78
chr11-72439223-DEL-75
chr11-72452695-DEL-148
chr11-72554972-DEL-763
chr11-72583842-DEL-77
chr11-72608611-DEL-227
chr11-72626305-DEL-292
chr11-72626751-DEL-400
chr11-72652800-DEL-206
chr11-72656282-DEL-246
chr11-72656344-DEL-1407
chr11-72671390-DEL-193
chr11-72677296-DEL-546
chr11-72692620-DEL-207
chr11-72699177-DEL-1561
chr11-72726976-DEL-228
chr11-72728954-DEL-223
chr11-72740666-DEL-98
chr11-72793838-DEL-234
chr11-72812490-DEL-198
chr11-72823064-DEL-1439
chr11-72828707-DEL-1442
chr11-72942385-DEL-274
chr11-72994027-DEL-80
chr11-73036585-DEL-75
chr11-73052925-DEL-153
chr11-73091471-DEL-53
chr11-73095294-DEL-88
chr11-73100112-DEL-59
chr11-73107634-DEL-76
chr11-73115394-DEL-1104
chr11-73150162-DEL-56
chr11-73195613-DEL-92
chr11-73199149-DEL-258
chr11-73205997-DEL-552
chr11-73216210-DEL-127
chr11-73251919-DEL-8425
chr11-73260712-DEL-497
chr11-73263980-DEL-1146
chr11-73369625-DEL-177
chr11-73392248-DEL-57
chr11-73429677-DEL-40987
chr11-73476755-DEL-55
chr11-73492270-DEL-257
chr11-73496664-DEL-400
chr11-73510360-DEL-5503
chr11-73582340-DEL-558
chr11-73598733-DEL-413
chr11-73619402-DEL-6567
chr11-73633889-DEL-409
chr11-73657885-DEL-127
chr11-73662310-DEL-111
chr11-73675790-DEL-3384
chr11-73688467-DEL-61
chr11-73709113-DEL-388
chr11-73747226-DEL-27356
chr11-73750850-DEL-5993
chr11-73757563-DEL-5511
chr11-73796119-DEL-51
chr11-73812545-DEL-59
chr11-73823092-DEL-179
chr11-73862409-DEL-778
chr11-73863489-DEL-197
chr11-73865666-DEL-549
chr11-73866411-DEL-3049
chr11-73873650-DEL-547
chr11-73876342-DEL-517
chr11-73884355-DEL-3814
chr11-73888476-DEL-116
chr11-73896694-DEL-4713
chr11-73921039-DEL-235
chr11-73924423-DEL-1000
chr11-73930182-DEL-200
chr11-73988844-DEL-11574
chr11-73996707-DEL-1201

chr11-74038716-DEL-107302
chr11-74060758-DEL-417
chr11-74067632-DEL-172
chr11-74073539-DEL-63
chr11-74090609-DEL-2519
chr11-74118134-DEL-484
chr11-74123018-DEL-958
chr11-74125232-DEL-824
chr11-74150057-DEL-136
chr11-74209118-DEL-1350
chr11-74230329-DEL-550
chr11-74281311-DEL-123
chr11-74321417-DEL-362
chr11-74351904-DEL-345
chr11-74496163-DEL-54
chr11-74503133-DEL-65
chr11-74518147-DEL-179
chr11-74535020-DEL-57
chr11-74537803-DEL-64
chr11-74569511-DEL-55
chr11-74574942-DEL-56
chr11-74577474-DEL-156
chr11-74633919-DEL-60
chr11-74648424-DEL-200
chr11-74652773-DEL-82
chr11-74692206-DEL-221
chr11-74701545-DEL-333
chr11-74714283-DEL-71
chr11-74718634-DEL-62
chr11-74738367-DEL-174
chr11-74745854-DEL-210
chr11-74786285-DEL-73
chr11-74824454-DEL-856
chr11-74862997-DEL-197
chr11-74891205-DEL-54
chr11-74910175-DEL-262
chr11-74928250-DEL-74
chr11-74956680-DEL-995
chr11-74978283-DEL-1140
chr11-74978949-DEL-600
chr11-74982341-DEL-52
chr11-74996449-DEL-1066
chr11-75015539-DEL-58
chr11-75042997-DEL-389
chr11-75078158-DEL-183
chr11-75078898-DEL-198
chr11-75112006-DEL-82
chr11-75114549-DEL-66
chr11-75208245-DEL-66
chr11-75229505-DEL-63
chr11-75346640-DEL-77
chr11-75379643-DEL-57
chr11-75398710-DEL-153
chr11-75418295-DEL-1038
chr11-75456992-DEL-77
chr11-75467166-DEL-52
chr11-75472857-DEL-94
chr11-75529115-DEL-276
chr11-75555760-DEL-181
chr11-75577716-DEL-254
chr11-75599809-DEL-193
chr11-75614333-DEL-451
chr11-75626598-DEL-249
chr11-75633386-DEL-201
chr11-75635842-DEL-59
chr11-75680568-DEL-56
chr11-75691345-DEL-174
chr11-75696931-DEL-197
chr11-75702153-DEL-221
chr11-75710195-DEL-110
chr11-75716259-DEL-271
chr11-75717927-DEL-917
chr11-75719827-DEL-216
chr11-75766134-DEL-51
chr11-75768388-DEL-62
chr11-75807711-DEL-61
chr11-75831074-DEL-90

chr11-75834762-DEL-176
chr11-75839784-DEL-74
chr11-75870070-DEL-53
chr11-75872644-DEL-240
chr11-75874806-DEL-471
chr11-75898915-DEL-97
chr11-75947761-DEL-182
chr11-76008885-DEL-133
chr11-76017797-DEL-903
chr11-76018981-DEL-205
chr11-76065621-DEL-51
chr11-76068593-DEL-202
chr11-76084287-DEL-238
chr11-76094382-DEL-72
chr11-76111455-DEL-216
chr11-76116206-DEL-72
chr11-76154970-DEL-1130
chr11-76161287-DEL-301
chr11-76162539-DEL-52
chr11-76165784-DEL-102
chr11-76171142-DEL-72
chr11-76202643-DEL-6050
chr11-76205363-DEL-1512
chr11-76268958-DEL-66
chr11-76291773-DEL-206
chr11-76295758-DEL-520
chr11-76305431-DEL-100
chr11-76335052-DEL-157
chr11-76344134-DEL-201
chr11-76344894-DEL-69
chr11-76353477-DEL-112
chr11-76394184-DEL-146
chr11-76407581-DEL-67
chr11-76477347-DEL-363
chr11-76499734-DEL-857
chr11-76518767-DEL-630
chr11-76521011-DEL-91
chr11-76536333-DEL-76
chr11-76569220-DEL-58
chr11-76573938-DEL-650
chr11-76579718-DEL-60
chr11-76592297-DEL-169
chr11-76610197-DEL-121
chr11-76621424-DEL-84
chr11-76631313-DEL-68
chr11-76673541-DEL-260
chr11-76764478-DEL-58
chr11-76802471-DEL-97
chr11-76815500-DEL-90
chr11-76843558-DEL-51
chr11-76848711-DEL-165
chr11-76852327-DEL-1935
chr11-76854082-DEL-198
chr11-76868118-DEL-242
chr11-76884792-DEL-1699
chr11-76916847-DEL-122
chr11-76925724-DEL-86
chr11-76928411-DEL-1719
chr11-76929666-DEL-251
chr11-76963500-DEL-59
chr11-77004811-DEL-209
chr11-77017647-DEL-1242
chr11-77025104-DEL-466
chr11-77058906-DEL-280
chr11-77068786-DEL-206
chr11-77147527-DEL-333
chr11-77211208-DEL-400
chr11-77213955-DEL-804
chr11-77220968-DEL-86
chr11-77280593-DEL-206
chr11-77321983-DEL-512
chr11-77336723-DEL-118
chr11-77458796-DEL-1019
chr11-77484532-DEL-516
chr11-77485201-DEL-204
chr11-77503235-DEL-178
chr11-77506957-DEL-293

chr11-77533421-DEL-76
chr11-77534874-DEL-679
chr11-77540421-DEL-341
chr11-77540562-DEL-622
chr11-77549900-DEL-90
chr11-77554155-DEL-223
chr11-77556687-DEL-4512
chr11-77563132-DEL-191
chr11-77567926-DEL-144
chr11-77570122-DEL-209
chr11-77590585-DEL-416
chr11-77593862-DEL-1435
chr11-77597953-DEL-384
chr11-77617223-DEL-70
chr11-77618914-DEL-56
chr11-77628733-DEL-53
chr11-77636076-DEL-1932
chr11-77665843-DEL-184
chr11-77669394-DEL-348
chr11-77678267-DEL-513
chr11-77685514-DEL-201
chr11-77688388-DEL-87
chr11-77694601-DEL-60
chr11-77722822-DEL-110
chr11-77737994-DEL-72
chr11-77768528-DEL-63
chr11-77809245-DEL-204
chr11-77809818-DEL-96
chr11-77826557-DEL-72
chr11-77839886-DEL-103
chr11-77902567-DEL-212
chr11-77917456-DEL-72
chr11-77934106-DEL-173
chr11-77973433-DEL-172
chr11-77975824-DEL-546
chr11-77976125-DEL-174
chr11-77994091-DEL-64
chr11-77997837-DEL-161
chr11-78002324-DEL-56
chr11-78003950-DEL-189
chr11-78075332-DEL-205
chr11-78090410-DEL-214
chr11-78101006-DEL-64
chr11-78105921-DEL-141
chr11-78108095-DEL-103
chr11-78132347-DEL-2095
chr11-78134378-DEL-136
chr11-78140532-DEL-197
chr11-78150781-DEL-186
chr11-78172465-DEL-55
chr11-78193945-DEL-830
chr11-78206544-DEL-102
chr11-78275183-DEL-125
chr11-78319987-DEL-213
chr11-78351902-DEL-247
chr11-78366804-DEL-58
chr11-78372819-DEL-51
chr11-78384639-DEL-68
chr11-78539217-DEL-168
chr11-78548775-DEL-285
chr11-78559884-DEL-85
chr11-78561471-DEL-77
chr11-78605231-DEL-204
chr11-78607942-DEL-205
chr11-78622566-DEL-137
chr11-78633451-DEL-4103
chr11-78645516-DEL-57
chr11-78664675-DEL-70
chr11-78699741-DEL-66
chr11-78742200-DEL-92
chr11-78744088-DEL-548
chr11-78749042-DEL-64
chr11-78750298-DEL-81
chr11-78788233-DEL-336
chr11-78817863-DEL-102
chr11-78885990-DEL-186
chr11-78892358-DEL-74

chr11-78897952-DEL-213
chr11-78912785-DEL-220
chr11-78920411-DEL-100
chr11-78930792-DEL-108
chr11-78946345-DEL-51
chr11-78978127-DEL-110
chr11-78995698-DEL-221
chr11-79001929-DEL-260
chr11-79022678-DEL-54
chr11-79045603-DEL-108
chr11-79063568-DEL-70
chr11-79068520-DEL-316
chr11-79071083-DEL-72
chr11-79080586-DEL-181
chr11-79092769-DEL-407
chr11-79171722-DEL-90
chr11-79173783-DEL-84
chr11-79190815-DEL-69
chr11-79216038-DEL-201
chr11-79243841-DEL-124
chr11-79294481-DEL-89
chr11-79315114-DEL-680
chr11-79318076-DEL-405
chr11-79351338-DEL-55
chr11-79352809-DEL-125
chr11-79398997-DEL-1117
chr11-79429999-DEL-96
chr11-79435823-DEL-182
chr11-79476060-DEL-310
chr11-79478606-DEL-51
chr11-79487276-DEL-60
chr11-79492578-DEL-76
chr11-79502937-DEL-108
chr11-79524111-DEL-1983
chr11-79533031-DEL-83
chr11-79533868-DEL-86
chr11-79541889-DEL-750
chr11-79612563-DEL-182
chr11-79623175-DEL-80
chr11-79682173-DEL-423
chr11-79689875-DEL-227
chr11-79703193-DEL-54
chr11-79709932-DEL-110
chr11-79727317-DEL-343
chr11-79734623-DEL-197
chr11-79739445-DEL-2802
chr11-79773212-DEL-261
chr11-79817296-DEL-204
chr11-79818645-DEL-205
chr11-79830634-DEL-186
chr11-79840032-DEL-602
chr11-79860204-DEL-541
chr11-79862189-DEL-87
chr11-79871262-DEL-62
chr11-79891207-DEL-133
chr11-79897401-DEL-255
chr11-79901160-DEL-200
chr11-79907158-DEL-2836
chr11-79918855-DEL-359
chr11-79920203-DEL-206
chr11-79928707-DEL-92
chr11-79957101-DEL-370
chr11-79960962-DEL-271
chr11-79962081-DEL-84
chr11-79974929-DEL-60
chr11-79975413-DEL-162
chr11-79977219-DEL-197
chr11-79987394-DEL-914
chr11-80015549-DEL-206
chr11-80037233-DEL-198
chr11-80039619-DEL-65
chr11-80072762-DEL-1141
chr11-80073672-DEL-489
chr11-80095406-DEL-162
chr11-80099081-DEL-58
chr11-80112089-DEL-78
chr11-80118339-DEL-76

chr11-80149781-DEL-203
chr11-80158040-DEL-352
chr11-80169408-DEL-179
chr11-80170925-DEL-111
chr11-80179714-DEL-311
chr11-80181984-DEL-951
chr11-80223403-DEL-66
chr11-80234737-DEL-174
chr11-80235743-DEL-487
chr11-80239862-DEL-213
chr11-80247462-DEL-292
chr11-80257700-DEL-192
chr11-80262508-DEL-86
chr11-80298014-DEL-243
chr11-80314392-DEL-55
chr11-80315871-DEL-94
chr11-80321708-DEL-200
chr11-80322368-DEL-114
chr11-80343787-DEL-181
chr11-80356399-DEL-80
chr11-80357260-DEL-249
chr11-80376240-DEL-99
chr11-80385223-DEL-362
chr11-80429829-DEL-55
chr11-80440082-DEL-103
chr11-80443435-DEL-56
chr11-80476584-DEL-120
chr11-80488690-DEL-80
chr11-80490016-DEL-430
chr11-80493749-DEL-120
chr11-80501496-DEL-143
chr11-80526517-DEL-216
chr11-80574606-DEL-91
chr11-80616085-DEL-87
chr11-80625335-DEL-329
chr11-80627842-DEL-3365
chr11-80631084-DEL-105
chr11-80639299-DEL-66
chr11-80694943-DEL-594
chr11-80720383-DEL-148
chr11-80753513-DEL-112
chr11-80828408-DEL-60
chr11-80836468-DEL-64
chr11-80885652-DEL-684
chr11-80901286-DEL-52
chr11-80906500-DEL-340
chr11-80921039-DEL-483
chr11-80972097-DEL-68
chr11-81007553-DEL-54
chr11-81029191-DEL-2211
chr11-81041863-DEL-65
chr11-81048283-DEL-54
chr11-81070382-DEL-120
chr11-81073082-DEL-179
chr11-81127363-DEL-68
chr11-81147344-DEL-56
chr11-81150991-DEL-69
chr11-81163563-DEL-139
chr11-81193665-DEL-1837
chr11-81219695-DEL-51
chr11-81240042-DEL-1865
chr11-81243336-DEL-71
chr11-81280607-DEL-80
chr11-81302854-DEL-72
chr11-81329454-DEL-65
chr11-81339172-DEL-256
chr11-81353414-DEL-55
chr11-81376567-DEL-217
chr11-81397677-DEL-505
chr11-81413527-DEL-54
chr11-81434900-DEL-63
chr11-81452155-DEL-137
chr11-81453953-DEL-86
chr11-81458475-DEL-54
chr11-81459798-DEL-62
chr11-81474042-DEL-79
chr11-81481789-DEL-58

chr11-81506845-DEL-152
chr11-81511333-DEL-855
chr11-81543131-DEL-852
chr11-81566357-DEL-146
chr11-81608871-DEL-1300
chr11-81619415-DEL-51
chr11-81661711-DEL-53
chr11-81666821-DEL-177
chr11-81722054-DEL-70
chr11-81737418-DEL-755
chr11-81738136-DEL-138
chr11-81767607-DEL-60
chr11-81793661-DEL-89
chr11-81801205-DEL-293
chr11-81831893-DEL-1212
chr11-81851268-DEL-382
chr11-81875881-DEL-190
chr11-81893260-DEL-51
chr11-81904495-DEL-82
chr11-81931684-DEL-247
chr11-81949851-DEL-392
chr11-81964378-DEL-61
chr11-81970828-DEL-257
chr11-81996500-DEL-70
chr11-82002260-DEL-109
chr11-82012802-DEL-216
chr11-82015564-DEL-4086
chr11-82024713-DEL-1342
chr11-82031114-DEL-123
chr11-82032691-DEL-2555
chr11-82044278-DEL-864
chr11-82119726-DEL-86
chr11-82132201-DEL-2757
chr11-82168071-DEL-65
chr11-82177903-DEL-688
chr11-82264482-DEL-98
chr11-82267458-DEL-57
chr11-82272956-DEL-339
chr11-82359010-DEL-67
chr11-82407254-DEL-58
chr11-82448775-DEL-4392
chr11-82462301-DEL-66
chr11-82490626-DEL-75
chr11-82549084-DEL-455
chr11-82563642-DEL-59
chr11-82591450-DEL-332
chr11-82620404-DEL-60
chr11-82621961-DEL-71
chr11-82635173-DEL-87
chr11-82658359-DEL-221
chr11-82671078-DEL-430
chr11-82678730-DEL-182
chr11-82710645-DEL-97
chr11-82723166-DEL-199
chr11-82757124-DEL-61
chr11-82759261-DEL-139
chr11-82782435-DEL-65
chr11-82822408-DEL-153
chr11-82823691-DEL-51
chr11-82835115-DEL-78
chr11-82845569-DEL-57
chr11-82888481-DEL-335
chr11-82892757-DEL-66
chr11-82905576-DEL-359
chr11-82988239-DEL-6973
chr11-82988757-DEL-6973
chr11-83023042-DEL-64
chr11-83051701-DEL-74
chr11-83058081-DEL-58
chr11-83210848-DEL-1225
chr11-83381728-DEL-197
chr11-83385458-DEL-305
chr11-83397593-DEL-52
chr11-83408274-DEL-106
chr11-83456597-DEL-2864
chr11-83471065-DEL-58
chr11-83476134-DEL-55

chr11-83482053-DEL-1172
chr11-83501146-DEL-398
chr11-83510364-DEL-513
chr11-83583795-DEL-1543
chr11-83591701-DEL-2516
chr11-83597601-DEL-352
chr11-83603520-DEL-55
chr11-83720286-DEL-76
chr11-83728892-DEL-104
chr11-83730926-DEL-62
chr11-83740348-DEL-80
chr11-83753870-DEL-108
chr11-83767197-DEL-88
chr11-83767852-DEL-58
chr11-83808254-DEL-87
chr11-83821202-DEL-561
chr11-83868576-DEL-65
chr11-83887913-DEL-94
chr11-83932710-DEL-203
chr11-83936243-DEL-512
chr11-83944193-DEL-191
chr11-83949906-DEL-590
chr11-83960606-DEL-68
chr11-83966303-DEL-61
chr11-83972168-DEL-66
chr11-83978462-DEL-84
chr11-84042868-DEL-264
chr11-84053272-DEL-84
chr11-84065061-DEL-468
chr11-84088175-DEL-71
chr11-84102931-DEL-91
chr11-84130054-DEL-688
chr11-84131093-DEL-168
chr11-84148443-DEL-67
chr11-84184910-DEL-199
chr11-84333945-DEL-51
chr11-84346592-DEL-299
chr11-84366955-DEL-196
chr11-84422106-DEL-134
chr11-84436216-DEL-116
chr11-84463002-DEL-156
chr11-84478983-DEL-68
chr11-84525758-DEL-76
chr11-84636814-DEL-1244
chr11-84639940-DEL-125
chr11-84720591-DEL-104
chr11-84726616-DEL-273
chr11-84752188-DEL-54
chr11-84766090-DEL-720
chr11-84784362-DEL-479
chr11-84794157-DEL-1289
chr11-84806583-DEL-81
chr11-84815260-DEL-160
chr11-84829615-DEL-72
chr11-84866529-DEL-195
chr11-84867473-DEL-63
chr11-84867912-DEL-195
chr11-84869581-DEL-174
chr11-84882287-DEL-201
chr11-84890135-DEL-195
chr11-84932455-DEL-281
chr11-84943172-DEL-688
chr11-84948208-DEL-72
chr11-84950315-DEL-696
chr11-84964660-DEL-58
chr11-84976405-DEL-172
chr11-85023400-DEL-1255
chr11-85052559-DEL-865
chr11-85066446-DEL-52
chr11-85114379-DEL-210
chr11-85138520-DEL-91
chr11-85147395-DEL-187
chr11-85153004-DEL-939
chr11-85156435-DEL-124
chr11-85189203-DEL-64
chr11-85191910-DEL-196
chr11-85197727-DEL-66

chr11-85198704-DEL-364
chr11-85200028-DEL-535
chr11-85210261-DEL-68
chr11-85219086-DEL-216
chr11-85296991-DEL-81
chr11-85306914-DEL-4570
chr11-85328926-DEL-5970
chr11-85330263-DEL-2837
chr11-85336552-DEL-60
chr11-85338282-DEL-246
chr11-85349998-DEL-112
chr11-85356499-DEL-203
chr11-85393966-DEL-232
chr11-85395083-DEL-246
chr11-85395422-DEL-336
chr11-85409790-DEL-80
chr11-85482750-DEL-186
chr11-85486317-DEL-262
chr11-85493137-DEL-52
chr11-85494604-DEL-164
chr11-85498899-DEL-84
chr11-85499990-DEL-108
chr11-85539515-DEL-78
chr11-85564879-DEL-141
chr11-85567199-DEL-197
chr11-85607705-DEL-185
chr11-85618037-DEL-237
chr11-85662570-DEL-64
chr11-85693362-DEL-467
chr11-85719114-DEL-56
chr11-85722355-DEL-84
chr11-85758589-DEL-170
chr11-85762298-DEL-379
chr11-85810737-DEL-60
chr11-85823313-DEL-57
chr11-85833408-DEL-559
chr11-85913833-DEL-102
chr11-85920039-DEL-58
chr11-85935007-DEL-192
chr11-85960843-DEL-55
chr11-86004055-DEL-77
chr11-86009654-DEL-4793
chr11-86011070-DEL-85
chr11-86024484-DEL-1086
chr11-86056396-DEL-187
chr11-86068604-DEL-208
chr11-86152851-DEL-58
chr11-86208539-DEL-203
chr11-86211844-DEL-305
chr11-86239170-DEL-57
chr11-86330541-DEL-66
chr11-86340471-DEL-77
chr11-86346150-DEL-124
chr11-86364909-DEL-235
chr11-86373878-DEL-206
chr11-86397006-DEL-55
chr11-86397997-DEL-2376
chr11-86449501-DEL-165
chr11-86476922-DEL-66
chr11-86484630-DEL-1979
chr11-86509902-DEL-53
chr11-86511158-DEL-128
chr11-86521909-DEL-417
chr11-86564882-DEL-53
chr11-86598789-DEL-65
chr11-86614693-DEL-167
chr11-86617869-DEL-186
chr11-86623127-DEL-79
chr11-86635364-DEL-159
chr11-86665618-DEL-210
chr11-86668372-DEL-206
chr11-86673301-DEL-158
chr11-86690868-DEL-235
chr11-86694402-DEL-71
chr11-86715022-DEL-190
chr11-86746226-DEL-135
chr11-86755403-DEL-66

chr11-86756093-DEL-163
chr11-86771371-DEL-1771
chr11-86773384-DEL-199
chr11-86874877-DEL-54
chr11-86902490-DEL-380
chr11-86946463-DEL-106
chr11-86946758-DEL-706
chr11-86974201-DEL-200
chr11-87060118-DEL-700
chr11-87112384-DEL-52
chr11-87175041-DEL-57
chr11-87185409-DEL-11162
chr11-87202826-DEL-314
chr11-87210611-DEL-84
chr11-87221184-DEL-105
chr11-87243327-DEL-103
chr11-87253480-DEL-253
chr11-87260851-DEL-51
chr11-87268287-DEL-142
chr11-87274965-DEL-201
chr11-87275848-DEL-224
chr11-87283396-DEL-156
chr11-87301140-DEL-76
chr11-87314641-DEL-182
chr11-87322977-DEL-250
chr11-87326351-DEL-147
chr11-87343330-DEL-411
chr11-87345167-DEL-63
chr11-87355100-DEL-197
chr11-87365827-DEL-283
chr11-87368733-DEL-5003
chr11-87392284-DEL-615
chr11-87437136-DEL-760
chr11-87448782-DEL-895
chr11-87450433-DEL-732
chr11-87463652-DEL-81
chr11-87546367-DEL-325
chr11-87579559-DEL-60
chr11-87584210-DEL-2978
chr11-87610769-DEL-124
chr11-87629695-DEL-1596
chr11-87662638-DEL-159
chr11-87687655-DEL-75
chr11-87704185-DEL-375
chr11-87741233-DEL-83
chr11-87757400-DEL-456
chr11-87781004-DEL-51
chr11-87786144-DEL-335
chr11-87788757-DEL-69
chr11-87791334-DEL-202
chr11-87807623-DEL-93
chr11-87819236-DEL-82
chr11-87870885-DEL-74
chr11-87901433-DEL-142
chr11-87906355-DEL-2519
chr11-87934246-DEL-197
chr11-87952284-DEL-65
chr11-87993698-DEL-77
chr11-88003691-DEL-602
chr11-88025813-DEL-57
chr11-88052383-DEL-52
chr11-88054892-DEL-208
chr11-88076448-DEL-60
chr11-88081663-DEL-125
chr11-88099977-DEL-460
chr11-88120306-DEL-115
chr11-88128446-DEL-75
chr11-88145805-DEL-64
chr11-88149213-DEL-467
chr11-88170078-DEL-56
chr11-88177533-DEL-52
chr11-88227695-DEL-75
chr11-88300082-DEL-196
chr11-88303247-DEL-60
chr11-88386392-DEL-165
chr11-88398625-DEL-93
chr11-88430919-DEL-60

chr11-88454471-DEL-127
chr11-88485983-DEL-114
chr11-88549481-DEL-170
chr11-88562919-DEL-106
chr11-88579397-DEL-52
chr11-88631373-DEL-169
chr11-88671376-DEL-67
chr11-88673818-DEL-445
chr11-88677180-DEL-190
chr11-88691727-DEL-80
chr11-88700765-DEL-68
chr11-88709381-DEL-210
chr11-88756453-DEL-184
chr11-88764847-DEL-192
chr11-88768946-DEL-86
chr11-88797694-DEL-864
chr11-88801178-DEL-74
chr11-88806655-DEL-968
chr11-88811591-DEL-227
chr11-88816837-DEL-64
chr11-88828658-DEL-250
chr11-88833478-DEL-99
chr11-88838556-DEL-75
chr11-88848424-DEL-54
chr11-88860443-DEL-82
chr11-88873990-DEL-1041
chr11-88876428-DEL-99
chr11-88880813-DEL-198
chr11-88892726-DEL-217
chr11-88897332-DEL-64
chr11-88902501-DEL-62
chr11-88928297-DEL-164
chr11-88952941-DEL-115
chr11-88974123-DEL-107
chr11-88983334-DEL-95
chr11-89101387-DEL-70
chr11-89105228-DEL-58
chr11-89113349-DEL-83
chr11-89123025-DEL-92
chr11-89138549-DEL-58
chr11-89169932-DEL-98
chr11-89234768-DEL-84
chr11-89244620-DEL-399
chr11-89323649-DEL-52
chr11-89331118-DEL-52
chr11-89355109-DEL-76
chr11-89362961-DEL-186
chr11-89539579-DEL-201
chr11-89559959-DEL-184
chr11-89568999-DEL-83
chr11-89577305-DEL-52
chr11-89591785-DEL-51
chr11-89604279-DEL-1576
chr11-89606054-DEL-176
chr11-89652719-DEL-810
chr11-89677258-DEL-2279
chr11-89701729-DEL-66
chr11-89721468-DEL-906
chr11-89756694-DEL-300
chr11-89757951-DEL-68
chr11-89771276-DEL-56
chr11-89783890-DEL-57
chr11-89842938-DEL-2382
chr11-89851638-DEL-6243
chr11-89888403-DEL-166
chr11-89910803-DEL-68
chr11-89912926-DEL-57
chr11-89929647-DEL-449
chr11-89942166-DEL-75
chr11-89951506-DEL-52
chr11-89976380-DEL-126
chr11-89982628-DEL-72
chr11-89994831-DEL-133
chr11-90038577-DEL-376
chr11-90047610-DEL-1156
chr11-90052752-DEL-206
chr11-90064985-DEL-52

chr11-90074526-DEL-373
chr11-90096306-DEL-57
chr11-90104147-DEL-84
chr11-90108223-DEL-114
chr11-90108859-DEL-1180
chr11-90109826-DEL-487
chr11-90146207-DEL-62
chr11-90161950-DEL-74
chr11-90164495-DEL-80
chr11-90169576-DEL-67
chr11-90170803-DEL-125
chr11-90180371-DEL-62
chr11-90192874-DEL-236
chr11-90205800-DEL-76
chr11-90206269-DEL-181
chr11-90215277-DEL-57
chr11-90316486-DEL-295
chr11-90319726-DEL-1615
chr11-90346376-DEL-116
chr11-90384434-DEL-55
chr11-90389718-DEL-106
chr11-90398770-DEL-2404
chr11-90544786-DEL-1489
chr11-90554021-DEL-53
chr11-90562823-DEL-276
chr11-90571673-DEL-2143
chr11-90575142-DEL-73
chr11-90583070-DEL-199
chr11-90592663-DEL-493
chr11-90595881-DEL-60
chr11-90600965-DEL-54
chr11-90620987-DEL-52
chr11-90655562-DEL-126
chr11-90704439-DEL-368
chr11-90711565-DEL-940
chr11-90726847-DEL-139
chr11-90753593-DEL-220
chr11-90776715-DEL-537
chr11-90802640-DEL-1009
chr11-90803750-DEL-362
chr11-90804379-DEL-970
chr11-90850903-DEL-111
chr11-90861428-DEL-93
chr11-90869438-DEL-124
chr11-90881039-DEL-134
chr11-90903950-DEL-2559
chr11-90918287-DEL-362
chr11-90936665-DEL-52
chr11-90937067-DEL-3099
chr11-90940190-DEL-1345
chr11-90966251-DEL-102
chr11-90983708-DEL-217
chr11-90995892-DEL-886
chr11-91003034-DEL-53
chr11-91010513-DEL-437
chr11-91034904-DEL-502
chr11-91038293-DEL-86
chr11-91083948-DEL-240
chr11-91090261-DEL-155
chr11-91092169-DEL-60
chr11-91145443-DEL-52
chr11-91156656-DEL-788
chr11-91164823-DEL-66
chr11-91191003-DEL-65
chr11-91196555-DEL-54
chr11-91211849-DEL-64
chr11-91219420-DEL-74
chr11-91234172-DEL-1320
chr11-91238203-DEL-93
chr11-91291788-DEL-1602
chr11-91294035-DEL-249
chr11-91297634-DEL-1043
chr11-91320356-DEL-59
chr11-91340387-DEL-1192
chr11-91349514-DEL-167
chr11-91374133-DEL-74
chr11-91385283-DEL-781

chr11-91400772-DEL-2549
chr11-91409251-DEL-149
chr11-91410358-DEL-52
chr11-91428238-DEL-67
chr11-91468716-DEL-141
chr11-91494729-DEL-2590
chr11-91537379-DEL-3378
chr11-91560567-DEL-52
chr11-91579770-DEL-63
chr11-91585366-DEL-2054
chr11-91589891-DEL-76
chr11-91591190-DEL-328
chr11-91595372-DEL-66139
chr11-91607359-DEL-288
chr11-91634465-DEL-2215
chr11-91653324-DEL-646
chr11-91668470-DEL-365
chr11-91680349-DEL-426
chr11-91689955-DEL-109
chr11-91692902-DEL-189
chr11-91695083-DEL-475
chr11-91700737-DEL-206
chr11-91703537-DEL-698
chr11-91730506-DEL-85
chr11-91739954-DEL-1255
chr11-91751332-DEL-1417
chr11-91754062-DEL-184
chr11-91762409-DEL-64
chr11-91762691-DEL-83
chr11-91788520-DEL-79
chr11-91796042-DEL-1574
chr11-91806023-DEL-417
chr11-91807592-DEL-153
chr11-91860261-DEL-495
chr11-91891439-DEL-2374
chr11-91909050-DEL-781
chr11-91915978-DEL-196
chr11-91920973-DEL-171
chr11-91929197-DEL-133
chr11-91983840-DEL-135
chr11-92018743-DEL-123
chr11-92023391-DEL-1113
chr11-92027799-DEL-203
chr11-92055858-DEL-52
chr11-92058160-DEL-51
chr11-92074812-DEL-971
chr11-92093737-DEL-878
chr11-92134485-DEL-6244
chr11-92192319-DEL-879
chr11-92205574-DEL-175
chr11-92215980-DEL-156
chr11-92217514-DEL-68
chr11-92217520-DEL-944
chr11-92227572-DEL-1890
chr11-92235066-DEL-168
chr11-92276115-DEL-171
chr11-92277534-DEL-93
chr11-92278560-DEL-426
chr11-92296867-DEL-4351
chr11-92300983-DEL-466
chr11-92302598-DEL-105
chr11-92315827-DEL-1995
chr11-92327014-DEL-54
chr11-92369815-DEL-386
chr11-92406749-DEL-258
chr11-92407103-DEL-726
chr11-92415958-DEL-764
chr11-92422990-DEL-57
chr11-92452939-DEL-491
chr11-92455044-DEL-616
chr11-92465869-DEL-111
chr11-92485406-DEL-7040
chr11-92485839-DEL-624
chr11-92502847-DEL-79
chr11-92535345-DEL-1965
chr11-92571904-DEL-488
chr11-92598815-DEL-209

chr11-92606540-DEL-66
chr11-92611164-DEL-500
chr11-92628380-DEL-88
chr11-92682131-DEL-298
chr11-92819367-DEL-56
chr11-92868771-DEL-84
chr11-92870239-DEL-68
chr11-92888297-DEL-3083
chr11-92891347-DEL-509
chr11-92896066-DEL-72
chr11-92914490-DEL-269
chr11-92976273-DEL-158
chr11-92978240-DEL-207
chr11-93006426-DEL-280
chr11-93018256-DEL-110
chr11-93036488-DEL-62
chr11-93048192-DEL-5653
chr11-93110859-DEL-108
chr11-93129031-DEL-69
chr11-93202281-DEL-108
chr11-93262661-DEL-860
chr11-93310589-DEL-53
chr11-93318571-DEL-1886
chr11-93323145-DEL-69
chr11-93362174-DEL-84
chr11-93395530-DEL-225
chr11-93396394-DEL-106
chr11-93396847-DEL-375
chr11-93408194-DEL-108
chr11-93421371-DEL-474
chr11-93452343-DEL-90
chr11-93464325-DEL-270
chr11-93468259-DEL-51
chr11-93577887-DEL-92
chr11-93583506-DEL-369
chr11-93595286-DEL-60
chr11-93628889-DEL-6198
chr11-93721994-DEL-936
chr11-93753905-DEL-1565
chr11-93763565-DEL-813
chr11-93780791-DEL-88
chr11-93793266-DEL-119
chr11-93892369-DEL-79
chr11-93892552-DEL-641
chr11-93902965-DEL-55
chr11-93915919-DEL-63
chr11-93939586-DEL-2028
chr11-93985193-DEL-488
chr11-93994363-DEL-250
chr11-94000496-DEL-198
chr11-94001958-DEL-187
chr11-94008571-DEL-189
chr11-94044781-DEL-265
chr11-94065100-DEL-110
chr11-94074884-DEL-78
chr11-94093673-DEL-198
chr11-94123321-DEL-58
chr11-94131792-DEL-154
chr11-94162117-DEL-115
chr11-94168687-DEL-59
chr11-94187444-DEL-180
chr11-94213163-DEL-77
chr11-94216747-DEL-163
chr11-94260356-DEL-106
chr11-94282825-DEL-65
chr11-94326274-DEL-256
chr11-94380651-DEL-138
chr11-94393783-DEL-59
chr11-94401542-DEL-307
chr11-94404167-DEL-73
chr11-94416084-DEL-230
chr11-94467424-DEL-411
chr11-94483216-DEL-118
chr11-94485507-DEL-704
chr11-94500857-DEL-171
chr11-94503504-DEL-119
chr11-94506630-DEL-682

chr11-94507396-DEL-259
chr11-94539637-DEL-217
chr11-94548669-DEL-68
chr11-94584735-DEL-211
chr11-94641357-DEL-1032
chr11-94656863-DEL-1567
chr11-94691890-DEL-177
chr11-94707494-DEL-173
chr11-94761214-DEL-994
chr11-94784700-DEL-622
chr11-94790701-DEL-179
chr11-94802798-DEL-78
chr11-94816882-DEL-243
chr11-94855749-DEL-65
chr11-94871329-DEL-124
chr11-94874101-DEL-88
chr11-94902185-DEL-66
chr11-94904352-DEL-51
chr11-94921220-DEL-140
chr11-94923949-DEL-1256
chr11-94940636-DEL-198
chr11-94951277-DEL-71
chr11-94971166-DEL-77
chr11-95001780-DEL-196
chr11-95005476-DEL-102
chr11-95006923-DEL-147
chr11-95038052-DEL-63
chr11-95069860-DEL-65
chr11-95195714-DEL-752
chr11-95218800-DEL-170
chr11-95223874-DEL-102
chr11-95232962-DEL-55
chr11-95281855-DEL-56
chr11-95288277-DEL-70
chr11-95311054-DEL-170
chr11-95357884-DEL-339
chr11-95369051-DEL-197
chr11-95425652-DEL-65
chr11-95427925-DEL-118
chr11-95479100-DEL-63
chr11-95496385-DEL-918
chr11-95546395-DEL-731
chr11-95591976-DEL-204
chr11-95597823-DEL-89
chr11-95648341-DEL-209
chr11-95667161-DEL-139
chr11-95693633-DEL-186
chr11-95704332-DEL-60
chr11-95745433-DEL-427
chr11-95746743-DEL-170
chr11-95767723-DEL-112
chr11-95778934-DEL-84
chr11-95803404-DEL-121
chr11-95807182-DEL-55
chr11-95807700-DEL-84
chr11-95817749-DEL-688
chr11-95820446-DEL-192
chr11-95825288-DEL-965
chr11-95839915-DEL-2767
chr11-95844495-DEL-79
chr11-95904843-DEL-184
chr11-95913736-DEL-273
chr11-95920511-DEL-60
chr11-95922564-DEL-220
chr11-95931057-DEL-119
chr11-95943674-DEL-359
chr11-95946202-DEL-210
chr11-95957708-DEL-3081
chr11-95973233-DEL-195
chr11-95980181-DEL-96
chr11-95985337-DEL-174
chr11-95988897-DEL-134
chr11-95990337-DEL-386
chr11-96010040-DEL-223
chr11-96010623-DEL-163
chr11-96013210-DEL-185
chr11-96017763-DEL-110

chr11-96023921-DEL-61
chr11-96037215-DEL-143
chr11-96038911-DEL-71
chr11-96046531-DEL-176
chr11-96047058-DEL-83
chr11-96064547-DEL-315
chr11-96083805-DEL-60
chr11-96111491-DEL-1467
chr11-96120612-DEL-62
chr11-96125990-DEL-54
chr11-96131966-DEL-57
chr11-96284430-DEL-164
chr11-96294286-DEL-94
chr11-96309671-DEL-159
chr11-96328826-DEL-55
chr11-96393031-DEL-74
chr11-96414133-DEL-68
chr11-96463741-DEL-72
chr11-96478562-DEL-52
chr11-96488725-DEL-437
chr11-96669294-DEL-199
chr11-96701625-DEL-197
chr11-96756069-DEL-345
chr11-96757954-DEL-67
chr11-96822145-DEL-115
chr11-96837912-DEL-76
chr11-96847899-DEL-125
chr11-96906251-DEL-87
chr11-96982461-DEL-80
chr11-96984872-DEL-102
chr11-97002693-DEL-197
chr11-97003370-DEL-66
chr11-97033534-DEL-183
chr11-97073865-DEL-261
chr11-97085576-DEL-82
chr11-97127771-DEL-223
chr11-97129363-DEL-821
chr11-97138256-DEL-175
chr11-97180545-DEL-109
chr11-97191698-DEL-151
chr11-97205197-DEL-202
chr11-97212583-DEL-853
chr11-97230290-DEL-52
chr11-97245804-DEL-51
chr11-97267021-DEL-95
chr11-97269548-DEL-207
chr11-97273610-DEL-56
chr11-97304372-DEL-165
chr11-97321431-DEL-730
chr11-97371822-DEL-2512
chr11-97373860-DEL-113
chr11-97374642-DEL-63
chr11-97377364-DEL-221
chr11-97397055-DEL-109
chr11-97436471-DEL-71
chr11-97481757-DEL-132
chr11-97486877-DEL-105
chr11-97491537-DEL-82
chr11-97507627-DEL-77
chr11-97512188-DEL-168
chr11-97530141-DEL-298
chr11-97530743-DEL-296
chr11-97532042-DEL-74
chr11-97537576-DEL-222
chr11-97544067-DEL-435
chr11-97587996-DEL-203
chr11-97599600-DEL-71
chr11-97600592-DEL-57
chr11-97605281-DEL-233
chr11-97610099-DEL-356
chr11-97642743-DEL-110
chr11-97658840-DEL-477
chr11-97664636-DEL-168
chr11-97715135-DEL-75
chr11-97735413-DEL-100
chr11-97735989-DEL-350
chr11-97749378-DEL-82

chr11-97752127-DEL-170
chr11-97761458-DEL-989
chr11-97771635-DEL-424
chr11-97794515-DEL-60
chr11-97800528-DEL-754
chr11-97809256-DEL-77
chr11-97840192-DEL-102
chr11-97850504-DEL-51
chr11-97854021-DEL-145
chr11-97856891-DEL-62
chr11-97866621-DEL-58
chr11-97882608-DEL-60
chr11-97914991-DEL-181
chr11-97920439-DEL-74
chr11-97931556-DEL-172
chr11-97939098-DEL-117
chr11-97947188-DEL-187
chr11-97971698-DEL-77
chr11-98007193-DEL-116
chr11-98010350-DEL-54
chr11-98011502-DEL-106
chr11-98020282-DEL-150
chr11-98022825-DEL-486
chr11-98062259-DEL-52
chr11-98073205-DEL-439
chr11-98082366-DEL-208
chr11-98087940-DEL-179
chr11-98100719-DEL-197
chr11-98112027-DEL-189
chr11-98133839-DEL-212
chr11-98154750-DEL-191
chr11-98192703-DEL-62
chr11-98199854-DEL-108
chr11-98204218-DEL-138
chr11-98255925-DEL-170
chr11-98295100-DEL-201
chr11-98307870-DEL-75
chr11-98324366-DEL-75
chr11-98351432-DEL-90
chr11-98365340-DEL-92
chr11-98383467-DEL-57
chr11-98399196-DEL-172
chr11-98420825-DEL-1343
chr11-98461259-DEL-1514
chr11-98463853-DEL-177
chr11-98495487-DEL-51
chr11-98538717-DEL-199
chr11-98570151-DEL-60
chr11-98575478-DEL-215
chr11-98705683-DEL-73
chr11-98721669-DEL-170
chr11-98732593-DEL-207
chr11-98788778-DEL-258
chr11-98802557-DEL-74
chr11-98837514-DEL-552
chr11-98878855-DEL-1055
chr11-98881304-DEL-162
chr11-98897364-DEL-193
chr11-98898898-DEL-193
chr11-98906604-DEL-107
chr11-98911321-DEL-190
chr11-98920557-DEL-196
chr11-98926785-DEL-102
chr11-98933049-DEL-65
chr11-98949440-DEL-134
chr11-99006133-DEL-72
chr11-99021833-DEL-52
chr11-99050446-DEL-66
chr11-99070641-DEL-72
chr11-99076138-DEL-340
chr11-99092759-DEL-131
chr11-99132538-DEL-58
chr11-99149262-DEL-68
chr11-99178311-DEL-68
chr11-99231528-DEL-1610
chr11-99233963-DEL-971
chr11-99258940-DEL-258

chr11-99259518-DEL-123
chr11-99295509-DEL-156
chr11-99298783-DEL-204
chr11-99312288-DEL-125
chr11-99318099-DEL-134
chr11-99324296-DEL-871
chr11-99333740-DEL-302
chr11-99334987-DEL-3289
chr11-99375674-DEL-82
chr11-99419632-DEL-82
chr11-99428773-DEL-58
chr11-99435640-DEL-69
chr11-99444041-DEL-50
chr11-99465033-DEL-91
chr11-99487778-DEL-57
chr11-99489407-DEL-52
chr11-99491716-DEL-120
chr11-99525473-DEL-60
chr11-99529377-DEL-2307
chr11-99567927-DEL-783
chr11-99568879-DEL-1367
chr11-99587116-DEL-51224
chr11-99680083-DEL-207
chr11-99686279-DEL-5768
chr11-99687345-DEL-299
chr11-99710135-DEL-946
chr11-99711225-DEL-788
chr11-99712140-DEL-539
chr11-99712774-DEL-14753
chr11-99716660-DEL-636
chr11-99728053-DEL-177
chr11-99735572-DEL-102
chr11-99741968-DEL-57
chr11-99747118-DEL-912
chr11-99748289-DEL-66
chr11-99760927-DEL-254
chr11-99780333-DEL-208
chr11-99799759-DEL-390
chr11-99823707-DEL-1007
chr11-99839973-DEL-72
chr11-99981474-DEL-100
chr11-100001542-DEL-93
chr11-100076577-DEL-310
chr11-100079687-DEL-89
chr11-100084841-DEL-175
chr11-100102035-DEL-198
chr11-100107017-DEL-65
chr11-100121001-DEL-5080
chr11-100136252-DEL-60
chr11-100162310-DEL-120
chr11-100217473-DEL-247
chr11-100222609-DEL-78
chr11-100222816-DEL-7490
chr11-100237918-DEL-126
chr11-100255792-DEL-89
chr11-100263011-DEL-58
chr11-100272093-DEL-173
chr11-100309010-DEL-236
chr11-100310198-DEL-203
chr11-100335744-DEL-249
chr11-100364510-DEL-186
chr11-100378483-DEL-2504
chr11-100420287-DEL-116
chr11-100422277-DEL-80
chr11-100431047-DEL-206
chr11-100431602-DEL-52
chr11-100432855-DEL-180
chr11-100445817-DEL-100
chr11-100451972-DEL-83
chr11-100456479-DEL-60
chr11-100463942-DEL-62
chr11-100482709-DEL-181
chr11-100486089-DEL-68
chr11-100503479-DEL-106
chr11-100507660-DEL-173
chr11-100540503-DEL-162
chr11-100573512-DEL-113

chr11-100574818-DEL-81
chr11-100616750-DEL-237
chr11-100654675-DEL-52
chr11-100655590-DEL-87
chr11-100657086-DEL-64
chr11-100700139-DEL-52
chr11-100782784-DEL-66
chr11-100785759-DEL-71
chr11-100791848-DEL-52
chr11-100803336-DEL-204
chr11-100818676-DEL-184
chr11-100824412-DEL-134
chr11-100835579-DEL-52
chr11-100839495-DEL-198
chr11-100876350-DEL-83
chr11-100878140-DEL-254
chr11-100880377-DEL-111
chr11-100881101-DEL-590
chr11-100884135-DEL-171
chr11-100896075-DEL-70
chr11-100906581-DEL-147
chr11-100917678-DEL-101
chr11-100920257-DEL-54
chr11-100948179-DEL-153
chr11-100953304-DEL-52
chr11-100959820-DEL-57
chr11-101000916-DEL-1085
chr11-101004611-DEL-1444
chr11-101018871-DEL-185
chr11-101035095-DEL-224
chr11-101036034-DEL-57
chr11-101093503-DEL-92
chr11-101105463-DEL-63
chr11-101131258-DEL-58
chr11-101132419-DEL-169
chr11-101166645-DEL-76
chr11-101187561-DEL-950
chr11-101237699-DEL-1364
chr11-101247422-DEL-200
chr11-101253781-DEL-56
chr11-101274119-DEL-213
chr11-101275833-DEL-90
chr11-101277218-DEL-55
chr11-101284735-DEL-129
chr11-101299789-DEL-76
chr11-101305280-DEL-202
chr11-101309270-DEL-70
chr11-101349357-DEL-82
chr11-101372793-DEL-208
chr11-101377560-DEL-188
chr11-101392248-DEL-67
chr11-101428272-DEL-69
chr11-101431812-DEL-461
chr11-101461173-DEL-68
chr11-101464570-DEL-709
chr11-101476117-DEL-400
chr11-101479459-DEL-84
chr11-101495915-DEL-141
chr11-101502712-DEL-88
chr11-101511912-DEL-152
chr11-101524480-DEL-68
chr11-101532997-DEL-56
chr11-101549284-DEL-393
chr11-101556767-DEL-90
chr11-101576138-DEL-83
chr11-101593621-DEL-159
chr11-101609604-DEL-238
chr11-101614007-DEL-1662
chr11-101630735-DEL-699
chr11-101665510-DEL-376
chr11-101680795-DEL-99
chr11-101715388-DEL-109
chr11-101743987-DEL-54
chr11-101747597-DEL-51
chr11-101754046-DEL-92
chr11-101755850-DEL-747
chr11-101777985-DEL-57

chr11-101790170-DEL-53
chr11-101808822-DEL-84
chr11-101849238-DEL-125
chr11-101869993-DEL-68
chr11-101873581-DEL-126
chr11-101904558-DEL-86
chr11-101924524-DEL-104
chr11-101955692-DEL-72
chr11-101965419-DEL-1422
chr11-101987760-DEL-155
chr11-102004529-DEL-154
chr11-102005358-DEL-335
chr11-102006359-DEL-168
chr11-102008071-DEL-94
chr11-102019025-DEL-56
chr11-102026906-DEL-184
chr11-102028867-DEL-126
chr11-102049520-DEL-204
chr11-102051020-DEL-192
chr11-102061020-DEL-72
chr11-102148201-DEL-60
chr11-102189217-DEL-77
chr11-102192640-DEL-190
chr11-102219963-DEL-184
chr11-102223088-DEL-196
chr11-102231422-DEL-1262
chr11-102245562-DEL-162
chr11-102258021-DEL-207
chr11-102264193-DEL-62
chr11-102271697-DEL-64
chr11-102301346-DEL-204
chr11-102303840-DEL-64
chr11-102314520-DEL-55
chr11-102327713-DEL-84
chr11-102374315-DEL-213
chr11-102384844-DEL-201
chr11-102394789-DEL-60
chr11-102430523-DEL-82
chr11-102450498-DEL-67
chr11-102490097-DEL-70
chr11-102517029-DEL-188
chr11-102519805-DEL-205
chr11-102523952-DEL-602
chr11-102524577-DEL-1324
chr11-102548788-DEL-61
chr11-102549562-DEL-230
chr11-102583401-DEL-184
chr11-102590677-DEL-196
chr11-102592848-DEL-814
chr11-102605813-DEL-641
chr11-102609702-DEL-68
chr11-102699105-DEL-55
chr11-102738440-DEL-133
chr11-102745122-DEL-118
chr11-102754128-DEL-88
chr11-102763742-DEL-127
chr11-102763883-DEL-548
chr11-102781433-DEL-395
chr11-102788843-DEL-58
chr11-102806121-DEL-172
chr11-102816868-DEL-4837
chr11-102858060-DEL-59
chr11-102867112-DEL-368
chr11-102895836-DEL-81
chr11-102904471-DEL-811
chr11-102905046-DEL-475
chr11-102911156-DEL-560
chr11-102927439-DEL-222
chr11-102932432-DEL-206
chr11-102934775-DEL-71
chr11-102939396-DEL-195
chr11-102940177-DEL-308
chr11-102962772-DEL-77
chr11-103041683-DEL-105
chr11-103074295-DEL-64
chr11-103121451-DEL-74
chr11-103202245-DEL-104

chr11-103205513-DEL-68
chr11-103354850-DEL-169
chr11-103358748-DEL-934
chr11-103369531-DEL-6724
chr11-103399750-DEL-20459
chr11-103402276-DEL-181
chr11-103408335-DEL-340
chr11-103418467-DEL-57
chr11-103427770-DEL-145
chr11-103443861-DEL-159
chr11-103467471-DEL-63
chr11-103513484-DEL-273
chr11-103553981-DEL-54
chr11-103560926-DEL-78
chr11-103606526-DEL-189
chr11-103616720-DEL-374
chr11-103645569-DEL-155
chr11-103648124-DEL-201
chr11-103655339-DEL-52
chr11-103657041-DEL-64
chr11-103702030-DEL-67
chr11-103723874-DEL-62
chr11-103753652-DEL-86
chr11-103758195-DEL-90
chr11-103762869-DEL-62
chr11-103768263-DEL-208
chr11-103770530-DEL-210
chr11-103779202-DEL-209
chr11-103808121-DEL-843
chr11-103818862-DEL-141
chr11-103835443-DEL-66
chr11-103846369-DEL-219
chr11-103847901-DEL-186
chr11-103854500-DEL-66
chr11-103855628-DEL-222
chr11-103858927-DEL-196
chr11-103904688-DEL-186
chr11-103914649-DEL-65
chr11-103921232-DEL-59
chr11-103932692-DEL-112
chr11-104014980-DEL-180
chr11-104058804-DEL-83
chr11-104187007-DEL-82
chr11-104204775-DEL-132
chr11-104216385-DEL-119
chr11-104218670-DEL-52
chr11-104240680-DEL-204
chr11-104248954-DEL-128
chr11-104261596-DEL-199
chr11-104291898-DEL-266
chr11-104292216-DEL-1126
chr11-104299373-DEL-2951
chr11-104319314-DEL-187
chr11-104341848-DEL-196
chr11-104344128-DEL-66
chr11-104385781-DEL-262
chr11-104421087-DEL-162
chr11-104427049-DEL-64
chr11-104445685-DEL-423
chr11-104466637-DEL-55
chr11-104470968-DEL-51
chr11-104497509-DEL-122
chr11-104558968-DEL-66
chr11-104565395-DEL-3797
chr11-104578777-DEL-158
chr11-104598911-DEL-119
chr11-104621316-DEL-162
chr11-104638889-DEL-67
chr11-104639597-DEL-105
chr11-104705869-DEL-90
chr11-104724437-DEL-886
chr11-104763678-DEL-71
chr11-104808505-DEL-1706
chr11-104816763-DEL-196
chr11-104832045-DEL-253
chr11-104911495-DEL-443
chr11-104915542-DEL-180

chr11-104963976-DEL-1101
chr11-104978606-DEL-118
chr11-104993751-DEL-81
chr11-105012840-DEL-198
chr11-105045118-DEL-258
chr11-105051287-DEL-123
chr11-105053069-DEL-187
chr11-105053592-DEL-243
chr11-105068667-DEL-162
chr11-105074432-DEL-200
chr11-105094445-DEL-93
chr11-105161456-DEL-54
chr11-105162669-DEL-910
chr11-105166164-DEL-207
chr11-105180621-DEL-175
chr11-105206029-DEL-111
chr11-105217135-DEL-63
chr11-105230686-DEL-136
chr11-105264570-DEL-167
chr11-105271825-DEL-173
chr11-105316084-DEL-105
chr11-105352904-DEL-227
chr11-105392774-DEL-112
chr11-105393868-DEL-86
chr11-105445223-DEL-64
chr11-105446224-DEL-187
chr11-105498070-DEL-583
chr11-105542896-DEL-147
chr11-105545039-DEL-863
chr11-105594309-DEL-8846
chr11-105654512-DEL-1892
chr11-105699462-DEL-614
chr11-105720759-DEL-83
chr11-105729247-DEL-135
chr11-105730733-DEL-107
chr11-105738706-DEL-6475
chr11-105754979-DEL-314
chr11-105785879-DEL-54
chr11-105837272-DEL-108
chr11-105844922-DEL-646
chr11-105853485-DEL-487
chr11-105917026-DEL-196
chr11-105933934-DEL-68
chr11-105959473-DEL-263
chr11-105961207-DEL-150
chr11-105990519-DEL-61
chr11-106026508-DEL-740
chr11-106029447-DEL-133
chr11-106033646-DEL-2175
chr11-106037944-DEL-207
chr11-106079657-DEL-229
chr11-106101606-DEL-198
chr11-106128181-DEL-700
chr11-106193995-DEL-286
chr11-106200043-DEL-71
chr11-106224517-DEL-122
chr11-106245434-DEL-129
chr11-106270283-DEL-59
chr11-106327097-DEL-198
chr11-106335812-DEL-210
chr11-106364671-DEL-405
chr11-106388470-DEL-206
chr11-106390077-DEL-86
chr11-106398523-DEL-132
chr11-106399548-DEL-93
chr11-106407241-DEL-183
chr11-106409361-DEL-79
chr11-106428603-DEL-77
chr11-106430370-DEL-103
chr11-106440977-DEL-121
chr11-106469262-DEL-70
chr11-106474568-DEL-257
chr11-106498550-DEL-172
chr11-106504836-DEL-52
chr11-106509807-DEL-198
chr11-106514090-DEL-64
chr11-106516575-DEL-359

chr11-106518338-DEL-199
chr11-106521803-DEL-1927
chr11-106531894-DEL-183
chr11-106546996-DEL-189
chr11-106555937-DEL-247
chr11-106570785-DEL-64
chr11-106580308-DEL-70
chr11-106599615-DEL-83
chr11-106609796-DEL-224
chr11-106616301-DEL-86
chr11-106616451-DEL-614
chr11-106622834-DEL-59
chr11-106639779-DEL-201
chr11-106646098-DEL-69
chr11-106656583-DEL-790
chr11-106659338-DEL-999
chr11-106669388-DEL-76
chr11-106694374-DEL-791
chr11-106697438-DEL-108
chr11-106741884-DEL-201
chr11-106760218-DEL-91
chr11-106774499-DEL-72
chr11-106787994-DEL-360
chr11-106792672-DEL-207
chr11-106834527-DEL-4287
chr11-106834556-DEL-79
chr11-106834721-DEL-6033
chr11-106835800-DEL-1345
chr11-106838082-DEL-106
chr11-106838816-DEL-169
chr11-106839362-DEL-265
chr11-106841139-DEL-174
chr11-106854151-DEL-51
chr11-106861953-DEL-1851
chr11-106876475-DEL-210
chr11-106910132-DEL-310
chr11-106918199-DEL-72
chr11-106931994-DEL-352
chr11-106939280-DEL-2731
chr11-106939281-DEL-1677
chr11-106939438-DEL-731
chr11-106940359-DEL-105
chr11-106940856-DEL-843
chr11-106941749-DEL-159
chr11-106942296-DEL-56
chr11-106950576-DEL-223
chr11-106980404-DEL-1108
chr11-106981661-DEL-56
chr11-106983993-DEL-206
chr11-106987521-DEL-203
chr11-106995813-DEL-167
chr11-106996718-DEL-217
chr11-106997607-DEL-352
chr11-107029869-DEL-204
chr11-107032623-DEL-221
chr11-107042259-DEL-196
chr11-107049337-DEL-158
chr11-107063779-DEL-56
chr11-107087892-DEL-74
chr11-107088351-DEL-331
chr11-107153003-DEL-65
chr11-107156699-DEL-150
chr11-107170555-DEL-74
chr11-107186122-DEL-145
chr11-107220908-DEL-208
chr11-107237634-DEL-64
chr11-107243194-DEL-124
chr11-107287691-DEL-330
chr11-107318613-DEL-147
chr11-107334808-DEL-185
chr11-107386919-DEL-59
chr11-107396763-DEL-56
chr11-107397875-DEL-200
chr11-107400815-DEL-85
chr11-107418223-DEL-155
chr11-107444152-DEL-54
chr11-107448251-DEL-94

chr11-107455095-DEL-214
chr11-107457819-DEL-53
chr11-107463379-DEL-385
chr11-107492633-DEL-54
chr11-107502097-DEL-132
chr11-107511992-DEL-1243
chr11-107548508-DEL-114
chr11-107561115-DEL-63
chr11-107566424-DEL-280
chr11-107591885-DEL-89
chr11-107624630-DEL-59
chr11-107639799-DEL-136
chr11-107641021-DEL-165
chr11-107645424-DEL-75
chr11-107661851-DEL-55
chr11-107671952-DEL-196
chr11-107682523-DEL-93
chr11-107727863-DEL-689
chr11-107733425-DEL-1007
chr11-107747182-DEL-96
chr11-107749720-DEL-243
chr11-107770394-DEL-196
chr11-107791041-DEL-51
chr11-107821327-DEL-157
chr11-107907533-DEL-493
chr11-107919129-DEL-99
chr11-107955679-DEL-52
chr11-107961253-DEL-163
chr11-107992784-DEL-1087
chr11-108040639-DEL-105
chr11-108044000-DEL-72
chr11-108051460-DEL-57
chr11-108054331-DEL-56
chr11-108066034-DEL-151
chr11-108089007-DEL-90
chr11-108105984-DEL-91
chr11-108108679-DEL-317
chr11-108128642-DEL-84
chr11-108148269-DEL-147
chr11-108153130-DEL-304
chr11-108160806-DEL-1826
chr11-108194664-DEL-869
chr11-108206412-DEL-66
chr11-108364858-DEL-115
chr11-108406474-DEL-1084
chr11-108416452-DEL-1380
chr11-108417052-DEL-420
chr11-108417613-DEL-120
chr11-108432920-DEL-663
chr11-108455372-DEL-525
chr11-108485324-DEL-63
chr11-108487469-DEL-61
chr11-108522980-DEL-705
chr11-108571631-DEL-2638
chr11-108589402-DEL-54
chr11-108629247-DEL-1206
chr11-108660328-DEL-1701
chr11-108702359-DEL-105
chr11-108714471-DEL-660
chr11-108729763-DEL-53
chr11-108748466-DEL-128
chr11-108764327-DEL-89
chr11-108804976-DEL-52
chr11-108806498-DEL-69
chr11-108822547-DEL-117
chr11-108834150-DEL-204
chr11-108859643-DEL-105
chr11-108913708-DEL-2188
chr11-108919569-DEL-68
chr11-108961030-DEL-81
chr11-108966921-DEL-52
chr11-108980322-DEL-346
chr11-109000301-DEL-56
chr11-109040564-DEL-52
chr11-109055847-DEL-593
chr11-109055873-DEL-1167
chr11-109056533-DEL-65

chr11-109061391-DEL-108
chr11-109066021-DEL-89
chr11-109105335-DEL-51
chr11-109127623-DEL-155
chr11-109152424-DEL-86
chr11-109162085-DEL-867
chr11-109163004-DEL-116
chr11-109173052-DEL-135
chr11-109186285-DEL-100
chr11-109202416-DEL-399
chr11-109226351-DEL-58
chr11-109230109-DEL-52
chr11-109293604-DEL-69
chr11-109295158-DEL-199
chr11-109322198-DEL-231
chr11-109345591-DEL-52
chr11-109347617-DEL-250
chr11-109390346-DEL-108
chr11-109392809-DEL-51
chr11-109406692-DEL-455
chr11-109418353-DEL-59
chr11-109428651-DEL-68
chr11-109444698-DEL-54
chr11-109445926-DEL-88
chr11-109446617-DEL-113
chr11-109456022-DEL-170
chr11-109456302-DEL-350
chr11-109491259-DEL-560
chr11-109530002-DEL-763
chr11-109547337-DEL-65
chr11-109581633-DEL-243
chr11-109597900-DEL-108
chr11-109660401-DEL-241
chr11-109705765-DEL-52
chr11-109731072-DEL-109
chr11-109760558-DEL-215
chr11-109761878-DEL-872
chr11-109768950-DEL-144
chr11-109784342-DEL-279
chr11-109797550-DEL-64
chr11-109802666-DEL-401
chr11-109850705-DEL-446
chr11-109876915-DEL-107
chr11-109877217-DEL-412
chr11-109878683-DEL-386
chr11-109889120-DEL-1438
chr11-109898572-DEL-198
chr11-109910469-DEL-4401
chr11-109923470-DEL-336
chr11-109984514-DEL-105
chr11-109987327-DEL-69
chr11-110032656-DEL-667
chr11-110060143-DEL-379
chr11-110086269-DEL-101
chr11-110103526-DEL-132
chr11-110112689-DEL-67
chr11-110126597-DEL-515
chr11-110150369-DEL-164
chr11-110155712-DEL-86
chr11-110171609-DEL-1247
chr11-110213206-DEL-72
chr11-110224845-DEL-93
chr11-110235116-DEL-188
chr11-110248081-DEL-83
chr11-110269016-DEL-412
chr11-110274581-DEL-155
chr11-110277759-DEL-53
chr11-110279211-DEL-317
chr11-110310536-DEL-56
chr11-110313264-DEL-76
chr11-110338260-DEL-103
chr11-110345790-DEL-88
chr11-110358378-DEL-1176
chr11-110358704-DEL-742
chr11-110359295-DEL-60
chr11-110427662-DEL-600
chr11-110429675-DEL-79

chr11-110432325-DEL-152
chr11-110436806-DEL-227
chr11-110444043-DEL-1602
chr11-110450208-DEL-57
chr11-110467001-DEL-56
chr11-110474868-DEL-298
chr11-110502506-DEL-207
chr11-110519672-DEL-84
chr11-110525121-DEL-83
chr11-110537318-DEL-1056
chr11-110582006-DEL-248
chr11-110607515-DEL-64
chr11-110612992-DEL-220
chr11-110615979-DEL-86
chr11-110634414-DEL-6408
chr11-110680746-DEL-6139
chr11-110715800-DEL-118
chr11-110723333-DEL-101
chr11-110738115-DEL-95
chr11-110759096-DEL-127
chr11-110772655-DEL-98
chr11-110863404-DEL-449
chr11-110866684-DEL-730
chr11-110874725-DEL-118
chr11-110874776-DEL-1257
chr11-110897299-DEL-150
chr11-110909927-DEL-97
chr11-110930593-DEL-56
chr11-110949960-DEL-174
chr11-110974319-DEL-639
chr11-110999401-DEL-70
chr11-111006482-DEL-71
chr11-111094965-DEL-22247
chr11-111095926-DEL-580
chr11-111097310-DEL-166
chr11-111128448-DEL-128
chr11-111148963-DEL-179
chr11-111160923-DEL-182
chr11-111180410-DEL-124
chr11-111186790-DEL-95
chr11-111193535-DEL-399
chr11-111271514-DEL-58
chr11-111273359-DEL-135
chr11-111279254-DEL-768
chr11-111292152-DEL-428
chr11-111312722-DEL-583
chr11-111364870-DEL-517
chr11-111365568-DEL-5751
chr11-111398131-DEL-483
chr11-111417773-DEL-122
chr11-111434805-DEL-187
chr11-111482453-DEL-1134
chr11-111505350-DEL-119
chr11-111573453-DEL-67
chr11-111620358-DEL-56
chr11-111628307-DEL-74
chr11-111664190-DEL-394
chr11-111677904-DEL-766
chr11-111746520-DEL-171
chr11-111772253-DEL-61
chr11-111805755-DEL-79
chr11-111851377-DEL-112
chr11-111879670-DEL-73
chr11-111895315-DEL-744
chr11-111924699-DEL-53
chr11-111955128-DEL-54
chr11-111987627-DEL-63
chr11-112009613-DEL-229
chr11-112013474-DEL-52
chr11-112045057-DEL-200
chr11-112057222-DEL-213
chr11-112079530-DEL-210
chr11-112127883-DEL-111
chr11-112154344-DEL-94
chr11-112178165-DEL-278
chr11-112185224-DEL-51
chr11-112202430-DEL-206

chr11-112230250-DEL-96
chr11-112333790-DEL-130
chr11-112390331-DEL-803
chr11-112413258-DEL-101
chr11-112443003-DEL-140
chr11-112480552-DEL-605
chr11-112539234-DEL-57
chr11-112565379-DEL-182
chr11-112571630-DEL-105
chr11-112577108-DEL-579
chr11-112580558-DEL-92
chr11-112585086-DEL-89
chr11-112589515-DEL-70
chr11-112625678-DEL-306
chr11-112642144-DEL-903
chr11-112690698-DEL-191
chr11-112695465-DEL-1195
chr11-112720997-DEL-124
chr11-112753019-DEL-982
chr11-112777595-DEL-66
chr11-112778110-DEL-214
chr11-112838536-DEL-213
chr11-112891376-DEL-77
chr11-112898203-DEL-74
chr11-112909604-DEL-92
chr11-112914600-DEL-154
chr11-112920510-DEL-108
chr11-112939770-DEL-58
chr11-112957965-DEL-63
chr11-112985498-DEL-58
chr11-112988940-DEL-86
chr11-113003425-DEL-182
chr11-113067808-DEL-144
chr11-113083477-DEL-146
chr11-113132858-DEL-524
chr11-113139333-DEL-55
chr11-113142992-DEL-344
chr11-113156166-DEL-57
chr11-113191971-DEL-752
chr11-113261069-DEL-70
chr11-113267927-DEL-106
chr11-113282853-DEL-102
chr11-113286415-DEL-436
chr11-113298977-DEL-52
chr11-113312798-DEL-798
chr11-113314874-DEL-85
chr11-113324697-DEL-90
chr11-113332776-DEL-55
chr11-113383924-DEL-206
chr11-113400871-DEL-98
chr11-113403055-DEL-143
chr11-113405758-DEL-91
chr11-113413115-DEL-54
chr11-113415288-DEL-75
chr11-113434794-DEL-140
chr11-113438403-DEL-52
chr11-113527050-DEL-584
chr11-113561646-DEL-207
chr11-113565981-DEL-785
chr11-113569631-DEL-236
chr11-113681100-DEL-283
chr11-113705730-DEL-59
chr11-113735929-DEL-152
chr11-113761500-DEL-56
chr11-113770869-DEL-167
chr11-113869403-DEL-174
chr11-113896560-DEL-121
chr11-113936761-DEL-296
chr11-113939505-DEL-175
chr11-113946658-DEL-109
chr11-113972800-DEL-92
chr11-114016421-DEL-101
chr11-114038934-DEL-686
chr11-114045032-DEL-2561
chr11-114111376-DEL-67
chr11-114172981-DEL-96
chr11-114194272-DEL-51

chr11-114230297-DEL-78
chr11-114234271-DEL-51
chr11-114239725-DEL-96
chr11-114260444-DEL-118
chr11-114262719-DEL-78
chr11-114271020-DEL-52
chr11-114272064-DEL-142
chr11-114272984-DEL-225
chr11-114302909-DEL-99
chr11-114312512-DEL-399
chr11-114374430-DEL-218
chr11-114383513-DEL-359
chr11-114392720-DEL-110
chr11-114393718-DEL-55
chr11-114418680-DEL-205
chr11-114420943-DEL-99
chr11-114456485-DEL-738
chr11-114473611-DEL-399
chr11-114520367-DEL-63
chr11-114529553-DEL-697
chr11-114534523-DEL-184
chr11-114569075-DEL-90
chr11-114594520-DEL-76
chr11-114609727-DEL-126
chr11-114621033-DEL-72
chr11-114633367-DEL-54
chr11-114647203-DEL-205
chr11-114658567-DEL-137
chr11-114660561-DEL-254
chr11-114689848-DEL-203
chr11-114693499-DEL-52
chr11-114712962-DEL-67
chr11-114718580-DEL-87
chr11-114729731-DEL-81
chr11-114765406-DEL-934
chr11-114774702-DEL-52
chr11-114798078-DEL-63
chr11-114854902-DEL-397
chr11-114861464-DEL-1818
chr11-114864175-DEL-7303
chr11-114875459-DEL-89
chr11-114903194-DEL-59
chr11-114908205-DEL-151
chr11-114909699-DEL-58
chr11-114924726-DEL-505
chr11-114929761-DEL-232
chr11-114935363-DEL-505
chr11-114948925-DEL-144
chr11-114956959-DEL-230
chr11-114963409-DEL-226
chr11-114975997-DEL-1777
chr11-114988626-DEL-77
chr11-114998279-DEL-6551
chr11-115022367-DEL-51
chr11-115033285-DEL-73
chr11-115065317-DEL-160
chr11-115076059-DEL-80
chr11-115125782-DEL-158
chr11-115137159-DEL-55
chr11-115163264-DEL-90
chr11-115191240-DEL-80
chr11-115195316-DEL-2029
chr11-115204621-DEL-284
chr11-115258944-DEL-111
chr11-115267394-DEL-60
chr11-115271065-DEL-56
chr11-115314153-DEL-425
chr11-115329965-DEL-206
chr11-115362228-DEL-211
chr11-115368547-DEL-310
chr11-115377341-DEL-64
chr11-115378352-DEL-388
chr11-115384526-DEL-187
chr11-115386346-DEL-894
chr11-115407350-DEL-205
chr11-115412367-DEL-194
chr11-115421471-DEL-178

chr11-115430090-DEL-222
chr11-115432044-DEL-189
chr11-115438476-DEL-966
chr11-115448096-DEL-407
chr11-115449229-DEL-361
chr11-115452549-DEL-64
chr11-115462852-DEL-183
chr11-115466657-DEL-171
chr11-115472357-DEL-172
chr11-115480009-DEL-399
chr11-115512696-DEL-762
chr11-115513543-DEL-1030
chr11-115519488-DEL-217
chr11-115524424-DEL-56
chr11-115543481-DEL-127
chr11-115547254-DEL-60
chr11-115548534-DEL-416
chr11-115549161-DEL-80
chr11-115568588-DEL-427
chr11-115592497-DEL-201
chr11-115610967-DEL-96
chr11-115614892-DEL-129
chr11-115616158-DEL-193
chr11-115626132-DEL-206
chr11-115640800-DEL-99
chr11-115641830-DEL-188
chr11-115643220-DEL-86
chr11-115662324-DEL-214
chr11-115662353-DEL-2409
chr11-115666003-DEL-60
chr11-115681351-DEL-172
chr11-115732628-DEL-81
chr11-115769577-DEL-122
chr11-115811753-DEL-104
chr11-115833938-DEL-69
chr11-115847159-DEL-69
chr11-115892593-DEL-565
chr11-115910620-DEL-222
chr11-115919248-DEL-133
chr11-115934449-DEL-58
chr11-115981824-DEL-382
chr11-116006216-DEL-188
chr11-116014900-DEL-64
chr11-116028848-DEL-94
chr11-116033580-DEL-62
chr11-116049562-DEL-62
chr11-116087412-DEL-57
chr11-116098022-DEL-378
chr11-116107988-DEL-53
chr11-116115644-DEL-168
chr11-116126452-DEL-54
chr11-116129981-DEL-193
chr11-116149484-DEL-65
chr11-116224492-DEL-130
chr11-116248073-DEL-169
chr11-116317397-DEL-233
chr11-116331141-DEL-178
chr11-116340824-DEL-1796
chr11-116347656-DEL-180
chr11-116348127-DEL-240
chr11-116358554-DEL-70
chr11-116359167-DEL-222
chr11-116365549-DEL-198
chr11-116386544-DEL-203
chr11-116414348-DEL-207
chr11-116442016-DEL-221
chr11-116453720-DEL-638
chr11-116485761-DEL-96
chr11-116527584-DEL-195
chr11-116555992-DEL-88
chr11-116574324-DEL-200
chr11-116595154-DEL-102
chr11-116607569-DEL-585
chr11-116612082-DEL-78
chr11-116617521-DEL-54
chr11-116633911-DEL-1820
chr11-116638314-DEL-403

chr11-116676176-DEL-7881
chr11-116706707-DEL-92
chr11-116708069-DEL-626
chr11-116714962-DEL-211
chr11-116760404-DEL-85
chr11-116774493-DEL-52
chr11-116787958-DEL-3197
chr11-116797308-DEL-52
chr11-116820964-DEL-103
chr11-116831818-DEL-327
chr11-116835173-DEL-53
chr11-116838357-DEL-270
chr11-116840280-DEL-56
chr11-116842040-DEL-602
chr11-116852828-DEL-113
chr11-116857835-DEL-70
chr11-116859847-DEL-224
chr11-116862162-DEL-196
chr11-116869299-DEL-73
chr11-116880290-DEL-223
chr11-116907036-DEL-98
chr11-116911530-DEL-114
chr11-116934698-DEL-846
chr11-116943585-DEL-500
chr11-116944895-DEL-200
chr11-116958529-DEL-253
chr11-116964584-DEL-213
chr11-116966891-DEL-306
chr11-116973827-DEL-54
chr11-116986863-DEL-441
chr11-117001100-DEL-202
chr11-117003202-DEL-226
chr11-117004721-DEL-89
chr11-117033159-DEL-65
chr11-117036213-DEL-267
chr11-117044849-DEL-68
chr11-117048531-DEL-73
chr11-117056338-DEL-68
chr11-117083388-DEL-530
chr11-117087954-DEL-132
chr11-117092135-DEL-58
chr11-117103657-DEL-194
chr11-117123673-DEL-53
chr11-117165255-DEL-347
chr11-117191259-DEL-272
chr11-117198239-DEL-180
chr11-117226762-DEL-118
chr11-117231026-DEL-222
chr11-117232802-DEL-84
chr11-117272112-DEL-223
chr11-117283665-DEL-222
chr11-117287066-DEL-2770
chr11-117288186-DEL-196
chr11-117290551-DEL-56
chr11-117299837-DEL-2862
chr11-117315294-DEL-193
chr11-117323340-DEL-343
chr11-117329850-DEL-59
chr11-117338541-DEL-494
chr11-117342416-DEL-60
chr11-117342870-DEL-216
chr11-117365723-DEL-75
chr11-117379160-DEL-1848
chr11-117390626-DEL-5268
chr11-117408566-DEL-111
chr11-117427919-DEL-63
chr11-117430423-DEL-190
chr11-117435656-DEL-319
chr11-117458286-DEL-154
chr11-117499597-DEL-144
chr11-117511083-DEL-97
chr11-117520976-DEL-198
chr11-117521271-DEL-205
chr11-117522559-DEL-70
chr11-117525805-DEL-247
chr11-117526271-DEL-1988
chr11-117566815-DEL-197

chr11-117585245-DEL-194
chr11-117607152-DEL-57
chr11-117695192-DEL-60
chr11-117711346-DEL-1442
chr11-117722379-DEL-258
chr11-117734934-DEL-196
chr11-117745621-DEL-182
chr11-117748532-DEL-398
chr11-117788274-DEL-833
chr11-117789570-DEL-1861
chr11-117799459-DEL-179
chr11-117803989-DEL-86
chr11-117828018-DEL-207
chr11-117838265-DEL-67
chr11-117847721-DEL-99
chr11-117904437-DEL-501
chr11-117907877-DEL-77
chr11-117938212-DEL-1257
chr11-117950504-DEL-71
chr11-117956029-DEL-339
chr11-117980731-DEL-128
chr11-117993865-DEL-55
chr11-118020350-DEL-65
chr11-118028546-DEL-389
chr11-118034129-DEL-196
chr11-118035428-DEL-924
chr11-118046015-DEL-383
chr11-118050763-DEL-1379
chr11-118080778-DEL-66
chr11-118082078-DEL-360
chr11-118220080-DEL-441
chr11-118229885-DEL-777
chr11-118278672-DEL-124
chr11-118286279-DEL-86
chr11-118341167-DEL-116
chr11-118343007-DEL-51
chr11-118349339-DEL-51
chr11-118359098-DEL-136
chr11-118403075-DEL-52
chr11-118411617-DEL-153
chr11-118451538-DEL-58
chr11-118452731-DEL-104
chr11-118491799-DEL-151
chr11-118508888-DEL-112
chr11-118510901-DEL-124
chr11-118532234-DEL-187
chr11-118555737-DEL-109
chr11-118561730-DEL-56
chr11-118575049-DEL-65
chr11-118575692-DEL-172
chr11-118591517-DEL-55
chr11-118607218-DEL-276
chr11-118615118-DEL-144
chr11-118659152-DEL-73
chr11-118670655-DEL-66
chr11-118675677-DEL-242
chr11-118681213-DEL-557
chr11-118692012-DEL-231
chr11-118692310-DEL-297
chr11-118713826-DEL-199
chr11-118722040-DEL-59
chr11-118736194-DEL-1474
chr11-118753665-DEL-68
chr11-118754138-DEL-52
chr11-118777827-DEL-94
chr11-118785404-DEL-128
chr11-118822183-DEL-112
chr11-118861330-DEL-64
chr11-118903715-DEL-56
chr11-118904821-DEL-116
chr11-118905698-DEL-713
chr11-118908067-DEL-1194
chr11-118917415-DEL-103
chr11-118936344-DEL-58
chr11-118950743-DEL-234
chr11-118960435-DEL-166
chr11-118985835-DEL-72

chr11-119003450-DEL-165
chr11-119011076-DEL-58
chr11-119013773-DEL-69
chr11-119026718-DEL-152
chr11-119108602-DEL-109
chr11-119139104-DEL-177
chr11-119142444-DEL-104
chr11-119173272-DEL-353
chr11-119227935-DEL-64
chr11-119232559-DEL-198
chr11-119244471-DEL-120
chr11-119251271-DEL-182
chr11-119256590-DEL-192
chr11-119264127-DEL-179
chr11-119266110-DEL-96
chr11-119269457-DEL-140
chr11-119274143-DEL-190
chr11-119277659-DEL-81
chr11-119319617-DEL-172
chr11-119401530-DEL-126
chr11-119458297-DEL-96
chr11-119463673-DEL-579
chr11-119472946-DEL-370
chr11-119493109-DEL-70
chr11-119503978-DEL-256
chr11-119522756-DEL-100
chr11-119545144-DEL-188
chr11-119586551-DEL-2382
chr11-119606852-DEL-108
chr11-119642242-DEL-923
chr11-119654307-DEL-270
chr11-119654950-DEL-204
chr11-119669565-DEL-62
chr11-119670090-DEL-186
chr11-119704285-DEL-160
chr11-119750541-DEL-62
chr11-119755877-DEL-111
chr11-119778229-DEL-110
chr11-119802613-DEL-52
chr11-119818299-DEL-673
chr11-119903910-DEL-133
chr11-119919782-DEL-198
chr11-119922133-DEL-214
chr11-119925468-DEL-62
chr11-119980025-DEL-54
chr11-120022757-DEL-144
chr11-120046001-DEL-421
chr11-120046655-DEL-196
chr11-120053421-DEL-171
chr11-120068470-DEL-56
chr11-120073874-DEL-150
chr11-120095567-DEL-381
chr11-120099368-DEL-228
chr11-120103545-DEL-2084
chr11-120140075-DEL-975
chr11-120211367-DEL-440
chr11-120232806-DEL-420
chr11-120278180-DEL-808
chr11-120280336-DEL-51
chr11-120286308-DEL-840
chr11-120330744-DEL-222
chr11-120332926-DEL-401
chr11-120349658-DEL-60
chr11-120367469-DEL-169
chr11-120376948-DEL-153
chr11-120397448-DEL-208
chr11-120404816-DEL-70
chr11-120424586-DEL-338
chr11-120448689-DEL-144
chr11-120461229-DEL-149
chr11-120466904-DEL-61
chr11-120488085-DEL-60
chr11-120492579-DEL-120
chr11-120525461-DEL-866
chr11-120528924-DEL-55
chr11-120529753-DEL-64
chr11-120542499-DEL-79

chr11-120543524-DEL-153
chr11-120546427-DEL-54
chr11-120578830-DEL-78
chr11-120607112-DEL-199
chr11-120615145-DEL-62
chr11-120622370-DEL-85
chr11-120633760-DEL-377
chr11-120633937-DEL-611
chr11-120651723-DEL-438
chr11-120656313-DEL-75
chr11-120662427-DEL-367
chr11-120665475-DEL-738
chr11-120688859-DEL-745
chr11-120689848-DEL-63
chr11-120728565-DEL-205
chr11-120755150-DEL-220
chr11-120771047-DEL-378
chr11-120782196-DEL-1276
chr11-120783827-DEL-206
chr11-120787655-DEL-202
chr11-120788158-DEL-91
chr11-120827063-DEL-434
chr11-120830882-DEL-80
chr11-120872230-DEL-215
chr11-120915775-DEL-2180
chr11-120918079-DEL-91
chr11-120925402-DEL-99
chr11-120957818-DEL-241
chr11-121001356-DEL-74
chr11-121018149-DEL-666
chr11-121029740-DEL-57
chr11-121075468-DEL-338
chr11-121076065-DEL-75
chr11-121101766-DEL-55
chr11-121124908-DEL-879
chr11-121270893-DEL-57
chr11-121302671-DEL-1270
chr11-121360742-DEL-207
chr11-121444862-DEL-190
chr11-121456623-DEL-2736
chr11-121572657-DEL-410
chr11-121584252-DEL-137
chr11-121657691-DEL-60
chr11-121703833-DEL-400
chr11-121717527-DEL-65
chr11-121750617-DEL-420
chr11-121772305-DEL-107
chr12-3052332-DEL-7147
chr12-3083742-DEL-4574
chr12-3108209-DEL-10101
chr12-3108262-DEL-5117
chr12-3146604-DEL-53
chr12-3149367-DEL-107
chr12-3159909-DEL-203
chr12-3162431-DEL-65
chr12-3171125-DEL-53
chr12-3182259-DEL-112
chr12-3196851-DEL-169
chr12-3199244-DEL-73
chr12-3217358-DEL-158
chr12-3239598-DEL-90
chr12-3246550-DEL-343
chr12-3246919-DEL-7043
chr12-3257740-DEL-626
chr12-3275352-DEL-1892
chr12-3312456-DEL-129
chr12-3325801-DEL-95
chr12-3344000-DEL-53
chr12-3368872-DEL-70
chr12-3370287-DEL-123
chr12-3370859-DEL-484
chr12-3374246-DEL-66
chr12-3377632-DEL-83
chr12-3395899-DEL-52
chr12-3398231-DEL-123
chr12-3407405-DEL-183
chr12-3426091-DEL-159

chr12-3435905-DEL-61
chr12-3442350-DEL-51
chr12-3446042-DEL-209
chr12-3463877-DEL-63
chr12-3484868-DEL-205
chr12-3490690-DEL-53
chr12-3496768-DEL-56
chr12-3534997-DEL-470
chr12-3547186-DEL-275
chr12-3562451-DEL-53
chr12-3567516-DEL-166
chr12-3580159-DEL-87
chr12-3588190-DEL-811
chr12-3613318-DEL-1036
chr12-3619675-DEL-200
chr12-3628101-DEL-239
chr12-3632103-DEL-232
chr12-3633227-DEL-992
chr12-3639635-DEL-629
chr12-3643850-DEL-55
chr12-3645121-DEL-681
chr12-3678666-DEL-554
chr12-3805135-DEL-96
chr12-3808945-DEL-56
chr12-3862142-DEL-90
chr12-3879976-DEL-3078
chr12-3880784-DEL-210
chr12-3897077-DEL-156
chr12-3903342-DEL-96
chr12-3914460-DEL-700
chr12-3926397-DEL-116
chr12-3933062-DEL-108
chr12-3975123-DEL-111
chr12-3989058-DEL-1038
chr12-4022075-DEL-203
chr12-4023195-DEL-114
chr12-4057310-DEL-82
chr12-4108010-DEL-177
chr12-4113095-DEL-193
chr12-4113547-DEL-54
chr12-4121446-DEL-54
chr12-4123088-DEL-147
chr12-4139473-DEL-51
chr12-4151657-DEL-202
chr12-4159494-DEL-76
chr12-4160837-DEL-1468
chr12-4163371-DEL-725
chr12-4163415-DEL-167
chr12-4168261-DEL-231
chr12-4194672-DEL-28119
chr12-4202469-DEL-447
chr12-4205323-DEL-114
chr12-4212406-DEL-96
chr12-4218192-DEL-402
chr12-4222506-DEL-431
chr12-4236337-DEL-291
chr12-4237524-DEL-282
chr12-4254278-DEL-346
chr12-4257454-DEL-70
chr12-4269345-DEL-390
chr12-4269390-DEL-2672
chr12-4269429-DEL-1355
chr12-4270539-DEL-612
chr12-4271184-DEL-172
chr12-4272743-DEL-56
chr12-4277989-DEL-51
chr12-4322902-DEL-1036
chr12-4331597-DEL-77
chr12-4342465-DEL-457
chr12-4343249-DEL-293
chr12-4383621-DEL-141
chr12-4408549-DEL-191
chr12-4413641-DEL-200
chr12-4420454-DEL-199
chr12-4440596-DEL-139
chr12-4454498-DEL-413
chr12-4469029-DEL-67

chr12-4474494-DEL-78
chr12-4480932-DEL-52
chr12-4487226-DEL-96
chr12-4516710-DEL-204
chr12-4516743-DEL-815
chr12-4517256-DEL-510
chr12-4559129-DEL-54
chr12-4562124-DEL-56
chr12-4564623-DEL-54
chr12-4597304-DEL-242
chr12-4613488-DEL-61
chr12-4616932-DEL-60
chr12-4622384-DEL-201
chr12-4633483-DEL-212
chr12-4660830-DEL-64
chr12-4721880-DEL-55
chr12-4734319-DEL-730
chr12-4742999-DEL-97
chr12-4744956-DEL-323
chr12-4784145-DEL-422
chr12-4784840-DEL-77
chr12-4788005-DEL-109
chr12-4824741-DEL-104
chr12-4829396-DEL-587
chr12-4845260-DEL-264
chr12-4847610-DEL-58
chr12-4865507-DEL-150
chr12-4893471-DEL-93
chr12-4899697-DEL-87
chr12-4909067-DEL-204
chr12-4915463-DEL-81
chr12-4927639-DEL-63
chr12-4932694-DEL-70
chr12-4934559-DEL-60
chr12-4950833-DEL-51
chr12-4957568-DEL-53
chr12-4964158-DEL-122
chr12-4977249-DEL-179
chr12-5008257-DEL-148
chr12-5013403-DEL-209
chr12-5038748-DEL-248
chr12-5098365-DEL-70
chr12-5100828-DEL-124
chr12-5102040-DEL-2974
chr12-5105282-DEL-123
chr12-5108207-DEL-1239
chr12-5109374-DEL-155
chr12-5111123-DEL-342
chr12-5113411-DEL-351
chr12-5162638-DEL-188
chr12-5344455-DEL-357
chr12-5357118-DEL-62
chr12-5359527-DEL-55
chr12-5430095-DEL-626
chr12-5453355-DEL-79
chr12-5462639-DEL-54
chr12-5467552-DEL-66
chr12-5473494-DEL-57
chr12-5547702-DEL-81
chr12-5556540-DEL-144
chr12-5560726-DEL-1103
chr12-5560727-DEL-51
chr12-5569201-DEL-985
chr12-5579935-DEL-63
chr12-5618090-DEL-118
chr12-5624729-DEL-56
chr12-5635814-DEL-250
chr12-5729157-DEL-83
chr12-5740156-DEL-135
chr12-5754687-DEL-132
chr12-5773787-DEL-119
chr12-5790899-DEL-1137
chr12-5803143-DEL-332
chr12-5822436-DEL-73
chr12-5830430-DEL-73
chr12-5856229-DEL-61
chr12-5910919-DEL-554

chr12-5921743-DEL-90
chr12-5946975-DEL-106
chr12-5978207-DEL-206
chr12-6014494-DEL-795
chr12-6015384-DEL-442
chr12-6021732-DEL-1317
chr12-6071460-DEL-54
chr12-6118421-DEL-84
chr12-6224822-DEL-630
chr12-6226243-DEL-112
chr12-6244454-DEL-1473
chr12-6262378-DEL-1857
chr12-6273565-DEL-784
chr12-6284926-DEL-7617
chr12-6329289-DEL-400
chr12-6369925-DEL-127
chr12-6382689-DEL-6248
chr12-6386346-DEL-2383
chr12-6399496-DEL-351
chr12-6401203-DEL-86
chr12-6402130-DEL-60
chr12-6406666-DEL-64
chr12-6421863-DEL-88
chr12-6434347-DEL-58
chr12-6446888-DEL-348
chr12-6478541-DEL-2161
chr12-6482410-DEL-582
chr12-6497068-DEL-429
chr12-6562956-DEL-13481
chr12-6647951-DEL-180
chr12-6651064-DEL-54
chr12-6662599-DEL-59
chr12-6674420-DEL-583
chr12-6676423-DEL-109
chr12-6699206-DEL-2492
chr12-6708459-DEL-80
chr12-6712141-DEL-443
chr12-6750009-DEL-436
chr12-6804542-DEL-128
chr12-6835647-DEL-54
chr12-6844072-DEL-51
chr12-6847325-DEL-399
chr12-6893192-DEL-3991
chr12-6912350-DEL-68
chr12-6920634-DEL-86
chr12-6924257-DEL-53
chr12-6948332-DEL-209
chr12-6953501-DEL-66
chr12-6978292-DEL-114
chr12-7010156-DEL-72
chr12-7037487-DEL-993
chr12-7043767-DEL-54
chr12-7050986-DEL-310
chr12-7051727-DEL-75
chr12-7060500-DEL-611
chr12-7062295-DEL-359
chr12-7068113-DEL-359
chr12-7099756-DEL-225
chr12-7155185-DEL-101
chr12-7191140-DEL-1792
chr12-7192478-DEL-351
chr12-7193228-DEL-8007
chr12-7207784-DEL-785
chr12-7270437-DEL-516
chr12-7273768-DEL-51
chr12-7296093-DEL-303
chr12-7314883-DEL-74
chr12-7336537-DEL-383
chr12-7358934-DEL-198
chr12-7363466-DEL-133
chr12-7367507-DEL-839
chr12-7383330-DEL-62
chr12-7393772-DEL-1122
chr12-7410104-DEL-100
chr12-7446298-DEL-56
chr12-7548712-DEL-179
chr12-7569466-DEL-60

chr12-7577263-DEL-65
chr12-7695337-DEL-194
chr12-7715098-DEL-167
chr12-7722890-DEL-57
chr12-7723640-DEL-552
chr12-7770343-DEL-62
chr12-7804588-DEL-104
chr12-7852234-DEL-212
chr12-7868819-DEL-264
chr12-7888132-DEL-723
chr12-7899658-DEL-2331
chr12-7909674-DEL-2004
chr12-7915385-DEL-797
chr12-7915406-DEL-99
chr12-7940117-DEL-124
chr12-7949651-DEL-502
chr12-7951189-DEL-1553
chr12-7982931-DEL-479
chr12-7988284-DEL-1538
chr12-8001091-DEL-57
chr12-8138487-DEL-226
chr12-8151220-DEL-58
chr12-8153808-DEL-128
chr12-8183290-DEL-1933
chr12-8211342-DEL-105
chr12-8228859-DEL-227
chr12-8235871-DEL-324
chr12-8256929-DEL-66
chr12-8259664-DEL-204
chr12-8275261-DEL-58
chr12-8307407-DEL-1950
chr12-8343688-DEL-58
chr12-8354550-DEL-213
chr12-8401260-DEL-68
chr12-8418092-DEL-146
chr12-8426512-DEL-66
chr12-8431579-DEL-961
chr12-8449570-DEL-56
chr12-8471975-DEL-53
chr12-8483541-DEL-59
chr12-8497946-DEL-64
chr12-8509509-DEL-108
chr12-8519183-DEL-233
chr12-8536900-DEL-97
chr12-8539646-DEL-608
chr12-8539711-DEL-1727
chr12-8540329-DEL-1242
chr12-8541186-DEL-385
chr12-8553068-DEL-200
chr12-8556222-DEL-84
chr12-8561903-DEL-219
chr12-8564041-DEL-319
chr12-8593164-DEL-58
chr12-8601578-DEL-819
chr12-8615262-DEL-106
chr12-8645862-DEL-311
chr12-8648666-DEL-1049
chr12-8649857-DEL-116
chr12-8657293-DEL-118
chr12-8658234-DEL-358
chr12-8658522-DEL-769
chr12-8659604-DEL-6680
chr12-8667156-DEL-141
chr12-8703143-DEL-57
chr12-8707738-DEL-88
chr12-8714964-DEL-209
chr12-8718419-DEL-218
chr12-8755742-DEL-56
chr12-8795024-DEL-81
chr12-8805123-DEL-744
chr12-8865126-DEL-65
chr12-8871007-DEL-69
chr12-8892725-DEL-72
chr12-8900541-DEL-80
chr12-8910587-DEL-54
chr12-8942402-DEL-52
chr12-8951674-DEL-175

chr12-8961526-DEL-62
chr12-8965669-DEL-276
chr12-8980682-DEL-387
chr12-9005840-DEL-151
chr12-9009455-DEL-51
chr12-9014292-DEL-1206
chr12-9014575-DEL-759
chr12-9015169-DEL-113
chr12-9016067-DEL-130
chr12-9018221-DEL-342
chr12-9054806-DEL-280
chr12-9095431-DEL-66
chr12-9125909-DEL-60
chr12-9135602-DEL-90
chr12-9177100-DEL-79
chr12-9193064-DEL-56
chr12-9222560-DEL-164
chr12-9226129-DEL-54
chr12-9249780-DEL-727
chr12-9266599-DEL-541
chr12-9280400-DEL-170
chr12-9296278-DEL-64
chr12-9298057-DEL-110
chr12-9304799-DEL-106
chr12-9358970-DEL-170
chr12-9391807-DEL-102
chr12-9393905-DEL-52
chr12-9448501-DEL-197
chr12-9475252-DEL-581
chr12-9475513-DEL-1769
chr12-9475830-DEL-131
chr12-9476220-DEL-162
chr12-9476599-DEL-402
chr12-9493832-DEL-1315
chr12-9494412-DEL-345
chr12-9495338-DEL-168
chr12-9509411-DEL-156
chr12-9585464-DEL-198
chr12-9674846-DEL-8387
chr12-9683174-DEL-55
chr12-9689701-DEL-1286
chr12-9691795-DEL-323
chr12-9698604-DEL-1358
chr12-9733266-DEL-212
chr12-9740098-DEL-1654
chr12-9750294-DEL-4978
chr12-9769105-DEL-1183
chr12-9792422-DEL-61
chr12-9833141-DEL-2025
chr12-9873671-DEL-222
chr12-9877760-DEL-229
chr12-9884470-DEL-506
chr12-9886093-DEL-399
chr12-9914848-DEL-66
chr12-9924855-DEL-57
chr12-9925808-DEL-158
chr12-9962245-DEL-129
chr12-9964632-DEL-80
chr12-10001631-DEL-52
chr12-10012920-DEL-3951
chr12-10029992-DEL-72
chr12-10092269-DEL-71
chr12-10228850-DEL-107
chr12-10234703-DEL-1467
chr12-10240552-DEL-914
chr12-10324111-DEL-211
chr12-10341442-DEL-284
chr12-10368874-DEL-254
chr12-10443120-DEL-65
chr12-10464878-DEL-142
chr12-10465603-DEL-1926
chr12-10499589-DEL-304
chr12-10535229-DEL-1459
chr12-10596029-DEL-155
chr12-10747918-DEL-2634
chr12-10759847-DEL-1144
chr12-10784277-DEL-1341

chr12-10814539-DEL-56
chr12-10815164-DEL-223
chr12-10826470-DEL-918
chr12-10835410-DEL-92
chr12-10857595-DEL-558
chr12-10867279-DEL-80
chr12-10872819-DEL-56
chr12-10879588-DEL-64
chr12-10906249-DEL-55
chr12-10912680-DEL-742
chr12-10962099-DEL-14723
chr12-10979817-DEL-204
chr12-10982549-DEL-333
chr12-11007464-DEL-550
chr12-11021152-DEL-6640
chr12-11038621-DEL-6323
chr12-11053380-DEL-1389
chr12-11059888-DEL-97
chr12-11116248-DEL-160
chr12-11126870-DEL-89
chr12-11184826-DEL-85
chr12-11229105-DEL-1457
chr12-11284212-DEL-1388
chr12-11302131-DEL-221
chr12-11305410-DEL-189
chr12-11354991-DEL-180
chr12-11379245-DEL-136
chr12-11386245-DEL-185
chr12-11400846-DEL-72
chr12-11402975-DEL-228
chr12-11478894-DEL-404
chr12-11496441-DEL-235
chr12-11513081-DEL-115
chr12-11516380-DEL-259
chr12-11528513-DEL-414
chr12-11542064-DEL-6293
chr12-11556214-DEL-6478
chr12-11594666-DEL-4710
chr12-11598454-DEL-212
chr12-11624228-DEL-70
chr12-11631534-DEL-704
chr12-11637007-DEL-297
chr12-11639059-DEL-400
chr12-11642151-DEL-6249
chr12-11655098-DEL-595
chr12-11656905-DEL-111
chr12-11667676-DEL-138
chr12-11739792-DEL-80
chr12-11742757-DEL-95
chr12-11765713-DEL-134
chr12-11766497-DEL-395
chr12-11779190-DEL-233
chr12-11816123-DEL-1060
chr12-11827171-DEL-1448
chr12-11865777-DEL-218
chr12-11873616-DEL-1388
chr12-11873941-DEL-52
chr12-11883766-DEL-165
chr12-11900658-DEL-83
chr12-11909207-DEL-83
chr12-11914109-DEL-139
chr12-11917987-DEL-194
chr12-11922999-DEL-60
chr12-11941835-DEL-66
chr12-11969722-DEL-1297
chr12-11971617-DEL-1176
chr12-11974770-DEL-194
chr12-11992226-DEL-100
chr12-12000174-DEL-3124
chr12-12097261-DEL-177
chr12-12098105-DEL-102
chr12-12115436-DEL-111
chr12-12160717-DEL-279
chr12-12201274-DEL-1339
chr12-12232152-DEL-80
chr12-12237491-DEL-280
chr12-12299323-DEL-66

chr12-12310743-DEL-102
chr12-12349060-DEL-231
chr12-12375998-DEL-184
chr12-12380665-DEL-2964
chr12-12400706-DEL-83
chr12-12427777-DEL-338
chr12-12455396-DEL-146
chr12-12561231-DEL-94
chr12-12601628-DEL-66
chr12-12633782-DEL-56
chr12-12685336-DEL-57
chr12-12692023-DEL-88
chr12-12736447-DEL-952
chr12-12750024-DEL-678
chr12-12811764-DEL-365
chr12-12822934-DEL-83
chr12-12854661-DEL-78
chr12-12866138-DEL-70
chr12-12901261-DEL-60
chr12-12961302-DEL-1343
chr12-12963692-DEL-79
chr12-12994636-DEL-60
chr12-12997502-DEL-56
chr12-13007313-DEL-397
chr12-13014363-DEL-51
chr12-13021258-DEL-56
chr12-13026978-DEL-74
chr12-13032552-DEL-56
chr12-13071504-DEL-1092
chr12-13075778-DEL-232
chr12-13101841-DEL-82
chr12-13112484-DEL-1885
chr12-13202163-DEL-576
chr12-13241921-DEL-148
chr12-13263035-DEL-200
chr12-13302535-DEL-554
chr12-13309397-DEL-179
chr12-13313095-DEL-52
chr12-13342741-DEL-60
chr12-13352746-DEL-175
chr12-13386798-DEL-63
chr12-13388128-DEL-120
chr12-13426493-DEL-118
chr12-13438807-DEL-183
chr12-13475462-DEL-212
chr12-13504657-DEL-1353
chr12-13506090-DEL-668
chr12-13511125-DEL-56
chr12-13517777-DEL-85
chr12-13534477-DEL-1116
chr12-13562345-DEL-298
chr12-13576905-DEL-600
chr12-13585105-DEL-52
chr12-13598996-DEL-5321
chr12-13617586-DEL-84
chr12-13618852-DEL-4434
chr12-13647401-DEL-334
chr12-13651975-DEL-226
chr12-13668653-DEL-300
chr12-13669873-DEL-1467
chr12-13697020-DEL-57
chr12-13698602-DEL-65
chr12-13735553-DEL-52
chr12-13757113-DEL-111
chr12-13846935-DEL-1288
chr12-13867781-DEL-57
chr12-13907520-DEL-1655
chr12-13937726-DEL-869
chr12-13977985-DEL-849
chr12-13994263-DEL-139
chr12-14010108-DEL-72
chr12-14029310-DEL-87
chr12-14041000-DEL-52
chr12-14092124-DEL-171
chr12-14095527-DEL-77
chr12-14121916-DEL-2226
chr12-14138684-DEL-59

chr12-14142651-DEL-145
chr12-14146662-DEL-81
chr12-14160479-DEL-64
chr12-14203229-DEL-74
chr12-14207878-DEL-202
chr12-14290464-DEL-183
chr12-14404019-DEL-56
chr12-14418617-DEL-235
chr12-14419596-DEL-100
chr12-14485016-DEL-56
chr12-14488094-DEL-1791
chr12-14491072-DEL-153
chr12-14494847-DEL-79
chr12-14566149-DEL-57
chr12-14587309-DEL-137
chr12-14607813-DEL-64
chr12-14619047-DEL-86
chr12-14636407-DEL-52
chr12-14653973-DEL-1798
chr12-14660134-DEL-756
chr12-14693015-DEL-718
chr12-14709382-DEL-615
chr12-14736642-DEL-90
chr12-14767768-DEL-182
chr12-14779513-DEL-82
chr12-14798858-DEL-108
chr12-14833520-DEL-651
chr12-14834892-DEL-470
chr12-14841125-DEL-416
chr12-14869875-DEL-148
chr12-14901715-DEL-259
chr12-14924223-DEL-97
chr12-14949390-DEL-927
chr12-14984790-DEL-424
chr12-14988146-DEL-6326
chr12-14994294-DEL-4753
chr12-15003662-DEL-1832
chr12-15070329-DEL-529
chr12-15098243-DEL-54
chr12-15164647-DEL-2001
chr12-15165361-DEL-1501
chr12-15177025-DEL-62
chr12-15190829-DEL-100
chr12-15196010-DEL-749
chr12-15222815-DEL-380
chr12-15226531-DEL-69
chr12-15230086-DEL-270
chr12-15233398-DEL-71
chr12-15234781-DEL-76
chr12-15256026-DEL-88
chr12-15257895-DEL-86
chr12-15280676-DEL-133
chr12-15312779-DEL-62
chr12-15421730-DEL-6450
chr12-15442378-DEL-61
chr12-15457839-DEL-251
chr12-15467423-DEL-70
chr12-15489408-DEL-161
chr12-15507571-DEL-66
chr12-15521413-DEL-63
chr12-15545070-DEL-216
chr12-15646053-DEL-408
chr12-15718307-DEL-3638
chr12-15742925-DEL-60
chr12-15771699-DEL-482
chr12-15803671-DEL-594
chr12-15870674-DEL-402
chr12-15927576-DEL-161
chr12-15942582-DEL-56
chr12-15953409-DEL-98
chr12-15954672-DEL-373
chr12-16006438-DEL-66
chr12-16016356-DEL-68
chr12-16040837-DEL-60
chr12-16054242-DEL-699
chr12-16087147-DEL-258
chr12-16088212-DEL-2777

chr12-16091070-DEL-61
chr12-16133077-DEL-66
chr12-16152414-DEL-74
chr12-16161339-DEL-150
chr12-16170614-DEL-730
chr12-16197381-DEL-60
chr12-16210194-DEL-66
chr12-16253918-DEL-810
chr12-16292047-DEL-61
chr12-16315399-DEL-52
chr12-16330551-DEL-53
chr12-16360093-DEL-75
chr12-16366441-DEL-1457
chr12-16450867-DEL-66
chr12-16472712-DEL-77
chr12-16473222-DEL-53
chr12-16604183-DEL-62
chr12-16621934-DEL-733
chr12-16642112-DEL-187
chr12-16652676-DEL-52
chr12-16663540-DEL-755
chr12-16682748-DEL-64
chr12-16688582-DEL-214
chr12-16701320-DEL-262
chr12-16739422-DEL-55
chr12-16841745-DEL-56
chr12-16889148-DEL-1300
chr12-16904283-DEL-136
chr12-16905575-DEL-62
chr12-16922767-DEL-108
chr12-16962323-DEL-522
chr12-16999586-DEL-113
chr12-17070728-DEL-581
chr12-17089944-DEL-56
chr12-17095405-DEL-1113
chr12-17145538-DEL-485
chr12-17195784-DEL-55
chr12-17196475-DEL-315
chr12-17215290-DEL-550
chr12-17217160-DEL-56
chr12-17230309-DEL-58
chr12-17234883-DEL-92
chr12-17243895-DEL-126
chr12-17256893-DEL-980
chr12-17283591-DEL-500
chr12-17285713-DEL-58
chr12-17345455-DEL-219
chr12-17376105-DEL-71
chr12-17435208-DEL-99
chr12-17458414-DEL-3103
chr12-17469560-DEL-980
chr12-17478867-DEL-59
chr12-17488116-DEL-215
chr12-17537036-DEL-70
chr12-17555536-DEL-70
chr12-17562708-DEL-2269
chr12-17641841-DEL-147
chr12-17695323-DEL-60
chr12-17803253-DEL-54
chr12-17814509-DEL-5448
chr12-17921038-DEL-84
chr12-17953488-DEL-83
chr12-17974358-DEL-52
chr12-17978648-DEL-2510
chr12-17993381-DEL-163
chr12-18024073-DEL-615
chr12-18037917-DEL-55
chr12-18039491-DEL-113
chr12-18046181-DEL-52
chr12-18047089-DEL-461
chr12-18057339-DEL-1083
chr12-18060734-DEL-416
chr12-18063603-DEL-68
chr12-18145451-DEL-86
chr12-18180470-DEL-83
chr12-18256632-DEL-1362
chr12-18262599-DEL-77

chr12-18263173-DEL-91
chr12-18264197-DEL-244
chr12-18266279-DEL-57
chr12-18266528-DEL-1311
chr12-18267810-DEL-216
chr12-18303660-DEL-6199
chr12-18322965-DEL-10366
chr12-18325105-DEL-7852
chr12-18385503-DEL-62
chr12-18385676-DEL-439
chr12-18563572-DEL-209
chr12-18569028-DEL-59
chr12-18669355-DEL-1095
chr12-18670118-DEL-52
chr12-18698754-DEL-84
chr12-18706095-DEL-4768
chr12-18710634-DEL-252
chr12-18743780-DEL-95
chr12-18771743-DEL-65
chr12-18774800-DEL-382
chr12-18784494-DEL-5078
chr12-18788450-DEL-252
chr12-18829354-DEL-498
chr12-18902986-DEL-341
chr12-18908389-DEL-232
chr12-18922713-DEL-210
chr12-18923062-DEL-471
chr12-18973240-DEL-64
chr12-19039121-DEL-6666
chr12-19046772-DEL-74
chr12-19048205-DEL-143
chr12-19066845-DEL-174
chr12-19078416-DEL-1885
chr12-19144969-DEL-341
chr12-19200067-DEL-51
chr12-19208656-DEL-60
chr12-19224986-DEL-430
chr12-19248901-DEL-168
chr12-19291728-DEL-1421
chr12-19293147-DEL-6568
chr12-19294523-DEL-63
chr12-19296130-DEL-96
chr12-19296138-DEL-820
chr12-19298536-DEL-296
chr12-19308254-DEL-7129
chr12-19335112-DEL-113
chr12-19372076-DEL-532
chr12-19436313-DEL-203
chr12-19440557-DEL-210
chr12-19442823-DEL-58
chr12-19648344-DEL-280
chr12-19660449-DEL-484
chr12-19668730-DEL-3730
chr12-19676896-DEL-1771
chr12-19678977-DEL-1966
chr12-19680966-DEL-2691
chr12-19689848-DEL-161
chr12-19693961-DEL-342
chr12-19697678-DEL-102
chr12-19698499-DEL-800
chr12-19705755-DEL-72
chr12-19829907-DEL-118
chr12-19854704-DEL-136
chr12-19873534-DEL-153
chr12-19878779-DEL-255
chr12-19908910-DEL-239
chr12-19909188-DEL-583
chr12-19929568-DEL-630
chr12-19938459-DEL-598
chr12-19939994-DEL-92
chr12-19962119-DEL-15834
chr12-20008774-DEL-57
chr12-20042363-DEL-51
chr12-20055455-DEL-2697
chr12-20055515-DEL-67
chr12-20071175-DEL-163
chr12-20072621-DEL-190

chr12-20107926-DEL-7739
chr12-20169944-DEL-202
chr12-20203503-DEL-8134
chr12-20249444-DEL-934
chr12-20261147-DEL-2580
chr12-20276139-DEL-8811
chr12-20352036-DEL-64
chr12-20353970-DEL-243
chr12-20354557-DEL-3272
chr12-20384778-DEL-116
chr12-20402676-DEL-60
chr12-20441351-DEL-66
chr12-20462978-DEL-80
chr12-20508714-DEL-1101
chr12-20543465-DEL-3349
chr12-20616275-DEL-55
chr12-20624757-DEL-5576
chr12-20627467-DEL-76
chr12-20633531-DEL-462
chr12-20634092-DEL-1149
chr12-20636963-DEL-1893
chr12-20645454-DEL-221
chr12-20695162-DEL-80
chr12-20728524-DEL-771
chr12-20733445-DEL-53
chr12-20744352-DEL-12805
chr12-20758788-DEL-8734
chr12-20794870-DEL-2777
chr12-20797679-DEL-2995
chr12-20822076-DEL-61
chr12-20841536-DEL-201
chr12-20916131-DEL-64
chr12-20926402-DEL-52
chr12-20942006-DEL-133
chr12-20945690-DEL-3435
chr12-20963795-DEL-570
chr12-20965639-DEL-597
chr12-20978121-DEL-565
chr12-20991097-DEL-2353
chr12-20997600-DEL-258
chr12-21020700-DEL-221
chr12-21033314-DEL-319
chr12-21034514-DEL-2378
chr12-21038981-DEL-202
chr12-21043210-DEL-209
chr12-21045355-DEL-343
chr12-21045800-DEL-2661
chr12-21053189-DEL-65
chr12-21066058-DEL-168
chr12-21070916-DEL-388
chr12-21095647-DEL-74
chr12-21107745-DEL-2244
chr12-21232108-DEL-74
chr12-21232585-DEL-179
chr12-21271908-DEL-1419
chr12-21294259-DEL-225
chr12-21300086-DEL-78
chr12-21309131-DEL-102
chr12-21324235-DEL-84
chr12-21399211-DEL-136
chr12-21402877-DEL-58
chr12-21417831-DEL-426
chr12-21481814-DEL-555
chr12-21488437-DEL-314
chr12-21536780-DEL-161
chr12-21591807-DEL-53
chr12-21596198-DEL-59
chr12-21615116-DEL-502
chr12-21674990-DEL-111
chr12-21719477-DEL-1752
chr12-21779083-DEL-1156
chr12-21782549-DEL-52
chr12-21785376-DEL-6433
chr12-21809171-DEL-84
chr12-21811882-DEL-185
chr12-21830498-DEL-6152
chr12-21836421-DEL-177

chr12-21843460-DEL-433
chr12-21852437-DEL-1990
chr12-21853950-DEL-211
chr12-21943735-DEL-802
chr12-21946326-DEL-201
chr12-21952482-DEL-113
chr12-21964720-DEL-625
chr12-22003747-DEL-58
chr12-22009578-DEL-53
chr12-22012563-DEL-52
chr12-22016890-DEL-180
chr12-22027490-DEL-64
chr12-22057023-DEL-61
chr12-22075896-DEL-115
chr12-22088781-DEL-97
chr12-22100267-DEL-72
chr12-22133985-DEL-4809
chr12-22139302-DEL-56
chr12-22158648-DEL-163
chr12-22178166-DEL-252
chr12-22187874-DEL-6198
chr12-22211763-DEL-58
chr12-22243026-DEL-154
chr12-22370571-DEL-54
chr12-22373254-DEL-57
chr12-22395958-DEL-70
chr12-22442104-DEL-61
chr12-22458129-DEL-153
chr12-22473942-DEL-161
chr12-22497201-DEL-160
chr12-22497729-DEL-168
chr12-22497992-DEL-504
chr12-22507451-DEL-67
chr12-22511992-DEL-58
chr12-22529173-DEL-113
chr12-22533452-DEL-608
chr12-22539357-DEL-63
chr12-22545071-DEL-220
chr12-22579641-DEL-474
chr12-22661579-DEL-74
chr12-22681540-DEL-315
chr12-22704501-DEL-57
chr12-22706946-DEL-59
chr12-22709014-DEL-52
chr12-22740128-DEL-202
chr12-22775210-DEL-84
chr12-22801840-DEL-321
chr12-22802742-DEL-602
chr12-22821045-DEL-2808
chr12-22831229-DEL-4875
chr12-22834216-DEL-1092
chr12-22839200-DEL-61
chr12-22850973-DEL-66
chr12-22864132-DEL-201
chr12-22887014-DEL-135
chr12-22891975-DEL-710
chr12-22967758-DEL-55
chr12-22969745-DEL-264
chr12-22970420-DEL-1220
chr12-22978442-DEL-84
chr12-22979355-DEL-1062
chr12-22983912-DEL-60
chr12-22996756-DEL-312
chr12-23008337-DEL-219
chr12-23011142-DEL-494
chr12-23027181-DEL-53
chr12-23048417-DEL-238
chr12-23051136-DEL-78
chr12-23063729-DEL-112
chr12-23071839-DEL-5205
chr12-23071843-DEL-4620
chr12-23073473-DEL-63
chr12-23083915-DEL-6526
chr12-23114518-DEL-100
chr12-23125460-DEL-57
chr12-23126701-DEL-1663
chr12-23131345-DEL-64

chr12-23174779-DEL-63
chr12-23196965-DEL-84
chr12-23208059-DEL-5502
chr12-23216639-DEL-60
chr12-23247656-DEL-80
chr12-23262681-DEL-53
chr12-23336979-DEL-168
chr12-23346603-DEL-6182
chr12-23370473-DEL-58
chr12-23403296-DEL-60
chr12-23508684-DEL-94
chr12-23510297-DEL-6496
chr12-23527816-DEL-259
chr12-23554897-DEL-105
chr12-23593483-DEL-76
chr12-23601114-DEL-86
chr12-23603821-DEL-154
chr12-23606500-DEL-96
chr12-23608870-DEL-315
chr12-23632972-DEL-161
chr12-23647154-DEL-62
chr12-23647683-DEL-8326
chr12-23656010-DEL-3589
chr12-23660015-DEL-666
chr12-23662095-DEL-221
chr12-23665454-DEL-672
chr12-23679674-DEL-58
chr12-23712764-DEL-220
chr12-23731116-DEL-935
chr12-23734905-DEL-215
chr12-23738704-DEL-52
chr12-23739764-DEL-85
chr12-23747737-DEL-7421
chr12-23790211-DEL-72
chr12-23793900-DEL-1799
chr12-23797663-DEL-55
chr12-23801220-DEL-68
chr12-23806256-DEL-65
chr12-23845920-DEL-232
chr12-23851381-DEL-3469
chr12-23861250-DEL-321
chr12-23862451-DEL-326
chr12-23864088-DEL-3607
chr12-23864687-DEL-1740
chr12-23871996-DEL-10567
chr12-23890818-DEL-9232
chr12-23900052-DEL-568
chr12-23903659-DEL-153
chr12-23907432-DEL-51
chr12-23908520-DEL-99
chr12-23908695-DEL-1597
chr12-23913272-DEL-120
chr12-23930347-DEL-391
chr12-23934639-DEL-71
chr12-23948076-DEL-216
chr12-23978543-DEL-84
chr12-24007329-DEL-74
chr12-24014070-DEL-403
chr12-24015372-DEL-946
chr12-24019986-DEL-2330
chr12-24022309-DEL-144
chr12-24037154-DEL-200
chr12-24041577-DEL-118
chr12-24050336-DEL-107
chr12-24085872-DEL-204
chr12-24090997-DEL-210
chr12-24111798-DEL-445
chr12-24112893-DEL-81
chr12-24122137-DEL-168
chr12-24130505-DEL-5140
chr12-24142277-DEL-117
chr12-24143946-DEL-128
chr12-24154827-DEL-608
chr12-24155505-DEL-58
chr12-24161798-DEL-151
chr12-24170161-DEL-405
chr12-24176049-DEL-486

chr12-24186133-DEL-2556
chr12-24225167-DEL-70
chr12-24232884-DEL-167
chr12-24233061-DEL-1228
chr12-24248859-DEL-61
chr12-24264066-DEL-84
chr12-24264890-DEL-336
chr12-24265562-DEL-84
chr12-24271056-DEL-167
chr12-24274213-DEL-102
chr12-24315485-DEL-76
chr12-24340121-DEL-52
chr12-24342808-DEL-56
chr12-24395671-DEL-2359
chr12-24403566-DEL-202
chr12-24465682-DEL-1977
chr12-24485104-DEL-6262
chr12-24523674-DEL-1648
chr12-24533988-DEL-150
chr12-24612027-DEL-140
chr12-24613324-DEL-160
chr12-24663733-DEL-128
chr12-24680113-DEL-112
chr12-24690496-DEL-216
chr12-24719371-DEL-59
chr12-24721215-DEL-294
chr12-24726720-DEL-426
chr12-24754002-DEL-756
chr12-24762028-DEL-53
chr12-24770047-DEL-81
chr12-24795982-DEL-234
chr12-24805806-DEL-221
chr12-24809084-DEL-58
chr12-24828218-DEL-190
chr12-24851768-DEL-66
chr12-24854217-DEL-184
chr12-24872627-DEL-86
chr12-24898076-DEL-701
chr12-24911454-DEL-115
chr12-24926304-DEL-406
chr12-24938910-DEL-56
chr12-24947936-DEL-182
chr12-24954107-DEL-57
chr12-24972767-DEL-58
chr12-24979649-DEL-209
chr12-24999888-DEL-227
chr12-25032555-DEL-173
chr12-25034924-DEL-55
chr12-25061791-DEL-61
chr12-25094999-DEL-54
chr12-25157421-DEL-91
chr12-25191095-DEL-58
chr12-25224909-DEL-106
chr12-25229292-DEL-125
chr12-25321924-DEL-204
chr12-25359277-DEL-669
chr12-25417969-DEL-441
chr12-25418686-DEL-94
chr12-25419783-DEL-187
chr12-25463640-DEL-144
chr12-25472650-DEL-1673
chr12-25476118-DEL-140
chr12-25488028-DEL-99
chr12-25506210-DEL-620
chr12-25547609-DEL-546
chr12-25573827-DEL-199
chr12-25619386-DEL-1166
chr12-25620603-DEL-419
chr12-25629743-DEL-391
chr12-25641809-DEL-3423
chr12-25654374-DEL-285
chr12-25658795-DEL-58
chr12-25669944-DEL-60
chr12-25685950-DEL-1496
chr12-25745457-DEL-54
chr12-25750015-DEL-95
chr12-25751969-DEL-64

chr12-25761843-DEL-2145
chr12-25770391-DEL-54
chr12-25850862-DEL-253
chr12-25897817-DEL-913
chr12-25911818-DEL-604
chr12-25956651-DEL-66
chr12-25959514-DEL-1180
chr12-26023147-DEL-54
chr12-26024339-DEL-332
chr12-26027941-DEL-82
chr12-26034517-DEL-70
chr12-26040871-DEL-535
chr12-26046190-DEL-453
chr12-26050771-DEL-604
chr12-26058490-DEL-132
chr12-26074589-DEL-620
chr12-26083448-DEL-68
chr12-26084892-DEL-4146
chr12-26089056-DEL-2404
chr12-26091838-DEL-54
chr12-26106546-DEL-306
chr12-26235897-DEL-1685
chr12-26235899-DEL-773
chr12-26237334-DEL-219
chr12-26281030-DEL-76
chr12-26318308-DEL-382
chr12-26344212-DEL-957
chr12-26362497-DEL-59
chr12-26363859-DEL-119
chr12-26373319-DEL-143
chr12-26410221-DEL-297
chr12-26448168-DEL-62
chr12-26448999-DEL-1399
chr12-26456542-DEL-68
chr12-26466539-DEL-58
chr12-26509869-DEL-169
chr12-26521063-DEL-87
chr12-26649833-DEL-148
chr12-26672359-DEL-62
chr12-26680245-DEL-241
chr12-26691425-DEL-84
chr12-26699111-DEL-97
chr12-26749620-DEL-56
chr12-26769320-DEL-54
chr12-26814290-DEL-76
chr12-26847468-DEL-1076
chr12-26893834-DEL-350
chr12-26906044-DEL-54
chr12-26946676-DEL-70
chr12-26954505-DEL-435
chr12-26996306-DEL-155
chr12-27001810-DEL-51
chr12-27041650-DEL-1669
chr12-27044392-DEL-1643
chr12-27173933-DEL-60
chr12-27214049-DEL-302
chr12-27245302-DEL-60
chr12-27279564-DEL-188
chr12-27291769-DEL-2626
chr12-27291946-DEL-1375
chr12-27317797-DEL-74
chr12-27330413-DEL-56
chr12-27365445-DEL-60
chr12-27408001-DEL-62
chr12-27449915-DEL-6491
chr12-27465170-DEL-1604
chr12-27468429-DEL-5484
chr12-27490193-DEL-212
chr12-27501771-DEL-74
chr12-27543007-DEL-111
chr12-27548256-DEL-93
chr12-27586129-DEL-236
chr12-27601650-DEL-2304
chr12-27650350-DEL-64
chr12-27661943-DEL-183
chr12-27719058-DEL-150
chr12-27728653-DEL-247

chr12-27740444-DEL-244
chr12-27748827-DEL-820
chr12-27765459-DEL-68
chr12-27789972-DEL-82
chr12-27798971-DEL-66
chr12-27803412-DEL-466
chr12-27902239-DEL-88
chr12-27910329-DEL-1447
chr12-27921445-DEL-87
chr12-27927010-DEL-177
chr12-27930426-DEL-53
chr12-27956532-DEL-429
chr12-28037628-DEL-65
chr12-28086188-DEL-261
chr12-28138863-DEL-70
chr12-28139411-DEL-54
chr12-28146450-DEL-194
chr12-28165595-DEL-631
chr12-28167617-DEL-339
chr12-28183172-DEL-70
chr12-28205035-DEL-605
chr12-28259053-DEL-268
chr12-28276944-DEL-50
chr12-28288834-DEL-68
chr12-28306179-DEL-1782
chr12-28376709-DEL-152
chr12-28385480-DEL-71
chr12-28386414-DEL-3263
chr12-28427114-DEL-104
chr12-28445777-DEL-108
chr12-28446151-DEL-253
chr12-28453313-DEL-664
chr12-28458774-DEL-149
chr12-28476863-DEL-56
chr12-28502722-DEL-72
chr12-28536365-DEL-63
chr12-28556336-DEL-61
chr12-28579572-DEL-54
chr12-28590367-DEL-58
chr12-28590803-DEL-200
chr12-28641828-DEL-2137
chr12-28660819-DEL-531
chr12-28688247-DEL-175
chr12-28711395-DEL-65
chr12-28711819-DEL-192
chr12-28766463-DEL-75
chr12-28789658-DEL-64
chr12-28794142-DEL-206
chr12-28804650-DEL-453
chr12-28813380-DEL-1110
chr12-28838313-DEL-239
chr12-28840541-DEL-136
chr12-28846382-DEL-109
chr12-28864560-DEL-106
chr12-28871766-DEL-366
chr12-28891915-DEL-58
chr12-28910983-DEL-161
chr12-28933250-DEL-1311
chr12-28937026-DEL-105
chr12-28986668-DEL-55
chr12-28995178-DEL-58
chr12-29007791-DEL-63
chr12-29033408-DEL-607
chr12-29050940-DEL-92
chr12-29098170-DEL-394
chr12-29115452-DEL-203
chr12-29136407-DEL-73
chr12-29160103-DEL-56
chr12-29196408-DEL-279
chr12-29214407-DEL-176
chr12-29229009-DEL-249
chr12-29281395-DEL-60
chr12-29291999-DEL-120
chr12-29315719-DEL-1144
chr12-29316172-DEL-63
chr12-29364493-DEL-86
chr12-29367592-DEL-334

chr12-29382816-DEL-485
chr12-29401626-DEL-62
chr12-29448833-DEL-1892
chr12-29476691-DEL-77
chr12-29506926-DEL-86
chr12-29539890-DEL-310
chr12-29544014-DEL-936
chr12-29554774-DEL-146
chr12-29559536-DEL-132
chr12-29589976-DEL-77
chr12-29641840-DEL-94
chr12-29657703-DEL-88
chr12-29706718-DEL-84
chr12-29750967-DEL-120
chr12-29762775-DEL-56
chr12-29829760-DEL-53
chr12-29892038-DEL-179
chr12-29895955-DEL-107
chr12-29914457-DEL-69
chr12-29960461-DEL-186
chr12-29978021-DEL-92
chr12-29996932-DEL-54
chr12-30112691-DEL-493
chr12-30122042-DEL-65
chr12-30184328-DEL-491
chr12-30196735-DEL-6197
chr12-30216424-DEL-53
chr12-30237010-DEL-58
chr12-30254647-DEL-352
chr12-30262633-DEL-276
chr12-30360617-DEL-183
chr12-30369921-DEL-82
chr12-30401004-DEL-56
chr12-30414651-DEL-54
chr12-30418721-DEL-64
chr12-30441990-DEL-119
chr12-30469876-DEL-236
chr12-30509023-DEL-100
chr12-30520809-DEL-66
chr12-30539541-DEL-76
chr12-30626704-DEL-442
chr12-30628622-DEL-208
chr12-30670218-DEL-62
chr12-30683420-DEL-180
chr12-30710593-DEL-1510
chr12-30726473-DEL-106
chr12-30741277-DEL-225
chr12-30743920-DEL-72
chr12-30747185-DEL-87
chr12-30748660-DEL-52
chr12-30781186-DEL-264
chr12-30793616-DEL-52
chr12-30814220-DEL-974
chr12-30822917-DEL-1219
chr12-30928288-DEL-70
chr12-30935837-DEL-53
chr12-30953166-DEL-331
chr12-30972477-DEL-51
chr12-30977385-DEL-155
chr12-30997877-DEL-78
chr12-31020779-DEL-170
chr12-31022544-DEL-2269
chr12-31040585-DEL-63
chr12-31050296-DEL-107
chr12-31056189-DEL-1884
chr12-31083803-DEL-169
chr12-31092841-DEL-9845
chr12-31124679-DEL-141
chr12-31134665-DEL-102
chr12-31136243-DEL-131
chr12-31142850-DEL-51
chr12-31148004-DEL-53
chr12-31202519-DEL-128
chr12-31212587-DEL-2647
chr12-31219350-DEL-132
chr12-31242964-DEL-59
chr12-31249869-DEL-973

chr12-31251923-DEL-445
chr12-31262066-DEL-66
chr12-31272137-DEL-176
chr12-31288507-DEL-217
chr12-31290566-DEL-66
chr12-31297268-DEL-226
chr12-31320181-DEL-236
chr12-31343699-DEL-100
chr12-31408159-DEL-576
chr12-31408763-DEL-6465
chr12-31420159-DEL-507
chr12-31426297-DEL-71
chr12-31431308-DEL-68
chr12-31439914-DEL-164
chr12-31443560-DEL-180
chr12-31501575-DEL-73
chr12-31516903-DEL-138
chr12-31522276-DEL-138
chr12-31528422-DEL-205
chr12-31561076-DEL-70
chr12-31602926-DEL-140
chr12-31607853-DEL-76
chr12-31607881-DEL-916
chr12-31611011-DEL-100
chr12-31633433-DEL-132
chr12-31655866-DEL-526
chr12-31665080-DEL-68
chr12-31745912-DEL-216
chr12-31768577-DEL-236
chr12-31774321-DEL-160
chr12-31820471-DEL-129
chr12-31827562-DEL-1194
chr12-31853625-DEL-70
chr12-31861074-DEL-198
chr12-31874158-DEL-67
chr12-31882751-DEL-102
chr12-31933626-DEL-201
chr12-32014198-DEL-61
chr12-32050478-DEL-1293
chr12-32062861-DEL-109
chr12-32070344-DEL-1099
chr12-32072149-DEL-834
chr12-32074842-DEL-754
chr12-32147616-DEL-83
chr12-32153360-DEL-141
chr12-32161307-DEL-58
chr12-32167238-DEL-4521
chr12-32206326-DEL-118
chr12-32214356-DEL-251
chr12-32237511-DEL-90
chr12-32259013-DEL-1208
chr12-32275623-DEL-142
chr12-32290799-DEL-74
chr12-32291291-DEL-136
chr12-32293095-DEL-76
chr12-32326752-DEL-1154
chr12-32366554-DEL-102
chr12-32385651-DEL-172
chr12-32393447-DEL-52
chr12-32463892-DEL-124
chr12-32468475-DEL-212
chr12-32469751-DEL-871
chr12-32501905-DEL-419
chr12-32518130-DEL-717
chr12-32611944-DEL-372
chr12-32633507-DEL-298
chr12-32636003-DEL-576
chr12-32636698-DEL-144
chr12-32639993-DEL-58
chr12-32662543-DEL-86
chr12-32781512-DEL-1101
chr12-32825504-DEL-60
chr12-32836346-DEL-64
chr12-32839581-DEL-659
chr12-32861983-DEL-923
chr12-32863767-DEL-63
chr12-32923683-DEL-439

chr12-32941335-DEL-288
chr12-32952327-DEL-4104
chr12-32952327-DEL-728
chr12-32992236-DEL-117
chr12-33032915-DEL-52
chr12-33056643-DEL-136
chr12-33066356-DEL-62
chr12-33089027-DEL-991
chr12-33110818-DEL-80
chr12-33156347-DEL-82
chr12-33181678-DEL-66
chr12-33331837-DEL-54
chr12-33369639-DEL-240
chr12-33378521-DEL-483
chr12-33420492-DEL-83
chr12-33427845-DEL-188
chr12-33431065-DEL-260
chr12-33443218-DEL-70
chr12-33445432-DEL-81
chr12-33457685-DEL-1573
chr12-33457992-DEL-254
chr12-33458203-DEL-897
chr12-33458957-DEL-138
chr12-33459512-DEL-713
chr12-33463582-DEL-398
chr12-33471170-DEL-172
chr12-33524314-DEL-189
chr12-33525037-DEL-647
chr12-33527775-DEL-469
chr12-33553264-DEL-4657
chr12-33559181-DEL-2257
chr12-33584845-DEL-1766
chr12-33601792-DEL-343
chr12-33635020-DEL-398
chr12-33693347-DEL-3443
chr12-33697026-DEL-2425
chr12-33707856-DEL-619
chr12-33712277-DEL-369
chr12-33816511-DEL-214
chr12-33830902-DEL-162
chr12-33873796-DEL-110
chr12-33926388-DEL-6169
chr12-33930111-DEL-394
chr12-33940466-DEL-115
chr12-33955065-DEL-1176
chr12-33963244-DEL-68
chr12-33973960-DEL-75
chr12-33981902-DEL-100
chr12-33986265-DEL-275
chr12-34004395-DEL-92
chr12-34070179-DEL-1500
chr12-34071489-DEL-1373
chr12-34072976-DEL-135
chr12-34140784-DEL-112
chr12-34178658-DEL-2624
chr12-34181323-DEL-696
chr12-34187148-DEL-185
chr12-34212640-DEL-626
chr12-34221515-DEL-92
chr12-34235030-DEL-84
chr12-34283982-DEL-80
chr12-34307590-DEL-54
chr12-34310216-DEL-106
chr12-34372033-DEL-786
chr12-34372782-DEL-170
chr12-34387297-DEL-80
chr12-34405186-DEL-64
chr12-34408124-DEL-174
chr12-34417869-DEL-280
chr12-34476324-DEL-751
chr12-34504596-DEL-110
chr12-34526963-DEL-171
chr12-34533050-DEL-174
chr12-34563363-DEL-76
chr12-34583779-DEL-79
chr12-34584163-DEL-212
chr12-34636384-DEL-62

chr12-34654192-DEL-16439
chr12-34659341-DEL-58
chr12-34663521-DEL-78
chr12-34701851-DEL-130
chr12-34719031-DEL-2761
chr12-34726897-DEL-69
chr12-34765380-DEL-67
chr12-34775878-DEL-2699
chr12-34814877-DEL-1151
chr12-34824818-DEL-52
chr12-34865576-DEL-221
chr12-34881031-DEL-56
chr12-34893977-DEL-70
chr12-34897101-DEL-66
chr12-34900588-DEL-56
chr12-34908352-DEL-62
chr12-34944289-DEL-98
chr12-34948597-DEL-129
chr12-34989899-DEL-1331
chr12-35027825-DEL-52
chr12-35046579-DEL-186
chr12-35060073-DEL-132
chr12-35074835-DEL-330
chr12-35082687-DEL-1208
chr12-35106800-DEL-586
chr12-35113451-DEL-64
chr12-35127752-DEL-183
chr12-35157064-DEL-60
chr12-35171379-DEL-984
chr12-35245285-DEL-326
chr12-35290915-DEL-479
chr12-35302468-DEL-777
chr12-35305159-DEL-103
chr12-35315519-DEL-56
chr12-35336699-DEL-69
chr12-35352582-DEL-229
chr12-35371842-DEL-370
chr12-35420919-DEL-395
chr12-35453157-DEL-54
chr12-35459030-DEL-1176
chr12-35471561-DEL-63
chr12-35474862-DEL-4081
chr12-35505162-DEL-165
chr12-35529702-DEL-494
chr12-35530482-DEL-199
chr12-35530781-DEL-1558
chr12-35567269-DEL-567
chr12-35567906-DEL-388
chr12-35605075-DEL-174
chr12-35625051-DEL-109
chr12-35688358-DEL-644
chr12-35793498-DEL-1499
chr12-35838889-DEL-1103
chr12-35851279-DEL-13095
chr12-35852417-DEL-6800
chr12-35870168-DEL-73
chr12-35888051-DEL-54
chr12-35914365-DEL-5999
chr12-35933296-DEL-76
chr12-35951437-DEL-293
chr12-36032186-DEL-85
chr12-36063174-DEL-84
chr12-36075993-DEL-1144
chr12-36110975-DEL-80
chr12-36137827-DEL-70
chr12-36154910-DEL-1998
chr12-36154952-DEL-74
chr12-36155499-DEL-1406
chr12-36156427-DEL-74
chr12-36195237-DEL-518
chr12-36198492-DEL-1009
chr12-36228311-DEL-141
chr12-36239081-DEL-240
chr12-36277201-DEL-65
chr12-36279119-DEL-238
chr12-36292736-DEL-101
chr12-36299702-DEL-52

chr12-36311697-DEL-237
chr12-36315119-DEL-532
chr12-36317221-DEL-120
chr12-36357588-DEL-444
chr12-36413478-DEL-602
chr12-36421795-DEL-149
chr12-36438726-DEL-83
chr12-36449611-DEL-111
chr12-36500190-DEL-150
chr12-36514542-DEL-218
chr12-36518445-DEL-149
chr12-36534569-DEL-509
chr12-36564613-DEL-107
chr12-36567341-DEL-480
chr12-36581994-DEL-399
chr12-36609950-DEL-63
chr12-36642294-DEL-70
chr12-36645099-DEL-56
chr12-36649692-DEL-60
chr12-36662055-DEL-69
chr12-36699836-DEL-586
chr12-36702177-DEL-67
chr12-36729109-DEL-1479
chr12-36732792-DEL-132
chr12-36745773-DEL-399
chr12-36813722-DEL-79
chr12-36835498-DEL-53
chr12-36866239-DEL-1602
chr12-36877928-DEL-109
chr12-36911708-DEL-53
chr12-36920099-DEL-69
chr12-36980026-DEL-1290
chr12-37009394-DEL-190
chr12-37010174-DEL-590
chr12-37055667-DEL-193
chr12-37070506-DEL-55
chr12-37091166-DEL-54
chr12-37094718-DEL-73
chr12-37113851-DEL-7735
chr12-37190919-DEL-154
chr12-37243469-DEL-474
chr12-37246259-DEL-804
chr12-37249532-DEL-2966
chr12-37254231-DEL-6233
chr12-37258389-DEL-65
chr12-37265049-DEL-131
chr12-37287787-DEL-65
chr12-37299577-DEL-85
chr12-37368063-DEL-65
chr12-37375318-DEL-425
chr12-37375321-DEL-1605
chr12-37389419-DEL-1305
chr12-37404278-DEL-757
chr12-37412912-DEL-6697
chr12-37417600-DEL-225
chr12-37457459-DEL-58
chr12-37485225-DEL-82
chr12-37514129-DEL-56
chr12-37535192-DEL-1512
chr12-37559277-DEL-447
chr12-37662607-DEL-282
chr12-37740847-DEL-226
chr12-37749530-DEL-150
chr12-37779991-DEL-654
chr12-37783445-DEL-668
chr12-37787682-DEL-126
chr12-37792192-DEL-533
chr12-37894462-DEL-117
chr12-37900853-DEL-72
chr12-37912546-DEL-136
chr12-37929427-DEL-61
chr12-37946686-DEL-64
chr12-37991332-DEL-866
chr12-37998615-DEL-56
chr12-38032184-DEL-110
chr12-38045594-DEL-540
chr12-38047632-DEL-1349

chr12-38073859-DEL-1369
chr12-38080353-DEL-2172
chr12-38087893-DEL-6310
chr12-38132331-DEL-386
chr12-38152964-DEL-176
chr12-38177053-DEL-53
chr12-38206708-DEL-1304
chr12-38210643-DEL-1407
chr12-38213348-DEL-53
chr12-38227252-DEL-51
chr12-38232092-DEL-139
chr12-38248736-DEL-2218
chr12-38257077-DEL-93
chr12-38271898-DEL-3739
chr12-38303945-DEL-58
chr12-38335931-DEL-51
chr12-38386539-DEL-125
chr12-38417398-DEL-87
chr12-38436465-DEL-84
chr12-38462303-DEL-122
chr12-38470186-DEL-424
chr12-38504011-DEL-215
chr12-38508407-DEL-1452
chr12-38541406-DEL-472
chr12-38552596-DEL-3424
chr12-38559108-DEL-6207
chr12-38573510-DEL-766
chr12-38574031-DEL-104
chr12-38577365-DEL-6470
chr12-38601296-DEL-1571
chr12-38639146-DEL-197
chr12-38664926-DEL-190
chr12-38671653-DEL-64
chr12-38696145-DEL-560
chr12-38704392-DEL-492
chr12-38726426-DEL-2012
chr12-38726833-DEL-81
chr12-38729029-DEL-56
chr12-38741115-DEL-261
chr12-38758626-DEL-56
chr12-38761750-DEL-52
chr12-38786564-DEL-66
chr12-38818763-DEL-130
chr12-38892597-DEL-579
chr12-38939883-DEL-1246
chr12-38945215-DEL-630
chr12-39005781-DEL-201
chr12-39057918-DEL-154
chr12-39087193-DEL-65
chr12-39129090-DEL-82
chr12-39162201-DEL-356
chr12-39183154-DEL-185
chr12-39183570-DEL-210
chr12-39196292-DEL-192
chr12-39219153-DEL-110
chr12-39253493-DEL-480
chr12-39263589-DEL-706
chr12-39288428-DEL-80
chr12-39288974-DEL-3205
chr12-39288978-DEL-501
chr12-39289532-DEL-1127
chr12-39290693-DEL-1481
chr12-39365483-DEL-122
chr12-39468588-DEL-2230
chr12-39491012-DEL-477
chr12-39494149-DEL-90
chr12-39497117-DEL-358
chr12-39504417-DEL-51
chr12-39535287-DEL-839
chr12-39578396-DEL-64
chr12-39578629-DEL-373
chr12-39585243-DEL-181
chr12-39610782-DEL-1933
chr12-39612442-DEL-688
chr12-39615530-DEL-3239
chr12-39618791-DEL-2584
chr12-39623706-DEL-767

chr12-39661864-DEL-853
chr12-39666523-DEL-1075
chr12-39700695-DEL-67
chr12-39719783-DEL-580
chr12-39721419-DEL-875
chr12-39728691-DEL-51
chr12-39746505-DEL-54
chr12-39765809-DEL-122
chr12-39809155-DEL-99
chr12-39837859-DEL-70
chr12-39864649-DEL-194
chr12-39867444-DEL-79
chr12-39884551-DEL-400
chr12-39893729-DEL-102
chr12-39913203-DEL-2034
chr12-39962612-DEL-334
chr12-39994785-DEL-52
chr12-40000926-DEL-58
chr12-40012085-DEL-1747
chr12-40015675-DEL-110
chr12-40016700-DEL-263
chr12-40029674-DEL-117
chr12-40034067-DEL-218
chr12-40038597-DEL-253
chr12-40067677-DEL-433
chr12-40076659-DEL-104
chr12-40079182-DEL-248
chr12-40103449-DEL-202
chr12-40149202-DEL-1164
chr12-40170398-DEL-429
chr12-40211126-DEL-98
chr12-40213307-DEL-1101
chr12-40215980-DEL-450
chr12-40220202-DEL-2012
chr12-40220268-DEL-102
chr12-40224988-DEL-74
chr12-40235223-DEL-55
chr12-40287374-DEL-53
chr12-40314541-DEL-71
chr12-40326978-DEL-60
chr12-40333287-DEL-73
chr12-40359450-DEL-1000
chr12-40361411-DEL-729
chr12-40369362-DEL-759
chr12-40386943-DEL-724
chr12-40407546-DEL-157
chr12-40417495-DEL-10016
chr12-40426933-DEL-205
chr12-40434684-DEL-91
chr12-40443831-DEL-639
chr12-40451049-DEL-121
chr12-40494647-DEL-80
chr12-40515376-DEL-67
chr12-40652404-DEL-352
chr12-40755066-DEL-192
chr12-40769190-DEL-112
chr12-40783682-DEL-62
chr12-40801281-DEL-2005
chr12-40813975-DEL-64
chr12-40815323-DEL-89
chr12-40852739-DEL-115
chr12-40908666-DEL-498
chr12-40923404-DEL-116
chr12-40961867-DEL-64
chr12-40964591-DEL-1254
chr12-40965013-DEL-113
chr12-41000186-DEL-196
chr12-41010092-DEL-77
chr12-41053337-DEL-60
chr12-41055892-DEL-1161
chr12-41138588-DEL-1091
chr12-41163270-DEL-719
chr12-41163886-DEL-200
chr12-41175384-DEL-103
chr12-41186876-DEL-6383
chr12-41212717-DEL-840
chr12-41233842-DEL-517

chr12-41234508-DEL-684
chr12-41244778-DEL-132
chr12-41267650-DEL-138
chr12-41300719-DEL-87
chr12-41332760-DEL-2073
chr12-41394685-DEL-74
chr12-41429235-DEL-6267
chr12-41437217-DEL-95
chr12-41487214-DEL-173
chr12-41525578-DEL-196
chr12-41547124-DEL-62
chr12-41561804-DEL-168
chr12-41574507-DEL-58
chr12-41579221-DEL-98
chr12-41582524-DEL-56
chr12-41585837-DEL-306
chr12-41617179-DEL-217
chr12-41617721-DEL-1383
chr12-41617927-DEL-941
chr12-41619213-DEL-1123
chr12-41620007-DEL-328
chr12-41639785-DEL-204
chr12-41653916-DEL-88
chr12-41666630-DEL-350
chr12-41700119-DEL-419
chr12-41737717-DEL-448
chr12-41788634-DEL-624
chr12-41858257-DEL-4950
chr12-41869686-DEL-1199
chr12-41870588-DEL-238
chr12-41873219-DEL-1375
chr12-41897438-DEL-58
chr12-41933984-DEL-51
chr12-41939745-DEL-152
chr12-41940354-DEL-715
chr12-41967912-DEL-113
chr12-41972743-DEL-72
chr12-41982196-DEL-574
chr12-41993242-DEL-1087
chr12-42001157-DEL-297
chr12-42068071-DEL-517
chr12-42090023-DEL-514
chr12-42195183-DEL-115
chr12-42275170-DEL-59
chr12-42276751-DEL-77
chr12-42283110-DEL-81
chr12-42288069-DEL-208
chr12-42310087-DEL-1281
chr12-42312295-DEL-71
chr12-42315613-DEL-188
chr12-42328607-DEL-69
chr12-42339578-DEL-784
chr12-42342787-DEL-156
chr12-42361693-DEL-54
chr12-42368231-DEL-106
chr12-42394223-DEL-67
chr12-42410744-DEL-322
chr12-42414675-DEL-381
chr12-42453443-DEL-62
chr12-42483896-DEL-6300
chr12-42502158-DEL-995
chr12-42581358-DEL-282
chr12-42585072-DEL-5107
chr12-42591324-DEL-364
chr12-42599972-DEL-599
chr12-42614240-DEL-111
chr12-42617096-DEL-309
chr12-42637875-DEL-63
chr12-42653938-DEL-118
chr12-42658751-DEL-158
chr12-42707854-DEL-912
chr12-42710566-DEL-2296
chr12-42730716-DEL-2246
chr12-42738581-DEL-54
chr12-42752630-DEL-103
chr12-42766574-DEL-96
chr12-42777418-DEL-690

chr12-42779197-DEL-567
chr12-42800531-DEL-173
chr12-42805522-DEL-58
chr12-42818003-DEL-226
chr12-42820866-DEL-58
chr12-42914225-DEL-55
chr12-42920177-DEL-81
chr12-42932420-DEL-55
chr12-42972481-DEL-62
chr12-42984769-DEL-454
chr12-43019347-DEL-621
chr12-43024860-DEL-131
chr12-43049584-DEL-132
chr12-43057142-DEL-306
chr12-43062426-DEL-499
chr12-43100225-DEL-2612
chr12-43129289-DEL-2364
chr12-43132197-DEL-61
chr12-43140984-DEL-538
chr12-43183369-DEL-2665
chr12-43186179-DEL-567
chr12-43186890-DEL-2137
chr12-43189335-DEL-201
chr12-43191013-DEL-383
chr12-43192004-DEL-521
chr12-43210539-DEL-71
chr12-43267043-DEL-2611
chr12-43267044-DEL-1153
chr12-43267044-DEL-441
chr12-43268173-DEL-300
chr12-43304286-DEL-1485
chr12-43313637-DEL-99
chr12-43336001-DEL-1724
chr12-43343092-DEL-407
chr12-43407086-DEL-1278
chr12-43424019-DEL-659
chr12-43440396-DEL-213
chr12-43448865-DEL-62
chr12-43459193-DEL-4398
chr12-43479834-DEL-546
chr12-43486028-DEL-182
chr12-43488209-DEL-152
chr12-43498262-DEL-56
chr12-43525841-DEL-281
chr12-43527542-DEL-1551
chr12-43538023-DEL-698
chr12-43554398-DEL-155
chr12-43558137-DEL-52
chr12-43563559-DEL-168
chr12-43571939-DEL-52
chr12-43572853-DEL-189
chr12-43603193-DEL-362
chr12-43607896-DEL-245
chr12-43629228-DEL-53
chr12-43704205-DEL-6907
chr12-43730502-DEL-268
chr12-43755346-DEL-440
chr12-43757268-DEL-1065
chr12-43767659-DEL-100
chr12-43790841-DEL-133
chr12-43792428-DEL-987
chr12-43794296-DEL-438
chr12-43805451-DEL-917
chr12-43861806-DEL-137
chr12-43864403-DEL-64
chr12-43868927-DEL-180
chr12-43870779-DEL-139
chr12-43942659-DEL-78
chr12-43944355-DEL-62
chr12-43952600-DEL-131
chr12-43975637-DEL-6432
chr12-44000241-DEL-8419
chr12-44011406-DEL-93
chr12-44013864-DEL-233
chr12-44015703-DEL-5083
chr12-44022394-DEL-742
chr12-44067852-DEL-53

chr12-44111088-DEL-6617
chr12-44140405-DEL-2265
chr12-44155705-DEL-1323
chr12-44172478-DEL-543
chr12-44234683-DEL-113
chr12-44239346-DEL-152
chr12-44268765-DEL-168
chr12-44288257-DEL-105
chr12-44298590-DEL-3833
chr12-44302285-DEL-278
chr12-44313070-DEL-339
chr12-44337149-DEL-164
chr12-44349617-DEL-64
chr12-44409008-DEL-180
chr12-44446265-DEL-503
chr12-44470390-DEL-93
chr12-44576052-DEL-141
chr12-44580461-DEL-258
chr12-44629001-DEL-196
chr12-44648152-DEL-262
chr12-44708235-DEL-641
chr12-44741236-DEL-10604
chr12-44759176-DEL-5504
chr12-44767300-DEL-1387
chr12-44768773-DEL-249
chr12-44770335-DEL-1864
chr12-44771568-DEL-203
chr12-44774717-DEL-291
chr12-44775527-DEL-414
chr12-44777842-DEL-7023
chr12-44788571-DEL-163
chr12-44799471-DEL-2038
chr12-44812498-DEL-6555
chr12-44820559-DEL-1875
chr12-44822830-DEL-694
chr12-44827452-DEL-5666
chr12-44827510-DEL-4884
chr12-44836294-DEL-6514
chr12-44846681-DEL-369
chr12-44870187-DEL-67
chr12-44873172-DEL-1411
chr12-44878003-DEL-74
chr12-44892446-DEL-53
chr12-44893915-DEL-126
chr12-44907607-DEL-322
chr12-44913532-DEL-721
chr12-44926605-DEL-780
chr12-44955651-DEL-294
chr12-44961098-DEL-2702
chr12-44971485-DEL-350
chr12-44981583-DEL-359
chr12-45010793-DEL-93
chr12-45023073-DEL-56
chr12-45086099-DEL-3240
chr12-45133561-DEL-176
chr12-45136556-DEL-1045
chr12-45146652-DEL-122
chr12-45159008-DEL-942
chr12-45175689-DEL-2158
chr12-45242391-DEL-211
chr12-45243427-DEL-524
chr12-45244705-DEL-204
chr12-45250368-DEL-5473
chr12-45260546-DEL-66
chr12-45268021-DEL-2746
chr12-45271157-DEL-820
chr12-45297606-DEL-1723
chr12-45300754-DEL-59
chr12-45319647-DEL-1052
chr12-45327693-DEL-75
chr12-45342217-DEL-577
chr12-45343414-DEL-7432
chr12-45355404-DEL-59
chr12-45366633-DEL-210
chr12-45408680-DEL-201
chr12-45410368-DEL-1107
chr12-45411605-DEL-53

chr12-45416862-DEL-62
chr12-45449563-DEL-111
chr12-45461705-DEL-626
chr12-45464314-DEL-649
chr12-45465483-DEL-283
chr12-45472410-DEL-730
chr12-45472617-DEL-284
chr12-45475913-DEL-299
chr12-45477812-DEL-75
chr12-45497642-DEL-6441
chr12-45507781-DEL-184
chr12-45516640-DEL-68
chr12-45616770-DEL-1370
chr12-45662886-DEL-56
chr12-45690338-DEL-2967
chr12-45707334-DEL-349
chr12-45720868-DEL-716
chr12-45734213-DEL-3751
chr12-45748723-DEL-2013
chr12-45774672-DEL-1150
chr12-45779439-DEL-556
chr12-45783579-DEL-58
chr12-45786693-DEL-57
chr12-45797443-DEL-107
chr12-45799245-DEL-3336
chr12-45825557-DEL-118
chr12-45835560-DEL-9209
chr12-45858237-DEL-135
chr12-45861669-DEL-980
chr12-45873182-DEL-60
chr12-45901251-DEL-536
chr12-45985167-DEL-54
chr12-45993065-DEL-96
chr12-45996330-DEL-4899
chr12-46009197-DEL-64
chr12-46030178-DEL-11338
chr12-46045842-DEL-80
chr12-46048417-DEL-72
chr12-46057657-DEL-262
chr12-46058413-DEL-999
chr12-46065028-DEL-51
chr12-46139412-DEL-90
chr12-46148877-DEL-54
chr12-46155982-DEL-57
chr12-46170934-DEL-64
chr12-46172410-DEL-53
chr12-46208234-DEL-78
chr12-46231920-DEL-94
chr12-46266270-DEL-319
chr12-46283572-DEL-1924
chr12-46287982-DEL-1088
chr12-46308597-DEL-397
chr12-46311537-DEL-677
chr12-46316180-DEL-13910
chr12-46382299-DEL-352
chr12-46391568-DEL-1490
chr12-46476366-DEL-195
chr12-46501555-DEL-1168
chr12-46584634-DEL-60
chr12-46587965-DEL-52
chr12-46596266-DEL-306
chr12-46597471-DEL-190
chr12-46605687-DEL-2189
chr12-46605777-DEL-624
chr12-46605778-DEL-1272
chr12-46606453-DEL-601
chr12-46607146-DEL-311
chr12-46668590-DEL-84
chr12-46761696-DEL-72
chr12-46794770-DEL-95
chr12-46797929-DEL-256
chr12-46798912-DEL-54
chr12-46840253-DEL-1549
chr12-46871472-DEL-54
chr12-46888689-DEL-207
chr12-46898848-DEL-2349
chr12-46921283-DEL-92

chr12-46924005-DEL-766
chr12-46925592-DEL-150
chr12-46943471-DEL-384
chr12-46954137-DEL-55
chr12-46980501-DEL-408
chr12-46987339-DEL-409
chr12-47031647-DEL-519
chr12-47033483-DEL-77
chr12-47043911-DEL-469
chr12-47048089-DEL-1579
chr12-47055846-DEL-73
chr12-47068837-DEL-584
chr12-47073915-DEL-6033
chr12-47085077-DEL-1184
chr12-47088873-DEL-100
chr12-47116130-DEL-204
chr12-47147190-DEL-348
chr12-47169644-DEL-52
chr12-47174690-DEL-96
chr12-47188341-DEL-161
chr12-47193231-DEL-594
chr12-47193971-DEL-1261
chr12-47202357-DEL-485
chr12-47210680-DEL-56
chr12-47213582-DEL-640
chr12-47215332-DEL-58
chr12-47237642-DEL-59
chr12-47243122-DEL-52
chr12-47259107-DEL-2603
chr12-47281774-DEL-807
chr12-47318762-DEL-259
chr12-47354302-DEL-77
chr12-47361467-DEL-399
chr12-47373435-DEL-328
chr12-47379350-DEL-315
chr12-47383867-DEL-678
chr12-47401369-DEL-729
chr12-47443888-DEL-156
chr12-47456776-DEL-55
chr12-47481111-DEL-313
chr12-47492490-DEL-774
chr12-47503263-DEL-170
chr12-47509109-DEL-147
chr12-47535006-DEL-2131
chr12-47546229-DEL-226
chr12-47549985-DEL-82
chr12-47555010-DEL-1894
chr12-47556887-DEL-60
chr12-47567637-DEL-257
chr12-47568635-DEL-1028
chr12-47581003-DEL-198
chr12-47587716-DEL-122
chr12-47592734-DEL-3490
chr12-47599461-DEL-326
chr12-47645315-DEL-72
chr12-47649047-DEL-551
chr12-47671995-DEL-415
chr12-47676769-DEL-60
chr12-47692730-DEL-701
chr12-47710988-DEL-1489
chr12-47717501-DEL-970
chr12-47732955-DEL-358
chr12-47745310-DEL-725
chr12-47749934-DEL-1374
chr12-47751682-DEL-72
chr12-47822530-DEL-650
chr12-47841031-DEL-109
chr12-47853846-DEL-60
chr12-47853949-DEL-325
chr12-47861254-DEL-513
chr12-47869430-DEL-152
chr12-47872002-DEL-103
chr12-47874144-DEL-73
chr12-47877736-DEL-552
chr12-47879357-DEL-194
chr12-47880704-DEL-198
chr12-47884220-DEL-2182

chr12-47896701-DEL-96
chr12-47906208-DEL-6834
chr12-47912751-DEL-208
chr12-47913926-DEL-1979
chr12-47930191-DEL-389
chr12-47955270-DEL-914
chr12-47975958-DEL-780
chr12-48005437-DEL-619
chr12-48029510-DEL-1746
chr12-48040861-DEL-198
chr12-48048955-DEL-498
chr12-48051700-DEL-596
chr12-48054309-DEL-105
chr12-48076274-DEL-17292
chr12-48095088-DEL-6234
chr12-48112911-DEL-139
chr12-48117875-DEL-57
chr12-48137599-DEL-2314
chr12-48184439-DEL-226
chr12-48194299-DEL-1895
chr12-48205653-DEL-546
chr12-48221970-DEL-155
chr12-48229373-DEL-72
chr12-48260445-DEL-92
chr12-48292703-DEL-6408
chr12-48306824-DEL-64
chr12-48308201-DEL-56
chr12-48325847-DEL-1224
chr12-48333010-DEL-82
chr12-48340145-DEL-1748
chr12-48342775-DEL-62
chr12-48357223-DEL-1054
chr12-48360270-DEL-80
chr12-48366210-DEL-404
chr12-48371445-DEL-163
chr12-48404400-DEL-1511
chr12-48405804-DEL-437
chr12-48407958-DEL-112
chr12-48424603-DEL-1987
chr12-48461731-DEL-2103
chr12-48463866-DEL-2625
chr12-48465942-DEL-6327
chr12-48472653-DEL-1784
chr12-48478738-DEL-196
chr12-48480621-DEL-75
chr12-48502500-DEL-95
chr12-48518320-DEL-78
chr12-48561193-DEL-6299
chr12-48569722-DEL-2168
chr12-48571540-DEL-1800
chr12-48588216-DEL-121
chr12-48591771-DEL-60
chr12-48603115-DEL-3391
chr12-48606576-DEL-642
chr12-48607682-DEL-1017
chr12-48621255-DEL-64
chr12-48664980-DEL-124
chr12-48669778-DEL-94
chr12-48695382-DEL-60
chr12-48701763-DEL-6090
chr12-48726417-DEL-116
chr12-48731718-DEL-62
chr12-48748442-DEL-2369
chr12-48757888-DEL-65
chr12-48761797-DEL-164
chr12-48769049-DEL-507
chr12-48772492-DEL-2040
chr12-48811495-DEL-52
chr12-48824513-DEL-180
chr12-48834248-DEL-239
chr12-48848653-DEL-226
chr12-48851968-DEL-130
chr12-48857879-DEL-99
chr12-48907204-DEL-422
chr12-48928693-DEL-711
chr12-48929565-DEL-2628
chr12-48941888-DEL-538

chr12-48943128-DEL-941
chr12-48957586-DEL-6044
chr12-48991705-DEL-2039
chr12-49001094-DEL-231
chr12-49010772-DEL-100
chr12-49017189-DEL-2031
chr12-49031203-DEL-90
chr12-49049730-DEL-110
chr12-49050164-DEL-340
chr12-49060516-DEL-1991
chr12-49064073-DEL-1193
chr12-49065769-DEL-168
chr12-49077326-DEL-2616
chr12-49088630-DEL-59
chr12-49104464-DEL-212
chr12-49119378-DEL-102
chr12-49123086-DEL-6427
chr12-49164629-DEL-326
chr12-49171662-DEL-7606
chr12-49176930-DEL-1195
chr12-49202816-DEL-6279
chr12-49206657-DEL-71
chr12-49223022-DEL-1368
chr12-49274311-DEL-121
chr12-49280199-DEL-80
chr12-49283961-DEL-435
chr12-49291867-DEL-6765
chr12-49303059-DEL-21120
chr12-49307521-DEL-6208
chr12-49343588-DEL-96
chr12-49350996-DEL-65
chr12-49354621-DEL-380
chr12-49374633-DEL-126
chr12-49386833-DEL-4578
chr12-49398427-DEL-939
chr12-49479038-DEL-74
chr12-49481405-DEL-1114
chr12-49498865-DEL-1678
chr12-49518830-DEL-404
chr12-49521577-DEL-123
chr12-49566715-DEL-107
chr12-49568490-DEL-12554
chr12-49611543-DEL-101
chr12-49621088-DEL-519
chr12-49690873-DEL-431
chr12-49692861-DEL-400
chr12-49696177-DEL-71
chr12-49710954-DEL-62
chr12-49734068-DEL-90
chr12-49742910-DEL-12379
chr12-49756180-DEL-887
chr12-49774174-DEL-55
chr12-49792728-DEL-84
chr12-49795453-DEL-146
chr12-49798896-DEL-86
chr12-49822811-DEL-791
chr12-49828489-DEL-1052
chr12-49835090-DEL-739
chr12-49842361-DEL-265
chr12-49845285-DEL-378
chr12-49903927-DEL-3040
chr12-49961224-DEL-66
chr12-50022145-DEL-129
chr12-50032820-DEL-739
chr12-50050475-DEL-122
chr12-50062779-DEL-14295
chr12-50077257-DEL-81
chr12-50085003-DEL-607
chr12-50101099-DEL-676
chr12-50139743-DEL-133
chr12-50147595-DEL-62
chr12-50150659-DEL-220
chr12-50174704-DEL-1036
chr12-50183042-DEL-178
chr12-50220240-DEL-1797
chr12-50264748-DEL-58
chr12-50265756-DEL-364

chr12-50295231-DEL-208
chr12-50309260-DEL-106
chr12-50310928-DEL-56
chr12-50358426-DEL-70
chr12-50375541-DEL-941
chr12-50378656-DEL-381
chr12-50394180-DEL-79
chr12-50399782-DEL-183
chr12-50414390-DEL-345
chr12-50449410-DEL-2985
chr12-50452435-DEL-2600
chr12-50455058-DEL-911
chr12-50457329-DEL-902
chr12-50571504-DEL-500
chr12-50594566-DEL-80
chr12-50620235-DEL-55
chr12-50627809-DEL-1193
chr12-50629020-DEL-5166
chr12-50642776-DEL-80
chr12-50680508-DEL-96
chr12-50688448-DEL-97
chr12-50705526-DEL-119
chr12-50749810-DEL-63
chr12-50766701-DEL-68
chr12-50822018-DEL-4829
chr12-50839637-DEL-84
chr12-50849853-DEL-58
chr12-50871075-DEL-77
chr12-50907246-DEL-88
chr12-50918877-DEL-230
chr12-50953242-DEL-6350
chr12-50997860-DEL-52
chr12-51052181-DEL-64
chr12-51089859-DEL-62
chr12-51097722-DEL-60
chr12-51198609-DEL-1155
chr12-51205943-DEL-406
chr12-51274100-DEL-151
chr12-51304401-DEL-137
chr12-51323409-DEL-189
chr12-51327033-DEL-105
chr12-51330991-DEL-82
chr12-51333000-DEL-57
chr12-51337838-DEL-57
chr12-51339781-DEL-372
chr12-51347640-DEL-8869
chr12-51354349-DEL-403
chr12-51357393-DEL-152
chr12-51366990-DEL-64
chr12-51386519-DEL-250
chr12-51398955-DEL-254
chr12-51401375-DEL-261
chr12-51443212-DEL-71
chr12-51447654-DEL-68
chr12-51449180-DEL-1287
chr12-51466352-DEL-53
chr12-51492026-DEL-68
chr12-51516465-DEL-70
chr12-51520903-DEL-234
chr12-51556377-DEL-60
chr12-51576076-DEL-54
chr12-51582837-DEL-102
chr12-51582873-DEL-743
chr12-51590990-DEL-101
chr12-51596639-DEL-391
chr12-51597021-DEL-2229
chr12-51598982-DEL-650
chr12-51599859-DEL-1550
chr12-51621573-DEL-51
chr12-51627954-DEL-332
chr12-51635908-DEL-221
chr12-51662612-DEL-204
chr12-51699131-DEL-230
chr12-51700093-DEL-69
chr12-51700902-DEL-205
chr12-51714123-DEL-133
chr12-51761212-DEL-436

chr12-51779832-DEL-68
chr12-51799455-DEL-85
chr12-51802886-DEL-58
chr12-51827081-DEL-65
chr12-51834131-DEL-134
chr12-51840391-DEL-215
chr12-51846046-DEL-807
chr12-51868607-DEL-280
chr12-51878993-DEL-58
chr12-51894254-DEL-166
chr12-51902604-DEL-866
chr12-51902712-DEL-3598
chr12-51905654-DEL-80
chr12-51916999-DEL-412
chr12-51934633-DEL-132
chr12-51938748-DEL-112
chr12-51947939-DEL-168
chr12-51949168-DEL-398
chr12-51951226-DEL-54
chr12-51975377-DEL-228
chr12-51985911-DEL-202
chr12-52021338-DEL-60
chr12-52027194-DEL-1260
chr12-52043315-DEL-148
chr12-52050577-DEL-68
chr12-52106947-DEL-197
chr12-52132504-DEL-1186
chr12-52152662-DEL-783
chr12-52154531-DEL-293
chr12-52189257-DEL-577
chr12-52197649-DEL-184
chr12-52204670-DEL-6247
chr12-52212377-DEL-4100
chr12-52222432-DEL-75
chr12-52229323-DEL-1379
chr12-52231009-DEL-787
chr12-52246730-DEL-202
chr12-52247206-DEL-7339
chr12-52260110-DEL-58
chr12-52262301-DEL-7788
chr12-52291699-DEL-69
chr12-52295808-DEL-492
chr12-52298004-DEL-302
chr12-52455598-DEL-108
chr12-52463715-DEL-186
chr12-52490837-DEL-83
chr12-52508349-DEL-792
chr12-52530420-DEL-214
chr12-52550132-DEL-73
chr12-52587421-DEL-1131
chr12-52713830-DEL-205
chr12-52716071-DEL-503
chr12-52723927-DEL-315
chr12-52728834-DEL-56
chr12-52756227-DEL-56
chr12-52758777-DEL-201
chr12-52762029-DEL-60
chr12-52769203-DEL-95
chr12-52774667-DEL-125
chr12-52776109-DEL-521
chr12-52817008-DEL-314
chr12-52895367-DEL-1991
chr12-52905901-DEL-998
chr12-52943093-DEL-206
chr12-52949850-DEL-124
chr12-52953620-DEL-273
chr12-52973427-DEL-740
chr12-52973464-DEL-5261
chr12-52974418-DEL-299
chr12-52975064-DEL-691
chr12-52975845-DEL-57
chr12-52976517-DEL-2387
chr12-52977473-DEL-1737
chr12-52978873-DEL-396
chr12-52983297-DEL-106
chr12-52996351-DEL-60
chr12-53037892-DEL-201

chr12-53039216-DEL-152
chr12-53048421-DEL-805
chr12-53058704-DEL-268
chr12-53170303-DEL-69
chr12-53216301-DEL-2096
chr12-53224643-DEL-57
chr12-53229011-DEL-1982
chr12-53233303-DEL-207
chr12-53252816-DEL-220
chr12-53304570-DEL-185
chr12-53317315-DEL-264
chr12-53320840-DEL-2236
chr12-53351974-DEL-81
chr12-53375669-DEL-51
chr12-53376428-DEL-60
chr12-53395377-DEL-181
chr12-53396245-DEL-55
chr12-53428842-DEL-55
chr12-53471989-DEL-64
chr12-53577326-DEL-1898
chr12-53674914-DEL-812
chr12-53737233-DEL-74
chr12-53754127-DEL-369
chr12-53762505-DEL-58
chr12-53777664-DEL-93
chr12-53820606-DEL-51
chr12-53832063-DEL-87
chr12-53847724-DEL-240
chr12-53903677-DEL-333
chr12-53905793-DEL-135
chr12-54003471-DEL-54
chr12-54078205-DEL-51
chr12-54093285-DEL-59
chr12-54121369-DEL-413
chr12-54129297-DEL-186
chr12-54173573-DEL-431
chr12-54192890-DEL-51
chr12-54193523-DEL-58
chr12-54210417-DEL-69
chr12-54218294-DEL-71
chr12-54218947-DEL-78
chr12-54262068-DEL-55
chr12-54266770-DEL-55
chr12-54271035-DEL-115
chr12-54282767-DEL-788
chr12-54306578-DEL-94
chr12-54307218-DEL-109
chr12-54325786-DEL-58
chr12-54338255-DEL-1056
chr12-54364504-DEL-75
chr12-54376591-DEL-198
chr12-54380218-DEL-56
chr12-54391210-DEL-52
chr12-54395160-DEL-405
chr12-54420186-DEL-120
chr12-54438105-DEL-643
chr12-54438458-DEL-2977
chr12-54440142-DEL-882
chr12-54450254-DEL-57
chr12-54514540-DEL-60
chr12-54551681-DEL-270
chr12-54557148-DEL-107
chr12-54560857-DEL-56
chr12-54567106-DEL-133
chr12-54568064-DEL-60
chr12-54575266-DEL-88
chr12-54583014-DEL-73
chr12-54589491-DEL-109
chr12-54607136-DEL-306
chr12-54607928-DEL-63
chr12-54611899-DEL-75
chr12-54614039-DEL-72
chr12-54621346-DEL-544
chr12-54626873-DEL-442
chr12-54630981-DEL-55
chr12-54670044-DEL-244
chr12-54671656-DEL-5715

chr12-54685045-DEL-117
chr12-54687371-DEL-130
chr12-54695971-DEL-131
chr12-54709814-DEL-365
chr12-54728055-DEL-2016
chr12-54728341-DEL-714
chr12-54729420-DEL-672
chr12-54732097-DEL-180
chr12-54741416-DEL-195
chr12-54751796-DEL-62
chr12-54759487-DEL-183
chr12-54764145-DEL-606
chr12-54777726-DEL-616
chr12-54782820-DEL-182
chr12-54790639-DEL-63
chr12-54795948-DEL-2354
chr12-54807113-DEL-60
chr12-54817751-DEL-384
chr12-54838609-DEL-336
chr12-54850329-DEL-352
chr12-54851987-DEL-199
chr12-54853772-DEL-223
chr12-54858159-DEL-104
chr12-54878883-DEL-114
chr12-54888276-DEL-1250
chr12-54919259-DEL-62
chr12-54928745-DEL-4247
chr12-54943143-DEL-156
chr12-54946543-DEL-51
chr12-54951365-DEL-64
chr12-54999312-DEL-102
chr12-55002536-DEL-175
chr12-55016793-DEL-314
chr12-55026492-DEL-202
chr12-55056762-DEL-65
chr12-55074608-DEL-296
chr12-55075006-DEL-407
chr12-55075895-DEL-1725
chr12-55080694-DEL-88
chr12-55088586-DEL-205
chr12-55112937-DEL-59
chr12-55114084-DEL-67
chr12-55124635-DEL-749
chr12-55203783-DEL-131
chr12-55207343-DEL-222
chr12-55243398-DEL-258
chr12-55278847-DEL-131
chr12-55282408-DEL-222
chr12-55287456-DEL-13034
chr12-55297101-DEL-213
chr12-55305774-DEL-390
chr12-55317601-DEL-86
chr12-55341777-DEL-79
chr12-55378374-DEL-85
chr12-55385845-DEL-60
chr12-55389084-DEL-235
chr12-55392763-DEL-492
chr12-55402405-DEL-60
chr12-55403543-DEL-114
chr12-55451509-DEL-1147
chr12-55462143-DEL-65
chr12-55478005-DEL-56
chr12-55488636-DEL-626
chr12-55547324-DEL-93
chr12-55603280-DEL-275
chr12-55637880-DEL-56
chr12-55665523-DEL-459
chr12-55749692-DEL-155
chr12-55762248-DEL-74
chr12-55803804-DEL-74
chr12-55810907-DEL-181
chr12-55863260-DEL-89
chr12-55868626-DEL-136
chr12-55872196-DEL-96
chr12-55921201-DEL-485
chr12-55924808-DEL-63
chr12-55938151-DEL-1954

chr12-55945792-DEL-1582
chr12-55955798-DEL-188
chr12-55976237-DEL-55
chr12-55983366-DEL-714
chr12-55987319-DEL-815
chr12-55992870-DEL-207
chr12-55999294-DEL-1552
chr12-56019245-DEL-212
chr12-56020446-DEL-495
chr12-56059184-DEL-278
chr12-56099073-DEL-58
chr12-56108878-DEL-364
chr12-56149873-DEL-640
chr12-56218167-DEL-60
chr12-56227346-DEL-821
chr12-56227374-DEL-1757
chr12-56228227-DEL-2013
chr12-56229166-DEL-1072
chr12-56248744-DEL-66
chr12-56250504-DEL-224
chr12-56310288-DEL-83
chr12-56319755-DEL-62
chr12-56327646-DEL-159
chr12-56353446-DEL-67
chr12-56425165-DEL-195
chr12-56440110-DEL-86
chr12-56456323-DEL-150
chr12-56460024-DEL-170
chr12-56468768-DEL-441
chr12-56470507-DEL-1203
chr12-56500686-DEL-81
chr12-56514218-DEL-106
chr12-56525343-DEL-85
chr12-56551554-DEL-103
chr12-56554287-DEL-102
chr12-56563160-DEL-153
chr12-56614358-DEL-106
chr12-56619973-DEL-391
chr12-56639762-DEL-594
chr12-56641737-DEL-198
chr12-56663146-DEL-195
chr12-56681404-DEL-300
chr12-56688443-DEL-1647
chr12-56703924-DEL-392
chr12-56751717-DEL-96
chr12-56754143-DEL-98
chr12-56791213-DEL-300
chr12-56834203-DEL-94
chr12-56839894-DEL-110
chr12-56851975-DEL-88
chr12-56892488-DEL-339
chr12-56920167-DEL-75
chr12-56936736-DEL-332
chr12-56958178-DEL-2364
chr12-57013021-DEL-420
chr12-57039531-DEL-125
chr12-57048212-DEL-1319
chr12-57052989-DEL-143
chr12-57067620-DEL-91
chr12-57070377-DEL-59
chr12-57083618-DEL-55
chr12-57092091-DEL-6098
chr12-57147782-DEL-81
chr12-57148323-DEL-71
chr12-57177882-DEL-57
chr12-57198996-DEL-1028
chr12-57214610-DEL-57
chr12-57228251-DEL-289
chr12-57246961-DEL-61
chr12-57251506-DEL-77
chr12-57257001-DEL-193
chr12-57260634-DEL-122
chr12-57272831-DEL-73
chr12-57303903-DEL-52
chr12-57306385-DEL-195
chr12-57310745-DEL-61
chr12-57316938-DEL-154

chr12-57351418-DEL-60
chr12-57351627-DEL-686
chr12-57355617-DEL-270
chr12-57356155-DEL-2198
chr12-57382363-DEL-96
chr12-57385663-DEL-2443
chr12-57391223-DEL-2296
chr12-57397296-DEL-140
chr12-57411927-DEL-79
chr12-57419133-DEL-105
chr12-57431559-DEL-203
chr12-57436140-DEL-249
chr12-57458028-DEL-70
chr12-57472661-DEL-134
chr12-57476365-DEL-1010
chr12-57522981-DEL-462
chr12-57568116-DEL-55
chr12-57569344-DEL-1573
chr12-57571587-DEL-653
chr12-57635358-DEL-82
chr12-57682480-DEL-1072
chr12-57690010-DEL-85
chr12-57723052-DEL-386
chr12-57745180-DEL-808
chr12-57764751-DEL-6911
chr12-57771903-DEL-109
chr12-57791592-DEL-1165
chr12-57792936-DEL-806
chr12-57799036-DEL-351
chr12-57826542-DEL-60
chr12-57829108-DEL-456
chr12-57831615-DEL-196
chr12-57843832-DEL-812
chr12-57848586-DEL-6168
chr12-57848803-DEL-497
chr12-57858372-DEL-2671
chr12-57859716-DEL-95
chr12-57891405-DEL-53
chr12-57918827-DEL-160
chr12-57920666-DEL-149
chr12-57924861-DEL-72
chr12-57927584-DEL-6039
chr12-57966684-DEL-1768
chr12-58005834-DEL-626
chr12-58026930-DEL-80
chr12-58027945-DEL-175
chr12-58028655-DEL-1161
chr12-58043127-DEL-158
chr12-58054093-DEL-165
chr12-58087349-DEL-54
chr12-58151131-DEL-400
chr12-58167123-DEL-623
chr12-58167856-DEL-201
chr12-58232325-DEL-392
chr12-58234837-DEL-6304
chr12-58266077-DEL-51
chr12-58273078-DEL-199
chr12-58281980-DEL-764
chr12-58284872-DEL-72
chr12-58290359-DEL-95
chr12-58293767-DEL-861
chr12-58363883-DEL-129
chr12-58393424-DEL-2971
chr12-58420745-DEL-56
chr12-58421294-DEL-375
chr12-58432956-DEL-59
chr12-58438152-DEL-568
chr12-58452159-DEL-389
chr12-58464626-DEL-118
chr12-58465883-DEL-80
chr12-58488546-DEL-82
chr12-58521286-DEL-53
chr12-58534117-DEL-6284
chr12-58557854-DEL-54
chr12-58567877-DEL-246
chr12-58622423-DEL-54
chr12-58626144-DEL-502

chr12-58638663-DEL-1205
chr12-58655233-DEL-61
chr12-58664749-DEL-254
chr12-58672636-DEL-537
chr12-58689742-DEL-76
chr12-58719435-DEL-105
chr12-58733645-DEL-6144
chr12-58756253-DEL-202
chr12-58769486-DEL-815
chr12-58782000-DEL-739
chr12-58816055-DEL-12048
chr12-58816776-DEL-212
chr12-58823028-DEL-1317
chr12-58870573-DEL-53
chr12-58908095-DEL-5476
chr12-58968215-DEL-182
chr12-58971810-DEL-95
chr12-58983660-DEL-192
chr12-58986748-DEL-318
chr12-58987970-DEL-97
chr12-59012026-DEL-96
chr12-59022955-DEL-64
chr12-59028105-DEL-54
chr12-59081758-DEL-1995
chr12-59081758-DEL-2945
chr12-59081758-DEL-855
chr12-59081760-DEL-95
chr12-59081831-DEL-4084
chr12-59082294-DEL-2850
chr12-59082858-DEL-3040
chr12-59083722-DEL-1900
chr12-59084291-DEL-1330
chr12-59085423-DEL-237
chr12-59087388-DEL-340
chr12-59092913-DEL-258
chr12-59101284-DEL-177
chr12-59130185-DEL-102
chr12-59131715-DEL-3459
chr12-59138786-DEL-1684
chr12-59145854-DEL-140
chr12-59160195-DEL-1045
chr12-59161666-DEL-348
chr12-59167259-DEL-439
chr12-59174180-DEL-250
chr12-59183371-DEL-111
chr12-59184822-DEL-72
chr12-59200773-DEL-170
chr12-59214581-DEL-66
chr12-59227653-DEL-77
chr12-59233281-DEL-182
chr12-59247020-DEL-314
chr12-59257084-DEL-106
chr12-59269408-DEL-3918
chr12-59310677-DEL-200
chr12-59316263-DEL-449
chr12-59340285-DEL-1129
chr12-59374537-DEL-2543
chr12-59432294-DEL-114
chr12-59447197-DEL-1303
chr12-59466612-DEL-385
chr12-59469277-DEL-1248
chr12-59486360-DEL-550
chr12-59523855-DEL-81
chr12-59554428-DEL-171
chr12-59618165-DEL-502
chr12-59700951-DEL-134
chr12-59750920-DEL-518
chr12-59751563-DEL-71
chr12-59752205-DEL-127
chr12-59783993-DEL-389
chr12-59799342-DEL-69
chr12-59800856-DEL-1333
chr12-59801102-DEL-817
chr12-59816162-DEL-446
chr12-59825609-DEL-198
chr12-59828926-DEL-153
chr12-59877470-DEL-833

chr12-59903342-DEL-155
chr12-59906467-DEL-288
chr12-59932006-DEL-74
chr12-59933108-DEL-4196
chr12-59950171-DEL-80
chr12-59952668-DEL-2661
chr12-59955777-DEL-77
chr12-59962810-DEL-8216
chr12-59987872-DEL-559
chr12-60000395-DEL-2593
chr12-60021668-DEL-239
chr12-60029510-DEL-519
chr12-60045246-DEL-315
chr12-60060439-DEL-1207
chr12-60065565-DEL-6001
chr12-60083021-DEL-57
chr12-60114945-DEL-665
chr12-60118651-DEL-189
chr12-60135767-DEL-364
chr12-60148677-DEL-54
chr12-60152370-DEL-1711
chr12-60171997-DEL-2836
chr12-60180072-DEL-528
chr12-60192584-DEL-124
chr12-60240201-DEL-57
chr12-60240805-DEL-52
chr12-60242434-DEL-70
chr12-60271906-DEL-4208
chr12-60276579-DEL-116
chr12-60297711-DEL-52
chr12-60307038-DEL-598
chr12-60315885-DEL-610
chr12-60318674-DEL-262
chr12-60339902-DEL-90
chr12-60345444-DEL-444
chr12-60381064-DEL-350
chr12-60422664-DEL-62
chr12-60433094-DEL-486
chr12-60438757-DEL-66
chr12-60443075-DEL-134
chr12-60455174-DEL-252
chr12-60457611-DEL-70
chr12-60483386-DEL-84
chr12-60490088-DEL-208
chr12-60502327-DEL-288
chr12-60512734-DEL-87
chr12-60526868-DEL-589
chr12-60541987-DEL-1846
chr12-60544180-DEL-832
chr12-60562731-DEL-459
chr12-60582486-DEL-78
chr12-60612855-DEL-602
chr12-60615525-DEL-392
chr12-60631303-DEL-807
chr12-60637079-DEL-622
chr12-60638518-DEL-292
chr12-60685971-DEL-83
chr12-60693989-DEL-71
chr12-60704596-DEL-1888
chr12-60730399-DEL-3029
chr12-60740551-DEL-4008
chr12-60748537-DEL-13341
chr12-60768244-DEL-383
chr12-60796872-DEL-8139
chr12-60806457-DEL-65
chr12-60827284-DEL-602
chr12-60831426-DEL-79
chr12-60874354-DEL-1354
chr12-60875794-DEL-137
chr12-60902343-DEL-56
chr12-60902892-DEL-217
chr12-60928185-DEL-59
chr12-60939088-DEL-338
chr12-60955826-DEL-1347
chr12-60959394-DEL-2212
chr12-60980525-DEL-66
chr12-60995628-DEL-67

chr12-61014849-DEL-105
chr12-61016118-DEL-122
chr12-61028534-DEL-237
chr12-61048413-DEL-51
chr12-61079395-DEL-1524
chr12-61081613-DEL-91
chr12-61104221-DEL-54
chr12-61126038-DEL-58
chr12-61134731-DEL-1287
chr12-61185345-DEL-148
chr12-61207958-DEL-74
chr12-61222724-DEL-1610
chr12-61225089-DEL-6159
chr12-61226956-DEL-1508
chr12-61229389-DEL-1593
chr12-61229907-DEL-1076
chr12-61232965-DEL-72
chr12-61236674-DEL-5447
chr12-61249585-DEL-131
chr12-61257593-DEL-595
chr12-61261631-DEL-75
chr12-61310285-DEL-54
chr12-61317006-DEL-154
chr12-61367406-DEL-1149
chr12-61367460-DEL-210
chr12-61399525-DEL-58
chr12-61416412-DEL-423
chr12-61419687-DEL-1261
chr12-61421006-DEL-4940
chr12-61429963-DEL-237
chr12-61448444-DEL-57
chr12-61485156-DEL-3104
chr12-61490288-DEL-935
chr12-61537235-DEL-61
chr12-61557103-DEL-55
chr12-61566811-DEL-161
chr12-61582075-DEL-5455
chr12-61587958-DEL-52
chr12-61597231-DEL-54
chr12-61598010-DEL-135
chr12-61614929-DEL-127
chr12-61637165-DEL-59
chr12-61642999-DEL-186
chr12-61650701-DEL-329
chr12-61666556-DEL-263
chr12-61671774-DEL-193
chr12-61677181-DEL-406
chr12-61679771-DEL-6665
chr12-61703183-DEL-431
chr12-61715036-DEL-105
chr12-61726947-DEL-228
chr12-61733055-DEL-447
chr12-61735227-DEL-120
chr12-61756592-DEL-663
chr12-61759777-DEL-239
chr12-61760912-DEL-52
chr12-61767233-DEL-181
chr12-61769968-DEL-261
chr12-61770942-DEL-79
chr12-61778123-DEL-80
chr12-61789397-DEL-553
chr12-61801314-DEL-60
chr12-61803172-DEL-2664
chr12-61805855-DEL-4319
chr12-61821290-DEL-197
chr12-61830283-DEL-63
chr12-61858280-DEL-65
chr12-61869694-DEL-400
chr12-61874270-DEL-591
chr12-61874994-DEL-162
chr12-61875058-DEL-3117
chr12-61927051-DEL-663
chr12-62003182-DEL-5266
chr12-62007356-DEL-86
chr12-62011328-DEL-1093
chr12-62012466-DEL-5246
chr12-62078336-DEL-6724

chr12-62102038-DEL-6264
chr12-62112612-DEL-51
chr12-62115040-DEL-398
chr12-62124084-DEL-150
chr12-62124248-DEL-3355
chr12-62149892-DEL-60
chr12-62194042-DEL-1824
chr12-62209627-DEL-6240
chr12-62241725-DEL-131
chr12-62260184-DEL-52
chr12-62289670-DEL-55
chr12-62290791-DEL-6840
chr12-62345490-DEL-265
chr12-62347308-DEL-74
chr12-62378117-DEL-140
chr12-62383918-DEL-880
chr12-62397684-DEL-1257
chr12-62404328-DEL-135
chr12-62423006-DEL-439
chr12-62436405-DEL-62
chr12-62471493-DEL-611
chr12-62471500-DEL-1714
chr12-62472220-DEL-300
chr12-62472965-DEL-915
chr12-62473773-DEL-280
chr12-62477265-DEL-2740
chr12-62503709-DEL-73
chr12-62524293-DEL-600
chr12-62536312-DEL-194
chr12-62558198-DEL-80
chr12-62574369-DEL-76
chr12-62594526-DEL-1484
chr12-62594660-DEL-212
chr12-62622697-DEL-334
chr12-62627895-DEL-62
chr12-62628588-DEL-3191
chr12-62632845-DEL-390
chr12-62666449-DEL-3057
chr12-62682995-DEL-272
chr12-62697670-DEL-96
chr12-62702207-DEL-245
chr12-62721035-DEL-2266
chr12-62753717-DEL-567
chr12-62769412-DEL-161
chr12-62774339-DEL-122
chr12-62793587-DEL-249
chr12-62799560-DEL-3077
chr12-62811424-DEL-134
chr12-62820459-DEL-1048
chr12-62828112-DEL-87
chr12-62847326-DEL-58
chr12-62857860-DEL-87
chr12-62858613-DEL-60
chr12-62861624-DEL-1004
chr12-62863694-DEL-1623
chr12-62869748-DEL-6215
chr12-62892342-DEL-502
chr12-62901500-DEL-231
chr12-62902007-DEL-1808
chr12-62930384-DEL-197
chr12-62932101-DEL-52
chr12-62996876-DEL-57
chr12-63017875-DEL-55
chr12-63023114-DEL-5859
chr12-63031792-DEL-6409
chr12-63031901-DEL-212
chr12-63044983-DEL-1280
chr12-63058482-DEL-62
chr12-63086625-DEL-1092
chr12-63189126-DEL-106
chr12-63191678-DEL-5488
chr12-63200045-DEL-64
chr12-63204508-DEL-1335
chr12-63210453-DEL-1177
chr12-63227131-DEL-920
chr12-63228592-DEL-5398
chr12-63243128-DEL-2680

chr12-63366872-DEL-2495
chr12-63375481-DEL-56
chr12-63395818-DEL-106
chr12-63421141-DEL-204
chr12-63440816-DEL-212
chr12-63469676-DEL-4805
chr12-63477876-DEL-53
chr12-63479703-DEL-357
chr12-63484105-DEL-694
chr12-63545559-DEL-56
chr12-63560774-DEL-5137
chr12-63568624-DEL-472
chr12-63582709-DEL-394
chr12-63589865-DEL-330
chr12-63610073-DEL-1126
chr12-63612064-DEL-185
chr12-63616930-DEL-10431
chr12-63619398-DEL-4220
chr12-63652962-DEL-721
chr12-63677726-DEL-1844
chr12-63694254-DEL-257
chr12-63711454-DEL-1047
chr12-63712371-DEL-294
chr12-63721204-DEL-64
chr12-63723372-DEL-75
chr12-63736142-DEL-1028
chr12-63764652-DEL-62
chr12-63770511-DEL-6133
chr12-63788240-DEL-6254
chr12-63797062-DEL-68
chr12-63811581-DEL-377
chr12-63833146-DEL-243
chr12-63844849-DEL-697
chr12-63847324-DEL-123
chr12-63849133-DEL-71
chr12-63864045-DEL-238
chr12-63867655-DEL-95
chr12-63936992-DEL-2586
chr12-63939294-DEL-592
chr12-63943090-DEL-81
chr12-63945857-DEL-167
chr12-63955662-DEL-6379
chr12-63972595-DEL-1227
chr12-63974584-DEL-1813
chr12-63976839-DEL-875
chr12-64033401-DEL-7879
chr12-64041405-DEL-3663
chr12-64047876-DEL-192
chr12-64081690-DEL-307
chr12-64086973-DEL-54
chr12-64102523-DEL-139
chr12-64104271-DEL-14973
chr12-64107464-DEL-68
chr12-64107641-DEL-1461
chr12-64142268-DEL-2041
chr12-64170700-DEL-167
chr12-64177843-DEL-236
chr12-64193620-DEL-6401
chr12-64261595-DEL-344
chr12-64268890-DEL-108
chr12-64275312-DEL-505
chr12-64288251-DEL-303
chr12-64297458-DEL-64
chr12-64300307-DEL-529
chr12-64330578-DEL-100
chr12-64335931-DEL-91
chr12-64399266-DEL-340
chr12-64401119-DEL-621
chr12-64417014-DEL-91
chr12-64471653-DEL-367
chr12-64473478-DEL-5800
chr12-64489209-DEL-1058
chr12-64494073-DEL-132
chr12-64495544-DEL-82
chr12-64505942-DEL-4205
chr12-64513775-DEL-664
chr12-64534172-DEL-912

chr12-64534354-DEL-190
chr12-64543649-DEL-602
chr12-64550681-DEL-129
chr12-64556486-DEL-531
chr12-64585180-DEL-790
chr12-64646296-DEL-58
chr12-64679909-DEL-126
chr12-64695509-DEL-392
chr12-64696887-DEL-1710
chr12-64699257-DEL-591
chr12-64710935-DEL-394
chr12-64711344-DEL-2813
chr12-64714169-DEL-876
chr12-64725329-DEL-1893
chr12-64737030-DEL-1221
chr12-64740371-DEL-67
chr12-64779774-DEL-288
chr12-64784270-DEL-3713
chr12-64796387-DEL-162
chr12-64819923-DEL-51
chr12-64841688-DEL-2720
chr12-64890759-DEL-74
chr12-64891772-DEL-4183
chr12-64905731-DEL-52
chr12-64930094-DEL-70
chr12-64931394-DEL-86
chr12-64933479-DEL-7028
chr12-64946005-DEL-406
chr12-64982732-DEL-62
chr12-65017050-DEL-172
chr12-65020659-DEL-111
chr12-65021425-DEL-1656
chr12-65037241-DEL-90
chr12-65043865-DEL-95
chr12-65045114-DEL-58
chr12-65049665-DEL-54
chr12-65066284-DEL-89
chr12-65103719-DEL-177
chr12-65125220-DEL-52
chr12-65128589-DEL-656
chr12-65128597-DEL-75
chr12-65138237-DEL-240
chr12-65170241-DEL-81
chr12-65184178-DEL-310
chr12-65194744-DEL-167
chr12-65223147-DEL-482
chr12-65242811-DEL-54
chr12-65254161-DEL-97
chr12-65259391-DEL-236
chr12-65261013-DEL-226
chr12-65284523-DEL-81
chr12-65292465-DEL-1677
chr12-65322133-DEL-72
chr12-65348963-DEL-173
chr12-65485037-DEL-67
chr12-65495112-DEL-57
chr12-65501238-DEL-1177
chr12-65570813-DEL-277
chr12-65581818-DEL-70
chr12-65594112-DEL-226
chr12-65603784-DEL-596
chr12-65618202-DEL-97
chr12-65656673-DEL-1152
chr12-65658860-DEL-851
chr12-65758243-DEL-148
chr12-65761647-DEL-167
chr12-65762299-DEL-84
chr12-65767472-DEL-270
chr12-65777290-DEL-494
chr12-65800727-DEL-73
chr12-65823307-DEL-56
chr12-65831238-DEL-2827
chr12-65855924-DEL-1502
chr12-65887829-DEL-298
chr12-65896787-DEL-56
chr12-65957919-DEL-1699
chr12-65962244-DEL-212

chr12-65982665-DEL-862
chr12-65997089-DEL-160
chr12-66017770-DEL-82
chr12-66110676-DEL-103
chr12-66111012-DEL-1353
chr12-66179461-DEL-58
chr12-66185961-DEL-56
chr12-66206457-DEL-57
chr12-66218511-DEL-106
chr12-66219246-DEL-62
chr12-66255353-DEL-162
chr12-66287702-DEL-168
chr12-66293301-DEL-106
chr12-66297451-DEL-1018
chr12-66333433-DEL-325
chr12-66341904-DEL-228
chr12-66371622-DEL-4702
chr12-66387269-DEL-140
chr12-66397692-DEL-388
chr12-66400911-DEL-297
chr12-66405380-DEL-99
chr12-66408612-DEL-487
chr12-66472130-DEL-277
chr12-66473107-DEL-250
chr12-66473950-DEL-518
chr12-66476232-DEL-315
chr12-66490381-DEL-851
chr12-66576602-DEL-56
chr12-66577731-DEL-703
chr12-66578719-DEL-1497
chr12-66587524-DEL-3349
chr12-66592893-DEL-724
chr12-66633308-DEL-1710
chr12-66640792-DEL-158
chr12-66685006-DEL-80
chr12-66692933-DEL-1576
chr12-66729495-DEL-88
chr12-66758582-DEL-78
chr12-66767186-DEL-1631
chr12-66787952-DEL-630
chr12-66800993-DEL-92
chr12-66866633-DEL-3784
chr12-66883227-DEL-60
chr12-66898961-DEL-98
chr12-66911029-DEL-373
chr12-66923962-DEL-57
chr12-66931296-DEL-1169
chr12-66955388-DEL-1093
chr12-66970123-DEL-1594
chr12-67031748-DEL-496
chr12-67038985-DEL-88
chr12-67040480-DEL-58
chr12-67043790-DEL-171
chr12-67084639-DEL-165
chr12-67094057-DEL-290
chr12-67098877-DEL-73
chr12-67105936-DEL-791
chr12-67107593-DEL-347
chr12-67133147-DEL-108
chr12-67138889-DEL-105
chr12-67156346-DEL-153
chr12-67193815-DEL-1252
chr12-67209957-DEL-54
chr12-67210284-DEL-427
chr12-67290449-DEL-52
chr12-67322437-DEL-6386
chr12-67333493-DEL-79
chr12-67345809-DEL-109
chr12-67354062-DEL-90
chr12-67390631-DEL-51
chr12-67406397-DEL-881
chr12-67410634-DEL-1495
chr12-67425969-DEL-60
chr12-67433346-DEL-90
chr12-67441831-DEL-119
chr12-67458586-DEL-1079
chr12-67486786-DEL-200

chr12-67497520-DEL-64
chr12-67499055-DEL-6970
chr12-67529600-DEL-1436
chr12-67531971-DEL-595
chr12-67533138-DEL-92
chr12-67563964-DEL-114
chr12-67570927-DEL-297
chr12-67579916-DEL-8172
chr12-67634516-DEL-60
chr12-67733312-DEL-5782
chr12-67740336-DEL-77
chr12-67745242-DEL-94
chr12-67774030-DEL-218
chr12-67793047-DEL-1378
chr12-67796297-DEL-200
chr12-67810056-DEL-3605
chr12-67845232-DEL-864
chr12-67847649-DEL-443
chr12-67865922-DEL-60
chr12-67869161-DEL-351
chr12-67875629-DEL-7827
chr12-67881985-DEL-415
chr12-67882723-DEL-208
chr12-67886539-DEL-171
chr12-67914502-DEL-88
chr12-67949795-DEL-98
chr12-67971750-DEL-875
chr12-67979166-DEL-57
chr12-68023009-DEL-54
chr12-68023816-DEL-2315
chr12-68029574-DEL-60
chr12-68032151-DEL-337
chr12-68047951-DEL-1250
chr12-68073593-DEL-619
chr12-68074939-DEL-1789
chr12-68116589-DEL-667
chr12-68139347-DEL-1228
chr12-68165557-DEL-879
chr12-68169235-DEL-95
chr12-68217456-DEL-74
chr12-68291122-DEL-56
chr12-68293636-DEL-915
chr12-68305253-DEL-1212
chr12-68313909-DEL-936
chr12-68335402-DEL-1412
chr12-68414746-DEL-1445
chr12-68428042-DEL-362
chr12-68440461-DEL-74
chr12-68445931-DEL-2119
chr12-68454572-DEL-361
chr12-68490979-DEL-16667
chr12-68507432-DEL-1127
chr12-68546302-DEL-1352
chr12-68570082-DEL-66
chr12-68573122-DEL-56
chr12-68580846-DEL-61
chr12-68590265-DEL-164
chr12-68613768-DEL-70
chr12-68619681-DEL-56
chr12-68624097-DEL-68
chr12-68664590-DEL-70
chr12-68728715-DEL-325
chr12-68771487-DEL-439
chr12-68810878-DEL-72
chr12-68819042-DEL-105
chr12-68832605-DEL-150
chr12-68834237-DEL-74
chr12-68850474-DEL-160
chr12-68856191-DEL-1253
chr12-68883866-DEL-125
chr12-68914187-DEL-3853
chr12-68934801-DEL-60
chr12-68948231-DEL-186
chr12-68949985-DEL-59
chr12-68950522-DEL-1592
chr12-68960619-DEL-8795
chr12-68974534-DEL-7638

chr12-68997695-DEL-196
chr12-69040393-DEL-112
chr12-69074886-DEL-100
chr12-69080017-DEL-401
chr12-69091721-DEL-53
chr12-69109163-DEL-2155
chr12-69118280-DEL-211
chr12-69146663-DEL-59
chr12-69159929-DEL-435
chr12-69183504-DEL-84
chr12-69185466-DEL-199
chr12-69210887-DEL-1438
chr12-69220788-DEL-71
chr12-69226149-DEL-56
chr12-69240264-DEL-404
chr12-69257129-DEL-189
chr12-69271269-DEL-70
chr12-69276015-DEL-260
chr12-69281721-DEL-184
chr12-69286289-DEL-69
chr12-69301947-DEL-170
chr12-69323006-DEL-538
chr12-69333395-DEL-234
chr12-69340427-DEL-119
chr12-69370018-DEL-87
chr12-69381850-DEL-56
chr12-69386838-DEL-214
chr12-69409314-DEL-135
chr12-69433032-DEL-259
chr12-69448425-DEL-76
chr12-69477907-DEL-148
chr12-69485753-DEL-516
chr12-69503660-DEL-168
chr12-69506250-DEL-308
chr12-69520018-DEL-90
chr12-69528306-DEL-51
chr12-69535561-DEL-159
chr12-69551217-DEL-102
chr12-69556510-DEL-254
chr12-69567178-DEL-209
chr12-69577627-DEL-537
chr12-69592357-DEL-55
chr12-69616131-DEL-80
chr12-69616802-DEL-1587
chr12-69623777-DEL-75
chr12-69647137-DEL-130
chr12-69655265-DEL-228
chr12-69671016-DEL-234
chr12-69677215-DEL-143
chr12-69699298-DEL-182
chr12-69701618-DEL-2967
chr12-69712634-DEL-6079
chr12-69749225-DEL-232
chr12-69781674-DEL-59
chr12-69790996-DEL-2452
chr12-69799734-DEL-89
chr12-69801460-DEL-255
chr12-69809018-DEL-73
chr12-69817935-DEL-59
chr12-69831183-DEL-63
chr12-69832681-DEL-66
chr12-69845151-DEL-192
chr12-69886099-DEL-75
chr12-69898396-DEL-59
chr12-69902415-DEL-284
chr12-69920402-DEL-92
chr12-69958380-DEL-228
chr12-69963608-DEL-60
chr12-69968745-DEL-74
chr12-70002587-DEL-649
chr12-70003162-DEL-155
chr12-70075498-DEL-70
chr12-70076519-DEL-102
chr12-70082080-DEL-542
chr12-70093288-DEL-90
chr12-70111221-DEL-261
chr12-70112348-DEL-948

chr12-70121370-DEL-59
chr12-70125240-DEL-107
chr12-70133815-DEL-64
chr12-70168687-DEL-1286
chr12-70230418-DEL-6192
chr12-70250243-DEL-475
chr12-70252134-DEL-189
chr12-70270601-DEL-178
chr12-70285157-DEL-55
chr12-70315631-DEL-52
chr12-70327997-DEL-400
chr12-70333929-DEL-69
chr12-70360811-DEL-70
chr12-70366119-DEL-688
chr12-70368802-DEL-611
chr12-70424827-DEL-70
chr12-70449495-DEL-112
chr12-70495293-DEL-59
chr12-70516863-DEL-123
chr12-70577288-DEL-509
chr12-70595666-DEL-64
chr12-70600370-DEL-393
chr12-70600968-DEL-1063
chr12-70600978-DEL-74
chr12-70642634-DEL-80
chr12-70667082-DEL-67
chr12-70692369-DEL-54
chr12-70696973-DEL-165
chr12-70706166-DEL-202
chr12-70730402-DEL-84
chr12-70734732-DEL-76
chr12-70744001-DEL-73
chr12-70775791-DEL-81
chr12-70821881-DEL-91
chr12-70868460-DEL-52
chr12-70889115-DEL-167
chr12-70944781-DEL-53
chr12-70958326-DEL-186
chr12-70975297-DEL-70
chr12-70984411-DEL-99
chr12-70989440-DEL-512
chr12-70992316-DEL-1493
chr12-71002274-DEL-170
chr12-71016988-DEL-146
chr12-71023306-DEL-835
chr12-71075029-DEL-147
chr12-71092489-DEL-228
chr12-71101678-DEL-560
chr12-71134493-DEL-567
chr12-71134965-DEL-6096
chr12-71135155-DEL-1213
chr12-71135353-DEL-1744
chr12-71135577-DEL-63
chr12-71136250-DEL-91
chr12-71136294-DEL-1083
chr12-71136545-DEL-639
chr12-71137117-DEL-3944
chr12-71137255-DEL-2078
chr12-71137744-DEL-619
chr12-71138348-DEL-546
chr12-71139109-DEL-283
chr12-71139386-DEL-1322
chr12-71139907-DEL-861
chr12-71141006-DEL-55
chr12-71146141-DEL-201
chr12-71149056-DEL-84
chr12-71152894-DEL-264
chr12-71160584-DEL-361
chr12-71164005-DEL-163
chr12-71165399-DEL-86
chr12-71166522-DEL-424
chr12-71175151-DEL-54
chr12-71181702-DEL-629
chr12-71184900-DEL-124
chr12-71240413-DEL-150
chr12-71249692-DEL-74
chr12-71312773-DEL-63

chr12-71360757-DEL-197
chr12-71376925-DEL-208
chr12-71393083-DEL-444
chr12-71426423-DEL-613
chr12-71430324-DEL-390
chr12-71431369-DEL-66
chr12-71456025-DEL-349
chr12-71492722-DEL-148
chr12-71513209-DEL-235
chr12-71517278-DEL-310
chr12-71602474-DEL-206
chr12-71620868-DEL-54
chr12-71663450-DEL-519
chr12-71698517-DEL-287
chr12-71710349-DEL-210
chr12-71744073-DEL-218
chr12-71746947-DEL-1167
chr12-71753023-DEL-984
chr12-71755318-DEL-268
chr12-71760449-DEL-327
chr12-71798437-DEL-467
chr12-71806828-DEL-399
chr12-71841527-DEL-234
chr12-71851709-DEL-6707
chr12-71892782-DEL-51
chr12-71894297-DEL-54
chr12-71896366-DEL-167
chr12-71900609-DEL-75
chr12-71923751-DEL-58
chr12-71938176-DEL-62
chr12-71971073-DEL-57
chr12-71988402-DEL-126
chr12-72015334-DEL-218
chr12-72021023-DEL-100
chr12-72026843-DEL-59
chr12-72044162-DEL-280
chr12-72102212-DEL-1115
chr12-72106394-DEL-99
chr12-72145392-DEL-574
chr12-72154793-DEL-60
chr12-72174402-DEL-282
chr12-72186092-DEL-140
chr12-72193903-DEL-520
chr12-72196079-DEL-275
chr12-72212182-DEL-853
chr12-72228051-DEL-85
chr12-72246650-DEL-61
chr12-72289304-DEL-81
chr12-72294401-DEL-55
chr12-72304085-DEL-112
chr12-72306781-DEL-189
chr12-72352867-DEL-1774
chr12-72371759-DEL-295
chr12-72415001-DEL-90
chr12-72441615-DEL-106
chr12-72461817-DEL-219
chr12-72463697-DEL-193
chr12-72486148-DEL-61
chr12-72498243-DEL-146
chr12-72508276-DEL-70
chr12-72546588-DEL-712
chr12-72556938-DEL-71
chr12-72576890-DEL-853
chr12-72582232-DEL-175
chr12-72595717-DEL-74
chr12-72603767-DEL-9520
chr12-72655134-DEL-190
chr12-72661795-DEL-318
chr12-72670662-DEL-805
chr12-72681628-DEL-274
chr12-72681960-DEL-1248
chr12-72682055-DEL-213
chr12-72682937-DEL-96
chr12-72685416-DEL-3698
chr12-72695638-DEL-56
chr12-72731523-DEL-210
chr12-72732943-DEL-465

chr12-72746041-DEL-149
chr12-72747364-DEL-200
chr12-72773601-DEL-196
chr12-72777779-DEL-156
chr12-72790037-DEL-443
chr12-72800281-DEL-279
chr12-72867521-DEL-103
chr12-72881163-DEL-78
chr12-72898356-DEL-280
chr12-72922454-DEL-196
chr12-72937438-DEL-92
chr12-72944850-DEL-72
chr12-72951197-DEL-214
chr12-72962303-DEL-110
chr12-72964285-DEL-159
chr12-72993292-DEL-78
chr12-73034364-DEL-145
chr12-73037474-DEL-202
chr12-73047331-DEL-95
chr12-73049150-DEL-54
chr12-73070018-DEL-370
chr12-73070758-DEL-494
chr12-73072826-DEL-256
chr12-73076506-DEL-66
chr12-73078927-DEL-67
chr12-73108953-DEL-1690
chr12-73110055-DEL-214
chr12-73110615-DEL-258
chr12-73113294-DEL-276
chr12-73118190-DEL-803
chr12-73118423-DEL-76
chr12-73161555-DEL-82
chr12-73163659-DEL-60
chr12-73168272-DEL-162
chr12-73171600-DEL-257
chr12-73179249-DEL-190
chr12-73180629-DEL-126
chr12-73182083-DEL-416
chr12-73183264-DEL-204
chr12-73186241-DEL-51
chr12-73198298-DEL-201
chr12-73202016-DEL-1547
chr12-73207741-DEL-76
chr12-73227130-DEL-553
chr12-73240897-DEL-245
chr12-73280605-DEL-578
chr12-73284170-DEL-87
chr12-73294897-DEL-216
chr12-73304061-DEL-916
chr12-73338476-DEL-59
chr12-73369619-DEL-57
chr12-73395515-DEL-151
chr12-73403081-DEL-55
chr12-73416740-DEL-52
chr12-73440104-DEL-52
chr12-73445733-DEL-200
chr12-73460868-DEL-122
chr12-73473046-DEL-86
chr12-73476557-DEL-500
chr12-73520933-DEL-60
chr12-73522227-DEL-57
chr12-73539766-DEL-81
chr12-73575041-DEL-148
chr12-73603074-DEL-67
chr12-73608032-DEL-80
chr12-73614028-DEL-88
chr12-73627891-DEL-105
chr12-73629075-DEL-60
chr12-73633544-DEL-56
chr12-73665346-DEL-118
chr12-73681737-DEL-65
chr12-73682528-DEL-52
chr12-73686083-DEL-76
chr12-73789049-DEL-194
chr12-73792576-DEL-56
chr12-73807550-DEL-402
chr12-73830767-DEL-60

chr12-73909242-DEL-237
chr12-73925342-DEL-670
chr12-73949729-DEL-677
chr12-74001056-DEL-76
chr12-74008026-DEL-1930
chr12-74058884-DEL-76
chr12-74071509-DEL-67
chr12-74081379-DEL-156
chr12-74126788-DEL-521
chr12-74140316-DEL-78
chr12-74146880-DEL-154
chr12-74148872-DEL-101
chr12-74158888-DEL-115
chr12-74162109-DEL-52
chr12-74166145-DEL-58
chr12-74190512-DEL-323
chr12-74265790-DEL-110
chr12-74266860-DEL-769
chr12-74277357-DEL-54
chr12-74325152-DEL-87
chr12-74339693-DEL-329
chr12-74377977-DEL-69
chr12-74383977-DEL-947
chr12-74404475-DEL-173
chr12-74426818-DEL-63
chr12-74462263-DEL-77
chr12-74485217-DEL-1132
chr12-74486873-DEL-1502
chr12-74492664-DEL-132
chr12-74500659-DEL-76
chr12-74508282-DEL-68
chr12-74510263-DEL-804
chr12-74530156-DEL-55
chr12-74534471-DEL-60
chr12-74536031-DEL-197
chr12-74589427-DEL-77
chr12-74636248-DEL-224
chr12-74646114-DEL-558
chr12-74657699-DEL-717
chr12-74661497-DEL-170
chr12-74678735-DEL-104
chr12-74692627-DEL-76
chr12-74705723-DEL-554
chr12-74740952-DEL-68
chr12-74842829-DEL-53
chr12-74850752-DEL-116
chr12-74854110-DEL-1081
chr12-74878441-DEL-55
chr12-74899625-DEL-319
chr12-74903251-DEL-6222
chr12-74930484-DEL-207
chr12-74949582-DEL-52
chr12-74955316-DEL-4241
chr12-75019877-DEL-81
chr12-75035153-DEL-250
chr12-75083898-DEL-52
chr12-75093229-DEL-122
chr12-75101075-DEL-65
chr12-75136096-DEL-59
chr12-75165105-DEL-70
chr12-75181991-DEL-132
chr12-75263440-DEL-87
chr12-75289143-DEL-6400
chr12-75326783-DEL-759
chr12-75336978-DEL-263
chr12-75351521-DEL-54
chr12-75364604-DEL-363
chr12-75383965-DEL-124
chr12-75413240-DEL-125
chr12-75451821-DEL-227
chr12-75453463-DEL-700
chr12-75454388-DEL-100
chr12-75460260-DEL-164
chr12-75484696-DEL-628
chr12-75502509-DEL-335
chr12-75555567-DEL-51
chr12-75580044-DEL-52

chr12-75596692-DEL-151
chr12-75601755-DEL-76
chr12-75625606-DEL-72
chr12-75627332-DEL-169
chr12-75671009-DEL-57
chr12-75713330-DEL-165
chr12-75745322-DEL-100
chr12-75759727-DEL-61
chr12-75764892-DEL-88
chr12-75797071-DEL-319
chr12-75802910-DEL-56
chr12-75811455-DEL-316
chr12-75817449-DEL-75
chr12-75829741-DEL-438
chr12-75843541-DEL-1081
chr12-75859484-DEL-99
chr12-75860322-DEL-84
chr12-75875789-DEL-107
chr12-75887099-DEL-235
chr12-75889724-DEL-1431
chr12-75890107-DEL-53
chr12-75895983-DEL-693
chr12-75904795-DEL-57
chr12-75931919-DEL-71
chr12-75940535-DEL-131
chr12-75964799-DEL-61
chr12-75974569-DEL-79
chr12-75976257-DEL-68
chr12-75988218-DEL-71
chr12-75996588-DEL-142
chr12-76009975-DEL-63
chr12-76086419-DEL-67
chr12-76094498-DEL-253
chr12-76117376-DEL-121
chr12-76138058-DEL-53
chr12-76146513-DEL-54
chr12-76195134-DEL-91
chr12-76197076-DEL-999
chr12-76203348-DEL-331
chr12-76207578-DEL-94
chr12-76211281-DEL-197
chr12-76218345-DEL-95
chr12-76226998-DEL-136
chr12-76230308-DEL-183
chr12-76236229-DEL-666
chr12-76283300-DEL-192
chr12-76298959-DEL-69
chr12-76304827-DEL-223
chr12-76305443-DEL-360
chr12-76306922-DEL-1757
chr12-76331685-DEL-86
chr12-76336725-DEL-169
chr12-76352793-DEL-138
chr12-76360782-DEL-89
chr12-76366932-DEL-60
chr12-76376363-DEL-72
chr12-76384074-DEL-876
chr12-76386061-DEL-56
chr12-76429875-DEL-61
chr12-76443901-DEL-238
chr12-76447615-DEL-68
chr12-76458726-DEL-1266
chr12-76462127-DEL-206
chr12-76468420-DEL-240
chr12-76508941-DEL-143
chr12-76538851-DEL-255
chr12-76545281-DEL-56
chr12-76615664-DEL-142
chr12-76632153-DEL-63
chr12-76648111-DEL-59
chr12-76660290-DEL-486
chr12-76683653-DEL-52
chr12-76715169-DEL-162
chr12-76762132-DEL-420
chr12-76770727-DEL-201
chr12-76773488-DEL-80
chr12-76775474-DEL-225

chr12-76782548-DEL-8758
chr12-76791381-DEL-4375
chr12-76798549-DEL-337
chr12-76824131-DEL-431
chr12-76826249-DEL-206
chr12-76827991-DEL-98
chr12-76838889-DEL-59
chr12-76862327-DEL-88
chr12-76881276-DEL-174
chr12-76886861-DEL-419
chr12-76893041-DEL-131
chr12-76904694-DEL-84
chr12-76911165-DEL-181
chr12-76929934-DEL-119
chr12-76965354-DEL-194
chr12-76971106-DEL-184
chr12-76976410-DEL-61
chr12-77001288-DEL-181
chr12-77007208-DEL-66
chr12-77049046-DEL-116
chr12-77057327-DEL-54
chr12-77079523-DEL-63
chr12-77083739-DEL-72
chr12-77103380-DEL-53
chr12-77131065-DEL-71
chr12-77172337-DEL-2228
chr12-77174763-DEL-463
chr12-77228254-DEL-301
chr12-77234357-DEL-225
chr12-77253026-DEL-344
chr12-77273304-DEL-54
chr12-77276868-DEL-74
chr12-77280378-DEL-201
chr12-77293680-DEL-66
chr12-77309980-DEL-54
chr12-77327377-DEL-579
chr12-77340713-DEL-446
chr12-77354817-DEL-561
chr12-77391823-DEL-916
chr12-77419469-DEL-146
chr12-77485970-DEL-102
chr12-77490762-DEL-61
chr12-77524583-DEL-157
chr12-77532108-DEL-3279
chr12-77554689-DEL-170
chr12-77567506-DEL-6199
chr12-77599942-DEL-212
chr12-77659701-DEL-65
chr12-77663179-DEL-126
chr12-77674526-DEL-60
chr12-77683095-DEL-60
chr12-77705383-DEL-52
chr12-77734261-DEL-84
chr12-77746178-DEL-55
chr12-77752332-DEL-51
chr12-77798715-DEL-53
chr12-77829122-DEL-230
chr12-77895620-DEL-104
chr12-77917983-DEL-1832
chr12-77929823-DEL-2191
chr12-77946013-DEL-96
chr12-77946816-DEL-104
chr12-77974653-DEL-82
chr12-77989575-DEL-80
chr12-78039414-DEL-124
chr12-78085294-DEL-668
chr12-78105137-DEL-66
chr12-78137258-DEL-96
chr12-78144771-DEL-81
chr12-78245675-DEL-824
chr12-78306381-DEL-1772
chr12-78319942-DEL-1969
chr12-78334523-DEL-2817
chr12-78339754-DEL-788
chr12-78354824-DEL-2193
chr12-78377093-DEL-136
chr12-78381882-DEL-2154

chr12-78393781-DEL-1281
chr12-78397642-DEL-302
chr12-78452897-DEL-5541
chr12-78466403-DEL-53
chr12-78483852-DEL-105
chr12-78509113-DEL-6189
chr12-78558246-DEL-100
chr12-78607936-DEL-223
chr12-78624211-DEL-650
chr12-78625397-DEL-74
chr12-78646788-DEL-387
chr12-78682168-DEL-1621
chr12-78696385-DEL-1267
chr12-78700583-DEL-388
chr12-78708557-DEL-344
chr12-78709972-DEL-322
chr12-78740860-DEL-80
chr12-78746799-DEL-5034
chr12-78763460-DEL-379
chr12-78786913-DEL-108
chr12-78813387-DEL-473
chr12-78831620-DEL-107
chr12-78837127-DEL-59
chr12-78845058-DEL-844
chr12-78845810-DEL-92
chr12-78866479-DEL-1052
chr12-78891100-DEL-237
chr12-78917799-DEL-188
chr12-78945323-DEL-205
chr12-78949577-DEL-264
chr12-78972523-DEL-221
chr12-78977373-DEL-9747
chr12-78987998-DEL-240
chr12-78998563-DEL-68
chr12-79013875-DEL-2941
chr12-79014311-DEL-124
chr12-79022169-DEL-56
chr12-79118499-DEL-116
chr12-79136889-DEL-232
chr12-79154217-DEL-308
chr12-79156482-DEL-374
chr12-79181120-DEL-64
chr12-79217908-DEL-130
chr12-79262369-DEL-234
chr12-79272234-DEL-345
chr12-79300065-DEL-60
chr12-79303789-DEL-244
chr12-79306150-DEL-74
chr12-79333359-DEL-360
chr12-79348965-DEL-176
chr12-79367881-DEL-1320
chr12-79486055-DEL-191
chr12-79499447-DEL-67
chr12-79521175-DEL-51
chr12-79530143-DEL-183
chr12-79530589-DEL-2885
chr12-79541805-DEL-150
chr12-79555429-DEL-1284
chr12-79564570-DEL-1414
chr12-79575186-DEL-88
chr12-79587560-DEL-132
chr12-79601699-DEL-105
chr12-79608352-DEL-7398
chr12-79641657-DEL-110
chr12-79646672-DEL-71
chr12-79654402-DEL-221
chr12-79661996-DEL-170
chr12-79724221-DEL-426
chr12-79759164-DEL-846
chr12-79817147-DEL-66
chr12-79828421-DEL-391
chr12-79866336-DEL-75
chr12-79968839-DEL-72
chr12-79993808-DEL-319
chr12-80002785-DEL-464
chr12-80012708-DEL-114
chr12-80025217-DEL-52

chr12-80175276-DEL-188
chr12-80186969-DEL-437
chr12-80193318-DEL-52
chr12-80200175-DEL-102
chr12-80202344-DEL-77
chr12-80219328-DEL-90
chr12-80232758-DEL-86
chr12-80303057-DEL-152
chr12-80368273-DEL-186
chr12-80425201-DEL-211
chr12-80427010-DEL-434
chr12-80464029-DEL-7156
chr12-80530177-DEL-292
chr12-80562839-DEL-200
chr12-80669437-DEL-169
chr12-80670395-DEL-149
chr12-80679454-DEL-195
chr12-80723528-DEL-164
chr12-80747686-DEL-51
chr12-80789128-DEL-159
chr12-80795991-DEL-80
chr12-80813254-DEL-137
chr12-80814572-DEL-176
chr12-80820902-DEL-1553
chr12-80835708-DEL-167
chr12-80929496-DEL-55
chr12-80939890-DEL-181
chr12-81009763-DEL-83
chr12-81057414-DEL-75
chr12-81099709-DEL-76
chr12-81105901-DEL-1811
chr12-81108098-DEL-64
chr12-81111157-DEL-52
chr12-81112254-DEL-88
chr12-81118717-DEL-80
chr12-81153789-DEL-451
chr12-81157085-DEL-208
chr12-81160460-DEL-79
chr12-81193536-DEL-85
chr12-81358967-DEL-69
chr12-81485543-DEL-507
chr12-81535523-DEL-2622
chr12-81544538-DEL-52
chr12-81550686-DEL-188
chr12-81577580-DEL-371
chr12-81588358-DEL-370
chr12-81589106-DEL-91
chr12-81622801-DEL-84
chr12-81623032-DEL-1476
chr12-81654421-DEL-197
chr12-81675807-DEL-892
chr12-81684588-DEL-2283
chr12-81697341-DEL-96
chr12-81701122-DEL-56
chr12-81705840-DEL-52
chr12-81737214-DEL-59
chr12-81753017-DEL-1964
chr12-81765529-DEL-53
chr12-81776722-DEL-52
chr12-81817876-DEL-63
chr12-81841895-DEL-998
chr12-81844690-DEL-133
chr12-81855698-DEL-168
chr12-81863140-DEL-52
chr12-81880145-DEL-66
chr12-81909709-DEL-1897
chr12-81932425-DEL-87
chr12-81985150-DEL-102
chr12-82009767-DEL-325
chr12-82032682-DEL-390
chr12-82063943-DEL-996
chr12-82092460-DEL-1003
chr12-82094766-DEL-996
chr12-82108830-DEL-4365
chr12-82178692-DEL-169
chr12-82183122-DEL-1471
chr12-82201792-DEL-52

chr12-82210500-DEL-418
chr12-82320518-DEL-124
chr12-82327997-DEL-1864
chr12-82328397-DEL-294
chr12-82328546-DEL-713
chr12-82328964-DEL-844
chr12-82329496-DEL-171
chr12-82331941-DEL-86
chr12-82351697-DEL-128
chr12-82364667-DEL-1529
chr12-82367563-DEL-72
chr12-82396908-DEL-54
chr12-82403277-DEL-128
chr12-82439020-DEL-175
chr12-82474967-DEL-1954
chr12-82486180-DEL-424
chr12-82522399-DEL-192
chr12-82524076-DEL-95
chr12-82551656-DEL-464
chr12-82556798-DEL-372
chr12-82564878-DEL-60
chr12-82579407-DEL-1126
chr12-82609289-DEL-2905
chr12-82609885-DEL-88
chr12-82654500-DEL-165
chr12-82686503-DEL-424
chr12-82688744-DEL-1364
chr12-82692632-DEL-1698
chr12-82703713-DEL-57
chr12-82744630-DEL-97
chr12-82758083-DEL-285
chr12-82770374-DEL-105
chr12-82781017-DEL-331
chr12-82785674-DEL-82
chr12-82796613-DEL-83
chr12-82817924-DEL-156
chr12-82894090-DEL-141
chr12-82912242-DEL-52
chr12-82930317-DEL-196
chr12-82938951-DEL-834
chr12-82941261-DEL-96
chr12-82965287-DEL-282
chr12-82974798-DEL-76
chr12-83003909-DEL-183
chr12-83058978-DEL-190
chr12-83081729-DEL-950
chr12-83092072-DEL-294
chr12-83096981-DEL-64
chr12-83139800-DEL-1917
chr12-83143990-DEL-90
chr12-83148410-DEL-102
chr12-83153802-DEL-129
chr12-83159116-DEL-68
chr12-83165252-DEL-204
chr12-83165820-DEL-70
chr12-83169103-DEL-1539
chr12-83179263-DEL-64
chr12-83185629-DEL-325
chr12-83203863-DEL-60
chr12-83242566-DEL-268
chr12-83245338-DEL-163
chr12-83256832-DEL-52
chr12-83351154-DEL-66
chr12-83440053-DEL-56
chr12-83476025-DEL-271
chr12-83503143-DEL-196
chr12-83517514-DEL-57
chr12-83521568-DEL-204
chr12-83551112-DEL-62
chr12-83568795-DEL-187
chr12-83631871-DEL-96
chr12-83637000-DEL-206
chr12-83637662-DEL-86
chr12-83650781-DEL-72
chr12-83659486-DEL-88
chr12-83664495-DEL-171
chr12-83711490-DEL-534

chr12-83712047-DEL-1271
chr12-83731761-DEL-181
chr12-83757257-DEL-52
chr12-83763209-DEL-498
chr12-83774446-DEL-56
chr12-83777647-DEL-54
chr12-83798154-DEL-230
chr12-83806902-DEL-735
chr12-83851663-DEL-205
chr12-83888639-DEL-336
chr12-83911862-DEL-58
chr12-83940107-DEL-276
chr12-83943395-DEL-159
chr12-83979963-DEL-58
chr12-83995103-DEL-215
chr12-84012248-DEL-365
chr12-84040417-DEL-66
chr12-84053806-DEL-56
chr12-84054755-DEL-428
chr12-84061091-DEL-51
chr12-84071362-DEL-206
chr12-84078619-DEL-155
chr12-84093153-DEL-93
chr12-84096217-DEL-453
chr12-84099354-DEL-1210
chr12-84107226-DEL-380
chr12-84111130-DEL-52
chr12-84133242-DEL-92
chr12-84135183-DEL-53
chr12-84153750-DEL-138
chr12-84170147-DEL-131
chr12-84205965-DEL-311
chr12-84299934-DEL-755
chr12-84304604-DEL-142
chr12-84310529-DEL-56
chr12-84316615-DEL-690
chr12-84316929-DEL-104
chr12-84319646-DEL-214
chr12-84323235-DEL-709
chr12-84326750-DEL-200
chr12-84329763-DEL-172
chr12-84331433-DEL-99
chr12-84358034-DEL-79
chr12-84381842-DEL-1062
chr12-84412451-DEL-112
chr12-84441605-DEL-263
chr12-84502719-DEL-71
chr12-84517043-DEL-64
chr12-84555810-DEL-217
chr12-84603088-DEL-52
chr12-84622849-DEL-56
chr12-84631447-DEL-66
chr12-84642548-DEL-73
chr12-84646646-DEL-2479
chr12-84665006-DEL-196
chr12-84669330-DEL-96
chr12-84676429-DEL-285
chr12-84689814-DEL-1794
chr12-84696964-DEL-54
chr12-84702703-DEL-222
chr12-84769322-DEL-149
chr12-84776469-DEL-107
chr12-84784724-DEL-222
chr12-84792877-DEL-56
chr12-84826430-DEL-257
chr12-84857158-DEL-146
chr12-84885857-DEL-182
chr12-84892806-DEL-83
chr12-84903380-DEL-179
chr12-84930028-DEL-915
chr12-84956110-DEL-430
chr12-84971886-DEL-77
chr12-84984344-DEL-650
chr12-85018100-DEL-54
chr12-85039700-DEL-204
chr12-85041003-DEL-432
chr12-85062804-DEL-161

chr12-85115415-DEL-240
chr12-85122732-DEL-174
chr12-85177916-DEL-89
chr12-85188831-DEL-226
chr12-85190234-DEL-842
chr12-85210102-DEL-91
chr12-85220160-DEL-211
chr12-85235547-DEL-105
chr12-85237399-DEL-51
chr12-85238118-DEL-263
chr12-85263824-DEL-52
chr12-85363360-DEL-132
chr12-85365009-DEL-881
chr12-85373839-DEL-96
chr12-85378930-DEL-79
chr12-85438372-DEL-215
chr12-85474233-DEL-69
chr12-85519649-DEL-60
chr12-85565941-DEL-59
chr12-85567549-DEL-83
chr12-85574138-DEL-126
chr12-85589555-DEL-224
chr12-85596417-DEL-410
chr12-85623967-DEL-130
chr12-85639147-DEL-127
chr12-85665061-DEL-51
chr12-85668281-DEL-68
chr12-85680851-DEL-233
chr12-85709373-DEL-94
chr12-85715185-DEL-52
chr12-85742044-DEL-169
chr12-85772102-DEL-52
chr12-85780836-DEL-63
chr12-85785415-DEL-72
chr12-85796607-DEL-94
chr12-85821910-DEL-117
chr12-85839019-DEL-81
chr12-85882474-DEL-1303
chr12-85911669-DEL-116
chr12-85914603-DEL-1086
chr12-85915190-DEL-96
chr12-85988804-DEL-965
chr12-86011483-DEL-1723
chr12-86112738-DEL-199
chr12-86142133-DEL-1675
chr12-86143616-DEL-311
chr12-86157901-DEL-57
chr12-86164132-DEL-79
chr12-86172320-DEL-208
chr12-86273241-DEL-63
chr12-86295057-DEL-466
chr12-86353884-DEL-931
chr12-86357481-DEL-176
chr12-86374051-DEL-91
chr12-86395864-DEL-101
chr12-86398641-DEL-65
chr12-86400103-DEL-70
chr12-86430262-DEL-187
chr12-86487439-DEL-283
chr12-86491114-DEL-222
chr12-86499033-DEL-205
chr12-86504764-DEL-176
chr12-86512323-DEL-127
chr12-86521012-DEL-232
chr12-86534509-DEL-151
chr12-86546072-DEL-56
chr12-86549300-DEL-189
chr12-86562457-DEL-241
chr12-86569408-DEL-189
chr12-86572204-DEL-308
chr12-86579880-DEL-67
chr12-86582498-DEL-727
chr12-86598646-DEL-114
chr12-86608047-DEL-54
chr12-86608623-DEL-245
chr12-86614971-DEL-83
chr12-86628438-DEL-88

chr12-86670946-DEL-56
chr12-86684384-DEL-118
chr12-86691143-DEL-56
chr12-86719836-DEL-142
chr12-86740540-DEL-72
chr12-86771224-DEL-83
chr12-86812404-DEL-138
chr12-86819207-DEL-103
chr12-86820755-DEL-428
chr12-86829302-DEL-435
chr12-86833753-DEL-92
chr12-86866563-DEL-396
chr12-86867270-DEL-169
chr12-86966562-DEL-53
chr12-86972866-DEL-158
chr12-86976364-DEL-61
chr12-87016394-DEL-60
chr12-87026271-DEL-118
chr12-87083344-DEL-141
chr12-87087292-DEL-71
chr12-87088472-DEL-614
chr12-87097396-DEL-281
chr12-87139144-DEL-807
chr12-87160075-DEL-68
chr12-87200552-DEL-85
chr12-87228993-DEL-102
chr12-87256737-DEL-170
chr12-87263888-DEL-136
chr12-87274095-DEL-205
chr12-87277250-DEL-79
chr12-87296378-DEL-100
chr12-87315142-DEL-70
chr12-87318726-DEL-56
chr12-87322457-DEL-2360
chr12-87324165-DEL-669
chr12-87337696-DEL-108
chr12-87339985-DEL-184
chr12-87380313-DEL-97
chr12-87383560-DEL-415
chr12-87385716-DEL-1752
chr12-87404997-DEL-485
chr12-87421826-DEL-87
chr12-87454114-DEL-150
chr12-87505099-DEL-444
chr12-87520752-DEL-159
chr12-87531924-DEL-8553
chr12-87531926-DEL-11319
chr12-87538209-DEL-5050
chr12-87566776-DEL-251
chr12-87569342-DEL-319
chr12-87574616-DEL-101
chr12-87576116-DEL-6246
chr12-87593448-DEL-395542
chr12-87599752-DEL-660
chr12-87600503-DEL-13969
chr12-87603821-DEL-63
chr12-87736244-DEL-2102
chr12-87755310-DEL-70
chr12-87780735-DEL-1530
chr12-87847933-DEL-2602
chr12-87855693-DEL-72
chr12-87868276-DEL-4478
chr12-87873298-DEL-56
chr12-87880684-DEL-4941
chr12-87897244-DEL-58
chr12-87915809-DEL-51
chr12-87921789-DEL-69
chr12-87935427-DEL-306
chr12-87935498-DEL-772
chr12-87937879-DEL-81
chr12-87939280-DEL-216
chr12-87940517-DEL-6262
chr12-87959214-DEL-824
chr12-87988260-DEL-164
chr12-87990609-DEL-329
chr12-88008601-DEL-1564
chr12-88019114-DEL-55

chr12-88054594-DEL-236
chr12-88056899-DEL-16878
chr12-88085765-DEL-70
chr12-88109649-DEL-3733
chr12-88114738-DEL-23759
chr12-88116443-DEL-20207
chr12-88196300-DEL-6277
chr12-88245217-DEL-70
chr12-88261522-DEL-326
chr12-88298733-DEL-109
chr12-88302641-DEL-60
chr12-88306624-DEL-292
chr12-88313359-DEL-328
chr12-88349255-DEL-556
chr12-88368449-DEL-126
chr12-88448804-DEL-62
chr12-88463678-DEL-87
chr12-88490039-DEL-132
chr12-88496456-DEL-488
chr12-88529697-DEL-55
chr12-88537103-DEL-307
chr12-88566867-DEL-1523
chr12-88583030-DEL-391
chr12-88613126-DEL-965
chr12-88638974-DEL-137
chr12-88667508-DEL-88
chr12-88681939-DEL-90
chr12-88711932-DEL-56
chr12-88753454-DEL-58
chr12-88765668-DEL-78
chr12-88771406-DEL-55
chr12-88836069-DEL-84
chr12-88881436-DEL-131
chr12-88962052-DEL-157
chr12-88980186-DEL-114
chr12-88997349-DEL-738
chr12-89011073-DEL-1161
chr12-89041052-DEL-52
chr12-89080973-DEL-75
chr12-89130875-DEL-2004
chr12-89131410-DEL-156
chr12-89134003-DEL-144
chr12-89189411-DEL-106
chr12-89233037-DEL-710
chr12-89235177-DEL-108
chr12-89248042-DEL-71
chr12-89425973-DEL-87
chr12-89459629-DEL-224
chr12-89496784-DEL-122
chr12-89508012-DEL-60
chr12-89524408-DEL-55
chr12-89618443-DEL-136
chr12-89627797-DEL-354
chr12-89628527-DEL-141
chr12-89634474-DEL-912
chr12-89676900-DEL-157
chr12-89700539-DEL-3039
chr12-89710559-DEL-501
chr12-89751774-DEL-925
chr12-89766159-DEL-1539
chr12-89831765-DEL-303
chr12-89873040-DEL-60
chr12-89896595-DEL-188
chr12-89909106-DEL-60
chr12-89911705-DEL-53
chr12-89956286-DEL-196
chr12-89975515-DEL-56
chr12-89999823-DEL-486
chr12-90005721-DEL-552
chr12-90048352-DEL-204
chr12-90049199-DEL-872
chr12-90053272-DEL-895
chr12-90114643-DEL-66
chr12-90116094-DEL-74
chr12-90161795-DEL-151
chr12-90163382-DEL-689
chr12-90170329-DEL-59

chr12-90226904-DEL-159
chr12-90232172-DEL-55
chr12-90238245-DEL-56
chr12-90251728-DEL-424
chr12-90305877-DEL-122
chr12-90321333-DEL-2079
chr12-90325696-DEL-6271
chr12-90354013-DEL-408
chr12-90394398-DEL-175
chr12-90414521-DEL-323
chr12-90422351-DEL-134
chr12-90441552-DEL-298
chr12-90485041-DEL-1309
chr12-90495542-DEL-68
chr12-90525206-DEL-481
chr12-90526294-DEL-94
chr12-90543821-DEL-152
chr12-90549844-DEL-231
chr12-90567039-DEL-60
chr12-90582794-DEL-1833
chr12-90628081-DEL-168
chr12-90635321-DEL-4955
chr12-90638401-DEL-1111
chr12-90662102-DEL-51
chr12-90665387-DEL-61
chr12-90669735-DEL-72
chr12-90709252-DEL-80
chr12-90719569-DEL-64
chr12-90722482-DEL-52
chr12-90741910-DEL-63
chr12-90759398-DEL-143
chr12-90782710-DEL-231
chr12-90807740-DEL-212
chr12-90818994-DEL-489
chr12-90845879-DEL-80
chr12-90887626-DEL-98
chr12-90903874-DEL-88
chr12-91011552-DEL-406
chr12-91033898-DEL-2632
chr12-91043178-DEL-2466
chr12-91072958-DEL-379
chr12-91089164-DEL-1498
chr12-91099387-DEL-99
chr12-91114657-DEL-166
chr12-91150532-DEL-1006
chr12-91201717-DEL-1229
chr12-91239712-DEL-200
chr12-91242244-DEL-51
chr12-91251872-DEL-149
chr12-91259836-DEL-501
chr12-91291754-DEL-57
chr12-91297916-DEL-593
chr12-91304098-DEL-105
chr12-91323657-DEL-7421
chr12-91381102-DEL-115
chr12-91412067-DEL-17391
chr12-91422827-DEL-6601
chr12-91423885-DEL-5569
chr12-91433356-DEL-15436
chr12-91434184-DEL-7557
chr12-91434199-DEL-8539
chr12-91451295-DEL-72
chr12-91454264-DEL-61
chr12-91457825-DEL-222
chr12-91464848-DEL-349
chr12-91469861-DEL-65
chr12-91483920-DEL-358
chr12-91512979-DEL-211
chr12-91538870-DEL-55
chr12-91562702-DEL-1449
chr12-91593099-DEL-4825
chr12-91602958-DEL-60
chr12-91606669-DEL-60
chr12-91638510-DEL-242
chr12-91683855-DEL-220
chr12-91686766-DEL-124
chr12-91709782-DEL-133

chr12-91727477-DEL-462
chr12-91764408-DEL-201
chr12-91797070-DEL-52
chr12-91866006-DEL-76
chr12-91885275-DEL-152
chr12-91885908-DEL-1396
chr12-91887711-DEL-70
chr12-91909916-DEL-424
chr12-91925744-DEL-202
chr12-91931769-DEL-106
chr12-91932770-DEL-2605
chr12-91967812-DEL-104
chr12-91979732-DEL-65
chr12-92024746-DEL-498
chr12-92025635-DEL-314
chr12-92077008-DEL-3107
chr12-92086608-DEL-56
chr12-92097237-DEL-2973
chr12-92106559-DEL-297
chr12-92115079-DEL-61
chr12-92149637-DEL-89
chr12-92161475-DEL-532
chr12-92174550-DEL-79
chr12-92206305-DEL-200
chr12-92278003-DEL-80
chr12-92291847-DEL-52
chr12-92308821-DEL-1679
chr12-92324349-DEL-104
chr12-92358518-DEL-2001
chr12-92389387-DEL-241
chr12-92533936-DEL-208
chr12-92569258-DEL-800
chr12-92596175-DEL-54
chr12-92617253-DEL-6353
chr12-92640489-DEL-1238
chr12-92687361-DEL-158
chr12-92701428-DEL-77
chr12-92744826-DEL-54
chr12-92748485-DEL-135
chr12-92769907-DEL-65
chr12-92782233-DEL-90
chr12-92805227-DEL-6121
chr12-92814454-DEL-81
chr12-92846893-DEL-54
chr12-92869438-DEL-1347
chr12-92881570-DEL-66
chr12-92894890-DEL-70
chr12-92910311-DEL-2030
chr12-92940387-DEL-1357
chr12-92946436-DEL-2365
chr12-92983802-DEL-437
chr12-92992058-DEL-82
chr12-92999591-DEL-68
chr12-93057325-DEL-94
chr12-93059260-DEL-1486
chr12-93088538-DEL-294
chr12-93112721-DEL-283
chr12-93173707-DEL-77
chr12-93184428-DEL-159
chr12-93200272-DEL-6182
chr12-93234954-DEL-527
chr12-93247785-DEL-74
chr12-93250761-DEL-721
chr12-93272221-DEL-284
chr12-93276838-DEL-60
chr12-93307261-DEL-1431
chr12-93342808-DEL-355
chr12-93352068-DEL-4347
chr12-93356595-DEL-448
chr12-93358206-DEL-181
chr12-93358498-DEL-2460
chr12-93360673-DEL-64
chr12-93369151-DEL-161
chr12-93369384-DEL-1349
chr12-93371213-DEL-836
chr12-93411829-DEL-389
chr12-93433675-DEL-164

chr12-93436648-DEL-335
chr12-93454654-DEL-5665
chr12-93462525-DEL-144
chr12-93487444-DEL-772
chr12-93494810-DEL-74
chr12-93494899-DEL-1176
chr12-93543789-DEL-66
chr12-93560345-DEL-506
chr12-93579244-DEL-102
chr12-93579831-DEL-409
chr12-93599205-DEL-94
chr12-93603702-DEL-1854
chr12-93607135-DEL-290
chr12-93609015-DEL-416
chr12-93615161-DEL-53
chr12-93626635-DEL-178
chr12-93630189-DEL-196
chr12-93667824-DEL-753
chr12-93738579-DEL-60
chr12-93817624-DEL-161
chr12-93826587-DEL-300
chr12-93873332-DEL-82
chr12-93884272-DEL-799
chr12-93887490-DEL-52
chr12-93903642-DEL-104
chr12-93910541-DEL-55
chr12-93934060-DEL-112
chr12-93986508-DEL-3544
chr12-94009409-DEL-63
chr12-94045890-DEL-2255
chr12-94055732-DEL-6501
chr12-94062483-DEL-1318
chr12-94092299-DEL-145
chr12-94098254-DEL-73
chr12-94126946-DEL-63
chr12-94156133-DEL-62
chr12-94165394-DEL-205
chr12-94175194-DEL-1000
chr12-94200700-DEL-2541
chr12-94205622-DEL-89
chr12-94245213-DEL-312
chr12-94245921-DEL-404
chr12-94277168-DEL-2330
chr12-94280304-DEL-5038
chr12-94306431-DEL-6035
chr12-94312309-DEL-1151
chr12-94314863-DEL-51
chr12-94345104-DEL-79
chr12-94347674-DEL-150
chr12-94380098-DEL-574
chr12-94398818-DEL-63
chr12-94422264-DEL-54
chr12-94444501-DEL-58
chr12-94454435-DEL-64
chr12-94487327-DEL-560
chr12-94488099-DEL-70
chr12-94516790-DEL-6196
chr12-94591223-DEL-587
chr12-94595040-DEL-165
chr12-94612385-DEL-99
chr12-94655444-DEL-387
chr12-94661398-DEL-3071
chr12-94661946-DEL-3066
chr12-94664179-DEL-4600
chr12-94707931-DEL-78
chr12-94710204-DEL-586
chr12-94710952-DEL-2211
chr12-94732455-DEL-58
chr12-94734165-DEL-82
chr12-94754194-DEL-779
chr12-94782949-DEL-227
chr12-94845228-DEL-98
chr12-94852550-DEL-658
chr12-94863690-DEL-79
chr12-94880548-DEL-60
chr12-94886037-DEL-3923
chr12-94890201-DEL-75

chr12-94921638-DEL-390
chr12-94924225-DEL-193
chr12-94946596-DEL-178
chr12-94956568-DEL-4367
chr12-94978679-DEL-124
chr12-95006157-DEL-76
chr12-95008576-DEL-68
chr12-95018981-DEL-4517
chr12-95040941-DEL-324
chr12-95053114-DEL-55
chr12-95124604-DEL-832
chr12-95130200-DEL-4035
chr12-95130800-DEL-499
chr12-95190502-DEL-404
chr12-95212007-DEL-101
chr12-95225192-DEL-6180
chr12-95242564-DEL-105
chr12-95248347-DEL-145
chr12-95264687-DEL-112
chr12-95276792-DEL-122
chr12-95294331-DEL-52
chr12-95335734-DEL-68
chr12-95337849-DEL-245
chr12-95373254-DEL-86
chr12-95379779-DEL-93
chr12-95406255-DEL-53
chr12-95408150-DEL-219
chr12-95418662-DEL-515
chr12-95534193-DEL-67
chr12-95539246-DEL-1211
chr12-95552283-DEL-212
chr12-95560633-DEL-488
chr12-95563017-DEL-1554
chr12-95613381-DEL-1068
chr12-95617351-DEL-52
chr12-95635507-DEL-598
chr12-95668695-DEL-80
chr12-95684716-DEL-109
chr12-95685445-DEL-60
chr12-95707236-DEL-511
chr12-95777425-DEL-1140
chr12-95860076-DEL-425
chr12-95878902-DEL-115
chr12-95895658-DEL-154
chr12-95901573-DEL-1124
chr12-95959626-DEL-155
chr12-95973275-DEL-402
chr12-95982225-DEL-6008
chr12-96018117-DEL-1851
chr12-96074164-DEL-5463
chr12-96094681-DEL-74
chr12-96126525-DEL-183
chr12-96145827-DEL-1204
chr12-96147267-DEL-168
chr12-96158646-DEL-93
chr12-96174587-DEL-382
chr12-96183864-DEL-1018
chr12-96233772-DEL-57
chr12-96248202-DEL-171
chr12-96269881-DEL-149
chr12-96328793-DEL-122
chr12-96340872-DEL-168
chr12-96345192-DEL-2078
chr12-96346934-DEL-554
chr12-96397055-DEL-2619
chr12-96401591-DEL-76
chr12-96426816-DEL-319
chr12-96429507-DEL-52
chr12-96478176-DEL-212
chr12-96490457-DEL-238
chr12-96499729-DEL-183
chr12-96532271-DEL-179
chr12-96559640-DEL-56
chr12-96576213-DEL-1155
chr12-96582566-DEL-2854
chr12-96616659-DEL-1809
chr12-96635799-DEL-440

chr12-96642484-DEL-61
chr12-96653976-DEL-1834
chr12-96671130-DEL-408
chr12-96709035-DEL-1706
chr12-96768711-DEL-159
chr12-96769540-DEL-145
chr12-96774765-DEL-237
chr12-96787555-DEL-121
chr12-96801987-DEL-118
chr12-96816294-DEL-66
chr12-96816923-DEL-182
chr12-96842776-DEL-74
chr12-96852549-DEL-404
chr12-96882892-DEL-927
chr12-96894568-DEL-56
chr12-96895779-DEL-123
chr12-96903451-DEL-3745
chr12-96915046-DEL-356
chr12-96938544-DEL-1148
chr12-96952625-DEL-124
chr12-96959698-DEL-174
chr12-96974959-DEL-6172
chr12-96993635-DEL-501
chr12-97000757-DEL-62
chr12-97004454-DEL-424
chr12-97035487-DEL-460
chr12-97039266-DEL-108
chr12-97041868-DEL-79
chr12-97042934-DEL-540
chr12-97045688-DEL-54
chr12-97074109-DEL-162
chr12-97098200-DEL-57
chr12-97138485-DEL-171
chr12-97150690-DEL-439
chr12-97156947-DEL-545
chr12-97160268-DEL-64
chr12-97171688-DEL-54
chr12-97264901-DEL-866
chr12-97279640-DEL-54
chr12-97295191-DEL-860
chr12-97299099-DEL-451
chr12-97321444-DEL-130
chr12-97351041-DEL-456
chr12-97353816-DEL-2365
chr12-97360427-DEL-70
chr12-97380786-DEL-70
chr12-97381929-DEL-240
chr12-97407564-DEL-88
chr12-97415662-DEL-275
chr12-97511405-DEL-341
chr12-97533040-DEL-6006
chr12-97561754-DEL-61
chr12-97581262-DEL-215
chr12-97582507-DEL-6176
chr12-97586870-DEL-1823
chr12-97597212-DEL-248
chr12-97600298-DEL-1140
chr12-97602728-DEL-702
chr12-97666484-DEL-149
chr12-97676024-DEL-67
chr12-97691845-DEL-3556
chr12-97707681-DEL-105
chr12-97708680-DEL-128
chr12-97719283-DEL-53
chr12-97735800-DEL-325
chr12-97757571-DEL-59
chr12-97768524-DEL-87
chr12-97786101-DEL-6230
chr12-97814889-DEL-54
chr12-97829617-DEL-77
chr12-97830661-DEL-84
chr12-97832544-DEL-870
chr12-97846021-DEL-153
chr12-97901144-DEL-400
chr12-97931477-DEL-51
chr12-97957463-DEL-861
chr12-97975408-DEL-70

chr12-97992141-DEL-164
chr12-98000024-DEL-1135
chr12-98001257-DEL-1446
chr12-98007790-DEL-1306
chr12-98023629-DEL-399
chr12-98027943-DEL-90
chr12-98055345-DEL-60
chr12-98060476-DEL-342
chr12-98123409-DEL-92
chr12-98143162-DEL-230
chr12-98146693-DEL-410
chr12-98165825-DEL-169
chr12-98191081-DEL-60
chr12-98209843-DEL-66
chr12-98226695-DEL-124
chr12-98250099-DEL-747
chr12-98288909-DEL-721
chr12-98293957-DEL-60
chr12-98294489-DEL-65
chr12-98319345-DEL-924
chr12-98330979-DEL-170
chr12-98340269-DEL-52
chr12-98379043-DEL-289
chr12-98404855-DEL-57
chr12-98427863-DEL-53
chr12-98449371-DEL-54
chr12-98465920-DEL-397
chr12-98467112-DEL-191
chr12-98471190-DEL-118
chr12-98522462-DEL-89
chr12-98570481-DEL-958
chr12-98583414-DEL-1184
chr12-98587116-DEL-206
chr12-98589690-DEL-345
chr12-98598580-DEL-143
chr12-98610311-DEL-88
chr12-98631656-DEL-63
chr12-98632435-DEL-1467
chr12-98639333-DEL-101
chr12-98666898-DEL-737
chr12-98672264-DEL-70
chr12-98682215-DEL-52
chr12-98688679-DEL-1153
chr12-98694253-DEL-180
chr12-98729756-DEL-210
chr12-98772182-DEL-1834
chr12-98796446-DEL-77
chr12-98809816-DEL-52
chr12-98824226-DEL-205
chr12-98871546-DEL-61
chr12-98877197-DEL-122
chr12-98895923-DEL-75
chr12-98964968-DEL-461
chr12-98973409-DEL-99
chr12-98981795-DEL-202
chr12-98984740-DEL-190
chr12-99012286-DEL-53
chr12-99024504-DEL-549
chr12-99065693-DEL-136
chr12-99092486-DEL-58
chr12-99097328-DEL-138
chr12-99099864-DEL-6343
chr12-99099877-DEL-70
chr12-99107319-DEL-136
chr12-99130599-DEL-77
chr12-99147910-DEL-80
chr12-99155174-DEL-155
chr12-99155968-DEL-60
chr12-99211891-DEL-76
chr12-99267754-DEL-72
chr12-99293904-DEL-1428
chr12-99317001-DEL-217
chr12-99329007-DEL-95
chr12-99366795-DEL-73
chr12-99370114-DEL-193
chr12-99372284-DEL-66
chr12-99376204-DEL-304

chr12-99404436-DEL-186
chr12-99457811-DEL-52
chr12-99471022-DEL-485
chr12-99471389-DEL-4355
chr12-99471601-DEL-675
chr12-99471936-DEL-1040
chr12-99473052-DEL-215
chr12-99473451-DEL-1536
chr12-99474061-DEL-243
chr12-99474765-DEL-261
chr12-99475414-DEL-190
chr12-99490033-DEL-116
chr12-99521125-DEL-126
chr12-99551730-DEL-140
chr12-99598965-DEL-132
chr12-99607563-DEL-137
chr12-99622433-DEL-264
chr12-99626603-DEL-51
chr12-99671521-DEL-396
chr12-99721123-DEL-261
chr12-99725162-DEL-2452
chr12-99761226-DEL-62
chr12-99791729-DEL-77
chr12-99805124-DEL-121
chr12-99805348-DEL-432
chr12-99806361-DEL-117
chr12-99862979-DEL-166
chr12-99877286-DEL-66
chr12-99885604-DEL-1086
chr12-99896747-DEL-237
chr12-99900758-DEL-171
chr12-99937974-DEL-103
chr12-99970043-DEL-73
chr12-99974773-DEL-76
chr12-99991838-DEL-73
chr12-100014970-DEL-115
chr12-100047648-DEL-200
chr12-100062338-DEL-789
chr12-100088766-DEL-81
chr12-100103142-DEL-481
chr12-100123842-DEL-61
chr12-100128508-DEL-189
chr12-100130177-DEL-56
chr12-100191591-DEL-2903
chr12-100208533-DEL-62
chr12-100232744-DEL-82
chr12-100254103-DEL-74
chr12-100259084-DEL-68
chr12-100273711-DEL-54
chr12-100308864-DEL-210
chr12-100309627-DEL-166
chr12-100323238-DEL-77
chr12-100343822-DEL-145
chr12-100352848-DEL-215
chr12-100360812-DEL-65
chr12-100425100-DEL-1138
chr12-100429703-DEL-307
chr12-100429804-DEL-1660
chr12-100457242-DEL-980
chr12-100491653-DEL-185
chr12-100506695-DEL-204
chr12-100510170-DEL-69
chr12-100680969-DEL-70
chr12-100694682-DEL-79
chr12-100717918-DEL-56
chr12-100733069-DEL-2318
chr12-100758433-DEL-117
chr12-100763594-DEL-6365
chr12-100784744-DEL-457
chr12-100816781-DEL-200
chr12-100828257-DEL-54
chr12-100829177-DEL-1129
chr12-100830859-DEL-148
chr12-100832451-DEL-207
chr12-100835038-DEL-429
chr12-100850390-DEL-53
chr12-100886046-DEL-182

chr12-100900455-DEL-63
chr12-100913159-DEL-99
chr12-100952363-DEL-79
chr12-100967005-DEL-71
chr12-100977667-DEL-133
chr12-101032236-DEL-159
chr12-101037302-DEL-608
chr12-101071524-DEL-51
chr12-101098516-DEL-62
chr12-101138792-DEL-68
chr12-101154421-DEL-408
chr12-101169666-DEL-4456
chr12-101175375-DEL-1861
chr12-101187958-DEL-1999
chr12-101193126-DEL-15782
chr12-101204641-DEL-4120
chr12-101211589-DEL-189
chr12-101213817-DEL-448
chr12-101220391-DEL-114
chr12-101223997-DEL-1271
chr12-101241983-DEL-51
chr12-101255516-DEL-62
chr12-101257642-DEL-92
chr12-101260896-DEL-817
chr12-101267819-DEL-62
chr12-101308498-DEL-390
chr12-101311531-DEL-7126
chr12-101343403-DEL-4591
chr12-101362398-DEL-439
chr12-101372690-DEL-2625
chr12-101374629-DEL-721
chr12-101394143-DEL-6235
chr12-101405222-DEL-2000
chr12-101436816-DEL-766
chr12-101439342-DEL-59
chr12-101457924-DEL-55
chr12-101458594-DEL-135
chr12-101494839-DEL-779
chr12-101505354-DEL-554
chr12-101506180-DEL-154
chr12-101537334-DEL-460
chr12-101541300-DEL-204
chr12-101546133-DEL-416
chr12-101556172-DEL-70
chr12-101571541-DEL-322
chr12-101598577-DEL-673
chr12-101613683-DEL-202
chr12-101645464-DEL-224
chr12-101684985-DEL-61
chr12-101695870-DEL-7036
chr12-101723074-DEL-108
chr12-101757101-DEL-52
chr12-101765442-DEL-62
chr12-101806415-DEL-63
chr12-101807172-DEL-131
chr12-101817494-DEL-560
chr12-101843213-DEL-489
chr12-101873805-DEL-537
chr12-101902713-DEL-99
chr12-101907028-DEL-126
chr12-101920152-DEL-564
chr12-101930186-DEL-2991
chr12-101992863-DEL-1793
chr12-102013420-DEL-1643
chr12-102033880-DEL-52
chr12-102035565-DEL-75
chr12-102039061-DEL-1856
chr12-102100123-DEL-331
chr12-102101220-DEL-73
chr12-102141674-DEL-401
chr12-102155363-DEL-51
chr12-102184637-DEL-112
chr12-102192284-DEL-51
chr12-102207595-DEL-227
chr12-102235800-DEL-57
chr12-102262465-DEL-113
chr12-102290770-DEL-54

chr12-102310776-DEL-181
chr12-102320163-DEL-183
chr12-102322586-DEL-138
chr12-102343640-DEL-248
chr12-102413209-DEL-533
chr12-102417337-DEL-91
chr12-102431877-DEL-102
chr12-102456298-DEL-475
chr12-102461152-DEL-132
chr12-102480568-DEL-101
chr12-102496631-DEL-1414
chr12-102516146-DEL-51
chr12-102531790-DEL-1891
chr12-102555359-DEL-55
chr12-102617575-DEL-60
chr12-102622490-DEL-131
chr12-102625350-DEL-1537
chr12-102636173-DEL-882
chr12-102675304-DEL-212
chr12-102680981-DEL-55
chr12-102687709-DEL-171
chr12-102736796-DEL-104
chr12-102739529-DEL-82
chr12-102757822-DEL-103
chr12-102780917-DEL-243
chr12-102789621-DEL-729
chr12-102791175-DEL-2194
chr12-102832812-DEL-969
chr12-102834035-DEL-589
chr12-102847524-DEL-1494
chr12-102887742-DEL-179
chr12-102904808-DEL-59
chr12-102933545-DEL-54
chr12-102959895-DEL-95
chr12-102983106-DEL-425
chr12-103042919-DEL-117
chr12-103081857-DEL-448
chr12-103082589-DEL-66
chr12-103083033-DEL-457
chr12-103117737-DEL-70
chr12-103152159-DEL-89
chr12-103189476-DEL-83
chr12-103225426-DEL-330
chr12-103266469-DEL-96
chr12-103279716-DEL-60
chr12-103296866-DEL-99
chr12-103348150-DEL-285
chr12-103360729-DEL-314
chr12-103407137-DEL-7999
chr12-103418893-DEL-4527
chr12-103423780-DEL-13799
chr12-103428286-DEL-6201
chr12-103437991-DEL-1653
chr12-103444858-DEL-371
chr12-103505874-DEL-62
chr12-103521390-DEL-68
chr12-103544420-DEL-152
chr12-103560341-DEL-405
chr12-103563696-DEL-129
chr12-103564548-DEL-51
chr12-103572674-DEL-217
chr12-103575334-DEL-465
chr12-103583731-DEL-6187
chr12-103605457-DEL-1894
chr12-103608253-DEL-1520
chr12-103627248-DEL-700
chr12-103629646-DEL-420
chr12-103630737-DEL-628
chr12-103665026-DEL-89
chr12-103678229-DEL-70
chr12-103710122-DEL-1732
chr12-103716118-DEL-52
chr12-103719308-DEL-53
chr12-103727173-DEL-102
chr12-103739244-DEL-78
chr12-103742402-DEL-6633
chr12-103748506-DEL-213

chr12-103750566-DEL-1651
chr12-103756165-DEL-1760
chr12-103758559-DEL-1992
chr12-103760725-DEL-6948
chr12-103761602-DEL-7325
chr12-103779745-DEL-96
chr12-103793402-DEL-205
chr12-103799913-DEL-357
chr12-103800470-DEL-613
chr12-103801675-DEL-77
chr12-103805283-DEL-3987
chr12-103815568-DEL-96
chr12-103817115-DEL-104
chr12-103839256-DEL-7217
chr12-103898673-DEL-3843
chr12-103952909-DEL-76
chr12-103957232-DEL-840
chr12-103970599-DEL-1537
chr12-103974375-DEL-80
chr12-103979532-DEL-690
chr12-104019523-DEL-303
chr12-104024417-DEL-126
chr12-104028857-DEL-61
chr12-104031558-DEL-664
chr12-104032976-DEL-553
chr12-104034362-DEL-192
chr12-104041401-DEL-84
chr12-104052327-DEL-408
chr12-104057087-DEL-68
chr12-104074548-DEL-93
chr12-104079539-DEL-2839
chr12-104086401-DEL-110
chr12-104120554-DEL-397
chr12-104121789-DEL-312
chr12-104151997-DEL-70
chr12-104194086-DEL-83
chr12-104227907-DEL-68
chr12-104248439-DEL-190
chr12-104251796-DEL-1187
chr12-104253164-DEL-1497
chr12-104254729-DEL-114
chr12-104260015-DEL-56
chr12-104278230-DEL-684
chr12-104314519-DEL-53
chr12-104323365-DEL-4470
chr12-104382100-DEL-131
chr12-104385260-DEL-54
chr12-104387686-DEL-198
chr12-104391507-DEL-112
chr12-104403148-DEL-52
chr12-104408740-DEL-233
chr12-104413276-DEL-86
chr12-104418355-DEL-84
chr12-104448179-DEL-54
chr12-104478212-DEL-356
chr12-104555165-DEL-59
chr12-104584212-DEL-169
chr12-104611016-DEL-117
chr12-104633462-DEL-57
chr12-104643179-DEL-59
chr12-104658155-DEL-52
chr12-104661746-DEL-303
chr12-104745075-DEL-135
chr12-104748688-DEL-81
chr12-104754895-DEL-1035
chr12-104762906-DEL-73
chr12-104787267-DEL-132
chr12-104799408-DEL-182
chr12-104840476-DEL-336
chr12-104849393-DEL-465
chr12-104856541-DEL-52
chr12-104877450-DEL-168
chr12-104891039-DEL-224
chr12-104892305-DEL-102
chr12-104895574-DEL-64
chr12-104903146-DEL-84
chr12-104907432-DEL-129

chr12-104909953-DEL-75
chr12-104914503-DEL-65
chr12-104932243-DEL-61
chr12-104933011-DEL-78
chr12-104984641-DEL-100
chr12-104989827-DEL-445
chr12-105008706-DEL-73
chr12-105025601-DEL-178
chr12-105030336-DEL-71
chr12-105038840-DEL-59
chr12-105046562-DEL-2939
chr12-105071690-DEL-150
chr12-105073520-DEL-185
chr12-105078168-DEL-57
chr12-105108166-DEL-77
chr12-105115252-DEL-140
chr12-105124092-DEL-111
chr12-105127740-DEL-52
chr12-105143713-DEL-58
chr12-105148048-DEL-160
chr12-105148338-DEL-440
chr12-105150092-DEL-322
chr12-105196485-DEL-152
chr12-105220268-DEL-241
chr12-105225840-DEL-91
chr12-105236192-DEL-119
chr12-105241853-DEL-56
chr12-105284017-DEL-1145
chr12-105299642-DEL-105
chr12-105312730-DEL-222
chr12-105328835-DEL-396
chr12-105365886-DEL-52
chr12-105386486-DEL-66
chr12-105389030-DEL-74
chr12-105390784-DEL-407
chr12-105402365-DEL-87
chr12-105410331-DEL-660
chr12-105432761-DEL-108
chr12-105440193-DEL-1105
chr12-105510760-DEL-119
chr12-105511632-DEL-62
chr12-105536035-DEL-482
chr12-105537740-DEL-59
chr12-105572530-DEL-1514
chr12-105594283-DEL-2742
chr12-105597361-DEL-168
chr12-105635283-DEL-53
chr12-105686847-DEL-150
chr12-105721541-DEL-58
chr12-105723319-DEL-90
chr12-105733422-DEL-886
chr12-105749933-DEL-113
chr12-105792631-DEL-54
chr12-105817844-DEL-56
chr12-105826182-DEL-416
chr12-105840509-DEL-68
chr12-105846481-DEL-69
chr12-105872461-DEL-64
chr12-105873058-DEL-70
chr12-105876755-DEL-462
chr12-105886006-DEL-84
chr12-105913359-DEL-192
chr12-105914468-DEL-139
chr12-105920135-DEL-293
chr12-105925764-DEL-182
chr12-105948870-DEL-55
chr12-105961935-DEL-54
chr12-106025031-DEL-190
chr12-106072152-DEL-70
chr12-106077730-DEL-387
chr12-106081862-DEL-1267
chr12-106108603-DEL-117
chr12-106124911-DEL-105
chr12-106154302-DEL-150
chr12-106162691-DEL-54
chr12-106175507-DEL-282
chr12-106195177-DEL-71

chr12-106217593-DEL-170
chr12-106256121-DEL-54
chr12-106256818-DEL-199
chr12-106267783-DEL-68
chr12-106290060-DEL-522
chr12-106321247-DEL-244
chr12-106327742-DEL-901
chr12-106331543-DEL-260
chr12-106334082-DEL-137
chr12-106368950-DEL-55
chr12-106372838-DEL-1180
chr12-106374118-DEL-1124
chr12-106375027-DEL-239
chr12-106375696-DEL-336
chr12-106393817-DEL-54
chr12-106394859-DEL-56
chr12-106426305-DEL-57
chr12-106447368-DEL-146
chr12-106469033-DEL-226
chr12-106486400-DEL-54
chr12-106498354-DEL-78
chr12-106521866-DEL-1833
chr12-106543218-DEL-96
chr12-106571685-DEL-326
chr12-106583156-DEL-245
chr12-106592608-DEL-52
chr12-106597834-DEL-106
chr12-106600788-DEL-75
chr12-106604138-DEL-63
chr12-106629879-DEL-56
chr12-106630662-DEL-114
chr12-106643712-DEL-253
chr12-106648272-DEL-53
chr12-106655369-DEL-2152
chr12-106718584-DEL-66
chr12-106745758-DEL-51
chr12-106753827-DEL-61
chr12-106757484-DEL-56
chr12-106777471-DEL-1125
chr12-106780407-DEL-56
chr12-106823004-DEL-68
chr12-106841027-DEL-213
chr12-106844239-DEL-164
chr12-106852023-DEL-110
chr12-106862389-DEL-118
chr12-106925928-DEL-60
chr12-106974188-DEL-90
chr12-107042321-DEL-82
chr12-107047417-DEL-370
chr12-107055043-DEL-67
chr12-107077434-DEL-390
chr12-107111526-DEL-54
chr12-107132763-DEL-684
chr12-107135369-DEL-2342
chr12-107164662-DEL-151
chr12-107174004-DEL-224
chr12-107221557-DEL-975
chr12-107236296-DEL-232
chr12-107296591-DEL-57
chr12-107320122-DEL-607
chr12-107330323-DEL-114
chr12-107333860-DEL-2364
chr12-107359919-DEL-97
chr12-107415463-DEL-184
chr12-107451903-DEL-108
chr12-107454394-DEL-276
chr12-107460807-DEL-410
chr12-107486572-DEL-70
chr12-107507275-DEL-126
chr12-107522354-DEL-284
chr12-107522397-DEL-462
chr12-107543764-DEL-233
chr12-107546487-DEL-69
chr12-107549717-DEL-1414
chr12-107566096-DEL-342
chr12-107573138-DEL-59
chr12-107579855-DEL-70

chr12-107613644-DEL-144
chr12-107646926-DEL-55
chr12-107669568-DEL-356
chr12-107755725-DEL-84
chr12-107793020-DEL-116
chr12-107805822-DEL-310
chr12-107808836-DEL-224
chr12-107812976-DEL-746
chr12-107829120-DEL-1467
chr12-107830702-DEL-468
chr12-107843010-DEL-1374
chr12-107862541-DEL-209
chr12-107995341-DEL-129
chr12-108003173-DEL-66
chr12-108012844-DEL-270
chr12-108045520-DEL-4231
chr12-108049902-DEL-673
chr12-108059295-DEL-167
chr12-108174840-DEL-87
chr12-108181865-DEL-681
chr12-108189989-DEL-54
chr12-108216487-DEL-919
chr12-108225054-DEL-513
chr12-108225899-DEL-175
chr12-108239043-DEL-106
chr12-108248862-DEL-118
chr12-108254234-DEL-208
chr12-108265174-DEL-938
chr12-108289606-DEL-114
chr12-108303935-DEL-155
chr12-108304978-DEL-61
chr12-108315598-DEL-369
chr12-108324787-DEL-581
chr12-108332005-DEL-52
chr12-108350862-DEL-288
chr12-108351410-DEL-305
chr12-108354680-DEL-97
chr12-108370927-DEL-82
chr12-108418806-DEL-522
chr12-108420782-DEL-78
chr12-108443519-DEL-101
chr12-108467541-DEL-66
chr12-108486505-DEL-110
chr12-108492508-DEL-59
chr12-108568105-DEL-172
chr12-108590671-DEL-960
chr12-108603173-DEL-60
chr12-108621979-DEL-64
chr12-108664207-DEL-98
chr12-108676139-DEL-191
chr12-108695040-DEL-69
chr12-108724483-DEL-69
chr12-108735544-DEL-199
chr12-108751828-DEL-94
chr12-108754906-DEL-112
chr12-108804580-DEL-274
chr12-108838742-DEL-427
chr12-108846404-DEL-57
chr12-108894995-DEL-357
chr12-108898974-DEL-288
chr12-108901341-DEL-56
chr12-108925757-DEL-54
chr12-108944749-DEL-72
chr12-108971444-DEL-79
chr12-108991461-DEL-93
chr12-108993866-DEL-54
chr12-109040583-DEL-65
chr12-109071654-DEL-1021
chr12-109101680-DEL-2078
chr12-109107976-DEL-116
chr12-109143740-DEL-6901
chr12-109150919-DEL-4938
chr12-109182363-DEL-771
chr12-109183445-DEL-78
chr12-109196899-DEL-64
chr12-109197546-DEL-6486
chr12-109206242-DEL-6342

chr12-109215089-DEL-2184
chr12-109218744-DEL-153
chr12-109240014-DEL-1519
chr12-109288684-DEL-82
chr12-109290255-DEL-288
chr12-109308644-DEL-197
chr12-109324956-DEL-193
chr12-109396733-DEL-128
chr12-109397721-DEL-81
chr12-109445517-DEL-135
chr12-109451970-DEL-153
chr12-109453121-DEL-53
chr12-109458849-DEL-210
chr12-109503824-DEL-63
chr12-109561377-DEL-66
chr12-109596954-DEL-129
chr12-109756934-DEL-248
chr12-109766881-DEL-117
chr12-109769376-DEL-145
chr12-109770168-DEL-58
chr12-109800189-DEL-51
chr12-109823408-DEL-422
chr12-109829753-DEL-96
chr12-109842657-DEL-574
chr12-109931708-DEL-63
chr12-109932193-DEL-63
chr12-109977291-DEL-1248
chr12-109977299-DEL-1059
chr12-109977301-DEL-2716
chr12-109978532-DEL-206
chr12-109978670-DEL-776
chr12-109979642-DEL-188
chr12-109980869-DEL-97
chr12-109983530-DEL-806
chr12-110007986-DEL-238
chr12-110008499-DEL-1230
chr12-110018934-DEL-110
chr12-110020984-DEL-59
chr12-110040982-DEL-240
chr12-110046353-DEL-131
chr12-110057841-DEL-176
chr12-110080582-DEL-54
chr12-110096560-DEL-74
chr12-110100011-DEL-152
chr12-110100352-DEL-307
chr12-110115684-DEL-132
chr12-110116708-DEL-252
chr12-110129044-DEL-122
chr12-110146509-DEL-732
chr12-110159537-DEL-63
chr12-110183604-DEL-121
chr12-110228834-DEL-298
chr12-110255753-DEL-63
chr12-110298915-DEL-535
chr12-110306431-DEL-218
chr12-110328913-DEL-315
chr12-110335773-DEL-170
chr12-110337439-DEL-135
chr12-110337934-DEL-68
chr12-110360915-DEL-80
chr12-110372368-DEL-245
chr12-110399131-DEL-74
chr12-110448525-DEL-92
chr12-110471133-DEL-113
chr12-110476556-DEL-106
chr12-110530313-DEL-712
chr12-110571114-DEL-569
chr12-110581677-DEL-215
chr12-110619127-DEL-121
chr12-110630057-DEL-153
chr12-110638176-DEL-199
chr12-110642366-DEL-189
chr12-110646704-DEL-310
chr12-110667914-DEL-136
chr12-110676070-DEL-289
chr12-110686136-DEL-360
chr12-110694864-DEL-63

chr12-110700902-DEL-55
chr12-110733027-DEL-203
chr12-110754589-DEL-205
chr12-110849718-DEL-348
chr12-110851965-DEL-199
chr12-110865847-DEL-73
chr12-110890004-DEL-606
chr12-110897282-DEL-204
chr12-110918199-DEL-426
chr12-110920740-DEL-95
chr12-110980957-DEL-58
chr12-110981763-DEL-54
chr12-111035783-DEL-274
chr12-111036924-DEL-138
chr12-111039583-DEL-340
chr12-111042478-DEL-202
chr12-111070834-DEL-83
chr12-111113974-DEL-689
chr12-111122117-DEL-60
chr12-111144748-DEL-124
chr12-111146179-DEL-76
chr12-111180572-DEL-122
chr12-111202354-DEL-126
chr12-111226262-DEL-614
chr12-111249394-DEL-276
chr12-111256634-DEL-57
chr12-111281948-DEL-53
chr12-111293347-DEL-70
chr12-111307569-DEL-277
chr12-111311713-DEL-108
chr12-111314606-DEL-89
chr12-111318863-DEL-89
chr12-111364298-DEL-2269
chr12-111374734-DEL-75
chr12-111377198-DEL-201
chr12-111400160-DEL-60
chr12-111419224-DEL-128
chr12-111432864-DEL-76
chr12-111440920-DEL-180
chr12-111442537-DEL-57
chr12-111448207-DEL-196
chr12-111464291-DEL-82
chr12-111467660-DEL-355
chr12-111472318-DEL-602
chr12-111476561-DEL-74
chr12-111477614-DEL-72
chr12-111513573-DEL-811
chr12-111530147-DEL-76
chr12-111530867-DEL-184
chr12-111582893-DEL-199
chr12-111587659-DEL-252
chr12-111596523-DEL-51
chr12-111657285-DEL-88
chr12-111668428-DEL-512
chr12-111686819-DEL-133
chr12-111696797-DEL-224
chr12-111697645-DEL-211
chr12-111699156-DEL-78
chr12-111704657-DEL-82
chr12-111716095-DEL-254
chr12-111754351-DEL-57
chr12-111762689-DEL-982
chr12-111834203-DEL-560
chr12-111834219-DEL-200
chr12-111866594-DEL-56
chr12-111895298-DEL-66
chr12-111916450-DEL-62
chr12-111925421-DEL-431
chr12-111969760-DEL-88
chr12-111971372-DEL-486
chr12-112037475-DEL-94
chr12-112051277-DEL-663
chr12-112109540-DEL-51
chr12-112132518-DEL-53
chr12-112158508-DEL-1257
chr12-112173189-DEL-118
chr12-112181584-DEL-179

chr12-112196148-DEL-91
chr12-112236840-DEL-196
chr12-112251056-DEL-52
chr12-112263882-DEL-111
chr12-112292209-DEL-216
chr12-112347218-DEL-198
chr12-112358310-DEL-73
chr12-112380836-DEL-240
chr12-112386679-DEL-714
chr12-112402792-DEL-108
chr12-112411434-DEL-875
chr12-112452946-DEL-471
chr12-112505358-DEL-213
chr12-112513574-DEL-351
chr12-112514033-DEL-712
chr12-112527427-DEL-169
chr12-112538168-DEL-108
chr12-112544472-DEL-92
chr12-112548611-DEL-106
chr12-112553388-DEL-69
chr12-112573076-DEL-230
chr12-112603333-DEL-744
chr12-112607975-DEL-51
chr12-112628090-DEL-249
chr12-112637826-DEL-52
chr12-112653530-DEL-224
chr12-112666078-DEL-91
chr12-112751060-DEL-97
chr12-112774049-DEL-65
chr12-112807873-DEL-124
chr12-112834887-DEL-337
chr12-112840737-DEL-59
chr12-112842952-DEL-198
chr12-112856730-DEL-68
chr12-112922704-DEL-167
chr12-112957539-DEL-58
chr12-112986305-DEL-138
chr12-113012049-DEL-62
chr12-113041995-DEL-70
chr12-113043472-DEL-189
chr12-113046507-DEL-806
chr12-113068371-DEL-69
chr12-113073038-DEL-63
chr12-113131146-DEL-171
chr12-113137259-DEL-833
chr12-113143666-DEL-1378
chr12-113163018-DEL-695
chr12-113164482-DEL-60
chr12-113199622-DEL-285
chr12-113211332-DEL-2941
chr12-113274314-DEL-534
chr12-113276376-DEL-147
chr12-113298700-DEL-59
chr12-113300337-DEL-97
chr12-113403372-DEL-3655
chr12-113403801-DEL-792
chr12-113414166-DEL-5869
chr12-113414482-DEL-5860
chr12-113442878-DEL-52
chr12-113447878-DEL-243
chr12-113450566-DEL-188
chr12-113482178-DEL-100
chr12-113487143-DEL-60
chr12-113527263-DEL-173
chr12-113558976-DEL-964
chr12-113576848-DEL-76
chr12-113586557-DEL-2447
chr12-113604353-DEL-244
chr12-113611627-DEL-571
chr12-113619449-DEL-65
chr12-113620781-DEL-98
chr12-113629603-DEL-1879
chr12-113644206-DEL-369
chr12-113647557-DEL-186
chr12-113653615-DEL-2294
chr12-113653953-DEL-746
chr12-113660706-DEL-62

chr12-113670455-DEL-6438
chr12-113694513-DEL-143
chr12-113701821-DEL-299
chr12-113739397-DEL-68
chr12-113740194-DEL-418
chr12-113745164-DEL-85
chr12-113756326-DEL-227
chr12-113804124-DEL-1032
chr12-113805175-DEL-265
chr12-113829943-DEL-6460
chr12-113851997-DEL-371
chr12-113870577-DEL-13626
chr12-113884514-DEL-3019
chr12-113893298-DEL-51
chr12-113904877-DEL-375
chr12-113907164-DEL-126
chr12-113933539-DEL-3170
chr12-113934946-DEL-360
chr12-113944488-DEL-64
chr12-113946156-DEL-2507
chr12-113947745-DEL-745
chr12-113959929-DEL-185
chr12-113960476-DEL-213
chr12-113979210-DEL-1138
chr12-113983729-DEL-1068
chr12-114006291-DEL-503
chr12-114012520-DEL-1756
chr12-114036568-DEL-6049
chr12-114059645-DEL-204
chr12-114070533-DEL-166
chr12-114074677-DEL-202
chr12-114076873-DEL-6165
chr12-114087217-DEL-1024
chr12-114100285-DEL-529
chr12-114109630-DEL-68
chr12-114115664-DEL-16309
chr12-114173760-DEL-414
chr12-114198588-DEL-120
chr12-114228415-DEL-82
chr12-114248908-DEL-52
chr12-114266678-DEL-487
chr12-114267718-DEL-8161
chr12-114288238-DEL-7005
chr12-114290548-DEL-4971
chr12-114298200-DEL-59
chr12-114305842-DEL-119
chr12-114375796-DEL-273
chr12-114376867-DEL-8858
chr12-114393855-DEL-127
chr12-114413739-DEL-1429
chr12-114413742-DEL-115
chr12-114421002-DEL-78
chr12-114437524-DEL-13535
chr12-114445724-DEL-3135
chr12-114455027-DEL-1035
chr12-114457611-DEL-10666
chr12-114466044-DEL-1086
chr12-114497163-DEL-1925
chr12-114500133-DEL-1635
chr12-114545765-DEL-52
chr12-114594423-DEL-1251
chr12-114611072-DEL-401
chr12-114611552-DEL-53
chr12-114617580-DEL-293
chr12-114631494-DEL-52
chr12-114638884-DEL-188
chr12-114661591-DEL-5221
chr12-114662633-DEL-115
chr12-114671984-DEL-60
chr12-114677561-DEL-224
chr12-114699236-DEL-60
chr12-114702525-DEL-2310
chr12-114713869-DEL-1429
chr12-114716831-DEL-188
chr12-114732092-DEL-70
chr12-114735907-DEL-107
chr12-114738528-DEL-70

chr12-114748314-DEL-54
chr12-114759914-DEL-2031
chr12-114787123-DEL-150
chr12-114789125-DEL-5927
chr12-114812958-DEL-1706
chr12-114820817-DEL-58
chr12-114821567-DEL-1426
chr12-114825341-DEL-2015
chr12-114838387-DEL-1812
chr12-114840772-DEL-293
chr12-114843225-DEL-59
chr12-114929526-DEL-72
chr12-115063530-DEL-7379
chr12-115071192-DEL-333
chr12-115080397-DEL-2572
chr12-115095352-DEL-102
chr12-115097565-DEL-79
chr12-115160577-DEL-52
chr12-115169733-DEL-1072
chr12-115208879-DEL-1522
chr12-115211877-DEL-62
chr12-115213260-DEL-291
chr12-115217385-DEL-70
chr12-115219052-DEL-2898
chr12-115227140-DEL-140
chr12-115231482-DEL-799
chr12-115232326-DEL-11610
chr12-115233314-DEL-7096
chr12-115246831-DEL-4751
chr12-115260486-DEL-160
chr12-115268194-DEL-1419
chr12-115271714-DEL-366
chr12-115299638-DEL-466
chr12-115308514-DEL-102
chr12-115310745-DEL-62
chr12-115340558-DEL-300
chr12-115343406-DEL-59
chr12-115352357-DEL-1561
chr12-115355810-DEL-260
chr12-115356790-DEL-291
chr12-115360931-DEL-67
chr12-115363452-DEL-1207
chr12-115381974-DEL-6549
chr12-115400627-DEL-6329
chr12-115439683-DEL-350
chr12-115444569-DEL-616
chr12-115474260-DEL-86
chr12-115475069-DEL-6027
chr12-115488800-DEL-511
chr12-115514669-DEL-93
chr12-115518884-DEL-822
chr12-115521630-DEL-7352
chr12-115531401-DEL-1797
chr12-115534229-DEL-1991
chr12-115543196-DEL-6388
chr12-115554366-DEL-303
chr12-115588828-DEL-806
chr12-115590158-DEL-376
chr12-115598723-DEL-181
chr12-115606037-DEL-203
chr12-115612911-DEL-206
chr12-115629209-DEL-6081
chr12-115631211-DEL-10478
chr12-115635376-DEL-84
chr12-115636843-DEL-1702
chr12-115645834-DEL-114
chr12-115648871-DEL-6362
chr12-115656618-DEL-59
chr12-115677209-DEL-100
chr12-115689318-DEL-6263
chr12-115696262-DEL-6295
chr12-115701415-DEL-59
chr12-115711191-DEL-1170
chr12-115717693-DEL-100
chr12-115737739-DEL-245
chr12-115742634-DEL-616
chr12-115761144-DEL-51

chr12-115769366-DEL-86
chr12-115782248-DEL-1604
chr12-115812331-DEL-166
chr12-115828583-DEL-92
chr12-115831464-DEL-282
chr12-115835707-DEL-52
chr12-115836299-DEL-253
chr12-115842558-DEL-895
chr12-115842562-DEL-19566
chr12-115843805-DEL-124
chr12-115849768-DEL-6227
chr12-115854340-DEL-62
chr12-115867019-DEL-8409
chr12-115885007-DEL-56
chr12-115900187-DEL-194
chr12-115911636-DEL-2756
chr12-115942052-DEL-320
chr12-115944312-DEL-12952
chr12-115946641-DEL-6921
chr12-115952713-DEL-428
chr12-115962359-DEL-723
chr12-115962471-DEL-723
chr12-115966262-DEL-51
chr12-115980855-DEL-128
chr12-115996891-DEL-1850
chr12-116001318-DEL-250
chr12-116025836-DEL-374
chr12-116027465-DEL-423
chr12-116069889-DEL-1343
chr12-116151134-DEL-311
chr12-116177515-DEL-155
chr12-116192009-DEL-126
chr12-116211984-DEL-249
chr12-116212960-DEL-91
chr12-116213061-DEL-717
chr12-116213482-DEL-907
chr12-116214086-DEL-660
chr12-116214266-DEL-97
chr12-116214603-DEL-133
chr12-116214961-DEL-435
chr12-116235685-DEL-51
chr12-116239960-DEL-52
chr12-116340542-DEL-501
chr12-116347905-DEL-1058
chr12-116349044-DEL-4550
chr12-116355741-DEL-171
chr12-116395217-DEL-61
chr12-116441991-DEL-224
chr12-116447624-DEL-58
chr12-116455932-DEL-86
chr12-116478518-DEL-68
chr12-116481650-DEL-73
chr12-116504605-DEL-6614
chr12-116520894-DEL-6380
chr12-116540652-DEL-850
chr12-116546608-DEL-252
chr12-116564603-DEL-55
chr12-116602638-DEL-67
chr12-116607379-DEL-215
chr12-116613188-DEL-52
chr12-116622635-DEL-231
chr12-116646176-DEL-206
chr12-116655496-DEL-167
chr12-116658154-DEL-677
chr12-116663341-DEL-6378
chr12-116671844-DEL-93
chr12-116695802-DEL-510
chr12-116717505-DEL-53
chr12-116721831-DEL-187
chr12-116751389-DEL-206
chr12-116786916-DEL-53
chr12-116841829-DEL-62
chr12-116851844-DEL-130
chr12-116944955-DEL-54
chr12-116970855-DEL-554
chr12-117023031-DEL-55
chr12-117025087-DEL-244

chr12-117134571-DEL-60
chr12-117176348-DEL-52
chr12-117203047-DEL-162
chr12-117253450-DEL-116
chr12-117258040-DEL-2317
chr12-117298303-DEL-62
chr12-117331465-DEL-2420
chr12-117342107-DEL-247
chr12-117349181-DEL-78
chr12-117351783-DEL-183
chr12-117355241-DEL-449
chr12-117357949-DEL-180
chr12-117358565-DEL-84
chr12-117358834-DEL-432
chr12-117399115-DEL-57
chr12-117409428-DEL-129
chr12-117427789-DEL-78
chr12-117446296-DEL-52
chr12-117484949-DEL-120
chr12-117486358-DEL-60
chr12-117556920-DEL-87
chr12-117561350-DEL-54
chr12-117578204-DEL-102
chr12-117579705-DEL-137
chr12-117627833-DEL-272
chr12-117664881-DEL-97
chr12-117701617-DEL-349
chr12-117711278-DEL-51
chr12-117724177-DEL-88
chr12-117753596-DEL-2489
chr12-117760556-DEL-192
chr12-117804991-DEL-71
chr12-117812854-DEL-315
chr12-117825160-DEL-626
chr12-117869588-DEL-118
chr12-117899362-DEL-83
chr12-117932634-DEL-81
chr12-117937930-DEL-183
chr12-118072251-DEL-120
chr12-118108680-DEL-174
chr12-118137263-DEL-397
chr12-118144353-DEL-51
chr12-118266730-DEL-137
chr12-118275717-DEL-52
chr12-118280009-DEL-94
chr12-118304721-DEL-6332
chr12-118304739-DEL-4005
chr12-118309536-DEL-1517
chr12-118317992-DEL-4617
chr12-118326970-DEL-1630
chr12-118385733-DEL-78
chr12-118396869-DEL-112
chr12-118486938-DEL-116
chr12-118486944-DEL-1008
chr12-118494694-DEL-74
chr12-118514234-DEL-55
chr12-118546134-DEL-56
chr12-118610585-DEL-214
chr12-118631888-DEL-123
chr12-118633493-DEL-54
chr12-118635447-DEL-58
chr12-118647373-DEL-426
chr12-118725320-DEL-201
chr12-118878639-DEL-90
chr12-118918629-DEL-5574
chr12-119043939-DEL-214
chr12-119082505-DEL-854
chr12-119085890-DEL-6415
chr12-119094659-DEL-175
chr12-119213836-DEL-2557
chr12-119309933-DEL-7531
chr12-119357508-DEL-247
chr12-119420133-DEL-65
chr12-119460341-DEL-440
chr12-119596349-DEL-75
chr12-119602263-DEL-558
chr12-119607945-DEL-104

chr12-119622176-DEL-12804
chr12-119637268-DEL-208
chr12-119649413-DEL-70
chr12-119716268-DEL-198
chr12-119736844-DEL-71
chr12-119750318-DEL-272
chr12-119765671-DEL-398
chr12-119768344-DEL-1223
chr12-119773464-DEL-3229
chr12-119817343-DEL-1689
chr12-119819082-DEL-3629
chr12-119827969-DEL-347
chr12-119831222-DEL-84
chr12-119853582-DEL-94
chr12-119854835-DEL-822
chr12-119870743-DEL-2589
chr12-119907307-DEL-846
chr12-119931041-DEL-9062
chr12-119950492-DEL-6563
chr12-119966341-DEL-778
chr13-3057061-DEL-4281
chr13-3068862-DEL-15423
chr13-3087273-DEL-215
chr13-3378290-DEL-3106
chr13-3384159-DEL-1044
chr13-3398478-DEL-3271
chr13-3433703-DEL-3548
chr13-3473437-DEL-195
chr13-3489015-DEL-84
chr13-3523542-DEL-115
chr13-3534051-DEL-4841
chr13-3550519-DEL-82
chr13-3556855-DEL-3936
chr13-3572126-DEL-55
chr13-3587577-DEL-55
chr13-3605123-DEL-338
chr13-3627991-DEL-673
chr13-3637300-DEL-230
chr13-3646978-DEL-207
chr13-3671055-DEL-256
chr13-3695055-DEL-501
chr13-3714622-DEL-65
chr13-3728432-DEL-57
chr13-3772510-DEL-100
chr13-3782666-DEL-6243
chr13-3790689-DEL-52
chr13-3817628-DEL-53
chr13-3830262-DEL-183
chr13-3865469-DEL-207
chr13-3890057-DEL-54
chr13-3901565-DEL-351
chr13-3906763-DEL-110
chr13-4009468-DEL-912
chr13-4012546-DEL-408
chr13-4014777-DEL-102
chr13-4108500-DEL-63
chr13-4119909-DEL-121
chr13-4149244-DEL-198
chr13-4161087-DEL-59
chr13-4197855-DEL-53
chr13-4207530-DEL-1943
chr13-4209566-DEL-152
chr13-4211123-DEL-52
chr13-4234553-DEL-443
chr13-4256938-DEL-169
chr13-4307016-DEL-75
chr13-4338496-DEL-1848
chr13-4342159-DEL-850
chr13-4344334-DEL-188
chr13-4346233-DEL-719
chr13-4361343-DEL-486
chr13-4401859-DEL-89
chr13-4412299-DEL-84
chr13-4458698-DEL-675
chr13-4491758-DEL-3133
chr13-4508724-DEL-336
chr13-4613142-DEL-525

chr13-4623708-DEL-78
chr13-4654633-DEL-68
chr13-4655239-DEL-114
chr13-4655898-DEL-230
chr13-4656492-DEL-446
chr13-4686062-DEL-1234
chr13-4714571-DEL-53
chr13-4750964-DEL-638
chr13-4765485-DEL-149
chr13-4811971-DEL-3472
chr13-4829724-DEL-86
chr13-4850245-DEL-287
chr13-4863976-DEL-6607
chr13-4877503-DEL-3005
chr13-4890873-DEL-1574
chr13-4892767-DEL-617
chr13-4900387-DEL-4601
chr13-4911197-DEL-229
chr13-4938274-DEL-593
chr13-4951734-DEL-3276
chr13-4989015-DEL-1482
chr13-5001557-DEL-201
chr13-5009220-DEL-55
chr13-5029958-DEL-10787
chr13-5031789-DEL-75
chr13-5045539-DEL-1501
chr13-5055264-DEL-824
chr13-5081993-DEL-8708
chr13-5124876-DEL-71
chr13-5130616-DEL-56
chr13-5139436-DEL-51
chr13-5149794-DEL-132
chr13-5175319-DEL-1520
chr13-5176969-DEL-71
chr13-5177484-DEL-7977
chr13-5199019-DEL-510
chr13-5200344-DEL-77
chr13-5237090-DEL-284
chr13-5241414-DEL-406
chr13-5256947-DEL-1440
chr13-5285042-DEL-74
chr13-5315851-DEL-225
chr13-5327864-DEL-139
chr13-5340828-DEL-82
chr13-5372623-DEL-492
chr13-5413027-DEL-248
chr13-5426409-DEL-569
chr13-5435724-DEL-1127
chr13-5437299-DEL-255
chr13-5439504-DEL-232
chr13-5439989-DEL-401
chr13-5464115-DEL-332
chr13-5480577-DEL-123
chr13-5510516-DEL-66
chr13-5529169-DEL-202
chr13-5532180-DEL-52
chr13-5585540-DEL-71
chr13-5613008-DEL-127
chr13-5621292-DEL-743
chr13-5652827-DEL-106
chr13-5685452-DEL-84
chr13-5723709-DEL-85
chr13-5769332-DEL-944
chr13-5772235-DEL-89
chr13-5804062-DEL-581
chr13-5856369-DEL-84
chr13-5903605-DEL-1658
chr13-5909513-DEL-51
chr13-5958214-DEL-653
chr13-5976833-DEL-72
chr13-5982619-DEL-229
chr13-6005238-DEL-100
chr13-6013528-DEL-601
chr13-6043399-DEL-79
chr13-6047052-DEL-186
chr13-6059110-DEL-150
chr13-6095583-DEL-77

chr13-6118279-DEL-75
chr13-6126719-DEL-274
chr13-6150442-DEL-51
chr13-6200150-DEL-222
chr13-6210508-DEL-1747
chr13-6274373-DEL-89
chr13-6291596-DEL-1066
chr13-6299346-DEL-463
chr13-6332473-DEL-52
chr13-6340687-DEL-1373
chr13-6365497-DEL-58
chr13-6368941-DEL-56
chr13-6372341-DEL-71
chr13-6377426-DEL-98
chr13-6381686-DEL-62
chr13-6384321-DEL-583
chr13-6448597-DEL-129
chr13-6480552-DEL-135
chr13-6502051-DEL-991
chr13-6502094-DEL-315
chr13-6502744-DEL-79
chr13-6514430-DEL-93
chr13-6527957-DEL-252
chr13-6554118-DEL-56
chr13-6557460-DEL-64
chr13-6561098-DEL-4482
chr13-6572125-DEL-6394
chr13-6595986-DEL-113
chr13-6601177-DEL-156
chr13-6604059-DEL-53
chr13-6634134-DEL-81
chr13-6642364-DEL-97
chr13-6655816-DEL-7080
chr13-6656318-DEL-2100
chr13-6656440-DEL-720
chr13-6659502-DEL-60
chr13-6659868-DEL-147
chr13-6659919-DEL-2520
chr13-6661657-DEL-900
chr13-6677160-DEL-223
chr13-6690492-DEL-102
chr13-6708502-DEL-64
chr13-6711224-DEL-135
chr13-6726882-DEL-241
chr13-6730502-DEL-97
chr13-6742251-DEL-3597
chr13-6747046-DEL-6215
chr13-6760422-DEL-63
chr13-6761501-DEL-5319
chr13-6772502-DEL-6591
chr13-6779485-DEL-1221
chr13-6779677-DEL-6447
chr13-6828619-DEL-212
chr13-6841392-DEL-52
chr13-6883665-DEL-93
chr13-6907332-DEL-90
chr13-6918764-DEL-395
chr13-6988339-DEL-133
chr13-7001907-DEL-1306
chr13-7014100-DEL-735
chr13-7015892-DEL-3373
chr13-7022201-DEL-1773
chr13-7031432-DEL-134
chr13-7049027-DEL-736
chr13-7062179-DEL-446
chr13-7078387-DEL-1347
chr13-7097940-DEL-52
chr13-7117000-DEL-9406
chr13-7128872-DEL-509
chr13-7139614-DEL-173
chr13-7180445-DEL-212
chr13-7193275-DEL-77
chr13-7263633-DEL-6176
chr13-7272110-DEL-6237
chr13-7314728-DEL-96
chr13-7338314-DEL-1070
chr13-7340106-DEL-537

chr13-7346559-DEL-1135
chr13-7347366-DEL-85
chr13-7348296-DEL-1615
chr13-7349194-DEL-215
chr13-7355662-DEL-52
chr13-7367911-DEL-2640
chr13-7371916-DEL-51
chr13-7382103-DEL-2524
chr13-7384778-DEL-491
chr13-7389455-DEL-6197
chr13-7419824-DEL-2100
chr13-7450540-DEL-5122
chr13-7458534-DEL-441
chr13-7511151-DEL-477
chr13-7527323-DEL-3535
chr13-7547381-DEL-2188
chr13-7550770-DEL-2202
chr13-7651810-DEL-3646
chr13-7672861-DEL-55
chr13-7681283-DEL-377
chr13-7683933-DEL-446
chr13-7698030-DEL-440
chr13-7746839-DEL-76
chr13-7771846-DEL-426
chr13-7779659-DEL-1288
chr13-7784156-DEL-58
chr13-7792490-DEL-52
chr13-7799595-DEL-199
chr13-7844496-DEL-257
chr13-7856006-DEL-6241
chr13-7887410-DEL-79
chr13-7891983-DEL-682
chr13-7894966-DEL-167
chr13-7919535-DEL-349
chr13-7936541-DEL-482
chr13-7943421-DEL-6343
chr13-7950173-DEL-420
chr13-7959937-DEL-59
chr13-7970786-DEL-52
chr13-7972154-DEL-820
chr13-7974123-DEL-1329
chr13-8013771-DEL-1611
chr13-8015403-DEL-1046
chr13-8023855-DEL-132
chr13-8035612-DEL-4272
chr13-8040578-DEL-1162
chr13-8057106-DEL-732
chr13-8141848-DEL-126
chr13-8157285-DEL-109
chr13-8203414-DEL-1220
chr13-8217044-DEL-6250
chr13-8250169-DEL-174
chr13-8252151-DEL-71
chr13-8269527-DEL-950
chr13-8288393-DEL-151
chr13-8289684-DEL-146
chr13-8340368-DEL-319
chr13-8360143-DEL-190
chr13-8380808-DEL-65
chr13-8440951-DEL-164
chr13-8445036-DEL-55
chr13-8446815-DEL-53
chr13-8456941-DEL-158
chr13-8475201-DEL-829
chr13-8495808-DEL-241
chr13-8603335-DEL-78
chr13-8627635-DEL-349
chr13-8692694-DEL-251
chr13-8739361-DEL-78
chr13-8746700-DEL-148
chr13-8769780-DEL-123
chr13-8804862-DEL-1053
chr13-8805265-DEL-124
chr13-8819790-DEL-344
chr13-8822401-DEL-334
chr13-8836428-DEL-74
chr13-8849789-DEL-55

chr13-8859538-DEL-207
chr13-8879553-DEL-54
chr13-8927332-DEL-1217
chr13-8934127-DEL-203
chr13-8936077-DEL-68
chr13-8971578-DEL-55
chr13-8973644-DEL-504
chr13-8991371-DEL-119
chr13-8993044-DEL-66
chr13-9000302-DEL-740
chr13-9040457-DEL-171
chr13-9047611-DEL-291
chr13-9051376-DEL-163
chr13-9059173-DEL-327
chr13-9061961-DEL-169
chr13-9082179-DEL-186
chr13-9104595-DEL-118
chr13-9120070-DEL-54
chr13-9203773-DEL-62
chr13-9215634-DEL-515
chr13-9239003-DEL-110
chr13-9240730-DEL-88
chr13-9245931-DEL-590
chr13-9269207-DEL-64
chr13-9308777-DEL-504
chr13-9371340-DEL-109
chr13-9401209-DEL-53
chr13-9409966-DEL-199
chr13-9417353-DEL-84
chr13-9442683-DEL-194
chr13-9449866-DEL-1254
chr13-9453733-DEL-384
chr13-9471488-DEL-5226
chr13-9489314-DEL-79
chr13-9509416-DEL-55
chr13-9562803-DEL-16294
chr13-9568129-DEL-349
chr13-9572159-DEL-839
chr13-9574605-DEL-3404
chr13-9637978-DEL-2201
chr13-9679702-DEL-750
chr13-9703043-DEL-78
chr13-9730515-DEL-159
chr13-9782053-DEL-140
chr13-9789183-DEL-56
chr13-9790694-DEL-91
chr13-9797809-DEL-203
chr13-9798936-DEL-275
chr13-9805980-DEL-84
chr13-9818328-DEL-61
chr13-9847882-DEL-61
chr13-9856317-DEL-1154
chr13-9863404-DEL-135
chr13-9870749-DEL-138
chr13-9881028-DEL-2731
chr13-9882052-DEL-6650
chr13-9882428-DEL-292
chr13-9886976-DEL-1724
chr13-9891793-DEL-519
chr13-9895454-DEL-55
chr13-9902813-DEL-7404
chr13-9933608-DEL-825
chr13-9934063-DEL-207
chr13-9939787-DEL-174
chr13-9972926-DEL-819
chr13-10001740-DEL-65
chr13-10037772-DEL-65
chr13-10099636-DEL-180
chr13-10133762-DEL-686
chr13-10156448-DEL-1522
chr13-10176145-DEL-58
chr13-10199625-DEL-61
chr13-10204902-DEL-804
chr13-10206678-DEL-82
chr13-10210096-DEL-77
chr13-10252995-DEL-66
chr13-10268772-DEL-174

chr13-10349559-DEL-1038
chr13-10351910-DEL-6232
chr13-10362686-DEL-92
chr13-10367786-DEL-75
chr13-10372113-DEL-67
chr13-10425443-DEL-767
chr13-10467723-DEL-132
chr13-10473073-DEL-54
chr13-10474039-DEL-203
chr13-10479079-DEL-3665
chr13-10499913-DEL-330
chr13-10503764-DEL-118
chr13-10524309-DEL-55
chr13-10533822-DEL-56
chr13-10558834-DEL-55
chr13-10559035-DEL-1127
chr13-10566006-DEL-1889
chr13-10569171-DEL-709
chr13-10618762-DEL-54
chr13-10619919-DEL-80
chr13-10637995-DEL-253
chr13-10648208-DEL-813
chr13-10670149-DEL-96
chr13-10691691-DEL-59
chr13-10700839-DEL-112
chr13-10715827-DEL-111
chr13-10733139-DEL-346
chr13-10746672-DEL-89
chr13-10767788-DEL-367
chr13-10810704-DEL-358
chr13-10830293-DEL-60
chr13-10886843-DEL-2501
chr13-10901707-DEL-351
chr13-10905807-DEL-7569
chr13-10923903-DEL-92
chr13-10960061-DEL-154
chr13-11000658-DEL-571
chr13-11008595-DEL-8488
chr13-11008597-DEL-1534
chr13-11016413-DEL-670
chr13-11022904-DEL-67
chr13-11093070-DEL-1206
chr13-11122665-DEL-172
chr13-11126477-DEL-1849
chr13-11128461-DEL-60
chr13-11142212-DEL-2022
chr13-11149834-DEL-6164
chr13-11162956-DEL-68
chr13-11209145-DEL-150
chr13-11210358-DEL-197
chr13-11226084-DEL-1290
chr13-11232608-DEL-129
chr13-11235955-DEL-817
chr13-11258879-DEL-1324
chr13-11353964-DEL-1447
chr13-11358874-DEL-76
chr13-11394279-DEL-6197
chr13-11446748-DEL-51
chr13-11474262-DEL-64
chr13-11482252-DEL-1250
chr13-11510528-DEL-222
chr13-11535638-DEL-203
chr13-11578993-DEL-104
chr13-11586363-DEL-400
chr13-11622986-DEL-304
chr13-11622997-DEL-1474
chr13-11703209-DEL-1849
chr13-11727757-DEL-169
chr13-11788914-DEL-1486
chr13-11854016-DEL-90
chr13-11896568-DEL-237
chr13-11901434-DEL-522
chr13-11904829-DEL-132
chr13-11913845-DEL-164
chr13-11926885-DEL-334
chr13-11937889-DEL-866
chr13-12000350-DEL-924

chr13-12006759-DEL-63
chr13-12007490-DEL-596
chr13-12008416-DEL-108
chr13-12014989-DEL-1679
chr13-12075325-DEL-66
chr13-12080604-DEL-303
chr13-12081364-DEL-399
chr13-12097077-DEL-222
chr13-12112543-DEL-97
chr13-12133399-DEL-1399
chr13-12137468-DEL-52
chr13-12138516-DEL-53
chr13-12152400-DEL-722
chr13-12180271-DEL-51
chr13-12184915-DEL-103
chr13-12202217-DEL-317
chr13-12202506-DEL-317
chr13-12216661-DEL-52
chr13-12238009-DEL-1519
chr13-12239938-DEL-252
chr13-12241895-DEL-220
chr13-12250969-DEL-197
chr13-12252464-DEL-78
chr13-12276922-DEL-308
chr13-12282529-DEL-193
chr13-12296888-DEL-204
chr13-12297886-DEL-533
chr13-12321268-DEL-61
chr13-12346351-DEL-199
chr13-12369652-DEL-399
chr13-12405243-DEL-1039
chr13-12414496-DEL-194
chr13-12451150-DEL-377
chr13-12451703-DEL-56
chr13-12508898-DEL-92
chr13-12528583-DEL-225
chr13-12541191-DEL-314
chr13-12541557-DEL-1151
chr13-12551952-DEL-393
chr13-12555645-DEL-204
chr13-12576383-DEL-60
chr13-12576997-DEL-62
chr13-12611302-DEL-52
chr13-12612088-DEL-659
chr13-12617505-DEL-1141
chr13-12619029-DEL-51
chr13-12639335-DEL-61
chr13-12645661-DEL-522
chr13-12668058-DEL-6439
chr13-12684979-DEL-55
chr13-12686471-DEL-58
chr13-12693188-DEL-1252
chr13-12694520-DEL-768
chr13-12695333-DEL-119
chr13-12705316-DEL-180
chr13-12709279-DEL-216
chr13-12735994-DEL-76
chr13-13145632-DEL-79
chr13-13196180-DEL-201
chr13-13431554-DEL-2303
chr13-13434012-DEL-433
chr13-13498800-DEL-53
chr13-13511205-DEL-128
chr13-13512295-DEL-3127
chr13-13520792-DEL-1133
chr13-13522080-DEL-178
chr13-13537028-DEL-133
chr13-13542747-DEL-61
chr13-13552075-DEL-65
chr13-13581462-DEL-130
chr13-13583065-DEL-52
chr13-13589831-DEL-129
chr13-13600179-DEL-182
chr13-13601654-DEL-52
chr13-13626289-DEL-58
chr13-13667959-DEL-170
chr13-13692293-DEL-584

chr13-13696107-DEL-1131
chr13-13696348-DEL-849
chr13-13702086-DEL-1078
chr13-13702214-DEL-438
chr13-13750187-DEL-96
chr13-13757406-DEL-77
chr13-13770667-DEL-436
chr13-13843801-DEL-60
chr13-13887593-DEL-8735
chr13-13927735-DEL-986
chr13-13927953-DEL-237
chr13-13928918-DEL-1125
chr13-13928934-DEL-852
chr13-13929593-DEL-324
chr13-13940592-DEL-95
chr13-13993656-DEL-400
chr13-14016150-DEL-2418
chr13-14035219-DEL-354
chr13-14043682-DEL-144
chr13-14046375-DEL-64
chr13-14064383-DEL-158
chr13-14076873-DEL-7208
chr13-14085317-DEL-628
chr13-14088866-DEL-55
chr13-14108241-DEL-333
chr13-14108574-DEL-9886
chr13-14123689-DEL-3180
chr13-14152663-DEL-178
chr13-14166703-DEL-119
chr13-14211815-DEL-440
chr13-14226421-DEL-51
chr13-14228071-DEL-68
chr13-14239571-DEL-54
chr13-14251327-DEL-112
chr13-14262018-DEL-127
chr13-14265485-DEL-1690
chr13-14272109-DEL-119
chr13-14278841-DEL-544
chr13-14295087-DEL-1285
chr13-14298166-DEL-223
chr13-14305277-DEL-117
chr13-14356122-DEL-278
chr13-14391202-DEL-223
chr13-14413615-DEL-316
chr13-14445379-DEL-60
chr13-14492700-DEL-114
chr13-14517276-DEL-52
chr13-14522471-DEL-106
chr13-14524406-DEL-232
chr13-14547244-DEL-78
chr13-14550174-DEL-87
chr13-14572052-DEL-1570
chr13-14597236-DEL-249
chr13-14603506-DEL-134
chr13-14624751-DEL-603
chr13-14629073-DEL-788
chr13-14630033-DEL-490
chr13-14640184-DEL-1672
chr13-14669965-DEL-522
chr13-14697871-DEL-59
chr13-14701443-DEL-225
chr13-14702330-DEL-718
chr13-14707172-DEL-146
chr13-14738439-DEL-7321
chr13-14759636-DEL-249
chr13-14763595-DEL-173
chr13-14765281-DEL-321
chr13-14766960-DEL-2929
chr13-14781107-DEL-632
chr13-14787173-DEL-86
chr13-14792024-DEL-518
chr13-14837061-DEL-1154
chr13-14848928-DEL-134
chr13-14874487-DEL-57
chr13-14877234-DEL-201
chr13-14899005-DEL-89
chr13-14903724-DEL-1412

chr13-14916341-DEL-6930
chr13-14938102-DEL-6229
chr13-14946013-DEL-6196
chr13-14955536-DEL-2717
chr13-14997596-DEL-349
chr13-15000035-DEL-839
chr13-15016588-DEL-774
chr13-15024144-DEL-377
chr13-15026396-DEL-6959
chr13-15033384-DEL-2131
chr13-15041621-DEL-4568
chr13-15048052-DEL-3930
chr13-15064028-DEL-56
chr13-15080088-DEL-331
chr13-15156430-DEL-78
chr13-15163516-DEL-2496
chr13-15174048-DEL-116
chr13-15178775-DEL-852
chr13-15185368-DEL-329
chr13-15337088-DEL-96
chr13-15351724-DEL-1787
chr13-15410815-DEL-6267
chr13-15421417-DEL-12863
chr13-15441222-DEL-6710
chr13-15464089-DEL-174
chr13-15498951-DEL-351
chr13-15522437-DEL-372
chr13-15525214-DEL-400
chr13-15528416-DEL-53
chr13-15544701-DEL-70
chr13-15559815-DEL-440
chr13-15563227-DEL-82
chr13-15571715-DEL-359
chr13-15586885-DEL-60
chr13-15607205-DEL-123
chr13-15642645-DEL-81
chr13-15653403-DEL-71
chr13-15674052-DEL-63
chr13-15683723-DEL-350
chr13-15685120-DEL-107
chr13-15689951-DEL-52
chr13-15694363-DEL-572
chr13-15735530-DEL-91
chr13-15741576-DEL-109
chr13-15747177-DEL-52
chr13-15768729-DEL-877
chr13-15769225-DEL-1534
chr13-15837840-DEL-145
chr13-15847202-DEL-370
chr13-15932819-DEL-73
chr13-16006147-DEL-5695
chr13-16008714-DEL-177
chr13-16057735-DEL-6066
chr13-16070487-DEL-1301
chr13-16077392-DEL-74
chr13-16103266-DEL-225
chr13-16129119-DEL-2776
chr13-16137241-DEL-6452
chr13-16211858-DEL-135
chr13-16222734-DEL-140
chr13-16232907-DEL-1445
chr13-16244095-DEL-92
chr13-16281494-DEL-62
chr13-16419575-DEL-58
chr13-16425621-DEL-1298
chr13-16451923-DEL-182
chr13-16508383-DEL-2143
chr13-16510901-DEL-56
chr13-16540185-DEL-515
chr13-16551802-DEL-400
chr13-16556828-DEL-75
chr13-16570741-DEL-178
chr13-16571612-DEL-277
chr13-16604904-DEL-1011
chr13-16604904-DEL-824
chr13-16605453-DEL-444
chr13-16606241-DEL-174

chr13-16645092-DEL-53
chr13-16662839-DEL-122
chr13-16713747-DEL-118
chr13-16721893-DEL-388
chr13-16742833-DEL-880
chr13-16743796-DEL-454
chr13-16750611-DEL-401
chr13-16785374-DEL-69
chr13-16799198-DEL-76
chr13-16860044-DEL-60
chr13-16869853-DEL-235
chr13-16908916-DEL-131
chr13-16959493-DEL-112
chr13-16981481-DEL-537
chr13-17021744-DEL-54
chr13-17037915-DEL-862
chr13-17039224-DEL-90
chr13-17047983-DEL-806
chr13-17055528-DEL-2873
chr13-17080487-DEL-112
chr13-17086128-DEL-75
chr13-17110781-DEL-157
chr13-17124537-DEL-257
chr13-17171337-DEL-138
chr13-17204824-DEL-54
chr13-17225247-DEL-207
chr13-17282118-DEL-52
chr13-17317449-DEL-2250
chr13-17353203-DEL-296
chr13-17369115-DEL-120
chr13-17403519-DEL-3825
chr13-17409485-DEL-586
chr13-17418914-DEL-99
chr13-17433845-DEL-898
chr13-17500781-DEL-254
chr13-17516917-DEL-75
chr13-17517155-DEL-377
chr13-17702237-DEL-1831
chr13-17704353-DEL-1074
chr13-17713901-DEL-2621
chr13-17715920-DEL-607
chr13-17730601-DEL-951
chr13-17738067-DEL-6261
chr13-17763836-DEL-6103
chr13-17770097-DEL-1143
chr13-17826330-DEL-2282
chr13-17834957-DEL-430
chr13-17861971-DEL-79
chr13-17907407-DEL-646
chr13-17923378-DEL-135
chr13-17958009-DEL-344
chr13-17994864-DEL-250
chr13-18003249-DEL-57
chr13-18025230-DEL-879
chr13-18042027-DEL-9848
chr13-18057927-DEL-51
chr13-18069358-DEL-52
chr13-18084921-DEL-62
chr13-18091477-DEL-7251
chr13-18091543-DEL-6798
chr13-18148880-DEL-272
chr13-18152736-DEL-83
chr13-18165913-DEL-198
chr13-18168951-DEL-806
chr13-18221866-DEL-2933
chr13-18240226-DEL-82
chr13-18258421-DEL-386
chr13-18261917-DEL-4037
chr13-18275476-DEL-683
chr13-18282004-DEL-243
chr13-18311627-DEL-52
chr13-18340116-DEL-84
chr13-18357044-DEL-80
chr13-18385718-DEL-66
chr13-18397082-DEL-192
chr13-18412766-DEL-64
chr13-18434425-DEL-62

chr13-18446375-DEL-52
chr13-18520456-DEL-63
chr13-18530046-DEL-75
chr13-18564505-DEL-61
chr13-18565340-DEL-130
chr13-18664066-DEL-132
chr13-18720631-DEL-5968
chr13-18723186-DEL-163
chr13-18726643-DEL-2077
chr13-18747610-DEL-154
chr13-18748339-DEL-688
chr13-18762433-DEL-1212
chr13-18781409-DEL-909
chr13-18837573-DEL-2219
chr13-18842902-DEL-54
chr13-18871807-DEL-6302
chr13-18890516-DEL-4862
chr13-18953801-DEL-246
chr13-18960557-DEL-426
chr13-19040916-DEL-85
chr13-19063055-DEL-109
chr13-19115292-DEL-14306
chr13-19214205-DEL-133
chr13-19272602-DEL-137
chr13-19273570-DEL-240
chr13-19320899-DEL-550
chr13-19328656-DEL-195
chr13-19331884-DEL-123
chr13-19340089-DEL-52
chr13-19386606-DEL-2821
chr13-19415516-DEL-4334
chr13-19420765-DEL-1157
chr13-19422038-DEL-1218
chr13-19445891-DEL-996
chr13-19488240-DEL-74
chr13-19515917-DEL-2415
chr13-19562348-DEL-73
chr13-19568137-DEL-435
chr13-19591104-DEL-98
chr13-19619556-DEL-118
chr13-19632836-DEL-68
chr13-19645530-DEL-444
chr13-19648654-DEL-490
chr13-19682466-DEL-545
chr13-19686885-DEL-467
chr13-19702407-DEL-53
chr13-19749921-DEL-62
chr13-19759623-DEL-95
chr13-19833773-DEL-62
chr13-19837878-DEL-62
chr13-19865057-DEL-2079
chr13-19869171-DEL-212
chr13-19896580-DEL-924
chr13-19919545-DEL-192
chr13-19923240-DEL-681
chr13-19972044-DEL-1012
chr13-19993919-DEL-1505
chr13-20008770-DEL-54
chr13-20013179-DEL-76
chr13-20018451-DEL-860
chr13-20042095-DEL-344
chr13-20042754-DEL-99
chr13-20053541-DEL-1265
chr13-20062359-DEL-112
chr13-20062434-DEL-699
chr13-20067752-DEL-117
chr13-20081882-DEL-159
chr13-20099031-DEL-1846
chr13-20104345-DEL-113
chr13-20138299-DEL-9676
chr13-20149150-DEL-124
chr13-20187592-DEL-1357
chr13-20192274-DEL-255
chr13-20206672-DEL-74
chr13-20212479-DEL-334
chr13-20214162-DEL-64
chr13-20240319-DEL-56

chr13-20315447-DEL-114
chr13-20328908-DEL-320
chr13-20329573-DEL-214
chr13-20463237-DEL-55
chr13-20481672-DEL-572
chr13-20492567-DEL-58
chr13-20510841-DEL-196
chr13-20528651-DEL-325
chr13-20536020-DEL-73
chr13-20548056-DEL-139
chr13-20548339-DEL-1539
chr13-20616314-DEL-57
chr13-20662023-DEL-132
chr13-20671788-DEL-173
chr13-20694240-DEL-494
chr13-20706269-DEL-91
chr13-20730410-DEL-52
chr13-20752165-DEL-150
chr13-20762814-DEL-318
chr13-20834147-DEL-8824
chr13-20846510-DEL-134
chr13-20848879-DEL-896
chr13-20851566-DEL-606
chr13-20852481-DEL-8593
chr13-20861340-DEL-90
chr13-20926104-DEL-1117
chr13-20935278-DEL-3462
chr13-20968921-DEL-111
chr13-20974198-DEL-52
chr13-20994272-DEL-224
chr13-21044851-DEL-6473
chr13-21051855-DEL-15134
chr13-21062678-DEL-2153
chr13-21083901-DEL-77
chr13-21091283-DEL-182
chr13-21107778-DEL-84
chr13-21108593-DEL-57
chr13-21119542-DEL-53
chr13-21144528-DEL-1538
chr13-21198705-DEL-62
chr13-21201712-DEL-1870
chr13-21206100-DEL-523
chr13-21215518-DEL-54
chr13-21224028-DEL-350
chr13-21238654-DEL-4077
chr13-21238944-DEL-203
chr13-21242986-DEL-1914
chr13-21265574-DEL-102
chr13-21268082-DEL-53
chr13-21355682-DEL-93
chr13-21367337-DEL-503
chr13-21368533-DEL-545
chr13-21373131-DEL-79
chr13-21388449-DEL-2880
chr13-21394493-DEL-815
chr13-21398511-DEL-52
chr13-21415200-DEL-161
chr13-21416469-DEL-155
chr13-21431475-DEL-135
chr13-21458043-DEL-439
chr13-21458715-DEL-150
chr13-21470003-DEL-397
chr13-21471095-DEL-61
chr13-21482181-DEL-52
chr13-21485072-DEL-777
chr13-21513940-DEL-6332
chr13-21540349-DEL-56
chr13-21554668-DEL-712
chr13-21564479-DEL-84
chr13-21611599-DEL-63
chr13-21615610-DEL-88
chr13-21625965-DEL-192
chr13-21644529-DEL-66
chr13-21657205-DEL-2552
chr13-21738732-DEL-3545
chr13-21766794-DEL-285
chr13-21772277-DEL-210

chr13-21843609-DEL-76
chr13-21845506-DEL-63
chr13-21871549-DEL-1024
chr13-21902329-DEL-73
chr13-21916074-DEL-275
chr13-21930307-DEL-149
chr13-21946964-DEL-4796
chr13-21949473-DEL-99
chr13-21952538-DEL-3614
chr13-21987178-DEL-152
chr13-21996577-DEL-7858
chr13-22006097-DEL-84
chr13-22010895-DEL-771
chr13-22014921-DEL-621
chr13-22015763-DEL-59
chr13-22024952-DEL-152
chr13-22027053-DEL-120
chr13-22073863-DEL-168
chr13-22103996-DEL-131
chr13-22137940-DEL-498
chr13-22155251-DEL-57
chr13-22190697-DEL-173
chr13-22211516-DEL-75
chr13-22217626-DEL-60
chr13-22224768-DEL-86
chr13-22268070-DEL-1792
chr13-22270040-DEL-3472
chr13-22273171-DEL-287
chr13-22280420-DEL-128
chr13-22341463-DEL-6338
chr13-22344304-DEL-106
chr13-22357978-DEL-312
chr13-22380241-DEL-335
chr13-22388513-DEL-8244
chr13-22420429-DEL-528
chr13-22430966-DEL-4868
chr13-22467222-DEL-282
chr13-22472811-DEL-322
chr13-22478282-DEL-1395
chr13-22486473-DEL-120
chr13-22555784-DEL-283
chr13-22605519-DEL-199
chr13-22607638-DEL-212
chr13-22630824-DEL-111
chr13-22641079-DEL-6448
chr13-22682810-DEL-432
chr13-22756627-DEL-54
chr13-22758773-DEL-944
chr13-22760995-DEL-5297
chr13-22767704-DEL-126
chr13-22777933-DEL-77
chr13-22782472-DEL-8619
chr13-22828670-DEL-6000
chr13-22848673-DEL-95
chr13-22909626-DEL-94
chr13-22915862-DEL-2003
chr13-22929818-DEL-203
chr13-22958771-DEL-881
chr13-22979000-DEL-102
chr13-23000187-DEL-7392
chr13-23044672-DEL-959
chr13-23064254-DEL-76296
chr13-23079557-DEL-76
chr13-23143585-DEL-241
chr13-23217802-DEL-120
chr13-23257459-DEL-211
chr13-23262119-DEL-2366
chr13-23268665-DEL-410
chr13-23272215-DEL-405
chr13-23274923-DEL-2931
chr13-23278569-DEL-352
chr13-23295570-DEL-980
chr13-23309006-DEL-565
chr13-23318507-DEL-54
chr13-23325663-DEL-3604
chr13-23350940-DEL-10473
chr13-23362257-DEL-1179

chr13-23408105-DEL-616
chr13-23429848-DEL-141
chr13-23444676-DEL-7152
chr13-23444881-DEL-69
chr13-23452929-DEL-983
chr13-23453911-DEL-507
chr13-23465094-DEL-440
chr13-23491184-DEL-59
chr13-23514345-DEL-66
chr13-23516931-DEL-103
chr13-23535179-DEL-182
chr13-23571765-DEL-125
chr13-23572214-DEL-891
chr13-23589443-DEL-230
chr13-23590337-DEL-1198
chr13-23592639-DEL-113
chr13-23601588-DEL-336
chr13-23609697-DEL-119
chr13-23631093-DEL-487
chr13-23638643-DEL-157
chr13-23673371-DEL-231
chr13-23724206-DEL-203
chr13-23794787-DEL-1586
chr13-23802051-DEL-54
chr13-23822687-DEL-9046
chr13-23859598-DEL-3327
chr13-23862900-DEL-58
chr13-23868911-DEL-113
chr13-23869948-DEL-91
chr13-23886826-DEL-400
chr13-23941511-DEL-93
chr13-23979233-DEL-1348
chr13-23983535-DEL-103
chr13-23993723-DEL-66
chr13-24096133-DEL-100
chr13-24137068-DEL-54
chr13-24182178-DEL-976
chr13-24247204-DEL-64
chr13-24366541-DEL-61
chr13-24493548-DEL-145
chr13-24500478-DEL-1167
chr13-24503738-DEL-58
chr13-24526795-DEL-3710
chr13-24535736-DEL-639
chr13-24547970-DEL-56
chr13-24566144-DEL-137
chr13-24573815-DEL-763
chr13-24592173-DEL-498
chr13-24595215-DEL-136
chr13-24605831-DEL-103
chr13-24617841-DEL-457
chr13-24629473-DEL-1085
chr13-24634667-DEL-1798
chr13-24653877-DEL-438
chr13-24655326-DEL-5284
chr13-24684617-DEL-58
chr13-24694947-DEL-60
chr13-24713619-DEL-2951
chr13-24740272-DEL-53
chr13-24742557-DEL-1340
chr13-24747152-DEL-74
chr13-24767856-DEL-129
chr13-24837183-DEL-57
chr13-24841516-DEL-781
chr13-24887542-DEL-62
chr13-24900334-DEL-266
chr13-24907516-DEL-129
chr13-24947983-DEL-10719
chr13-24960430-DEL-94
chr13-24974892-DEL-357
chr13-24988211-DEL-62
chr13-25072745-DEL-966
chr13-25075113-DEL-296
chr13-25118182-DEL-136
chr13-25140603-DEL-203
chr13-25172955-DEL-276
chr13-25224708-DEL-6111

chr13-25256157-DEL-645
chr13-25274039-DEL-118
chr13-25290010-DEL-123
chr13-25330480-DEL-147
chr13-25376823-DEL-516
chr13-25381412-DEL-466
chr13-25409767-DEL-6227
chr13-25419586-DEL-1110
chr13-25420717-DEL-5465
chr13-25477497-DEL-200
chr13-25482807-DEL-600
chr13-25491568-DEL-217
chr13-25502957-DEL-4028
chr13-25513850-DEL-141
chr13-25518747-DEL-753
chr13-25583742-DEL-476
chr13-25584326-DEL-75
chr13-25608901-DEL-82
chr13-25611049-DEL-388
chr13-25634357-DEL-221
chr13-25661052-DEL-4741
chr13-25674476-DEL-402
chr13-25734117-DEL-194
chr13-25745523-DEL-308
chr13-25787117-DEL-661
chr13-25797025-DEL-54
chr13-25815854-DEL-1256
chr13-25824693-DEL-436
chr13-25825456-DEL-169
chr13-25834471-DEL-2752
chr13-25838030-DEL-587
chr13-25839612-DEL-1814
chr13-25865740-DEL-117
chr13-25873449-DEL-1189
chr13-25883506-DEL-95
chr13-25889090-DEL-51
chr13-25916426-DEL-149
chr13-25934216-DEL-127
chr13-25942092-DEL-92
chr13-26012374-DEL-71
chr13-26089237-DEL-234
chr13-26091256-DEL-314
chr13-26092477-DEL-1192
chr13-26113480-DEL-123
chr13-26156603-DEL-58
chr13-26174640-DEL-170
chr13-26211664-DEL-2192
chr13-26224282-DEL-56
chr13-26244738-DEL-54
chr13-26254991-DEL-99
chr13-26289698-DEL-1051
chr13-26297615-DEL-240
chr13-26367846-DEL-71
chr13-26383313-DEL-267
chr13-26413620-DEL-312
chr13-26416928-DEL-616
chr13-26448874-DEL-638
chr13-26451796-DEL-94
chr13-26497696-DEL-434
chr13-26516708-DEL-303
chr13-26517375-DEL-202
chr13-26531002-DEL-51
chr13-26584407-DEL-63
chr13-26599747-DEL-337
chr13-26643997-DEL-86
chr13-26705853-DEL-6513
chr13-26714465-DEL-711
chr13-26725437-DEL-6255
chr13-26760884-DEL-1150
chr13-26764654-DEL-66
chr13-26784262-DEL-1551
chr13-26803684-DEL-629
chr13-26844994-DEL-439
chr13-26854838-DEL-6211
chr13-26865790-DEL-93
chr13-26866645-DEL-9417
chr13-26897595-DEL-222

chr13-26919419-DEL-2337
chr13-26987888-DEL-113
chr13-27001752-DEL-6206
chr13-27011891-DEL-123
chr13-27013311-DEL-52
chr13-27014867-DEL-6724
chr13-27020949-DEL-212
chr13-27023739-DEL-87
chr13-27023867-DEL-6561
chr13-27032153-DEL-1872
chr13-27037606-DEL-204
chr13-27072771-DEL-214
chr13-27080797-DEL-1015
chr13-27104067-DEL-70
chr13-27120178-DEL-279
chr13-27173994-DEL-352
chr13-27183518-DEL-87
chr13-27190981-DEL-1961
chr13-27192999-DEL-187
chr13-27206467-DEL-64
chr13-27213366-DEL-90
chr13-27230970-DEL-68
chr13-27234382-DEL-1277
chr13-27255862-DEL-1436
chr13-27268323-DEL-5197
chr13-27273432-DEL-264
chr13-27283797-DEL-62
chr13-27286085-DEL-392
chr13-27287301-DEL-157
chr13-27321653-DEL-313
chr13-27355224-DEL-104
chr13-27372148-DEL-122
chr13-27376842-DEL-105
chr13-27377872-DEL-124
chr13-27381530-DEL-102
chr13-27391290-DEL-5979
chr13-27393653-DEL-1977
chr13-27410314-DEL-126
chr13-27438562-DEL-86
chr13-27440662-DEL-251
chr13-27445390-DEL-6271
chr13-27486183-DEL-2640
chr13-27506628-DEL-1517
chr13-27531122-DEL-372
chr13-27533692-DEL-54
chr13-27543167-DEL-6340
chr13-27551525-DEL-6259
chr13-27568965-DEL-60
chr13-27634580-DEL-200
chr13-27644803-DEL-1234
chr13-27659242-DEL-2105
chr13-27694612-DEL-517
chr13-27703838-DEL-3212
chr13-27707434-DEL-129
chr13-27742483-DEL-214
chr13-27752080-DEL-84
chr13-27754387-DEL-204
chr13-27758269-DEL-1208
chr13-27773587-DEL-326
chr13-27796379-DEL-124
chr13-27813565-DEL-950
chr13-27826836-DEL-109
chr13-27828750-DEL-544
chr13-27850284-DEL-1303
chr13-27860871-DEL-1757
chr13-27877931-DEL-569
chr13-27961717-DEL-54
chr13-27977286-DEL-90
chr13-28016684-DEL-3943
chr13-28059793-DEL-6407
chr13-28066540-DEL-7670
chr13-28066966-DEL-6303
chr13-28109602-DEL-431
chr13-28118112-DEL-4275
chr13-28135493-DEL-1126
chr13-28139015-DEL-993
chr13-28167156-DEL-6435

chr13-28185855-DEL-538
chr13-28196784-DEL-64
chr13-28213300-DEL-1336
chr13-28231752-DEL-175
chr13-28232425-DEL-256
chr13-28233493-DEL-847
chr13-28273441-DEL-871
chr13-28277644-DEL-64
chr13-28293327-DEL-429
chr13-28316132-DEL-976
chr13-28319499-DEL-66
chr13-28339397-DEL-168
chr13-28438867-DEL-647
chr13-28441565-DEL-563
chr13-28458230-DEL-58
chr13-28460390-DEL-501
chr13-28484414-DEL-560
chr13-28517835-DEL-407
chr13-28521511-DEL-195
chr13-28543164-DEL-1899
chr13-28552477-DEL-3282
chr13-28585358-DEL-93
chr13-28596560-DEL-100
chr13-28640351-DEL-100
chr13-28729949-DEL-65
chr13-28733563-DEL-81
chr13-28750194-DEL-209
chr13-28756724-DEL-360
chr13-28798834-DEL-84
chr13-28815623-DEL-71
chr13-28831588-DEL-193
chr13-28861561-DEL-300
chr13-28910695-DEL-101
chr13-28968124-DEL-122
chr13-28974394-DEL-52
chr13-29001226-DEL-54
chr13-29003612-DEL-499
chr13-29020896-DEL-88
chr13-29037242-DEL-219
chr13-29053923-DEL-58
chr13-29056206-DEL-61
chr13-29056815-DEL-137
chr13-29085607-DEL-178
chr13-29094694-DEL-343
chr13-29116322-DEL-55
chr13-29146622-DEL-62
chr13-29155093-DEL-188
chr13-29160218-DEL-372
chr13-29165787-DEL-188
chr13-29192298-DEL-70
chr13-29196197-DEL-618
chr13-29217495-DEL-357
chr13-29219241-DEL-519
chr13-29222634-DEL-233
chr13-29226526-DEL-905
chr13-29254812-DEL-636
chr13-29266062-DEL-67
chr13-29281744-DEL-303
chr13-29366042-DEL-71
chr13-29367294-DEL-60
chr13-29398563-DEL-719
chr13-29404055-DEL-53
chr13-29420854-DEL-6074
chr13-29429550-DEL-327
chr13-29430947-DEL-60
chr13-29470362-DEL-583
chr13-29476329-DEL-155
chr13-29500899-DEL-374
chr13-29502549-DEL-73
chr13-29503909-DEL-167
chr13-29513736-DEL-94
chr13-29522207-DEL-83
chr13-29546708-DEL-103
chr13-29560573-DEL-2283
chr13-29606504-DEL-101
chr13-29659436-DEL-311
chr13-29667499-DEL-2185

chr13-29695826-DEL-56
chr13-29712451-DEL-53
chr13-29715991-DEL-950
chr13-29782032-DEL-96
chr13-29797386-DEL-53
chr13-29799783-DEL-71
chr13-29803108-DEL-64
chr13-29812953-DEL-81
chr13-29873124-DEL-1301
chr13-29873310-DEL-496
chr13-29934179-DEL-127
chr13-29988349-DEL-63
chr13-30059084-DEL-67
chr13-30104910-DEL-171
chr13-30175972-DEL-106
chr13-30189016-DEL-87
chr13-30255347-DEL-889
chr13-30265897-DEL-71
chr13-30273774-DEL-68
chr13-30274771-DEL-77
chr13-30276475-DEL-149
chr13-30278979-DEL-51
chr13-30300897-DEL-2131
chr13-30303582-DEL-1427
chr13-30305056-DEL-2072
chr13-30312366-DEL-691
chr13-30313178-DEL-294
chr13-30313857-DEL-756
chr13-30314543-DEL-139
chr13-30327803-DEL-60
chr13-30333909-DEL-136
chr13-30398605-DEL-118
chr13-30434198-DEL-62
chr13-30445693-DEL-1059
chr13-30510481-DEL-2084
chr13-30517049-DEL-54
chr13-30531456-DEL-386
chr13-30537131-DEL-1101
chr13-30579340-DEL-604
chr13-30587039-DEL-56
chr13-30619101-DEL-199
chr13-30622270-DEL-214
chr13-30639928-DEL-5624
chr13-30645440-DEL-255
chr13-30757589-DEL-76
chr13-30760156-DEL-58
chr13-30804158-DEL-73
chr13-30810031-DEL-87
chr13-30877720-DEL-518
chr13-30883137-DEL-331
chr13-30964412-DEL-88
chr13-30971727-DEL-62
chr13-31011786-DEL-57
chr13-31175840-DEL-80
chr13-31182526-DEL-128
chr13-31276759-DEL-138
chr13-31308511-DEL-5982
chr13-31308663-DEL-120
chr13-31339500-DEL-135
chr13-31389074-DEL-128
chr13-31395390-DEL-123
chr13-31399754-DEL-835
chr13-31408682-DEL-207
chr13-31430738-DEL-135
chr13-31465581-DEL-88
chr13-31512331-DEL-60
chr13-31515462-DEL-18701
chr13-31518279-DEL-85
chr13-31557514-DEL-531
chr13-31566773-DEL-666
chr13-31580951-DEL-118
chr13-31698405-DEL-58
chr13-31720720-DEL-399
chr13-31734975-DEL-87
chr13-31801086-DEL-52
chr13-31819450-DEL-75
chr13-31820138-DEL-128

chr13-31840163-DEL-118
chr13-31846134-DEL-70
chr13-31855315-DEL-537
chr13-31860148-DEL-83
chr13-31893687-DEL-212
chr13-31896783-DEL-315
chr13-31934666-DEL-206
chr13-31965438-DEL-164
chr13-32007158-DEL-95
chr13-32025741-DEL-1135
chr13-32027501-DEL-172
chr13-32057949-DEL-52
chr13-32085484-DEL-121
chr13-32237992-DEL-182
chr13-32240510-DEL-2760
chr13-32250804-DEL-67
chr13-32297074-DEL-890
chr13-32308616-DEL-97
chr13-32324712-DEL-78
chr13-32337445-DEL-168
chr13-32369280-DEL-894
chr13-32371750-DEL-818
chr13-32375803-DEL-446
chr13-32395387-DEL-142
chr13-32398521-DEL-60
chr13-32405335-DEL-243
chr13-32408922-DEL-51
chr13-32417223-DEL-187
chr13-32429537-DEL-5501
chr13-32434617-DEL-857
chr13-32450133-DEL-58
chr13-32486093-DEL-3986
chr13-32559165-DEL-220
chr13-32584898-DEL-401
chr13-32592796-DEL-72
chr13-32594352-DEL-70
chr13-32596133-DEL-457
chr13-32596267-DEL-1392
chr13-32607295-DEL-102
chr13-32616204-DEL-196
chr13-32617202-DEL-424
chr13-32644945-DEL-598
chr13-32646708-DEL-868
chr13-32653904-DEL-611
chr13-32712272-DEL-234
chr13-32712650-DEL-502
chr13-32716693-DEL-468
chr13-32719123-DEL-371
chr13-32724947-DEL-423
chr13-32741401-DEL-204
chr13-32743027-DEL-88
chr13-32744410-DEL-115
chr13-32774385-DEL-887
chr13-32779875-DEL-252
chr13-32805580-DEL-66
chr13-32824620-DEL-177
chr13-32830176-DEL-114
chr13-32840401-DEL-751
chr13-32842104-DEL-687
chr13-32843094-DEL-60
chr13-32847649-DEL-386
chr13-32860563-DEL-88
chr13-32868622-DEL-720
chr13-32888556-DEL-54
chr13-32897160-DEL-237
chr13-32930713-DEL-425
chr13-32952957-DEL-52
chr13-32979875-DEL-591
chr13-32982151-DEL-209
chr13-32984133-DEL-59
chr13-33027694-DEL-67
chr13-33031778-DEL-63
chr13-33048997-DEL-56
chr13-33069582-DEL-743
chr13-33072661-DEL-91
chr13-33073190-DEL-6392
chr13-33084565-DEL-124

chr13-33094138-DEL-2184
chr13-33143908-DEL-66
chr13-33149329-DEL-70
chr13-33164170-DEL-9210
chr13-33220015-DEL-79
chr13-33226378-DEL-377
chr13-33240599-DEL-252
chr13-33260726-DEL-79
chr13-33272064-DEL-670
chr13-33272861-DEL-286
chr13-33282900-DEL-2198
chr13-33314751-DEL-4029
chr13-33324585-DEL-6563
chr13-33366502-DEL-2143
chr13-33373234-DEL-59
chr13-33414387-DEL-6211
chr13-33454713-DEL-173
chr13-33530339-DEL-63
chr13-33557153-DEL-7260
chr13-33566678-DEL-565
chr13-33613282-DEL-84
chr13-33621642-DEL-7089
chr13-33629012-DEL-53
chr13-33644562-DEL-568
chr13-33648179-DEL-2493
chr13-33652214-DEL-2505
chr13-33717976-DEL-6327
chr13-33728087-DEL-1951
chr13-33752683-DEL-776
chr13-33788524-DEL-10705
chr13-33791351-DEL-6773
chr13-33799091-DEL-124
chr13-33832725-DEL-102
chr13-33839248-DEL-112
chr13-33974825-DEL-2129
chr13-33977142-DEL-1963
chr13-33981204-DEL-201
chr13-33991217-DEL-1061
chr13-33992436-DEL-89
chr13-33992733-DEL-5594
chr13-33998424-DEL-2819
chr13-34027872-DEL-53
chr13-34043573-DEL-112
chr13-34044393-DEL-61
chr13-34072183-DEL-104
chr13-34076361-DEL-493
chr13-34090757-DEL-52
chr13-34247038-DEL-273
chr13-34267162-DEL-491
chr13-34287521-DEL-220
chr13-34313593-DEL-67
chr13-34344810-DEL-73
chr13-34349597-DEL-174
chr13-34425860-DEL-108
chr13-34494777-DEL-52
chr13-34506311-DEL-701
chr13-34540336-DEL-836
chr13-34580029-DEL-108
chr13-34603552-DEL-386
chr13-34642253-DEL-171
chr13-34647624-DEL-56
chr13-34670350-DEL-1039
chr13-34671620-DEL-51
chr13-34691160-DEL-101
chr13-34719348-DEL-270
chr13-34726479-DEL-409
chr13-34769564-DEL-442
chr13-34774617-DEL-188
chr13-34778670-DEL-60
chr13-34782293-DEL-642
chr13-34797082-DEL-5407
chr13-34821656-DEL-308
chr13-34854997-DEL-53
chr13-34876324-DEL-161
chr13-34903370-DEL-148
chr13-34919696-DEL-104
chr13-34922622-DEL-3112

chr13-34941891-DEL-111
chr13-34945524-DEL-431
chr13-35015589-DEL-1086
chr13-35047209-DEL-91
chr13-35068484-DEL-86
chr13-35121047-DEL-6292
chr13-35127568-DEL-256
chr13-35153706-DEL-108
chr13-35169587-DEL-60
chr13-35189114-DEL-8705
chr13-35208637-DEL-1838
chr13-35223034-DEL-151
chr13-35228230-DEL-12757
chr13-35284672-DEL-106
chr13-35292713-DEL-381
chr13-35377888-DEL-373
chr13-35382487-DEL-1210
chr13-35412125-DEL-83
chr13-35546130-DEL-128
chr13-35583754-DEL-101
chr13-35653851-DEL-153
chr13-35658388-DEL-104
chr13-35666881-DEL-448
chr13-35695208-DEL-399
chr13-35700792-DEL-79
chr13-35729884-DEL-66
chr13-35737469-DEL-1176
chr13-35751411-DEL-212
chr13-35758549-DEL-6108
chr13-35762503-DEL-1933
chr13-35768799-DEL-227
chr13-35770296-DEL-220
chr13-35773714-DEL-178
chr13-35808136-DEL-81
chr13-35814311-DEL-6399
chr13-35814364-DEL-199
chr13-35871319-DEL-1536
chr13-35873749-DEL-509
chr13-35883464-DEL-329
chr13-35899861-DEL-547
chr13-35900513-DEL-65
chr13-35912151-DEL-145
chr13-35921453-DEL-56
chr13-35948268-DEL-401
chr13-36079262-DEL-126
chr13-36089227-DEL-172
chr13-36091042-DEL-474
chr13-36103224-DEL-325
chr13-36108314-DEL-3107
chr13-36124337-DEL-52
chr13-36157203-DEL-77
chr13-36194364-DEL-51
chr13-36196791-DEL-684
chr13-36203766-DEL-196
chr13-36207941-DEL-56
chr13-36263735-DEL-67
chr13-36279214-DEL-799
chr13-36289122-DEL-97
chr13-36372918-DEL-81
chr13-36387389-DEL-176
chr13-36390271-DEL-54
chr13-36401637-DEL-81
chr13-36432164-DEL-1196
chr13-36470177-DEL-2555
chr13-36473260-DEL-88
chr13-36474944-DEL-899
chr13-36478232-DEL-51
chr13-36478976-DEL-54
chr13-36484909-DEL-709
chr13-36515566-DEL-1656
chr13-36517149-DEL-402
chr13-36517623-DEL-2105
chr13-36538236-DEL-3119
chr13-36549889-DEL-91
chr13-36659256-DEL-160
chr13-36749721-DEL-76
chr13-36837144-DEL-52

chr13-36839117-DEL-250
chr13-36847615-DEL-6297
chr13-36856015-DEL-267
chr13-36913529-DEL-109
chr13-36922898-DEL-1265
chr13-36944695-DEL-56
chr13-36971997-DEL-113
chr13-36980191-DEL-172
chr13-37052984-DEL-828
chr13-37066215-DEL-524
chr13-37090190-DEL-52
chr13-37092434-DEL-82
chr13-37106836-DEL-2862
chr13-37118802-DEL-54
chr13-37123657-DEL-106
chr13-37133003-DEL-63
chr13-37139472-DEL-1023
chr13-37149082-DEL-60
chr13-37162377-DEL-254
chr13-37170968-DEL-59
chr13-37198134-DEL-58
chr13-37206443-DEL-77
chr13-37289160-DEL-672
chr13-37305061-DEL-550
chr13-37325548-DEL-73
chr13-37351656-DEL-415
chr13-37433134-DEL-201
chr13-37438642-DEL-77
chr13-37444136-DEL-1154
chr13-37471667-DEL-52
chr13-37518168-DEL-511
chr13-37539208-DEL-60
chr13-37558370-DEL-87
chr13-37559441-DEL-330
chr13-37566879-DEL-1224
chr13-37616323-DEL-69
chr13-37647263-DEL-64
chr13-37680566-DEL-74
chr13-37683742-DEL-181
chr13-37704616-DEL-125
chr13-37747057-DEL-147
chr13-37802312-DEL-976
chr13-37841685-DEL-62
chr13-37874239-DEL-274
chr13-37892516-DEL-71
chr13-37903487-DEL-613
chr13-37927507-DEL-617
chr13-37928766-DEL-51
chr13-37938037-DEL-198
chr13-37952398-DEL-311
chr13-37958783-DEL-123
chr13-37969642-DEL-88
chr13-38024822-DEL-87
chr13-38043253-DEL-122
chr13-38075520-DEL-683
chr13-38142337-DEL-66
chr13-38148452-DEL-64
chr13-38157941-DEL-216
chr13-38173698-DEL-72
chr13-38188178-DEL-123
chr13-38188345-DEL-876
chr13-38243436-DEL-63
chr13-38257554-DEL-164
chr13-38314094-DEL-63
chr13-38323804-DEL-58
chr13-38343914-DEL-268
chr13-38345482-DEL-72
chr13-38350468-DEL-194
chr13-38390546-DEL-203
chr13-38391259-DEL-73
chr13-38417136-DEL-73
chr13-38425480-DEL-78
chr13-38453125-DEL-57
chr13-38474989-DEL-239
chr13-38493013-DEL-625
chr13-38499078-DEL-116
chr13-38505468-DEL-138

chr13-38509718-DEL-187
chr13-38518711-DEL-623
chr13-38522106-DEL-535
chr13-38527508-DEL-201
chr13-38554194-DEL-79
chr13-38566763-DEL-664
chr13-38577100-DEL-101
chr13-38596083-DEL-109
chr13-38599651-DEL-65
chr13-38624066-DEL-52
chr13-38627692-DEL-79
chr13-38640790-DEL-90
chr13-38755717-DEL-810
chr13-38759482-DEL-585
chr13-38760241-DEL-237
chr13-38804108-DEL-303
chr13-38837041-DEL-1309
chr13-38853233-DEL-79
chr13-38892145-DEL-99
chr13-38908587-DEL-396
chr13-38914017-DEL-61
chr13-38929031-DEL-77
chr13-38951286-DEL-1286
chr13-38956136-DEL-87
chr13-38966521-DEL-58
chr13-38969800-DEL-396
chr13-38971578-DEL-103
chr13-39033711-DEL-103
chr13-39036210-DEL-120
chr13-39037321-DEL-122
chr13-39045318-DEL-160
chr13-39050482-DEL-1609
chr13-39067658-DEL-6912
chr13-39100127-DEL-70
chr13-39105197-DEL-59
chr13-39147406-DEL-6161
chr13-39169102-DEL-56
chr13-39203495-DEL-434
chr13-39206583-DEL-66
chr13-39207304-DEL-109
chr13-39243339-DEL-64
chr13-39282503-DEL-112
chr13-39299254-DEL-245
chr13-39314127-DEL-5988
chr13-39334520-DEL-129
chr13-39397991-DEL-773
chr13-39487551-DEL-103
chr13-39541726-DEL-91
chr13-39571056-DEL-545
chr13-39583938-DEL-84
chr13-39586753-DEL-57
chr13-39593497-DEL-52
chr13-39617016-DEL-62
chr13-39644324-DEL-150
chr13-39659578-DEL-102
chr13-39667078-DEL-169
chr13-39708392-DEL-52
chr13-39718850-DEL-98
chr13-39723565-DEL-116
chr13-39757900-DEL-84
chr13-39765278-DEL-81
chr13-39777907-DEL-7249
chr13-39791452-DEL-1103
chr13-39809803-DEL-127
chr13-39820408-DEL-204
chr13-39867634-DEL-624
chr13-39915565-DEL-543
chr13-39931373-DEL-333
chr13-39938094-DEL-56
chr13-39968342-DEL-113
chr13-39971268-DEL-82
chr13-39990284-DEL-69
chr13-39995532-DEL-54
chr13-40088612-DEL-60
chr13-40093007-DEL-2979
chr13-40093340-DEL-211
chr13-40097113-DEL-2645

chr13-40140202-DEL-655
chr13-40145915-DEL-87
chr13-40238399-DEL-106
chr13-40264948-DEL-57
chr13-40267307-DEL-151
chr13-40269243-DEL-68
chr13-40287188-DEL-52
chr13-40327004-DEL-92
chr13-40368538-DEL-67
chr13-40387281-DEL-384
chr13-40388576-DEL-140
chr13-40427771-DEL-76
chr13-40443076-DEL-80
chr13-40449573-DEL-58
chr13-40506852-DEL-58
chr13-40510427-DEL-587
chr13-40520974-DEL-85
chr13-40539571-DEL-76
chr13-40551309-DEL-57
chr13-40555268-DEL-200
chr13-40562468-DEL-625
chr13-40593547-DEL-469
chr13-40617411-DEL-1031
chr13-40635032-DEL-132
chr13-40639459-DEL-70
chr13-40668817-DEL-58
chr13-40686594-DEL-128
chr13-40701801-DEL-65
chr13-40750545-DEL-66
chr13-40770787-DEL-153
chr13-40800700-DEL-76
chr13-40832931-DEL-2766
chr13-40846027-DEL-60
chr13-40852059-DEL-394
chr13-40928240-DEL-300
chr13-40983464-DEL-8059
chr13-40985033-DEL-6488
chr13-40991281-DEL-533
chr13-41009653-DEL-170
chr13-41016874-DEL-60
chr13-41021094-DEL-59
chr13-41022288-DEL-51
chr13-41047094-DEL-2988
chr13-41051575-DEL-189
chr13-41056850-DEL-451
chr13-41063212-DEL-605
chr13-41074587-DEL-54
chr13-41076149-DEL-87
chr13-41107369-DEL-87
chr13-41114660-DEL-2062
chr13-41115711-DEL-3093
chr13-41136122-DEL-65
chr13-41136880-DEL-315
chr13-41141099-DEL-56
chr13-41141672-DEL-1010
chr13-41149226-DEL-188
chr13-41152477-DEL-76
chr13-41175784-DEL-63
chr13-41189833-DEL-64
chr13-41201870-DEL-301
chr13-41208029-DEL-195
chr13-41208644-DEL-59
chr13-41236053-DEL-68
chr13-41246379-DEL-129
chr13-41273759-DEL-358
chr13-41280893-DEL-720
chr13-41321489-DEL-2080
chr13-41340209-DEL-58
chr13-41361240-DEL-172
chr13-41461363-DEL-66
chr13-41518583-DEL-608
chr13-41526022-DEL-78
chr13-41560544-DEL-364
chr13-41571405-DEL-108
chr13-41623915-DEL-51
chr13-41661305-DEL-67
chr13-41782725-DEL-64

chr13-41787055-DEL-122
chr13-41808346-DEL-60
chr13-41863596-DEL-247
chr13-41896541-DEL-1040
chr13-41897268-DEL-67
chr13-41965298-DEL-224
chr13-41967153-DEL-165
chr13-41996562-DEL-389
chr13-42025189-DEL-95
chr13-42040124-DEL-74
chr13-42060378-DEL-78
chr13-42131540-DEL-51
chr13-42157148-DEL-349
chr13-42171050-DEL-538
chr13-42174669-DEL-115
chr13-42216365-DEL-395
chr13-42231034-DEL-68
chr13-42237912-DEL-2398
chr13-42240020-DEL-286
chr13-42293484-DEL-384
chr13-42328786-DEL-52
chr13-42347415-DEL-54
chr13-42348632-DEL-1228
chr13-42357682-DEL-687
chr13-42366064-DEL-78
chr13-42383870-DEL-199
chr13-42388066-DEL-156
chr13-42415147-DEL-60
chr13-42444275-DEL-67
chr13-42512704-DEL-422
chr13-42513532-DEL-72
chr13-42517429-DEL-60
chr13-42532714-DEL-78
chr13-42536972-DEL-1174
chr13-42543795-DEL-91
chr13-42561750-DEL-94
chr13-42571705-DEL-328
chr13-42592911-DEL-283
chr13-42619078-DEL-397
chr13-42626853-DEL-74
chr13-42721749-DEL-178
chr13-42745328-DEL-61
chr13-42764154-DEL-87
chr13-42778448-DEL-98
chr13-42841045-DEL-70
chr13-42869683-DEL-113
chr13-42870435-DEL-60
chr13-42879707-DEL-134
chr13-42886482-DEL-3736
chr13-42923105-DEL-67
chr13-42948812-DEL-93
chr13-42953614-DEL-178
chr13-42987957-DEL-79
chr13-42993881-DEL-1164
chr13-43000046-DEL-482
chr13-43113825-DEL-380
chr13-43121538-DEL-64
chr13-43137987-DEL-327
chr13-43157905-DEL-131
chr13-43173760-DEL-64
chr13-43181342-DEL-64
chr13-43234247-DEL-90
chr13-43283205-DEL-64
chr13-43313709-DEL-56
chr13-43315344-DEL-422
chr13-43329779-DEL-201
chr13-43332668-DEL-78
chr13-43334525-DEL-92
chr13-43391549-DEL-54
chr13-43395048-DEL-61
chr13-43398136-DEL-472
chr13-43411070-DEL-123
chr13-43430733-DEL-79
chr13-43443343-DEL-59
chr13-43476133-DEL-587
chr13-43510584-DEL-72
chr13-43513160-DEL-66

chr13-43517777-DEL-257
chr13-43522682-DEL-110
chr13-43525541-DEL-204
chr13-43534065-DEL-110
chr13-43535657-DEL-55
chr13-43664890-DEL-65
chr13-43677014-DEL-183
chr13-43715111-DEL-75
chr13-43727024-DEL-59
chr13-43739848-DEL-357
chr13-43793754-DEL-176
chr13-43798850-DEL-62
chr13-43799474-DEL-241
chr13-43813326-DEL-188
chr13-43815813-DEL-262
chr13-43830642-DEL-64
chr13-43846535-DEL-67
chr13-43870018-DEL-63
chr13-43905632-DEL-832
chr13-43912416-DEL-554
chr13-43941031-DEL-103
chr13-43969650-DEL-101
chr13-44038279-DEL-960
chr13-44094805-DEL-194
chr13-44109298-DEL-146
chr13-44164335-DEL-72
chr13-44168275-DEL-52
chr13-44215540-DEL-2283
chr13-44256126-DEL-531
chr13-44275343-DEL-1411
chr13-44298491-DEL-118
chr13-44325708-DEL-658
chr13-44360380-DEL-54
chr13-44388576-DEL-111
chr13-44403728-DEL-86
chr13-44429679-DEL-268
chr13-44450147-DEL-87
chr13-44460555-DEL-255
chr13-44467304-DEL-191
chr13-44515082-DEL-146
chr13-44525384-DEL-171
chr13-44541698-DEL-438
chr13-44548741-DEL-307
chr13-44598434-DEL-60
chr13-44629074-DEL-546
chr13-44632298-DEL-515
chr13-44647802-DEL-53
chr13-44698824-DEL-183
chr13-44699920-DEL-56
chr13-44721817-DEL-59
chr13-44739587-DEL-81
chr13-44746895-DEL-390
chr13-44772827-DEL-127
chr13-44777999-DEL-115
chr13-44790628-DEL-536
chr13-44791343-DEL-400
chr13-44801761-DEL-91
chr13-44806903-DEL-200
chr13-44809053-DEL-231
chr13-44815760-DEL-109
chr13-44847456-DEL-53
chr13-44870033-DEL-1586
chr13-44871967-DEL-1131
chr13-44892145-DEL-107
chr13-44893182-DEL-199
chr13-44895619-DEL-86
chr13-44915087-DEL-196
chr13-44922183-DEL-63
chr13-44936175-DEL-65
chr13-44944669-DEL-607
chr13-44954297-DEL-297
chr13-44963226-DEL-4999
chr13-44974953-DEL-599
chr13-44999961-DEL-853
chr13-45004935-DEL-639
chr13-45016208-DEL-305
chr13-45022737-DEL-916

chr13-45028140-DEL-194
chr13-45034792-DEL-191
chr13-45061921-DEL-505
chr13-45082133-DEL-117
chr13-45082679-DEL-211
chr13-45114566-DEL-94
chr13-45164148-DEL-80
chr13-45180983-DEL-58
chr13-45195326-DEL-323
chr13-45203879-DEL-444
chr13-45208441-DEL-243
chr13-45214835-DEL-938
chr13-45273012-DEL-612
chr13-45278695-DEL-119
chr13-45285858-DEL-68
chr13-45300441-DEL-154
chr13-45307637-DEL-98
chr13-45329211-DEL-2598
chr13-45357139-DEL-569
chr13-45380054-DEL-93
chr13-45395608-DEL-55
chr13-45413053-DEL-120
chr13-45413931-DEL-62
chr13-45447201-DEL-139
chr13-45482153-DEL-299
chr13-45486491-DEL-84
chr13-45522761-DEL-932
chr13-45552483-DEL-52
chr13-45555038-DEL-165
chr13-45573305-DEL-229
chr13-45610455-DEL-72
chr13-45621526-DEL-68
chr13-45648132-DEL-76
chr13-45685444-DEL-54
chr13-45728249-DEL-185
chr13-45737776-DEL-135
chr13-45764295-DEL-133
chr13-45799759-DEL-3040
chr13-45800913-DEL-671
chr13-45832999-DEL-136
chr13-45873390-DEL-226
chr13-45876699-DEL-87
chr13-45916389-DEL-287
chr13-45918453-DEL-894
chr13-45921372-DEL-59
chr13-45926103-DEL-55
chr13-45943449-DEL-153
chr13-45952783-DEL-1949
chr13-45973500-DEL-73
chr13-45985542-DEL-90
chr13-45987312-DEL-112
chr13-45987504-DEL-1476
chr13-45992528-DEL-78
chr13-46008894-DEL-206
chr13-46031560-DEL-192
chr13-46032287-DEL-155
chr13-46040634-DEL-232
chr13-46074993-DEL-60
chr13-46080680-DEL-280
chr13-46091123-DEL-180
chr13-46109693-DEL-67
chr13-46123178-DEL-52
chr13-46164888-DEL-62
chr13-46190516-DEL-215
chr13-46211087-DEL-244
chr13-46219841-DEL-100
chr13-46261169-DEL-55
chr13-46268711-DEL-135
chr13-46278821-DEL-802
chr13-46285130-DEL-112
chr13-46295122-DEL-115
chr13-46304738-DEL-159
chr13-46328289-DEL-76
chr13-46335723-DEL-2034
chr13-46382313-DEL-114
chr13-46423554-DEL-59
chr13-46428801-DEL-140

chr13-46450667-DEL-376
chr13-46459538-DEL-155
chr13-46461380-DEL-106
chr13-46470427-DEL-1495
chr13-46470440-DEL-200
chr13-46471920-DEL-211
chr13-46489462-DEL-96
chr13-46508128-DEL-78
chr13-46562473-DEL-393
chr13-46613458-DEL-638
chr13-46619362-DEL-2545
chr13-46643148-DEL-181
chr13-46665395-DEL-92
chr13-46672343-DEL-310
chr13-46713705-DEL-56
chr13-46729414-DEL-6540
chr13-46729642-DEL-199
chr13-46730392-DEL-2354
chr13-46757317-DEL-75
chr13-46791584-DEL-70
chr13-46818914-DEL-248
chr13-46856601-DEL-301
chr13-46862395-DEL-352
chr13-46873963-DEL-82
chr13-46878752-DEL-72
chr13-46900271-DEL-659
chr13-46919603-DEL-55
chr13-46935100-DEL-51
chr13-46947714-DEL-778
chr13-46956876-DEL-114
chr13-46958735-DEL-1814
chr13-46980895-DEL-207
chr13-46992391-DEL-61
chr13-47005071-DEL-62
chr13-47073099-DEL-224
chr13-47074145-DEL-70
chr13-47091646-DEL-56
chr13-47122344-DEL-549
chr13-47142787-DEL-620
chr13-47148578-DEL-178
chr13-47149669-DEL-174
chr13-47159463-DEL-108
chr13-47215552-DEL-149
chr13-47216694-DEL-54
chr13-47228663-DEL-239
chr13-47296892-DEL-77
chr13-47302333-DEL-2170
chr13-47314369-DEL-77
chr13-47329180-DEL-59
chr13-47340344-DEL-67
chr13-47348223-DEL-60
chr13-47375913-DEL-399
chr13-47384075-DEL-245
chr13-47399685-DEL-51
chr13-47422863-DEL-1489
chr13-47438171-DEL-715
chr13-47445608-DEL-98
chr13-47448006-DEL-52
chr13-47493770-DEL-52
chr13-47520893-DEL-92
chr13-47587431-DEL-62
chr13-47594273-DEL-406
chr13-47627872-DEL-234
chr13-47631233-DEL-1039
chr13-47631874-DEL-1031
chr13-47633572-DEL-59
chr13-47639741-DEL-60
chr13-47640576-DEL-63
chr13-47647781-DEL-93
chr13-47670674-DEL-1675
chr13-47685195-DEL-161
chr13-47723635-DEL-68
chr13-47728218-DEL-70
chr13-47772834-DEL-64
chr13-47822693-DEL-361
chr13-47836991-DEL-108
chr13-47869161-DEL-58

chr13-47882337-DEL-801
chr13-47912534-DEL-563
chr13-47921787-DEL-59
chr13-47923553-DEL-89
chr13-47937480-DEL-136
chr13-47944550-DEL-997
chr13-47957829-DEL-1677
chr13-47969791-DEL-58
chr13-47999745-DEL-55
chr13-48041581-DEL-250
chr13-48084829-DEL-178
chr13-48106784-DEL-126
chr13-48120290-DEL-514
chr13-48203843-DEL-52
chr13-48238661-DEL-91
chr13-48263217-DEL-80
chr13-48270694-DEL-1682
chr13-48293838-DEL-113
chr13-48302854-DEL-893
chr13-48327679-DEL-115
chr13-48335991-DEL-214
chr13-48367132-DEL-66
chr13-48444768-DEL-162
chr13-48476128-DEL-401
chr13-48476710-DEL-55
chr13-48527163-DEL-65
chr13-48541603-DEL-59
chr13-48567417-DEL-323
chr13-48586126-DEL-89
chr13-48587510-DEL-139
chr13-48618408-DEL-249
chr13-48668235-DEL-209
chr13-48677328-DEL-76
chr13-48680203-DEL-76
chr13-48697965-DEL-85
chr13-48718150-DEL-196
chr13-48721883-DEL-148
chr13-48801803-DEL-83
chr13-48861349-DEL-68
chr13-48867681-DEL-96
chr13-48898099-DEL-86
chr13-48901065-DEL-60
chr13-48913890-DEL-69
chr13-48917817-DEL-101
chr13-48932549-DEL-89
chr13-48988043-DEL-76
chr13-48994819-DEL-189
chr13-48999119-DEL-248
chr13-49004308-DEL-72
chr13-49019266-DEL-154
chr13-49036008-DEL-615
chr13-49037877-DEL-126
chr13-49063091-DEL-67
chr13-49079492-DEL-147
chr13-49080988-DEL-86
chr13-49108303-DEL-341
chr13-49138762-DEL-219
chr13-49139055-DEL-1857
chr13-49141465-DEL-95
chr13-49145339-DEL-219
chr13-49159690-DEL-155
chr13-49183540-DEL-214
chr13-49207516-DEL-340
chr13-49208861-DEL-118
chr13-49218984-DEL-1159
chr13-49228513-DEL-71
chr13-49229154-DEL-89
chr13-49251494-DEL-207
chr13-49302992-DEL-238
chr13-49305581-DEL-1400
chr13-49365643-DEL-134
chr13-49375338-DEL-68
chr13-49466202-DEL-212
chr13-49524059-DEL-198
chr13-49545833-DEL-107
chr13-49552797-DEL-774
chr13-49592676-DEL-56

chr13-49647061-DEL-2812
chr13-49651389-DEL-498
chr13-49713882-DEL-58
chr13-49721160-DEL-87
chr13-49722545-DEL-190
chr13-49741091-DEL-233
chr13-49744365-DEL-1008
chr13-49802795-DEL-221
chr13-49823939-DEL-74
chr13-49879417-DEL-52
chr13-49889107-DEL-51
chr13-49898286-DEL-58
chr13-49902854-DEL-1142
chr13-49907102-DEL-326
chr13-49934337-DEL-132
chr13-49946064-DEL-321
chr13-49951451-DEL-308
chr13-49959653-DEL-241
chr13-49975919-DEL-81
chr13-49981546-DEL-474
chr13-49985607-DEL-1029
chr13-49987328-DEL-753
chr13-50005845-DEL-77
chr13-50007250-DEL-140
chr13-50009519-DEL-375
chr13-50036105-DEL-179
chr13-50036967-DEL-483
chr13-50076439-DEL-95
chr13-50078661-DEL-5487
chr13-50088464-DEL-1142
chr13-50120280-DEL-1709
chr13-50128317-DEL-113
chr13-50134939-DEL-6402
chr13-50146885-DEL-193
chr13-50147778-DEL-78
chr13-50149492-DEL-1318
chr13-50169191-DEL-6572
chr13-50191099-DEL-5830
chr13-50231754-DEL-6422
chr13-50273352-DEL-7114
chr13-50283083-DEL-6386
chr13-50324953-DEL-730
chr13-50344567-DEL-57
chr13-50366188-DEL-607
chr13-50374903-DEL-538
chr13-50391341-DEL-328
chr13-50411018-DEL-894
chr13-50447168-DEL-7699
chr13-50456561-DEL-2304
chr13-50483727-DEL-433
chr13-50491961-DEL-9280
chr13-50505916-DEL-3383
chr13-50631814-DEL-128
chr13-50635814-DEL-1341
chr13-50679964-DEL-94
chr13-50681194-DEL-288
chr13-50682157-DEL-6571
chr13-50682440-DEL-199
chr13-50767037-DEL-166
chr13-50814743-DEL-4157
chr13-50831898-DEL-8854
chr13-50843824-DEL-6568
chr13-50865546-DEL-8753
chr13-50882067-DEL-498
chr13-50884547-DEL-11352
chr13-50884679-DEL-208
chr13-50909938-DEL-565
chr13-50920752-DEL-70
chr13-50942763-DEL-52
chr13-50945360-DEL-1318
chr13-50948697-DEL-135
chr13-50957051-DEL-6206
chr13-50964576-DEL-1793
chr13-50966338-DEL-403
chr13-50968676-DEL-115
chr13-50974104-DEL-2133
chr13-51013733-DEL-483

chr13-51014896-DEL-179
chr13-51018759-DEL-395
chr13-51023978-DEL-6168
chr13-51100190-DEL-134
chr13-51157155-DEL-86
chr13-51163453-DEL-802
chr13-51209274-DEL-6475
chr13-51219499-DEL-655
chr13-51229948-DEL-1286
chr13-51231308-DEL-116
chr13-51290673-DEL-429
chr13-51309937-DEL-56
chr13-51315423-DEL-117
chr13-51317534-DEL-1118
chr13-51328001-DEL-100
chr13-51331368-DEL-1028
chr13-51355064-DEL-2728
chr13-51366554-DEL-125
chr13-51388395-DEL-351
chr13-51447456-DEL-80
chr13-51447821-DEL-821
chr13-51452052-DEL-59
chr13-51458311-DEL-239
chr13-51465891-DEL-63
chr13-51466875-DEL-344
chr13-51582818-DEL-170
chr13-51586953-DEL-60
chr13-51594771-DEL-1283
chr13-51604870-DEL-69
chr13-51609795-DEL-55
chr13-51633374-DEL-117
chr13-51638166-DEL-989
chr13-51661117-DEL-73
chr13-51709446-DEL-58
chr13-51712513-DEL-79
chr13-51714258-DEL-86
chr13-51725301-DEL-214
chr13-51776415-DEL-51
chr13-51787938-DEL-5197
chr13-51787970-DEL-3011
chr13-51791012-DEL-1945
chr13-51818144-DEL-165
chr13-51871465-DEL-243
chr13-51994605-DEL-59
chr13-52006442-DEL-51
chr13-52028792-DEL-85
chr13-52044094-DEL-1782
chr13-52131386-DEL-567
chr13-52154586-DEL-63
chr13-52166830-DEL-56
chr13-52224311-DEL-62
chr13-52257108-DEL-148
chr13-52296894-DEL-695
chr13-52319930-DEL-221
chr13-52320540-DEL-674
chr13-52323460-DEL-56
chr13-52379152-DEL-397
chr13-52395768-DEL-59
chr13-52403500-DEL-94
chr13-52436859-DEL-161
chr13-52450084-DEL-59
chr13-52495547-DEL-53
chr13-52498135-DEL-180
chr13-52510376-DEL-1045
chr13-52511552-DEL-70
chr13-52522808-DEL-1792
chr13-52544171-DEL-345
chr13-52547664-DEL-60
chr13-52571476-DEL-2033
chr13-52572643-DEL-240
chr13-52579394-DEL-901
chr13-52580630-DEL-356
chr13-52586208-DEL-460
chr13-52591429-DEL-1469
chr13-52598469-DEL-337
chr13-52679902-DEL-85
chr13-52691067-DEL-1379

chr13-52703742-DEL-1020
chr13-52725039-DEL-68
chr13-52726873-DEL-68
chr13-52823082-DEL-117
chr13-52825521-DEL-203
chr13-52860895-DEL-2315
chr13-52869725-DEL-58
chr13-52896695-DEL-72
chr13-52901104-DEL-352
chr13-52914153-DEL-175
chr13-52942430-DEL-81
chr13-53042184-DEL-305
chr13-53061092-DEL-1418
chr13-53062844-DEL-1349
chr13-53074616-DEL-150
chr13-53085202-DEL-72
chr13-53116685-DEL-1333
chr13-53158476-DEL-111
chr13-53176983-DEL-324
chr13-53193145-DEL-86
chr13-53203628-DEL-162
chr13-53213798-DEL-129
chr13-53226348-DEL-88
chr13-53231166-DEL-57
chr13-53238063-DEL-479
chr13-53277430-DEL-79
chr13-53278688-DEL-113
chr13-53283404-DEL-76
chr13-53293536-DEL-128
chr13-53340178-DEL-128
chr13-53342374-DEL-107
chr13-53344253-DEL-198
chr13-53382313-DEL-325
chr13-53423707-DEL-102
chr13-53427092-DEL-334
chr13-53464917-DEL-54
chr13-53471271-DEL-422
chr13-53480075-DEL-199
chr13-53483967-DEL-199
chr13-53536663-DEL-319
chr13-53546144-DEL-118
chr13-53569803-DEL-94
chr13-53573411-DEL-75
chr13-53611403-DEL-201
chr13-53637928-DEL-240
chr13-53644757-DEL-52
chr13-53651652-DEL-56
chr13-53660202-DEL-139
chr13-53690703-DEL-100
chr13-53766154-DEL-2500
chr13-53789812-DEL-63
chr13-53798798-DEL-66
chr13-53820897-DEL-60
chr13-53823099-DEL-128
chr13-53843394-DEL-66
chr13-53846484-DEL-66
chr13-53896633-DEL-207
chr13-53982371-DEL-101
chr13-54013356-DEL-107
chr13-54044660-DEL-3266
chr13-54071657-DEL-481
chr13-54093979-DEL-651
chr13-54097046-DEL-642
chr13-54129853-DEL-938
chr13-54135553-DEL-51
chr13-54165333-DEL-1596
chr13-54173906-DEL-460
chr13-54174112-DEL-465
chr13-54271209-DEL-491
chr13-54290134-DEL-59
chr13-54292594-DEL-849
chr13-54293722-DEL-382
chr13-54316670-DEL-91
chr13-54327284-DEL-108
chr13-54349665-DEL-246
chr13-54462529-DEL-139
chr13-54497011-DEL-107

chr13-54516337-DEL-54
chr13-54527795-DEL-99
chr13-54552698-DEL-107
chr13-54553047-DEL-742
chr13-54600550-DEL-1406
chr13-54601908-DEL-182
chr13-54610882-DEL-168
chr13-54644305-DEL-58
chr13-54653684-DEL-163
chr13-54673080-DEL-63
chr13-54683852-DEL-234
chr13-54696749-DEL-55
chr13-54708764-DEL-594
chr13-54718818-DEL-65
chr13-54753006-DEL-90
chr13-54760251-DEL-94
chr13-54760906-DEL-161
chr13-54782202-DEL-103
chr13-54808151-DEL-205
chr13-54820948-DEL-1251
chr13-54833631-DEL-53
chr13-54844961-DEL-108
chr13-54890245-DEL-78
chr13-54915736-DEL-118
chr13-54957084-DEL-58
chr13-54960743-DEL-79
chr13-54978708-DEL-82
chr13-55016552-DEL-72
chr13-55050540-DEL-387
chr13-55076917-DEL-56
chr13-55109041-DEL-71
chr13-55155637-DEL-10876
chr13-55169493-DEL-284
chr13-55178483-DEL-58
chr13-55215877-DEL-225
chr13-55218243-DEL-205
chr13-55231321-DEL-102
chr13-55234788-DEL-365
chr13-55290251-DEL-90
chr13-55310477-DEL-52
chr13-55323661-DEL-169
chr13-55329431-DEL-198
chr13-55339386-DEL-520
chr13-55340658-DEL-130
chr13-55344386-DEL-165
chr13-55352417-DEL-175
chr13-55368431-DEL-54
chr13-55401261-DEL-838
chr13-55421947-DEL-147
chr13-55439720-DEL-1420
chr13-55447975-DEL-211
chr13-55475020-DEL-114
chr13-55481327-DEL-303
chr13-55489470-DEL-190
chr13-55497163-DEL-213
chr13-55507435-DEL-61
chr13-55512271-DEL-186
chr13-55535004-DEL-217
chr13-55564617-DEL-211
chr13-55572286-DEL-91
chr13-55576214-DEL-56
chr13-55616315-DEL-169
chr13-55638363-DEL-188
chr13-55641249-DEL-139
chr13-55678721-DEL-144
chr13-55733239-DEL-196
chr13-55735558-DEL-52
chr13-55753396-DEL-197
chr13-55781506-DEL-188
chr13-55808736-DEL-56
chr13-55819736-DEL-722
chr13-55834628-DEL-199
chr13-55849612-DEL-53
chr13-55862000-DEL-98
chr13-55876609-DEL-116
chr13-55905852-DEL-62
chr13-55912520-DEL-132

chr13-55921807-DEL-249
chr13-55926880-DEL-1370
chr13-55928267-DEL-3292
chr13-55960316-DEL-68
chr13-55966229-DEL-67
chr13-55967387-DEL-224
chr13-56047226-DEL-74
chr13-56079340-DEL-239
chr13-56122696-DEL-313
chr13-56185324-DEL-76
chr13-56207894-DEL-202
chr13-56245783-DEL-87
chr13-56253375-DEL-177
chr13-56341276-DEL-841
chr13-56389771-DEL-140
chr13-56396507-DEL-60
chr13-56430786-DEL-575
chr13-56562315-DEL-61
chr13-56579718-DEL-95
chr13-56614452-DEL-404
chr13-56634475-DEL-436
chr13-56647440-DEL-846
chr13-56672483-DEL-588
chr13-56684208-DEL-73
chr13-56725420-DEL-897
chr13-56727451-DEL-428
chr13-56763369-DEL-54
chr13-56781159-DEL-188
chr13-56801128-DEL-309
chr13-56817791-DEL-175
chr13-56818740-DEL-195
chr13-56836607-DEL-119
chr13-56892625-DEL-468
chr13-56892865-DEL-110
chr13-56919023-DEL-252
chr13-56921840-DEL-121
chr13-56935648-DEL-56
chr13-56948906-DEL-827
chr13-56949195-DEL-176
chr13-56984143-DEL-115
chr13-57012774-DEL-60
chr13-57024926-DEL-152
chr13-57036272-DEL-436
chr13-57070760-DEL-78
chr13-57079476-DEL-54
chr13-57090421-DEL-385
chr13-57097048-DEL-1057
chr13-57103726-DEL-338
chr13-57112958-DEL-2289
chr13-57134894-DEL-70
chr13-57140410-DEL-3175
chr13-57142463-DEL-95
chr13-57148711-DEL-291
chr13-57152237-DEL-69
chr13-57152867-DEL-51
chr13-57166179-DEL-60
chr13-57193769-DEL-339
chr13-57213215-DEL-1586
chr13-57228579-DEL-56
chr13-57229890-DEL-204
chr13-57259537-DEL-60
chr13-57282786-DEL-355
chr13-57296969-DEL-54
chr13-57329917-DEL-131
chr13-57349683-DEL-2503
chr13-57349687-DEL-65
chr13-57352197-DEL-3835
chr13-57357055-DEL-2070
chr13-57381678-DEL-100
chr13-57420393-DEL-58
chr13-57422326-DEL-68
chr13-57443073-DEL-117
chr13-57461489-DEL-70
chr13-57463144-DEL-83
chr13-57468056-DEL-138
chr13-57500347-DEL-345
chr13-57524861-DEL-217

chr13-57531253-DEL-131
chr13-57533753-DEL-72
chr13-57563876-DEL-180
chr13-57588761-DEL-73
chr13-57608663-DEL-3266
chr13-57614458-DEL-85
chr13-57667551-DEL-197
chr13-57707737-DEL-61
chr13-57720260-DEL-71
chr13-57740919-DEL-56
chr13-57763965-DEL-70
chr13-57765120-DEL-133
chr13-57780905-DEL-1448
chr13-57797335-DEL-163
chr13-57818735-DEL-248
chr13-57821144-DEL-148
chr13-57826499-DEL-469
chr13-57858257-DEL-274
chr13-57889926-DEL-17434
chr13-57916565-DEL-444
chr13-57932874-DEL-299
chr13-57983472-DEL-96
chr13-58046500-DEL-151
chr13-58108036-DEL-146
chr13-58119186-DEL-179
chr13-58119448-DEL-120
chr13-58125853-DEL-80
chr13-58191802-DEL-192
chr13-58192277-DEL-2600
chr13-58222338-DEL-52
chr13-58233904-DEL-59
chr13-58248767-DEL-381
chr13-58299238-DEL-856
chr13-58332438-DEL-194
chr13-58341543-DEL-178
chr13-58383261-DEL-56
chr13-58390974-DEL-202
chr13-58410475-DEL-154
chr13-58415473-DEL-204
chr13-58425247-DEL-93
chr13-58436502-DEL-71
chr13-58453546-DEL-311
chr13-58483497-DEL-474
chr13-58489162-DEL-767
chr13-58494251-DEL-3437
chr13-58494270-DEL-2333
chr13-58494650-DEL-244
chr13-58494952-DEL-2367
chr13-58495932-DEL-158
chr13-58496277-DEL-317
chr13-58496886-DEL-758
chr13-58497564-DEL-62
chr13-58511015-DEL-936
chr13-58537965-DEL-57
chr13-58568556-DEL-2263
chr13-58649466-DEL-51
chr13-58650056-DEL-1493
chr13-58679340-DEL-173
chr13-58700614-DEL-57
chr13-58702297-DEL-82
chr13-58714849-DEL-358
chr13-58726770-DEL-100
chr13-58785103-DEL-165
chr13-58790111-DEL-329
chr13-58794625-DEL-1500
chr13-58801883-DEL-287
chr13-58809692-DEL-212
chr13-58810442-DEL-52
chr13-58857695-DEL-129
chr13-58859859-DEL-89
chr13-58864544-DEL-123
chr13-58873222-DEL-3265
chr13-58880711-DEL-100
chr13-58899148-DEL-242
chr13-58922177-DEL-50
chr13-58973453-DEL-51
chr13-58989032-DEL-53

chr13-59083471-DEL-76
chr13-59096146-DEL-123
chr13-59098587-DEL-60
chr13-59180083-DEL-120
chr13-59202458-DEL-85
chr13-59209309-DEL-90
chr13-59212837-DEL-582
chr13-59237445-DEL-68
chr13-59285946-DEL-84
chr13-59287030-DEL-324
chr13-59287508-DEL-72
chr13-59353646-DEL-78
chr13-59355718-DEL-127
chr13-59356993-DEL-180
chr13-59374323-DEL-64
chr13-59381955-DEL-152
chr13-59398106-DEL-294
chr13-59400642-DEL-61
chr13-59440298-DEL-52
chr13-59467523-DEL-54
chr13-59536582-DEL-2358
chr13-59564913-DEL-51
chr13-59571364-DEL-92
chr13-59576342-DEL-62
chr13-59589887-DEL-276
chr13-59593915-DEL-108
chr13-59594546-DEL-164
chr13-59611292-DEL-255
chr13-59619059-DEL-1366
chr13-59625686-DEL-314
chr13-59635650-DEL-62
chr13-59660679-DEL-170
chr13-59673591-DEL-377
chr13-59709510-DEL-210
chr13-59740807-DEL-676
chr13-59765187-DEL-57
chr13-59787142-DEL-58
chr13-59799401-DEL-70
chr13-59809920-DEL-102
chr13-59819543-DEL-51
chr13-59820573-DEL-83
chr13-59824399-DEL-58
chr13-59825933-DEL-93
chr13-59832399-DEL-2329
chr13-59847525-DEL-215
chr13-59869728-DEL-259
chr13-59886101-DEL-151
chr13-59918193-DEL-434
chr13-59988726-DEL-73
chr13-59996242-DEL-220
chr13-60010399-DEL-148
chr13-60041615-DEL-78
chr13-60044632-DEL-2273
chr13-60046971-DEL-85
chr13-60063204-DEL-55
chr13-60102746-DEL-272
chr13-60110118-DEL-275
chr13-60130211-DEL-329
chr13-60133349-DEL-76
chr13-60137424-DEL-55
chr13-60157188-DEL-213
chr13-60164764-DEL-54
chr13-60218173-DEL-90
chr13-60222254-DEL-402
chr13-60224707-DEL-87
chr13-60244571-DEL-810
chr13-60245189-DEL-72
chr13-60285873-DEL-173
chr13-60307774-DEL-125
chr13-60317721-DEL-64
chr13-60327169-DEL-242
chr13-60331786-DEL-326
chr13-60354160-DEL-546
chr13-60381172-DEL-56
chr13-60432163-DEL-53
chr13-60452209-DEL-61
chr13-60533999-DEL-54

chr13-60537275-DEL-6647
chr13-60544788-DEL-1690
chr13-60552575-DEL-55
chr13-60576549-DEL-52
chr13-60583799-DEL-318
chr13-60590623-DEL-70
chr13-60594356-DEL-57
chr13-60630119-DEL-73
chr13-60639379-DEL-194
chr13-60650606-DEL-98
chr13-60663781-DEL-425
chr13-60681455-DEL-54
chr13-60687674-DEL-192
chr13-60714593-DEL-556
chr13-60738556-DEL-245
chr13-60747061-DEL-164
chr13-60753560-DEL-139
chr13-60757314-DEL-66
chr13-60789004-DEL-76
chr13-60802740-DEL-760
chr13-60814750-DEL-73
chr13-60828914-DEL-725
chr13-60829561-DEL-130
chr13-60850059-DEL-61
chr13-60851685-DEL-61
chr13-60922140-DEL-136
chr13-60936326-DEL-140
chr13-60943967-DEL-61
chr13-60965878-DEL-64
chr13-60966909-DEL-395
chr13-60971665-DEL-1851
chr13-60991988-DEL-54
chr13-60992706-DEL-67
chr13-61017212-DEL-60
chr13-61018276-DEL-125
chr13-61044808-DEL-172
chr13-61049013-DEL-106
chr13-61055141-DEL-733
chr13-61071323-DEL-2501
chr13-61079264-DEL-562
chr13-61079805-DEL-68
chr13-61081086-DEL-490
chr13-61088608-DEL-193
chr13-61108310-DEL-472
chr13-61156799-DEL-476
chr13-61169930-DEL-505
chr13-61195070-DEL-1371
chr13-61249923-DEL-13653
chr13-61259948-DEL-201
chr13-61276558-DEL-116
chr13-61282281-DEL-354
chr13-61284039-DEL-2551
chr13-61305477-DEL-204
chr13-61320911-DEL-67
chr13-61335574-DEL-1012
chr13-61345004-DEL-231
chr13-61391988-DEL-280
chr13-61459358-DEL-1065
chr13-61466963-DEL-7728
chr13-61478126-DEL-6426
chr13-61520574-DEL-1170
chr13-61527475-DEL-5346
chr13-61538094-DEL-65
chr13-61545112-DEL-6421
chr13-61557199-DEL-9470
chr13-61557483-DEL-1178
chr13-61559400-DEL-630
chr13-61566275-DEL-96
chr13-61569830-DEL-6852
chr13-61611222-DEL-54
chr13-61611334-DEL-2361
chr13-61611334-DEL-6923
chr13-61614039-DEL-4219
chr13-61614054-DEL-519
chr13-61638732-DEL-267
chr13-61663932-DEL-6071
chr13-61682720-DEL-122

chr13-61695613-DEL-1909
chr13-61738381-DEL-76
chr13-61779146-DEL-4605
chr13-61783987-DEL-84
chr13-61794598-DEL-185
chr13-61803469-DEL-1538
chr13-61875705-DEL-544
chr13-61892865-DEL-168
chr13-61924573-DEL-93
chr13-61939966-DEL-60
chr13-61942449-DEL-2039
chr13-61950068-DEL-206
chr13-61952827-DEL-9227
chr13-61954111-DEL-215
chr13-61971134-DEL-430
chr13-61975440-DEL-6448
chr13-61998283-DEL-3949
chr13-62009048-DEL-168
chr13-62010548-DEL-197
chr13-62012673-DEL-936
chr13-62018636-DEL-218
chr13-62039881-DEL-82
chr13-62050756-DEL-212
chr13-62068335-DEL-51
chr13-62070291-DEL-247
chr13-62074623-DEL-99
chr13-62075656-DEL-216
chr13-62082717-DEL-115
chr13-62085154-DEL-6936
chr13-62121911-DEL-123
chr13-62138322-DEL-200
chr13-62140772-DEL-3541
chr13-62143115-DEL-182
chr13-62144338-DEL-54
chr13-62146528-DEL-332
chr13-62156079-DEL-4568
chr13-62213145-DEL-156
chr13-62226736-DEL-526
chr13-62229888-DEL-11543
chr13-62250962-DEL-162
chr13-62259423-DEL-3959
chr13-62286476-DEL-996
chr13-62289184-DEL-141
chr13-62308979-DEL-206
chr13-62325357-DEL-197
chr13-62385765-DEL-72
chr13-62433217-DEL-539
chr13-62435225-DEL-398
chr13-62436845-DEL-201
chr13-62439491-DEL-5231
chr13-62439861-DEL-245
chr13-62443216-DEL-68
chr13-62554119-DEL-215
chr13-62562102-DEL-207
chr13-62566529-DEL-975
chr13-62576560-DEL-821
chr13-62582895-DEL-59
chr13-62584237-DEL-178
chr13-62591075-DEL-84
chr13-62598411-DEL-90
chr13-62600851-DEL-166
chr13-62618164-DEL-209
chr13-62628918-DEL-205
chr13-62646894-DEL-155
chr13-62699858-DEL-5670
chr13-62711221-DEL-140
chr13-62734446-DEL-66
chr13-62753980-DEL-680
chr13-62756094-DEL-149
chr13-62764577-DEL-260
chr13-62765031-DEL-1079
chr13-62785422-DEL-74
chr13-62800501-DEL-4095
chr13-62827182-DEL-208
chr13-62837774-DEL-9238
chr13-62839038-DEL-2515
chr13-62848079-DEL-84

chr13-62849225-DEL-1247
chr13-62859881-DEL-60
chr13-62862071-DEL-8943
chr13-62862214-DEL-8001
chr13-62898555-DEL-155
chr13-62906095-DEL-213
chr13-62920705-DEL-220
chr13-62923626-DEL-205
chr13-62934497-DEL-60
chr13-62939310-DEL-850
chr13-62942585-DEL-2955
chr13-62951263-DEL-263
chr13-62952401-DEL-1538
chr13-62959869-DEL-12410
chr13-62963417-DEL-132
chr13-62964946-DEL-400
chr13-62965426-DEL-1178
chr13-62973180-DEL-2617
chr13-62979374-DEL-1098
chr13-62980538-DEL-913
chr13-62986463-DEL-140
chr13-62987493-DEL-198
chr13-63018130-DEL-54
chr13-63038314-DEL-69
chr13-63077199-DEL-193
chr13-63079611-DEL-377
chr13-63135925-DEL-3136
chr13-63156606-DEL-506
chr13-63174849-DEL-60
chr13-63191332-DEL-78
chr13-63210895-DEL-56
chr13-63256991-DEL-51
chr13-63311901-DEL-530
chr13-63313136-DEL-559
chr13-63313306-DEL-147
chr13-63340920-DEL-58
chr13-63343691-DEL-74
chr13-63348849-DEL-68
chr13-63354256-DEL-603
chr13-63399953-DEL-111
chr13-63508461-DEL-78
chr13-63527219-DEL-152
chr13-63527231-DEL-647
chr13-63527743-DEL-380
chr13-63571752-DEL-486
chr13-63584236-DEL-8116
chr13-63605277-DEL-199
chr13-63639069-DEL-59
chr13-63725536-DEL-240
chr13-63730461-DEL-57
chr13-63738191-DEL-68
chr13-63763034-DEL-196
chr13-63763983-DEL-70
chr13-63767678-DEL-56
chr13-63788652-DEL-166
chr13-63834831-DEL-104
chr13-63844601-DEL-113
chr13-63896598-DEL-435
chr13-63901903-DEL-2093
chr13-63910174-DEL-421
chr13-63931024-DEL-52
chr13-63943475-DEL-74
chr13-64023784-DEL-54
chr13-64043805-DEL-76
chr13-64056592-DEL-114
chr13-64057487-DEL-67
chr13-64062180-DEL-57
chr13-64080807-DEL-79
chr13-64106978-DEL-136
chr13-64120408-DEL-897
chr13-64192556-DEL-106
chr13-64205432-DEL-317
chr13-64216515-DEL-467
chr13-64226587-DEL-263
chr13-64314981-DEL-59
chr13-64317708-DEL-72
chr13-64354697-DEL-98

chr13-64389604-DEL-123
chr13-64420170-DEL-88
chr13-64435187-DEL-583
chr13-64442241-DEL-897
chr13-64444579-DEL-201
chr13-64453498-DEL-352
chr13-64476242-DEL-206
chr13-64495984-DEL-98
chr13-64518837-DEL-561
chr13-64550776-DEL-194
chr13-64554566-DEL-58
chr13-64561878-DEL-59
chr13-64596211-DEL-133
chr13-64643775-DEL-116
chr13-64687696-DEL-6272
chr13-64710745-DEL-90
chr13-64711717-DEL-610
chr13-64722980-DEL-2060
chr13-64730214-DEL-75
chr13-64769471-DEL-75
chr13-64771954-DEL-85
chr13-64805669-DEL-7389
chr13-64822794-DEL-1325
chr13-64831458-DEL-569
chr13-64846444-DEL-218
chr13-64858176-DEL-3727
chr13-64861622-DEL-112
chr13-64963203-DEL-992
chr13-64966299-DEL-573
chr13-64993530-DEL-536
chr13-64994563-DEL-552
chr13-64999537-DEL-838
chr13-65052112-DEL-54
chr13-65080487-DEL-65188
chr13-65086030-DEL-1023
chr13-65132315-DEL-1049
chr13-65156633-DEL-75
chr13-65157447-DEL-400
chr13-65170031-DEL-2253
chr13-65187858-DEL-56
chr13-65194688-DEL-52
chr13-65209915-DEL-257
chr13-65218498-DEL-565
chr13-65230035-DEL-62
chr13-65306286-DEL-56
chr13-65327730-DEL-52
chr13-65340432-DEL-173
chr13-65348263-DEL-52
chr13-65360192-DEL-72
chr13-65366661-DEL-518
chr13-65372911-DEL-210
chr13-65374004-DEL-462
chr13-65448024-DEL-68
chr13-65450897-DEL-67
chr13-65477705-DEL-199
chr13-65478762-DEL-387
chr13-65486486-DEL-168
chr13-65533285-DEL-606
chr13-65534340-DEL-1887
chr13-65543539-DEL-1460
chr13-65545386-DEL-252
chr13-65546492-DEL-84
chr13-65549192-DEL-195
chr13-65549859-DEL-6796
chr13-65558729-DEL-261
chr13-65560552-DEL-213
chr13-65562144-DEL-207
chr13-65563588-DEL-1834
chr13-65587731-DEL-52
chr13-65590904-DEL-7365
chr13-65615548-DEL-130
chr13-65666550-DEL-2673
chr13-65694876-DEL-294
chr13-65695052-DEL-165
chr13-65701298-DEL-261
chr13-65704714-DEL-207
chr13-65706282-DEL-1833

chr13-65727975-DEL-60
chr13-65746692-DEL-2838
chr13-65767478-DEL-270
chr13-65770471-DEL-196
chr13-65780618-DEL-103
chr13-65784929-DEL-222
chr13-65789021-DEL-2432
chr13-65813452-DEL-12559
chr13-65847479-DEL-129
chr13-65856136-DEL-400
chr13-65894286-DEL-68
chr13-65906137-DEL-1346
chr13-65913322-DEL-195
chr13-65951699-DEL-84
chr13-65952420-DEL-84
chr13-65960922-DEL-4606
chr13-65979184-DEL-84
chr13-66045245-DEL-1205
chr13-66051269-DEL-194
chr13-66055455-DEL-168
chr13-66064333-DEL-9391
chr13-66065000-DEL-800
chr13-66065874-DEL-5437
chr13-66115073-DEL-244
chr13-66120065-DEL-90
chr13-66133985-DEL-188
chr13-66135214-DEL-3143
chr13-66142417-DEL-84
chr13-66148087-DEL-243
chr13-66155343-DEL-4505
chr13-66160107-DEL-7099
chr13-66170470-DEL-400
chr13-66174059-DEL-5075
chr13-66179412-DEL-129
chr13-66186918-DEL-54
chr13-66195446-DEL-498
chr13-66209949-DEL-72
chr13-66242898-DEL-62
chr13-66249449-DEL-95
chr13-66269383-DEL-252
chr13-66273176-DEL-52
chr13-66311170-DEL-210
chr13-66314331-DEL-143
chr13-66338280-DEL-183
chr13-66340811-DEL-3845
chr13-66357512-DEL-262
chr13-66375101-DEL-125
chr13-66377029-DEL-7109
chr13-66393154-DEL-13656
chr13-66429913-DEL-167
chr13-66468478-DEL-484
chr13-66472283-DEL-135
chr13-66473797-DEL-556
chr13-66474706-DEL-291
chr13-66485221-DEL-843
chr13-66487695-DEL-168
chr13-66488636-DEL-168
chr13-66517087-DEL-117
chr13-66522482-DEL-6962
chr13-66530899-DEL-5109
chr13-66565779-DEL-55
chr13-66573030-DEL-195
chr13-66578515-DEL-2172
chr13-66581003-DEL-88
chr13-66590668-DEL-15188
chr13-66624589-DEL-2829
chr13-66628372-DEL-302
chr13-66629506-DEL-83
chr13-66636792-DEL-56
chr13-66639844-DEL-200
chr13-66640341-DEL-375
chr13-66647810-DEL-139
chr13-66673071-DEL-195
chr13-66676080-DEL-168
chr13-66676780-DEL-335
chr13-66681851-DEL-168
chr13-66685518-DEL-693

chr13-66689597-DEL-519
chr13-66690503-DEL-347
chr13-66691994-DEL-69
chr13-66692275-DEL-191
chr13-66693527-DEL-3469
chr13-66698363-DEL-137
chr13-66700052-DEL-70
chr13-66704956-DEL-30917
chr13-66707153-DEL-51
chr13-66722788-DEL-83
chr13-66748215-DEL-173
chr13-66753276-DEL-170
chr13-66755643-DEL-251
chr13-66759497-DEL-399
chr13-66771422-DEL-657
chr13-66857408-DEL-101
chr13-66862727-DEL-142
chr13-66885540-DEL-245
chr13-66893444-DEL-6282
chr13-66895748-DEL-116
chr13-66915149-DEL-5094
chr13-66940992-DEL-126
chr13-66943660-DEL-209
chr13-66951866-DEL-14938
chr13-66953469-DEL-1481
chr13-66958805-DEL-3843
chr13-66975529-DEL-83
chr13-66982957-DEL-6470
chr13-66999582-DEL-85
chr13-66999958-DEL-168
chr13-67000377-DEL-252
chr13-67000629-DEL-381
chr13-67012491-DEL-1739
chr13-67030290-DEL-1009
chr13-67037939-DEL-3984
chr13-67054009-DEL-1571
chr13-67055398-DEL-96
chr13-67069500-DEL-933
chr13-67083839-DEL-1568
chr13-67115297-DEL-750
chr13-67115299-DEL-152
chr13-67115315-DEL-1153
chr13-67116161-DEL-139
chr13-67131892-DEL-72
chr13-67137984-DEL-314
chr13-67181312-DEL-56
chr13-67196507-DEL-712
chr13-67201341-DEL-2065
chr13-67201344-DEL-591
chr13-67201888-DEL-1883
chr13-67233203-DEL-302
chr13-67246855-DEL-84
chr13-67263544-DEL-58
chr13-67264866-DEL-484
chr13-67296997-DEL-554
chr13-67317826-DEL-3131
chr13-67329898-DEL-420
chr13-67371459-DEL-3322
chr13-67402186-DEL-52
chr13-67419468-DEL-7727
chr13-67426229-DEL-57
chr13-67466100-DEL-60
chr13-67476285-DEL-51
chr13-67490889-DEL-99
chr13-67502735-DEL-156
chr13-67503496-DEL-327
chr13-67520409-DEL-1974
chr13-67537506-DEL-314
chr13-67538280-DEL-252
chr13-67541177-DEL-2257
chr13-67543623-DEL-160
chr13-67545129-DEL-117
chr13-67547115-DEL-165
chr13-67553704-DEL-5708
chr13-67563150-DEL-133
chr13-67564589-DEL-226
chr13-67571393-DEL-73

chr13-67576146-DEL-52
chr13-67580243-DEL-161
chr13-67584879-DEL-210
chr13-67586309-DEL-404
chr13-67598624-DEL-448
chr13-67655692-DEL-76
chr13-67702436-DEL-1944
chr13-67709995-DEL-1017
chr13-67718929-DEL-492
chr13-67721874-DEL-502
chr13-67725494-DEL-84
chr13-67746488-DEL-69
chr13-67751950-DEL-473
chr13-67774621-DEL-68
chr13-67779790-DEL-513
chr13-67780702-DEL-396
chr13-67827777-DEL-1446
chr13-67833317-DEL-1091
chr13-67838752-DEL-327
chr13-67873248-DEL-84
chr13-67903581-DEL-348
chr13-67910804-DEL-7524
chr13-67946247-DEL-9693
chr13-67958284-DEL-1118
chr13-67959592-DEL-741
chr13-67960530-DEL-678
chr13-67988652-DEL-5267
chr13-67988947-DEL-1677
chr13-67990305-DEL-96
chr13-68002814-DEL-85
chr13-68008436-DEL-1256
chr13-68010237-DEL-52
chr13-68013680-DEL-978
chr13-68015202-DEL-395
chr13-68032382-DEL-144
chr13-68045784-DEL-85
chr13-68049476-DEL-128
chr13-68051680-DEL-595
chr13-68058848-DEL-425
chr13-68072753-DEL-581
chr13-68100134-DEL-116
chr13-68212493-DEL-7602
chr13-68226405-DEL-498
chr13-68237879-DEL-14011
chr13-68298454-DEL-1827
chr13-68303366-DEL-69
chr13-68307333-DEL-1548
chr13-68309073-DEL-196
chr13-68310764-DEL-66
chr13-68337080-DEL-8013
chr13-68343493-DEL-59
chr13-68362907-DEL-59
chr13-68382589-DEL-58
chr13-68385897-DEL-105
chr13-68397705-DEL-6901
chr13-68403998-DEL-2979
chr13-68425097-DEL-116
chr13-68444210-DEL-5070
chr13-68457115-DEL-69
chr13-68464015-DEL-53
chr13-68480247-DEL-1105
chr13-68491970-DEL-564
chr13-68502614-DEL-78
chr13-68505902-DEL-1477
chr13-68506016-DEL-397
chr13-68524300-DEL-220
chr13-68531230-DEL-69
chr13-68568905-DEL-6615
chr13-68584412-DEL-270
chr13-68590114-DEL-8740
chr13-68601882-DEL-346
chr13-68603478-DEL-1512
chr13-68605884-DEL-293
chr13-68610401-DEL-337
chr13-68614018-DEL-196
chr13-68619111-DEL-232
chr13-68693003-DEL-486

chr13-68694920-DEL-554
chr13-68706118-DEL-343
chr13-68707034-DEL-361
chr13-68717199-DEL-151
chr13-68753928-DEL-75
chr13-68792464-DEL-4862
chr13-68839065-DEL-497
chr13-68854709-DEL-234
chr13-68866392-DEL-611
chr13-68878103-DEL-88
chr13-68898575-DEL-62
chr13-68928155-DEL-235
chr13-68930925-DEL-160
chr13-68931663-DEL-71
chr13-68953885-DEL-67
chr13-69004584-DEL-746
chr13-69078628-DEL-210
chr13-69113234-DEL-66
chr13-69183676-DEL-72
chr13-69190404-DEL-57
chr13-69220271-DEL-470
chr13-69243514-DEL-6193
chr13-69266038-DEL-76
chr13-69267010-DEL-110
chr13-69268289-DEL-52
chr13-69289576-DEL-1004
chr13-69346502-DEL-80
chr13-69355739-DEL-593
chr13-69371405-DEL-58
chr13-69392594-DEL-101
chr13-69393429-DEL-2887
chr13-69407293-DEL-55
chr13-69418805-DEL-184
chr13-69423140-DEL-72
chr13-69447583-DEL-56
chr13-69485181-DEL-151
chr13-69515744-DEL-81
chr13-69517710-DEL-79
chr13-69532876-DEL-101
chr13-69540616-DEL-56
chr13-69545754-DEL-477
chr13-69553788-DEL-63
chr13-69579675-DEL-158
chr13-69582340-DEL-227
chr13-69585090-DEL-263
chr13-69605530-DEL-63
chr13-69620513-DEL-300
chr13-69690265-DEL-120
chr13-69697876-DEL-53
chr13-69719887-DEL-141
chr13-69723461-DEL-54
chr13-69739779-DEL-85
chr13-69773147-DEL-104
chr13-69794873-DEL-217
chr13-69808501-DEL-184
chr13-69817594-DEL-80
chr13-69867370-DEL-282
chr13-69891852-DEL-280
chr13-69911343-DEL-150
chr13-69913955-DEL-78
chr13-69923608-DEL-503
chr13-69983582-DEL-52
chr13-69989926-DEL-61
chr13-70010683-DEL-90
chr13-70017686-DEL-445
chr13-70023725-DEL-1012
chr13-70026025-DEL-414
chr13-70026060-DEL-1927
chr13-70052087-DEL-635
chr13-70052110-DEL-159
chr13-70070103-DEL-51
chr13-70139139-DEL-898
chr13-70145434-DEL-486
chr13-70151098-DEL-390
chr13-70168024-DEL-93
chr13-70206649-DEL-229
chr13-70207139-DEL-2577

chr13-70243038-DEL-246
chr13-70255770-DEL-63
chr13-70287257-DEL-111
chr13-70314960-DEL-235
chr13-70349090-DEL-3448
chr13-70349816-DEL-122
chr13-70350826-DEL-498
chr13-70356116-DEL-124
chr13-70366958-DEL-150
chr13-70372749-DEL-58
chr13-70380061-DEL-1653
chr13-70389807-DEL-56
chr13-70411741-DEL-6148
chr13-70412525-DEL-2494
chr13-70419515-DEL-1153
chr13-70438208-DEL-562
chr13-70449946-DEL-76
chr13-70464645-DEL-149
chr13-70493116-DEL-184
chr13-70497831-DEL-138
chr13-70498473-DEL-1366
chr13-70500038-DEL-1057
chr13-70508184-DEL-614
chr13-70512129-DEL-54
chr13-70518364-DEL-332
chr13-70527541-DEL-100
chr13-70537545-DEL-6368
chr13-70548821-DEL-2058
chr13-70614033-DEL-3456
chr13-70630475-DEL-4703
chr13-70636310-DEL-1018
chr13-70641344-DEL-83
chr13-70662686-DEL-376
chr13-70677998-DEL-232
chr13-70685699-DEL-1782
chr13-70691083-DEL-385
chr13-70693756-DEL-77
chr13-70704678-DEL-52
chr13-70807427-DEL-112
chr13-70852456-DEL-60
chr13-70884173-DEL-59
chr13-70895186-DEL-175
chr13-70912812-DEL-500
chr13-70923043-DEL-130
chr13-70925005-DEL-62
chr13-70944032-DEL-75
chr13-70962247-DEL-200
chr13-70975492-DEL-76
chr13-70991927-DEL-78
chr13-71002349-DEL-118
chr13-71006876-DEL-53
chr13-71010998-DEL-70
chr13-71042559-DEL-431
chr13-71056247-DEL-51
chr13-71063408-DEL-198
chr13-71081575-DEL-87
chr13-71105253-DEL-1835
chr13-71124705-DEL-1159
chr13-71144741-DEL-65
chr13-71157159-DEL-1222
chr13-71162180-DEL-72
chr13-71166411-DEL-660
chr13-71167278-DEL-1567
chr13-71171297-DEL-105
chr13-71232849-DEL-70
chr13-71274145-DEL-109
chr13-71295558-DEL-338
chr13-71302229-DEL-198
chr13-71315163-DEL-97
chr13-71334124-DEL-55
chr13-71342242-DEL-177
chr13-71354906-DEL-56
chr13-71358442-DEL-425
chr13-71370055-DEL-261
chr13-71406849-DEL-66
chr13-71410221-DEL-95
chr13-71449294-DEL-64

chr13-71453932-DEL-58
chr13-71454338-DEL-1482
chr13-71460783-DEL-6699
chr13-71460905-DEL-198
chr13-71472336-DEL-359
chr13-71529521-DEL-6481
chr13-71529870-DEL-154
chr13-71540324-DEL-77
chr13-71544092-DEL-78
chr13-71554532-DEL-60
chr13-71567633-DEL-80
chr13-71648155-DEL-55
chr13-71662288-DEL-97
chr13-71690624-DEL-129
chr13-71699971-DEL-4616
chr13-71715573-DEL-63
chr13-71777025-DEL-71
chr13-71788732-DEL-2582
chr13-71820103-DEL-667
chr13-71861928-DEL-338
chr13-71886031-DEL-194
chr13-71942562-DEL-515
chr13-72024964-DEL-72
chr13-72054042-DEL-178
chr13-72093321-DEL-1219
chr13-72093857-DEL-367
chr13-72150173-DEL-1423
chr13-72151898-DEL-180
chr13-72153083-DEL-59
chr13-72153535-DEL-2616
chr13-72154584-DEL-1540
chr13-72168382-DEL-1823
chr13-72192293-DEL-58
chr13-72228959-DEL-87
chr13-72271314-DEL-1636
chr13-72313995-DEL-203
chr13-72334742-DEL-122
chr13-72401492-DEL-578
chr13-72459930-DEL-289
chr13-72474403-DEL-89
chr13-72481929-DEL-72
chr13-72499508-DEL-73
chr13-72559601-DEL-210
chr13-72587436-DEL-182
chr13-72601395-DEL-1118
chr13-72601409-DEL-1998
chr13-72602685-DEL-1560
chr13-72602761-DEL-723
chr13-72603611-DEL-646
chr13-72635975-DEL-802
chr13-72636664-DEL-546
chr13-72647969-DEL-178
chr13-72650587-DEL-90
chr13-72653768-DEL-91
chr13-72670471-DEL-4058
chr13-72676877-DEL-1643
chr13-72691593-DEL-1722
chr13-72753192-DEL-2144
chr13-72785287-DEL-252
chr13-72825661-DEL-200
chr13-72840196-DEL-64
chr13-72857202-DEL-66
chr13-72858075-DEL-52
chr13-72867634-DEL-67
chr13-72882669-DEL-734
chr13-72896530-DEL-608
chr13-72942498-DEL-199
chr13-72959707-DEL-663
chr13-72971154-DEL-55
chr13-72972141-DEL-61
chr13-73063595-DEL-102
chr13-73074664-DEL-66
chr13-73128638-DEL-452
chr13-73129869-DEL-125
chr13-73133279-DEL-6231
chr13-73147421-DEL-87
chr13-73154274-DEL-137

chr13-73188839-DEL-62
chr13-73254486-DEL-58
chr13-73290852-DEL-3457
chr13-73290929-DEL-54
chr13-73313530-DEL-2019
chr13-73355041-DEL-70
chr13-73428736-DEL-64
chr13-73472417-DEL-480
chr13-73516734-DEL-347
chr13-73522240-DEL-526
chr13-73534368-DEL-60
chr13-73549891-DEL-758
chr13-73569855-DEL-72
chr13-73591114-DEL-51
chr13-73610560-DEL-185
chr13-73668922-DEL-222
chr13-73678092-DEL-55
chr13-73695014-DEL-3943
chr13-73695106-DEL-2512
chr13-73702046-DEL-284
chr13-73714578-DEL-708
chr13-73748974-DEL-235
chr13-73767323-DEL-52
chr13-73783983-DEL-248
chr13-73788708-DEL-171
chr13-73800485-DEL-206
chr13-73863624-DEL-189
chr13-73874665-DEL-231
chr13-73906376-DEL-57
chr13-73977977-DEL-736
chr13-73992058-DEL-81
chr13-74007525-DEL-58
chr13-74123695-DEL-101
chr13-74195437-DEL-54
chr13-74203973-DEL-66
chr13-74228665-DEL-109
chr13-74254892-DEL-278
chr13-74256085-DEL-432
chr13-74257190-DEL-305
chr13-74302467-DEL-306
chr13-74333871-DEL-165
chr13-74379206-DEL-1753
chr13-74434886-DEL-228
chr13-74532387-DEL-331
chr13-74579278-DEL-1035
chr13-74613824-DEL-113
chr13-74623876-DEL-354
chr13-74656010-DEL-332
chr13-74660309-DEL-91
chr13-74676553-DEL-2655
chr13-74677347-DEL-140
chr13-74678063-DEL-286
chr13-74681646-DEL-1219
chr13-74685498-DEL-7842
chr13-74696905-DEL-503
chr13-74706583-DEL-204
chr13-74709751-DEL-206
chr13-74716546-DEL-400
chr13-74721317-DEL-423
chr13-74860289-DEL-82
chr13-74878109-DEL-659
chr13-74888805-DEL-156
chr13-74900190-DEL-130
chr13-74925921-DEL-191
chr13-74926334-DEL-962
chr13-74927215-DEL-285
chr13-74937060-DEL-75
chr13-74940150-DEL-72
chr13-74967687-DEL-318
chr13-74983977-DEL-6620
chr13-74997930-DEL-202
chr13-75000811-DEL-106
chr13-75025679-DEL-6189
chr13-75035154-DEL-198
chr13-75070885-DEL-217
chr13-75075707-DEL-202
chr13-75084463-DEL-120

chr13-75089926-DEL-568
chr13-75093477-DEL-4847
chr13-75127569-DEL-51
chr13-75137729-DEL-474
chr13-75145239-DEL-902
chr13-75150080-DEL-470
chr13-75171046-DEL-1223
chr13-75172486-DEL-2609
chr13-75211601-DEL-144
chr13-75255204-DEL-155
chr13-75272913-DEL-56
chr13-75296319-DEL-1443
chr13-75301351-DEL-64
chr13-75314503-DEL-2124
chr13-75354143-DEL-200
chr13-75356807-DEL-2525
chr13-75363964-DEL-161
chr13-75369851-DEL-103
chr13-75397605-DEL-180
chr13-75503424-DEL-63
chr13-75526335-DEL-712
chr13-75548030-DEL-108
chr13-75557666-DEL-1636
chr13-75576024-DEL-52
chr13-75585434-DEL-599
chr13-75594899-DEL-767
chr13-75618667-DEL-74
chr13-75626967-DEL-84
chr13-75629367-DEL-130
chr13-75660746-DEL-1065
chr13-75697299-DEL-398
chr13-75732408-DEL-429
chr13-75762589-DEL-107
chr13-75812973-DEL-220
chr13-75821868-DEL-580
chr13-75850643-DEL-182
chr13-75851649-DEL-458
chr13-75879632-DEL-1234
chr13-75882656-DEL-264
chr13-75908994-DEL-72
chr13-75918015-DEL-81
chr13-75968598-DEL-2120
chr13-75970323-DEL-1095
chr13-75979020-DEL-1509
chr13-76003811-DEL-373
chr13-76013633-DEL-72
chr13-76019673-DEL-52
chr13-76023472-DEL-218
chr13-76042653-DEL-5334
chr13-76052665-DEL-81
chr13-76053794-DEL-63
chr13-76058190-DEL-108
chr13-76062522-DEL-57
chr13-76092928-DEL-120
chr13-76112192-DEL-219
chr13-76142066-DEL-156
chr13-76142723-DEL-68
chr13-76172286-DEL-172
chr13-76187051-DEL-53
chr13-76215872-DEL-2330
chr13-76215897-DEL-61
chr13-76262827-DEL-67
chr13-76270642-DEL-70
chr13-76293066-DEL-72
chr13-76314219-DEL-5360
chr13-76346787-DEL-366
chr13-76351926-DEL-4589
chr13-76356531-DEL-1790
chr13-76372564-DEL-67
chr13-76374583-DEL-761
chr13-76386899-DEL-565
chr13-76403910-DEL-55
chr13-76451449-DEL-148
chr13-76479461-DEL-72
chr13-76483165-DEL-178
chr13-76487314-DEL-560
chr13-76524645-DEL-132

chr13-76529681-DEL-191
chr13-76530291-DEL-52
chr13-76581758-DEL-4349
chr13-76598211-DEL-5614
chr13-76612679-DEL-79
chr13-76622235-DEL-764
chr13-76622236-DEL-62
chr13-76623273-DEL-2479
chr13-76625818-DEL-694
chr13-76641139-DEL-12867
chr13-76641156-DEL-7965
chr13-76657138-DEL-1452
chr13-76699808-DEL-108
chr13-76738971-DEL-204
chr13-76777780-DEL-531
chr13-76779021-DEL-94
chr13-76791163-DEL-171
chr13-76805051-DEL-1068
chr13-76831928-DEL-56
chr13-76862554-DEL-56
chr13-76871118-DEL-181
chr13-76896162-DEL-894
chr13-76899567-DEL-58
chr13-76923290-DEL-3473
chr13-76963200-DEL-123
chr13-76979906-DEL-53
chr13-77014347-DEL-129
chr13-77040614-DEL-267
chr13-77080089-DEL-84
chr13-77088477-DEL-215
chr13-77136938-DEL-8812
chr13-77154565-DEL-54
chr13-77175119-DEL-195
chr13-77202549-DEL-4638
chr13-77237255-DEL-71
chr13-77257638-DEL-195
chr13-77266748-DEL-85
chr13-77271722-DEL-189
chr13-77277879-DEL-85
chr13-77286813-DEL-286
chr13-77294459-DEL-641
chr13-77334960-DEL-4258
chr13-77403372-DEL-553
chr13-77431473-DEL-156
chr13-77454013-DEL-445
chr13-77454508-DEL-300
chr13-77461274-DEL-101
chr13-77482723-DEL-68
chr13-77507071-DEL-136
chr13-77507167-DEL-1571
chr13-77509129-DEL-5643
chr13-77514777-DEL-6464
chr13-77515799-DEL-4149
chr13-77519967-DEL-1272
chr13-77523590-DEL-6084
chr13-77564807-DEL-396
chr13-77566370-DEL-267
chr13-77569963-DEL-1367
chr13-77576292-DEL-6590
chr13-77583336-DEL-1392
chr13-77588036-DEL-93
chr13-77590695-DEL-1357
chr13-77604491-DEL-57
chr13-77606595-DEL-186
chr13-77613170-DEL-254
chr13-77668171-DEL-137
chr13-77711717-DEL-13502
chr13-77717401-DEL-6901
chr13-77724964-DEL-208
chr13-77729148-DEL-89
chr13-77733582-DEL-849
chr13-77746323-DEL-7114
chr13-77802006-DEL-306
chr13-77805702-DEL-291
chr13-77817718-DEL-499
chr13-77818815-DEL-4516
chr13-77836043-DEL-1025

chr13-77837038-DEL-62
chr13-77853306-DEL-488
chr13-77864793-DEL-4857
chr13-77872417-DEL-700
chr13-77875564-DEL-184
chr13-77885396-DEL-125
chr13-77915111-DEL-9370
chr13-77923380-DEL-78
chr13-77929208-DEL-1099
chr13-77937222-DEL-5386
chr13-77937567-DEL-16870
chr13-77952776-DEL-96
chr13-77960022-DEL-110
chr13-77962654-DEL-218
chr13-78001798-DEL-357
chr13-78054564-DEL-558
chr13-78069412-DEL-133
chr13-78077722-DEL-268
chr13-78078325-DEL-1163
chr13-78079564-DEL-435
chr13-78080996-DEL-1996
chr13-78122880-DEL-51
chr13-78138410-DEL-461
chr13-78179426-DEL-242
chr13-78187589-DEL-476
chr13-78194685-DEL-79
chr13-78198170-DEL-6613
chr13-78202949-DEL-5577
chr13-78212776-DEL-138
chr13-78217675-DEL-5214
chr13-78225623-DEL-2487
chr13-78252873-DEL-1641
chr13-78299141-DEL-58
chr13-78307983-DEL-413
chr13-78373400-DEL-784
chr13-78384372-DEL-1526
chr13-78396500-DEL-198
chr13-78423051-DEL-50
chr13-78427643-DEL-349
chr13-78469427-DEL-1358
chr13-78500913-DEL-6404
chr13-78507040-DEL-199
chr13-78534953-DEL-164
chr13-78535909-DEL-1887
chr13-78679658-DEL-4038
chr13-78708601-DEL-96
chr13-78711776-DEL-913
chr13-78757974-DEL-3532
chr13-78814764-DEL-51
chr13-78835918-DEL-252
chr13-78867222-DEL-342
chr13-78986035-DEL-6630
chr13-78986675-DEL-3361
chr13-78991446-DEL-136
chr13-79044086-DEL-1227
chr13-79045393-DEL-172
chr13-79060267-DEL-386
chr13-79072972-DEL-64
chr13-79088503-DEL-191
chr13-79105448-DEL-536
chr13-79131940-DEL-181
chr13-79161941-DEL-621
chr13-79163009-DEL-289
chr13-79221857-DEL-114
chr13-79234314-DEL-521
chr13-79238854-DEL-855
chr13-79251122-DEL-756
chr13-79265161-DEL-1417
chr13-79310822-DEL-136
chr13-79334157-DEL-501
chr13-79335026-DEL-329
chr13-79336676-DEL-1520
chr13-79337608-DEL-66
chr13-79377297-DEL-246
chr13-79395950-DEL-155
chr13-79452282-DEL-204
chr13-79456731-DEL-1264

chr13-79474937-DEL-144
chr13-79478704-DEL-4616
chr13-79478802-DEL-199
chr13-79545973-DEL-55
chr13-79584719-DEL-76
chr13-79590319-DEL-118
chr13-79610546-DEL-431
chr13-79621654-DEL-1473
chr13-79632711-DEL-695
chr13-79664122-DEL-496
chr13-79671518-DEL-2981
chr13-79700465-DEL-150
chr13-79702454-DEL-318
chr13-79728444-DEL-2349
chr13-79746684-DEL-481
chr13-79752019-DEL-53
chr13-79771354-DEL-683
chr13-79785852-DEL-116
chr13-79799323-DEL-158
chr13-79816004-DEL-326
chr13-79820758-DEL-52
chr13-79842016-DEL-2917
chr13-79848966-DEL-90
chr13-79863331-DEL-6704
chr13-79864908-DEL-876
chr13-79880458-DEL-88
chr13-79886036-DEL-64
chr13-79894753-DEL-387
chr13-79902870-DEL-6033
chr13-79919931-DEL-705
chr13-79938979-DEL-1074
chr13-79940584-DEL-70
chr13-79942768-DEL-52
chr13-79963300-DEL-59
chr13-79995078-DEL-1033
chr13-80041678-DEL-358
chr13-80042386-DEL-570
chr13-80043303-DEL-6957
chr13-80058168-DEL-206
chr13-80059304-DEL-87
chr13-80067239-DEL-4860
chr13-80075155-DEL-688
chr13-80095925-DEL-597
chr13-80106222-DEL-267
chr13-80157086-DEL-1722
chr13-80159677-DEL-70
chr13-80162375-DEL-56
chr13-80167323-DEL-108
chr13-80197122-DEL-68
chr13-80197509-DEL-6277
chr13-80209218-DEL-77
chr13-80219404-DEL-1309
chr13-80229698-DEL-78
chr13-80238520-DEL-55
chr13-80241481-DEL-71
chr13-80247472-DEL-82
chr13-80248495-DEL-190
chr13-80280383-DEL-51
chr13-80308102-DEL-52
chr13-80308555-DEL-162
chr13-80337375-DEL-80
chr13-80351669-DEL-208
chr13-80360670-DEL-438
chr13-80362187-DEL-98
chr13-80386161-DEL-60
chr13-80401476-DEL-4308
chr13-80458690-DEL-53
chr13-80459601-DEL-67
chr13-80473891-DEL-440
chr13-80487651-DEL-66
chr13-80520102-DEL-243
chr13-80554085-DEL-115
chr13-80560412-DEL-485
chr13-80616699-DEL-129
chr13-80654908-DEL-1550
chr13-80656653-DEL-1361
chr13-80675471-DEL-64

chr13-80684858-DEL-1887
chr13-80706326-DEL-6313
chr13-80745485-DEL-360
chr13-80800600-DEL-142
chr13-80902178-DEL-64
chr13-80915081-DEL-258
chr13-80962436-DEL-260
chr13-80976983-DEL-77
chr13-81000160-DEL-1835
chr13-81006717-DEL-930
chr13-81020074-DEL-75
chr13-81047730-DEL-125
chr13-81060235-DEL-343
chr13-81071901-DEL-86
chr13-81072065-DEL-582
chr13-81087940-DEL-87
chr13-81095423-DEL-66
chr13-81104830-DEL-59
chr13-81108482-DEL-187
chr13-81115587-DEL-200
chr13-81137265-DEL-192
chr13-81140357-DEL-140
chr13-81165674-DEL-249
chr13-81181180-DEL-138
chr13-81198308-DEL-66
chr13-81211467-DEL-435
chr13-81266008-DEL-81
chr13-81290020-DEL-98
chr13-81342131-DEL-516
chr13-81355551-DEL-81
chr13-81376080-DEL-209
chr13-81380048-DEL-63
chr13-81394589-DEL-183
chr13-81403442-DEL-476
chr13-81404339-DEL-741
chr13-81416072-DEL-92
chr13-81420740-DEL-77
chr13-81444857-DEL-700
chr13-81457769-DEL-403
chr13-81466212-DEL-55
chr13-81469462-DEL-195
chr13-81477589-DEL-224
chr13-81563194-DEL-184
chr13-81617767-DEL-117
chr13-81621761-DEL-264
chr13-81633891-DEL-181
chr13-81653085-DEL-58
chr13-81654993-DEL-573
chr13-81715848-DEL-58
chr13-81749344-DEL-411
chr13-81750930-DEL-392
chr13-81784033-DEL-184
chr13-81797951-DEL-870
chr13-81829309-DEL-350
chr13-81856115-DEL-61
chr13-81858343-DEL-74
chr13-81875483-DEL-816
chr13-81887678-DEL-492
chr13-81896088-DEL-99
chr13-81966340-DEL-3868
chr13-82009414-DEL-81
chr13-82032292-DEL-910
chr13-82065679-DEL-295
chr13-82121744-DEL-2157
chr13-82125390-DEL-218
chr13-82127393-DEL-69
chr13-82140305-DEL-63
chr13-82163306-DEL-8874
chr13-82163689-DEL-64
chr13-82166676-DEL-5370
chr13-82168409-DEL-1909
chr13-82205408-DEL-72
chr13-82215599-DEL-51
chr13-82259773-DEL-81
chr13-82269110-DEL-209
chr13-82313967-DEL-266
chr13-82315027-DEL-346

chr13-82415323-DEL-84
chr13-82473134-DEL-1707
chr13-82483780-DEL-214
chr13-82512133-DEL-79
chr13-82531552-DEL-1993
chr13-82543209-DEL-56
chr13-82582136-DEL-146
chr13-82622871-DEL-295
chr13-82629682-DEL-55
chr13-82637239-DEL-400
chr13-82671101-DEL-2621
chr13-82673862-DEL-3643
chr13-82689078-DEL-78
chr13-82704058-DEL-799
chr13-82748522-DEL-309
chr13-82749906-DEL-895
chr13-82790600-DEL-764
chr13-82796270-DEL-52
chr13-82828838-DEL-77
chr13-82860847-DEL-69
chr13-82887557-DEL-591
chr13-82928123-DEL-54
chr13-82976960-DEL-58
chr13-83006987-DEL-76
chr13-83014924-DEL-829
chr13-83017348-DEL-60
chr13-83024604-DEL-70
chr13-83090576-DEL-84
chr13-83121378-DEL-71
chr13-83130634-DEL-192
chr13-83137020-DEL-90
chr13-83149866-DEL-303
chr13-83190186-DEL-182
chr13-83197852-DEL-2234
chr13-83233788-DEL-86
chr13-83235485-DEL-68
chr13-83236020-DEL-650
chr13-83245498-DEL-131
chr13-83292444-DEL-70
chr13-83316387-DEL-73
chr13-83318890-DEL-57
chr13-83341490-DEL-7393
chr13-83353493-DEL-609
chr13-83361192-DEL-176
chr13-83401018-DEL-118
chr13-83427183-DEL-208
chr13-83457195-DEL-915
chr13-83487644-DEL-185
chr13-83521711-DEL-1041
chr13-83544919-DEL-3462
chr13-83567089-DEL-1556
chr13-83568935-DEL-1868
chr13-83571801-DEL-2292
chr13-83574172-DEL-74
chr13-83575044-DEL-69
chr13-83618803-DEL-875
chr13-83625410-DEL-226
chr13-83661981-DEL-53
chr13-83945030-DEL-82
chr13-83983851-DEL-1489
chr13-83989402-DEL-135
chr13-84008134-DEL-60
chr13-84040860-DEL-1356
chr13-84047370-DEL-95
chr13-84048207-DEL-210
chr13-84067787-DEL-370
chr13-84069168-DEL-400
chr13-84085145-DEL-427
chr13-84110939-DEL-708
chr13-84112677-DEL-1269
chr13-84117210-DEL-4114
chr13-84144150-DEL-165
chr13-84162455-DEL-370
chr13-84184634-DEL-103
chr13-84185397-DEL-52
chr13-84190148-DEL-212
chr13-84226234-DEL-60

chr13-84236611-DEL-204
chr13-84246258-DEL-322
chr13-84274949-DEL-122
chr13-84285588-DEL-84
chr13-84309052-DEL-225
chr13-84313012-DEL-54
chr13-84382979-DEL-3092
chr13-84391545-DEL-72
chr13-84394683-DEL-454
chr13-84397283-DEL-71
chr13-84413333-DEL-234
chr13-84507826-DEL-330
chr13-84510341-DEL-1284
chr13-84524923-DEL-2417
chr13-84540514-DEL-60
chr13-84550499-DEL-51
chr13-84555443-DEL-156
chr13-84555541-DEL-849
chr13-84566123-DEL-69
chr13-84573550-DEL-147
chr13-84637748-DEL-212
chr13-84667162-DEL-157
chr13-84679725-DEL-83
chr13-84680143-DEL-225
chr13-84691713-DEL-55
chr13-84730201-DEL-661
chr13-84751301-DEL-102
chr13-84762402-DEL-64
chr13-84763706-DEL-396
chr13-84778517-DEL-458
chr13-84799204-DEL-1282
chr13-84841041-DEL-187
chr13-84841070-DEL-1274
chr13-84845909-DEL-211
chr13-84862796-DEL-1791
chr13-84880433-DEL-1200
chr13-84881941-DEL-143
chr13-84882579-DEL-460
chr13-84887781-DEL-441
chr13-84939069-DEL-405
chr13-84958047-DEL-22631
chr13-84959493-DEL-6231
chr13-84970043-DEL-314
chr13-84976906-DEL-281
chr13-84996055-DEL-6135
chr13-85003255-DEL-2696
chr13-85004135-DEL-556
chr13-85012159-DEL-79
chr13-85024936-DEL-257
chr13-85050277-DEL-385
chr13-85076410-DEL-100
chr13-85077215-DEL-156
chr13-85081715-DEL-896
chr13-85100472-DEL-66
chr13-85137551-DEL-2591
chr13-85160350-DEL-287
chr13-85162616-DEL-2200
chr13-85193925-DEL-74
chr13-85243730-DEL-843
chr13-85260731-DEL-642
chr13-85274293-DEL-298
chr13-85291282-DEL-915
chr13-85292228-DEL-1815
chr13-85317431-DEL-1621
chr13-85461034-DEL-77
chr13-85485863-DEL-164
chr13-85502120-DEL-262
chr13-85538727-DEL-271
chr13-85570906-DEL-937
chr13-85574147-DEL-341
chr13-85575966-DEL-121
chr13-85699379-DEL-1193
chr13-85724159-DEL-403
chr13-85724748-DEL-62
chr13-85736024-DEL-55
chr13-85742383-DEL-1482
chr13-85758630-DEL-67

chr13-85761262-DEL-1052
chr13-85763959-DEL-79
chr13-85813564-DEL-174
chr13-85826895-DEL-368
chr13-85850394-DEL-455
chr13-85853282-DEL-63
chr13-85870475-DEL-182
chr13-85874752-DEL-6336
chr13-85881814-DEL-61
chr13-85916175-DEL-80
chr13-85938329-DEL-110
chr13-85964290-DEL-61
chr13-86006858-DEL-194
chr13-86024113-DEL-6224
chr13-86037976-DEL-182
chr13-86062364-DEL-329
chr13-86077478-DEL-6405
chr13-86092261-DEL-485
chr13-86094585-DEL-587
chr13-86115493-DEL-65
chr13-86171200-DEL-406
chr13-86171739-DEL-717
chr13-86194078-DEL-205
chr13-86206483-DEL-524
chr13-86213837-DEL-95
chr13-86241776-DEL-721
chr13-86268852-DEL-7290
chr13-86284531-DEL-2529
chr13-86298134-DEL-952
chr13-86318120-DEL-1869
chr13-86322349-DEL-179
chr13-86334297-DEL-15001
chr13-86335359-DEL-8811
chr13-86355178-DEL-102
chr13-86387496-DEL-93
chr13-86389611-DEL-1339
chr13-86404278-DEL-944
chr13-86406418-DEL-6334
chr13-86412537-DEL-194
chr13-86420491-DEL-52
chr13-86435992-DEL-6672
chr13-86436232-DEL-214
chr13-86443820-DEL-4038
chr13-86445605-DEL-8422
chr13-86454213-DEL-61
chr13-86458267-DEL-2037
chr13-86460346-DEL-2512
chr13-86466064-DEL-220
chr13-86470119-DEL-4371
chr13-86474927-DEL-167
chr13-86492027-DEL-183
chr13-86504520-DEL-666
chr13-86510868-DEL-226
chr13-86538450-DEL-1640
chr13-86538887-DEL-151
chr13-86554894-DEL-438
chr13-86564093-DEL-127
chr13-86570984-DEL-74
chr13-86574734-DEL-2734
chr13-86614040-DEL-1120
chr13-86614773-DEL-53
chr13-86624837-DEL-1114
chr13-86641307-DEL-582
chr13-86665610-DEL-578
chr13-86674963-DEL-244
chr13-86676763-DEL-521
chr13-86701369-DEL-400
chr13-86707020-DEL-334
chr13-86709702-DEL-202
chr13-86724100-DEL-6210
chr13-86762379-DEL-426
chr13-86820544-DEL-60
chr13-86832485-DEL-1341
chr13-86832486-DEL-6212
chr13-86880482-DEL-2149
chr13-86889368-DEL-3551
chr13-86896930-DEL-884

chr13-86901281-DEL-6258
chr13-86908497-DEL-115
chr13-86910138-DEL-680
chr13-86913149-DEL-2791
chr13-86914025-DEL-197
chr13-86926680-DEL-717
chr13-86937374-DEL-121
chr13-86940410-DEL-480
chr13-86953675-DEL-536
chr13-86961229-DEL-473
chr13-86961922-DEL-130
chr13-86999218-DEL-67
chr13-87022626-DEL-177
chr13-87049054-DEL-1303
chr13-87070331-DEL-2809
chr13-87093938-DEL-74
chr13-87094175-DEL-1230
chr13-87098816-DEL-520
chr13-87126225-DEL-219
chr13-87155112-DEL-66
chr13-87166079-DEL-144
chr13-87190861-DEL-3554
chr13-87207645-DEL-400
chr13-87209838-DEL-341
chr13-87226939-DEL-816
chr13-87235766-DEL-1204
chr13-87237125-DEL-69
chr13-87278548-DEL-536
chr13-87285464-DEL-6434
chr13-87287767-DEL-19307
chr13-87291912-DEL-1104
chr13-87293061-DEL-1302
chr13-87294385-DEL-656
chr13-87315282-DEL-8863
chr13-87318067-DEL-4516
chr13-87324876-DEL-677
chr13-87333557-DEL-6035
chr13-87333924-DEL-199
chr13-87346567-DEL-790
chr13-87354503-DEL-90
chr13-87360500-DEL-196
chr13-87393647-DEL-140
chr13-87397057-DEL-52
chr13-87401283-DEL-144
chr13-87464140-DEL-204
chr13-87476922-DEL-3608
chr13-87495844-DEL-347
chr13-87504122-DEL-6312
chr13-87516352-DEL-8484
chr13-87518883-DEL-4431
chr13-87539969-DEL-1770
chr13-87543365-DEL-577
chr13-87553776-DEL-205
chr13-87573955-DEL-460
chr13-87589115-DEL-6329
chr13-87644286-DEL-196
chr13-87669728-DEL-6150
chr13-87685324-DEL-383
chr13-87692393-DEL-93
chr13-87694698-DEL-249
chr13-87704419-DEL-5742
chr13-87756835-DEL-1518
chr13-87768558-DEL-5810
chr13-87777556-DEL-61
chr13-87784221-DEL-288
chr13-87856383-DEL-60
chr13-87881197-DEL-52
chr13-87899275-DEL-113
chr13-87900933-DEL-54
chr13-87903573-DEL-205
chr13-87910479-DEL-161
chr13-87911424-DEL-112
chr13-87917736-DEL-2859
chr13-87950972-DEL-3338
chr13-87959890-DEL-1194
chr13-87961406-DEL-849
chr13-87964460-DEL-8649

chr13-87984341-DEL-7828
chr13-87986798-DEL-2263
chr13-88020955-DEL-1156
chr13-88022394-DEL-1029
chr13-88025649-DEL-139
chr13-88026403-DEL-155
chr13-88036692-DEL-167
chr13-88056421-DEL-265
chr13-88067796-DEL-51
chr13-88069558-DEL-86
chr13-88081629-DEL-5362
chr13-88087022-DEL-840
chr13-88087939-DEL-188
chr13-88091341-DEL-64
chr13-88094954-DEL-51
chr13-88105847-DEL-101
chr13-88134350-DEL-53
chr13-88146544-DEL-97
chr13-88151380-DEL-82
chr13-88182795-DEL-79
chr13-88206491-DEL-4802
chr13-88220239-DEL-96
chr13-88260000-DEL-51
chr13-88262693-DEL-200
chr13-88265218-DEL-130
chr13-88314114-DEL-258
chr13-88321955-DEL-1969
chr13-88326580-DEL-606
chr13-88329354-DEL-110
chr13-88330752-DEL-498
chr13-88344942-DEL-468
chr13-88350421-DEL-1029
chr13-88377127-DEL-2464
chr13-88400953-DEL-108
chr13-88424289-DEL-326
chr13-88439592-DEL-208
chr13-88465127-DEL-1873
chr13-88471863-DEL-112
chr13-88488310-DEL-357
chr13-88495328-DEL-641
chr13-88499204-DEL-52
chr13-88500196-DEL-2630
chr13-88502844-DEL-934
chr13-88504351-DEL-71
chr13-88515341-DEL-1387
chr13-88517001-DEL-77
chr13-88518582-DEL-1344
chr13-88531833-DEL-1044
chr13-88538923-DEL-60
chr13-88554580-DEL-5385
chr13-88561332-DEL-826
chr13-88564520-DEL-275
chr13-88567048-DEL-588
chr13-88568327-DEL-144
chr13-88571543-DEL-94
chr13-88596326-DEL-72
chr13-88616766-DEL-230
chr13-88621861-DEL-1182
chr13-88625988-DEL-640
chr13-88632238-DEL-2443
chr13-88669036-DEL-290
chr13-88685766-DEL-538
chr13-88720520-DEL-71
chr13-88722828-DEL-754
chr13-88747159-DEL-64
chr13-88785818-DEL-6415
chr13-88802677-DEL-268
chr13-88814635-DEL-2437
chr13-88836770-DEL-571
chr13-88854780-DEL-83
chr13-88876898-DEL-277
chr13-88881373-DEL-67
chr13-88895109-DEL-51
chr13-88912160-DEL-193
chr13-88918967-DEL-72
chr13-88925369-DEL-387
chr13-88948672-DEL-390

chr13-88954855-DEL-121
chr13-88980647-DEL-54
chr13-88984308-DEL-420
chr13-88989320-DEL-75
chr13-89002454-DEL-140
chr13-89002641-DEL-2956
chr13-89081722-DEL-126
chr13-89085252-DEL-426
chr13-89103775-DEL-279
chr13-89131698-DEL-187
chr13-89141300-DEL-1880
chr13-89147502-DEL-311
chr13-89150062-DEL-3351
chr13-89150288-DEL-829
chr13-89171394-DEL-601
chr13-89240534-DEL-698
chr13-89271438-DEL-1057
chr13-89275301-DEL-80
chr13-89275861-DEL-281
chr13-89323855-DEL-57
chr13-89373573-DEL-375
chr13-89408109-DEL-366
chr13-89493724-DEL-72
chr13-89547045-DEL-55
chr13-89556647-DEL-107
chr13-89567039-DEL-59
chr13-89595702-DEL-71
chr13-89615331-DEL-106
chr13-89617481-DEL-744
chr13-89648789-DEL-2460
chr13-89682989-DEL-69
chr13-89691945-DEL-255
chr13-89698884-DEL-516
chr13-89709988-DEL-59
chr13-89716006-DEL-59
chr13-89723000-DEL-252
chr13-89723928-DEL-2519
chr13-89736231-DEL-128
chr13-89785610-DEL-73
chr13-89806912-DEL-103
chr13-89857405-DEL-482
chr13-89881946-DEL-57
chr13-89910312-DEL-66
chr13-89920787-DEL-247
chr13-89928916-DEL-84
chr13-89943363-DEL-372
chr13-89971007-DEL-70
chr13-89994421-DEL-243
chr13-90009847-DEL-5153
chr13-90015112-DEL-1819
chr13-90041294-DEL-84
chr13-90056829-DEL-68
chr13-90067821-DEL-154
chr13-90104595-DEL-445
chr13-90107618-DEL-760
chr13-90124651-DEL-98
chr13-90154536-DEL-134
chr13-90155973-DEL-129
chr13-90183775-DEL-133
chr13-90216657-DEL-55
chr13-90223373-DEL-219
chr13-90233858-DEL-226
chr13-90242956-DEL-100
chr13-90278788-DEL-222
chr13-90302834-DEL-82
chr13-90319268-DEL-53
chr13-90348085-DEL-461
chr13-90348790-DEL-50
chr13-90357139-DEL-60
chr13-90446433-DEL-77
chr13-90451239-DEL-71
chr13-90479671-DEL-177
chr13-90511779-DEL-120
chr13-90523967-DEL-59
chr13-90551199-DEL-58
chr13-90565375-DEL-55
chr13-90575853-DEL-359

chr13-90591531-DEL-624
chr13-90624478-DEL-68
chr13-90716086-DEL-99
chr13-90761839-DEL-66
chr13-90762215-DEL-1139
chr13-90766578-DEL-688
chr13-90775894-DEL-62
chr13-90785627-DEL-154
chr13-90788170-DEL-162
chr13-90895278-DEL-482
chr13-90902284-DEL-2260
chr13-90909305-DEL-178
chr13-90928803-DEL-899
chr13-90936131-DEL-649
chr13-90953287-DEL-308
chr13-90996875-DEL-797
chr13-91005216-DEL-545
chr13-91025316-DEL-203
chr13-91026918-DEL-335
chr13-91039311-DEL-54
chr13-91041341-DEL-58
chr13-91048715-DEL-64
chr13-91138389-DEL-1470
chr13-91157724-DEL-103
chr13-91164895-DEL-296
chr13-91172136-DEL-286
chr13-91174782-DEL-1947
chr13-91217794-DEL-218
chr13-91232685-DEL-928
chr13-91233014-DEL-638
chr13-91251886-DEL-1250
chr13-91270912-DEL-62
chr13-91321609-DEL-83
chr13-91322255-DEL-534
chr13-91322400-DEL-284
chr13-91328698-DEL-97
chr13-91366246-DEL-138
chr13-91374802-DEL-341
chr13-91402235-DEL-498
chr13-91455864-DEL-54
chr13-91473178-DEL-60
chr13-91506355-DEL-138
chr13-91507043-DEL-52
chr13-91549073-DEL-161
chr13-91552507-DEL-117
chr13-91564639-DEL-136
chr13-91571507-DEL-51
chr13-91626380-DEL-114
chr13-91730236-DEL-423
chr13-91747574-DEL-316
chr13-91748118-DEL-762
chr13-91771698-DEL-366
chr13-91786643-DEL-153
chr13-91890266-DEL-105
chr13-91928522-DEL-138
chr13-91937913-DEL-1203
chr13-91939722-DEL-1297
chr13-91956146-DEL-57
chr13-91957137-DEL-580
chr13-91969945-DEL-117
chr13-92001824-DEL-362
chr13-92004861-DEL-335
chr13-92011877-DEL-104
chr13-92012619-DEL-112
chr13-92015207-DEL-333
chr13-92051605-DEL-120
chr13-92053453-DEL-105
chr13-92092104-DEL-3650
chr13-92102588-DEL-104
chr13-92109101-DEL-3935
chr13-92118012-DEL-307
chr13-92136284-DEL-54
chr13-92146284-DEL-132
chr13-92180499-DEL-72
chr13-92187961-DEL-54
chr13-92190911-DEL-1923
chr13-92212400-DEL-118

chr13-92227356-DEL-1270
chr13-92287934-DEL-144
chr13-92326750-DEL-495
chr13-92328535-DEL-161
chr13-92344634-DEL-87
chr13-92352734-DEL-2435
chr13-92366213-DEL-596
chr13-92390129-DEL-78
chr13-92402799-DEL-82
chr13-92460291-DEL-1532
chr13-92501293-DEL-180
chr13-92505942-DEL-263
chr13-92513906-DEL-129
chr13-92539013-DEL-143
chr13-92580941-DEL-88
chr13-92596864-DEL-465
chr13-92606877-DEL-146
chr13-92625665-DEL-1018
chr13-92633013-DEL-52
chr13-92656911-DEL-241
chr13-92668142-DEL-68
chr13-92687622-DEL-106
chr13-92692633-DEL-78
chr13-92705873-DEL-64
chr13-92752595-DEL-292
chr13-92762724-DEL-184
chr13-92783018-DEL-289
chr13-92790128-DEL-442
chr13-92790697-DEL-379
chr13-92805761-DEL-385
chr13-92821926-DEL-54
chr13-92845596-DEL-218
chr13-92864629-DEL-280
chr13-92939994-DEL-84
chr13-92948545-DEL-54
chr13-93009408-DEL-6230
chr13-93039522-DEL-1525
chr13-93046570-DEL-146
chr13-93070604-DEL-301
chr13-93082492-DEL-56
chr13-93090106-DEL-2047
chr13-93104930-DEL-54
chr13-93131985-DEL-57
chr13-93134389-DEL-96
chr13-93145840-DEL-53
chr13-93149192-DEL-63
chr13-93162536-DEL-68
chr13-93188308-DEL-54
chr13-93191612-DEL-52
chr13-93237225-DEL-74
chr13-93263149-DEL-65
chr13-93287576-DEL-3186
chr13-93330373-DEL-749
chr13-93354596-DEL-185
chr13-93394373-DEL-64
chr13-93408189-DEL-80
chr13-93491463-DEL-72
chr13-93508099-DEL-408
chr13-93529197-DEL-220
chr13-93545208-DEL-59
chr13-93576866-DEL-150
chr13-93580901-DEL-5109
chr13-93587630-DEL-400
chr13-93610417-DEL-139
chr13-93630613-DEL-115
chr13-93637150-DEL-98
chr13-93656023-DEL-146
chr13-93666394-DEL-4187
chr13-93666618-DEL-587
chr13-93696255-DEL-135
chr13-93706997-DEL-196
chr13-93711168-DEL-84
chr13-93735035-DEL-52
chr13-93742038-DEL-114
chr13-93745688-DEL-186
chr13-93763963-DEL-103
chr13-93772679-DEL-89

chr13-93778152-DEL-64
chr13-93786053-DEL-1862
chr13-93801012-DEL-595
chr13-93863056-DEL-179
chr13-93875891-DEL-148
chr13-93890457-DEL-196
chr13-93912046-DEL-1070
chr13-93918910-DEL-108
chr13-93952107-DEL-485
chr13-93966123-DEL-90
chr13-93978171-DEL-1113
chr13-93978235-DEL-834
chr13-93979366-DEL-118
chr13-93985772-DEL-58
chr13-93996703-DEL-73
chr13-94028671-DEL-719
chr13-94073869-DEL-309
chr13-94099463-DEL-778
chr13-94120739-DEL-90
chr13-94162452-DEL-71
chr13-94217231-DEL-186
chr13-94217832-DEL-444
chr13-94232257-DEL-155
chr13-94244337-DEL-66
chr13-94260741-DEL-202
chr13-94293880-DEL-54
chr13-94298590-DEL-5451
chr13-94312314-DEL-283
chr13-94359145-DEL-118
chr13-94365314-DEL-131
chr13-94379797-DEL-95
chr13-94417246-DEL-111
chr13-94436103-DEL-60
chr13-94478365-DEL-71
chr13-94481188-DEL-317
chr13-94525574-DEL-179
chr13-94530590-DEL-122
chr13-94542258-DEL-73
chr13-94565459-DEL-62
chr13-94579802-DEL-107
chr13-94580756-DEL-126
chr13-94614318-DEL-58
chr13-94643871-DEL-225
chr13-94650486-DEL-599
chr13-94700258-DEL-90
chr13-94706321-DEL-56
chr13-94731210-DEL-120
chr13-94741967-DEL-60
chr13-94764491-DEL-293
chr13-94767024-DEL-2085
chr13-94793520-DEL-1178
chr13-94815708-DEL-698
chr13-94825821-DEL-54
chr13-94863251-DEL-77
chr13-94867885-DEL-126
chr13-94895797-DEL-63
chr13-94897313-DEL-82
chr13-94899145-DEL-2693
chr13-94952379-DEL-103
chr13-95030159-DEL-78
chr13-95044505-DEL-295
chr13-95058110-DEL-677
chr13-95095104-DEL-120
chr13-95109134-DEL-85
chr13-95175704-DEL-85
chr13-95240533-DEL-289
chr13-95249923-DEL-109
chr13-95279955-DEL-876
chr13-95280397-DEL-1615
chr13-95280839-DEL-634
chr13-95282120-DEL-1129
chr13-95306491-DEL-167
chr13-95317634-DEL-973
chr13-95338192-DEL-57
chr13-95352397-DEL-118
chr13-95381991-DEL-63
chr13-95396018-DEL-54

chr13-95406281-DEL-125
chr13-95411554-DEL-72
chr13-95422884-DEL-280
chr13-95449615-DEL-162
chr13-95495723-DEL-88
chr13-95514238-DEL-267
chr13-95539043-DEL-83
chr13-95545083-DEL-66
chr13-95556647-DEL-56
chr13-95569925-DEL-216
chr13-95599128-DEL-55
chr13-95602843-DEL-400
chr13-95630346-DEL-223
chr13-95635530-DEL-133
chr13-95646895-DEL-253
chr13-95680052-DEL-51
chr13-95697010-DEL-82
chr13-95702676-DEL-87
chr13-95704555-DEL-280
chr13-95811192-DEL-143
chr13-95886976-DEL-267
chr13-95930020-DEL-60
chr13-95934340-DEL-58
chr13-95951118-DEL-60
chr13-95962058-DEL-146
chr13-96004945-DEL-119
chr13-96035007-DEL-76
chr13-96038781-DEL-52
chr13-96043794-DEL-79
chr13-96045446-DEL-179
chr13-96052955-DEL-275
chr13-96065176-DEL-5011
chr13-96067422-DEL-1224
chr13-96078006-DEL-74
chr13-96095993-DEL-55
chr13-96142012-DEL-1217
chr13-96157588-DEL-1433
chr13-96187561-DEL-1325
chr13-96213300-DEL-582
chr13-96213924-DEL-1169
chr13-96257832-DEL-418
chr13-96304222-DEL-56
chr13-96311121-DEL-811
chr13-96348794-DEL-120
chr13-96362165-DEL-382
chr13-96366148-DEL-6417
chr13-96412762-DEL-1139
chr13-96418099-DEL-135
chr13-96433203-DEL-328
chr13-96440707-DEL-65
chr13-96452488-DEL-95
chr13-96466652-DEL-105
chr13-96561615-DEL-449
chr13-96598019-DEL-203
chr13-96611813-DEL-1455
chr13-96625185-DEL-96
chr13-96652064-DEL-402
chr13-96657232-DEL-2611
chr13-96697253-DEL-2871
chr13-96712516-DEL-221
chr13-96723882-DEL-660
chr13-96750521-DEL-350
chr13-96781401-DEL-140
chr13-96804592-DEL-67
chr13-96809764-DEL-116
chr13-96841873-DEL-761
chr13-96857373-DEL-120
chr13-96869256-DEL-100
chr13-96896756-DEL-622
chr13-96897629-DEL-211
chr13-96943773-DEL-3109
chr13-96971375-DEL-77
chr13-97008105-DEL-86
chr13-97025012-DEL-2472
chr13-97033587-DEL-1964
chr13-97059859-DEL-274
chr13-97079714-DEL-54

chr13-97100724-DEL-95
chr13-97113215-DEL-86
chr13-97134154-DEL-226
chr13-97173386-DEL-6440
chr13-97180676-DEL-204
chr13-97192463-DEL-52
chr13-97202695-DEL-194
chr13-97238776-DEL-161
chr13-97262570-DEL-54
chr13-97298729-DEL-142
chr13-97302609-DEL-165
chr13-97322439-DEL-127
chr13-97327054-DEL-78
chr13-97336807-DEL-197
chr13-97365218-DEL-209
chr13-97369340-DEL-209
chr13-97407986-DEL-501
chr13-97445079-DEL-116
chr13-97457225-DEL-92
chr13-97459102-DEL-1519
chr13-97510989-DEL-233
chr13-97528086-DEL-65
chr13-97554283-DEL-157
chr13-97562807-DEL-97
chr13-97563465-DEL-92
chr13-97585062-DEL-203
chr13-97597164-DEL-98
chr13-97603356-DEL-141
chr13-97632090-DEL-72
chr13-97665102-DEL-193
chr13-97714862-DEL-160
chr13-97736116-DEL-78
chr13-97834952-DEL-1142
chr13-97837629-DEL-876
chr13-97846939-DEL-76
chr13-97851383-DEL-680
chr13-97852764-DEL-6442
chr13-97863006-DEL-6466
chr13-97889144-DEL-84
chr13-97899175-DEL-120
chr13-97906494-DEL-4653
chr13-97933074-DEL-267
chr13-97948898-DEL-190
chr13-97955893-DEL-690
chr13-97984537-DEL-5295
chr13-98038056-DEL-181
chr13-98044883-DEL-136
chr13-98085144-DEL-98
chr13-98087133-DEL-569
chr13-98108005-DEL-58
chr13-98115302-DEL-142
chr13-98152006-DEL-53
chr13-98245479-DEL-54
chr13-98279069-DEL-77
chr13-98294777-DEL-1524
chr13-98300091-DEL-441
chr13-98303757-DEL-131
chr13-98304464-DEL-310
chr13-98325384-DEL-1061
chr13-98392646-DEL-76
chr13-98482831-DEL-376
chr13-98502046-DEL-68
chr13-98507599-DEL-257
chr13-98562276-DEL-366
chr13-98569744-DEL-52
chr13-98592987-DEL-365
chr13-98609198-DEL-99
chr13-98617202-DEL-56
chr13-98631497-DEL-504
chr13-98645513-DEL-167
chr13-98694347-DEL-302
chr13-98696920-DEL-94
chr13-98792872-DEL-101
chr13-98804658-DEL-122
chr13-98821888-DEL-462
chr13-98840612-DEL-71
chr13-98852425-DEL-224

chr13-98871317-DEL-72
chr13-98915996-DEL-183
chr13-98948784-DEL-53
chr13-98953508-DEL-86
chr13-98962876-DEL-239
chr13-98964621-DEL-271
chr13-98967479-DEL-180
chr13-98969949-DEL-99
chr13-98971244-DEL-89
chr13-99044318-DEL-77
chr13-99053312-DEL-896
chr13-99061303-DEL-66
chr13-99069830-DEL-313
chr13-99081841-DEL-73
chr13-99098096-DEL-3362
chr13-99110883-DEL-176
chr13-99117219-DEL-53
chr13-99136194-DEL-202
chr13-99137747-DEL-801
chr13-99148022-DEL-120
chr13-99172211-DEL-495
chr13-99205670-DEL-164
chr13-99221326-DEL-504
chr13-99222192-DEL-186
chr13-99236374-DEL-54
chr13-99250739-DEL-182
chr13-99267333-DEL-7237
chr13-99316650-DEL-255
chr13-99320283-DEL-58
chr13-99337936-DEL-51
chr13-99360887-DEL-781
chr13-99361672-DEL-177
chr13-99384420-DEL-230
chr13-99386353-DEL-123
chr13-99393693-DEL-545
chr13-99395680-DEL-78
chr13-99459288-DEL-237
chr13-99485864-DEL-66
chr13-99497103-DEL-409
chr13-99544914-DEL-52
chr13-99575306-DEL-56
chr13-99616828-DEL-69
chr13-99636084-DEL-68
chr13-99654659-DEL-400
chr13-99680868-DEL-284
chr13-99688685-DEL-112
chr13-99692709-DEL-400
chr13-99707604-DEL-68
chr13-99735254-DEL-701
chr13-99747591-DEL-61
chr13-99767041-DEL-108
chr13-99786420-DEL-124
chr13-99883398-DEL-61
chr13-99888973-DEL-257
chr13-99891983-DEL-123
chr13-99939346-DEL-384
chr13-99946674-DEL-1342
chr13-99986456-DEL-215
chr13-99995356-DEL-96
chr13-100004246-DEL-2809
chr13-100038104-DEL-1879
chr13-100038159-DEL-1040
chr13-100038483-DEL-987
chr13-100070945-DEL-377
chr13-100084769-DEL-139
chr13-100097056-DEL-299
chr13-100106941-DEL-73
chr13-100131900-DEL-268
chr13-100145807-DEL-270
chr13-100147913-DEL-196
chr13-100194428-DEL-101
chr13-100231431-DEL-618
chr13-100266555-DEL-205
chr13-100279739-DEL-142
chr13-100304971-DEL-104
chr13-100314985-DEL-76
chr13-100354570-DEL-149

chr13-100360960-DEL-94
chr13-100370410-DEL-8257
chr13-100370429-DEL-5575
chr13-100370511-DEL-415
chr13-100377227-DEL-1470
chr13-100390743-DEL-264
chr13-100434314-DEL-86
chr13-100526780-DEL-9453
chr13-100529665-DEL-1132
chr13-100558740-DEL-161
chr13-100569974-DEL-65
chr13-100620941-DEL-156
chr13-100668725-DEL-148
chr13-100699801-DEL-610
chr13-100706131-DEL-189
chr13-100754634-DEL-195
chr13-100755844-DEL-795
chr13-100757759-DEL-232
chr13-100764842-DEL-195
chr13-100771927-DEL-188
chr13-100773998-DEL-198
chr13-100776479-DEL-200
chr13-100790074-DEL-236
chr13-100801788-DEL-221
chr13-100808719-DEL-193
chr13-100822958-DEL-197
chr13-100843345-DEL-514
chr13-100850747-DEL-1916
chr13-100870486-DEL-228
chr13-100885049-DEL-88
chr13-100906902-DEL-449
chr13-100924538-DEL-182
chr13-100926334-DEL-113
chr13-100930343-DEL-75
chr13-100938822-DEL-140
chr13-100948261-DEL-278
chr13-100948917-DEL-209
chr13-100952568-DEL-881
chr13-100957308-DEL-195
chr13-100964687-DEL-76
chr13-100981122-DEL-95
chr13-101010273-DEL-57
chr13-101010900-DEL-156
chr13-101044154-DEL-2660
chr13-101070415-DEL-1793
chr13-101081634-DEL-3660
chr13-101081821-DEL-224
chr13-101083437-DEL-1237
chr13-101109086-DEL-98
chr13-101133075-DEL-154
chr13-101133381-DEL-94
chr13-101158116-DEL-619
chr13-101192267-DEL-100
chr13-101200263-DEL-336
chr13-101217149-DEL-248
chr13-101263025-DEL-54
chr13-101264483-DEL-101
chr13-101289469-DEL-68
chr13-101310005-DEL-51
chr13-101328085-DEL-1504
chr13-101330215-DEL-416
chr13-101332510-DEL-58
chr13-101361376-DEL-77
chr13-101416387-DEL-849
chr13-101430403-DEL-75
chr13-101468321-DEL-573
chr13-101472078-DEL-440
chr13-101531622-DEL-329
chr13-101539118-DEL-198
chr13-101571693-DEL-121
chr13-101581553-DEL-58
chr13-101582820-DEL-111
chr13-101686564-DEL-59
chr13-101702581-DEL-60
chr13-101703262-DEL-2172
chr13-101709385-DEL-217
chr13-101719439-DEL-1502

chr13-101755208-DEL-196
chr13-101813653-DEL-120
chr13-101880025-DEL-68
chr13-101892202-DEL-58
chr13-101906012-DEL-187
chr13-101937717-DEL-124
chr13-101957272-DEL-3154
chr13-101985662-DEL-283
chr13-102019181-DEL-199
chr13-102080090-DEL-53
chr13-102088215-DEL-133
chr13-102102010-DEL-62
chr13-102111297-DEL-63
chr13-102121818-DEL-182
chr13-102135986-DEL-59
chr13-102147423-DEL-273
chr13-102159684-DEL-129
chr13-102164093-DEL-97
chr13-102184656-DEL-1791
chr13-102191747-DEL-70
chr13-102193151-DEL-105
chr13-102221869-DEL-678
chr13-102231409-DEL-52
chr13-102243506-DEL-55
chr13-102285133-DEL-72
chr13-102287423-DEL-66
chr13-102327446-DEL-1712
chr13-102420951-DEL-94
chr13-102453639-DEL-200
chr13-102460165-DEL-123
chr13-102483712-DEL-172
chr13-102493585-DEL-226
chr13-102495760-DEL-262
chr13-102500017-DEL-77
chr13-102509693-DEL-80
chr13-102525467-DEL-1161
chr13-102550308-DEL-6467
chr13-102557823-DEL-6663
chr13-102580662-DEL-70
chr13-102585102-DEL-76
chr13-102593717-DEL-743
chr13-102693664-DEL-130
chr13-102717544-DEL-180
chr13-102768195-DEL-68
chr13-102823688-DEL-657
chr13-102831790-DEL-59
chr13-102867940-DEL-262
chr13-102880857-DEL-967
chr13-102883652-DEL-72
chr13-102899672-DEL-143
chr13-102932296-DEL-74
chr13-102949165-DEL-86
chr13-102973622-DEL-54
chr13-102993075-DEL-195
chr13-103006097-DEL-4899
chr13-103112934-DEL-1825
chr13-103115701-DEL-70
chr13-103175051-DEL-52
chr13-103177096-DEL-57
chr13-103182302-DEL-52
chr13-103223065-DEL-212
chr13-103253898-DEL-570
chr13-103253968-DEL-148
chr13-103321900-DEL-159
chr13-103400652-DEL-152
chr13-103416816-DEL-55
chr13-103457218-DEL-80
chr13-103520883-DEL-768
chr13-103521612-DEL-56
chr13-103527074-DEL-444
chr13-103537249-DEL-401
chr13-103563120-DEL-55
chr13-103607465-DEL-196
chr13-103634017-DEL-73
chr13-103666768-DEL-161
chr13-103718680-DEL-6319
chr13-103718746-DEL-213

chr13-103727788-DEL-341
chr13-103759204-DEL-56
chr13-103778421-DEL-428
chr13-103812168-DEL-168
chr13-103821054-DEL-1200
chr13-103826028-DEL-4634
chr13-103831187-DEL-86
chr13-103842311-DEL-73
chr13-103883664-DEL-795
chr13-103924203-DEL-2038
chr13-103973877-DEL-61
chr13-103999496-DEL-92
chr13-104001559-DEL-68
chr13-104066450-DEL-7496
chr13-104078340-DEL-559
chr13-104107501-DEL-77
chr13-104119138-DEL-884
chr13-104187707-DEL-429
chr13-104190387-DEL-1945
chr13-104206951-DEL-65
chr13-104212572-DEL-146
chr13-104223383-DEL-219
chr13-104230050-DEL-119
chr13-104274110-DEL-112
chr13-104285701-DEL-131
chr13-104297965-DEL-63
chr13-104358801-DEL-98
chr13-104375761-DEL-180
chr13-104382369-DEL-113
chr13-104389386-DEL-510
chr13-104390472-DEL-1661
chr13-104403872-DEL-83
chr13-104410721-DEL-186
chr13-104411893-DEL-4845
chr13-104416942-DEL-1299
chr13-104418672-DEL-198
chr13-104459659-DEL-3623
chr13-104487307-DEL-106
chr13-104531255-DEL-117
chr13-104547340-DEL-121
chr13-104548784-DEL-432
chr13-104592619-DEL-262
chr13-104660547-DEL-150
chr13-104727573-DEL-257
chr13-104732246-DEL-333
chr13-104753717-DEL-144
chr13-104797348-DEL-350
chr13-104812863-DEL-3193
chr13-104847175-DEL-68
chr13-104868673-DEL-69
chr13-104895728-DEL-195
chr13-104911334-DEL-1222
chr13-104979741-DEL-198
chr13-104981148-DEL-83
chr13-105028523-DEL-144
chr13-105037108-DEL-6518
chr13-105057710-DEL-113
chr13-105058382-DEL-352
chr13-105099144-DEL-51
chr13-105135281-DEL-97
chr13-105157385-DEL-94
chr13-105163943-DEL-271
chr13-105207574-DEL-54
chr13-105221665-DEL-77
chr13-105251661-DEL-62
chr13-105262090-DEL-2377
chr13-105285983-DEL-102
chr13-105287940-DEL-600
chr13-105292542-DEL-520
chr13-105316587-DEL-108
chr13-105348213-DEL-177
chr13-105362858-DEL-203
chr13-105374894-DEL-308
chr13-105392441-DEL-66
chr13-105405376-DEL-1445
chr13-105407758-DEL-184
chr13-105427279-DEL-260

chr13-105447135-DEL-66
chr13-105458945-DEL-971
chr13-105540490-DEL-66
chr13-105549577-DEL-66
chr13-105555397-DEL-63
chr13-105560039-DEL-1728
chr13-105594567-DEL-392
chr13-105631021-DEL-61
chr13-105632906-DEL-776
chr13-105665468-DEL-1936
chr13-105696085-DEL-54
chr13-105709680-DEL-76
chr13-105711179-DEL-81
chr13-105752832-DEL-1343
chr13-105753921-DEL-387
chr13-105766658-DEL-64
chr13-105776594-DEL-79
chr13-105809249-DEL-3792
chr13-105835310-DEL-123
chr13-105844370-DEL-350
chr13-105868094-DEL-321
chr13-105894661-DEL-916
chr13-105899452-DEL-60
chr13-105916445-DEL-3886
chr13-105916839-DEL-640
chr13-105920590-DEL-3444
chr13-105920660-DEL-1469
chr13-105939126-DEL-126
chr13-105940515-DEL-133
chr13-105949022-DEL-1120
chr13-105951334-DEL-64
chr13-106020750-DEL-108
chr13-106029410-DEL-83
chr13-106032988-DEL-992
chr13-106035810-DEL-2458
chr13-106042827-DEL-69
chr13-106088374-DEL-3316
chr13-106106637-DEL-537
chr13-106192212-DEL-6175
chr13-106210237-DEL-332
chr13-106218602-DEL-56
chr13-106235860-DEL-1892
chr13-106251086-DEL-167
chr13-106258406-DEL-1334
chr13-106272781-DEL-180
chr13-106274880-DEL-2774
chr13-106286034-DEL-56
chr13-106303308-DEL-358
chr13-106314817-DEL-252
chr13-106322212-DEL-512
chr13-106329183-DEL-68
chr13-106353509-DEL-112
chr13-106381133-DEL-647
chr13-106407839-DEL-2758
chr13-106422182-DEL-208
chr13-106437880-DEL-94
chr13-106441739-DEL-4726
chr13-106462538-DEL-525
chr13-106516813-DEL-585
chr13-106538993-DEL-72
chr13-106539623-DEL-133
chr13-106540254-DEL-52
chr13-106562631-DEL-740
chr13-106566401-DEL-60
chr13-106580916-DEL-60
chr13-106623191-DEL-491
chr13-106636975-DEL-234
chr13-106641029-DEL-132
chr13-106643324-DEL-3466
chr13-106655925-DEL-237
chr13-106720352-DEL-6267
chr13-106733968-DEL-1456
chr13-106750417-DEL-1288
chr13-106752344-DEL-1703
chr13-106754532-DEL-3320
chr13-106763222-DEL-6416
chr13-106774771-DEL-54

chr13-106809867-DEL-62
chr13-106816127-DEL-6503
chr13-106840486-DEL-53
chr13-106867818-DEL-590
chr13-106872577-DEL-805
chr13-106880647-DEL-918
chr13-106901409-DEL-347
chr13-106915583-DEL-128
chr13-106923987-DEL-811
chr13-106929326-DEL-52
chr13-106942111-DEL-75
chr13-106956933-DEL-182
chr13-106964779-DEL-114
chr13-106976180-DEL-111
chr13-106983215-DEL-115
chr13-106989653-DEL-96
chr13-106996153-DEL-80
chr13-106999393-DEL-479
chr13-107049352-DEL-576
chr13-107061766-DEL-77
chr13-107062528-DEL-94
chr13-107070151-DEL-63
chr13-107076385-DEL-54
chr13-107078586-DEL-68
chr13-107104470-DEL-196
chr13-107108074-DEL-68
chr13-107134307-DEL-172
chr13-107194861-DEL-400
chr13-107220864-DEL-91
chr13-107221347-DEL-185
chr13-107242382-DEL-77
chr13-107256184-DEL-357
chr13-107258204-DEL-377
chr13-107266177-DEL-1526
chr13-107302200-DEL-115
chr13-107340886-DEL-158
chr13-107405340-DEL-59
chr13-107408135-DEL-227
chr13-107443936-DEL-282
chr13-107459211-DEL-333
chr13-107486195-DEL-411
chr13-107511543-DEL-135
chr13-107548705-DEL-360
chr13-107565325-DEL-303
chr13-107588949-DEL-70
chr13-107603345-DEL-69
chr13-107619617-DEL-74
chr13-107623024-DEL-79
chr13-107683569-DEL-58
chr13-107689998-DEL-75
chr13-107723999-DEL-52
chr13-107737448-DEL-52
chr13-107763281-DEL-81
chr13-107768958-DEL-54
chr13-107788175-DEL-789
chr13-107801543-DEL-51
chr13-107813934-DEL-298
chr13-107869317-DEL-60
chr13-107909339-DEL-65
chr13-107910636-DEL-2086
chr13-107930327-DEL-112
chr13-107931461-DEL-58
chr13-107971651-DEL-287
chr13-107994789-DEL-1326
chr13-107994789-DEL-936
chr13-108014072-DEL-102
chr13-108047578-DEL-550
chr13-108066769-DEL-496
chr13-108087647-DEL-52
chr13-108097984-DEL-57
chr13-108118267-DEL-107
chr13-108170414-DEL-71
chr13-108192768-DEL-404
chr13-108203432-DEL-252
chr13-108214620-DEL-80
chr13-108263934-DEL-168
chr13-108275835-DEL-212

chr13-108287226-DEL-115
chr13-108321077-DEL-1946
chr13-108324724-DEL-850
chr13-108326195-DEL-206
chr13-108329007-DEL-573
chr13-108360210-DEL-51
chr13-108371619-DEL-54
chr13-108381549-DEL-83
chr13-108392599-DEL-60
chr13-108402814-DEL-476
chr13-108521655-DEL-106
chr13-108555799-DEL-595
chr13-108557838-DEL-71
chr13-108568826-DEL-344
chr13-108575158-DEL-333
chr13-108581860-DEL-56
chr13-108615128-DEL-53
chr13-108615640-DEL-1222
chr13-108636125-DEL-61
chr13-108645749-DEL-424
chr13-108683328-DEL-291
chr13-108700702-DEL-164
chr13-108763851-DEL-397
chr13-108814988-DEL-685
chr13-108858684-DEL-128
chr13-108898755-DEL-86
chr13-108907511-DEL-269
chr13-108954654-DEL-497
chr13-108985813-DEL-55
chr13-109014526-DEL-185
chr13-109017679-DEL-52
chr13-109052227-DEL-65
chr13-109072669-DEL-194
chr13-109119901-DEL-100
chr13-109185663-DEL-140
chr13-109240768-DEL-2704
chr13-109243183-DEL-609
chr13-109331254-DEL-60
chr13-109335345-DEL-102
chr13-109354278-DEL-284
chr13-109475452-DEL-127
chr13-109518005-DEL-60
chr13-109525122-DEL-51
chr13-109544524-DEL-96
chr13-109558488-DEL-211
chr13-109650596-DEL-198
chr13-109670182-DEL-56
chr13-109684020-DEL-140
chr13-109694195-DEL-176
chr13-109699736-DEL-872
chr13-109702163-DEL-73
chr13-109712477-DEL-61
chr13-109719639-DEL-128
chr13-109738255-DEL-81
chr13-109760332-DEL-393
chr13-109807494-DEL-166
chr13-109814936-DEL-75
chr13-109820096-DEL-269
chr13-109879637-DEL-1866
chr13-109891101-DEL-248
chr13-109914755-DEL-176
chr13-109947647-DEL-949
chr13-109957243-DEL-73
chr13-109998392-DEL-53
chr13-110013416-DEL-482
chr13-110030536-DEL-358
chr13-110118291-DEL-78
chr13-110135680-DEL-64
chr13-110137435-DEL-1087
chr13-110171147-DEL-80
chr13-110172153-DEL-2521
chr13-110196728-DEL-57
chr13-110206633-DEL-404
chr13-110233571-DEL-60
chr13-110245883-DEL-220
chr13-110285524-DEL-190
chr13-110297363-DEL-635

chr13-110356368-DEL-74
chr13-110400674-DEL-62
chr13-110415788-DEL-71
chr13-110446059-DEL-428
chr13-110458585-DEL-1236
chr13-110465787-DEL-429
chr13-110467200-DEL-84
chr13-110467806-DEL-402
chr13-110486661-DEL-891
chr13-110508823-DEL-480
chr13-110521464-DEL-126
chr13-110605369-DEL-156
chr13-110619740-DEL-83
chr13-110635413-DEL-94
chr13-110639734-DEL-247
chr13-110643132-DEL-205
chr13-110650651-DEL-111
chr13-110660853-DEL-82
chr13-110671597-DEL-307
chr13-110739583-DEL-66
chr13-110767463-DEL-672
chr13-110777713-DEL-114
chr13-110860326-DEL-72
chr13-110873295-DEL-68
chr13-110880538-DEL-244
chr13-110934113-DEL-399
chr13-110956797-DEL-905
chr13-110970953-DEL-400
chr13-110984127-DEL-886
chr13-110999206-DEL-65
chr13-111004026-DEL-79
chr13-111014514-DEL-628
chr13-111095266-DEL-217
chr13-111169299-DEL-2430
chr13-111193273-DEL-106
chr13-111210632-DEL-212
chr13-111217585-DEL-1239
chr13-111221943-DEL-1441
chr13-111250513-DEL-1298
chr13-111344145-DEL-1366
chr13-111366752-DEL-91
chr13-111444772-DEL-843
chr13-111474567-DEL-541
chr13-111477050-DEL-200
chr13-111479139-DEL-109
chr13-111494293-DEL-199
chr13-111511572-DEL-52
chr13-111516606-DEL-106
chr13-111521530-DEL-1653
chr13-111526762-DEL-715
chr13-111541100-DEL-246
chr13-111543803-DEL-1072
chr13-111553097-DEL-132
chr13-111554435-DEL-57
chr13-111555952-DEL-62
chr13-111564093-DEL-727
chr13-111594848-DEL-214
chr13-111608285-DEL-181
chr13-111609071-DEL-244
chr13-111630207-DEL-788
chr13-111673750-DEL-575
chr13-111696456-DEL-68
chr13-111704961-DEL-56
chr13-111707092-DEL-195
chr13-111708480-DEL-476
chr13-111709111-DEL-58
chr13-111712033-DEL-223
chr13-111722720-DEL-195
chr13-111733788-DEL-75
chr13-111736512-DEL-62
chr13-111738465-DEL-186
chr13-111740463-DEL-321
chr13-111742085-DEL-127
chr13-111772273-DEL-215
chr13-111806221-DEL-460
chr13-111880784-DEL-325
chr13-111905666-DEL-68

chr13-111911206-DEL-176
chr13-111950927-DEL-189
chr13-111965163-DEL-77
chr13-112013885-DEL-100
chr13-112022980-DEL-82
chr13-112036619-DEL-52
chr13-112045595-DEL-148
chr13-112056101-DEL-359
chr13-112088440-DEL-433
chr13-112160846-DEL-75
chr13-112228647-DEL-1997
chr13-112247458-DEL-1414
chr13-112311519-DEL-2139
chr13-112341623-DEL-220
chr13-112352229-DEL-92
chr13-112365197-DEL-55
chr13-112377513-DEL-96
chr13-112391375-DEL-55
chr13-112400030-DEL-56
chr13-112430988-DEL-62
chr13-112465351-DEL-88
chr13-112515613-DEL-1387
chr13-112515614-DEL-949
chr13-112535061-DEL-121
chr13-112539050-DEL-329
chr13-112556278-DEL-58
chr13-112567094-DEL-1580
chr13-112568315-DEL-280
chr13-112588976-DEL-94
chr13-112595817-DEL-64
chr13-112604994-DEL-115
chr13-112614530-DEL-136
chr13-112622557-DEL-1157
chr13-112658415-DEL-70
chr13-112661618-DEL-230
chr13-112676163-DEL-152
chr13-112703057-DEL-67
chr13-112711997-DEL-1137
chr13-112771641-DEL-137
chr13-112773003-DEL-132
chr13-112807115-DEL-78
chr13-112815517-DEL-66
chr13-112844187-DEL-134
chr13-112845190-DEL-486
chr13-112853505-DEL-1671
chr13-112861196-DEL-272
chr13-112861427-DEL-685
chr13-112866943-DEL-132
chr13-112870586-DEL-220
chr13-112880932-DEL-118
chr13-112923298-DEL-56
chr13-112949133-DEL-128
chr13-112979239-DEL-65
chr13-112980764-DEL-595
chr13-112991468-DEL-52
chr13-112992275-DEL-51
chr13-113024637-DEL-2695
chr13-113025092-DEL-139
chr13-113029520-DEL-1488
chr13-113055797-DEL-56
chr13-113082658-DEL-173
chr13-113147454-DEL-75
chr13-113151849-DEL-163
chr13-113154279-DEL-63
chr13-113160165-DEL-106
chr13-113189065-DEL-228
chr13-113220089-DEL-68
chr13-113242336-DEL-88
chr13-113242871-DEL-190
chr13-113243501-DEL-1687
chr13-113249893-DEL-795
chr13-113255654-DEL-81
chr13-113289278-DEL-202
chr13-113322422-DEL-259
chr13-113335159-DEL-1325
chr13-113348160-DEL-173
chr13-113360390-DEL-60

chr13-113361223-DEL-60
chr13-113386645-DEL-53
chr13-113405065-DEL-382
chr13-113409471-DEL-7144
chr13-113440862-DEL-61
chr13-113452592-DEL-4331
chr13-113484083-DEL-251
chr13-113501519-DEL-123
chr13-113508488-DEL-134
chr13-113513511-DEL-1883
chr13-113515853-DEL-128
chr13-113534456-DEL-138
chr13-113536401-DEL-67
chr13-113550125-DEL-1208
chr13-113643931-DEL-505
chr13-113658880-DEL-202
chr13-113669498-DEL-278
chr13-113683002-DEL-66
chr13-113692612-DEL-199
chr13-113726477-DEL-198
chr13-113757407-DEL-147
chr13-113762929-DEL-99
chr13-113776857-DEL-125
chr13-113809734-DEL-231
chr13-113816276-DEL-350
chr13-113829199-DEL-164
chr13-113847572-DEL-67
chr13-113854953-DEL-2146
chr13-113857151-DEL-3334
chr13-113875284-DEL-126
chr13-113897382-DEL-313
chr13-113904681-DEL-196
chr13-113913900-DEL-115
chr13-113926916-DEL-472
chr13-113935396-DEL-193
chr13-113939863-DEL-528
chr13-113941054-DEL-264
chr13-113942244-DEL-351
chr13-113943166-DEL-286
chr13-113954885-DEL-169
chr13-113995035-DEL-148
chr13-113996104-DEL-256
chr13-113998190-DEL-56
chr13-114006439-DEL-384
chr13-114022754-DEL-60
chr13-114030289-DEL-51
chr13-114068081-DEL-95
chr13-114146755-DEL-108
chr13-114161157-DEL-58
chr13-114163602-DEL-123
chr13-114174309-DEL-60
chr13-114210190-DEL-170
chr13-114248100-DEL-149
chr13-114264850-DEL-946
chr13-114289570-DEL-64
chr13-114305080-DEL-755
chr13-114307792-DEL-66
chr13-114329200-DEL-54
chr13-114331013-DEL-1695
chr13-114341250-DEL-201
chr13-114357822-DEL-129
chr13-114376189-DEL-73
chr13-114406340-DEL-400
chr13-114439205-DEL-92
chr13-114449072-DEL-114
chr13-114461751-DEL-562
chr13-114470908-DEL-51
chr13-114474982-DEL-72
chr13-114501112-DEL-97
chr13-114514989-DEL-80
chr13-114530409-DEL-70
chr13-114562849-DEL-165
chr13-114567654-DEL-68
chr13-114605044-DEL-62
chr13-114618977-DEL-300
chr13-114619534-DEL-57
chr13-114627802-DEL-949

chr13-114643362-DEL-60
chr13-114652302-DEL-651
chr13-114661183-DEL-606
chr13-114667036-DEL-60
chr13-114675521-DEL-1135
chr13-114713296-DEL-91
chr13-114718640-DEL-50
chr13-114724936-DEL-776
chr13-114751717-DEL-113
chr13-114790454-DEL-86
chr13-114822430-DEL-52
chr13-114825214-DEL-209
chr13-114918481-DEL-87
chr13-114920688-DEL-51
chr13-114938124-DEL-133
chr13-114940623-DEL-459
chr13-114981463-DEL-86
chr13-115005246-DEL-396
chr13-115016516-DEL-78
chr13-115031161-DEL-230
chr13-115033160-DEL-76
chr13-115080040-DEL-90
chr13-115098860-DEL-1700
chr13-115105212-DEL-86
chr13-115134917-DEL-208
chr13-115142431-DEL-62
chr13-115151081-DEL-292
chr13-115151840-DEL-70
chr13-115224667-DEL-79
chr13-115239142-DEL-75
chr13-115255769-DEL-200
chr13-115275429-DEL-61
chr13-115298194-DEL-109
chr13-115298678-DEL-776
chr13-115312113-DEL-150
chr13-115321184-DEL-222
chr13-115330754-DEL-1431
chr13-115342865-DEL-80
chr13-115353397-DEL-91
chr13-115356784-DEL-148
chr13-115364965-DEL-454
chr13-115377882-DEL-121
chr13-115381113-DEL-403
chr13-115418602-DEL-607
chr13-115504308-DEL-327
chr13-115522191-DEL-92
chr13-115556979-DEL-442
chr13-115567099-DEL-54
chr13-115574679-DEL-619
chr13-115590010-DEL-1352
chr13-115591387-DEL-2022
chr13-115599142-DEL-127
chr13-115613316-DEL-404
chr13-115627088-DEL-148
chr13-115633313-DEL-88
chr13-115634634-DEL-6411
chr13-115638958-DEL-305
chr13-115640522-DEL-200
chr13-115665014-DEL-1321
chr13-115684533-DEL-58
chr13-115702474-DEL-106
chr13-115717672-DEL-150
chr13-115735779-DEL-3880
chr13-115791026-DEL-110
chr13-115794638-DEL-121
chr13-115838402-DEL-492
chr13-115854867-DEL-79
chr13-115860078-DEL-87
chr13-115910387-DEL-51
chr13-115917805-DEL-127
chr13-115926585-DEL-1390
chr13-115938664-DEL-693
chr13-116000403-DEL-1142
chr13-116010250-DEL-6920
chr13-116014711-DEL-217
chr13-116016004-DEL-253
chr13-116067493-DEL-559

chr13-116086620-DEL-60
chr13-116109591-DEL-778
chr13-116121559-DEL-404
chr13-116135054-DEL-1021
chr13-116141249-DEL-512
chr13-116181317-DEL-76
chr13-116221711-DEL-170
chr13-116259201-DEL-284
chr13-116268889-DEL-1149
chr13-116294698-DEL-594
chr13-116317936-DEL-378
chr13-116344945-DEL-1644
chr13-116354212-DEL-73
chr13-116364913-DEL-258
chr13-116427484-DEL-1083
chr13-116441208-DEL-51
chr13-116457861-DEL-99
chr13-116461693-DEL-114
chr13-116466602-DEL-85
chr13-116505390-DEL-51
chr13-116508606-DEL-249
chr13-116511842-DEL-680
chr13-116546362-DEL-85
chr13-116571023-DEL-70
chr13-116572608-DEL-62
chr13-116619472-DEL-162
chr13-116627837-DEL-60
chr13-116640655-DEL-3411
chr13-116648813-DEL-106
chr13-116661250-DEL-2996
chr13-116704753-DEL-1213
chr13-116753093-DEL-66
chr13-116800614-DEL-401
chr13-116815225-DEL-55
chr13-116817929-DEL-453
chr13-116838897-DEL-81
chr13-116925751-DEL-2560
chr13-116968947-DEL-382
chr13-116983417-DEL-60
chr13-116994216-DEL-359
chr13-117002030-DEL-220
chr13-117005189-DEL-132
chr13-117022012-DEL-479
chr13-117026318-DEL-241
chr13-117065649-DEL-70
chr13-117104579-DEL-381
chr13-117112724-DEL-74
chr13-117137297-DEL-52
chr13-117139146-DEL-179
chr13-117152778-DEL-189
chr13-117172301-DEL-300
chr13-117176657-DEL-53
chr13-117198565-DEL-137
chr13-117237796-DEL-185
chr13-117289972-DEL-683
chr13-117319052-DEL-1128
chr13-117362314-DEL-282
chr13-117364721-DEL-218
chr13-117365878-DEL-801
chr13-117384582-DEL-198
chr13-117387456-DEL-3108
chr13-117423398-DEL-2365
chr13-117423421-DEL-100
chr13-117444074-DEL-70
chr13-117453959-DEL-75
chr13-117456313-DEL-77
chr13-117466756-DEL-148
chr13-117483788-DEL-486
chr13-117509344-DEL-5388
chr13-117525224-DEL-59
chr13-117577511-DEL-4216
chr13-117637847-DEL-97
chr13-117743186-DEL-114
chr13-117773059-DEL-56
chr13-117815041-DEL-59
chr13-117815829-DEL-322
chr13-117837172-DEL-61

chr13-117866562-DEL-82
chr13-117945323-DEL-54
chr13-117955352-DEL-989
chr13-117956790-DEL-55
chr13-117979761-DEL-5827
chr13-117992418-DEL-443
chr13-118018751-DEL-391
chr13-118020423-DEL-96
chr13-118022326-DEL-140
chr13-118046025-DEL-95
chr13-118056794-DEL-91
chr13-118094493-DEL-53
chr13-118099408-DEL-96
chr13-118131949-DEL-170
chr13-118191814-DEL-200
chr13-118245274-DEL-80
chr13-118250833-DEL-862
chr13-118256754-DEL-629
chr13-118257508-DEL-96
chr13-118329689-DEL-66
chr13-118375117-DEL-866
chr13-118412009-DEL-90
chr13-118412655-DEL-225
chr13-118430597-DEL-57
chr13-118472889-DEL-224
chr13-118479676-DEL-288
chr13-118488935-DEL-436
chr13-118513363-DEL-114
chr13-118548963-DEL-693
chr13-118566513-DEL-69
chr13-118602595-DEL-648
chr13-118603754-DEL-400
chr13-118632134-DEL-341
chr13-118656514-DEL-720
chr13-118705703-DEL-235
chr13-118767132-DEL-52
chr13-118768318-DEL-504
chr13-118770164-DEL-77
chr13-118781094-DEL-79
chr13-118782087-DEL-300
chr13-118802212-DEL-145
chr13-118835489-DEL-181
chr13-118844187-DEL-51
chr13-118888097-DEL-678
chr13-118969195-DEL-96
chr13-119001833-DEL-329
chr13-119036007-DEL-57
chr13-119095713-DEL-90
chr13-119101060-DEL-184
chr13-119117117-DEL-140
chr13-119125337-DEL-6181
chr13-119128610-DEL-179
chr13-119128810-DEL-4806
chr13-119151473-DEL-51
chr13-119167495-DEL-52
chr13-119182575-DEL-717
chr13-119185110-DEL-55
chr13-119207142-DEL-64
chr13-119215319-DEL-1637
chr13-119244400-DEL-204
chr13-119251247-DEL-876
chr13-119260545-DEL-65
chr13-119284868-DEL-208
chr13-119297306-DEL-1129
chr13-119306486-DEL-457
chr13-119324471-DEL-7005
chr13-119333338-DEL-290
chr13-119338730-DEL-321
chr13-119348033-DEL-6195
chr13-119358100-DEL-6202
chr13-119402051-DEL-204
chr13-119453200-DEL-540
chr13-119480242-DEL-382
chr13-119480807-DEL-51
chr13-119489340-DEL-92
chr13-119518105-DEL-67
chr13-119527796-DEL-139

chr13-119529107-DEL-69
chr13-119529766-DEL-656
chr13-119539893-DEL-285
chr13-119541589-DEL-86
chr13-119555757-DEL-937
chr13-119557632-DEL-90
chr13-119564710-DEL-68
chr13-119566290-DEL-94
chr13-119607007-DEL-148
chr13-119624151-DEL-257
chr13-119625843-DEL-230
chr13-120057226-DEL-91
chr13-120061419-DEL-91
chr13-120063867-DEL-91
chr13-120072886-DEL-1526
chr13-120074254-DEL-138
chr13-120078477-DEL-52
chr13-120079573-DEL-66
chr13-120092170-DEL-4548
chr13-120119523-DEL-73
chr13-120120646-DEL-78
chr13-120123135-DEL-8326
chr13-120138585-DEL-65
chr13-120159749-DEL-81
chr13-120202669-DEL-76
chr13-120209440-DEL-369
chr13-120209847-DEL-732
chr13-120212936-DEL-53
chr13-120224212-DEL-159
chr13-120233833-DEL-175
chr13-120268882-DEL-230
chr13-120276559-DEL-4823
chr13-120276861-DEL-1751
chr13-120311189-DEL-602
chr13-120326435-DEL-6401
chr13-120335245-DEL-115
chr13-120363298-DEL-65
chr13-120367892-DEL-240
chr13-120383352-DEL-225
chr13-120390618-DEL-6936
chr13-120390851-DEL-53
chr13-120392006-DEL-566
chr13-120407609-DEL-87
chr13-120411490-DEL-4247
chr13-120415828-DEL-2640
chr13-120418001-DEL-548
chr13-120419533-DEL-565
chr13-120419681-DEL-3373
chr13-120420581-DEL-1064
chr13-120423411-DEL-3060
chr13-120423567-DEL-112
chr13-120437912-DEL-3795
chr13-120459392-DEL-62
chr13-120469752-DEL-239
chr13-120475103-DEL-2526
chr13-120481727-DEL-53
chr13-120490678-DEL-241
chr13-120494225-DEL-339
chr13-120504227-DEL-172
chr13-120507985-DEL-2533
chr13-120531513-DEL-113
chr13-120543896-DEL-3795
chr13-120556311-DEL-335
chr13-120559166-DEL-2949
chr13-120562482-DEL-3373
chr13-120563886-DEL-1064
chr13-120565440-DEL-565
chr13-120567012-DEL-622
chr13-120570894-DEL-7006
chr13-120595163-DEL-53
chr13-120600016-DEL-565
chr13-120601684-DEL-53
chr13-120630952-DEL-397
chr13-120631845-DEL-4829
chr13-120646749-DEL-172
chr13-120655299-DEL-115
chr13-120656649-DEL-1099

chr13-120661156-DEL-72
chr13-120662108-DEL-1009
chr13-120664486-DEL-5538
chr13-120664931-DEL-10669
chr13-120672548-DEL-602
chr13-120686031-DEL-7252
chr13-120716023-DEL-224
chr13-120736678-DEL-284
chr13-120742596-DEL-498
chr13-120751545-DEL-65
chr13-120751803-DEL-6651
chr13-120762345-DEL-56
chr13-120765662-DEL-554
chr13-120773275-DEL-252
chr14-3088438-DEL-566
chr14-3108226-DEL-53
chr14-3119099-DEL-3482
chr14-3155732-DEL-84
chr14-3158923-DEL-86
chr14-3159319-DEL-6437
chr14-3258999-DEL-1094
chr14-3272299-DEL-1761
chr14-3332627-DEL-121
chr14-3366560-DEL-125
chr14-3368720-DEL-6506
chr14-3382904-DEL-51
chr14-3404004-DEL-86
chr14-3406140-DEL-186
chr14-3425385-DEL-74
chr14-3431961-DEL-301
chr14-3451788-DEL-486
chr14-3478088-DEL-55
chr14-3535747-DEL-77
chr14-3555509-DEL-62
chr14-3559361-DEL-548
chr14-3563000-DEL-2202
chr14-3568019-DEL-117
chr14-3572559-DEL-513
chr14-3615157-DEL-174
chr14-3716015-DEL-84
chr14-3728598-DEL-71
chr14-3739007-DEL-1981
chr14-3744063-DEL-1517
chr14-3757557-DEL-1686
chr14-3760341-DEL-100
chr14-3765729-DEL-2407
chr14-3765812-DEL-263
chr14-3810634-DEL-338
chr14-3817202-DEL-252
chr14-3829043-DEL-871
chr14-3875230-DEL-2400
chr14-3878874-DEL-72
chr14-3890352-DEL-236
chr14-3900675-DEL-81
chr14-3902124-DEL-451
chr14-3919140-DEL-56
chr14-3962138-DEL-212
chr14-4029101-DEL-194
chr14-4053509-DEL-3837
chr14-4064792-DEL-185
chr14-4085925-DEL-1211
chr14-4160336-DEL-71
chr14-4233521-DEL-209
chr14-4248766-DEL-80
chr14-4281855-DEL-13864
chr14-4373284-DEL-77
chr14-4374691-DEL-52
chr14-4390395-DEL-6170
chr14-4412524-DEL-1243
chr14-4436467-DEL-108
chr14-4450967-DEL-225
chr14-4493869-DEL-789
chr14-4516889-DEL-245
chr14-4544502-DEL-194
chr14-4545195-DEL-291
chr14-4609905-DEL-101
chr14-4625529-DEL-4818

chr14-4630714-DEL-90
chr14-4649933-DEL-227
chr14-4667980-DEL-80
chr14-4678558-DEL-433
chr14-4689748-DEL-1221
chr14-4714029-DEL-79
chr14-4720972-DEL-182
chr14-4745377-DEL-92
chr14-4782979-DEL-68
chr14-4857736-DEL-1367
chr14-4893168-DEL-735
chr14-4895190-DEL-103
chr14-4909848-DEL-580
chr14-4956824-DEL-185
chr14-4966221-DEL-445
chr14-4971568-DEL-6147
chr14-5001320-DEL-96
chr14-5012525-DEL-105
chr14-5013070-DEL-20436
chr14-5013601-DEL-2606
chr14-5020383-DEL-3021
chr14-5025530-DEL-401
chr14-5079296-DEL-1868
chr14-5113028-DEL-130
chr14-5118727-DEL-83
chr14-5142987-DEL-12429
chr14-5162297-DEL-2403
chr14-5176886-DEL-294
chr14-5181050-DEL-6378
chr14-5189723-DEL-1889
chr14-5218433-DEL-2408
chr14-5219663-DEL-103
chr14-5266921-DEL-113
chr14-5281622-DEL-1828
chr14-5285835-DEL-3713
chr14-5303658-DEL-366
chr14-5323170-DEL-58
chr14-5430469-DEL-63
chr14-5431457-DEL-53
chr14-5450760-DEL-755
chr14-5487812-DEL-57
chr14-5497798-DEL-1934
chr14-5609069-DEL-990
chr14-5613407-DEL-270
chr14-5614362-DEL-1140
chr14-5623812-DEL-208
chr14-5631283-DEL-6754
chr14-5656202-DEL-222
chr14-5751926-DEL-313
chr14-5803187-DEL-6014
chr14-5820120-DEL-199
chr14-5848967-DEL-115
chr14-5893345-DEL-52
chr14-5942609-DEL-60
chr14-5965833-DEL-590
chr14-5987665-DEL-61
chr14-6003311-DEL-4438
chr14-6010254-DEL-83
chr14-6059373-DEL-57
chr14-6063763-DEL-65
chr14-6073428-DEL-52
chr14-6080454-DEL-799
chr14-6109955-DEL-1556
chr14-6119015-DEL-183
chr14-6126667-DEL-513
chr14-6239030-DEL-384
chr14-6247208-DEL-1674
chr14-6272333-DEL-283
chr14-6297345-DEL-1970
chr14-6335154-DEL-310
chr14-6340151-DEL-1383
chr14-6352501-DEL-6659
chr14-6379572-DEL-84
chr14-6391137-DEL-449
chr14-6397073-DEL-4475
chr14-6419718-DEL-451
chr14-6430093-DEL-519

chr14-6431614-DEL-884
chr14-6466351-DEL-166
chr14-6467552-DEL-233
chr14-6470007-DEL-52
chr14-6474036-DEL-275
chr14-6494381-DEL-1039
chr14-6527384-DEL-458
chr14-6558177-DEL-330
chr14-6563478-DEL-57
chr14-6611618-DEL-9187
chr14-6626673-DEL-62
chr14-6628439-DEL-5987
chr14-6638641-DEL-55
chr14-6645130-DEL-7259
chr14-6655603-DEL-154
chr14-6659623-DEL-1014
chr14-6671185-DEL-52
chr14-6685744-DEL-1215
chr14-6740904-DEL-55
chr14-6759285-DEL-408
chr14-6784415-DEL-66
chr14-6822930-DEL-64
chr14-6844700-DEL-730
chr14-6864269-DEL-680
chr14-6893582-DEL-96
chr14-6897142-DEL-204
chr14-6909574-DEL-198
chr14-6916484-DEL-5526
chr14-6925466-DEL-316
chr14-6959699-DEL-6458
chr14-6968179-DEL-195
chr14-6997534-DEL-299
chr14-7067352-DEL-181
chr14-7070027-DEL-399
chr14-7120039-DEL-220
chr14-7176865-DEL-171
chr14-7213308-DEL-92
chr14-7267260-DEL-111
chr14-7297957-DEL-55
chr14-7301345-DEL-670
chr14-7302435-DEL-207
chr14-7417751-DEL-76
chr14-7426332-DEL-1584
chr14-7464197-DEL-174
chr14-7465624-DEL-178
chr14-7481542-DEL-686
chr14-7486890-DEL-76
chr14-7521409-DEL-155
chr14-7556328-DEL-157
chr14-7568182-DEL-420
chr14-7609313-DEL-83
chr14-7651713-DEL-492
chr14-7662752-DEL-826
chr14-7674101-DEL-239
chr14-7677097-DEL-425
chr14-7754052-DEL-85
chr14-7785952-DEL-408
chr14-7791424-DEL-2311
chr14-7849881-DEL-447
chr14-7899710-DEL-7668
chr14-7935746-DEL-56
chr14-7942033-DEL-452
chr14-7967529-DEL-116
chr14-7998969-DEL-259
chr14-8000383-DEL-2673
chr14-8022601-DEL-57
chr14-8058223-DEL-909
chr14-8081673-DEL-6265
chr14-8104787-DEL-190
chr14-8108234-DEL-609
chr14-8108988-DEL-333
chr14-8152503-DEL-198
chr14-8157956-DEL-156
chr14-8160006-DEL-6560
chr14-8264156-DEL-251
chr14-8265336-DEL-250
chr14-8267841-DEL-100

chr14-8271397-DEL-1319
chr14-8283704-DEL-88
chr14-8307662-DEL-2145
chr14-8315892-DEL-3453
chr14-8340573-DEL-1512
chr14-8355161-DEL-231
chr14-8475167-DEL-109
chr14-8512992-DEL-292
chr14-8532900-DEL-4426
chr14-8549798-DEL-258
chr14-8556151-DEL-193
chr14-8627765-DEL-248
chr14-8670438-DEL-855
chr14-8697010-DEL-54
chr14-8732078-DEL-58
chr14-8755773-DEL-57
chr14-8756657-DEL-7476
chr14-8758369-DEL-2552
chr14-8839122-DEL-57
chr14-8842404-DEL-1085
chr14-8866755-DEL-447
chr14-8967225-DEL-53
chr14-8986289-DEL-1522
chr14-8986331-DEL-245
chr14-9041622-DEL-89
chr14-9049813-DEL-1044
chr14-9077094-DEL-92
chr14-9089613-DEL-168
chr14-9157069-DEL-9185
chr14-9195356-DEL-324
chr14-9195462-DEL-1169
chr14-9203609-DEL-61
chr14-9214119-DEL-83
chr14-9221050-DEL-2687
chr14-9238672-DEL-100
chr14-9266975-DEL-56
chr14-9285108-DEL-1520
chr14-9325768-DEL-1134
chr14-9328737-DEL-112
chr14-9329662-DEL-1578
chr14-9331650-DEL-52
chr14-9338635-DEL-376
chr14-9382503-DEL-286
chr14-9399038-DEL-356
chr14-9453022-DEL-506
chr14-9471987-DEL-73
chr14-9479937-DEL-377
chr14-9518377-DEL-63
chr14-9535808-DEL-72
chr14-9559657-DEL-519
chr14-9561333-DEL-127
chr14-9602249-DEL-280
chr14-9614628-DEL-5499
chr14-9653989-DEL-116
chr14-9711580-DEL-163
chr14-9732175-DEL-824
chr14-9756190-DEL-4813
chr14-9810939-DEL-72
chr14-9817543-DEL-74
chr14-9820923-DEL-80
chr14-9823077-DEL-59
chr14-9830327-DEL-89
chr14-9835485-DEL-445
chr14-9841350-DEL-52
chr14-9854720-DEL-558
chr14-9871091-DEL-134
chr14-9873745-DEL-128
chr14-9875504-DEL-62
chr14-9915184-DEL-111
chr14-9916078-DEL-1978
chr14-9945420-DEL-138
chr14-9987605-DEL-174
chr14-10025782-DEL-162
chr14-10050324-DEL-57
chr14-10088025-DEL-78
chr14-10108113-DEL-2861
chr14-10110905-DEL-115

chr14-10134112-DEL-64
chr14-10155164-DEL-78
chr14-10215693-DEL-415
chr14-10267250-DEL-663
chr14-10327460-DEL-493
chr14-10331070-DEL-195
chr14-10381762-DEL-64
chr14-10390879-DEL-110
chr14-10405243-DEL-212
chr14-10430804-DEL-335
chr14-10444903-DEL-87
chr14-10447482-DEL-444
chr14-10484827-DEL-309
chr14-10514561-DEL-84
chr14-10528471-DEL-83
chr14-10534393-DEL-146
chr14-10554723-DEL-1475
chr14-10558993-DEL-79
chr14-10582346-DEL-55
chr14-10588716-DEL-57
chr14-10642755-DEL-383
chr14-10652160-DEL-67
chr14-10655856-DEL-3228
chr14-10665770-DEL-73
chr14-10671438-DEL-270
chr14-10717786-DEL-61
chr14-10725722-DEL-64
chr14-10754501-DEL-52
chr14-10810572-DEL-188
chr14-10825269-DEL-56
chr14-10837440-DEL-201
chr14-10851883-DEL-145
chr14-10883469-DEL-64
chr14-10888563-DEL-617
chr14-10896995-DEL-58
chr14-10918800-DEL-155
chr14-10927912-DEL-231
chr14-10929062-DEL-100
chr14-10971435-DEL-1079
chr14-10989398-DEL-959
chr14-10990081-DEL-201
chr14-11005522-DEL-113
chr14-11043009-DEL-306
chr14-11069686-DEL-74
chr14-11073715-DEL-99
chr14-11098813-DEL-93
chr14-11125419-DEL-2137
chr14-11197188-DEL-1420
chr14-11226370-DEL-54
chr14-11242590-DEL-62
chr14-11327017-DEL-52
chr14-11351167-DEL-602
chr14-11372410-DEL-1159
chr14-11419956-DEL-1602
chr14-11421180-DEL-432
chr14-11422450-DEL-395
chr14-11449714-DEL-56
chr14-11453128-DEL-3822
chr14-11492390-DEL-260
chr14-11512468-DEL-246
chr14-11514355-DEL-342
chr14-11545092-DEL-179
chr14-11568359-DEL-217
chr14-11570662-DEL-1236
chr14-11617298-DEL-58
chr14-11629396-DEL-68
chr14-11633208-DEL-58
chr14-11639929-DEL-1137
chr14-11643056-DEL-66
chr14-11686171-DEL-282
chr14-11741299-DEL-171
chr14-11774303-DEL-482
chr14-11778401-DEL-6222
chr14-11790158-DEL-915
chr14-11802616-DEL-101
chr14-11805963-DEL-52
chr14-11849902-DEL-1729

chr14-11884035-DEL-70
chr14-11905189-DEL-293
chr14-11965417-DEL-4090
chr14-12065878-DEL-60
chr14-12068345-DEL-59
chr14-12116029-DEL-81
chr14-12146849-DEL-56
chr14-12173521-DEL-77
chr14-12197561-DEL-2118
chr14-12254325-DEL-56
chr14-12258583-DEL-241
chr14-12303477-DEL-543
chr14-12310488-DEL-64
chr14-12338530-DEL-52
chr14-12340051-DEL-596
chr14-12351283-DEL-6213
chr14-12361863-DEL-6456
chr14-12376830-DEL-6529
chr14-12391825-DEL-5293
chr14-12395452-DEL-384
chr14-12459762-DEL-1081
chr14-12524945-DEL-72
chr14-12527116-DEL-54
chr14-12577799-DEL-88
chr14-12582991-DEL-838
chr14-12587274-DEL-1676
chr14-12603698-DEL-2077
chr14-12618188-DEL-643
chr14-12622863-DEL-391
chr14-12690911-DEL-1248
chr14-12696264-DEL-57
chr14-12722180-DEL-176
chr14-12763900-DEL-74
chr14-12788193-DEL-503
chr14-12809329-DEL-97
chr14-12821404-DEL-67
chr14-12899429-DEL-66
chr14-12914949-DEL-106
chr14-12920032-DEL-70
chr14-12939934-DEL-51
chr14-12964496-DEL-427
chr14-12986004-DEL-93
chr14-13001682-DEL-83
chr14-13043062-DEL-12453
chr14-13054884-DEL-905
chr14-13071699-DEL-52
chr14-13079703-DEL-280
chr14-13099246-DEL-90
chr14-13116667-DEL-99
chr14-13147996-DEL-104
chr14-13188965-DEL-159
chr14-13196960-DEL-81
chr14-13216126-DEL-125
chr14-13306195-DEL-159
chr14-13330330-DEL-4102
chr14-13348929-DEL-85
chr14-13438098-DEL-484
chr14-13449793-DEL-1493
chr14-13485769-DEL-69
chr14-13505142-DEL-285
chr14-13507223-DEL-108
chr14-13582904-DEL-6586
chr14-13592976-DEL-64
chr14-13608237-DEL-3948
chr14-13631010-DEL-75
chr14-13636964-DEL-87
chr14-13649745-DEL-55
chr14-13679191-DEL-278
chr14-13689791-DEL-82
chr14-13703071-DEL-67
chr14-13711130-DEL-134
chr14-13718973-DEL-7496
chr14-13731129-DEL-344
chr14-13744490-DEL-1165
chr14-13745800-DEL-1148
chr14-13820334-DEL-66
chr14-13829574-DEL-176

chr14-13872145-DEL-1077
chr14-13886847-DEL-72
chr14-13891215-DEL-62
chr14-13892373-DEL-245
chr14-13902339-DEL-234
chr14-13923199-DEL-153
chr14-13980524-DEL-53
chr14-13982116-DEL-1248
chr14-13990110-DEL-124
chr14-14005880-DEL-120
chr14-14013279-DEL-134
chr14-14018412-DEL-720
chr14-14019160-DEL-1163
chr14-14020376-DEL-149
chr14-14044866-DEL-112
chr14-14055753-DEL-942
chr14-14109611-DEL-664
chr14-14129026-DEL-63
chr14-14133717-DEL-480
chr14-14195009-DEL-5998
chr14-14197280-DEL-58
chr14-14237497-DEL-55
chr14-14256758-DEL-793
chr14-14261410-DEL-213
chr14-14271423-DEL-175
chr14-14284644-DEL-106
chr14-14310838-DEL-53
chr14-14361029-DEL-191
chr14-14383205-DEL-218
chr14-14395832-DEL-5890
chr14-14404480-DEL-84
chr14-14409579-DEL-804
chr14-14424924-DEL-292
chr14-14426148-DEL-54
chr14-14436317-DEL-74
chr14-14450869-DEL-957
chr14-14451269-DEL-1056
chr14-14455270-DEL-196
chr14-14463606-DEL-84
chr14-14471281-DEL-218
chr14-14477234-DEL-2850
chr14-14481324-DEL-698
chr14-14488852-DEL-210
chr14-14520423-DEL-64
chr14-14522313-DEL-51
chr14-14534223-DEL-97
chr14-14571889-DEL-79
chr14-14640650-DEL-224
chr14-14681929-DEL-499
chr14-14683064-DEL-165
chr14-14696962-DEL-175
chr14-14719375-DEL-63
chr14-14723829-DEL-75
chr14-14749465-DEL-64
chr14-14758204-DEL-272
chr14-14778604-DEL-4258
chr14-14783824-DEL-1264
chr14-14804658-DEL-182
chr14-14808183-DEL-1304
chr14-14808607-DEL-59
chr14-14809585-DEL-4089
chr14-14875784-DEL-6538
chr14-14890091-DEL-227
chr14-14891061-DEL-57
chr14-14897912-DEL-76
chr14-14898164-DEL-1355
chr14-14922341-DEL-957
chr14-14945398-DEL-786
chr14-14949274-DEL-1371
chr14-14952571-DEL-142
chr14-14979153-DEL-96
chr14-14999809-DEL-2234
chr14-15011373-DEL-75
chr14-15029785-DEL-466
chr14-15104038-DEL-68
chr14-15111965-DEL-5359
chr14-15194801-DEL-60

chr14-15226499-DEL-137
chr14-15253335-DEL-1946
chr14-15263570-DEL-132
chr14-15290858-DEL-62
chr14-15293559-DEL-4130
chr14-15300311-DEL-634
chr14-15322326-DEL-1946
chr14-15332461-DEL-153
chr14-15359989-DEL-62
chr14-15511099-DEL-3793
chr14-15544800-DEL-1708
chr14-15552829-DEL-7102
chr14-15569346-DEL-5138
chr14-15593982-DEL-64
chr14-15604146-DEL-52
chr14-15610413-DEL-65
chr14-15617643-DEL-209
chr14-15620846-DEL-231
chr14-15633547-DEL-1419
chr14-15646919-DEL-113
chr14-15662222-DEL-111
chr14-15668962-DEL-1503
chr14-15677285-DEL-1709
chr14-15683061-DEL-62
chr14-15683315-DEL-273
chr14-15834881-DEL-284
chr14-15839580-DEL-6527
chr14-15846057-DEL-444
chr14-15855532-DEL-190
chr14-15886876-DEL-705
chr14-15926549-DEL-410
chr14-15988782-DEL-498
chr14-16088735-DEL-315
chr14-16186647-DEL-208
chr14-16218992-DEL-85
chr14-16236947-DEL-1883
chr14-16244573-DEL-412
chr14-16249959-DEL-51
chr14-16291810-DEL-409
chr14-16332188-DEL-494
chr14-16340200-DEL-536
chr14-16352391-DEL-92
chr14-16366390-DEL-1437
chr14-16375135-DEL-60
chr14-16460187-DEL-142
chr14-16537914-DEL-191
chr14-16555139-DEL-90
chr14-16557735-DEL-60
chr14-16561772-DEL-95
chr14-16578683-DEL-2154
chr14-16586140-DEL-96
chr14-16587839-DEL-5161
chr14-16653788-DEL-67
chr14-16767540-DEL-3400
chr14-16922232-DEL-337
chr14-16941618-DEL-64
chr14-16979767-DEL-1689
chr14-16982733-DEL-359
chr14-17008990-DEL-127
chr14-17057447-DEL-74
chr14-17208532-DEL-68
chr14-17350626-DEL-54
chr14-17384699-DEL-60
chr14-17410652-DEL-3007
chr14-17443791-DEL-272
chr14-17451447-DEL-3077
chr14-17470061-DEL-73
chr14-17492477-DEL-6452
chr14-17521352-DEL-4118
chr14-17541496-DEL-2478
chr14-17627633-DEL-370
chr14-17700208-DEL-195
chr14-17707241-DEL-84
chr14-17772887-DEL-60
chr14-17984263-DEL-2645
chr14-18057139-DEL-84
chr14-18097814-DEL-98

chr14-18136756-DEL-103
chr14-18163804-DEL-522
chr14-18202319-DEL-138
chr14-18225719-DEL-412
chr14-18290049-DEL-78
chr14-18314514-DEL-61
chr14-18371640-DEL-6383
chr14-18381545-DEL-373
chr14-18391943-DEL-108
chr14-18446884-DEL-203
chr14-18452686-DEL-1229
chr14-18459543-DEL-416
chr14-18474676-DEL-52
chr14-18483035-DEL-51
chr14-18484371-DEL-158
chr14-18489985-DEL-58
chr14-18522328-DEL-82
chr14-18538526-DEL-107
chr14-18543694-DEL-284
chr14-18677808-DEL-70
chr14-18682954-DEL-284
chr14-18701088-DEL-107
chr14-18702102-DEL-250
chr14-18791219-DEL-92
chr14-18853948-DEL-2151
chr14-18923785-DEL-90
chr14-18940482-DEL-107
chr14-18941496-DEL-250
chr14-18973356-DEL-16625
chr14-18974453-DEL-16629
chr14-19088379-DEL-60
chr14-19101116-DEL-1350
chr14-19155199-DEL-82
chr14-19160359-DEL-284
chr14-19285693-DEL-122
chr14-19292512-DEL-6229
chr14-19329473-DEL-957
chr14-19338716-DEL-5242
chr14-19341960-DEL-108
chr14-19352568-DEL-130
chr14-19353262-DEL-112
chr14-19368357-DEL-56
chr14-19386297-DEL-51
chr14-19404025-DEL-455
chr14-19417235-DEL-1016
chr14-19427725-DEL-2164
chr14-19451904-DEL-77
chr14-19591261-DEL-7083
chr14-19653157-DEL-57
chr14-19695819-DEL-331
chr14-19775691-DEL-1025
chr14-19776581-DEL-1059
chr14-19819118-DEL-186
chr14-19840596-DEL-340
chr14-19899596-DEL-1888
chr14-19927746-DEL-310
chr14-19975935-DEL-1339
chr14-19979460-DEL-90
chr14-20015008-DEL-111
chr14-20022832-DEL-72
chr14-20031662-DEL-164
chr14-20045581-DEL-65
chr14-20050379-DEL-477
chr14-20057290-DEL-206
chr14-20064513-DEL-94
chr14-20070835-DEL-193
chr14-20077756-DEL-1793
chr14-20083004-DEL-56
chr14-20105638-DEL-287
chr14-20137213-DEL-60
chr14-20167846-DEL-365
chr14-20203679-DEL-55
chr14-20216149-DEL-157
chr14-20223556-DEL-113
chr14-20237972-DEL-56
chr14-20247141-DEL-111
chr14-20259081-DEL-105

chr14-20270530-DEL-100
chr14-20289736-DEL-75
chr14-20302124-DEL-140
chr14-20303744-DEL-3345
chr14-20306115-DEL-719
chr14-20342728-DEL-108
chr14-20361324-DEL-67
chr14-20368102-DEL-99
chr14-20403078-DEL-134
chr14-20428961-DEL-69
chr14-20453545-DEL-64
chr14-20456059-DEL-384
chr14-20469009-DEL-101
chr14-20469518-DEL-560
chr14-20499054-DEL-74
chr14-20509767-DEL-118
chr14-20524412-DEL-206
chr14-20525149-DEL-251
chr14-20560574-DEL-94
chr14-20589852-DEL-963
chr14-20607476-DEL-535
chr14-20610141-DEL-71
chr14-20618959-DEL-60
chr14-20625516-DEL-200
chr14-20648158-DEL-71
chr14-20651883-DEL-59
chr14-20664672-DEL-304
chr14-20709124-DEL-85
chr14-20720359-DEL-190
chr14-20728623-DEL-80
chr14-20762599-DEL-159
chr14-20819786-DEL-220
chr14-20826486-DEL-351
chr14-20829656-DEL-203
chr14-20857157-DEL-90
chr14-20951845-DEL-194
chr14-20953120-DEL-88
chr14-20967869-DEL-104
chr14-20994729-DEL-212
chr14-20999850-DEL-58
chr14-21016746-DEL-54
chr14-21019002-DEL-91
chr14-21054386-DEL-62
chr14-21062459-DEL-56
chr14-21072249-DEL-65
chr14-21114908-DEL-57
chr14-21135160-DEL-80
chr14-21157010-DEL-58
chr14-21171328-DEL-73
chr14-21199936-DEL-56
chr14-21203623-DEL-248
chr14-21226779-DEL-2856
chr14-21329963-DEL-84
chr14-21341984-DEL-2558
chr14-21350156-DEL-614
chr14-21386139-DEL-177
chr14-21463981-DEL-97
chr14-21486931-DEL-98
chr14-21526288-DEL-58
chr14-21535161-DEL-131
chr14-21605950-DEL-124
chr14-21610198-DEL-55
chr14-21656815-DEL-295
chr14-21668485-DEL-164
chr14-21692703-DEL-66
chr14-21733033-DEL-70
chr14-21742398-DEL-580
chr14-21769011-DEL-65
chr14-21781567-DEL-161
chr14-21806920-DEL-346
chr14-21811780-DEL-112
chr14-21838676-DEL-201
chr14-21853216-DEL-151
chr14-21857397-DEL-70
chr14-21861075-DEL-314
chr14-21889968-DEL-72
chr14-21963564-DEL-117

chr14-22019663-DEL-72
chr14-22022269-DEL-74
chr14-22059563-DEL-84
chr14-22082490-DEL-96
chr14-22115758-DEL-93
chr14-22190540-DEL-79
chr14-22219917-DEL-356
chr14-22264903-DEL-210
chr14-22348884-DEL-56
chr14-22360919-DEL-91
chr14-22385760-DEL-956
chr14-22474555-DEL-51
chr14-22494653-DEL-73860
chr14-22528956-DEL-2165
chr14-22536885-DEL-52
chr14-22568833-DEL-67
chr14-22611764-DEL-740
chr14-22672999-DEL-1387
chr14-22678093-DEL-157
chr14-22735074-DEL-532
chr14-22747142-DEL-85
chr14-22772634-DEL-89
chr14-22776958-DEL-65
chr14-22781079-DEL-95
chr14-22805375-DEL-96
chr14-22849024-DEL-91
chr14-22852008-DEL-72
chr14-22856620-DEL-58
chr14-22964877-DEL-82
chr14-22973356-DEL-71
chr14-22995464-DEL-52
chr14-23012291-DEL-55
chr14-23027591-DEL-313
chr14-23037537-DEL-399
chr14-23038877-DEL-54
chr14-23042178-DEL-86
chr14-23064551-DEL-52
chr14-23083773-DEL-343
chr14-23099885-DEL-202
chr14-23127556-DEL-444
chr14-23150209-DEL-77
chr14-23171874-DEL-205
chr14-23178876-DEL-91
chr14-23207668-DEL-80
chr14-23225840-DEL-374
chr14-23239909-DEL-257
chr14-23300964-DEL-98
chr14-23307978-DEL-578
chr14-23419057-DEL-104
chr14-23437777-DEL-204
chr14-23545725-DEL-56
chr14-23566741-DEL-86
chr14-23571903-DEL-72
chr14-23585068-DEL-3506
chr14-23596293-DEL-65
chr14-23603971-DEL-257
chr14-23667876-DEL-140
chr14-23710882-DEL-2006
chr14-23760014-DEL-126
chr14-23771114-DEL-66
chr14-23815406-DEL-56
chr14-23839049-DEL-77
chr14-23884366-DEL-417
chr14-23900972-DEL-217
chr14-23909981-DEL-60
chr14-23967516-DEL-68
chr14-23969261-DEL-81
chr14-24027593-DEL-698
chr14-24030276-DEL-59
chr14-24077212-DEL-72
chr14-24078208-DEL-980
chr14-24119791-DEL-1193
chr14-24121569-DEL-131
chr14-24125798-DEL-568
chr14-24191188-DEL-210
chr14-24210851-DEL-65
chr14-24238258-DEL-52

chr14-24260417-DEL-81
chr14-24270007-DEL-52
chr14-24274458-DEL-156
chr14-24309793-DEL-70
chr14-24310655-DEL-868
chr14-24360221-DEL-449
chr14-24361337-DEL-100
chr14-24400324-DEL-487
chr14-24401410-DEL-1710
chr14-24449825-DEL-225
chr14-24528065-DEL-174
chr14-24537433-DEL-368
chr14-24538256-DEL-63
chr14-24547543-DEL-151
chr14-24593744-DEL-102
chr14-24602494-DEL-53
chr14-24691609-DEL-61
chr14-24743604-DEL-83
chr14-24764975-DEL-55
chr14-24796431-DEL-1188
chr14-24873204-DEL-74
chr14-24894991-DEL-52
chr14-24906593-DEL-139
chr14-24944698-DEL-69
chr14-24954205-DEL-245
chr14-24993756-DEL-60
chr14-25024764-DEL-72
chr14-25116631-DEL-91
chr14-25122764-DEL-145
chr14-25144404-DEL-519
chr14-25153820-DEL-56
chr14-25201582-DEL-58
chr14-25213063-DEL-156
chr14-25227904-DEL-126
chr14-25248590-DEL-134
chr14-25293588-DEL-388
chr14-25308914-DEL-275
chr14-25330610-DEL-207
chr14-25330752-DEL-162
chr14-25331799-DEL-298
chr14-25335394-DEL-115
chr14-25345092-DEL-206
chr14-25346990-DEL-52
chr14-25381200-DEL-208
chr14-25387989-DEL-72
chr14-25417161-DEL-56
chr14-25445337-DEL-76
chr14-25467664-DEL-132
chr14-25553206-DEL-56
chr14-25556974-DEL-195
chr14-25568158-DEL-293
chr14-25612707-DEL-62
chr14-25619219-DEL-52
chr14-25675409-DEL-448
chr14-25690570-DEL-113
chr14-25744655-DEL-85
chr14-25754614-DEL-76
chr14-25761026-DEL-142
chr14-25790347-DEL-60
chr14-25798355-DEL-68
chr14-25809075-DEL-559
chr14-25820781-DEL-2312
chr14-25827844-DEL-197
chr14-25827992-DEL-739
chr14-25912138-DEL-97
chr14-25912256-DEL-1733
chr14-25915180-DEL-197
chr14-25915781-DEL-103
chr14-25921436-DEL-550
chr14-25922116-DEL-255
chr14-25933775-DEL-79
chr14-25947836-DEL-63
chr14-25959600-DEL-513
chr14-25963068-DEL-1071
chr14-25993531-DEL-54
chr14-26007341-DEL-179
chr14-26016233-DEL-176

chr14-26022005-DEL-930
chr14-26026876-DEL-138
chr14-26071413-DEL-1399
chr14-26072968-DEL-1100
chr14-26074186-DEL-1095
chr14-26103319-DEL-172
chr14-26110599-DEL-587
chr14-26112461-DEL-199
chr14-26119108-DEL-184
chr14-26166144-DEL-56
chr14-26168191-DEL-174
chr14-26188337-DEL-78
chr14-26192438-DEL-58
chr14-26208478-DEL-60
chr14-26217375-DEL-66
chr14-26231219-DEL-58
chr14-26245239-DEL-68
chr14-26289988-DEL-92
chr14-26297022-DEL-268
chr14-26393282-DEL-341
chr14-26403603-DEL-180
chr14-26462474-DEL-239
chr14-26467128-DEL-69
chr14-26468603-DEL-101
chr14-26472354-DEL-81
chr14-26476194-DEL-54
chr14-26479600-DEL-3885
chr14-26525216-DEL-1397
chr14-26533290-DEL-51
chr14-26562381-DEL-1879
chr14-26581564-DEL-52
chr14-26615832-DEL-62
chr14-26625603-DEL-176
chr14-26638697-DEL-123
chr14-26648518-DEL-321
chr14-26666485-DEL-60
chr14-26695710-DEL-144
chr14-26708063-DEL-190
chr14-26717094-DEL-226
chr14-26720580-DEL-700
chr14-26730240-DEL-174
chr14-26735327-DEL-51
chr14-26736942-DEL-78
chr14-26745193-DEL-96
chr14-26746596-DEL-338
chr14-26776642-DEL-113
chr14-26779830-DEL-658
chr14-26782355-DEL-496
chr14-26800108-DEL-535
chr14-26808286-DEL-135
chr14-26819251-DEL-202
chr14-26857645-DEL-216
chr14-26939234-DEL-56
chr14-26945375-DEL-72
chr14-27002681-DEL-508
chr14-27014549-DEL-1000
chr14-27022161-DEL-51
chr14-27029383-DEL-7035
chr14-27078047-DEL-54
chr14-27126251-DEL-205
chr14-27135462-DEL-71
chr14-27143456-DEL-83
chr14-27181051-DEL-561
chr14-27183998-DEL-1530
chr14-27250456-DEL-178
chr14-27273492-DEL-65
chr14-27307535-DEL-1780
chr14-27311591-DEL-70
chr14-27313752-DEL-112
chr14-27352708-DEL-134
chr14-27357507-DEL-60
chr14-27366152-DEL-56
chr14-27386625-DEL-58
chr14-27423248-DEL-52
chr14-27450952-DEL-127
chr14-27455182-DEL-60
chr14-27546917-DEL-83

chr14-27548184-DEL-128
chr14-27579808-DEL-181
chr14-27618003-DEL-114
chr14-27717352-DEL-420
chr14-27719933-DEL-54
chr14-27745249-DEL-472
chr14-27753590-DEL-56
chr14-27807190-DEL-53
chr14-27813465-DEL-58
chr14-27867580-DEL-339
chr14-27922848-DEL-896
chr14-27976001-DEL-130
chr14-27977996-DEL-90
chr14-27983426-DEL-1313
chr14-27999807-DEL-86
chr14-28082061-DEL-1414
chr14-28130320-DEL-1015
chr14-28166000-DEL-52
chr14-28196412-DEL-98
chr14-28217465-DEL-52
chr14-28219362-DEL-268
chr14-28240971-DEL-64
chr14-28261001-DEL-248
chr14-28326315-DEL-6235
chr14-28335915-DEL-91
chr14-28336333-DEL-6836
chr14-28355109-DEL-1175
chr14-28388659-DEL-62
chr14-28389706-DEL-70
chr14-28421471-DEL-88
chr14-28431122-DEL-1865
chr14-28435836-DEL-73
chr14-28439320-DEL-73
chr14-28510739-DEL-1646
chr14-28522352-DEL-167
chr14-28541300-DEL-205
chr14-28561786-DEL-83
chr14-28583705-DEL-83
chr14-28589850-DEL-52
chr14-28617929-DEL-119
chr14-28622117-DEL-296
chr14-28662753-DEL-134
chr14-28667815-DEL-63
chr14-28672079-DEL-64
chr14-28687759-DEL-109
chr14-28776418-DEL-142
chr14-28790511-DEL-232
chr14-28916162-DEL-1244
chr14-28921901-DEL-62
chr14-28923702-DEL-250
chr14-28949174-DEL-336
chr14-28983657-DEL-64
chr14-28986272-DEL-59
chr14-29000324-DEL-78
chr14-29002493-DEL-775
chr14-29005976-DEL-1852
chr14-29008551-DEL-164
chr14-29031832-DEL-165
chr14-29047294-DEL-1198
chr14-29133607-DEL-63
chr14-29135648-DEL-54
chr14-29196237-DEL-113
chr14-29204659-DEL-108
chr14-29230516-DEL-269
chr14-29295356-DEL-208
chr14-29311273-DEL-192
chr14-29328521-DEL-265
chr14-29336503-DEL-64
chr14-29430364-DEL-302
chr14-29459501-DEL-221
chr14-29464286-DEL-166
chr14-29482235-DEL-104
chr14-29492395-DEL-475
chr14-29494115-DEL-242
chr14-29497616-DEL-152
chr14-29515163-DEL-81
chr14-29553904-DEL-392

chr14-29568325-DEL-66
chr14-29593770-DEL-660
chr14-29625662-DEL-72
chr14-29633658-DEL-67
chr14-29652818-DEL-108
chr14-29671594-DEL-350
chr14-29747196-DEL-63
chr14-29867987-DEL-81
chr14-29904111-DEL-175
chr14-29948913-DEL-1995
chr14-29959383-DEL-72
chr14-30117414-DEL-914
chr14-30119539-DEL-290
chr14-30151720-DEL-122
chr14-30187708-DEL-103
chr14-30256408-DEL-59
chr14-30285305-DEL-66
chr14-30310718-DEL-80
chr14-30311264-DEL-201
chr14-30319943-DEL-85
chr14-30328482-DEL-166
chr14-30342528-DEL-72
chr14-30349247-DEL-200
chr14-30389325-DEL-83
chr14-30420636-DEL-60
chr14-30451034-DEL-2569
chr14-30455836-DEL-244
chr14-30466535-DEL-76
chr14-30470731-DEL-6036
chr14-30498631-DEL-370
chr14-30523338-DEL-58
chr14-30545321-DEL-100
chr14-30561611-DEL-57
chr14-30629295-DEL-72
chr14-30632421-DEL-82
chr14-30680029-DEL-629
chr14-30683936-DEL-208
chr14-30684509-DEL-143
chr14-30719731-DEL-215
chr14-30729456-DEL-196
chr14-30733969-DEL-72
chr14-30737685-DEL-191
chr14-30756624-DEL-296
chr14-30757331-DEL-194
chr14-30779893-DEL-216
chr14-30794558-DEL-934
chr14-30797921-DEL-80
chr14-30798431-DEL-136
chr14-30900161-DEL-256
chr14-30940479-DEL-84
chr14-31048339-DEL-195
chr14-31072989-DEL-287
chr14-31123487-DEL-181
chr14-31147793-DEL-76
chr14-31171605-DEL-179
chr14-31174735-DEL-1467
chr14-31177260-DEL-1268
chr14-31183981-DEL-64
chr14-31199421-DEL-400
chr14-31248679-DEL-206
chr14-31275069-DEL-68
chr14-31276617-DEL-218
chr14-31282074-DEL-127
chr14-31303135-DEL-615
chr14-31304940-DEL-148
chr14-31306450-DEL-57
chr14-31309410-DEL-871
chr14-31315299-DEL-711
chr14-31317272-DEL-51
chr14-31366037-DEL-197
chr14-31367888-DEL-116
chr14-31401967-DEL-205
chr14-31405495-DEL-183
chr14-31431702-DEL-346
chr14-31513155-DEL-60
chr14-31515321-DEL-254
chr14-31517560-DEL-2482

chr14-31532745-DEL-152
chr14-31534209-DEL-89
chr14-31569633-DEL-197
chr14-31581145-DEL-52
chr14-31581868-DEL-958
chr14-31591012-DEL-58
chr14-31617314-DEL-55
chr14-31619429-DEL-234
chr14-31625897-DEL-52
chr14-31636752-DEL-55
chr14-31639352-DEL-482
chr14-31668707-DEL-245
chr14-31676704-DEL-105
chr14-31702249-DEL-701
chr14-31713430-DEL-249
chr14-31723324-DEL-1292
chr14-31735502-DEL-187
chr14-31761032-DEL-280
chr14-31763659-DEL-122
chr14-31794627-DEL-1654
chr14-31819981-DEL-1038
chr14-31845176-DEL-907
chr14-31847632-DEL-52
chr14-31853491-DEL-70
chr14-31865110-DEL-51
chr14-31865560-DEL-137
chr14-31868479-DEL-229
chr14-31869079-DEL-130
chr14-31872638-DEL-54
chr14-31878234-DEL-60
chr14-31906990-DEL-138
chr14-31909980-DEL-65
chr14-31921526-DEL-209
chr14-31981344-DEL-376
chr14-32069704-DEL-224
chr14-32077528-DEL-52
chr14-32079960-DEL-3767
chr14-32083288-DEL-2010
chr14-32083348-DEL-270
chr14-32084153-DEL-180
chr14-32084318-DEL-555
chr14-32096167-DEL-84
chr14-32099121-DEL-69
chr14-32131175-DEL-63
chr14-32164238-DEL-98
chr14-32171943-DEL-434
chr14-32173582-DEL-160
chr14-32178394-DEL-523
chr14-32214251-DEL-67
chr14-32220376-DEL-557
chr14-32310004-DEL-104
chr14-32348810-DEL-58
chr14-32379663-DEL-61
chr14-32444942-DEL-108
chr14-32497531-DEL-246
chr14-32600466-DEL-73
chr14-32600759-DEL-350
chr14-32621902-DEL-75
chr14-32648689-DEL-54
chr14-32666817-DEL-54
chr14-32666820-DEL-662
chr14-32734131-DEL-63
chr14-32835843-DEL-62
chr14-32892831-DEL-72
chr14-32932238-DEL-61
chr14-32939703-DEL-144
chr14-32940340-DEL-93
chr14-32965690-DEL-54
chr14-32974068-DEL-1014
chr14-33008230-DEL-80
chr14-33008980-DEL-54
chr14-33024405-DEL-62
chr14-33028809-DEL-61
chr14-33041140-DEL-56
chr14-33135634-DEL-60
chr14-33142507-DEL-58
chr14-33150471-DEL-752

chr14-33192089-DEL-163
chr14-33278016-DEL-191
chr14-33288591-DEL-64
chr14-33294342-DEL-104
chr14-33296524-DEL-272
chr14-33313649-DEL-556
chr14-33321878-DEL-116
chr14-33330083-DEL-950
chr14-33403627-DEL-569
chr14-33423776-DEL-56
chr14-33425775-DEL-2584
chr14-33428371-DEL-4398
chr14-33444154-DEL-540
chr14-33466116-DEL-1150
chr14-33487640-DEL-108
chr14-33505044-DEL-121
chr14-33508275-DEL-1947
chr14-33521537-DEL-84
chr14-33545581-DEL-205
chr14-33566423-DEL-828
chr14-33568714-DEL-56
chr14-33582469-DEL-80
chr14-33583208-DEL-808
chr14-33594943-DEL-2854
chr14-33629409-DEL-94
chr14-33662042-DEL-81
chr14-33673054-DEL-60
chr14-33689322-DEL-962
chr14-33698648-DEL-53
chr14-33733004-DEL-3084
chr14-33770962-DEL-1248
chr14-33786677-DEL-82
chr14-33801534-DEL-479
chr14-33802822-DEL-312
chr14-33810620-DEL-381
chr14-33812787-DEL-58
chr14-33851091-DEL-65
chr14-33930939-DEL-53
chr14-33956258-DEL-63
chr14-33970305-DEL-62
chr14-33971733-DEL-168
chr14-33994107-DEL-168
chr14-34053338-DEL-206
chr14-34061061-DEL-193
chr14-34105597-DEL-57
chr14-34127309-DEL-53
chr14-34132076-DEL-121
chr14-34171142-DEL-55
chr14-34187305-DEL-74
chr14-34229639-DEL-130
chr14-34257124-DEL-62
chr14-34278412-DEL-208
chr14-34328320-DEL-591
chr14-34372121-DEL-452
chr14-34401759-DEL-191
chr14-34409692-DEL-1155
chr14-34418208-DEL-60
chr14-34422966-DEL-602
chr14-34424176-DEL-7302
chr14-34440974-DEL-238
chr14-34457876-DEL-199
chr14-34475157-DEL-75
chr14-34502841-DEL-338
chr14-34565752-DEL-71
chr14-34582436-DEL-126
chr14-34590438-DEL-72
chr14-34617367-DEL-626
chr14-34647947-DEL-52
chr14-34687831-DEL-445
chr14-34689722-DEL-168
chr14-34699743-DEL-65
chr14-34708877-DEL-158
chr14-34779548-DEL-90
chr14-34797884-DEL-76
chr14-34799884-DEL-241
chr14-34808596-DEL-359
chr14-34855042-DEL-102

chr14-34909344-DEL-55
chr14-34919394-DEL-54
chr14-34952102-DEL-69
chr14-35005941-DEL-1169
chr14-35082295-DEL-602
chr14-35098519-DEL-88
chr14-35103240-DEL-131
chr14-35186274-DEL-125
chr14-35204255-DEL-92
chr14-35229611-DEL-1977
chr14-35244976-DEL-65
chr14-35255491-DEL-208
chr14-35279862-DEL-56
chr14-35315710-DEL-58
chr14-35316674-DEL-51
chr14-35318549-DEL-133
chr14-35319216-DEL-654
chr14-35340785-DEL-131
chr14-35388370-DEL-110
chr14-35390189-DEL-96
chr14-35392344-DEL-1404
chr14-35437151-DEL-201
chr14-35490625-DEL-107
chr14-35491475-DEL-132
chr14-35515755-DEL-62
chr14-35526970-DEL-1614
chr14-35534249-DEL-56
chr14-35535895-DEL-2462
chr14-35543476-DEL-6318
chr14-35543489-DEL-201
chr14-35561777-DEL-76
chr14-35571890-DEL-58
chr14-35650132-DEL-106
chr14-35686887-DEL-499
chr14-35704883-DEL-56
chr14-35736534-DEL-56
chr14-35750309-DEL-431
chr14-35763976-DEL-57
chr14-35773662-DEL-114
chr14-35789791-DEL-92
chr14-35799554-DEL-194
chr14-35801695-DEL-6109
chr14-35830207-DEL-142
chr14-35859912-DEL-1281
chr14-35867474-DEL-127
chr14-35871212-DEL-51
chr14-35918946-DEL-257
chr14-35959799-DEL-56
chr14-35964690-DEL-438
chr14-35968680-DEL-331
chr14-36019538-DEL-51
chr14-36031146-DEL-16685
chr14-36033139-DEL-61
chr14-36077747-DEL-52
chr14-36091006-DEL-1235
chr14-36098181-DEL-6155
chr14-36126173-DEL-56
chr14-36134140-DEL-465
chr14-36141917-DEL-5197
chr14-36156536-DEL-629
chr14-36165726-DEL-1211
chr14-36181787-DEL-1088
chr14-36237247-DEL-635
chr14-36282631-DEL-280
chr14-36286716-DEL-1010
chr14-36292329-DEL-357
chr14-36318389-DEL-208
chr14-36391387-DEL-2303
chr14-36432698-DEL-94
chr14-36442739-DEL-213
chr14-36445966-DEL-58
chr14-36466269-DEL-579
chr14-36472675-DEL-134
chr14-36501898-DEL-519
chr14-36552862-DEL-351
chr14-36568372-DEL-98
chr14-36627423-DEL-94

chr14-36628470-DEL-111
chr14-36669449-DEL-416
chr14-36694618-DEL-60
chr14-36696620-DEL-318
chr14-36730341-DEL-55
chr14-36740013-DEL-61
chr14-36745761-DEL-102
chr14-36752876-DEL-212
chr14-36755020-DEL-297
chr14-36777622-DEL-140
chr14-36809931-DEL-66
chr14-36826607-DEL-82
chr14-36899905-DEL-615
chr14-36909214-DEL-1130
chr14-36994693-DEL-59
chr14-37056281-DEL-91
chr14-37079831-DEL-2403
chr14-37105844-DEL-92
chr14-37110883-DEL-56
chr14-37248337-DEL-1778
chr14-37259140-DEL-72
chr14-37313328-DEL-6316
chr14-37333039-DEL-526
chr14-37334619-DEL-2259
chr14-37334656-DEL-324
chr14-37341472-DEL-7087
chr14-37350693-DEL-6477
chr14-37414583-DEL-925
chr14-37439360-DEL-540
chr14-37442218-DEL-123
chr14-37446749-DEL-2795
chr14-37461474-DEL-62
chr14-37487870-DEL-439
chr14-37541895-DEL-161
chr14-37544194-DEL-165
chr14-37552092-DEL-520
chr14-37557271-DEL-424
chr14-37577932-DEL-61
chr14-37583683-DEL-1331
chr14-37588200-DEL-737
chr14-37622950-DEL-90
chr14-37643689-DEL-82
chr14-37664113-DEL-661
chr14-37719094-DEL-800
chr14-37730924-DEL-67
chr14-37742282-DEL-788
chr14-37755177-DEL-1158
chr14-37762778-DEL-368
chr14-37773401-DEL-602
chr14-37782752-DEL-1376
chr14-37786264-DEL-1394
chr14-37871132-DEL-154
chr14-37878358-DEL-235
chr14-37901493-DEL-398
chr14-37943653-DEL-1057
chr14-37949414-DEL-58
chr14-38017719-DEL-54
chr14-38188668-DEL-3250
chr14-38197884-DEL-410
chr14-38199326-DEL-289
chr14-38215615-DEL-207
chr14-38250651-DEL-82
chr14-38265767-DEL-75
chr14-38267134-DEL-54
chr14-38276437-DEL-85
chr14-38313591-DEL-1638
chr14-38356362-DEL-116
chr14-38357734-DEL-525
chr14-38360757-DEL-69
chr14-38404259-DEL-83
chr14-38429932-DEL-61
chr14-38439185-DEL-251
chr14-38460689-DEL-461
chr14-38474730-DEL-71
chr14-38479075-DEL-72
chr14-38512091-DEL-97
chr14-38558183-DEL-115

chr14-38567589-DEL-156
chr14-38576644-DEL-124
chr14-38589444-DEL-676
chr14-38602755-DEL-54
chr14-38606161-DEL-71
chr14-38621731-DEL-222
chr14-38754107-DEL-59
chr14-38781300-DEL-155210
chr14-38792338-DEL-1866
chr14-38808372-DEL-2034
chr14-38850602-DEL-1461
chr14-38862521-DEL-120
chr14-38897600-DEL-1314
chr14-38899964-DEL-51
chr14-38900457-DEL-86
chr14-38922125-DEL-426
chr14-39033620-DEL-1304
chr14-39036794-DEL-255
chr14-39132308-DEL-120
chr14-39187629-DEL-142
chr14-39247463-DEL-85
chr14-39276662-DEL-602
chr14-39294393-DEL-758
chr14-39353482-DEL-60
chr14-39444003-DEL-79
chr14-39460906-DEL-66
chr14-39463407-DEL-81
chr14-39516310-DEL-5379
chr14-39533305-DEL-483
chr14-39536318-DEL-161
chr14-39596607-DEL-663
chr14-39637823-DEL-103
chr14-39642020-DEL-628
chr14-39692535-DEL-180
chr14-39695518-DEL-56
chr14-39746489-DEL-57
chr14-39759467-DEL-52
chr14-39784729-DEL-63
chr14-39787174-DEL-574
chr14-39790543-DEL-2084
chr14-39802823-DEL-55
chr14-39816150-DEL-64
chr14-39831864-DEL-298
chr14-39862307-DEL-73
chr14-39863792-DEL-491
chr14-39878624-DEL-1321
chr14-39936882-DEL-527
chr14-39940167-DEL-4490
chr14-39951237-DEL-71
chr14-39960115-DEL-86
chr14-39962172-DEL-241
chr14-39966214-DEL-52
chr14-39967672-DEL-6523
chr14-39986921-DEL-54
chr14-39999220-DEL-1235
chr14-40002150-DEL-583
chr14-40002811-DEL-1010
chr14-40043026-DEL-50
chr14-40048143-DEL-526
chr14-40053335-DEL-3485
chr14-40056947-DEL-1936
chr14-40059823-DEL-332
chr14-40064782-DEL-5698
chr14-40091074-DEL-203
chr14-40117468-DEL-400
chr14-40133070-DEL-70
chr14-40171796-DEL-444
chr14-40173892-DEL-621
chr14-40185299-DEL-510
chr14-40198192-DEL-842
chr14-40206842-DEL-54
chr14-40215513-DEL-304
chr14-40234658-DEL-147
chr14-40245958-DEL-1270
chr14-40250297-DEL-122
chr14-40254500-DEL-77
chr14-40269353-DEL-424

chr14-40282753-DEL-106
chr14-40370525-DEL-794
chr14-40379819-DEL-6292
chr14-40398726-DEL-578
chr14-40439424-DEL-782
chr14-40448029-DEL-66
chr14-40475638-DEL-102
chr14-40518846-DEL-130
chr14-40525702-DEL-131
chr14-40527128-DEL-335
chr14-40549477-DEL-63
chr14-40566368-DEL-416
chr14-40600583-DEL-148
chr14-40605194-DEL-182
chr14-40613820-DEL-521
chr14-40621174-DEL-452
chr14-40626391-DEL-1821
chr14-40634087-DEL-624
chr14-40683316-DEL-165
chr14-40709690-DEL-190
chr14-40746939-DEL-172
chr14-40754166-DEL-100
chr14-40755342-DEL-91
chr14-40756386-DEL-151
chr14-40788740-DEL-773
chr14-40806727-DEL-424
chr14-40810076-DEL-177
chr14-40822030-DEL-54
chr14-40836982-DEL-79
chr14-40858689-DEL-1189
chr14-40872060-DEL-51
chr14-40901005-DEL-292
chr14-40936912-DEL-104
chr14-40969627-DEL-4200
chr14-40974782-DEL-598
chr14-40976423-DEL-8195
chr14-40990472-DEL-2412
chr14-40997688-DEL-71
chr14-41004742-DEL-1884
chr14-41014521-DEL-6474
chr14-41050194-DEL-777
chr14-41064721-DEL-647
chr14-41073568-DEL-4587
chr14-41139591-DEL-1420
chr14-41141057-DEL-2382
chr14-41171625-DEL-60
chr14-41179692-DEL-7005
chr14-41200818-DEL-338
chr14-41203240-DEL-936
chr14-41260378-DEL-522
chr14-41396654-DEL-1076
chr14-41415679-DEL-5036
chr14-41430541-DEL-1272
chr14-41743441-DEL-223
chr14-41747740-DEL-1064
chr14-41760009-DEL-1058
chr14-41761452-DEL-94
chr14-41765156-DEL-63
chr14-41775225-DEL-4843
chr14-41865587-DEL-199
chr14-41889954-DEL-1078
chr14-41949595-DEL-4820
chr14-42006215-DEL-1453
chr14-42047300-DEL-8085
chr14-42081397-DEL-164
chr14-42091946-DEL-100
chr14-42096435-DEL-5923
chr14-42102418-DEL-53
chr14-42145327-DEL-523
chr14-42222626-DEL-790
chr14-42226412-DEL-3170
chr14-42234344-DEL-1079
chr14-42267636-DEL-4566
chr14-42268687-DEL-1271
chr14-42277764-DEL-3188
chr14-42279994-DEL-647
chr14-42292755-DEL-168

chr14-42302261-DEL-1162
chr14-42323662-DEL-201
chr14-42353052-DEL-597
chr14-42354691-DEL-8194
chr14-42368726-DEL-2413
chr14-42375917-DEL-71
chr14-42443918-DEL-52
chr14-42525343-DEL-199
chr14-42551166-DEL-1078
chr14-42559530-DEL-6603
chr14-42570709-DEL-69
chr14-42577857-DEL-143
chr14-42597467-DEL-7198
chr14-42648267-DEL-1077
chr14-42674789-DEL-202
chr14-42898039-DEL-53
chr14-42917148-DEL-337
chr14-42927401-DEL-1124
chr14-42959796-DEL-6451
chr14-43023623-DEL-684
chr14-43024849-DEL-3401
chr14-43050984-DEL-4986
chr14-43056206-DEL-6402
chr14-43057112-DEL-183
chr14-43067878-DEL-158
chr14-43075747-DEL-6426
chr14-43075759-DEL-212
chr14-43101717-DEL-2370
chr14-43162100-DEL-58
chr14-43183148-DEL-1770
chr14-43752735-DEL-79
chr14-43762147-DEL-1421
chr14-43763610-DEL-2370
chr14-43775326-DEL-553
chr14-43823282-DEL-6925
chr14-43858637-DEL-228
chr14-43897975-DEL-623
chr14-43899720-DEL-7039
chr14-43931066-DEL-203
chr14-43955909-DEL-405
chr14-43957057-DEL-179
chr14-44040678-DEL-54
chr14-44041824-DEL-8321
chr14-44055053-DEL-236
chr14-44055646-DEL-494
chr14-44082256-DEL-5960
chr14-44125853-DEL-130106
chr14-44180242-DEL-51
chr14-44186046-DEL-122
chr14-44191790-DEL-56
chr14-44202847-DEL-103
chr14-44230078-DEL-3614
chr14-44248628-DEL-1388
chr14-44252049-DEL-114
chr14-44258059-DEL-66
chr14-44366501-DEL-52
chr14-44378906-DEL-60
chr14-44383733-DEL-6248
chr14-44403722-DEL-99
chr14-44413968-DEL-652
chr14-44418797-DEL-768
chr14-44421090-DEL-2379
chr14-44423511-DEL-1419
chr14-44427144-DEL-183
chr14-44429292-DEL-195
chr14-44470757-DEL-6701
chr14-44502473-DEL-3546
chr14-44514971-DEL-4075
chr14-44537102-DEL-79
chr14-44587563-DEL-4384
chr14-44651479-DEL-1806
chr14-44663115-DEL-2612
chr14-44693117-DEL-97
chr14-44693591-DEL-1054
chr14-44694945-DEL-8903
chr14-44752079-DEL-1886
chr14-44868496-DEL-1055

chr14-44869893-DEL-6027
chr14-44879155-DEL-7925
chr14-44931065-DEL-779
chr14-44947681-DEL-103
chr14-44969480-DEL-183
chr14-44972003-DEL-1493
chr14-44974634-DEL-364
chr14-44986756-DEL-14189
chr14-45015271-DEL-134
chr14-45031420-DEL-162
chr14-45035373-DEL-305
chr14-45043003-DEL-3372
chr14-45060751-DEL-63
chr14-45066182-DEL-438
chr14-45080299-DEL-108
chr14-45093568-DEL-121
chr14-45106878-DEL-109
chr14-45113025-DEL-152
chr14-45114199-DEL-1113
chr14-45131472-DEL-365
chr14-45151437-DEL-57
chr14-45163769-DEL-118
chr14-45171646-DEL-815
chr14-45176164-DEL-131
chr14-45199481-DEL-9237
chr14-45245550-DEL-155
chr14-45250994-DEL-8280
chr14-45262124-DEL-316
chr14-45263086-DEL-55
chr14-45264491-DEL-6215
chr14-45325824-DEL-477
chr14-45343971-DEL-310
chr14-45366419-DEL-1561
chr14-45378739-DEL-144
chr14-45380419-DEL-252
chr14-45399198-DEL-161
chr14-45405130-DEL-109
chr14-45414616-DEL-103
chr14-45435997-DEL-95
chr14-45511636-DEL-94
chr14-45523659-DEL-239
chr14-45574133-DEL-116
chr14-45604417-DEL-96
chr14-45620091-DEL-83
chr14-45631485-DEL-1968
chr14-45642568-DEL-52
chr14-45646510-DEL-51
chr14-45650173-DEL-177
chr14-45657082-DEL-336
chr14-45677268-DEL-197
chr14-45679448-DEL-406
chr14-45698866-DEL-265
chr14-45716179-DEL-91
chr14-45717365-DEL-177
chr14-45717956-DEL-1544
chr14-45740449-DEL-170
chr14-45761487-DEL-364
chr14-45774677-DEL-201
chr14-45781849-DEL-180
chr14-45805118-DEL-117
chr14-45806904-DEL-210
chr14-45809264-DEL-57
chr14-45827390-DEL-772
chr14-45854677-DEL-532
chr14-45926443-DEL-51
chr14-46000354-DEL-167
chr14-46009089-DEL-112
chr14-46036292-DEL-153
chr14-46042463-DEL-63
chr14-46054064-DEL-343
chr14-46075474-DEL-2485
chr14-46078169-DEL-2026
chr14-46083214-DEL-183
chr14-46086559-DEL-144
chr14-46105050-DEL-64
chr14-46108645-DEL-987
chr14-46108960-DEL-131

chr14-46113118-DEL-293
chr14-46117104-DEL-58
chr14-46126581-DEL-868
chr14-46127539-DEL-5583
chr14-46133563-DEL-215
chr14-46165276-DEL-102
chr14-46194948-DEL-75
chr14-46204611-DEL-1430
chr14-46281492-DEL-58
chr14-46286748-DEL-102
chr14-46307397-DEL-216
chr14-46338016-DEL-230
chr14-46399400-DEL-52
chr14-46405476-DEL-373
chr14-46436836-DEL-76
chr14-46446564-DEL-938
chr14-46456420-DEL-773
chr14-46487042-DEL-190
chr14-46488467-DEL-1442
chr14-46490128-DEL-994
chr14-46497244-DEL-161
chr14-46505679-DEL-6218
chr14-46513771-DEL-144
chr14-46546973-DEL-192
chr14-46584084-DEL-64
chr14-46587765-DEL-64
chr14-46605891-DEL-51
chr14-46610251-DEL-87
chr14-46701235-DEL-87
chr14-46703993-DEL-996
chr14-46763774-DEL-2083
chr14-46768695-DEL-340
chr14-46776850-DEL-196
chr14-46789583-DEL-975
chr14-46807326-DEL-51
chr14-46826328-DEL-76
chr14-46859612-DEL-439
chr14-46865239-DEL-70
chr14-46903917-DEL-60
chr14-46922028-DEL-478
chr14-46928341-DEL-118
chr14-46937688-DEL-52
chr14-46951321-DEL-398
chr14-46975908-DEL-157
chr14-46976649-DEL-292
chr14-46991954-DEL-341
chr14-47024082-DEL-169
chr14-47032289-DEL-4395
chr14-47058328-DEL-72
chr14-47067648-DEL-172
chr14-47079482-DEL-119
chr14-47106151-DEL-97
chr14-47131825-DEL-75
chr14-47137388-DEL-159
chr14-47152317-DEL-52
chr14-47168962-DEL-88
chr14-47205582-DEL-67
chr14-47223301-DEL-64
chr14-47250305-DEL-1085
chr14-47269338-DEL-119
chr14-47294664-DEL-78
chr14-47299054-DEL-69
chr14-47305018-DEL-300
chr14-47359837-DEL-70
chr14-47369815-DEL-75
chr14-47384056-DEL-195
chr14-47412072-DEL-168
chr14-47412848-DEL-68
chr14-47414893-DEL-52
chr14-47430421-DEL-139
chr14-47464396-DEL-83
chr14-47490913-DEL-54
chr14-47492141-DEL-104
chr14-47495332-DEL-244
chr14-47501225-DEL-207
chr14-47506757-DEL-95
chr14-47540103-DEL-68

chr14-47564666-DEL-200
chr14-47568582-DEL-249
chr14-47569448-DEL-89
chr14-47576519-DEL-156
chr14-47580299-DEL-765
chr14-47581152-DEL-562
chr14-47587008-DEL-193
chr14-47593700-DEL-111
chr14-47596302-DEL-66
chr14-47630482-DEL-378
chr14-47639226-DEL-180
chr14-47674121-DEL-351
chr14-47675143-DEL-351
chr14-47678695-DEL-431
chr14-47684812-DEL-120
chr14-47705669-DEL-201
chr14-47730198-DEL-85
chr14-47733282-DEL-203
chr14-47764136-DEL-124
chr14-47797777-DEL-167
chr14-47799004-DEL-135
chr14-47808280-DEL-161
chr14-47811503-DEL-205
chr14-47812731-DEL-203
chr14-47814545-DEL-108
chr14-47815286-DEL-123
chr14-47817745-DEL-471
chr14-47824162-DEL-367
chr14-47827489-DEL-178
chr14-47835086-DEL-286
chr14-47849339-DEL-122
chr14-47854892-DEL-83
chr14-47857423-DEL-93
chr14-47866538-DEL-162
chr14-47880763-DEL-66
chr14-47909451-DEL-72
chr14-47916361-DEL-219
chr14-47923490-DEL-201
chr14-47947589-DEL-198
chr14-47978235-DEL-513
chr14-48014493-DEL-271
chr14-48016203-DEL-54
chr14-48048084-DEL-73
chr14-48051271-DEL-434
chr14-48057666-DEL-81
chr14-48077748-DEL-205
chr14-48167625-DEL-83
chr14-48200723-DEL-210
chr14-48207906-DEL-183
chr14-48215067-DEL-2419
chr14-48230024-DEL-54
chr14-48239556-DEL-213
chr14-48244187-DEL-545
chr14-48251695-DEL-53
chr14-48259042-DEL-1741
chr14-48270688-DEL-3540
chr14-48310302-DEL-84
chr14-48383747-DEL-98
chr14-48385096-DEL-204
chr14-48393331-DEL-134
chr14-48403190-DEL-189
chr14-48441778-DEL-56
chr14-48452666-DEL-522
chr14-48453323-DEL-228
chr14-48474906-DEL-74
chr14-48483035-DEL-64
chr14-48535206-DEL-190
chr14-48575118-DEL-407
chr14-48619050-DEL-59
chr14-48641932-DEL-101
chr14-48645996-DEL-1272
chr14-48663773-DEL-232
chr14-48671023-DEL-602
chr14-48680929-DEL-201
chr14-48705681-DEL-183
chr14-48713036-DEL-100
chr14-48740201-DEL-114

chr14-48759612-DEL-641
chr14-48766018-DEL-329
chr14-48814738-DEL-195
chr14-48847894-DEL-198
chr14-48850851-DEL-60
chr14-48864668-DEL-574
chr14-48955724-DEL-345
chr14-48964986-DEL-231
chr14-48992704-DEL-195
chr14-49001551-DEL-55
chr14-49007441-DEL-84
chr14-49007799-DEL-408
chr14-49008989-DEL-146
chr14-49010970-DEL-1334
chr14-49016176-DEL-55
chr14-49019219-DEL-71
chr14-49022238-DEL-2145
chr14-49028283-DEL-462
chr14-49039306-DEL-62
chr14-49045712-DEL-149
chr14-49062520-DEL-60
chr14-49066042-DEL-1525
chr14-49071120-DEL-59
chr14-49079416-DEL-111
chr14-49125538-DEL-55
chr14-49146656-DEL-139
chr14-49160264-DEL-1190
chr14-49170246-DEL-68
chr14-49186578-DEL-573
chr14-49192710-DEL-67
chr14-49202810-DEL-835
chr14-49209772-DEL-104
chr14-49222528-DEL-102
chr14-49262119-DEL-2577
chr14-49274080-DEL-90
chr14-49281632-DEL-1540
chr14-49284393-DEL-52
chr14-49286015-DEL-58
chr14-49325253-DEL-70
chr14-49331798-DEL-1603
chr14-49334434-DEL-103
chr14-49351697-DEL-1065
chr14-49373601-DEL-174
chr14-49456542-DEL-699
chr14-49467399-DEL-370
chr14-49505233-DEL-1401
chr14-49511305-DEL-62
chr14-49618549-DEL-1484
chr14-49628252-DEL-463
chr14-49632392-DEL-265
chr14-49670225-DEL-2840
chr14-49685533-DEL-51
chr14-49686846-DEL-84
chr14-49696414-DEL-1391
chr14-49707191-DEL-4247
chr14-49727211-DEL-2717
chr14-49767199-DEL-3983
chr14-49791341-DEL-62
chr14-49801848-DEL-715
chr14-49802850-DEL-108
chr14-49815254-DEL-1890
chr14-49860172-DEL-63
chr14-49876465-DEL-583
chr14-49894440-DEL-397
chr14-49901433-DEL-108
chr14-49968428-DEL-118
chr14-49996555-DEL-362
chr14-50042990-DEL-81
chr14-50066993-DEL-151
chr14-50081628-DEL-2337
chr14-50085425-DEL-60
chr14-50087193-DEL-6374
chr14-50098315-DEL-52
chr14-50122005-DEL-214
chr14-50123617-DEL-70
chr14-50142925-DEL-5543
chr14-50162123-DEL-200

chr14-50190366-DEL-72
chr14-50194883-DEL-167
chr14-50239380-DEL-154
chr14-50241349-DEL-145
chr14-50250052-DEL-6306
chr14-50257817-DEL-291
chr14-50262934-DEL-85
chr14-50291142-DEL-383
chr14-50298591-DEL-7111
chr14-50310876-DEL-184
chr14-50322620-DEL-124
chr14-50322656-DEL-912
chr14-50326483-DEL-51
chr14-50329804-DEL-448
chr14-50335215-DEL-542
chr14-50380817-DEL-1564
chr14-50430076-DEL-2187
chr14-50439018-DEL-7401
chr14-50455106-DEL-565
chr14-50460198-DEL-402
chr14-50464289-DEL-761
chr14-50486824-DEL-208
chr14-50495846-DEL-1163
chr14-50505037-DEL-352
chr14-50512503-DEL-1048
chr14-50514363-DEL-954
chr14-50523816-DEL-93
chr14-50529081-DEL-123
chr14-50667815-DEL-481
chr14-50690042-DEL-67
chr14-50702334-DEL-59
chr14-50703523-DEL-118
chr14-50708119-DEL-119
chr14-50713304-DEL-76
chr14-50745875-DEL-85
chr14-50772725-DEL-10016
chr14-50795240-DEL-6715
chr14-50810785-DEL-187
chr14-50820365-DEL-177
chr14-50833235-DEL-67
chr14-50837270-DEL-849
chr14-50857394-DEL-80
chr14-50860791-DEL-210
chr14-50862158-DEL-10780
chr14-50882561-DEL-170
chr14-50989471-DEL-308
chr14-51001816-DEL-83
chr14-51033225-DEL-347
chr14-51050500-DEL-54
chr14-51069798-DEL-56
chr14-51072717-DEL-104
chr14-51085945-DEL-68
chr14-51093414-DEL-585
chr14-51099659-DEL-180
chr14-51135949-DEL-199
chr14-51142148-DEL-208
chr14-51151295-DEL-368
chr14-51200015-DEL-86
chr14-51201918-DEL-415
chr14-51213317-DEL-965
chr14-51237996-DEL-51
chr14-51251385-DEL-194
chr14-51274933-DEL-109
chr14-51300128-DEL-60
chr14-51336628-DEL-69
chr14-51410509-DEL-68
chr14-51414813-DEL-90
chr14-51448215-DEL-2656
chr14-51449804-DEL-1068
chr14-51487056-DEL-554
chr14-51504415-DEL-126
chr14-51534355-DEL-885
chr14-51561531-DEL-150
chr14-51578469-DEL-104
chr14-51674911-DEL-144
chr14-51725350-DEL-6528
chr14-51744950-DEL-9183

chr14-51747879-DEL-6312
chr14-51755867-DEL-391
chr14-51756725-DEL-52
chr14-51794662-DEL-61
chr14-51905472-DEL-166
chr14-51921338-DEL-177
chr14-51957325-DEL-447
chr14-51959424-DEL-592
chr14-51964176-DEL-6200
chr14-51970352-DEL-220
chr14-51988148-DEL-2715
chr14-51988946-DEL-73
chr14-51995119-DEL-3906
chr14-52000677-DEL-199
chr14-52002648-DEL-3101
chr14-52013972-DEL-1087
chr14-52051888-DEL-1490
chr14-52067424-DEL-18960
chr14-52068486-DEL-6391
chr14-52078228-DEL-489
chr14-52097528-DEL-163
chr14-52103183-DEL-193
chr14-52103844-DEL-959
chr14-52112423-DEL-423
chr14-52125221-DEL-181
chr14-52137736-DEL-199
chr14-52170485-DEL-85
chr14-52181263-DEL-232
chr14-52203274-DEL-88
chr14-52270114-DEL-586
chr14-52274314-DEL-271
chr14-52293004-DEL-258
chr14-52298089-DEL-191
chr14-52333307-DEL-66
chr14-52342229-DEL-215
chr14-52347222-DEL-242
chr14-52351859-DEL-129
chr14-52359818-DEL-199
chr14-52372531-DEL-232
chr14-52393163-DEL-59
chr14-52394133-DEL-104
chr14-52446558-DEL-100
chr14-52471381-DEL-151
chr14-52488476-DEL-196
chr14-52518720-DEL-98
chr14-52541185-DEL-199
chr14-52545602-DEL-440
chr14-52573317-DEL-846
chr14-52592246-DEL-218
chr14-52594134-DEL-2852
chr14-52607792-DEL-314
chr14-52613760-DEL-58
chr14-52621314-DEL-409
chr14-52663265-DEL-143
chr14-52709372-DEL-5959
chr14-52717447-DEL-62
chr14-52735260-DEL-6147
chr14-52747697-DEL-2152
chr14-52754682-DEL-202
chr14-52758361-DEL-59
chr14-52764546-DEL-1617
chr14-52787703-DEL-1062
chr14-52793915-DEL-6394
chr14-52813708-DEL-924
chr14-52881318-DEL-60
chr14-52889844-DEL-808
chr14-52894570-DEL-313
chr14-52895893-DEL-814
chr14-52898226-DEL-968
chr14-52904580-DEL-139
chr14-52910222-DEL-105
chr14-52910476-DEL-6683
chr14-52924671-DEL-574
chr14-52926252-DEL-235
chr14-52940908-DEL-903
chr14-52950984-DEL-54
chr14-52958919-DEL-1605

chr14-52962279-DEL-240
chr14-52975003-DEL-3194
chr14-53012165-DEL-1433
chr14-53030100-DEL-2327
chr14-53049405-DEL-1789
chr14-53076818-DEL-56
chr14-53090332-DEL-161
chr14-53095612-DEL-112
chr14-53098434-DEL-1779
chr14-53140209-DEL-540
chr14-53144800-DEL-54
chr14-53145513-DEL-357
chr14-53157579-DEL-8586
chr14-53212099-DEL-973
chr14-53216805-DEL-4183
chr14-53234234-DEL-4560
chr14-53248876-DEL-94
chr14-53249865-DEL-6904
chr14-53257637-DEL-70
chr14-53262050-DEL-83
chr14-53268578-DEL-101
chr14-53271090-DEL-274
chr14-53284239-DEL-233
chr14-53284770-DEL-9719
chr14-53286463-DEL-705
chr14-53297181-DEL-12132
chr14-53347579-DEL-221
chr14-53348889-DEL-2922
chr14-53349400-DEL-75
chr14-53355482-DEL-374
chr14-53363056-DEL-2362
chr14-53372005-DEL-54
chr14-53376741-DEL-272
chr14-53382101-DEL-170
chr14-53383133-DEL-3753
chr14-53401376-DEL-161
chr14-53409394-DEL-1779
chr14-53451137-DEL-535
chr14-53455729-DEL-56
chr14-53458876-DEL-126
chr14-53504209-DEL-96
chr14-53505227-DEL-58
chr14-53518511-DEL-125
chr14-53523234-DEL-973
chr14-53545405-DEL-4560
chr14-53552505-DEL-252
chr14-53561821-DEL-102
chr14-53583961-DEL-267
chr14-53590408-DEL-56
chr14-53590753-DEL-290
chr14-53597607-DEL-741
chr14-53600053-DEL-1227
chr14-53608166-DEL-430
chr14-53621815-DEL-97
chr14-53623397-DEL-203
chr14-53632942-DEL-234
chr14-53640191-DEL-2308
chr14-53647111-DEL-6744
chr14-53654228-DEL-52
chr14-53687487-DEL-204
chr14-53703367-DEL-1436
chr14-53708067-DEL-51
chr14-53711517-DEL-53
chr14-53751413-DEL-226
chr14-53766081-DEL-69
chr14-53783076-DEL-163
chr14-53785285-DEL-482
chr14-53785981-DEL-4180
chr14-53798619-DEL-2125
chr14-53801095-DEL-7947
chr14-53823053-DEL-66
chr14-53826214-DEL-1870
chr14-53835887-DEL-212
chr14-53854440-DEL-59
chr14-53860752-DEL-64
chr14-53871538-DEL-105
chr14-53886680-DEL-56

chr14-53897463-DEL-166
chr14-53898908-DEL-1076
chr14-53909278-DEL-4450
chr14-53909823-DEL-2603
chr14-53910120-DEL-203
chr14-53922876-DEL-8815
chr14-53944591-DEL-63
chr14-53951162-DEL-51
chr14-53953145-DEL-115
chr14-53956684-DEL-371
chr14-53958519-DEL-389
chr14-53965234-DEL-101
chr14-53967844-DEL-70
chr14-53982967-DEL-2176
chr14-53991824-DEL-1558
chr14-54016023-DEL-54
chr14-54119163-DEL-71
chr14-54165206-DEL-71
chr14-54166598-DEL-7276
chr14-54174211-DEL-7205
chr14-54175022-DEL-66
chr14-54184612-DEL-85
chr14-54192158-DEL-521
chr14-54277118-DEL-398
chr14-54332529-DEL-60
chr14-54337914-DEL-72
chr14-54340370-DEL-88
chr14-54426246-DEL-64
chr14-54444105-DEL-220
chr14-54447417-DEL-64
chr14-54500970-DEL-72
chr14-54502268-DEL-104
chr14-54542703-DEL-352
chr14-54543259-DEL-1999
chr14-54556375-DEL-197
chr14-54567290-DEL-67
chr14-54575435-DEL-105
chr14-54576896-DEL-197
chr14-54580358-DEL-212
chr14-54594277-DEL-81
chr14-54599596-DEL-173
chr14-54622828-DEL-53
chr14-54630211-DEL-1336
chr14-54636972-DEL-206
chr14-54648757-DEL-80
chr14-54654564-DEL-52
chr14-54658266-DEL-246
chr14-54748065-DEL-172
chr14-54763295-DEL-97
chr14-54776830-DEL-226
chr14-54781193-DEL-100
chr14-54781577-DEL-62
chr14-54790080-DEL-86
chr14-54803002-DEL-209
chr14-54829304-DEL-215
chr14-54895882-DEL-419
chr14-54928796-DEL-1271
chr14-54946253-DEL-76
chr14-54951016-DEL-70
chr14-54955550-DEL-55
chr14-55030436-DEL-58
chr14-55071259-DEL-335
chr14-55073334-DEL-199
chr14-55095071-DEL-66
chr14-55102371-DEL-194
chr14-55104912-DEL-54
chr14-55137310-DEL-52
chr14-55209272-DEL-244
chr14-55214942-DEL-81
chr14-55221398-DEL-591
chr14-55228810-DEL-242
chr14-55240534-DEL-143
chr14-55260375-DEL-72
chr14-55320565-DEL-89
chr14-55372668-DEL-2462
chr14-55404547-DEL-88
chr14-55415012-DEL-75

chr14-55418894-DEL-87
chr14-55420206-DEL-76
chr14-55461226-DEL-55
chr14-55498190-DEL-416
chr14-55515587-DEL-53
chr14-55518453-DEL-1169
chr14-55522235-DEL-56
chr14-55524629-DEL-161
chr14-55525019-DEL-911
chr14-55532488-DEL-115
chr14-55544742-DEL-106
chr14-55547832-DEL-52
chr14-55583792-DEL-72
chr14-55604491-DEL-86
chr14-55605333-DEL-74
chr14-55613327-DEL-140
chr14-55699154-DEL-114
chr14-55710513-DEL-647
chr14-55722655-DEL-145
chr14-55769941-DEL-445
chr14-55773351-DEL-52
chr14-55789841-DEL-174
chr14-55820256-DEL-117
chr14-55837799-DEL-304
chr14-55838332-DEL-75
chr14-55845782-DEL-212
chr14-55848732-DEL-1993
chr14-55851639-DEL-62
chr14-55882825-DEL-188
chr14-55891255-DEL-103
chr14-55891416-DEL-685
chr14-55901416-DEL-121
chr14-55925395-DEL-228
chr14-55978440-DEL-168
chr14-55993345-DEL-574
chr14-55997507-DEL-332
chr14-56042149-DEL-6147
chr14-56049482-DEL-221
chr14-56074009-DEL-201
chr14-56078819-DEL-162
chr14-56148916-DEL-710
chr14-56160759-DEL-63
chr14-56186788-DEL-256
chr14-56193209-DEL-65
chr14-56198724-DEL-200
chr14-56214975-DEL-93
chr14-56235224-DEL-703
chr14-56246577-DEL-338
chr14-56305170-DEL-3913
chr14-56381779-DEL-415
chr14-56425718-DEL-13999
chr14-56432705-DEL-129
chr14-56438208-DEL-731
chr14-56464712-DEL-1588
chr14-56490543-DEL-53
chr14-56513219-DEL-266
chr14-56513679-DEL-542
chr14-56526710-DEL-157
chr14-56542838-DEL-111
chr14-56547654-DEL-586
chr14-56549598-DEL-57
chr14-56557511-DEL-894
chr14-56564948-DEL-72
chr14-56580749-DEL-4140
chr14-56594679-DEL-640
chr14-56620385-DEL-100
chr14-56630148-DEL-519
chr14-56649606-DEL-1037
chr14-56693865-DEL-720
chr14-56702562-DEL-328
chr14-56711424-DEL-169
chr14-56763582-DEL-394
chr14-56773443-DEL-2020
chr14-56775265-DEL-1990
chr14-56782231-DEL-1351
chr14-56783726-DEL-2367
chr14-56794130-DEL-56

chr14-56797751-DEL-440
chr14-56803171-DEL-135
chr14-56826312-DEL-85
chr14-56826982-DEL-228
chr14-56840382-DEL-64
chr14-56846976-DEL-198
chr14-56854377-DEL-78
chr14-56874215-DEL-877
chr14-56881777-DEL-197
chr14-56883717-DEL-524
chr14-56918692-DEL-2344
chr14-56971388-DEL-140
chr14-57024786-DEL-681
chr14-57025807-DEL-453
chr14-57026561-DEL-1620
chr14-57077186-DEL-164
chr14-57077530-DEL-445
chr14-57122312-DEL-199
chr14-57134339-DEL-212
chr14-57143783-DEL-185
chr14-57186009-DEL-141
chr14-57217214-DEL-78
chr14-57236293-DEL-2329
chr14-57241941-DEL-58
chr14-57264287-DEL-466
chr14-57282722-DEL-61
chr14-57284471-DEL-61
chr14-57286054-DEL-78
chr14-57320732-DEL-77
chr14-57410122-DEL-537
chr14-57434283-DEL-66
chr14-57447531-DEL-66
chr14-57460881-DEL-437
chr14-57490287-DEL-1370
chr14-57507373-DEL-3823
chr14-57515972-DEL-58
chr14-57536477-DEL-418
chr14-57559170-DEL-54
chr14-57595594-DEL-182
chr14-57618230-DEL-220
chr14-57625138-DEL-98
chr14-57625976-DEL-64
chr14-57640119-DEL-79
chr14-57652558-DEL-5246
chr14-57658758-DEL-609
chr14-57665834-DEL-772
chr14-57731750-DEL-51
chr14-57738759-DEL-220
chr14-57742718-DEL-87
chr14-57789269-DEL-162
chr14-57794796-DEL-179
chr14-57796584-DEL-767
chr14-57796586-DEL-123
chr14-57797409-DEL-72
chr14-57842077-DEL-56
chr14-57859926-DEL-471
chr14-57890859-DEL-197
chr14-57894327-DEL-65
chr14-57896506-DEL-226
chr14-57919098-DEL-68
chr14-57924820-DEL-72
chr14-57938385-DEL-402
chr14-57954897-DEL-55
chr14-57959346-DEL-58
chr14-57965943-DEL-518
chr14-57989056-DEL-62
chr14-57992507-DEL-230
chr14-57996598-DEL-178
chr14-58019197-DEL-76
chr14-58031187-DEL-72
chr14-58075980-DEL-78
chr14-58084947-DEL-76
chr14-58107626-DEL-109
chr14-58122201-DEL-161
chr14-58125987-DEL-171
chr14-58129164-DEL-622
chr14-58149408-DEL-1225

chr14-58149861-DEL-75
chr14-58157044-DEL-2229
chr14-58184181-DEL-569
chr14-58192563-DEL-1048
chr14-58196285-DEL-222
chr14-58205399-DEL-79
chr14-58224848-DEL-60
chr14-58230980-DEL-144
chr14-58247011-DEL-70
chr14-58265642-DEL-184
chr14-58266792-DEL-729
chr14-58267675-DEL-76
chr14-58271588-DEL-168
chr14-58325959-DEL-53
chr14-58327811-DEL-162
chr14-58348364-DEL-64
chr14-58357370-DEL-150
chr14-58361093-DEL-146
chr14-58365784-DEL-226
chr14-58376294-DEL-82
chr14-58419124-DEL-205
chr14-58434282-DEL-5096
chr14-58439910-DEL-69
chr14-58459580-DEL-1729
chr14-58495370-DEL-74
chr14-58499081-DEL-92
chr14-58517556-DEL-96
chr14-58555577-DEL-59
chr14-58562142-DEL-464
chr14-58587439-DEL-200
chr14-58660985-DEL-92
chr14-58687402-DEL-703
chr14-58772917-DEL-895
chr14-58795610-DEL-120
chr14-58803127-DEL-92
chr14-58805699-DEL-1698
chr14-58823793-DEL-1157
chr14-58824197-DEL-399
chr14-58837575-DEL-1645
chr14-58849229-DEL-341
chr14-58859072-DEL-400
chr14-58863215-DEL-259
chr14-58899677-DEL-78
chr14-58905769-DEL-82
chr14-58933385-DEL-804
chr14-58938017-DEL-502
chr14-58962580-DEL-2156
chr14-59044708-DEL-2090
chr14-59051521-DEL-193
chr14-59055508-DEL-191
chr14-59067738-DEL-3794
chr14-59109067-DEL-630
chr14-59128502-DEL-268
chr14-59130900-DEL-724
chr14-59139481-DEL-88
chr14-59158728-DEL-1841
chr14-59185574-DEL-6280
chr14-59192558-DEL-1019
chr14-59193590-DEL-1386
chr14-59239389-DEL-1659
chr14-59258663-DEL-1299
chr14-59288675-DEL-117
chr14-59293393-DEL-66
chr14-59312831-DEL-167
chr14-59316946-DEL-54
chr14-59336733-DEL-271
chr14-59338481-DEL-414
chr14-59394297-DEL-139
chr14-59402229-DEL-6516
chr14-59415461-DEL-54
chr14-59421485-DEL-100
chr14-59433214-DEL-86
chr14-59454223-DEL-64
chr14-59500195-DEL-7384
chr14-59537430-DEL-392
chr14-59539370-DEL-6408
chr14-59627831-DEL-85

chr14-59664041-DEL-418
chr14-59683306-DEL-178
chr14-59692284-DEL-62
chr14-59720159-DEL-56
chr14-59726056-DEL-2103
chr14-59742331-DEL-100
chr14-59745780-DEL-846
chr14-59762394-DEL-54
chr14-59767479-DEL-526
chr14-59775996-DEL-198
chr14-59821453-DEL-52
chr14-59830902-DEL-69
chr14-59856203-DEL-165
chr14-59889734-DEL-106
chr14-59892676-DEL-52
chr14-59905708-DEL-175
chr14-59916481-DEL-61
chr14-59916962-DEL-182
chr14-59926141-DEL-158
chr14-59946188-DEL-766
chr14-59953995-DEL-144
chr14-59958436-DEL-88
chr14-60007558-DEL-185
chr14-60025437-DEL-180
chr14-60033259-DEL-101
chr14-60041832-DEL-115
chr14-60048037-DEL-97
chr14-60070687-DEL-1929
chr14-60075215-DEL-468
chr14-60092424-DEL-758
chr14-60128044-DEL-163
chr14-60164007-DEL-2812
chr14-60188072-DEL-53
chr14-60217139-DEL-8316
chr14-60229199-DEL-3336
chr14-60229293-DEL-359
chr14-60231146-DEL-240
chr14-60231997-DEL-156
chr14-60247898-DEL-353
chr14-60256611-DEL-416
chr14-60261401-DEL-112
chr14-60263016-DEL-71
chr14-60278002-DEL-971
chr14-60278195-DEL-307
chr14-60279087-DEL-109
chr14-60282812-DEL-176
chr14-60314640-DEL-490
chr14-60343091-DEL-88
chr14-60356627-DEL-71
chr14-60369694-DEL-530
chr14-60376073-DEL-238
chr14-60419322-DEL-72
chr14-60424577-DEL-56
chr14-60429255-DEL-69
chr14-60445171-DEL-1191
chr14-60486985-DEL-347
chr14-60498098-DEL-1283
chr14-60525820-DEL-724
chr14-60552009-DEL-1687
chr14-60564580-DEL-12637
chr14-60587674-DEL-55
chr14-60596789-DEL-54
chr14-60599916-DEL-54
chr14-60673456-DEL-836
chr14-60675145-DEL-243
chr14-60684289-DEL-72
chr14-60695758-DEL-145
chr14-60696960-DEL-64
chr14-60703568-DEL-385
chr14-60713845-DEL-3923
chr14-60731859-DEL-95
chr14-60735127-DEL-160
chr14-60801293-DEL-72
chr14-60807487-DEL-142
chr14-60817135-DEL-542
chr14-60820539-DEL-1560
chr14-60832494-DEL-108

chr14-60833838-DEL-214
chr14-60852922-DEL-89
chr14-60905440-DEL-344
chr14-60906542-DEL-338
chr14-60938903-DEL-70
chr14-60951526-DEL-761
chr14-60976668-DEL-118
chr14-60985991-DEL-52
chr14-60989919-DEL-235
chr14-60997595-DEL-63
chr14-61019117-DEL-176
chr14-61020184-DEL-181
chr14-61024983-DEL-157
chr14-61073549-DEL-194
chr14-61075184-DEL-174
chr14-61120105-DEL-1709
chr14-61125899-DEL-60
chr14-61156440-DEL-62
chr14-61166306-DEL-1248
chr14-61186552-DEL-227
chr14-61191270-DEL-2233
chr14-61196807-DEL-3717
chr14-61212400-DEL-101
chr14-61239001-DEL-58
chr14-61254169-DEL-88
chr14-61261373-DEL-97
chr14-61265337-DEL-125
chr14-61291845-DEL-261
chr14-61307921-DEL-54
chr14-61344694-DEL-125
chr14-61352951-DEL-350
chr14-61367329-DEL-150
chr14-61374553-DEL-58
chr14-61382951-DEL-68
chr14-61453855-DEL-203
chr14-61459343-DEL-192
chr14-61464203-DEL-266
chr14-61478091-DEL-1649
chr14-61499325-DEL-80
chr14-61499453-DEL-1106
chr14-61511877-DEL-87
chr14-61521627-DEL-1931
chr14-61536361-DEL-82
chr14-61550678-DEL-1287
chr14-61560983-DEL-164
chr14-61594638-DEL-134
chr14-61606090-DEL-179
chr14-61697372-DEL-66
chr14-61707202-DEL-163
chr14-61708731-DEL-160
chr14-61723075-DEL-257
chr14-61726316-DEL-96
chr14-61741789-DEL-53
chr14-61755004-DEL-770
chr14-61788380-DEL-69
chr14-61806266-DEL-195
chr14-61812203-DEL-90
chr14-61814725-DEL-52
chr14-61815942-DEL-159
chr14-61817918-DEL-6275
chr14-61840939-DEL-280
chr14-61843559-DEL-135
chr14-61855550-DEL-60
chr14-61856887-DEL-67
chr14-61858400-DEL-51
chr14-61891423-DEL-641
chr14-61892978-DEL-152
chr14-61896058-DEL-341
chr14-62019739-DEL-626
chr14-62040199-DEL-82
chr14-62043831-DEL-76
chr14-62072674-DEL-74
chr14-62084662-DEL-198
chr14-62088092-DEL-139
chr14-62092509-DEL-1630
chr14-62105548-DEL-170
chr14-62109612-DEL-223

chr14-62131685-DEL-134
chr14-62189172-DEL-1330
chr14-62194754-DEL-56
chr14-62234984-DEL-108
chr14-62245238-DEL-57
chr14-62257167-DEL-98
chr14-62272552-DEL-54
chr14-62287842-DEL-89
chr14-62310291-DEL-755
chr14-62337596-DEL-664
chr14-62342203-DEL-213
chr14-62344902-DEL-5630
chr14-62404279-DEL-763
chr14-62418085-DEL-90
chr14-62419312-DEL-100
chr14-62449996-DEL-1387
chr14-62454732-DEL-237
chr14-62465865-DEL-58
chr14-62470038-DEL-184
chr14-62509510-DEL-400
chr14-62515579-DEL-109
chr14-62605156-DEL-128
chr14-62619673-DEL-91
chr14-62641714-DEL-81
chr14-62664302-DEL-104
chr14-62666006-DEL-1439
chr14-62724974-DEL-212
chr14-62765904-DEL-52
chr14-62798358-DEL-52
chr14-62818535-DEL-67
chr14-62822780-DEL-60
chr14-62827644-DEL-872
chr14-62906017-DEL-64
chr14-62969976-DEL-102
chr14-63022500-DEL-104
chr14-63045059-DEL-173
chr14-63051943-DEL-69
chr14-63097434-DEL-62
chr14-63154030-DEL-119
chr14-63224354-DEL-233
chr14-63261242-DEL-524
chr14-63312090-DEL-151
chr14-63321619-DEL-201
chr14-63376738-DEL-54
chr14-63378395-DEL-97
chr14-63388785-DEL-54
chr14-63395232-DEL-301
chr14-63406444-DEL-2115
chr14-63411704-DEL-201
chr14-63495910-DEL-58
chr14-63534390-DEL-175
chr14-63570892-DEL-654
chr14-63630606-DEL-3503
chr14-63639127-DEL-94
chr14-63663686-DEL-74
chr14-63698099-DEL-58
chr14-63701080-DEL-400
chr14-63753583-DEL-54
chr14-63762540-DEL-146
chr14-63769956-DEL-53
chr14-63774014-DEL-202
chr14-63786377-DEL-1115
chr14-63797396-DEL-89
chr14-63804835-DEL-199
chr14-63840798-DEL-191
chr14-63860886-DEL-100
chr14-63920112-DEL-119
chr14-63933939-DEL-69
chr14-63939715-DEL-54
chr14-63996258-DEL-160
chr14-64054729-DEL-52
chr14-64212580-DEL-70
chr14-64215500-DEL-73
chr14-64351614-DEL-81
chr14-64353487-DEL-90
chr14-64354974-DEL-546
chr14-64392961-DEL-76

chr14-64425485-DEL-356
chr14-64441979-DEL-98
chr14-64477604-DEL-72
chr14-64524253-DEL-128
chr14-64550384-DEL-60
chr14-64648334-DEL-65
chr14-64752994-DEL-1102
chr14-64775953-DEL-59
chr14-64841441-DEL-195
chr14-64850077-DEL-203
chr14-64882817-DEL-364
chr14-64904968-DEL-64
chr14-65118077-DEL-80
chr14-65175572-DEL-118
chr14-65175896-DEL-837
chr14-65192214-DEL-174
chr14-65224156-DEL-259
chr14-65224243-DEL-1220
chr14-65224714-DEL-739
chr14-65259354-DEL-132
chr14-65273378-DEL-192
chr14-65277773-DEL-177
chr14-65281820-DEL-164
chr14-65284818-DEL-101
chr14-65304707-DEL-209
chr14-65339629-DEL-103
chr14-65384389-DEL-59
chr14-65393049-DEL-607
chr14-65403997-DEL-51
chr14-65405863-DEL-289
chr14-65410307-DEL-557
chr14-65427408-DEL-236
chr14-65429241-DEL-52
chr14-65535433-DEL-81
chr14-65544555-DEL-158
chr14-65575821-DEL-52
chr14-65634324-DEL-52
chr14-65650266-DEL-84
chr14-65708091-DEL-5962
chr14-65711745-DEL-901
chr14-65724986-DEL-1137
chr14-65742948-DEL-69
chr14-65778478-DEL-272
chr14-65811363-DEL-261
chr14-65835846-DEL-67
chr14-65862567-DEL-240
chr14-65863667-DEL-383
chr14-65867801-DEL-145
chr14-65890067-DEL-3516
chr14-65926233-DEL-108
chr14-65947157-DEL-116
chr14-65960624-DEL-66
chr14-65995343-DEL-417
chr14-66005084-DEL-914
chr14-66006390-DEL-512
chr14-66006971-DEL-225
chr14-66023510-DEL-93
chr14-66063006-DEL-74
chr14-66110690-DEL-238
chr14-66129858-DEL-1377
chr14-66228196-DEL-65
chr14-66258061-DEL-344
chr14-66285504-DEL-610
chr14-66318672-DEL-403
chr14-66320534-DEL-145
chr14-66337036-DEL-74
chr14-66338317-DEL-73
chr14-66372113-DEL-185
chr14-66374626-DEL-94
chr14-66376214-DEL-202
chr14-66404804-DEL-56
chr14-66412070-DEL-91
chr14-66417900-DEL-96
chr14-66421774-DEL-122
chr14-66461715-DEL-75
chr14-66471947-DEL-81
chr14-66487555-DEL-676

chr14-66504601-DEL-68
chr14-66610150-DEL-78
chr14-66614233-DEL-195
chr14-66620292-DEL-72
chr14-66628665-DEL-397
chr14-66647840-DEL-62
chr14-66649891-DEL-137
chr14-66661155-DEL-62
chr14-66661667-DEL-811
chr14-66662264-DEL-226
chr14-66671504-DEL-79
chr14-66733040-DEL-86
chr14-66749852-DEL-102
chr14-66806323-DEL-265
chr14-66814020-DEL-934
chr14-66833745-DEL-58
chr14-66837796-DEL-1731
chr14-66899852-DEL-54
chr14-66911048-DEL-1220
chr14-66924872-DEL-82
chr14-66925478-DEL-994
chr14-66941880-DEL-236
chr14-66953322-DEL-1555
chr14-66978333-DEL-147
chr14-67000248-DEL-180
chr14-67003810-DEL-106
chr14-67013089-DEL-343
chr14-67050897-DEL-50
chr14-67055238-DEL-67
chr14-67065476-DEL-251
chr14-67086783-DEL-80
chr14-67109879-DEL-52
chr14-67141448-DEL-986
chr14-67151194-DEL-160
chr14-67209425-DEL-136
chr14-67216559-DEL-60
chr14-67240989-DEL-55
chr14-67254739-DEL-181
chr14-67278374-DEL-911
chr14-67282156-DEL-97
chr14-67294626-DEL-53
chr14-67325258-DEL-56
chr14-67331739-DEL-63
chr14-67341008-DEL-295
chr14-67399807-DEL-192
chr14-67428271-DEL-52
chr14-67441290-DEL-72
chr14-67442971-DEL-347
chr14-67449444-DEL-61
chr14-67465165-DEL-70
chr14-67479293-DEL-52
chr14-67488634-DEL-69
chr14-67594172-DEL-71
chr14-67636473-DEL-158
chr14-67676495-DEL-110
chr14-67688013-DEL-272
chr14-67701102-DEL-61
chr14-67705013-DEL-206
chr14-67745990-DEL-191
chr14-67754306-DEL-614
chr14-67755059-DEL-115
chr14-67793049-DEL-52
chr14-67810481-DEL-56
chr14-67835904-DEL-197
chr14-67864247-DEL-281
chr14-67882617-DEL-53
chr14-67886588-DEL-60
chr14-67887211-DEL-2636
chr14-67894012-DEL-85
chr14-67911432-DEL-805
chr14-67929499-DEL-242
chr14-67935869-DEL-74
chr14-67937688-DEL-60
chr14-67941890-DEL-71
chr14-67991615-DEL-190
chr14-67999371-DEL-108
chr14-68002737-DEL-83

chr14-68042058-DEL-108
chr14-68046772-DEL-202
chr14-68053138-DEL-60
chr14-68061007-DEL-123
chr14-68088636-DEL-90
chr14-68090006-DEL-60
chr14-68134326-DEL-83
chr14-68178267-DEL-60
chr14-68196556-DEL-52
chr14-68197808-DEL-59
chr14-68202366-DEL-413
chr14-68209111-DEL-56
chr14-68232737-DEL-60
chr14-68239917-DEL-352
chr14-68240641-DEL-172
chr14-68268871-DEL-55
chr14-68282850-DEL-151
chr14-68287849-DEL-268
chr14-68297389-DEL-121
chr14-68329257-DEL-444
chr14-68335083-DEL-65
chr14-68344615-DEL-1099
chr14-68345837-DEL-1187
chr14-68377049-DEL-300
chr14-68404230-DEL-155
chr14-68405305-DEL-155
chr14-68407765-DEL-263
chr14-68416226-DEL-324
chr14-68451785-DEL-1094
chr14-68470690-DEL-223
chr14-68491942-DEL-6433
chr14-68505189-DEL-131
chr14-68508530-DEL-7194
chr14-68521608-DEL-128
chr14-68522175-DEL-2517
chr14-68547854-DEL-1410
chr14-68568834-DEL-91
chr14-68612078-DEL-181
chr14-68615728-DEL-295
chr14-68713316-DEL-1400
chr14-68713316-DEL-797
chr14-68713447-DEL-2010
chr14-68730378-DEL-202
chr14-68760478-DEL-222
chr14-68778515-DEL-317
chr14-68779499-DEL-80
chr14-68788454-DEL-2812
chr14-68791381-DEL-1801
chr14-68960093-DEL-57
chr14-68975698-DEL-238
chr14-68978083-DEL-52
chr14-68981377-DEL-60
chr14-69009619-DEL-1136
chr14-69055901-DEL-79
chr14-69079128-DEL-154
chr14-69109270-DEL-4492
chr14-69127485-DEL-171
chr14-69130493-DEL-100
chr14-69207990-DEL-436
chr14-69212293-DEL-56
chr14-69257576-DEL-53
chr14-69275018-DEL-186
chr14-69311284-DEL-3208
chr14-69347228-DEL-416
chr14-69358902-DEL-61
chr14-69369990-DEL-74
chr14-69399458-DEL-260
chr14-69407235-DEL-107
chr14-69440772-DEL-238
chr14-69443600-DEL-54
chr14-69453294-DEL-68
chr14-69455839-DEL-106
chr14-69465013-DEL-57
chr14-69498656-DEL-145
chr14-69504381-DEL-276
chr14-69506518-DEL-72
chr14-69997082-DEL-184

chr14-70153436-DEL-388
chr14-70167106-DEL-145
chr14-70182779-DEL-67
chr14-70204835-DEL-81
chr14-70230170-DEL-51
chr14-70288522-DEL-63
chr14-70290532-DEL-110
chr14-70342471-DEL-69
chr14-70401231-DEL-143
chr14-70420239-DEL-53
chr14-70427578-DEL-199
chr14-70450928-DEL-150
chr14-70473580-DEL-145
chr14-70502782-DEL-279
chr14-70576137-DEL-212
chr14-70661724-DEL-401
chr14-70670249-DEL-60
chr14-70674778-DEL-64
chr14-70819010-DEL-74
chr14-70838993-DEL-78
chr14-70870069-DEL-71
chr14-70891640-DEL-187
chr14-70934450-DEL-209
chr14-70937513-DEL-72
chr14-70962814-DEL-96
chr14-70963979-DEL-270
chr14-70964319-DEL-261
chr14-70982291-DEL-5081
chr14-70994084-DEL-98
chr14-70998276-DEL-201
chr14-71023409-DEL-232
chr14-71048080-DEL-1038
chr14-71094253-DEL-96
chr14-71106842-DEL-672
chr14-71120449-DEL-353
chr14-71146941-DEL-103
chr14-71151159-DEL-401
chr14-71181231-DEL-492
chr14-71221634-DEL-71
chr14-71272548-DEL-396
chr14-71279016-DEL-60
chr14-71302202-DEL-65
chr14-71328595-DEL-247
chr14-71346147-DEL-92
chr14-71371779-DEL-106
chr14-71374076-DEL-142
chr14-71383048-DEL-62
chr14-71406924-DEL-2748
chr14-71418904-DEL-97
chr14-71517765-DEL-120
chr14-71544569-DEL-56
chr14-71563733-DEL-107
chr14-71577558-DEL-137
chr14-71623737-DEL-1579
chr14-71628866-DEL-13165
chr14-71647792-DEL-215
chr14-71647904-DEL-963
chr14-71655463-DEL-90
chr14-71671491-DEL-53
chr14-71680837-DEL-844
chr14-71681735-DEL-3017
chr14-71712433-DEL-59
chr14-71743253-DEL-79
chr14-71819105-DEL-421
chr14-71827014-DEL-52
chr14-71838723-DEL-153
chr14-71854442-DEL-485
chr14-71873631-DEL-177
chr14-71983971-DEL-67
chr14-72018417-DEL-104
chr14-72020880-DEL-8653
chr14-72021929-DEL-1055
chr14-72028600-DEL-208
chr14-72032956-DEL-63
chr14-72074254-DEL-54
chr14-72096424-DEL-53
chr14-72139370-DEL-3144

chr14-72178311-DEL-267
chr14-72224693-DEL-256
chr14-72225695-DEL-395
chr14-72226328-DEL-881
chr14-72275067-DEL-354
chr14-72316595-DEL-922
chr14-72383391-DEL-107
chr14-72393311-DEL-87
chr14-72394311-DEL-310
chr14-72403039-DEL-317
chr14-72416064-DEL-70
chr14-72447384-DEL-1486
chr14-72488198-DEL-1490
chr14-72501007-DEL-54
chr14-72506962-DEL-191
chr14-72560389-DEL-86
chr14-72593713-DEL-1138
chr14-72623076-DEL-180
chr14-72624929-DEL-102
chr14-72628022-DEL-75
chr14-72645378-DEL-324
chr14-72649630-DEL-628
chr14-72658682-DEL-4248
chr14-72675931-DEL-546
chr14-72686604-DEL-90
chr14-72706278-DEL-84
chr14-72739398-DEL-64
chr14-72750655-DEL-68
chr14-72757685-DEL-6563
chr14-72762959-DEL-1119
chr14-72883574-DEL-799
chr14-72899755-DEL-140
chr14-72910171-DEL-112
chr14-72920020-DEL-300
chr14-72926031-DEL-68
chr14-72966761-DEL-60
chr14-72981134-DEL-64
chr14-73015900-DEL-120
chr14-73018427-DEL-143
chr14-73076297-DEL-114
chr14-73095126-DEL-72
chr14-73128904-DEL-400
chr14-73143724-DEL-56
chr14-73146766-DEL-3401
chr14-73150192-DEL-4217
chr14-73164009-DEL-78
chr14-73165191-DEL-59
chr14-73202097-DEL-74
chr14-73204004-DEL-1368
chr14-73205460-DEL-6220
chr14-73214405-DEL-496
chr14-73216784-DEL-55
chr14-73235494-DEL-62
chr14-73250835-DEL-116
chr14-73264281-DEL-70
chr14-73274613-DEL-208
chr14-73290963-DEL-59
chr14-73321813-DEL-245
chr14-73321957-DEL-648
chr14-73327185-DEL-428
chr14-73378885-DEL-54
chr14-73381375-DEL-163
chr14-73392945-DEL-269
chr14-73412598-DEL-168
chr14-73416319-DEL-66
chr14-73478291-DEL-58
chr14-73490253-DEL-154
chr14-73491722-DEL-53
chr14-73503823-DEL-122
chr14-73538908-DEL-69
chr14-73566340-DEL-234
chr14-73594169-DEL-98
chr14-73598641-DEL-472
chr14-73602633-DEL-192
chr14-73642586-DEL-227
chr14-73645323-DEL-53
chr14-73669881-DEL-56

chr14-73674231-DEL-1200
chr14-73681838-DEL-221
chr14-73686436-DEL-52
chr14-73705779-DEL-191
chr14-73719950-DEL-65
chr14-73745790-DEL-68
chr14-73748877-DEL-58
chr14-73758525-DEL-245
chr14-73772227-DEL-64
chr14-73820288-DEL-211
chr14-73838315-DEL-56
chr14-73853650-DEL-399
chr14-73921967-DEL-197
chr14-73963416-DEL-432
chr14-73996982-DEL-179
chr14-74000989-DEL-4429
chr14-74022346-DEL-101
chr14-74055751-DEL-1534
chr14-74085006-DEL-2281
chr14-74089177-DEL-341
chr14-74093264-DEL-1350
chr14-74096846-DEL-1907
chr14-74136001-DEL-134
chr14-74156468-DEL-115
chr14-74163485-DEL-58
chr14-74188604-DEL-101
chr14-74199448-DEL-53
chr14-74202235-DEL-111
chr14-74221729-DEL-979
chr14-74238958-DEL-5106
chr14-74268809-DEL-63
chr14-74278266-DEL-512
chr14-74279766-DEL-478
chr14-74287898-DEL-1368
chr14-74292669-DEL-208
chr14-74315864-DEL-55
chr14-74318561-DEL-84
chr14-74324257-DEL-3414
chr14-74348578-DEL-266
chr14-74352134-DEL-393
chr14-74388058-DEL-104
chr14-74392371-DEL-74
chr14-74392976-DEL-75
chr14-74401183-DEL-1352
chr14-74448577-DEL-150
chr14-74499244-DEL-997
chr14-74525530-DEL-702
chr14-74528760-DEL-575
chr14-74570208-DEL-77
chr14-74571712-DEL-62
chr14-74590123-DEL-114
chr14-74611120-DEL-3381
chr14-74658279-DEL-56
chr14-74668343-DEL-65
chr14-74672059-DEL-1565
chr14-74672151-DEL-6109
chr14-74673668-DEL-4584
chr14-74707053-DEL-301
chr14-74709051-DEL-2233
chr14-74728290-DEL-240
chr14-74785513-DEL-1891
chr14-74803441-DEL-60
chr14-74860173-DEL-278
chr14-74890587-DEL-52
chr14-74912769-DEL-477
chr14-74948133-DEL-114
chr14-74988768-DEL-620
chr14-75013379-DEL-7549
chr14-75013829-DEL-374
chr14-75074646-DEL-55
chr14-75091967-DEL-403
chr14-75101305-DEL-268
chr14-75133286-DEL-2313
chr14-75154488-DEL-296
chr14-75163123-DEL-208
chr14-75172503-DEL-977
chr14-75189328-DEL-5302

chr14-75190014-DEL-57
chr14-75214167-DEL-249
chr14-75224170-DEL-127
chr14-75226648-DEL-55
chr14-75246534-DEL-64
chr14-75253023-DEL-218
chr14-75261763-DEL-61
chr14-75285900-DEL-52
chr14-75295041-DEL-91
chr14-75320161-DEL-368
chr14-75362246-DEL-172
chr14-75380934-DEL-79
chr14-75394300-DEL-342
chr14-75399131-DEL-432
chr14-75426843-DEL-78
chr14-75456351-DEL-195
chr14-75460288-DEL-57
chr14-75494473-DEL-80
chr14-75574548-DEL-176
chr14-75595511-DEL-387
chr14-75608134-DEL-1592
chr14-75611616-DEL-84
chr14-75648149-DEL-180
chr14-75672254-DEL-7787
chr14-75672653-DEL-6184
chr14-75704457-DEL-141
chr14-75706546-DEL-69
chr14-75710921-DEL-96
chr14-75717841-DEL-70
chr14-75735096-DEL-1191
chr14-75761088-DEL-105
chr14-75776126-DEL-224
chr14-75819421-DEL-70
chr14-75822778-DEL-1182
chr14-75833344-DEL-1350
chr14-75855439-DEL-60
chr14-75860130-DEL-52
chr14-75872200-DEL-70
chr14-75890953-DEL-157
chr14-75893432-DEL-57
chr14-75896462-DEL-144
chr14-75900228-DEL-150
chr14-75930970-DEL-74
chr14-75938089-DEL-400
chr14-75943538-DEL-70
chr14-75948201-DEL-111
chr14-76003273-DEL-86
chr14-76016547-DEL-88
chr14-76020792-DEL-77
chr14-76034913-DEL-55
chr14-76041993-DEL-112
chr14-76051272-DEL-363
chr14-76060436-DEL-1091
chr14-76060653-DEL-641
chr14-76062190-DEL-473
chr14-76063301-DEL-72
chr14-76067890-DEL-660
chr14-76068888-DEL-64
chr14-76070938-DEL-2008
chr14-76075983-DEL-193
chr14-76086845-DEL-154
chr14-76094131-DEL-373
chr14-76096717-DEL-241
chr14-76097973-DEL-207
chr14-76103258-DEL-85
chr14-76122419-DEL-183
chr14-76139097-DEL-52
chr14-76159871-DEL-417
chr14-76170441-DEL-2585
chr14-76172276-DEL-2623
chr14-76198558-DEL-163
chr14-76233611-DEL-203
chr14-76260292-DEL-56
chr14-76269434-DEL-184
chr14-76280683-DEL-294
chr14-76285982-DEL-142
chr14-76287967-DEL-872

chr14-76300733-DEL-57
chr14-76338511-DEL-1214
chr14-76342736-DEL-54
chr14-76346680-DEL-655
chr14-76350261-DEL-192
chr14-76365582-DEL-218
chr14-76371896-DEL-153
chr14-76372869-DEL-120
chr14-76376959-DEL-674
chr14-76377558-DEL-190
chr14-76403440-DEL-533
chr14-76425556-DEL-78
chr14-76426286-DEL-342
chr14-76456726-DEL-112
chr14-76472249-DEL-58
chr14-76473396-DEL-1861
chr14-76525078-DEL-65
chr14-76537346-DEL-56
chr14-76589718-DEL-95
chr14-76613185-DEL-149
chr14-76620014-DEL-58
chr14-76622829-DEL-52
chr14-76647194-DEL-108
chr14-76700031-DEL-169
chr14-76701808-DEL-84
chr14-76708750-DEL-1416
chr14-76722371-DEL-1837
chr14-76726419-DEL-86
chr14-76790006-DEL-115
chr14-76808224-DEL-2677
chr14-76828982-DEL-435
chr14-76842253-DEL-56
chr14-76922215-DEL-53
chr14-76977372-DEL-96
chr14-76982649-DEL-110
chr14-76987991-DEL-273
chr14-77006468-DEL-514
chr14-77014967-DEL-145
chr14-77043898-DEL-152
chr14-77045411-DEL-146
chr14-77096291-DEL-472
chr14-77119085-DEL-64
chr14-77128079-DEL-177
chr14-77225393-DEL-128
chr14-77245345-DEL-52
chr14-77248276-DEL-213
chr14-77273379-DEL-124
chr14-77285856-DEL-2950
chr14-77285882-DEL-2700
chr14-77292589-DEL-2108
chr14-77298833-DEL-1117
chr14-77301242-DEL-250
chr14-77342439-DEL-542
chr14-77348161-DEL-70
chr14-77353750-DEL-5994
chr14-77364969-DEL-65
chr14-77368673-DEL-127
chr14-77391274-DEL-182
chr14-77400619-DEL-476
chr14-77453198-DEL-423
chr14-77456098-DEL-815
chr14-77508579-DEL-54
chr14-77557634-DEL-54
chr14-77561081-DEL-102
chr14-77574941-DEL-1279
chr14-77583010-DEL-60
chr14-77594832-DEL-84
chr14-77670593-DEL-387
chr14-77720570-DEL-113
chr14-77837575-DEL-57
chr14-77856956-DEL-61
chr14-77904715-DEL-58
chr14-77937477-DEL-202
chr14-77953513-DEL-311
chr14-77967227-DEL-54
chr14-77987337-DEL-104
chr14-78017891-DEL-122

chr14-78020699-DEL-1193
chr14-78026191-DEL-106
chr14-78050020-DEL-111
chr14-78067869-DEL-760
chr14-78142906-DEL-1178
chr14-78159079-DEL-55
chr14-78179523-DEL-112
chr14-78225109-DEL-56
chr14-78230274-DEL-73
chr14-78288115-DEL-57
chr14-78290801-DEL-63
chr14-78296855-DEL-53
chr14-78311378-DEL-54
chr14-78324250-DEL-537
chr14-78377613-DEL-400
chr14-78435398-DEL-357
chr14-78475838-DEL-75
chr14-78483791-DEL-155
chr14-78494421-DEL-470
chr14-78504798-DEL-148
chr14-78517984-DEL-84
chr14-78541285-DEL-98
chr14-78554538-DEL-259
chr14-78616624-DEL-87
chr14-78627287-DEL-182
chr14-78639107-DEL-1586
chr14-78681012-DEL-54
chr14-78685004-DEL-275
chr14-78712079-DEL-58
chr14-78725047-DEL-93
chr14-78771988-DEL-204
chr14-78779666-DEL-54
chr14-78798215-DEL-62
chr14-78799968-DEL-61
chr14-78815263-DEL-425
chr14-78852073-DEL-158
chr14-78860181-DEL-730
chr14-78944388-DEL-177
chr14-78977347-DEL-6543
chr14-78994803-DEL-74
chr14-79001231-DEL-228
chr14-79032746-DEL-70
chr14-79042048-DEL-111
chr14-79049960-DEL-73
chr14-79051713-DEL-1742
chr14-79056725-DEL-1393
chr14-79066163-DEL-164
chr14-79122279-DEL-238
chr14-79166341-DEL-1791
chr14-79202716-DEL-206
chr14-79203269-DEL-761
chr14-79222416-DEL-1478
chr14-79222467-DEL-180
chr14-79222521-DEL-809
chr14-79223500-DEL-311
chr14-79230649-DEL-93
chr14-79243622-DEL-73
chr14-79254249-DEL-1015
chr14-79255307-DEL-105
chr14-79305659-DEL-62
chr14-79328615-DEL-608
chr14-79346830-DEL-81
chr14-79351046-DEL-329
chr14-79370146-DEL-201
chr14-79373434-DEL-853
chr14-79405430-DEL-6194
chr14-79436768-DEL-121
chr14-79450869-DEL-128
chr14-79455152-DEL-60
chr14-79455908-DEL-84
chr14-79487211-DEL-300
chr14-79494706-DEL-332
chr14-79520049-DEL-1914
chr14-79522141-DEL-129
chr14-79533020-DEL-57
chr14-79549610-DEL-89
chr14-79629665-DEL-486

chr14-79652645-DEL-400
chr14-79659069-DEL-162
chr14-79679783-DEL-90
chr14-79706666-DEL-55
chr14-79711739-DEL-196
chr14-79713629-DEL-267
chr14-79723043-DEL-96
chr14-79728144-DEL-63
chr14-79731474-DEL-60
chr14-79732105-DEL-94
chr14-79741538-DEL-534
chr14-79780410-DEL-353
chr14-79796132-DEL-259
chr14-79797580-DEL-55
chr14-79815643-DEL-179
chr14-79821190-DEL-146
chr14-79828583-DEL-651
chr14-79828891-DEL-795
chr14-79829658-DEL-967
chr14-79830574-DEL-165
chr14-79846785-DEL-59
chr14-79857496-DEL-57
chr14-79870475-DEL-149
chr14-79894327-DEL-112
chr14-79900919-DEL-306
chr14-79900924-DEL-1028
chr14-79901428-DEL-825
chr14-79906078-DEL-52
chr14-79920529-DEL-259
chr14-79933419-DEL-463
chr14-79968458-DEL-56
chr14-79999181-DEL-273
chr14-80073082-DEL-424
chr14-80076460-DEL-119
chr14-80104304-DEL-122
chr14-80106702-DEL-871
chr14-80123380-DEL-73
chr14-80129368-DEL-292
chr14-80133354-DEL-945
chr14-80147991-DEL-82
chr14-80175569-DEL-425
chr14-80203696-DEL-217
chr14-80228704-DEL-100
chr14-80285912-DEL-2596
chr14-80301897-DEL-79
chr14-80314092-DEL-219
chr14-80326675-DEL-212
chr14-80334608-DEL-98
chr14-80380148-DEL-847
chr14-80397927-DEL-79
chr14-80409866-DEL-92
chr14-80416488-DEL-7967
chr14-80424600-DEL-416
chr14-80452382-DEL-243
chr14-80473956-DEL-468
chr14-80512290-DEL-2417
chr14-80549883-DEL-663
chr14-80560392-DEL-93
chr14-80592575-DEL-848
chr14-80596655-DEL-60
chr14-80630744-DEL-329
chr14-80652360-DEL-149
chr14-80656839-DEL-67
chr14-80659044-DEL-6241
chr14-80679682-DEL-165
chr14-80680950-DEL-60
chr14-80697803-DEL-945
chr14-80800818-DEL-54
chr14-80803140-DEL-475
chr14-80814995-DEL-7475
chr14-80889656-DEL-75
chr14-80892997-DEL-97
chr14-80896937-DEL-67
chr14-80902352-DEL-142
chr14-80924109-DEL-51
chr14-80941431-DEL-6164
chr14-80960979-DEL-65

chr14-80967964-DEL-715
chr14-80997839-DEL-61
chr14-81000502-DEL-1525
chr14-81026731-DEL-73
chr14-81037838-DEL-292
chr14-81043881-DEL-2316
chr14-81044941-DEL-477
chr14-81056079-DEL-89
chr14-81090573-DEL-72
chr14-81100180-DEL-199
chr14-81102537-DEL-155
chr14-81109347-DEL-847
chr14-81118631-DEL-64
chr14-81125896-DEL-675
chr14-81161201-DEL-208
chr14-81198497-DEL-67
chr14-81211513-DEL-215
chr14-81239996-DEL-52
chr14-81269797-DEL-52
chr14-81315453-DEL-70
chr14-81315776-DEL-722
chr14-81325282-DEL-7190
chr14-81344058-DEL-990
chr14-81348128-DEL-637
chr14-81353713-DEL-159
chr14-81363112-DEL-126
chr14-81387199-DEL-1569
chr14-81388105-DEL-3540
chr14-81389043-DEL-102
chr14-81452211-DEL-179
chr14-81479223-DEL-685
chr14-81491201-DEL-180
chr14-81514184-DEL-396
chr14-81517665-DEL-386
chr14-81524090-DEL-68
chr14-81537402-DEL-111
chr14-81551141-DEL-80
chr14-81557788-DEL-164
chr14-81577691-DEL-53
chr14-81600313-DEL-62
chr14-81633670-DEL-190
chr14-81654442-DEL-721
chr14-81658104-DEL-52
chr14-81721875-DEL-58
chr14-81731844-DEL-485
chr14-81741745-DEL-129
chr14-81744674-DEL-133
chr14-81759490-DEL-72
chr14-81763341-DEL-3338
chr14-81772379-DEL-1369
chr14-81784155-DEL-1450
chr14-81840276-DEL-65
chr14-81859422-DEL-54
chr14-81864828-DEL-1745
chr14-81866682-DEL-162
chr14-81870698-DEL-913
chr14-81890090-DEL-5325
chr14-81894851-DEL-4574
chr14-81895864-DEL-604
chr14-81898139-DEL-4939
chr14-81911924-DEL-430
chr14-81921309-DEL-463
chr14-81925805-DEL-12464
chr14-81933694-DEL-4127
chr14-81949835-DEL-1188
chr14-81955956-DEL-1674
chr14-81995141-DEL-51
chr14-82028266-DEL-7708
chr14-82028329-DEL-6936
chr14-82043573-DEL-328
chr14-82050448-DEL-173
chr14-82086429-DEL-53
chr14-82087478-DEL-64
chr14-82111953-DEL-5855
chr14-82297885-DEL-611
chr14-82310640-DEL-250
chr14-82313830-DEL-160

chr14-82321853-DEL-212
chr14-82339590-DEL-103
chr14-82401771-DEL-3096
chr14-82404516-DEL-2437
chr14-82408849-DEL-414
chr14-82426632-DEL-390
chr14-82436404-DEL-190
chr14-82484040-DEL-552
chr14-82523062-DEL-159
chr14-82538933-DEL-5881
chr14-82550075-DEL-377
chr14-82576090-DEL-1116
chr14-82613990-DEL-2901
chr14-82630410-DEL-293
chr14-82643785-DEL-486
chr14-82644509-DEL-229
chr14-82694725-DEL-67
chr14-82722520-DEL-133
chr14-82726022-DEL-4759
chr14-82738958-DEL-131
chr14-82742970-DEL-2666
chr14-82769940-DEL-756
chr14-82779950-DEL-6636
chr14-82779964-DEL-194
chr14-82787736-DEL-195
chr14-82859110-DEL-465
chr14-82860731-DEL-1981
chr14-82899799-DEL-6950
chr14-82899914-DEL-555
chr14-82985010-DEL-55
chr14-83005017-DEL-57
chr14-83019740-DEL-56
chr14-83034673-DEL-131
chr14-83035964-DEL-2576
chr14-83061431-DEL-86
chr14-83092009-DEL-396
chr14-83098783-DEL-5974
chr14-83105365-DEL-108
chr14-83124904-DEL-6402
chr14-83144943-DEL-64
chr14-83145594-DEL-1003
chr14-83170778-DEL-63
chr14-83204272-DEL-657
chr14-83234422-DEL-54
chr14-83263954-DEL-1477
chr14-83269419-DEL-99
chr14-83270538-DEL-621
chr14-83281215-DEL-13205
chr14-83342120-DEL-242
chr14-83358415-DEL-2222
chr14-83361921-DEL-535
chr14-83368311-DEL-1263
chr14-83371468-DEL-1081
chr14-83381278-DEL-3478
chr14-83389629-DEL-314
chr14-83392528-DEL-79
chr14-83467540-DEL-51
chr14-83469242-DEL-72
chr14-83494123-DEL-71
chr14-83500910-DEL-1970
chr14-83517651-DEL-63
chr14-83523104-DEL-125
chr14-83529340-DEL-451
chr14-83530251-DEL-6249
chr14-83539091-DEL-486
chr14-83540645-DEL-103
chr14-83541632-DEL-100
chr14-83542540-DEL-1489
chr14-83573777-DEL-197
chr14-83588566-DEL-126
chr14-83591807-DEL-739
chr14-83609641-DEL-60
chr14-83615176-DEL-310
chr14-83626935-DEL-2489
chr14-83743874-DEL-6214
chr14-83771865-DEL-220
chr14-83777074-DEL-166

chr14-83787695-DEL-2629
chr14-83819784-DEL-190
chr14-83822771-DEL-1239
chr14-83836928-DEL-207
chr14-83892130-DEL-82
chr14-83916982-DEL-890
chr14-83953294-DEL-2709
chr14-84093378-DEL-1795
chr14-84121398-DEL-231
chr14-84127242-DEL-667
chr14-84132006-DEL-472
chr14-84147319-DEL-620
chr14-84148276-DEL-874
chr14-84194686-DEL-150
chr14-84206839-DEL-66
chr14-84219404-DEL-70
chr14-84245221-DEL-261
chr14-84246024-DEL-2485
chr14-84246822-DEL-84
chr14-84251935-DEL-80
chr14-84260191-DEL-728
chr14-84312671-DEL-268
chr14-84319950-DEL-653
chr14-84399068-DEL-1977
chr14-84401548-DEL-780
chr14-84406954-DEL-1373
chr14-84429910-DEL-2047
chr14-84432934-DEL-51
chr14-84452625-DEL-1449
chr14-84454102-DEL-5380
chr14-84457506-DEL-54
chr14-84468232-DEL-2953
chr14-84493978-DEL-51
chr14-84507288-DEL-60
chr14-84569683-DEL-95
chr14-84598111-DEL-793
chr14-84603086-DEL-1096
chr14-84603463-DEL-243
chr14-84605577-DEL-6610
chr14-84605592-DEL-214
chr14-84615419-DEL-949
chr14-84623371-DEL-6300
chr14-84629312-DEL-126
chr14-84641233-DEL-6380
chr14-84677639-DEL-139
chr14-84720731-DEL-322
chr14-84733467-DEL-218
chr14-84847843-DEL-118
chr14-84900046-DEL-349
chr14-84952122-DEL-675
chr14-84962212-DEL-895
chr14-84964624-DEL-75
chr14-84973746-DEL-75
chr14-84989616-DEL-7880
chr14-85003063-DEL-862
chr14-85008674-DEL-1978
chr14-85012047-DEL-1130
chr14-85013939-DEL-7803
chr14-85027612-DEL-1225
chr14-85049502-DEL-102
chr14-85051223-DEL-479
chr14-85064210-DEL-4583
chr14-85081152-DEL-1645
chr14-85130275-DEL-52
chr14-85150064-DEL-198
chr14-85175216-DEL-208
chr14-85175658-DEL-415
chr14-85192831-DEL-91
chr14-85199052-DEL-57
chr14-85215608-DEL-477
chr14-85268453-DEL-59
chr14-85275891-DEL-145
chr14-85276763-DEL-260
chr14-85290668-DEL-536
chr14-85313005-DEL-1210
chr14-85391981-DEL-54
chr14-85397257-DEL-65

chr14-85404824-DEL-159
chr14-85407397-DEL-62
chr14-85418996-DEL-75
chr14-85432433-DEL-914
chr14-85433459-DEL-7601
chr14-85434440-DEL-6409
chr14-85451618-DEL-74
chr14-85484055-DEL-6271
chr14-85525848-DEL-336
chr14-85527999-DEL-78
chr14-85547671-DEL-64
chr14-85553919-DEL-437
chr14-85565746-DEL-543
chr14-85618844-DEL-370
chr14-85641672-DEL-83
chr14-85676393-DEL-66
chr14-85715655-DEL-89
chr14-85742197-DEL-55
chr14-85742728-DEL-84
chr14-85790591-DEL-70
chr14-85798095-DEL-113
chr14-85798604-DEL-1272
chr14-85820630-DEL-1985
chr14-85839717-DEL-2854
chr14-85871370-DEL-547
chr14-85895168-DEL-820
chr14-85916200-DEL-55
chr14-85928445-DEL-1250
chr14-85937768-DEL-385
chr14-85968390-DEL-415
chr14-85998105-DEL-208
chr14-86023552-DEL-139
chr14-86026278-DEL-348
chr14-86029968-DEL-103
chr14-86051800-DEL-86
chr14-86059477-DEL-5286
chr14-86073217-DEL-400
chr14-86095191-DEL-54
chr14-86101290-DEL-6691
chr14-86114529-DEL-6433
chr14-86138179-DEL-796
chr14-86177246-DEL-57
chr14-86190736-DEL-5007
chr14-86196097-DEL-1265
chr14-86199269-DEL-61
chr14-86212294-DEL-124
chr14-86225470-DEL-100
chr14-86238598-DEL-305
chr14-86241412-DEL-183
chr14-86244842-DEL-68
chr14-86256365-DEL-58
chr14-86260539-DEL-145
chr14-86269635-DEL-6188
chr14-86280406-DEL-51
chr14-86320810-DEL-830
chr14-86363758-DEL-151
chr14-86378511-DEL-100
chr14-86383302-DEL-62
chr14-86424548-DEL-124
chr14-86442076-DEL-274
chr14-86458149-DEL-675
chr14-86477954-DEL-354
chr14-86497492-DEL-128
chr14-86541917-DEL-160
chr14-86556605-DEL-193
chr14-86611332-DEL-1245
chr14-86612634-DEL-632
chr14-86619514-DEL-1002
chr14-86676197-DEL-75
chr14-86734985-DEL-477
chr14-86735864-DEL-377
chr14-86759653-DEL-338
chr14-86764680-DEL-1101
chr14-86799121-DEL-208
chr14-86817083-DEL-552
chr14-86845849-DEL-6190
chr14-86853136-DEL-114

chr14-86862904-DEL-515
chr14-86895989-DEL-173
chr14-86948510-DEL-66
chr14-86957533-DEL-401
chr14-87037360-DEL-52
chr14-87061730-DEL-128
chr14-87062292-DEL-474
chr14-87081889-DEL-145
chr14-87084171-DEL-86
chr14-87091153-DEL-66
chr14-87135070-DEL-798
chr14-87141154-DEL-250
chr14-87169378-DEL-64
chr14-87176959-DEL-96
chr14-87189017-DEL-343
chr14-87189777-DEL-766
chr14-87192178-DEL-668
chr14-87243598-DEL-80
chr14-87247546-DEL-4874
chr14-87250069-DEL-716
chr14-87275523-DEL-117
chr14-87278912-DEL-293
chr14-87280429-DEL-999
chr14-87323931-DEL-387
chr14-87352372-DEL-54
chr14-87369255-DEL-52
chr14-87435607-DEL-686
chr14-87456980-DEL-60
chr14-87458109-DEL-180
chr14-87462012-DEL-108
chr14-87479988-DEL-1791
chr14-87480112-DEL-325
chr14-87487145-DEL-579
chr14-87520181-DEL-1084
chr14-87524425-DEL-89
chr14-87535566-DEL-399
chr14-87556427-DEL-104
chr14-87576634-DEL-60
chr14-87645126-DEL-179
chr14-87675695-DEL-144
chr14-87692570-DEL-54
chr14-87700508-DEL-239
chr14-87715803-DEL-278
chr14-87716142-DEL-1288
chr14-87732866-DEL-859
chr14-87745157-DEL-211
chr14-87753353-DEL-76
chr14-87758362-DEL-81
chr14-87770521-DEL-125
chr14-87789255-DEL-64
chr14-87818091-DEL-843
chr14-87823551-DEL-1495
chr14-87834497-DEL-60
chr14-87861741-DEL-852
chr14-87891341-DEL-140
chr14-87897714-DEL-876
chr14-87957587-DEL-80
chr14-87965575-DEL-2165
chr14-87976470-DEL-279
chr14-88027896-DEL-6387
chr14-88033920-DEL-200
chr14-88061055-DEL-715
chr14-88063701-DEL-6021
chr14-88099782-DEL-166
chr14-88118776-DEL-588
chr14-88124256-DEL-153
chr14-88172613-DEL-6158
chr14-88214628-DEL-796
chr14-88217153-DEL-1396
chr14-88259587-DEL-176
chr14-88285608-DEL-65
chr14-88289306-DEL-1490
chr14-88291258-DEL-400
chr14-88293894-DEL-478
chr14-88299772-DEL-52
chr14-88308528-DEL-892
chr14-88313490-DEL-288

chr14-88315166-DEL-216
chr14-88370128-DEL-66
chr14-88388109-DEL-976
chr14-88402475-DEL-3044
chr14-88405733-DEL-233
chr14-88406914-DEL-6000
chr14-88419578-DEL-52
chr14-88423546-DEL-87
chr14-88438490-DEL-1822
chr14-88449301-DEL-79
chr14-88460594-DEL-234
chr14-88470268-DEL-118
chr14-88473717-DEL-52
chr14-88489970-DEL-53
chr14-88492093-DEL-217
chr14-88503570-DEL-1193
chr14-88508994-DEL-66
chr14-88527804-DEL-73
chr14-88528585-DEL-387
chr14-88538222-DEL-673
chr14-88546475-DEL-155
chr14-88572903-DEL-106
chr14-88587242-DEL-5856
chr14-88595443-DEL-55
chr14-88619570-DEL-686
chr14-88620449-DEL-254
chr14-88650766-DEL-81
chr14-88667285-DEL-558
chr14-88679502-DEL-1025
chr14-88731378-DEL-400
chr14-88734867-DEL-75
chr14-88739226-DEL-432
chr14-88744177-DEL-63
chr14-88747561-DEL-79
chr14-88753895-DEL-1385
chr14-88767161-DEL-79
chr14-88785975-DEL-971
chr14-88810925-DEL-1629
chr14-88810925-DEL-756
chr14-88811960-DEL-578
chr14-88812598-DEL-1019
chr14-88812870-DEL-3125
chr14-88813843-DEL-300
chr14-88814665-DEL-1330
chr14-88827821-DEL-1688
chr14-88837018-DEL-164
chr14-88882647-DEL-1652
chr14-88885781-DEL-2143
chr14-88892789-DEL-6189
chr14-88925980-DEL-160
chr14-88927449-DEL-6303
chr14-88991407-DEL-2391
chr14-89001799-DEL-1639
chr14-89047613-DEL-1250
chr14-89059932-DEL-125
chr14-89061972-DEL-52
chr14-89064755-DEL-377
chr14-89069771-DEL-296
chr14-89091596-DEL-3190
chr14-89098052-DEL-103
chr14-89117732-DEL-1032
chr14-89131937-DEL-145
chr14-89151939-DEL-51
chr14-89214975-DEL-52
chr14-89278601-DEL-7708
chr14-89281025-DEL-1163
chr14-89282648-DEL-535
chr14-89282951-DEL-1400
chr14-89291059-DEL-1982
chr14-89296070-DEL-1289
chr14-89297381-DEL-78
chr14-89356090-DEL-75
chr14-89365353-DEL-524
chr14-89380981-DEL-89
chr14-89382052-DEL-297
chr14-89385333-DEL-711
chr14-89396931-DEL-52

chr14-89400776-DEL-1337
chr14-89407893-DEL-205
chr14-89430786-DEL-57
chr14-89433521-DEL-548
chr14-89441616-DEL-94
chr14-89442952-DEL-53
chr14-89491083-DEL-112
chr14-89493570-DEL-3314
chr14-89494221-DEL-3657
chr14-89515441-DEL-2065
chr14-89525459-DEL-6468
chr14-89539898-DEL-92
chr14-89550876-DEL-64
chr14-89553716-DEL-64
chr14-89568165-DEL-6060
chr14-89584974-DEL-62
chr14-89609280-DEL-83
chr14-89654536-DEL-147
chr14-89690816-DEL-463
chr14-89691350-DEL-327
chr14-89698149-DEL-97
chr14-89698657-DEL-3959
chr14-89762385-DEL-87
chr14-89804824-DEL-1172
chr14-89815385-DEL-316
chr14-89818613-DEL-3320
chr14-89836532-DEL-213
chr14-89852637-DEL-2420
chr14-89852667-DEL-6252
chr14-89855077-DEL-3843
chr14-89896069-DEL-73
chr14-89916896-DEL-60
chr14-89917374-DEL-778
chr14-89923555-DEL-121
chr14-89955166-DEL-67
chr14-89966452-DEL-472
chr14-89967004-DEL-1727
chr14-89968793-DEL-344
chr14-89976284-DEL-156
chr14-90059614-DEL-56
chr14-90080400-DEL-1265
chr14-90102156-DEL-73
chr14-90154936-DEL-485
chr14-90161193-DEL-154
chr14-90171542-DEL-97
chr14-90178829-DEL-115
chr14-90201793-DEL-375
chr14-90240116-DEL-1007
chr14-90270243-DEL-345
chr14-90307378-DEL-2509
chr14-90312505-DEL-105
chr14-90372703-DEL-60
chr14-90376087-DEL-80
chr14-90381529-DEL-95
chr14-90384404-DEL-456
chr14-90386617-DEL-3538
chr14-90391750-DEL-211
chr14-90398036-DEL-59
chr14-90455792-DEL-2106
chr14-90467991-DEL-52
chr14-90475811-DEL-242
chr14-90483118-DEL-62
chr14-90487430-DEL-127
chr14-90499584-DEL-57
chr14-90504819-DEL-923
chr14-90512619-DEL-4410
chr14-90567690-DEL-351
chr14-90569559-DEL-127
chr14-90593307-DEL-61
chr14-90594271-DEL-140
chr14-90603097-DEL-103
chr14-90645463-DEL-356
chr14-90709188-DEL-908
chr14-90711667-DEL-983
chr14-90735300-DEL-1298
chr14-90737095-DEL-711
chr14-90737235-DEL-342

chr14-90755640-DEL-423
chr14-90794227-DEL-56
chr14-90807947-DEL-978
chr14-90836541-DEL-58
chr14-90873805-DEL-8579
chr14-90882626-DEL-392
chr14-90897299-DEL-189
chr14-90919554-DEL-8733
chr14-90930239-DEL-3580
chr14-90950004-DEL-53
chr14-90950894-DEL-403
chr14-90954485-DEL-56
chr14-90965297-DEL-1986
chr14-90967960-DEL-56
chr14-91013767-DEL-3563
chr14-91026706-DEL-504
chr14-91031726-DEL-62
chr14-91071732-DEL-182
chr14-91080097-DEL-58
chr14-91085507-DEL-100
chr14-91094047-DEL-1392
chr14-91100661-DEL-570
chr14-91134087-DEL-62
chr14-91140824-DEL-650
chr14-91206976-DEL-54
chr14-91211815-DEL-6642
chr14-91234403-DEL-3709
chr14-91243987-DEL-712
chr14-91247661-DEL-375
chr14-91255955-DEL-56
chr14-91269050-DEL-54
chr14-91272581-DEL-4173
chr14-91280238-DEL-708
chr14-91294333-DEL-252
chr14-91294421-DEL-964
chr14-91294989-DEL-183
chr14-91313166-DEL-859
chr14-91339130-DEL-96
chr14-91356818-DEL-90
chr14-91390392-DEL-84
chr14-91394139-DEL-1179
chr14-91397241-DEL-5424
chr14-91442328-DEL-1254
chr14-91446774-DEL-142
chr14-91459001-DEL-230
chr14-91473765-DEL-53
chr14-91498299-DEL-258
chr14-91509353-DEL-1041
chr14-91513953-DEL-77
chr14-91514445-DEL-1162
chr14-91548806-DEL-269
chr14-91559790-DEL-52
chr14-91565140-DEL-179
chr14-91580353-DEL-347
chr14-91597032-DEL-52
chr14-91620845-DEL-109
chr14-91634255-DEL-7476
chr14-91651485-DEL-6646
chr14-91651489-DEL-10523
chr14-91659088-DEL-334
chr14-91665682-DEL-329
chr14-91676637-DEL-3699
chr14-91681036-DEL-1650
chr14-91685578-DEL-308
chr14-91693128-DEL-66
chr14-91694041-DEL-791
chr14-91810921-DEL-472
chr14-91816997-DEL-610
chr14-91822868-DEL-60
chr14-91847368-DEL-52
chr14-91882304-DEL-113
chr14-91886891-DEL-1398
chr14-91915293-DEL-1151
chr14-91935775-DEL-364
chr14-91936479-DEL-151
chr14-91942368-DEL-137
chr14-91967853-DEL-549

chr14-91970918-DEL-181
chr14-91985898-DEL-88
chr14-91987656-DEL-13836
chr14-91991164-DEL-456
chr14-91992074-DEL-6220
chr14-91993545-DEL-2895
chr14-92017480-DEL-704
chr14-92018998-DEL-360
chr14-92019665-DEL-69
chr14-92028616-DEL-446
chr14-92035096-DEL-64
chr14-92037657-DEL-6934
chr14-92043904-DEL-400
chr14-92046700-DEL-358
chr14-92056598-DEL-9980
chr14-92074835-DEL-1835
chr14-92098873-DEL-76
chr14-92124648-DEL-76
chr14-92140517-DEL-60
chr14-92141252-DEL-318
chr14-92155811-DEL-176
chr14-92158745-DEL-256
chr14-92170904-DEL-6120
chr14-92177970-DEL-386
chr14-92186904-DEL-4974
chr14-92217039-DEL-53
chr14-92217050-DEL-2689
chr14-92222164-DEL-483
chr14-92242222-DEL-1146
chr14-92267250-DEL-124
chr14-92327249-DEL-161
chr14-92328192-DEL-5956
chr14-92334505-DEL-3503
chr14-92334672-DEL-5208
chr14-92338195-DEL-1686
chr14-92349973-DEL-594
chr14-92356658-DEL-1972
chr14-92372098-DEL-53
chr14-92393345-DEL-505
chr14-92403663-DEL-61
chr14-92453490-DEL-193
chr14-92490189-DEL-60
chr14-92500600-DEL-110
chr14-92528919-DEL-54
chr14-92553266-DEL-55
chr14-92559960-DEL-1036
chr14-92572001-DEL-1127
chr14-92599698-DEL-591
chr14-92608443-DEL-58
chr14-92617138-DEL-224
chr14-92628855-DEL-233
chr14-92647457-DEL-2953
chr14-92652368-DEL-406
chr14-92658731-DEL-122
chr14-92663676-DEL-52
chr14-92680021-DEL-69
chr14-92692568-DEL-218
chr14-92719140-DEL-356
chr14-92723167-DEL-66
chr14-92731177-DEL-1148
chr14-92738357-DEL-358
chr14-92746194-DEL-53
chr14-92755047-DEL-112
chr14-92760410-DEL-94
chr14-92773405-DEL-7191
chr14-92781794-DEL-84
chr14-92828274-DEL-371
chr14-92841805-DEL-1672
chr14-92856790-DEL-3955
chr14-92862281-DEL-344
chr14-92876647-DEL-130
chr14-92877736-DEL-59
chr14-92965000-DEL-96
chr14-92982497-DEL-2307
chr14-93009985-DEL-100
chr14-93031748-DEL-408
chr14-93039139-DEL-300

chr14-93076454-DEL-88
chr14-93077226-DEL-53
chr14-93081028-DEL-226
chr14-93097716-DEL-73
chr14-93104211-DEL-436
chr14-93121486-DEL-1520
chr14-93132894-DEL-2495
chr14-93143734-DEL-58
chr14-93145476-DEL-73
chr14-93158159-DEL-6396
chr14-93175037-DEL-6459
chr14-93189042-DEL-1967
chr14-93209327-DEL-385
chr14-93223886-DEL-109
chr14-93242758-DEL-235
chr14-93244333-DEL-59
chr14-93269519-DEL-58
chr14-93307200-DEL-54
chr14-93308934-DEL-68
chr14-93317364-DEL-263
chr14-93324326-DEL-58
chr14-93340664-DEL-52
chr14-93359576-DEL-112
chr14-93379363-DEL-356
chr14-93396062-DEL-141
chr14-93410314-DEL-96
chr14-93510466-DEL-74
chr14-93529484-DEL-52
chr14-93532578-DEL-97
chr14-93592467-DEL-6892
chr14-93625598-DEL-252
chr14-93645066-DEL-162
chr14-93647412-DEL-153
chr14-93673317-DEL-98
chr14-93686606-DEL-200
chr14-93702915-DEL-1257
chr14-93725746-DEL-1610
chr14-93728646-DEL-847
chr14-93778278-DEL-81
chr14-93829871-DEL-126
chr14-93846614-DEL-446
chr14-93879908-DEL-176
chr14-93882032-DEL-75
chr14-93897112-DEL-175
chr14-93929571-DEL-164
chr14-93950439-DEL-185
chr14-93954754-DEL-949
chr14-93987656-DEL-83
chr14-94002606-DEL-1331
chr14-94015556-DEL-56
chr14-94049528-DEL-134
chr14-94066707-DEL-475
chr14-94096457-DEL-85
chr14-94140800-DEL-68
chr14-94194538-DEL-87
chr14-94226962-DEL-688
chr14-94247978-DEL-56
chr14-94251334-DEL-73
chr14-94271882-DEL-86
chr14-94288058-DEL-7204
chr14-94295486-DEL-1163
chr14-94304486-DEL-55
chr14-94308835-DEL-106
chr14-94318897-DEL-609
chr14-94326555-DEL-51
chr14-94389554-DEL-81
chr14-94401219-DEL-61
chr14-94423240-DEL-3587
chr14-94432404-DEL-546
chr14-94436155-DEL-154
chr14-94441880-DEL-124
chr14-94468885-DEL-892
chr14-94479273-DEL-1177
chr14-94502345-DEL-382
chr14-94557047-DEL-1180
chr14-94562915-DEL-2371
chr14-94580356-DEL-681

chr14-94587127-DEL-230
chr14-94594472-DEL-58
chr14-94594691-DEL-356
chr14-94626032-DEL-1227
chr14-94638629-DEL-150
chr14-94639801-DEL-97
chr14-94699729-DEL-1908
chr14-94740624-DEL-72
chr14-94743071-DEL-1189
chr14-94753130-DEL-523
chr14-94765366-DEL-61
chr14-94766186-DEL-2194
chr14-94773326-DEL-54
chr14-94776533-DEL-156
chr14-94801865-DEL-477
chr14-94819797-DEL-248
chr14-94836910-DEL-326
chr14-94847931-DEL-87
chr14-94851397-DEL-333
chr14-94878813-DEL-8466
chr14-94896436-DEL-771
chr14-94906168-DEL-106
chr14-94910485-DEL-178
chr14-94914254-DEL-243
chr14-94916522-DEL-53
chr14-94924618-DEL-236
chr14-94930094-DEL-72
chr14-94934255-DEL-1621
chr14-94937550-DEL-63
chr14-94939789-DEL-54
chr14-94977488-DEL-284
chr14-95005625-DEL-2645
chr14-95011330-DEL-104
chr14-95034595-DEL-584
chr14-95044957-DEL-178
chr14-95047532-DEL-3041
chr14-95067013-DEL-109
chr14-95113985-DEL-323
chr14-95144823-DEL-3064
chr14-95179651-DEL-127
chr14-95182620-DEL-51
chr14-95200474-DEL-1040
chr14-95225846-DEL-88
chr14-95245351-DEL-2544
chr14-95245534-DEL-346
chr14-95246007-DEL-809
chr14-95246193-DEL-1154
chr14-95246829-DEL-142
chr14-95247223-DEL-309
chr14-95260541-DEL-827
chr14-95282833-DEL-2287
chr14-95303301-DEL-3198
chr14-95341829-DEL-143
chr14-95355765-DEL-86
chr14-95391248-DEL-55
chr14-95399913-DEL-2115
chr14-95403711-DEL-4770
chr14-95408929-DEL-64
chr14-95423944-DEL-1316
chr14-95446848-DEL-1690
chr14-95462969-DEL-73
chr14-95463985-DEL-60
chr14-95493186-DEL-60
chr14-95495410-DEL-376
chr14-95542364-DEL-80
chr14-95542818-DEL-302
chr14-95565645-DEL-175
chr14-95568417-DEL-6706
chr14-95579293-DEL-352
chr14-95618222-DEL-74
chr14-95620007-DEL-1012
chr14-95655300-DEL-197
chr14-95665836-DEL-407
chr14-95680255-DEL-56
chr14-95681644-DEL-62
chr14-95685021-DEL-268
chr14-95688375-DEL-366

chr14-95703975-DEL-468
chr14-95708719-DEL-6225
chr14-95740575-DEL-52
chr14-95788867-DEL-194
chr14-95796007-DEL-67
chr14-95796722-DEL-507
chr14-95812755-DEL-614
chr14-95814932-DEL-1321
chr14-95822092-DEL-1268
chr14-95842334-DEL-371
chr14-95855408-DEL-801
chr14-95860038-DEL-155
chr14-95899131-DEL-1891
chr14-95904374-DEL-60
chr14-95915674-DEL-1927
chr14-95952482-DEL-51
chr14-95970397-DEL-1204
chr14-95971125-DEL-133
chr14-95975169-DEL-168
chr14-95990097-DEL-582
chr14-96025236-DEL-465
chr14-96032594-DEL-53
chr14-96040958-DEL-54
chr14-96042324-DEL-69
chr14-96065270-DEL-52
chr14-96071588-DEL-755
chr14-96084120-DEL-593
chr14-96093374-DEL-887
chr14-96128737-DEL-334
chr14-96135250-DEL-341
chr14-96143036-DEL-53
chr14-96143831-DEL-6051
chr14-96161277-DEL-571
chr14-96170774-DEL-426
chr14-96201881-DEL-225
chr14-96210170-DEL-184
chr14-96240795-DEL-56
chr14-96243296-DEL-140
chr14-96269303-DEL-211
chr14-96301131-DEL-69
chr14-96321142-DEL-212
chr14-96323678-DEL-59
chr14-96333215-DEL-2004
chr14-96336129-DEL-771
chr14-96344300-DEL-73
chr14-96345163-DEL-9172
chr14-96390083-DEL-83
chr14-96393351-DEL-51
chr14-96402779-DEL-610
chr14-96406746-DEL-310
chr14-96425272-DEL-200
chr14-96435646-DEL-517
chr14-96442519-DEL-1280
chr14-96471633-DEL-81
chr14-96500895-DEL-68
chr14-96538854-DEL-427
chr14-96544551-DEL-99
chr14-96588977-DEL-100
chr14-96617237-DEL-130
chr14-96631941-DEL-238
chr14-96632606-DEL-635
chr14-96658773-DEL-64
chr14-96730610-DEL-1433
chr14-96734805-DEL-2582
chr14-96749889-DEL-328
chr14-96765573-DEL-98
chr14-96803870-DEL-3648
chr14-96809363-DEL-1834
chr14-96813508-DEL-60
chr14-96814342-DEL-6725
chr14-96834319-DEL-490
chr14-96842260-DEL-373
chr14-96870709-DEL-440
chr14-96884351-DEL-621
chr14-96889563-DEL-54
chr14-96891751-DEL-626
chr14-96945736-DEL-1966

chr14-96973418-DEL-167
chr14-97000738-DEL-119
chr14-97003115-DEL-875
chr14-97022699-DEL-4318
chr14-97035514-DEL-4182
chr14-97039995-DEL-189
chr14-97043714-DEL-174
chr14-97053593-DEL-174
chr14-97054977-DEL-141
chr14-97061144-DEL-52
chr14-97076012-DEL-306
chr14-97077094-DEL-64
chr14-97082586-DEL-138
chr14-97100063-DEL-116
chr14-97126188-DEL-56
chr14-97126903-DEL-729
chr14-97158536-DEL-55
chr14-97208425-DEL-151
chr14-97224849-DEL-111
chr14-97269278-DEL-5905
chr14-97302172-DEL-494
chr14-97341732-DEL-318
chr14-97347177-DEL-3942
chr14-97361360-DEL-3235
chr14-97361904-DEL-80
chr14-97370904-DEL-1202
chr14-97374122-DEL-903
chr14-97392677-DEL-3219
chr14-97395919-DEL-3185
chr14-97437604-DEL-315
chr14-97442478-DEL-144
chr14-97444526-DEL-121
chr14-97448885-DEL-856
chr14-97564739-DEL-419
chr14-97575384-DEL-5962
chr14-97593545-DEL-1344
chr14-97614627-DEL-291
chr14-97629848-DEL-813
chr14-97635669-DEL-315
chr14-97655837-DEL-749
chr14-97662035-DEL-60
chr14-97689934-DEL-4400
chr14-97695983-DEL-157
chr14-97736140-DEL-373
chr14-97754748-DEL-2315
chr14-97784252-DEL-72
chr14-97822684-DEL-444
chr14-97828059-DEL-4291
chr14-97855841-DEL-54
chr14-97861881-DEL-215
chr14-97891679-DEL-2201
chr14-97901654-DEL-2711
chr14-97945549-DEL-400
chr14-97960919-DEL-6549
chr14-98037584-DEL-56
chr14-98049015-DEL-177
chr14-98054928-DEL-849
chr14-98061583-DEL-51
chr14-98064770-DEL-55
chr14-98169858-DEL-367
chr14-98238327-DEL-64
chr14-98250983-DEL-261
chr14-98293194-DEL-67
chr14-98299303-DEL-382
chr14-98306546-DEL-70
chr14-98324275-DEL-76
chr14-98338183-DEL-182
chr14-98374863-DEL-188
chr14-98456136-DEL-96
chr14-98461252-DEL-738
chr14-98489083-DEL-193
chr14-98498906-DEL-52
chr14-98501921-DEL-89
chr14-98503346-DEL-61
chr14-98553126-DEL-120
chr14-98601737-DEL-396
chr14-98621147-DEL-198

chr14-98622581-DEL-88
chr14-98629233-DEL-103
chr14-98649084-DEL-66
chr14-98682162-DEL-381
chr14-98694504-DEL-515
chr14-98696207-DEL-1156
chr14-98702063-DEL-58
chr14-98724320-DEL-115
chr14-98727512-DEL-6050
chr14-98736225-DEL-56
chr14-98753752-DEL-71
chr14-98763699-DEL-166
chr14-98765267-DEL-254
chr14-98769453-DEL-887
chr14-98786574-DEL-110
chr14-98854780-DEL-605
chr14-98856650-DEL-52
chr14-98881997-DEL-57
chr14-98897362-DEL-295
chr14-98906996-DEL-1747
chr14-98916208-DEL-51
chr14-98990490-DEL-89
chr14-99027430-DEL-96
chr14-99031750-DEL-72
chr14-99038079-DEL-2141
chr14-99204763-DEL-75
chr14-99220023-DEL-184
chr14-99231305-DEL-490
chr14-99232921-DEL-219
chr14-99278698-DEL-1970
chr14-99297754-DEL-74
chr14-99307980-DEL-152
chr14-99312965-DEL-640
chr14-99355924-DEL-209
chr14-99423609-DEL-238
chr14-99457869-DEL-156
chr14-99470009-DEL-57
chr14-99520936-DEL-1130
chr14-99523590-DEL-128
chr14-99525611-DEL-59
chr14-99543849-DEL-187
chr14-99549876-DEL-365
chr14-99560610-DEL-67
chr14-99573385-DEL-218
chr14-99613997-DEL-120
chr14-99622555-DEL-58
chr14-99643953-DEL-861
chr14-99674638-DEL-58
chr14-99694662-DEL-256
chr14-99702218-DEL-222
chr14-99705082-DEL-328
chr14-99714222-DEL-114
chr14-99728147-DEL-55
chr14-99766890-DEL-68
chr14-99782427-DEL-195
chr14-99784471-DEL-52
chr14-99815919-DEL-55
chr14-99819678-DEL-76
chr14-99821108-DEL-77
chr14-99853358-DEL-390
chr14-99893332-DEL-68
chr14-99906895-DEL-134
chr14-99937899-DEL-101
chr14-99963384-DEL-365
chr14-99967263-DEL-196
chr14-99970202-DEL-674
chr14-100018754-DEL-54
chr14-100028537-DEL-138
chr14-100031561-DEL-69
chr14-100046501-DEL-56
chr14-100076623-DEL-579
chr14-100151631-DEL-81
chr14-100171340-DEL-174
chr14-100208366-DEL-558
chr14-100209288-DEL-309
chr14-100222055-DEL-1572
chr14-100275020-DEL-277

chr14-100304169-DEL-134
chr14-100326710-DEL-55
chr14-100332745-DEL-190
chr14-100400789-DEL-1166
chr14-100412511-DEL-129
chr14-100425926-DEL-92
chr14-100451951-DEL-802
chr14-100453332-DEL-70
chr14-100457418-DEL-52
chr14-100494863-DEL-52
chr14-100503528-DEL-356
chr14-100513942-DEL-158
chr14-100526936-DEL-54
chr14-100536003-DEL-75
chr14-100554370-DEL-175
chr14-100579554-DEL-365
chr14-100613596-DEL-53
chr14-100636607-DEL-132
chr14-100637775-DEL-1047
chr14-100645323-DEL-119
chr14-100669616-DEL-293
chr14-100706346-DEL-65
chr14-100722070-DEL-249
chr14-100723651-DEL-84
chr14-100769513-DEL-87
chr14-100770809-DEL-1004
chr14-100776102-DEL-960
chr14-100781284-DEL-57
chr14-100799665-DEL-86
chr14-100800312-DEL-191
chr14-100805805-DEL-234
chr14-100806961-DEL-274
chr14-100849452-DEL-144
chr14-100864290-DEL-383
chr14-100915860-DEL-5111
chr14-100923844-DEL-62
chr14-100933747-DEL-314
chr14-100935374-DEL-113
chr14-100959328-DEL-219
chr14-100974775-DEL-208
chr14-100986269-DEL-300
chr14-100998730-DEL-134
chr14-101025657-DEL-637
chr14-101039187-DEL-423
chr14-101043336-DEL-406
chr14-101072054-DEL-1154
chr14-101154523-DEL-52
chr14-101166535-DEL-447
chr14-101167266-DEL-629
chr14-101189122-DEL-55
chr14-101226470-DEL-130
chr14-101228781-DEL-303
chr14-101247617-DEL-2795
chr14-101267091-DEL-396
chr14-101273840-DEL-176
chr14-101289719-DEL-565
chr14-101323154-DEL-96
chr14-101325210-DEL-3151
chr14-101342730-DEL-955
chr14-101345805-DEL-254
chr14-101354696-DEL-56
chr14-101365949-DEL-2263
chr14-101399166-DEL-1632
chr14-101400827-DEL-3222
chr14-101405304-DEL-549
chr14-101424545-DEL-80
chr14-101424745-DEL-852
chr14-101427379-DEL-60
chr14-101431846-DEL-1340
chr14-101540915-DEL-583
chr14-101551973-DEL-721
chr14-101571824-DEL-130
chr14-101579968-DEL-111
chr14-101645248-DEL-197
chr14-101646403-DEL-6269
chr14-101672415-DEL-62
chr14-101715588-DEL-110

chr14-101717793-DEL-86
chr14-101732764-DEL-193
chr14-101733021-DEL-639
chr14-101747383-DEL-632
chr14-101758173-DEL-145
chr14-101772246-DEL-160
chr14-101789855-DEL-90
chr14-101812380-DEL-68
chr14-101837339-DEL-94
chr14-101842511-DEL-62
chr14-101882162-DEL-61
chr14-101886491-DEL-93
chr14-101924384-DEL-59
chr14-101925523-DEL-200
chr14-101935652-DEL-244
chr14-101941300-DEL-198
chr14-101968941-DEL-72
chr14-101989507-DEL-224
chr14-101994467-DEL-58
chr14-102005704-DEL-212
chr14-102013135-DEL-65
chr14-102063147-DEL-1089
chr14-102140310-DEL-861
chr14-102187163-DEL-98
chr14-102190975-DEL-120
chr14-102203241-DEL-243
chr14-102214850-DEL-432
chr14-102234251-DEL-112
chr14-102240417-DEL-126
chr14-102247246-DEL-197
chr14-102255501-DEL-310
chr14-102261819-DEL-754
chr14-102263434-DEL-120
chr14-102264811-DEL-1150
chr14-102269764-DEL-68
chr14-102272061-DEL-2524
chr14-102280751-DEL-565
chr14-102281708-DEL-1290
chr14-102298070-DEL-697
chr14-102304004-DEL-56
chr14-102322964-DEL-218
chr14-102338813-DEL-237
chr14-102359428-DEL-3170
chr14-102361309-DEL-674
chr14-102362587-DEL-6211
chr14-102365165-DEL-7069
chr14-102370955-DEL-111
chr14-102380684-DEL-6808
chr14-102380846-DEL-208
chr14-102391623-DEL-74
chr14-102406291-DEL-86
chr14-102408004-DEL-463
chr14-102417110-DEL-517
chr14-102428162-DEL-56
chr14-102435083-DEL-174
chr14-102438785-DEL-1158
chr14-102438785-DEL-854
chr14-102439673-DEL-1060
chr14-102439967-DEL-682
chr14-102440713-DEL-110
chr14-102457717-DEL-70
chr14-102458947-DEL-2126
chr14-102460815-DEL-80
chr14-102496855-DEL-439
chr14-102520434-DEL-2847
chr14-102525465-DEL-204
chr14-102542642-DEL-1006
chr14-102543981-DEL-2555
chr14-102559253-DEL-6157
chr14-102577703-DEL-58
chr14-102579796-DEL-222
chr14-102604341-DEL-116
chr14-102612348-DEL-5443
chr14-102730792-DEL-1689
chr14-102768575-DEL-1469
chr14-102773000-DEL-743
chr14-102785954-DEL-4065

chr14-102792521-DEL-518
chr14-102793450-DEL-76
chr14-102796725-DEL-6488
chr14-102820969-DEL-955
chr14-102824307-DEL-6031
chr14-102825387-DEL-4373
chr14-102840978-DEL-1394
chr14-102859923-DEL-916
chr14-102863670-DEL-57
chr14-102865032-DEL-94
chr14-102872456-DEL-5256
chr14-102890960-DEL-393
chr14-102905552-DEL-661
chr14-102912158-DEL-54
chr14-102941634-DEL-52
chr14-102965410-DEL-51
chr14-102971376-DEL-80
chr14-102993150-DEL-6926
chr14-103040793-DEL-873
chr14-103050991-DEL-64
chr14-103051093-DEL-1425
chr14-103052909-DEL-100
chr14-103054016-DEL-1120
chr14-103054303-DEL-574
chr14-103079251-DEL-1992
chr14-103104276-DEL-938
chr14-103110951-DEL-3150
chr14-103153192-DEL-93
chr14-103155067-DEL-219
chr14-103179111-DEL-97
chr14-103192034-DEL-54
chr14-103209111-DEL-72
chr14-103241282-DEL-60
chr14-103269023-DEL-808
chr14-103290655-DEL-162
chr14-103344082-DEL-345
chr14-103345078-DEL-1886
chr14-103353176-DEL-67
chr14-103395528-DEL-64
chr14-103439537-DEL-230
chr14-103475681-DEL-412
chr14-103492248-DEL-253
chr14-103537758-DEL-2358
chr14-103545651-DEL-80
chr14-103612180-DEL-623
chr14-103630950-DEL-932
chr14-103634337-DEL-75
chr14-103664941-DEL-6226
chr14-103667358-DEL-115
chr14-103678598-DEL-56
chr14-103681364-DEL-499
chr14-103696456-DEL-56
chr14-103702033-DEL-56
chr14-103736668-DEL-71
chr14-103741943-DEL-122
chr14-103743225-DEL-88
chr14-103780392-DEL-75
chr14-103801438-DEL-598
chr14-103819677-DEL-114
chr14-103846199-DEL-65
chr14-103856039-DEL-635
chr14-103857000-DEL-59
chr14-103857735-DEL-66
chr14-103871609-DEL-59
chr14-103904016-DEL-94
chr14-103910632-DEL-60
chr14-103914552-DEL-731
chr14-103915935-DEL-73
chr14-103945364-DEL-104
chr14-103970916-DEL-485
chr14-104006476-DEL-64
chr14-104043050-DEL-176
chr14-104044642-DEL-235
chr14-104045912-DEL-64
chr14-104047147-DEL-116
chr14-104067511-DEL-82
chr14-104095968-DEL-5991

chr14-104105779-DEL-6000
chr14-104114865-DEL-300
chr14-104139482-DEL-54
chr14-104153171-DEL-54
chr14-104165916-DEL-74
chr14-104178146-DEL-203
chr14-104209278-DEL-66
chr14-104219466-DEL-82
chr14-104221200-DEL-104
chr14-104228510-DEL-54
chr14-104228534-DEL-694
chr14-104255828-DEL-54
chr14-104266306-DEL-83
chr14-104269284-DEL-480
chr14-104308822-DEL-212
chr14-104342259-DEL-7143
chr14-104365560-DEL-72
chr14-104414362-DEL-52
chr14-104416847-DEL-79
chr14-104427268-DEL-65
chr14-104446649-DEL-121
chr14-104455798-DEL-79
chr14-104468982-DEL-85
chr14-104482961-DEL-111
chr14-104489474-DEL-1525
chr14-104514235-DEL-369
chr14-104529856-DEL-51
chr14-104549065-DEL-93
chr14-104553842-DEL-248
chr14-104557069-DEL-216
chr14-104603805-DEL-119
chr14-104611085-DEL-475
chr14-104657347-DEL-54
chr14-104709637-DEL-65
chr14-104763361-DEL-60
chr14-104786225-DEL-58
chr14-104798233-DEL-424
chr14-104843993-DEL-92
chr14-104854953-DEL-3094
chr14-104863727-DEL-244
chr14-104891627-DEL-92
chr14-104894441-DEL-85
chr14-104895315-DEL-1742
chr14-104905454-DEL-75
chr14-104909552-DEL-51
chr14-104917279-DEL-60
chr14-104936209-DEL-60
chr14-104941366-DEL-87
chr14-104952530-DEL-145
chr14-105001165-DEL-451
chr14-105008377-DEL-543
chr14-105030454-DEL-62
chr14-105061149-DEL-82
chr14-105076612-DEL-53
chr14-105088327-DEL-170
chr14-105129975-DEL-66423
chr14-105133190-DEL-88
chr14-105144956-DEL-4038
chr14-105153740-DEL-96
chr14-105179785-DEL-203
chr14-105185388-DEL-1735
chr14-105189438-DEL-138
chr14-105223476-DEL-62
chr14-105241293-DEL-2542
chr14-105277091-DEL-51
chr14-105277397-DEL-538
chr14-105283606-DEL-51
chr14-105297285-DEL-237
chr14-105303511-DEL-576
chr14-105318573-DEL-235
chr14-105341389-DEL-96
chr14-105374331-DEL-149
chr14-105381248-DEL-93
chr14-105382447-DEL-67
chr14-105385396-DEL-1823
chr14-105393590-DEL-136
chr14-105425863-DEL-130

chr14-105442203-DEL-868
chr14-105471353-DEL-224
chr14-105474720-DEL-96
chr14-105482021-DEL-219
chr14-105484746-DEL-87
chr14-105501710-DEL-56
chr14-105507367-DEL-317
chr14-105526381-DEL-1287
chr14-105526775-DEL-82
chr14-105527373-DEL-183
chr14-105588146-DEL-106
chr14-105661533-DEL-93
chr14-105704644-DEL-162
chr14-105714357-DEL-52
chr14-105720051-DEL-122
chr14-105737664-DEL-92
chr14-105764374-DEL-98
chr14-105768310-DEL-154
chr14-105773488-DEL-180
chr14-105784746-DEL-1139
chr14-105801459-DEL-765
chr14-105819753-DEL-88
chr14-105821961-DEL-79
chr14-105838005-DEL-189
chr14-105855234-DEL-56
chr14-105867549-DEL-146
chr14-105883652-DEL-84
chr14-105910201-DEL-1062
chr14-105918465-DEL-54
chr14-105987361-DEL-120
chr14-106008549-DEL-69
chr14-106019793-DEL-86
chr14-106031768-DEL-205
chr14-106034229-DEL-284
chr14-106035071-DEL-102
chr14-106070792-DEL-170
chr14-106079909-DEL-195
chr14-106106560-DEL-435
chr14-106121999-DEL-56
chr14-106143363-DEL-684
chr14-106148279-DEL-58
chr14-106156350-DEL-63
chr14-106164918-DEL-6339
chr14-106180674-DEL-6440
chr14-106189799-DEL-77
chr14-106197381-DEL-10109
chr14-106198142-DEL-6401
chr14-106208519-DEL-7092
chr14-106223480-DEL-85
chr14-106251637-DEL-932
chr14-106262498-DEL-401
chr14-106301781-DEL-287
chr14-106307614-DEL-447
chr14-106308356-DEL-79
chr14-106349489-DEL-5892
chr14-106363072-DEL-289
chr14-106387526-DEL-60
chr14-106401499-DEL-112
chr14-106406318-DEL-79
chr14-106412296-DEL-281
chr14-106421646-DEL-134
chr14-106441421-DEL-265
chr14-106442102-DEL-1372
chr14-106452224-DEL-750
chr14-106455856-DEL-866
chr14-106486018-DEL-645
chr14-106492137-DEL-413
chr14-106494191-DEL-174
chr14-106548594-DEL-1729
chr14-106566506-DEL-137
chr14-106587887-DEL-2452
chr14-106599676-DEL-634
chr14-106604138-DEL-60
chr14-106618646-DEL-279
chr14-106624240-DEL-327
chr14-106642564-DEL-639
chr14-106681365-DEL-579

chr14-106684852-DEL-3362
chr14-106690782-DEL-4434
chr14-106695222-DEL-13416
chr14-106724541-DEL-356
chr14-106734824-DEL-133
chr14-106763586-DEL-54
chr14-106780207-DEL-118
chr14-106829149-DEL-86
chr14-106835924-DEL-61
chr14-106865326-DEL-819
chr14-106866189-DEL-265
chr14-106872909-DEL-355
chr14-106883590-DEL-90
chr14-106889896-DEL-140
chr14-106909424-DEL-80
chr14-106913225-DEL-94
chr14-106936592-DEL-114
chr14-106939712-DEL-123
chr14-106942600-DEL-773
chr14-106950241-DEL-133
chr14-106955720-DEL-60
chr14-106974848-DEL-93
chr14-107019817-DEL-170
chr14-107040445-DEL-1530
chr14-107068258-DEL-227
chr14-107071201-DEL-1951
chr14-107071327-DEL-54
chr14-107098259-DEL-1277
chr14-107104461-DEL-1163
chr14-107113790-DEL-121
chr14-107125296-DEL-74
chr14-107138121-DEL-100
chr14-107139116-DEL-1192
chr14-107142538-DEL-930
chr14-107201379-DEL-7371
chr14-107226795-DEL-61
chr14-107244695-DEL-64
chr14-107272559-DEL-52
chr14-107273594-DEL-869
chr14-107275886-DEL-277
chr14-107277809-DEL-244
chr14-107281132-DEL-55
chr14-107296159-DEL-152
chr14-107306329-DEL-221
chr14-107311824-DEL-873
chr14-107312773-DEL-63
chr14-107326354-DEL-57
chr14-107334323-DEL-126
chr14-107336434-DEL-425
chr14-107352679-DEL-341
chr14-107355272-DEL-306
chr14-107381773-DEL-53
chr14-107402749-DEL-532
chr14-107403541-DEL-208
chr14-107430955-DEL-56
chr14-107432187-DEL-304
chr14-107432931-DEL-75
chr14-107444477-DEL-6358
chr14-107445350-DEL-94
chr14-107472792-DEL-1545
chr14-107473991-DEL-347
chr14-107477291-DEL-63
chr14-107481935-DEL-223
chr14-107506426-DEL-78
chr14-107516314-DEL-529
chr14-107548343-DEL-103
chr14-107556819-DEL-137
chr14-107557086-DEL-494
chr14-107596089-DEL-166
chr14-107599389-DEL-66
chr14-107608368-DEL-860
chr14-107611202-DEL-83
chr14-107646901-DEL-120
chr14-107655894-DEL-82
chr14-107682358-DEL-279
chr14-107683601-DEL-153
chr14-107693174-DEL-288

chr14-107699461-DEL-51
chr14-107701304-DEL-107
chr14-107702991-DEL-381
chr14-107719001-DEL-71
chr14-107725149-DEL-516
chr14-107728284-DEL-1787
chr14-107736267-DEL-106
chr14-107756939-DEL-252
chr14-107761289-DEL-56
chr14-107771657-DEL-54
chr14-107800621-DEL-295
chr14-107808688-DEL-3426
chr14-107812311-DEL-614
chr14-107823571-DEL-209
chr14-107832888-DEL-169
chr14-107833806-DEL-2936
chr14-107855543-DEL-441
chr14-107862693-DEL-832
chr14-107864978-DEL-115
chr14-107875397-DEL-144
chr14-107909639-DEL-310
chr14-107921432-DEL-643
chr14-107932892-DEL-62
chr14-107943481-DEL-1010
chr14-107947993-DEL-57
chr14-107952533-DEL-79
chr14-107968870-DEL-1110
chr14-107972834-DEL-1751
chr14-107977652-DEL-64
chr14-108001823-DEL-54
chr14-108095937-DEL-129
chr14-108123837-DEL-914
chr14-108146647-DEL-3509
chr14-108151828-DEL-65
chr14-108159333-DEL-109
chr14-108174235-DEL-120
chr14-108191440-DEL-100
chr14-108226688-DEL-159
chr14-108252103-DEL-1503
chr14-108299220-DEL-1007
chr14-108313287-DEL-328
chr14-108325276-DEL-283
chr14-108340499-DEL-83
chr14-108342626-DEL-52
chr14-108345427-DEL-4622
chr14-108352468-DEL-297
chr14-108365333-DEL-3452
chr14-108392867-DEL-950
chr14-108417230-DEL-671
chr14-108421142-DEL-378
chr14-108446389-DEL-113
chr14-108451867-DEL-4198
chr14-108485694-DEL-140
chr14-108495158-DEL-156
chr14-108516879-DEL-713
chr14-108520377-DEL-3313
chr14-108525858-DEL-54
chr14-108550293-DEL-52
chr14-108558022-DEL-232
chr14-108570855-DEL-56
chr14-108586208-DEL-157
chr14-108599533-DEL-6180
chr14-108616475-DEL-1834
chr14-108616480-DEL-1317
chr14-108618015-DEL-288
chr14-108635385-DEL-112
chr14-108641130-DEL-88
chr14-108645063-DEL-89
chr14-108646363-DEL-219
chr14-108671364-DEL-56
chr14-108671662-DEL-624
chr14-108691259-DEL-2702
chr14-108702656-DEL-1044
chr14-108703930-DEL-282
chr14-108731501-DEL-297
chr14-108749733-DEL-2150
chr14-108765801-DEL-93

chr14-108804554-DEL-1137
chr14-108832708-DEL-167
chr14-108846007-DEL-505
chr14-108855155-DEL-84
chr14-108885117-DEL-2209
chr14-108889285-DEL-241
chr14-108906761-DEL-401
chr14-108920661-DEL-148
chr14-108956677-DEL-6594
chr14-108971075-DEL-70
chr14-108991169-DEL-54
chr14-109017555-DEL-99
chr14-109031988-DEL-604
chr14-109070471-DEL-447
chr14-109076643-DEL-315
chr14-109097968-DEL-58
chr14-109120760-DEL-52
chr14-109179333-DEL-64
chr14-109196703-DEL-918
chr14-109214702-DEL-85
chr14-109225988-DEL-475
chr14-109234755-DEL-54
chr14-109239961-DEL-159
chr14-109248327-DEL-4014
chr14-109278212-DEL-203
chr14-109292150-DEL-4872
chr14-109311170-DEL-82
chr14-109313837-DEL-130
chr14-109336417-DEL-666
chr14-109336936-DEL-189
chr14-109345701-DEL-688
chr14-109349972-DEL-70
chr14-109355659-DEL-960
chr14-109366487-DEL-492
chr14-109370023-DEL-4323
chr14-109370852-DEL-60
chr14-109378961-DEL-534
chr14-109383639-DEL-1029
chr14-109393442-DEL-912
chr14-109394413-DEL-315
chr14-109439365-DEL-113
chr14-109454535-DEL-108
chr14-109467199-DEL-162
chr14-109481401-DEL-55
chr14-109511701-DEL-325
chr14-109515336-DEL-257
chr14-109526552-DEL-2389
chr14-109526558-DEL-1316
chr14-109526735-DEL-51
chr14-109528246-DEL-742
chr14-109557456-DEL-2340
chr14-109559150-DEL-89
chr14-109572443-DEL-223
chr14-109573776-DEL-137
chr14-109578841-DEL-608
chr14-109579786-DEL-7793
chr14-109581267-DEL-203
chr14-109581895-DEL-1697
chr14-109583153-DEL-119
chr14-109598702-DEL-937
chr14-109609552-DEL-180
chr14-109611641-DEL-718
chr14-109623952-DEL-112
chr14-109632588-DEL-1168
chr14-109639695-DEL-2446
chr14-109643075-DEL-791
chr14-109648063-DEL-93
chr14-109675399-DEL-76
chr14-109686164-DEL-53
chr14-109715246-DEL-353
chr14-109719221-DEL-109
chr14-109730049-DEL-63
chr14-109738325-DEL-534
chr14-109774489-DEL-210
chr14-109781640-DEL-766
chr14-109808746-DEL-3749
chr14-109911660-DEL-73

chr14-109915153-DEL-52
chr14-109932859-DEL-60
chr14-109934486-DEL-266
chr14-109947330-DEL-52
chr14-109951425-DEL-189
chr14-109997100-DEL-50
chr14-110035993-DEL-1947
chr14-110072067-DEL-137
chr14-110092656-DEL-6296
chr14-110113541-DEL-229
chr14-110115147-DEL-73
chr14-110123173-DEL-962
chr14-110160866-DEL-296
chr14-110165066-DEL-325
chr14-110175524-DEL-3021
chr14-110184045-DEL-55
chr14-110196339-DEL-7069
chr14-110212037-DEL-134
chr14-110242155-DEL-82
chr14-110242976-DEL-101
chr14-110252032-DEL-96
chr14-110263789-DEL-165
chr14-110275257-DEL-5170
chr14-110275953-DEL-54
chr14-110292473-DEL-483
chr14-110293115-DEL-6372
chr14-110305422-DEL-51
chr14-110308696-DEL-58
chr14-110329960-DEL-1091
chr14-110337755-DEL-315
chr14-110355117-DEL-63
chr14-110361873-DEL-562
chr14-110370430-DEL-223
chr14-110395334-DEL-59
chr14-110420381-DEL-58
chr14-110431178-DEL-827
chr14-110436942-DEL-1083
chr14-110439990-DEL-59
chr14-110446055-DEL-1182
chr14-110455058-DEL-442
chr14-110475869-DEL-57
chr14-110480155-DEL-1195
chr14-110484312-DEL-623
chr14-110510106-DEL-76
chr14-110512673-DEL-678
chr14-110545271-DEL-54
chr14-110545778-DEL-691
chr14-110546484-DEL-1695
chr14-110548233-DEL-251
chr14-110571720-DEL-4783
chr14-110581130-DEL-3698
chr14-110624643-DEL-519
chr14-110669181-DEL-98
chr14-110690588-DEL-315
chr14-110719314-DEL-838
chr14-110737404-DEL-130
chr14-110751090-DEL-586
chr14-110797593-DEL-142
chr14-110810388-DEL-60
chr14-110831064-DEL-5115
chr14-110843866-DEL-1156
chr14-110848444-DEL-1617
chr14-110880846-DEL-581
chr14-110888444-DEL-133
chr14-110901780-DEL-68
chr14-110927298-DEL-6164
chr14-110958701-DEL-425
chr14-110969644-DEL-65
chr14-111040554-DEL-400
chr14-111071034-DEL-1274
chr14-111082980-DEL-130
chr14-111096609-DEL-53
chr14-111097889-DEL-111
chr14-111098371-DEL-7145
chr14-111115798-DEL-10276
chr14-111139571-DEL-227
chr14-111140361-DEL-701

chr14-111180079-DEL-78
chr14-111181499-DEL-474
chr14-111221392-DEL-296
chr14-111223109-DEL-302
chr14-111230152-DEL-1894
chr14-111233891-DEL-323
chr14-111241124-DEL-83
chr14-111256782-DEL-200
chr14-111299814-DEL-843
chr14-111309953-DEL-529
chr14-111326825-DEL-2724
chr14-111343087-DEL-1129
chr14-111392223-DEL-133
chr14-111417514-DEL-59
chr14-111451136-DEL-476
chr14-111476421-DEL-315
chr14-111480915-DEL-243
chr14-111482944-DEL-67
chr14-111496336-DEL-139
chr14-111513631-DEL-53
chr14-111518749-DEL-58
chr14-111519490-DEL-72
chr14-111521830-DEL-454
chr14-111526818-DEL-493
chr14-111528732-DEL-463
chr14-111540498-DEL-1002
chr14-111545550-DEL-2392
chr14-111570045-DEL-492
chr14-111571880-DEL-182
chr14-111576677-DEL-5860
chr14-111598101-DEL-56
chr14-111611594-DEL-1091
chr14-111614790-DEL-200
chr14-111615618-DEL-99
chr14-111616929-DEL-202
chr14-111735946-DEL-1456
chr14-111737534-DEL-6219
chr14-111827012-DEL-54
chr14-111830926-DEL-138
chr14-111831342-DEL-1647
chr14-111839697-DEL-121
chr14-111873217-DEL-869
chr14-111888143-DEL-145
chr14-111927742-DEL-145
chr14-111954840-DEL-1304
chr14-112008904-DEL-57
chr14-112057277-DEL-195
chr14-112065203-DEL-160
chr14-112071724-DEL-3995
chr14-112118798-DEL-408
chr14-112137898-DEL-8625
chr14-112147236-DEL-1046
chr14-112186429-DEL-426
chr14-112194747-DEL-506
chr14-112204297-DEL-1208
chr14-112207331-DEL-62
chr14-112209368-DEL-68
chr14-112217434-DEL-238
chr14-112223002-DEL-498
chr14-112237485-DEL-1477
chr14-112241777-DEL-164
chr14-112255049-DEL-202
chr14-112259020-DEL-395
chr14-112268920-DEL-93
chr14-112296956-DEL-630
chr14-112327166-DEL-3075
chr14-112339837-DEL-731
chr14-112375201-DEL-405
chr14-112398926-DEL-242
chr14-112404557-DEL-280
chr14-112421745-DEL-87
chr14-112427397-DEL-81
chr14-112440768-DEL-106
chr14-112475281-DEL-215
chr14-112502902-DEL-71
chr14-112537101-DEL-390
chr14-112548860-DEL-52

chr14-112561915-DEL-59
chr14-112566853-DEL-60
chr14-112591842-DEL-63
chr14-112596751-DEL-626
chr14-112626979-DEL-53
chr14-112645787-DEL-1000
chr14-112646557-DEL-231
chr14-112649970-DEL-124
chr14-112652451-DEL-7325
chr14-112662163-DEL-158
chr14-112670169-DEL-525
chr14-112671437-DEL-629
chr14-112683243-DEL-59
chr14-112704727-DEL-56
chr14-112730288-DEL-2238
chr14-112748352-DEL-61
chr14-112752666-DEL-108
chr14-112772115-DEL-1207
chr14-112775914-DEL-1502
chr14-112789388-DEL-61
chr14-112801247-DEL-3226
chr14-112804704-DEL-56
chr14-112807463-DEL-290
chr14-112827542-DEL-51
chr14-112828740-DEL-369
chr14-112829208-DEL-1193
chr14-112832145-DEL-245
chr14-112835698-DEL-1102
chr14-112899042-DEL-1495
chr14-112903435-DEL-96
chr14-112922246-DEL-66
chr14-112932205-DEL-446
chr14-112934606-DEL-924
chr14-112946319-DEL-76
chr14-112955001-DEL-72
chr14-112963147-DEL-1491
chr14-112973775-DEL-52
chr14-112989048-DEL-72
chr14-112990249-DEL-889
chr14-112999500-DEL-1666
chr14-113008928-DEL-229
chr14-113018533-DEL-74
chr14-113024436-DEL-70
chr14-113025291-DEL-62
chr14-113046128-DEL-498
chr14-113092355-DEL-1863
chr14-113094365-DEL-77
chr14-113101569-DEL-106
chr14-113108282-DEL-173
chr14-113116291-DEL-139
chr14-113147215-DEL-322
chr14-113162109-DEL-108
chr14-113164615-DEL-350
chr14-113179331-DEL-618
chr14-113182832-DEL-138
chr14-113192082-DEL-84
chr14-113200986-DEL-91
chr14-113215475-DEL-67
chr14-113223847-DEL-126
chr14-113231179-DEL-58
chr14-113239603-DEL-88
chr14-113273516-DEL-1326
chr14-113294585-DEL-150
chr14-113297093-DEL-486
chr14-113308667-DEL-475
chr14-113312776-DEL-306
chr14-113313581-DEL-210
chr14-113329369-DEL-235
chr14-113331625-DEL-3348
chr14-113350810-DEL-66
chr14-113352261-DEL-1181
chr14-113370863-DEL-62
chr14-113375242-DEL-2012
chr14-113392502-DEL-5155
chr14-113399279-DEL-685
chr14-113402460-DEL-1704
chr14-113446651-DEL-313

chr14-113447411-DEL-339
chr14-113463497-DEL-70
chr14-113498864-DEL-71
chr14-113516182-DEL-208
chr14-113522826-DEL-2157
chr14-113524394-DEL-7169
chr14-113538125-DEL-51
chr14-113556084-DEL-177
chr14-113567018-DEL-2144
chr14-113572374-DEL-396
chr14-113584205-DEL-91
chr14-113587656-DEL-2490
chr14-113594152-DEL-550
chr14-113602938-DEL-256
chr14-113607607-DEL-66
chr14-113610371-DEL-6976
chr14-113610697-DEL-118
chr14-113613790-DEL-614
chr14-113622358-DEL-445
chr14-113625413-DEL-879
chr14-113649261-DEL-231
chr14-113658684-DEL-218
chr14-113662762-DEL-54
chr14-113759892-DEL-56
chr14-113773586-DEL-74
chr14-113784417-DEL-2524
chr14-113793470-DEL-316
chr14-113823313-DEL-201
chr14-113831031-DEL-842
chr14-113845071-DEL-3724
chr14-113850141-DEL-5832
chr14-113865093-DEL-612
chr14-113866773-DEL-1564
chr14-113872836-DEL-100
chr14-113887403-DEL-212
chr14-113895771-DEL-977
chr14-113899707-DEL-823
chr14-113900655-DEL-3589
chr14-113915850-DEL-2550
chr14-113945263-DEL-774
chr14-113974401-DEL-7344
chr14-113980052-DEL-467
chr14-114007710-DEL-2373
chr14-114010983-DEL-1233
chr14-114030711-DEL-6597
chr14-114047123-DEL-160
chr14-114048020-DEL-5004
chr14-114058524-DEL-162
chr14-114083692-DEL-56
chr14-114085619-DEL-139
chr14-114099961-DEL-1196
chr14-114101579-DEL-676
chr14-114125466-DEL-1164
chr14-114146781-DEL-80
chr14-114164649-DEL-305
chr14-114175270-DEL-1155
chr14-114178592-DEL-129
chr14-114184352-DEL-213
chr14-114205607-DEL-2861
chr14-114225873-DEL-104
chr14-114233393-DEL-779
chr14-114233875-DEL-1852
chr14-114244378-DEL-516
chr14-114264029-DEL-73
chr14-114295980-DEL-53
chr14-114317758-DEL-350
chr14-114327896-DEL-98
chr14-114343147-DEL-73
chr14-114344280-DEL-81
chr14-114352044-DEL-187
chr14-114374332-DEL-121
chr14-114379115-DEL-79
chr14-114425320-DEL-3222
chr14-114425364-DEL-2658
chr14-114428095-DEL-70
chr14-114431946-DEL-57
chr14-114434207-DEL-399

chr14-114440081-DEL-232
chr14-114453810-DEL-96
chr14-114455361-DEL-1132
chr14-114503027-DEL-207
chr14-114506422-DEL-674
chr14-114508483-DEL-66
chr14-114512470-DEL-517
chr14-114515749-DEL-90659
chr14-114529875-DEL-243
chr14-114530648-DEL-152
chr14-114537065-DEL-106
chr14-114570718-DEL-745
chr14-114581278-DEL-87
chr14-114624838-DEL-115
chr14-114643966-DEL-54
chr14-114671823-DEL-90
chr14-114672987-DEL-100
chr14-114689266-DEL-1212
chr14-114704069-DEL-1189
chr14-114723359-DEL-53
chr14-114760426-DEL-259
chr14-114772004-DEL-202
chr14-114793286-DEL-2132
chr14-114811127-DEL-61
chr14-114826591-DEL-73
chr14-114829124-DEL-421
chr14-114834153-DEL-176
chr14-114850174-DEL-578
chr14-114852529-DEL-2405
chr14-114874484-DEL-479
chr14-114878916-DEL-53
chr14-114887707-DEL-63
chr14-114902942-DEL-197
chr14-114930477-DEL-51
chr14-114942415-DEL-6208
chr14-114949192-DEL-66
chr14-114960059-DEL-210
chr14-114974095-DEL-203
chr14-114987878-DEL-98
chr14-115001258-DEL-234
chr14-115026993-DEL-51
chr14-115064930-DEL-456
chr14-115095979-DEL-54
chr14-115096407-DEL-845
chr14-115117145-DEL-3468
chr14-115124531-DEL-68
chr14-115140499-DEL-636
chr14-115178795-DEL-52
chr14-115208470-DEL-745
chr14-115216578-DEL-60
chr14-115220114-DEL-812
chr14-115243280-DEL-53
chr14-115250414-DEL-307
chr14-115272207-DEL-730
chr14-115297309-DEL-1443
chr14-115353482-DEL-832
chr14-115354104-DEL-207
chr14-115358581-DEL-171
chr14-115359834-DEL-86
chr14-115369352-DEL-74
chr14-115410624-DEL-156
chr14-115430821-DEL-759
chr14-115450442-DEL-302
chr14-115451738-DEL-356
chr14-115479056-DEL-86
chr14-115510982-DEL-6726
chr14-115563021-DEL-206
chr14-115573010-DEL-466
chr14-115595723-DEL-59
chr14-115642775-DEL-560
chr14-115665194-DEL-61
chr14-115678133-DEL-1450
chr14-115706692-DEL-104
chr14-115728748-DEL-61
chr14-115741994-DEL-70
chr14-115757157-DEL-632
chr14-115792374-DEL-318

chr14-115813886-DEL-257
chr14-115826214-DEL-3070
chr14-115909206-DEL-180
chr14-115915588-DEL-135
chr14-115933932-DEL-454
chr14-115941719-DEL-1144
chr14-115954867-DEL-267
chr14-115955829-DEL-53
chr14-115970953-DEL-256
chr14-115984860-DEL-68
chr14-115987087-DEL-312
chr14-115989793-DEL-51
chr14-116000908-DEL-507
chr14-116034405-DEL-62
chr14-116036052-DEL-94
chr14-116046810-DEL-517
chr14-116050860-DEL-2138
chr14-116052468-DEL-425
chr14-116099104-DEL-602
chr14-116125270-DEL-165
chr14-116136719-DEL-60
chr14-116144274-DEL-4298
chr14-116144277-DEL-1218
chr14-116152401-DEL-159
chr14-116164471-DEL-73
chr14-116174361-DEL-60
chr14-116180866-DEL-1093
chr14-116189591-DEL-534
chr14-116215515-DEL-161
chr14-116229962-DEL-923
chr14-116234347-DEL-136
chr14-116245560-DEL-304
chr14-116252490-DEL-68
chr14-116252929-DEL-949
chr14-116265667-DEL-74
chr14-116294338-DEL-329
chr14-116312122-DEL-318
chr14-116330314-DEL-442
chr14-116350145-DEL-51
chr14-116382219-DEL-291
chr14-116395797-DEL-3350
chr14-116439379-DEL-681
chr14-116440985-DEL-408
chr14-116448699-DEL-86
chr14-116459314-DEL-438
chr14-116470472-DEL-206
chr14-116495821-DEL-83
chr14-116498372-DEL-258
chr14-116509669-DEL-102
chr14-116521836-DEL-2388
chr14-116531278-DEL-242
chr14-116535460-DEL-56
chr14-116535664-DEL-2538
chr14-116592854-DEL-203
chr14-116603665-DEL-1841
chr14-116613913-DEL-3065
chr14-116618397-DEL-159
chr14-116645991-DEL-99
chr14-116665189-DEL-70
chr14-116678332-DEL-125
chr14-116684009-DEL-156
chr14-116684233-DEL-1290
chr14-116702841-DEL-6502
chr14-116738459-DEL-56
chr14-116742343-DEL-113
chr14-116742631-DEL-1163
chr14-116764353-DEL-54
chr14-116807438-DEL-595
chr14-116812960-DEL-1418
chr14-116825951-DEL-1000
chr14-116852952-DEL-3619
chr14-116860818-DEL-74
chr14-116862183-DEL-56
chr14-116880352-DEL-122
chr14-116883534-DEL-79
chr14-116898026-DEL-278
chr14-116921987-DEL-1268

chr14-116933776-DEL-95
chr14-116957362-DEL-64
chr14-116973077-DEL-668
chr14-116974199-DEL-57
chr14-116976253-DEL-105
chr14-116982618-DEL-55
chr14-117034137-DEL-60
chr14-117049312-DEL-85
chr14-117054320-DEL-5987
chr14-117061736-DEL-343
chr14-117064423-DEL-117
chr14-117070595-DEL-502
chr14-117073468-DEL-400
chr14-117123531-DEL-66
chr14-117155019-DEL-441
chr14-117170191-DEL-217
chr14-117177494-DEL-71
chr14-117201002-DEL-190
chr14-117205963-DEL-108
chr14-117207971-DEL-207
chr14-117223329-DEL-193
chr14-117225210-DEL-5346
chr14-117258921-DEL-1348
chr14-117261141-DEL-723
chr14-117272649-DEL-88
chr14-117293069-DEL-650
chr14-117299955-DEL-107
chr14-117306072-DEL-87
chr14-117322133-DEL-522
chr14-117340945-DEL-105
chr14-117350935-DEL-58
chr14-117397044-DEL-246
chr14-117399801-DEL-572
chr14-117450383-DEL-52
chr14-117480174-DEL-74
chr14-117480573-DEL-592
chr14-117491109-DEL-344
chr14-117506584-DEL-220
chr14-117520299-DEL-310
chr14-117556365-DEL-990
chr14-117587459-DEL-188
chr14-117600921-DEL-426
chr14-117613275-DEL-53
chr14-117616667-DEL-178
chr14-117623576-DEL-561
chr14-117628020-DEL-483
chr14-117640031-DEL-1495
chr14-117641101-DEL-426
chr14-117662290-DEL-407
chr14-117664854-DEL-327
chr14-117678939-DEL-137
chr14-117692504-DEL-140
chr14-117716815-DEL-164
chr14-117717986-DEL-58
chr14-117739356-DEL-84
chr14-117765196-DEL-178
chr14-117783773-DEL-809
chr14-117787803-DEL-56
chr14-117822462-DEL-732
chr14-117866292-DEL-55
chr14-117870611-DEL-231
chr14-117884631-DEL-84
chr14-117897467-DEL-3623
chr14-117932638-DEL-1873
chr14-117939973-DEL-227
chr14-117954930-DEL-52
chr14-117958119-DEL-126
chr14-117979944-DEL-220
chr14-118006271-DEL-2480
chr14-118034482-DEL-729
chr14-118044797-DEL-102
chr14-118056438-DEL-731
chr14-118069452-DEL-191
chr14-118076062-DEL-417
chr14-118110104-DEL-54
chr14-118169885-DEL-124
chr14-118170970-DEL-265

chr14-118171957-DEL-225
chr14-118205790-DEL-555
chr14-118238671-DEL-54
chr14-118265997-DEL-118
chr14-118279174-DEL-54
chr14-118294924-DEL-69
chr14-118297634-DEL-150
chr14-118382996-DEL-59
chr14-118387728-DEL-215
chr14-118389333-DEL-83
chr14-118393388-DEL-308
chr14-118412105-DEL-403
chr14-118435982-DEL-504
chr14-118443594-DEL-133
chr14-118466575-DEL-54
chr14-118500829-DEL-60
chr14-118509004-DEL-102
chr14-118532470-DEL-164
chr14-118574490-DEL-66
chr14-118589978-DEL-56
chr14-118602791-DEL-166
chr14-118614024-DEL-53
chr14-118691021-DEL-123
chr14-118769563-DEL-544
chr14-118784427-DEL-170
chr14-118785004-DEL-170
chr14-118785508-DEL-242
chr14-118803387-DEL-83
chr14-118807746-DEL-274
chr14-118819627-DEL-213
chr14-118844327-DEL-52
chr14-118847031-DEL-221
chr14-118855716-DEL-119
chr14-118863343-DEL-60
chr14-118878192-DEL-112
chr14-118890828-DEL-163
chr14-118913600-DEL-56
chr14-118938026-DEL-73
chr14-118958178-DEL-127
chr14-119000948-DEL-197
chr14-119030079-DEL-123
chr14-119039376-DEL-570
chr14-119049051-DEL-810
chr14-119049773-DEL-135
chr14-119053885-DEL-113
chr14-119066030-DEL-54
chr14-119087082-DEL-181
chr14-119093391-DEL-59
chr14-119099629-DEL-78
chr14-119107872-DEL-247
chr14-119118646-DEL-349
chr14-119132051-DEL-64
chr14-119143167-DEL-53
chr14-119179776-DEL-80
chr14-119210723-DEL-228
chr14-119214927-DEL-85
chr14-119239664-DEL-435
chr14-119271416-DEL-106
chr14-119284037-DEL-242
chr14-119295377-DEL-52
chr14-119301909-DEL-336
chr14-119308507-DEL-52
chr14-119347512-DEL-90
chr14-119359179-DEL-53
chr14-119379694-DEL-100
chr14-119382348-DEL-71
chr14-119384540-DEL-550
chr14-119419343-DEL-142
chr14-119421485-DEL-282
chr14-119421625-DEL-478
chr14-119426719-DEL-183
chr14-119441824-DEL-116
chr14-119462629-DEL-56
chr14-119497093-DEL-61
chr14-119535757-DEL-68
chr14-119585754-DEL-52
chr14-119596554-DEL-82

chr14-119611996-DEL-59
chr14-119662380-DEL-75
chr14-119697836-DEL-65
chr14-119729135-DEL-59
chr14-119746759-DEL-1233
chr14-119757960-DEL-122
chr14-119847852-DEL-856
chr14-119848977-DEL-155
chr14-119905496-DEL-304
chr14-119911163-DEL-184
chr14-119911369-DEL-1917
chr14-119914076-DEL-365
chr14-119921500-DEL-62
chr14-119928746-DEL-71
chr14-119964143-DEL-67
chr14-120011527-DEL-83
chr14-120039789-DEL-145
chr14-120092462-DEL-122
chr14-120099987-DEL-64
chr14-120102386-DEL-1206
chr14-120112018-DEL-183
chr14-120143326-DEL-88
chr14-120211557-DEL-148
chr14-120217434-DEL-930
chr14-120223413-DEL-83
chr14-120249037-DEL-218
chr14-120269734-DEL-278
chr14-120288285-DEL-55
chr14-120295899-DEL-216
chr14-120308834-DEL-205
chr14-120335377-DEL-883
chr14-120336736-DEL-56
chr14-120358083-DEL-780
chr14-120363536-DEL-62
chr14-120365312-DEL-115
chr14-120369834-DEL-62
chr14-120379669-DEL-61
chr14-120388968-DEL-88
chr14-120389164-DEL-1080
chr14-120431062-DEL-55
chr14-120433441-DEL-6660
chr14-120441004-DEL-200
chr14-120489443-DEL-1248
chr14-120507034-DEL-317
chr14-120527574-DEL-80
chr14-120546097-DEL-74
chr14-120587747-DEL-441
chr14-120609299-DEL-68
chr14-120638291-DEL-283
chr14-120648946-DEL-85
chr14-120659666-DEL-57
chr14-120674882-DEL-96
chr14-120677600-DEL-56
chr14-120695369-DEL-170
chr14-120701805-DEL-52
chr14-120718299-DEL-305
chr14-120740186-DEL-267
chr14-120765538-DEL-1086
chr14-120795007-DEL-125
chr14-120811696-DEL-338
chr14-120832890-DEL-56
chr14-120835853-DEL-62
chr14-120865818-DEL-884
chr14-120872218-DEL-2214
chr14-120876479-DEL-184
chr14-120895455-DEL-106
chr14-120900895-DEL-60
chr14-120910346-DEL-263
chr14-120930411-DEL-99
chr14-120933576-DEL-226
chr14-120934586-DEL-142
chr14-120955198-DEL-183
chr14-121000435-DEL-330
chr14-121003715-DEL-51
chr14-121026166-DEL-328
chr14-121038248-DEL-1492
chr14-121041596-DEL-1168

chr14-121042810-DEL-2472
chr14-121049880-DEL-500
chr14-121054548-DEL-205
chr14-121066269-DEL-762
chr14-121067948-DEL-165
chr14-121082213-DEL-128
chr14-121101547-DEL-150
chr14-121102538-DEL-1766
chr14-121108879-DEL-76
chr14-121145117-DEL-54
chr14-121165237-DEL-63
chr14-121188880-DEL-366
chr14-121205876-DEL-777
chr14-121214370-DEL-514
chr14-121216335-DEL-889
chr14-121233403-DEL-524
chr14-121253114-DEL-101
chr14-121256285-DEL-409
chr14-121269679-DEL-66
chr14-121282253-DEL-147
chr14-121294328-DEL-86
chr14-121295004-DEL-80
chr14-121310982-DEL-123
chr14-121317413-DEL-1321
chr14-121322317-DEL-147
chr14-121341666-DEL-98
chr14-121366791-DEL-6354
chr14-121385926-DEL-1476
chr14-121388541-DEL-93
chr14-121396051-DEL-802
chr14-121406563-DEL-655
chr14-121457620-DEL-81
chr14-121465522-DEL-145
chr14-121485352-DEL-82
chr14-121506551-DEL-64
chr14-121527504-DEL-581
chr14-121528522-DEL-53
chr14-121533152-DEL-199
chr14-121541385-DEL-56
chr14-121543592-DEL-54
chr14-121545850-DEL-86
chr14-121555218-DEL-88
chr14-121561694-DEL-86
chr14-121563965-DEL-56
chr14-121602044-DEL-53
chr14-121610207-DEL-114
chr14-121625623-DEL-56
chr14-121632968-DEL-51
chr14-121640972-DEL-52
chr14-121653843-DEL-177
chr14-121662703-DEL-61
chr14-121664176-DEL-51
chr14-121676196-DEL-291
chr14-121685956-DEL-56
chr14-121720704-DEL-192
chr14-121721288-DEL-84
chr14-121731520-DEL-53
chr14-121787369-DEL-140
chr14-121798525-DEL-197
chr14-121817719-DEL-129
chr14-121831330-DEL-59
chr14-121862269-DEL-158
chr14-121876267-DEL-246
chr14-121934532-DEL-359
chr14-121960516-DEL-69
chr14-122010480-DEL-316
chr14-122022525-DEL-57
chr14-122037180-DEL-95
chr14-122048035-DEL-265
chr14-122105737-DEL-171
chr14-122113545-DEL-214
chr14-122134930-DEL-83
chr14-122154717-DEL-1161
chr14-122157957-DEL-80
chr14-122205727-DEL-158
chr14-122248782-DEL-163
chr14-122259120-DEL-86

chr14-122263169-DEL-78
chr14-122283088-DEL-347
chr14-122288927-DEL-165
chr14-122291779-DEL-181
chr14-122309568-DEL-52
chr14-122355797-DEL-3694
chr14-122420458-DEL-151
chr14-122424103-DEL-377
chr14-122426231-DEL-80
chr14-122430831-DEL-105
chr14-122475840-DEL-192
chr14-122497590-DEL-209
chr14-122531678-DEL-202
chr14-122538201-DEL-473
chr14-122560380-DEL-216
chr14-122595575-DEL-90
chr14-122597876-DEL-144
chr14-122632684-DEL-226
chr14-122635729-DEL-52
chr14-122657432-DEL-298
chr14-122681900-DEL-99
chr14-122689724-DEL-123
chr14-122709391-DEL-90
chr14-122725427-DEL-68
chr14-122733389-DEL-53
chr14-122745962-DEL-219
chr14-122754275-DEL-9434
chr14-122763025-DEL-203
chr14-122768776-DEL-59
chr14-122780272-DEL-4793
chr14-122811167-DEL-85
chr14-122832501-DEL-423
chr14-122839718-DEL-53
chr14-122842005-DEL-161
chr14-122870462-DEL-84
chr14-122898476-DEL-54
chr14-122901866-DEL-270
chr14-122909304-DEL-54
chr14-122936002-DEL-60
chr14-122967038-DEL-69
chr14-123038002-DEL-627
chr14-123071894-DEL-113
chr14-123072913-DEL-118
chr14-123098597-DEL-80
chr14-123100892-DEL-457
chr14-123108582-DEL-162
chr14-123151092-DEL-529
chr14-123182619-DEL-232
chr14-123237395-DEL-214
chr14-123273642-DEL-1566
chr14-123279704-DEL-677
chr14-123304330-DEL-6454
chr14-123320602-DEL-706
chr14-123419735-DEL-438
chr14-123564147-DEL-2274
chr14-123582346-DEL-398
chr14-123609786-DEL-248
chr14-123614782-DEL-430
chr14-123628472-DEL-323
chr14-123657956-DEL-131
chr14-123664193-DEL-7337
chr14-123674496-DEL-360
chr14-123677095-DEL-52
chr14-123684866-DEL-607
chr14-123696288-DEL-1263
chr14-123704162-DEL-779
chr14-123704933-DEL-61
chr14-123708426-DEL-519
chr14-123711875-DEL-72
chr14-123759638-DEL-183
chr14-123792197-DEL-71
chr14-123839658-DEL-182
chr14-123859305-DEL-284
chr14-123871317-DEL-155
chr14-123882767-DEL-392
chr14-123894897-DEL-65
chr14-123912194-DEL-133

chr14-123953521-DEL-61
chr14-123959241-DEL-18898
chr14-123982178-DEL-119
chr14-124030518-DEL-208
chr14-124053120-DEL-440
chr14-124058511-DEL-6325
chr14-124067951-DEL-6147
chr14-124103053-DEL-1008
chr14-124126088-DEL-2248
chr14-124131771-DEL-65
chr14-124141984-DEL-3977
chr14-124163530-DEL-466
chr14-124172541-DEL-1354
chr14-124239388-DEL-1722
chr14-124246716-DEL-1534
chr14-124250086-DEL-6201
chr14-124256394-DEL-1978
chr14-124257476-DEL-229
chr14-124265216-DEL-1483
chr14-124278288-DEL-59
chr14-124339469-DEL-64
chr14-124355045-DEL-3156
chr14-124385219-DEL-59
chr14-124389854-DEL-253
chr14-124392463-DEL-474
chr14-124399666-DEL-602
chr14-124404130-DEL-1079
chr14-124410477-DEL-626
chr14-124465390-DEL-77
chr14-124484709-DEL-210
chr14-124503168-DEL-345
chr14-124506549-DEL-81
chr14-124527004-DEL-74
chr14-124535923-DEL-111
chr14-124541922-DEL-55
chr14-124552378-DEL-172
chr14-124555511-DEL-201
chr14-124571228-DEL-6399
chr14-124582767-DEL-767
chr14-124615778-DEL-592
chr14-124620594-DEL-200
chr14-124630411-DEL-204
chr14-124652304-DEL-7917
chr14-124652928-DEL-409
chr14-124659537-DEL-680
chr14-124670454-DEL-281
chr14-124671616-DEL-2278
chr14-124738579-DEL-52
chr14-124762324-DEL-195
chr14-124782494-DEL-2576
chr14-124803100-DEL-746
chr14-124863025-DEL-95
chr14-124920266-DEL-1493
chr14-124922111-DEL-56
chr14-124965620-DEL-814
chr14-124988686-DEL-6339
chr14-124995266-DEL-5461
chr14-125016594-DEL-19984
chr14-125023490-DEL-3232
chr14-125034890-DEL-797
chr14-125038889-DEL-73
chr15-3070412-DEL-6369
chr15-3111884-DEL-664
chr15-3131049-DEL-5735
chr15-3162640-DEL-77
chr15-3180298-DEL-6141
chr15-3188914-DEL-1360
chr15-3195385-DEL-4375
chr15-3213013-DEL-1519
chr15-3235429-DEL-328
chr15-3248508-DEL-154
chr15-3274591-DEL-339
chr15-3277832-DEL-84
chr15-3285145-DEL-536
chr15-3363783-DEL-69
chr15-3364022-DEL-1752
chr15-3368694-DEL-166

chr15-3409037-DEL-212
chr15-3442282-DEL-112
chr15-3492319-DEL-603
chr15-3563427-DEL-145
chr15-3573539-DEL-97
chr15-3676955-DEL-383
chr15-3709958-DEL-77
chr15-3757353-DEL-6154
chr15-3800636-DEL-95
chr15-3801845-DEL-3420
chr15-3808128-DEL-906
chr15-3815307-DEL-276
chr15-3817070-DEL-919
chr15-3836507-DEL-6825
chr15-3836862-DEL-203
chr15-3851490-DEL-2779
chr15-3904755-DEL-243
chr15-3938054-DEL-79
chr15-3972689-DEL-7883
chr15-4041850-DEL-51
chr15-4047165-DEL-168
chr15-4106410-DEL-1524
chr15-4113621-DEL-106
chr15-4120992-DEL-61
chr15-4125697-DEL-75
chr15-4133021-DEL-140
chr15-4142734-DEL-64
chr15-4147682-DEL-53
chr15-4185276-DEL-6156
chr15-4194104-DEL-506
chr15-4196874-DEL-60
chr15-4213030-DEL-62
chr15-4274575-DEL-180
chr15-4321885-DEL-95
chr15-4340291-DEL-146
chr15-4409482-DEL-67
chr15-4411927-DEL-155
chr15-4430444-DEL-60
chr15-4466321-DEL-64
chr15-4493001-DEL-282
chr15-4499576-DEL-210
chr15-4519520-DEL-212
chr15-4545396-DEL-51
chr15-4596281-DEL-104
chr15-4621353-DEL-62
chr15-4624982-DEL-3927
chr15-4637388-DEL-6157
chr15-4638225-DEL-4607
chr15-4647511-DEL-764
chr15-4679060-DEL-53
chr15-4679232-DEL-846
chr15-4687948-DEL-101
chr15-4738614-DEL-410
chr15-4747475-DEL-3652
chr15-4749869-DEL-1868
chr15-4793489-DEL-10817
chr15-4804829-DEL-3340
chr15-4809056-DEL-276
chr15-4823650-DEL-54
chr15-4824130-DEL-297
chr15-4831613-DEL-1945
chr15-4856198-DEL-65
chr15-4860528-DEL-63
chr15-4881516-DEL-7183
chr15-4881525-DEL-19304
chr15-4891571-DEL-5768
chr15-4910216-DEL-56
chr15-4968517-DEL-489
chr15-4972548-DEL-82
chr15-4975208-DEL-458
chr15-5022458-DEL-354
chr15-5055382-DEL-1518
chr15-5081834-DEL-2775
chr15-5120358-DEL-107
chr15-5128053-DEL-120
chr15-5143513-DEL-80
chr15-5157856-DEL-55

chr15-5162359-DEL-229
chr15-5163266-DEL-54
chr15-5164023-DEL-3210
chr15-5190227-DEL-1263
chr15-5190434-DEL-609
chr15-5191468-DEL-478
chr15-5191608-DEL-793
chr15-5198991-DEL-350
chr15-5233250-DEL-946
chr15-5241209-DEL-81
chr15-5248509-DEL-119
chr15-5257575-DEL-778
chr15-5260117-DEL-56
chr15-5283248-DEL-236
chr15-5294949-DEL-101
chr15-5304552-DEL-1054
chr15-5333880-DEL-138
chr15-5341862-DEL-1892
chr15-5351450-DEL-78
chr15-5360342-DEL-123
chr15-5366654-DEL-1681
chr15-5396555-DEL-1684
chr15-5398419-DEL-6791
chr15-5398429-DEL-4850
chr15-5402974-DEL-1961
chr15-5407730-DEL-7223
chr15-5450210-DEL-466
chr15-5470783-DEL-291
chr15-5498946-DEL-742
chr15-5504617-DEL-1075
chr15-5506128-DEL-52
chr15-5516292-DEL-636
chr15-5592764-DEL-922
chr15-5605429-DEL-4457
chr15-5605953-DEL-91
chr15-5610527-DEL-52
chr15-5613380-DEL-365
chr15-5631166-DEL-400
chr15-5673473-DEL-244
chr15-5691452-DEL-62
chr15-5696550-DEL-200
chr15-5699471-DEL-2626
chr15-5701195-DEL-59
chr15-5726551-DEL-731
chr15-5744743-DEL-258
chr15-5750221-DEL-67
chr15-5757743-DEL-200
chr15-5762200-DEL-1577
chr15-5776059-DEL-136
chr15-5782427-DEL-83
chr15-5804152-DEL-268
chr15-5806423-DEL-53
chr15-5847902-DEL-306
chr15-5864742-DEL-245
chr15-5869959-DEL-184
chr15-5873219-DEL-1226
chr15-5880811-DEL-380
chr15-5991476-DEL-1423
chr15-5998897-DEL-326
chr15-6008922-DEL-2773
chr15-6009357-DEL-124
chr15-6029602-DEL-402
chr15-6033913-DEL-438
chr15-6047331-DEL-51
chr15-6065754-DEL-56
chr15-6118339-DEL-76
chr15-6138558-DEL-316
chr15-6157572-DEL-105
chr15-6168457-DEL-1070
chr15-6197637-DEL-197
chr15-6200106-DEL-315
chr15-6203397-DEL-674
chr15-6206976-DEL-441
chr15-6232529-DEL-2121
chr15-6235217-DEL-204
chr15-6242731-DEL-92
chr15-6280055-DEL-2369

chr15-6313364-DEL-1189
chr15-6329397-DEL-1541
chr15-6378837-DEL-186
chr15-6381536-DEL-220
chr15-6383713-DEL-79
chr15-6392148-DEL-976
chr15-6426765-DEL-156
chr15-6450287-DEL-61
chr15-6541296-DEL-938
chr15-6545515-DEL-196
chr15-6548324-DEL-185
chr15-6557825-DEL-54
chr15-6558666-DEL-2285
chr15-6558862-DEL-155
chr15-6559276-DEL-184
chr15-6560955-DEL-168
chr15-6561993-DEL-352
chr15-6575328-DEL-498
chr15-6577453-DEL-56
chr15-6597355-DEL-63
chr15-6616023-DEL-90
chr15-6637810-DEL-263
chr15-6640453-DEL-522
chr15-6647300-DEL-1466
chr15-6707874-DEL-66
chr15-6709077-DEL-132
chr15-6709285-DEL-5431
chr15-6710026-DEL-124
chr15-6718011-DEL-781
chr15-6723629-DEL-7169
chr15-6757965-DEL-54
chr15-6852192-DEL-56
chr15-6855192-DEL-116
chr15-6902537-DEL-130
chr15-6912285-DEL-61
chr15-6929167-DEL-58
chr15-6953032-DEL-2208
chr15-6962340-DEL-62
chr15-6986621-DEL-620
chr15-6987265-DEL-111
chr15-6987433-DEL-1750
chr15-7011882-DEL-58
chr15-7020232-DEL-1514
chr15-7022227-DEL-215
chr15-7036129-DEL-106
chr15-7081710-DEL-63
chr15-7083574-DEL-1486
chr15-7164457-DEL-537
chr15-7242947-DEL-53
chr15-7271587-DEL-95
chr15-7282286-DEL-79
chr15-7314935-DEL-58
chr15-7339083-DEL-127
chr15-7379323-DEL-92
chr15-7389967-DEL-250
chr15-7390319-DEL-1055
chr15-7391466-DEL-5289
chr15-7448823-DEL-142
chr15-7451833-DEL-429
chr15-7451862-DEL-69
chr15-7452804-DEL-153
chr15-7470183-DEL-53
chr15-7474632-DEL-80
chr15-7476883-DEL-54
chr15-7495165-DEL-258
chr15-7506907-DEL-52
chr15-7535587-DEL-1815
chr15-7537124-DEL-54
chr15-7558778-DEL-60
chr15-7589419-DEL-1240
chr15-7594879-DEL-59
chr15-7598912-DEL-1395
chr15-7602737-DEL-67
chr15-7607096-DEL-84
chr15-7634901-DEL-51
chr15-7677914-DEL-657
chr15-7691667-DEL-54

chr15-7729236-DEL-97
chr15-7785865-DEL-65
chr15-7793567-DEL-298
chr15-7809297-DEL-53
chr15-7817468-DEL-176
chr15-7830587-DEL-318
chr15-7831375-DEL-391
chr15-7875617-DEL-140
chr15-7927347-DEL-589
chr15-7932598-DEL-55
chr15-7947910-DEL-412
chr15-7948559-DEL-51
chr15-7967205-DEL-124
chr15-8003133-DEL-102
chr15-8004516-DEL-738
chr15-8021383-DEL-192
chr15-8028434-DEL-109
chr15-8068279-DEL-816
chr15-8083814-DEL-56
chr15-8097039-DEL-80
chr15-8101025-DEL-3855
chr15-8102455-DEL-3515
chr15-8149658-DEL-53
chr15-8157194-DEL-308
chr15-8205165-DEL-203
chr15-8206001-DEL-112
chr15-8209663-DEL-163
chr15-8222221-DEL-386
chr15-8225726-DEL-202
chr15-8238184-DEL-122
chr15-8281989-DEL-60
chr15-8282481-DEL-1512
chr15-8282814-DEL-2242
chr15-8296273-DEL-1043
chr15-8297555-DEL-1323
chr15-8298124-DEL-197
chr15-8345417-DEL-54
chr15-8381359-DEL-5014
chr15-8444371-DEL-205
chr15-8450807-DEL-203
chr15-8463022-DEL-84
chr15-8509946-DEL-166
chr15-8516193-DEL-1537
chr15-8524013-DEL-106
chr15-8547737-DEL-1748
chr15-8550770-DEL-302
chr15-8571113-DEL-837
chr15-8586545-DEL-495
chr15-8594350-DEL-90
chr15-8596857-DEL-95
chr15-8599600-DEL-179
chr15-8610357-DEL-123
chr15-8630232-DEL-132
chr15-8699265-DEL-352
chr15-8704140-DEL-994
chr15-8708291-DEL-460
chr15-8723542-DEL-241
chr15-8724279-DEL-54
chr15-8742677-DEL-136
chr15-8747142-DEL-182
chr15-8753418-DEL-467
chr15-8760894-DEL-575
chr15-8761579-DEL-463
chr15-8762144-DEL-678
chr15-8770124-DEL-201
chr15-8780856-DEL-256
chr15-8782250-DEL-85
chr15-8790785-DEL-65
chr15-8795817-DEL-2860
chr15-8834739-DEL-427
chr15-8842260-DEL-201
chr15-8876285-DEL-82
chr15-8878420-DEL-162
chr15-8885789-DEL-115
chr15-8904972-DEL-253
chr15-8932038-DEL-132
chr15-8947437-DEL-10355

chr15-8959516-DEL-439
chr15-8977956-DEL-60
chr15-9004641-DEL-58
chr15-9036034-DEL-127
chr15-9045517-DEL-7383
chr15-9045520-DEL-4278
chr15-9045645-DEL-759
chr15-9049304-DEL-1380
chr15-9049787-DEL-552
chr15-9052346-DEL-138
chr15-9101505-DEL-88
chr15-9103151-DEL-156
chr15-9120860-DEL-104
chr15-9132518-DEL-84
chr15-9154520-DEL-63
chr15-9189208-DEL-93
chr15-9202967-DEL-342
chr15-9212817-DEL-52
chr15-9217336-DEL-10699
chr15-9217539-DEL-1036
chr15-9227294-DEL-326
chr15-9236585-DEL-51
chr15-9265134-DEL-1727
chr15-9268068-DEL-365
chr15-9293019-DEL-52
chr15-9299771-DEL-197
chr15-9327509-DEL-59
chr15-9352140-DEL-58
chr15-9354711-DEL-2644
chr15-9376402-DEL-361
chr15-9386535-DEL-6322
chr15-9422411-DEL-56
chr15-9436376-DEL-113
chr15-9446299-DEL-1659
chr15-9448788-DEL-337
chr15-9461954-DEL-135
chr15-9467627-DEL-132
chr15-9468718-DEL-547
chr15-9578052-DEL-105
chr15-9615737-DEL-466
chr15-9619361-DEL-371
chr15-9682880-DEL-640
chr15-9706133-DEL-90
chr15-9707086-DEL-53
chr15-9737716-DEL-139
chr15-9766380-DEL-2167
chr15-9772870-DEL-6422
chr15-9780142-DEL-379
chr15-9830877-DEL-7562
chr15-9866887-DEL-57
chr15-9888217-DEL-165
chr15-9901890-DEL-2540
chr15-9911622-DEL-127
chr15-9917665-DEL-207
chr15-9992041-DEL-208
chr15-10012242-DEL-1387
chr15-10019594-DEL-5951
chr15-10026996-DEL-273
chr15-10052988-DEL-96
chr15-10060705-DEL-5490
chr15-10103379-DEL-943
chr15-10174667-DEL-112
chr15-10244611-DEL-123
chr15-10261337-DEL-456
chr15-10323676-DEL-1313
chr15-10401493-DEL-153
chr15-10460618-DEL-97
chr15-10465945-DEL-299
chr15-10494199-DEL-166
chr15-10498016-DEL-144
chr15-10510316-DEL-551
chr15-10533541-DEL-70
chr15-10536827-DEL-64
chr15-10543257-DEL-964
chr15-10561541-DEL-128
chr15-10573514-DEL-52
chr15-10578280-DEL-78

chr15-10579660-DEL-1427
chr15-10579675-DEL-755
chr15-10580693-DEL-1428
chr15-10581912-DEL-82
chr15-10583669-DEL-59
chr15-10590414-DEL-840
chr15-10619206-DEL-96
chr15-10619724-DEL-83
chr15-10658202-DEL-75
chr15-10766199-DEL-142
chr15-10778103-DEL-624
chr15-10781265-DEL-6204
chr15-10849680-DEL-1339
chr15-10853241-DEL-10599
chr15-10858096-DEL-531
chr15-10874838-DEL-54
chr15-10886025-DEL-443
chr15-10906914-DEL-251
chr15-10939008-DEL-338
chr15-10946338-DEL-54
chr15-10968929-DEL-54
chr15-10978880-DEL-104
chr15-10991667-DEL-187
chr15-10998865-DEL-188
chr15-11004333-DEL-529
chr15-11011348-DEL-98
chr15-11032842-DEL-62
chr15-11060636-DEL-64
chr15-11063697-DEL-89
chr15-11073671-DEL-68
chr15-11085321-DEL-168
chr15-11092737-DEL-77
chr15-11095079-DEL-55
chr15-11100776-DEL-740
chr15-11142295-DEL-78
chr15-11142947-DEL-1913
chr15-11165568-DEL-170
chr15-11175903-DEL-200
chr15-11182488-DEL-1890
chr15-11189346-DEL-159
chr15-11204132-DEL-64
chr15-11204164-DEL-760
chr15-11246142-DEL-1729
chr15-11252718-DEL-278
chr15-11267260-DEL-128
chr15-11284294-DEL-62
chr15-11312229-DEL-192
chr15-11314974-DEL-105
chr15-11325581-DEL-58
chr15-11330055-DEL-102
chr15-11350373-DEL-72
chr15-11366661-DEL-881
chr15-11402919-DEL-79
chr15-11403965-DEL-807
chr15-11480113-DEL-244
chr15-11535987-DEL-56
chr15-11567469-DEL-399
chr15-11579408-DEL-117
chr15-11580374-DEL-2515
chr15-11580375-DEL-373
chr15-11594357-DEL-61
chr15-11600692-DEL-4223
chr15-11610672-DEL-73
chr15-11611366-DEL-72
chr15-11611509-DEL-148
chr15-11618163-DEL-6244
chr15-11659100-DEL-1443
chr15-11660072-DEL-620
chr15-11727374-DEL-54
chr15-11749609-DEL-1793
chr15-11762659-DEL-313
chr15-11828490-DEL-2082
chr15-11849363-DEL-90
chr15-11857821-DEL-228
chr15-11892698-DEL-474
chr15-11916678-DEL-7996
chr15-11951627-DEL-209

chr15-11971694-DEL-200
chr15-11972972-DEL-684
chr15-11975217-DEL-88
chr15-11997503-DEL-1512
chr15-12008540-DEL-112
chr15-12033668-DEL-499
chr15-12036299-DEL-60
chr15-12043533-DEL-400
chr15-12045905-DEL-460
chr15-12065455-DEL-128
chr15-12074969-DEL-248
chr15-12080171-DEL-283
chr15-12086873-DEL-75
chr15-12096078-DEL-86
chr15-12098498-DEL-258
chr15-12098509-DEL-738
chr15-12098513-DEL-1275
chr15-12164437-DEL-572
chr15-12173241-DEL-203
chr15-12181155-DEL-721
chr15-12200404-DEL-344
chr15-12224425-DEL-145
chr15-12225129-DEL-52
chr15-12232355-DEL-56
chr15-12235224-DEL-88
chr15-12249796-DEL-682
chr15-12256303-DEL-205
chr15-12259989-DEL-401
chr15-12293294-DEL-3076
chr15-12300157-DEL-8977
chr15-12332881-DEL-354
chr15-12336339-DEL-118
chr15-12346584-DEL-1846
chr15-12412335-DEL-200
chr15-12418237-DEL-52
chr15-12460574-DEL-79
chr15-12463168-DEL-88
chr15-12473278-DEL-138
chr15-12495367-DEL-144
chr15-12531967-DEL-53
chr15-12537311-DEL-386
chr15-12570010-DEL-141
chr15-12580329-DEL-290
chr15-12612891-DEL-112
chr15-12666219-DEL-60
chr15-12686630-DEL-56
chr15-12711059-DEL-1189
chr15-12724389-DEL-52
chr15-12748897-DEL-52
chr15-12787151-DEL-61
chr15-12795457-DEL-65
chr15-12803551-DEL-77
chr15-12835282-DEL-1598
chr15-12869081-DEL-56
chr15-12869747-DEL-350
chr15-12872785-DEL-683
chr15-12874065-DEL-202
chr15-12876210-DEL-1410
chr15-12885401-DEL-1505
chr15-12886765-DEL-142
chr15-12888864-DEL-92
chr15-12899074-DEL-61
chr15-12901671-DEL-52
chr15-12908919-DEL-51
chr15-12923594-DEL-123
chr15-12968341-DEL-82
chr15-13001457-DEL-52
chr15-13002844-DEL-1564
chr15-13003435-DEL-543
chr15-13004221-DEL-51
chr15-13046975-DEL-434
chr15-13110884-DEL-2035
chr15-13145288-DEL-55
chr15-13220702-DEL-589
chr15-13226604-DEL-282
chr15-13233877-DEL-106
chr15-13237728-DEL-184

chr15-13249714-DEL-66
chr15-13264274-DEL-66
chr15-13282659-DEL-65
chr15-13334506-DEL-54
chr15-13383509-DEL-368
chr15-13412087-DEL-86
chr15-13556460-DEL-55
chr15-13645213-DEL-202
chr15-13659792-DEL-356
chr15-13673322-DEL-67
chr15-13678865-DEL-219
chr15-13695365-DEL-135
chr15-13704451-DEL-53
chr15-13720824-DEL-792
chr15-13742212-DEL-738
chr15-13770662-DEL-76
chr15-13787217-DEL-429
chr15-13789143-DEL-173
chr15-13798143-DEL-504
chr15-13798427-DEL-1005
chr15-13798991-DEL-72
chr15-13843443-DEL-330
chr15-13846511-DEL-3723
chr15-13868673-DEL-258
chr15-13887508-DEL-1715
chr15-13922172-DEL-8469
chr15-13926406-DEL-475
chr15-13942002-DEL-54
chr15-13965347-DEL-55
chr15-13967854-DEL-72
chr15-13987695-DEL-879
chr15-13990812-DEL-910
chr15-13991916-DEL-87
chr15-13994414-DEL-62
chr15-14029854-DEL-61
chr15-14046479-DEL-74
chr15-14067295-DEL-2119
chr15-14086658-DEL-268
chr15-14096284-DEL-119
chr15-14100782-DEL-59
chr15-14108392-DEL-6203
chr15-14125733-DEL-629
chr15-14200522-DEL-406
chr15-14232959-DEL-804
chr15-14239390-DEL-55
chr15-14241833-DEL-88
chr15-14255474-DEL-117
chr15-14279402-DEL-76
chr15-14294944-DEL-250
chr15-14306454-DEL-82
chr15-14308554-DEL-54
chr15-14329315-DEL-61
chr15-14330985-DEL-88
chr15-14336787-DEL-97
chr15-14384068-DEL-102
chr15-14394989-DEL-1051
chr15-14396737-DEL-56
chr15-14397936-DEL-442
chr15-14398437-DEL-1294
chr15-14420636-DEL-83
chr15-14424536-DEL-299
chr15-14445795-DEL-138
chr15-14466826-DEL-90
chr15-14471352-DEL-94
chr15-14475052-DEL-63
chr15-14476678-DEL-8628
chr15-14480550-DEL-4436
chr15-14497652-DEL-293
chr15-14506819-DEL-1296
chr15-14510492-DEL-957
chr15-14513144-DEL-2064
chr15-14524443-DEL-383
chr15-14531498-DEL-6246
chr15-14564375-DEL-62
chr15-14567920-DEL-5501
chr15-14568834-DEL-96
chr15-14573385-DEL-348

chr15-14574099-DEL-1224
chr15-14596791-DEL-61
chr15-14607274-DEL-1260
chr15-14618232-DEL-862
chr15-14621390-DEL-724
chr15-14644320-DEL-61
chr15-14646263-DEL-91
chr15-14677676-DEL-373
chr15-14681904-DEL-73
chr15-14687150-DEL-764
chr15-14689642-DEL-112
chr15-14708048-DEL-734
chr15-14716140-DEL-413
chr15-14738678-DEL-78
chr15-14754497-DEL-2705
chr15-14764555-DEL-2860
chr15-14777892-DEL-54
chr15-14799334-DEL-64
chr15-14857428-DEL-51
chr15-14870822-DEL-921
chr15-14885366-DEL-138
chr15-14891251-DEL-11226
chr15-14896255-DEL-5007
chr15-14935454-DEL-289
chr15-14937069-DEL-1295
chr15-14946300-DEL-75
chr15-14959770-DEL-1692
chr15-14963141-DEL-1739
chr15-14978641-DEL-2411
chr15-15002203-DEL-81
chr15-15043513-DEL-1218
chr15-15048690-DEL-12835
chr15-15071025-DEL-110
chr15-15073918-DEL-328
chr15-15076808-DEL-7210
chr15-15084222-DEL-79
chr15-15086172-DEL-1270
chr15-15100867-DEL-963
chr15-15112093-DEL-501
chr15-15115868-DEL-76
chr15-15123404-DEL-1603
chr15-15136202-DEL-2339
chr15-15143383-DEL-1648
chr15-15152053-DEL-121
chr15-15152727-DEL-1106
chr15-15199017-DEL-75
chr15-15221078-DEL-61
chr15-15237799-DEL-144
chr15-15250136-DEL-76
chr15-15254409-DEL-114
chr15-15260376-DEL-428
chr15-15261437-DEL-1878
chr15-15265443-DEL-6310
chr15-15273000-DEL-6021
chr15-15282572-DEL-60
chr15-15300652-DEL-263
chr15-15304696-DEL-146
chr15-15324254-DEL-99
chr15-15326833-DEL-2817
chr15-15342899-DEL-1142
chr15-15360549-DEL-57
chr15-15366168-DEL-111
chr15-15372766-DEL-77
chr15-15417406-DEL-3375
chr15-15429726-DEL-52
chr15-15478941-DEL-611
chr15-15490519-DEL-1488
chr15-15500241-DEL-219
chr15-15525483-DEL-5305
chr15-15555364-DEL-442
chr15-15557785-DEL-56
chr15-15592063-DEL-65
chr15-15610057-DEL-52
chr15-15628665-DEL-364
chr15-15676915-DEL-532
chr15-15681111-DEL-194
chr15-15692415-DEL-150

chr15-15698145-DEL-899
chr15-15712071-DEL-206
chr15-15717772-DEL-58
chr15-15721437-DEL-1753
chr15-15760119-DEL-94
chr15-15771532-DEL-1217
chr15-15789895-DEL-1390
chr15-15796297-DEL-54
chr15-15828560-DEL-81
chr15-15835884-DEL-56
chr15-15879110-DEL-327
chr15-15901958-DEL-390
chr15-15914608-DEL-6208
chr15-15935708-DEL-558
chr15-15937263-DEL-613
chr15-15947289-DEL-210
chr15-15948721-DEL-222
chr15-15961395-DEL-861
chr15-15987495-DEL-688
chr15-15989323-DEL-961
chr15-15998341-DEL-52
chr15-16003688-DEL-242
chr15-16022498-DEL-452
chr15-16050013-DEL-2415
chr15-16064215-DEL-155
chr15-16068290-DEL-58
chr15-16072852-DEL-710
chr15-16078068-DEL-64
chr15-16080499-DEL-60
chr15-16087465-DEL-836
chr15-16125621-DEL-237
chr15-16149547-DEL-51
chr15-16150452-DEL-283
chr15-16155539-DEL-54
chr15-16168235-DEL-1683
chr15-16171232-DEL-430
chr15-16172191-DEL-1379
chr15-16174074-DEL-378
chr15-16175580-DEL-1477
chr15-16213909-DEL-56
chr15-16241434-DEL-967
chr15-16259223-DEL-931
chr15-16271032-DEL-54
chr15-16283095-DEL-262
chr15-16321581-DEL-194
chr15-16345859-DEL-58
chr15-16373752-DEL-637
chr15-16374673-DEL-418
chr15-16382875-DEL-583
chr15-16415594-DEL-347
chr15-16440312-DEL-60
chr15-16447029-DEL-107
chr15-16460218-DEL-58
chr15-16461535-DEL-87
chr15-16461893-DEL-2010
chr15-16464518-DEL-533
chr15-16477183-DEL-53
chr15-16507472-DEL-91
chr15-16573781-DEL-782
chr15-16587797-DEL-333
chr15-16588225-DEL-1651
chr15-16619209-DEL-166
chr15-16626439-DEL-106
chr15-16647351-DEL-544
chr15-16649203-DEL-459
chr15-16651088-DEL-98
chr15-16691596-DEL-292
chr15-16704368-DEL-2143
chr15-16708230-DEL-52
chr15-16709637-DEL-751
chr15-16716429-DEL-1189
chr15-16718805-DEL-66
chr15-16804146-DEL-105
chr15-16812393-DEL-54
chr15-16845051-DEL-1250
chr15-16866286-DEL-350
chr15-16870103-DEL-1195

chr15-16873007-DEL-768
chr15-16899050-DEL-24077
chr15-16901412-DEL-3864
chr15-16905302-DEL-2305
chr15-16939792-DEL-1087
chr15-16939793-DEL-888
chr15-16940701-DEL-378
chr15-17011584-DEL-291
chr15-17022717-DEL-196
chr15-17027608-DEL-55
chr15-17049631-DEL-57
chr15-17050285-DEL-246
chr15-17083961-DEL-1056
chr15-17088867-DEL-199
chr15-17088867-DEL-6607
chr15-17110411-DEL-54
chr15-17131705-DEL-169
chr15-17132367-DEL-1391
chr15-17133091-DEL-109
chr15-17136534-DEL-336
chr15-17140619-DEL-601
chr15-17143221-DEL-112
chr15-17169193-DEL-416
chr15-17171099-DEL-143
chr15-17189663-DEL-2386
chr15-17210508-DEL-399
chr15-17229910-DEL-647
chr15-17237814-DEL-1889
chr15-17248115-DEL-116
chr15-17258921-DEL-62
chr15-17266008-DEL-52
chr15-17311303-DEL-54
chr15-17314479-DEL-149
chr15-17339201-DEL-1770
chr15-17346555-DEL-57
chr15-17358910-DEL-671
chr15-17359832-DEL-1161
chr15-17378080-DEL-196
chr15-17388754-DEL-74
chr15-17394666-DEL-55
chr15-17432710-DEL-706
chr15-17456040-DEL-193
chr15-17469811-DEL-121
chr15-17470450-DEL-123
chr15-17478908-DEL-1652
chr15-17489156-DEL-130
chr15-17510684-DEL-74
chr15-17515069-DEL-439
chr15-17527128-DEL-93
chr15-17550149-DEL-3821
chr15-17577958-DEL-237
chr15-17587878-DEL-97
chr15-17615705-DEL-184
chr15-17619874-DEL-116
chr15-17681756-DEL-83
chr15-17682524-DEL-6413
chr15-17701202-DEL-844
chr15-17701756-DEL-444
chr15-17704276-DEL-104
chr15-17711598-DEL-86
chr15-17721826-DEL-2234
chr15-17737045-DEL-1843
chr15-17747264-DEL-411
chr15-17750475-DEL-6175
chr15-17795156-DEL-112
chr15-17856803-DEL-107
chr15-17896404-DEL-1109
chr15-17903200-DEL-6258
chr15-17918893-DEL-6657
chr15-17927527-DEL-238
chr15-17944249-DEL-72
chr15-17958715-DEL-105
chr15-17959669-DEL-964
chr15-17969262-DEL-3062
chr15-17971005-DEL-110
chr15-17989139-DEL-76
chr15-18009120-DEL-334

chr15-18044475-DEL-394
chr15-18045488-DEL-68
chr15-18093720-DEL-63
chr15-18107070-DEL-4636
chr15-18118610-DEL-791
chr15-18138983-DEL-85
chr15-18151575-DEL-1913
chr15-18175006-DEL-857
chr15-18177048-DEL-423
chr15-18183777-DEL-292
chr15-18191807-DEL-438
chr15-18192941-DEL-62
chr15-18218244-DEL-117
chr15-18223361-DEL-54
chr15-18238332-DEL-1758
chr15-18248496-DEL-51
chr15-18265761-DEL-90
chr15-18267483-DEL-6237
chr15-18276496-DEL-537
chr15-18309787-DEL-2099
chr15-18312061-DEL-100
chr15-18325717-DEL-109
chr15-18329464-DEL-103
chr15-18354962-DEL-1108
chr15-18389049-DEL-67
chr15-18404211-DEL-556
chr15-18411387-DEL-77
chr15-18433634-DEL-177
chr15-18441829-DEL-316
chr15-18444914-DEL-2514
chr15-18459934-DEL-315
chr15-18461203-DEL-408
chr15-18474609-DEL-495
chr15-18570353-DEL-172
chr15-18599174-DEL-558
chr15-18615364-DEL-53
chr15-18630849-DEL-212
chr15-18632020-DEL-461
chr15-18635690-DEL-79
chr15-18640940-DEL-60
chr15-18641681-DEL-495
chr15-18662297-DEL-2011
chr15-18670549-DEL-204
chr15-18675174-DEL-107
chr15-18676910-DEL-115
chr15-18683979-DEL-111
chr15-18712340-DEL-64
chr15-18725190-DEL-6405
chr15-18735675-DEL-77
chr15-18794117-DEL-2483
chr15-18803639-DEL-131
chr15-18814532-DEL-469
chr15-18856796-DEL-282
chr15-18860891-DEL-52
chr15-18863510-DEL-1945
chr15-18871235-DEL-4780
chr15-18924348-DEL-56
chr15-18944621-DEL-60
chr15-18982242-DEL-551
chr15-18988482-DEL-787
chr15-18999273-DEL-258
chr15-19005423-DEL-1549
chr15-19020315-DEL-118
chr15-19075350-DEL-8047
chr15-19080958-DEL-284
chr15-19095686-DEL-6446
chr15-19103097-DEL-130
chr15-19135082-DEL-1946
chr15-19146040-DEL-52
chr15-19158678-DEL-838
chr15-19159297-DEL-831
chr15-19175929-DEL-98
chr15-19176798-DEL-2279
chr15-19199596-DEL-1727
chr15-19218555-DEL-1992
chr15-19238658-DEL-59
chr15-19282741-DEL-81

chr15-19291126-DEL-2319
chr15-19294828-DEL-112
chr15-19302906-DEL-112
chr15-19338169-DEL-53
chr15-19365498-DEL-996
chr15-19368923-DEL-482
chr15-19388149-DEL-465
chr15-19390855-DEL-545
chr15-19395456-DEL-240
chr15-19448153-DEL-6562
chr15-19452158-DEL-1490
chr15-19456186-DEL-86
chr15-19465954-DEL-175
chr15-19474187-DEL-2962
chr15-19495443-DEL-4039
chr15-19507280-DEL-918
chr15-19509720-DEL-335
chr15-19514929-DEL-81
chr15-19515193-DEL-2105
chr15-19520564-DEL-6475
chr15-19523973-DEL-7694
chr15-19530420-DEL-56
chr15-19556637-DEL-1788
chr15-19558794-DEL-6089
chr15-19567903-DEL-6422
chr15-19567959-DEL-12655
chr15-19569798-DEL-11233
chr15-19570051-DEL-1311
chr15-19571261-DEL-343
chr15-19571770-DEL-2528
chr15-19574400-DEL-6216
chr15-19576294-DEL-4736
chr15-19592735-DEL-1835
chr15-19602431-DEL-88
chr15-19609436-DEL-1468
chr15-19626159-DEL-199
chr15-19636930-DEL-1421
chr15-19639655-DEL-6195
chr15-19659517-DEL-80
chr15-19679080-DEL-208
chr15-19695201-DEL-2149
chr15-19697487-DEL-115
chr15-19709932-DEL-80
chr15-19718135-DEL-294
chr15-19737047-DEL-126
chr15-19745208-DEL-1436
chr15-19752819-DEL-1552
chr15-19776057-DEL-457
chr15-19802987-DEL-68
chr15-19806995-DEL-62
chr15-19825666-DEL-75
chr15-19843684-DEL-6248
chr15-19851683-DEL-52
chr15-19858079-DEL-9924
chr15-19861075-DEL-72
chr15-19875087-DEL-184
chr15-19884272-DEL-633
chr15-19899723-DEL-1527
chr15-19901351-DEL-753
chr15-19942660-DEL-55
chr15-19945211-DEL-1190
chr15-19951172-DEL-3549
chr15-19954287-DEL-6198
chr15-19960679-DEL-1002
chr15-19963873-DEL-81
chr15-19974581-DEL-939
chr15-19991450-DEL-565
chr15-19997428-DEL-8410
chr15-20009134-DEL-4148
chr15-20013491-DEL-5016
chr15-20017019-DEL-4897
chr15-20046447-DEL-193
chr15-20049518-DEL-190
chr15-20054471-DEL-1843
chr15-20060186-DEL-7624
chr15-20078256-DEL-72
chr15-20099134-DEL-436

chr15-20107993-DEL-210
chr15-20109606-DEL-433
chr15-20123284-DEL-184
chr15-20150098-DEL-51
chr15-20151131-DEL-62
chr15-20154789-DEL-5085
chr15-20160763-DEL-78
chr15-20168700-DEL-6107
chr15-20175051-DEL-54
chr15-20176600-DEL-1626
chr15-20178344-DEL-585
chr15-20202122-DEL-6471
chr15-20204540-DEL-398
chr15-20220037-DEL-84
chr15-20227227-DEL-1050
chr15-20250502-DEL-325
chr15-20254885-DEL-140
chr15-20257240-DEL-199
chr15-20259034-DEL-142
chr15-20259873-DEL-421
chr15-20261227-DEL-261
chr15-20274906-DEL-2844
chr15-20284808-DEL-1533
chr15-20292275-DEL-70
chr15-20292478-DEL-1011
chr15-20309538-DEL-112
chr15-20343228-DEL-3158
chr15-20348790-DEL-5687
chr15-20355050-DEL-894
chr15-20363187-DEL-293
chr15-20377935-DEL-685
chr15-20427547-DEL-64
chr15-20453601-DEL-67
chr15-20482747-DEL-177
chr15-20489739-DEL-90
chr15-20497963-DEL-9447
chr15-20509056-DEL-1210
chr15-20584204-DEL-146
chr15-20611726-DEL-112
chr15-20624548-DEL-482
chr15-20644768-DEL-2490
chr15-20644924-DEL-120
chr15-20670005-DEL-208
chr15-20681525-DEL-1100
chr15-20699537-DEL-931
chr15-20701231-DEL-1654
chr15-20701754-DEL-52
chr15-20710224-DEL-1563
chr15-20745994-DEL-397
chr15-20782771-DEL-2218
chr15-20805147-DEL-1369
chr15-20841783-DEL-64
chr15-20857578-DEL-58
chr15-20870333-DEL-1288
chr15-20899459-DEL-2081
chr15-20966844-DEL-138
chr15-20992087-DEL-68
chr15-20993578-DEL-202
chr15-21032439-DEL-1254
chr15-21035672-DEL-358
chr15-21042725-DEL-71
chr15-21051116-DEL-208
chr15-21057902-DEL-874
chr15-21074292-DEL-81
chr15-21077268-DEL-109
chr15-21092279-DEL-492
chr15-21101034-DEL-500
chr15-21114438-DEL-681
chr15-21145826-DEL-78
chr15-21150570-DEL-94
chr15-21152524-DEL-73
chr15-21163565-DEL-797
chr15-21197797-DEL-854
chr15-21214574-DEL-193
chr15-21249300-DEL-56
chr15-21274122-DEL-502
chr15-21304005-DEL-94

chr15-21318498-DEL-1454
chr15-21334707-DEL-217
chr15-21361643-DEL-2594
chr15-21364328-DEL-106
chr15-21423009-DEL-2524
chr15-21425825-DEL-64
chr15-21437066-DEL-943
chr15-21443693-DEL-10953
chr15-21443704-DEL-416
chr15-21457322-DEL-1139
chr15-21472157-DEL-6150
chr15-21480939-DEL-56
chr15-21485191-DEL-2626
chr15-21489628-DEL-214
chr15-21496589-DEL-629
chr15-21498231-DEL-62
chr15-21522075-DEL-112
chr15-21524267-DEL-60
chr15-21551696-DEL-856
chr15-21554389-DEL-117
chr15-21565293-DEL-340
chr15-21580995-DEL-266
chr15-21621627-DEL-945
chr15-21632764-DEL-491
chr15-21636511-DEL-61
chr15-21649754-DEL-61
chr15-21680981-DEL-354
chr15-21705342-DEL-1450
chr15-21706983-DEL-286
chr15-21728371-DEL-80
chr15-21740038-DEL-1246
chr15-21740589-DEL-473
chr15-21750906-DEL-468
chr15-21755279-DEL-179
chr15-21755730-DEL-831
chr15-21777315-DEL-358
chr15-21784794-DEL-53
chr15-21806382-DEL-1266
chr15-21814917-DEL-6193
chr15-21823712-DEL-51
chr15-21846618-DEL-78
chr15-21852834-DEL-2585
chr15-21861030-DEL-19258
chr15-21861922-DEL-6186
chr15-21869970-DEL-6675
chr15-21902467-DEL-945
chr15-21917714-DEL-7355
chr15-21928319-DEL-1361
chr15-21931869-DEL-322
chr15-21937837-DEL-64
chr15-21942640-DEL-94
chr15-21944665-DEL-2764
chr15-21952636-DEL-102
chr15-21958360-DEL-1570
chr15-21964348-DEL-1041
chr15-21966360-DEL-329
chr15-21967396-DEL-101
chr15-21978350-DEL-503
chr15-21981595-DEL-722
chr15-21982454-DEL-216
chr15-21985773-DEL-424
chr15-21987779-DEL-102
chr15-21992570-DEL-60
chr15-22006663-DEL-1945
chr15-22012539-DEL-401
chr15-22037965-DEL-1972
chr15-22049605-DEL-231
chr15-22051632-DEL-281
chr15-22067679-DEL-4346
chr15-22084223-DEL-431
chr15-22097907-DEL-1298
chr15-22172053-DEL-168
chr15-22178331-DEL-939
chr15-22182605-DEL-1386
chr15-22185058-DEL-844
chr15-22202170-DEL-141
chr15-22211536-DEL-660

chr15-22211656-DEL-142
chr15-22291296-DEL-6191
chr15-22303993-DEL-2938
chr15-22320166-DEL-342
chr15-22333181-DEL-245
chr15-22381173-DEL-1617
chr15-22383924-DEL-337
chr15-22395924-DEL-191
chr15-22410755-DEL-219
chr15-22416035-DEL-118
chr15-22421465-DEL-2911
chr15-22445132-DEL-135
chr15-22445999-DEL-751
chr15-22448435-DEL-68
chr15-22475051-DEL-106
chr15-22487938-DEL-6498
chr15-22492846-DEL-1178
chr15-22499871-DEL-251
chr15-22515355-DEL-60
chr15-22578550-DEL-147
chr15-22603891-DEL-145
chr15-22615085-DEL-56
chr15-22629479-DEL-58
chr15-22656516-DEL-182
chr15-22660518-DEL-1273
chr15-22676901-DEL-1861
chr15-22684189-DEL-6260
chr15-22693154-DEL-6362
chr15-22699732-DEL-5738
chr15-22713219-DEL-1758
chr15-22715149-DEL-525
chr15-22762533-DEL-570
chr15-22763279-DEL-361
chr15-22766889-DEL-53
chr15-22771251-DEL-3836
chr15-22777610-DEL-58
chr15-22783675-DEL-183
chr15-22799717-DEL-110
chr15-22799717-DEL-1210
chr15-22799718-DEL-605
chr15-22805795-DEL-6214
chr15-22868782-DEL-218
chr15-22939716-DEL-669
chr15-22953231-DEL-252
chr15-22964502-DEL-69
chr15-22978321-DEL-64
chr15-22982233-DEL-597
chr15-22987367-DEL-73
chr15-22988974-DEL-74
chr15-23000024-DEL-74
chr15-23001083-DEL-100
chr15-23022136-DEL-52
chr15-23048067-DEL-340
chr15-23062027-DEL-249
chr15-23085042-DEL-98
chr15-23127514-DEL-61
chr15-23140156-DEL-653
chr15-23143249-DEL-161
chr15-23145997-DEL-3095
chr15-23168387-DEL-401
chr15-23169341-DEL-53
chr15-23193193-DEL-103
chr15-23215365-DEL-898
chr15-23237412-DEL-881
chr15-23238529-DEL-56
chr15-23259941-DEL-94
chr15-23263859-DEL-51
chr15-23275595-DEL-64
chr15-23280641-DEL-286
chr15-23281187-DEL-1862
chr15-23285108-DEL-106
chr15-23297477-DEL-212
chr15-23344275-DEL-2188
chr15-23356479-DEL-342
chr15-23361149-DEL-72
chr15-23362689-DEL-1266
chr15-23382151-DEL-278

chr15-23384521-DEL-263
chr15-23395107-DEL-327
chr15-23402975-DEL-1115
chr15-23419149-DEL-53
chr15-23439913-DEL-199
chr15-23462562-DEL-6141
chr15-23490967-DEL-257
chr15-23507956-DEL-428
chr15-23513466-DEL-65
chr15-23520402-DEL-299
chr15-23529991-DEL-1829
chr15-23534065-DEL-172
chr15-23538615-DEL-680
chr15-23540320-DEL-491
chr15-23561197-DEL-85
chr15-23566165-DEL-108
chr15-23574279-DEL-1360
chr15-23615857-DEL-62
chr15-23625988-DEL-595
chr15-23639851-DEL-500
chr15-23643176-DEL-438
chr15-23651224-DEL-120
chr15-23668419-DEL-110
chr15-23674276-DEL-64
chr15-23683563-DEL-286
chr15-23693640-DEL-361
chr15-23700212-DEL-2606
chr15-23703226-DEL-1070
chr15-23710641-DEL-1470
chr15-23717094-DEL-429
chr15-23719727-DEL-74
chr15-23725446-DEL-56
chr15-23738100-DEL-4716
chr15-23761364-DEL-58
chr15-23770551-DEL-6331
chr15-23786803-DEL-2045
chr15-23793128-DEL-2724
chr15-23806171-DEL-8337
chr15-23820720-DEL-54
chr15-23861243-DEL-80
chr15-23862220-DEL-311
chr15-23893946-DEL-83
chr15-23897429-DEL-611
chr15-23906834-DEL-331
chr15-23913283-DEL-923
chr15-23919706-DEL-199
chr15-23926951-DEL-3033
chr15-23944053-DEL-52
chr15-23958428-DEL-3405
chr15-23963452-DEL-1328
chr15-23977431-DEL-85
chr15-23994918-DEL-128
chr15-23997606-DEL-512
chr15-24023381-DEL-706
chr15-24064357-DEL-6216
chr15-24120748-DEL-322
chr15-24129229-DEL-100
chr15-24145276-DEL-60
chr15-24176991-DEL-652
chr15-24180597-DEL-103
chr15-24210717-DEL-1358
chr15-24214865-DEL-6243
chr15-24247426-DEL-1009
chr15-24252595-DEL-52
chr15-24269064-DEL-60
chr15-24278315-DEL-340
chr15-24285003-DEL-118
chr15-24294951-DEL-2264
chr15-24297865-DEL-401
chr15-24299724-DEL-400
chr15-24309347-DEL-7699
chr15-24347873-DEL-245
chr15-24355342-DEL-51
chr15-24366238-DEL-6020
chr15-24376617-DEL-2809
chr15-24379182-DEL-545
chr15-24384867-DEL-163

chr15-24387570-DEL-61
chr15-24396237-DEL-437
chr15-24400331-DEL-890
chr15-24411878-DEL-208
chr15-24425510-DEL-76
chr15-24433861-DEL-453
chr15-24440467-DEL-253
chr15-24447034-DEL-1260
chr15-24467721-DEL-56
chr15-24476025-DEL-62
chr15-24477018-DEL-4402
chr15-24479178-DEL-106
chr15-24540305-DEL-82
chr15-24551589-DEL-147
chr15-24587921-DEL-66
chr15-24588909-DEL-166
chr15-24637047-DEL-1726
chr15-24642739-DEL-6210
chr15-24653712-DEL-6517
chr15-24679751-DEL-68
chr15-24714991-DEL-198
chr15-24715588-DEL-447
chr15-24722152-DEL-96
chr15-24781700-DEL-2297
chr15-24788033-DEL-721
chr15-24801596-DEL-7722
chr15-24822729-DEL-62
chr15-24889543-DEL-70
chr15-24904747-DEL-109
chr15-24914138-DEL-67
chr15-24930400-DEL-87
chr15-24942406-DEL-603
chr15-24945009-DEL-641
chr15-24948233-DEL-918
chr15-24968197-DEL-85
chr15-24975876-DEL-76
chr15-24977788-DEL-58
chr15-25036830-DEL-502
chr15-25044838-DEL-204
chr15-25070795-DEL-107
chr15-25084376-DEL-152
chr15-25209529-DEL-74
chr15-25211285-DEL-375
chr15-25266253-DEL-145
chr15-25347654-DEL-81
chr15-25372061-DEL-163
chr15-25385894-DEL-55
chr15-25416411-DEL-95
chr15-25437063-DEL-184
chr15-25446992-DEL-55
chr15-25480319-DEL-360
chr15-25484533-DEL-99
chr15-25500654-DEL-910
chr15-25508882-DEL-89
chr15-25520889-DEL-497
chr15-25570216-DEL-90
chr15-25592886-DEL-58
chr15-25604647-DEL-2291
chr15-25611496-DEL-891
chr15-25613883-DEL-260
chr15-25617746-DEL-231
chr15-25629201-DEL-118
chr15-25635230-DEL-75
chr15-25695120-DEL-196
chr15-25699953-DEL-190
chr15-25745849-DEL-57
chr15-25834482-DEL-1767
chr15-25852687-DEL-55
chr15-25853914-DEL-69
chr15-25862368-DEL-110
chr15-25874046-DEL-75
chr15-25901592-DEL-84
chr15-25903579-DEL-70
chr15-25917360-DEL-157
chr15-25940216-DEL-65
chr15-25944774-DEL-181
chr15-25954395-DEL-160

chr15-25975774-DEL-105
chr15-25980598-DEL-75
chr15-25998998-DEL-96
chr15-26000721-DEL-236
chr15-26002797-DEL-195
chr15-26023314-DEL-418
chr15-26035754-DEL-110
chr15-26111503-DEL-54
chr15-26112620-DEL-6798
chr15-26112729-DEL-212
chr15-26123532-DEL-111
chr15-26125873-DEL-309
chr15-26168603-DEL-869
chr15-26186934-DEL-137
chr15-26219509-DEL-144
chr15-26233898-DEL-762
chr15-26255849-DEL-51
chr15-26256668-DEL-3222
chr15-26279667-DEL-100
chr15-26288177-DEL-302
chr15-26288860-DEL-57
chr15-26294014-DEL-5679
chr15-26331530-DEL-1094
chr15-26335304-DEL-408
chr15-26421468-DEL-1845
chr15-26428491-DEL-72
chr15-26433867-DEL-59
chr15-26478056-DEL-6319
chr15-26490581-DEL-1078
chr15-26495332-DEL-4077
chr15-26545058-DEL-193
chr15-26606245-DEL-128
chr15-26616573-DEL-177
chr15-26624886-DEL-114
chr15-26647018-DEL-1736
chr15-26665143-DEL-96
chr15-26670457-DEL-65
chr15-26672755-DEL-271
chr15-26681389-DEL-212
chr15-26709679-DEL-180
chr15-26913791-DEL-1129
chr15-26917997-DEL-890
chr15-26940798-DEL-77
chr15-26966461-DEL-72
chr15-26975894-DEL-81
chr15-27007097-DEL-1580
chr15-27025371-DEL-997
chr15-27082631-DEL-203
chr15-27106213-DEL-88
chr15-27177513-DEL-784
chr15-27180207-DEL-4692
chr15-27203841-DEL-2308
chr15-27206248-DEL-50
chr15-27262977-DEL-126
chr15-27273838-DEL-5565
chr15-27286237-DEL-7220
chr15-27304504-DEL-63
chr15-27319019-DEL-992
chr15-27341456-DEL-202
chr15-27347222-DEL-174
chr15-27349760-DEL-2614
chr15-27389017-DEL-81
chr15-27393362-DEL-821
chr15-27394891-DEL-2205
chr15-27397201-DEL-206
chr15-27399037-DEL-400
chr15-27414406-DEL-66
chr15-27420201-DEL-158
chr15-27488314-DEL-52
chr15-27497785-DEL-3983
chr15-27514908-DEL-353
chr15-27560938-DEL-180
chr15-27565336-DEL-211
chr15-27578474-DEL-97
chr15-27601591-DEL-2243
chr15-27646640-DEL-164
chr15-27657900-DEL-92

chr15-27663235-DEL-81
chr15-27682352-DEL-242
chr15-27728635-DEL-123
chr15-27767184-DEL-66
chr15-27807849-DEL-70
chr15-27868180-DEL-89
chr15-27885893-DEL-122
chr15-27893719-DEL-104
chr15-27914397-DEL-407
chr15-27920281-DEL-189
chr15-27935484-DEL-106
chr15-27948553-DEL-52
chr15-27956942-DEL-105
chr15-27971127-DEL-137
chr15-27971209-DEL-1428
chr15-27971716-DEL-330
chr15-27972399-DEL-314
chr15-27977373-DEL-60
chr15-27992877-DEL-151
chr15-28011370-DEL-255
chr15-28083056-DEL-67
chr15-28106595-DEL-51
chr15-28117609-DEL-594
chr15-28134652-DEL-143
chr15-28137261-DEL-60
chr15-28138493-DEL-54
chr15-28169122-DEL-909
chr15-28190503-DEL-101
chr15-28193949-DEL-1149
chr15-28224809-DEL-82
chr15-28250126-DEL-54
chr15-28274885-DEL-3319
chr15-28280442-DEL-215
chr15-28319651-DEL-943
chr15-28393581-DEL-60
chr15-28458163-DEL-56
chr15-28468484-DEL-74
chr15-28482914-DEL-57
chr15-28484325-DEL-354
chr15-28493906-DEL-78
chr15-28500764-DEL-897
chr15-28512693-DEL-75
chr15-28518875-DEL-599
chr15-28527820-DEL-1357
chr15-28544740-DEL-64
chr15-28564336-DEL-6037
chr15-28588855-DEL-157
chr15-28601090-DEL-60
chr15-28604998-DEL-156
chr15-28614089-DEL-115
chr15-28621440-DEL-1828
chr15-28628932-DEL-57
chr15-28633265-DEL-869
chr15-28656953-DEL-78
chr15-28672167-DEL-57
chr15-28675658-DEL-82
chr15-28746438-DEL-195
chr15-28767765-DEL-114
chr15-28805035-DEL-79
chr15-28805545-DEL-167
chr15-28816510-DEL-1192
chr15-28844447-DEL-682
chr15-28894527-DEL-736
chr15-28941156-DEL-102
chr15-28965574-DEL-63
chr15-28966921-DEL-75
chr15-28971186-DEL-51
chr15-28988412-DEL-65
chr15-28996181-DEL-194
chr15-29034072-DEL-178
chr15-29058044-DEL-240
chr15-29082936-DEL-414
chr15-29121815-DEL-1888
chr15-29140727-DEL-381
chr15-29178808-DEL-132
chr15-29185317-DEL-140
chr15-29197603-DEL-70

chr15-29201822-DEL-1934
chr15-29215622-DEL-76
chr15-29227767-DEL-421
chr15-29262647-DEL-155
chr15-29262947-DEL-485
chr15-29268421-DEL-859
chr15-29271720-DEL-619
chr15-29284884-DEL-6677
chr15-29293086-DEL-293
chr15-29331672-DEL-516
chr15-29332890-DEL-249
chr15-29343450-DEL-234
chr15-29345639-DEL-7843
chr15-29350477-DEL-630
chr15-29366300-DEL-6283
chr15-29375858-DEL-176
chr15-29381838-DEL-60
chr15-29388403-DEL-97
chr15-29440612-DEL-60
chr15-29456874-DEL-59
chr15-29462827-DEL-69
chr15-29486597-DEL-374
chr15-29490414-DEL-202
chr15-29527058-DEL-2067
chr15-29530704-DEL-147
chr15-29533427-DEL-6200
chr15-29541679-DEL-2509
chr15-29545083-DEL-6228
chr15-29556110-DEL-269
chr15-29558594-DEL-2892
chr15-29570104-DEL-229
chr15-29575520-DEL-107
chr15-29605740-DEL-103
chr15-29644562-DEL-84
chr15-29648326-DEL-6196
chr15-29663337-DEL-62
chr15-29675831-DEL-1226
chr15-29748376-DEL-61
chr15-29751358-DEL-258
chr15-29752138-DEL-1764
chr15-29760291-DEL-135
chr15-29764899-DEL-62
chr15-29781898-DEL-2134
chr15-29862264-DEL-683
chr15-29866850-DEL-671
chr15-29867827-DEL-727
chr15-29886113-DEL-57
chr15-29895372-DEL-368
chr15-29908296-DEL-197
chr15-29928289-DEL-1473
chr15-29931067-DEL-56
chr15-29939362-DEL-1506
chr15-29944077-DEL-1477
chr15-29951337-DEL-721
chr15-29954170-DEL-1493
chr15-29963066-DEL-64
chr15-29967994-DEL-53
chr15-29968939-DEL-131
chr15-29995380-DEL-7793
chr15-30038183-DEL-427
chr15-30041061-DEL-329
chr15-30042792-DEL-207
chr15-30054537-DEL-741
chr15-30062312-DEL-417
chr15-30102761-DEL-91
chr15-30106348-DEL-99
chr15-30114124-DEL-1439
chr15-30114135-DEL-514
chr15-30115007-DEL-150
chr15-30129447-DEL-52
chr15-30140240-DEL-1441
chr15-30144554-DEL-410
chr15-30147828-DEL-57
chr15-30148540-DEL-973
chr15-30169372-DEL-110
chr15-30301307-DEL-1297
chr15-30333392-DEL-498

chr15-30368844-DEL-608
chr15-30380603-DEL-489
chr15-30422267-DEL-93
chr15-30518905-DEL-703
chr15-30539942-DEL-66
chr15-30649035-DEL-113
chr15-30672396-DEL-152
chr15-30684883-DEL-76
chr15-30693661-DEL-56
chr15-30704303-DEL-53
chr15-30730379-DEL-104
chr15-30733402-DEL-503
chr15-30754587-DEL-212
chr15-30769186-DEL-154
chr15-30820757-DEL-80
chr15-30841042-DEL-130
chr15-30841837-DEL-831
chr15-30881558-DEL-1037
chr15-30934063-DEL-313
chr15-30948978-DEL-195
chr15-30996792-DEL-6474
chr15-31009942-DEL-6440
chr15-31056853-DEL-385
chr15-31072348-DEL-3331
chr15-31088131-DEL-52
chr15-31089889-DEL-136
chr15-31095369-DEL-983
chr15-31117665-DEL-75
chr15-31176143-DEL-363
chr15-31221369-DEL-62
chr15-31229937-DEL-368
chr15-31310118-DEL-353
chr15-31317411-DEL-196
chr15-31374992-DEL-105
chr15-31380671-DEL-58
chr15-31415111-DEL-141
chr15-31440545-DEL-555
chr15-31444916-DEL-1480
chr15-31450536-DEL-131
chr15-31464447-DEL-70
chr15-31472078-DEL-70
chr15-31504739-DEL-299
chr15-31507092-DEL-396
chr15-31516436-DEL-213
chr15-31532973-DEL-63
chr15-31567228-DEL-349
chr15-31605311-DEL-179
chr15-31629453-DEL-77
chr15-31652352-DEL-192
chr15-31656396-DEL-3264
chr15-31696978-DEL-96
chr15-31714400-DEL-84
chr15-31745971-DEL-66
chr15-31748019-DEL-366
chr15-31754788-DEL-579
chr15-31770934-DEL-185
chr15-31810767-DEL-209
chr15-31833529-DEL-88
chr15-31833837-DEL-834
chr15-31837259-DEL-115
chr15-31848382-DEL-103
chr15-31888040-DEL-6236
chr15-31892825-DEL-1451
chr15-31930762-DEL-1605
chr15-32035771-DEL-70
chr15-32054591-DEL-90
chr15-32077174-DEL-114
chr15-32081454-DEL-404
chr15-32109456-DEL-65
chr15-32110526-DEL-87
chr15-32155988-DEL-10925
chr15-32224341-DEL-73
chr15-32229102-DEL-161
chr15-32285988-DEL-110
chr15-32287008-DEL-567
chr15-32296350-DEL-88
chr15-32318847-DEL-412

chr15-32322966-DEL-107
chr15-32336156-DEL-1983
chr15-32357663-DEL-670
chr15-32418345-DEL-598
chr15-32427180-DEL-52
chr15-32454086-DEL-66
chr15-32464256-DEL-1091
chr15-32490361-DEL-581
chr15-32500748-DEL-81
chr15-32585370-DEL-74
chr15-32642608-DEL-77
chr15-32697620-DEL-76
chr15-32709997-DEL-56
chr15-32712534-DEL-935
chr15-32747928-DEL-524
chr15-32748582-DEL-57
chr15-32749379-DEL-1450
chr15-32769333-DEL-621
chr15-32788845-DEL-939
chr15-32800235-DEL-82
chr15-32812142-DEL-4478
chr15-32833110-DEL-6646
chr15-32841008-DEL-6402
chr15-32856873-DEL-6307
chr15-32869767-DEL-8687
chr15-32880070-DEL-205
chr15-32925039-DEL-73
chr15-32959417-DEL-400
chr15-33012238-DEL-433
chr15-33016782-DEL-411
chr15-33045260-DEL-298
chr15-33065452-DEL-117
chr15-33096802-DEL-90
chr15-33148939-DEL-58
chr15-33155331-DEL-152
chr15-33160779-DEL-141
chr15-33165013-DEL-8535
chr15-33222575-DEL-1241
chr15-33246424-DEL-55
chr15-33286710-DEL-1706
chr15-33294396-DEL-2477
chr15-33386537-DEL-66
chr15-33388037-DEL-235
chr15-33417453-DEL-3951
chr15-33437810-DEL-88
chr15-33444085-DEL-1907
chr15-33461469-DEL-64
chr15-33468101-DEL-478
chr15-33485004-DEL-88
chr15-33493138-DEL-212
chr15-33508138-DEL-106
chr15-33536711-DEL-478
chr15-33575994-DEL-56
chr15-33614580-DEL-299
chr15-33700354-DEL-71
chr15-33704903-DEL-277
chr15-33705475-DEL-823
chr15-33711957-DEL-1015
chr15-33761964-DEL-505
chr15-33778087-DEL-54
chr15-33786764-DEL-428
chr15-33789219-DEL-6169
chr15-33808252-DEL-268
chr15-33814307-DEL-2524
chr15-33822743-DEL-55
chr15-33823750-DEL-104
chr15-33839698-DEL-6399
chr15-33865869-DEL-198
chr15-33867664-DEL-6001
chr15-33879958-DEL-7709
chr15-33880302-DEL-6338
chr15-33910060-DEL-69
chr15-33916953-DEL-543
chr15-33921061-DEL-625
chr15-33954477-DEL-3375
chr15-33995817-DEL-135
chr15-33996178-DEL-2347

chr15-33998632-DEL-76
chr15-34027464-DEL-53
chr15-34044568-DEL-206
chr15-34049862-DEL-51
chr15-34055882-DEL-183
chr15-34057094-DEL-84
chr15-34075566-DEL-52
chr15-34103123-DEL-185
chr15-34113575-DEL-114
chr15-34150183-DEL-454
chr15-34155820-DEL-77
chr15-34175589-DEL-84
chr15-34191202-DEL-828
chr15-34201488-DEL-460
chr15-34207564-DEL-3179
chr15-34214901-DEL-154
chr15-34217838-DEL-2579
chr15-34267947-DEL-155
chr15-34310210-DEL-202
chr15-34326386-DEL-57
chr15-34354117-DEL-54
chr15-34362990-DEL-74
chr15-34376029-DEL-175
chr15-34379776-DEL-171
chr15-34413623-DEL-75
chr15-34416896-DEL-56
chr15-34431220-DEL-200
chr15-34469980-DEL-89
chr15-34473808-DEL-326
chr15-34501128-DEL-302
chr15-34501695-DEL-296
chr15-34505538-DEL-225
chr15-34510596-DEL-374
chr15-34522665-DEL-2134
chr15-34576840-DEL-54
chr15-34611596-DEL-543
chr15-34670476-DEL-1011
chr15-34675449-DEL-77
chr15-34693038-DEL-54
chr15-34722127-DEL-56
chr15-34733267-DEL-1253
chr15-34749598-DEL-230
chr15-34772033-DEL-1877
chr15-34798102-DEL-6198
chr15-34812344-DEL-628
chr15-34858909-DEL-1887
chr15-34862556-DEL-51
chr15-34956918-DEL-126
chr15-35013803-DEL-631
chr15-35014365-DEL-177
chr15-35028339-DEL-13879
chr15-35130066-DEL-341
chr15-35160549-DEL-6919
chr15-35173754-DEL-254
chr15-35189370-DEL-212
chr15-35204410-DEL-121
chr15-35222024-DEL-56
chr15-35278170-DEL-72
chr15-35281534-DEL-205
chr15-35312181-DEL-81
chr15-35316802-DEL-432
chr15-35329001-DEL-788
chr15-35381647-DEL-6165
chr15-35391257-DEL-7740
chr15-35403992-DEL-4294
chr15-35410914-DEL-505
chr15-35439878-DEL-173
chr15-35459069-DEL-59
chr15-35475527-DEL-58
chr15-35491646-DEL-1228
chr15-35534769-DEL-76
chr15-35536110-DEL-301
chr15-35543103-DEL-199
chr15-35548550-DEL-57
chr15-35553862-DEL-1227
chr15-35598432-DEL-550
chr15-35616829-DEL-56

chr15-35629771-DEL-6044
chr15-35646878-DEL-1023
chr15-35664970-DEL-57
chr15-35684808-DEL-2189
chr15-35684900-DEL-1046
chr15-35685627-DEL-144
chr15-35686497-DEL-212
chr15-35737313-DEL-1725
chr15-35752294-DEL-714
chr15-35758614-DEL-101
chr15-35790438-DEL-68
chr15-35791042-DEL-79
chr15-35796816-DEL-349
chr15-35817975-DEL-521
chr15-35818191-DEL-928
chr15-35818898-DEL-290
chr15-35820002-DEL-933
chr15-35833366-DEL-222
chr15-35836919-DEL-307
chr15-35855687-DEL-4838
chr15-35908672-DEL-364
chr15-35909134-DEL-760
chr15-35941889-DEL-1097
chr15-35962706-DEL-8042
chr15-35967050-DEL-239
chr15-35968339-DEL-71
chr15-35977368-DEL-138
chr15-36002398-DEL-138
chr15-36012258-DEL-96
chr15-36027304-DEL-114
chr15-36047057-DEL-67
chr15-36081565-DEL-192
chr15-36092081-DEL-705
chr15-36124018-DEL-90
chr15-36145754-DEL-111
chr15-36158285-DEL-105
chr15-36165133-DEL-72
chr15-36170949-DEL-212
chr15-36202329-DEL-1133
chr15-36202401-DEL-335
chr15-36202687-DEL-713
chr15-36203234-DEL-253
chr15-36203608-DEL-744
chr15-36214742-DEL-102
chr15-36216909-DEL-80
chr15-36223487-DEL-320
chr15-36243954-DEL-82
chr15-36249955-DEL-64
chr15-36278788-DEL-167
chr15-36308218-DEL-894
chr15-36314393-DEL-1303
chr15-36356095-DEL-192
chr15-36362288-DEL-64
chr15-36371388-DEL-54
chr15-36377136-DEL-921
chr15-36383398-DEL-123
chr15-36407480-DEL-64
chr15-36424499-DEL-64
chr15-36426934-DEL-55
chr15-36431440-DEL-524
chr15-36433893-DEL-64
chr15-36457711-DEL-309
chr15-36459319-DEL-4190
chr15-36472394-DEL-315
chr15-36519114-DEL-875
chr15-36534405-DEL-52
chr15-36567957-DEL-75
chr15-36568588-DEL-988
chr15-36570907-DEL-75
chr15-36575258-DEL-271
chr15-36575530-DEL-623
chr15-36585341-DEL-1011
chr15-36586956-DEL-98
chr15-36603755-DEL-97
chr15-36614742-DEL-109
chr15-36642341-DEL-171
chr15-36655896-DEL-170

chr15-36656580-DEL-185
chr15-36663841-DEL-101
chr15-36665758-DEL-52
chr15-36690695-DEL-75
chr15-36735584-DEL-559
chr15-36736084-DEL-207
chr15-36749164-DEL-119
chr15-36804834-DEL-294
chr15-36809074-DEL-100
chr15-36837296-DEL-68
chr15-36842294-DEL-56
chr15-36854858-DEL-68
chr15-36882930-DEL-110
chr15-36887075-DEL-55
chr15-36892227-DEL-52
chr15-36909011-DEL-176
chr15-36941109-DEL-459
chr15-36962975-DEL-164
chr15-36969237-DEL-178
chr15-36974396-DEL-1125
chr15-36978550-DEL-109
chr15-37013884-DEL-52
chr15-37021630-DEL-257
chr15-37042388-DEL-312
chr15-37059439-DEL-62
chr15-37059784-DEL-894
chr15-37088397-DEL-2755
chr15-37105302-DEL-201
chr15-37108208-DEL-630
chr15-37110332-DEL-2579
chr15-37112060-DEL-63
chr15-37113615-DEL-298
chr15-37124977-DEL-220
chr15-37125547-DEL-102
chr15-37142271-DEL-88
chr15-37143396-DEL-78
chr15-37152990-DEL-90
chr15-37156163-DEL-132
chr15-37175529-DEL-1208
chr15-37175530-DEL-639
chr15-37175530-DEL-973
chr15-37176570-DEL-151
chr15-37239923-DEL-125
chr15-37259075-DEL-312
chr15-37261380-DEL-110
chr15-37263178-DEL-144
chr15-37272193-DEL-142
chr15-37300103-DEL-66
chr15-37312525-DEL-53
chr15-37317000-DEL-74
chr15-37326319-DEL-77
chr15-37338704-DEL-3696
chr15-37347798-DEL-88
chr15-37360386-DEL-82
chr15-37380929-DEL-51
chr15-37389172-DEL-64
chr15-37420003-DEL-419
chr15-37446427-DEL-52
chr15-37490154-DEL-93
chr15-37518242-DEL-58
chr15-37526395-DEL-64
chr15-37530629-DEL-1269
chr15-37541920-DEL-304
chr15-37569618-DEL-296
chr15-37576809-DEL-64
chr15-37586440-DEL-64
chr15-37607832-DEL-65
chr15-37629509-DEL-72
chr15-37668696-DEL-63
chr15-37677823-DEL-66
chr15-37692583-DEL-60
chr15-37705854-DEL-400
chr15-37715688-DEL-74
chr15-37763941-DEL-105
chr15-37764306-DEL-98
chr15-37780125-DEL-107
chr15-37820628-DEL-66

chr15-37830419-DEL-135
chr15-37852622-DEL-94
chr15-37860617-DEL-6427
chr15-37872459-DEL-1379
chr15-37889734-DEL-172
chr15-37951083-DEL-107
chr15-37954715-DEL-202
chr15-37963530-DEL-51
chr15-37982033-DEL-161
chr15-37988068-DEL-311
chr15-38048885-DEL-157
chr15-38090355-DEL-95
chr15-38094183-DEL-243
chr15-38104895-DEL-164
chr15-38142310-DEL-180
chr15-38148436-DEL-51
chr15-38162104-DEL-78
chr15-38186536-DEL-168
chr15-38190183-DEL-98
chr15-38193202-DEL-2012
chr15-38198707-DEL-77
chr15-38204028-DEL-537
chr15-38205333-DEL-167
chr15-38206969-DEL-102
chr15-38212399-DEL-64
chr15-38216856-DEL-1598
chr15-38224555-DEL-68
chr15-38274583-DEL-64
chr15-38279291-DEL-51
chr15-38291893-DEL-160
chr15-38308796-DEL-62
chr15-38315754-DEL-126
chr15-38357984-DEL-52
chr15-38394644-DEL-208
chr15-38396929-DEL-66
chr15-38397335-DEL-1733
chr15-38425329-DEL-71
chr15-38449246-DEL-123
chr15-38455150-DEL-394
chr15-38466640-DEL-51
chr15-38522442-DEL-220
chr15-38545935-DEL-58
chr15-38564314-DEL-51
chr15-38574961-DEL-1123
chr15-38597456-DEL-206
chr15-38608515-DEL-62
chr15-38631615-DEL-193
chr15-38633814-DEL-186
chr15-38638053-DEL-1468
chr15-38648404-DEL-76
chr15-38665464-DEL-85
chr15-38679067-DEL-182
chr15-38703964-DEL-72
chr15-38708983-DEL-146
chr15-38719200-DEL-82
chr15-38720500-DEL-376
chr15-38734602-DEL-82
chr15-38735142-DEL-160
chr15-38738443-DEL-77
chr15-38750546-DEL-1403
chr15-38802754-DEL-54
chr15-38837965-DEL-459
chr15-38848278-DEL-1686
chr15-38863584-DEL-1097
chr15-38865056-DEL-2089
chr15-38912213-DEL-109
chr15-38932053-DEL-207
chr15-38999416-DEL-66
chr15-39001303-DEL-62
chr15-39007300-DEL-828
chr15-39018052-DEL-202
chr15-39027525-DEL-153
chr15-39057304-DEL-57
chr15-39077197-DEL-939
chr15-39102781-DEL-64
chr15-39105236-DEL-5683
chr15-39170835-DEL-8015

chr15-39194318-DEL-1204
chr15-39199095-DEL-690
chr15-39199102-DEL-972
chr15-39199370-DEL-1048
chr15-39200516-DEL-2731
chr15-39205383-DEL-105
chr15-39249496-DEL-2357
chr15-39285651-DEL-68
chr15-39307640-DEL-53
chr15-39309230-DEL-70
chr15-39323899-DEL-72
chr15-39337615-DEL-82
chr15-39341291-DEL-59
chr15-39358382-DEL-83
chr15-39372318-DEL-60
chr15-39382960-DEL-546
chr15-39386913-DEL-325
chr15-39413554-DEL-335
chr15-39444322-DEL-346
chr15-39474213-DEL-56
chr15-39534847-DEL-201
chr15-39561116-DEL-5945
chr15-39567119-DEL-247
chr15-39576696-DEL-165
chr15-39605193-DEL-69
chr15-39613276-DEL-277
chr15-39617309-DEL-53
chr15-39642982-DEL-2476
chr15-39682103-DEL-156
chr15-39692518-DEL-174
chr15-39727318-DEL-171
chr15-39786919-DEL-1886
chr15-39787440-DEL-332
chr15-39817687-DEL-211
chr15-39860738-DEL-69
chr15-39888921-DEL-64
chr15-39919431-DEL-363
chr15-40000387-DEL-2143
chr15-40000959-DEL-148
chr15-40002049-DEL-980
chr15-40024794-DEL-73
chr15-40028036-DEL-302
chr15-40034531-DEL-60
chr15-40043091-DEL-87
chr15-40132811-DEL-382
chr15-40143337-DEL-1305
chr15-40212910-DEL-87
chr15-40248801-DEL-80
chr15-40265032-DEL-136
chr15-40274513-DEL-1986
chr15-40288533-DEL-124
chr15-40301206-DEL-799
chr15-40301891-DEL-128
chr15-40331304-DEL-95
chr15-40359186-DEL-288
chr15-40360248-DEL-82
chr15-40438057-DEL-56
chr15-40440293-DEL-765
chr15-40455173-DEL-148
chr15-40500299-DEL-865
chr15-40536009-DEL-64
chr15-40545224-DEL-58
chr15-40605260-DEL-269
chr15-40639900-DEL-199
chr15-40643513-DEL-91
chr15-40699513-DEL-86
chr15-40708152-DEL-201
chr15-40782554-DEL-1461
chr15-40811645-DEL-109
chr15-40869830-DEL-275
chr15-40902683-DEL-117
chr15-40905071-DEL-267
chr15-40910728-DEL-265
chr15-40938637-DEL-60
chr15-40944452-DEL-75
chr15-41002210-DEL-81
chr15-41021593-DEL-165

chr15-41039054-DEL-52
chr15-41048868-DEL-4285
chr15-41054036-DEL-243
chr15-41058524-DEL-1170
chr15-41071650-DEL-259
chr15-41094460-DEL-431
chr15-41107718-DEL-6671
chr15-41107722-DEL-213
chr15-41114697-DEL-324
chr15-41115988-DEL-932
chr15-41143562-DEL-1202
chr15-41158870-DEL-178
chr15-41177031-DEL-54
chr15-41200786-DEL-755
chr15-41233477-DEL-60
chr15-41255048-DEL-80
chr15-41288076-DEL-2213
chr15-41394671-DEL-1957
chr15-41494655-DEL-2297
chr15-41519566-DEL-287
chr15-41521315-DEL-78
chr15-41536345-DEL-2717
chr15-41565175-DEL-2364
chr15-41598571-DEL-242
chr15-41700598-DEL-55
chr15-41734988-DEL-1076
chr15-41748974-DEL-56
chr15-41750955-DEL-784
chr15-41772834-DEL-1572
chr15-41781112-DEL-274
chr15-41796135-DEL-212
chr15-41832801-DEL-76
chr15-41840760-DEL-57
chr15-41842396-DEL-1669
chr15-41850190-DEL-68
chr15-41850896-DEL-452
chr15-41853554-DEL-110
chr15-41855169-DEL-180
chr15-41872185-DEL-116
chr15-41872403-DEL-1224
chr15-41873299-DEL-58
chr15-41877249-DEL-383
chr15-41880315-DEL-548
chr15-41887302-DEL-146
chr15-41929465-DEL-356
chr15-41961525-DEL-2152
chr15-41979863-DEL-51
chr15-41999881-DEL-6447
chr15-42008307-DEL-190
chr15-42025757-DEL-409
chr15-42041045-DEL-2233
chr15-42047535-DEL-307
chr15-42073316-DEL-144
chr15-42094608-DEL-77
chr15-42096388-DEL-804
chr15-42149933-DEL-57
chr15-42158247-DEL-64
chr15-42161429-DEL-56
chr15-42200949-DEL-2974
chr15-42211385-DEL-82
chr15-42212375-DEL-85
chr15-42214135-DEL-4127
chr15-42214137-DEL-141
chr15-42214626-DEL-919
chr15-42215591-DEL-2666
chr15-42218247-DEL-6283
chr15-42218723-DEL-4469
chr15-42221958-DEL-760
chr15-42243262-DEL-269
chr15-42264020-DEL-2622
chr15-42275706-DEL-68
chr15-42276430-DEL-1171
chr15-42317281-DEL-1496
chr15-42346806-DEL-2947
chr15-42352406-DEL-269
chr15-42356379-DEL-336
chr15-42361413-DEL-72

chr15-42404039-DEL-58
chr15-42466198-DEL-348
chr15-42497765-DEL-223
chr15-42515067-DEL-246
chr15-42529482-DEL-63
chr15-42587217-DEL-112
chr15-42587718-DEL-631
chr15-42610707-DEL-1017
chr15-42635019-DEL-92
chr15-42660636-DEL-2113
chr15-42668302-DEL-548
chr15-42698240-DEL-51
chr15-42717170-DEL-366
chr15-42720771-DEL-248
chr15-42739172-DEL-57
chr15-42748999-DEL-53
chr15-42752309-DEL-266
chr15-42764303-DEL-1015
chr15-42782414-DEL-1252
chr15-42788813-DEL-2359
chr15-42793025-DEL-12737
chr15-42808423-DEL-53
chr15-42822282-DEL-95
chr15-42826614-DEL-621
chr15-42835231-DEL-325
chr15-42845088-DEL-608
chr15-42863837-DEL-100
chr15-42895068-DEL-99
chr15-42896097-DEL-159
chr15-42932878-DEL-115
chr15-42970607-DEL-56
chr15-43047516-DEL-1231
chr15-43096061-DEL-207
chr15-43150338-DEL-6760
chr15-43156820-DEL-82
chr15-43173351-DEL-1439
chr15-43181734-DEL-158
chr15-43240887-DEL-399
chr15-43251696-DEL-137
chr15-43350874-DEL-1356
chr15-43365189-DEL-379
chr15-43365214-DEL-742
chr15-43365446-DEL-1303
chr15-43365895-DEL-931
chr15-43366094-DEL-228
chr15-43378365-DEL-56
chr15-43411255-DEL-92
chr15-43413649-DEL-1519
chr15-43419621-DEL-89
chr15-43420294-DEL-730
chr15-43425719-DEL-102
chr15-43456493-DEL-212
chr15-43463742-DEL-70
chr15-43496087-DEL-527
chr15-43502372-DEL-629
chr15-43510771-DEL-339
chr15-43523787-DEL-54
chr15-43534695-DEL-297
chr15-43542370-DEL-221
chr15-43593235-DEL-546
chr15-43598375-DEL-354
chr15-43608215-DEL-4587
chr15-43665684-DEL-566
chr15-43679118-DEL-5411
chr15-43687499-DEL-79
chr15-43691855-DEL-1451
chr15-43702996-DEL-269
chr15-43726418-DEL-102
chr15-43748842-DEL-342
chr15-43770840-DEL-59
chr15-43810983-DEL-1218
chr15-43811290-DEL-3185
chr15-43831462-DEL-102
chr15-43840677-DEL-932
chr15-43849670-DEL-196
chr15-43854326-DEL-111
chr15-43904477-DEL-1490

chr15-43942240-DEL-3818
chr15-43993463-DEL-495
chr15-44013570-DEL-86
chr15-44055925-DEL-1045
chr15-44070100-DEL-1286
chr15-44093484-DEL-185
chr15-44106091-DEL-397
chr15-44107785-DEL-6107
chr15-44135528-DEL-365
chr15-44145447-DEL-853
chr15-44199023-DEL-3661
chr15-44202933-DEL-217
chr15-44231650-DEL-53
chr15-44288674-DEL-72
chr15-44298643-DEL-146
chr15-44309424-DEL-373
chr15-44341154-DEL-396
chr15-44346022-DEL-104
chr15-44403236-DEL-217
chr15-44456728-DEL-185
chr15-44470800-DEL-71
chr15-44512240-DEL-11167
chr15-44532467-DEL-96
chr15-44564011-DEL-66
chr15-44596977-DEL-56
chr15-44628587-DEL-153
chr15-44629407-DEL-364
chr15-44659175-DEL-214
chr15-44662432-DEL-64
chr15-44695274-DEL-70
chr15-44700923-DEL-56
chr15-44770110-DEL-1830
chr15-44779168-DEL-198
chr15-44784528-DEL-405
chr15-44789266-DEL-58
chr15-44822901-DEL-630
chr15-44848341-DEL-5930
chr15-44853140-DEL-66
chr15-44854025-DEL-200
chr15-44872890-DEL-51
chr15-44894006-DEL-2109
chr15-44918437-DEL-55
chr15-44936009-DEL-58
chr15-44993227-DEL-422
chr15-45055216-DEL-92
chr15-45080439-DEL-1779
chr15-45080603-DEL-57
chr15-45082418-DEL-987
chr15-45101548-DEL-105
chr15-45162794-DEL-1054
chr15-45176031-DEL-1227
chr15-45199824-DEL-58
chr15-45203379-DEL-191
chr15-45211216-DEL-202
chr15-45218798-DEL-299
chr15-45234344-DEL-714
chr15-45254399-DEL-840
chr15-45285741-DEL-2993
chr15-45312293-DEL-928
chr15-45331582-DEL-1283
chr15-45340432-DEL-1432
chr15-45354278-DEL-201
chr15-45363236-DEL-81
chr15-45376547-DEL-72
chr15-45378890-DEL-76
chr15-45379818-DEL-1928
chr15-45389216-DEL-294
chr15-45389562-DEL-2383
chr15-45398103-DEL-55
chr15-45411418-DEL-380
chr15-45446122-DEL-8579
chr15-45446902-DEL-7102
chr15-45470819-DEL-229
chr15-45495596-DEL-1265
chr15-45498904-DEL-6760
chr15-45512818-DEL-1403
chr15-45515046-DEL-203

chr15-45518854-DEL-176
chr15-45530994-DEL-107
chr15-45534180-DEL-162
chr15-45539299-DEL-4730
chr15-45544068-DEL-994
chr15-45547694-DEL-811
chr15-45551510-DEL-1743
chr15-45553772-DEL-2137
chr15-45586718-DEL-68
chr15-45589075-DEL-2739
chr15-45606376-DEL-133
chr15-45618879-DEL-7255
chr15-45620006-DEL-6071
chr15-45629821-DEL-667
chr15-45675241-DEL-1203
chr15-45714103-DEL-6170
chr15-45737837-DEL-52
chr15-45770760-DEL-3622
chr15-45799087-DEL-311
chr15-45822828-DEL-723
chr15-45855245-DEL-6199
chr15-45896487-DEL-1205
chr15-45918601-DEL-96
chr15-45936679-DEL-4693
chr15-45961836-DEL-1675
chr15-45990463-DEL-71
chr15-46046702-DEL-51
chr15-46061383-DEL-1748
chr15-46063133-DEL-1930
chr15-46067853-DEL-11688
chr15-46073035-DEL-5297
chr15-46137765-DEL-5040
chr15-46156109-DEL-1399
chr15-46189372-DEL-568
chr15-46214113-DEL-66
chr15-46222329-DEL-723
chr15-46260013-DEL-1065
chr15-46264871-DEL-53
chr15-46280765-DEL-176
chr15-46290793-DEL-898
chr15-46327426-DEL-146
chr15-46333329-DEL-579
chr15-46363976-DEL-99
chr15-46377747-DEL-2151
chr15-46383820-DEL-390
chr15-46456841-DEL-103
chr15-46481242-DEL-64
chr15-46505513-DEL-3037
chr15-46512698-DEL-362
chr15-46528782-DEL-842
chr15-46569745-DEL-5239
chr15-46577827-DEL-2313
chr15-46601504-DEL-232
chr15-46606131-DEL-312
chr15-46609419-DEL-108
chr15-46641648-DEL-11567
chr15-46655529-DEL-939
chr15-46656417-DEL-2114
chr15-46658681-DEL-2833
chr15-46691539-DEL-404
chr15-46694899-DEL-2101
chr15-46696791-DEL-407
chr15-46704780-DEL-785
chr15-46722939-DEL-88
chr15-46745031-DEL-65
chr15-46752049-DEL-208
chr15-46841845-DEL-6353
chr15-46883933-DEL-473
chr15-46885259-DEL-530
chr15-46885988-DEL-6497
chr15-46895009-DEL-1177
chr15-46899150-DEL-60
chr15-46917494-DEL-164
chr15-46949073-DEL-538
chr15-46971941-DEL-7566
chr15-46984559-DEL-6241
chr15-47003178-DEL-64

chr15-47053446-DEL-1121
chr15-47112613-DEL-98
chr15-47139549-DEL-56
chr15-47142033-DEL-759
chr15-47150835-DEL-325
chr15-47231140-DEL-52
chr15-47232468-DEL-7806
chr15-47250095-DEL-83
chr15-47251594-DEL-237
chr15-47279089-DEL-1296
chr15-47290312-DEL-305
chr15-47321831-DEL-97
chr15-47338028-DEL-57
chr15-47339880-DEL-203
chr15-47363949-DEL-2319
chr15-47385011-DEL-5791
chr15-47394680-DEL-75
chr15-47402741-DEL-181
chr15-47411854-DEL-412
chr15-47419235-DEL-55
chr15-47425598-DEL-6248
chr15-47443298-DEL-76
chr15-47453489-DEL-451
chr15-47461700-DEL-1087
chr15-47474390-DEL-389
chr15-47489386-DEL-137
chr15-47495168-DEL-68
chr15-47516382-DEL-140
chr15-47579778-DEL-51
chr15-47584319-DEL-55
chr15-47607042-DEL-290
chr15-47639250-DEL-72
chr15-47648427-DEL-341
chr15-47650483-DEL-1356
chr15-47665070-DEL-2459
chr15-47720367-DEL-156
chr15-47727234-DEL-305
chr15-47734914-DEL-135
chr15-47749605-DEL-689
chr15-47768546-DEL-282
chr15-47820077-DEL-66
chr15-47827990-DEL-54
chr15-47830351-DEL-411
chr15-47832833-DEL-62
chr15-47840612-DEL-75
chr15-47876420-DEL-966
chr15-47895613-DEL-74
chr15-47921192-DEL-505
chr15-47928251-DEL-1253
chr15-47930950-DEL-603
chr15-47937065-DEL-690
chr15-47945566-DEL-1076
chr15-47946691-DEL-1258
chr15-47952975-DEL-1314
chr15-47969373-DEL-171
chr15-47986980-DEL-4101
chr15-48046947-DEL-57
chr15-48050479-DEL-77
chr15-48055789-DEL-196
chr15-48065676-DEL-1357
chr15-48069671-DEL-172
chr15-48070070-DEL-2113
chr15-48071279-DEL-343
chr15-48095107-DEL-234
chr15-48118318-DEL-552
chr15-48134647-DEL-220
chr15-48164087-DEL-56
chr15-48184544-DEL-665
chr15-48188353-DEL-270
chr15-48193856-DEL-58
chr15-48214133-DEL-52
chr15-48235838-DEL-194
chr15-48256583-DEL-400
chr15-48257771-DEL-88
chr15-48258218-DEL-145
chr15-48259197-DEL-5493
chr15-48266697-DEL-1889

chr15-48301755-DEL-203
chr15-48308409-DEL-191
chr15-48353008-DEL-810
chr15-48358134-DEL-67
chr15-48390818-DEL-84
chr15-48429749-DEL-90
chr15-48430181-DEL-277
chr15-48430663-DEL-355
chr15-48436025-DEL-52
chr15-48445826-DEL-96
chr15-48456022-DEL-663
chr15-48462799-DEL-124
chr15-48463795-DEL-540
chr15-48477493-DEL-56
chr15-48518888-DEL-53
chr15-48521148-DEL-923
chr15-48591778-DEL-88
chr15-48601211-DEL-382
chr15-48623937-DEL-110
chr15-48643183-DEL-60
chr15-48660353-DEL-250
chr15-48673280-DEL-175
chr15-48677112-DEL-262
chr15-48681255-DEL-245
chr15-48710788-DEL-478
chr15-48744488-DEL-126
chr15-48744951-DEL-6141
chr15-48787529-DEL-77
chr15-48798072-DEL-58
chr15-48804013-DEL-408
chr15-48841518-DEL-605
chr15-48850698-DEL-61
chr15-48851486-DEL-2105
chr15-48854046-DEL-6192
chr15-48871936-DEL-9801
chr15-48893339-DEL-636
chr15-48902374-DEL-1941
chr15-48914493-DEL-289
chr15-48928405-DEL-60
chr15-48931576-DEL-401
chr15-48933691-DEL-51
chr15-48957537-DEL-6418
chr15-48966779-DEL-880
chr15-48972080-DEL-6605
chr15-48991388-DEL-554
chr15-48993805-DEL-156
chr15-49030267-DEL-56
chr15-49030816-DEL-746
chr15-49045457-DEL-235
chr15-49062504-DEL-61
chr15-49092199-DEL-509
chr15-49117443-DEL-214
chr15-49132109-DEL-278
chr15-49145718-DEL-7015
chr15-49156857-DEL-6207
chr15-49168440-DEL-194
chr15-49191316-DEL-506
chr15-49194366-DEL-1260
chr15-49207458-DEL-196
chr15-49220359-DEL-67
chr15-49222964-DEL-407
chr15-49224834-DEL-6208
chr15-49249512-DEL-82
chr15-49250440-DEL-958
chr15-49251698-DEL-613
chr15-49254555-DEL-371
chr15-49268272-DEL-501
chr15-49277804-DEL-6180
chr15-49285076-DEL-349
chr15-49288049-DEL-63
chr15-49311976-DEL-60
chr15-49329328-DEL-550
chr15-49339343-DEL-52
chr15-49377712-DEL-1270
chr15-49387531-DEL-189
chr15-49417991-DEL-153
chr15-49450335-DEL-857

chr15-49458709-DEL-1787
chr15-49474379-DEL-439
chr15-49488202-DEL-323
chr15-49492636-DEL-158
chr15-49518589-DEL-6193
chr15-49559503-DEL-63
chr15-49563473-DEL-204
chr15-49592602-DEL-58
chr15-49632330-DEL-97
chr15-49636707-DEL-3581
chr15-49640697-DEL-542
chr15-49647550-DEL-4882
chr15-49654564-DEL-1186
chr15-49693998-DEL-1168
chr15-49695422-DEL-591
chr15-49724858-DEL-64
chr15-49773182-DEL-80
chr15-49783153-DEL-53
chr15-49788152-DEL-154
chr15-49821994-DEL-160
chr15-49829625-DEL-4757
chr15-49833854-DEL-731
chr15-49881573-DEL-751
chr15-49882399-DEL-1742
chr15-49899951-DEL-56
chr15-49904779-DEL-420
chr15-49930425-DEL-315
chr15-49934382-DEL-4147
chr15-49945685-DEL-66
chr15-49951125-DEL-51
chr15-49974952-DEL-56
chr15-49980280-DEL-62
chr15-50051746-DEL-132
chr15-50053548-DEL-113
chr15-50070320-DEL-606
chr15-50076291-DEL-214
chr15-50083497-DEL-853
chr15-50095993-DEL-65
chr15-50096947-DEL-225
chr15-50137072-DEL-587
chr15-50153073-DEL-638
chr15-50159557-DEL-1122
chr15-50210705-DEL-7417
chr15-50234466-DEL-62
chr15-50256246-DEL-374
chr15-50270952-DEL-58
chr15-50289992-DEL-154
chr15-50312578-DEL-60
chr15-50355225-DEL-573
chr15-50391281-DEL-1862
chr15-50418424-DEL-6026
chr15-50430213-DEL-615
chr15-50501197-DEL-6232
chr15-50536941-DEL-70
chr15-50554832-DEL-106
chr15-50564541-DEL-52
chr15-50583548-DEL-84
chr15-50595462-DEL-206
chr15-50704092-DEL-80
chr15-50720889-DEL-93
chr15-50759931-DEL-107
chr15-50771173-DEL-486
chr15-50773155-DEL-171
chr15-50866377-DEL-66
chr15-50888760-DEL-63
chr15-50894214-DEL-6610
chr15-50911646-DEL-6796
chr15-50911911-DEL-207
chr15-50921871-DEL-425
chr15-51033754-DEL-1369
chr15-51046543-DEL-75
chr15-51066265-DEL-336
chr15-51078062-DEL-66
chr15-51079558-DEL-75
chr15-51087334-DEL-199
chr15-51151803-DEL-53
chr15-51175824-DEL-66

chr15-51178578-DEL-85
chr15-51183061-DEL-182
chr15-51207817-DEL-344
chr15-51210377-DEL-91
chr15-51347054-DEL-204
chr15-51349593-DEL-1198
chr15-51353147-DEL-75
chr15-51358662-DEL-550
chr15-51363256-DEL-310
chr15-51369920-DEL-124
chr15-51372711-DEL-55
chr15-51376610-DEL-1000
chr15-51383977-DEL-61
chr15-51386986-DEL-54
chr15-51394982-DEL-75
chr15-51412814-DEL-65
chr15-51431473-DEL-88
chr15-51488195-DEL-194
chr15-51489325-DEL-295
chr15-51494511-DEL-54
chr15-51504813-DEL-52
chr15-51519356-DEL-170
chr15-51522590-DEL-157
chr15-51524359-DEL-1547
chr15-51546110-DEL-73
chr15-51560833-DEL-118
chr15-51583768-DEL-69
chr15-51601626-DEL-486
chr15-51626813-DEL-92
chr15-51672541-DEL-583
chr15-51679655-DEL-283
chr15-51686700-DEL-67
chr15-51692321-DEL-814
chr15-51710836-DEL-114
chr15-51711859-DEL-114
chr15-51712828-DEL-99
chr15-51714101-DEL-71
chr15-51747028-DEL-59
chr15-51781335-DEL-92
chr15-51842385-DEL-2394
chr15-51846605-DEL-163
chr15-51851841-DEL-176
chr15-51853343-DEL-192
chr15-51859699-DEL-169
chr15-51861532-DEL-194
chr15-51867757-DEL-152
chr15-51877719-DEL-2857
chr15-51891302-DEL-214
chr15-51937046-DEL-86
chr15-51969095-DEL-121
chr15-51982332-DEL-57
chr15-51986602-DEL-335
chr15-51988991-DEL-1316
chr15-51997617-DEL-82
chr15-52033837-DEL-190
chr15-52054235-DEL-185
chr15-52073150-DEL-52
chr15-52079783-DEL-120
chr15-52092755-DEL-5164
chr15-52110847-DEL-2677
chr15-52129251-DEL-2117
chr15-52143558-DEL-695
chr15-52151908-DEL-898
chr15-52156117-DEL-168
chr15-52159434-DEL-338
chr15-52190561-DEL-52
chr15-52192260-DEL-60
chr15-52215545-DEL-1070
chr15-52218186-DEL-2514
chr15-52244208-DEL-863
chr15-52273068-DEL-2348
chr15-52305803-DEL-131
chr15-52334936-DEL-1248
chr15-52352556-DEL-555
chr15-52359286-DEL-177
chr15-52361092-DEL-211
chr15-52387379-DEL-172

chr15-52392357-DEL-1045
chr15-52407305-DEL-514
chr15-52425548-DEL-54
chr15-52437403-DEL-872
chr15-52490741-DEL-51
chr15-52512012-DEL-6587
chr15-52523321-DEL-6659
chr15-52546717-DEL-580
chr15-52549127-DEL-658
chr15-52550294-DEL-779
chr15-52634910-DEL-1564
chr15-52651926-DEL-2029
chr15-52706972-DEL-424
chr15-52712596-DEL-259
chr15-52725501-DEL-54
chr15-52737118-DEL-7706
chr15-52769460-DEL-62
chr15-52781463-DEL-869
chr15-52865622-DEL-2623
chr15-52871507-DEL-292
chr15-52874662-DEL-66
chr15-52888656-DEL-1415
chr15-52891131-DEL-398
chr15-52901342-DEL-181
chr15-52917062-DEL-66
chr15-52921623-DEL-2041
chr15-52976371-DEL-323
chr15-53093026-DEL-119
chr15-53140344-DEL-64
chr15-53169672-DEL-359
chr15-53180326-DEL-84
chr15-53190293-DEL-167
chr15-53246417-DEL-306
chr15-53248001-DEL-87
chr15-53336979-DEL-60
chr15-53340622-DEL-198
chr15-53353560-DEL-597
chr15-53420833-DEL-56
chr15-53447166-DEL-51
chr15-53462677-DEL-401
chr15-53463684-DEL-135
chr15-53485546-DEL-234
chr15-53498168-DEL-1709
chr15-53584843-DEL-77
chr15-53613439-DEL-52
chr15-53623433-DEL-75
chr15-53634683-DEL-98
chr15-53650129-DEL-52
chr15-53664736-DEL-211
chr15-53670896-DEL-182
chr15-53676714-DEL-210
chr15-53677922-DEL-714
chr15-53713492-DEL-328
chr15-53716823-DEL-78
chr15-53752059-DEL-1398
chr15-53815463-DEL-6038
chr15-53827602-DEL-2001
chr15-53829666-DEL-117
chr15-53835030-DEL-54
chr15-53840128-DEL-7625
chr15-53840177-DEL-1486
chr15-53973331-DEL-84
chr15-54123987-DEL-588
chr15-54148228-DEL-75
chr15-54178072-DEL-1891
chr15-54181357-DEL-64
chr15-54221077-DEL-673
chr15-54239558-DEL-827
chr15-54264043-DEL-1293
chr15-54285972-DEL-128
chr15-54290353-DEL-632
chr15-54332242-DEL-54
chr15-54351800-DEL-172
chr15-54358810-DEL-68
chr15-54371929-DEL-95
chr15-54383014-DEL-89
chr15-54395138-DEL-639

chr15-54405458-DEL-808
chr15-54416983-DEL-5870
chr15-54431741-DEL-98
chr15-54440752-DEL-55
chr15-54446163-DEL-231
chr15-54488496-DEL-270
chr15-54493110-DEL-55
chr15-54560574-DEL-1497
chr15-54646322-DEL-63
chr15-54653948-DEL-144
chr15-54665660-DEL-97
chr15-54677378-DEL-146
chr15-54694141-DEL-2730
chr15-54740088-DEL-64
chr15-54768519-DEL-197
chr15-54812587-DEL-173
chr15-54832808-DEL-232
chr15-54834223-DEL-3128
chr15-54875152-DEL-473
chr15-54884779-DEL-203
chr15-54913817-DEL-161
chr15-54961114-DEL-216
chr15-54999610-DEL-91
chr15-55013550-DEL-235
chr15-55022534-DEL-2581
chr15-55031885-DEL-199
chr15-55033238-DEL-81
chr15-55037294-DEL-91
chr15-55045069-DEL-163
chr15-55070890-DEL-124
chr15-55079408-DEL-202
chr15-55091977-DEL-319
chr15-55093039-DEL-162
chr15-55114133-DEL-408
chr15-55142112-DEL-359
chr15-55142825-DEL-99
chr15-55154225-DEL-52
chr15-55157216-DEL-92
chr15-55158295-DEL-62
chr15-55178312-DEL-311
chr15-55186160-DEL-62
chr15-55191103-DEL-59
chr15-55195189-DEL-616
chr15-55196043-DEL-67
chr15-55213583-DEL-61
chr15-55238059-DEL-62
chr15-55246455-DEL-51
chr15-55247592-DEL-57
chr15-55303318-DEL-129
chr15-55343863-DEL-1435
chr15-55392532-DEL-54
chr15-55414395-DEL-1065
chr15-55434025-DEL-353
chr15-55445074-DEL-88
chr15-55463436-DEL-159
chr15-55475777-DEL-69
chr15-55477124-DEL-205
chr15-55483414-DEL-82
chr15-55521341-DEL-61
chr15-55539935-DEL-138
chr15-55547560-DEL-212
chr15-55560212-DEL-160
chr15-55630010-DEL-103
chr15-55639611-DEL-101
chr15-55671942-DEL-362
chr15-55678052-DEL-55
chr15-55683061-DEL-375
chr15-55686032-DEL-58
chr15-55719416-DEL-52
chr15-55722962-DEL-278
chr15-55723885-DEL-806
chr15-55744089-DEL-54
chr15-55757662-DEL-2462
chr15-55767360-DEL-67
chr15-55770707-DEL-113
chr15-55791579-DEL-1820
chr15-55810680-DEL-1586

chr15-55839703-DEL-90
chr15-55847316-DEL-399
chr15-55853793-DEL-722
chr15-55855621-DEL-2115
chr15-55864833-DEL-453
chr15-55873709-DEL-985
chr15-55880835-DEL-107
chr15-55930110-DEL-108
chr15-55936750-DEL-777
chr15-55957082-DEL-2319
chr15-55964915-DEL-106
chr15-55969129-DEL-59
chr15-55983313-DEL-55
chr15-55994071-DEL-1336
chr15-56003494-DEL-62
chr15-56009143-DEL-6171
chr15-56031412-DEL-65
chr15-56068336-DEL-1522
chr15-56071084-DEL-6220
chr15-56117606-DEL-395
chr15-56118049-DEL-1294
chr15-56122093-DEL-2194
chr15-56131845-DEL-198
chr15-56148811-DEL-1744
chr15-56168074-DEL-4394
chr15-56170856-DEL-1141
chr15-56185791-DEL-401
chr15-56199384-DEL-336
chr15-56215596-DEL-62
chr15-56225183-DEL-6198
chr15-56238124-DEL-54
chr15-56238899-DEL-90
chr15-56258949-DEL-62
chr15-56287042-DEL-365
chr15-56296281-DEL-453
chr15-56319858-DEL-253
chr15-56321743-DEL-1017
chr15-56365940-DEL-1691
chr15-56383825-DEL-6465
chr15-56383972-DEL-214
chr15-56415631-DEL-53
chr15-56415950-DEL-319
chr15-56421487-DEL-79
chr15-56453136-DEL-1064
chr15-56454800-DEL-4149
chr15-56455880-DEL-501
chr15-56456172-DEL-1487
chr15-56465084-DEL-1071
chr15-56473717-DEL-144
chr15-56538978-DEL-115
chr15-56543853-DEL-323
chr15-56547300-DEL-462
chr15-56549791-DEL-214
chr15-56559079-DEL-59
chr15-56581859-DEL-520
chr15-56587612-DEL-202
chr15-56613329-DEL-872
chr15-56690908-DEL-104
chr15-56701015-DEL-56
chr15-56716326-DEL-100
chr15-56730006-DEL-92
chr15-56731959-DEL-296
chr15-56732192-DEL-576
chr15-56737206-DEL-2610
chr15-56748513-DEL-58
chr15-56748912-DEL-65
chr15-56748927-DEL-6638
chr15-56748988-DEL-1129
chr15-56758829-DEL-208
chr15-56765388-DEL-6613
chr15-56776751-DEL-6812
chr15-56776856-DEL-710
chr15-56792521-DEL-54
chr15-56793047-DEL-759
chr15-56828237-DEL-348
chr15-56831979-DEL-59
chr15-56845257-DEL-6434

chr15-56850913-DEL-2534
chr15-56852668-DEL-4266
chr15-56875662-DEL-77
chr15-56921051-DEL-86
chr15-56951078-DEL-2040
chr15-56965678-DEL-196
chr15-56971718-DEL-58
chr15-56999961-DEL-1515
chr15-57023741-DEL-57
chr15-57069293-DEL-902
chr15-57073966-DEL-107
chr15-57081683-DEL-404
chr15-57092090-DEL-117
chr15-57096779-DEL-51
chr15-57107617-DEL-485
chr15-57142502-DEL-214
chr15-57155199-DEL-230
chr15-57172718-DEL-5285
chr15-57194522-DEL-52
chr15-57210366-DEL-295
chr15-57211809-DEL-184
chr15-57225411-DEL-1800
chr15-57239955-DEL-69
chr15-57257377-DEL-56
chr15-57273682-DEL-60
chr15-57287238-DEL-212
chr15-57301846-DEL-58
chr15-57318016-DEL-66
chr15-57347108-DEL-897
chr15-57418762-DEL-56
chr15-57427879-DEL-564
chr15-57451401-DEL-69
chr15-57474246-DEL-997
chr15-57502018-DEL-98
chr15-57505851-DEL-147
chr15-57508514-DEL-146
chr15-57518965-DEL-126
chr15-57528707-DEL-876
chr15-57538024-DEL-1296
chr15-57584444-DEL-326
chr15-57604562-DEL-78
chr15-57607060-DEL-304
chr15-57619365-DEL-74
chr15-57640123-DEL-5331
chr15-57648956-DEL-153
chr15-57671674-DEL-52
chr15-57678951-DEL-151
chr15-57687312-DEL-54
chr15-57697334-DEL-78
chr15-57698745-DEL-1177
chr15-57784250-DEL-460
chr15-57800986-DEL-257
chr15-57803412-DEL-218
chr15-57806157-DEL-1000
chr15-57820641-DEL-174
chr15-57860786-DEL-1052
chr15-57860811-DEL-356
chr15-57861396-DEL-692
chr15-57874566-DEL-73
chr15-57907324-DEL-196
chr15-57930244-DEL-516
chr15-57986720-DEL-206
chr15-58002353-DEL-261
chr15-58029300-DEL-558
chr15-58070557-DEL-55
chr15-58133172-DEL-92
chr15-58173375-DEL-62
chr15-58180589-DEL-119
chr15-58193282-DEL-103
chr15-58242429-DEL-1082
chr15-58258301-DEL-55
chr15-58258864-DEL-277
chr15-58309636-DEL-256
chr15-58309656-DEL-2943
chr15-58310848-DEL-1280
chr15-58336683-DEL-817
chr15-58339975-DEL-1544

chr15-58354103-DEL-5757
chr15-58386337-DEL-252
chr15-58397922-DEL-95
chr15-58422144-DEL-198
chr15-58422730-DEL-105
chr15-58451639-DEL-95
chr15-58470150-DEL-212
chr15-58475084-DEL-135
chr15-58513355-DEL-66
chr15-58521508-DEL-7583
chr15-58527702-DEL-136
chr15-58536973-DEL-94
chr15-58607952-DEL-66
chr15-58639864-DEL-123
chr15-58781808-DEL-3526
chr15-58781936-DEL-259
chr15-58818981-DEL-90
chr15-58833478-DEL-228
chr15-58872299-DEL-1921
chr15-58895254-DEL-354
chr15-58914708-DEL-364
chr15-58945577-DEL-324
chr15-58963375-DEL-113
chr15-58991681-DEL-74
chr15-59005712-DEL-145
chr15-59010239-DEL-432
chr15-59023041-DEL-420
chr15-59033542-DEL-1326
chr15-59034896-DEL-4408
chr15-59054144-DEL-403
chr15-59080624-DEL-217
chr15-59092782-DEL-185
chr15-59110295-DEL-587
chr15-59115633-DEL-1635
chr15-59115928-DEL-219
chr15-59120751-DEL-78
chr15-59149541-DEL-170
chr15-59152013-DEL-70
chr15-59175845-DEL-152
chr15-59181625-DEL-4411
chr15-59195838-DEL-106
chr15-59219226-DEL-630
chr15-59266659-DEL-334
chr15-59303751-DEL-2929
chr15-59303799-DEL-1079
chr15-59303839-DEL-2244
chr15-59305161-DEL-682
chr15-59306266-DEL-456
chr15-59339645-DEL-55
chr15-59346211-DEL-64
chr15-59350827-DEL-59
chr15-59476615-DEL-66
chr15-59562668-DEL-112
chr15-59583509-DEL-152
chr15-59585109-DEL-130
chr15-59604763-DEL-81
chr15-59658449-DEL-71
chr15-59684635-DEL-208
chr15-59720918-DEL-105
chr15-59721850-DEL-958
chr15-59742539-DEL-2460
chr15-59745015-DEL-1972
chr15-59760766-DEL-52
chr15-59761286-DEL-153
chr15-59775698-DEL-68
chr15-59820737-DEL-845
chr15-59831439-DEL-145
chr15-59843838-DEL-656
chr15-59848481-DEL-770
chr15-59908514-DEL-65
chr15-59940435-DEL-454
chr15-59941386-DEL-183
chr15-59975621-DEL-225
chr15-59997696-DEL-199
chr15-60035945-DEL-2092
chr15-60045850-DEL-2892
chr15-60048386-DEL-1006

chr15-60091745-DEL-62
chr15-60101018-DEL-72
chr15-60114137-DEL-56
chr15-60124828-DEL-1354
chr15-60129261-DEL-58
chr15-60168135-DEL-190
chr15-60169700-DEL-143
chr15-60195083-DEL-84
chr15-60196250-DEL-400
chr15-60236965-DEL-606
chr15-60267182-DEL-917
chr15-60267886-DEL-55
chr15-60281367-DEL-6579
chr15-60290370-DEL-107
chr15-60317886-DEL-1825
chr15-60341446-DEL-98
chr15-60368217-DEL-148
chr15-60392163-DEL-741
chr15-60456135-DEL-368
chr15-60458734-DEL-397
chr15-60469650-DEL-192
chr15-60478752-DEL-150
chr15-60479633-DEL-56
chr15-60517600-DEL-179
chr15-60533563-DEL-1648
chr15-60538383-DEL-59
chr15-60541080-DEL-98
chr15-60543082-DEL-175
chr15-60551604-DEL-99
chr15-60568163-DEL-60
chr15-60622512-DEL-76
chr15-60647237-DEL-143
chr15-60653300-DEL-807
chr15-60655596-DEL-52
chr15-60659761-DEL-53
chr15-60660375-DEL-1269
chr15-60742997-DEL-60
chr15-60763581-DEL-1582
chr15-60765400-DEL-76
chr15-60775318-DEL-340
chr15-60790389-DEL-267
chr15-60811076-DEL-9102
chr15-60813499-DEL-6370
chr15-60824550-DEL-367
chr15-60828925-DEL-2646
chr15-60831965-DEL-6027
chr15-60845220-DEL-6698
chr15-60907974-DEL-447
chr15-60909042-DEL-361
chr15-60931831-DEL-725
chr15-60940513-DEL-895
chr15-60975148-DEL-106
chr15-61046391-DEL-1539
chr15-61050585-DEL-99
chr15-61067217-DEL-351
chr15-61068282-DEL-152
chr15-61098413-DEL-142
chr15-61124431-DEL-57
chr15-61150073-DEL-7529
chr15-61156715-DEL-173
chr15-61239721-DEL-455
chr15-61245159-DEL-3773
chr15-61289341-DEL-51
chr15-61333345-DEL-204
chr15-61353259-DEL-72
chr15-61360885-DEL-602
chr15-61400623-DEL-214
chr15-61401975-DEL-66
chr15-61413282-DEL-108
chr15-61415728-DEL-107
chr15-61437835-DEL-6218
chr15-61461292-DEL-372
chr15-61507714-DEL-1280
chr15-61528058-DEL-221
chr15-61532024-DEL-146
chr15-61558887-DEL-219
chr15-61567655-DEL-57

chr15-61579596-DEL-54
chr15-61613034-DEL-213
chr15-61633406-DEL-54
chr15-61685418-DEL-441
chr15-61701955-DEL-525
chr15-61717638-DEL-2598
chr15-61729971-DEL-91
chr15-61739341-DEL-57
chr15-61750299-DEL-792
chr15-61769912-DEL-3898
chr15-61793336-DEL-52
chr15-61808041-DEL-56
chr15-61817296-DEL-199
chr15-61838178-DEL-55
chr15-61845753-DEL-103
chr15-61870970-DEL-526
chr15-61880226-DEL-441
chr15-61890176-DEL-1040
chr15-61891898-DEL-1933
chr15-61894305-DEL-1754
chr15-61902854-DEL-186
chr15-61908411-DEL-492
chr15-61949950-DEL-150
chr15-61954304-DEL-755
chr15-61965984-DEL-93
chr15-61992412-DEL-68
chr15-62036403-DEL-63
chr15-62048960-DEL-421
chr15-62058710-DEL-102
chr15-62098135-DEL-475
chr15-62098637-DEL-1969
chr15-62137122-DEL-126
chr15-62143936-DEL-120
chr15-62188677-DEL-618
chr15-62199848-DEL-68
chr15-62204759-DEL-60
chr15-62239254-DEL-2677
chr15-62248836-DEL-64
chr15-62284250-DEL-91
chr15-62312563-DEL-674
chr15-62314913-DEL-139
chr15-62320136-DEL-70
chr15-62370710-DEL-204
chr15-62386578-DEL-267
chr15-62441409-DEL-132
chr15-62470896-DEL-83
chr15-62507818-DEL-734
chr15-62524506-DEL-2197
chr15-62533436-DEL-60
chr15-62541539-DEL-56
chr15-62548900-DEL-265
chr15-62548993-DEL-1224
chr15-62549961-DEL-711
chr15-62642066-DEL-568
chr15-62714499-DEL-216
chr15-62725835-DEL-908
chr15-62728020-DEL-55
chr15-62735211-DEL-235
chr15-62735214-DEL-678
chr15-62735943-DEL-693
chr15-62737095-DEL-207
chr15-62785108-DEL-682
chr15-62821435-DEL-268
chr15-62888208-DEL-190
chr15-62900426-DEL-2209
chr15-62901529-DEL-1469
chr15-62911366-DEL-80
chr15-62974288-DEL-205
chr15-62999701-DEL-208
chr15-63025255-DEL-171
chr15-63053561-DEL-2163
chr15-63071097-DEL-1183
chr15-63074028-DEL-105
chr15-63076244-DEL-192
chr15-63084404-DEL-258
chr15-63111939-DEL-488
chr15-63118757-DEL-64

chr15-63125368-DEL-61
chr15-63133520-DEL-93
chr15-63135280-DEL-90
chr15-63139759-DEL-148
chr15-63168588-DEL-60
chr15-63188344-DEL-132
chr15-63192795-DEL-981
chr15-63200475-DEL-6479
chr15-63210313-DEL-99
chr15-63257038-DEL-489
chr15-63262149-DEL-80
chr15-63270230-DEL-85
chr15-63285193-DEL-64
chr15-63319935-DEL-191
chr15-63328482-DEL-71
chr15-63385230-DEL-528
chr15-63386008-DEL-1637
chr15-63386041-DEL-262
chr15-63386851-DEL-181
chr15-63407347-DEL-534
chr15-63420215-DEL-744
chr15-63444686-DEL-230
chr15-63470098-DEL-190
chr15-63470427-DEL-958
chr15-63544742-DEL-183
chr15-63562495-DEL-69
chr15-63575658-DEL-66
chr15-63586775-DEL-732
chr15-63633332-DEL-316
chr15-63661220-DEL-1478
chr15-63700937-DEL-77
chr15-63713462-DEL-698
chr15-63722892-DEL-1665
chr15-63724648-DEL-687
chr15-63741754-DEL-69
chr15-63746843-DEL-697
chr15-63780492-DEL-763
chr15-63781186-DEL-64
chr15-63807523-DEL-365
chr15-63811959-DEL-213
chr15-63830745-DEL-58
chr15-63855848-DEL-97
chr15-63883376-DEL-238
chr15-63937773-DEL-196
chr15-63941130-DEL-204
chr15-63941900-DEL-51
chr15-63968803-DEL-195
chr15-64014216-DEL-54
chr15-64033928-DEL-410
chr15-64093059-DEL-54
chr15-64101745-DEL-123
chr15-64120356-DEL-4260
chr15-64161470-DEL-54
chr15-64239002-DEL-75
chr15-64246076-DEL-914
chr15-64264010-DEL-72
chr15-64306216-DEL-99
chr15-64333851-DEL-444
chr15-64395171-DEL-6019
chr15-64409595-DEL-67
chr15-64410117-DEL-714
chr15-64423563-DEL-88
chr15-64475049-DEL-66
chr15-64480287-DEL-2880
chr15-64497049-DEL-52
chr15-64501970-DEL-76
chr15-64518486-DEL-63
chr15-64518959-DEL-2929
chr15-64519458-DEL-98
chr15-64529021-DEL-285
chr15-64547813-DEL-7275
chr15-64565721-DEL-1642
chr15-64569402-DEL-224
chr15-64570177-DEL-59
chr15-64621810-DEL-55
chr15-64623650-DEL-123
chr15-64635663-DEL-264

chr15-64677549-DEL-688
chr15-64681776-DEL-334
chr15-64690491-DEL-1558
chr15-64692969-DEL-290
chr15-64731828-DEL-84
chr15-64758304-DEL-203
chr15-64812190-DEL-137
chr15-64813174-DEL-108
chr15-64817116-DEL-113
chr15-64825923-DEL-73
chr15-64839612-DEL-117
chr15-64843964-DEL-55
chr15-64879625-DEL-147
chr15-64926028-DEL-1331
chr15-64976981-DEL-258
chr15-64993959-DEL-52
chr15-64994263-DEL-1106
chr15-64999261-DEL-102
chr15-65024200-DEL-6213
chr15-65046087-DEL-2205
chr15-65061650-DEL-205
chr15-65081890-DEL-60
chr15-65083080-DEL-152
chr15-65096603-DEL-2139
chr15-65101275-DEL-73
chr15-65128515-DEL-332
chr15-65131267-DEL-159
chr15-65132785-DEL-51
chr15-65140231-DEL-150
chr15-65145154-DEL-212
chr15-65186400-DEL-407
chr15-65267864-DEL-54
chr15-65271666-DEL-6421
chr15-65271740-DEL-4342
chr15-65277826-DEL-207
chr15-65279345-DEL-162
chr15-65284550-DEL-66
chr15-65287418-DEL-62
chr15-65308062-DEL-58
chr15-65313709-DEL-909
chr15-65328704-DEL-6475
chr15-65334911-DEL-64
chr15-65351890-DEL-211
chr15-65360464-DEL-454
chr15-65364000-DEL-55
chr15-65390904-DEL-56
chr15-65400621-DEL-1213
chr15-65433231-DEL-3281
chr15-65476626-DEL-881
chr15-65495252-DEL-1245
chr15-65514790-DEL-331
chr15-65534922-DEL-75
chr15-65585897-DEL-11192
chr15-65609419-DEL-212
chr15-65609985-DEL-212
chr15-65624389-DEL-972
chr15-65701988-DEL-4345
chr15-65730884-DEL-1053
chr15-65739895-DEL-136
chr15-65791930-DEL-553
chr15-65794003-DEL-61
chr15-65800089-DEL-757
chr15-65805822-DEL-207
chr15-65807905-DEL-68
chr15-65821503-DEL-6214
chr15-65830054-DEL-6165
chr15-65896545-DEL-73
chr15-65900681-DEL-1210
chr15-65925573-DEL-1156
chr15-65928697-DEL-9058
chr15-65966485-DEL-64
chr15-66016433-DEL-136
chr15-66023632-DEL-85
chr15-66038317-DEL-166
chr15-66042729-DEL-166
chr15-66045942-DEL-438
chr15-66051036-DEL-52

chr15-66059815-DEL-714
chr15-66083512-DEL-151
chr15-66088256-DEL-3242
chr15-66093409-DEL-87
chr15-66122830-DEL-60
chr15-66131056-DEL-52
chr15-66151157-DEL-815
chr15-66171124-DEL-61
chr15-66240855-DEL-65
chr15-66306513-DEL-53
chr15-66507360-DEL-276
chr15-66521958-DEL-296
chr15-66524672-DEL-56
chr15-66526393-DEL-73
chr15-66527267-DEL-261
chr15-66527877-DEL-155
chr15-66535487-DEL-430
chr15-66547025-DEL-72
chr15-66591917-DEL-92
chr15-66603480-DEL-52
chr15-66604006-DEL-246
chr15-66646040-DEL-1094
chr15-66707466-DEL-509
chr15-66741020-DEL-322
chr15-66846215-DEL-1099
chr15-66935400-DEL-76
chr15-66951581-DEL-141
chr15-66976623-DEL-161
chr15-66985089-DEL-98
chr15-67002090-DEL-701
chr15-67034070-DEL-52
chr15-67067535-DEL-325
chr15-67068279-DEL-84
chr15-67073287-DEL-1074
chr15-67104578-DEL-52
chr15-67127127-DEL-56
chr15-67161397-DEL-168
chr15-67177565-DEL-178
chr15-67190314-DEL-58
chr15-67196932-DEL-605
chr15-67274435-DEL-117
chr15-67328755-DEL-1051
chr15-67337122-DEL-1001
chr15-67340566-DEL-204
chr15-67360204-DEL-132
chr15-67368706-DEL-5584
chr15-67388867-DEL-236
chr15-67389039-DEL-424
chr15-67394950-DEL-414
chr15-67467827-DEL-202
chr15-67473878-DEL-97
chr15-67480767-DEL-859
chr15-67487089-DEL-921
chr15-67524766-DEL-650
chr15-67550318-DEL-55
chr15-67566842-DEL-2211
chr15-67638061-DEL-1076
chr15-67678665-DEL-532
chr15-67708910-DEL-226
chr15-67722281-DEL-367
chr15-67737198-DEL-120
chr15-67738959-DEL-204
chr15-67765373-DEL-151
chr15-67793837-DEL-76
chr15-67795548-DEL-316
chr15-67817468-DEL-1345
chr15-67838966-DEL-654
chr15-67847633-DEL-86
chr15-67952625-DEL-56
chr15-67998172-DEL-54
chr15-68008377-DEL-782
chr15-68017892-DEL-110
chr15-68083995-DEL-434
chr15-68106468-DEL-53
chr15-68115795-DEL-477
chr15-68125619-DEL-70
chr15-68134881-DEL-144

chr15-68136804-DEL-141
chr15-68139929-DEL-105
chr15-68186482-DEL-108
chr15-68324620-DEL-2539
chr15-68330116-DEL-145
chr15-68378849-DEL-51
chr15-68379720-DEL-61
chr15-68391753-DEL-156
chr15-68392543-DEL-54
chr15-68415657-DEL-1132
chr15-68429748-DEL-443
chr15-68505266-DEL-201
chr15-68512214-DEL-1262
chr15-68513803-DEL-1108
chr15-68521069-DEL-64
chr15-68575713-DEL-355
chr15-68576547-DEL-773
chr15-68577094-DEL-194
chr15-68772333-DEL-104
chr15-68772899-DEL-53
chr15-68830441-DEL-60
chr15-68841357-DEL-1217
chr15-68857839-DEL-212
chr15-68875450-DEL-4554
chr15-68889698-DEL-97
chr15-68892597-DEL-58
chr15-68903745-DEL-62
chr15-68971317-DEL-53
chr15-69030653-DEL-889
chr15-69034533-DEL-1886
chr15-69042230-DEL-75
chr15-69049759-DEL-59
chr15-69051267-DEL-115
chr15-69058193-DEL-54
chr15-69063520-DEL-347
chr15-69080101-DEL-537
chr15-69107850-DEL-399
chr15-69108927-DEL-827
chr15-69127782-DEL-58
chr15-69136594-DEL-6560
chr15-69169318-DEL-55
chr15-69219686-DEL-100
chr15-69223420-DEL-487
chr15-69229076-DEL-54
chr15-69237964-DEL-484
chr15-69262051-DEL-6497
chr15-69271603-DEL-59
chr15-69282994-DEL-106
chr15-69286044-DEL-52
chr15-69293590-DEL-57
chr15-69308884-DEL-54
chr15-69343004-DEL-1730
chr15-69362781-DEL-487
chr15-69365180-DEL-577
chr15-69406583-DEL-531
chr15-69420492-DEL-177
chr15-69433973-DEL-168
chr15-69465311-DEL-142
chr15-69472642-DEL-418
chr15-69488954-DEL-793
chr15-69498136-DEL-84
chr15-69506189-DEL-197
chr15-69529761-DEL-695
chr15-69538770-DEL-775
chr15-69605288-DEL-86
chr15-69615184-DEL-729
chr15-69655043-DEL-112
chr15-69672394-DEL-169
chr15-69688817-DEL-341
chr15-69705442-DEL-537
chr15-69708088-DEL-69
chr15-69753054-DEL-108
chr15-69754208-DEL-604
chr15-69818524-DEL-5947
chr15-69837638-DEL-1452
chr15-69840407-DEL-615
chr15-69862423-DEL-337

chr15-69885501-DEL-59
chr15-69919504-DEL-281
chr15-69954939-DEL-213
chr15-69970940-DEL-1526
chr15-69997520-DEL-455
chr15-69999142-DEL-774
chr15-70040419-DEL-371
chr15-70046078-DEL-194
chr15-70078985-DEL-487
chr15-70081664-DEL-54
chr15-70087464-DEL-923
chr15-70101650-DEL-65
chr15-70112356-DEL-441
chr15-70123425-DEL-932
chr15-70132694-DEL-64
chr15-70141800-DEL-1014
chr15-70148047-DEL-100
chr15-70201103-DEL-620
chr15-70218719-DEL-70
chr15-70261748-DEL-105
chr15-70263654-DEL-80
chr15-70275751-DEL-142
chr15-70303768-DEL-177
chr15-70323682-DEL-350
chr15-70331088-DEL-418
chr15-70384193-DEL-555
chr15-70386512-DEL-201
chr15-70397141-DEL-537
chr15-70403853-DEL-5986
chr15-70437981-DEL-362
chr15-70447909-DEL-1132
chr15-70460148-DEL-1407
chr15-70473434-DEL-72
chr15-70484295-DEL-204
chr15-70505757-DEL-109
chr15-70524253-DEL-197
chr15-70538575-DEL-503
chr15-70570246-DEL-1648
chr15-70582862-DEL-172
chr15-70587488-DEL-68
chr15-70592736-DEL-68
chr15-70595608-DEL-83
chr15-70603931-DEL-87
chr15-70611393-DEL-387
chr15-70616592-DEL-75
chr15-70637788-DEL-63
chr15-70639513-DEL-403
chr15-70658141-DEL-64
chr15-70675159-DEL-66
chr15-70715184-DEL-66
chr15-70716353-DEL-198
chr15-70728905-DEL-212
chr15-70743625-DEL-86
chr15-70752585-DEL-57
chr15-70761844-DEL-501
chr15-70763769-DEL-83
chr15-70766130-DEL-116
chr15-70809573-DEL-92
chr15-70819104-DEL-126
chr15-70821841-DEL-195
chr15-70829557-DEL-2166
chr15-70843487-DEL-90
chr15-70857077-DEL-809
chr15-70861352-DEL-7004
chr15-70874875-DEL-190
chr15-70897373-DEL-55
chr15-70912323-DEL-1104
chr15-70937360-DEL-107
chr15-70956509-DEL-123
chr15-70956978-DEL-837
chr15-70972403-DEL-302
chr15-70974218-DEL-682
chr15-70982582-DEL-1527
chr15-70986668-DEL-110
chr15-70993243-DEL-96
chr15-71001174-DEL-62
chr15-71011427-DEL-171

chr15-71039436-DEL-568
chr15-71052621-DEL-1174
chr15-71082540-DEL-235
chr15-71089815-DEL-463
chr15-71122450-DEL-194
chr15-71128415-DEL-3340
chr15-71134777-DEL-6349
chr15-71135614-DEL-67
chr15-71136541-DEL-1565
chr15-71149851-DEL-55
chr15-71168154-DEL-70
chr15-71177500-DEL-256
chr15-71180780-DEL-92
chr15-71194085-DEL-65
chr15-71218857-DEL-868
chr15-71222164-DEL-169
chr15-71240530-DEL-56
chr15-71252987-DEL-1010
chr15-71261012-DEL-2006
chr15-71261062-DEL-226
chr15-71275131-DEL-751
chr15-71279222-DEL-211
chr15-71281815-DEL-4148
chr15-71282215-DEL-3276
chr15-71348231-DEL-719
chr15-71367885-DEL-78
chr15-71436620-DEL-7621
chr15-71443045-DEL-112
chr15-71456999-DEL-6273
chr15-71463565-DEL-179
chr15-71469749-DEL-7235
chr15-71494314-DEL-95
chr15-71521248-DEL-208
chr15-71522187-DEL-6789
chr15-71559112-DEL-505
chr15-71590413-DEL-60
chr15-71608673-DEL-1193
chr15-71608748-DEL-482
chr15-71609592-DEL-67
chr15-71668569-DEL-175
chr15-71728454-DEL-131
chr15-71761850-DEL-52
chr15-71776183-DEL-54
chr15-71783066-DEL-81
chr15-71790902-DEL-192
chr15-71793247-DEL-198
chr15-71797775-DEL-331
chr15-71817395-DEL-321
chr15-71817764-DEL-614
chr15-71920001-DEL-751
chr15-71920701-DEL-529
chr15-71926778-DEL-97
chr15-71968463-DEL-66
chr15-71975571-DEL-541
chr15-71984759-DEL-2491
chr15-72061107-DEL-144
chr15-72062475-DEL-106
chr15-72089909-DEL-720
chr15-72092421-DEL-199
chr15-72108857-DEL-2146
chr15-72118093-DEL-81
chr15-72121284-DEL-3394
chr15-72121421-DEL-1697
chr15-72122654-DEL-3409
chr15-72126438-DEL-867
chr15-72172983-DEL-55
chr15-72216777-DEL-148
chr15-72245380-DEL-260
chr15-72327211-DEL-658
chr15-72331735-DEL-5173
chr15-72347066-DEL-65
chr15-72432646-DEL-87
chr15-72482513-DEL-116
chr15-72595976-DEL-108
chr15-72655172-DEL-245
chr15-72688023-DEL-61
chr15-72710437-DEL-235

chr15-72742314-DEL-185
chr15-72785148-DEL-510
chr15-72798844-DEL-144
chr15-72803918-DEL-114
chr15-72821938-DEL-53
chr15-72848568-DEL-1023
chr15-72849340-DEL-105
chr15-72850736-DEL-173
chr15-72868833-DEL-288
chr15-72869718-DEL-160
chr15-72918530-DEL-200
chr15-72935858-DEL-194
chr15-73038032-DEL-85
chr15-73102909-DEL-56
chr15-73119867-DEL-1949
chr15-73139670-DEL-158
chr15-73201809-DEL-247
chr15-73205267-DEL-69
chr15-73217720-DEL-140
chr15-73222982-DEL-123
chr15-73273546-DEL-9535
chr15-73283589-DEL-175
chr15-73290164-DEL-686
chr15-73299886-DEL-280
chr15-73310871-DEL-159
chr15-73338911-DEL-63
chr15-73352628-DEL-1009
chr15-73354477-DEL-307
chr15-73367982-DEL-67
chr15-73380813-DEL-231
chr15-73409600-DEL-69
chr15-73464306-DEL-669
chr15-73487433-DEL-128
chr15-73512808-DEL-51
chr15-73514001-DEL-118
chr15-73554000-DEL-138
chr15-73572756-DEL-912
chr15-73646667-DEL-58
chr15-73656058-DEL-254
chr15-73695473-DEL-1068
chr15-73755237-DEL-829
chr15-73761984-DEL-178
chr15-73770659-DEL-102
chr15-73781333-DEL-56
chr15-73846341-DEL-110
chr15-73852085-DEL-804
chr15-73853511-DEL-101
chr15-73891394-DEL-89
chr15-73892966-DEL-60
chr15-73906228-DEL-57
chr15-73977194-DEL-613
chr15-74073683-DEL-55
chr15-74102515-DEL-95
chr15-74132684-DEL-65
chr15-74138312-DEL-136
chr15-74159718-DEL-135
chr15-74187591-DEL-87
chr15-74211325-DEL-1438
chr15-74231078-DEL-52
chr15-74240312-DEL-185
chr15-74249199-DEL-55
chr15-74260951-DEL-81
chr15-74282407-DEL-595
chr15-74287010-DEL-59
chr15-74292094-DEL-56
chr15-74424586-DEL-183
chr15-74449877-DEL-101
chr15-74496119-DEL-126
chr15-74521289-DEL-1432
chr15-74525815-DEL-1435
chr15-74538845-DEL-114
chr15-74585826-DEL-171
chr15-74605256-DEL-52
chr15-74669821-DEL-71
chr15-74696914-DEL-4751
chr15-74720460-DEL-179
chr15-74728964-DEL-103

chr15-74762289-DEL-92
chr15-74801940-DEL-113
chr15-74878033-DEL-72
chr15-74896267-DEL-1403
chr15-74906333-DEL-55
chr15-74957243-DEL-1649
chr15-75010780-DEL-338
chr15-75014208-DEL-423
chr15-75036362-DEL-279
chr15-75036685-DEL-10903
chr15-75039794-DEL-6649
chr15-75064210-DEL-1055
chr15-75088228-DEL-183
chr15-75091351-DEL-104
chr15-75095776-DEL-97
chr15-75113218-DEL-255
chr15-75125162-DEL-64
chr15-75142133-DEL-1236
chr15-75144636-DEL-1397
chr15-75156015-DEL-826
chr15-75162925-DEL-62
chr15-75169650-DEL-6006
chr15-75203494-DEL-52
chr15-75215240-DEL-62
chr15-75363176-DEL-54
chr15-75409358-DEL-4790
chr15-75450724-DEL-439
chr15-75510550-DEL-396
chr15-75521944-DEL-253
chr15-75526905-DEL-104
chr15-75593829-DEL-325
chr15-75598934-DEL-99
chr15-75608456-DEL-54
chr15-75622308-DEL-59
chr15-75652381-DEL-315
chr15-75701429-DEL-62
chr15-75713110-DEL-95
chr15-75723634-DEL-218
chr15-75753254-DEL-190
chr15-75770331-DEL-281
chr15-75792785-DEL-63
chr15-75822839-DEL-9385
chr15-75851423-DEL-153
chr15-75871246-DEL-98
chr15-75874612-DEL-66
chr15-75898450-DEL-167
chr15-75900513-DEL-67
chr15-75915683-DEL-53
chr15-75968818-DEL-189
chr15-75975007-DEL-13896
chr15-76014013-DEL-149
chr15-76144872-DEL-101
chr15-76158700-DEL-5925
chr15-76166953-DEL-551
chr15-76171406-DEL-54
chr15-76173502-DEL-98
chr15-76178214-DEL-112
chr15-76221768-DEL-863
chr15-76249403-DEL-180
chr15-76250484-DEL-215
chr15-76251211-DEL-227
chr15-76271454-DEL-156
chr15-76281509-DEL-58
chr15-76296934-DEL-132
chr15-76307654-DEL-95
chr15-76322211-DEL-86
chr15-76398631-DEL-82
chr15-76414702-DEL-54
chr15-76454469-DEL-270
chr15-76456022-DEL-216
chr15-76464634-DEL-222
chr15-76465430-DEL-104
chr15-76492313-DEL-245
chr15-76493922-DEL-85
chr15-76519894-DEL-63
chr15-76570313-DEL-195
chr15-76599399-DEL-57

chr15-76616102-DEL-107
chr15-76622497-DEL-177
chr15-76643178-DEL-450
chr15-76693287-DEL-161
chr15-76742646-DEL-216
chr15-76745858-DEL-439
chr15-76762618-DEL-366
chr15-76766473-DEL-397
chr15-76777229-DEL-160
chr15-76777894-DEL-199
chr15-76779751-DEL-418
chr15-76780643-DEL-1344
chr15-76785419-DEL-317
chr15-76800528-DEL-72
chr15-76806847-DEL-201
chr15-76846753-DEL-204
chr15-76857271-DEL-175
chr15-76885513-DEL-649
chr15-76917550-DEL-78
chr15-76926648-DEL-436
chr15-76932682-DEL-574
chr15-76936361-DEL-9717
chr15-77005523-DEL-988
chr15-77060689-DEL-2597
chr15-77101421-DEL-55
chr15-77163397-DEL-108
chr15-77198582-DEL-156
chr15-77214931-DEL-64
chr15-77217187-DEL-554
chr15-77239043-DEL-3536
chr15-77254292-DEL-94
chr15-77256768-DEL-402
chr15-77274790-DEL-162
chr15-77276119-DEL-168
chr15-77314616-DEL-6423
chr15-77382599-DEL-88
chr15-77388692-DEL-201
chr15-77389860-DEL-863
chr15-77398836-DEL-663
chr15-77413199-DEL-82
chr15-77418676-DEL-572
chr15-77424123-DEL-210
chr15-77433682-DEL-6425
chr15-77441813-DEL-117
chr15-77443997-DEL-60
chr15-77453652-DEL-2539
chr15-77488539-DEL-600
chr15-77490847-DEL-395
chr15-77529238-DEL-894
chr15-77568069-DEL-8271
chr15-77580847-DEL-1107
chr15-77581936-DEL-301
chr15-77591024-DEL-121
chr15-77631138-DEL-58
chr15-77638132-DEL-52
chr15-77684535-DEL-100
chr15-77709126-DEL-54
chr15-77710941-DEL-101
chr15-77723525-DEL-57
chr15-77731345-DEL-204
chr15-77737327-DEL-430
chr15-77750311-DEL-82
chr15-77755572-DEL-3571
chr15-77770877-DEL-82
chr15-77771557-DEL-97
chr15-77800092-DEL-253
chr15-77822498-DEL-60
chr15-77859580-DEL-60
chr15-77880079-DEL-162
chr15-77923153-DEL-138
chr15-77963931-DEL-100
chr15-78009964-DEL-97
chr15-78083132-DEL-68
chr15-78090996-DEL-1126
chr15-78093159-DEL-234
chr15-78095210-DEL-76
chr15-78110953-DEL-76

chr15-78121607-DEL-119
chr15-78126641-DEL-405
chr15-78132512-DEL-88
chr15-78153378-DEL-382
chr15-78154455-DEL-270
chr15-78160009-DEL-137
chr15-78168154-DEL-76
chr15-78179833-DEL-147
chr15-78201841-DEL-1427
chr15-78213999-DEL-63
chr15-78255046-DEL-78
chr15-78273274-DEL-97
chr15-78307170-DEL-377
chr15-78364406-DEL-70
chr15-78375412-DEL-56
chr15-78397069-DEL-1086
chr15-78419481-DEL-58
chr15-78463425-DEL-51
chr15-78464507-DEL-1234
chr15-78534348-DEL-64
chr15-78593672-DEL-121
chr15-78610530-DEL-304
chr15-78632053-DEL-149
chr15-78637287-DEL-85
chr15-78644678-DEL-95
chr15-78673259-DEL-60
chr15-78684069-DEL-117
chr15-78694920-DEL-882
chr15-78705710-DEL-204
chr15-78706698-DEL-171
chr15-78713379-DEL-283
chr15-78714193-DEL-51
chr15-78794824-DEL-569
chr15-78805322-DEL-181
chr15-78813253-DEL-154
chr15-78817868-DEL-98
chr15-78818468-DEL-51
chr15-78821798-DEL-69
chr15-78828188-DEL-147
chr15-78864977-DEL-142
chr15-78895042-DEL-445
chr15-78901965-DEL-80
chr15-78907839-DEL-1879
chr15-78930550-DEL-213
chr15-78932392-DEL-189
chr15-78962918-DEL-60
chr15-78987170-DEL-82
chr15-78996541-DEL-84
chr15-79045593-DEL-90
chr15-79087419-DEL-1364
chr15-79093920-DEL-126
chr15-79122993-DEL-76
chr15-79156728-DEL-59
chr15-79176731-DEL-1220
chr15-79187387-DEL-63
chr15-79197511-DEL-550
chr15-79200029-DEL-121
chr15-79202519-DEL-116
chr15-79222813-DEL-214
chr15-79224867-DEL-286
chr15-79229701-DEL-54
chr15-79239881-DEL-173
chr15-79248732-DEL-59
chr15-79318695-DEL-69
chr15-79346518-DEL-643
chr15-79346886-DEL-187
chr15-79397673-DEL-51
chr15-79405613-DEL-172
chr15-79410550-DEL-464
chr15-79450376-DEL-2203
chr15-79480183-DEL-205
chr15-79498142-DEL-209
chr15-79502189-DEL-196
chr15-79503719-DEL-57
chr15-79523820-DEL-105
chr15-79529119-DEL-189
chr15-79538805-DEL-186

chr15-79552087-DEL-219
chr15-79605970-DEL-424
chr15-79606121-DEL-292
chr15-79614903-DEL-76
chr15-79627441-DEL-53
chr15-79631192-DEL-126
chr15-79666830-DEL-689
chr15-79667228-DEL-255
chr15-79684965-DEL-62
chr15-79735787-DEL-196
chr15-79793580-DEL-241
chr15-79796127-DEL-113
chr15-80040942-DEL-80
chr15-80050249-DEL-383
chr15-80061577-DEL-93
chr15-80062716-DEL-203
chr15-80068361-DEL-135
chr15-80090885-DEL-55
chr15-80116095-DEL-206
chr15-80124354-DEL-327
chr15-80125035-DEL-85
chr15-80125196-DEL-1310
chr15-80125441-DEL-287
chr15-80126056-DEL-452
chr15-80185042-DEL-52
chr15-80186423-DEL-176
chr15-80192601-DEL-161
chr15-80197863-DEL-64
chr15-80217748-DEL-251
chr15-80222720-DEL-473
chr15-80224221-DEL-137
chr15-80232445-DEL-82
chr15-80235252-DEL-145
chr15-80245323-DEL-186
chr15-80278374-DEL-411
chr15-80307733-DEL-82
chr15-80319077-DEL-215
chr15-80328334-DEL-559
chr15-80372509-DEL-171
chr15-80399067-DEL-127
chr15-80412949-DEL-68
chr15-80419541-DEL-100
chr15-80424137-DEL-1464
chr15-80442907-DEL-54
chr15-80496749-DEL-96
chr15-80523910-DEL-60
chr15-80552897-DEL-288
chr15-80593823-DEL-102
chr15-80636033-DEL-82
chr15-80658045-DEL-1675
chr15-80672645-DEL-670
chr15-80696309-DEL-92
chr15-80697573-DEL-1033
chr15-80732791-DEL-72
chr15-80739319-DEL-664
chr15-80752392-DEL-66
chr15-80753013-DEL-204
chr15-80789915-DEL-403
chr15-80791149-DEL-171
chr15-80794611-DEL-102
chr15-80804182-DEL-124
chr15-80805236-DEL-324
chr15-80826375-DEL-389
chr15-80831535-DEL-91
chr15-80835374-DEL-358
chr15-80845299-DEL-97
chr15-80849120-DEL-200
chr15-80868306-DEL-258
chr15-80875668-DEL-223
chr15-80888424-DEL-204
chr15-80895938-DEL-56
chr15-80910983-DEL-88
chr15-80913705-DEL-275
chr15-81022985-DEL-300
chr15-81056159-DEL-252
chr15-81060187-DEL-165
chr15-81070028-DEL-139

chr15-81100403-DEL-306
chr15-81167599-DEL-250
chr15-81170325-DEL-55
chr15-81194063-DEL-340
chr15-81272084-DEL-76
chr15-81275745-DEL-222
chr15-81330743-DEL-192
chr15-81331625-DEL-282
chr15-81334880-DEL-55
chr15-81358483-DEL-213
chr15-81364717-DEL-86
chr15-81369624-DEL-181
chr15-81386305-DEL-198
chr15-81399053-DEL-1995
chr15-81404647-DEL-66
chr15-81431285-DEL-179
chr15-81432641-DEL-59
chr15-81437459-DEL-164
chr15-81438197-DEL-183
chr15-81445081-DEL-53
chr15-81463515-DEL-243
chr15-81497706-DEL-2343
chr15-81500688-DEL-122
chr15-81508202-DEL-87
chr15-81542358-DEL-238
chr15-81549668-DEL-89
chr15-81554862-DEL-219
chr15-81562207-DEL-64
chr15-81563077-DEL-2290
chr15-81580682-DEL-119
chr15-81609744-DEL-245
chr15-81614862-DEL-749
chr15-81631277-DEL-81
chr15-81649528-DEL-195
chr15-81654244-DEL-173
chr15-81673947-DEL-1677
chr15-81714495-DEL-53
chr15-81760785-DEL-154
chr15-81782223-DEL-72
chr15-81807253-DEL-63
chr15-81826049-DEL-174
chr15-81845910-DEL-58
chr15-81886411-DEL-77
chr15-81902399-DEL-267
chr15-81904890-DEL-659
chr15-81941294-DEL-550
chr15-81951293-DEL-706
chr15-81974461-DEL-52
chr15-81982838-DEL-296
chr15-81997397-DEL-51
chr15-82003987-DEL-72
chr15-82012339-DEL-89
chr15-82033989-DEL-217
chr15-82042231-DEL-1249
chr15-82093803-DEL-57
chr15-82097390-DEL-163
chr15-82099329-DEL-72
chr15-82101878-DEL-151
chr15-82115355-DEL-355
chr15-82120241-DEL-154
chr15-82126150-DEL-519
chr15-82126817-DEL-203
chr15-82158384-DEL-99
chr15-82210355-DEL-77
chr15-82232272-DEL-67
chr15-82243280-DEL-3454
chr15-82246234-DEL-248
chr15-82271896-DEL-115
chr15-82296178-DEL-66
chr15-82311137-DEL-66
chr15-82372378-DEL-6387
chr15-82384392-DEL-198
chr15-82400443-DEL-3044
chr15-82406848-DEL-1253
chr15-82407694-DEL-6664
chr15-82413429-DEL-97
chr15-82447429-DEL-727

chr15-82455397-DEL-283
chr15-82503425-DEL-237
chr15-82507665-DEL-84
chr15-82580041-DEL-508
chr15-82597513-DEL-60
chr15-82616689-DEL-607
chr15-82730142-DEL-967
chr15-82758488-DEL-6762
chr15-82786187-DEL-52
chr15-82904622-DEL-244
chr15-82911121-DEL-76
chr15-82964729-DEL-384
chr15-82996087-DEL-59
chr15-83009537-DEL-206
chr15-83039125-DEL-96
chr15-83048375-DEL-327
chr15-83068560-DEL-106
chr15-83077955-DEL-8419
chr15-83090933-DEL-193
chr15-83133355-DEL-207
chr15-83142649-DEL-464
chr15-83154938-DEL-250
chr15-83157268-DEL-98
chr15-83161104-DEL-310
chr15-83175756-DEL-91
chr15-83193407-DEL-586
chr15-83205466-DEL-1012
chr15-83207977-DEL-248
chr15-83210443-DEL-167
chr15-83214385-DEL-166
chr15-83252542-DEL-52
chr15-83275285-DEL-185
chr15-83304643-DEL-455
chr15-83311826-DEL-202
chr15-83314868-DEL-54
chr15-83368966-DEL-176
chr15-83372847-DEL-64
chr15-83385772-DEL-52
chr15-83415445-DEL-73
chr15-83506446-DEL-97
chr15-83510101-DEL-1098
chr15-83512390-DEL-87
chr15-83516354-DEL-87
chr15-83525717-DEL-255
chr15-83532443-DEL-71
chr15-83591453-DEL-541
chr15-83607593-DEL-239
chr15-83623021-DEL-94
chr15-83628354-DEL-178
chr15-83629080-DEL-814
chr15-83632412-DEL-872
chr15-83641785-DEL-820
chr15-83644085-DEL-79
chr15-83650794-DEL-1544
chr15-83651442-DEL-84
chr15-83654624-DEL-121
chr15-83693996-DEL-88
chr15-83695353-DEL-61
chr15-83710841-DEL-110
chr15-83731552-DEL-51
chr15-83804744-DEL-517
chr15-83821044-DEL-1452
chr15-83822552-DEL-74
chr15-83834329-DEL-1358
chr15-83842112-DEL-542
chr15-83846366-DEL-52
chr15-83854355-DEL-130
chr15-83862839-DEL-83
chr15-83864304-DEL-206
chr15-83883810-DEL-198
chr15-83884759-DEL-76
chr15-83914948-DEL-68
chr15-83916269-DEL-468
chr15-83919389-DEL-83
chr15-83920102-DEL-315
chr15-83922681-DEL-565
chr15-83923792-DEL-194

chr15-83925464-DEL-61
chr15-83993678-DEL-106
chr15-83994863-DEL-153
chr15-84028360-DEL-1988
chr15-84030734-DEL-2936
chr15-84037159-DEL-221
chr15-84047668-DEL-599
chr15-84054346-DEL-443
chr15-84078487-DEL-906
chr15-84107270-DEL-787
chr15-84120163-DEL-90
chr15-84125133-DEL-79
chr15-84159388-DEL-117
chr15-84201430-DEL-60
chr15-84203853-DEL-140
chr15-84226994-DEL-2898
chr15-84243770-DEL-86
chr15-84246680-DEL-210
chr15-84258986-DEL-448
chr15-84272159-DEL-219
chr15-84281138-DEL-77
chr15-84286420-DEL-280
chr15-84295313-DEL-76
chr15-84311141-DEL-146
chr15-84355973-DEL-582
chr15-84361881-DEL-235
chr15-84398821-DEL-353
chr15-84449050-DEL-299
chr15-84483283-DEL-251
chr15-84491192-DEL-52
chr15-84519674-DEL-249
chr15-84539194-DEL-272
chr15-84561161-DEL-67
chr15-84618456-DEL-154
chr15-84621883-DEL-944
chr15-84623971-DEL-465
chr15-84666305-DEL-53
chr15-84677612-DEL-300
chr15-84699556-DEL-65
chr15-84700277-DEL-237
chr15-84743416-DEL-1953
chr15-84769524-DEL-215
chr15-84796817-DEL-348
chr15-84797842-DEL-1280
chr15-84802942-DEL-52
chr15-84854724-DEL-1140
chr15-84854837-DEL-540
chr15-84854993-DEL-240
chr15-84855417-DEL-320
chr15-84865498-DEL-377
chr15-84869074-DEL-191
chr15-84895740-DEL-176
chr15-84907136-DEL-133
chr15-84917298-DEL-905
chr15-84986109-DEL-448
chr15-85018136-DEL-92
chr15-85048875-DEL-58
chr15-85053575-DEL-290
chr15-85058179-DEL-1525
chr15-85061946-DEL-55
chr15-85086286-DEL-184
chr15-85096727-DEL-78
chr15-85141214-DEL-198
chr15-85158852-DEL-526
chr15-85177606-DEL-54
chr15-85179497-DEL-68
chr15-85185223-DEL-113
chr15-85191042-DEL-158
chr15-85204927-DEL-136
chr15-85227078-DEL-123
chr15-85242586-DEL-779
chr15-85257644-DEL-694
chr15-85258900-DEL-322
chr15-85301585-DEL-60
chr15-85313190-DEL-116
chr15-85356175-DEL-216
chr15-85385030-DEL-66

chr15-85393712-DEL-112
chr15-85399499-DEL-377
chr15-85416934-DEL-280
chr15-85474972-DEL-276
chr15-85521921-DEL-288
chr15-85532226-DEL-192
chr15-85533862-DEL-232
chr15-85629799-DEL-66
chr15-85648235-DEL-73
chr15-85649795-DEL-61
chr15-85662944-DEL-194
chr15-85669517-DEL-186
chr15-85674332-DEL-284
chr15-85696676-DEL-52
chr15-85712820-DEL-51
chr15-85722045-DEL-692
chr15-85735958-DEL-91
chr15-85748777-DEL-169
chr15-85787889-DEL-135
chr15-85797323-DEL-56
chr15-85797824-DEL-89
chr15-85800404-DEL-85
chr15-85807409-DEL-735
chr15-85838842-DEL-72
chr15-85864341-DEL-620
chr15-85872490-DEL-82
chr15-85873125-DEL-84
chr15-85884157-DEL-380
chr15-85929763-DEL-188
chr15-85932324-DEL-104
chr15-85937076-DEL-225
chr15-85939377-DEL-60
chr15-85968764-DEL-750
chr15-86001452-DEL-428
chr15-86039493-DEL-166
chr15-86041960-DEL-201
chr15-86082365-DEL-62
chr15-86099994-DEL-94
chr15-86100034-DEL-3856
chr15-86106179-DEL-8402
chr15-86107979-DEL-116
chr15-86117476-DEL-100
chr15-86129554-DEL-225
chr15-86129858-DEL-295
chr15-86144792-DEL-103
chr15-86158366-DEL-420
chr15-86191978-DEL-134
chr15-86209505-DEL-1021
chr15-86209687-DEL-821
chr15-86221802-DEL-588
chr15-86231610-DEL-697
chr15-86247887-DEL-109
chr15-86287298-DEL-1137
chr15-86405977-DEL-581
chr15-86424295-DEL-436
chr15-86435698-DEL-86
chr15-86459449-DEL-173
chr15-86470760-DEL-52
chr15-86496783-DEL-79
chr15-86502966-DEL-78
chr15-86504613-DEL-53
chr15-86534647-DEL-138
chr15-86549287-DEL-90
chr15-86624135-DEL-60
chr15-86628239-DEL-592
chr15-86637780-DEL-170
chr15-86644153-DEL-2229
chr15-86646475-DEL-66
chr15-86655381-DEL-128
chr15-86666394-DEL-52
chr15-86717037-DEL-57
chr15-86740718-DEL-752
chr15-86753767-DEL-452
chr15-86755458-DEL-105
chr15-86777821-DEL-65
chr15-86789391-DEL-524
chr15-86816242-DEL-1955

chr15-86824328-DEL-70
chr15-86832205-DEL-120
chr15-86842840-DEL-78
chr15-86902884-DEL-134
chr15-86918768-DEL-72
chr15-86921225-DEL-139
chr15-86933105-DEL-363
chr15-86953220-DEL-64
chr15-86977633-DEL-225
chr15-87003123-DEL-54
chr15-87061931-DEL-61
chr15-87092721-DEL-52
chr15-87105252-DEL-81
chr15-87120786-DEL-160
chr15-87276054-DEL-52
chr15-87280882-DEL-60
chr15-87310068-DEL-63
chr15-87325468-DEL-54
chr15-87329655-DEL-286
chr15-87355041-DEL-338
chr15-87368914-DEL-85
chr15-87371791-DEL-52
chr15-87373056-DEL-90
chr15-87385767-DEL-82
chr15-87433266-DEL-283
chr15-87461907-DEL-134
chr15-87498493-DEL-276
chr15-87548964-DEL-294
chr15-87551992-DEL-58
chr15-87575316-DEL-238
chr15-87615020-DEL-132
chr15-87633260-DEL-153
chr15-87657151-DEL-109
chr15-87658504-DEL-483
chr15-87690535-DEL-68
chr15-87706837-DEL-1415
chr15-87711650-DEL-158
chr15-87725408-DEL-67
chr15-87837259-DEL-53
chr15-87842873-DEL-51
chr15-87845423-DEL-864
chr15-87845429-DEL-1475
chr15-87845866-DEL-274
chr15-87846206-DEL-848
chr15-87852069-DEL-229
chr15-87865762-DEL-162
chr15-87905362-DEL-87
chr15-87928850-DEL-107
chr15-87939585-DEL-422
chr15-87944529-DEL-80
chr15-87946013-DEL-56
chr15-87979450-DEL-473
chr15-87995104-DEL-266
chr15-88031249-DEL-85
chr15-88062441-DEL-386
chr15-88071167-DEL-75
chr15-88085279-DEL-69
chr15-88119109-DEL-1008
chr15-88140465-DEL-145
chr15-88166481-DEL-1345
chr15-88192217-DEL-69
chr15-88216439-DEL-1025
chr15-88228897-DEL-189
chr15-88240716-DEL-176
chr15-88279823-DEL-243
chr15-88284667-DEL-58
chr15-88308777-DEL-1269
chr15-88308778-DEL-826
chr15-88309101-DEL-430
chr15-88309595-DEL-457
chr15-88330926-DEL-420
chr15-88348850-DEL-67
chr15-88394550-DEL-547
chr15-88406654-DEL-486
chr15-88447560-DEL-63
chr15-88460008-DEL-942
chr15-88483544-DEL-249

chr15-88516517-DEL-96
chr15-88519711-DEL-184
chr15-88524642-DEL-335
chr15-88530278-DEL-51
chr15-88543861-DEL-86
chr15-88555286-DEL-208
chr15-88566298-DEL-518
chr15-88570124-DEL-199
chr15-88612191-DEL-837
chr15-88648514-DEL-561
chr15-88649326-DEL-1186
chr15-88657498-DEL-2392
chr15-88670874-DEL-202
chr15-88678888-DEL-1406
chr15-88681727-DEL-219
chr15-88687933-DEL-51
chr15-88692342-DEL-2025
chr15-88698109-DEL-215
chr15-88702429-DEL-98
chr15-88711077-DEL-171
chr15-88713542-DEL-56
chr15-88722738-DEL-122
chr15-88724846-DEL-208
chr15-88760972-DEL-58
chr15-88768221-DEL-433
chr15-88781384-DEL-52
chr15-88789180-DEL-66
chr15-88790717-DEL-1866
chr15-88825321-DEL-79
chr15-88826601-DEL-284
chr15-88835207-DEL-90
chr15-88836002-DEL-57
chr15-88844977-DEL-1215
chr15-88851205-DEL-60
chr15-88853089-DEL-55
chr15-88859179-DEL-1260
chr15-88859197-DEL-168
chr15-88859862-DEL-545
chr15-88878648-DEL-252
chr15-88912616-DEL-151
chr15-88913934-DEL-2346
chr15-88976225-DEL-259
chr15-88987342-DEL-81
chr15-88991338-DEL-64
chr15-88996264-DEL-65
chr15-88997084-DEL-1672
chr15-89003569-DEL-94
chr15-89037410-DEL-207
chr15-89093630-DEL-75
chr15-89103185-DEL-844
chr15-89108171-DEL-89
chr15-89108401-DEL-1190
chr15-89118818-DEL-196
chr15-89119514-DEL-53
chr15-89131421-DEL-316
chr15-89132464-DEL-2559
chr15-89133082-DEL-90
chr15-89182781-DEL-233
chr15-89228722-DEL-407
chr15-89243147-DEL-79
chr15-89247394-DEL-55
chr15-89282157-DEL-57
chr15-89286792-DEL-140
chr15-89287381-DEL-109
chr15-89290126-DEL-198
chr15-89297539-DEL-60
chr15-89299121-DEL-219
chr15-89305980-DEL-54
chr15-89318362-DEL-458
chr15-89321215-DEL-342
chr15-89344427-DEL-84
chr15-89345197-DEL-169
chr15-89354773-DEL-408
chr15-89369953-DEL-108
chr15-89376491-DEL-104
chr15-89517956-DEL-61
chr15-89538434-DEL-2997

chr15-89541214-DEL-134
chr15-89578557-DEL-748
chr15-89600143-DEL-76
chr15-89614387-DEL-1685
chr15-89646984-DEL-1156
chr15-89675576-DEL-187
chr15-89692109-DEL-1656
chr15-89705267-DEL-92
chr15-89710044-DEL-251
chr15-89712403-DEL-79
chr15-89756678-DEL-1121
chr15-89790432-DEL-335
chr15-89838481-DEL-221
chr15-89840994-DEL-55
chr15-89841981-DEL-6972
chr15-89858641-DEL-108
chr15-89900500-DEL-66
chr15-89905165-DEL-54
chr15-89923643-DEL-1383
chr15-89932949-DEL-137
chr15-89933640-DEL-295
chr15-89934595-DEL-1399
chr15-89935873-DEL-1424
chr15-89973244-DEL-254
chr15-89977761-DEL-437
chr15-89981667-DEL-115
chr15-90022667-DEL-77
chr15-90065769-DEL-794
chr15-90100502-DEL-273
chr15-90127012-DEL-379
chr15-90144703-DEL-675
chr15-90178560-DEL-76
chr15-90188763-DEL-208
chr15-90193872-DEL-65
chr15-90202376-DEL-477
chr15-90242904-DEL-173
chr15-90254814-DEL-70
chr15-90287319-DEL-283
chr15-90294109-DEL-1054
chr15-90311333-DEL-2340
chr15-90319202-DEL-65
chr15-90328026-DEL-165
chr15-90333860-DEL-61
chr15-90397305-DEL-71
chr15-90417831-DEL-67
chr15-90470847-DEL-66
chr15-90472357-DEL-308
chr15-90475100-DEL-383
chr15-90476513-DEL-150
chr15-90500739-DEL-52
chr15-90503517-DEL-78
chr15-90506104-DEL-52
chr15-90513430-DEL-338
chr15-90572610-DEL-89
chr15-90608302-DEL-352
chr15-90616364-DEL-56
chr15-90643336-DEL-398
chr15-90676855-DEL-131
chr15-90683428-DEL-270
chr15-90734983-DEL-56
chr15-90740058-DEL-57
chr15-90749110-DEL-70
chr15-90763270-DEL-78
chr15-90766475-DEL-88
chr15-90766680-DEL-1450
chr15-90769033-DEL-1289
chr15-90778329-DEL-599
chr15-90785841-DEL-152
chr15-90788047-DEL-97
chr15-90827340-DEL-222
chr15-90845763-DEL-428
chr15-90874754-DEL-69
chr15-90879474-DEL-70
chr15-90911766-DEL-215
chr15-90991989-DEL-1056
chr15-91006938-DEL-355
chr15-91017788-DEL-1165

chr15-91059931-DEL-992
chr15-91081078-DEL-433
chr15-91084118-DEL-62
chr15-91088282-DEL-127
chr15-91094999-DEL-298
chr15-91118276-DEL-178
chr15-91130733-DEL-129
chr15-91156710-DEL-85
chr15-91172340-DEL-583
chr15-91226417-DEL-79
chr15-91277768-DEL-52
chr15-91288075-DEL-438
chr15-91292366-DEL-109
chr15-91363171-DEL-80
chr15-91392212-DEL-90
chr15-91407760-DEL-336
chr15-91409563-DEL-52
chr15-91451725-DEL-162
chr15-91462586-DEL-228
chr15-91485816-DEL-433
chr15-91496114-DEL-939
chr15-91504363-DEL-123
chr15-91519062-DEL-95
chr15-91520601-DEL-70
chr15-91525586-DEL-108
chr15-91540793-DEL-55
chr15-91570804-DEL-104
chr15-91582954-DEL-523
chr15-91603173-DEL-72
chr15-91605046-DEL-1873
chr15-91619415-DEL-66
chr15-91670498-DEL-56
chr15-91682473-DEL-101
chr15-91730865-DEL-3514
chr15-91826852-DEL-99
chr15-91830818-DEL-565
chr15-91834871-DEL-113
chr15-91853172-DEL-84
chr15-91865075-DEL-55
chr15-91874465-DEL-5897
chr15-91894737-DEL-77
chr15-91913257-DEL-6214
chr15-91941276-DEL-64
chr15-91957493-DEL-72
chr15-91964371-DEL-290
chr15-92025834-DEL-114
chr15-92045800-DEL-793
chr15-92051832-DEL-172
chr15-92055109-DEL-517
chr15-92069792-DEL-182
chr15-92094235-DEL-51
chr15-92136780-DEL-104
chr15-92172794-DEL-56
chr15-92175935-DEL-103
chr15-92200370-DEL-160
chr15-92201770-DEL-72
chr15-92205323-DEL-134
chr15-92217323-DEL-217
chr15-92249663-DEL-122
chr15-92255289-DEL-78
chr15-92264795-DEL-433
chr15-92269733-DEL-62
chr15-92300931-DEL-396
chr15-92429854-DEL-157
chr15-92442576-DEL-910
chr15-92454784-DEL-104
chr15-92486044-DEL-140
chr15-92511644-DEL-4529
chr15-92521053-DEL-100
chr15-92524794-DEL-231
chr15-92528937-DEL-224
chr15-92547586-DEL-248
chr15-92659629-DEL-122
chr15-92663932-DEL-175
chr15-92665082-DEL-571
chr15-92680532-DEL-82
chr15-92684033-DEL-193

chr15-92698474-DEL-108
chr15-92726444-DEL-682
chr15-92743118-DEL-925
chr15-92746768-DEL-820
chr15-92808382-DEL-80
chr15-92811795-DEL-105
chr15-92840024-DEL-56
chr15-92841826-DEL-75
chr15-92885453-DEL-1249
chr15-92892458-DEL-2660
chr15-92900552-DEL-53
chr15-92912888-DEL-288
chr15-92918570-DEL-55
chr15-92953231-DEL-6451
chr15-92988901-DEL-621
chr15-93006390-DEL-240
chr15-93006427-DEL-1967
chr15-93029691-DEL-67
chr15-93044950-DEL-88
chr15-93077387-DEL-69
chr15-93083844-DEL-902
chr15-93111182-DEL-51
chr15-93140726-DEL-169
chr15-93158034-DEL-70
chr15-93198756-DEL-63
chr15-93222151-DEL-74
chr15-93228065-DEL-92
chr15-93236296-DEL-202
chr15-93238163-DEL-202
chr15-93244735-DEL-3668
chr15-93251685-DEL-179
chr15-93252304-DEL-191
chr15-93270494-DEL-122
chr15-93272957-DEL-190
chr15-93274858-DEL-55
chr15-93305458-DEL-914
chr15-93328261-DEL-51
chr15-93330635-DEL-72
chr15-93334975-DEL-171
chr15-93341634-DEL-106
chr15-93347573-DEL-61
chr15-93365754-DEL-924
chr15-93365794-DEL-377
chr15-93365797-DEL-1294
chr15-93376141-DEL-1024
chr15-93401958-DEL-207
chr15-93407598-DEL-125
chr15-93410186-DEL-130
chr15-93420500-DEL-96
chr15-93434650-DEL-60
chr15-93448150-DEL-95
chr15-93475791-DEL-1501
chr15-93501450-DEL-271
chr15-93563372-DEL-137
chr15-93570082-DEL-56
chr15-93583001-DEL-156
chr15-93615141-DEL-151
chr15-93615907-DEL-62
chr15-93650300-DEL-114
chr15-93657606-DEL-882
chr15-93662168-DEL-80
chr15-93664036-DEL-735
chr15-93676005-DEL-69
chr15-93717372-DEL-121
chr15-93732045-DEL-399
chr15-93762872-DEL-62
chr15-93772486-DEL-99
chr15-93785439-DEL-139
chr15-93829115-DEL-198
chr15-93843116-DEL-314
chr15-93893266-DEL-120
chr15-93899918-DEL-138
chr15-93908329-DEL-7855
chr15-93927500-DEL-62
chr15-93932163-DEL-54
chr15-93933348-DEL-224
chr15-93945920-DEL-92

chr15-93962304-DEL-63
chr15-93982378-DEL-59
chr15-93986343-DEL-1022
chr15-93986891-DEL-63
chr15-94005548-DEL-114
chr15-94016075-DEL-6159
chr15-94024950-DEL-56
chr15-94094985-DEL-53
chr15-94114731-DEL-62
chr15-94117617-DEL-134
chr15-94126905-DEL-175
chr15-94153282-DEL-71
chr15-94231018-DEL-402
chr15-94238366-DEL-96
chr15-94239120-DEL-51
chr15-94253674-DEL-60
chr15-94281485-DEL-1802
chr15-94282347-DEL-283
chr15-94298420-DEL-96
chr15-94312372-DEL-668
chr15-94338988-DEL-71
chr15-94343230-DEL-226
chr15-94350061-DEL-56
chr15-94366766-DEL-51
chr15-94388115-DEL-270
chr15-94394894-DEL-70
chr15-94396733-DEL-65
chr15-94411935-DEL-148
chr15-94417097-DEL-52
chr15-94419527-DEL-1613
chr15-94443000-DEL-57
chr15-94453157-DEL-52
chr15-94459787-DEL-345
chr15-94474699-DEL-167
chr15-94491332-DEL-74
chr15-94494958-DEL-90
chr15-94505329-DEL-100
chr15-94552820-DEL-52
chr15-94565045-DEL-60
chr15-94583409-DEL-66
chr15-94590902-DEL-2120
chr15-94595315-DEL-204
chr15-94607544-DEL-2936
chr15-94622475-DEL-577
chr15-94623953-DEL-616
chr15-94647752-DEL-171
chr15-94655289-DEL-3014
chr15-94668400-DEL-165
chr15-94685596-DEL-212
chr15-94724357-DEL-1098
chr15-94743754-DEL-375
chr15-94748191-DEL-1313
chr15-94763228-DEL-65
chr15-94767835-DEL-60
chr15-94798557-DEL-137
chr15-94808996-DEL-72
chr15-94817876-DEL-55
chr15-94828647-DEL-216
chr15-94845677-DEL-126
chr15-94884419-DEL-321
chr15-94890421-DEL-231
chr15-94893630-DEL-169
chr15-94896985-DEL-297
chr15-94915387-DEL-74
chr15-94916330-DEL-214
chr15-94928739-DEL-392
chr15-94963250-DEL-62
chr15-94968746-DEL-189
chr15-95076644-DEL-467
chr15-95077156-DEL-1552
chr15-95089763-DEL-79
chr15-95091369-DEL-172
chr15-95099752-DEL-412
chr15-95166289-DEL-658
chr15-95189595-DEL-53
chr15-95220106-DEL-76
chr15-95310891-DEL-65

chr15-95337314-DEL-119
chr15-95345611-DEL-2287
chr15-95352382-DEL-388
chr15-95353815-DEL-84
chr15-95362024-DEL-54
chr15-95367577-DEL-159
chr15-95382600-DEL-400
chr15-95420298-DEL-126
chr15-95459839-DEL-207
chr15-95467675-DEL-67
chr15-95474773-DEL-187
chr15-95477383-DEL-64
chr15-95477858-DEL-1156
chr15-95533776-DEL-94
chr15-95546834-DEL-51
chr15-95564165-DEL-59
chr15-95566021-DEL-246
chr15-95577166-DEL-248
chr15-95583182-DEL-775
chr15-95623377-DEL-74
chr15-95635171-DEL-101
chr15-95652175-DEL-1357
chr15-95659102-DEL-75
chr15-95664252-DEL-54
chr15-95744420-DEL-2126
chr15-95744955-DEL-51
chr15-95745294-DEL-76
chr15-95750594-DEL-156
chr15-95750982-DEL-854
chr15-95764843-DEL-245
chr15-95790575-DEL-75
chr15-95816556-DEL-110
chr15-95839264-DEL-85
chr15-95841770-DEL-219
chr15-95866927-DEL-56
chr15-95891364-DEL-302
chr15-95908145-DEL-125
chr15-95911139-DEL-211
chr15-95929747-DEL-87
chr15-95932843-DEL-72
chr15-95954362-DEL-55
chr15-95968525-DEL-743
chr15-95981816-DEL-64
chr15-95999116-DEL-69
chr15-96012354-DEL-124
chr15-96043037-DEL-98
chr15-96064580-DEL-290
chr15-96070970-DEL-192
chr15-96126552-DEL-536
chr15-96131017-DEL-94
chr15-96136515-DEL-400
chr15-96141942-DEL-183
chr15-96151556-DEL-76
chr15-96201857-DEL-69
chr15-96203659-DEL-207
chr15-96208888-DEL-638
chr15-96214518-DEL-201
chr15-96222801-DEL-793
chr15-96226545-DEL-212
chr15-96251218-DEL-68
chr15-96293865-DEL-199
chr15-96337389-DEL-91
chr15-96339486-DEL-344
chr15-96391835-DEL-75
chr15-96495484-DEL-127
chr15-96547388-DEL-104
chr15-96552143-DEL-105
chr15-96553913-DEL-314
chr15-96567086-DEL-1104
chr15-96567262-DEL-51
chr15-96570834-DEL-72
chr15-96571193-DEL-244
chr15-96578021-DEL-70
chr15-96578313-DEL-594
chr15-96619976-DEL-121
chr15-96685967-DEL-157
chr15-96693781-DEL-109

chr15-96721722-DEL-52
chr15-96800454-DEL-53
chr15-96820284-DEL-85
chr15-96832578-DEL-390
chr15-96833188-DEL-52
chr15-96840052-DEL-367
chr15-96840730-DEL-841
chr15-96841591-DEL-610
chr15-96886256-DEL-343
chr15-96928471-DEL-51
chr15-96974473-DEL-76
chr15-96976291-DEL-168
chr15-96979686-DEL-604
chr15-97030816-DEL-158
chr15-97043271-DEL-55
chr15-97051052-DEL-328
chr15-97052787-DEL-108
chr15-97082612-DEL-349
chr15-97113075-DEL-111
chr15-97116847-DEL-121
chr15-97123779-DEL-66
chr15-97154055-DEL-60
chr15-97172145-DEL-83
chr15-97202398-DEL-89
chr15-97208691-DEL-65
chr15-97220417-DEL-381
chr15-97227096-DEL-54
chr15-97231804-DEL-1106
chr15-97233201-DEL-225
chr15-97243000-DEL-89
chr15-97243765-DEL-51
chr15-97264552-DEL-91
chr15-97269046-DEL-74
chr15-97293103-DEL-127
chr15-97301279-DEL-212
chr15-97303297-DEL-200
chr15-97319993-DEL-175
chr15-97324509-DEL-198
chr15-97328362-DEL-76
chr15-97328371-DEL-1401
chr15-97335164-DEL-232
chr15-97340778-DEL-347
chr15-97343227-DEL-61
chr15-97346229-DEL-56
chr15-97359577-DEL-217
chr15-97364668-DEL-1843
chr15-97394666-DEL-126
chr15-97397581-DEL-83
chr15-97419597-DEL-105
chr15-97464434-DEL-542
chr15-97482268-DEL-116
chr15-97495378-DEL-396
chr15-97506806-DEL-322
chr15-97525864-DEL-274
chr15-97526282-DEL-5819
chr15-97549262-DEL-2866
chr15-97549353-DEL-90
chr15-97560388-DEL-336
chr15-97566940-DEL-213
chr15-97570973-DEL-147
chr15-97572337-DEL-55
chr15-97635995-DEL-73
chr15-97692809-DEL-568
chr15-97708448-DEL-148
chr15-97714178-DEL-59
chr15-97725191-DEL-76
chr15-97762899-DEL-171
chr15-97818459-DEL-229
chr15-97819007-DEL-95
chr15-97824166-DEL-84
chr15-97826645-DEL-198
chr15-97844613-DEL-945
chr15-97845616-DEL-121
chr15-97856810-DEL-273
chr15-97865475-DEL-55
chr15-97920207-DEL-289
chr15-97953137-DEL-63

chr15-97972178-DEL-65
chr15-97998345-DEL-583
chr15-98022185-DEL-98
chr15-98035200-DEL-435
chr15-98053639-DEL-229
chr15-98114971-DEL-1587
chr15-98140236-DEL-1487
chr15-98147820-DEL-102
chr15-98163079-DEL-76
chr15-98221386-DEL-4433
chr15-98251139-DEL-53
chr15-98251828-DEL-66
chr15-98264992-DEL-1374
chr15-98266464-DEL-162
chr15-98274905-DEL-694
chr15-98280202-DEL-2403
chr15-98285523-DEL-219
chr15-98300532-DEL-104
chr15-98344664-DEL-221
chr15-98370284-DEL-54
chr15-98390962-DEL-90
chr15-98394357-DEL-354
chr15-98399499-DEL-60
chr15-98401574-DEL-64
chr15-98456444-DEL-53
chr15-98458489-DEL-193
chr15-98482528-DEL-64
chr15-98546986-DEL-191
chr15-98585833-DEL-54
chr15-98591036-DEL-202
chr15-98596329-DEL-200
chr15-98598358-DEL-1630
chr15-98603423-DEL-52
chr15-98620006-DEL-62
chr15-98623989-DEL-73
chr15-98630727-DEL-80
chr15-98636347-DEL-160
chr15-98649162-DEL-147
chr15-98653995-DEL-68
chr15-98664344-DEL-200
chr15-98739438-DEL-110
chr15-98756944-DEL-52
chr15-98804494-DEL-231
chr15-98835725-DEL-72
chr15-98869591-DEL-77
chr15-98894867-DEL-70
chr15-98901255-DEL-218
chr15-98904171-DEL-88
chr15-98924489-DEL-182
chr15-98932528-DEL-484
chr15-98933440-DEL-1462
chr15-98960296-DEL-203
chr15-98961027-DEL-199
chr15-98964942-DEL-930
chr15-98971096-DEL-286
chr15-98977624-DEL-65
chr15-99000131-DEL-538
chr15-99005620-DEL-355
chr15-99033648-DEL-73
chr15-99035970-DEL-76
chr15-99037205-DEL-140
chr15-99059384-DEL-3161
chr15-99067826-DEL-118
chr15-99068531-DEL-750
chr15-99074202-DEL-61
chr15-99079346-DEL-182
chr15-99087582-DEL-329
chr15-99087977-DEL-4327
chr15-99090732-DEL-122
chr15-99097579-DEL-79
chr15-99106374-DEL-70
chr15-99114316-DEL-318
chr15-99134126-DEL-70
chr15-99166684-DEL-82
chr15-99278665-DEL-368
chr15-99279693-DEL-52
chr15-99287323-DEL-51

chr15-99313626-DEL-105
chr15-99315350-DEL-71
chr15-99345838-DEL-154
chr15-99346458-DEL-437
chr15-99377986-DEL-190
chr15-99388302-DEL-1504
chr15-99405351-DEL-135
chr15-99450105-DEL-1104
chr15-99450765-DEL-262
chr15-99516302-DEL-262
chr15-99520009-DEL-54
chr15-99533039-DEL-195
chr15-99536778-DEL-280
chr15-99566043-DEL-205
chr15-99629377-DEL-192
chr15-99640744-DEL-345
chr15-99642219-DEL-80
chr15-99663829-DEL-80
chr15-99670781-DEL-57
chr15-99679234-DEL-113
chr15-99693541-DEL-458
chr15-99707270-DEL-607
chr15-99729931-DEL-221
chr15-99739013-DEL-89
chr15-99749122-DEL-199
chr15-99762379-DEL-184
chr15-99769232-DEL-178
chr15-99796931-DEL-417
chr15-99820994-DEL-1041
chr15-99834565-DEL-241
chr15-99842051-DEL-114
chr15-99849794-DEL-60
chr15-99854456-DEL-53
chr15-99875804-DEL-210
chr15-99884643-DEL-197
chr15-99896114-DEL-130
chr15-99896434-DEL-215
chr15-99906993-DEL-947
chr15-99914411-DEL-78
chr15-99920031-DEL-121
chr15-99925660-DEL-293
chr15-99928618-DEL-1704
chr15-99933903-DEL-1831
chr15-99954294-DEL-53
chr15-99972786-DEL-144
chr15-99989013-DEL-55
chr15-99999802-DEL-71
chr15-100027692-DEL-204
chr15-100046707-DEL-196
chr15-100063063-DEL-925
chr15-100063173-DEL-756
chr15-100068091-DEL-258
chr15-100090911-DEL-163
chr15-100096397-DEL-73
chr15-100098321-DEL-550
chr15-100144885-DEL-96
chr15-100165314-DEL-55
chr15-100167485-DEL-73
chr15-100176039-DEL-54
chr15-100204863-DEL-103
chr15-100205755-DEL-321
chr15-100209516-DEL-232
chr15-100210151-DEL-274
chr15-100217718-DEL-53
chr15-100234506-DEL-195
chr15-100252971-DEL-95
chr15-100254379-DEL-191
chr15-100266044-DEL-9287
chr15-100268241-DEL-136
chr15-100294109-DEL-51
chr15-100309106-DEL-69
chr15-100314691-DEL-174
chr15-100323170-DEL-59
chr15-100325001-DEL-60
chr15-100333758-DEL-97
chr15-100340600-DEL-120
chr15-100375048-DEL-68

chr15-100396009-DEL-120
chr15-100413692-DEL-72
chr15-100416107-DEL-176
chr15-100423594-DEL-184
chr15-100429225-DEL-366
chr15-100454354-DEL-87
chr15-100470486-DEL-202
chr15-100482214-DEL-106
chr15-100482611-DEL-483
chr15-100488738-DEL-196
chr15-100502300-DEL-58
chr15-100504586-DEL-88
chr15-100508655-DEL-153
chr15-100521327-DEL-100
chr15-100526065-DEL-349
chr15-100527725-DEL-52
chr15-100539153-DEL-51
chr15-100539697-DEL-64
chr15-100540577-DEL-144
chr15-100543948-DEL-175
chr15-100545012-DEL-1371
chr15-100553714-DEL-137
chr15-100594311-DEL-54
chr15-100625280-DEL-384
chr15-100632650-DEL-167
chr15-100634714-DEL-77
chr15-100642391-DEL-275
chr15-100643482-DEL-94
chr15-100667003-DEL-81
chr15-100677335-DEL-1507
chr15-100720354-DEL-171
chr15-100747135-DEL-548
chr15-100762320-DEL-291
chr15-100764617-DEL-932
chr15-100764917-DEL-612
chr15-100774492-DEL-134
chr15-100783623-DEL-103
chr15-100796680-DEL-59
chr15-100802385-DEL-176
chr15-100823032-DEL-116
chr15-100824875-DEL-393
chr15-100829175-DEL-100
chr15-100835846-DEL-162
chr15-100863177-DEL-216
chr15-100868804-DEL-189
chr15-100876298-DEL-151
chr15-100891714-DEL-290
chr15-100930287-DEL-82
chr15-100961380-DEL-168
chr15-101021646-DEL-261
chr15-101053558-DEL-96
chr15-101094258-DEL-188
chr15-101097480-DEL-106
chr15-101101193-DEL-468
chr15-101103485-DEL-228
chr15-101115828-DEL-220
chr15-101124509-DEL-196
chr15-101142824-DEL-51
chr15-101157487-DEL-179
chr15-101196853-DEL-192
chr15-101199050-DEL-84
chr15-101201037-DEL-159
chr15-101203374-DEL-403
chr15-101205017-DEL-109
chr15-101218145-DEL-98
chr15-101222398-DEL-221
chr15-101224012-DEL-182
chr15-101244446-DEL-440
chr15-101254667-DEL-273
chr15-101264969-DEL-98
chr15-101299456-DEL-331
chr15-101315489-DEL-66
chr15-101321782-DEL-739
chr15-101341231-DEL-512
chr15-101365453-DEL-54
chr15-101415345-DEL-54
chr15-101419521-DEL-74

chr15-101425678-DEL-440
chr15-101430897-DEL-57
chr15-101458315-DEL-72
chr15-101459782-DEL-64
chr15-101496321-DEL-65
chr15-101498205-DEL-2520
chr15-101505346-DEL-138
chr15-101506699-DEL-51
chr15-101523839-DEL-93
chr15-101526910-DEL-783
chr15-101547860-DEL-90
chr15-101564073-DEL-6478
chr15-101572343-DEL-87
chr15-101581857-DEL-586
chr15-101581928-DEL-1464
chr15-101637246-DEL-2505
chr15-101658626-DEL-655
chr15-101663375-DEL-65
chr15-101674314-DEL-559
chr15-101676531-DEL-735
chr15-101688393-DEL-129
chr15-101698209-DEL-89
chr15-101728283-DEL-84
chr15-101762533-DEL-72
chr15-101809456-DEL-66
chr15-101836972-DEL-224
chr15-101855079-DEL-66
chr15-101855640-DEL-99
chr15-101898304-DEL-72
chr15-101962722-DEL-66
chr15-102058083-DEL-85
chr15-102064005-DEL-55
chr15-102089840-DEL-72
chr15-102107203-DEL-96
chr15-102119598-DEL-219
chr15-102184332-DEL-237
chr15-102203175-DEL-202
chr15-102208871-DEL-54
chr15-102210662-DEL-217
chr15-102227429-DEL-208
chr15-102235984-DEL-384
chr15-102238035-DEL-77
chr15-102241285-DEL-61
chr15-102256869-DEL-906
chr15-102259558-DEL-89
chr15-102263811-DEL-117
chr15-102296009-DEL-211
chr15-102297210-DEL-60
chr15-102306109-DEL-1017
chr15-102319758-DEL-103
chr15-102323565-DEL-204
chr15-102325214-DEL-847
chr15-102366239-DEL-370
chr15-102371468-DEL-84
chr15-102375838-DEL-208
chr15-102376370-DEL-118
chr15-102389153-DEL-374
chr15-102389724-DEL-96
chr15-102405589-DEL-197
chr15-102415628-DEL-257
chr15-102416336-DEL-62
chr15-102458775-DEL-198
chr15-102467689-DEL-201
chr15-102469594-DEL-573
chr15-102496196-DEL-218
chr15-102519126-DEL-375
chr15-102536554-DEL-214
chr15-102547875-DEL-175
chr15-102562132-DEL-83
chr15-102592701-DEL-475
chr15-102597044-DEL-99
chr15-102607680-DEL-58
chr15-102618581-DEL-207
chr15-102639909-DEL-432
chr15-102666064-DEL-57
chr15-102673276-DEL-366
chr15-102696343-DEL-54

chr15-102768116-DEL-389
chr15-102815172-DEL-70
chr15-102897583-DEL-62
chr15-102981751-DEL-197
chr15-103050692-DEL-489
chr15-103059418-DEL-82
chr15-103083930-DEL-59
chr15-103108400-DEL-52
chr15-103114919-DEL-189
chr15-103126035-DEL-737
chr15-103134100-DEL-58
chr15-103146213-DEL-158
chr15-103155473-DEL-252
chr15-103158374-DEL-108
chr15-103190913-DEL-324
chr15-103219608-DEL-193
chr15-103283131-DEL-66
chr15-103297419-DEL-170
chr15-103301876-DEL-194
chr15-103308259-DEL-132
chr15-103319060-DEL-1783
chr15-103320931-DEL-563
chr15-103325308-DEL-522
chr15-103326967-DEL-233
chr15-103352217-DEL-112
chr15-103488081-DEL-54
chr15-103500623-DEL-1211
chr15-103551771-DEL-201
chr15-103555744-DEL-201
chr15-103583422-DEL-53
chr15-103590333-DEL-53
chr15-103604737-DEL-2970
chr15-103610019-DEL-407
chr15-103646711-DEL-2914
chr15-103652188-DEL-277
chr15-103654761-DEL-1183
chr15-103656834-DEL-14319
chr15-103658478-DEL-413
chr15-103678726-DEL-138
chr15-103686470-DEL-6222
chr15-103706705-DEL-7968
chr15-103727408-DEL-857
chr15-103729406-DEL-135
chr15-103735773-DEL-80
chr15-103742293-DEL-367
chr15-103747421-DEL-205
chr15-103759871-DEL-95
chr15-103783727-DEL-161
chr15-103797268-DEL-332
chr15-103797273-DEL-6552
chr15-103810423-DEL-7120
chr15-103842438-DEL-3688
chr15-103846177-DEL-6467
chr15-103864065-DEL-64
chr15-103876348-DEL-75
chr15-103892033-DEL-60
chr15-103933457-DEL-80
chr16-3051731-DEL-62
chr16-3054925-DEL-64
chr16-3056541-DEL-230
chr16-3058316-DEL-186
chr16-3059722-DEL-63
chr16-3063498-DEL-310
chr16-3068019-DEL-603
chr16-3068705-DEL-1660
chr16-3108284-DEL-2717
chr16-3134645-DEL-4395
chr16-3146833-DEL-8770
chr16-3147182-DEL-425
chr16-3158320-DEL-812
chr16-3201639-DEL-13290
chr16-3215827-DEL-252
chr16-3222073-DEL-1153
chr16-3292119-DEL-53
chr16-3292435-DEL-1076
chr16-3322120-DEL-931
chr16-3334064-DEL-102

chr16-3355176-DEL-1729
chr16-3361103-DEL-8710
chr16-3380630-DEL-223
chr16-3395629-DEL-908
chr16-3423315-DEL-198
chr16-3431017-DEL-101
chr16-3433740-DEL-1979
chr16-3449193-DEL-140
chr16-3467945-DEL-3052
chr16-3472143-DEL-150
chr16-3474962-DEL-100
chr16-3485035-DEL-32664
chr16-3548110-DEL-2459
chr16-3554490-DEL-113
chr16-3556403-DEL-384
chr16-3607774-DEL-214
chr16-3623383-DEL-106
chr16-3629295-DEL-606
chr16-3631883-DEL-73
chr16-3654174-DEL-81
chr16-3668240-DEL-76
chr16-3669404-DEL-81
chr16-3755588-DEL-2638
chr16-3759423-DEL-368
chr16-3763467-DEL-187
chr16-3784669-DEL-238
chr16-3789693-DEL-286
chr16-3814424-DEL-205
chr16-3832828-DEL-53
chr16-3835092-DEL-1831
chr16-3838245-DEL-111
chr16-3872922-DEL-212
chr16-3896122-DEL-195
chr16-3914627-DEL-1127
chr16-4035071-DEL-62
chr16-4057010-DEL-169
chr16-4070131-DEL-180
chr16-4100599-DEL-62
chr16-4102173-DEL-212
chr16-4118752-DEL-151
chr16-4157208-DEL-200
chr16-4180151-DEL-4604
chr16-4192091-DEL-78
chr16-4193405-DEL-66
chr16-4202785-DEL-61
chr16-4210292-DEL-97
chr16-4281770-DEL-55
chr16-4304695-DEL-62
chr16-4343402-DEL-79
chr16-4347496-DEL-336
chr16-4357807-DEL-99
chr16-4361758-DEL-119
chr16-4402790-DEL-202
chr16-4408031-DEL-157
chr16-4427398-DEL-60
chr16-4450744-DEL-197
chr16-4455919-DEL-209
chr16-4497942-DEL-57
chr16-4506633-DEL-207
chr16-4541906-DEL-1531
chr16-4547562-DEL-97
chr16-4592147-DEL-96
chr16-4626931-DEL-529
chr16-4632773-DEL-822
chr16-4679905-DEL-390
chr16-4685027-DEL-52
chr16-4711209-DEL-188
chr16-4713389-DEL-233
chr16-4722125-DEL-550
chr16-4730225-DEL-80
chr16-4753326-DEL-345
chr16-4791796-DEL-76
chr16-4813497-DEL-147
chr16-4818013-DEL-152
chr16-4831415-DEL-192
chr16-4874253-DEL-355
chr16-4896909-DEL-54

chr16-4912524-DEL-220
chr16-4917333-DEL-154
chr16-4952317-DEL-260
chr16-4955468-DEL-58
chr16-4962770-DEL-553
chr16-4995446-DEL-84
chr16-5024438-DEL-386
chr16-5026119-DEL-101
chr16-5033381-DEL-119
chr16-5041177-DEL-4556
chr16-5074175-DEL-58
chr16-5091128-DEL-68
chr16-5100382-DEL-440
chr16-5119954-DEL-65
chr16-5129108-DEL-4558
chr16-5135151-DEL-80
chr16-5170146-DEL-195
chr16-5178444-DEL-2131
chr16-5193706-DEL-181
chr16-5212717-DEL-77
chr16-5215838-DEL-2601
chr16-5223915-DEL-97
chr16-5278622-DEL-79
chr16-5294343-DEL-151
chr16-5329657-DEL-65
chr16-5346610-DEL-72
chr16-5358063-DEL-113
chr16-5371322-DEL-80
chr16-5375182-DEL-6158
chr16-5386163-DEL-198
chr16-5398424-DEL-2177
chr16-5418946-DEL-81
chr16-5432332-DEL-66
chr16-5479041-DEL-80
chr16-5513393-DEL-113
chr16-5525791-DEL-230
chr16-5527041-DEL-71
chr16-5536213-DEL-66
chr16-5547975-DEL-206
chr16-5588600-DEL-141
chr16-5595794-DEL-1839
chr16-5601225-DEL-161
chr16-5662209-DEL-334
chr16-5699615-DEL-57
chr16-5734012-DEL-81
chr16-5752228-DEL-59
chr16-5758683-DEL-52
chr16-5839519-DEL-141
chr16-5859503-DEL-56
chr16-5978253-DEL-56
chr16-6010035-DEL-172
chr16-6021032-DEL-1770
chr16-6048574-DEL-86
chr16-6050380-DEL-377
chr16-6051815-DEL-464
chr16-6058429-DEL-837
chr16-6089985-DEL-122
chr16-6159170-DEL-915
chr16-6193658-DEL-596
chr16-6262375-DEL-65
chr16-6264523-DEL-127
chr16-6273945-DEL-91
chr16-6275370-DEL-243
chr16-6418020-DEL-1069
chr16-6434741-DEL-62
chr16-6448675-DEL-105
chr16-6453452-DEL-82
chr16-6482434-DEL-70
chr16-6484441-DEL-830
chr16-6486113-DEL-994
chr16-6498786-DEL-51
chr16-6633815-DEL-72
chr16-6638820-DEL-192
chr16-6669962-DEL-59
chr16-6706270-DEL-279
chr16-6717365-DEL-75
chr16-6722659-DEL-56

chr16-6730041-DEL-60
chr16-6746680-DEL-57
chr16-6758793-DEL-592
chr16-6778417-DEL-594
chr16-6825801-DEL-62
chr16-6838515-DEL-90
chr16-6844665-DEL-809
chr16-6872259-DEL-96
chr16-6890645-DEL-74
chr16-6900697-DEL-120
chr16-6962686-DEL-52
chr16-7022907-DEL-692
chr16-7048202-DEL-127
chr16-7050207-DEL-77
chr16-7066517-DEL-298
chr16-7096217-DEL-92
chr16-7102254-DEL-124
chr16-7121444-DEL-127
chr16-7132592-DEL-591
chr16-7145714-DEL-63
chr16-7152132-DEL-1221
chr16-7186489-DEL-66
chr16-7219568-DEL-453
chr16-7231220-DEL-170
chr16-7254833-DEL-575
chr16-7261837-DEL-74
chr16-7313479-DEL-1531
chr16-7330300-DEL-54
chr16-7364428-DEL-265
chr16-7390013-DEL-147
chr16-7434763-DEL-105
chr16-7462576-DEL-923
chr16-7471729-DEL-159
chr16-7476998-DEL-815
chr16-7502937-DEL-345
chr16-7530641-DEL-67
chr16-7531363-DEL-111
chr16-7561313-DEL-446
chr16-7584210-DEL-436
chr16-7629940-DEL-3127
chr16-7642297-DEL-91
chr16-7643188-DEL-67
chr16-7687139-DEL-53
chr16-7702556-DEL-62
chr16-7713938-DEL-70
chr16-7718184-DEL-76
chr16-7730781-DEL-2852
chr16-7755626-DEL-116
chr16-7780203-DEL-67
chr16-7795883-DEL-238
chr16-7798900-DEL-53
chr16-7801844-DEL-550
chr16-7802499-DEL-1737
chr16-7802741-DEL-117
chr16-7828315-DEL-1596
chr16-7835217-DEL-89
chr16-7848997-DEL-12570
chr16-7860808-DEL-60
chr16-7894251-DEL-51
chr16-7901987-DEL-301
chr16-7919244-DEL-646
chr16-7920035-DEL-369
chr16-7927578-DEL-70
chr16-7928874-DEL-102
chr16-7979828-DEL-1073
chr16-8019670-DEL-59
chr16-8077663-DEL-1374
chr16-8088945-DEL-220
chr16-8123458-DEL-56
chr16-8165684-DEL-51
chr16-8171536-DEL-83
chr16-8263367-DEL-1361
chr16-8270391-DEL-117
chr16-8299640-DEL-116
chr16-8303547-DEL-64
chr16-8305404-DEL-254
chr16-8310416-DEL-175

chr16-8317917-DEL-3551
chr16-8337484-DEL-54
chr16-8338273-DEL-90
chr16-8345033-DEL-512
chr16-8356317-DEL-54
chr16-8370069-DEL-209
chr16-8388671-DEL-161
chr16-8393051-DEL-54
chr16-8405842-DEL-792
chr16-8408310-DEL-1698
chr16-8419677-DEL-146
chr16-8422356-DEL-123
chr16-8424088-DEL-249
chr16-8425284-DEL-207
chr16-8448486-DEL-2459
chr16-8520995-DEL-212
chr16-8527405-DEL-55
chr16-8575841-DEL-235
chr16-8578702-DEL-223
chr16-8583185-DEL-1073
chr16-8587633-DEL-144
chr16-8591740-DEL-72
chr16-8603219-DEL-1107
chr16-8607357-DEL-218
chr16-8616589-DEL-757
chr16-8652662-DEL-1652
chr16-8715921-DEL-172
chr16-8717066-DEL-551
chr16-8725940-DEL-279
chr16-8742494-DEL-88
chr16-8825020-DEL-342
chr16-8870443-DEL-400
chr16-8919531-DEL-1053
chr16-8919599-DEL-985
chr16-8927319-DEL-646
chr16-8929200-DEL-57
chr16-8948500-DEL-114
chr16-8990718-DEL-2971
chr16-9013775-DEL-56
chr16-9041551-DEL-477
chr16-9070774-DEL-185
chr16-9089418-DEL-289
chr16-9117344-DEL-54
chr16-9118324-DEL-1048
chr16-9120296-DEL-4208
chr16-9147254-DEL-63
chr16-9148805-DEL-266
chr16-9161244-DEL-55
chr16-9168940-DEL-867
chr16-9172919-DEL-481
chr16-9185928-DEL-78
chr16-9215552-DEL-58
chr16-9263898-DEL-55
chr16-9283699-DEL-292
chr16-9298320-DEL-93
chr16-9300911-DEL-62
chr16-9311775-DEL-122
chr16-9342637-DEL-54
chr16-9346410-DEL-121
chr16-9362287-DEL-255
chr16-9366321-DEL-106
chr16-9401382-DEL-360
chr16-9404123-DEL-216
chr16-9406926-DEL-272
chr16-9437231-DEL-63
chr16-9443067-DEL-106
chr16-9524765-DEL-59
chr16-9540884-DEL-701
chr16-9541553-DEL-128
chr16-9559381-DEL-407
chr16-9560646-DEL-72
chr16-9564762-DEL-54
chr16-9603020-DEL-100
chr16-9607813-DEL-138
chr16-9620385-DEL-58
chr16-9632827-DEL-117
chr16-9633564-DEL-60

chr16-9638613-DEL-131
chr16-9670647-DEL-97
chr16-9691480-DEL-51
chr16-9697171-DEL-247
chr16-9714053-DEL-97
chr16-9732513-DEL-1490
chr16-9737430-DEL-100
chr16-9763384-DEL-173
chr16-9826511-DEL-192
chr16-9838435-DEL-167
chr16-9854778-DEL-69
chr16-9893074-DEL-1155
chr16-9904909-DEL-390
chr16-9941406-DEL-346
chr16-9973048-DEL-4451
chr16-9979485-DEL-660
chr16-10032917-DEL-136
chr16-10040290-DEL-5002
chr16-10083869-DEL-184
chr16-10088302-DEL-390
chr16-10092076-DEL-219
chr16-10094589-DEL-198
chr16-10097043-DEL-832
chr16-10102097-DEL-51
chr16-10132454-DEL-128
chr16-10135894-DEL-162
chr16-10142007-DEL-163
chr16-10148278-DEL-105
chr16-10154642-DEL-160
chr16-10189482-DEL-178
chr16-10205835-DEL-418
chr16-10226108-DEL-200
chr16-10246730-DEL-81
chr16-10335193-DEL-68
chr16-10369804-DEL-248
chr16-10379623-DEL-1209
chr16-10382011-DEL-159
chr16-10406963-DEL-221
chr16-10417391-DEL-59
chr16-10455928-DEL-77
chr16-10500772-DEL-97
chr16-10506809-DEL-64
chr16-10524608-DEL-62
chr16-10539562-DEL-75
chr16-10555685-DEL-92
chr16-10578575-DEL-84
chr16-10597174-DEL-229
chr16-10611818-DEL-78
chr16-10616349-DEL-169
chr16-10654062-DEL-67
chr16-10673712-DEL-58
chr16-10674355-DEL-68
chr16-10702071-DEL-65
chr16-10705371-DEL-153
chr16-10712568-DEL-82
chr16-10716073-DEL-98
chr16-10717632-DEL-205
chr16-10750468-DEL-389
chr16-10766047-DEL-84
chr16-10813021-DEL-278
chr16-10833538-DEL-92
chr16-10840252-DEL-147
chr16-10850072-DEL-58
chr16-10856575-DEL-1562
chr16-10923922-DEL-563
chr16-10936577-DEL-54
chr16-10939112-DEL-56
chr16-10945391-DEL-156
chr16-10985162-DEL-5311
chr16-10994899-DEL-242
chr16-11027291-DEL-164
chr16-11031007-DEL-2408
chr16-11080575-DEL-409
chr16-11099892-DEL-202
chr16-11108759-DEL-142
chr16-11113972-DEL-76
chr16-11116017-DEL-2807

chr16-11129215-DEL-194
chr16-11150376-DEL-92
chr16-11161668-DEL-1590
chr16-11169307-DEL-80
chr16-11171590-DEL-178
chr16-11179012-DEL-84
chr16-11182763-DEL-95
chr16-11237059-DEL-677
chr16-11241621-DEL-228
chr16-11271204-DEL-63
chr16-11283997-DEL-142
chr16-11351384-DEL-1340
chr16-11379381-DEL-114
chr16-11413850-DEL-94
chr16-11452383-DEL-1413
chr16-11452688-DEL-228
chr16-11455460-DEL-138
chr16-11474438-DEL-204
chr16-11478068-DEL-54
chr16-11488213-DEL-84
chr16-11495685-DEL-427
chr16-11504333-DEL-887
chr16-11504578-DEL-1231
chr16-11505348-DEL-205
chr16-11561592-DEL-239
chr16-11575115-DEL-72
chr16-11585734-DEL-64
chr16-11590889-DEL-56
chr16-11638345-DEL-569
chr16-11656151-DEL-341
chr16-11662010-DEL-83
chr16-11677639-DEL-212
chr16-11691019-DEL-69
chr16-11715029-DEL-62
chr16-11731801-DEL-226
chr16-11742184-DEL-106
chr16-11757237-DEL-500
chr16-11760769-DEL-94
chr16-11784086-DEL-65
chr16-11784907-DEL-298
chr16-11811085-DEL-55
chr16-11854102-DEL-102
chr16-11955584-DEL-2170
chr16-12012167-DEL-51
chr16-12026371-DEL-452
chr16-12031794-DEL-53
chr16-12083030-DEL-96
chr16-12123453-DEL-93
chr16-12150826-DEL-4734
chr16-12157581-DEL-4504
chr16-12166380-DEL-1051
chr16-12170342-DEL-91
chr16-12172755-DEL-641
chr16-12173898-DEL-734
chr16-12176416-DEL-491
chr16-12177546-DEL-327
chr16-12195288-DEL-1202
chr16-12201456-DEL-68
chr16-12233050-DEL-8898
chr16-12292955-DEL-77
chr16-12316965-DEL-51
chr16-12356280-DEL-180
chr16-12387498-DEL-498
chr16-12391836-DEL-51
chr16-12395230-DEL-127
chr16-12439709-DEL-5615
chr16-12461345-DEL-224
chr16-12516478-DEL-196
chr16-12529272-DEL-11263
chr16-12573895-DEL-63
chr16-12575645-DEL-54
chr16-12579618-DEL-170
chr16-12589723-DEL-68
chr16-12598033-DEL-1272
chr16-12605920-DEL-434
chr16-12646657-DEL-272
chr16-12648652-DEL-133

chr16-12669902-DEL-125
chr16-12676348-DEL-82
chr16-12782381-DEL-145
chr16-12831241-DEL-328
chr16-12836689-DEL-351
chr16-12842788-DEL-64
chr16-12845800-DEL-281
chr16-12893831-DEL-550
chr16-12900509-DEL-209
chr16-12901888-DEL-62
chr16-12905569-DEL-6413
chr16-12913834-DEL-155
chr16-12917621-DEL-7152
chr16-12983377-DEL-777
chr16-12984790-DEL-101
chr16-13001593-DEL-302
chr16-13022603-DEL-68
chr16-13026715-DEL-56
chr16-13046675-DEL-272
chr16-13105124-DEL-2155
chr16-13119536-DEL-383
chr16-13123092-DEL-57
chr16-13162278-DEL-209
chr16-13206422-DEL-197
chr16-13224595-DEL-133
chr16-13242461-DEL-163
chr16-13247339-DEL-607
chr16-13299552-DEL-2425
chr16-13307296-DEL-68
chr16-13331157-DEL-561
chr16-13363483-DEL-234
chr16-13381009-DEL-180
chr16-13417549-DEL-893
chr16-13475639-DEL-426
chr16-13505814-DEL-145
chr16-13510711-DEL-56
chr16-13531280-DEL-196
chr16-13544026-DEL-1031
chr16-13555654-DEL-256
chr16-13560649-DEL-537
chr16-13561523-DEL-238
chr16-13564703-DEL-170
chr16-13567096-DEL-174
chr16-13568778-DEL-132
chr16-13572137-DEL-117
chr16-13573660-DEL-173
chr16-13602172-DEL-56
chr16-13604548-DEL-187
chr16-13614148-DEL-109
chr16-13630495-DEL-148
chr16-13635831-DEL-89
chr16-13643145-DEL-223
chr16-13658734-DEL-60
chr16-13671120-DEL-64
chr16-13679442-DEL-73
chr16-13725778-DEL-127
chr16-13755331-DEL-398
chr16-13761830-DEL-78
chr16-13768569-DEL-122
chr16-13808762-DEL-217
chr16-13812551-DEL-91
chr16-13822241-DEL-51
chr16-13882505-DEL-51
chr16-13906284-DEL-80
chr16-13908749-DEL-199
chr16-13912418-DEL-876
chr16-13914538-DEL-117
chr16-13915438-DEL-197
chr16-13937381-DEL-58
chr16-13973764-DEL-486
chr16-13979306-DEL-111
chr16-14006536-DEL-160
chr16-14011007-DEL-58
chr16-14030652-DEL-106
chr16-14066862-DEL-145
chr16-14076178-DEL-116
chr16-14093441-DEL-122

chr16-14103577-DEL-403
chr16-14104456-DEL-60
chr16-14109328-DEL-104
chr16-14116991-DEL-60
chr16-14121733-DEL-269
chr16-14161981-DEL-105
chr16-14164583-DEL-254
chr16-14166898-DEL-782
chr16-14243911-DEL-249
chr16-14247260-DEL-645
chr16-14260181-DEL-102
chr16-14287373-DEL-95
chr16-14302414-DEL-199
chr16-14305603-DEL-523
chr16-14320796-DEL-56
chr16-14328101-DEL-194
chr16-14344939-DEL-4524
chr16-14364436-DEL-132
chr16-14366846-DEL-335
chr16-14376702-DEL-383
chr16-14386554-DEL-531
chr16-14401877-DEL-70
chr16-14409955-DEL-977
chr16-14413456-DEL-607
chr16-14417016-DEL-128
chr16-14425687-DEL-66
chr16-14427319-DEL-59
chr16-14436824-DEL-123
chr16-14448610-DEL-622
chr16-14460013-DEL-75
chr16-14463025-DEL-632
chr16-14472577-DEL-57
chr16-14474455-DEL-9590
chr16-14474995-DEL-65
chr16-14484343-DEL-323
chr16-14491726-DEL-20819
chr16-14542903-DEL-6151
chr16-14550187-DEL-1047
chr16-14560413-DEL-293
chr16-14590087-DEL-101
chr16-14591346-DEL-156
chr16-14595972-DEL-68
chr16-14601912-DEL-272
chr16-14646009-DEL-274
chr16-14648557-DEL-86
chr16-14693323-DEL-4478
chr16-14699117-DEL-2744
chr16-14728256-DEL-114
chr16-14781009-DEL-52
chr16-14860377-DEL-4615
chr16-14876180-DEL-1555
chr16-14876749-DEL-51
chr16-14878402-DEL-290
chr16-14941392-DEL-76
chr16-14958079-DEL-191
chr16-14965423-DEL-52
chr16-15044617-DEL-6376
chr16-15065589-DEL-54
chr16-15082387-DEL-6263
chr16-15096382-DEL-244
chr16-15097217-DEL-7102
chr16-15107946-DEL-76
chr16-15161986-DEL-79
chr16-15174583-DEL-179
chr16-15180305-DEL-614
chr16-15212085-DEL-166
chr16-15274328-DEL-4913
chr16-15292728-DEL-184
chr16-15311062-DEL-114
chr16-15337606-DEL-85
chr16-15346096-DEL-328
chr16-15377120-DEL-156
chr16-15424459-DEL-183
chr16-15457879-DEL-2988
chr16-15459868-DEL-185
chr16-15462386-DEL-198
chr16-15498748-DEL-480

chr16-15522340-DEL-378
chr16-15535350-DEL-85
chr16-15536007-DEL-118
chr16-15574380-DEL-1202
chr16-15578481-DEL-472
chr16-15608786-DEL-73
chr16-15678008-DEL-75
chr16-15717012-DEL-181
chr16-15755839-DEL-369
chr16-15825418-DEL-1446
chr16-15846419-DEL-198
chr16-15849399-DEL-66
chr16-15864061-DEL-577
chr16-15882582-DEL-345
chr16-15909281-DEL-597
chr16-15940620-DEL-52
chr16-15940946-DEL-1521
chr16-15948062-DEL-6189
chr16-15986836-DEL-80
chr16-15989412-DEL-14966
chr16-15995274-DEL-3612
chr16-16086097-DEL-55
chr16-16127620-DEL-654
chr16-16128727-DEL-422
chr16-16144070-DEL-2350
chr16-16150202-DEL-348
chr16-16164503-DEL-2917
chr16-16170435-DEL-169
chr16-16185345-DEL-546
chr16-16270640-DEL-181
chr16-16278763-DEL-191
chr16-16282493-DEL-204
chr16-16311224-DEL-104
chr16-16319045-DEL-68
chr16-16324456-DEL-803
chr16-16325641-DEL-455
chr16-16333533-DEL-1136
chr16-16343074-DEL-66
chr16-16344193-DEL-156
chr16-16378634-DEL-1345
chr16-16385010-DEL-112
chr16-16392484-DEL-63
chr16-16422011-DEL-2629
chr16-16430829-DEL-1201
chr16-16437802-DEL-160
chr16-16454762-DEL-878
chr16-16460979-DEL-217
chr16-16476638-DEL-7580
chr16-16530012-DEL-4867
chr16-16566632-DEL-127
chr16-16579016-DEL-448
chr16-16600913-DEL-74
chr16-16602551-DEL-835
chr16-16615971-DEL-9939
chr16-16616123-DEL-431
chr16-16616941-DEL-6067
chr16-16650190-DEL-213
chr16-16658708-DEL-1982
chr16-16684965-DEL-64
chr16-16716003-DEL-309
chr16-16719295-DEL-134
chr16-16734349-DEL-201
chr16-16757816-DEL-99
chr16-16767718-DEL-186
chr16-16772598-DEL-1463
chr16-16816948-DEL-161
chr16-16825379-DEL-383
chr16-16828332-DEL-527
chr16-16846879-DEL-205
chr16-16849746-DEL-211
chr16-16858880-DEL-577
chr16-16877084-DEL-403
chr16-16921498-DEL-565
chr16-16924480-DEL-197
chr16-16964309-DEL-194
chr16-16973108-DEL-2597
chr16-17010025-DEL-244

chr16-17016243-DEL-193
chr16-17022529-DEL-111
chr16-17025949-DEL-59
chr16-17048829-DEL-494
chr16-17058946-DEL-506
chr16-17059634-DEL-211
chr16-17072651-DEL-135
chr16-17074461-DEL-179
chr16-17106162-DEL-224
chr16-17110011-DEL-96
chr16-17137374-DEL-449
chr16-17138616-DEL-61
chr16-17146826-DEL-139
chr16-17214112-DEL-190
chr16-17235187-DEL-401
chr16-17238018-DEL-59
chr16-17238529-DEL-137
chr16-17243967-DEL-73
chr16-17250040-DEL-872
chr16-17276538-DEL-8517
chr16-17299609-DEL-55
chr16-17311789-DEL-76
chr16-17357878-DEL-3233
chr16-17376267-DEL-118
chr16-17378195-DEL-250
chr16-17402056-DEL-197
chr16-17419257-DEL-545
chr16-17422581-DEL-52
chr16-17428356-DEL-62
chr16-17432904-DEL-240
chr16-17449694-DEL-125
chr16-17528933-DEL-210
chr16-17551646-DEL-329
chr16-17556231-DEL-412
chr16-17573898-DEL-515
chr16-17590871-DEL-198
chr16-17629106-DEL-228
chr16-17642089-DEL-839
chr16-17685251-DEL-199
chr16-17696423-DEL-144
chr16-17727507-DEL-192
chr16-17737052-DEL-195
chr16-17739303-DEL-71
chr16-17758864-DEL-73
chr16-17765135-DEL-480
chr16-17806846-DEL-909
chr16-17829236-DEL-5701
chr16-17829860-DEL-94
chr16-17848316-DEL-438
chr16-17868267-DEL-96
chr16-17876569-DEL-176
chr16-17889820-DEL-58
chr16-17938908-DEL-437
chr16-17940274-DEL-102
chr16-18028145-DEL-90
chr16-18064810-DEL-148
chr16-18078568-DEL-207
chr16-18108170-DEL-197
chr16-18113028-DEL-202
chr16-18131011-DEL-256
chr16-18145893-DEL-5728
chr16-18150527-DEL-314
chr16-18164000-DEL-64
chr16-18172909-DEL-195
chr16-18177393-DEL-206
chr16-18181349-DEL-202
chr16-18207426-DEL-59
chr16-18215947-DEL-414
chr16-18252841-DEL-284
chr16-18299649-DEL-121
chr16-18309040-DEL-88
chr16-18313083-DEL-213
chr16-18329860-DEL-204
chr16-18342025-DEL-265
chr16-18372278-DEL-54
chr16-18393436-DEL-208
chr16-18411074-DEL-147

chr16-18434624-DEL-171
chr16-18470762-DEL-355
chr16-18507198-DEL-68
chr16-18516956-DEL-174
chr16-18532851-DEL-155
chr16-18544553-DEL-232
chr16-18545150-DEL-198
chr16-18556472-DEL-68
chr16-18559147-DEL-64
chr16-18582514-DEL-1856
chr16-18634488-DEL-65
chr16-18659209-DEL-285
chr16-18663292-DEL-226
chr16-18665100-DEL-349
chr16-18665625-DEL-372
chr16-18666513-DEL-7775
chr16-18679469-DEL-1889
chr16-18703074-DEL-186
chr16-18705253-DEL-204
chr16-18710181-DEL-1454
chr16-18718233-DEL-163
chr16-18749188-DEL-100
chr16-18762226-DEL-73
chr16-18794434-DEL-9022
chr16-18821284-DEL-135
chr16-18926904-DEL-12788
chr16-18927103-DEL-16327
chr16-18951746-DEL-51
chr16-18952818-DEL-244
chr16-18955754-DEL-962
chr16-18975316-DEL-947
chr16-18994970-DEL-1116
chr16-19002944-DEL-6337
chr16-19021206-DEL-129
chr16-19035838-DEL-762
chr16-19037622-DEL-62
chr16-19060610-DEL-5947
chr16-19067195-DEL-88
chr16-19102492-DEL-60
chr16-19116034-DEL-6742
chr16-19122628-DEL-352
chr16-19160663-DEL-7095
chr16-19170866-DEL-6176
chr16-19188892-DEL-6188
chr16-19215412-DEL-2306
chr16-19219550-DEL-123
chr16-19220400-DEL-1006
chr16-19225206-DEL-430
chr16-19231086-DEL-158
chr16-19244612-DEL-239
chr16-19251904-DEL-134
chr16-19259326-DEL-95
chr16-19272551-DEL-4515
chr16-19362028-DEL-6170
chr16-19449346-DEL-1259
chr16-19464019-DEL-4165
chr16-19467097-DEL-3734
chr16-19470308-DEL-901
chr16-19473774-DEL-72
chr16-19558946-DEL-119
chr16-19582613-DEL-202
chr16-19608131-DEL-234
chr16-19610483-DEL-68
chr16-19711651-DEL-113
chr16-19747051-DEL-6643
chr16-19757136-DEL-962
chr16-19758336-DEL-6591
chr16-19768070-DEL-101
chr16-19769074-DEL-1747
chr16-19794428-DEL-59
chr16-19819124-DEL-53
chr16-19819782-DEL-8731
chr16-19838599-DEL-146
chr16-19843310-DEL-6393
chr16-19873241-DEL-11011
chr16-19893788-DEL-112
chr16-19928288-DEL-69

chr16-19945091-DEL-180
chr16-19948265-DEL-590
chr16-19986680-DEL-169
chr16-20031419-DEL-306
chr16-20061589-DEL-119
chr16-20065296-DEL-460
chr16-20072536-DEL-213
chr16-20084631-DEL-318
chr16-20085054-DEL-1166
chr16-20086410-DEL-144
chr16-20091806-DEL-115
chr16-20136526-DEL-860
chr16-20155652-DEL-55
chr16-20160056-DEL-1326
chr16-20162992-DEL-142
chr16-20181925-DEL-697
chr16-20184200-DEL-89
chr16-20217177-DEL-280
chr16-20239216-DEL-152
chr16-20252406-DEL-91
chr16-20333078-DEL-222
chr16-20333192-DEL-1867
chr16-20386373-DEL-171
chr16-20388703-DEL-178
chr16-20465742-DEL-77
chr16-20467081-DEL-198
chr16-20467680-DEL-379
chr16-20486429-DEL-68
chr16-20559050-DEL-94
chr16-20566510-DEL-185
chr16-20572976-DEL-204
chr16-20585618-DEL-60
chr16-20594757-DEL-214
chr16-20599010-DEL-229
chr16-20634735-DEL-522
chr16-20640836-DEL-603
chr16-20641479-DEL-534
chr16-20653325-DEL-70
chr16-20660614-DEL-352
chr16-20667477-DEL-173
chr16-20668601-DEL-1076
chr16-20672791-DEL-339
chr16-20689590-DEL-4556
chr16-20689720-DEL-10705
chr16-20703027-DEL-498
chr16-20745343-DEL-293
chr16-20774964-DEL-1553
chr16-20800988-DEL-7134
chr16-20823015-DEL-401
chr16-20831811-DEL-238
chr16-20835683-DEL-135
chr16-20890777-DEL-1963
chr16-20898980-DEL-294
chr16-20910202-DEL-1254
chr16-20921324-DEL-408
chr16-20921919-DEL-902
chr16-20923664-DEL-4327
chr16-20972790-DEL-113
chr16-20985394-DEL-1322
chr16-21032121-DEL-212
chr16-21079058-DEL-1573
chr16-21094106-DEL-84
chr16-21160896-DEL-203
chr16-21187645-DEL-86
chr16-21203125-DEL-116
chr16-21255060-DEL-1929
chr16-21275345-DEL-107
chr16-21281764-DEL-179
chr16-21300794-DEL-130
chr16-21325946-DEL-74
chr16-21332728-DEL-998
chr16-21367172-DEL-87
chr16-21390445-DEL-156
chr16-21415861-DEL-204
chr16-21451930-DEL-332
chr16-21475325-DEL-1567
chr16-21477282-DEL-52

chr16-21487655-DEL-57
chr16-21497390-DEL-264
chr16-21530943-DEL-164
chr16-21531439-DEL-789
chr16-21540748-DEL-52
chr16-21545229-DEL-152
chr16-21563558-DEL-7162
chr16-21575927-DEL-56
chr16-21630913-DEL-98
chr16-21648225-DEL-177
chr16-21696170-DEL-2090
chr16-21788725-DEL-178
chr16-21801468-DEL-523
chr16-21810859-DEL-197
chr16-21824264-DEL-3892
chr16-21912777-DEL-235
chr16-21932963-DEL-199
chr16-21951097-DEL-82
chr16-21954869-DEL-55
chr16-21961418-DEL-57
chr16-22004059-DEL-5353
chr16-22013124-DEL-305
chr16-22028998-DEL-100
chr16-22032842-DEL-55
chr16-22037058-DEL-5092
chr16-22049084-DEL-1323
chr16-22054369-DEL-58
chr16-22066775-DEL-199
chr16-22091554-DEL-3057
chr16-22101448-DEL-58
chr16-22108093-DEL-86
chr16-22148828-DEL-400
chr16-22168641-DEL-107
chr16-22171617-DEL-162
chr16-22172841-DEL-235
chr16-22187108-DEL-185
chr16-22194305-DEL-177
chr16-22196185-DEL-134
chr16-22213908-DEL-74
chr16-22255413-DEL-56
chr16-22269443-DEL-334
chr16-22278822-DEL-88
chr16-22296148-DEL-807
chr16-22296334-DEL-909
chr16-22301579-DEL-58
chr16-22330525-DEL-51
chr16-22348218-DEL-265
chr16-22360711-DEL-71
chr16-22365196-DEL-80
chr16-22381009-DEL-75
chr16-22381622-DEL-106
chr16-22389433-DEL-138
chr16-22421852-DEL-81
chr16-22457084-DEL-203
chr16-22478901-DEL-181
chr16-22528654-DEL-322
chr16-22568583-DEL-74
chr16-22578112-DEL-4286
chr16-22600154-DEL-76
chr16-22604455-DEL-291
chr16-22629214-DEL-62
chr16-22632808-DEL-53
chr16-22649450-DEL-215
chr16-22672396-DEL-66
chr16-22737612-DEL-77
chr16-22738632-DEL-172
chr16-22759403-DEL-969
chr16-22760976-DEL-538
chr16-22808699-DEL-123
chr16-22857277-DEL-114
chr16-22890545-DEL-193
chr16-22904255-DEL-399
chr16-22914528-DEL-171
chr16-22917478-DEL-408
chr16-22922060-DEL-186
chr16-22981171-DEL-198
chr16-23029395-DEL-1500

chr16-23040890-DEL-127
chr16-23050944-DEL-52
chr16-23051995-DEL-65
chr16-23053556-DEL-156
chr16-23078713-DEL-79
chr16-23081446-DEL-144
chr16-23085359-DEL-3245
chr16-23128599-DEL-60
chr16-23156725-DEL-54
chr16-23159465-DEL-1562
chr16-23161451-DEL-2338
chr16-23166780-DEL-484
chr16-23168786-DEL-148
chr16-23219291-DEL-527
chr16-23232586-DEL-138
chr16-23255576-DEL-76
chr16-23272624-DEL-61
chr16-23277057-DEL-320
chr16-23300600-DEL-401
chr16-23335827-DEL-71
chr16-23354559-DEL-2589
chr16-23365579-DEL-108
chr16-23378632-DEL-958
chr16-23379257-DEL-52
chr16-23381076-DEL-72
chr16-23418625-DEL-128
chr16-23434059-DEL-223
chr16-23438995-DEL-221
chr16-23440632-DEL-1002
chr16-23442541-DEL-60
chr16-23465485-DEL-104
chr16-23468707-DEL-130
chr16-23473317-DEL-287
chr16-23482173-DEL-118
chr16-23483587-DEL-207
chr16-23526618-DEL-84
chr16-23545971-DEL-215
chr16-23547873-DEL-1152
chr16-23574287-DEL-398
chr16-23616371-DEL-147
chr16-23621751-DEL-191
chr16-23681895-DEL-1042
chr16-23734280-DEL-242
chr16-23767371-DEL-278
chr16-23794798-DEL-157
chr16-23821939-DEL-126
chr16-23857372-DEL-63
chr16-23860861-DEL-86
chr16-23934925-DEL-1101
chr16-23944532-DEL-1693
chr16-23986630-DEL-60
chr16-24011179-DEL-85
chr16-24092302-DEL-773
chr16-24093156-DEL-304
chr16-24130800-DEL-429
chr16-24154904-DEL-160
chr16-24170502-DEL-172
chr16-24180132-DEL-51
chr16-24187684-DEL-66
chr16-24208249-DEL-407
chr16-24259502-DEL-90
chr16-24276874-DEL-132
chr16-24282319-DEL-75
chr16-24298669-DEL-97
chr16-24366382-DEL-64
chr16-24369719-DEL-106
chr16-24402332-DEL-198
chr16-24424224-DEL-237
chr16-24442210-DEL-110
chr16-24467439-DEL-53
chr16-24483160-DEL-3796
chr16-24526330-DEL-62
chr16-24551087-DEL-60
chr16-24561095-DEL-734
chr16-24587407-DEL-208
chr16-24609784-DEL-228
chr16-24613368-DEL-224

chr16-24632178-DEL-165
chr16-24643854-DEL-206
chr16-24650603-DEL-214
chr16-24684869-DEL-69
chr16-24694055-DEL-189
chr16-24754994-DEL-595
chr16-24843974-DEL-265
chr16-24855276-DEL-225
chr16-24875051-DEL-561
chr16-24924970-DEL-397
chr16-24926396-DEL-57
chr16-24943044-DEL-336
chr16-25005913-DEL-121
chr16-25016712-DEL-126
chr16-25027548-DEL-172
chr16-25034905-DEL-1822
chr16-25045036-DEL-128
chr16-25052495-DEL-267
chr16-25130458-DEL-58
chr16-25166147-DEL-85
chr16-25171796-DEL-62
chr16-25181459-DEL-3476
chr16-25189929-DEL-542
chr16-25207636-DEL-150
chr16-25229721-DEL-76
chr16-25239138-DEL-80
chr16-25253286-DEL-464
chr16-25271485-DEL-87
chr16-25374236-DEL-1176
chr16-25392703-DEL-322
chr16-25417575-DEL-58
chr16-25423706-DEL-1175
chr16-25455883-DEL-1088
chr16-25459777-DEL-2276
chr16-25461649-DEL-649
chr16-25472510-DEL-182
chr16-25473518-DEL-535
chr16-25494871-DEL-52
chr16-25500664-DEL-61
chr16-25533718-DEL-54
chr16-25547645-DEL-618
chr16-25568099-DEL-328
chr16-25589843-DEL-1665
chr16-25629009-DEL-209
chr16-25674151-DEL-106
chr16-25720927-DEL-72
chr16-25726881-DEL-58
chr16-25741631-DEL-1231
chr16-25743476-DEL-2688
chr16-25748253-DEL-66
chr16-25776232-DEL-594
chr16-25782075-DEL-78
chr16-25790657-DEL-207
chr16-25793239-DEL-806
chr16-25833223-DEL-486
chr16-25847783-DEL-120
chr16-25884218-DEL-69
chr16-25934192-DEL-56
chr16-25994588-DEL-56
chr16-25997338-DEL-363
chr16-26011111-DEL-638
chr16-26013828-DEL-544
chr16-26028734-DEL-1062
chr16-26050553-DEL-782
chr16-26088431-DEL-1492
chr16-26092864-DEL-168
chr16-26097781-DEL-214
chr16-26103978-DEL-190
chr16-26116869-DEL-747
chr16-26136032-DEL-6489
chr16-26148501-DEL-189
chr16-26158468-DEL-205
chr16-26160873-DEL-346
chr16-26169871-DEL-229
chr16-26202443-DEL-70
chr16-26229204-DEL-102
chr16-26230623-DEL-1552

chr16-26232832-DEL-103
chr16-26233956-DEL-950
chr16-26300936-DEL-54
chr16-26315626-DEL-841
chr16-26317591-DEL-768
chr16-26333518-DEL-56
chr16-26381694-DEL-470
chr16-26383355-DEL-439
chr16-26392975-DEL-1080
chr16-26419497-DEL-833
chr16-26420879-DEL-66
chr16-26436654-DEL-72
chr16-26440247-DEL-578
chr16-26447357-DEL-58
chr16-26505207-DEL-99
chr16-26506573-DEL-163
chr16-26527606-DEL-1121
chr16-26567985-DEL-86
chr16-26572396-DEL-179
chr16-26627204-DEL-6533
chr16-26638033-DEL-142
chr16-26656160-DEL-907
chr16-26657237-DEL-1667
chr16-26659670-DEL-277
chr16-26664603-DEL-105
chr16-26673010-DEL-84
chr16-26681305-DEL-204
chr16-26685123-DEL-4702
chr16-26696531-DEL-7514
chr16-26709258-DEL-871
chr16-26755545-DEL-7726
chr16-26769550-DEL-68
chr16-26789014-DEL-120
chr16-26805912-DEL-515
chr16-26816031-DEL-213
chr16-26821312-DEL-1533
chr16-26846636-DEL-1293
chr16-26848180-DEL-265
chr16-26883457-DEL-2753
chr16-26890628-DEL-7384
chr16-26926952-DEL-1864
chr16-26928596-DEL-487
chr16-26937320-DEL-104
chr16-26944222-DEL-80
chr16-26961825-DEL-52
chr16-26990489-DEL-19315
chr16-27013225-DEL-144
chr16-27020942-DEL-584
chr16-27030885-DEL-8374
chr16-27035064-DEL-2144
chr16-27086283-DEL-191
chr16-27089704-DEL-138
chr16-27104006-DEL-664
chr16-27158626-DEL-94
chr16-27252201-DEL-51
chr16-27269138-DEL-70
chr16-27278616-DEL-2222
chr16-27282727-DEL-2436
chr16-27314904-DEL-943
chr16-27386357-DEL-64
chr16-27401749-DEL-610
chr16-27410400-DEL-6220
chr16-27431746-DEL-117
chr16-27438278-DEL-3261
chr16-27471417-DEL-7473
chr16-27483310-DEL-98
chr16-27483806-DEL-506
chr16-27484946-DEL-6222
chr16-27492338-DEL-3960
chr16-27521017-DEL-193
chr16-27537153-DEL-750
chr16-27567623-DEL-960
chr16-27619759-DEL-1246
chr16-27675713-DEL-259
chr16-27676286-DEL-78
chr16-27730973-DEL-803
chr16-27770854-DEL-231

chr16-27787282-DEL-1337
chr16-27794039-DEL-73
chr16-27819646-DEL-900
chr16-27820641-DEL-71
chr16-27835987-DEL-74
chr16-27836744-DEL-53
chr16-27853521-DEL-55
chr16-27856562-DEL-56
chr16-27865073-DEL-819
chr16-27870358-DEL-152
chr16-27876144-DEL-56
chr16-27889980-DEL-1695
chr16-27893371-DEL-170
chr16-27895022-DEL-198
chr16-27943929-DEL-79
chr16-27952081-DEL-276
chr16-27953045-DEL-249
chr16-27957384-DEL-75
chr16-27965498-DEL-389
chr16-28003370-DEL-71
chr16-28041634-DEL-922
chr16-28043084-DEL-82
chr16-28091423-DEL-161
chr16-28100553-DEL-322
chr16-28112043-DEL-416
chr16-28152380-DEL-195
chr16-28284989-DEL-62
chr16-28296542-DEL-538
chr16-28319249-DEL-208
chr16-28333660-DEL-67
chr16-28339124-DEL-55
chr16-28343439-DEL-62
chr16-28344442-DEL-230
chr16-28393902-DEL-94
chr16-28397384-DEL-80
chr16-28408174-DEL-510
chr16-28409120-DEL-168
chr16-28462226-DEL-80
chr16-28464581-DEL-379
chr16-28486890-DEL-343
chr16-28530629-DEL-96
chr16-28548843-DEL-325
chr16-28600461-DEL-1490
chr16-28639167-DEL-354
chr16-28671556-DEL-217
chr16-28672162-DEL-78
chr16-28685436-DEL-53
chr16-28707745-DEL-538
chr16-28715484-DEL-124
chr16-28727165-DEL-52
chr16-28770176-DEL-1503
chr16-28785934-DEL-87
chr16-28794743-DEL-440
chr16-28797912-DEL-80
chr16-28835284-DEL-202
chr16-28855812-DEL-59
chr16-28930967-DEL-118
chr16-28947182-DEL-68
chr16-28968859-DEL-233
chr16-29000352-DEL-84
chr16-29002036-DEL-90
chr16-29122604-DEL-132
chr16-29132416-DEL-498
chr16-29152459-DEL-73
chr16-29156459-DEL-409
chr16-29202737-DEL-63
chr16-29208546-DEL-866
chr16-29209317-DEL-96
chr16-29226707-DEL-107
chr16-29236754-DEL-324
chr16-29303420-DEL-323
chr16-29304816-DEL-52
chr16-29334249-DEL-70
chr16-29334738-DEL-361
chr16-29375625-DEL-878
chr16-29401606-DEL-220
chr16-29401705-DEL-908

chr16-29402209-DEL-221
chr16-29402812-DEL-130
chr16-29403078-DEL-377
chr16-29412251-DEL-170
chr16-29496102-DEL-341
chr16-29503978-DEL-737
chr16-29510901-DEL-52
chr16-29527686-DEL-53
chr16-29537208-DEL-56
chr16-29546831-DEL-103
chr16-29566215-DEL-122
chr16-29572287-DEL-834
chr16-29592401-DEL-140
chr16-29596727-DEL-62
chr16-29600145-DEL-294
chr16-29613773-DEL-51
chr16-29624283-DEL-92
chr16-29696885-DEL-73
chr16-29704702-DEL-117
chr16-29728555-DEL-53
chr16-29731625-DEL-56
chr16-29732738-DEL-92
chr16-29788207-DEL-77
chr16-29794019-DEL-59
chr16-29802669-DEL-198
chr16-29851605-DEL-54
chr16-29863317-DEL-74
chr16-29871791-DEL-51
chr16-29897745-DEL-53
chr16-29902703-DEL-137
chr16-29908102-DEL-222
chr16-29920418-DEL-76
chr16-29923147-DEL-85
chr16-29956327-DEL-140
chr16-29958180-DEL-521
chr16-29960484-DEL-179
chr16-29985718-DEL-191
chr16-29995427-DEL-58
chr16-30042481-DEL-126
chr16-30043631-DEL-82
chr16-30047234-DEL-219
chr16-30051605-DEL-98
chr16-30104856-DEL-60
chr16-30107534-DEL-343
chr16-30119016-DEL-798
chr16-30148577-DEL-2050
chr16-30161224-DEL-933
chr16-30183856-DEL-115
chr16-30191564-DEL-141
chr16-30192345-DEL-363
chr16-30214941-DEL-231
chr16-30223645-DEL-2203
chr16-30240133-DEL-57
chr16-30275531-DEL-56
chr16-30278350-DEL-202
chr16-30281371-DEL-531
chr16-30295543-DEL-86
chr16-30327457-DEL-78
chr16-30344481-DEL-75
chr16-30366302-DEL-202
chr16-30377025-DEL-202
chr16-30378590-DEL-1211
chr16-30412586-DEL-164
chr16-30447700-DEL-72
chr16-30457532-DEL-595
chr16-30459932-DEL-365
chr16-30462684-DEL-84
chr16-30503709-DEL-56
chr16-30532333-DEL-401
chr16-30534411-DEL-284
chr16-30536306-DEL-155
chr16-30547834-DEL-539
chr16-30549082-DEL-69
chr16-30552432-DEL-93
chr16-30556567-DEL-73
chr16-30586137-DEL-182
chr16-30632070-DEL-99

chr16-30663031-DEL-69
chr16-30692320-DEL-90
chr16-30713640-DEL-94
chr16-30736073-DEL-71
chr16-30747614-DEL-128
chr16-30768725-DEL-132
chr16-30771675-DEL-320
chr16-30807669-DEL-63
chr16-30833580-DEL-94
chr16-30842132-DEL-246
chr16-30845189-DEL-621
chr16-30845256-DEL-122
chr16-30872232-DEL-177
chr16-30873046-DEL-85
chr16-30879710-DEL-134
chr16-30891699-DEL-62
chr16-30903408-DEL-5625
chr16-30916036-DEL-64
chr16-30917909-DEL-130
chr16-30944035-DEL-111
chr16-30969297-DEL-72
chr16-30977179-DEL-51
chr16-30999358-DEL-198
chr16-31006105-DEL-101
chr16-31012434-DEL-243
chr16-31024819-DEL-205
chr16-31053512-DEL-71
chr16-31055274-DEL-772
chr16-31096517-DEL-82
chr16-31098172-DEL-1291
chr16-31141090-DEL-137
chr16-31175793-DEL-1742
chr16-31180855-DEL-115
chr16-31181691-DEL-116
chr16-31198755-DEL-5929
chr16-31203775-DEL-232
chr16-31220510-DEL-481
chr16-31227513-DEL-57
chr16-31253777-DEL-91
chr16-31254438-DEL-191
chr16-31255439-DEL-400
chr16-31258632-DEL-53
chr16-31262019-DEL-196
chr16-31309497-DEL-211
chr16-31327028-DEL-128
chr16-31331181-DEL-140
chr16-31344347-DEL-183
chr16-31349910-DEL-96
chr16-31357897-DEL-56
chr16-31363142-DEL-125
chr16-31383281-DEL-265
chr16-31436515-DEL-72
chr16-31439679-DEL-72
chr16-31448885-DEL-202
chr16-31462737-DEL-667
chr16-31501554-DEL-230
chr16-31506297-DEL-198
chr16-31522034-DEL-1108
chr16-31605220-DEL-423
chr16-31607139-DEL-340
chr16-31613759-DEL-52
chr16-31633946-DEL-116
chr16-31723604-DEL-54
chr16-31728980-DEL-175
chr16-31732081-DEL-317
chr16-31761244-DEL-51
chr16-31768956-DEL-94
chr16-31770346-DEL-1736
chr16-31793634-DEL-83
chr16-31800886-DEL-202
chr16-31804953-DEL-136
chr16-31812808-DEL-83
chr16-31813441-DEL-212
chr16-31824020-DEL-176
chr16-31849861-DEL-192
chr16-31864340-DEL-60
chr16-31891539-DEL-455

chr16-31899097-DEL-399
chr16-31907271-DEL-452
chr16-31910144-DEL-179
chr16-31931142-DEL-176
chr16-31976876-DEL-191
chr16-32037819-DEL-56
chr16-32041590-DEL-62
chr16-32072461-DEL-212
chr16-32087538-DEL-156
chr16-32089382-DEL-122
chr16-32094164-DEL-167
chr16-32125984-DEL-1270
chr16-32134537-DEL-98
chr16-32137721-DEL-194
chr16-32141945-DEL-184
chr16-32144341-DEL-197
chr16-32159092-DEL-88
chr16-32230245-DEL-1453
chr16-32237429-DEL-151
chr16-32241888-DEL-247
chr16-32309240-DEL-815
chr16-32313267-DEL-57
chr16-32333535-DEL-65
chr16-32338253-DEL-87
chr16-32353703-DEL-185
chr16-32367853-DEL-349
chr16-32368909-DEL-1444
chr16-32370322-DEL-93
chr16-32371016-DEL-52
chr16-32372033-DEL-1662
chr16-32386278-DEL-119
chr16-32391144-DEL-59
chr16-32393662-DEL-74
chr16-32428676-DEL-67
chr16-32432111-DEL-197
chr16-32441004-DEL-208
chr16-32448519-DEL-142
chr16-32462019-DEL-91
chr16-32468923-DEL-202
chr16-32509866-DEL-493
chr16-32534444-DEL-1193
chr16-32578350-DEL-52
chr16-32591791-DEL-53
chr16-32604034-DEL-51
chr16-32614734-DEL-92
chr16-32658546-DEL-51
chr16-32676868-DEL-65
chr16-32691091-DEL-1240
chr16-32725126-DEL-61
chr16-32781489-DEL-97
chr16-32820431-DEL-177
chr16-32823342-DEL-706
chr16-32867946-DEL-274
chr16-32872543-DEL-581
chr16-32908068-DEL-66
chr16-32927654-DEL-191
chr16-32953448-DEL-81
chr16-32955895-DEL-1420
chr16-32986899-DEL-177
chr16-33046050-DEL-200
chr16-33054543-DEL-83
chr16-33091670-DEL-386
chr16-33092553-DEL-931
chr16-33094161-DEL-59
chr16-33096256-DEL-98
chr16-33107855-DEL-54
chr16-33140895-DEL-164
chr16-33145447-DEL-51
chr16-33147317-DEL-1091
chr16-33152834-DEL-160
chr16-33172814-DEL-875
chr16-33193450-DEL-2157
chr16-33208219-DEL-198
chr16-33210477-DEL-699
chr16-33217025-DEL-221
chr16-33222871-DEL-979
chr16-33280459-DEL-206

chr16-33301162-DEL-64
chr16-33302408-DEL-77
chr16-33314376-DEL-203
chr16-33359425-DEL-69
chr16-33367037-DEL-355
chr16-33368664-DEL-52
chr16-33372262-DEL-220
chr16-33379804-DEL-114
chr16-33396748-DEL-102
chr16-33479240-DEL-54
chr16-33509487-DEL-199
chr16-33527577-DEL-248
chr16-33541518-DEL-120
chr16-33545269-DEL-56
chr16-33601354-DEL-101
chr16-33625398-DEL-157
chr16-33655510-DEL-1242
chr16-33713630-DEL-487
chr16-33730080-DEL-57
chr16-33739049-DEL-54
chr16-33741357-DEL-65
chr16-33744289-DEL-1220
chr16-33749215-DEL-2552
chr16-33761678-DEL-52
chr16-33767703-DEL-95
chr16-33796997-DEL-66
chr16-33802667-DEL-199
chr16-33804117-DEL-343
chr16-33823865-DEL-91
chr16-33826997-DEL-990
chr16-33835910-DEL-144
chr16-33839090-DEL-7775
chr16-33846895-DEL-110
chr16-33895658-DEL-236
chr16-33919361-DEL-66
chr16-33925341-DEL-114
chr16-33951879-DEL-1399
chr16-33953892-DEL-55
chr16-33956750-DEL-80
chr16-33983117-DEL-72
chr16-33988360-DEL-98
chr16-34014085-DEL-178
chr16-34028093-DEL-959
chr16-34043462-DEL-57
chr16-34066965-DEL-210
chr16-34153540-DEL-61
chr16-34177059-DEL-152
chr16-34206769-DEL-224
chr16-34211230-DEL-67
chr16-34223082-DEL-79
chr16-34229142-DEL-126
chr16-34250977-DEL-608
chr16-34268932-DEL-52
chr16-34273096-DEL-53
chr16-34295189-DEL-77
chr16-34327653-DEL-199
chr16-34386801-DEL-102
chr16-34400824-DEL-52
chr16-34401317-DEL-60
chr16-34401476-DEL-10355
chr16-34463057-DEL-65
chr16-34463305-DEL-58
chr16-34475598-DEL-82
chr16-34477571-DEL-1214
chr16-34501046-DEL-183
chr16-34506523-DEL-107
chr16-34508513-DEL-79
chr16-34530281-DEL-68
chr16-34568590-DEL-90
chr16-34592080-DEL-375
chr16-34594478-DEL-113
chr16-34595451-DEL-1691
chr16-34597829-DEL-54
chr16-34632033-DEL-57
chr16-34639349-DEL-818
chr16-34704081-DEL-134
chr16-34758097-DEL-71

chr16-34761325-DEL-95
chr16-34784683-DEL-107
chr16-34819323-DEL-71
chr16-34911514-DEL-149
chr16-34950198-DEL-87
chr16-34977904-DEL-248
chr16-34979080-DEL-73
chr16-35015634-DEL-401
chr16-35017917-DEL-166
chr16-35019010-DEL-137
chr16-35037755-DEL-60
chr16-35078942-DEL-54
chr16-35107846-DEL-64
chr16-35143331-DEL-311
chr16-35151884-DEL-87
chr16-35225033-DEL-162
chr16-35239645-DEL-53
chr16-35253328-DEL-142
chr16-35260417-DEL-525
chr16-35282238-DEL-748
chr16-35321428-DEL-52
chr16-35384446-DEL-262
chr16-35394144-DEL-93
chr16-35427321-DEL-78
chr16-35432005-DEL-2942
chr16-35464176-DEL-909
chr16-35472417-DEL-57
chr16-35475388-DEL-69
chr16-35521238-DEL-126
chr16-35562064-DEL-66
chr16-35563251-DEL-86
chr16-35564742-DEL-52
chr16-35619974-DEL-331
chr16-35621525-DEL-58
chr16-35641741-DEL-1673
chr16-35726182-DEL-106
chr16-35740128-DEL-82
chr16-35745674-DEL-419
chr16-35770871-DEL-772
chr16-35804809-DEL-58
chr16-35824855-DEL-120
chr16-35830657-DEL-960
chr16-35831356-DEL-381
chr16-35866143-DEL-332
chr16-35872444-DEL-95
chr16-35882695-DEL-275
chr16-35939267-DEL-227
chr16-35960710-DEL-96
chr16-35981696-DEL-93
chr16-35982563-DEL-72
chr16-35983615-DEL-2098
chr16-35985771-DEL-200
chr16-36000838-DEL-77
chr16-36005857-DEL-80
chr16-36011694-DEL-564
chr16-36019730-DEL-70
chr16-36021849-DEL-59
chr16-36039988-DEL-193
chr16-36076676-DEL-103
chr16-36078594-DEL-156
chr16-36143718-DEL-198
chr16-36144715-DEL-7138
chr16-36153011-DEL-827
chr16-36183190-DEL-101
chr16-36205877-DEL-56
chr16-36223942-DEL-72
chr16-36238207-DEL-99
chr16-36264941-DEL-3245
chr16-36298981-DEL-276
chr16-36308820-DEL-1254
chr16-36340497-DEL-74
chr16-36348431-DEL-300
chr16-36401396-DEL-377
chr16-36430462-DEL-681
chr16-36437862-DEL-67
chr16-36439132-DEL-1212
chr16-36450592-DEL-1765

chr16-36480470-DEL-409
chr16-36507530-DEL-161
chr16-36520289-DEL-68
chr16-36533898-DEL-298
chr16-36537406-DEL-168
chr16-36558590-DEL-56
chr16-36568467-DEL-205
chr16-36622425-DEL-82
chr16-36624949-DEL-79
chr16-36630358-DEL-63
chr16-36664583-DEL-67
chr16-36667206-DEL-196
chr16-36742232-DEL-92
chr16-36765981-DEL-193
chr16-36811607-DEL-144
chr16-36822136-DEL-746
chr16-36823022-DEL-212
chr16-36828113-DEL-109
chr16-36844497-DEL-1331
chr16-36876342-DEL-75
chr16-37016931-DEL-286
chr16-37043652-DEL-335
chr16-37044873-DEL-58
chr16-37047210-DEL-205
chr16-37065195-DEL-222
chr16-37077701-DEL-236
chr16-37116364-DEL-1324
chr16-37125949-DEL-4787
chr16-37138959-DEL-143
chr16-37150064-DEL-64
chr16-37216077-DEL-453
chr16-37228442-DEL-52
chr16-37239350-DEL-68
chr16-37243979-DEL-59
chr16-37249704-DEL-61
chr16-37297450-DEL-382
chr16-37323619-DEL-280
chr16-37329007-DEL-64
chr16-37357246-DEL-85
chr16-37414231-DEL-936
chr16-37432826-DEL-135
chr16-37460672-DEL-92
chr16-37480158-DEL-118
chr16-37494562-DEL-58
chr16-37510592-DEL-227
chr16-37512124-DEL-79
chr16-37518993-DEL-1246
chr16-37522344-DEL-1581
chr16-37523169-DEL-145
chr16-37543525-DEL-864
chr16-37602026-DEL-443
chr16-37615609-DEL-57
chr16-37637705-DEL-254
chr16-37685973-DEL-172
chr16-37696064-DEL-192
chr16-37701056-DEL-55
chr16-37710106-DEL-154
chr16-37711398-DEL-53
chr16-37753924-DEL-101
chr16-37774250-DEL-137
chr16-37810445-DEL-247
chr16-37814791-DEL-65
chr16-37859082-DEL-830
chr16-37894664-DEL-575
chr16-37939971-DEL-200
chr16-37959804-DEL-57
chr16-37971964-DEL-986
chr16-38011028-DEL-53
chr16-38046257-DEL-150
chr16-38052430-DEL-215
chr16-38053800-DEL-2902
chr16-38063413-DEL-66
chr16-38086808-DEL-120
chr16-38102542-DEL-52
chr16-38120453-DEL-219
chr16-38129626-DEL-1401
chr16-38154730-DEL-63

chr16-38158299-DEL-77
chr16-38162800-DEL-544
chr16-38173592-DEL-256
chr16-38203688-DEL-176
chr16-38213540-DEL-100
chr16-38218070-DEL-503
chr16-38232916-DEL-1895
chr16-38257811-DEL-63
chr16-38265009-DEL-52
chr16-38268155-DEL-326
chr16-38276718-DEL-52
chr16-38285987-DEL-53
chr16-38320339-DEL-291
chr16-38356075-DEL-278
chr16-38373698-DEL-365
chr16-38435629-DEL-239
chr16-38476662-DEL-760
chr16-38499233-DEL-65
chr16-38616733-DEL-1480
chr16-38622336-DEL-54
chr16-38638972-DEL-88
chr16-38651387-DEL-5206
chr16-38672477-DEL-2601
chr16-38697517-DEL-5518
chr16-38755752-DEL-59
chr16-38821692-DEL-65
chr16-38823591-DEL-75
chr16-38853512-DEL-650
chr16-38861741-DEL-1900
chr16-38925026-DEL-107
chr16-38999318-DEL-1588
chr16-39001707-DEL-207
chr16-39016550-DEL-598
chr16-39024371-DEL-2037
chr16-39034470-DEL-129
chr16-39036609-DEL-52
chr16-39039754-DEL-1150
chr16-39089089-DEL-112
chr16-39101330-DEL-63
chr16-39119843-DEL-225
chr16-39139298-DEL-82
chr16-39165000-DEL-648
chr16-39169257-DEL-107
chr16-39169490-DEL-445
chr16-39173730-DEL-139
chr16-39176475-DEL-60
chr16-39194206-DEL-52
chr16-39214328-DEL-86
chr16-39216209-DEL-149
chr16-39230345-DEL-55
chr16-39234803-DEL-123
chr16-39241309-DEL-51
chr16-39246259-DEL-51
chr16-39270360-DEL-94
chr16-39275116-DEL-494
chr16-39296657-DEL-1455
chr16-39313815-DEL-5998
chr16-39354462-DEL-58
chr16-39371977-DEL-234
chr16-39406564-DEL-1699
chr16-39417671-DEL-55
chr16-39431037-DEL-308
chr16-39522327-DEL-922
chr16-39530816-DEL-76
chr16-39542611-DEL-485
chr16-39543515-DEL-68
chr16-39561142-DEL-194
chr16-39604356-DEL-60
chr16-39604971-DEL-6117
chr16-39623221-DEL-63
chr16-39681545-DEL-65
chr16-39693498-DEL-188
chr16-39761106-DEL-119
chr16-39770607-DEL-221
chr16-39788468-DEL-739
chr16-39792608-DEL-125
chr16-39797173-DEL-329

chr16-39811318-DEL-52
chr16-39828555-DEL-71
chr16-39830201-DEL-75
chr16-39838095-DEL-68
chr16-39901553-DEL-135
chr16-39908060-DEL-80
chr16-39921708-DEL-94
chr16-39934984-DEL-60
chr16-39998440-DEL-435
chr16-40033259-DEL-73
chr16-40068112-DEL-340
chr16-40068403-DEL-637
chr16-40093723-DEL-73
chr16-40097128-DEL-301
chr16-40106843-DEL-199
chr16-40140307-DEL-84
chr16-40228946-DEL-1053
chr16-40234994-DEL-52
chr16-40280970-DEL-1837
chr16-40292331-DEL-72
chr16-40353082-DEL-56
chr16-40384906-DEL-669
chr16-40448560-DEL-698
chr16-40487396-DEL-54
chr16-40488427-DEL-1787
chr16-40493371-DEL-51
chr16-40521312-DEL-64
chr16-40536416-DEL-9155
chr16-40537788-DEL-5711
chr16-40538703-DEL-6291
chr16-40544495-DEL-578
chr16-40549089-DEL-78
chr16-40556163-DEL-53
chr16-40579784-DEL-1516
chr16-40619775-DEL-199
chr16-40632364-DEL-9607
chr16-40673226-DEL-589
chr16-40676546-DEL-170
chr16-40682557-DEL-53
chr16-40731538-DEL-308
chr16-40770473-DEL-1365
chr16-40806403-DEL-324
chr16-40807720-DEL-627
chr16-40810417-DEL-370
chr16-40834289-DEL-61
chr16-40840399-DEL-96
chr16-40848742-DEL-136
chr16-40854482-DEL-217
chr16-40869446-DEL-135
chr16-40880760-DEL-212
chr16-40889784-DEL-52
chr16-40895599-DEL-64
chr16-40918412-DEL-65
chr16-40927168-DEL-75
chr16-40940713-DEL-2176
chr16-40962358-DEL-56
chr16-40963922-DEL-178
chr16-40968090-DEL-724
chr16-40987909-DEL-77
chr16-41000935-DEL-61
chr16-41015527-DEL-1109
chr16-41015528-DEL-3219
chr16-41015528-DEL-7141
chr16-41016680-DEL-6002
chr16-41018772-DEL-3909
chr16-41061085-DEL-121
chr16-41085072-DEL-75
chr16-41088511-DEL-6214
chr16-41133067-DEL-223
chr16-41164182-DEL-151
chr16-41181684-DEL-76
chr16-41195265-DEL-138
chr16-41205870-DEL-1225
chr16-41210632-DEL-415
chr16-41213286-DEL-6334
chr16-41236406-DEL-75
chr16-41275547-DEL-150

chr16-41310364-DEL-100
chr16-41333001-DEL-69
chr16-41395417-DEL-1068
chr16-41396080-DEL-212
chr16-41478366-DEL-63
chr16-41486785-DEL-1779
chr16-41509080-DEL-57
chr16-41535989-DEL-51
chr16-41542870-DEL-161
chr16-41559934-DEL-2377
chr16-41651801-DEL-183
chr16-41665056-DEL-69
chr16-41668711-DEL-56
chr16-41691243-DEL-60
chr16-41711999-DEL-166
chr16-41800478-DEL-728
chr16-41818658-DEL-202
chr16-41846294-DEL-443
chr16-41865083-DEL-671
chr16-42013289-DEL-94
chr16-42013780-DEL-174
chr16-42014493-DEL-72
chr16-42020136-DEL-449
chr16-42054630-DEL-67
chr16-42064173-DEL-142
chr16-42067495-DEL-73
chr16-42079204-DEL-117
chr16-42087583-DEL-178
chr16-42094214-DEL-138
chr16-42121878-DEL-592
chr16-42124951-DEL-573
chr16-42130753-DEL-76
chr16-42134049-DEL-63
chr16-42134950-DEL-126
chr16-42143686-DEL-100
chr16-42194059-DEL-168
chr16-42195450-DEL-76
chr16-42226689-DEL-161
chr16-42229885-DEL-377
chr16-42232845-DEL-52
chr16-42261867-DEL-59
chr16-42262057-DEL-481
chr16-42315327-DEL-95
chr16-42345858-DEL-820
chr16-42378996-DEL-4232
chr16-42410306-DEL-225
chr16-42477496-DEL-4157
chr16-42507412-DEL-125
chr16-42539595-DEL-569
chr16-42545731-DEL-170
chr16-42550818-DEL-300
chr16-42583630-DEL-67
chr16-42588241-DEL-89
chr16-42620921-DEL-64
chr16-42639928-DEL-106
chr16-42665322-DEL-3335
chr16-42669301-DEL-3317
chr16-42684165-DEL-643
chr16-42798083-DEL-76
chr16-42805549-DEL-208
chr16-42811330-DEL-500
chr16-42881225-DEL-106
chr16-42953194-DEL-51
chr16-42995263-DEL-985
chr16-43022988-DEL-413
chr16-43060693-DEL-60
chr16-43154120-DEL-103
chr16-43157949-DEL-66
chr16-43267407-DEL-81
chr16-43288891-DEL-88
chr16-43315828-DEL-77
chr16-43339077-DEL-251
chr16-43389138-DEL-51
chr16-43485327-DEL-1241
chr16-43488269-DEL-1011
chr16-43493320-DEL-250
chr16-43504686-DEL-56

chr16-43513772-DEL-618
chr16-43532761-DEL-133
chr16-43558054-DEL-59
chr16-43564056-DEL-189
chr16-43567335-DEL-162
chr16-43568179-DEL-78
chr16-43575383-DEL-323
chr16-43596368-DEL-204
chr16-43598902-DEL-1890
chr16-43606895-DEL-84
chr16-43608782-DEL-217
chr16-43650187-DEL-123
chr16-43654519-DEL-863
chr16-43732596-DEL-157
chr16-43733132-DEL-1664
chr16-43734899-DEL-155
chr16-43743593-DEL-702
chr16-43767574-DEL-55
chr16-43802632-DEL-52
chr16-43867931-DEL-1517
chr16-43873843-DEL-126
chr16-43886701-DEL-56
chr16-43889996-DEL-146
chr16-43913284-DEL-3696
chr16-43970761-DEL-391
chr16-43971925-DEL-1892
chr16-43988376-DEL-257
chr16-44015207-DEL-190
chr16-44025450-DEL-66
chr16-44055835-DEL-54
chr16-44072570-DEL-393
chr16-44083534-DEL-58
chr16-44140811-DEL-134
chr16-44158683-DEL-69
chr16-44164250-DEL-332
chr16-44180340-DEL-77
chr16-44185075-DEL-75
chr16-44191567-DEL-199
chr16-44216738-DEL-638
chr16-44219493-DEL-108
chr16-44262059-DEL-467
chr16-44295083-DEL-63
chr16-44306645-DEL-102
chr16-44337444-DEL-59
chr16-44345991-DEL-88
chr16-44348497-DEL-60
chr16-44362033-DEL-115
chr16-44381775-DEL-114
chr16-44389746-DEL-55
chr16-44407808-DEL-222
chr16-44448794-DEL-70
chr16-44472196-DEL-111
chr16-44495436-DEL-473
chr16-44520126-DEL-191
chr16-44526965-DEL-131
chr16-44555165-DEL-260
chr16-44568966-DEL-84
chr16-44626848-DEL-124
chr16-44629853-DEL-449
chr16-44641684-DEL-70
chr16-44642704-DEL-61
chr16-44642991-DEL-3393
chr16-44659503-DEL-2594
chr16-44725931-DEL-1652
chr16-44780337-DEL-315
chr16-44791281-DEL-930
chr16-44819216-DEL-136
chr16-44859167-DEL-2544
chr16-44881598-DEL-7727
chr16-44882100-DEL-2922
chr16-44901765-DEL-117
chr16-44953221-DEL-6185
chr16-45019057-DEL-161
chr16-45038805-DEL-62
chr16-45056712-DEL-1410
chr16-45091279-DEL-929
chr16-45092804-DEL-1637

chr16-45115276-DEL-1663
chr16-45119904-DEL-1356
chr16-45129815-DEL-867
chr16-45142462-DEL-85
chr16-45153403-DEL-151
chr16-45175895-DEL-163
chr16-45185576-DEL-51
chr16-45226169-DEL-503
chr16-45229745-DEL-121
chr16-45254748-DEL-78
chr16-45255543-DEL-2000
chr16-45280105-DEL-85
chr16-45281900-DEL-6218
chr16-45312778-DEL-811
chr16-45317521-DEL-1656
chr16-45350425-DEL-1447
chr16-45393520-DEL-112
chr16-45421454-DEL-61
chr16-45515497-DEL-124
chr16-45518988-DEL-110
chr16-45528383-DEL-106
chr16-45548110-DEL-1316
chr16-45591861-DEL-61
chr16-45599854-DEL-518
chr16-45666453-DEL-1274
chr16-45708615-DEL-1054
chr16-45730450-DEL-306
chr16-45835580-DEL-104
chr16-45845848-DEL-84
chr16-45910702-DEL-600
chr16-45923834-DEL-1185
chr16-45925043-DEL-1214
chr16-45935303-DEL-811
chr16-45964197-DEL-104
chr16-45978950-DEL-141
chr16-45992413-DEL-428
chr16-45995870-DEL-480
chr16-46017758-DEL-126
chr16-46028420-DEL-259
chr16-46028740-DEL-631
chr16-46074486-DEL-888
chr16-46132534-DEL-5613
chr16-46155113-DEL-54
chr16-46194799-DEL-1073
chr16-46245767-DEL-382
chr16-46285598-DEL-55
chr16-46312956-DEL-52
chr16-46328779-DEL-8104
chr16-46350044-DEL-66
chr16-46354293-DEL-422
chr16-46374928-DEL-225
chr16-46448384-DEL-1082
chr16-46464294-DEL-211
chr16-46476072-DEL-51
chr16-46501397-DEL-64
chr16-46511574-DEL-3138
chr16-46511895-DEL-1663
chr16-46525377-DEL-84
chr16-46549662-DEL-197
chr16-46572913-DEL-96
chr16-46573886-DEL-12477
chr16-46594810-DEL-808
chr16-46641115-DEL-290
chr16-46725019-DEL-172
chr16-46746523-DEL-261
chr16-46761088-DEL-1140
chr16-46804491-DEL-88
chr16-46841019-DEL-180
chr16-46841943-DEL-1171
chr16-46856334-DEL-1202
chr16-46919392-DEL-1890
chr16-46937150-DEL-172
chr16-46951564-DEL-6575
chr16-46963390-DEL-5843
chr16-46986551-DEL-2278
chr16-46992299-DEL-2619
chr16-47018969-DEL-3593

chr16-47022848-DEL-76
chr16-47070014-DEL-1161
chr16-47074439-DEL-139
chr16-47083756-DEL-115
chr16-47085678-DEL-561
chr16-47099954-DEL-502
chr16-47133600-DEL-123
chr16-47170082-DEL-52
chr16-47194686-DEL-58
chr16-47212241-DEL-205
chr16-47222333-DEL-260
chr16-47299121-DEL-447
chr16-47328872-DEL-115
chr16-47342960-DEL-109
chr16-47344452-DEL-110
chr16-47353111-DEL-359
chr16-47372534-DEL-2663
chr16-47415589-DEL-528
chr16-47434581-DEL-53
chr16-47458267-DEL-56
chr16-47460942-DEL-198
chr16-47468407-DEL-216
chr16-47477785-DEL-1394
chr16-47517239-DEL-3396
chr16-47523639-DEL-1595
chr16-47547405-DEL-408
chr16-47548513-DEL-231
chr16-47561917-DEL-2726
chr16-47576690-DEL-56
chr16-47579528-DEL-226
chr16-47589746-DEL-56
chr16-47597401-DEL-672
chr16-47621666-DEL-6022
chr16-47640478-DEL-3287
chr16-47646989-DEL-110
chr16-47649995-DEL-7292
chr16-47692841-DEL-431
chr16-47714632-DEL-73
chr16-47783840-DEL-1646
chr16-47791301-DEL-665
chr16-47801818-DEL-208
chr16-47828529-DEL-7374
chr16-47832854-DEL-1232
chr16-47836077-DEL-336
chr16-47837556-DEL-325
chr16-47841994-DEL-642
chr16-47902781-DEL-8404
chr16-47904294-DEL-6204
chr16-47915934-DEL-8651
chr16-47916139-DEL-2314
chr16-47921590-DEL-2277
chr16-47923145-DEL-1414
chr16-47945444-DEL-674
chr16-47950656-DEL-303
chr16-48068226-DEL-438
chr16-48087881-DEL-598
chr16-48091449-DEL-142
chr16-48101807-DEL-440
chr16-48102640-DEL-975
chr16-48106866-DEL-58
chr16-48112621-DEL-237
chr16-48129150-DEL-184
chr16-48133380-DEL-186
chr16-48175733-DEL-247
chr16-48182388-DEL-92
chr16-48198960-DEL-52
chr16-48199967-DEL-51
chr16-48211474-DEL-401
chr16-48213626-DEL-56
chr16-48287199-DEL-56
chr16-48317846-DEL-212
chr16-48329911-DEL-182
chr16-48348227-DEL-7230
chr16-48358947-DEL-9555
chr16-48400270-DEL-949
chr16-48400299-DEL-1397
chr16-48401173-DEL-521

chr16-48409040-DEL-249
chr16-48425154-DEL-343
chr16-48446293-DEL-52
chr16-48486051-DEL-51
chr16-48509158-DEL-5067
chr16-48516368-DEL-211
chr16-48518704-DEL-83
chr16-48536431-DEL-3821
chr16-48539032-DEL-51
chr16-48546576-DEL-137
chr16-48548789-DEL-66
chr16-48601763-DEL-85
chr16-48642204-DEL-52
chr16-48673967-DEL-80
chr16-48686582-DEL-3140
chr16-48726818-DEL-192
chr16-48731850-DEL-273
chr16-48808967-DEL-57
chr16-48816426-DEL-77
chr16-48851334-DEL-66
chr16-48862243-DEL-461
chr16-48899098-DEL-1114
chr16-48903894-DEL-1457
chr16-48906151-DEL-625
chr16-48926113-DEL-58
chr16-48968752-DEL-1891
chr16-48968870-DEL-385
chr16-49006993-DEL-55
chr16-49011799-DEL-62
chr16-49067422-DEL-1375
chr16-49086352-DEL-54
chr16-49151966-DEL-84
chr16-49204310-DEL-181
chr16-49224068-DEL-790
chr16-49231270-DEL-52
chr16-49233395-DEL-61
chr16-49240340-DEL-588
chr16-49241003-DEL-1289
chr16-49256308-DEL-56
chr16-49284478-DEL-66
chr16-49311332-DEL-1629
chr16-49314060-DEL-1132
chr16-49315060-DEL-277
chr16-49335517-DEL-60
chr16-49381119-DEL-191
chr16-49414424-DEL-1275
chr16-49432830-DEL-61
chr16-49439828-DEL-488
chr16-49452484-DEL-1044
chr16-49477811-DEL-473
chr16-49496341-DEL-400
chr16-49506264-DEL-108
chr16-49507438-DEL-397
chr16-49508607-DEL-55
chr16-49510336-DEL-92
chr16-49615106-DEL-54
chr16-49701595-DEL-66
chr16-49746171-DEL-83
chr16-49777574-DEL-56
chr16-49792963-DEL-2132
chr16-49821532-DEL-62
chr16-49939230-DEL-158
chr16-49960927-DEL-2785
chr16-49970619-DEL-403
chr16-49984834-DEL-2088
chr16-50091215-DEL-63
chr16-50106137-DEL-2158
chr16-50109144-DEL-1528
chr16-50110732-DEL-1366
chr16-50116405-DEL-51
chr16-50147057-DEL-98
chr16-50188015-DEL-94
chr16-50239249-DEL-125
chr16-50262041-DEL-51
chr16-50296920-DEL-102
chr16-50327923-DEL-148
chr16-50341590-DEL-82

chr16-50342683-DEL-324
chr16-50360709-DEL-1916
chr16-50421705-DEL-117
chr16-50458657-DEL-93
chr16-50495796-DEL-185
chr16-50499923-DEL-2382
chr16-50505632-DEL-195
chr16-50523619-DEL-58
chr16-50526667-DEL-838
chr16-50561798-DEL-134
chr16-50586567-DEL-112
chr16-50597515-DEL-400
chr16-50602468-DEL-400
chr16-50723183-DEL-1283
chr16-50729412-DEL-663
chr16-50745111-DEL-2140
chr16-50752665-DEL-172
chr16-50757528-DEL-901
chr16-50802978-DEL-814
chr16-50814936-DEL-168
chr16-50816475-DEL-342
chr16-50825622-DEL-112
chr16-50881721-DEL-1498
chr16-50895291-DEL-272
chr16-50907533-DEL-7077
chr16-50919327-DEL-1035
chr16-51005519-DEL-1747
chr16-51024552-DEL-141
chr16-51027379-DEL-60
chr16-51055723-DEL-451
chr16-51056663-DEL-1201
chr16-51058441-DEL-3926
chr16-51072362-DEL-6667
chr16-51086327-DEL-313
chr16-51088216-DEL-7844
chr16-51123158-DEL-97
chr16-51131623-DEL-510
chr16-51137631-DEL-52
chr16-51160026-DEL-534
chr16-51164218-DEL-690
chr16-51165061-DEL-139
chr16-51193357-DEL-1467
chr16-51198208-DEL-520
chr16-51220777-DEL-399
chr16-51248781-DEL-474
chr16-51263542-DEL-5960
chr16-51269483-DEL-2388
chr16-51307013-DEL-6120
chr16-51344271-DEL-679
chr16-51351887-DEL-754
chr16-51388355-DEL-84
chr16-51388670-DEL-297
chr16-51425601-DEL-2239
chr16-51436911-DEL-637
chr16-51445406-DEL-57
chr16-51459736-DEL-1154
chr16-51475123-DEL-105
chr16-51475191-DEL-1030
chr16-51476515-DEL-139
chr16-51482820-DEL-372
chr16-51484276-DEL-184
chr16-51494409-DEL-106
chr16-51563048-DEL-715
chr16-51565212-DEL-51
chr16-51570958-DEL-635
chr16-51593878-DEL-203
chr16-51595639-DEL-1371
chr16-51604492-DEL-52
chr16-51617986-DEL-6179
chr16-51643404-DEL-1104
chr16-51662211-DEL-132
chr16-51672598-DEL-64
chr16-51686744-DEL-230
chr16-51716478-DEL-235
chr16-51718210-DEL-115
chr16-51723061-DEL-1339
chr16-51741822-DEL-183

chr16-51751340-DEL-60
chr16-51866728-DEL-86
chr16-51874369-DEL-246
chr16-51878568-DEL-308
chr16-51883123-DEL-520
chr16-51903629-DEL-89
chr16-51940937-DEL-567
chr16-51981065-DEL-56
chr16-52022687-DEL-57
chr16-52032207-DEL-180
chr16-52060530-DEL-76
chr16-52060921-DEL-853
chr16-52075089-DEL-1199
chr16-52126921-DEL-90
chr16-52134924-DEL-79
chr16-52136574-DEL-1279
chr16-52138229-DEL-58
chr16-52168949-DEL-53
chr16-52169848-DEL-90
chr16-52225091-DEL-69
chr16-52267993-DEL-257
chr16-52355662-DEL-107
chr16-52359731-DEL-852
chr16-52362466-DEL-445
chr16-52440590-DEL-490
chr16-52477070-DEL-88
chr16-52484883-DEL-2305
chr16-52510191-DEL-379
chr16-52518398-DEL-1254
chr16-52642360-DEL-330
chr16-52643786-DEL-3211
chr16-52648106-DEL-2480
chr16-52666387-DEL-269
chr16-52678500-DEL-60
chr16-52689458-DEL-150
chr16-52695721-DEL-112
chr16-52701754-DEL-250
chr16-52711649-DEL-124
chr16-52741036-DEL-900
chr16-52744822-DEL-70
chr16-52763060-DEL-912
chr16-52772950-DEL-101
chr16-52774611-DEL-596
chr16-52831437-DEL-385
chr16-52840940-DEL-68
chr16-52842765-DEL-205
chr16-52864635-DEL-64
chr16-52898294-DEL-223
chr16-52903072-DEL-52
chr16-52929913-DEL-271
chr16-52937895-DEL-548
chr16-52954411-DEL-6134
chr16-52963980-DEL-5770
chr16-52983307-DEL-197
chr16-53012877-DEL-5734
chr16-53020582-DEL-87
chr16-53024864-DEL-2465
chr16-53036254-DEL-56
chr16-53059821-DEL-596
chr16-53073874-DEL-91
chr16-53074829-DEL-2189
chr16-53077419-DEL-221
chr16-53084536-DEL-98
chr16-53084833-DEL-363
chr16-53120323-DEL-111
chr16-53168179-DEL-2677
chr16-53172338-DEL-2166
chr16-53176508-DEL-6215
chr16-53186939-DEL-3086
chr16-53200203-DEL-451
chr16-53209239-DEL-417
chr16-53210270-DEL-1057
chr16-53214560-DEL-129
chr16-53215383-DEL-237
chr16-53217475-DEL-440
chr16-53232370-DEL-3761
chr16-53290220-DEL-264

chr16-53308876-DEL-76
chr16-53315404-DEL-628
chr16-53339858-DEL-1044
chr16-53386841-DEL-428
chr16-53391668-DEL-94
chr16-53391839-DEL-5098
chr16-53400018-DEL-3919
chr16-53403342-DEL-78
chr16-53409054-DEL-1099
chr16-53432234-DEL-100
chr16-53503317-DEL-76
chr16-53503429-DEL-2838
chr16-53537581-DEL-1089
chr16-53561032-DEL-546
chr16-53571326-DEL-2361
chr16-53573954-DEL-275
chr16-53581344-DEL-1904
chr16-53601483-DEL-209
chr16-53622707-DEL-367
chr16-53638871-DEL-54
chr16-53659226-DEL-1120
chr16-53661014-DEL-168
chr16-53694752-DEL-194
chr16-53698767-DEL-6147
chr16-53786196-DEL-874
chr16-53826773-DEL-240
chr16-53868564-DEL-822
chr16-53887164-DEL-2422
chr16-53942266-DEL-58
chr16-53944495-DEL-2869
chr16-53975712-DEL-52
chr16-53983274-DEL-56
chr16-53992939-DEL-61
chr16-54069026-DEL-57
chr16-54071993-DEL-1209
chr16-54079186-DEL-66
chr16-54084238-DEL-58
chr16-54100439-DEL-72
chr16-54104406-DEL-513
chr16-54106734-DEL-105
chr16-54110656-DEL-537
chr16-54142383-DEL-58
chr16-54145617-DEL-56
chr16-54156607-DEL-322
chr16-54183763-DEL-4574
chr16-54196230-DEL-2704
chr16-54197036-DEL-2542
chr16-54200597-DEL-1285
chr16-54205207-DEL-1840
chr16-54246241-DEL-6699
chr16-54264716-DEL-79
chr16-54285027-DEL-6613
chr16-54321869-DEL-2401
chr16-54360157-DEL-587
chr16-54365317-DEL-52
chr16-54391197-DEL-2045
chr16-54497441-DEL-1002
chr16-54502124-DEL-91
chr16-54521826-DEL-179
chr16-54551747-DEL-87
chr16-54563137-DEL-108
chr16-54574343-DEL-179
chr16-54584169-DEL-60
chr16-54606879-DEL-71
chr16-54620528-DEL-231
chr16-54636027-DEL-169
chr16-54667585-DEL-468
chr16-54679599-DEL-1045
chr16-54681847-DEL-6826
chr16-54698108-DEL-96
chr16-54712962-DEL-730
chr16-54716296-DEL-118
chr16-54728095-DEL-1404
chr16-54732293-DEL-87
chr16-54771190-DEL-250
chr16-54794688-DEL-142
chr16-54828349-DEL-324

chr16-54861406-DEL-94
chr16-54867487-DEL-699
chr16-54884798-DEL-53
chr16-54909935-DEL-64
chr16-54916341-DEL-1097
chr16-54916797-DEL-72
chr16-54919974-DEL-18523
chr16-54957794-DEL-149
chr16-54986000-DEL-55
chr16-55037412-DEL-406
chr16-55043549-DEL-470
chr16-55072454-DEL-711
chr16-55103369-DEL-94
chr16-55126252-DEL-291
chr16-55151129-DEL-235
chr16-55153664-DEL-314
chr16-55161994-DEL-1757
chr16-55165211-DEL-107
chr16-55166809-DEL-992
chr16-55199171-DEL-60
chr16-55228666-DEL-113
chr16-55258030-DEL-312
chr16-55318251-DEL-54
chr16-55358970-DEL-1486
chr16-55359131-DEL-106
chr16-55410565-DEL-65
chr16-55422955-DEL-897
chr16-55484988-DEL-55
chr16-55494770-DEL-7174
chr16-55509467-DEL-53
chr16-55510588-DEL-101
chr16-55524275-DEL-757
chr16-55526327-DEL-122
chr16-55537798-DEL-367
chr16-55571394-DEL-174
chr16-55575536-DEL-51
chr16-55588260-DEL-62
chr16-55597709-DEL-167
chr16-55627514-DEL-133
chr16-55670671-DEL-519
chr16-55674655-DEL-64
chr16-55675152-DEL-314
chr16-55676723-DEL-53
chr16-55694473-DEL-254
chr16-55727041-DEL-185
chr16-55730502-DEL-564
chr16-55753471-DEL-202
chr16-55758881-DEL-4712
chr16-55766603-DEL-54
chr16-55768615-DEL-127
chr16-55770475-DEL-177
chr16-55771283-DEL-947
chr16-55773098-DEL-126
chr16-55813040-DEL-559
chr16-55823821-DEL-1357
chr16-55834201-DEL-376
chr16-55835703-DEL-61
chr16-55837859-DEL-88
chr16-55875445-DEL-272
chr16-55880193-DEL-170
chr16-55916817-DEL-84
chr16-55965027-DEL-190
chr16-55973568-DEL-82
chr16-55979282-DEL-294
chr16-55985239-DEL-415
chr16-55988660-DEL-159
chr16-56021138-DEL-194
chr16-56029665-DEL-280
chr16-56035233-DEL-269
chr16-56040431-DEL-72
chr16-56047835-DEL-53
chr16-56064802-DEL-333
chr16-56065503-DEL-66
chr16-56102561-DEL-79
chr16-56124355-DEL-138
chr16-56129485-DEL-205
chr16-56133246-DEL-1043

chr16-56138989-DEL-441
chr16-56142527-DEL-13056
chr16-56145070-DEL-6159
chr16-56171562-DEL-300
chr16-56254182-DEL-271
chr16-56258328-DEL-1727
chr16-56265588-DEL-778
chr16-56277507-DEL-281
chr16-56297262-DEL-343
chr16-56326636-DEL-572
chr16-56333070-DEL-52
chr16-56360901-DEL-156
chr16-56371353-DEL-182
chr16-56385957-DEL-316
chr16-56401816-DEL-60
chr16-56414374-DEL-235
chr16-56423488-DEL-512
chr16-56435068-DEL-923
chr16-56468200-DEL-157
chr16-56493741-DEL-433
chr16-56496962-DEL-556
chr16-56571541-DEL-1129
chr16-56613235-DEL-546
chr16-56622234-DEL-439
chr16-56656622-DEL-2476
chr16-56674323-DEL-981
chr16-56741518-DEL-433
chr16-56746810-DEL-271
chr16-56754508-DEL-6784
chr16-56760932-DEL-204
chr16-56770321-DEL-6273
chr16-56781597-DEL-102
chr16-56788727-DEL-52
chr16-56796438-DEL-93
chr16-56808559-DEL-10410
chr16-56820632-DEL-236
chr16-56821021-DEL-1459
chr16-56839207-DEL-2074
chr16-56865872-DEL-208
chr16-56903412-DEL-282
chr16-56914508-DEL-1795
chr16-56936038-DEL-74
chr16-56994025-DEL-53
chr16-57066579-DEL-58
chr16-57068599-DEL-64
chr16-57083037-DEL-64
chr16-57083782-DEL-57
chr16-57084279-DEL-1358
chr16-57084903-DEL-162
chr16-57085922-DEL-102
chr16-57090698-DEL-2247
chr16-57119216-DEL-54
chr16-57126286-DEL-861
chr16-57152252-DEL-300
chr16-57163619-DEL-53
chr16-57200455-DEL-352
chr16-57260007-DEL-134
chr16-57274896-DEL-67
chr16-57279057-DEL-83
chr16-57279897-DEL-115
chr16-57297128-DEL-96
chr16-57378959-DEL-81
chr16-57387322-DEL-441
chr16-57400537-DEL-67
chr16-57428081-DEL-127
chr16-57509854-DEL-98
chr16-57512370-DEL-132
chr16-57542710-DEL-58
chr16-57557229-DEL-8280
chr16-57566377-DEL-881
chr16-57572357-DEL-215
chr16-57588153-DEL-1081
chr16-57595919-DEL-1926
chr16-57603900-DEL-119
chr16-57604769-DEL-141
chr16-57607434-DEL-54
chr16-57637182-DEL-794

chr16-57638331-DEL-131
chr16-57662520-DEL-53
chr16-57668369-DEL-341
chr16-57680416-DEL-124
chr16-57696056-DEL-72
chr16-57725938-DEL-88
chr16-57726916-DEL-58
chr16-57730173-DEL-150
chr16-57779877-DEL-312
chr16-57806042-DEL-1049
chr16-57808077-DEL-1096
chr16-57815193-DEL-252
chr16-57818478-DEL-156
chr16-57825205-DEL-212
chr16-57869510-DEL-728
chr16-57915860-DEL-3104
chr16-57919754-DEL-64
chr16-57946015-DEL-131
chr16-57961440-DEL-1036
chr16-57963393-DEL-69
chr16-58000941-DEL-761
chr16-58003443-DEL-3641
chr16-58021858-DEL-6483
chr16-58034085-DEL-6423
chr16-58053796-DEL-132
chr16-58059664-DEL-6324
chr16-58066790-DEL-321
chr16-58085138-DEL-591
chr16-58085752-DEL-142
chr16-58101567-DEL-56
chr16-58114212-DEL-60
chr16-58119394-DEL-541
chr16-58127288-DEL-361
chr16-58128583-DEL-6565
chr16-58139703-DEL-6439
chr16-58146569-DEL-923
chr16-58148460-DEL-105
chr16-58204106-DEL-674
chr16-58211877-DEL-75
chr16-58258160-DEL-4065
chr16-58282652-DEL-62
chr16-58283979-DEL-109
chr16-58322791-DEL-253
chr16-58348674-DEL-1833
chr16-58359016-DEL-489
chr16-58359535-DEL-2972
chr16-58372969-DEL-65
chr16-58383238-DEL-6188
chr16-58392410-DEL-1492
chr16-58431049-DEL-61
chr16-58448794-DEL-65
chr16-58458385-DEL-176
chr16-58461400-DEL-1002
chr16-58503684-DEL-288
chr16-58533544-DEL-1758
chr16-58566226-DEL-64
chr16-58579778-DEL-210
chr16-58584456-DEL-3126
chr16-58589598-DEL-3049
chr16-58592052-DEL-72
chr16-58601531-DEL-79
chr16-58609550-DEL-350
chr16-58637141-DEL-2018
chr16-58663767-DEL-13441
chr16-58671736-DEL-991
chr16-58682002-DEL-8476
chr16-58697915-DEL-6254
chr16-58706276-DEL-64
chr16-58709158-DEL-54
chr16-58751836-DEL-209
chr16-58778616-DEL-56
chr16-58782591-DEL-707
chr16-58787846-DEL-71
chr16-58797125-DEL-254
chr16-58797440-DEL-7117
chr16-58804385-DEL-88
chr16-58811533-DEL-68

chr16-58822366-DEL-2353
chr16-58869574-DEL-701
chr16-58870114-DEL-7853
chr16-58877105-DEL-178
chr16-58878903-DEL-85
chr16-59015988-DEL-193
chr16-59034018-DEL-110
chr16-59038942-DEL-84
chr16-59106654-DEL-137
chr16-59124596-DEL-89
chr16-59130021-DEL-201
chr16-59153849-DEL-56
chr16-59177839-DEL-56
chr16-59192866-DEL-6339
chr16-59219030-DEL-847
chr16-59251195-DEL-98
chr16-59259627-DEL-615
chr16-59288901-DEL-486
chr16-59291632-DEL-164
chr16-59335191-DEL-1852
chr16-59346584-DEL-76
chr16-59366633-DEL-802
chr16-59370137-DEL-80
chr16-59384933-DEL-60
chr16-59411856-DEL-180
chr16-59415898-DEL-111
chr16-59427983-DEL-723
chr16-59429220-DEL-385
chr16-59440557-DEL-68
chr16-59445205-DEL-140
chr16-59449443-DEL-1069
chr16-59489517-DEL-108
chr16-59547373-DEL-315
chr16-59553521-DEL-71
chr16-59610224-DEL-96
chr16-59678101-DEL-911
chr16-59712271-DEL-13090
chr16-59764426-DEL-494
chr16-59770619-DEL-558
chr16-59831735-DEL-72
chr16-59844712-DEL-121
chr16-59845298-DEL-243
chr16-59845808-DEL-136
chr16-59892590-DEL-85
chr16-59928154-DEL-95
chr16-59941744-DEL-58
chr16-59971703-DEL-376
chr16-59991915-DEL-2726
chr16-60014720-DEL-1861
chr16-60095118-DEL-1116
chr16-60101176-DEL-1070
chr16-60105060-DEL-183
chr16-60115279-DEL-274
chr16-60124259-DEL-105
chr16-60138457-DEL-91
chr16-60146739-DEL-993
chr16-60164212-DEL-128
chr16-60210578-DEL-58
chr16-60213702-DEL-70
chr16-60236994-DEL-92
chr16-60280158-DEL-2223
chr16-60285104-DEL-183
chr16-60290813-DEL-873
chr16-60320261-DEL-61
chr16-60322978-DEL-77
chr16-60388839-DEL-890
chr16-60405973-DEL-117
chr16-60504657-DEL-1509
chr16-60510371-DEL-230
chr16-60517082-DEL-103
chr16-60534788-DEL-64
chr16-60544678-DEL-7184
chr16-60618651-DEL-71
chr16-60619389-DEL-58
chr16-60664410-DEL-54
chr16-60665609-DEL-998
chr16-60667913-DEL-93

chr16-60673088-DEL-356
chr16-60673852-DEL-2427
chr16-60677049-DEL-375
chr16-60698108-DEL-63
chr16-60703717-DEL-2206
chr16-60741459-DEL-777
chr16-60742308-DEL-1761
chr16-60754232-DEL-1106
chr16-60760610-DEL-1638
chr16-60762540-DEL-236
chr16-60762976-DEL-365
chr16-60786036-DEL-4968
chr16-60799851-DEL-1694
chr16-60822728-DEL-55
chr16-60825848-DEL-59
chr16-60837259-DEL-113
chr16-60851977-DEL-64
chr16-60853068-DEL-131
chr16-60859751-DEL-56
chr16-60904212-DEL-80
chr16-60916031-DEL-6215
chr16-60925040-DEL-75
chr16-60954404-DEL-75
chr16-60955561-DEL-965
chr16-60957779-DEL-71
chr16-60966769-DEL-596
chr16-60970032-DEL-8294
chr16-61025347-DEL-1146
chr16-61033359-DEL-140
chr16-61034902-DEL-1654
chr16-61045409-DEL-3779
chr16-61054351-DEL-1816
chr16-61057140-DEL-340
chr16-61070398-DEL-9512
chr16-61117456-DEL-1851
chr16-61120775-DEL-232
chr16-61120803-DEL-6580
chr16-61125003-DEL-6072
chr16-61140171-DEL-88
chr16-61225655-DEL-180
chr16-61289510-DEL-352
chr16-61290867-DEL-53
chr16-61342504-DEL-181
chr16-61344189-DEL-2039
chr16-61348732-DEL-893
chr16-61357758-DEL-229
chr16-61379008-DEL-2928
chr16-61393035-DEL-121
chr16-61395910-DEL-662
chr16-61408927-DEL-7460
chr16-61422525-DEL-6199
chr16-61452717-DEL-2293
chr16-61476918-DEL-735
chr16-61481295-DEL-73
chr16-61490797-DEL-346
chr16-61503826-DEL-60
chr16-61504984-DEL-210
chr16-61508249-DEL-327
chr16-61543296-DEL-2772
chr16-61547269-DEL-60
chr16-61574747-DEL-82
chr16-61583760-DEL-70
chr16-61611373-DEL-65
chr16-61621457-DEL-77
chr16-61633382-DEL-8122
chr16-61643728-DEL-156
chr16-61648208-DEL-240
chr16-61649544-DEL-174
chr16-61663236-DEL-56
chr16-61683527-DEL-245
chr16-61709278-DEL-63
chr16-61717365-DEL-2093
chr16-61727361-DEL-73
chr16-61737714-DEL-368
chr16-61742977-DEL-129
chr16-61766177-DEL-353
chr16-61779741-DEL-186

chr16-61781140-DEL-564
chr16-61782542-DEL-72
chr16-61786449-DEL-77
chr16-61793651-DEL-829
chr16-61811481-DEL-2906
chr16-61819030-DEL-657
chr16-61826254-DEL-431
chr16-61839539-DEL-2427
chr16-61848098-DEL-191
chr16-61852727-DEL-731
chr16-61870793-DEL-97
chr16-61886178-DEL-930
chr16-61914412-DEL-2259
chr16-61916173-DEL-81
chr16-61926039-DEL-57
chr16-61927372-DEL-208
chr16-61975905-DEL-1145
chr16-62024773-DEL-207
chr16-62030424-DEL-61
chr16-62033071-DEL-2986
chr16-62044143-DEL-89
chr16-62061544-DEL-839
chr16-62095679-DEL-54
chr16-62100330-DEL-52
chr16-62130654-DEL-144
chr16-62182409-DEL-185
chr16-62183611-DEL-429
chr16-62215031-DEL-5585
chr16-62233385-DEL-70
chr16-62240218-DEL-2319
chr16-62254924-DEL-70
chr16-62257785-DEL-445
chr16-62294636-DEL-66
chr16-62320183-DEL-52
chr16-62324338-DEL-457
chr16-62337198-DEL-1813
chr16-62396085-DEL-58
chr16-62396540-DEL-1899
chr16-62418506-DEL-1058
chr16-62439499-DEL-271
chr16-62457939-DEL-3753
chr16-62484576-DEL-6738
chr16-62495426-DEL-1187
chr16-62508668-DEL-182
chr16-62514070-DEL-100
chr16-62535653-DEL-82
chr16-62618581-DEL-285
chr16-62648185-DEL-60
chr16-62653564-DEL-338
chr16-62654737-DEL-188
chr16-62702788-DEL-378
chr16-62711514-DEL-84
chr16-62752027-DEL-64
chr16-62759508-DEL-371
chr16-62796226-DEL-379
chr16-62804443-DEL-115
chr16-62813342-DEL-65
chr16-62826949-DEL-1114
chr16-62844731-DEL-4637
chr16-62859621-DEL-149
chr16-62903431-DEL-109
chr16-62910671-DEL-180
chr16-62933399-DEL-3862
chr16-62942024-DEL-2003
chr16-62948558-DEL-208
chr16-62963795-DEL-53
chr16-62966382-DEL-6295
chr16-63038761-DEL-675
chr16-63061460-DEL-113
chr16-63062031-DEL-297
chr16-63096312-DEL-166
chr16-63103055-DEL-930
chr16-63121619-DEL-1368
chr16-63148915-DEL-66
chr16-63159943-DEL-136
chr16-63168158-DEL-82
chr16-63217033-DEL-839

chr16-63217095-DEL-141
chr16-63219126-DEL-187
chr16-63219526-DEL-3834
chr16-63226263-DEL-132
chr16-63231684-DEL-68
chr16-63243785-DEL-653
chr16-63254037-DEL-2293
chr16-63259290-DEL-2904
chr16-63265961-DEL-98
chr16-63279309-DEL-814
chr16-63280687-DEL-3243
chr16-63280733-DEL-297
chr16-63282018-DEL-277
chr16-63282587-DEL-307
chr16-63282928-DEL-885
chr16-63382778-DEL-51
chr16-63395764-DEL-403
chr16-63426132-DEL-204
chr16-63426994-DEL-606
chr16-63478271-DEL-60
chr16-63479809-DEL-289
chr16-63521972-DEL-157
chr16-63550805-DEL-527
chr16-63552185-DEL-500
chr16-63561329-DEL-217
chr16-63583462-DEL-67
chr16-63601074-DEL-357
chr16-63616404-DEL-94
chr16-63653332-DEL-1397
chr16-63653931-DEL-747
chr16-63704935-DEL-243
chr16-63727343-DEL-120
chr16-63732682-DEL-75
chr16-63739666-DEL-3680
chr16-63745955-DEL-946
chr16-63747813-DEL-869
chr16-63759189-DEL-417
chr16-63760231-DEL-198
chr16-63761467-DEL-110
chr16-63777556-DEL-82
chr16-63788922-DEL-6498
chr16-63788939-DEL-2149
chr16-63845924-DEL-416
chr16-63859076-DEL-77
chr16-63877374-DEL-1536
chr16-63892511-DEL-338
chr16-63900437-DEL-397
chr16-63951338-DEL-145
chr16-63953237-DEL-1661
chr16-63954645-DEL-258
chr16-63955245-DEL-1370
chr16-63968473-DEL-51
chr16-63972241-DEL-222
chr16-63992657-DEL-70
chr16-63998315-DEL-226
chr16-64001191-DEL-54
chr16-64012149-DEL-3124
chr16-64026039-DEL-74
chr16-64035734-DEL-63
chr16-64048542-DEL-51
chr16-64069471-DEL-68
chr16-64122712-DEL-6486
chr16-64140723-DEL-1053
chr16-64155558-DEL-75
chr16-64164790-DEL-82
chr16-64175608-DEL-97
chr16-64200774-DEL-92
chr16-64229787-DEL-241
chr16-64230783-DEL-6318
chr16-64238251-DEL-926
chr16-64270615-DEL-6421
chr16-64270747-DEL-212
chr16-64285840-DEL-68
chr16-64311784-DEL-128
chr16-64329819-DEL-1316
chr16-64402599-DEL-537
chr16-64436185-DEL-570

chr16-64494695-DEL-60
chr16-64500117-DEL-348
chr16-64506574-DEL-74
chr16-64514656-DEL-213
chr16-64535649-DEL-607
chr16-64550790-DEL-84
chr16-64559390-DEL-93
chr16-64597157-DEL-74
chr16-64657277-DEL-92
chr16-64665558-DEL-91
chr16-64674592-DEL-119
chr16-64684548-DEL-204
chr16-64712430-DEL-66
chr16-64730709-DEL-90
chr16-64751385-DEL-614
chr16-64766793-DEL-137
chr16-64770299-DEL-296
chr16-64774978-DEL-1142
chr16-64784904-DEL-208
chr16-64793294-DEL-177
chr16-64796035-DEL-11535
chr16-64799728-DEL-507
chr16-64836995-DEL-138
chr16-64850681-DEL-2899
chr16-64854841-DEL-1015
chr16-64854842-DEL-615
chr16-64855136-DEL-478
chr16-64857063-DEL-501
chr16-64868503-DEL-76
chr16-64876215-DEL-6667
chr16-64928680-DEL-57
chr16-64932355-DEL-54
chr16-64946206-DEL-95
chr16-64965431-DEL-320
chr16-64983830-DEL-87
chr16-64984623-DEL-224
chr16-65033200-DEL-95
chr16-65040375-DEL-2957
chr16-65047252-DEL-6137
chr16-65055625-DEL-1196
chr16-65057063-DEL-6130
chr16-65062505-DEL-245
chr16-65071756-DEL-93
chr16-65072030-DEL-438
chr16-65075495-DEL-89
chr16-65108898-DEL-117
chr16-65112706-DEL-6257
chr16-65119344-DEL-255
chr16-65125384-DEL-762
chr16-65150175-DEL-103
chr16-65158841-DEL-269
chr16-65162066-DEL-52
chr16-65164839-DEL-421
chr16-65168701-DEL-206
chr16-65169317-DEL-439
chr16-65228814-DEL-523
chr16-65242538-DEL-66
chr16-65256274-DEL-1878
chr16-65260111-DEL-64
chr16-65270638-DEL-82
chr16-65272373-DEL-96
chr16-65287445-DEL-122
chr16-65290023-DEL-73
chr16-65298037-DEL-51
chr16-65352815-DEL-70
chr16-65379003-DEL-57
chr16-65385076-DEL-173
chr16-65394091-DEL-62
chr16-65420106-DEL-82
chr16-65438903-DEL-91
chr16-65444138-DEL-67
chr16-65464825-DEL-53
chr16-65473000-DEL-217
chr16-65477898-DEL-679
chr16-65494252-DEL-681
chr16-65495761-DEL-518
chr16-65504983-DEL-146

chr16-65529117-DEL-56
chr16-65557762-DEL-1521
chr16-65671791-DEL-104
chr16-65692035-DEL-72
chr16-65694889-DEL-2118
chr16-65706740-DEL-186
chr16-65707127-DEL-378
chr16-65708501-DEL-1003
chr16-65710871-DEL-62
chr16-65725949-DEL-5742
chr16-65736590-DEL-205
chr16-65793603-DEL-64
chr16-65799614-DEL-588
chr16-65803457-DEL-62
chr16-65821868-DEL-62
chr16-65826436-DEL-455
chr16-65854265-DEL-3956
chr16-65876560-DEL-2768
chr16-65921332-DEL-53
chr16-65954853-DEL-825
chr16-65972921-DEL-650
chr16-65977461-DEL-1628
chr16-65987945-DEL-56
chr16-66003375-DEL-625
chr16-66006034-DEL-347
chr16-66016674-DEL-77
chr16-66018223-DEL-172
chr16-66026046-DEL-6497
chr16-66032918-DEL-74
chr16-66033475-DEL-1103
chr16-66037338-DEL-170
chr16-66038935-DEL-5480
chr16-66118224-DEL-775
chr16-66157605-DEL-214
chr16-66159973-DEL-1008
chr16-66164601-DEL-53
chr16-66167921-DEL-78
chr16-66181063-DEL-54
chr16-66191192-DEL-5734
chr16-66215035-DEL-93
chr16-66223221-DEL-6246
chr16-66250562-DEL-172
chr16-66300231-DEL-61
chr16-66320447-DEL-228
chr16-66321241-DEL-212
chr16-66358601-DEL-60
chr16-66365591-DEL-320
chr16-66372301-DEL-418
chr16-66405073-DEL-60
chr16-66406672-DEL-95
chr16-66467062-DEL-154
chr16-66502461-DEL-3398
chr16-66504837-DEL-370
chr16-66535110-DEL-642
chr16-66546437-DEL-537
chr16-66576804-DEL-119
chr16-66655218-DEL-52
chr16-66708185-DEL-87
chr16-66738902-DEL-255
chr16-66745609-DEL-419
chr16-66751196-DEL-72
chr16-66779242-DEL-58
chr16-66789891-DEL-60
chr16-66828661-DEL-60
chr16-66856811-DEL-216
chr16-66874148-DEL-82
chr16-66882752-DEL-51
chr16-66883963-DEL-310
chr16-66892409-DEL-234
chr16-66931253-DEL-73
chr16-66962595-DEL-2029
chr16-67006431-DEL-52
chr16-67033950-DEL-1115
chr16-67117888-DEL-1578
chr16-67120912-DEL-53
chr16-67123485-DEL-72
chr16-67153150-DEL-93

chr16-67165312-DEL-60
chr16-67181358-DEL-58
chr16-67194021-DEL-118
chr16-67205706-DEL-190
chr16-67213099-DEL-297
chr16-67217282-DEL-426
chr16-67225628-DEL-115
chr16-67241792-DEL-482
chr16-67259865-DEL-330
chr16-67294993-DEL-77
chr16-67355956-DEL-597
chr16-67450458-DEL-88
chr16-67453822-DEL-244
chr16-67456977-DEL-63
chr16-67536576-DEL-385
chr16-67550757-DEL-70
chr16-67574168-DEL-63
chr16-67579207-DEL-421
chr16-67630581-DEL-748
chr16-67662372-DEL-177
chr16-67662888-DEL-53
chr16-67664820-DEL-2979
chr16-67688712-DEL-185
chr16-67710190-DEL-359
chr16-67718025-DEL-55
chr16-67753671-DEL-619
chr16-67770239-DEL-1211
chr16-67798653-DEL-52
chr16-67800009-DEL-944
chr16-67805823-DEL-401
chr16-67822814-DEL-563
chr16-67829357-DEL-1654
chr16-67855975-DEL-247
chr16-67871182-DEL-194
chr16-67897336-DEL-217
chr16-67904775-DEL-72
chr16-67924616-DEL-373
chr16-67936615-DEL-64
chr16-67971048-DEL-99
chr16-67987637-DEL-456
chr16-67998256-DEL-4339
chr16-68009790-DEL-66
chr16-68018071-DEL-69
chr16-68060232-DEL-343
chr16-68071221-DEL-191
chr16-68094062-DEL-79
chr16-68097098-DEL-5947
chr16-68104160-DEL-178
chr16-68109651-DEL-173
chr16-68114645-DEL-203
chr16-68124887-DEL-140
chr16-68136005-DEL-6595
chr16-68172036-DEL-3238
chr16-68181608-DEL-391
chr16-68227943-DEL-54
chr16-68231357-DEL-63
chr16-68283847-DEL-107
chr16-68284944-DEL-54
chr16-68286630-DEL-5046
chr16-68298855-DEL-361
chr16-68307676-DEL-899
chr16-68326230-DEL-6327
chr16-68332465-DEL-163
chr16-68332788-DEL-1945
chr16-68334758-DEL-142
chr16-68338528-DEL-6532
chr16-68344744-DEL-214
chr16-68369855-DEL-53
chr16-68370296-DEL-2815
chr16-68382991-DEL-77
chr16-68389820-DEL-710
chr16-68390669-DEL-1000
chr16-68391747-DEL-424
chr16-68392244-DEL-1204
chr16-68396814-DEL-52
chr16-68407641-DEL-348
chr16-68413078-DEL-2382

chr16-68428016-DEL-129
chr16-68443833-DEL-73
chr16-68478224-DEL-56
chr16-68479387-DEL-328
chr16-68482949-DEL-92
chr16-68514826-DEL-157
chr16-68519715-DEL-692
chr16-68530689-DEL-89
chr16-68534375-DEL-7586
chr16-68542539-DEL-1738
chr16-68566885-DEL-111
chr16-68569523-DEL-1235
chr16-68620173-DEL-63
chr16-68632114-DEL-1485
chr16-68682185-DEL-221
chr16-68703224-DEL-1462
chr16-68710206-DEL-58
chr16-68712593-DEL-6386
chr16-68756210-DEL-111
chr16-68802280-DEL-105
chr16-68835075-DEL-1512
chr16-68910626-DEL-1466
chr16-68929218-DEL-1465
chr16-68947429-DEL-1725
chr16-68948720-DEL-61
chr16-68970912-DEL-442
chr16-69032329-DEL-56
chr16-69040934-DEL-705
chr16-69051036-DEL-162
chr16-69054307-DEL-68
chr16-69058810-DEL-63
chr16-69063363-DEL-462
chr16-69077093-DEL-1712
chr16-69091345-DEL-101
chr16-69091518-DEL-2175
chr16-69108722-DEL-58
chr16-69110107-DEL-549
chr16-69120160-DEL-3574
chr16-69125119-DEL-82
chr16-69125995-DEL-6205
chr16-69147154-DEL-152
chr16-69170284-DEL-55
chr16-69171473-DEL-66
chr16-69188579-DEL-6722
chr16-69188625-DEL-4816
chr16-69195479-DEL-60
chr16-69217877-DEL-2072
chr16-69240941-DEL-2393
chr16-69262829-DEL-99
chr16-69265426-DEL-70
chr16-69268398-DEL-53
chr16-69275679-DEL-93
chr16-69286689-DEL-1317
chr16-69295484-DEL-125
chr16-69313932-DEL-251
chr16-69317470-DEL-482
chr16-69327959-DEL-521
chr16-69328595-DEL-355
chr16-69358027-DEL-442
chr16-69382489-DEL-393
chr16-69400807-DEL-99
chr16-69401090-DEL-522
chr16-69402853-DEL-710
chr16-69416223-DEL-436
chr16-69426557-DEL-64
chr16-69436635-DEL-803
chr16-69476459-DEL-64
chr16-69485403-DEL-69
chr16-69485974-DEL-643
chr16-69494043-DEL-56
chr16-69502896-DEL-214
chr16-69515651-DEL-286
chr16-69546582-DEL-282
chr16-69580598-DEL-57
chr16-69584578-DEL-95
chr16-69613165-DEL-83
chr16-69619064-DEL-1654

chr16-69643970-DEL-55
chr16-69684203-DEL-56
chr16-69718024-DEL-93
chr16-69728331-DEL-241
chr16-69734952-DEL-1891
chr16-69741235-DEL-119
chr16-69751009-DEL-51
chr16-69779868-DEL-61
chr16-69786102-DEL-60
chr16-69798104-DEL-92
chr16-69825442-DEL-329
chr16-69827668-DEL-193
chr16-69915594-DEL-174
chr16-69926684-DEL-138
chr16-69931969-DEL-537
chr16-69941748-DEL-1071
chr16-69942411-DEL-672
chr16-69942989-DEL-146
chr16-69994829-DEL-181
chr16-70003852-DEL-55
chr16-70020714-DEL-163
chr16-70042174-DEL-520
chr16-70067563-DEL-5024
chr16-70078185-DEL-153
chr16-70084941-DEL-734
chr16-70097965-DEL-122
chr16-70118280-DEL-146
chr16-70271957-DEL-87
chr16-70276996-DEL-299
chr16-70279918-DEL-421
chr16-70304405-DEL-144
chr16-70349180-DEL-6784
chr16-70372897-DEL-399
chr16-70408367-DEL-3581
chr16-70415104-DEL-67
chr16-70430873-DEL-2251
chr16-70430873-DEL-91
chr16-70433145-DEL-3417
chr16-70464384-DEL-207
chr16-70480472-DEL-1283
chr16-70527985-DEL-55
chr16-70641571-DEL-196
chr16-70648839-DEL-62
chr16-70682323-DEL-172
chr16-70687277-DEL-1036
chr16-70688107-DEL-160
chr16-70700952-DEL-831
chr16-70743744-DEL-290
chr16-70748283-DEL-158
chr16-70757952-DEL-120
chr16-70763389-DEL-579
chr16-70779851-DEL-214
chr16-70799184-DEL-595
chr16-70815290-DEL-74
chr16-70819835-DEL-2192
chr16-70824679-DEL-2421
chr16-70831151-DEL-271
chr16-70831995-DEL-77
chr16-70835404-DEL-1029
chr16-70848736-DEL-1585
chr16-70858834-DEL-85
chr16-70897875-DEL-165
chr16-70898767-DEL-310
chr16-70925755-DEL-59
chr16-70941094-DEL-158
chr16-70960631-DEL-1157
chr16-70986759-DEL-1132
chr16-71042233-DEL-52
chr16-71072108-DEL-485
chr16-71072847-DEL-118
chr16-71080449-DEL-57
chr16-71104693-DEL-2237
chr16-71132200-DEL-58
chr16-71154978-DEL-2148
chr16-71172368-DEL-55
chr16-71182107-DEL-232
chr16-71190769-DEL-7391

chr16-71206825-DEL-54
chr16-71208152-DEL-6141
chr16-71219090-DEL-52
chr16-71224750-DEL-541
chr16-71249621-DEL-96
chr16-71252777-DEL-419
chr16-71278295-DEL-6905
chr16-71278680-DEL-208
chr16-71290770-DEL-334
chr16-71301599-DEL-68
chr16-71315780-DEL-753
chr16-71332379-DEL-285
chr16-71336837-DEL-56
chr16-71344466-DEL-142
chr16-71371082-DEL-161
chr16-71381524-DEL-146
chr16-71386026-DEL-135
chr16-71388412-DEL-3287
chr16-71402887-DEL-354
chr16-71406652-DEL-150
chr16-71421246-DEL-217
chr16-71442882-DEL-105
chr16-71445012-DEL-53
chr16-71453632-DEL-71
chr16-71463091-DEL-74
chr16-71469585-DEL-788
chr16-71504805-DEL-74
chr16-71510441-DEL-1822
chr16-71542019-DEL-142
chr16-71555120-DEL-466
chr16-71575685-DEL-167
chr16-71616685-DEL-140
chr16-71656868-DEL-62
chr16-71679821-DEL-1343
chr16-71691514-DEL-166
chr16-71696881-DEL-87
chr16-71734597-DEL-105
chr16-71760236-DEL-123
chr16-71764122-DEL-51
chr16-71769864-DEL-226
chr16-71772720-DEL-4183
chr16-71789964-DEL-53
chr16-71837899-DEL-180
chr16-71846020-DEL-522
chr16-71890044-DEL-1753
chr16-71893725-DEL-421
chr16-71894551-DEL-204
chr16-71903512-DEL-178
chr16-71920837-DEL-2233
chr16-71965537-DEL-192
chr16-71968700-DEL-1433
chr16-71988983-DEL-2205
chr16-72008616-DEL-85
chr16-72019889-DEL-58
chr16-72031816-DEL-95
chr16-72040427-DEL-60
chr16-72108936-DEL-163
chr16-72109775-DEL-720
chr16-72165386-DEL-6668
chr16-72172790-DEL-137
chr16-72174806-DEL-71
chr16-72238938-DEL-58
chr16-72262981-DEL-76
chr16-72277363-DEL-135
chr16-72296384-DEL-695
chr16-72313978-DEL-129
chr16-72356389-DEL-104
chr16-72377025-DEL-92
chr16-72440001-DEL-1919
chr16-72440451-DEL-234
chr16-72449139-DEL-56
chr16-72468645-DEL-109
chr16-72484876-DEL-73
chr16-72545775-DEL-4793
chr16-72610297-DEL-100
chr16-72642584-DEL-1367
chr16-72657479-DEL-78

chr16-72752340-DEL-122
chr16-72766391-DEL-64
chr16-72767677-DEL-117
chr16-72796441-DEL-182
chr16-72808216-DEL-105
chr16-72856466-DEL-92
chr16-72871480-DEL-57
chr16-72871787-DEL-257
chr16-72873789-DEL-71
chr16-72876940-DEL-79
chr16-72943727-DEL-1277
chr16-72955767-DEL-117
chr16-72986837-DEL-51
chr16-73036959-DEL-603
chr16-73049452-DEL-81
chr16-73059255-DEL-90
chr16-73064878-DEL-55
chr16-73081748-DEL-55
chr16-73102561-DEL-560
chr16-73129990-DEL-198
chr16-73145022-DEL-60
chr16-73151240-DEL-6295
chr16-73151336-DEL-214
chr16-73153911-DEL-3669
chr16-73161765-DEL-153
chr16-73323188-DEL-185
chr16-73387498-DEL-533
chr16-73402038-DEL-105
chr16-73431257-DEL-674
chr16-73447452-DEL-3079
chr16-73463127-DEL-66
chr16-73501397-DEL-372
chr16-73504706-DEL-471
chr16-73517705-DEL-244
chr16-73573568-DEL-463
chr16-73595465-DEL-2126
chr16-73630177-DEL-93
chr16-73638379-DEL-56
chr16-73665303-DEL-200
chr16-73680269-DEL-52
chr16-73695472-DEL-145
chr16-73714649-DEL-58
chr16-73755530-DEL-229
chr16-73862655-DEL-164
chr16-73931109-DEL-72
chr16-73971669-DEL-328
chr16-73992184-DEL-56
chr16-74036734-DEL-553
chr16-74046867-DEL-94
chr16-74067873-DEL-79
chr16-74068477-DEL-119
chr16-74094794-DEL-348
chr16-74099254-DEL-241
chr16-74122795-DEL-376
chr16-74140796-DEL-251
chr16-74173517-DEL-200
chr16-74191448-DEL-385
chr16-74213656-DEL-297
chr16-74219118-DEL-109
chr16-74258396-DEL-60
chr16-74269129-DEL-155
chr16-74304609-DEL-760
chr16-74321694-DEL-65
chr16-74356853-DEL-53
chr16-74386011-DEL-107
chr16-74407412-DEL-95
chr16-74422472-DEL-55
chr16-74429807-DEL-74
chr16-74468879-DEL-331
chr16-74474952-DEL-54
chr16-74483403-DEL-909
chr16-74484740-DEL-824
chr16-74526319-DEL-459
chr16-74532884-DEL-97
chr16-74556991-DEL-183
chr16-74591121-DEL-96
chr16-74598823-DEL-786

chr16-74613499-DEL-132
chr16-74631614-DEL-370
chr16-74650884-DEL-262
chr16-74655295-DEL-66
chr16-74661869-DEL-52
chr16-74689541-DEL-402
chr16-74699363-DEL-172
chr16-74706206-DEL-653
chr16-74712673-DEL-74
chr16-74720322-DEL-116
chr16-74722731-DEL-235
chr16-74751644-DEL-54
chr16-74756494-DEL-119
chr16-74784168-DEL-132
chr16-74784838-DEL-258
chr16-74800328-DEL-3035
chr16-74833694-DEL-56
chr16-74851142-DEL-868
chr16-74861437-DEL-60
chr16-74870950-DEL-461
chr16-74907430-DEL-1425
chr16-74912617-DEL-660
chr16-74923171-DEL-56
chr16-74935445-DEL-78
chr16-74946026-DEL-158
chr16-74956298-DEL-973
chr16-74986313-DEL-82
chr16-74987224-DEL-60
chr16-74993386-DEL-390
chr16-75021318-DEL-2439
chr16-75033792-DEL-236
chr16-75036454-DEL-477
chr16-75043519-DEL-614
chr16-75065246-DEL-173
chr16-75070610-DEL-126
chr16-75087003-DEL-52
chr16-75088760-DEL-184
chr16-75102091-DEL-2013
chr16-75102241-DEL-171
chr16-75102550-DEL-245
chr16-75103917-DEL-342
chr16-75113515-DEL-87
chr16-75143722-DEL-332
chr16-75179900-DEL-848
chr16-75192355-DEL-141
chr16-75231385-DEL-404
chr16-75256547-DEL-104
chr16-75278479-DEL-801
chr16-75305515-DEL-52
chr16-75306708-DEL-62
chr16-75349794-DEL-234
chr16-75385831-DEL-1474
chr16-75417091-DEL-1914
chr16-75424421-DEL-55
chr16-75452710-DEL-84
chr16-75461091-DEL-2301
chr16-75462853-DEL-1183
chr16-75476348-DEL-59
chr16-75524537-DEL-6436
chr16-75533641-DEL-6598
chr16-75566696-DEL-174
chr16-75595693-DEL-6299
chr16-75643838-DEL-220
chr16-75652957-DEL-124
chr16-75692913-DEL-138
chr16-75696687-DEL-84
chr16-75743307-DEL-91
chr16-75780496-DEL-87
chr16-75785761-DEL-710
chr16-75807599-DEL-94
chr16-75816442-DEL-55
chr16-75832164-DEL-72
chr16-75841715-DEL-60
chr16-75867557-DEL-60
chr16-75884919-DEL-100
chr16-75887320-DEL-61
chr16-75917194-DEL-53

chr16-75922026-DEL-121
chr16-75943993-DEL-733
chr16-75986911-DEL-181
chr16-75988528-DEL-174
chr16-76002101-DEL-76
chr16-76060575-DEL-51
chr16-76064759-DEL-63
chr16-76066744-DEL-108
chr16-76083832-DEL-72
chr16-76099296-DEL-67
chr16-76188783-DEL-789
chr16-76253467-DEL-83
chr16-76277245-DEL-69
chr16-76277863-DEL-55
chr16-76302862-DEL-73
chr16-76341494-DEL-126
chr16-76357750-DEL-671
chr16-76360093-DEL-56
chr16-76404592-DEL-132
chr16-76411007-DEL-57
chr16-76414162-DEL-52
chr16-76442269-DEL-3449
chr16-76444752-DEL-650
chr16-76459735-DEL-148
chr16-76474754-DEL-54
chr16-76483632-DEL-357
chr16-76544901-DEL-456
chr16-76547618-DEL-117
chr16-76584446-DEL-175
chr16-76609452-DEL-58
chr16-76614504-DEL-52
chr16-76628986-DEL-61
chr16-76694017-DEL-52
chr16-76710875-DEL-77
chr16-76739170-DEL-23367
chr16-76739800-DEL-400
chr16-76772375-DEL-399
chr16-76776261-DEL-243
chr16-76779308-DEL-73
chr16-76780498-DEL-65
chr16-76785531-DEL-3212
chr16-76946810-DEL-133
chr16-76958806-DEL-54
chr16-77004384-DEL-891
chr16-77015637-DEL-194
chr16-77019437-DEL-3131
chr16-77025784-DEL-1921
chr16-77041061-DEL-493
chr16-77054796-DEL-142
chr16-77098138-DEL-1218
chr16-77099367-DEL-1175
chr16-77138473-DEL-60
chr16-77162713-DEL-74
chr16-77172801-DEL-2130
chr16-77175054-DEL-67
chr16-77219779-DEL-4980
chr16-77377549-DEL-328
chr16-77417899-DEL-72
chr16-77477878-DEL-686
chr16-77480446-DEL-430
chr16-77499276-DEL-81
chr16-77540424-DEL-67
chr16-77598294-DEL-77
chr16-77611902-DEL-213
chr16-77620071-DEL-80
chr16-77622181-DEL-2126
chr16-77633606-DEL-542
chr16-77638638-DEL-669
chr16-77652587-DEL-281
chr16-77680128-DEL-61
chr16-77725274-DEL-210
chr16-77752393-DEL-57
chr16-77763922-DEL-394
chr16-77771033-DEL-73
chr16-77772742-DEL-136
chr16-77773333-DEL-513
chr16-77779674-DEL-72

chr16-77782290-DEL-196
chr16-77792623-DEL-939
chr16-77793573-DEL-313
chr16-77840821-DEL-235
chr16-77846720-DEL-70
chr16-77850960-DEL-789
chr16-77856954-DEL-93
chr16-77870725-DEL-58
chr16-77871666-DEL-112
chr16-77874571-DEL-501
chr16-77875158-DEL-56
chr16-77905434-DEL-119
chr16-77996601-DEL-359
chr16-77997903-DEL-190
chr16-78014243-DEL-60
chr16-78046646-DEL-295
chr16-78047225-DEL-153
chr16-78049671-DEL-226
chr16-78067954-DEL-86
chr16-78089044-DEL-87
chr16-78104492-DEL-137
chr16-78117892-DEL-4075
chr16-78196797-DEL-63
chr16-78208199-DEL-61
chr16-78233737-DEL-88
chr16-78247990-DEL-115
chr16-78253876-DEL-79
chr16-78257553-DEL-3790
chr16-78280145-DEL-363
chr16-78298333-DEL-1192
chr16-78301376-DEL-61
chr16-78331387-DEL-53
chr16-78392453-DEL-2935
chr16-78436433-DEL-58
chr16-78440061-DEL-147
chr16-78458569-DEL-60
chr16-78482931-DEL-126
chr16-78484048-DEL-596
chr16-78489651-DEL-951
chr16-78495583-DEL-150
chr16-78515717-DEL-53
chr16-78550187-DEL-576
chr16-78589570-DEL-234
chr16-78600961-DEL-458
chr16-78610198-DEL-220
chr16-78613316-DEL-450
chr16-78653510-DEL-754
chr16-78658293-DEL-69
chr16-78659412-DEL-65
chr16-78666736-DEL-1389
chr16-78683889-DEL-2838
chr16-78689470-DEL-84
chr16-78696471-DEL-360
chr16-78708866-DEL-265
chr16-78755255-DEL-66
chr16-78768930-DEL-67
chr16-78781925-DEL-71
chr16-78784137-DEL-61
chr16-78837463-DEL-199
chr16-78845036-DEL-2101
chr16-78866572-DEL-83
chr16-78893593-DEL-256
chr16-78898499-DEL-77
chr16-78912442-DEL-5208
chr16-78921043-DEL-503
chr16-78943165-DEL-125
chr16-78950462-DEL-84
chr16-78952427-DEL-56
chr16-78954854-DEL-409
chr16-78963036-DEL-1084
chr16-79021863-DEL-200
chr16-79025064-DEL-163
chr16-79072529-DEL-354
chr16-79119697-DEL-201
chr16-79202246-DEL-160
chr16-79207238-DEL-118
chr16-79209477-DEL-161

chr16-79210843-DEL-782
chr16-79221797-DEL-693
chr16-79255097-DEL-2313
chr16-79257475-DEL-2826
chr16-79280044-DEL-818
chr16-79312251-DEL-346
chr16-79333682-DEL-419
chr16-79365610-DEL-96
chr16-79378670-DEL-279
chr16-79405913-DEL-434
chr16-79408638-DEL-1575
chr16-79419949-DEL-54
chr16-79421001-DEL-51
chr16-79425878-DEL-73
chr16-79433057-DEL-184
chr16-79446751-DEL-1080
chr16-79512869-DEL-63
chr16-79552911-DEL-547
chr16-79566621-DEL-115
chr16-79573973-DEL-197
chr16-79600956-DEL-155
chr16-79605449-DEL-93
chr16-79621469-DEL-599
chr16-79628885-DEL-327
chr16-79633826-DEL-240
chr16-79642879-DEL-260
chr16-79660768-DEL-84
chr16-79669016-DEL-4651
chr16-79678206-DEL-402
chr16-79703893-DEL-8343
chr16-79719475-DEL-970
chr16-79728034-DEL-355
chr16-79736005-DEL-68
chr16-79748688-DEL-916
chr16-79755020-DEL-151
chr16-79772675-DEL-787
chr16-79788272-DEL-831
chr16-79795726-DEL-2897
chr16-79801256-DEL-3081
chr16-79807204-DEL-91
chr16-79828756-DEL-64
chr16-79837280-DEL-441
chr16-79852276-DEL-62
chr16-79868681-DEL-378
chr16-79889119-DEL-85
chr16-79890371-DEL-953
chr16-79914591-DEL-115
chr16-79917169-DEL-1045
chr16-79925731-DEL-109
chr16-79960316-DEL-71
chr16-79967211-DEL-773
chr16-79982529-DEL-282
chr16-79994968-DEL-117
chr16-80003399-DEL-1402
chr16-80033614-DEL-312
chr16-80038727-DEL-571
chr16-80049185-DEL-75
chr16-80061208-DEL-126
chr16-80064977-DEL-2648
chr16-80084708-DEL-7115
chr16-80095827-DEL-185
chr16-80101274-DEL-5304
chr16-80118747-DEL-417
chr16-80151846-DEL-511
chr16-80155160-DEL-408
chr16-80161608-DEL-657
chr16-80165167-DEL-54
chr16-80175131-DEL-68
chr16-80176898-DEL-412
chr16-80236247-DEL-644
chr16-80238243-DEL-529
chr16-80246530-DEL-1364
chr16-80253689-DEL-52
chr16-80288351-DEL-283
chr16-80343772-DEL-359
chr16-80383644-DEL-79
chr16-80390865-DEL-60

chr16-80397759-DEL-859
chr16-80411808-DEL-6040
chr16-80420350-DEL-114
chr16-80429055-DEL-76
chr16-80474197-DEL-51
chr16-80499930-DEL-74
chr16-80521998-DEL-2609
chr16-80536070-DEL-200
chr16-80540879-DEL-232
chr16-80612481-DEL-60
chr16-80651083-DEL-61
chr16-80669573-DEL-1177
chr16-80675324-DEL-303
chr16-80681815-DEL-139
chr16-80706670-DEL-1089
chr16-80745033-DEL-52
chr16-80788060-DEL-752
chr16-80799916-DEL-152
chr16-80802877-DEL-1349
chr16-80810941-DEL-125
chr16-80826254-DEL-6969
chr16-80832440-DEL-214
chr16-80839172-DEL-3029
chr16-80848074-DEL-56
chr16-80849875-DEL-720
chr16-80868808-DEL-127
chr16-80870462-DEL-1888
chr16-80875763-DEL-127
chr16-80905090-DEL-54
chr16-80917302-DEL-299
chr16-80948513-DEL-2532
chr16-80953149-DEL-80
chr16-80958456-DEL-109
chr16-80978295-DEL-52
chr16-81007651-DEL-168
chr16-81022873-DEL-357
chr16-81064158-DEL-224
chr16-81078796-DEL-690
chr16-81084392-DEL-833
chr16-81142509-DEL-301
chr16-81165779-DEL-60
chr16-81177070-DEL-2508
chr16-81179618-DEL-1634
chr16-81185241-DEL-290
chr16-81186179-DEL-474
chr16-81222356-DEL-3111
chr16-81291880-DEL-452
chr16-81295151-DEL-251
chr16-81313442-DEL-403
chr16-81338310-DEL-4298
chr16-81408432-DEL-281
chr16-81447471-DEL-558
chr16-81455560-DEL-6467
chr16-81461993-DEL-138
chr16-81480598-DEL-572
chr16-81483272-DEL-6194
chr16-81503610-DEL-202
chr16-81522375-DEL-133
chr16-81533958-DEL-118
chr16-81536962-DEL-4285
chr16-81609156-DEL-264
chr16-81625746-DEL-62
chr16-81637537-DEL-873
chr16-81671060-DEL-56
chr16-81696381-DEL-152
chr16-81730313-DEL-61
chr16-81796588-DEL-975
chr16-81799574-DEL-183
chr16-81838753-DEL-6377
chr16-81852529-DEL-92
chr16-81858287-DEL-60
chr16-81898340-DEL-119
chr16-81908926-DEL-55
chr16-81921816-DEL-85
chr16-81938671-DEL-79
chr16-81958671-DEL-943
chr16-81959859-DEL-271

chr16-81962413-DEL-856
chr16-81963906-DEL-205
chr16-81985120-DEL-948
chr16-81987334-DEL-130
chr16-81998939-DEL-224
chr16-82002474-DEL-54
chr16-82026281-DEL-55
chr16-82042008-DEL-78
chr16-82053771-DEL-505
chr16-82073553-DEL-3195
chr16-82075269-DEL-944
chr16-82083033-DEL-58
chr16-82084541-DEL-582
chr16-82097651-DEL-3077
chr16-82123241-DEL-1219
chr16-82128189-DEL-74
chr16-82146489-DEL-575
chr16-82195655-DEL-2453
chr16-82206814-DEL-208
chr16-82211033-DEL-197
chr16-82216907-DEL-150
chr16-82218440-DEL-236
chr16-82220847-DEL-350
chr16-82226621-DEL-130
chr16-82230736-DEL-95
chr16-82242575-DEL-478
chr16-82262323-DEL-111
chr16-82265075-DEL-919
chr16-82268606-DEL-116
chr16-82274027-DEL-82
chr16-82278528-DEL-549
chr16-82303313-DEL-54
chr16-82304515-DEL-212
chr16-82311038-DEL-102124
chr16-82343565-DEL-938
chr16-82360752-DEL-436
chr16-82400956-DEL-181
chr16-82411041-DEL-60
chr16-82415719-DEL-374
chr16-82422003-DEL-53
chr16-82429812-DEL-9377
chr16-82480294-DEL-908
chr16-82482650-DEL-131
chr16-82508963-DEL-132
chr16-82518475-DEL-660
chr16-82521789-DEL-2793
chr16-82523920-DEL-81
chr16-82538409-DEL-3020
chr16-82545657-DEL-182
chr16-82559373-DEL-209
chr16-82565828-DEL-3807
chr16-82575606-DEL-860
chr16-82587975-DEL-76
chr16-82601180-DEL-65
chr16-82605309-DEL-60
chr16-82632064-DEL-121
chr16-82640798-DEL-1784
chr16-82660074-DEL-554
chr16-82664541-DEL-387
chr16-82673290-DEL-684
chr16-82699490-DEL-315
chr16-82702732-DEL-58
chr16-82703717-DEL-76
chr16-82715576-DEL-204
chr16-82716925-DEL-271
chr16-82742554-DEL-118
chr16-82743745-DEL-395
chr16-82760490-DEL-97
chr16-82763752-DEL-251
chr16-82778545-DEL-745
chr16-82783224-DEL-1273
chr16-82791786-DEL-58
chr16-82804967-DEL-185
chr16-82809068-DEL-1792
chr16-82847200-DEL-4975
chr16-82855164-DEL-132
chr16-82869234-DEL-2843

chr16-82869538-DEL-273
chr16-82872132-DEL-2005
chr16-82887527-DEL-1570
chr16-82897001-DEL-227
chr16-82906507-DEL-114
chr16-82910697-DEL-438
chr16-82923800-DEL-748
chr16-82934201-DEL-6136
chr16-82955004-DEL-3194
chr16-82996619-DEL-324
chr16-83009240-DEL-951
chr16-83038844-DEL-240
chr16-83039472-DEL-75
chr16-83093370-DEL-65
chr16-83106993-DEL-256
chr16-83111339-DEL-109
chr16-83129136-DEL-1877
chr16-83150566-DEL-1699
chr16-83150760-DEL-836
chr16-83184575-DEL-326
chr16-83189753-DEL-463
chr16-83206718-DEL-315
chr16-83226938-DEL-388
chr16-83254565-DEL-60
chr16-83257254-DEL-918
chr16-83293090-DEL-333
chr16-83295883-DEL-6104
chr16-83302445-DEL-57
chr16-83338047-DEL-71
chr16-83340704-DEL-63
chr16-83347514-DEL-302
chr16-83359186-DEL-287
chr16-83365631-DEL-56
chr16-83375042-DEL-1416
chr16-83457317-DEL-103
chr16-83487250-DEL-2092
chr16-83502707-DEL-75
chr16-83514970-DEL-112
chr16-83520912-DEL-209
chr16-83527409-DEL-68
chr16-83635031-DEL-1760
chr16-83649899-DEL-469
chr16-83667413-DEL-59
chr16-83683414-DEL-403
chr16-83747195-DEL-6443
chr16-83762294-DEL-196
chr16-83763306-DEL-60
chr16-83779881-DEL-1104
chr16-83794015-DEL-56
chr16-83795923-DEL-390
chr16-83801409-DEL-51
chr16-83801662-DEL-7586
chr16-83821886-DEL-96
chr16-83826181-DEL-68
chr16-83846173-DEL-71
chr16-83863404-DEL-57
chr16-83864153-DEL-546
chr16-83867782-DEL-304
chr16-83877629-DEL-4400
chr16-83878893-DEL-2652
chr16-83881574-DEL-921
chr16-83885057-DEL-493
chr16-83888368-DEL-76
chr16-83892450-DEL-2455
chr16-83892454-DEL-1614
chr16-83894140-DEL-763
chr16-83952288-DEL-217
chr16-83989080-DEL-84
chr16-83996668-DEL-370
chr16-84004101-DEL-55
chr16-84010481-DEL-213
chr16-84037311-DEL-58
chr16-84040716-DEL-907
chr16-84042762-DEL-124
chr16-84093310-DEL-469
chr16-84162001-DEL-68
chr16-84165200-DEL-74

chr16-84174693-DEL-168
chr16-84193415-DEL-62
chr16-84202989-DEL-1277
chr16-84203059-DEL-335
chr16-84203857-DEL-538
chr16-84203961-DEL-236
chr16-84263225-DEL-2262
chr16-84287105-DEL-571
chr16-84328854-DEL-262
chr16-84341691-DEL-898
chr16-84360246-DEL-139
chr16-84361285-DEL-51
chr16-84370298-DEL-264
chr16-84395800-DEL-3357
chr16-84399767-DEL-146
chr16-84408115-DEL-1394
chr16-84445888-DEL-1081
chr16-84453585-DEL-499
chr16-84483060-DEL-436
chr16-84485696-DEL-247
chr16-84490832-DEL-134
chr16-84521192-DEL-557
chr16-84523113-DEL-57
chr16-84541042-DEL-209
chr16-84565506-DEL-231
chr16-84594536-DEL-201
chr16-84600860-DEL-60
chr16-84604740-DEL-164
chr16-84608842-DEL-85
chr16-84612366-DEL-545
chr16-84648078-DEL-160
chr16-84710773-DEL-2533
chr16-84722377-DEL-51
chr16-84729256-DEL-353
chr16-84739199-DEL-424
chr16-84743268-DEL-385
chr16-84744711-DEL-382
chr16-84755351-DEL-552
chr16-84762469-DEL-70
chr16-84790400-DEL-58
chr16-84792896-DEL-82
chr16-84794047-DEL-90
chr16-84804640-DEL-566
chr16-84805613-DEL-75
chr16-84858031-DEL-96
chr16-84869644-DEL-785
chr16-84877046-DEL-125
chr16-84883634-DEL-135
chr16-84935296-DEL-153
chr16-84936245-DEL-121
chr16-84948177-DEL-109
chr16-84958600-DEL-353
chr16-84959043-DEL-63
chr16-84986037-DEL-306
chr16-85019887-DEL-78
chr16-85024376-DEL-514
chr16-85031275-DEL-12633
chr16-85043834-DEL-221
chr16-85051961-DEL-54
chr16-85053137-DEL-54
chr16-85054295-DEL-52
chr16-85063813-DEL-3230
chr16-85110551-DEL-200
chr16-85112174-DEL-346
chr16-85117458-DEL-100
chr16-85124718-DEL-418
chr16-85160102-DEL-141
chr16-85160975-DEL-265
chr16-85163858-DEL-397
chr16-85179467-DEL-114
chr16-85240759-DEL-420
chr16-85242210-DEL-82
chr16-85246539-DEL-52
chr16-85247030-DEL-2495
chr16-85247082-DEL-2462
chr16-85253508-DEL-65
chr16-85267865-DEL-168

chr16-85358748-DEL-401
chr16-85361053-DEL-140
chr16-85422775-DEL-78
chr16-85424604-DEL-132
chr16-85454051-DEL-1156
chr16-85455260-DEL-857
chr16-85476392-DEL-530
chr16-85486086-DEL-97
chr16-85497425-DEL-215
chr16-85545452-DEL-534
chr16-85568413-DEL-52
chr16-85569699-DEL-59
chr16-85578750-DEL-1652
chr16-85585182-DEL-117
chr16-85603163-DEL-60
chr16-85605475-DEL-1553
chr16-85610342-DEL-77
chr16-85620231-DEL-68
chr16-85641670-DEL-89
chr16-85690483-DEL-60
chr16-85868591-DEL-51
chr16-85891113-DEL-53
chr16-85969996-DEL-5402
chr16-86055767-DEL-1195
chr16-86070275-DEL-119
chr16-86083817-DEL-1423
chr16-86096854-DEL-221
chr16-86100681-DEL-2380
chr16-86101123-DEL-267
chr16-86109239-DEL-166
chr16-86110638-DEL-2255
chr16-86123887-DEL-365
chr16-86143761-DEL-76
chr16-86162750-DEL-2252
chr16-86188930-DEL-708
chr16-86191542-DEL-304
chr16-86205206-DEL-2788
chr16-86213589-DEL-193
chr16-86236069-DEL-72
chr16-86242952-DEL-569
chr16-86266856-DEL-194
chr16-86268597-DEL-768
chr16-86273171-DEL-51
chr16-86278783-DEL-186
chr16-86294908-DEL-91
chr16-86320436-DEL-295
chr16-86321806-DEL-970
chr16-86326812-DEL-163
chr16-86346776-DEL-130
chr16-86412565-DEL-393
chr16-86459245-DEL-184
chr16-86474733-DEL-5991
chr16-86513728-DEL-1328
chr16-86544419-DEL-110
chr16-86558474-DEL-391
chr16-86559367-DEL-99
chr16-86570618-DEL-648
chr16-86575469-DEL-70
chr16-86584521-DEL-80
chr16-86586374-DEL-1437
chr16-86598421-DEL-757
chr16-86599669-DEL-63
chr16-86605071-DEL-271
chr16-86655227-DEL-606
chr16-86660077-DEL-2770
chr16-86705218-DEL-388
chr16-86707063-DEL-52
chr16-86782790-DEL-1087
chr16-86808793-DEL-541
chr16-86867772-DEL-693
chr16-86909038-DEL-212
chr16-86924861-DEL-127
chr16-86929539-DEL-57
chr16-86944241-DEL-1688
chr16-86962068-DEL-204
chr16-87027843-DEL-307
chr16-87040284-DEL-87

chr16-87147713-DEL-1182
chr16-87168095-DEL-90
chr16-87169348-DEL-444
chr16-87170265-DEL-1666
chr16-87170284-DEL-378
chr16-87171980-DEL-1120
chr16-87183011-DEL-535
chr16-87183750-DEL-84
chr16-87246657-DEL-486
chr16-87253687-DEL-342
chr16-87286756-DEL-430
chr16-87289343-DEL-75
chr16-87302652-DEL-72
chr16-87305506-DEL-776
chr16-87305522-DEL-1670
chr16-87306099-DEL-355
chr16-87306752-DEL-455
chr16-87389484-DEL-161
chr16-87397214-DEL-625
chr16-87431619-DEL-150
chr16-87445988-DEL-107
chr16-87539249-DEL-85
chr16-87547761-DEL-2011
chr16-87592205-DEL-349
chr16-87600607-DEL-51
chr16-87618369-DEL-53
chr16-87635985-DEL-784
chr16-87638979-DEL-399
chr16-87651591-DEL-109
chr16-87677697-DEL-130
chr16-87711781-DEL-72
chr16-87745155-DEL-739
chr16-87777079-DEL-66
chr16-87798948-DEL-56
chr16-87802282-DEL-419
chr16-87803446-DEL-55
chr16-87833571-DEL-57
chr16-87908505-DEL-52
chr16-87910861-DEL-74
chr16-88003138-DEL-68
chr16-88029346-DEL-60
chr16-88043337-DEL-60
chr16-88076358-DEL-824
chr16-88088628-DEL-63
chr16-88150844-DEL-65
chr16-88164727-DEL-52
chr16-88185395-DEL-148
chr16-88186593-DEL-84
chr16-88210485-DEL-64
chr16-88215352-DEL-2454
chr16-88236324-DEL-120
chr16-88246846-DEL-2481
chr16-88251569-DEL-1126
chr16-88255073-DEL-183
chr16-88264542-DEL-131
chr16-88265979-DEL-527
chr16-88316672-DEL-237
chr16-88362892-DEL-54
chr16-88414104-DEL-86
chr16-88420230-DEL-2159
chr16-88433149-DEL-82
chr16-88442052-DEL-516
chr16-88445719-DEL-458
chr16-88446594-DEL-136
chr16-88448243-DEL-63
chr16-88483212-DEL-1184
chr16-88505442-DEL-135
chr16-88522250-DEL-68
chr16-88530587-DEL-53
chr16-88547177-DEL-66
chr16-88563180-DEL-1155
chr16-88593458-DEL-111
chr16-88646023-DEL-93
chr16-88646727-DEL-54
chr16-88668617-DEL-1058
chr16-88686873-DEL-356
chr16-88687034-DEL-1346

chr16-88696979-DEL-374
chr16-88709407-DEL-172
chr16-88723913-DEL-659
chr16-88783249-DEL-107
chr16-88784297-DEL-284
chr16-88806234-DEL-165
chr16-88809220-DEL-578
chr16-88821065-DEL-2402
chr16-88826842-DEL-151
chr16-88830171-DEL-137
chr16-88844958-DEL-646
chr16-88880333-DEL-204
chr16-88883972-DEL-1262
chr16-88885259-DEL-4155
chr16-88890600-DEL-1369
chr16-88896903-DEL-87
chr16-88900779-DEL-1472
chr16-88905262-DEL-2368
chr16-88923792-DEL-54
chr16-88952877-DEL-400
chr16-88973748-DEL-1896
chr16-88978470-DEL-183
chr16-88981249-DEL-2629
chr16-88986578-DEL-6474
chr16-89004497-DEL-55
chr16-89005553-DEL-106
chr16-89006700-DEL-69
chr16-89021149-DEL-643
chr16-89038695-DEL-55
chr16-89076317-DEL-587
chr16-89108906-DEL-918
chr16-89108907-DEL-160
chr16-89126033-DEL-5263
chr16-89131439-DEL-962
chr16-89133475-DEL-178
chr16-89133972-DEL-895
chr16-89150587-DEL-999
chr16-89153364-DEL-834
chr16-89159169-DEL-369
chr16-89161117-DEL-126
chr16-89192960-DEL-79
chr16-89207414-DEL-302
chr16-89215377-DEL-220
chr16-89222850-DEL-566
chr16-89224652-DEL-131
chr16-89230002-DEL-2174
chr16-89254215-DEL-998
chr16-89264561-DEL-72
chr16-89281099-DEL-51
chr16-89285330-DEL-58
chr16-89314543-DEL-97
chr16-89338442-DEL-58
chr16-89350368-DEL-89
chr16-89352985-DEL-487
chr16-89358189-DEL-95
chr16-89388430-DEL-95
chr16-89399876-DEL-63
chr16-89432862-DEL-7046
chr16-89432897-DEL-856
chr16-89432941-DEL-214
chr16-89441005-DEL-60
chr16-89442147-DEL-93
chr16-89446856-DEL-133
chr16-89448540-DEL-66
chr16-89453583-DEL-57
chr16-89473100-DEL-58
chr16-89484992-DEL-173
chr16-89528591-DEL-51
chr16-89538164-DEL-108
chr16-89554914-DEL-84
chr16-89557963-DEL-84
chr16-89574045-DEL-131
chr16-89634991-DEL-80
chr16-89675092-DEL-400
chr16-89686780-DEL-204
chr16-89701998-DEL-941
chr16-89702336-DEL-293

chr16-89714715-DEL-320
chr16-89730190-DEL-92
chr16-89735452-DEL-216
chr16-89748447-DEL-84
chr16-89759196-DEL-2626
chr16-89760375-DEL-200
chr16-89760861-DEL-550
chr16-89763142-DEL-1094
chr16-89767586-DEL-424
chr16-89799665-DEL-70
chr16-89879072-DEL-71
chr16-89893538-DEL-70
chr16-89899571-DEL-79
chr16-89907885-DEL-164
chr16-89912432-DEL-206
chr16-89920970-DEL-370
chr16-89929060-DEL-89
chr16-89930308-DEL-97
chr16-89946024-DEL-112
chr16-89956178-DEL-58
chr16-89958668-DEL-64
chr16-89985830-DEL-480
chr16-89991204-DEL-559
chr16-90008216-DEL-70
chr16-90040258-DEL-111
chr16-90057754-DEL-317
chr16-90066806-DEL-213
chr16-90082824-DEL-207
chr16-90095201-DEL-52
chr16-90108230-DEL-164
chr16-90123032-DEL-207
chr16-90163267-DEL-162
chr16-90177708-DEL-108
chr16-90179724-DEL-99
chr16-90211237-DEL-214
chr16-90211823-DEL-110
chr16-90212371-DEL-70
chr16-90221834-DEL-169
chr16-90224962-DEL-130
chr16-90226555-DEL-122
chr16-90230159-DEL-70
chr16-90244294-DEL-69
chr16-90254490-DEL-69
chr16-90257756-DEL-290
chr16-90274854-DEL-422
chr16-90286346-DEL-261
chr16-90311437-DEL-80
chr16-90312082-DEL-140
chr16-90323434-DEL-3178
chr16-90369269-DEL-3809
chr16-90399788-DEL-96
chr16-90400443-DEL-100
chr16-90403380-DEL-642
chr16-90410044-DEL-124
chr16-90420311-DEL-60
chr16-90450918-DEL-62
chr16-90454551-DEL-64
chr16-90476541-DEL-56
chr16-90482511-DEL-743
chr16-90486705-DEL-160
chr16-90511260-DEL-834
chr16-90529027-DEL-51
chr16-90536021-DEL-77
chr16-90536837-DEL-63
chr16-90541336-DEL-60
chr16-90542041-DEL-398
chr16-90555684-DEL-57
chr16-90566786-DEL-660
chr16-90591098-DEL-639
chr16-90603934-DEL-190
chr16-90604982-DEL-137
chr16-90616803-DEL-225
chr16-90628606-DEL-74
chr16-90661401-DEL-63
chr16-90687883-DEL-301
chr16-90696104-DEL-90
chr16-90699368-DEL-64

chr16-90710490-DEL-65
chr16-90712672-DEL-56
chr16-90723812-DEL-93
chr16-90749108-DEL-59
chr16-90817725-DEL-96
chr16-90836683-DEL-338
chr16-90865302-DEL-151
chr16-90923704-DEL-60
chr16-90927538-DEL-220
chr16-90928456-DEL-63
chr16-90968493-DEL-169
chr16-90974313-DEL-315
chr16-90979234-DEL-263
chr16-90993176-DEL-501
chr16-91031893-DEL-75
chr16-91036916-DEL-130
chr16-91037905-DEL-99
chr16-91043797-DEL-62
chr16-91054299-DEL-247
chr16-91061943-DEL-407
chr16-91077238-DEL-51
chr16-91104662-DEL-173
chr16-91109806-DEL-72
chr16-91139240-DEL-448
chr16-91140361-DEL-161
chr16-91146598-DEL-71
chr16-91150723-DEL-788
chr16-91152029-DEL-995
chr16-91170142-DEL-624
chr16-91213219-DEL-134
chr16-91214827-DEL-107
chr16-91245595-DEL-153
chr16-91251336-DEL-224
chr16-91251836-DEL-767
chr16-91264392-DEL-2748
chr16-91268604-DEL-656
chr16-91279855-DEL-203
chr16-91284599-DEL-200
chr16-91289187-DEL-218
chr16-91293163-DEL-119
chr16-91298900-DEL-896
chr16-91309532-DEL-127
chr16-91327347-DEL-486
chr16-91335340-DEL-151
chr16-91355521-DEL-56
chr16-91364330-DEL-80
chr16-91371242-DEL-79
chr16-91385377-DEL-455
chr16-91411898-DEL-153
chr16-91485888-DEL-105
chr16-91504678-DEL-196
chr16-91518285-DEL-92
chr16-91541690-DEL-443
chr16-91547324-DEL-143
chr16-91593048-DEL-2199
chr16-91595546-DEL-385
chr16-91616034-DEL-260
chr16-91625376-DEL-1142
chr16-91714190-DEL-62
chr16-91730866-DEL-185
chr16-91745847-DEL-92
chr16-91749989-DEL-56
chr16-91761093-DEL-1275
chr16-91782592-DEL-379
chr16-91787324-DEL-96
chr16-91791529-DEL-58
chr16-91795548-DEL-63
chr16-91801083-DEL-233
chr16-91802037-DEL-232
chr16-91805210-DEL-79
chr16-91807762-DEL-220
chr16-91808341-DEL-61
chr16-91809438-DEL-78
chr16-91811157-DEL-65
chr16-91818999-DEL-84
chr16-91825771-DEL-273
chr16-91828632-DEL-228

chr16-91839388-DEL-134
chr16-91850729-DEL-521
chr16-91890832-DEL-178
chr16-91938226-DEL-61
chr16-91952913-DEL-334
chr16-91961559-DEL-135
chr16-91979604-DEL-133
chr16-91992282-DEL-832
chr16-91996627-DEL-136
chr16-92007619-DEL-1969
chr16-92042403-DEL-154
chr16-92045620-DEL-102
chr16-92046502-DEL-69
chr16-92049199-DEL-135
chr16-92051580-DEL-2061
chr16-92051655-DEL-1427
chr16-92052341-DEL-422
chr16-92053030-DEL-94
chr16-92087149-DEL-148
chr16-92108277-DEL-76
chr16-92120365-DEL-128
chr16-92128443-DEL-409
chr16-92131920-DEL-80
chr16-92137245-DEL-67
chr16-92142596-DEL-174
chr16-92144410-DEL-384
chr16-92147068-DEL-63
chr16-92150154-DEL-54
chr16-92163926-DEL-162
chr16-92174948-DEL-60
chr16-92192546-DEL-86
chr16-92201221-DEL-969
chr16-92207672-DEL-306
chr16-92210699-DEL-706
chr16-92255345-DEL-51
chr16-92273975-DEL-81
chr16-92306671-DEL-193
chr16-92310497-DEL-137
chr16-92326022-DEL-72
chr16-92328007-DEL-73
chr16-92332365-DEL-88
chr16-92358821-DEL-94
chr16-92366567-DEL-51
chr16-92403665-DEL-53
chr16-92466154-DEL-421
chr16-92511032-DEL-143
chr16-92514065-DEL-52
chr16-92527285-DEL-1142
chr16-92555377-DEL-64
chr16-92556062-DEL-63
chr16-92562836-DEL-150
chr16-92569132-DEL-895
chr16-92571202-DEL-273
chr16-92571906-DEL-58
chr16-92595367-DEL-236
chr16-92597071-DEL-91
chr16-92639606-DEL-208
chr16-92675907-DEL-286
chr16-92713359-DEL-4290
chr16-92722439-DEL-658
chr16-92735907-DEL-75
chr16-92756211-DEL-523
chr16-92761343-DEL-86
chr16-92798633-DEL-181
chr16-92837428-DEL-144
chr16-92880557-DEL-54
chr16-92906926-DEL-82
chr16-92908409-DEL-53
chr16-92914710-DEL-64
chr16-92922012-DEL-52
chr16-92933448-DEL-385
chr16-92961560-DEL-66
chr16-92962815-DEL-568
chr16-92966480-DEL-717
chr16-92969742-DEL-169
chr16-92984648-DEL-53
chr16-93013723-DEL-54

chr16-93024391-DEL-538
chr16-93025261-DEL-60
chr16-93075239-DEL-113
chr16-93079805-DEL-321
chr16-93133011-DEL-828
chr16-93177369-DEL-139
chr16-93186470-DEL-260
chr16-93199619-DEL-58
chr16-93204673-DEL-52
chr16-93205711-DEL-320
chr16-93241470-DEL-86
chr16-93243833-DEL-113
chr16-93252980-DEL-210
chr16-93254980-DEL-225
chr16-93258371-DEL-52
chr16-93261004-DEL-453
chr16-93265030-DEL-79
chr16-93274399-DEL-2056
chr16-93275338-DEL-53
chr16-93282721-DEL-612
chr16-93291208-DEL-56
chr16-93299829-DEL-470
chr16-93305291-DEL-748
chr16-93319747-DEL-1218
chr16-93321226-DEL-104
chr16-93322737-DEL-186
chr16-93335092-DEL-220
chr16-93338518-DEL-58
chr16-93358605-DEL-3805
chr16-93360772-DEL-124
chr16-93363731-DEL-100
chr16-93370148-DEL-101
chr16-93376730-DEL-90
chr16-93385602-DEL-195
chr16-93388447-DEL-97
chr16-93396914-DEL-58
chr16-93402008-DEL-610
chr16-93404236-DEL-52
chr16-93410815-DEL-57
chr16-93411015-DEL-440
chr16-93420107-DEL-18386
chr16-93429364-DEL-2485
chr16-93430290-DEL-2226
chr16-93432953-DEL-902
chr16-93437052-DEL-894
chr16-93437071-DEL-1100
chr16-93443614-DEL-147
chr16-93451575-DEL-139
chr16-93467628-DEL-158
chr16-93494339-DEL-8985
chr16-93508164-DEL-145
chr16-93510375-DEL-430
chr16-93523676-DEL-348
chr16-93544099-DEL-51
chr16-93569150-DEL-518
chr16-93575217-DEL-202
chr16-93580513-DEL-584
chr16-93589392-DEL-56
chr16-93592157-DEL-82
chr16-93599695-DEL-436
chr16-93601316-DEL-188
chr16-93610363-DEL-153
chr16-93619067-DEL-643
chr16-93651519-DEL-192
chr16-93671375-DEL-54
chr16-93676461-DEL-271
chr16-93686387-DEL-1195
chr16-93710098-DEL-200
chr16-93712629-DEL-70
chr16-93720469-DEL-114
chr16-93724370-DEL-78
chr16-93742959-DEL-494
chr16-93743596-DEL-104
chr16-93771601-DEL-92
chr16-93774667-DEL-66
chr16-93789271-DEL-1121
chr16-93790046-DEL-171

chr16-93790092-DEL-310
chr16-93790554-DEL-3666
chr16-93791107-DEL-3015
chr16-93791526-DEL-375
chr16-93791769-DEL-678
chr16-93792458-DEL-845
chr16-93792788-DEL-153
chr16-93793472-DEL-754
chr16-93796620-DEL-1283
chr16-93796862-DEL-260
chr16-93797582-DEL-492
chr16-93802411-DEL-51
chr16-93804738-DEL-52
chr16-93811649-DEL-52
chr16-93816193-DEL-209
chr16-93820923-DEL-56
chr16-93833946-DEL-81
chr16-93842711-DEL-93
chr16-93855750-DEL-54
chr16-93870074-DEL-52
chr16-93871128-DEL-93
chr16-93881969-DEL-209
chr16-93887273-DEL-53
chr16-93901847-DEL-96
chr16-93916540-DEL-129
chr16-93955821-DEL-53
chr16-93998271-DEL-255
chr16-94016885-DEL-94
chr16-94038444-DEL-81
chr16-94058921-DEL-66
chr16-94067496-DEL-287
chr16-94072569-DEL-388
chr16-94096338-DEL-74
chr16-94097294-DEL-1476
chr16-94100431-DEL-345
chr16-94134801-DEL-174
chr16-94152354-DEL-1497
chr16-94176502-DEL-594
chr16-94231310-DEL-119
chr16-94233737-DEL-58
chr16-94267382-DEL-152
chr16-94277504-DEL-312
chr16-94285319-DEL-4392
chr16-94293537-DEL-64
chr16-94315409-DEL-347
chr16-94324131-DEL-194
chr16-94347379-DEL-68
chr16-94358386-DEL-90
chr16-94366037-DEL-202
chr16-94393701-DEL-545
chr16-94418451-DEL-252
chr16-94431100-DEL-104
chr16-94444819-DEL-1487
chr16-94449874-DEL-57
chr16-94468066-DEL-71
chr16-94476729-DEL-288
chr16-94499144-DEL-174
chr16-94499748-DEL-144
chr16-94526521-DEL-145
chr16-94541433-DEL-72
chr16-94542156-DEL-99
chr16-94549489-DEL-54
chr16-94557787-DEL-175
chr16-94576353-DEL-116
chr16-94586121-DEL-70
chr16-94610430-DEL-122
chr16-94611061-DEL-1102
chr16-94619403-DEL-168
chr16-94630935-DEL-115
chr16-94657769-DEL-161
chr16-94661490-DEL-53
chr16-94671088-DEL-160
chr16-94684529-DEL-58
chr16-94706328-DEL-98
chr16-94724820-DEL-448
chr16-94725370-DEL-139
chr16-94729098-DEL-589

chr16-94746167-DEL-90
chr16-94802513-DEL-53
chr16-94838008-DEL-66
chr16-94850054-DEL-224
chr16-94870806-DEL-60
chr16-94891391-DEL-148
chr16-94931558-DEL-61
chr16-94939360-DEL-98
chr16-94966428-DEL-84
chr16-94966962-DEL-85
chr16-95012826-DEL-1408
chr16-95046070-DEL-1359
chr16-95056074-DEL-301
chr16-95078168-DEL-69
chr16-95088160-DEL-81
chr16-95151035-DEL-52
chr16-95156872-DEL-120
chr16-95165041-DEL-55
chr16-95200842-DEL-103
chr16-95215945-DEL-60
chr16-95229047-DEL-58
chr16-95273395-DEL-130
chr16-95274357-DEL-80
chr16-95277556-DEL-657
chr16-95289322-DEL-109
chr16-95292565-DEL-70
chr16-95301440-DEL-1387
chr16-95308998-DEL-52
chr16-95309819-DEL-2061
chr16-95338536-DEL-86
chr16-95344563-DEL-60
chr16-95360691-DEL-55
chr16-95407297-DEL-572
chr16-95409907-DEL-55
chr16-95413634-DEL-249
chr16-95435534-DEL-171
chr16-95448016-DEL-337
chr16-95498910-DEL-93
chr16-95525331-DEL-92
chr16-95532962-DEL-133
chr16-95533743-DEL-542
chr16-95542988-DEL-55
chr16-95545471-DEL-157
chr16-95575407-DEL-76
chr16-95596651-DEL-65
chr16-95601253-DEL-61
chr16-95607057-DEL-181
chr16-95611018-DEL-215
chr16-95618097-DEL-203
chr16-95630782-DEL-119
chr16-95642761-DEL-123
chr16-95653838-DEL-191
chr16-95674120-DEL-91
chr16-95700748-DEL-621
chr16-95731972-DEL-118
chr16-95735259-DEL-54
chr16-95776640-DEL-70
chr16-95777694-DEL-608
chr16-95780748-DEL-117
chr16-95783628-DEL-200
chr16-95807299-DEL-1161
chr16-95828088-DEL-205
chr16-95894551-DEL-1188
chr16-95942835-DEL-55
chr16-95951820-DEL-197
chr16-95963958-DEL-140
chr16-95972829-DEL-96
chr16-95987437-DEL-75
chr16-96004272-DEL-66
chr16-96005848-DEL-119
chr16-96007969-DEL-232
chr16-96026579-DEL-118
chr16-96048316-DEL-225
chr16-96059523-DEL-68
chr16-96079557-DEL-69
chr16-96098368-DEL-110
chr16-96102499-DEL-165

chr16-96106664-DEL-70
chr16-96128362-DEL-322
chr16-96148259-DEL-61
chr16-96149691-DEL-97
chr16-96152841-DEL-54
chr16-96177080-DEL-490
chr16-96192938-DEL-560
chr16-96193613-DEL-167
chr16-96194223-DEL-553
chr16-96210484-DEL-453
chr16-96243848-DEL-393
chr16-96258231-DEL-80
chr16-96304649-DEL-938
chr16-96326351-DEL-188
chr16-96343604-DEL-185
chr16-96382389-DEL-310
chr16-96424411-DEL-53
chr16-96436631-DEL-108
chr16-96464130-DEL-102
chr16-96476632-DEL-289
chr16-96480351-DEL-461
chr16-96521381-DEL-190
chr16-96524578-DEL-107
chr16-96630524-DEL-88
chr16-96659901-DEL-64
chr16-96666581-DEL-372
chr16-96717866-DEL-448
chr16-96757826-DEL-196
chr16-96814430-DEL-57
chr16-96840607-DEL-163
chr16-96855488-DEL-556
chr16-96871137-DEL-158
chr16-96875403-DEL-65
chr16-96879079-DEL-76
chr16-96886224-DEL-650
chr16-96930669-DEL-395
chr16-97006741-DEL-51
chr16-97007052-DEL-1083
chr16-97035162-DEL-141
chr16-97112023-DEL-649
chr16-97165265-DEL-128
chr16-97165800-DEL-80
chr16-97214984-DEL-310
chr16-97266294-DEL-175
chr16-97314700-DEL-133
chr16-97379262-DEL-421
chr16-97422386-DEL-117
chr16-97437647-DEL-74
chr16-97441615-DEL-122
chr16-97451308-DEL-53
chr16-97461242-DEL-705
chr16-97469425-DEL-118
chr16-97505146-DEL-203
chr16-97560345-DEL-68
chr16-97625870-DEL-52
chr16-97672189-DEL-63
chr16-97744508-DEL-85
chr16-97748041-DEL-160
chr16-97813108-DEL-84
chr16-97817754-DEL-1321
chr16-97854228-DEL-54
chr16-97870316-DEL-51
chr16-97871929-DEL-74
chr16-97874526-DEL-52
chr16-97887955-DEL-164
chr16-97901539-DEL-3304
chr16-97908218-DEL-154
chr17-3052919-DEL-181
chr17-3056293-DEL-105
chr17-3067423-DEL-3381
chr17-3069497-DEL-60
chr17-3074418-DEL-145
chr17-3085829-DEL-502
chr17-3086743-DEL-125
chr17-3088054-DEL-480
chr17-3094116-DEL-929
chr17-3102080-DEL-337

chr17-3103789-DEL-511
chr17-3106845-DEL-313
chr17-3111665-DEL-158
chr17-3112605-DEL-6821
chr17-3121320-DEL-54
chr17-3125610-DEL-87
chr17-3126737-DEL-1645
chr17-3139558-DEL-172
chr17-3149191-DEL-579
chr17-3156413-DEL-1412
chr17-3163338-DEL-855
chr17-3239233-DEL-265
chr17-3270362-DEL-2829
chr17-3276346-DEL-62
chr17-3328168-DEL-257
chr17-3338542-DEL-57
chr17-3353875-DEL-68
chr17-3375540-DEL-118
chr17-3434664-DEL-162
chr17-3436426-DEL-6111
chr17-3493845-DEL-69
chr17-3494404-DEL-1290
chr17-3499765-DEL-244
chr17-3531010-DEL-435
chr17-3544816-DEL-70
chr17-3556751-DEL-142
chr17-3561969-DEL-513
chr17-3579798-DEL-52
chr17-3589794-DEL-51
chr17-3622143-DEL-1500
chr17-3639216-DEL-125
chr17-3643063-DEL-53
chr17-3655230-DEL-144
chr17-3735206-DEL-273
chr17-3747156-DEL-125
chr17-3771972-DEL-400
chr17-3796575-DEL-1405
chr17-3824794-DEL-64
chr17-3828561-DEL-58
chr17-3830070-DEL-384
chr17-3843674-DEL-58
chr17-3916566-DEL-130
chr17-3918192-DEL-67
chr17-3961126-DEL-150
chr17-3981526-DEL-54
chr17-4046479-DEL-194
chr17-4068401-DEL-6039
chr17-4088213-DEL-71
chr17-4091686-DEL-134
chr17-4101266-DEL-54
chr17-4109022-DEL-200
chr17-4139247-DEL-57
chr17-4188859-DEL-433
chr17-4232889-DEL-54
chr17-4233490-DEL-57
chr17-4290369-DEL-1465
chr17-4309589-DEL-991
chr17-4330371-DEL-78
chr17-4349880-DEL-68
chr17-4427030-DEL-990
chr17-4427038-DEL-282
chr17-4427423-DEL-124
chr17-4444558-DEL-676
chr17-4456538-DEL-94
chr17-4460553-DEL-1331
chr17-4484175-DEL-108
chr17-4493144-DEL-114
chr17-4510144-DEL-100
chr17-4556922-DEL-197
chr17-4609496-DEL-61
chr17-4623261-DEL-383
chr17-4637708-DEL-1710
chr17-4702708-DEL-522
chr17-4730215-DEL-159
chr17-4767871-DEL-117
chr17-4770850-DEL-615
chr17-4782401-DEL-56

chr17-4788765-DEL-243
chr17-4791200-DEL-82
chr17-4836526-DEL-497
chr17-4844795-DEL-156
chr17-4845964-DEL-195
chr17-4860304-DEL-70
chr17-4861521-DEL-611
chr17-4865752-DEL-282
chr17-4903811-DEL-178
chr17-4911941-DEL-220
chr17-4925019-DEL-153
chr17-4930843-DEL-750
chr17-4987390-DEL-388
chr17-4993761-DEL-63
chr17-5014262-DEL-79
chr17-5035989-DEL-60
chr17-5054294-DEL-115
chr17-5061921-DEL-105
chr17-5069759-DEL-234
chr17-5085897-DEL-53
chr17-5088622-DEL-1239
chr17-5105622-DEL-375
chr17-5121867-DEL-565
chr17-5143692-DEL-275
chr17-5153997-DEL-166
chr17-5168819-DEL-869
chr17-5191612-DEL-1009
chr17-5192720-DEL-1240
chr17-5196360-DEL-2094
chr17-5196419-DEL-1204
chr17-5196661-DEL-120
chr17-5197321-DEL-1703
chr17-5197386-DEL-681
chr17-5197802-DEL-1223
chr17-5198245-DEL-182
chr17-5198308-DEL-319
chr17-5208839-DEL-552
chr17-5208839-DEL-92
chr17-5245290-DEL-83
chr17-5263962-DEL-58
chr17-5266304-DEL-1008
chr17-5275090-DEL-120
chr17-5291546-DEL-429
chr17-5305114-DEL-72
chr17-5322516-DEL-167
chr17-5365211-DEL-70
chr17-5419585-DEL-64
chr17-5426264-DEL-679
chr17-5482494-DEL-859
chr17-5487054-DEL-438
chr17-5494589-DEL-551
chr17-5504971-DEL-538
chr17-5508774-DEL-110
chr17-5548608-DEL-399
chr17-5554646-DEL-68
chr17-5571445-DEL-801
chr17-5572841-DEL-181
chr17-5590737-DEL-148
chr17-5598184-DEL-102
chr17-5606517-DEL-154
chr17-5629397-DEL-102
chr17-5637013-DEL-172
chr17-5643161-DEL-165
chr17-5651861-DEL-201
chr17-5670055-DEL-947
chr17-5686114-DEL-56
chr17-5691366-DEL-59
chr17-5701542-DEL-201
chr17-5725369-DEL-163
chr17-5733696-DEL-51
chr17-5741133-DEL-838
chr17-5766826-DEL-1785
chr17-5768516-DEL-60
chr17-5789550-DEL-77
chr17-5791623-DEL-72
chr17-5795970-DEL-314
chr17-5809289-DEL-61

chr17-5812045-DEL-210
chr17-5818913-DEL-544
chr17-5823708-DEL-5348
chr17-5838921-DEL-1556
chr17-5841000-DEL-479
chr17-5861692-DEL-98
chr17-5901457-DEL-108
chr17-5902618-DEL-369
chr17-5912664-DEL-307
chr17-5912688-DEL-1366
chr17-5913880-DEL-54
chr17-5914308-DEL-2107
chr17-5914432-DEL-495
chr17-5914619-DEL-1006
chr17-5915052-DEL-217
chr17-5915659-DEL-76
chr17-5916249-DEL-1123
chr17-5916724-DEL-111
chr17-5917248-DEL-98
chr17-5929872-DEL-2239
chr17-5930559-DEL-205
chr17-5930635-DEL-2933
chr17-5931176-DEL-402
chr17-5931179-DEL-1946
chr17-5932202-DEL-122
chr17-5933285-DEL-266
chr17-5933616-DEL-729
chr17-5934016-DEL-1249
chr17-5934405-DEL-362
chr17-5934948-DEL-117
chr17-5938481-DEL-93
chr17-5942317-DEL-56
chr17-5945425-DEL-1724
chr17-5965381-DEL-54
chr17-5975033-DEL-368
chr17-5978593-DEL-200
chr17-5999680-DEL-130
chr17-6011599-DEL-154
chr17-6021511-DEL-1303
chr17-6025101-DEL-134
chr17-6057121-DEL-214
chr17-6060161-DEL-153
chr17-6064583-DEL-108
chr17-6068802-DEL-82
chr17-6077210-DEL-56
chr17-6086391-DEL-72
chr17-6107401-DEL-74
chr17-6126710-DEL-200
chr17-6129756-DEL-72
chr17-6141675-DEL-4672
chr17-6150177-DEL-753
chr17-6171324-DEL-1183
chr17-6174207-DEL-58
chr17-6175532-DEL-148
chr17-6185675-DEL-196
chr17-6204199-DEL-245
chr17-6218575-DEL-232
chr17-6234891-DEL-111
chr17-6249275-DEL-107
chr17-6259625-DEL-130
chr17-6273444-DEL-64
chr17-6280780-DEL-112
chr17-6291636-DEL-54
chr17-6323838-DEL-127
chr17-6383944-DEL-203
chr17-6395131-DEL-5174
chr17-6406127-DEL-87
chr17-6408128-DEL-64
chr17-6413643-DEL-200
chr17-6434907-DEL-223
chr17-6445398-DEL-165
chr17-6449092-DEL-1498
chr17-6451224-DEL-473
chr17-6455827-DEL-830
chr17-6652875-DEL-223
chr17-6662332-DEL-350
chr17-6672952-DEL-830

chr17-6680411-DEL-181
chr17-6682846-DEL-196
chr17-6688640-DEL-2050
chr17-6696762-DEL-135
chr17-6741329-DEL-126
chr17-6745743-DEL-80
chr17-6759552-DEL-2169
chr17-6764108-DEL-537
chr17-6767265-DEL-55
chr17-6774043-DEL-54
chr17-6776282-DEL-12123
chr17-6829314-DEL-181
chr17-6838240-DEL-830
chr17-6843111-DEL-5873
chr17-6851848-DEL-305
chr17-6852640-DEL-1455
chr17-6860177-DEL-2052
chr17-6865357-DEL-78
chr17-6882325-DEL-140
chr17-6884234-DEL-89
chr17-6922849-DEL-204
chr17-6923912-DEL-135
chr17-6946434-DEL-396
chr17-6977949-DEL-1027
chr17-7012617-DEL-234
chr17-7038713-DEL-60
chr17-7098287-DEL-1044
chr17-7105897-DEL-1438
chr17-7107003-DEL-745
chr17-7109531-DEL-153
chr17-7109591-DEL-1064
chr17-7118084-DEL-195
chr17-7188182-DEL-5582
chr17-7198475-DEL-7354
chr17-7199312-DEL-1896
chr17-7199401-DEL-924
chr17-7199750-DEL-4044
chr17-7199839-DEL-1116
chr17-7201585-DEL-3048
chr17-7202173-DEL-474
chr17-7202223-DEL-1148
chr17-7202300-DEL-2785
chr17-7203998-DEL-212
chr17-7204668-DEL-625
chr17-7208007-DEL-224
chr17-7210840-DEL-652
chr17-7211903-DEL-58
chr17-7212469-DEL-99
chr17-7221654-DEL-77
chr17-7225677-DEL-60
chr17-7257857-DEL-72
chr17-7259530-DEL-55
chr17-7331211-DEL-55
chr17-7340122-DEL-54
chr17-7356971-DEL-101
chr17-7404988-DEL-226
chr17-7411927-DEL-167
chr17-7423599-DEL-53
chr17-7434260-DEL-1066
chr17-7440317-DEL-356
chr17-7539562-DEL-205
chr17-7564593-DEL-80
chr17-7601398-DEL-51
chr17-7602258-DEL-2859
chr17-7635846-DEL-735
chr17-7742447-DEL-177
chr17-7743457-DEL-347
chr17-7754222-DEL-54
chr17-7757014-DEL-184
chr17-7768146-DEL-196
chr17-7799994-DEL-180
chr17-7823286-DEL-54
chr17-7840693-DEL-655
chr17-7890020-DEL-76
chr17-7901655-DEL-57
chr17-7903683-DEL-306
chr17-7917629-DEL-439

chr17-7919030-DEL-446
chr17-7928750-DEL-109
chr17-7945305-DEL-60
chr17-7957075-DEL-62
chr17-7993669-DEL-104
chr17-7997748-DEL-218
chr17-8004302-DEL-180
chr17-8029090-DEL-216
chr17-8032927-DEL-111
chr17-8039184-DEL-134
chr17-8053974-DEL-66
chr17-8063428-DEL-344
chr17-8086425-DEL-59
chr17-8088174-DEL-219
chr17-8089575-DEL-1947
chr17-8113949-DEL-88
chr17-8120377-DEL-51
chr17-8125782-DEL-167
chr17-8126126-DEL-133
chr17-8126148-DEL-2067
chr17-8126989-DEL-186
chr17-8128013-DEL-64
chr17-8128640-DEL-174
chr17-8129045-DEL-396
chr17-8130974-DEL-129
chr17-8131565-DEL-152
chr17-8132525-DEL-52
chr17-8168011-DEL-99
chr17-8169048-DEL-646
chr17-8172286-DEL-224
chr17-8175236-DEL-888
chr17-8175508-DEL-375
chr17-8181722-DEL-489
chr17-8188349-DEL-64
chr17-8243940-DEL-1292
chr17-8253417-DEL-69
chr17-8255200-DEL-197
chr17-8262285-DEL-65
chr17-8272175-DEL-54
chr17-8305258-DEL-74
chr17-8310037-DEL-51
chr17-8333392-DEL-415
chr17-8353332-DEL-55
chr17-8387207-DEL-60
chr17-8427114-DEL-96
chr17-8445712-DEL-587
chr17-8497171-DEL-891
chr17-8516655-DEL-150
chr17-8525620-DEL-95
chr17-8545293-DEL-209
chr17-8565999-DEL-90
chr17-8572511-DEL-1357
chr17-8575566-DEL-75
chr17-8603924-DEL-208
chr17-8604831-DEL-100
chr17-8607543-DEL-59
chr17-8617185-DEL-54
chr17-8622910-DEL-299
chr17-8716886-DEL-823
chr17-8736138-DEL-68
chr17-8762764-DEL-137
chr17-8779217-DEL-7070
chr17-8793380-DEL-602
chr17-8842770-DEL-66
chr17-8847256-DEL-61
chr17-8890008-DEL-505
chr17-8908213-DEL-78
chr17-8919212-DEL-178
chr17-8926757-DEL-1983
chr17-8955768-DEL-192
chr17-8979728-DEL-215
chr17-8983388-DEL-132
chr17-8983433-DEL-1254
chr17-8991766-DEL-68
chr17-9012534-DEL-192
chr17-9014574-DEL-61
chr17-9023636-DEL-120

chr17-9044063-DEL-245
chr17-9071165-DEL-324
chr17-9108066-DEL-1199
chr17-9124999-DEL-745
chr17-9131265-DEL-1442
chr17-9154814-DEL-336
chr17-9249853-DEL-233
chr17-9252151-DEL-133
chr17-9297895-DEL-101
chr17-9316316-DEL-93
chr17-9316712-DEL-362
chr17-9326086-DEL-413
chr17-9333476-DEL-59
chr17-9334200-DEL-70
chr17-9362656-DEL-69
chr17-9422473-DEL-56
chr17-9445881-DEL-56
chr17-9457250-DEL-432
chr17-9479299-DEL-101
chr17-9480331-DEL-53
chr17-9503766-DEL-89
chr17-9577484-DEL-8129
chr17-9580692-DEL-6186
chr17-9649211-DEL-495
chr17-9707212-DEL-53
chr17-9741386-DEL-54
chr17-9759496-DEL-1038
chr17-9806404-DEL-1876
chr17-9850422-DEL-122
chr17-9859360-DEL-1874
chr17-9898825-DEL-353
chr17-9903966-DEL-120
chr17-9942859-DEL-267
chr17-9958543-DEL-107
chr17-9963732-DEL-74
chr17-9970037-DEL-60
chr17-9990934-DEL-104
chr17-10013456-DEL-105
chr17-10016494-DEL-503
chr17-10044550-DEL-88
chr17-10049947-DEL-228
chr17-10059102-DEL-538
chr17-10074254-DEL-58
chr17-10078641-DEL-90
chr17-10085872-DEL-235
chr17-10104779-DEL-233
chr17-10127326-DEL-60
chr17-10159383-DEL-236
chr17-10214973-DEL-219
chr17-10233557-DEL-220
chr17-10248936-DEL-56
chr17-10251926-DEL-65
chr17-10304516-DEL-56
chr17-10341635-DEL-54
chr17-10394157-DEL-58
chr17-10405118-DEL-86
chr17-10442672-DEL-1493
chr17-10461895-DEL-2492
chr17-10485392-DEL-556
chr17-10542923-DEL-280
chr17-10559059-DEL-102
chr17-10570045-DEL-86
chr17-10583096-DEL-197
chr17-10584785-DEL-54
chr17-10611995-DEL-104
chr17-10643962-DEL-56
chr17-10663923-DEL-97
chr17-10664097-DEL-304
chr17-10684944-DEL-60
chr17-10706640-DEL-279
chr17-10718518-DEL-1253
chr17-10721665-DEL-53
chr17-10727643-DEL-178
chr17-10743984-DEL-155
chr17-10760290-DEL-53
chr17-10822436-DEL-74
chr17-10939035-DEL-66

chr17-10968786-DEL-1104
chr17-11011714-DEL-1982
chr17-11012848-DEL-62
chr17-11062957-DEL-136
chr17-11126482-DEL-65
chr17-11167073-DEL-1015
chr17-11170029-DEL-6062
chr17-11229888-DEL-122
chr17-11235668-DEL-54
chr17-11268042-DEL-158
chr17-11276420-DEL-881
chr17-11282570-DEL-66
chr17-11300118-DEL-74
chr17-11350876-DEL-153
chr17-11355339-DEL-1300
chr17-11356672-DEL-301
chr17-11382670-DEL-808
chr17-11386186-DEL-86
chr17-11424496-DEL-1141
chr17-11445510-DEL-768
chr17-11511502-DEL-6569
chr17-11513772-DEL-3947
chr17-11544110-DEL-198
chr17-11546972-DEL-425
chr17-11564544-DEL-120
chr17-11574070-DEL-160
chr17-11597167-DEL-63
chr17-11603923-DEL-51
chr17-11657898-DEL-59
chr17-11665308-DEL-1282
chr17-11669703-DEL-751
chr17-11680916-DEL-57
chr17-11743629-DEL-126
chr17-11817930-DEL-169
chr17-11834456-DEL-70
chr17-11847631-DEL-74
chr17-11851829-DEL-392
chr17-11971897-DEL-61
chr17-12009510-DEL-632
chr17-12044647-DEL-172
chr17-12048349-DEL-55
chr17-12067922-DEL-122
chr17-12094610-DEL-438
chr17-12102144-DEL-53
chr17-12107377-DEL-113
chr17-12112100-DEL-197
chr17-12137339-DEL-159
chr17-12150535-DEL-96
chr17-12150860-DEL-778
chr17-12179803-DEL-98
chr17-12219566-DEL-1811
chr17-12223112-DEL-56
chr17-12228386-DEL-219
chr17-12239662-DEL-201
chr17-12243960-DEL-739
chr17-12304930-DEL-58
chr17-12310243-DEL-74
chr17-12359119-DEL-2448
chr17-12362505-DEL-57
chr17-12418784-DEL-1376
chr17-12431427-DEL-69
chr17-12447922-DEL-171
chr17-12464907-DEL-504
chr17-12485895-DEL-94
chr17-12498937-DEL-94
chr17-12502579-DEL-4718
chr17-12508493-DEL-57
chr17-12576248-DEL-821
chr17-12640716-DEL-75
chr17-12651706-DEL-224
chr17-12661277-DEL-368
chr17-12664304-DEL-74
chr17-12666467-DEL-271
chr17-12690888-DEL-216
chr17-12730897-DEL-71
chr17-12745912-DEL-1074
chr17-12773033-DEL-63

chr17-12776191-DEL-408
chr17-12807520-DEL-51
chr17-12808762-DEL-55
chr17-12811069-DEL-109
chr17-12825379-DEL-58
chr17-12854930-DEL-512
chr17-12855021-DEL-1430
chr17-12856170-DEL-281
chr17-12868225-DEL-61
chr17-12873760-DEL-61
chr17-12894855-DEL-52
chr17-12906923-DEL-143
chr17-12947445-DEL-444
chr17-13018866-DEL-53
chr17-13027628-DEL-75
chr17-13078840-DEL-57
chr17-13086671-DEL-92
chr17-13115690-DEL-204
chr17-13116976-DEL-150
chr17-13118019-DEL-51
chr17-13127503-DEL-411
chr17-13133245-DEL-429
chr17-13151343-DEL-187
chr17-13175418-DEL-186
chr17-13190584-DEL-1060
chr17-13195788-DEL-167
chr17-13199534-DEL-54
chr17-13204123-DEL-139
chr17-13208662-DEL-219
chr17-13215821-DEL-340
chr17-13223109-DEL-66
chr17-13241232-DEL-116
chr17-13259293-DEL-263
chr17-13285456-DEL-104
chr17-13289499-DEL-52
chr17-13302923-DEL-6833
chr17-13309319-DEL-212
chr17-13361517-DEL-56
chr17-13368474-DEL-18302
chr17-13371991-DEL-7160
chr17-13461232-DEL-7112
chr17-13534993-DEL-71
chr17-13580338-DEL-1111
chr17-13618361-DEL-506
chr17-13628784-DEL-12553
chr17-13633677-DEL-5423
chr17-13700421-DEL-85
chr17-13720294-DEL-70
chr17-13762360-DEL-150
chr17-13770953-DEL-2332
chr17-13773181-DEL-288
chr17-13777756-DEL-209
chr17-13796451-DEL-197
chr17-13801279-DEL-86
chr17-13805953-DEL-69504
chr17-13887744-DEL-173
chr17-13891276-DEL-66
chr17-13899180-DEL-53
chr17-13916787-DEL-560
chr17-13927496-DEL-445
chr17-13957438-DEL-69
chr17-14007431-DEL-1692
chr17-14031406-DEL-67
chr17-14058272-DEL-7082
chr17-14135520-DEL-416
chr17-14136107-DEL-1003
chr17-14190593-DEL-6212
chr17-14219242-DEL-284
chr17-14234035-DEL-7758
chr17-14252061-DEL-67
chr17-14277328-DEL-72
chr17-14314647-DEL-5503
chr17-14398202-DEL-55
chr17-14428794-DEL-61
chr17-14432635-DEL-137
chr17-14445108-DEL-208
chr17-14445179-DEL-1130

chr17-14469191-DEL-57
chr17-14505663-DEL-70
chr17-14523744-DEL-74
chr17-14528635-DEL-67
chr17-14538515-DEL-1851
chr17-14570429-DEL-122
chr17-14585715-DEL-1239
chr17-14618048-DEL-67
chr17-14636212-DEL-82
chr17-14644220-DEL-340
chr17-14665713-DEL-64
chr17-14668419-DEL-254
chr17-14669103-DEL-431
chr17-14674879-DEL-63
chr17-14678233-DEL-77
chr17-14692211-DEL-80
chr17-14720651-DEL-75
chr17-14736675-DEL-70
chr17-14752393-DEL-51
chr17-14782960-DEL-328
chr17-14786438-DEL-3625
chr17-14792672-DEL-98
chr17-14862970-DEL-56
chr17-14865584-DEL-1321
chr17-14903944-DEL-237
chr17-14925170-DEL-590
chr17-14929229-DEL-1282
chr17-14933915-DEL-140
chr17-14982373-DEL-1748
chr17-15013467-DEL-72
chr17-15015729-DEL-74
chr17-15020087-DEL-455
chr17-15082068-DEL-156
chr17-15098754-DEL-58
chr17-15107915-DEL-55
chr17-15112100-DEL-174
chr17-15115197-DEL-152
chr17-15154520-DEL-70
chr17-15156799-DEL-87
chr17-15176636-DEL-54
chr17-15193151-DEL-219
chr17-15194780-DEL-203
chr17-15205422-DEL-117
chr17-15213065-DEL-77
chr17-15238818-DEL-75
chr17-15244431-DEL-77
chr17-15246278-DEL-57
chr17-15257501-DEL-203
chr17-15268169-DEL-116
chr17-15362020-DEL-129
chr17-15380052-DEL-1352
chr17-15398869-DEL-160
chr17-15472243-DEL-55
chr17-15481288-DEL-60
chr17-15482780-DEL-263
chr17-15539430-DEL-2650
chr17-15591766-DEL-62
chr17-15606548-DEL-115
chr17-15613366-DEL-65
chr17-15639198-DEL-568
chr17-15656434-DEL-380
chr17-15658248-DEL-399
chr17-15667881-DEL-252
chr17-15684106-DEL-1191
chr17-15695468-DEL-71
chr17-15706138-DEL-2890
chr17-15706139-DEL-1808
chr17-15706157-DEL-362
chr17-15707014-DEL-866
chr17-15707840-DEL-249
chr17-15708830-DEL-424
chr17-15709479-DEL-52
chr17-15750769-DEL-101
chr17-15751391-DEL-308
chr17-15752856-DEL-5556
chr17-15764871-DEL-86
chr17-15769750-DEL-212

chr17-15770284-DEL-2523
chr17-15772521-DEL-303
chr17-15777039-DEL-104
chr17-15785170-DEL-358
chr17-15795208-DEL-79
chr17-15798599-DEL-202
chr17-15806450-DEL-64
chr17-15816096-DEL-305
chr17-15834290-DEL-205
chr17-15835749-DEL-78
chr17-15849229-DEL-192
chr17-15859699-DEL-122
chr17-15866776-DEL-237
chr17-15878357-DEL-213
chr17-15892596-DEL-216
chr17-15894331-DEL-80
chr17-15911637-DEL-204
chr17-15943181-DEL-122
chr17-15984552-DEL-56
chr17-15996590-DEL-59
chr17-15999332-DEL-77
chr17-16001310-DEL-4181
chr17-16023351-DEL-73
chr17-16042965-DEL-63
chr17-16051843-DEL-229
chr17-16065594-DEL-145
chr17-16126229-DEL-57
chr17-16134533-DEL-669
chr17-16152658-DEL-2503
chr17-16165552-DEL-402
chr17-16181768-DEL-112
chr17-16218753-DEL-206
chr17-16237511-DEL-250
chr17-16254054-DEL-58
chr17-16286212-DEL-1153
chr17-16311812-DEL-158
chr17-16339181-DEL-12318
chr17-16357267-DEL-6464
chr17-16364623-DEL-72
chr17-16365856-DEL-3163
chr17-16378530-DEL-51
chr17-16385154-DEL-109
chr17-16429476-DEL-66
chr17-16446164-DEL-67
chr17-16456059-DEL-462
chr17-16475550-DEL-107
chr17-16509153-DEL-226
chr17-16509747-DEL-398
chr17-16521148-DEL-497
chr17-16544062-DEL-306
chr17-16553120-DEL-798
chr17-16562243-DEL-76
chr17-16571287-DEL-1955
chr17-16577020-DEL-808
chr17-16590673-DEL-1165
chr17-16597148-DEL-62
chr17-16608651-DEL-6496
chr17-16617941-DEL-358
chr17-16624438-DEL-53
chr17-16625105-DEL-6308
chr17-16664609-DEL-52
chr17-16698306-DEL-314
chr17-16739432-DEL-76
chr17-16746677-DEL-209
chr17-16767685-DEL-4360
chr17-16785557-DEL-482
chr17-16813304-DEL-69
chr17-16825898-DEL-2932
chr17-16845269-DEL-890
chr17-16853402-DEL-52
chr17-16879363-DEL-113
chr17-16901961-DEL-54
chr17-16909474-DEL-73
chr17-16917006-DEL-3565
chr17-16930282-DEL-2377
chr17-16932990-DEL-364
chr17-16944762-DEL-400

chr17-16949253-DEL-1379
chr17-16962410-DEL-6258
chr17-16996717-DEL-176
chr17-17001268-DEL-4070
chr17-17040035-DEL-431
chr17-17055065-DEL-59
chr17-17102943-DEL-882
chr17-17110099-DEL-285
chr17-17125995-DEL-173
chr17-17138740-DEL-844
chr17-17150245-DEL-209
chr17-17167119-DEL-60
chr17-17194861-DEL-424
chr17-17199746-DEL-78
chr17-17213450-DEL-72
chr17-17241635-DEL-560
chr17-17248307-DEL-79
chr17-17253927-DEL-687
chr17-17279067-DEL-94
chr17-17308746-DEL-84
chr17-17309627-DEL-342
chr17-17325624-DEL-193
chr17-17329903-DEL-322
chr17-17332376-DEL-306
chr17-17333698-DEL-6226
chr17-17366495-DEL-253
chr17-17435083-DEL-6209
chr17-17494274-DEL-385
chr17-17548872-DEL-462
chr17-17560376-DEL-134
chr17-17584972-DEL-207
chr17-17603308-DEL-201
chr17-17606026-DEL-55
chr17-17629072-DEL-58
chr17-17648825-DEL-8011
chr17-17657717-DEL-781
chr17-17683114-DEL-143
chr17-17701965-DEL-739
chr17-17724638-DEL-4812
chr17-17725589-DEL-188
chr17-17725678-DEL-610
chr17-17759071-DEL-555
chr17-17796376-DEL-109
chr17-17814950-DEL-610
chr17-17834658-DEL-252
chr17-17886975-DEL-153
chr17-17920243-DEL-252
chr17-17958084-DEL-197
chr17-17961048-DEL-588
chr17-17967691-DEL-2953
chr17-17973296-DEL-1937
chr17-18024325-DEL-61
chr17-18029871-DEL-105
chr17-18040769-DEL-82
chr17-18043164-DEL-90
chr17-18053607-DEL-159
chr17-18069350-DEL-226
chr17-18156187-DEL-996
chr17-18179565-DEL-1173
chr17-18217512-DEL-1885
chr17-18244331-DEL-1124
chr17-18278884-DEL-1741
chr17-18296288-DEL-1060
chr17-18303338-DEL-231
chr17-18335742-DEL-2204
chr17-18338643-DEL-4311
chr17-18355000-DEL-193
chr17-18360178-DEL-142
chr17-18396514-DEL-1604
chr17-18410762-DEL-4364
chr17-18413542-DEL-1511
chr17-18432086-DEL-7502
chr17-18446036-DEL-327
chr17-18456892-DEL-397
chr17-18505551-DEL-52
chr17-18552554-DEL-56
chr17-18561881-DEL-399

chr17-18652070-DEL-524
chr17-18657547-DEL-961
chr17-18657575-DEL-356
chr17-18662603-DEL-302
chr17-18666730-DEL-513
chr17-18805292-DEL-123
chr17-18811289-DEL-62
chr17-18814713-DEL-57
chr17-18836694-DEL-1683
chr17-18850600-DEL-61
chr17-18890808-DEL-212
chr17-18943082-DEL-90
chr17-18952026-DEL-80
chr17-18954989-DEL-1821
chr17-18958025-DEL-208
chr17-19031793-DEL-154
chr17-19040863-DEL-343
chr17-19092707-DEL-327
chr17-19101022-DEL-1920
chr17-19111732-DEL-11401
chr17-19138873-DEL-2779
chr17-19192276-DEL-90
chr17-19194821-DEL-555
chr17-19316374-DEL-195
chr17-19316647-DEL-492
chr17-19319059-DEL-84
chr17-19339176-DEL-296
chr17-19353381-DEL-108
chr17-19357273-DEL-109
chr17-19391662-DEL-3120
chr17-19430855-DEL-12860
chr17-19448135-DEL-64
chr17-19466174-DEL-202
chr17-19490555-DEL-1666
chr17-19513147-DEL-124
chr17-19619004-DEL-118
chr17-19643154-DEL-37834
chr17-19695633-DEL-839
chr17-19702259-DEL-1249
chr17-19709638-DEL-961
chr17-19734129-DEL-16430
chr17-19775315-DEL-3494
chr17-19784247-DEL-348
chr17-19827619-DEL-67
chr17-19837570-DEL-71
chr17-19901805-DEL-943
chr17-19924819-DEL-5984
chr17-19936374-DEL-4225
chr17-19946302-DEL-336
chr17-19954870-DEL-5582
chr17-19987885-DEL-61
chr17-19994384-DEL-186
chr17-19996207-DEL-90
chr17-20030500-DEL-58
chr17-20054686-DEL-151
chr17-20064271-DEL-80
chr17-20088055-DEL-557
chr17-20203361-DEL-136
chr17-20204900-DEL-54
chr17-20252483-DEL-3580
chr17-20264013-DEL-325
chr17-20265719-DEL-187
chr17-20274192-DEL-80
chr17-20294466-DEL-520
chr17-20319177-DEL-8241
chr17-20340240-DEL-75
chr17-20348634-DEL-1079
chr17-20354417-DEL-58
chr17-20362878-DEL-52
chr17-20364512-DEL-6555
chr17-20421131-DEL-1272
chr17-20426022-DEL-388
chr17-20432203-DEL-1552
chr17-20434627-DEL-58
chr17-20453299-DEL-77
chr17-20455801-DEL-127
chr17-20473334-DEL-77

chr17-20486579-DEL-13739
chr17-20500753-DEL-254
chr17-20507542-DEL-6234
chr17-20516827-DEL-408
chr17-20524936-DEL-1395
chr17-20563599-DEL-119
chr17-20588944-DEL-149
chr17-20604096-DEL-328
chr17-20610831-DEL-6227
chr17-20621759-DEL-287
chr17-20624521-DEL-66
chr17-20650625-DEL-463
chr17-20676931-DEL-120
chr17-20710409-DEL-147
chr17-20726506-DEL-133
chr17-20729988-DEL-4268
chr17-20805208-DEL-314
chr17-20840785-DEL-593
chr17-20847996-DEL-6699
chr17-20848255-DEL-186
chr17-20889516-DEL-146
chr17-20922402-DEL-6044
chr17-20932221-DEL-7375
chr17-20933270-DEL-1020
chr17-20939346-DEL-196
chr17-20940478-DEL-9400
chr17-20940716-DEL-72
chr17-20941711-DEL-404
chr17-20964065-DEL-882
chr17-20967339-DEL-8683
chr17-21045091-DEL-115
chr17-21074269-DEL-771
chr17-21078252-DEL-52
chr17-21105767-DEL-2122
chr17-21116824-DEL-158
chr17-21128992-DEL-305
chr17-21132184-DEL-68
chr17-21148821-DEL-55
chr17-21155711-DEL-619
chr17-21213501-DEL-72
chr17-21220360-DEL-100
chr17-21237069-DEL-200
chr17-21255484-DEL-5544
chr17-21264023-DEL-76
chr17-21266272-DEL-62
chr17-21267329-DEL-59
chr17-21319275-DEL-113
chr17-21365252-DEL-66
chr17-21432162-DEL-845
chr17-21438479-DEL-2729
chr17-21442183-DEL-188
chr17-21454812-DEL-2891
chr17-21460621-DEL-114
chr17-21463589-DEL-1213
chr17-21535760-DEL-72
chr17-21585742-DEL-3248
chr17-21632474-DEL-1594
chr17-21635676-DEL-2666
chr17-21637800-DEL-973
chr17-21645833-DEL-206
chr17-21656046-DEL-401
chr17-21657918-DEL-175
chr17-21659477-DEL-201
chr17-21661025-DEL-1353
chr17-21662630-DEL-498
chr17-21663421-DEL-1332
chr17-21734760-DEL-326
chr17-21742996-DEL-51
chr17-21760178-DEL-116
chr17-21774482-DEL-193
chr17-21799772-DEL-61
chr17-21827437-DEL-6157
chr17-21853008-DEL-4466
chr17-21915481-DEL-94
chr17-21918101-DEL-5412
chr17-21955415-DEL-195
chr17-21976506-DEL-4546

chr17-21981755-DEL-83
chr17-21983485-DEL-56
chr17-21985657-DEL-1007
chr17-21991204-DEL-518
chr17-22013256-DEL-52
chr17-22020057-DEL-7446
chr17-22035275-DEL-188
chr17-22037626-DEL-84
chr17-22045036-DEL-141
chr17-22115422-DEL-174
chr17-22154541-DEL-1010
chr17-22159811-DEL-1098
chr17-22300927-DEL-927
chr17-22328959-DEL-1875
chr17-22333251-DEL-1870
chr17-22390216-DEL-659
chr17-22395344-DEL-4450
chr17-22401472-DEL-97
chr17-22402363-DEL-4762
chr17-22436580-DEL-73
chr17-22441312-DEL-710
chr17-22449385-DEL-7447
chr17-22474418-DEL-118
chr17-22509788-DEL-409
chr17-22513373-DEL-117
chr17-22516808-DEL-1007
chr17-22519425-DEL-2875
chr17-22540668-DEL-6179
chr17-22566892-DEL-198
chr17-22609950-DEL-253
chr17-22684742-DEL-12613
chr17-22702007-DEL-11802
chr17-22706490-DEL-7106
chr17-22715933-DEL-255
chr17-22740870-DEL-170
chr17-22791561-DEL-56
chr17-22815279-DEL-62
chr17-22846551-DEL-201
chr17-22867415-DEL-6790
chr17-22875320-DEL-205
chr17-22916668-DEL-135
chr17-22947992-DEL-76
chr17-22952948-DEL-150
chr17-22972407-DEL-61
chr17-22972914-DEL-7262
chr17-23013528-DEL-1283
chr17-23022131-DEL-67
chr17-23057978-DEL-166
chr17-23072838-DEL-104
chr17-23117830-DEL-2111
chr17-23137210-DEL-152
chr17-23156158-DEL-11268
chr17-23156758-DEL-89
chr17-23159666-DEL-79
chr17-23173650-DEL-60
chr17-23185175-DEL-346
chr17-23189372-DEL-6423
chr17-23202033-DEL-3197
chr17-23223586-DEL-61
chr17-23243114-DEL-2391
chr17-23248534-DEL-343
chr17-23268965-DEL-122
chr17-23314839-DEL-2389
chr17-23323945-DEL-196
chr17-23332843-DEL-114
chr17-23338201-DEL-4507
chr17-23350110-DEL-5213
chr17-23400042-DEL-170
chr17-23415050-DEL-75
chr17-23421046-DEL-10558
chr17-23442077-DEL-1682
chr17-23463957-DEL-6527
chr17-23473213-DEL-6406
chr17-23481440-DEL-1445
chr17-23490951-DEL-1927
chr17-23533303-DEL-176
chr17-23631122-DEL-6920

chr17-23639877-DEL-848
chr17-23653060-DEL-841
chr17-23661813-DEL-796
chr17-23682351-DEL-196
chr17-23683347-DEL-885
chr17-23689375-DEL-211
chr17-23723387-DEL-6423
chr17-23734766-DEL-4735
chr17-23741922-DEL-819
chr17-23790827-DEL-207
chr17-23801325-DEL-278
chr17-23806723-DEL-501
chr17-23807848-DEL-68
chr17-23813364-DEL-53
chr17-23822416-DEL-52
chr17-23851424-DEL-88
chr17-23855897-DEL-103
chr17-23874311-DEL-171
chr17-23877770-DEL-61
chr17-23915627-DEL-96
chr17-23920442-DEL-177
chr17-23943192-DEL-197
chr17-24014155-DEL-130
chr17-24032267-DEL-333
chr17-24033896-DEL-83
chr17-24056659-DEL-195
chr17-24080947-DEL-62
chr17-24110987-DEL-3118
chr17-24125271-DEL-79
chr17-24127957-DEL-661
chr17-24129763-DEL-3134
chr17-24188225-DEL-55
chr17-24249810-DEL-68
chr17-24254600-DEL-93
chr17-24261163-DEL-132
chr17-24262657-DEL-127
chr17-24305628-DEL-124
chr17-24315240-DEL-367
chr17-24315771-DEL-283
chr17-24337618-DEL-196
chr17-24348612-DEL-1046
chr17-24351124-DEL-201
chr17-24381487-DEL-119
chr17-24384988-DEL-154
chr17-24391568-DEL-168
chr17-24399068-DEL-146
chr17-24405879-DEL-196
chr17-24412415-DEL-321
chr17-24413847-DEL-155
chr17-24443610-DEL-60
chr17-24459156-DEL-329
chr17-24474679-DEL-601
chr17-24474680-DEL-1061
chr17-24474951-DEL-1523
chr17-24476393-DEL-82
chr17-24478833-DEL-233
chr17-24479021-DEL-1037
chr17-24480029-DEL-181
chr17-24480112-DEL-625
chr17-24481285-DEL-189
chr17-24505656-DEL-3138
chr17-24509136-DEL-4917
chr17-24511515-DEL-74
chr17-24512716-DEL-88
chr17-24518511-DEL-79
chr17-24523383-DEL-492
chr17-24541903-DEL-167
chr17-24591409-DEL-683
chr17-24613177-DEL-182
chr17-24676232-DEL-78
chr17-24678712-DEL-1892
chr17-24743594-DEL-52
chr17-24758697-DEL-215
chr17-24763867-DEL-82
chr17-24781184-DEL-710
chr17-24843983-DEL-129
chr17-24881199-DEL-55

chr17-24886146-DEL-150
chr17-24910812-DEL-161
chr17-24922348-DEL-200
chr17-24943674-DEL-54
chr17-24948558-DEL-858
chr17-24959534-DEL-217
chr17-24968595-DEL-214
chr17-24999494-DEL-399
chr17-25006517-DEL-198
chr17-25080923-DEL-82
chr17-25152976-DEL-61
chr17-25154578-DEL-114
chr17-25159038-DEL-504
chr17-25165407-DEL-1411
chr17-25167440-DEL-85
chr17-25173253-DEL-58
chr17-25177416-DEL-172
chr17-25220733-DEL-487
chr17-25227263-DEL-72
chr17-25240613-DEL-200
chr17-25243145-DEL-67
chr17-25269968-DEL-369
chr17-25272114-DEL-151
chr17-25277900-DEL-1687
chr17-25287635-DEL-60
chr17-25290989-DEL-96
chr17-25302160-DEL-64
chr17-25307973-DEL-58
chr17-25314822-DEL-55
chr17-25377920-DEL-70
chr17-25414146-DEL-98
chr17-25445238-DEL-51
chr17-25496159-DEL-1253
chr17-25544864-DEL-167
chr17-25561695-DEL-64
chr17-25566823-DEL-121
chr17-25571270-DEL-3859
chr17-25583660-DEL-71
chr17-25686101-DEL-126
chr17-25689742-DEL-61
chr17-25717089-DEL-60
chr17-25723245-DEL-56
chr17-25779448-DEL-56
chr17-25837321-DEL-86
chr17-25847733-DEL-97
chr17-25870642-DEL-500
chr17-25877064-DEL-561
chr17-25877568-DEL-380
chr17-25902243-DEL-3757
chr17-25919206-DEL-202
chr17-25983166-DEL-146
chr17-26018259-DEL-537
chr17-26023682-DEL-68
chr17-26068985-DEL-79
chr17-26096367-DEL-85
chr17-26120577-DEL-568
chr17-26151338-DEL-54
chr17-26210089-DEL-141
chr17-26252026-DEL-881
chr17-26257009-DEL-118
chr17-26265022-DEL-246
chr17-26284544-DEL-96
chr17-26287759-DEL-69
chr17-26293836-DEL-52
chr17-26298706-DEL-296
chr17-26317298-DEL-124
chr17-26323019-DEL-84
chr17-26328772-DEL-209
chr17-26386546-DEL-113
chr17-26389436-DEL-201
chr17-26433903-DEL-99
chr17-26437439-DEL-72
chr17-26441577-DEL-184
chr17-26446046-DEL-195
chr17-26456433-DEL-189
chr17-26457182-DEL-91
chr17-26474217-DEL-55

chr17-26487824-DEL-180
chr17-26488316-DEL-1511
chr17-26492273-DEL-77
chr17-26510423-DEL-6853
chr17-26527260-DEL-229
chr17-26537544-DEL-60
chr17-26538198-DEL-281
chr17-26538916-DEL-91
chr17-26539549-DEL-268
chr17-26541058-DEL-84
chr17-26541789-DEL-289
chr17-26543193-DEL-129
chr17-26545605-DEL-175
chr17-26549153-DEL-62
chr17-26549432-DEL-479
chr17-26549652-DEL-2020
chr17-26638543-DEL-281
chr17-26645883-DEL-51
chr17-26691233-DEL-106
chr17-26698877-DEL-2029
chr17-26712738-DEL-506
chr17-26718994-DEL-56
chr17-26724326-DEL-61
chr17-26737175-DEL-784
chr17-26767239-DEL-176
chr17-26777534-DEL-133
chr17-26779178-DEL-197
chr17-26784681-DEL-253
chr17-26796412-DEL-279
chr17-26820134-DEL-67
chr17-26832834-DEL-86
chr17-26852061-DEL-116
chr17-26880068-DEL-214
chr17-26894590-DEL-203
chr17-26902075-DEL-1291
chr17-26902286-DEL-128
chr17-26902819-DEL-4419
chr17-26917517-DEL-61
chr17-26983292-DEL-597
chr17-27003979-DEL-968
chr17-27020252-DEL-500
chr17-27020910-DEL-1030
chr17-27045491-DEL-354
chr17-27046284-DEL-223
chr17-27068029-DEL-67
chr17-27077332-DEL-77
chr17-27090270-DEL-197
chr17-27114747-DEL-60
chr17-27127982-DEL-432
chr17-27148211-DEL-70
chr17-27216075-DEL-200
chr17-27244610-DEL-57
chr17-27250932-DEL-229
chr17-27256383-DEL-212
chr17-27265330-DEL-89
chr17-27299277-DEL-428
chr17-27311479-DEL-160
chr17-27381062-DEL-250
chr17-27384997-DEL-204
chr17-27396185-DEL-176
chr17-27402052-DEL-343
chr17-27456496-DEL-78
chr17-27481796-DEL-1629
chr17-27507232-DEL-2094
chr17-27523735-DEL-114
chr17-27541033-DEL-154
chr17-27543010-DEL-219
chr17-27544922-DEL-59
chr17-27569940-DEL-4081
chr17-27702491-DEL-81
chr17-27708602-DEL-68
chr17-27709449-DEL-54
chr17-27722534-DEL-74
chr17-27748576-DEL-347
chr17-27754933-DEL-393
chr17-27759634-DEL-51
chr17-27830849-DEL-2803

chr17-27834037-DEL-52
chr17-27836383-DEL-138
chr17-27838312-DEL-190
chr17-27877619-DEL-66
chr17-27892577-DEL-190
chr17-27912162-DEL-179
chr17-27918881-DEL-165
chr17-27948348-DEL-102
chr17-27951223-DEL-63
chr17-27963106-DEL-98
chr17-27965042-DEL-193
chr17-27966043-DEL-1351
chr17-28000888-DEL-761
chr17-28005816-DEL-194
chr17-28028127-DEL-429
chr17-28033879-DEL-400
chr17-28045112-DEL-127
chr17-28060563-DEL-176
chr17-28084912-DEL-72
chr17-28087903-DEL-1419
chr17-28115637-DEL-107
chr17-28123397-DEL-185
chr17-28134034-DEL-208
chr17-28141921-DEL-521
chr17-28161928-DEL-406
chr17-28165372-DEL-647
chr17-28166577-DEL-207
chr17-28168858-DEL-206
chr17-28179829-DEL-120
chr17-28180887-DEL-407
chr17-28181717-DEL-467
chr17-28211358-DEL-63
chr17-28282241-DEL-88
chr17-28294086-DEL-1287
chr17-28313094-DEL-576
chr17-28321518-DEL-54
chr17-28338530-DEL-165
chr17-28371500-DEL-223
chr17-28429742-DEL-385
chr17-28433650-DEL-204
chr17-28438357-DEL-227
chr17-28440496-DEL-63
chr17-28452802-DEL-204
chr17-28454276-DEL-413
chr17-28460729-DEL-597
chr17-28487795-DEL-616
chr17-28507891-DEL-478
chr17-28511026-DEL-1689
chr17-28519320-DEL-58
chr17-28530063-DEL-172
chr17-28540687-DEL-923
chr17-28542074-DEL-132
chr17-28577843-DEL-54
chr17-28585662-DEL-199
chr17-28586988-DEL-160
chr17-28598910-DEL-73
chr17-28606065-DEL-219
chr17-28627615-DEL-225
chr17-28637294-DEL-98
chr17-28645603-DEL-253
chr17-28654745-DEL-75
chr17-28656387-DEL-528
chr17-28667984-DEL-204
chr17-28673417-DEL-55
chr17-28684389-DEL-76
chr17-28693159-DEL-610
chr17-28721042-DEL-64
chr17-28728398-DEL-61
chr17-28745733-DEL-588
chr17-28750481-DEL-376
chr17-28752873-DEL-183
chr17-28760290-DEL-239
chr17-28766989-DEL-164
chr17-28805300-DEL-198
chr17-28814406-DEL-237
chr17-28828296-DEL-58
chr17-28845536-DEL-69

chr17-28858283-DEL-1119
chr17-28859460-DEL-58
chr17-28866738-DEL-55
chr17-28897335-DEL-186
chr17-28904441-DEL-219
chr17-28925663-DEL-66
chr17-28956742-DEL-54
chr17-28969407-DEL-81
chr17-28973694-DEL-104
chr17-28978905-DEL-225
chr17-28981614-DEL-275
chr17-28985461-DEL-117
chr17-28991828-DEL-61
chr17-28992490-DEL-191
chr17-28993517-DEL-189
chr17-29007936-DEL-68
chr17-29009570-DEL-403
chr17-29010296-DEL-152
chr17-29040624-DEL-108
chr17-29058681-DEL-436
chr17-29061304-DEL-56
chr17-29067527-DEL-104
chr17-29096736-DEL-100
chr17-29104182-DEL-279
chr17-29140357-DEL-377
chr17-29152429-DEL-174
chr17-29160065-DEL-51
chr17-29199343-DEL-187
chr17-29199697-DEL-330
chr17-29219744-DEL-109
chr17-29220441-DEL-149
chr17-29222282-DEL-190
chr17-29227813-DEL-51
chr17-29248702-DEL-83
chr17-29264231-DEL-58
chr17-29265645-DEL-215
chr17-29273871-DEL-124
chr17-29290382-DEL-191
chr17-29292057-DEL-77
chr17-29296729-DEL-60
chr17-29302128-DEL-188
chr17-29304903-DEL-173
chr17-29320026-DEL-870
chr17-29320066-DEL-643
chr17-29320440-DEL-7968
chr17-29320621-DEL-202
chr17-29321244-DEL-169
chr17-29321873-DEL-210
chr17-29322439-DEL-104
chr17-29323106-DEL-1536
chr17-29325535-DEL-223
chr17-29325856-DEL-266
chr17-29325861-DEL-1533
chr17-29326111-DEL-74
chr17-29326818-DEL-502
chr17-29327619-DEL-141
chr17-29332108-DEL-194
chr17-29337574-DEL-51
chr17-29340442-DEL-395
chr17-29351469-DEL-128
chr17-29364623-DEL-127
chr17-29391079-DEL-79
chr17-29393044-DEL-135
chr17-29412452-DEL-74
chr17-29417057-DEL-676
chr17-29417152-DEL-916
chr17-29421190-DEL-156
chr17-29426880-DEL-54
chr17-29437997-DEL-446
chr17-29461654-DEL-82
chr17-29497896-DEL-90
chr17-29519527-DEL-205
chr17-29531130-DEL-151
chr17-29536353-DEL-443
chr17-29549960-DEL-71
chr17-29553771-DEL-86
chr17-29572925-DEL-212

chr17-29585677-DEL-93
chr17-29587673-DEL-57
chr17-29589873-DEL-57
chr17-29633385-DEL-160
chr17-29655279-DEL-235
chr17-29681991-DEL-170
chr17-29734996-DEL-134
chr17-29740681-DEL-182
chr17-29748618-DEL-7291
chr17-29771790-DEL-198
chr17-29791380-DEL-167
chr17-29793033-DEL-211
chr17-29813871-DEL-58
chr17-29827444-DEL-193
chr17-29842636-DEL-167
chr17-29849336-DEL-195
chr17-29875105-DEL-176
chr17-29876433-DEL-179
chr17-29882414-DEL-141
chr17-29947631-DEL-855
chr17-29976720-DEL-205
chr17-29989067-DEL-117
chr17-30009959-DEL-69
chr17-30021687-DEL-623
chr17-30030468-DEL-57
chr17-30076014-DEL-104
chr17-30081739-DEL-222
chr17-30149259-DEL-67
chr17-30186577-DEL-64
chr17-30216209-DEL-60
chr17-30245360-DEL-196
chr17-30252804-DEL-215
chr17-30302055-DEL-762
chr17-30308035-DEL-59
chr17-30312954-DEL-176
chr17-30357019-DEL-170
chr17-30359155-DEL-278
chr17-30359254-DEL-763
chr17-30391418-DEL-54
chr17-30440551-DEL-152
chr17-30476164-DEL-68
chr17-30479554-DEL-169
chr17-30489143-DEL-179
chr17-30491163-DEL-150
chr17-30495211-DEL-94
chr17-30521031-DEL-136
chr17-30521497-DEL-596
chr17-30551055-DEL-192
chr17-30614514-DEL-515
chr17-30623390-DEL-70
chr17-30624841-DEL-128
chr17-30641669-DEL-183
chr17-30683998-DEL-1502
chr17-30761662-DEL-154
chr17-30774739-DEL-158
chr17-30781430-DEL-1219
chr17-30782975-DEL-426
chr17-30793127-DEL-153
chr17-30810680-DEL-201
chr17-30820148-DEL-1262
chr17-30827609-DEL-84
chr17-30834350-DEL-2919
chr17-30844034-DEL-2467
chr17-30846562-DEL-1100
chr17-30847548-DEL-195
chr17-30850076-DEL-104
chr17-30852970-DEL-90
chr17-30877360-DEL-346
chr17-30878918-DEL-2016
chr17-30881468-DEL-55
chr17-30889230-DEL-129
chr17-30895295-DEL-61
chr17-30904491-DEL-146
chr17-30905802-DEL-387
chr17-30936883-DEL-307
chr17-30975028-DEL-200
chr17-30978437-DEL-57

chr17-30986597-DEL-58
chr17-30992988-DEL-79
chr17-31005710-DEL-87
chr17-31006288-DEL-84
chr17-31016884-DEL-1191
chr17-31017471-DEL-251
chr17-31045527-DEL-92
chr17-31087352-DEL-81
chr17-31098772-DEL-62
chr17-31115345-DEL-140
chr17-31115640-DEL-485
chr17-31136788-DEL-167
chr17-31137581-DEL-111
chr17-31144740-DEL-58
chr17-31171183-DEL-170
chr17-31182359-DEL-99
chr17-31211973-DEL-118
chr17-31256585-DEL-74
chr17-31275314-DEL-455
chr17-31311811-DEL-54
chr17-31362494-DEL-167
chr17-31370404-DEL-56
chr17-31424240-DEL-81
chr17-31425960-DEL-134
chr17-31435040-DEL-276
chr17-31448764-DEL-157
chr17-31449354-DEL-138
chr17-31459517-DEL-190
chr17-31469261-DEL-379
chr17-31481333-DEL-51
chr17-31483914-DEL-155
chr17-31503306-DEL-1223
chr17-31510614-DEL-230
chr17-31513515-DEL-109
chr17-31528695-DEL-472
chr17-31554802-DEL-56
chr17-31560773-DEL-56
chr17-31578234-DEL-139
chr17-31591045-DEL-5218
chr17-31614290-DEL-65
chr17-31616179-DEL-59
chr17-31657068-DEL-94
chr17-31665821-DEL-73
chr17-31672884-DEL-801
chr17-31674453-DEL-480
chr17-31682403-DEL-86
chr17-31683494-DEL-97
chr17-31692560-DEL-62
chr17-31696731-DEL-1738
chr17-31710324-DEL-91
chr17-31710973-DEL-68
chr17-31712421-DEL-154
chr17-31742892-DEL-195
chr17-31756871-DEL-52
chr17-31761717-DEL-64
chr17-31780109-DEL-130
chr17-31782124-DEL-183
chr17-31800310-DEL-56
chr17-31804430-DEL-233
chr17-31817082-DEL-186
chr17-31828623-DEL-64
chr17-31844285-DEL-62
chr17-31848707-DEL-136
chr17-31852352-DEL-482
chr17-31859680-DEL-61
chr17-31860379-DEL-256
chr17-31862812-DEL-968
chr17-31920473-DEL-285
chr17-31930461-DEL-112
chr17-31994299-DEL-172
chr17-32016904-DEL-2020
chr17-32023662-DEL-912
chr17-32041825-DEL-115
chr17-32085668-DEL-276
chr17-32093715-DEL-87
chr17-32119848-DEL-90
chr17-32143979-DEL-74

chr17-32156910-DEL-1878
chr17-32232796-DEL-100
chr17-32306969-DEL-290
chr17-32342453-DEL-51
chr17-32347265-DEL-53
chr17-32354897-DEL-1038
chr17-32354995-DEL-420
chr17-32378749-DEL-60
chr17-32402971-DEL-106
chr17-32434318-DEL-140
chr17-32447997-DEL-221
chr17-32493418-DEL-61
chr17-32507948-DEL-185
chr17-32508096-DEL-7436
chr17-32511380-DEL-171
chr17-32517800-DEL-223
chr17-32542972-DEL-324
chr17-32545606-DEL-4106
chr17-32547222-DEL-236
chr17-32618716-DEL-68
chr17-32619382-DEL-90
chr17-32621725-DEL-80
chr17-32624961-DEL-1633
chr17-32626663-DEL-279
chr17-32635391-DEL-142
chr17-32640907-DEL-253
chr17-32646796-DEL-68
chr17-32648319-DEL-511
chr17-32648988-DEL-404
chr17-32676173-DEL-82
chr17-32677098-DEL-56
chr17-32731452-DEL-130
chr17-32737432-DEL-64
chr17-32745436-DEL-118
chr17-32746551-DEL-68
chr17-32754958-DEL-89
chr17-32757482-DEL-57
chr17-32767103-DEL-188
chr17-32774267-DEL-24994
chr17-32802996-DEL-1034
chr17-32814297-DEL-3743
chr17-32822074-DEL-396
chr17-32823976-DEL-202
chr17-32894187-DEL-103
chr17-32908037-DEL-1353
chr17-32913474-DEL-181
chr17-32921989-DEL-5514
chr17-32935465-DEL-3630
chr17-32943564-DEL-1714
chr17-32951362-DEL-195
chr17-32960993-DEL-71
chr17-32968514-DEL-625
chr17-32970182-DEL-177
chr17-32982309-DEL-208
chr17-33005005-DEL-141
chr17-33008877-DEL-112
chr17-33029331-DEL-72
chr17-33035550-DEL-60
chr17-33044178-DEL-1071
chr17-33047528-DEL-96
chr17-33064943-DEL-54
chr17-33066048-DEL-1168
chr17-33067579-DEL-2009
chr17-33080271-DEL-476
chr17-33090117-DEL-152
chr17-33093750-DEL-183
chr17-33098388-DEL-222
chr17-33118689-DEL-70
chr17-33125883-DEL-55
chr17-33129185-DEL-89
chr17-33137385-DEL-713
chr17-33143096-DEL-91
chr17-33149352-DEL-92
chr17-33157247-DEL-72
chr17-33193281-DEL-1058
chr17-33199675-DEL-72
chr17-33227282-DEL-1280

chr17-33234744-DEL-102
chr17-33239396-DEL-76
chr17-33242448-DEL-172
chr17-33245924-DEL-434
chr17-33254958-DEL-311
chr17-33289356-DEL-141
chr17-33295847-DEL-2193
chr17-33298035-DEL-2090
chr17-33301640-DEL-430
chr17-33305134-DEL-160
chr17-33317340-DEL-84
chr17-33329796-DEL-119
chr17-33340645-DEL-353
chr17-33360530-DEL-79
chr17-33364768-DEL-73
chr17-33377021-DEL-406
chr17-33377817-DEL-220
chr17-33378124-DEL-1844
chr17-33382889-DEL-2643
chr17-33405567-DEL-2643
chr17-33414309-DEL-2267
chr17-33422177-DEL-566
chr17-33424453-DEL-536
chr17-33433743-DEL-58
chr17-33461235-DEL-95
chr17-33492124-DEL-212
chr17-33497596-DEL-6951
chr17-33508899-DEL-62
chr17-33524255-DEL-55
chr17-33529067-DEL-2009
chr17-33560227-DEL-151
chr17-33560971-DEL-179
chr17-33561555-DEL-379
chr17-33563277-DEL-3932
chr17-33572786-DEL-65
chr17-33578884-DEL-9866
chr17-33581162-DEL-7385
chr17-33619982-DEL-54
chr17-33630221-DEL-1644
chr17-33653519-DEL-68
chr17-33655493-DEL-990
chr17-33659450-DEL-58
chr17-33661628-DEL-71
chr17-33670643-DEL-3201
chr17-33676559-DEL-940
chr17-33698426-DEL-53
chr17-33709612-DEL-77
chr17-33719071-DEL-112
chr17-33720896-DEL-56
chr17-33760845-DEL-62
chr17-33779829-DEL-94
chr17-33800168-DEL-217
chr17-33807456-DEL-149
chr17-33813558-DEL-54
chr17-33822794-DEL-61
chr17-33825846-DEL-87
chr17-33880890-DEL-1093
chr17-33912691-DEL-110
chr17-33915902-DEL-68
chr17-33916722-DEL-58
chr17-33921856-DEL-70
chr17-33924023-DEL-191
chr17-33942532-DEL-59
chr17-33944693-DEL-603
chr17-33945314-DEL-3461
chr17-33961191-DEL-144
chr17-33986727-DEL-1782
chr17-33991978-DEL-191
chr17-34009862-DEL-126
chr17-34014782-DEL-121
chr17-34018406-DEL-96
chr17-34019088-DEL-205
chr17-34025386-DEL-123
chr17-34026734-DEL-74
chr17-34027914-DEL-106
chr17-34058562-DEL-187
chr17-34060258-DEL-184

chr17-34085219-DEL-393
chr17-34111964-DEL-5890
chr17-34131900-DEL-72
chr17-34171284-DEL-90
chr17-34182553-DEL-390
chr17-34183381-DEL-454
chr17-34190284-DEL-795
chr17-34209814-DEL-109
chr17-34309974-DEL-67
chr17-34376890-DEL-75
chr17-34387902-DEL-56
chr17-34398681-DEL-57
chr17-34413595-DEL-79
chr17-34421115-DEL-285
chr17-34435596-DEL-106
chr17-34437670-DEL-70
chr17-34474839-DEL-326
chr17-34484744-DEL-945
chr17-34490485-DEL-54
chr17-34491734-DEL-252
chr17-34499605-DEL-398
chr17-34508482-DEL-55
chr17-34549548-DEL-182
chr17-34571986-DEL-55
chr17-34572712-DEL-125
chr17-34591558-DEL-55
chr17-34662494-DEL-57
chr17-34681559-DEL-2982
chr17-34681574-DEL-1491
chr17-34682640-DEL-1490
chr17-34714877-DEL-266
chr17-34717804-DEL-223
chr17-34724042-DEL-159
chr17-34732036-DEL-111
chr17-34741397-DEL-5531
chr17-34741400-DEL-3524
chr17-34741402-DEL-1407
chr17-34742880-DEL-4042
chr17-34744976-DEL-1946
chr17-34746707-DEL-197
chr17-34786847-DEL-56
chr17-34788689-DEL-170
chr17-34798402-DEL-152
chr17-34828468-DEL-54
chr17-34840872-DEL-388
chr17-34843680-DEL-363
chr17-34854038-DEL-64
chr17-34882183-DEL-1374
chr17-34885403-DEL-203
chr17-34957632-DEL-134
chr17-35100383-DEL-54
chr17-35128159-DEL-478
chr17-35131173-DEL-100
chr17-35155100-DEL-125
chr17-35179054-DEL-60
chr17-35180004-DEL-54
chr17-35195441-DEL-207
chr17-35206281-DEL-73
chr17-35206653-DEL-957
chr17-35215151-DEL-341
chr17-35253122-DEL-122
chr17-35275046-DEL-121
chr17-35303466-DEL-75
chr17-35304940-DEL-206
chr17-35306519-DEL-205
chr17-35311089-DEL-153
chr17-35323374-DEL-742
chr17-35335775-DEL-211
chr17-35344842-DEL-64
chr17-35351238-DEL-228
chr17-35388359-DEL-217
chr17-35416754-DEL-75
chr17-35423855-DEL-56
chr17-35443904-DEL-2456
chr17-35475814-DEL-188
chr17-35488869-DEL-2723
chr17-35493210-DEL-207

chr17-35497854-DEL-1589
chr17-35506594-DEL-59
chr17-35519936-DEL-68
chr17-35525054-DEL-51
chr17-35526624-DEL-4321
chr17-35528933-DEL-54
chr17-35533461-DEL-153
chr17-35559285-DEL-192
chr17-35570503-DEL-113
chr17-35572898-DEL-614
chr17-35575433-DEL-263
chr17-35586318-DEL-7092
chr17-35615608-DEL-389
chr17-35641542-DEL-141
chr17-35659948-DEL-104
chr17-35673428-DEL-1919
chr17-35679164-DEL-85
chr17-35680857-DEL-208
chr17-35733130-DEL-126
chr17-35735522-DEL-72
chr17-35753878-DEL-86
chr17-35771124-DEL-66
chr17-35789952-DEL-76
chr17-35796441-DEL-56
chr17-35805990-DEL-203
chr17-35807165-DEL-206
chr17-35842686-DEL-101
chr17-35847342-DEL-1100
chr17-35858656-DEL-56
chr17-35890199-DEL-190
chr17-35903667-DEL-181
chr17-35914055-DEL-5886
chr17-35924211-DEL-232
chr17-35931199-DEL-90
chr17-35934072-DEL-208
chr17-35949688-DEL-125
chr17-35953560-DEL-85
chr17-35955933-DEL-52
chr17-36014396-DEL-116
chr17-36020764-DEL-261
chr17-36030832-DEL-450
chr17-36034305-DEL-68
chr17-36039169-DEL-66
chr17-36040666-DEL-200
chr17-36057203-DEL-1043
chr17-36062340-DEL-81
chr17-36065569-DEL-1489
chr17-36072396-DEL-228
chr17-36080802-DEL-203
chr17-36086246-DEL-74
chr17-36091474-DEL-55
chr17-36097603-DEL-366
chr17-36140178-DEL-175
chr17-36142190-DEL-74
chr17-36155552-DEL-113
chr17-36214331-DEL-416
chr17-36246271-DEL-72
chr17-36251281-DEL-56
chr17-36267169-DEL-59
chr17-36282893-DEL-189
chr17-36293650-DEL-90
chr17-36296755-DEL-1518
chr17-36303733-DEL-130
chr17-36306786-DEL-96
chr17-36314722-DEL-171
chr17-36347637-DEL-202
chr17-36348370-DEL-86
chr17-36354953-DEL-79
chr17-36356049-DEL-58
chr17-36378644-DEL-8706
chr17-36388883-DEL-181
chr17-36392428-DEL-196
chr17-36396520-DEL-365
chr17-36404377-DEL-478
chr17-36411439-DEL-6633
chr17-36450117-DEL-223
chr17-36451397-DEL-199

chr17-36464734-DEL-792
chr17-36466063-DEL-81
chr17-36468741-DEL-2244
chr17-36472203-DEL-766
chr17-36483020-DEL-1745
chr17-36507047-DEL-123
chr17-36513048-DEL-673
chr17-36518880-DEL-3030
chr17-36532572-DEL-117
chr17-36534524-DEL-91
chr17-36536157-DEL-53
chr17-36538863-DEL-53
chr17-36539934-DEL-137
chr17-36542210-DEL-174
chr17-36544095-DEL-212
chr17-36546321-DEL-344
chr17-36548017-DEL-169
chr17-36556234-DEL-159
chr17-36557726-DEL-667
chr17-36559287-DEL-4016
chr17-36563707-DEL-116
chr17-36612766-DEL-299
chr17-36620153-DEL-1488
chr17-36676099-DEL-769
chr17-36689396-DEL-2870
chr17-36716885-DEL-607
chr17-36740246-DEL-327
chr17-36752313-DEL-1440
chr17-36778072-DEL-202
chr17-36798311-DEL-6399
chr17-36801704-DEL-827
chr17-36808536-DEL-84
chr17-36865017-DEL-171
chr17-36877132-DEL-6398
chr17-36883188-DEL-199
chr17-36893178-DEL-1175
chr17-36898720-DEL-19112
chr17-36916662-DEL-506
chr17-36928902-DEL-74
chr17-36963133-DEL-117
chr17-36966545-DEL-183
chr17-36983885-DEL-4975
chr17-36994382-DEL-102
chr17-37044979-DEL-26036
chr17-37045158-DEL-13075
chr17-37100469-DEL-929
chr17-37105208-DEL-56
chr17-37187090-DEL-122
chr17-37189725-DEL-255
chr17-37191685-DEL-106
chr17-37201352-DEL-4241
chr17-37254924-DEL-145
chr17-37278885-DEL-133
chr17-37282748-DEL-212
chr17-37301251-DEL-243
chr17-37324549-DEL-61
chr17-37330494-DEL-361
chr17-37332489-DEL-307
chr17-37336981-DEL-203
chr17-37340393-DEL-144
chr17-37347795-DEL-56
chr17-37410550-DEL-6424
chr17-37419711-DEL-433
chr17-37448378-DEL-198
chr17-37463376-DEL-55
chr17-37496921-DEL-69
chr17-37520847-DEL-113
chr17-37570484-DEL-435
chr17-37570922-DEL-1843
chr17-37614086-DEL-78
chr17-37621645-DEL-387
chr17-37622212-DEL-66
chr17-37652916-DEL-57
chr17-37660322-DEL-6206
chr17-37666936-DEL-1924
chr17-37677862-DEL-123
chr17-37711044-DEL-498

chr17-37732843-DEL-7639
chr17-37746203-DEL-556
chr17-37748787-DEL-198
chr17-37783532-DEL-6335
chr17-37798011-DEL-328
chr17-37798844-DEL-66
chr17-37849145-DEL-5309
chr17-37857039-DEL-504
chr17-37883016-DEL-437
chr17-37912054-DEL-5935
chr17-37924658-DEL-59
chr17-37940377-DEL-389
chr17-37941811-DEL-346
chr17-37946186-DEL-2705
chr17-37951597-DEL-2739
chr17-37960148-DEL-204
chr17-37962907-DEL-221
chr17-37964653-DEL-926
chr17-37978567-DEL-494
chr17-38000090-DEL-556
chr17-38023034-DEL-612
chr17-38023717-DEL-1109
chr17-38041891-DEL-6165
chr17-38081974-DEL-69
chr17-38099104-DEL-4011
chr17-38101839-DEL-66
chr17-38109563-DEL-421
chr17-38122035-DEL-6628
chr17-38133637-DEL-6684
chr17-38135153-DEL-57
chr17-38138755-DEL-58
chr17-38175218-DEL-1406
chr17-38176305-DEL-274
chr17-38195544-DEL-477
chr17-38200236-DEL-938
chr17-38227146-DEL-175
chr17-38251205-DEL-643
chr17-38291135-DEL-223
chr17-38302116-DEL-697
chr17-38303365-DEL-565
chr17-38312842-DEL-186
chr17-38318462-DEL-74
chr17-38322795-DEL-83
chr17-38325460-DEL-320
chr17-38336039-DEL-487
chr17-38345106-DEL-552
chr17-38346008-DEL-1481
chr17-38351887-DEL-51
chr17-38358055-DEL-2966
chr17-38358741-DEL-81
chr17-38364194-DEL-249
chr17-38373024-DEL-87
chr17-38377244-DEL-405
chr17-38381596-DEL-899
chr17-38389642-DEL-159
chr17-38392420-DEL-183
chr17-38413968-DEL-196
chr17-38431596-DEL-520
chr17-38434765-DEL-685
chr17-38446414-DEL-3351
chr17-38516617-DEL-202
chr17-38535955-DEL-6376
chr17-38535962-DEL-216
chr17-38551547-DEL-6209
chr17-38560681-DEL-82
chr17-38568043-DEL-1922
chr17-38577053-DEL-795
chr17-38614972-DEL-73176
chr17-38625477-DEL-313
chr17-38628649-DEL-130
chr17-38630095-DEL-368
chr17-38636835-DEL-142
chr17-38648144-DEL-69
chr17-38658079-DEL-108
chr17-38663926-DEL-600
chr17-38668725-DEL-7645
chr17-38670257-DEL-698

chr17-38686464-DEL-196
chr17-38696176-DEL-516
chr17-38697050-DEL-2630
chr17-38699746-DEL-192
chr17-38700142-DEL-2006
chr17-38706581-DEL-78
chr17-38715763-DEL-2178
chr17-38732712-DEL-660
chr17-38737917-DEL-82
chr17-38739444-DEL-2307
chr17-38741150-DEL-1085
chr17-38742148-DEL-359
chr17-38742991-DEL-1026
chr17-38757078-DEL-119
chr17-38761900-DEL-95
chr17-38771285-DEL-845
chr17-38809156-DEL-3398
chr17-38817389-DEL-190
chr17-38825871-DEL-6202
chr17-38841514-DEL-252
chr17-38843676-DEL-1050
chr17-38847171-DEL-7830
chr17-38855095-DEL-56
chr17-38863796-DEL-680
chr17-38908854-DEL-958
chr17-38910518-DEL-3021
chr17-38915637-DEL-62
chr17-38918409-DEL-74
chr17-38921913-DEL-1174
chr17-38927593-DEL-56
chr17-38930755-DEL-61
chr17-38934366-DEL-200
chr17-38947454-DEL-95
chr17-38952597-DEL-201
chr17-38972523-DEL-200
chr17-38975353-DEL-85
chr17-38986302-DEL-59
chr17-38987921-DEL-6590
chr17-38998409-DEL-75
chr17-39051801-DEL-58
chr17-39066929-DEL-1882
chr17-39073094-DEL-5715
chr17-39083873-DEL-4027
chr17-39113513-DEL-1257
chr17-39115924-DEL-504
chr17-39118447-DEL-248
chr17-39175894-DEL-310
chr17-39201598-DEL-6703
chr17-39208354-DEL-6490
chr17-39211779-DEL-4889
chr17-39239674-DEL-74
chr17-39255650-DEL-67
chr17-39269514-DEL-54
chr17-39287423-DEL-6381
chr17-39310228-DEL-6231
chr17-39321286-DEL-8965
chr17-39321573-DEL-664
chr17-39331592-DEL-98
chr17-39332514-DEL-88
chr17-39345095-DEL-676
chr17-39347439-DEL-53
chr17-39355043-DEL-66
chr17-39364864-DEL-2901
chr17-39369942-DEL-158
chr17-39371300-DEL-55
chr17-39381097-DEL-2208
chr17-39387031-DEL-1954
chr17-39437163-DEL-3609
chr17-39442661-DEL-2323
chr17-39449946-DEL-75
chr17-39460313-DEL-61
chr17-39498105-DEL-54
chr17-39503878-DEL-583
chr17-39535735-DEL-57
chr17-39536091-DEL-6200
chr17-39549056-DEL-98
chr17-39554449-DEL-248

chr17-39562785-DEL-52
chr17-39573671-DEL-3006
chr17-39573765-DEL-159
chr17-39576741-DEL-1547
chr17-39580548-DEL-382
chr17-39588903-DEL-221
chr17-39591199-DEL-1135
chr17-39591553-DEL-566
chr17-39601311-DEL-401
chr17-39637240-DEL-292
chr17-39653036-DEL-1839
chr17-39670229-DEL-1167
chr17-39710214-DEL-126
chr17-39712307-DEL-4123
chr17-39737218-DEL-814
chr17-39740348-DEL-158
chr17-39741387-DEL-64
chr17-39769047-DEL-166
chr17-39772270-DEL-4496
chr17-39776797-DEL-1614
chr17-39778906-DEL-2111
chr17-39782383-DEL-1918
chr17-39786499-DEL-86
chr17-39808733-DEL-2362
chr17-39817998-DEL-326
chr17-39861082-DEL-807
chr17-39918037-DEL-66
chr17-39921702-DEL-56
chr17-39928925-DEL-1089
chr17-39929584-DEL-73
chr17-39941588-DEL-64
chr17-39944105-DEL-222
chr17-39947521-DEL-525
chr17-39957745-DEL-359
chr17-39960528-DEL-3903
chr17-39960531-DEL-24985
chr17-39967339-DEL-68
chr17-39975209-DEL-60
chr17-39979804-DEL-52
chr17-39989661-DEL-2120
chr17-40035251-DEL-74
chr17-40042652-DEL-68
chr17-40046962-DEL-1131
chr17-40079706-DEL-213
chr17-40164737-DEL-124
chr17-40216634-DEL-307
chr17-40227349-DEL-241
chr17-40243823-DEL-129
chr17-40246113-DEL-169
chr17-40247558-DEL-436
chr17-40279935-DEL-1781
chr17-40285747-DEL-358
chr17-40314679-DEL-132
chr17-40339255-DEL-936
chr17-40345689-DEL-70
chr17-40354224-DEL-1487
chr17-40387405-DEL-62
chr17-40401410-DEL-4129
chr17-40420497-DEL-812
chr17-40426093-DEL-2864
chr17-40481417-DEL-54
chr17-40484250-DEL-568
chr17-40489308-DEL-5619
chr17-40503102-DEL-220
chr17-40507860-DEL-1449
chr17-40514751-DEL-185
chr17-40522689-DEL-6388
chr17-40556347-DEL-76
chr17-40565857-DEL-4378
chr17-40573021-DEL-52
chr17-40579178-DEL-54
chr17-40582472-DEL-355
chr17-40587960-DEL-6011
chr17-40595851-DEL-434
chr17-40607025-DEL-177
chr17-40609252-DEL-58
chr17-40649011-DEL-1293

chr17-40650611-DEL-135
chr17-40660644-DEL-423
chr17-40661451-DEL-2130
chr17-40667307-DEL-789
chr17-40673719-DEL-200
chr17-40675154-DEL-518
chr17-40676673-DEL-6412
chr17-40685153-DEL-6244
chr17-40689761-DEL-2556
chr17-40701490-DEL-162
chr17-40703690-DEL-1255
chr17-40743601-DEL-605
chr17-40744910-DEL-656
chr17-40748382-DEL-68
chr17-40750642-DEL-4503
chr17-40768999-DEL-992
chr17-40775094-DEL-6238
chr17-40794785-DEL-594
chr17-40814682-DEL-60
chr17-40838465-DEL-68
chr17-40842893-DEL-178
chr17-40846007-DEL-7152
chr17-40852201-DEL-642
chr17-40852467-DEL-427
chr17-40868082-DEL-952
chr17-40880729-DEL-59
chr17-40883910-DEL-2508
chr17-40889324-DEL-10854
chr17-40992804-DEL-447
chr17-41005869-DEL-9417
chr17-41034848-DEL-2205
chr17-41057474-DEL-216
chr17-41061497-DEL-161
chr17-41061793-DEL-1300
chr17-41065181-DEL-82
chr17-41092869-DEL-108
chr17-41105386-DEL-53
chr17-41113328-DEL-424
chr17-41126903-DEL-54
chr17-41130778-DEL-606
chr17-41140554-DEL-1114
chr17-41160355-DEL-1592
chr17-41162397-DEL-1813
chr17-41183724-DEL-52
chr17-41206173-DEL-2964
chr17-41217945-DEL-262
chr17-41223962-DEL-586
chr17-41229097-DEL-52
chr17-41240536-DEL-140
chr17-41276226-DEL-6709
chr17-41290353-DEL-6741
chr17-41318451-DEL-1873
chr17-41339325-DEL-8594
chr17-41341197-DEL-1765
chr17-41347269-DEL-204
chr17-41363402-DEL-240
chr17-41371044-DEL-1084
chr17-41386974-DEL-1624
chr17-41388799-DEL-214
chr17-41395777-DEL-121
chr17-41420737-DEL-52
chr17-41431406-DEL-167
chr17-41435973-DEL-591
chr17-41463920-DEL-52
chr17-41465746-DEL-1842
chr17-41473165-DEL-55
chr17-41477211-DEL-59
chr17-41505628-DEL-86
chr17-41508320-DEL-62
chr17-41520286-DEL-1364
chr17-41523269-DEL-203
chr17-41533392-DEL-507
chr17-41550467-DEL-1649
chr17-41583577-DEL-2350
chr17-41606981-DEL-1960
chr17-41614626-DEL-15472
chr17-41625573-DEL-723

chr17-41629295-DEL-94
chr17-41643922-DEL-92
chr17-41701931-DEL-1476
chr17-41728296-DEL-805
chr17-41741423-DEL-105
chr17-41748244-DEL-1622
chr17-41751626-DEL-1020
chr17-41773181-DEL-186
chr17-41779960-DEL-4764
chr17-41787112-DEL-618
chr17-41805564-DEL-863
chr17-41807370-DEL-768
chr17-41813495-DEL-204
chr17-41823312-DEL-653
chr17-41864405-DEL-67
chr17-41871835-DEL-357
chr17-41893320-DEL-956
chr17-41895097-DEL-67
chr17-41897369-DEL-2946
chr17-41901675-DEL-63
chr17-41904493-DEL-651
chr17-41914559-DEL-6466
chr17-41953763-DEL-1070
chr17-41979558-DEL-2602
chr17-41985721-DEL-2570
chr17-41998158-DEL-6021
chr17-42006840-DEL-67
chr17-42007883-DEL-52
chr17-42013067-DEL-2215
chr17-42015185-DEL-120
chr17-42015761-DEL-580
chr17-42067649-DEL-500
chr17-42078538-DEL-6204
chr17-42090371-DEL-94
chr17-42105720-DEL-366
chr17-42108142-DEL-760
chr17-42111924-DEL-221
chr17-42118050-DEL-1011
chr17-42122224-DEL-6202
chr17-42132467-DEL-90
chr17-42136414-DEL-1377
chr17-42138882-DEL-87
chr17-42150370-DEL-89
chr17-42153286-DEL-2000
chr17-42160615-DEL-635
chr17-42168909-DEL-717
chr17-42170697-DEL-6183
chr17-42182879-DEL-641
chr17-42185011-DEL-1505
chr17-42186575-DEL-189
chr17-42186829-DEL-1415
chr17-42196739-DEL-197
chr17-42197829-DEL-114
chr17-42222373-DEL-129
chr17-42225238-DEL-6363
chr17-42241595-DEL-392
chr17-42242786-DEL-1983
chr17-42258139-DEL-812
chr17-42267185-DEL-228
chr17-42297387-DEL-142
chr17-42325590-DEL-6234
chr17-42362970-DEL-2226
chr17-42365771-DEL-130
chr17-42397127-DEL-863
chr17-42405553-DEL-82
chr17-42408800-DEL-491
chr17-42411349-DEL-55
chr17-42413752-DEL-337
chr17-42416500-DEL-816
chr17-42416676-DEL-177
chr17-42419746-DEL-178
chr17-42438553-DEL-234
chr17-42439732-DEL-875
chr17-42450468-DEL-119
chr17-42454312-DEL-332
chr17-42462973-DEL-187
chr17-42513100-DEL-114

chr17-42535718-DEL-367
chr17-42536818-DEL-888
chr17-42542018-DEL-1473
chr17-42555950-DEL-631
chr17-42571893-DEL-399
chr17-42576607-DEL-703
chr17-42620405-DEL-124
chr17-42663469-DEL-292
chr17-42678160-DEL-56
chr17-42705272-DEL-169
chr17-42758611-DEL-171
chr17-42764299-DEL-218
chr17-42770449-DEL-132
chr17-42785109-DEL-1376
chr17-42785689-DEL-52
chr17-42801834-DEL-1217
chr17-42836144-DEL-66
chr17-42846547-DEL-72
chr17-42857009-DEL-56
chr17-42865767-DEL-544
chr17-42889435-DEL-875
chr17-42899666-DEL-2112
chr17-42902211-DEL-52
chr17-42910378-DEL-188
chr17-42915559-DEL-85
chr17-42998201-DEL-89
chr17-43005652-DEL-153
chr17-43016471-DEL-188
chr17-43027254-DEL-64
chr17-43031461-DEL-104
chr17-43038937-DEL-80
chr17-43041398-DEL-98
chr17-43068140-DEL-204
chr17-43068142-DEL-20295
chr17-43074800-DEL-5398
chr17-43076048-DEL-485
chr17-43083162-DEL-1758
chr17-43085543-DEL-119
chr17-43109216-DEL-57
chr17-43125393-DEL-88
chr17-43132184-DEL-189
chr17-43153637-DEL-196
chr17-43174368-DEL-70
chr17-43192322-DEL-207
chr17-43201710-DEL-76
chr17-43203510-DEL-99
chr17-43206311-DEL-276
chr17-43211711-DEL-2120
chr17-43216685-DEL-53
chr17-43220295-DEL-54
chr17-43242485-DEL-167
chr17-43243606-DEL-115
chr17-43278322-DEL-52
chr17-43297561-DEL-52
chr17-43317099-DEL-167
chr17-43361187-DEL-116
chr17-43362468-DEL-102
chr17-43406012-DEL-56
chr17-43409107-DEL-878
chr17-43409754-DEL-1356
chr17-43432958-DEL-399
chr17-43452201-DEL-453
chr17-43476455-DEL-84
chr17-43484500-DEL-86
chr17-43503915-DEL-142
chr17-43521711-DEL-899
chr17-43522609-DEL-79
chr17-43532926-DEL-69
chr17-43595971-DEL-88
chr17-43603666-DEL-96
chr17-43614988-DEL-356
chr17-43616959-DEL-208
chr17-43626558-DEL-174
chr17-43629947-DEL-52
chr17-43630691-DEL-859
chr17-43641331-DEL-60
chr17-43664188-DEL-229

chr17-43682657-DEL-346
chr17-43690506-DEL-877
chr17-43708045-DEL-58
chr17-43789379-DEL-52
chr17-43794045-DEL-249
chr17-43794649-DEL-581
chr17-43800979-DEL-174
chr17-43801245-DEL-608
chr17-43805503-DEL-90
chr17-43820490-DEL-318
chr17-43853188-DEL-59
chr17-43885076-DEL-4960
chr17-43901707-DEL-811
chr17-43909454-DEL-64
chr17-43918727-DEL-174
chr17-43925860-DEL-99
chr17-43950516-DEL-1265
chr17-43959154-DEL-80
chr17-43972704-DEL-62
chr17-43989262-DEL-76
chr17-44015338-DEL-72
chr17-44030360-DEL-2075
chr17-44033313-DEL-130
chr17-44101211-DEL-106
chr17-44106337-DEL-810
chr17-44153997-DEL-97
chr17-44162224-DEL-1083
chr17-44163360-DEL-910
chr17-44185502-DEL-69
chr17-44195833-DEL-63
chr17-44198009-DEL-75
chr17-44220040-DEL-230
chr17-44271380-DEL-99
chr17-44290638-DEL-96
chr17-44296050-DEL-139
chr17-44350770-DEL-145
chr17-44358679-DEL-110
chr17-44376795-DEL-54
chr17-44399512-DEL-56
chr17-44419655-DEL-6418
chr17-44445971-DEL-8917
chr17-44493489-DEL-439
chr17-44498468-DEL-78
chr17-44502282-DEL-65
chr17-44508307-DEL-106
chr17-44511169-DEL-1556
chr17-44563462-DEL-103
chr17-44572976-DEL-120
chr17-44597766-DEL-5772
chr17-44609425-DEL-68
chr17-44624303-DEL-378
chr17-44625438-DEL-1461
chr17-44625808-DEL-82
chr17-44628108-DEL-118
chr17-44641965-DEL-84
chr17-44650090-DEL-121
chr17-44667369-DEL-149
chr17-44680799-DEL-72
chr17-44711298-DEL-56
chr17-44714940-DEL-154
chr17-44725965-DEL-52
chr17-44732498-DEL-224
chr17-44751553-DEL-60
chr17-44791398-DEL-156
chr17-44812083-DEL-418
chr17-44813066-DEL-172
chr17-44825581-DEL-67
chr17-44882005-DEL-110
chr17-44885980-DEL-59
chr17-44893979-DEL-51
chr17-44896334-DEL-529
chr17-44913380-DEL-147
chr17-44959802-DEL-152
chr17-44979553-DEL-84
chr17-44989846-DEL-63
chr17-45051195-DEL-60
chr17-45065897-DEL-1413

chr17-45076033-DEL-53
chr17-45132972-DEL-66
chr17-45155233-DEL-157
chr17-45163737-DEL-692
chr17-45164448-DEL-199
chr17-45164757-DEL-2560
chr17-45174647-DEL-74
chr17-45189692-DEL-84
chr17-45201410-DEL-510
chr17-45229785-DEL-350
chr17-45239439-DEL-1526
chr17-45257847-DEL-67
chr17-45271541-DEL-60
chr17-45275847-DEL-115
chr17-45285929-DEL-85
chr17-45305514-DEL-52
chr17-45326151-DEL-359
chr17-45380742-DEL-64
chr17-45383038-DEL-125
chr17-45468811-DEL-119
chr17-45472042-DEL-79
chr17-45475759-DEL-78
chr17-45500981-DEL-407
chr17-45517515-DEL-77
chr17-45534836-DEL-56
chr17-45539232-DEL-99
chr17-45542658-DEL-68
chr17-45673876-DEL-177
chr17-45676787-DEL-64
chr17-45685820-DEL-166
chr17-45697336-DEL-123
chr17-45701544-DEL-67
chr17-45727091-DEL-71
chr17-45766343-DEL-824
chr17-45770024-DEL-64
chr17-45773031-DEL-1906
chr17-45775907-DEL-62
chr17-45777983-DEL-185
chr17-45791183-DEL-149
chr17-45794607-DEL-100
chr17-45795247-DEL-73
chr17-45822032-DEL-127
chr17-45891081-DEL-810
chr17-45893065-DEL-61
chr17-45908490-DEL-85
chr17-45934862-DEL-227
chr17-45949332-DEL-57
chr17-46018722-DEL-1007
chr17-46082094-DEL-63
chr17-46090958-DEL-228
chr17-46119904-DEL-128
chr17-46230166-DEL-70
chr17-46231794-DEL-60
chr17-46294234-DEL-116
chr17-46306612-DEL-66
chr17-46321950-DEL-94
chr17-46322903-DEL-88
chr17-46325179-DEL-68
chr17-46355463-DEL-52
chr17-46421344-DEL-298
chr17-46431957-DEL-329
chr17-46444956-DEL-168
chr17-46447015-DEL-432
chr17-46448273-DEL-88
chr17-46496072-DEL-211
chr17-46545883-DEL-375
chr17-46598362-DEL-162
chr17-46601282-DEL-164
chr17-46603745-DEL-105
chr17-46607223-DEL-116
chr17-46615486-DEL-75
chr17-46622194-DEL-846
chr17-46640657-DEL-132
chr17-46646112-DEL-68
chr17-46650590-DEL-70
chr17-46655691-DEL-12080
chr17-46669234-DEL-1115

chr17-46692461-DEL-262
chr17-46694076-DEL-163
chr17-46697163-DEL-141
chr17-46776329-DEL-179
chr17-46787022-DEL-79
chr17-46842117-DEL-1362
chr17-46844608-DEL-191
chr17-46894998-DEL-86
chr17-46948623-DEL-80
chr17-46950185-DEL-94
chr17-46966156-DEL-101
chr17-47030074-DEL-112
chr17-47033683-DEL-72
chr17-47038468-DEL-202
chr17-47041231-DEL-1379
chr17-47060112-DEL-67
chr17-47078138-DEL-85
chr17-47094662-DEL-227
chr17-47095417-DEL-342
chr17-47096686-DEL-172
chr17-47130855-DEL-55
chr17-47164640-DEL-192
chr17-47182589-DEL-170
chr17-47206260-DEL-161
chr17-47216099-DEL-1282
chr17-47261191-DEL-512
chr17-47261360-DEL-1255
chr17-47275786-DEL-101
chr17-47336675-DEL-4489
chr17-47403594-DEL-3861
chr17-47417406-DEL-51
chr17-47447192-DEL-72
chr17-47499444-DEL-56
chr17-47504687-DEL-61
chr17-47518114-DEL-375
chr17-47527270-DEL-122
chr17-47582780-DEL-57
chr17-47649442-DEL-59
chr17-47662813-DEL-169
chr17-47721157-DEL-1559
chr17-47725928-DEL-135
chr17-47727571-DEL-64
chr17-47740694-DEL-412
chr17-47767205-DEL-601
chr17-47817158-DEL-259
chr17-47820701-DEL-50
chr17-47833861-DEL-87
chr17-47836156-DEL-170
chr17-47864664-DEL-79
chr17-47876212-DEL-709
chr17-47877750-DEL-185
chr17-47890878-DEL-210
chr17-47892323-DEL-187
chr17-47953318-DEL-143
chr17-47958139-DEL-219
chr17-47972583-DEL-2788
chr17-48028041-DEL-417
chr17-48051607-DEL-1929
chr17-48055149-DEL-74
chr17-48057285-DEL-66
chr17-48059120-DEL-82
chr17-48073287-DEL-96
chr17-48085306-DEL-93
chr17-48092869-DEL-137
chr17-48096026-DEL-63
chr17-48134942-DEL-72
chr17-48151455-DEL-102
chr17-48176213-DEL-94
chr17-48206195-DEL-60
chr17-48213415-DEL-361
chr17-48216630-DEL-112
chr17-48224602-DEL-97
chr17-48253606-DEL-166
chr17-48258854-DEL-172
chr17-48274370-DEL-141
chr17-48276812-DEL-80
chr17-48282007-DEL-232

chr17-48305825-DEL-196
chr17-48316472-DEL-74
chr17-48356321-DEL-94
chr17-48361194-DEL-107
chr17-48397146-DEL-139
chr17-48407648-DEL-189
chr17-48413682-DEL-103
chr17-48422581-DEL-104
chr17-48429710-DEL-51
chr17-48433587-DEL-2986
chr17-48444841-DEL-268
chr17-48450979-DEL-580
chr17-48460709-DEL-666
chr17-48461487-DEL-67
chr17-48466933-DEL-905
chr17-48475394-DEL-80
chr17-48481177-DEL-7089
chr17-48501000-DEL-364
chr17-48501735-DEL-55
chr17-48503952-DEL-58
chr17-48504188-DEL-231
chr17-48506637-DEL-118
chr17-48508957-DEL-52
chr17-48514523-DEL-4826
chr17-48523083-DEL-134
chr17-48526226-DEL-378
chr17-48528409-DEL-72
chr17-48528681-DEL-6822
chr17-48528975-DEL-2445
chr17-48531484-DEL-4021
chr17-48535866-DEL-56
chr17-48571097-DEL-91
chr17-48600297-DEL-108
chr17-48652033-DEL-250
chr17-48662344-DEL-336
chr17-48664315-DEL-192
chr17-48689008-DEL-1089
chr17-48691671-DEL-1120
chr17-48712503-DEL-1586
chr17-48783029-DEL-128
chr17-48792629-DEL-72
chr17-48801356-DEL-105
chr17-48810556-DEL-445
chr17-48828597-DEL-74
chr17-48831280-DEL-185
chr17-48833829-DEL-88
chr17-48842554-DEL-52
chr17-48866532-DEL-62
chr17-48909039-DEL-196
chr17-48917500-DEL-131
chr17-48928111-DEL-468
chr17-48929889-DEL-534
chr17-48944425-DEL-52
chr17-48948489-DEL-71
chr17-48971686-DEL-56
chr17-48972918-DEL-90
chr17-49037377-DEL-65
chr17-49044495-DEL-180
chr17-49093844-DEL-115
chr17-49107236-DEL-84
chr17-49123303-DEL-72
chr17-49130126-DEL-65
chr17-49163136-DEL-90
chr17-49182310-DEL-114
chr17-49187896-DEL-77
chr17-49190993-DEL-62
chr17-49194504-DEL-57
chr17-49218835-DEL-144
chr17-49260774-DEL-54
chr17-49268285-DEL-118
chr17-49335302-DEL-820
chr17-49348608-DEL-52
chr17-49451840-DEL-294
chr17-49456201-DEL-75
chr17-49507062-DEL-54
chr17-49526470-DEL-70
chr17-49531520-DEL-108

chr17-49532764-DEL-1504
chr17-49551266-DEL-52
chr17-49562489-DEL-74
chr17-49564139-DEL-72
chr17-49570915-DEL-99
chr17-49576594-DEL-75
chr17-49577306-DEL-56
chr17-49578955-DEL-66
chr17-49584916-DEL-135
chr17-49587190-DEL-273
chr17-49591667-DEL-91
chr17-49593168-DEL-93
chr17-49598160-DEL-60
chr17-49614174-DEL-853
chr17-49614370-DEL-74
chr17-49614956-DEL-202
chr17-49659579-DEL-546
chr17-49673185-DEL-54
chr17-49681841-DEL-160
chr17-49683212-DEL-104
chr17-49686311-DEL-109
chr17-49706472-DEL-54
chr17-49780726-DEL-64
chr17-49788117-DEL-68
chr17-49801863-DEL-61
chr17-49812903-DEL-3094
chr17-49824458-DEL-66
chr17-49834507-DEL-51
chr17-49842362-DEL-54
chr17-49843992-DEL-573
chr17-49899252-DEL-173
chr17-49904899-DEL-190
chr17-49941561-DEL-399
chr17-49952223-DEL-64
chr17-49953259-DEL-54
chr17-49957224-DEL-55
chr17-49983048-DEL-54
chr17-49999964-DEL-55
chr17-50022696-DEL-448
chr17-50024848-DEL-256
chr17-50035930-DEL-173
chr17-50036532-DEL-67
chr17-50087507-DEL-70
chr17-50089787-DEL-62
chr17-50097910-DEL-59
chr17-50129668-DEL-709
chr17-50149414-DEL-104
chr17-50155008-DEL-80
chr17-50166827-DEL-73
chr17-50172364-DEL-1478
chr17-50193490-DEL-275
chr17-50218613-DEL-3377
chr17-50223009-DEL-5351
chr17-50224330-DEL-3995
chr17-50232286-DEL-180
chr17-50249498-DEL-634
chr17-50277949-DEL-180
chr17-50280742-DEL-81
chr17-50304832-DEL-93
chr17-50308200-DEL-78
chr17-50318401-DEL-109
chr17-50325046-DEL-58
chr17-50330504-DEL-3755
chr17-50334457-DEL-824
chr17-50336594-DEL-99
chr17-50339088-DEL-156
chr17-50345130-DEL-163
chr17-50351181-DEL-87
chr17-50368953-DEL-67
chr17-50371598-DEL-402
chr17-50380930-DEL-5642
chr17-50387255-DEL-1212
chr17-50422692-DEL-128
chr17-50422845-DEL-703
chr17-50423642-DEL-207
chr17-50429169-DEL-214
chr17-50448673-DEL-215

chr17-50460770-DEL-922
chr17-50488052-DEL-56
chr17-50490784-DEL-54
chr17-50521375-DEL-194
chr17-50534949-DEL-146
chr17-50539235-DEL-196
chr17-50577018-DEL-54
chr17-50602821-DEL-194
chr17-50608016-DEL-1467
chr17-50631354-DEL-267
chr17-50641053-DEL-1691
chr17-50652587-DEL-396
chr17-50656324-DEL-67
chr17-50687826-DEL-68
chr17-50693264-DEL-705
chr17-50706262-DEL-112
chr17-50778195-DEL-57
chr17-50778956-DEL-112
chr17-50791512-DEL-82
chr17-50836362-DEL-366
chr17-50870406-DEL-1161
chr17-50938228-DEL-600
chr17-50951299-DEL-56
chr17-51013338-DEL-321
chr17-51022792-DEL-52
chr17-51035029-DEL-791
chr17-51035758-DEL-51
chr17-51046371-DEL-194
chr17-51078411-DEL-1911
chr17-51083919-DEL-118
chr17-51113659-DEL-1400
chr17-51119568-DEL-53
chr17-51156679-DEL-251
chr17-51163814-DEL-1195
chr17-51164368-DEL-54
chr17-51185121-DEL-79
chr17-51191590-DEL-1091
chr17-51210477-DEL-316
chr17-51255964-DEL-73
chr17-51266705-DEL-636
chr17-51267383-DEL-109
chr17-51271374-DEL-171
chr17-51295694-DEL-567
chr17-51308605-DEL-331
chr17-51313130-DEL-80
chr17-51363780-DEL-109
chr17-51375615-DEL-324
chr17-51406548-DEL-4732
chr17-51434616-DEL-57
chr17-51497561-DEL-404
chr17-51540098-DEL-780
chr17-51555958-DEL-56
chr17-51571435-DEL-4633
chr17-51612151-DEL-79
chr17-51621255-DEL-195
chr17-51626697-DEL-186
chr17-51636739-DEL-754
chr17-51645712-DEL-2145
chr17-51657535-DEL-418
chr17-51660392-DEL-60
chr17-51755198-DEL-298
chr17-51764975-DEL-52
chr17-51783746-DEL-399
chr17-51860755-DEL-80
chr17-51868189-DEL-400
chr17-51871301-DEL-70
chr17-51874864-DEL-60
chr17-51888081-DEL-66
chr17-51962513-DEL-480
chr17-51969067-DEL-70
chr17-51987708-DEL-898
chr17-52000086-DEL-182
chr17-52019287-DEL-74
chr17-52021875-DEL-52
chr17-52022834-DEL-6159
chr17-52030297-DEL-206
chr17-52051377-DEL-51

chr17-52073968-DEL-124
chr17-52125731-DEL-54
chr17-52165543-DEL-56
chr17-52209628-DEL-137
chr17-52217253-DEL-204
chr17-52264079-DEL-2888
chr17-52289226-DEL-145
chr17-52334157-DEL-85
chr17-52376609-DEL-171
chr17-52411513-DEL-74
chr17-52439925-DEL-66
chr17-52449468-DEL-1593
chr17-52477032-DEL-476
chr17-52513748-DEL-64
chr17-52534158-DEL-706
chr17-52537056-DEL-69
chr17-52544979-DEL-58
chr17-52554701-DEL-51
chr17-52560462-DEL-149
chr17-52560662-DEL-781
chr17-52610422-DEL-66
chr17-52629044-DEL-1800
chr17-52633289-DEL-2785
chr17-52648056-DEL-293
chr17-52679947-DEL-54
chr17-52719093-DEL-467
chr17-52721546-DEL-455
chr17-52730856-DEL-430
chr17-52809239-DEL-1657
chr17-52809938-DEL-114
chr17-52819350-DEL-75
chr17-52823884-DEL-928
chr17-52825039-DEL-1581
chr17-52827062-DEL-292
chr17-52835262-DEL-140
chr17-52846595-DEL-1017
chr17-52849845-DEL-6397
chr17-52857331-DEL-426
chr17-52857865-DEL-174
chr17-52872414-DEL-119
chr17-52893782-DEL-186
chr17-52902857-DEL-196
chr17-52937400-DEL-591
chr17-52942016-DEL-1089
chr17-53022352-DEL-98
chr17-53043632-DEL-513
chr17-53050454-DEL-374
chr17-53079873-DEL-123
chr17-53080516-DEL-1955
chr17-53097344-DEL-2384
chr17-53102918-DEL-643
chr17-53114493-DEL-128
chr17-53169070-DEL-157
chr17-53191978-DEL-64
chr17-53247385-DEL-2872
chr17-53277493-DEL-3168
chr17-53339888-DEL-6411
chr17-53373782-DEL-544
chr17-53397444-DEL-280
chr17-53405321-DEL-794
chr17-53413540-DEL-22423
chr17-53414696-DEL-6314
chr17-53421479-DEL-224
chr17-53423942-DEL-6710
chr17-53442155-DEL-91
chr17-53464670-DEL-196
chr17-53468330-DEL-1122
chr17-53517716-DEL-1865
chr17-53519814-DEL-73
chr17-53554444-DEL-63
chr17-53563875-DEL-52
chr17-53589844-DEL-1115
chr17-53593329-DEL-82
chr17-53609800-DEL-2011
chr17-53626556-DEL-111
chr17-53639293-DEL-80
chr17-53646413-DEL-73

chr17-53697858-DEL-178
chr17-53716699-DEL-127
chr17-53731450-DEL-74
chr17-53732116-DEL-52
chr17-53738173-DEL-928
chr17-53771329-DEL-64
chr17-53818521-DEL-204
chr17-53830696-DEL-460
chr17-53837683-DEL-1569
chr17-53839728-DEL-65
chr17-53853137-DEL-310
chr17-53859756-DEL-535
chr17-53865214-DEL-261
chr17-53869507-DEL-211
chr17-53877822-DEL-122
chr17-53909591-DEL-208
chr17-53909592-DEL-5903
chr17-53911767-DEL-156
chr17-53912047-DEL-5634
chr17-53915286-DEL-208
chr17-53917501-DEL-225
chr17-53964629-DEL-164
chr17-53981998-DEL-236
chr17-53984167-DEL-1639
chr17-54023215-DEL-76
chr17-54043691-DEL-213
chr17-54060862-DEL-52
chr17-54067422-DEL-8789
chr17-54088741-DEL-233
chr17-54098662-DEL-348
chr17-54110534-DEL-342
chr17-54111069-DEL-6336
chr17-54150028-DEL-81
chr17-54160277-DEL-250
chr17-54169336-DEL-69
chr17-54204291-DEL-665
chr17-54214992-DEL-218
chr17-54243076-DEL-212
chr17-54267672-DEL-110
chr17-54288215-DEL-73
chr17-54312817-DEL-1771
chr17-54318673-DEL-222
chr17-54345252-DEL-120
chr17-54350865-DEL-6634
chr17-54367330-DEL-78
chr17-54377042-DEL-212
chr17-54391407-DEL-58
chr17-54407357-DEL-1296
chr17-54419747-DEL-416
chr17-54454236-DEL-9068
chr17-54466364-DEL-625
chr17-54504758-DEL-312
chr17-54540490-DEL-110
chr17-54540614-DEL-3303
chr17-54541095-DEL-1435
chr17-54542588-DEL-1187
chr17-54627092-DEL-332
chr17-54637882-DEL-1158
chr17-54648424-DEL-236
chr17-54651218-DEL-55
chr17-54715200-DEL-105
chr17-54736940-DEL-708
chr17-54751975-DEL-71
chr17-54761558-DEL-82
chr17-54764250-DEL-90
chr17-54781813-DEL-62
chr17-54784148-DEL-587
chr17-54845189-DEL-334
chr17-54853203-DEL-6196
chr17-54860107-DEL-618
chr17-54861106-DEL-62
chr17-54886813-DEL-64
chr17-54892729-DEL-129
chr17-54920684-DEL-1096
chr17-54952942-DEL-2561
chr17-54960387-DEL-56
chr17-54975319-DEL-82

chr17-55066741-DEL-2458
chr17-55099383-DEL-80
chr17-55105424-DEL-3501
chr17-55117916-DEL-1041
chr17-55126965-DEL-98
chr17-55139148-DEL-62
chr17-55144956-DEL-190
chr17-55147851-DEL-187
chr17-55206200-DEL-1628
chr17-55229892-DEL-1988
chr17-55273936-DEL-85
chr17-55316645-DEL-366
chr17-55326873-DEL-2243
chr17-55354316-DEL-389
chr17-55380352-DEL-95
chr17-55421572-DEL-540
chr17-55426039-DEL-353
chr17-55442099-DEL-54
chr17-55465832-DEL-1307
chr17-55477196-DEL-558
chr17-55495051-DEL-151
chr17-55498835-DEL-936
chr17-55553310-DEL-2950
chr17-55553339-DEL-1603
chr17-55555200-DEL-607
chr17-55577712-DEL-5182
chr17-55606761-DEL-2552
chr17-55606768-DEL-4312
chr17-55609352-DEL-3659
chr17-55611104-DEL-1908
chr17-55613724-DEL-378
chr17-55628626-DEL-952
chr17-55640809-DEL-271
chr17-55653004-DEL-189
chr17-55654819-DEL-2141
chr17-55676510-DEL-850
chr17-55683165-DEL-393
chr17-55684488-DEL-6779
chr17-55691261-DEL-299
chr17-55693740-DEL-3439
chr17-55704841-DEL-56
chr17-55732513-DEL-1302
chr17-55762416-DEL-51
chr17-55826236-DEL-105
chr17-55842213-DEL-91
chr17-55844700-DEL-6261
chr17-55854254-DEL-66
chr17-55855331-DEL-74
chr17-55858898-DEL-6490
chr17-55876679-DEL-77
chr17-55882135-DEL-6133
chr17-55943090-DEL-435
chr17-55979683-DEL-118
chr17-55987470-DEL-145
chr17-56055919-DEL-198
chr17-56077454-DEL-1547
chr17-56083928-DEL-578
chr17-56119320-DEL-311
chr17-56135111-DEL-75
chr17-56187781-DEL-96
chr17-56222312-DEL-593
chr17-56244735-DEL-51
chr17-56271347-DEL-94
chr17-56328448-DEL-96
chr17-56355570-DEL-167
chr17-56359325-DEL-352
chr17-56372834-DEL-201
chr17-56398780-DEL-298
chr17-56401786-DEL-199
chr17-56402287-DEL-244
chr17-56406592-DEL-206
chr17-56411965-DEL-990
chr17-56412786-DEL-198
chr17-56435813-DEL-115
chr17-56462380-DEL-222
chr17-56473466-DEL-57
chr17-56489995-DEL-52

chr17-56506956-DEL-100
chr17-56532327-DEL-195
chr17-56558828-DEL-190
chr17-56560062-DEL-61
chr17-56570708-DEL-189
chr17-56594473-DEL-176
chr17-56652705-DEL-582
chr17-56673055-DEL-114
chr17-56690810-DEL-123
chr17-56730041-DEL-72
chr17-56808588-DEL-68
chr17-56812095-DEL-51
chr17-56814102-DEL-9900
chr17-56867674-DEL-79
chr17-56882589-DEL-83
chr17-56886320-DEL-68
chr17-56949782-DEL-119
chr17-56979317-DEL-61
chr17-57009447-DEL-64
chr17-57010641-DEL-131
chr17-57048776-DEL-194
chr17-57073177-DEL-194
chr17-57093167-DEL-52
chr17-57114635-DEL-239
chr17-57138614-DEL-75
chr17-57214623-DEL-55
chr17-57216374-DEL-56
chr17-57232684-DEL-259
chr17-57258869-DEL-52
chr17-57263508-DEL-56
chr17-57311888-DEL-52
chr17-57314346-DEL-53
chr17-57324353-DEL-192
chr17-57336192-DEL-163
chr17-57391838-DEL-396
chr17-57408781-DEL-92
chr17-57425599-DEL-52
chr17-57462852-DEL-719
chr17-57463107-DEL-83
chr17-57475229-DEL-61
chr17-57516884-DEL-149
chr17-57539358-DEL-110
chr17-57540355-DEL-97
chr17-57541141-DEL-54
chr17-57548457-DEL-351
chr17-57549582-DEL-60
chr17-57593756-DEL-51
chr17-57610267-DEL-52
chr17-57650139-DEL-58
chr17-57671544-DEL-995
chr17-57711317-DEL-476
chr17-57723247-DEL-86
chr17-57732033-DEL-10020
chr17-57748848-DEL-1674
chr17-57770586-DEL-498
chr17-57775763-DEL-488
chr17-57781046-DEL-951
chr17-57789170-DEL-123
chr17-57795824-DEL-59
chr17-57796812-DEL-715
chr17-57799958-DEL-6159
chr17-57841245-DEL-203
chr17-57890656-DEL-54
chr17-57893678-DEL-15864
chr17-57899683-DEL-9812
chr17-57909413-DEL-127
chr17-57914682-DEL-62
chr17-57925517-DEL-56
chr17-57946508-DEL-1749
chr17-57950893-DEL-837
chr17-57965917-DEL-212
chr17-57969876-DEL-135
chr17-57977684-DEL-2097
chr17-57979833-DEL-56
chr17-57993907-DEL-6206
chr17-58003237-DEL-1407
chr17-58012988-DEL-1194

chr17-58033607-DEL-117
chr17-58035135-DEL-122
chr17-58037407-DEL-1370
chr17-58041377-DEL-5791
chr17-58084769-DEL-494
chr17-58088311-DEL-805
chr17-58101253-DEL-205
chr17-58104590-DEL-51
chr17-58107302-DEL-322
chr17-58134525-DEL-59
chr17-58135697-DEL-289
chr17-58140089-DEL-65
chr17-58188564-DEL-69
chr17-58193077-DEL-1130
chr17-58237555-DEL-131
chr17-58240613-DEL-168
chr17-58249874-DEL-2081
chr17-58261075-DEL-57
chr17-58261813-DEL-81
chr17-58262235-DEL-3708
chr17-58318378-DEL-77
chr17-58324426-DEL-3490
chr17-58327941-DEL-1451
chr17-58332500-DEL-82
chr17-58363806-DEL-4720
chr17-58393562-DEL-83
chr17-58405876-DEL-300
chr17-58416571-DEL-2701
chr17-58443059-DEL-2464
chr17-58452823-DEL-55
chr17-58472437-DEL-124
chr17-58478080-DEL-6501
chr17-58478094-DEL-3946
chr17-58482061-DEL-2580
chr17-58518858-DEL-11280
chr17-58519762-DEL-6283
chr17-58530753-DEL-99
chr17-58532969-DEL-62
chr17-58534568-DEL-6084
chr17-58580110-DEL-1269
chr17-58641557-DEL-5689
chr17-58664379-DEL-186
chr17-58708648-DEL-806
chr17-58736438-DEL-1457
chr17-58737927-DEL-1809
chr17-58744303-DEL-57
chr17-58754516-DEL-56
chr17-58775035-DEL-240
chr17-58790379-DEL-78
chr17-58792315-DEL-5610
chr17-58811660-DEL-144
chr17-58834668-DEL-913
chr17-58836959-DEL-607
chr17-58841661-DEL-1932
chr17-58844480-DEL-1439
chr17-58848541-DEL-1211
chr17-58851023-DEL-15608
chr17-58851324-DEL-77
chr17-58851587-DEL-1824
chr17-58858454-DEL-6483
chr17-58866131-DEL-108
chr17-58874330-DEL-146
chr17-58879039-DEL-638
chr17-58883364-DEL-176
chr17-58886018-DEL-2194
chr17-58890822-DEL-470
chr17-58894168-DEL-102
chr17-58900601-DEL-3730
chr17-58923968-DEL-52
chr17-58955525-DEL-588
chr17-58957058-DEL-66
chr17-58971986-DEL-142
chr17-58993659-DEL-371
chr17-59008465-DEL-76
chr17-59011556-DEL-1348
chr17-59014181-DEL-327
chr17-59024387-DEL-6255

chr17-59035799-DEL-8037
chr17-59043127-DEL-726
chr17-59044400-DEL-53
chr17-59062270-DEL-76
chr17-59079430-DEL-1034
chr17-59089008-DEL-79
chr17-59100682-DEL-401
chr17-59119822-DEL-100
chr17-59138946-DEL-96
chr17-59147116-DEL-8829
chr17-59160527-DEL-914
chr17-59211250-DEL-110
chr17-59288180-DEL-252
chr17-59288703-DEL-748
chr17-59321805-DEL-968
chr17-59378762-DEL-791
chr17-59392050-DEL-831
chr17-59417403-DEL-299
chr17-59439361-DEL-437
chr17-59449943-DEL-207
chr17-59466716-DEL-954
chr17-59499006-DEL-131
chr17-59522570-DEL-4069
chr17-59540791-DEL-1053
chr17-59554679-DEL-231
chr17-59602534-DEL-62
chr17-59621605-DEL-55
chr17-59626629-DEL-194
chr17-59658128-DEL-103
chr17-59662959-DEL-80
chr17-59674148-DEL-69
chr17-59680412-DEL-53
chr17-59691288-DEL-333
chr17-59694239-DEL-811
chr17-59702368-DEL-423
chr17-59719856-DEL-4067
chr17-59762407-DEL-56
chr17-59763176-DEL-154
chr17-59792411-DEL-1941
chr17-59811272-DEL-855
chr17-59812201-DEL-398
chr17-59818582-DEL-82
chr17-59819520-DEL-213
chr17-59830357-DEL-498
chr17-59836361-DEL-6179
chr17-59868500-DEL-6712
chr17-59882429-DEL-333
chr17-59883762-DEL-6672
chr17-59892049-DEL-726
chr17-59931188-DEL-54
chr17-59974785-DEL-86
chr17-59977860-DEL-1049
chr17-59984463-DEL-529
chr17-59986504-DEL-53
chr17-59988635-DEL-60
chr17-60001331-DEL-833
chr17-60109346-DEL-527
chr17-60117512-DEL-185
chr17-60162369-DEL-147
chr17-60163150-DEL-85
chr17-60195959-DEL-56
chr17-60235879-DEL-390
chr17-60277596-DEL-159
chr17-60318941-DEL-368
chr17-60347519-DEL-255
chr17-60375414-DEL-56
chr17-60393252-DEL-322
chr17-60477582-DEL-184
chr17-60485303-DEL-330
chr17-60507095-DEL-301
chr17-60514309-DEL-299
chr17-60518715-DEL-778
chr17-60526777-DEL-297
chr17-60535751-DEL-440
chr17-60576764-DEL-6225
chr17-60592445-DEL-256
chr17-60595539-DEL-4821

chr17-60610832-DEL-104
chr17-60618422-DEL-240
chr17-60634925-DEL-894
chr17-60657675-DEL-153
chr17-60666613-DEL-55
chr17-60680210-DEL-163
chr17-60687966-DEL-239
chr17-60705028-DEL-118
chr17-60710226-DEL-58
chr17-60723000-DEL-352
chr17-60783825-DEL-435
chr17-60784265-DEL-1311
chr17-60784874-DEL-720
chr17-60795762-DEL-1569
chr17-60806175-DEL-95
chr17-60815340-DEL-303
chr17-60882114-DEL-485
chr17-60894512-DEL-59
chr17-60908935-DEL-105
chr17-60924080-DEL-214
chr17-60927170-DEL-704
chr17-60931967-DEL-1817
chr17-60935019-DEL-62
chr17-60957303-DEL-64
chr17-60958015-DEL-1583
chr17-60991627-DEL-185
chr17-61004275-DEL-269
chr17-61011493-DEL-52
chr17-61039070-DEL-82
chr17-61056601-DEL-71
chr17-61107163-DEL-86
chr17-61115356-DEL-5058
chr17-61164250-DEL-64
chr17-61167604-DEL-1651
chr17-61179797-DEL-271
chr17-61184889-DEL-518
chr17-61241587-DEL-893
chr17-61249632-DEL-53
chr17-61259955-DEL-432
chr17-61324715-DEL-264
chr17-61368537-DEL-132
chr17-61370693-DEL-69
chr17-61412741-DEL-67
chr17-61417709-DEL-69
chr17-61433432-DEL-82
chr17-61488396-DEL-2803
chr17-61493075-DEL-218
chr17-61500923-DEL-1439
chr17-61506250-DEL-57
chr17-61529761-DEL-56
chr17-61533179-DEL-767
chr17-61536569-DEL-128
chr17-61556590-DEL-135
chr17-61571121-DEL-68
chr17-61581147-DEL-791
chr17-61591282-DEL-320
chr17-61605542-DEL-2101
chr17-61624670-DEL-1142
chr17-61633621-DEL-454
chr17-61668842-DEL-223
chr17-61673248-DEL-174
chr17-61707464-DEL-73
chr17-61712605-DEL-834
chr17-61723158-DEL-62
chr17-61739335-DEL-771
chr17-61783556-DEL-775
chr17-61787774-DEL-350
chr17-61790083-DEL-4070
chr17-61796566-DEL-121
chr17-61805586-DEL-329
chr17-61807401-DEL-4078
chr17-61834394-DEL-1628
chr17-61836964-DEL-338
chr17-61847035-DEL-1394
chr17-61848394-DEL-196
chr17-61880299-DEL-60
chr17-61907883-DEL-65

chr17-61917621-DEL-89
chr17-61929015-DEL-51
chr17-61951799-DEL-522
chr17-61953118-DEL-218
chr17-61961580-DEL-53
chr17-61992347-DEL-90
chr17-62010859-DEL-1005
chr17-62011701-DEL-399
chr17-62021225-DEL-92
chr17-62021719-DEL-2813
chr17-62034778-DEL-208
chr17-62087707-DEL-1254
chr17-62100671-DEL-72
chr17-62103375-DEL-608
chr17-62136105-DEL-1072
chr17-62165741-DEL-71
chr17-62177489-DEL-550
chr17-62224042-DEL-1145
chr17-62234751-DEL-92
chr17-62235641-DEL-818
chr17-62246256-DEL-76
chr17-62279223-DEL-139
chr17-62309379-DEL-170
chr17-62321820-DEL-1322
chr17-62333080-DEL-212
chr17-62380366-DEL-66
chr17-62555838-DEL-738
chr17-62560372-DEL-77
chr17-62598240-DEL-78
chr17-62608960-DEL-408
chr17-62622059-DEL-240
chr17-62622167-DEL-606
chr17-62631083-DEL-1884
chr17-62649881-DEL-1344
chr17-62657927-DEL-369
chr17-62674641-DEL-134
chr17-62677671-DEL-6617
chr17-62703448-DEL-6287
chr17-62747372-DEL-52
chr17-62800467-DEL-51
chr17-62818893-DEL-68
chr17-62859113-DEL-64
chr17-62872445-DEL-224
chr17-62883990-DEL-367
chr17-62889735-DEL-78
chr17-62897234-DEL-82
chr17-62975032-DEL-145
chr17-62978105-DEL-52
chr17-63045454-DEL-51
chr17-63059004-DEL-143
chr17-63087020-DEL-68
chr17-63243158-DEL-76
chr17-63273597-DEL-248
chr17-63461360-DEL-1984
chr17-63488900-DEL-249
chr17-63547708-DEL-30564
chr17-63549584-DEL-61
chr17-63550567-DEL-242
chr17-63576033-DEL-177
chr17-63587444-DEL-106
chr17-63588332-DEL-132
chr17-63598757-DEL-261
chr17-63626621-DEL-96
chr17-63701208-DEL-53
chr17-63713669-DEL-158
chr17-63720289-DEL-317
chr17-63722249-DEL-133
chr17-63814507-DEL-54
chr17-63819080-DEL-175
chr17-63821983-DEL-157
chr17-63824342-DEL-61
chr17-63828753-DEL-109
chr17-63870007-DEL-391
chr17-63885495-DEL-1466
chr17-63889499-DEL-73
chr17-63907276-DEL-295
chr17-63909710-DEL-65

chr17-63915563-DEL-77
chr17-63916794-DEL-8594
chr17-63937922-DEL-96
chr17-63959937-DEL-179
chr17-63961815-DEL-215
chr17-63999610-DEL-251
chr17-64010112-DEL-55
chr17-64089499-DEL-558
chr17-64090504-DEL-172
chr17-64121214-DEL-105
chr17-64122347-DEL-178
chr17-64137440-DEL-280
chr17-64141884-DEL-140
chr17-64151847-DEL-181
chr17-64182196-DEL-73
chr17-64184731-DEL-54
chr17-64191445-DEL-4551
chr17-64206931-DEL-98
chr17-64219524-DEL-194
chr17-64235488-DEL-2105
chr17-64262385-DEL-466
chr17-64274817-DEL-928
chr17-64282018-DEL-85
chr17-64289702-DEL-4641
chr17-64306894-DEL-52
chr17-64307821-DEL-398
chr17-64354273-DEL-143
chr17-64368860-DEL-1256
chr17-64375373-DEL-117
chr17-64393284-DEL-222
chr17-64394353-DEL-2492
chr17-64396893-DEL-153
chr17-64402686-DEL-868
chr17-64458006-DEL-387
chr17-64497155-DEL-84
chr17-64505054-DEL-423
chr17-64521354-DEL-249
chr17-64540262-DEL-395
chr17-64541192-DEL-127
chr17-64546983-DEL-345
chr17-64576769-DEL-51
chr17-64586806-DEL-371
chr17-64652871-DEL-83
chr17-64682399-DEL-90
chr17-64707328-DEL-214
chr17-64718945-DEL-75
chr17-64731412-DEL-220
chr17-64746579-DEL-132
chr17-64759479-DEL-399
chr17-64769433-DEL-61
chr17-64779895-DEL-103
chr17-64781841-DEL-526
chr17-64824878-DEL-74
chr17-64831063-DEL-126
chr17-64832223-DEL-314
chr17-64850075-DEL-3059
chr17-64861181-DEL-72
chr17-64874849-DEL-70
chr17-64893254-DEL-54
chr17-64905879-DEL-68
chr17-64910890-DEL-81
chr17-64923653-DEL-160
chr17-64929386-DEL-334
chr17-64945520-DEL-1695
chr17-64946601-DEL-519
chr17-64982363-DEL-53
chr17-64988173-DEL-56
chr17-65033659-DEL-1480
chr17-65036263-DEL-54
chr17-65064169-DEL-730
chr17-65149444-DEL-88
chr17-65171922-DEL-56
chr17-65181859-DEL-129
chr17-65185840-DEL-198
chr17-65188101-DEL-1364
chr17-65227434-DEL-54
chr17-65306142-DEL-1243

chr17-65376318-DEL-911
chr17-65443409-DEL-5472
chr17-65449613-DEL-115
chr17-65552158-DEL-77
chr17-65597874-DEL-6225
chr17-65614572-DEL-129
chr17-65669284-DEL-2208
chr17-65685486-DEL-239
chr17-65686107-DEL-117
chr17-65731398-DEL-62
chr17-65737027-DEL-75
chr17-65749088-DEL-413
chr17-65760032-DEL-328
chr17-65764003-DEL-11945
chr17-65775982-DEL-6256
chr17-65938859-DEL-195
chr17-65939687-DEL-888
chr17-65945262-DEL-135
chr17-65963881-DEL-58
chr17-65992209-DEL-2754
chr17-66028491-DEL-57
chr17-66035457-DEL-1103
chr17-66068236-DEL-73
chr17-66085933-DEL-212
chr17-66101367-DEL-99
chr17-66154229-DEL-322
chr17-66172548-DEL-167
chr17-66184506-DEL-3346
chr17-66216942-DEL-1026
chr17-66222336-DEL-182
chr17-66227656-DEL-220
chr17-66235215-DEL-196
chr17-66249948-DEL-172
chr17-66283946-DEL-158
chr17-66353009-DEL-113
chr17-66410539-DEL-329
chr17-66429116-DEL-132
chr17-66432966-DEL-107
chr17-66441348-DEL-215
chr17-66448017-DEL-299
chr17-66461994-DEL-57
chr17-66465055-DEL-65
chr17-66467004-DEL-1744
chr17-66469484-DEL-6703
chr17-66494044-DEL-61
chr17-66498600-DEL-89
chr17-66504939-DEL-415
chr17-66516588-DEL-72
chr17-66532265-DEL-115
chr17-66546500-DEL-200
chr17-66560342-DEL-1016
chr17-66571444-DEL-152
chr17-66591269-DEL-123
chr17-66598048-DEL-99
chr17-66609811-DEL-6153
chr17-66611579-DEL-6440
chr17-66615977-DEL-2158
chr17-66625662-DEL-1542
chr17-66636768-DEL-101
chr17-66672466-DEL-107
chr17-66679173-DEL-172
chr17-66747344-DEL-52
chr17-66748071-DEL-829
chr17-66761071-DEL-125
chr17-66767073-DEL-866
chr17-66772374-DEL-64
chr17-66799733-DEL-81
chr17-66816061-DEL-333
chr17-66838946-DEL-51
chr17-66840880-DEL-51
chr17-66842336-DEL-140
chr17-66853256-DEL-705
chr17-66855261-DEL-170
chr17-66858584-DEL-242
chr17-66859789-DEL-457
chr17-66875345-DEL-1341
chr17-66881451-DEL-64

chr17-66922071-DEL-149
chr17-66932298-DEL-1103
chr17-66953759-DEL-274
chr17-66985909-DEL-57
chr17-67017966-DEL-103
chr17-67032032-DEL-598
chr17-67033695-DEL-94
chr17-67065446-DEL-2086
chr17-67072947-DEL-123
chr17-67079382-DEL-53
chr17-67098358-DEL-126
chr17-67109260-DEL-96
chr17-67123611-DEL-370
chr17-67147980-DEL-161
chr17-67158714-DEL-687
chr17-67245870-DEL-92
chr17-67271189-DEL-181
chr17-67271577-DEL-794
chr17-67281014-DEL-491
chr17-67289492-DEL-55
chr17-67316903-DEL-2104
chr17-67322707-DEL-287
chr17-67383215-DEL-66
chr17-67396060-DEL-83
chr17-67404614-DEL-174
chr17-67446528-DEL-54
chr17-67461062-DEL-1142
chr17-67475552-DEL-93
chr17-67487311-DEL-219
chr17-67522304-DEL-64
chr17-67532576-DEL-188
chr17-67537474-DEL-580
chr17-67557268-DEL-200
chr17-67564249-DEL-435
chr17-67565488-DEL-67
chr17-67570163-DEL-64
chr17-67597530-DEL-72
chr17-67640250-DEL-93
chr17-67663944-DEL-60
chr17-67666572-DEL-231
chr17-67670313-DEL-141
chr17-67683766-DEL-136
chr17-67711372-DEL-293
chr17-67726826-DEL-161
chr17-67741664-DEL-108
chr17-67822905-DEL-52
chr17-67853935-DEL-336
chr17-67889403-DEL-702
chr17-67907085-DEL-145
chr17-67908492-DEL-228
chr17-67919354-DEL-67
chr17-67967300-DEL-666
chr17-67974696-DEL-102
chr17-68003210-DEL-116
chr17-68027253-DEL-121
chr17-68048603-DEL-54
chr17-68055916-DEL-103
chr17-68057303-DEL-128
chr17-68079012-DEL-292
chr17-68142921-DEL-180
chr17-68148294-DEL-94
chr17-68148559-DEL-336
chr17-68174628-DEL-62
chr17-68217990-DEL-67
chr17-68232552-DEL-110
chr17-68236441-DEL-52
chr17-68258733-DEL-73
chr17-68271617-DEL-840
chr17-68278832-DEL-113
chr17-68284741-DEL-340
chr17-68300568-DEL-215
chr17-68330658-DEL-51
chr17-68370610-DEL-135
chr17-68425385-DEL-108
chr17-68441332-DEL-58
chr17-68515788-DEL-63
chr17-68521402-DEL-190

chr17-68531508-DEL-330
chr17-68547726-DEL-96
chr17-68550872-DEL-84
chr17-68574538-DEL-60
chr17-68627137-DEL-81
chr17-68656666-DEL-6217
chr17-68677088-DEL-804
chr17-68802440-DEL-101
chr17-68813145-DEL-2924
chr17-68847493-DEL-81
chr17-68857088-DEL-125
chr17-68861283-DEL-142
chr17-68868081-DEL-55
chr17-68884218-DEL-101
chr17-68905130-DEL-109
chr17-68938283-DEL-52
chr17-68975198-DEL-798
chr17-68976672-DEL-203
chr17-68994383-DEL-7999
chr17-69005637-DEL-914
chr17-69007203-DEL-114
chr17-69021565-DEL-415
chr17-69029450-DEL-486
chr17-69044545-DEL-66
chr17-69050731-DEL-215
chr17-69051885-DEL-350
chr17-69079228-DEL-52
chr17-69122495-DEL-69
chr17-69135114-DEL-780
chr17-69162038-DEL-150
chr17-69191101-DEL-148
chr17-69204340-DEL-320
chr17-69204749-DEL-440
chr17-69207175-DEL-147
chr17-69247035-DEL-65
chr17-69248622-DEL-7164
chr17-69255787-DEL-65
chr17-69256384-DEL-956
chr17-69259079-DEL-79
chr17-69317356-DEL-380
chr17-69327905-DEL-716
chr17-69328990-DEL-142
chr17-69331637-DEL-61
chr17-69371964-DEL-331
chr17-69373260-DEL-290
chr17-69399172-DEL-1146
chr17-69401920-DEL-459
chr17-69413002-DEL-140
chr17-69450876-DEL-121
chr17-69460876-DEL-111
chr17-69486064-DEL-75
chr17-69495622-DEL-67
chr17-69539925-DEL-143
chr17-69564748-DEL-830
chr17-69583209-DEL-204
chr17-69605195-DEL-432
chr17-69605205-DEL-954
chr17-69605834-DEL-227
chr17-69611626-DEL-167
chr17-69668829-DEL-991
chr17-69710757-DEL-4504
chr17-69723252-DEL-125
chr17-69776673-DEL-63
chr17-69801329-DEL-68
chr17-69825738-DEL-101
chr17-69826057-DEL-6597
chr17-69841000-DEL-2960
chr17-69850740-DEL-290
chr17-69854058-DEL-6242
chr17-69863216-DEL-410
chr17-69913490-DEL-99
chr17-69915282-DEL-61
chr17-69915729-DEL-3942
chr17-69921488-DEL-1656
chr17-69933648-DEL-101
chr17-69938078-DEL-132
chr17-69941318-DEL-73

chr17-69962585-DEL-956
chr17-69968690-DEL-368
chr17-69994278-DEL-5956
chr17-70009544-DEL-6593
chr17-70033546-DEL-209
chr17-70096621-DEL-309
chr17-70097501-DEL-619
chr17-70112659-DEL-62
chr17-70121636-DEL-5999
chr17-70142210-DEL-290
chr17-70143213-DEL-1065
chr17-70186963-DEL-112
chr17-70209382-DEL-214
chr17-70215329-DEL-86
chr17-70282474-DEL-93
chr17-70344181-DEL-2925
chr17-70366844-DEL-525
chr17-70373416-DEL-276
chr17-70378050-DEL-311
chr17-70382427-DEL-702
chr17-70401491-DEL-862
chr17-70410298-DEL-116
chr17-70430585-DEL-1470
chr17-70455258-DEL-9110
chr17-70483433-DEL-1798
chr17-70497347-DEL-68
chr17-70503065-DEL-3456
chr17-70526994-DEL-2160
chr17-70535665-DEL-597
chr17-70536318-DEL-424
chr17-70580534-DEL-319
chr17-70689253-DEL-56
chr17-70692767-DEL-91
chr17-70705375-DEL-502
chr17-70710106-DEL-643
chr17-70750498-DEL-108
chr17-70751489-DEL-409
chr17-70771155-DEL-105
chr17-70783099-DEL-74
chr17-70836342-DEL-100
chr17-70837228-DEL-149
chr17-70849048-DEL-279
chr17-70898842-DEL-62
chr17-70927366-DEL-52
chr17-70968033-DEL-216
chr17-71012333-DEL-76
chr17-71032150-DEL-2024
chr17-71040965-DEL-200
chr17-71070807-DEL-582
chr17-71081490-DEL-162
chr17-71110449-DEL-81
chr17-71117315-DEL-206
chr17-71129878-DEL-67
chr17-71131276-DEL-198
chr17-71144700-DEL-320
chr17-71149003-DEL-127
chr17-71161710-DEL-2169
chr17-71178109-DEL-174
chr17-71182402-DEL-85
chr17-71182806-DEL-532
chr17-71188756-DEL-537
chr17-71191527-DEL-64
chr17-71208865-DEL-94
chr17-71214426-DEL-51
chr17-71226591-DEL-187
chr17-71250596-DEL-54
chr17-71254131-DEL-1740
chr17-71262779-DEL-295
chr17-71266510-DEL-62
chr17-71335830-DEL-105
chr17-71355016-DEL-3497
chr17-71355125-DEL-1733
chr17-71356878-DEL-1762
chr17-71358177-DEL-475
chr17-71358878-DEL-178
chr17-71360415-DEL-104
chr17-71408478-DEL-69

chr17-71425549-DEL-51
chr17-71463670-DEL-896
chr17-71541021-DEL-204
chr17-71614707-DEL-57
chr17-71626580-DEL-3042
chr17-71632092-DEL-1581
chr17-71650134-DEL-174
chr17-71667344-DEL-532
chr17-71675828-DEL-168
chr17-71682182-DEL-176
chr17-71723790-DEL-77
chr17-71729316-DEL-376
chr17-71745115-DEL-133
chr17-71746093-DEL-155
chr17-71746561-DEL-144
chr17-71764586-DEL-199
chr17-71779399-DEL-205
chr17-71785589-DEL-140
chr17-71791078-DEL-208
chr17-71826072-DEL-176
chr17-71845134-DEL-2029
chr17-71881589-DEL-119
chr17-71907999-DEL-133
chr17-71916084-DEL-2062
chr17-71938624-DEL-100
chr17-71971878-DEL-166
chr17-71984174-DEL-176
chr17-71985165-DEL-60
chr17-72039055-DEL-54
chr17-72168397-DEL-1456
chr17-72190079-DEL-61
chr17-72203650-DEL-80
chr17-72212326-DEL-192
chr17-72214703-DEL-66
chr17-72220614-DEL-245
chr17-72225960-DEL-84
chr17-72241091-DEL-190
chr17-72266551-DEL-89
chr17-72386213-DEL-146
chr17-72397971-DEL-1481
chr17-72421287-DEL-123
chr17-72454114-DEL-384
chr17-72489604-DEL-72
chr17-72496420-DEL-143
chr17-72627013-DEL-63
chr17-72628933-DEL-51
chr17-72638825-DEL-466
chr17-72647900-DEL-55
chr17-72668406-DEL-72
chr17-72689006-DEL-56
chr17-72692721-DEL-66
chr17-72698685-DEL-122
chr17-72740469-DEL-6540
chr17-72756319-DEL-7381
chr17-72767586-DEL-1335
chr17-72771936-DEL-1194
chr17-72865425-DEL-286
chr17-72896775-DEL-102
chr17-72900189-DEL-174
chr17-72922256-DEL-833
chr17-72923124-DEL-3619
chr17-72942356-DEL-212
chr17-72992473-DEL-83
chr17-73013227-DEL-90
chr17-73028874-DEL-65
chr17-73030359-DEL-1025
chr17-73033359-DEL-174
chr17-73073334-DEL-58
chr17-73126095-DEL-170
chr17-73128411-DEL-54
chr17-73137167-DEL-81
chr17-73158267-DEL-1112
chr17-73198523-DEL-787
chr17-73206927-DEL-70
chr17-73265141-DEL-127
chr17-73308284-DEL-83
chr17-73380143-DEL-350

chr17-73381002-DEL-54
chr17-73387154-DEL-131
chr17-73422599-DEL-389
chr17-73450378-DEL-1141
chr17-73457687-DEL-145
chr17-73459401-DEL-62
chr17-73484556-DEL-4900
chr17-73484883-DEL-56
chr17-73502076-DEL-53
chr17-73505759-DEL-3312
chr17-73525597-DEL-318
chr17-73558973-DEL-65
chr17-73563444-DEL-70
chr17-73569726-DEL-55
chr17-73586626-DEL-1633
chr17-73599511-DEL-110
chr17-73607958-DEL-56
chr17-73613248-DEL-93
chr17-73652372-DEL-302
chr17-73663997-DEL-2118
chr17-73682570-DEL-84
chr17-73710702-DEL-57
chr17-73719851-DEL-118
chr17-73729060-DEL-672
chr17-73730767-DEL-1775
chr17-73760080-DEL-67
chr17-73892590-DEL-105
chr17-73918507-DEL-99
chr17-73951388-DEL-84
chr17-73996292-DEL-1204
chr17-74011707-DEL-1942
chr17-74072759-DEL-328
chr17-74096362-DEL-460
chr17-74157157-DEL-1383
chr17-74163136-DEL-1772
chr17-74192519-DEL-81
chr17-74209752-DEL-374
chr17-74215793-DEL-56
chr17-74274132-DEL-2058
chr17-74279229-DEL-151
chr17-74326110-DEL-1984
chr17-74333168-DEL-120
chr17-74336999-DEL-131
chr17-74376258-DEL-134
chr17-74384446-DEL-100
chr17-74392219-DEL-245
chr17-74394137-DEL-219
chr17-74413210-DEL-137
chr17-74447812-DEL-219
chr17-74456960-DEL-212
chr17-74486292-DEL-1165
chr17-74489220-DEL-204
chr17-74498650-DEL-137
chr17-74558028-DEL-56
chr17-74565194-DEL-72
chr17-74583516-DEL-68
chr17-74586573-DEL-91
chr17-74603695-DEL-193
chr17-74627454-DEL-54
chr17-74636177-DEL-936
chr17-74655772-DEL-61
chr17-74677729-DEL-298
chr17-74720001-DEL-1750
chr17-74745188-DEL-474
chr17-74746280-DEL-206
chr17-74770909-DEL-226
chr17-74780647-DEL-80
chr17-74784566-DEL-301
chr17-74785439-DEL-109
chr17-74792763-DEL-230
chr17-74801487-DEL-128
chr17-74818753-DEL-211
chr17-74822134-DEL-63
chr17-74842052-DEL-2209
chr17-74847829-DEL-848
chr17-74848713-DEL-610
chr17-74849495-DEL-115

chr17-74858727-DEL-1568
chr17-74862839-DEL-247
chr17-74866320-DEL-118
chr17-74869333-DEL-1461
chr17-74874007-DEL-201
chr17-74875796-DEL-383
chr17-74878501-DEL-175
chr17-74911705-DEL-796
chr17-74919482-DEL-90
chr17-74920892-DEL-202
chr17-74951977-DEL-189
chr17-74962825-DEL-196
chr17-74987051-DEL-264
chr17-75000183-DEL-87
chr17-75033311-DEL-181
chr17-75038174-DEL-206
chr17-75059714-DEL-466
chr17-75062393-DEL-87
chr17-75078705-DEL-2647
chr17-75096885-DEL-2014
chr17-75102620-DEL-54
chr17-75129689-DEL-106
chr17-75159645-DEL-697
chr17-75162619-DEL-51
chr17-75181194-DEL-69
chr17-75199038-DEL-6916
chr17-75212528-DEL-7215
chr17-75227596-DEL-158
chr17-75241263-DEL-196
chr17-75242389-DEL-269
chr17-75245550-DEL-52
chr17-75306801-DEL-96
chr17-75317060-DEL-234
chr17-75336086-DEL-643
chr17-75369720-DEL-128
chr17-75383965-DEL-55
chr17-75405861-DEL-2927
chr17-75420698-DEL-139
chr17-75445699-DEL-3750
chr17-75461794-DEL-54
chr17-75508992-DEL-53
chr17-75537277-DEL-51
chr17-75622663-DEL-1343
chr17-75642252-DEL-189
chr17-75684522-DEL-54
chr17-75723598-DEL-974
chr17-75725796-DEL-155
chr17-75753842-DEL-130
chr17-75781134-DEL-423
chr17-75806380-DEL-52
chr17-75861976-DEL-350
chr17-75862539-DEL-417
chr17-75883613-DEL-83
chr17-75886033-DEL-81
chr17-75896988-DEL-90
chr17-75899024-DEL-56
chr17-75903584-DEL-3289
chr17-75915795-DEL-1604
chr17-75924049-DEL-122
chr17-75971077-DEL-146
chr17-75971759-DEL-520
chr17-76017223-DEL-54
chr17-76018601-DEL-60
chr17-76113625-DEL-175
chr17-76137603-DEL-199
chr17-76169750-DEL-52
chr17-76228848-DEL-680
chr17-76233167-DEL-596
chr17-76237755-DEL-322
chr17-76263065-DEL-85
chr17-76272806-DEL-113
chr17-76282922-DEL-60
chr17-76292141-DEL-84
chr17-76305051-DEL-72
chr17-76308111-DEL-91
chr17-76332719-DEL-2725
chr17-76340411-DEL-60

chr17-76390004-DEL-641
chr17-76402422-DEL-138
chr17-76421309-DEL-117
chr17-76446536-DEL-340
chr17-76497369-DEL-58
chr17-76501278-DEL-202
chr17-76502908-DEL-1762
chr17-76507600-DEL-297
chr17-76518617-DEL-2497
chr17-76534729-DEL-51
chr17-76546917-DEL-704
chr17-76590048-DEL-2612
chr17-76616633-DEL-171
chr17-76658347-DEL-267
chr17-76688067-DEL-265
chr17-76706286-DEL-198
chr17-76707367-DEL-1017
chr17-76731808-DEL-943
chr17-76789403-DEL-92
chr17-76841144-DEL-120
chr17-76853390-DEL-81
chr17-76864401-DEL-795
chr17-76873331-DEL-295
chr17-76886444-DEL-987
chr17-76887377-DEL-51
chr17-76901341-DEL-473
chr17-76905323-DEL-79
chr17-76915397-DEL-6385
chr17-76930416-DEL-2520
chr17-76946552-DEL-765
chr17-76963421-DEL-747
chr17-76970682-DEL-532
chr17-76973866-DEL-223
chr17-76985786-DEL-62
chr17-77012437-DEL-77
chr17-77018537-DEL-3106
chr17-77036837-DEL-56
chr17-77043016-DEL-123
chr17-77045772-DEL-60
chr17-77053730-DEL-3770
chr17-77070889-DEL-313
chr17-77088442-DEL-55
chr17-77098204-DEL-2566
chr17-77105036-DEL-298
chr17-77123609-DEL-442
chr17-77129206-DEL-320
chr17-77173859-DEL-1139
chr17-77179459-DEL-58
chr17-77232524-DEL-193
chr17-77265591-DEL-742
chr17-77331797-DEL-60
chr17-77332996-DEL-2012
chr17-77344700-DEL-155
chr17-77348000-DEL-3477
chr17-77376627-DEL-121
chr17-77386527-DEL-213
chr17-77396524-DEL-88
chr17-77407287-DEL-195
chr17-77424657-DEL-55
chr17-77453181-DEL-58
chr17-77466369-DEL-137
chr17-77466676-DEL-925
chr17-77468589-DEL-554
chr17-77470568-DEL-889
chr17-77508115-DEL-70
chr17-77530319-DEL-1462
chr17-77578907-DEL-203
chr17-77647216-DEL-552
chr17-77650054-DEL-525
chr17-77663806-DEL-2823
chr17-77666976-DEL-448
chr17-77666979-DEL-1924
chr17-77687073-DEL-21666
chr17-77698655-DEL-79
chr17-77715260-DEL-661
chr17-77746840-DEL-58
chr17-77755835-DEL-1647

chr17-77790838-DEL-301
chr17-77802544-DEL-53
chr17-77818175-DEL-183
chr17-77820681-DEL-1257
chr17-77824868-DEL-963
chr17-77849081-DEL-1399
chr17-77851053-DEL-383
chr17-77861588-DEL-854
chr17-77895080-DEL-723
chr17-77921690-DEL-64
chr17-77930898-DEL-349
chr17-77940861-DEL-102
chr17-77959630-DEL-3436
chr17-77964968-DEL-55
chr17-77970719-DEL-168
chr17-78019904-DEL-662
chr17-78031281-DEL-749
chr17-78039396-DEL-330
chr17-78045683-DEL-384
chr17-78058274-DEL-194
chr17-78062284-DEL-491
chr17-78066956-DEL-4169
chr17-78077862-DEL-340
chr17-78100833-DEL-102
chr17-78101872-DEL-399
chr17-78152199-DEL-228
chr17-78159683-DEL-60
chr17-78167363-DEL-73
chr17-78171513-DEL-107
chr17-78201371-DEL-86
chr17-78205388-DEL-71
chr17-78209155-DEL-154
chr17-78222672-DEL-338
chr17-78244519-DEL-238
chr17-78263823-DEL-196
chr17-78265731-DEL-178
chr17-78297339-DEL-82
chr17-78312819-DEL-112
chr17-78340429-DEL-135
chr17-78343663-DEL-58
chr17-78344700-DEL-293
chr17-78369466-DEL-68
chr17-78382617-DEL-428
chr17-78394884-DEL-62
chr17-78403806-DEL-418
chr17-78408380-DEL-111
chr17-78422636-DEL-177
chr17-78441755-DEL-93
chr17-78484735-DEL-72
chr17-78486625-DEL-283
chr17-78492161-DEL-321
chr17-78515984-DEL-65
chr17-78631427-DEL-231
chr17-78723651-DEL-424
chr17-78731726-DEL-142
chr17-78736245-DEL-1674
chr17-78746679-DEL-69
chr17-78765026-DEL-61
chr17-78769598-DEL-94
chr17-78776120-DEL-94
chr17-78794819-DEL-1491
chr17-78829422-DEL-58
chr17-78836446-DEL-78
chr17-78846361-DEL-120
chr17-78858673-DEL-80
chr17-78886702-DEL-52
chr17-78906474-DEL-254
chr17-78968972-DEL-192
chr17-79021377-DEL-1015
chr17-79055717-DEL-122
chr17-79074879-DEL-242
chr17-79112362-DEL-73
chr17-79116825-DEL-115
chr17-79153733-DEL-205
chr17-79173045-DEL-123
chr17-79184109-DEL-297
chr17-79199337-DEL-144

chr17-79211291-DEL-90
chr17-79232441-DEL-53
chr17-79288759-DEL-88
chr17-79316590-DEL-80
chr17-79372810-DEL-96
chr17-79391075-DEL-106
chr17-79394972-DEL-450
chr17-79408248-DEL-224
chr17-79413895-DEL-666
chr17-79414260-DEL-763
chr17-79417514-DEL-85
chr17-79436740-DEL-72
chr17-79462391-DEL-1736
chr17-79481855-DEL-242
chr17-79491966-DEL-146
chr17-79519750-DEL-246
chr17-79523615-DEL-514
chr17-79524290-DEL-1908
chr17-79529324-DEL-114
chr17-79574438-DEL-119
chr17-79582288-DEL-60
chr17-79610301-DEL-361
chr17-79626458-DEL-102
chr17-79628680-DEL-115
chr17-79686419-DEL-113
chr17-79703323-DEL-132
chr17-79735945-DEL-996
chr17-79768559-DEL-51
chr17-79781543-DEL-97
chr17-79796407-DEL-409
chr17-79816094-DEL-72
chr17-79819639-DEL-59
chr17-79820524-DEL-81
chr17-79826930-DEL-173
chr17-79828276-DEL-102
chr17-79836814-DEL-58
chr17-79841729-DEL-2066
chr17-79846741-DEL-221
chr17-79848604-DEL-301
chr17-79857450-DEL-151
chr17-79867337-DEL-2431
chr17-79879634-DEL-56
chr17-79900337-DEL-394
chr17-79921293-DEL-167
chr17-79933543-DEL-170
chr17-79942754-DEL-69
chr17-79943551-DEL-68
chr17-79968140-DEL-98
chr17-80048750-DEL-211
chr17-80078188-DEL-103
chr17-80088454-DEL-51
chr17-80101347-DEL-219
chr17-80109159-DEL-82
chr17-80116281-DEL-159
chr17-80128295-DEL-264
chr17-80154599-DEL-69
chr17-80178323-DEL-938
chr17-80185745-DEL-173
chr17-80187858-DEL-287
chr17-80205268-DEL-56
chr17-80210602-DEL-788
chr17-80242774-DEL-113
chr17-80259523-DEL-109
chr17-80277631-DEL-852
chr17-80280303-DEL-96
chr17-80286626-DEL-70
chr17-80327242-DEL-67
chr17-80398670-DEL-71
chr17-80417871-DEL-7022
chr17-80426177-DEL-116
chr17-80458247-DEL-252
chr17-80477287-DEL-830
chr17-80483228-DEL-226
chr17-80498389-DEL-202
chr17-80516923-DEL-215
chr17-80530998-DEL-175
chr17-80538022-DEL-110

chr17-80547032-DEL-57
chr17-80548858-DEL-66
chr17-80556017-DEL-425
chr17-80561944-DEL-509
chr17-80571487-DEL-106
chr17-80592441-DEL-1354
chr17-80594415-DEL-901
chr17-80628178-DEL-58
chr17-80634707-DEL-53
chr17-80635268-DEL-196
chr17-80639201-DEL-66
chr17-80645963-DEL-104
chr17-80653432-DEL-154
chr17-80666464-DEL-541
chr17-80709167-DEL-92
chr17-80725529-DEL-395
chr17-80728730-DEL-874
chr17-80732357-DEL-52
chr17-80743238-DEL-4188
chr17-80746594-DEL-131
chr17-80776018-DEL-89
chr17-80811512-DEL-64
chr17-80815119-DEL-208
chr17-80819180-DEL-272
chr17-80827732-DEL-189
chr17-80861508-DEL-106
chr17-80863880-DEL-1864
chr17-80864072-DEL-170
chr17-80868826-DEL-56
chr17-80906369-DEL-328
chr17-80909744-DEL-1399
chr17-80929705-DEL-57
chr17-80930408-DEL-878
chr17-80942218-DEL-57
chr17-80944261-DEL-202
chr17-80951105-DEL-192
chr17-80981671-DEL-329
chr17-80992111-DEL-148
chr17-80995028-DEL-56
chr17-81040263-DEL-72
chr17-81044448-DEL-56
chr17-81052042-DEL-260
chr17-81069588-DEL-137
chr17-81073046-DEL-603
chr17-81074835-DEL-208
chr17-81077799-DEL-1210
chr17-81085256-DEL-1414
chr17-81088703-DEL-53
chr17-81159035-DEL-134
chr17-81161335-DEL-7171
chr17-81161404-DEL-1565
chr17-81163053-DEL-1999
chr17-81165507-DEL-2996
chr17-81171876-DEL-247
chr17-81234253-DEL-432
chr17-81234944-DEL-116
chr17-81262349-DEL-527
chr17-81317643-DEL-70
chr17-81338227-DEL-104
chr17-81354921-DEL-58
chr17-81355823-DEL-104
chr17-81382768-DEL-204
chr17-81384897-DEL-528
chr17-81410708-DEL-65
chr17-81412358-DEL-82
chr17-81421430-DEL-1302
chr17-81491251-DEL-116
chr17-81522756-DEL-149
chr17-81545673-DEL-228
chr17-81548511-DEL-4840
chr17-81564583-DEL-77
chr17-81583894-DEL-104
chr17-81590589-DEL-60
chr17-81600880-DEL-523
chr17-81621621-DEL-1570
chr17-81624776-DEL-151
chr17-81652092-DEL-265

chr17-81747788-DEL-74
chr17-81755137-DEL-55
chr17-81847330-DEL-51
chr17-81865987-DEL-885
chr17-81888481-DEL-118
chr17-81910728-DEL-56
chr17-81933626-DEL-534
chr17-81935081-DEL-56
chr17-81941491-DEL-84
chr17-81972698-DEL-205
chr17-82011356-DEL-72
chr17-82054387-DEL-962
chr17-82056122-DEL-56
chr17-82074043-DEL-91
chr17-82081167-DEL-2629
chr17-82084058-DEL-196
chr17-82102830-DEL-368
chr17-82141480-DEL-746
chr17-82144959-DEL-787
chr17-82150157-DEL-114
chr17-82198388-DEL-421
chr17-82199225-DEL-1899
chr17-82211238-DEL-2601
chr17-82222178-DEL-68
chr17-82238965-DEL-354
chr17-82241255-DEL-242
chr17-82289432-DEL-91
chr17-82297129-DEL-660
chr17-82305402-DEL-310
chr17-82306087-DEL-6446
chr17-82315069-DEL-324
chr17-82344727-DEL-72
chr17-82369784-DEL-112
chr17-82384706-DEL-96
chr17-82429846-DEL-66
chr17-82434222-DEL-62
chr17-82458091-DEL-962
chr17-82492216-DEL-116
chr17-82500969-DEL-525
chr17-82502121-DEL-87
chr17-82521248-DEL-624
chr17-82523972-DEL-875
chr17-82531839-DEL-520
chr17-82532624-DEL-871
chr17-82542603-DEL-565
chr17-82543238-DEL-400
chr17-82544525-DEL-217
chr17-82553375-DEL-1552
chr17-82563025-DEL-214
chr17-82572698-DEL-275
chr17-82581956-DEL-92
chr17-82599898-DEL-888
chr17-82615680-DEL-136
chr17-82646259-DEL-71
chr17-82651894-DEL-52
chr17-82663430-DEL-1402
chr17-82668595-DEL-210
chr17-82686779-DEL-554
chr17-82696745-DEL-102
chr17-82712323-DEL-399
chr17-82733898-DEL-1493
chr17-82739009-DEL-341
chr17-82750030-DEL-147
chr17-82758812-DEL-205
chr17-82785958-DEL-130
chr17-82794243-DEL-287
chr17-82827197-DEL-284
chr17-82831341-DEL-6223
chr17-82843248-DEL-203
chr17-82844632-DEL-108
chr17-82859919-DEL-143
chr17-82891486-DEL-56
chr17-82905657-DEL-92
chr17-82920032-DEL-853
chr17-82931900-DEL-87
chr17-82975974-DEL-222
chr17-82990163-DEL-6080

chr17-83005802-DEL-6957
chr17-83007139-DEL-485
chr17-83010277-DEL-17893
chr17-83023770-DEL-6540
chr17-83042569-DEL-221
chr17-83063303-DEL-58
chr17-83081556-DEL-1870
chr17-83083091-DEL-462
chr17-83101060-DEL-235
chr17-83118515-DEL-345
chr17-83133370-DEL-1558
chr17-83169469-DEL-164
chr17-83192202-DEL-2378
chr17-83196968-DEL-631
chr17-83207012-DEL-273
chr17-83241106-DEL-111
chr17-83256843-DEL-151
chr17-83294407-DEL-56
chr17-83295391-DEL-3628
chr17-83350621-DEL-188
chr17-83352952-DEL-837
chr17-83354591-DEL-106
chr17-83369984-DEL-682
chr17-83370074-DEL-641
chr17-83399266-DEL-1107
chr17-83434562-DEL-132
chr17-83475362-DEL-2291
chr17-83475725-DEL-199
chr17-83475876-DEL-787
chr17-83476486-DEL-994
chr17-83477272-DEL-192
chr17-83496928-DEL-89
chr17-83519911-DEL-90
chr17-83540559-DEL-71
chr17-83562539-DEL-89
chr17-83570811-DEL-102
chr17-83571952-DEL-90
chr17-83573448-DEL-206
chr17-83590217-DEL-62
chr17-83607396-DEL-208
chr17-83626104-DEL-54
chr17-83629246-DEL-200
chr17-83651068-DEL-142
chr17-83654952-DEL-232
chr17-83664792-DEL-1658
chr17-83708701-DEL-3409
chr17-83710415-DEL-128
chr17-83714075-DEL-1928
chr17-83720336-DEL-331
chr17-83735091-DEL-55
chr17-83743363-DEL-80
chr17-83749073-DEL-184
chr17-83813042-DEL-202
chr17-83825895-DEL-68
chr17-83866113-DEL-111
chr17-83867875-DEL-166
chr17-83898375-DEL-62
chr17-83910077-DEL-68
chr17-83917062-DEL-105
chr17-83926397-DEL-173
chr17-83973495-DEL-290
chr17-83983655-DEL-250
chr17-83984716-DEL-74
chr17-83989133-DEL-195
chr17-83996197-DEL-63
chr17-84011264-DEL-84
chr17-84018686-DEL-342
chr17-84028698-DEL-120
chr17-84046795-DEL-344
chr17-84065013-DEL-243
chr17-84069783-DEL-112
chr17-84076155-DEL-170
chr17-84101587-DEL-161
chr17-84124815-DEL-791
chr17-84133823-DEL-108
chr17-84163120-DEL-246
chr17-84173722-DEL-52

chr17-84181706-DEL-64
chr17-84186098-DEL-189
chr17-84256053-DEL-122
chr17-84433253-DEL-52
chr17-84434637-DEL-219
chr17-84442616-DEL-87
chr17-84447062-DEL-400
chr17-84458397-DEL-201
chr17-84461098-DEL-61
chr17-84462926-DEL-429
chr17-84517378-DEL-201
chr17-84520399-DEL-81
chr17-84529806-DEL-67
chr17-84542492-DEL-146
chr17-84553848-DEL-64
chr17-84554270-DEL-347
chr17-84568051-DEL-118
chr17-84569306-DEL-140
chr17-84593384-DEL-163
chr17-84599862-DEL-385
chr17-84607999-DEL-884
chr17-84610305-DEL-695
chr17-84611236-DEL-119
chr17-84618867-DEL-62
chr17-84658257-DEL-73
chr17-84685133-DEL-1065
chr17-84694801-DEL-60
chr17-84694881-DEL-1500
chr17-84695295-DEL-138
chr17-84762760-DEL-117
chr17-84766524-DEL-5051
chr17-84767453-DEL-250
chr17-84768300-DEL-221
chr17-84769520-DEL-330
chr17-84771120-DEL-276
chr17-84823300-DEL-52
chr17-84846007-DEL-101
chr17-84846462-DEL-382
chr17-84855067-DEL-78
chr17-84861601-DEL-60
chr17-84872011-DEL-454
chr17-84887684-DEL-81
chr17-84895319-DEL-188
chr17-84921657-DEL-340
chr17-84930110-DEL-140
chr17-84946088-DEL-191
chr17-84960967-DEL-110
chr17-84967005-DEL-296
chr17-84970554-DEL-84
chr17-84974079-DEL-738
chr17-84987012-DEL-93
chr17-85066424-DEL-61
chr17-85088709-DEL-2008
chr17-85093935-DEL-126
chr17-85095152-DEL-518
chr17-85099895-DEL-146
chr17-85109480-DEL-67
chr17-85146312-DEL-80
chr17-85149410-DEL-59
chr17-85189072-DEL-207
chr17-85207545-DEL-433
chr17-85208045-DEL-1067
chr17-85216354-DEL-73
chr17-85227149-DEL-178
chr17-85228283-DEL-880
chr17-85242257-DEL-68
chr17-85276403-DEL-1163
chr17-85295581-DEL-72
chr17-85303979-DEL-140
chr17-85308308-DEL-1184
chr17-85308534-DEL-230
chr17-85341941-DEL-285
chr17-85343582-DEL-72
chr17-85347231-DEL-87
chr17-85366214-DEL-79
chr17-85412559-DEL-348
chr17-85413677-DEL-75

chr17-85450901-DEL-135
chr17-85460167-DEL-199
chr17-85487028-DEL-61
chr17-85518721-DEL-51
chr17-85561395-DEL-108
chr17-85562419-DEL-586
chr17-85584756-DEL-98
chr17-85609583-DEL-979
chr17-85612682-DEL-56
chr17-85662134-DEL-116
chr17-85664993-DEL-140
chr17-85672638-DEL-80
chr17-85676565-DEL-140
chr17-85677337-DEL-111
chr17-85687732-DEL-365
chr17-85691322-DEL-67
chr17-85692513-DEL-75
chr17-85693529-DEL-54
chr17-85791400-DEL-58
chr17-85798078-DEL-53
chr17-85806034-DEL-480
chr17-85812551-DEL-92
chr17-85830352-DEL-99
chr17-85832394-DEL-59
chr17-85853132-DEL-1482
chr17-85854720-DEL-71
chr17-85872232-DEL-56
chr17-85889950-DEL-75
chr17-85891729-DEL-59
chr17-85945872-DEL-101
chr17-86013178-DEL-308
chr17-86017633-DEL-1389
chr17-86019010-DEL-73
chr17-86019904-DEL-381
chr17-86038313-DEL-97
chr17-86062403-DEL-107
chr17-86079698-DEL-60
chr17-86080170-DEL-235
chr17-86097708-DEL-68
chr17-86124236-DEL-60
chr17-86128618-DEL-881
chr17-86208346-DEL-593
chr17-86211483-DEL-109
chr17-86218774-DEL-645
chr17-86241865-DEL-120
chr17-86262364-DEL-82
chr17-86266068-DEL-369
chr17-86280789-DEL-175
chr17-86281576-DEL-741
chr17-86285180-DEL-989
chr17-86288091-DEL-74
chr17-86323683-DEL-281
chr17-86370855-DEL-162
chr17-86371697-DEL-419
chr17-86392434-DEL-528
chr17-86394920-DEL-86
chr17-86431213-DEL-389
chr17-86444138-DEL-74
chr17-86447795-DEL-610
chr17-86466275-DEL-66
chr17-86470505-DEL-74
chr17-86481760-DEL-58
chr17-86486282-DEL-111
chr17-86496046-DEL-1289
chr17-86515263-DEL-78
chr17-86531351-DEL-64
chr17-86535269-DEL-85
chr17-86535860-DEL-292
chr17-86537857-DEL-66
chr17-86555826-DEL-416
chr17-86565121-DEL-147
chr17-86589463-DEL-114
chr17-86597582-DEL-59
chr17-86603524-DEL-80
chr17-86607744-DEL-54
chr17-86615164-DEL-1019
chr17-86617746-DEL-68

chr17-86629405-DEL-51
chr17-86664624-DEL-575
chr17-86668047-DEL-102
chr17-86694081-DEL-1476
chr17-86710025-DEL-65
chr17-86742939-DEL-415
chr17-86743730-DEL-56
chr17-86750397-DEL-85
chr17-86784933-DEL-116
chr17-86793886-DEL-214
chr17-86827158-DEL-66
chr17-86853078-DEL-55
chr17-86864042-DEL-65
chr17-86869848-DEL-222
chr17-86875607-DEL-116
chr17-86909089-DEL-133
chr17-86935081-DEL-203
chr17-86952665-DEL-55
chr17-86991958-DEL-787
chr17-86998127-DEL-60
chr17-87040668-DEL-61
chr17-87053044-DEL-155
chr17-87118663-DEL-66
chr17-87134132-DEL-72
chr17-87139068-DEL-56
chr17-87143208-DEL-1954
chr17-87147401-DEL-1584
chr17-87149742-DEL-1525
chr17-87151326-DEL-266
chr17-87154999-DEL-178
chr17-87165034-DEL-440
chr17-87181073-DEL-147
chr17-87204838-DEL-61
chr17-87220318-DEL-58
chr17-87229996-DEL-482
chr17-87255908-DEL-63
chr17-87256907-DEL-2218
chr17-87278161-DEL-239
chr17-87288039-DEL-264
chr17-87288671-DEL-669
chr17-87289092-DEL-112
chr17-87290748-DEL-94
chr17-87319798-DEL-1914
chr17-87319826-DEL-362
chr17-87320453-DEL-544
chr17-87320651-DEL-924
chr17-87339046-DEL-59
chr17-87355658-DEL-80
chr17-87359320-DEL-151
chr17-87379681-DEL-92
chr17-87405784-DEL-107
chr17-87409056-DEL-83
chr17-87412916-DEL-364
chr17-87423956-DEL-293
chr17-87439368-DEL-78
chr17-87458768-DEL-158
chr17-87491418-DEL-474
chr17-87509258-DEL-395
chr17-87516297-DEL-1003
chr17-87516689-DEL-531
chr17-87516962-DEL-215
chr17-87523703-DEL-241
chr17-87529419-DEL-1176
chr17-87532511-DEL-102
chr17-87550405-DEL-56
chr17-87557677-DEL-100
chr17-87604567-DEL-802
chr17-87605265-DEL-105
chr17-87608649-DEL-117
chr17-87625335-DEL-79
chr17-87626442-DEL-690
chr17-87635454-DEL-159
chr17-87683947-DEL-73
chr17-87702305-DEL-206
chr17-87707508-DEL-599
chr17-87723089-DEL-180
chr17-87724966-DEL-344

chr17-87728917-DEL-179
chr17-87729576-DEL-54
chr17-87731573-DEL-193
chr17-87733892-DEL-112
chr17-87736318-DEL-369
chr17-87737344-DEL-88
chr17-87741785-DEL-121
chr17-87745014-DEL-736
chr17-87748468-DEL-72
chr17-87789239-DEL-163
chr17-87791710-DEL-536
chr17-87796520-DEL-57
chr17-87802461-DEL-2241
chr17-87811196-DEL-214
chr17-87813853-DEL-67
chr17-87823132-DEL-153
chr17-87828119-DEL-114
chr17-87856633-DEL-62
chr17-87858210-DEL-178
chr17-87864052-DEL-2581
chr17-87888956-DEL-51
chr17-87906026-DEL-1298
chr17-87930484-DEL-66
chr17-87938747-DEL-173
chr17-87941159-DEL-117
chr17-87955759-DEL-268
chr17-87969618-DEL-200
chr17-87970859-DEL-940
chr17-87987657-DEL-98
chr17-88020652-DEL-412
chr17-88021386-DEL-276
chr17-88042471-DEL-215
chr17-88048480-DEL-78
chr17-88056251-DEL-5312
chr17-88061965-DEL-168
chr17-88062861-DEL-59
chr17-88069414-DEL-59
chr17-88069987-DEL-57
chr17-88081341-DEL-60
chr17-88084887-DEL-630
chr17-88086783-DEL-228
chr17-88120598-DEL-155
chr17-88133816-DEL-280
chr17-88143690-DEL-375
chr17-88147086-DEL-89
chr17-88153893-DEL-246
chr17-88162334-DEL-5776
chr17-88175817-DEL-763
chr17-88176644-DEL-412
chr17-88199236-DEL-354
chr17-88202784-DEL-117
chr17-88208242-DEL-59
chr17-88209605-DEL-85
chr17-88216143-DEL-71
chr17-88218058-DEL-203
chr17-88226771-DEL-154
chr17-88229956-DEL-89
chr17-88234549-DEL-63
chr17-88241824-DEL-567
chr17-88245365-DEL-52
chr17-88247476-DEL-59
chr17-88249485-DEL-57
chr17-88252648-DEL-901
chr17-88261935-DEL-585
chr17-88262814-DEL-154
chr17-88309707-DEL-66
chr17-88311466-DEL-72
chr17-88317814-DEL-164
chr17-88329311-DEL-91
chr17-88335675-DEL-159
chr17-88348874-DEL-102
chr17-88354181-DEL-468
chr17-88355990-DEL-54
chr17-88367105-DEL-57
chr17-88368984-DEL-66
chr17-88386084-DEL-177
chr17-88389793-DEL-1260

chr17-88408978-DEL-218
chr17-88411390-DEL-172
chr17-88427741-DEL-105
chr17-88432262-DEL-177
chr17-88447829-DEL-54
chr17-88452624-DEL-130
chr17-88455524-DEL-400
chr17-88473901-DEL-496
chr17-88485201-DEL-119
chr17-88486623-DEL-13320
chr17-88496023-DEL-338
chr17-88535595-DEL-155
chr17-88547146-DEL-104
chr17-88575204-DEL-703
chr17-88585775-DEL-430
chr17-88590265-DEL-100
chr17-88619747-DEL-61
chr17-88625894-DEL-84
chr17-88632257-DEL-64
chr17-88673575-DEL-71
chr17-88687677-DEL-58
chr17-88691433-DEL-165
chr17-88700017-DEL-57
chr17-88705012-DEL-398
chr17-88712962-DEL-198
chr17-88716077-DEL-175
chr17-88741339-DEL-53
chr17-88761918-DEL-98
chr17-88772571-DEL-1080
chr17-88773897-DEL-1077
chr17-88808222-DEL-105
chr17-88808386-DEL-2066
chr17-88810519-DEL-818
chr17-88811178-DEL-419
chr17-88818949-DEL-212
chr17-88820353-DEL-264
chr17-88832148-DEL-60
chr17-88839710-DEL-155
chr17-88840949-DEL-73
chr17-88919868-DEL-1173
chr17-88920655-DEL-112
chr17-88921101-DEL-2767
chr17-88921157-DEL-1436
chr17-88932232-DEL-99
chr17-88950914-DEL-290
chr17-88980292-DEL-101
chr17-88984604-DEL-90
chr17-88994790-DEL-229
chr17-88998004-DEL-51
chr17-89017383-DEL-261
chr17-89024514-DEL-312
chr17-89025320-DEL-117
chr17-89058986-DEL-57
chr17-89082654-DEL-312
chr17-89102758-DEL-62
chr17-89108548-DEL-753
chr17-89120428-DEL-160
chr17-89127070-DEL-76
chr17-89127607-DEL-340
chr17-89140063-DEL-62
chr17-89144042-DEL-100
chr17-89146428-DEL-458
chr17-89153466-DEL-123
chr17-89174958-DEL-77
chr17-89175488-DEL-966
chr17-89182778-DEL-92
chr17-89190025-DEL-432
chr17-89224149-DEL-202
chr17-89246371-DEL-82
chr17-89264233-DEL-58
chr17-89308370-DEL-248
chr17-89317781-DEL-115
chr17-89352749-DEL-161
chr17-89393873-DEL-334
chr17-89400329-DEL-123
chr17-89415082-DEL-1529
chr17-89418551-DEL-168

chr17-89420786-DEL-60
chr17-89423811-DEL-62
chr17-89429465-DEL-221
chr17-89451262-DEL-106
chr17-89461484-DEL-201
chr17-89470499-DEL-447
chr17-89475350-DEL-70
chr17-89479046-DEL-1481
chr17-89483941-DEL-79
chr17-89492678-DEL-1742
chr17-89498216-DEL-116
chr17-89510977-DEL-897
chr17-89513852-DEL-707
chr17-89583819-DEL-143
chr17-89612247-DEL-182
chr17-89616297-DEL-1696
chr17-89629634-DEL-97
chr17-89655905-DEL-71
chr17-89662717-DEL-134
chr17-89684064-DEL-667
chr17-89685381-DEL-190
chr17-89690950-DEL-54
chr17-89728622-DEL-746
chr17-89740202-DEL-748
chr17-89740707-DEL-626
chr17-89747992-DEL-6415
chr17-89754954-DEL-408
chr17-89848412-DEL-54
chr17-89854270-DEL-64
chr17-89886571-DEL-982
chr17-89887714-DEL-252
chr17-89951605-DEL-161
chr17-89963515-DEL-53
chr17-90021969-DEL-230
chr17-90044274-DEL-94
chr17-90059166-DEL-6211
chr17-90070386-DEL-578
chr17-90084796-DEL-1504
chr17-90098267-DEL-66
chr17-90147453-DEL-212
chr17-90164409-DEL-202
chr17-90171106-DEL-227
chr17-90181961-DEL-6648
chr17-90212242-DEL-497
chr17-90250527-DEL-57
chr17-90259093-DEL-53
chr17-90268513-DEL-86
chr17-90278062-DEL-399
chr17-90292126-DEL-105
chr17-90313869-DEL-95
chr17-90323281-DEL-51
chr17-90480310-DEL-73
chr17-90483923-DEL-76
chr17-90484572-DEL-921
chr17-90491991-DEL-238
chr17-90501352-DEL-269
chr17-90528022-DEL-334
chr17-90533030-DEL-52
chr17-90542287-DEL-411
chr17-90548478-DEL-52
chr17-90564708-DEL-115
chr17-90591477-DEL-1541
chr17-90612057-DEL-600
chr17-90612963-DEL-177
chr17-90641451-DEL-255
chr17-90644845-DEL-220
chr17-90653228-DEL-92
chr17-90699439-DEL-144
chr17-90710652-DEL-106
chr17-90736652-DEL-613
chr17-90752173-DEL-170
chr17-90766350-DEL-61
chr17-90772914-DEL-345
chr17-90785109-DEL-419
chr17-90785687-DEL-122
chr17-90786450-DEL-382
chr17-90787745-DEL-51

chr17-90815640-DEL-89
chr17-90818714-DEL-126
chr17-90825364-DEL-2265
chr17-90833952-DEL-98
chr17-90899986-DEL-1587
chr17-90901318-DEL-118
chr17-90947583-DEL-968
chr17-90968864-DEL-258
chr17-90970379-DEL-106
chr17-90981021-DEL-70
chr17-90986375-DEL-246
chr17-90989617-DEL-556
chr17-90997285-DEL-114
chr17-91022840-DEL-62
chr17-91054444-DEL-69
chr17-91143422-DEL-52
chr17-91148913-DEL-101
chr17-91215553-DEL-66
chr17-91215738-DEL-1368
chr17-91240425-DEL-98
chr17-91268909-DEL-53
chr17-91333255-DEL-283
chr17-91341107-DEL-63
chr17-91369096-DEL-128
chr17-91371545-DEL-962
chr17-91375855-DEL-611
chr17-91388763-DEL-446
chr17-91409204-DEL-1347
chr17-91426335-DEL-180
chr17-91438443-DEL-771
chr17-91445721-DEL-77
chr17-91457174-DEL-58
chr17-91494837-DEL-195
chr17-91499592-DEL-90
chr17-91522031-DEL-1763
chr17-91527432-DEL-204
chr17-91540101-DEL-142
chr17-91542080-DEL-641
chr17-91568784-DEL-62
chr17-91578267-DEL-3870
chr17-91601766-DEL-407
chr17-91605263-DEL-381
chr17-91606308-DEL-166
chr17-91677381-DEL-192
chr17-91678027-DEL-350
chr17-91682908-DEL-559
chr17-91692846-DEL-528
chr17-91731409-DEL-70
chr17-91751978-DEL-315
chr17-91793879-DEL-3080
chr17-91796708-DEL-508
chr17-91797216-DEL-6264
chr17-91864252-DEL-474
chr17-91869247-DEL-2508
chr17-91871975-DEL-186
chr17-91872219-DEL-1230
chr17-91873367-DEL-461
chr17-91880052-DEL-6411
chr17-91897942-DEL-76
chr17-91928266-DEL-665
chr17-91929737-DEL-214
chr17-91945132-DEL-58
chr17-91948581-DEL-6598
chr17-91975588-DEL-275
chr17-91976379-DEL-847
chr17-91981234-DEL-107
chr17-91982075-DEL-56
chr17-92012238-DEL-124
chr17-92028354-DEL-655
chr17-92044594-DEL-1041
chr17-92057209-DEL-320
chr17-92102538-DEL-1347
chr17-92104432-DEL-1124
chr17-92107834-DEL-134
chr17-92110739-DEL-229
chr17-92120418-DEL-74
chr17-92120504-DEL-10747

chr17-92127383-DEL-72
chr17-92171654-DEL-86
chr17-92183893-DEL-170
chr17-92191417-DEL-504
chr17-92209306-DEL-510
chr17-92232145-DEL-8469
chr17-92245081-DEL-811
chr17-92258412-DEL-380
chr17-92263051-DEL-152
chr17-92281182-DEL-167
chr17-92302095-DEL-238
chr17-92305053-DEL-90
chr17-92309839-DEL-53
chr17-92320822-DEL-325
chr17-92335818-DEL-65
chr17-92346002-DEL-2840
chr17-92357996-DEL-56
chr17-92377321-DEL-1010
chr17-92378323-DEL-101
chr17-92384401-DEL-57
chr17-92402824-DEL-611
chr17-92407849-DEL-91
chr17-92408358-DEL-162
chr17-92423776-DEL-2168
chr17-92431076-DEL-180
chr17-92440043-DEL-65
chr17-92442682-DEL-68
chr17-92444457-DEL-553
chr17-92459674-DEL-395
chr17-92482924-DEL-9039
chr17-92494770-DEL-568
chr17-92500230-DEL-91
chr17-92515278-DEL-431
chr17-92561867-DEL-1274
chr17-92568969-DEL-340
chr17-92570091-DEL-2018
chr17-92577064-DEL-1123
chr17-92593357-DEL-265
chr17-92613093-DEL-416
chr17-92621792-DEL-62
chr17-92626911-DEL-6267
chr17-92628928-DEL-115
chr17-92651847-DEL-3607
chr17-92662529-DEL-4092
chr17-92672094-DEL-202
chr17-92687694-DEL-186
chr17-92696444-DEL-60
chr17-92702639-DEL-1443
chr17-92703499-DEL-61
chr17-92706484-DEL-1364
chr17-92708319-DEL-1584
chr17-92709244-DEL-355
chr17-92741853-DEL-178
chr17-92742566-DEL-1218
chr17-92743582-DEL-2435
chr17-92745053-DEL-1129
chr17-92759351-DEL-7260
chr17-92793197-DEL-147
chr17-92793362-DEL-1984
chr17-92796080-DEL-63
chr17-92798198-DEL-58
chr17-92803234-DEL-64
chr17-92809788-DEL-393
chr17-92810200-DEL-2529
chr17-92824247-DEL-73
chr17-92835554-DEL-276
chr17-92836417-DEL-401
chr17-92874847-DEL-375
chr17-92886023-DEL-342
chr17-92893329-DEL-428
chr17-92910094-DEL-60
chr17-92955851-DEL-54
chr17-92961954-DEL-1196
chr17-92982872-DEL-1108
chr17-93043309-DEL-315
chr17-93048788-DEL-1458
chr17-93059492-DEL-73

chr17-93063642-DEL-129
chr17-93067375-DEL-762
chr17-93068333-DEL-65
chr17-93075010-DEL-52
chr17-93088252-DEL-1331
chr17-93097252-DEL-1109
chr17-93120833-DEL-1461
chr17-93126664-DEL-54
chr17-93140753-DEL-104
chr17-93142198-DEL-6516
chr17-93187028-DEL-1046
chr17-93216305-DEL-639
chr17-93225876-DEL-84
chr17-93245476-DEL-1478
chr17-93250317-DEL-6524
chr17-93292117-DEL-2477
chr17-93292964-DEL-6418
chr17-93296939-DEL-2372
chr17-93318310-DEL-1279
chr17-93340677-DEL-6316
chr17-93347174-DEL-1433
chr17-93352256-DEL-60
chr17-93356859-DEL-74
chr17-93358928-DEL-56
chr17-93392842-DEL-356
chr17-93400359-DEL-133
chr17-93415599-DEL-350
chr17-93423837-DEL-2994
chr17-93427840-DEL-100
chr17-93464311-DEL-1203
chr17-93498777-DEL-104
chr17-93535204-DEL-238
chr17-93572198-DEL-1558
chr17-93579788-DEL-688
chr17-93586669-DEL-599
chr17-93588048-DEL-424
chr17-93659714-DEL-56
chr17-93680943-DEL-98
chr17-93707225-DEL-314
chr17-93709513-DEL-3596
chr17-93742041-DEL-506
chr17-93770144-DEL-95
chr17-93775472-DEL-204
chr17-93780529-DEL-135
chr17-93781715-DEL-201
chr17-93808783-DEL-198
chr17-93818136-DEL-68
chr17-93820496-DEL-62
chr17-93853623-DEL-111
chr17-93883752-DEL-76
chr17-93930946-DEL-6448
chr17-93990708-DEL-103
chr17-94077683-DEL-190
chr17-94092360-DEL-1132
chr17-94107097-DEL-52
chr17-94151958-DEL-115
chr17-94166137-DEL-511
chr17-94174901-DEL-126
chr17-94183625-DEL-179
chr17-94203464-DEL-358
chr17-94205336-DEL-517
chr17-94213462-DEL-1574
chr17-94215934-DEL-520
chr17-94222123-DEL-136
chr17-94242258-DEL-795
chr17-94255653-DEL-62
chr17-94273918-DEL-2473
chr17-94291943-DEL-51
chr17-94321944-DEL-533
chr17-94327065-DEL-6079
chr17-94382339-DEL-1266
chr17-94414787-DEL-64
chr17-94417704-DEL-207
chr17-94430240-DEL-158
chr17-94441004-DEL-8776
chr17-94448845-DEL-1811
chr17-94464655-DEL-60

chr17-94490767-DEL-74
chr17-94500559-DEL-146
chr17-94534804-DEL-3333
chr17-94547710-DEL-59
chr17-94558372-DEL-844
chr17-94579000-DEL-52
chr17-94600275-DEL-261
chr17-94614436-DEL-314
chr17-94619938-DEL-159
chr17-94635336-DEL-356
chr17-94659838-DEL-396
chr17-94747400-DEL-2290
chr17-94762887-DEL-70
chr17-94763266-DEL-208
chr17-94768305-DEL-69
chr17-94834533-DEL-929
chr17-94843322-DEL-6555
chr17-94843349-DEL-214
chr17-94864037-DEL-4659
chr17-94915559-DEL-430
chr17-94917375-DEL-6108
chr17-94927795-DEL-127
chr17-94963931-DEL-78
chr17-94971320-DEL-8513
chr17-94980466-DEL-149
chr17-94986863-DEL-94
chr17-94990112-DEL-5217
chr17-94998715-DEL-78
chr17-95079357-DEL-3245
chr17-95089420-DEL-64
chr17-95125729-DEL-168
chr17-95143338-DEL-143
chr17-95145046-DEL-57
chr17-95149215-DEL-444
chr17-95163601-DEL-359
chr17-95165247-DEL-161
chr17-95176882-DEL-173
chr18-3011496-DEL-575
chr18-3021782-DEL-5176
chr18-3028865-DEL-399
chr18-3031018-DEL-6686
chr18-3044190-DEL-23084
chr18-3044784-DEL-57
chr18-3047622-DEL-4211
chr18-3053247-DEL-6473
chr18-3061502-DEL-1871
chr18-3071979-DEL-448
chr18-3104589-DEL-196
chr18-3125235-DEL-108
chr18-3163708-DEL-815
chr18-3166513-DEL-85
chr18-3178409-DEL-26799
chr18-3220722-DEL-178
chr18-3225840-DEL-263
chr18-3251983-DEL-3435
chr18-3288894-DEL-70
chr18-3290275-DEL-64
chr18-3354123-DEL-328
chr18-3355730-DEL-53
chr18-3377159-DEL-95
chr18-3387585-DEL-91
chr18-3394773-DEL-90
chr18-3443622-DEL-358
chr18-3452924-DEL-13616
chr18-3536705-DEL-205
chr18-3582661-DEL-410
chr18-3594507-DEL-58
chr18-3614308-DEL-1553
chr18-3672547-DEL-3906
chr18-3689825-DEL-64
chr18-3698633-DEL-4464
chr18-3702360-DEL-79
chr18-3705224-DEL-6246
chr18-3735530-DEL-4054
chr18-3748250-DEL-1283
chr18-3799505-DEL-273
chr18-3804593-DEL-67

chr18-3880091-DEL-1891
chr18-3883644-DEL-136
chr18-3897895-DEL-79
chr18-3909115-DEL-97
chr18-3911351-DEL-190
chr18-3913064-DEL-79
chr18-3939858-DEL-2379
chr18-3960276-DEL-150
chr18-3962556-DEL-219
chr18-3982932-DEL-175
chr18-4011346-DEL-235
chr18-4065280-DEL-216
chr18-4110148-DEL-1354
chr18-4144611-DEL-86
chr18-4147300-DEL-129
chr18-4152466-DEL-152
chr18-4165507-DEL-60
chr18-4166437-DEL-1014
chr18-4177179-DEL-209
chr18-4222485-DEL-6405
chr18-4245689-DEL-1006
chr18-4246733-DEL-872
chr18-4255040-DEL-209
chr18-4293649-DEL-332
chr18-4301251-DEL-444
chr18-4307316-DEL-657
chr18-4371212-DEL-203
chr18-4384080-DEL-160
chr18-4406377-DEL-428
chr18-4418037-DEL-1433
chr18-4426761-DEL-5305
chr18-4435179-DEL-261
chr18-4441433-DEL-3965
chr18-4458081-DEL-77
chr18-4467444-DEL-54
chr18-4490526-DEL-3952
chr18-4494439-DEL-198
chr18-4503719-DEL-835
chr18-4553926-DEL-406
chr18-4714937-DEL-56
chr18-4727616-DEL-846
chr18-4734660-DEL-176
chr18-4767983-DEL-513
chr18-4777080-DEL-3476
chr18-4784703-DEL-2573
chr18-4803459-DEL-75
chr18-4901926-DEL-173
chr18-4916585-DEL-109
chr18-4976246-DEL-110
chr18-4982153-DEL-96
chr18-4982930-DEL-96
chr18-4984416-DEL-198
chr18-5026269-DEL-59
chr18-5035694-DEL-54
chr18-5062676-DEL-151
chr18-5131003-DEL-69
chr18-5144258-DEL-355
chr18-5158749-DEL-74
chr18-5187619-DEL-216
chr18-5188532-DEL-2328
chr18-5195939-DEL-155
chr18-5204992-DEL-662
chr18-5219460-DEL-164
chr18-5241498-DEL-68
chr18-5249620-DEL-288
chr18-5264032-DEL-414
chr18-5278451-DEL-823
chr18-5282227-DEL-157
chr18-5314079-DEL-78
chr18-5325616-DEL-130
chr18-5341344-DEL-83
chr18-5363243-DEL-182
chr18-5378506-DEL-1711
chr18-5389384-DEL-53
chr18-5433111-DEL-74
chr18-5448499-DEL-148
chr18-5471069-DEL-101

chr18-5503481-DEL-86
chr18-5528039-DEL-51
chr18-5531311-DEL-1874
chr18-5550749-DEL-547
chr18-5570541-DEL-58
chr18-5575822-DEL-5528
chr18-5576830-DEL-195
chr18-5581936-DEL-305
chr18-5584348-DEL-114
chr18-5603236-DEL-176
chr18-5630470-DEL-72
chr18-5661111-DEL-177
chr18-5671360-DEL-56
chr18-5688956-DEL-373
chr18-5747974-DEL-120
chr18-5788457-DEL-296
chr18-5788811-DEL-1434
chr18-5789718-DEL-661
chr18-5883550-DEL-64
chr18-5894181-DEL-6188
chr18-5911005-DEL-103
chr18-5925377-DEL-124
chr18-5947846-DEL-383
chr18-5981932-DEL-60
chr18-6068291-DEL-200
chr18-6081545-DEL-971
chr18-6081693-DEL-216
chr18-6090289-DEL-60
chr18-6091141-DEL-54
chr18-6094325-DEL-400
chr18-6112693-DEL-249
chr18-6156970-DEL-465
chr18-6157127-DEL-1544
chr18-6157899-DEL-185
chr18-6158064-DEL-743
chr18-6158473-DEL-85
chr18-6252219-DEL-3581
chr18-6282752-DEL-63
chr18-6291425-DEL-1094
chr18-6292742-DEL-252
chr18-6293930-DEL-98
chr18-6326959-DEL-717
chr18-6332573-DEL-513
chr18-6349430-DEL-225
chr18-6364391-DEL-1495
chr18-6377713-DEL-80
chr18-6389800-DEL-1283
chr18-6395878-DEL-6476
chr18-6403918-DEL-117
chr18-6421043-DEL-492
chr18-6497364-DEL-96
chr18-6505751-DEL-414
chr18-6542134-DEL-12214
chr18-6545859-DEL-5625
chr18-6556179-DEL-249
chr18-6559930-DEL-52
chr18-6561941-DEL-1416
chr18-6564182-DEL-6757
chr18-6643832-DEL-198
chr18-6675741-DEL-52
chr18-6701260-DEL-119
chr18-6702419-DEL-68
chr18-6709344-DEL-1258
chr18-6721596-DEL-424
chr18-6732137-DEL-124
chr18-6746030-DEL-193
chr18-6749310-DEL-2271
chr18-6759574-DEL-305
chr18-6760074-DEL-880
chr18-6765780-DEL-178
chr18-6796381-DEL-6706
chr18-6848231-DEL-107
chr18-6924076-DEL-255
chr18-6939868-DEL-214
chr18-6948929-DEL-206
chr18-6973748-DEL-841
chr18-7013648-DEL-661

chr18-7033729-DEL-71
chr18-7044912-DEL-113
chr18-7057840-DEL-162
chr18-7058647-DEL-4493
chr18-7070107-DEL-4433
chr18-7074043-DEL-529
chr18-7088752-DEL-1049
chr18-7092997-DEL-104
chr18-7113431-DEL-57
chr18-7113680-DEL-356
chr18-7127618-DEL-60
chr18-7142579-DEL-83
chr18-7145800-DEL-701
chr18-7151715-DEL-52
chr18-7170156-DEL-198
chr18-7174496-DEL-144
chr18-7208859-DEL-1030
chr18-7228473-DEL-176
chr18-7255213-DEL-539
chr18-7265796-DEL-399
chr18-7274435-DEL-7540
chr18-7358778-DEL-61
chr18-7401638-DEL-79
chr18-7415459-DEL-163
chr18-7416674-DEL-222
chr18-7425877-DEL-107
chr18-7437139-DEL-128
chr18-7486672-DEL-1314
chr18-7500742-DEL-158
chr18-7506072-DEL-75
chr18-7515849-DEL-101
chr18-7546184-DEL-168
chr18-7547472-DEL-119
chr18-7556369-DEL-356
chr18-7569734-DEL-421
chr18-7573207-DEL-57
chr18-7608038-DEL-109
chr18-7608947-DEL-360
chr18-7608950-DEL-948
chr18-7612037-DEL-57
chr18-7701510-DEL-124
chr18-7708983-DEL-147
chr18-7713859-DEL-7652
chr18-7724772-DEL-3455
chr18-7730402-DEL-55
chr18-7732117-DEL-488
chr18-7755119-DEL-132
chr18-7760926-DEL-3021
chr18-7779498-DEL-63
chr18-7781302-DEL-245
chr18-7790405-DEL-61
chr18-7791196-DEL-78
chr18-7818093-DEL-1444
chr18-7818180-DEL-528
chr18-7824019-DEL-294
chr18-7836083-DEL-50
chr18-7858684-DEL-66
chr18-7859137-DEL-2037
chr18-7859225-DEL-240
chr18-7859773-DEL-840
chr18-7860581-DEL-160
chr18-7860797-DEL-120
chr18-7865401-DEL-2812
chr18-7909622-DEL-94
chr18-7932168-DEL-169
chr18-7943054-DEL-55
chr18-7964306-DEL-7423
chr18-7979864-DEL-920
chr18-8004094-DEL-344
chr18-8020217-DEL-332
chr18-8028033-DEL-92
chr18-8106501-DEL-1437
chr18-8184332-DEL-83
chr18-8222021-DEL-1688
chr18-8223409-DEL-2489
chr18-8225706-DEL-2462
chr18-8229566-DEL-581

chr18-8232216-DEL-44855
chr18-8233234-DEL-278
chr18-8242821-DEL-9386
chr18-8268618-DEL-2463
chr18-8283889-DEL-65
chr18-8285990-DEL-219
chr18-8299652-DEL-322
chr18-8314325-DEL-934
chr18-8319971-DEL-94
chr18-8382954-DEL-468
chr18-8386529-DEL-205
chr18-8437713-DEL-893
chr18-8462106-DEL-2260
chr18-8491185-DEL-56
chr18-8515139-DEL-130
chr18-8538556-DEL-132
chr18-8573880-DEL-140
chr18-8626740-DEL-546
chr18-8627144-DEL-1397
chr18-8631317-DEL-60
chr18-8702190-DEL-491
chr18-8708745-DEL-368
chr18-8721860-DEL-5257
chr18-8731742-DEL-4721
chr18-8745176-DEL-1894
chr18-8876505-DEL-86
chr18-8942033-DEL-3025
chr18-8954012-DEL-926
chr18-8972748-DEL-84
chr18-8974429-DEL-1338
chr18-8978666-DEL-784
chr18-8980810-DEL-70
chr18-9014719-DEL-388
chr18-9035658-DEL-146
chr18-9042527-DEL-93
chr18-9061041-DEL-1488
chr18-9099111-DEL-73
chr18-9121707-DEL-411
chr18-9122288-DEL-84
chr18-9181943-DEL-400
chr18-9181945-DEL-1500
chr18-9200634-DEL-551
chr18-9225225-DEL-1884
chr18-9286011-DEL-2204
chr18-9288368-DEL-77
chr18-9291217-DEL-462
chr18-9328292-DEL-66
chr18-9349029-DEL-85
chr18-9352408-DEL-52
chr18-9419827-DEL-6304
chr18-9437334-DEL-51
chr18-9454714-DEL-91
chr18-9461632-DEL-1240
chr18-9462762-DEL-329
chr18-9464625-DEL-57
chr18-9487309-DEL-520
chr18-9489123-DEL-247
chr18-9505182-DEL-616
chr18-9515677-DEL-197
chr18-9520222-DEL-201
chr18-9526726-DEL-1086
chr18-9571152-DEL-570
chr18-9585641-DEL-191
chr18-9592386-DEL-5104
chr18-9602151-DEL-66
chr18-9635375-DEL-10058
chr18-9649597-DEL-1115
chr18-9668480-DEL-244
chr18-9678383-DEL-553
chr18-9680150-DEL-11375
chr18-9689052-DEL-1271
chr18-9711388-DEL-68
chr18-9733060-DEL-816
chr18-9733076-DEL-1189
chr18-9741592-DEL-795
chr18-9781201-DEL-883
chr18-9789715-DEL-81

chr18-9797213-DEL-62
chr18-9802685-DEL-60
chr18-9832112-DEL-63
chr18-9839013-DEL-96
chr18-9889165-DEL-5515
chr18-9905720-DEL-2216
chr18-9908733-DEL-670
chr18-9909442-DEL-1390
chr18-9909902-DEL-214
chr18-9911681-DEL-4268
chr18-9912020-DEL-142
chr18-9917101-DEL-1037
chr18-9924615-DEL-158
chr18-9927511-DEL-79
chr18-9936618-DEL-3336
chr18-9945766-DEL-332
chr18-9980652-DEL-55
chr18-10014908-DEL-256
chr18-10015731-DEL-485
chr18-10026300-DEL-1456
chr18-10035030-DEL-1896
chr18-10148304-DEL-449
chr18-10152877-DEL-462
chr18-10204304-DEL-210
chr18-10206723-DEL-783
chr18-10274262-DEL-403
chr18-10297704-DEL-260
chr18-10331773-DEL-96
chr18-10334529-DEL-83
chr18-10349700-DEL-52
chr18-10362597-DEL-9407
chr18-10372806-DEL-1400
chr18-10428913-DEL-76
chr18-10431148-DEL-260
chr18-10442825-DEL-182
chr18-10478566-DEL-273
chr18-10508397-DEL-82
chr18-10527402-DEL-152
chr18-10538700-DEL-87
chr18-10545693-DEL-72
chr18-10548129-DEL-899
chr18-10574464-DEL-203
chr18-10596583-DEL-63
chr18-10598444-DEL-210
chr18-10639639-DEL-70
chr18-10641081-DEL-52
chr18-10642457-DEL-164
chr18-10653212-DEL-58
chr18-10663853-DEL-264
chr18-10687261-DEL-1430
chr18-10687266-DEL-4722
chr18-10691944-DEL-523
chr18-10692049-DEL-1444
chr18-10692222-DEL-246
chr18-10693834-DEL-391
chr18-10727088-DEL-60
chr18-10834954-DEL-57
chr18-10888904-DEL-317
chr18-10916048-DEL-799
chr18-10934490-DEL-352
chr18-10955097-DEL-74
chr18-10963438-DEL-74
chr18-10967131-DEL-56
chr18-10991784-DEL-63
chr18-11010770-DEL-72
chr18-11019978-DEL-1316
chr18-11034609-DEL-51
chr18-11168493-DEL-347
chr18-11177114-DEL-6169
chr18-11198752-DEL-54
chr18-11228856-DEL-186
chr18-11249098-DEL-331
chr18-11252234-DEL-1373
chr18-11302672-DEL-875
chr18-11322781-DEL-60
chr18-11334613-DEL-2622
chr18-11378116-DEL-1838

chr18-11378977-DEL-310
chr18-11380794-DEL-4880
chr18-11385885-DEL-1507
chr18-11394633-DEL-204
chr18-11400657-DEL-1141
chr18-11411064-DEL-212
chr18-11413240-DEL-95
chr18-11417353-DEL-66
chr18-11432799-DEL-148
chr18-11449359-DEL-55
chr18-11496700-DEL-273
chr18-11500265-DEL-536
chr18-11544629-DEL-93
chr18-11548050-DEL-167
chr18-11555119-DEL-124
chr18-11559111-DEL-64
chr18-11594437-DEL-381
chr18-11607896-DEL-135
chr18-11618929-DEL-73
chr18-11621229-DEL-85
chr18-11622568-DEL-108
chr18-11627850-DEL-361
chr18-11644630-DEL-87
chr18-11655410-DEL-2473
chr18-11675641-DEL-7562
chr18-11707187-DEL-225
chr18-11710460-DEL-308
chr18-11728859-DEL-110
chr18-11734212-DEL-400
chr18-11756460-DEL-167
chr18-11803291-DEL-85
chr18-11816678-DEL-52
chr18-11992488-DEL-62
chr18-12057066-DEL-58
chr18-12102573-DEL-654
chr18-12120256-DEL-68
chr18-12145609-DEL-185
chr18-12150203-DEL-819
chr18-12150898-DEL-126
chr18-12166802-DEL-788
chr18-12167327-DEL-175
chr18-12188877-DEL-381
chr18-12218585-DEL-4677
chr18-12224748-DEL-2459
chr18-12224929-DEL-877
chr18-12227233-DEL-2035
chr18-12299550-DEL-339
chr18-12305689-DEL-168
chr18-12319331-DEL-209
chr18-12336676-DEL-52
chr18-12342293-DEL-205
chr18-12393753-DEL-72
chr18-12401114-DEL-61
chr18-12428551-DEL-60
chr18-12516926-DEL-300
chr18-12518196-DEL-58
chr18-12544099-DEL-93
chr18-12566651-DEL-325
chr18-12584323-DEL-221
chr18-12585279-DEL-237
chr18-12585332-DEL-746
chr18-12585903-DEL-172
chr18-12587419-DEL-580
chr18-12591267-DEL-341
chr18-12609135-DEL-60
chr18-12613030-DEL-264
chr18-12623088-DEL-123
chr18-12642527-DEL-102
chr18-12651886-DEL-60
chr18-12675751-DEL-196
chr18-12743106-DEL-358
chr18-12752953-DEL-1398
chr18-12752956-DEL-722
chr18-12754237-DEL-302
chr18-12754466-DEL-114
chr18-12771874-DEL-84
chr18-12816383-DEL-237

chr18-12823458-DEL-269
chr18-12834725-DEL-92
chr18-12840385-DEL-101
chr18-12847207-DEL-63
chr18-12849513-DEL-156
chr18-12851423-DEL-87
chr18-12856926-DEL-413
chr18-12864343-DEL-81
chr18-12876819-DEL-55
chr18-12878823-DEL-150
chr18-12907923-DEL-392
chr18-12934799-DEL-416
chr18-12939586-DEL-288
chr18-12963706-DEL-55
chr18-12964607-DEL-388
chr18-12988918-DEL-347
chr18-12993040-DEL-72
chr18-12998962-DEL-1538
chr18-13039093-DEL-683
chr18-13044450-DEL-73
chr18-13055806-DEL-53
chr18-13077638-DEL-90
chr18-13083726-DEL-972
chr18-13086211-DEL-62
chr18-13116000-DEL-137
chr18-13122316-DEL-69
chr18-13156107-DEL-81
chr18-13160150-DEL-106
chr18-13162529-DEL-295
chr18-13173036-DEL-1990
chr18-13175852-DEL-612
chr18-13176326-DEL-306
chr18-13208725-DEL-638
chr18-13228291-DEL-223
chr18-13237207-DEL-1119
chr18-13255669-DEL-84
chr18-13263480-DEL-120
chr18-13279799-DEL-74
chr18-13336539-DEL-1238
chr18-13338203-DEL-4662
chr18-13348173-DEL-1219
chr18-13361482-DEL-61
chr18-13372594-DEL-6170
chr18-13382207-DEL-213
chr18-13382824-DEL-1136
chr18-13383592-DEL-142
chr18-13402934-DEL-93
chr18-13424858-DEL-2182
chr18-13429799-DEL-180
chr18-13430299-DEL-130
chr18-13431007-DEL-810
chr18-13431933-DEL-170
chr18-13432384-DEL-251
chr18-13516985-DEL-54
chr18-13533713-DEL-54
chr18-13554723-DEL-54
chr18-13555778-DEL-94
chr18-13564215-DEL-57
chr18-13591761-DEL-91
chr18-13597183-DEL-170
chr18-13621317-DEL-60
chr18-13658926-DEL-764
chr18-13669567-DEL-62
chr18-13760143-DEL-1374
chr18-13778444-DEL-1296
chr18-13808806-DEL-51
chr18-13810222-DEL-87
chr18-13846532-DEL-66
chr18-13903879-DEL-506
chr18-14054860-DEL-136
chr18-14073197-DEL-124
chr18-14091846-DEL-573
chr18-14141945-DEL-396
chr18-14159847-DEL-420
chr18-14225436-DEL-643
chr18-14234014-DEL-73
chr18-14240766-DEL-51

chr18-14268472-DEL-400
chr18-14296997-DEL-447
chr18-14307529-DEL-75
chr18-14338641-DEL-100
chr18-14352813-DEL-3024
chr18-14383687-DEL-482
chr18-14402670-DEL-52
chr18-14411646-DEL-74
chr18-14436327-DEL-139
chr18-14442706-DEL-54
chr18-14450434-DEL-57
chr18-14451517-DEL-184
chr18-14470865-DEL-135
chr18-14480445-DEL-54
chr18-14509902-DEL-895
chr18-14526656-DEL-244
chr18-14532282-DEL-100
chr18-14542458-DEL-58
chr18-14547060-DEL-134
chr18-14551010-DEL-88
chr18-14568800-DEL-175
chr18-14572422-DEL-2680
chr18-14655217-DEL-106
chr18-14693467-DEL-1149
chr18-14695328-DEL-53
chr18-14696259-DEL-160
chr18-14700726-DEL-59
chr18-14754305-DEL-203
chr18-14780786-DEL-200
chr18-14793465-DEL-346
chr18-14808317-DEL-64
chr18-14811303-DEL-106
chr18-14829460-DEL-300
chr18-14833994-DEL-62
chr18-14837946-DEL-65
chr18-14861848-DEL-345
chr18-14877903-DEL-284
chr18-14881371-DEL-1917
chr18-14891230-DEL-563
chr18-14900007-DEL-228
chr18-14905480-DEL-623
chr18-14965032-DEL-1025
chr18-14973829-DEL-66
chr18-14985003-DEL-2243
chr18-14998029-DEL-107
chr18-15004480-DEL-2620
chr18-15017577-DEL-108
chr18-15046749-DEL-356
chr18-15051080-DEL-367
chr18-15055575-DEL-602
chr18-15058792-DEL-97
chr18-15082267-DEL-401
chr18-15144246-DEL-797
chr18-15146176-DEL-251
chr18-15150626-DEL-74
chr18-15153598-DEL-63
chr18-15155447-DEL-72
chr18-15160277-DEL-189
chr18-15186875-DEL-191
chr18-15187898-DEL-86
chr18-15191802-DEL-221
chr18-15247010-DEL-1204
chr18-15247012-DEL-236
chr18-15248081-DEL-153
chr18-15264202-DEL-392
chr18-15310218-DEL-84
chr18-15331494-DEL-72
chr18-15338395-DEL-60
chr18-15354646-DEL-152
chr18-15355928-DEL-237
chr18-15378085-DEL-108
chr18-15409569-DEL-400
chr18-15422829-DEL-114
chr18-15429467-DEL-57
chr18-15441160-DEL-300
chr18-15477185-DEL-1040
chr18-15491450-DEL-78

chr18-15514693-DEL-1332
chr18-15520758-DEL-56
chr18-15564664-DEL-199
chr18-15567408-DEL-312
chr18-15574280-DEL-153
chr18-15577154-DEL-221
chr18-15595876-DEL-64
chr18-15626254-DEL-51
chr18-15630561-DEL-278
chr18-15634468-DEL-494
chr18-15647467-DEL-6195
chr18-15652610-DEL-117
chr18-15690659-DEL-1847
chr18-15723466-DEL-70
chr18-15725532-DEL-126
chr18-15730300-DEL-5116
chr18-15743037-DEL-64
chr18-15745458-DEL-56
chr18-15792186-DEL-597
chr18-15797066-DEL-52
chr18-15828144-DEL-6806
chr18-15836584-DEL-140
chr18-15863454-DEL-208
chr18-15864079-DEL-208
chr18-15872946-DEL-162
chr18-15902993-DEL-403
chr18-15914281-DEL-158
chr18-15941498-DEL-574
chr18-15979179-DEL-1797
chr18-15983035-DEL-71
chr18-15995780-DEL-143
chr18-16012446-DEL-53
chr18-16016682-DEL-65
chr18-16023607-DEL-1164
chr18-16023962-DEL-73
chr18-16027852-DEL-378
chr18-16046845-DEL-228
chr18-16070784-DEL-1706
chr18-16077166-DEL-6438
chr18-16088978-DEL-7252
chr18-16114055-DEL-9199
chr18-16123734-DEL-290
chr18-16126150-DEL-251
chr18-16128189-DEL-156
chr18-16135138-DEL-80
chr18-16186397-DEL-243
chr18-16187495-DEL-58
chr18-16189649-DEL-960
chr18-16221189-DEL-350
chr18-16229279-DEL-58
chr18-16294044-DEL-883
chr18-16294951-DEL-1550
chr18-16321233-DEL-666
chr18-16337883-DEL-230
chr18-16376131-DEL-68
chr18-16378318-DEL-131
chr18-16446882-DEL-262
chr18-16484660-DEL-404
chr18-16486392-DEL-126
chr18-16490001-DEL-116
chr18-16527741-DEL-243
chr18-16555759-DEL-1702
chr18-16563862-DEL-399
chr18-16585828-DEL-88
chr18-16586538-DEL-73
chr18-16603142-DEL-206
chr18-16643221-DEL-162
chr18-16651698-DEL-299
chr18-16656075-DEL-156
chr18-16682080-DEL-1300
chr18-16753098-DEL-1311
chr18-16754927-DEL-219
chr18-16768028-DEL-130
chr18-16827514-DEL-427
chr18-16843962-DEL-191
chr18-16868420-DEL-76
chr18-16878949-DEL-64

chr18-16897339-DEL-393
chr18-16911222-DEL-373
chr18-16958618-DEL-1189
chr18-16961464-DEL-261
chr18-17000314-DEL-1346
chr18-17001087-DEL-64
chr18-17042039-DEL-1447
chr18-17069701-DEL-258
chr18-17104514-DEL-146
chr18-17121412-DEL-248
chr18-17123155-DEL-356
chr18-17149558-DEL-1557
chr18-17188867-DEL-640
chr18-17201570-DEL-956
chr18-17216774-DEL-67
chr18-17235745-DEL-1086
chr18-17242598-DEL-86
chr18-17262201-DEL-867
chr18-17294499-DEL-265
chr18-17303253-DEL-739
chr18-17325947-DEL-350
chr18-17332512-DEL-1297
chr18-17359484-DEL-180
chr18-17360295-DEL-96
chr18-17365387-DEL-51
chr18-17386000-DEL-55
chr18-17392686-DEL-135
chr18-17404918-DEL-2012
chr18-17408455-DEL-446
chr18-17413647-DEL-55
chr18-17440852-DEL-61
chr18-17449598-DEL-514
chr18-17510852-DEL-390
chr18-17557456-DEL-545
chr18-17558145-DEL-6410
chr18-17566947-DEL-6589
chr18-17587145-DEL-6396
chr18-17594012-DEL-55
chr18-17597827-DEL-5493
chr18-17606484-DEL-71
chr18-17607563-DEL-55
chr18-17612339-DEL-6229
chr18-17641069-DEL-60
chr18-17641595-DEL-180
chr18-17666964-DEL-103
chr18-17668714-DEL-102
chr18-17689655-DEL-997
chr18-17700984-DEL-77
chr18-17708735-DEL-1194
chr18-17709766-DEL-210
chr18-17714291-DEL-956
chr18-17753704-DEL-444
chr18-17827448-DEL-65
chr18-17872396-DEL-57
chr18-17874742-DEL-59
chr18-17887510-DEL-166
chr18-17945422-DEL-130
chr18-17983200-DEL-506
chr18-17992209-DEL-2960
chr18-18018173-DEL-497
chr18-18024266-DEL-2575
chr18-18047244-DEL-54
chr18-18151267-DEL-82
chr18-18235053-DEL-399
chr18-18242328-DEL-151
chr18-18265854-DEL-1149
chr18-18274979-DEL-54
chr18-18363449-DEL-180
chr18-18382725-DEL-352
chr18-18388262-DEL-6202
chr18-18411275-DEL-94
chr18-18418201-DEL-64
chr18-18423074-DEL-541
chr18-18433332-DEL-1350
chr18-18434724-DEL-2106
chr18-18447704-DEL-993
chr18-18452826-DEL-258

chr18-18519320-DEL-77
chr18-18534858-DEL-1225
chr18-18557112-DEL-58
chr18-18563275-DEL-781
chr18-18568175-DEL-2309
chr18-18582476-DEL-318
chr18-18596497-DEL-173
chr18-18598286-DEL-2061
chr18-18605163-DEL-150
chr18-18606942-DEL-354
chr18-18608198-DEL-104
chr18-18616872-DEL-379
chr18-18648844-DEL-167
chr18-18670995-DEL-1608
chr18-18677249-DEL-213
chr18-18681310-DEL-214
chr18-18688297-DEL-440
chr18-18689717-DEL-226
chr18-18715186-DEL-354
chr18-18716928-DEL-1699
chr18-18722058-DEL-60
chr18-18742931-DEL-94
chr18-18773252-DEL-221
chr18-18812886-DEL-196
chr18-18835757-DEL-4083
chr18-18854260-DEL-327
chr18-18867896-DEL-260
chr18-18883093-DEL-61
chr18-18887342-DEL-211
chr18-18899826-DEL-312
chr18-18908691-DEL-61
chr18-18915579-DEL-899
chr18-18916558-DEL-88
chr18-18924584-DEL-108
chr18-18949579-DEL-7325
chr18-18956584-DEL-214
chr18-18961947-DEL-7430
chr18-18974701-DEL-652
chr18-18976981-DEL-74
chr18-18984686-DEL-639
chr18-18991484-DEL-1878
chr18-19016932-DEL-2126
chr18-19029200-DEL-377
chr18-19070224-DEL-54
chr18-19080781-DEL-704
chr18-19087675-DEL-10602
chr18-19103872-DEL-60
chr18-19108385-DEL-9966
chr18-19110058-DEL-7115
chr18-19121036-DEL-3359
chr18-19121040-DEL-5137
chr18-19124451-DEL-1726
chr18-19127215-DEL-108
chr18-19138459-DEL-55
chr18-19154100-DEL-148
chr18-19159106-DEL-67
chr18-19267868-DEL-108
chr18-19282228-DEL-2397
chr18-19308416-DEL-1454
chr18-19325368-DEL-182
chr18-19358112-DEL-358
chr18-19393837-DEL-6571
chr18-19393901-DEL-205
chr18-19406976-DEL-6571
chr18-19450972-DEL-1207
chr18-19459304-DEL-428
chr18-19464390-DEL-71
chr18-19475055-DEL-401
chr18-19499074-DEL-319
chr18-19513850-DEL-573
chr18-19550172-DEL-7096
chr18-19572812-DEL-5175
chr18-19625917-DEL-460
chr18-19629090-DEL-108
chr18-19653887-DEL-264
chr18-19659196-DEL-151
chr18-19733338-DEL-2474

chr18-19845444-DEL-72
chr18-19888835-DEL-649
chr18-19901415-DEL-148
chr18-19916281-DEL-1481
chr18-19957197-DEL-341
chr18-19960646-DEL-53
chr18-19967297-DEL-54
chr18-19973770-DEL-154
chr18-19974869-DEL-1569
chr18-19991490-DEL-59
chr18-20032555-DEL-215
chr18-20034801-DEL-7940
chr18-20077556-DEL-1492
chr18-20081412-DEL-113
chr18-20107894-DEL-71
chr18-20119216-DEL-56
chr18-20130071-DEL-237
chr18-20133010-DEL-119
chr18-20144032-DEL-270
chr18-20274770-DEL-62
chr18-20298572-DEL-8264
chr18-20316847-DEL-213
chr18-20362911-DEL-226
chr18-20367787-DEL-52
chr18-20372282-DEL-303
chr18-20377605-DEL-214
chr18-20418399-DEL-126
chr18-20422431-DEL-52
chr18-20435365-DEL-6156
chr18-20453628-DEL-84
chr18-20463560-DEL-364
chr18-20504155-DEL-1435
chr18-20521465-DEL-106
chr18-20602505-DEL-63
chr18-20621562-DEL-986
chr18-20644389-DEL-96
chr18-20659453-DEL-75
chr18-20696751-DEL-54
chr18-20720969-DEL-76
chr18-20722762-DEL-92
chr18-20728403-DEL-52
chr18-20745125-DEL-229
chr18-20748066-DEL-155
chr18-20749156-DEL-129
chr18-20750036-DEL-437
chr18-20750578-DEL-373
chr18-20758752-DEL-152
chr18-20761780-DEL-461
chr18-20770045-DEL-60
chr18-20777611-DEL-98
chr18-20781219-DEL-201
chr18-20802205-DEL-127
chr18-20812298-DEL-301
chr18-20850153-DEL-214
chr18-20859532-DEL-259
chr18-20866432-DEL-196
chr18-20885112-DEL-54
chr18-20927955-DEL-51
chr18-20932155-DEL-105
chr18-20932789-DEL-132
chr18-21004825-DEL-122
chr18-21007154-DEL-98
chr18-21008142-DEL-208
chr18-21011621-DEL-131
chr18-21045850-DEL-95
chr18-21047082-DEL-563
chr18-21051732-DEL-197
chr18-21054280-DEL-216
chr18-21082556-DEL-321
chr18-21108508-DEL-198
chr18-21139199-DEL-63
chr18-21140237-DEL-255
chr18-21148934-DEL-182
chr18-21171370-DEL-145
chr18-21173119-DEL-197
chr18-21178191-DEL-232
chr18-21193415-DEL-4246

chr18-21223509-DEL-69
chr18-21232197-DEL-694
chr18-21255842-DEL-128
chr18-21290328-DEL-53
chr18-21303807-DEL-99
chr18-21309152-DEL-161
chr18-21315346-DEL-138
chr18-21322013-DEL-57
chr18-21333751-DEL-119
chr18-21359229-DEL-130
chr18-21364253-DEL-62
chr18-21375856-DEL-2764
chr18-21439439-DEL-150
chr18-21455276-DEL-53
chr18-21474273-DEL-1232
chr18-21522949-DEL-207
chr18-21543418-DEL-6348
chr18-21558897-DEL-202
chr18-21570323-DEL-89
chr18-21586218-DEL-473
chr18-21588260-DEL-1492
chr18-21601662-DEL-616
chr18-21604084-DEL-361
chr18-21614545-DEL-151
chr18-21663498-DEL-2659
chr18-21703796-DEL-58
chr18-21707964-DEL-57
chr18-21714374-DEL-64
chr18-21722788-DEL-548
chr18-21733162-DEL-587
chr18-21882115-DEL-745
chr18-21899002-DEL-64
chr18-21971281-DEL-91
chr18-21976128-DEL-141
chr18-22030636-DEL-1220
chr18-22099105-DEL-1424
chr18-22100578-DEL-344
chr18-22104839-DEL-52
chr18-22111899-DEL-148
chr18-22142745-DEL-58
chr18-22162182-DEL-227
chr18-22223316-DEL-190
chr18-22225316-DEL-361
chr18-22238103-DEL-59
chr18-22242263-DEL-360
chr18-22246937-DEL-3035
chr18-22249094-DEL-907
chr18-22250715-DEL-121
chr18-22273884-DEL-502
chr18-22289075-DEL-66
chr18-22311621-DEL-71
chr18-22320942-DEL-58
chr18-22338415-DEL-72
chr18-22342517-DEL-8328
chr18-22355003-DEL-92
chr18-22375315-DEL-6209
chr18-22384372-DEL-60
chr18-22392346-DEL-389
chr18-22421982-DEL-70
chr18-22449735-DEL-149
chr18-22461027-DEL-299
chr18-22465814-DEL-414
chr18-22466569-DEL-96
chr18-22503643-DEL-95
chr18-22515926-DEL-52
chr18-22549535-DEL-142
chr18-22565758-DEL-52
chr18-22671983-DEL-3372
chr18-22688248-DEL-57
chr18-22717736-DEL-444
chr18-22731806-DEL-237
chr18-22733621-DEL-401
chr18-22777003-DEL-52
chr18-22812462-DEL-732
chr18-22841618-DEL-173
chr18-22842492-DEL-65
chr18-22849654-DEL-72

chr18-22884797-DEL-159
chr18-22910973-DEL-65
chr18-22922303-DEL-1106
chr18-22946055-DEL-364
chr18-22972381-DEL-84
chr18-23010832-DEL-212
chr18-23067851-DEL-63
chr18-23068437-DEL-258
chr18-23076518-DEL-1190
chr18-23081931-DEL-109
chr18-23104499-DEL-224
chr18-23143008-DEL-52
chr18-23146835-DEL-279
chr18-23160147-DEL-84
chr18-23181687-DEL-209
chr18-23236351-DEL-795
chr18-23236524-DEL-427
chr18-23236550-DEL-853
chr18-23248885-DEL-75
chr18-23252999-DEL-171
chr18-23254171-DEL-3493
chr18-23261959-DEL-64
chr18-23270829-DEL-66
chr18-23287868-DEL-5978
chr18-23294353-DEL-86
chr18-23298539-DEL-1311
chr18-23337986-DEL-2758
chr18-23342457-DEL-6137
chr18-23365875-DEL-11469
chr18-23371874-DEL-2375
chr18-23381994-DEL-1488
chr18-23396046-DEL-6012
chr18-23412319-DEL-1740
chr18-23427390-DEL-114
chr18-23451541-DEL-271
chr18-23455693-DEL-565
chr18-23466927-DEL-56
chr18-23500887-DEL-113
chr18-23516516-DEL-78
chr18-23537918-DEL-3654
chr18-23541243-DEL-716
chr18-23551582-DEL-321
chr18-23556650-DEL-91
chr18-23557202-DEL-128
chr18-23595839-DEL-129
chr18-23621715-DEL-85
chr18-23627142-DEL-55
chr18-23629138-DEL-59
chr18-23633293-DEL-343
chr18-23635901-DEL-63
chr18-23638011-DEL-1247
chr18-23642890-DEL-57
chr18-23679664-DEL-71
chr18-23683081-DEL-91
chr18-23687449-DEL-2136
chr18-23699805-DEL-69
chr18-23725144-DEL-75
chr18-23758964-DEL-230
chr18-23761954-DEL-1091
chr18-23814096-DEL-53
chr18-23819213-DEL-2155
chr18-23868696-DEL-3963
chr18-23868926-DEL-467
chr18-23877688-DEL-73
chr18-23907958-DEL-158
chr18-23952122-DEL-108
chr18-23999152-DEL-1391
chr18-24015823-DEL-217
chr18-24018962-DEL-59
chr18-24030337-DEL-483
chr18-24053143-DEL-59
chr18-24055551-DEL-150
chr18-24098169-DEL-84
chr18-24114959-DEL-321
chr18-24117829-DEL-261
chr18-24134590-DEL-246
chr18-24169671-DEL-90

chr18-24230609-DEL-75
chr18-24248701-DEL-219
chr18-24279818-DEL-120
chr18-24280294-DEL-780
chr18-24297797-DEL-79
chr18-24314955-DEL-63
chr18-24322570-DEL-201
chr18-24327513-DEL-183
chr18-24333495-DEL-56
chr18-24352155-DEL-327
chr18-24363275-DEL-115
chr18-24367874-DEL-93
chr18-24378877-DEL-178
chr18-24381136-DEL-311
chr18-24384928-DEL-321
chr18-24387123-DEL-387
chr18-24392221-DEL-184
chr18-24421297-DEL-64
chr18-24437796-DEL-284
chr18-24438976-DEL-65
chr18-24556427-DEL-66
chr18-24563075-DEL-79
chr18-24588078-DEL-371
chr18-24614252-DEL-162
chr18-24615396-DEL-2273
chr18-24615573-DEL-78
chr18-24629280-DEL-53
chr18-24665197-DEL-8348
chr18-24670127-DEL-61
chr18-24676655-DEL-222
chr18-24678983-DEL-207
chr18-24682439-DEL-1760
chr18-24685009-DEL-573
chr18-24686298-DEL-5594
chr18-24699242-DEL-590
chr18-24728595-DEL-80
chr18-24758901-DEL-62
chr18-24760850-DEL-82
chr18-24778686-DEL-843
chr18-24790510-DEL-242
chr18-24810351-DEL-295
chr18-24831658-DEL-52
chr18-24861822-DEL-107
chr18-24876558-DEL-52
chr18-24877268-DEL-116
chr18-24881407-DEL-71
chr18-24947602-DEL-80
chr18-24980015-DEL-160
chr18-25038902-DEL-498
chr18-25053017-DEL-93
chr18-25080543-DEL-58
chr18-25099385-DEL-907
chr18-25129649-DEL-219
chr18-25188000-DEL-100
chr18-25243434-DEL-304
chr18-25278789-DEL-295
chr18-25300676-DEL-52
chr18-25371160-DEL-567
chr18-25385538-DEL-400
chr18-25386063-DEL-6598
chr18-25392361-DEL-192
chr18-25416771-DEL-74
chr18-25428759-DEL-98
chr18-25449232-DEL-11380
chr18-25452308-DEL-2399
chr18-25468785-DEL-2176
chr18-25488873-DEL-87
chr18-25519634-DEL-179
chr18-25532033-DEL-737
chr18-25546300-DEL-3254
chr18-25546973-DEL-88
chr18-25569102-DEL-2835
chr18-25593706-DEL-426
chr18-25640327-DEL-71
chr18-25715077-DEL-175
chr18-25759921-DEL-205
chr18-25866867-DEL-75

chr18-25886997-DEL-60
chr18-25895229-DEL-90
chr18-25922587-DEL-75
chr18-25931255-DEL-78
chr18-25936760-DEL-57
chr18-25951122-DEL-52
chr18-26002610-DEL-245
chr18-26055022-DEL-71
chr18-26058740-DEL-2244
chr18-26059577-DEL-2253
chr18-26135007-DEL-278
chr18-26136104-DEL-2869
chr18-26138997-DEL-2223
chr18-26143580-DEL-96
chr18-26189424-DEL-244
chr18-26194676-DEL-88
chr18-26208071-DEL-74
chr18-26250017-DEL-913
chr18-26278937-DEL-231
chr18-26349800-DEL-74
chr18-26373561-DEL-503
chr18-26419478-DEL-68
chr18-26446950-DEL-2489
chr18-26489012-DEL-110
chr18-26520455-DEL-657
chr18-26524537-DEL-118
chr18-26557843-DEL-90
chr18-26561419-DEL-431
chr18-26577548-DEL-464
chr18-26598179-DEL-51
chr18-26605940-DEL-73
chr18-26606847-DEL-197
chr18-26609301-DEL-1003
chr18-26623876-DEL-115
chr18-26665305-DEL-1255
chr18-26721970-DEL-3037
chr18-26735120-DEL-253
chr18-26748715-DEL-127
chr18-26759178-DEL-907
chr18-26829814-DEL-214
chr18-26838628-DEL-114
chr18-26854696-DEL-199
chr18-26899727-DEL-69
chr18-26912124-DEL-916
chr18-27002232-DEL-138
chr18-27036540-DEL-108
chr18-27065309-DEL-512
chr18-27084281-DEL-183
chr18-27128833-DEL-52
chr18-27139813-DEL-1187
chr18-27142747-DEL-428
chr18-27153311-DEL-74
chr18-27188500-DEL-383
chr18-27201166-DEL-214
chr18-27209633-DEL-145
chr18-27224596-DEL-73
chr18-27254717-DEL-128
chr18-27266201-DEL-204
chr18-27274082-DEL-83
chr18-27280143-DEL-154
chr18-27283099-DEL-1114
chr18-27310587-DEL-6271
chr18-27327003-DEL-127
chr18-27334304-DEL-102
chr18-27354365-DEL-93
chr18-27356093-DEL-64
chr18-27367251-DEL-214
chr18-27408296-DEL-454
chr18-27441238-DEL-238
chr18-27441589-DEL-1228
chr18-27454929-DEL-63
chr18-27462901-DEL-101
chr18-27480857-DEL-122
chr18-27512763-DEL-141
chr18-27513619-DEL-394
chr18-27519707-DEL-138
chr18-27528628-DEL-54

chr18-27564636-DEL-52
chr18-27565410-DEL-365
chr18-27606910-DEL-489
chr18-27617469-DEL-378
chr18-27656231-DEL-1163
chr18-27722858-DEL-565
chr18-27734222-DEL-5773
chr18-27734366-DEL-3173
chr18-27734891-DEL-3142
chr18-27748651-DEL-56
chr18-27809014-DEL-201
chr18-27842657-DEL-309
chr18-27848460-DEL-362
chr18-27849399-DEL-200
chr18-27875897-DEL-637
chr18-27896185-DEL-6036
chr18-27908913-DEL-1491
chr18-27910406-DEL-10318
chr18-27936715-DEL-783
chr18-27973718-DEL-105
chr18-28011061-DEL-2316
chr18-28021076-DEL-115
chr18-28036409-DEL-1160
chr18-28051482-DEL-2968
chr18-28136491-DEL-109
chr18-28140605-DEL-1470
chr18-28171387-DEL-281
chr18-28205573-DEL-995
chr18-28217865-DEL-159
chr18-28221443-DEL-74
chr18-28224370-DEL-403
chr18-28236699-DEL-2331
chr18-28264932-DEL-846
chr18-28272916-DEL-1851
chr18-28286088-DEL-1796
chr18-28299288-DEL-1893
chr18-28308941-DEL-725
chr18-28335720-DEL-51
chr18-28342793-DEL-1290
chr18-28354423-DEL-922
chr18-28377914-DEL-13205
chr18-28453951-DEL-88
chr18-28465175-DEL-1286
chr18-28476388-DEL-244
chr18-28477752-DEL-1108
chr18-28510459-DEL-180
chr18-28511273-DEL-195
chr18-28524242-DEL-247
chr18-28538580-DEL-152
chr18-28582830-DEL-87
chr18-28583951-DEL-364
chr18-28589455-DEL-4193
chr18-28606960-DEL-93
chr18-28611010-DEL-199
chr18-28684606-DEL-1166
chr18-28688471-DEL-90
chr18-28690536-DEL-68
chr18-28703528-DEL-93
chr18-28711380-DEL-232
chr18-28719833-DEL-2256
chr18-28726885-DEL-1438
chr18-28730552-DEL-5979
chr18-28743935-DEL-6427
chr18-28756025-DEL-7456
chr18-28763053-DEL-208
chr18-28784310-DEL-360
chr18-28794557-DEL-467
chr18-28821567-DEL-120
chr18-28869562-DEL-908
chr18-28870819-DEL-114
chr18-28885502-DEL-111
chr18-28933000-DEL-75
chr18-28949908-DEL-79
chr18-29037156-DEL-3075
chr18-29066247-DEL-2648
chr18-29076552-DEL-188
chr18-29147823-DEL-7109

chr18-29164396-DEL-170
chr18-29170769-DEL-77
chr18-29176352-DEL-123
chr18-29201334-DEL-315
chr18-29211465-DEL-108
chr18-29312322-DEL-56
chr18-29313333-DEL-5827
chr18-29313449-DEL-70
chr18-29313724-DEL-1120
chr18-29327682-DEL-7086
chr18-29327713-DEL-212
chr18-29327882-DEL-636
chr18-29355743-DEL-114
chr18-29358155-DEL-556
chr18-29358788-DEL-4236
chr18-29373461-DEL-91
chr18-29415762-DEL-144
chr18-29418860-DEL-60
chr18-29463476-DEL-461
chr18-29479907-DEL-1285
chr18-29516234-DEL-102
chr18-29553066-DEL-3072
chr18-29585440-DEL-666
chr18-29605410-DEL-1241
chr18-29632187-DEL-58
chr18-29661112-DEL-4492
chr18-29669808-DEL-87
chr18-29684523-DEL-2227
chr18-29686545-DEL-6346
chr18-29714645-DEL-1845
chr18-29732498-DEL-138
chr18-29754056-DEL-73
chr18-29782874-DEL-105
chr18-29820979-DEL-200
chr18-29905121-DEL-642
chr18-30021618-DEL-3214
chr18-30033076-DEL-299
chr18-30073212-DEL-241
chr18-30079373-DEL-64
chr18-30080545-DEL-102
chr18-30123094-DEL-1200
chr18-30169791-DEL-201
chr18-30176013-DEL-74
chr18-30189927-DEL-73
chr18-30201221-DEL-188
chr18-30209021-DEL-54
chr18-30246623-DEL-52
chr18-30272210-DEL-231
chr18-30293746-DEL-479
chr18-30301805-DEL-249
chr18-30380157-DEL-187
chr18-30393907-DEL-570
chr18-30415421-DEL-179
chr18-30428334-DEL-210
chr18-30431862-DEL-179
chr18-30457683-DEL-104
chr18-30459809-DEL-609
chr18-30492291-DEL-60
chr18-30507603-DEL-233
chr18-30517327-DEL-181
chr18-30529640-DEL-93
chr18-30540716-DEL-76
chr18-30559354-DEL-225
chr18-30560204-DEL-2133
chr18-30562130-DEL-64
chr18-30562375-DEL-1526
chr18-30604871-DEL-114
chr18-30657486-DEL-83
chr18-30682423-DEL-84
chr18-30705949-DEL-51
chr18-30722172-DEL-64
chr18-30740247-DEL-344
chr18-30765135-DEL-67
chr18-30803493-DEL-134
chr18-30807813-DEL-55
chr18-30809064-DEL-376
chr18-30809504-DEL-1827

chr18-30829377-DEL-201
chr18-30841663-DEL-62
chr18-30891201-DEL-98
chr18-30907930-DEL-64
chr18-30920411-DEL-6487
chr18-30927396-DEL-77
chr18-30935299-DEL-6363
chr18-30950350-DEL-228
chr18-30974531-DEL-60
chr18-31003898-DEL-2507
chr18-31004661-DEL-828
chr18-31009004-DEL-311
chr18-31024227-DEL-409
chr18-31073705-DEL-1866
chr18-31081187-DEL-84
chr18-31096378-DEL-68
chr18-31154029-DEL-3935
chr18-31177443-DEL-458
chr18-31194228-DEL-387
chr18-31217088-DEL-56
chr18-31245338-DEL-565
chr18-31246374-DEL-111
chr18-31274533-DEL-960
chr18-31317347-DEL-207
chr18-31319873-DEL-456
chr18-31413922-DEL-6226
chr18-31426268-DEL-1152
chr18-31451325-DEL-304
chr18-31475513-DEL-188
chr18-31482635-DEL-359
chr18-31483876-DEL-630
chr18-31515655-DEL-63
chr18-31532617-DEL-1567
chr18-31538964-DEL-52
chr18-31558354-DEL-56
chr18-31559904-DEL-2489
chr18-31560825-DEL-60
chr18-31587258-DEL-135
chr18-31603114-DEL-153
chr18-31614289-DEL-52
chr18-31626356-DEL-65
chr18-31735438-DEL-104
chr18-31752726-DEL-2087
chr18-31752747-DEL-63
chr18-31753607-DEL-400
chr18-31802880-DEL-74
chr18-31819470-DEL-577
chr18-31835848-DEL-116
chr18-31837394-DEL-3114
chr18-31898672-DEL-77
chr18-31919779-DEL-74
chr18-31933947-DEL-189
chr18-31941061-DEL-332
chr18-31946612-DEL-167
chr18-31952016-DEL-58
chr18-31958560-DEL-204
chr18-31966123-DEL-104
chr18-31975332-DEL-350
chr18-31990148-DEL-71
chr18-31994691-DEL-57
chr18-32000774-DEL-228
chr18-32027499-DEL-54
chr18-32098744-DEL-80
chr18-32141978-DEL-1113
chr18-32144573-DEL-83
chr18-32160173-DEL-189
chr18-32176552-DEL-6436
chr18-32196026-DEL-651
chr18-32204747-DEL-227
chr18-32245630-DEL-71
chr18-32249412-DEL-531
chr18-32258958-DEL-827
chr18-32261171-DEL-102
chr18-32262477-DEL-70
chr18-32279126-DEL-140
chr18-32295607-DEL-127
chr18-32311690-DEL-2468

chr18-32317823-DEL-285
chr18-32327264-DEL-93
chr18-32334731-DEL-561
chr18-32354052-DEL-154
chr18-32370795-DEL-128
chr18-32375820-DEL-77
chr18-32445999-DEL-60
chr18-32463225-DEL-166
chr18-32473992-DEL-84
chr18-32496348-DEL-74
chr18-32497856-DEL-553
chr18-32500028-DEL-577
chr18-32527687-DEL-84
chr18-32550121-DEL-150
chr18-32557677-DEL-63
chr18-32600150-DEL-1222
chr18-32601276-DEL-144
chr18-32603482-DEL-1423
chr18-32609396-DEL-863
chr18-32611247-DEL-342
chr18-32625408-DEL-295
chr18-32632230-DEL-163
chr18-32654644-DEL-309
chr18-32668924-DEL-89
chr18-32670755-DEL-259
chr18-32677023-DEL-63
chr18-32686100-DEL-248
chr18-32695907-DEL-326
chr18-32697098-DEL-555
chr18-32713980-DEL-2098
chr18-32723204-DEL-64
chr18-32730617-DEL-1218
chr18-32735121-DEL-2834
chr18-32739457-DEL-301
chr18-32741652-DEL-197
chr18-32742370-DEL-4025
chr18-32750148-DEL-182
chr18-32757814-DEL-62
chr18-32765523-DEL-70
chr18-32792417-DEL-1284
chr18-32815816-DEL-6448
chr18-32815820-DEL-4291
chr18-32822268-DEL-275
chr18-32880467-DEL-64
chr18-32903610-DEL-11819
chr18-32904588-DEL-90
chr18-32912364-DEL-927
chr18-32916421-DEL-90
chr18-33074914-DEL-70
chr18-33144464-DEL-6883
chr18-33153745-DEL-15861
chr18-33162846-DEL-6760
chr18-33170721-DEL-68
chr18-33177119-DEL-6465
chr18-33201003-DEL-123
chr18-33203362-DEL-241
chr18-33238453-DEL-51
chr18-33245875-DEL-92
chr18-33274190-DEL-797
chr18-33294597-DEL-131
chr18-33315521-DEL-97
chr18-33329587-DEL-856
chr18-33343272-DEL-196
chr18-33351471-DEL-151
chr18-33356419-DEL-643
chr18-33393967-DEL-1074
chr18-33455593-DEL-8269
chr18-33463097-DEL-213
chr18-33513747-DEL-112
chr18-33516925-DEL-1136
chr18-33522070-DEL-1473
chr18-33527965-DEL-424
chr18-33561369-DEL-68
chr18-33599789-DEL-70
chr18-33608951-DEL-60
chr18-33628833-DEL-505
chr18-33631944-DEL-920

chr18-33645230-DEL-446
chr18-33648327-DEL-52
chr18-33650182-DEL-754
chr18-33655129-DEL-82
chr18-33668748-DEL-390
chr18-33696699-DEL-534
chr18-33725512-DEL-104
chr18-33776017-DEL-68
chr18-33784759-DEL-383
chr18-33806469-DEL-447
chr18-33814804-DEL-2650
chr18-33896647-DEL-1015
chr18-33898074-DEL-58
chr18-33899212-DEL-689
chr18-33900132-DEL-1193
chr18-33924255-DEL-188
chr18-33926134-DEL-52
chr18-33934865-DEL-76
chr18-33938527-DEL-67
chr18-33943557-DEL-352
chr18-33954690-DEL-1001
chr18-33973281-DEL-3274
chr18-33973706-DEL-1536
chr18-33979858-DEL-1393
chr18-33991185-DEL-306
chr18-33997424-DEL-62
chr18-34002889-DEL-54
chr18-34026560-DEL-78
chr18-34030046-DEL-222
chr18-34031140-DEL-6128
chr18-34041775-DEL-105
chr18-34090014-DEL-231
chr18-34091445-DEL-71
chr18-34093639-DEL-1478
chr18-34121138-DEL-66
chr18-34173547-DEL-275
chr18-34177178-DEL-275
chr18-34208398-DEL-126
chr18-34216376-DEL-130
chr18-34220989-DEL-230
chr18-34228389-DEL-402
chr18-34261824-DEL-90
chr18-34266187-DEL-71
chr18-34307992-DEL-398
chr18-34327945-DEL-89
chr18-34341219-DEL-56
chr18-34347681-DEL-1946
chr18-34363379-DEL-184
chr18-34373126-DEL-104
chr18-34386303-DEL-79
chr18-34402303-DEL-106
chr18-34410045-DEL-2056
chr18-34456993-DEL-84
chr18-34457045-DEL-4117
chr18-34459421-DEL-1741
chr18-34472843-DEL-69
chr18-34479271-DEL-101
chr18-34497040-DEL-71
chr18-34536525-DEL-105
chr18-34544234-DEL-198
chr18-34559456-DEL-175
chr18-34566654-DEL-809
chr18-34609589-DEL-181
chr18-34634172-DEL-160
chr18-34652627-DEL-57
chr18-34691978-DEL-178
chr18-34703872-DEL-135
chr18-34713669-DEL-171
chr18-34714383-DEL-203
chr18-34726444-DEL-104
chr18-34748002-DEL-994
chr18-34777243-DEL-249
chr18-34795414-DEL-247
chr18-34827487-DEL-227
chr18-34834127-DEL-281
chr18-34839503-DEL-94
chr18-34842622-DEL-210

chr18-34848367-DEL-3325
chr18-34861423-DEL-190
chr18-34864988-DEL-177
chr18-34872865-DEL-81
chr18-34923237-DEL-264
chr18-34944774-DEL-115
chr18-34953928-DEL-57
chr18-34959102-DEL-59
chr18-34968494-DEL-125
chr18-34970082-DEL-55
chr18-35011961-DEL-122
chr18-35019299-DEL-65
chr18-35022522-DEL-201
chr18-35092637-DEL-766
chr18-35102689-DEL-1199
chr18-35106738-DEL-206
chr18-35112741-DEL-326
chr18-35183918-DEL-78
chr18-35200905-DEL-119
chr18-35246272-DEL-357
chr18-35259864-DEL-60
chr18-35300767-DEL-211
chr18-35334502-DEL-380
chr18-35337336-DEL-139
chr18-35338678-DEL-169
chr18-35380657-DEL-268
chr18-35388647-DEL-234
chr18-35391487-DEL-87
chr18-35410237-DEL-89
chr18-35416209-DEL-108
chr18-35428608-DEL-163
chr18-35472770-DEL-740
chr18-35511771-DEL-329
chr18-35570269-DEL-350
chr18-35647039-DEL-353
chr18-35652945-DEL-343
chr18-35669958-DEL-328
chr18-35683233-DEL-72
chr18-35714407-DEL-66
chr18-35717139-DEL-171
chr18-35728619-DEL-193
chr18-35737608-DEL-764
chr18-35766584-DEL-226
chr18-35771030-DEL-182
chr18-35777087-DEL-186
chr18-35779708-DEL-66
chr18-35796626-DEL-62
chr18-35812209-DEL-103
chr18-35842972-DEL-56
chr18-35852135-DEL-415
chr18-35870016-DEL-476
chr18-35891097-DEL-197
chr18-35893107-DEL-184
chr18-35894028-DEL-111
chr18-35911564-DEL-130
chr18-35914676-DEL-122
chr18-36018347-DEL-188
chr18-36029811-DEL-198
chr18-36056603-DEL-155
chr18-36132591-DEL-475
chr18-36145363-DEL-82
chr18-36184041-DEL-58
chr18-36193710-DEL-74
chr18-36194054-DEL-825
chr18-36278423-DEL-156
chr18-36295118-DEL-91
chr18-36304630-DEL-211
chr18-36339734-DEL-239
chr18-36344680-DEL-524
chr18-36353645-DEL-139
chr18-36374332-DEL-176
chr18-36376294-DEL-3481
chr18-36380375-DEL-859
chr18-36380766-DEL-7321
chr18-36382658-DEL-272
chr18-36382986-DEL-1757
chr18-36407816-DEL-270

chr18-36436702-DEL-73
chr18-36437863-DEL-539
chr18-36441779-DEL-431
chr18-36457496-DEL-164
chr18-36470261-DEL-407
chr18-36506279-DEL-70
chr18-36523890-DEL-76
chr18-36570092-DEL-107
chr18-36593521-DEL-59
chr18-36595239-DEL-84
chr18-36600163-DEL-204
chr18-36604174-DEL-53
chr18-36621239-DEL-60
chr18-36678641-DEL-226
chr18-36681230-DEL-51
chr18-36720279-DEL-1246
chr18-36722969-DEL-160
chr18-36728693-DEL-224
chr18-36740207-DEL-2009
chr18-36745068-DEL-159
chr18-36748834-DEL-128
chr18-36782689-DEL-109
chr18-36827526-DEL-91
chr18-36830017-DEL-108
chr18-36842528-DEL-471
chr18-36846862-DEL-190
chr18-36857464-DEL-369
chr18-36862457-DEL-183
chr18-36908821-DEL-170
chr18-36910906-DEL-259
chr18-36943734-DEL-525
chr18-36948903-DEL-204
chr18-36949384-DEL-2885
chr18-36950462-DEL-250
chr18-36950842-DEL-991
chr18-36963012-DEL-2802
chr18-36968010-DEL-202
chr18-36974155-DEL-66
chr18-36987174-DEL-2463
chr18-37000031-DEL-908
chr18-37008216-DEL-351
chr18-37013951-DEL-2591
chr18-37021601-DEL-3630
chr18-37028563-DEL-458
chr18-37031823-DEL-1708
chr18-37090782-DEL-64
chr18-37155982-DEL-86
chr18-37157211-DEL-255
chr18-37162637-DEL-109
chr18-37172430-DEL-6900
chr18-37180136-DEL-69
chr18-37259353-DEL-119
chr18-37262477-DEL-1139
chr18-37304606-DEL-417
chr18-37341775-DEL-183
chr18-37361424-DEL-202
chr18-37362879-DEL-95
chr18-37364549-DEL-1631
chr18-37387043-DEL-1977
chr18-37418946-DEL-66
chr18-37431575-DEL-76
chr18-37473945-DEL-131
chr18-37491737-DEL-399
chr18-37496719-DEL-8219
chr18-37506624-DEL-86
chr18-37521199-DEL-6401
chr18-37530328-DEL-88
chr18-37532266-DEL-934
chr18-37593420-DEL-144
chr18-37598421-DEL-336
chr18-37672856-DEL-1258
chr18-37676628-DEL-6388
chr18-37705677-DEL-1021
chr18-37708821-DEL-7612
chr18-37720710-DEL-6273
chr18-37727262-DEL-299
chr18-37729187-DEL-62

chr18-37733595-DEL-6165
chr18-37733778-DEL-1914
chr18-37735928-DEL-224
chr18-37744289-DEL-1141
chr18-37773060-DEL-106
chr18-37831424-DEL-85
chr18-37835862-DEL-189
chr18-37838264-DEL-89
chr18-37845120-DEL-195
chr18-37846324-DEL-64
chr18-37916096-DEL-119
chr18-37922337-DEL-902
chr18-37924113-DEL-869
chr18-37992766-DEL-201
chr18-38029951-DEL-78
chr18-38054025-DEL-71
chr18-38115412-DEL-72
chr18-38149924-DEL-110
chr18-38167822-DEL-52
chr18-38239716-DEL-413
chr18-38240320-DEL-306
chr18-38260922-DEL-1499
chr18-38262786-DEL-675
chr18-38268847-DEL-129
chr18-38283713-DEL-59
chr18-38351390-DEL-83
chr18-38371467-DEL-73
chr18-38374944-DEL-180
chr18-38377845-DEL-640
chr18-38444299-DEL-201
chr18-38452478-DEL-93
chr18-38463059-DEL-101
chr18-38481182-DEL-142
chr18-38484394-DEL-333
chr18-38489192-DEL-58
chr18-38506842-DEL-58
chr18-38515517-DEL-210
chr18-38530599-DEL-350
chr18-38556139-DEL-196
chr18-38578456-DEL-1051
chr18-38582012-DEL-115
chr18-38598463-DEL-82
chr18-38622045-DEL-66
chr18-38630405-DEL-1645
chr18-38648291-DEL-173
chr18-38664276-DEL-187
chr18-38687561-DEL-1727
chr18-38704177-DEL-209
chr18-38704764-DEL-71
chr18-38736636-DEL-186
chr18-38739856-DEL-171
chr18-38742496-DEL-52
chr18-38768636-DEL-586
chr18-38792857-DEL-224
chr18-38843341-DEL-532
chr18-38884008-DEL-143
chr18-38905886-DEL-56
chr18-38906771-DEL-60
chr18-38917639-DEL-73
chr18-38984534-DEL-238
chr18-39025367-DEL-155
chr18-39027301-DEL-94
chr18-39035098-DEL-195
chr18-39066231-DEL-83
chr18-39069019-DEL-481
chr18-39080993-DEL-69
chr18-39114856-DEL-509
chr18-39170710-DEL-123
chr18-39173447-DEL-163
chr18-39173908-DEL-227
chr18-39308070-DEL-77
chr18-39313607-DEL-148
chr18-39348543-DEL-256
chr18-39438541-DEL-96
chr18-39469226-DEL-76
chr18-39519536-DEL-62
chr18-39528576-DEL-123

chr18-39551128-DEL-815
chr18-39660151-DEL-76
chr18-39706528-DEL-98
chr18-39721767-DEL-155
chr18-39726045-DEL-1252
chr18-39763638-DEL-145
chr18-39772527-DEL-113
chr18-39777053-DEL-56
chr18-39803793-DEL-730
chr18-39815347-DEL-83
chr18-39841534-DEL-72
chr18-39900440-DEL-265
chr18-39922303-DEL-129
chr18-39997888-DEL-70
chr18-40018681-DEL-69
chr18-40064918-DEL-109
chr18-40066036-DEL-2877
chr18-40066773-DEL-96
chr18-40130321-DEL-73
chr18-40133767-DEL-226
chr18-40142761-DEL-1731
chr18-40181640-DEL-97
chr18-40228102-DEL-69
chr18-40283245-DEL-759
chr18-40336898-DEL-328
chr18-40360968-DEL-107
chr18-40365142-DEL-55
chr18-40369424-DEL-1259
chr18-40375794-DEL-209
chr18-40394259-DEL-86
chr18-40420585-DEL-325
chr18-40439498-DEL-59
chr18-40470402-DEL-64
chr18-40553088-DEL-155
chr18-40557935-DEL-276
chr18-40558615-DEL-260
chr18-40579484-DEL-70
chr18-40582456-DEL-56
chr18-40592704-DEL-6198
chr18-40616778-DEL-360
chr18-40625124-DEL-197
chr18-40642182-DEL-429
chr18-40650663-DEL-118
chr18-40709482-DEL-2398
chr18-40718942-DEL-357
chr18-40734614-DEL-8621
chr18-40743539-DEL-5733
chr18-40772499-DEL-227
chr18-40780776-DEL-6204
chr18-40819189-DEL-57
chr18-40858467-DEL-182
chr18-40872408-DEL-92
chr18-40875058-DEL-80
chr18-40877496-DEL-432
chr18-40879091-DEL-209
chr18-40917205-DEL-1241
chr18-40932160-DEL-56
chr18-40967038-DEL-134
chr18-40988935-DEL-1026
chr18-41072428-DEL-542
chr18-41075995-DEL-69
chr18-41078228-DEL-4221
chr18-41135894-DEL-301
chr18-41217531-DEL-1170
chr18-41225400-DEL-52
chr18-41245258-DEL-2541
chr18-41254722-DEL-59
chr18-41271395-DEL-51
chr18-41296823-DEL-58
chr18-41312809-DEL-950
chr18-41335643-DEL-605
chr18-41340212-DEL-9834
chr18-41367271-DEL-948
chr18-41423156-DEL-342
chr18-41444766-DEL-60
chr18-41468592-DEL-1217
chr18-41469826-DEL-1597

chr18-41483717-DEL-71
chr18-41493152-DEL-86
chr18-41521124-DEL-371
chr18-41523895-DEL-72
chr18-41553218-DEL-211
chr18-41559748-DEL-57
chr18-41575703-DEL-82
chr18-41643221-DEL-200
chr18-41653838-DEL-798
chr18-41752820-DEL-86
chr18-41766093-DEL-586
chr18-41790546-DEL-54
chr18-41805303-DEL-60
chr18-41837493-DEL-127
chr18-41869042-DEL-1000
chr18-41871666-DEL-64
chr18-41889254-DEL-788
chr18-41970147-DEL-1367
chr18-41976917-DEL-277
chr18-41980666-DEL-77
chr18-42015276-DEL-84
chr18-42050135-DEL-118
chr18-42073571-DEL-224
chr18-42086574-DEL-181
chr18-42105701-DEL-195
chr18-42128846-DEL-356
chr18-42129654-DEL-494
chr18-42145969-DEL-71
chr18-42176624-DEL-56
chr18-42218007-DEL-1766
chr18-42231058-DEL-171
chr18-42249512-DEL-200
chr18-42298160-DEL-81
chr18-42302337-DEL-168
chr18-42312484-DEL-56
chr18-42339052-DEL-55
chr18-42383641-DEL-51
chr18-42384759-DEL-233
chr18-42385421-DEL-511
chr18-42402544-DEL-74
chr18-42440225-DEL-189
chr18-42442549-DEL-216
chr18-42464879-DEL-111
chr18-42487570-DEL-69
chr18-42499044-DEL-56
chr18-42505988-DEL-264
chr18-42517948-DEL-90
chr18-42559171-DEL-142
chr18-42565164-DEL-67
chr18-42579872-DEL-64
chr18-42618139-DEL-474
chr18-42649032-DEL-227
chr18-42652974-DEL-198
chr18-42653484-DEL-244
chr18-42677496-DEL-410
chr18-42714137-DEL-221
chr18-42717790-DEL-144
chr18-42729605-DEL-1032
chr18-42736829-DEL-412
chr18-42765326-DEL-486
chr18-42780386-DEL-105
chr18-42808978-DEL-75
chr18-42842986-DEL-356
chr18-42848900-DEL-457
chr18-42852288-DEL-708
chr18-42897167-DEL-54
chr18-42898076-DEL-309
chr18-42920726-DEL-64
chr18-42991796-DEL-1146
chr18-43054984-DEL-62
chr18-43059994-DEL-148
chr18-43085727-DEL-390
chr18-43091136-DEL-316
chr18-43104493-DEL-69
chr18-43164667-DEL-60
chr18-43218106-DEL-68
chr18-43224069-DEL-99

chr18-43228371-DEL-69
chr18-43276730-DEL-78
chr18-43304153-DEL-3387
chr18-43323713-DEL-465
chr18-43325162-DEL-54
chr18-43325621-DEL-166
chr18-43328276-DEL-56
chr18-43338319-DEL-179
chr18-43361015-DEL-70
chr18-43403036-DEL-844
chr18-43405373-DEL-51
chr18-43441834-DEL-381
chr18-43453103-DEL-53
chr18-43502485-DEL-58
chr18-43600574-DEL-77
chr18-43620317-DEL-265
chr18-43636552-DEL-60
chr18-43637844-DEL-483
chr18-43686725-DEL-723
chr18-43727197-DEL-157
chr18-43751825-DEL-106
chr18-43755907-DEL-2805
chr18-43759463-DEL-197
chr18-43764082-DEL-110
chr18-43790047-DEL-170
chr18-43825325-DEL-108
chr18-43837999-DEL-100
chr18-43855874-DEL-165
chr18-43864422-DEL-487
chr18-43890582-DEL-116
chr18-43921168-DEL-398
chr18-43937246-DEL-114
chr18-43938631-DEL-50
chr18-43948105-DEL-51
chr18-43968900-DEL-198
chr18-44033158-DEL-201
chr18-44062401-DEL-76
chr18-44082801-DEL-304
chr18-44112916-DEL-452
chr18-44180254-DEL-6238
chr18-44214545-DEL-339
chr18-44218395-DEL-71
chr18-44239084-DEL-75
chr18-44261805-DEL-516
chr18-44272829-DEL-536
chr18-44276042-DEL-255
chr18-44302867-DEL-55
chr18-44309122-DEL-63
chr18-44317162-DEL-3640
chr18-44326099-DEL-379
chr18-44329521-DEL-606
chr18-44339292-DEL-4641
chr18-44345938-DEL-4249
chr18-44352427-DEL-4754
chr18-44353493-DEL-77
chr18-44360553-DEL-6344
chr18-44375169-DEL-6250
chr18-44389106-DEL-51
chr18-44389285-DEL-963
chr18-44407942-DEL-59
chr18-44422939-DEL-6546
chr18-44460483-DEL-900
chr18-44470608-DEL-87
chr18-44493788-DEL-2507
chr18-44509956-DEL-242
chr18-44610827-DEL-147
chr18-44652454-DEL-71
chr18-44681287-DEL-184
chr18-44683951-DEL-129
chr18-44736682-DEL-192
chr18-44760885-DEL-52
chr18-44811434-DEL-346
chr18-44835152-DEL-1420
chr18-44852415-DEL-214
chr18-44875480-DEL-92
chr18-44913107-DEL-449
chr18-44925077-DEL-57

chr18-44931725-DEL-132
chr18-44946536-DEL-52
chr18-44970900-DEL-958
chr18-45004792-DEL-59
chr18-45020011-DEL-91
chr18-45027091-DEL-221
chr18-45048719-DEL-52
chr18-45063793-DEL-170
chr18-45073229-DEL-342
chr18-45107366-DEL-231
chr18-45146864-DEL-51
chr18-45183172-DEL-31519
chr18-45206900-DEL-69
chr18-45227178-DEL-56
chr18-45245285-DEL-62
chr18-45278767-DEL-1427
chr18-45307352-DEL-969
chr18-45322661-DEL-676
chr18-45333193-DEL-92
chr18-45338463-DEL-16564
chr18-45340492-DEL-6295
chr18-45356915-DEL-318
chr18-45367670-DEL-2877
chr18-45424123-DEL-5486
chr18-45480481-DEL-60
chr18-45481941-DEL-532
chr18-45490418-DEL-7429
chr18-45530906-DEL-52
chr18-45540349-DEL-60
chr18-45541971-DEL-2525
chr18-45547331-DEL-1431
chr18-45586213-DEL-2709
chr18-45589267-DEL-1793
chr18-45595292-DEL-716
chr18-45651226-DEL-78
chr18-45685783-DEL-222
chr18-45698592-DEL-64
chr18-45720067-DEL-100
chr18-45761274-DEL-52
chr18-45819867-DEL-289
chr18-45846238-DEL-264
chr18-45855584-DEL-54
chr18-45859089-DEL-1891
chr18-45893443-DEL-1090
chr18-45894320-DEL-78
chr18-45926022-DEL-95
chr18-45963649-DEL-604
chr18-45963987-DEL-899
chr18-45983203-DEL-52
chr18-45995917-DEL-1846
chr18-46003077-DEL-5876
chr18-46056375-DEL-2456
chr18-46064886-DEL-147
chr18-46090857-DEL-199
chr18-46130842-DEL-1449
chr18-46144485-DEL-68
chr18-46154353-DEL-6372
chr18-46162191-DEL-8923
chr18-46163016-DEL-382
chr18-46163929-DEL-926
chr18-46172075-DEL-201
chr18-46173569-DEL-106
chr18-46174940-DEL-1219
chr18-46214586-DEL-6435
chr18-46222949-DEL-1397
chr18-46283460-DEL-175
chr18-46283942-DEL-1041
chr18-46294347-DEL-78
chr18-46340648-DEL-52
chr18-46392012-DEL-66
chr18-46400070-DEL-103
chr18-46406066-DEL-69
chr18-46417579-DEL-125
chr18-46447015-DEL-539
chr18-46462432-DEL-317
chr18-46463230-DEL-1491
chr18-46464945-DEL-408

chr18-46466358-DEL-70
chr18-46485197-DEL-68
chr18-46513146-DEL-223
chr18-46519197-DEL-290
chr18-46536499-DEL-367
chr18-46546226-DEL-2109
chr18-46596344-DEL-59
chr18-46602539-DEL-406
chr18-46603662-DEL-163
chr18-46626773-DEL-70
chr18-46628626-DEL-76
chr18-46648885-DEL-885
chr18-46655282-DEL-72
chr18-46661206-DEL-725
chr18-46664644-DEL-190
chr18-46689035-DEL-146
chr18-46689855-DEL-206
chr18-46711928-DEL-3550
chr18-46723634-DEL-104
chr18-46754550-DEL-60
chr18-46765236-DEL-321
chr18-46874231-DEL-114
chr18-46883239-DEL-496
chr18-46924504-DEL-1888
chr18-46942165-DEL-164
chr18-46964691-DEL-89
chr18-46973541-DEL-59
chr18-46976943-DEL-78
chr18-47005877-DEL-78
chr18-47029705-DEL-80
chr18-47076612-DEL-65
chr18-47086663-DEL-197
chr18-47090085-DEL-187
chr18-47094838-DEL-237
chr18-47131127-DEL-99
chr18-47174090-DEL-69
chr18-47216384-DEL-90
chr18-47218631-DEL-93
chr18-47239887-DEL-52
chr18-47271826-DEL-1075
chr18-47315429-DEL-2004
chr18-47315432-DEL-8263
chr18-47317445-DEL-6254
chr18-47321441-DEL-616
chr18-47397265-DEL-440
chr18-47431476-DEL-57
chr18-47461367-DEL-206
chr18-47507126-DEL-54
chr18-47518993-DEL-442
chr18-47529155-DEL-2198
chr18-47536833-DEL-104
chr18-47540298-DEL-1023
chr18-47547331-DEL-245
chr18-47552389-DEL-199
chr18-47573565-DEL-138
chr18-47578024-DEL-950
chr18-47618724-DEL-231
chr18-47642628-DEL-215
chr18-47652034-DEL-66
chr18-47737828-DEL-149
chr18-47744864-DEL-196
chr18-47766442-DEL-1710
chr18-47766449-DEL-478
chr18-47793503-DEL-82
chr18-47813102-DEL-159
chr18-47823366-DEL-108
chr18-47919757-DEL-55
chr18-47931349-DEL-56
chr18-47957366-DEL-384
chr18-48008394-DEL-3060
chr18-48015335-DEL-52
chr18-48032669-DEL-60
chr18-48035335-DEL-502
chr18-48051224-DEL-125
chr18-48086098-DEL-81
chr18-48106850-DEL-530
chr18-48182329-DEL-347

chr18-48228929-DEL-70
chr18-48269235-DEL-52
chr18-48286264-DEL-72
chr18-48309040-DEL-77
chr18-48328515-DEL-1004
chr18-48332527-DEL-65
chr18-48333828-DEL-20135
chr18-48339249-DEL-14787
chr18-48350046-DEL-428
chr18-48359170-DEL-64
chr18-48377599-DEL-52
chr18-48415109-DEL-71
chr18-48418358-DEL-160
chr18-48425575-DEL-917
chr18-48464038-DEL-80
chr18-48476583-DEL-1370
chr18-48492692-DEL-70
chr18-48507528-DEL-4795
chr18-48518017-DEL-138
chr18-48571093-DEL-132
chr18-48613635-DEL-280
chr18-48620055-DEL-1159
chr18-48623706-DEL-439
chr18-48626631-DEL-594
chr18-48630465-DEL-137
chr18-48643745-DEL-4080
chr18-48650836-DEL-84
chr18-48711234-DEL-129
chr18-48715529-DEL-51
chr18-48722984-DEL-743
chr18-48732489-DEL-51
chr18-48743018-DEL-180
chr18-48751603-DEL-324
chr18-48773987-DEL-68
chr18-48777641-DEL-170
chr18-48783146-DEL-67
chr18-48790545-DEL-150
chr18-48882918-DEL-385
chr18-48891417-DEL-6267
chr18-48899969-DEL-180
chr18-48906392-DEL-2723
chr18-48913086-DEL-195
chr18-48950369-DEL-1108
chr18-48950509-DEL-901
chr18-48963092-DEL-6197
chr18-49012163-DEL-87
chr18-49013988-DEL-706
chr18-49020230-DEL-766
chr18-49092297-DEL-102
chr18-49094292-DEL-1984
chr18-49106224-DEL-71
chr18-49115719-DEL-484
chr18-49147520-DEL-1169
chr18-49151210-DEL-750
chr18-49165534-DEL-558
chr18-49169823-DEL-102
chr18-49175504-DEL-156
chr18-49195528-DEL-726
chr18-49222299-DEL-57
chr18-49311828-DEL-413
chr18-49343616-DEL-79
chr18-49344593-DEL-1450
chr18-49348593-DEL-329
chr18-49482037-DEL-526
chr18-49549125-DEL-82
chr18-49553320-DEL-576
chr18-49579506-DEL-151
chr18-49612544-DEL-67
chr18-49619030-DEL-155
chr18-49640085-DEL-1111
chr18-49665228-DEL-351
chr18-49730053-DEL-52
chr18-49741942-DEL-115
chr18-49744671-DEL-192
chr18-49757223-DEL-104
chr18-49771220-DEL-56
chr18-49819845-DEL-152

chr18-49822379-DEL-7119
chr18-49868052-DEL-51
chr18-49875274-DEL-57
chr18-49876396-DEL-129
chr18-49877502-DEL-217
chr18-49901192-DEL-385
chr18-49908300-DEL-247
chr18-49926552-DEL-144
chr18-49962229-DEL-89
chr18-49985588-DEL-221
chr18-49992758-DEL-1306
chr18-50009790-DEL-153
chr18-50030519-DEL-109
chr18-50051287-DEL-1284
chr18-50063497-DEL-623
chr18-50065719-DEL-427
chr18-50076772-DEL-164
chr18-50104854-DEL-130
chr18-50121002-DEL-340
chr18-50122880-DEL-236
chr18-50135514-DEL-58
chr18-50235322-DEL-401
chr18-50240808-DEL-174
chr18-50276050-DEL-196
chr18-50279639-DEL-115
chr18-50304815-DEL-81
chr18-50341794-DEL-430
chr18-50353089-DEL-330
chr18-50354400-DEL-486
chr18-50360877-DEL-1246
chr18-50365808-DEL-61
chr18-50372510-DEL-201
chr18-50379097-DEL-57
chr18-50385301-DEL-94
chr18-50385552-DEL-5859
chr18-50386377-DEL-6563
chr18-50391966-DEL-1149
chr18-50439372-DEL-737
chr18-50440842-DEL-158
chr18-50441880-DEL-349
chr18-50442963-DEL-204
chr18-50444148-DEL-301
chr18-50455496-DEL-312
chr18-50457302-DEL-73
chr18-50458508-DEL-6215
chr18-50475161-DEL-3020
chr18-50480197-DEL-1381
chr18-50486648-DEL-68
chr18-50511951-DEL-607
chr18-50513298-DEL-56
chr18-50515036-DEL-415
chr18-50570179-DEL-270
chr18-50580164-DEL-466
chr18-50588840-DEL-989
chr18-50600770-DEL-1609
chr18-50615251-DEL-96
chr18-50636282-DEL-985
chr18-50636339-DEL-1052
chr18-50644508-DEL-94
chr18-50646132-DEL-428
chr18-50656653-DEL-726
chr18-50671549-DEL-208
chr18-50674029-DEL-959
chr18-50727491-DEL-383
chr18-50745185-DEL-82
chr18-50746258-DEL-381
chr18-50756741-DEL-872
chr18-50817395-DEL-83
chr18-50823156-DEL-9534
chr18-50832588-DEL-238
chr18-50863440-DEL-407
chr18-50867213-DEL-1396
chr18-50873694-DEL-415
chr18-50896552-DEL-70
chr18-51020302-DEL-68
chr18-51066826-DEL-1652
chr18-51097934-DEL-79

chr18-51149423-DEL-96
chr18-51151701-DEL-85
chr18-51188960-DEL-171
chr18-51190375-DEL-1872
chr18-51197006-DEL-299
chr18-51198122-DEL-303
chr18-51198794-DEL-230
chr18-51211640-DEL-333
chr18-51246498-DEL-724
chr18-51272447-DEL-84
chr18-51274191-DEL-290
chr18-51276456-DEL-86
chr18-51280230-DEL-97
chr18-51283121-DEL-99
chr18-51298555-DEL-179
chr18-51305942-DEL-71
chr18-51325421-DEL-109
chr18-51336962-DEL-54
chr18-51341598-DEL-675
chr18-51343788-DEL-60
chr18-51364764-DEL-83
chr18-51369460-DEL-130
chr18-51400394-DEL-181
chr18-51404085-DEL-84
chr18-51406011-DEL-278
chr18-51425317-DEL-166
chr18-51444145-DEL-64
chr18-51456561-DEL-291
chr18-51460109-DEL-145
chr18-51469471-DEL-84
chr18-51486691-DEL-53
chr18-51513952-DEL-166
chr18-51537905-DEL-75
chr18-51542601-DEL-764
chr18-51553859-DEL-1757
chr18-51559562-DEL-88
chr18-51574153-DEL-56
chr18-51579483-DEL-334
chr18-51595155-DEL-90
chr18-51617454-DEL-173
chr18-51626161-DEL-205
chr18-51632476-DEL-1607
chr18-51634448-DEL-1679
chr18-51649229-DEL-935
chr18-51698303-DEL-451
chr18-51699976-DEL-56
chr18-51706005-DEL-135
chr18-51714465-DEL-122
chr18-51721049-DEL-574
chr18-51765524-DEL-901
chr18-51779265-DEL-81
chr18-51790539-DEL-505
chr18-51791016-DEL-1718
chr18-51792830-DEL-730
chr18-51793574-DEL-1652
chr18-51805679-DEL-2701
chr18-51839287-DEL-222
chr18-51841646-DEL-75
chr18-51878526-DEL-369
chr18-51934243-DEL-534
chr18-52044120-DEL-56
chr18-52153102-DEL-6261
chr18-52167769-DEL-6327
chr18-52179707-DEL-1603
chr18-52250330-DEL-95
chr18-52262321-DEL-57
chr18-52272966-DEL-165
chr18-52320717-DEL-58
chr18-52356692-DEL-51
chr18-52375469-DEL-126
chr18-52401150-DEL-807
chr18-52411057-DEL-130
chr18-52452442-DEL-127
chr18-52455659-DEL-113
chr18-52456558-DEL-106
chr18-52485048-DEL-1153
chr18-52489071-DEL-172

chr18-52490600-DEL-160
chr18-52519746-DEL-760
chr18-52520816-DEL-67
chr18-52611213-DEL-70
chr18-52627698-DEL-51
chr18-52630171-DEL-128
chr18-52658663-DEL-105
chr18-52702425-DEL-334
chr18-52712835-DEL-2595
chr18-52716989-DEL-495
chr18-52802806-DEL-1406
chr18-52802812-DEL-163
chr18-52814406-DEL-216
chr18-52829009-DEL-817
chr18-52881395-DEL-60
chr18-52889205-DEL-84
chr18-52990322-DEL-3413
chr18-53009731-DEL-212
chr18-53021388-DEL-110
chr18-53036073-DEL-187
chr18-53059434-DEL-319
chr18-53118752-DEL-94
chr18-53131392-DEL-374
chr18-53143622-DEL-59
chr18-53147933-DEL-66
chr18-53162112-DEL-166
chr18-53189740-DEL-51
chr18-53195271-DEL-1548
chr18-53195579-DEL-144
chr18-53195763-DEL-564
chr18-53196581-DEL-618
chr18-53216230-DEL-461
chr18-53217960-DEL-131
chr18-53251726-DEL-55
chr18-53279592-DEL-553
chr18-53307914-DEL-52
chr18-53333873-DEL-284
chr18-53346744-DEL-1811
chr18-53346951-DEL-897
chr18-53347649-DEL-265
chr18-53363326-DEL-121
chr18-53388333-DEL-129
chr18-53393833-DEL-203
chr18-53397544-DEL-178
chr18-53402848-DEL-1387
chr18-53405744-DEL-169
chr18-53437040-DEL-108
chr18-53437656-DEL-134
chr18-53437782-DEL-95
chr18-53488055-DEL-120
chr18-53488416-DEL-319
chr18-53492509-DEL-65
chr18-53533614-DEL-200
chr18-53540690-DEL-95
chr18-53636711-DEL-105
chr18-53637630-DEL-306
chr18-53637652-DEL-1120
chr18-53638142-DEL-202
chr18-53638844-DEL-80
chr18-53641704-DEL-62
chr18-53651713-DEL-67
chr18-53709773-DEL-175
chr18-53727614-DEL-100
chr18-53757269-DEL-212
chr18-53795366-DEL-68
chr18-53856018-DEL-61
chr18-53934026-DEL-126
chr18-53954780-DEL-170
chr18-53956432-DEL-966
chr18-53972959-DEL-218
chr18-54019683-DEL-73
chr18-54043899-DEL-2702
chr18-54093561-DEL-355
chr18-54107824-DEL-52
chr18-54170816-DEL-69
chr18-54172151-DEL-217
chr18-54181488-DEL-668

chr18-54193650-DEL-202
chr18-54196667-DEL-188
chr18-54200912-DEL-561
chr18-54211467-DEL-64
chr18-54230932-DEL-631
chr18-54265572-DEL-570
chr18-54288589-DEL-3948
chr18-54385205-DEL-58
chr18-54420028-DEL-953
chr18-54421088-DEL-2534
chr18-54431640-DEL-63
chr18-54440348-DEL-115
chr18-54476707-DEL-53
chr18-54534041-DEL-135
chr18-54618610-DEL-81
chr18-54671792-DEL-134
chr18-54736341-DEL-68
chr18-54742873-DEL-58
chr18-54774035-DEL-306
chr18-54781443-DEL-199
chr18-54813372-DEL-448
chr18-54815026-DEL-70
chr18-54839267-DEL-159
chr18-54861529-DEL-64
chr18-54867869-DEL-56
chr18-54896140-DEL-83
chr18-54904181-DEL-58
chr18-54913268-DEL-116
chr18-54999138-DEL-68
chr18-55036141-DEL-439
chr18-55090173-DEL-52
chr18-55106675-DEL-78
chr18-55130209-DEL-58
chr18-55144109-DEL-611
chr18-55204499-DEL-59
chr18-55233412-DEL-160
chr18-55244389-DEL-80
chr18-55264007-DEL-65
chr18-55305295-DEL-75
chr18-55307468-DEL-395
chr18-55312501-DEL-61
chr18-55364735-DEL-93
chr18-55368385-DEL-290
chr18-55400932-DEL-59
chr18-55439825-DEL-96
chr18-55465027-DEL-100
chr18-55470520-DEL-409
chr18-55482366-DEL-74
chr18-55500158-DEL-313
chr18-55504535-DEL-224
chr18-55588724-DEL-66
chr18-55647295-DEL-59
chr18-55667976-DEL-98
chr18-55678374-DEL-56
chr18-55688420-DEL-994
chr18-55702358-DEL-185
chr18-55738117-DEL-60
chr18-55741139-DEL-195
chr18-55744721-DEL-326
chr18-55751376-DEL-71
chr18-55778587-DEL-52
chr18-55789805-DEL-117
chr18-55805077-DEL-881
chr18-55843122-DEL-462
chr18-55870777-DEL-53
chr18-55880320-DEL-1039
chr18-55881735-DEL-72
chr18-55884004-DEL-56
chr18-55921554-DEL-113
chr18-55946582-DEL-604
chr18-55981206-DEL-95
chr18-55986343-DEL-978
chr18-55987287-DEL-1031
chr18-55988312-DEL-1049
chr18-56009039-DEL-104
chr18-56056693-DEL-91
chr18-56072541-DEL-187

chr18-56079009-DEL-82
chr18-56081068-DEL-6016
chr18-56117265-DEL-232
chr18-56132122-DEL-477
chr18-56187725-DEL-54
chr18-56276880-DEL-557
chr18-56294391-DEL-95
chr18-56316209-DEL-164
chr18-56348175-DEL-486
chr18-56396012-DEL-245
chr18-56405983-DEL-6280
chr18-56434713-DEL-832
chr18-56458384-DEL-112
chr18-56509397-DEL-235
chr18-56545164-DEL-80
chr18-56562571-DEL-564
chr18-56570192-DEL-71
chr18-56577886-DEL-66
chr18-56579339-DEL-264
chr18-56582858-DEL-627
chr18-56584222-DEL-54
chr18-56602925-DEL-260
chr18-56625462-DEL-1762
chr18-56630066-DEL-109
chr18-56642147-DEL-53
chr18-56660649-DEL-83
chr18-56669076-DEL-205
chr18-56686188-DEL-946
chr18-56697273-DEL-192
chr18-56712623-DEL-147
chr18-56726818-DEL-85
chr18-56787749-DEL-66
chr18-56788604-DEL-85
chr18-56810321-DEL-70
chr18-56813053-DEL-92
chr18-56815964-DEL-53
chr18-56817358-DEL-84
chr18-56854385-DEL-381
chr18-56860849-DEL-202
chr18-56866556-DEL-251
chr18-56876405-DEL-209
chr18-56897707-DEL-222
chr18-56979511-DEL-93
chr18-56986726-DEL-331
chr18-57027388-DEL-102
chr18-57037698-DEL-19441
chr18-57049909-DEL-111
chr18-57051937-DEL-59
chr18-57066964-DEL-83
chr18-57080420-DEL-86
chr18-57102164-DEL-153
chr18-57149358-DEL-942
chr18-57168536-DEL-54
chr18-57174488-DEL-62
chr18-57201579-DEL-100
chr18-57251351-DEL-1291
chr18-57251386-DEL-815
chr18-57252236-DEL-347
chr18-57298421-DEL-1000
chr18-57304880-DEL-301
chr18-57318091-DEL-1956
chr18-57320439-DEL-152
chr18-57346656-DEL-114
chr18-57377257-DEL-100
chr18-57381451-DEL-211
chr18-57402786-DEL-517
chr18-57443479-DEL-69
chr18-57455123-DEL-54
chr18-57457110-DEL-56
chr18-57460986-DEL-59
chr18-57463812-DEL-100
chr18-57480978-DEL-56
chr18-57497551-DEL-312
chr18-57498655-DEL-58
chr18-57501704-DEL-579
chr18-57502514-DEL-97
chr18-57506091-DEL-76

chr18-57545200-DEL-798
chr18-57562996-DEL-624
chr18-57638065-DEL-1455
chr18-57703175-DEL-155
chr18-57708653-DEL-825
chr18-57715078-DEL-84
chr18-57723072-DEL-62
chr18-57771124-DEL-82
chr18-57791241-DEL-119
chr18-57826055-DEL-576
chr18-57829439-DEL-301
chr18-57845355-DEL-946
chr18-57855374-DEL-51
chr18-57916039-DEL-192
chr18-57922791-DEL-93
chr18-57932827-DEL-103
chr18-57977946-DEL-74
chr18-57999549-DEL-163
chr18-58018532-DEL-1874
chr18-58090301-DEL-240
chr18-58092085-DEL-51
chr18-58104297-DEL-2666
chr18-58107697-DEL-1094
chr18-58161792-DEL-138
chr18-58163268-DEL-53
chr18-58171014-DEL-156
chr18-58185777-DEL-174
chr18-58227121-DEL-194
chr18-58228383-DEL-53
chr18-58284343-DEL-127
chr18-58300265-DEL-161
chr18-58327504-DEL-438
chr18-58329242-DEL-108
chr18-58329891-DEL-93
chr18-58335418-DEL-97
chr18-58338177-DEL-58
chr18-58358004-DEL-93
chr18-58385095-DEL-122
chr18-58398849-DEL-171
chr18-58458478-DEL-187
chr18-58466585-DEL-412
chr18-58473271-DEL-90
chr18-58486366-DEL-52
chr18-58532475-DEL-161
chr18-58552996-DEL-391
chr18-58559981-DEL-449
chr18-58567255-DEL-338
chr18-58570611-DEL-204
chr18-58585585-DEL-60
chr18-58590679-DEL-58
chr18-58591429-DEL-640
chr18-58616851-DEL-52
chr18-58643532-DEL-81
chr18-58645554-DEL-469
chr18-58649929-DEL-55
chr18-58677614-DEL-90
chr18-58699137-DEL-67
chr18-58772199-DEL-368
chr18-58808665-DEL-566
chr18-58815282-DEL-2534
chr18-58827586-DEL-132
chr18-58844858-DEL-562
chr18-58847650-DEL-759
chr18-58860579-DEL-446
chr18-58874868-DEL-124
chr18-58885609-DEL-253
chr18-58916820-DEL-1831
chr18-58920028-DEL-133
chr18-58947034-DEL-427
chr18-58948037-DEL-518
chr18-58948763-DEL-112
chr18-58967844-DEL-87
chr18-59027948-DEL-1132
chr18-59048417-DEL-112
chr18-59057217-DEL-71
chr18-59067650-DEL-1782
chr18-59088289-DEL-1582

chr18-59091229-DEL-433
chr18-59100287-DEL-88
chr18-59108227-DEL-286
chr18-59114929-DEL-96
chr18-59115081-DEL-958
chr18-59116060-DEL-1883
chr18-59138135-DEL-130
chr18-59143755-DEL-256
chr18-59149908-DEL-262
chr18-59150428-DEL-1622
chr18-59175606-DEL-150
chr18-59230937-DEL-160
chr18-59231675-DEL-1156
chr18-59234108-DEL-53
chr18-59247860-DEL-91
chr18-59323717-DEL-56
chr18-59338107-DEL-125
chr18-59351490-DEL-201
chr18-59371488-DEL-679
chr18-59392959-DEL-69
chr18-59420214-DEL-118
chr18-59441598-DEL-70
chr18-59443478-DEL-100
chr18-59455446-DEL-922
chr18-59466452-DEL-56
chr18-59468875-DEL-230
chr18-59488434-DEL-92
chr18-59506645-DEL-312
chr18-59513092-DEL-267
chr18-59519715-DEL-66
chr18-59577733-DEL-95
chr18-59583031-DEL-69
chr18-59587380-DEL-486
chr18-59606510-DEL-1982
chr18-59611253-DEL-351
chr18-59628919-DEL-5341
chr18-59629113-DEL-212
chr18-59636421-DEL-151
chr18-59637610-DEL-6473
chr18-59637744-DEL-200
chr18-59666998-DEL-340
chr18-59688420-DEL-94
chr18-59755577-DEL-631
chr18-59763124-DEL-60
chr18-59777939-DEL-70
chr18-59938486-DEL-208
chr18-59952874-DEL-89
chr18-59982637-DEL-53
chr18-59986280-DEL-954
chr18-60000050-DEL-112
chr18-60004920-DEL-276
chr18-60062822-DEL-6203
chr18-60062848-DEL-2444
chr18-60065321-DEL-3679
chr18-60069296-DEL-195
chr18-60080341-DEL-986
chr18-60081587-DEL-7112
chr18-60087113-DEL-143
chr18-60122447-DEL-6560
chr18-60150217-DEL-50
chr18-60187651-DEL-597
chr18-60194418-DEL-1064
chr18-60202636-DEL-60
chr18-60218730-DEL-386
chr18-60227808-DEL-767
chr18-60228989-DEL-51
chr18-60236247-DEL-65
chr18-60241673-DEL-717
chr18-60270080-DEL-1149
chr18-60273351-DEL-3108
chr18-60286021-DEL-804
chr18-60300165-DEL-3401
chr18-60323112-DEL-366
chr18-60351424-DEL-2141
chr18-60370248-DEL-688
chr18-60386196-DEL-640
chr18-60390288-DEL-124

chr18-60395918-DEL-508
chr18-60402190-DEL-183
chr18-60463079-DEL-16552
chr18-60529991-DEL-2400
chr18-60539616-DEL-5236
chr18-60548068-DEL-2945
chr18-60573814-DEL-544
chr18-60584266-DEL-439
chr18-60594188-DEL-68
chr18-60605359-DEL-103
chr18-60622937-DEL-76
chr18-60661438-DEL-2071
chr18-60663598-DEL-3194
chr18-60671927-DEL-69
chr18-60679590-DEL-205
chr18-60729726-DEL-62
chr18-60745546-DEL-70
chr18-60747377-DEL-64
chr18-60774946-DEL-52
chr18-60798916-DEL-144
chr18-60824616-DEL-120
chr18-60863948-DEL-196
chr18-60902091-DEL-80
chr18-60906433-DEL-168
chr18-60922896-DEL-1386
chr18-60926362-DEL-452
chr18-60966188-DEL-90
chr18-60991071-DEL-206
chr18-61029113-DEL-441
chr18-61060139-DEL-160
chr18-61079370-DEL-167
chr18-61100620-DEL-137
chr18-61112243-DEL-504
chr18-61113835-DEL-92
chr18-61123246-DEL-274
chr18-61136009-DEL-119
chr18-61195202-DEL-84
chr18-61207062-DEL-68
chr18-61243706-DEL-63
chr18-61267264-DEL-156
chr18-61279053-DEL-75
chr18-61327044-DEL-52
chr18-61372249-DEL-99
chr18-61395014-DEL-74
chr18-61415855-DEL-150
chr18-61446801-DEL-1644
chr18-61453646-DEL-125
chr18-61469443-DEL-180
chr18-61489642-DEL-180
chr18-61497129-DEL-1843
chr18-61551882-DEL-71
chr18-61558370-DEL-62
chr18-61565049-DEL-293
chr18-61575756-DEL-55
chr18-61588711-DEL-136
chr18-61633306-DEL-948
chr18-61641271-DEL-114
chr18-61646720-DEL-205
chr18-61671450-DEL-77
chr18-61671534-DEL-510
chr18-61733019-DEL-442
chr18-61741818-DEL-400
chr18-61746052-DEL-156
chr18-61803196-DEL-195
chr18-61804053-DEL-203
chr18-61855173-DEL-206
chr18-61857317-DEL-803
chr18-61866134-DEL-129
chr18-61902259-DEL-125
chr18-61908059-DEL-60
chr18-61917807-DEL-77
chr18-61927863-DEL-245
chr18-61942781-DEL-851
chr18-61969413-DEL-235
chr18-61999292-DEL-701
chr18-62050137-DEL-69
chr18-62052647-DEL-105

chr18-62073133-DEL-290
chr18-62094703-DEL-1759
chr18-62127058-DEL-260
chr18-62133593-DEL-219
chr18-62137622-DEL-1554
chr18-62156525-DEL-293
chr18-62189076-DEL-1770
chr18-62210527-DEL-55
chr18-62278830-DEL-59
chr18-62301326-DEL-86
chr18-62366306-DEL-677
chr18-62440655-DEL-128
chr18-62451287-DEL-319
chr18-62453096-DEL-54
chr18-62527540-DEL-158
chr18-62550263-DEL-68
chr18-62582183-DEL-401
chr18-62626575-DEL-82
chr18-62722225-DEL-546
chr18-62750557-DEL-650
chr18-62752745-DEL-2431
chr18-62760325-DEL-190
chr18-62794470-DEL-1592
chr18-62814962-DEL-180
chr18-62815959-DEL-62
chr18-62829439-DEL-2888
chr18-62852767-DEL-95
chr18-62887261-DEL-70
chr18-62896828-DEL-110
chr18-62923482-DEL-370
chr18-62976712-DEL-727
chr18-63026108-DEL-627
chr18-63047703-DEL-56
chr18-63052925-DEL-91
chr18-63121289-DEL-2868
chr18-63135319-DEL-89
chr18-63137841-DEL-653
chr18-63240861-DEL-53
chr18-63286877-DEL-57
chr18-63290550-DEL-111
chr18-63320875-DEL-112
chr18-63331133-DEL-344
chr18-63343666-DEL-1505
chr18-63412570-DEL-547
chr18-63435981-DEL-416
chr18-63440802-DEL-51
chr18-63453394-DEL-52
chr18-63492589-DEL-224
chr18-63498874-DEL-133
chr18-63515926-DEL-67
chr18-63530448-DEL-52
chr18-63532698-DEL-51
chr18-63533039-DEL-50
chr18-63539543-DEL-129
chr18-63592453-DEL-429
chr18-63632933-DEL-88
chr18-63657968-DEL-322
chr18-63660655-DEL-157
chr18-63663538-DEL-174
chr18-63673860-DEL-624
chr18-63676479-DEL-1797
chr18-63686155-DEL-181
chr18-63699291-DEL-51
chr18-63702794-DEL-51
chr18-63743254-DEL-55
chr18-63744719-DEL-156
chr18-63756043-DEL-154
chr18-63758008-DEL-68
chr18-63800539-DEL-89
chr18-63811791-DEL-584
chr18-63816453-DEL-221
chr18-63832355-DEL-3145
chr18-63844316-DEL-72
chr18-63909532-DEL-144
chr18-63946265-DEL-221
chr18-63948864-DEL-80
chr18-63955217-DEL-54

chr18-64059283-DEL-96
chr18-64099913-DEL-90
chr18-64148989-DEL-60
chr18-64213278-DEL-1325
chr18-64259304-DEL-361
chr18-64264249-DEL-3153
chr18-64264503-DEL-514
chr18-64265409-DEL-143
chr18-64329413-DEL-198
chr18-64334837-DEL-552
chr18-64359096-DEL-1592
chr18-64362877-DEL-56
chr18-64369905-DEL-1365
chr18-64421399-DEL-523
chr18-64438485-DEL-99
chr18-64455867-DEL-54
chr18-64475012-DEL-77
chr18-64491493-DEL-208
chr18-64516475-DEL-1304
chr18-64517066-DEL-214
chr18-64607249-DEL-292
chr18-64612579-DEL-117
chr18-64624993-DEL-78
chr18-64636151-DEL-88
chr18-64640445-DEL-88
chr18-64680334-DEL-2582
chr18-64685199-DEL-79
chr18-64701900-DEL-152
chr18-64703358-DEL-169
chr18-64707057-DEL-118
chr18-64709118-DEL-108
chr18-64722274-DEL-75
chr18-64731854-DEL-128
chr18-64735104-DEL-70
chr18-64781727-DEL-2781
chr18-64784981-DEL-93
chr18-64865277-DEL-244
chr18-64901622-DEL-172
chr18-64943270-DEL-134
chr18-64960736-DEL-277
chr18-64968824-DEL-3320
chr18-64981477-DEL-87
chr18-65059460-DEL-91
chr18-65083341-DEL-55
chr18-65099598-DEL-198
chr18-65114335-DEL-134
chr18-65119086-DEL-207
chr18-65138105-DEL-116
chr18-65144574-DEL-191
chr18-65147989-DEL-568
chr18-65199569-DEL-109
chr18-65206783-DEL-129
chr18-65245381-DEL-71
chr18-65271468-DEL-286
chr18-65340043-DEL-94
chr18-65360209-DEL-513
chr18-65382423-DEL-849
chr18-65385881-DEL-132
chr18-65409079-DEL-74
chr18-65469675-DEL-68
chr18-65519518-DEL-139
chr18-65520829-DEL-175
chr18-65541766-DEL-3625
chr18-65553852-DEL-698
chr18-65588418-DEL-198
chr18-65598181-DEL-149
chr18-65610664-DEL-136
chr18-65623257-DEL-223
chr18-65631366-DEL-197
chr18-65641049-DEL-57
chr18-65667866-DEL-2864
chr18-65749918-DEL-489
chr18-65755805-DEL-69
chr18-65775816-DEL-60
chr18-65776862-DEL-110
chr18-65816185-DEL-161
chr18-65825255-DEL-83

chr18-65825874-DEL-183
chr18-65854302-DEL-231
chr18-65888737-DEL-51
chr18-65949115-DEL-78
chr18-65965197-DEL-130
chr18-65986780-DEL-208
chr18-66011855-DEL-197
chr18-66024155-DEL-60
chr18-66040731-DEL-116
chr18-66049024-DEL-103
chr18-66055045-DEL-94
chr18-66098439-DEL-171
chr18-66099255-DEL-57
chr18-66119406-DEL-59
chr18-66163021-DEL-488
chr18-66214484-DEL-75
chr18-66215869-DEL-95
chr18-66220147-DEL-128
chr18-66224483-DEL-246
chr18-66254981-DEL-506
chr18-66257476-DEL-52
chr18-66260316-DEL-3874
chr18-66281583-DEL-90
chr18-66289573-DEL-96
chr18-66308864-DEL-200
chr18-66315488-DEL-52
chr18-66319072-DEL-114
chr18-66333327-DEL-120
chr18-66334452-DEL-130
chr18-66345731-DEL-1509
chr18-66349155-DEL-154
chr18-66371000-DEL-1235
chr18-66372224-DEL-75
chr18-66373619-DEL-647
chr18-66397520-DEL-347
chr18-66431605-DEL-59
chr18-66470467-DEL-109
chr18-66514520-DEL-181
chr18-66515428-DEL-77
chr18-66523955-DEL-1549
chr18-66540714-DEL-80
chr18-66559763-DEL-438
chr18-66576555-DEL-263
chr18-66620979-DEL-173
chr18-66634023-DEL-87
chr18-66640285-DEL-476
chr18-66651384-DEL-78
chr18-66665598-DEL-173
chr18-66668425-DEL-52
chr18-66675700-DEL-6258
chr18-66722161-DEL-1746
chr18-66723930-DEL-1107
chr18-66739989-DEL-719
chr18-66766520-DEL-529
chr18-66774003-DEL-72
chr18-66794467-DEL-6197
chr18-66802179-DEL-129
chr18-66804409-DEL-137
chr18-66818779-DEL-100
chr18-66823746-DEL-128
chr18-66834562-DEL-403
chr18-66835746-DEL-324
chr18-66838912-DEL-54
chr18-66850875-DEL-80
chr18-66891996-DEL-54
chr18-66894770-DEL-178
chr18-66905586-DEL-122
chr18-66912372-DEL-303
chr18-66932518-DEL-6269
chr18-66952090-DEL-248
chr18-66965105-DEL-58
chr18-66979958-DEL-732
chr18-67007871-DEL-53
chr18-67013901-DEL-6392
chr18-67037296-DEL-68
chr18-67051309-DEL-66
chr18-67082113-DEL-3526

chr18-67084335-DEL-1650
chr18-67092165-DEL-403
chr18-67107689-DEL-224
chr18-67108637-DEL-904
chr18-67158459-DEL-4207
chr18-67163536-DEL-115
chr18-67182264-DEL-93
chr18-67214604-DEL-80
chr18-67224675-DEL-101
chr18-67230012-DEL-85
chr18-67256749-DEL-95
chr18-67259810-DEL-58
chr18-67265455-DEL-128
chr18-67271888-DEL-64
chr18-67307315-DEL-747
chr18-67309494-DEL-481
chr18-67319642-DEL-155
chr18-67320706-DEL-108
chr18-67322777-DEL-299
chr18-67371343-DEL-64
chr18-67371899-DEL-1074
chr18-67410819-DEL-70
chr18-67414931-DEL-151
chr18-67417985-DEL-190
chr18-67446206-DEL-61
chr18-67466545-DEL-914
chr18-67469072-DEL-57
chr18-67472529-DEL-87
chr18-67481371-DEL-92
chr18-67483654-DEL-94
chr18-67485461-DEL-1026
chr18-67486370-DEL-109
chr18-67488513-DEL-184
chr18-67519938-DEL-278
chr18-67528084-DEL-62
chr18-67590323-DEL-57
chr18-67616233-DEL-177
chr18-67624815-DEL-241
chr18-67627418-DEL-211
chr18-67654205-DEL-612
chr18-67677262-DEL-68
chr18-67694801-DEL-144
chr18-67718440-DEL-77
chr18-67746899-DEL-1116
chr18-67749000-DEL-639
chr18-67786448-DEL-81
chr18-67799690-DEL-238
chr18-67810384-DEL-196
chr18-67831207-DEL-206
chr18-67833398-DEL-798
chr18-67850786-DEL-362
chr18-67860997-DEL-577
chr18-67889834-DEL-60
chr18-67895831-DEL-1279
chr18-67899416-DEL-65
chr18-67924397-DEL-407
chr18-67932297-DEL-60
chr18-67938925-DEL-398
chr18-67943186-DEL-60
chr18-67969877-DEL-86
chr18-68005163-DEL-333
chr18-68022128-DEL-324
chr18-68023434-DEL-150
chr18-68032785-DEL-213
chr18-68058040-DEL-80
chr18-68073094-DEL-63
chr18-68083120-DEL-86
chr18-68116081-DEL-52
chr18-68156652-DEL-58
chr18-68249845-DEL-130
chr18-68297649-DEL-50
chr18-68373947-DEL-158
chr18-68443160-DEL-155
chr18-68460723-DEL-199
chr18-68492490-DEL-51
chr18-68498291-DEL-4965
chr18-68514656-DEL-904

chr18-68520230-DEL-208
chr18-68521958-DEL-98
chr18-68527801-DEL-52
chr18-68555496-DEL-52
chr18-68568442-DEL-154
chr18-68580615-DEL-71
chr18-68605581-DEL-202
chr18-68639645-DEL-76
chr18-68652100-DEL-186
chr18-68658398-DEL-77
chr18-68664849-DEL-140
chr18-68678914-DEL-86
chr18-68703371-DEL-212
chr18-68705802-DEL-2813
chr18-68729123-DEL-1061
chr18-68754518-DEL-110
chr18-68769538-DEL-54
chr18-68797850-DEL-52
chr18-68813114-DEL-114
chr18-68839876-DEL-375
chr18-68851521-DEL-402
chr18-68897015-DEL-90
chr18-68900648-DEL-287
chr18-68911692-DEL-3819
chr18-68949145-DEL-87
chr18-68958818-DEL-51
chr18-68983369-DEL-60
chr18-68990598-DEL-266
chr18-69058321-DEL-119
chr18-69066817-DEL-56
chr18-69071742-DEL-55
chr18-69120711-DEL-68
chr18-69126336-DEL-697
chr18-69201268-DEL-524
chr18-69236849-DEL-1571
chr18-69242834-DEL-83
chr18-69352807-DEL-55
chr18-69362917-DEL-798
chr18-69365025-DEL-90
chr18-69365934-DEL-75
chr18-69373638-DEL-323
chr18-69375005-DEL-405
chr18-69391092-DEL-63
chr18-69395300-DEL-1350
chr18-69405249-DEL-535
chr18-69409267-DEL-153
chr18-69475548-DEL-358
chr18-69492685-DEL-153
chr18-69500575-DEL-72
chr18-69515447-DEL-212
chr18-69590755-DEL-148
chr18-69593618-DEL-100
chr18-69619934-DEL-64
chr18-69644779-DEL-75
chr18-69678393-DEL-56
chr18-69737450-DEL-58
chr18-69758489-DEL-181
chr18-69771228-DEL-152
chr18-69803051-DEL-104
chr18-69804515-DEL-51
chr18-69832488-DEL-144
chr18-69855378-DEL-109
chr18-69859450-DEL-1473
chr18-69880935-DEL-601
chr18-69919726-DEL-52
chr18-69936967-DEL-249
chr18-69944010-DEL-490
chr18-69944538-DEL-56
chr18-69962848-DEL-412
chr18-70000489-DEL-1912
chr18-70008490-DEL-52
chr18-70031008-DEL-55
chr18-70060156-DEL-97
chr18-70131325-DEL-55
chr18-70175020-DEL-1531
chr18-70193739-DEL-102
chr18-70217748-DEL-54

chr18-70227671-DEL-771
chr18-70262143-DEL-75
chr18-70266241-DEL-1185
chr18-70289792-DEL-2854
chr18-70305021-DEL-814
chr18-70329734-DEL-241
chr18-70330928-DEL-284
chr18-70353877-DEL-91
chr18-70360404-DEL-830
chr18-70367270-DEL-62
chr18-70383182-DEL-65
chr18-70415277-DEL-494
chr18-70420543-DEL-1330
chr18-70428462-DEL-116
chr18-70435034-DEL-72
chr18-70456505-DEL-9335
chr18-70457186-DEL-2746
chr18-70495822-DEL-167
chr18-70508246-DEL-84
chr18-70539952-DEL-93
chr18-70572674-DEL-429
chr18-70592579-DEL-85
chr18-70594262-DEL-181
chr18-70614767-DEL-104
chr18-70671185-DEL-91
chr18-70678509-DEL-52
chr18-70747468-DEL-100
chr18-70750681-DEL-62
chr18-70764821-DEL-148
chr18-70768516-DEL-1118
chr18-70769775-DEL-2698
chr18-70800698-DEL-4492
chr18-70806292-DEL-552
chr18-70860083-DEL-1230
chr18-70904027-DEL-554
chr18-70908372-DEL-6573
chr18-70921453-DEL-6162
chr18-70967195-DEL-146
chr18-70968811-DEL-138
chr18-70970196-DEL-148
chr18-70970908-DEL-148
chr18-71001668-DEL-76
chr18-71035611-DEL-67
chr18-71073731-DEL-70
chr18-71131830-DEL-109
chr18-71138827-DEL-135
chr18-71147277-DEL-68
chr18-71158429-DEL-417
chr18-71176146-DEL-327
chr18-71191823-DEL-54
chr18-71223533-DEL-2932
chr18-71241162-DEL-633
chr18-71242857-DEL-51
chr18-71249802-DEL-1285
chr18-71272180-DEL-214
chr18-71295455-DEL-60
chr18-71323846-DEL-838
chr18-71339949-DEL-208
chr18-71356365-DEL-383
chr18-71358056-DEL-1564
chr18-71362950-DEL-6099
chr18-71363518-DEL-568
chr18-71367593-DEL-541
chr18-71378300-DEL-1315
chr18-71384890-DEL-54
chr18-71398761-DEL-401
chr18-71414886-DEL-1158
chr18-71424613-DEL-316
chr18-71523161-DEL-166
chr18-71546478-DEL-72
chr18-71555086-DEL-61
chr18-71555515-DEL-377
chr18-71574421-DEL-52
chr18-71577511-DEL-383
chr18-71585756-DEL-84
chr18-71604489-DEL-60
chr18-71617824-DEL-54

chr18-71623277-DEL-195
chr18-71627074-DEL-58
chr18-71634719-DEL-1120
chr18-71653715-DEL-1142
chr18-71692313-DEL-336
chr18-71821909-DEL-1067
chr18-71827978-DEL-297
chr18-71834017-DEL-369
chr18-71886968-DEL-112
chr18-71887162-DEL-1143
chr18-71907451-DEL-1521
chr18-71949355-DEL-103
chr18-71972993-DEL-169
chr18-72001612-DEL-179
chr18-72006313-DEL-281
chr18-72015408-DEL-492
chr18-72019915-DEL-180
chr18-72020806-DEL-654
chr18-72023627-DEL-86
chr18-72031758-DEL-209
chr18-72042651-DEL-114
chr18-72043307-DEL-179
chr18-72066022-DEL-1860
chr18-72074876-DEL-59
chr18-72131146-DEL-66
chr18-72173347-DEL-324
chr18-72176457-DEL-103
chr18-72180118-DEL-930
chr18-72257308-DEL-124
chr18-72282034-DEL-314
chr18-72288761-DEL-172
chr18-72294780-DEL-228
chr18-72296925-DEL-150
chr18-72352625-DEL-57
chr18-72367238-DEL-66
chr18-72379654-DEL-68
chr18-72443332-DEL-81
chr18-72448576-DEL-64
chr18-72467443-DEL-383
chr18-72522664-DEL-106
chr18-72530093-DEL-53
chr18-72541088-DEL-350
chr18-72548161-DEL-83
chr18-72590216-DEL-1800
chr18-72590218-DEL-71
chr18-72592028-DEL-1183
chr18-72592940-DEL-277
chr18-72600069-DEL-338
chr18-72604983-DEL-321
chr18-72623440-DEL-9073
chr18-72689614-DEL-63
chr18-72691314-DEL-2855
chr18-72709896-DEL-424
chr18-72730240-DEL-460
chr18-72735105-DEL-564
chr18-72760847-DEL-7402
chr18-72763228-DEL-641
chr18-72763928-DEL-1194
chr18-72768592-DEL-1603
chr18-72812408-DEL-1031
chr18-72864352-DEL-315
chr18-72894353-DEL-849
chr18-72905057-DEL-1969
chr18-72905231-DEL-555
chr18-72906030-DEL-1039
chr18-72906240-DEL-148
chr18-72906389-DEL-525
chr18-72938751-DEL-261
chr18-72983722-DEL-51
chr18-72988219-DEL-104
chr18-73010613-DEL-5620
chr18-73039712-DEL-64
chr18-73042611-DEL-1380
chr18-73088161-DEL-65
chr18-73124340-DEL-52
chr18-73187898-DEL-529
chr18-73190402-DEL-534

chr18-73202334-DEL-1460
chr18-73219424-DEL-116
chr18-73268376-DEL-54
chr18-73273253-DEL-2710
chr18-73364076-DEL-6214
chr18-73364080-DEL-383
chr18-73369095-DEL-464
chr18-73389778-DEL-1180
chr18-73391958-DEL-337
chr18-73403090-DEL-579
chr18-73404852-DEL-64
chr18-73414195-DEL-70
chr18-73418634-DEL-66
chr18-73419921-DEL-62
chr18-73421504-DEL-1717
chr18-73421505-DEL-139
chr18-73422019-DEL-129
chr18-73428929-DEL-68
chr18-73435696-DEL-151
chr18-73439085-DEL-686
chr18-73448850-DEL-105
chr18-73464284-DEL-6257
chr18-73556650-DEL-1560
chr18-73557010-DEL-1259
chr18-73557010-DEL-210
chr18-73557148-DEL-902
chr18-73558084-DEL-180
chr18-73558933-DEL-90
chr18-73573131-DEL-104
chr18-73574193-DEL-309
chr18-73580751-DEL-528
chr18-73584515-DEL-215
chr18-73616325-DEL-76
chr18-73623636-DEL-341
chr18-73628821-DEL-308
chr18-73668917-DEL-246
chr18-73677446-DEL-307
chr18-73685293-DEL-142
chr18-73689179-DEL-99
chr18-73689657-DEL-1137
chr18-73712835-DEL-163
chr18-73747535-DEL-653
chr18-73761210-DEL-575
chr18-73774265-DEL-53
chr18-73785541-DEL-145
chr18-73822632-DEL-136
chr18-73822808-DEL-1310
chr18-73822869-DEL-1959
chr18-73823424-DEL-334
chr18-73823861-DEL-66
chr18-73823888-DEL-2200
chr18-73823889-DEL-1036
chr18-73824658-DEL-188
chr18-73824929-DEL-1552
chr18-73825239-DEL-198
chr18-73825763-DEL-229
chr18-73862578-DEL-59
chr18-73904930-DEL-491
chr18-73922680-DEL-108
chr18-73929072-DEL-59
chr18-73936458-DEL-600
chr18-73944709-DEL-119
chr18-73959002-DEL-901
chr18-73960102-DEL-295
chr18-73965616-DEL-51
chr18-73978580-DEL-1328
chr18-73986226-DEL-654
chr18-73999095-DEL-63
chr18-74008087-DEL-138
chr18-74024667-DEL-405
chr18-74027832-DEL-390
chr18-74054324-DEL-62
chr18-74070996-DEL-150
chr18-74081534-DEL-162
chr18-74082413-DEL-51
chr18-74126130-DEL-109
chr18-74133747-DEL-71

chr18-74145159-DEL-54
chr18-74149438-DEL-72
chr18-74163612-DEL-353
chr18-74170941-DEL-112
chr18-74182180-DEL-67
chr18-74205971-DEL-67
chr18-74272999-DEL-401
chr18-74284370-DEL-338
chr18-74307601-DEL-82
chr18-74311917-DEL-81
chr18-74346739-DEL-51
chr18-74347563-DEL-59
chr18-74363520-DEL-298
chr18-74376084-DEL-96
chr18-74387853-DEL-108
chr18-74428348-DEL-59
chr18-74436356-DEL-697
chr18-74442044-DEL-179
chr18-74445236-DEL-58
chr18-74469697-DEL-53
chr18-74478277-DEL-2324
chr18-74486305-DEL-70
chr18-74526896-DEL-125
chr18-74527367-DEL-4999
chr18-74546112-DEL-158
chr18-74551491-DEL-559
chr18-74553428-DEL-445
chr18-74560450-DEL-265
chr18-74592991-DEL-95
chr18-74608633-DEL-59
chr18-74623027-DEL-130
chr18-74626419-DEL-154
chr18-74637374-DEL-62
chr18-74650206-DEL-53
chr18-74686130-DEL-776
chr18-74687007-DEL-412
chr18-74689405-DEL-80
chr18-74694467-DEL-68
chr18-74711616-DEL-125
chr18-74720607-DEL-108
chr18-74773198-DEL-92
chr18-74818800-DEL-111
chr18-74835279-DEL-195
chr18-74871833-DEL-59
chr18-74882974-DEL-6440
chr18-74895966-DEL-195
chr18-74896164-DEL-507
chr18-74924571-DEL-550
chr18-74938669-DEL-57
chr18-74950076-DEL-402
chr18-74961131-DEL-681
chr18-74975112-DEL-399
chr18-74975663-DEL-324
chr18-74988155-DEL-182
chr18-75004928-DEL-61
chr18-75013649-DEL-460
chr18-75014008-DEL-347
chr18-75039337-DEL-75
chr18-75050639-DEL-54
chr18-75053785-DEL-60
chr18-75055266-DEL-81
chr18-75086827-DEL-129
chr18-75108435-DEL-66
chr18-75125422-DEL-197
chr18-75149192-DEL-53
chr18-75170621-DEL-57
chr18-75172038-DEL-55
chr18-75178783-DEL-261
chr18-75192935-DEL-84
chr18-75211274-DEL-88
chr18-75219527-DEL-308
chr18-75223809-DEL-60
chr18-75274536-DEL-98
chr18-75275922-DEL-171
chr18-75300405-DEL-151
chr18-75306357-DEL-84
chr18-75325001-DEL-849

chr18-75334682-DEL-67
chr18-75353005-DEL-200
chr18-75358022-DEL-1818
chr18-75375148-DEL-52
chr18-75388765-DEL-56
chr18-75401355-DEL-1251
chr18-75401477-DEL-154
chr18-75403648-DEL-1042
chr18-75425719-DEL-109
chr18-75434683-DEL-100
chr18-75444565-DEL-56
chr18-75470571-DEL-61
chr18-75481813-DEL-113
chr18-75539708-DEL-75
chr18-75542114-DEL-113
chr18-75565904-DEL-62
chr18-75584735-DEL-806
chr18-75594593-DEL-52
chr18-75613523-DEL-118
chr18-75650766-DEL-51
chr18-75703352-DEL-55
chr18-75715302-DEL-109
chr18-75721327-DEL-850
chr18-75733649-DEL-99
chr18-75742481-DEL-65
chr18-75748305-DEL-576
chr18-75765939-DEL-135
chr18-75782955-DEL-407
chr18-75840924-DEL-80
chr18-75852556-DEL-785
chr18-75877401-DEL-62
chr18-75936987-DEL-76
chr18-75944008-DEL-183
chr18-75968843-DEL-3129
chr18-75987530-DEL-588
chr18-76001691-DEL-1594
chr18-76009801-DEL-65
chr18-76029625-DEL-56
chr18-76083270-DEL-71
chr18-76102355-DEL-71
chr18-76103069-DEL-74
chr18-76151372-DEL-369
chr18-76174409-DEL-215
chr18-76239876-DEL-1750
chr18-76239881-DEL-180
chr18-76276349-DEL-67
chr18-76284585-DEL-56
chr18-76287165-DEL-52
chr18-76318184-DEL-56
chr18-76377092-DEL-215
chr18-76401602-DEL-509
chr18-76412637-DEL-81
chr18-76443115-DEL-61
chr18-76486679-DEL-153
chr18-76493788-DEL-115
chr18-76495983-DEL-66
chr18-76516653-DEL-136
chr18-76523534-DEL-88
chr18-76593193-DEL-52
chr18-76614844-DEL-128
chr18-76618245-DEL-645
chr18-76626932-DEL-70
chr18-76655491-DEL-697
chr18-76655503-DEL-249
chr18-76662719-DEL-155
chr18-76672053-DEL-1809
chr18-76677466-DEL-86
chr18-76685487-DEL-326
chr18-76689646-DEL-335
chr18-76724788-DEL-945
chr18-76736687-DEL-56
chr18-76743280-DEL-160
chr18-76752135-DEL-296
chr18-76759409-DEL-71
chr18-76778170-DEL-51
chr18-76786862-DEL-82
chr18-76799705-DEL-689

chr18-76827380-DEL-721
chr18-76838275-DEL-63
chr18-76844916-DEL-55
chr18-76928816-DEL-157
chr18-76932798-DEL-94
chr18-76941942-DEL-93
chr18-76987133-DEL-144
chr18-76989936-DEL-214
chr18-77090522-DEL-574
chr18-77101769-DEL-247
chr18-77102439-DEL-1208
chr18-77105832-DEL-204
chr18-77108747-DEL-215
chr18-77122229-DEL-62
chr18-77123022-DEL-205
chr18-77131812-DEL-72
chr18-77174807-DEL-196
chr18-77175177-DEL-5887
chr18-77189775-DEL-69
chr18-77204958-DEL-386
chr18-77226722-DEL-164
chr18-77253984-DEL-227
chr18-77261044-DEL-83
chr18-77261774-DEL-822
chr18-77270364-DEL-158
chr18-77276999-DEL-447
chr18-77289242-DEL-1921
chr18-77309177-DEL-158
chr18-77323887-DEL-739
chr18-77333004-DEL-78
chr18-77334109-DEL-51
chr18-77348499-DEL-177
chr18-77349498-DEL-9306
chr18-77349805-DEL-84
chr18-77385735-DEL-120
chr18-77420828-DEL-249
chr18-77436834-DEL-97
chr18-77458102-DEL-625
chr18-77464785-DEL-60
chr18-77478372-DEL-150
chr18-77524643-DEL-55
chr18-77526376-DEL-63
chr18-77527300-DEL-225
chr18-77545201-DEL-183
chr18-77552286-DEL-96
chr18-77591516-DEL-125
chr18-77618592-DEL-66
chr18-77622890-DEL-58
chr18-77680901-DEL-92
chr18-77692488-DEL-69
chr18-77707040-DEL-105
chr18-77710048-DEL-158
chr18-77716643-DEL-179
chr18-77765409-DEL-220
chr18-77775207-DEL-826
chr18-77778439-DEL-231
chr18-77785363-DEL-105
chr18-77792957-DEL-63
chr18-77805596-DEL-182
chr18-77806252-DEL-200
chr18-77810007-DEL-179
chr18-77810205-DEL-1341
chr18-77836317-DEL-205
chr18-77839779-DEL-94
chr18-77841319-DEL-119
chr18-77858969-DEL-359
chr18-77878507-DEL-127
chr18-77892101-DEL-163
chr18-77892888-DEL-198
chr18-77910959-DEL-54
chr18-77946635-DEL-247
chr18-77961502-DEL-200
chr18-77989479-DEL-56
chr18-78006346-DEL-75
chr18-78008270-DEL-242
chr18-78010646-DEL-600
chr18-78030647-DEL-86

chr18-78037128-DEL-335
chr18-78039227-DEL-1012
chr18-78059633-DEL-1173
chr18-78059819-DEL-344
chr18-78059851-DEL-896
chr18-78110829-DEL-58
chr18-78127741-DEL-126
chr18-78137343-DEL-82
chr18-78174364-DEL-164
chr18-78196427-DEL-357
chr18-78208703-DEL-92
chr18-78211846-DEL-379
chr18-78241497-DEL-78
chr18-78248221-DEL-110
chr18-78315026-DEL-1009
chr18-78374270-DEL-229
chr18-78379086-DEL-233
chr18-78413077-DEL-275
chr18-78432480-DEL-60
chr18-78443824-DEL-61
chr18-78455098-DEL-695
chr18-78456602-DEL-7529
chr18-78502903-DEL-492
chr18-78541805-DEL-265
chr18-78542379-DEL-240
chr18-78554137-DEL-388
chr18-78560270-DEL-629
chr18-78706087-DEL-428
chr18-78774869-DEL-71
chr18-78782549-DEL-96
chr18-78796040-DEL-90
chr18-78864067-DEL-58
chr18-78898969-DEL-69
chr18-78947770-DEL-7543
chr18-78950907-DEL-69
chr18-78976319-DEL-137
chr18-78984943-DEL-153
chr18-79004979-DEL-56
chr18-79028576-DEL-110
chr18-79112072-DEL-130
chr18-79113338-DEL-101
chr18-79125134-DEL-57
chr18-79159317-DEL-186
chr18-79184074-DEL-217
chr18-79202854-DEL-3291
chr18-79238664-DEL-501
chr18-79243502-DEL-68
chr18-79268069-DEL-256
chr18-79274625-DEL-124
chr18-79294059-DEL-253
chr18-79300410-DEL-78
chr18-79386198-DEL-766
chr18-79391233-DEL-370
chr18-79402918-DEL-6280
chr18-79435122-DEL-389
chr18-79446916-DEL-962
chr18-79449002-DEL-6712
chr18-79457306-DEL-124
chr18-79481070-DEL-1560
chr18-79481268-DEL-912
chr18-79484969-DEL-1266
chr18-79485223-DEL-472
chr18-79496651-DEL-195
chr18-79522060-DEL-464
chr18-79544742-DEL-64
chr18-79551044-DEL-212
chr18-79572103-DEL-129
chr18-79583045-DEL-238
chr18-79591304-DEL-82
chr18-79597257-DEL-80
chr18-79598180-DEL-466
chr18-79650177-DEL-281
chr18-79682225-DEL-954
chr18-79694307-DEL-58
chr18-79716720-DEL-56
chr18-79719592-DEL-1569
chr18-79722930-DEL-1937

chr18-79755354-DEL-427
chr18-79762929-DEL-294
chr18-79763468-DEL-862
chr18-79767484-DEL-1640
chr18-79768412-DEL-196
chr18-79786247-DEL-8019
chr18-79794189-DEL-253
chr18-79797007-DEL-63
chr18-79815885-DEL-81
chr18-79864753-DEL-1278
chr18-79892205-DEL-62
chr18-79922672-DEL-83
chr18-79954706-DEL-1372
chr18-79970553-DEL-295
chr18-79973323-DEL-111
chr18-79983623-DEL-176
chr18-80012743-DEL-206
chr18-80014996-DEL-53
chr18-80015568-DEL-180
chr18-80017423-DEL-59
chr18-80021905-DEL-615
chr18-80029688-DEL-1211
chr18-80031144-DEL-118
chr18-80055108-DEL-6919
chr18-80055160-DEL-3271
chr18-80055475-DEL-399
chr18-80058650-DEL-389
chr18-80059472-DEL-3269
chr18-80064829-DEL-531
chr18-80086849-DEL-1160
chr18-80088711-DEL-70
chr18-80132996-DEL-162
chr18-80148004-DEL-146
chr18-80149563-DEL-702
chr18-80151198-DEL-51
chr18-80168035-DEL-263
chr18-80177321-DEL-88
chr18-80204366-DEL-74
chr18-80211831-DEL-201
chr18-80214293-DEL-81
chr18-80259624-DEL-78
chr18-80268717-DEL-94
chr18-80285915-DEL-99
chr18-80297256-DEL-249
chr18-80307697-DEL-303
chr18-80344221-DEL-372
chr18-80385590-DEL-194
chr18-80455147-DEL-87
chr18-80500047-DEL-98
chr18-80500166-DEL-1003
chr18-80514335-DEL-2115
chr18-80534211-DEL-72
chr18-80546027-DEL-97
chr18-80645952-DEL-782
chr18-80647705-DEL-124
chr18-80674188-DEL-220
chr18-80755202-DEL-53
chr18-80763573-DEL-181
chr18-80785770-DEL-51
chr18-80790043-DEL-1213
chr18-80809324-DEL-81
chr18-80850899-DEL-188
chr18-80939933-DEL-207
chr18-80964528-DEL-352
chr18-80980205-DEL-76
chr18-81055031-DEL-65
chr18-81064530-DEL-87
chr18-81065467-DEL-55
chr18-81070552-DEL-60
chr18-81072468-DEL-52
chr18-81098401-DEL-179
chr18-81111067-DEL-533
chr18-81111897-DEL-6466
chr18-81136146-DEL-140
chr18-81139401-DEL-1404
chr18-81139811-DEL-364
chr18-81144143-DEL-513

chr18-81159397-DEL-97
chr18-81201071-DEL-439
chr18-81236089-DEL-599
chr18-81274247-DEL-62
chr18-81299743-DEL-60
chr18-81310459-DEL-78
chr18-81335484-DEL-422
chr18-81336447-DEL-171
chr18-81372713-DEL-80
chr18-81380225-DEL-60
chr18-81396652-DEL-109
chr18-81447614-DEL-161
chr18-81487885-DEL-104
chr18-81489452-DEL-208
chr18-81523772-DEL-825
chr18-81533974-DEL-60
chr18-81557887-DEL-135
chr18-81568553-DEL-73
chr18-81611422-DEL-98
chr18-81643734-DEL-248
chr18-81697965-DEL-1262
chr18-81714479-DEL-242
chr18-81728189-DEL-1402
chr18-81740262-DEL-238
chr18-81750231-DEL-88
chr18-81755524-DEL-299
chr18-81755845-DEL-520
chr18-81774303-DEL-86
chr18-81805358-DEL-57
chr18-81807850-DEL-66
chr18-81829987-DEL-84
chr18-81838202-DEL-340
chr18-81866129-DEL-83
chr18-81883373-DEL-663
chr18-81915964-DEL-352
chr18-81929526-DEL-88
chr18-81943146-DEL-52
chr18-81998308-DEL-118
chr18-82004637-DEL-74
chr18-82012680-DEL-4533
chr18-82019620-DEL-168
chr18-82024038-DEL-87
chr18-82040257-DEL-260
chr18-82048324-DEL-725
chr18-82079649-DEL-86
chr18-82095313-DEL-6197
chr18-82198868-DEL-2162
chr18-82204121-DEL-35252
chr18-82204747-DEL-201
chr18-82206033-DEL-366
chr18-82206063-DEL-160
chr18-82207007-DEL-653
chr18-82208581-DEL-199
chr18-82210348-DEL-290
chr18-82211297-DEL-366
chr18-82212473-DEL-240
chr18-82214100-DEL-15228
chr18-82232332-DEL-452
chr18-82234074-DEL-202
chr18-82235384-DEL-527
chr18-82236671-DEL-694
chr18-82238267-DEL-161
chr18-82238991-DEL-82
chr18-82269231-DEL-159
chr18-82269253-DEL-1970
chr18-82281355-DEL-54
chr18-82290106-DEL-83
chr18-82291853-DEL-84
chr18-82320947-DEL-92
chr18-82333837-DEL-78
chr18-82338732-DEL-151
chr18-82372450-DEL-3640
chr18-82378394-DEL-77
chr18-82392930-DEL-270
chr18-82397234-DEL-654
chr18-82401784-DEL-1549
chr18-82426844-DEL-740

chr18-82429964-DEL-334
chr18-82437854-DEL-132
chr18-82451605-DEL-67
chr18-82454287-DEL-760
chr18-82489223-DEL-683
chr18-82490713-DEL-1130
chr18-82496825-DEL-664
chr18-82504907-DEL-210
chr18-82509111-DEL-119
chr18-82520745-DEL-950
chr18-82526707-DEL-109
chr18-82533308-DEL-202
chr18-82548050-DEL-98
chr18-82561401-DEL-100
chr18-82562669-DEL-125
chr18-82566599-DEL-82
chr18-82587126-DEL-62
chr18-82604888-DEL-62
chr18-82608180-DEL-648
chr18-82622633-DEL-274
chr18-82633668-DEL-62
chr18-82637159-DEL-492
chr18-82637863-DEL-358
chr18-82650168-DEL-128
chr18-82651502-DEL-86
chr18-82694472-DEL-94
chr18-82723342-DEL-1477
chr18-82726586-DEL-128
chr18-82737920-DEL-63
chr18-82743245-DEL-67
chr18-82749298-DEL-1933
chr18-82764901-DEL-116
chr18-82793834-DEL-64
chr18-82796316-DEL-211
chr18-82805782-DEL-71
chr18-82815487-DEL-57
chr18-82818149-DEL-61
chr18-82858339-DEL-82
chr18-82882056-DEL-147
chr18-82893373-DEL-288
chr18-82895480-DEL-1426
chr18-82941522-DEL-114
chr18-82942086-DEL-192
chr18-82948737-DEL-122
chr18-82966111-DEL-2149
chr18-82971077-DEL-145
chr18-82994475-DEL-86
chr18-83002045-DEL-204
chr18-83003698-DEL-286
chr18-83009422-DEL-261
chr18-83025547-DEL-298
chr18-83036541-DEL-96
chr18-83053986-DEL-89
chr18-83056741-DEL-142
chr18-83064040-DEL-231
chr18-83066859-DEL-106
chr18-83075063-DEL-200
chr18-83089213-DEL-113
chr18-83105330-DEL-314
chr18-83111015-DEL-81
chr18-83120831-DEL-213
chr18-83137266-DEL-65
chr18-83156500-DEL-286
chr18-83157062-DEL-145
chr18-83174074-DEL-63
chr18-83203125-DEL-60
chr18-83205604-DEL-2925
chr18-83213335-DEL-96
chr18-83215583-DEL-64
chr18-83274022-DEL-76
chr18-83317034-DEL-439
chr18-83334359-DEL-77
chr18-83344058-DEL-116
chr18-83352624-DEL-68
chr18-83388265-DEL-208
chr18-83390773-DEL-69
chr18-83441443-DEL-120

chr18-83443569-DEL-64
chr18-83449734-DEL-134
chr18-83451693-DEL-68
chr18-83491559-DEL-88
chr18-83503067-DEL-236
chr18-83515133-DEL-77
chr18-83520596-DEL-57
chr18-83526429-DEL-121
chr18-83532519-DEL-1269
chr18-83556256-DEL-54
chr18-83559421-DEL-280
chr18-83561771-DEL-330
chr18-83570517-DEL-286
chr18-83586482-DEL-270
chr18-83607001-DEL-160
chr18-83635234-DEL-148
chr18-83654622-DEL-74
chr18-83661017-DEL-1216
chr18-83678539-DEL-114
chr18-83686259-DEL-81
chr18-83692298-DEL-73
chr18-83705885-DEL-78
chr18-83785676-DEL-334
chr18-83799248-DEL-221
chr18-83803998-DEL-128
chr18-83817531-DEL-1662
chr18-83862307-DEL-200
chr18-83873432-DEL-58
chr18-83875971-DEL-72
chr18-83889347-DEL-60
chr18-83905329-DEL-138
chr18-83910874-DEL-79
chr18-83920789-DEL-1882
chr18-83926551-DEL-54
chr18-83943269-DEL-234
chr18-83943750-DEL-1321
chr18-84000613-DEL-118
chr18-84017142-DEL-62
chr18-84040388-DEL-336
chr18-84044587-DEL-122
chr18-84056687-DEL-80
chr18-84059111-DEL-127
chr18-84081550-DEL-55
chr18-84084199-DEL-243
chr18-84109349-DEL-51
chr18-84122702-DEL-54
chr18-84200809-DEL-115
chr18-84204540-DEL-478
chr18-84212903-DEL-842
chr18-84221226-DEL-69
chr18-84223569-DEL-150
chr18-84244284-DEL-2457
chr18-84247041-DEL-195
chr18-84289469-DEL-77
chr18-84290480-DEL-196
chr18-84309443-DEL-206
chr18-84376252-DEL-61
chr18-84399580-DEL-221
chr18-84407939-DEL-252
chr18-84442785-DEL-253
chr18-84466238-DEL-102
chr18-84511333-DEL-58
chr18-84519722-DEL-82
chr18-84553393-DEL-55
chr18-84582066-DEL-460
chr18-84582323-DEL-1096
chr18-84582798-DEL-440
chr18-84596075-DEL-51
chr18-84604440-DEL-348
chr18-84606745-DEL-69
chr18-84661148-DEL-99
chr18-84672427-DEL-82
chr18-84675894-DEL-64
chr18-84676693-DEL-1373
chr18-84680769-DEL-86
chr18-84683537-DEL-60
chr18-84689105-DEL-122

chr18-84690499-DEL-167
chr18-84704683-DEL-118
chr18-84705528-DEL-183
chr18-84729462-DEL-226
chr18-84747001-DEL-58
chr18-84752136-DEL-79
chr18-84784110-DEL-1982
chr18-84787803-DEL-245
chr18-84792268-DEL-1454
chr18-84800333-DEL-53
chr18-84848902-DEL-2736
chr18-84849673-DEL-63
chr18-84849887-DEL-99
chr18-84850344-DEL-129
chr18-84851123-DEL-309
chr18-84851174-DEL-90
chr18-84892074-DEL-81
chr18-84893017-DEL-703
chr18-84905176-DEL-2127
chr18-84925034-DEL-2625
chr18-84945403-DEL-187
chr18-84956214-DEL-2110
chr18-84956908-DEL-1900
chr18-84971070-DEL-344
chr18-84972903-DEL-374
chr18-84992052-DEL-2396
chr18-84995864-DEL-278
chr18-85003519-DEL-150
chr18-85006720-DEL-2231
chr18-85026993-DEL-55
chr18-85042053-DEL-79
chr18-85044720-DEL-502
chr18-85051212-DEL-197
chr18-85052828-DEL-16200
chr18-85052828-DEL-6439
chr18-85062590-DEL-9375
chr18-85140814-DEL-217
chr18-85177576-DEL-80
chr18-85197311-DEL-96
chr18-85215178-DEL-55
chr18-85232254-DEL-337
chr18-85254293-DEL-79
chr18-85261916-DEL-72
chr18-85266449-DEL-166
chr18-85278039-DEL-117
chr18-85281293-DEL-1988
chr18-85298633-DEL-70
chr18-85365613-DEL-52
chr18-85367069-DEL-173
chr18-85386424-DEL-58
chr18-85405411-DEL-861
chr18-85406575-DEL-965
chr18-85455940-DEL-399
chr18-85457984-DEL-6148
chr18-85471601-DEL-57
chr18-85518699-DEL-121
chr18-85560247-DEL-1007
chr18-85620263-DEL-1453
chr18-85624452-DEL-462
chr18-85625157-DEL-57
chr18-85639762-DEL-68
chr18-85643595-DEL-192
chr18-85668828-DEL-2651
chr18-85671372-DEL-806
chr18-85708398-DEL-718
chr18-85760428-DEL-53
chr18-85775220-DEL-902
chr18-85780070-DEL-125
chr18-85820485-DEL-65
chr18-85824352-DEL-970
chr18-85848874-DEL-129
chr18-85863926-DEL-2339
chr18-85866079-DEL-224
chr18-85889882-DEL-963
chr18-85896438-DEL-696
chr18-85898859-DEL-357
chr18-85899338-DEL-1652

chr18-85901687-DEL-6261
chr18-85911804-DEL-57
chr18-85915524-DEL-6587
chr18-85935333-DEL-211
chr18-85938415-DEL-4254
chr18-85939952-DEL-2126
chr18-85961773-DEL-571
chr18-85993720-DEL-488
chr18-86005159-DEL-107
chr18-86037456-DEL-71
chr18-86044665-DEL-71
chr18-86049794-DEL-104
chr18-86062226-DEL-5647
chr18-86068929-DEL-223
chr18-86069149-DEL-195
chr18-86084337-DEL-6828
chr18-86114771-DEL-141
chr18-86119616-DEL-920
chr18-86134761-DEL-204
chr18-86174728-DEL-284
chr18-86194902-DEL-86
chr18-86233361-DEL-99
chr18-86270022-DEL-399
chr18-86287424-DEL-223
chr18-86297915-DEL-275
chr18-86315341-DEL-130
chr18-86320888-DEL-189
chr18-86324112-DEL-118
chr18-86327043-DEL-55
chr18-86333078-DEL-64
chr18-86335103-DEL-879
chr18-86384140-DEL-64
chr18-86389561-DEL-63
chr18-86391719-DEL-61
chr18-86444420-DEL-368
chr18-86499358-DEL-172
chr18-86506244-DEL-64
chr18-86507839-DEL-95
chr18-86537013-DEL-89
chr18-86648208-DEL-1776
chr18-86658578-DEL-241
chr18-86690896-DEL-301
chr18-86695196-DEL-790
chr18-86711497-DEL-905
chr18-86718943-DEL-144
chr18-86748290-DEL-52
chr18-86749442-DEL-74
chr18-86759961-DEL-68
chr18-86763890-DEL-410
chr18-86787590-DEL-67
chr18-86801422-DEL-944
chr18-86814209-DEL-102
chr18-86846481-DEL-5980
chr18-86873358-DEL-100
chr18-86889488-DEL-2048
chr18-86892396-DEL-523
chr18-86893913-DEL-700
chr18-86944322-DEL-72
chr18-86953783-DEL-68
chr18-86978634-DEL-260
chr18-86995069-DEL-69
chr18-87007812-DEL-52
chr18-87023709-DEL-100
chr18-87024328-DEL-3304
chr18-87028475-DEL-100
chr18-87028890-DEL-974
chr18-87029997-DEL-134
chr18-87075139-DEL-334
chr18-87079599-DEL-166
chr18-87100001-DEL-3538
chr18-87155284-DEL-94
chr18-87162136-DEL-6405
chr18-87171039-DEL-58
chr18-87172762-DEL-6606
chr18-87180507-DEL-52
chr18-87213493-DEL-438
chr18-87240862-DEL-2527

chr18-87250621-DEL-187
chr18-87254576-DEL-88
chr18-87259166-DEL-1076
chr18-87262020-DEL-7403
chr18-87288376-DEL-148
chr18-87289863-DEL-6270
chr18-87313222-DEL-332
chr18-87332282-DEL-251
chr18-87335813-DEL-403
chr18-87344799-DEL-58
chr18-87346475-DEL-152
chr18-87351135-DEL-78
chr18-87351549-DEL-1822
chr18-87359775-DEL-75
chr18-87386586-DEL-1058
chr18-87395868-DEL-904
chr18-87404745-DEL-185
chr18-87409631-DEL-233
chr18-87446649-DEL-2156
chr18-87448846-DEL-1919
chr18-87451590-DEL-1656
chr18-87488573-DEL-527
chr18-87493435-DEL-472
chr18-87497056-DEL-400
chr18-87520688-DEL-461
chr18-87536582-DEL-86
chr18-87536965-DEL-764
chr18-87545297-DEL-1232
chr18-87558592-DEL-1810
chr18-87589327-DEL-369
chr18-87618049-DEL-58
chr18-87659313-DEL-70
chr18-87701036-DEL-207
chr18-87741917-DEL-608
chr18-87747380-DEL-64
chr18-87757237-DEL-333
chr18-87773569-DEL-3935
chr18-87781689-DEL-199
chr18-87784630-DEL-769
chr18-87795172-DEL-6285
chr18-87857963-DEL-927
chr18-87897216-DEL-10151
chr18-87911392-DEL-56
chr18-87931544-DEL-133
chr18-87951096-DEL-5510
chr18-87958551-DEL-612
chr18-87960006-DEL-557
chr18-87966259-DEL-2544
chr18-87971406-DEL-8713
chr18-87986679-DEL-672
chr18-87991030-DEL-229
chr18-88038566-DEL-948
chr18-88041182-DEL-99
chr18-88064287-DEL-71
chr18-88128958-DEL-91
chr18-88136802-DEL-916
chr18-88137954-DEL-85
chr18-88147627-DEL-499
chr18-88154057-DEL-62
chr18-88174125-DEL-67
chr18-88178886-DEL-205
chr18-88190997-DEL-611
chr18-88193974-DEL-125
chr18-88206140-DEL-1477
chr18-88219968-DEL-90
chr18-88232702-DEL-5451
chr18-88279340-DEL-1019
chr18-88280495-DEL-1397
chr18-88281352-DEL-894
chr18-88283879-DEL-1008
chr18-88321419-DEL-208
chr18-88326814-DEL-86
chr18-88328581-DEL-253
chr18-88422114-DEL-917
chr18-88436805-DEL-798
chr18-88456791-DEL-273
chr18-88457452-DEL-687

chr18-88459026-DEL-353
chr18-88470480-DEL-181
chr18-88481220-DEL-92
chr18-88507186-DEL-1361
chr18-88524643-DEL-84
chr18-88546791-DEL-119
chr18-88547916-DEL-2249
chr18-88591135-DEL-679
chr18-88596598-DEL-561
chr18-88602499-DEL-332
chr18-88630400-DEL-156
chr18-88634369-DEL-154
chr18-88637607-DEL-6271
chr18-88645592-DEL-278
chr18-88689076-DEL-94
chr18-88715003-DEL-201
chr18-88807985-DEL-53
chr18-88919739-DEL-58
chr18-89023902-DEL-54
chr18-89026354-DEL-62
chr18-89050538-DEL-87
chr18-89052755-DEL-1890
chr18-89056133-DEL-1511
chr18-89104264-DEL-65
chr18-89114861-DEL-1546
chr18-89234468-DEL-71
chr18-89304417-DEL-611
chr18-89329865-DEL-125
chr18-89338731-DEL-99
chr18-89359594-DEL-635
chr18-89379175-DEL-373
chr18-89425760-DEL-6207
chr18-89447920-DEL-55
chr18-89474065-DEL-52
chr18-89492281-DEL-132
chr18-89515145-DEL-108
chr18-89532765-DEL-2013
chr18-89534837-DEL-2128
chr18-89587277-DEL-81
chr18-89629841-DEL-9364
chr18-89644654-DEL-223
chr18-89652723-DEL-1340
chr18-89681873-DEL-10857
chr18-89740243-DEL-1796
chr18-89784243-DEL-102
chr18-89785325-DEL-84
chr18-89791120-DEL-2651
chr18-89808445-DEL-640
chr18-89854604-DEL-76
chr18-89877359-DEL-298
chr18-89884480-DEL-147
chr18-89889223-DEL-1096
chr18-89898027-DEL-770
chr18-89899923-DEL-95
chr18-89925287-DEL-774
chr18-89941242-DEL-6095
chr18-89984461-DEL-336
chr18-90041441-DEL-1559
chr18-90073791-DEL-1856
chr18-90104368-DEL-5988
chr18-90154203-DEL-126
chr18-90171116-DEL-804
chr18-90208993-DEL-2067
chr18-90267700-DEL-236
chr18-90312955-DEL-231
chr18-90332338-DEL-40494
chr18-90414879-DEL-252
chr18-90430252-DEL-124
chr18-90442147-DEL-316
chr18-90442162-DEL-6668
chr18-90529831-DEL-453
chr18-90554807-DEL-102
chr18-90562284-DEL-3163
chr18-90568630-DEL-2162
chr18-90571338-DEL-7103
chr18-90594517-DEL-2098
chr18-90620071-DEL-65

chr19-3052378-DEL-212
chr19-3052833-DEL-3763
chr19-3083269-DEL-5414
chr19-3090047-DEL-208
chr19-3175080-DEL-928
chr19-3195820-DEL-436
chr19-3210859-DEL-7516
chr19-3223195-DEL-4849
chr19-3253675-DEL-2365
chr19-3256521-DEL-229
chr19-3288181-DEL-2795
chr19-3337324-DEL-66
chr19-3348120-DEL-194
chr19-3368224-DEL-169
chr19-3422385-DEL-418
chr19-3432887-DEL-722
chr19-3439427-DEL-82
chr19-3443286-DEL-175
chr19-3445270-DEL-217
chr19-3470270-DEL-401
chr19-3480780-DEL-118
chr19-3491802-DEL-74
chr19-3497470-DEL-95
chr19-3521830-DEL-91
chr19-3541827-DEL-209
chr19-3586840-DEL-199
chr19-3587990-DEL-169
chr19-3599010-DEL-625
chr19-3602629-DEL-206
chr19-3638052-DEL-98
chr19-3638580-DEL-204
chr19-3645412-DEL-203
chr19-3684398-DEL-288
chr19-3729707-DEL-66
chr19-3739036-DEL-208
chr19-3740421-DEL-157
chr19-3757090-DEL-60
chr19-3785637-DEL-195
chr19-3787095-DEL-207
chr19-3792275-DEL-192
chr19-3796550-DEL-448
chr19-3806054-DEL-519
chr19-3819472-DEL-196
chr19-3837483-DEL-98
chr19-3843569-DEL-118
chr19-3857462-DEL-66
chr19-3880423-DEL-57
chr19-3882417-DEL-168
chr19-3902944-DEL-58
chr19-3903501-DEL-130
chr19-3911477-DEL-124
chr19-3927092-DEL-194
chr19-3966101-DEL-58
chr19-3967029-DEL-183
chr19-4008123-DEL-200
chr19-4041493-DEL-74
chr19-4073870-DEL-83
chr19-4123852-DEL-179
chr19-4156082-DEL-72
chr19-4175421-DEL-179
chr19-4200950-DEL-56
chr19-4216846-DEL-104
chr19-4280893-DEL-162
chr19-4291428-DEL-64
chr19-4305168-DEL-124
chr19-4309014-DEL-208
chr19-4328888-DEL-168
chr19-4349333-DEL-60
chr19-4358876-DEL-336
chr19-4403509-DEL-51
chr19-4410964-DEL-198
chr19-4433795-DEL-205
chr19-4434617-DEL-79
chr19-4504175-DEL-212
chr19-4513014-DEL-139
chr19-4521644-DEL-131
chr19-4562501-DEL-78

chr19-4611538-DEL-178
chr19-4636882-DEL-140
chr19-4649982-DEL-97
chr19-4687036-DEL-64
chr19-4696033-DEL-60
chr19-4706436-DEL-223
chr19-4713006-DEL-135
chr19-4734461-DEL-200
chr19-4736260-DEL-104
chr19-4817801-DEL-76
chr19-4819134-DEL-197
chr19-4823156-DEL-211
chr19-4831622-DEL-164
chr19-4858855-DEL-56
chr19-4863259-DEL-55
chr19-4868321-DEL-1811
chr19-4868321-DEL-788
chr19-4868797-DEL-1412
chr19-4869504-DEL-235
chr19-4878018-DEL-198
chr19-4888057-DEL-53
chr19-4898968-DEL-197
chr19-5004095-DEL-1345
chr19-5015000-DEL-751
chr19-5019723-DEL-189
chr19-5045107-DEL-471
chr19-5058452-DEL-67
chr19-5059610-DEL-168
chr19-5062897-DEL-74
chr19-5064564-DEL-73
chr19-5207540-DEL-1826
chr19-5212999-DEL-713
chr19-5213001-DEL-151
chr19-5242082-DEL-299
chr19-5251334-DEL-58
chr19-5256036-DEL-371
chr19-5277368-DEL-161
chr19-5291912-DEL-98
chr19-5306876-DEL-53
chr19-5318954-DEL-2934
chr19-5325686-DEL-52
chr19-5330370-DEL-92
chr19-5344924-DEL-51
chr19-5354888-DEL-100
chr19-5382365-DEL-57
chr19-5391780-DEL-144
chr19-5392851-DEL-174
chr19-5457437-DEL-66
chr19-5470307-DEL-61
chr19-5526847-DEL-56
chr19-5547230-DEL-58
chr19-5585571-DEL-303
chr19-5586985-DEL-227
chr19-5591119-DEL-446
chr19-5591777-DEL-237
chr19-5592669-DEL-255
chr19-5625188-DEL-153
chr19-5637501-DEL-74
chr19-5648696-DEL-248
chr19-5650470-DEL-226
chr19-5668856-DEL-90
chr19-5676271-DEL-87
chr19-5720925-DEL-156
chr19-5736740-DEL-336
chr19-5759996-DEL-75
chr19-5787734-DEL-795
chr19-5812717-DEL-199
chr19-5832583-DEL-188
chr19-5836183-DEL-189
chr19-5863132-DEL-1970
chr19-5865922-DEL-92
chr19-5872793-DEL-249
chr19-5873845-DEL-94
chr19-5876596-DEL-66
chr19-5911093-DEL-531
chr19-5911125-DEL-1064
chr19-5937752-DEL-71

chr19-5945158-DEL-184
chr19-5950553-DEL-198
chr19-5976691-DEL-192
chr19-5988895-DEL-376
chr19-6017696-DEL-52
chr19-6021017-DEL-62
chr19-6028729-DEL-200
chr19-6029548-DEL-185
chr19-6033803-DEL-313
chr19-6045369-DEL-1193
chr19-6045493-DEL-193
chr19-6052107-DEL-202
chr19-6052937-DEL-65
chr19-6054283-DEL-294
chr19-6059644-DEL-209
chr19-6087171-DEL-186
chr19-6089000-DEL-1050
chr19-6110169-DEL-192
chr19-6142731-DEL-358
chr19-6147072-DEL-93
chr19-6160652-DEL-682
chr19-6182717-DEL-609
chr19-6204166-DEL-337
chr19-6211355-DEL-604
chr19-6225502-DEL-241
chr19-6228457-DEL-69
chr19-6252733-DEL-366
chr19-6277230-DEL-76
chr19-6298476-DEL-773
chr19-6298986-DEL-122
chr19-6309850-DEL-69
chr19-6319215-DEL-195
chr19-6320354-DEL-171
chr19-6333549-DEL-143
chr19-6334274-DEL-52
chr19-6377837-DEL-237
chr19-6379332-DEL-292
chr19-6409443-DEL-60
chr19-6485653-DEL-3523
chr19-6508211-DEL-51
chr19-6530174-DEL-62
chr19-6543951-DEL-76
chr19-6555564-DEL-104
chr19-6558990-DEL-262
chr19-6584874-DEL-54
chr19-6605592-DEL-52
chr19-6608402-DEL-141
chr19-6608572-DEL-450
chr19-6609417-DEL-1601
chr19-6610666-DEL-736
chr19-6624554-DEL-79
chr19-6625285-DEL-962
chr19-6628614-DEL-174
chr19-6671811-DEL-288
chr19-6683984-DEL-627
chr19-6692221-DEL-70
chr19-6723544-DEL-510
chr19-6733570-DEL-152
chr19-6738091-DEL-257
chr19-6739433-DEL-51011
chr19-6739909-DEL-50059
chr19-6836347-DEL-95
chr19-6847215-DEL-224
chr19-6867532-DEL-206
chr19-6868956-DEL-201
chr19-6883469-DEL-145
chr19-6907929-DEL-64
chr19-6960537-DEL-197
chr19-6988638-DEL-201
chr19-6988669-DEL-412
chr19-6993882-DEL-196
chr19-7039803-DEL-962
chr19-7093660-DEL-154
chr19-7095545-DEL-62
chr19-7101159-DEL-614
chr19-7109161-DEL-52
chr19-7117526-DEL-66

chr19-7150242-DEL-200
chr19-7179359-DEL-93
chr19-7188308-DEL-134
chr19-7195576-DEL-170
chr19-7199705-DEL-290
chr19-7228107-DEL-198
chr19-7312319-DEL-198
chr19-7341829-DEL-166
chr19-7346771-DEL-828
chr19-7349511-DEL-53
chr19-7373107-DEL-63
chr19-7429730-DEL-778
chr19-7451066-DEL-52
chr19-7456352-DEL-367
chr19-7476508-DEL-200
chr19-7485536-DEL-528
chr19-7547893-DEL-190
chr19-7583480-DEL-59
chr19-7604298-DEL-243
chr19-7615140-DEL-144
chr19-7625617-DEL-4720
chr19-7652006-DEL-6449
chr19-7665040-DEL-1649
chr19-7711573-DEL-4642
chr19-7751777-DEL-208
chr19-7787373-DEL-6730
chr19-7849825-DEL-334
chr19-7877433-DEL-158
chr19-7877923-DEL-2864
chr19-7890556-DEL-416
chr19-7891100-DEL-208
chr19-7897521-DEL-179
chr19-7897527-DEL-1573
chr19-7912061-DEL-22743
chr19-7924699-DEL-267
chr19-7934587-DEL-1895
chr19-7974182-DEL-446
chr19-8010852-DEL-3589
chr19-8033071-DEL-720
chr19-8052760-DEL-58
chr19-8067380-DEL-3695
chr19-8071256-DEL-948
chr19-8085888-DEL-1122
chr19-8098332-DEL-7397
chr19-8120437-DEL-88
chr19-8149166-DEL-91
chr19-8175800-DEL-13110
chr19-8190891-DEL-1865
chr19-8215972-DEL-14916
chr19-8220904-DEL-6803
chr19-8232110-DEL-55
chr19-8269470-DEL-143
chr19-8323009-DEL-1361
chr19-8328229-DEL-54
chr19-8378492-DEL-2563
chr19-8405977-DEL-72
chr19-8462297-DEL-136
chr19-8479966-DEL-15860
chr19-8496891-DEL-4756
chr19-8551764-DEL-312
chr19-8600475-DEL-60
chr19-8609187-DEL-1186
chr19-8611455-DEL-2210
chr19-8611648-DEL-441
chr19-8613706-DEL-1026
chr19-8613706-DEL-998
chr19-8615238-DEL-156
chr19-8630849-DEL-135
chr19-8635646-DEL-63
chr19-8653813-DEL-203
chr19-8691988-DEL-78
chr19-8693108-DEL-178
chr19-8699821-DEL-52
chr19-8704720-DEL-242
chr19-8729232-DEL-236
chr19-8763992-DEL-191
chr19-8769305-DEL-401

chr19-8827080-DEL-224
chr19-8830583-DEL-76
chr19-8834823-DEL-1078
chr19-8839201-DEL-147
chr19-8845623-DEL-147
chr19-8860819-DEL-166
chr19-8881778-DEL-56
chr19-8894881-DEL-138
chr19-8896092-DEL-100
chr19-8924191-DEL-89
chr19-8927018-DEL-578
chr19-8934846-DEL-1217
chr19-8946989-DEL-125
chr19-9004891-DEL-57
chr19-9017939-DEL-186
chr19-9019891-DEL-486
chr19-9029210-DEL-131
chr19-9057956-DEL-72
chr19-9064423-DEL-139
chr19-9091009-DEL-103
chr19-9129625-DEL-9466
chr19-9173999-DEL-578
chr19-9255172-DEL-145
chr19-9258151-DEL-190
chr19-9259679-DEL-3124
chr19-9260585-DEL-1287
chr19-9328321-DEL-52
chr19-9334140-DEL-1278
chr19-9339555-DEL-434
chr19-9339862-DEL-1447
chr19-9362079-DEL-191
chr19-9367499-DEL-416
chr19-9368333-DEL-1971
chr19-9368420-DEL-110
chr19-9393667-DEL-377
chr19-9429353-DEL-2709
chr19-9455619-DEL-5091
chr19-9465109-DEL-54
chr19-9479475-DEL-5755
chr19-9498155-DEL-311
chr19-9558512-DEL-1438
chr19-9571436-DEL-54
chr19-9584951-DEL-84
chr19-9633683-DEL-188
chr19-9634678-DEL-90
chr19-9636831-DEL-145
chr19-9644020-DEL-112
chr19-9646933-DEL-90
chr19-9682546-DEL-2440
chr19-9689177-DEL-6186
chr19-9708854-DEL-432
chr19-9731095-DEL-6689
chr19-9742615-DEL-465
chr19-9762878-DEL-53
chr19-9774493-DEL-6361
chr19-9775232-DEL-56037
chr19-9785556-DEL-2902
chr19-9796796-DEL-750
chr19-9806205-DEL-8323
chr19-9843243-DEL-216
chr19-9851101-DEL-160
chr19-9909392-DEL-68
chr19-9915188-DEL-290
chr19-9954983-DEL-74
chr19-9967306-DEL-173
chr19-9972986-DEL-751
chr19-9977231-DEL-72
chr19-9982811-DEL-199
chr19-9989498-DEL-211
chr19-10028982-DEL-200
chr19-10037574-DEL-83
chr19-10054403-DEL-288
chr19-10063536-DEL-891
chr19-10065163-DEL-142
chr19-10084144-DEL-7841
chr19-10109116-DEL-87
chr19-10116569-DEL-52

chr19-10149765-DEL-191
chr19-10155424-DEL-176
chr19-10194957-DEL-169
chr19-10197380-DEL-373
chr19-10199414-DEL-56
chr19-10231368-DEL-206
chr19-10251929-DEL-497
chr19-10283667-DEL-104
chr19-10292254-DEL-625
chr19-10305267-DEL-1159
chr19-10335908-DEL-211
chr19-10349103-DEL-201
chr19-10364305-DEL-992
chr19-10372008-DEL-63
chr19-10376746-DEL-108
chr19-10444628-DEL-66
chr19-10445762-DEL-184
chr19-10457408-DEL-177
chr19-10466378-DEL-52
chr19-10485561-DEL-1545
chr19-10485587-DEL-166
chr19-10486091-DEL-242
chr19-10486612-DEL-190
chr19-10490139-DEL-1999
chr19-10538032-DEL-371
chr19-10543104-DEL-62
chr19-10543972-DEL-76
chr19-10601808-DEL-145
chr19-10622947-DEL-87
chr19-10675089-DEL-56
chr19-10681919-DEL-54
chr19-10695008-DEL-288
chr19-10723804-DEL-440
chr19-10728221-DEL-94
chr19-10730413-DEL-83
chr19-10752761-DEL-56
chr19-10770533-DEL-74
chr19-10786410-DEL-418
chr19-10793322-DEL-163
chr19-10863336-DEL-822
chr19-10885769-DEL-59
chr19-10926166-DEL-180
chr19-10992057-DEL-196
chr19-10997224-DEL-69
chr19-11001054-DEL-2745
chr19-11006665-DEL-325
chr19-11086018-DEL-640
chr19-11094041-DEL-367
chr19-11097206-DEL-195
chr19-11097533-DEL-1353
chr19-11107040-DEL-430
chr19-11120612-DEL-193
chr19-11130028-DEL-1673
chr19-11152707-DEL-5202
chr19-11154170-DEL-111
chr19-11165198-DEL-60
chr19-11176941-DEL-6383
chr19-11183944-DEL-169
chr19-11186999-DEL-2297
chr19-11210566-DEL-60
chr19-11256461-DEL-122
chr19-11265145-DEL-846
chr19-11309706-DEL-254
chr19-11313713-DEL-4768
chr19-11334521-DEL-6496
chr19-11344209-DEL-521
chr19-11370237-DEL-118
chr19-11374880-DEL-93
chr19-11398290-DEL-159
chr19-11410108-DEL-68
chr19-11421804-DEL-98
chr19-11429294-DEL-74
chr19-11433010-DEL-1829
chr19-11433868-DEL-572
chr19-11433991-DEL-2270
chr19-11439875-DEL-881
chr19-11441000-DEL-54

chr19-11457033-DEL-80
chr19-11484000-DEL-4022
chr19-11499675-DEL-113
chr19-11508479-DEL-970
chr19-11514612-DEL-138
chr19-11551056-DEL-204
chr19-11629110-DEL-551
chr19-11641064-DEL-356
chr19-11674539-DEL-120
chr19-11678948-DEL-60
chr19-11708374-DEL-3873
chr19-11775295-DEL-677
chr19-11776554-DEL-194
chr19-11781241-DEL-82
chr19-11794216-DEL-58
chr19-11817007-DEL-559
chr19-11819797-DEL-279
chr19-11863706-DEL-505
chr19-11985201-DEL-122
chr19-11994430-DEL-662
chr19-12001094-DEL-110
chr19-12010977-DEL-196
chr19-12015466-DEL-56
chr19-12022290-DEL-152
chr19-12057799-DEL-55
chr19-12142010-DEL-689
chr19-12149223-DEL-3614
chr19-12166188-DEL-8163
chr19-12182423-DEL-86
chr19-12191410-DEL-280
chr19-12200033-DEL-1030
chr19-12210999-DEL-425
chr19-12227332-DEL-851
chr19-12243414-DEL-494
chr19-12318604-DEL-80
chr19-12459140-DEL-53
chr19-12494808-DEL-58
chr19-12496115-DEL-1722
chr19-12504540-DEL-95
chr19-12600396-DEL-467
chr19-12608617-DEL-382
chr19-12614586-DEL-84
chr19-12664458-DEL-393
chr19-12666904-DEL-1144
chr19-12702510-DEL-51
chr19-12708560-DEL-51
chr19-12727346-DEL-2127
chr19-12730517-DEL-63
chr19-12734112-DEL-70
chr19-12791682-DEL-1305
chr19-12826379-DEL-221
chr19-12829149-DEL-187
chr19-12830045-DEL-128
chr19-12832033-DEL-806
chr19-12845773-DEL-87
chr19-12873126-DEL-3556
chr19-12886985-DEL-83
chr19-12890877-DEL-754
chr19-12924164-DEL-258
chr19-12941695-DEL-407
chr19-12955534-DEL-8470
chr19-12980647-DEL-825
chr19-12996047-DEL-1290
chr19-13007319-DEL-609
chr19-13009519-DEL-105
chr19-13015814-DEL-15103
chr19-13033300-DEL-295
chr19-13081008-DEL-56
chr19-13084488-DEL-405
chr19-13093754-DEL-13247
chr19-13097807-DEL-134
chr19-13115263-DEL-7448
chr19-13173524-DEL-83
chr19-13196369-DEL-94
chr19-13204263-DEL-155
chr19-13219333-DEL-1140
chr19-13219688-DEL-102

chr19-13221498-DEL-1489
chr19-13237919-DEL-1629
chr19-13240561-DEL-389
chr19-13246611-DEL-75
chr19-13265013-DEL-204
chr19-13282067-DEL-245
chr19-13291918-DEL-102
chr19-13301584-DEL-56
chr19-13307944-DEL-1199
chr19-13309495-DEL-3950
chr19-13329734-DEL-6156
chr19-13344520-DEL-93
chr19-13348663-DEL-95
chr19-13354632-DEL-976
chr19-13377070-DEL-2385
chr19-13396633-DEL-748
chr19-13404572-DEL-55
chr19-13511698-DEL-407
chr19-13546861-DEL-1463
chr19-13602194-DEL-1623
chr19-13631521-DEL-152
chr19-13650306-DEL-69
chr19-13662693-DEL-5518
chr19-13665310-DEL-1723
chr19-13682734-DEL-66
chr19-13709220-DEL-730
chr19-13715460-DEL-377
chr19-13736880-DEL-227
chr19-13737585-DEL-2597
chr19-13747922-DEL-212
chr19-13762110-DEL-498
chr19-13793597-DEL-178
chr19-13829698-DEL-70
chr19-13914267-DEL-56
chr19-13951187-DEL-174
chr19-13969804-DEL-62
chr19-13970970-DEL-265
chr19-13976970-DEL-402
chr19-14008066-DEL-68
chr19-14008589-DEL-432
chr19-14026705-DEL-53
chr19-14035149-DEL-696
chr19-14060712-DEL-409
chr19-14083308-DEL-372
chr19-14114154-DEL-136
chr19-14159714-DEL-77
chr19-14174141-DEL-584
chr19-14206166-DEL-201
chr19-14245857-DEL-1304
chr19-14264437-DEL-720
chr19-14271461-DEL-288
chr19-14282487-DEL-1561
chr19-14289948-DEL-394
chr19-14334853-DEL-75
chr19-14335971-DEL-154
chr19-14343364-DEL-306
chr19-14354131-DEL-93
chr19-14374345-DEL-310
chr19-14386616-DEL-98
chr19-14418629-DEL-353
chr19-14456238-DEL-108
chr19-14505622-DEL-243
chr19-14510779-DEL-212
chr19-14519854-DEL-98
chr19-14536297-DEL-214
chr19-14597565-DEL-67
chr19-14602509-DEL-59
chr19-14646908-DEL-111
chr19-14759279-DEL-132
chr19-14766227-DEL-89
chr19-14842406-DEL-60
chr19-14844140-DEL-268
chr19-14858625-DEL-516
chr19-14895963-DEL-84
chr19-14911688-DEL-1369
chr19-14914023-DEL-53
chr19-15004594-DEL-224

chr19-15036749-DEL-287
chr19-15041942-DEL-75
chr19-15043015-DEL-664
chr19-15049760-DEL-74
chr19-15095056-DEL-57
chr19-15125967-DEL-1930
chr19-15127914-DEL-1477
chr19-15146068-DEL-57
chr19-15160487-DEL-58
chr19-15166737-DEL-458
chr19-15225043-DEL-122
chr19-15242712-DEL-1000
chr19-15243313-DEL-309
chr19-15273317-DEL-61
chr19-15285657-DEL-8420
chr19-15295773-DEL-57
chr19-15321585-DEL-1149
chr19-15335895-DEL-812
chr19-15346453-DEL-632
chr19-15347822-DEL-242
chr19-15352432-DEL-633
chr19-15377476-DEL-78
chr19-15392703-DEL-74
chr19-15449453-DEL-213
chr19-15489512-DEL-89
chr19-15498919-DEL-172
chr19-15514207-DEL-58
chr19-15564273-DEL-83
chr19-15603107-DEL-181
chr19-15649492-DEL-1557
chr19-15669241-DEL-52
chr19-15684891-DEL-366
chr19-15685806-DEL-172
chr19-15701945-DEL-55
chr19-15720532-DEL-541
chr19-15758753-DEL-756
chr19-15776956-DEL-308
chr19-15782670-DEL-3253
chr19-15790239-DEL-125
chr19-15796393-DEL-157
chr19-15838452-DEL-53
chr19-15858241-DEL-1056
chr19-15858805-DEL-52
chr19-15860587-DEL-1682
chr19-15880108-DEL-61
chr19-15913853-DEL-289
chr19-15920637-DEL-51
chr19-15931764-DEL-452
chr19-15949790-DEL-85
chr19-15952888-DEL-282
chr19-15953925-DEL-106
chr19-15956506-DEL-285
chr19-15975350-DEL-117
chr19-15979813-DEL-54
chr19-16007192-DEL-516
chr19-16012609-DEL-202
chr19-16017374-DEL-59
chr19-16036887-DEL-217
chr19-16053207-DEL-105
chr19-16057982-DEL-3726
chr19-16058597-DEL-75
chr19-16083509-DEL-71
chr19-16086163-DEL-746
chr19-16093860-DEL-60
chr19-16094155-DEL-2904
chr19-16115569-DEL-120
chr19-16128665-DEL-2087
chr19-16132469-DEL-94
chr19-16135440-DEL-191
chr19-16162624-DEL-54
chr19-16168005-DEL-1744
chr19-16204769-DEL-69
chr19-16208822-DEL-72
chr19-16243786-DEL-1214
chr19-16408054-DEL-54
chr19-16424455-DEL-291
chr19-16482473-DEL-129

chr19-16526923-DEL-1966
chr19-16551374-DEL-328
chr19-16567084-DEL-107
chr19-16638076-DEL-55
chr19-16639402-DEL-374
chr19-16647454-DEL-200
chr19-16691882-DEL-283
chr19-16721468-DEL-71
chr19-16755039-DEL-202
chr19-16772007-DEL-142
chr19-16832008-DEL-2653
chr19-16837928-DEL-2548
chr19-16864025-DEL-80
chr19-16869329-DEL-603
chr19-16871781-DEL-55
chr19-16881854-DEL-53
chr19-16903056-DEL-432
chr19-17016258-DEL-156
chr19-17022347-DEL-291
chr19-17023248-DEL-172
chr19-17023521-DEL-278
chr19-17069599-DEL-52
chr19-17116466-DEL-119
chr19-17123059-DEL-523
chr19-17140086-DEL-268
chr19-17153512-DEL-76
chr19-17168369-DEL-166
chr19-17254610-DEL-210
chr19-17273222-DEL-55
chr19-17294463-DEL-159
chr19-17315279-DEL-654
chr19-17329061-DEL-964
chr19-17344950-DEL-83
chr19-17354760-DEL-198
chr19-17362346-DEL-56
chr19-17387255-DEL-210
chr19-17407142-DEL-189
chr19-17414286-DEL-110
chr19-17441179-DEL-1669
chr19-17501902-DEL-93
chr19-17574619-DEL-67
chr19-17618309-DEL-84
chr19-17645563-DEL-1451
chr19-17650491-DEL-57
chr19-17658114-DEL-653
chr19-17678453-DEL-63
chr19-17685022-DEL-64
chr19-17693444-DEL-111
chr19-17775417-DEL-60
chr19-17786870-DEL-54
chr19-17795351-DEL-70
chr19-17803123-DEL-52
chr19-17828714-DEL-139
chr19-17830406-DEL-519
chr19-17840024-DEL-145
chr19-17845121-DEL-2050
chr19-17878522-DEL-57
chr19-17878839-DEL-631
chr19-17885952-DEL-918
chr19-17890138-DEL-78
chr19-17898341-DEL-82
chr19-17900500-DEL-1249
chr19-17929498-DEL-114
chr19-18069712-DEL-59
chr19-18142300-DEL-57
chr19-18147958-DEL-134
chr19-18156632-DEL-119
chr19-18191467-DEL-106
chr19-18227751-DEL-118
chr19-18238897-DEL-554
chr19-18254967-DEL-171
chr19-18257279-DEL-476
chr19-18335281-DEL-200
chr19-18363046-DEL-76
chr19-18422873-DEL-6310
chr19-18429617-DEL-1565
chr19-18485116-DEL-84

chr19-18486238-DEL-92
chr19-18507110-DEL-58
chr19-18511643-DEL-1955
chr19-18517303-DEL-130
chr19-18536526-DEL-186
chr19-18566137-DEL-482
chr19-18602764-DEL-245
chr19-18678249-DEL-99
chr19-18693315-DEL-354
chr19-18697318-DEL-250
chr19-18722950-DEL-63
chr19-18728900-DEL-108
chr19-18758563-DEL-1299
chr19-18771279-DEL-153
chr19-18816089-DEL-76
chr19-18823575-DEL-328
chr19-18828279-DEL-54
chr19-18868360-DEL-78
chr19-18890270-DEL-60
chr19-18891592-DEL-2902
chr19-18895887-DEL-979
chr19-18998241-DEL-1480
chr19-19001671-DEL-64
chr19-19111939-DEL-74
chr19-19134208-DEL-6208
chr19-19147780-DEL-70
chr19-19171948-DEL-333
chr19-19182729-DEL-88
chr19-19212823-DEL-6203
chr19-19246595-DEL-372
chr19-19252347-DEL-63
chr19-19274332-DEL-68
chr19-19290084-DEL-58
chr19-19299304-DEL-118
chr19-19320048-DEL-96
chr19-19327394-DEL-893
chr19-19331949-DEL-192
chr19-19412927-DEL-312
chr19-19419849-DEL-63
chr19-19438229-DEL-311
chr19-19440486-DEL-399
chr19-19444161-DEL-294
chr19-19447320-DEL-140
chr19-19530539-DEL-308
chr19-19534194-DEL-88
chr19-19535758-DEL-389
chr19-19536231-DEL-1439
chr19-19561287-DEL-51
chr19-19592340-DEL-1230
chr19-19594524-DEL-461
chr19-19652616-DEL-128
chr19-19660607-DEL-916
chr19-19671044-DEL-233
chr19-19673615-DEL-196
chr19-19679784-DEL-382
chr19-19680981-DEL-340
chr19-19699052-DEL-216
chr19-19718880-DEL-85
chr19-19727722-DEL-135
chr19-19763500-DEL-1850
chr19-19805764-DEL-2061
chr19-19832225-DEL-1140
chr19-19869549-DEL-68
chr19-19878833-DEL-224
chr19-19939514-DEL-52
chr19-19959504-DEL-1670
chr19-20003922-DEL-134
chr19-20009865-DEL-290
chr19-20044995-DEL-198
chr19-20057380-DEL-1478
chr19-20077626-DEL-170
chr19-20082503-DEL-2319
chr19-20098467-DEL-64
chr19-20126790-DEL-198
chr19-20151606-DEL-627
chr19-20164246-DEL-453
chr19-20181120-DEL-875

chr19-20181756-DEL-231
chr19-20236540-DEL-2378
chr19-20334628-DEL-54
chr19-20341046-DEL-629
chr19-20343678-DEL-107
chr19-20351183-DEL-588
chr19-20400306-DEL-400
chr19-20400857-DEL-129
chr19-20432491-DEL-7757
chr19-20442317-DEL-1351
chr19-20476432-DEL-2638
chr19-20492653-DEL-480
chr19-20531446-DEL-248
chr19-20568104-DEL-1185
chr19-20593513-DEL-9181
chr19-20598532-DEL-54
chr19-20672589-DEL-317
chr19-20681756-DEL-540
chr19-20689606-DEL-2303
chr19-20708467-DEL-108
chr19-20711995-DEL-2597
chr19-20743882-DEL-8191
chr19-20783796-DEL-62
chr19-20789319-DEL-55
chr19-20790886-DEL-56
chr19-20830959-DEL-73
chr19-20844644-DEL-293
chr19-20851442-DEL-52
chr19-20856348-DEL-140
chr19-20896825-DEL-73
chr19-20920460-DEL-1017
chr19-20927367-DEL-92
chr19-20969246-DEL-62
chr19-20975775-DEL-425
chr19-20990209-DEL-3549
chr19-21032958-DEL-252
chr19-21043606-DEL-209
chr19-21088516-DEL-161
chr19-21095670-DEL-146
chr19-21111173-DEL-129
chr19-21180737-DEL-74
chr19-21187053-DEL-100
chr19-21221621-DEL-92
chr19-21280003-DEL-421
chr19-21321125-DEL-585
chr19-21338932-DEL-242
chr19-21340813-DEL-230
chr19-21370892-DEL-56
chr19-21445740-DEL-52
chr19-21452595-DEL-62
chr19-21481118-DEL-78
chr19-21496063-DEL-169
chr19-21505676-DEL-272
chr19-21508973-DEL-313
chr19-21544556-DEL-356
chr19-21581081-DEL-209
chr19-21586309-DEL-64
chr19-21605744-DEL-150
chr19-21613767-DEL-93
chr19-21634692-DEL-357
chr19-21700893-DEL-58
chr19-21752029-DEL-165
chr19-21781333-DEL-58
chr19-21810090-DEL-73
chr19-21837357-DEL-92
chr19-21852282-DEL-374
chr19-21886189-DEL-84
chr19-21893935-DEL-80
chr19-21897720-DEL-14149
chr19-21899451-DEL-7522
chr19-21921899-DEL-502
chr19-21941513-DEL-10981
chr19-21962918-DEL-162
chr19-21974565-DEL-394
chr19-21979543-DEL-6152
chr19-21986421-DEL-366
chr19-21990668-DEL-78

chr19-22024220-DEL-52
chr19-22028731-DEL-243
chr19-22031181-DEL-85
chr19-22072497-DEL-68
chr19-22131769-DEL-2833
chr19-22176755-DEL-1004
chr19-22213114-DEL-979
chr19-22276990-DEL-140
chr19-22299315-DEL-102
chr19-22313820-DEL-84
chr19-22392540-DEL-481
chr19-22501282-DEL-53
chr19-22587426-DEL-82
chr19-22597904-DEL-495
chr19-22612286-DEL-210
chr19-22664503-DEL-110
chr19-22668136-DEL-435
chr19-22669229-DEL-58
chr19-22756025-DEL-87
chr19-22761828-DEL-641
chr19-22763631-DEL-665
chr19-22831427-DEL-126
chr19-22840885-DEL-255
chr19-22843361-DEL-684
chr19-22857452-DEL-796
chr19-22864530-DEL-172
chr19-22867880-DEL-91
chr19-22875963-DEL-88
chr19-22889993-DEL-266
chr19-22930898-DEL-1451
chr19-22933126-DEL-1189
chr19-23008931-DEL-60
chr19-23010695-DEL-55
chr19-23014898-DEL-1518
chr19-23020549-DEL-82
chr19-23021679-DEL-141
chr19-23031458-DEL-51
chr19-23048421-DEL-102
chr19-23057106-DEL-86
chr19-23066952-DEL-72
chr19-23068797-DEL-66
chr19-23079681-DEL-94
chr19-23093340-DEL-82
chr19-23094911-DEL-176
chr19-23100521-DEL-138
chr19-23106716-DEL-387
chr19-23134916-DEL-1953
chr19-23178080-DEL-191
chr19-23190478-DEL-134
chr19-23207663-DEL-240
chr19-23209999-DEL-347
chr19-23220314-DEL-94
chr19-23222952-DEL-3679
chr19-23227546-DEL-831
chr19-23230650-DEL-179
chr19-23251358-DEL-299
chr19-23283575-DEL-54
chr19-23295810-DEL-170
chr19-23334793-DEL-154
chr19-23353130-DEL-127
chr19-23367271-DEL-132
chr19-23404490-DEL-74
chr19-23410197-DEL-58
chr19-23411001-DEL-126
chr19-23413188-DEL-155
chr19-23432679-DEL-136
chr19-23433735-DEL-112
chr19-23447314-DEL-3718
chr19-23460185-DEL-552
chr19-23481010-DEL-54
chr19-23549066-DEL-173
chr19-23575115-DEL-89
chr19-23580283-DEL-51
chr19-23580493-DEL-396
chr19-23592506-DEL-127
chr19-23594533-DEL-3550
chr19-23600498-DEL-66

chr19-23640079-DEL-499
chr19-23642023-DEL-82
chr19-23719928-DEL-285
chr19-23723478-DEL-792
chr19-23880148-DEL-94
chr19-23929706-DEL-52
chr19-23985047-DEL-1299
chr19-23992497-DEL-62
chr19-24036499-DEL-99
chr19-24043884-DEL-116
chr19-24051369-DEL-1425
chr19-24070475-DEL-202
chr19-24093531-DEL-593
chr19-24122793-DEL-63
chr19-24127604-DEL-75
chr19-24148725-DEL-213
chr19-24159304-DEL-55
chr19-24166359-DEL-84
chr19-24194442-DEL-196
chr19-24195160-DEL-675
chr19-24197133-DEL-158
chr19-24217475-DEL-399
chr19-24240480-DEL-114
chr19-24267285-DEL-153
chr19-24273794-DEL-322
chr19-24296461-DEL-160
chr19-24312173-DEL-51
chr19-24343698-DEL-99
chr19-24355606-DEL-127
chr19-24359315-DEL-701
chr19-24409828-DEL-68
chr19-24492946-DEL-399
chr19-24537273-DEL-54
chr19-24551333-DEL-191
chr19-24563669-DEL-51
chr19-24564417-DEL-139
chr19-24602468-DEL-84
chr19-24616857-DEL-233
chr19-24638074-DEL-77
chr19-24642398-DEL-429
chr19-24688741-DEL-62
chr19-24746194-DEL-1227
chr19-24758528-DEL-177
chr19-24777333-DEL-60
chr19-24786565-DEL-646
chr19-24815582-DEL-83
chr19-24835324-DEL-159
chr19-24849389-DEL-51
chr19-24859001-DEL-818
chr19-24867692-DEL-2080
chr19-24869234-DEL-54
chr19-24873710-DEL-392
chr19-24889508-DEL-7019
chr19-24889802-DEL-6307
chr19-24945586-DEL-191
chr19-24979973-DEL-218
chr19-24987285-DEL-274
chr19-24995072-DEL-209
chr19-25063085-DEL-114
chr19-25102241-DEL-135
chr19-25123261-DEL-51
chr19-25133817-DEL-237
chr19-25157229-DEL-134
chr19-25192497-DEL-386
chr19-25195880-DEL-172
chr19-25200763-DEL-51
chr19-25202750-DEL-69
chr19-25210806-DEL-119
chr19-25228421-DEL-128
chr19-25242730-DEL-205
chr19-25257416-DEL-829
chr19-25273925-DEL-135
chr19-25287141-DEL-68
chr19-25310805-DEL-66
chr19-25320262-DEL-194
chr19-25333859-DEL-52
chr19-25348355-DEL-476

chr19-25348699-DEL-900
chr19-25349346-DEL-224
chr19-25384811-DEL-348
chr19-25434824-DEL-387
chr19-25438241-DEL-59
chr19-25442204-DEL-54
chr19-25449346-DEL-89
chr19-25460850-DEL-463
chr19-25476100-DEL-133
chr19-25489575-DEL-60
chr19-25490795-DEL-196
chr19-25517783-DEL-135
chr19-25537540-DEL-74
chr19-25566406-DEL-1872
chr19-25566433-DEL-63
chr19-25571885-DEL-397
chr19-25594735-DEL-81
chr19-25614123-DEL-86
chr19-25631225-DEL-97
chr19-25636824-DEL-141
chr19-25677206-DEL-421
chr19-25680758-DEL-56
chr19-25684102-DEL-164
chr19-25688997-DEL-321
chr19-25721540-DEL-1213
chr19-25723602-DEL-751
chr19-25724467-DEL-190
chr19-25732520-DEL-5999
chr19-25754558-DEL-75
chr19-25760517-DEL-59
chr19-25808623-DEL-66
chr19-25831357-DEL-66
chr19-25839117-DEL-623
chr19-25847756-DEL-201
chr19-25869426-DEL-124
chr19-25889730-DEL-52916
chr19-25954626-DEL-93
chr19-25963269-DEL-98
chr19-25981050-DEL-224
chr19-26002951-DEL-58
chr19-26019994-DEL-309
chr19-26041201-DEL-188
chr19-26051724-DEL-273
chr19-26071494-DEL-1282
chr19-26083665-DEL-59
chr19-26086962-DEL-462
chr19-26088937-DEL-55
chr19-26107768-DEL-143
chr19-26115539-DEL-85
chr19-26133994-DEL-650
chr19-26136049-DEL-458
chr19-26142903-DEL-2663
chr19-26176032-DEL-65
chr19-26177908-DEL-399
chr19-26182947-DEL-62
chr19-26189246-DEL-1206
chr19-26204022-DEL-1808
chr19-26270732-DEL-566
chr19-26282413-DEL-53
chr19-26324254-DEL-72
chr19-26341299-DEL-70
chr19-26369020-DEL-96
chr19-26416052-DEL-1732
chr19-26440740-DEL-487
chr19-26518366-DEL-83
chr19-26534806-DEL-106
chr19-26615002-DEL-172
chr19-26621586-DEL-104
chr19-26703025-DEL-347
chr19-26774038-DEL-64
chr19-26785076-DEL-862
chr19-26795828-DEL-180
chr19-26798872-DEL-352
chr19-26815742-DEL-380
chr19-26823685-DEL-262
chr19-26869596-DEL-100
chr19-26872022-DEL-160

chr19-26874856-DEL-328
chr19-26893326-DEL-2842
chr19-26901611-DEL-66
chr19-26903843-DEL-70
chr19-26909691-DEL-3262
chr19-26914980-DEL-528
chr19-26923410-DEL-54
chr19-26971045-DEL-134
chr19-26973072-DEL-2000
chr19-27018053-DEL-52
chr19-27032259-DEL-1916
chr19-27060271-DEL-52
chr19-27070012-DEL-112
chr19-27091284-DEL-52
chr19-27119388-DEL-68
chr19-27137147-DEL-206
chr19-27148165-DEL-98
chr19-27159502-DEL-199
chr19-27219662-DEL-75
chr19-27251059-DEL-2154
chr19-27263467-DEL-184
chr19-27275063-DEL-69
chr19-27370035-DEL-589
chr19-27398983-DEL-98
chr19-27426412-DEL-623
chr19-27428313-DEL-235
chr19-27470867-DEL-75
chr19-27539353-DEL-129
chr19-27550140-DEL-847
chr19-27627516-DEL-351
chr19-27630912-DEL-51
chr19-27644969-DEL-54
chr19-27647821-DEL-54
chr19-27658366-DEL-74
chr19-27729031-DEL-212
chr19-27747114-DEL-204
chr19-27773889-DEL-113
chr19-27877672-DEL-257
chr19-27889716-DEL-103
chr19-27933145-DEL-363
chr19-27948423-DEL-1151
chr19-27951245-DEL-438
chr19-27958884-DEL-54
chr19-27964464-DEL-56
chr19-28002192-DEL-2544
chr19-28016067-DEL-1393
chr19-28019639-DEL-76
chr19-28032197-DEL-53
chr19-28047714-DEL-57
chr19-28071519-DEL-77
chr19-28109109-DEL-55
chr19-28219459-DEL-293
chr19-28220569-DEL-75
chr19-28246244-DEL-67
chr19-28255178-DEL-133
chr19-28288680-DEL-61
chr19-28307779-DEL-60
chr19-28348080-DEL-167
chr19-28378002-DEL-157
chr19-28382705-DEL-178
chr19-28456394-DEL-74
chr19-28463704-DEL-1019
chr19-28503994-DEL-88
chr19-28605319-DEL-64
chr19-28637841-DEL-78
chr19-28667832-DEL-61
chr19-28675940-DEL-90
chr19-28677314-DEL-94
chr19-28683794-DEL-347
chr19-28688731-DEL-196
chr19-28713470-DEL-95
chr19-28723615-DEL-59
chr19-28742443-DEL-458
chr19-28848376-DEL-53
chr19-28860602-DEL-51
chr19-28938228-DEL-237
chr19-28980530-DEL-54

chr19-29012493-DEL-54
chr19-29036965-DEL-263
chr19-29080398-DEL-201
chr19-29084579-DEL-104
chr19-29168776-DEL-1173
chr19-29182452-DEL-96
chr19-29196781-DEL-1880
chr19-29207168-DEL-147
chr19-29233879-DEL-260
chr19-29238196-DEL-366
chr19-29291287-DEL-240
chr19-29297987-DEL-372
chr19-29304190-DEL-361
chr19-29318912-DEL-204
chr19-29353739-DEL-72
chr19-29364426-DEL-56
chr19-29383165-DEL-54
chr19-29390154-DEL-245
chr19-29392323-DEL-89
chr19-29403803-DEL-79
chr19-29410757-DEL-2524
chr19-29428468-DEL-486
chr19-29430637-DEL-69
chr19-29449999-DEL-347
chr19-29461634-DEL-271
chr19-29484026-DEL-397
chr19-29493595-DEL-56
chr19-29500333-DEL-69
chr19-29507702-DEL-60
chr19-29513862-DEL-166
chr19-29547683-DEL-65
chr19-29576194-DEL-100
chr19-29591158-DEL-392
chr19-29617901-DEL-116
chr19-29619075-DEL-93
chr19-29640098-DEL-562
chr19-29648451-DEL-68
chr19-29652405-DEL-224
chr19-29654656-DEL-187
chr19-29660818-DEL-689
chr19-29676552-DEL-93
chr19-29691252-DEL-75
chr19-29738429-DEL-191
chr19-29829627-DEL-454
chr19-29832077-DEL-170
chr19-29843076-DEL-1625
chr19-29870971-DEL-101
chr19-29901635-DEL-172
chr19-29909259-DEL-146
chr19-29947095-DEL-345
chr19-30006858-DEL-115
chr19-30020538-DEL-101
chr19-30050246-DEL-204
chr19-30112018-DEL-63
chr19-30118246-DEL-1741
chr19-30125250-DEL-57
chr19-30133328-DEL-267
chr19-30160537-DEL-89
chr19-30170949-DEL-167
chr19-30185202-DEL-157
chr19-30188509-DEL-101
chr19-30191413-DEL-236
chr19-30193568-DEL-262
chr19-30199108-DEL-205
chr19-30201850-DEL-3141
chr19-30205843-DEL-110
chr19-30213101-DEL-311
chr19-30222929-DEL-72
chr19-30228625-DEL-152
chr19-30229471-DEL-99
chr19-30267304-DEL-59
chr19-30317325-DEL-221
chr19-30320070-DEL-810
chr19-30325487-DEL-402
chr19-30346887-DEL-281
chr19-30350642-DEL-1557
chr19-30353707-DEL-373

chr19-30360604-DEL-8427
chr19-30362974-DEL-289
chr19-30365186-DEL-2372
chr19-30365353-DEL-1488
chr19-30375459-DEL-102
chr19-30384835-DEL-111
chr19-30401845-DEL-59
chr19-30455733-DEL-163
chr19-30468006-DEL-141
chr19-30478068-DEL-1520
chr19-30478478-DEL-545
chr19-30490136-DEL-113
chr19-30514821-DEL-2455
chr19-30538876-DEL-1764
chr19-30598300-DEL-55
chr19-30625893-DEL-203
chr19-30648724-DEL-229
chr19-30710755-DEL-63
chr19-30778174-DEL-54
chr19-30817537-DEL-173
chr19-30832006-DEL-57
chr19-30869432-DEL-157
chr19-30881120-DEL-54
chr19-30884776-DEL-2080
chr19-30896039-DEL-378
chr19-30898944-DEL-2149
chr19-30957996-DEL-241
chr19-30977089-DEL-133
chr19-31011090-DEL-57
chr19-31015499-DEL-59
chr19-31037518-DEL-133
chr19-31062514-DEL-212
chr19-31077558-DEL-2587
chr19-31100920-DEL-57
chr19-31117062-DEL-76
chr19-31126756-DEL-62
chr19-31152710-DEL-92
chr19-31172052-DEL-66
chr19-31183492-DEL-444
chr19-31223687-DEL-98
chr19-31255983-DEL-90
chr19-31260840-DEL-114
chr19-31298327-DEL-74
chr19-31379327-DEL-51
chr19-31396861-DEL-82
chr19-31440526-DEL-277
chr19-31461948-DEL-265
chr19-31465882-DEL-87
chr19-31470098-DEL-93
chr19-31488591-DEL-83
chr19-31529843-DEL-58
chr19-31549982-DEL-180
chr19-31568471-DEL-101
chr19-31589463-DEL-1818
chr19-31595222-DEL-183
chr19-31667990-DEL-53
chr19-31686195-DEL-79
chr19-31709054-DEL-230
chr19-31709770-DEL-1425
chr19-31714234-DEL-1583
chr19-31745649-DEL-71
chr19-31763506-DEL-69
chr19-31766984-DEL-145
chr19-31795665-DEL-82
chr19-31829579-DEL-170
chr19-31831047-DEL-53
chr19-31869640-DEL-108
chr19-31907013-DEL-1002
chr19-31914573-DEL-212
chr19-31939381-DEL-56
chr19-31951206-DEL-72
chr19-31955896-DEL-61
chr19-31980849-DEL-111
chr19-31998083-DEL-126
chr19-32017340-DEL-54
chr19-32022852-DEL-149
chr19-32024632-DEL-1741

chr19-32025947-DEL-1327
chr19-32027589-DEL-498
chr19-32045588-DEL-67
chr19-32057825-DEL-359
chr19-32077604-DEL-78
chr19-32088515-DEL-52
chr19-32109266-DEL-445
chr19-32123751-DEL-56
chr19-32132582-DEL-58
chr19-32162547-DEL-94
chr19-32213428-DEL-83
chr19-32281408-DEL-56
chr19-32284425-DEL-539
chr19-32341028-DEL-102
chr19-32372615-DEL-179
chr19-32392648-DEL-82
chr19-32432839-DEL-400
chr19-32479476-DEL-892
chr19-32497301-DEL-72
chr19-32500842-DEL-274
chr19-32540399-DEL-744
chr19-32547422-DEL-51
chr19-32564131-DEL-224
chr19-32570736-DEL-98
chr19-32605174-DEL-1207
chr19-32607218-DEL-289
chr19-32632209-DEL-154
chr19-32651989-DEL-178
chr19-32655627-DEL-214
chr19-32663858-DEL-195
chr19-32674594-DEL-51
chr19-32676035-DEL-68
chr19-32699555-DEL-94
chr19-32722023-DEL-53
chr19-32725930-DEL-263
chr19-32747873-DEL-195
chr19-32763394-DEL-145
chr19-32850902-DEL-214
chr19-32852524-DEL-129
chr19-32897410-DEL-54
chr19-32947968-DEL-581
chr19-32960681-DEL-81
chr19-32982730-DEL-92
chr19-32989144-DEL-65
chr19-32991951-DEL-508
chr19-33013958-DEL-54
chr19-33022762-DEL-587
chr19-33024330-DEL-61
chr19-33036481-DEL-309
chr19-33052868-DEL-82
chr19-33100009-DEL-62
chr19-33104364-DEL-60
chr19-33109290-DEL-56
chr19-33121960-DEL-285
chr19-33213045-DEL-712
chr19-33232348-DEL-99
chr19-33258704-DEL-233
chr19-33416733-DEL-6529
chr19-33460164-DEL-10252
chr19-33471253-DEL-2319
chr19-33473050-DEL-105
chr19-33489299-DEL-3048
chr19-33514785-DEL-2599
chr19-33588847-DEL-56
chr19-33631437-DEL-81
chr19-33635679-DEL-119
chr19-33637444-DEL-336
chr19-33683854-DEL-6193
chr19-33739791-DEL-6438
chr19-33763573-DEL-199
chr19-33788361-DEL-6358
chr19-33796213-DEL-124
chr19-33800287-DEL-4168
chr19-33808419-DEL-4187
chr19-33833052-DEL-73
chr19-33840050-DEL-2450
chr19-33842695-DEL-347

chr19-33843204-DEL-1110
chr19-33844344-DEL-1665
chr19-33852773-DEL-2627
chr19-33867341-DEL-1746
chr19-33873995-DEL-326
chr19-33897324-DEL-1593
chr19-33902614-DEL-72
chr19-33926395-DEL-10437
chr19-33940640-DEL-105
chr19-34002798-DEL-6447
chr19-34009151-DEL-278
chr19-34021601-DEL-86
chr19-34029998-DEL-59
chr19-34046349-DEL-95
chr19-34059707-DEL-68
chr19-34060577-DEL-87
chr19-34074697-DEL-54
chr19-34101013-DEL-65
chr19-34121483-DEL-144
chr19-34123251-DEL-123
chr19-34129812-DEL-1124
chr19-34146026-DEL-112
chr19-34158685-DEL-98
chr19-34179173-DEL-206
chr19-34251391-DEL-341
chr19-34274047-DEL-134
chr19-34347892-DEL-169
chr19-34373039-DEL-57
chr19-34399674-DEL-938
chr19-34424342-DEL-526
chr19-34436241-DEL-92
chr19-34459744-DEL-1008
chr19-34467918-DEL-1489
chr19-34475258-DEL-52
chr19-34538287-DEL-1167
chr19-34576083-DEL-2623
chr19-34797564-DEL-198
chr19-34806132-DEL-70
chr19-34830618-DEL-729
chr19-34911886-DEL-176
chr19-34934980-DEL-65
chr19-34941110-DEL-128
chr19-34952105-DEL-438
chr19-34956718-DEL-538
chr19-35003678-DEL-149
chr19-35006249-DEL-252
chr19-35006259-DEL-1395
chr19-35012127-DEL-948
chr19-35012765-DEL-94
chr19-35033682-DEL-84
chr19-35038651-DEL-626
chr19-35042990-DEL-1801
chr19-35073148-DEL-92
chr19-35093129-DEL-60
chr19-35131397-DEL-84
chr19-35132826-DEL-54
chr19-35155459-DEL-1004
chr19-35168568-DEL-68
chr19-35171976-DEL-474
chr19-35199858-DEL-137
chr19-35240235-DEL-108
chr19-35250091-DEL-1259
chr19-35269645-DEL-66
chr19-35301214-DEL-191
chr19-35302534-DEL-121
chr19-35310414-DEL-6625
chr19-35316835-DEL-7654
chr19-35317733-DEL-54
chr19-35334160-DEL-69
chr19-35341726-DEL-109
chr19-35361380-DEL-6156
chr19-35364034-DEL-1866
chr19-35370921-DEL-151
chr19-35372358-DEL-63
chr19-35399829-DEL-299
chr19-35447983-DEL-208
chr19-35488373-DEL-58

chr19-35537125-DEL-1895
chr19-35562158-DEL-6647
chr19-35574164-DEL-5172
chr19-35576791-DEL-416
chr19-35582332-DEL-52
chr19-35612443-DEL-181
chr19-35624266-DEL-308
chr19-35640365-DEL-815
chr19-35643920-DEL-6975
chr19-35644259-DEL-204
chr19-35679745-DEL-58
chr19-35691687-DEL-73
chr19-35739130-DEL-469
chr19-35777136-DEL-61
chr19-35826009-DEL-183
chr19-35862683-DEL-133
chr19-35866261-DEL-894
chr19-35881401-DEL-636
chr19-35914216-DEL-3537
chr19-35933965-DEL-709
chr19-35939284-DEL-135
chr19-35958317-DEL-862
chr19-35964580-DEL-103
chr19-35967265-DEL-81
chr19-35969667-DEL-185
chr19-35991394-DEL-2023
chr19-36025520-DEL-82
chr19-36047790-DEL-2932
chr19-36047794-DEL-289
chr19-36050488-DEL-234
chr19-36065198-DEL-117
chr19-36070645-DEL-85
chr19-36079480-DEL-53
chr19-36082506-DEL-78
chr19-36085990-DEL-630
chr19-36150163-DEL-73
chr19-36189523-DEL-51
chr19-36281876-DEL-385
chr19-36304862-DEL-191
chr19-36320742-DEL-400
chr19-36333934-DEL-59
chr19-36379932-DEL-399
chr19-36383404-DEL-59
chr19-36490282-DEL-66
chr19-36495423-DEL-7633
chr19-36502794-DEL-111
chr19-36562527-DEL-96
chr19-36585755-DEL-52
chr19-36605602-DEL-88
chr19-36607457-DEL-1952
chr19-36624993-DEL-293
chr19-36651846-DEL-172
chr19-36661902-DEL-104
chr19-36671265-DEL-63
chr19-36676566-DEL-231
chr19-36684158-DEL-93
chr19-36684972-DEL-58
chr19-36727840-DEL-198
chr19-36750143-DEL-3249
chr19-36764240-DEL-1633
chr19-36781419-DEL-72
chr19-36783539-DEL-102
chr19-36796137-DEL-596
chr19-36805737-DEL-70
chr19-36848715-DEL-182
chr19-36866555-DEL-146
chr19-36890161-DEL-166
chr19-36902059-DEL-80
chr19-36919686-DEL-205
chr19-36941213-DEL-1772
chr19-37012653-DEL-135
chr19-37054350-DEL-287
chr19-37085448-DEL-160
chr19-37128059-DEL-172
chr19-37130397-DEL-206
chr19-37160509-DEL-209
chr19-37168073-DEL-60

chr19-37174338-DEL-210
chr19-37200374-DEL-199
chr19-37238818-DEL-176
chr19-37276167-DEL-754
chr19-37276180-DEL-2238
chr19-37276180-DEL-2934
chr19-37276453-DEL-1421
chr19-37277904-DEL-174
chr19-37284299-DEL-94
chr19-37302989-DEL-55
chr19-37308301-DEL-96
chr19-37310393-DEL-2611
chr19-37319340-DEL-328
chr19-37347599-DEL-203
chr19-37350817-DEL-61
chr19-37352397-DEL-65
chr19-37374341-DEL-156
chr19-37383352-DEL-180
chr19-37396214-DEL-59
chr19-37412769-DEL-94
chr19-37432975-DEL-420
chr19-37438327-DEL-64
chr19-37465882-DEL-1244
chr19-37471655-DEL-70
chr19-37493502-DEL-159
chr19-37512755-DEL-208
chr19-37532703-DEL-64
chr19-37534750-DEL-62
chr19-37555536-DEL-57
chr19-37571343-DEL-225
chr19-37592721-DEL-227
chr19-37616097-DEL-86
chr19-37619096-DEL-136
chr19-37633516-DEL-130
chr19-37650491-DEL-1398
chr19-37656611-DEL-60
chr19-37663581-DEL-176
chr19-37665987-DEL-302
chr19-37710846-DEL-306
chr19-37714736-DEL-136
chr19-37722066-DEL-210
chr19-37730033-DEL-54
chr19-37745888-DEL-1224
chr19-37775091-DEL-1106
chr19-37780175-DEL-70
chr19-37792342-DEL-67
chr19-37812687-DEL-51
chr19-37817041-DEL-1344
chr19-37818942-DEL-131
chr19-37819525-DEL-69
chr19-37838321-DEL-1226
chr19-37856726-DEL-784
chr19-37864912-DEL-149
chr19-37935475-DEL-145
chr19-37981485-DEL-102
chr19-38017607-DEL-336
chr19-38017755-DEL-945
chr19-38021351-DEL-56
chr19-38040585-DEL-384
chr19-38047294-DEL-442
chr19-38069880-DEL-329
chr19-38075018-DEL-53
chr19-38077744-DEL-61
chr19-38091060-DEL-228
chr19-38111340-DEL-60
chr19-38117194-DEL-849
chr19-38182828-DEL-373
chr19-38185564-DEL-449
chr19-38195998-DEL-265
chr19-38199945-DEL-1129
chr19-38209314-DEL-64
chr19-38231310-DEL-134
chr19-38245787-DEL-206
chr19-38270109-DEL-199
chr19-38277381-DEL-114
chr19-38306590-DEL-109
chr19-38329273-DEL-114

chr19-38351322-DEL-66
chr19-38408601-DEL-2720
chr19-38410816-DEL-931
chr19-38412033-DEL-56
chr19-38414318-DEL-152
chr19-38428747-DEL-52
chr19-38438217-DEL-58
chr19-38446055-DEL-99
chr19-38450613-DEL-226
chr19-38453779-DEL-446
chr19-38463436-DEL-105
chr19-38478749-DEL-4881
chr19-38484779-DEL-195
chr19-38490177-DEL-262
chr19-38494335-DEL-233
chr19-38511878-DEL-82
chr19-38547932-DEL-195
chr19-38552931-DEL-60
chr19-38632971-DEL-78
chr19-38635854-DEL-361
chr19-38645303-DEL-129
chr19-38653170-DEL-80
chr19-38673409-DEL-310
chr19-38694965-DEL-113
chr19-38718077-DEL-91
chr19-38744749-DEL-202
chr19-38775505-DEL-68
chr19-38794400-DEL-61
chr19-38840733-DEL-264
chr19-38842488-DEL-375
chr19-38866409-DEL-1533
chr19-38926179-DEL-151
chr19-38957416-DEL-85
chr19-38966126-DEL-1143
chr19-38966353-DEL-679
chr19-38990089-DEL-82
chr19-38996168-DEL-593
chr19-39007913-DEL-68
chr19-39019923-DEL-204
chr19-39041205-DEL-480
chr19-39041831-DEL-1428
chr19-39043046-DEL-381
chr19-39066304-DEL-1605
chr19-39126723-DEL-2256
chr19-39133130-DEL-455
chr19-39145251-DEL-6358
chr19-39145346-DEL-311
chr19-39171566-DEL-200
chr19-39209974-DEL-3270
chr19-39230937-DEL-6765
chr19-39244762-DEL-1477
chr19-39293206-DEL-1248
chr19-39336205-DEL-10166
chr19-39341123-DEL-1981
chr19-39365436-DEL-89
chr19-39380865-DEL-6238
chr19-39425491-DEL-56
chr19-39453212-DEL-6392
chr19-39467387-DEL-3146
chr19-39480686-DEL-1894
chr19-39490875-DEL-378
chr19-39492402-DEL-58
chr19-39560447-DEL-489
chr19-39564912-DEL-1383
chr19-39614942-DEL-56
chr19-39633365-DEL-54
chr19-39635310-DEL-1381
chr19-39643854-DEL-383
chr19-39644373-DEL-63
chr19-39645455-DEL-6257
chr19-39728722-DEL-214
chr19-39769189-DEL-5937
chr19-39797334-DEL-54
chr19-39831624-DEL-5526
chr19-39842313-DEL-5870
chr19-39860842-DEL-2343
chr19-39874376-DEL-54

chr19-39879803-DEL-1992
chr19-39886931-DEL-386
chr19-39899477-DEL-2185
chr19-39922697-DEL-59
chr19-39944903-DEL-16760
chr19-39950518-DEL-8796
chr19-39951382-DEL-114
chr19-40005527-DEL-400
chr19-40094851-DEL-184
chr19-40110345-DEL-59
chr19-40123248-DEL-59
chr19-40163890-DEL-655
chr19-40181740-DEL-16565
chr19-40197248-DEL-1421
chr19-40215615-DEL-56
chr19-40236913-DEL-62
chr19-40242972-DEL-779
chr19-40243912-DEL-104
chr19-40250175-DEL-1810
chr19-40317071-DEL-901
chr19-40317097-DEL-1065
chr19-40317356-DEL-451
chr19-40356877-DEL-98
chr19-40447100-DEL-2000
chr19-40490168-DEL-293
chr19-40507093-DEL-125
chr19-40510428-DEL-126
chr19-40519637-DEL-82
chr19-40528484-DEL-124
chr19-40558471-DEL-78
chr19-40696324-DEL-283
chr19-40697440-DEL-169
chr19-40746615-DEL-59
chr19-40751532-DEL-740
chr19-40752323-DEL-231
chr19-40804757-DEL-260
chr19-40830743-DEL-462
chr19-40852510-DEL-113
chr19-40864573-DEL-577
chr19-40916331-DEL-459
chr19-40936138-DEL-96
chr19-40938061-DEL-404
chr19-41002686-DEL-4843
chr19-41040849-DEL-104
chr19-41046483-DEL-55
chr19-41065262-DEL-67
chr19-41065944-DEL-1692
chr19-41082505-DEL-129
chr19-41090257-DEL-211
chr19-41092631-DEL-71
chr19-41103790-DEL-97
chr19-41115287-DEL-1149
chr19-41125593-DEL-132
chr19-41160776-DEL-115
chr19-41161410-DEL-161
chr19-41174689-DEL-88
chr19-41230343-DEL-60
chr19-41231456-DEL-208
chr19-41257543-DEL-84
chr19-41280396-DEL-109
chr19-41330030-DEL-79
chr19-41340129-DEL-107
chr19-41350770-DEL-88
chr19-41376819-DEL-175
chr19-41437432-DEL-110
chr19-41449247-DEL-689
chr19-41452030-DEL-183
chr19-41456335-DEL-89
chr19-41464112-DEL-246
chr19-41474335-DEL-93
chr19-41519147-DEL-291
chr19-41521086-DEL-198
chr19-41522344-DEL-2043
chr19-41528773-DEL-147
chr19-41619617-DEL-64
chr19-41621667-DEL-271
chr19-41624474-DEL-730

chr19-41643203-DEL-197
chr19-41757715-DEL-66
chr19-41778498-DEL-112
chr19-41793900-DEL-125
chr19-41796883-DEL-1560
chr19-41804902-DEL-327
chr19-41812922-DEL-86
chr19-41815390-DEL-92
chr19-41837822-DEL-175
chr19-41861501-DEL-140
chr19-41877715-DEL-208
chr19-41907979-DEL-52
chr19-41939556-DEL-295
chr19-41948078-DEL-64
chr19-41965198-DEL-243
chr19-41978856-DEL-197
chr19-41983327-DEL-130
chr19-42000825-DEL-103
chr19-42028071-DEL-77
chr19-42045477-DEL-101
chr19-42072236-DEL-52
chr19-42096979-DEL-69
chr19-42099701-DEL-840
chr19-42121889-DEL-215
chr19-42142848-DEL-160
chr19-42151188-DEL-209
chr19-42155573-DEL-941
chr19-42216864-DEL-457
chr19-42250127-DEL-64
chr19-42390420-DEL-216
chr19-42411534-DEL-1426
chr19-42449333-DEL-52
chr19-42480638-DEL-64
chr19-42579431-DEL-68
chr19-42605657-DEL-116
chr19-42663506-DEL-166
chr19-42679493-DEL-117
chr19-42687854-DEL-53
chr19-42704027-DEL-209
chr19-42770693-DEL-431
chr19-42806282-DEL-81
chr19-42879913-DEL-64
chr19-42893945-DEL-72
chr19-42945092-DEL-128
chr19-42954532-DEL-74
chr19-42998171-DEL-151
chr19-43012503-DEL-61
chr19-43018283-DEL-81
chr19-43062620-DEL-148
chr19-43072798-DEL-261
chr19-43131159-DEL-62
chr19-43162257-DEL-54
chr19-43198824-DEL-1027
chr19-43224473-DEL-1875
chr19-43237616-DEL-615
chr19-43238480-DEL-242
chr19-43242035-DEL-346
chr19-43274003-DEL-70
chr19-43303702-DEL-4664
chr19-43308346-DEL-225
chr19-43335949-DEL-3659
chr19-43388885-DEL-327
chr19-43412472-DEL-63
chr19-43413189-DEL-980
chr19-43424007-DEL-79
chr19-43469821-DEL-62
chr19-43481885-DEL-65
chr19-43516529-DEL-56
chr19-43525069-DEL-731
chr19-43528229-DEL-431
chr19-43529938-DEL-56
chr19-43541975-DEL-58
chr19-43561314-DEL-102
chr19-43620321-DEL-142
chr19-43637313-DEL-236
chr19-43683007-DEL-250
chr19-43684570-DEL-65

chr19-43685098-DEL-77
chr19-43696161-DEL-78
chr19-43775209-DEL-58
chr19-43785597-DEL-224
chr19-43807165-DEL-56
chr19-43808476-DEL-227
chr19-43829294-DEL-486
chr19-43832626-DEL-208
chr19-43869858-DEL-194
chr19-43880037-DEL-344
chr19-43886422-DEL-1317
chr19-43895742-DEL-86
chr19-43896554-DEL-127
chr19-43910403-DEL-60
chr19-43913187-DEL-64
chr19-43925975-DEL-288
chr19-44008961-DEL-164
chr19-44009536-DEL-212
chr19-44012453-DEL-66
chr19-44015819-DEL-300
chr19-44114297-DEL-112
chr19-44132717-DEL-67
chr19-44139456-DEL-200
chr19-44151443-DEL-216
chr19-44177015-DEL-61
chr19-44193258-DEL-77
chr19-44204390-DEL-283
chr19-44240085-DEL-78
chr19-44254186-DEL-89
chr19-44257474-DEL-88
chr19-44273923-DEL-174
chr19-44279557-DEL-100
chr19-44285202-DEL-53
chr19-44305784-DEL-753
chr19-44305795-DEL-177
chr19-44310677-DEL-445
chr19-44311502-DEL-102
chr19-44336852-DEL-373
chr19-44340874-DEL-121
chr19-44347505-DEL-2090
chr19-44379113-DEL-242
chr19-44395352-DEL-146
chr19-44405408-DEL-779
chr19-44416431-DEL-93
chr19-44427187-DEL-247
chr19-44441902-DEL-188
chr19-44476485-DEL-111
chr19-44477263-DEL-347
chr19-44486454-DEL-347
chr19-44515344-DEL-60
chr19-44515792-DEL-273
chr19-44597899-DEL-113
chr19-44630798-DEL-52
chr19-44648854-DEL-120
chr19-44695658-DEL-63
chr19-44756912-DEL-53
chr19-44790782-DEL-54
chr19-44832219-DEL-78
chr19-44901985-DEL-58
chr19-44904748-DEL-298
chr19-44911498-DEL-2146
chr19-44938313-DEL-411
chr19-44943542-DEL-445
chr19-45006207-DEL-65
chr19-45022541-DEL-216
chr19-45030561-DEL-121
chr19-45051440-DEL-182
chr19-45051993-DEL-170
chr19-45159670-DEL-1392
chr19-45181727-DEL-320
chr19-45189960-DEL-204
chr19-45191302-DEL-1733
chr19-45193158-DEL-876
chr19-45198127-DEL-163
chr19-45243856-DEL-200
chr19-45248408-DEL-405
chr19-45261926-DEL-206

chr19-45286743-DEL-53
chr19-45308082-DEL-108
chr19-45312422-DEL-332
chr19-45328038-DEL-177
chr19-45381280-DEL-56
chr19-45388092-DEL-51
chr19-45403521-DEL-619
chr19-45409713-DEL-101
chr19-45471258-DEL-142
chr19-45545333-DEL-142
chr19-45549726-DEL-223
chr19-45582107-DEL-68
chr19-45597699-DEL-203
chr19-45603147-DEL-66
chr19-45607189-DEL-218
chr19-45627354-DEL-51
chr19-45639497-DEL-110
chr19-45651887-DEL-257
chr19-45658525-DEL-273
chr19-45696210-DEL-278
chr19-45756767-DEL-1080
chr19-45774607-DEL-731
chr19-45778538-DEL-176
chr19-45794460-DEL-150
chr19-45817309-DEL-111
chr19-45833184-DEL-63
chr19-45852888-DEL-146
chr19-45878961-DEL-172
chr19-45892933-DEL-76
chr19-45896191-DEL-485
chr19-45929356-DEL-66
chr19-45956686-DEL-1201
chr19-45973842-DEL-200
chr19-45973844-DEL-113
chr19-45978927-DEL-200
chr19-45984179-DEL-205
chr19-46000357-DEL-204
chr19-46006443-DEL-744
chr19-46007417-DEL-1050
chr19-46010735-DEL-200
chr19-46012705-DEL-189
chr19-46013918-DEL-222
chr19-46040401-DEL-56
chr19-46041315-DEL-369
chr19-46066109-DEL-257
chr19-46075387-DEL-352
chr19-46077938-DEL-149
chr19-46079033-DEL-74
chr19-46084211-DEL-78
chr19-46088295-DEL-164
chr19-46091441-DEL-217
chr19-46095991-DEL-133
chr19-46096556-DEL-197
chr19-46108766-DEL-170
chr19-46117634-DEL-79
chr19-46120063-DEL-54
chr19-46155270-DEL-114
chr19-46192669-DEL-345
chr19-46202985-DEL-63
chr19-46220032-DEL-329
chr19-46228954-DEL-247
chr19-46288488-DEL-63
chr19-46289615-DEL-1136
chr19-46331838-DEL-198
chr19-46375120-DEL-515
chr19-46415982-DEL-150
chr19-46418912-DEL-219
chr19-46427894-DEL-198
chr19-46430876-DEL-177
chr19-46450529-DEL-202
chr19-46452622-DEL-150
chr19-46482686-DEL-498
chr19-46545479-DEL-903
chr19-46548324-DEL-971
chr19-46559955-DEL-213
chr19-46568479-DEL-571
chr19-46569689-DEL-997

chr19-46586972-DEL-106
chr19-46599860-DEL-95
chr19-46615181-DEL-169
chr19-46620948-DEL-66
chr19-46623130-DEL-159
chr19-46665575-DEL-82
chr19-46684990-DEL-570
chr19-46685928-DEL-75
chr19-46701123-DEL-60
chr19-46701634-DEL-312
chr19-46738885-DEL-189
chr19-46740184-DEL-54
chr19-46742331-DEL-143
chr19-46748608-DEL-193
chr19-46761614-DEL-66
chr19-46769258-DEL-897
chr19-46778197-DEL-287
chr19-46799802-DEL-72
chr19-46816223-DEL-66
chr19-46841156-DEL-60
chr19-46856093-DEL-53
chr19-46958710-DEL-51
chr19-46975907-DEL-127
chr19-46985051-DEL-503
chr19-46989065-DEL-208
chr19-46992821-DEL-89
chr19-46995189-DEL-68
chr19-47035681-DEL-144
chr19-47062036-DEL-202
chr19-47090124-DEL-95
chr19-47108929-DEL-62
chr19-47150043-DEL-52
chr19-47162711-DEL-81
chr19-47200729-DEL-2283
chr19-47265568-DEL-54
chr19-47283221-DEL-67
chr19-47296333-DEL-183
chr19-47320527-DEL-178
chr19-47324378-DEL-186
chr19-47357752-DEL-152
chr19-47389986-DEL-734
chr19-47405299-DEL-55
chr19-47408777-DEL-131
chr19-47419645-DEL-68
chr19-47464647-DEL-95
chr19-47470066-DEL-659
chr19-47470855-DEL-60
chr19-47484040-DEL-410
chr19-47484945-DEL-599
chr19-47518408-DEL-151
chr19-47547116-DEL-65
chr19-47574363-DEL-58
chr19-47579782-DEL-11291
chr19-47622455-DEL-819
chr19-47641175-DEL-168
chr19-47643486-DEL-104
chr19-47649631-DEL-75
chr19-47658020-DEL-54
chr19-47708115-DEL-83
chr19-47709323-DEL-147
chr19-47719370-DEL-68
chr19-47731784-DEL-54
chr19-47735647-DEL-88
chr19-47736901-DEL-79
chr19-47782010-DEL-751
chr19-47807229-DEL-113
chr19-47810565-DEL-71
chr19-47814866-DEL-78
chr19-47822837-DEL-178
chr19-47849682-DEL-88
chr19-47923121-DEL-297
chr19-47942313-DEL-56
chr19-47945187-DEL-57
chr19-47968354-DEL-75
chr19-48025495-DEL-52
chr19-48039977-DEL-324
chr19-48042359-DEL-168

chr19-48044351-DEL-878
chr19-48053719-DEL-56
chr19-48082914-DEL-1970
chr19-48102440-DEL-78
chr19-48165991-DEL-405
chr19-48232185-DEL-164
chr19-48234535-DEL-65
chr19-48244721-DEL-137
chr19-48253823-DEL-806
chr19-48256514-DEL-134
chr19-48397452-DEL-112
chr19-48408944-DEL-56
chr19-48419972-DEL-58
chr19-48420853-DEL-140
chr19-48443625-DEL-2480
chr19-48453149-DEL-120
chr19-48458204-DEL-59
chr19-48465347-DEL-95
chr19-48473062-DEL-969
chr19-48480147-DEL-337
chr19-48507069-DEL-547
chr19-48537362-DEL-99
chr19-48543170-DEL-97
chr19-48545733-DEL-164
chr19-48546123-DEL-195
chr19-48556887-DEL-53
chr19-48659251-DEL-122
chr19-48682996-DEL-63
chr19-48684264-DEL-72
chr19-48713880-DEL-54
chr19-48748830-DEL-972
chr19-48765145-DEL-51
chr19-48820200-DEL-56
chr19-48821288-DEL-80
chr19-48869048-DEL-303
chr19-48899473-DEL-61
chr19-48929614-DEL-1146
chr19-48950257-DEL-607
chr19-48992589-DEL-74
chr19-48998123-DEL-113
chr19-49011732-DEL-242
chr19-49024754-DEL-383
chr19-49032119-DEL-572
chr19-49063538-DEL-120
chr19-49070785-DEL-56
chr19-49082870-DEL-208
chr19-49087429-DEL-55
chr19-49090232-DEL-85
chr19-49206151-DEL-588
chr19-49215262-DEL-121
chr19-49217888-DEL-133
chr19-49224412-DEL-8401
chr19-49306768-DEL-141
chr19-49323255-DEL-13427
chr19-49329263-DEL-212
chr19-49370722-DEL-2215
chr19-49384078-DEL-56
chr19-49434289-DEL-257
chr19-49484844-DEL-137
chr19-49490589-DEL-219
chr19-49518686-DEL-676
chr19-49543476-DEL-1134
chr19-49548483-DEL-142
chr19-49554438-DEL-300
chr19-49583775-DEL-63
chr19-49593274-DEL-332
chr19-49599575-DEL-377
chr19-49604544-DEL-409
chr19-49621413-DEL-101
chr19-49651696-DEL-2830
chr19-49656362-DEL-181
chr19-49680605-DEL-52
chr19-49697425-DEL-57
chr19-49702693-DEL-66
chr19-49702969-DEL-755
chr19-49710594-DEL-65
chr19-49718183-DEL-86

chr19-49754713-DEL-132
chr19-49793735-DEL-128
chr19-49811905-DEL-900
chr19-49876039-DEL-240
chr19-49879050-DEL-57
chr19-49892692-DEL-51
chr19-49906475-DEL-74
chr19-49913848-DEL-72
chr19-49922019-DEL-207
chr19-49927040-DEL-65
chr19-49929761-DEL-550
chr19-49935077-DEL-99
chr19-49940479-DEL-356
chr19-49945011-DEL-131
chr19-49965144-DEL-55
chr19-49966344-DEL-433
chr19-49968658-DEL-95
chr19-49991077-DEL-80
chr19-50017080-DEL-70
chr19-50021096-DEL-78
chr19-50036079-DEL-85
chr19-50039235-DEL-98
chr19-50145625-DEL-98
chr19-50172991-DEL-134
chr19-50175552-DEL-92
chr19-50182646-DEL-60
chr19-50188625-DEL-342
chr19-50200565-DEL-52
chr19-50239867-DEL-340
chr19-50249132-DEL-154
chr19-50251817-DEL-63
chr19-50265236-DEL-54
chr19-50282473-DEL-62
chr19-50302063-DEL-608
chr19-50312321-DEL-180
chr19-50334655-DEL-436
chr19-50355404-DEL-179
chr19-50366240-DEL-56
chr19-50395035-DEL-101
chr19-50450629-DEL-62
chr19-50462740-DEL-303
chr19-50468853-DEL-196
chr19-50484623-DEL-83
chr19-50488773-DEL-175
chr19-50511272-DEL-72
chr19-50602814-DEL-1413
chr19-50621749-DEL-84
chr19-50678498-DEL-52
chr19-50688566-DEL-1453
chr19-50741934-DEL-193
chr19-50757714-DEL-339
chr19-50767675-DEL-463
chr19-50776235-DEL-86
chr19-50776926-DEL-68
chr19-50781815-DEL-1409
chr19-50815518-DEL-86
chr19-50824873-DEL-98
chr19-50836144-DEL-65
chr19-50890700-DEL-250
chr19-50949380-DEL-2323
chr19-50986427-DEL-54
chr19-51063083-DEL-217
chr19-51069124-DEL-149
chr19-51086166-DEL-142
chr19-51087157-DEL-70
chr19-51103949-DEL-427
chr19-51119611-DEL-55
chr19-51139775-DEL-181
chr19-51144225-DEL-2364
chr19-51151077-DEL-384
chr19-51153941-DEL-263
chr19-51157761-DEL-97
chr19-51158355-DEL-52
chr19-51161794-DEL-1765
chr19-51167241-DEL-61
chr19-51175743-DEL-290
chr19-51185375-DEL-1995

chr19-51220303-DEL-66
chr19-51236639-DEL-3000
chr19-51283405-DEL-1281
chr19-51288895-DEL-280
chr19-51290639-DEL-146
chr19-51301263-DEL-1093
chr19-51353499-DEL-94
chr19-51368502-DEL-116
chr19-51400865-DEL-77
chr19-51409127-DEL-99
chr19-51427451-DEL-72
chr19-51448300-DEL-6363
chr19-51458004-DEL-6454
chr19-51464341-DEL-335
chr19-51470154-DEL-141
chr19-51484120-DEL-163
chr19-51496997-DEL-138
chr19-51514329-DEL-687
chr19-51565699-DEL-236
chr19-51591585-DEL-398
chr19-51610478-DEL-147
chr19-51636681-DEL-9338
chr19-51674843-DEL-88
chr19-51684035-DEL-6107
chr19-51727015-DEL-299
chr19-51757304-DEL-180
chr19-51791535-DEL-1534
chr19-51830395-DEL-2422
chr19-51833583-DEL-147
chr19-51887532-DEL-352
chr19-51902286-DEL-78
chr19-51940411-DEL-2954
chr19-51950525-DEL-56
chr19-51954883-DEL-52
chr19-51975143-DEL-403
chr19-51975427-DEL-805
chr19-52013719-DEL-73
chr19-52023471-DEL-222
chr19-52046375-DEL-68
chr19-52053527-DEL-92
chr19-52055392-DEL-64
chr19-52066097-DEL-117
chr19-52077465-DEL-1503
chr19-52082606-DEL-92
chr19-52097830-DEL-408
chr19-52119516-DEL-898
chr19-52153163-DEL-435
chr19-52168579-DEL-217
chr19-52193422-DEL-53
chr19-52196690-DEL-500
chr19-52207438-DEL-185
chr19-52211090-DEL-102
chr19-52218876-DEL-1411
chr19-52231678-DEL-176
chr19-52232106-DEL-4117
chr19-52312832-DEL-114
chr19-52348759-DEL-1140
chr19-52350472-DEL-639
chr19-52376650-DEL-131
chr19-52380905-DEL-60
chr19-52393198-DEL-86
chr19-52398954-DEL-64
chr19-52413003-DEL-270
chr19-52415805-DEL-205
chr19-52450866-DEL-1306
chr19-52459869-DEL-155
chr19-52485183-DEL-1428
chr19-52501169-DEL-2945
chr19-52504710-DEL-106
chr19-52510856-DEL-69
chr19-52530100-DEL-62
chr19-52533988-DEL-555
chr19-52685751-DEL-87
chr19-52709023-DEL-415
chr19-52823280-DEL-158
chr19-52844314-DEL-268
chr19-52859597-DEL-77

chr19-52865384-DEL-56
chr19-52891399-DEL-80
chr19-52904056-DEL-152
chr19-52956368-DEL-1489
chr19-52968261-DEL-190
chr19-52971781-DEL-160
chr19-53022583-DEL-763
chr19-53028183-DEL-66
chr19-53048850-DEL-62
chr19-53084612-DEL-76
chr19-53105158-DEL-448
chr19-53139882-DEL-62
chr19-53203744-DEL-267
chr19-53214248-DEL-167
chr19-53254743-DEL-404
chr19-53256741-DEL-206
chr19-53260584-DEL-76
chr19-53271431-DEL-452
chr19-53289325-DEL-332
chr19-53308089-DEL-54
chr19-53311615-DEL-192
chr19-53315366-DEL-306
chr19-53317364-DEL-76
chr19-53350850-DEL-67
chr19-53414856-DEL-52
chr19-53486157-DEL-113
chr19-53511231-DEL-3529
chr19-53512540-DEL-81
chr19-53513468-DEL-1463
chr19-53513850-DEL-91
chr19-53514752-DEL-208
chr19-53528282-DEL-257
chr19-53551890-DEL-730
chr19-53571968-DEL-204
chr19-53573157-DEL-409
chr19-53575528-DEL-289
chr19-53581608-DEL-100
chr19-53582510-DEL-5483
chr19-53587569-DEL-425
chr19-53592937-DEL-1474
chr19-53620570-DEL-196
chr19-53621096-DEL-135
chr19-53623712-DEL-52
chr19-53631094-DEL-424
chr19-53651474-DEL-97
chr19-53652250-DEL-97
chr19-53657760-DEL-64
chr19-53677638-DEL-929
chr19-53681228-DEL-125
chr19-53681833-DEL-3439
chr19-53706534-DEL-72
chr19-53729222-DEL-163
chr19-53763911-DEL-2069
chr19-53763913-DEL-90
chr19-53765298-DEL-315
chr19-53770482-DEL-148
chr19-53795222-DEL-116
chr19-53804296-DEL-126
chr19-53810985-DEL-146
chr19-53826445-DEL-419
chr19-53856204-DEL-90
chr19-53868206-DEL-139
chr19-53891118-DEL-85
chr19-53920797-DEL-173
chr19-53947737-DEL-166
chr19-53959943-DEL-59
chr19-53962902-DEL-128
chr19-53967575-DEL-1431
chr19-53980494-DEL-52
chr19-53991933-DEL-220
chr19-54007576-DEL-208
chr19-54010475-DEL-68
chr19-54056507-DEL-78
chr19-54058596-DEL-76
chr19-54078522-DEL-52
chr19-54114148-DEL-129
chr19-54123090-DEL-55

chr19-54124804-DEL-6199
chr19-54136005-DEL-2843
chr19-54136985-DEL-464
chr19-54139095-DEL-549
chr19-54140041-DEL-88
chr19-54140341-DEL-130
chr19-54143540-DEL-99
chr19-54174133-DEL-140
chr19-54185029-DEL-548
chr19-54195597-DEL-84
chr19-54238929-DEL-126
chr19-54263052-DEL-240
chr19-54297851-DEL-163
chr19-54312653-DEL-60
chr19-54324554-DEL-2001
chr19-54342392-DEL-1864
chr19-54347788-DEL-54
chr19-54354762-DEL-1828
chr19-54378296-DEL-152
chr19-54470688-DEL-116
chr19-54476555-DEL-87
chr19-54477044-DEL-188
chr19-54490853-DEL-84
chr19-54546259-DEL-56
chr19-54602584-DEL-983
chr19-54608301-DEL-452
chr19-54641658-DEL-146
chr19-54673683-DEL-302
chr19-54694664-DEL-60
chr19-54700075-DEL-59
chr19-54706682-DEL-150
chr19-54712088-DEL-102
chr19-54721439-DEL-450
chr19-54724223-DEL-261
chr19-54727515-DEL-1587
chr19-54734715-DEL-401
chr19-54757071-DEL-64
chr19-54760024-DEL-134
chr19-54845393-DEL-385
chr19-54849223-DEL-68
chr19-54864285-DEL-400
chr19-54898327-DEL-80
chr19-54944437-DEL-72
chr19-54967810-DEL-94
chr19-55002376-DEL-2810
chr19-55006496-DEL-52
chr19-55020452-DEL-206
chr19-55029611-DEL-69
chr19-55040313-DEL-90
chr19-55042504-DEL-202
chr19-55044923-DEL-51
chr19-55051534-DEL-80
chr19-55068296-DEL-53
chr19-55093445-DEL-144
chr19-55099912-DEL-168
chr19-55118779-DEL-199
chr19-55120920-DEL-608
chr19-55124102-DEL-75
chr19-55126542-DEL-82
chr19-55131437-DEL-311
chr19-55151119-DEL-61
chr19-55153226-DEL-340
chr19-55230945-DEL-124
chr19-55251849-DEL-62
chr19-55290437-DEL-54
chr19-55357003-DEL-252
chr19-55361872-DEL-583
chr19-55451488-DEL-123
chr19-55462534-DEL-108
chr19-55488081-DEL-223
chr19-55498528-DEL-110
chr19-55535187-DEL-94
chr19-55553769-DEL-144
chr19-55594126-DEL-54
chr19-55616594-DEL-57
chr19-55623178-DEL-66
chr19-55624542-DEL-104

chr19-55643994-DEL-84
chr19-55671145-DEL-56
chr19-55673507-DEL-75
chr19-55674266-DEL-60
chr19-55675981-DEL-261
chr19-55684293-DEL-145
chr19-55717014-DEL-55
chr19-55719707-DEL-160
chr19-55758960-DEL-384
chr19-55775993-DEL-64
chr19-55799647-DEL-988
chr19-55809347-DEL-154
chr19-55810939-DEL-74
chr19-55831417-DEL-126
chr19-55834447-DEL-80
chr19-55841754-DEL-59
chr19-55871126-DEL-114
chr19-55874726-DEL-59
chr19-55906361-DEL-174
chr19-55906927-DEL-210
chr19-55909635-DEL-121
chr19-55911231-DEL-477
chr19-56004210-DEL-131
chr19-56026611-DEL-101
chr19-56077825-DEL-55
chr19-56094560-DEL-93
chr19-56102849-DEL-52
chr19-56131691-DEL-136
chr19-56141196-DEL-919
chr19-56151909-DEL-1486
chr19-56161851-DEL-59
chr19-56190225-DEL-241
chr19-56203214-DEL-546
chr19-56264534-DEL-134
chr19-56272650-DEL-628
chr19-56311166-DEL-168
chr19-56317680-DEL-81
chr19-56339450-DEL-316
chr19-56344752-DEL-110
chr19-56366046-DEL-170
chr19-56367865-DEL-116
chr19-56373329-DEL-55
chr19-56398750-DEL-6276
chr19-56415858-DEL-135
chr19-56436160-DEL-535
chr19-56463090-DEL-52
chr19-56477412-DEL-447
chr19-56511774-DEL-123
chr19-56544069-DEL-1028
chr19-56557828-DEL-66
chr19-56561295-DEL-68
chr19-56590327-DEL-106
chr19-56594899-DEL-197
chr19-56731983-DEL-88
chr19-56737036-DEL-71
chr19-56746954-DEL-122
chr19-56767126-DEL-120
chr19-56782252-DEL-51
chr19-56785581-DEL-68
chr19-56834416-DEL-61
chr19-56836004-DEL-631
chr19-56883160-DEL-134
chr19-56885159-DEL-144
chr19-56940867-DEL-72
chr19-56960718-DEL-264
chr19-56967601-DEL-86
chr19-56968626-DEL-147
chr19-56981295-DEL-68
chr19-57059106-DEL-104
chr19-57067158-DEL-76
chr19-57104464-DEL-102
chr19-57109787-DEL-134
chr19-57121470-DEL-268
chr19-57128009-DEL-85
chr19-57131408-DEL-203
chr19-57147785-DEL-54
chr19-57181603-DEL-68

chr19-57220484-DEL-120
chr19-57247558-DEL-75
chr19-57288688-DEL-98
chr19-57313474-DEL-54
chr19-57315303-DEL-81
chr19-57334021-DEL-476
chr19-57334906-DEL-76
chr19-57336568-DEL-55
chr19-57345170-DEL-150
chr19-57352491-DEL-94
chr19-57374134-DEL-58
chr19-57383475-DEL-76
chr19-57389045-DEL-220
chr19-57398157-DEL-96
chr19-57399625-DEL-151
chr19-57405290-DEL-73
chr19-57422676-DEL-1278
chr19-57437324-DEL-108
chr19-57437623-DEL-163
chr19-57439733-DEL-65
chr19-57441067-DEL-143
chr19-57448498-DEL-115
chr19-57509680-DEL-94
chr19-57566651-DEL-117
chr19-57583720-DEL-610
chr19-57585747-DEL-68
chr19-57595105-DEL-105
chr19-57597831-DEL-73
chr19-57613318-DEL-239
chr19-57630557-DEL-62
chr19-57631143-DEL-63
chr19-57659319-DEL-240
chr19-57659491-DEL-742
chr19-57660140-DEL-260
chr19-57668946-DEL-118
chr19-57697509-DEL-169
chr19-57698035-DEL-68
chr19-57703729-DEL-64
chr19-57727269-DEL-153
chr19-57730377-DEL-283
chr19-57747148-DEL-1460
chr19-57747515-DEL-120
chr19-57760140-DEL-60
chr19-57765572-DEL-711
chr19-57792650-DEL-78
chr19-57802437-DEL-77
chr19-57804545-DEL-136
chr19-57828676-DEL-56
chr19-57847483-DEL-55
chr19-57848422-DEL-79
chr19-57855617-DEL-139
chr19-57868678-DEL-131
chr19-57877281-DEL-173
chr19-57917798-DEL-125
chr19-57925963-DEL-66
chr19-57942642-DEL-62
chr19-57968884-DEL-222
chr19-57972450-DEL-87
chr19-57982951-DEL-98
chr19-58000240-DEL-65
chr19-58003832-DEL-562
chr19-58006472-DEL-78
chr19-58009148-DEL-72
chr19-58037631-DEL-91
chr19-58059505-DEL-65
chr19-58092676-DEL-391
chr19-58101635-DEL-492
chr19-58111031-DEL-62
chr19-58122317-DEL-51
chr19-58131924-DEL-86
chr19-58132825-DEL-72
chr19-58148851-DEL-262
chr19-58187148-DEL-52
chr19-58200475-DEL-52
chr19-58210141-DEL-807
chr19-58220789-DEL-51
chr19-58228662-DEL-64

chr19-58263679-DEL-55
chr19-58271830-DEL-532
chr19-58272680-DEL-82
chr19-58280446-DEL-63
chr19-58281704-DEL-108
chr19-58297416-DEL-80
chr19-58321421-DEL-67
chr19-58334623-DEL-454
chr19-58423616-DEL-58
chr19-58479088-DEL-52
chr19-58483398-DEL-85
chr19-58489923-DEL-51
chr19-58526300-DEL-575
chr19-58529349-DEL-224
chr19-58534081-DEL-55
chr19-58540671-DEL-62
chr19-58574353-DEL-1135
chr19-58601874-DEL-52
chr19-58643991-DEL-52
chr19-58662557-DEL-163
chr19-58664141-DEL-65
chr19-58694140-DEL-926
chr19-58711989-DEL-193
chr19-58727663-DEL-431
chr19-58739461-DEL-228
chr19-58742499-DEL-833
chr19-58760407-DEL-160
chr19-58776413-DEL-374
chr19-58778636-DEL-2044
chr19-58809038-DEL-78
chr19-58821303-DEL-87
chr19-58870801-DEL-364
chr19-58884482-DEL-130
chr19-58938376-DEL-394
chr19-58939212-DEL-146
chr19-58948825-DEL-60
chr19-58954155-DEL-1257
chr19-58955693-DEL-136
chr19-58967281-DEL-63
chr19-59034150-DEL-51
chr19-59039892-DEL-93
chr19-59070875-DEL-82
chr19-59072001-DEL-100
chr19-59072840-DEL-148
chr19-59096541-DEL-110
chr19-59129772-DEL-116
chr19-59130423-DEL-69
chr19-59136355-DEL-82
chr19-59137156-DEL-72
chr19-59140507-DEL-355
chr19-59144399-DEL-717
chr19-59167122-DEL-71
chr19-59168740-DEL-181
chr19-59170131-DEL-56
chr19-59171658-DEL-199
chr19-59188609-DEL-768
chr19-59201638-DEL-828
chr19-59224417-DEL-79
chr19-59238927-DEL-78
chr19-59326432-DEL-57
chr19-59381799-DEL-69
chr19-59404647-DEL-94
chr19-59432256-DEL-127
chr19-59434285-DEL-58
chr19-59472008-DEL-959
chr19-59486236-DEL-61
chr19-59519526-DEL-294
chr19-59537359-DEL-68
chr19-59541184-DEL-75
chr19-59544457-DEL-100
chr19-59546898-DEL-52
chr19-59560378-DEL-79
chr19-59575110-DEL-51
chr19-59591865-DEL-66
chr19-59599509-DEL-644
chr19-59621534-DEL-379
chr19-59653411-DEL-183

chr19-59655389-DEL-158
chr19-59660140-DEL-456
chr19-59661928-DEL-68
chr19-59672054-DEL-81
chr19-59703680-DEL-798
chr19-59724367-DEL-592
chr19-59801889-DEL-195
chr19-59846709-DEL-1120
chr19-59869933-DEL-330
chr19-59873562-DEL-82
chr19-59897658-DEL-405
chr19-59972520-DEL-73
chr19-59989470-DEL-78
chr19-60072232-DEL-6074
chr19-60075683-DEL-5410
chr19-60091938-DEL-103
chr19-60190672-DEL-61
chr19-60224011-DEL-648
chr19-60259119-DEL-2287
chr19-60264649-DEL-176
chr19-60266679-DEL-3924
chr19-60305148-DEL-159
chr19-60325317-DEL-201
chr19-60334869-DEL-51
chr19-60343779-DEL-989
chr19-60355268-DEL-69
chr19-60355376-DEL-2093
chr19-60398912-DEL-104
chr19-60470472-DEL-68
chr19-60475955-DEL-52
chr19-60488893-DEL-631
chr19-60493274-DEL-517
chr19-60505008-DEL-74
chr19-60506507-DEL-505
chr19-60508665-DEL-52
chr19-60522286-DEL-223
chr19-60552321-DEL-140
chr19-60644184-DEL-56
chr19-60671884-DEL-112
chr19-60714887-DEL-350
chr19-60721458-DEL-51
chr19-60734930-DEL-57
chr19-60745537-DEL-132
chr19-60784499-DEL-389
chr19-60794600-DEL-56
chr19-60815767-DEL-134
chr19-60841429-DEL-134
chr19-60851447-DEL-56
chr19-60854535-DEL-1207
chr19-60856549-DEL-241
chr19-60864402-DEL-220
chr19-60870552-DEL-182
chr19-60913102-DEL-64
chr19-60931591-DEL-706
chr19-60936558-DEL-139
chr19-60943734-DEL-61
chr19-61014550-DEL-66
chr19-61054530-DEL-74
chr19-61088024-DEL-222
chr19-61170701-DEL-142
chr19-61176975-DEL-198
chr19-61193893-DEL-1487
chr19-61227318-DEL-4027
chr19-61236903-DEL-3245
chr19-61256150-DEL-974
chr19-61259405-DEL-207
chr19-61260318-DEL-9963
chr2-3070644-DEL-5808
chr2-3085678-DEL-54
chr2-3138804-DEL-53
chr2-3153056-DEL-344
chr2-3165494-DEL-82
chr2-3189454-DEL-970
chr2-3196899-DEL-133
chr2-3223168-DEL-56
chr2-3233997-DEL-74
chr2-3241370-DEL-191

chr2-3271814-DEL-1555
chr2-3288945-DEL-187
chr2-3339065-DEL-99
chr2-3350990-DEL-513
chr2-3380355-DEL-81
chr2-3398216-DEL-408
chr2-3433837-DEL-908
chr2-3442066-DEL-81
chr2-3446507-DEL-416
chr2-3453753-DEL-83
chr2-3477446-DEL-1198
chr2-3529619-DEL-68
chr2-3530113-DEL-312
chr2-3530391-DEL-129
chr2-3544534-DEL-60
chr2-3551071-DEL-8413
chr2-3593668-DEL-119
chr2-3597159-DEL-51
chr2-3612345-DEL-1145
chr2-3620158-DEL-1364
chr2-3626167-DEL-252
chr2-3627118-DEL-221
chr2-3635138-DEL-304
chr2-3636597-DEL-372
chr2-3674686-DEL-199
chr2-3685551-DEL-445
chr2-3686681-DEL-141
chr2-3687244-DEL-174
chr2-3704219-DEL-147
chr2-3734675-DEL-160
chr2-3743033-DEL-166
chr2-3768204-DEL-53
chr2-3775500-DEL-78
chr2-3785018-DEL-192
chr2-3786263-DEL-240
chr2-3789710-DEL-1637
chr2-3793599-DEL-64
chr2-3798665-DEL-90
chr2-3799871-DEL-1178
chr2-3807655-DEL-203
chr2-3835197-DEL-318
chr2-3835215-DEL-4772
chr2-3836248-DEL-1092
chr2-3852035-DEL-77
chr2-3875335-DEL-63
chr2-3904478-DEL-115
chr2-3909841-DEL-53
chr2-3913869-DEL-2178
chr2-3921531-DEL-183
chr2-3943116-DEL-199
chr2-3958817-DEL-74
chr2-3966269-DEL-2767
chr2-3967831-DEL-376
chr2-3987049-DEL-115
chr2-3991556-DEL-913
chr2-3997467-DEL-214
chr2-4002958-DEL-431
chr2-4043947-DEL-183
chr2-4050523-DEL-286
chr2-4141716-DEL-70
chr2-4149114-DEL-77
chr2-4184627-DEL-592
chr2-4214578-DEL-76
chr2-4224854-DEL-114
chr2-4255570-DEL-80
chr2-4321438-DEL-60
chr2-4323696-DEL-108
chr2-4378597-DEL-67
chr2-4388789-DEL-197
chr2-4406554-DEL-53
chr2-4498707-DEL-240
chr2-4529151-DEL-73
chr2-4544410-DEL-91
chr2-4569263-DEL-102
chr2-4600430-DEL-196
chr2-4602857-DEL-115
chr2-4619209-DEL-166

chr2-4623446-DEL-1081
chr2-4623526-DEL-452
chr2-4627844-DEL-91
chr2-4630716-DEL-1961
chr2-4646929-DEL-616
chr2-4658320-DEL-90
chr2-4668259-DEL-122
chr2-4670673-DEL-168
chr2-4683225-DEL-68
chr2-4705851-DEL-405
chr2-4728781-DEL-158
chr2-4738026-DEL-114
chr2-4749738-DEL-52
chr2-4749842-DEL-936
chr2-4821182-DEL-588
chr2-4821923-DEL-169
chr2-4847135-DEL-291
chr2-4849989-DEL-60
chr2-4866540-DEL-60
chr2-4872061-DEL-58
chr2-4877723-DEL-156
chr2-4896731-DEL-199
chr2-4949457-DEL-5562
chr2-4966030-DEL-195
chr2-4966882-DEL-137
chr2-4982047-DEL-203
chr2-4984998-DEL-203
chr2-4986971-DEL-93
chr2-5014577-DEL-525
chr2-5030045-DEL-51
chr2-5043648-DEL-211
chr2-5060131-DEL-156
chr2-5063928-DEL-54
chr2-5080994-DEL-52
chr2-5088446-DEL-76
chr2-5097461-DEL-57
chr2-5098525-DEL-158
chr2-5124252-DEL-124
chr2-5135868-DEL-54
chr2-5144035-DEL-51
chr2-5149868-DEL-149
chr2-5150470-DEL-472
chr2-5163480-DEL-173
chr2-5211864-DEL-877
chr2-5220046-DEL-176
chr2-5230729-DEL-682
chr2-5243747-DEL-70
chr2-5248862-DEL-70
chr2-5261339-DEL-400
chr2-5298128-DEL-58
chr2-5323489-DEL-172
chr2-5340398-DEL-901
chr2-5344395-DEL-160
chr2-5354404-DEL-56
chr2-5355914-DEL-68
chr2-5376853-DEL-63
chr2-5381030-DEL-480
chr2-5407054-DEL-144
chr2-5423671-DEL-288
chr2-5425914-DEL-108
chr2-5437105-DEL-53
chr2-5458572-DEL-66
chr2-5481197-DEL-102
chr2-5487027-DEL-96
chr2-5495849-DEL-134
chr2-5496748-DEL-187
chr2-5503858-DEL-83
chr2-5522149-DEL-313
chr2-5539224-DEL-756
chr2-5540041-DEL-94
chr2-5547046-DEL-420
chr2-5554115-DEL-79
chr2-5572040-DEL-272
chr2-5574753-DEL-77
chr2-5606110-DEL-110
chr2-5621176-DEL-51
chr2-5638672-DEL-209

chr2-5643830-DEL-235
chr2-5648881-DEL-107
chr2-5650880-DEL-81
chr2-5657971-DEL-238
chr2-5660503-DEL-134
chr2-5682190-DEL-63
chr2-5710576-DEL-54
chr2-5723292-DEL-93
chr2-5742272-DEL-56
chr2-5743478-DEL-59
chr2-5760910-DEL-86
chr2-5817611-DEL-107
chr2-5842340-DEL-200
chr2-5846388-DEL-104
chr2-5856437-DEL-147
chr2-5861177-DEL-205
chr2-5891223-DEL-54
chr2-5905839-DEL-254
chr2-5920007-DEL-399
chr2-5929250-DEL-1041
chr2-5958394-DEL-182
chr2-5984046-DEL-87
chr2-6004119-DEL-201
chr2-6037565-DEL-82
chr2-6043335-DEL-334
chr2-6058754-DEL-204
chr2-6063307-DEL-888
chr2-6066781-DEL-157
chr2-6083292-DEL-1025
chr2-6088351-DEL-111
chr2-6108432-DEL-62
chr2-6109454-DEL-201
chr2-6126302-DEL-330
chr2-6151828-DEL-75
chr2-6160389-DEL-66
chr2-6163739-DEL-244
chr2-6184113-DEL-272
chr2-6255447-DEL-77
chr2-6275390-DEL-345
chr2-6279932-DEL-74
chr2-6342522-DEL-59
chr2-6368283-DEL-527
chr2-6370192-DEL-392
chr2-6385988-DEL-195
chr2-6438636-DEL-126
chr2-6443081-DEL-250
chr2-6470539-DEL-426
chr2-6494096-DEL-155
chr2-6500391-DEL-196
chr2-6503059-DEL-340
chr2-6503704-DEL-152
chr2-6532095-DEL-66
chr2-6571173-DEL-62
chr2-6598258-DEL-74
chr2-6605267-DEL-224
chr2-6632737-DEL-218
chr2-6753111-DEL-212
chr2-6777871-DEL-4408
chr2-6780655-DEL-95
chr2-6802527-DEL-3265
chr2-6818008-DEL-136
chr2-6878947-DEL-60
chr2-6895904-DEL-807
chr2-6992451-DEL-145
chr2-7035421-DEL-68
chr2-7109181-DEL-67
chr2-7130455-DEL-57
chr2-7149192-DEL-188
chr2-7208381-DEL-771
chr2-7245021-DEL-167
chr2-7278971-DEL-3204
chr2-7309143-DEL-57
chr2-7335645-DEL-254
chr2-7341942-DEL-57
chr2-7369885-DEL-262
chr2-7390752-DEL-502
chr2-7391355-DEL-154

chr2-7419895-DEL-6445
chr2-7435881-DEL-3625
chr2-7441555-DEL-70
chr2-7446920-DEL-1809
chr2-7449041-DEL-4545
chr2-7463272-DEL-114
chr2-7555414-DEL-66
chr2-7563516-DEL-82
chr2-7568024-DEL-699
chr2-7576515-DEL-70
chr2-7576620-DEL-6197
chr2-7585353-DEL-6637
chr2-7593700-DEL-12373
chr2-7598759-DEL-6072
chr2-7605836-DEL-201
chr2-7612968-DEL-2289
chr2-7615090-DEL-143
chr2-7629233-DEL-102
chr2-7653731-DEL-206
chr2-7692154-DEL-73
chr2-7756933-DEL-5246
chr2-7757340-DEL-6204
chr2-7762643-DEL-958
chr2-7763725-DEL-63
chr2-7770774-DEL-131
chr2-7780525-DEL-199
chr2-7784184-DEL-84
chr2-7799941-DEL-98
chr2-7826938-DEL-80
chr2-7829354-DEL-58
chr2-7852607-DEL-82
chr2-7876679-DEL-136
chr2-7882965-DEL-201
chr2-7896627-DEL-56
chr2-7935401-DEL-1040
chr2-7936997-DEL-1059
chr2-7956713-DEL-216
chr2-7979222-DEL-69
chr2-8007543-DEL-1065
chr2-8009773-DEL-6261
chr2-8018855-DEL-168
chr2-8025159-DEL-347
chr2-8025590-DEL-432
chr2-8039341-DEL-10403
chr2-8179626-DEL-54
chr2-8187726-DEL-16005
chr2-8206358-DEL-71
chr2-8207838-DEL-1400
chr2-8222986-DEL-2064
chr2-8274234-DEL-6188
chr2-8285837-DEL-90
chr2-8333737-DEL-132
chr2-8356735-DEL-226
chr2-8378826-DEL-312
chr2-8380605-DEL-2802
chr2-8399385-DEL-78
chr2-8409002-DEL-1086
chr2-8424033-DEL-54
chr2-8439647-DEL-60
chr2-8454597-DEL-503
chr2-8464806-DEL-87
chr2-8493837-DEL-2535
chr2-8498477-DEL-124
chr2-8528995-DEL-57
chr2-8545381-DEL-51
chr2-8571240-DEL-89
chr2-8613651-DEL-53
chr2-8618282-DEL-197
chr2-8626839-DEL-762
chr2-8639163-DEL-1652
chr2-8643894-DEL-184
chr2-8664572-DEL-238
chr2-8700020-DEL-56
chr2-8706391-DEL-75
chr2-8721494-DEL-137
chr2-8734681-DEL-59
chr2-8789705-DEL-57

chr2-8790846-DEL-717
chr2-8823871-DEL-2764
chr2-8833206-DEL-315
chr2-8833729-DEL-52
chr2-8837193-DEL-1156
chr2-8849727-DEL-444
chr2-8907374-DEL-205
chr2-8940987-DEL-112
chr2-8990682-DEL-76
chr2-8992764-DEL-76
chr2-8993428-DEL-532
chr2-9004134-DEL-356
chr2-9024438-DEL-643
chr2-9025648-DEL-89
chr2-9029232-DEL-3033
chr2-9031285-DEL-75
chr2-9072554-DEL-123
chr2-9084308-DEL-1488
chr2-9113764-DEL-249
chr2-9120288-DEL-130
chr2-9167952-DEL-142
chr2-9201115-DEL-218
chr2-9215073-DEL-63
chr2-9217565-DEL-212
chr2-9228603-DEL-96
chr2-9239803-DEL-2625
chr2-9285622-DEL-150
chr2-9299526-DEL-2218
chr2-9302314-DEL-60
chr2-9409109-DEL-170
chr2-9409918-DEL-76
chr2-9470473-DEL-82
chr2-9538185-DEL-172
chr2-9583431-DEL-66
chr2-9641839-DEL-64
chr2-9665958-DEL-99
chr2-9689238-DEL-66
chr2-9696593-DEL-190
chr2-9778400-DEL-260
chr2-9780284-DEL-857
chr2-9782065-DEL-58
chr2-9784285-DEL-2198
chr2-9802252-DEL-358
chr2-9811912-DEL-164
chr2-9817153-DEL-412
chr2-9823281-DEL-406
chr2-9850301-DEL-72
chr2-9868805-DEL-55
chr2-9896379-DEL-232
chr2-9909751-DEL-84
chr2-9963804-DEL-483
chr2-9997456-DEL-114
chr2-10007924-DEL-209
chr2-10023413-DEL-51
chr2-10031568-DEL-80
chr2-10035324-DEL-122
chr2-10049330-DEL-78
chr2-10078875-DEL-1531
chr2-10087039-DEL-201
chr2-10089627-DEL-140
chr2-10141967-DEL-2413
chr2-10176650-DEL-98
chr2-10206028-DEL-51
chr2-10219190-DEL-1265
chr2-10234875-DEL-112
chr2-10252901-DEL-296
chr2-10273998-DEL-366
chr2-10286954-DEL-178
chr2-10314318-DEL-65
chr2-10342424-DEL-107
chr2-10363941-DEL-68
chr2-10373445-DEL-60
chr2-10381495-DEL-56
chr2-10392792-DEL-674
chr2-10400418-DEL-646
chr2-10405283-DEL-52
chr2-10411517-DEL-130

chr2-10417964-DEL-2319
chr2-10423090-DEL-52
chr2-10466970-DEL-102
chr2-10487302-DEL-2439
chr2-10488841-DEL-924
chr2-10490816-DEL-2465
chr2-10492084-DEL-2464
chr2-10494105-DEL-1636
chr2-10499015-DEL-1657
chr2-10501486-DEL-1600
chr2-10503905-DEL-1616
chr2-10506326-DEL-1639
chr2-10517939-DEL-58
chr2-10534875-DEL-56
chr2-10541722-DEL-54
chr2-10558887-DEL-623
chr2-10565628-DEL-59
chr2-10576293-DEL-60
chr2-10640799-DEL-351
chr2-10648358-DEL-1512
chr2-10653367-DEL-272
chr2-10661227-DEL-66
chr2-10683782-DEL-2355
chr2-10708116-DEL-606
chr2-10737983-DEL-121
chr2-10738710-DEL-393
chr2-10765480-DEL-64
chr2-10776299-DEL-54
chr2-10787143-DEL-160
chr2-10796724-DEL-466
chr2-10799416-DEL-56
chr2-10800153-DEL-161
chr2-10811644-DEL-219
chr2-10814687-DEL-100
chr2-10820571-DEL-65
chr2-10829032-DEL-273
chr2-10835415-DEL-56
chr2-10846768-DEL-115
chr2-10860573-DEL-266
chr2-10872032-DEL-929
chr2-10872962-DEL-297
chr2-10883328-DEL-52
chr2-10909743-DEL-6200
chr2-10916072-DEL-112
chr2-10917915-DEL-2700
chr2-10925334-DEL-55
chr2-10957763-DEL-449
chr2-10973281-DEL-123
chr2-11008193-DEL-215
chr2-11038537-DEL-77
chr2-11053908-DEL-86
chr2-11067252-DEL-60
chr2-11079356-DEL-688
chr2-11101321-DEL-1055
chr2-11102772-DEL-2410
chr2-11102819-DEL-572
chr2-11115295-DEL-58
chr2-11121806-DEL-60
chr2-11126470-DEL-636
chr2-11144215-DEL-273
chr2-11168127-DEL-1982
chr2-11184875-DEL-530
chr2-11203036-DEL-1916
chr2-11218278-DEL-322
chr2-11229300-DEL-51
chr2-11283189-DEL-58
chr2-11284091-DEL-93
chr2-11322108-DEL-76
chr2-11322535-DEL-226
chr2-11325869-DEL-699
chr2-11336152-DEL-51
chr2-11339996-DEL-560
chr2-11350317-DEL-111
chr2-11391542-DEL-310
chr2-11392085-DEL-103
chr2-11404668-DEL-1403
chr2-11417174-DEL-97

chr2-11438079-DEL-100
chr2-11466686-DEL-140
chr2-11468869-DEL-58
chr2-11501018-DEL-169
chr2-11509964-DEL-86
chr2-11515665-DEL-1330
chr2-11535976-DEL-96
chr2-11537685-DEL-70
chr2-11550392-DEL-55
chr2-11552305-DEL-86
chr2-11588242-DEL-187
chr2-11598534-DEL-287
chr2-11619676-DEL-178
chr2-11632378-DEL-69
chr2-11643980-DEL-460
chr2-11666582-DEL-64
chr2-11691161-DEL-95
chr2-11704406-DEL-51
chr2-11728967-DEL-625
chr2-11730006-DEL-146
chr2-11749547-DEL-247
chr2-11791321-DEL-87
chr2-11811094-DEL-1685
chr2-11816843-DEL-599
chr2-11819542-DEL-6794
chr2-11829441-DEL-700
chr2-11873505-DEL-77
chr2-11890137-DEL-92
chr2-11900854-DEL-981
chr2-11910245-DEL-490
chr2-11923082-DEL-255
chr2-11937399-DEL-2583
chr2-11942172-DEL-6278
chr2-11950155-DEL-4003
chr2-11973212-DEL-221
chr2-11976950-DEL-120
chr2-11980284-DEL-271
chr2-11982576-DEL-6390
chr2-11992207-DEL-538
chr2-12031004-DEL-70
chr2-12044491-DEL-923
chr2-12058671-DEL-91
chr2-12074187-DEL-332
chr2-12121435-DEL-94
chr2-12140607-DEL-53
chr2-12157552-DEL-545
chr2-12180717-DEL-942
chr2-12229641-DEL-78
chr2-12235120-DEL-177
chr2-12269023-DEL-92
chr2-12274276-DEL-587
chr2-12288692-DEL-93
chr2-12312785-DEL-87
chr2-12325309-DEL-59
chr2-12337864-DEL-64
chr2-12341065-DEL-539
chr2-12392337-DEL-104
chr2-12411454-DEL-145
chr2-12415118-DEL-54
chr2-12427818-DEL-507
chr2-12432248-DEL-58
chr2-12435064-DEL-57
chr2-12436329-DEL-380
chr2-12437321-DEL-55
chr2-12442035-DEL-105
chr2-12501379-DEL-68
chr2-12515769-DEL-1570
chr2-12533458-DEL-1289
chr2-12536735-DEL-247
chr2-12550455-DEL-170
chr2-12553560-DEL-72
chr2-12610167-DEL-3110
chr2-12612846-DEL-60
chr2-12626596-DEL-1066
chr2-12645923-DEL-3264
chr2-12645936-DEL-6801
chr2-12663086-DEL-82

chr2-12664722-DEL-1639
chr2-12761303-DEL-259
chr2-12833868-DEL-676
chr2-12840046-DEL-51
chr2-12911595-DEL-522
chr2-12950133-DEL-204
chr2-12953737-DEL-201
chr2-12977114-DEL-399
chr2-13036174-DEL-56
chr2-13050293-DEL-377
chr2-13076344-DEL-199
chr2-13110295-DEL-76
chr2-13115235-DEL-82
chr2-13127448-DEL-98
chr2-13142069-DEL-72
chr2-13171443-DEL-54
chr2-13172290-DEL-74
chr2-13198956-DEL-954
chr2-13199817-DEL-502
chr2-13216489-DEL-1199
chr2-13222923-DEL-70
chr2-13240041-DEL-53
chr2-13262603-DEL-162
chr2-13266013-DEL-1756
chr2-13291032-DEL-64
chr2-13348518-DEL-481
chr2-13414079-DEL-131
chr2-13450455-DEL-58
chr2-13455022-DEL-70
chr2-13464356-DEL-56
chr2-13471158-DEL-70
chr2-13479015-DEL-1049
chr2-13488812-DEL-64
chr2-13501900-DEL-102
chr2-13513806-DEL-138
chr2-13546721-DEL-52
chr2-13625336-DEL-585
chr2-13626752-DEL-480
chr2-13628324-DEL-197
chr2-13640326-DEL-289
chr2-13650994-DEL-62
chr2-13663194-DEL-424
chr2-13671281-DEL-74
chr2-13678314-DEL-112
chr2-13689913-DEL-505
chr2-13702515-DEL-215
chr2-13750031-DEL-277
chr2-13769292-DEL-130
chr2-13795354-DEL-137
chr2-13808800-DEL-189
chr2-13812322-DEL-71
chr2-13844700-DEL-171
chr2-13861329-DEL-314
chr2-13868272-DEL-2841
chr2-13897506-DEL-55
chr2-13900185-DEL-376
chr2-13920938-DEL-68
chr2-13932712-DEL-159
chr2-13941413-DEL-1006
chr2-13955085-DEL-213
chr2-13972944-DEL-186
chr2-13979331-DEL-274
chr2-13986529-DEL-58
chr2-14001900-DEL-140
chr2-14016650-DEL-122
chr2-14017142-DEL-3625
chr2-14018886-DEL-1735
chr2-14021823-DEL-55
chr2-14029327-DEL-83
chr2-14041821-DEL-52
chr2-14059419-DEL-163
chr2-14090846-DEL-204
chr2-14115992-DEL-113
chr2-14172948-DEL-60
chr2-14195677-DEL-62
chr2-14200175-DEL-231
chr2-14227094-DEL-65

chr2-14255412-DEL-417
chr2-14287005-DEL-6668
chr2-14299450-DEL-95
chr2-14300566-DEL-7712
chr2-14319408-DEL-66
chr2-14337652-DEL-84
chr2-14339785-DEL-95
chr2-14350099-DEL-6787
chr2-14350175-DEL-5568
chr2-14363885-DEL-6273
chr2-14375203-DEL-6780
chr2-14387853-DEL-3517
chr2-14395140-DEL-51
chr2-14402937-DEL-84
chr2-14430175-DEL-843
chr2-14458636-DEL-751
chr2-14459686-DEL-108
chr2-14479877-DEL-114
chr2-14481580-DEL-1067
chr2-14493878-DEL-823
chr2-14523523-DEL-381
chr2-14532471-DEL-52
chr2-14569880-DEL-133
chr2-14601979-DEL-74
chr2-14654453-DEL-292
chr2-14731798-DEL-214
chr2-14763676-DEL-184
chr2-14817382-DEL-68
chr2-14825199-DEL-92
chr2-14831009-DEL-162
chr2-14839357-DEL-59
chr2-14881077-DEL-56
chr2-14891755-DEL-74
chr2-14981684-DEL-83
chr2-15005168-DEL-579
chr2-15006300-DEL-3910
chr2-15024893-DEL-5838
chr2-15047259-DEL-62
chr2-15093272-DEL-277
chr2-15119465-DEL-199
chr2-15127687-DEL-111
chr2-15130737-DEL-527
chr2-15247802-DEL-393
chr2-15249351-DEL-6183
chr2-15258706-DEL-1482
chr2-15277169-DEL-358
chr2-15279925-DEL-434
chr2-15291526-DEL-2331
chr2-15305133-DEL-111
chr2-15305774-DEL-14777
chr2-15321754-DEL-1599
chr2-15369948-DEL-58
chr2-15376123-DEL-63
chr2-15380458-DEL-973
chr2-15406435-DEL-709
chr2-15443295-DEL-1047
chr2-15466326-DEL-213
chr2-15485513-DEL-392
chr2-15495786-DEL-599
chr2-15502610-DEL-166
chr2-15530953-DEL-135
chr2-15534130-DEL-61
chr2-15537939-DEL-119
chr2-15551195-DEL-121
chr2-15585364-DEL-1571
chr2-15589045-DEL-1413
chr2-15595499-DEL-129
chr2-15659446-DEL-72
chr2-15735661-DEL-94
chr2-15765089-DEL-1137
chr2-15775689-DEL-327
chr2-15801274-DEL-250
chr2-15812828-DEL-878
chr2-15830309-DEL-56
chr2-15859320-DEL-654
chr2-15878672-DEL-498
chr2-15892615-DEL-180

chr2-15918549-DEL-9260
chr2-15927080-DEL-260
chr2-15938798-DEL-173
chr2-15939345-DEL-149
chr2-15942107-DEL-1042
chr2-15943762-DEL-534
chr2-15956503-DEL-98
chr2-15960913-DEL-76
chr2-15981105-DEL-54
chr2-15985824-DEL-542
chr2-15994082-DEL-259
chr2-16010242-DEL-88
chr2-16057923-DEL-58
chr2-16068985-DEL-1946
chr2-16070875-DEL-6187
chr2-16097452-DEL-82
chr2-16119317-DEL-143
chr2-16183112-DEL-139
chr2-16197049-DEL-6368
chr2-16217689-DEL-422
chr2-16218225-DEL-53
chr2-16238426-DEL-1912
chr2-16243205-DEL-74
chr2-16247161-DEL-91
chr2-16265424-DEL-7011
chr2-16265589-DEL-400
chr2-16274529-DEL-515
chr2-16285460-DEL-2728
chr2-16298574-DEL-63
chr2-16302314-DEL-128
chr2-16346904-DEL-136
chr2-16348939-DEL-60
chr2-16386038-DEL-424
chr2-16400778-DEL-533
chr2-16449222-DEL-90
chr2-16469057-DEL-131
chr2-16500386-DEL-222
chr2-16520492-DEL-121
chr2-16547830-DEL-400
chr2-16550364-DEL-68
chr2-16566308-DEL-2051
chr2-16614803-DEL-2749
chr2-16632450-DEL-743
chr2-16643791-DEL-1673
chr2-16700317-DEL-276
chr2-16731296-DEL-56
chr2-16733460-DEL-108
chr2-16750734-DEL-1980
chr2-16764073-DEL-380
chr2-16786964-DEL-2721
chr2-16789811-DEL-3917
chr2-16861381-DEL-74
chr2-16864337-DEL-120
chr2-16866510-DEL-1293
chr2-16873542-DEL-67
chr2-16877647-DEL-1757
chr2-16899514-DEL-6279
chr2-16916331-DEL-89
chr2-16922242-DEL-430
chr2-16927211-DEL-426
chr2-16931108-DEL-68
chr2-16938634-DEL-389
chr2-16999068-DEL-390
chr2-17004444-DEL-220
chr2-17035382-DEL-23778
chr2-17037984-DEL-4326
chr2-17047401-DEL-6245
chr2-17064108-DEL-6808
chr2-17089248-DEL-115
chr2-17103176-DEL-340
chr2-17112899-DEL-451
chr2-17128967-DEL-1617
chr2-17131989-DEL-2217
chr2-17136137-DEL-156
chr2-17152217-DEL-651
chr2-17158519-DEL-757
chr2-17160517-DEL-890

chr2-17172236-DEL-55
chr2-17180708-DEL-2498
chr2-17185429-DEL-593
chr2-17190970-DEL-222
chr2-17198872-DEL-84
chr2-17204792-DEL-567
chr2-17209231-DEL-162
chr2-17214085-DEL-742
chr2-17235925-DEL-1731
chr2-17268090-DEL-100
chr2-17289805-DEL-1075
chr2-17387786-DEL-2162
chr2-17399680-DEL-851
chr2-17413400-DEL-173
chr2-17417219-DEL-113
chr2-17432592-DEL-52
chr2-17444447-DEL-60
chr2-17450280-DEL-57
chr2-17462527-DEL-1419
chr2-17476117-DEL-709
chr2-17478257-DEL-2205
chr2-17480932-DEL-249
chr2-17483128-DEL-54
chr2-17486001-DEL-264
chr2-17491123-DEL-2543
chr2-17491136-DEL-6246
chr2-17511911-DEL-104
chr2-17533652-DEL-175
chr2-17537581-DEL-806
chr2-17546323-DEL-763
chr2-17547333-DEL-76
chr2-17559622-DEL-100
chr2-17577640-DEL-228
chr2-17580123-DEL-1118
chr2-17592940-DEL-60
chr2-17625770-DEL-82
chr2-17632532-DEL-342
chr2-17745412-DEL-291
chr2-17777672-DEL-78
chr2-17793700-DEL-117
chr2-17799479-DEL-55
chr2-17811477-DEL-470
chr2-17812601-DEL-85
chr2-17827849-DEL-81
chr2-17852002-DEL-72
chr2-17861314-DEL-317
chr2-17869112-DEL-55
chr2-17878821-DEL-277
chr2-17893322-DEL-187
chr2-17901168-DEL-65
chr2-17904318-DEL-82
chr2-17912106-DEL-91
chr2-17923255-DEL-396
chr2-17931798-DEL-72
chr2-17966607-DEL-287
chr2-17972235-DEL-52
chr2-17985803-DEL-167
chr2-18102660-DEL-175
chr2-18109582-DEL-145
chr2-18123122-DEL-68
chr2-18132998-DEL-445
chr2-18135242-DEL-88
chr2-18174154-DEL-86
chr2-18230969-DEL-300
chr2-18341135-DEL-209
chr2-18342850-DEL-1752
chr2-18358965-DEL-107
chr2-18367039-DEL-154
chr2-18380063-DEL-139
chr2-18399098-DEL-251
chr2-18403029-DEL-104
chr2-18426899-DEL-2235
chr2-18435918-DEL-86
chr2-18446290-DEL-254
chr2-18458864-DEL-1460
chr2-18463890-DEL-514
chr2-18484107-DEL-4156

chr2-18501118-DEL-69
chr2-18540525-DEL-231
chr2-18544294-DEL-173
chr2-18550939-DEL-135
chr2-18579482-DEL-111
chr2-18644848-DEL-115
chr2-18651711-DEL-186
chr2-18723909-DEL-353
chr2-18724265-DEL-1037
chr2-18725343-DEL-153
chr2-18740602-DEL-52
chr2-18756442-DEL-885
chr2-18783631-DEL-78
chr2-18788978-DEL-75
chr2-18791476-DEL-106
chr2-18793154-DEL-2535
chr2-18809682-DEL-192
chr2-18811643-DEL-346
chr2-18837680-DEL-1142
chr2-18839693-DEL-393
chr2-18873269-DEL-2277
chr2-18898259-DEL-69
chr2-18907301-DEL-63
chr2-18956403-DEL-132
chr2-18972151-DEL-612
chr2-18994254-DEL-1638
chr2-19024424-DEL-84
chr2-19025031-DEL-500
chr2-19092228-DEL-106
chr2-19102346-DEL-54
chr2-19128006-DEL-6192
chr2-19135658-DEL-845
chr2-19210213-DEL-11490
chr2-19254478-DEL-71
chr2-19261674-DEL-135
chr2-19262417-DEL-52
chr2-19266877-DEL-576
chr2-19267172-DEL-691
chr2-19286697-DEL-130
chr2-19288786-DEL-251
chr2-19291401-DEL-462
chr2-19295052-DEL-1839
chr2-19305894-DEL-67
chr2-19318914-DEL-166
chr2-19369148-DEL-809
chr2-19394281-DEL-75
chr2-19409436-DEL-404
chr2-19421030-DEL-276
chr2-19441355-DEL-60
chr2-19444183-DEL-198
chr2-19503373-DEL-673
chr2-19527589-DEL-62
chr2-19562317-DEL-68
chr2-19597708-DEL-241
chr2-19615117-DEL-78
chr2-19615793-DEL-94
chr2-19620540-DEL-385
chr2-19637991-DEL-253
chr2-19648065-DEL-822
chr2-19658457-DEL-989
chr2-19677433-DEL-100
chr2-19692112-DEL-131
chr2-19698972-DEL-55
chr2-19701992-DEL-152
chr2-19710311-DEL-401
chr2-19712343-DEL-6255
chr2-19713343-DEL-70
chr2-19718286-DEL-189
chr2-19733858-DEL-57
chr2-19736635-DEL-134
chr2-19830633-DEL-53
chr2-19854175-DEL-60
chr2-19855190-DEL-99
chr2-19873202-DEL-443
chr2-19923072-DEL-54
chr2-19965695-DEL-1774
chr2-19967335-DEL-107

chr2-19970402-DEL-1949
chr2-19972842-DEL-358
chr2-19989012-DEL-840
chr2-20011158-DEL-66
chr2-20016495-DEL-316
chr2-20034276-DEL-90
chr2-20050962-DEL-1302
chr2-20058009-DEL-401
chr2-20061217-DEL-61
chr2-20071045-DEL-4314
chr2-20096604-DEL-1484
chr2-20098114-DEL-2321
chr2-20102397-DEL-117
chr2-20116849-DEL-67
chr2-20117316-DEL-228
chr2-20141270-DEL-90
chr2-20150742-DEL-220
chr2-20179013-DEL-5994
chr2-20196404-DEL-201
chr2-20202625-DEL-155
chr2-20210104-DEL-576
chr2-20212618-DEL-333
chr2-20230077-DEL-362
chr2-20353129-DEL-64
chr2-20392592-DEL-1755
chr2-20409058-DEL-67
chr2-20442868-DEL-72
chr2-20460197-DEL-290
chr2-20470682-DEL-64
chr2-20484199-DEL-1216
chr2-20488006-DEL-76
chr2-20490197-DEL-471
chr2-20514482-DEL-341
chr2-20516063-DEL-50
chr2-20532140-DEL-229
chr2-20540831-DEL-435
chr2-20543418-DEL-142
chr2-20570736-DEL-1122
chr2-20582517-DEL-96
chr2-20596925-DEL-436
chr2-20611957-DEL-71
chr2-20621763-DEL-112
chr2-20643324-DEL-164
chr2-20655441-DEL-57
chr2-20659463-DEL-76
chr2-20675921-DEL-185
chr2-20820437-DEL-393
chr2-20821816-DEL-209
chr2-20825559-DEL-118
chr2-20962330-DEL-177
chr2-20992225-DEL-66
chr2-21022153-DEL-84
chr2-21050396-DEL-173
chr2-21068130-DEL-169
chr2-21120600-DEL-78
chr2-21148452-DEL-53
chr2-21159071-DEL-168
chr2-21173472-DEL-431
chr2-21177117-DEL-104
chr2-21270819-DEL-670
chr2-21272814-DEL-83
chr2-21333982-DEL-4344
chr2-21338647-DEL-605
chr2-21343360-DEL-64
chr2-21356087-DEL-136
chr2-21371103-DEL-90
chr2-21374676-DEL-109
chr2-21380806-DEL-56
chr2-21397184-DEL-421
chr2-21411645-DEL-149
chr2-21469674-DEL-72
chr2-21486979-DEL-165
chr2-21551070-DEL-477
chr2-21557086-DEL-97
chr2-21565941-DEL-66
chr2-21574629-DEL-94
chr2-21582929-DEL-3447

chr2-21588253-DEL-4868
chr2-21590157-DEL-212
chr2-21601585-DEL-6408
chr2-21607656-DEL-200
chr2-21615041-DEL-160
chr2-21615958-DEL-71
chr2-21626148-DEL-2698
chr2-21629912-DEL-199
chr2-21655336-DEL-55
chr2-21667977-DEL-69
chr2-21679938-DEL-470
chr2-21692790-DEL-104
chr2-21701069-DEL-75
chr2-21720224-DEL-101
chr2-21722234-DEL-1384
chr2-21737164-DEL-364
chr2-21738717-DEL-2214
chr2-21745207-DEL-87
chr2-21774740-DEL-165
chr2-21776061-DEL-560
chr2-21786231-DEL-180
chr2-21791133-DEL-503
chr2-21793194-DEL-2208
chr2-21806848-DEL-2204
chr2-21828749-DEL-272
chr2-21845144-DEL-66
chr2-21883005-DEL-663
chr2-21891337-DEL-4501
chr2-21925979-DEL-68
chr2-21928681-DEL-1824
chr2-21947213-DEL-101
chr2-21952167-DEL-963
chr2-21955926-DEL-75
chr2-22036803-DEL-89
chr2-22074932-DEL-4668
chr2-22080158-DEL-77
chr2-22081535-DEL-6575
chr2-22096746-DEL-295
chr2-22102327-DEL-64
chr2-22104844-DEL-113
chr2-22115251-DEL-1894
chr2-22129437-DEL-2069
chr2-22169326-DEL-636
chr2-22174655-DEL-175
chr2-22189901-DEL-119
chr2-22192234-DEL-1531
chr2-22196359-DEL-5346
chr2-22235764-DEL-1599
chr2-22263105-DEL-1542
chr2-22283497-DEL-65
chr2-22293912-DEL-165
chr2-22341796-DEL-66
chr2-22362608-DEL-765
chr2-22389899-DEL-133
chr2-22390699-DEL-6311
chr2-22432599-DEL-1000
chr2-22432600-DEL-1002
chr2-22471001-DEL-214
chr2-22477285-DEL-3268
chr2-22494423-DEL-187
chr2-22536942-DEL-91
chr2-22556674-DEL-69
chr2-22558069-DEL-54
chr2-22566118-DEL-59
chr2-22621146-DEL-92
chr2-22641529-DEL-91
chr2-22726248-DEL-3456
chr2-22732282-DEL-152
chr2-22756626-DEL-67
chr2-22768079-DEL-276
chr2-22798966-DEL-1574
chr2-22800286-DEL-1295
chr2-22810762-DEL-934
chr2-22842645-DEL-53
chr2-22845177-DEL-188
chr2-22873989-DEL-52
chr2-22883111-DEL-75

chr2-22894656-DEL-183
chr2-22917552-DEL-180
chr2-22936279-DEL-3975
chr2-22948825-DEL-411
chr2-22950118-DEL-3076
chr2-22953582-DEL-72
chr2-22956495-DEL-276
chr2-22962500-DEL-123
chr2-22974831-DEL-198
chr2-23080120-DEL-83
chr2-23094657-DEL-291
chr2-23117190-DEL-178
chr2-23139336-DEL-292
chr2-23160921-DEL-134
chr2-23163905-DEL-2493
chr2-23185934-DEL-6059
chr2-23192010-DEL-5498
chr2-23210372-DEL-57
chr2-23333039-DEL-2203
chr2-23339221-DEL-51
chr2-23344597-DEL-182
chr2-23353914-DEL-1871
chr2-23375062-DEL-53
chr2-23442263-DEL-74
chr2-23476324-DEL-62
chr2-23481957-DEL-2100
chr2-23513781-DEL-2159
chr2-23522245-DEL-65
chr2-23534567-DEL-1199
chr2-23552572-DEL-141
chr2-23557940-DEL-60
chr2-23561284-DEL-3909
chr2-23568454-DEL-445
chr2-23583665-DEL-1512
chr2-23592736-DEL-6669
chr2-23599119-DEL-199
chr2-23599738-DEL-683
chr2-23600822-DEL-85
chr2-23622950-DEL-3762
chr2-23639135-DEL-104
chr2-23641928-DEL-150
chr2-23656524-DEL-810
chr2-23666636-DEL-260
chr2-23672995-DEL-1981
chr2-23701404-DEL-86
chr2-23742293-DEL-60
chr2-23750087-DEL-1151
chr2-23752330-DEL-4228
chr2-23766149-DEL-1499
chr2-23772905-DEL-172
chr2-23806707-DEL-100
chr2-23808760-DEL-7643
chr2-23809745-DEL-102
chr2-23820246-DEL-2263
chr2-23827993-DEL-18782
chr2-23828242-DEL-1157
chr2-23829570-DEL-135
chr2-23836075-DEL-10386
chr2-23850695-DEL-388
chr2-23877881-DEL-175
chr2-23879735-DEL-892
chr2-23882751-DEL-321
chr2-23901374-DEL-75
chr2-23916574-DEL-267
chr2-23924011-DEL-4952
chr2-23974626-DEL-53
chr2-23982417-DEL-56
chr2-24016521-DEL-965
chr2-24079664-DEL-508
chr2-24126225-DEL-1026
chr2-24143830-DEL-440
chr2-24191260-DEL-832
chr2-24267305-DEL-56
chr2-24278232-DEL-62
chr2-24279275-DEL-182
chr2-24308509-DEL-71
chr2-24317037-DEL-101

chr2-24371276-DEL-51
chr2-24395856-DEL-76
chr2-24406341-DEL-72
chr2-24426390-DEL-358
chr2-24479739-DEL-1892
chr2-24524248-DEL-332
chr2-24556955-DEL-79
chr2-24624980-DEL-73
chr2-24630173-DEL-91
chr2-24633707-DEL-86
chr2-24657345-DEL-78
chr2-24659501-DEL-161
chr2-24669460-DEL-208
chr2-24700797-DEL-120
chr2-24708646-DEL-186
chr2-24724937-DEL-98
chr2-24744264-DEL-209
chr2-24746155-DEL-157
chr2-24788701-DEL-164
chr2-24811183-DEL-327
chr2-24822666-DEL-52
chr2-24826898-DEL-933
chr2-24828441-DEL-156
chr2-24845320-DEL-2414
chr2-24856211-DEL-84
chr2-24870407-DEL-237
chr2-24883433-DEL-56
chr2-24925186-DEL-71
chr2-24932243-DEL-57
chr2-24937193-DEL-280
chr2-24958954-DEL-62
chr2-24959921-DEL-245
chr2-25020060-DEL-86
chr2-25048700-DEL-428
chr2-25051356-DEL-68
chr2-25099614-DEL-64
chr2-25128484-DEL-108
chr2-25191434-DEL-219
chr2-25191504-DEL-56
chr2-25207314-DEL-71
chr2-25249294-DEL-157
chr2-25250435-DEL-157
chr2-25257959-DEL-110
chr2-25300560-DEL-136
chr2-25408515-DEL-95
chr2-25415309-DEL-52
chr2-25419169-DEL-63
chr2-25443447-DEL-223
chr2-25461695-DEL-799
chr2-25500078-DEL-70
chr2-25509006-DEL-600
chr2-25512775-DEL-141
chr2-25575342-DEL-51
chr2-25624623-DEL-55
chr2-25630754-DEL-882
chr2-25678417-DEL-262
chr2-25696786-DEL-2976
chr2-25728654-DEL-164
chr2-25747725-DEL-423
chr2-25794456-DEL-68
chr2-25805443-DEL-68
chr2-25836176-DEL-134
chr2-25899561-DEL-75
chr2-25926554-DEL-89
chr2-25961571-DEL-119
chr2-25974471-DEL-134
chr2-25986651-DEL-156
chr2-26024635-DEL-628
chr2-26041333-DEL-1653
chr2-26108038-DEL-74
chr2-26150877-DEL-56
chr2-26176984-DEL-174
chr2-26195259-DEL-97
chr2-26272415-DEL-76
chr2-26294241-DEL-415
chr2-26326656-DEL-57
chr2-26340628-DEL-248

chr2-26448283-DEL-137
chr2-26449298-DEL-64
chr2-26491312-DEL-63
chr2-26491938-DEL-95
chr2-26532638-DEL-6156
chr2-26538272-DEL-57
chr2-26546390-DEL-62
chr2-26565739-DEL-2792
chr2-26573822-DEL-1466
chr2-26588038-DEL-7415
chr2-26588039-DEL-617
chr2-26588942-DEL-6798
chr2-26600369-DEL-208
chr2-26600997-DEL-6184
chr2-26608323-DEL-56
chr2-26613130-DEL-4509
chr2-26617128-DEL-318
chr2-26625053-DEL-97
chr2-26673790-DEL-84
chr2-26694260-DEL-194
chr2-26698482-DEL-440
chr2-26723354-DEL-598
chr2-26725177-DEL-60
chr2-26741195-DEL-238
chr2-26742748-DEL-573
chr2-26755674-DEL-178
chr2-26783265-DEL-204
chr2-26834732-DEL-60
chr2-26849153-DEL-408
chr2-26859817-DEL-58
chr2-26877923-DEL-100
chr2-26884371-DEL-56
chr2-26909674-DEL-137
chr2-26996035-DEL-191
chr2-27005487-DEL-64
chr2-27007742-DEL-114
chr2-27019312-DEL-924
chr2-27051454-DEL-54
chr2-27091737-DEL-282
chr2-27098951-DEL-58
chr2-27169582-DEL-71
chr2-27207931-DEL-162
chr2-27209070-DEL-929
chr2-27210418-DEL-52
chr2-27263900-DEL-793
chr2-27272780-DEL-608
chr2-27328432-DEL-64
chr2-27329258-DEL-144
chr2-27375806-DEL-393
chr2-27376511-DEL-179
chr2-27377146-DEL-93
chr2-27399540-DEL-52
chr2-27407816-DEL-202
chr2-27411668-DEL-1760
chr2-27413469-DEL-232
chr2-27428631-DEL-501
chr2-27429680-DEL-107
chr2-27456025-DEL-82
chr2-27468539-DEL-1463
chr2-27476279-DEL-76
chr2-27505685-DEL-52
chr2-27538892-DEL-204
chr2-27656466-DEL-86
chr2-27659218-DEL-117
chr2-27661030-DEL-76
chr2-27680189-DEL-486
chr2-27709371-DEL-112
chr2-27711347-DEL-169
chr2-27728612-DEL-59
chr2-27761494-DEL-176
chr2-27783008-DEL-142
chr2-27783919-DEL-163
chr2-27796309-DEL-221
chr2-27828820-DEL-84
chr2-27833505-DEL-568
chr2-27851276-DEL-248
chr2-27882859-DEL-74

chr2-27901907-DEL-298
chr2-27953383-DEL-88
chr2-27963490-DEL-435
chr2-27979806-DEL-52
chr2-28007394-DEL-88
chr2-28010613-DEL-51
chr2-28092434-DEL-81
chr2-28149305-DEL-606
chr2-28159647-DEL-86
chr2-28194952-DEL-2115
chr2-28207330-DEL-2078
chr2-28222820-DEL-844
chr2-28223894-DEL-829
chr2-28224938-DEL-266
chr2-28226150-DEL-90
chr2-28229206-DEL-140
chr2-28246993-DEL-1048
chr2-28261944-DEL-84
chr2-28281569-DEL-1791
chr2-28283738-DEL-429
chr2-28300292-DEL-54
chr2-28321642-DEL-234
chr2-28333978-DEL-66
chr2-28338945-DEL-204
chr2-28349352-DEL-188
chr2-28367106-DEL-541
chr2-28383982-DEL-166
chr2-28390002-DEL-1594
chr2-28426074-DEL-54
chr2-28453763-DEL-112
chr2-28455558-DEL-526
chr2-28485889-DEL-136
chr2-28490629-DEL-109
chr2-28491091-DEL-366
chr2-28492834-DEL-348
chr2-28520889-DEL-659
chr2-28545437-DEL-110
chr2-28568485-DEL-54
chr2-28616773-DEL-756
chr2-28634460-DEL-98
chr2-28652633-DEL-417
chr2-28656434-DEL-76
chr2-28657538-DEL-773
chr2-28695057-DEL-133
chr2-28698545-DEL-131
chr2-28772455-DEL-259
chr2-28788212-DEL-82
chr2-28851109-DEL-59
chr2-28862368-DEL-68
chr2-28874939-DEL-118
chr2-28898435-DEL-415
chr2-28945768-DEL-212
chr2-28952111-DEL-68
chr2-28993499-DEL-12919
chr2-29011452-DEL-135
chr2-29024871-DEL-178
chr2-29044558-DEL-61
chr2-29046113-DEL-201
chr2-29057761-DEL-205
chr2-29059342-DEL-80
chr2-29076377-DEL-122
chr2-29112993-DEL-181
chr2-29148441-DEL-58
chr2-29224963-DEL-619
chr2-29291190-DEL-503
chr2-29325065-DEL-58
chr2-29388618-DEL-54
chr2-29391518-DEL-83
chr2-29401611-DEL-357
chr2-29417753-DEL-240
chr2-29421154-DEL-76
chr2-29431073-DEL-146
chr2-29435252-DEL-56
chr2-29439670-DEL-64
chr2-29461541-DEL-250
chr2-29473271-DEL-318
chr2-29483972-DEL-134

chr2-29486093-DEL-168
chr2-29516088-DEL-1078
chr2-29530215-DEL-205
chr2-29535389-DEL-96
chr2-29540416-DEL-205
chr2-29557739-DEL-113
chr2-29644322-DEL-536
chr2-29645138-DEL-199
chr2-29646283-DEL-178
chr2-29647844-DEL-478
chr2-29651956-DEL-961
chr2-29659154-DEL-106
chr2-29666261-DEL-429
chr2-29675829-DEL-171
chr2-29691251-DEL-543
chr2-29713261-DEL-169
chr2-29751049-DEL-571
chr2-29751277-DEL-897
chr2-29756073-DEL-89
chr2-29767354-DEL-106
chr2-29778447-DEL-423
chr2-29785821-DEL-212
chr2-29799504-DEL-210
chr2-29801061-DEL-209
chr2-29803080-DEL-330
chr2-29813812-DEL-83
chr2-29826534-DEL-208
chr2-29831661-DEL-227
chr2-29837107-DEL-153
chr2-29844895-DEL-202
chr2-29869595-DEL-252
chr2-29940394-DEL-96
chr2-29941172-DEL-135
chr2-30010612-DEL-54
chr2-30017391-DEL-76
chr2-30020908-DEL-495
chr2-30035706-DEL-138
chr2-30044999-DEL-69
chr2-30048378-DEL-187
chr2-30049543-DEL-119
chr2-30056474-DEL-2908
chr2-30059916-DEL-205
chr2-30072884-DEL-160
chr2-30106850-DEL-172
chr2-30121845-DEL-206
chr2-30143230-DEL-706
chr2-30154397-DEL-57
chr2-30161075-DEL-202
chr2-30165808-DEL-160
chr2-30177693-DEL-331
chr2-30181150-DEL-228
chr2-30185593-DEL-151
chr2-30186357-DEL-158
chr2-30197068-DEL-161
chr2-30208400-DEL-741
chr2-30244463-DEL-80
chr2-30309718-DEL-205
chr2-30315395-DEL-223
chr2-30345255-DEL-243
chr2-30349933-DEL-54
chr2-30377388-DEL-163
chr2-30384055-DEL-250
chr2-30389029-DEL-1030
chr2-30396912-DEL-58
chr2-30470444-DEL-90
chr2-30511576-DEL-60
chr2-30525954-DEL-96
chr2-30534172-DEL-342
chr2-30545358-DEL-122
chr2-30562691-DEL-132
chr2-30578012-DEL-51
chr2-30580376-DEL-51
chr2-30626064-DEL-135
chr2-30630465-DEL-142
chr2-30630785-DEL-539
chr2-30638953-DEL-172
chr2-30655301-DEL-152

chr2-30678373-DEL-83
chr2-30679233-DEL-190
chr2-30683577-DEL-181
chr2-30698500-DEL-64
chr2-30717760-DEL-134
chr2-30744530-DEL-56
chr2-30749244-DEL-107
chr2-30762619-DEL-56
chr2-30763433-DEL-443
chr2-30784758-DEL-92
chr2-30785350-DEL-72
chr2-30813480-DEL-96
chr2-30820322-DEL-1164
chr2-30861748-DEL-264
chr2-30864625-DEL-582
chr2-30866624-DEL-80
chr2-30887709-DEL-77
chr2-30896779-DEL-598
chr2-30924728-DEL-500
chr2-30945871-DEL-270
chr2-30947424-DEL-60
chr2-30988518-DEL-97
chr2-30990024-DEL-129
chr2-31002747-DEL-171
chr2-31086646-DEL-56
chr2-31106901-DEL-60
chr2-31119287-DEL-281
chr2-31126737-DEL-52
chr2-31178906-DEL-95
chr2-31211442-DEL-110
chr2-31218866-DEL-54
chr2-31237318-DEL-77
chr2-31252610-DEL-311
chr2-31255878-DEL-122
chr2-31263979-DEL-52
chr2-31269746-DEL-204
chr2-31280374-DEL-92
chr2-31286306-DEL-94
chr2-31287934-DEL-89
chr2-31322520-DEL-52
chr2-31338185-DEL-219
chr2-31340244-DEL-74
chr2-31356000-DEL-219
chr2-31365533-DEL-199
chr2-31434049-DEL-80
chr2-31441625-DEL-194
chr2-31446762-DEL-134
chr2-31457476-DEL-53
chr2-31543513-DEL-108
chr2-31561702-DEL-205
chr2-31563014-DEL-52
chr2-31564417-DEL-56
chr2-31576666-DEL-379
chr2-31586547-DEL-69
chr2-31615218-DEL-167
chr2-31619572-DEL-170
chr2-31627021-DEL-182
chr2-31630153-DEL-190
chr2-31645289-DEL-78
chr2-31648157-DEL-338
chr2-31652227-DEL-67
chr2-31684258-DEL-56
chr2-31685014-DEL-200
chr2-31695017-DEL-363
chr2-31712199-DEL-82
chr2-31743188-DEL-479
chr2-31757228-DEL-94
chr2-31763088-DEL-285
chr2-31765202-DEL-75
chr2-31767815-DEL-93
chr2-31799698-DEL-51
chr2-31830188-DEL-103
chr2-31834608-DEL-59
chr2-31897624-DEL-436
chr2-31919227-DEL-235
chr2-31931633-DEL-140
chr2-31944235-DEL-204

chr2-31978699-DEL-61
chr2-31988164-DEL-77
chr2-31999309-DEL-164
chr2-32000583-DEL-57
chr2-32011010-DEL-90
chr2-32013294-DEL-94
chr2-32022764-DEL-69
chr2-32028295-DEL-54
chr2-32031196-DEL-169
chr2-32032939-DEL-233
chr2-32059690-DEL-94
chr2-32066455-DEL-188
chr2-32075772-DEL-115
chr2-32158590-DEL-99
chr2-32160812-DEL-79
chr2-32166917-DEL-1057
chr2-32214039-DEL-3041
chr2-32263569-DEL-51
chr2-32283878-DEL-438
chr2-32284675-DEL-172
chr2-32300060-DEL-169
chr2-32314835-DEL-51
chr2-32350535-DEL-200
chr2-32355771-DEL-190
chr2-32363373-DEL-4457
chr2-32371837-DEL-291
chr2-32394218-DEL-121
chr2-32394976-DEL-443
chr2-32403178-DEL-583
chr2-32406917-DEL-279
chr2-32410893-DEL-184
chr2-32451281-DEL-206
chr2-32470601-DEL-85
chr2-32502754-DEL-167
chr2-32521214-DEL-325
chr2-32552821-DEL-377
chr2-32556893-DEL-618
chr2-32565569-DEL-848
chr2-32566559-DEL-147
chr2-32604836-DEL-63
chr2-32605614-DEL-145
chr2-32628846-DEL-210
chr2-32652030-DEL-74
chr2-32677130-DEL-186
chr2-32706465-DEL-112
chr2-32714993-DEL-71
chr2-32757024-DEL-207
chr2-32793002-DEL-199
chr2-32796166-DEL-91
chr2-32823562-DEL-53
chr2-32824656-DEL-87
chr2-32867555-DEL-79
chr2-32873330-DEL-315
chr2-32889658-DEL-183
chr2-32898915-DEL-229
chr2-32932379-DEL-263
chr2-32938136-DEL-96
chr2-32951309-DEL-251
chr2-32960146-DEL-171
chr2-32987287-DEL-180
chr2-33007975-DEL-635
chr2-33011075-DEL-58
chr2-33045790-DEL-144
chr2-33102087-DEL-205
chr2-33103534-DEL-174
chr2-33142867-DEL-63
chr2-33182820-DEL-66
chr2-33189902-DEL-197
chr2-33222390-DEL-91
chr2-33274998-DEL-400
chr2-33278506-DEL-114
chr2-33298086-DEL-126
chr2-33310375-DEL-167
chr2-33330201-DEL-132
chr2-33368892-DEL-96
chr2-33376524-DEL-96
chr2-33394148-DEL-56

chr2-33396204-DEL-60
chr2-33399767-DEL-70
chr2-33405043-DEL-55
chr2-33455363-DEL-75
chr2-33473058-DEL-51
chr2-33511106-DEL-332
chr2-33569748-DEL-56
chr2-33617564-DEL-83
chr2-33629962-DEL-70
chr2-33635156-DEL-96
chr2-33667473-DEL-200
chr2-33735766-DEL-51
chr2-33800286-DEL-53
chr2-33812580-DEL-68
chr2-33815945-DEL-86
chr2-33818263-DEL-71
chr2-33840718-DEL-99
chr2-33902718-DEL-56
chr2-33935887-DEL-1976
chr2-33940146-DEL-638
chr2-33940985-DEL-198
chr2-33977871-DEL-66
chr2-34015379-DEL-941
chr2-34029434-DEL-65
chr2-34044775-DEL-132
chr2-34046341-DEL-109
chr2-34093840-DEL-184
chr2-34158101-DEL-57
chr2-34185941-DEL-208
chr2-34190716-DEL-1445
chr2-34277453-DEL-216
chr2-34307627-DEL-214
chr2-34384071-DEL-914
chr2-34454971-DEL-891
chr2-34470243-DEL-238
chr2-34500712-DEL-207
chr2-34525238-DEL-64
chr2-34558874-DEL-78
chr2-34562512-DEL-52
chr2-34612653-DEL-67
chr2-34634149-DEL-189
chr2-34634730-DEL-214
chr2-34639121-DEL-204
chr2-34648469-DEL-2917
chr2-34654471-DEL-434
chr2-34659738-DEL-204
chr2-34663402-DEL-199
chr2-34723311-DEL-1317
chr2-34734815-DEL-2165
chr2-34774305-DEL-121
chr2-34886336-DEL-160
chr2-34894068-DEL-116
chr2-34989208-DEL-272
chr2-34994840-DEL-93
chr2-35012089-DEL-306
chr2-35064593-DEL-107
chr2-35112366-DEL-136
chr2-35134033-DEL-127
chr2-35214794-DEL-58
chr2-35247926-DEL-420
chr2-35248887-DEL-180
chr2-35274084-DEL-106
chr2-35319925-DEL-75
chr2-35356013-DEL-51
chr2-35377601-DEL-198
chr2-35430354-DEL-146
chr2-35433745-DEL-367
chr2-35535348-DEL-126
chr2-35626421-DEL-65
chr2-35647300-DEL-856
chr2-35668385-DEL-122
chr2-35683000-DEL-198
chr2-35692446-DEL-181
chr2-35698493-DEL-180
chr2-35703192-DEL-391
chr2-35703740-DEL-1197
chr2-35709078-DEL-70

chr2-35751927-DEL-215
chr2-35807867-DEL-473
chr2-35810272-DEL-185
chr2-35812744-DEL-82
chr2-35825660-DEL-54
chr2-35829622-DEL-127
chr2-35840981-DEL-400
chr2-35874742-DEL-160
chr2-35886600-DEL-149
chr2-35887105-DEL-1098
chr2-35892960-DEL-72
chr2-35935248-DEL-512
chr2-35999076-DEL-81
chr2-36008940-DEL-282
chr2-36082659-DEL-108
chr2-36095958-DEL-384
chr2-36122557-DEL-125
chr2-36159053-DEL-54
chr2-36159929-DEL-93
chr2-36182899-DEL-100
chr2-36236115-DEL-11611
chr2-36238691-DEL-226
chr2-36239375-DEL-7342
chr2-36284033-DEL-20201
chr2-36325014-DEL-119
chr2-36333354-DEL-624
chr2-36353311-DEL-6404
chr2-36383091-DEL-3129
chr2-36392427-DEL-1275
chr2-36401085-DEL-7314
chr2-36423869-DEL-3267
chr2-36432749-DEL-7775
chr2-36441334-DEL-1801
chr2-36453606-DEL-2377
chr2-36463902-DEL-533
chr2-36465888-DEL-124
chr2-36485363-DEL-204
chr2-36486035-DEL-203
chr2-36493126-DEL-5852
chr2-36493336-DEL-4085
chr2-36502981-DEL-92
chr2-36513444-DEL-117
chr2-36515097-DEL-54
chr2-36523169-DEL-1049
chr2-36524120-DEL-498
chr2-36538023-DEL-129
chr2-36561264-DEL-16705
chr2-36584762-DEL-127
chr2-36585340-DEL-3134
chr2-36618128-DEL-1893
chr2-36647351-DEL-121
chr2-36648097-DEL-58
chr2-36675021-DEL-1279
chr2-36697780-DEL-660
chr2-36709034-DEL-51
chr2-36717796-DEL-167
chr2-36727224-DEL-106
chr2-36751560-DEL-80
chr2-36801329-DEL-792
chr2-36820626-DEL-59
chr2-36822117-DEL-73
chr2-36824734-DEL-248
chr2-36841089-DEL-22875
chr2-36842399-DEL-859
chr2-36848537-DEL-3315
chr2-36854486-DEL-6321
chr2-36891959-DEL-51
chr2-36912729-DEL-3835
chr2-36927391-DEL-79
chr2-36957630-DEL-62
chr2-36961764-DEL-106
chr2-36988138-DEL-1490
chr2-37006093-DEL-57
chr2-37032283-DEL-99
chr2-37040770-DEL-417
chr2-37042517-DEL-409
chr2-37045900-DEL-208

chr2-37057334-DEL-123
chr2-37059892-DEL-6181
chr2-37072620-DEL-1322
chr2-37086773-DEL-449
chr2-37088663-DEL-1344
chr2-37094642-DEL-88
chr2-37097796-DEL-751
chr2-37112089-DEL-60
chr2-37159300-DEL-1037
chr2-37172048-DEL-4707
chr2-37194454-DEL-861
chr2-37226142-DEL-165
chr2-37271309-DEL-181
chr2-37280413-DEL-1503
chr2-37308024-DEL-231
chr2-37338945-DEL-115
chr2-37345218-DEL-219
chr2-37355085-DEL-205
chr2-37355454-DEL-517
chr2-37395346-DEL-55
chr2-37438463-DEL-151
chr2-37480341-DEL-109
chr2-37511206-DEL-104
chr2-37597124-DEL-62
chr2-37608235-DEL-107
chr2-37628446-DEL-224
chr2-37654768-DEL-320
chr2-37662291-DEL-219
chr2-37760401-DEL-604
chr2-37766897-DEL-54
chr2-37775206-DEL-65
chr2-37808629-DEL-59
chr2-37877785-DEL-160
chr2-37943795-DEL-212
chr2-37970499-DEL-1142
chr2-37978305-DEL-1458
chr2-38023148-DEL-114
chr2-38096438-DEL-3234
chr2-38102036-DEL-436
chr2-38122610-DEL-196
chr2-38147271-DEL-2276
chr2-38150649-DEL-511
chr2-38295348-DEL-209
chr2-38317900-DEL-455
chr2-38325796-DEL-193
chr2-38335992-DEL-223
chr2-38343290-DEL-204
chr2-38374245-DEL-74
chr2-38397838-DEL-63
chr2-38474307-DEL-103
chr2-38545417-DEL-355
chr2-38579752-DEL-52
chr2-38580200-DEL-539
chr2-38622790-DEL-98
chr2-38628272-DEL-117
chr2-38635131-DEL-2540
chr2-38636713-DEL-1630
chr2-38646068-DEL-401
chr2-38647866-DEL-135
chr2-38669109-DEL-74
chr2-38678435-DEL-200
chr2-38694248-DEL-427
chr2-38705195-DEL-243
chr2-38705958-DEL-1004
chr2-38711129-DEL-98
chr2-38727025-DEL-71
chr2-38759603-DEL-186
chr2-38787149-DEL-3297
chr2-38790730-DEL-1985
chr2-38818679-DEL-333
chr2-38821303-DEL-55
chr2-38829418-DEL-205
chr2-38830125-DEL-206
chr2-38842301-DEL-122
chr2-38866024-DEL-57
chr2-38876098-DEL-69
chr2-38933575-DEL-76

chr2-39071103-DEL-401
chr2-39103207-DEL-509
chr2-39124550-DEL-298
chr2-39130488-DEL-57
chr2-39143527-DEL-97
chr2-39157129-DEL-410
chr2-39173878-DEL-69
chr2-39177539-DEL-6187
chr2-39218164-DEL-69
chr2-39223430-DEL-69
chr2-39229923-DEL-51
chr2-39234452-DEL-1801
chr2-39250338-DEL-123
chr2-39272226-DEL-132
chr2-39292963-DEL-334
chr2-39297516-DEL-401
chr2-39318796-DEL-204
chr2-39328183-DEL-1735
chr2-39379224-DEL-168
chr2-39404315-DEL-8722
chr2-39407293-DEL-14309
chr2-39429819-DEL-1224
chr2-39442821-DEL-1648
chr2-39468884-DEL-169
chr2-39473649-DEL-51
chr2-39486269-DEL-151
chr2-39499970-DEL-888
chr2-39524947-DEL-76
chr2-39525383-DEL-12834
chr2-39535353-DEL-214
chr2-39567848-DEL-2817
chr2-39580361-DEL-3735
chr2-39654353-DEL-170
chr2-39681123-DEL-69
chr2-39681317-DEL-1733
chr2-39686951-DEL-245
chr2-39692537-DEL-116
chr2-39696374-DEL-3671
chr2-39701635-DEL-62
chr2-39717260-DEL-171
chr2-39737577-DEL-611
chr2-39746675-DEL-233
chr2-39777472-DEL-1540
chr2-39781774-DEL-51
chr2-39792014-DEL-153
chr2-39796276-DEL-127
chr2-39798103-DEL-8963
chr2-39810246-DEL-2003
chr2-39812649-DEL-2956
chr2-39828450-DEL-537
chr2-39880120-DEL-327
chr2-39892698-DEL-678
chr2-39907890-DEL-57
chr2-39910874-DEL-65
chr2-39916209-DEL-52
chr2-39918484-DEL-4033
chr2-39931724-DEL-52
chr2-39950503-DEL-1025
chr2-39957873-DEL-51
chr2-39977027-DEL-1821
chr2-39994569-DEL-77
chr2-39995686-DEL-50904
chr2-40049311-DEL-493
chr2-40068769-DEL-54
chr2-40090886-DEL-203
chr2-40103973-DEL-1250
chr2-40136969-DEL-504
chr2-40152558-DEL-6342
chr2-40176389-DEL-60
chr2-40212928-DEL-256
chr2-40237357-DEL-86
chr2-40256841-DEL-317
chr2-40258277-DEL-6407
chr2-40317210-DEL-2067
chr2-40340573-DEL-69
chr2-40370445-DEL-54
chr2-40421662-DEL-54

chr2-40434436-DEL-945
chr2-40444817-DEL-403
chr2-40445693-DEL-1778
chr2-40460955-DEL-259
chr2-40465615-DEL-2803
chr2-40471157-DEL-385
chr2-40482365-DEL-134
chr2-40498946-DEL-55
chr2-40511509-DEL-288
chr2-40523665-DEL-1328
chr2-40524901-DEL-84
chr2-40534695-DEL-110
chr2-40541357-DEL-1163
chr2-40542428-DEL-166
chr2-40546737-DEL-131
chr2-40587811-DEL-138
chr2-40597988-DEL-155
chr2-40599524-DEL-2006
chr2-40616401-DEL-231
chr2-40643527-DEL-849
chr2-40646773-DEL-131
chr2-40648212-DEL-554
chr2-40653853-DEL-6126
chr2-40672374-DEL-74
chr2-40677796-DEL-6240
chr2-40698431-DEL-1466
chr2-40709065-DEL-115
chr2-40713162-DEL-53
chr2-40718529-DEL-4788
chr2-40731257-DEL-2193
chr2-40741205-DEL-52
chr2-40741996-DEL-227
chr2-40753378-DEL-200
chr2-40754164-DEL-52
chr2-40759805-DEL-259
chr2-40794457-DEL-51
chr2-40841343-DEL-60
chr2-40857361-DEL-2500
chr2-40866363-DEL-912
chr2-40897929-DEL-476
chr2-40924503-DEL-800
chr2-40937266-DEL-26755
chr2-40945004-DEL-1600
chr2-40967124-DEL-694
chr2-40991250-DEL-9262
chr2-41009063-DEL-80
chr2-41016037-DEL-4592
chr2-41051269-DEL-261
chr2-41071806-DEL-539
chr2-41074712-DEL-54
chr2-41081314-DEL-125
chr2-41125860-DEL-52
chr2-41163746-DEL-595
chr2-41164488-DEL-963
chr2-41166462-DEL-345
chr2-41180958-DEL-199
chr2-41181601-DEL-2004
chr2-41194155-DEL-63
chr2-41201621-DEL-154
chr2-41323842-DEL-121
chr2-41326956-DEL-177
chr2-41341272-DEL-87
chr2-41345870-DEL-57
chr2-41357324-DEL-84
chr2-41368257-DEL-170
chr2-41405273-DEL-577
chr2-41418018-DEL-858
chr2-41418167-DEL-90
chr2-41432933-DEL-2935
chr2-41448088-DEL-71
chr2-41449400-DEL-113
chr2-41471246-DEL-206
chr2-41478708-DEL-387
chr2-41498222-DEL-843
chr2-41555813-DEL-1088
chr2-41558807-DEL-59
chr2-41566613-DEL-3361

chr2-41567137-DEL-346
chr2-41584476-DEL-4298
chr2-41599186-DEL-78
chr2-41599383-DEL-1637
chr2-41633143-DEL-239
chr2-41647664-DEL-72
chr2-41649863-DEL-60
chr2-41662752-DEL-291
chr2-41664484-DEL-801
chr2-41672176-DEL-108
chr2-41677013-DEL-75
chr2-41679785-DEL-55
chr2-41700687-DEL-82
chr2-41746463-DEL-144
chr2-41786148-DEL-78
chr2-41814987-DEL-629
chr2-41822594-DEL-733
chr2-41837767-DEL-386
chr2-41880964-DEL-607
chr2-41891255-DEL-488
chr2-41924197-DEL-108
chr2-41925351-DEL-1205
chr2-41943722-DEL-58
chr2-41948475-DEL-86
chr2-41956321-DEL-200
chr2-41963506-DEL-6174
chr2-41973452-DEL-6879
chr2-42010152-DEL-120
chr2-42011251-DEL-279
chr2-42012375-DEL-1693
chr2-42075049-DEL-362
chr2-42089418-DEL-150
chr2-42090170-DEL-1058
chr2-42091246-DEL-1755
chr2-42096316-DEL-69
chr2-42115994-DEL-414
chr2-42120552-DEL-117
chr2-42120762-DEL-777
chr2-42141933-DEL-66
chr2-42177762-DEL-149
chr2-42215662-DEL-74
chr2-42235127-DEL-1001
chr2-42246375-DEL-52
chr2-42294723-DEL-59
chr2-42317908-DEL-52
chr2-42318189-DEL-499
chr2-42343130-DEL-2422
chr2-42369085-DEL-136
chr2-42374692-DEL-134
chr2-42391050-DEL-1317
chr2-42396240-DEL-651
chr2-42436718-DEL-431
chr2-42468812-DEL-66
chr2-42473846-DEL-137
chr2-42497765-DEL-235
chr2-42512950-DEL-56
chr2-42543639-DEL-68
chr2-42563835-DEL-52
chr2-42581967-DEL-75
chr2-42594114-DEL-6969
chr2-42602651-DEL-85
chr2-42626943-DEL-752
chr2-42643096-DEL-2490
chr2-42657375-DEL-355
chr2-42677008-DEL-720
chr2-42695859-DEL-404
chr2-42719880-DEL-51
chr2-42725911-DEL-57
chr2-42741386-DEL-144
chr2-42767541-DEL-56
chr2-42779199-DEL-86
chr2-42847712-DEL-2788
chr2-42892073-DEL-154
chr2-42901999-DEL-51
chr2-42954780-DEL-83
chr2-42958416-DEL-599
chr2-42988563-DEL-206

chr2-43016383-DEL-554
chr2-43017292-DEL-65
chr2-43025524-DEL-1406
chr2-43109468-DEL-277
chr2-43114361-DEL-1024
chr2-43190189-DEL-824
chr2-43197034-DEL-123
chr2-43204974-DEL-862
chr2-43216987-DEL-3379
chr2-43217949-DEL-147
chr2-43219448-DEL-360
chr2-43235333-DEL-68
chr2-43237773-DEL-8954
chr2-43281179-DEL-326
chr2-43286355-DEL-117
chr2-43340978-DEL-81
chr2-43341380-DEL-1374
chr2-43388681-DEL-726
chr2-43389978-DEL-6251
chr2-43400217-DEL-69
chr2-43428662-DEL-78
chr2-43434343-DEL-56
chr2-43443181-DEL-55
chr2-43449720-DEL-320
chr2-43450175-DEL-417
chr2-43488354-DEL-353
chr2-43497616-DEL-55
chr2-43534828-DEL-69
chr2-43547748-DEL-2409
chr2-43597727-DEL-288
chr2-43603422-DEL-940
chr2-43641760-DEL-121
chr2-43668898-DEL-76
chr2-43732783-DEL-82
chr2-43740719-DEL-126
chr2-43741957-DEL-120
chr2-43744970-DEL-277
chr2-43812322-DEL-99
chr2-43935148-DEL-111
chr2-43938312-DEL-401
chr2-44000762-DEL-120
chr2-44064229-DEL-214
chr2-44068078-DEL-83
chr2-44091978-DEL-401
chr2-44110109-DEL-2754
chr2-44204067-DEL-211
chr2-44207761-DEL-258
chr2-44214173-DEL-73
chr2-44226005-DEL-99
chr2-44280498-DEL-382
chr2-44349641-DEL-1443
chr2-44385538-DEL-1120
chr2-44386940-DEL-102
chr2-44402311-DEL-93
chr2-44406732-DEL-644
chr2-44457192-DEL-219
chr2-44472834-DEL-56
chr2-44480507-DEL-325
chr2-44530397-DEL-59
chr2-44564363-DEL-1480
chr2-44613421-DEL-73
chr2-44622137-DEL-354
chr2-44669898-DEL-519
chr2-44670596-DEL-3414
chr2-44687271-DEL-275
chr2-44721265-DEL-152
chr2-44756541-DEL-1310
chr2-44796863-DEL-52
chr2-44826024-DEL-6482
chr2-44834665-DEL-61
chr2-44923982-DEL-53
chr2-44933299-DEL-80
chr2-44934042-DEL-71
chr2-44935610-DEL-262
chr2-45029810-DEL-730
chr2-45054992-DEL-52
chr2-45080887-DEL-255

chr2-45132082-DEL-82
chr2-45185361-DEL-399
chr2-45188027-DEL-72
chr2-45209824-DEL-3129
chr2-45211715-DEL-1079
chr2-45228357-DEL-112
chr2-45242177-DEL-62
chr2-45264276-DEL-95
chr2-45284627-DEL-702
chr2-45310412-DEL-186
chr2-45347681-DEL-57
chr2-45358289-DEL-112
chr2-45372212-DEL-576
chr2-45407673-DEL-218
chr2-45473208-DEL-56
chr2-45480167-DEL-215
chr2-45488733-DEL-228
chr2-45516629-DEL-372
chr2-45539054-DEL-199
chr2-45555392-DEL-54
chr2-45579436-DEL-251
chr2-45581071-DEL-994
chr2-45610262-DEL-217
chr2-45623201-DEL-392
chr2-45631853-DEL-922
chr2-45656667-DEL-135
chr2-45660291-DEL-176
chr2-45689538-DEL-210
chr2-45714563-DEL-744
chr2-45741336-DEL-65
chr2-45754383-DEL-1998
chr2-45823799-DEL-6372
chr2-45854124-DEL-2542
chr2-45915140-DEL-53
chr2-45937139-DEL-78
chr2-45941992-DEL-1492
chr2-45973435-DEL-59
chr2-46027643-DEL-64
chr2-46039924-DEL-148
chr2-46047963-DEL-393
chr2-46056530-DEL-55
chr2-46072124-DEL-141
chr2-46089020-DEL-159
chr2-46116998-DEL-59
chr2-46117686-DEL-67
chr2-46122810-DEL-1718
chr2-46137791-DEL-140
chr2-46165447-DEL-599
chr2-46171968-DEL-329
chr2-46225324-DEL-206
chr2-46233251-DEL-1497
chr2-46233365-DEL-4491
chr2-46233949-DEL-2997
chr2-46319403-DEL-52
chr2-46324053-DEL-56
chr2-46348622-DEL-70
chr2-46353576-DEL-210
chr2-46365966-DEL-76
chr2-46366863-DEL-62
chr2-46381310-DEL-3206
chr2-46394314-DEL-80
chr2-46404465-DEL-455
chr2-46405374-DEL-2401
chr2-46518360-DEL-202
chr2-46527524-DEL-80
chr2-46574668-DEL-425
chr2-46592855-DEL-540
chr2-46651233-DEL-442
chr2-46656246-DEL-557
chr2-46661010-DEL-472
chr2-46666972-DEL-58
chr2-46676600-DEL-202
chr2-46678595-DEL-400
chr2-46696453-DEL-70
chr2-46724378-DEL-7363
chr2-46738580-DEL-128
chr2-46798350-DEL-126

chr2-46807121-DEL-128
chr2-46837224-DEL-222
chr2-46843355-DEL-138
chr2-46866053-DEL-3047
chr2-46898684-DEL-1247
chr2-46915784-DEL-550
chr2-46928553-DEL-293
chr2-46931828-DEL-354
chr2-46958697-DEL-284
chr2-46971704-DEL-76
chr2-46980321-DEL-1158
chr2-46997322-DEL-132
chr2-47036604-DEL-10864
chr2-47084727-DEL-7068
chr2-47129801-DEL-115
chr2-47156757-DEL-208
chr2-47208683-DEL-59
chr2-47246865-DEL-399
chr2-47286282-DEL-132
chr2-47334388-DEL-725
chr2-47346007-DEL-1180
chr2-47347167-DEL-168
chr2-47367513-DEL-1081
chr2-47407123-DEL-182
chr2-47414958-DEL-283
chr2-47419330-DEL-392
chr2-47420202-DEL-5057
chr2-47429234-DEL-58
chr2-47435835-DEL-7080
chr2-47475459-DEL-1937
chr2-47478537-DEL-318
chr2-47494266-DEL-1063
chr2-47500804-DEL-1512
chr2-47513402-DEL-775
chr2-47514502-DEL-99
chr2-47517518-DEL-1058
chr2-47523697-DEL-374
chr2-47537299-DEL-144
chr2-47560658-DEL-4267
chr2-47564161-DEL-358
chr2-47566359-DEL-225
chr2-47576734-DEL-54
chr2-47582166-DEL-2879
chr2-47631595-DEL-150
chr2-47653995-DEL-1319
chr2-47695126-DEL-386
chr2-47699379-DEL-1716
chr2-47705146-DEL-2361
chr2-47750685-DEL-718
chr2-47791605-DEL-4174
chr2-47821212-DEL-1497
chr2-47827408-DEL-14613
chr2-47832382-DEL-1445
chr2-47837275-DEL-57
chr2-47854773-DEL-2624
chr2-47860154-DEL-85
chr2-47873716-DEL-102
chr2-47874397-DEL-284
chr2-47948682-DEL-372
chr2-47962362-DEL-3146
chr2-47978429-DEL-678
chr2-47986063-DEL-447
chr2-47996103-DEL-145
chr2-48021750-DEL-93
chr2-48103505-DEL-3860
chr2-48108464-DEL-62
chr2-48121096-DEL-307
chr2-48133468-DEL-384
chr2-48144365-DEL-261
chr2-48148626-DEL-55
chr2-48177201-DEL-410
chr2-48181540-DEL-53
chr2-48199797-DEL-843
chr2-48204899-DEL-78
chr2-48228006-DEL-3041
chr2-48249699-DEL-125
chr2-48258736-DEL-1460

chr2-48262201-DEL-1751
chr2-48273228-DEL-820
chr2-48304327-DEL-51
chr2-48329374-DEL-116
chr2-48372022-DEL-8997
chr2-48421091-DEL-760
chr2-48432550-DEL-784
chr2-48441864-DEL-86
chr2-48450830-DEL-393
chr2-48501643-DEL-108
chr2-48619895-DEL-69
chr2-48626958-DEL-79
chr2-48681060-DEL-2809
chr2-48690091-DEL-77
chr2-48726384-DEL-567
chr2-48796600-DEL-64
chr2-48810154-DEL-73
chr2-48810757-DEL-210
chr2-48838725-DEL-99
chr2-48865945-DEL-2946
chr2-48870272-DEL-219
chr2-48880732-DEL-166
chr2-48912118-DEL-1825
chr2-48946811-DEL-625
chr2-48947592-DEL-157
chr2-48952276-DEL-1451
chr2-49020316-DEL-81
chr2-49041944-DEL-90
chr2-49043077-DEL-101
chr2-49044030-DEL-3183
chr2-49069914-DEL-498
chr2-49072418-DEL-876
chr2-49098130-DEL-65
chr2-49101210-DEL-5276
chr2-49105517-DEL-1007
chr2-49115845-DEL-374
chr2-49130109-DEL-242
chr2-49138050-DEL-132
chr2-49138347-DEL-184
chr2-49260054-DEL-242
chr2-49263447-DEL-102
chr2-49291673-DEL-1112
chr2-49346116-DEL-699
chr2-49357864-DEL-206
chr2-49363425-DEL-52
chr2-49454454-DEL-2793
chr2-49497961-DEL-718
chr2-49522120-DEL-72
chr2-49572586-DEL-197
chr2-49650638-DEL-407
chr2-49669173-DEL-55
chr2-49675027-DEL-118
chr2-49729427-DEL-58
chr2-49731535-DEL-356
chr2-49756054-DEL-52
chr2-49764260-DEL-62
chr2-49819281-DEL-4205
chr2-49850375-DEL-54
chr2-49912582-DEL-70
chr2-49942726-DEL-453
chr2-49950294-DEL-197
chr2-50018174-DEL-56
chr2-50038168-DEL-925
chr2-50049262-DEL-125
chr2-50093628-DEL-747
chr2-50124169-DEL-199
chr2-50143649-DEL-728
chr2-50180965-DEL-92
chr2-50242231-DEL-447
chr2-50249199-DEL-137
chr2-50253296-DEL-56
chr2-50262897-DEL-52
chr2-50277768-DEL-6395
chr2-50279950-DEL-238
chr2-50358230-DEL-62
chr2-50384492-DEL-214
chr2-50446584-DEL-90

chr2-50455861-DEL-64
chr2-50490474-DEL-214
chr2-50523787-DEL-1783
chr2-50587259-DEL-80
chr2-50594031-DEL-65
chr2-50595547-DEL-203
chr2-50608871-DEL-2552
chr2-50662670-DEL-435
chr2-50662672-DEL-1463
chr2-50710035-DEL-120
chr2-50788944-DEL-382
chr2-50844104-DEL-101
chr2-50866985-DEL-64
chr2-50874574-DEL-65
chr2-50899088-DEL-54
chr2-50930768-DEL-54
chr2-50932380-DEL-61
chr2-50945121-DEL-184
chr2-50978523-DEL-139
chr2-50991666-DEL-194
chr2-50993416-DEL-58
chr2-51010608-DEL-52
chr2-51011480-DEL-178
chr2-51026316-DEL-113
chr2-51042935-DEL-51
chr2-51047625-DEL-122
chr2-51090538-DEL-152
chr2-51092725-DEL-220
chr2-51123480-DEL-92
chr2-51194223-DEL-103
chr2-51213767-DEL-56
chr2-51234068-DEL-55
chr2-51256777-DEL-481
chr2-51261419-DEL-97
chr2-51293596-DEL-52
chr2-51315023-DEL-141
chr2-51441807-DEL-68
chr2-51442867-DEL-2304
chr2-51478338-DEL-52
chr2-51488291-DEL-313
chr2-51493157-DEL-499
chr2-51515957-DEL-51
chr2-51516718-DEL-88
chr2-51519393-DEL-54
chr2-51550690-DEL-134
chr2-51557466-DEL-83
chr2-51579834-DEL-2344
chr2-51580238-DEL-252
chr2-51580455-DEL-1389
chr2-51581849-DEL-151
chr2-51596669-DEL-426
chr2-51601740-DEL-597
chr2-51681736-DEL-57
chr2-51696655-DEL-198
chr2-51706911-DEL-52
chr2-51730087-DEL-162
chr2-51781618-DEL-53
chr2-51783479-DEL-84
chr2-51806983-DEL-54
chr2-51836313-DEL-1656
chr2-51837572-DEL-423
chr2-51885084-DEL-249
chr2-51908575-DEL-93
chr2-51923594-DEL-66
chr2-51952532-DEL-216
chr2-51957349-DEL-268
chr2-51992460-DEL-79
chr2-52003852-DEL-257
chr2-52048987-DEL-91
chr2-52069626-DEL-82
chr2-52070161-DEL-140
chr2-52070685-DEL-333
chr2-52081154-DEL-73
chr2-52112059-DEL-161
chr2-52156618-DEL-58
chr2-52168654-DEL-57
chr2-52190200-DEL-719

chr2-52196959-DEL-54
chr2-52224620-DEL-89
chr2-52236119-DEL-146
chr2-52236929-DEL-2503
chr2-52238629-DEL-1487
chr2-52252308-DEL-68
chr2-52258279-DEL-86
chr2-52296312-DEL-149
chr2-52306024-DEL-68
chr2-52412105-DEL-59
chr2-52421653-DEL-207
chr2-52429100-DEL-67
chr2-52436949-DEL-86
chr2-52438193-DEL-60
chr2-52441375-DEL-188
chr2-52450089-DEL-488
chr2-52471642-DEL-354
chr2-52476072-DEL-350
chr2-52482010-DEL-106
chr2-52498798-DEL-1532
chr2-52509735-DEL-903
chr2-52519523-DEL-75
chr2-52549356-DEL-488
chr2-52562152-DEL-76
chr2-52586591-DEL-54
chr2-52597102-DEL-96
chr2-52607491-DEL-392
chr2-52610037-DEL-53
chr2-52625094-DEL-106
chr2-52627904-DEL-492
chr2-52697552-DEL-57
chr2-52727668-DEL-66
chr2-52728912-DEL-192
chr2-52734186-DEL-55
chr2-52815682-DEL-261
chr2-52838368-DEL-56
chr2-52850474-DEL-2089
chr2-52873804-DEL-58
chr2-52877315-DEL-73
chr2-52894840-DEL-204
chr2-52929924-DEL-80
chr2-52952579-DEL-67
chr2-52982970-DEL-265
chr2-53014112-DEL-72
chr2-53032764-DEL-1763
chr2-53037544-DEL-119
chr2-53056290-DEL-150
chr2-53060871-DEL-135
chr2-53063756-DEL-233
chr2-53086909-DEL-56
chr2-53101877-DEL-52
chr2-53125821-DEL-97
chr2-53131947-DEL-68
chr2-53150749-DEL-68
chr2-53161247-DEL-533
chr2-53177143-DEL-64
chr2-53207686-DEL-6219
chr2-53213789-DEL-6646
chr2-53220716-DEL-56
chr2-53237766-DEL-6634
chr2-53245251-DEL-65
chr2-53251279-DEL-6345
chr2-53262171-DEL-95
chr2-53270349-DEL-56
chr2-53275627-DEL-198
chr2-53277296-DEL-63
chr2-53280338-DEL-78
chr2-53297903-DEL-81
chr2-53310640-DEL-1553
chr2-53317479-DEL-547
chr2-53319252-DEL-60
chr2-53321226-DEL-405
chr2-53330528-DEL-2499
chr2-53345418-DEL-428
chr2-53354665-DEL-148
chr2-53366988-DEL-212
chr2-53414912-DEL-981

chr2-53417914-DEL-61
chr2-53419829-DEL-149
chr2-53424163-DEL-268
chr2-53443989-DEL-6172
chr2-53484077-DEL-777
chr2-53505802-DEL-652
chr2-53506624-DEL-1271
chr2-53549802-DEL-1208
chr2-53556253-DEL-213
chr2-53574415-DEL-78
chr2-53583563-DEL-224
chr2-53610621-DEL-54
chr2-53618643-DEL-62
chr2-53634991-DEL-6001
chr2-53665941-DEL-6382
chr2-53665943-DEL-12950
chr2-53679201-DEL-6022
chr2-53722855-DEL-165
chr2-53739327-DEL-133
chr2-53760167-DEL-285
chr2-53784187-DEL-88
chr2-53794144-DEL-181
chr2-53816951-DEL-98
chr2-53850363-DEL-308
chr2-53860463-DEL-196
chr2-53876979-DEL-1402
chr2-53881303-DEL-6367
chr2-53896745-DEL-1308
chr2-53898663-DEL-5242
chr2-53904607-DEL-111
chr2-53914135-DEL-352
chr2-53961359-DEL-200
chr2-54007353-DEL-62
chr2-54038940-DEL-620
chr2-54059055-DEL-3187
chr2-54102373-DEL-739
chr2-54117385-DEL-79
chr2-54132237-DEL-192
chr2-54159995-DEL-82
chr2-54199350-DEL-6267
chr2-54233784-DEL-138
chr2-54234322-DEL-176
chr2-54262043-DEL-747
chr2-54267709-DEL-154
chr2-54289397-DEL-5163
chr2-54334245-DEL-951
chr2-54368309-DEL-52
chr2-54403794-DEL-133
chr2-54419233-DEL-129
chr2-54443725-DEL-232
chr2-54457496-DEL-2367
chr2-54458837-DEL-6290
chr2-54460008-DEL-6898
chr2-54496960-DEL-796
chr2-54497821-DEL-985
chr2-54512554-DEL-532
chr2-54518744-DEL-100
chr2-54533215-DEL-902
chr2-54545796-DEL-528
chr2-54579679-DEL-116
chr2-54585097-DEL-441
chr2-54587980-DEL-26315
chr2-54587981-DEL-24172
chr2-54605358-DEL-146
chr2-54664340-DEL-907
chr2-54667186-DEL-117
chr2-54720287-DEL-69
chr2-54721902-DEL-133
chr2-54739331-DEL-831
chr2-54745761-DEL-83
chr2-54764050-DEL-60
chr2-54764704-DEL-92
chr2-54769363-DEL-131
chr2-54775645-DEL-207
chr2-54782046-DEL-2453
chr2-54795278-DEL-240
chr2-54804605-DEL-62

chr2-54827696-DEL-169
chr2-54843679-DEL-53
chr2-54858383-DEL-104
chr2-54962093-DEL-51
chr2-54981066-DEL-55
chr2-54993061-DEL-3631
chr2-55024485-DEL-296
chr2-55034733-DEL-3276
chr2-55142259-DEL-124
chr2-55162195-DEL-316
chr2-55251720-DEL-239
chr2-55259068-DEL-602
chr2-55285085-DEL-4050
chr2-55294510-DEL-63
chr2-55299736-DEL-88
chr2-55353655-DEL-75
chr2-55371295-DEL-65
chr2-55379652-DEL-400
chr2-55404084-DEL-60
chr2-55423256-DEL-54
chr2-55426478-DEL-467
chr2-55443263-DEL-534
chr2-55445252-DEL-586
chr2-55452582-DEL-618
chr2-55470831-DEL-70
chr2-55478092-DEL-60
chr2-55478093-DEL-2170
chr2-55512548-DEL-6750
chr2-55521669-DEL-92
chr2-55523333-DEL-265
chr2-55533090-DEL-157
chr2-55536853-DEL-334
chr2-55586263-DEL-1362
chr2-55594795-DEL-4978
chr2-55663609-DEL-98
chr2-55670372-DEL-73
chr2-55689397-DEL-608
chr2-55703774-DEL-1024
chr2-55705517-DEL-324
chr2-55706360-DEL-114
chr2-55713746-DEL-1795
chr2-55713746-DEL-261
chr2-55713746-DEL-4772
chr2-55715579-DEL-4512
chr2-55718553-DEL-1534
chr2-55725255-DEL-63
chr2-55744697-DEL-94
chr2-55745523-DEL-671
chr2-55773854-DEL-1284
chr2-55804514-DEL-52
chr2-55806775-DEL-645
chr2-55826695-DEL-352
chr2-55884756-DEL-419
chr2-55911875-DEL-782
chr2-55935589-DEL-10982
chr2-55935698-DEL-10224
chr2-55947091-DEL-271
chr2-55948213-DEL-184
chr2-55948905-DEL-231
chr2-55973389-DEL-65
chr2-55974664-DEL-50
chr2-55986912-DEL-2868
chr2-56016193-DEL-233
chr2-56048286-DEL-726
chr2-56059345-DEL-53
chr2-56063412-DEL-67
chr2-56082836-DEL-1844
chr2-56099212-DEL-598
chr2-56118858-DEL-179
chr2-56124311-DEL-27004
chr2-56131593-DEL-248
chr2-56132508-DEL-777
chr2-56137003-DEL-52
chr2-56138244-DEL-350
chr2-56141334-DEL-108
chr2-56151122-DEL-1377
chr2-56289595-DEL-415

chr2-56297790-DEL-64
chr2-56300576-DEL-486
chr2-56302321-DEL-52
chr2-56320119-DEL-228
chr2-56322240-DEL-51
chr2-56324843-DEL-364
chr2-56330943-DEL-173
chr2-56338459-DEL-140
chr2-56390370-DEL-400
chr2-56395858-DEL-114
chr2-56396547-DEL-102
chr2-56399169-DEL-831
chr2-56401442-DEL-52
chr2-56415015-DEL-52
chr2-56434124-DEL-78
chr2-56488056-DEL-114
chr2-56542777-DEL-663
chr2-56556721-DEL-71
chr2-56587522-DEL-139
chr2-56589865-DEL-457
chr2-56600417-DEL-198
chr2-56647346-DEL-300
chr2-56666623-DEL-148
chr2-56678541-DEL-84
chr2-56679590-DEL-124
chr2-56689340-DEL-423
chr2-56700253-DEL-436
chr2-56729791-DEL-6229
chr2-56750164-DEL-192
chr2-56795825-DEL-1331
chr2-56817777-DEL-604
chr2-56839522-DEL-351
chr2-56852232-DEL-1700
chr2-56857498-DEL-594
chr2-56865514-DEL-229
chr2-56928176-DEL-108
chr2-56939759-DEL-1399
chr2-56967794-DEL-70
chr2-56980551-DEL-67
chr2-57026806-DEL-423
chr2-57070307-DEL-62
chr2-57083985-DEL-753
chr2-57102323-DEL-92
chr2-57167522-DEL-1113
chr2-57183641-DEL-67
chr2-57216811-DEL-64
chr2-57275737-DEL-320
chr2-57284675-DEL-92
chr2-57306342-DEL-186
chr2-57319838-DEL-2058
chr2-57375728-DEL-83
chr2-57401477-DEL-1264
chr2-57474591-DEL-97
chr2-57475974-DEL-2305
chr2-57518355-DEL-7747
chr2-57535425-DEL-379
chr2-57565766-DEL-3618
chr2-57577081-DEL-283
chr2-57616038-DEL-198
chr2-57628441-DEL-97
chr2-57630459-DEL-258
chr2-57673483-DEL-628
chr2-57681523-DEL-58
chr2-57689591-DEL-1585
chr2-57704443-DEL-1179
chr2-57706613-DEL-85
chr2-57718265-DEL-59
chr2-57752668-DEL-1316
chr2-57760654-DEL-119
chr2-57775614-DEL-73
chr2-57777451-DEL-124
chr2-57786825-DEL-80
chr2-57833298-DEL-82
chr2-57884082-DEL-396
chr2-57953319-DEL-130
chr2-57957711-DEL-6226
chr2-58006182-DEL-886

chr2-58014394-DEL-5267
chr2-58025915-DEL-54
chr2-58041563-DEL-64
chr2-58051865-DEL-154
chr2-58059555-DEL-142
chr2-58070390-DEL-407
chr2-58096918-DEL-320
chr2-58101323-DEL-4488
chr2-58170016-DEL-179
chr2-58180696-DEL-130
chr2-58181945-DEL-53
chr2-58193779-DEL-82
chr2-58230587-DEL-77
chr2-58261244-DEL-134
chr2-58280860-DEL-51
chr2-58308479-DEL-61
chr2-58311408-DEL-248
chr2-58315903-DEL-249
chr2-58328503-DEL-793
chr2-58330059-DEL-177
chr2-58376402-DEL-84
chr2-58382241-DEL-87
chr2-58384547-DEL-132
chr2-58448016-DEL-202
chr2-58456868-DEL-106
chr2-58469567-DEL-154
chr2-58480124-DEL-57
chr2-58514057-DEL-198
chr2-58537713-DEL-6594
chr2-58537812-DEL-201
chr2-58549324-DEL-119
chr2-58588211-DEL-74
chr2-58602618-DEL-223
chr2-58607313-DEL-478
chr2-58769147-DEL-187
chr2-58771240-DEL-6777
chr2-58777505-DEL-208
chr2-58778552-DEL-208
chr2-58784324-DEL-1797
chr2-58786591-DEL-5971
chr2-58799974-DEL-146
chr2-58805216-DEL-1824
chr2-58851739-DEL-51
chr2-58873616-DEL-60
chr2-58875346-DEL-81
chr2-58915584-DEL-51
chr2-58920000-DEL-74
chr2-58946624-DEL-554
chr2-58947118-DEL-386
chr2-58947843-DEL-411
chr2-58948754-DEL-219
chr2-58949653-DEL-128
chr2-58958266-DEL-1447
chr2-59018810-DEL-412
chr2-59037731-DEL-6737
chr2-59052956-DEL-915
chr2-59116071-DEL-64
chr2-59130340-DEL-1193
chr2-59149533-DEL-68
chr2-59189295-DEL-188
chr2-59220049-DEL-184
chr2-59223080-DEL-54
chr2-59227665-DEL-55
chr2-59250580-DEL-90
chr2-59274550-DEL-2595
chr2-59291229-DEL-71
chr2-59308677-DEL-52
chr2-59382258-DEL-452
chr2-59545738-DEL-60
chr2-59554101-DEL-240
chr2-59581370-DEL-92
chr2-59611708-DEL-110
chr2-59612587-DEL-82
chr2-59620539-DEL-91
chr2-59642550-DEL-67
chr2-59642875-DEL-2477
chr2-59646339-DEL-56

chr2-59649187-DEL-95
chr2-59658465-DEL-81
chr2-59666958-DEL-56
chr2-59671967-DEL-70
chr2-59687126-DEL-1294
chr2-59700411-DEL-173
chr2-59735773-DEL-63
chr2-59759266-DEL-1085
chr2-59759526-DEL-318
chr2-59801891-DEL-1071
chr2-59834303-DEL-192
chr2-59839284-DEL-453
chr2-59847380-DEL-200
chr2-59871437-DEL-175
chr2-59900945-DEL-231
chr2-59940342-DEL-232
chr2-59967389-DEL-200
chr2-59987736-DEL-1989
chr2-60000329-DEL-77
chr2-60006839-DEL-1143
chr2-60009672-DEL-766
chr2-60018966-DEL-309
chr2-60047752-DEL-1651
chr2-60057478-DEL-64
chr2-60090326-DEL-174
chr2-60095027-DEL-498
chr2-60100746-DEL-330
chr2-60119574-DEL-73
chr2-60134770-DEL-202
chr2-60166066-DEL-61
chr2-60168791-DEL-82
chr2-60169412-DEL-455
chr2-60219905-DEL-61
chr2-60231314-DEL-396
chr2-60267823-DEL-61
chr2-60278133-DEL-60
chr2-60298993-DEL-108
chr2-60337698-DEL-882
chr2-60351931-DEL-792
chr2-60396050-DEL-323
chr2-60404121-DEL-194
chr2-60417310-DEL-6124
chr2-60446048-DEL-337
chr2-60457432-DEL-80
chr2-60462155-DEL-868
chr2-60483017-DEL-191
chr2-60488594-DEL-52
chr2-60536721-DEL-85
chr2-60626433-DEL-60
chr2-60637226-DEL-97
chr2-60754515-DEL-73
chr2-60759034-DEL-65
chr2-60783011-DEL-76
chr2-60845207-DEL-54
chr2-60846742-DEL-327
chr2-60879382-DEL-60
chr2-60908585-DEL-178
chr2-60921611-DEL-244
chr2-60941841-DEL-598
chr2-60961207-DEL-174
chr2-60994561-DEL-76
chr2-61026563-DEL-53
chr2-61047566-DEL-645
chr2-61057039-DEL-54
chr2-61140089-DEL-52
chr2-61196505-DEL-52
chr2-61237829-DEL-412
chr2-61249723-DEL-378
chr2-61255906-DEL-182
chr2-61265624-DEL-108
chr2-61283646-DEL-71
chr2-61330718-DEL-505
chr2-61358673-DEL-61
chr2-61389565-DEL-1429
chr2-61402932-DEL-921
chr2-61415995-DEL-1417
chr2-61438794-DEL-62

chr2-61439629-DEL-2846
chr2-61504187-DEL-75
chr2-61507421-DEL-60
chr2-61509319-DEL-779
chr2-61545259-DEL-534
chr2-61607378-DEL-58
chr2-61620929-DEL-892
chr2-61622291-DEL-109
chr2-61668116-DEL-460
chr2-61676999-DEL-6879
chr2-61685837-DEL-1269
chr2-61691878-DEL-95
chr2-61693575-DEL-5749
chr2-61734082-DEL-502
chr2-61746223-DEL-168
chr2-61748008-DEL-1133
chr2-61801353-DEL-1697
chr2-61804049-DEL-2227
chr2-61823011-DEL-88
chr2-61830602-DEL-1821
chr2-61906099-DEL-75
chr2-61922336-DEL-53
chr2-61950902-DEL-190
chr2-61962543-DEL-1058
chr2-61998111-DEL-753
chr2-62042549-DEL-82
chr2-62113578-DEL-53
chr2-62158525-DEL-57
chr2-62202459-DEL-71
chr2-62219743-DEL-868
chr2-62246470-DEL-340
chr2-62272708-DEL-236
chr2-62301427-DEL-158
chr2-62303253-DEL-148
chr2-62316664-DEL-60
chr2-62324379-DEL-65
chr2-62327515-DEL-64
chr2-62369939-DEL-434
chr2-62394888-DEL-66
chr2-62486813-DEL-394
chr2-62506415-DEL-68
chr2-62540570-DEL-72
chr2-62603586-DEL-66
chr2-62615925-DEL-86
chr2-62641788-DEL-199
chr2-62659454-DEL-75
chr2-62704809-DEL-169
chr2-62710835-DEL-92
chr2-62714123-DEL-54
chr2-62719892-DEL-1594
chr2-62731838-DEL-80
chr2-62816731-DEL-70
chr2-62861178-DEL-55
chr2-62880854-DEL-58
chr2-62942132-DEL-69
chr2-62950506-DEL-533
chr2-62976499-DEL-66
chr2-62991312-DEL-105
chr2-63055698-DEL-138
chr2-63071149-DEL-2136
chr2-63073304-DEL-1105
chr2-63081009-DEL-178
chr2-63171143-DEL-110
chr2-63214160-DEL-645
chr2-63217048-DEL-263
chr2-63225852-DEL-259
chr2-63259690-DEL-396
chr2-63270737-DEL-52
chr2-63289192-DEL-114
chr2-63294790-DEL-76
chr2-63298675-DEL-53
chr2-63339110-DEL-415
chr2-63385347-DEL-203
chr2-63429506-DEL-391
chr2-63477896-DEL-427
chr2-63503789-DEL-251
chr2-63540583-DEL-377

chr2-63544666-DEL-81
chr2-63612049-DEL-202
chr2-63635852-DEL-408
chr2-63720540-DEL-52
chr2-63721125-DEL-64
chr2-63734455-DEL-386
chr2-63783190-DEL-329
chr2-63833646-DEL-124
chr2-63844573-DEL-62
chr2-63936517-DEL-485
chr2-63940747-DEL-51
chr2-63955173-DEL-196
chr2-63967214-DEL-320
chr2-64010943-DEL-68
chr2-64019957-DEL-820
chr2-64042262-DEL-450
chr2-64058679-DEL-3763
chr2-64062525-DEL-68
chr2-64086296-DEL-840
chr2-64104273-DEL-778
chr2-64105266-DEL-641
chr2-64111565-DEL-78
chr2-64139265-DEL-1690
chr2-64166996-DEL-94
chr2-64217499-DEL-346
chr2-64232234-DEL-890
chr2-64298486-DEL-60
chr2-64307732-DEL-126
chr2-64329077-DEL-1707
chr2-64335206-DEL-438
chr2-64373031-DEL-76
chr2-64389133-DEL-54
chr2-64491943-DEL-174
chr2-64508357-DEL-2447
chr2-64526144-DEL-80
chr2-64545286-DEL-152
chr2-64550740-DEL-396
chr2-64563170-DEL-51
chr2-64573713-DEL-6412
chr2-64581018-DEL-511
chr2-64585149-DEL-809
chr2-64602785-DEL-112
chr2-64710957-DEL-68
chr2-64714461-DEL-105
chr2-64772079-DEL-164
chr2-64829388-DEL-118
chr2-64846508-DEL-72
chr2-64906672-DEL-323
chr2-64939593-DEL-70
chr2-64967041-DEL-837
chr2-64972269-DEL-250
chr2-65016630-DEL-62
chr2-65020410-DEL-239
chr2-65024407-DEL-75
chr2-65026900-DEL-1631
chr2-65033106-DEL-385
chr2-65073387-DEL-82
chr2-65095756-DEL-111
chr2-65131999-DEL-88
chr2-65170209-DEL-210
chr2-65187684-DEL-204
chr2-65203662-DEL-1935
chr2-65214106-DEL-99
chr2-65216848-DEL-53
chr2-65261290-DEL-397
chr2-65271136-DEL-53
chr2-65281560-DEL-101
chr2-65303754-DEL-312
chr2-65328585-DEL-59
chr2-65396609-DEL-57
chr2-65410137-DEL-2495
chr2-65411462-DEL-162
chr2-65435471-DEL-55
chr2-65460488-DEL-94
chr2-65463096-DEL-165
chr2-65475662-DEL-299
chr2-65496675-DEL-112

chr2-65527044-DEL-331
chr2-65533662-DEL-991
chr2-65593459-DEL-147
chr2-65619781-DEL-62
chr2-65625510-DEL-1192
chr2-65625707-DEL-765
chr2-65626370-DEL-864
chr2-65626877-DEL-297
chr2-65632144-DEL-294
chr2-65645275-DEL-67
chr2-65673127-DEL-106
chr2-65692397-DEL-59
chr2-65705534-DEL-678
chr2-65714888-DEL-51
chr2-65729385-DEL-78
chr2-65766111-DEL-252
chr2-65814720-DEL-1891
chr2-65838989-DEL-215
chr2-65875678-DEL-1310
chr2-65875876-DEL-161
chr2-65886078-DEL-6365
chr2-65959908-DEL-316
chr2-66005653-DEL-75
chr2-66009358-DEL-436
chr2-66045056-DEL-64
chr2-66090815-DEL-498
chr2-66117489-DEL-116
chr2-66136357-DEL-6360
chr2-66144396-DEL-2275
chr2-66151462-DEL-60
chr2-66212522-DEL-498
chr2-66216255-DEL-239
chr2-66242656-DEL-66
chr2-66292873-DEL-65
chr2-66318248-DEL-61
chr2-66336280-DEL-270
chr2-66377087-DEL-70
chr2-66420919-DEL-1137
chr2-66422557-DEL-78
chr2-66436519-DEL-174
chr2-66459109-DEL-60
chr2-66467305-DEL-55
chr2-66471988-DEL-56
chr2-66483265-DEL-1576
chr2-66488064-DEL-96
chr2-66495272-DEL-231
chr2-66529370-DEL-354
chr2-66548129-DEL-112
chr2-66561284-DEL-6184
chr2-66574535-DEL-6246
chr2-66721748-DEL-104
chr2-66738477-DEL-155
chr2-66738985-DEL-790
chr2-66748167-DEL-337
chr2-66760487-DEL-102
chr2-66775629-DEL-88
chr2-66789654-DEL-56
chr2-66836684-DEL-6304
chr2-66843419-DEL-52
chr2-66846686-DEL-78
chr2-66858341-DEL-63
chr2-66864087-DEL-55
chr2-66887374-DEL-6175
chr2-66920214-DEL-206
chr2-66945264-DEL-3016
chr2-66981593-DEL-313
chr2-66989382-DEL-141
chr2-67020745-DEL-64
chr2-67073538-DEL-161
chr2-67076859-DEL-65
chr2-67082906-DEL-2785
chr2-67085007-DEL-1384
chr2-67093829-DEL-67
chr2-67127587-DEL-56
chr2-67149580-DEL-199
chr2-67172666-DEL-161
chr2-67179203-DEL-120

chr2-67210287-DEL-63
chr2-67222021-DEL-139
chr2-67238667-DEL-6280
chr2-67246782-DEL-149
chr2-67278199-DEL-98
chr2-67297009-DEL-54
chr2-67319612-DEL-59
chr2-67360819-DEL-230
chr2-67365025-DEL-95
chr2-67366699-DEL-197
chr2-67399380-DEL-104
chr2-67414747-DEL-52
chr2-67428876-DEL-571
chr2-67434105-DEL-780
chr2-67439847-DEL-66
chr2-67474922-DEL-149
chr2-67489649-DEL-579
chr2-67490122-DEL-386
chr2-67550658-DEL-836
chr2-67552161-DEL-62
chr2-67559358-DEL-86
chr2-67651965-DEL-82
chr2-67686872-DEL-459
chr2-67690256-DEL-7329
chr2-67704320-DEL-57
chr2-67717821-DEL-776
chr2-67860999-DEL-742
chr2-67898826-DEL-87
chr2-67906932-DEL-115
chr2-67915309-DEL-107
chr2-67933021-DEL-68
chr2-67960084-DEL-8162
chr2-67981787-DEL-3688
chr2-68033148-DEL-871
chr2-68059099-DEL-62
chr2-68098889-DEL-104
chr2-68128683-DEL-1994
chr2-68214754-DEL-62
chr2-68224807-DEL-231
chr2-68227835-DEL-246
chr2-68239250-DEL-113
chr2-68252702-DEL-444
chr2-68263789-DEL-717
chr2-68267155-DEL-63
chr2-68275988-DEL-97
chr2-68278832-DEL-85
chr2-68284615-DEL-93
chr2-68291129-DEL-1171
chr2-68345427-DEL-64
chr2-68346696-DEL-89
chr2-68347601-DEL-678
chr2-68376795-DEL-10108
chr2-68394148-DEL-159
chr2-68404017-DEL-54
chr2-68440060-DEL-145
chr2-68465956-DEL-75
chr2-68485554-DEL-1621
chr2-68488985-DEL-766
chr2-68504889-DEL-204
chr2-68556748-DEL-239
chr2-68584231-DEL-310
chr2-68643360-DEL-143
chr2-68695954-DEL-170
chr2-68760261-DEL-52
chr2-68835452-DEL-199
chr2-68838396-DEL-56
chr2-68851089-DEL-138
chr2-68854251-DEL-132
chr2-68870495-DEL-66
chr2-68904300-DEL-146
chr2-68943367-DEL-207
chr2-68977291-DEL-317
chr2-69063326-DEL-74
chr2-69124619-DEL-882
chr2-69133249-DEL-77
chr2-69134810-DEL-54
chr2-69150610-DEL-56

chr2-69177591-DEL-690
chr2-69178480-DEL-278
chr2-69188145-DEL-55
chr2-69198321-DEL-51
chr2-69221419-DEL-73
chr2-69252468-DEL-57
chr2-69258079-DEL-68
chr2-69287407-DEL-1092
chr2-69361660-DEL-161
chr2-69392034-DEL-234
chr2-69406446-DEL-59
chr2-69407750-DEL-64
chr2-69410033-DEL-59
chr2-69426605-DEL-182
chr2-69443752-DEL-57
chr2-69473981-DEL-175
chr2-69486458-DEL-62
chr2-69544132-DEL-83
chr2-69556368-DEL-525
chr2-69559209-DEL-504
chr2-69567194-DEL-1358
chr2-69596719-DEL-902
chr2-69601563-DEL-180
chr2-69635212-DEL-184
chr2-69639593-DEL-59
chr2-69640092-DEL-404
chr2-69667812-DEL-225
chr2-69718488-DEL-214
chr2-69723079-DEL-194
chr2-69732036-DEL-2344
chr2-69745537-DEL-193
chr2-69745983-DEL-536
chr2-69748870-DEL-58
chr2-69760865-DEL-842
chr2-69793758-DEL-105
chr2-69820991-DEL-62
chr2-69831678-DEL-112
chr2-69842187-DEL-173
chr2-69856937-DEL-54
chr2-69859112-DEL-209
chr2-69865670-DEL-52
chr2-69873835-DEL-206
chr2-69896539-DEL-104
chr2-69918476-DEL-1893
chr2-69924260-DEL-241
chr2-69973927-DEL-123
chr2-69995690-DEL-105
chr2-70052220-DEL-114
chr2-70055651-DEL-64
chr2-70092388-DEL-69
chr2-70106762-DEL-165
chr2-70182288-DEL-2201
chr2-70206528-DEL-54
chr2-70223517-DEL-480
chr2-70227615-DEL-126
chr2-70240752-DEL-57
chr2-70243199-DEL-80
chr2-70250650-DEL-95
chr2-70253233-DEL-63
chr2-70258615-DEL-148
chr2-70284094-DEL-80
chr2-70295313-DEL-118
chr2-70317702-DEL-251
chr2-70320216-DEL-1822
chr2-70323914-DEL-408
chr2-70326734-DEL-211
chr2-70336899-DEL-158
chr2-70357532-DEL-78
chr2-70368685-DEL-86
chr2-70409638-DEL-62
chr2-70412842-DEL-74
chr2-70416462-DEL-101
chr2-70467680-DEL-336
chr2-70470476-DEL-444
chr2-70502063-DEL-175
chr2-70524264-DEL-405
chr2-70529944-DEL-207

chr2-70531946-DEL-612
chr2-70550995-DEL-63
chr2-70554780-DEL-207
chr2-70584242-DEL-79
chr2-70627774-DEL-187
chr2-70634421-DEL-52
chr2-70636705-DEL-72
chr2-70638276-DEL-208
chr2-70690372-DEL-200
chr2-70692842-DEL-56
chr2-70697457-DEL-55
chr2-70700775-DEL-59
chr2-70721580-DEL-85
chr2-70721793-DEL-1586
chr2-70728414-DEL-113
chr2-70729626-DEL-522
chr2-70760598-DEL-435
chr2-70842924-DEL-68
chr2-70873001-DEL-428
chr2-70894462-DEL-245
chr2-70924537-DEL-198
chr2-70927722-DEL-183
chr2-70936824-DEL-2564
chr2-70992084-DEL-361
chr2-71003775-DEL-178
chr2-71016294-DEL-1266
chr2-71024305-DEL-382
chr2-71040366-DEL-51
chr2-71043190-DEL-227
chr2-71049514-DEL-69
chr2-71094971-DEL-111
chr2-71116454-DEL-170
chr2-71118508-DEL-54
chr2-71134131-DEL-731
chr2-71172889-DEL-56
chr2-71180726-DEL-323
chr2-71203237-DEL-179
chr2-71207332-DEL-763
chr2-71225767-DEL-211
chr2-71240631-DEL-212
chr2-71257687-DEL-320
chr2-71270602-DEL-52
chr2-71278820-DEL-90
chr2-71290350-DEL-1416
chr2-71311801-DEL-91
chr2-71386387-DEL-1122
chr2-71389579-DEL-71
chr2-71413981-DEL-286
chr2-71440158-DEL-121
chr2-71463293-DEL-55
chr2-71491620-DEL-179
chr2-71516257-DEL-284
chr2-71524614-DEL-110
chr2-71548121-DEL-58
chr2-71552099-DEL-76
chr2-71567660-DEL-305
chr2-71643647-DEL-54
chr2-71734652-DEL-59
chr2-71738377-DEL-330
chr2-71772482-DEL-3672
chr2-71776187-DEL-3448
chr2-71789711-DEL-192
chr2-71801320-DEL-89
chr2-71818373-DEL-68
chr2-71848958-DEL-90
chr2-71857192-DEL-58
chr2-71873761-DEL-68
chr2-71904548-DEL-68
chr2-71939951-DEL-167
chr2-71970292-DEL-72
chr2-72024107-DEL-123
chr2-72098555-DEL-120
chr2-72147141-DEL-71
chr2-72217618-DEL-165
chr2-72242575-DEL-839
chr2-72244489-DEL-109
chr2-72285018-DEL-204

chr2-72315665-DEL-120
chr2-72331820-DEL-102
chr2-72366364-DEL-95
chr2-72396291-DEL-159
chr2-72407885-DEL-52
chr2-72494125-DEL-74
chr2-72536391-DEL-502
chr2-72549465-DEL-1224
chr2-72550609-DEL-476
chr2-72633505-DEL-56
chr2-72634826-DEL-118
chr2-72638114-DEL-65
chr2-72650445-DEL-53
chr2-72666779-DEL-63
chr2-72673185-DEL-87
chr2-72688732-DEL-433
chr2-72720710-DEL-114
chr2-72745031-DEL-54
chr2-72775218-DEL-190
chr2-72825270-DEL-155
chr2-72827826-DEL-57
chr2-72835358-DEL-62
chr2-72853709-DEL-112
chr2-72876221-DEL-60
chr2-72936438-DEL-170
chr2-72953013-DEL-72
chr2-72966227-DEL-384
chr2-72976206-DEL-94
chr2-73011400-DEL-719
chr2-73056600-DEL-167
chr2-73087283-DEL-188
chr2-73091758-DEL-205
chr2-73095521-DEL-53
chr2-73126858-DEL-176
chr2-73144487-DEL-93
chr2-73223725-DEL-332
chr2-73248614-DEL-69
chr2-73255852-DEL-312
chr2-73285822-DEL-130
chr2-73306861-DEL-61
chr2-73323075-DEL-58
chr2-73367295-DEL-93
chr2-73397916-DEL-121
chr2-73399323-DEL-75
chr2-73425994-DEL-55
chr2-73456828-DEL-68
chr2-73460845-DEL-90
chr2-73480899-DEL-88
chr2-73499377-DEL-3910
chr2-73502652-DEL-104
chr2-73516929-DEL-296
chr2-73539425-DEL-125
chr2-73541824-DEL-51
chr2-73543590-DEL-83
chr2-73547789-DEL-90
chr2-73556167-DEL-547
chr2-73562213-DEL-198
chr2-73759111-DEL-1579
chr2-73824061-DEL-258
chr2-73828342-DEL-60
chr2-73838234-DEL-78
chr2-73840371-DEL-72
chr2-73868883-DEL-299
chr2-73872736-DEL-3218
chr2-73935269-DEL-90
chr2-73945681-DEL-70
chr2-73967486-DEL-169
chr2-73970914-DEL-58
chr2-74027519-DEL-72
chr2-74051624-DEL-613
chr2-74069403-DEL-216
chr2-74081701-DEL-60
chr2-74097474-DEL-68
chr2-74132029-DEL-90
chr2-74137233-DEL-1858
chr2-74151540-DEL-66
chr2-74245357-DEL-167

chr2-74292619-DEL-75
chr2-74293013-DEL-346
chr2-74304911-DEL-1112
chr2-74339686-DEL-114
chr2-74392339-DEL-52
chr2-74416510-DEL-5326
chr2-74424180-DEL-1045
chr2-74448547-DEL-67
chr2-74469099-DEL-55
chr2-74600857-DEL-3547
chr2-74621182-DEL-84
chr2-74635877-DEL-504
chr2-74690598-DEL-91
chr2-74701615-DEL-241
chr2-74772489-DEL-63
chr2-74780377-DEL-183
chr2-74789543-DEL-56
chr2-74791892-DEL-184
chr2-74802490-DEL-98
chr2-74834697-DEL-52
chr2-74864247-DEL-272
chr2-74877094-DEL-60
chr2-75012294-DEL-739
chr2-75017447-DEL-274
chr2-75020372-DEL-202
chr2-75035567-DEL-59
chr2-75065796-DEL-172
chr2-75161716-DEL-160
chr2-75243205-DEL-65
chr2-75265231-DEL-152
chr2-75268115-DEL-179
chr2-75286213-DEL-172
chr2-75360628-DEL-1490
chr2-75369436-DEL-84
chr2-75371393-DEL-161
chr2-75384404-DEL-114
chr2-75405257-DEL-181
chr2-75412090-DEL-54
chr2-75456362-DEL-54
chr2-75466209-DEL-86
chr2-75486829-DEL-179
chr2-75500198-DEL-256
chr2-75501955-DEL-204
chr2-75503447-DEL-213
chr2-75578483-DEL-130
chr2-75579572-DEL-61
chr2-75608207-DEL-347
chr2-75633738-DEL-482
chr2-75638580-DEL-1132
chr2-75650680-DEL-829
chr2-75678958-DEL-545
chr2-75692548-DEL-118
chr2-75705019-DEL-497
chr2-75716783-DEL-252
chr2-75802370-DEL-64
chr2-75805881-DEL-63
chr2-75873206-DEL-844
chr2-75912051-DEL-67
chr2-75949176-DEL-203
chr2-75971883-DEL-2881
chr2-75999117-DEL-265
chr2-76025848-DEL-381
chr2-76078843-DEL-52
chr2-76098061-DEL-104
chr2-76141957-DEL-483
chr2-76209946-DEL-673
chr2-76243737-DEL-60
chr2-76274053-DEL-1283
chr2-76294516-DEL-63
chr2-76309411-DEL-226
chr2-76348834-DEL-88
chr2-76357273-DEL-194
chr2-76368000-DEL-73
chr2-76413360-DEL-57
chr2-76463007-DEL-396
chr2-76487408-DEL-54
chr2-76510003-DEL-58

chr2-76525839-DEL-87
chr2-76633894-DEL-63
chr2-76675738-DEL-318
chr2-76740543-DEL-161
chr2-76752503-DEL-66
chr2-76758991-DEL-64
chr2-76824391-DEL-142
chr2-76833000-DEL-56
chr2-76900680-DEL-1219
chr2-77033234-DEL-115
chr2-77045693-DEL-86
chr2-77046788-DEL-70
chr2-77105733-DEL-58
chr2-77116491-DEL-76
chr2-77168977-DEL-102
chr2-77188301-DEL-542
chr2-77195288-DEL-63
chr2-77238248-DEL-93
chr2-77326853-DEL-128
chr2-77355256-DEL-215
chr2-77378211-DEL-120
chr2-77382527-DEL-100
chr2-77440232-DEL-51
chr2-77473413-DEL-918
chr2-77479193-DEL-2018
chr2-77479419-DEL-66
chr2-77484478-DEL-197
chr2-77536939-DEL-61
chr2-77564332-DEL-64
chr2-77581905-DEL-84
chr2-77716949-DEL-467
chr2-77782210-DEL-7251
chr2-77782273-DEL-1265
chr2-77785945-DEL-3571
chr2-77788764-DEL-416
chr2-77794177-DEL-6236
chr2-77826178-DEL-1062
chr2-77827983-DEL-6273
chr2-77869008-DEL-52
chr2-77901055-DEL-498
chr2-77902232-DEL-671
chr2-77927892-DEL-2323
chr2-77953824-DEL-3043
chr2-77953874-DEL-6780
chr2-77956936-DEL-3719
chr2-77992560-DEL-212
chr2-78041795-DEL-55
chr2-78096125-DEL-62
chr2-78211413-DEL-873
chr2-78231214-DEL-1078
chr2-78306389-DEL-715
chr2-78342685-DEL-1047
chr2-78356887-DEL-65
chr2-78374039-DEL-968
chr2-78419926-DEL-57
chr2-78444712-DEL-108
chr2-78482914-DEL-280
chr2-78491005-DEL-8082
chr2-78505736-DEL-235
chr2-78519965-DEL-415
chr2-78524306-DEL-271
chr2-78550249-DEL-71
chr2-78567195-DEL-82
chr2-78577564-DEL-56
chr2-78581076-DEL-60
chr2-78597250-DEL-305
chr2-78604535-DEL-58
chr2-78605109-DEL-314
chr2-78620504-DEL-70
chr2-78628965-DEL-198
chr2-78635410-DEL-111
chr2-78651625-DEL-1709
chr2-78669192-DEL-534
chr2-78673803-DEL-89
chr2-78675291-DEL-73
chr2-78698426-DEL-123
chr2-78712232-DEL-176

chr2-78721215-DEL-59
chr2-78766097-DEL-517
chr2-78787643-DEL-200
chr2-78843352-DEL-589
chr2-78872181-DEL-102
chr2-78898493-DEL-1152
chr2-78900790-DEL-943
chr2-78936477-DEL-852
chr2-78938924-DEL-200
chr2-78950433-DEL-210
chr2-78979671-DEL-798
chr2-78985541-DEL-258
chr2-78987857-DEL-211
chr2-79000771-DEL-101
chr2-79010684-DEL-228
chr2-79030269-DEL-347
chr2-79030848-DEL-52
chr2-79038730-DEL-1504
chr2-79090681-DEL-985
chr2-79124029-DEL-120
chr2-79178545-DEL-1653
chr2-79207950-DEL-94
chr2-79282175-DEL-59
chr2-79386559-DEL-1544
chr2-79445654-DEL-1914
chr2-79509444-DEL-198
chr2-79515721-DEL-167
chr2-79594993-DEL-92
chr2-79596034-DEL-287
chr2-79606325-DEL-307
chr2-79627404-DEL-109
chr2-79672000-DEL-6250
chr2-79679905-DEL-65
chr2-79729985-DEL-602
chr2-79730920-DEL-431
chr2-79730934-DEL-580
chr2-79834350-DEL-107
chr2-79835782-DEL-454
chr2-79873222-DEL-804
chr2-79954787-DEL-57
chr2-80044981-DEL-995
chr2-80046707-DEL-328
chr2-80050410-DEL-60
chr2-80079334-DEL-2019
chr2-80094619-DEL-135
chr2-80134682-DEL-79
chr2-80196675-DEL-71
chr2-80233684-DEL-478
chr2-80246310-DEL-87
chr2-80262359-DEL-62
chr2-80282859-DEL-136
chr2-80288454-DEL-60
chr2-80321376-DEL-175
chr2-80323625-DEL-641
chr2-80324420-DEL-77
chr2-80336124-DEL-115
chr2-80351013-DEL-187
chr2-80388980-DEL-192
chr2-80398457-DEL-204
chr2-80409182-DEL-64
chr2-80427162-DEL-102
chr2-80441947-DEL-167
chr2-80490128-DEL-112
chr2-80498509-DEL-66
chr2-80508467-DEL-73
chr2-80535816-DEL-148
chr2-80540376-DEL-342
chr2-80585495-DEL-910
chr2-80591960-DEL-60
chr2-80613055-DEL-212
chr2-80622753-DEL-361
chr2-80636189-DEL-11595
chr2-80642185-DEL-624
chr2-80644374-DEL-624
chr2-80651288-DEL-13144
chr2-80656176-DEL-7031
chr2-80689845-DEL-378

chr2-80702728-DEL-55
chr2-80728151-DEL-777
chr2-80740874-DEL-605
chr2-80779592-DEL-821
chr2-80802405-DEL-63
chr2-80807155-DEL-1404
chr2-80855369-DEL-443
chr2-80862285-DEL-51
chr2-80869167-DEL-41515
chr2-80883206-DEL-196
chr2-80891710-DEL-3521
chr2-80899124-DEL-6427
chr2-80918432-DEL-53
chr2-80923269-DEL-173
chr2-80971064-DEL-62
chr2-80975570-DEL-328
chr2-80982747-DEL-5498
chr2-81038677-DEL-110
chr2-81040101-DEL-6606
chr2-81042101-DEL-56
chr2-81043189-DEL-199
chr2-81065569-DEL-1162
chr2-81066429-DEL-56
chr2-81071070-DEL-2137
chr2-81090467-DEL-93
chr2-81094210-DEL-406
chr2-81095402-DEL-213
chr2-81173943-DEL-252
chr2-81181798-DEL-1636
chr2-81196363-DEL-1260
chr2-81206046-DEL-72
chr2-81231103-DEL-55
chr2-81234053-DEL-356
chr2-81260561-DEL-236
chr2-81269033-DEL-62
chr2-81290282-DEL-1346
chr2-81290647-DEL-6766
chr2-81291258-DEL-4085
chr2-81294069-DEL-1902
chr2-81295022-DEL-1370
chr2-81296457-DEL-1250
chr2-81298000-DEL-438
chr2-81301694-DEL-70
chr2-81304609-DEL-566
chr2-81316024-DEL-143
chr2-81347791-DEL-22477
chr2-81353889-DEL-2153
chr2-81356201-DEL-129
chr2-81377485-DEL-54
chr2-81404332-DEL-248
chr2-81406729-DEL-57
chr2-81419486-DEL-6236
chr2-81538494-DEL-102
chr2-81550651-DEL-582
chr2-81580922-DEL-6351
chr2-81599395-DEL-910
chr2-81606688-DEL-342
chr2-81617236-DEL-477
chr2-81628102-DEL-3337
chr2-81632223-DEL-54
chr2-81658952-DEL-378
chr2-81718545-DEL-519
chr2-81728605-DEL-69
chr2-81731098-DEL-244
chr2-81733335-DEL-178
chr2-81744426-DEL-183
chr2-81756282-DEL-71
chr2-81772611-DEL-1838
chr2-81776759-DEL-174
chr2-81805643-DEL-375
chr2-81807084-DEL-1299
chr2-81816532-DEL-8977
chr2-81819187-DEL-5033
chr2-81919306-DEL-796
chr2-81937068-DEL-4731
chr2-81949758-DEL-108
chr2-81954114-DEL-2511

chr2-81958575-DEL-1494
chr2-81968465-DEL-70
chr2-82025463-DEL-84
chr2-82055845-DEL-484
chr2-82055941-DEL-987
chr2-82058277-DEL-668
chr2-82065440-DEL-76
chr2-82073784-DEL-558
chr2-82095758-DEL-80
chr2-82126827-DEL-166
chr2-82127529-DEL-1793
chr2-82144379-DEL-691
chr2-82151753-DEL-262
chr2-82155055-DEL-98
chr2-82164827-DEL-5923
chr2-82198391-DEL-102
chr2-82201474-DEL-55
chr2-82204702-DEL-537
chr2-82210577-DEL-84
chr2-82217821-DEL-71
chr2-82222056-DEL-755
chr2-82229274-DEL-54
chr2-82233808-DEL-380
chr2-82237240-DEL-5241
chr2-82255110-DEL-6615
chr2-82261241-DEL-416
chr2-82278050-DEL-2891
chr2-82296668-DEL-420
chr2-82310170-DEL-378
chr2-82312941-DEL-1115
chr2-82345029-DEL-60
chr2-82379438-DEL-515
chr2-82382543-DEL-3444
chr2-82386003-DEL-180
chr2-82391389-DEL-83
chr2-82398791-DEL-540
chr2-82461950-DEL-52
chr2-82469682-DEL-142
chr2-82484289-DEL-52
chr2-82487222-DEL-854
chr2-82526636-DEL-52
chr2-82534746-DEL-165
chr2-82590010-DEL-378
chr2-82591306-DEL-213
chr2-82599243-DEL-346
chr2-82613804-DEL-6178
chr2-82613966-DEL-1029
chr2-82640492-DEL-172
chr2-82647127-DEL-854
chr2-82699007-DEL-6064
chr2-82706542-DEL-400
chr2-82744844-DEL-1008
chr2-82761260-DEL-75
chr2-82761851-DEL-6074
chr2-82786935-DEL-949
chr2-82797007-DEL-258
chr2-82820867-DEL-108
chr2-82864571-DEL-157
chr2-82870532-DEL-379
chr2-82871537-DEL-6400
chr2-82871825-DEL-199
chr2-82888442-DEL-112
chr2-82902783-DEL-6243
chr2-82917238-DEL-415
chr2-82932920-DEL-60
chr2-82938203-DEL-118
chr2-82945342-DEL-794
chr2-82970926-DEL-6206
chr2-82998461-DEL-60
chr2-83009176-DEL-2415
chr2-83022228-DEL-5617
chr2-83028017-DEL-3129
chr2-83034554-DEL-257
chr2-83085105-DEL-52
chr2-83095148-DEL-375
chr2-83122783-DEL-310
chr2-83124812-DEL-418

chr2-83154484-DEL-4340
chr2-83176834-DEL-56
chr2-83179231-DEL-62
chr2-83192678-DEL-6188
chr2-83203826-DEL-6380
chr2-83240496-DEL-52
chr2-83242710-DEL-198
chr2-83243828-DEL-7079
chr2-83251759-DEL-343
chr2-83260276-DEL-7025
chr2-83272215-DEL-105
chr2-83275311-DEL-401
chr2-83282645-DEL-5635
chr2-83286652-DEL-850
chr2-83301707-DEL-6503
chr2-83327519-DEL-222
chr2-83329626-DEL-130
chr2-83337789-DEL-145
chr2-83347320-DEL-85
chr2-83360311-DEL-7533
chr2-83371237-DEL-6542
chr2-83386888-DEL-7088
chr2-83387418-DEL-212
chr2-83396919-DEL-61
chr2-83400990-DEL-7781
chr2-83401113-DEL-6170
chr2-83408873-DEL-568
chr2-83408875-DEL-10970
chr2-83409605-DEL-10232
chr2-83429235-DEL-60
chr2-83533073-DEL-90
chr2-83671180-DEL-462
chr2-83777663-DEL-6454
chr2-83777667-DEL-5950
chr2-84024446-DEL-196
chr2-84025234-DEL-6541
chr2-84034213-DEL-7492
chr2-84182756-DEL-779
chr2-84187952-DEL-113
chr2-84216780-DEL-107
chr2-84225600-DEL-277
chr2-84243770-DEL-679
chr2-84362377-DEL-2785
chr2-84384686-DEL-387
chr2-84403187-DEL-381
chr2-84416796-DEL-93
chr2-84424099-DEL-113
chr2-84451745-DEL-238
chr2-84458453-DEL-201
chr2-84492638-DEL-51
chr2-84509788-DEL-2622
chr2-84514518-DEL-176
chr2-84528453-DEL-110
chr2-84531468-DEL-202
chr2-84577337-DEL-629
chr2-84619857-DEL-53
chr2-84627497-DEL-203
chr2-84630309-DEL-68
chr2-84639918-DEL-510
chr2-84664984-DEL-52
chr2-84684337-DEL-315
chr2-84718956-DEL-64
chr2-84726697-DEL-190
chr2-84751043-DEL-175
chr2-84752035-DEL-128
chr2-84753055-DEL-750
chr2-84763162-DEL-55
chr2-84766071-DEL-786
chr2-84786063-DEL-72
chr2-84800589-DEL-173
chr2-84860104-DEL-74
chr2-84915971-DEL-119
chr2-84923653-DEL-365
chr2-84942291-DEL-51
chr2-84945757-DEL-570
chr2-84950571-DEL-238
chr2-84999884-DEL-171

chr2-85001807-DEL-56
chr2-85004394-DEL-265
chr2-85030241-DEL-72
chr2-85033287-DEL-720
chr2-85048296-DEL-440
chr2-85054476-DEL-78
chr2-85071095-DEL-307
chr2-85084940-DEL-206
chr2-85090866-DEL-56
chr2-85153564-DEL-762
chr2-85183699-DEL-11542
chr2-85183702-DEL-17814
chr2-85192621-DEL-644
chr2-85195265-DEL-6591
chr2-85201842-DEL-888
chr2-85214121-DEL-205
chr2-85220757-DEL-975
chr2-85223985-DEL-1089
chr2-85274672-DEL-1135
chr2-85276603-DEL-221
chr2-85288124-DEL-56
chr2-85349655-DEL-153
chr2-85406163-DEL-80
chr2-85429613-DEL-1163
chr2-85448588-DEL-263
chr2-85495443-DEL-645
chr2-85507680-DEL-1063
chr2-85510024-DEL-75
chr2-85511166-DEL-78
chr2-85523342-DEL-135
chr2-85557213-DEL-92
chr2-85568887-DEL-6012
chr2-85618767-DEL-66
chr2-85621987-DEL-2362
chr2-85622387-DEL-142
chr2-85624473-DEL-52
chr2-85685209-DEL-68
chr2-85704147-DEL-2355
chr2-85710157-DEL-822
chr2-85723891-DEL-2173
chr2-85747846-DEL-74
chr2-85761223-DEL-6170
chr2-85764804-DEL-3261
chr2-85777686-DEL-278
chr2-85780557-DEL-84
chr2-85788400-DEL-6972
chr2-85799008-DEL-1192
chr2-85847233-DEL-10015
chr2-85863794-DEL-366
chr2-85873591-DEL-169
chr2-85883520-DEL-470
chr2-85903924-DEL-54
chr2-85909424-DEL-1104
chr2-85922364-DEL-2500
chr2-85939323-DEL-58
chr2-85946169-DEL-63
chr2-85960032-DEL-143
chr2-85962726-DEL-877
chr2-85999316-DEL-178
chr2-86024697-DEL-1117
chr2-86025357-DEL-51
chr2-86050968-DEL-1915
chr2-86055636-DEL-94
chr2-86060923-DEL-102
chr2-86112096-DEL-1361
chr2-86121339-DEL-101
chr2-86123690-DEL-91
chr2-86133582-DEL-6185
chr2-86146980-DEL-1359
chr2-86162185-DEL-241
chr2-86187638-DEL-118
chr2-86214731-DEL-1509
chr2-86216112-DEL-165
chr2-86217777-DEL-27728
chr2-86223318-DEL-982
chr2-86226523-DEL-107
chr2-86228589-DEL-2401

chr2-86239670-DEL-62
chr2-86248934-DEL-60
chr2-86254344-DEL-61
chr2-86258795-DEL-436
chr2-86259610-DEL-186
chr2-86263526-DEL-1502
chr2-86264765-DEL-519
chr2-86281642-DEL-253
chr2-86296399-DEL-58
chr2-86315191-DEL-156
chr2-86331446-DEL-693
chr2-86341188-DEL-91
chr2-86346138-DEL-124
chr2-86376917-DEL-71
chr2-86377777-DEL-1061
chr2-86384640-DEL-90
chr2-86395063-DEL-1780
chr2-86405398-DEL-60
chr2-86412426-DEL-3765
chr2-86420904-DEL-722
chr2-86424224-DEL-22469
chr2-86475324-DEL-6158
chr2-86482832-DEL-929
chr2-86486959-DEL-51
chr2-86504310-DEL-51
chr2-86527563-DEL-463
chr2-86532100-DEL-2755
chr2-86536077-DEL-9736
chr2-86543823-DEL-198
chr2-86548400-DEL-435
chr2-86549114-DEL-51
chr2-86561920-DEL-2020
chr2-86576556-DEL-102
chr2-86585096-DEL-54
chr2-86592101-DEL-233
chr2-86599571-DEL-66
chr2-86607410-DEL-211
chr2-86644267-DEL-1142
chr2-86660735-DEL-56
chr2-86689698-DEL-2663
chr2-86726688-DEL-53
chr2-86729731-DEL-313
chr2-86793355-DEL-68
chr2-86807882-DEL-93
chr2-86812398-DEL-4210
chr2-86826499-DEL-52
chr2-86834350-DEL-67
chr2-86845965-DEL-83
chr2-86863132-DEL-125
chr2-86950822-DEL-2298
chr2-86958688-DEL-221
chr2-86967609-DEL-371
chr2-86975531-DEL-2607
chr2-87021196-DEL-107
chr2-87042619-DEL-75
chr2-87049195-DEL-95
chr2-87052006-DEL-6400
chr2-87086156-DEL-501
chr2-87086702-DEL-1118
chr2-87093315-DEL-108
chr2-87094833-DEL-181
chr2-87108840-DEL-192
chr2-87111081-DEL-56
chr2-87111990-DEL-1261
chr2-87119640-DEL-89
chr2-87140880-DEL-70
chr2-87151940-DEL-53
chr2-87156419-DEL-3830
chr2-87231541-DEL-277
chr2-87233592-DEL-201
chr2-87239887-DEL-377
chr2-87270919-DEL-5765
chr2-87277582-DEL-6052
chr2-87292740-DEL-677
chr2-87293459-DEL-267
chr2-87293804-DEL-625
chr2-87294493-DEL-236

chr2-87305894-DEL-1914
chr2-87307887-DEL-803
chr2-87319616-DEL-87
chr2-87331841-DEL-219
chr2-87350019-DEL-213
chr2-87373771-DEL-76
chr2-87378775-DEL-1220
chr2-87395701-DEL-215
chr2-87409546-DEL-403
chr2-87468902-DEL-52
chr2-87490337-DEL-188
chr2-87493143-DEL-2628
chr2-87495930-DEL-516
chr2-87506059-DEL-6587
chr2-87531187-DEL-6227
chr2-87538046-DEL-1037
chr2-87546975-DEL-5781
chr2-87555495-DEL-5748
chr2-87564722-DEL-3103
chr2-87566820-DEL-243
chr2-87581761-DEL-303
chr2-87587447-DEL-60
chr2-87621025-DEL-6180
chr2-87635923-DEL-116
chr2-87644956-DEL-1381
chr2-87648136-DEL-484
chr2-87652369-DEL-4231
chr2-87675641-DEL-639
chr2-87691436-DEL-184
chr2-87712452-DEL-863
chr2-87713438-DEL-669
chr2-87714122-DEL-211
chr2-87714313-DEL-8245
chr2-87745457-DEL-3875
chr2-87756596-DEL-5117
chr2-87782053-DEL-103
chr2-87783865-DEL-117
chr2-87823928-DEL-109
chr2-87840496-DEL-90
chr2-87851450-DEL-59
chr2-87947054-DEL-97
chr2-87958895-DEL-307
chr2-87959097-DEL-1868
chr2-87961011-DEL-337
chr2-87961986-DEL-184
chr2-88010032-DEL-795
chr2-88010053-DEL-202
chr2-88013081-DEL-817
chr2-88020464-DEL-1889
chr2-88054906-DEL-2133
chr2-88062444-DEL-862
chr2-88067217-DEL-120
chr2-88087695-DEL-4129
chr2-88136473-DEL-171
chr2-88147628-DEL-4995
chr2-88161911-DEL-599
chr2-88180759-DEL-52
chr2-88191306-DEL-487
chr2-88226127-DEL-460
chr2-88244298-DEL-943
chr2-88252494-DEL-217
chr2-88256775-DEL-92
chr2-88351904-DEL-55
chr2-88421128-DEL-232
chr2-88461696-DEL-263
chr2-88472033-DEL-55
chr2-88494517-DEL-6461
chr2-88499910-DEL-1069
chr2-88500654-DEL-212
chr2-88505052-DEL-103
chr2-88507133-DEL-55
chr2-88511725-DEL-61
chr2-88515999-DEL-115
chr2-88520166-DEL-2733
chr2-88526074-DEL-223
chr2-88533661-DEL-1228
chr2-88548478-DEL-151

chr2-88551677-DEL-434
chr2-88574696-DEL-400
chr2-88627809-DEL-243
chr2-88635627-DEL-70
chr2-88643913-DEL-196
chr2-88650440-DEL-211
chr2-88708915-DEL-60
chr2-88725726-DEL-54
chr2-88736711-DEL-1197
chr2-88743113-DEL-1253
chr2-88748539-DEL-345
chr2-88749780-DEL-186
chr2-88754245-DEL-354
chr2-88756784-DEL-283
chr2-88757583-DEL-684
chr2-88758265-DEL-374
chr2-88762570-DEL-747
chr2-88771110-DEL-1388
chr2-88774422-DEL-52
chr2-88778090-DEL-181
chr2-88779678-DEL-828
chr2-88793722-DEL-6382
chr2-88816485-DEL-64
chr2-88835947-DEL-900
chr2-88850825-DEL-52
chr2-88857549-DEL-350
chr2-88893052-DEL-59
chr2-88909909-DEL-996
chr2-88916134-DEL-303
chr2-88948637-DEL-66
chr2-88969982-DEL-105
chr2-89012976-DEL-1938
chr2-89015805-DEL-104
chr2-89034398-DEL-71
chr2-89040615-DEL-2049
chr2-89043847-DEL-55
chr2-89072666-DEL-2052
chr2-89075905-DEL-55
chr2-89103986-DEL-837
chr2-89111141-DEL-494
chr2-89115660-DEL-52
chr2-89121187-DEL-92
chr2-89122438-DEL-55
chr2-89139063-DEL-644
chr2-89174729-DEL-6010
chr2-89184131-DEL-397
chr2-89187420-DEL-326
chr2-89189236-DEL-227
chr2-89203682-DEL-88
chr2-89203693-DEL-4149
chr2-89213143-DEL-801
chr2-89216086-DEL-268
chr2-89226189-DEL-352
chr2-89231683-DEL-213
chr2-89232230-DEL-304
chr2-89251965-DEL-1455
chr2-89255055-DEL-2458
chr2-89276806-DEL-64
chr2-89278618-DEL-446
chr2-89288455-DEL-184
chr2-89298264-DEL-6535
chr2-89300004-DEL-6259
chr2-89319751-DEL-126
chr2-89374731-DEL-228
chr2-89387751-DEL-447
chr2-89396943-DEL-335
chr2-89424842-DEL-201
chr2-89475683-DEL-151
chr2-89481313-DEL-621
chr2-89484609-DEL-1015
chr2-89504011-DEL-108
chr2-89505723-DEL-6222
chr2-89524233-DEL-4089
chr2-89533595-DEL-618
chr2-89547908-DEL-155
chr2-89560615-DEL-7002
chr2-89569973-DEL-4405

chr2-89579310-DEL-92
chr2-89599446-DEL-6036
chr2-89607317-DEL-341
chr2-89618151-DEL-123
chr2-89700923-DEL-6396
chr2-89717474-DEL-84
chr2-89719850-DEL-13976
chr2-89733542-DEL-66
chr2-89750837-DEL-769
chr2-89796517-DEL-92
chr2-89875134-DEL-4440
chr2-89904674-DEL-319
chr2-89925938-DEL-6243
chr2-89932447-DEL-2857
chr2-89985750-DEL-105
chr2-89997115-DEL-82432
chr2-90036266-DEL-1440
chr2-90044637-DEL-763
chr2-90097867-DEL-351
chr2-90102948-DEL-6491
chr2-90120855-DEL-180
chr2-90130156-DEL-5095
chr2-90130379-DEL-89
chr2-90143635-DEL-6215
chr2-90145647-DEL-144
chr2-90155775-DEL-499
chr2-90161977-DEL-763
chr2-90191511-DEL-7056
chr2-90200057-DEL-1241
chr2-90203509-DEL-6755
chr2-90206086-DEL-501
chr2-90206605-DEL-4859
chr2-90214924-DEL-358
chr2-90219591-DEL-6735
chr2-90221602-DEL-144
chr2-90225474-DEL-517
chr2-90226379-DEL-761
chr2-90246864-DEL-89
chr2-90256180-DEL-51
chr2-90307146-DEL-150
chr2-90326886-DEL-382
chr2-90361223-DEL-1198
chr2-90394659-DEL-460
chr2-90402168-DEL-95
chr2-90414854-DEL-64
chr2-90420316-DEL-1221
chr2-90434944-DEL-200
chr2-90449889-DEL-64
chr2-90452157-DEL-2103
chr2-90452947-DEL-219
chr2-90472552-DEL-66
chr2-90484247-DEL-196
chr2-90493940-DEL-211
chr2-90541132-DEL-193
chr2-90545957-DEL-99
chr2-90560273-DEL-82
chr2-90617216-DEL-200
chr2-90621297-DEL-158
chr2-90634006-DEL-1017
chr2-90636049-DEL-53
chr2-90644727-DEL-92
chr2-90659139-DEL-2002
chr2-90661461-DEL-204
chr2-90673104-DEL-2369
chr2-90712796-DEL-377
chr2-90729868-DEL-557
chr2-90758835-DEL-59
chr2-90801899-DEL-54
chr2-90810152-DEL-1309
chr2-90813470-DEL-629
chr2-90832028-DEL-78
chr2-90856322-DEL-245
chr2-90880479-DEL-78
chr2-90947067-DEL-63
chr2-90973599-DEL-207
chr2-91019033-DEL-64
chr2-91039331-DEL-144

chr2-91052912-DEL-183
chr2-91069936-DEL-66
chr2-91070247-DEL-200
chr2-91132749-DEL-55
chr2-91140467-DEL-184
chr2-91179631-DEL-76
chr2-91183416-DEL-73
chr2-91205008-DEL-792
chr2-91209588-DEL-832
chr2-91212317-DEL-55
chr2-91213520-DEL-167
chr2-91221824-DEL-88
chr2-91224022-DEL-82
chr2-91249553-DEL-211
chr2-91262005-DEL-67
chr2-91263816-DEL-87
chr2-91314351-DEL-89
chr2-91314892-DEL-204
chr2-91405156-DEL-1031
chr2-91413501-DEL-485
chr2-91428783-DEL-183
chr2-91432203-DEL-69
chr2-91447608-DEL-57
chr2-91455363-DEL-67
chr2-91494905-DEL-846
chr2-91506900-DEL-488
chr2-91527846-DEL-67
chr2-91534044-DEL-521
chr2-91562223-DEL-985
chr2-91563567-DEL-205
chr2-91591138-DEL-227
chr2-91592100-DEL-2341
chr2-91594558-DEL-206
chr2-91594860-DEL-1531
chr2-91597619-DEL-57
chr2-91600666-DEL-99
chr2-91638376-DEL-59
chr2-91647758-DEL-475
chr2-91648685-DEL-507
chr2-91657892-DEL-72
chr2-91661088-DEL-107
chr2-91667023-DEL-222
chr2-91694197-DEL-241
chr2-91724445-DEL-73
chr2-91725703-DEL-129
chr2-91731933-DEL-77
chr2-91791305-DEL-165
chr2-91812553-DEL-96
chr2-91884001-DEL-91
chr2-91914752-DEL-239
chr2-91923939-DEL-156
chr2-91940361-DEL-188
chr2-91995649-DEL-60
chr2-91996854-DEL-153
chr2-92008866-DEL-195
chr2-92010903-DEL-78
chr2-92036445-DEL-58
chr2-92041717-DEL-194
chr2-92100843-DEL-1818
chr2-92112320-DEL-412
chr2-92168733-DEL-414
chr2-92211273-DEL-51
chr2-92224531-DEL-222
chr2-92248081-DEL-51
chr2-92267308-DEL-228
chr2-92273363-DEL-56
chr2-92275010-DEL-3628
chr2-92301721-DEL-138
chr2-92391980-DEL-1398
chr2-92413864-DEL-145
chr2-92433555-DEL-76
chr2-92479806-DEL-158
chr2-92487452-DEL-59
chr2-92491343-DEL-74
chr2-92563127-DEL-485
chr2-92602681-DEL-88
chr2-92671214-DEL-159

chr2-92681672-DEL-1241
chr2-92686873-DEL-1911
chr2-92747318-DEL-77
chr2-92770542-DEL-358
chr2-92780309-DEL-110
chr2-92790471-DEL-129
chr2-92800815-DEL-73
chr2-92864134-DEL-92
chr2-92872344-DEL-52
chr2-92891759-DEL-195
chr2-92893721-DEL-69
chr2-92895024-DEL-67
chr2-92898808-DEL-195
chr2-92901506-DEL-98
chr2-92916491-DEL-113
chr2-92926147-DEL-99
chr2-92934391-DEL-56
chr2-92947155-DEL-56
chr2-92970336-DEL-100
chr2-92983839-DEL-631
chr2-93052461-DEL-84
chr2-93065566-DEL-471
chr2-93070419-DEL-301
chr2-93089873-DEL-481
chr2-93117448-DEL-58
chr2-93120131-DEL-63
chr2-93132068-DEL-265
chr2-93135675-DEL-252
chr2-93136745-DEL-63
chr2-93146909-DEL-138
chr2-93157126-DEL-160
chr2-93177823-DEL-54
chr2-93180723-DEL-574
chr2-93181550-DEL-604
chr2-93202906-DEL-78
chr2-93203544-DEL-135
chr2-93216380-DEL-510
chr2-93268047-DEL-1442
chr2-93279765-DEL-89
chr2-93314686-DEL-110
chr2-93316921-DEL-52
chr2-93327498-DEL-68
chr2-93343886-DEL-498
chr2-93353201-DEL-54
chr2-93362032-DEL-178
chr2-93390427-DEL-382
chr2-93408822-DEL-78
chr2-93410773-DEL-54
chr2-93439605-DEL-175
chr2-93545934-DEL-84
chr2-93551669-DEL-137
chr2-93581989-DEL-139
chr2-93602566-DEL-64
chr2-93603795-DEL-140
chr2-93604404-DEL-96
chr2-93620769-DEL-342
chr2-93633407-DEL-114
chr2-93645111-DEL-64
chr2-93650663-DEL-60
chr2-93673910-DEL-194
chr2-93678907-DEL-567
chr2-93703449-DEL-58
chr2-93720491-DEL-87
chr2-93770073-DEL-436
chr2-93819797-DEL-588
chr2-93867681-DEL-59
chr2-93869156-DEL-1475
chr2-93914603-DEL-2250
chr2-93917889-DEL-192
chr2-93940272-DEL-606
chr2-93965789-DEL-237
chr2-93997281-DEL-7038
chr2-94020069-DEL-139
chr2-94028071-DEL-124
chr2-94092488-DEL-53
chr2-94114175-DEL-60
chr2-94169616-DEL-59

chr2-94179104-DEL-351
chr2-94179720-DEL-117
chr2-94184621-DEL-202
chr2-94198484-DEL-233
chr2-94224520-DEL-205
chr2-94265325-DEL-65
chr2-94269162-DEL-149
chr2-94292326-DEL-405
chr2-94299160-DEL-102
chr2-94387033-DEL-54
chr2-94400923-DEL-52
chr2-94414217-DEL-60
chr2-94454106-DEL-553
chr2-94463758-DEL-6083
chr2-94474626-DEL-576
chr2-94480432-DEL-6608
chr2-94521435-DEL-291
chr2-94548177-DEL-75
chr2-94568330-DEL-988
chr2-94630592-DEL-288
chr2-94679697-DEL-6118
chr2-94696100-DEL-1188
chr2-94751012-DEL-1244
chr2-94760378-DEL-52
chr2-94766354-DEL-1217
chr2-94796701-DEL-113
chr2-94798286-DEL-527
chr2-94804127-DEL-866
chr2-94820848-DEL-65
chr2-94870467-DEL-448
chr2-94871536-DEL-786
chr2-94872406-DEL-1255
chr2-94879109-DEL-834
chr2-94922853-DEL-561
chr2-94924681-DEL-758
chr2-94938199-DEL-54
chr2-94987963-DEL-1172
chr2-95014269-DEL-70
chr2-95018357-DEL-105
chr2-95037057-DEL-1603
chr2-95055658-DEL-56
chr2-95058300-DEL-191
chr2-95065184-DEL-1838
chr2-95067406-DEL-835
chr2-95076800-DEL-891
chr2-95096118-DEL-124
chr2-95119100-DEL-101
chr2-95129634-DEL-2488
chr2-95132013-DEL-108
chr2-95137620-DEL-354
chr2-95138900-DEL-223
chr2-95144211-DEL-11625
chr2-95147480-DEL-2317
chr2-95164203-DEL-3529
chr2-95168798-DEL-71
chr2-95174527-DEL-825
chr2-95180824-DEL-65
chr2-95181331-DEL-1481
chr2-95208239-DEL-53
chr2-95211892-DEL-234
chr2-95216693-DEL-58
chr2-95244409-DEL-8093
chr2-95251836-DEL-84
chr2-95321809-DEL-719
chr2-95361671-DEL-60
chr2-95370527-DEL-1567
chr2-95371139-DEL-59
chr2-95389741-DEL-338
chr2-95390986-DEL-841
chr2-95414288-DEL-150
chr2-95414357-DEL-1308
chr2-95418310-DEL-1559
chr2-95440760-DEL-53
chr2-95442833-DEL-787
chr2-95460753-DEL-430
chr2-95466817-DEL-188
chr2-95469046-DEL-714

chr2-95513242-DEL-365
chr2-95526468-DEL-5134
chr2-95527322-DEL-54
chr2-95534331-DEL-4860
chr2-95549988-DEL-106
chr2-95553448-DEL-3099
chr2-95559183-DEL-273
chr2-95578125-DEL-96
chr2-95594428-DEL-82
chr2-95629602-DEL-400
chr2-95632536-DEL-66
chr2-95636709-DEL-583
chr2-95643742-DEL-358
chr2-95676079-DEL-52
chr2-95679203-DEL-91
chr2-95705286-DEL-2802
chr2-95710671-DEL-479
chr2-95727186-DEL-92
chr2-95772428-DEL-132
chr2-95773369-DEL-237
chr2-95791962-DEL-404
chr2-95797861-DEL-2923
chr2-95818816-DEL-461
chr2-95820585-DEL-828
chr2-95827429-DEL-60
chr2-95840275-DEL-196
chr2-95845071-DEL-401
chr2-95859279-DEL-213
chr2-95870250-DEL-101
chr2-95875453-DEL-56
chr2-95907238-DEL-76
chr2-95917070-DEL-56
chr2-95920131-DEL-271
chr2-95940920-DEL-59
chr2-95947532-DEL-957
chr2-95948638-DEL-656
chr2-95977045-DEL-1463
chr2-95978917-DEL-72
chr2-96017271-DEL-654
chr2-96033385-DEL-5917
chr2-96040776-DEL-420
chr2-96041970-DEL-220
chr2-96046723-DEL-99
chr2-96073007-DEL-797
chr2-96075137-DEL-53
chr2-96078405-DEL-58
chr2-96087456-DEL-1519
chr2-96106859-DEL-195
chr2-96121146-DEL-55
chr2-96140577-DEL-59
chr2-96189538-DEL-719
chr2-96203548-DEL-6072
chr2-96223373-DEL-3637
chr2-96230595-DEL-250
chr2-96238636-DEL-2681
chr2-96322446-DEL-227
chr2-96370351-DEL-989
chr2-96384698-DEL-84
chr2-96387874-DEL-811
chr2-96391245-DEL-6245
chr2-96423361-DEL-59
chr2-96467262-DEL-925
chr2-96474890-DEL-71
chr2-96477881-DEL-62
chr2-96517946-DEL-739
chr2-96541326-DEL-1443
chr2-96568366-DEL-99
chr2-96580172-DEL-4344
chr2-96618287-DEL-1013
chr2-96638468-DEL-171
chr2-96644946-DEL-2980
chr2-96666295-DEL-89
chr2-96670980-DEL-2274
chr2-96800497-DEL-8067
chr2-96800640-DEL-7142
chr2-96818153-DEL-170
chr2-96842099-DEL-51

chr2-96866545-DEL-52
chr2-96877363-DEL-1168
chr2-96892123-DEL-400
chr2-96893842-DEL-156
chr2-96896367-DEL-2786
chr2-96902960-DEL-902
chr2-96954019-DEL-116
chr2-96959021-DEL-83
chr2-96960844-DEL-1265
chr2-96998349-DEL-498
chr2-97010153-DEL-5017
chr2-97030834-DEL-51
chr2-97040745-DEL-2629
chr2-97053082-DEL-99
chr2-97055566-DEL-1303
chr2-97080719-DEL-10297
chr2-97093017-DEL-84
chr2-97121204-DEL-734
chr2-97124359-DEL-1815
chr2-97138228-DEL-6189
chr2-97145860-DEL-88
chr2-97157706-DEL-208
chr2-97229460-DEL-64
chr2-97240049-DEL-2537
chr2-97284075-DEL-63
chr2-97304692-DEL-95
chr2-97343284-DEL-703
chr2-97351783-DEL-621
chr2-97369818-DEL-97
chr2-97387001-DEL-92
chr2-97443288-DEL-85
chr2-97447708-DEL-99
chr2-97452992-DEL-678
chr2-97472586-DEL-97
chr2-97479513-DEL-439
chr2-97488913-DEL-167
chr2-97497561-DEL-1457
chr2-97543580-DEL-215
chr2-97544497-DEL-523
chr2-97554686-DEL-261
chr2-97583883-DEL-322
chr2-97588125-DEL-80
chr2-97590075-DEL-114
chr2-97590923-DEL-5240
chr2-97597486-DEL-493
chr2-97610607-DEL-513
chr2-97621520-DEL-831
chr2-97636329-DEL-365
chr2-97638221-DEL-442
chr2-97644179-DEL-85
chr2-97647452-DEL-389
chr2-97664034-DEL-12638
chr2-97685835-DEL-83
chr2-97693333-DEL-204
chr2-97702544-DEL-4699
chr2-97709042-DEL-720
chr2-97721872-DEL-194
chr2-97724173-DEL-233
chr2-97746929-DEL-52
chr2-97749542-DEL-294
chr2-97764356-DEL-1493
chr2-97778112-DEL-1988
chr2-97798456-DEL-290
chr2-97829746-DEL-856
chr2-97830863-DEL-108
chr2-97866502-DEL-112
chr2-97886320-DEL-1679
chr2-97906282-DEL-940
chr2-97909294-DEL-1577
chr2-97961589-DEL-58906
chr2-97977111-DEL-62
chr2-97982141-DEL-428
chr2-97991246-DEL-613
chr2-98001158-DEL-5279
chr2-98038050-DEL-1011
chr2-98043288-DEL-665
chr2-98047345-DEL-832

chr2-98091339-DEL-231
chr2-98093053-DEL-354
chr2-98097817-DEL-63
chr2-98126174-DEL-4020
chr2-98159962-DEL-8299
chr2-98187754-DEL-653
chr2-98207624-DEL-84
chr2-98239758-DEL-6273
chr2-98257477-DEL-251
chr2-98275788-DEL-2152
chr2-98279348-DEL-582
chr2-98292486-DEL-66
chr2-98293346-DEL-744
chr2-98311563-DEL-78
chr2-98317126-DEL-161
chr2-98331868-DEL-2632
chr2-98352903-DEL-1577
chr2-98436725-DEL-5977
chr2-98456449-DEL-3938
chr2-98462486-DEL-416
chr2-98490859-DEL-11580
chr2-98516816-DEL-6637
chr2-98555071-DEL-1022
chr2-98595705-DEL-6117
chr2-98642078-DEL-152
chr2-98673412-DEL-851
chr2-98677332-DEL-226
chr2-98679372-DEL-1239
chr2-98694904-DEL-1118
chr2-98699081-DEL-829
chr2-98721490-DEL-768
chr2-98728730-DEL-2590
chr2-98746966-DEL-2558
chr2-98753836-DEL-66
chr2-98755341-DEL-686
chr2-98760480-DEL-753
chr2-98761458-DEL-840
chr2-98767620-DEL-697
chr2-98782020-DEL-502
chr2-98782290-DEL-2099
chr2-98784042-DEL-194
chr2-98802551-DEL-1478
chr2-98826448-DEL-86
chr2-98853580-DEL-52
chr2-98864610-DEL-2461
chr2-98880252-DEL-8743
chr2-98888609-DEL-385
chr2-98914943-DEL-273
chr2-98942888-DEL-244
chr2-98943266-DEL-554
chr2-99025983-DEL-54
chr2-99037471-DEL-6172
chr2-99037838-DEL-670
chr2-99054793-DEL-74
chr2-99113553-DEL-450
chr2-99119593-DEL-133
chr2-99149418-DEL-1460
chr2-99154643-DEL-689
chr2-99166506-DEL-74
chr2-99170367-DEL-2869
chr2-99186575-DEL-448
chr2-99196039-DEL-6177
chr2-99202413-DEL-1216
chr2-99213638-DEL-628
chr2-99282075-DEL-2434
chr2-99284775-DEL-251
chr2-99346639-DEL-1491
chr2-99362986-DEL-241
chr2-99365965-DEL-157
chr2-99369674-DEL-122
chr2-99409573-DEL-625
chr2-99419670-DEL-196
chr2-99422165-DEL-63
chr2-99430637-DEL-487
chr2-99443904-DEL-110
chr2-99450407-DEL-3554
chr2-99466514-DEL-111

chr2-99481112-DEL-288
chr2-99487319-DEL-96
chr2-99494213-DEL-345
chr2-99506211-DEL-60
chr2-99539587-DEL-4635
chr2-99552827-DEL-1336
chr2-99585547-DEL-91
chr2-99618296-DEL-307
chr2-99657691-DEL-333
chr2-99658754-DEL-75
chr2-99669545-DEL-122
chr2-99689147-DEL-82
chr2-99708654-DEL-68
chr2-99709705-DEL-158
chr2-99724265-DEL-608
chr2-99731623-DEL-69
chr2-99733438-DEL-6667
chr2-99756156-DEL-139
chr2-99764084-DEL-76
chr2-99774711-DEL-6038
chr2-99785118-DEL-118
chr2-99789690-DEL-1205
chr2-99795806-DEL-76
chr2-99797148-DEL-96
chr2-99798373-DEL-99
chr2-99799338-DEL-1962
chr2-99811463-DEL-492
chr2-99850838-DEL-192
chr2-99854742-DEL-250
chr2-99858392-DEL-1350
chr2-99872210-DEL-2323
chr2-99912679-DEL-51
chr2-99923750-DEL-71
chr2-99935725-DEL-416
chr2-100012481-DEL-64
chr2-100037885-DEL-109
chr2-100054245-DEL-1229
chr2-100066733-DEL-2109
chr2-100088465-DEL-400
chr2-100159437-DEL-349
chr2-100181801-DEL-6155
chr2-100196182-DEL-437
chr2-100212512-DEL-203
chr2-100228306-DEL-1622
chr2-100244797-DEL-222
chr2-100245921-DEL-103
chr2-100267674-DEL-538
chr2-100369159-DEL-59
chr2-100370354-DEL-7705
chr2-100394188-DEL-77
chr2-100423584-DEL-113
chr2-100426246-DEL-302
chr2-100440729-DEL-1772
chr2-100445772-DEL-84
chr2-100628420-DEL-103
chr2-100662584-DEL-157
chr2-100676146-DEL-2810
chr2-100702228-DEL-51
chr2-100713196-DEL-1041
chr2-100714637-DEL-1003
chr2-100719667-DEL-397
chr2-100727610-DEL-6456
chr2-100727813-DEL-54
chr2-100747623-DEL-92
chr2-100755912-DEL-57
chr2-100759149-DEL-229
chr2-100764789-DEL-3030
chr2-100803340-DEL-523
chr2-100881321-DEL-400
chr2-100887044-DEL-826
chr2-100896822-DEL-58
chr2-100922004-DEL-83
chr2-100927280-DEL-181
chr2-101071036-DEL-338
chr2-101093968-DEL-2326
chr2-101170076-DEL-131
chr2-101181409-DEL-282

chr2-101208593-DEL-738
chr2-101231653-DEL-181
chr2-101234438-DEL-58
chr2-101236109-DEL-201
chr2-101236847-DEL-60
chr2-101279318-DEL-106
chr2-101283279-DEL-299
chr2-101285168-DEL-63
chr2-101297798-DEL-1171
chr2-101299366-DEL-1393
chr2-101305575-DEL-200
chr2-101307618-DEL-320
chr2-101311881-DEL-6426
chr2-101336614-DEL-194
chr2-101354831-DEL-300
chr2-101399397-DEL-295
chr2-101488060-DEL-165
chr2-101497528-DEL-486
chr2-101532625-DEL-1376
chr2-101539115-DEL-81
chr2-101544080-DEL-80
chr2-101604049-DEL-130
chr2-101680547-DEL-60
chr2-101683020-DEL-1066
chr2-101699562-DEL-891
chr2-101706464-DEL-212
chr2-101732216-DEL-97
chr2-101741929-DEL-2534
chr2-101775868-DEL-535
chr2-101780221-DEL-60
chr2-101790190-DEL-445
chr2-101792601-DEL-134
chr2-101819533-DEL-56
chr2-101837576-DEL-88
chr2-101888800-DEL-257
chr2-101897002-DEL-880
chr2-101998959-DEL-1811
chr2-102003595-DEL-494
chr2-102037652-DEL-90
chr2-102082252-DEL-136
chr2-102095518-DEL-61
chr2-102161438-DEL-54
chr2-102198976-DEL-446
chr2-102206038-DEL-60
chr2-102261445-DEL-56
chr2-102320216-DEL-607
chr2-102323753-DEL-52
chr2-102325373-DEL-365
chr2-102332337-DEL-75
chr2-102363189-DEL-73
chr2-102398263-DEL-96
chr2-102430948-DEL-587
chr2-102454363-DEL-82
chr2-102456889-DEL-52
chr2-102510709-DEL-59
chr2-102605835-DEL-69
chr2-102635098-DEL-210
chr2-102682860-DEL-84
chr2-102701170-DEL-170
chr2-102719577-DEL-57
chr2-102733825-DEL-178
chr2-102767868-DEL-373
chr2-102791267-DEL-1686
chr2-102841547-DEL-66
chr2-102879208-DEL-92
chr2-102879717-DEL-488
chr2-102936670-DEL-90
chr2-102953929-DEL-1044
chr2-102981316-DEL-122
chr2-103011319-DEL-54
chr2-103050742-DEL-54
chr2-103086604-DEL-75
chr2-103141668-DEL-68
chr2-103222667-DEL-748
chr2-103308418-DEL-137
chr2-103378874-DEL-260
chr2-103379280-DEL-2239

chr2-103464333-DEL-62
chr2-103465175-DEL-175
chr2-103467499-DEL-742
chr2-103648618-DEL-329
chr2-103651637-DEL-687
chr2-103681921-DEL-52
chr2-103682288-DEL-5660
chr2-103682490-DEL-7835
chr2-103688863-DEL-547
chr2-103689855-DEL-885
chr2-103696411-DEL-196
chr2-103705503-DEL-99
chr2-103709288-DEL-124
chr2-103728009-DEL-168
chr2-103774053-DEL-226
chr2-103793716-DEL-1702
chr2-103794099-DEL-614
chr2-103798338-DEL-96
chr2-103828120-DEL-131
chr2-103842486-DEL-122
chr2-103854441-DEL-86
chr2-103864873-DEL-202
chr2-103877585-DEL-51
chr2-103900492-DEL-63
chr2-103902354-DEL-66
chr2-103911953-DEL-1668
chr2-103913266-DEL-64
chr2-103917474-DEL-4976
chr2-103918225-DEL-3426
chr2-103925438-DEL-186
chr2-103928954-DEL-94
chr2-103992300-DEL-58
chr2-104016794-DEL-353
chr2-104121832-DEL-1278
chr2-104132332-DEL-105
chr2-104133713-DEL-53
chr2-104149040-DEL-62
chr2-104194167-DEL-140
chr2-104196392-DEL-163
chr2-104242078-DEL-58
chr2-104270077-DEL-56
chr2-104278518-DEL-80
chr2-104315954-DEL-264
chr2-104346036-DEL-844
chr2-104359937-DEL-68
chr2-104365539-DEL-54
chr2-104377821-DEL-53
chr2-104393990-DEL-802
chr2-104395607-DEL-98
chr2-104397959-DEL-836
chr2-104412858-DEL-70
chr2-104429581-DEL-74
chr2-104448082-DEL-667
chr2-104475441-DEL-498
chr2-104482962-DEL-84
chr2-104524667-DEL-221
chr2-104524914-DEL-1666
chr2-104526594-DEL-1107
chr2-104581771-DEL-163
chr2-104595292-DEL-228
chr2-104677975-DEL-61
chr2-104691574-DEL-55
chr2-104701340-DEL-98
chr2-104704703-DEL-343
chr2-104708047-DEL-74
chr2-104710044-DEL-51
chr2-104717890-DEL-190
chr2-104751943-DEL-100
chr2-104758283-DEL-200
chr2-104778626-DEL-1220
chr2-104789100-DEL-178
chr2-104803285-DEL-72
chr2-104855088-DEL-77
chr2-104876610-DEL-57
chr2-104878875-DEL-3413
chr2-104889964-DEL-226
chr2-104898303-DEL-518

chr2-104922472-DEL-86
chr2-105005245-DEL-225
chr2-105036585-DEL-60
chr2-105050607-DEL-60
chr2-105088135-DEL-195
chr2-105088183-DEL-819
chr2-105099605-DEL-836
chr2-105108616-DEL-63
chr2-105119880-DEL-180
chr2-105144414-DEL-56
chr2-105150271-DEL-258
chr2-105181872-DEL-72
chr2-105182794-DEL-139
chr2-105210876-DEL-109
chr2-105251029-DEL-61
chr2-105274973-DEL-99
chr2-105291020-DEL-72
chr2-105361335-DEL-706
chr2-105362513-DEL-1152
chr2-105377486-DEL-243
chr2-105387428-DEL-6126
chr2-105394022-DEL-316
chr2-105395016-DEL-64
chr2-105409158-DEL-77
chr2-105420775-DEL-173
chr2-105472302-DEL-51
chr2-105475375-DEL-202
chr2-105475948-DEL-80
chr2-105481762-DEL-2085
chr2-105579422-DEL-21176
chr2-105611353-DEL-69
chr2-105624123-DEL-64
chr2-105652567-DEL-86
chr2-105671173-DEL-543
chr2-105675435-DEL-6999
chr2-105688843-DEL-6621
chr2-105719683-DEL-1072
chr2-105760043-DEL-90
chr2-105778726-DEL-208
chr2-105812124-DEL-74
chr2-105830236-DEL-90
chr2-105836201-DEL-429
chr2-105884447-DEL-113
chr2-105895375-DEL-66
chr2-105898329-DEL-188
chr2-105900175-DEL-55
chr2-105943677-DEL-259
chr2-105955179-DEL-257
chr2-105958447-DEL-81
chr2-105977597-DEL-351
chr2-105991343-DEL-78
chr2-106001956-DEL-78
chr2-106011767-DEL-134
chr2-106045391-DEL-325
chr2-106067644-DEL-142
chr2-106098109-DEL-1671
chr2-106100402-DEL-474
chr2-106108191-DEL-547
chr2-106121255-DEL-2999
chr2-106126035-DEL-400
chr2-106181025-DEL-289
chr2-106201638-DEL-79
chr2-106260790-DEL-187
chr2-106268798-DEL-638
chr2-106295728-DEL-208
chr2-106312750-DEL-1769
chr2-106313044-DEL-62
chr2-106313441-DEL-157
chr2-106313489-DEL-3855
chr2-106330711-DEL-56
chr2-106352136-DEL-136
chr2-106367482-DEL-924
chr2-106385560-DEL-681
chr2-106397259-DEL-2164
chr2-106455800-DEL-422
chr2-106504170-DEL-91
chr2-106607553-DEL-1143

chr2-106616648-DEL-55
chr2-106646397-DEL-2799
chr2-106647151-DEL-68
chr2-106654538-DEL-103
chr2-106657740-DEL-52
chr2-106662900-DEL-67
chr2-106669766-DEL-216
chr2-106672362-DEL-145
chr2-106745369-DEL-89
chr2-106750057-DEL-67
chr2-106750530-DEL-6234
chr2-106764662-DEL-72
chr2-106810402-DEL-357
chr2-106838288-DEL-1163
chr2-106850655-DEL-201
chr2-106863680-DEL-1495
chr2-106907973-DEL-60
chr2-106989178-DEL-352
chr2-107002129-DEL-496
chr2-107056889-DEL-509
chr2-107140504-DEL-455
chr2-107147800-DEL-148
chr2-107154927-DEL-153
chr2-107165317-DEL-121
chr2-107166339-DEL-73
chr2-107167665-DEL-105
chr2-107280608-DEL-145
chr2-107341651-DEL-1815
chr2-107359325-DEL-58
chr2-107365683-DEL-332
chr2-107375123-DEL-1254
chr2-107391272-DEL-554
chr2-107415358-DEL-2848
chr2-107421877-DEL-460
chr2-107483116-DEL-824
chr2-107494427-DEL-116
chr2-107512736-DEL-67
chr2-107521748-DEL-64
chr2-107536669-DEL-6194
chr2-107549658-DEL-53
chr2-107562912-DEL-66
chr2-107585707-DEL-9602
chr2-107589678-DEL-60
chr2-107612061-DEL-75
chr2-107652643-DEL-1507
chr2-107676820-DEL-656
chr2-107692016-DEL-468
chr2-107695219-DEL-139
chr2-107719386-DEL-52
chr2-107723820-DEL-1293
chr2-107725760-DEL-52
chr2-107736919-DEL-64
chr2-107742927-DEL-8583
chr2-107804728-DEL-117
chr2-107813965-DEL-52
chr2-107845632-DEL-256
chr2-107944835-DEL-64
chr2-107956939-DEL-401
chr2-107965731-DEL-1335
chr2-107973267-DEL-76
chr2-107981805-DEL-805
chr2-108037292-DEL-60
chr2-108040033-DEL-2855
chr2-108055808-DEL-605
chr2-108074372-DEL-573
chr2-108091598-DEL-1464
chr2-108140712-DEL-68
chr2-108147366-DEL-1560
chr2-108166717-DEL-184
chr2-108173732-DEL-678
chr2-108184152-DEL-1443
chr2-108202760-DEL-1205
chr2-108205970-DEL-780
chr2-108220856-DEL-133
chr2-108228258-DEL-80
chr2-108288173-DEL-158
chr2-108292940-DEL-54

chr2-108293959-DEL-9975
chr2-108309723-DEL-276
chr2-108319733-DEL-6317
chr2-108351210-DEL-63
chr2-108433911-DEL-64
chr2-108438744-DEL-6176
chr2-108483269-DEL-216
chr2-108523570-DEL-237
chr2-108566441-DEL-81
chr2-108611038-DEL-837
chr2-108660863-DEL-150
chr2-108669819-DEL-1532
chr2-108677550-DEL-57
chr2-108688249-DEL-578
chr2-108700112-DEL-101
chr2-108752576-DEL-85
chr2-108769926-DEL-141
chr2-108772637-DEL-83
chr2-108774624-DEL-7779
chr2-108806646-DEL-66
chr2-108836017-DEL-64
chr2-108844700-DEL-341
chr2-108869487-DEL-395
chr2-108870930-DEL-718
chr2-108877601-DEL-299
chr2-108887412-DEL-2473
chr2-108902823-DEL-2268
chr2-108941453-DEL-141
chr2-108953285-DEL-118
chr2-108978727-DEL-526
chr2-109011820-DEL-1587
chr2-109022491-DEL-6971
chr2-109093719-DEL-270
chr2-109122951-DEL-185
chr2-109142835-DEL-1992
chr2-109157883-DEL-1322
chr2-109218057-DEL-62
chr2-109252847-DEL-84
chr2-109296275-DEL-1096
chr2-109324574-DEL-117
chr2-109333277-DEL-66
chr2-109365768-DEL-52
chr2-109367708-DEL-68
chr2-109410605-DEL-155
chr2-109454888-DEL-54
chr2-109558567-DEL-116
chr2-109583087-DEL-63
chr2-109588718-DEL-2237
chr2-109621580-DEL-80
chr2-109632588-DEL-520
chr2-109633391-DEL-86
chr2-109666421-DEL-578
chr2-109678366-DEL-76
chr2-109692302-DEL-52
chr2-109699211-DEL-207
chr2-109735883-DEL-162
chr2-109771022-DEL-525
chr2-109779180-DEL-401
chr2-109782368-DEL-188
chr2-109839591-DEL-136
chr2-109844861-DEL-84
chr2-109921544-DEL-456
chr2-109934933-DEL-6257
chr2-109946616-DEL-12514
chr2-109950728-DEL-6516
chr2-109957267-DEL-221
chr2-109972467-DEL-153
chr2-109973210-DEL-1758
chr2-110025946-DEL-54
chr2-110065383-DEL-64
chr2-110082919-DEL-2097
chr2-110110746-DEL-399
chr2-110127648-DEL-664
chr2-110165897-DEL-2257
chr2-110211309-DEL-178
chr2-110235031-DEL-1887
chr2-110246048-DEL-948

chr2-110251077-DEL-4432
chr2-110255813-DEL-135
chr2-110292505-DEL-5183
chr2-110368944-DEL-317
chr2-110396848-DEL-64
chr2-110433430-DEL-55
chr2-110436935-DEL-58
chr2-110448453-DEL-106
chr2-110458566-DEL-1735
chr2-110494297-DEL-688
chr2-110495520-DEL-80
chr2-110499811-DEL-238
chr2-110525489-DEL-58
chr2-110532204-DEL-98
chr2-110533755-DEL-398
chr2-110553225-DEL-1132
chr2-110565441-DEL-533
chr2-110586360-DEL-67
chr2-110592762-DEL-3005
chr2-110658851-DEL-68
chr2-110681355-DEL-60
chr2-110692446-DEL-58
chr2-110772493-DEL-623
chr2-110811604-DEL-206
chr2-110835866-DEL-12259
chr2-110839981-DEL-342
chr2-110842411-DEL-3870
chr2-110850428-DEL-4451
chr2-110871227-DEL-6189
chr2-110911799-DEL-80
chr2-110947206-DEL-4210
chr2-110951438-DEL-2075
chr2-110953255-DEL-214
chr2-111002913-DEL-52
chr2-111019810-DEL-361
chr2-111028508-DEL-108
chr2-111091609-DEL-93
chr2-111155546-DEL-210
chr2-111167206-DEL-82
chr2-111175005-DEL-8598
chr2-111183496-DEL-723
chr2-111215003-DEL-66
chr2-111269936-DEL-309
chr2-111272118-DEL-139
chr2-111288123-DEL-356
chr2-111288659-DEL-4806
chr2-111333163-DEL-321
chr2-111339678-DEL-150
chr2-111345223-DEL-293
chr2-111350640-DEL-452
chr2-111372540-DEL-4203
chr2-111403836-DEL-400
chr2-111414103-DEL-72
chr2-111452602-DEL-97
chr2-111467097-DEL-164
chr2-111472814-DEL-73
chr2-111480241-DEL-201
chr2-111487873-DEL-220
chr2-111496614-DEL-74
chr2-111497677-DEL-99
chr2-111498756-DEL-879
chr2-111503652-DEL-6185
chr2-111517367-DEL-200
chr2-111523154-DEL-326
chr2-111538322-DEL-312
chr2-111566609-DEL-55
chr2-111570696-DEL-350
chr2-111575275-DEL-946
chr2-111578491-DEL-97
chr2-111582786-DEL-66
chr2-111606089-DEL-86
chr2-111624832-DEL-96
chr2-111654109-DEL-82
chr2-111659601-DEL-143
chr2-111660420-DEL-592
chr2-111667409-DEL-3898
chr2-111675272-DEL-6638

chr2-111710991-DEL-238
chr2-111714252-DEL-91
chr2-111720860-DEL-6438
chr2-111752230-DEL-1281
chr2-111758402-DEL-1066
chr2-111763685-DEL-654
chr2-111766569-DEL-440
chr2-111770120-DEL-237
chr2-111780934-DEL-113
chr2-111782462-DEL-1817
chr2-111797413-DEL-7384
chr2-111855503-DEL-445
chr2-111862506-DEL-58
chr2-111883874-DEL-6474
chr2-111897384-DEL-597
chr2-111903334-DEL-51
chr2-111905579-DEL-289
chr2-111909876-DEL-61
chr2-111926903-DEL-78
chr2-111933263-DEL-517
chr2-111944432-DEL-149
chr2-111959503-DEL-309
chr2-111961659-DEL-64
chr2-111975838-DEL-335
chr2-111983990-DEL-185
chr2-111990707-DEL-6762
chr2-111990839-DEL-204
chr2-111998859-DEL-124
chr2-112014709-DEL-6490
chr2-112021606-DEL-225
chr2-112037383-DEL-82
chr2-112047401-DEL-4007
chr2-112056220-DEL-1444
chr2-112227387-DEL-106
chr2-112265313-DEL-806
chr2-112266705-DEL-75
chr2-112278346-DEL-188
chr2-112291529-DEL-67
chr2-112292440-DEL-215
chr2-112293855-DEL-222
chr2-112296207-DEL-166
chr2-112298818-DEL-79
chr2-112305151-DEL-57
chr2-112322651-DEL-54
chr2-112337954-DEL-73
chr2-112341119-DEL-2348
chr2-112353643-DEL-643
chr2-112390949-DEL-61
chr2-112397937-DEL-3620
chr2-112434096-DEL-255
chr2-112434222-DEL-147
chr2-112489677-DEL-84
chr2-112544756-DEL-290
chr2-112548132-DEL-170
chr2-112550601-DEL-148
chr2-112554689-DEL-287
chr2-112582854-DEL-204
chr2-112602475-DEL-55
chr2-112639125-DEL-377
chr2-112659514-DEL-490
chr2-112688291-DEL-84
chr2-112700017-DEL-628
chr2-112720146-DEL-158
chr2-112783743-DEL-327
chr2-112833368-DEL-56
chr2-112835569-DEL-390
chr2-112875717-DEL-83
chr2-112886679-DEL-102
chr2-112892857-DEL-184
chr2-112910772-DEL-128
chr2-112988978-DEL-590
chr2-113018380-DEL-417
chr2-113022213-DEL-66
chr2-113035628-DEL-97
chr2-113090437-DEL-60
chr2-113098695-DEL-360
chr2-113115676-DEL-56

chr2-113126490-DEL-102
chr2-113127834-DEL-4315
chr2-113133315-DEL-79
chr2-113140011-DEL-55
chr2-113142351-DEL-391
chr2-113153029-DEL-252
chr2-113164849-DEL-508
chr2-113203238-DEL-301
chr2-113307530-DEL-82
chr2-113313648-DEL-140
chr2-113356491-DEL-1441
chr2-113421949-DEL-636
chr2-113483346-DEL-147
chr2-113511348-DEL-384
chr2-113578657-DEL-65
chr2-113602209-DEL-134
chr2-113629899-DEL-160
chr2-113648941-DEL-132
chr2-113674276-DEL-213
chr2-113682482-DEL-6329
chr2-113689999-DEL-196
chr2-113694269-DEL-919
chr2-113707867-DEL-80
chr2-113766349-DEL-51
chr2-113782385-DEL-66
chr2-113794010-DEL-139
chr2-113867888-DEL-478
chr2-113881148-DEL-56
chr2-114059904-DEL-96
chr2-114083256-DEL-80
chr2-114098969-DEL-136304
chr2-114116924-DEL-649
chr2-114208651-DEL-58
chr2-114229954-DEL-148
chr2-114308017-DEL-1799
chr2-114310611-DEL-176
chr2-114379950-DEL-6312
chr2-114400988-DEL-1996
chr2-114408981-DEL-73
chr2-114415448-DEL-1352
chr2-114469744-DEL-56
chr2-114481037-DEL-199
chr2-114481616-DEL-655
chr2-114498882-DEL-93
chr2-114574415-DEL-53
chr2-114594519-DEL-552
chr2-114619190-DEL-114
chr2-114630707-DEL-119
chr2-114632346-DEL-76
chr2-114658353-DEL-117
chr2-114686885-DEL-60
chr2-114705994-DEL-64
chr2-114770656-DEL-82
chr2-114778321-DEL-4787
chr2-114783556-DEL-213
chr2-114810477-DEL-731
chr2-114811568-DEL-60
chr2-114811895-DEL-1073
chr2-114836179-DEL-2163
chr2-114848679-DEL-172
chr2-114849864-DEL-265
chr2-114866326-DEL-51
chr2-114905146-DEL-59
chr2-114910533-DEL-59
chr2-114972329-DEL-1237
chr2-115011015-DEL-65
chr2-115039978-DEL-496
chr2-115056181-DEL-302
chr2-115073834-DEL-52
chr2-115076368-DEL-87
chr2-115131101-DEL-76
chr2-115151907-DEL-64
chr2-115162780-DEL-104
chr2-115197325-DEL-206
chr2-115204702-DEL-51
chr2-115219241-DEL-80
chr2-115251752-DEL-223

chr2-115272237-DEL-486
chr2-115300702-DEL-110
chr2-115304929-DEL-142
chr2-115310103-DEL-56
chr2-115316001-DEL-750
chr2-115366737-DEL-173
chr2-115375648-DEL-164
chr2-115445285-DEL-173
chr2-115454731-DEL-779
chr2-115485723-DEL-53
chr2-115616507-DEL-65
chr2-115653474-DEL-143
chr2-115712682-DEL-70
chr2-115920430-DEL-114
chr2-115921556-DEL-95
chr2-115948877-DEL-1651
chr2-115972659-DEL-56
chr2-115975938-DEL-86
chr2-116007258-DEL-190
chr2-116021844-DEL-57
chr2-116035184-DEL-53
chr2-116061620-DEL-330
chr2-116092088-DEL-134
chr2-116096630-DEL-10876
chr2-116098785-DEL-2174
chr2-116102770-DEL-1483
chr2-116104449-DEL-784
chr2-116124618-DEL-261
chr2-116158326-DEL-3245
chr2-116175339-DEL-62
chr2-116244700-DEL-56
chr2-116257855-DEL-1256
chr2-116319409-DEL-13536
chr2-116365706-DEL-449
chr2-116376962-DEL-749
chr2-116377633-DEL-174
chr2-116378972-DEL-277
chr2-116388024-DEL-3050
chr2-116388656-DEL-87
chr2-116391427-DEL-394
chr2-116392004-DEL-1071
chr2-116405098-DEL-78
chr2-116430366-DEL-156
chr2-116449309-DEL-799
chr2-116457287-DEL-64
chr2-116463567-DEL-79
chr2-116493432-DEL-86
chr2-116524929-DEL-296
chr2-116545452-DEL-958
chr2-116594751-DEL-248
chr2-116604800-DEL-316
chr2-116616681-DEL-148
chr2-116622276-DEL-54
chr2-116625955-DEL-228
chr2-116628690-DEL-70
chr2-116658821-DEL-400
chr2-116697080-DEL-98
chr2-116702571-DEL-663
chr2-116705892-DEL-401
chr2-116724737-DEL-57
chr2-116751983-DEL-147
chr2-116905175-DEL-224
chr2-116912472-DEL-168
chr2-116975750-DEL-52
chr2-116996169-DEL-514
chr2-116998569-DEL-3352
chr2-117032109-DEL-3574
chr2-117063393-DEL-684
chr2-117063607-DEL-1557
chr2-117064202-DEL-912
chr2-117093080-DEL-92
chr2-117098446-DEL-74
chr2-117200769-DEL-85
chr2-117218269-DEL-125
chr2-117222014-DEL-83
chr2-117242186-DEL-143
chr2-117276119-DEL-268

chr2-117278654-DEL-178
chr2-117330396-DEL-302
chr2-117399748-DEL-59
chr2-117435666-DEL-136
chr2-117438639-DEL-51
chr2-117443032-DEL-757
chr2-117449961-DEL-52
chr2-117512422-DEL-489
chr2-117518897-DEL-65
chr2-117528449-DEL-825
chr2-117586892-DEL-565
chr2-117591195-DEL-64
chr2-117602012-DEL-314
chr2-117610219-DEL-261
chr2-117615234-DEL-424
chr2-117639390-DEL-66
chr2-117729540-DEL-218
chr2-117731636-DEL-76
chr2-117735512-DEL-387
chr2-117752970-DEL-168
chr2-117770494-DEL-58
chr2-117791702-DEL-148
chr2-117808939-DEL-52
chr2-117824904-DEL-65
chr2-117844011-DEL-239
chr2-117920038-DEL-60
chr2-117934492-DEL-85
chr2-117968596-DEL-119
chr2-117970167-DEL-348
chr2-117971269-DEL-189
chr2-117971952-DEL-63
chr2-117982991-DEL-1510
chr2-118000596-DEL-219
chr2-118011890-DEL-61
chr2-118031590-DEL-112
chr2-118042261-DEL-288
chr2-118044692-DEL-137
chr2-118056677-DEL-174
chr2-118065393-DEL-365
chr2-118132461-DEL-1187
chr2-118135039-DEL-100
chr2-118145238-DEL-65
chr2-118147944-DEL-62
chr2-118207729-DEL-580
chr2-118211969-DEL-3545
chr2-118248103-DEL-672
chr2-118257161-DEL-299
chr2-118274810-DEL-892
chr2-118275771-DEL-66
chr2-118279705-DEL-95
chr2-118312599-DEL-159
chr2-118324516-DEL-208
chr2-118338878-DEL-282
chr2-118352187-DEL-93
chr2-118429891-DEL-67
chr2-118481559-DEL-302
chr2-118483741-DEL-1111
chr2-118514530-DEL-68
chr2-118521922-DEL-211
chr2-118529080-DEL-194
chr2-118551741-DEL-67
chr2-118552289-DEL-51
chr2-118607727-DEL-53
chr2-118609353-DEL-56
chr2-118643087-DEL-52
chr2-118685979-DEL-1121
chr2-118687119-DEL-1441
chr2-118702089-DEL-208
chr2-118730245-DEL-84
chr2-118765665-DEL-200
chr2-118767666-DEL-50
chr2-118777675-DEL-124
chr2-118782812-DEL-84
chr2-118784937-DEL-453
chr2-118791291-DEL-322
chr2-118819236-DEL-162
chr2-118824105-DEL-95

chr2-118862714-DEL-706
chr2-118882902-DEL-71
chr2-118914670-DEL-54
chr2-118922827-DEL-327
chr2-118923444-DEL-146
chr2-118937554-DEL-229
chr2-118938584-DEL-303
chr2-118955099-DEL-484
chr2-118959278-DEL-1869
chr2-118963720-DEL-288
chr2-118990121-DEL-97
chr2-119015119-DEL-203
chr2-119020941-DEL-198
chr2-119030974-DEL-349
chr2-119037835-DEL-52
chr2-119058679-DEL-57
chr2-119074288-DEL-577
chr2-119084486-DEL-96
chr2-119084998-DEL-55
chr2-119111535-DEL-239
chr2-119117018-DEL-90
chr2-119122733-DEL-55
chr2-119138328-DEL-187
chr2-119149606-DEL-530
chr2-119160321-DEL-140
chr2-119171211-DEL-409
chr2-119188042-DEL-74
chr2-119191042-DEL-931
chr2-119214292-DEL-149
chr2-119226745-DEL-770
chr2-119252102-DEL-177
chr2-119266907-DEL-161
chr2-119287724-DEL-144
chr2-119291009-DEL-226
chr2-119301099-DEL-1242
chr2-119316285-DEL-134
chr2-119327967-DEL-197
chr2-119345415-DEL-55
chr2-119373324-DEL-54
chr2-119381347-DEL-194
chr2-119387930-DEL-74
chr2-119391532-DEL-54
chr2-119398074-DEL-209
chr2-119419554-DEL-163
chr2-119442843-DEL-103
chr2-119446526-DEL-211
chr2-119456866-DEL-231
chr2-119472587-DEL-88
chr2-119495569-DEL-1064
chr2-119498599-DEL-4776
chr2-119532635-DEL-363
chr2-119535414-DEL-475
chr2-119554201-DEL-178
chr2-119569628-DEL-64
chr2-119598970-DEL-106
chr2-119606834-DEL-56
chr2-119613244-DEL-72
chr2-119667372-DEL-55
chr2-119690502-DEL-52
chr2-119698045-DEL-56
chr2-119710872-DEL-66
chr2-119740255-DEL-53
chr2-119768054-DEL-139
chr2-119785548-DEL-542
chr2-119789530-DEL-63
chr2-119817833-DEL-812
chr2-119824495-DEL-133
chr2-119859448-DEL-280
chr2-119873832-DEL-64
chr2-119926605-DEL-830
chr2-119943196-DEL-113
chr2-119947125-DEL-83
chr2-119986152-DEL-132
chr2-119996259-DEL-181
chr2-120008545-DEL-53
chr2-120023316-DEL-52
chr2-120077185-DEL-2691

chr2-120078496-DEL-125
chr2-120090097-DEL-205
chr2-120091602-DEL-87
chr2-120098269-DEL-191
chr2-120101859-DEL-126
chr2-120135947-DEL-74
chr2-120138127-DEL-199
chr2-120185040-DEL-86
chr2-120237367-DEL-4456
chr2-120281374-DEL-59
chr2-120292904-DEL-73
chr2-120301669-DEL-720
chr2-120306123-DEL-72
chr2-120329540-DEL-164
chr2-120349978-DEL-133
chr2-120358439-DEL-138
chr2-120399525-DEL-124
chr2-120452796-DEL-394
chr2-120453844-DEL-999
chr2-120457642-DEL-418
chr2-120478737-DEL-1971
chr2-120481519-DEL-846
chr2-120492615-DEL-263
chr2-120505282-DEL-2799
chr2-120514025-DEL-83
chr2-120540355-DEL-123
chr2-120590377-DEL-60
chr2-120625779-DEL-204
chr2-120626797-DEL-5484
chr2-120632342-DEL-200
chr2-120641979-DEL-245
chr2-120644399-DEL-7125
chr2-120659740-DEL-51
chr2-120695389-DEL-74
chr2-120700035-DEL-110
chr2-120713648-DEL-182
chr2-120767512-DEL-1630
chr2-120804249-DEL-1534
chr2-120805394-DEL-357
chr2-120821846-DEL-63
chr2-120914360-DEL-72
chr2-120944205-DEL-51
chr2-120964817-DEL-111
chr2-120976071-DEL-153
chr2-120987463-DEL-561
chr2-120995537-DEL-55
chr2-121088954-DEL-400
chr2-121093113-DEL-1390
chr2-121108921-DEL-210
chr2-121110099-DEL-1903
chr2-121115210-DEL-88
chr2-121148164-DEL-177
chr2-121155669-DEL-205
chr2-121221261-DEL-51
chr2-121226725-DEL-84
chr2-121239602-DEL-194
chr2-121245980-DEL-56
chr2-121249714-DEL-52
chr2-121257394-DEL-204
chr2-121267809-DEL-145
chr2-121297607-DEL-152
chr2-121308435-DEL-63
chr2-121309046-DEL-183
chr2-121311447-DEL-668
chr2-121314244-DEL-77
chr2-121318668-DEL-1263
chr2-121330158-DEL-154
chr2-121336063-DEL-184
chr2-121370798-DEL-943
chr2-121386681-DEL-147
chr2-121465814-DEL-69
chr2-121508974-DEL-290
chr2-121550494-DEL-477
chr2-121551726-DEL-258
chr2-121553764-DEL-130
chr2-121560387-DEL-217
chr2-121608245-DEL-210

chr2-121618210-DEL-53
chr2-121647380-DEL-69
chr2-121660135-DEL-285
chr2-121664605-DEL-130
chr2-121720855-DEL-58
chr2-121750255-DEL-268
chr2-121778549-DEL-505
chr2-121789196-DEL-208
chr2-121791500-DEL-758
chr2-121804257-DEL-182
chr2-121820019-DEL-3324
chr2-121832803-DEL-74
chr2-121835490-DEL-198
chr2-121850219-DEL-52
chr2-121853258-DEL-195
chr2-121868100-DEL-203
chr2-121898929-DEL-57
chr2-121909473-DEL-403
chr2-121910142-DEL-312
chr2-121925709-DEL-64
chr2-121964754-DEL-511
chr2-121970718-DEL-401
chr2-121995097-DEL-410
chr2-122038634-DEL-112
chr2-122046959-DEL-1465
chr2-122056508-DEL-159
chr2-122057320-DEL-53
chr2-122075077-DEL-66
chr2-122084261-DEL-936
chr2-122119354-DEL-185
chr2-122127576-DEL-72
chr2-122155628-DEL-150
chr2-122161019-DEL-199
chr2-122191211-DEL-53
chr2-122216189-DEL-102
chr2-122225324-DEL-124
chr2-122237261-DEL-196
chr2-122248330-DEL-60
chr2-122287415-DEL-139
chr2-122349039-DEL-213
chr2-122349964-DEL-460
chr2-122389085-DEL-5144
chr2-122443889-DEL-1451
chr2-122470194-DEL-78
chr2-122476085-DEL-69
chr2-122506739-DEL-153
chr2-122512436-DEL-343
chr2-122517388-DEL-81
chr2-122533630-DEL-85
chr2-122574739-DEL-489
chr2-122593419-DEL-113
chr2-122612202-DEL-53
chr2-122620035-DEL-60
chr2-122753066-DEL-1678
chr2-122774019-DEL-2010
chr2-122822220-DEL-2052
chr2-122853022-DEL-107
chr2-122932440-DEL-237
chr2-122974580-DEL-357
chr2-123037004-DEL-134
chr2-123113895-DEL-2973
chr2-123128533-DEL-148
chr2-123149057-DEL-314
chr2-123164430-DEL-82
chr2-123207254-DEL-886
chr2-123225503-DEL-524
chr2-123233497-DEL-112
chr2-123273375-DEL-351
chr2-123292151-DEL-57
chr2-123334297-DEL-235
chr2-123337966-DEL-159
chr2-123338455-DEL-7052
chr2-123405454-DEL-6208
chr2-123412132-DEL-101
chr2-123415299-DEL-56
chr2-123462943-DEL-603
chr2-123486421-DEL-61

chr2-123513269-DEL-134
chr2-123580701-DEL-332
chr2-123598472-DEL-85
chr2-123619051-DEL-975
chr2-123619141-DEL-486
chr2-123626022-DEL-52
chr2-123662414-DEL-65
chr2-123668698-DEL-80
chr2-123690886-DEL-496
chr2-123692274-DEL-266
chr2-123696807-DEL-87
chr2-123698024-DEL-368
chr2-123703498-DEL-758
chr2-123734509-DEL-579
chr2-123747156-DEL-5903
chr2-123758869-DEL-77
chr2-123760483-DEL-237
chr2-123774325-DEL-216
chr2-123796627-DEL-456
chr2-123802982-DEL-621
chr2-123842959-DEL-56
chr2-123849210-DEL-810
chr2-123899072-DEL-1795
chr2-124012412-DEL-201
chr2-124163965-DEL-344
chr2-124192006-DEL-2909
chr2-124207428-DEL-144
chr2-124229252-DEL-302
chr2-124233434-DEL-216
chr2-124266306-DEL-921
chr2-124280620-DEL-108
chr2-124314532-DEL-59
chr2-124324195-DEL-329
chr2-124336003-DEL-75
chr2-124435477-DEL-54
chr2-124522402-DEL-54
chr2-124555432-DEL-1807
chr2-124570495-DEL-66
chr2-124579543-DEL-1330
chr2-124585646-DEL-67
chr2-124592359-DEL-382
chr2-124593444-DEL-71
chr2-124598758-DEL-4948
chr2-124633075-DEL-75
chr2-124664141-DEL-1221
chr2-124698172-DEL-1207
chr2-124699146-DEL-639
chr2-124709796-DEL-58
chr2-124741081-DEL-104
chr2-124763210-DEL-60
chr2-124785522-DEL-150
chr2-124821384-DEL-6417
chr2-124831149-DEL-183
chr2-124900741-DEL-258
chr2-124900913-DEL-132
chr2-124901481-DEL-995
chr2-124902415-DEL-397
chr2-124946913-DEL-359
chr2-124987429-DEL-421
chr2-125081894-DEL-647
chr2-125162503-DEL-51
chr2-125174007-DEL-129
chr2-125174729-DEL-68
chr2-125268452-DEL-746
chr2-125313734-DEL-65
chr2-125328539-DEL-276
chr2-125335396-DEL-56
chr2-125396020-DEL-60
chr2-125403337-DEL-52
chr2-125417557-DEL-70
chr2-125430071-DEL-123
chr2-125437118-DEL-1110
chr2-125443732-DEL-505
chr2-125445793-DEL-291
chr2-125459552-DEL-128
chr2-125467935-DEL-1283
chr2-125472082-DEL-87

chr2-125526779-DEL-56
chr2-125608707-DEL-153
chr2-125615458-DEL-111
chr2-125658215-DEL-199
chr2-125659792-DEL-180
chr2-125661191-DEL-465
chr2-125679525-DEL-211
chr2-125694045-DEL-4572
chr2-125740841-DEL-165
chr2-125781878-DEL-104
chr2-125787351-DEL-308
chr2-125794552-DEL-1602
chr2-125799092-DEL-4548
chr2-125803870-DEL-1058
chr2-125845639-DEL-57
chr2-125853628-DEL-957
chr2-125858182-DEL-954
chr2-125869821-DEL-166
chr2-125870832-DEL-100
chr2-125890294-DEL-74
chr2-125890827-DEL-520
chr2-125974322-DEL-213
chr2-125981519-DEL-3062
chr2-125993631-DEL-68
chr2-126008766-DEL-116
chr2-126037307-DEL-54
chr2-126068033-DEL-218
chr2-126115330-DEL-223
chr2-126165754-DEL-166
chr2-126208273-DEL-310
chr2-126221410-DEL-67
chr2-126236460-DEL-155
chr2-126252247-DEL-3384
chr2-126272504-DEL-160
chr2-126280702-DEL-573
chr2-126287383-DEL-1882
chr2-126295333-DEL-429
chr2-126307836-DEL-87
chr2-126326314-DEL-93
chr2-126431340-DEL-224
chr2-126465440-DEL-330
chr2-126536974-DEL-200
chr2-126555087-DEL-280
chr2-126568224-DEL-53
chr2-126586303-DEL-176
chr2-126595350-DEL-127
chr2-126605674-DEL-72
chr2-126635814-DEL-54
chr2-126691636-DEL-55
chr2-126767122-DEL-52
chr2-126771491-DEL-761
chr2-126794789-DEL-79
chr2-126846607-DEL-65
chr2-126873331-DEL-199
chr2-126880897-DEL-99
chr2-126914248-DEL-4761
chr2-126932193-DEL-173
chr2-126939832-DEL-100
chr2-126941901-DEL-174
chr2-126942006-DEL-849
chr2-126967861-DEL-86
chr2-126978833-DEL-87
chr2-126993882-DEL-197
chr2-127010427-DEL-1143
chr2-127011595-DEL-1175
chr2-127026411-DEL-1281
chr2-127048550-DEL-274
chr2-127091850-DEL-204
chr2-127111259-DEL-67
chr2-127145365-DEL-545
chr2-127145829-DEL-212
chr2-127240117-DEL-57
chr2-127311938-DEL-92
chr2-127317937-DEL-190
chr2-127319769-DEL-241
chr2-127408240-DEL-58
chr2-127442927-DEL-55

chr2-127459575-DEL-317
chr2-127462877-DEL-106
chr2-127528825-DEL-60
chr2-127607393-DEL-52
chr2-127737363-DEL-623
chr2-127738215-DEL-57
chr2-127774649-DEL-161
chr2-127808459-DEL-1354
chr2-127818401-DEL-86
chr2-127871207-DEL-71
chr2-127885995-DEL-52
chr2-127888352-DEL-259
chr2-127911345-DEL-78
chr2-127944930-DEL-75
chr2-127950224-DEL-62
chr2-127952862-DEL-185
chr2-127990766-DEL-87
chr2-128026898-DEL-94
chr2-128034069-DEL-173
chr2-128049967-DEL-58
chr2-128069074-DEL-1644
chr2-128090382-DEL-134
chr2-128115077-DEL-56
chr2-128124591-DEL-57
chr2-128148353-DEL-60
chr2-128164835-DEL-322
chr2-128251841-DEL-128
chr2-128263430-DEL-208
chr2-128265416-DEL-505
chr2-128267932-DEL-113
chr2-128291027-DEL-57
chr2-128390086-DEL-132
chr2-128393369-DEL-60
chr2-128395677-DEL-577
chr2-128420671-DEL-55
chr2-128427339-DEL-373
chr2-128434931-DEL-403
chr2-128522516-DEL-126
chr2-128533119-DEL-1389
chr2-128566836-DEL-216
chr2-128572582-DEL-123
chr2-128575146-DEL-144
chr2-128588770-DEL-58
chr2-128598900-DEL-55
chr2-128611797-DEL-157
chr2-128622021-DEL-924
chr2-128627784-DEL-58
chr2-128634402-DEL-81
chr2-128646821-DEL-167
chr2-128652647-DEL-56
chr2-128655598-DEL-56
chr2-128656780-DEL-114
chr2-128665230-DEL-205
chr2-128715487-DEL-727
chr2-128719633-DEL-722
chr2-128726464-DEL-63
chr2-128752582-DEL-771
chr2-128752636-DEL-1819
chr2-128798052-DEL-77
chr2-128798785-DEL-6721
chr2-128802250-DEL-506
chr2-128831524-DEL-165
chr2-128853269-DEL-105
chr2-128896282-DEL-150
chr2-128903103-DEL-775
chr2-128914385-DEL-102
chr2-128916936-DEL-463
chr2-128920268-DEL-127
chr2-128923685-DEL-64
chr2-128929544-DEL-336
chr2-128962778-DEL-100
chr2-128987414-DEL-1030
chr2-128994330-DEL-176
chr2-129057036-DEL-132
chr2-129075762-DEL-55
chr2-129078867-DEL-205
chr2-129086393-DEL-59

chr2-129093023-DEL-51
chr2-129123492-DEL-217
chr2-129158881-DEL-923
chr2-129174037-DEL-132
chr2-129204528-DEL-105
chr2-129212773-DEL-125
chr2-129274746-DEL-1714
chr2-129302855-DEL-103
chr2-129469361-DEL-128
chr2-129564672-DEL-142
chr2-129566923-DEL-158
chr2-129568574-DEL-105
chr2-129620118-DEL-84
chr2-129631961-DEL-1483
chr2-129638552-DEL-126
chr2-129684148-DEL-1413
chr2-129685574-DEL-3209
chr2-129717661-DEL-388
chr2-129759071-DEL-715
chr2-129770176-DEL-62
chr2-129773809-DEL-440
chr2-129782857-DEL-273
chr2-129788704-DEL-4789
chr2-129821942-DEL-186
chr2-129827399-DEL-85
chr2-129938283-DEL-167
chr2-129950607-DEL-63
chr2-129969232-DEL-64
chr2-129977563-DEL-55
chr2-130050446-DEL-428
chr2-130050995-DEL-301
chr2-130075257-DEL-99
chr2-130089141-DEL-175
chr2-130089652-DEL-70
chr2-130093869-DEL-128
chr2-130102743-DEL-93
chr2-130172977-DEL-510
chr2-130186161-DEL-64
chr2-130230554-DEL-1057
chr2-130267586-DEL-194
chr2-130290586-DEL-771
chr2-130318789-DEL-56
chr2-130329221-DEL-207
chr2-130348738-DEL-56
chr2-130368008-DEL-54
chr2-130372602-DEL-112
chr2-130458907-DEL-61
chr2-130459687-DEL-451
chr2-130470678-DEL-73
chr2-130475437-DEL-54
chr2-130510434-DEL-609
chr2-130534606-DEL-207
chr2-130555574-DEL-104
chr2-130594225-DEL-59
chr2-130597229-DEL-85
chr2-130598985-DEL-217
chr2-130632704-DEL-713
chr2-130657791-DEL-271
chr2-130667524-DEL-678
chr2-130716515-DEL-573
chr2-130730347-DEL-66
chr2-130730361-DEL-850
chr2-130744808-DEL-125
chr2-130758725-DEL-65
chr2-130771479-DEL-331
chr2-130781509-DEL-172
chr2-130811478-DEL-364
chr2-130833953-DEL-262
chr2-130846769-DEL-165
chr2-130908062-DEL-398
chr2-130913951-DEL-201
chr2-130941887-DEL-180
chr2-131073004-DEL-55
chr2-131099775-DEL-124
chr2-131102635-DEL-53
chr2-131146790-DEL-306
chr2-131151815-DEL-52

chr2-131160490-DEL-116
chr2-131163149-DEL-102
chr2-131163842-DEL-445
chr2-131200343-DEL-53
chr2-131210024-DEL-70
chr2-131230970-DEL-3615
chr2-131236716-DEL-171
chr2-131250060-DEL-88
chr2-131251266-DEL-444
chr2-131259997-DEL-196
chr2-131291606-DEL-82
chr2-131313649-DEL-2592
chr2-131316192-DEL-67
chr2-131380311-DEL-272
chr2-131400648-DEL-125
chr2-131406435-DEL-135
chr2-131410061-DEL-153
chr2-131506087-DEL-232
chr2-131564341-DEL-2867
chr2-131581468-DEL-78
chr2-131584023-DEL-432
chr2-131625448-DEL-1516
chr2-131652837-DEL-58
chr2-131672710-DEL-544
chr2-131691265-DEL-80
chr2-131719643-DEL-52
chr2-131726078-DEL-88
chr2-131729915-DEL-191
chr2-131776794-DEL-541
chr2-131791410-DEL-93
chr2-131804605-DEL-56
chr2-131816742-DEL-295
chr2-131840824-DEL-297
chr2-131855518-DEL-56
chr2-131857640-DEL-208
chr2-131910313-DEL-124
chr2-131917981-DEL-367
chr2-131938789-DEL-93
chr2-131971640-DEL-206
chr2-131979586-DEL-95
chr2-132006705-DEL-166
chr2-132010799-DEL-1762
chr2-132012206-DEL-174
chr2-132012788-DEL-189
chr2-132015309-DEL-60
chr2-132023337-DEL-62
chr2-132054982-DEL-64
chr2-132169147-DEL-82
chr2-132183235-DEL-6635
chr2-132185270-DEL-158
chr2-132192366-DEL-549
chr2-132193046-DEL-1108
chr2-132193306-DEL-824
chr2-132216287-DEL-138
chr2-132319161-DEL-64
chr2-132320895-DEL-57
chr2-132344350-DEL-1139
chr2-132349959-DEL-1000
chr2-132353506-DEL-115
chr2-132365068-DEL-164
chr2-132491415-DEL-77
chr2-132494161-DEL-52
chr2-132502092-DEL-666
chr2-132553483-DEL-245
chr2-132569877-DEL-247
chr2-132580078-DEL-112
chr2-132580797-DEL-436
chr2-132607851-DEL-58
chr2-132611702-DEL-121
chr2-132641828-DEL-82
chr2-132643930-DEL-52
chr2-132727517-DEL-6475
chr2-132749730-DEL-1712
chr2-132774218-DEL-152
chr2-132802516-DEL-998
chr2-132850643-DEL-88
chr2-132852695-DEL-84

chr2-132865935-DEL-2097
chr2-132889702-DEL-59
chr2-132898898-DEL-644
chr2-132917268-DEL-650
chr2-132943805-DEL-5566
chr2-132954206-DEL-183
chr2-132966574-DEL-199
chr2-132998027-DEL-56
chr2-133039117-DEL-3616
chr2-133043788-DEL-1818
chr2-133074432-DEL-84
chr2-133075710-DEL-159
chr2-133076400-DEL-186
chr2-133114940-DEL-93
chr2-133117145-DEL-86
chr2-133139141-DEL-66
chr2-133175517-DEL-401
chr2-133205728-DEL-147
chr2-133314267-DEL-61
chr2-133318517-DEL-61
chr2-133319108-DEL-109
chr2-133333432-DEL-56
chr2-133357385-DEL-52
chr2-133423183-DEL-50
chr2-133525785-DEL-52
chr2-133579691-DEL-54
chr2-133606593-DEL-154
chr2-133621143-DEL-97
chr2-133680606-DEL-105
chr2-133687601-DEL-78
chr2-133719648-DEL-109
chr2-133720227-DEL-649
chr2-133755284-DEL-968
chr2-133873950-DEL-58
chr2-133885511-DEL-51
chr2-133897788-DEL-660
chr2-133928862-DEL-64
chr2-133971366-DEL-1394
chr2-133991081-DEL-99
chr2-134010838-DEL-1928
chr2-134035030-DEL-337
chr2-134072681-DEL-806
chr2-134075806-DEL-8269
chr2-134119316-DEL-1986
chr2-134129580-DEL-106
chr2-134176906-DEL-349
chr2-134180563-DEL-153
chr2-134208107-DEL-196
chr2-134221035-DEL-110
chr2-134244985-DEL-108
chr2-134256929-DEL-1608
chr2-134267012-DEL-6063
chr2-134293472-DEL-85
chr2-134305891-DEL-81
chr2-134317423-DEL-275
chr2-134344233-DEL-72
chr2-134347939-DEL-636
chr2-134369041-DEL-62
chr2-134371663-DEL-202
chr2-134425536-DEL-128
chr2-134432463-DEL-506
chr2-134451813-DEL-1777
chr2-134463430-DEL-5144
chr2-134477423-DEL-65
chr2-134508801-DEL-213
chr2-134522921-DEL-383
chr2-134524648-DEL-292
chr2-134569401-DEL-1173
chr2-134569577-DEL-99
chr2-134579723-DEL-218
chr2-134619380-DEL-108
chr2-134642066-DEL-104
chr2-134647847-DEL-212
chr2-134665831-DEL-1172
chr2-134679663-DEL-236
chr2-134762000-DEL-60
chr2-134808564-DEL-1101

chr2-134838607-DEL-72
chr2-134877045-DEL-307
chr2-134925306-DEL-628
chr2-134961248-DEL-57
chr2-134987720-DEL-6151
chr2-134988198-DEL-986
chr2-135000304-DEL-732
chr2-135009187-DEL-354
chr2-135019845-DEL-499
chr2-135058069-DEL-401
chr2-135078094-DEL-103
chr2-135089376-DEL-1893
chr2-135104266-DEL-101
chr2-135125371-DEL-56
chr2-135171292-DEL-95
chr2-135178393-DEL-140
chr2-135193242-DEL-903
chr2-135204363-DEL-782
chr2-135216251-DEL-68
chr2-135320238-DEL-3875
chr2-135343392-DEL-1970
chr2-135349647-DEL-603
chr2-135437868-DEL-52
chr2-135472073-DEL-585
chr2-135480611-DEL-176
chr2-135492442-DEL-351
chr2-135520026-DEL-965
chr2-135520029-DEL-275
chr2-135524141-DEL-124
chr2-135528184-DEL-150
chr2-135529083-DEL-174
chr2-135551613-DEL-217
chr2-135559170-DEL-73
chr2-135577305-DEL-118
chr2-135603777-DEL-189
chr2-135609873-DEL-53
chr2-135624574-DEL-51
chr2-135629822-DEL-187
chr2-135642118-DEL-88
chr2-135666015-DEL-52
chr2-135692328-DEL-662
chr2-135732664-DEL-417
chr2-135746104-DEL-845
chr2-135751438-DEL-220
chr2-135758162-DEL-57
chr2-135767278-DEL-111
chr2-135769403-DEL-134
chr2-135777799-DEL-108
chr2-135801089-DEL-101
chr2-135822726-DEL-68
chr2-135827534-DEL-1082
chr2-135830473-DEL-232
chr2-135941712-DEL-57
chr2-135970888-DEL-96
chr2-135992149-DEL-131
chr2-135994271-DEL-128
chr2-136022177-DEL-87
chr2-136025014-DEL-1559
chr2-136041810-DEL-264
chr2-136041825-DEL-806
chr2-136046284-DEL-1528
chr2-136060235-DEL-106
chr2-136071425-DEL-1897
chr2-136073964-DEL-104
chr2-136090093-DEL-99
chr2-136096809-DEL-840
chr2-136096909-DEL-113
chr2-136107042-DEL-60
chr2-136132008-DEL-75
chr2-136171583-DEL-68
chr2-136184682-DEL-58
chr2-136204557-DEL-90
chr2-136264167-DEL-270
chr2-136296352-DEL-991
chr2-136302264-DEL-175
chr2-136305762-DEL-605
chr2-136318035-DEL-84

chr2-136419362-DEL-60
chr2-136420608-DEL-77
chr2-136421779-DEL-187
chr2-136454781-DEL-506
chr2-136489629-DEL-358
chr2-136507752-DEL-203
chr2-136525262-DEL-580
chr2-136549535-DEL-847
chr2-136568395-DEL-64
chr2-136592994-DEL-53
chr2-136678953-DEL-746
chr2-136711753-DEL-206
chr2-136727093-DEL-66
chr2-136728370-DEL-85
chr2-136741069-DEL-404
chr2-136742983-DEL-2456
chr2-136745855-DEL-78
chr2-136800964-DEL-64
chr2-136806367-DEL-3049
chr2-136807790-DEL-6196
chr2-136823011-DEL-52
chr2-136851787-DEL-131
chr2-136852551-DEL-425
chr2-136857625-DEL-65
chr2-136859592-DEL-167
chr2-136870040-DEL-6279
chr2-136917256-DEL-158
chr2-136966163-DEL-173
chr2-137027594-DEL-113
chr2-137029607-DEL-162
chr2-137041201-DEL-204
chr2-137050731-DEL-441
chr2-137092788-DEL-63
chr2-137105417-DEL-51
chr2-137116157-DEL-2292
chr2-137174867-DEL-155
chr2-137187706-DEL-7226
chr2-137205373-DEL-715
chr2-137205394-DEL-6541
chr2-137206116-DEL-5813
chr2-137209872-DEL-1608
chr2-137215246-DEL-58
chr2-137231776-DEL-155
chr2-137264002-DEL-412
chr2-137371153-DEL-212
chr2-137420823-DEL-200
chr2-137424975-DEL-2468
chr2-137431117-DEL-989
chr2-137459823-DEL-1954
chr2-137480781-DEL-3881
chr2-137483247-DEL-466
chr2-137484169-DEL-214
chr2-137487324-DEL-68
chr2-137497572-DEL-164
chr2-137564433-DEL-115
chr2-137577147-DEL-250
chr2-137634656-DEL-745
chr2-137641179-DEL-593
chr2-137653853-DEL-64
chr2-137680812-DEL-54
chr2-137697764-DEL-502
chr2-137705466-DEL-258
chr2-137727209-DEL-538
chr2-137728238-DEL-66
chr2-137730759-DEL-6184
chr2-137737854-DEL-96
chr2-137770280-DEL-141
chr2-137803015-DEL-56
chr2-137812818-DEL-3070
chr2-137853166-DEL-52
chr2-137857013-DEL-80
chr2-137867443-DEL-503
chr2-137894636-DEL-55
chr2-137903626-DEL-5786
chr2-137918829-DEL-327
chr2-137919644-DEL-1007
chr2-137959991-DEL-1040

chr2-138010587-DEL-6026
chr2-138071594-DEL-482
chr2-138085196-DEL-138
chr2-138217287-DEL-901
chr2-138237180-DEL-212
chr2-138330303-DEL-134
chr2-138392645-DEL-52
chr2-138404295-DEL-8847
chr2-138415728-DEL-6578
chr2-138467788-DEL-2826
chr2-138467873-DEL-84
chr2-138468906-DEL-668
chr2-138600969-DEL-83
chr2-138605170-DEL-66
chr2-138609977-DEL-62
chr2-138617571-DEL-698
chr2-138636548-DEL-231
chr2-138640621-DEL-57
chr2-138664804-DEL-155
chr2-138709673-DEL-63
chr2-138809201-DEL-1739
chr2-138824313-DEL-71
chr2-138852098-DEL-52
chr2-138932448-DEL-141
chr2-138938797-DEL-2540
chr2-138958661-DEL-98
chr2-138962533-DEL-78
chr2-138967490-DEL-708
chr2-139000337-DEL-2958
chr2-139030154-DEL-361
chr2-139051242-DEL-777
chr2-139064086-DEL-201
chr2-139080099-DEL-909
chr2-139080686-DEL-2924
chr2-139081027-DEL-3168
chr2-139096794-DEL-173
chr2-139102951-DEL-1579
chr2-139133762-DEL-139
chr2-139148810-DEL-402
chr2-139226715-DEL-51
chr2-139253240-DEL-685
chr2-139260279-DEL-218
chr2-139262588-DEL-72
chr2-139296774-DEL-8497
chr2-139346074-DEL-403
chr2-139364203-DEL-208
chr2-139381367-DEL-1889
chr2-139439023-DEL-73
chr2-139506215-DEL-67
chr2-139509002-DEL-53
chr2-139542428-DEL-85
chr2-139636434-DEL-131
chr2-139660223-DEL-60
chr2-139689129-DEL-1075
chr2-139730725-DEL-133
chr2-139753875-DEL-73
chr2-139803285-DEL-74
chr2-139831606-DEL-354
chr2-139901335-DEL-77
chr2-139918151-DEL-90
chr2-139938122-DEL-53
chr2-139953373-DEL-96
chr2-139958200-DEL-196
chr2-139973099-DEL-1686
chr2-139993118-DEL-3247
chr2-140000770-DEL-214
chr2-140012020-DEL-185
chr2-140140827-DEL-569
chr2-140153952-DEL-215
chr2-140156330-DEL-2744
chr2-140161073-DEL-59
chr2-140173490-DEL-58
chr2-140199014-DEL-2166
chr2-140227919-DEL-1107
chr2-140300139-DEL-977
chr2-140301661-DEL-285
chr2-140308286-DEL-5414

chr2-140313250-DEL-60
chr2-140371210-DEL-1107
chr2-140382047-DEL-6476
chr2-140382112-DEL-214
chr2-140394039-DEL-57
chr2-140407269-DEL-164
chr2-140413919-DEL-291
chr2-140415422-DEL-2040
chr2-140473892-DEL-259
chr2-140492413-DEL-138
chr2-140516552-DEL-244
chr2-140523300-DEL-146
chr2-140559005-DEL-225
chr2-140586069-DEL-358
chr2-140660542-DEL-201
chr2-140707542-DEL-256
chr2-140761960-DEL-128
chr2-140923671-DEL-139
chr2-140940582-DEL-103
chr2-140963320-DEL-55
chr2-140974654-DEL-783
chr2-140975244-DEL-188
chr2-140998197-DEL-70
chr2-141022048-DEL-73
chr2-141043980-DEL-66
chr2-141050257-DEL-171
chr2-141070639-DEL-66
chr2-141076031-DEL-107
chr2-141089927-DEL-81
chr2-141176568-DEL-1604
chr2-141195739-DEL-823
chr2-141213875-DEL-82
chr2-141227233-DEL-112
chr2-141235591-DEL-74
chr2-141242429-DEL-720
chr2-141251607-DEL-1425
chr2-141306972-DEL-467
chr2-141405963-DEL-107270
chr2-141515857-DEL-51
chr2-141603982-DEL-840
chr2-141612765-DEL-553
chr2-141629348-DEL-1702
chr2-141756312-DEL-54
chr2-141772858-DEL-90
chr2-141798697-DEL-366
chr2-141801189-DEL-88
chr2-141853548-DEL-1690
chr2-141871886-DEL-201
chr2-141884394-DEL-714
chr2-141894635-DEL-52
chr2-141930299-DEL-1210
chr2-141985155-DEL-63
chr2-142047077-DEL-248
chr2-142079019-DEL-133
chr2-142091127-DEL-341
chr2-142181615-DEL-112
chr2-142220224-DEL-922
chr2-142220886-DEL-274
chr2-142264096-DEL-1611
chr2-142271432-DEL-391
chr2-142274291-DEL-222
chr2-142306167-DEL-6415
chr2-142315402-DEL-2565
chr2-142318793-DEL-6452
chr2-142328424-DEL-331
chr2-142374050-DEL-80
chr2-142387378-DEL-69
chr2-142394777-DEL-117
chr2-142484822-DEL-1596
chr2-142516740-DEL-378
chr2-142517751-DEL-63
chr2-142613544-DEL-87
chr2-142618067-DEL-1439
chr2-142643807-DEL-72
chr2-142666131-DEL-180
chr2-142698313-DEL-64
chr2-142720301-DEL-60

chr2-142727212-DEL-70
chr2-142744094-DEL-60
chr2-142767964-DEL-455
chr2-142779594-DEL-80
chr2-142817407-DEL-51
chr2-142832708-DEL-201
chr2-142854155-DEL-286
chr2-142870371-DEL-76
chr2-142880505-DEL-57
chr2-142884649-DEL-201
chr2-142946719-DEL-101
chr2-142947574-DEL-508
chr2-142993635-DEL-56
chr2-142998677-DEL-685
chr2-143003729-DEL-1008
chr2-143083106-DEL-174
chr2-143088416-DEL-477
chr2-143100635-DEL-1505
chr2-143102755-DEL-208
chr2-143122467-DEL-205
chr2-143130312-DEL-6629
chr2-143136518-DEL-333
chr2-143168202-DEL-192
chr2-143172000-DEL-138
chr2-143230432-DEL-64
chr2-143239023-DEL-615
chr2-143242830-DEL-272
chr2-143253634-DEL-686
chr2-143263606-DEL-75
chr2-143268704-DEL-6884
chr2-143287392-DEL-152
chr2-143293484-DEL-87
chr2-143333712-DEL-52
chr2-143357505-DEL-99
chr2-143407420-DEL-853
chr2-143414654-DEL-88
chr2-143433024-DEL-56
chr2-143459388-DEL-454
chr2-143475995-DEL-54
chr2-143547793-DEL-112
chr2-143562371-DEL-194
chr2-143564552-DEL-55
chr2-143578207-DEL-64
chr2-143588493-DEL-89
chr2-143617958-DEL-521
chr2-143646618-DEL-54
chr2-143674531-DEL-80
chr2-143675516-DEL-391
chr2-143702264-DEL-66
chr2-143715520-DEL-1270
chr2-143715685-DEL-977
chr2-143715731-DEL-237
chr2-143729518-DEL-562
chr2-143767302-DEL-441
chr2-143772174-DEL-113
chr2-143775091-DEL-74
chr2-143797613-DEL-59
chr2-143821596-DEL-53
chr2-143829892-DEL-52
chr2-143835833-DEL-78
chr2-143860647-DEL-171
chr2-143880457-DEL-240
chr2-143903131-DEL-51
chr2-143916305-DEL-93
chr2-143973546-DEL-387
chr2-143982489-DEL-734
chr2-144062133-DEL-127
chr2-144066620-DEL-70
chr2-144074093-DEL-155
chr2-144074429-DEL-649
chr2-144090028-DEL-1357
chr2-144093986-DEL-200
chr2-144164321-DEL-98
chr2-144167291-DEL-359
chr2-144177544-DEL-566
chr2-144189699-DEL-293
chr2-144206669-DEL-209

chr2-144233214-DEL-210
chr2-144237754-DEL-182
chr2-144240554-DEL-102
chr2-144251611-DEL-163
chr2-144253683-DEL-85
chr2-144254173-DEL-201
chr2-144254960-DEL-888
chr2-144286867-DEL-68
chr2-144288869-DEL-66
chr2-144304671-DEL-76
chr2-144311011-DEL-270
chr2-144349606-DEL-305
chr2-144356984-DEL-414
chr2-144447455-DEL-172
chr2-144468701-DEL-110
chr2-144529092-DEL-230
chr2-144532281-DEL-479
chr2-144584066-DEL-120
chr2-144658179-DEL-140
chr2-144668262-DEL-193
chr2-144696682-DEL-286
chr2-144717070-DEL-1182
chr2-144746945-DEL-66
chr2-144758865-DEL-80
chr2-144776934-DEL-1029
chr2-144782513-DEL-51
chr2-144787357-DEL-66
chr2-144791710-DEL-342
chr2-144815156-DEL-872
chr2-144835001-DEL-67
chr2-144847910-DEL-72
chr2-144857234-DEL-107
chr2-144903797-DEL-942
chr2-144929729-DEL-131
chr2-144934053-DEL-292
chr2-144966923-DEL-182
chr2-144981943-DEL-6412
chr2-144988442-DEL-61
chr2-145019821-DEL-56
chr2-145027340-DEL-434
chr2-145041432-DEL-115
chr2-145092352-DEL-69
chr2-145109327-DEL-68
chr2-145174994-DEL-60
chr2-145202050-DEL-79
chr2-145235529-DEL-640
chr2-145249477-DEL-468
chr2-145257150-DEL-204
chr2-145307430-DEL-518
chr2-145332394-DEL-197
chr2-145335942-DEL-2720
chr2-145384036-DEL-86
chr2-145440785-DEL-99
chr2-145503069-DEL-142
chr2-145503442-DEL-344
chr2-145521239-DEL-58
chr2-145650690-DEL-1792
chr2-145654177-DEL-541
chr2-145710896-DEL-1239
chr2-145713185-DEL-66
chr2-145731639-DEL-385
chr2-145742309-DEL-409
chr2-145764574-DEL-171
chr2-145771080-DEL-192
chr2-145780480-DEL-434
chr2-145826322-DEL-62
chr2-145826982-DEL-59
chr2-145844379-DEL-6184
chr2-145859737-DEL-4005
chr2-145899916-DEL-410
chr2-146017047-DEL-1825
chr2-146047168-DEL-226
chr2-146051412-DEL-53
chr2-146071719-DEL-279
chr2-146084900-DEL-172
chr2-146128747-DEL-701
chr2-146137478-DEL-75

chr2-146235598-DEL-202
chr2-146241058-DEL-62
chr2-146245566-DEL-486
chr2-146251055-DEL-66
chr2-146270058-DEL-56
chr2-146275928-DEL-238
chr2-146331067-DEL-507
chr2-146334561-DEL-58
chr2-146365029-DEL-1240
chr2-146371691-DEL-247
chr2-146394487-DEL-76
chr2-146414715-DEL-1436
chr2-146505628-DEL-594
chr2-146510396-DEL-56
chr2-146574949-DEL-140
chr2-146576211-DEL-53
chr2-146611251-DEL-2493
chr2-146638253-DEL-295
chr2-146647157-DEL-87
chr2-146734801-DEL-316
chr2-146738434-DEL-2215
chr2-146747423-DEL-1182
chr2-146755314-DEL-59
chr2-146765758-DEL-51
chr2-146781437-DEL-169
chr2-146790922-DEL-490
chr2-146795347-DEL-490
chr2-146802120-DEL-242
chr2-146850859-DEL-61
chr2-146975207-DEL-61
chr2-146976701-DEL-786
chr2-146989230-DEL-1332
chr2-147007604-DEL-1174
chr2-147014395-DEL-70
chr2-147018045-DEL-210
chr2-147037848-DEL-113
chr2-147043452-DEL-73
chr2-147052731-DEL-126
chr2-147056445-DEL-52
chr2-147067533-DEL-391
chr2-147093047-DEL-262
chr2-147108380-DEL-57
chr2-147152621-DEL-588
chr2-147170025-DEL-378
chr2-147183695-DEL-238
chr2-147226873-DEL-1379
chr2-147229490-DEL-1547
chr2-147242763-DEL-666
chr2-147260326-DEL-167
chr2-147278387-DEL-139
chr2-147287086-DEL-529
chr2-147351319-DEL-146
chr2-147393211-DEL-134
chr2-147403543-DEL-71
chr2-147429493-DEL-333
chr2-147435205-DEL-2798
chr2-147438166-DEL-52
chr2-147472517-DEL-112
chr2-147497398-DEL-1197
chr2-147540263-DEL-1796
chr2-147621060-DEL-197
chr2-147639663-DEL-85
chr2-147643384-DEL-495
chr2-147664429-DEL-329
chr2-147673611-DEL-90
chr2-147706616-DEL-6467
chr2-147801036-DEL-152
chr2-147807221-DEL-170
chr2-147833293-DEL-70
chr2-147926568-DEL-92
chr2-147981195-DEL-111
chr2-148023108-DEL-290
chr2-148049120-DEL-54
chr2-148072904-DEL-55
chr2-148091182-DEL-634
chr2-148096308-DEL-994
chr2-148157859-DEL-56

chr2-148172319-DEL-352
chr2-148181165-DEL-84
chr2-148182333-DEL-1331
chr2-148190474-DEL-77
chr2-148208642-DEL-127
chr2-148277640-DEL-61
chr2-148287509-DEL-90
chr2-148302358-DEL-211
chr2-148314538-DEL-1342
chr2-148323324-DEL-175
chr2-148324544-DEL-69
chr2-148331690-DEL-172
chr2-148364633-DEL-106
chr2-148390207-DEL-393
chr2-148392515-DEL-185
chr2-148489014-DEL-98
chr2-148490243-DEL-210
chr2-148583720-DEL-69
chr2-148597871-DEL-71
chr2-148607512-DEL-96
chr2-148613561-DEL-72
chr2-148657228-DEL-52
chr2-148659603-DEL-115
chr2-148683730-DEL-496
chr2-148685437-DEL-668
chr2-148686752-DEL-264
chr2-148695296-DEL-319
chr2-148699065-DEL-1541
chr2-148749526-DEL-822
chr2-148755917-DEL-2575
chr2-148776914-DEL-63
chr2-148780103-DEL-2225
chr2-148783221-DEL-562
chr2-148822077-DEL-6195
chr2-148846868-DEL-83
chr2-148881900-DEL-1991
chr2-148888183-DEL-103
chr2-148891611-DEL-6389
chr2-148909263-DEL-51
chr2-148932872-DEL-89
chr2-148936395-DEL-2345
chr2-148950962-DEL-22796
chr2-148982329-DEL-87
chr2-149025912-DEL-42212
chr2-149086370-DEL-1366
chr2-149088584-DEL-51
chr2-149110515-DEL-6120
chr2-149124195-DEL-52
chr2-149127928-DEL-6759
chr2-149161610-DEL-10341
chr2-149171236-DEL-1631
chr2-149179198-DEL-302
chr2-149188172-DEL-53
chr2-149192396-DEL-1118
chr2-149198295-DEL-1433
chr2-149199008-DEL-8743
chr2-149207411-DEL-62
chr2-149211002-DEL-3056
chr2-149228436-DEL-4175
chr2-149270474-DEL-2813
chr2-149285060-DEL-1953
chr2-149355501-DEL-6211
chr2-149362133-DEL-2823
chr2-149394188-DEL-1997
chr2-149403079-DEL-61
chr2-149404352-DEL-2183
chr2-149426999-DEL-182
chr2-149430774-DEL-286
chr2-149464618-DEL-881
chr2-149465381-DEL-1667
chr2-149482840-DEL-55
chr2-149490184-DEL-505
chr2-149494340-DEL-95
chr2-149530200-DEL-736
chr2-149531987-DEL-400
chr2-149566648-DEL-651
chr2-149630371-DEL-699

chr2-149758290-DEL-95
chr2-149777993-DEL-80
chr2-149810598-DEL-812
chr2-149812335-DEL-571
chr2-149814676-DEL-87
chr2-149856385-DEL-51
chr2-149861791-DEL-88
chr2-149865098-DEL-129
chr2-149910830-DEL-6147
chr2-149920468-DEL-12122
chr2-149928908-DEL-4090
chr2-149929470-DEL-810
chr2-149947331-DEL-5723
chr2-149949569-DEL-604
chr2-150057827-DEL-361
chr2-150136844-DEL-206
chr2-150141250-DEL-77
chr2-150154201-DEL-221
chr2-150203548-DEL-229
chr2-150215421-DEL-401
chr2-150237820-DEL-194
chr2-150241467-DEL-74
chr2-150253941-DEL-6942
chr2-150266199-DEL-6190
chr2-150276965-DEL-1359
chr2-150298506-DEL-96
chr2-150311130-DEL-1082
chr2-150330061-DEL-78
chr2-150339866-DEL-2967
chr2-150350649-DEL-196
chr2-150357763-DEL-31858
chr2-150360824-DEL-137
chr2-150386497-DEL-1314
chr2-150391455-DEL-486
chr2-150438197-DEL-1629
chr2-150445281-DEL-72
chr2-150483949-DEL-60
chr2-150490887-DEL-51
chr2-150508396-DEL-206
chr2-150595534-DEL-995
chr2-150637922-DEL-1875
chr2-150647496-DEL-68
chr2-150663735-DEL-370
chr2-150691609-DEL-101
chr2-150709240-DEL-111
chr2-150721216-DEL-1465
chr2-150725325-DEL-68
chr2-150735308-DEL-566
chr2-150741602-DEL-405
chr2-150745288-DEL-107
chr2-150764530-DEL-56
chr2-150826450-DEL-64
chr2-150844213-DEL-97
chr2-150874318-DEL-55
chr2-150885805-DEL-6228
chr2-150894040-DEL-1168
chr2-150898781-DEL-296
chr2-150900480-DEL-1424
chr2-150924509-DEL-66
chr2-150926159-DEL-12616
chr2-150936711-DEL-7822
chr2-150952374-DEL-4795
chr2-150966696-DEL-146
chr2-150969041-DEL-71
chr2-150971126-DEL-3586
chr2-150971681-DEL-6765
chr2-151001650-DEL-7263
chr2-151001794-DEL-416
chr2-151015207-DEL-66
chr2-151021136-DEL-4817
chr2-151060625-DEL-53
chr2-151064292-DEL-272
chr2-151066473-DEL-6142
chr2-151072412-DEL-64
chr2-151163343-DEL-900
chr2-151175746-DEL-7377
chr2-151176316-DEL-79

chr2-151184680-DEL-6601
chr2-151225091-DEL-969
chr2-151229634-DEL-990
chr2-151232807-DEL-1654
chr2-151236083-DEL-66
chr2-151265499-DEL-79
chr2-151295373-DEL-1129
chr2-151303047-DEL-1668
chr2-151304737-DEL-5619
chr2-151311971-DEL-61
chr2-151312873-DEL-61
chr2-151320445-DEL-1669
chr2-151378029-DEL-133
chr2-151392820-DEL-60
chr2-151412881-DEL-194
chr2-151458047-DEL-240
chr2-151459461-DEL-561
chr2-151482113-DEL-1717
chr2-151492175-DEL-170
chr2-151499020-DEL-54
chr2-151574746-DEL-54
chr2-151588461-DEL-222
chr2-151605559-DEL-120
chr2-151611410-DEL-214
chr2-151612420-DEL-1035
chr2-151618061-DEL-55
chr2-151618329-DEL-1677
chr2-151641006-DEL-51
chr2-151664697-DEL-65
chr2-151667499-DEL-68
chr2-151722012-DEL-7977
chr2-151723943-DEL-481
chr2-151756680-DEL-88
chr2-151798519-DEL-1892
chr2-151864550-DEL-820
chr2-151868074-DEL-662
chr2-151874928-DEL-849
chr2-151893192-DEL-55
chr2-151896049-DEL-286
chr2-151960878-DEL-128
chr2-151964182-DEL-95
chr2-151967843-DEL-87
chr2-151968644-DEL-74
chr2-152036565-DEL-856
chr2-152041543-DEL-60
chr2-152054298-DEL-329
chr2-152057716-DEL-64
chr2-152060233-DEL-60
chr2-152095793-DEL-100
chr2-152131486-DEL-197
chr2-152195451-DEL-1273
chr2-152231249-DEL-175
chr2-152233923-DEL-209
chr2-152235086-DEL-175
chr2-152258599-DEL-249
chr2-152262433-DEL-88
chr2-152287985-DEL-836
chr2-152292114-DEL-107
chr2-152294705-DEL-71
chr2-152302708-DEL-61
chr2-152305003-DEL-92
chr2-152327766-DEL-90
chr2-152339659-DEL-63
chr2-152349833-DEL-86
chr2-152350677-DEL-78
chr2-152400825-DEL-53
chr2-152467745-DEL-56
chr2-152494253-DEL-148
chr2-152505662-DEL-62
chr2-152515551-DEL-209
chr2-152556039-DEL-881
chr2-152556234-DEL-170
chr2-152594935-DEL-59
chr2-152628472-DEL-57
chr2-152649962-DEL-200
chr2-152665021-DEL-182
chr2-152685751-DEL-203

chr2-152687071-DEL-255
chr2-152688912-DEL-56
chr2-152693660-DEL-1080
chr2-152744538-DEL-84
chr2-152808046-DEL-1380
chr2-152812169-DEL-419
chr2-152846705-DEL-68
chr2-152880959-DEL-62
chr2-152921794-DEL-240
chr2-152935421-DEL-71
chr2-152935982-DEL-2304
chr2-152936007-DEL-1344
chr2-152936052-DEL-226
chr2-152936085-DEL-760
chr2-152936954-DEL-1532
chr2-152936976-DEL-110
chr2-152937539-DEL-794
chr2-152938576-DEL-154
chr2-152938800-DEL-2004
chr2-152939621-DEL-1036
chr2-152941051-DEL-484
chr2-152941052-DEL-1408
chr2-152941995-DEL-418
chr2-152958570-DEL-199
chr2-153013052-DEL-208
chr2-153038589-DEL-355
chr2-153085971-DEL-398
chr2-153122660-DEL-388
chr2-153124101-DEL-286
chr2-153201132-DEL-213
chr2-153201930-DEL-738
chr2-153230786-DEL-1011
chr2-153268512-DEL-305
chr2-153294118-DEL-195
chr2-153295774-DEL-202
chr2-153307967-DEL-1083
chr2-153341501-DEL-196
chr2-153349675-DEL-201
chr2-153382937-DEL-79
chr2-153397876-DEL-52
chr2-153413589-DEL-55
chr2-153426986-DEL-81
chr2-153459322-DEL-261
chr2-153510216-DEL-68
chr2-153550539-DEL-51
chr2-153551696-DEL-52
chr2-153570161-DEL-56
chr2-153647757-DEL-144
chr2-153657117-DEL-51
chr2-153667262-DEL-202
chr2-153683904-DEL-59
chr2-153687531-DEL-68
chr2-153692983-DEL-464
chr2-153693655-DEL-391
chr2-153699275-DEL-125
chr2-153715800-DEL-362
chr2-153742645-DEL-844
chr2-153772680-DEL-84
chr2-153782327-DEL-172
chr2-153793010-DEL-73
chr2-153845093-DEL-69
chr2-153865925-DEL-7657
chr2-153879201-DEL-1594
chr2-153892048-DEL-17221
chr2-153894782-DEL-662
chr2-153911045-DEL-717
chr2-153920359-DEL-265
chr2-153925584-DEL-88
chr2-153925780-DEL-1024
chr2-153926891-DEL-440
chr2-153929843-DEL-6589
chr2-154130437-DEL-71
chr2-154131720-DEL-263
chr2-154136578-DEL-333
chr2-154145562-DEL-87
chr2-154169773-DEL-98
chr2-154209003-DEL-372

chr2-154222238-DEL-134
chr2-154230447-DEL-198
chr2-154248454-DEL-93
chr2-154274935-DEL-89
chr2-154285469-DEL-1437
chr2-154289481-DEL-160
chr2-154304566-DEL-52
chr2-154311954-DEL-216
chr2-154317332-DEL-83
chr2-154383983-DEL-91
chr2-154414680-DEL-66
chr2-154440729-DEL-121
chr2-154447803-DEL-457
chr2-154456691-DEL-52
chr2-154494295-DEL-78
chr2-154497752-DEL-88
chr2-154512560-DEL-779
chr2-154520594-DEL-205
chr2-154533348-DEL-200
chr2-154539323-DEL-1051
chr2-154545672-DEL-1013
chr2-154546864-DEL-212
chr2-154570550-DEL-231
chr2-154596313-DEL-202
chr2-154608657-DEL-204
chr2-154694367-DEL-243
chr2-154700832-DEL-61
chr2-154732819-DEL-174
chr2-154743727-DEL-188
chr2-154755438-DEL-86
chr2-154764933-DEL-234
chr2-154769477-DEL-189
chr2-154843944-DEL-756
chr2-154845695-DEL-54
chr2-154859783-DEL-8816
chr2-154876780-DEL-1039
chr2-154881105-DEL-219
chr2-154900467-DEL-403
chr2-154918695-DEL-408
chr2-154939149-DEL-436
chr2-154947804-DEL-399
chr2-154949114-DEL-492
chr2-154961421-DEL-256
chr2-154961879-DEL-9146
chr2-154994189-DEL-68
chr2-155055524-DEL-197
chr2-155069528-DEL-822
chr2-155071851-DEL-498
chr2-155094065-DEL-520
chr2-155096869-DEL-471
chr2-155112123-DEL-202
chr2-155128272-DEL-68
chr2-155134973-DEL-201
chr2-155141958-DEL-147
chr2-155167627-DEL-170
chr2-155202696-DEL-1016
chr2-155207953-DEL-224
chr2-155209185-DEL-203
chr2-155240215-DEL-52
chr2-155255146-DEL-196
chr2-155259090-DEL-815
chr2-155273983-DEL-54
chr2-155319824-DEL-75
chr2-155353594-DEL-144
chr2-155357718-DEL-63
chr2-155387839-DEL-194
chr2-155458154-DEL-52
chr2-155486715-DEL-111
chr2-155489408-DEL-78
chr2-155496592-DEL-55
chr2-155518724-DEL-184
chr2-155523885-DEL-63
chr2-155527301-DEL-76
chr2-155536006-DEL-197
chr2-155548649-DEL-57
chr2-155561205-DEL-52
chr2-155583462-DEL-175

chr2-155608373-DEL-68
chr2-155609023-DEL-77
chr2-155612448-DEL-93
chr2-155620910-DEL-101
chr2-155627261-DEL-60
chr2-155679646-DEL-58
chr2-155683231-DEL-364
chr2-155735247-DEL-179
chr2-155743645-DEL-200
chr2-155762249-DEL-94
chr2-155788769-DEL-471
chr2-155790760-DEL-65
chr2-155797371-DEL-258
chr2-155804446-DEL-51
chr2-155807542-DEL-109
chr2-155814400-DEL-383
chr2-155827473-DEL-138
chr2-155868031-DEL-356
chr2-155880041-DEL-404
chr2-155944578-DEL-69
chr2-155954312-DEL-948
chr2-155971128-DEL-2143
chr2-155974006-DEL-202
chr2-155978607-DEL-1374
chr2-155982919-DEL-198
chr2-156001895-DEL-243
chr2-156005180-DEL-274
chr2-156037220-DEL-404
chr2-156047519-DEL-207
chr2-156056610-DEL-812
chr2-156058436-DEL-210
chr2-156072910-DEL-144
chr2-156074322-DEL-60
chr2-156116792-DEL-112
chr2-156117373-DEL-491
chr2-156124299-DEL-68
chr2-156164186-DEL-103
chr2-156177669-DEL-74
chr2-156179019-DEL-84
chr2-156185862-DEL-218
chr2-156187270-DEL-279
chr2-156190206-DEL-157
chr2-156195410-DEL-54
chr2-156206514-DEL-232
chr2-156208388-DEL-63
chr2-156253224-DEL-214
chr2-156254194-DEL-74
chr2-156257050-DEL-54
chr2-156323334-DEL-208
chr2-156326011-DEL-795
chr2-156329178-DEL-560
chr2-156358637-DEL-195
chr2-156370622-DEL-68
chr2-156388145-DEL-87
chr2-156432483-DEL-67
chr2-156443180-DEL-73
chr2-156444275-DEL-68
chr2-156478930-DEL-741
chr2-156503660-DEL-213
chr2-156543698-DEL-210
chr2-156551959-DEL-124
chr2-156580892-DEL-144
chr2-156583583-DEL-195
chr2-156618416-DEL-149
chr2-156641184-DEL-10249
chr2-156646497-DEL-81
chr2-156659291-DEL-301
chr2-156666964-DEL-204
chr2-156679339-DEL-54
chr2-156702381-DEL-383
chr2-156702786-DEL-376
chr2-156709653-DEL-206
chr2-156730584-DEL-300
chr2-156737200-DEL-180
chr2-156737698-DEL-1226
chr2-156740642-DEL-96
chr2-156742533-DEL-56

chr2-156758859-DEL-6189
chr2-156765398-DEL-63
chr2-156767045-DEL-79
chr2-156775075-DEL-309
chr2-156785173-DEL-94
chr2-156835199-DEL-200
chr2-156851041-DEL-710
chr2-156895309-DEL-102
chr2-156898433-DEL-201
chr2-156912999-DEL-107
chr2-156951732-DEL-104
chr2-156990981-DEL-161
chr2-157007807-DEL-440
chr2-157019790-DEL-477
chr2-157040085-DEL-506
chr2-157045225-DEL-270
chr2-157054871-DEL-201
chr2-157073821-DEL-219
chr2-157075265-DEL-196
chr2-157077917-DEL-186
chr2-157146968-DEL-52
chr2-157149100-DEL-108
chr2-157153162-DEL-171
chr2-157181234-DEL-77
chr2-157203058-DEL-203
chr2-157216570-DEL-520
chr2-157233766-DEL-57
chr2-157250584-DEL-160
chr2-157255787-DEL-185
chr2-157287006-DEL-398
chr2-157291191-DEL-415
chr2-157327023-DEL-346
chr2-157347531-DEL-84
chr2-157363387-DEL-56
chr2-157365831-DEL-80
chr2-157422014-DEL-225
chr2-157427501-DEL-2564
chr2-157430106-DEL-1495
chr2-157431650-DEL-1060
chr2-157435279-DEL-857
chr2-157440650-DEL-192
chr2-157458999-DEL-126
chr2-157462143-DEL-4732
chr2-157466421-DEL-65
chr2-157479759-DEL-169
chr2-157515595-DEL-224
chr2-157563689-DEL-361
chr2-157565400-DEL-426
chr2-157590394-DEL-116
chr2-157621035-DEL-554
chr2-157622122-DEL-152
chr2-157653958-DEL-939
chr2-157743213-DEL-54
chr2-157801861-DEL-52
chr2-157802518-DEL-147
chr2-157825553-DEL-62
chr2-157891015-DEL-69
chr2-157892541-DEL-452
chr2-157898656-DEL-143
chr2-157905613-DEL-806
chr2-157906886-DEL-541
chr2-157934611-DEL-67
chr2-157948454-DEL-130
chr2-157952603-DEL-862
chr2-157990579-DEL-132
chr2-158025770-DEL-331
chr2-158092239-DEL-140
chr2-158098037-DEL-52
chr2-158115612-DEL-51
chr2-158122134-DEL-58
chr2-158124500-DEL-119
chr2-158153342-DEL-520
chr2-158154603-DEL-155
chr2-158187764-DEL-54
chr2-158194522-DEL-59
chr2-158271044-DEL-500
chr2-158288531-DEL-344

chr2-158306680-DEL-347
chr2-158325882-DEL-108
chr2-158358397-DEL-165
chr2-158361545-DEL-78
chr2-158396620-DEL-580
chr2-158407428-DEL-868
chr2-158411472-DEL-853
chr2-158472859-DEL-60
chr2-158530242-DEL-51
chr2-158533880-DEL-58
chr2-158536096-DEL-130
chr2-158546039-DEL-115
chr2-158577454-DEL-4600
chr2-158579202-DEL-116
chr2-158589355-DEL-126
chr2-158591832-DEL-1089
chr2-158632113-DEL-400
chr2-158666343-DEL-1693
chr2-158681176-DEL-216
chr2-158689526-DEL-70
chr2-158692903-DEL-101
chr2-158709276-DEL-165
chr2-158736205-DEL-454
chr2-158767642-DEL-52
chr2-158803941-DEL-58
chr2-158820536-DEL-579
chr2-158893502-DEL-51
chr2-158898323-DEL-661
chr2-158901823-DEL-126
chr2-158920715-DEL-185
chr2-158958369-DEL-408
chr2-158991494-DEL-173
chr2-159064791-DEL-51
chr2-159066634-DEL-2036
chr2-159230714-DEL-748
chr2-159267389-DEL-132
chr2-159344326-DEL-186
chr2-159360894-DEL-59
chr2-159363344-DEL-62
chr2-159384878-DEL-136
chr2-159447715-DEL-52
chr2-159475215-DEL-125
chr2-159476414-DEL-3048
chr2-159478281-DEL-52
chr2-159515381-DEL-3051
chr2-159535211-DEL-52
chr2-159599176-DEL-2304
chr2-159619169-DEL-1676
chr2-159622611-DEL-103
chr2-159655450-DEL-83
chr2-159679048-DEL-1011
chr2-159709807-DEL-94
chr2-159713881-DEL-64
chr2-159714169-DEL-404
chr2-159714246-DEL-744
chr2-159718985-DEL-58
chr2-159727388-DEL-107
chr2-159737951-DEL-337
chr2-159744154-DEL-63
chr2-159768187-DEL-478
chr2-159788207-DEL-90
chr2-159825054-DEL-158
chr2-159933495-DEL-58
chr2-159933980-DEL-81
chr2-159941227-DEL-654
chr2-159944671-DEL-402
chr2-159981321-DEL-174
chr2-160037435-DEL-2108
chr2-160157455-DEL-1268
chr2-160179131-DEL-64
chr2-160185909-DEL-1518
chr2-160193964-DEL-89
chr2-160237035-DEL-51
chr2-160251801-DEL-113
chr2-160284138-DEL-68
chr2-160309339-DEL-60
chr2-160312001-DEL-320

chr2-160354362-DEL-245
chr2-160383240-DEL-119
chr2-160386391-DEL-64
chr2-160397881-DEL-2536
chr2-160408665-DEL-395
chr2-160410478-DEL-373
chr2-160424827-DEL-840
chr2-160431294-DEL-54
chr2-160431837-DEL-53
chr2-160443355-DEL-345
chr2-160452571-DEL-415
chr2-160453194-DEL-103
chr2-160473735-DEL-5755
chr2-160479513-DEL-857
chr2-160551103-DEL-56
chr2-160559764-DEL-146
chr2-160570182-DEL-680
chr2-160585189-DEL-157
chr2-160650541-DEL-1669
chr2-160761687-DEL-100
chr2-160799637-DEL-210
chr2-160827932-DEL-101
chr2-160841328-DEL-902
chr2-160860911-DEL-134
chr2-160883422-DEL-54
chr2-160925620-DEL-1025
chr2-160931519-DEL-65
chr2-160960802-DEL-102
chr2-160980154-DEL-64
chr2-160997980-DEL-1989
chr2-160999319-DEL-170
chr2-161014686-DEL-500
chr2-161016022-DEL-80
chr2-161039747-DEL-401
chr2-161044161-DEL-52
chr2-161110471-DEL-178
chr2-161112881-DEL-113
chr2-161161547-DEL-494
chr2-161161573-DEL-69
chr2-161199451-DEL-64
chr2-161206809-DEL-90
chr2-161243374-DEL-473
chr2-161255981-DEL-54
chr2-161324323-DEL-688
chr2-161346609-DEL-496
chr2-161373506-DEL-128
chr2-161374380-DEL-397
chr2-161403903-DEL-95
chr2-161404546-DEL-59
chr2-161406428-DEL-91
chr2-161438279-DEL-400
chr2-161523305-DEL-57
chr2-161524792-DEL-1263
chr2-161570053-DEL-519
chr2-161575292-DEL-4433
chr2-161591386-DEL-6189
chr2-161620182-DEL-59
chr2-161639587-DEL-88
chr2-161641343-DEL-63
chr2-161643822-DEL-108
chr2-161666126-DEL-55
chr2-161690696-DEL-6159
chr2-161732903-DEL-138
chr2-161737914-DEL-69
chr2-161750702-DEL-1992
chr2-161775278-DEL-96
chr2-161776704-DEL-1076
chr2-161797497-DEL-68
chr2-161823353-DEL-151
chr2-161883846-DEL-201
chr2-161914299-DEL-62
chr2-161929735-DEL-126
chr2-161953794-DEL-468
chr2-161961771-DEL-54
chr2-161987949-DEL-6849
chr2-162002408-DEL-6233
chr2-162013216-DEL-63

chr2-162016741-DEL-540
chr2-162071397-DEL-706
chr2-162095811-DEL-235
chr2-162107518-DEL-831
chr2-162110370-DEL-69
chr2-162119208-DEL-87
chr2-162150485-DEL-518
chr2-162163756-DEL-147
chr2-162290713-DEL-507
chr2-162291343-DEL-222
chr2-162310773-DEL-126
chr2-162315532-DEL-288
chr2-162330919-DEL-67
chr2-162336752-DEL-160
chr2-162405091-DEL-98
chr2-162437093-DEL-1248
chr2-162457442-DEL-81
chr2-162459338-DEL-88
chr2-162461374-DEL-107
chr2-162461898-DEL-1017
chr2-162469987-DEL-54
chr2-162488486-DEL-217
chr2-162494755-DEL-62
chr2-162521744-DEL-110
chr2-162524158-DEL-171
chr2-162526800-DEL-200
chr2-162627477-DEL-68
chr2-162661316-DEL-82
chr2-162669382-DEL-104
chr2-162705711-DEL-99
chr2-162745180-DEL-3521
chr2-162797468-DEL-108
chr2-162804030-DEL-62
chr2-162842784-DEL-213
chr2-162848272-DEL-86
chr2-162849963-DEL-200
chr2-162860350-DEL-201
chr2-162870029-DEL-836
chr2-162879672-DEL-56
chr2-162907250-DEL-196
chr2-162927963-DEL-131
chr2-162931026-DEL-71
chr2-162943867-DEL-112
chr2-162960098-DEL-116
chr2-162980384-DEL-60
chr2-163007038-DEL-80
chr2-163057487-DEL-179
chr2-163076435-DEL-149
chr2-163105113-DEL-117
chr2-163161583-DEL-107
chr2-163165519-DEL-250
chr2-163170636-DEL-567
chr2-163173159-DEL-189
chr2-163222430-DEL-71
chr2-163224825-DEL-3603
chr2-163228161-DEL-121
chr2-163232007-DEL-84
chr2-163238392-DEL-79
chr2-163241064-DEL-72
chr2-163254519-DEL-169
chr2-163273671-DEL-189
chr2-163274651-DEL-64
chr2-163275264-DEL-484
chr2-163277707-DEL-54
chr2-163290174-DEL-464
chr2-163293229-DEL-70
chr2-163317209-DEL-2861
chr2-163342049-DEL-69
chr2-163358973-DEL-188
chr2-163398410-DEL-66
chr2-163409077-DEL-254
chr2-163418230-DEL-93
chr2-163419292-DEL-70
chr2-163419876-DEL-56
chr2-163427723-DEL-88
chr2-163439521-DEL-107
chr2-163484803-DEL-409

chr2-163529259-DEL-684
chr2-163529271-DEL-1489
chr2-163530966-DEL-59
chr2-163532895-DEL-407
chr2-163543587-DEL-500
chr2-163569927-DEL-58
chr2-163607329-DEL-438
chr2-163646517-DEL-108
chr2-163665528-DEL-133
chr2-163667790-DEL-55
chr2-163682109-DEL-233
chr2-163690925-DEL-54
chr2-163694653-DEL-55
chr2-163780926-DEL-146
chr2-163814229-DEL-3711
chr2-163815218-DEL-390
chr2-163835251-DEL-229
chr2-163849134-DEL-235
chr2-163883430-DEL-153
chr2-163890363-DEL-155
chr2-163899897-DEL-117
chr2-163947653-DEL-57
chr2-163971678-DEL-623
chr2-163972759-DEL-674
chr2-163986355-DEL-67
chr2-164001642-DEL-95
chr2-164016177-DEL-63
chr2-164049735-DEL-625
chr2-164052962-DEL-842
chr2-164079460-DEL-69
chr2-164082045-DEL-416
chr2-164100893-DEL-12704
chr2-164118351-DEL-70
chr2-164120962-DEL-188
chr2-164123184-DEL-191
chr2-164206254-DEL-54
chr2-164239921-DEL-126
chr2-164275568-DEL-91
chr2-164299386-DEL-120
chr2-164349615-DEL-193
chr2-164387682-DEL-400
chr2-164414153-DEL-200
chr2-164429828-DEL-321
chr2-164446765-DEL-68
chr2-164450335-DEL-95
chr2-164461657-DEL-58
chr2-164468809-DEL-52
chr2-164475776-DEL-68
chr2-164496470-DEL-60
chr2-164500010-DEL-207
chr2-164506003-DEL-89
chr2-164506812-DEL-152
chr2-164509965-DEL-53
chr2-164535061-DEL-543
chr2-164552218-DEL-1390
chr2-164556203-DEL-62
chr2-164557493-DEL-137
chr2-164571312-DEL-190
chr2-164575337-DEL-167
chr2-164592725-DEL-2702
chr2-164616581-DEL-75
chr2-164624748-DEL-51
chr2-164640370-DEL-196
chr2-164642551-DEL-102
chr2-164690221-DEL-119
chr2-164706205-DEL-56
chr2-164760530-DEL-201
chr2-164777319-DEL-885
chr2-164788581-DEL-54
chr2-164820206-DEL-80
chr2-164835245-DEL-52
chr2-164905846-DEL-79
chr2-164934130-DEL-53
chr2-164964828-DEL-76
chr2-164967698-DEL-83
chr2-164996495-DEL-59
chr2-165015937-DEL-100

chr2-165021902-DEL-62
chr2-165027930-DEL-76
chr2-165044971-DEL-119
chr2-165078003-DEL-419
chr2-165107592-DEL-201
chr2-165109944-DEL-169
chr2-165143332-DEL-143
chr2-165145692-DEL-67
chr2-165197190-DEL-394
chr2-165198908-DEL-186
chr2-165212025-DEL-1916
chr2-165224993-DEL-144
chr2-165226471-DEL-110
chr2-165230749-DEL-360
chr2-165240334-DEL-174
chr2-165244552-DEL-68
chr2-165294439-DEL-1205
chr2-165321597-DEL-92
chr2-165323508-DEL-162
chr2-165344606-DEL-291
chr2-165347856-DEL-93
chr2-165351238-DEL-945
chr2-165353006-DEL-396
chr2-165355840-DEL-112
chr2-165404249-DEL-239
chr2-165422518-DEL-1224
chr2-165456858-DEL-220
chr2-165457692-DEL-240
chr2-165488836-DEL-63
chr2-165536096-DEL-69
chr2-165537967-DEL-71
chr2-165539006-DEL-687
chr2-165543448-DEL-140
chr2-165549658-DEL-73
chr2-165570526-DEL-1012
chr2-165597273-DEL-114
chr2-165598989-DEL-92
chr2-165615706-DEL-122
chr2-165621895-DEL-53
chr2-165638148-DEL-140
chr2-165644021-DEL-107
chr2-165645786-DEL-69
chr2-165703643-DEL-69
chr2-165715722-DEL-53
chr2-165733103-DEL-1674
chr2-165735470-DEL-201
chr2-165738955-DEL-58
chr2-165750136-DEL-129
chr2-165752404-DEL-159
chr2-165767847-DEL-259
chr2-165776979-DEL-1024
chr2-165780828-DEL-98
chr2-165789997-DEL-139
chr2-165791153-DEL-222
chr2-165802701-DEL-83
chr2-165804753-DEL-125
chr2-165825880-DEL-59
chr2-165833601-DEL-193
chr2-165839554-DEL-460
chr2-165854657-DEL-180
chr2-165855861-DEL-175
chr2-165861702-DEL-100
chr2-165862428-DEL-137
chr2-165868178-DEL-86
chr2-165885314-DEL-51
chr2-165919818-DEL-468
chr2-165926667-DEL-222
chr2-165953092-DEL-74
chr2-165960488-DEL-174
chr2-165970158-DEL-200
chr2-166003054-DEL-199
chr2-166015047-DEL-1317
chr2-166034591-DEL-131
chr2-166045249-DEL-236
chr2-166053240-DEL-1056
chr2-166065829-DEL-95
chr2-166075122-DEL-129

chr2-166082323-DEL-140
chr2-166098388-DEL-1201
chr2-166105733-DEL-386
chr2-166127883-DEL-54
chr2-166159020-DEL-66
chr2-166182198-DEL-81
chr2-166231614-DEL-293
chr2-166273384-DEL-54
chr2-166274126-DEL-72
chr2-166295334-DEL-133
chr2-166303877-DEL-75
chr2-166332070-DEL-54
chr2-166335927-DEL-54
chr2-166347749-DEL-72
chr2-166364603-DEL-196
chr2-166393747-DEL-69
chr2-166446006-DEL-217
chr2-166456668-DEL-78
chr2-166473243-DEL-51
chr2-166496484-DEL-53
chr2-166503446-DEL-148
chr2-166512648-DEL-193
chr2-166544736-DEL-81
chr2-166558464-DEL-617
chr2-166563157-DEL-302
chr2-166574253-DEL-69
chr2-166575721-DEL-156
chr2-166579884-DEL-805
chr2-166594208-DEL-256
chr2-166608842-DEL-64
chr2-166616870-DEL-508
chr2-166621459-DEL-242
chr2-166624609-DEL-636
chr2-166625295-DEL-117
chr2-166627253-DEL-149
chr2-166630114-DEL-187
chr2-166672678-DEL-1040
chr2-166672886-DEL-320
chr2-166673619-DEL-100
chr2-166674607-DEL-1607
chr2-166676900-DEL-64
chr2-166681943-DEL-329
chr2-166685033-DEL-51
chr2-166689320-DEL-133
chr2-166696980-DEL-558
chr2-166712351-DEL-198
chr2-166718204-DEL-161
chr2-166719330-DEL-177
chr2-166722254-DEL-205
chr2-166725528-DEL-451
chr2-166725544-DEL-140
chr2-166760527-DEL-75
chr2-166773250-DEL-204
chr2-166803174-DEL-203
chr2-166809919-DEL-311
chr2-166811387-DEL-188
chr2-166815017-DEL-726
chr2-166832167-DEL-102
chr2-166840370-DEL-205
chr2-166860741-DEL-193
chr2-166871848-DEL-176
chr2-166873580-DEL-89
chr2-166874490-DEL-83
chr2-166887538-DEL-108
chr2-166927482-DEL-114
chr2-166935022-DEL-56
chr2-166972345-DEL-53
chr2-167000095-DEL-90
chr2-167001246-DEL-135
chr2-167020710-DEL-204
chr2-167040254-DEL-120
chr2-167054186-DEL-80
chr2-167059169-DEL-56
chr2-167070962-DEL-204
chr2-167105171-DEL-103
chr2-167119560-DEL-202
chr2-167124389-DEL-7969

chr2-167124411-DEL-105
chr2-167124656-DEL-7081
chr2-167132026-DEL-89
chr2-167135557-DEL-72
chr2-167207265-DEL-208
chr2-167209568-DEL-73
chr2-167210545-DEL-68
chr2-167212591-DEL-57
chr2-167249818-DEL-6122
chr2-167295314-DEL-86
chr2-167312904-DEL-70
chr2-167329711-DEL-158
chr2-167331935-DEL-77
chr2-167355614-DEL-155
chr2-167362437-DEL-4813
chr2-167421037-DEL-60
chr2-167433015-DEL-53
chr2-167475608-DEL-530
chr2-167511097-DEL-52
chr2-167511783-DEL-53
chr2-167514001-DEL-133
chr2-167519030-DEL-108
chr2-167519721-DEL-154
chr2-167521757-DEL-187
chr2-167523755-DEL-183
chr2-167524431-DEL-226
chr2-167586510-DEL-61
chr2-167592252-DEL-814
chr2-167592908-DEL-265
chr2-167638215-DEL-493
chr2-167660038-DEL-66
chr2-167674102-DEL-340
chr2-167701163-DEL-1074
chr2-167712369-DEL-81
chr2-167718670-DEL-494
chr2-167743704-DEL-217
chr2-167763339-DEL-144
chr2-167832666-DEL-196
chr2-167840337-DEL-107
chr2-167851259-DEL-52
chr2-167856011-DEL-151
chr2-167866735-DEL-198
chr2-167879987-DEL-72
chr2-167895527-DEL-200
chr2-167900136-DEL-119
chr2-167945534-DEL-53
chr2-167952853-DEL-68
chr2-167964761-DEL-516
chr2-167984777-DEL-132
chr2-168012871-DEL-52
chr2-168015058-DEL-261
chr2-168016243-DEL-194
chr2-168043185-DEL-71
chr2-168057445-DEL-181
chr2-168078362-DEL-149
chr2-168079354-DEL-85
chr2-168109756-DEL-77
chr2-168139440-DEL-480
chr2-168159010-DEL-124
chr2-168203646-DEL-360
chr2-168218815-DEL-144
chr2-168226089-DEL-194
chr2-168244799-DEL-76
chr2-168252559-DEL-237
chr2-168255729-DEL-61
chr2-168269435-DEL-60
chr2-168273013-DEL-203
chr2-168288686-DEL-157
chr2-168337982-DEL-134
chr2-168338879-DEL-185
chr2-168356826-DEL-130
chr2-168370393-DEL-57
chr2-168382493-DEL-155
chr2-168399190-DEL-60
chr2-168441975-DEL-53
chr2-168461103-DEL-99
chr2-168487292-DEL-105

chr2-168518524-DEL-213
chr2-168519349-DEL-464
chr2-168519839-DEL-1543
chr2-168521312-DEL-101
chr2-168527516-DEL-386
chr2-168535226-DEL-124
chr2-168589358-DEL-343
chr2-168595070-DEL-162
chr2-168620341-DEL-197
chr2-168632007-DEL-174
chr2-168633776-DEL-70
chr2-168639662-DEL-178
chr2-168644081-DEL-60
chr2-168660050-DEL-199
chr2-168667295-DEL-61
chr2-168669110-DEL-146
chr2-168684712-DEL-60
chr2-168699288-DEL-164
chr2-168710275-DEL-79
chr2-168719506-DEL-52
chr2-168738724-DEL-52
chr2-168751607-DEL-74
chr2-168756086-DEL-1604
chr2-168757401-DEL-662
chr2-168766378-DEL-56
chr2-168769332-DEL-252
chr2-168776103-DEL-167
chr2-168780434-DEL-106
chr2-168790295-DEL-70
chr2-168818418-DEL-88
chr2-168822346-DEL-63
chr2-168837431-DEL-160
chr2-168841692-DEL-106
chr2-168849070-DEL-63
chr2-168854298-DEL-50
chr2-168865525-DEL-427
chr2-168898627-DEL-58
chr2-168905592-DEL-1109
chr2-168914578-DEL-83
chr2-168947372-DEL-64
chr2-168949343-DEL-54
chr2-168957610-DEL-207
chr2-168970605-DEL-175
chr2-168986790-DEL-105
chr2-169004726-DEL-286
chr2-169012411-DEL-189
chr2-169060701-DEL-52
chr2-169104967-DEL-414
chr2-169152685-DEL-52
chr2-169204901-DEL-60
chr2-169235873-DEL-84
chr2-169263399-DEL-192
chr2-169313255-DEL-115
chr2-169319979-DEL-128
chr2-169339315-DEL-96
chr2-169405729-DEL-143
chr2-169409258-DEL-213
chr2-169423881-DEL-68
chr2-169429474-DEL-406
chr2-169433792-DEL-54
chr2-169460062-DEL-162
chr2-169472555-DEL-73
chr2-169477926-DEL-57
chr2-169487027-DEL-73
chr2-169524925-DEL-186
chr2-169568451-DEL-611
chr2-169598686-DEL-230
chr2-169634439-DEL-53
chr2-169644906-DEL-101
chr2-169665945-DEL-213
chr2-169677679-DEL-100
chr2-169684794-DEL-148
chr2-169692194-DEL-107
chr2-169695378-DEL-96
chr2-169696302-DEL-80
chr2-169746351-DEL-119
chr2-169776446-DEL-58

chr2-169780965-DEL-816
chr2-169786718-DEL-152
chr2-169794292-DEL-529
chr2-169799429-DEL-78
chr2-169818859-DEL-72
chr2-169820098-DEL-68
chr2-169822191-DEL-175
chr2-169828444-DEL-190
chr2-169866175-DEL-212
chr2-169868506-DEL-140
chr2-169880367-DEL-65
chr2-169891667-DEL-132
chr2-169896937-DEL-76
chr2-169905168-DEL-185
chr2-169935076-DEL-128
chr2-169944164-DEL-237
chr2-169977064-DEL-131
chr2-169988328-DEL-116
chr2-170005164-DEL-209
chr2-170010525-DEL-210
chr2-170011481-DEL-1373
chr2-170020216-DEL-63
chr2-170021974-DEL-70
chr2-170022507-DEL-253
chr2-170025053-DEL-384
chr2-170032411-DEL-115
chr2-170053297-DEL-51
chr2-170058170-DEL-77
chr2-170064914-DEL-64
chr2-170066401-DEL-191
chr2-170066446-DEL-2262
chr2-170075094-DEL-243
chr2-170083171-DEL-115
chr2-170085185-DEL-106
chr2-170089454-DEL-108
chr2-170099840-DEL-177
chr2-170114625-DEL-57
chr2-170115192-DEL-1557
chr2-170116029-DEL-153
chr2-170120523-DEL-52
chr2-170125186-DEL-53
chr2-170141637-DEL-399
chr2-170156113-DEL-57
chr2-170159145-DEL-65
chr2-170161019-DEL-141
chr2-170162845-DEL-259
chr2-170170943-DEL-66
chr2-170177487-DEL-111
chr2-170184884-DEL-204
chr2-170191771-DEL-59
chr2-170193855-DEL-167
chr2-170202509-DEL-854
chr2-170202673-DEL-153
chr2-170203269-DEL-115
chr2-170250902-DEL-402
chr2-170276306-DEL-58
chr2-170277505-DEL-166
chr2-170306570-DEL-177
chr2-170307627-DEL-60
chr2-170346416-DEL-661
chr2-170346440-DEL-103
chr2-170365762-DEL-86
chr2-170373315-DEL-64
chr2-170384631-DEL-66
chr2-170426670-DEL-96
chr2-170443293-DEL-66
chr2-170445650-DEL-54
chr2-170451467-DEL-441
chr2-170452240-DEL-136
chr2-170454158-DEL-350
chr2-170459650-DEL-52
chr2-170465079-DEL-457
chr2-170468453-DEL-66
chr2-170495113-DEL-66
chr2-170506169-DEL-99
chr2-170507250-DEL-52
chr2-170510404-DEL-82

chr2-170512831-DEL-1103
chr2-170519262-DEL-220
chr2-170521853-DEL-294
chr2-170530309-DEL-170
chr2-170541552-DEL-146
chr2-170560417-DEL-117
chr2-170561619-DEL-206
chr2-170563338-DEL-58
chr2-170571278-DEL-140
chr2-170584415-DEL-81
chr2-170593258-DEL-94
chr2-170601042-DEL-134
chr2-170614832-DEL-56
chr2-170628296-DEL-60
chr2-170645516-DEL-59
chr2-170662599-DEL-545
chr2-170666649-DEL-142
chr2-170678123-DEL-58
chr2-170679298-DEL-53
chr2-170680887-DEL-164
chr2-170684443-DEL-111
chr2-170707004-DEL-74
chr2-170716895-DEL-81
chr2-170717971-DEL-74
chr2-170722753-DEL-701
chr2-170734186-DEL-810
chr2-170740472-DEL-94
chr2-170758201-DEL-408
chr2-170765786-DEL-74
chr2-170795865-DEL-88
chr2-170797893-DEL-112
chr2-170801805-DEL-187
chr2-170815613-DEL-74
chr2-170833029-DEL-135
chr2-170839683-DEL-270
chr2-170864276-DEL-224
chr2-170877024-DEL-50
chr2-170910008-DEL-52
chr2-170913834-DEL-321
chr2-170920417-DEL-88
chr2-170952562-DEL-143
chr2-170984294-DEL-1325
chr2-170987822-DEL-72
chr2-170997278-DEL-140
chr2-171016032-DEL-127
chr2-171031280-DEL-54
chr2-171052480-DEL-407
chr2-171090974-DEL-2187
chr2-171092004-DEL-647
chr2-171092595-DEL-243
chr2-171096121-DEL-96
chr2-171118514-DEL-140
chr2-171132595-DEL-1220
chr2-171132662-DEL-619
chr2-171132896-DEL-973
chr2-171138426-DEL-97
chr2-171143444-DEL-55
chr2-171148901-DEL-66
chr2-171152085-DEL-231
chr2-171157960-DEL-203
chr2-171181917-DEL-78
chr2-171192905-DEL-429
chr2-171204179-DEL-167
chr2-171232757-DEL-220
chr2-171236016-DEL-131
chr2-171253994-DEL-435
chr2-171257317-DEL-150
chr2-171266375-DEL-121
chr2-171267884-DEL-89
chr2-171280779-DEL-138
chr2-171317619-DEL-72
chr2-171318363-DEL-117
chr2-171333766-DEL-52
chr2-171335438-DEL-87
chr2-171347382-DEL-957
chr2-171347599-DEL-792
chr2-171359327-DEL-74

chr2-171382520-DEL-64
chr2-171383194-DEL-5815
chr2-171389043-DEL-740
chr2-171389807-DEL-264
chr2-171390103-DEL-1018
chr2-171400472-DEL-123
chr2-171402858-DEL-692
chr2-171410086-DEL-180
chr2-171427660-DEL-80
chr2-171437628-DEL-72
chr2-171453679-DEL-56
chr2-171467282-DEL-2949
chr2-171504845-DEL-61
chr2-171507163-DEL-58
chr2-171516000-DEL-1922
chr2-171550324-DEL-372
chr2-171552956-DEL-70
chr2-171560976-DEL-91
chr2-171565790-DEL-77
chr2-171578095-DEL-61
chr2-171633982-DEL-108
chr2-171645168-DEL-159
chr2-171651861-DEL-51
chr2-171657004-DEL-59
chr2-171674659-DEL-127
chr2-171704913-DEL-182
chr2-171704955-DEL-308
chr2-171724860-DEL-162
chr2-171730077-DEL-66
chr2-171734865-DEL-53
chr2-171741219-DEL-54
chr2-171756247-DEL-105
chr2-171787949-DEL-552
chr2-171799629-DEL-52
chr2-171801248-DEL-52
chr2-171802113-DEL-74
chr2-171811433-DEL-75
chr2-171821565-DEL-59
chr2-171836441-DEL-219
chr2-171841759-DEL-77
chr2-171843224-DEL-820
chr2-171871404-DEL-71
chr2-171907812-DEL-278
chr2-171909611-DEL-57
chr2-171927062-DEL-415
chr2-171930222-DEL-315
chr2-171943337-DEL-69
chr2-171970258-DEL-2220
chr2-171975866-DEL-423
chr2-171982914-DEL-56
chr2-171988813-DEL-329
chr2-172000364-DEL-157
chr2-172006149-DEL-852
chr2-172043616-DEL-376
chr2-172046280-DEL-527
chr2-172046466-DEL-290
chr2-172047588-DEL-156
chr2-172056800-DEL-97
chr2-172061394-DEL-341
chr2-172064672-DEL-158
chr2-172066271-DEL-96
chr2-172069292-DEL-1243
chr2-172094218-DEL-1230
chr2-172095452-DEL-57
chr2-172116593-DEL-57
chr2-172131898-DEL-320
chr2-172143775-DEL-598
chr2-172145525-DEL-87
chr2-172169733-DEL-154
chr2-172180750-DEL-76
chr2-172219315-DEL-120
chr2-172229135-DEL-317
chr2-172233286-DEL-82
chr2-172297387-DEL-51
chr2-172308743-DEL-217
chr2-172309469-DEL-553
chr2-172325666-DEL-99

chr2-172359541-DEL-102
chr2-172366986-DEL-811
chr2-172370454-DEL-160
chr2-172406715-DEL-254
chr2-172436990-DEL-492
chr2-172466714-DEL-237
chr2-172468280-DEL-82
chr2-172468876-DEL-58
chr2-172484073-DEL-64
chr2-172490340-DEL-52
chr2-172540603-DEL-78
chr2-172556186-DEL-252
chr2-172560164-DEL-90
chr2-172644367-DEL-219
chr2-172645554-DEL-196
chr2-172660590-DEL-63
chr2-172683625-DEL-292
chr2-172716039-DEL-517
chr2-172757338-DEL-56
chr2-172763526-DEL-52
chr2-172767548-DEL-70
chr2-172798246-DEL-59
chr2-172806644-DEL-194
chr2-172835914-DEL-74
chr2-172844686-DEL-70
chr2-172847734-DEL-203
chr2-172878389-DEL-87
chr2-172908362-DEL-167
chr2-172940621-DEL-1707
chr2-172940774-DEL-349
chr2-172941529-DEL-112
chr2-172953507-DEL-454
chr2-172959823-DEL-66
chr2-172961440-DEL-154
chr2-172963577-DEL-79
chr2-172991979-DEL-245
chr2-173004833-DEL-171
chr2-173017011-DEL-90
chr2-173051354-DEL-180
chr2-173078072-DEL-112
chr2-173081282-DEL-177
chr2-173184185-DEL-109
chr2-173205523-DEL-52
chr2-173211463-DEL-56
chr2-173214812-DEL-75
chr2-173242175-DEL-167
chr2-173242691-DEL-480
chr2-173258118-DEL-63
chr2-173268843-DEL-663
chr2-173291895-DEL-124
chr2-173303026-DEL-83
chr2-173330702-DEL-601
chr2-173335172-DEL-66
chr2-173336803-DEL-97
chr2-173348818-DEL-86
chr2-173351685-DEL-61
chr2-173356044-DEL-264
chr2-173372057-DEL-140
chr2-173380867-DEL-61
chr2-173446470-DEL-489
chr2-173469194-DEL-107
chr2-173473428-DEL-147
chr2-173477499-DEL-468
chr2-173478334-DEL-145
chr2-173521416-DEL-56
chr2-173549429-DEL-290
chr2-173555141-DEL-100
chr2-173580961-DEL-203
chr2-173587527-DEL-260
chr2-173594475-DEL-568
chr2-173595543-DEL-155
chr2-173610465-DEL-85
chr2-173611579-DEL-670
chr2-173611597-DEL-225
chr2-173619652-DEL-71
chr2-173639058-DEL-345
chr2-173676307-DEL-146

chr2-173717009-DEL-160
chr2-173760924-DEL-53
chr2-173800671-DEL-52
chr2-173804847-DEL-69
chr2-173809719-DEL-144
chr2-173938325-DEL-179
chr2-173946751-DEL-52
chr2-173984880-DEL-79
chr2-173985444-DEL-68
chr2-173997870-DEL-128
chr2-174010613-DEL-157
chr2-174018292-DEL-68
chr2-174062811-DEL-111
chr2-174089109-DEL-4196
chr2-174131583-DEL-74
chr2-174198449-DEL-643
chr2-174202575-DEL-68
chr2-174244555-DEL-52
chr2-174257234-DEL-169
chr2-174329652-DEL-131
chr2-174332112-DEL-470
chr2-174348936-DEL-146
chr2-174387211-DEL-1614
chr2-174425749-DEL-164
chr2-174511795-DEL-226
chr2-174556286-DEL-77
chr2-174679646-DEL-61
chr2-174783716-DEL-175
chr2-174794489-DEL-54
chr2-174811185-DEL-142
chr2-174825795-DEL-7115
chr2-174834646-DEL-1530
chr2-174841945-DEL-208
chr2-174842426-DEL-935
chr2-174844069-DEL-320
chr2-174844627-DEL-618
chr2-174855301-DEL-718
chr2-174859483-DEL-8794
chr2-174860309-DEL-84
chr2-174860739-DEL-84
chr2-174881729-DEL-84
chr2-174883729-DEL-2883
chr2-174947363-DEL-104
chr2-174954961-DEL-144
chr2-174956256-DEL-1224
chr2-174970547-DEL-208
chr2-174972834-DEL-757
chr2-174978710-DEL-216
chr2-174990064-DEL-849
chr2-175004700-DEL-1161
chr2-175011022-DEL-168
chr2-175011035-DEL-504
chr2-175018394-DEL-166
chr2-175020998-DEL-1223
chr2-175046165-DEL-207
chr2-175264014-DEL-419
chr2-175663363-DEL-207
chr2-175763454-DEL-1894
chr2-175987589-DEL-59
chr2-175989868-DEL-92
chr2-176020228-DEL-202
chr2-176029253-DEL-200
chr2-176032343-DEL-290
chr2-176033610-DEL-180
chr2-176036147-DEL-953
chr2-176039599-DEL-362
chr2-176372294-DEL-252
chr2-176373431-DEL-165
chr2-176406447-DEL-1894
chr2-176519637-DEL-144
chr2-176527274-DEL-124
chr2-176809939-DEL-8511
chr2-176815361-DEL-234
chr2-176832417-DEL-84
chr2-176835311-DEL-345
chr2-176848382-DEL-7159
chr2-176848836-DEL-79

chr2-176875028-DEL-169
chr2-176896612-DEL-360
chr2-176918362-DEL-252
chr2-176923229-DEL-1027
chr2-176939843-DEL-205
chr2-176942708-DEL-5498
chr2-176945813-DEL-370
chr2-176946641-DEL-83
chr2-176949086-DEL-166
chr2-176962590-DEL-476
chr2-176997697-DEL-143
chr2-177020556-DEL-226
chr2-177054153-DEL-2612
chr2-177084627-DEL-1212
chr2-177163917-DEL-556
chr2-177167206-DEL-126
chr2-177178410-DEL-1455
chr2-177193281-DEL-84
chr2-177201178-DEL-84
chr2-177223854-DEL-196
chr2-177266507-DEL-197
chr2-177279870-DEL-204
chr2-177291152-DEL-205
chr2-177291624-DEL-1432
chr2-177296262-DEL-5316
chr2-177376702-DEL-68
chr2-177451365-DEL-94
chr2-177457530-DEL-185
chr2-177464455-DEL-241
chr2-177488851-DEL-231
chr2-177548045-DEL-84
chr2-177609249-DEL-4352
chr2-177620239-DEL-808
chr2-177636395-DEL-51
chr2-177648178-DEL-6744
chr2-177674913-DEL-84
chr2-177693330-DEL-604
chr2-177714502-DEL-430
chr2-177728472-DEL-289
chr2-177735589-DEL-64
chr2-177742559-DEL-125
chr2-177745716-DEL-233
chr2-177750555-DEL-397
chr2-177769135-DEL-174
chr2-177792153-DEL-284
chr2-177820092-DEL-101
chr2-177822278-DEL-125
chr2-177828765-DEL-597
chr2-177838289-DEL-188
chr2-177851079-DEL-54
chr2-177863168-DEL-88
chr2-177899891-DEL-151
chr2-177907591-DEL-186
chr2-177912567-DEL-60
chr2-177913375-DEL-398
chr2-177932334-DEL-56
chr2-177940665-DEL-164
chr2-177940712-DEL-128
chr2-177963630-DEL-130
chr2-177983142-DEL-1211
chr2-178034214-DEL-177
chr2-178071979-DEL-52
chr2-178104038-DEL-126
chr2-178156827-DEL-676
chr2-178170013-DEL-377
chr2-178170477-DEL-116
chr2-178180973-DEL-53
chr2-178192137-DEL-55
chr2-178220340-DEL-65
chr2-178220797-DEL-856
chr2-178240705-DEL-2482
chr2-178249471-DEL-262
chr2-178268289-DEL-128
chr2-178283461-DEL-208
chr2-178287024-DEL-1807
chr2-178295175-DEL-570
chr2-178317667-DEL-56

chr2-178352226-DEL-757
chr2-178365846-DEL-63
chr2-178379242-DEL-628
chr2-178381402-DEL-56
chr2-178382636-DEL-112
chr2-178395146-DEL-56
chr2-178411497-DEL-62
chr2-178422771-DEL-332
chr2-178439221-DEL-62
chr2-178449327-DEL-92
chr2-178458724-DEL-101
chr2-178492210-DEL-70
chr2-178511406-DEL-54
chr2-178525409-DEL-54
chr2-178535493-DEL-63
chr2-178552914-DEL-81
chr2-178578843-DEL-121
chr2-178584612-DEL-77
chr2-178603274-DEL-85
chr2-178646501-DEL-52
chr2-178659038-DEL-79
chr2-178672547-DEL-91
chr2-178688276-DEL-120
chr2-178744903-DEL-54
chr2-178775232-DEL-70
chr2-178779160-DEL-95
chr2-178783363-DEL-398
chr2-178785427-DEL-92
chr2-178837536-DEL-135
chr2-178840475-DEL-65
chr2-178859842-DEL-62
chr2-178892773-DEL-83
chr2-178931003-DEL-61
chr2-178951271-DEL-263
chr2-178951933-DEL-66
chr2-178952557-DEL-99
chr2-178957013-DEL-372
chr2-179021428-DEL-65
chr2-179033799-DEL-88
chr2-179049938-DEL-52
chr2-179052500-DEL-1078
chr2-179085780-DEL-52
chr2-179089756-DEL-321
chr2-179113199-DEL-51
chr2-179121997-DEL-572
chr2-179134205-DEL-198
chr2-179155391-DEL-108
chr2-179168162-DEL-865
chr2-179251187-DEL-54
chr2-179259312-DEL-63
chr2-179270864-DEL-74
chr2-179296972-DEL-180
chr2-179303779-DEL-463
chr2-179316869-DEL-66
chr2-179331353-DEL-76
chr2-179341473-DEL-148
chr2-179345675-DEL-269
chr2-179350070-DEL-84
chr2-179361193-DEL-139
chr2-179364870-DEL-558
chr2-179368992-DEL-51
chr2-179378421-DEL-51
chr2-179380342-DEL-204
chr2-179384389-DEL-83
chr2-179387469-DEL-250
chr2-179413665-DEL-163
chr2-179414754-DEL-146
chr2-179430048-DEL-59
chr2-179467208-DEL-781
chr2-179468996-DEL-56
chr2-179529673-DEL-55
chr2-179550606-DEL-102
chr2-179569628-DEL-142
chr2-179612129-DEL-126
chr2-179643392-DEL-3056
chr2-179706354-DEL-2582
chr2-179706406-DEL-388

chr2-179707144-DEL-494
chr2-179707801-DEL-240
chr2-179708200-DEL-591
chr2-179716775-DEL-102
chr2-179724737-DEL-65
chr2-179795849-DEL-465
chr2-179797249-DEL-171
chr2-179799941-DEL-200
chr2-179868769-DEL-89
chr2-179886153-DEL-1304
chr2-179888251-DEL-516
chr2-179910999-DEL-208
chr2-179915362-DEL-98
chr2-179932522-DEL-705
chr2-179937076-DEL-202
chr2-179940762-DEL-179
chr2-179954262-DEL-56
chr2-179966678-DEL-61
chr2-179982072-DEL-59
chr2-180033697-DEL-74
chr2-180082078-DEL-84
chr2-180126266-DEL-85
chr2-180202461-DEL-55
chr2-180223539-DEL-295
chr2-180229574-DEL-180
chr2-180260593-DEL-56
chr2-180260990-DEL-222
chr2-180269615-DEL-53
chr2-180345753-DEL-94
chr2-180372595-DEL-352
chr2-180385580-DEL-192
chr2-180391794-DEL-460
chr2-180392358-DEL-222
chr2-180392757-DEL-236
chr2-180415450-DEL-72
chr2-180422989-DEL-355
chr2-180469032-DEL-60
chr2-180474525-DEL-1495
chr2-180489942-DEL-209
chr2-180537639-DEL-68
chr2-180540320-DEL-64
chr2-180549569-DEL-204
chr2-180557187-DEL-315
chr2-180603570-DEL-451
chr2-180668434-DEL-804
chr2-180708546-DEL-102
chr2-180770676-DEL-87
chr2-180781260-DEL-53
chr2-180824385-DEL-451
chr2-180839676-DEL-62
chr2-180851887-DEL-180
chr2-180914320-DEL-185
chr2-180918219-DEL-284
chr2-181059361-DEL-138
chr2-181089657-DEL-193
chr2-181091932-DEL-68
chr2-181177051-DEL-208
chr2-181179468-DEL-1201
chr2-181180715-DEL-1514
chr2-181185419-DEL-311
chr2-181200625-DEL-208
chr2-181201163-DEL-199
chr2-181208329-DEL-164
chr2-181247832-DEL-243
chr2-181250178-DEL-69
chr2-181267330-DEL-199
chr2-181279049-DEL-201
chr2-181295996-DEL-89
chr2-181392521-DEL-200
chr2-181420527-DEL-198
chr2-181443428-DEL-51
chr2-181459080-DEL-100
chr2-181485421-DEL-615
chr2-181569405-DEL-69
chr2-181570480-DEL-52
chr2-181645558-DEL-4426
chr3-3011286-DEL-105

chr3-3082604-DEL-3596
chr3-3119373-DEL-77
chr3-3124452-DEL-57
chr3-3148985-DEL-6184
chr3-3165472-DEL-12655
chr3-3190866-DEL-116
chr3-3196395-DEL-180
chr3-3222400-DEL-86
chr3-3228727-DEL-1533
chr3-3236744-DEL-4217
chr3-3261940-DEL-374
chr3-3263243-DEL-1864
chr3-3273732-DEL-63
chr3-3315066-DEL-15208
chr3-3315246-DEL-201
chr3-3320771-DEL-8262
chr3-3344123-DEL-846
chr3-3347726-DEL-1688
chr3-3357651-DEL-51
chr3-3359314-DEL-56
chr3-3369617-DEL-1694
chr3-3391731-DEL-5507
chr3-3406667-DEL-98
chr3-3431904-DEL-1712
chr3-3457847-DEL-56
chr3-3463573-DEL-171
chr3-3464450-DEL-402
chr3-3480917-DEL-6311
chr3-3489308-DEL-7472
chr3-3497235-DEL-4399
chr3-3504356-DEL-7851
chr3-3513292-DEL-2136
chr3-3520310-DEL-157
chr3-3552289-DEL-1082
chr3-3557493-DEL-1745
chr3-3560298-DEL-62
chr3-3570044-DEL-60
chr3-3627562-DEL-90
chr3-3629963-DEL-149
chr3-3636531-DEL-101
chr3-3660445-DEL-104
chr3-3665571-DEL-654
chr3-3671840-DEL-66
chr3-3679988-DEL-69
chr3-3685137-DEL-1997
chr3-3690643-DEL-68
chr3-3814370-DEL-82
chr3-3818433-DEL-1134
chr3-3831255-DEL-128
chr3-3831607-DEL-2464
chr3-3894684-DEL-471
chr3-3904510-DEL-6199
chr3-3922961-DEL-6236
chr3-3952204-DEL-697
chr3-3964951-DEL-62
chr3-3985806-DEL-215
chr3-4000268-DEL-216
chr3-4023107-DEL-81
chr3-4027243-DEL-514
chr3-4027990-DEL-1639
chr3-4031134-DEL-5627
chr3-4050211-DEL-98
chr3-4052098-DEL-1092
chr3-4063406-DEL-1009
chr3-4072296-DEL-109
chr3-4093368-DEL-1315
chr3-4108438-DEL-326
chr3-4143773-DEL-704
chr3-4147719-DEL-54
chr3-4149598-DEL-5697
chr3-4150808-DEL-456
chr3-4151325-DEL-3451
chr3-4169531-DEL-406
chr3-4173929-DEL-6189
chr3-4190379-DEL-428
chr3-4192973-DEL-86
chr3-4210995-DEL-2093

chr3-4216417-DEL-339
chr3-4255248-DEL-208
chr3-4266424-DEL-2313
chr3-4279482-DEL-109
chr3-4306638-DEL-1218
chr3-4317563-DEL-137
chr3-4319955-DEL-6143
chr3-4327459-DEL-1544
chr3-4330689-DEL-60
chr3-4353417-DEL-3420
chr3-4367303-DEL-498
chr3-4370115-DEL-1478
chr3-4400692-DEL-614
chr3-4401760-DEL-3761
chr3-4403473-DEL-3433
chr3-4415595-DEL-105
chr3-4445898-DEL-783
chr3-4451458-DEL-7470
chr3-4517334-DEL-249
chr3-4520870-DEL-2092
chr3-4547100-DEL-51
chr3-4549761-DEL-78
chr3-4550947-DEL-470
chr3-4552036-DEL-494
chr3-4556623-DEL-583
chr3-4561862-DEL-70
chr3-4566434-DEL-64
chr3-4583292-DEL-1748
chr3-4588775-DEL-115
chr3-4589509-DEL-389
chr3-4600647-DEL-1881
chr3-4603616-DEL-504
chr3-4630764-DEL-374
chr3-4684344-DEL-497
chr3-4685214-DEL-91
chr3-4685425-DEL-6187
chr3-4686391-DEL-4234
chr3-4686469-DEL-3239
chr3-4690424-DEL-481
chr3-4694104-DEL-75
chr3-4700718-DEL-592
chr3-4705191-DEL-199
chr3-4716974-DEL-94
chr3-4763757-DEL-2102
chr3-4770424-DEL-10929
chr3-4778339-DEL-3028
chr3-4812798-DEL-481
chr3-4829315-DEL-871
chr3-4853410-DEL-281
chr3-4855611-DEL-217
chr3-4870225-DEL-126
chr3-4918480-DEL-97
chr3-4935313-DEL-330
chr3-4936129-DEL-1106
chr3-4955132-DEL-136
chr3-4957476-DEL-534
chr3-4985477-DEL-521
chr3-4987182-DEL-3825
chr3-4995951-DEL-1626
chr3-5035325-DEL-1606
chr3-5047414-DEL-1672
chr3-5048760-DEL-326
chr3-5066490-DEL-1872
chr3-5078138-DEL-2233
chr3-5087785-DEL-110
chr3-5098948-DEL-1029
chr3-5110113-DEL-83
chr3-5145765-DEL-1836
chr3-5178422-DEL-200
chr3-5179975-DEL-1546
chr3-5208732-DEL-51
chr3-5235640-DEL-430
chr3-5240454-DEL-84
chr3-5254088-DEL-147
chr3-5262650-DEL-86
chr3-5281004-DEL-53
chr3-5304288-DEL-706

chr3-5320944-DEL-61
chr3-5326940-DEL-90
chr3-5355907-DEL-169
chr3-5386284-DEL-851
chr3-5426452-DEL-199
chr3-5434752-DEL-656
chr3-5471862-DEL-75
chr3-5519766-DEL-64
chr3-5542430-DEL-143
chr3-5548025-DEL-604
chr3-5549054-DEL-923
chr3-5600869-DEL-792
chr3-5646456-DEL-6114
chr3-5713087-DEL-2329
chr3-5715446-DEL-3997
chr3-5725319-DEL-1637
chr3-5729586-DEL-933
chr3-5731716-DEL-147
chr3-5732147-DEL-1948
chr3-5734079-DEL-846
chr3-5737458-DEL-523
chr3-5769789-DEL-2704
chr3-5780773-DEL-1028
chr3-5789934-DEL-79
chr3-5815849-DEL-501
chr3-5842834-DEL-976
chr3-5864456-DEL-6219
chr3-5875284-DEL-88
chr3-5877312-DEL-63
chr3-5913773-DEL-350
chr3-5952083-DEL-268
chr3-5960046-DEL-51
chr3-6028578-DEL-96
chr3-6043224-DEL-328
chr3-6066260-DEL-582
chr3-6094391-DEL-67
chr3-6095615-DEL-54
chr3-6096876-DEL-64
chr3-6126402-DEL-3031
chr3-6127587-DEL-461
chr3-6145311-DEL-1221
chr3-6163589-DEL-81
chr3-6209780-DEL-246
chr3-6235669-DEL-155
chr3-6282359-DEL-52
chr3-6330063-DEL-1485
chr3-6362580-DEL-831
chr3-6362664-DEL-208
chr3-6369720-DEL-417
chr3-6381032-DEL-3328
chr3-6411315-DEL-8780
chr3-6452578-DEL-54
chr3-6475383-DEL-56
chr3-6475923-DEL-449
chr3-6481834-DEL-6286
chr3-6489015-DEL-927
chr3-6490824-DEL-537
chr3-6494463-DEL-65
chr3-6537362-DEL-6541
chr3-6558886-DEL-55
chr3-6559910-DEL-1748
chr3-6567369-DEL-1361
chr3-6582395-DEL-705
chr3-6604445-DEL-843
chr3-6620853-DEL-3256
chr3-6629435-DEL-66
chr3-6659847-DEL-54
chr3-6663675-DEL-482
chr3-6694151-DEL-913
chr3-6721771-DEL-3856
chr3-6761260-DEL-6469
chr3-6771431-DEL-2726
chr3-6775015-DEL-1795
chr3-6818937-DEL-55
chr3-6823577-DEL-84
chr3-6825683-DEL-253
chr3-6837255-DEL-118

chr3-6844828-DEL-657
chr3-6860237-DEL-699
chr3-6882287-DEL-330
chr3-6886706-DEL-59
chr3-6892463-DEL-75
chr3-6903548-DEL-60
chr3-6913011-DEL-483
chr3-6928775-DEL-491
chr3-6941938-DEL-597
chr3-6958306-DEL-1054
chr3-6964605-DEL-6222
chr3-7012760-DEL-132
chr3-7013952-DEL-512
chr3-7050768-DEL-77
chr3-7057538-DEL-400
chr3-7093796-DEL-69
chr3-7111225-DEL-60
chr3-7117847-DEL-75
chr3-7125983-DEL-501
chr3-7128648-DEL-353
chr3-7147905-DEL-6712
chr3-7147952-DEL-2724
chr3-7150708-DEL-3735
chr3-7154620-DEL-200
chr3-7156651-DEL-6256
chr3-7179182-DEL-215
chr3-7203440-DEL-1786
chr3-7219053-DEL-51
chr3-7224293-DEL-150
chr3-7228763-DEL-424
chr3-7230565-DEL-1485
chr3-7236494-DEL-132
chr3-7237990-DEL-1089
chr3-7242839-DEL-26797
chr3-7242851-DEL-6517
chr3-7253720-DEL-351
chr3-7269784-DEL-80
chr3-7281823-DEL-8302
chr3-7292659-DEL-104
chr3-7296616-DEL-2070
chr3-7298482-DEL-104
chr3-7299707-DEL-1181
chr3-7308480-DEL-393
chr3-7314140-DEL-135
chr3-7394918-DEL-214
chr3-7399742-DEL-85
chr3-7438157-DEL-50
chr3-7488972-DEL-517
chr3-7495935-DEL-492
chr3-7501896-DEL-66
chr3-7514998-DEL-237
chr3-7522412-DEL-67
chr3-7525415-DEL-63
chr3-7539075-DEL-2509
chr3-7542447-DEL-6383
chr3-7573499-DEL-98
chr3-7584341-DEL-811
chr3-7589336-DEL-727
chr3-7637474-DEL-101
chr3-7646101-DEL-72
chr3-7657150-DEL-388
chr3-7732084-DEL-63
chr3-7748052-DEL-764
chr3-7748970-DEL-469
chr3-7768351-DEL-75
chr3-7789768-DEL-147
chr3-7805266-DEL-6522
chr3-7816733-DEL-69
chr3-7821862-DEL-69
chr3-7836554-DEL-562
chr3-7855747-DEL-5187
chr3-7868426-DEL-54
chr3-7902507-DEL-3090
chr3-7983401-DEL-234
chr3-8006750-DEL-80
chr3-8010115-DEL-4173
chr3-8030361-DEL-629

chr3-8046052-DEL-3135
chr3-8049557-DEL-272
chr3-8051848-DEL-141
chr3-8052925-DEL-325
chr3-8089837-DEL-53
chr3-8115691-DEL-64
chr3-8157620-DEL-1074
chr3-8209525-DEL-315
chr3-8224318-DEL-111
chr3-8246482-DEL-288
chr3-8258047-DEL-303
chr3-8262668-DEL-1422
chr3-8279079-DEL-53
chr3-8283992-DEL-682
chr3-8290083-DEL-2460
chr3-8293538-DEL-97
chr3-8300999-DEL-1400
chr3-8306705-DEL-2202
chr3-8309310-DEL-2348
chr3-8325862-DEL-2182
chr3-8361171-DEL-1119
chr3-8373387-DEL-910
chr3-8377594-DEL-62
chr3-8395253-DEL-451
chr3-8418897-DEL-510
chr3-8438300-DEL-70
chr3-8505759-DEL-55
chr3-8539782-DEL-173
chr3-8541827-DEL-6207
chr3-8565052-DEL-137
chr3-8579975-DEL-73
chr3-8594423-DEL-84
chr3-8598316-DEL-95
chr3-8631409-DEL-208
chr3-8660437-DEL-57
chr3-8679998-DEL-381
chr3-8685355-DEL-1544
chr3-8714625-DEL-361
chr3-8719050-DEL-3341
chr3-8734484-DEL-54
chr3-8766774-DEL-56
chr3-8768063-DEL-66
chr3-8772088-DEL-122
chr3-8811149-DEL-68
chr3-8825495-DEL-141
chr3-8828885-DEL-408
chr3-8835840-DEL-202
chr3-8841361-DEL-194
chr3-8842933-DEL-345
chr3-8859043-DEL-162
chr3-8863604-DEL-476
chr3-8864159-DEL-153
chr3-8887748-DEL-60
chr3-8896872-DEL-7007
chr3-8897092-DEL-416
chr3-8911835-DEL-100
chr3-8942463-DEL-2089
chr3-8971687-DEL-5985
chr3-8980885-DEL-203
chr3-8983634-DEL-343
chr3-9006862-DEL-53
chr3-9012948-DEL-372
chr3-9029180-DEL-68
chr3-9050033-DEL-678
chr3-9050054-DEL-89
chr3-9051305-DEL-60
chr3-9053426-DEL-203
chr3-9073253-DEL-1208
chr3-9075404-DEL-560
chr3-9076549-DEL-146
chr3-9079030-DEL-74
chr3-9091809-DEL-87
chr3-9098316-DEL-143
chr3-9111896-DEL-71
chr3-9122824-DEL-420
chr3-9127671-DEL-156
chr3-9127836-DEL-1159

chr3-9135190-DEL-349
chr3-9198191-DEL-2393
chr3-9214382-DEL-221
chr3-9237252-DEL-805
chr3-9252219-DEL-210
chr3-9271056-DEL-2890
chr3-9282475-DEL-70
chr3-9296116-DEL-79
chr3-9305110-DEL-64
chr3-9325435-DEL-915
chr3-9358261-DEL-73
chr3-9375650-DEL-6370
chr3-9384977-DEL-72
chr3-9386983-DEL-91
chr3-9393142-DEL-55
chr3-9394887-DEL-295
chr3-9395730-DEL-82
chr3-9409453-DEL-60
chr3-9430490-DEL-70
chr3-9450659-DEL-761
chr3-9452152-DEL-6931
chr3-9461028-DEL-695
chr3-9481066-DEL-67
chr3-9484849-DEL-1346
chr3-9501713-DEL-166
chr3-9511975-DEL-148
chr3-9516192-DEL-137
chr3-9516446-DEL-120
chr3-9532850-DEL-110
chr3-9537300-DEL-199
chr3-9547267-DEL-52
chr3-9551345-DEL-1909
chr3-9566539-DEL-1104
chr3-9566967-DEL-340
chr3-9593033-DEL-102
chr3-9593538-DEL-75
chr3-9605588-DEL-156
chr3-9629104-DEL-272
chr3-9636126-DEL-105
chr3-9667302-DEL-350
chr3-9682413-DEL-88
chr3-9683804-DEL-80
chr3-9696771-DEL-207
chr3-9701051-DEL-302
chr3-9707950-DEL-160
chr3-9767880-DEL-109
chr3-9768257-DEL-1245
chr3-9803909-DEL-84
chr3-9809871-DEL-289
chr3-9824190-DEL-258
chr3-9828437-DEL-445
chr3-9834914-DEL-76
chr3-9840725-DEL-79
chr3-9842846-DEL-431
chr3-9867998-DEL-700
chr3-9877619-DEL-55
chr3-9887171-DEL-120
chr3-9909877-DEL-101
chr3-9913747-DEL-270
chr3-9921734-DEL-84
chr3-9926727-DEL-191
chr3-9932710-DEL-348
chr3-9953887-DEL-66
chr3-9962643-DEL-81
chr3-9990257-DEL-60
chr3-9992907-DEL-77
chr3-10009143-DEL-58
chr3-10015013-DEL-5896
chr3-10037272-DEL-175
chr3-10038171-DEL-74
chr3-10039454-DEL-71
chr3-10041326-DEL-1488
chr3-10043668-DEL-112
chr3-10048114-DEL-509
chr3-10100615-DEL-256
chr3-10131849-DEL-1170
chr3-10133898-DEL-560

chr3-10137347-DEL-1477
chr3-10181738-DEL-569
chr3-10187848-DEL-734
chr3-10199138-DEL-186
chr3-10212007-DEL-1405
chr3-10214632-DEL-306
chr3-10248886-DEL-53
chr3-10249543-DEL-110
chr3-10282752-DEL-535
chr3-10285014-DEL-53
chr3-10286015-DEL-2678
chr3-10316177-DEL-120
chr3-10329232-DEL-67
chr3-10353223-DEL-197
chr3-10360984-DEL-96
chr3-10364912-DEL-82
chr3-10384874-DEL-64
chr3-10389343-DEL-120
chr3-10402547-DEL-148
chr3-10435191-DEL-86
chr3-10471680-DEL-63
chr3-10473737-DEL-62
chr3-10489140-DEL-62
chr3-10494472-DEL-103
chr3-10496824-DEL-131
chr3-10498680-DEL-82
chr3-10501483-DEL-177
chr3-10547296-DEL-6409
chr3-10576921-DEL-2054
chr3-10590477-DEL-3400
chr3-10593803-DEL-1048
chr3-10594242-DEL-2411
chr3-10595192-DEL-1874
chr3-10596600-DEL-145
chr3-10599878-DEL-76
chr3-10610018-DEL-433
chr3-10624850-DEL-57
chr3-10683643-DEL-1807
chr3-10688765-DEL-308
chr3-10717111-DEL-1620
chr3-10777693-DEL-1578
chr3-10777837-DEL-2151
chr3-10777967-DEL-943
chr3-10778601-DEL-1331
chr3-10793153-DEL-70
chr3-10802426-DEL-152
chr3-10837816-DEL-73
chr3-10855376-DEL-234
chr3-10859996-DEL-317
chr3-10875686-DEL-207
chr3-10879020-DEL-263
chr3-10963170-DEL-752
chr3-10977855-DEL-121
chr3-10998013-DEL-62
chr3-11009660-DEL-1065
chr3-11033171-DEL-208
chr3-11038101-DEL-174
chr3-11056203-DEL-60
chr3-11073635-DEL-299
chr3-11077772-DEL-399
chr3-11109952-DEL-196
chr3-11112232-DEL-12662
chr3-11149453-DEL-79
chr3-11168259-DEL-1295
chr3-11173619-DEL-118
chr3-11185875-DEL-13855
chr3-11205694-DEL-8699
chr3-11215147-DEL-130
chr3-11239499-DEL-205
chr3-11251322-DEL-680
chr3-11256446-DEL-56
chr3-11310643-DEL-147
chr3-11311096-DEL-5499
chr3-11323905-DEL-861
chr3-11332591-DEL-215
chr3-11334144-DEL-1681
chr3-11338123-DEL-469

chr3-11395366-DEL-5178
chr3-11417232-DEL-329
chr3-11438090-DEL-1225
chr3-11462998-DEL-358
chr3-11464365-DEL-85
chr3-11471600-DEL-65
chr3-11480043-DEL-6197
chr3-11516157-DEL-8205
chr3-11541967-DEL-113
chr3-11576562-DEL-5338
chr3-11579621-DEL-111
chr3-11583666-DEL-6275
chr3-11600190-DEL-588
chr3-11605042-DEL-2844
chr3-11610155-DEL-1337
chr3-11641416-DEL-362
chr3-11692877-DEL-401
chr3-11735297-DEL-58
chr3-11738784-DEL-1187
chr3-11760979-DEL-2106
chr3-11771824-DEL-223
chr3-11772346-DEL-727
chr3-11787361-DEL-141
chr3-11788834-DEL-5021
chr3-11789305-DEL-2597
chr3-11795140-DEL-320
chr3-11817755-DEL-77
chr3-11837678-DEL-379
chr3-11838903-DEL-365
chr3-11879358-DEL-178
chr3-11882885-DEL-239
chr3-11883037-DEL-1370
chr3-11888215-DEL-250
chr3-11912347-DEL-99
chr3-11915904-DEL-9024
chr3-11940909-DEL-73
chr3-11943950-DEL-1138
chr3-11945228-DEL-6208
chr3-11973607-DEL-1014
chr3-11982877-DEL-67
chr3-11985520-DEL-473
chr3-11998299-DEL-6461
chr3-12007713-DEL-60
chr3-12026519-DEL-1805
chr3-12068225-DEL-228
chr3-12088530-DEL-340
chr3-12110383-DEL-194
chr3-12111701-DEL-2163
chr3-12125197-DEL-693
chr3-12146590-DEL-181
chr3-12163604-DEL-784
chr3-12182039-DEL-664
chr3-12190291-DEL-6194
chr3-12215384-DEL-73
chr3-12240231-DEL-71
chr3-12241075-DEL-569
chr3-12244302-DEL-369
chr3-12257116-DEL-146
chr3-12295415-DEL-6495
chr3-12309863-DEL-847
chr3-12313400-DEL-6390
chr3-12341359-DEL-456
chr3-12343783-DEL-57
chr3-12366565-DEL-64
chr3-12388584-DEL-1246
chr3-12400401-DEL-6501
chr3-12406441-DEL-201
chr3-12416794-DEL-353
chr3-12420936-DEL-85
chr3-12434241-DEL-10504
chr3-12449020-DEL-215
chr3-12457750-DEL-85
chr3-12464327-DEL-59
chr3-12469615-DEL-84
chr3-12487055-DEL-167
chr3-12489255-DEL-6063
chr3-12498439-DEL-194

chr3-12501565-DEL-1434
chr3-12505004-DEL-101
chr3-12510579-DEL-66
chr3-12532846-DEL-13366
chr3-12577318-DEL-54
chr3-12605217-DEL-6398
chr3-12622518-DEL-1243
chr3-12638731-DEL-6339
chr3-12648178-DEL-1177
chr3-12659885-DEL-72
chr3-12684891-DEL-204
chr3-12710504-DEL-5648
chr3-12729994-DEL-75
chr3-12739166-DEL-77
chr3-12742071-DEL-78
chr3-12744388-DEL-302
chr3-12747097-DEL-2232
chr3-12775303-DEL-3887
chr3-12783188-DEL-155
chr3-12786710-DEL-6206
chr3-12807639-DEL-2965
chr3-12844257-DEL-92
chr3-12848971-DEL-77
chr3-12853235-DEL-6352
chr3-12864453-DEL-1795
chr3-12878542-DEL-125
chr3-12885782-DEL-58
chr3-12915332-DEL-60
chr3-12924838-DEL-99
chr3-12928762-DEL-194
chr3-12939424-DEL-829
chr3-12965878-DEL-1089
chr3-12985434-DEL-2367
chr3-13006816-DEL-63
chr3-13014760-DEL-72
chr3-13044313-DEL-192
chr3-13049664-DEL-2930
chr3-13069095-DEL-546
chr3-13084111-DEL-171
chr3-13089332-DEL-92
chr3-13122918-DEL-804
chr3-13123739-DEL-9995
chr3-13123796-DEL-8465
chr3-13160814-DEL-210
chr3-13171107-DEL-6210
chr3-13172745-DEL-1903
chr3-13173636-DEL-1461
chr3-13175139-DEL-1446
chr3-13177696-DEL-189
chr3-13182511-DEL-119
chr3-13194339-DEL-133
chr3-13195531-DEL-2106
chr3-13196008-DEL-60
chr3-13204479-DEL-221
chr3-13252916-DEL-1899
chr3-13255362-DEL-211
chr3-13271353-DEL-186
chr3-13308548-DEL-68
chr3-13348804-DEL-599
chr3-13354332-DEL-131
chr3-13372297-DEL-67
chr3-13380341-DEL-587
chr3-13382773-DEL-514
chr3-13406715-DEL-197
chr3-13417634-DEL-6186
chr3-13417670-DEL-1394
chr3-13421838-DEL-1998
chr3-13426481-DEL-450
chr3-13436784-DEL-96
chr3-13438474-DEL-6513
chr3-13444559-DEL-402
chr3-13448001-DEL-71
chr3-13452546-DEL-8304
chr3-13456390-DEL-1503
chr3-13494246-DEL-178
chr3-13528132-DEL-60
chr3-13541806-DEL-228

chr3-13598615-DEL-146
chr3-13621281-DEL-107
chr3-13646330-DEL-456
chr3-13654066-DEL-52
chr3-13669428-DEL-358
chr3-13684970-DEL-804
chr3-13686507-DEL-85
chr3-13688743-DEL-167
chr3-13688791-DEL-1588
chr3-13695325-DEL-61
chr3-13720880-DEL-145
chr3-13735897-DEL-672
chr3-13777267-DEL-56
chr3-13826399-DEL-96
chr3-13855688-DEL-5460
chr3-13861235-DEL-572
chr3-13930507-DEL-56
chr3-13944008-DEL-60
chr3-13958052-DEL-442
chr3-13963308-DEL-437
chr3-13971706-DEL-282
chr3-13980372-DEL-3644
chr3-14024609-DEL-102
chr3-14046997-DEL-55
chr3-14064140-DEL-58
chr3-14082298-DEL-87
chr3-14119240-DEL-998
chr3-14121163-DEL-183
chr3-14224790-DEL-2474
chr3-14249200-DEL-244
chr3-14254366-DEL-504
chr3-14282610-DEL-1749
chr3-14294257-DEL-3244
chr3-14334638-DEL-6085
chr3-14341830-DEL-926
chr3-14355248-DEL-728
chr3-14356521-DEL-3974
chr3-14369646-DEL-6161
chr3-14376320-DEL-58
chr3-14378398-DEL-537
chr3-14384930-DEL-62
chr3-14385685-DEL-6272
chr3-14393034-DEL-1523
chr3-14422296-DEL-5922
chr3-14453951-DEL-80
chr3-14454827-DEL-692
chr3-14457164-DEL-270
chr3-14483983-DEL-251
chr3-14494612-DEL-5141
chr3-14512332-DEL-5777
chr3-14558208-DEL-66
chr3-14558671-DEL-870
chr3-14593954-DEL-1667
chr3-14647988-DEL-62
chr3-14657224-DEL-131
chr3-14658371-DEL-4189
chr3-14663054-DEL-445
chr3-14700976-DEL-166
chr3-14766809-DEL-187
chr3-14770057-DEL-17275
chr3-14787762-DEL-1021
chr3-14796406-DEL-372
chr3-14804706-DEL-6427
chr3-14812966-DEL-269
chr3-14815626-DEL-588
chr3-14846835-DEL-2693
chr3-14849617-DEL-298
chr3-14910706-DEL-67
chr3-14920881-DEL-849
chr3-14923894-DEL-54
chr3-14940760-DEL-213
chr3-14987543-DEL-6253
chr3-15009384-DEL-65
chr3-15041408-DEL-221
chr3-15041948-DEL-57
chr3-15045792-DEL-6847
chr3-15071862-DEL-493

chr3-15103950-DEL-113
chr3-15108121-DEL-2185
chr3-15113169-DEL-68
chr3-15139300-DEL-241
chr3-15201043-DEL-468
chr3-15294380-DEL-1065
chr3-15297916-DEL-382
chr3-15298490-DEL-535
chr3-15299659-DEL-130
chr3-15303911-DEL-983
chr3-15343740-DEL-395
chr3-15371521-DEL-8294
chr3-15408334-DEL-1574
chr3-15431769-DEL-3886
chr3-15462005-DEL-702
chr3-15479153-DEL-357
chr3-15514679-DEL-75
chr3-15533849-DEL-54
chr3-15549841-DEL-60
chr3-15567466-DEL-3450
chr3-15577981-DEL-6728
chr3-15578024-DEL-8946
chr3-15584880-DEL-2195
chr3-15610108-DEL-224
chr3-15612895-DEL-267
chr3-15628740-DEL-413
chr3-15657720-DEL-1200
chr3-15670193-DEL-408
chr3-15676064-DEL-728
chr3-15703352-DEL-1573
chr3-15820670-DEL-1569
chr3-15822927-DEL-52
chr3-15829887-DEL-351
chr3-15838258-DEL-60
chr3-15862176-DEL-1724
chr3-15865822-DEL-1771
chr3-15900870-DEL-152
chr3-15908283-DEL-64
chr3-15920593-DEL-72
chr3-15922636-DEL-6168
chr3-15946000-DEL-6309
chr3-15975071-DEL-13963
chr3-15989691-DEL-402
chr3-16011838-DEL-1455
chr3-16032859-DEL-1761
chr3-16046349-DEL-98
chr3-16059013-DEL-64
chr3-16067769-DEL-857
chr3-16088707-DEL-65
chr3-16093154-DEL-143
chr3-16105859-DEL-11514
chr3-16106193-DEL-234
chr3-16108856-DEL-483
chr3-16112662-DEL-138
chr3-16130097-DEL-102
chr3-16133320-DEL-604
chr3-16141189-DEL-218
chr3-16272514-DEL-205
chr3-16299158-DEL-152
chr3-16304045-DEL-514
chr3-16313767-DEL-51
chr3-16321678-DEL-154
chr3-16393653-DEL-1008
chr3-16414970-DEL-62
chr3-16434544-DEL-2797
chr3-16456230-DEL-127
chr3-16472287-DEL-1184
chr3-16472835-DEL-6269
chr3-16506101-DEL-52
chr3-16527704-DEL-6201
chr3-16538137-DEL-2456
chr3-16540744-DEL-117
chr3-16554501-DEL-210
chr3-16557927-DEL-341
chr3-16560223-DEL-564
chr3-16572992-DEL-74
chr3-16578513-DEL-5317

chr3-16593962-DEL-7230
chr3-16599823-DEL-717
chr3-16620914-DEL-6430
chr3-16678650-DEL-16218
chr3-16731867-DEL-1001
chr3-16799892-DEL-1019
chr3-16807788-DEL-1340
chr3-16815984-DEL-406
chr3-16825919-DEL-1273
chr3-16843399-DEL-1814
chr3-16876817-DEL-67
chr3-16891553-DEL-555
chr3-16893521-DEL-395
chr3-16894333-DEL-592
chr3-16928820-DEL-206
chr3-16944392-DEL-56
chr3-16962633-DEL-102
chr3-16965072-DEL-1377
chr3-16973997-DEL-1202
chr3-17013320-DEL-4027
chr3-17021193-DEL-411
chr3-17034226-DEL-597
chr3-17048599-DEL-643
chr3-17090855-DEL-78
chr3-17103801-DEL-2878
chr3-17150393-DEL-362
chr3-17152530-DEL-1133
chr3-17153939-DEL-442
chr3-17159621-DEL-576
chr3-17163176-DEL-3293
chr3-17164685-DEL-429
chr3-17186736-DEL-93
chr3-17200834-DEL-4179
chr3-17239556-DEL-2454
chr3-17242770-DEL-2858
chr3-17245700-DEL-6570
chr3-17251907-DEL-283
chr3-17295097-DEL-1501
chr3-17297386-DEL-846
chr3-17340151-DEL-675
chr3-17357322-DEL-249
chr3-17364754-DEL-1242
chr3-17369030-DEL-201
chr3-17374468-DEL-637
chr3-17396773-DEL-54
chr3-17398945-DEL-177
chr3-17401910-DEL-107
chr3-17417479-DEL-342
chr3-17418054-DEL-359
chr3-17432904-DEL-7797
chr3-17473667-DEL-6183
chr3-17484044-DEL-2664
chr3-17508225-DEL-1129
chr3-17509997-DEL-1218
chr3-17518014-DEL-149
chr3-17539760-DEL-65
chr3-17546001-DEL-1285
chr3-17555232-DEL-470
chr3-17598883-DEL-60
chr3-17625170-DEL-397
chr3-17649912-DEL-673
chr3-17656035-DEL-334
chr3-17666043-DEL-462
chr3-17668027-DEL-200
chr3-17692569-DEL-162
chr3-17758281-DEL-202
chr3-17761787-DEL-4099
chr3-17776064-DEL-495
chr3-17805409-DEL-1230
chr3-17857331-DEL-64
chr3-17866749-DEL-372
chr3-17872784-DEL-112
chr3-17883640-DEL-334
chr3-17886275-DEL-984
chr3-17923144-DEL-1339
chr3-17933762-DEL-398
chr3-17979392-DEL-558

chr3-17981833-DEL-58
chr3-18025731-DEL-59
chr3-18028853-DEL-6901
chr3-18039333-DEL-327
chr3-18043899-DEL-526
chr3-18045476-DEL-6160
chr3-18070938-DEL-204
chr3-18071180-DEL-5363
chr3-18076769-DEL-794
chr3-18077768-DEL-94
chr3-18105802-DEL-162
chr3-18138493-DEL-86
chr3-18156253-DEL-80
chr3-18164297-DEL-988
chr3-18180554-DEL-81
chr3-18197587-DEL-98
chr3-18201337-DEL-52
chr3-18255584-DEL-105
chr3-18276100-DEL-666
chr3-18276953-DEL-616
chr3-18277860-DEL-337
chr3-18313265-DEL-56
chr3-18367778-DEL-82
chr3-18375669-DEL-445
chr3-18375683-DEL-1886
chr3-18380129-DEL-240
chr3-18462982-DEL-108
chr3-18495230-DEL-74
chr3-18551162-DEL-78
chr3-18592778-DEL-6253
chr3-18598899-DEL-51
chr3-18612827-DEL-56
chr3-18616737-DEL-485
chr3-18677539-DEL-67
chr3-18709287-DEL-140
chr3-18735916-DEL-56
chr3-18745892-DEL-91
chr3-18817702-DEL-123
chr3-18819927-DEL-182
chr3-18853543-DEL-2033
chr3-18888236-DEL-1349
chr3-18973151-DEL-92
chr3-19008928-DEL-1716
chr3-19031689-DEL-339
chr3-19033882-DEL-1559
chr3-19118141-DEL-1129
chr3-19125184-DEL-66
chr3-19132030-DEL-171
chr3-19148833-DEL-6553
chr3-19149114-DEL-65
chr3-19169379-DEL-449
chr3-19200298-DEL-160
chr3-19201377-DEL-52
chr3-19206663-DEL-58
chr3-19224071-DEL-70
chr3-19261628-DEL-80
chr3-19292379-DEL-60
chr3-19307538-DEL-240
chr3-19350743-DEL-63
chr3-19354582-DEL-501
chr3-19356158-DEL-72
chr3-19389848-DEL-140
chr3-19392534-DEL-5784
chr3-19393300-DEL-220
chr3-19403497-DEL-1098
chr3-19458529-DEL-126
chr3-19468960-DEL-56
chr3-19478458-DEL-131
chr3-19488284-DEL-577
chr3-19550635-DEL-129
chr3-19597233-DEL-2285
chr3-19629966-DEL-749
chr3-19641842-DEL-193
chr3-19649002-DEL-3170
chr3-19649070-DEL-6246
chr3-19652617-DEL-901
chr3-19656041-DEL-543

chr3-19687865-DEL-65
chr3-19709573-DEL-359
chr3-19713865-DEL-76
chr3-19721279-DEL-268
chr3-19736663-DEL-1046
chr3-19737233-DEL-476
chr3-19772601-DEL-70
chr3-19797836-DEL-513
chr3-19828291-DEL-91
chr3-19833809-DEL-454
chr3-19847870-DEL-184
chr3-19867849-DEL-7394
chr3-19937180-DEL-331
chr3-19938113-DEL-307
chr3-19953171-DEL-182
chr3-19968372-DEL-1143
chr3-19971926-DEL-3816
chr3-19977162-DEL-6122
chr3-19988089-DEL-2517
chr3-20055288-DEL-258
chr3-20060199-DEL-51
chr3-20062311-DEL-85
chr3-20078496-DEL-140
chr3-20102425-DEL-513
chr3-20130277-DEL-200
chr3-20134116-DEL-418
chr3-20175901-DEL-161
chr3-20240613-DEL-130
chr3-20243563-DEL-1418
chr3-20246135-DEL-207
chr3-20260426-DEL-154
chr3-20276029-DEL-71
chr3-20310155-DEL-86
chr3-20313687-DEL-1000
chr3-20322062-DEL-194
chr3-20337545-DEL-80
chr3-20347097-DEL-498
chr3-20348745-DEL-84
chr3-20404863-DEL-940
chr3-20405505-DEL-190
chr3-20437320-DEL-4485
chr3-20443306-DEL-1128
chr3-20456133-DEL-250
chr3-20469207-DEL-4314
chr3-20501357-DEL-6389
chr3-20552924-DEL-122
chr3-20563833-DEL-56
chr3-20567434-DEL-4987
chr3-20573040-DEL-1041
chr3-20577472-DEL-272
chr3-20579133-DEL-57
chr3-20611526-DEL-406
chr3-20640208-DEL-1241
chr3-20645621-DEL-52
chr3-20667262-DEL-60
chr3-20671406-DEL-6807
chr3-20681276-DEL-756
chr3-20682349-DEL-198
chr3-20704630-DEL-630
chr3-20740620-DEL-54
chr3-20761803-DEL-694
chr3-20765997-DEL-577
chr3-20767446-DEL-108
chr3-20773480-DEL-6226
chr3-20819643-DEL-74
chr3-20847317-DEL-59
chr3-20861666-DEL-1889
chr3-20863718-DEL-69
chr3-20887681-DEL-6500
chr3-20897927-DEL-10771
chr3-20930351-DEL-82
chr3-20932763-DEL-1265
chr3-20935616-DEL-60
chr3-20939107-DEL-1262
chr3-20986062-DEL-6336
chr3-20987274-DEL-132
chr3-21014632-DEL-2260

chr3-21055873-DEL-6265
chr3-21093887-DEL-70
chr3-21100584-DEL-262
chr3-21132922-DEL-2007
chr3-21163888-DEL-167
chr3-21175237-DEL-1456
chr3-21209780-DEL-4825
chr3-21212299-DEL-833
chr3-21213819-DEL-84
chr3-21221169-DEL-1862
chr3-21222066-DEL-145
chr3-21282518-DEL-584
chr3-21291671-DEL-1097
chr3-21297123-DEL-2826
chr3-21316475-DEL-397
chr3-21334635-DEL-301
chr3-21349368-DEL-83
chr3-21371685-DEL-52
chr3-21380459-DEL-93
chr3-21402477-DEL-438
chr3-21423391-DEL-109
chr3-21424907-DEL-83
chr3-21429402-DEL-83
chr3-21500050-DEL-328
chr3-21504430-DEL-8362
chr3-21513788-DEL-320
chr3-21524495-DEL-79
chr3-21553663-DEL-341
chr3-21566949-DEL-6194
chr3-21609483-DEL-233
chr3-21609811-DEL-542
chr3-21635760-DEL-65
chr3-21700914-DEL-566
chr3-21748509-DEL-77
chr3-21762570-DEL-539
chr3-21769662-DEL-56
chr3-21779663-DEL-995
chr3-21828251-DEL-54
chr3-21836792-DEL-1551
chr3-21860914-DEL-458
chr3-21881019-DEL-56
chr3-21902278-DEL-57
chr3-21941477-DEL-63
chr3-21943021-DEL-790
chr3-21944680-DEL-400
chr3-21949371-DEL-210
chr3-21985013-DEL-89
chr3-21986783-DEL-343
chr3-22003844-DEL-116
chr3-22007489-DEL-53
chr3-22050778-DEL-140
chr3-22079892-DEL-7791
chr3-22123412-DEL-1331
chr3-22136674-DEL-60
chr3-22141874-DEL-58
chr3-22154531-DEL-54
chr3-22215642-DEL-78
chr3-22272214-DEL-133
chr3-22294917-DEL-6359
chr3-22300948-DEL-142
chr3-22301311-DEL-11096
chr3-22303083-DEL-4310
chr3-22312239-DEL-3008
chr3-22318079-DEL-3779
chr3-22326026-DEL-276
chr3-22347941-DEL-4375
chr3-22360482-DEL-330
chr3-22373247-DEL-200
chr3-22390385-DEL-199
chr3-22403279-DEL-401
chr3-22407643-DEL-101
chr3-22415632-DEL-10801
chr3-22457383-DEL-70
chr3-22459994-DEL-1328
chr3-22462768-DEL-1281
chr3-22467587-DEL-3492
chr3-22477489-DEL-132

chr3-22483910-DEL-1251
chr3-22502327-DEL-12929
chr3-22517372-DEL-888
chr3-22523612-DEL-98
chr3-22538049-DEL-4215
chr3-22553211-DEL-222
chr3-22560099-DEL-58
chr3-22569061-DEL-6199
chr3-22580115-DEL-60
chr3-22580871-DEL-81
chr3-22584622-DEL-74
chr3-22586154-DEL-260
chr3-22607016-DEL-403
chr3-22609832-DEL-1151
chr3-22636383-DEL-962
chr3-22641458-DEL-137
chr3-22666156-DEL-6206
chr3-22675535-DEL-6589
chr3-22697750-DEL-3714
chr3-22706567-DEL-2042
chr3-22706677-DEL-5425
chr3-22712126-DEL-1365
chr3-22713100-DEL-214
chr3-22717206-DEL-465
chr3-22718571-DEL-75
chr3-22719014-DEL-637
chr3-22723761-DEL-6484
chr3-22736592-DEL-130
chr3-22741101-DEL-289
chr3-22743967-DEL-100
chr3-22744285-DEL-5988
chr3-22756062-DEL-7405
chr3-22769454-DEL-6586
chr3-22784842-DEL-1456
chr3-22789191-DEL-199
chr3-22805538-DEL-1701
chr3-22806937-DEL-282
chr3-22834350-DEL-76
chr3-22855388-DEL-101
chr3-22872163-DEL-324
chr3-22880553-DEL-260
chr3-22883521-DEL-904
chr3-22884525-DEL-1950
chr3-22948730-DEL-51
chr3-22954820-DEL-18163
chr3-22973899-DEL-388
chr3-22986783-DEL-6457
chr3-22988224-DEL-2737
chr3-23003185-DEL-6399
chr3-23015627-DEL-6304
chr3-23035720-DEL-128
chr3-23040605-DEL-1105
chr3-23063003-DEL-6243
chr3-23082505-DEL-2104
chr3-23090159-DEL-883
chr3-23092480-DEL-1183
chr3-23101358-DEL-71
chr3-23105180-DEL-202
chr3-23121536-DEL-92
chr3-23152644-DEL-1411
chr3-23153265-DEL-1803
chr3-23199035-DEL-516
chr3-23209452-DEL-563
chr3-23210054-DEL-1213
chr3-23309969-DEL-7278
chr3-23326381-DEL-5426
chr3-23332202-DEL-63
chr3-23344667-DEL-365
chr3-23352630-DEL-1639
chr3-23356196-DEL-12753
chr3-23373038-DEL-6221
chr3-23376006-DEL-145
chr3-23393174-DEL-222
chr3-23399259-DEL-63
chr3-23405402-DEL-920
chr3-23424345-DEL-133
chr3-23448404-DEL-252

chr3-23475211-DEL-80
chr3-23487296-DEL-67
chr3-23514224-DEL-1089
chr3-23516999-DEL-6555
chr3-23539189-DEL-2981
chr3-23548961-DEL-4037
chr3-23633644-DEL-6235
chr3-23651889-DEL-839
chr3-23653439-DEL-58
chr3-23665155-DEL-53
chr3-23681176-DEL-52
chr3-23693858-DEL-56
chr3-23701255-DEL-1043
chr3-23728307-DEL-1276
chr3-23792032-DEL-54
chr3-23836451-DEL-999
chr3-23837355-DEL-96
chr3-23842494-DEL-895
chr3-23867035-DEL-1956
chr3-23899351-DEL-71
chr3-23951321-DEL-204
chr3-23974682-DEL-80
chr3-24024459-DEL-1297
chr3-24042212-DEL-915
chr3-24043767-DEL-228
chr3-24060098-DEL-128
chr3-24085021-DEL-711
chr3-24121812-DEL-59
chr3-24200618-DEL-117
chr3-24209567-DEL-458
chr3-24233861-DEL-3022
chr3-24238771-DEL-137
chr3-24249365-DEL-60
chr3-24256407-DEL-56
chr3-24293400-DEL-206
chr3-24306057-DEL-1756
chr3-24312789-DEL-2134
chr3-24317683-DEL-572
chr3-24404185-DEL-401
chr3-24410536-DEL-80
chr3-24410923-DEL-390
chr3-24420591-DEL-649
chr3-24431726-DEL-352
chr3-24466576-DEL-52
chr3-24472728-DEL-100
chr3-24478278-DEL-59
chr3-24482137-DEL-695
chr3-24493187-DEL-126
chr3-24498772-DEL-486
chr3-24564271-DEL-277
chr3-24588792-DEL-92
chr3-24629301-DEL-176
chr3-24634074-DEL-132
chr3-24661858-DEL-399
chr3-24676912-DEL-6115
chr3-24703439-DEL-99
chr3-24708688-DEL-6853
chr3-24716733-DEL-87
chr3-24718967-DEL-263
chr3-24724314-DEL-56
chr3-24733535-DEL-200
chr3-24758659-DEL-80
chr3-24777966-DEL-2747
chr3-24807948-DEL-1169
chr3-24813485-DEL-2038
chr3-24814466-DEL-124
chr3-24839829-DEL-580
chr3-24898421-DEL-102
chr3-24942364-DEL-244
chr3-24951143-DEL-60
chr3-25002197-DEL-1374
chr3-25002200-DEL-5009
chr3-25002200-DEL-6185
chr3-25003607-DEL-4777
chr3-25007237-DEL-1145
chr3-25038108-DEL-54
chr3-25057291-DEL-496

chr3-25121790-DEL-54
chr3-25144943-DEL-669
chr3-25146368-DEL-334
chr3-25151628-DEL-75
chr3-25170605-DEL-715
chr3-25187353-DEL-65
chr3-25201833-DEL-116
chr3-25202623-DEL-538
chr3-25247713-DEL-69
chr3-25257960-DEL-506
chr3-25268924-DEL-54
chr3-25276742-DEL-6490
chr3-25296398-DEL-320
chr3-25304177-DEL-6873
chr3-25317194-DEL-204
chr3-25323710-DEL-221
chr3-25330107-DEL-3066
chr3-25332886-DEL-52
chr3-25341744-DEL-8995
chr3-25351383-DEL-243
chr3-25354292-DEL-580
chr3-25360005-DEL-6066
chr3-25363919-DEL-182
chr3-25397146-DEL-1460
chr3-25400049-DEL-2157
chr3-25411703-DEL-73
chr3-25421179-DEL-124
chr3-25452633-DEL-65
chr3-25465247-DEL-830
chr3-25469215-DEL-509
chr3-25512365-DEL-208
chr3-25517946-DEL-2015
chr3-25541731-DEL-103
chr3-25547373-DEL-373
chr3-25604280-DEL-55
chr3-25620513-DEL-92
chr3-25636341-DEL-113
chr3-25642207-DEL-676
chr3-25643972-DEL-55
chr3-25728674-DEL-76
chr3-25802891-DEL-122
chr3-25804065-DEL-856
chr3-25810427-DEL-72
chr3-25841362-DEL-436
chr3-25843863-DEL-147
chr3-25941214-DEL-55
chr3-26025543-DEL-2225
chr3-26030324-DEL-56
chr3-26034867-DEL-191
chr3-26036818-DEL-87
chr3-26145866-DEL-2489
chr3-26192852-DEL-118
chr3-26201928-DEL-54
chr3-26258371-DEL-493
chr3-26261149-DEL-1605
chr3-26274453-DEL-597
chr3-26287563-DEL-218
chr3-26333911-DEL-80
chr3-26337723-DEL-108
chr3-26362067-DEL-59
chr3-26399979-DEL-88
chr3-26422663-DEL-1461
chr3-26474161-DEL-4290
chr3-26478211-DEL-630
chr3-26487011-DEL-79
chr3-26493176-DEL-401
chr3-26512038-DEL-89
chr3-26558548-DEL-397
chr3-26570415-DEL-388
chr3-26596634-DEL-1027
chr3-26618394-DEL-1121
chr3-26640326-DEL-181
chr3-26648756-DEL-54
chr3-26655771-DEL-103
chr3-26655796-DEL-676
chr3-26657919-DEL-1049
chr3-26664250-DEL-72

chr3-26685495-DEL-147
chr3-26719593-DEL-229
chr3-26745602-DEL-762
chr3-26763649-DEL-482
chr3-26771639-DEL-55
chr3-26772035-DEL-701
chr3-26779072-DEL-409
chr3-26825243-DEL-148
chr3-26845324-DEL-1055
chr3-26859873-DEL-2060
chr3-26864960-DEL-56
chr3-26876965-DEL-66
chr3-26947353-DEL-4674
chr3-26949377-DEL-58
chr3-26972292-DEL-56
chr3-27011713-DEL-310
chr3-27016790-DEL-1240
chr3-27040288-DEL-335
chr3-27045486-DEL-402
chr3-27046361-DEL-312
chr3-27046942-DEL-1156
chr3-27051519-DEL-250
chr3-27052644-DEL-458
chr3-27088198-DEL-59
chr3-27095192-DEL-54
chr3-27115895-DEL-104
chr3-27121298-DEL-143
chr3-27137462-DEL-185
chr3-27148090-DEL-1415
chr3-27149300-DEL-627
chr3-27211941-DEL-1197
chr3-27271640-DEL-335
chr3-27291997-DEL-124
chr3-27306227-DEL-54
chr3-27314620-DEL-246
chr3-27333562-DEL-420
chr3-27337895-DEL-1152
chr3-27351575-DEL-53
chr3-27354130-DEL-1407
chr3-27355711-DEL-82
chr3-27358760-DEL-1455
chr3-27374416-DEL-54
chr3-27391878-DEL-204
chr3-27400287-DEL-204
chr3-27420176-DEL-53
chr3-27480711-DEL-125
chr3-27494958-DEL-245
chr3-27516819-DEL-195
chr3-27520397-DEL-63
chr3-27529155-DEL-265
chr3-27606515-DEL-66
chr3-27651237-DEL-167
chr3-27694589-DEL-51
chr3-27699756-DEL-65
chr3-27715293-DEL-119
chr3-27725753-DEL-176
chr3-27735516-DEL-84
chr3-27794361-DEL-52
chr3-27795132-DEL-531
chr3-27805163-DEL-77
chr3-27825954-DEL-73
chr3-27831197-DEL-100
chr3-27845329-DEL-95
chr3-27850502-DEL-153
chr3-27857112-DEL-73
chr3-27884902-DEL-51
chr3-27887473-DEL-519
chr3-27911311-DEL-667
chr3-27970364-DEL-135
chr3-27977574-DEL-255
chr3-27981560-DEL-124
chr3-27987370-DEL-2206
chr3-28002138-DEL-366
chr3-28009515-DEL-121
chr3-28032569-DEL-63
chr3-28034086-DEL-2263
chr3-28049279-DEL-68

chr3-28060162-DEL-465
chr3-28063618-DEL-475
chr3-28076846-DEL-115
chr3-28111310-DEL-198
chr3-28133680-DEL-73
chr3-28147499-DEL-206
chr3-28168106-DEL-216
chr3-28170833-DEL-406
chr3-28193129-DEL-1594
chr3-28216303-DEL-165
chr3-28223192-DEL-58
chr3-28228660-DEL-171
chr3-28315636-DEL-74
chr3-28342360-DEL-134
chr3-28369960-DEL-51
chr3-28373936-DEL-63
chr3-28393702-DEL-137
chr3-28398964-DEL-53
chr3-28428792-DEL-73
chr3-28435150-DEL-184
chr3-28450590-DEL-93
chr3-28532514-DEL-204
chr3-28548721-DEL-76
chr3-28553857-DEL-68
chr3-28562357-DEL-430
chr3-28598380-DEL-285
chr3-28601339-DEL-210
chr3-28628067-DEL-197
chr3-28669180-DEL-316
chr3-28689334-DEL-67
chr3-28694328-DEL-60
chr3-28698328-DEL-83
chr3-28714963-DEL-66
chr3-28791644-DEL-940
chr3-28793701-DEL-308
chr3-28830225-DEL-327
chr3-28871799-DEL-1835
chr3-28886917-DEL-78
chr3-28887608-DEL-1876
chr3-28906501-DEL-5512
chr3-28950510-DEL-153
chr3-28981780-DEL-374
chr3-28992818-DEL-124
chr3-29021702-DEL-53
chr3-29045073-DEL-6499
chr3-29062192-DEL-6205
chr3-29093102-DEL-820
chr3-29099645-DEL-314
chr3-29115874-DEL-247
chr3-29175939-DEL-61
chr3-29203945-DEL-64
chr3-29229200-DEL-65
chr3-29288715-DEL-86
chr3-29331347-DEL-616
chr3-29344338-DEL-64
chr3-29368928-DEL-459
chr3-29427582-DEL-2646
chr3-29445117-DEL-73
chr3-29448110-DEL-86
chr3-29449350-DEL-57
chr3-29506049-DEL-94
chr3-29535215-DEL-313
chr3-29535648-DEL-1057
chr3-29536738-DEL-693
chr3-29537549-DEL-291
chr3-29541730-DEL-6221
chr3-29551852-DEL-302
chr3-29558970-DEL-65
chr3-29604165-DEL-301
chr3-29606105-DEL-5380
chr3-29644133-DEL-64
chr3-29658163-DEL-54
chr3-29674783-DEL-1966
chr3-29706305-DEL-55
chr3-29707906-DEL-142
chr3-29747116-DEL-52
chr3-29750673-DEL-59

chr3-29808178-DEL-93
chr3-29837225-DEL-3076
chr3-29850591-DEL-109
chr3-29886140-DEL-62
chr3-29896646-DEL-104
chr3-29897708-DEL-297
chr3-29926609-DEL-212
chr3-29953121-DEL-140
chr3-29959543-DEL-61
chr3-29961395-DEL-244
chr3-29962298-DEL-7919
chr3-29965202-DEL-6940
chr3-29981135-DEL-583
chr3-29993541-DEL-65
chr3-30044498-DEL-182
chr3-30044842-DEL-645
chr3-30152534-DEL-171
chr3-30178760-DEL-56
chr3-30247442-DEL-65
chr3-30247952-DEL-207
chr3-30321085-DEL-80
chr3-30378234-DEL-54
chr3-30382631-DEL-413
chr3-30417871-DEL-67
chr3-30437059-DEL-54
chr3-30439694-DEL-3950
chr3-30475562-DEL-59
chr3-30484158-DEL-109
chr3-30510002-DEL-67
chr3-30534340-DEL-210
chr3-30549841-DEL-54
chr3-30579985-DEL-197
chr3-30584811-DEL-522
chr3-30645011-DEL-167
chr3-30647716-DEL-58
chr3-30655511-DEL-59
chr3-30660927-DEL-62
chr3-30667897-DEL-1248
chr3-30676574-DEL-200
chr3-30679426-DEL-78
chr3-30692478-DEL-726
chr3-30717980-DEL-64
chr3-30719438-DEL-2351
chr3-30723489-DEL-1207
chr3-30757367-DEL-1889
chr3-30758245-DEL-156
chr3-30781672-DEL-1384
chr3-30787225-DEL-753
chr3-30832854-DEL-511
chr3-30877915-DEL-62
chr3-30893604-DEL-1436
chr3-30937658-DEL-1124
chr3-30940911-DEL-56
chr3-30956908-DEL-54
chr3-30971114-DEL-176
chr3-30984653-DEL-80
chr3-30994798-DEL-101
chr3-30995360-DEL-56
chr3-31009485-DEL-197
chr3-31038888-DEL-2662
chr3-31063131-DEL-3587
chr3-31079962-DEL-181
chr3-31090551-DEL-61
chr3-31107102-DEL-119
chr3-31110168-DEL-200
chr3-31111679-DEL-2073
chr3-31117494-DEL-182
chr3-31124235-DEL-528
chr3-31137742-DEL-54
chr3-31158646-DEL-81
chr3-31160646-DEL-77
chr3-31178390-DEL-236
chr3-31186423-DEL-169
chr3-31189520-DEL-91
chr3-31191950-DEL-99
chr3-31210711-DEL-330
chr3-31223611-DEL-1613

chr3-31230477-DEL-1476
chr3-31235298-DEL-54
chr3-31246267-DEL-109
chr3-31265922-DEL-282
chr3-31268441-DEL-94
chr3-31278853-DEL-1584
chr3-31294954-DEL-237
chr3-31328798-DEL-306
chr3-31348685-DEL-272
chr3-31352886-DEL-222
chr3-31387817-DEL-449
chr3-31400044-DEL-53
chr3-31404792-DEL-204
chr3-31410852-DEL-1269
chr3-31419412-DEL-54
chr3-31433465-DEL-4297
chr3-31446462-DEL-128
chr3-31449445-DEL-70
chr3-31482186-DEL-1518
chr3-31490556-DEL-231
chr3-31491710-DEL-132
chr3-31521675-DEL-61
chr3-31523512-DEL-8735
chr3-31534202-DEL-53
chr3-31608286-DEL-60
chr3-31621843-DEL-968
chr3-31651603-DEL-1431
chr3-31654526-DEL-613
chr3-31678835-DEL-880
chr3-31686952-DEL-70
chr3-31690425-DEL-225
chr3-31697654-DEL-366
chr3-31705256-DEL-6472
chr3-31722380-DEL-59
chr3-31729768-DEL-6263
chr3-31784588-DEL-653
chr3-31827311-DEL-6075
chr3-31885455-DEL-253
chr3-31891173-DEL-162
chr3-31933563-DEL-1450
chr3-31935524-DEL-97
chr3-32016754-DEL-62
chr3-32021509-DEL-72
chr3-32040958-DEL-261
chr3-32046826-DEL-764
chr3-32068523-DEL-121
chr3-32126757-DEL-104
chr3-32192329-DEL-389
chr3-32193767-DEL-1888
chr3-32196740-DEL-66
chr3-32225365-DEL-795
chr3-32269366-DEL-475
chr3-32277659-DEL-1478
chr3-32291829-DEL-54
chr3-32311900-DEL-252
chr3-32312283-DEL-448
chr3-32319236-DEL-90
chr3-32333369-DEL-1023
chr3-32334561-DEL-249
chr3-32338069-DEL-61
chr3-32368662-DEL-400
chr3-32399123-DEL-235
chr3-32475476-DEL-106
chr3-32504951-DEL-60
chr3-32544041-DEL-139
chr3-32552042-DEL-120
chr3-32582437-DEL-124
chr3-32630465-DEL-90
chr3-32663418-DEL-61
chr3-32674218-DEL-68
chr3-32699800-DEL-65
chr3-32731465-DEL-355
chr3-32751846-DEL-2014
chr3-32772279-DEL-131
chr3-32777112-DEL-684
chr3-32788797-DEL-92
chr3-32791884-DEL-123

chr3-32799859-DEL-82
chr3-32806345-DEL-58
chr3-32816449-DEL-3543
chr3-32822380-DEL-70
chr3-32870384-DEL-58
chr3-32900306-DEL-51
chr3-32918260-DEL-56
chr3-32922955-DEL-89
chr3-32931872-DEL-66
chr3-32937121-DEL-125
chr3-32990625-DEL-96
chr3-33038738-DEL-66
chr3-33054696-DEL-56
chr3-33093469-DEL-68
chr3-33148872-DEL-748
chr3-33242561-DEL-136
chr3-33246164-DEL-130
chr3-33266215-DEL-90
chr3-33296973-DEL-9466
chr3-33296991-DEL-615
chr3-33298162-DEL-6282
chr3-33307169-DEL-54
chr3-33330554-DEL-397
chr3-33336939-DEL-3247
chr3-33337149-DEL-75
chr3-33382950-DEL-4431
chr3-33391589-DEL-93
chr3-33438786-DEL-455
chr3-33444805-DEL-3948
chr3-33508359-DEL-56
chr3-33514542-DEL-523
chr3-33515810-DEL-4202
chr3-33533884-DEL-237
chr3-33582373-DEL-6213
chr3-33600019-DEL-186
chr3-33651078-DEL-12422
chr3-33668235-DEL-131
chr3-33673101-DEL-9518
chr3-33683649-DEL-242
chr3-33686161-DEL-174
chr3-33687881-DEL-181
chr3-33812225-DEL-85
chr3-33849214-DEL-52
chr3-33882129-DEL-176
chr3-33895489-DEL-186
chr3-33897257-DEL-55
chr3-33901154-DEL-51
chr3-33919169-DEL-106
chr3-33946725-DEL-111
chr3-33976788-DEL-434
chr3-33983928-DEL-6182
chr3-33993226-DEL-762
chr3-34034939-DEL-137
chr3-34036021-DEL-487
chr3-34039526-DEL-509
chr3-34069606-DEL-51
chr3-34079962-DEL-176
chr3-34081897-DEL-220
chr3-34086365-DEL-173
chr3-34110672-DEL-312
chr3-34165673-DEL-60
chr3-34177304-DEL-1170
chr3-34182534-DEL-61
chr3-34187773-DEL-109
chr3-34221345-DEL-350
chr3-34257090-DEL-55
chr3-34287541-DEL-156
chr3-34313324-DEL-405
chr3-34317780-DEL-52
chr3-34350742-DEL-61
chr3-34430046-DEL-65
chr3-34444098-DEL-163
chr3-34479115-DEL-64
chr3-34508370-DEL-174
chr3-34543863-DEL-1008
chr3-34634039-DEL-52
chr3-34642330-DEL-52

chr3-34656160-DEL-68
chr3-34657309-DEL-117
chr3-34661429-DEL-112
chr3-34670469-DEL-197
chr3-34735975-DEL-93
chr3-34738325-DEL-140
chr3-34743790-DEL-861
chr3-34752799-DEL-62
chr3-34754311-DEL-92
chr3-34782918-DEL-212
chr3-34821979-DEL-57
chr3-34826304-DEL-298
chr3-34857798-DEL-353
chr3-34896065-DEL-381
chr3-34921014-DEL-336
chr3-34948665-DEL-88
chr3-35014476-DEL-94
chr3-35033277-DEL-246
chr3-35037926-DEL-740
chr3-35067655-DEL-457
chr3-35087139-DEL-81
chr3-35090100-DEL-754
chr3-35105370-DEL-2192
chr3-35106801-DEL-62
chr3-35118318-DEL-1320
chr3-35189086-DEL-51
chr3-35197355-DEL-403
chr3-35212050-DEL-175
chr3-35239269-DEL-310
chr3-35264023-DEL-96
chr3-35330134-DEL-68
chr3-35338413-DEL-291
chr3-35363413-DEL-76
chr3-35402046-DEL-98
chr3-35425065-DEL-335
chr3-35445785-DEL-113
chr3-35467716-DEL-5548
chr3-35497068-DEL-3584
chr3-35506638-DEL-276
chr3-35531093-DEL-52
chr3-35533547-DEL-334
chr3-35542926-DEL-504
chr3-35558568-DEL-938
chr3-35565995-DEL-85
chr3-35580722-DEL-56
chr3-35592594-DEL-1609
chr3-35642517-DEL-2121
chr3-35658140-DEL-401
chr3-35661263-DEL-6877
chr3-35670316-DEL-471
chr3-35692584-DEL-897
chr3-35697539-DEL-2578
chr3-35706709-DEL-1857
chr3-35719953-DEL-51
chr3-35729997-DEL-170
chr3-35734829-DEL-144
chr3-35744105-DEL-1137
chr3-35767864-DEL-174
chr3-35776876-DEL-610
chr3-35785450-DEL-60
chr3-35792778-DEL-2160
chr3-35816000-DEL-203
chr3-35849732-DEL-70
chr3-35882174-DEL-74
chr3-35950716-DEL-115
chr3-35962522-DEL-358
chr3-35971977-DEL-72
chr3-35993041-DEL-205
chr3-36035528-DEL-1138
chr3-36054184-DEL-60
chr3-36058275-DEL-104
chr3-36079802-DEL-143
chr3-36082212-DEL-1762
chr3-36122479-DEL-104
chr3-36125864-DEL-95
chr3-36148940-DEL-139
chr3-36155194-DEL-451

chr3-36185319-DEL-4715
chr3-36207056-DEL-144
chr3-36246039-DEL-594
chr3-36253073-DEL-57
chr3-36260379-DEL-1335
chr3-36276867-DEL-100
chr3-36278581-DEL-141
chr3-36312453-DEL-292
chr3-36322500-DEL-1205
chr3-36382509-DEL-138
chr3-36417254-DEL-2192
chr3-36443854-DEL-885
chr3-36448020-DEL-66
chr3-36469133-DEL-226
chr3-36469654-DEL-4451
chr3-36483342-DEL-116
chr3-36528296-DEL-118
chr3-36531491-DEL-88
chr3-36542867-DEL-3407
chr3-36558890-DEL-179
chr3-36559608-DEL-68
chr3-36572214-DEL-198
chr3-36634324-DEL-74
chr3-36641640-DEL-2478
chr3-36653835-DEL-1577
chr3-36681516-DEL-70
chr3-36707481-DEL-301
chr3-36778996-DEL-61
chr3-36799919-DEL-920
chr3-36816561-DEL-93
chr3-36839505-DEL-208
chr3-36852633-DEL-65
chr3-36856507-DEL-5419
chr3-36877307-DEL-62
chr3-36890423-DEL-3770
chr3-36901356-DEL-6432
chr3-37030092-DEL-86
chr3-37036024-DEL-75
chr3-37045129-DEL-396
chr3-37049932-DEL-60
chr3-37057023-DEL-115
chr3-37168611-DEL-58
chr3-37194845-DEL-670
chr3-37207537-DEL-323
chr3-37225291-DEL-139
chr3-37231603-DEL-58
chr3-37236251-DEL-64
chr3-37237077-DEL-65
chr3-37251465-DEL-55
chr3-37262985-DEL-203
chr3-37274744-DEL-553
chr3-37296467-DEL-809
chr3-37298753-DEL-55
chr3-37344229-DEL-2181
chr3-37428611-DEL-633
chr3-37438079-DEL-2637
chr3-37476563-DEL-87
chr3-37476824-DEL-625
chr3-37501429-DEL-684
chr3-37521315-DEL-198
chr3-37568951-DEL-792
chr3-37576056-DEL-1215
chr3-37590099-DEL-116
chr3-37621704-DEL-56
chr3-37638662-DEL-230
chr3-37709473-DEL-83
chr3-37739559-DEL-81
chr3-37759117-DEL-60
chr3-37827110-DEL-57
chr3-37855462-DEL-256
chr3-37856507-DEL-59
chr3-37867236-DEL-1501
chr3-37930959-DEL-196
chr3-37951555-DEL-60
chr3-38009332-DEL-106
chr3-38020659-DEL-88
chr3-38037274-DEL-1200

chr3-38043316-DEL-55
chr3-38073695-DEL-61
chr3-38074444-DEL-10274
chr3-38078034-DEL-2166
chr3-38086576-DEL-244
chr3-38089609-DEL-197
chr3-38095326-DEL-67
chr3-38110653-DEL-76
chr3-38135759-DEL-101
chr3-38162288-DEL-106
chr3-38187831-DEL-146
chr3-38191225-DEL-89
chr3-38218388-DEL-2250
chr3-38221752-DEL-1033
chr3-38235603-DEL-425
chr3-38237179-DEL-51
chr3-38245346-DEL-249
chr3-38264877-DEL-6685
chr3-38278288-DEL-75
chr3-38279702-DEL-76
chr3-38302741-DEL-185
chr3-38346931-DEL-82
chr3-38352292-DEL-69
chr3-38363374-DEL-1167
chr3-38392935-DEL-291
chr3-38404849-DEL-89
chr3-38424549-DEL-677
chr3-38445073-DEL-1423
chr3-38467223-DEL-54
chr3-38552136-DEL-68
chr3-38559707-DEL-403
chr3-38579915-DEL-72
chr3-38587425-DEL-79
chr3-38615041-DEL-71
chr3-38626910-DEL-194
chr3-38648923-DEL-178
chr3-38658764-DEL-136
chr3-38701576-DEL-100
chr3-38706449-DEL-400
chr3-38715711-DEL-51
chr3-38717012-DEL-215
chr3-38730486-DEL-53
chr3-38734313-DEL-845
chr3-38744072-DEL-842
chr3-38767906-DEL-52
chr3-38801013-DEL-377
chr3-38832791-DEL-1254
chr3-38834561-DEL-154
chr3-38841618-DEL-341
chr3-38915331-DEL-420
chr3-38918781-DEL-67
chr3-38920322-DEL-1571
chr3-38980606-DEL-130
chr3-39052768-DEL-54
chr3-39079947-DEL-6139
chr3-39118316-DEL-374
chr3-39123713-DEL-896
chr3-39177492-DEL-394
chr3-39215885-DEL-1470
chr3-39289950-DEL-214
chr3-39320648-DEL-182
chr3-39331274-DEL-72
chr3-39335796-DEL-1671
chr3-39368390-DEL-110
chr3-39389554-DEL-81
chr3-39406435-DEL-96
chr3-39416680-DEL-51
chr3-39476268-DEL-391
chr3-39511143-DEL-176
chr3-39530978-DEL-56
chr3-39598221-DEL-12429
chr3-39617845-DEL-266
chr3-39623552-DEL-210
chr3-39652284-DEL-76
chr3-39693743-DEL-779
chr3-39694585-DEL-2326
chr3-39696955-DEL-2524

chr3-39699527-DEL-1342
chr3-39716378-DEL-995
chr3-39746918-DEL-96
chr3-39748615-DEL-53
chr3-39756963-DEL-1370
chr3-39769299-DEL-58
chr3-39782141-DEL-139
chr3-39803283-DEL-57
chr3-39822524-DEL-64
chr3-39876730-DEL-66
chr3-39906933-DEL-1423
chr3-39911860-DEL-6362
chr3-39922327-DEL-2005
chr3-39927204-DEL-53
chr3-39931260-DEL-6488
chr3-39980823-DEL-536
chr3-40010646-DEL-263
chr3-40013544-DEL-78
chr3-40074894-DEL-297
chr3-40077630-DEL-58
chr3-40090713-DEL-67
chr3-40091836-DEL-57
chr3-40102308-DEL-56
chr3-40105631-DEL-64
chr3-40140528-DEL-64
chr3-40142421-DEL-78
chr3-40167196-DEL-166
chr3-40174167-DEL-56
chr3-40227356-DEL-54
chr3-40231647-DEL-840
chr3-40274176-DEL-553
chr3-40306200-DEL-551
chr3-40348250-DEL-709
chr3-40354579-DEL-51
chr3-40365637-DEL-120
chr3-40366120-DEL-6169
chr3-40408577-DEL-666
chr3-40442347-DEL-364
chr3-40483327-DEL-71
chr3-40504378-DEL-268
chr3-40506403-DEL-60
chr3-40527856-DEL-69
chr3-40550769-DEL-72
chr3-40569837-DEL-175
chr3-40576481-DEL-56
chr3-40589228-DEL-150
chr3-40603873-DEL-2008
chr3-40604592-DEL-197
chr3-40605117-DEL-573
chr3-40605235-DEL-143
chr3-40616312-DEL-657
chr3-40627496-DEL-72
chr3-40634339-DEL-2485
chr3-40638168-DEL-903
chr3-40644003-DEL-106
chr3-40652789-DEL-125
chr3-40691233-DEL-170
chr3-40715829-DEL-2563
chr3-40758498-DEL-117
chr3-40811324-DEL-670
chr3-40812681-DEL-1896
chr3-40819099-DEL-161
chr3-40845179-DEL-401
chr3-40869721-DEL-60
chr3-40880499-DEL-277
chr3-40908171-DEL-1094
chr3-40952597-DEL-68
chr3-40987336-DEL-54
chr3-41008686-DEL-51
chr3-41025717-DEL-74
chr3-41033707-DEL-161
chr3-41043187-DEL-126
chr3-41080390-DEL-94
chr3-41081883-DEL-60
chr3-41091459-DEL-168
chr3-41175754-DEL-72
chr3-41180238-DEL-72

chr3-41247928-DEL-218
chr3-41262602-DEL-156
chr3-41273537-DEL-70
chr3-41343592-DEL-76
chr3-41353146-DEL-90
chr3-41354912-DEL-53
chr3-41357626-DEL-77
chr3-41386112-DEL-109
chr3-41396086-DEL-211
chr3-41399458-DEL-54
chr3-41404512-DEL-115
chr3-41429240-DEL-87
chr3-41456179-DEL-100
chr3-41456765-DEL-98
chr3-41477960-DEL-1480
chr3-41483409-DEL-90
chr3-41492892-DEL-142
chr3-41512608-DEL-187
chr3-41513669-DEL-109
chr3-41549705-DEL-64
chr3-41619362-DEL-107
chr3-41626477-DEL-62
chr3-41662483-DEL-251
chr3-41686765-DEL-66
chr3-41735031-DEL-56
chr3-41735187-DEL-4766
chr3-41735729-DEL-23308
chr3-41749137-DEL-3698
chr3-41772591-DEL-353
chr3-41804926-DEL-1487
chr3-41824672-DEL-965
chr3-41824732-DEL-5216
chr3-41829772-DEL-421
chr3-41834730-DEL-929
chr3-41837387-DEL-9049
chr3-41847551-DEL-552
chr3-41857194-DEL-52
chr3-41891631-DEL-605
chr3-41940059-DEL-56
chr3-41947802-DEL-6141
chr3-42015516-DEL-405
chr3-42039109-DEL-68
chr3-42040788-DEL-1984
chr3-42051054-DEL-79
chr3-42052363-DEL-78
chr3-42090322-DEL-381
chr3-42135252-DEL-198
chr3-42162579-DEL-303
chr3-42181352-DEL-247
chr3-42196604-DEL-220
chr3-42215959-DEL-106
chr3-42227500-DEL-1168
chr3-42238356-DEL-90
chr3-42286750-DEL-560
chr3-42293831-DEL-1654
chr3-42306456-DEL-105
chr3-42351447-DEL-52
chr3-42365929-DEL-310
chr3-42368116-DEL-120
chr3-42384471-DEL-450
chr3-42386718-DEL-2087
chr3-42412048-DEL-664
chr3-42430734-DEL-1721
chr3-42448622-DEL-91
chr3-42475978-DEL-70
chr3-42499022-DEL-130
chr3-42555511-DEL-110
chr3-42560084-DEL-6040
chr3-42571922-DEL-125
chr3-42574943-DEL-359
chr3-42581901-DEL-191
chr3-42593023-DEL-1491
chr3-42598986-DEL-100
chr3-42636763-DEL-1200
chr3-42638227-DEL-175
chr3-42652487-DEL-72
chr3-42656404-DEL-938

chr3-42663288-DEL-214
chr3-42668593-DEL-153
chr3-42678492-DEL-620
chr3-42718145-DEL-266
chr3-42735306-DEL-557
chr3-42747850-DEL-1294
chr3-42800381-DEL-771
chr3-42815649-DEL-53
chr3-42833555-DEL-140
chr3-42834070-DEL-159
chr3-42914249-DEL-280
chr3-42925658-DEL-558
chr3-42958005-DEL-400
chr3-42986847-DEL-55
chr3-43001639-DEL-313
chr3-43016474-DEL-65
chr3-43041723-DEL-240
chr3-43044605-DEL-2214
chr3-43051267-DEL-86
chr3-43077340-DEL-169
chr3-43129968-DEL-198
chr3-43149429-DEL-525
chr3-43161844-DEL-1143
chr3-43163424-DEL-492
chr3-43171457-DEL-852
chr3-43173584-DEL-147
chr3-43176245-DEL-64
chr3-43186831-DEL-1030
chr3-43192601-DEL-114
chr3-43249103-DEL-54
chr3-43264477-DEL-970
chr3-43282043-DEL-515
chr3-43290367-DEL-91
chr3-43317780-DEL-100
chr3-43358439-DEL-123
chr3-43391286-DEL-54
chr3-43414416-DEL-730
chr3-43420913-DEL-4819
chr3-43448515-DEL-3201
chr3-43492654-DEL-232
chr3-43498424-DEL-1543
chr3-43521586-DEL-904
chr3-43530209-DEL-569
chr3-43563431-DEL-872
chr3-43571065-DEL-571
chr3-43579512-DEL-56
chr3-43586441-DEL-54
chr3-43613073-DEL-224
chr3-43618151-DEL-58
chr3-43656791-DEL-203
chr3-43658491-DEL-357
chr3-43685192-DEL-89
chr3-43695515-DEL-1076
chr3-43713007-DEL-1704
chr3-43716948-DEL-382
chr3-43722621-DEL-3988
chr3-43734535-DEL-74
chr3-43750882-DEL-479
chr3-43751796-DEL-85
chr3-43768916-DEL-204
chr3-43780566-DEL-6061
chr3-43808754-DEL-7097
chr3-43817682-DEL-225
chr3-43817715-DEL-832
chr3-43819107-DEL-3313
chr3-43832764-DEL-6417
chr3-43838802-DEL-214
chr3-43846895-DEL-962
chr3-43856278-DEL-2396
chr3-43866658-DEL-62
chr3-43876242-DEL-314
chr3-43899915-DEL-52
chr3-43908658-DEL-6215
chr3-43922612-DEL-11029
chr3-43930977-DEL-54
chr3-43959035-DEL-6787
chr3-43972506-DEL-176

chr3-43975525-DEL-645
chr3-43978772-DEL-5332
chr3-43998157-DEL-337
chr3-43998158-DEL-2205
chr3-44006064-DEL-1141
chr3-44025674-DEL-1668
chr3-44028971-DEL-268
chr3-44032975-DEL-1042
chr3-44114380-DEL-74
chr3-44115672-DEL-660
chr3-44146594-DEL-214
chr3-44148014-DEL-78
chr3-44153200-DEL-1328
chr3-44172351-DEL-66
chr3-44201896-DEL-208
chr3-44211380-DEL-8618
chr3-44221238-DEL-396
chr3-44253967-DEL-542
chr3-44275895-DEL-58
chr3-44298824-DEL-2134
chr3-44314454-DEL-1227
chr3-44322494-DEL-138
chr3-44324364-DEL-392
chr3-44351546-DEL-108
chr3-44366544-DEL-98
chr3-44371686-DEL-960
chr3-44372798-DEL-405
chr3-44399525-DEL-184
chr3-44403022-DEL-684
chr3-44411717-DEL-6300
chr3-44419329-DEL-56
chr3-44446357-DEL-62
chr3-44447122-DEL-668
chr3-44452077-DEL-174
chr3-44458171-DEL-6329
chr3-44467314-DEL-1586
chr3-44476147-DEL-165
chr3-44479587-DEL-345
chr3-44491416-DEL-1401
chr3-44496909-DEL-2881
chr3-44501809-DEL-560
chr3-44545258-DEL-55
chr3-44549501-DEL-5871
chr3-44567777-DEL-3544
chr3-44601415-DEL-6069
chr3-44647480-DEL-499
chr3-44652175-DEL-164
chr3-44682636-DEL-62
chr3-44687467-DEL-345
chr3-44690629-DEL-797
chr3-44693496-DEL-967
chr3-44700165-DEL-334
chr3-44702852-DEL-93
chr3-44724372-DEL-80
chr3-44768236-DEL-653
chr3-44786319-DEL-56
chr3-44796927-DEL-12815
chr3-44799624-DEL-115
chr3-44799951-DEL-573
chr3-44838875-DEL-6304
chr3-44871194-DEL-74
chr3-44877625-DEL-76
chr3-44885053-DEL-829
chr3-44901687-DEL-55
chr3-44905471-DEL-2085
chr3-44917586-DEL-54
chr3-44927261-DEL-454
chr3-44928616-DEL-534
chr3-44945514-DEL-969
chr3-44948286-DEL-80
chr3-44951027-DEL-55
chr3-44956495-DEL-459
chr3-44964078-DEL-100
chr3-44966105-DEL-323
chr3-44968669-DEL-191
chr3-44969842-DEL-198
chr3-44972284-DEL-51

chr3-44973306-DEL-3512
chr3-44983591-DEL-782
chr3-45019384-DEL-100
chr3-45022967-DEL-249
chr3-45027727-DEL-191
chr3-45039943-DEL-1137
chr3-45044151-DEL-54
chr3-45078497-DEL-2006
chr3-45078912-DEL-2869
chr3-45103533-DEL-191
chr3-45108797-DEL-7261
chr3-45114928-DEL-601
chr3-45139979-DEL-598
chr3-45144254-DEL-114
chr3-45144695-DEL-390
chr3-45198710-DEL-720
chr3-45199468-DEL-314
chr3-45233527-DEL-109
chr3-45241454-DEL-710
chr3-45287130-DEL-124
chr3-45416986-DEL-58
chr3-45421047-DEL-1857
chr3-45452031-DEL-404
chr3-45455934-DEL-578
chr3-45457138-DEL-469
chr3-45466483-DEL-54
chr3-45475866-DEL-58
chr3-45479189-DEL-88
chr3-45484407-DEL-67
chr3-45502660-DEL-74
chr3-45526477-DEL-51
chr3-45532652-DEL-13748
chr3-45533278-DEL-424
chr3-45540042-DEL-270
chr3-45555050-DEL-91
chr3-45586819-DEL-81
chr3-45617675-DEL-5500
chr3-45628481-DEL-55
chr3-45644488-DEL-162
chr3-45684789-DEL-80
chr3-45721552-DEL-54
chr3-45747658-DEL-614
chr3-45753175-DEL-127
chr3-45754743-DEL-65
chr3-45774048-DEL-1625
chr3-45784505-DEL-1347
chr3-45802743-DEL-2286
chr3-45855932-DEL-369
chr3-45857492-DEL-182
chr3-45876659-DEL-77
chr3-45900844-DEL-70
chr3-45902157-DEL-316
chr3-45922587-DEL-291
chr3-45926022-DEL-733
chr3-45942265-DEL-1870
chr3-45979973-DEL-110
chr3-45989779-DEL-84
chr3-46007386-DEL-76
chr3-46014176-DEL-127
chr3-46020365-DEL-1810
chr3-46030876-DEL-206
chr3-46034797-DEL-358
chr3-46062628-DEL-327
chr3-46072266-DEL-399
chr3-46119644-DEL-441
chr3-46161687-DEL-12392
chr3-46169779-DEL-163
chr3-46180232-DEL-190
chr3-46190228-DEL-445
chr3-46194107-DEL-5372
chr3-46197490-DEL-706
chr3-46197799-DEL-2649
chr3-46198536-DEL-279
chr3-46200516-DEL-2687
chr3-46202157-DEL-846
chr3-46218921-DEL-1672
chr3-46224341-DEL-54

chr3-46226289-DEL-58
chr3-46228877-DEL-6450
chr3-46241545-DEL-580
chr3-46276675-DEL-521
chr3-46296054-DEL-51
chr3-46309161-DEL-52
chr3-46319566-DEL-774
chr3-46342295-DEL-406
chr3-46405383-DEL-75
chr3-46408762-DEL-6007
chr3-46468528-DEL-59
chr3-46531891-DEL-129
chr3-46537196-DEL-104
chr3-46578681-DEL-232
chr3-46589902-DEL-255
chr3-46605314-DEL-81
chr3-46635700-DEL-1489
chr3-46650277-DEL-60
chr3-46668701-DEL-401
chr3-46677526-DEL-73
chr3-46710506-DEL-65
chr3-46772804-DEL-56
chr3-46784153-DEL-519
chr3-46805107-DEL-102
chr3-46810228-DEL-762
chr3-46811144-DEL-143
chr3-46811535-DEL-716
chr3-46821135-DEL-57
chr3-46875497-DEL-6469
chr3-46884337-DEL-6403
chr3-46902008-DEL-97
chr3-46945777-DEL-1620
chr3-46960082-DEL-152
chr3-46963111-DEL-77
chr3-46968731-DEL-581
chr3-46988012-DEL-57
chr3-47006851-DEL-2680
chr3-47008374-DEL-621
chr3-47009428-DEL-214
chr3-47034382-DEL-495
chr3-47049138-DEL-75
chr3-47058264-DEL-219
chr3-47067677-DEL-59
chr3-47084749-DEL-83
chr3-47097346-DEL-279
chr3-47103381-DEL-1322
chr3-47121181-DEL-82
chr3-47141116-DEL-2621
chr3-47148696-DEL-470
chr3-47191395-DEL-491
chr3-47198922-DEL-1102
chr3-47200430-DEL-668
chr3-47208010-DEL-3247
chr3-47237376-DEL-6407
chr3-47268239-DEL-1040
chr3-47287851-DEL-60
chr3-47315843-DEL-1684
chr3-47316034-DEL-841
chr3-47317531-DEL-4570
chr3-47317585-DEL-3883
chr3-47321774-DEL-328
chr3-47323430-DEL-9525
chr3-47326390-DEL-2907
chr3-47329327-DEL-3313
chr3-47336258-DEL-204
chr3-47349244-DEL-158
chr3-47434061-DEL-140
chr3-47458752-DEL-72
chr3-47488228-DEL-609
chr3-47489820-DEL-6217
chr3-47496819-DEL-76
chr3-47545051-DEL-858
chr3-47554714-DEL-90
chr3-47581877-DEL-1433
chr3-47585041-DEL-339
chr3-47604106-DEL-190
chr3-47616172-DEL-51

chr3-47620239-DEL-59
chr3-47633129-DEL-1644
chr3-47639139-DEL-92
chr3-47665820-DEL-1966
chr3-47670879-DEL-52
chr3-47700768-DEL-210
chr3-47705013-DEL-59
chr3-47719300-DEL-226
chr3-47721280-DEL-392
chr3-47733620-DEL-184
chr3-47750829-DEL-344
chr3-47752160-DEL-1889
chr3-47772703-DEL-1312
chr3-47779128-DEL-368
chr3-47794989-DEL-51
chr3-47806692-DEL-6192
chr3-47813560-DEL-206
chr3-47815237-DEL-56
chr3-47826808-DEL-4846
chr3-47837328-DEL-1368
chr3-47841542-DEL-435
chr3-47866947-DEL-60
chr3-47867744-DEL-269
chr3-47890058-DEL-810
chr3-47896961-DEL-59
chr3-47897707-DEL-312
chr3-47917244-DEL-208
chr3-47920158-DEL-2941
chr3-47931438-DEL-1300
chr3-47954609-DEL-2500
chr3-47981275-DEL-6028
chr3-47995426-DEL-701
chr3-47999492-DEL-67
chr3-48026589-DEL-55
chr3-48033010-DEL-82
chr3-48082023-DEL-384
chr3-48098808-DEL-114
chr3-48106746-DEL-149
chr3-48108632-DEL-92
chr3-48135658-DEL-96
chr3-48142923-DEL-109
chr3-48163075-DEL-509
chr3-48177530-DEL-123
chr3-48188370-DEL-982
chr3-48204622-DEL-104
chr3-48236104-DEL-94
chr3-48237848-DEL-1007
chr3-48248968-DEL-210
chr3-48253791-DEL-5918
chr3-48263400-DEL-65
chr3-48269516-DEL-6480
chr3-48291925-DEL-57
chr3-48295132-DEL-65
chr3-48302305-DEL-4132
chr3-48337401-DEL-481
chr3-48388250-DEL-7032
chr3-48399534-DEL-6237
chr3-48413582-DEL-2925
chr3-48436392-DEL-59
chr3-48442325-DEL-68
chr3-48454501-DEL-380
chr3-48538500-DEL-267
chr3-48586696-DEL-242
chr3-48631245-DEL-72
chr3-48660023-DEL-231
chr3-48673711-DEL-56
chr3-48694426-DEL-189
chr3-48716984-DEL-111
chr3-48723388-DEL-124
chr3-48746328-DEL-62
chr3-48748329-DEL-1371
chr3-48757392-DEL-402
chr3-48769500-DEL-538
chr3-48775267-DEL-88
chr3-48791946-DEL-76
chr3-48818981-DEL-242
chr3-48819439-DEL-356

chr3-48834753-DEL-202
chr3-48849979-DEL-3704
chr3-48909081-DEL-68
chr3-48915884-DEL-1045
chr3-48938775-DEL-52
chr3-48943080-DEL-94
chr3-48950690-DEL-365
chr3-48955600-DEL-5928
chr3-48969019-DEL-2505
chr3-48981476-DEL-3068
chr3-48986907-DEL-114
chr3-49001191-DEL-2926
chr3-49008265-DEL-3011
chr3-49016987-DEL-65
chr3-49020805-DEL-294
chr3-49067814-DEL-72
chr3-49109821-DEL-81
chr3-49134889-DEL-3411
chr3-49139155-DEL-1394
chr3-49144910-DEL-209
chr3-49145526-DEL-91
chr3-49155951-DEL-70
chr3-49161167-DEL-97
chr3-49200141-DEL-307
chr3-49209634-DEL-59
chr3-49232330-DEL-205
chr3-49250968-DEL-78
chr3-49261956-DEL-232
chr3-49299702-DEL-114
chr3-49300882-DEL-117
chr3-49318828-DEL-571
chr3-49360599-DEL-13580
chr3-49384284-DEL-100
chr3-49385070-DEL-5173
chr3-49389952-DEL-12020
chr3-49398006-DEL-6305
chr3-49408942-DEL-638
chr3-49412748-DEL-6408
chr3-49420245-DEL-56
chr3-49424481-DEL-2518
chr3-49429049-DEL-1397
chr3-49430543-DEL-6784
chr3-49445511-DEL-59
chr3-49486225-DEL-63
chr3-49494234-DEL-199
chr3-49496724-DEL-63
chr3-49501285-DEL-1878
chr3-49522896-DEL-80
chr3-49532365-DEL-1551
chr3-49574197-DEL-1289
chr3-49582057-DEL-90
chr3-49582597-DEL-1749
chr3-49657429-DEL-58
chr3-49662129-DEL-116
chr3-49681745-DEL-6669
chr3-49714648-DEL-60
chr3-49767005-DEL-730
chr3-49771316-DEL-12336
chr3-49816031-DEL-86
chr3-49878414-DEL-871
chr3-49881722-DEL-139
chr3-49897388-DEL-57
chr3-49918850-DEL-52
chr3-49947277-DEL-1118
chr3-49993834-DEL-310
chr3-50054880-DEL-420
chr3-50069099-DEL-134
chr3-50079473-DEL-984
chr3-50082904-DEL-157
chr3-50087831-DEL-1150
chr3-50088622-DEL-62
chr3-50096614-DEL-697
chr3-50141503-DEL-108
chr3-50159147-DEL-143
chr3-50194389-DEL-65
chr3-50210486-DEL-337
chr3-50308500-DEL-81

chr3-50396603-DEL-57
chr3-50415249-DEL-94
chr3-50431527-DEL-86
chr3-50442765-DEL-1697
chr3-50445083-DEL-1270
chr3-50465456-DEL-62
chr3-50506273-DEL-93
chr3-50511403-DEL-6035
chr3-50534071-DEL-100
chr3-50571455-DEL-55
chr3-50617019-DEL-64
chr3-50617083-DEL-608
chr3-50696757-DEL-155
chr3-50711527-DEL-759
chr3-50719830-DEL-1637
chr3-50720693-DEL-735
chr3-50736165-DEL-1340
chr3-50787912-DEL-70
chr3-50795272-DEL-62
chr3-50814277-DEL-156
chr3-50826045-DEL-504
chr3-50829188-DEL-65
chr3-50897254-DEL-68
chr3-50922242-DEL-69
chr3-50970916-DEL-173
chr3-51033215-DEL-178
chr3-51060304-DEL-169
chr3-51078486-DEL-256
chr3-51097425-DEL-138
chr3-51118113-DEL-117
chr3-51142721-DEL-770
chr3-51180174-DEL-160
chr3-51205497-DEL-74
chr3-51206139-DEL-225
chr3-51236678-DEL-94
chr3-51237853-DEL-451
chr3-51283355-DEL-271
chr3-51297202-DEL-135
chr3-51304810-DEL-208
chr3-51311421-DEL-513
chr3-51333797-DEL-238
chr3-51336582-DEL-69
chr3-51341408-DEL-227
chr3-51349206-DEL-53
chr3-51375518-DEL-141
chr3-51394593-DEL-105
chr3-51397477-DEL-56
chr3-51403447-DEL-211
chr3-51419956-DEL-419
chr3-51420852-DEL-394
chr3-51431528-DEL-137
chr3-51444831-DEL-196
chr3-51449445-DEL-79
chr3-51461984-DEL-280
chr3-51477465-DEL-179
chr3-51480903-DEL-200
chr3-51494381-DEL-56
chr3-51515172-DEL-80
chr3-51533028-DEL-59
chr3-51554494-DEL-71
chr3-51561769-DEL-191
chr3-51568336-DEL-77
chr3-51576652-DEL-1161
chr3-51600249-DEL-72
chr3-51615022-DEL-145
chr3-51619737-DEL-99
chr3-51673516-DEL-724
chr3-51691144-DEL-480
chr3-51740144-DEL-90
chr3-51783420-DEL-92
chr3-51790003-DEL-96
chr3-51804971-DEL-238
chr3-51818106-DEL-167
chr3-51829590-DEL-103
chr3-51881188-DEL-1016
chr3-51915969-DEL-163
chr3-51964611-DEL-821

chr3-51969011-DEL-71
chr3-52017470-DEL-208
chr3-52032383-DEL-181
chr3-52051895-DEL-70
chr3-52059840-DEL-74
chr3-52085713-DEL-105
chr3-52096221-DEL-103
chr3-52158150-DEL-76
chr3-52191044-DEL-75
chr3-52192755-DEL-54
chr3-52204983-DEL-71
chr3-52215400-DEL-701
chr3-52235314-DEL-1031
chr3-52235314-DEL-945
chr3-52236775-DEL-86
chr3-52261389-DEL-98
chr3-52274068-DEL-294
chr3-52323630-DEL-54
chr3-52333320-DEL-83
chr3-52337474-DEL-71
chr3-52354715-DEL-58
chr3-52356799-DEL-192
chr3-52365143-DEL-60
chr3-52392341-DEL-176
chr3-52394850-DEL-122
chr3-52396263-DEL-790
chr3-52425240-DEL-57
chr3-52458650-DEL-1031
chr3-52527453-DEL-67
chr3-52553289-DEL-98
chr3-52556060-DEL-66
chr3-52642531-DEL-115
chr3-52645433-DEL-51
chr3-52688377-DEL-51
chr3-52694989-DEL-211
chr3-52704018-DEL-151
chr3-52730933-DEL-81
chr3-52765039-DEL-365
chr3-52775805-DEL-81
chr3-52837470-DEL-60
chr3-52949910-DEL-74
chr3-52961358-DEL-767
chr3-52968090-DEL-410
chr3-53035506-DEL-65
chr3-53036843-DEL-6789
chr3-53043767-DEL-1604
chr3-53077683-DEL-2233
chr3-53124104-DEL-66
chr3-53127238-DEL-66
chr3-53140920-DEL-214
chr3-53162211-DEL-425
chr3-53163715-DEL-52
chr3-53174276-DEL-128
chr3-53179449-DEL-61
chr3-53181632-DEL-96
chr3-53195805-DEL-64
chr3-53231266-DEL-85
chr3-53279901-DEL-195
chr3-53281935-DEL-79
chr3-53284062-DEL-140
chr3-53305965-DEL-66
chr3-53317199-DEL-88
chr3-53334161-DEL-6310
chr3-53358003-DEL-442
chr3-53370032-DEL-63
chr3-53375579-DEL-850
chr3-53400794-DEL-264
chr3-53403158-DEL-99
chr3-53406392-DEL-130
chr3-53406874-DEL-477
chr3-53409443-DEL-130
chr3-53419089-DEL-59
chr3-53422161-DEL-227
chr3-53426391-DEL-77
chr3-53442298-DEL-60
chr3-53451555-DEL-95
chr3-53456420-DEL-283

chr3-53461371-DEL-127
chr3-53489552-DEL-199
chr3-53520229-DEL-4303
chr3-53536714-DEL-195
chr3-53548774-DEL-2233
chr3-53569673-DEL-64
chr3-53590969-DEL-136
chr3-53592866-DEL-892
chr3-53603773-DEL-780
chr3-53616008-DEL-723
chr3-53619498-DEL-1346
chr3-53653876-DEL-102
chr3-53675366-DEL-70
chr3-53680833-DEL-90
chr3-53693344-DEL-202
chr3-53710115-DEL-3866
chr3-53743079-DEL-709
chr3-53743601-DEL-6423
chr3-53759913-DEL-469
chr3-53768104-DEL-155
chr3-53785448-DEL-317
chr3-53795549-DEL-59
chr3-53800936-DEL-62
chr3-53806223-DEL-212
chr3-53830281-DEL-58
chr3-53832778-DEL-250
chr3-53859686-DEL-122
chr3-53903361-DEL-84
chr3-53929309-DEL-125
chr3-53966106-DEL-487
chr3-54006741-DEL-148
chr3-54010809-DEL-83
chr3-54029471-DEL-621
chr3-54030074-DEL-6425
chr3-54036215-DEL-213
chr3-54066986-DEL-156
chr3-54079044-DEL-136
chr3-54093321-DEL-58
chr3-54098507-DEL-57
chr3-54126390-DEL-819
chr3-54137504-DEL-2073
chr3-54183232-DEL-342
chr3-54183881-DEL-60
chr3-54238731-DEL-576
chr3-54246511-DEL-166
chr3-54249138-DEL-1206
chr3-54261180-DEL-92
chr3-54263925-DEL-744
chr3-54283655-DEL-52
chr3-54327392-DEL-104
chr3-54345158-DEL-188
chr3-54390657-DEL-508
chr3-54399796-DEL-571
chr3-54404719-DEL-2696
chr3-54440210-DEL-51
chr3-54441012-DEL-125
chr3-54458271-DEL-55
chr3-54474978-DEL-103
chr3-54479880-DEL-72
chr3-54484545-DEL-65
chr3-54498103-DEL-909
chr3-54503792-DEL-450
chr3-54522944-DEL-871
chr3-54545143-DEL-1194
chr3-54585991-DEL-441
chr3-54601594-DEL-73
chr3-54611034-DEL-187
chr3-54642530-DEL-101
chr3-54684188-DEL-54
chr3-54695413-DEL-82
chr3-54716420-DEL-160
chr3-54829093-DEL-51
chr3-54837809-DEL-74
chr3-54858960-DEL-1002
chr3-54903875-DEL-900
chr3-54906370-DEL-2494
chr3-54915799-DEL-116

chr3-54934471-DEL-56
chr3-54941042-DEL-69
chr3-54948781-DEL-66
chr3-54964157-DEL-919
chr3-54967906-DEL-61
chr3-54995628-DEL-56
chr3-55009554-DEL-398
chr3-55057663-DEL-69
chr3-55069616-DEL-492
chr3-55111431-DEL-192
chr3-55118189-DEL-878
chr3-55142239-DEL-112
chr3-55146104-DEL-69
chr3-55156935-DEL-75
chr3-55177907-DEL-74
chr3-55192221-DEL-51
chr3-55240632-DEL-79
chr3-55248400-DEL-68
chr3-55251356-DEL-223
chr3-55275156-DEL-75
chr3-55276463-DEL-127
chr3-55289425-DEL-94
chr3-55314400-DEL-56
chr3-55315838-DEL-131
chr3-55323652-DEL-252
chr3-55335090-DEL-80
chr3-55388521-DEL-141
chr3-55494669-DEL-4091
chr3-55517317-DEL-212
chr3-55518816-DEL-58
chr3-55527645-DEL-2989
chr3-55541857-DEL-148
chr3-55609688-DEL-290
chr3-55628346-DEL-55
chr3-55629728-DEL-78
chr3-55633678-DEL-2457
chr3-55729783-DEL-118
chr3-55751995-DEL-3763
chr3-55757185-DEL-134
chr3-55817082-DEL-1575
chr3-55832354-DEL-66
chr3-55844864-DEL-62
chr3-55878338-DEL-6708
chr3-55897226-DEL-55
chr3-55902026-DEL-61
chr3-55966424-DEL-920
chr3-56004545-DEL-122
chr3-56042987-DEL-104
chr3-56070548-DEL-165
chr3-56101503-DEL-74
chr3-56116912-DEL-214
chr3-56140080-DEL-62
chr3-56154345-DEL-208
chr3-56167964-DEL-159
chr3-56233133-DEL-402
chr3-56234880-DEL-162
chr3-56241650-DEL-2002
chr3-56260156-DEL-244
chr3-56270609-DEL-1062
chr3-56272147-DEL-401
chr3-56276621-DEL-134
chr3-56286188-DEL-75
chr3-56333059-DEL-641
chr3-56369198-DEL-378
chr3-56386545-DEL-286
chr3-56416310-DEL-100
chr3-56449620-DEL-519
chr3-56452386-DEL-214
chr3-56454658-DEL-289
chr3-56455177-DEL-597
chr3-56457013-DEL-1166
chr3-56458217-DEL-2135
chr3-56469093-DEL-1240
chr3-56498166-DEL-129
chr3-56501179-DEL-220
chr3-56521979-DEL-311
chr3-56531547-DEL-82

chr3-56532390-DEL-643
chr3-56538475-DEL-1118
chr3-56552666-DEL-192
chr3-56555907-DEL-61
chr3-56573857-DEL-282
chr3-56578921-DEL-1097
chr3-56607202-DEL-52
chr3-56649738-DEL-9354
chr3-56706455-DEL-528
chr3-56797839-DEL-76
chr3-56810196-DEL-109
chr3-56811298-DEL-6456
chr3-56818270-DEL-1256
chr3-56821749-DEL-415
chr3-56882301-DEL-2118
chr3-56905138-DEL-82
chr3-56922473-DEL-779
chr3-56928083-DEL-2585
chr3-56947913-DEL-100
chr3-56966797-DEL-10833
chr3-56980380-DEL-478
chr3-56983158-DEL-54
chr3-57027973-DEL-1322
chr3-57063852-DEL-355
chr3-57159584-DEL-85
chr3-57162384-DEL-2770
chr3-57167881-DEL-227
chr3-57187962-DEL-630
chr3-57199822-DEL-108
chr3-57246999-DEL-891
chr3-57279352-DEL-7438
chr3-57300521-DEL-83
chr3-57315828-DEL-58
chr3-57321932-DEL-1095
chr3-57324613-DEL-93
chr3-57335691-DEL-666
chr3-57352414-DEL-227
chr3-57353890-DEL-133
chr3-57531635-DEL-63
chr3-57535511-DEL-142
chr3-57541178-DEL-112
chr3-57586412-DEL-56
chr3-57597240-DEL-172
chr3-57605194-DEL-160
chr3-57605845-DEL-85
chr3-57606410-DEL-600
chr3-57607765-DEL-6218
chr3-57614114-DEL-105
chr3-57651473-DEL-81
chr3-57664833-DEL-601
chr3-57672510-DEL-158
chr3-57718883-DEL-404
chr3-57722248-DEL-493
chr3-57723830-DEL-62
chr3-57736546-DEL-3297
chr3-57743992-DEL-56
chr3-57775699-DEL-880
chr3-57794802-DEL-2507
chr3-57799126-DEL-144
chr3-57834178-DEL-1373
chr3-57834207-DEL-2465
chr3-57836482-DEL-1207
chr3-57849225-DEL-338
chr3-57898542-DEL-86
chr3-57916192-DEL-172
chr3-57950874-DEL-58
chr3-57958533-DEL-1880
chr3-57973487-DEL-2039
chr3-57977490-DEL-183
chr3-57981992-DEL-86
chr3-58049468-DEL-113
chr3-58081102-DEL-98
chr3-58087825-DEL-58
chr3-58094263-DEL-208
chr3-58136698-DEL-518
chr3-58159290-DEL-242
chr3-58160208-DEL-548

chr3-58188187-DEL-165
chr3-58236720-DEL-83
chr3-58319588-DEL-1247
chr3-58356143-DEL-54
chr3-58381787-DEL-257
chr3-58384306-DEL-1937
chr3-58386930-DEL-66
chr3-58397243-DEL-271
chr3-58436654-DEL-525
chr3-58449542-DEL-842
chr3-58458373-DEL-868
chr3-58469619-DEL-106
chr3-58473860-DEL-196
chr3-58474854-DEL-6706
chr3-58478364-DEL-341
chr3-58481271-DEL-376
chr3-58482824-DEL-121
chr3-58486434-DEL-154
chr3-58488050-DEL-306
chr3-58509347-DEL-164
chr3-58522217-DEL-107
chr3-58529220-DEL-161
chr3-58601743-DEL-100
chr3-58641645-DEL-58
chr3-58649109-DEL-86
chr3-58651006-DEL-814
chr3-58663712-DEL-62
chr3-58677989-DEL-63
chr3-58679380-DEL-922
chr3-58684781-DEL-56
chr3-58750418-DEL-103
chr3-58768282-DEL-119
chr3-58891616-DEL-51
chr3-58896420-DEL-378
chr3-58903604-DEL-115
chr3-58923296-DEL-60
chr3-58939901-DEL-276
chr3-58953724-DEL-463
chr3-58988337-DEL-779
chr3-59001527-DEL-1386
chr3-59011838-DEL-141
chr3-59017079-DEL-157
chr3-59025253-DEL-62
chr3-59030495-DEL-65
chr3-59062171-DEL-171
chr3-59133347-DEL-144
chr3-59150157-DEL-334
chr3-59179395-DEL-478
chr3-59266717-DEL-517
chr3-59301759-DEL-6449
chr3-59326382-DEL-6067
chr3-59341345-DEL-66
chr3-59382133-DEL-58
chr3-59390894-DEL-66
chr3-59396365-DEL-6228
chr3-59419162-DEL-230
chr3-59419529-DEL-7068
chr3-59515015-DEL-578
chr3-59516574-DEL-6094
chr3-59529201-DEL-105
chr3-59532367-DEL-9957
chr3-59563268-DEL-6205
chr3-59590619-DEL-7796
chr3-59644157-DEL-586
chr3-59649523-DEL-725
chr3-59662704-DEL-216
chr3-59670841-DEL-9960
chr3-59674630-DEL-1706
chr3-59736418-DEL-250
chr3-59740477-DEL-58
chr3-59759766-DEL-400
chr3-59782215-DEL-144
chr3-59782798-DEL-183
chr3-59821019-DEL-163
chr3-59825651-DEL-60
chr3-59857838-DEL-52
chr3-59861520-DEL-493

chr3-59866859-DEL-499
chr3-59939441-DEL-114
chr3-59952005-DEL-54
chr3-59952472-DEL-1008
chr3-59977574-DEL-146
chr3-59979849-DEL-69
chr3-60000715-DEL-127
chr3-60026738-DEL-68
chr3-60033479-DEL-1267
chr3-60035765-DEL-606
chr3-60053668-DEL-109
chr3-60085306-DEL-139
chr3-60098378-DEL-337
chr3-60102928-DEL-74
chr3-60124078-DEL-55
chr3-60134983-DEL-1293
chr3-60149842-DEL-3419
chr3-60153268-DEL-697
chr3-60165173-DEL-625
chr3-60192869-DEL-194
chr3-60232702-DEL-65
chr3-60289767-DEL-62
chr3-60298093-DEL-90
chr3-60300588-DEL-60
chr3-60302674-DEL-435
chr3-60305234-DEL-57
chr3-60337940-DEL-6647
chr3-60372265-DEL-117
chr3-60373003-DEL-529
chr3-60374247-DEL-518
chr3-60376436-DEL-455
chr3-60382909-DEL-58
chr3-60388712-DEL-176
chr3-60418752-DEL-383
chr3-60431858-DEL-201
chr3-60458462-DEL-141
chr3-60501324-DEL-170
chr3-60509567-DEL-97
chr3-60561573-DEL-1902
chr3-60571562-DEL-6490
chr3-60588701-DEL-72
chr3-60592058-DEL-3174
chr3-60611021-DEL-938
chr3-60615721-DEL-170
chr3-60620101-DEL-237
chr3-60625463-DEL-344
chr3-60643559-DEL-132
chr3-60656061-DEL-578
chr3-60676863-DEL-117
chr3-60717161-DEL-62
chr3-60735036-DEL-86
chr3-60736130-DEL-664
chr3-60768912-DEL-66
chr3-60772847-DEL-243
chr3-60775540-DEL-240
chr3-60814304-DEL-115
chr3-60843616-DEL-56
chr3-60850484-DEL-1931
chr3-60902881-DEL-581
chr3-60921306-DEL-57
chr3-60961477-DEL-256
chr3-60995299-DEL-289
chr3-61072167-DEL-1270
chr3-61073601-DEL-199
chr3-61087614-DEL-3380
chr3-61113944-DEL-8021
chr3-61164063-DEL-3136
chr3-61168105-DEL-6452
chr3-61206014-DEL-183
chr3-61227225-DEL-206
chr3-61234524-DEL-326
chr3-61240952-DEL-1860
chr3-61286994-DEL-79
chr3-61383821-DEL-1750
chr3-61388196-DEL-1619
chr3-61395752-DEL-61
chr3-61408672-DEL-199

chr3-61414667-DEL-677
chr3-61440313-DEL-197
chr3-61462414-DEL-2827
chr3-61474901-DEL-74
chr3-61484154-DEL-959
chr3-61511224-DEL-88
chr3-61514459-DEL-123
chr3-61514680-DEL-1269
chr3-61516092-DEL-238
chr3-61526400-DEL-1985
chr3-61526404-DEL-164
chr3-61537522-DEL-2733
chr3-61561124-DEL-698
chr3-61570709-DEL-65
chr3-61579618-DEL-2579
chr3-61580863-DEL-1295
chr3-61582525-DEL-3166
chr3-61583142-DEL-203
chr3-61588672-DEL-64
chr3-61600925-DEL-297
chr3-61614997-DEL-6646
chr3-61627501-DEL-6447
chr3-61640080-DEL-438
chr3-61641945-DEL-498
chr3-61653442-DEL-3867
chr3-61681347-DEL-6026
chr3-61697210-DEL-456
chr3-61755190-DEL-607
chr3-61756370-DEL-64
chr3-61777836-DEL-52
chr3-61796631-DEL-182
chr3-61810614-DEL-6160
chr3-61825507-DEL-1037
chr3-61841188-DEL-55
chr3-61843364-DEL-423
chr3-61852011-DEL-66
chr3-61861888-DEL-56
chr3-61863200-DEL-656
chr3-61864073-DEL-1632
chr3-61883037-DEL-66
chr3-61900743-DEL-2746
chr3-61907622-DEL-1215
chr3-61913309-DEL-115
chr3-61925870-DEL-127
chr3-61934070-DEL-485
chr3-61935785-DEL-290
chr3-61939603-DEL-6413
chr3-61945579-DEL-200
chr3-61960686-DEL-70
chr3-61968835-DEL-135
chr3-61996558-DEL-170
chr3-62008027-DEL-178
chr3-62015552-DEL-25437
chr3-62028946-DEL-4310
chr3-62029654-DEL-6226
chr3-62033952-DEL-1927
chr3-62037818-DEL-10737
chr3-62053247-DEL-599
chr3-62054772-DEL-6210
chr3-62062513-DEL-2305
chr3-62065046-DEL-891
chr3-62066368-DEL-219
chr3-62096326-DEL-303
chr3-62098276-DEL-594
chr3-62102415-DEL-69
chr3-62166707-DEL-351
chr3-62168874-DEL-72
chr3-62176933-DEL-373
chr3-62218590-DEL-189
chr3-62283127-DEL-54
chr3-62292134-DEL-203
chr3-62357788-DEL-136
chr3-62376688-DEL-273
chr3-62410192-DEL-1571
chr3-62411077-DEL-815
chr3-62418925-DEL-6029
chr3-62425765-DEL-6235

chr3-62505542-DEL-291
chr3-62516014-DEL-936
chr3-62521933-DEL-1029
chr3-62532166-DEL-88
chr3-62539656-DEL-732
chr3-62553178-DEL-1720
chr3-62575618-DEL-82
chr3-62584736-DEL-86
chr3-62586681-DEL-423
chr3-62616675-DEL-2443
chr3-62619878-DEL-292
chr3-62621227-DEL-483
chr3-62649809-DEL-212
chr3-62656350-DEL-120
chr3-62658851-DEL-6252
chr3-62695408-DEL-63
chr3-62718687-DEL-79
chr3-62724511-DEL-375
chr3-62786623-DEL-428
chr3-62794837-DEL-204
chr3-62809582-DEL-1278
chr3-62811315-DEL-401
chr3-62822918-DEL-199
chr3-62840188-DEL-213
chr3-62898977-DEL-208
chr3-62901670-DEL-58
chr3-62903813-DEL-417
chr3-62915214-DEL-10532
chr3-62928256-DEL-126
chr3-62945451-DEL-206
chr3-62994194-DEL-840
chr3-63015625-DEL-5464
chr3-63030203-DEL-2012
chr3-63109977-DEL-5355
chr3-63132664-DEL-71
chr3-63149294-DEL-252
chr3-63172628-DEL-112
chr3-63196582-DEL-175
chr3-63199588-DEL-508
chr3-63236194-DEL-58
chr3-63239221-DEL-685
chr3-63245631-DEL-201
chr3-63246732-DEL-315
chr3-63249989-DEL-116
chr3-63262018-DEL-50
chr3-63271575-DEL-403
chr3-63286407-DEL-163
chr3-63307623-DEL-257
chr3-63314731-DEL-56
chr3-63362638-DEL-540
chr3-63417579-DEL-821
chr3-63437608-DEL-83
chr3-63459593-DEL-60
chr3-63461090-DEL-1219
chr3-63462596-DEL-61
chr3-63471885-DEL-90
chr3-63510311-DEL-74
chr3-63540596-DEL-3282
chr3-63573809-DEL-136
chr3-63584603-DEL-61
chr3-63588373-DEL-84
chr3-63618154-DEL-84
chr3-63654000-DEL-732
chr3-63654752-DEL-2141
chr3-63718463-DEL-117
chr3-63732844-DEL-356
chr3-63744218-DEL-149
chr3-63792180-DEL-1109
chr3-63808951-DEL-584
chr3-63814304-DEL-65
chr3-63815708-DEL-72
chr3-63826524-DEL-110
chr3-63841519-DEL-3040
chr3-63854352-DEL-481
chr3-63882520-DEL-174
chr3-63889167-DEL-76
chr3-63891373-DEL-77

chr3-63905263-DEL-201
chr3-63919294-DEL-215
chr3-63931271-DEL-274
chr3-63931430-DEL-1814
chr3-63961684-DEL-52
chr3-63975218-DEL-176
chr3-64016125-DEL-891
chr3-64017281-DEL-537
chr3-64019241-DEL-65
chr3-64026821-DEL-1815
chr3-64031974-DEL-102
chr3-64034442-DEL-3303
chr3-64043429-DEL-164
chr3-64078655-DEL-289
chr3-64079384-DEL-235
chr3-64121826-DEL-948
chr3-64126425-DEL-60
chr3-64135349-DEL-64
chr3-64146543-DEL-159
chr3-64158963-DEL-140
chr3-64184904-DEL-208
chr3-64207851-DEL-305
chr3-64211351-DEL-189
chr3-64226916-DEL-152
chr3-64271139-DEL-76
chr3-64314783-DEL-171
chr3-64316216-DEL-102
chr3-64321837-DEL-55
chr3-64342210-DEL-2277
chr3-64345878-DEL-336
chr3-64421499-DEL-576
chr3-64438405-DEL-387
chr3-64469743-DEL-520
chr3-64476805-DEL-967
chr3-64557688-DEL-185
chr3-64564972-DEL-4554
chr3-64582088-DEL-5970
chr3-64586035-DEL-521
chr3-64589144-DEL-427
chr3-64591369-DEL-112
chr3-64629519-DEL-461
chr3-64692958-DEL-68
chr3-64731030-DEL-204
chr3-64857245-DEL-65
chr3-64919569-DEL-69
chr3-64933118-DEL-72
chr3-64936914-DEL-307
chr3-64954559-DEL-207
chr3-64964712-DEL-88
chr3-64982888-DEL-1707
chr3-64988271-DEL-107
chr3-65030441-DEL-98
chr3-65047278-DEL-51
chr3-65066185-DEL-2338
chr3-65096048-DEL-72
chr3-65103878-DEL-644
chr3-65212236-DEL-51
chr3-65254554-DEL-148
chr3-65260037-DEL-1144
chr3-65268304-DEL-516
chr3-65297740-DEL-754
chr3-65329191-DEL-460
chr3-65366278-DEL-52
chr3-65374283-DEL-98
chr3-65404512-DEL-112
chr3-65409197-DEL-1266
chr3-65577703-DEL-62
chr3-65601432-DEL-207
chr3-65604257-DEL-63
chr3-65662184-DEL-8696
chr3-65676759-DEL-468
chr3-65703076-DEL-102
chr3-65723989-DEL-401
chr3-65773024-DEL-76
chr3-65786111-DEL-74
chr3-65801637-DEL-681
chr3-65813721-DEL-213

chr3-65825794-DEL-307
chr3-65840401-DEL-108
chr3-65887370-DEL-434
chr3-65907343-DEL-55
chr3-65930120-DEL-58
chr3-65949313-DEL-103
chr3-65951251-DEL-131
chr3-65978824-DEL-6652
chr3-65984682-DEL-780
chr3-65987666-DEL-106
chr3-66035888-DEL-262
chr3-66067514-DEL-200
chr3-66112713-DEL-63
chr3-66141339-DEL-293
chr3-66190278-DEL-237
chr3-66218605-DEL-80
chr3-66226057-DEL-131
chr3-66236507-DEL-53
chr3-66244059-DEL-175
chr3-66273995-DEL-283
chr3-66278625-DEL-688
chr3-66471486-DEL-104
chr3-66474665-DEL-82
chr3-66523562-DEL-63
chr3-66527493-DEL-61
chr3-66556804-DEL-76
chr3-66578002-DEL-205
chr3-66601295-DEL-64
chr3-66630160-DEL-426
chr3-66714584-DEL-204
chr3-66722325-DEL-146
chr3-66758318-DEL-105
chr3-66796963-DEL-130
chr3-66809578-DEL-8203
chr3-66821550-DEL-6224
chr3-66963275-DEL-3020
chr3-66976118-DEL-212
chr3-66981627-DEL-252
chr3-66992971-DEL-919
chr3-67015086-DEL-83
chr3-67050307-DEL-149
chr3-67069118-DEL-208
chr3-67081955-DEL-154
chr3-67111549-DEL-708
chr3-67149004-DEL-237
chr3-67170510-DEL-596
chr3-67235076-DEL-568
chr3-67254949-DEL-856
chr3-67268338-DEL-159
chr3-67277179-DEL-69
chr3-67282410-DEL-51
chr3-67297584-DEL-93
chr3-67299494-DEL-68
chr3-67314729-DEL-1245
chr3-67461350-DEL-58
chr3-67463523-DEL-69
chr3-67516017-DEL-1231
chr3-67516288-DEL-54
chr3-67529399-DEL-51
chr3-67535351-DEL-437
chr3-67544698-DEL-59
chr3-67572768-DEL-594
chr3-67586842-DEL-6103
chr3-67593434-DEL-1550
chr3-67599499-DEL-416
chr3-67609811-DEL-138
chr3-67611164-DEL-248
chr3-67660709-DEL-214
chr3-67685051-DEL-68
chr3-67703607-DEL-78
chr3-67707242-DEL-740
chr3-67739108-DEL-411
chr3-67753116-DEL-409
chr3-67766400-DEL-1579
chr3-67770370-DEL-249
chr3-67771789-DEL-281
chr3-67773546-DEL-488

chr3-67795353-DEL-1339
chr3-67806658-DEL-188
chr3-67845614-DEL-101
chr3-67956324-DEL-94
chr3-67990356-DEL-1590
chr3-68033247-DEL-136
chr3-68042010-DEL-90
chr3-68068687-DEL-3936
chr3-68074410-DEL-232
chr3-68107151-DEL-58
chr3-68115332-DEL-83
chr3-68116744-DEL-59
chr3-68138173-DEL-442
chr3-68172355-DEL-260
chr3-68208034-DEL-53
chr3-68232460-DEL-118
chr3-68262127-DEL-127
chr3-68271170-DEL-543
chr3-68277150-DEL-412
chr3-68298263-DEL-1003
chr3-68298880-DEL-379
chr3-68304802-DEL-101
chr3-68322236-DEL-1723
chr3-68353228-DEL-220
chr3-68355243-DEL-259
chr3-68356761-DEL-473
chr3-68360976-DEL-52
chr3-68409844-DEL-89
chr3-68425863-DEL-71
chr3-68474460-DEL-301
chr3-68476743-DEL-468
chr3-68506220-DEL-194
chr3-68509838-DEL-186
chr3-68526579-DEL-68
chr3-68530744-DEL-57
chr3-68534542-DEL-129
chr3-68562288-DEL-280
chr3-68566765-DEL-100
chr3-68587396-DEL-2798
chr3-68609372-DEL-56
chr3-68639882-DEL-5978
chr3-68639892-DEL-7091
chr3-68642324-DEL-4657
chr3-68645889-DEL-1095
chr3-68653615-DEL-245
chr3-68655051-DEL-6872
chr3-68661511-DEL-214
chr3-68681797-DEL-87
chr3-68693040-DEL-169
chr3-68701601-DEL-94
chr3-68712000-DEL-53
chr3-68727104-DEL-205
chr3-68737661-DEL-60
chr3-68739675-DEL-1882
chr3-68794770-DEL-3507
chr3-68796635-DEL-239
chr3-68848157-DEL-53
chr3-68891232-DEL-93
chr3-68901700-DEL-62
chr3-68958527-DEL-61
chr3-68960381-DEL-602
chr3-68964966-DEL-127
chr3-69012979-DEL-55
chr3-69051891-DEL-3066
chr3-69055026-DEL-1552
chr3-69059070-DEL-55
chr3-69060633-DEL-175
chr3-69061113-DEL-158
chr3-69078926-DEL-169
chr3-69090154-DEL-59
chr3-69102829-DEL-291
chr3-69110346-DEL-1648
chr3-69117513-DEL-130
chr3-69124115-DEL-913
chr3-69124975-DEL-298
chr3-69161818-DEL-83
chr3-69176975-DEL-302

chr3-69182615-DEL-55
chr3-69191492-DEL-196
chr3-69194647-DEL-52
chr3-69202619-DEL-78
chr3-69230657-DEL-144
chr3-69234087-DEL-3450
chr3-69245397-DEL-64
chr3-69251794-DEL-113
chr3-69252327-DEL-553
chr3-69253121-DEL-183
chr3-69273973-DEL-189
chr3-69305513-DEL-55
chr3-69309544-DEL-90
chr3-69320214-DEL-100
chr3-69328329-DEL-77
chr3-69363290-DEL-1703
chr3-69396676-DEL-52
chr3-69407181-DEL-106
chr3-69421915-DEL-82
chr3-69448074-DEL-64
chr3-69450421-DEL-1147
chr3-69451469-DEL-88
chr3-69472071-DEL-56
chr3-69473051-DEL-200
chr3-69476977-DEL-228
chr3-69495068-DEL-327
chr3-69506414-DEL-123
chr3-69513589-DEL-58
chr3-69530350-DEL-991
chr3-69541651-DEL-85
chr3-69542934-DEL-72
chr3-69548916-DEL-1898
chr3-69552408-DEL-297
chr3-69566988-DEL-645
chr3-69570457-DEL-219
chr3-69575223-DEL-1015
chr3-69579242-DEL-59
chr3-69588412-DEL-93
chr3-69591233-DEL-159
chr3-69594171-DEL-63
chr3-69603237-DEL-98
chr3-69617051-DEL-72
chr3-69622622-DEL-485
chr3-69627811-DEL-642
chr3-69638059-DEL-55
chr3-69648001-DEL-196
chr3-69651808-DEL-52
chr3-69666933-DEL-54
chr3-69702744-DEL-750
chr3-69707048-DEL-66
chr3-69735889-DEL-984
chr3-69742995-DEL-622
chr3-69752026-DEL-91
chr3-69755209-DEL-192
chr3-69782935-DEL-159
chr3-69792024-DEL-461
chr3-69817782-DEL-62
chr3-69846320-DEL-311
chr3-69847478-DEL-63
chr3-69851001-DEL-384
chr3-69851640-DEL-6485
chr3-69857808-DEL-272
chr3-69867298-DEL-332
chr3-69873054-DEL-87
chr3-69878015-DEL-276
chr3-69889212-DEL-119
chr3-69922413-DEL-214
chr3-69928389-DEL-280
chr3-69953513-DEL-6753
chr3-69962399-DEL-262
chr3-69971229-DEL-5060
chr3-69978707-DEL-718
chr3-69981531-DEL-75
chr3-70035635-DEL-76
chr3-70116764-DEL-52
chr3-70139449-DEL-1001
chr3-70173457-DEL-56

chr3-70183526-DEL-481
chr3-70196341-DEL-193
chr3-70218054-DEL-114
chr3-70242324-DEL-58
chr3-70285211-DEL-354
chr3-70296066-DEL-186
chr3-70305563-DEL-84
chr3-70319132-DEL-1696
chr3-70324931-DEL-105
chr3-70326763-DEL-492
chr3-70328891-DEL-65
chr3-70349499-DEL-67
chr3-70355862-DEL-136
chr3-70356943-DEL-647
chr3-70359093-DEL-97
chr3-70363205-DEL-325
chr3-70372194-DEL-68
chr3-70382970-DEL-155
chr3-70384724-DEL-162
chr3-70400881-DEL-236
chr3-70409275-DEL-74
chr3-70417983-DEL-560
chr3-70427952-DEL-1797
chr3-70464522-DEL-96
chr3-70466335-DEL-72
chr3-70487752-DEL-51
chr3-70489333-DEL-338
chr3-70536870-DEL-54
chr3-70541126-DEL-252
chr3-70550474-DEL-191
chr3-70551575-DEL-177
chr3-70553823-DEL-623
chr3-70561665-DEL-3628
chr3-70574865-DEL-56
chr3-70577780-DEL-9520
chr3-70589134-DEL-214
chr3-70595043-DEL-7059
chr3-70595506-DEL-208
chr3-70602447-DEL-542
chr3-70611497-DEL-4235
chr3-70616140-DEL-819
chr3-70617237-DEL-199
chr3-70642933-DEL-259
chr3-70672334-DEL-187
chr3-70688553-DEL-401
chr3-70699656-DEL-51
chr3-70700693-DEL-75
chr3-70723653-DEL-2239
chr3-70725905-DEL-1513
chr3-70734544-DEL-6432
chr3-70743908-DEL-2968
chr3-70743987-DEL-5532
chr3-70755870-DEL-6389
chr3-70762871-DEL-1933
chr3-70770073-DEL-52
chr3-70787957-DEL-115
chr3-70808770-DEL-1523
chr3-70819017-DEL-1581
chr3-70826919-DEL-153
chr3-70838189-DEL-119
chr3-70895633-DEL-613
chr3-70904409-DEL-170
chr3-70964203-DEL-52
chr3-70975262-DEL-3865
chr3-71024627-DEL-6527
chr3-71030747-DEL-200
chr3-71076538-DEL-64
chr3-71081266-DEL-327
chr3-71091131-DEL-142
chr3-71092549-DEL-297
chr3-71129064-DEL-567
chr3-71130659-DEL-57
chr3-71162134-DEL-55
chr3-71185290-DEL-54
chr3-71227492-DEL-73
chr3-71228211-DEL-7648
chr3-71228212-DEL-2708

chr3-71240528-DEL-96
chr3-71270788-DEL-700
chr3-71309311-DEL-3209
chr3-71315882-DEL-756
chr3-71346368-DEL-309
chr3-71377902-DEL-3227
chr3-71406705-DEL-2209
chr3-71426597-DEL-1259
chr3-71438423-DEL-1368
chr3-71446693-DEL-1041
chr3-71459665-DEL-834
chr3-71477976-DEL-101
chr3-71478250-DEL-1017
chr3-71491899-DEL-3406
chr3-71536350-DEL-101
chr3-71549630-DEL-749
chr3-71550912-DEL-459
chr3-71572050-DEL-56
chr3-71584436-DEL-6209
chr3-71599726-DEL-98
chr3-71606347-DEL-93
chr3-71617795-DEL-1847
chr3-71621438-DEL-1056
chr3-71622887-DEL-70
chr3-71661874-DEL-266
chr3-71693415-DEL-103
chr3-71710530-DEL-83
chr3-71730935-DEL-213
chr3-71739546-DEL-128
chr3-71758388-DEL-1230
chr3-71793393-DEL-63
chr3-71845626-DEL-70
chr3-71848928-DEL-6631
chr3-71851766-DEL-101
chr3-71891296-DEL-310
chr3-71896877-DEL-200
chr3-71927693-DEL-58
chr3-71958661-DEL-539
chr3-71966132-DEL-71
chr3-71978021-DEL-161
chr3-72020308-DEL-65
chr3-72030267-DEL-92
chr3-72035079-DEL-133
chr3-72042030-DEL-103
chr3-72093666-DEL-420
chr3-72126719-DEL-144
chr3-72131490-DEL-570
chr3-72134610-DEL-191
chr3-72138694-DEL-72
chr3-72163829-DEL-6056
chr3-72170268-DEL-68
chr3-72173417-DEL-59
chr3-72180813-DEL-6267
chr3-72189960-DEL-15094
chr3-72189984-DEL-162
chr3-72215608-DEL-203
chr3-72219145-DEL-373
chr3-72234145-DEL-76
chr3-72247608-DEL-206
chr3-72254162-DEL-6124
chr3-72272824-DEL-213
chr3-72274737-DEL-104
chr3-72282243-DEL-58
chr3-72283141-DEL-1702
chr3-72287064-DEL-2563
chr3-72305681-DEL-366
chr3-72311510-DEL-248
chr3-72339359-DEL-758
chr3-72351704-DEL-158
chr3-72359463-DEL-389
chr3-72361179-DEL-126
chr3-72424993-DEL-563
chr3-72431456-DEL-2137
chr3-72444074-DEL-1353
chr3-72451414-DEL-1751
chr3-72470592-DEL-412
chr3-72482287-DEL-307

chr3-72482871-DEL-168
chr3-72491004-DEL-449
chr3-72504399-DEL-613
chr3-72517868-DEL-375
chr3-72521531-DEL-1635
chr3-72539484-DEL-2812
chr3-72543621-DEL-1293
chr3-72553189-DEL-612
chr3-72562045-DEL-2537
chr3-72576414-DEL-65
chr3-72637080-DEL-68
chr3-72639178-DEL-51
chr3-72647255-DEL-543
chr3-72654338-DEL-99
chr3-72657288-DEL-431
chr3-72683392-DEL-83
chr3-72787788-DEL-78
chr3-72789117-DEL-369
chr3-72816743-DEL-206
chr3-72831435-DEL-130
chr3-72847796-DEL-223
chr3-72861030-DEL-154
chr3-72887677-DEL-660
chr3-72908234-DEL-922
chr3-72916761-DEL-8353
chr3-72955109-DEL-64
chr3-72970192-DEL-253
chr3-73013283-DEL-51
chr3-73029313-DEL-201
chr3-73029905-DEL-951
chr3-73034393-DEL-94
chr3-73050835-DEL-638
chr3-73056400-DEL-51
chr3-73097034-DEL-381
chr3-73101048-DEL-188
chr3-73109345-DEL-661
chr3-73126478-DEL-54
chr3-73133197-DEL-3356
chr3-73146003-DEL-52
chr3-73167422-DEL-337
chr3-73182373-DEL-1560
chr3-73187611-DEL-994
chr3-73235296-DEL-833
chr3-73241040-DEL-51
chr3-73249023-DEL-219
chr3-73296371-DEL-296
chr3-73307126-DEL-76
chr3-73310692-DEL-837
chr3-73311954-DEL-482
chr3-73315706-DEL-380
chr3-73320479-DEL-392
chr3-73358988-DEL-1499
chr3-73383045-DEL-77
chr3-73383753-DEL-63
chr3-73393976-DEL-1889
chr3-73403193-DEL-266
chr3-73405090-DEL-200
chr3-73413776-DEL-1894
chr3-73440802-DEL-306
chr3-73466896-DEL-486
chr3-73478817-DEL-81
chr3-73488527-DEL-830
chr3-73541025-DEL-416
chr3-73560027-DEL-94
chr3-73567182-DEL-1913
chr3-73576257-DEL-332
chr3-73585318-DEL-2172
chr3-73636587-DEL-106
chr3-73665694-DEL-123
chr3-73680384-DEL-338
chr3-73682004-DEL-62
chr3-73685151-DEL-52
chr3-73685898-DEL-1553
chr3-73706143-DEL-3419
chr3-73731074-DEL-61
chr3-73738475-DEL-81
chr3-73740665-DEL-65

chr3-73762788-DEL-7721
chr3-73763877-DEL-5646
chr3-73788881-DEL-66
chr3-73816002-DEL-401
chr3-73829345-DEL-317
chr3-73862308-DEL-1572
chr3-73865067-DEL-59
chr3-73873008-DEL-323
chr3-73892960-DEL-82
chr3-73928861-DEL-81
chr3-73938239-DEL-90
chr3-73939217-DEL-437
chr3-73947568-DEL-901
chr3-73948962-DEL-65
chr3-73963153-DEL-264
chr3-73982406-DEL-328
chr3-74039020-DEL-3708
chr3-74067939-DEL-79
chr3-74148659-DEL-249
chr3-74172012-DEL-7872
chr3-74239625-DEL-2334
chr3-74290585-DEL-192
chr3-74300045-DEL-2876
chr3-74312248-DEL-83
chr3-74313864-DEL-1226
chr3-74336469-DEL-58
chr3-74359353-DEL-6573
chr3-74369626-DEL-69
chr3-74370435-DEL-113
chr3-74377909-DEL-6695
chr3-74387086-DEL-3578
chr3-74488777-DEL-6004
chr3-74525988-DEL-57
chr3-74545011-DEL-181
chr3-74572217-DEL-56
chr3-74584728-DEL-300
chr3-74588463-DEL-1563
chr3-74642708-DEL-85
chr3-74650820-DEL-752
chr3-74659261-DEL-3139
chr3-74682955-DEL-92
chr3-74692297-DEL-80
chr3-74711462-DEL-254
chr3-74714069-DEL-288
chr3-74724610-DEL-2078
chr3-74771484-DEL-7970
chr3-74772318-DEL-6178
chr3-74782978-DEL-6351
chr3-74790470-DEL-2535
chr3-74811265-DEL-66
chr3-74815905-DEL-188
chr3-74819371-DEL-260
chr3-74822381-DEL-351
chr3-74848389-DEL-629
chr3-74871560-DEL-8936
chr3-74908180-DEL-552
chr3-74914673-DEL-202
chr3-74918497-DEL-74
chr3-74922994-DEL-413
chr3-74926730-DEL-353
chr3-74975237-DEL-1687
chr3-74996686-DEL-1142
chr3-75006078-DEL-2786
chr3-75042659-DEL-80
chr3-75069118-DEL-679
chr3-75121902-DEL-462
chr3-75160907-DEL-99
chr3-75188065-DEL-6311
chr3-75190676-DEL-1433
chr3-75243563-DEL-63
chr3-75255756-DEL-741
chr3-75256513-DEL-4062
chr3-75261569-DEL-523
chr3-75262923-DEL-329
chr3-75298090-DEL-66
chr3-75325978-DEL-5342
chr3-75345562-DEL-454

chr3-75353523-DEL-2647
chr3-75390667-DEL-73
chr3-75400928-DEL-359
chr3-75519781-DEL-52
chr3-75573478-DEL-1879
chr3-75585784-DEL-189
chr3-75612070-DEL-97
chr3-75625054-DEL-384
chr3-75633349-DEL-9678
chr3-75658993-DEL-491
chr3-75666164-DEL-686
chr3-75680266-DEL-109
chr3-75742671-DEL-101
chr3-75785812-DEL-76
chr3-75793259-DEL-152
chr3-75814890-DEL-68
chr3-75854064-DEL-239
chr3-75878916-DEL-104
chr3-75886909-DEL-84
chr3-75888399-DEL-300
chr3-75913631-DEL-70
chr3-75920871-DEL-539
chr3-75928987-DEL-1167
chr3-75979712-DEL-106
chr3-76008582-DEL-1254
chr3-76027689-DEL-79
chr3-76029472-DEL-109
chr3-76036884-DEL-112
chr3-76037526-DEL-215
chr3-76049435-DEL-54
chr3-76070470-DEL-513
chr3-76080937-DEL-66
chr3-76103024-DEL-195
chr3-76105286-DEL-239
chr3-76108907-DEL-225
chr3-76139282-DEL-389
chr3-76176135-DEL-219
chr3-76202476-DEL-1247
chr3-76206956-DEL-204
chr3-76225055-DEL-529
chr3-76240744-DEL-640
chr3-76248500-DEL-332
chr3-76249345-DEL-1471
chr3-76270304-DEL-51
chr3-76301570-DEL-3353
chr3-76309759-DEL-108
chr3-76317425-DEL-91
chr3-76347926-DEL-269
chr3-76360230-DEL-112
chr3-76363589-DEL-313
chr3-76373915-DEL-2418
chr3-76376556-DEL-1410
chr3-76378020-DEL-1246
chr3-76392089-DEL-55
chr3-76403601-DEL-59
chr3-76408970-DEL-60
chr3-76422003-DEL-4763
chr3-76428135-DEL-274
chr3-76431460-DEL-125
chr3-76437121-DEL-2899
chr3-76451365-DEL-843
chr3-76456428-DEL-169
chr3-76459835-DEL-283
chr3-76476093-DEL-559
chr3-76479271-DEL-604
chr3-76483408-DEL-1091
chr3-76484505-DEL-472
chr3-76490070-DEL-174
chr3-76580642-DEL-53
chr3-76584870-DEL-1335
chr3-76599151-DEL-748
chr3-76607657-DEL-1896
chr3-76609771-DEL-432
chr3-76655004-DEL-66
chr3-76663845-DEL-6413
chr3-76671449-DEL-507
chr3-76713322-DEL-72

chr3-76722621-DEL-429
chr3-76730645-DEL-119
chr3-76736149-DEL-508
chr3-76751089-DEL-870
chr3-76754019-DEL-74
chr3-76806618-DEL-739
chr3-76828884-DEL-61
chr3-76828995-DEL-3048
chr3-76843002-DEL-351
chr3-76845853-DEL-452
chr3-76875664-DEL-159
chr3-76879468-DEL-82
chr3-76882023-DEL-62
chr3-76898206-DEL-188
chr3-76900382-DEL-1007
chr3-76901404-DEL-1148
chr3-76924661-DEL-168
chr3-76935591-DEL-226
chr3-76943868-DEL-62
chr3-76950335-DEL-158
chr3-76956367-DEL-152
chr3-76966749-DEL-160
chr3-76988432-DEL-1012
chr3-76995757-DEL-520
chr3-76997732-DEL-93
chr3-77026794-DEL-1270
chr3-77028904-DEL-87
chr3-77033648-DEL-288
chr3-77061758-DEL-66
chr3-77109787-DEL-76
chr3-77136130-DEL-134
chr3-77152220-DEL-804
chr3-77152403-DEL-52
chr3-77187991-DEL-89
chr3-77205754-DEL-139
chr3-77206898-DEL-58
chr3-77216877-DEL-706
chr3-77224088-DEL-585
chr3-77246924-DEL-72
chr3-77261994-DEL-67
chr3-77266386-DEL-168
chr3-77298707-DEL-434
chr3-77301342-DEL-148
chr3-77328784-DEL-436
chr3-77338435-DEL-1494
chr3-77370448-DEL-102
chr3-77390964-DEL-991
chr3-77439620-DEL-1868
chr3-77447003-DEL-68
chr3-77478903-DEL-325
chr3-77480288-DEL-2512
chr3-77496519-DEL-2151
chr3-77504347-DEL-199
chr3-77534104-DEL-98
chr3-77574577-DEL-6298
chr3-77610243-DEL-1783
chr3-77618968-DEL-4699
chr3-77623687-DEL-1414
chr3-77633983-DEL-259
chr3-77719377-DEL-6792
chr3-77729593-DEL-401
chr3-77760272-DEL-70
chr3-77772232-DEL-274
chr3-77816461-DEL-60
chr3-77833245-DEL-855
chr3-77844081-DEL-56
chr3-77858917-DEL-1893
chr3-77859644-DEL-1133
chr3-77872185-DEL-61
chr3-77874888-DEL-3390
chr3-77891176-DEL-1797
chr3-77910773-DEL-80
chr3-77929535-DEL-538
chr3-77933293-DEL-6309
chr3-77939268-DEL-128
chr3-77949809-DEL-186
chr3-77975647-DEL-132

chr3-77979212-DEL-418
chr3-77999802-DEL-516
chr3-78001967-DEL-64
chr3-78012649-DEL-663
chr3-78029138-DEL-1645
chr3-78032969-DEL-51
chr3-78066529-DEL-118
chr3-78099795-DEL-66
chr3-78122765-DEL-70
chr3-78151774-DEL-5805
chr3-78166339-DEL-152
chr3-78169490-DEL-57
chr3-78201627-DEL-54
chr3-78207557-DEL-124
chr3-78222360-DEL-61
chr3-78232480-DEL-71
chr3-78245863-DEL-198
chr3-78246394-DEL-199
chr3-78293070-DEL-487
chr3-78333751-DEL-130
chr3-78366069-DEL-208
chr3-78393555-DEL-14956
chr3-78393671-DEL-268
chr3-78393672-DEL-7397
chr3-78401009-DEL-176
chr3-78401241-DEL-7271
chr3-78450694-DEL-75
chr3-78526786-DEL-63
chr3-78532274-DEL-106
chr3-78552968-DEL-623
chr3-78571526-DEL-853
chr3-78573645-DEL-77
chr3-78622981-DEL-116
chr3-78648680-DEL-4361
chr3-78655238-DEL-1580
chr3-78658017-DEL-787
chr3-78672501-DEL-86
chr3-78710777-DEL-778
chr3-78747012-DEL-74
chr3-78760560-DEL-1773
chr3-78763286-DEL-533
chr3-78793411-DEL-1482
chr3-78824476-DEL-1310
chr3-78830077-DEL-252
chr3-78840132-DEL-266
chr3-78849411-DEL-275
chr3-78853290-DEL-759
chr3-78883786-DEL-357
chr3-78926074-DEL-107
chr3-78948184-DEL-1999
chr3-78953530-DEL-6488
chr3-78960133-DEL-64
chr3-78983193-DEL-342
chr3-78997500-DEL-1317
chr3-79058808-DEL-422
chr3-79059700-DEL-57
chr3-79134398-DEL-125
chr3-79140918-DEL-138
chr3-79150089-DEL-193
chr3-79213019-DEL-86
chr3-79222179-DEL-108
chr3-79244081-DEL-507
chr3-79245026-DEL-538
chr3-79249769-DEL-221
chr3-79273182-DEL-104
chr3-79275575-DEL-152
chr3-79287804-DEL-1316
chr3-79304017-DEL-207
chr3-79332379-DEL-392
chr3-79347492-DEL-360
chr3-79360901-DEL-218
chr3-79365317-DEL-110
chr3-79402943-DEL-388
chr3-79435983-DEL-184
chr3-79480923-DEL-392
chr3-79573478-DEL-56
chr3-79578733-DEL-151

chr3-79586196-DEL-56
chr3-79587774-DEL-129
chr3-79625368-DEL-345
chr3-79633561-DEL-106
chr3-79662658-DEL-52
chr3-79669609-DEL-8575
chr3-79676204-DEL-313
chr3-79688726-DEL-152
chr3-79693432-DEL-141
chr3-79735356-DEL-282
chr3-79736535-DEL-69
chr3-79751159-DEL-96
chr3-79761166-DEL-108
chr3-79764586-DEL-53
chr3-79775906-DEL-58
chr3-79779005-DEL-292
chr3-79796000-DEL-58
chr3-79821090-DEL-186
chr3-79857006-DEL-318
chr3-79863599-DEL-103
chr3-79906526-DEL-288
chr3-79917551-DEL-9386
chr3-79920396-DEL-2270
chr3-80001090-DEL-56
chr3-80012870-DEL-232
chr3-80029245-DEL-878
chr3-80032644-DEL-489
chr3-80068543-DEL-60
chr3-80118201-DEL-60
chr3-80125795-DEL-1188
chr3-80156905-DEL-94
chr3-80171584-DEL-2623
chr3-80176158-DEL-530
chr3-80189163-DEL-866
chr3-80190294-DEL-177
chr3-80213023-DEL-111
chr3-80224977-DEL-117
chr3-80233666-DEL-80
chr3-80246370-DEL-69
chr3-80247735-DEL-66
chr3-80264097-DEL-117
chr3-80286007-DEL-596
chr3-80291827-DEL-1117
chr3-80310169-DEL-51
chr3-80335984-DEL-52
chr3-80403051-DEL-355
chr3-80409968-DEL-175
chr3-80432194-DEL-4705
chr3-80470769-DEL-64
chr3-80494996-DEL-58
chr3-80527983-DEL-1840
chr3-80529617-DEL-6194
chr3-80543038-DEL-3297
chr3-80565370-DEL-51
chr3-80566397-DEL-1066
chr3-80611925-DEL-115
chr3-80624781-DEL-161
chr3-80625656-DEL-1855
chr3-80630204-DEL-196
chr3-80634450-DEL-1270
chr3-80649905-DEL-212
chr3-80685507-DEL-202
chr3-80741171-DEL-182
chr3-80757211-DEL-283
chr3-80774282-DEL-81
chr3-80793267-DEL-331
chr3-80805566-DEL-572
chr3-80848450-DEL-6219
chr3-80902889-DEL-133
chr3-80925474-DEL-58
chr3-81075204-DEL-155
chr3-81096961-DEL-148
chr3-81109576-DEL-56
chr3-81145731-DEL-3343
chr3-81149108-DEL-1162
chr3-81160321-DEL-722
chr3-81161080-DEL-5451

chr3-81178397-DEL-76
chr3-81191790-DEL-54
chr3-81222015-DEL-200
chr3-81223794-DEL-330
chr3-81224884-DEL-185
chr3-81237525-DEL-164
chr3-81291737-DEL-52
chr3-81318867-DEL-69
chr3-81340441-DEL-108
chr3-81356182-DEL-310
chr3-81447282-DEL-56
chr3-81502546-DEL-1086
chr3-81523072-DEL-65
chr3-81523722-DEL-6496
chr3-81530214-DEL-636
chr3-81557176-DEL-51
chr3-81592673-DEL-242
chr3-81612865-DEL-6831
chr3-81632054-DEL-51
chr3-81632376-DEL-799
chr3-81640823-DEL-90
chr3-81665269-DEL-90
chr3-81673271-DEL-3153
chr3-81682132-DEL-6414
chr3-81749756-DEL-6419
chr3-81764504-DEL-443
chr3-81779723-DEL-651
chr3-81782099-DEL-703
chr3-81782909-DEL-1176
chr3-81814115-DEL-107
chr3-81826809-DEL-76
chr3-81878345-DEL-66
chr3-81886730-DEL-196
chr3-81902536-DEL-103
chr3-81912359-DEL-974
chr3-81957754-DEL-121
chr3-81993405-DEL-533
chr3-82011411-DEL-93
chr3-82057065-DEL-82
chr3-82106908-DEL-623
chr3-82109925-DEL-970
chr3-82114313-DEL-401
chr3-82115182-DEL-129
chr3-82127402-DEL-61
chr3-82135068-DEL-66
chr3-82200625-DEL-190
chr3-82215572-DEL-61
chr3-82227975-DEL-124
chr3-82241559-DEL-1235
chr3-82242841-DEL-544
chr3-82244043-DEL-139
chr3-82322101-DEL-64
chr3-82329834-DEL-608
chr3-82347543-DEL-111
chr3-82364517-DEL-56
chr3-82365837-DEL-637
chr3-82433803-DEL-198
chr3-82450631-DEL-1151
chr3-82458263-DEL-58
chr3-82464091-DEL-56
chr3-82470574-DEL-104
chr3-82502658-DEL-400
chr3-82506302-DEL-1181
chr3-82539248-DEL-108
chr3-82597991-DEL-1246
chr3-82608934-DEL-2758
chr3-82649470-DEL-146
chr3-82686020-DEL-85
chr3-82698026-DEL-688
chr3-82701510-DEL-125
chr3-82726512-DEL-225
chr3-82738018-DEL-206
chr3-82812281-DEL-158
chr3-82827068-DEL-1126
chr3-82837777-DEL-665
chr3-82845803-DEL-904
chr3-82861964-DEL-184

chr3-82943623-DEL-192
chr3-82971389-DEL-248
chr3-82992893-DEL-101
chr3-82999024-DEL-245
chr3-83056280-DEL-103
chr3-83060174-DEL-73
chr3-83064891-DEL-51
chr3-83070420-DEL-191
chr3-83072725-DEL-1403
chr3-83104258-DEL-83
chr3-83113194-DEL-60
chr3-83213389-DEL-375
chr3-83215271-DEL-69
chr3-83242805-DEL-1809
chr3-83243887-DEL-6581
chr3-83244640-DEL-2443
chr3-83247570-DEL-3684
chr3-83248918-DEL-2336
chr3-83285531-DEL-7859
chr3-83357373-DEL-495
chr3-83378693-DEL-308
chr3-83382646-DEL-1747
chr3-83440387-DEL-70
chr3-83468702-DEL-56
chr3-83477352-DEL-141
chr3-83479263-DEL-364
chr3-83494451-DEL-63
chr3-83499826-DEL-112
chr3-83525141-DEL-107
chr3-83538886-DEL-363
chr3-83553516-DEL-690
chr3-83572642-DEL-1450
chr3-83581797-DEL-737
chr3-83609543-DEL-1993
chr3-83633270-DEL-259
chr3-83676894-DEL-1593
chr3-83732297-DEL-194
chr3-83738738-DEL-306
chr3-83752957-DEL-2678
chr3-83754125-DEL-2243
chr3-83754624-DEL-203
chr3-83755535-DEL-91
chr3-83755858-DEL-481
chr3-83763130-DEL-164
chr3-83767341-DEL-70
chr3-83796172-DEL-58
chr3-83811162-DEL-768
chr3-83815121-DEL-212
chr3-83825341-DEL-509
chr3-83847181-DEL-150
chr3-83870188-DEL-68
chr3-83888276-DEL-114
chr3-83958505-DEL-554
chr3-83961026-DEL-3874
chr3-83989977-DEL-268
chr3-84007013-DEL-577
chr3-84020690-DEL-729
chr3-84064134-DEL-71
chr3-84079854-DEL-75
chr3-84110967-DEL-133
chr3-84114249-DEL-64
chr3-84124849-DEL-85
chr3-84131475-DEL-1111
chr3-84191880-DEL-69
chr3-84201091-DEL-96
chr3-84230591-DEL-346
chr3-84237058-DEL-52
chr3-84237940-DEL-205
chr3-84238506-DEL-229
chr3-84295469-DEL-187
chr3-84314928-DEL-587
chr3-84325251-DEL-83
chr3-84366965-DEL-51
chr3-84391972-DEL-56
chr3-84499128-DEL-153
chr3-84516525-DEL-79
chr3-84526905-DEL-1404

chr3-84530566-DEL-612
chr3-84532633-DEL-81
chr3-84571094-DEL-73
chr3-84582348-DEL-84
chr3-84627422-DEL-92
chr3-84680906-DEL-110
chr3-84751898-DEL-59
chr3-84752018-DEL-767
chr3-84801652-DEL-206
chr3-84819916-DEL-93
chr3-84864543-DEL-93
chr3-84866333-DEL-2311
chr3-84888110-DEL-241
chr3-84894730-DEL-402
chr3-84948117-DEL-95
chr3-84961625-DEL-1084
chr3-84971917-DEL-480
chr3-84978297-DEL-88
chr3-85037308-DEL-82
chr3-85037939-DEL-147
chr3-85040463-DEL-600
chr3-85048583-DEL-138
chr3-85136255-DEL-10611
chr3-85147930-DEL-112
chr3-85185410-DEL-858
chr3-85185538-DEL-1065
chr3-85188954-DEL-165
chr3-85206403-DEL-320
chr3-85312310-DEL-52
chr3-85322236-DEL-78
chr3-85327765-DEL-53
chr3-85344543-DEL-1214
chr3-85356021-DEL-53
chr3-85398716-DEL-73
chr3-85504517-DEL-474
chr3-85559109-DEL-67
chr3-85580065-DEL-63
chr3-85617102-DEL-362
chr3-85621525-DEL-59
chr3-85661487-DEL-529
chr3-85688539-DEL-174
chr3-85689969-DEL-783
chr3-85695321-DEL-320
chr3-85763704-DEL-66
chr3-85799930-DEL-223
chr3-85806778-DEL-56
chr3-85819122-DEL-1689
chr3-85822995-DEL-497
chr3-85834121-DEL-184
chr3-85848488-DEL-72
chr3-85864172-DEL-74
chr3-85927190-DEL-93
chr3-85949285-DEL-292
chr3-85979212-DEL-83
chr3-85994842-DEL-59
chr3-85997507-DEL-1542
chr3-86017758-DEL-84
chr3-86044720-DEL-72
chr3-86053440-DEL-3696
chr3-86084441-DEL-60
chr3-86085205-DEL-209
chr3-86100940-DEL-1179
chr3-86102948-DEL-55
chr3-86108810-DEL-198
chr3-86153931-DEL-98
chr3-86163968-DEL-390
chr3-86174168-DEL-876
chr3-86229141-DEL-1115
chr3-86278654-DEL-51
chr3-86329632-DEL-286
chr3-86351342-DEL-60
chr3-86371907-DEL-58
chr3-86387518-DEL-2967
chr3-86462960-DEL-173
chr3-86534623-DEL-74
chr3-86544230-DEL-279
chr3-86545593-DEL-72

chr3-86562803-DEL-120
chr3-86579030-DEL-521
chr3-86597647-DEL-883
chr3-86603289-DEL-117
chr3-86633464-DEL-1609
chr3-86677685-DEL-54
chr3-86686856-DEL-78
chr3-86711108-DEL-58
chr3-86713232-DEL-148
chr3-86723375-DEL-72
chr3-86732900-DEL-402
chr3-86733280-DEL-457
chr3-86748563-DEL-505
chr3-86771101-DEL-58
chr3-86784965-DEL-207
chr3-86855261-DEL-235
chr3-86900598-DEL-89
chr3-86902648-DEL-66
chr3-86923156-DEL-403
chr3-86926798-DEL-58
chr3-86983583-DEL-60
chr3-87071946-DEL-51
chr3-87085238-DEL-228
chr3-87114610-DEL-727
chr3-87137597-DEL-146
chr3-87176266-DEL-1291
chr3-87210225-DEL-251
chr3-87210528-DEL-1101
chr3-87210550-DEL-863
chr3-87211534-DEL-198
chr3-87212910-DEL-495
chr3-87220139-DEL-6474
chr3-87226206-DEL-204
chr3-87260211-DEL-1681
chr3-87274611-DEL-72
chr3-87312626-DEL-176
chr3-87317712-DEL-80
chr3-87319937-DEL-108
chr3-87366327-DEL-26405
chr3-87374859-DEL-11397
chr3-87403137-DEL-52
chr3-87412426-DEL-880
chr3-87475838-DEL-52
chr3-87485754-DEL-1012
chr3-87489504-DEL-752
chr3-87493557-DEL-104
chr3-87543348-DEL-62
chr3-87564416-DEL-767
chr3-87587377-DEL-168
chr3-87593729-DEL-207
chr3-87658064-DEL-96
chr3-87667808-DEL-132
chr3-87727658-DEL-57
chr3-87760125-DEL-154
chr3-87786751-DEL-207
chr3-87796215-DEL-156
chr3-87799259-DEL-60
chr3-87850174-DEL-57
chr3-87851497-DEL-98
chr3-87864754-DEL-207
chr3-87875153-DEL-1212
chr3-87915117-DEL-6554
chr3-87919248-DEL-73
chr3-87926838-DEL-195
chr3-87933010-DEL-224
chr3-87946408-DEL-59
chr3-87969691-DEL-1155
chr3-88003625-DEL-115
chr3-88037897-DEL-210
chr3-88044063-DEL-231
chr3-88060153-DEL-74
chr3-88083561-DEL-460
chr3-88103308-DEL-60
chr3-88177638-DEL-129
chr3-88179646-DEL-2588
chr3-88184619-DEL-98
chr3-88224164-DEL-1218

chr3-88234722-DEL-88
chr3-88292823-DEL-80
chr3-88296570-DEL-221
chr3-88311731-DEL-289
chr3-88321184-DEL-51
chr3-88335429-DEL-202
chr3-88336196-DEL-569
chr3-88340692-DEL-51
chr3-88350896-DEL-102
chr3-88354661-DEL-138
chr3-88369808-DEL-240
chr3-88370752-DEL-459
chr3-88379021-DEL-259
chr3-88379763-DEL-94
chr3-88382675-DEL-534
chr3-88432893-DEL-63
chr3-88463468-DEL-204
chr3-88481992-DEL-274
chr3-88483509-DEL-1001
chr3-88488738-DEL-1667
chr3-88488992-DEL-200
chr3-88503277-DEL-229
chr3-88503963-DEL-93
chr3-88519852-DEL-298
chr3-88569438-DEL-66
chr3-88590386-DEL-201
chr3-88597232-DEL-2591
chr3-88606370-DEL-79
chr3-88614949-DEL-411
chr3-88631453-DEL-67
chr3-88631975-DEL-53
chr3-88638724-DEL-59
chr3-88644659-DEL-95
chr3-88653810-DEL-195
chr3-88670907-DEL-1431
chr3-88698384-DEL-199
chr3-88710208-DEL-181
chr3-88722389-DEL-4031
chr3-88724404-DEL-4038
chr3-88725268-DEL-181
chr3-88730129-DEL-87
chr3-88747098-DEL-212
chr3-88790478-DEL-93
chr3-88842764-DEL-167
chr3-88843732-DEL-166
chr3-88886409-DEL-764
chr3-88940904-DEL-255
chr3-88946956-DEL-202
chr3-88957186-DEL-132
chr3-88965196-DEL-65
chr3-89029356-DEL-241
chr3-89050807-DEL-75
chr3-89106531-DEL-331
chr3-89108874-DEL-88
chr3-89111000-DEL-194
chr3-89135078-DEL-232
chr3-89161805-DEL-194
chr3-89167119-DEL-117
chr3-89190240-DEL-156
chr3-89202121-DEL-183
chr3-89277012-DEL-57
chr3-89382026-DEL-72
chr3-89397627-DEL-123
chr3-89405097-DEL-231
chr3-89424715-DEL-200
chr3-89438866-DEL-134
chr3-89453036-DEL-69
chr3-89554551-DEL-110
chr3-89569497-DEL-126
chr3-89593254-DEL-846
chr3-89599926-DEL-56
chr3-89602296-DEL-192
chr3-89611139-DEL-150
chr3-89613618-DEL-301
chr3-89630575-DEL-52
chr3-89638932-DEL-407
chr3-89655911-DEL-208

chr3-89673420-DEL-180
chr3-89674627-DEL-84
chr3-89676079-DEL-170
chr3-89697343-DEL-469
chr3-89697864-DEL-234
chr3-89713236-DEL-87
chr3-89714039-DEL-838
chr3-89743590-DEL-247
chr3-89745748-DEL-57
chr3-89770274-DEL-534
chr3-89775194-DEL-436
chr3-89779173-DEL-72
chr3-89787016-DEL-170
chr3-89789385-DEL-109
chr3-89790637-DEL-61
chr3-89796893-DEL-215
chr3-89800908-DEL-210
chr3-89807716-DEL-197
chr3-89810000-DEL-157
chr3-89826908-DEL-498
chr3-89832429-DEL-106
chr3-89836990-DEL-162
chr3-89849483-DEL-70
chr3-89862112-DEL-156
chr3-89875087-DEL-379
chr3-89888380-DEL-197
chr3-89890435-DEL-1603
chr3-89896363-DEL-162
chr3-90018702-DEL-195
chr3-90021272-DEL-178
chr3-90023627-DEL-196
chr3-90024600-DEL-176
chr3-90033827-DEL-594
chr3-90038155-DEL-192
chr3-90048748-DEL-1577
chr3-90071017-DEL-89
chr3-90074553-DEL-228
chr3-90101670-DEL-200
chr3-90118390-DEL-55
chr3-90141710-DEL-171
chr3-90157349-DEL-329
chr3-90160975-DEL-67
chr3-90183100-DEL-203
chr3-90203597-DEL-55
chr3-90275400-DEL-428
chr3-90288285-DEL-170
chr3-90302932-DEL-67
chr3-90313152-DEL-112
chr3-90329199-DEL-207
chr3-90350203-DEL-84
chr3-90360444-DEL-63
chr3-90399248-DEL-608
chr3-90425976-DEL-205
chr3-90428892-DEL-123
chr3-90440843-DEL-687
chr3-90514292-DEL-109
chr3-90544709-DEL-53
chr3-90565422-DEL-163
chr3-90574603-DEL-96
chr3-90582152-DEL-1899
chr3-90587546-DEL-63
chr3-90599474-DEL-80
chr3-90607554-DEL-109
chr3-90622612-DEL-822
chr3-90642119-DEL-60
chr3-90654147-DEL-129
chr3-90670835-DEL-6305
chr3-90683057-DEL-1867
chr3-90691125-DEL-261
chr3-90701872-DEL-209
chr3-90830061-DEL-526
chr3-90854687-DEL-7966
chr3-90870616-DEL-6709
chr3-90907300-DEL-831
chr3-90988322-DEL-70
chr3-91003821-DEL-416
chr3-91005077-DEL-1143

chr3-91006191-DEL-623
chr3-91019116-DEL-68
chr3-91032428-DEL-10951
chr3-91033921-DEL-7498
chr3-91035376-DEL-99016
chr3-91065909-DEL-835
chr3-91098816-DEL-296
chr3-91174126-DEL-309
chr3-91176049-DEL-100
chr3-91194277-DEL-1442
chr3-91202225-DEL-4045
chr3-91208982-DEL-78
chr3-91216130-DEL-680
chr3-91313754-DEL-64
chr3-91324050-DEL-4827
chr3-91361189-DEL-2399
chr3-91398538-DEL-656
chr3-91399546-DEL-3295
chr3-91400038-DEL-430
chr3-91402460-DEL-863
chr3-91412916-DEL-1042
chr3-91439799-DEL-56
chr3-91444961-DEL-52
chr3-91473621-DEL-1876
chr3-91480668-DEL-120
chr3-91482306-DEL-95
chr3-91482843-DEL-5006
chr3-91484868-DEL-200
chr3-91491637-DEL-376
chr3-91500118-DEL-81
chr3-91506793-DEL-403
chr3-91517979-DEL-60
chr3-91527431-DEL-404
chr3-91546504-DEL-5114
chr3-91556735-DEL-888
chr3-91583492-DEL-6314
chr3-91609265-DEL-6675
chr3-91621269-DEL-6314
chr3-91654896-DEL-1121
chr3-91657561-DEL-1476
chr3-91679358-DEL-1070
chr3-91681219-DEL-64
chr3-91682426-DEL-258
chr3-91745179-DEL-1169
chr3-91751929-DEL-1235
chr3-91763916-DEL-6194
chr3-91771867-DEL-1163
chr3-91788862-DEL-208
chr3-91840757-DEL-78
chr3-91860045-DEL-1880
chr3-91862230-DEL-1658
chr3-91886805-DEL-1761
chr3-91896957-DEL-1466
chr3-91922970-DEL-6117
chr3-91949782-DEL-2400
chr3-91967738-DEL-61
chr3-91978582-DEL-114
chr3-91999791-DEL-115
chr3-92027189-DEL-51
chr3-92046022-DEL-170
chr3-92053097-DEL-63
chr3-92055045-DEL-81
chr3-92078633-DEL-87
chr3-92087103-DEL-77
chr3-92107841-DEL-2664
chr3-92121654-DEL-334
chr3-92123051-DEL-116
chr3-92123592-DEL-2426
chr3-92149139-DEL-111
chr3-92158405-DEL-2427
chr3-92179136-DEL-891
chr3-92182080-DEL-1082
chr3-92183930-DEL-111
chr3-92191152-DEL-334
chr3-92192548-DEL-116
chr3-92193089-DEL-2427
chr3-92216307-DEL-332

chr3-92227207-DEL-574
chr3-92227956-DEL-196
chr3-92231098-DEL-64
chr3-92280087-DEL-2252
chr3-92294699-DEL-141
chr3-92325388-DEL-55
chr3-92326206-DEL-55
chr3-92360945-DEL-110
chr3-92383161-DEL-2050
chr3-92388840-DEL-76
chr3-92397819-DEL-84
chr3-92433585-DEL-980
chr3-92472533-DEL-104
chr3-92613738-DEL-145
chr3-92626239-DEL-60
chr3-92644975-DEL-2231
chr3-92658507-DEL-321
chr3-92681724-DEL-885
chr3-92689076-DEL-103
chr3-92770488-DEL-92
chr3-92848841-DEL-55
chr3-92859608-DEL-128
chr3-92860056-DEL-2949
chr3-92881248-DEL-2298
chr3-92895594-DEL-674
chr3-92914297-DEL-54
chr3-92925573-DEL-51
chr3-92966403-DEL-3086
chr3-93011480-DEL-59
chr3-93111167-DEL-1176
chr3-93111393-DEL-1644
chr3-93113695-DEL-234
chr3-93114954-DEL-702
chr3-93115214-DEL-234
chr3-93116827-DEL-240
chr3-93117681-DEL-690
chr3-93119062-DEL-708
chr3-93119787-DEL-228
chr3-93120807-DEL-708
chr3-93121209-DEL-228
chr3-93123541-DEL-228
chr3-93123631-DEL-1404
chr3-93124455-DEL-233
chr3-93124925-DEL-462
chr3-93126106-DEL-455
chr3-93132996-DEL-231
chr3-93150915-DEL-608
chr3-93168071-DEL-90
chr3-93188348-DEL-1501
chr3-93189441-DEL-2248
chr3-93189493-DEL-739
chr3-93189614-DEL-1477
chr3-93192468-DEL-771
chr3-93193957-DEL-765
chr3-93197839-DEL-750
chr3-93198146-DEL-750
chr3-93232064-DEL-1044
chr3-93233062-DEL-530
chr3-93234188-DEL-787
chr3-93235828-DEL-513
chr3-93243583-DEL-126
chr3-93258251-DEL-101
chr3-93281152-DEL-59
chr3-93303697-DEL-72
chr3-93304757-DEL-72
chr3-93317505-DEL-53
chr3-93356562-DEL-183
chr3-93366189-DEL-102
chr3-93381203-DEL-393
chr3-93386950-DEL-1716
chr3-93394059-DEL-323
chr3-93418639-DEL-129
chr3-93432448-DEL-212
chr3-93455365-DEL-135
chr3-93506381-DEL-544
chr3-93528183-DEL-56
chr3-93544853-DEL-59

chr3-93565733-DEL-52
chr3-93619344-DEL-7496
chr3-93628502-DEL-1833
chr3-93631025-DEL-504
chr3-93639570-DEL-892
chr3-93642637-DEL-231
chr3-93662756-DEL-5515
chr3-93698243-DEL-56
chr3-93744648-DEL-12776
chr3-93750302-DEL-1305
chr3-93767299-DEL-3508
chr3-93787754-DEL-486
chr3-93791151-DEL-6260
chr3-93800269-DEL-7476
chr3-93807151-DEL-5036
chr3-93820947-DEL-504
chr3-93829549-DEL-624
chr3-93875249-DEL-1299
chr3-93881111-DEL-187
chr3-93893660-DEL-323
chr3-93897874-DEL-6403
chr3-93910498-DEL-486
chr3-93947182-DEL-1206
chr3-93974663-DEL-10529
chr3-93980018-DEL-171
chr3-93985377-DEL-613
chr3-94014319-DEL-485
chr3-94016011-DEL-6299
chr3-94039331-DEL-3252
chr3-94040463-DEL-1906
chr3-94053586-DEL-6214
chr3-94060257-DEL-72
chr3-94157485-DEL-274
chr3-94159499-DEL-60
chr3-94190714-DEL-6392
chr3-94201469-DEL-202
chr3-94204335-DEL-68
chr3-94241877-DEL-115
chr3-94262985-DEL-325
chr3-94272291-DEL-67
chr3-94276339-DEL-579
chr3-94315705-DEL-66
chr3-94341229-DEL-67
chr3-94346931-DEL-57
chr3-94456106-DEL-164
chr3-94465306-DEL-400
chr3-94482275-DEL-66
chr3-94506182-DEL-785
chr3-94511655-DEL-500
chr3-94513305-DEL-195
chr3-94536972-DEL-334
chr3-94544294-DEL-129
chr3-94554892-DEL-100
chr3-94583433-DEL-199
chr3-94594822-DEL-180
chr3-94595241-DEL-1551
chr3-94598270-DEL-65
chr3-94608298-DEL-176
chr3-94616095-DEL-7652
chr3-94619192-DEL-1001
chr3-94629748-DEL-134
chr3-94638589-DEL-199
chr3-94648269-DEL-805
chr3-94666052-DEL-197
chr3-94696222-DEL-60
chr3-94712771-DEL-55
chr3-94722587-DEL-324
chr3-94737649-DEL-74
chr3-94809171-DEL-198
chr3-94822586-DEL-53
chr3-94830647-DEL-167
chr3-94841253-DEL-152
chr3-94856858-DEL-813
chr3-94874798-DEL-317
chr3-94899032-DEL-231
chr3-94924601-DEL-570
chr3-94927390-DEL-1018

chr3-94928481-DEL-200
chr3-94939467-DEL-206
chr3-94969574-DEL-150
chr3-94979424-DEL-178
chr3-94992925-DEL-197
chr3-94997545-DEL-194
chr3-94999559-DEL-579
chr3-95052454-DEL-93
chr3-95057403-DEL-241
chr3-95058320-DEL-91
chr3-95083449-DEL-201
chr3-95179108-DEL-51
chr3-95218762-DEL-53
chr3-95241982-DEL-2170
chr3-95243238-DEL-115
chr3-95258507-DEL-632
chr3-95264986-DEL-201
chr3-95268095-DEL-933
chr3-95286729-DEL-58
chr3-95303344-DEL-1396
chr3-95303345-DEL-5146
chr3-95304790-DEL-475
chr3-95304819-DEL-4059
chr3-95308139-DEL-719
chr3-95313990-DEL-329
chr3-95320964-DEL-180
chr3-95323072-DEL-506
chr3-95330455-DEL-250
chr3-95331965-DEL-169
chr3-95342762-DEL-106
chr3-95350698-DEL-568
chr3-95352422-DEL-188
chr3-95378799-DEL-996
chr3-95425855-DEL-257
chr3-95430673-DEL-127
chr3-95447699-DEL-87
chr3-95479374-DEL-2995
chr3-95486179-DEL-56
chr3-95503157-DEL-177
chr3-95503425-DEL-1531
chr3-95547677-DEL-91
chr3-95552664-DEL-57
chr3-95555200-DEL-214
chr3-95577540-DEL-270
chr3-95579838-DEL-110
chr3-95598145-DEL-206
chr3-95606628-DEL-158
chr3-95610322-DEL-52
chr3-95708278-DEL-53
chr3-95730860-DEL-219
chr3-95773039-DEL-56
chr3-95823343-DEL-76
chr3-95833050-DEL-203
chr3-95838949-DEL-377
chr3-95841513-DEL-55
chr3-95842538-DEL-228
chr3-95864347-DEL-116
chr3-95906812-DEL-86
chr3-95952165-DEL-68
chr3-95993999-DEL-631
chr3-96002033-DEL-493
chr3-96007988-DEL-100
chr3-96028337-DEL-1265
chr3-96139502-DEL-57
chr3-96177630-DEL-1489
chr3-96194830-DEL-116
chr3-96229010-DEL-110
chr3-96230213-DEL-215
chr3-96230771-DEL-215
chr3-96231324-DEL-142
chr3-96238459-DEL-358
chr3-96266814-DEL-368
chr3-96271379-DEL-8772
chr3-96279219-DEL-76
chr3-96281585-DEL-364
chr3-96296448-DEL-76
chr3-96313941-DEL-166

chr3-96315198-DEL-440
chr3-96361784-DEL-1179
chr3-96373248-DEL-202
chr3-96386092-DEL-60
chr3-96390122-DEL-265
chr3-96396558-DEL-5298
chr3-96423087-DEL-56
chr3-96442411-DEL-204
chr3-96456164-DEL-153
chr3-96498179-DEL-236
chr3-96502017-DEL-152
chr3-96513297-DEL-202
chr3-96513805-DEL-60
chr3-96526714-DEL-415
chr3-96557500-DEL-59
chr3-96561105-DEL-222
chr3-96568633-DEL-53
chr3-96572395-DEL-245
chr3-96579865-DEL-236
chr3-96593586-DEL-197
chr3-96594568-DEL-60
chr3-96618347-DEL-167
chr3-96632591-DEL-168
chr3-96642339-DEL-144
chr3-96647905-DEL-844
chr3-96658475-DEL-336
chr3-96683459-DEL-54
chr3-96726990-DEL-103
chr3-96747958-DEL-4262
chr3-96748061-DEL-1195
chr3-96748321-DEL-3068
chr3-96748455-DEL-727
chr3-96748752-DEL-337
chr3-96748891-DEL-2966
chr3-96749442-DEL-260
chr3-96750200-DEL-1463
chr3-96773937-DEL-247
chr3-96829642-DEL-481
chr3-96853189-DEL-142
chr3-96878306-DEL-55
chr3-96887748-DEL-81
chr3-96891348-DEL-862
chr3-96893165-DEL-524
chr3-96946485-DEL-242
chr3-96970900-DEL-76
chr3-96987011-DEL-67
chr3-97007435-DEL-1665
chr3-97011875-DEL-194
chr3-97015830-DEL-6168
chr3-97021545-DEL-4181
chr3-97032286-DEL-759
chr3-97049074-DEL-71
chr3-97067891-DEL-63
chr3-97068645-DEL-128
chr3-97091589-DEL-57
chr3-97094650-DEL-359
chr3-97098077-DEL-116
chr3-97168246-DEL-319
chr3-97190341-DEL-77
chr3-97193839-DEL-100
chr3-97222627-DEL-617
chr3-97228606-DEL-71
chr3-97238881-DEL-923
chr3-97259759-DEL-179
chr3-97262713-DEL-1730
chr3-97314586-DEL-64
chr3-97315738-DEL-292
chr3-97340749-DEL-497
chr3-97360709-DEL-139
chr3-97369588-DEL-284
chr3-97372245-DEL-95
chr3-97376906-DEL-474
chr3-97385953-DEL-5362
chr3-97391907-DEL-80
chr3-97402459-DEL-120
chr3-97405761-DEL-1558
chr3-97419332-DEL-116

chr3-97423084-DEL-1309
chr3-97444650-DEL-546
chr3-97450835-DEL-82
chr3-97451414-DEL-59
chr3-97458212-DEL-131
chr3-97486253-DEL-68
chr3-97520390-DEL-1034
chr3-97551956-DEL-788
chr3-97553410-DEL-150
chr3-97588888-DEL-69
chr3-97590405-DEL-140
chr3-97596421-DEL-65
chr3-97642588-DEL-134
chr3-97644956-DEL-63
chr3-97686321-DEL-102
chr3-97783628-DEL-76
chr3-97790181-DEL-54
chr3-97792814-DEL-268
chr3-97829562-DEL-121
chr3-97844226-DEL-124
chr3-97874967-DEL-51
chr3-97877740-DEL-227
chr3-97887136-DEL-52
chr3-97897207-DEL-136
chr3-97908435-DEL-210
chr3-97951726-DEL-222
chr3-97969856-DEL-74
chr3-97976125-DEL-131
chr3-97986265-DEL-1395
chr3-98013054-DEL-54
chr3-98034473-DEL-96
chr3-98039156-DEL-172
chr3-98062973-DEL-3376
chr3-98079022-DEL-3494
chr3-98084721-DEL-90
chr3-98101318-DEL-191
chr3-98103489-DEL-499
chr3-98115288-DEL-72
chr3-98121599-DEL-116
chr3-98164413-DEL-400
chr3-98165246-DEL-246
chr3-98181534-DEL-273
chr3-98232224-DEL-136
chr3-98238058-DEL-58
chr3-98264099-DEL-1197
chr3-98265779-DEL-183
chr3-98268703-DEL-535
chr3-98270581-DEL-436
chr3-98303333-DEL-15666
chr3-98313991-DEL-63
chr3-98330601-DEL-75
chr3-98355016-DEL-1286
chr3-98376917-DEL-541
chr3-98453517-DEL-185
chr3-98483997-DEL-65
chr3-98501293-DEL-3519
chr3-98540320-DEL-1575
chr3-98575463-DEL-384
chr3-98576042-DEL-1290
chr3-98616560-DEL-1073
chr3-98634045-DEL-6316
chr3-98664568-DEL-2461
chr3-98670087-DEL-761
chr3-98707128-DEL-220
chr3-98714170-DEL-161
chr3-98728256-DEL-1618
chr3-98729983-DEL-343
chr3-98757115-DEL-590
chr3-98757739-DEL-1388
chr3-98766518-DEL-1468
chr3-98785456-DEL-74
chr3-98806080-DEL-79
chr3-98806672-DEL-6396
chr3-98815489-DEL-1858
chr3-98852158-DEL-479
chr3-98871743-DEL-902
chr3-98887233-DEL-630

chr3-98893285-DEL-1478
chr3-98903006-DEL-370
chr3-98939198-DEL-3430
chr3-98943605-DEL-168
chr3-98953868-DEL-51
chr3-98955262-DEL-424
chr3-98981193-DEL-57
chr3-98987304-DEL-327
chr3-98997291-DEL-151
chr3-99034309-DEL-175
chr3-99040068-DEL-730
chr3-99047155-DEL-103
chr3-99067014-DEL-198
chr3-99070517-DEL-221
chr3-99102149-DEL-57
chr3-99102891-DEL-74
chr3-99113774-DEL-117
chr3-99181066-DEL-60
chr3-99258071-DEL-499
chr3-99282827-DEL-219
chr3-99290857-DEL-779
chr3-99313413-DEL-7056
chr3-99319775-DEL-720
chr3-99322027-DEL-468
chr3-99336035-DEL-657
chr3-99336680-DEL-6174
chr3-99445469-DEL-56
chr3-99445606-DEL-2495
chr3-99456945-DEL-66
chr3-99457773-DEL-463
chr3-99461600-DEL-281
chr3-99510999-DEL-107
chr3-99515595-DEL-200
chr3-99529458-DEL-215
chr3-99565531-DEL-973
chr3-99579877-DEL-401
chr3-99589478-DEL-1090
chr3-99614978-DEL-79
chr3-99625269-DEL-96
chr3-99751067-DEL-52
chr3-99752855-DEL-69
chr3-99757382-DEL-85
chr3-99766521-DEL-2741
chr3-99893450-DEL-60
chr3-99904438-DEL-65
chr3-99908469-DEL-72
chr3-99913140-DEL-109
chr3-99927882-DEL-51
chr3-99990342-DEL-79
chr3-99997336-DEL-1455
chr3-100004722-DEL-1338
chr3-100011802-DEL-80
chr3-100066084-DEL-207
chr3-100067367-DEL-112
chr3-100096181-DEL-77
chr3-100131117-DEL-60
chr3-100136943-DEL-750
chr3-100139115-DEL-5699
chr3-100140112-DEL-1500
chr3-100172909-DEL-57
chr3-100181424-DEL-272
chr3-100197014-DEL-111
chr3-100212852-DEL-80
chr3-100221663-DEL-600
chr3-100236574-DEL-203
chr3-100291367-DEL-2136
chr3-100299450-DEL-674
chr3-100307101-DEL-323
chr3-100332752-DEL-70
chr3-100333373-DEL-445
chr3-100361016-DEL-120
chr3-100391059-DEL-58
chr3-100397293-DEL-92
chr3-100408373-DEL-67
chr3-100425685-DEL-196
chr3-100436554-DEL-961
chr3-100456204-DEL-341

chr3-100458319-DEL-90
chr3-100499768-DEL-51
chr3-100516733-DEL-345
chr3-100567793-DEL-86
chr3-100574336-DEL-65
chr3-100580564-DEL-61
chr3-100653980-DEL-75
chr3-100661015-DEL-892
chr3-100676099-DEL-51
chr3-100683846-DEL-233
chr3-100685536-DEL-100
chr3-100714439-DEL-832
chr3-100717235-DEL-132
chr3-100731783-DEL-99
chr3-100779213-DEL-102
chr3-100826186-DEL-59
chr3-100839236-DEL-736
chr3-100842144-DEL-55
chr3-100931737-DEL-144
chr3-100937644-DEL-61
chr3-100943692-DEL-53
chr3-100961721-DEL-96
chr3-100989860-DEL-53
chr3-101002624-DEL-78
chr3-101005950-DEL-72
chr3-101022648-DEL-72
chr3-101040968-DEL-60
chr3-101082649-DEL-277
chr3-101133858-DEL-83
chr3-101151295-DEL-560
chr3-101159561-DEL-2604
chr3-101169892-DEL-56
chr3-101180795-DEL-243
chr3-101203773-DEL-430
chr3-101217877-DEL-122
chr3-101271538-DEL-76
chr3-101301156-DEL-54
chr3-101358612-DEL-123
chr3-101376802-DEL-69
chr3-101379671-DEL-86
chr3-101406095-DEL-51
chr3-101408141-DEL-96
chr3-101408732-DEL-75
chr3-101431444-DEL-236
chr3-101442677-DEL-160
chr3-101450763-DEL-53
chr3-101459852-DEL-113
chr3-101472980-DEL-210
chr3-101479924-DEL-138
chr3-101546979-DEL-76
chr3-101566122-DEL-72
chr3-101575794-DEL-81
chr3-101602264-DEL-348
chr3-101622495-DEL-114
chr3-101628711-DEL-190
chr3-101646469-DEL-187
chr3-101694344-DEL-892
chr3-101707594-DEL-148
chr3-101731729-DEL-640
chr3-101756048-DEL-132
chr3-101793401-DEL-72
chr3-101803553-DEL-79
chr3-101857465-DEL-790
chr3-101866254-DEL-104
chr3-101889772-DEL-56
chr3-101900263-DEL-62
chr3-101905851-DEL-786
chr3-101952125-DEL-93
chr3-101978748-DEL-54
chr3-101990815-DEL-291
chr3-102059986-DEL-162
chr3-102060602-DEL-54
chr3-102061634-DEL-80
chr3-102062488-DEL-232
chr3-102083016-DEL-107
chr3-102096280-DEL-53
chr3-102133514-DEL-52

chr3-102146982-DEL-296
chr3-102171274-DEL-57
chr3-102196829-DEL-447
chr3-102197528-DEL-438
chr3-102212544-DEL-111
chr3-102218383-DEL-70
chr3-102278331-DEL-78
chr3-102300678-DEL-214
chr3-102309582-DEL-129
chr3-102318574-DEL-247
chr3-102319467-DEL-380
chr3-102321539-DEL-54
chr3-102333741-DEL-54
chr3-102438324-DEL-1665
chr3-102444629-DEL-1071
chr3-102462982-DEL-58
chr3-102502038-DEL-176
chr3-102518595-DEL-62
chr3-102573784-DEL-189
chr3-102589236-DEL-67
chr3-102637472-DEL-110
chr3-102656716-DEL-314
chr3-102719282-DEL-6226
chr3-102737798-DEL-721
chr3-102740350-DEL-212
chr3-102773482-DEL-6494
chr3-102794044-DEL-6320
chr3-102802703-DEL-401
chr3-102850781-DEL-149
chr3-102867102-DEL-60
chr3-102867709-DEL-183
chr3-102917195-DEL-69
chr3-102945162-DEL-88
chr3-102949700-DEL-158
chr3-102968026-DEL-98
chr3-102978325-DEL-204
chr3-103003831-DEL-285
chr3-103012834-DEL-228
chr3-103020415-DEL-170
chr3-103042008-DEL-1307
chr3-103047962-DEL-194
chr3-103097442-DEL-204
chr3-103105971-DEL-117
chr3-103112653-DEL-174
chr3-103113414-DEL-412
chr3-103118359-DEL-64
chr3-103127021-DEL-53
chr3-103162212-DEL-194
chr3-103199779-DEL-258
chr3-103201698-DEL-90
chr3-103204564-DEL-207
chr3-103207610-DEL-202
chr3-103240129-DEL-205
chr3-103279337-DEL-518
chr3-103282064-DEL-125
chr3-103284686-DEL-183
chr3-103286118-DEL-68
chr3-103302634-DEL-416
chr3-103302879-DEL-464
chr3-103324011-DEL-130
chr3-103333116-DEL-100
chr3-103354345-DEL-77
chr3-103379082-DEL-108
chr3-103383578-DEL-59
chr3-103470505-DEL-81
chr3-103528681-DEL-385
chr3-103576287-DEL-909
chr3-103596449-DEL-80
chr3-103597991-DEL-198
chr3-103605534-DEL-6159
chr3-103668418-DEL-89
chr3-103672578-DEL-198
chr3-103692980-DEL-304
chr3-103723395-DEL-74
chr3-103729601-DEL-547
chr3-103735245-DEL-2618
chr3-103756283-DEL-72

chr3-103771687-DEL-208
chr3-103825521-DEL-605
chr3-103841360-DEL-61
chr3-103848523-DEL-1102
chr3-103850787-DEL-78
chr3-103863071-DEL-300
chr3-103883626-DEL-51
chr3-103916864-DEL-76
chr3-103933403-DEL-69
chr3-103936371-DEL-226
chr3-103987559-DEL-1203
chr3-104000872-DEL-124
chr3-104008973-DEL-69
chr3-104021726-DEL-309
chr3-104024082-DEL-6257
chr3-104036054-DEL-103
chr3-104069864-DEL-1017
chr3-104080287-DEL-325
chr3-104101373-DEL-6584
chr3-104107664-DEL-191
chr3-104120417-DEL-249
chr3-104149061-DEL-71
chr3-104152334-DEL-120
chr3-104165787-DEL-847
chr3-104190569-DEL-52
chr3-104212993-DEL-85
chr3-104233408-DEL-60
chr3-104236159-DEL-103
chr3-104254469-DEL-75
chr3-104263248-DEL-52
chr3-104308005-DEL-76
chr3-104313616-DEL-445
chr3-104354649-DEL-70
chr3-104367325-DEL-849
chr3-104377695-DEL-51
chr3-104400317-DEL-64
chr3-104434133-DEL-130
chr3-104439627-DEL-184
chr3-104454520-DEL-52
chr3-104488531-DEL-457
chr3-104489047-DEL-195
chr3-104494810-DEL-52
chr3-104511134-DEL-549
chr3-104551554-DEL-102
chr3-104584728-DEL-172
chr3-104592036-DEL-4529
chr3-104598738-DEL-51
chr3-104606677-DEL-714
chr3-104618615-DEL-329
chr3-104649864-DEL-177
chr3-104652317-DEL-61
chr3-104652749-DEL-150
chr3-104657955-DEL-238
chr3-104659686-DEL-937
chr3-104672880-DEL-245
chr3-104701059-DEL-68
chr3-104705067-DEL-116
chr3-104737274-DEL-66
chr3-104745736-DEL-55
chr3-104782454-DEL-250
chr3-104817076-DEL-69
chr3-104820067-DEL-229
chr3-104827369-DEL-93
chr3-104829723-DEL-54
chr3-104839322-DEL-270
chr3-104881980-DEL-83
chr3-104891260-DEL-76
chr3-104935327-DEL-71
chr3-104935704-DEL-1756
chr3-104937021-DEL-70
chr3-104951489-DEL-52
chr3-104964208-DEL-628
chr3-104977434-DEL-52
chr3-104982081-DEL-81
chr3-105037734-DEL-121
chr3-105068912-DEL-167
chr3-105100972-DEL-123

chr3-105110107-DEL-55
chr3-105168921-DEL-120
chr3-105202853-DEL-101
chr3-105239399-DEL-399
chr3-105305563-DEL-119
chr3-105368693-DEL-70
chr3-105415453-DEL-73
chr3-105455918-DEL-79
chr3-105466056-DEL-115
chr3-105479296-DEL-58
chr3-105485109-DEL-74
chr3-105494159-DEL-1118
chr3-105501510-DEL-2615
chr3-105554238-DEL-67
chr3-105637127-DEL-52
chr3-105655192-DEL-184
chr3-105658656-DEL-404
chr3-105741072-DEL-415
chr3-105792954-DEL-1386
chr3-105794371-DEL-1787
chr3-105800480-DEL-71
chr3-105815552-DEL-559
chr3-105816508-DEL-593
chr3-105824747-DEL-115
chr3-105941816-DEL-237
chr3-105948451-DEL-782
chr3-105953369-DEL-6313
chr3-105960967-DEL-16477
chr3-105976607-DEL-438
chr3-106017438-DEL-496
chr3-106050212-DEL-204
chr3-106120668-DEL-581
chr3-106151170-DEL-50678
chr3-106185522-DEL-1524
chr3-106213929-DEL-239
chr3-106261937-DEL-247
chr3-106318564-DEL-551
chr3-106338442-DEL-1026
chr3-106339625-DEL-58
chr3-106344156-DEL-411
chr3-106345134-DEL-1111
chr3-106384750-DEL-98
chr3-106405901-DEL-65
chr3-106431631-DEL-838
chr3-106450830-DEL-84
chr3-106511377-DEL-208
chr3-106542954-DEL-1106
chr3-106562808-DEL-87
chr3-106579991-DEL-6528
chr3-106580199-DEL-190
chr3-106612174-DEL-5255
chr3-106620301-DEL-71
chr3-106656045-DEL-170
chr3-106718781-DEL-935
chr3-106719895-DEL-731
chr3-106721534-DEL-263
chr3-106722593-DEL-6280
chr3-106744041-DEL-4764
chr3-106761060-DEL-4704
chr3-106772672-DEL-2112
chr3-106808826-DEL-1780
chr3-106817796-DEL-71
chr3-106829313-DEL-405
chr3-106829956-DEL-3146
chr3-106829971-DEL-6970
chr3-106833152-DEL-3816
chr3-106844229-DEL-1693
chr3-106987461-DEL-268
chr3-106989972-DEL-270
chr3-107110485-DEL-61
chr3-107131813-DEL-60
chr3-107174734-DEL-158
chr3-107180092-DEL-82
chr3-107180670-DEL-194
chr3-107222012-DEL-51
chr3-107225731-DEL-99
chr3-107249658-DEL-60

chr3-107264938-DEL-119
chr3-107289486-DEL-61
chr3-107302854-DEL-3387
chr3-107355911-DEL-284
chr3-107385912-DEL-399
chr3-107386949-DEL-182
chr3-107393150-DEL-58
chr3-107418554-DEL-249
chr3-107481897-DEL-1835
chr3-107487510-DEL-53
chr3-107515808-DEL-51
chr3-107669059-DEL-342
chr3-107696828-DEL-1055
chr3-107721494-DEL-199
chr3-107730725-DEL-66
chr3-107774934-DEL-225
chr3-107831541-DEL-1438
chr3-107907493-DEL-486
chr3-107915459-DEL-74
chr3-107919051-DEL-52
chr3-107953173-DEL-712
chr3-107961965-DEL-987
chr3-108021787-DEL-321
chr3-108069088-DEL-87
chr3-108099972-DEL-118
chr3-108125879-DEL-90
chr3-108146596-DEL-712
chr3-108179085-DEL-74
chr3-108183146-DEL-276
chr3-108187662-DEL-66
chr3-108255224-DEL-176
chr3-108275648-DEL-64
chr3-108288357-DEL-62
chr3-108328891-DEL-80
chr3-108345411-DEL-66
chr3-108349850-DEL-52
chr3-108371116-DEL-56
chr3-108391283-DEL-53
chr3-108400454-DEL-87
chr3-108405028-DEL-433
chr3-108424152-DEL-562
chr3-108439738-DEL-67
chr3-108459375-DEL-1217
chr3-108476874-DEL-210
chr3-108484033-DEL-529
chr3-108530195-DEL-182
chr3-108589203-DEL-94
chr3-108595655-DEL-81
chr3-108619875-DEL-658
chr3-108663358-DEL-52
chr3-108682783-DEL-56
chr3-108683264-DEL-1847
chr3-108714697-DEL-291
chr3-108715078-DEL-469
chr3-108723949-DEL-296
chr3-108724212-DEL-320
chr3-108724534-DEL-606
chr3-108731233-DEL-527
chr3-108762046-DEL-162
chr3-108774691-DEL-58
chr3-108834193-DEL-177
chr3-108839218-DEL-191
chr3-108921180-DEL-375
chr3-108949938-DEL-130
chr3-108956392-DEL-66
chr3-109041512-DEL-84
chr3-109043051-DEL-83
chr3-109044052-DEL-783
chr3-109059129-DEL-92
chr3-109071146-DEL-321
chr3-109156942-DEL-552
chr3-109172792-DEL-252
chr3-109179310-DEL-382
chr3-109182736-DEL-435
chr3-109189174-DEL-668
chr3-109196440-DEL-222
chr3-109224678-DEL-110

chr3-109236004-DEL-139
chr3-109236858-DEL-177
chr3-109242760-DEL-56
chr3-109256439-DEL-85
chr3-109257144-DEL-415
chr3-109260922-DEL-767
chr3-109265959-DEL-189
chr3-109267180-DEL-87
chr3-109288347-DEL-159
chr3-109294310-DEL-67
chr3-109302037-DEL-111
chr3-109317401-DEL-58
chr3-109333162-DEL-138
chr3-109389052-DEL-58
chr3-109504741-DEL-399
chr3-109649231-DEL-62
chr3-109678283-DEL-58
chr3-109779409-DEL-92
chr3-109821097-DEL-394
chr3-109849913-DEL-138
chr3-109894913-DEL-6182
chr3-109905865-DEL-68
chr3-109919361-DEL-64
chr3-109940172-DEL-68
chr3-109947609-DEL-56
chr3-109965024-DEL-212
chr3-109982256-DEL-54
chr3-109987924-DEL-124
chr3-110033483-DEL-60
chr3-110062630-DEL-78
chr3-110063563-DEL-55
chr3-110065965-DEL-54
chr3-110080619-DEL-1790
chr3-110088875-DEL-163
chr3-110142784-DEL-208
chr3-110175017-DEL-60
chr3-110182725-DEL-908
chr3-110213931-DEL-107
chr3-110233410-DEL-214
chr3-110256177-DEL-162
chr3-110284134-DEL-404
chr3-110299029-DEL-321
chr3-110331093-DEL-118
chr3-110342989-DEL-300
chr3-110363579-DEL-882
chr3-110371078-DEL-640
chr3-110414099-DEL-87
chr3-110414453-DEL-1654
chr3-110416969-DEL-64
chr3-110435651-DEL-1299
chr3-110460198-DEL-288
chr3-110505859-DEL-224
chr3-110534477-DEL-1166
chr3-110545204-DEL-60
chr3-110610142-DEL-3305
chr3-110627008-DEL-76
chr3-110646869-DEL-1060
chr3-110693993-DEL-53
chr3-110701442-DEL-2136
chr3-110755439-DEL-3576
chr3-110769006-DEL-473
chr3-110833638-DEL-670
chr3-110842120-DEL-58
chr3-110852892-DEL-68
chr3-110875603-DEL-62
chr3-110903172-DEL-52
chr3-111019567-DEL-1858
chr3-111036902-DEL-114
chr3-111065721-DEL-135
chr3-111069211-DEL-59
chr3-111085685-DEL-52
chr3-111087716-DEL-2864
chr3-111090887-DEL-3541
chr3-111097888-DEL-132
chr3-111142723-DEL-613
chr3-111145664-DEL-94
chr3-111153517-DEL-610

chr3-111163391-DEL-6057
chr3-111163409-DEL-12578
chr3-111232410-DEL-331
chr3-111235595-DEL-64
chr3-111265306-DEL-549
chr3-111268386-DEL-162
chr3-111295785-DEL-60
chr3-111302814-DEL-175
chr3-111322011-DEL-86
chr3-111325057-DEL-126
chr3-111345823-DEL-369
chr3-111370839-DEL-102
chr3-111396152-DEL-60
chr3-111425517-DEL-206
chr3-111444654-DEL-103
chr3-111456763-DEL-397
chr3-111479569-DEL-383
chr3-111484230-DEL-9976
chr3-111489685-DEL-1111
chr3-111504729-DEL-9995
chr3-111528772-DEL-181
chr3-111531162-DEL-71
chr3-111536840-DEL-54
chr3-111544270-DEL-108
chr3-111546488-DEL-939
chr3-111559984-DEL-334
chr3-111560926-DEL-1244
chr3-111566228-DEL-81
chr3-111575377-DEL-248
chr3-111582689-DEL-83
chr3-111593430-DEL-7388
chr3-111619704-DEL-899
chr3-111655308-DEL-3977
chr3-111684913-DEL-76
chr3-111690918-DEL-89
chr3-111693894-DEL-80
chr3-111701266-DEL-4989
chr3-111713388-DEL-105
chr3-111724147-DEL-51
chr3-111748534-DEL-869
chr3-111762664-DEL-54
chr3-111769841-DEL-3272
chr3-111794165-DEL-273
chr3-111799323-DEL-457
chr3-111809372-DEL-6294
chr3-111827656-DEL-68
chr3-111829746-DEL-281
chr3-111843105-DEL-63
chr3-111904599-DEL-368
chr3-111923169-DEL-614
chr3-111930657-DEL-113
chr3-111934764-DEL-970
chr3-111935877-DEL-4164
chr3-111947738-DEL-2549
chr3-111956920-DEL-98
chr3-111981362-DEL-179
chr3-111983954-DEL-65
chr3-111997664-DEL-401
chr3-112023905-DEL-1886
chr3-112033294-DEL-540
chr3-112055976-DEL-1275
chr3-112058534-DEL-351
chr3-112067921-DEL-999
chr3-112068938-DEL-141
chr3-112075678-DEL-1961
chr3-112080420-DEL-1358
chr3-112099628-DEL-214
chr3-112100486-DEL-588
chr3-112133105-DEL-1130
chr3-112149708-DEL-128
chr3-112164408-DEL-171
chr3-112172206-DEL-1022
chr3-112188403-DEL-6247
chr3-112222588-DEL-6404
chr3-112236251-DEL-59
chr3-112254325-DEL-5613
chr3-112272820-DEL-244

chr3-112275013-DEL-618
chr3-112312703-DEL-187
chr3-112315808-DEL-194
chr3-112348456-DEL-1150
chr3-112370953-DEL-64
chr3-112371260-DEL-1230
chr3-112373109-DEL-448
chr3-112384249-DEL-99
chr3-112426564-DEL-488
chr3-112432547-DEL-136
chr3-112438116-DEL-63
chr3-112440401-DEL-190
chr3-112475172-DEL-69
chr3-112479647-DEL-68
chr3-112566085-DEL-534
chr3-112570024-DEL-56
chr3-112581351-DEL-815
chr3-112616900-DEL-439
chr3-112654196-DEL-324
chr3-112672031-DEL-52
chr3-112718277-DEL-315
chr3-112736659-DEL-110
chr3-112747323-DEL-212
chr3-112752315-DEL-214
chr3-112769668-DEL-2780
chr3-112772635-DEL-297
chr3-112778256-DEL-118
chr3-112820822-DEL-8086
chr3-112844532-DEL-298
chr3-112856021-DEL-139
chr3-112858942-DEL-844
chr3-112863983-DEL-1437
chr3-112905182-DEL-68
chr3-112932500-DEL-834
chr3-112934164-DEL-1685
chr3-112941457-DEL-1358
chr3-112944481-DEL-2344
chr3-112949298-DEL-1277
chr3-112953609-DEL-90
chr3-112961848-DEL-67
chr3-113016766-DEL-59
chr3-113021037-DEL-186
chr3-113119030-DEL-8247
chr3-113126822-DEL-657
chr3-113134705-DEL-6626
chr3-113143354-DEL-2947
chr3-113148911-DEL-100
chr3-113164685-DEL-486
chr3-113170371-DEL-1498
chr3-113202983-DEL-1498
chr3-113324629-DEL-487
chr3-113348420-DEL-77
chr3-113380468-DEL-78
chr3-113460960-DEL-9203
chr3-113460964-DEL-10614
chr3-113463273-DEL-6298
chr3-113470205-DEL-1174
chr3-113470652-DEL-645
chr3-113471440-DEL-110
chr3-113502682-DEL-2004
chr3-113505824-DEL-880
chr3-113507058-DEL-6599
chr3-113513773-DEL-3192
chr3-113520183-DEL-223
chr3-113521553-DEL-506
chr3-113534787-DEL-90
chr3-113554131-DEL-733
chr3-113558812-DEL-60
chr3-113581912-DEL-1782
chr3-113601465-DEL-358
chr3-113640360-DEL-6358
chr3-113654891-DEL-66
chr3-113656986-DEL-658
chr3-113656997-DEL-6413
chr3-113673230-DEL-65
chr3-113679339-DEL-61
chr3-113739754-DEL-787

chr3-113751613-DEL-295
chr3-113771885-DEL-7669
chr3-113779873-DEL-1176
chr3-113801003-DEL-95
chr3-113802626-DEL-63
chr3-113807481-DEL-847
chr3-113809572-DEL-377
chr3-113813913-DEL-1557
chr3-113832760-DEL-2360
chr3-113832802-DEL-98
chr3-113833870-DEL-64
chr3-113837104-DEL-67
chr3-113844726-DEL-216
chr3-113877923-DEL-179
chr3-113883979-DEL-61
chr3-113908649-DEL-158
chr3-113990264-DEL-839
chr3-113996633-DEL-452
chr3-114016996-DEL-54
chr3-114040391-DEL-198
chr3-114042414-DEL-797
chr3-114072462-DEL-208
chr3-114134744-DEL-147
chr3-114138965-DEL-5567
chr3-114210201-DEL-75
chr3-114233661-DEL-1517
chr3-114256282-DEL-152
chr3-114274014-DEL-89
chr3-114278369-DEL-124
chr3-114296959-DEL-72
chr3-114302463-DEL-145
chr3-114309314-DEL-59
chr3-114312636-DEL-228
chr3-114326344-DEL-324
chr3-114333693-DEL-630
chr3-114352980-DEL-489
chr3-114362383-DEL-1544
chr3-114395634-DEL-1598
chr3-114439182-DEL-168
chr3-114450284-DEL-6014
chr3-114451544-DEL-658
chr3-114455670-DEL-2467
chr3-114463086-DEL-56
chr3-114463243-DEL-6261
chr3-114469714-DEL-302
chr3-114486368-DEL-56
chr3-114504308-DEL-59
chr3-114510032-DEL-2469
chr3-114527608-DEL-2528
chr3-114577478-DEL-591
chr3-114590665-DEL-58
chr3-114622986-DEL-109
chr3-114644142-DEL-6184
chr3-114700803-DEL-76
chr3-114704817-DEL-113
chr3-114900446-DEL-55
chr3-114901423-DEL-72
chr3-114902214-DEL-603
chr3-114905389-DEL-445
chr3-114969126-DEL-1404
chr3-114975456-DEL-129
chr3-114975996-DEL-14961
chr3-115023095-DEL-98
chr3-115043294-DEL-101
chr3-115045041-DEL-62
chr3-115061027-DEL-344
chr3-115061801-DEL-68
chr3-115108423-DEL-60
chr3-115142636-DEL-238
chr3-115152011-DEL-705
chr3-115173669-DEL-1072
chr3-115176874-DEL-1067
chr3-115185021-DEL-8617
chr3-115185058-DEL-1430
chr3-115188400-DEL-1626
chr3-115202523-DEL-369
chr3-115214163-DEL-6382

chr3-115216837-DEL-154
chr3-115226895-DEL-79
chr3-115227969-DEL-11813
chr3-115230320-DEL-6445
chr3-115236397-DEL-214
chr3-115240346-DEL-839
chr3-115241973-DEL-144
chr3-115259167-DEL-179
chr3-115276054-DEL-1931
chr3-115285070-DEL-393
chr3-115314548-DEL-68
chr3-115320150-DEL-52
chr3-115373158-DEL-111
chr3-115388228-DEL-1281
chr3-115437143-DEL-1071
chr3-115438271-DEL-902
chr3-115438624-DEL-1232
chr3-115463233-DEL-784
chr3-115470486-DEL-58
chr3-115477932-DEL-213
chr3-115487335-DEL-584
chr3-115497534-DEL-57
chr3-115521749-DEL-63
chr3-115537518-DEL-76
chr3-115581483-DEL-101
chr3-115582524-DEL-141
chr3-115583234-DEL-108
chr3-115598170-DEL-154
chr3-115606425-DEL-89
chr3-115640475-DEL-291
chr3-115669684-DEL-59
chr3-115700006-DEL-373
chr3-115700808-DEL-52
chr3-115709936-DEL-424
chr3-115721324-DEL-743
chr3-115729106-DEL-606
chr3-115749988-DEL-64
chr3-115767172-DEL-622
chr3-115781845-DEL-127
chr3-115817069-DEL-79
chr3-115880235-DEL-68
chr3-115886853-DEL-123
chr3-115895138-DEL-240
chr3-115921354-DEL-76
chr3-115932424-DEL-192
chr3-115936056-DEL-57
chr3-115943990-DEL-377
chr3-115952671-DEL-300
chr3-115975859-DEL-1600
chr3-115986675-DEL-66
chr3-116003602-DEL-56
chr3-116053991-DEL-97
chr3-116078765-DEL-57
chr3-116123434-DEL-64
chr3-116124686-DEL-67
chr3-116147710-DEL-101
chr3-116148372-DEL-66
chr3-116179008-DEL-70
chr3-116203674-DEL-919
chr3-116206257-DEL-123
chr3-116260352-DEL-86
chr3-116266333-DEL-224
chr3-116268026-DEL-77
chr3-116278368-DEL-788
chr3-116278928-DEL-80
chr3-116330647-DEL-1779
chr3-116332506-DEL-112
chr3-116335695-DEL-169
chr3-116349710-DEL-3572
chr3-116387653-DEL-242
chr3-116406591-DEL-714
chr3-116408540-DEL-1802
chr3-116447928-DEL-168
chr3-116470177-DEL-51
chr3-116473911-DEL-184
chr3-116475947-DEL-384
chr3-116488402-DEL-1297

chr3-116488439-DEL-212
chr3-116513877-DEL-212
chr3-116525249-DEL-1170
chr3-116527817-DEL-439
chr3-116529495-DEL-3023
chr3-116560724-DEL-297
chr3-116562158-DEL-166
chr3-116606543-DEL-56
chr3-116611402-DEL-69
chr3-116634753-DEL-514
chr3-116636841-DEL-87
chr3-116639528-DEL-89
chr3-116644688-DEL-171
chr3-116656938-DEL-80
chr3-116660041-DEL-120
chr3-116663058-DEL-323
chr3-116667068-DEL-736
chr3-116685173-DEL-93
chr3-116710758-DEL-1042
chr3-116723617-DEL-389
chr3-116724466-DEL-135
chr3-116733922-DEL-73
chr3-116743961-DEL-224
chr3-116748204-DEL-65
chr3-116786974-DEL-650
chr3-116852864-DEL-467
chr3-116852866-DEL-84
chr3-116892622-DEL-1062
chr3-116904471-DEL-342
chr3-116912530-DEL-293
chr3-116980832-DEL-63
chr3-116991028-DEL-62
chr3-117009248-DEL-65
chr3-117029888-DEL-2213
chr3-117052431-DEL-361
chr3-117063958-DEL-199
chr3-117095792-DEL-1526
chr3-117102274-DEL-390
chr3-117172389-DEL-249
chr3-117184881-DEL-58
chr3-117196315-DEL-175
chr3-117208412-DEL-58
chr3-117223983-DEL-814
chr3-117264659-DEL-3957
chr3-117266215-DEL-129
chr3-117298493-DEL-194
chr3-117301049-DEL-1208
chr3-117358607-DEL-332
chr3-117385063-DEL-54
chr3-117404789-DEL-51
chr3-117415756-DEL-136
chr3-117427478-DEL-52
chr3-117452820-DEL-1960
chr3-117462371-DEL-92
chr3-117466563-DEL-2035
chr3-117560546-DEL-4415
chr3-117591745-DEL-280
chr3-117596803-DEL-244
chr3-117612823-DEL-180
chr3-117642204-DEL-112
chr3-117665075-DEL-705
chr3-117676796-DEL-53
chr3-117688104-DEL-212
chr3-117689196-DEL-483
chr3-117705670-DEL-1251
chr3-117707695-DEL-125
chr3-117717586-DEL-65
chr3-117743737-DEL-492
chr3-117769852-DEL-66
chr3-117795662-DEL-5041
chr3-117795808-DEL-2519
chr3-117795809-DEL-2519
chr3-117797148-DEL-2522
chr3-117810566-DEL-280
chr3-117832929-DEL-165
chr3-117872649-DEL-65
chr3-117879721-DEL-472

chr3-117944027-DEL-710
chr3-117948533-DEL-210
chr3-117962892-DEL-183
chr3-118007364-DEL-70
chr3-118009241-DEL-206
chr3-118038617-DEL-90
chr3-118048334-DEL-314
chr3-118052834-DEL-863
chr3-118095997-DEL-282
chr3-118098017-DEL-88
chr3-118135726-DEL-51
chr3-118146714-DEL-110
chr3-118185494-DEL-686
chr3-118186812-DEL-106
chr3-118188612-DEL-523
chr3-118289005-DEL-320
chr3-118294921-DEL-296
chr3-118325348-DEL-188
chr3-118387966-DEL-52
chr3-118462746-DEL-123
chr3-118471665-DEL-187
chr3-118494416-DEL-91
chr3-118529733-DEL-4061
chr3-118542348-DEL-927
chr3-118562918-DEL-1089
chr3-118588894-DEL-221
chr3-118605328-DEL-84
chr3-118611399-DEL-4997
chr3-118618141-DEL-97
chr3-118625701-DEL-52
chr3-118662918-DEL-272
chr3-118678306-DEL-58
chr3-118733664-DEL-52
chr3-118740189-DEL-951
chr3-118754556-DEL-58
chr3-118785281-DEL-167
chr3-118803273-DEL-3811
chr3-118820109-DEL-252
chr3-118826012-DEL-884
chr3-118831058-DEL-117
chr3-118835218-DEL-1771
chr3-118878623-DEL-72
chr3-118879090-DEL-487
chr3-118881798-DEL-924
chr3-118917189-DEL-337
chr3-118928345-DEL-154
chr3-118937639-DEL-263
chr3-118984583-DEL-73
chr3-119001463-DEL-1056
chr3-119003358-DEL-3260
chr3-119010004-DEL-692
chr3-119028341-DEL-1016
chr3-119037566-DEL-382
chr3-119074239-DEL-177
chr3-119105829-DEL-358
chr3-119107090-DEL-105
chr3-119110844-DEL-293
chr3-119143274-DEL-324
chr3-119174318-DEL-290
chr3-119177524-DEL-60
chr3-119219079-DEL-102
chr3-119254493-DEL-354
chr3-119283685-DEL-60
chr3-119295575-DEL-97
chr3-119296140-DEL-52
chr3-119312646-DEL-327
chr3-119317980-DEL-52
chr3-119381063-DEL-306
chr3-119398804-DEL-99
chr3-119408174-DEL-191
chr3-119420838-DEL-53
chr3-119425358-DEL-51
chr3-119566714-DEL-806
chr3-119592160-DEL-1104
chr3-119642279-DEL-310
chr3-119644316-DEL-103
chr3-119651294-DEL-118

chr3-119652281-DEL-126
chr3-119664376-DEL-63
chr3-119673049-DEL-478
chr3-119772900-DEL-4730
chr3-119810970-DEL-96
chr3-119823269-DEL-921
chr3-119838652-DEL-1779
chr3-119852637-DEL-58
chr3-119920596-DEL-52
chr3-119943166-DEL-62
chr3-119947207-DEL-100
chr3-119954046-DEL-134
chr3-119954305-DEL-457
chr3-119969172-DEL-72
chr3-119975737-DEL-1737
chr3-120045124-DEL-56
chr3-120047364-DEL-334
chr3-120047467-DEL-753
chr3-120052196-DEL-110
chr3-120053600-DEL-399
chr3-120100911-DEL-3025
chr3-120104237-DEL-677
chr3-120118998-DEL-90
chr3-120154416-DEL-52
chr3-120154790-DEL-2301
chr3-120162189-DEL-899
chr3-120194209-DEL-51
chr3-120199778-DEL-81
chr3-120202313-DEL-447
chr3-120213948-DEL-54
chr3-120260738-DEL-141
chr3-120298772-DEL-416
chr3-120301714-DEL-163
chr3-120311908-DEL-161
chr3-120373766-DEL-62
chr3-120379453-DEL-60
chr3-120429492-DEL-217
chr3-120497885-DEL-1320
chr3-120504441-DEL-100
chr3-120506305-DEL-52
chr3-120565712-DEL-571
chr3-120596534-DEL-207
chr3-120656719-DEL-119
chr3-120660959-DEL-92
chr3-120684700-DEL-95
chr3-120715230-DEL-6856
chr3-120723631-DEL-346
chr3-120727316-DEL-53
chr3-120732200-DEL-6625
chr3-120769735-DEL-58
chr3-120770750-DEL-469
chr3-120779644-DEL-290
chr3-120792214-DEL-621
chr3-120815193-DEL-162
chr3-120821382-DEL-51
chr3-120838244-DEL-74
chr3-120890527-DEL-197
chr3-120956263-DEL-57
chr3-120965512-DEL-75
chr3-120986329-DEL-62
chr3-121001170-DEL-94
chr3-121043427-DEL-236
chr3-121044192-DEL-66
chr3-121048181-DEL-467
chr3-121093136-DEL-521
chr3-121168336-DEL-102
chr3-121171094-DEL-80
chr3-121179291-DEL-522
chr3-121202761-DEL-275
chr3-121233142-DEL-416
chr3-121235610-DEL-183
chr3-121247146-DEL-312
chr3-121259120-DEL-550
chr3-121259863-DEL-60
chr3-121284293-DEL-54
chr3-121290351-DEL-53
chr3-121326470-DEL-148

chr3-121362315-DEL-957
chr3-121366464-DEL-546
chr3-121379910-DEL-96
chr3-121384926-DEL-60
chr3-121386428-DEL-58
chr3-121387205-DEL-51
chr3-121401570-DEL-176
chr3-121422150-DEL-289
chr3-121434066-DEL-197
chr3-121449078-DEL-651
chr3-121490103-DEL-75
chr3-121520536-DEL-433
chr3-121523321-DEL-461
chr3-121567172-DEL-137
chr3-121584007-DEL-70
chr3-121607207-DEL-169
chr3-121613021-DEL-160
chr3-121623445-DEL-52
chr3-121650978-DEL-51
chr3-121655068-DEL-64
chr3-121675621-DEL-290
chr3-121676138-DEL-291
chr3-121676899-DEL-108
chr3-121684511-DEL-170
chr3-121699082-DEL-54
chr3-121706743-DEL-717
chr3-121715083-DEL-52
chr3-121717929-DEL-117
chr3-121725782-DEL-64
chr3-121734412-DEL-230
chr3-121755998-DEL-60
chr3-121757022-DEL-206
chr3-121798229-DEL-196
chr3-121814461-DEL-83
chr3-121817059-DEL-180
chr3-121823233-DEL-90
chr3-121826470-DEL-241
chr3-121874437-DEL-68
chr3-121879404-DEL-52
chr3-121902574-DEL-504
chr3-121907601-DEL-1244
chr3-121929003-DEL-123
chr3-121940122-DEL-52
chr3-121945704-DEL-70
chr3-121947310-DEL-341
chr3-121965674-DEL-60
chr3-122000704-DEL-203
chr3-122021410-DEL-2024
chr3-122023445-DEL-9501
chr3-122044460-DEL-51
chr3-122050322-DEL-764
chr3-122064074-DEL-2436
chr3-122083921-DEL-54
chr3-122100043-DEL-608
chr3-122124834-DEL-78
chr3-122168334-DEL-63
chr3-122180484-DEL-186
chr3-122209520-DEL-249
chr3-122241498-DEL-204
chr3-122307152-DEL-159
chr3-122326508-DEL-122
chr3-122404992-DEL-218
chr3-122415092-DEL-681
chr3-122439805-DEL-140
chr3-122442953-DEL-214
chr3-122526302-DEL-61
chr3-122543900-DEL-62
chr3-122554932-DEL-166
chr3-122573022-DEL-1881
chr3-122598252-DEL-386
chr3-122598797-DEL-616
chr3-122606084-DEL-98
chr3-122619179-DEL-53
chr3-122624482-DEL-809
chr3-122637774-DEL-485
chr3-122721815-DEL-220
chr3-122728741-DEL-541

chr3-122730396-DEL-2081
chr3-122757152-DEL-409
chr3-122770405-DEL-76
chr3-122856926-DEL-351
chr3-122893066-DEL-91
chr3-122917254-DEL-64
chr3-122919063-DEL-148
chr3-122933583-DEL-58
chr3-122951284-DEL-82
chr3-122975508-DEL-66
chr3-122985828-DEL-106
chr3-123003777-DEL-102
chr3-123014732-DEL-68
chr3-123039699-DEL-1320
chr3-123041810-DEL-51
chr3-123063602-DEL-567
chr3-123077357-DEL-56
chr3-123080532-DEL-74
chr3-123134394-DEL-490
chr3-123160988-DEL-140
chr3-123166955-DEL-366
chr3-123177649-DEL-981
chr3-123180359-DEL-250
chr3-123183218-DEL-52
chr3-123203504-DEL-2761
chr3-123233658-DEL-378
chr3-123243139-DEL-154
chr3-123265406-DEL-146
chr3-123265513-DEL-2305
chr3-123267872-DEL-1061
chr3-123269019-DEL-410
chr3-123276767-DEL-54
chr3-123290530-DEL-79
chr3-123300348-DEL-508
chr3-123304874-DEL-268
chr3-123306897-DEL-72
chr3-123316179-DEL-668
chr3-123341311-DEL-232
chr3-123346538-DEL-146
chr3-123368611-DEL-108
chr3-123371343-DEL-217
chr3-123376547-DEL-84
chr3-123388573-DEL-61
chr3-123413487-DEL-65
chr3-123463249-DEL-1245
chr3-123471038-DEL-637
chr3-123471783-DEL-585
chr3-123472550-DEL-259
chr3-123472964-DEL-1623
chr3-123473370-DEL-117
chr3-123474059-DEL-85
chr3-123542333-DEL-1050
chr3-123635226-DEL-167
chr3-123673857-DEL-5206
chr3-123702990-DEL-2622
chr3-123718389-DEL-13162
chr3-123735284-DEL-6174
chr3-123750065-DEL-1337
chr3-123764505-DEL-1157
chr3-123790566-DEL-1562
chr3-123791425-DEL-108
chr3-123792441-DEL-467
chr3-123826345-DEL-208
chr3-123847575-DEL-148
chr3-123857481-DEL-689
chr3-123887299-DEL-129
chr3-123978082-DEL-275
chr3-123987204-DEL-7170
chr3-123995990-DEL-51
chr3-123997884-DEL-3756
chr3-124003950-DEL-461
chr3-124004603-DEL-57
chr3-124046357-DEL-180
chr3-124049460-DEL-524
chr3-124128432-DEL-2053
chr3-124140934-DEL-448
chr3-124184309-DEL-210

chr3-124197733-DEL-72
chr3-124227154-DEL-318
chr3-124310324-DEL-287
chr3-124324456-DEL-118
chr3-124328980-DEL-52
chr3-124336406-DEL-878
chr3-124363794-DEL-431
chr3-124369836-DEL-244
chr3-124380049-DEL-575
chr3-124392018-DEL-400
chr3-124404245-DEL-204
chr3-124435252-DEL-1778
chr3-124476702-DEL-120
chr3-124485999-DEL-1207
chr3-124504886-DEL-145
chr3-124518929-DEL-319
chr3-124525251-DEL-79
chr3-124525712-DEL-122
chr3-124595937-DEL-2496
chr3-124620404-DEL-12209
chr3-124622057-DEL-6473
chr3-124642493-DEL-373
chr3-124653046-DEL-55
chr3-124695700-DEL-55
chr3-124696722-DEL-214
chr3-124737535-DEL-1364
chr3-124751200-DEL-203
chr3-124759118-DEL-6275
chr3-124772658-DEL-3307
chr3-124786655-DEL-6181
chr3-124817418-DEL-73
chr3-124864518-DEL-54
chr3-124869519-DEL-162
chr3-124869925-DEL-4453
chr3-124874370-DEL-1290
chr3-124875507-DEL-446
chr3-124875864-DEL-378
chr3-124901363-DEL-347
chr3-124918598-DEL-186
chr3-124928426-DEL-154
chr3-124966509-DEL-53
chr3-124966927-DEL-2111
chr3-124978102-DEL-263
chr3-124987439-DEL-122
chr3-125046144-DEL-140
chr3-125058580-DEL-469
chr3-125062062-DEL-427
chr3-125078392-DEL-214
chr3-125093523-DEL-105
chr3-125126361-DEL-70
chr3-125155524-DEL-137
chr3-125217512-DEL-385
chr3-125218730-DEL-265
chr3-125223862-DEL-75
chr3-125225304-DEL-66
chr3-125250073-DEL-2601
chr3-125264930-DEL-283
chr3-125294752-DEL-432
chr3-125346409-DEL-1439
chr3-125346425-DEL-958
chr3-125347216-DEL-409
chr3-125364084-DEL-106
chr3-125369053-DEL-84
chr3-125380839-DEL-70
chr3-125442618-DEL-55
chr3-125443543-DEL-104
chr3-125497071-DEL-464
chr3-125528494-DEL-1385
chr3-125575949-DEL-523
chr3-125576796-DEL-418
chr3-125588771-DEL-1366
chr3-125602124-DEL-126
chr3-125603684-DEL-1221
chr3-125604978-DEL-497
chr3-125615786-DEL-665
chr3-125643797-DEL-110
chr3-125718239-DEL-86

chr3-125723110-DEL-55
chr3-125743657-DEL-68
chr3-125778566-DEL-245
chr3-125790056-DEL-52
chr3-125814613-DEL-52
chr3-125823765-DEL-5494
chr3-125839653-DEL-51
chr3-125848705-DEL-100
chr3-125854428-DEL-6139
chr3-125862007-DEL-100
chr3-125877004-DEL-627
chr3-125915060-DEL-153
chr3-125933414-DEL-998
chr3-125934155-DEL-196
chr3-125952721-DEL-68
chr3-125961078-DEL-1148
chr3-125974040-DEL-3965
chr3-126005992-DEL-220
chr3-126014210-DEL-5388
chr3-126046779-DEL-83
chr3-126067014-DEL-92
chr3-126070490-DEL-253
chr3-126079862-DEL-150
chr3-126091808-DEL-56
chr3-126107149-DEL-62
chr3-126111976-DEL-121
chr3-126160374-DEL-185
chr3-126179794-DEL-56
chr3-126184004-DEL-216
chr3-126334731-DEL-68
chr3-126336395-DEL-538
chr3-126344999-DEL-1934
chr3-126374403-DEL-87
chr3-126403850-DEL-94
chr3-126448577-DEL-616
chr3-126451317-DEL-101
chr3-126464590-DEL-279
chr3-126475126-DEL-56
chr3-126501796-DEL-181
chr3-126514023-DEL-164
chr3-126540664-DEL-102
chr3-126550266-DEL-120
chr3-126562650-DEL-262
chr3-126572942-DEL-149
chr3-126577391-DEL-68
chr3-126626037-DEL-56
chr3-126646240-DEL-4292
chr3-126653375-DEL-6741
chr3-126704688-DEL-224
chr3-126839348-DEL-64
chr3-126852486-DEL-734
chr3-126874414-DEL-115
chr3-126970434-DEL-57
chr3-126971969-DEL-90
chr3-126980620-DEL-67
chr3-127007198-DEL-342
chr3-127024141-DEL-54
chr3-127048137-DEL-826
chr3-127049986-DEL-798
chr3-127055521-DEL-120
chr3-127056787-DEL-726
chr3-127057574-DEL-233
chr3-127083365-DEL-85
chr3-127098336-DEL-56
chr3-127118607-DEL-55
chr3-127129244-DEL-88
chr3-127151476-DEL-176
chr3-127165356-DEL-995
chr3-127193668-DEL-59
chr3-127198048-DEL-338
chr3-127216901-DEL-216
chr3-127237387-DEL-219
chr3-127238045-DEL-330
chr3-127266288-DEL-210
chr3-127274061-DEL-478
chr3-127305399-DEL-64
chr3-127310828-DEL-917

chr3-127315371-DEL-288
chr3-127319983-DEL-91
chr3-127352409-DEL-213
chr3-127356829-DEL-81
chr3-127387128-DEL-183
chr3-127423088-DEL-86
chr3-127459933-DEL-61
chr3-127461534-DEL-111
chr3-127469098-DEL-56
chr3-127475278-DEL-95
chr3-127483367-DEL-444
chr3-127512395-DEL-439
chr3-127553733-DEL-66
chr3-127581254-DEL-1756
chr3-127597471-DEL-1426
chr3-127621363-DEL-709
chr3-127663169-DEL-136
chr3-127676816-DEL-55
chr3-127702426-DEL-672
chr3-127704187-DEL-491
chr3-127758999-DEL-53
chr3-127764177-DEL-69
chr3-127790346-DEL-91
chr3-127793605-DEL-83
chr3-127800120-DEL-1123
chr3-127813453-DEL-221
chr3-127815212-DEL-68
chr3-127899026-DEL-910
chr3-127919178-DEL-77
chr3-127921784-DEL-97
chr3-127997186-DEL-1492
chr3-128005322-DEL-104
chr3-128143025-DEL-275
chr3-128160969-DEL-72
chr3-128183700-DEL-173
chr3-128184532-DEL-61
chr3-128198640-DEL-102
chr3-128211262-DEL-336
chr3-128263763-DEL-490
chr3-128281883-DEL-1018
chr3-128283464-DEL-674
chr3-128288435-DEL-1468
chr3-128344251-DEL-1413
chr3-128360534-DEL-122
chr3-128393956-DEL-57
chr3-128438646-DEL-1652
chr3-128467234-DEL-68
chr3-128481347-DEL-1410
chr3-128495793-DEL-74
chr3-128505937-DEL-165
chr3-128562045-DEL-142
chr3-128566428-DEL-425
chr3-128570687-DEL-7615
chr3-128571536-DEL-462
chr3-128627687-DEL-426
chr3-128668380-DEL-220
chr3-128699301-DEL-644
chr3-128711359-DEL-681
chr3-128733284-DEL-554
chr3-128789365-DEL-417
chr3-128792213-DEL-1208
chr3-128797461-DEL-132
chr3-128815285-DEL-400
chr3-128850453-DEL-158
chr3-128850891-DEL-2123
chr3-128904784-DEL-1740
chr3-128908919-DEL-60
chr3-128921087-DEL-72
chr3-128929347-DEL-134
chr3-128940832-DEL-169
chr3-129030975-DEL-70
chr3-129099839-DEL-195
chr3-129116747-DEL-62
chr3-129118888-DEL-105
chr3-129119377-DEL-60
chr3-129129547-DEL-174
chr3-129146801-DEL-203

chr3-129178528-DEL-389
chr3-129179132-DEL-268
chr3-129203259-DEL-69
chr3-129219189-DEL-720
chr3-129220167-DEL-132
chr3-129222340-DEL-140
chr3-129230564-DEL-97
chr3-129241797-DEL-61
chr3-129261334-DEL-436
chr3-129262139-DEL-96
chr3-129285326-DEL-77
chr3-129301899-DEL-70
chr3-129319096-DEL-210
chr3-129344483-DEL-147
chr3-129354211-DEL-143
chr3-129358011-DEL-56
chr3-129389893-DEL-74
chr3-129397937-DEL-60
chr3-129413837-DEL-211
chr3-129452856-DEL-210
chr3-129455747-DEL-72
chr3-129477474-DEL-245
chr3-129483123-DEL-201
chr3-129485130-DEL-228
chr3-129552975-DEL-77
chr3-129578031-DEL-68
chr3-129598132-DEL-616
chr3-129608273-DEL-304
chr3-129624065-DEL-61
chr3-129627792-DEL-57
chr3-129643964-DEL-62
chr3-129666239-DEL-222
chr3-129673701-DEL-110
chr3-129685185-DEL-401
chr3-129685804-DEL-251
chr3-129719221-DEL-78
chr3-129725269-DEL-479
chr3-129728543-DEL-100
chr3-129733432-DEL-122
chr3-129743883-DEL-73
chr3-129751558-DEL-148
chr3-129771021-DEL-137
chr3-129771776-DEL-84
chr3-129878234-DEL-71
chr3-129890902-DEL-439
chr3-129904485-DEL-52
chr3-129915864-DEL-70
chr3-129923943-DEL-675
chr3-129924646-DEL-1351
chr3-129959593-DEL-508
chr3-130020611-DEL-173
chr3-130026302-DEL-490
chr3-130041496-DEL-56
chr3-130072996-DEL-435
chr3-130114304-DEL-4358
chr3-130149376-DEL-355
chr3-130170322-DEL-240
chr3-130173559-DEL-53
chr3-130189917-DEL-654
chr3-130213677-DEL-54
chr3-130276999-DEL-1324
chr3-130321804-DEL-137
chr3-130402722-DEL-111
chr3-130404142-DEL-52
chr3-130445989-DEL-58
chr3-130457752-DEL-6288
chr3-130470360-DEL-328
chr3-130494594-DEL-777
chr3-130529141-DEL-52
chr3-130596384-DEL-4762
chr3-130643240-DEL-511
chr3-130684693-DEL-64
chr3-130719567-DEL-52
chr3-130724553-DEL-90
chr3-130753224-DEL-96
chr3-130797608-DEL-2663
chr3-130801789-DEL-400

chr3-130817279-DEL-107
chr3-130829909-DEL-186
chr3-130856061-DEL-208
chr3-130857865-DEL-101
chr3-130864758-DEL-215
chr3-130865283-DEL-72
chr3-130865794-DEL-160
chr3-130870519-DEL-243
chr3-130873820-DEL-73
chr3-130919861-DEL-63
chr3-130924330-DEL-207
chr3-130956570-DEL-800
chr3-130960692-DEL-102
chr3-130964765-DEL-82
chr3-131004657-DEL-293
chr3-131098244-DEL-255
chr3-131117561-DEL-78
chr3-131123623-DEL-102
chr3-131145663-DEL-108
chr3-131169028-DEL-69
chr3-131214493-DEL-95
chr3-131301122-DEL-363
chr3-131308860-DEL-162
chr3-131346435-DEL-56
chr3-131402667-DEL-469
chr3-131416269-DEL-5887
chr3-131432609-DEL-66
chr3-131440805-DEL-765
chr3-131479857-DEL-261
chr3-131493785-DEL-395
chr3-131530883-DEL-51
chr3-131531027-DEL-335
chr3-131531358-DEL-934
chr3-131531390-DEL-1108
chr3-131555600-DEL-52
chr3-131563853-DEL-213
chr3-131572974-DEL-71
chr3-131580740-DEL-6605
chr3-131609785-DEL-486
chr3-131631647-DEL-123
chr3-131634309-DEL-6442
chr3-131641458-DEL-215
chr3-131648310-DEL-74
chr3-131675421-DEL-734
chr3-131715756-DEL-65
chr3-131795511-DEL-70
chr3-131812912-DEL-62
chr3-131815682-DEL-55
chr3-131826071-DEL-352
chr3-131846136-DEL-70
chr3-131850159-DEL-128
chr3-131890268-DEL-59
chr3-131928149-DEL-2029
chr3-131990357-DEL-150
chr3-131993934-DEL-476
chr3-132000287-DEL-392
chr3-132003797-DEL-12974
chr3-132045374-DEL-102
chr3-132067820-DEL-53
chr3-132078513-DEL-619
chr3-132092739-DEL-57
chr3-132099433-DEL-113
chr3-132225549-DEL-66
chr3-132231027-DEL-388
chr3-132265515-DEL-72
chr3-132344074-DEL-336
chr3-132352348-DEL-77
chr3-132392283-DEL-175
chr3-132397147-DEL-57
chr3-132416836-DEL-961
chr3-132424499-DEL-652
chr3-132425749-DEL-169
chr3-132441546-DEL-80
chr3-132447166-DEL-84
chr3-132460464-DEL-1693
chr3-132490239-DEL-109
chr3-132505081-DEL-250

chr3-132520384-DEL-57
chr3-132522127-DEL-1898
chr3-132551111-DEL-526
chr3-132571931-DEL-60
chr3-132576369-DEL-305
chr3-132584099-DEL-992
chr3-132685181-DEL-1075
chr3-132690270-DEL-233
chr3-132722450-DEL-222
chr3-132733815-DEL-274
chr3-132742743-DEL-73
chr3-132745500-DEL-227
chr3-132759524-DEL-81
chr3-132769124-DEL-6199
chr3-132841334-DEL-55
chr3-132854264-DEL-203
chr3-132862179-DEL-120
chr3-132895644-DEL-6279
chr3-132954720-DEL-5204
chr3-132969834-DEL-6397
chr3-133012998-DEL-59
chr3-133023779-DEL-54
chr3-133043369-DEL-106
chr3-133057132-DEL-1477
chr3-133062837-DEL-107
chr3-133084407-DEL-74
chr3-133092560-DEL-369
chr3-133106927-DEL-1493
chr3-133145351-DEL-123
chr3-133158974-DEL-1531
chr3-133160969-DEL-60
chr3-133166955-DEL-317
chr3-133244208-DEL-223
chr3-133283215-DEL-101
chr3-133285921-DEL-128
chr3-133336526-DEL-136
chr3-133347956-DEL-212
chr3-133364425-DEL-72
chr3-133384750-DEL-209
chr3-133436882-DEL-75
chr3-133453379-DEL-626
chr3-133459549-DEL-70
chr3-133485654-DEL-94
chr3-133488007-DEL-583
chr3-133497427-DEL-52
chr3-133503048-DEL-201
chr3-133559876-DEL-6365
chr3-133568460-DEL-207
chr3-133623659-DEL-80
chr3-133627049-DEL-545
chr3-133633229-DEL-712
chr3-133645999-DEL-68
chr3-133696963-DEL-205
chr3-133730165-DEL-1517
chr3-133731761-DEL-3611
chr3-133750854-DEL-111
chr3-133751813-DEL-84
chr3-133792395-DEL-718
chr3-133793291-DEL-338
chr3-133802155-DEL-467
chr3-133806477-DEL-108
chr3-133820451-DEL-63
chr3-133847765-DEL-97
chr3-133859864-DEL-874
chr3-133902262-DEL-1196
chr3-133911506-DEL-77
chr3-133930354-DEL-1443
chr3-133969029-DEL-58
chr3-133996070-DEL-276
chr3-134019172-DEL-56
chr3-134031583-DEL-80
chr3-134045134-DEL-224
chr3-134045405-DEL-1759
chr3-134056007-DEL-62
chr3-134058516-DEL-62
chr3-134060348-DEL-81
chr3-134078232-DEL-288

chr3-134092319-DEL-344
chr3-134101954-DEL-74
chr3-134109596-DEL-171
chr3-134150669-DEL-209
chr3-134185269-DEL-81
chr3-134190961-DEL-120
chr3-134211891-DEL-94
chr3-134228374-DEL-186
chr3-134235152-DEL-6927
chr3-134239596-DEL-794
chr3-134241413-DEL-570
chr3-134242586-DEL-659
chr3-134246152-DEL-396
chr3-134251516-DEL-6395
chr3-134268026-DEL-381
chr3-134284515-DEL-62
chr3-134285198-DEL-1369
chr3-134325089-DEL-172
chr3-134345708-DEL-94
chr3-134362948-DEL-59
chr3-134378515-DEL-3444
chr3-134390348-DEL-6201
chr3-134397495-DEL-185
chr3-134400087-DEL-489
chr3-134420898-DEL-116
chr3-134458186-DEL-146
chr3-134469601-DEL-16755
chr3-134473896-DEL-97
chr3-134478099-DEL-1673
chr3-134479954-DEL-3246
chr3-134481832-DEL-65
chr3-134486252-DEL-123
chr3-134506039-DEL-62
chr3-134533860-DEL-177
chr3-134551398-DEL-54
chr3-134554573-DEL-202
chr3-134567975-DEL-6404
chr3-134584835-DEL-406
chr3-134605243-DEL-358
chr3-134647257-DEL-4778
chr3-134659031-DEL-208
chr3-134718004-DEL-1026
chr3-134763510-DEL-142
chr3-134815044-DEL-297
chr3-134823146-DEL-283
chr3-134826500-DEL-64
chr3-134827368-DEL-64
chr3-134832702-DEL-4082
chr3-134837213-DEL-81
chr3-134853206-DEL-67
chr3-134858862-DEL-87
chr3-134907456-DEL-1022
chr3-134968420-DEL-650
chr3-134981238-DEL-182
chr3-134991838-DEL-407
chr3-134992816-DEL-734
chr3-135015103-DEL-54
chr3-135040877-DEL-76
chr3-135048617-DEL-280
chr3-135062178-DEL-51
chr3-135078664-DEL-55
chr3-135081129-DEL-52
chr3-135081991-DEL-2226
chr3-135096863-DEL-147
chr3-135130403-DEL-109
chr3-135131151-DEL-65
chr3-135178459-DEL-497
chr3-135181377-DEL-59
chr3-135212879-DEL-185
chr3-135213740-DEL-108
chr3-135221023-DEL-972
chr3-135227000-DEL-118
chr3-135236387-DEL-610
chr3-135242158-DEL-2891
chr3-135245142-DEL-1006
chr3-135259604-DEL-542
chr3-135263660-DEL-1911

chr3-135282905-DEL-272
chr3-135305286-DEL-67
chr3-135323683-DEL-51
chr3-135338231-DEL-79
chr3-135354450-DEL-58
chr3-135372555-DEL-56
chr3-135386259-DEL-76
chr3-135390688-DEL-76
chr3-135399448-DEL-221
chr3-135429974-DEL-568
chr3-135434152-DEL-259
chr3-135457704-DEL-66
chr3-135463761-DEL-769
chr3-135466948-DEL-220
chr3-135475398-DEL-5284
chr3-135484049-DEL-357
chr3-135492623-DEL-1162
chr3-135513354-DEL-120
chr3-135523209-DEL-86
chr3-135557081-DEL-116
chr3-135564663-DEL-90
chr3-135569456-DEL-1347
chr3-135579703-DEL-475
chr3-135588480-DEL-887
chr3-135620372-DEL-148
chr3-135631310-DEL-56
chr3-135671107-DEL-57
chr3-135683474-DEL-2897
chr3-135720213-DEL-622
chr3-135727434-DEL-1609
chr3-135767509-DEL-72
chr3-135768225-DEL-714
chr3-135778114-DEL-397
chr3-135778878-DEL-710
chr3-135812691-DEL-221
chr3-135830985-DEL-1136
chr3-135841185-DEL-534
chr3-135848585-DEL-137
chr3-135863030-DEL-62
chr3-135896767-DEL-54
chr3-135897658-DEL-739
chr3-135902092-DEL-52
chr3-135907168-DEL-53
chr3-135912971-DEL-831
chr3-135934674-DEL-1162
chr3-135954777-DEL-132
chr3-135994960-DEL-917
chr3-136032953-DEL-389
chr3-136070520-DEL-2327
chr3-136094957-DEL-1835
chr3-136117745-DEL-2753
chr3-136125244-DEL-132
chr3-136146294-DEL-214
chr3-136149309-DEL-222
chr3-136170036-DEL-197
chr3-136174840-DEL-796
chr3-136177854-DEL-522
chr3-136179018-DEL-248
chr3-136189333-DEL-152
chr3-136240025-DEL-575
chr3-136285053-DEL-234
chr3-136294949-DEL-162
chr3-136300836-DEL-723
chr3-136306092-DEL-453
chr3-136341154-DEL-435
chr3-136383086-DEL-216
chr3-136419755-DEL-51
chr3-136443679-DEL-110
chr3-136465708-DEL-68
chr3-136484584-DEL-60
chr3-136495649-DEL-52
chr3-136500393-DEL-102
chr3-136536315-DEL-207
chr3-136556593-DEL-177
chr3-136598597-DEL-579
chr3-136608736-DEL-90
chr3-136648161-DEL-180

chr3-136728606-DEL-108
chr3-136747607-DEL-92
chr3-136782902-DEL-92
chr3-136785296-DEL-74
chr3-136803791-DEL-79
chr3-136837733-DEL-3262
chr3-136850371-DEL-812
chr3-136866130-DEL-166
chr3-136872336-DEL-327
chr3-136878431-DEL-92
chr3-136945576-DEL-54
chr3-136946579-DEL-121
chr3-136949466-DEL-297
chr3-136963617-DEL-688
chr3-136993741-DEL-55
chr3-137016849-DEL-162
chr3-137113125-DEL-6203
chr3-137133494-DEL-83
chr3-137139353-DEL-5302
chr3-137141939-DEL-6482
chr3-137149427-DEL-11412
chr3-137149432-DEL-13605
chr3-137151142-DEL-70
chr3-137166532-DEL-2562
chr3-137177069-DEL-53
chr3-137197281-DEL-90
chr3-137219222-DEL-413
chr3-137244030-DEL-505
chr3-137261347-DEL-85
chr3-137264315-DEL-58
chr3-137277509-DEL-159
chr3-137295088-DEL-148
chr3-137296680-DEL-667
chr3-137316456-DEL-100
chr3-137340144-DEL-1893
chr3-137364451-DEL-6379
chr3-137381456-DEL-1526
chr3-137416928-DEL-176
chr3-137421406-DEL-130
chr3-137440893-DEL-133
chr3-137451339-DEL-159
chr3-137505578-DEL-177
chr3-137526276-DEL-308
chr3-137540999-DEL-114
chr3-137546458-DEL-52
chr3-137554355-DEL-1272
chr3-137555716-DEL-54
chr3-137562477-DEL-124
chr3-137615460-DEL-78
chr3-137619803-DEL-265
chr3-137670104-DEL-52
chr3-137714717-DEL-78
chr3-137776388-DEL-73
chr3-137800131-DEL-127
chr3-137807815-DEL-222
chr3-137812104-DEL-103
chr3-137816776-DEL-362
chr3-137844081-DEL-55
chr3-137867896-DEL-174
chr3-137870051-DEL-66
chr3-137897977-DEL-51
chr3-137930550-DEL-134
chr3-137966992-DEL-83
chr3-137973295-DEL-196
chr3-137995426-DEL-56
chr3-138002465-DEL-95
chr3-138004745-DEL-232
chr3-138046329-DEL-56
chr3-138052359-DEL-58
chr3-138060299-DEL-51
chr3-138110391-DEL-66
chr3-138144038-DEL-320
chr3-138146947-DEL-55
chr3-138160245-DEL-174
chr3-138168818-DEL-305
chr3-138191085-DEL-61
chr3-138242411-DEL-308

chr3-138243006-DEL-1124
chr3-138250596-DEL-54
chr3-138294485-DEL-154
chr3-138306799-DEL-149
chr3-138319561-DEL-95
chr3-138339838-DEL-120
chr3-138347903-DEL-55
chr3-138353262-DEL-77
chr3-138355441-DEL-632
chr3-138379810-DEL-103
chr3-138387626-DEL-380
chr3-138401015-DEL-79
chr3-138456180-DEL-160
chr3-138456464-DEL-910
chr3-138457639-DEL-64
chr3-138458448-DEL-94
chr3-138501031-DEL-124
chr3-138511434-DEL-105
chr3-138596199-DEL-60
chr3-138642417-DEL-144
chr3-138642418-DEL-2586
chr3-138709195-DEL-100
chr3-138744457-DEL-184
chr3-138764297-DEL-248
chr3-138799637-DEL-74
chr3-138819557-DEL-860
chr3-138821912-DEL-159
chr3-138882494-DEL-108
chr3-138884608-DEL-335
chr3-138891177-DEL-218
chr3-138891637-DEL-2448
chr3-138925749-DEL-679
chr3-138965190-DEL-2829
chr3-138971646-DEL-387
chr3-138972762-DEL-1547
chr3-138984045-DEL-1481
chr3-139031529-DEL-6408
chr3-139049815-DEL-241
chr3-139071258-DEL-9512
chr3-139073068-DEL-6267
chr3-139085450-DEL-1070
chr3-139087054-DEL-1251
chr3-139093768-DEL-6222
chr3-139102505-DEL-96
chr3-139151628-DEL-80
chr3-139153452-DEL-68
chr3-139158392-DEL-210
chr3-139181166-DEL-57
chr3-139181699-DEL-2158
chr3-139212942-DEL-59
chr3-139225606-DEL-988
chr3-139270789-DEL-636
chr3-139300859-DEL-68
chr3-139355059-DEL-1276
chr3-139385348-DEL-228
chr3-139388989-DEL-1039
chr3-139398470-DEL-911
chr3-139400816-DEL-74
chr3-139420051-DEL-90
chr3-139424543-DEL-64
chr3-139464782-DEL-524
chr3-139477593-DEL-60
chr3-139518199-DEL-624
chr3-139521572-DEL-6223
chr3-139583863-DEL-59
chr3-139649676-DEL-83
chr3-139652618-DEL-143
chr3-139721707-DEL-84
chr3-139738598-DEL-87
chr3-139749757-DEL-157
chr3-139763101-DEL-163
chr3-139773212-DEL-1460
chr3-139809509-DEL-63
chr3-139843437-DEL-177
chr3-139846322-DEL-296
chr3-139876579-DEL-67
chr3-139880456-DEL-107

chr3-139909116-DEL-162
chr3-139926317-DEL-138
chr3-139936165-DEL-1024
chr3-139943368-DEL-55
chr3-139953404-DEL-757
chr3-139956123-DEL-172
chr3-139967031-DEL-84
chr3-139977939-DEL-1124
chr3-139986904-DEL-96
chr3-140037448-DEL-51
chr3-140052836-DEL-168
chr3-140091986-DEL-56
chr3-140098436-DEL-702
chr3-140105613-DEL-437
chr3-140115952-DEL-1294
chr3-140118972-DEL-83
chr3-140148883-DEL-734
chr3-140161315-DEL-2555
chr3-140180419-DEL-841
chr3-140203320-DEL-149
chr3-140230535-DEL-1831
chr3-140257080-DEL-65
chr3-140335402-DEL-91
chr3-140343646-DEL-245
chr3-140353056-DEL-555
chr3-140364968-DEL-96
chr3-140379483-DEL-123
chr3-140395679-DEL-136
chr3-140396372-DEL-646
chr3-140409805-DEL-56
chr3-140410365-DEL-60
chr3-140449354-DEL-107
chr3-140466362-DEL-296
chr3-140466851-DEL-12810
chr3-140478818-DEL-208
chr3-140495990-DEL-133
chr3-140503251-DEL-93
chr3-140526832-DEL-64
chr3-140546441-DEL-928
chr3-140550842-DEL-289
chr3-140557120-DEL-1936
chr3-140569623-DEL-783
chr3-140594491-DEL-671
chr3-140657773-DEL-290
chr3-140676930-DEL-54
chr3-140732641-DEL-60
chr3-140773391-DEL-16174
chr3-140779094-DEL-76
chr3-140852575-DEL-8182
chr3-140861146-DEL-1111
chr3-140868383-DEL-912
chr3-140960120-DEL-2178
chr3-140963731-DEL-54
chr3-140975219-DEL-3890
chr3-140996229-DEL-52
chr3-141014240-DEL-598
chr3-141090868-DEL-187
chr3-141131311-DEL-90
chr3-141216194-DEL-121
chr3-141227443-DEL-559
chr3-141249020-DEL-540
chr3-141389263-DEL-90
chr3-141395673-DEL-138
chr3-141401387-DEL-106
chr3-141411384-DEL-83
chr3-141424793-DEL-127
chr3-141458583-DEL-1024
chr3-141471775-DEL-212
chr3-141552879-DEL-399
chr3-141565152-DEL-86
chr3-141565878-DEL-2143
chr3-141595747-DEL-86
chr3-141597050-DEL-66
chr3-141598949-DEL-109
chr3-141628198-DEL-76
chr3-141668216-DEL-98
chr3-141715899-DEL-85

chr3-141733837-DEL-801
chr3-141825715-DEL-52
chr3-141854942-DEL-62
chr3-141917130-DEL-208
chr3-141965927-DEL-482
chr3-141973373-DEL-445
chr3-141990931-DEL-195
chr3-142003433-DEL-125
chr3-142028256-DEL-96
chr3-142101122-DEL-52
chr3-142138930-DEL-88
chr3-142163588-DEL-285
chr3-142268019-DEL-219
chr3-142284815-DEL-611
chr3-142314684-DEL-829
chr3-142347314-DEL-92
chr3-142353715-DEL-349
chr3-142367107-DEL-139
chr3-142377938-DEL-56
chr3-142382685-DEL-197
chr3-142389088-DEL-146
chr3-142402499-DEL-65
chr3-142413682-DEL-520
chr3-142418870-DEL-59
chr3-142434688-DEL-500
chr3-142479296-DEL-51
chr3-142513382-DEL-276
chr3-142554232-DEL-203
chr3-142567400-DEL-81
chr3-142587200-DEL-80
chr3-142610440-DEL-204
chr3-142621961-DEL-64
chr3-142654275-DEL-52
chr3-142737432-DEL-76
chr3-142737874-DEL-576
chr3-142767772-DEL-2026
chr3-142819394-DEL-516
chr3-142850393-DEL-60
chr3-142874269-DEL-141
chr3-142876606-DEL-831
chr3-142882257-DEL-1718
chr3-142889765-DEL-782
chr3-142894475-DEL-226
chr3-142906673-DEL-145
chr3-142908525-DEL-64
chr3-142916132-DEL-2100
chr3-142944909-DEL-72
chr3-143016670-DEL-330
chr3-143044870-DEL-72
chr3-143147308-DEL-51
chr3-143191176-DEL-1785
chr3-143217539-DEL-52
chr3-143263020-DEL-328
chr3-143306995-DEL-264
chr3-143351984-DEL-55
chr3-143364942-DEL-246
chr3-143378063-DEL-58
chr3-143382949-DEL-934
chr3-143394198-DEL-113
chr3-143395226-DEL-341
chr3-143432537-DEL-115
chr3-143536364-DEL-72
chr3-143556037-DEL-7340
chr3-143581189-DEL-1256
chr3-143587758-DEL-58
chr3-143617345-DEL-64
chr3-143680670-DEL-126
chr3-143692494-DEL-56
chr3-143736029-DEL-6219
chr3-143756823-DEL-82
chr3-143764739-DEL-96
chr3-143814867-DEL-141
chr3-143850147-DEL-109
chr3-143900588-DEL-79
chr3-143934374-DEL-346
chr3-143949392-DEL-78
chr3-143955212-DEL-131

chr3-143955601-DEL-836
chr3-144037722-DEL-68
chr3-144039042-DEL-70
chr3-144051967-DEL-331
chr3-144079860-DEL-85
chr3-144108879-DEL-66
chr3-144129229-DEL-508
chr3-144132887-DEL-584
chr3-144187361-DEL-214
chr3-144199599-DEL-111
chr3-144232722-DEL-144
chr3-144252665-DEL-1254
chr3-144252822-DEL-358
chr3-144252937-DEL-806
chr3-144260447-DEL-2010
chr3-144268939-DEL-202
chr3-144277303-DEL-57
chr3-144311848-DEL-181
chr3-144338787-DEL-195
chr3-144344473-DEL-150
chr3-144345115-DEL-71
chr3-144357300-DEL-420
chr3-144358099-DEL-9751
chr3-144384472-DEL-199
chr3-144420918-DEL-62
chr3-144437942-DEL-1477
chr3-144477429-DEL-2774
chr3-144509685-DEL-1024
chr3-144509956-DEL-165
chr3-144510583-DEL-66
chr3-144528603-DEL-162
chr3-144533601-DEL-196
chr3-144616438-DEL-281
chr3-144622874-DEL-506
chr3-144693310-DEL-98
chr3-144739319-DEL-142
chr3-144747885-DEL-91
chr3-144755123-DEL-317
chr3-144758343-DEL-227
chr3-144795998-DEL-234
chr3-144801711-DEL-53
chr3-144868875-DEL-291
chr3-144877366-DEL-68
chr3-144893668-DEL-2700
chr3-144894632-DEL-242
chr3-144928626-DEL-607
chr3-144981853-DEL-55
chr3-145007088-DEL-102
chr3-145017083-DEL-83
chr3-145026576-DEL-140
chr3-145027342-DEL-57
chr3-145037550-DEL-66
chr3-145052081-DEL-90
chr3-145057871-DEL-432
chr3-145089651-DEL-633
chr3-145092564-DEL-51
chr3-145127070-DEL-150
chr3-145140069-DEL-101
chr3-145184829-DEL-66
chr3-145235761-DEL-53
chr3-145249800-DEL-318
chr3-145251964-DEL-149
chr3-145266763-DEL-57
chr3-145268744-DEL-96
chr3-145292816-DEL-290
chr3-145311904-DEL-125
chr3-145349843-DEL-147
chr3-145392203-DEL-740
chr3-145405115-DEL-462
chr3-145415493-DEL-280
chr3-145416784-DEL-1177
chr3-145431751-DEL-174
chr3-145450303-DEL-73
chr3-145484607-DEL-527
chr3-145502065-DEL-147
chr3-145547380-DEL-64
chr3-145554282-DEL-454

chr3-145554835-DEL-288
chr3-145556409-DEL-256
chr3-145566230-DEL-156
chr3-145570856-DEL-416
chr3-145579384-DEL-172
chr3-145592722-DEL-52
chr3-145604322-DEL-102
chr3-145634842-DEL-172
chr3-145658176-DEL-224
chr3-145660053-DEL-102
chr3-145664998-DEL-101
chr3-145673805-DEL-197
chr3-145717260-DEL-61
chr3-145719397-DEL-114
chr3-145732199-DEL-97
chr3-145734289-DEL-59
chr3-145741447-DEL-57
chr3-145744594-DEL-151
chr3-145767799-DEL-106
chr3-145778918-DEL-6310
chr3-145780650-DEL-102
chr3-145785605-DEL-152
chr3-145802429-DEL-60
chr3-145806973-DEL-481
chr3-145828173-DEL-52
chr3-145829926-DEL-93
chr3-145834733-DEL-54
chr3-145839759-DEL-132
chr3-145850421-DEL-56
chr3-145866617-DEL-484
chr3-145870520-DEL-81
chr3-145906455-DEL-56
chr3-145916086-DEL-257
chr3-145960793-DEL-105
chr3-145974446-DEL-66
chr3-145977103-DEL-59
chr3-146028663-DEL-1680
chr3-146035272-DEL-53
chr3-146039525-DEL-60
chr3-146054819-DEL-62
chr3-146072507-DEL-54
chr3-146075840-DEL-152
chr3-146082920-DEL-1479
chr3-146103283-DEL-260
chr3-146104884-DEL-56
chr3-146120988-DEL-927
chr3-146133415-DEL-78
chr3-146138924-DEL-168
chr3-146146521-DEL-208
chr3-146173070-DEL-55
chr3-146182524-DEL-1779
chr3-146182569-DEL-214
chr3-146183407-DEL-596
chr3-146208070-DEL-439
chr3-146214279-DEL-56
chr3-146214939-DEL-409
chr3-146221476-DEL-51
chr3-146228136-DEL-185
chr3-146237026-DEL-201
chr3-146238403-DEL-1181
chr3-146258408-DEL-863
chr3-146265740-DEL-318
chr3-146273376-DEL-58
chr3-146277393-DEL-104
chr3-146279071-DEL-350
chr3-146320262-DEL-54
chr3-146366493-DEL-64
chr3-146372833-DEL-465
chr3-146377286-DEL-708
chr3-146401521-DEL-107
chr3-146417719-DEL-66
chr3-146442477-DEL-117
chr3-146460642-DEL-566
chr3-146542944-DEL-138
chr3-146608460-DEL-58
chr3-146654688-DEL-54
chr3-146672050-DEL-719

chr3-146724182-DEL-69
chr3-146729091-DEL-1521
chr3-146744661-DEL-207
chr3-146754640-DEL-600
chr3-146770976-DEL-213
chr3-146809851-DEL-63
chr3-146811744-DEL-8332
chr3-146814879-DEL-1888
chr3-146846127-DEL-60
chr3-146853710-DEL-252
chr3-146856027-DEL-463
chr3-146933536-DEL-2330
chr3-146937573-DEL-56
chr3-146940411-DEL-478
chr3-146983122-DEL-78
chr3-146984009-DEL-74
chr3-147096919-DEL-53
chr3-147100391-DEL-73
chr3-147131003-DEL-525
chr3-147134440-DEL-60
chr3-147140460-DEL-88
chr3-147154136-DEL-211
chr3-147155088-DEL-123
chr3-147199484-DEL-81
chr3-147214978-DEL-54
chr3-147225076-DEL-126
chr3-147245816-DEL-127
chr3-147247338-DEL-576
chr3-147253926-DEL-68
chr3-147331245-DEL-122
chr3-147334757-DEL-90
chr3-147335274-DEL-444
chr3-147365690-DEL-420
chr3-147372259-DEL-316
chr3-147372580-DEL-1314
chr3-147373501-DEL-239
chr3-147375709-DEL-172
chr3-147387799-DEL-97
chr3-147487557-DEL-65
chr3-147497439-DEL-1263
chr3-147520159-DEL-458
chr3-147521924-DEL-107
chr3-147627985-DEL-621
chr3-147631333-DEL-68
chr3-147646280-DEL-56
chr3-147678906-DEL-241
chr3-147691599-DEL-228
chr3-147727052-DEL-129
chr3-147768621-DEL-61
chr3-147769392-DEL-169
chr3-147785499-DEL-430
chr3-147800772-DEL-84
chr3-147807196-DEL-62
chr3-147851212-DEL-587
chr3-147857012-DEL-1244
chr3-147873917-DEL-143
chr3-147888975-DEL-64
chr3-147916235-DEL-55
chr3-147924991-DEL-132
chr3-147928915-DEL-105
chr3-147935657-DEL-53
chr3-147936393-DEL-54
chr3-147949566-DEL-856
chr3-147998506-DEL-313
chr3-148026680-DEL-83
chr3-148056321-DEL-76
chr3-148060293-DEL-76
chr3-148089187-DEL-51
chr3-148106368-DEL-56
chr3-148140319-DEL-80
chr3-148146062-DEL-124
chr3-148171557-DEL-92
chr3-148191849-DEL-260
chr3-148210882-DEL-60
chr3-148218104-DEL-86
chr3-148248363-DEL-68
chr3-148273200-DEL-62

chr3-148281387-DEL-114
chr3-148290699-DEL-294
chr3-148303552-DEL-62
chr3-148332438-DEL-1478
chr3-148345402-DEL-53
chr3-148360755-DEL-136
chr3-148365653-DEL-602
chr3-148377911-DEL-93
chr3-148409878-DEL-6226
chr3-148431985-DEL-54
chr3-148468993-DEL-654
chr3-148474153-DEL-57
chr3-148505209-DEL-62
chr3-148514094-DEL-81
chr3-148517175-DEL-60
chr3-148533964-DEL-65
chr3-148584758-DEL-60
chr3-148695859-DEL-92
chr3-148764441-DEL-317
chr3-148772896-DEL-72
chr3-148779053-DEL-53
chr3-148782335-DEL-53
chr3-148792050-DEL-70
chr3-148792862-DEL-84
chr3-148803350-DEL-102
chr3-148814063-DEL-54
chr3-148888563-DEL-56
chr3-148892408-DEL-52
chr3-148893169-DEL-120
chr3-148912134-DEL-88
chr3-148922330-DEL-66
chr3-148928613-DEL-364
chr3-148948330-DEL-1250
chr3-148982380-DEL-631
chr3-149023564-DEL-1075
chr3-149028385-DEL-872
chr3-149032983-DEL-287
chr3-149051517-DEL-51
chr3-149143445-DEL-65
chr3-149143959-DEL-104
chr3-149153539-DEL-1331
chr3-149177633-DEL-971
chr3-149178735-DEL-1639
chr3-149211082-DEL-242
chr3-149212311-DEL-205
chr3-149218993-DEL-165
chr3-149232305-DEL-168
chr3-149233645-DEL-353
chr3-149253349-DEL-154
chr3-149297533-DEL-52
chr3-149301962-DEL-402
chr3-149371881-DEL-401
chr3-149373358-DEL-193
chr3-149400769-DEL-76
chr3-149443268-DEL-52
chr3-149445691-DEL-192
chr3-149450484-DEL-160
chr3-149481870-DEL-4140
chr3-149488039-DEL-358
chr3-149490251-DEL-316
chr3-149506529-DEL-65
chr3-149556787-DEL-612
chr3-149569345-DEL-677
chr3-149576835-DEL-59
chr3-149581137-DEL-313
chr3-149586379-DEL-65
chr3-149600490-DEL-169
chr3-149630984-DEL-107
chr3-149644458-DEL-425
chr3-149686379-DEL-167
chr3-149693020-DEL-186
chr3-149741127-DEL-203
chr3-149743244-DEL-65
chr3-149795404-DEL-73
chr3-149798147-DEL-58
chr3-149800721-DEL-304
chr3-149814250-DEL-51

chr3-149825975-DEL-155
chr3-149833140-DEL-2129
chr3-149835932-DEL-740
chr3-149836802-DEL-242
chr3-149846997-DEL-83
chr3-149852142-DEL-87
chr3-149887648-DEL-191
chr3-149931436-DEL-168
chr3-149938283-DEL-52
chr3-149980948-DEL-173
chr3-149995096-DEL-146
chr3-150006951-DEL-176
chr3-150055024-DEL-877
chr3-150065634-DEL-183
chr3-150067822-DEL-58
chr3-150077608-DEL-180
chr3-150079780-DEL-51
chr3-150126392-DEL-933
chr3-150145757-DEL-341
chr3-150146491-DEL-121
chr3-150179880-DEL-168
chr3-150208412-DEL-126
chr3-150217019-DEL-471
chr3-150226309-DEL-78
chr3-150229431-DEL-197
chr3-150277171-DEL-1459
chr3-150279169-DEL-53
chr3-150301156-DEL-2426
chr3-150344147-DEL-584
chr3-150426774-DEL-6165
chr3-150447365-DEL-3883
chr3-150461897-DEL-63
chr3-150468400-DEL-72
chr3-150469797-DEL-58
chr3-150539403-DEL-3651
chr3-150568225-DEL-979
chr3-150613215-DEL-172
chr3-150618797-DEL-52
chr3-150632412-DEL-433
chr3-150660288-DEL-52
chr3-150664876-DEL-1794
chr3-150701055-DEL-100
chr3-150718730-DEL-67
chr3-150763651-DEL-62
chr3-150799254-DEL-143
chr3-150813558-DEL-111
chr3-150851861-DEL-250
chr3-150858795-DEL-205
chr3-150900334-DEL-3661
chr3-150918152-DEL-88
chr3-150951760-DEL-93
chr3-150962032-DEL-918
chr3-150974871-DEL-72
chr3-151014911-DEL-211
chr3-151051493-DEL-70
chr3-151074250-DEL-425
chr3-151150646-DEL-595
chr3-151167354-DEL-115
chr3-151183476-DEL-54
chr3-151238205-DEL-52
chr3-151245112-DEL-180
chr3-151246858-DEL-52
chr3-151304237-DEL-99
chr3-151307063-DEL-313
chr3-151317918-DEL-312
chr3-151372483-DEL-105
chr3-151374748-DEL-164
chr3-151407934-DEL-528
chr3-151448106-DEL-1874
chr3-151456513-DEL-130
chr3-151478132-DEL-1671
chr3-151491915-DEL-60
chr3-151495492-DEL-204
chr3-151497084-DEL-85
chr3-151508964-DEL-286
chr3-151524257-DEL-186
chr3-151529370-DEL-117

chr3-151554316-DEL-454
chr3-151560659-DEL-388
chr3-151572378-DEL-59
chr3-151579632-DEL-80
chr3-151663749-DEL-1555
chr3-151705478-DEL-81
chr3-151709349-DEL-85
chr3-151736618-DEL-888
chr3-151740739-DEL-369
chr3-151745135-DEL-129
chr3-151754241-DEL-1689
chr3-151761887-DEL-262
chr3-151790243-DEL-159
chr3-151793613-DEL-54
chr3-151845263-DEL-123
chr3-151864485-DEL-80
chr3-151914874-DEL-95
chr3-151918411-DEL-76
chr3-151945437-DEL-129
chr3-151955642-DEL-79
chr3-151977368-DEL-154
chr3-151985336-DEL-97
chr3-152022126-DEL-77
chr3-152038191-DEL-99
chr3-152053489-DEL-170
chr3-152075450-DEL-160
chr3-152105239-DEL-4359
chr3-152108655-DEL-157
chr3-152117256-DEL-141
chr3-152126766-DEL-207
chr3-152182033-DEL-93
chr3-152184671-DEL-89
chr3-152185107-DEL-226
chr3-152191756-DEL-360
chr3-152196858-DEL-96
chr3-152205712-DEL-73
chr3-152208562-DEL-115
chr3-152224959-DEL-177
chr3-152227380-DEL-77
chr3-152239444-DEL-319
chr3-152252037-DEL-162
chr3-152261698-DEL-96
chr3-152268823-DEL-194
chr3-152269767-DEL-99
chr3-152275847-DEL-6205
chr3-152295172-DEL-66
chr3-152304597-DEL-238
chr3-152355693-DEL-94
chr3-152360136-DEL-183
chr3-152388672-DEL-421
chr3-152389163-DEL-980
chr3-152389472-DEL-78
chr3-152421554-DEL-53
chr3-152422967-DEL-91
chr3-152427346-DEL-118
chr3-152445996-DEL-58
chr3-152447713-DEL-315
chr3-152449458-DEL-155
chr3-152457274-DEL-330
chr3-152461260-DEL-52
chr3-152482369-DEL-1672
chr3-152484749-DEL-184
chr3-152490145-DEL-206
chr3-152550785-DEL-160
chr3-152556798-DEL-113
chr3-152568007-DEL-70
chr3-152569266-DEL-107
chr3-152596855-DEL-132
chr3-152614794-DEL-96
chr3-152621805-DEL-126
chr3-152663167-DEL-94
chr3-152706791-DEL-153
chr3-152709840-DEL-60
chr3-152710927-DEL-358
chr3-152712369-DEL-63
chr3-152730735-DEL-184
chr3-152738849-DEL-112

chr3-152748239-DEL-135
chr3-152758770-DEL-62
chr3-152763001-DEL-1660
chr3-152765272-DEL-73
chr3-152774359-DEL-65
chr3-152805665-DEL-57
chr3-152814228-DEL-129
chr3-152819197-DEL-77
chr3-152845024-DEL-63
chr3-152883597-DEL-1132
chr3-152931035-DEL-60
chr3-152931864-DEL-145
chr3-152967796-DEL-67
chr3-152972839-DEL-52
chr3-152992759-DEL-145
chr3-153017754-DEL-134
chr3-153024940-DEL-81
chr3-153035470-DEL-72
chr3-153038944-DEL-586
chr3-153059544-DEL-65
chr3-153061874-DEL-57
chr3-153074674-DEL-58
chr3-153076603-DEL-67
chr3-153112650-DEL-98
chr3-153119679-DEL-125
chr3-153132424-DEL-325
chr3-153133094-DEL-395
chr3-153134245-DEL-196
chr3-153139779-DEL-154
chr3-153152649-DEL-205
chr3-153186559-DEL-831
chr3-153212171-DEL-288
chr3-153229801-DEL-554
chr3-153249473-DEL-295
chr3-153272058-DEL-253
chr3-153278325-DEL-118
chr3-153313864-DEL-69
chr3-153321790-DEL-368
chr3-153330315-DEL-383
chr3-153332527-DEL-77
chr3-153364821-DEL-90
chr3-153371738-DEL-554
chr3-153377245-DEL-143
chr3-153410776-DEL-142
chr3-153415989-DEL-210
chr3-153427427-DEL-1239
chr3-153436463-DEL-56
chr3-153461639-DEL-53
chr3-153465617-DEL-120
chr3-153475588-DEL-307
chr3-153477322-DEL-213
chr3-153494301-DEL-181
chr3-153516229-DEL-106
chr3-153525120-DEL-209
chr3-153525927-DEL-830
chr3-153556009-DEL-123
chr3-153587467-DEL-137
chr3-153600324-DEL-116
chr3-153604420-DEL-865
chr3-153605522-DEL-56
chr3-153624897-DEL-191
chr3-153626529-DEL-199
chr3-153627743-DEL-123
chr3-153672784-DEL-59
chr3-153677316-DEL-173
chr3-153685701-DEL-87
chr3-153700166-DEL-172
chr3-153724138-DEL-398
chr3-153735205-DEL-201
chr3-153772051-DEL-322
chr3-153772052-DEL-919
chr3-153772236-DEL-831
chr3-153779817-DEL-271
chr3-153811355-DEL-52
chr3-153837020-DEL-80
chr3-153860938-DEL-2874
chr3-153874476-DEL-55

chr3-153897548-DEL-56
chr3-153944252-DEL-79
chr3-153959961-DEL-160
chr3-153978070-DEL-61
chr3-153987270-DEL-83
chr3-154021542-DEL-181
chr3-154040501-DEL-63
chr3-154054736-DEL-361
chr3-154056517-DEL-52
chr3-154061904-DEL-4143
chr3-154069736-DEL-131
chr3-154099916-DEL-56
chr3-154102020-DEL-56
chr3-154110838-DEL-630
chr3-154120580-DEL-1458
chr3-154121237-DEL-801
chr3-154128228-DEL-1202
chr3-154128316-DEL-280
chr3-154160822-DEL-96
chr3-154178465-DEL-640
chr3-154179487-DEL-160
chr3-154191963-DEL-54
chr3-154209749-DEL-94
chr3-154223231-DEL-60
chr3-154247707-DEL-116
chr3-154283946-DEL-506
chr3-154286533-DEL-52
chr3-154305687-DEL-72
chr3-154305975-DEL-996
chr3-154311480-DEL-228
chr3-154321849-DEL-59
chr3-154363259-DEL-64
chr3-154365082-DEL-75
chr3-154368573-DEL-136
chr3-154381809-DEL-54
chr3-154385634-DEL-3377
chr3-154386881-DEL-167
chr3-154438430-DEL-57
chr3-154503660-DEL-196
chr3-154513632-DEL-103
chr3-154529550-DEL-87
chr3-154532776-DEL-224
chr3-154549904-DEL-59
chr3-154551888-DEL-878
chr3-154552615-DEL-134
chr3-154579324-DEL-304
chr3-154590432-DEL-164
chr3-154599045-DEL-68
chr3-154630697-DEL-96
chr3-154684066-DEL-205
chr3-154685096-DEL-52
chr3-154688699-DEL-160
chr3-154706397-DEL-533
chr3-154711347-DEL-51
chr3-154712013-DEL-216
chr3-154750828-DEL-71
chr3-154770306-DEL-52
chr3-154776764-DEL-62
chr3-154797246-DEL-156
chr3-154808417-DEL-274
chr3-154847317-DEL-292
chr3-154851576-DEL-6414
chr3-154868851-DEL-56
chr3-154872137-DEL-845
chr3-154888553-DEL-194
chr3-154890972-DEL-221
chr3-154894381-DEL-1036
chr3-154922779-DEL-130
chr3-154947241-DEL-102
chr3-154950430-DEL-99
chr3-154994188-DEL-641
chr3-155063053-DEL-193
chr3-155082472-DEL-305
chr3-155098810-DEL-192
chr3-155108929-DEL-120
chr3-155138758-DEL-92
chr3-155156294-DEL-69

chr3-155170519-DEL-1161
chr3-155172612-DEL-52
chr3-155230223-DEL-1506
chr3-155246206-DEL-1167
chr3-155252202-DEL-77
chr3-155255949-DEL-299
chr3-155277051-DEL-203
chr3-155288834-DEL-987
chr3-155290462-DEL-575
chr3-155298736-DEL-216
chr3-155308221-DEL-127
chr3-155312716-DEL-144
chr3-155318388-DEL-54
chr3-155322479-DEL-482
chr3-155351907-DEL-379
chr3-155355929-DEL-140
chr3-155376208-DEL-95
chr3-155407712-DEL-4223
chr3-155413666-DEL-55
chr3-155461889-DEL-422
chr3-155500905-DEL-1025
chr3-155531563-DEL-1655
chr3-155550052-DEL-91
chr3-155554323-DEL-170
chr3-155591136-DEL-618
chr3-155608542-DEL-6168
chr3-155643596-DEL-537
chr3-155650762-DEL-848
chr3-155667123-DEL-73
chr3-155683253-DEL-1233
chr3-155716873-DEL-104
chr3-155740956-DEL-1638
chr3-155751122-DEL-154
chr3-155754320-DEL-2624
chr3-155760186-DEL-53
chr3-155796557-DEL-603
chr3-155817814-DEL-1546
chr3-155838769-DEL-92
chr3-155849024-DEL-66
chr3-155849355-DEL-369
chr3-155853742-DEL-1043
chr3-155895297-DEL-1387
chr3-155902382-DEL-243
chr3-155922618-DEL-483
chr3-155933827-DEL-74
chr3-155960374-DEL-466
chr3-155962654-DEL-5741
chr3-155984042-DEL-52
chr3-156011860-DEL-446
chr3-156082691-DEL-438
chr3-156158655-DEL-178
chr3-156180174-DEL-582
chr3-156182502-DEL-286
chr3-156194087-DEL-4096
chr3-156243839-DEL-321
chr3-156286567-DEL-114
chr3-156306741-DEL-352
chr3-156375261-DEL-746
chr3-156407550-DEL-62
chr3-156408809-DEL-71
chr3-156428830-DEL-54
chr3-156460784-DEL-52
chr3-156467202-DEL-56
chr3-156489000-DEL-74
chr3-156533568-DEL-108
chr3-156580753-DEL-102
chr3-156599691-DEL-167
chr3-156622122-DEL-88
chr3-156666289-DEL-59
chr3-156677454-DEL-73
chr3-156711624-DEL-307
chr3-156723583-DEL-143
chr3-156788128-DEL-59
chr3-156798788-DEL-152
chr3-156800225-DEL-142
chr3-156856523-DEL-64
chr3-156939585-DEL-91

chr3-156954374-DEL-70
chr3-157033768-DEL-132
chr3-157051041-DEL-2954
chr3-157088898-DEL-51
chr3-157117705-DEL-6895
chr3-157146634-DEL-1384
chr3-157187936-DEL-195
chr3-157226804-DEL-78
chr3-157231188-DEL-227
chr3-157256757-DEL-1271
chr3-157319065-DEL-64
chr3-157338666-DEL-131
chr3-157351195-DEL-2428
chr3-157385208-DEL-123
chr3-157412329-DEL-59
chr3-157440158-DEL-97
chr3-157477754-DEL-5512
chr3-157498740-DEL-81
chr3-157506570-DEL-66
chr3-157529255-DEL-125
chr3-157530115-DEL-1143
chr3-157564287-DEL-77
chr3-157565438-DEL-174
chr3-157576757-DEL-56
chr3-157591699-DEL-127
chr3-157606466-DEL-3602
chr3-157636051-DEL-96
chr3-157645972-DEL-2228
chr3-157647135-DEL-486
chr3-157659111-DEL-201
chr3-157669080-DEL-140
chr3-157670523-DEL-57
chr3-157671299-DEL-184
chr3-157673025-DEL-411
chr3-157677468-DEL-118
chr3-157688050-DEL-107
chr3-157701718-DEL-79
chr3-157767937-DEL-58
chr3-157800904-DEL-6922
chr3-157818182-DEL-6380
chr3-157842549-DEL-300
chr3-157843163-DEL-208
chr3-158006565-DEL-83
chr3-158063795-DEL-2109
chr3-158078767-DEL-127
chr3-158087904-DEL-303
chr3-158138929-DEL-2312
chr3-158141641-DEL-58
chr3-158155903-DEL-138
chr3-158160914-DEL-770
chr3-158168391-DEL-174
chr3-158235025-DEL-514
chr3-158240130-DEL-392
chr3-158243461-DEL-510
chr3-158277367-DEL-326
chr3-158279151-DEL-1657
chr3-158285368-DEL-59
chr3-158308624-DEL-216
chr3-158323918-DEL-1104
chr3-158342838-DEL-607
chr3-158345204-DEL-86
chr3-158354753-DEL-304
chr3-158357841-DEL-82
chr3-158388530-DEL-259
chr3-158392011-DEL-68
chr3-158393050-DEL-689
chr3-158395889-DEL-885
chr3-158442969-DEL-1872
chr3-158443772-DEL-208
chr3-158445048-DEL-208
chr3-158470129-DEL-68
chr3-158518279-DEL-138
chr3-158536753-DEL-53
chr3-158539469-DEL-838
chr3-158543759-DEL-81
chr3-158561821-DEL-6194
chr3-158574734-DEL-52

chr3-158656760-DEL-304
chr3-158688152-DEL-139
chr3-158705604-DEL-1166
chr3-158707044-DEL-1174
chr3-158766018-DEL-2585
chr3-158769109-DEL-54
chr3-158777948-DEL-190
chr3-158786641-DEL-595
chr3-158802575-DEL-160
chr3-158839065-DEL-8138
chr3-158862431-DEL-429
chr3-158869005-DEL-68
chr3-158871389-DEL-6607
chr3-158879409-DEL-6037
chr3-158904180-DEL-191
chr3-158910244-DEL-80
chr3-158956834-DEL-58
chr3-158998276-DEL-6033
chr3-159060109-DEL-181
chr3-159090888-DEL-6283
chr3-159116274-DEL-2024
chr3-159116464-DEL-166
chr3-159162598-DEL-206
chr3-159170534-DEL-22550
chr3-159229987-DEL-74
chr3-159246974-DEL-209
chr3-159257405-DEL-5864
chr3-159290191-DEL-363
chr3-159299188-DEL-367
chr3-159374382-DEL-6669
chr3-159452801-DEL-54
chr3-159454777-DEL-2010
chr3-159484814-DEL-65
chr3-159529931-DEL-57
chr3-159530474-DEL-2445
chr3-159579985-DEL-198
chr3-159626826-DEL-62
chr4-3053669-DEL-62
chr4-3060087-DEL-833
chr4-3061177-DEL-105
chr4-3072866-DEL-6199
chr4-3079731-DEL-108
chr4-3085534-DEL-63
chr4-3088551-DEL-565
chr4-3090247-DEL-63
chr4-3101251-DEL-76
chr4-3114898-DEL-214
chr4-3161203-DEL-121
chr4-3175202-DEL-570
chr4-3180466-DEL-175
chr4-3230528-DEL-1148
chr4-3233728-DEL-948
chr4-3240074-DEL-244
chr4-3269739-DEL-76
chr4-3278392-DEL-443
chr4-3300395-DEL-67
chr4-3330532-DEL-8285
chr4-3358032-DEL-2732
chr4-3371165-DEL-366
chr4-3382759-DEL-55
chr4-3386365-DEL-474
chr4-3422268-DEL-81
chr4-3512366-DEL-131
chr4-3540976-DEL-3324
chr4-3543623-DEL-4282
chr4-3595830-DEL-55
chr4-3613368-DEL-576
chr4-3614057-DEL-350
chr4-3627441-DEL-4932
chr4-3661522-DEL-254
chr4-3669186-DEL-60
chr4-3692176-DEL-78
chr4-3697194-DEL-125
chr4-3755470-DEL-242
chr4-3760101-DEL-121
chr4-3791919-DEL-60
chr4-3796832-DEL-162

chr4-3798115-DEL-358
chr4-3801358-DEL-75
chr4-3806583-DEL-3102
chr4-3811465-DEL-401
chr4-3818245-DEL-465
chr4-3889303-DEL-73
chr4-3889494-DEL-1852
chr4-3967092-DEL-55
chr4-4004734-DEL-59
chr4-4014003-DEL-298
chr4-4028773-DEL-6172
chr4-4040653-DEL-5825
chr4-4049109-DEL-214
chr4-4082941-DEL-92
chr4-4091042-DEL-5610
chr4-4120067-DEL-3293
chr4-4149227-DEL-118
chr4-4160963-DEL-6278
chr4-4188984-DEL-489
chr4-4211503-DEL-70
chr4-4223534-DEL-1295
chr4-4248572-DEL-675
chr4-4268458-DEL-237
chr4-4275774-DEL-10161
chr4-4288489-DEL-130
chr4-4301174-DEL-55
chr4-4312241-DEL-1758
chr4-4357502-DEL-2960
chr4-4389779-DEL-6047
chr4-4405157-DEL-51
chr4-4424539-DEL-1205
chr4-4460548-DEL-467
chr4-4470085-DEL-99
chr4-4481286-DEL-186
chr4-4536001-DEL-3289
chr4-4562192-DEL-85
chr4-4604962-DEL-221
chr4-4623137-DEL-64
chr4-4627740-DEL-825
chr4-4649588-DEL-122
chr4-4686862-DEL-52
chr4-4713141-DEL-136
chr4-4724881-DEL-3287
chr4-4729403-DEL-153
chr4-4744386-DEL-78
chr4-4799798-DEL-12357
chr4-4809009-DEL-55
chr4-4820219-DEL-62
chr4-4828572-DEL-66
chr4-4843715-DEL-3513
chr4-4858818-DEL-3508
chr4-4862362-DEL-160
chr4-4881528-DEL-71
chr4-4895970-DEL-6221
chr4-4909139-DEL-6527
chr4-4933429-DEL-593
chr4-4950370-DEL-197
chr4-5019082-DEL-199
chr4-5024170-DEL-52
chr4-5039269-DEL-6605
chr4-5045479-DEL-201
chr4-5050841-DEL-1129
chr4-5066506-DEL-77
chr4-5078068-DEL-400
chr4-5090705-DEL-6412
chr4-5105769-DEL-2453
chr4-5115768-DEL-722
chr4-5234751-DEL-1482
chr4-5248511-DEL-1013
chr4-5265772-DEL-470
chr4-5272129-DEL-871
chr4-5294438-DEL-97
chr4-5357622-DEL-80
chr4-5369355-DEL-6414
chr4-5407926-DEL-58
chr4-5411210-DEL-90
chr4-5419822-DEL-67

chr4-5442730-DEL-162
chr4-5467158-DEL-59
chr4-5470449-DEL-73
chr4-5503493-DEL-150
chr4-5524623-DEL-2061
chr4-5526670-DEL-738
chr4-5535663-DEL-249
chr4-5541404-DEL-190
chr4-5549111-DEL-68
chr4-5567357-DEL-62
chr4-5635613-DEL-401
chr4-5647494-DEL-716
chr4-5659498-DEL-436
chr4-5748573-DEL-350
chr4-5765373-DEL-189
chr4-5849594-DEL-439
chr4-5862283-DEL-117
chr4-5867208-DEL-62
chr4-5874330-DEL-784
chr4-5877293-DEL-7329
chr4-5927544-DEL-5641
chr4-5952964-DEL-836
chr4-6031269-DEL-116
chr4-6050097-DEL-80
chr4-6051377-DEL-175
chr4-6067958-DEL-62
chr4-6155825-DEL-277
chr4-6194994-DEL-63
chr4-6264670-DEL-108
chr4-6283869-DEL-334
chr4-6314928-DEL-206
chr4-6319370-DEL-9812
chr4-6360383-DEL-1694
chr4-6363006-DEL-303
chr4-6431522-DEL-1073
chr4-6431701-DEL-359
chr4-6442221-DEL-371
chr4-6447814-DEL-628
chr4-6465810-DEL-172
chr4-6467368-DEL-106
chr4-6470282-DEL-52
chr4-6482306-DEL-92
chr4-6510507-DEL-181
chr4-6556681-DEL-60
chr4-6557745-DEL-147
chr4-6560046-DEL-463
chr4-6565044-DEL-69
chr4-6576256-DEL-309
chr4-6586304-DEL-383
chr4-6591344-DEL-57
chr4-6603461-DEL-6401
chr4-6624877-DEL-1864
chr4-6629304-DEL-77
chr4-6635854-DEL-200
chr4-6638572-DEL-561
chr4-6685615-DEL-268
chr4-6705086-DEL-93
chr4-6721827-DEL-108
chr4-6725779-DEL-222
chr4-6737791-DEL-72
chr4-6750921-DEL-86
chr4-6759999-DEL-89
chr4-6850808-DEL-757
chr4-6871859-DEL-115
chr4-6976539-DEL-56
chr4-6992135-DEL-86
chr4-7008248-DEL-1357
chr4-7020068-DEL-69
chr4-7024532-DEL-74
chr4-7033363-DEL-196
chr4-7043663-DEL-472
chr4-7054300-DEL-62
chr4-7066473-DEL-106
chr4-7108793-DEL-59
chr4-7113255-DEL-56
chr4-7113828-DEL-61
chr4-7133530-DEL-487

chr4-7219890-DEL-348
chr4-7249200-DEL-144
chr4-7268038-DEL-87
chr4-7284654-DEL-61
chr4-7345327-DEL-52
chr4-7390434-DEL-623
chr4-7391753-DEL-79
chr4-7412801-DEL-66
chr4-7454064-DEL-158
chr4-7511638-DEL-53
chr4-7518601-DEL-485
chr4-7519608-DEL-1476
chr4-7523360-DEL-2112
chr4-7526227-DEL-1050
chr4-7530336-DEL-320
chr4-7535548-DEL-357
chr4-7569399-DEL-141
chr4-7597357-DEL-52
chr4-7617728-DEL-54
chr4-7627614-DEL-12781
chr4-7627623-DEL-2557
chr4-7628273-DEL-6975
chr4-7651836-DEL-1072
chr4-7659454-DEL-2666
chr4-7673341-DEL-93
chr4-7692274-DEL-3448
chr4-7695312-DEL-424
chr4-7702743-DEL-6603
chr4-7713047-DEL-1497
chr4-7720631-DEL-64
chr4-7721289-DEL-142
chr4-7733675-DEL-6475
chr4-7800914-DEL-865
chr4-7858080-DEL-201
chr4-7869600-DEL-6228
chr4-7878168-DEL-55
chr4-7879265-DEL-11501
chr4-7914119-DEL-765
chr4-7936375-DEL-2580
chr4-7972280-DEL-56
chr4-7982666-DEL-92
chr4-7984458-DEL-52
chr4-8006506-DEL-232
chr4-8014683-DEL-116
chr4-8018817-DEL-519
chr4-8081069-DEL-310
chr4-8104372-DEL-178
chr4-8116959-DEL-7518
chr4-8133258-DEL-3734
chr4-8144069-DEL-74
chr4-8152122-DEL-278
chr4-8155511-DEL-419
chr4-8160515-DEL-2404
chr4-8173984-DEL-320
chr4-8267769-DEL-59
chr4-8269008-DEL-108
chr4-8273985-DEL-52
chr4-8294929-DEL-94
chr4-8316555-DEL-54
chr4-8388125-DEL-494
chr4-8424894-DEL-373
chr4-8438582-DEL-2017
chr4-8440739-DEL-683
chr4-8457376-DEL-2245
chr4-8462078-DEL-972
chr4-8473283-DEL-54
chr4-8479469-DEL-6289
chr4-8500165-DEL-64
chr4-8512975-DEL-1894
chr4-8555472-DEL-1191
chr4-8599349-DEL-329
chr4-8652304-DEL-321
chr4-8664934-DEL-1615
chr4-8714450-DEL-71
chr4-8715229-DEL-56
chr4-8730938-DEL-374
chr4-8755831-DEL-135

chr4-8758667-DEL-318
chr4-8765779-DEL-60
chr4-8780278-DEL-146
chr4-8817041-DEL-389
chr4-8837949-DEL-52
chr4-8880278-DEL-81
chr4-8884812-DEL-2102
chr4-8888664-DEL-646
chr4-8894640-DEL-86
chr4-8900337-DEL-67
chr4-8908896-DEL-70
chr4-8937067-DEL-409
chr4-8947109-DEL-71
chr4-8981502-DEL-244
chr4-9020420-DEL-66
chr4-9021567-DEL-320
chr4-9025820-DEL-60
chr4-9034823-DEL-98
chr4-9063847-DEL-190
chr4-9086642-DEL-110
chr4-9124191-DEL-141
chr4-9167766-DEL-166
chr4-9201548-DEL-69
chr4-9214545-DEL-758
chr4-9261251-DEL-502
chr4-9287148-DEL-61
chr4-9306065-DEL-68
chr4-9309278-DEL-805
chr4-9309313-DEL-1049
chr4-9346492-DEL-298
chr4-9348911-DEL-228
chr4-9400224-DEL-193
chr4-9426119-DEL-399
chr4-9432463-DEL-188
chr4-9434866-DEL-70
chr4-9438047-DEL-1564
chr4-9442110-DEL-51
chr4-9455167-DEL-76
chr4-9481351-DEL-2122
chr4-9487803-DEL-182
chr4-9502641-DEL-396
chr4-9519589-DEL-344
chr4-9534068-DEL-51
chr4-9585009-DEL-9547
chr4-9604925-DEL-277
chr4-9606896-DEL-361
chr4-9620218-DEL-61
chr4-9633592-DEL-56
chr4-9650358-DEL-54
chr4-9677367-DEL-889
chr4-9696433-DEL-72
chr4-9698352-DEL-1941
chr4-9717182-DEL-6832
chr4-9723463-DEL-408
chr4-9727956-DEL-7109
chr4-9742721-DEL-59
chr4-9768571-DEL-134
chr4-9799283-DEL-332
chr4-9824267-DEL-391
chr4-9827016-DEL-661
chr4-9903402-DEL-5265
chr4-9931483-DEL-224
chr4-9935045-DEL-78
chr4-9989804-DEL-203
chr4-10045057-DEL-239
chr4-10045919-DEL-439
chr4-10048722-DEL-99
chr4-10093417-DEL-6443
chr4-10112107-DEL-53
chr4-10136845-DEL-710
chr4-10136872-DEL-1079
chr4-10137766-DEL-183
chr4-10159569-DEL-6183
chr4-10166530-DEL-208
chr4-10177543-DEL-280
chr4-10199405-DEL-120
chr4-10232682-DEL-1022

chr4-10238089-DEL-86
chr4-10276410-DEL-60
chr4-10289479-DEL-62
chr4-10299443-DEL-2640
chr4-10349596-DEL-794
chr4-10360557-DEL-199
chr4-10363447-DEL-339
chr4-10412243-DEL-309
chr4-10444255-DEL-1913
chr4-10465319-DEL-63
chr4-10472172-DEL-6261
chr4-10483863-DEL-206
chr4-10543918-DEL-279
chr4-10549772-DEL-826
chr4-10582595-DEL-1130
chr4-10603678-DEL-66
chr4-10610692-DEL-97
chr4-10622199-DEL-91
chr4-10632464-DEL-1056
chr4-10663084-DEL-646
chr4-10663915-DEL-55
chr4-10666074-DEL-65
chr4-10686302-DEL-63
chr4-10716048-DEL-315
chr4-10717084-DEL-56
chr4-10734606-DEL-409
chr4-10742204-DEL-761
chr4-10754488-DEL-169
chr4-10759823-DEL-554
chr4-10825866-DEL-56
chr4-10885780-DEL-4174
chr4-10892069-DEL-56
chr4-10917721-DEL-236
chr4-10939066-DEL-315
chr4-10949761-DEL-206
chr4-10951981-DEL-1626
chr4-10972334-DEL-74
chr4-11005076-DEL-612
chr4-11015685-DEL-204
chr4-11023417-DEL-79
chr4-11029911-DEL-226
chr4-11034032-DEL-55
chr4-11035633-DEL-70
chr4-11045938-DEL-452
chr4-11053928-DEL-2115
chr4-11053981-DEL-1710
chr4-11054923-DEL-407
chr4-11055656-DEL-225
chr4-11059291-DEL-57
chr4-11065537-DEL-78
chr4-11069472-DEL-271
chr4-11117319-DEL-1453
chr4-11125953-DEL-54
chr4-11138226-DEL-113
chr4-11160016-DEL-221
chr4-11176293-DEL-7991
chr4-11226385-DEL-193
chr4-11241804-DEL-110
chr4-11270542-DEL-212
chr4-11273531-DEL-123
chr4-11278489-DEL-936
chr4-11304967-DEL-197
chr4-11306387-DEL-70
chr4-11317910-DEL-2267
chr4-11317912-DEL-104
chr4-11396446-DEL-195
chr4-11397344-DEL-124
chr4-11400053-DEL-1800
chr4-11413285-DEL-116
chr4-11427167-DEL-140
chr4-11434391-DEL-13509
chr4-11441734-DEL-467
chr4-11466083-DEL-168
chr4-11479145-DEL-127
chr4-11511339-DEL-260
chr4-11531783-DEL-620
chr4-11542497-DEL-196

chr4-11543832-DEL-57
chr4-11545887-DEL-74
chr4-11553277-DEL-56
chr4-11559528-DEL-357
chr4-11571143-DEL-186
chr4-11621436-DEL-57
chr4-11636200-DEL-202
chr4-11658190-DEL-717
chr4-11718828-DEL-316
chr4-11741037-DEL-1414
chr4-11775962-DEL-340
chr4-11777757-DEL-83
chr4-11794524-DEL-152
chr4-11815284-DEL-73
chr4-11831327-DEL-53
chr4-11832414-DEL-75
chr4-11994403-DEL-917
chr4-12024136-DEL-2095
chr4-12027994-DEL-71
chr4-12031742-DEL-112
chr4-12048559-DEL-499
chr4-12066809-DEL-1611
chr4-12071220-DEL-997
chr4-12113869-DEL-11006
chr4-12152521-DEL-116
chr4-12174713-DEL-1536
chr4-12176298-DEL-228
chr4-12188636-DEL-401
chr4-12189191-DEL-67
chr4-12196680-DEL-1045
chr4-12200404-DEL-2835
chr4-12227483-DEL-51
chr4-12245409-DEL-87
chr4-12309151-DEL-839
chr4-12314371-DEL-506
chr4-12331429-DEL-64
chr4-12336032-DEL-72
chr4-12357691-DEL-215
chr4-12358456-DEL-106
chr4-12386888-DEL-87
chr4-12395849-DEL-122
chr4-12427633-DEL-545
chr4-12445660-DEL-4182
chr4-12484400-DEL-974
chr4-12517450-DEL-98
chr4-12551209-DEL-115
chr4-12594392-DEL-54
chr4-12625731-DEL-2665
chr4-12630344-DEL-53
chr4-12638111-DEL-137
chr4-12701180-DEL-143
chr4-12702494-DEL-1233
chr4-12707441-DEL-1174
chr4-12715392-DEL-495
chr4-12818105-DEL-101
chr4-12854543-DEL-957
chr4-12873569-DEL-195
chr4-12884088-DEL-394
chr4-12903105-DEL-62
chr4-12922337-DEL-162
chr4-12934535-DEL-64
chr4-12945960-DEL-144
chr4-12951571-DEL-431
chr4-12985295-DEL-565
chr4-12988470-DEL-3760
chr4-12997372-DEL-6296
chr4-13003386-DEL-64
chr4-13006034-DEL-272
chr4-13014663-DEL-2455
chr4-13017723-DEL-732
chr4-13064301-DEL-156
chr4-13106758-DEL-159
chr4-13129976-DEL-1058
chr4-13165971-DEL-199
chr4-13174878-DEL-69
chr4-13178778-DEL-2293
chr4-13205567-DEL-78

chr4-13233106-DEL-608
chr4-13263140-DEL-1088
chr4-13306338-DEL-506
chr4-13328319-DEL-668
chr4-13330698-DEL-204
chr4-13354443-DEL-1848
chr4-13375432-DEL-219
chr4-13394752-DEL-343
chr4-13395370-DEL-299
chr4-13450514-DEL-67
chr4-13488254-DEL-221
chr4-13533763-DEL-927
chr4-13575437-DEL-179
chr4-13684176-DEL-74
chr4-13705899-DEL-1271
chr4-13722032-DEL-377
chr4-13731941-DEL-401
chr4-13793019-DEL-207
chr4-13819066-DEL-152
chr4-13865529-DEL-98
chr4-13897543-DEL-188
chr4-13924635-DEL-322
chr4-13928540-DEL-54
chr4-13930189-DEL-6426
chr4-13931096-DEL-66
chr4-13985862-DEL-312
chr4-14010551-DEL-70
chr4-14012267-DEL-411
chr4-14063633-DEL-270
chr4-14073362-DEL-1495
chr4-14085917-DEL-1425
chr4-14105157-DEL-237
chr4-14112938-DEL-635
chr4-14131371-DEL-428
chr4-14181987-DEL-80
chr4-14210091-DEL-732
chr4-14210755-DEL-64
chr4-14229684-DEL-328
chr4-14238339-DEL-92
chr4-14258442-DEL-98
chr4-14282849-DEL-493
chr4-14304276-DEL-6208
chr4-14312053-DEL-1160
chr4-14320772-DEL-1959
chr4-14346887-DEL-1115
chr4-14355940-DEL-371
chr4-14361085-DEL-95
chr4-14386127-DEL-85
chr4-14394525-DEL-221
chr4-14446627-DEL-612
chr4-14459586-DEL-73
chr4-14464017-DEL-2152
chr4-14500822-DEL-53
chr4-14532900-DEL-477
chr4-14548153-DEL-86
chr4-14643462-DEL-1217
chr4-14692909-DEL-551
chr4-14741160-DEL-244
chr4-14753396-DEL-280
chr4-14757058-DEL-212
chr4-14758850-DEL-6267
chr4-14776766-DEL-65
chr4-14789268-DEL-397
chr4-14790860-DEL-2102
chr4-14793065-DEL-62
chr4-14823875-DEL-239
chr4-14827112-DEL-102
chr4-14834063-DEL-1454
chr4-14866012-DEL-384
chr4-14872734-DEL-284
chr4-14886009-DEL-78
chr4-14888969-DEL-261
chr4-14920061-DEL-287
chr4-14943036-DEL-1021
chr4-14957090-DEL-51
chr4-14984362-DEL-332
chr4-14985185-DEL-53

chr4-15035498-DEL-419
chr4-15073561-DEL-535
chr4-15113497-DEL-277
chr4-15133857-DEL-2116
chr4-15135778-DEL-356
chr4-15169676-DEL-57
chr4-15170313-DEL-51
chr4-15203392-DEL-70
chr4-15223807-DEL-1313
chr4-15243031-DEL-319
chr4-15292535-DEL-71
chr4-15317067-DEL-69
chr4-15320563-DEL-576
chr4-15353181-DEL-103
chr4-15388007-DEL-1865
chr4-15392191-DEL-66
chr4-15424652-DEL-155
chr4-15443359-DEL-56
chr4-15461777-DEL-6243
chr4-15468780-DEL-2527
chr4-15503837-DEL-69
chr4-15546006-DEL-827
chr4-15547928-DEL-127
chr4-15571641-DEL-1054
chr4-15572892-DEL-269
chr4-15597923-DEL-191
chr4-15674639-DEL-2114
chr4-15696143-DEL-52
chr4-15733122-DEL-621
chr4-15740673-DEL-789
chr4-15747037-DEL-4363
chr4-15747038-DEL-1326
chr4-15781017-DEL-137
chr4-15859696-DEL-669
chr4-15861825-DEL-6662
chr4-15868842-DEL-351
chr4-15889923-DEL-99
chr4-15912672-DEL-4290
chr4-15937522-DEL-55
chr4-15954392-DEL-174
chr4-15960500-DEL-201
chr4-15971945-DEL-1356
chr4-15976407-DEL-132
chr4-15982004-DEL-53
chr4-15987590-DEL-107
chr4-16021868-DEL-130
chr4-16040263-DEL-129
chr4-16041256-DEL-109
chr4-16077055-DEL-294
chr4-16088464-DEL-90
chr4-16104540-DEL-915
chr4-16125318-DEL-832
chr4-16129561-DEL-82
chr4-16147136-DEL-995
chr4-16148139-DEL-1830
chr4-16186314-DEL-4447
chr4-16198148-DEL-59
chr4-16211926-DEL-76
chr4-16227274-DEL-2398
chr4-16246944-DEL-2029
chr4-16252278-DEL-422
chr4-16263366-DEL-764
chr4-16293740-DEL-716
chr4-16302997-DEL-7203
chr4-16310874-DEL-207
chr4-16316679-DEL-100
chr4-16319790-DEL-7363
chr4-16357410-DEL-54
chr4-16367745-DEL-727
chr4-16373614-DEL-716
chr4-16404197-DEL-1033
chr4-16408669-DEL-121
chr4-16428913-DEL-475
chr4-16445440-DEL-82
chr4-16450903-DEL-978
chr4-16453836-DEL-67
chr4-16462106-DEL-78

chr4-16463816-DEL-243
chr4-16471410-DEL-53
chr4-16472022-DEL-946
chr4-16478716-DEL-52
chr4-16490793-DEL-209
chr4-16508022-DEL-52
chr4-16525894-DEL-173
chr4-16529472-DEL-56
chr4-16622543-DEL-66
chr4-16689271-DEL-782
chr4-16713661-DEL-58
chr4-16727785-DEL-72
chr4-16760653-DEL-3107
chr4-16763870-DEL-57
chr4-16823573-DEL-1484
chr4-16827147-DEL-93
chr4-16835201-DEL-6931
chr4-16868674-DEL-131
chr4-16869239-DEL-362
chr4-16913346-DEL-988
chr4-16919670-DEL-428
chr4-16928092-DEL-1366
chr4-16953657-DEL-202
chr4-16964502-DEL-2007
chr4-16973881-DEL-834
chr4-16984908-DEL-333
chr4-16990084-DEL-4023
chr4-16991677-DEL-191
chr4-16998319-DEL-371
chr4-17002238-DEL-161
chr4-17011899-DEL-6396
chr4-17025594-DEL-64
chr4-17046602-DEL-184
chr4-17082014-DEL-83
chr4-17135803-DEL-83
chr4-17143673-DEL-400
chr4-17154111-DEL-634
chr4-17157441-DEL-58
chr4-17171690-DEL-6205
chr4-17180174-DEL-619
chr4-17196161-DEL-64
chr4-17203559-DEL-508
chr4-17204902-DEL-381
chr4-17207195-DEL-10522
chr4-17207443-DEL-66
chr4-17215191-DEL-97
chr4-17216727-DEL-135
chr4-17237795-DEL-61
chr4-17255169-DEL-54
chr4-17334617-DEL-702
chr4-17346574-DEL-355
chr4-17353552-DEL-359
chr4-17354625-DEL-678
chr4-17357420-DEL-55
chr4-17372581-DEL-688
chr4-17375544-DEL-335
chr4-17377369-DEL-562
chr4-17383812-DEL-542
chr4-17384207-DEL-5556
chr4-17384397-DEL-4292
chr4-17391827-DEL-9379
chr4-17393896-DEL-3484
chr4-17397529-DEL-529
chr4-17402302-DEL-893
chr4-17402992-DEL-203
chr4-17403486-DEL-1341
chr4-17408680-DEL-8528
chr4-17425280-DEL-151
chr4-17442252-DEL-174
chr4-17466195-DEL-1151
chr4-17474966-DEL-877
chr4-17488020-DEL-108
chr4-17533896-DEL-71
chr4-17538701-DEL-3829
chr4-17563672-DEL-67
chr4-17618888-DEL-168
chr4-17659492-DEL-935

chr4-17673658-DEL-1176
chr4-17677921-DEL-721
chr4-17688011-DEL-3157
chr4-17694374-DEL-73
chr4-17701471-DEL-77
chr4-17707132-DEL-212
chr4-17716790-DEL-603
chr4-17732677-DEL-381
chr4-17743796-DEL-128
chr4-17744540-DEL-6187
chr4-17770431-DEL-372
chr4-17823124-DEL-63
chr4-17859065-DEL-167
chr4-17886842-DEL-175
chr4-17895481-DEL-66
chr4-17907261-DEL-10129
chr4-17915175-DEL-1006
chr4-17925101-DEL-898
chr4-17932918-DEL-1879
chr4-17941309-DEL-494
chr4-17989834-DEL-533
chr4-18002806-DEL-186
chr4-18031017-DEL-275
chr4-18055326-DEL-86
chr4-18069023-DEL-509
chr4-18088458-DEL-461
chr4-18140536-DEL-409
chr4-18151752-DEL-2062
chr4-18165076-DEL-213
chr4-18198982-DEL-313
chr4-18202419-DEL-2184
chr4-18204234-DEL-559
chr4-18204819-DEL-1743
chr4-18206615-DEL-298
chr4-18210785-DEL-527
chr4-18218184-DEL-390
chr4-18229924-DEL-95
chr4-18248284-DEL-529
chr4-18263644-DEL-146
chr4-18279924-DEL-110
chr4-18283646-DEL-203
chr4-18317527-DEL-378
chr4-18323393-DEL-112
chr4-18333363-DEL-2588
chr4-18343686-DEL-938
chr4-18349061-DEL-73
chr4-18391503-DEL-59
chr4-18396626-DEL-64
chr4-18437349-DEL-203
chr4-18454189-DEL-366
chr4-18522347-DEL-1883
chr4-18528773-DEL-367
chr4-18547942-DEL-516
chr4-18551315-DEL-121
chr4-18607206-DEL-212
chr4-18641698-DEL-2914
chr4-18690185-DEL-425
chr4-18698868-DEL-6289
chr4-18708051-DEL-15355
chr4-18732607-DEL-14315
chr4-18764826-DEL-2711
chr4-18771366-DEL-6043
chr4-18802519-DEL-62
chr4-18819002-DEL-318
chr4-18828568-DEL-123
chr4-18848512-DEL-406
chr4-18901879-DEL-110
chr4-18905029-DEL-228
chr4-18919631-DEL-53
chr4-18952974-DEL-96
chr4-18957113-DEL-1139
chr4-19007232-DEL-81
chr4-19043265-DEL-60
chr4-19061362-DEL-8188
chr4-19083787-DEL-51
chr4-19098055-DEL-74
chr4-19125918-DEL-163

chr4-19128739-DEL-285
chr4-19133479-DEL-325
chr4-19134110-DEL-629
chr4-19143705-DEL-155
chr4-19148916-DEL-211
chr4-19149443-DEL-7489
chr4-19188264-DEL-63
chr4-19188519-DEL-12883
chr4-19207309-DEL-1931
chr4-19229616-DEL-1377
chr4-19238387-DEL-114
chr4-19243638-DEL-235
chr4-19252899-DEL-75
chr4-19274320-DEL-5058
chr4-19279272-DEL-132
chr4-19289455-DEL-199
chr4-19294167-DEL-3071
chr4-19305493-DEL-58
chr4-19307249-DEL-124
chr4-19334678-DEL-108
chr4-19346853-DEL-290
chr4-19348491-DEL-199
chr4-19350089-DEL-676
chr4-19352140-DEL-95
chr4-19377661-DEL-197
chr4-19382489-DEL-5674
chr4-19388206-DEL-2488
chr4-19391247-DEL-2359
chr4-19401334-DEL-519
chr4-19546065-DEL-1250
chr4-19561347-DEL-1914
chr4-19563585-DEL-220
chr4-19688201-DEL-260
chr4-19695437-DEL-1102
chr4-19729774-DEL-120
chr4-19755743-DEL-157
chr4-19769152-DEL-2086
chr4-19787343-DEL-601
chr4-19800260-DEL-669
chr4-19810910-DEL-100
chr4-19828922-DEL-228
chr4-19908290-DEL-75
chr4-19927654-DEL-238
chr4-19965293-DEL-272
chr4-19985089-DEL-54
chr4-20005886-DEL-96
chr4-20034062-DEL-75
chr4-20047667-DEL-144
chr4-20078192-DEL-519
chr4-20127078-DEL-72
chr4-20141534-DEL-87
chr4-20168639-DEL-6611
chr4-20182301-DEL-403
chr4-20194128-DEL-75
chr4-20196621-DEL-76
chr4-20233999-DEL-74
chr4-20256268-DEL-470
chr4-20263074-DEL-822
chr4-20265486-DEL-592
chr4-20272042-DEL-252
chr4-20278397-DEL-602
chr4-20284235-DEL-130
chr4-20299216-DEL-302
chr4-20324877-DEL-480
chr4-20334674-DEL-128
chr4-20409117-DEL-2715
chr4-20415860-DEL-54
chr4-20454155-DEL-78
chr4-20462602-DEL-2047
chr4-20491819-DEL-66
chr4-20512142-DEL-79
chr4-20615488-DEL-56
chr4-20663582-DEL-61
chr4-20675710-DEL-63
chr4-20713209-DEL-877
chr4-20737613-DEL-63
chr4-20753943-DEL-65

chr4-20771940-DEL-74
chr4-20779869-DEL-1935
chr4-20826142-DEL-1732
chr4-20855843-DEL-5426
chr4-20862444-DEL-374
chr4-20913989-DEL-6210
chr4-20946076-DEL-211
chr4-20955009-DEL-56
chr4-20955979-DEL-759
chr4-20959541-DEL-54
chr4-21001631-DEL-62
chr4-21024495-DEL-247
chr4-21027599-DEL-713
chr4-21030783-DEL-6185
chr4-21039314-DEL-204
chr4-21060871-DEL-196
chr4-21107538-DEL-328
chr4-21109374-DEL-753
chr4-21120354-DEL-253
chr4-21121617-DEL-516
chr4-21139319-DEL-68
chr4-21139902-DEL-1917
chr4-21147211-DEL-3454
chr4-21164454-DEL-970
chr4-21172599-DEL-80
chr4-21197141-DEL-132
chr4-21242751-DEL-2692
chr4-21284631-DEL-1300
chr4-21287457-DEL-424
chr4-21291204-DEL-1763
chr4-21293032-DEL-4947
chr4-21308909-DEL-60
chr4-21309542-DEL-56
chr4-21347239-DEL-118
chr4-21362982-DEL-406
chr4-21367763-DEL-123
chr4-21369905-DEL-20977
chr4-21381349-DEL-8224
chr4-21382037-DEL-417
chr4-21395220-DEL-100
chr4-21399530-DEL-6737
chr4-21408290-DEL-184
chr4-21415683-DEL-310
chr4-21422740-DEL-1015
chr4-21424293-DEL-3064
chr4-21431815-DEL-800
chr4-21442706-DEL-110
chr4-21449172-DEL-1918
chr4-21455400-DEL-533
chr4-21466249-DEL-55
chr4-21469473-DEL-6882
chr4-21480607-DEL-193
chr4-21482163-DEL-3754
chr4-21488565-DEL-3147
chr4-21498507-DEL-886
chr4-21503309-DEL-96
chr4-21505265-DEL-14899
chr4-21509540-DEL-2775
chr4-21513667-DEL-2775
chr4-21533279-DEL-66
chr4-21540674-DEL-721
chr4-21579072-DEL-7875
chr4-21587003-DEL-77
chr4-21624950-DEL-380
chr4-21681820-DEL-52
chr4-21729883-DEL-115
chr4-21758566-DEL-66
chr4-21771451-DEL-1246
chr4-21788120-DEL-1306
chr4-21789773-DEL-192
chr4-21800872-DEL-535
chr4-21829982-DEL-176
chr4-21843119-DEL-137
chr4-21848747-DEL-118
chr4-21880506-DEL-71
chr4-21885611-DEL-171
chr4-21913232-DEL-167

chr4-21922952-DEL-1469
chr4-21924462-DEL-2332
chr4-21938227-DEL-420
chr4-21938374-DEL-1531
chr4-21938485-DEL-393
chr4-21961507-DEL-85
chr4-21963849-DEL-116
chr4-21975549-DEL-72
chr4-22003860-DEL-233
chr4-22026504-DEL-768
chr4-22053668-DEL-336
chr4-22150712-DEL-89
chr4-22171364-DEL-80
chr4-22177798-DEL-334
chr4-22178961-DEL-68
chr4-22202299-DEL-3647
chr4-22211769-DEL-846
chr4-22230714-DEL-52
chr4-22231982-DEL-1017
chr4-22233471-DEL-51
chr4-22235115-DEL-493
chr4-22240437-DEL-513
chr4-22259690-DEL-61
chr4-22308751-DEL-60
chr4-22312712-DEL-213
chr4-22324164-DEL-67
chr4-22334816-DEL-58
chr4-22347111-DEL-243
chr4-22349334-DEL-4383
chr4-22399372-DEL-1079
chr4-22404060-DEL-179
chr4-22411821-DEL-4793
chr4-22460210-DEL-12964
chr4-22503123-DEL-68
chr4-22511428-DEL-54
chr4-22535875-DEL-61
chr4-22542330-DEL-1310
chr4-22588544-DEL-124
chr4-22591760-DEL-7269
chr4-22594662-DEL-210
chr4-22596210-DEL-6199
chr4-22609352-DEL-2176
chr4-22611695-DEL-1950
chr4-22626611-DEL-6243
chr4-22673455-DEL-336
chr4-22675013-DEL-119
chr4-22690727-DEL-6237
chr4-22739079-DEL-3073
chr4-22790662-DEL-109
chr4-22804794-DEL-145
chr4-22814804-DEL-362
chr4-22820365-DEL-76
chr4-22841956-DEL-188
chr4-22875063-DEL-56
chr4-22889484-DEL-60
chr4-22897326-DEL-51
chr4-22901268-DEL-6462
chr4-22953909-DEL-422
chr4-22959676-DEL-96
chr4-23003638-DEL-103960
chr4-23006949-DEL-758
chr4-23008063-DEL-362
chr4-23010393-DEL-2549
chr4-23012026-DEL-5610
chr4-23069918-DEL-71
chr4-23086489-DEL-2996
chr4-23087210-DEL-60
chr4-23093430-DEL-8237
chr4-23107548-DEL-6614
chr4-23127340-DEL-237
chr4-23128579-DEL-214
chr4-23134803-DEL-94
chr4-23137911-DEL-1719
chr4-23141730-DEL-408
chr4-23151481-DEL-3334
chr4-23201143-DEL-6583
chr4-23218819-DEL-186

chr4-23228313-DEL-58
chr4-23229929-DEL-618
chr4-23230917-DEL-62
chr4-23236782-DEL-2290
chr4-23276622-DEL-227
chr4-23283099-DEL-55
chr4-23289158-DEL-424
chr4-23296201-DEL-400
chr4-23297281-DEL-520
chr4-23298110-DEL-1787
chr4-23334781-DEL-77
chr4-23348690-DEL-286
chr4-23374400-DEL-67
chr4-23425061-DEL-67
chr4-23438864-DEL-95
chr4-23448894-DEL-146
chr4-23472302-DEL-291
chr4-23481576-DEL-505
chr4-23519224-DEL-62
chr4-23531202-DEL-481
chr4-23541632-DEL-1353
chr4-23625363-DEL-827
chr4-23659323-DEL-471
chr4-23694729-DEL-193
chr4-23701057-DEL-239
chr4-23706266-DEL-624
chr4-23771958-DEL-63
chr4-23824284-DEL-53
chr4-23836506-DEL-64
chr4-23883488-DEL-1623
chr4-23884272-DEL-204
chr4-23885821-DEL-124
chr4-23888186-DEL-1081
chr4-23890995-DEL-773
chr4-23900810-DEL-72
chr4-23936138-DEL-57
chr4-23949414-DEL-632
chr4-23963552-DEL-106
chr4-23963697-DEL-508
chr4-23977104-DEL-60
chr4-23981057-DEL-199
chr4-24014570-DEL-182
chr4-24025418-DEL-1445
chr4-24045143-DEL-727
chr4-24048324-DEL-555
chr4-24065883-DEL-530
chr4-24069178-DEL-698
chr4-24071306-DEL-213
chr4-24078934-DEL-341
chr4-24103014-DEL-62
chr4-24115684-DEL-280
chr4-24138669-DEL-56
chr4-24145787-DEL-76
chr4-24160412-DEL-189
chr4-24164706-DEL-574
chr4-24177617-DEL-263
chr4-24194147-DEL-424
chr4-24204750-DEL-89
chr4-24226345-DEL-344
chr4-24231269-DEL-1144
chr4-24256727-DEL-188
chr4-24258833-DEL-70
chr4-24269691-DEL-6239
chr4-24269724-DEL-205
chr4-24290182-DEL-169
chr4-24315888-DEL-2116
chr4-24338238-DEL-64
chr4-24349123-DEL-58
chr4-24350498-DEL-166
chr4-24351248-DEL-63
chr4-24363139-DEL-119
chr4-24371058-DEL-179
chr4-24434477-DEL-457
chr4-24435960-DEL-406
chr4-24438101-DEL-2154
chr4-24448835-DEL-423
chr4-24449386-DEL-232

chr4-24450953-DEL-97
chr4-24545885-DEL-90
chr4-24624998-DEL-93
chr4-24668287-DEL-55
chr4-24726221-DEL-52
chr4-24731820-DEL-286
chr4-24757216-DEL-53
chr4-24769026-DEL-84
chr4-24858582-DEL-141
chr4-24875869-DEL-54
chr4-24883525-DEL-58
chr4-24899008-DEL-451
chr4-24903264-DEL-165
chr4-24910463-DEL-2872
chr4-24913568-DEL-303
chr4-24926139-DEL-1211
chr4-24934035-DEL-567
chr4-24945646-DEL-340
chr4-24974409-DEL-99
chr4-25004797-DEL-846
chr4-25009431-DEL-134
chr4-25030808-DEL-250
chr4-25032129-DEL-238
chr4-25047959-DEL-143
chr4-25048281-DEL-513
chr4-25050364-DEL-1239
chr4-25060461-DEL-1032
chr4-25061364-DEL-245
chr4-25069380-DEL-898
chr4-25070106-DEL-2188
chr4-25070923-DEL-636
chr4-25085381-DEL-660
chr4-25098319-DEL-90
chr4-25107980-DEL-6564
chr4-25118492-DEL-3381
chr4-25118957-DEL-1114
chr4-25120014-DEL-125
chr4-25139741-DEL-69
chr4-25147042-DEL-54
chr4-25148039-DEL-1121
chr4-25172046-DEL-2835
chr4-25181196-DEL-83
chr4-25188867-DEL-212
chr4-25192610-DEL-243
chr4-25205643-DEL-81
chr4-25261189-DEL-151
chr4-25263679-DEL-60
chr4-25312293-DEL-561
chr4-25331096-DEL-467
chr4-25377022-DEL-4694
chr4-25423055-DEL-132
chr4-25425218-DEL-67
chr4-25442033-DEL-54
chr4-25456179-DEL-1072
chr4-25458508-DEL-52
chr4-25479172-DEL-447
chr4-25493979-DEL-8355
chr4-25532438-DEL-52
chr4-25554644-DEL-82
chr4-25555532-DEL-2662
chr4-25560120-DEL-6764
chr4-25574402-DEL-6390
chr4-25594364-DEL-1192
chr4-25596749-DEL-1097
chr4-25618782-DEL-321
chr4-25687946-DEL-232
chr4-25705418-DEL-56
chr4-25737766-DEL-55
chr4-25749772-DEL-194
chr4-25771893-DEL-3855
chr4-25801230-DEL-263
chr4-25834822-DEL-6392
chr4-25848212-DEL-5707
chr4-25894082-DEL-142
chr4-25895146-DEL-118
chr4-25900399-DEL-187
chr4-25927441-DEL-291

chr4-25945071-DEL-331
chr4-26004634-DEL-200
chr4-26008872-DEL-428
chr4-26009507-DEL-65
chr4-26032139-DEL-3017
chr4-26047861-DEL-1375
chr4-26050035-DEL-53
chr4-26051852-DEL-6602
chr4-26061344-DEL-1612
chr4-26077704-DEL-117
chr4-26085567-DEL-1240
chr4-26092172-DEL-3657
chr4-26099521-DEL-64
chr4-26102902-DEL-851
chr4-26109498-DEL-7152
chr4-26128464-DEL-741
chr4-26136341-DEL-125
chr4-26137718-DEL-100
chr4-26148631-DEL-1065
chr4-26202163-DEL-1258
chr4-26207957-DEL-336
chr4-26219759-DEL-2573
chr4-26220448-DEL-90
chr4-26241265-DEL-884
chr4-26262077-DEL-1290
chr4-26309604-DEL-11918
chr4-26339738-DEL-154
chr4-26355499-DEL-122
chr4-26376475-DEL-772
chr4-26400492-DEL-2114
chr4-26410181-DEL-1671
chr4-26411146-DEL-805
chr4-26439252-DEL-903
chr4-26493379-DEL-80
chr4-26495372-DEL-399
chr4-26497660-DEL-1012
chr4-26498744-DEL-2069
chr4-26503599-DEL-832
chr4-26509177-DEL-328
chr4-26552126-DEL-17600
chr4-26622665-DEL-118
chr4-26624858-DEL-944
chr4-26640821-DEL-470
chr4-26650665-DEL-2472
chr4-26672165-DEL-65
chr4-26673568-DEL-73
chr4-26674592-DEL-242
chr4-26675977-DEL-371
chr4-26687333-DEL-126
chr4-26695210-DEL-52
chr4-26703704-DEL-1774
chr4-26712327-DEL-239
chr4-26718295-DEL-117
chr4-26732440-DEL-203
chr4-26782753-DEL-52
chr4-26845066-DEL-165
chr4-26872829-DEL-77
chr4-26876432-DEL-253
chr4-26890544-DEL-627
chr4-26905443-DEL-2531
chr4-26909156-DEL-2224
chr4-26952514-DEL-1039
chr4-26970469-DEL-1555
chr4-27004260-DEL-1404
chr4-27043085-DEL-71
chr4-27065227-DEL-6199
chr4-27137497-DEL-72
chr4-27152272-DEL-574
chr4-27155772-DEL-162
chr4-27162187-DEL-4425
chr4-27194827-DEL-62
chr4-27212955-DEL-6200
chr4-27237094-DEL-768
chr4-27262276-DEL-57
chr4-27265826-DEL-108
chr4-27283200-DEL-1776
chr4-27300941-DEL-6019

chr4-27311328-DEL-2041
chr4-27314143-DEL-74
chr4-27380175-DEL-274
chr4-27439810-DEL-56
chr4-27456239-DEL-156
chr4-27511893-DEL-65
chr4-27513088-DEL-3371
chr4-27519956-DEL-104
chr4-27548370-DEL-746
chr4-27589993-DEL-240
chr4-27590890-DEL-70
chr4-27598277-DEL-461
chr4-27618121-DEL-69
chr4-27670580-DEL-65
chr4-27682188-DEL-204
chr4-27683642-DEL-209
chr4-27740533-DEL-283
chr4-27774380-DEL-393
chr4-27775271-DEL-1583
chr4-27825728-DEL-364
chr4-27850357-DEL-688
chr4-27859165-DEL-1824
chr4-27860993-DEL-255
chr4-27861108-DEL-1640
chr4-27862597-DEL-5158
chr4-27884710-DEL-117
chr4-27920357-DEL-1374
chr4-27946652-DEL-118
chr4-27953544-DEL-510
chr4-28012097-DEL-72
chr4-28022064-DEL-64
chr4-28033126-DEL-5322
chr4-28041111-DEL-159
chr4-28060667-DEL-580
chr4-28074872-DEL-1242
chr4-28103313-DEL-1932
chr4-28129768-DEL-1075
chr4-28163750-DEL-62
chr4-28176773-DEL-443
chr4-28208421-DEL-67
chr4-28246581-DEL-52
chr4-28255551-DEL-343
chr4-28261796-DEL-800
chr4-28263811-DEL-534
chr4-28278478-DEL-62
chr4-28282450-DEL-160
chr4-28301515-DEL-60
chr4-28307618-DEL-241
chr4-28321197-DEL-568
chr4-28331512-DEL-900
chr4-28376699-DEL-95
chr4-28380163-DEL-64
chr4-28424047-DEL-52
chr4-28446443-DEL-215
chr4-28470551-DEL-193
chr4-28471850-DEL-149
chr4-28491111-DEL-2039
chr4-28493281-DEL-2181
chr4-28514611-DEL-328
chr4-28516510-DEL-72
chr4-28521840-DEL-61
chr4-28533866-DEL-1276
chr4-28550788-DEL-98
chr4-28552289-DEL-66
chr4-28555371-DEL-51
chr4-28596155-DEL-1606
chr4-28597991-DEL-262
chr4-28670595-DEL-430
chr4-28691713-DEL-2458
chr4-28737559-DEL-110
chr4-28759122-DEL-434
chr4-28764460-DEL-180
chr4-28765634-DEL-3107
chr4-28774097-DEL-267
chr4-28806917-DEL-924
chr4-28842225-DEL-59
chr4-28845333-DEL-1782

chr4-28888469-DEL-136
chr4-28908914-DEL-76
chr4-28918710-DEL-291
chr4-28954226-DEL-111
chr4-28967980-DEL-52
chr4-28995367-DEL-219
chr4-29025438-DEL-204
chr4-29027064-DEL-2688
chr4-29085919-DEL-2557
chr4-29095925-DEL-173
chr4-29141671-DEL-455
chr4-29158457-DEL-56
chr4-29198730-DEL-88
chr4-29221214-DEL-632
chr4-29224397-DEL-256
chr4-29233512-DEL-1494
chr4-29257553-DEL-1588
chr4-29272339-DEL-897
chr4-29297210-DEL-133
chr4-29298166-DEL-94
chr4-29322941-DEL-83
chr4-29333188-DEL-68
chr4-29341909-DEL-258
chr4-29354989-DEL-486
chr4-29374318-DEL-53
chr4-29405036-DEL-901
chr4-29420485-DEL-71
chr4-29422493-DEL-227
chr4-29475281-DEL-135
chr4-29478034-DEL-1648
chr4-29538755-DEL-65
chr4-29570498-DEL-469
chr4-29600995-DEL-2059
chr4-29626043-DEL-164
chr4-29626371-DEL-705
chr4-29692815-DEL-51
chr4-29696865-DEL-186
chr4-29740277-DEL-231
chr4-29745617-DEL-6442
chr4-29751620-DEL-198
chr4-29759100-DEL-3523
chr4-29790970-DEL-176
chr4-29830793-DEL-60
chr4-29834695-DEL-192
chr4-29848591-DEL-72
chr4-29853293-DEL-59
chr4-29886794-DEL-2080
chr4-29891264-DEL-76
chr4-29932206-DEL-1880
chr4-29952114-DEL-57
chr4-29975518-DEL-55
chr4-29993085-DEL-2232
chr4-30018498-DEL-56
chr4-30047902-DEL-636
chr4-30062573-DEL-80
chr4-30068752-DEL-712
chr4-30114658-DEL-122
chr4-30119908-DEL-740
chr4-30137224-DEL-140
chr4-30152645-DEL-104
chr4-30157730-DEL-71
chr4-30169975-DEL-2807
chr4-30169984-DEL-196
chr4-30178069-DEL-2607
chr4-30188848-DEL-143
chr4-30197015-DEL-52
chr4-30200863-DEL-59
chr4-30215212-DEL-143
chr4-30228575-DEL-4034
chr4-30233004-DEL-2951
chr4-30260062-DEL-71
chr4-30326854-DEL-208
chr4-30347594-DEL-96
chr4-30357768-DEL-8289
chr4-30374030-DEL-69
chr4-30376137-DEL-679
chr4-30384845-DEL-213

chr4-30392337-DEL-1144
chr4-30397559-DEL-126
chr4-30401369-DEL-137
chr4-30419949-DEL-163
chr4-30442515-DEL-57
chr4-30452951-DEL-70
chr4-30453880-DEL-83
chr4-30461879-DEL-603
chr4-30479373-DEL-135
chr4-30505618-DEL-346
chr4-30525032-DEL-1533
chr4-30532973-DEL-1169
chr4-30534163-DEL-384
chr4-30536034-DEL-2354
chr4-30537539-DEL-849
chr4-30604458-DEL-52
chr4-30631192-DEL-2374
chr4-30631503-DEL-2373
chr4-30637751-DEL-134
chr4-30684009-DEL-70
chr4-30746502-DEL-69
chr4-30748663-DEL-55
chr4-30749814-DEL-83
chr4-30752264-DEL-51
chr4-30758175-DEL-578
chr4-30758834-DEL-221
chr4-30813276-DEL-1269
chr4-30837775-DEL-1858
chr4-30844936-DEL-431
chr4-30845703-DEL-819
chr4-30877402-DEL-1359
chr4-30879835-DEL-6228
chr4-30904563-DEL-55
chr4-30956799-DEL-2353
chr4-30960281-DEL-79
chr4-30964879-DEL-157
chr4-31036078-DEL-428
chr4-31062726-DEL-58
chr4-31077679-DEL-399
chr4-31086695-DEL-115
chr4-31090853-DEL-457
chr4-31123529-DEL-452
chr4-31132280-DEL-652
chr4-31133065-DEL-1240
chr4-31179651-DEL-63
chr4-31215779-DEL-299
chr4-31222392-DEL-137
chr4-31230070-DEL-238
chr4-31230924-DEL-3087
chr4-31245241-DEL-886
chr4-31284351-DEL-74
chr4-31302961-DEL-432
chr4-31355757-DEL-1867
chr4-31378818-DEL-82
chr4-31421819-DEL-210
chr4-31477540-DEL-163
chr4-31501516-DEL-423
chr4-31599305-DEL-168
chr4-31645453-DEL-308
chr4-31772372-DEL-77
chr4-31786800-DEL-1250
chr4-31794313-DEL-840
chr4-31804668-DEL-516
chr4-31826773-DEL-999
chr4-31855128-DEL-60
chr4-31913895-DEL-114
chr4-31917063-DEL-206
chr4-31934694-DEL-372
chr4-31987104-DEL-667
chr4-32045107-DEL-157
chr4-32052217-DEL-384
chr4-32053522-DEL-263
chr4-32058907-DEL-446
chr4-32089923-DEL-175
chr4-32117215-DEL-51
chr4-32119312-DEL-100
chr4-32132390-DEL-463

chr4-32137887-DEL-100
chr4-32145849-DEL-116
chr4-32179543-DEL-122
chr4-32185501-DEL-191
chr4-32249741-DEL-70
chr4-32294719-DEL-378
chr4-32304126-DEL-74
chr4-32306029-DEL-126
chr4-32315026-DEL-88
chr4-32321189-DEL-1008
chr4-32323129-DEL-384
chr4-32339865-DEL-197
chr4-32341382-DEL-216
chr4-32344377-DEL-1407
chr4-32361234-DEL-74
chr4-32386170-DEL-67
chr4-32416853-DEL-123
chr4-32418984-DEL-184
chr4-32465167-DEL-129
chr4-32484358-DEL-58
chr4-32506400-DEL-571
chr4-32516745-DEL-134
chr4-32527189-DEL-221
chr4-32545743-DEL-1780
chr4-32552997-DEL-338
chr4-32587777-DEL-65
chr4-32593227-DEL-57
chr4-32606193-DEL-204
chr4-32612950-DEL-169
chr4-32624672-DEL-63
chr4-32631920-DEL-89
chr4-32726169-DEL-66
chr4-32734584-DEL-198
chr4-32744256-DEL-128
chr4-32754734-DEL-210
chr4-32755942-DEL-98
chr4-32783257-DEL-185
chr4-32784508-DEL-1313
chr4-32816457-DEL-80
chr4-32841619-DEL-208
chr4-32902780-DEL-61
chr4-32929781-DEL-1296
chr4-32931589-DEL-396
chr4-32934184-DEL-156
chr4-32941975-DEL-207
chr4-32945889-DEL-231
chr4-32969765-DEL-51
chr4-32970036-DEL-1245
chr4-32991536-DEL-236
chr4-33002578-DEL-128
chr4-33008965-DEL-96
chr4-33011422-DEL-204
chr4-33018252-DEL-208
chr4-33053361-DEL-1042
chr4-33071703-DEL-73
chr4-33073525-DEL-257
chr4-33083489-DEL-75
chr4-33097048-DEL-202
chr4-33108113-DEL-64
chr4-33144086-DEL-183
chr4-33168718-DEL-904
chr4-33171422-DEL-350
chr4-33191468-DEL-341
chr4-33211518-DEL-206
chr4-33234255-DEL-53
chr4-33239103-DEL-60
chr4-33240860-DEL-85
chr4-33251911-DEL-243
chr4-33260868-DEL-78
chr4-33286176-DEL-54
chr4-33348593-DEL-51
chr4-33349655-DEL-118
chr4-33404806-DEL-2004
chr4-33419000-DEL-216
chr4-33445222-DEL-205
chr4-33469957-DEL-55
chr4-33496265-DEL-185

chr4-33514242-DEL-63
chr4-33517503-DEL-247
chr4-33551901-DEL-65
chr4-33606526-DEL-401
chr4-33628722-DEL-489
chr4-33632160-DEL-164
chr4-33636760-DEL-220
chr4-33648651-DEL-178
chr4-33672393-DEL-160
chr4-33673900-DEL-136
chr4-33681859-DEL-1362
chr4-33695709-DEL-161
chr4-33714980-DEL-56
chr4-33728761-DEL-400
chr4-33729638-DEL-243
chr4-33753868-DEL-138
chr4-33789820-DEL-178
chr4-33808130-DEL-1312
chr4-33815238-DEL-344
chr4-33873757-DEL-55
chr4-33950667-DEL-68
chr4-33955470-DEL-1810
chr4-34000465-DEL-93
chr4-34012019-DEL-51
chr4-34031053-DEL-93
chr4-34085647-DEL-60
chr4-34139729-DEL-123
chr4-34206068-DEL-74
chr4-34208987-DEL-65
chr4-34242470-DEL-124
chr4-34250881-DEL-1190
chr4-34270207-DEL-183
chr4-34278046-DEL-446
chr4-34290318-DEL-1867
chr4-34299673-DEL-538
chr4-34309590-DEL-280
chr4-34335136-DEL-191
chr4-34350811-DEL-2441
chr4-34357678-DEL-402
chr4-34384825-DEL-102
chr4-34395960-DEL-1197
chr4-34416512-DEL-210
chr4-34426609-DEL-1153
chr4-34434523-DEL-381
chr4-34465285-DEL-236
chr4-34497481-DEL-103
chr4-34500505-DEL-7267
chr4-34575840-DEL-51
chr4-34617190-DEL-176
chr4-34640396-DEL-1636
chr4-34649203-DEL-136
chr4-34758002-DEL-436
chr4-34800050-DEL-722
chr4-34812084-DEL-146
chr4-34816092-DEL-171
chr4-34829791-DEL-410
chr4-34843509-DEL-3308
chr4-34856884-DEL-174
chr4-34865689-DEL-63
chr4-34876179-DEL-204
chr4-34891638-DEL-105
chr4-34908167-DEL-404
chr4-34916262-DEL-698
chr4-34918657-DEL-368
chr4-34925949-DEL-194
chr4-34972659-DEL-387
chr4-35000994-DEL-1242
chr4-35025869-DEL-82
chr4-35062238-DEL-60
chr4-35062829-DEL-54
chr4-35081689-DEL-126
chr4-35087965-DEL-378
chr4-35113073-DEL-70
chr4-35141771-DEL-176
chr4-35189253-DEL-54
chr4-35199412-DEL-273
chr4-35204782-DEL-291

chr4-35237467-DEL-74
chr4-35257353-DEL-57
chr4-35268642-DEL-1013
chr4-35269708-DEL-101
chr4-35296563-DEL-447
chr4-35325642-DEL-98
chr4-35352013-DEL-66
chr4-35360382-DEL-771
chr4-35377121-DEL-8291
chr4-35425774-DEL-6343
chr4-35435334-DEL-682
chr4-35436937-DEL-90
chr4-35437819-DEL-401
chr4-35440965-DEL-368
chr4-35465252-DEL-430
chr4-35523280-DEL-218
chr4-35527027-DEL-2921
chr4-35598175-DEL-6401
chr4-35613090-DEL-1617
chr4-35615940-DEL-3567
chr4-35620500-DEL-2049
chr4-35655230-DEL-594
chr4-35656016-DEL-9366
chr4-35739724-DEL-106
chr4-35748765-DEL-1150
chr4-35756238-DEL-433
chr4-35810581-DEL-772
chr4-35820540-DEL-96
chr4-35833565-DEL-2468
chr4-35839835-DEL-185
chr4-35866911-DEL-208
chr4-35953789-DEL-328
chr4-35973244-DEL-6057
chr4-35989384-DEL-57
chr4-35996688-DEL-2102
chr4-35997479-DEL-422
chr4-36004890-DEL-189
chr4-36023158-DEL-6192
chr4-36046332-DEL-78
chr4-36097927-DEL-218
chr4-36109146-DEL-244
chr4-36142970-DEL-861
chr4-36149499-DEL-70
chr4-36185371-DEL-64
chr4-36186812-DEL-323
chr4-36188358-DEL-72
chr4-36220152-DEL-2504
chr4-36229178-DEL-1454
chr4-36235293-DEL-384
chr4-36239038-DEL-464
chr4-36254799-DEL-458
chr4-36267638-DEL-380
chr4-36280528-DEL-1005
chr4-36302552-DEL-913
chr4-36305789-DEL-103
chr4-36314490-DEL-79
chr4-36319067-DEL-352
chr4-36351572-DEL-938
chr4-36354349-DEL-82
chr4-36369537-DEL-543
chr4-36373976-DEL-95
chr4-36403003-DEL-167
chr4-36414105-DEL-11053
chr4-36420695-DEL-377
chr4-36421321-DEL-132
chr4-36456314-DEL-170
chr4-36465337-DEL-126276
chr4-36475385-DEL-58
chr4-36484063-DEL-77
chr4-36497150-DEL-1692
chr4-36504323-DEL-995
chr4-36520266-DEL-190
chr4-36524351-DEL-319
chr4-36582241-DEL-52
chr4-36586999-DEL-191
chr4-36596297-DEL-297
chr4-36605927-DEL-2901

chr4-36608863-DEL-2577
chr4-36611488-DEL-514
chr4-36620324-DEL-68
chr4-36670226-DEL-69
chr4-36672118-DEL-147
chr4-36682379-DEL-82
chr4-36686473-DEL-54
chr4-36693190-DEL-457
chr4-36696017-DEL-311
chr4-36697205-DEL-52
chr4-36720767-DEL-146
chr4-36721289-DEL-368
chr4-36723364-DEL-51
chr4-36733894-DEL-1258
chr4-36756682-DEL-204
chr4-36778719-DEL-454
chr4-36783933-DEL-107
chr4-36787923-DEL-723
chr4-36792453-DEL-1650
chr4-36805774-DEL-211
chr4-36817664-DEL-89
chr4-36826430-DEL-85
chr4-36829995-DEL-324
chr4-36833959-DEL-204
chr4-36847290-DEL-1492
chr4-36860728-DEL-1265
chr4-36880783-DEL-506
chr4-36881840-DEL-57
chr4-36892738-DEL-1212
chr4-36917526-DEL-284
chr4-36926008-DEL-1858
chr4-36940704-DEL-213
chr4-36957165-DEL-74
chr4-36958049-DEL-204
chr4-36988413-DEL-383
chr4-37012006-DEL-101
chr4-37020016-DEL-99
chr4-37086579-DEL-92
chr4-37115997-DEL-60
chr4-37116763-DEL-3120
chr4-37134504-DEL-1583
chr4-37138872-DEL-994
chr4-37172505-DEL-57
chr4-37176177-DEL-347
chr4-37179431-DEL-4595
chr4-37186720-DEL-307
chr4-37194571-DEL-1428
chr4-37215145-DEL-58
chr4-37216419-DEL-532
chr4-37240568-DEL-166
chr4-37248631-DEL-1089
chr4-37271006-DEL-157
chr4-37272374-DEL-514
chr4-37301292-DEL-78
chr4-37338682-DEL-80
chr4-37381828-DEL-820
chr4-37407805-DEL-459
chr4-37411121-DEL-59
chr4-37418799-DEL-68
chr4-37433165-DEL-912
chr4-37436661-DEL-53
chr4-37439565-DEL-83
chr4-37443950-DEL-5438
chr4-37450537-DEL-52
chr4-37454923-DEL-580
chr4-37456743-DEL-69
chr4-37504024-DEL-267
chr4-37546746-DEL-2186
chr4-37552686-DEL-387
chr4-37562217-DEL-622
chr4-37565954-DEL-582
chr4-37576093-DEL-538
chr4-37581696-DEL-336
chr4-37590129-DEL-2366
chr4-37595061-DEL-6140
chr4-37628610-DEL-855
chr4-37669225-DEL-187

chr4-37680831-DEL-186
chr4-37681921-DEL-279
chr4-37715636-DEL-404
chr4-37726053-DEL-302
chr4-37746355-DEL-646
chr4-37757415-DEL-6481
chr4-37789392-DEL-1824
chr4-37792968-DEL-1408
chr4-37826978-DEL-512
chr4-37827673-DEL-933
chr4-37846041-DEL-80
chr4-37871121-DEL-52
chr4-37887464-DEL-53
chr4-37900467-DEL-115
chr4-37901831-DEL-838
chr4-37930820-DEL-2691
chr4-37934214-DEL-1186
chr4-37937568-DEL-5089
chr4-37949954-DEL-70
chr4-37968708-DEL-2377
chr4-37970309-DEL-217
chr4-37984044-DEL-2406
chr4-37997208-DEL-164
chr4-38023016-DEL-3388
chr4-38023700-DEL-1498
chr4-38039530-DEL-481
chr4-38054032-DEL-359
chr4-38054664-DEL-1235
chr4-38055925-DEL-137
chr4-38058732-DEL-58
chr4-38079246-DEL-865
chr4-38094824-DEL-1075
chr4-38107326-DEL-4014
chr4-38107735-DEL-683
chr4-38114644-DEL-52
chr4-38116392-DEL-2475
chr4-38123353-DEL-380
chr4-38125862-DEL-61
chr4-38126762-DEL-1039
chr4-38155129-DEL-60
chr4-38157652-DEL-312
chr4-38168180-DEL-60
chr4-38168409-DEL-737
chr4-38171506-DEL-363
chr4-38172967-DEL-153
chr4-38184561-DEL-753
chr4-38190082-DEL-118
chr4-38205966-DEL-108
chr4-38214477-DEL-6404
chr4-38225415-DEL-5993
chr4-38239300-DEL-5759
chr4-38248225-DEL-360
chr4-38265260-DEL-107
chr4-38276519-DEL-119
chr4-38305408-DEL-3849
chr4-38307306-DEL-1675
chr4-38351701-DEL-864
chr4-38359018-DEL-68
chr4-38412635-DEL-562
chr4-38418493-DEL-7427
chr4-38447711-DEL-106
chr4-38455384-DEL-266
chr4-38458164-DEL-297
chr4-38485185-DEL-64
chr4-38499156-DEL-80
chr4-38500991-DEL-176
chr4-38517840-DEL-416
chr4-38597765-DEL-296
chr4-38599638-DEL-335
chr4-38605137-DEL-318
chr4-38638569-DEL-344
chr4-38639182-DEL-571
chr4-38652053-DEL-2989
chr4-38671048-DEL-51
chr4-38717846-DEL-3138
chr4-38724816-DEL-401
chr4-38855458-DEL-8854

chr4-38916549-DEL-3512
chr4-38924569-DEL-1423
chr4-38959700-DEL-6054
chr4-38982625-DEL-1687
chr4-38986053-DEL-222
chr4-38991429-DEL-196
chr4-38998641-DEL-1307
chr4-39031466-DEL-440
chr4-39049797-DEL-203
chr4-39059544-DEL-208
chr4-39113140-DEL-548
chr4-39153753-DEL-322
chr4-39167811-DEL-85
chr4-39190692-DEL-768
chr4-39222296-DEL-84
chr4-39273994-DEL-108
chr4-39277497-DEL-5267
chr4-39285607-DEL-502
chr4-39310694-DEL-71
chr4-39327311-DEL-87
chr4-39375943-DEL-314
chr4-39392278-DEL-478
chr4-39406063-DEL-2059
chr4-39414216-DEL-270
chr4-39416682-DEL-74
chr4-39421695-DEL-64
chr4-39466978-DEL-211
chr4-39478862-DEL-135
chr4-39495267-DEL-52
chr4-39535755-DEL-754
chr4-39538272-DEL-80
chr4-39548243-DEL-138
chr4-39564893-DEL-209
chr4-39572527-DEL-65
chr4-39589367-DEL-660
chr4-39593845-DEL-101
chr4-39623919-DEL-117
chr4-39632673-DEL-321
chr4-39663073-DEL-54
chr4-39669107-DEL-538
chr4-39673704-DEL-74
chr4-39677801-DEL-110
chr4-39688376-DEL-190
chr4-39700390-DEL-352
chr4-39709589-DEL-374
chr4-39738003-DEL-1166
chr4-39789385-DEL-618
chr4-39799203-DEL-477
chr4-39840620-DEL-385
chr4-39850021-DEL-2366
chr4-39856078-DEL-6244
chr4-39885172-DEL-7881
chr4-39908385-DEL-55
chr4-39921410-DEL-232
chr4-39923444-DEL-1889
chr4-39929064-DEL-118
chr4-39944307-DEL-64
chr4-39945144-DEL-86
chr4-39949917-DEL-52
chr4-39958158-DEL-206
chr4-39969768-DEL-231
chr4-40002291-DEL-52
chr4-40018638-DEL-128
chr4-40029874-DEL-60
chr4-40033110-DEL-3103
chr4-40045718-DEL-79
chr4-40102305-DEL-278
chr4-40114982-DEL-331
chr4-40115974-DEL-60
chr4-40146578-DEL-463
chr4-40153585-DEL-71
chr4-40169693-DEL-198
chr4-40179441-DEL-66
chr4-40195798-DEL-109
chr4-40215773-DEL-51
chr4-40235535-DEL-1114
chr4-40294801-DEL-3629

chr4-40311899-DEL-64
chr4-40312856-DEL-92
chr4-40315113-DEL-69
chr4-40355158-DEL-58
chr4-40356304-DEL-336
chr4-40385027-DEL-97
chr4-40412398-DEL-177
chr4-40414554-DEL-3553
chr4-40424436-DEL-2978
chr4-40435312-DEL-8672
chr4-40447149-DEL-574
chr4-40458067-DEL-51
chr4-40458209-DEL-1375
chr4-40470792-DEL-99
chr4-40528848-DEL-57
chr4-40582085-DEL-252
chr4-40589499-DEL-54
chr4-40591254-DEL-1052
chr4-40613821-DEL-60
chr4-40627417-DEL-292
chr4-40633421-DEL-69
chr4-40635934-DEL-431
chr4-40661018-DEL-115
chr4-40677571-DEL-293
chr4-40685609-DEL-321
chr4-40688439-DEL-76
chr4-40714569-DEL-188
chr4-40744292-DEL-195
chr4-40747066-DEL-165
chr4-40748206-DEL-202
chr4-40749073-DEL-445
chr4-40764786-DEL-1260
chr4-40775078-DEL-696
chr4-40799207-DEL-196
chr4-40819269-DEL-100
chr4-40824713-DEL-280
chr4-40858105-DEL-855
chr4-40860756-DEL-114
chr4-40861569-DEL-244
chr4-40922554-DEL-122
chr4-40928050-DEL-70
chr4-40934622-DEL-351
chr4-40961220-DEL-630
chr4-40965576-DEL-262
chr4-40970023-DEL-144
chr4-40981401-DEL-105
chr4-41003893-DEL-388
chr4-41006762-DEL-229
chr4-41033339-DEL-54
chr4-41046319-DEL-54
chr4-41058560-DEL-796
chr4-41082230-DEL-322
chr4-41146195-DEL-191
chr4-41148532-DEL-78
chr4-41153502-DEL-186
chr4-41235889-DEL-210
chr4-41246035-DEL-76
chr4-41267309-DEL-164
chr4-41268884-DEL-213
chr4-41274238-DEL-53
chr4-41297934-DEL-201
chr4-41309470-DEL-201
chr4-41326398-DEL-113
chr4-41328660-DEL-298
chr4-41345295-DEL-67
chr4-41355312-DEL-182
chr4-41364336-DEL-85
chr4-41373819-DEL-79
chr4-41381519-DEL-60
chr4-41420631-DEL-91
chr4-41433022-DEL-78
chr4-41452061-DEL-2153
chr4-41454959-DEL-63
chr4-41485024-DEL-206
chr4-41567655-DEL-60
chr4-41588520-DEL-210
chr4-41593014-DEL-613

chr4-41608640-DEL-193
chr4-41611178-DEL-146
chr4-41643377-DEL-1019
chr4-41781874-DEL-51
chr4-41877330-DEL-5127
chr4-41883854-DEL-191
chr4-41886726-DEL-191
chr4-41889596-DEL-191
chr4-41892470-DEL-191
chr4-42028081-DEL-4346
chr4-42092094-DEL-172
chr4-42232294-DEL-2873
chr4-42380293-DEL-4346
chr4-42462168-DEL-3969
chr4-42530514-DEL-6469
chr4-42589424-DEL-172
chr4-42643669-DEL-467
chr4-42644713-DEL-54
chr4-42652418-DEL-635
chr4-42691932-DEL-90
chr4-42718123-DEL-402
chr4-42738105-DEL-57
chr4-42744016-DEL-498
chr4-42768998-DEL-324
chr4-42786801-DEL-163
chr4-42803779-DEL-144
chr4-42844252-DEL-426
chr4-42908280-DEL-379
chr4-42914830-DEL-54
chr4-42936576-DEL-182
chr4-42974527-DEL-262
chr4-43012559-DEL-309
chr4-43026725-DEL-278
chr4-43046563-DEL-82
chr4-43079708-DEL-622
chr4-43162742-DEL-60
chr4-43262016-DEL-840
chr4-43285556-DEL-941
chr4-43379339-DEL-54
chr4-43433622-DEL-766
chr4-43434435-DEL-506
chr4-43501026-DEL-56
chr4-43599517-DEL-138
chr4-43683622-DEL-116
chr4-43698457-DEL-62
chr4-43734216-DEL-56
chr4-43766846-DEL-983
chr4-43804766-DEL-120
chr4-43812993-DEL-205
chr4-43848422-DEL-136
chr4-43946831-DEL-399
chr4-43948480-DEL-398
chr4-44007026-DEL-80
chr4-44011669-DEL-205
chr4-44038054-DEL-256
chr4-44063281-DEL-1890
chr4-44079609-DEL-125
chr4-44091480-DEL-397
chr4-44115413-DEL-59
chr4-44123226-DEL-1195
chr4-44169721-DEL-68
chr4-44176737-DEL-353
chr4-44196517-DEL-211
chr4-44238189-DEL-555
chr4-44246735-DEL-191
chr4-44259052-DEL-78
chr4-44271022-DEL-152
chr4-44290191-DEL-201
chr4-44297675-DEL-514
chr4-44303723-DEL-457
chr4-44385679-DEL-400
chr4-44393770-DEL-142
chr4-44397910-DEL-52
chr4-44409338-DEL-119
chr4-44456039-DEL-52
chr4-44465674-DEL-84
chr4-44473367-DEL-6163

chr4-44499327-DEL-96
chr4-44540737-DEL-60
chr4-44592641-DEL-131
chr4-44598592-DEL-191
chr4-44605992-DEL-1468
chr4-44617078-DEL-59
chr4-44619986-DEL-134
chr4-44640377-DEL-58
chr4-44675348-DEL-55
chr4-44678586-DEL-122
chr4-44683932-DEL-58
chr4-44733580-DEL-108
chr4-44736058-DEL-243
chr4-44738123-DEL-88
chr4-44745490-DEL-63
chr4-44792058-DEL-278
chr4-44816377-DEL-643
chr4-44832204-DEL-223
chr4-44861872-DEL-196
chr4-44865257-DEL-78
chr4-44867944-DEL-1564
chr4-44881829-DEL-59
chr4-44908907-DEL-82
chr4-45020637-DEL-229
chr4-45038909-DEL-229
chr4-45051988-DEL-89
chr4-45053303-DEL-52
chr4-45063984-DEL-88
chr4-45076186-DEL-570
chr4-45081212-DEL-218
chr4-45093121-DEL-1607
chr4-45098697-DEL-1411
chr4-45113168-DEL-1857
chr4-45115109-DEL-1866
chr4-45139893-DEL-163
chr4-45154831-DEL-58
chr4-45219031-DEL-55
chr4-45220822-DEL-72
chr4-45222899-DEL-131
chr4-45232165-DEL-126
chr4-45264319-DEL-1721
chr4-45289404-DEL-184
chr4-45312339-DEL-118
chr4-45313254-DEL-746
chr4-45330083-DEL-70
chr4-45336048-DEL-70
chr4-45395691-DEL-69
chr4-45404326-DEL-81
chr4-45417823-DEL-57
chr4-45463993-DEL-209
chr4-45510884-DEL-179
chr4-45524774-DEL-519
chr4-45533230-DEL-204
chr4-45564811-DEL-131
chr4-45569462-DEL-754
chr4-45578334-DEL-206
chr4-45614107-DEL-65
chr4-45643422-DEL-245
chr4-45681984-DEL-301
chr4-45698154-DEL-981
chr4-45704490-DEL-1737
chr4-45709147-DEL-252
chr4-45709745-DEL-246
chr4-45722036-DEL-54
chr4-45730669-DEL-82
chr4-45773205-DEL-222
chr4-45773441-DEL-660
chr4-45883558-DEL-310
chr4-45891721-DEL-119
chr4-45942148-DEL-87
chr4-45999534-DEL-58
chr4-46019173-DEL-52
chr4-46029261-DEL-166
chr4-46031579-DEL-103
chr4-46037660-DEL-151
chr4-46045434-DEL-394
chr4-46050940-DEL-981

chr4-46067898-DEL-100
chr4-46119472-DEL-56
chr4-46145825-DEL-140
chr4-46169435-DEL-327
chr4-46179381-DEL-666
chr4-46187018-DEL-285
chr4-46200058-DEL-1255
chr4-46215942-DEL-90
chr4-46240822-DEL-3115
chr4-46246978-DEL-204
chr4-46254162-DEL-545
chr4-46256266-DEL-67
chr4-46271469-DEL-1661
chr4-46286525-DEL-134
chr4-46289244-DEL-82
chr4-46329317-DEL-132
chr4-46335402-DEL-53
chr4-46348489-DEL-169
chr4-46367284-DEL-52
chr4-46412586-DEL-139
chr4-46423357-DEL-1616
chr4-46428047-DEL-212
chr4-46429907-DEL-56
chr4-46439265-DEL-1561
chr4-46481926-DEL-751
chr4-46482275-DEL-120
chr4-46519603-DEL-153
chr4-46525893-DEL-62
chr4-46560918-DEL-204
chr4-46570441-DEL-201
chr4-46591010-DEL-387
chr4-46608427-DEL-343
chr4-46631854-DEL-1071
chr4-46634058-DEL-115
chr4-46637210-DEL-164
chr4-46704154-DEL-391
chr4-46705592-DEL-195
chr4-46722994-DEL-57
chr4-46745965-DEL-88
chr4-46773089-DEL-543
chr4-46778655-DEL-142
chr4-46796574-DEL-62
chr4-46812650-DEL-323
chr4-46817017-DEL-84
chr4-46831653-DEL-3293
chr4-46840792-DEL-400
chr4-46893476-DEL-60
chr4-46917265-DEL-1650
chr4-46924163-DEL-315
chr4-46925746-DEL-62
chr4-46948910-DEL-58
chr4-46954032-DEL-101
chr4-46962590-DEL-84
chr4-46985256-DEL-131
chr4-47062380-DEL-59
chr4-47074170-DEL-56
chr4-47105990-DEL-56
chr4-47142889-DEL-51
chr4-47165086-DEL-659
chr4-47188768-DEL-136
chr4-47216386-DEL-119
chr4-47242654-DEL-198
chr4-47261420-DEL-493
chr4-47313537-DEL-180
chr4-47377927-DEL-204
chr4-47390748-DEL-208
chr4-47428942-DEL-68
chr4-47430622-DEL-2249
chr4-47437070-DEL-78
chr4-47494498-DEL-91
chr4-47501053-DEL-225
chr4-47577887-DEL-72
chr4-47581438-DEL-62
chr4-47608339-DEL-78
chr4-47678173-DEL-53
chr4-47736878-DEL-76
chr4-47743422-DEL-151

chr4-47749947-DEL-2428
chr4-47787492-DEL-1377
chr4-47790601-DEL-58
chr4-47811555-DEL-59
chr4-47827183-DEL-344
chr4-47857689-DEL-671
chr4-47905153-DEL-296
chr4-47912505-DEL-52
chr4-47931195-DEL-84
chr4-47969020-DEL-850
chr4-47969052-DEL-119
chr4-47969182-DEL-295
chr4-48031730-DEL-92
chr4-48035923-DEL-424
chr4-48071398-DEL-186
chr4-48086812-DEL-61
chr4-48179147-DEL-92
chr4-48227905-DEL-84
chr4-48246136-DEL-3525
chr4-48250941-DEL-1022
chr4-48272865-DEL-54
chr4-48306232-DEL-379
chr4-48313526-DEL-1516
chr4-48315312-DEL-56
chr4-48347383-DEL-343
chr4-48347930-DEL-225
chr4-48351579-DEL-752
chr4-48356596-DEL-53
chr4-48385563-DEL-190
chr4-48394961-DEL-541
chr4-48398923-DEL-70
chr4-48408990-DEL-54
chr4-48500256-DEL-147
chr4-48507352-DEL-104
chr4-48508096-DEL-260
chr4-48513775-DEL-5179
chr4-48521640-DEL-194
chr4-48525876-DEL-1068
chr4-48534209-DEL-194
chr4-48550218-DEL-177
chr4-48588003-DEL-70
chr4-48614688-DEL-200
chr4-48678530-DEL-426
chr4-48686211-DEL-118
chr4-48687566-DEL-558
chr4-48700755-DEL-5988
chr4-48715568-DEL-76
chr4-48739370-DEL-195
chr4-48763038-DEL-198
chr4-48764799-DEL-105
chr4-48794621-DEL-525
chr4-48802083-DEL-119
chr4-48805372-DEL-2845
chr4-48812736-DEL-84
chr4-48887303-DEL-386
chr4-48904376-DEL-64
chr4-48911884-DEL-581
chr4-48971812-DEL-192
chr4-48976448-DEL-1160
chr4-48976869-DEL-209
chr4-49024526-DEL-84
chr4-49027360-DEL-6167
chr4-49043415-DEL-60
chr4-49082561-DEL-61
chr4-49083826-DEL-518
chr4-49100219-DEL-7863
chr4-49112643-DEL-960
chr4-49130085-DEL-2253
chr4-49135032-DEL-84
chr4-49136749-DEL-6366
chr4-49146416-DEL-51
chr4-49168425-DEL-261
chr4-49170830-DEL-109
chr4-49173038-DEL-271
chr4-49177133-DEL-52
chr4-49191192-DEL-132
chr4-49202847-DEL-843

chr4-49227287-DEL-66
chr4-49228705-DEL-654
chr4-49230812-DEL-204
chr4-49257764-DEL-2142
chr4-49260266-DEL-898
chr4-49290254-DEL-61
chr4-49323257-DEL-67
chr4-49347883-DEL-134
chr4-49387526-DEL-83
chr4-49420404-DEL-9184
chr4-49449934-DEL-190
chr4-49460295-DEL-179
chr4-49466877-DEL-212
chr4-49481535-DEL-86
chr4-49496166-DEL-438
chr4-49515412-DEL-1809
chr4-49542459-DEL-64
chr4-49588623-DEL-133
chr4-49605351-DEL-1488
chr4-49619686-DEL-122
chr4-49620541-DEL-637
chr4-49639498-DEL-60
chr4-49640373-DEL-52
chr4-49687608-DEL-166
chr4-49740185-DEL-133
chr4-49764072-DEL-1474
chr4-49790349-DEL-213
chr4-49868518-DEL-71
chr4-49908854-DEL-551
chr4-49950569-DEL-357
chr4-50002101-DEL-1011
chr4-50032142-DEL-219
chr4-50052633-DEL-190
chr4-50071532-DEL-428
chr4-50085907-DEL-347
chr4-50088488-DEL-403
chr4-50091297-DEL-1885
chr4-50093039-DEL-137
chr4-50150476-DEL-99
chr4-50164070-DEL-55
chr4-50210357-DEL-477
chr4-50213645-DEL-79
chr4-50270286-DEL-381
chr4-50341491-DEL-2781
chr4-50345983-DEL-56
chr4-50355624-DEL-61
chr4-50365516-DEL-61
chr4-50365830-DEL-2128
chr4-50373486-DEL-202
chr4-50392681-DEL-924
chr4-50398909-DEL-103
chr4-50448224-DEL-1646
chr4-50459825-DEL-15853
chr4-50464519-DEL-405
chr4-50466630-DEL-1521
chr4-50466656-DEL-10411
chr4-50488986-DEL-138
chr4-50489614-DEL-400
chr4-50502081-DEL-80
chr4-50518050-DEL-7052
chr4-50539561-DEL-60
chr4-50587322-DEL-2784
chr4-50601026-DEL-87
chr4-50615328-DEL-843
chr4-50619153-DEL-286
chr4-50631263-DEL-131
chr4-50634555-DEL-100
chr4-50643036-DEL-69
chr4-50644676-DEL-1135
chr4-50677908-DEL-275
chr4-50696983-DEL-180
chr4-50700046-DEL-6432
chr4-50700061-DEL-200
chr4-50723881-DEL-298
chr4-50741325-DEL-2170
chr4-50759508-DEL-101
chr4-50804142-DEL-1380

chr4-50820861-DEL-135
chr4-50885947-DEL-1918
chr4-50890203-DEL-4127
chr4-50894505-DEL-434
chr4-50899298-DEL-136
chr4-50922437-DEL-105
chr4-50923470-DEL-1243
chr4-50945929-DEL-6373
chr4-50969387-DEL-831
chr4-50977789-DEL-841
chr4-50983794-DEL-147
chr4-50991989-DEL-392
chr4-50992675-DEL-2832
chr4-51007631-DEL-447
chr4-51022511-DEL-562
chr4-51060080-DEL-6133
chr4-51087700-DEL-1592
chr4-51149468-DEL-390
chr4-51171951-DEL-3804
chr4-51176688-DEL-99
chr4-51178875-DEL-1616
chr4-51223539-DEL-84
chr4-51226393-DEL-1591
chr4-51232575-DEL-716
chr4-51236229-DEL-1279
chr4-51259528-DEL-52
chr4-51260795-DEL-96
chr4-51261683-DEL-1057
chr4-51276440-DEL-1324
chr4-51283070-DEL-116
chr4-51286623-DEL-76
chr4-51305866-DEL-305
chr4-51309217-DEL-1120
chr4-51313432-DEL-65
chr4-51316923-DEL-1143
chr4-51319525-DEL-13808
chr4-51344361-DEL-732
chr4-51354904-DEL-1191
chr4-51379749-DEL-392
chr4-51385033-DEL-577
chr4-51385674-DEL-6012
chr4-51386895-DEL-4025
chr4-51398434-DEL-54
chr4-51405192-DEL-88
chr4-51413321-DEL-1767
chr4-51427822-DEL-556
chr4-51447395-DEL-147
chr4-51488947-DEL-135
chr4-51492258-DEL-710
chr4-51494282-DEL-6240
chr4-51500911-DEL-63
chr4-51523766-DEL-184
chr4-51532811-DEL-89
chr4-51558482-DEL-78
chr4-51559296-DEL-184
chr4-51606550-DEL-3165
chr4-51610841-DEL-95
chr4-51617841-DEL-872
chr4-51619781-DEL-62
chr4-51630981-DEL-1367
chr4-51645882-DEL-1118
chr4-51648053-DEL-181
chr4-51662261-DEL-52
chr4-51694087-DEL-56
chr4-51702253-DEL-2719
chr4-51702775-DEL-6269
chr4-51730627-DEL-5521
chr4-51736733-DEL-556
chr4-51749317-DEL-300
chr4-51751846-DEL-213
chr4-51822434-DEL-393
chr4-51857059-DEL-668
chr4-51860670-DEL-158
chr4-51867222-DEL-2089
chr4-51869389-DEL-946
chr4-51925161-DEL-311
chr4-51956357-DEL-1170

chr4-51957021-DEL-4634
chr4-51959160-DEL-61
chr4-51969084-DEL-61
chr4-51994970-DEL-335
chr4-52025150-DEL-719
chr4-52041682-DEL-666
chr4-52063618-DEL-6276
chr4-52069952-DEL-58
chr4-52071722-DEL-204
chr4-52072859-DEL-69
chr4-52079676-DEL-7938
chr4-52086014-DEL-847
chr4-52086555-DEL-848
chr4-52086699-DEL-847
chr4-52160483-DEL-498
chr4-52192831-DEL-899
chr4-52242532-DEL-5245
chr4-52253733-DEL-349
chr4-52287042-DEL-6274
chr4-52299418-DEL-1474
chr4-52351194-DEL-1003
chr4-52368015-DEL-65
chr4-52432547-DEL-1787
chr4-52440724-DEL-59
chr4-52448901-DEL-192
chr4-52472325-DEL-1774
chr4-52535816-DEL-6490
chr4-52542079-DEL-7930
chr4-52558878-DEL-123
chr4-52600697-DEL-875
chr4-52641314-DEL-55
chr4-52710412-DEL-86
chr4-52736429-DEL-247
chr4-52810567-DEL-68
chr4-52852925-DEL-55
chr4-52867584-DEL-387
chr4-52880247-DEL-140
chr4-52895834-DEL-1928
chr4-52900840-DEL-2386
chr4-52933202-DEL-111
chr4-52943233-DEL-801
chr4-52969588-DEL-168
chr4-52983226-DEL-195
chr4-53013031-DEL-224
chr4-53037857-DEL-141
chr4-53058196-DEL-636
chr4-53101198-DEL-446
chr4-53127852-DEL-61
chr4-53135374-DEL-55
chr4-53158424-DEL-143
chr4-53164934-DEL-105
chr4-53190075-DEL-71
chr4-53201415-DEL-83
chr4-53214982-DEL-215
chr4-53243621-DEL-429
chr4-53286717-DEL-89
chr4-53294798-DEL-393
chr4-53313046-DEL-56
chr4-53348251-DEL-85
chr4-53349391-DEL-118
chr4-53362928-DEL-3137
chr4-53364105-DEL-233
chr4-53375431-DEL-90
chr4-53448214-DEL-1685
chr4-53456661-DEL-1112
chr4-53457353-DEL-338
chr4-53463748-DEL-315
chr4-53508827-DEL-72
chr4-53515274-DEL-259
chr4-53533606-DEL-394
chr4-53549981-DEL-197
chr4-53560078-DEL-64
chr4-53601847-DEL-1360
chr4-53604118-DEL-728
chr4-53668534-DEL-211
chr4-53668933-DEL-424
chr4-53680886-DEL-78

chr4-53741839-DEL-1388
chr4-53765854-DEL-3286
chr4-53781147-DEL-186
chr4-53791654-DEL-193
chr4-53792148-DEL-979
chr4-53793950-DEL-212
chr4-53796092-DEL-129
chr4-53801052-DEL-119
chr4-53810960-DEL-1788
chr4-53814217-DEL-1954
chr4-53816212-DEL-1015
chr4-53840327-DEL-71
chr4-53853928-DEL-57
chr4-53877646-DEL-57
chr4-53885738-DEL-262
chr4-53909949-DEL-211
chr4-53912921-DEL-93
chr4-53932943-DEL-63
chr4-53933631-DEL-140
chr4-53936806-DEL-68
chr4-53939145-DEL-8606
chr4-53939683-DEL-185
chr4-53953825-DEL-1024
chr4-53956135-DEL-253
chr4-53966253-DEL-495
chr4-53983901-DEL-195
chr4-53986672-DEL-147
chr4-53992572-DEL-124
chr4-54012964-DEL-330
chr4-54068045-DEL-76
chr4-54114411-DEL-56
chr4-54166638-DEL-82
chr4-54191587-DEL-87
chr4-54209000-DEL-347
chr4-54241108-DEL-523
chr4-54281389-DEL-72
chr4-54320799-DEL-55
chr4-54340408-DEL-55
chr4-54364237-DEL-56
chr4-54395263-DEL-6025
chr4-54429878-DEL-52
chr4-54440929-DEL-339
chr4-54498101-DEL-54
chr4-54598400-DEL-1628
chr4-54648372-DEL-55
chr4-54695713-DEL-6219
chr4-54701646-DEL-230
chr4-54706527-DEL-108
chr4-54748774-DEL-259
chr4-54767958-DEL-84
chr4-54814888-DEL-58
chr4-54883253-DEL-1400
chr4-54888115-DEL-222
chr4-54927844-DEL-133
chr4-54929340-DEL-511
chr4-54939345-DEL-61
chr4-54942466-DEL-63
chr4-55048528-DEL-67
chr4-55087694-DEL-68
chr4-55107106-DEL-90
chr4-55170702-DEL-64
chr4-55213617-DEL-52
chr4-55234087-DEL-382
chr4-55306443-DEL-69
chr4-55307658-DEL-226
chr4-55321577-DEL-328
chr4-55323597-DEL-51
chr4-55324089-DEL-194
chr4-55330976-DEL-173
chr4-55343236-DEL-74
chr4-55368203-DEL-71
chr4-55370553-DEL-102
chr4-55393339-DEL-210
chr4-55396785-DEL-473
chr4-55449628-DEL-85
chr4-55461121-DEL-186
chr4-55493247-DEL-62

chr4-55549808-DEL-61
chr4-55557508-DEL-1120
chr4-55559478-DEL-1047
chr4-55578507-DEL-115
chr4-55591538-DEL-151
chr4-55689418-DEL-54
chr4-55701906-DEL-276
chr4-55734690-DEL-166
chr4-55747714-DEL-189
chr4-55788211-DEL-137
chr4-55792721-DEL-63
chr4-55803369-DEL-64
chr4-55822409-DEL-52
chr4-55829398-DEL-1145
chr4-55877188-DEL-346
chr4-55885564-DEL-55
chr4-55897691-DEL-418
chr4-55977288-DEL-57
chr4-55978770-DEL-440
chr4-55999443-DEL-98
chr4-56054599-DEL-88
chr4-56078675-DEL-160
chr4-56079247-DEL-55
chr4-56092424-DEL-52
chr4-56110196-DEL-51
chr4-56143377-DEL-313
chr4-56173453-DEL-52
chr4-56201702-DEL-62
chr4-56227569-DEL-55
chr4-56234522-DEL-124
chr4-56243166-DEL-64
chr4-56262460-DEL-81
chr4-56293582-DEL-116
chr4-56352929-DEL-60
chr4-56394675-DEL-6351
chr4-56402217-DEL-99
chr4-56416943-DEL-326
chr4-56428956-DEL-116
chr4-56434603-DEL-7574
chr4-56435029-DEL-200
chr4-56461884-DEL-152
chr4-56498096-DEL-223
chr4-56537343-DEL-69
chr4-56540395-DEL-1039
chr4-56554059-DEL-6172
chr4-56561558-DEL-1124
chr4-56567520-DEL-73
chr4-56570140-DEL-6245
chr4-56625233-DEL-151
chr4-56636287-DEL-72
chr4-56651056-DEL-182
chr4-56653338-DEL-449
chr4-56653922-DEL-86
chr4-56667829-DEL-181
chr4-56668880-DEL-1105
chr4-56688644-DEL-187
chr4-56689013-DEL-711
chr4-56693221-DEL-315
chr4-56698573-DEL-56
chr4-56702484-DEL-976
chr4-56736155-DEL-109
chr4-56738728-DEL-525
chr4-56788779-DEL-157
chr4-56815583-DEL-118
chr4-56830431-DEL-251
chr4-56842794-DEL-228
chr4-56853239-DEL-60
chr4-56914156-DEL-737
chr4-56955382-DEL-111
chr4-56963490-DEL-64
chr4-56964072-DEL-204
chr4-56970558-DEL-114
chr4-57034044-DEL-96
chr4-57074860-DEL-179
chr4-57075540-DEL-88
chr4-57124004-DEL-230
chr4-57150447-DEL-55

chr4-57179202-DEL-127
chr4-57181927-DEL-124
chr4-57198929-DEL-425
chr4-57226104-DEL-51
chr4-57236107-DEL-200
chr4-57272406-DEL-84
chr4-57315028-DEL-56
chr4-57317673-DEL-966
chr4-57353210-DEL-57
chr4-57410205-DEL-190
chr4-57414792-DEL-56
chr4-57473742-DEL-55
chr4-57486980-DEL-52
chr4-57509475-DEL-1337
chr4-57532417-DEL-388
chr4-57541652-DEL-151
chr4-57589966-DEL-486
chr4-57605495-DEL-118
chr4-57623821-DEL-121
chr4-57697095-DEL-116
chr4-57721368-DEL-65
chr4-57767457-DEL-62
chr4-57774494-DEL-93
chr4-57811427-DEL-220
chr4-57820311-DEL-117
chr4-57839023-DEL-81
chr4-57869928-DEL-210
chr4-57906462-DEL-148
chr4-57929717-DEL-60
chr4-57930621-DEL-156
chr4-57934637-DEL-52
chr4-57964803-DEL-304
chr4-57971527-DEL-270
chr4-57996438-DEL-61
chr4-58005459-DEL-66
chr4-58022472-DEL-59
chr4-58026033-DEL-60
chr4-58058758-DEL-704
chr4-58087275-DEL-327
chr4-58099456-DEL-394
chr4-58118316-DEL-129
chr4-58169749-DEL-181
chr4-58173828-DEL-52
chr4-58210135-DEL-90
chr4-58222802-DEL-85
chr4-58239775-DEL-151
chr4-58252016-DEL-65
chr4-58254977-DEL-6220
chr4-58273964-DEL-234
chr4-58322820-DEL-177
chr4-58337474-DEL-123
chr4-58379806-DEL-1040
chr4-58396013-DEL-71
chr4-58406481-DEL-6408
chr4-58432731-DEL-375
chr4-58448187-DEL-120
chr4-58457524-DEL-65
chr4-58458232-DEL-100
chr4-58559739-DEL-131
chr4-58561389-DEL-162
chr4-58575071-DEL-64
chr4-58579750-DEL-246
chr4-58593750-DEL-66
chr4-58599997-DEL-129
chr4-58649592-DEL-57
chr4-58651588-DEL-3885
chr4-58654784-DEL-481
chr4-58698216-DEL-68
chr4-58717305-DEL-84
chr4-58719053-DEL-186
chr4-58767404-DEL-51
chr4-58775560-DEL-74
chr4-58810213-DEL-239
chr4-58845229-DEL-690
chr4-58866772-DEL-95
chr4-58885605-DEL-56
chr4-58885738-DEL-1079

chr4-58950192-DEL-435
chr4-58951294-DEL-106
chr4-58976192-DEL-79
chr4-58994310-DEL-74
chr4-59001033-DEL-210
chr4-59007425-DEL-61
chr4-59020905-DEL-159
chr4-59045451-DEL-173
chr4-59058069-DEL-1150
chr4-59091910-DEL-83
chr4-59098577-DEL-63
chr4-59100237-DEL-3633
chr4-59133295-DEL-226
chr4-59133451-DEL-107
chr4-59166324-DEL-100
chr4-59180973-DEL-77
chr4-59219987-DEL-135
chr4-59242971-DEL-53
chr4-59246759-DEL-7506
chr4-59249153-DEL-222
chr4-59270534-DEL-156
chr4-59272528-DEL-58
chr4-59274349-DEL-220
chr4-59295882-DEL-116
chr4-59300702-DEL-167
chr4-59302917-DEL-4435
chr4-59341470-DEL-369
chr4-59342811-DEL-200
chr4-59396006-DEL-215
chr4-59403785-DEL-54
chr4-59406289-DEL-121
chr4-59433867-DEL-122
chr4-59516876-DEL-176
chr4-59541711-DEL-174
chr4-59555997-DEL-64
chr4-59556231-DEL-580
chr4-59557294-DEL-188
chr4-59563077-DEL-71
chr4-59571702-DEL-182
chr4-59573247-DEL-209
chr4-59573603-DEL-362
chr4-59601932-DEL-1807
chr4-59610572-DEL-53
chr4-59651208-DEL-57
chr4-59659313-DEL-210
chr4-59678525-DEL-1447
chr4-59678617-DEL-219
chr4-59678861-DEL-569
chr4-59679705-DEL-78
chr4-59732274-DEL-319
chr4-59742199-DEL-304
chr4-59753706-DEL-95
chr4-59764238-DEL-59
chr4-59797811-DEL-1404
chr4-59827949-DEL-53
chr4-59836356-DEL-187
chr4-59956085-DEL-54
chr4-59960904-DEL-64
chr4-59971808-DEL-6630
chr4-59972266-DEL-179
chr4-60011173-DEL-4772
chr4-60018055-DEL-133
chr4-60037673-DEL-3242
chr4-60037677-DEL-66
chr4-60041377-DEL-53
chr4-60049127-DEL-7473
chr4-60070878-DEL-114
chr4-60077730-DEL-787
chr4-60185233-DEL-2384
chr4-60218185-DEL-3440
chr4-60220913-DEL-51
chr4-60223412-DEL-175
chr4-60227076-DEL-563
chr4-60227820-DEL-76
chr4-60233960-DEL-60
chr4-60279011-DEL-1127
chr4-60280044-DEL-205

chr4-60296861-DEL-192
chr4-60300785-DEL-66
chr4-60308760-DEL-55
chr4-60333091-DEL-885
chr4-60338477-DEL-1310
chr4-60363113-DEL-123
chr4-60407915-DEL-6866
chr4-60507726-DEL-6684
chr4-60519928-DEL-4818
chr4-60527605-DEL-5326
chr4-60549369-DEL-102
chr4-60557313-DEL-56
chr4-60575761-DEL-665
chr4-60660876-DEL-424
chr4-61111029-DEL-779
chr4-61134288-DEL-7585
chr4-61229445-DEL-4204
chr4-61338456-DEL-1893
chr4-61344192-DEL-1210
chr4-61347358-DEL-5339
chr4-61362950-DEL-65
chr4-61377175-DEL-394
chr4-61378448-DEL-364
chr4-61433396-DEL-3419
chr4-61472101-DEL-1303
chr4-61534478-DEL-364
chr4-61566863-DEL-5692
chr4-61574506-DEL-1519
chr4-61596885-DEL-9937
chr4-61601583-DEL-7168
chr4-61603223-DEL-3318
chr4-61604011-DEL-406
chr4-61605917-DEL-624
chr4-61608647-DEL-4685
chr4-61612939-DEL-72
chr4-61613246-DEL-338
chr4-61613653-DEL-1473
chr4-61615147-DEL-5320
chr4-61685403-DEL-7544
chr4-61728232-DEL-487
chr4-61738621-DEL-58
chr4-61753688-DEL-63
chr4-61812857-DEL-64
chr4-61837076-DEL-3530
chr4-61848846-DEL-465
chr4-61852749-DEL-2022
chr4-61860259-DEL-1322
chr4-61863803-DEL-200
chr4-61884278-DEL-203
chr4-61886164-DEL-551
chr4-61894638-DEL-290
chr4-61910586-DEL-197
chr4-61913221-DEL-2023
chr4-61913536-DEL-1012
chr4-61914404-DEL-1012
chr4-61916049-DEL-563
chr4-61921270-DEL-3177
chr4-61922256-DEL-515
chr4-61937090-DEL-585
chr4-61938539-DEL-190
chr4-61939298-DEL-196
chr4-61948595-DEL-92
chr4-61953986-DEL-2020
chr4-61965054-DEL-200
chr4-61970525-DEL-91
chr4-62065812-DEL-2869
chr4-62072071-DEL-336
chr4-62091791-DEL-330
chr4-62138543-DEL-165
chr4-62171820-DEL-53
chr4-62182534-DEL-579
chr4-62186200-DEL-5286
chr4-62202331-DEL-329
chr4-62243069-DEL-972
chr4-62298603-DEL-209
chr4-62299594-DEL-60
chr4-62318981-DEL-68

chr4-62357511-DEL-165
chr4-62434620-DEL-176
chr4-62508732-DEL-52
chr4-62519196-DEL-495
chr4-62519841-DEL-57
chr4-62555318-DEL-357
chr4-62563257-DEL-370
chr4-62638386-DEL-78
chr4-62680432-DEL-226
chr4-62760496-DEL-401
chr4-62776981-DEL-56
chr4-62802132-DEL-76
chr4-62813750-DEL-850
chr4-62824199-DEL-60
chr4-62876246-DEL-64
chr4-62895319-DEL-81
chr4-62909147-DEL-261
chr4-62950040-DEL-71
chr4-62981016-DEL-107
chr4-63025304-DEL-62
chr4-63049589-DEL-143
chr4-63076109-DEL-68
chr4-63110899-DEL-54
chr4-63177764-DEL-108
chr4-63183059-DEL-85
chr4-63186922-DEL-3440
chr4-63247012-DEL-208
chr4-63250613-DEL-84
chr4-63253620-DEL-86
chr4-63258480-DEL-426
chr4-63290915-DEL-68
chr4-63340935-DEL-56
chr4-63374595-DEL-74
chr4-63420020-DEL-88
chr4-63426777-DEL-116
chr4-63431009-DEL-58
chr4-63439218-DEL-103
chr4-63448210-DEL-101
chr4-63460889-DEL-140
chr4-63498325-DEL-72
chr4-63507224-DEL-73
chr4-63520280-DEL-362
chr4-63535240-DEL-65
chr4-63538584-DEL-53
chr4-63559678-DEL-1335
chr4-63582106-DEL-380
chr4-63582808-DEL-266
chr4-63583325-DEL-292
chr4-63610196-DEL-70
chr4-63619982-DEL-120
chr4-63637288-DEL-74
chr4-63667814-DEL-115
chr4-63698145-DEL-1777
chr4-63706607-DEL-843
chr4-63720381-DEL-3338
chr4-63735196-DEL-56
chr4-63766194-DEL-51
chr4-63800967-DEL-51
chr4-63813728-DEL-634
chr4-63818262-DEL-208
chr4-63851990-DEL-1483
chr4-63869418-DEL-1431
chr4-63915078-DEL-115
chr4-63916500-DEL-84
chr4-63926659-DEL-109
chr4-63928050-DEL-200
chr4-63945385-DEL-98
chr4-63960569-DEL-73
chr4-63968670-DEL-183
chr4-63970550-DEL-104
chr4-64015872-DEL-52
chr4-64048868-DEL-70
chr4-64075133-DEL-60
chr4-64091207-DEL-71
chr4-64095672-DEL-298
chr4-64101725-DEL-2343
chr4-64104350-DEL-144

chr4-64105858-DEL-279
chr4-64119007-DEL-55
chr4-64141468-DEL-116
chr4-64158111-DEL-64
chr4-64172171-DEL-543
chr4-64185066-DEL-76
chr4-64193968-DEL-125
chr4-64268953-DEL-52
chr4-64273761-DEL-115
chr4-64280171-DEL-70
chr4-64297620-DEL-467
chr4-64302561-DEL-52
chr4-64305002-DEL-51
chr4-64305675-DEL-54
chr4-64309532-DEL-75
chr4-64312763-DEL-55
chr4-64319271-DEL-185
chr4-64325162-DEL-129
chr4-64387445-DEL-198
chr4-64394329-DEL-1617
chr4-64415924-DEL-494
chr4-64443760-DEL-194
chr4-64450138-DEL-1109
chr4-64535241-DEL-125
chr4-64564810-DEL-92
chr4-64585011-DEL-153
chr4-64623635-DEL-1332
chr4-64627024-DEL-1477
chr4-64667466-DEL-1190
chr4-64677311-DEL-910
chr4-64681095-DEL-354
chr4-64683377-DEL-137
chr4-64709270-DEL-132
chr4-64714347-DEL-1499
chr4-64715342-DEL-719
chr4-64733545-DEL-1411
chr4-64736755-DEL-53
chr4-64744940-DEL-219
chr4-64753924-DEL-95
chr4-64764190-DEL-60
chr4-64778973-DEL-754
chr4-64797506-DEL-141
chr4-64800843-DEL-140
chr4-64809249-DEL-1717
chr4-64835093-DEL-1693
chr4-64839929-DEL-266
chr4-64859553-DEL-56
chr4-64867471-DEL-1397
chr4-64887579-DEL-85
chr4-64889986-DEL-401
chr4-64916935-DEL-54
chr4-64935482-DEL-72
chr4-64944239-DEL-92
chr4-64957945-DEL-60
chr4-64974492-DEL-4391
chr4-64978180-DEL-1886
chr4-65018140-DEL-53
chr4-65092543-DEL-51
chr4-65112533-DEL-52
chr4-65129894-DEL-387
chr4-65160285-DEL-82
chr4-65182152-DEL-107
chr4-65233137-DEL-72
chr4-65257972-DEL-472
chr4-65338036-DEL-55
chr4-65369499-DEL-1451
chr4-65374741-DEL-78
chr4-65378030-DEL-2290
chr4-65385693-DEL-166
chr4-65386644-DEL-1356
chr4-65407145-DEL-127
chr4-65409578-DEL-64
chr4-65431571-DEL-87
chr4-65459853-DEL-806
chr4-65472543-DEL-62
chr4-65485658-DEL-6276
chr4-65509492-DEL-183

chr4-65517852-DEL-111
chr4-65571802-DEL-54
chr4-65578795-DEL-77
chr4-65614700-DEL-101
chr4-65692883-DEL-101
chr4-65709136-DEL-182
chr4-65713892-DEL-392
chr4-65734428-DEL-51
chr4-65759951-DEL-67
chr4-65772497-DEL-768
chr4-65775831-DEL-56
chr4-65794628-DEL-60
chr4-65818740-DEL-191
chr4-65881181-DEL-803
chr4-65886541-DEL-95
chr4-65914582-DEL-2247
chr4-65937170-DEL-65
chr4-65948266-DEL-242
chr4-65974541-DEL-171
chr4-65982864-DEL-6656
chr4-65996483-DEL-6462
chr4-66004638-DEL-141
chr4-66014704-DEL-108
chr4-66055582-DEL-164
chr4-66064880-DEL-161
chr4-66068075-DEL-90
chr4-66068919-DEL-1718
chr4-66084601-DEL-57
chr4-66091625-DEL-52
chr4-66115888-DEL-975
chr4-66129299-DEL-89
chr4-66136878-DEL-133
chr4-66140020-DEL-68
chr4-66162683-DEL-660
chr4-66165734-DEL-128
chr4-66183357-DEL-70
chr4-66184977-DEL-464
chr4-66195105-DEL-152
chr4-66205545-DEL-236
chr4-66248745-DEL-62
chr4-66298922-DEL-271
chr4-66306506-DEL-212
chr4-66334396-DEL-54
chr4-66359710-DEL-150
chr4-66407499-DEL-1514
chr4-66410040-DEL-1867
chr4-66411524-DEL-879
chr4-66424936-DEL-61
chr4-66425950-DEL-6210
chr4-66467049-DEL-81
chr4-66555740-DEL-212
chr4-66559761-DEL-6292
chr4-66562704-DEL-205
chr4-66574178-DEL-6341
chr4-66581268-DEL-173
chr4-66602061-DEL-277
chr4-66622832-DEL-6260
chr4-66650415-DEL-430
chr4-66656452-DEL-339
chr4-66657685-DEL-4411
chr4-66693258-DEL-8111
chr4-66701631-DEL-468
chr4-66713505-DEL-496
chr4-66727596-DEL-70
chr4-66768954-DEL-170
chr4-66786218-DEL-318
chr4-66787614-DEL-142
chr4-66806717-DEL-732
chr4-66807383-DEL-181
chr4-66820290-DEL-5089
chr4-66850284-DEL-55
chr4-66861690-DEL-56
chr4-66877925-DEL-147
chr4-66880744-DEL-72
chr4-66935333-DEL-82
chr4-66956197-DEL-191
chr4-66966931-DEL-1040

chr4-66985477-DEL-73
chr4-66996028-DEL-74
chr4-67009068-DEL-58
chr4-67011044-DEL-1105
chr4-67037688-DEL-2575
chr4-67050297-DEL-550
chr4-67055025-DEL-667
chr4-67063160-DEL-53
chr4-67069903-DEL-350
chr4-67091831-DEL-644
chr4-67098105-DEL-1306
chr4-67126771-DEL-57
chr4-67150430-DEL-104
chr4-67151852-DEL-214
chr4-67161643-DEL-65
chr4-67169786-DEL-103
chr4-67190383-DEL-118
chr4-67193917-DEL-5805
chr4-67203836-DEL-54
chr4-67207633-DEL-76
chr4-67258756-DEL-550
chr4-67262614-DEL-54
chr4-67265520-DEL-2420
chr4-67274357-DEL-73
chr4-67315714-DEL-166
chr4-67327390-DEL-65
chr4-67348975-DEL-85
chr4-67365123-DEL-1447
chr4-67413671-DEL-79
chr4-67419543-DEL-154
chr4-67427261-DEL-108
chr4-67428213-DEL-1175
chr4-67428317-DEL-586
chr4-67494616-DEL-51
chr4-67510157-DEL-572
chr4-67530070-DEL-57
chr4-67542235-DEL-116
chr4-67565619-DEL-82
chr4-67579712-DEL-98
chr4-67582402-DEL-61
chr4-67585323-DEL-318
chr4-67607462-DEL-138
chr4-67624525-DEL-273
chr4-67660944-DEL-52
chr4-67667081-DEL-112
chr4-67672839-DEL-458
chr4-67733869-DEL-164
chr4-67766805-DEL-1483
chr4-67785857-DEL-429
chr4-67791485-DEL-89
chr4-67813977-DEL-829
chr4-67815211-DEL-59
chr4-67834246-DEL-424
chr4-67873922-DEL-51
chr4-67881078-DEL-127
chr4-67890125-DEL-98
chr4-67892219-DEL-102
chr4-67911977-DEL-93
chr4-67917661-DEL-220
chr4-67941718-DEL-1284
chr4-67974791-DEL-51
chr4-68005484-DEL-127
chr4-68017666-DEL-2387
chr4-68023310-DEL-58
chr4-68024924-DEL-168
chr4-68029327-DEL-498
chr4-68030489-DEL-358
chr4-68032994-DEL-94
chr4-68052699-DEL-985
chr4-68055825-DEL-116
chr4-68069839-DEL-60
chr4-68091292-DEL-155
chr4-68107230-DEL-51
chr4-68110235-DEL-1615
chr4-68166164-DEL-574
chr4-68223876-DEL-5116
chr4-68244248-DEL-689

chr4-68248411-DEL-56
chr4-68252131-DEL-72
chr4-68263589-DEL-521
chr4-68281793-DEL-541
chr4-68293811-DEL-456
chr4-68339536-DEL-55
chr4-68344309-DEL-6374
chr4-68344523-DEL-64
chr4-68356752-DEL-574
chr4-68364392-DEL-1940
chr4-68407886-DEL-197
chr4-68411736-DEL-146
chr4-68413878-DEL-52
chr4-68423098-DEL-59
chr4-68481210-DEL-162
chr4-68493543-DEL-75
chr4-68506470-DEL-1873
chr4-68576056-DEL-55
chr4-68576565-DEL-717
chr4-68648190-DEL-66
chr4-68702051-DEL-948
chr4-68745966-DEL-132
chr4-68777334-DEL-52
chr4-68799982-DEL-115
chr4-68816662-DEL-252
chr4-68842732-DEL-83
chr4-68869752-DEL-68
chr4-68906025-DEL-470
chr4-68912365-DEL-91
chr4-68913771-DEL-6175
chr4-68932787-DEL-117
chr4-68978718-DEL-183
chr4-68979412-DEL-514
chr4-68985054-DEL-54
chr4-69009907-DEL-516
chr4-69030231-DEL-69
chr4-69055150-DEL-72
chr4-69064534-DEL-1600
chr4-69084128-DEL-376
chr4-69089404-DEL-6290
chr4-69134687-DEL-3401
chr4-69134713-DEL-6461
chr4-69141205-DEL-114
chr4-69178886-DEL-234
chr4-69187822-DEL-208
chr4-69195842-DEL-76
chr4-69216201-DEL-200
chr4-69222809-DEL-1235
chr4-69225048-DEL-143
chr4-69228280-DEL-4834
chr4-69249973-DEL-88
chr4-69277770-DEL-95
chr4-69278350-DEL-841
chr4-69279215-DEL-288
chr4-69300524-DEL-581
chr4-69303390-DEL-645
chr4-69328580-DEL-331
chr4-69333403-DEL-6244
chr4-69352516-DEL-5131
chr4-69359886-DEL-65
chr4-69372892-DEL-168
chr4-69379213-DEL-521
chr4-69381804-DEL-304
chr4-69383203-DEL-857
chr4-69390495-DEL-177
chr4-69445506-DEL-60
chr4-69459754-DEL-79
chr4-69482207-DEL-6072
chr4-69493468-DEL-60
chr4-69538610-DEL-52
chr4-69556102-DEL-70
chr4-69570878-DEL-445
chr4-69573259-DEL-109
chr4-69607204-DEL-56
chr4-69610170-DEL-725
chr4-69695516-DEL-64
chr4-69700782-DEL-2308

chr4-69715683-DEL-181
chr4-69741387-DEL-54
chr4-69745451-DEL-554
chr4-69807107-DEL-458
chr4-69860931-DEL-223
chr4-69886130-DEL-495
chr4-69892848-DEL-364
chr4-69918902-DEL-559
chr4-69986468-DEL-60
chr4-69986934-DEL-199
chr4-69999213-DEL-6492
chr4-70035175-DEL-105
chr4-70073346-DEL-922
chr4-70100837-DEL-358
chr4-70125634-DEL-928
chr4-70231142-DEL-1026
chr4-70252239-DEL-172
chr4-70285368-DEL-5615
chr4-70296354-DEL-131
chr4-70312854-DEL-89
chr4-70338216-DEL-79
chr4-70339949-DEL-329
chr4-70356764-DEL-605
chr4-70361476-DEL-1087
chr4-70443926-DEL-552
chr4-70484039-DEL-1124
chr4-70495575-DEL-828
chr4-70513210-DEL-83
chr4-70531397-DEL-153
chr4-70531579-DEL-814
chr4-70548609-DEL-57
chr4-70554513-DEL-2054
chr4-70556663-DEL-53
chr4-70559231-DEL-78
chr4-70559467-DEL-1966
chr4-70584752-DEL-1214
chr4-70630628-DEL-636
chr4-70649494-DEL-366
chr4-70658788-DEL-254
chr4-70663104-DEL-198
chr4-70673879-DEL-6360
chr4-70696709-DEL-1893
chr4-70737208-DEL-168
chr4-70740626-DEL-57
chr4-70770301-DEL-140
chr4-70830041-DEL-110
chr4-70846508-DEL-213
chr4-70877154-DEL-55
chr4-70877523-DEL-252
chr4-70881274-DEL-59
chr4-70888825-DEL-53
chr4-70897581-DEL-214
chr4-70967413-DEL-156
chr4-70990209-DEL-2779
chr4-71039643-DEL-885
chr4-71059186-DEL-2904
chr4-71100415-DEL-2556
chr4-71150779-DEL-219
chr4-71154553-DEL-1998
chr4-71211458-DEL-2094
chr4-71270154-DEL-216
chr4-71281486-DEL-54
chr4-71294848-DEL-52
chr4-71304987-DEL-3766
chr4-71327764-DEL-385
chr4-71332655-DEL-1067
chr4-71343740-DEL-795
chr4-71383023-DEL-174
chr4-71412300-DEL-6181
chr4-71453455-DEL-105
chr4-71458688-DEL-62
chr4-71463972-DEL-108
chr4-71474475-DEL-736
chr4-71483559-DEL-942
chr4-71500820-DEL-361
chr4-71505041-DEL-936
chr4-71536152-DEL-3691

chr4-71560728-DEL-60
chr4-71563710-DEL-117
chr4-71588232-DEL-132
chr4-71594065-DEL-88
chr4-71597215-DEL-51
chr4-71604776-DEL-304
chr4-71619638-DEL-131
chr4-71621047-DEL-7239
chr4-71665428-DEL-70
chr4-71687256-DEL-149
chr4-71695993-DEL-116
chr4-71697319-DEL-11330
chr4-71721362-DEL-69
chr4-71725746-DEL-182
chr4-71751463-DEL-76
chr4-71800512-DEL-1130
chr4-71813312-DEL-197
chr4-71816408-DEL-110
chr4-71821361-DEL-6454
chr4-71827543-DEL-214
chr4-71899260-DEL-92
chr4-71911852-DEL-6292
chr4-71927799-DEL-942
chr4-71933442-DEL-647
chr4-71944934-DEL-123
chr4-71966363-DEL-78
chr4-71977147-DEL-486
chr4-71978557-DEL-147
chr4-71998210-DEL-424
chr4-72006371-DEL-6181
chr4-72038949-DEL-62
chr4-72182889-DEL-2087
chr4-72185192-DEL-892
chr4-72194693-DEL-75
chr4-72198024-DEL-1192
chr4-72201216-DEL-865
chr4-72209247-DEL-1755
chr4-72211245-DEL-1328
chr4-72215896-DEL-58
chr4-72218501-DEL-206
chr4-72227308-DEL-356
chr4-72227986-DEL-1081
chr4-72272802-DEL-6408
chr4-72319085-DEL-414
chr4-72335160-DEL-629
chr4-72400498-DEL-1841
chr4-72402761-DEL-1184
chr4-72409241-DEL-1522
chr4-72428914-DEL-436
chr4-72437511-DEL-1997
chr4-72441935-DEL-60
chr4-72443249-DEL-476
chr4-72470601-DEL-54
chr4-72476857-DEL-69
chr4-72519145-DEL-467
chr4-72538980-DEL-3646
chr4-72571310-DEL-217
chr4-72610894-DEL-524
chr4-72613918-DEL-56
chr4-72631359-DEL-212
chr4-72640189-DEL-138
chr4-72652230-DEL-166
chr4-72658160-DEL-1287
chr4-72660355-DEL-54
chr4-72662376-DEL-669
chr4-72665717-DEL-375
chr4-72672570-DEL-352
chr4-72695032-DEL-378
chr4-72707706-DEL-99
chr4-72715880-DEL-401
chr4-72779226-DEL-66
chr4-72797939-DEL-1397
chr4-72808034-DEL-214
chr4-72812640-DEL-58
chr4-72813666-DEL-72
chr4-72823194-DEL-62
chr4-72823333-DEL-1268

chr4-72839998-DEL-150
chr4-72842328-DEL-56
chr4-72847213-DEL-150
chr4-72851318-DEL-853
chr4-72852398-DEL-302
chr4-72855425-DEL-1283
chr4-72866935-DEL-19025
chr4-72874092-DEL-6094
chr4-72881772-DEL-4535
chr4-72884231-DEL-1787
chr4-72897588-DEL-2486
chr4-72902403-DEL-457
chr4-72904316-DEL-510
chr4-72908024-DEL-1903
chr4-72923195-DEL-590
chr4-72951985-DEL-233
chr4-72952515-DEL-56
chr4-72971745-DEL-81
chr4-72982769-DEL-2717
chr4-72990770-DEL-444
chr4-72992544-DEL-87
chr4-72994464-DEL-936
chr4-72995984-DEL-90
chr4-73037902-DEL-5971
chr4-73062642-DEL-66
chr4-73093043-DEL-54
chr4-73101305-DEL-1436
chr4-73102871-DEL-13621
chr4-73102888-DEL-606
chr4-73104047-DEL-7429
chr4-73112420-DEL-169
chr4-73114199-DEL-1289
chr4-73120474-DEL-7088
chr4-73129458-DEL-4032
chr4-73141476-DEL-71
chr4-73143561-DEL-1280
chr4-73146879-DEL-464
chr4-73151551-DEL-6219
chr4-73175663-DEL-14572
chr4-73225813-DEL-6352
chr4-73283710-DEL-246
chr4-73325762-DEL-267
chr4-73328318-DEL-73
chr4-73336557-DEL-2847
chr4-73346428-DEL-1313
chr4-73356740-DEL-139
chr4-73382722-DEL-1183
chr4-73396383-DEL-60
chr4-73478185-DEL-1489
chr4-73489927-DEL-214
chr4-73491282-DEL-197
chr4-73522674-DEL-5075
chr4-73523383-DEL-259
chr4-73533275-DEL-54
chr4-73583855-DEL-262
chr4-73593753-DEL-58
chr4-73599031-DEL-257
chr4-73608910-DEL-53
chr4-73628123-DEL-77
chr4-73630434-DEL-838
chr4-73638311-DEL-278
chr4-73665442-DEL-502
chr4-73668677-DEL-6278
chr4-73703258-DEL-1585
chr4-73716629-DEL-6197
chr4-73746997-DEL-6269
chr4-73771741-DEL-415
chr4-73773314-DEL-6450
chr4-73782421-DEL-3411
chr4-73788723-DEL-7842
chr4-73791544-DEL-1844
chr4-73831646-DEL-21642
chr4-73856508-DEL-154
chr4-73880634-DEL-56
chr4-73897904-DEL-546
chr4-73952868-DEL-204
chr4-73964227-DEL-74

chr4-73992846-DEL-784
chr4-73999246-DEL-619
chr4-74057723-DEL-387
chr4-74146290-DEL-1917
chr4-74148440-DEL-61
chr4-74156620-DEL-85
chr4-74202200-DEL-699
chr4-74218605-DEL-128
chr4-74224847-DEL-213
chr4-74243407-DEL-214
chr4-74284153-DEL-127
chr4-74301973-DEL-474
chr4-74314384-DEL-388
chr4-74329057-DEL-54
chr4-74336796-DEL-195
chr4-74362456-DEL-125
chr4-74404438-DEL-6185
chr4-74414585-DEL-3008
chr4-74417521-DEL-6898
chr4-74423511-DEL-85
chr4-74424190-DEL-272
chr4-74424902-DEL-177
chr4-74443562-DEL-71
chr4-74457283-DEL-117
chr4-74507857-DEL-99
chr4-74536273-DEL-820
chr4-74537704-DEL-1201
chr4-74547430-DEL-2287
chr4-74555818-DEL-149
chr4-74563504-DEL-3933
chr4-74575370-DEL-262
chr4-74577975-DEL-65
chr4-74584061-DEL-6233
chr4-74606280-DEL-1494
chr4-74711579-DEL-326
chr4-74719972-DEL-6014
chr4-74725433-DEL-1834
chr4-74746046-DEL-8393
chr4-74749588-DEL-92
chr4-74772477-DEL-213
chr4-74773661-DEL-145
chr4-74778947-DEL-72
chr4-74798472-DEL-3492
chr4-74810769-DEL-843
chr4-74832334-DEL-2856
chr4-74838439-DEL-63
chr4-74865213-DEL-1016
chr4-74876341-DEL-1383
chr4-74880607-DEL-1389
chr4-74893095-DEL-72
chr4-74898339-DEL-644
chr4-74905871-DEL-305
chr4-74906318-DEL-1628
chr4-74922297-DEL-62
chr4-74935474-DEL-764
chr4-74963156-DEL-6299
chr4-75000371-DEL-1550
chr4-75005029-DEL-108
chr4-75080440-DEL-10481
chr4-75100411-DEL-1063
chr4-75115924-DEL-1279
chr4-75120838-DEL-56
chr4-75128617-DEL-138
chr4-75160019-DEL-57
chr4-75160872-DEL-400
chr4-75173561-DEL-1206
chr4-75185464-DEL-300
chr4-75202868-DEL-56
chr4-75213133-DEL-2312
chr4-75216886-DEL-482
chr4-75236005-DEL-1892
chr4-75251580-DEL-53
chr4-75290775-DEL-490
chr4-75291705-DEL-1082
chr4-75302407-DEL-310
chr4-75318216-DEL-81
chr4-75319106-DEL-2252

chr4-75328558-DEL-1014
chr4-75349997-DEL-199
chr4-75392229-DEL-66
chr4-75404391-DEL-6309
chr4-75413595-DEL-55
chr4-75413909-DEL-6126
chr4-75422930-DEL-6900
chr4-75456117-DEL-746
chr4-75483270-DEL-51
chr4-75513937-DEL-122
chr4-75570879-DEL-315
chr4-75573541-DEL-401
chr4-75578631-DEL-518
chr4-75596122-DEL-283
chr4-75626490-DEL-2459
chr4-75630698-DEL-135
chr4-75632051-DEL-293
chr4-75645956-DEL-66
chr4-75653894-DEL-2103
chr4-75732311-DEL-1085
chr4-75735210-DEL-92
chr4-75754415-DEL-59
chr4-75825449-DEL-65
chr4-75857915-DEL-146
chr4-75938280-DEL-75
chr4-75955760-DEL-72
chr4-76027811-DEL-462
chr4-76048867-DEL-655
chr4-76052324-DEL-82
chr4-76079259-DEL-70
chr4-76093083-DEL-52
chr4-76137328-DEL-209
chr4-76151616-DEL-83
chr4-76154334-DEL-112
chr4-76200744-DEL-440
chr4-76240110-DEL-59
chr4-76254124-DEL-610
chr4-76290311-DEL-2022
chr4-76325537-DEL-235
chr4-76381082-DEL-52
chr4-76458014-DEL-1482
chr4-76488538-DEL-493
chr4-76546021-DEL-65
chr4-76585549-DEL-52
chr4-76617254-DEL-77
chr4-76621860-DEL-7326
chr4-76629879-DEL-1078
chr4-76638961-DEL-625
chr4-76655680-DEL-1095
chr4-76665268-DEL-326
chr4-76666387-DEL-343
chr4-76709337-DEL-69
chr4-76735407-DEL-863
chr4-76745823-DEL-151
chr4-76747903-DEL-201
chr4-76750060-DEL-86
chr4-76774474-DEL-195
chr4-76780399-DEL-1709
chr4-76781569-DEL-741
chr4-76801463-DEL-64
chr4-76804121-DEL-616
chr4-76810136-DEL-338
chr4-76814985-DEL-1527
chr4-76844804-DEL-3173
chr4-76853824-DEL-181
chr4-76868114-DEL-414
chr4-76877188-DEL-60
chr4-76913303-DEL-332
chr4-76924837-DEL-712
chr4-76931068-DEL-222
chr4-76944951-DEL-52
chr4-76993327-DEL-60
chr4-76999094-DEL-659
chr4-77040510-DEL-88
chr4-77056054-DEL-142
chr4-77099845-DEL-117
chr4-77153904-DEL-593

chr4-77161866-DEL-61
chr4-77228329-DEL-162
chr4-77268473-DEL-94
chr4-77281680-DEL-54
chr4-77293677-DEL-142
chr4-77332408-DEL-6551
chr4-77358573-DEL-60
chr4-77361586-DEL-270
chr4-77362509-DEL-62
chr4-77365292-DEL-928
chr4-77390663-DEL-398
chr4-77399962-DEL-77
chr4-77411614-DEL-72
chr4-77417812-DEL-6202
chr4-77471665-DEL-1364
chr4-77479784-DEL-260
chr4-77492953-DEL-5518
chr4-77500376-DEL-3766
chr4-77501871-DEL-958
chr4-77506660-DEL-6531
chr4-77514283-DEL-1034
chr4-77535114-DEL-855
chr4-77543900-DEL-63
chr4-77551911-DEL-573
chr4-77584308-DEL-599
chr4-77607099-DEL-52
chr4-77607900-DEL-60
chr4-77614348-DEL-218
chr4-77641492-DEL-174
chr4-77654938-DEL-114
chr4-77659663-DEL-157
chr4-77663740-DEL-6003
chr4-77663750-DEL-11065
chr4-77669887-DEL-4928
chr4-77712929-DEL-217
chr4-77731750-DEL-53237
chr4-77733431-DEL-63
chr4-77742027-DEL-55
chr4-77769204-DEL-88
chr4-77823069-DEL-65
chr4-77874928-DEL-366
chr4-77877836-DEL-52
chr4-77886070-DEL-1950
chr4-77940004-DEL-1493
chr4-77957645-DEL-76
chr4-77959286-DEL-5072
chr4-77987622-DEL-61
chr4-78034248-DEL-54
chr4-78039430-DEL-4563
chr4-78044237-DEL-4056
chr4-78052219-DEL-1711
chr4-78064587-DEL-652
chr4-78095810-DEL-54
chr4-78098668-DEL-88
chr4-78103172-DEL-211
chr4-78146424-DEL-213
chr4-78153395-DEL-52
chr4-78163725-DEL-258
chr4-78218692-DEL-100
chr4-78250164-DEL-134
chr4-78273747-DEL-860
chr4-78273755-DEL-1030
chr4-78274702-DEL-432
chr4-78297174-DEL-157
chr4-78311637-DEL-201
chr4-78369697-DEL-137
chr4-78397692-DEL-2153
chr4-78422800-DEL-69
chr4-78444521-DEL-176
chr4-78506796-DEL-101
chr4-78520288-DEL-66
chr4-78524763-DEL-192
chr4-78531912-DEL-796
chr4-78536058-DEL-60
chr4-78536267-DEL-1050
chr4-78536645-DEL-62
chr4-78537816-DEL-190

chr4-78538908-DEL-55
chr4-78557385-DEL-1047
chr4-78564114-DEL-385
chr4-78571303-DEL-136
chr4-78574377-DEL-602
chr4-78589837-DEL-219
chr4-78621511-DEL-237
chr4-78630311-DEL-1157
chr4-78638155-DEL-127
chr4-78639102-DEL-84
chr4-78641304-DEL-1104
chr4-78654880-DEL-539
chr4-78655434-DEL-1621
chr4-78656546-DEL-509
chr4-78722704-DEL-956
chr4-78794275-DEL-75
chr4-78846445-DEL-198
chr4-78846748-DEL-744
chr4-78847992-DEL-2127
chr4-78848520-DEL-1466
chr4-78848927-DEL-860
chr4-78878321-DEL-437
chr4-78906250-DEL-1625
chr4-78913963-DEL-7438
chr4-78921649-DEL-52
chr4-78940522-DEL-537
chr4-78953723-DEL-301
chr4-78984993-DEL-1104
chr4-78987379-DEL-830
chr4-78989257-DEL-778
chr4-79000788-DEL-1592
chr4-79016464-DEL-3555
chr4-79025335-DEL-200
chr4-79029886-DEL-1363
chr4-79075747-DEL-268
chr4-79096667-DEL-1051
chr4-79100378-DEL-200
chr4-79104185-DEL-358
chr4-79106753-DEL-77
chr4-79114504-DEL-636
chr4-79129622-DEL-196
chr4-79151495-DEL-244
chr4-79169233-DEL-55
chr4-79175499-DEL-6233
chr4-79211578-DEL-15313
chr4-79237772-DEL-97
chr4-79259480-DEL-99
chr4-79261023-DEL-52
chr4-79265541-DEL-1769
chr4-79294245-DEL-611
chr4-79301966-DEL-138
chr4-79324369-DEL-71
chr4-79324804-DEL-844
chr4-79333924-DEL-2648
chr4-79338081-DEL-109
chr4-79339996-DEL-306
chr4-79341774-DEL-1648
chr4-79350484-DEL-121
chr4-79360058-DEL-1332
chr4-79377490-DEL-146
chr4-79382312-DEL-431
chr4-79403172-DEL-6091
chr4-79423969-DEL-672
chr4-79444587-DEL-3355
chr4-79459179-DEL-279
chr4-79465495-DEL-204
chr4-79472563-DEL-208
chr4-79490005-DEL-1708
chr4-79502219-DEL-3518
chr4-79518599-DEL-51
chr4-79521605-DEL-59
chr4-79572887-DEL-175
chr4-79574819-DEL-82
chr4-79577181-DEL-154
chr4-79586295-DEL-779
chr4-79656621-DEL-267
chr4-79664514-DEL-164

chr4-79667193-DEL-1075
chr4-79689370-DEL-883
chr4-79691639-DEL-325
chr4-79695125-DEL-524
chr4-79697718-DEL-1389
chr4-79714785-DEL-716
chr4-79727764-DEL-66
chr4-79741429-DEL-58
chr4-79769160-DEL-194
chr4-79819628-DEL-182
chr4-79827584-DEL-334
chr4-79852708-DEL-6401
chr4-79855229-DEL-2060
chr4-79873051-DEL-1511
chr4-79882671-DEL-997
chr4-79890937-DEL-1700
chr4-79932316-DEL-912
chr4-79938563-DEL-62
chr4-79948127-DEL-51
chr4-79958263-DEL-59
chr4-79991504-DEL-319
chr4-79992277-DEL-2452
chr4-80007623-DEL-65
chr4-80008990-DEL-609
chr4-80031877-DEL-88
chr4-80032189-DEL-491
chr4-80036239-DEL-2173
chr4-80040726-DEL-72
chr4-80075070-DEL-268
chr4-80081992-DEL-81
chr4-80090765-DEL-241
chr4-80096578-DEL-61
chr4-80102725-DEL-58
chr4-80139431-DEL-191
chr4-80145507-DEL-6130
chr4-80153616-DEL-121
chr4-80159386-DEL-6359
chr4-80170589-DEL-651
chr4-80171689-DEL-223
chr4-80201774-DEL-921
chr4-80212435-DEL-212
chr4-80263696-DEL-156
chr4-80270989-DEL-984
chr4-80287666-DEL-1356
chr4-80293435-DEL-187
chr4-80303311-DEL-125
chr4-80312158-DEL-148
chr4-80318630-DEL-156
chr4-80390531-DEL-152
chr4-80439339-DEL-54
chr4-80447994-DEL-488
chr4-80453146-DEL-508
chr4-80488897-DEL-68
chr4-80492139-DEL-1504
chr4-80520345-DEL-955
chr4-80525542-DEL-213
chr4-80622446-DEL-114
chr4-80623689-DEL-94
chr4-80629023-DEL-177
chr4-80694521-DEL-155
chr4-80718244-DEL-617
chr4-80719453-DEL-9191
chr4-80754498-DEL-209
chr4-80763755-DEL-81
chr4-80767407-DEL-137
chr4-80822352-DEL-1869
chr4-80852610-DEL-347
chr4-80861962-DEL-58
chr4-80885635-DEL-420
chr4-80892005-DEL-70
chr4-80911096-DEL-278
chr4-80992776-DEL-2495
chr4-81055421-DEL-153
chr4-81056424-DEL-66
chr4-81061454-DEL-6674
chr4-81067570-DEL-224
chr4-81071585-DEL-84

chr4-81080018-DEL-6194
chr4-81129592-DEL-57
chr4-81142701-DEL-772
chr4-81147277-DEL-1683
chr4-81148692-DEL-4662
chr4-81167991-DEL-66
chr4-81175429-DEL-6413
chr4-81177711-DEL-83
chr4-81241765-DEL-89
chr4-81247901-DEL-822
chr4-81247901-DEL-86
chr4-81254535-DEL-654
chr4-81257479-DEL-3404
chr4-81258536-DEL-1304
chr4-81265788-DEL-413
chr4-81275354-DEL-225
chr4-81314796-DEL-111
chr4-81389477-DEL-95
chr4-81424359-DEL-162
chr4-81444854-DEL-86
chr4-81476588-DEL-53
chr4-81505634-DEL-74
chr4-81576095-DEL-324
chr4-81601932-DEL-610
chr4-81636171-DEL-4333
chr4-81655770-DEL-400
chr4-81689223-DEL-114
chr4-81717541-DEL-6163
chr4-81734887-DEL-54
chr4-81757185-DEL-162
chr4-81791632-DEL-8136
chr4-81792907-DEL-5830
chr4-81794179-DEL-4558
chr4-81833849-DEL-144
chr4-81837682-DEL-2120
chr4-81869574-DEL-120
chr4-81889535-DEL-118
chr4-81929839-DEL-1266
chr4-81931411-DEL-201
chr4-81942397-DEL-134
chr4-82028516-DEL-59
chr4-82063009-DEL-351
chr4-82128743-DEL-6270
chr4-82153931-DEL-242
chr4-82317272-DEL-62
chr4-82338509-DEL-361
chr4-82346684-DEL-173
chr4-82354762-DEL-58
chr4-82475123-DEL-1317
chr4-82475309-DEL-815
chr4-82571603-DEL-584
chr4-82577895-DEL-50
chr4-82579037-DEL-949
chr4-82583703-DEL-103
chr4-82600887-DEL-79
chr4-82638807-DEL-54
chr4-82643017-DEL-91
chr4-82673977-DEL-183
chr4-82689989-DEL-104
chr4-82690635-DEL-5990
chr4-82741200-DEL-824
chr4-82754045-DEL-237
chr4-82794947-DEL-1385
chr4-82806483-DEL-89
chr4-82813938-DEL-197
chr4-82849699-DEL-76
chr4-82865985-DEL-105
chr4-82878407-DEL-84
chr4-82887222-DEL-203
chr4-82893154-DEL-94
chr4-82893365-DEL-1902
chr4-82898079-DEL-1838
chr4-82905560-DEL-873
chr4-82917358-DEL-84
chr4-82919083-DEL-62
chr4-82923117-DEL-156
chr4-82948919-DEL-87

chr4-82951662-DEL-165
chr4-82955854-DEL-333
chr4-82967517-DEL-56
chr4-82977628-DEL-401
chr4-82984303-DEL-624
chr4-82993025-DEL-414
chr4-82993609-DEL-414
chr4-83005359-DEL-85
chr4-83022206-DEL-69
chr4-83036253-DEL-114
chr4-83037206-DEL-72
chr4-83050457-DEL-1981
chr4-83053579-DEL-111
chr4-83105890-DEL-51
chr4-83208025-DEL-161
chr4-83211333-DEL-1818
chr4-83228544-DEL-467
chr4-83274242-DEL-171
chr4-83279862-DEL-62
chr4-83302019-DEL-86
chr4-83309640-DEL-200
chr4-83395586-DEL-235
chr4-83420183-DEL-88
chr4-83423078-DEL-287
chr4-83423839-DEL-1165
chr4-83441371-DEL-183
chr4-83452475-DEL-63
chr4-83473638-DEL-429
chr4-83515947-DEL-1067
chr4-83541572-DEL-143
chr4-83561456-DEL-212
chr4-83572539-DEL-61
chr4-83582585-DEL-124
chr4-83592760-DEL-62
chr4-83593517-DEL-86
chr4-83606223-DEL-468
chr4-83619676-DEL-120
chr4-83654615-DEL-51
chr4-83660074-DEL-105
chr4-83674579-DEL-6199
chr4-83682825-DEL-69
chr4-83707366-DEL-407
chr4-83709009-DEL-147
chr4-83730957-DEL-55
chr4-83759982-DEL-1296
chr4-83759994-DEL-80
chr4-83760071-DEL-767
chr4-83760720-DEL-304
chr4-83798871-DEL-52
chr4-83813138-DEL-153
chr4-83818199-DEL-90
chr4-83832701-DEL-249
chr4-83927728-DEL-70
chr4-83961882-DEL-633
chr4-83969535-DEL-1241
chr4-84003299-DEL-57
chr4-84085986-DEL-335
chr4-84112562-DEL-1174
chr4-84177084-DEL-200
chr4-84177387-DEL-948
chr4-84303263-DEL-157
chr4-84327052-DEL-698
chr4-84371345-DEL-186
chr4-84403751-DEL-353
chr4-84408925-DEL-54
chr4-84429230-DEL-66
chr4-84432026-DEL-53
chr4-84435202-DEL-55
chr4-84436806-DEL-62
chr4-84452722-DEL-161
chr4-84506301-DEL-490
chr4-84508695-DEL-57
chr4-84536077-DEL-63
chr4-84537325-DEL-180
chr4-84552232-DEL-62
chr4-84559901-DEL-671
chr4-84583035-DEL-71

chr4-84635310-DEL-53
chr4-84657678-DEL-174
chr4-84715656-DEL-123
chr4-84774506-DEL-167
chr4-84784175-DEL-52
chr4-84839882-DEL-379
chr4-84890314-DEL-175
chr4-84930905-DEL-54
chr4-85034472-DEL-57
chr4-85071567-DEL-291
chr4-85105264-DEL-264
chr4-85107270-DEL-5304
chr4-85117555-DEL-64
chr4-85129022-DEL-52
chr4-85140367-DEL-691
chr4-85142079-DEL-51
chr4-85150139-DEL-578
chr4-85159889-DEL-189
chr4-85170978-DEL-94
chr4-85191804-DEL-106
chr4-85207043-DEL-507
chr4-85222180-DEL-51
chr4-85228529-DEL-56
chr4-85234823-DEL-1317
chr4-85286059-DEL-55
chr4-85333129-DEL-1200
chr4-85335423-DEL-972
chr4-85362322-DEL-319
chr4-85372760-DEL-303
chr4-85429407-DEL-263
chr4-85458743-DEL-100
chr4-85514067-DEL-1780
chr4-85517918-DEL-2888
chr4-85520415-DEL-195
chr4-85535967-DEL-930
chr4-85621390-DEL-558
chr4-85650496-DEL-60
chr4-85656336-DEL-80
chr4-85667074-DEL-129
chr4-85675991-DEL-191
chr4-85677540-DEL-157
chr4-85701425-DEL-77
chr4-85715225-DEL-578
chr4-85718493-DEL-77
chr4-85721090-DEL-1730
chr4-85724838-DEL-60
chr4-85799932-DEL-54
chr4-85869470-DEL-62
chr4-85887133-DEL-499
chr4-85902092-DEL-134
chr4-85917358-DEL-208
chr4-85927525-DEL-159
chr4-85991609-DEL-77
chr4-85992565-DEL-94
chr4-86001974-DEL-66
chr4-86063168-DEL-1666
chr4-86079552-DEL-182
chr4-86083093-DEL-744
chr4-86116847-DEL-88
chr4-86148935-DEL-172
chr4-86191310-DEL-74
chr4-86228292-DEL-131
chr4-86234503-DEL-555
chr4-86239898-DEL-181
chr4-86262846-DEL-96
chr4-86273876-DEL-439
chr4-86376558-DEL-4870
chr4-86376692-DEL-2979
chr4-86396770-DEL-99
chr4-86429406-DEL-111
chr4-86458880-DEL-229
chr4-86476005-DEL-57
chr4-86516529-DEL-76
chr4-86534635-DEL-106
chr4-86539105-DEL-56
chr4-86556066-DEL-820
chr4-86558311-DEL-445

chr4-86573051-DEL-102
chr4-86593447-DEL-70
chr4-86594448-DEL-397
chr4-86596898-DEL-53
chr4-86600014-DEL-69
chr4-86606471-DEL-167
chr4-86608766-DEL-54
chr4-86610547-DEL-253
chr4-86644772-DEL-186
chr4-86652753-DEL-2275
chr4-86663301-DEL-65
chr4-86714856-DEL-185
chr4-86749318-DEL-1626
chr4-86750602-DEL-361
chr4-86752283-DEL-1811
chr4-86762282-DEL-103
chr4-86763294-DEL-56
chr4-86808087-DEL-434
chr4-86808417-DEL-744
chr4-86810268-DEL-141
chr4-86823724-DEL-90
chr4-86837690-DEL-70
chr4-86844280-DEL-150
chr4-86849089-DEL-177
chr4-86871650-DEL-1947
chr4-86884585-DEL-311
chr4-86910579-DEL-52
chr4-86921521-DEL-70
chr4-86922182-DEL-66
chr4-86934739-DEL-510
chr4-86978550-DEL-287
chr4-86982601-DEL-81
chr4-87108347-DEL-69
chr4-87111323-DEL-175
chr4-87121270-DEL-2085
chr4-87159505-DEL-56
chr4-87167344-DEL-3970
chr4-87175825-DEL-444
chr4-87180977-DEL-95
chr4-87225012-DEL-11797
chr4-87232819-DEL-996
chr4-87267509-DEL-51
chr4-87269694-DEL-1214
chr4-87285176-DEL-296
chr4-87355277-DEL-1361
chr4-87359996-DEL-78
chr4-87407204-DEL-101
chr4-87417928-DEL-80
chr4-87421228-DEL-208
chr4-87434596-DEL-836
chr4-87436154-DEL-128
chr4-87450668-DEL-64
chr4-87451912-DEL-229
chr4-87453139-DEL-52
chr4-87453940-DEL-630
chr4-87479962-DEL-362
chr4-87513583-DEL-132
chr4-87553508-DEL-114
chr4-87559390-DEL-401
chr4-87564396-DEL-52
chr4-87605514-DEL-696
chr4-87619245-DEL-81
chr4-87621992-DEL-91
chr4-87677816-DEL-366
chr4-87705746-DEL-74
chr4-87789688-DEL-239
chr4-87792500-DEL-57
chr4-87794285-DEL-52
chr4-87889845-DEL-81
chr4-87891813-DEL-56
chr4-87903945-DEL-101
chr4-87909908-DEL-74
chr4-87960575-DEL-492
chr4-87991528-DEL-1468
chr4-88031365-DEL-1488
chr4-88035385-DEL-126
chr4-88048442-DEL-99

chr4-88107260-DEL-3942
chr4-88146705-DEL-54
chr4-88165985-DEL-198
chr4-88177678-DEL-707
chr4-88210648-DEL-228
chr4-88223925-DEL-61
chr4-88237736-DEL-425
chr4-88279060-DEL-6607
chr4-88307003-DEL-970
chr4-88319671-DEL-406
chr4-88399607-DEL-6048
chr4-88472228-DEL-104
chr4-88481679-DEL-6981
chr4-88483028-DEL-3552
chr4-88493299-DEL-3566
chr4-88515926-DEL-176
chr4-88533415-DEL-6009
chr4-88543106-DEL-330
chr4-88549438-DEL-665
chr4-88558495-DEL-496
chr4-88560994-DEL-92
chr4-88562758-DEL-469
chr4-88564180-DEL-229
chr4-88577748-DEL-51
chr4-88616778-DEL-75
chr4-88625388-DEL-111
chr4-88631517-DEL-82
chr4-88642442-DEL-194
chr4-88673191-DEL-2976
chr4-88682442-DEL-7346
chr4-88682652-DEL-66
chr4-88689570-DEL-66
chr4-88750834-DEL-52
chr4-88772138-DEL-2383
chr4-88788259-DEL-150
chr4-88802114-DEL-126
chr4-88814947-DEL-264
chr4-88822736-DEL-69
chr4-88823417-DEL-709
chr4-88843509-DEL-199
chr4-88876291-DEL-205
chr4-88882826-DEL-582
chr4-88892130-DEL-92
chr4-88894615-DEL-1780
chr4-88898669-DEL-60
chr4-88911649-DEL-104
chr4-88921516-DEL-2954
chr4-88937099-DEL-1043
chr4-88987029-DEL-121
chr4-88991511-DEL-83
chr4-89014881-DEL-95
chr4-89048560-DEL-78
chr4-89050797-DEL-298
chr4-89081093-DEL-1651
chr4-89087164-DEL-54
chr4-89093053-DEL-440
chr4-89103234-DEL-7735
chr4-89118107-DEL-8338
chr4-89126744-DEL-52
chr4-89135349-DEL-55
chr4-89138480-DEL-432
chr4-89143226-DEL-192
chr4-89155765-DEL-346
chr4-89179714-DEL-2127
chr4-89180125-DEL-2126
chr4-89181235-DEL-197
chr4-89191402-DEL-241
chr4-89203310-DEL-305
chr4-89240709-DEL-100
chr4-89269079-DEL-52
chr4-89275808-DEL-288
chr4-89289370-DEL-70
chr4-89293148-DEL-275
chr4-89305744-DEL-63
chr4-89308215-DEL-922
chr4-89331158-DEL-514
chr4-89338865-DEL-86

chr4-89348952-DEL-56
chr4-89354630-DEL-54
chr4-89376452-DEL-121
chr4-89428426-DEL-4596
chr4-89434854-DEL-102
chr4-89462161-DEL-507
chr4-89463141-DEL-102
chr4-89484478-DEL-134
chr4-89487938-DEL-696
chr4-89497812-DEL-52
chr4-89553926-DEL-189
chr4-89590513-DEL-166
chr4-89599661-DEL-12582
chr4-89614571-DEL-970
chr4-89615675-DEL-463
chr4-89678361-DEL-13692
chr4-89681132-DEL-211
chr4-89699022-DEL-71
chr4-89762841-DEL-58
chr4-89765691-DEL-3087
chr4-89785975-DEL-93
chr4-89819539-DEL-83
chr4-89850390-DEL-826
chr4-89862380-DEL-56
chr4-89881794-DEL-1838
chr4-89884676-DEL-66
chr4-89902030-DEL-100
chr4-89907474-DEL-1509
chr4-89918204-DEL-1131
chr4-89944611-DEL-58
chr4-89952287-DEL-1428
chr4-89954217-DEL-73
chr4-89986214-DEL-72
chr4-90022595-DEL-62
chr4-90063386-DEL-315
chr4-90081085-DEL-113
chr4-90092188-DEL-1257
chr4-90096705-DEL-424
chr4-90106787-DEL-124
chr4-90109246-DEL-180
chr4-90128704-DEL-94
chr4-90269090-DEL-56
chr4-90275593-DEL-90
chr4-90280773-DEL-508
chr4-90281023-DEL-2002
chr4-90281359-DEL-3923
chr4-90341021-DEL-764
chr4-90394208-DEL-400
chr4-90407564-DEL-230
chr4-90411151-DEL-53
chr4-90462431-DEL-337
chr4-90471339-DEL-634
chr4-90479272-DEL-58
chr4-90486047-DEL-78
chr4-90508000-DEL-669
chr4-90538980-DEL-7260
chr4-90546966-DEL-6254
chr4-90640824-DEL-886
chr4-90658366-DEL-2548
chr4-90661109-DEL-553
chr4-90662039-DEL-1758
chr4-90688245-DEL-730
chr4-90714615-DEL-902
chr4-90719780-DEL-789
chr4-90775762-DEL-338
chr4-90867420-DEL-13542
chr4-90867840-DEL-423
chr4-90868611-DEL-411
chr4-90883427-DEL-173
chr4-90887588-DEL-6220
chr4-90901945-DEL-352
chr4-90926839-DEL-108
chr4-90935427-DEL-1437
chr4-90944381-DEL-128
chr4-90951226-DEL-6239
chr4-91033933-DEL-67
chr4-91046261-DEL-85

chr4-91051803-DEL-114
chr4-91057049-DEL-6637
chr4-91103083-DEL-67
chr4-91124071-DEL-247
chr4-91229839-DEL-57
chr4-91247731-DEL-215
chr4-91279204-DEL-74
chr4-91303341-DEL-57
chr4-91318148-DEL-1576
chr4-91327896-DEL-165
chr4-91355276-DEL-6188
chr4-91372401-DEL-142
chr4-91415109-DEL-67
chr4-91430004-DEL-241
chr4-91493907-DEL-3768
chr4-91497581-DEL-310
chr4-91570380-DEL-3983
chr4-91583999-DEL-4403
chr4-91590403-DEL-61
chr4-91599715-DEL-65
chr4-91609342-DEL-1863
chr4-91642853-DEL-3532
chr4-91647805-DEL-239
chr4-91686919-DEL-8743
chr4-91708521-DEL-109
chr4-91724923-DEL-6067
chr4-91802848-DEL-58
chr4-91831436-DEL-194
chr4-91841806-DEL-86
chr4-91846445-DEL-424
chr4-91858725-DEL-9904
chr4-91860666-DEL-7223
chr4-91867931-DEL-188
chr4-91871947-DEL-183
chr4-91880330-DEL-820
chr4-91882581-DEL-319
chr4-91890641-DEL-52
chr4-91919527-DEL-148
chr4-91932373-DEL-72
chr4-91937685-DEL-326
chr4-91963581-DEL-1490
chr4-91981596-DEL-90
chr4-92003064-DEL-269
chr4-92027670-DEL-52
chr4-92030133-DEL-91
chr4-92038171-DEL-1357
chr4-92048685-DEL-369
chr4-92128128-DEL-113
chr4-92139998-DEL-65
chr4-92143357-DEL-432
chr4-92149910-DEL-1309
chr4-92167724-DEL-4718
chr4-92200128-DEL-52
chr4-92223322-DEL-1565
chr4-92224918-DEL-2483
chr4-92233890-DEL-994
chr4-92241782-DEL-90
chr4-92254862-DEL-175
chr4-92292279-DEL-6199
chr4-92304009-DEL-401
chr4-92316882-DEL-72
chr4-92323206-DEL-119
chr4-92346056-DEL-67
chr4-92363395-DEL-542
chr4-92386039-DEL-70
chr4-92426804-DEL-204
chr4-92466212-DEL-55
chr4-92475287-DEL-70
chr4-92494484-DEL-1243
chr4-92514079-DEL-952
chr4-92515112-DEL-565
chr4-92516792-DEL-1855
chr4-92521936-DEL-1026
chr4-92550646-DEL-986
chr4-92661852-DEL-176
chr4-92682757-DEL-935
chr4-92691736-DEL-3286

chr4-92699168-DEL-214
chr4-92734736-DEL-52
chr4-92745993-DEL-984
chr4-92791555-DEL-464
chr4-92811017-DEL-161
chr4-92816282-DEL-64
chr4-92832463-DEL-415
chr4-92846930-DEL-173
chr4-92873856-DEL-7163
chr4-92884695-DEL-102
chr4-92898768-DEL-3554
chr4-92915059-DEL-83
chr4-92926384-DEL-6205
chr4-92933350-DEL-1473
chr4-92939593-DEL-914
chr4-92950327-DEL-138
chr4-92967705-DEL-408
chr4-92980016-DEL-126
chr4-92999893-DEL-260
chr4-93003906-DEL-132
chr4-93018153-DEL-414
chr4-93035085-DEL-93
chr4-93036751-DEL-1522
chr4-93045138-DEL-3370
chr4-93053075-DEL-1310
chr4-93112229-DEL-568
chr4-93151385-DEL-271
chr4-93268780-DEL-750
chr4-93269629-DEL-309
chr4-93312355-DEL-487
chr4-93314688-DEL-89
chr4-93318139-DEL-2736
chr4-93322540-DEL-1378
chr4-93329534-DEL-75
chr4-93426533-DEL-53
chr4-93459592-DEL-163
chr4-93469179-DEL-385
chr4-93474689-DEL-2544
chr4-93495996-DEL-750
chr4-93544219-DEL-145
chr4-93584556-DEL-184
chr4-93633917-DEL-524
chr4-93635093-DEL-2037
chr4-93636991-DEL-144
chr4-93670562-DEL-1982
chr4-93704542-DEL-767
chr4-93723556-DEL-72
chr4-93732643-DEL-95
chr4-93738875-DEL-2536
chr4-93839670-DEL-55
chr4-93859132-DEL-62
chr4-93865360-DEL-3606
chr4-93886148-DEL-113
chr4-93902738-DEL-64
chr4-93923612-DEL-375
chr4-93958846-DEL-114
chr4-94013671-DEL-855
chr4-94024177-DEL-988
chr4-94055875-DEL-1821
chr4-94059087-DEL-78
chr4-94078759-DEL-175
chr4-94147578-DEL-267
chr4-94195674-DEL-355
chr4-94197948-DEL-448
chr4-94223534-DEL-133
chr4-94243736-DEL-70
chr4-94252767-DEL-79
chr4-94255078-DEL-53
chr4-94289247-DEL-365
chr4-94313726-DEL-302
chr4-94326984-DEL-90
chr4-94351403-DEL-157
chr4-94442104-DEL-2151
chr4-94466113-DEL-97
chr4-94487598-DEL-213
chr4-94504008-DEL-250
chr4-94509759-DEL-170

chr4-94523057-DEL-66
chr4-94567596-DEL-360
chr4-94611239-DEL-60
chr4-94613059-DEL-73
chr4-94670013-DEL-1136
chr4-94755123-DEL-68
chr4-94765798-DEL-198
chr4-94768313-DEL-6191
chr4-94779956-DEL-423
chr4-94843550-DEL-108
chr4-94862872-DEL-186
chr4-94868319-DEL-87
chr4-94870239-DEL-1224
chr4-94872080-DEL-882
chr4-94877653-DEL-342
chr4-94879043-DEL-1253
chr4-94892202-DEL-138
chr4-94896835-DEL-242
chr4-94900002-DEL-435
chr4-94907969-DEL-51
chr4-94930081-DEL-66
chr4-94941201-DEL-63
chr4-94994022-DEL-64
chr4-94995518-DEL-52
chr4-94997949-DEL-93
chr4-95022882-DEL-457
chr4-95023863-DEL-63
chr4-95033251-DEL-80
chr4-95062071-DEL-55
chr4-95064207-DEL-55
chr4-95099112-DEL-87
chr4-95103769-DEL-51
chr4-95108057-DEL-154
chr4-95140887-DEL-1539
chr4-95187261-DEL-78
chr4-95203345-DEL-196
chr4-95218400-DEL-501
chr4-95345733-DEL-1843
chr4-95362296-DEL-70
chr4-95368676-DEL-130
chr4-95398453-DEL-416
chr4-95422790-DEL-837
chr4-95449207-DEL-67
chr4-95477270-DEL-473
chr4-95479535-DEL-2173
chr4-95502934-DEL-68
chr4-95530875-DEL-55
chr4-95539336-DEL-265
chr4-95562144-DEL-79
chr4-95582936-DEL-1143
chr4-95632492-DEL-110
chr4-95682629-DEL-55
chr4-95706657-DEL-258
chr4-95739318-DEL-156
chr4-95741774-DEL-58
chr4-95761106-DEL-1626
chr4-95776880-DEL-971
chr4-95778280-DEL-75
chr4-95783145-DEL-260
chr4-95812629-DEL-371
chr4-95838722-DEL-53
chr4-95864541-DEL-6284
chr4-95874073-DEL-172
chr4-95874887-DEL-1386
chr4-95883316-DEL-575
chr4-95894881-DEL-75
chr4-95918881-DEL-760
chr4-95925415-DEL-90
chr4-95931804-DEL-73
chr4-95934115-DEL-59
chr4-95947421-DEL-202
chr4-95992457-DEL-1887
chr4-96008505-DEL-125
chr4-96010128-DEL-149
chr4-96092851-DEL-8516
chr4-96095620-DEL-2866
chr4-96096742-DEL-2866

chr4-96137263-DEL-88
chr4-96188969-DEL-6448
chr4-96217612-DEL-59
chr4-96238994-DEL-56
chr4-96243891-DEL-81
chr4-96265793-DEL-1489
chr4-96267785-DEL-79
chr4-96268847-DEL-3509
chr4-96282738-DEL-248
chr4-96283517-DEL-77
chr4-96285366-DEL-381
chr4-96288646-DEL-631
chr4-96302941-DEL-245
chr4-96306846-DEL-1037
chr4-96307829-DEL-62
chr4-96327144-DEL-693
chr4-96333629-DEL-118
chr4-96346263-DEL-6150
chr4-96374112-DEL-81
chr4-96398412-DEL-1049
chr4-96401401-DEL-73
chr4-96415888-DEL-993
chr4-96417934-DEL-1521
chr4-96491524-DEL-1891
chr4-96494172-DEL-1293
chr4-96509344-DEL-81
chr4-96515312-DEL-62
chr4-96565181-DEL-639
chr4-96567044-DEL-240
chr4-96569101-DEL-67
chr4-96605601-DEL-55
chr4-96618279-DEL-65
chr4-96647466-DEL-54
chr4-96660396-DEL-307
chr4-96670271-DEL-1689
chr4-96683198-DEL-170
chr4-96696804-DEL-59
chr4-96707574-DEL-1667
chr4-96718711-DEL-1747
chr4-96727577-DEL-499
chr4-96756388-DEL-7141
chr4-96768074-DEL-487
chr4-96833633-DEL-9326
chr4-96877129-DEL-778
chr4-96905649-DEL-97
chr4-97033134-DEL-130
chr4-97061242-DEL-135
chr4-97065671-DEL-395
chr4-97103816-DEL-69
chr4-97194394-DEL-61
chr4-97210064-DEL-1094
chr4-97229934-DEL-84
chr4-97236762-DEL-275
chr4-97258216-DEL-5492
chr4-97299517-DEL-752
chr4-97345182-DEL-299
chr4-97345750-DEL-1097
chr4-97351188-DEL-908
chr4-97386465-DEL-54
chr4-97422046-DEL-1247
chr4-97447240-DEL-63
chr4-97477305-DEL-125
chr4-97531141-DEL-53
chr4-97532266-DEL-298
chr4-97546132-DEL-54
chr4-97577111-DEL-155
chr4-97593779-DEL-124
chr4-97605615-DEL-76
chr4-97608321-DEL-950
chr4-97635203-DEL-82
chr4-97637958-DEL-239
chr4-97695778-DEL-52
chr4-97700659-DEL-76
chr4-97705937-DEL-84
chr4-97780143-DEL-131
chr4-97833580-DEL-118
chr4-97853919-DEL-54

chr4-97866530-DEL-62
chr4-97913961-DEL-63
chr4-97925866-DEL-1024
chr4-97926503-DEL-143
chr4-97957858-DEL-1388
chr4-97976404-DEL-51
chr4-97977181-DEL-58
chr4-97979267-DEL-172
chr4-98020193-DEL-1301
chr4-98094444-DEL-59
chr4-98102270-DEL-447
chr4-98108018-DEL-144
chr4-98130778-DEL-182
chr4-98168081-DEL-393
chr4-98175953-DEL-5925
chr4-98183112-DEL-75
chr4-98187753-DEL-62
chr4-98188764-DEL-342
chr4-98213994-DEL-1927
chr4-98230420-DEL-113
chr4-98277780-DEL-724
chr4-98288077-DEL-77
chr4-98291538-DEL-57
chr4-98294605-DEL-193
chr4-98303641-DEL-125
chr4-98313756-DEL-58
chr4-98372667-DEL-1072
chr4-98375553-DEL-52
chr4-98386008-DEL-1140
chr4-98388240-DEL-89
chr4-98444469-DEL-169
chr4-98472911-DEL-342
chr4-98505106-DEL-53
chr4-98512703-DEL-2734
chr4-98519928-DEL-62
chr4-98546188-DEL-86
chr4-98559330-DEL-479
chr4-98561027-DEL-66
chr4-98569610-DEL-110
chr4-98584573-DEL-70
chr4-98597265-DEL-174
chr4-98603970-DEL-70
chr4-98609777-DEL-51
chr4-98635155-DEL-5320
chr4-98698365-DEL-176
chr4-98714610-DEL-115
chr4-98738849-DEL-52
chr4-98739369-DEL-167
chr4-98790098-DEL-51
chr4-98825831-DEL-109
chr4-98845147-DEL-195
chr4-98848935-DEL-506
chr4-98866732-DEL-180
chr4-98873478-DEL-1204
chr4-98885123-DEL-352
chr4-98890774-DEL-594
chr4-98898162-DEL-64
chr4-98912504-DEL-178
chr4-98968467-DEL-143
chr4-98986163-DEL-68
chr4-98988919-DEL-63
chr4-99022347-DEL-58
chr4-99029313-DEL-288
chr4-99053641-DEL-188
chr4-99060619-DEL-1653
chr4-99074356-DEL-58
chr4-99089239-DEL-57
chr4-99112822-DEL-443
chr4-99136684-DEL-140
chr4-99141649-DEL-276
chr4-99142742-DEL-399
chr4-99181807-DEL-58
chr4-99245074-DEL-139
chr4-99280030-DEL-64
chr4-99297381-DEL-92
chr4-99321752-DEL-220
chr4-99402710-DEL-585

chr4-99412435-DEL-259
chr4-99420661-DEL-192
chr4-99433553-DEL-434
chr4-99444199-DEL-372
chr4-99459160-DEL-669
chr4-99498162-DEL-5270
chr4-99511468-DEL-60
chr4-99515979-DEL-80
chr4-99552328-DEL-255
chr4-99584026-DEL-173
chr4-99588621-DEL-66
chr4-99592217-DEL-52
chr4-99599320-DEL-74
chr4-99604841-DEL-401
chr4-99629409-DEL-110
chr4-99633847-DEL-65
chr4-99662442-DEL-66
chr4-99717712-DEL-1531
chr4-99733202-DEL-1434
chr4-99736191-DEL-76
chr4-99825740-DEL-75
chr4-99844976-DEL-1019
chr4-99850720-DEL-2369
chr4-99870442-DEL-948
chr4-99873751-DEL-6533
chr4-99899844-DEL-64
chr4-99911472-DEL-748
chr4-99914522-DEL-1326
chr4-99935787-DEL-66
chr4-99936821-DEL-118
chr4-99946216-DEL-4069
chr4-99968620-DEL-166
chr4-99995276-DEL-1495
chr4-100010676-DEL-106
chr4-100038421-DEL-122
chr4-100052131-DEL-54
chr4-100065955-DEL-2052
chr4-100067778-DEL-69
chr4-100076816-DEL-179
chr4-100096439-DEL-90
chr4-100122840-DEL-58
chr4-100170319-DEL-1261
chr4-100170321-DEL-68
chr4-100170985-DEL-464
chr4-100175234-DEL-60
chr4-100197398-DEL-300
chr4-100200869-DEL-403
chr4-100218702-DEL-205
chr4-100244428-DEL-92
chr4-100260495-DEL-135
chr4-100263414-DEL-52
chr4-100288074-DEL-63
chr4-100294931-DEL-283
chr4-100313410-DEL-1520
chr4-100314712-DEL-250
chr4-100314955-DEL-3731
chr4-100340310-DEL-1779
chr4-100349906-DEL-83
chr4-100366082-DEL-170
chr4-100413833-DEL-86
chr4-100414812-DEL-1234
chr4-100446839-DEL-56
chr4-100474036-DEL-2255
chr4-100481758-DEL-77
chr4-100482874-DEL-3735
chr4-100482993-DEL-130
chr4-100486178-DEL-102
chr4-100487414-DEL-400
chr4-100489654-DEL-629
chr4-100515678-DEL-416
chr4-100528589-DEL-608
chr4-100553087-DEL-952
chr4-100553822-DEL-417
chr4-100591784-DEL-523
chr4-100662248-DEL-64
chr4-100718200-DEL-57
chr4-100719535-DEL-116

chr4-100752948-DEL-231
chr4-100772875-DEL-332
chr4-100788838-DEL-6721
chr4-100864647-DEL-240
chr4-100878945-DEL-419
chr4-100889181-DEL-153
chr4-100901682-DEL-76
chr4-100904416-DEL-210
chr4-100909571-DEL-312
chr4-100919938-DEL-72
chr4-100931799-DEL-246
chr4-100931934-DEL-1457
chr4-100932531-DEL-531
chr4-100933152-DEL-207
chr4-101084968-DEL-982
chr4-101092275-DEL-178
chr4-101094602-DEL-1793
chr4-101104470-DEL-129
chr4-101122788-DEL-74
chr4-101129273-DEL-479
chr4-101140086-DEL-74
chr4-101152387-DEL-1970
chr4-101160993-DEL-96
chr4-101169822-DEL-143
chr4-101169822-DEL-2002
chr4-101170587-DEL-1121
chr4-101170719-DEL-64
chr4-101171106-DEL-81
chr4-101171412-DEL-104
chr4-101180599-DEL-117
chr4-101220472-DEL-61
chr4-101246542-DEL-380
chr4-101250082-DEL-75
chr4-101285971-DEL-74
chr4-101300360-DEL-64
chr4-101325358-DEL-65
chr4-101338535-DEL-52
chr4-101368170-DEL-343
chr4-101393981-DEL-264
chr4-101400940-DEL-86
chr4-101402002-DEL-204
chr4-101416444-DEL-526
chr4-101437112-DEL-138
chr4-101437749-DEL-77
chr4-101453760-DEL-84
chr4-101458009-DEL-228
chr4-101458538-DEL-151
chr4-101518614-DEL-245
chr4-101538889-DEL-131
chr4-101547401-DEL-700
chr4-101570297-DEL-63
chr4-101585698-DEL-322
chr4-101586923-DEL-71
chr4-101597003-DEL-58
chr4-101608486-DEL-577
chr4-101609613-DEL-125
chr4-101626522-DEL-1008
chr4-101640152-DEL-99
chr4-101661190-DEL-214
chr4-101665255-DEL-245
chr4-101696729-DEL-76
chr4-101697242-DEL-57
chr4-101701018-DEL-478
chr4-101714780-DEL-129
chr4-101762191-DEL-76
chr4-101770476-DEL-219
chr4-101803357-DEL-2488
chr4-101811376-DEL-75
chr4-101824978-DEL-146
chr4-101850147-DEL-54
chr4-101857112-DEL-51
chr4-101874470-DEL-100
chr4-101875542-DEL-202
chr4-101892603-DEL-199
chr4-101895270-DEL-830
chr4-101896121-DEL-435
chr4-101905266-DEL-2911

chr4-101912572-DEL-621
chr4-101920040-DEL-68
chr4-101960282-DEL-98
chr4-102002100-DEL-889
chr4-102002807-DEL-303
chr4-102054114-DEL-223
chr4-102073751-DEL-1192
chr4-102074279-DEL-127
chr4-102099279-DEL-636
chr4-102105865-DEL-66
chr4-102175313-DEL-618
chr4-102199078-DEL-537
chr4-102236550-DEL-151
chr4-102322694-DEL-2277
chr4-102346649-DEL-178
chr4-102346746-DEL-1908
chr4-102351385-DEL-238
chr4-102370555-DEL-24479
chr4-102401445-DEL-70
chr4-102432086-DEL-1301
chr4-102464607-DEL-87
chr4-102486904-DEL-1136
chr4-102495570-DEL-96
chr4-102522205-DEL-75
chr4-102556814-DEL-6551
chr4-102591157-DEL-59
chr4-102612511-DEL-51
chr4-102654355-DEL-1823
chr4-102687742-DEL-154
chr4-102715903-DEL-560
chr4-102738794-DEL-312
chr4-102815748-DEL-751
chr4-102818005-DEL-1237
chr4-102818007-DEL-142
chr4-102841270-DEL-456
chr4-102869479-DEL-130
chr4-102885148-DEL-479
chr4-102911738-DEL-492
chr4-102915631-DEL-506
chr4-102927209-DEL-236
chr4-102932999-DEL-146
chr4-102945815-DEL-77
chr4-103004622-DEL-763
chr4-103009149-DEL-325
chr4-103010380-DEL-132
chr4-103026243-DEL-502
chr4-103042575-DEL-3362
chr4-103049830-DEL-182
chr4-103063026-DEL-66
chr4-103080250-DEL-344
chr4-103088963-DEL-3366
chr4-103114356-DEL-1116
chr4-103128778-DEL-73
chr4-103130513-DEL-212
chr4-103185289-DEL-52
chr4-103188703-DEL-106
chr4-103193558-DEL-2217
chr4-103216301-DEL-91
chr4-103217900-DEL-126
chr4-103218919-DEL-102
chr4-103230911-DEL-6407
chr4-103298085-DEL-56
chr4-103303545-DEL-70
chr4-103310580-DEL-1699
chr4-103353632-DEL-119
chr4-103382880-DEL-522
chr4-103422659-DEL-126
chr4-103440480-DEL-81
chr4-103444454-DEL-55
chr4-103447369-DEL-4622
chr4-103485789-DEL-320
chr4-103531668-DEL-163
chr4-103533786-DEL-152
chr4-103543673-DEL-60
chr4-103613764-DEL-147
chr4-103616573-DEL-268
chr4-103674438-DEL-53

chr4-103682618-DEL-91
chr4-103699645-DEL-399
chr4-103737635-DEL-85
chr4-103741035-DEL-60
chr4-103801865-DEL-240
chr4-103831368-DEL-1423
chr4-103837010-DEL-11379
chr4-103860234-DEL-200
chr4-103937882-DEL-60
chr4-103946932-DEL-212
chr4-103973073-DEL-662
chr4-103975135-DEL-57
chr4-103992591-DEL-52
chr4-104032479-DEL-79
chr4-104118117-DEL-53
chr4-104133148-DEL-88
chr4-104183990-DEL-64
chr4-104236843-DEL-197
chr4-104308674-DEL-90
chr4-104310975-DEL-56
chr4-104319507-DEL-189
chr4-104323606-DEL-61
chr4-104332173-DEL-198
chr4-104341655-DEL-71
chr4-104382755-DEL-112
chr4-104465864-DEL-232
chr4-104502485-DEL-194
chr4-104505935-DEL-157
chr4-104517161-DEL-104
chr4-104541555-DEL-160
chr4-104555620-DEL-84
chr4-104613912-DEL-57
chr4-104617219-DEL-88
chr4-104625044-DEL-60
chr4-104632708-DEL-58
chr4-104647930-DEL-575
chr4-104664675-DEL-1606
chr4-104712354-DEL-166
chr4-104720208-DEL-60
chr4-104737974-DEL-353
chr4-104780165-DEL-1062
chr4-104783591-DEL-277
chr4-104785362-DEL-2419
chr4-104823284-DEL-133
chr4-104838283-DEL-152
chr4-104930994-DEL-320
chr4-104939912-DEL-122
chr4-105015967-DEL-65
chr4-105066832-DEL-127
chr4-105094405-DEL-54
chr4-105098146-DEL-63
chr4-105160460-DEL-73
chr4-105161938-DEL-69
chr4-105167188-DEL-438
chr4-105184748-DEL-64
chr4-105200862-DEL-294
chr4-105230337-DEL-93
chr4-105360406-DEL-427
chr4-105385210-DEL-325
chr4-105453908-DEL-3156
chr4-105462219-DEL-1531
chr4-105491903-DEL-3697
chr4-105528421-DEL-59
chr4-105532204-DEL-3633
chr4-105582826-DEL-318
chr4-105592679-DEL-212
chr4-105599918-DEL-447
chr4-105645094-DEL-396
chr4-105662861-DEL-53
chr4-105704106-DEL-91
chr4-105781304-DEL-1159
chr4-105786976-DEL-110
chr4-105788790-DEL-617
chr4-105813892-DEL-652
chr4-105817756-DEL-58
chr4-105917651-DEL-215
chr4-105940577-DEL-157

chr4-105977134-DEL-112
chr4-105989465-DEL-168
chr4-105992282-DEL-85
chr4-105995722-DEL-674
chr4-106010211-DEL-227
chr4-106027450-DEL-52
chr4-106034440-DEL-400
chr4-106045439-DEL-80
chr4-106058128-DEL-1828
chr4-106093401-DEL-3671
chr4-106101241-DEL-149
chr4-106115699-DEL-68
chr4-106125945-DEL-3026
chr4-106126126-DEL-51
chr4-106146406-DEL-85
chr4-106152569-DEL-115
chr4-106184976-DEL-74
chr4-106213546-DEL-1419
chr4-106273596-DEL-78
chr4-106317069-DEL-67
chr4-106365614-DEL-92
chr4-106374121-DEL-200
chr4-106376314-DEL-153
chr4-106427011-DEL-62
chr4-106461259-DEL-415
chr4-106462729-DEL-1535
chr4-106464839-DEL-459
chr4-106480465-DEL-90
chr4-106538587-DEL-197
chr4-106542226-DEL-1064
chr4-106542260-DEL-1520
chr4-106542261-DEL-874
chr4-106560723-DEL-73
chr4-106575449-DEL-62
chr4-106588704-DEL-10574
chr4-106594962-DEL-164
chr4-106615877-DEL-195
chr4-106623479-DEL-55
chr4-106625505-DEL-161
chr4-106633288-DEL-67
chr4-106647841-DEL-96
chr4-106667568-DEL-84
chr4-106682911-DEL-139
chr4-106686734-DEL-134
chr4-106691309-DEL-932
chr4-106735791-DEL-349
chr4-106741940-DEL-235
chr4-106751963-DEL-132
chr4-106757082-DEL-86
chr4-106831694-DEL-61
chr4-106899987-DEL-104
chr4-106907609-DEL-129
chr4-106947805-DEL-58
chr4-106966166-DEL-180
chr4-106971213-DEL-76
chr4-106989059-DEL-178
chr4-107032518-DEL-71
chr4-107079307-DEL-129
chr4-107153174-DEL-87
chr4-107178167-DEL-143
chr4-107181064-DEL-476
chr4-107198928-DEL-65
chr4-107205032-DEL-96
chr4-107215063-DEL-389
chr4-107234639-DEL-145
chr4-107245340-DEL-213
chr4-107249241-DEL-209
chr4-107253066-DEL-90
chr4-107287005-DEL-212
chr4-107338579-DEL-51
chr4-107338973-DEL-189
chr4-107411481-DEL-79
chr4-107432356-DEL-204
chr4-107443336-DEL-52
chr4-107453297-DEL-55
chr4-107455972-DEL-178
chr4-107467917-DEL-421

chr4-107490597-DEL-59
chr4-107525126-DEL-238
chr4-107546218-DEL-52
chr4-107554912-DEL-492
chr4-107633531-DEL-76
chr4-107651553-DEL-1253
chr4-107658369-DEL-73
chr4-107673572-DEL-67
chr4-107682550-DEL-992
chr4-107684423-DEL-885
chr4-107685358-DEL-152
chr4-107720090-DEL-213
chr4-107737659-DEL-80
chr4-107758735-DEL-61
chr4-107793546-DEL-560
chr4-107803845-DEL-138
chr4-107804404-DEL-875
chr4-107824635-DEL-268
chr4-107849905-DEL-135
chr4-107862240-DEL-62
chr4-107871194-DEL-436
chr4-107886972-DEL-73
chr4-107893286-DEL-203
chr4-107897586-DEL-77
chr4-107960815-DEL-193
chr4-107975817-DEL-60
chr4-107982770-DEL-2779
chr4-107995505-DEL-4622
chr4-108019659-DEL-214
chr4-108042409-DEL-268
chr4-108046996-DEL-895
chr4-108056961-DEL-401
chr4-108059354-DEL-167
chr4-108079282-DEL-501
chr4-108085383-DEL-103
chr4-108095428-DEL-174
chr4-108126968-DEL-75
chr4-108140160-DEL-445
chr4-108153588-DEL-208
chr4-108163750-DEL-201
chr4-108164299-DEL-383
chr4-108176653-DEL-1575
chr4-108178631-DEL-201
chr4-108181257-DEL-87
chr4-108200659-DEL-198
chr4-108204408-DEL-932
chr4-108229258-DEL-88
chr4-108240950-DEL-121
chr4-108255429-DEL-57
chr4-108267176-DEL-203
chr4-108307799-DEL-54
chr4-108324076-DEL-180
chr4-108351120-DEL-272
chr4-108362108-DEL-80
chr4-108412073-DEL-129
chr4-108432028-DEL-90
chr4-108435776-DEL-181
chr4-108440298-DEL-209
chr4-108459332-DEL-829
chr4-108492097-DEL-104
chr4-108515993-DEL-329
chr4-108563484-DEL-809
chr4-108580072-DEL-66
chr4-108586502-DEL-202
chr4-108590826-DEL-638
chr4-108591643-DEL-621
chr4-108598101-DEL-157
chr4-108622094-DEL-420
chr4-108663936-DEL-72
chr4-108677453-DEL-195
chr4-108685166-DEL-1532
chr4-108686626-DEL-106
chr4-108693042-DEL-82
chr4-108698227-DEL-141
chr4-108737463-DEL-54
chr4-108758400-DEL-65
chr4-108802417-DEL-65

chr4-108803262-DEL-66
chr4-108824424-DEL-117
chr4-108830945-DEL-183
chr4-108836529-DEL-879
chr4-108846510-DEL-50
chr4-108865604-DEL-649
chr4-108991783-DEL-53
chr4-109016839-DEL-67
chr4-109080230-DEL-202
chr4-109144566-DEL-80
chr4-109156779-DEL-228
chr4-109157198-DEL-428
chr4-109158840-DEL-61
chr4-109171087-DEL-130
chr4-109175731-DEL-160
chr4-109183782-DEL-109
chr4-109185547-DEL-206
chr4-109188888-DEL-505
chr4-109197656-DEL-116
chr4-109205327-DEL-584
chr4-109205993-DEL-5664
chr4-109215099-DEL-181
chr4-109236768-DEL-122
chr4-109247863-DEL-52
chr4-109256775-DEL-64
chr4-109260454-DEL-320
chr4-109271404-DEL-137
chr4-109286377-DEL-253
chr4-109290056-DEL-107
chr4-109292585-DEL-229
chr4-109314367-DEL-1155
chr4-109348796-DEL-76
chr4-109352298-DEL-89
chr4-109368215-DEL-399
chr4-109404616-DEL-1422
chr4-109414761-DEL-1537
chr4-109421480-DEL-60
chr4-109439221-DEL-164
chr4-109444926-DEL-62
chr4-109490040-DEL-700
chr4-109501448-DEL-256
chr4-109528177-DEL-149
chr4-109536029-DEL-73
chr4-109565889-DEL-110
chr4-109589555-DEL-147
chr4-109651052-DEL-90
chr4-109688056-DEL-67
chr4-109688248-DEL-2660
chr4-109858456-DEL-61
chr4-109878043-DEL-703
chr4-109886419-DEL-126
chr4-109896856-DEL-578
chr4-109911002-DEL-115
chr4-109912420-DEL-7131
chr4-109937424-DEL-82
chr4-109960924-DEL-424
chr4-109967355-DEL-52
chr4-109983489-DEL-71
chr4-110000694-DEL-170
chr4-110003225-DEL-122
chr4-110039180-DEL-82
chr4-110057333-DEL-59
chr4-110091379-DEL-241
chr4-110092200-DEL-201
chr4-110107589-DEL-102
chr4-110107627-DEL-901
chr4-110115634-DEL-219
chr4-110155099-DEL-300
chr4-110204155-DEL-58
chr4-110272733-DEL-254
chr4-110274958-DEL-62
chr4-110296109-DEL-226
chr4-110303232-DEL-1903
chr4-110326747-DEL-6596
chr4-110339902-DEL-299
chr4-110344360-DEL-6585
chr4-110357735-DEL-6197

chr4-110364517-DEL-62
chr4-110375694-DEL-83
chr4-110380730-DEL-66
chr4-110393693-DEL-180
chr4-110408704-DEL-88
chr4-110408965-DEL-622
chr4-110423436-DEL-1622
chr4-110425316-DEL-395
chr4-110426312-DEL-52
chr4-110431849-DEL-111
chr4-110432437-DEL-2801
chr4-110441052-DEL-59
chr4-110455927-DEL-2073
chr4-110462427-DEL-52
chr4-110469542-DEL-6159
chr4-110472717-DEL-187
chr4-110477472-DEL-12065
chr4-110480406-DEL-6230
chr4-110483626-DEL-187
chr4-110523426-DEL-361
chr4-110526511-DEL-704
chr4-110527523-DEL-178
chr4-110577308-DEL-191
chr4-110595467-DEL-2687
chr4-110595678-DEL-364
chr4-110618468-DEL-2397
chr4-110620896-DEL-3892
chr4-110627668-DEL-129
chr4-110629028-DEL-523
chr4-110659505-DEL-89
chr4-110662490-DEL-56
chr4-110674816-DEL-214
chr4-110703859-DEL-638
chr4-110716233-DEL-2453
chr4-110722121-DEL-7101
chr4-110732135-DEL-1035
chr4-110752066-DEL-943
chr4-110787485-DEL-1746
chr4-110787490-DEL-70
chr4-110816730-DEL-6576
chr4-110816789-DEL-201
chr4-110823833-DEL-409
chr4-110872160-DEL-557
chr4-110884098-DEL-387
chr4-110906582-DEL-1492
chr4-110908335-DEL-1065
chr4-110931673-DEL-52
chr4-110941274-DEL-911
chr4-110959440-DEL-1490
chr4-110963478-DEL-369
chr4-111008557-DEL-103
chr4-111011556-DEL-604
chr4-111023578-DEL-8278
chr4-111023863-DEL-1295
chr4-111050635-DEL-439
chr4-111076479-DEL-555
chr4-111080338-DEL-159
chr4-111090594-DEL-70
chr4-111094061-DEL-68
chr4-111123286-DEL-1582
chr4-111144620-DEL-1376
chr4-111158075-DEL-52
chr4-111181906-DEL-1165
chr4-111219221-DEL-1329
chr4-111241974-DEL-268
chr4-111295404-DEL-438
chr4-111295436-DEL-82
chr4-111318942-DEL-287
chr4-111396344-DEL-564
chr4-111412043-DEL-96
chr4-111415873-DEL-1106
chr4-111427933-DEL-97
chr4-111442427-DEL-195
chr4-111447856-DEL-96
chr4-111449302-DEL-72
chr4-111451053-DEL-79
chr4-111490636-DEL-357

chr4-111545740-DEL-84
chr4-111553171-DEL-148
chr4-111561073-DEL-108
chr4-111618761-DEL-2692
chr4-111640769-DEL-57
chr4-111642840-DEL-53
chr4-111649845-DEL-963
chr4-111650683-DEL-277
chr4-111680493-DEL-3353
chr4-111685162-DEL-169
chr4-111689546-DEL-112
chr4-111702213-DEL-3022
chr4-111702820-DEL-65
chr4-111711312-DEL-318
chr4-111730789-DEL-3535
chr4-111740068-DEL-53
chr4-111768778-DEL-52
chr4-111777813-DEL-60
chr4-111827525-DEL-64
chr4-111833786-DEL-58
chr4-111847066-DEL-3055
chr4-111855969-DEL-506
chr4-111896229-DEL-3148
chr4-111896996-DEL-481
chr4-111903521-DEL-861
chr4-111904529-DEL-408
chr4-111905010-DEL-478
chr4-111908145-DEL-1304
chr4-111908372-DEL-247
chr4-111915047-DEL-6223
chr4-111923812-DEL-1388
chr4-111925386-DEL-1554
chr4-111928990-DEL-328
chr4-111946728-DEL-57
chr4-111960637-DEL-6189
chr4-111981361-DEL-184
chr4-111988162-DEL-1068
chr4-111990659-DEL-81
chr4-112014741-DEL-62
chr4-112025682-DEL-2621
chr4-112033792-DEL-2048
chr4-112036855-DEL-4876
chr4-112043911-DEL-1935
chr4-112087679-DEL-498
chr4-112091907-DEL-417
chr4-112097143-DEL-64
chr4-112128659-DEL-510
chr4-112132673-DEL-336
chr4-112143839-DEL-66
chr4-112163842-DEL-1107
chr4-112181453-DEL-14833
chr4-112200558-DEL-6188
chr4-112229855-DEL-551
chr4-112232265-DEL-1885
chr4-112242791-DEL-97
chr4-112244674-DEL-379
chr4-112245113-DEL-379
chr4-112251641-DEL-910
chr4-112253722-DEL-74
chr4-112284420-DEL-63
chr4-112308046-DEL-115
chr4-112308378-DEL-6156
chr4-112310909-DEL-1453
chr4-112354976-DEL-1222
chr4-112367637-DEL-84
chr4-112371513-DEL-366
chr4-112376563-DEL-9328
chr4-112379737-DEL-1882
chr4-112399805-DEL-217
chr4-112402417-DEL-127
chr4-112403057-DEL-159
chr4-112425658-DEL-6370
chr4-112436120-DEL-103
chr4-112459075-DEL-69
chr4-112463260-DEL-4380
chr4-112474666-DEL-1449
chr4-112475275-DEL-1140

chr4-112493729-DEL-2618
chr4-112520219-DEL-3423
chr4-112556469-DEL-155
chr4-112629491-DEL-163
chr4-112634471-DEL-186
chr4-112649081-DEL-412
chr4-112681655-DEL-256
chr4-112694989-DEL-81
chr4-112701369-DEL-51
chr4-112704698-DEL-408
chr4-112712637-DEL-154
chr4-112715767-DEL-192
chr4-112727021-DEL-401
chr4-112728396-DEL-532
chr4-112734699-DEL-141
chr4-112738273-DEL-183
chr4-112743857-DEL-834
chr4-112744773-DEL-2858
chr4-112747763-DEL-342
chr4-112753223-DEL-230
chr4-112761408-DEL-67
chr4-112764875-DEL-72
chr4-112800418-DEL-111
chr4-112801963-DEL-1662
chr4-112805221-DEL-498
chr4-112849635-DEL-305
chr4-112855383-DEL-565
chr4-112860304-DEL-238
chr4-112863232-DEL-78
chr4-112864385-DEL-146
chr4-112943347-DEL-184
chr4-112948302-DEL-307
chr4-112952997-DEL-471
chr4-112955409-DEL-1244
chr4-112961654-DEL-144
chr4-112966169-DEL-690
chr4-112968120-DEL-120
chr4-112969498-DEL-2196
chr4-112969617-DEL-340
chr4-112970219-DEL-266
chr4-112970611-DEL-1542
chr4-112976968-DEL-108
chr4-112983200-DEL-144
chr4-112986215-DEL-215
chr4-112987648-DEL-91
chr4-112993906-DEL-89
chr4-113011912-DEL-71
chr4-113013916-DEL-89
chr4-113032412-DEL-74
chr4-113034415-DEL-88
chr4-113072456-DEL-67
chr4-113074694-DEL-191
chr4-113076663-DEL-587
chr4-113089098-DEL-111
chr4-113115253-DEL-400
chr4-113153654-DEL-184
chr4-113160290-DEL-347
chr4-113233654-DEL-3391
chr4-113252181-DEL-6351
chr4-113263581-DEL-475
chr4-113271228-DEL-7004
chr4-113282049-DEL-231
chr4-113292021-DEL-2755
chr4-113305929-DEL-808
chr4-113321261-DEL-8160
chr4-113330466-DEL-1626
chr4-113337443-DEL-52
chr4-113339452-DEL-97
chr4-113340086-DEL-858
chr4-113348129-DEL-6183
chr4-113354670-DEL-437
chr4-113357140-DEL-2565
chr4-113357229-DEL-65
chr4-113367408-DEL-57
chr4-113381696-DEL-328
chr4-113385114-DEL-80
chr4-113395076-DEL-54

chr4-113397055-DEL-57
chr4-113416835-DEL-65
chr4-113422288-DEL-865
chr4-113434130-DEL-265
chr4-113437857-DEL-6183
chr4-113445181-DEL-2816
chr4-113453180-DEL-446
chr4-113456802-DEL-6623
chr4-113499257-DEL-153
chr4-113502025-DEL-60
chr4-113504956-DEL-3195
chr4-113518178-DEL-197
chr4-113519942-DEL-1300
chr4-113521297-DEL-84
chr4-113549289-DEL-332
chr4-113550450-DEL-438
chr4-113554277-DEL-55
chr4-113558157-DEL-54
chr4-113565318-DEL-1961
chr4-113569701-DEL-2855
chr4-113573838-DEL-51
chr4-113577072-DEL-97
chr4-113578244-DEL-437
chr4-113590465-DEL-767
chr4-113593485-DEL-870
chr4-113602034-DEL-540
chr4-113602729-DEL-124
chr4-113603911-DEL-342
chr4-113631079-DEL-763
chr4-113639554-DEL-299
chr4-113640566-DEL-8535
chr4-113652299-DEL-82
chr4-113655220-DEL-498
chr4-113660709-DEL-144
chr4-113661954-DEL-292
chr4-113664965-DEL-66
chr4-113675485-DEL-65
chr4-113681729-DEL-56
chr4-113701684-DEL-61
chr4-113717113-DEL-52
chr4-113718628-DEL-223
chr4-113719981-DEL-53
chr4-113731441-DEL-81
chr4-113736331-DEL-1011
chr4-113749174-DEL-67
chr4-113753346-DEL-894
chr4-113762135-DEL-96
chr4-113765199-DEL-79
chr4-113765482-DEL-3776
chr4-113766908-DEL-208
chr4-113770206-DEL-71
chr4-113782647-DEL-1320
chr4-113796929-DEL-645
chr4-113806364-DEL-1634
chr4-113814970-DEL-53
chr4-113817542-DEL-88
chr4-113828614-DEL-154
chr4-113831536-DEL-104
chr4-113855305-DEL-66
chr4-113858829-DEL-110
chr4-113867599-DEL-4947
chr4-113874269-DEL-602
chr4-113876804-DEL-55
chr4-113879987-DEL-57
chr4-113911970-DEL-147
chr4-113913356-DEL-1146
chr4-113914597-DEL-3647
chr4-113915879-DEL-365
chr4-113925498-DEL-10951
chr4-113945834-DEL-613
chr4-113960875-DEL-3649
chr4-113962761-DEL-151
chr4-113964102-DEL-288
chr4-113968349-DEL-2917
chr4-113970576-DEL-401
chr4-113972130-DEL-652
chr4-113986350-DEL-14468

chr4-113990438-DEL-8212
chr4-114021790-DEL-77
chr4-114022094-DEL-837
chr4-114022990-DEL-374
chr4-114064216-DEL-58
chr4-114068790-DEL-7101
chr4-114078024-DEL-204
chr4-114085368-DEL-201
chr4-114089703-DEL-4706
chr4-114095088-DEL-1886
chr4-114096972-DEL-499
chr4-114121955-DEL-75
chr4-114135506-DEL-6242
chr4-114285808-DEL-180
chr4-114287845-DEL-97
chr4-114365173-DEL-56
chr4-114383851-DEL-3879
chr4-114405920-DEL-424
chr4-114417888-DEL-56
chr4-114434016-DEL-68
chr4-114462805-DEL-89
chr4-114522108-DEL-105
chr4-114555999-DEL-110
chr4-114579744-DEL-760
chr4-114591641-DEL-309
chr4-114597911-DEL-856
chr4-114609760-DEL-289
chr4-114618145-DEL-198
chr4-114664500-DEL-143
chr4-114689511-DEL-281
chr4-114705701-DEL-1871
chr4-114746224-DEL-979
chr4-114762590-DEL-65
chr4-114811264-DEL-749
chr4-114828184-DEL-557
chr4-114830409-DEL-557
chr4-114872793-DEL-190
chr4-114890206-DEL-1501
chr4-114962538-DEL-217
chr4-114965355-DEL-172
chr4-114970483-DEL-2545
chr4-114973308-DEL-269
chr4-114992420-DEL-92
chr4-114993514-DEL-63
chr4-114998503-DEL-77
chr4-115000802-DEL-6208
chr4-115000926-DEL-199
chr4-115021814-DEL-671
chr4-115029213-DEL-329
chr4-115045970-DEL-1493
chr4-115050569-DEL-107
chr4-115125665-DEL-6213
chr4-115164753-DEL-8512
chr4-115173361-DEL-102
chr4-115175018-DEL-2222
chr4-115178840-DEL-185
chr4-115182685-DEL-55
chr4-115198145-DEL-12353
chr4-115236884-DEL-2111
chr4-115272045-DEL-92
chr4-115282367-DEL-198
chr4-115298167-DEL-86
chr4-115321361-DEL-3211
chr4-115342855-DEL-65
chr4-115383921-DEL-1710
chr4-115429388-DEL-1709
chr4-115500989-DEL-59
chr4-115507256-DEL-309
chr4-115523638-DEL-2436
chr4-115529836-DEL-223
chr4-115550521-DEL-636
chr4-115568677-DEL-311
chr4-115574651-DEL-199
chr4-115621089-DEL-112
chr4-115627866-DEL-305
chr4-115645635-DEL-118
chr4-115671611-DEL-61

chr4-115679735-DEL-53
chr4-115693578-DEL-436
chr4-115695639-DEL-121
chr4-115708520-DEL-54
chr4-115797801-DEL-146
chr4-115836217-DEL-69
chr4-115848640-DEL-84
chr4-115863461-DEL-103
chr4-115866769-DEL-191
chr4-115893587-DEL-199
chr4-115898291-DEL-574
chr4-115902213-DEL-192
chr4-115902247-DEL-1280
chr4-115902536-DEL-511
chr4-115902947-DEL-192
chr4-115938849-DEL-209
chr4-115951772-DEL-74
chr4-115958655-DEL-72
chr4-115985890-DEL-116
chr4-116004107-DEL-191
chr4-116005557-DEL-356
chr4-116035526-DEL-125
chr4-116050624-DEL-102
chr4-116051912-DEL-90
chr4-116052812-DEL-992
chr4-116057211-DEL-260
chr4-116057450-DEL-747
chr4-116059634-DEL-726
chr4-116082656-DEL-221
chr4-116087581-DEL-58
chr4-116094151-DEL-107
chr4-116099011-DEL-1932
chr4-116102626-DEL-56
chr4-116109459-DEL-56
chr4-116125184-DEL-141
chr4-116155380-DEL-384
chr4-116178545-DEL-191
chr4-116202929-DEL-184
chr4-116250947-DEL-478
chr4-116284045-DEL-102
chr4-116337653-DEL-68
chr4-116345202-DEL-73
chr4-116373824-DEL-3448
chr4-116419344-DEL-864
chr4-116421559-DEL-167
chr4-116477072-DEL-56
chr4-116487079-DEL-214
chr4-116497509-DEL-242
chr4-116523129-DEL-1938
chr4-116526566-DEL-368
chr4-116529659-DEL-123
chr4-116598248-DEL-104
chr4-116635051-DEL-261
chr4-116641408-DEL-223
chr4-116643120-DEL-197
chr4-116644669-DEL-86
chr4-116653169-DEL-153
chr4-116654599-DEL-165
chr4-116665655-DEL-174
chr4-116680998-DEL-62
chr4-116689096-DEL-133
chr4-116695615-DEL-73
chr4-116700443-DEL-64
chr4-116708554-DEL-407
chr4-116749736-DEL-174
chr4-116752670-DEL-54
chr4-116760450-DEL-188
chr4-116791520-DEL-58
chr4-116805759-DEL-220
chr4-116808812-DEL-59
chr4-116823886-DEL-199
chr4-116835312-DEL-51
chr4-116837257-DEL-69
chr4-116866992-DEL-94
chr4-116900919-DEL-157
chr4-116950403-DEL-210
chr4-116951641-DEL-54

chr4-116956760-DEL-413
chr4-116994529-DEL-204
chr4-116995062-DEL-600
chr4-116999438-DEL-58
chr4-117027832-DEL-331
chr4-117030367-DEL-172
chr4-117036232-DEL-208
chr4-117038214-DEL-149
chr4-117044368-DEL-449
chr4-117045050-DEL-194
chr4-117062065-DEL-430
chr4-117063057-DEL-218
chr4-117064232-DEL-173
chr4-117077563-DEL-102
chr4-117078418-DEL-215
chr4-117094266-DEL-120
chr4-117095674-DEL-54
chr4-117101530-DEL-86
chr4-117102174-DEL-266
chr4-117106052-DEL-168
chr4-117111718-DEL-688
chr4-117158850-DEL-156
chr4-117235090-DEL-185
chr4-117256538-DEL-128
chr4-117261078-DEL-95
chr4-117282188-DEL-96
chr4-117289328-DEL-52
chr4-117305519-DEL-117
chr4-117317067-DEL-58
chr4-117320305-DEL-736
chr4-117328459-DEL-139
chr4-117364102-DEL-444
chr4-117366420-DEL-129
chr4-117372961-DEL-259
chr4-117401297-DEL-106
chr4-117564366-DEL-98
chr4-117580625-DEL-93
chr4-117605394-DEL-861
chr4-117606245-DEL-206
chr4-117628609-DEL-120
chr4-117654340-DEL-1138
chr4-117667056-DEL-65
chr4-117795151-DEL-159
chr4-117821049-DEL-79
chr4-117829429-DEL-125
chr4-117838751-DEL-87
chr4-117840320-DEL-194
chr4-117856464-DEL-60
chr4-117865085-DEL-62
chr4-117869881-DEL-103
chr4-117879323-DEL-53
chr4-117912723-DEL-399
chr4-117925524-DEL-69
chr4-117952806-DEL-87
chr4-117954019-DEL-266
chr4-117959098-DEL-689
chr4-118034872-DEL-210
chr4-118040549-DEL-1587
chr4-118040549-DEL-7231
chr4-118040550-DEL-1286
chr4-118041892-DEL-1002
chr4-118042949-DEL-1308
chr4-118044382-DEL-3399
chr4-118053842-DEL-399
chr4-118057552-DEL-197
chr4-118064253-DEL-210
chr4-118228506-DEL-140
chr4-118239521-DEL-54
chr4-118261180-DEL-120
chr4-118261213-DEL-606
chr4-118279655-DEL-236
chr4-118294728-DEL-344
chr4-118296658-DEL-222
chr4-118306544-DEL-72
chr4-118310472-DEL-54
chr4-118325316-DEL-97
chr4-118345196-DEL-147

chr4-118371346-DEL-265
chr4-118375789-DEL-258
chr4-118376946-DEL-78
chr4-118383517-DEL-148
chr4-118404254-DEL-1630
chr4-118412950-DEL-527
chr4-118412978-DEL-1255
chr4-118413718-DEL-356
chr4-118432525-DEL-148
chr4-118435205-DEL-66
chr4-118444167-DEL-60
chr4-118454430-DEL-280
chr4-118459192-DEL-153
chr4-118460299-DEL-338
chr4-118465509-DEL-691
chr4-118479755-DEL-187
chr4-118490828-DEL-151
chr4-118500024-DEL-268
chr4-118508500-DEL-5661
chr4-118520399-DEL-207
chr4-118546184-DEL-267
chr4-118548389-DEL-84
chr4-118578306-DEL-599
chr4-118579866-DEL-58
chr4-118615672-DEL-67501
chr4-118630014-DEL-842
chr4-118690590-DEL-88
chr4-118691789-DEL-1380
chr4-118696966-DEL-654
chr4-118699787-DEL-65
chr4-118728438-DEL-98
chr4-118730966-DEL-207
chr4-118736657-DEL-5388
chr4-118743308-DEL-126
chr4-118749593-DEL-84
chr4-118765425-DEL-105
chr4-118798943-DEL-1928
chr4-118807665-DEL-1369
chr4-118850299-DEL-279
chr4-118851727-DEL-2807
chr4-118859943-DEL-148
chr4-118880339-DEL-52
chr4-118902600-DEL-189
chr4-118903707-DEL-220
chr4-118922466-DEL-397
chr4-119003605-DEL-354
chr4-119071504-DEL-121
chr4-119077470-DEL-186
chr4-119124593-DEL-203
chr4-119125751-DEL-223
chr4-119130761-DEL-80
chr4-119146445-DEL-264
chr4-119157459-DEL-200
chr4-119212341-DEL-60
chr4-119213621-DEL-176
chr4-119239940-DEL-162
chr4-119242846-DEL-58
chr4-119246643-DEL-89
chr4-119249866-DEL-233
chr4-119273252-DEL-58
chr4-119288613-DEL-99
chr4-119294588-DEL-428
chr4-119311376-DEL-182
chr4-119341241-DEL-650
chr4-119343256-DEL-183
chr4-119360190-DEL-105
chr4-119365912-DEL-827
chr4-119399736-DEL-839
chr4-119401103-DEL-52
chr4-119405280-DEL-181
chr4-119421139-DEL-198
chr4-119474073-DEL-478
chr4-119491673-DEL-456
chr4-119530450-DEL-54
chr4-119539489-DEL-656
chr4-119570828-DEL-92
chr4-119580795-DEL-202

chr4-119614661-DEL-262
chr4-119615015-DEL-578
chr4-119620792-DEL-1032
chr4-119657361-DEL-74
chr4-119695678-DEL-55
chr4-119698691-DEL-200
chr4-119748055-DEL-84
chr4-119788500-DEL-108
chr4-119811345-DEL-860
chr4-119813676-DEL-51
chr4-119815578-DEL-200
chr4-119853086-DEL-710
chr4-119853135-DEL-82
chr4-119886671-DEL-82
chr4-119950296-DEL-148
chr4-119975339-DEL-91
chr4-119982229-DEL-119
chr4-120004375-DEL-152
chr4-120034972-DEL-100
chr4-120077636-DEL-98
chr4-120113328-DEL-680
chr4-120152118-DEL-160
chr4-120175354-DEL-303
chr4-120180414-DEL-71
chr4-120247530-DEL-1112
chr4-120256539-DEL-425
chr4-120272024-DEL-410
chr4-120289081-DEL-207
chr4-120297405-DEL-66
chr4-120350562-DEL-66
chr4-120350994-DEL-749
chr4-120388461-DEL-154
chr4-120417972-DEL-540
chr4-120442877-DEL-275
chr4-120464066-DEL-116
chr4-120485730-DEL-54
chr4-120502571-DEL-122
chr4-120510990-DEL-820
chr4-120521101-DEL-499
chr4-120534534-DEL-420
chr4-120548143-DEL-224
chr4-120576998-DEL-51
chr4-120607806-DEL-61
chr4-120611746-DEL-52
chr4-120631614-DEL-4310
chr4-120676439-DEL-216
chr4-120721562-DEL-405
chr4-120745426-DEL-58
chr4-120755109-DEL-193
chr4-120756896-DEL-76
chr4-120761466-DEL-150
chr4-120780685-DEL-74
chr4-120804505-DEL-78
chr4-120833701-DEL-59
chr4-120837094-DEL-277
chr4-120878794-DEL-162
chr4-120892655-DEL-129
chr4-120903863-DEL-65
chr4-120965273-DEL-112
chr4-120972282-DEL-1592
chr4-120977142-DEL-105
chr4-121038755-DEL-553
chr4-121043579-DEL-193
chr4-121062933-DEL-202
chr4-121077326-DEL-161
chr4-121111851-DEL-408
chr4-121125109-DEL-2462
chr4-121128229-DEL-529
chr4-121132617-DEL-504
chr4-121135812-DEL-383
chr4-121146106-DEL-6261
chr4-121175929-DEL-352
chr4-121193116-DEL-64
chr4-121230124-DEL-973
chr4-121292069-DEL-105
chr4-121305409-DEL-54
chr4-121313633-DEL-68

chr4-121325028-DEL-197
chr4-121335441-DEL-131
chr4-121346434-DEL-92
chr4-121382510-DEL-6669
chr4-121409250-DEL-2714
chr4-121416018-DEL-89
chr4-121423437-DEL-60
chr4-121480077-DEL-362
chr4-121492679-DEL-53
chr4-121506574-DEL-72
chr4-121524363-DEL-2772
chr4-121538557-DEL-136
chr4-121541592-DEL-271
chr4-121546187-DEL-72
chr4-121547498-DEL-54
chr4-121594991-DEL-85
chr4-121642976-DEL-55
chr4-121644315-DEL-1461
chr4-121650472-DEL-341
chr4-121678259-DEL-84
chr4-121960860-DEL-208
chr4-122046277-DEL-80
chr4-122380007-DEL-1697
chr4-122407875-DEL-54
chr4-122411177-DEL-138
chr4-122417051-DEL-223
chr4-122447297-DEL-89
chr4-122449699-DEL-69
chr4-122483506-DEL-506
chr4-122545336-DEL-590
chr4-122547267-DEL-52
chr4-122548767-DEL-115
chr4-122551891-DEL-678
chr4-122556206-DEL-4985
chr4-122606563-DEL-212
chr4-122718772-DEL-75
chr4-122720840-DEL-136
chr4-122773848-DEL-205
chr4-122793711-DEL-157
chr4-122818862-DEL-159
chr4-122836701-DEL-68
chr4-122851778-DEL-122
chr4-122883442-DEL-78
chr4-122895415-DEL-137
chr4-122917942-DEL-400
chr4-122929303-DEL-68
chr4-122934339-DEL-1221
chr4-122958017-DEL-464
chr4-122972631-DEL-60
chr4-122973594-DEL-63
chr4-122974485-DEL-52
chr4-123012679-DEL-59
chr4-123023103-DEL-98
chr4-123029656-DEL-58
chr4-123032357-DEL-121
chr4-123052078-DEL-69
chr4-123106470-DEL-116
chr4-123163791-DEL-502
chr4-123170097-DEL-167
chr4-123181937-DEL-62
chr4-123187876-DEL-193
chr4-123195806-DEL-52
chr4-123207483-DEL-564
chr4-123221808-DEL-56
chr4-123223018-DEL-110
chr4-123233625-DEL-142
chr4-123240046-DEL-324
chr4-123263075-DEL-51
chr4-123267505-DEL-60
chr4-123314711-DEL-180
chr4-123350781-DEL-52
chr4-123351311-DEL-85
chr4-123353843-DEL-979
chr4-123361391-DEL-510
chr4-123378857-DEL-207
chr4-123401542-DEL-63
chr4-123413807-DEL-84

chr4-123430556-DEL-126
chr4-123434495-DEL-493
chr4-123439321-DEL-2112
chr4-123477260-DEL-195
chr4-123504912-DEL-188
chr4-123506755-DEL-103
chr4-123520327-DEL-402
chr4-123522511-DEL-91
chr4-123548139-DEL-204
chr4-123561719-DEL-93
chr4-123566521-DEL-52
chr4-123568974-DEL-88
chr4-123576035-DEL-57
chr4-123594244-DEL-1335
chr4-123600217-DEL-54
chr4-123600821-DEL-305
chr4-123603688-DEL-736
chr4-123605725-DEL-189
chr4-123614000-DEL-90
chr4-123625204-DEL-221
chr4-123647359-DEL-311
chr4-123660388-DEL-266
chr4-123674748-DEL-6291
chr4-123700962-DEL-277
chr4-123703103-DEL-355
chr4-123737300-DEL-8776
chr4-123746581-DEL-2377
chr4-123756395-DEL-7285
chr4-123819736-DEL-60
chr4-123826626-DEL-92
chr4-123869712-DEL-173
chr4-123879156-DEL-52
chr4-123914215-DEL-155
chr4-123949610-DEL-277
chr4-123968853-DEL-171
chr4-124019753-DEL-17634
chr4-124033649-DEL-63
chr4-124046191-DEL-52
chr4-124049146-DEL-455
chr4-124112525-DEL-55
chr4-124114196-DEL-54
chr4-124157043-DEL-154
chr4-124183202-DEL-56
chr4-124230290-DEL-56
chr4-124255857-DEL-83
chr4-124276175-DEL-80
chr4-124317539-DEL-1064
chr4-124326672-DEL-136
chr4-124348234-DEL-79
chr4-124354373-DEL-314
chr4-124395626-DEL-169
chr4-124424673-DEL-117
chr4-124443185-DEL-120
chr4-124504222-DEL-631
chr4-124536252-DEL-61
chr4-124539338-DEL-71
chr4-124581508-DEL-1085
chr4-124615217-DEL-145
chr4-124617722-DEL-116
chr4-124629423-DEL-524
chr4-124643131-DEL-98
chr4-124656269-DEL-205
chr4-124719580-DEL-188
chr4-124724372-DEL-93
chr4-124730140-DEL-100
chr4-124746126-DEL-67
chr4-124760739-DEL-135
chr4-124764596-DEL-74
chr4-124769705-DEL-1247
chr4-124770773-DEL-186
chr4-124794843-DEL-209
chr4-124797644-DEL-93
chr4-124800911-DEL-115
chr4-124802288-DEL-68
chr4-124826670-DEL-1712
chr4-124827117-DEL-1717
chr4-124832043-DEL-97

chr4-124834430-DEL-324
chr4-124857425-DEL-110
chr4-124870229-DEL-153
chr4-124906560-DEL-565
chr4-124907978-DEL-118
chr4-124916047-DEL-89
chr4-124919837-DEL-84
chr4-124922025-DEL-71
chr4-124929812-DEL-464
chr4-124956873-DEL-191
chr4-124977025-DEL-95
chr4-124985774-DEL-68
chr4-124991661-DEL-259
chr4-124991740-DEL-933
chr4-125040732-DEL-316
chr4-125048628-DEL-176
chr4-125103714-DEL-126
chr4-125108937-DEL-98
chr4-125139806-DEL-155
chr4-125152618-DEL-66
chr4-125154123-DEL-152
chr4-125165621-DEL-208
chr4-125166366-DEL-220
chr4-125184260-DEL-72
chr4-125213990-DEL-144
chr4-125224807-DEL-83
chr4-125228305-DEL-210
chr4-125231172-DEL-66
chr4-125283677-DEL-108
chr4-125332281-DEL-757
chr4-125407815-DEL-55
chr4-125427029-DEL-223
chr4-125432866-DEL-81
chr4-125475514-DEL-279
chr4-125478956-DEL-210
chr4-125486105-DEL-273
chr4-125559107-DEL-56
chr4-125561076-DEL-2639
chr4-125565433-DEL-211
chr4-125587277-DEL-93
chr4-125589256-DEL-51
chr4-125597609-DEL-98
chr4-125604594-DEL-52
chr4-125624218-DEL-1684
chr4-125650941-DEL-154
chr4-125666752-DEL-800
chr4-125699370-DEL-187
chr4-125762503-DEL-346
chr4-125763642-DEL-518
chr4-125797975-DEL-98
chr4-125879454-DEL-106
chr4-125887648-DEL-109
chr4-125944564-DEL-88
chr4-125945571-DEL-402
chr4-125953654-DEL-402
chr4-125967041-DEL-54
chr4-126049701-DEL-108
chr4-126117084-DEL-58
chr4-126168285-DEL-364
chr4-126200322-DEL-115
chr4-126280674-DEL-62
chr4-126308178-DEL-376
chr4-126341450-DEL-61
chr4-126345676-DEL-92
chr4-126369819-DEL-1353
chr4-126407118-DEL-233
chr4-126414433-DEL-58
chr4-126478859-DEL-206
chr4-126482733-DEL-142
chr4-126482783-DEL-811
chr4-126490839-DEL-59
chr4-126543900-DEL-61
chr4-126545420-DEL-503
chr4-126570725-DEL-51
chr4-126634498-DEL-195
chr4-126657122-DEL-60
chr4-126684772-DEL-183

chr4-126689804-DEL-55
chr4-126720555-DEL-198
chr4-126740710-DEL-86
chr4-126807178-DEL-204
chr4-126827209-DEL-235
chr4-126836387-DEL-53
chr4-126843136-DEL-1520
chr4-126845691-DEL-77
chr4-126870959-DEL-187
chr4-126872140-DEL-165
chr4-126878851-DEL-190
chr4-126882834-DEL-201
chr4-126884861-DEL-202
chr4-126895335-DEL-197
chr4-126897263-DEL-203
chr4-126918605-DEL-145
chr4-126931746-DEL-3505
chr4-126961181-DEL-289
chr4-126964359-DEL-376
chr4-126996894-DEL-147
chr4-127015233-DEL-649
chr4-127024281-DEL-189
chr4-127052229-DEL-96
chr4-127056096-DEL-336
chr4-127057116-DEL-250
chr4-127065765-DEL-80
chr4-127076155-DEL-108
chr4-127102289-DEL-67
chr4-127106537-DEL-503
chr4-127149418-DEL-958
chr4-127167272-DEL-66
chr4-127169801-DEL-201
chr4-127200393-DEL-114
chr4-127202785-DEL-180
chr4-127237613-DEL-271
chr4-127257041-DEL-551
chr4-127271465-DEL-63
chr4-127271974-DEL-78
chr4-127284389-DEL-62
chr4-127318941-DEL-57
chr4-127352436-DEL-210
chr4-127375464-DEL-141
chr4-127383635-DEL-63
chr4-127430791-DEL-57
chr4-127431880-DEL-108
chr4-127436274-DEL-140
chr4-127442367-DEL-54
chr4-127444039-DEL-768
chr4-127503050-DEL-98
chr4-127525859-DEL-59
chr4-127532995-DEL-87
chr4-127543972-DEL-75
chr4-127608420-DEL-101
chr4-127618536-DEL-209
chr4-127629886-DEL-64
chr4-127671544-DEL-56
chr4-127676887-DEL-75
chr4-127678055-DEL-88
chr4-127694875-DEL-1003
chr4-127756687-DEL-79
chr4-127762311-DEL-66
chr4-127784209-DEL-235
chr4-127788868-DEL-58
chr4-127837199-DEL-87
chr4-127876659-DEL-104
chr4-127901531-DEL-54
chr4-127908260-DEL-259
chr4-127911869-DEL-189
chr4-127918706-DEL-189
chr4-127949167-DEL-195
chr4-127963231-DEL-95
chr4-127987891-DEL-174
chr4-128002528-DEL-136
chr4-128050891-DEL-70
chr4-128062724-DEL-83
chr4-128063638-DEL-52
chr4-128083480-DEL-370

chr4-128086080-DEL-63
chr4-128102628-DEL-66
chr4-128106208-DEL-60
chr4-128115603-DEL-252
chr4-128132474-DEL-54
chr4-128160076-DEL-441
chr4-128201034-DEL-363
chr4-128208034-DEL-60
chr4-128217726-DEL-52
chr4-128235785-DEL-236
chr4-128270388-DEL-67
chr4-128280050-DEL-151
chr4-128283031-DEL-62
chr4-128348877-DEL-97
chr4-128384374-DEL-74
chr4-128419187-DEL-55
chr4-128426443-DEL-120
chr4-128452542-DEL-70
chr4-128457426-DEL-144
chr4-128463505-DEL-586
chr4-128494908-DEL-1520
chr4-128505752-DEL-203
chr4-128524096-DEL-79
chr4-128527204-DEL-250
chr4-128542035-DEL-200
chr4-128560562-DEL-726
chr4-128566064-DEL-86
chr4-128596889-DEL-69
chr4-128658975-DEL-202
chr4-128675426-DEL-72
chr4-128692545-DEL-629
chr4-128722787-DEL-64
chr4-128732443-DEL-64
chr4-128758976-DEL-184
chr4-128799583-DEL-167
chr4-128802417-DEL-69
chr4-128822225-DEL-405
chr4-128832211-DEL-52
chr4-128832808-DEL-96
chr4-128846868-DEL-80
chr4-128857313-DEL-174
chr4-128866934-DEL-550
chr4-128869411-DEL-169
chr4-128872099-DEL-187
chr4-128881807-DEL-853
chr4-128885879-DEL-55
chr4-128921767-DEL-1837
chr4-128922668-DEL-177
chr4-128922781-DEL-203
chr4-128923686-DEL-395
chr4-128924085-DEL-553
chr4-128945262-DEL-110
chr4-128958258-DEL-53
chr4-128961600-DEL-221
chr4-128973130-DEL-282
chr4-128979675-DEL-145
chr4-128990635-DEL-76
chr4-129040945-DEL-233
chr4-129091528-DEL-77
chr4-129106601-DEL-223
chr4-129109537-DEL-158
chr4-129123401-DEL-86
chr4-129125616-DEL-119
chr4-129127724-DEL-52
chr4-129146206-DEL-159
chr4-129146907-DEL-116
chr4-129170133-DEL-101
chr4-129176427-DEL-58
chr4-129177871-DEL-200
chr4-129199777-DEL-206
chr4-129200885-DEL-287
chr4-129204233-DEL-127
chr4-129207942-DEL-265
chr4-129210508-DEL-142
chr4-129230421-DEL-187
chr4-129233859-DEL-80
chr4-129242086-DEL-190

chr4-129242940-DEL-127
chr4-129259411-DEL-2192
chr4-129266053-DEL-198
chr4-129266978-DEL-52
chr4-129270047-DEL-51
chr4-129281082-DEL-277
chr4-129297097-DEL-177
chr4-129307948-DEL-138
chr4-129357302-DEL-84
chr4-129414398-DEL-170
chr4-129422964-DEL-296
chr4-129425132-DEL-55
chr4-129432820-DEL-181
chr4-129437191-DEL-517
chr4-129446394-DEL-511
chr4-129454967-DEL-202
chr4-129488160-DEL-195
chr4-129529335-DEL-173
chr4-129562177-DEL-217
chr4-129574569-DEL-170
chr4-129588399-DEL-208
chr4-129619808-DEL-72
chr4-129631494-DEL-203
chr4-129643820-DEL-200
chr4-129645192-DEL-200
chr4-129646698-DEL-79
chr4-129651697-DEL-450
chr4-129656789-DEL-207
chr4-129664743-DEL-185
chr4-129674879-DEL-209
chr4-129678424-DEL-282
chr4-129679396-DEL-52
chr4-129680474-DEL-84
chr4-129707300-DEL-426
chr4-129731754-DEL-150
chr4-129737729-DEL-216
chr4-129756821-DEL-622
chr4-129778991-DEL-207
chr4-129788014-DEL-61
chr4-129792733-DEL-1506
chr4-129805180-DEL-93
chr4-129810507-DEL-143
chr4-129831757-DEL-59
chr4-129832472-DEL-114
chr4-129834437-DEL-197
chr4-129835038-DEL-231
chr4-129837810-DEL-76
chr4-129844202-DEL-143
chr4-129851456-DEL-278
chr4-129904404-DEL-197
chr4-129952612-DEL-89
chr4-129961785-DEL-58
chr4-129962855-DEL-54
chr4-129965274-DEL-114
chr4-129978856-DEL-60
chr4-130002139-DEL-90
chr4-130015779-DEL-99
chr4-130017202-DEL-188
chr4-130034448-DEL-56
chr4-130036698-DEL-236
chr4-130044409-DEL-143
chr4-130046528-DEL-326
chr4-130094529-DEL-57
chr4-130095100-DEL-54
chr4-130099184-DEL-144
chr4-130099996-DEL-63
chr4-130102069-DEL-119
chr4-130109681-DEL-219
chr4-130114760-DEL-220
chr4-130124101-DEL-62
chr4-130125193-DEL-63
chr4-130135250-DEL-356
chr4-130154918-DEL-101
chr4-130185671-DEL-80
chr4-130215148-DEL-252
chr4-130228740-DEL-54
chr4-130241087-DEL-77

chr4-130268148-DEL-147
chr4-130270375-DEL-481
chr4-130285741-DEL-160
chr4-130289161-DEL-58
chr4-130303894-DEL-195
chr4-130312458-DEL-226
chr4-130321899-DEL-53
chr4-130330734-DEL-56
chr4-130342950-DEL-116
chr4-130344338-DEL-120
chr4-130345556-DEL-1358
chr4-130352222-DEL-166
chr4-130352732-DEL-73
chr4-130368980-DEL-885
chr4-130371182-DEL-193
chr4-130417852-DEL-387
chr4-130462113-DEL-94
chr4-130504752-DEL-77
chr4-130509295-DEL-86
chr4-130514355-DEL-356
chr4-130534982-DEL-349
chr4-130539336-DEL-67
chr4-130573100-DEL-195
chr4-130577430-DEL-175
chr4-130613954-DEL-60
chr4-130616321-DEL-180
chr4-130623547-DEL-66
chr4-130635241-DEL-260
chr4-130662440-DEL-324
chr4-130667641-DEL-94
chr4-130675095-DEL-1891
chr4-130681220-DEL-92
chr4-130692407-DEL-62
chr4-130702917-DEL-355
chr4-130739844-DEL-324
chr4-130760442-DEL-67
chr4-130763900-DEL-84
chr4-130765296-DEL-63
chr4-130781934-DEL-54
chr4-130820017-DEL-833
chr4-130821173-DEL-257
chr4-130823503-DEL-326
chr4-130826508-DEL-119
chr4-130826686-DEL-536
chr4-130844413-DEL-80
chr4-130854829-DEL-82
chr4-130857013-DEL-58
chr4-130859172-DEL-160
chr4-130949427-DEL-6679
chr4-131023957-DEL-2203
chr4-131053262-DEL-120
chr4-131070147-DEL-72
chr4-131074926-DEL-216
chr4-131090948-DEL-1730
chr4-131109898-DEL-92
chr4-131111755-DEL-93
chr4-131112281-DEL-1312
chr4-131129000-DEL-1938
chr4-131130305-DEL-58
chr4-131136862-DEL-83
chr4-131141398-DEL-205
chr4-131236686-DEL-207
chr4-131248592-DEL-51
chr4-131270515-DEL-77
chr4-131271136-DEL-53
chr4-131302499-DEL-55
chr4-131310559-DEL-67
chr4-131346075-DEL-64
chr4-131365120-DEL-83
chr4-131406729-DEL-127
chr4-131417568-DEL-2139
chr4-131421174-DEL-51
chr4-131433413-DEL-68
chr4-131504864-DEL-54
chr4-131521087-DEL-102
chr4-131532045-DEL-269
chr4-131541831-DEL-93

chr4-131555260-DEL-134
chr4-131572381-DEL-57
chr4-131585739-DEL-195
chr4-131622658-DEL-1875
chr4-131625450-DEL-203
chr4-131630098-DEL-527
chr4-131660453-DEL-236
chr4-131667854-DEL-199
chr4-131683427-DEL-52
chr4-131741782-DEL-309
chr4-131789129-DEL-171
chr4-131833503-DEL-204
chr4-131842123-DEL-661
chr4-131852745-DEL-62
chr4-131855680-DEL-155
chr4-131857260-DEL-90
chr4-131866425-DEL-233
chr4-131875675-DEL-559
chr4-131876243-DEL-1155
chr4-131888085-DEL-183
chr4-131891160-DEL-340
chr4-131902960-DEL-80
chr4-131918471-DEL-790
chr4-131919622-DEL-172
chr4-131926759-DEL-617
chr4-131941545-DEL-67
chr4-131949572-DEL-204
chr4-131956650-DEL-109
chr4-131973105-DEL-61
chr4-131982508-DEL-300
chr4-132004027-DEL-195
chr4-132011639-DEL-64
chr4-132025203-DEL-209
chr4-132031192-DEL-366
chr4-132038339-DEL-180
chr4-132044932-DEL-756
chr4-132062897-DEL-86
chr4-132064125-DEL-98
chr4-132160903-DEL-104
chr4-132163139-DEL-56
chr4-132165717-DEL-505
chr4-132171203-DEL-369
chr4-132172171-DEL-1863
chr4-132177484-DEL-109
chr4-132198607-DEL-52
chr4-132201107-DEL-73
chr4-132204636-DEL-154
chr4-132212841-DEL-200
chr4-132222972-DEL-460
chr4-132243155-DEL-989
chr4-132256342-DEL-1105
chr4-132265671-DEL-125
chr4-132324629-DEL-177
chr4-132331297-DEL-193
chr4-132339688-DEL-200
chr4-132354149-DEL-202
chr4-132355360-DEL-1284
chr4-132358491-DEL-199
chr4-132386912-DEL-63
chr4-132424475-DEL-69
chr4-132425964-DEL-122
chr4-132441657-DEL-773
chr4-132449981-DEL-62
chr4-132486330-DEL-168
chr4-132487310-DEL-167
chr4-132497309-DEL-129
chr4-132513693-DEL-202
chr4-132514290-DEL-259
chr4-132520883-DEL-207
chr4-132524528-DEL-73
chr4-132526245-DEL-210
chr4-132530847-DEL-208
chr4-132550376-DEL-942
chr4-132589386-DEL-187
chr4-132614920-DEL-175
chr4-132622575-DEL-88
chr4-132624724-DEL-398

chr4-132632084-DEL-2304
chr4-132635801-DEL-633
chr4-132646644-DEL-152
chr4-132655140-DEL-76
chr4-132659154-DEL-52
chr4-132664712-DEL-61
chr4-132678019-DEL-253
chr4-132681190-DEL-97
chr4-132683060-DEL-875
chr4-132685514-DEL-182
chr4-132687491-DEL-78
chr4-132692545-DEL-407
chr4-132696221-DEL-141
chr4-132708055-DEL-308
chr4-132719947-DEL-224
chr4-132759597-DEL-55
chr4-132829770-DEL-60
chr4-132833289-DEL-533
chr4-132851197-DEL-206
chr4-132855895-DEL-57
chr4-132865898-DEL-54
chr4-132870188-DEL-129
chr4-132877753-DEL-203
chr4-132890767-DEL-68
chr4-132907918-DEL-113
chr4-132928991-DEL-76
chr4-132941301-DEL-81
chr4-132952415-DEL-220
chr4-132962270-DEL-260
chr4-132999546-DEL-965
chr4-133015167-DEL-85
chr4-133017789-DEL-144
chr4-133034365-DEL-1634
chr4-133045773-DEL-193
chr4-133055256-DEL-1599
chr4-133072583-DEL-141
chr4-133078780-DEL-409
chr4-133080489-DEL-52
chr4-133110219-DEL-321
chr4-133129985-DEL-604
chr4-133132271-DEL-53
chr4-133143977-DEL-101
chr4-133146384-DEL-119
chr4-133152951-DEL-54
chr4-133158404-DEL-156
chr4-133168648-DEL-173
chr4-133194803-DEL-171
chr4-133201201-DEL-562
chr4-133239969-DEL-230
chr4-133242114-DEL-92
chr4-133250511-DEL-181
chr4-133254054-DEL-144
chr4-133284209-DEL-208
chr4-133287703-DEL-443
chr4-133292482-DEL-197
chr4-133298692-DEL-54
chr4-133299208-DEL-51
chr4-133300647-DEL-198
chr4-133304302-DEL-229
chr4-133313755-DEL-579
chr4-133315169-DEL-488
chr4-133319883-DEL-64
chr4-133343722-DEL-76
chr4-133357282-DEL-211
chr4-133368493-DEL-244
chr4-133370479-DEL-72
chr4-133424769-DEL-68
chr4-133463698-DEL-4029
chr4-133508349-DEL-57
chr4-133517503-DEL-1202
chr4-133519180-DEL-57
chr4-133529410-DEL-236
chr4-133546751-DEL-178
chr4-133551082-DEL-52
chr4-133582818-DEL-93
chr4-133616832-DEL-214
chr4-133658490-DEL-56

chr4-133669841-DEL-57
chr4-133683637-DEL-205
chr4-133689447-DEL-120
chr4-133690207-DEL-520
chr4-133693465-DEL-102
chr4-133706597-DEL-209
chr4-133715561-DEL-166
chr4-133759538-DEL-229
chr4-133763750-DEL-252
chr4-133764730-DEL-93
chr4-133773927-DEL-174
chr4-133828557-DEL-164
chr4-133831091-DEL-209
chr4-133859314-DEL-199
chr4-133916003-DEL-171
chr4-133925326-DEL-181
chr4-133929522-DEL-249
chr4-133938771-DEL-185
chr4-133944522-DEL-114
chr4-133952379-DEL-70
chr4-133953499-DEL-85
chr4-133970173-DEL-83
chr4-133990241-DEL-313
chr4-134000572-DEL-77
chr4-134011060-DEL-66
chr4-134018509-DEL-198
chr4-134088066-DEL-186
chr4-134094415-DEL-54
chr4-134108029-DEL-642
chr4-134108048-DEL-809
chr4-134127930-DEL-103
chr4-134129140-DEL-109
chr4-134134838-DEL-84
chr4-134139797-DEL-387
chr4-134162523-DEL-387
chr4-134188576-DEL-59
chr4-134216314-DEL-102
chr4-134218247-DEL-325
chr4-134221434-DEL-179
chr4-134280577-DEL-77
chr4-134312080-DEL-485
chr4-134314345-DEL-66
chr4-134316841-DEL-57
chr4-134317754-DEL-780
chr4-134322177-DEL-93
chr4-134433384-DEL-95
chr4-134443834-DEL-204
chr4-134458005-DEL-52
chr4-134497895-DEL-241
chr4-134522647-DEL-371
chr4-134537598-DEL-60
chr4-134542995-DEL-53
chr4-134553136-DEL-195
chr4-134555955-DEL-162
chr4-134558299-DEL-197
chr4-134572169-DEL-94
chr4-134572643-DEL-438
chr4-134608251-DEL-220
chr4-134614848-DEL-211
chr4-134637586-DEL-51
chr4-134659251-DEL-1039
chr4-134660669-DEL-907
chr4-134679909-DEL-101
chr4-134685875-DEL-385
chr4-134691109-DEL-68
chr4-134697934-DEL-169
chr4-134700300-DEL-359
chr4-134720627-DEL-51
chr4-134903538-DEL-137
chr4-134904198-DEL-130
chr4-134924356-DEL-733
chr4-134926540-DEL-60
chr4-134944730-DEL-51
chr4-134952939-DEL-211
chr4-134955127-DEL-252
chr4-134979815-DEL-252
chr4-135006252-DEL-187

chr4-135026752-DEL-147
chr4-135028007-DEL-493
chr4-135039132-DEL-155
chr4-135059927-DEL-52
chr4-135062025-DEL-176
chr4-135088096-DEL-202
chr4-135089836-DEL-83
chr4-135098596-DEL-189
chr4-135114919-DEL-61
chr4-135116384-DEL-3947
chr4-135136511-DEL-73
chr4-135149815-DEL-205
chr4-135157690-DEL-62
chr4-135214642-DEL-134
chr4-135224612-DEL-2168
chr4-135226588-DEL-186
chr4-135236309-DEL-97
chr4-135239839-DEL-51
chr4-135246832-DEL-180
chr4-135248956-DEL-83
chr4-135250272-DEL-78
chr4-135268233-DEL-70
chr4-135279528-DEL-90
chr4-135358665-DEL-270
chr4-135370172-DEL-1270
chr4-135371412-DEL-162
chr4-135375172-DEL-166
chr4-135395510-DEL-219
chr4-135405935-DEL-503
chr4-135418052-DEL-77
chr4-135481253-DEL-70
chr4-135511543-DEL-250
chr4-135526513-DEL-75
chr4-135529446-DEL-358
chr4-135532927-DEL-117
chr4-135550236-DEL-182
chr4-135609625-DEL-651
chr4-135613733-DEL-138
chr4-135615493-DEL-67
chr4-135617705-DEL-62
chr4-135619214-DEL-395
chr4-135626256-DEL-206
chr4-135636324-DEL-124
chr4-135661196-DEL-669
chr4-135662538-DEL-77
chr4-135684390-DEL-155
chr4-135705382-DEL-389
chr4-135725413-DEL-1071
chr4-135766790-DEL-204
chr4-135768331-DEL-79
chr4-135785681-DEL-204
chr4-135787173-DEL-96
chr4-135826515-DEL-315
chr4-135829278-DEL-196
chr4-135851987-DEL-56
chr4-135857199-DEL-52
chr4-135859780-DEL-892
chr4-135864572-DEL-135
chr4-135879363-DEL-63
chr4-135880872-DEL-89
chr4-135882936-DEL-62
chr4-135892883-DEL-80
chr4-135900632-DEL-2010
chr4-135924477-DEL-400
chr4-135929412-DEL-83
chr4-135935158-DEL-95
chr4-135941425-DEL-205
chr4-135985652-DEL-168
chr4-135999255-DEL-71
chr4-136031787-DEL-199
chr4-136044822-DEL-184
chr4-136045742-DEL-166
chr4-136055875-DEL-216
chr4-136077881-DEL-868
chr4-136078852-DEL-131
chr4-136087678-DEL-503
chr4-136107382-DEL-86

chr4-136115070-DEL-1947
chr4-136117262-DEL-116
chr4-136117747-DEL-240
chr4-136120359-DEL-1022
chr4-136126213-DEL-158
chr4-136127252-DEL-400
chr4-136130955-DEL-180
chr4-136159726-DEL-230
chr4-136167268-DEL-365
chr4-136174266-DEL-352
chr4-136175243-DEL-69
chr4-136191922-DEL-55
chr4-136233713-DEL-404
chr4-136357770-DEL-208
chr4-136371797-DEL-52
chr4-136418721-DEL-54
chr4-136422637-DEL-100
chr4-136431003-DEL-53
chr4-136434418-DEL-60
chr4-136515321-DEL-279
chr4-136526440-DEL-58
chr4-136536904-DEL-172
chr4-136542300-DEL-53
chr4-136546710-DEL-51
chr4-136596366-DEL-162
chr4-136606920-DEL-102
chr4-136645618-DEL-402
chr4-136685336-DEL-158
chr4-136686874-DEL-58
chr4-136739549-DEL-757
chr4-136752122-DEL-504
chr4-136773065-DEL-81
chr4-136774466-DEL-236
chr4-136777025-DEL-93
chr4-136780804-DEL-53
chr4-136801139-DEL-163
chr4-136848122-DEL-187
chr4-136849556-DEL-1792
chr4-136902147-DEL-111
chr4-136979322-DEL-62
chr4-136985675-DEL-65
chr4-137001449-DEL-54
chr4-137041412-DEL-82
chr4-137071967-DEL-89
chr4-137101901-DEL-654
chr4-137107567-DEL-176
chr4-137117113-DEL-592
chr4-137118168-DEL-193
chr4-137118357-DEL-517
chr4-137122523-DEL-514
chr4-137137930-DEL-198
chr4-137142574-DEL-205
chr4-137154002-DEL-189
chr4-137156992-DEL-193
chr4-137218788-DEL-233
chr4-137227743-DEL-80
chr4-137242277-DEL-289
chr4-137253094-DEL-71
chr4-137253890-DEL-77
chr4-137301102-DEL-172
chr4-137306639-DEL-101
chr4-137308456-DEL-103
chr4-137309946-DEL-955
chr4-137311491-DEL-178
chr4-137315835-DEL-156
chr4-137320077-DEL-95
chr4-137327344-DEL-54
chr4-137345194-DEL-211
chr4-137362877-DEL-193
chr4-137412001-DEL-72
chr4-137455203-DEL-91
chr4-137491443-DEL-56
chr4-137495857-DEL-542
chr4-137508162-DEL-172
chr4-137519545-DEL-197
chr4-137542454-DEL-233
chr4-137559010-DEL-60

chr4-137570913-DEL-53
chr4-137620813-DEL-83
chr4-137621748-DEL-248
chr4-137657672-DEL-139
chr4-137661128-DEL-294
chr4-137693725-DEL-375
chr4-137698059-DEL-8448
chr4-137714360-DEL-89
chr4-137716543-DEL-54
chr4-137728543-DEL-208
chr4-137729826-DEL-1199
chr4-137740031-DEL-79
chr4-137770697-DEL-544
chr4-137771415-DEL-73
chr4-137774371-DEL-174
chr4-137784798-DEL-73
chr4-137789207-DEL-158
chr4-137795471-DEL-66
chr4-137817892-DEL-61
chr4-137841115-DEL-128
chr4-137875960-DEL-52
chr4-137898011-DEL-76
chr4-137911438-DEL-151
chr4-137940668-DEL-235
chr4-137947856-DEL-66
chr4-137976465-DEL-53
chr4-137980068-DEL-177
chr4-138004579-DEL-57
chr4-138007087-DEL-90
chr4-138007819-DEL-455
chr4-138019654-DEL-964
chr4-138022345-DEL-204
chr4-138029616-DEL-82
chr4-138052267-DEL-102
chr4-138085934-DEL-214
chr4-138116636-DEL-51
chr4-138125911-DEL-860
chr4-138129617-DEL-1683
chr4-138136047-DEL-96
chr4-138140880-DEL-360
chr4-138163331-DEL-1470
chr4-138177291-DEL-826
chr4-138193707-DEL-124
chr4-138201651-DEL-85
chr4-138203189-DEL-63
chr4-138204399-DEL-90
chr4-138208778-DEL-182
chr4-138230066-DEL-802
chr4-138237981-DEL-138
chr4-138274513-DEL-85
chr4-138309526-DEL-70
chr4-138330422-DEL-61
chr4-138378937-DEL-92
chr4-138382113-DEL-1121
chr4-138383242-DEL-1421
chr4-138388479-DEL-206
chr4-138390057-DEL-59
chr4-138403387-DEL-89
chr4-138445419-DEL-144
chr4-138475408-DEL-94
chr4-138558725-DEL-93
chr4-138587994-DEL-160
chr4-138602486-DEL-126
chr4-138611338-DEL-157
chr4-138624572-DEL-69
chr4-138651803-DEL-110
chr4-138657018-DEL-1407
chr4-138659604-DEL-99
chr4-138663370-DEL-287
chr4-138664123-DEL-88
chr4-138667476-DEL-77
chr4-138672289-DEL-198
chr4-138707107-DEL-802
chr4-138745486-DEL-188
chr4-138748544-DEL-60
chr4-138752393-DEL-191
chr4-138753217-DEL-268

chr4-138753420-DEL-170
chr4-138753830-DEL-1154
chr4-138754282-DEL-106
chr4-138769687-DEL-124
chr4-138804481-DEL-75
chr4-138805198-DEL-112
chr4-138864765-DEL-110
chr4-138865059-DEL-501
chr4-138887221-DEL-938
chr4-138900027-DEL-381
chr4-138905752-DEL-72
chr4-138937281-DEL-278
chr4-138997774-DEL-52
chr4-139030210-DEL-55
chr4-139043694-DEL-219
chr4-139050649-DEL-102
chr4-139063530-DEL-853
chr4-139064418-DEL-1931
chr4-139066309-DEL-174
chr4-139070869-DEL-60
chr4-139072420-DEL-70
chr4-139112088-DEL-54
chr4-139126658-DEL-54
chr4-139144741-DEL-176
chr4-139183117-DEL-73
chr4-139211654-DEL-894
chr4-139224122-DEL-132
chr4-139230808-DEL-78
chr4-139260616-DEL-108
chr4-139316604-DEL-59
chr4-139321030-DEL-130
chr4-139326818-DEL-63
chr4-139371050-DEL-133
chr4-139379565-DEL-78
chr4-139430891-DEL-406
chr4-139446029-DEL-313
chr4-139448485-DEL-90
chr4-139456180-DEL-63
chr4-139502158-DEL-74
chr4-139509268-DEL-97
chr4-139512946-DEL-111
chr4-139520234-DEL-75
chr4-139530701-DEL-75
chr4-139600749-DEL-78
chr4-139606267-DEL-302
chr4-139611014-DEL-51
chr4-139621371-DEL-76
chr4-139638475-DEL-197
chr4-139657971-DEL-127
chr4-139699915-DEL-80
chr4-139713001-DEL-56
chr4-139718390-DEL-159
chr4-139759684-DEL-78
chr4-139774819-DEL-73
chr4-139791720-DEL-54
chr4-139798081-DEL-315
chr4-139824719-DEL-727
chr4-139844084-DEL-74
chr4-139856870-DEL-52
chr4-139860333-DEL-88
chr4-139868034-DEL-1269
chr4-139898844-DEL-69
chr4-139910658-DEL-797
chr4-139928233-DEL-203
chr4-139942827-DEL-124
chr4-139978688-DEL-73
chr4-139986424-DEL-106
chr4-139987442-DEL-82
chr4-139992868-DEL-63
chr4-140007261-DEL-122
chr4-140015983-DEL-56
chr4-140026093-DEL-58
chr4-140029427-DEL-141
chr4-140038167-DEL-115
chr4-140044396-DEL-342
chr4-140075122-DEL-242
chr4-140096873-DEL-170

chr4-140099832-DEL-52
chr4-140120422-DEL-129
chr4-140131381-DEL-1242
chr4-140134922-DEL-1548
chr4-140191615-DEL-64
chr4-140211900-DEL-88
chr4-140217209-DEL-156
chr4-140218391-DEL-84
chr4-140230928-DEL-125
chr4-140236458-DEL-64
chr4-140236580-DEL-96
chr4-140246967-DEL-58
chr4-140272849-DEL-200
chr4-140272927-DEL-667
chr4-140287835-DEL-876
chr4-140291041-DEL-52
chr4-140314910-DEL-100
chr4-140318004-DEL-212
chr4-140324204-DEL-163
chr4-140327301-DEL-128
chr4-140350519-DEL-52
chr4-140403627-DEL-101
chr4-140405223-DEL-458
chr4-140414918-DEL-200
chr4-140459339-DEL-90
chr4-140474096-DEL-618
chr4-140491258-DEL-179
chr4-140494160-DEL-78
chr4-140511576-DEL-81
chr4-140535397-DEL-52
chr4-140538568-DEL-117
chr4-140567626-DEL-146
chr4-140587543-DEL-110
chr4-140608190-DEL-59
chr4-140612958-DEL-51
chr4-140620430-DEL-291
chr4-140624379-DEL-811
chr4-140627122-DEL-52
chr4-140641731-DEL-187
chr4-140650625-DEL-137
chr4-140666197-DEL-851
chr4-140670316-DEL-1011
chr4-140681908-DEL-221
chr4-140708312-DEL-73
chr4-140720317-DEL-143
chr4-140736670-DEL-105
chr4-140782921-DEL-203
chr4-140783193-DEL-732
chr4-140785775-DEL-156
chr4-140820721-DEL-1100
chr4-140825589-DEL-214
chr4-140836900-DEL-902
chr4-140837941-DEL-609
chr4-140838794-DEL-97
chr4-140865306-DEL-201
chr4-140891572-DEL-60
chr4-140895921-DEL-102
chr4-140907271-DEL-185
chr4-140912213-DEL-156
chr4-140916939-DEL-390
chr4-140918092-DEL-1024
chr4-140923534-DEL-899
chr4-140939468-DEL-232
chr4-140947957-DEL-461
chr4-140951550-DEL-69
chr4-140991609-DEL-1806
chr4-141003723-DEL-307
chr4-141008245-DEL-52
chr4-141022284-DEL-57
chr4-141082031-DEL-623
chr4-141082865-DEL-102
chr4-141094214-DEL-70
chr4-141128118-DEL-212
chr4-141142741-DEL-120
chr4-141146072-DEL-96
chr4-141161787-DEL-2000
chr4-141165017-DEL-51

chr4-141167363-DEL-589
chr4-141209704-DEL-76
chr4-141213925-DEL-91
chr4-141233383-DEL-176
chr4-141289198-DEL-695
chr4-141292246-DEL-200
chr4-141299025-DEL-318
chr4-141307619-DEL-147
chr4-141366730-DEL-160
chr4-141382803-DEL-177
chr4-141384914-DEL-59
chr4-141387941-DEL-92
chr4-141393712-DEL-121
chr4-141396909-DEL-66
chr4-141444513-DEL-2148
chr4-141458177-DEL-80
chr4-141461044-DEL-204
chr4-141470578-DEL-513
chr4-141477987-DEL-214
chr4-141481485-DEL-190
chr4-141500755-DEL-114
chr4-141535145-DEL-1704
chr4-141614032-DEL-1261
chr4-141615971-DEL-64
chr4-141692651-DEL-6246
chr4-141695957-DEL-1593
chr4-141704436-DEL-356
chr4-141726461-DEL-95
chr4-141734130-DEL-74
chr4-141739599-DEL-92
chr4-141742588-DEL-87
chr4-141743872-DEL-1320
chr4-141747798-DEL-61
chr4-141748682-DEL-1324
chr4-141753389-DEL-86
chr4-141785636-DEL-549
chr4-141816607-DEL-81
chr4-141826962-DEL-377
chr4-141851100-DEL-66
chr4-141865197-DEL-164
chr4-141907217-DEL-52
chr4-141925187-DEL-384
chr4-141927313-DEL-83
chr4-141943034-DEL-136
chr4-141967893-DEL-158
chr4-141970811-DEL-144
chr4-141973057-DEL-169
chr4-142000085-DEL-108
chr4-142022688-DEL-76
chr4-142081874-DEL-78
chr4-142092984-DEL-371
chr4-142132656-DEL-190
chr4-142136404-DEL-53
chr4-142157101-DEL-62
chr4-142187077-DEL-181
chr4-142208509-DEL-119
chr4-142236597-DEL-115
chr4-142294493-DEL-324
chr4-142350059-DEL-61
chr4-142362413-DEL-64
chr4-142367794-DEL-66
chr4-142433343-DEL-1359
chr4-142447516-DEL-1082
chr4-142464248-DEL-1296
chr4-142467086-DEL-1569
chr4-142469106-DEL-132
chr4-142493724-DEL-176
chr4-142495522-DEL-60
chr4-142502847-DEL-54
chr4-142520630-DEL-165
chr4-142524689-DEL-357
chr4-142559935-DEL-64
chr4-142566320-DEL-348
chr4-142567869-DEL-81
chr4-142571461-DEL-168
chr4-142577790-DEL-132
chr4-142585019-DEL-4138

chr4-142596315-DEL-206
chr4-142633561-DEL-80
chr4-142645100-DEL-67
chr4-142646119-DEL-61
chr4-142687064-DEL-271
chr4-142689373-DEL-53
chr4-142699367-DEL-1458
chr4-142721549-DEL-148
chr4-142750336-DEL-54
chr4-142828511-DEL-195
chr4-142863968-DEL-1466
chr4-142868095-DEL-105
chr4-142874234-DEL-51
chr4-142879754-DEL-75
chr4-142909244-DEL-54
chr4-142922370-DEL-1223
chr4-142924265-DEL-2747
chr4-142949945-DEL-95
chr4-142958600-DEL-99
chr4-142961144-DEL-57
chr4-142962694-DEL-580
chr4-143000741-DEL-146
chr4-143002789-DEL-168
chr4-143037661-DEL-185
chr4-143044157-DEL-171
chr4-143057744-DEL-176
chr4-143090880-DEL-68
chr4-143112673-DEL-454
chr4-143133189-DEL-316
chr4-143170210-DEL-103
chr4-143183673-DEL-62
chr4-143222483-DEL-90
chr4-143241559-DEL-9058
chr4-143293579-DEL-1924
chr4-143296284-DEL-325
chr4-143296901-DEL-60
chr4-143311551-DEL-6476
chr4-143318724-DEL-56
chr4-143336357-DEL-113
chr4-143393633-DEL-409
chr4-143396742-DEL-576
chr4-143401446-DEL-22417
chr4-143440120-DEL-728
chr4-143442053-DEL-71
chr4-143491744-DEL-117
chr4-143493379-DEL-1311
chr4-143524643-DEL-1610
chr4-143527249-DEL-863
chr4-143546904-DEL-1464
chr4-143554044-DEL-6213
chr4-143562732-DEL-829
chr4-143565613-DEL-51
chr4-143573439-DEL-54
chr4-143616195-DEL-141
chr4-143644893-DEL-477
chr4-143668500-DEL-15396
chr4-143702098-DEL-4947
chr4-143707799-DEL-1014
chr4-143710517-DEL-4883
chr4-143710629-DEL-60
chr4-143715082-DEL-60
chr4-143729037-DEL-2541
chr4-143740463-DEL-209
chr4-143767342-DEL-191
chr4-143775978-DEL-212
chr4-143777288-DEL-342
chr4-143784010-DEL-1397
chr4-143809647-DEL-53
chr4-143820483-DEL-178
chr4-143838560-DEL-9838
chr4-143851181-DEL-101
chr4-143852435-DEL-8803
chr4-143864547-DEL-57
chr4-143867800-DEL-13738
chr4-143906832-DEL-473
chr4-143939328-DEL-70
chr4-144028273-DEL-102

chr4-144029197-DEL-114
chr4-144044814-DEL-3877
chr4-144046708-DEL-193
chr4-144048953-DEL-227
chr4-144075988-DEL-353
chr4-144076484-DEL-1526
chr4-144078907-DEL-566
chr4-144092090-DEL-973
chr4-144111500-DEL-787
chr4-144115272-DEL-62
chr4-144126938-DEL-226
chr4-144128783-DEL-879
chr4-144137146-DEL-54
chr4-144167942-DEL-84
chr4-144256883-DEL-656
chr4-144296449-DEL-1408
chr4-144327621-DEL-410
chr4-144463638-DEL-1084
chr4-144483204-DEL-93
chr4-144516752-DEL-2577
chr4-144561542-DEL-1791
chr4-144561561-DEL-962
chr4-144562036-DEL-844
chr4-144562180-DEL-130
chr4-144562288-DEL-1010
chr4-144563030-DEL-231
chr4-144565406-DEL-10274
chr4-144575347-DEL-53
chr4-144577927-DEL-4310
chr4-144612352-DEL-52
chr4-144648513-DEL-62
chr4-144650858-DEL-134
chr4-144669524-DEL-645
chr4-144686694-DEL-129
chr4-144717609-DEL-336
chr4-144733888-DEL-79
chr4-144807392-DEL-60
chr4-144855353-DEL-94
chr4-144909066-DEL-200
chr4-144917499-DEL-216
chr4-144953681-DEL-154
chr4-144981842-DEL-102
chr4-145009069-DEL-76
chr4-145020422-DEL-188
chr4-145075458-DEL-1007
chr4-145075489-DEL-984
chr4-145092774-DEL-63
chr4-145104127-DEL-24968
chr4-145131552-DEL-105
chr4-145151892-DEL-64
chr4-145199693-DEL-2296
chr4-145211998-DEL-307
chr4-145255186-DEL-225
chr4-145276574-DEL-569
chr4-145295285-DEL-5228
chr4-145303387-DEL-326
chr4-145310663-DEL-54
chr4-145317994-DEL-10202
chr4-145327978-DEL-437
chr4-145349304-DEL-756
chr4-145350086-DEL-252
chr4-145428536-DEL-168
chr4-145442985-DEL-8151
chr4-145454280-DEL-102
chr4-145458570-DEL-400
chr4-145491885-DEL-52
chr4-145495676-DEL-181
chr4-145499254-DEL-340
chr4-145500519-DEL-261
chr4-145506824-DEL-227
chr4-145507812-DEL-171
chr4-145509188-DEL-79
chr4-145511554-DEL-164
chr4-145523813-DEL-61
chr4-145532473-DEL-424
chr4-145538734-DEL-83
chr4-145558901-DEL-194

chr4-145563536-DEL-159
chr4-145568612-DEL-929
chr4-145580288-DEL-831
chr4-145585098-DEL-197
chr4-145590345-DEL-223
chr4-145594578-DEL-136
chr4-145601737-DEL-2089
chr4-145608323-DEL-7344
chr4-145618823-DEL-101
chr4-146030826-DEL-51
chr4-146034070-DEL-231
chr4-146050102-DEL-486
chr4-146057870-DEL-54
chr4-146064507-DEL-790
chr4-146118463-DEL-5457
chr4-146132721-DEL-84
chr4-146133509-DEL-84
chr4-146158737-DEL-122
chr4-146180291-DEL-227
chr4-146194564-DEL-187
chr4-146198148-DEL-340
chr4-146199419-DEL-178
chr4-146205765-DEL-129
chr4-146207012-DEL-171
chr4-146222706-DEL-58
chr4-146237643-DEL-83
chr4-146257802-DEL-193
chr4-146262453-DEL-159
chr4-146267158-DEL-165
chr4-146267515-DEL-930
chr4-146284014-DEL-197
chr4-146324480-DEL-3939
chr4-146325944-DEL-177
chr4-146336338-DEL-4812
chr4-146362018-DEL-52
chr4-146379855-DEL-484
chr4-146383828-DEL-89
chr4-146384149-DEL-57
chr4-146388153-DEL-194
chr4-146390904-DEL-414
chr4-146392179-DEL-168
chr4-146416386-DEL-8887
chr4-146458694-DEL-51
chr4-146461692-DEL-1114
chr4-146465515-DEL-1275
chr4-146498990-DEL-74
chr4-146586563-DEL-10746
chr4-146587535-DEL-8763
chr4-146614648-DEL-69
chr4-146630396-DEL-331
chr4-146647063-DEL-118
chr4-146661181-DEL-213
chr4-146664379-DEL-51
chr4-146665557-DEL-256
chr4-146674474-DEL-195
chr4-146687890-DEL-262
chr4-146687955-DEL-1469
chr4-146694358-DEL-171
chr4-146695376-DEL-109
chr4-146696565-DEL-79
chr4-146698910-DEL-169
chr4-146702887-DEL-188
chr4-146802584-DEL-84
chr4-146804288-DEL-168
chr4-146811976-DEL-259
chr4-146825431-DEL-215
chr4-146829663-DEL-5773
chr4-146832027-DEL-86
chr4-146892978-DEL-54
chr4-146897721-DEL-327
chr4-146913258-DEL-84
chr4-146915015-DEL-1047
chr4-146921194-DEL-64
chr4-146987256-DEL-1274
chr4-146990296-DEL-1675
chr4-146996165-DEL-7607
chr4-147009059-DEL-197

chr4-147010245-DEL-108
chr4-147017411-DEL-1718
chr4-147021284-DEL-4425
chr4-147025987-DEL-4532
chr4-147028938-DEL-199
chr4-147067023-DEL-54
chr4-147088865-DEL-119
chr4-147095074-DEL-502
chr4-147104968-DEL-60
chr4-147128665-DEL-5480
chr4-147142352-DEL-84
chr4-147164173-DEL-107
chr4-147214434-DEL-1512
chr4-147216710-DEL-171
chr4-147247107-DEL-58
chr4-147267425-DEL-197
chr4-147276214-DEL-115
chr4-147276978-DEL-930
chr4-147285036-DEL-169
chr4-147287032-DEL-7180
chr4-147295836-DEL-831
chr4-147300642-DEL-198
chr4-147313607-DEL-51
chr4-147319009-DEL-208
chr4-147333244-DEL-87
chr4-147382181-DEL-601
chr4-147383274-DEL-80
chr4-147385540-DEL-427
chr4-147390769-DEL-494
chr4-147392554-DEL-320
chr4-147395706-DEL-1256
chr4-147395819-DEL-169
chr4-147405842-DEL-170
chr4-147412717-DEL-194
chr4-147414570-DEL-1044
chr4-147460035-DEL-196
chr4-147497307-DEL-141
chr4-147498605-DEL-82
chr4-147499272-DEL-1283
chr4-147508013-DEL-58
chr4-147519213-DEL-1921
chr4-147531690-DEL-1404
chr4-147532495-DEL-317
chr4-147584478-DEL-7223
chr4-147597069-DEL-504
chr4-147597619-DEL-83
chr4-147597759-DEL-670
chr4-147598622-DEL-209
chr4-147628973-DEL-5565
chr4-147655898-DEL-881
chr4-147705156-DEL-65
chr4-147738357-DEL-510
chr4-147743430-DEL-327
chr4-147744975-DEL-60
chr4-147751839-DEL-77
chr4-147760621-DEL-1046
chr4-147760806-DEL-676
chr4-147763235-DEL-195
chr4-147766885-DEL-75
chr4-147770587-DEL-480
chr4-147786392-DEL-221
chr4-147790039-DEL-72
chr4-147791355-DEL-194
chr4-147807850-DEL-6438
chr4-147824640-DEL-185
chr4-147839599-DEL-336
chr4-147880884-DEL-6160
chr4-147887764-DEL-121
chr4-147893750-DEL-349
chr4-147926585-DEL-95
chr4-147943630-DEL-290
chr4-147951413-DEL-116
chr4-147958724-DEL-64
chr4-147992377-DEL-86
chr4-148009168-DEL-108
chr4-148023964-DEL-172
chr4-148034061-DEL-88

chr4-148049308-DEL-146
chr4-148057121-DEL-225
chr4-148061538-DEL-85
chr4-148063276-DEL-245
chr4-148081879-DEL-328
chr4-148117157-DEL-76
chr4-148152242-DEL-749
chr4-148158162-DEL-433
chr4-148198204-DEL-138
chr4-148200733-DEL-106
chr4-148226624-DEL-185
chr4-148238964-DEL-67
chr4-148253610-DEL-164
chr4-148261315-DEL-195
chr4-148275559-DEL-92
chr4-148278953-DEL-52
chr4-148290352-DEL-253
chr4-148303701-DEL-121
chr4-148305487-DEL-252
chr4-148320678-DEL-144
chr4-148326084-DEL-178
chr4-148332766-DEL-408
chr4-148349456-DEL-84
chr4-148364589-DEL-91
chr4-148408300-DEL-446
chr4-148417388-DEL-129
chr4-148450200-DEL-232
chr4-148451494-DEL-74
chr4-148455560-DEL-112
chr4-148471670-DEL-174
chr4-148506953-DEL-67
chr4-148519572-DEL-132
chr4-148534336-DEL-60
chr4-148544950-DEL-112
chr4-148574568-DEL-200
chr4-148600525-DEL-85
chr4-148612626-DEL-202
chr4-148613382-DEL-199
chr4-148632956-DEL-206
chr4-148669020-DEL-108
chr4-148669544-DEL-2042
chr4-148679636-DEL-596
chr4-148715874-DEL-140
chr4-148734756-DEL-51
chr4-148737128-DEL-206
chr4-148738157-DEL-913
chr4-148743978-DEL-755
chr4-148749560-DEL-85
chr4-148765337-DEL-60
chr4-148781636-DEL-55
chr4-148801467-DEL-830
chr4-148801494-DEL-128
chr4-148804684-DEL-158
chr4-148807286-DEL-1350
chr4-148836646-DEL-108
chr4-148852032-DEL-121
chr4-148873343-DEL-80
chr4-148911766-DEL-131
chr4-148934410-DEL-70
chr4-148940567-DEL-60
chr4-148953146-DEL-245
chr4-148996188-DEL-488
chr4-149094237-DEL-743
chr4-149094931-DEL-187
chr4-149179609-DEL-355
chr4-149196371-DEL-424
chr4-149209366-DEL-90
chr4-149219539-DEL-220
chr4-149223709-DEL-112
chr4-149228351-DEL-664
chr4-149260196-DEL-226
chr4-149269001-DEL-59
chr4-149274869-DEL-209
chr4-149275903-DEL-82
chr4-149281275-DEL-97
chr4-149291673-DEL-66
chr4-149292483-DEL-128

chr4-149293241-DEL-204
chr4-149295948-DEL-195
chr4-149312522-DEL-77
chr4-149344687-DEL-167
chr4-149352672-DEL-161
chr4-149356112-DEL-676
chr4-149365788-DEL-76
chr4-149368403-DEL-474
chr4-149397494-DEL-72
chr4-149401215-DEL-209
chr4-149434842-DEL-252
chr4-149436388-DEL-120
chr4-149443014-DEL-117
chr4-149460712-DEL-81
chr4-149464779-DEL-96
chr4-149516273-DEL-53
chr4-149517882-DEL-56
chr4-149532637-DEL-181
chr4-149535677-DEL-109
chr4-149540327-DEL-784
chr4-149552719-DEL-204
chr4-149557350-DEL-66
chr4-149562806-DEL-177
chr4-149567071-DEL-261
chr4-149575080-DEL-216
chr4-149581515-DEL-216
chr4-149583143-DEL-54
chr4-149594884-DEL-353
chr4-149597917-DEL-84
chr4-149613006-DEL-272
chr4-149668053-DEL-65
chr4-149669127-DEL-392
chr4-149677541-DEL-76
chr4-149688087-DEL-66
chr4-149693190-DEL-78
chr4-149707318-DEL-284
chr4-149724147-DEL-185
chr4-149747166-DEL-126
chr4-149750676-DEL-65
chr4-149755489-DEL-183
chr4-149778640-DEL-190
chr4-149793337-DEL-61
chr4-149804401-DEL-1141
chr4-149850931-DEL-73
chr4-149853319-DEL-327
chr4-149854025-DEL-61
chr4-149894689-DEL-481
chr4-149902732-DEL-1047
chr4-149902895-DEL-237
chr4-149906454-DEL-54
chr4-149907992-DEL-458
chr4-149908933-DEL-63
chr4-149911628-DEL-138
chr4-149913606-DEL-60
chr4-149916278-DEL-58
chr4-149938322-DEL-86
chr4-149945364-DEL-160
chr4-149956529-DEL-52
chr4-149957763-DEL-316
chr4-149984728-DEL-112
chr4-150015153-DEL-73
chr4-150029307-DEL-304
chr4-150060989-DEL-51
chr4-150094026-DEL-648
chr4-150095973-DEL-203
chr4-150101781-DEL-169
chr4-150108612-DEL-323
chr4-150120264-DEL-89
chr4-150130718-DEL-198
chr4-150147364-DEL-70
chr4-150200849-DEL-78
chr4-150218199-DEL-52
chr4-150236035-DEL-60
chr4-150254795-DEL-153
chr4-150264745-DEL-457
chr4-150272811-DEL-53
chr4-150288521-DEL-52

chr4-150296239-DEL-197
chr4-150358300-DEL-159
chr4-150370554-DEL-54
chr4-150384350-DEL-206
chr4-150397627-DEL-213
chr4-150409227-DEL-82
chr4-150433221-DEL-114
chr4-150443015-DEL-1008
chr4-150447637-DEL-340
chr4-150448335-DEL-804
chr4-150460670-DEL-157
chr4-150465611-DEL-210
chr4-150503028-DEL-202
chr4-150508545-DEL-59
chr4-150556364-DEL-120
chr4-150564774-DEL-86
chr4-150574407-DEL-303
chr4-150595608-DEL-324
chr4-150596275-DEL-63
chr4-150598190-DEL-342
chr4-150598929-DEL-304
chr4-150605189-DEL-204
chr4-150631900-DEL-264
chr4-150667082-DEL-366
chr4-150672892-DEL-6300
chr4-150673071-DEL-208
chr4-150683441-DEL-218
chr4-150687082-DEL-228
chr4-150726069-DEL-54
chr4-150730759-DEL-89
chr4-150738658-DEL-62
chr4-150763357-DEL-80
chr4-150768415-DEL-419
chr4-150771289-DEL-98
chr4-150784599-DEL-444
chr4-150807050-DEL-64
chr4-150814184-DEL-214
chr4-150816584-DEL-186
chr4-150828582-DEL-76
chr4-150831500-DEL-56
chr4-150838774-DEL-204
chr4-150848439-DEL-290
chr4-150849648-DEL-54
chr4-150863571-DEL-165
chr4-150895068-DEL-98
chr4-150926116-DEL-156
chr4-150932623-DEL-81
chr4-150988891-DEL-650
chr4-150995459-DEL-183
chr4-151028347-DEL-193
chr4-151060067-DEL-362
chr4-151072349-DEL-257
chr4-151108130-DEL-235
chr4-151134855-DEL-201
chr4-151165386-DEL-79
chr4-151203372-DEL-101
chr4-151230771-DEL-54
chr4-151237037-DEL-56
chr4-151256276-DEL-306
chr4-151299205-DEL-356
chr4-151300625-DEL-209
chr4-151315723-DEL-596
chr4-151331140-DEL-66
chr4-151335418-DEL-254
chr4-151360793-DEL-217
chr4-151387395-DEL-619
chr4-151392137-DEL-71
chr4-151393130-DEL-800
chr4-151393156-DEL-248
chr4-151410067-DEL-163
chr4-151415460-DEL-65
chr4-151462763-DEL-369
chr4-151475704-DEL-99
chr4-151501172-DEL-82
chr4-151512465-DEL-78
chr4-151515288-DEL-217
chr4-151561097-DEL-125

chr4-151566116-DEL-132
chr4-151569483-DEL-75
chr4-151591486-DEL-57
chr4-151594732-DEL-68
chr4-151616791-DEL-88
chr4-151619541-DEL-198
chr4-151633131-DEL-198
chr4-151660125-DEL-72
chr4-151660749-DEL-266
chr4-151667357-DEL-58
chr4-151706243-DEL-552
chr4-151708696-DEL-98
chr4-151714197-DEL-127
chr4-151734897-DEL-63
chr4-151736024-DEL-434
chr4-151738920-DEL-53
chr4-151773769-DEL-124
chr4-151789282-DEL-90
chr4-151815788-DEL-148
chr4-151816533-DEL-146
chr4-151844545-DEL-51
chr4-151852428-DEL-54
chr4-151862344-DEL-64
chr4-151888511-DEL-149
chr4-151902772-DEL-53
chr4-151924850-DEL-472
chr4-151930005-DEL-136
chr4-151968488-DEL-184
chr4-151971672-DEL-83
chr4-151975585-DEL-125
chr4-152011940-DEL-301
chr4-152017892-DEL-115
chr4-152044557-DEL-85
chr4-152054046-DEL-180
chr4-152056165-DEL-82
chr4-152063061-DEL-194
chr4-152125201-DEL-61
chr4-152129361-DEL-82
chr4-152135409-DEL-57
chr4-152154954-DEL-62
chr4-152178451-DEL-208
chr4-152257086-DEL-67
chr4-152259907-DEL-54
chr4-152269510-DEL-427
chr4-152276378-DEL-60
chr4-152282934-DEL-72
chr4-152288558-DEL-58
chr4-152314835-DEL-271
chr4-152318660-DEL-157
chr4-152320046-DEL-57
chr4-152362223-DEL-206
chr4-152373433-DEL-699
chr4-152379469-DEL-52
chr4-152403860-DEL-196
chr4-152408112-DEL-181
chr4-152448127-DEL-52
chr4-152454227-DEL-276
chr4-152458978-DEL-107
chr4-152463715-DEL-77
chr4-152507735-DEL-116
chr4-152531970-DEL-60
chr4-152570450-DEL-56
chr4-152570939-DEL-246
chr4-152586391-DEL-169
chr4-152609612-DEL-57
chr4-152641963-DEL-89
chr4-152651718-DEL-88
chr4-152659408-DEL-65
chr4-152673096-DEL-52
chr4-152702499-DEL-1363
chr4-152703016-DEL-349
chr4-152729863-DEL-51
chr4-152730391-DEL-52
chr4-152776670-DEL-705
chr4-152794808-DEL-62
chr4-152824761-DEL-85
chr4-152862633-DEL-55

chr4-152868072-DEL-1322
chr4-152874084-DEL-71
chr4-152898205-DEL-466
chr4-152915025-DEL-197
chr4-152938144-DEL-239
chr4-152949357-DEL-58
chr4-152953142-DEL-110
chr4-152978195-DEL-336
chr4-152990420-DEL-76
chr4-152994771-DEL-54
chr4-153000173-DEL-58
chr4-153001841-DEL-268
chr4-153038385-DEL-190
chr4-153071226-DEL-2527
chr4-153075980-DEL-60
chr4-153096623-DEL-64
chr4-153097245-DEL-68
chr4-153133869-DEL-143
chr4-153169652-DEL-112
chr4-153171868-DEL-179
chr4-153182255-DEL-77
chr4-153205911-DEL-120
chr4-153222689-DEL-89
chr4-153224797-DEL-122
chr4-153234593-DEL-58
chr4-153237551-DEL-271
chr4-153242740-DEL-67
chr4-153271958-DEL-66
chr4-153274602-DEL-56
chr4-153279621-DEL-528
chr4-153288170-DEL-724
chr4-153299071-DEL-583
chr4-153326614-DEL-54
chr4-153337569-DEL-200
chr4-153338379-DEL-366
chr4-153365109-DEL-60
chr4-153385281-DEL-56
chr4-153422964-DEL-54
chr4-153446573-DEL-54
chr4-153451826-DEL-237
chr4-153474673-DEL-288
chr4-153475032-DEL-256
chr4-153481997-DEL-117
chr4-153488552-DEL-95
chr4-153499390-DEL-625
chr4-153537685-DEL-76
chr4-153557203-DEL-80
chr4-153559481-DEL-159
chr4-153587280-DEL-300
chr4-153595041-DEL-131
chr4-153605848-DEL-1763
chr4-153651781-DEL-96
chr4-153690584-DEL-60
chr4-153713206-DEL-400
chr4-153751330-DEL-98
chr4-153754734-DEL-166
chr4-153757506-DEL-66
chr4-153758253-DEL-431
chr4-153769854-DEL-286
chr4-153791969-DEL-94
chr4-153805494-DEL-92
chr4-153826848-DEL-66
chr4-153841212-DEL-72
chr4-153842571-DEL-242
chr4-153859663-DEL-53
chr4-153892072-DEL-317
chr4-153901508-DEL-90
chr4-153907342-DEL-118
chr4-153915215-DEL-83
chr4-153915414-DEL-501
chr4-153930513-DEL-116
chr4-153964836-DEL-348
chr4-153977896-DEL-3522
chr4-153996373-DEL-137
chr4-154004171-DEL-192
chr4-154009732-DEL-142
chr4-154066652-DEL-87

chr4-154070519-DEL-96
chr4-154078980-DEL-84
chr4-154084750-DEL-478
chr4-154084771-DEL-1438
chr4-154087275-DEL-135
chr4-154089548-DEL-93
chr4-154104689-DEL-125
chr4-154116679-DEL-70
chr4-154134551-DEL-52
chr4-154144280-DEL-510
chr4-154147633-DEL-102
chr4-154159976-DEL-197
chr4-154176643-DEL-181
chr4-154177242-DEL-72
chr4-154204589-DEL-85
chr4-154211066-DEL-134
chr4-154228723-DEL-100
chr4-154232643-DEL-66
chr4-154233520-DEL-197
chr4-154259064-DEL-70
chr4-154264440-DEL-428
chr4-154293066-DEL-232
chr4-154295037-DEL-56
chr4-154434373-DEL-59
chr4-154444653-DEL-61
chr4-154458633-DEL-167
chr4-154461951-DEL-795
chr4-154484466-DEL-102
chr4-154489456-DEL-241
chr4-154498962-DEL-60
chr4-154501756-DEL-330
chr4-154520831-DEL-104
chr4-154530628-DEL-59
chr4-154571343-DEL-103
chr4-154602406-DEL-51
chr4-154610618-DEL-124
chr4-154619620-DEL-76
chr4-154644995-DEL-113
chr4-154709879-DEL-56
chr4-154723437-DEL-71
chr4-154754733-DEL-113
chr4-154757187-DEL-52
chr4-154760831-DEL-52
chr4-154765180-DEL-127
chr4-154771485-DEL-146
chr4-154772347-DEL-2052
chr4-154779930-DEL-52
chr4-154852425-DEL-825
chr4-154861122-DEL-428
chr4-154922268-DEL-296
chr4-154925160-DEL-2908
chr4-154925450-DEL-3202
chr4-154925798-DEL-71
chr4-154927477-DEL-2346
chr4-154928205-DEL-447
chr4-154928937-DEL-959
chr4-154930057-DEL-5421
chr4-154930243-DEL-2524
chr4-154930441-DEL-1134
chr4-154931712-DEL-3723
chr4-154931738-DEL-487
chr4-154932938-DEL-540
chr4-154934238-DEL-222
chr4-154935797-DEL-1258
chr4-154935797-DEL-936
chr4-154946863-DEL-101
chr4-154956990-DEL-188
chr4-154960564-DEL-1032
chr4-154974482-DEL-80
chr4-154988896-DEL-208
chr4-154993650-DEL-432
chr4-154996056-DEL-174
chr4-154996647-DEL-143
chr4-155014891-DEL-1880
chr4-155029310-DEL-189
chr4-155031306-DEL-280
chr4-155032388-DEL-207

chr4-155035140-DEL-346
chr4-155081032-DEL-75
chr4-155137366-DEL-300
chr4-155210861-DEL-114
chr4-155221144-DEL-287
chr4-155231979-DEL-64
chr4-155282748-DEL-372
chr4-155284311-DEL-723
chr4-155297122-DEL-100
chr4-155313433-DEL-140
chr4-155316845-DEL-204
chr4-155341466-DEL-195
chr4-155364129-DEL-150
chr4-155394480-DEL-203
chr4-155395347-DEL-63
chr4-155411214-DEL-215
chr4-155434094-DEL-179
chr4-155434277-DEL-855
chr4-155452674-DEL-239
chr4-155453533-DEL-137
chr4-155476837-DEL-112
chr4-155486553-DEL-202
chr4-155497580-DEL-107
chr4-155508817-DEL-73
chr4-155527770-DEL-79
chr4-155530061-DEL-57
chr4-155533674-DEL-390
chr4-155537632-DEL-150
chr4-155538964-DEL-90
chr4-155543640-DEL-60
chr4-155551095-DEL-56
chr4-155559585-DEL-58
chr4-155561145-DEL-322
chr4-155563402-DEL-227
chr4-155601743-DEL-214
chr4-155615121-DEL-53
chr4-155627248-DEL-56
chr4-155631605-DEL-78
chr4-155671392-DEL-236
chr4-155690219-DEL-159
chr4-155707934-DEL-58
chr4-155712649-DEL-198
chr4-155717731-DEL-59
chr4-155736283-DEL-63
chr4-155746913-DEL-213
chr4-155773161-DEL-224
chr4-155787642-DEL-140
chr4-155797794-DEL-737
chr4-155798968-DEL-62
chr4-155836962-DEL-56
chr4-155849221-DEL-1262
chr4-155880180-DEL-133
chr4-155887602-DEL-204
chr4-155890254-DEL-93
chr4-155900408-DEL-353
chr4-155968244-DEL-446
chr4-155969132-DEL-61
chr4-155981809-DEL-921
chr4-155982437-DEL-290
chr4-155995774-DEL-2023
chr4-156007936-DEL-1818
chr4-156012792-DEL-202
chr4-156030163-DEL-185
chr4-156054050-DEL-205
chr4-156060847-DEL-308
chr4-156062408-DEL-176
chr4-156069237-DEL-238
chr4-156077226-DEL-192
chr4-156085808-DEL-615
chr4-156101818-DEL-86
chr4-156121637-DEL-185
chr4-156126047-DEL-70
chr4-156130367-DEL-215
chr4-156195559-DEL-183
chr4-156199817-DEL-1356
chr4-156221537-DEL-1150
chr4-156221608-DEL-126

chr4-156222283-DEL-135
chr4-156237792-DEL-114
chr4-156247741-DEL-52
chr4-156268575-DEL-179
chr4-156281957-DEL-97
chr4-156286131-DEL-87
chr4-156321062-DEL-100
chr4-156326310-DEL-82
chr4-156330655-DEL-206
chr4-156337395-DEL-1451
chr4-156370937-DEL-88
chr4-156372925-DEL-211
chr4-156392038-DEL-104
chr4-156440573-DEL-5848
chr4-156496467-DEL-72
chr4-156496725-DEL-60
chr4-156497779-DEL-486
chr4-156498013-DEL-664
chr4-156498782-DEL-903
chr4-156499791-DEL-264
chr4-156501447-DEL-190
chr4-156502753-DEL-203
chr4-156509316-DEL-7218
chr4-156521614-DEL-1270
chr4-156556848-DEL-53
chr4-156564229-DEL-166
chr4-156565728-DEL-119
chr4-156566454-DEL-106
chr4-156567529-DEL-96
chr4-156590104-DEL-202
chr4-156593953-DEL-5690
chr4-156597628-DEL-293
chr4-156679718-DEL-330
chr4-156680653-DEL-151
chr5-3134300-DEL-921
chr5-3134509-DEL-3608
chr5-3149550-DEL-287
chr5-3178853-DEL-341
chr5-3184391-DEL-358
chr5-3190709-DEL-343
chr5-3229968-DEL-7047
chr5-3230444-DEL-5234
chr5-3236576-DEL-225
chr5-3239022-DEL-194
chr5-3241178-DEL-453
chr5-3256591-DEL-54
chr5-3271549-DEL-115
chr5-3290000-DEL-2991
chr5-3293591-DEL-114
chr5-3307698-DEL-85
chr5-3333724-DEL-157
chr5-3333904-DEL-1896
chr5-3337190-DEL-1496
chr5-3367555-DEL-201
chr5-3429682-DEL-291
chr5-3445902-DEL-55
chr5-3446340-DEL-2192
chr5-3473254-DEL-2434
chr5-3486815-DEL-204
chr5-3487817-DEL-179
chr5-3559395-DEL-118
chr5-3561861-DEL-70
chr5-3575950-DEL-830
chr5-3656764-DEL-125
chr5-3660600-DEL-216
chr5-3660928-DEL-216
chr5-3661268-DEL-378
chr5-3663781-DEL-528
chr5-3679190-DEL-75
chr5-3733230-DEL-113
chr5-3737277-DEL-106
chr5-3763691-DEL-186
chr5-3778157-DEL-1146
chr5-3783120-DEL-84
chr5-3806113-DEL-635
chr5-3810646-DEL-6303
chr5-3819975-DEL-188

chr5-3898903-DEL-54
chr5-3904646-DEL-480
chr5-3933636-DEL-72
chr5-3943591-DEL-53
chr5-3971508-DEL-55
chr5-3981896-DEL-60
chr5-3995136-DEL-1308
chr5-4007448-DEL-106
chr5-4012460-DEL-300
chr5-4012614-DEL-180
chr5-4030704-DEL-52
chr5-4089602-DEL-75
chr5-4101317-DEL-443
chr5-4102214-DEL-86
chr5-4102259-DEL-1693
chr5-4102543-DEL-193
chr5-4102696-DEL-118
chr5-4103517-DEL-899
chr5-4103561-DEL-85
chr5-4104164-DEL-106
chr5-4107491-DEL-97
chr5-4116765-DEL-141
chr5-4118223-DEL-205
chr5-4121344-DEL-54
chr5-4123377-DEL-64
chr5-4128947-DEL-84
chr5-4147369-DEL-275
chr5-4167637-DEL-274
chr5-4184374-DEL-59
chr5-4206023-DEL-66
chr5-4211576-DEL-114
chr5-4214107-DEL-197
chr5-4239112-DEL-340
chr5-4239895-DEL-67
chr5-4242648-DEL-53
chr5-4244527-DEL-232
chr5-4263952-DEL-288
chr5-4282872-DEL-168
chr5-4313396-DEL-66
chr5-4315064-DEL-504
chr5-4316252-DEL-454
chr5-4369625-DEL-80
chr5-4402347-DEL-1474
chr5-4452690-DEL-1064
chr5-4542004-DEL-52
chr5-4543591-DEL-54
chr5-4551407-DEL-108
chr5-4588422-DEL-51
chr5-4597816-DEL-331
chr5-4606914-DEL-54
chr5-4666292-DEL-52
chr5-4720376-DEL-449
chr5-4729961-DEL-2738
chr5-4749078-DEL-201
chr5-4774592-DEL-6236
chr5-4813298-DEL-59
chr5-4825134-DEL-54
chr5-4827973-DEL-66
chr5-4846406-DEL-2138
chr5-4873350-DEL-74
chr5-4917191-DEL-81
chr5-4963625-DEL-81
chr5-4971239-DEL-421
chr5-4986694-DEL-91
chr5-5000142-DEL-58
chr5-5021962-DEL-52
chr5-5038216-DEL-102
chr5-5059974-DEL-122
chr5-5061745-DEL-108
chr5-5142216-DEL-167
chr5-5179331-DEL-106
chr5-5201689-DEL-70
chr5-5223881-DEL-204
chr5-5263856-DEL-95
chr5-5277913-DEL-1286
chr5-5278326-DEL-150
chr5-5323338-DEL-171

chr5-5339718-DEL-218
chr5-5349730-DEL-334
chr5-5402789-DEL-204
chr5-5449304-DEL-79
chr5-5537750-DEL-93
chr5-5549173-DEL-260
chr5-5568056-DEL-170
chr5-5570537-DEL-6441
chr5-5578307-DEL-135
chr5-5621533-DEL-633
chr5-5647175-DEL-6156
chr5-5691094-DEL-191
chr5-5710050-DEL-147
chr5-5714547-DEL-54
chr5-5721319-DEL-59
chr5-5752373-DEL-371
chr5-5770855-DEL-3174
chr5-5791202-DEL-141
chr5-5803727-DEL-850
chr5-5810988-DEL-483
chr5-5831499-DEL-2147
chr5-5844272-DEL-103
chr5-5849482-DEL-99
chr5-5849884-DEL-7021
chr5-5860094-DEL-3485
chr5-5861657-DEL-63
chr5-5902507-DEL-377
chr5-5903624-DEL-486
chr5-5904114-DEL-3041
chr5-5907523-DEL-191
chr5-5907755-DEL-2160
chr5-5909933-DEL-1300
chr5-5913337-DEL-449
chr5-5915822-DEL-52
chr5-5930082-DEL-52
chr5-5942939-DEL-109
chr5-5953555-DEL-1479
chr5-5960876-DEL-204
chr5-5968281-DEL-311
chr5-6008747-DEL-102
chr5-6033096-DEL-161
chr5-6082336-DEL-6220
chr5-6099047-DEL-51
chr5-6104103-DEL-400
chr5-6104859-DEL-171
chr5-6130671-DEL-11763
chr5-6134170-DEL-2021
chr5-6139456-DEL-52
chr5-6150334-DEL-846
chr5-6153154-DEL-80
chr5-6232546-DEL-145
chr5-6244836-DEL-2517
chr5-6259930-DEL-6357
chr5-6268602-DEL-1244
chr5-6270171-DEL-137
chr5-6272106-DEL-239
chr5-6272421-DEL-10767
chr5-6285554-DEL-367
chr5-6285890-DEL-604
chr5-6286301-DEL-1345
chr5-6288089-DEL-82
chr5-6321727-DEL-3751
chr5-6359472-DEL-6481
chr5-6436530-DEL-102
chr5-6441969-DEL-1087
chr5-6454418-DEL-89
chr5-6465686-DEL-6165
chr5-6497146-DEL-62
chr5-6498718-DEL-170
chr5-6503512-DEL-96
chr5-6506371-DEL-329
chr5-6515671-DEL-6427
chr5-6523648-DEL-73
chr5-6550332-DEL-56
chr5-6560153-DEL-6168
chr5-6561459-DEL-458
chr5-6561899-DEL-458

chr5-6562501-DEL-592
chr5-6572384-DEL-1158
chr5-6587977-DEL-176
chr5-6612127-DEL-4514
chr5-6621086-DEL-184
chr5-6669663-DEL-2045
chr5-6671060-DEL-1705
chr5-6678154-DEL-88
chr5-6721966-DEL-51
chr5-6734386-DEL-4539
chr5-6739382-DEL-1443
chr5-6750057-DEL-175
chr5-6753458-DEL-176
chr5-6770969-DEL-841
chr5-6771880-DEL-157
chr5-6773839-DEL-461
chr5-6805157-DEL-1042
chr5-6834550-DEL-62
chr5-6850841-DEL-2526
chr5-6866129-DEL-901
chr5-6867665-DEL-162
chr5-6894759-DEL-2004
chr5-6900643-DEL-1338
chr5-6905896-DEL-413
chr5-6928460-DEL-151
chr5-6967949-DEL-59
chr5-6986555-DEL-543
chr5-7055034-DEL-219
chr5-7069104-DEL-54
chr5-7079907-DEL-2964
chr5-7113060-DEL-118
chr5-7135469-DEL-424
chr5-7154703-DEL-520
chr5-7229286-DEL-1564
chr5-7250069-DEL-7132
chr5-7277649-DEL-875
chr5-7326490-DEL-2867
chr5-7345555-DEL-148
chr5-7358131-DEL-12032
chr5-7371329-DEL-640
chr5-7372126-DEL-52
chr5-7413192-DEL-65
chr5-7454454-DEL-17197
chr5-7460526-DEL-623
chr5-7462959-DEL-142
chr5-7472986-DEL-59
chr5-7495582-DEL-609
chr5-7515580-DEL-2649
chr5-7548142-DEL-897
chr5-7551279-DEL-8444
chr5-7572079-DEL-103
chr5-7575488-DEL-6438
chr5-7623177-DEL-1636
chr5-7707764-DEL-150
chr5-7734507-DEL-1171
chr5-7737252-DEL-1654
chr5-7739848-DEL-324
chr5-7742972-DEL-51
chr5-7751057-DEL-1828
chr5-7759999-DEL-355
chr5-7766001-DEL-131
chr5-7769587-DEL-72
chr5-7774541-DEL-56
chr5-7790268-DEL-838
chr5-7834968-DEL-72
chr5-7841616-DEL-61
chr5-7857177-DEL-3977
chr5-7869897-DEL-1339
chr5-7936355-DEL-161
chr5-7943183-DEL-6221
chr5-7950967-DEL-64
chr5-7971945-DEL-54
chr5-7987265-DEL-13636
chr5-8042386-DEL-58
chr5-8067762-DEL-7951
chr5-8080854-DEL-594
chr5-8092846-DEL-312

chr5-8110775-DEL-200
chr5-8115241-DEL-338
chr5-8115255-DEL-510
chr5-8117357-DEL-99
chr5-8133379-DEL-288
chr5-8146845-DEL-494
chr5-8155286-DEL-70
chr5-8161444-DEL-84
chr5-8176693-DEL-276
chr5-8191930-DEL-161
chr5-8197308-DEL-229
chr5-8218077-DEL-7941
chr5-8228218-DEL-151
chr5-8229041-DEL-260
chr5-8242634-DEL-565
chr5-8256414-DEL-357
chr5-8297124-DEL-274
chr5-8301321-DEL-878
chr5-8359070-DEL-119
chr5-8421781-DEL-60
chr5-8451108-DEL-61
chr5-8470699-DEL-56
chr5-8483692-DEL-2778
chr5-8488097-DEL-198
chr5-8491593-DEL-653
chr5-8492186-DEL-98
chr5-8534613-DEL-68
chr5-8534918-DEL-418
chr5-8538983-DEL-220
chr5-8548131-DEL-200
chr5-8555596-DEL-60
chr5-8590108-DEL-94
chr5-8593054-DEL-863
chr5-8649425-DEL-215
chr5-8703041-DEL-208
chr5-8727262-DEL-74
chr5-8764203-DEL-70
chr5-8791837-DEL-2858
chr5-8805115-DEL-209
chr5-8813789-DEL-106
chr5-8814503-DEL-6233
chr5-8834048-DEL-1072
chr5-8837396-DEL-174
chr5-8839618-DEL-288
chr5-8841219-DEL-55
chr5-8846334-DEL-54
chr5-8870390-DEL-246
chr5-8880065-DEL-598
chr5-8957715-DEL-53
chr5-8987053-DEL-203
chr5-8994544-DEL-150
chr5-8994619-DEL-650
chr5-9021897-DEL-525
chr5-9029514-DEL-60
chr5-9030110-DEL-261
chr5-9044415-DEL-183
chr5-9047340-DEL-73
chr5-9060053-DEL-98
chr5-9066399-DEL-159
chr5-9069362-DEL-70
chr5-9073466-DEL-2175
chr5-9074780-DEL-2038
chr5-9089041-DEL-246
chr5-9096838-DEL-182
chr5-9128516-DEL-166
chr5-9139137-DEL-87
chr5-9146786-DEL-848
chr5-9205217-DEL-136
chr5-9215432-DEL-165
chr5-9222825-DEL-98
chr5-9289218-DEL-1067
chr5-9294881-DEL-56
chr5-9303399-DEL-242
chr5-9312883-DEL-175
chr5-9339603-DEL-4006
chr5-9341070-DEL-4109
chr5-9370353-DEL-896

chr5-9392395-DEL-1518
chr5-9393920-DEL-72
chr5-9431364-DEL-286
chr5-9451845-DEL-6440
chr5-9461356-DEL-82
chr5-9502475-DEL-730
chr5-9523912-DEL-243
chr5-9590498-DEL-51
chr5-9617706-DEL-54
chr5-9655073-DEL-2547
chr5-9666260-DEL-74
chr5-9667754-DEL-152
chr5-9668829-DEL-466
chr5-9675597-DEL-1315
chr5-9735321-DEL-160
chr5-9778329-DEL-53
chr5-9798853-DEL-54
chr5-9809094-DEL-115
chr5-9812737-DEL-54
chr5-9846852-DEL-77
chr5-9871876-DEL-402
chr5-9898650-DEL-8812
chr5-9918797-DEL-55
chr5-9971885-DEL-826
chr5-9987031-DEL-60
chr5-10054520-DEL-1467
chr5-10075461-DEL-51
chr5-10101571-DEL-310
chr5-10114899-DEL-67
chr5-10118063-DEL-78
chr5-10139746-DEL-55
chr5-10155308-DEL-400
chr5-10155340-DEL-74
chr5-10157325-DEL-62
chr5-10159894-DEL-56
chr5-10182511-DEL-428
chr5-10198788-DEL-633
chr5-10208567-DEL-98
chr5-10235599-DEL-2445
chr5-10250447-DEL-330
chr5-10251763-DEL-100
chr5-10256438-DEL-113
chr5-10260924-DEL-195
chr5-10261112-DEL-4939
chr5-10296958-DEL-1623
chr5-10298037-DEL-811
chr5-10310572-DEL-3222
chr5-10319628-DEL-65
chr5-10337567-DEL-64
chr5-10345919-DEL-2165
chr5-10353629-DEL-57
chr5-10381634-DEL-494
chr5-10389599-DEL-61
chr5-10405053-DEL-9718
chr5-10442309-DEL-58
chr5-10445992-DEL-90
chr5-10446232-DEL-2584
chr5-10516268-DEL-67
chr5-10529872-DEL-1037
chr5-10531490-DEL-68
chr5-10543921-DEL-87
chr5-10576087-DEL-357
chr5-10577363-DEL-1461
chr5-10589503-DEL-26525
chr5-10596386-DEL-417
chr5-10626430-DEL-1336
chr5-10630411-DEL-1259
chr5-10637136-DEL-311
chr5-10645484-DEL-1433
chr5-10660569-DEL-122
chr5-10681274-DEL-886
chr5-10684861-DEL-120
chr5-10702091-DEL-591
chr5-10717350-DEL-469
chr5-10750643-DEL-2638
chr5-10767752-DEL-1965
chr5-10782015-DEL-332

chr5-10792042-DEL-772
chr5-10797545-DEL-51
chr5-10821210-DEL-398
chr5-10839486-DEL-527
chr5-10900315-DEL-100
chr5-10934105-DEL-56
chr5-10957975-DEL-6350
chr5-10980369-DEL-1093
chr5-10998655-DEL-67
chr5-11007421-DEL-4526
chr5-11021677-DEL-3397
chr5-11036524-DEL-1051
chr5-11051078-DEL-304
chr5-11051697-DEL-1043
chr5-11052692-DEL-766
chr5-11066771-DEL-2684
chr5-11074526-DEL-220
chr5-11114193-DEL-208
chr5-11178528-DEL-71
chr5-11196485-DEL-1041
chr5-11198274-DEL-137
chr5-11205359-DEL-487
chr5-11208114-DEL-59
chr5-11219191-DEL-3897
chr5-11258650-DEL-192
chr5-11267523-DEL-1657
chr5-11282751-DEL-125
chr5-11297464-DEL-60
chr5-11306812-DEL-70
chr5-11326753-DEL-529
chr5-11330444-DEL-430
chr5-11375762-DEL-188
chr5-11384428-DEL-173
chr5-11411145-DEL-1378
chr5-11418844-DEL-2139
chr5-11421150-DEL-9717
chr5-11467253-DEL-105
chr5-11505460-DEL-1045
chr5-11508368-DEL-6554
chr5-11508431-DEL-204
chr5-11591372-DEL-1051
chr5-11645867-DEL-70
chr5-11668942-DEL-6402
chr5-11717483-DEL-74
chr5-11732864-DEL-70
chr5-11766102-DEL-115
chr5-11804786-DEL-116
chr5-11813416-DEL-174
chr5-11830149-DEL-4508
chr5-11848181-DEL-1714
chr5-11861212-DEL-1041
chr5-11873145-DEL-1239
chr5-11896373-DEL-105
chr5-11905103-DEL-4508
chr5-11916489-DEL-90
chr5-11955064-DEL-168
chr5-12027888-DEL-1434
chr5-12028109-DEL-4393
chr5-12080523-DEL-77
chr5-12084541-DEL-72
chr5-12102851-DEL-60
chr5-12114306-DEL-104
chr5-12140875-DEL-138
chr5-12162993-DEL-208
chr5-12170052-DEL-114
chr5-12206543-DEL-280
chr5-12208446-DEL-84
chr5-12234612-DEL-298
chr5-12251989-DEL-417
chr5-12261186-DEL-211
chr5-12269079-DEL-70
chr5-12313806-DEL-272
chr5-12314602-DEL-591
chr5-12397605-DEL-1494
chr5-12404632-DEL-1937
chr5-12409549-DEL-2833
chr5-12426676-DEL-74

chr5-12458147-DEL-1444
chr5-12514935-DEL-57
chr5-12531410-DEL-297
chr5-12551347-DEL-59
chr5-12558431-DEL-430
chr5-12578617-DEL-578
chr5-12603919-DEL-670
chr5-12614265-DEL-157
chr5-12634512-DEL-70
chr5-12649907-DEL-52
chr5-12672835-DEL-155
chr5-12717226-DEL-1982
chr5-12729876-DEL-313
chr5-12734152-DEL-238
chr5-12766299-DEL-62
chr5-12773663-DEL-1929
chr5-12810807-DEL-171
chr5-12831463-DEL-84
chr5-12842199-DEL-378
chr5-12864030-DEL-3443
chr5-12881780-DEL-564
chr5-12883524-DEL-54
chr5-12913258-DEL-441
chr5-12913833-DEL-74
chr5-12919492-DEL-126
chr5-12927732-DEL-53
chr5-12943430-DEL-130
chr5-12944179-DEL-384
chr5-12945890-DEL-189
chr5-12957550-DEL-66
chr5-12961064-DEL-441
chr5-12984328-DEL-507
chr5-13022122-DEL-76
chr5-13035174-DEL-206
chr5-13078641-DEL-798
chr5-13092103-DEL-56
chr5-13092141-DEL-1333
chr5-13100318-DEL-76
chr5-13101728-DEL-6209
chr5-13128117-DEL-77
chr5-13139987-DEL-336
chr5-13144617-DEL-56
chr5-13198850-DEL-84
chr5-13215108-DEL-72
chr5-13241832-DEL-105
chr5-13242499-DEL-495
chr5-13243617-DEL-351
chr5-13255167-DEL-57
chr5-13297187-DEL-121
chr5-13317961-DEL-85
chr5-13354614-DEL-51
chr5-13405124-DEL-633
chr5-13453393-DEL-830
chr5-13501719-DEL-94
chr5-13513163-DEL-107
chr5-13553592-DEL-67
chr5-13574694-DEL-262
chr5-13608084-DEL-53
chr5-13616080-DEL-506
chr5-13622658-DEL-174
chr5-13670052-DEL-68
chr5-13724655-DEL-6070
chr5-13732025-DEL-1900
chr5-13752736-DEL-1813
chr5-13841902-DEL-400
chr5-13868583-DEL-178
chr5-13916964-DEL-1555
chr5-13920007-DEL-1787
chr5-13924133-DEL-74
chr5-13930470-DEL-408
chr5-13964662-DEL-281
chr5-13965655-DEL-75
chr5-13987207-DEL-78
chr5-13999291-DEL-52
chr5-14000193-DEL-72
chr5-14013999-DEL-459
chr5-14100666-DEL-193

chr5-14140931-DEL-55
chr5-14187965-DEL-52
chr5-14195993-DEL-68
chr5-14200664-DEL-64
chr5-14206754-DEL-93
chr5-14256497-DEL-129
chr5-14266510-DEL-678
chr5-14267129-DEL-285
chr5-14286516-DEL-121
chr5-14319174-DEL-7907
chr5-14335704-DEL-397
chr5-14341391-DEL-1617
chr5-14345783-DEL-6997
chr5-14353981-DEL-58
chr5-14354877-DEL-1097
chr5-14370000-DEL-58
chr5-14378275-DEL-123
chr5-14386896-DEL-60
chr5-14393896-DEL-58
chr5-14414990-DEL-1510
chr5-14439022-DEL-305
chr5-14457709-DEL-2523
chr5-14493597-DEL-2125
chr5-14503036-DEL-2627
chr5-14506273-DEL-1042
chr5-14507347-DEL-1167
chr5-14605408-DEL-415
chr5-14638632-DEL-52
chr5-14683818-DEL-2322
chr5-14686281-DEL-182
chr5-14687653-DEL-801
chr5-14689128-DEL-78
chr5-14690511-DEL-1343
chr5-14706295-DEL-444
chr5-14708715-DEL-199
chr5-14710575-DEL-1724
chr5-14715692-DEL-881
chr5-14718145-DEL-108
chr5-14790175-DEL-128
chr5-14833908-DEL-82
chr5-14957444-DEL-2649
chr5-15014121-DEL-6358
chr5-15096720-DEL-1607
chr5-15100000-DEL-929
chr5-15696690-DEL-5550
chr5-15704189-DEL-4059
chr5-15711722-DEL-512
chr5-15747289-DEL-4971
chr5-15750315-DEL-2014
chr5-15769275-DEL-1574
chr5-15773211-DEL-141
chr5-15775511-DEL-985
chr5-15863073-DEL-66
chr5-15922805-DEL-67
chr5-15986342-DEL-241
chr5-16001690-DEL-452
chr5-16017060-DEL-1338
chr5-16029193-DEL-566
chr5-16064834-DEL-255
chr5-16186857-DEL-311
chr5-16225376-DEL-315
chr5-16241176-DEL-583
chr5-16243296-DEL-296
chr5-16251493-DEL-268
chr5-16284253-DEL-394
chr5-16333532-DEL-752
chr5-16347498-DEL-72
chr5-16362955-DEL-62
chr5-16384889-DEL-361
chr5-16404330-DEL-370
chr5-16420802-DEL-55
chr5-16455595-DEL-745
chr5-16477730-DEL-561
chr5-16501609-DEL-117
chr5-16505447-DEL-88
chr5-16552266-DEL-217
chr5-16571567-DEL-780

chr5-16571928-DEL-114
chr5-16580510-DEL-511
chr5-16581974-DEL-1172
chr5-16582192-DEL-724
chr5-16583085-DEL-63
chr5-16594430-DEL-416
chr5-16609087-DEL-5188
chr5-16645670-DEL-72
chr5-16679372-DEL-322
chr5-16691072-DEL-1871
chr5-16698795-DEL-61
chr5-16725379-DEL-450
chr5-16726055-DEL-72
chr5-16726735-DEL-59
chr5-16749313-DEL-59
chr5-16770796-DEL-266
chr5-16773295-DEL-139
chr5-16839864-DEL-95
chr5-16899758-DEL-1721
chr5-16912588-DEL-69
chr5-16922639-DEL-5456
chr5-16943627-DEL-6292
chr5-16953002-DEL-192
chr5-17019631-DEL-89
chr5-17054323-DEL-82
chr5-17080456-DEL-2188
chr5-17101309-DEL-490
chr5-17102090-DEL-254
chr5-17124920-DEL-201
chr5-17145433-DEL-17569
chr5-17182017-DEL-5699
chr5-17194757-DEL-406
chr5-17220398-DEL-114
chr5-17228565-DEL-920
chr5-17231642-DEL-176
chr5-17237834-DEL-176
chr5-17247791-DEL-6461
chr5-17259087-DEL-1939
chr5-17274884-DEL-81
chr5-17338440-DEL-129
chr5-17352334-DEL-427
chr5-17402165-DEL-725
chr5-17451501-DEL-212
chr5-17468241-DEL-1335
chr5-17479932-DEL-212
chr5-17517842-DEL-6311
chr5-17585614-DEL-511
chr5-17627461-DEL-228
chr5-17662768-DEL-932
chr5-17696177-DEL-105
chr5-17711623-DEL-58
chr5-17737969-DEL-2548
chr5-17743231-DEL-120
chr5-17812462-DEL-9243
chr5-17812980-DEL-8823
chr5-17858252-DEL-70
chr5-17863060-DEL-74
chr5-17881859-DEL-407
chr5-17938072-DEL-756
chr5-17946280-DEL-143
chr5-17954299-DEL-90
chr5-17975330-DEL-78
chr5-17980919-DEL-952
chr5-17991182-DEL-63
chr5-18004329-DEL-212
chr5-18017918-DEL-64
chr5-18028270-DEL-56
chr5-18031246-DEL-86
chr5-18064959-DEL-94
chr5-18083402-DEL-1465
chr5-18124591-DEL-2077
chr5-18140954-DEL-157
chr5-18145306-DEL-996
chr5-18165362-DEL-62
chr5-18194819-DEL-251
chr5-18220806-DEL-58
chr5-18226529-DEL-71

chr5-18230955-DEL-134
chr5-18262406-DEL-78
chr5-18263669-DEL-68
chr5-18306938-DEL-54
chr5-18358499-DEL-180
chr5-18361835-DEL-67
chr5-18367147-DEL-66
chr5-18369806-DEL-494
chr5-18377624-DEL-1721
chr5-18409147-DEL-99
chr5-18474180-DEL-188
chr5-18546983-DEL-98
chr5-18588473-DEL-895
chr5-18589450-DEL-846
chr5-18592506-DEL-1920
chr5-18602914-DEL-6195
chr5-18611003-DEL-226
chr5-18612795-DEL-70
chr5-18620823-DEL-156
chr5-18624037-DEL-105
chr5-18636014-DEL-89
chr5-18645413-DEL-85
chr5-18688766-DEL-1959
chr5-18701300-DEL-1568
chr5-18719947-DEL-213
chr5-18728884-DEL-228
chr5-18733382-DEL-254
chr5-18759327-DEL-534
chr5-18777108-DEL-366
chr5-18857347-DEL-522
chr5-18857436-DEL-522
chr5-18908518-DEL-6213
chr5-18991857-DEL-51
chr5-19007772-DEL-734
chr5-19016324-DEL-184
chr5-19053755-DEL-476
chr5-19055087-DEL-682
chr5-19070541-DEL-55
chr5-19078796-DEL-3375
chr5-19091785-DEL-1063
chr5-19098920-DEL-64
chr5-19109402-DEL-53
chr5-19111276-DEL-75
chr5-19156201-DEL-565
chr5-19165436-DEL-85
chr5-19204539-DEL-147
chr5-19234477-DEL-56
chr5-19248980-DEL-54
chr5-19254272-DEL-148
chr5-19260155-DEL-57
chr5-19277061-DEL-59
chr5-19282647-DEL-90
chr5-19286151-DEL-57
chr5-19300152-DEL-958
chr5-19323156-DEL-371
chr5-19328992-DEL-4069
chr5-19370535-DEL-126
chr5-19373484-DEL-2016
chr5-19376936-DEL-1571
chr5-19400322-DEL-73
chr5-19413677-DEL-194
chr5-19422085-DEL-2777
chr5-19428192-DEL-68
chr5-19438575-DEL-72
chr5-19448453-DEL-52
chr5-19456409-DEL-424
chr5-19466097-DEL-646
chr5-19484434-DEL-91
chr5-19492507-DEL-333
chr5-19495334-DEL-115
chr5-19529833-DEL-85
chr5-19610951-DEL-5946
chr5-19671582-DEL-110
chr5-19807602-DEL-234
chr5-19808327-DEL-214
chr5-19828074-DEL-293
chr5-19854128-DEL-206

chr5-19862330-DEL-1283
chr5-19944003-DEL-124
chr5-19951434-DEL-80
chr5-19960602-DEL-256
chr5-19981437-DEL-90
chr5-20016902-DEL-199
chr5-20022896-DEL-127
chr5-20042430-DEL-141
chr5-20047666-DEL-204
chr5-20056440-DEL-62
chr5-20064738-DEL-127
chr5-20087457-DEL-2424
chr5-20088463-DEL-2415
chr5-20108326-DEL-100
chr5-20146081-DEL-112
chr5-20153441-DEL-78
chr5-20167608-DEL-214
chr5-20170429-DEL-390
chr5-20179672-DEL-135
chr5-20235232-DEL-262
chr5-20236298-DEL-1391
chr5-20264481-DEL-69
chr5-20290654-DEL-61
chr5-20292794-DEL-287
chr5-20296305-DEL-197
chr5-20300220-DEL-80
chr5-20316334-DEL-90
chr5-20326203-DEL-6634
chr5-20344151-DEL-383
chr5-20415700-DEL-335
chr5-20443275-DEL-60
chr5-20444936-DEL-61
chr5-20463024-DEL-750
chr5-20478120-DEL-1217
chr5-20514593-DEL-130
chr5-20519644-DEL-444
chr5-20542396-DEL-1548
chr5-20576114-DEL-230
chr5-20599863-DEL-124
chr5-20630176-DEL-8239
chr5-20632799-DEL-1890
chr5-20636500-DEL-165
chr5-20639889-DEL-60
chr5-20652880-DEL-235
chr5-20660862-DEL-100
chr5-20716628-DEL-443
chr5-20731805-DEL-872
chr5-20834892-DEL-208
chr5-20848393-DEL-92
chr5-20850934-DEL-269
chr5-20910946-DEL-53
chr5-20911781-DEL-548
chr5-20942415-DEL-153
chr5-20954632-DEL-98
chr5-20991714-DEL-304
chr5-20994279-DEL-126
chr5-21014894-DEL-205
chr5-21020638-DEL-170
chr5-21027511-DEL-66
chr5-21063008-DEL-187
chr5-21064686-DEL-97
chr5-21075248-DEL-853
chr5-21084499-DEL-72
chr5-21133574-DEL-52
chr5-21164680-DEL-679
chr5-21175707-DEL-55
chr5-21177012-DEL-85
chr5-21183495-DEL-1026
chr5-21183823-DEL-693
chr5-21228475-DEL-153
chr5-21249196-DEL-364
chr5-21250194-DEL-65
chr5-21277325-DEL-337
chr5-21282679-DEL-193
chr5-21325768-DEL-63
chr5-21390130-DEL-128
chr5-21392811-DEL-84

chr5-21396469-DEL-55
chr5-21402696-DEL-235
chr5-21423485-DEL-211
chr5-21434982-DEL-434
chr5-21471852-DEL-163
chr5-21473098-DEL-694
chr5-21497136-DEL-52
chr5-21527881-DEL-138
chr5-21532393-DEL-248
chr5-21535363-DEL-97
chr5-21540675-DEL-5423
chr5-21542975-DEL-129
chr5-21556421-DEL-314
chr5-21560892-DEL-440
chr5-21585156-DEL-89
chr5-21608037-DEL-4249
chr5-21621712-DEL-77
chr5-21623456-DEL-4569
chr5-21693363-DEL-66
chr5-21734922-DEL-568
chr5-21801039-DEL-711
chr5-21801470-DEL-1922
chr5-21803445-DEL-245
chr5-21804939-DEL-64
chr5-21816588-DEL-2961
chr5-21847690-DEL-192
chr5-21859651-DEL-1885
chr5-21871700-DEL-470
chr5-21872471-DEL-56
chr5-21888983-DEL-68
chr5-21890470-DEL-173
chr5-21900403-DEL-294
chr5-21922723-DEL-82
chr5-21957165-DEL-160
chr5-21957944-DEL-219
chr5-21971371-DEL-62
chr5-21995119-DEL-1381
chr5-22001302-DEL-79
chr5-22015019-DEL-105
chr5-22020183-DEL-88
chr5-22024958-DEL-63
chr5-22091098-DEL-52
chr5-22092468-DEL-66
chr5-22101129-DEL-70
chr5-22111276-DEL-171
chr5-22111687-DEL-240
chr5-22112437-DEL-109
chr5-22113772-DEL-332
chr5-22115430-DEL-170
chr5-22116164-DEL-92
chr5-22127535-DEL-52
chr5-22133465-DEL-544
chr5-22134327-DEL-61
chr5-22174729-DEL-117
chr5-22198032-DEL-1865
chr5-22203214-DEL-115
chr5-22228818-DEL-51
chr5-22241063-DEL-182
chr5-22267908-DEL-66
chr5-22271724-DEL-406
chr5-22275115-DEL-1840
chr5-22319216-DEL-599
chr5-22325671-DEL-60
chr5-22411546-DEL-97
chr5-22419012-DEL-6412
chr5-22513086-DEL-658
chr5-22558812-DEL-54
chr5-22561662-DEL-576
chr5-22563552-DEL-5880
chr5-22563733-DEL-244
chr5-22569458-DEL-4947
chr5-22583755-DEL-125
chr5-22586171-DEL-3121
chr5-22621712-DEL-70
chr5-22635814-DEL-2174
chr5-22647850-DEL-121
chr5-22649533-DEL-270

chr5-22650169-DEL-366
chr5-22651417-DEL-316
chr5-22652345-DEL-404
chr5-22652663-DEL-458
chr5-22696972-DEL-5388
chr5-22696974-DEL-77
chr5-22697073-DEL-2824
chr5-22697404-DEL-1613
chr5-22697826-DEL-76
chr5-22698341-DEL-96
chr5-22699092-DEL-806
chr5-22700226-DEL-2100
chr5-22702011-DEL-220
chr5-22717801-DEL-1570
chr5-22723288-DEL-812
chr5-22738473-DEL-59
chr5-22753100-DEL-109
chr5-22812911-DEL-430
chr5-22887327-DEL-486
chr5-22900082-DEL-6101
chr5-22947110-DEL-117
chr5-23024912-DEL-1637
chr5-23027299-DEL-94
chr5-23033597-DEL-98
chr5-23038429-DEL-1398
chr5-23066977-DEL-809
chr5-23123981-DEL-71
chr5-23125074-DEL-11976
chr5-23173315-DEL-880
chr5-23174084-DEL-397
chr5-23179668-DEL-854
chr5-23205491-DEL-277
chr5-23228169-DEL-242
chr5-23242757-DEL-108
chr5-23264406-DEL-399
chr5-23276837-DEL-54
chr5-23277774-DEL-198
chr5-23305909-DEL-57
chr5-23316699-DEL-62
chr5-23340180-DEL-3237
chr5-23360787-DEL-671
chr5-23374276-DEL-249
chr5-23379445-DEL-56
chr5-23427881-DEL-64
chr5-23446018-DEL-99
chr5-23446637-DEL-2055
chr5-23464425-DEL-107
chr5-23483304-DEL-62
chr5-23523793-DEL-4962
chr5-23543816-DEL-144
chr5-23589452-DEL-651
chr5-23591282-DEL-66
chr5-23627454-DEL-120
chr5-23682157-DEL-51
chr5-23813021-DEL-54
chr5-23844349-DEL-63
chr5-23851737-DEL-1193
chr5-23877719-DEL-94
chr5-23888837-DEL-59
chr5-23895234-DEL-249
chr5-23899902-DEL-672
chr5-23929554-DEL-808
chr5-23931407-DEL-3827
chr5-23939179-DEL-192
chr5-23942857-DEL-339
chr5-23984658-DEL-57
chr5-23991940-DEL-164
chr5-24019124-DEL-1359
chr5-24039537-DEL-68
chr5-24042937-DEL-311
chr5-24050700-DEL-120
chr5-24059732-DEL-201
chr5-24071418-DEL-897
chr5-24129972-DEL-960
chr5-24202587-DEL-163
chr5-24208691-DEL-51
chr5-24219519-DEL-97

chr5-24228761-DEL-459
chr5-24238575-DEL-55
chr5-24266677-DEL-1864
chr5-24272753-DEL-281
chr5-24287811-DEL-92
chr5-24289531-DEL-1165
chr5-24314831-DEL-274
chr5-24315731-DEL-563
chr5-24319121-DEL-82
chr5-24341274-DEL-204
chr5-24348682-DEL-112
chr5-24362112-DEL-874
chr5-24397111-DEL-152
chr5-24406471-DEL-449
chr5-24410674-DEL-60
chr5-24415348-DEL-55
chr5-24428459-DEL-412
chr5-24511521-DEL-54
chr5-24515093-DEL-187
chr5-24580167-DEL-66
chr5-24601363-DEL-204
chr5-24617604-DEL-68
chr5-24645750-DEL-57
chr5-24698865-DEL-80
chr5-24712061-DEL-58
chr5-24713204-DEL-221
chr5-24752570-DEL-58
chr5-24754087-DEL-52
chr5-24767733-DEL-59
chr5-24769402-DEL-490
chr5-24819379-DEL-182
chr5-24831866-DEL-1010
chr5-24832144-DEL-76
chr5-24846574-DEL-156
chr5-24889879-DEL-56
chr5-24898990-DEL-69
chr5-24909134-DEL-2256
chr5-24913308-DEL-54
chr5-24921046-DEL-117
chr5-24929517-DEL-54
chr5-24932698-DEL-184
chr5-24937569-DEL-104
chr5-24957383-DEL-429
chr5-24977003-DEL-117
chr5-24982258-DEL-68
chr5-25055268-DEL-571
chr5-25073449-DEL-210
chr5-25101723-DEL-97
chr5-25110510-DEL-295
chr5-25130187-DEL-133
chr5-25143802-DEL-96
chr5-25185573-DEL-165
chr5-25194637-DEL-52
chr5-25216383-DEL-63
chr5-25244457-DEL-59
chr5-25258006-DEL-540
chr5-25270854-DEL-327
chr5-25274072-DEL-77
chr5-25282374-DEL-150
chr5-25288466-DEL-461
chr5-25296042-DEL-82
chr5-25307607-DEL-134
chr5-25317521-DEL-164
chr5-25322107-DEL-394
chr5-25331479-DEL-53
chr5-25335245-DEL-424
chr5-25343435-DEL-52
chr5-25382419-DEL-66
chr5-25401264-DEL-372
chr5-25421064-DEL-59
chr5-25472761-DEL-217
chr5-25490352-DEL-1529
chr5-25506941-DEL-108
chr5-25524377-DEL-62
chr5-25587115-DEL-52
chr5-25591009-DEL-85
chr5-25605128-DEL-79

chr5-25630954-DEL-75
chr5-25631518-DEL-141
chr5-25689257-DEL-69
chr5-25694776-DEL-149
chr5-25710218-DEL-138
chr5-25747865-DEL-68
chr5-25752139-DEL-52
chr5-25823859-DEL-269
chr5-25830791-DEL-72
chr5-25831282-DEL-1941
chr5-25832490-DEL-54
chr5-25833250-DEL-1492
chr5-25842291-DEL-263
chr5-25856304-DEL-135
chr5-25866725-DEL-644
chr5-25883707-DEL-1130
chr5-25890521-DEL-695
chr5-25895639-DEL-184
chr5-25905435-DEL-128
chr5-25909111-DEL-108
chr5-25970923-DEL-186
chr5-25973845-DEL-92
chr5-25976599-DEL-107
chr5-26021816-DEL-559
chr5-26022922-DEL-59
chr5-26024028-DEL-90
chr5-26030427-DEL-64
chr5-26036112-DEL-88
chr5-26038310-DEL-182
chr5-26040162-DEL-145
chr5-26048655-DEL-76
chr5-26050235-DEL-91
chr5-26050854-DEL-93
chr5-26060216-DEL-202
chr5-26081046-DEL-145
chr5-26088859-DEL-75
chr5-26114508-DEL-72
chr5-26125743-DEL-506
chr5-26145330-DEL-132
chr5-26157412-DEL-86
chr5-26166046-DEL-16101
chr5-26175632-DEL-132
chr5-26237023-DEL-497
chr5-26249332-DEL-2617
chr5-26265452-DEL-698
chr5-26273962-DEL-198
chr5-26276472-DEL-3725
chr5-26285225-DEL-1641
chr5-26292286-DEL-64
chr5-26312235-DEL-698
chr5-26313136-DEL-2239
chr5-26320634-DEL-5991
chr5-26352433-DEL-94
chr5-26374969-DEL-70
chr5-26378029-DEL-105
chr5-26378514-DEL-504
chr5-26401506-DEL-52
chr5-26490793-DEL-119
chr5-26512547-DEL-713
chr5-26519538-DEL-1848
chr5-26529462-DEL-776
chr5-26535953-DEL-402
chr5-26538138-DEL-764
chr5-26538972-DEL-1023
chr5-26583270-DEL-11007
chr5-26584404-DEL-7290
chr5-26584419-DEL-212
chr5-26647453-DEL-723
chr5-26718596-DEL-204
chr5-26719547-DEL-303
chr5-26747633-DEL-186
chr5-26800152-DEL-736
chr5-26829689-DEL-3742
chr5-26832762-DEL-695
chr5-26853908-DEL-448
chr5-26865110-DEL-804
chr5-26881852-DEL-3710

chr5-26889911-DEL-129
chr5-26902639-DEL-136
chr5-26947360-DEL-63
chr5-26964514-DEL-307
chr5-26996991-DEL-227
chr5-27020511-DEL-109
chr5-27031241-DEL-92
chr5-27038275-DEL-82
chr5-27053542-DEL-52
chr5-27075499-DEL-103
chr5-27086502-DEL-312
chr5-27094149-DEL-1766
chr5-27119531-DEL-66
chr5-27154959-DEL-165
chr5-27213855-DEL-80
chr5-27303649-DEL-328
chr5-27343778-DEL-5065
chr5-27451181-DEL-73
chr5-27475418-DEL-155
chr5-27478173-DEL-318
chr5-27510668-DEL-53
chr5-27514409-DEL-61
chr5-27520936-DEL-74
chr5-27526246-DEL-237
chr5-27563131-DEL-88
chr5-27632697-DEL-124
chr5-27675645-DEL-158
chr5-27676553-DEL-688
chr5-27726117-DEL-390
chr5-27730777-DEL-275
chr5-27731145-DEL-5445
chr5-27803361-DEL-54
chr5-27829926-DEL-195
chr5-27853612-DEL-210
chr5-27856873-DEL-94
chr5-27871981-DEL-75
chr5-27878606-DEL-300
chr5-27886428-DEL-164
chr5-27904076-DEL-549
chr5-27955768-DEL-63
chr5-27962893-DEL-108
chr5-27990924-DEL-517
chr5-28018840-DEL-183
chr5-28024549-DEL-201
chr5-28036108-DEL-52
chr5-28054125-DEL-107
chr5-28070959-DEL-2421
chr5-28073444-DEL-3174
chr5-28076441-DEL-423
chr5-28077559-DEL-112
chr5-28104375-DEL-89
chr5-28116900-DEL-3966
chr5-28121113-DEL-694
chr5-28125015-DEL-4597
chr5-28136703-DEL-219
chr5-28141091-DEL-635
chr5-28190187-DEL-6084
chr5-28220752-DEL-87
chr5-28229219-DEL-1034
chr5-28241464-DEL-68
chr5-28254177-DEL-2812
chr5-28288409-DEL-54
chr5-28302366-DEL-607
chr5-28303386-DEL-1185
chr5-28304184-DEL-181
chr5-28305101-DEL-189
chr5-28325285-DEL-59
chr5-28352575-DEL-66
chr5-28359923-DEL-84
chr5-28387108-DEL-141
chr5-28405292-DEL-55
chr5-28412635-DEL-62
chr5-28420030-DEL-100
chr5-28426509-DEL-2978
chr5-28461174-DEL-125
chr5-28462002-DEL-388
chr5-28473107-DEL-58

chr5-28480600-DEL-174
chr5-28507649-DEL-80
chr5-28516082-DEL-104
chr5-28601969-DEL-70
chr5-28645347-DEL-56
chr5-28656604-DEL-58
chr5-28789458-DEL-10424
chr5-28792530-DEL-6641
chr5-28806271-DEL-2774
chr5-28809064-DEL-6785
chr5-28859390-DEL-66
chr5-28860480-DEL-214
chr5-28864395-DEL-62
chr5-28874363-DEL-2656
chr5-28875161-DEL-320
chr5-28879342-DEL-58
chr5-28902413-DEL-152
chr5-28934460-DEL-99
chr5-28935861-DEL-81
chr5-28962614-DEL-833
chr5-29000414-DEL-102
chr5-29093140-DEL-51
chr5-29099887-DEL-93
chr5-29132956-DEL-311
chr5-29133103-DEL-431
chr5-29157680-DEL-1892
chr5-29181179-DEL-370
chr5-29209764-DEL-60
chr5-29211904-DEL-73
chr5-29213107-DEL-100
chr5-29243837-DEL-205
chr5-29252471-DEL-514
chr5-29267004-DEL-214
chr5-29305861-DEL-185
chr5-29361490-DEL-56
chr5-29368212-DEL-926
chr5-29377151-DEL-86
chr5-29415931-DEL-220
chr5-29439626-DEL-56
chr5-29449191-DEL-1698
chr5-29486841-DEL-149
chr5-29505854-DEL-53
chr5-29510382-DEL-437
chr5-29511049-DEL-1309
chr5-29512984-DEL-594
chr5-29548883-DEL-743
chr5-29560622-DEL-147
chr5-29578062-DEL-73
chr5-29589773-DEL-54
chr5-29592988-DEL-74
chr5-29657216-DEL-810
chr5-29687511-DEL-201
chr5-29703008-DEL-800
chr5-29723374-DEL-69
chr5-29728652-DEL-52
chr5-29766528-DEL-373
chr5-29810066-DEL-175
chr5-29817579-DEL-198
chr5-29823265-DEL-254
chr5-29889633-DEL-209
chr5-29902751-DEL-195
chr5-29909554-DEL-196
chr5-29924888-DEL-56
chr5-29933502-DEL-203
chr5-29994912-DEL-103
chr5-30004852-DEL-918
chr5-30017912-DEL-57
chr5-30033182-DEL-427
chr5-30145518-DEL-66
chr5-30167358-DEL-200
chr5-30191929-DEL-100
chr5-30230796-DEL-2659
chr5-30246471-DEL-182
chr5-30254754-DEL-6257
chr5-30293787-DEL-235
chr5-30372593-DEL-201
chr5-30378060-DEL-216

chr5-30398968-DEL-160
chr5-30414820-DEL-366
chr5-30418660-DEL-207
chr5-30424113-DEL-78
chr5-30428036-DEL-7119
chr5-30467157-DEL-124
chr5-30478286-DEL-272
chr5-30479591-DEL-4322
chr5-30489802-DEL-4433
chr5-30495065-DEL-63
chr5-30495093-DEL-1066
chr5-30495905-DEL-1132
chr5-30495951-DEL-402
chr5-30504840-DEL-129
chr5-30509432-DEL-133
chr5-30516238-DEL-131
chr5-30519164-DEL-61
chr5-30530467-DEL-89
chr5-30575026-DEL-202
chr5-30604013-DEL-138
chr5-30643138-DEL-117
chr5-30739882-DEL-287
chr5-30753261-DEL-112
chr5-30759704-DEL-60
chr5-30799143-DEL-128
chr5-30821623-DEL-69
chr5-30827271-DEL-136
chr5-30856140-DEL-68
chr5-30893407-DEL-90
chr5-30939019-DEL-129
chr5-30986976-DEL-200
chr5-31000267-DEL-195
chr5-31014345-DEL-202
chr5-31034274-DEL-79
chr5-31035820-DEL-105
chr5-31041244-DEL-347
chr5-31041749-DEL-120
chr5-31113865-DEL-816
chr5-31114715-DEL-252
chr5-31139155-DEL-512
chr5-31146953-DEL-367
chr5-31158701-DEL-186
chr5-31165867-DEL-1253
chr5-31169521-DEL-96
chr5-31222827-DEL-210
chr5-31307635-DEL-220
chr5-31321601-DEL-375
chr5-31329112-DEL-210
chr5-31343959-DEL-92
chr5-31346466-DEL-178
chr5-31349220-DEL-106
chr5-31376527-DEL-66
chr5-31414447-DEL-71
chr5-31422504-DEL-72
chr5-31430954-DEL-183
chr5-31504088-DEL-222
chr5-31525741-DEL-189
chr5-31554935-DEL-6786
chr5-31561294-DEL-464
chr5-31592180-DEL-504
chr5-31595301-DEL-791
chr5-31616140-DEL-795
chr5-31616778-DEL-161
chr5-31618210-DEL-169
chr5-31640899-DEL-206
chr5-31648555-DEL-213
chr5-31654846-DEL-186
chr5-31704979-DEL-257
chr5-31711480-DEL-117
chr5-31713680-DEL-201
chr5-31724678-DEL-67
chr5-31778659-DEL-219
chr5-31787245-DEL-53
chr5-31801876-DEL-54
chr5-31804141-DEL-52
chr5-31846534-DEL-352
chr5-31864897-DEL-102

chr5-31899755-DEL-83
chr5-31904971-DEL-274
chr5-31991614-DEL-69
chr5-32008082-DEL-327
chr5-32062159-DEL-207
chr5-32069855-DEL-87
chr5-32101459-DEL-112
chr5-32105170-DEL-1230
chr5-32105171-DEL-246
chr5-32105696-DEL-492
chr5-32130099-DEL-103
chr5-32135169-DEL-1138
chr5-32205590-DEL-196
chr5-32247560-DEL-1821
chr5-32343297-DEL-60
chr5-32365367-DEL-157
chr5-32385460-DEL-89
chr5-32391313-DEL-165
chr5-32395484-DEL-94
chr5-32407538-DEL-1419
chr5-32467506-DEL-68
chr5-32556731-DEL-57
chr5-32577875-DEL-80
chr5-32623310-DEL-1021
chr5-32627479-DEL-215
chr5-32703705-DEL-230
chr5-32708774-DEL-609
chr5-32721649-DEL-233
chr5-32729153-DEL-1733
chr5-32736350-DEL-250
chr5-32738053-DEL-548
chr5-32743773-DEL-182
chr5-32753369-DEL-53
chr5-32753792-DEL-571
chr5-32802458-DEL-204
chr5-32884040-DEL-736
chr5-32913727-DEL-3869
chr5-32952025-DEL-244
chr5-32981668-DEL-216
chr5-33005114-DEL-60
chr5-33033769-DEL-181
chr5-33077002-DEL-216
chr5-33078246-DEL-313
chr5-33119205-DEL-401
chr5-33133824-DEL-71
chr5-33134494-DEL-215
chr5-33189405-DEL-573
chr5-33194693-DEL-114
chr5-33198280-DEL-141
chr5-33236694-DEL-164
chr5-33257156-DEL-65
chr5-33273005-DEL-52
chr5-33275751-DEL-175
chr5-33312452-DEL-86
chr5-33360506-DEL-158
chr5-33394503-DEL-378
chr5-33404127-DEL-59
chr5-33438405-DEL-174
chr5-33439550-DEL-206
chr5-33458277-DEL-232
chr5-33475408-DEL-247
chr5-33481815-DEL-246
chr5-33511227-DEL-187
chr5-33512867-DEL-204
chr5-33537240-DEL-67
chr5-33547855-DEL-185
chr5-33574941-DEL-57
chr5-33593307-DEL-54
chr5-33603259-DEL-72
chr5-33613105-DEL-150
chr5-33614121-DEL-51
chr5-33617582-DEL-66
chr5-33687711-DEL-54
chr5-33710638-DEL-1182
chr5-33744331-DEL-88
chr5-33772493-DEL-215
chr5-33774306-DEL-217

chr5-33785004-DEL-89
chr5-33817982-DEL-244
chr5-33838058-DEL-169
chr5-33847785-DEL-198
chr5-33896265-DEL-210
chr5-33908351-DEL-90
chr5-33916117-DEL-81
chr5-33942910-DEL-1455
chr5-34004390-DEL-166
chr5-34040736-DEL-437
chr5-34094291-DEL-1893
chr5-34166734-DEL-143
chr5-34219820-DEL-1672
chr5-34246173-DEL-104
chr5-34301102-DEL-925
chr5-34302198-DEL-1965
chr5-34347673-DEL-53
chr5-34379116-DEL-80
chr5-34379692-DEL-62
chr5-34384114-DEL-80
chr5-34415344-DEL-60
chr5-34418483-DEL-123
chr5-34455665-DEL-414
chr5-34475657-DEL-931
chr5-34487951-DEL-438
chr5-34489821-DEL-190
chr5-34521733-DEL-216
chr5-34523275-DEL-201
chr5-34581674-DEL-198
chr5-34586162-DEL-116
chr5-34630853-DEL-1430
chr5-34668969-DEL-87
chr5-34674042-DEL-215
chr5-34677024-DEL-59
chr5-34724371-DEL-203
chr5-34778892-DEL-186
chr5-34814383-DEL-478
chr5-34829363-DEL-906
chr5-34830922-DEL-64
chr5-34884227-DEL-115
chr5-34920922-DEL-944
chr5-34935579-DEL-66
chr5-34938764-DEL-396
chr5-35022786-DEL-72
chr5-35159508-DEL-54
chr5-35192849-DEL-132
chr5-35193654-DEL-227
chr5-35206995-DEL-89
chr5-35233705-DEL-75
chr5-35235084-DEL-59
chr5-35248371-DEL-210
chr5-35285165-DEL-59
chr5-35324292-DEL-60
chr5-35332508-DEL-158
chr5-35345568-DEL-116
chr5-35348800-DEL-114
chr5-35375935-DEL-118
chr5-35416751-DEL-131
chr5-35419667-DEL-98
chr5-35430342-DEL-378
chr5-35476866-DEL-209
chr5-35486490-DEL-64
chr5-35496265-DEL-55
chr5-35533687-DEL-58
chr5-35541349-DEL-238
chr5-35569146-DEL-63
chr5-35670462-DEL-92
chr5-35678083-DEL-95
chr5-35678208-DEL-197
chr5-35684121-DEL-737
chr5-35701394-DEL-138
chr5-35713378-DEL-69
chr5-35750737-DEL-137
chr5-35761627-DEL-74
chr5-35803231-DEL-64
chr5-35806643-DEL-2318
chr5-35856968-DEL-52

chr5-36017128-DEL-78
chr5-36024900-DEL-57
chr5-36090095-DEL-1065
chr5-36096808-DEL-80
chr5-36112974-DEL-473
chr5-36116267-DEL-894
chr5-36136784-DEL-64
chr5-36193967-DEL-83
chr5-36200632-DEL-76
chr5-36220838-DEL-1152
chr5-36340864-DEL-204
chr5-36354642-DEL-97
chr5-36394844-DEL-63
chr5-36395757-DEL-212
chr5-36415207-DEL-100
chr5-36419751-DEL-99
chr5-36429628-DEL-154
chr5-36438806-DEL-52
chr5-36499021-DEL-123
chr5-36514749-DEL-82
chr5-36575493-DEL-2625
chr5-36678494-DEL-127
chr5-36681780-DEL-198
chr5-36688480-DEL-372
chr5-36706445-DEL-66
chr5-36712736-DEL-199
chr5-36714500-DEL-92
chr5-36753113-DEL-294
chr5-36859370-DEL-204
chr5-36865805-DEL-350
chr5-36890414-DEL-199
chr5-36892731-DEL-164
chr5-36902038-DEL-790
chr5-36906475-DEL-309
chr5-36925767-DEL-62
chr5-36931902-DEL-2438
chr5-36964836-DEL-106
chr5-36983047-DEL-210
chr5-36990465-DEL-109
chr5-37029724-DEL-407
chr5-37061450-DEL-65
chr5-37064575-DEL-413
chr5-37075328-DEL-84
chr5-37108820-DEL-232
chr5-37128638-DEL-59
chr5-37167326-DEL-162
chr5-37197320-DEL-298
chr5-37212212-DEL-142
chr5-37246134-DEL-258
chr5-37259328-DEL-51
chr5-37260257-DEL-61
chr5-37339138-DEL-109
chr5-37357907-DEL-116
chr5-37360873-DEL-111
chr5-37388749-DEL-369
chr5-37412662-DEL-1122
chr5-37421951-DEL-209
chr5-37435979-DEL-103
chr5-37453405-DEL-66
chr5-37459860-DEL-138
chr5-37511788-DEL-1342
chr5-37524147-DEL-228
chr5-37546571-DEL-88
chr5-37559728-DEL-1157
chr5-37564635-DEL-114
chr5-37648516-DEL-100
chr5-37704175-DEL-223
chr5-37728637-DEL-133
chr5-37746345-DEL-749
chr5-37753718-DEL-146
chr5-37887278-DEL-58
chr5-37897462-DEL-799
chr5-37969278-DEL-58
chr5-37979061-DEL-91
chr5-37992918-DEL-85
chr5-38014100-DEL-188
chr5-38110147-DEL-859

chr5-38112157-DEL-53
chr5-38118198-DEL-82
chr5-38160991-DEL-64
chr5-38180320-DEL-192
chr5-38259577-DEL-432
chr5-38277067-DEL-279
chr5-38384981-DEL-112
chr5-38396539-DEL-66
chr5-38402818-DEL-125
chr5-38470540-DEL-441
chr5-38527069-DEL-2257
chr5-38539624-DEL-96
chr5-38577543-DEL-194
chr5-38580604-DEL-51
chr5-38587192-DEL-72
chr5-38601308-DEL-1357
chr5-38605951-DEL-776
chr5-38627006-DEL-53
chr5-38636866-DEL-74
chr5-38674580-DEL-61
chr5-38746465-DEL-1846
chr5-38814184-DEL-105
chr5-38816278-DEL-132
chr5-38820433-DEL-200
chr5-38843334-DEL-411
chr5-38847760-DEL-103
chr5-38858229-DEL-165
chr5-38867271-DEL-79
chr5-38882083-DEL-287
chr5-38894110-DEL-208
chr5-38900421-DEL-332
chr5-38909328-DEL-200
chr5-39002016-DEL-70
chr5-39010677-DEL-178
chr5-39013505-DEL-103
chr5-39023641-DEL-71
chr5-39032112-DEL-89
chr5-39045338-DEL-516
chr5-39064574-DEL-707
chr5-39078962-DEL-249
chr5-39105950-DEL-79
chr5-39138699-DEL-981
chr5-39162883-DEL-62
chr5-39164188-DEL-58
chr5-39173362-DEL-1536
chr5-39188413-DEL-8032
chr5-39221327-DEL-924
chr5-39224607-DEL-89
chr5-39241183-DEL-189
chr5-39255981-DEL-115
chr5-39265256-DEL-126
chr5-39269108-DEL-254
chr5-39276376-DEL-124
chr5-39325923-DEL-61
chr5-39396598-DEL-60
chr5-39428120-DEL-96
chr5-39452524-DEL-1440
chr5-39495908-DEL-97
chr5-39522886-DEL-376
chr5-39577552-DEL-54
chr5-39580281-DEL-106
chr5-39591985-DEL-206
chr5-39607634-DEL-74
chr5-39639071-DEL-76
chr5-39679007-DEL-256
chr5-39749075-DEL-79
chr5-39750456-DEL-608
chr5-39757517-DEL-93
chr5-39829217-DEL-142
chr5-39874758-DEL-55
chr5-39877652-DEL-6876
chr5-39884983-DEL-1112
chr5-39893656-DEL-181
chr5-39898478-DEL-469
chr5-39914572-DEL-335
chr5-39924919-DEL-72
chr5-39969154-DEL-1133

chr5-39972140-DEL-1746
chr5-39985606-DEL-482
chr5-40019885-DEL-57
chr5-40074345-DEL-6447
chr5-40098325-DEL-59
chr5-40106497-DEL-715
chr5-40115179-DEL-750
chr5-40140441-DEL-644
chr5-40149132-DEL-327
chr5-40159798-DEL-87
chr5-40185655-DEL-55
chr5-40197324-DEL-1403
chr5-40215982-DEL-76
chr5-40233093-DEL-208
chr5-40234792-DEL-547
chr5-40235727-DEL-5075
chr5-40239583-DEL-2968
chr5-40245658-DEL-356
chr5-40246882-DEL-559
chr5-40260199-DEL-91
chr5-40266206-DEL-177
chr5-40272084-DEL-451
chr5-40283674-DEL-438
chr5-40301660-DEL-62
chr5-40307654-DEL-200
chr5-40308750-DEL-74
chr5-40315576-DEL-1666
chr5-40323291-DEL-60
chr5-40324399-DEL-205
chr5-40326261-DEL-258
chr5-40326843-DEL-965
chr5-40333477-DEL-379
chr5-40339247-DEL-283
chr5-40346352-DEL-390
chr5-40394197-DEL-56
chr5-40412723-DEL-251
chr5-40442336-DEL-52
chr5-40455725-DEL-1159
chr5-40463480-DEL-599
chr5-40470122-DEL-327
chr5-40471346-DEL-72
chr5-40515714-DEL-787
chr5-40544829-DEL-57
chr5-40603104-DEL-293
chr5-40634644-DEL-81
chr5-40637879-DEL-2089
chr5-40717928-DEL-2152
chr5-40768368-DEL-109
chr5-40786128-DEL-69
chr5-40791388-DEL-807
chr5-40800851-DEL-68
chr5-40824351-DEL-3487
chr5-40850520-DEL-1274
chr5-40855388-DEL-66
chr5-40876423-DEL-775
chr5-40882715-DEL-80
chr5-40901870-DEL-54
chr5-40926441-DEL-1620
chr5-40936414-DEL-461
chr5-40950362-DEL-2536
chr5-40965192-DEL-83
chr5-41023755-DEL-202
chr5-41028408-DEL-467
chr5-41051485-DEL-1130
chr5-41061021-DEL-289
chr5-41125458-DEL-854
chr5-41148959-DEL-83
chr5-41155125-DEL-86
chr5-41156354-DEL-82
chr5-41173227-DEL-1099
chr5-41194063-DEL-65
chr5-41214598-DEL-315
chr5-41230467-DEL-1935
chr5-41366919-DEL-4772
chr5-41450029-DEL-198
chr5-41462820-DEL-190
chr5-41474502-DEL-80

chr5-41475462-DEL-1188
chr5-41477165-DEL-997
chr5-41489171-DEL-116
chr5-41491468-DEL-958
chr5-41492667-DEL-1412
chr5-41507971-DEL-69
chr5-41596281-DEL-1189
chr5-41610654-DEL-1940
chr5-41621047-DEL-1222
chr5-41622556-DEL-6413
chr5-41628922-DEL-659
chr5-41629269-DEL-1972
chr5-41645645-DEL-1663
chr5-41646805-DEL-6932
chr5-41652173-DEL-1556
chr5-41675883-DEL-53
chr5-41709722-DEL-54
chr5-41731788-DEL-152
chr5-41769437-DEL-52
chr5-41801082-DEL-287
chr5-41817085-DEL-2570
chr5-41868996-DEL-2139
chr5-41876786-DEL-101
chr5-42017639-DEL-920
chr5-42022178-DEL-2230
chr5-42037962-DEL-280
chr5-42039879-DEL-82
chr5-42054840-DEL-54
chr5-42063886-DEL-199
chr5-42130449-DEL-60
chr5-42155727-DEL-19478
chr5-42170726-DEL-212
chr5-42195767-DEL-6186
chr5-42217427-DEL-701
chr5-42227646-DEL-399
chr5-42231884-DEL-68
chr5-42240296-DEL-102
chr5-42275103-DEL-51
chr5-42305783-DEL-62
chr5-42378942-DEL-99
chr5-42382645-DEL-743
chr5-42452855-DEL-4555
chr5-42479036-DEL-74
chr5-42498000-DEL-53
chr5-42515842-DEL-1200
chr5-42578449-DEL-64
chr5-42593471-DEL-2001
chr5-42596226-DEL-1777
chr5-42604295-DEL-576
chr5-42645972-DEL-6348
chr5-42664951-DEL-1205
chr5-42665847-DEL-353
chr5-42691694-DEL-68
chr5-42706719-DEL-56
chr5-42715852-DEL-332
chr5-42721539-DEL-91
chr5-42734368-DEL-520
chr5-42743536-DEL-1464
chr5-42760413-DEL-387
chr5-42774463-DEL-126
chr5-42781268-DEL-71
chr5-42805248-DEL-60
chr5-42810873-DEL-1314
chr5-42888429-DEL-7799
chr5-42912003-DEL-1478
chr5-42922808-DEL-1321
chr5-42951659-DEL-1848
chr5-42959698-DEL-55
chr5-42973511-DEL-664
chr5-43011719-DEL-71
chr5-43051999-DEL-299
chr5-43054739-DEL-115
chr5-43067728-DEL-311
chr5-43069696-DEL-2526
chr5-43106470-DEL-1332
chr5-43153274-DEL-10752
chr5-43167382-DEL-7397

chr5-43167409-DEL-6181
chr5-43210748-DEL-75
chr5-43218950-DEL-71
chr5-43277373-DEL-90
chr5-43331413-DEL-212
chr5-43334573-DEL-124
chr5-43409341-DEL-58
chr5-43462032-DEL-358
chr5-43466200-DEL-121
chr5-43470262-DEL-148
chr5-43478061-DEL-2015
chr5-43538440-DEL-424
chr5-43539782-DEL-623
chr5-43549659-DEL-302
chr5-43583511-DEL-6260
chr5-43603567-DEL-60
chr5-43649151-DEL-6308
chr5-43716018-DEL-1368
chr5-43793206-DEL-210
chr5-43796966-DEL-1971
chr5-43799002-DEL-1287
chr5-43807229-DEL-3805
chr5-43814179-DEL-1388
chr5-43837240-DEL-550
chr5-43843294-DEL-883
chr5-43958417-DEL-51
chr5-43973515-DEL-211
chr5-43999502-DEL-64
chr5-44001093-DEL-182
chr5-44008545-DEL-83
chr5-44013589-DEL-904
chr5-44028045-DEL-179
chr5-44053849-DEL-108
chr5-44070475-DEL-804
chr5-44104486-DEL-66
chr5-44116302-DEL-1036
chr5-44126808-DEL-52
chr5-44157696-DEL-52
chr5-44168678-DEL-880
chr5-44196098-DEL-654
chr5-44206238-DEL-1287
chr5-44222467-DEL-291
chr5-44227831-DEL-134
chr5-44289837-DEL-942
chr5-44302231-DEL-93
chr5-44344530-DEL-138
chr5-44388903-DEL-58
chr5-44418943-DEL-534
chr5-44425312-DEL-401
chr5-44429094-DEL-63
chr5-44455904-DEL-54
chr5-44457758-DEL-148
chr5-44470269-DEL-2999
chr5-44471734-DEL-1760
chr5-44518050-DEL-1436
chr5-44519157-DEL-356
chr5-44641262-DEL-65
chr5-44661258-DEL-875
chr5-44689002-DEL-467
chr5-44719236-DEL-114
chr5-44758425-DEL-301
chr5-44799167-DEL-105
chr5-44840720-DEL-218
chr5-44902090-DEL-158
chr5-44910878-DEL-184
chr5-44913420-DEL-590
chr5-44920236-DEL-55
chr5-44939139-DEL-400
chr5-44971348-DEL-450
chr5-44974420-DEL-62
chr5-45003324-DEL-68
chr5-45025606-DEL-805
chr5-45060747-DEL-235
chr5-45086051-DEL-8679
chr5-45087731-DEL-204
chr5-45115614-DEL-647
chr5-45115846-DEL-2860

chr5-45116506-DEL-198
chr5-45118278-DEL-6203
chr5-45146916-DEL-207
chr5-45204842-DEL-150
chr5-45219712-DEL-82
chr5-45221287-DEL-226
chr5-45242669-DEL-483
chr5-45243125-DEL-760
chr5-45309255-DEL-321
chr5-45372370-DEL-92
chr5-45392672-DEL-82
chr5-45408799-DEL-99
chr5-45438533-DEL-242
chr5-45449381-DEL-117
chr5-45462853-DEL-2723
chr5-45466207-DEL-97
chr5-45467484-DEL-66
chr5-45467965-DEL-781
chr5-45487783-DEL-974
chr5-45501036-DEL-85
chr5-45510923-DEL-94
chr5-45526197-DEL-426
chr5-45557926-DEL-180
chr5-45561089-DEL-55
chr5-45579301-DEL-501
chr5-45600954-DEL-109
chr5-45603853-DEL-239
chr5-45609749-DEL-137
chr5-45619167-DEL-614
chr5-45657407-DEL-103
chr5-45659732-DEL-70
chr5-45681306-DEL-187
chr5-45684419-DEL-298
chr5-45868014-DEL-87
chr5-45958618-DEL-278
chr5-45969680-DEL-474
chr5-45973383-DEL-6010
chr5-45979940-DEL-2524
chr5-46003442-DEL-60
chr5-46047222-DEL-111
chr5-46062438-DEL-234
chr5-46098779-DEL-1149
chr5-46121223-DEL-1067
chr5-46131518-DEL-69
chr5-46135451-DEL-6215
chr5-46152047-DEL-792
chr5-46157346-DEL-725
chr5-46182577-DEL-57
chr5-46201159-DEL-688
chr5-46207534-DEL-163
chr5-46215416-DEL-274
chr5-46231914-DEL-304
chr5-46232958-DEL-52
chr5-46234767-DEL-712
chr5-46401051-DEL-6235
chr5-46438136-DEL-627
chr5-46503274-DEL-95
chr5-46517833-DEL-62
chr5-46530237-DEL-6617
chr5-46628398-DEL-302
chr5-46657204-DEL-5175
chr5-46701753-DEL-68
chr5-46703597-DEL-349
chr5-46721632-DEL-103
chr5-46756505-DEL-160
chr5-46773802-DEL-1463
chr5-46776659-DEL-65
chr5-46816442-DEL-184
chr5-46841343-DEL-338
chr5-46852507-DEL-788
chr5-46855245-DEL-8437
chr5-46896670-DEL-463
chr5-46918863-DEL-6437
chr5-46918873-DEL-3759
chr5-46922662-DEL-135
chr5-46922827-DEL-2133
chr5-46925189-DEL-122

chr5-46978554-DEL-2512
chr5-47022577-DEL-105
chr5-47027053-DEL-115
chr5-47038205-DEL-185
chr5-47044029-DEL-551
chr5-47046669-DEL-203
chr5-47049928-DEL-174
chr5-47054178-DEL-76
chr5-47058289-DEL-82
chr5-47062522-DEL-53
chr5-47072828-DEL-604
chr5-47111538-DEL-2483
chr5-47136819-DEL-237
chr5-47140537-DEL-429
chr5-47176728-DEL-2058
chr5-47176729-DEL-1776
chr5-47178525-DEL-449
chr5-47212865-DEL-962
chr5-47282084-DEL-6250
chr5-47295460-DEL-1925
chr5-47302779-DEL-61
chr5-47305665-DEL-321
chr5-47329102-DEL-57
chr5-47335955-DEL-6313
chr5-47355567-DEL-94
chr5-47381867-DEL-388
chr5-47432178-DEL-56
chr5-47463942-DEL-1145
chr5-47498068-DEL-100
chr5-47537104-DEL-865
chr5-47572475-DEL-190
chr5-47639938-DEL-79
chr5-47673808-DEL-913
chr5-47681176-DEL-70
chr5-47689104-DEL-3568
chr5-47725483-DEL-68
chr5-47733302-DEL-82
chr5-47756860-DEL-404
chr5-47765468-DEL-70
chr5-47767379-DEL-72
chr5-47776314-DEL-2228
chr5-47847781-DEL-574
chr5-47862914-DEL-2604
chr5-47880433-DEL-100
chr5-47915419-DEL-6342
chr5-47915565-DEL-65
chr5-47949482-DEL-831
chr5-47974877-DEL-137
chr5-47985498-DEL-675
chr5-48036143-DEL-278
chr5-48043936-DEL-315
chr5-48045003-DEL-135
chr5-48045544-DEL-400
chr5-48087191-DEL-192
chr5-48105223-DEL-197
chr5-48159081-DEL-86
chr5-48198010-DEL-808
chr5-48216885-DEL-57
chr5-48225893-DEL-385
chr5-48234702-DEL-562
chr5-48248074-DEL-136
chr5-48255800-DEL-57
chr5-48281707-DEL-64
chr5-48366495-DEL-643
chr5-48422269-DEL-579
chr5-48425568-DEL-332
chr5-48435485-DEL-74
chr5-48450026-DEL-1117
chr5-48464241-DEL-69
chr5-48510477-DEL-53
chr5-48519136-DEL-4139
chr5-48524963-DEL-452
chr5-48570000-DEL-211
chr5-48605648-DEL-155
chr5-48607352-DEL-57
chr5-48645703-DEL-770
chr5-48668431-DEL-67

chr5-48688985-DEL-164
chr5-48775080-DEL-57
chr5-48783466-DEL-56
chr5-48815882-DEL-673
chr5-48839268-DEL-448
chr5-48913066-DEL-214
chr5-48938052-DEL-95
chr5-48944153-DEL-1116
chr5-48968162-DEL-70
chr5-48976637-DEL-69
chr5-49016961-DEL-75
chr5-49091565-DEL-70
chr5-49156826-DEL-1354
chr5-49181231-DEL-57
chr5-49213802-DEL-90
chr5-49223585-DEL-526
chr5-49225938-DEL-63
chr5-49229953-DEL-560
chr5-49256040-DEL-707
chr5-49261938-DEL-80
chr5-49288411-DEL-67
chr5-49288958-DEL-52
chr5-49322434-DEL-977
chr5-49348841-DEL-5028
chr5-49361130-DEL-83
chr5-49369235-DEL-58
chr5-49393776-DEL-760
chr5-49480218-DEL-167
chr5-49589712-DEL-581
chr5-49595070-DEL-447
chr5-49644834-DEL-84
chr5-49666444-DEL-78
chr5-49713286-DEL-53
chr5-49734531-DEL-497
chr5-49744334-DEL-137
chr5-49745417-DEL-63
chr5-49750215-DEL-61
chr5-49766536-DEL-52
chr5-49798465-DEL-64
chr5-49844909-DEL-76
chr5-49879123-DEL-6297
chr5-49932016-DEL-1570
chr5-49935354-DEL-269
chr5-49936857-DEL-1468
chr5-49997306-DEL-132
chr5-50026953-DEL-509
chr5-50038605-DEL-76
chr5-50070779-DEL-54
chr5-50077163-DEL-59
chr5-50091426-DEL-893
chr5-50094968-DEL-1483
chr5-50105264-DEL-102
chr5-50109400-DEL-2406
chr5-50228496-DEL-90
chr5-50253201-DEL-72
chr5-50258636-DEL-3178
chr5-50264679-DEL-1292
chr5-50266678-DEL-4786
chr5-50272773-DEL-1776
chr5-50337018-DEL-115
chr5-50344702-DEL-65
chr5-50349705-DEL-71
chr5-50361349-DEL-1845
chr5-50362879-DEL-1873
chr5-50368128-DEL-302
chr5-50418752-DEL-72
chr5-50421745-DEL-1977
chr5-50424012-DEL-205
chr5-50433685-DEL-89
chr5-50476364-DEL-212
chr5-50494397-DEL-703
chr5-50533505-DEL-58
chr5-50537851-DEL-100
chr5-50545777-DEL-60
chr5-50582988-DEL-106
chr5-50592966-DEL-806
chr5-50605380-DEL-400

chr5-50664170-DEL-56
chr5-50735623-DEL-65
chr5-50743167-DEL-315
chr5-50762286-DEL-1412
chr5-50779142-DEL-66
chr5-50784855-DEL-52
chr5-50787008-DEL-56
chr5-50868677-DEL-71
chr5-50984793-DEL-464
chr5-51000822-DEL-56
chr5-51049082-DEL-557
chr5-51075451-DEL-129
chr5-51134344-DEL-74
chr5-51138784-DEL-108
chr5-51187813-DEL-103
chr5-51200061-DEL-101
chr5-51207987-DEL-56
chr5-51209532-DEL-412
chr5-51264795-DEL-57
chr5-51307874-DEL-60
chr5-51313805-DEL-513
chr5-51316913-DEL-102
chr5-51347035-DEL-125
chr5-51353786-DEL-120
chr5-51358605-DEL-778
chr5-51428273-DEL-117
chr5-51482736-DEL-1690
chr5-51485098-DEL-638
chr5-51502149-DEL-570
chr5-51533008-DEL-5521
chr5-51580095-DEL-172
chr5-51587177-DEL-810
chr5-51596373-DEL-3339
chr5-51598180-DEL-167
chr5-51783585-DEL-595
chr5-51806767-DEL-220
chr5-51807852-DEL-400
chr5-51813067-DEL-302
chr5-51841633-DEL-55
chr5-51879890-DEL-89
chr5-51888869-DEL-172
chr5-51901705-DEL-478
chr5-51943000-DEL-78
chr5-51978556-DEL-56
chr5-51983409-DEL-236
chr5-52009801-DEL-66
chr5-52021165-DEL-237
chr5-52024267-DEL-288
chr5-52112114-DEL-203
chr5-52116502-DEL-204
chr5-52145343-DEL-226
chr5-52148082-DEL-53
chr5-52195582-DEL-88
chr5-52252386-DEL-60
chr5-52278282-DEL-60
chr5-52304055-DEL-52
chr5-52311191-DEL-131
chr5-52390806-DEL-140
chr5-52418019-DEL-52
chr5-52419160-DEL-513
chr5-52462087-DEL-539
chr5-52472914-DEL-816
chr5-52507381-DEL-72
chr5-52536211-DEL-52
chr5-52542928-DEL-54
chr5-52549656-DEL-56
chr5-52553746-DEL-538
chr5-52578821-DEL-243
chr5-52614895-DEL-1705
chr5-52620180-DEL-908
chr5-52668081-DEL-128
chr5-52707172-DEL-149
chr5-52708941-DEL-93
chr5-52738395-DEL-2822
chr5-52756071-DEL-1465
chr5-52828773-DEL-92
chr5-52853628-DEL-69

chr5-52858923-DEL-480
chr5-52911006-DEL-58
chr5-52916761-DEL-1211
chr5-52928812-DEL-57
chr5-52949307-DEL-309
chr5-52952487-DEL-153
chr5-52959719-DEL-72
chr5-52975147-DEL-170
chr5-52983211-DEL-208
chr5-53037872-DEL-77
chr5-53045300-DEL-51
chr5-53059142-DEL-416
chr5-53075559-DEL-752
chr5-53103982-DEL-391
chr5-53108504-DEL-102
chr5-53147354-DEL-202
chr5-53148800-DEL-86
chr5-53150233-DEL-223
chr5-53154418-DEL-315
chr5-53171612-DEL-75
chr5-53181213-DEL-89
chr5-53185904-DEL-516
chr5-53187360-DEL-855
chr5-53191646-DEL-1710
chr5-53192075-DEL-228
chr5-53195168-DEL-129
chr5-53223644-DEL-870
chr5-53235019-DEL-86
chr5-53240892-DEL-143
chr5-53241137-DEL-1954
chr5-53247709-DEL-109
chr5-53251794-DEL-87
chr5-53277975-DEL-60
chr5-53310234-DEL-61
chr5-53348108-DEL-63
chr5-53361481-DEL-96
chr5-53403543-DEL-57
chr5-53404884-DEL-55
chr5-53421611-DEL-54
chr5-53442823-DEL-65
chr5-53449057-DEL-223
chr5-53459752-DEL-288
chr5-53465480-DEL-3780
chr5-53469329-DEL-1631
chr5-53471227-DEL-224
chr5-53513073-DEL-81
chr5-53515328-DEL-98
chr5-53530451-DEL-52
chr5-53563159-DEL-63
chr5-53572256-DEL-426
chr5-53597921-DEL-100
chr5-53616996-DEL-177
chr5-53618660-DEL-1548
chr5-53628718-DEL-56
chr5-53706394-DEL-452
chr5-53707796-DEL-60
chr5-53713447-DEL-67
chr5-53725150-DEL-475
chr5-53730644-DEL-164
chr5-53735781-DEL-53
chr5-53740069-DEL-1341
chr5-53758093-DEL-211
chr5-53781745-DEL-115
chr5-53818180-DEL-162
chr5-53827156-DEL-453
chr5-53828379-DEL-152
chr5-53833379-DEL-164
chr5-53886064-DEL-750
chr5-53894293-DEL-60
chr5-53923531-DEL-89
chr5-53932061-DEL-278
chr5-53934277-DEL-1453
chr5-53971396-DEL-71
chr5-54062616-DEL-327
chr5-54136775-DEL-80
chr5-54147930-DEL-212
chr5-54184084-DEL-3421

chr5-54193784-DEL-71
chr5-54197847-DEL-245
chr5-54211436-DEL-666
chr5-54251286-DEL-1912
chr5-54269670-DEL-63
chr5-54278843-DEL-422
chr5-54288873-DEL-332
chr5-54316183-DEL-220
chr5-54330497-DEL-82
chr5-54400014-DEL-100
chr5-54444658-DEL-56
chr5-54454534-DEL-320
chr5-54466300-DEL-72
chr5-54471943-DEL-664
chr5-54478389-DEL-490
chr5-54489053-DEL-962
chr5-54490938-DEL-113
chr5-54500715-DEL-133
chr5-54501951-DEL-1425
chr5-54536341-DEL-53
chr5-54585294-DEL-85
chr5-54590094-DEL-54
chr5-54590534-DEL-217
chr5-54695277-DEL-194
chr5-54702194-DEL-55
chr5-54705903-DEL-5960
chr5-54715043-DEL-10340
chr5-54716112-DEL-113
chr5-54716982-DEL-7992
chr5-54732438-DEL-2277
chr5-54777614-DEL-868
chr5-54784989-DEL-58
chr5-54808899-DEL-649
chr5-54858125-DEL-332
chr5-54875091-DEL-372
chr5-54881867-DEL-200
chr5-54899218-DEL-389
chr5-54903289-DEL-75
chr5-54917388-DEL-345
chr5-54959693-DEL-68
chr5-54961402-DEL-87
chr5-55035257-DEL-1852
chr5-55063040-DEL-62
chr5-55095875-DEL-93
chr5-55108018-DEL-315
chr5-55125620-DEL-399
chr5-55129434-DEL-2676
chr5-55147502-DEL-398
chr5-55154978-DEL-51
chr5-55174473-DEL-213
chr5-55183969-DEL-71
chr5-55189220-DEL-51
chr5-55197431-DEL-448
chr5-55207626-DEL-56
chr5-55209643-DEL-72
chr5-55236988-DEL-1565
chr5-55278306-DEL-86
chr5-55306149-DEL-6167
chr5-55326139-DEL-294
chr5-55335332-DEL-3079
chr5-55346707-DEL-200
chr5-55381454-DEL-56
chr5-55385940-DEL-213
chr5-55424085-DEL-61
chr5-55424540-DEL-8508
chr5-55454239-DEL-757
chr5-55495496-DEL-684
chr5-55522727-DEL-56
chr5-55524632-DEL-51
chr5-55549150-DEL-1615
chr5-55563585-DEL-148
chr5-55590910-DEL-3016
chr5-55610495-DEL-654
chr5-55613589-DEL-142
chr5-55630543-DEL-1320
chr5-55641001-DEL-3647
chr5-55655086-DEL-1009

chr5-55689635-DEL-54
chr5-55702295-DEL-1420
chr5-55712721-DEL-1427
chr5-55721065-DEL-108
chr5-55721793-DEL-53
chr5-55725993-DEL-107
chr5-55749957-DEL-279
chr5-55753999-DEL-2005
chr5-55780339-DEL-64
chr5-55796402-DEL-77
chr5-55804215-DEL-11499
chr5-55838820-DEL-1132
chr5-55855630-DEL-56
chr5-55865227-DEL-92
chr5-55899393-DEL-4427
chr5-55934658-DEL-471
chr5-55935552-DEL-162
chr5-55959730-DEL-784
chr5-55964352-DEL-84
chr5-56007805-DEL-67
chr5-56010175-DEL-90
chr5-56021461-DEL-84
chr5-56023703-DEL-93
chr5-56028404-DEL-371
chr5-56035781-DEL-52
chr5-56055243-DEL-178
chr5-56084456-DEL-208
chr5-56089248-DEL-188
chr5-56098330-DEL-143
chr5-56101725-DEL-8576
chr5-56110323-DEL-1663
chr5-56129508-DEL-1583
chr5-56132152-DEL-65
chr5-56135450-DEL-205
chr5-56141584-DEL-432
chr5-56142122-DEL-64
chr5-56162465-DEL-162
chr5-56170593-DEL-1190
chr5-56182903-DEL-69
chr5-56205450-DEL-3923
chr5-56224381-DEL-171
chr5-56225635-DEL-5013
chr5-56256394-DEL-147
chr5-56261344-DEL-2732
chr5-56279773-DEL-231
chr5-56328083-DEL-342
chr5-56376971-DEL-80
chr5-56387622-DEL-383
chr5-56424181-DEL-225
chr5-56428367-DEL-151
chr5-56457521-DEL-2092
chr5-56495658-DEL-66
chr5-56518441-DEL-76
chr5-56522075-DEL-1607
chr5-56539164-DEL-57
chr5-56580270-DEL-792
chr5-56595820-DEL-68
chr5-56601881-DEL-73
chr5-56612289-DEL-113
chr5-56627775-DEL-7986
chr5-56628875-DEL-212
chr5-56635664-DEL-54
chr5-56637988-DEL-100
chr5-56647476-DEL-317
chr5-56682133-DEL-52
chr5-56684263-DEL-208
chr5-56686595-DEL-492
chr5-56692266-DEL-296
chr5-56695854-DEL-2020
chr5-56698074-DEL-1643
chr5-56729946-DEL-89
chr5-56741930-DEL-155
chr5-56747305-DEL-66
chr5-56754258-DEL-75
chr5-56793531-DEL-197
chr5-56794498-DEL-1402
chr5-56814230-DEL-2542

chr5-56823343-DEL-60
chr5-56852010-DEL-146
chr5-56853087-DEL-56
chr5-56854084-DEL-1292
chr5-56872731-DEL-52
chr5-56970325-DEL-1037
chr5-56975062-DEL-54
chr5-56983232-DEL-64
chr5-56992747-DEL-104
chr5-57002500-DEL-2703
chr5-57016489-DEL-86
chr5-57024767-DEL-619
chr5-57038498-DEL-275
chr5-57069579-DEL-82
chr5-57077289-DEL-151
chr5-57082036-DEL-357
chr5-57086470-DEL-63
chr5-57089861-DEL-388
chr5-57111293-DEL-2030
chr5-57122017-DEL-11558
chr5-57137732-DEL-93
chr5-57166329-DEL-68
chr5-57168693-DEL-56
chr5-57188859-DEL-1801
chr5-57218537-DEL-200
chr5-57221876-DEL-169
chr5-57232903-DEL-79
chr5-57279532-DEL-670
chr5-57280357-DEL-86
chr5-57280496-DEL-695
chr5-57320032-DEL-59
chr5-57323551-DEL-452
chr5-57325046-DEL-69
chr5-57326995-DEL-179
chr5-57329837-DEL-556
chr5-57374310-DEL-167
chr5-57379480-DEL-111
chr5-57396306-DEL-66
chr5-57406855-DEL-344
chr5-57429293-DEL-566
chr5-57433440-DEL-110
chr5-57514801-DEL-1026
chr5-57518000-DEL-325
chr5-57563330-DEL-119
chr5-57565465-DEL-1235
chr5-57576812-DEL-386
chr5-57577302-DEL-1266
chr5-57629840-DEL-215
chr5-57633560-DEL-1116
chr5-57636467-DEL-87
chr5-57642728-DEL-740
chr5-57648473-DEL-1560
chr5-57655625-DEL-6192
chr5-57743853-DEL-356
chr5-57765708-DEL-800
chr5-57780246-DEL-116
chr5-57782154-DEL-2735
chr5-57784932-DEL-285
chr5-57802663-DEL-68
chr5-57805033-DEL-10998
chr5-57823691-DEL-6688
chr5-57839432-DEL-364
chr5-57844013-DEL-60
chr5-57916904-DEL-286
chr5-57946831-DEL-400
chr5-58000850-DEL-58
chr5-58014159-DEL-72
chr5-58046199-DEL-220
chr5-58052470-DEL-90
chr5-58068549-DEL-76
chr5-58085890-DEL-68
chr5-58090855-DEL-140
chr5-58127043-DEL-56
chr5-58129518-DEL-1833
chr5-58172346-DEL-135
chr5-58185303-DEL-66
chr5-58199612-DEL-180

chr5-58236992-DEL-732
chr5-58302385-DEL-54
chr5-58323460-DEL-1019
chr5-58344007-DEL-88
chr5-58360322-DEL-315
chr5-58363082-DEL-72
chr5-58391377-DEL-3491
chr5-58430897-DEL-1459
chr5-58432866-DEL-1992
chr5-58460079-DEL-62
chr5-58461944-DEL-8886
chr5-58463047-DEL-6221
chr5-58475309-DEL-8186
chr5-58543624-DEL-243
chr5-58573363-DEL-331
chr5-58584483-DEL-878
chr5-58608407-DEL-399
chr5-58631362-DEL-1985
chr5-58655393-DEL-1598
chr5-58703877-DEL-97
chr5-58743794-DEL-98
chr5-58762795-DEL-2923
chr5-58772030-DEL-56
chr5-58798654-DEL-1442
chr5-58804318-DEL-3236
chr5-58821579-DEL-3492
chr5-58847102-DEL-2937
chr5-58857890-DEL-430
chr5-58870522-DEL-6198
chr5-58895921-DEL-270
chr5-58913091-DEL-550
chr5-58926481-DEL-96
chr5-58948968-DEL-235
chr5-58950217-DEL-110
chr5-58985021-DEL-141
chr5-59030858-DEL-2482
chr5-59033370-DEL-3899
chr5-59051322-DEL-58
chr5-59052410-DEL-4517
chr5-59068102-DEL-149
chr5-59069940-DEL-53
chr5-59078338-DEL-72
chr5-59103378-DEL-1825
chr5-59107200-DEL-91
chr5-59113439-DEL-1658
chr5-59113862-DEL-289
chr5-59170962-DEL-721
chr5-59197448-DEL-1102
chr5-59208671-DEL-150
chr5-59218014-DEL-126
chr5-59222968-DEL-273
chr5-59242696-DEL-1428
chr5-59371728-DEL-6876
chr5-59388296-DEL-91
chr5-59418349-DEL-199
chr5-59442452-DEL-844
chr5-59478737-DEL-549
chr5-59496565-DEL-1385
chr5-59504601-DEL-158
chr5-59505288-DEL-54
chr5-59505652-DEL-229
chr5-59512190-DEL-58
chr5-59517098-DEL-55
chr5-59529594-DEL-3819
chr5-59531697-DEL-936
chr5-59550508-DEL-180
chr5-59554742-DEL-354
chr5-59555642-DEL-490
chr5-59572014-DEL-1022
chr5-59610284-DEL-438
chr5-59612178-DEL-648
chr5-59623212-DEL-277
chr5-59630053-DEL-196
chr5-59670769-DEL-77
chr5-59686794-DEL-241
chr5-59692367-DEL-507
chr5-59696862-DEL-144

chr5-59778936-DEL-1118
chr5-59792662-DEL-2262
chr5-59794908-DEL-141
chr5-59816969-DEL-384
chr5-59837849-DEL-1650
chr5-59863110-DEL-139
chr5-59864382-DEL-4465
chr5-59874473-DEL-11583
chr5-59905656-DEL-206
chr5-59922375-DEL-506
chr5-59936476-DEL-198
chr5-59961003-DEL-52
chr5-59988808-DEL-183
chr5-59995377-DEL-637
chr5-60002622-DEL-176
chr5-60022521-DEL-647
chr5-60023598-DEL-490
chr5-60066499-DEL-118
chr5-60087397-DEL-214
chr5-60106293-DEL-7140
chr5-60115354-DEL-165
chr5-60115951-DEL-421
chr5-60131253-DEL-93
chr5-60157265-DEL-124
chr5-60174169-DEL-808
chr5-60176427-DEL-1878
chr5-60192390-DEL-63
chr5-60192637-DEL-975
chr5-60226958-DEL-1761
chr5-60265003-DEL-223
chr5-60265578-DEL-2571
chr5-60376695-DEL-406
chr5-60407259-DEL-145
chr5-60442269-DEL-226
chr5-60462656-DEL-594
chr5-60466404-DEL-673
chr5-60488853-DEL-2423
chr5-60509803-DEL-2265
chr5-60515812-DEL-466
chr5-60529552-DEL-56
chr5-60533596-DEL-190
chr5-60562941-DEL-265
chr5-60592848-DEL-621
chr5-60607163-DEL-60
chr5-60611268-DEL-979
chr5-60615170-DEL-93
chr5-60652014-DEL-6181
chr5-60664310-DEL-61
chr5-60783726-DEL-57
chr5-60786359-DEL-239
chr5-60803604-DEL-447
chr5-60804512-DEL-99
chr5-60828625-DEL-94
chr5-60831803-DEL-1215
chr5-60835976-DEL-2700
chr5-60873710-DEL-379
chr5-60902447-DEL-2716
chr5-60903019-DEL-2168
chr5-60909473-DEL-434
chr5-60918638-DEL-151
chr5-60974850-DEL-1651
chr5-60984813-DEL-1313
chr5-60995300-DEL-135
chr5-61005464-DEL-57
chr5-61006095-DEL-348
chr5-61015611-DEL-62
chr5-61022852-DEL-101
chr5-61069737-DEL-143
chr5-61071289-DEL-66
chr5-61074590-DEL-4254
chr5-61078234-DEL-1098
chr5-61089134-DEL-66
chr5-61109004-DEL-470
chr5-61115436-DEL-6239
chr5-61155404-DEL-64
chr5-61166767-DEL-421
chr5-61176387-DEL-98

chr5-61201208-DEL-987
chr5-61218364-DEL-104
chr5-61221563-DEL-63
chr5-61226191-DEL-62
chr5-61236479-DEL-51
chr5-61238766-DEL-63
chr5-61278353-DEL-338
chr5-61286300-DEL-75
chr5-61302002-DEL-60
chr5-61325543-DEL-511
chr5-61337597-DEL-716
chr5-61341583-DEL-241
chr5-61348723-DEL-85
chr5-61369114-DEL-573
chr5-61370290-DEL-1804
chr5-61375623-DEL-1077
chr5-61389625-DEL-6468
chr5-61389841-DEL-220
chr5-61398629-DEL-388
chr5-61442708-DEL-3511
chr5-61444171-DEL-454
chr5-61450128-DEL-7064
chr5-61450750-DEL-212
chr5-61465101-DEL-6205
chr5-61484239-DEL-400
chr5-61494578-DEL-222
chr5-61513547-DEL-211
chr5-61527943-DEL-68
chr5-61620326-DEL-210
chr5-61639647-DEL-56
chr5-61677451-DEL-627
chr5-61678285-DEL-699
chr5-61679227-DEL-246
chr5-61683370-DEL-51
chr5-61709214-DEL-479
chr5-61713205-DEL-109
chr5-61741665-DEL-787
chr5-61743865-DEL-7816
chr5-61745468-DEL-57
chr5-61747207-DEL-861
chr5-61747992-DEL-224
chr5-61792762-DEL-233
chr5-61809342-DEL-390
chr5-61811867-DEL-64
chr5-61839654-DEL-1293
chr5-61863096-DEL-74
chr5-61864754-DEL-923
chr5-61933330-DEL-26511
chr5-61936034-DEL-212
chr5-61953079-DEL-1572
chr5-61969469-DEL-247
chr5-61978801-DEL-58
chr5-61983169-DEL-2299
chr5-61992965-DEL-554
chr5-61994573-DEL-156
chr5-62009066-DEL-969
chr5-62016802-DEL-439
chr5-62021379-DEL-64
chr5-62035213-DEL-2968
chr5-62040178-DEL-197
chr5-62055819-DEL-295
chr5-62071332-DEL-6657
chr5-62103524-DEL-80
chr5-62122109-DEL-128
chr5-62124780-DEL-280
chr5-62136385-DEL-216
chr5-62144086-DEL-54
chr5-62157725-DEL-157
chr5-62162023-DEL-180
chr5-62167892-DEL-52
chr5-62168893-DEL-4517
chr5-62186143-DEL-95
chr5-62186849-DEL-6935
chr5-62186849-DEL-7509
chr5-62195305-DEL-62
chr5-62210475-DEL-78
chr5-62261714-DEL-911

chr5-62291294-DEL-1015
chr5-62295699-DEL-108
chr5-62324814-DEL-130
chr5-62327429-DEL-488
chr5-62331885-DEL-78
chr5-62366625-DEL-6419
chr5-62366721-DEL-212
chr5-62376921-DEL-6671
chr5-62383098-DEL-214
chr5-62385040-DEL-6869
chr5-62397995-DEL-6425
chr5-62419425-DEL-234
chr5-62429120-DEL-6401
chr5-62483414-DEL-6342
chr5-62529903-DEL-168
chr5-62543023-DEL-89
chr5-62544009-DEL-68
chr5-62550881-DEL-6274
chr5-62565470-DEL-2338
chr5-62575381-DEL-1423
chr5-62584858-DEL-1047
chr5-62589375-DEL-1719
chr5-62591325-DEL-224
chr5-62595263-DEL-51
chr5-62600810-DEL-1803
chr5-62623074-DEL-132
chr5-62626879-DEL-63
chr5-62672596-DEL-124
chr5-62674765-DEL-199
chr5-62681818-DEL-6341
chr5-62705156-DEL-1566
chr5-62717908-DEL-84
chr5-62720651-DEL-135
chr5-62732231-DEL-52
chr5-62755418-DEL-263
chr5-62789297-DEL-134
chr5-62818091-DEL-54
chr5-62839485-DEL-946
chr5-62876961-DEL-179
chr5-62886700-DEL-1052
chr5-62895001-DEL-76
chr5-62930173-DEL-2785
chr5-62932936-DEL-53
chr5-62939719-DEL-139
chr5-62947373-DEL-1466
chr5-62962283-DEL-56
chr5-62991714-DEL-1903
chr5-63079352-DEL-134
chr5-63094295-DEL-1875
chr5-63096990-DEL-1298
chr5-63134451-DEL-4170
chr5-63141694-DEL-86
chr5-63146669-DEL-759
chr5-63162851-DEL-978
chr5-63173261-DEL-66
chr5-63185074-DEL-854
chr5-63227723-DEL-877
chr5-63230477-DEL-100
chr5-63233358-DEL-650
chr5-63235127-DEL-85
chr5-63244790-DEL-5429
chr5-63269630-DEL-200
chr5-63281910-DEL-684
chr5-63292315-DEL-56
chr5-63295704-DEL-81
chr5-63343912-DEL-400
chr5-63347586-DEL-2450
chr5-63369368-DEL-107
chr5-63402350-DEL-272
chr5-63403207-DEL-407
chr5-63408399-DEL-1886
chr5-63433155-DEL-606
chr5-63451275-DEL-264
chr5-63478557-DEL-373
chr5-63525835-DEL-6218
chr5-63565090-DEL-251
chr5-63612506-DEL-2159

chr5-63628278-DEL-223
chr5-63677599-DEL-2529
chr5-63687233-DEL-224
chr5-63692221-DEL-137
chr5-63724799-DEL-52
chr5-63757713-DEL-6245
chr5-63767341-DEL-93
chr5-63782478-DEL-52
chr5-63797632-DEL-895
chr5-63840428-DEL-63
chr5-63843641-DEL-310
chr5-63846868-DEL-1039
chr5-63863684-DEL-583
chr5-63897283-DEL-582
chr5-63916494-DEL-731
chr5-63917345-DEL-1564
chr5-63931437-DEL-98
chr5-64039394-DEL-60
chr5-64054803-DEL-179
chr5-64108354-DEL-185
chr5-64110554-DEL-190
chr5-64129248-DEL-244
chr5-64153150-DEL-58
chr5-64158008-DEL-111
chr5-64170577-DEL-7626
chr5-64212948-DEL-1472
chr5-64218663-DEL-116
chr5-64228888-DEL-70
chr5-64235497-DEL-4472
chr5-64237174-DEL-387
chr5-64260872-DEL-121
chr5-64261909-DEL-953
chr5-64301608-DEL-312
chr5-64326171-DEL-233
chr5-64342800-DEL-676
chr5-64347728-DEL-57
chr5-64361289-DEL-165
chr5-64363068-DEL-87
chr5-64372175-DEL-829
chr5-64380022-DEL-217
chr5-64404928-DEL-64
chr5-64416414-DEL-90
chr5-64426213-DEL-75
chr5-64426623-DEL-595
chr5-64442508-DEL-55
chr5-64443598-DEL-93
chr5-64581617-DEL-1096
chr5-64592723-DEL-944
chr5-64607113-DEL-120
chr5-64650854-DEL-54
chr5-64666594-DEL-57
chr5-64670794-DEL-60
chr5-64675026-DEL-53
chr5-64702182-DEL-6345
chr5-64739452-DEL-66
chr5-64794583-DEL-190
chr5-64800389-DEL-65
chr5-64836143-DEL-209
chr5-64859962-DEL-57
chr5-64917296-DEL-510
chr5-64919989-DEL-73
chr5-65006457-DEL-73
chr5-65026445-DEL-1931
chr5-65029490-DEL-191
chr5-65039123-DEL-74
chr5-65041960-DEL-85
chr5-65058478-DEL-134
chr5-65090569-DEL-54
chr5-65093527-DEL-1237
chr5-65098682-DEL-1895
chr5-65106500-DEL-363
chr5-65123490-DEL-118
chr5-65138037-DEL-2494
chr5-65140564-DEL-1890
chr5-65160154-DEL-60
chr5-65170900-DEL-77
chr5-65199062-DEL-151

chr5-65201866-DEL-592
chr5-65272504-DEL-73
chr5-65321398-DEL-68
chr5-65389361-DEL-454
chr5-65393414-DEL-84
chr5-65447309-DEL-913
chr5-65454046-DEL-65
chr5-65471288-DEL-204
chr5-65490377-DEL-312
chr5-65496624-DEL-204
chr5-65513127-DEL-339
chr5-65537811-DEL-1282
chr5-65573781-DEL-71
chr5-65585425-DEL-102
chr5-65599899-DEL-163
chr5-65601049-DEL-1031
chr5-65604666-DEL-135
chr5-65605885-DEL-295
chr5-65619354-DEL-1145
chr5-65622756-DEL-145
chr5-65685256-DEL-2692
chr5-65709378-DEL-348
chr5-65710601-DEL-203
chr5-65719820-DEL-102
chr5-65746496-DEL-53
chr5-65780144-DEL-112
chr5-65818242-DEL-80
chr5-65820594-DEL-132
chr5-65827818-DEL-204
chr5-65830612-DEL-53
chr5-65864965-DEL-673
chr5-65871045-DEL-566
chr5-65874112-DEL-548
chr5-65875527-DEL-1065
chr5-65888026-DEL-56
chr5-65929631-DEL-1239
chr5-65953784-DEL-259
chr5-65994370-DEL-397
chr5-66047319-DEL-77
chr5-66069065-DEL-85
chr5-66073267-DEL-56
chr5-66081625-DEL-54
chr5-66120412-DEL-74
chr5-66134906-DEL-198
chr5-66141351-DEL-69
chr5-66149550-DEL-243
chr5-66231944-DEL-563
chr5-66232804-DEL-240
chr5-66243547-DEL-67
chr5-66245948-DEL-81
chr5-66248754-DEL-52
chr5-66315799-DEL-152
chr5-66346763-DEL-64
chr5-66359343-DEL-126
chr5-66370676-DEL-160
chr5-66392338-DEL-5610
chr5-66395907-DEL-84
chr5-66398612-DEL-210
chr5-66411464-DEL-1015
chr5-66415658-DEL-287
chr5-66416791-DEL-60
chr5-66422187-DEL-178
chr5-66425014-DEL-60
chr5-66432741-DEL-51
chr5-66441117-DEL-64
chr5-66447541-DEL-299
chr5-66521965-DEL-51
chr5-66559680-DEL-58
chr5-66701632-DEL-58
chr5-66707047-DEL-67
chr5-66761609-DEL-254
chr5-66762191-DEL-97
chr5-66768420-DEL-109
chr5-66786090-DEL-73
chr5-66794475-DEL-266
chr5-66818582-DEL-98
chr5-66831179-DEL-99

chr5-66870758-DEL-159
chr5-66884044-DEL-54
chr5-66913134-DEL-650
chr5-66918613-DEL-72
chr5-66962335-DEL-87
chr5-66969966-DEL-198
chr5-66985263-DEL-169
chr5-67014323-DEL-82
chr5-67043767-DEL-73
chr5-67067911-DEL-98
chr5-67098050-DEL-60
chr5-67122534-DEL-121
chr5-67153359-DEL-54
chr5-67160397-DEL-64
chr5-67161154-DEL-68
chr5-67219874-DEL-228
chr5-67254293-DEL-70
chr5-67275508-DEL-102
chr5-67313098-DEL-188
chr5-67358956-DEL-751
chr5-67382053-DEL-192
chr5-67396994-DEL-1173
chr5-67458764-DEL-318
chr5-67462614-DEL-191
chr5-67471614-DEL-54
chr5-67481467-DEL-162
chr5-67508137-DEL-56
chr5-67529503-DEL-2084
chr5-67537109-DEL-335
chr5-67579221-DEL-64
chr5-67615303-DEL-109
chr5-67648999-DEL-215
chr5-67671869-DEL-116
chr5-67674682-DEL-173
chr5-67717020-DEL-865
chr5-67719315-DEL-99
chr5-67734237-DEL-247
chr5-67747309-DEL-52
chr5-67792853-DEL-77
chr5-67823703-DEL-64
chr5-67857096-DEL-2091
chr5-67885735-DEL-72
chr5-67912485-DEL-120
chr5-67968077-DEL-52
chr5-68084634-DEL-1560
chr5-68094198-DEL-294
chr5-68098713-DEL-82
chr5-68109171-DEL-372
chr5-68116303-DEL-54
chr5-68154345-DEL-134
chr5-68173901-DEL-221
chr5-68175577-DEL-712
chr5-68209578-DEL-72
chr5-68225912-DEL-53
chr5-68252337-DEL-460
chr5-68256676-DEL-58
chr5-68263653-DEL-177
chr5-68321602-DEL-247
chr5-68325212-DEL-944
chr5-68344578-DEL-118
chr5-68363092-DEL-1019
chr5-68365569-DEL-110
chr5-68378190-DEL-208
chr5-68379925-DEL-887
chr5-68439025-DEL-1032
chr5-68440640-DEL-136
chr5-68469154-DEL-621
chr5-68513466-DEL-51
chr5-68529159-DEL-54
chr5-68560845-DEL-98
chr5-68565445-DEL-61
chr5-68568985-DEL-500
chr5-68596705-DEL-62
chr5-68604943-DEL-64
chr5-68606115-DEL-6042
chr5-68613974-DEL-1378
chr5-68624328-DEL-57

chr5-68628812-DEL-1333
chr5-68641239-DEL-112
chr5-68643159-DEL-554
chr5-68667260-DEL-143
chr5-68672858-DEL-204
chr5-68680461-DEL-429
chr5-68721288-DEL-790
chr5-68745596-DEL-207
chr5-68770355-DEL-95
chr5-68785313-DEL-577
chr5-68805608-DEL-58
chr5-68817650-DEL-976
chr5-68825313-DEL-1486
chr5-68828368-DEL-221
chr5-68829130-DEL-71
chr5-68834612-DEL-147
chr5-68835601-DEL-124
chr5-68874468-DEL-129
chr5-68882368-DEL-640
chr5-68891847-DEL-1958
chr5-68910617-DEL-226
chr5-68938623-DEL-82
chr5-68959686-DEL-1596
chr5-68995185-DEL-60
chr5-69005472-DEL-6189
chr5-69006647-DEL-1304
chr5-69030276-DEL-97
chr5-69073549-DEL-104
chr5-69100194-DEL-60
chr5-69104032-DEL-54
chr5-69154195-DEL-683
chr5-69160674-DEL-682
chr5-69228826-DEL-81
chr5-69229082-DEL-888
chr5-69249249-DEL-98
chr5-69280727-DEL-1255
chr5-69331673-DEL-61
chr5-69359967-DEL-62
chr5-69395801-DEL-68
chr5-69397770-DEL-1614
chr5-69431749-DEL-631
chr5-69461438-DEL-181
chr5-69508573-DEL-154
chr5-69537742-DEL-706
chr5-69571839-DEL-218
chr5-69647400-DEL-214
chr5-69647915-DEL-6248
chr5-69657461-DEL-332
chr5-69670736-DEL-895
chr5-69688806-DEL-6981
chr5-69736011-DEL-54
chr5-69766224-DEL-108
chr5-69772004-DEL-1034
chr5-69773192-DEL-96
chr5-69800378-DEL-1164
chr5-69814613-DEL-59
chr5-69849304-DEL-202
chr5-69858536-DEL-210
chr5-69859579-DEL-268
chr5-69894970-DEL-54
chr5-69918970-DEL-426
chr5-69921291-DEL-85
chr5-69977914-DEL-6139
chr5-69999662-DEL-53
chr5-70033668-DEL-131
chr5-70036964-DEL-91
chr5-70049730-DEL-520
chr5-70082175-DEL-52
chr5-70107017-DEL-145
chr5-70121290-DEL-9082
chr5-70140045-DEL-60
chr5-70153172-DEL-7674
chr5-70157217-DEL-1291
chr5-70159547-DEL-6182
chr5-70224879-DEL-92
chr5-70244350-DEL-2655
chr5-70256289-DEL-186

chr5-70273440-DEL-56
chr5-70274363-DEL-4197
chr5-70281666-DEL-813
chr5-70285931-DEL-5280
chr5-70291418-DEL-1275
chr5-70315369-DEL-55
chr5-70346627-DEL-96
chr5-70371225-DEL-60
chr5-70381211-DEL-112
chr5-70399426-DEL-2676
chr5-70416474-DEL-64
chr5-70454296-DEL-621
chr5-70494050-DEL-86
chr5-70522611-DEL-248
chr5-70530241-DEL-66
chr5-70535573-DEL-6585
chr5-70548843-DEL-67
chr5-70586832-DEL-199
chr5-70587210-DEL-4435
chr5-70607503-DEL-223
chr5-70616939-DEL-1086
chr5-70633294-DEL-93
chr5-70670467-DEL-2237
chr5-70673249-DEL-87
chr5-70674677-DEL-4284
chr5-70699056-DEL-419
chr5-70699632-DEL-6440
chr5-70714297-DEL-80
chr5-70715377-DEL-12475
chr5-70716463-DEL-556
chr5-70718659-DEL-7159
chr5-70743711-DEL-87
chr5-70745818-DEL-1735
chr5-70768997-DEL-51
chr5-70776418-DEL-133
chr5-70788053-DEL-365
chr5-70822161-DEL-99
chr5-70823596-DEL-133
chr5-70829146-DEL-334
chr5-70839333-DEL-6009
chr5-70867322-DEL-53
chr5-70872491-DEL-508
chr5-70881905-DEL-641
chr5-70918841-DEL-206
chr5-70962296-DEL-509
chr5-70968656-DEL-357
chr5-70990104-DEL-1221
chr5-71038660-DEL-55
chr5-71039298-DEL-718
chr5-71065468-DEL-72
chr5-71070417-DEL-449
chr5-71077813-DEL-162
chr5-71081652-DEL-718
chr5-71082966-DEL-118
chr5-71087456-DEL-670
chr5-71141632-DEL-340
chr5-71156692-DEL-1498
chr5-71195317-DEL-304
chr5-71202957-DEL-60
chr5-71207235-DEL-828
chr5-71225035-DEL-705
chr5-71296091-DEL-1320
chr5-71298213-DEL-86
chr5-71351173-DEL-403
chr5-71430349-DEL-1977
chr5-71435504-DEL-106
chr5-71462335-DEL-78
chr5-71464852-DEL-95
chr5-71496460-DEL-165
chr5-71499605-DEL-6595
chr5-71506930-DEL-689
chr5-71511337-DEL-841
chr5-71517832-DEL-870
chr5-71521043-DEL-679
chr5-71532973-DEL-706
chr5-71556598-DEL-4404
chr5-71556620-DEL-2784

chr5-71566573-DEL-83
chr5-71570193-DEL-303
chr5-71573509-DEL-1044
chr5-71590289-DEL-75
chr5-71603553-DEL-202
chr5-71615713-DEL-2369
chr5-71641779-DEL-66
chr5-71644319-DEL-109
chr5-71661248-DEL-91
chr5-71702910-DEL-74
chr5-71731255-DEL-75
chr5-71743494-DEL-96
chr5-71794966-DEL-52
chr5-71808231-DEL-386
chr5-71812160-DEL-96
chr5-71831251-DEL-107
chr5-71832036-DEL-2561
chr5-71846356-DEL-450
chr5-71861335-DEL-67
chr5-71898704-DEL-72
chr5-71925106-DEL-829
chr5-71977466-DEL-350
chr5-71978447-DEL-56
chr5-72020453-DEL-2760
chr5-72034370-DEL-1995
chr5-72037584-DEL-287
chr5-72054816-DEL-378
chr5-72087120-DEL-55
chr5-72118753-DEL-70
chr5-72160351-DEL-54
chr5-72171959-DEL-88
chr5-72189132-DEL-205
chr5-72207102-DEL-795
chr5-72215091-DEL-72
chr5-72215484-DEL-288
chr5-72238007-DEL-56
chr5-72244932-DEL-420
chr5-72265868-DEL-86
chr5-72269309-DEL-6451
chr5-72293525-DEL-88
chr5-72337638-DEL-1211
chr5-72372196-DEL-111
chr5-72390565-DEL-2371
chr5-72402250-DEL-185
chr5-72428408-DEL-692
chr5-72459664-DEL-203
chr5-72513922-DEL-61
chr5-72524324-DEL-535
chr5-72545650-DEL-88
chr5-72588444-DEL-54
chr5-72620197-DEL-1589
chr5-72626587-DEL-137
chr5-72627080-DEL-698
chr5-72649290-DEL-90
chr5-72692069-DEL-171
chr5-72708803-DEL-608
chr5-72742275-DEL-102
chr5-72789598-DEL-87
chr5-72800479-DEL-60
chr5-72843588-DEL-77
chr5-72848194-DEL-441
chr5-72872676-DEL-88
chr5-72949876-DEL-308
chr5-72983495-DEL-158
chr5-73000455-DEL-249
chr5-73003542-DEL-659
chr5-73004785-DEL-79
chr5-73014910-DEL-72
chr5-73056328-DEL-250
chr5-73060964-DEL-203
chr5-73064520-DEL-178
chr5-73081386-DEL-69
chr5-73088119-DEL-65
chr5-73102031-DEL-875
chr5-73119164-DEL-56
chr5-73146048-DEL-54
chr5-73146723-DEL-91

chr5-73150830-DEL-186
chr5-73155518-DEL-80
chr5-73159707-DEL-1311
chr5-73199106-DEL-2580
chr5-73215688-DEL-130
chr5-73276657-DEL-86
chr5-73316228-DEL-71
chr5-73323580-DEL-69
chr5-73355408-DEL-61
chr5-73371038-DEL-738
chr5-73394157-DEL-595
chr5-73396438-DEL-87
chr5-73424371-DEL-192
chr5-73482914-DEL-94
chr5-73497972-DEL-70
chr5-73503808-DEL-75
chr5-73508320-DEL-159
chr5-73550926-DEL-537
chr5-73554284-DEL-249
chr5-73556618-DEL-64
chr5-73559947-DEL-214
chr5-73562727-DEL-210
chr5-73569648-DEL-211
chr5-73591847-DEL-128
chr5-73601683-DEL-3704
chr5-73605407-DEL-113
chr5-73610825-DEL-295
chr5-73623121-DEL-158
chr5-73645914-DEL-343
chr5-73684836-DEL-228
chr5-73726696-DEL-58
chr5-73744898-DEL-6136
chr5-73756402-DEL-256
chr5-73757341-DEL-60
chr5-73759318-DEL-2791
chr5-73784372-DEL-152
chr5-73810235-DEL-1394
chr5-73811562-DEL-325
chr5-73819112-DEL-677
chr5-73853523-DEL-69
chr5-73856423-DEL-382
chr5-73888327-DEL-56
chr5-73916922-DEL-913
chr5-73958981-DEL-2580
chr5-73964272-DEL-198
chr5-73993907-DEL-6153
chr5-74018933-DEL-61
chr5-74041165-DEL-81
chr5-74099253-DEL-226
chr5-74109665-DEL-56
chr5-74155581-DEL-167
chr5-74156010-DEL-489
chr5-74168166-DEL-56
chr5-74169839-DEL-196
chr5-74183455-DEL-66
chr5-74191796-DEL-54
chr5-74192327-DEL-52
chr5-74195634-DEL-1693
chr5-74206612-DEL-152
chr5-74218003-DEL-414
chr5-74238408-DEL-137
chr5-74251119-DEL-163
chr5-74266753-DEL-1136
chr5-74274248-DEL-96
chr5-74301137-DEL-116
chr5-74432519-DEL-107
chr5-74441438-DEL-87
chr5-74447450-DEL-400
chr5-74459594-DEL-70
chr5-74469410-DEL-248
chr5-74477696-DEL-111
chr5-74500567-DEL-767
chr5-74510678-DEL-168
chr5-74512125-DEL-56
chr5-74565394-DEL-274
chr5-74580675-DEL-96
chr5-74621178-DEL-106

chr5-74629568-DEL-203
chr5-74646809-DEL-838
chr5-74647673-DEL-555
chr5-74655799-DEL-919
chr5-74656998-DEL-92
chr5-74659751-DEL-189
chr5-74695248-DEL-58
chr5-74710869-DEL-4176
chr5-74729325-DEL-110
chr5-74744034-DEL-306
chr5-74778946-DEL-55
chr5-74809677-DEL-3140
chr5-74827546-DEL-64
chr5-74832282-DEL-97
chr5-74879884-DEL-79
chr5-74906871-DEL-197
chr5-74913402-DEL-94
chr5-74932653-DEL-274
chr5-74942238-DEL-697
chr5-74942907-DEL-60
chr5-74951312-DEL-143
chr5-74956579-DEL-58
chr5-74996996-DEL-119
chr5-75010977-DEL-1611
chr5-75018432-DEL-712
chr5-75110943-DEL-87
chr5-75128053-DEL-52
chr5-75143844-DEL-128
chr5-75145376-DEL-57
chr5-75148642-DEL-2346
chr5-75149042-DEL-80
chr5-75194106-DEL-51
chr5-75194852-DEL-127
chr5-75259318-DEL-430
chr5-75271580-DEL-140
chr5-75281453-DEL-104
chr5-75285669-DEL-66
chr5-75388652-DEL-99
chr5-75397427-DEL-135
chr5-75425574-DEL-453
chr5-75455777-DEL-53
chr5-75457346-DEL-80
chr5-75463242-DEL-80
chr5-75488413-DEL-53
chr5-75517020-DEL-465
chr5-75540926-DEL-200
chr5-75551275-DEL-60
chr5-75594333-DEL-360
chr5-75608139-DEL-118
chr5-75612855-DEL-1043
chr5-75622728-DEL-262
chr5-75635325-DEL-317
chr5-75662188-DEL-112
chr5-75700695-DEL-151
chr5-75703334-DEL-341
chr5-75721140-DEL-727
chr5-75728232-DEL-54
chr5-75762739-DEL-320
chr5-75800932-DEL-60
chr5-75808210-DEL-129
chr5-75820973-DEL-490
chr5-75821780-DEL-249
chr5-75825967-DEL-51
chr5-75842635-DEL-62
chr5-75890157-DEL-204
chr5-75930976-DEL-132
chr5-75939890-DEL-62
chr5-75963567-DEL-84
chr5-76020134-DEL-89
chr5-76035445-DEL-62
chr5-76049647-DEL-52
chr5-76115470-DEL-53
chr5-76188579-DEL-124
chr5-76193969-DEL-68
chr5-76220550-DEL-914
chr5-76229130-DEL-381
chr5-76242894-DEL-290

chr5-76245321-DEL-1924
chr5-76254699-DEL-119
chr5-76267146-DEL-61
chr5-76299388-DEL-423
chr5-76305330-DEL-64
chr5-76310684-DEL-470
chr5-76314763-DEL-235
chr5-76335264-DEL-70
chr5-76376116-DEL-174
chr5-76385308-DEL-165
chr5-76394840-DEL-425
chr5-76436707-DEL-193
chr5-76476606-DEL-440
chr5-76574514-DEL-264
chr5-76583702-DEL-119
chr5-76615593-DEL-164
chr5-76646135-DEL-401
chr5-76651217-DEL-252
chr5-76666204-DEL-407
chr5-76690805-DEL-203
chr5-76699819-DEL-55
chr5-76713403-DEL-88
chr5-76721282-DEL-512
chr5-76728778-DEL-95
chr5-76760060-DEL-168
chr5-76767574-DEL-457
chr5-76786307-DEL-112
chr5-76826439-DEL-143
chr5-76829695-DEL-112
chr5-76838739-DEL-66
chr5-76871117-DEL-1267
chr5-76879924-DEL-70
chr5-76904736-DEL-128
chr5-76906030-DEL-134
chr5-76909773-DEL-191
chr5-76924177-DEL-94
chr5-76924203-DEL-1337
chr5-77043805-DEL-52
chr5-77050892-DEL-115
chr5-77056662-DEL-1922
chr5-77059462-DEL-203
chr5-77078156-DEL-10849
chr5-77103105-DEL-165
chr5-77106878-DEL-170
chr5-77135107-DEL-319
chr5-77186714-DEL-1498
chr5-77222621-DEL-1105
chr5-77226493-DEL-443
chr5-77272024-DEL-533
chr5-77291354-DEL-70
chr5-77295143-DEL-213
chr5-77313595-DEL-97
chr5-77318056-DEL-503
chr5-77351196-DEL-524
chr5-77353286-DEL-126
chr5-77355724-DEL-160
chr5-77376498-DEL-206
chr5-77391212-DEL-228
chr5-77406444-DEL-56
chr5-77410608-DEL-57
chr5-77429675-DEL-54
chr5-77437892-DEL-1848
chr5-77444237-DEL-226
chr5-77447772-DEL-257
chr5-77453063-DEL-210
chr5-77477501-DEL-54
chr5-77497562-DEL-52
chr5-77518672-DEL-408
chr5-77522110-DEL-56
chr5-77557452-DEL-6544
chr5-77569567-DEL-381
chr5-77571025-DEL-427
chr5-77602114-DEL-59
chr5-77617570-DEL-122
chr5-77618598-DEL-302
chr5-77643556-DEL-320
chr5-77654527-DEL-436

chr5-77662090-DEL-75
chr5-77720130-DEL-195
chr5-77744307-DEL-263
chr5-77753798-DEL-160
chr5-77762639-DEL-239
chr5-77789952-DEL-762
chr5-77800722-DEL-60
chr5-77811719-DEL-710
chr5-77884147-DEL-402
chr5-77899433-DEL-15239
chr5-77912908-DEL-10609
chr5-77918227-DEL-4521
chr5-77926041-DEL-58
chr5-77933399-DEL-66
chr5-77945711-DEL-200
chr5-77968381-DEL-83
chr5-77987705-DEL-100
chr5-78011491-DEL-991
chr5-78084737-DEL-2255
chr5-78136607-DEL-6614
chr5-78142702-DEL-427
chr5-78145176-DEL-6326
chr5-78168721-DEL-85
chr5-78172514-DEL-55
chr5-78173028-DEL-5452
chr5-78244972-DEL-196
chr5-78251191-DEL-104
chr5-78265444-DEL-66
chr5-78276149-DEL-6554
chr5-78276376-DEL-208
chr5-78284548-DEL-85
chr5-78288808-DEL-6490
chr5-78295367-DEL-91
chr5-78301805-DEL-6481
chr5-78309292-DEL-338
chr5-78323766-DEL-281
chr5-78381956-DEL-329
chr5-78385699-DEL-508
chr5-78386621-DEL-2297
chr5-78410139-DEL-624
chr5-78424482-DEL-642
chr5-78434717-DEL-353
chr5-78458560-DEL-117
chr5-78504989-DEL-155
chr5-78506002-DEL-1066
chr5-78511939-DEL-422
chr5-78560509-DEL-6236
chr5-78574628-DEL-184
chr5-78588741-DEL-1501
chr5-78612414-DEL-5172
chr5-78628719-DEL-74
chr5-78659115-DEL-1493
chr5-78694167-DEL-142
chr5-78695249-DEL-584
chr5-78716914-DEL-787
chr5-78730416-DEL-377
chr5-78754201-DEL-232
chr5-78799985-DEL-70
chr5-78839674-DEL-201
chr5-78843031-DEL-318
chr5-78875502-DEL-6212
chr5-78901405-DEL-231
chr5-78911885-DEL-133
chr5-78920722-DEL-325
chr5-78924438-DEL-311
chr5-78952260-DEL-100
chr5-79010383-DEL-66
chr5-79029010-DEL-72
chr5-79048164-DEL-125
chr5-79097416-DEL-154
chr5-79098712-DEL-141
chr5-79138158-DEL-428
chr5-79173775-DEL-2677
chr5-79174821-DEL-133
chr5-79193749-DEL-4224
chr5-79194369-DEL-3690
chr5-79204339-DEL-646

chr5-79238853-DEL-108
chr5-79256684-DEL-68
chr5-79269258-DEL-61
chr5-79296140-DEL-178
chr5-79327728-DEL-63
chr5-79333620-DEL-1141
chr5-79349966-DEL-3843
chr5-79362405-DEL-112
chr5-79362635-DEL-1005
chr5-79419328-DEL-256
chr5-79477300-DEL-1238
chr5-79490843-DEL-2330
chr5-79511674-DEL-55
chr5-79519473-DEL-117
chr5-79534320-DEL-96
chr5-79550780-DEL-1232
chr5-79553058-DEL-54
chr5-79579754-DEL-1525
chr5-79606335-DEL-444
chr5-79608287-DEL-1453
chr5-79613301-DEL-99
chr5-79643435-DEL-6217
chr5-79661266-DEL-153
chr5-79663440-DEL-183
chr5-79695292-DEL-54
chr5-79724790-DEL-3414
chr5-79728500-DEL-338
chr5-79729583-DEL-147
chr5-79744448-DEL-62
chr5-79762823-DEL-252
chr5-79785910-DEL-463
chr5-79841748-DEL-855
chr5-79846977-DEL-600
chr5-79894123-DEL-114
chr5-79898810-DEL-452
chr5-79929714-DEL-85
chr5-79981818-DEL-543
chr5-79982529-DEL-54
chr5-79989873-DEL-1592
chr5-79995157-DEL-344
chr5-80029976-DEL-52
chr5-80049458-DEL-265
chr5-80075636-DEL-288
chr5-80079451-DEL-204
chr5-80098931-DEL-98
chr5-80102369-DEL-6681
chr5-80117505-DEL-494
chr5-80135603-DEL-93
chr5-80140816-DEL-6445
chr5-80154727-DEL-358
chr5-80166534-DEL-88
chr5-80170387-DEL-1859
chr5-80183156-DEL-57
chr5-80201841-DEL-2253
chr5-80212382-DEL-186
chr5-80220790-DEL-338
chr5-80221171-DEL-2644
chr5-80261234-DEL-61
chr5-80279663-DEL-2778
chr5-80325360-DEL-157
chr5-80328934-DEL-52
chr5-80341542-DEL-53
chr5-80357219-DEL-2016
chr5-80473614-DEL-519
chr5-80482748-DEL-64
chr5-80488438-DEL-137
chr5-80498836-DEL-170
chr5-80535623-DEL-738
chr5-80554304-DEL-86
chr5-80565614-DEL-1176
chr5-80566963-DEL-829
chr5-80567849-DEL-244
chr5-80568162-DEL-1019
chr5-80594430-DEL-3321
chr5-80611292-DEL-834
chr5-80625429-DEL-211
chr5-80628533-DEL-405

chr5-80704672-DEL-6072
chr5-80735505-DEL-59
chr5-80739367-DEL-121
chr5-80785010-DEL-86
chr5-80803553-DEL-1684
chr5-80809265-DEL-58
chr5-80812140-DEL-311
chr5-80815506-DEL-439
chr5-80837303-DEL-916
chr5-80853173-DEL-52
chr5-80854220-DEL-97
chr5-80854955-DEL-1519
chr5-80856809-DEL-791
chr5-80879082-DEL-67
chr5-80881445-DEL-388
chr5-80913300-DEL-424
chr5-80928372-DEL-372
chr5-80929330-DEL-551
chr5-80929990-DEL-940
chr5-80969599-DEL-400
chr5-81008697-DEL-61
chr5-81059635-DEL-998
chr5-81105756-DEL-88
chr5-81119289-DEL-220
chr5-81124216-DEL-1342
chr5-81150756-DEL-80
chr5-81255787-DEL-737
chr5-81334535-DEL-103
chr5-81387396-DEL-2908
chr5-81412872-DEL-56
chr5-81420097-DEL-428
chr5-81423981-DEL-80
chr5-81522034-DEL-52
chr5-81545194-DEL-201
chr5-81547822-DEL-63
chr5-81569947-DEL-100
chr5-81571960-DEL-1739
chr5-81576626-DEL-388
chr5-81586558-DEL-2065
chr5-81587587-DEL-52
chr5-81599283-DEL-1173
chr5-81642865-DEL-2221
chr5-81663878-DEL-92
chr5-81676703-DEL-69
chr5-81679741-DEL-200
chr5-81703245-DEL-55
chr5-81710718-DEL-160
chr5-81739271-DEL-69
chr5-81814134-DEL-189
chr5-81845733-DEL-133
chr5-81867577-DEL-51
chr5-81876327-DEL-186
chr5-81953927-DEL-580
chr5-81966256-DEL-908
chr5-81969673-DEL-167
chr5-81986317-DEL-101
chr5-82001548-DEL-1623
chr5-82025467-DEL-744
chr5-82027321-DEL-79
chr5-82090611-DEL-442
chr5-82095015-DEL-55
chr5-82128346-DEL-53
chr5-82149097-DEL-1190
chr5-82211051-DEL-224
chr5-82212748-DEL-488
chr5-82215743-DEL-159
chr5-82234524-DEL-753
chr5-82242199-DEL-165
chr5-82251085-DEL-58
chr5-82252283-DEL-895
chr5-82268849-DEL-1414
chr5-82285257-DEL-2261
chr5-82292918-DEL-620
chr5-82308103-DEL-178
chr5-82317826-DEL-56
chr5-82327431-DEL-128
chr5-82327880-DEL-6255

chr5-82338680-DEL-57
chr5-82377560-DEL-448
chr5-82381369-DEL-674
chr5-82397567-DEL-373
chr5-82421510-DEL-717
chr5-82462564-DEL-506
chr5-82467900-DEL-152
chr5-82481631-DEL-147
chr5-82484679-DEL-890
chr5-82555786-DEL-1284
chr5-82574369-DEL-54
chr5-82617488-DEL-73
chr5-82623598-DEL-2561
chr5-82626389-DEL-2437
chr5-82630701-DEL-988
chr5-82666443-DEL-119
chr5-82695178-DEL-93
chr5-82698656-DEL-110
chr5-82741045-DEL-99
chr5-82755399-DEL-168
chr5-82758485-DEL-668
chr5-82761434-DEL-6177
chr5-82771626-DEL-304
chr5-82791389-DEL-172
chr5-82791803-DEL-6217
chr5-82801044-DEL-82
chr5-82801326-DEL-6678
chr5-82809638-DEL-1889
chr5-82813157-DEL-59
chr5-82813784-DEL-82
chr5-82832253-DEL-60
chr5-82834583-DEL-3077
chr5-82846026-DEL-1464
chr5-82849025-DEL-3429
chr5-82862094-DEL-93
chr5-82867204-DEL-261
chr5-82917840-DEL-2938
chr5-82924270-DEL-124
chr5-82961392-DEL-2933
chr5-82980485-DEL-73
chr5-82995194-DEL-80
chr5-83018319-DEL-352
chr5-83037110-DEL-5750
chr5-83043104-DEL-548
chr5-83050407-DEL-82
chr5-83134309-DEL-57
chr5-83205134-DEL-204
chr5-83218848-DEL-1753
chr5-83224946-DEL-3959
chr5-83251336-DEL-187
chr5-83290990-DEL-855
chr5-83310589-DEL-1467
chr5-83312778-DEL-108
chr5-83314006-DEL-69
chr5-83322849-DEL-64
chr5-83335803-DEL-1058
chr5-83380652-DEL-1212
chr5-83380894-DEL-52
chr5-83388069-DEL-218
chr5-83415885-DEL-6350
chr5-83438178-DEL-58
chr5-83470912-DEL-134
chr5-83473531-DEL-592
chr5-83482109-DEL-909
chr5-83510896-DEL-2116
chr5-83536876-DEL-424
chr5-83549657-DEL-6010
chr5-83573554-DEL-682
chr5-83620405-DEL-54
chr5-83627739-DEL-726
chr5-83630549-DEL-89
chr5-83639765-DEL-71
chr5-83661520-DEL-351
chr5-83668487-DEL-4169
chr5-83676083-DEL-395
chr5-83706611-DEL-2589
chr5-83708491-DEL-150

chr5-83718647-DEL-194
chr5-83750232-DEL-66
chr5-83782039-DEL-2027
chr5-83792717-DEL-783
chr5-83803641-DEL-237
chr5-83818873-DEL-924
chr5-83824724-DEL-185
chr5-83833146-DEL-1879
chr5-83847024-DEL-564
chr5-83856396-DEL-1045
chr5-83871445-DEL-115
chr5-83921648-DEL-1339
chr5-83941294-DEL-2457
chr5-83953701-DEL-139
chr5-83959830-DEL-102
chr5-83975263-DEL-2131
chr5-83999505-DEL-91
chr5-84008793-DEL-275
chr5-84032222-DEL-103
chr5-84040293-DEL-269
chr5-84136007-DEL-127
chr5-84143340-DEL-234
chr5-84187091-DEL-352
chr5-84309443-DEL-273
chr5-84319100-DEL-100
chr5-84319422-DEL-690
chr5-84325875-DEL-1376
chr5-84332698-DEL-432
chr5-84336033-DEL-186
chr5-84336466-DEL-1326
chr5-84341825-DEL-61
chr5-84352701-DEL-81
chr5-84401107-DEL-739
chr5-84406676-DEL-78
chr5-84435448-DEL-72
chr5-84450624-DEL-443
chr5-84484367-DEL-818
chr5-84502714-DEL-448
chr5-84525902-DEL-212
chr5-84536406-DEL-86
chr5-84537854-DEL-140
chr5-84583638-DEL-938
chr5-84620307-DEL-414
chr5-84625746-DEL-494
chr5-84647180-DEL-64
chr5-84647564-DEL-1339
chr5-84651216-DEL-553
chr5-84656302-DEL-1314
chr5-84706728-DEL-305
chr5-84757358-DEL-1066
chr5-84787650-DEL-78
chr5-84811517-DEL-179
chr5-84813063-DEL-155
chr5-84824438-DEL-150
chr5-84834910-DEL-1812
chr5-84861089-DEL-242
chr5-84865633-DEL-6036
chr5-84890560-DEL-1440
chr5-84901561-DEL-7038
chr5-84917540-DEL-6767
chr5-84935840-DEL-1636
chr5-84965277-DEL-8979
chr5-84977393-DEL-1311
chr5-84999649-DEL-175
chr5-85015003-DEL-79
chr5-85017179-DEL-397
chr5-85043753-DEL-68
chr5-85066297-DEL-61
chr5-85068416-DEL-484
chr5-85070501-DEL-165
chr5-85073612-DEL-51
chr5-85096211-DEL-62
chr5-85129016-DEL-154
chr5-85138535-DEL-89
chr5-85166613-DEL-260
chr5-85168833-DEL-101
chr5-85172213-DEL-90

chr5-85176846-DEL-1485
chr5-85185127-DEL-261
chr5-85189077-DEL-796
chr5-85194619-DEL-448
chr5-85208836-DEL-6266
chr5-85250460-DEL-1890
chr5-85257851-DEL-1294
chr5-85258523-DEL-1291
chr5-85310837-DEL-104
chr5-85311599-DEL-528
chr5-85321307-DEL-212
chr5-85346007-DEL-236
chr5-85444780-DEL-1883
chr5-85457595-DEL-59
chr5-85460852-DEL-66
chr5-85467728-DEL-254
chr5-85471449-DEL-56
chr5-85489812-DEL-992
chr5-85502901-DEL-5108
chr5-85529810-DEL-51
chr5-85533089-DEL-1250
chr5-85540074-DEL-52
chr5-85540103-DEL-1841
chr5-85540901-DEL-83
chr5-85541017-DEL-1673
chr5-85549214-DEL-56
chr5-85660309-DEL-9526
chr5-85671213-DEL-509
chr5-85678074-DEL-291
chr5-85697957-DEL-60
chr5-85721388-DEL-839
chr5-85750240-DEL-65
chr5-85753654-DEL-351
chr5-85767547-DEL-312
chr5-85788806-DEL-1050
chr5-85806217-DEL-116
chr5-85820557-DEL-52
chr5-85825492-DEL-1460
chr5-85827892-DEL-6176
chr5-85889579-DEL-606
chr5-85937110-DEL-122
chr5-85960562-DEL-53
chr5-85970518-DEL-6384
chr5-85984757-DEL-809
chr5-85986579-DEL-143
chr5-85996957-DEL-1625
chr5-86041226-DEL-65
chr5-86046598-DEL-184
chr5-86074820-DEL-1096
chr5-86076055-DEL-479
chr5-86093626-DEL-105
chr5-86101604-DEL-206
chr5-86107204-DEL-1498
chr5-86121789-DEL-198
chr5-86130930-DEL-819
chr5-86143824-DEL-82
chr5-86153564-DEL-6189
chr5-86167111-DEL-1432
chr5-86190994-DEL-187
chr5-86192954-DEL-94
chr5-86197036-DEL-698
chr5-86225465-DEL-63
chr5-86226362-DEL-51
chr5-86236524-DEL-2061
chr5-86239352-DEL-112
chr5-86252650-DEL-131
chr5-86348527-DEL-6800
chr5-86360089-DEL-9173
chr5-86375705-DEL-232
chr5-86383828-DEL-995
chr5-86398343-DEL-966
chr5-86429276-DEL-58
chr5-86460115-DEL-182
chr5-86467310-DEL-73
chr5-86468418-DEL-5046
chr5-86479663-DEL-1461
chr5-86508973-DEL-60

chr5-86516809-DEL-213
chr5-86518399-DEL-1100
chr5-86520727-DEL-183
chr5-86560658-DEL-129
chr5-86574351-DEL-234
chr5-86596786-DEL-373
chr5-86607415-DEL-262
chr5-86693882-DEL-60
chr5-86708072-DEL-1210
chr5-86718057-DEL-115
chr5-86725729-DEL-54
chr5-86737300-DEL-961
chr5-86738740-DEL-156
chr5-86741220-DEL-11124
chr5-86754891-DEL-59
chr5-86759985-DEL-139
chr5-86798617-DEL-98
chr5-86834543-DEL-3881
chr5-86836528-DEL-3485
chr5-86839750-DEL-226
chr5-86849115-DEL-51
chr5-86908421-DEL-54
chr5-86931465-DEL-62
chr5-86939179-DEL-461
chr5-86950428-DEL-258
chr5-86957351-DEL-219
chr5-87050656-DEL-186
chr5-87077596-DEL-3119
chr5-87093230-DEL-4458
chr5-87120265-DEL-98
chr5-87150821-DEL-210
chr5-87159502-DEL-70
chr5-87180255-DEL-16249
chr5-87223831-DEL-91
chr5-87232885-DEL-115
chr5-87248855-DEL-40169
chr5-87259899-DEL-412
chr5-87262817-DEL-54
chr5-87349444-DEL-588
chr5-87365970-DEL-477
chr5-87388948-DEL-166
chr5-87449312-DEL-250
chr5-87463414-DEL-112
chr5-87475643-DEL-222
chr5-87479424-DEL-282
chr5-87492554-DEL-203
chr5-87498460-DEL-61
chr5-87502458-DEL-221
chr5-87515951-DEL-2446
chr5-87528750-DEL-1547
chr5-87531880-DEL-6212
chr5-87555923-DEL-78
chr5-87641643-DEL-313
chr5-87677595-DEL-144
chr5-87686311-DEL-455
chr5-87687553-DEL-7019
chr5-87735281-DEL-68
chr5-87736698-DEL-315
chr5-87744330-DEL-57
chr5-87763274-DEL-66
chr5-87772395-DEL-3608
chr5-87800062-DEL-1610
chr5-87806339-DEL-71
chr5-87828082-DEL-132
chr5-87848238-DEL-505
chr5-87849942-DEL-66
chr5-87850512-DEL-2474
chr5-87850768-DEL-615
chr5-87852160-DEL-169
chr5-87878778-DEL-155
chr5-87888280-DEL-84
chr5-87892555-DEL-158
chr5-87942080-DEL-472
chr5-87944426-DEL-1013
chr5-87948014-DEL-63
chr5-88005905-DEL-181
chr5-88030236-DEL-1958

chr5-88078734-DEL-66
chr5-88095524-DEL-2497
chr5-88135817-DEL-55
chr5-88137742-DEL-3014
chr5-88141008-DEL-52
chr5-88142033-DEL-96
chr5-88198961-DEL-103
chr5-88203433-DEL-6309
chr5-88216329-DEL-131
chr5-88293782-DEL-7044
chr5-88294920-DEL-1244
chr5-88296202-DEL-603
chr5-88296841-DEL-3979
chr5-88310311-DEL-5549
chr5-88391536-DEL-6681
chr5-88401987-DEL-66
chr5-88407129-DEL-6286
chr5-88425767-DEL-996
chr5-88427373-DEL-493
chr5-88507491-DEL-3077
chr5-88512289-DEL-158
chr5-88553060-DEL-131
chr5-88554915-DEL-12269
chr5-88563776-DEL-25568
chr5-88576240-DEL-52
chr5-88577711-DEL-861
chr5-88582817-DEL-245
chr5-88622086-DEL-114
chr5-88624293-DEL-210
chr5-88630052-DEL-54
chr5-88668088-DEL-59
chr5-88728784-DEL-64
chr5-88759496-DEL-212
chr5-88768252-DEL-90
chr5-88770708-DEL-1546
chr5-88781011-DEL-246
chr5-88806753-DEL-203
chr5-88818813-DEL-144
chr5-88839020-DEL-82
chr5-88843514-DEL-1212
chr5-88883217-DEL-6456
chr5-88893150-DEL-104
chr5-88899027-DEL-172
chr5-88928811-DEL-103
chr5-88935924-DEL-1099
chr5-88948597-DEL-182
chr5-88952159-DEL-56
chr5-88963980-DEL-198
chr5-88984333-DEL-449
chr5-88992798-DEL-63
chr5-88993016-DEL-2595
chr5-89002840-DEL-282
chr5-89009664-DEL-61
chr5-89010988-DEL-372
chr5-89026297-DEL-68
chr5-89052417-DEL-83
chr5-89061207-DEL-128
chr5-89110866-DEL-66
chr5-89126165-DEL-714
chr5-89156278-DEL-141
chr5-89167728-DEL-139
chr5-89207160-DEL-53
chr5-89313467-DEL-59
chr5-89321994-DEL-115
chr5-89330150-DEL-428
chr5-89381702-DEL-62
chr5-89392848-DEL-500
chr5-89413700-DEL-53
chr5-89438027-DEL-61
chr5-89457604-DEL-6382
chr5-89466471-DEL-1163
chr5-89475698-DEL-5575
chr5-89475748-DEL-8159
chr5-89481473-DEL-2583
chr5-89481482-DEL-1291
chr5-89493645-DEL-53
chr5-89530273-DEL-1131

chr5-89530317-DEL-873
chr5-89530547-DEL-433
chr5-89567064-DEL-139
chr5-89571034-DEL-2529
chr5-89581791-DEL-424
chr5-89593076-DEL-52
chr5-89593850-DEL-135
chr5-89596150-DEL-355
chr5-89621464-DEL-317
chr5-89641167-DEL-1363
chr5-89655770-DEL-70
chr5-89659134-DEL-2146
chr5-89662619-DEL-540
chr5-89767670-DEL-2226
chr5-89767705-DEL-1428
chr5-89768081-DEL-167
chr5-89768526-DEL-714
chr5-89769393-DEL-167
chr5-89789982-DEL-167
chr5-89798473-DEL-55
chr5-89821470-DEL-87
chr5-89836542-DEL-147
chr5-89859881-DEL-950
chr5-89924686-DEL-52
chr5-89950269-DEL-1550
chr5-89981401-DEL-126
chr5-89993135-DEL-2634
chr5-90028234-DEL-72
chr5-90039200-DEL-403
chr5-90039952-DEL-4645
chr5-90064484-DEL-84
chr5-90081164-DEL-402
chr5-90096329-DEL-1550
chr5-90100781-DEL-8485
chr5-90115696-DEL-54
chr5-90116283-DEL-2475
chr5-90120206-DEL-494
chr5-90120869-DEL-262
chr5-90124393-DEL-54
chr5-90131967-DEL-2073
chr5-90146003-DEL-576
chr5-90150379-DEL-128
chr5-90171090-DEL-2130
chr5-90195388-DEL-52
chr5-90213237-DEL-87
chr5-90227675-DEL-228
chr5-90232696-DEL-260
chr5-90241827-DEL-336
chr5-90269067-DEL-62
chr5-90285935-DEL-59
chr5-90302473-DEL-122
chr5-90316597-DEL-7081
chr5-90336677-DEL-480
chr5-90398010-DEL-196
chr5-90401002-DEL-166
chr5-90407075-DEL-82
chr5-90424828-DEL-64
chr5-90450071-DEL-736
chr5-90456550-DEL-3688
chr5-90498553-DEL-2731
chr5-90527628-DEL-89
chr5-90533805-DEL-483
chr5-90536643-DEL-274
chr5-90539217-DEL-2202
chr5-90554504-DEL-56
chr5-90607511-DEL-67
chr5-90610022-DEL-57
chr5-90639774-DEL-168
chr5-90657369-DEL-142
chr5-90667278-DEL-238
chr5-90668607-DEL-301
chr5-90671163-DEL-235
chr5-90709733-DEL-85
chr5-90716099-DEL-60
chr5-90724484-DEL-187
chr5-90729369-DEL-80
chr5-90828335-DEL-462

chr5-90858801-DEL-1692
chr5-90860273-DEL-459
chr5-90864156-DEL-85
chr5-90874979-DEL-65
chr5-90878514-DEL-468
chr5-90882952-DEL-6826
chr5-90894217-DEL-414
chr5-90943823-DEL-100
chr5-90948826-DEL-55
chr5-90970069-DEL-7916
chr5-90975704-DEL-1487
chr5-90983821-DEL-1270
chr5-90986786-DEL-1420
chr5-90995514-DEL-2747
chr5-91000896-DEL-121
chr5-91029940-DEL-85
chr5-91045881-DEL-58
chr5-91057664-DEL-6481
chr5-91059154-DEL-8986
chr5-91070939-DEL-209
chr5-91086248-DEL-129
chr5-91139904-DEL-6207
chr5-91151602-DEL-136
chr5-91176422-DEL-110
chr5-91212783-DEL-1810
chr5-91256595-DEL-76
chr5-91278552-DEL-110
chr5-91279086-DEL-200
chr5-91301806-DEL-72
chr5-91351446-DEL-133
chr5-91373260-DEL-255
chr5-91395951-DEL-396
chr5-91396408-DEL-2455
chr5-91435268-DEL-59
chr5-91457357-DEL-1246
chr5-91476032-DEL-336
chr5-91495691-DEL-125
chr5-91527670-DEL-128
chr5-91565186-DEL-624
chr5-91566378-DEL-2483
chr5-91584301-DEL-56
chr5-91600008-DEL-2363
chr5-91621518-DEL-391
chr5-91632819-DEL-1137
chr5-91635711-DEL-1164
chr5-91638853-DEL-67
chr5-91668980-DEL-60
chr5-91778965-DEL-208
chr5-91793993-DEL-394
chr5-91834393-DEL-200
chr5-91835905-DEL-66
chr5-91866240-DEL-64
chr5-91878057-DEL-590
chr5-91889460-DEL-219
chr5-91900820-DEL-367
chr5-91942099-DEL-168
chr5-92022716-DEL-61
chr5-92026936-DEL-63
chr5-92040414-DEL-66
chr5-92043797-DEL-132
chr5-92055921-DEL-53
chr5-92071528-DEL-51
chr5-92081062-DEL-86
chr5-92112574-DEL-177
chr5-92130579-DEL-1844
chr5-92151157-DEL-75
chr5-92164308-DEL-64
chr5-92172079-DEL-58
chr5-92193175-DEL-651
chr5-92204400-DEL-75
chr5-92209036-DEL-67
chr5-92220792-DEL-82
chr5-92226434-DEL-101
chr5-92252779-DEL-212
chr5-92257493-DEL-67
chr5-92291315-DEL-1363
chr5-92301255-DEL-193

chr5-92316951-DEL-691
chr5-92317851-DEL-52
chr5-92328761-DEL-221
chr5-92370378-DEL-2828
chr5-92383293-DEL-83
chr5-92388255-DEL-234
chr5-92401455-DEL-2184
chr5-92409710-DEL-97
chr5-92445864-DEL-61
chr5-92458913-DEL-233
chr5-92489993-DEL-71
chr5-92493350-DEL-373
chr5-92551858-DEL-381
chr5-92562655-DEL-187
chr5-92607034-DEL-60
chr5-92630918-DEL-122
chr5-92641187-DEL-76
chr5-92649678-DEL-69
chr5-92675804-DEL-156
chr5-92678994-DEL-75
chr5-92687872-DEL-604
chr5-92735680-DEL-139
chr5-92741987-DEL-164
chr5-92745574-DEL-1362
chr5-92750251-DEL-50
chr5-92761762-DEL-1947
chr5-92762999-DEL-134
chr5-92763766-DEL-6531
chr5-92792757-DEL-65
chr5-92795068-DEL-72
chr5-92828026-DEL-54
chr5-92955728-DEL-75
chr5-92959400-DEL-84
chr5-92966250-DEL-277
chr5-92987337-DEL-1658
chr5-92989923-DEL-78
chr5-93011790-DEL-85
chr5-93020592-DEL-60
chr5-93033578-DEL-170
chr5-93060991-DEL-126
chr5-93078860-DEL-81
chr5-93104181-DEL-64
chr5-93111693-DEL-60
chr5-93116629-DEL-204
chr5-93119002-DEL-80
chr5-93151653-DEL-710
chr5-93203817-DEL-1172
chr5-93204100-DEL-63
chr5-93226875-DEL-208
chr5-93250189-DEL-629
chr5-93250200-DEL-164
chr5-93259227-DEL-160
chr5-93292041-DEL-245
chr5-93347187-DEL-132
chr5-93358899-DEL-84
chr5-93367023-DEL-382
chr5-93456076-DEL-172
chr5-93513243-DEL-3144
chr5-93523673-DEL-6399
chr5-93527819-DEL-6714
chr5-93530409-DEL-6040
chr5-93604209-DEL-143
chr5-93609698-DEL-163
chr5-93618815-DEL-71
chr5-93629322-DEL-84
chr5-93631042-DEL-81
chr5-93633512-DEL-468
chr5-93773045-DEL-8217
chr5-93856172-DEL-28929
chr5-93868531-DEL-53
chr5-93943347-DEL-72
chr5-93962701-DEL-374
chr5-93974933-DEL-76
chr5-94374389-DEL-56
chr5-94376874-DEL-6891
chr5-94495765-DEL-503
chr5-94518948-DEL-6436

chr5-94577399-DEL-1792
chr5-94639671-DEL-373
chr5-94790614-DEL-4758
chr5-95347339-DEL-56
chr5-95587954-DEL-2248
chr5-95593438-DEL-383
chr5-95604948-DEL-180
chr5-95622721-DEL-171
chr5-95624268-DEL-60
chr5-95625585-DEL-110
chr5-95633545-DEL-1024
chr5-95638388-DEL-56
chr5-95642344-DEL-6008
chr5-95656786-DEL-182
chr5-95659401-DEL-165
chr5-95667227-DEL-402
chr5-95859946-DEL-636
chr5-95969199-DEL-6618
chr5-95969330-DEL-201
chr5-96101980-DEL-810
chr5-96120657-DEL-70
chr5-96121099-DEL-2758
chr5-96171418-DEL-61
chr5-96192000-DEL-6267
chr5-96204277-DEL-7332
chr5-96244144-DEL-265
chr5-96257357-DEL-103
chr5-96332148-DEL-164
chr5-96354543-DEL-69
chr5-96364070-DEL-57
chr5-96375666-DEL-56
chr5-96399939-DEL-300
chr5-96434437-DEL-7215
chr5-96442983-DEL-197
chr5-96445325-DEL-2361
chr5-96473522-DEL-90
chr5-96482293-DEL-231
chr5-96486654-DEL-64
chr5-96491336-DEL-69
chr5-96507886-DEL-53
chr5-96531155-DEL-621
chr5-96541126-DEL-111
chr5-96550039-DEL-174
chr5-96588383-DEL-356
chr5-96590784-DEL-430
chr5-96594607-DEL-87
chr5-96656643-DEL-108
chr5-96774863-DEL-59
chr5-96950832-DEL-56
chr5-96962074-DEL-365
chr5-96972576-DEL-51
chr5-96981085-DEL-136
chr5-97009999-DEL-225
chr5-97046477-DEL-56
chr5-97053268-DEL-66
chr5-97060074-DEL-64
chr5-97086048-DEL-184
chr5-97090648-DEL-107
chr5-97094029-DEL-272
chr5-97095052-DEL-678
chr5-97138255-DEL-859
chr5-97169017-DEL-330
chr5-97174967-DEL-234
chr5-97176926-DEL-148
chr5-97191065-DEL-1262
chr5-97201355-DEL-221
chr5-97319363-DEL-56
chr5-97324080-DEL-193
chr5-97326283-DEL-164
chr5-97368551-DEL-134
chr5-97377714-DEL-344
chr5-97394714-DEL-849
chr5-97395801-DEL-220
chr5-97412511-DEL-167
chr5-97424045-DEL-100
chr5-97429081-DEL-914
chr5-97440345-DEL-76

chr5-97446443-DEL-128
chr5-97485712-DEL-59
chr5-97486340-DEL-84
chr5-97608786-DEL-196
chr5-97630750-DEL-89
chr5-97705211-DEL-59
chr5-97711169-DEL-236
chr5-97735518-DEL-56
chr5-97752135-DEL-111
chr5-97766765-DEL-67
chr5-97779063-DEL-102
chr5-97786808-DEL-245
chr5-97800249-DEL-56
chr5-97805977-DEL-221
chr5-97807290-DEL-959
chr5-97829659-DEL-801
chr5-97863902-DEL-802
chr5-97896871-DEL-6330
chr5-97903600-DEL-58
chr5-97931634-DEL-901
chr5-97965113-DEL-64
chr5-98019730-DEL-55
chr5-98037637-DEL-553
chr5-98043875-DEL-587
chr5-98045844-DEL-52
chr5-98052114-DEL-252
chr5-98125118-DEL-64
chr5-98132547-DEL-476
chr5-98136736-DEL-856
chr5-98153922-DEL-7207
chr5-98208175-DEL-66
chr5-98211383-DEL-1560
chr5-98213852-DEL-116
chr5-98249335-DEL-264
chr5-98266840-DEL-392
chr5-98322722-DEL-285
chr5-98347867-DEL-108
chr5-98370546-DEL-748
chr5-98381511-DEL-718
chr5-98383328-DEL-78
chr5-98443932-DEL-79
chr5-98462916-DEL-212
chr5-98478293-DEL-98
chr5-98503568-DEL-117
chr5-98515747-DEL-58
chr5-98520146-DEL-80
chr5-98526438-DEL-182
chr5-98542790-DEL-1117
chr5-98547323-DEL-915
chr5-98592505-DEL-52
chr5-98598401-DEL-280
chr5-98674545-DEL-268
chr5-98737411-DEL-65
chr5-98744826-DEL-70
chr5-98751125-DEL-96
chr5-98802113-DEL-969
chr5-98817148-DEL-107
chr5-98821942-DEL-352
chr5-98830843-DEL-155
chr5-98836494-DEL-400
chr5-98857149-DEL-192
chr5-98898384-DEL-2335
chr5-98906567-DEL-71
chr5-98925492-DEL-201
chr5-98961027-DEL-413
chr5-98984420-DEL-114
chr5-98995080-DEL-254
chr5-99013434-DEL-178
chr5-99033096-DEL-51
chr5-99040929-DEL-53
chr5-99129689-DEL-79
chr5-99135656-DEL-63
chr5-99146679-DEL-182
chr5-99203900-DEL-222
chr5-99210630-DEL-476
chr5-99246575-DEL-62
chr5-99254313-DEL-201

chr5-99280514-DEL-787
chr5-99298414-DEL-138
chr5-99306155-DEL-143
chr5-99309002-DEL-150
chr5-99335234-DEL-58
chr5-99335768-DEL-69
chr5-99340794-DEL-85
chr5-99348602-DEL-60
chr5-99353409-DEL-170
chr5-99362959-DEL-84
chr5-99413002-DEL-51
chr5-99457334-DEL-6051
chr5-99490419-DEL-54
chr5-99491980-DEL-1909
chr5-99494143-DEL-64
chr5-99517580-DEL-63
chr5-99599712-DEL-113
chr5-99640597-DEL-138
chr5-99676855-DEL-248
chr5-99682104-DEL-78
chr5-99696489-DEL-64
chr5-99698090-DEL-401
chr5-99699072-DEL-52
chr5-99714853-DEL-600
chr5-99724546-DEL-106
chr5-99732776-DEL-103
chr5-99779159-DEL-109
chr5-99791295-DEL-777
chr5-99791361-DEL-57
chr5-99792268-DEL-337
chr5-99806776-DEL-58
chr5-99841643-DEL-466
chr5-99878115-DEL-522
chr5-99896713-DEL-207
chr5-99902908-DEL-1813
chr5-99908231-DEL-82
chr5-99922302-DEL-61
chr5-99941613-DEL-229
chr5-99946854-DEL-114
chr5-99952861-DEL-68
chr5-99979166-DEL-220
chr5-99979923-DEL-252
chr5-100001259-DEL-55
chr5-100036551-DEL-64
chr5-100037456-DEL-228
chr5-100039176-DEL-84
chr5-100045914-DEL-51
chr5-100099990-DEL-54
chr5-100134385-DEL-931
chr5-100134387-DEL-7096
chr5-100135580-DEL-4554
chr5-100138836-DEL-4729
chr5-100140222-DEL-1262
chr5-100142247-DEL-1365
chr5-100146538-DEL-156
chr5-100148393-DEL-1921
chr5-100164277-DEL-122
chr5-100166253-DEL-950
chr5-100167716-DEL-70
chr5-100171200-DEL-57
chr5-100201352-DEL-598
chr5-100230553-DEL-121
chr5-100247561-DEL-130
chr5-100278052-DEL-6212
chr5-100291577-DEL-735
chr5-100315923-DEL-51
chr5-100320609-DEL-1259
chr5-100322866-DEL-1385
chr5-100330549-DEL-1514
chr5-100350789-DEL-107
chr5-100355494-DEL-582
chr5-100360124-DEL-128
chr5-100361033-DEL-570
chr5-100366784-DEL-76
chr5-100381276-DEL-550
chr5-100392040-DEL-174
chr5-100394659-DEL-123

chr5-100400399-DEL-69
chr5-100405768-DEL-66
chr5-100428471-DEL-73
chr5-100429718-DEL-2092
chr5-100429800-DEL-6795
chr5-100431900-DEL-4695
chr5-100459311-DEL-149
chr5-100460935-DEL-116
chr5-100461190-DEL-1034
chr5-100466620-DEL-84
chr5-100492115-DEL-196
chr5-100495813-DEL-95
chr5-100499127-DEL-53
chr5-100504608-DEL-83
chr5-100520573-DEL-138
chr5-100548987-DEL-199
chr5-100603528-DEL-200
chr5-100608853-DEL-122
chr5-100609299-DEL-211
chr5-100609300-DEL-1801
chr5-100610934-DEL-168
chr5-100615931-DEL-54
chr5-100643559-DEL-348
chr5-100678617-DEL-227
chr5-100686463-DEL-286
chr5-100692427-DEL-52
chr5-100700516-DEL-643
chr5-100797310-DEL-184
chr5-100844255-DEL-266
chr5-100900589-DEL-115
chr5-100906690-DEL-67
chr5-100909159-DEL-404
chr5-100948697-DEL-199
chr5-100961586-DEL-51
chr5-100964598-DEL-209
chr5-100975256-DEL-265
chr5-101014498-DEL-198
chr5-101038272-DEL-71
chr5-101066209-DEL-54
chr5-101073407-DEL-68
chr5-101082851-DEL-155
chr5-101089824-DEL-78
chr5-101111653-DEL-59
chr5-101117435-DEL-946
chr5-101126948-DEL-108
chr5-101134071-DEL-287
chr5-101140163-DEL-60
chr5-101170312-DEL-79
chr5-101171295-DEL-159
chr5-101183139-DEL-78
chr5-101187521-DEL-64
chr5-101198398-DEL-68
chr5-101241544-DEL-363
chr5-101276203-DEL-72
chr5-101284740-DEL-79
chr5-101306602-DEL-52
chr5-101307956-DEL-120
chr5-101317382-DEL-184
chr5-101355695-DEL-107
chr5-101398501-DEL-210
chr5-101428737-DEL-88
chr5-101484674-DEL-123
chr5-101492514-DEL-61
chr5-101519929-DEL-54
chr5-101527117-DEL-93
chr5-101533409-DEL-520
chr5-101544096-DEL-2985
chr5-101547949-DEL-522
chr5-101564916-DEL-180
chr5-101572243-DEL-717
chr5-101581935-DEL-93
chr5-101590170-DEL-86
chr5-101600164-DEL-144
chr5-101650355-DEL-134
chr5-101650875-DEL-1458
chr5-101666497-DEL-53
chr5-101667741-DEL-210

chr5-101682759-DEL-54
chr5-101692215-DEL-89
chr5-101729220-DEL-492
chr5-101735521-DEL-147
chr5-101747546-DEL-100
chr5-101749559-DEL-190
chr5-101782418-DEL-54
chr5-101783680-DEL-486
chr5-101865931-DEL-52
chr5-101872672-DEL-58
chr5-101875915-DEL-319
chr5-101887640-DEL-58
chr5-101895697-DEL-111
chr5-101897749-DEL-68
chr5-101911776-DEL-259
chr5-101922500-DEL-421
chr5-101961245-DEL-147
chr5-102005107-DEL-786
chr5-102008046-DEL-204
chr5-102009794-DEL-63
chr5-102045593-DEL-64
chr5-102064913-DEL-194
chr5-102083159-DEL-79
chr5-102103196-DEL-82
chr5-102112406-DEL-127
chr5-102120526-DEL-2040
chr5-102205580-DEL-104
chr5-102228856-DEL-98
chr5-102248061-DEL-72
chr5-102251798-DEL-810
chr5-102255098-DEL-52
chr5-102256837-DEL-76
chr5-102269215-DEL-74
chr5-102284396-DEL-65
chr5-102314107-DEL-60
chr5-102321607-DEL-69
chr5-102375892-DEL-72
chr5-102394786-DEL-122
chr5-102404544-DEL-297
chr5-102418055-DEL-4156
chr5-102424798-DEL-407
chr5-102445343-DEL-424
chr5-102473475-DEL-135
chr5-102477867-DEL-2722
chr5-102485472-DEL-93
chr5-102515301-DEL-276
chr5-102529562-DEL-123
chr5-102536111-DEL-201
chr5-102541600-DEL-352
chr5-102547328-DEL-54
chr5-102570125-DEL-100
chr5-102577361-DEL-65
chr5-102619701-DEL-179
chr5-102640594-DEL-104
chr5-102641528-DEL-192
chr5-102684946-DEL-76
chr5-102695463-DEL-77
chr5-102722782-DEL-91
chr5-102779886-DEL-1025
chr5-102782800-DEL-160
chr5-102805472-DEL-64
chr5-102833983-DEL-67
chr5-102839192-DEL-112
chr5-102867995-DEL-1986
chr5-102874802-DEL-55
chr5-102889990-DEL-51
chr5-102913601-DEL-68
chr5-102973486-DEL-53
chr5-102982064-DEL-150
chr5-103021197-DEL-192
chr5-103030932-DEL-468
chr5-103067426-DEL-273
chr5-103068169-DEL-465
chr5-103069357-DEL-1650
chr5-103082222-DEL-66
chr5-103115504-DEL-78
chr5-103119798-DEL-329

chr5-103122697-DEL-54
chr5-103214618-DEL-4349
chr5-103223488-DEL-400
chr5-103279494-DEL-343
chr5-103290603-DEL-107
chr5-103335222-DEL-53
chr5-103400514-DEL-1818
chr5-103426465-DEL-70
chr5-103441366-DEL-128
chr5-103454123-DEL-328
chr5-103459172-DEL-356
chr5-103461269-DEL-6082
chr5-103472421-DEL-55
chr5-103490102-DEL-270
chr5-103531899-DEL-927
chr5-103533217-DEL-317
chr5-103537810-DEL-70
chr5-103548158-DEL-1378
chr5-103581158-DEL-189
chr5-103583720-DEL-104
chr5-103614614-DEL-119
chr5-103618700-DEL-151
chr5-103619397-DEL-898
chr5-103625556-DEL-51
chr5-103692654-DEL-363
chr5-103713952-DEL-189
chr5-103717898-DEL-601
chr5-103723988-DEL-224
chr5-103748826-DEL-935
chr5-103786615-DEL-234
chr5-103789339-DEL-205
chr5-103800355-DEL-135
chr5-103808565-DEL-1395
chr5-103833997-DEL-294
chr5-103865546-DEL-67
chr5-103874398-DEL-222
chr5-103896539-DEL-100
chr5-103906802-DEL-139
chr5-103983022-DEL-1017
chr5-103985031-DEL-550
chr5-104005025-DEL-54
chr5-104006338-DEL-397
chr5-104018909-DEL-58
chr5-104030583-DEL-133
chr5-104040192-DEL-390
chr5-104049334-DEL-189
chr5-104053327-DEL-56
chr5-104059320-DEL-96
chr5-104062779-DEL-161
chr5-104075602-DEL-1023
chr5-104082945-DEL-1504
chr5-104084907-DEL-374
chr5-104090553-DEL-211
chr5-104094626-DEL-4142
chr5-104096448-DEL-78
chr5-104118885-DEL-826
chr5-104135700-DEL-1487
chr5-104141998-DEL-244
chr5-104146379-DEL-72
chr5-104165346-DEL-218
chr5-104175558-DEL-86
chr5-104189472-DEL-224
chr5-104213464-DEL-58
chr5-104221514-DEL-191
chr5-104250506-DEL-490
chr5-104252259-DEL-2688
chr5-104478980-DEL-1174
chr5-104479067-DEL-341
chr5-104479792-DEL-207
chr5-104488634-DEL-125
chr5-104558901-DEL-125
chr5-104572711-DEL-214
chr5-104576774-DEL-101
chr5-104626256-DEL-124
chr5-104750441-DEL-79
chr5-104755329-DEL-86
chr5-104766527-DEL-354

chr5-104767456-DEL-247
chr5-104790441-DEL-636
chr5-104819232-DEL-93
chr5-104826439-DEL-196
chr5-104838821-DEL-84
chr5-104839329-DEL-84
chr5-104839967-DEL-83
chr5-104841160-DEL-3583
chr5-104873128-DEL-91
chr5-104874829-DEL-194
chr5-104897966-DEL-252
chr5-104916213-DEL-96
chr5-104920839-DEL-111
chr5-104932525-DEL-165
chr5-104960277-DEL-60
chr5-105035102-DEL-7120
chr5-105041636-DEL-212
chr5-105068205-DEL-189
chr5-105101188-DEL-514
chr5-105127376-DEL-60
chr5-105240961-DEL-203
chr5-105331773-DEL-69
chr5-105344580-DEL-417
chr5-105480863-DEL-100
chr5-105484941-DEL-171
chr5-105520504-DEL-6380
chr5-105530664-DEL-6351
chr5-105610331-DEL-1631
chr5-105686941-DEL-113
chr5-105689076-DEL-920
chr5-105698979-DEL-54
chr5-105739358-DEL-59
chr5-105766478-DEL-131
chr5-105786419-DEL-57
chr5-105837011-DEL-58
chr5-105843018-DEL-204
chr5-105860619-DEL-58
chr5-105861877-DEL-120
chr5-105866617-DEL-184
chr5-105900224-DEL-221
chr5-105945153-DEL-454
chr5-105947188-DEL-176
chr5-105951924-DEL-2263
chr5-105957611-DEL-116
chr5-105977720-DEL-116
chr5-105986258-DEL-143
chr5-105998578-DEL-404
chr5-105999783-DEL-306
chr5-106006093-DEL-176
chr5-106012774-DEL-208
chr5-106032348-DEL-107
chr5-106077304-DEL-364
chr5-106077385-DEL-181
chr5-106113966-DEL-200
chr5-106122028-DEL-126
chr5-106133105-DEL-109
chr5-106137682-DEL-160
chr5-106155141-DEL-99
chr5-106165139-DEL-6317
chr5-106173876-DEL-411
chr5-106176467-DEL-51
chr5-106209011-DEL-125
chr5-106244202-DEL-155
chr5-106245743-DEL-78
chr5-106276374-DEL-184
chr5-106300876-DEL-868
chr5-106304087-DEL-547
chr5-106336831-DEL-1670
chr5-106347368-DEL-56
chr5-106377329-DEL-71
chr5-106389742-DEL-406
chr5-106409154-DEL-116
chr5-106516581-DEL-141
chr5-106519240-DEL-430
chr5-106551491-DEL-419
chr5-106555644-DEL-103
chr5-106556470-DEL-83

chr5-106578034-DEL-54
chr5-106737689-DEL-80
chr5-106816821-DEL-121
chr5-106824295-DEL-2666
chr5-106860485-DEL-111
chr5-106861177-DEL-8173
chr5-106914250-DEL-164
chr5-106956098-DEL-368
chr5-107001285-DEL-52
chr5-107044365-DEL-70
chr5-107090487-DEL-1451
chr5-107090489-DEL-1161
chr5-107091663-DEL-565
chr5-107115558-DEL-120
chr5-107125449-DEL-53
chr5-107132879-DEL-200
chr5-107180941-DEL-60
chr5-107226734-DEL-129
chr5-107241952-DEL-215
chr5-107259445-DEL-84
chr5-107322752-DEL-66
chr5-107377263-DEL-458
chr5-107378343-DEL-91
chr5-107392817-DEL-663
chr5-107457835-DEL-178
chr5-107458980-DEL-223
chr5-107496719-DEL-68
chr5-107534366-DEL-216
chr5-107537738-DEL-83
chr5-107599124-DEL-53
chr5-107694184-DEL-141
chr5-107697540-DEL-357
chr5-107706521-DEL-183
chr5-107723465-DEL-2249
chr5-107726770-DEL-7260
chr5-107737021-DEL-130
chr5-107739950-DEL-232
chr5-107762790-DEL-91
chr5-107767151-DEL-212
chr5-107817396-DEL-66
chr5-107837117-DEL-228
chr5-107839372-DEL-137
chr5-107850153-DEL-154
chr5-107889462-DEL-96
chr5-107899799-DEL-176
chr5-107914291-DEL-54
chr5-107915293-DEL-176
chr5-107917342-DEL-248
chr5-107922997-DEL-220
chr5-107929878-DEL-784
chr5-107952895-DEL-671
chr5-107967199-DEL-200
chr5-107998800-DEL-118
chr5-108041356-DEL-241
chr5-108047444-DEL-77
chr5-108058147-DEL-1313
chr5-108071412-DEL-138
chr5-108088846-DEL-117
chr5-108145436-DEL-112
chr5-108199193-DEL-130
chr5-108205451-DEL-197
chr5-108212042-DEL-158
chr5-108231258-DEL-537
chr5-108238851-DEL-193
chr5-108242218-DEL-52
chr5-108261391-DEL-90
chr5-108295402-DEL-338
chr5-108297170-DEL-622
chr5-108311928-DEL-86
chr5-108321123-DEL-143
chr5-108330271-DEL-219
chr5-108335019-DEL-1395
chr5-108431927-DEL-615
chr5-108434511-DEL-177
chr5-108436628-DEL-296
chr5-108440548-DEL-97
chr5-108442180-DEL-6241

chr5-108454596-DEL-410
chr5-108518214-DEL-52
chr5-108548366-DEL-211
chr5-108583929-DEL-60
chr5-108587311-DEL-1198
chr5-108592110-DEL-88
chr5-108626353-DEL-198
chr5-108657442-DEL-199
chr5-108664962-DEL-481
chr5-108722803-DEL-61
chr5-108735406-DEL-191
chr5-108791780-DEL-168
chr5-108830299-DEL-141
chr5-108834236-DEL-338
chr5-108838294-DEL-238
chr5-108912608-DEL-1103
chr5-108953013-DEL-7892
chr5-109023658-DEL-649
chr5-109028405-DEL-196
chr5-109030995-DEL-3911
chr5-109034217-DEL-78
chr5-109053562-DEL-1655
chr5-109059978-DEL-343
chr5-109069085-DEL-84
chr5-109069733-DEL-529
chr5-109081771-DEL-351
chr5-109096624-DEL-69
chr5-109099584-DEL-74
chr5-109107209-DEL-1785
chr5-109125605-DEL-2384
chr5-109131357-DEL-91
chr5-109140492-DEL-195
chr5-109146418-DEL-84
chr5-109173392-DEL-6278
chr5-109200571-DEL-110
chr5-109202170-DEL-60
chr5-109203335-DEL-60
chr5-109220927-DEL-6273
chr5-109233297-DEL-230
chr5-109236232-DEL-673
chr5-109254133-DEL-581
chr5-109265725-DEL-750
chr5-109279774-DEL-234
chr5-109280978-DEL-177
chr5-109293846-DEL-7002
chr5-109320592-DEL-13657
chr5-109324852-DEL-204
chr5-109401931-DEL-907
chr5-109427384-DEL-6215
chr5-109444623-DEL-66
chr5-109446849-DEL-71
chr5-109451427-DEL-400
chr5-109453648-DEL-124
chr5-109459176-DEL-67
chr5-109473571-DEL-321
chr5-109526219-DEL-814
chr5-109533836-DEL-92
chr5-109586400-DEL-4803
chr5-109591800-DEL-771
chr5-109602148-DEL-69
chr5-109603284-DEL-149
chr5-109623001-DEL-51
chr5-109651548-DEL-63
chr5-109677043-DEL-120
chr5-109689191-DEL-1959
chr5-109691497-DEL-185
chr5-109723510-DEL-494
chr5-109735241-DEL-7061
chr5-109766958-DEL-3529
chr5-109766963-DEL-69
chr5-109767004-DEL-6216
chr5-109770547-DEL-2673
chr5-109775847-DEL-6735
chr5-109858769-DEL-4090
chr5-109934544-DEL-1259
chr5-109936862-DEL-669
chr5-109971564-DEL-352

chr5-109972772-DEL-70
chr5-109985519-DEL-561
chr5-110006238-DEL-52
chr5-110021110-DEL-54
chr5-110022980-DEL-102
chr5-110047569-DEL-177
chr5-110081356-DEL-3873
chr5-110094119-DEL-524
chr5-110121742-DEL-6549
chr5-110133530-DEL-660
chr5-110172628-DEL-289
chr5-110175517-DEL-739
chr5-110176546-DEL-176
chr5-110186326-DEL-312
chr5-110187388-DEL-219
chr5-110235140-DEL-59
chr5-110257696-DEL-55
chr5-110273327-DEL-52
chr5-110289173-DEL-199
chr5-110297324-DEL-105
chr5-110320403-DEL-154
chr5-110331396-DEL-210
chr5-110335858-DEL-64
chr5-110396038-DEL-61
chr5-110400256-DEL-116
chr5-110404964-DEL-400
chr5-110429603-DEL-1888
chr5-110452083-DEL-536
chr5-110457338-DEL-57
chr5-110460172-DEL-186
chr5-110470198-DEL-84
chr5-110493933-DEL-73
chr5-110495024-DEL-185
chr5-110513667-DEL-358
chr5-110517936-DEL-168
chr5-110553187-DEL-638
chr5-110556251-DEL-327
chr5-110643486-DEL-9700
chr5-110687991-DEL-87
chr5-110715020-DEL-313
chr5-110747279-DEL-244
chr5-110756010-DEL-198
chr5-110766268-DEL-88
chr5-110779881-DEL-390
chr5-110852392-DEL-144
chr5-110869569-DEL-662
chr5-110921028-DEL-177
chr5-110951179-DEL-83
chr5-110959248-DEL-112
chr5-110969351-DEL-60
chr5-110984779-DEL-199
chr5-111009170-DEL-109
chr5-111022653-DEL-52
chr5-111043842-DEL-99
chr5-111047462-DEL-80
chr5-111049140-DEL-74
chr5-111061726-DEL-1487
chr5-111099643-DEL-3929
chr5-111101025-DEL-3132
chr5-111109796-DEL-135
chr5-111110036-DEL-2790
chr5-111111717-DEL-1109
chr5-111111861-DEL-191
chr5-111112421-DEL-270
chr5-111137670-DEL-96
chr5-111156009-DEL-51
chr5-111159062-DEL-57
chr5-111162368-DEL-154
chr5-111164109-DEL-219
chr5-111169664-DEL-61
chr5-111183783-DEL-168
chr5-111209983-DEL-512
chr5-111249133-DEL-200
chr5-111282778-DEL-155
chr5-111296770-DEL-324
chr5-111304580-DEL-136
chr5-111315881-DEL-69

chr5-111429338-DEL-260
chr5-111430414-DEL-79
chr5-111453293-DEL-191
chr5-111469691-DEL-233
chr5-111471852-DEL-320
chr5-111488750-DEL-148
chr5-111513297-DEL-1397
chr5-111586951-DEL-389
chr5-111588811-DEL-133
chr5-111594985-DEL-379
chr5-111606956-DEL-470
chr5-111617784-DEL-223
chr5-111655979-DEL-64
chr5-111692949-DEL-85
chr5-111700344-DEL-517
chr5-111706673-DEL-299
chr5-111723808-DEL-178
chr5-111756450-DEL-61
chr5-111778581-DEL-54
chr5-111787033-DEL-68
chr5-111817242-DEL-60
chr5-111828431-DEL-72
chr5-111840746-DEL-86
chr5-111849730-DEL-70
chr5-111874317-DEL-53
chr5-111882203-DEL-52
chr5-111883124-DEL-382
chr5-111899535-DEL-62
chr5-111900744-DEL-115
chr5-111905446-DEL-102
chr5-111924575-DEL-234
chr5-111929043-DEL-753
chr5-111934163-DEL-256
chr5-111963065-DEL-82
chr5-111965839-DEL-224
chr5-111994559-DEL-62
chr5-111996937-DEL-57
chr5-112145419-DEL-92
chr5-112181423-DEL-57
chr5-112191186-DEL-74
chr5-112236058-DEL-117
chr5-112238845-DEL-65
chr5-112254985-DEL-74
chr5-112326392-DEL-70
chr5-112408993-DEL-51
chr5-112415591-DEL-108
chr5-112446326-DEL-52
chr5-112448091-DEL-140
chr5-112499248-DEL-105
chr5-112513792-DEL-130
chr5-112533944-DEL-110
chr5-112603008-DEL-829
chr5-112605429-DEL-57
chr5-112611231-DEL-61
chr5-112698494-DEL-57
chr5-112710483-DEL-82
chr5-112751237-DEL-107
chr5-112805329-DEL-497
chr5-112828058-DEL-137
chr5-112873586-DEL-76
chr5-112889191-DEL-189
chr5-112909408-DEL-108
chr5-112919283-DEL-60
chr5-112926359-DEL-171
chr5-112937033-DEL-124
chr5-112948001-DEL-63
chr5-112955343-DEL-112
chr5-112966459-DEL-117
chr5-112983329-DEL-84
chr5-112994680-DEL-68
chr5-113015771-DEL-73
chr5-113031856-DEL-805
chr5-113034254-DEL-129
chr5-113072494-DEL-70
chr5-113075152-DEL-70
chr5-113113690-DEL-612
chr5-113145934-DEL-83

chr5-113170308-DEL-113
chr5-113199353-DEL-122
chr5-113223133-DEL-74
chr5-113231472-DEL-159
chr5-113251057-DEL-88
chr5-113254082-DEL-119
chr5-113262124-DEL-106
chr5-113311537-DEL-94
chr5-113312781-DEL-99
chr5-113320968-DEL-64
chr5-113357723-DEL-87
chr5-113359202-DEL-158
chr5-113360955-DEL-54
chr5-113364085-DEL-58
chr5-113377202-DEL-135
chr5-113409179-DEL-1830
chr5-113432437-DEL-148
chr5-113462531-DEL-273
chr5-113491389-DEL-614
chr5-113525917-DEL-191
chr5-113531547-DEL-56
chr5-113567527-DEL-194
chr5-113568988-DEL-53
chr5-113573849-DEL-56
chr5-113576537-DEL-110
chr5-113589206-DEL-79
chr5-113613599-DEL-120
chr5-113655649-DEL-53
chr5-113672380-DEL-90
chr5-113791673-DEL-166
chr5-113795648-DEL-204
chr5-113804004-DEL-79
chr5-113807532-DEL-78
chr5-113812221-DEL-123
chr5-113837165-DEL-947
chr5-113843988-DEL-59
chr5-113851454-DEL-78
chr5-113865943-DEL-62
chr5-113867951-DEL-227
chr5-113880421-DEL-54
chr5-113890618-DEL-73
chr5-113918127-DEL-180
chr5-113939314-DEL-344
chr5-113945043-DEL-83
chr5-113973537-DEL-213
chr5-113985101-DEL-355
chr5-114013379-DEL-217
chr5-114024953-DEL-228
chr5-114030843-DEL-1024
chr5-114085501-DEL-534
chr5-114140068-DEL-58
chr5-114151290-DEL-52
chr5-114196057-DEL-104
chr5-114216238-DEL-381
chr5-114226851-DEL-65
chr5-114233940-DEL-904
chr5-114249326-DEL-121
chr5-114251126-DEL-70
chr5-114255821-DEL-204
chr5-114280112-DEL-290
chr5-114288396-DEL-431
chr5-114295220-DEL-123
chr5-114298347-DEL-93
chr5-114305232-DEL-213
chr5-114307080-DEL-186
chr5-114310210-DEL-500
chr5-114336557-DEL-276
chr5-114395616-DEL-74
chr5-114410946-DEL-52
chr5-114419689-DEL-81
chr5-114438639-DEL-80
chr5-114453902-DEL-68
chr5-114461228-DEL-71
chr5-114466901-DEL-213
chr5-114470843-DEL-485
chr5-114478576-DEL-90
chr5-114478918-DEL-91

chr5-114488795-DEL-166
chr5-114517038-DEL-52
chr5-114547889-DEL-64
chr5-114557352-DEL-64
chr5-114563456-DEL-72
chr5-114589576-DEL-53
chr5-114592638-DEL-156
chr5-114638702-DEL-74
chr5-114639561-DEL-94
chr5-114667508-DEL-135
chr5-114686576-DEL-1726
chr5-114734065-DEL-187
chr5-114768287-DEL-80
chr5-114770086-DEL-73
chr5-114812029-DEL-72
chr5-114837530-DEL-458
chr5-114839728-DEL-79
chr5-114843086-DEL-879
chr5-114844701-DEL-406
chr5-114846571-DEL-77
chr5-114883967-DEL-524
chr5-114906016-DEL-136
chr5-114915342-DEL-196
chr5-114932270-DEL-66
chr5-114953245-DEL-10397
chr5-114967667-DEL-177
chr5-114989962-DEL-55
chr5-115017839-DEL-2679
chr5-115027983-DEL-5305
chr5-115068423-DEL-202
chr5-115076163-DEL-3386
chr5-115091191-DEL-138
chr5-115092582-DEL-51
chr5-115099008-DEL-204
chr5-115101270-DEL-62
chr5-115124691-DEL-227
chr5-115146840-DEL-235
chr5-115177006-DEL-120
chr5-115186790-DEL-2321
chr5-115220951-DEL-207
chr5-115245896-DEL-227
chr5-115247198-DEL-285
chr5-115257941-DEL-374
chr5-115259833-DEL-154
chr5-115278331-DEL-59
chr5-115304214-DEL-76
chr5-115345941-DEL-54
chr5-115389965-DEL-2735
chr5-115391694-DEL-1183
chr5-115458223-DEL-208
chr5-115487956-DEL-398
chr5-115516417-DEL-201
chr5-115517610-DEL-93
chr5-115518994-DEL-420
chr5-115525634-DEL-129
chr5-115530948-DEL-1023
chr5-115551708-DEL-233
chr5-115560170-DEL-196
chr5-115562281-DEL-887
chr5-115584446-DEL-219
chr5-115591287-DEL-61
chr5-115599573-DEL-756
chr5-115604559-DEL-75
chr5-115617184-DEL-93
chr5-115695237-DEL-55
chr5-115696172-DEL-54
chr5-115696901-DEL-53
chr5-115729209-DEL-728
chr5-115796902-DEL-68
chr5-115797151-DEL-713
chr5-115802488-DEL-210
chr5-115816128-DEL-56
chr5-115835590-DEL-180
chr5-115854059-DEL-705
chr5-115901279-DEL-66
chr5-115922570-DEL-349
chr5-115947501-DEL-156

chr5-115947501-DEL-984
chr5-115948210-DEL-131
chr5-115953881-DEL-61
chr5-115956292-DEL-260
chr5-115959341-DEL-70
chr5-115968601-DEL-159
chr5-115969819-DEL-160
chr5-116015536-DEL-86
chr5-116028764-DEL-251
chr5-116034264-DEL-1671
chr5-116080308-DEL-209
chr5-116095174-DEL-58
chr5-116098041-DEL-90
chr5-116129965-DEL-219
chr5-116136712-DEL-106
chr5-116160014-DEL-87
chr5-116180730-DEL-226
chr5-116191651-DEL-100
chr5-116193374-DEL-69
chr5-116196728-DEL-448
chr5-116202171-DEL-138
chr5-116237184-DEL-1260
chr5-116252863-DEL-296
chr5-116255528-DEL-66
chr5-116276211-DEL-478
chr5-116281498-DEL-136
chr5-116293748-DEL-95
chr5-116299201-DEL-1904
chr5-116304424-DEL-58
chr5-116317387-DEL-55
chr5-116329361-DEL-106
chr5-116355241-DEL-2036
chr5-116378614-DEL-64
chr5-116393002-DEL-150
chr5-116401307-DEL-1182
chr5-116423301-DEL-78
chr5-116424952-DEL-100
chr5-116430863-DEL-186
chr5-116454420-DEL-146
chr5-116467135-DEL-80
chr5-116481738-DEL-307
chr5-116484816-DEL-507
chr5-116485751-DEL-170
chr5-116552993-DEL-76
chr5-116556134-DEL-60
chr5-116557326-DEL-98
chr5-116559404-DEL-73
chr5-116562706-DEL-146
chr5-116563913-DEL-74
chr5-116594129-DEL-69
chr5-116604124-DEL-65
chr5-116608067-DEL-55
chr5-116611155-DEL-84
chr5-116629660-DEL-102
chr5-116664814-DEL-686
chr5-116690480-DEL-57
chr5-116705056-DEL-159
chr5-116706168-DEL-138
chr5-116725522-DEL-620
chr5-116739347-DEL-547
chr5-116772764-DEL-52
chr5-116828214-DEL-1386
chr5-116831765-DEL-71
chr5-116858868-DEL-487
chr5-116876545-DEL-890
chr5-116878497-DEL-195
chr5-116884358-DEL-244
chr5-116898839-DEL-58
chr5-116903891-DEL-256
chr5-116938318-DEL-229
chr5-116941655-DEL-270
chr5-116968279-DEL-57
chr5-116969629-DEL-65
chr5-116987395-DEL-1332
chr5-117043285-DEL-3469
chr5-117088664-DEL-52
chr5-117097554-DEL-107

chr5-117115359-DEL-52
chr5-117119482-DEL-162
chr5-117139653-DEL-74
chr5-117154681-DEL-75
chr5-117198457-DEL-135
chr5-117210974-DEL-61
chr5-117236634-DEL-135
chr5-117248344-DEL-195
chr5-117257429-DEL-65
chr5-117315587-DEL-67
chr5-117336319-DEL-266
chr5-117338672-DEL-185
chr5-117339873-DEL-84
chr5-117352591-DEL-61
chr5-117362798-DEL-71
chr5-117369505-DEL-684
chr5-117376146-DEL-524
chr5-117392935-DEL-523
chr5-117399156-DEL-86
chr5-117401930-DEL-51
chr5-117411517-DEL-91
chr5-117414766-DEL-70
chr5-117515029-DEL-77
chr5-117522717-DEL-62
chr5-117573230-DEL-87
chr5-117580874-DEL-108
chr5-117582701-DEL-264
chr5-117584621-DEL-52
chr5-117607255-DEL-175
chr5-117610693-DEL-406
chr5-117624639-DEL-164
chr5-117626943-DEL-82
chr5-117635719-DEL-103
chr5-117641440-DEL-205
chr5-117650863-DEL-1094
chr5-117665583-DEL-131
chr5-117670076-DEL-96
chr5-117676288-DEL-151
chr5-117690669-DEL-183
chr5-117704071-DEL-76
chr5-117724095-DEL-238
chr5-117732899-DEL-83
chr5-117763722-DEL-100
chr5-117772441-DEL-133
chr5-117802215-DEL-62
chr5-117823923-DEL-1016
chr5-117827062-DEL-57
chr5-117871425-DEL-527
chr5-117883416-DEL-60
chr5-117912338-DEL-63
chr5-117913534-DEL-387
chr5-117957886-DEL-60
chr5-118002361-DEL-53
chr5-118007782-DEL-341
chr5-118018310-DEL-56
chr5-118020411-DEL-637
chr5-118038064-DEL-212
chr5-118054842-DEL-51
chr5-118059672-DEL-199
chr5-118084349-DEL-270
chr5-118090364-DEL-55
chr5-118121883-DEL-544
chr5-118145169-DEL-96
chr5-118154553-DEL-207
chr5-118159655-DEL-224
chr5-118180281-DEL-186
chr5-118181972-DEL-133
chr5-118189937-DEL-86
chr5-118216289-DEL-69
chr5-118233093-DEL-241
chr5-118253683-DEL-55
chr5-118300860-DEL-78
chr5-118308654-DEL-66
chr5-118321410-DEL-56
chr5-118340411-DEL-105
chr5-118349265-DEL-265
chr5-118349862-DEL-1326

chr5-118374314-DEL-178
chr5-118387078-DEL-116
chr5-118392646-DEL-232
chr5-118423055-DEL-52
chr5-118452673-DEL-92
chr5-118458654-DEL-104
chr5-118464689-DEL-195
chr5-118474582-DEL-1103
chr5-118520187-DEL-227
chr5-118542884-DEL-1785
chr5-118566831-DEL-171
chr5-118570000-DEL-73
chr5-118570003-DEL-641
chr5-118637334-DEL-92
chr5-118644891-DEL-78
chr5-118689590-DEL-104
chr5-118709099-DEL-59
chr5-118719084-DEL-94
chr5-118726042-DEL-81
chr5-118735403-DEL-67
chr5-118764306-DEL-77
chr5-118768619-DEL-140
chr5-118778349-DEL-472
chr5-118812724-DEL-64
chr5-118815111-DEL-459
chr5-118824482-DEL-546
chr5-118825061-DEL-1344
chr5-118827480-DEL-931
chr5-118854980-DEL-72
chr5-118885959-DEL-60
chr5-118908576-DEL-329
chr5-118932145-DEL-690
chr5-118963815-DEL-333
chr5-118982010-DEL-56
chr5-118983006-DEL-92
chr5-119010193-DEL-82
chr5-119011175-DEL-89
chr5-119054534-DEL-59
chr5-119079721-DEL-82
chr5-119088493-DEL-107
chr5-119092711-DEL-84
chr5-119116357-DEL-72
chr5-119120929-DEL-102
chr5-119130767-DEL-280
chr5-119131759-DEL-206
chr5-119151903-DEL-104
chr5-119155413-DEL-72
chr5-119167947-DEL-60
chr5-119189891-DEL-76
chr5-119241547-DEL-70
chr5-119250864-DEL-135
chr5-119253075-DEL-416
chr5-119255159-DEL-52
chr5-119314501-DEL-63
chr5-119315691-DEL-70
chr5-119377503-DEL-139
chr5-119378294-DEL-127
chr5-119391714-DEL-55
chr5-119408473-DEL-121
chr5-119411935-DEL-384
chr5-119427167-DEL-138
chr5-119432341-DEL-454
chr5-119436657-DEL-134
chr5-119437833-DEL-99
chr5-119449548-DEL-691
chr5-119472254-DEL-225
chr5-119481860-DEL-139
chr5-119489238-DEL-347
chr5-119489386-DEL-315
chr5-119501257-DEL-52
chr5-119514014-DEL-51
chr5-119517645-DEL-143
chr5-119537468-DEL-74
chr5-119614931-DEL-366
chr5-119640201-DEL-1377
chr5-119659171-DEL-84
chr5-119661097-DEL-76

chr5-119661521-DEL-403
chr5-119672727-DEL-193
chr5-119699192-DEL-73
chr5-119703309-DEL-1061
chr5-119712731-DEL-51
chr5-119723554-DEL-205
chr5-119725898-DEL-114
chr5-119748588-DEL-150
chr5-119755047-DEL-78
chr5-119756925-DEL-51
chr5-119760777-DEL-60
chr5-119767272-DEL-53
chr5-119837946-DEL-78
chr5-119847660-DEL-199
chr5-119856146-DEL-398
chr5-119856770-DEL-90
chr5-119866702-DEL-277
chr5-119874396-DEL-341
chr5-119878011-DEL-66
chr5-119883620-DEL-83
chr5-119903975-DEL-234
chr5-119906709-DEL-200
chr5-119923296-DEL-182
chr5-119952130-DEL-202
chr5-119965937-DEL-54
chr5-119995994-DEL-57
chr5-120020167-DEL-62
chr5-120027545-DEL-68
chr5-120032903-DEL-240
chr5-120034004-DEL-82
chr5-120050868-DEL-65
chr5-120061891-DEL-168
chr5-120074490-DEL-81
chr5-120078950-DEL-154
chr5-120089146-DEL-55
chr5-120103579-DEL-122
chr5-120111926-DEL-1214
chr5-120114093-DEL-82
chr5-120164865-DEL-67
chr5-120196105-DEL-54
chr5-120227544-DEL-72
chr5-120237096-DEL-162
chr5-120245876-DEL-81
chr5-120253548-DEL-203
chr5-120290681-DEL-60
chr5-120323162-DEL-71
chr5-120325200-DEL-52
chr5-120337132-DEL-172
chr5-120349831-DEL-115
chr5-120413462-DEL-328
chr5-120446849-DEL-556
chr5-120448732-DEL-104
chr5-120462277-DEL-59
chr5-120486770-DEL-60
chr5-120510723-DEL-136
chr5-120513105-DEL-236
chr5-120542358-DEL-59
chr5-120545469-DEL-106
chr5-120578818-DEL-66
chr5-120583673-DEL-107
chr5-120587660-DEL-95
chr5-120603016-DEL-95
chr5-120606027-DEL-120
chr5-120637521-DEL-212
chr5-120644954-DEL-129
chr5-120656787-DEL-56
chr5-120718796-DEL-149
chr5-120729692-DEL-131
chr5-120752667-DEL-61
chr5-120775061-DEL-134
chr5-120775310-DEL-758
chr5-120777073-DEL-208
chr5-120783504-DEL-412
chr5-120804001-DEL-144
chr5-120859622-DEL-201
chr5-120862312-DEL-51
chr5-120866555-DEL-94

chr5-120883643-DEL-74
chr5-120892039-DEL-94
chr5-120916788-DEL-137
chr5-120921316-DEL-2963
chr5-120945900-DEL-54
chr5-121006441-DEL-689
chr5-121050920-DEL-292
chr5-121094296-DEL-64
chr5-121106892-DEL-118
chr5-121109693-DEL-64
chr5-121111264-DEL-54
chr5-121130377-DEL-56
chr5-121132360-DEL-951
chr5-121157241-DEL-70
chr5-121173664-DEL-366
chr5-121185122-DEL-70
chr5-121224641-DEL-96
chr5-121248205-DEL-59
chr5-121265094-DEL-181
chr5-121277624-DEL-913
chr5-121285789-DEL-1290
chr5-121286857-DEL-241
chr5-121313757-DEL-901
chr5-121327878-DEL-72
chr5-121333336-DEL-56
chr5-121347218-DEL-3105
chr5-121352304-DEL-463
chr5-121357185-DEL-89
chr5-121362057-DEL-203
chr5-121423367-DEL-53
chr5-121470594-DEL-124
chr5-121510790-DEL-80
chr5-121514979-DEL-210
chr5-121540506-DEL-163
chr5-121564250-DEL-60
chr5-121579260-DEL-185
chr5-121583044-DEL-53
chr5-121607590-DEL-2633
chr5-121611950-DEL-197
chr5-121620963-DEL-297
chr5-121647027-DEL-898
chr5-121660681-DEL-90
chr5-121665873-DEL-300
chr5-121668352-DEL-93
chr5-121677061-DEL-91
chr5-121701293-DEL-185
chr5-121720511-DEL-57
chr5-121751223-DEL-476
chr5-121836420-DEL-86
chr5-121864089-DEL-185
chr5-121871701-DEL-51
chr5-121881320-DEL-64
chr5-121890427-DEL-199
chr5-122029020-DEL-63
chr5-122044331-DEL-58
chr5-122077256-DEL-170
chr5-122102843-DEL-54
chr5-122125303-DEL-2332
chr5-122171751-DEL-108
chr5-122179486-DEL-179
chr5-122190696-DEL-52
chr5-122193596-DEL-57
chr5-122237806-DEL-51
chr5-122248395-DEL-80
chr5-122264244-DEL-185
chr5-122285224-DEL-3237
chr5-122314950-DEL-990
chr5-122316163-DEL-63
chr5-122321161-DEL-3313
chr5-122323126-DEL-77
chr5-122326758-DEL-189
chr5-122333646-DEL-1360
chr5-122365616-DEL-194
chr5-122369503-DEL-352
chr5-122375910-DEL-189
chr5-122419990-DEL-232
chr5-122498308-DEL-59

chr5-122534586-DEL-188
chr5-122542420-DEL-69
chr5-122567187-DEL-115
chr5-122568844-DEL-96
chr5-122570032-DEL-707
chr5-122608979-DEL-153
chr5-122623031-DEL-2028
chr5-122627620-DEL-566
chr5-122658546-DEL-171
chr5-122659633-DEL-832
chr5-122660344-DEL-57
chr5-122669743-DEL-1404
chr5-122671175-DEL-98
chr5-122695056-DEL-88
chr5-122700010-DEL-56
chr5-122711776-DEL-210
chr5-122761482-DEL-256
chr5-122779664-DEL-555
chr5-122785865-DEL-207
chr5-122787294-DEL-114
chr5-122830962-DEL-176
chr5-122832499-DEL-1026
chr5-122864475-DEL-86
chr5-122882808-DEL-138
chr5-122920567-DEL-299
chr5-122923343-DEL-69
chr5-122932849-DEL-358
chr5-122935719-DEL-187
chr5-122968518-DEL-163
chr5-122996272-DEL-389
chr5-123052827-DEL-439
chr5-123055903-DEL-570
chr5-123060620-DEL-56
chr5-123064712-DEL-55
chr5-123104894-DEL-80
chr5-123110977-DEL-52
chr5-123142836-DEL-198
chr5-123153129-DEL-61
chr5-123173268-DEL-78
chr5-123201471-DEL-204
chr5-123203176-DEL-217
chr5-123207667-DEL-79
chr5-123208184-DEL-116
chr5-123219756-DEL-189
chr5-123244244-DEL-84
chr5-123259658-DEL-132
chr5-123261829-DEL-611
chr5-123312138-DEL-105
chr5-123321993-DEL-138
chr5-123335970-DEL-6518
chr5-123360034-DEL-102
chr5-123380354-DEL-80
chr5-123383362-DEL-60
chr5-123384400-DEL-533
chr5-123384418-DEL-1271
chr5-123385565-DEL-81
chr5-123390547-DEL-80
chr5-123417120-DEL-288
chr5-123431243-DEL-198
chr5-123434071-DEL-51
chr5-123444966-DEL-54
chr5-123446305-DEL-58
chr5-123448972-DEL-107
chr5-123458638-DEL-140
chr5-123472065-DEL-64
chr5-123478902-DEL-53
chr5-123530397-DEL-215
chr5-123546508-DEL-187
chr5-123557697-DEL-160
chr5-123560568-DEL-207
chr5-123602987-DEL-60
chr5-123629837-DEL-135
chr5-123643751-DEL-311
chr5-123656928-DEL-261
chr5-123676786-DEL-7774
chr5-123692151-DEL-104
chr5-123706918-DEL-56

chr5-123718657-DEL-52
chr5-123736456-DEL-62
chr5-123737705-DEL-72
chr5-123748563-DEL-292
chr5-123758461-DEL-266
chr5-123759208-DEL-228
chr5-123779045-DEL-80
chr5-123795078-DEL-240
chr5-123853327-DEL-468
chr5-123876039-DEL-849
chr5-123917327-DEL-54
chr5-123921608-DEL-60
chr5-123938721-DEL-78
chr5-123963776-DEL-781
chr5-123967469-DEL-86
chr5-123993376-DEL-436
chr5-124041453-DEL-159
chr5-124043981-DEL-82
chr5-124055600-DEL-204
chr5-124061161-DEL-57
chr5-124077339-DEL-3985
chr5-124081882-DEL-52
chr5-124109754-DEL-124
chr5-124125933-DEL-178
chr5-124133045-DEL-75
chr5-124157059-DEL-105
chr5-124176265-DEL-154
chr5-124187774-DEL-118
chr5-124208804-DEL-515
chr5-124264936-DEL-64
chr5-124340432-DEL-98
chr5-124418837-DEL-203
chr5-124437464-DEL-367
chr5-124441205-DEL-159
chr5-124452723-DEL-177
chr5-124458167-DEL-150
chr5-124473165-DEL-217
chr5-124485872-DEL-223
chr5-124499098-DEL-196
chr5-124503674-DEL-1666
chr5-124505878-DEL-62
chr5-124517963-DEL-188
chr5-124539627-DEL-200
chr5-124540907-DEL-72
chr5-124557414-DEL-123
chr5-124565596-DEL-836
chr5-124593226-DEL-1175
chr5-124602417-DEL-207
chr5-124609356-DEL-56
chr5-124617496-DEL-126
chr5-124619146-DEL-1788
chr5-124619174-DEL-270
chr5-124620807-DEL-106
chr5-124662222-DEL-75
chr5-124701148-DEL-241
chr5-124701803-DEL-2210
chr5-124702123-DEL-615
chr5-124703100-DEL-156
chr5-124703751-DEL-78
chr5-124712902-DEL-405
chr5-124723946-DEL-721
chr5-124739355-DEL-220
chr5-124745109-DEL-69
chr5-124747533-DEL-207
chr5-124776073-DEL-174
chr5-124785764-DEL-324
chr5-124785827-DEL-1656
chr5-124787159-DEL-144
chr5-124788619-DEL-127
chr5-124806430-DEL-69
chr5-124813471-DEL-104
chr5-124822597-DEL-60
chr5-124855719-DEL-51
chr5-124856388-DEL-506
chr5-124877501-DEL-56
chr5-124881137-DEL-136
chr5-124924990-DEL-101

chr5-124946078-DEL-401
chr5-124976300-DEL-541
chr5-124984853-DEL-67
chr5-125026588-DEL-192
chr5-125049788-DEL-508
chr5-125050763-DEL-79
chr5-125071780-DEL-151
chr5-125146402-DEL-215
chr5-125212742-DEL-372
chr5-125261475-DEL-135
chr5-125263879-DEL-115
chr5-125265112-DEL-80
chr5-125282247-DEL-215
chr5-125291561-DEL-203
chr5-125318252-DEL-72
chr5-125320399-DEL-69
chr5-125335117-DEL-171
chr5-125337609-DEL-569
chr5-125444533-DEL-220
chr5-125464209-DEL-228
chr5-125474698-DEL-596
chr5-125475495-DEL-171
chr5-125528499-DEL-56
chr5-125535325-DEL-83
chr5-125538891-DEL-126
chr5-125549568-DEL-327
chr5-125573786-DEL-548
chr5-125579791-DEL-84
chr5-125582706-DEL-118
chr5-125601128-DEL-54
chr5-125638325-DEL-131
chr5-125658233-DEL-1057
chr5-125661748-DEL-111
chr5-125672909-DEL-77
chr5-125677561-DEL-179
chr5-125681471-DEL-121
chr5-125696160-DEL-140
chr5-125709300-DEL-198
chr5-125737307-DEL-162
chr5-125759128-DEL-202
chr5-125770508-DEL-1483
chr5-125772584-DEL-52
chr5-125774022-DEL-223
chr5-125779455-DEL-56
chr5-125788907-DEL-53
chr5-125800066-DEL-86
chr5-125816851-DEL-298
chr5-125829258-DEL-187
chr5-125887272-DEL-66
chr5-125947777-DEL-102
chr5-125962902-DEL-336
chr5-125976080-DEL-105
chr5-125976644-DEL-132
chr5-125982647-DEL-213
chr5-125988676-DEL-281
chr5-126013142-DEL-1105
chr5-126015803-DEL-89
chr5-126029445-DEL-88
chr5-126035895-DEL-2166
chr5-126044900-DEL-54
chr5-126058133-DEL-65
chr5-126067864-DEL-138
chr5-126094966-DEL-72
chr5-126098998-DEL-59
chr5-126112500-DEL-5028
chr5-126128708-DEL-328
chr5-126168999-DEL-156
chr5-126215885-DEL-72
chr5-126229429-DEL-70
chr5-126230067-DEL-54
chr5-126238610-DEL-82
chr5-126285373-DEL-1692
chr5-126295127-DEL-146
chr5-126403965-DEL-104
chr5-126409894-DEL-62
chr5-126411133-DEL-117
chr5-126415955-DEL-502

chr5-126475834-DEL-101
chr5-126549333-DEL-62
chr5-126575095-DEL-70
chr5-126626335-DEL-305
chr5-126644487-DEL-138
chr5-126654551-DEL-411
chr5-126713679-DEL-182
chr5-126714513-DEL-55
chr5-126723950-DEL-56
chr5-126733246-DEL-145
chr5-126760029-DEL-1135
chr5-126785723-DEL-65
chr5-126800085-DEL-64
chr5-126810893-DEL-51
chr5-126820421-DEL-161
chr5-126908127-DEL-79
chr5-126931784-DEL-995
chr5-126976555-DEL-298
chr5-126985276-DEL-253
chr5-126997408-DEL-288
chr5-127011083-DEL-247
chr5-127074130-DEL-52
chr5-127080862-DEL-66
chr5-127084693-DEL-6047
chr5-127153067-DEL-52
chr5-127225949-DEL-58
chr5-127236581-DEL-68
chr5-127244082-DEL-223
chr5-127254150-DEL-108
chr5-127258094-DEL-52
chr5-127262779-DEL-119
chr5-127283054-DEL-62
chr5-127383349-DEL-56
chr5-127387179-DEL-60
chr5-127423611-DEL-110
chr5-127450938-DEL-111
chr5-127504840-DEL-230
chr5-127512000-DEL-170
chr5-127588477-DEL-93
chr5-127599877-DEL-915
chr5-127653203-DEL-265
chr5-127670551-DEL-1710
chr5-127678167-DEL-206
chr5-127701824-DEL-132
chr5-127706373-DEL-51
chr5-127707927-DEL-54
chr5-127731877-DEL-706
chr5-127732397-DEL-496
chr5-127759252-DEL-52
chr5-127792363-DEL-55
chr5-127811948-DEL-225
chr5-127843119-DEL-56
chr5-127849972-DEL-81
chr5-127852504-DEL-100
chr5-127863764-DEL-61
chr5-127879302-DEL-52
chr5-127881765-DEL-62
chr5-127881897-DEL-394
chr5-127893252-DEL-53
chr5-127902537-DEL-74
chr5-127944404-DEL-2643
chr5-127951982-DEL-3964
chr5-127957133-DEL-3016
chr5-128047320-DEL-404
chr5-128066514-DEL-136
chr5-128097030-DEL-55
chr5-128123731-DEL-102
chr5-128142348-DEL-100
chr5-128146963-DEL-54
chr5-128173274-DEL-209
chr5-128175578-DEL-337
chr5-128208554-DEL-51
chr5-128228498-DEL-87
chr5-128234944-DEL-306
chr5-128241805-DEL-172
chr5-128286609-DEL-194
chr5-128291378-DEL-69

chr5-128305896-DEL-958
chr5-128391256-DEL-236
chr5-128392329-DEL-512
chr5-128422125-DEL-58
chr5-128530675-DEL-188
chr5-128531793-DEL-71
chr5-128532661-DEL-128
chr5-128548946-DEL-69
chr5-128567084-DEL-450
chr5-128597493-DEL-159
chr5-128699369-DEL-85
chr5-128758976-DEL-254
chr5-128768855-DEL-393
chr5-128798598-DEL-6431
chr5-128805279-DEL-283
chr5-128871145-DEL-64
chr5-128899587-DEL-53
chr5-128912291-DEL-221
chr5-128914160-DEL-289
chr5-128931657-DEL-51
chr5-128947178-DEL-267
chr5-129015802-DEL-141
chr5-129034217-DEL-220
chr5-129043602-DEL-190
chr5-129047051-DEL-1211
chr5-129048665-DEL-108
chr5-129049981-DEL-62
chr5-129062032-DEL-84
chr5-129120938-DEL-57
chr5-129130710-DEL-267
chr5-129131532-DEL-60
chr5-129165645-DEL-59
chr5-129181528-DEL-79
chr5-129182642-DEL-327
chr5-129212755-DEL-144
chr5-129234665-DEL-162
chr5-129265032-DEL-200
chr5-129283238-DEL-60
chr5-129287092-DEL-490
chr5-129287617-DEL-135
chr5-129305398-DEL-94
chr5-129330862-DEL-55
chr5-129331217-DEL-499
chr5-129337907-DEL-55
chr5-129350231-DEL-664
chr5-129355576-DEL-65
chr5-129361941-DEL-886
chr5-129402018-DEL-279
chr5-129411835-DEL-279
chr5-129426251-DEL-63
chr5-129439446-DEL-74
chr5-129446175-DEL-54
chr5-129463378-DEL-442
chr5-129479205-DEL-144
chr5-129495665-DEL-1308
chr5-129566226-DEL-166
chr5-129576018-DEL-165
chr5-129588180-DEL-56
chr5-129612212-DEL-972
chr5-129691182-DEL-650
chr5-129704144-DEL-208
chr5-129715388-DEL-182
chr5-129719241-DEL-431
chr5-129747595-DEL-54
chr5-129770673-DEL-201
chr5-129782578-DEL-1883
chr5-129799756-DEL-194
chr5-129871095-DEL-175
chr5-129871954-DEL-62
chr5-129879344-DEL-331
chr5-129892246-DEL-204
chr5-129892895-DEL-57
chr5-129894445-DEL-204
chr5-129904624-DEL-353
chr5-129921418-DEL-211
chr5-129928568-DEL-239
chr5-129930102-DEL-191

chr5-129958606-DEL-99
chr5-129982783-DEL-5662
chr5-129985014-DEL-88
chr5-129988037-DEL-87
chr5-130021030-DEL-226
chr5-130045934-DEL-72
chr5-130059751-DEL-108
chr5-130080089-DEL-240
chr5-130082888-DEL-109
chr5-130089770-DEL-495
chr5-130095172-DEL-199
chr5-130119422-DEL-1537
chr5-130121934-DEL-315
chr5-130148948-DEL-64
chr5-130151848-DEL-181
chr5-130185843-DEL-212
chr5-130197534-DEL-181
chr5-130220391-DEL-55
chr5-130231787-DEL-54
chr5-130241306-DEL-174
chr5-130259671-DEL-138
chr5-130276089-DEL-149
chr5-130277956-DEL-1684
chr5-130282603-DEL-82
chr5-130286962-DEL-58
chr5-130298589-DEL-2143
chr5-130337436-DEL-53
chr5-130345426-DEL-149
chr5-130348776-DEL-245
chr5-130356546-DEL-96
chr5-130360883-DEL-59
chr5-130365435-DEL-1067
chr5-130393997-DEL-106
chr5-130397824-DEL-53
chr5-130410917-DEL-154
chr5-130424544-DEL-56
chr5-130436575-DEL-195
chr5-130464661-DEL-244
chr5-130465443-DEL-197
chr5-130480394-DEL-56
chr5-130486161-DEL-57
chr5-130492978-DEL-53
chr5-130498420-DEL-142
chr5-130508445-DEL-79
chr5-130510737-DEL-121
chr5-130525515-DEL-1501
chr5-130533424-DEL-66
chr5-130541910-DEL-2045
chr5-130544666-DEL-177
chr5-130558377-DEL-81
chr5-130572780-DEL-51
chr5-130606905-DEL-145
chr5-130614887-DEL-1234
chr5-130615613-DEL-239
chr5-130637755-DEL-168
chr5-130656070-DEL-52
chr5-130681212-DEL-67
chr5-130712549-DEL-52
chr5-130769292-DEL-126
chr5-130793389-DEL-60
chr5-130814255-DEL-165
chr5-130817218-DEL-78
chr5-130822880-DEL-52
chr5-130830043-DEL-229
chr5-130858680-DEL-82
chr5-130863529-DEL-199
chr5-130864676-DEL-305
chr5-130917567-DEL-803
chr5-130950294-DEL-204
chr5-130989585-DEL-87
chr5-130999823-DEL-240
chr5-131001367-DEL-702
chr5-131027591-DEL-66
chr5-131028429-DEL-54
chr5-131039013-DEL-79
chr5-131056329-DEL-134
chr5-131064351-DEL-962

chr5-131078331-DEL-54
chr5-131134030-DEL-52
chr5-131137913-DEL-68
chr5-131158608-DEL-52
chr5-131177092-DEL-1166
chr5-131182857-DEL-123
chr5-131190642-DEL-221
chr5-131248481-DEL-53
chr5-131266780-DEL-937
chr5-131282204-DEL-203
chr5-131287198-DEL-52
chr5-131290692-DEL-60
chr5-131294280-DEL-58
chr5-131294831-DEL-110
chr5-131298559-DEL-308
chr5-131311654-DEL-55
chr5-131315417-DEL-106
chr5-131339270-DEL-159
chr5-131358393-DEL-116
chr5-131402993-DEL-152
chr5-131408345-DEL-66
chr5-131419562-DEL-120
chr5-131436722-DEL-156
chr5-131517802-DEL-109
chr5-131579746-DEL-72
chr5-131596931-DEL-88
chr5-131619952-DEL-77
chr5-131652302-DEL-159
chr5-131657638-DEL-334
chr5-131661534-DEL-1880
chr5-131661607-DEL-141
chr5-131661665-DEL-987
chr5-131661886-DEL-658
chr5-131662739-DEL-164
chr5-131667751-DEL-64
chr5-131677430-DEL-53
chr5-131686381-DEL-105
chr5-131690659-DEL-159
chr5-131742601-DEL-155
chr5-131765767-DEL-93
chr5-131784244-DEL-273
chr5-131821937-DEL-604
chr5-131826628-DEL-280
chr5-131831486-DEL-187
chr5-131839931-DEL-80
chr5-131860136-DEL-54
chr5-131882144-DEL-106
chr5-131900872-DEL-1040
chr5-131964710-DEL-63
chr5-131980510-DEL-1262
chr5-131989595-DEL-1516
chr5-132038237-DEL-139
chr5-132107379-DEL-149
chr5-132169394-DEL-130
chr5-132226472-DEL-575
chr5-132234969-DEL-59
chr5-132249955-DEL-54
chr5-132262695-DEL-72
chr5-132264174-DEL-62
chr5-132270878-DEL-78
chr5-132273407-DEL-76
chr5-132419299-DEL-296
chr5-132440467-DEL-132
chr5-132514123-DEL-59
chr5-132521294-DEL-238
chr5-132544857-DEL-102
chr5-132581078-DEL-90
chr5-132584070-DEL-74
chr5-132594518-DEL-72
chr5-132603158-DEL-58
chr5-132611343-DEL-61
chr5-132619876-DEL-216
chr5-132624548-DEL-159
chr5-132649936-DEL-160
chr5-132690547-DEL-1104
chr5-132694089-DEL-56
chr5-132700934-DEL-66

chr5-132735143-DEL-106
chr5-132750532-DEL-104
chr5-132754270-DEL-53
chr5-132755464-DEL-250
chr5-132756109-DEL-66
chr5-132765952-DEL-140
chr5-132777535-DEL-53
chr5-132780415-DEL-130
chr5-132786927-DEL-273
chr5-132795185-DEL-54
chr5-132802397-DEL-73
chr5-132807862-DEL-174
chr5-132815106-DEL-52
chr5-132841645-DEL-54
chr5-132855757-DEL-68
chr5-132859269-DEL-259
chr5-132863971-DEL-53
chr5-132876294-DEL-307
chr5-132886303-DEL-172
chr5-132908006-DEL-107
chr5-132912564-DEL-63
chr5-132933753-DEL-53
chr5-132938378-DEL-112
chr5-132943050-DEL-348
chr5-132944958-DEL-87
chr5-132949906-DEL-71
chr5-132956504-DEL-70
chr5-132989792-DEL-1298
chr5-132997550-DEL-53
chr5-133000327-DEL-175
chr5-133013882-DEL-265
chr5-133055997-DEL-67
chr5-133072161-DEL-60
chr5-133118064-DEL-590
chr5-133165944-DEL-71
chr5-133170528-DEL-167
chr5-133172681-DEL-104
chr5-133182776-DEL-83
chr5-133230968-DEL-60
chr5-133253750-DEL-126
chr5-133267086-DEL-51
chr5-133268258-DEL-74
chr5-133282626-DEL-430
chr5-133285897-DEL-201
chr5-133298194-DEL-59
chr5-133321346-DEL-60
chr5-133350001-DEL-52
chr5-133358087-DEL-58
chr5-133370380-DEL-251
chr5-133403510-DEL-58
chr5-133452335-DEL-208
chr5-133460068-DEL-62
chr5-133461632-DEL-64
chr5-133489593-DEL-7177
chr5-133496718-DEL-212
chr5-133515580-DEL-88
chr5-133524083-DEL-169
chr5-133524395-DEL-99
chr5-133538316-DEL-57
chr5-133565463-DEL-229
chr5-133567730-DEL-71
chr5-133569118-DEL-54
chr5-133606085-DEL-106
chr5-133643837-DEL-1306
chr5-133678894-DEL-58
chr5-133688642-DEL-264
chr5-133691329-DEL-55
chr5-133711132-DEL-96
chr5-133713536-DEL-64
chr5-133715807-DEL-52
chr5-133720999-DEL-195
chr5-133722718-DEL-74
chr5-133726493-DEL-6239
chr5-133755496-DEL-506
chr5-133756534-DEL-709
chr5-133757567-DEL-135
chr5-133764221-DEL-165

chr5-133772263-DEL-452
chr5-133797445-DEL-71
chr5-133810469-DEL-78
chr5-133822115-DEL-51
chr5-133857883-DEL-98
chr5-133857912-DEL-1137
chr5-133866671-DEL-338
chr5-133883169-DEL-596
chr5-133913392-DEL-96
chr5-133929128-DEL-106
chr5-133940313-DEL-137
chr5-133957754-DEL-280
chr5-133977745-DEL-107
chr5-133988393-DEL-424
chr5-133991238-DEL-94
chr5-134009715-DEL-115
chr5-134025172-DEL-58
chr5-134027826-DEL-210
chr5-134030349-DEL-106
chr5-134045604-DEL-477
chr5-134050317-DEL-74
chr5-134055716-DEL-385
chr5-134064663-DEL-324
chr5-134069034-DEL-126
chr5-134069817-DEL-165
chr5-134072458-DEL-160
chr5-134085702-DEL-57
chr5-134089249-DEL-62
chr5-134090301-DEL-52
chr5-134092072-DEL-67
chr5-134094095-DEL-199
chr5-134110730-DEL-702
chr5-134118902-DEL-132
chr5-134132827-DEL-264
chr5-134154286-DEL-57
chr5-134166705-DEL-55
chr5-134174440-DEL-58
chr5-134187935-DEL-1886
chr5-134191519-DEL-154
chr5-134193261-DEL-67
chr5-134197959-DEL-58
chr5-134198696-DEL-129
chr5-134217185-DEL-396
chr5-134221085-DEL-89
chr5-134223441-DEL-52
chr5-134224621-DEL-202
chr5-134226400-DEL-53
chr5-134257941-DEL-307
chr5-134321122-DEL-196
chr5-134324923-DEL-179
chr5-134326333-DEL-180
chr5-134329248-DEL-203
chr5-134337750-DEL-198
chr5-134362199-DEL-123
chr5-134363349-DEL-585
chr5-134364931-DEL-167
chr5-134368197-DEL-130
chr5-134370526-DEL-65
chr5-134376391-DEL-94
chr5-134384184-DEL-204
chr5-134388154-DEL-54
chr5-134407397-DEL-256
chr5-134417638-DEL-159
chr5-134424370-DEL-52
chr5-134428999-DEL-171
chr5-134450458-DEL-153
chr5-134478131-DEL-465
chr5-134491358-DEL-60
chr5-134492070-DEL-343
chr5-134501107-DEL-53
chr5-134512125-DEL-65
chr5-134529269-DEL-53
chr5-134537254-DEL-167
chr5-134543433-DEL-121
chr5-134556208-DEL-102
chr5-134564936-DEL-211
chr5-134624575-DEL-230

chr5-134637996-DEL-187
chr5-134639621-DEL-52
chr5-134669098-DEL-71
chr5-134672934-DEL-55
chr5-134677776-DEL-213
chr5-134736364-DEL-60
chr5-134750688-DEL-84
chr5-134752558-DEL-76
chr5-134754871-DEL-52
chr5-134767268-DEL-206
chr5-134767401-DEL-619
chr5-134782492-DEL-51
chr5-134791570-DEL-58
chr5-134802370-DEL-52
chr5-134808716-DEL-90
chr5-134809497-DEL-64
chr5-134812607-DEL-160
chr5-134834550-DEL-78
chr5-134836166-DEL-1078
chr5-134836395-DEL-168
chr5-134843636-DEL-53
chr5-134857322-DEL-1682
chr5-134872942-DEL-265
chr5-134889320-DEL-1066
chr5-134914807-DEL-53
chr5-134958040-DEL-113
chr5-134962516-DEL-127
chr5-134968467-DEL-622
chr5-134979127-DEL-302
chr5-134980998-DEL-86
chr5-134987329-DEL-108
chr5-134988542-DEL-121
chr5-134991139-DEL-209
chr5-135002933-DEL-58
chr5-135005441-DEL-53
chr5-135032175-DEL-409
chr5-135035323-DEL-367
chr5-135055744-DEL-430
chr5-135058927-DEL-53
chr5-135061159-DEL-117
chr5-135072351-DEL-194
chr5-135097632-DEL-170
chr5-135097633-DEL-1057
chr5-135109907-DEL-105
chr5-135118081-DEL-81
chr5-135122500-DEL-166
chr5-135125669-DEL-242
chr5-135148396-DEL-52
chr5-135198766-DEL-55
chr5-135232024-DEL-222
chr5-135286603-DEL-61
chr5-135294423-DEL-54
chr5-135295611-DEL-424
chr5-135296138-DEL-720
chr5-135298963-DEL-167
chr5-135299775-DEL-54
chr5-135306200-DEL-118
chr5-135314653-DEL-279
chr5-135316501-DEL-106
chr5-135318613-DEL-124
chr5-135345397-DEL-65
chr5-135379522-DEL-385
chr5-135388143-DEL-106
chr5-135422376-DEL-121
chr5-135441754-DEL-264
chr5-135464282-DEL-53
chr5-135475336-DEL-110
chr5-135480700-DEL-158
chr5-135497595-DEL-171
chr5-135498262-DEL-211
chr5-135512618-DEL-266
chr5-135525039-DEL-203
chr5-135527367-DEL-54
chr5-135529678-DEL-106
chr5-135587642-DEL-190
chr5-135589882-DEL-211
chr5-135603860-DEL-68

chr5-135606456-DEL-106
chr5-135613119-DEL-54
chr5-135615481-DEL-2699
chr5-135621578-DEL-52
chr5-135622992-DEL-388
chr5-135625437-DEL-94
chr5-135632340-DEL-2076
chr5-135633935-DEL-2070
chr5-135635910-DEL-207
chr5-135648794-DEL-115
chr5-135655205-DEL-99
chr5-135704689-DEL-87
chr5-135708428-DEL-65
chr5-135714373-DEL-317
chr5-135716291-DEL-282
chr5-135719024-DEL-60
chr5-135719660-DEL-212
chr5-135730068-DEL-376
chr5-135750310-DEL-265
chr5-135783319-DEL-147
chr5-135792622-DEL-138
chr5-135826976-DEL-53
chr5-135828155-DEL-62
chr5-135866439-DEL-381
chr5-135869715-DEL-53
chr5-135895139-DEL-59
chr5-135905869-DEL-214
chr5-135909767-DEL-112
chr5-135918561-DEL-217
chr5-135948959-DEL-71
chr5-135956594-DEL-173
chr5-135965739-DEL-75
chr5-135991115-DEL-55
chr5-135992541-DEL-319
chr5-135998619-DEL-178
chr5-136008325-DEL-53
chr5-136016969-DEL-54
chr5-136021396-DEL-194
chr5-136062459-DEL-101
chr5-136065728-DEL-52
chr5-136066733-DEL-55
chr5-136075995-DEL-209
chr5-136081972-DEL-53
chr5-136084762-DEL-158
chr5-136095570-DEL-163
chr5-136108845-DEL-325
chr5-136179981-DEL-52
chr5-136195265-DEL-395
chr5-136232881-DEL-52
chr5-136261286-DEL-77
chr5-136261741-DEL-175
chr5-136271864-DEL-71
chr5-136276582-DEL-52
chr5-136284140-DEL-212
chr5-136287409-DEL-105
chr5-136403069-DEL-58
chr5-136426429-DEL-105
chr5-136455170-DEL-104
chr5-136477301-DEL-61
chr5-136496084-DEL-217
chr5-136530579-DEL-81
chr5-136535487-DEL-52
chr5-136600632-DEL-84
chr5-136616793-DEL-69
chr5-136641441-DEL-85
chr5-136655461-DEL-53
chr5-136657250-DEL-80
chr5-136658633-DEL-55
chr5-136675396-DEL-106
chr5-136678957-DEL-52
chr5-136680623-DEL-60
chr5-136689722-DEL-54
chr5-136707475-DEL-106
chr5-136712932-DEL-66
chr5-136725935-DEL-53
chr5-136731401-DEL-99
chr5-136731990-DEL-159

chr5-136736294-DEL-54
chr5-136748476-DEL-69
chr5-136757075-DEL-278
chr5-136784030-DEL-104
chr5-136787327-DEL-204
chr5-136790088-DEL-84
chr5-136797336-DEL-264
chr5-136822654-DEL-80
chr5-136837038-DEL-70
chr5-136844400-DEL-340
chr5-136846716-DEL-237
chr5-136850070-DEL-423
chr5-136852961-DEL-158
chr5-136855822-DEL-739
chr5-136855930-DEL-132
chr5-136861339-DEL-264
chr5-136865934-DEL-54
chr5-136883241-DEL-149
chr5-136887708-DEL-104
chr5-136889581-DEL-54
chr5-136904039-DEL-213
chr5-136913632-DEL-53
chr5-136922996-DEL-146
chr5-136927207-DEL-51
chr5-136930973-DEL-53
chr5-136935677-DEL-202
chr5-136940345-DEL-65
chr5-136948970-DEL-67
chr5-136952152-DEL-53
chr5-136957298-DEL-270
chr5-136964467-DEL-207
chr5-136966937-DEL-105
chr5-136978372-DEL-315
chr5-137013160-DEL-68
chr5-137035117-DEL-65
chr5-137038084-DEL-78
chr5-137045730-DEL-57
chr5-137046849-DEL-53
chr5-137087291-DEL-60
chr5-137087709-DEL-159
chr5-137088299-DEL-182
chr5-137092608-DEL-108
chr5-137115865-DEL-137
chr5-137117590-DEL-51
chr5-137129604-DEL-332
chr5-137152965-DEL-173
chr5-137178240-DEL-270
chr5-137179302-DEL-108
chr5-137179896-DEL-213
chr5-137181846-DEL-90
chr5-137182489-DEL-144
chr5-137183055-DEL-108
chr5-137184075-DEL-108
chr5-137184375-DEL-1712
chr5-137184544-DEL-81
chr5-137186295-DEL-108
chr5-137186741-DEL-153
chr5-137201742-DEL-269
chr5-137217511-DEL-900
chr5-137229684-DEL-129
chr5-137231034-DEL-56
chr5-137248149-DEL-58
chr5-137256306-DEL-631
chr5-137270158-DEL-1361
chr5-137317600-DEL-212
chr5-137320175-DEL-53
chr5-137334079-DEL-272
chr5-137341850-DEL-138
chr5-137343511-DEL-281
chr5-137366463-DEL-53
chr5-137388829-DEL-58
chr5-137390394-DEL-428
chr5-137395229-DEL-263
chr5-137397655-DEL-196
chr5-137407438-DEL-80
chr5-137450159-DEL-64
chr5-137452966-DEL-319

chr5-137458777-DEL-200
chr5-137462707-DEL-176
chr5-137477773-DEL-905
chr5-137486921-DEL-118
chr5-137490944-DEL-84
chr5-137567324-DEL-54
chr5-137604080-DEL-138
chr5-137615497-DEL-160
chr5-137660897-DEL-190
chr5-137666157-DEL-211
chr5-137674261-DEL-266
chr5-137677378-DEL-199
chr5-137678216-DEL-97
chr5-137686687-DEL-188
chr5-137697580-DEL-214
chr5-137709251-DEL-56
chr5-137710709-DEL-204
chr5-137717991-DEL-2273
chr5-137720760-DEL-193
chr5-137721474-DEL-51
chr5-137753077-DEL-312
chr5-137772178-DEL-359
chr5-137810633-DEL-3478
chr5-137813309-DEL-75
chr5-137821250-DEL-227
chr5-137832653-DEL-67
chr5-137836114-DEL-6459
chr5-137843161-DEL-81
chr5-137856752-DEL-110
chr5-137857945-DEL-3209
chr5-137858680-DEL-198
chr5-137881178-DEL-655
chr5-137883779-DEL-64
chr5-137898130-DEL-149
chr5-137927089-DEL-267
chr5-137958364-DEL-696
chr5-137960440-DEL-85
chr5-137966946-DEL-322
chr5-137967513-DEL-111
chr5-137987051-DEL-254
chr5-138001134-DEL-2285
chr5-138022537-DEL-4494
chr5-138027094-DEL-831
chr5-138027961-DEL-1058
chr5-138038509-DEL-4494
chr5-138043932-DEL-1058
chr5-138072500-DEL-1047
chr5-138083189-DEL-80
chr5-138094503-DEL-137
chr5-138118980-DEL-2091
chr5-138119046-DEL-610
chr5-138119737-DEL-1330
chr5-138139693-DEL-109
chr5-138147382-DEL-1092
chr5-138147410-DEL-546
chr5-138158404-DEL-52
chr5-138162562-DEL-51
chr5-138234305-DEL-309
chr5-138235163-DEL-160
chr5-138244070-DEL-2465
chr5-138250795-DEL-71
chr5-138256528-DEL-53
chr5-138300535-DEL-64
chr5-138316135-DEL-497
chr5-138320632-DEL-460
chr5-138323053-DEL-440
chr5-138334932-DEL-52
chr5-138366318-DEL-6278
chr5-138374751-DEL-591
chr5-138375186-DEL-16941
chr5-138462288-DEL-917
chr5-138480203-DEL-14444
chr5-138544676-DEL-2730
chr5-138553571-DEL-727
chr5-138562214-DEL-69
chr5-138573750-DEL-1407
chr5-138608299-DEL-338

chr5-138613774-DEL-63
chr5-138674064-DEL-162
chr5-138701917-DEL-105
chr5-138728463-DEL-85
chr5-138737478-DEL-72
chr5-138767646-DEL-53
chr5-138812492-DEL-80
chr5-138821928-DEL-113
chr5-138833853-DEL-52
chr5-138862484-DEL-52
chr5-138898987-DEL-157
chr5-138913175-DEL-80
chr5-138919307-DEL-54
chr5-138940436-DEL-1232
chr5-138954345-DEL-177
chr5-139009704-DEL-199
chr5-139023319-DEL-157
chr5-139026844-DEL-54
chr5-139047834-DEL-212
chr5-139054254-DEL-144
chr5-139057992-DEL-53
chr5-139068893-DEL-159
chr5-139090111-DEL-285
chr5-139094793-DEL-108
chr5-139098839-DEL-202
chr5-139109341-DEL-52
chr5-139112909-DEL-159
chr5-139119583-DEL-316
chr5-139121414-DEL-52
chr5-139162465-DEL-100
chr5-139167349-DEL-70
chr5-139174354-DEL-481
chr5-139175054-DEL-108
chr5-139179958-DEL-68
chr5-139181902-DEL-407
chr5-139184701-DEL-735
chr5-139209250-DEL-52
chr5-139225015-DEL-733
chr5-139232979-DEL-53
chr5-139244915-DEL-665
chr5-139261841-DEL-136
chr5-139262376-DEL-87
chr5-139291073-DEL-402
chr5-139292231-DEL-260
chr5-139292894-DEL-62
chr5-139294098-DEL-52
chr5-139296790-DEL-212
chr5-139308045-DEL-52
chr5-139328902-DEL-124
chr5-139338832-DEL-52
chr5-139357666-DEL-69
chr5-139376802-DEL-56
chr5-139392506-DEL-61
chr5-139416152-DEL-386
chr5-139461129-DEL-108
chr5-139470794-DEL-236
chr5-139481304-DEL-140
chr5-139493121-DEL-314
chr5-139499100-DEL-66
chr5-139500005-DEL-65
chr5-139500754-DEL-73
chr5-139508695-DEL-368
chr5-139550552-DEL-53
chr5-139556844-DEL-57
chr5-139565166-DEL-180
chr5-139574249-DEL-53
chr5-139574960-DEL-56
chr5-139588312-DEL-51
chr5-139594375-DEL-60
chr5-139597512-DEL-313
chr5-139609848-DEL-204
chr5-139637222-DEL-81
chr5-139637453-DEL-1376
chr5-139638331-DEL-630
chr5-139641862-DEL-1580
chr5-139645609-DEL-52
chr5-139656253-DEL-104

chr5-139664477-DEL-75
chr5-139675531-DEL-216
chr5-139686913-DEL-118
chr5-139693512-DEL-265
chr5-139705740-DEL-78
chr5-139705938-DEL-1610
chr5-139726254-DEL-296
chr5-139758962-DEL-54
chr5-139763667-DEL-144
chr5-139768011-DEL-57
chr5-139789875-DEL-92
chr5-139813073-DEL-53
chr5-139823081-DEL-79
chr5-139828426-DEL-85
chr5-139862681-DEL-72
chr5-139886742-DEL-254
chr5-139891886-DEL-81
chr5-139922903-DEL-315
chr5-139925211-DEL-57
chr5-139928910-DEL-154
chr5-139934242-DEL-79
chr5-139940575-DEL-54
chr5-139979283-DEL-655
chr5-140052153-DEL-279
chr5-140071434-DEL-108
chr5-140084952-DEL-53
chr5-140107245-DEL-51
chr5-140183227-DEL-205
chr5-140199558-DEL-52
chr5-140204631-DEL-54
chr5-140206907-DEL-74
chr5-140267368-DEL-52
chr5-140285435-DEL-77
chr5-140296987-DEL-385
chr5-140304450-DEL-1344
chr5-140350786-DEL-165
chr5-140354128-DEL-53
chr5-140376085-DEL-75
chr5-140380774-DEL-208
chr5-140393687-DEL-480
chr5-140418996-DEL-281
chr5-140424433-DEL-528
chr5-140440675-DEL-120
chr5-140445562-DEL-169
chr5-140451422-DEL-643
chr5-140460156-DEL-184
chr5-140465357-DEL-101
chr5-140470030-DEL-105
chr5-140480446-DEL-90
chr5-140481595-DEL-553
chr5-140482623-DEL-2325
chr5-140487074-DEL-58
chr5-140498112-DEL-106
chr5-140501270-DEL-52
chr5-140504025-DEL-158
chr5-140515572-DEL-5276
chr5-140536675-DEL-170
chr5-140538584-DEL-60
chr5-140543875-DEL-52
chr5-140554854-DEL-82
chr5-140611585-DEL-71
chr5-140658550-DEL-65
chr5-140673059-DEL-53
chr5-140677619-DEL-126
chr5-140681624-DEL-146
chr5-140690003-DEL-171
chr5-140694890-DEL-203
chr5-140714219-DEL-202
chr5-140725242-DEL-103
chr5-140727561-DEL-51
chr5-140764911-DEL-617
chr5-140772120-DEL-57
chr5-140819817-DEL-76
chr5-140828318-DEL-165
chr5-140855066-DEL-56
chr5-140868692-DEL-99
chr5-140880089-DEL-87

chr5-140880527-DEL-208
chr5-140884475-DEL-104
chr5-140897791-DEL-52
chr5-140899657-DEL-160
chr5-140903979-DEL-93
chr5-140916981-DEL-142
chr5-140931542-DEL-94
chr5-140932410-DEL-74
chr5-140940500-DEL-336
chr5-140953324-DEL-128
chr5-140965436-DEL-103
chr5-140969517-DEL-52
chr5-140975587-DEL-88
chr5-141001391-DEL-100
chr5-141011260-DEL-74
chr5-141020734-DEL-53
chr5-141027098-DEL-71
chr5-141049309-DEL-105
chr5-141055793-DEL-476
chr5-141069202-DEL-60
chr5-141076440-DEL-99
chr5-141080246-DEL-1317
chr5-141094211-DEL-345
chr5-141106201-DEL-320
chr5-141130158-DEL-2441
chr5-141170483-DEL-262
chr5-141175477-DEL-71
chr5-141194572-DEL-98
chr5-141196198-DEL-220
chr5-141213472-DEL-95
chr5-141214536-DEL-105
chr5-141270359-DEL-157
chr5-141328082-DEL-75
chr5-141332409-DEL-60
chr5-141352848-DEL-333
chr5-141381521-DEL-108
chr5-141394029-DEL-174
chr5-141422898-DEL-80
chr5-141498428-DEL-205
chr5-141516658-DEL-926
chr5-141518737-DEL-479
chr5-141519729-DEL-613
chr5-141528079-DEL-53
chr5-141548756-DEL-102
chr5-141554032-DEL-98
chr5-141575355-DEL-263
chr5-141585006-DEL-6193
chr5-141598260-DEL-249
chr5-141622735-DEL-197
chr5-141633613-DEL-419
chr5-141642842-DEL-60
chr5-141700877-DEL-82
chr5-141718179-DEL-202
chr5-141723554-DEL-2476
chr5-141738580-DEL-82
chr5-141749307-DEL-52
chr5-141751932-DEL-140
chr5-141775139-DEL-112
chr5-141829485-DEL-160
chr5-141845096-DEL-114
chr5-141846992-DEL-96
chr5-141962778-DEL-63
chr5-141967892-DEL-125
chr5-141991313-DEL-192
chr5-142065979-DEL-222
chr5-142069215-DEL-106
chr5-142087538-DEL-67
chr5-142110493-DEL-141
chr5-142118512-DEL-775
chr5-142137665-DEL-105
chr5-142146377-DEL-57
chr5-142150521-DEL-110
chr5-142161620-DEL-78
chr5-142169462-DEL-53
chr5-142174284-DEL-53
chr5-142193005-DEL-53
chr5-142219940-DEL-58

chr5-142229329-DEL-371
chr5-142245050-DEL-53
chr5-142249450-DEL-80
chr5-142265990-DEL-103
chr5-142273441-DEL-469
chr5-142287902-DEL-173
chr5-142294648-DEL-59
chr5-142305986-DEL-90
chr5-142314482-DEL-369
chr5-142322912-DEL-84
chr5-142324933-DEL-125
chr5-142336637-DEL-58
chr5-142367878-DEL-69
chr5-142375907-DEL-144
chr5-142396654-DEL-51
chr5-142414151-DEL-102
chr5-142420506-DEL-225
chr5-142428941-DEL-72
chr5-142439024-DEL-221
chr5-142456598-DEL-424
chr5-142469777-DEL-164
chr5-142477024-DEL-436
chr5-142487399-DEL-108
chr5-142488413-DEL-84
chr5-142518995-DEL-220
chr5-142521518-DEL-210
chr5-142546174-DEL-64
chr5-142550987-DEL-161
chr5-142563926-DEL-159
chr5-142568969-DEL-52
chr5-142570554-DEL-69
chr5-142597402-DEL-743
chr5-142600013-DEL-105
chr5-142639358-DEL-882
chr5-142644402-DEL-120
chr5-142661521-DEL-163
chr5-142663350-DEL-152
chr5-142666799-DEL-96
chr5-142667404-DEL-70
chr5-142684832-DEL-94
chr5-142690806-DEL-51
chr5-142706399-DEL-365
chr5-142708761-DEL-116
chr5-142714176-DEL-317
chr5-142727513-DEL-50
chr5-142744760-DEL-112
chr5-142772293-DEL-197
chr5-142794563-DEL-422
chr5-142813488-DEL-83
chr5-142814298-DEL-268
chr5-142815303-DEL-86
chr5-142822482-DEL-82
chr5-142824045-DEL-1419
chr5-142825361-DEL-76
chr5-142825689-DEL-515
chr5-142844696-DEL-209
chr5-142868957-DEL-212
chr5-142883079-DEL-197
chr5-142905891-DEL-194
chr5-142909792-DEL-424
chr5-142910804-DEL-176
chr5-142914467-DEL-68
chr5-142919494-DEL-249
chr5-142924630-DEL-57
chr5-142937522-DEL-201
chr5-142960914-DEL-86
chr5-142970809-DEL-1907
chr5-142972615-DEL-66
chr5-142974604-DEL-190
chr5-142999937-DEL-385
chr5-143020534-DEL-188
chr5-143041243-DEL-95
chr5-143091847-DEL-219
chr5-143092335-DEL-359
chr5-143101704-DEL-258
chr5-143119617-DEL-760
chr5-143122012-DEL-67

chr5-143139001-DEL-689
chr5-143142264-DEL-175
chr5-143171626-DEL-53
chr5-143176763-DEL-53
chr5-143189064-DEL-55
chr5-143211197-DEL-66
chr5-143257102-DEL-56
chr5-143259137-DEL-178
chr5-143263305-DEL-980
chr5-143280708-DEL-52
chr5-143289973-DEL-62
chr5-143290997-DEL-165
chr5-143298796-DEL-63
chr5-143306595-DEL-185
chr5-143333102-DEL-52
chr5-143353006-DEL-72
chr5-143357376-DEL-316
chr5-143368880-DEL-171
chr5-143384533-DEL-136
chr5-143399769-DEL-810
chr5-143422507-DEL-219
chr5-143437533-DEL-1465
chr5-143439386-DEL-206
chr5-143442344-DEL-2311
chr5-143462355-DEL-278
chr5-143462874-DEL-56
chr5-143471943-DEL-70
chr5-143479184-DEL-62
chr5-143507136-DEL-206
chr5-143510538-DEL-93
chr5-143519274-DEL-108
chr5-143520317-DEL-210
chr5-143544835-DEL-150
chr5-143589554-DEL-368
chr5-143610237-DEL-134
chr5-143621087-DEL-162
chr5-143674548-DEL-320
chr5-143751473-DEL-152
chr5-143759077-DEL-8728
chr5-143771465-DEL-486
chr5-143789116-DEL-397
chr5-143802688-DEL-73
chr5-143807817-DEL-392
chr5-143825872-DEL-72
chr5-143830157-DEL-136
chr5-143851841-DEL-57
chr5-143869098-DEL-53
chr5-143876493-DEL-1052
chr5-143877765-DEL-855
chr5-143882178-DEL-55
chr5-143901607-DEL-105
chr5-143962923-DEL-78
chr5-143964813-DEL-262
chr5-143966058-DEL-448
chr5-143967462-DEL-151
chr5-143971141-DEL-160
chr5-143992349-DEL-192
chr5-143999118-DEL-59
chr5-144008628-DEL-112
chr5-144043100-DEL-1290
chr5-144050281-DEL-293
chr5-144091356-DEL-114
chr5-144098490-DEL-349
chr5-144104523-DEL-360
chr5-144105138-DEL-467
chr5-144117352-DEL-107
chr5-144118854-DEL-342
chr5-144145928-DEL-96
chr5-144150753-DEL-71
chr5-144152897-DEL-122
chr5-144172440-DEL-158
chr5-144174307-DEL-137
chr5-144177603-DEL-82
chr5-144182699-DEL-157
chr5-144188993-DEL-753
chr5-144209392-DEL-192
chr5-144214760-DEL-70

chr5-144228771-DEL-104
chr5-144229301-DEL-401
chr5-144237453-DEL-400
chr5-144245608-DEL-95
chr5-144252698-DEL-191
chr5-144276830-DEL-322
chr5-144303326-DEL-215
chr5-144314028-DEL-100
chr5-144323951-DEL-80
chr5-144324953-DEL-52
chr5-144330507-DEL-80
chr5-144337589-DEL-54
chr5-144340009-DEL-52
chr5-144351037-DEL-136
chr5-144355431-DEL-58
chr5-144358968-DEL-110
chr5-144361352-DEL-92
chr5-144367286-DEL-261
chr5-144368766-DEL-97
chr5-144376883-DEL-207
chr5-144378128-DEL-57
chr5-144387083-DEL-158
chr5-144392696-DEL-61
chr5-144400772-DEL-329
chr5-144412252-DEL-189
chr5-144420789-DEL-61
chr5-144431726-DEL-128
chr5-144437476-DEL-151
chr5-144447414-DEL-80
chr5-144453683-DEL-410
chr5-144457250-DEL-105
chr5-144469678-DEL-76
chr5-144503927-DEL-163
chr5-144515794-DEL-257
chr5-144528805-DEL-68
chr5-144538009-DEL-451
chr5-144553264-DEL-132
chr5-144556804-DEL-71
chr5-144570117-DEL-260
chr5-144574722-DEL-525
chr5-144575792-DEL-67
chr5-144583703-DEL-74
chr5-144585675-DEL-56
chr5-144589737-DEL-343
chr5-144590828-DEL-83
chr5-144594286-DEL-105
chr5-144595548-DEL-79
chr5-144636176-DEL-147
chr5-144645237-DEL-89
chr5-144659751-DEL-104
chr5-144673097-DEL-109
chr5-144682756-DEL-210
chr5-144701657-DEL-228
chr5-144703799-DEL-198
chr5-144732657-DEL-126
chr5-144793876-DEL-104
chr5-144798373-DEL-3886
chr5-144804201-DEL-205
chr5-144831559-DEL-499
chr5-144854965-DEL-80
chr5-144860946-DEL-140
chr5-144892135-DEL-158
chr5-144909355-DEL-72
chr5-144950289-DEL-79
chr5-144953798-DEL-324
chr5-144957527-DEL-298
chr5-144993524-DEL-86
chr5-145000272-DEL-385
chr5-145001516-DEL-62
chr5-145003734-DEL-92
chr5-145004480-DEL-811
chr5-145012303-DEL-204
chr5-145034438-DEL-140
chr5-145055782-DEL-70
chr5-145109554-DEL-51
chr5-145133333-DEL-113
chr5-145134786-DEL-470

chr5-145140115-DEL-438
chr5-145147526-DEL-264
chr5-145152966-DEL-180
chr5-145191237-DEL-4708
chr5-145212015-DEL-51
chr5-145230885-DEL-205
chr5-145238622-DEL-5957
chr5-145249549-DEL-204
chr5-145260284-DEL-3210
chr5-145266835-DEL-7127
chr5-145280982-DEL-4166
chr5-145289727-DEL-733
chr5-145290775-DEL-3045
chr5-145336389-DEL-475
chr5-145352402-DEL-97
chr5-145353236-DEL-654
chr5-145368982-DEL-226
chr5-145539475-DEL-120
chr5-145543240-DEL-1082
chr5-145543482-DEL-580
chr5-145578820-DEL-130
chr5-145580043-DEL-2677
chr5-145611517-DEL-2297
chr5-145666650-DEL-4214
chr5-145683162-DEL-580
chr5-145688461-DEL-3382
chr5-145692055-DEL-240
chr5-145704455-DEL-6316
chr5-145716707-DEL-6813
chr5-145763554-DEL-63
chr5-145825085-DEL-1885
chr5-145844752-DEL-219
chr5-145894406-DEL-7382
chr5-145909628-DEL-178
chr5-145933397-DEL-69
chr5-145962688-DEL-740
chr5-145974400-DEL-212
chr5-145984420-DEL-117
chr5-146014171-DEL-54
chr5-146028333-DEL-126
chr5-146029479-DEL-69
chr5-146032224-DEL-117
chr5-146089496-DEL-504
chr5-146096031-DEL-86
chr5-146153271-DEL-60
chr5-146156713-DEL-55
chr5-146200111-DEL-1160
chr5-146202984-DEL-172
chr5-146259740-DEL-120
chr5-146264868-DEL-284
chr5-146272592-DEL-63
chr5-146282817-DEL-216
chr5-146284937-DEL-234
chr5-146285936-DEL-54
chr5-146301974-DEL-142
chr5-146343677-DEL-75
chr5-146343722-DEL-777
chr5-146348380-DEL-498
chr5-146351296-DEL-73
chr5-146374246-DEL-440
chr5-146374756-DEL-152
chr5-146400470-DEL-585
chr5-146401076-DEL-60
chr5-146401404-DEL-399
chr5-146412649-DEL-153
chr5-146414581-DEL-2534
chr5-146469965-DEL-190
chr5-146478836-DEL-6592
chr5-146485724-DEL-611
chr5-146488455-DEL-62
chr5-146499568-DEL-80
chr5-146538051-DEL-148
chr5-146553188-DEL-84
chr5-146605925-DEL-68
chr5-146653583-DEL-327
chr5-146658181-DEL-128
chr5-146662793-DEL-81

chr5-146675258-DEL-80
chr5-146696277-DEL-659
chr5-146697199-DEL-1859
chr5-146735677-DEL-1124
chr5-146736167-DEL-60
chr5-146753820-DEL-63
chr5-146762008-DEL-74
chr5-146773217-DEL-1180
chr5-146774935-DEL-164
chr5-146792745-DEL-69
chr5-146818688-DEL-72
chr5-146826379-DEL-187
chr5-146829103-DEL-87
chr5-146832627-DEL-231
chr5-146859137-DEL-239
chr5-146882139-DEL-699
chr5-146884330-DEL-158
chr5-146892884-DEL-56
chr5-146902091-DEL-57
chr5-146918781-DEL-78
chr5-146957116-DEL-171
chr5-146982212-DEL-245
chr5-146990866-DEL-333
chr5-147024855-DEL-55
chr5-147028723-DEL-613
chr5-147030671-DEL-58
chr5-147049578-DEL-313
chr5-147079130-DEL-1214
chr5-147116532-DEL-158
chr5-147164732-DEL-1462
chr5-147176332-DEL-80
chr5-147178873-DEL-373
chr5-147193145-DEL-401
chr5-147213316-DEL-391
chr5-147215675-DEL-56
chr5-147232801-DEL-184
chr5-147268700-DEL-236
chr5-147270735-DEL-193
chr5-147289855-DEL-163
chr5-147320577-DEL-386
chr5-147331236-DEL-2279
chr5-147336437-DEL-53
chr5-147340640-DEL-488
chr5-147342213-DEL-224
chr5-147344596-DEL-190
chr5-147349842-DEL-307
chr5-147351088-DEL-103
chr5-147365623-DEL-278
chr5-147369847-DEL-76
chr5-147372881-DEL-211
chr5-147373704-DEL-176
chr5-147383975-DEL-898
chr5-147387153-DEL-55
chr5-147426252-DEL-53
chr5-147471199-DEL-747
chr5-147505040-DEL-202
chr5-147536694-DEL-559
chr5-147544992-DEL-449
chr5-147548286-DEL-170
chr5-147554273-DEL-220
chr5-147681209-DEL-387
chr5-147688977-DEL-296
chr5-147705893-DEL-153
chr5-147714092-DEL-190
chr5-147742092-DEL-82
chr5-147744769-DEL-128
chr5-147744886-DEL-2910
chr5-147747883-DEL-193
chr5-147767514-DEL-3090
chr5-147768825-DEL-88
chr5-147774400-DEL-528
chr5-147800064-DEL-167
chr5-147833129-DEL-79
chr5-147835358-DEL-51
chr5-147856117-DEL-62
chr5-147865353-DEL-56
chr5-147874165-DEL-227

chr5-147881654-DEL-237
chr5-147889118-DEL-160
chr5-147896212-DEL-374
chr5-147904034-DEL-250
chr5-147915607-DEL-191
chr5-147953716-DEL-226
chr5-147988800-DEL-513
chr5-148011666-DEL-60
chr5-148018753-DEL-93
chr5-148039887-DEL-141
chr5-148085032-DEL-1201
chr5-148095321-DEL-54
chr5-148105735-DEL-487
chr5-148114476-DEL-268
chr5-148121548-DEL-69
chr5-148157697-DEL-51
chr5-148170292-DEL-84
chr5-148182725-DEL-839
chr5-148189872-DEL-100
chr5-148214492-DEL-200
chr5-148235500-DEL-333
chr5-148244691-DEL-4515
chr5-148245533-DEL-364
chr5-148249381-DEL-65
chr5-148299500-DEL-873
chr5-148306775-DEL-59
chr5-148337586-DEL-79
chr5-148352391-DEL-119
chr5-148367650-DEL-105
chr5-148393190-DEL-2552
chr5-148412600-DEL-80
chr5-148428342-DEL-189
chr5-148549309-DEL-65
chr5-148554534-DEL-127
chr5-148593167-DEL-187
chr5-148596580-DEL-70
chr5-148602112-DEL-68
chr5-148651367-DEL-823
chr5-148672442-DEL-791
chr5-148674213-DEL-187
chr5-148685841-DEL-96
chr5-148708957-DEL-130
chr5-148709605-DEL-66
chr5-148716094-DEL-257
chr5-148737465-DEL-56
chr5-148755653-DEL-1358
chr5-148777578-DEL-67
chr5-148793326-DEL-66
chr5-148813301-DEL-338
chr5-148818079-DEL-164
chr5-148827435-DEL-77
chr5-148841087-DEL-96
chr5-148843925-DEL-278
chr5-148850956-DEL-189
chr5-148857072-DEL-460
chr5-148874102-DEL-2675
chr5-148970640-DEL-326
chr5-148975143-DEL-672
chr5-148996296-DEL-437
chr5-149004228-DEL-94
chr5-149009490-DEL-106
chr5-149019306-DEL-169
chr5-149034337-DEL-187
chr5-149041107-DEL-487
chr5-149086713-DEL-87
chr5-149095070-DEL-861
chr5-149108520-DEL-179
chr5-149117184-DEL-58
chr5-149152344-DEL-55
chr5-149159645-DEL-515
chr5-149189239-DEL-110
chr5-149202586-DEL-58
chr5-149210923-DEL-65
chr5-149225540-DEL-86
chr5-149235922-DEL-115
chr5-149253811-DEL-102
chr5-149254582-DEL-136

chr5-149260543-DEL-65
chr5-149282811-DEL-129
chr5-149297139-DEL-115
chr5-149306674-DEL-244
chr5-149312746-DEL-51
chr5-149357581-DEL-74
chr5-149358502-DEL-114
chr5-149367297-DEL-63
chr5-149400040-DEL-125
chr5-149415549-DEL-117
chr5-149434120-DEL-3208
chr5-149442643-DEL-139
chr5-149474580-DEL-104
chr5-149492650-DEL-1281
chr5-149498499-DEL-90
chr5-149499067-DEL-999
chr5-149501466-DEL-53
chr5-149503320-DEL-77
chr5-149542827-DEL-61
chr5-149566969-DEL-55
chr5-149571719-DEL-109
chr5-149578195-DEL-354
chr5-149607200-DEL-201
chr5-149620471-DEL-432
chr5-149660865-DEL-64
chr5-149663913-DEL-1444
chr5-149669190-DEL-101
chr5-149673282-DEL-55
chr5-149678854-DEL-126
chr5-149681470-DEL-248
chr5-149695331-DEL-99
chr5-149738115-DEL-66
chr5-149799114-DEL-874
chr5-149805841-DEL-87
chr5-149813092-DEL-100
chr5-149818302-DEL-53
chr5-149837305-DEL-70
chr5-149840444-DEL-233
chr5-149867568-DEL-160
chr5-149883656-DEL-58
chr5-149904552-DEL-146
chr5-149934046-DEL-64
chr5-149998458-DEL-51
chr5-150004276-DEL-77
chr5-150038837-DEL-108
chr5-150055400-DEL-58
chr5-150077984-DEL-52
chr5-150085999-DEL-204
chr5-150104270-DEL-60
chr5-150114036-DEL-196
chr5-150127334-DEL-116
chr5-150150923-DEL-57
chr5-150172432-DEL-508
chr5-150263988-DEL-56
chr5-150274642-DEL-104
chr5-150315916-DEL-333
chr5-150330959-DEL-64
chr5-150335234-DEL-888
chr5-150335377-DEL-712
chr5-150335911-DEL-213
chr5-150359449-DEL-58
chr5-150367305-DEL-471
chr5-150367810-DEL-1106
chr5-150367968-DEL-948
chr5-150376054-DEL-79
chr5-150392832-DEL-243
chr5-150395798-DEL-84
chr5-150418051-DEL-68
chr5-150422271-DEL-61
chr5-150431723-DEL-108
chr5-150457659-DEL-211
chr5-150512977-DEL-57
chr5-150521278-DEL-56
chr5-150572104-DEL-199
chr5-150619196-DEL-107
chr5-150655297-DEL-69
chr5-150664143-DEL-723

chr5-150720969-DEL-77
chr5-150756283-DEL-2023
chr5-150761138-DEL-177
chr5-150797628-DEL-357
chr5-150808568-DEL-149
chr5-150826065-DEL-78
chr5-150842695-DEL-491
chr5-150844370-DEL-1148
chr5-150848507-DEL-139
chr5-150917371-DEL-2272
chr5-150920287-DEL-127
chr5-150925583-DEL-51
chr5-150954436-DEL-519
chr5-151020358-DEL-696
chr5-151044178-DEL-132
chr5-151088763-DEL-132
chr5-151089667-DEL-88
chr5-151214091-DEL-60
chr5-151238699-DEL-64
chr5-151264300-DEL-198
chr5-151374327-DEL-7145
chr5-151386994-DEL-6354
chr5-151394053-DEL-5123
chr5-151416011-DEL-502
chr5-151418883-DEL-6257
chr5-151430723-DEL-3442
chr5-151435937-DEL-83
chr5-151448134-DEL-7084
chr5-151468410-DEL-52
chr5-151472961-DEL-123
chr5-151483668-DEL-198
chr5-151486581-DEL-2770
chr5-151487236-DEL-103
chr5-151488405-DEL-641
chr5-151516438-DEL-6167
chr5-151563019-DEL-774
chr5-151606944-DEL-85
chr5-151643210-DEL-468
chr5-151648096-DEL-2194
chr5-151653514-DEL-174
chr6-3057127-DEL-91
chr6-3069718-DEL-92
chr6-3094346-DEL-6431
chr6-3115443-DEL-1138
chr6-3116181-DEL-332
chr6-3122283-DEL-140
chr6-3129639-DEL-121
chr6-3132104-DEL-79
chr6-3133335-DEL-768
chr6-3155347-DEL-92
chr6-3158595-DEL-5497
chr6-3193791-DEL-91
chr6-3206455-DEL-1705
chr6-3216436-DEL-93
chr6-3224363-DEL-142
chr6-3224534-DEL-6216
chr6-3234216-DEL-620
chr6-3259396-DEL-65
chr6-3277456-DEL-1524
chr6-3291632-DEL-373
chr6-3305086-DEL-139
chr6-3312286-DEL-56
chr6-3320529-DEL-119
chr6-3365005-DEL-87
chr6-3388050-DEL-728
chr6-3388906-DEL-1618
chr6-3403659-DEL-486
chr6-3404703-DEL-322
chr6-3411388-DEL-476
chr6-3423923-DEL-669
chr6-3424744-DEL-86
chr6-3437155-DEL-106
chr6-3442911-DEL-66
chr6-3448230-DEL-1492
chr6-3460874-DEL-142
chr6-3526979-DEL-89
chr6-3558158-DEL-91

chr6-3560075-DEL-52
chr6-3579052-DEL-6193
chr6-3603810-DEL-66
chr6-3617200-DEL-1302
chr6-3622103-DEL-1384
chr6-3639632-DEL-578
chr6-3640763-DEL-6576
chr6-3648269-DEL-119
chr6-3650375-DEL-7082
chr6-3663650-DEL-1039
chr6-3666117-DEL-115
chr6-3814310-DEL-452
chr6-3819058-DEL-2955
chr6-3838291-DEL-1796
chr6-3853379-DEL-1241
chr6-3853380-DEL-883
chr6-3854189-DEL-88
chr6-3986005-DEL-51
chr6-4047179-DEL-95
chr6-4067914-DEL-209
chr6-4101123-DEL-105
chr6-4130875-DEL-116
chr6-4148061-DEL-247
chr6-4156242-DEL-50
chr6-4244339-DEL-346
chr6-4296224-DEL-290
chr6-4305365-DEL-69
chr6-4308090-DEL-96
chr6-4321998-DEL-207
chr6-4358504-DEL-71
chr6-4368386-DEL-303
chr6-4378870-DEL-60
chr6-4391345-DEL-52
chr6-4399854-DEL-335
chr6-4401425-DEL-78
chr6-4505038-DEL-64
chr6-4514526-DEL-159
chr6-4638614-DEL-117
chr6-4653707-DEL-65
chr6-4660145-DEL-58
chr6-4716791-DEL-120
chr6-4729032-DEL-61
chr6-4786644-DEL-71
chr6-4795781-DEL-148
chr6-4806684-DEL-130
chr6-4834069-DEL-327
chr6-4923773-DEL-229
chr6-4942757-DEL-74
chr6-4948918-DEL-195
chr6-4967614-DEL-97
chr6-4992719-DEL-127
chr6-4994043-DEL-174
chr6-5013666-DEL-208
chr6-5030795-DEL-397
chr6-5031168-DEL-1080
chr6-5047101-DEL-78
chr6-5047885-DEL-772
chr6-5051991-DEL-199
chr6-5079628-DEL-60
chr6-5086009-DEL-65
chr6-5106453-DEL-104
chr6-5107147-DEL-55
chr6-5190590-DEL-695
chr6-5193292-DEL-74
chr6-5195702-DEL-279
chr6-5208486-DEL-502
chr6-5231574-DEL-400
chr6-5235261-DEL-54
chr6-5245287-DEL-265
chr6-5250860-DEL-122
chr6-5252799-DEL-169
chr6-5295687-DEL-105
chr6-5317789-DEL-7256
chr6-5341086-DEL-184
chr6-5346413-DEL-87
chr6-5348750-DEL-161
chr6-5356835-DEL-329

chr6-5393212-DEL-54
chr6-5412006-DEL-595
chr6-5418119-DEL-65
chr6-5436185-DEL-95
chr6-5437945-DEL-3009
chr6-5461213-DEL-1507
chr6-5502932-DEL-54
chr6-5514426-DEL-161
chr6-5532493-DEL-110
chr6-5577986-DEL-71
chr6-5588532-DEL-76
chr6-5638477-DEL-446
chr6-5662149-DEL-160
chr6-5673367-DEL-60
chr6-5688551-DEL-216
chr6-5705054-DEL-258
chr6-5705317-DEL-608
chr6-5706037-DEL-209
chr6-5708754-DEL-61
chr6-5717800-DEL-58
chr6-5720513-DEL-202
chr6-5753925-DEL-1041
chr6-5795151-DEL-93
chr6-5866730-DEL-381
chr6-5912719-DEL-102
chr6-5915165-DEL-63
chr6-5934538-DEL-56
chr6-5948972-DEL-70
chr6-5954322-DEL-272
chr6-5955028-DEL-100
chr6-6029326-DEL-56
chr6-6126554-DEL-113
chr6-6170828-DEL-331
chr6-6179363-DEL-181
chr6-6213025-DEL-147
chr6-6223374-DEL-92
chr6-6229175-DEL-61
chr6-6233135-DEL-67
chr6-6237410-DEL-170
chr6-6244530-DEL-388
chr6-6274150-DEL-59
chr6-6274973-DEL-144
chr6-6275941-DEL-78
chr6-6282445-DEL-50
chr6-6283743-DEL-1440
chr6-6283748-DEL-176
chr6-6284937-DEL-256
chr6-6355427-DEL-98
chr6-6385471-DEL-533
chr6-6416629-DEL-673
chr6-6419623-DEL-486
chr6-6439357-DEL-58
chr6-6443096-DEL-1088
chr6-6459326-DEL-59
chr6-6463225-DEL-137
chr6-6481631-DEL-529
chr6-6489995-DEL-6072
chr6-6537859-DEL-201
chr6-6542892-DEL-141
chr6-6571050-DEL-969
chr6-6598206-DEL-190
chr6-6605538-DEL-196
chr6-6615030-DEL-2839
chr6-6646242-DEL-372
chr6-6720654-DEL-7453
chr6-6733659-DEL-208
chr6-6781387-DEL-58
chr6-6816644-DEL-88
chr6-6838940-DEL-56
chr6-6882241-DEL-108
chr6-6897309-DEL-91
chr6-6907150-DEL-54
chr6-6962186-DEL-255
chr6-6990881-DEL-1508
chr6-6990881-DEL-3672
chr6-6993094-DEL-1214
chr6-7002146-DEL-65

chr6-7004603-DEL-15667
chr6-7057473-DEL-398
chr6-7067233-DEL-84
chr6-7094150-DEL-1894
chr6-7139572-DEL-53
chr6-7161372-DEL-1492
chr6-7167886-DEL-894
chr6-7206811-DEL-229
chr6-7217713-DEL-175
chr6-7224916-DEL-6154
chr6-7231279-DEL-615
chr6-7259617-DEL-59
chr6-7282882-DEL-5389
chr6-7295688-DEL-97
chr6-7317735-DEL-61
chr6-7333041-DEL-208
chr6-7377422-DEL-64
chr6-7378508-DEL-106
chr6-7391783-DEL-193
chr6-7405854-DEL-176
chr6-7416258-DEL-546
chr6-7430859-DEL-752
chr6-7457306-DEL-64
chr6-7462901-DEL-396
chr6-7465603-DEL-873
chr6-7467166-DEL-127
chr6-7496063-DEL-63
chr6-7498974-DEL-101
chr6-7499243-DEL-6223
chr6-7514733-DEL-1507
chr6-7516458-DEL-3166
chr6-7554231-DEL-80
chr6-7566830-DEL-814
chr6-7567635-DEL-358
chr6-7608397-DEL-72
chr6-7613534-DEL-6988
chr6-7621356-DEL-354
chr6-7629009-DEL-97
chr6-7646453-DEL-121
chr6-7666607-DEL-804
chr6-7667767-DEL-136
chr6-7698314-DEL-418
chr6-7730733-DEL-1246
chr6-7757704-DEL-791
chr6-7763374-DEL-52
chr6-7769298-DEL-527
chr6-7775666-DEL-785
chr6-7780842-DEL-1393
chr6-7795149-DEL-2433
chr6-7798740-DEL-341
chr6-7813834-DEL-57
chr6-7821132-DEL-100
chr6-7827041-DEL-300
chr6-7852004-DEL-102
chr6-7880126-DEL-60
chr6-7883239-DEL-52
chr6-7901607-DEL-696
chr6-7910290-DEL-400
chr6-7916826-DEL-131
chr6-7924604-DEL-208
chr6-7928828-DEL-6262
chr6-7939064-DEL-2151
chr6-7953521-DEL-1499
chr6-7962914-DEL-62
chr6-7992844-DEL-51
chr6-8030560-DEL-52
chr6-8049789-DEL-157
chr6-8081735-DEL-1341
chr6-8088053-DEL-80
chr6-8099574-DEL-673
chr6-8109292-DEL-87
chr6-8112483-DEL-82
chr6-8156130-DEL-383
chr6-8165497-DEL-52
chr6-8217793-DEL-777
chr6-8251073-DEL-153
chr6-8307341-DEL-52

chr6-8381014-DEL-54
chr6-8414455-DEL-88
chr6-8484319-DEL-183
chr6-8487044-DEL-59
chr6-8547753-DEL-3154
chr6-8551000-DEL-2228
chr6-8573429-DEL-84
chr6-8581569-DEL-116
chr6-8606476-DEL-11739
chr6-8675557-DEL-78
chr6-8713107-DEL-209
chr6-8713710-DEL-60
chr6-8721029-DEL-152
chr6-8728725-DEL-97
chr6-8728766-DEL-636
chr6-8755625-DEL-61
chr6-8788670-DEL-914
chr6-8796906-DEL-58
chr6-8811064-DEL-3222
chr6-8819172-DEL-97
chr6-8825937-DEL-97
chr6-8846369-DEL-665
chr6-8874015-DEL-320
chr6-8903206-DEL-659
chr6-8971583-DEL-244
chr6-9015827-DEL-82
chr6-9034474-DEL-72
chr6-9073804-DEL-79
chr6-9085056-DEL-93
chr6-9088136-DEL-700
chr6-9125202-DEL-64
chr6-9126316-DEL-1881
chr6-9130239-DEL-53
chr6-9166871-DEL-77
chr6-9173615-DEL-101
chr6-9178030-DEL-65
chr6-9179826-DEL-53
chr6-9183749-DEL-71
chr6-9213112-DEL-1053
chr6-9219577-DEL-76
chr6-9245757-DEL-97
chr6-9257457-DEL-78
chr6-9272126-DEL-63
chr6-9283973-DEL-51
chr6-9289905-DEL-78
chr6-9317165-DEL-309
chr6-9360834-DEL-54
chr6-9367518-DEL-51
chr6-9392203-DEL-417
chr6-9414478-DEL-283
chr6-9430964-DEL-6380
chr6-9446339-DEL-6604
chr6-9462101-DEL-133
chr6-9486679-DEL-239
chr6-9490758-DEL-1567
chr6-9494066-DEL-219
chr6-9495563-DEL-121
chr6-9546952-DEL-1159
chr6-9554421-DEL-54
chr6-9575569-DEL-70
chr6-9576142-DEL-164
chr6-9601729-DEL-311
chr6-9632858-DEL-2163
chr6-9678716-DEL-848
chr6-9707951-DEL-441
chr6-9709070-DEL-156
chr6-9713315-DEL-3199
chr6-9716660-DEL-172
chr6-9717843-DEL-188
chr6-9720353-DEL-2715
chr6-9726590-DEL-102
chr6-9750405-DEL-1714
chr6-9758808-DEL-56
chr6-9779401-DEL-64
chr6-9804701-DEL-83
chr6-9815347-DEL-101
chr6-9850961-DEL-2562

chr6-9852949-DEL-74
chr6-9853539-DEL-2988
chr6-9859189-DEL-498
chr6-9860801-DEL-8157
chr6-9872304-DEL-94
chr6-9887946-DEL-1424
chr6-9889892-DEL-311
chr6-9892648-DEL-67
chr6-9893274-DEL-4513
chr6-9920734-DEL-905
chr6-9963911-DEL-1586
chr6-9974654-DEL-143
chr6-10001881-DEL-309
chr6-10016362-DEL-201
chr6-10063855-DEL-53
chr6-10073749-DEL-5983
chr6-10083715-DEL-1054
chr6-10100356-DEL-283
chr6-10146204-DEL-54
chr6-10165921-DEL-76
chr6-10190782-DEL-184
chr6-10206286-DEL-928
chr6-10224393-DEL-76
chr6-10225827-DEL-6127
chr6-10240565-DEL-6393
chr6-10249735-DEL-85
chr6-10251789-DEL-211
chr6-10256524-DEL-71
chr6-10268633-DEL-3177
chr6-10271566-DEL-248
chr6-10285248-DEL-527
chr6-10296081-DEL-230
chr6-10307000-DEL-6258
chr6-10313652-DEL-284
chr6-10329013-DEL-2879
chr6-10341050-DEL-2695
chr6-10360525-DEL-809
chr6-10361174-DEL-169
chr6-10367589-DEL-111
chr6-10372867-DEL-98
chr6-10407166-DEL-106
chr6-10432151-DEL-256
chr6-10436415-DEL-103
chr6-10442934-DEL-594
chr6-10470767-DEL-74
chr6-10477993-DEL-56
chr6-10478622-DEL-61
chr6-10482482-DEL-144
chr6-10487074-DEL-2852
chr6-10502733-DEL-366
chr6-10514604-DEL-803
chr6-10531972-DEL-52
chr6-10534785-DEL-844
chr6-10577998-DEL-6447
chr6-10578014-DEL-9035
chr6-10584379-DEL-2605
chr6-10584603-DEL-2604
chr6-10588425-DEL-372
chr6-10596346-DEL-54
chr6-10605725-DEL-225
chr6-10646358-DEL-854
chr6-10649178-DEL-5035
chr6-10657458-DEL-209
chr6-10672284-DEL-6211
chr6-10699344-DEL-6703
chr6-10709242-DEL-226
chr6-10711558-DEL-5193
chr6-10719468-DEL-1094
chr6-10742552-DEL-666
chr6-10744488-DEL-161
chr6-10748112-DEL-505
chr6-10753249-DEL-389
chr6-10755591-DEL-772
chr6-10771714-DEL-177
chr6-10774919-DEL-1471
chr6-10805233-DEL-2015
chr6-10808703-DEL-1030

chr6-10826734-DEL-197
chr6-10860612-DEL-548
chr6-10880520-DEL-848
chr6-10882802-DEL-3287
chr6-10903758-DEL-1658
chr6-10907395-DEL-337
chr6-10929349-DEL-968
chr6-10937781-DEL-160
chr6-10967264-DEL-671
chr6-10971588-DEL-258
chr6-11006616-DEL-104
chr6-11023962-DEL-123
chr6-11029006-DEL-1586
chr6-11037055-DEL-193
chr6-11039112-DEL-65
chr6-11044620-DEL-82
chr6-11046924-DEL-1613
chr6-11084209-DEL-3031
chr6-11087085-DEL-3014
chr6-11181846-DEL-8810
chr6-11189010-DEL-1596
chr6-11190194-DEL-362
chr6-11203098-DEL-967
chr6-11220830-DEL-53
chr6-11236936-DEL-54
chr6-11250868-DEL-6302
chr6-11276169-DEL-51
chr6-11341107-DEL-219
chr6-11358499-DEL-307
chr6-11376938-DEL-109
chr6-11378840-DEL-405
chr6-11380632-DEL-1447
chr6-11417096-DEL-4579
chr6-11428567-DEL-6406
chr6-11451055-DEL-208
chr6-11468722-DEL-647
chr6-11483228-DEL-101
chr6-11517212-DEL-187
chr6-11537708-DEL-402
chr6-11550238-DEL-66
chr6-11554438-DEL-1171
chr6-11570102-DEL-671
chr6-11570672-DEL-1759
chr6-11593510-DEL-204
chr6-11604770-DEL-57
chr6-11625042-DEL-1594
chr6-11646380-DEL-808
chr6-11669757-DEL-60
chr6-11674866-DEL-77
chr6-11680048-DEL-6454
chr6-11689493-DEL-274
chr6-11703174-DEL-6179
chr6-11759112-DEL-954
chr6-11774258-DEL-246
chr6-11797401-DEL-90
chr6-11800197-DEL-569
chr6-11822645-DEL-7521
chr6-11822853-DEL-41863
chr6-11848937-DEL-110
chr6-11854698-DEL-290
chr6-11875149-DEL-110
chr6-11886130-DEL-11323
chr6-11890900-DEL-2783
chr6-11890912-DEL-6241
chr6-11893703-DEL-3446
chr6-11901928-DEL-392
chr6-11938454-DEL-101
chr6-11948325-DEL-196
chr6-11965415-DEL-2084
chr6-11967524-DEL-872
chr6-11968442-DEL-1587
chr6-11970151-DEL-55
chr6-11974138-DEL-2131
chr6-11995418-DEL-187
chr6-12075611-DEL-69
chr6-12077752-DEL-176
chr6-12097594-DEL-339

chr6-12100550-DEL-312
chr6-12112829-DEL-1492
chr6-12135204-DEL-55
chr6-12139641-DEL-170
chr6-12145787-DEL-59
chr6-12157953-DEL-2831
chr6-12250588-DEL-121
chr6-12272182-DEL-688
chr6-12298652-DEL-6828
chr6-12322934-DEL-155
chr6-12348513-DEL-164
chr6-12360866-DEL-8762
chr6-12362528-DEL-2324
chr6-12374625-DEL-1454
chr6-12427744-DEL-70
chr6-12455496-DEL-2150
chr6-12459986-DEL-271
chr6-12460499-DEL-170
chr6-12469289-DEL-58
chr6-12474974-DEL-100
chr6-12486200-DEL-53
chr6-12497533-DEL-213
chr6-12505014-DEL-52
chr6-12512419-DEL-297
chr6-12537164-DEL-4030
chr6-12564978-DEL-60
chr6-12577341-DEL-427
chr6-12584227-DEL-56
chr6-12604685-DEL-108
chr6-12611417-DEL-55
chr6-12615343-DEL-163
chr6-12620841-DEL-5963
chr6-12630657-DEL-206
chr6-12659921-DEL-1249
chr6-12662495-DEL-168
chr6-12704336-DEL-994
chr6-12711712-DEL-55
chr6-12714819-DEL-585
chr6-12715695-DEL-311
chr6-12716696-DEL-166
chr6-12720050-DEL-1060
chr6-12722954-DEL-504
chr6-12738346-DEL-81
chr6-12768024-DEL-439
chr6-12780271-DEL-1508
chr6-12784397-DEL-458
chr6-12788112-DEL-1229
chr6-12799669-DEL-328
chr6-12813552-DEL-230
chr6-12819603-DEL-431
chr6-12914854-DEL-64
chr6-12950606-DEL-52
chr6-12974324-DEL-117
chr6-12978288-DEL-1489
chr6-13002100-DEL-112
chr6-13035063-DEL-2506
chr6-13061102-DEL-117
chr6-13062314-DEL-402
chr6-13064508-DEL-316
chr6-13090481-DEL-983
chr6-13091289-DEL-2270
chr6-13117201-DEL-2576
chr6-13121350-DEL-1683
chr6-13134791-DEL-105
chr6-13152591-DEL-104
chr6-13202184-DEL-1069
chr6-13213274-DEL-76
chr6-13220276-DEL-576
chr6-13221112-DEL-86
chr6-13229532-DEL-1018
chr6-13231384-DEL-155
chr6-13250491-DEL-1262
chr6-13284909-DEL-275
chr6-13320715-DEL-792
chr6-13341614-DEL-2278
chr6-13390341-DEL-6192
chr6-13398843-DEL-344

chr6-13417529-DEL-52
chr6-13426256-DEL-177
chr6-13431003-DEL-128
chr6-13468923-DEL-68
chr6-13504297-DEL-89
chr6-13519909-DEL-70
chr6-13523261-DEL-235
chr6-13543697-DEL-158
chr6-13549472-DEL-4721
chr6-13561761-DEL-62
chr6-13600465-DEL-248
chr6-13622543-DEL-138
chr6-13645000-DEL-181
chr6-13648089-DEL-1244
chr6-13656022-DEL-676
chr6-13660097-DEL-68
chr6-13679312-DEL-733
chr6-13723085-DEL-111
chr6-13780511-DEL-94
chr6-13783413-DEL-113
chr6-13786632-DEL-230
chr6-13793057-DEL-2289
chr6-13813650-DEL-396
chr6-13824332-DEL-468
chr6-13847005-DEL-77
chr6-13853591-DEL-815
chr6-13888054-DEL-828
chr6-13906161-DEL-328
chr6-13912239-DEL-2149
chr6-13929047-DEL-162
chr6-13934194-DEL-59
chr6-13948807-DEL-66
chr6-13949183-DEL-406
chr6-13956347-DEL-2086
chr6-13966785-DEL-68
chr6-14004040-DEL-761
chr6-14010504-DEL-6808
chr6-14048031-DEL-56
chr6-14058020-DEL-76
chr6-14069303-DEL-68
chr6-14072894-DEL-1913
chr6-14079896-DEL-76
chr6-14111269-DEL-564
chr6-14134229-DEL-2776
chr6-14198149-DEL-1593
chr6-14242825-DEL-8524
chr6-14244975-DEL-2153
chr6-14264657-DEL-80
chr6-14275750-DEL-60
chr6-14296229-DEL-79
chr6-14332865-DEL-52
chr6-14371461-DEL-6205
chr6-14380223-DEL-343
chr6-14394581-DEL-408
chr6-14419827-DEL-136
chr6-14440695-DEL-6181
chr6-14449290-DEL-6857
chr6-14566501-DEL-1301
chr6-14603536-DEL-2533
chr6-14612115-DEL-112
chr6-14629652-DEL-66
chr6-14639986-DEL-161
chr6-14640882-DEL-80
chr6-14641164-DEL-945
chr6-14667528-DEL-56
chr6-14675887-DEL-58
chr6-14681939-DEL-535
chr6-14703556-DEL-317
chr6-14704287-DEL-531
chr6-14751014-DEL-72
chr6-14768516-DEL-1204
chr6-14790985-DEL-19057
chr6-14794427-DEL-6493
chr6-14801438-DEL-4085
chr6-14805767-DEL-2137
chr6-14832073-DEL-56
chr6-14868206-DEL-1385

chr6-14944815-DEL-413
chr6-14956183-DEL-1967
chr6-14978175-DEL-132
chr6-14989427-DEL-233
chr6-14993981-DEL-239
chr6-15043212-DEL-58
chr6-15066817-DEL-331
chr6-15148170-DEL-325
chr6-15174615-DEL-652
chr6-15265212-DEL-67
chr6-15329904-DEL-230
chr6-15364590-DEL-56
chr6-15371967-DEL-58
chr6-15374302-DEL-58
chr6-15389382-DEL-1288
chr6-15418681-DEL-1777
chr6-15446780-DEL-64
chr6-15452175-DEL-6172
chr6-15472296-DEL-143
chr6-15511807-DEL-816
chr6-15527093-DEL-244
chr6-15531388-DEL-154
chr6-15556479-DEL-132
chr6-15568240-DEL-352
chr6-15594560-DEL-115
chr6-15616946-DEL-59
chr6-15667675-DEL-371
chr6-15669321-DEL-304
chr6-15714494-DEL-70
chr6-15755960-DEL-294
chr6-15756690-DEL-59
chr6-15775612-DEL-541
chr6-15781756-DEL-1182
chr6-15783299-DEL-54
chr6-15834594-DEL-509
chr6-15844105-DEL-154
chr6-15873858-DEL-73
chr6-15875697-DEL-312
chr6-15901596-DEL-446
chr6-15915933-DEL-152
chr6-15925906-DEL-3704
chr6-15936064-DEL-56
chr6-15983429-DEL-210
chr6-15994018-DEL-67
chr6-15997554-DEL-212
chr6-16007138-DEL-1481
chr6-16027254-DEL-2104
chr6-16029785-DEL-710
chr6-16032454-DEL-566
chr6-16081356-DEL-578
chr6-16102503-DEL-436
chr6-16112255-DEL-78
chr6-16115506-DEL-2546
chr6-16179733-DEL-315
chr6-16186082-DEL-426
chr6-16204517-DEL-52
chr6-16208452-DEL-53
chr6-16210072-DEL-363
chr6-16219203-DEL-181
chr6-16226018-DEL-181
chr6-16229654-DEL-127
chr6-16241575-DEL-6479
chr6-16243230-DEL-70
chr6-16259027-DEL-721
chr6-16261361-DEL-67
chr6-16339673-DEL-275
chr6-16367322-DEL-815
chr6-16368562-DEL-58
chr6-16377565-DEL-359
chr6-16382661-DEL-187
chr6-16391329-DEL-212
chr6-16399214-DEL-1221
chr6-16475276-DEL-590
chr6-16481647-DEL-96
chr6-16488119-DEL-1686
chr6-16512465-DEL-122
chr6-16525581-DEL-113

chr6-16563925-DEL-53
chr6-16616926-DEL-558
chr6-16627995-DEL-1667
chr6-16661715-DEL-434
chr6-16709417-DEL-2358
chr6-16784699-DEL-304
chr6-16798255-DEL-87
chr6-16799189-DEL-338
chr6-16821776-DEL-135
chr6-16837282-DEL-108
chr6-16875140-DEL-71
chr6-16879186-DEL-924
chr6-16922550-DEL-2224
chr6-16948170-DEL-158
chr6-16949736-DEL-312
chr6-16988016-DEL-288
chr6-17004029-DEL-7462
chr6-17004413-DEL-407
chr6-17029249-DEL-195
chr6-17032715-DEL-84
chr6-17064107-DEL-55
chr6-17069406-DEL-490
chr6-17094008-DEL-105
chr6-17108532-DEL-52
chr6-17123119-DEL-51
chr6-17168049-DEL-127
chr6-17188841-DEL-51
chr6-17225887-DEL-147
chr6-17277044-DEL-201
chr6-17278905-DEL-1053
chr6-17326249-DEL-91
chr6-17335416-DEL-796
chr6-17344951-DEL-58
chr6-17351223-DEL-78
chr6-17355160-DEL-164
chr6-17356238-DEL-122
chr6-17364326-DEL-60
chr6-17391695-DEL-1890
chr6-17408666-DEL-242
chr6-17454707-DEL-53
chr6-17492940-DEL-283
chr6-17570298-DEL-56
chr6-17602229-DEL-83
chr6-17609495-DEL-665
chr6-17658635-DEL-220
chr6-17671404-DEL-363
chr6-17706113-DEL-1522
chr6-17711431-DEL-117
chr6-17719322-DEL-427
chr6-17769306-DEL-445
chr6-17898927-DEL-75
chr6-17915171-DEL-1135
chr6-17935856-DEL-1080
chr6-17935904-DEL-594
chr6-17936855-DEL-297
chr6-17940721-DEL-155
chr6-17974202-DEL-188
chr6-17986537-DEL-350
chr6-17999953-DEL-250
chr6-18063113-DEL-7717
chr6-18094109-DEL-103
chr6-18132950-DEL-95
chr6-18144123-DEL-201
chr6-18152084-DEL-57
chr6-18156144-DEL-1097
chr6-18194959-DEL-66
chr6-18261879-DEL-3232
chr6-18264310-DEL-2909
chr6-18295627-DEL-104
chr6-18538030-DEL-60
chr6-18569597-DEL-200
chr6-18621360-DEL-75
chr6-18638470-DEL-688
chr6-18670299-DEL-1184
chr6-18675531-DEL-246
chr6-18675983-DEL-1313
chr6-18694172-DEL-226

chr6-18727285-DEL-5041
chr6-18736482-DEL-6241
chr6-18742915-DEL-174
chr6-18796963-DEL-238
chr6-18802323-DEL-126
chr6-18824502-DEL-1076
chr6-18842842-DEL-5293
chr6-18867668-DEL-59
chr6-18874925-DEL-2404
chr6-18880010-DEL-549
chr6-18936477-DEL-208
chr6-18944327-DEL-62
chr6-18960386-DEL-1776
chr6-18997520-DEL-470
chr6-18998029-DEL-105
chr6-19051880-DEL-53
chr6-19086074-DEL-1877
chr6-19092040-DEL-184
chr6-19096425-DEL-940
chr6-19099661-DEL-119
chr6-19108747-DEL-1041
chr6-19120687-DEL-150
chr6-19145466-DEL-7343
chr6-19146140-DEL-212
chr6-19159578-DEL-86
chr6-19174745-DEL-524
chr6-19197383-DEL-443
chr6-19204323-DEL-174
chr6-19217788-DEL-69
chr6-19221743-DEL-66
chr6-19238721-DEL-4016
chr6-19275528-DEL-953
chr6-19295628-DEL-80
chr6-19298297-DEL-793
chr6-19309700-DEL-5333
chr6-19319992-DEL-7411
chr6-19333230-DEL-470
chr6-19341153-DEL-1663
chr6-19388637-DEL-1437
chr6-19396649-DEL-56
chr6-19397733-DEL-342
chr6-19449002-DEL-6153
chr6-19459836-DEL-10500
chr6-19467238-DEL-3283
chr6-19474242-DEL-51
chr6-19478129-DEL-950
chr6-19480214-DEL-2385
chr6-19498578-DEL-2450
chr6-19500566-DEL-159
chr6-19505531-DEL-3298
chr6-19516261-DEL-91
chr6-19588888-DEL-58
chr6-19607802-DEL-3169
chr6-19665513-DEL-719
chr6-19671063-DEL-204
chr6-19687099-DEL-477
chr6-19688717-DEL-87
chr6-19712842-DEL-58
chr6-19734461-DEL-1313
chr6-19770089-DEL-2234
chr6-19785244-DEL-58
chr6-19801930-DEL-160
chr6-19803489-DEL-140
chr6-19827290-DEL-190
chr6-19838104-DEL-5263
chr6-19859321-DEL-3658
chr6-19865326-DEL-52
chr6-19885072-DEL-482
chr6-19893695-DEL-843
chr6-19899746-DEL-138
chr6-19907546-DEL-64
chr6-19926594-DEL-815
chr6-19959886-DEL-388
chr6-19994152-DEL-6142
chr6-20000878-DEL-104
chr6-20015206-DEL-479
chr6-20044634-DEL-206

chr6-20045990-DEL-466
chr6-20047089-DEL-1623
chr6-20049653-DEL-151
chr6-20056543-DEL-2783
chr6-20066301-DEL-357
chr6-20091520-DEL-285
chr6-20097883-DEL-2249
chr6-20103065-DEL-76
chr6-20113218-DEL-7167
chr6-20119017-DEL-865
chr6-20127882-DEL-6455
chr6-20149781-DEL-5925
chr6-20164280-DEL-6186
chr6-20200339-DEL-940
chr6-20210138-DEL-411
chr6-20261774-DEL-89
chr6-20276682-DEL-111
chr6-20278642-DEL-2766
chr6-20285226-DEL-1501
chr6-20294780-DEL-3969
chr6-20295604-DEL-199
chr6-20311817-DEL-98
chr6-20320185-DEL-225
chr6-20321448-DEL-2509
chr6-20337303-DEL-714
chr6-20357977-DEL-1377
chr6-20372853-DEL-134
chr6-20381334-DEL-69
chr6-20387314-DEL-116
chr6-20395482-DEL-60
chr6-20417851-DEL-524
chr6-20430767-DEL-110
chr6-20450912-DEL-83
chr6-20528156-DEL-8121
chr6-20528161-DEL-199
chr6-20536840-DEL-575
chr6-20577291-DEL-1126
chr6-20613950-DEL-884
chr6-20629056-DEL-184
chr6-20657923-DEL-63
chr6-20681867-DEL-70
chr6-20705003-DEL-62
chr6-20710718-DEL-7693
chr6-20750825-DEL-1971
chr6-20904171-DEL-78
chr6-20922635-DEL-2288
chr6-20935053-DEL-4691
chr6-20954478-DEL-121
chr6-20973138-DEL-70
chr6-21000024-DEL-1672
chr6-21010612-DEL-13802
chr6-21014212-DEL-6470
chr6-21040711-DEL-941
chr6-21047301-DEL-115
chr6-21056397-DEL-124
chr6-21061797-DEL-74
chr6-21064903-DEL-692
chr6-21084249-DEL-96
chr6-21090209-DEL-53
chr6-21114346-DEL-2110
chr6-21140186-DEL-102
chr6-21149520-DEL-941
chr6-21185591-DEL-235
chr6-21198163-DEL-268
chr6-21198543-DEL-2561
chr6-21206161-DEL-61
chr6-21209006-DEL-252
chr6-21222429-DEL-122
chr6-21252349-DEL-408
chr6-21255473-DEL-225
chr6-21263042-DEL-53
chr6-21285405-DEL-583
chr6-21296629-DEL-56
chr6-21318793-DEL-85
chr6-21330413-DEL-66
chr6-21331351-DEL-90
chr6-21349263-DEL-76

chr6-21359693-DEL-78
chr6-21422758-DEL-63
chr6-21438847-DEL-317
chr6-21439801-DEL-940
chr6-21441623-DEL-326
chr6-21448268-DEL-77
chr6-21485737-DEL-215
chr6-21492822-DEL-53
chr6-21495865-DEL-808
chr6-21520063-DEL-264
chr6-21539718-DEL-143
chr6-21540718-DEL-275
chr6-21553053-DEL-154
chr6-21583022-DEL-867
chr6-21604939-DEL-579
chr6-21640452-DEL-1862
chr6-21671477-DEL-174
chr6-21685591-DEL-111
chr6-21705050-DEL-57
chr6-21706710-DEL-104
chr6-21713560-DEL-293
chr6-21739183-DEL-126
chr6-21740895-DEL-102
chr6-21766535-DEL-236
chr6-21767213-DEL-99
chr6-21824534-DEL-317
chr6-21860011-DEL-345
chr6-21881409-DEL-1139
chr6-21882155-DEL-103
chr6-21888138-DEL-51
chr6-21904306-DEL-1525
chr6-21942702-DEL-61
chr6-21944605-DEL-523
chr6-21958832-DEL-66
chr6-21976316-DEL-6520
chr6-21993276-DEL-1408
chr6-22007156-DEL-560
chr6-22008305-DEL-51
chr6-22027430-DEL-1471
chr6-22052961-DEL-71
chr6-22065041-DEL-64
chr6-22075368-DEL-973
chr6-22086767-DEL-400
chr6-22101290-DEL-68
chr6-22101581-DEL-1020
chr6-22110535-DEL-523
chr6-22119664-DEL-354
chr6-22122034-DEL-62
chr6-22125095-DEL-6726
chr6-22132428-DEL-182
chr6-22144871-DEL-52
chr6-22154838-DEL-778
chr6-22173699-DEL-75
chr6-22190348-DEL-57
chr6-22199724-DEL-107
chr6-22269659-DEL-100
chr6-22275614-DEL-135
chr6-22301953-DEL-275
chr6-22305309-DEL-199
chr6-22308747-DEL-98
chr6-22312150-DEL-52
chr6-22318357-DEL-60
chr6-22386059-DEL-62
chr6-22388195-DEL-192
chr6-22390814-DEL-55
chr6-22400483-DEL-825
chr6-22421324-DEL-173
chr6-22513834-DEL-170
chr6-22515698-DEL-126
chr6-22543673-DEL-736
chr6-22551544-DEL-561
chr6-22617963-DEL-88
chr6-22643438-DEL-162
chr6-22653747-DEL-285
chr6-22657084-DEL-3476
chr6-22669514-DEL-115
chr6-22673216-DEL-73

chr6-22696006-DEL-77
chr6-22708350-DEL-75
chr6-22709645-DEL-3948
chr6-22732189-DEL-3634
chr6-22744415-DEL-278
chr6-22753885-DEL-355
chr6-22755649-DEL-52
chr6-22779802-DEL-6051
chr6-22780575-DEL-488
chr6-22784270-DEL-2074
chr6-22805533-DEL-1288
chr6-22840236-DEL-132
chr6-22861407-DEL-887
chr6-22888458-DEL-477
chr6-22913800-DEL-72
chr6-22924325-DEL-54
chr6-22935788-DEL-100
chr6-22939613-DEL-72
chr6-22975778-DEL-7002
chr6-23026252-DEL-58
chr6-23079478-DEL-167
chr6-23095669-DEL-208
chr6-23097696-DEL-160
chr6-23121803-DEL-204
chr6-23136902-DEL-358
chr6-23139359-DEL-1022
chr6-23178823-DEL-77
chr6-23288868-DEL-103
chr6-23326765-DEL-473
chr6-23333639-DEL-82
chr6-23341667-DEL-110
chr6-23348045-DEL-74
chr6-23378538-DEL-63
chr6-23388846-DEL-420
chr6-23432633-DEL-735
chr6-23453020-DEL-117
chr6-23459288-DEL-1332
chr6-23520815-DEL-55
chr6-23525143-DEL-505
chr6-23539085-DEL-320
chr6-23554681-DEL-10319
chr6-23570909-DEL-219
chr6-23604941-DEL-185
chr6-23604982-DEL-6403
chr6-23635434-DEL-63
chr6-23661016-DEL-1569
chr6-23662381-DEL-100
chr6-23676275-DEL-106
chr6-23686488-DEL-144
chr6-23687742-DEL-145
chr6-23691655-DEL-935
chr6-23711629-DEL-439
chr6-23722097-DEL-612
chr6-23755392-DEL-1247
chr6-23840706-DEL-60
chr6-23894492-DEL-54
chr6-23894600-DEL-4393
chr6-23949691-DEL-4143
chr6-23957185-DEL-3122
chr6-23978124-DEL-517
chr6-23983114-DEL-53
chr6-24015265-DEL-70
chr6-24051422-DEL-284
chr6-24060516-DEL-164
chr6-24119480-DEL-277
chr6-24140958-DEL-234
chr6-24161289-DEL-197
chr6-24180707-DEL-110
chr6-24227922-DEL-1281
chr6-24299171-DEL-58
chr6-24317232-DEL-794
chr6-24318025-DEL-122
chr6-24323030-DEL-120
chr6-24324149-DEL-80
chr6-24443682-DEL-422
chr6-24455384-DEL-410
chr6-24491931-DEL-424

chr6-24506915-DEL-2744
chr6-24511126-DEL-90
chr6-24553342-DEL-51
chr6-24568315-DEL-673
chr6-24641188-DEL-138
chr6-24702307-DEL-155
chr6-24717390-DEL-432
chr6-24732475-DEL-71
chr6-24737048-DEL-2084
chr6-24752637-DEL-189
chr6-24816222-DEL-60
chr6-24818223-DEL-368
chr6-24824082-DEL-570
chr6-24834439-DEL-90
chr6-24884857-DEL-903
chr6-24904924-DEL-56
chr6-24907192-DEL-103
chr6-24920410-DEL-205
chr6-24956846-DEL-107
chr6-24965009-DEL-72
chr6-25004667-DEL-276
chr6-25052027-DEL-89
chr6-25054672-DEL-342
chr6-25079153-DEL-124
chr6-25097619-DEL-671
chr6-25101330-DEL-71
chr6-25124441-DEL-1481
chr6-25167887-DEL-52
chr6-25207527-DEL-193
chr6-25208190-DEL-3547
chr6-25214263-DEL-181
chr6-25219096-DEL-267
chr6-25220921-DEL-6199
chr6-25233425-DEL-2128
chr6-25263287-DEL-189
chr6-25263993-DEL-351
chr6-25269361-DEL-52
chr6-25308344-DEL-144
chr6-25309472-DEL-3628
chr6-25331897-DEL-790
chr6-25337116-DEL-560
chr6-25347720-DEL-1008
chr6-25386109-DEL-86
chr6-25401732-DEL-481
chr6-25416713-DEL-180
chr6-25436793-DEL-165
chr6-25443113-DEL-87
chr6-25446210-DEL-813
chr6-25447966-DEL-185
chr6-25453771-DEL-89
chr6-25470133-DEL-1271
chr6-25475541-DEL-401
chr6-25492173-DEL-4306
chr6-25507777-DEL-13720
chr6-25520478-DEL-1020
chr6-25557274-DEL-212
chr6-25560561-DEL-5995
chr6-25652799-DEL-63
chr6-25661731-DEL-262
chr6-25663309-DEL-479
chr6-25711347-DEL-299
chr6-25723290-DEL-726
chr6-25737207-DEL-138
chr6-25758352-DEL-182
chr6-25764914-DEL-829
chr6-25767865-DEL-65
chr6-25779402-DEL-51
chr6-25808921-DEL-84
chr6-25837086-DEL-70
chr6-25891147-DEL-67
chr6-25892217-DEL-78
chr6-25918900-DEL-136
chr6-25934966-DEL-1667
chr6-25949258-DEL-70
chr6-25957038-DEL-724
chr6-26003718-DEL-355
chr6-26012217-DEL-3841

chr6-26016815-DEL-433
chr6-26030719-DEL-726
chr6-26039181-DEL-92
chr6-26040672-DEL-2185
chr6-26062609-DEL-135
chr6-26093420-DEL-105
chr6-26120075-DEL-1920
chr6-26147805-DEL-623
chr6-26149989-DEL-170
chr6-26153964-DEL-122
chr6-26175641-DEL-143
chr6-26247577-DEL-6222
chr6-26254519-DEL-1148
chr6-26304742-DEL-1656
chr6-26318850-DEL-440
chr6-26334582-DEL-51
chr6-26362196-DEL-739
chr6-26393149-DEL-1334
chr6-26415799-DEL-1040
chr6-26416594-DEL-468
chr6-26425912-DEL-268
chr6-26451275-DEL-681
chr6-26505241-DEL-6197
chr6-26537624-DEL-2051
chr6-26549081-DEL-386
chr6-26564191-DEL-255
chr6-26588309-DEL-76
chr6-26602097-DEL-1248
chr6-26604907-DEL-6412
chr6-26638461-DEL-72
chr6-26640557-DEL-186
chr6-26643437-DEL-166
chr6-26652169-DEL-6091
chr6-26668225-DEL-150
chr6-26680570-DEL-7188
chr6-26689900-DEL-58
chr6-26715726-DEL-170
chr6-26734680-DEL-141
chr6-26751832-DEL-2667
chr6-26779492-DEL-65
chr6-26779789-DEL-952
chr6-26785465-DEL-66
chr6-26786273-DEL-554
chr6-26834554-DEL-201
chr6-26849991-DEL-657
chr6-26859541-DEL-388
chr6-26861576-DEL-2026
chr6-26865641-DEL-1367
chr6-26867090-DEL-76
chr6-26868995-DEL-1359
chr6-26874924-DEL-6521
chr6-26932958-DEL-180
chr6-26939305-DEL-51
chr6-26960511-DEL-208
chr6-26968325-DEL-192
chr6-26971636-DEL-57
chr6-27006741-DEL-66
chr6-27015843-DEL-2112
chr6-27076010-DEL-52
chr6-27078316-DEL-6176
chr6-27091908-DEL-4455
chr6-27108024-DEL-118
chr6-27108118-DEL-6821
chr6-27108877-DEL-2937
chr6-27112634-DEL-2001
chr6-27117175-DEL-7076
chr6-27125418-DEL-4471
chr6-27140058-DEL-2592
chr6-27179420-DEL-61
chr6-27183749-DEL-90
chr6-27215410-DEL-486
chr6-27234385-DEL-892
chr6-27309133-DEL-70
chr6-27334369-DEL-535
chr6-27335427-DEL-371
chr6-27339635-DEL-84
chr6-27370095-DEL-72

chr6-27400730-DEL-59
chr6-27487585-DEL-93
chr6-27492484-DEL-643
chr6-27501686-DEL-214
chr6-27515011-DEL-187
chr6-27515977-DEL-56
chr6-27519924-DEL-6070
chr6-27550940-DEL-873
chr6-27602754-DEL-113
chr6-27667683-DEL-137
chr6-27688088-DEL-127
chr6-27738418-DEL-83
chr6-27767639-DEL-176
chr6-27786893-DEL-1319
chr6-27797384-DEL-293
chr6-27831501-DEL-124
chr6-27840608-DEL-852
chr6-27862013-DEL-648
chr6-27881225-DEL-62
chr6-27921185-DEL-63
chr6-27922464-DEL-407
chr6-28008594-DEL-172
chr6-28010962-DEL-207
chr6-28012400-DEL-78
chr6-28016207-DEL-314
chr6-28052827-DEL-52
chr6-28083484-DEL-164
chr6-28085704-DEL-2348
chr6-28146056-DEL-76
chr6-28161204-DEL-1461
chr6-28170034-DEL-455
chr6-28177055-DEL-56
chr6-28187592-DEL-126
chr6-28237268-DEL-56
chr6-28293675-DEL-101
chr6-28298413-DEL-128
chr6-28323111-DEL-820
chr6-28325939-DEL-190
chr6-28331330-DEL-51
chr6-28339682-DEL-801
chr6-28384315-DEL-237
chr6-28391811-DEL-269
chr6-28403117-DEL-2398
chr6-28414465-DEL-56
chr6-28476593-DEL-83
chr6-28488754-DEL-233
chr6-28532706-DEL-628
chr6-28544480-DEL-400
chr6-28570602-DEL-180
chr6-28575401-DEL-331
chr6-28595465-DEL-55
chr6-28685042-DEL-90
chr6-28742194-DEL-54
chr6-28754678-DEL-330
chr6-28807643-DEL-116
chr6-28845833-DEL-66
chr6-28911761-DEL-1741
chr6-28955186-DEL-177
chr6-28965058-DEL-191
chr6-29012440-DEL-604
chr6-29030868-DEL-126
chr6-29040463-DEL-138
chr6-29077644-DEL-117
chr6-29099317-DEL-83
chr6-29218196-DEL-52
chr6-29233474-DEL-218
chr6-29249332-DEL-155
chr6-29250773-DEL-70
chr6-29261755-DEL-131
chr6-29281902-DEL-559
chr6-29285036-DEL-108
chr6-29297775-DEL-1163
chr6-29300019-DEL-53
chr6-29330132-DEL-229
chr6-29349725-DEL-199
chr6-29382392-DEL-106
chr6-29383180-DEL-75

chr6-29386190-DEL-261
chr6-29390670-DEL-195
chr6-29449572-DEL-82
chr6-29519578-DEL-116
chr6-29542175-DEL-459
chr6-29606246-DEL-705
chr6-29610230-DEL-63
chr6-29623214-DEL-1620
chr6-29636075-DEL-58
chr6-29642854-DEL-103
chr6-29681198-DEL-141
chr6-29682831-DEL-91
chr6-29690074-DEL-2466
chr6-29690463-DEL-302
chr6-29716452-DEL-95
chr6-29726079-DEL-62
chr6-29747648-DEL-1542
chr6-29748857-DEL-333
chr6-29782418-DEL-56
chr6-29806237-DEL-688
chr6-29816201-DEL-209
chr6-29816576-DEL-375
chr6-29840873-DEL-203
chr6-29858565-DEL-183
chr6-29867612-DEL-193
chr6-29906323-DEL-64
chr6-29941539-DEL-176
chr6-29969048-DEL-622
chr6-29974390-DEL-385
chr6-29986718-DEL-1742
chr6-29991224-DEL-158
chr6-30011963-DEL-210
chr6-30024581-DEL-267
chr6-30027407-DEL-136
chr6-30031336-DEL-217
chr6-30038489-DEL-207
chr6-30061549-DEL-80
chr6-30111710-DEL-191
chr6-30124027-DEL-197
chr6-30153597-DEL-339
chr6-30184091-DEL-88
chr6-30186837-DEL-266
chr6-30194880-DEL-183
chr6-30206791-DEL-198
chr6-30246115-DEL-51
chr6-30294165-DEL-72
chr6-30359992-DEL-55
chr6-30381494-DEL-100
chr6-30396304-DEL-1027
chr6-30445556-DEL-84
chr6-30456576-DEL-442
chr6-30458635-DEL-272
chr6-30467750-DEL-302
chr6-30471718-DEL-77
chr6-30473848-DEL-200
chr6-30478450-DEL-938
chr6-30535815-DEL-187
chr6-30536671-DEL-178
chr6-30556077-DEL-98
chr6-30588487-DEL-76
chr6-30607688-DEL-84
chr6-30647563-DEL-64
chr6-30760317-DEL-52
chr6-30762303-DEL-166
chr6-30771245-DEL-261
chr6-30778126-DEL-159
chr6-30779929-DEL-1587
chr6-30858739-DEL-81
chr6-30903393-DEL-125
chr6-30917946-DEL-319
chr6-30921832-DEL-254
chr6-30953409-DEL-131
chr6-30969718-DEL-98
chr6-30973512-DEL-4059
chr6-31003718-DEL-1869
chr6-31022248-DEL-504
chr6-31028351-DEL-66

chr6-31044941-DEL-1721
chr6-31113208-DEL-807
chr6-31120832-DEL-382
chr6-31136595-DEL-63
chr6-31148064-DEL-285
chr6-31155909-DEL-113
chr6-31167010-DEL-53
chr6-31177863-DEL-470
chr6-31326781-DEL-166
chr6-31337592-DEL-123
chr6-31364254-DEL-1166
chr6-31365781-DEL-693
chr6-31405392-DEL-1340
chr6-31433296-DEL-53
chr6-31468298-DEL-373
chr6-31493641-DEL-60
chr6-31541086-DEL-59
chr6-31572717-DEL-539
chr6-31575105-DEL-72
chr6-31610757-DEL-90
chr6-31623049-DEL-60
chr6-31628813-DEL-191
chr6-31638807-DEL-129
chr6-31643272-DEL-120
chr6-31650683-DEL-53
chr6-31653511-DEL-148
chr6-31678699-DEL-261
chr6-31679528-DEL-63
chr6-31680848-DEL-82
chr6-31706203-DEL-75
chr6-31716085-DEL-757
chr6-31736805-DEL-1960
chr6-31741274-DEL-1235
chr6-31768363-DEL-185
chr6-31776305-DEL-5139
chr6-31776305-DEL-87
chr6-31781354-DEL-202
chr6-31793304-DEL-93
chr6-31797934-DEL-104
chr6-31808905-DEL-1316
chr6-31812850-DEL-112
chr6-31843476-DEL-186
chr6-31852359-DEL-225
chr6-31859233-DEL-1308
chr6-31862553-DEL-72
chr6-31863002-DEL-168
chr6-31915114-DEL-57
chr6-31934518-DEL-1454
chr6-32011993-DEL-1891
chr6-32070841-DEL-636
chr6-32105883-DEL-1358
chr6-32157786-DEL-408
chr6-32173598-DEL-106
chr6-32202385-DEL-112
chr6-32241984-DEL-138
chr6-32302889-DEL-250
chr6-32371730-DEL-355
chr6-32372631-DEL-486
chr6-32560196-DEL-82
chr6-32574868-DEL-329
chr6-32693258-DEL-1096
chr6-32696355-DEL-1138
chr6-32711200-DEL-145
chr6-32722183-DEL-105
chr6-32734390-DEL-54
chr6-32739715-DEL-80
chr6-32754264-DEL-120
chr6-32771990-DEL-55
chr6-32776104-DEL-72
chr6-32842732-DEL-195
chr6-32854484-DEL-59
chr6-32909734-DEL-114
chr6-32942894-DEL-55
chr6-33012892-DEL-57
chr6-33023586-DEL-76
chr6-33031999-DEL-763
chr6-33038961-DEL-164

chr6-33047183-DEL-602
chr6-33049807-DEL-61
chr6-33052714-DEL-457
chr6-33061742-DEL-156
chr6-33082336-DEL-62
chr6-33086998-DEL-100
chr6-33096352-DEL-217
chr6-33111470-DEL-1080
chr6-33118008-DEL-172
chr6-33125263-DEL-872
chr6-33139895-DEL-75
chr6-33150227-DEL-56
chr6-33177389-DEL-6268
chr6-33211306-DEL-83
chr6-33247749-DEL-191
chr6-33250829-DEL-560
chr6-33352587-DEL-144
chr6-33354704-DEL-1635
chr6-33362213-DEL-5476
chr6-33381990-DEL-75
chr6-33391643-DEL-66
chr6-33404365-DEL-1036
chr6-33411939-DEL-1301
chr6-33422292-DEL-58
chr6-33434537-DEL-84
chr6-33459798-DEL-1593
chr6-33467288-DEL-2660
chr6-33470028-DEL-2122
chr6-33471635-DEL-240
chr6-33518245-DEL-332
chr6-33520044-DEL-59
chr6-33595628-DEL-74
chr6-33602327-DEL-204
chr6-33643836-DEL-5926
chr6-33670568-DEL-197
chr6-33698142-DEL-60
chr6-33699760-DEL-107
chr6-33706495-DEL-119
chr6-33760287-DEL-571
chr6-33779549-DEL-408
chr6-33852231-DEL-166
chr6-33858108-DEL-1032
chr6-33875016-DEL-518
chr6-33892003-DEL-169
chr6-33910474-DEL-53
chr6-33920190-DEL-120
chr6-33967641-DEL-243
chr6-33995006-DEL-135
chr6-34062406-DEL-94
chr6-34118108-DEL-656
chr6-34165179-DEL-62
chr6-34177500-DEL-232
chr6-34184472-DEL-654
chr6-34186890-DEL-80
chr6-34226749-DEL-1565
chr6-34302348-DEL-1846
chr6-34307213-DEL-132
chr6-34348146-DEL-201
chr6-34351684-DEL-1040
chr6-34351775-DEL-123
chr6-34351813-DEL-592
chr6-34372642-DEL-128
chr6-34401191-DEL-59
chr6-34402730-DEL-789
chr6-34402829-DEL-6530
chr6-34404146-DEL-155
chr6-34406370-DEL-382
chr6-34406799-DEL-1394
chr6-34430886-DEL-658
chr6-34456000-DEL-71
chr6-34473692-DEL-79
chr6-34482141-DEL-52
chr6-34507839-DEL-97
chr6-34577556-DEL-207
chr6-34599082-DEL-228
chr6-34604649-DEL-467
chr6-34646065-DEL-61

chr6-34681110-DEL-64
chr6-34727049-DEL-88
chr6-34743266-DEL-66
chr6-34752450-DEL-342
chr6-34754696-DEL-180
chr6-34763597-DEL-892
chr6-34815627-DEL-113
chr6-34829530-DEL-91
chr6-34841192-DEL-98
chr6-34843405-DEL-297
chr6-34866627-DEL-52
chr6-34888939-DEL-167
chr6-34890003-DEL-66
chr6-34908280-DEL-70
chr6-34912852-DEL-198
chr6-34923929-DEL-63
chr6-34947036-DEL-326
chr6-35021145-DEL-167
chr6-35046719-DEL-256
chr6-35052747-DEL-115
chr6-35083629-DEL-1491
chr6-35098250-DEL-77
chr6-35102364-DEL-1070
chr6-35121323-DEL-64
chr6-35138907-DEL-54
chr6-35139777-DEL-432
chr6-35157392-DEL-305
chr6-35177658-DEL-205
chr6-35239774-DEL-1448
chr6-35260756-DEL-114
chr6-35268621-DEL-99
chr6-35346742-DEL-174
chr6-35364561-DEL-173
chr6-35403891-DEL-56
chr6-35410394-DEL-6400
chr6-35417623-DEL-63
chr6-35446182-DEL-6236
chr6-35464994-DEL-1650
chr6-35468305-DEL-199
chr6-35478009-DEL-61
chr6-35514262-DEL-88
chr6-35539666-DEL-54
chr6-35549193-DEL-6274
chr6-35582152-DEL-121
chr6-35654006-DEL-898
chr6-35675756-DEL-57
chr6-35676659-DEL-4340
chr6-35701481-DEL-6134
chr6-35745163-DEL-110
chr6-35762326-DEL-65
chr6-35762936-DEL-969
chr6-35788059-DEL-54
chr6-35867076-DEL-214
chr6-35910319-DEL-272
chr6-35910515-DEL-1143
chr6-35911703-DEL-1157
chr6-35970081-DEL-1202
chr6-35980435-DEL-84
chr6-35997562-DEL-56
chr6-36052714-DEL-62
chr6-36055867-DEL-62
chr6-36102673-DEL-267
chr6-36122001-DEL-821
chr6-36137254-DEL-442
chr6-36144967-DEL-82
chr6-36172780-DEL-100
chr6-36173189-DEL-2137
chr6-36295671-DEL-1179
chr6-36340617-DEL-523
chr6-36367919-DEL-140
chr6-36371770-DEL-121
chr6-36428312-DEL-51
chr6-36499157-DEL-51
chr6-36508791-DEL-199
chr6-36528546-DEL-1483
chr6-36551173-DEL-69
chr6-36552685-DEL-2180

chr6-36561008-DEL-3937
chr6-36588085-DEL-775
chr6-36596365-DEL-1644
chr6-36607498-DEL-565
chr6-36615287-DEL-56
chr6-36708836-DEL-119
chr6-36711633-DEL-595
chr6-36737100-DEL-101
chr6-36778645-DEL-225
chr6-36801360-DEL-430
chr6-36875231-DEL-187
chr6-36939755-DEL-169
chr6-37065124-DEL-117
chr6-37094340-DEL-70
chr6-37106883-DEL-191
chr6-37131782-DEL-65
chr6-37144338-DEL-167
chr6-37161816-DEL-95
chr6-37181667-DEL-63
chr6-37204464-DEL-58
chr6-37242462-DEL-1109
chr6-37268415-DEL-52
chr6-37300895-DEL-234
chr6-37325827-DEL-1535
chr6-37335759-DEL-134
chr6-37358616-DEL-104
chr6-37361934-DEL-587
chr6-37362569-DEL-1226
chr6-37381839-DEL-59
chr6-37436038-DEL-54
chr6-37475983-DEL-342
chr6-37503358-DEL-78
chr6-37538239-DEL-79
chr6-37544620-DEL-813
chr6-37547928-DEL-104
chr6-37551886-DEL-909
chr6-37555646-DEL-2084
chr6-37560401-DEL-59
chr6-37582330-DEL-54
chr6-37621840-DEL-60
chr6-37671895-DEL-74
chr6-37719491-DEL-58
chr6-37724308-DEL-565
chr6-37732317-DEL-54
chr6-37735019-DEL-858
chr6-37746891-DEL-52
chr6-37772544-DEL-68
chr6-37792500-DEL-210
chr6-37811585-DEL-225
chr6-37813164-DEL-467
chr6-37814694-DEL-221
chr6-37829332-DEL-540
chr6-37833484-DEL-106
chr6-37868224-DEL-157
chr6-37869965-DEL-260
chr6-37888249-DEL-1146
chr6-37899566-DEL-170
chr6-37919619-DEL-79
chr6-37945885-DEL-128
chr6-37993843-DEL-95
chr6-38041469-DEL-64
chr6-38043009-DEL-69
chr6-38071114-DEL-187
chr6-38110315-DEL-58
chr6-38114810-DEL-352
chr6-38135215-DEL-137
chr6-38167338-DEL-60
chr6-38185032-DEL-72
chr6-38188345-DEL-941
chr6-38189276-DEL-1340
chr6-38238442-DEL-59
chr6-38243747-DEL-103
chr6-38247101-DEL-54
chr6-38299092-DEL-93
chr6-38307153-DEL-86
chr6-38323458-DEL-1596
chr6-38336060-DEL-388

chr6-38342448-DEL-209
chr6-38344874-DEL-130
chr6-38348427-DEL-904
chr6-38348475-DEL-55
chr6-38350359-DEL-439
chr6-38395841-DEL-400
chr6-38421467-DEL-262
chr6-38457914-DEL-185
chr6-38502008-DEL-70
chr6-38504790-DEL-61
chr6-38520933-DEL-234
chr6-38538132-DEL-234
chr6-38551661-DEL-1270
chr6-38555792-DEL-290
chr6-38574995-DEL-227
chr6-38583065-DEL-232
chr6-38611308-DEL-84
chr6-38616272-DEL-86
chr6-38633494-DEL-191
chr6-38652043-DEL-207
chr6-38656860-DEL-96
chr6-38691851-DEL-102
chr6-38698904-DEL-138
chr6-38701632-DEL-87
chr6-38705987-DEL-115
chr6-38728570-DEL-54
chr6-38757344-DEL-903
chr6-38758205-DEL-180
chr6-38818654-DEL-78
chr6-38871820-DEL-117
chr6-38911302-DEL-382
chr6-38970098-DEL-70
chr6-38972580-DEL-532
chr6-38974279-DEL-95
chr6-38978442-DEL-69
chr6-38990220-DEL-80
chr6-39082355-DEL-56
chr6-39084161-DEL-54
chr6-39086830-DEL-59
chr6-39097050-DEL-520
chr6-39106684-DEL-867
chr6-39116718-DEL-174
chr6-39215370-DEL-207
chr6-39238336-DEL-78
chr6-39252424-DEL-60
chr6-39256445-DEL-62
chr6-39282423-DEL-109
chr6-39287283-DEL-96
chr6-39288054-DEL-206
chr6-39290121-DEL-225
chr6-39292808-DEL-60
chr6-39300258-DEL-130
chr6-39345856-DEL-78
chr6-39368744-DEL-210
chr6-39383683-DEL-86
chr6-39388055-DEL-219
chr6-39393943-DEL-697
chr6-39409613-DEL-229
chr6-39414530-DEL-495
chr6-39421473-DEL-206
chr6-39429428-DEL-549
chr6-39448643-DEL-80
chr6-39457978-DEL-642
chr6-39459307-DEL-125
chr6-39468716-DEL-51
chr6-39471072-DEL-462
chr6-39490221-DEL-190
chr6-39490915-DEL-204
chr6-39494444-DEL-413
chr6-39498312-DEL-70
chr6-39507977-DEL-2398
chr6-39535487-DEL-212
chr6-39566708-DEL-68
chr6-39567343-DEL-1803
chr6-39575950-DEL-54
chr6-39628053-DEL-145
chr6-39671464-DEL-945

chr6-39674497-DEL-409
chr6-39742273-DEL-224
chr6-39760837-DEL-100
chr6-39776659-DEL-410
chr6-39792045-DEL-1482
chr6-39812856-DEL-58
chr6-39858506-DEL-62
chr6-39863973-DEL-926
chr6-39925958-DEL-52
chr6-39998979-DEL-197
chr6-40076727-DEL-354
chr6-40085594-DEL-85
chr6-40139215-DEL-77
chr6-40168234-DEL-52
chr6-40209303-DEL-171
chr6-40244295-DEL-649
chr6-40267362-DEL-310
chr6-40282804-DEL-92
chr6-40284551-DEL-296
chr6-40287330-DEL-130
chr6-40332761-DEL-54
chr6-40384110-DEL-157
chr6-40442091-DEL-70
chr6-40492116-DEL-109
chr6-40523522-DEL-7586
chr6-40615151-DEL-92
chr6-40661082-DEL-81
chr6-40677357-DEL-92
chr6-40680576-DEL-460
chr6-40758575-DEL-61
chr6-40767336-DEL-65
chr6-40775293-DEL-213
chr6-40848985-DEL-96
chr6-40849917-DEL-67
chr6-40919606-DEL-4109
chr6-40929914-DEL-107
chr6-40948291-DEL-6594
chr6-40957899-DEL-100
chr6-40960123-DEL-100
chr6-40963242-DEL-6595
chr6-40980719-DEL-363
chr6-40984213-DEL-1092
chr6-40990715-DEL-636
chr6-41078589-DEL-76
chr6-41108144-DEL-62
chr6-41119395-DEL-77
chr6-41148439-DEL-598
chr6-41159830-DEL-377
chr6-41170053-DEL-52
chr6-41170901-DEL-827
chr6-41172689-DEL-106
chr6-41174079-DEL-4403
chr6-41190715-DEL-54
chr6-41207157-DEL-77
chr6-41228217-DEL-1122
chr6-41230265-DEL-896
chr6-41267558-DEL-285
chr6-41321804-DEL-315
chr6-41331672-DEL-83
chr6-41346050-DEL-374
chr6-41357849-DEL-724
chr6-41379889-DEL-733
chr6-41385193-DEL-128
chr6-41386315-DEL-150
chr6-41386891-DEL-1029
chr6-41414541-DEL-327
chr6-41452343-DEL-3634
chr6-41461242-DEL-1061
chr6-41478653-DEL-99
chr6-41506982-DEL-53
chr6-41515134-DEL-8512
chr6-41547637-DEL-71
chr6-41579194-DEL-53
chr6-41603226-DEL-99
chr6-41649721-DEL-52
chr6-41722742-DEL-70
chr6-41729489-DEL-1346

chr6-41747968-DEL-170
chr6-41778082-DEL-6824
chr6-41784226-DEL-208
chr6-41788175-DEL-287
chr6-41816941-DEL-59
chr6-41817846-DEL-6077
chr6-41832191-DEL-69
chr6-41849970-DEL-8254
chr6-41857065-DEL-416
chr6-41873689-DEL-61
chr6-41896833-DEL-156
chr6-41925952-DEL-62
chr6-41946399-DEL-37195
chr6-41960468-DEL-23572
chr6-42032025-DEL-148
chr6-42042378-DEL-3535
chr6-42068357-DEL-198
chr6-42070034-DEL-2483
chr6-42080631-DEL-203
chr6-42088850-DEL-78
chr6-42098222-DEL-9574
chr6-42122163-DEL-6353
chr6-42143176-DEL-8436
chr6-42147095-DEL-2048
chr6-42152230-DEL-852
chr6-42155836-DEL-99
chr6-42166101-DEL-1267
chr6-42182841-DEL-6821
chr6-42208745-DEL-153
chr6-42212375-DEL-724
chr6-42217274-DEL-58
chr6-42224229-DEL-87
chr6-42354921-DEL-92
chr6-42361326-DEL-158
chr6-42380946-DEL-68
chr6-42399084-DEL-98
chr6-42426330-DEL-63
chr6-42427685-DEL-6217
chr6-42551665-DEL-426
chr6-42594714-DEL-496
chr6-42594953-DEL-1307
chr6-42665342-DEL-476
chr6-42700914-DEL-104
chr6-42711116-DEL-89
chr6-42715035-DEL-640
chr6-42731325-DEL-935
chr6-42748416-DEL-4198
chr6-42785769-DEL-829
chr6-42802496-DEL-100
chr6-42808943-DEL-85
chr6-42817303-DEL-65
chr6-42822795-DEL-572
chr6-42840618-DEL-52
chr6-42850855-DEL-1840
chr6-42857554-DEL-104
chr6-42862690-DEL-4203
chr6-42867502-DEL-69
chr6-42876118-DEL-78
chr6-42911547-DEL-190
chr6-42913654-DEL-1512
chr6-42920644-DEL-2289
chr6-42927015-DEL-671
chr6-42943254-DEL-31350
chr6-42977119-DEL-10175
chr6-42978194-DEL-59
chr6-43007766-DEL-11308
chr6-43010168-DEL-82
chr6-43019422-DEL-97
chr6-43019767-DEL-439
chr6-43037796-DEL-60
chr6-43085633-DEL-2209
chr6-43102317-DEL-6264
chr6-43132250-DEL-358
chr6-43159484-DEL-799
chr6-43320690-DEL-84
chr6-43324547-DEL-798
chr6-43393933-DEL-1456

chr6-43395759-DEL-68
chr6-43432019-DEL-207
chr6-43475012-DEL-287
chr6-43475013-DEL-2895
chr6-43475357-DEL-1604
chr6-43476720-DEL-547
chr6-43477852-DEL-190
chr6-43491118-DEL-61
chr6-43516794-DEL-309
chr6-43689377-DEL-110
chr6-43700279-DEL-68
chr6-43705896-DEL-205
chr6-43729608-DEL-67
chr6-43743254-DEL-79
chr6-43761786-DEL-7150
chr6-43776302-DEL-6419
chr6-43828409-DEL-355
chr6-43843428-DEL-193
chr6-43867763-DEL-413
chr6-43876201-DEL-707
chr6-43881180-DEL-865
chr6-43882488-DEL-6280
chr6-43897585-DEL-5195
chr6-43899128-DEL-324
chr6-43918962-DEL-136
chr6-43928016-DEL-2716
chr6-43932908-DEL-83
chr6-43933423-DEL-6301
chr6-43949743-DEL-297
chr6-43955072-DEL-3005
chr6-44013080-DEL-89
chr6-44039448-DEL-1218
chr6-44110992-DEL-553
chr6-44147557-DEL-2254
chr6-44158238-DEL-395
chr6-44200506-DEL-53
chr6-44219132-DEL-324
chr6-44221273-DEL-75
chr6-44239888-DEL-212
chr6-44241797-DEL-6177
chr6-44287256-DEL-61
chr6-44363045-DEL-279
chr6-44410985-DEL-370
chr6-44416347-DEL-89
chr6-44449732-DEL-1157
chr6-44452262-DEL-99
chr6-44476866-DEL-1307
chr6-44484074-DEL-9928
chr6-44519229-DEL-75
chr6-44544133-DEL-56
chr6-44559743-DEL-4127
chr6-44564074-DEL-553
chr6-44564642-DEL-1178
chr6-44573788-DEL-63
chr6-44579334-DEL-1554
chr6-44581045-DEL-412
chr6-44597087-DEL-108
chr6-44601581-DEL-177
chr6-44613738-DEL-186
chr6-44636676-DEL-6213
chr6-44648416-DEL-160
chr6-44676729-DEL-59
chr6-44717754-DEL-6229
chr6-44727264-DEL-109
chr6-44760198-DEL-99
chr6-44772177-DEL-65
chr6-44813143-DEL-1230
chr6-44814996-DEL-198
chr6-44854476-DEL-606
chr6-44871182-DEL-297
chr6-44913021-DEL-1058
chr6-44948677-DEL-126
chr6-44961691-DEL-576
chr6-44966044-DEL-938
chr6-44967229-DEL-57
chr6-45021571-DEL-131
chr6-45025032-DEL-4234

chr6-45060744-DEL-2192
chr6-45113326-DEL-764
chr6-45244333-DEL-69
chr6-45315111-DEL-72
chr6-45327284-DEL-165
chr6-45333277-DEL-64
chr6-45362875-DEL-77
chr6-45402555-DEL-2450
chr6-45413790-DEL-54
chr6-45418388-DEL-128
chr6-45420603-DEL-354
chr6-45428560-DEL-113
chr6-45446892-DEL-291
chr6-45464193-DEL-162
chr6-45503509-DEL-158
chr6-45513520-DEL-193
chr6-45516246-DEL-52
chr6-45562850-DEL-1015
chr6-45679144-DEL-101
chr6-45699985-DEL-56
chr6-45732092-DEL-1309
chr6-45745149-DEL-225
chr6-45765149-DEL-84
chr6-45784181-DEL-134
chr6-45788108-DEL-422
chr6-45795657-DEL-84
chr6-45800455-DEL-2641
chr6-45824954-DEL-404
chr6-45869695-DEL-404
chr6-45881297-DEL-102
chr6-45898778-DEL-556
chr6-45900175-DEL-5601
chr6-45958583-DEL-405
chr6-45960886-DEL-477
chr6-45990470-DEL-134
chr6-45991065-DEL-200
chr6-46011105-DEL-341
chr6-46048186-DEL-61
chr6-46049592-DEL-678
chr6-46066260-DEL-908
chr6-46094884-DEL-280
chr6-46116749-DEL-210
chr6-46127083-DEL-85
chr6-46177055-DEL-172
chr6-46187453-DEL-309
chr6-46189207-DEL-206
chr6-46195114-DEL-182
chr6-46200272-DEL-3858
chr6-46243097-DEL-622
chr6-46303759-DEL-3724
chr6-46343788-DEL-130
chr6-46345395-DEL-62
chr6-46406406-DEL-214
chr6-46433746-DEL-173
chr6-46502046-DEL-163
chr6-46515222-DEL-161
chr6-46554062-DEL-6477
chr6-46563243-DEL-55
chr6-46565524-DEL-8811
chr6-46574516-DEL-177
chr6-46579544-DEL-51
chr6-46667975-DEL-54
chr6-46679546-DEL-110
chr6-46719339-DEL-89
chr6-46739267-DEL-65
chr6-46795257-DEL-4728
chr6-46796879-DEL-2257
chr6-46807542-DEL-1030
chr6-46809014-DEL-54
chr6-46811000-DEL-62
chr6-46821620-DEL-217
chr6-46826154-DEL-56
chr6-46841208-DEL-976
chr6-46842208-DEL-225
chr6-46848086-DEL-166
chr6-46852824-DEL-401
chr6-46983017-DEL-172

chr6-47009827-DEL-72
chr6-47083917-DEL-72
chr6-47096376-DEL-1835
chr6-47118246-DEL-113
chr6-47122436-DEL-161
chr6-47135428-DEL-502
chr6-47136525-DEL-394
chr6-47137411-DEL-285
chr6-47141362-DEL-161
chr6-47204829-DEL-250
chr6-47207693-DEL-72
chr6-47263397-DEL-102
chr6-47305819-DEL-87
chr6-47328132-DEL-406
chr6-47329643-DEL-150
chr6-47361367-DEL-68
chr6-47365153-DEL-69
chr6-47386889-DEL-725
chr6-47464418-DEL-93
chr6-47501027-DEL-60
chr6-47513281-DEL-1716
chr6-47538180-DEL-1652
chr6-47565583-DEL-54
chr6-47597540-DEL-112
chr6-47603894-DEL-194
chr6-47606970-DEL-203
chr6-47615491-DEL-2151
chr6-47624611-DEL-68
chr6-47730357-DEL-520
chr6-47738097-DEL-1263
chr6-47744130-DEL-95
chr6-47754741-DEL-95
chr6-47759619-DEL-498
chr6-47772586-DEL-168
chr6-47793175-DEL-53
chr6-47795999-DEL-176
chr6-47848946-DEL-93
chr6-47863668-DEL-105
chr6-47889320-DEL-765
chr6-47890587-DEL-948
chr6-47913171-DEL-158
chr6-47914723-DEL-505
chr6-47916325-DEL-899
chr6-47990587-DEL-2244
chr6-47992746-DEL-80
chr6-48008351-DEL-214
chr6-48063899-DEL-53
chr6-48086042-DEL-307
chr6-48103265-DEL-66
chr6-48128661-DEL-66
chr6-48134498-DEL-375
chr6-48143399-DEL-6459
chr6-48149469-DEL-214
chr6-48200706-DEL-96
chr6-48212249-DEL-6205
chr6-48231134-DEL-420
chr6-48244138-DEL-518
chr6-48247922-DEL-191
chr6-48261178-DEL-1103
chr6-48263530-DEL-1205
chr6-48266177-DEL-2792
chr6-48272989-DEL-5045
chr6-48296378-DEL-13348
chr6-48314485-DEL-1707
chr6-48338398-DEL-1548
chr6-48342024-DEL-295
chr6-48351526-DEL-4820
chr6-48356392-DEL-2233
chr6-48396601-DEL-351
chr6-48398522-DEL-109
chr6-48479106-DEL-202
chr6-48484154-DEL-1902
chr6-48494897-DEL-84
chr6-48523793-DEL-72
chr6-48600768-DEL-299
chr6-48611169-DEL-2168
chr6-48615876-DEL-713

chr6-48618301-DEL-93
chr6-48651480-DEL-483
chr6-48731487-DEL-562
chr6-48740280-DEL-1420
chr6-48764134-DEL-179
chr6-48764741-DEL-803
chr6-48769557-DEL-279
chr6-48793511-DEL-1289
chr6-48806488-DEL-72
chr6-48823198-DEL-56
chr6-48835502-DEL-63
chr6-48864904-DEL-4023
chr6-48869899-DEL-72
chr6-48886944-DEL-65
chr6-48915753-DEL-325
chr6-48917952-DEL-2996
chr6-48943608-DEL-54
chr6-48946653-DEL-93
chr6-48950926-DEL-807
chr6-48982059-DEL-1061
chr6-48989474-DEL-97
chr6-49001735-DEL-65
chr6-49002424-DEL-1030
chr6-49004297-DEL-70
chr6-49007015-DEL-735
chr6-49031390-DEL-188
chr6-49093186-DEL-58
chr6-49121170-DEL-156
chr6-49129348-DEL-97
chr6-49169283-DEL-214
chr6-49207101-DEL-983
chr6-49209962-DEL-204
chr6-49213379-DEL-170
chr6-49216412-DEL-614
chr6-49241338-DEL-183
chr6-49254398-DEL-53
chr6-49260707-DEL-171
chr6-49274790-DEL-1177
chr6-49288769-DEL-1653
chr6-49290464-DEL-2404
chr6-49302003-DEL-131
chr6-49314983-DEL-134
chr6-49323290-DEL-328
chr6-49334031-DEL-760
chr6-49339850-DEL-379
chr6-49350202-DEL-159
chr6-49354978-DEL-96
chr6-49373316-DEL-277
chr6-49383103-DEL-310
chr6-49385097-DEL-190
chr6-49418870-DEL-203
chr6-49421310-DEL-492
chr6-49424458-DEL-203
chr6-49457131-DEL-56
chr6-49465263-DEL-2216
chr6-49471921-DEL-5613
chr6-49486249-DEL-142
chr6-49502045-DEL-128
chr6-49512074-DEL-1171
chr6-49533008-DEL-1075
chr6-49536835-DEL-588
chr6-49566562-DEL-157
chr6-49567016-DEL-393
chr6-49591844-DEL-85
chr6-49697553-DEL-3002
chr6-49710948-DEL-68
chr6-49814838-DEL-1212
chr6-49889792-DEL-151
chr6-49919898-DEL-177
chr6-49920655-DEL-100
chr6-49979132-DEL-51
chr6-50022032-DEL-165
chr6-50023355-DEL-126
chr6-50042303-DEL-198
chr6-50044484-DEL-1047
chr6-50046671-DEL-66
chr6-50115153-DEL-544

chr6-50121356-DEL-63
chr6-50136191-DEL-129
chr6-50144122-DEL-57
chr6-50170691-DEL-60
chr6-50173099-DEL-203
chr6-50173882-DEL-350
chr6-50202263-DEL-189
chr6-50214152-DEL-52
chr6-50263885-DEL-59
chr6-50296325-DEL-371
chr6-50304687-DEL-109
chr6-50310996-DEL-154
chr6-50356467-DEL-280
chr6-50423202-DEL-443
chr6-50424973-DEL-55
chr6-50469032-DEL-90
chr6-50483868-DEL-1336
chr6-50489066-DEL-2086
chr6-50516366-DEL-1845
chr6-50524561-DEL-1065
chr6-50534246-DEL-597
chr6-50547424-DEL-874
chr6-50588870-DEL-3618
chr6-50595684-DEL-100
chr6-50600929-DEL-494
chr6-50612667-DEL-863
chr6-50671914-DEL-454
chr6-50775301-DEL-56
chr6-50879398-DEL-2215
chr6-50909894-DEL-312
chr6-50930711-DEL-50
chr6-50933175-DEL-70
chr6-50962333-DEL-64
chr6-50980976-DEL-149
chr6-51025173-DEL-431
chr6-51068948-DEL-118
chr6-51120564-DEL-437
chr6-51130965-DEL-120
chr6-51176110-DEL-799
chr6-51205899-DEL-6223
chr6-51257681-DEL-68
chr6-51259460-DEL-108
chr6-51261347-DEL-64
chr6-51273408-DEL-137
chr6-51315681-DEL-2899
chr6-51395060-DEL-80
chr6-51397395-DEL-779
chr6-51398049-DEL-66
chr6-51399312-DEL-927
chr6-51420025-DEL-65
chr6-51420397-DEL-206
chr6-51475591-DEL-105
chr6-51483450-DEL-171
chr6-51491140-DEL-1055
chr6-51491394-DEL-58
chr6-51510706-DEL-51
chr6-51521294-DEL-140
chr6-51532273-DEL-431
chr6-51539584-DEL-233
chr6-51540570-DEL-114
chr6-51580673-DEL-131
chr6-51603587-DEL-112
chr6-51690582-DEL-318
chr6-51707439-DEL-255
chr6-51753629-DEL-510
chr6-51764750-DEL-74
chr6-51797550-DEL-1020
chr6-51798316-DEL-242
chr6-51805314-DEL-695
chr6-51890070-DEL-120
chr6-51893228-DEL-96
chr6-51918360-DEL-482
chr6-51926237-DEL-4236
chr6-51931601-DEL-307
chr6-51932454-DEL-647
chr6-51950366-DEL-401
chr6-52047194-DEL-168

chr6-52057241-DEL-110
chr6-52057245-DEL-612
chr6-52057784-DEL-68
chr6-52074702-DEL-248
chr6-52096819-DEL-772
chr6-52109861-DEL-70
chr6-52119437-DEL-542
chr6-52128331-DEL-118
chr6-52256326-DEL-194
chr6-52259775-DEL-704
chr6-52271388-DEL-213
chr6-52322509-DEL-2386
chr6-52325391-DEL-1541
chr6-52326953-DEL-1084
chr6-52327083-DEL-954
chr6-52383825-DEL-356
chr6-52447709-DEL-178
chr6-52489710-DEL-100
chr6-52533831-DEL-52
chr6-52589825-DEL-4069
chr6-52715520-DEL-424
chr6-52757488-DEL-54
chr6-52761233-DEL-353
chr6-52761637-DEL-3198
chr6-52770637-DEL-200
chr6-52801145-DEL-432
chr6-52820266-DEL-54
chr6-52865546-DEL-96
chr6-52908910-DEL-112
chr6-52921940-DEL-286
chr6-52922627-DEL-422
chr6-52939484-DEL-290
chr6-52999671-DEL-137
chr6-53026436-DEL-350
chr6-53067248-DEL-415
chr6-53070776-DEL-51
chr6-53087234-DEL-52
chr6-53119706-DEL-62
chr6-53128860-DEL-114
chr6-53151188-DEL-8391
chr6-53153900-DEL-205
chr6-53194953-DEL-157
chr6-53210296-DEL-146
chr6-53214806-DEL-59
chr6-53226248-DEL-193
chr6-53379436-DEL-144
chr6-53510844-DEL-168
chr6-53519396-DEL-62
chr6-53520070-DEL-128
chr6-53527660-DEL-54
chr6-53536114-DEL-174
chr6-53543092-DEL-55
chr6-53565149-DEL-72
chr6-53632775-DEL-576
chr6-53667628-DEL-72
chr6-53688860-DEL-266
chr6-53712478-DEL-146
chr6-53716910-DEL-204
chr6-53722667-DEL-56
chr6-53723344-DEL-405
chr6-53775272-DEL-1121
chr6-53810680-DEL-91
chr6-53811979-DEL-71
chr6-53815231-DEL-762
chr6-53868873-DEL-58
chr6-53890975-DEL-82
chr6-53910608-DEL-136
chr6-53924816-DEL-97
chr6-53926151-DEL-61
chr6-53932177-DEL-78
chr6-54051047-DEL-89
chr6-54086433-DEL-184
chr6-54130456-DEL-355
chr6-54133053-DEL-57
chr6-54151033-DEL-120
chr6-54166040-DEL-328
chr6-54204694-DEL-51

chr6-54207138-DEL-54
chr6-54216225-DEL-60
chr6-54249164-DEL-81
chr6-54262180-DEL-92
chr6-54265496-DEL-233
chr6-54274561-DEL-97
chr6-54282215-DEL-51
chr6-54290452-DEL-56
chr6-54326900-DEL-1039
chr6-54339447-DEL-190
chr6-54351583-DEL-60
chr6-54369934-DEL-82
chr6-54372518-DEL-59
chr6-54374307-DEL-892
chr6-54379239-DEL-58
chr6-54419952-DEL-566
chr6-54433443-DEL-222
chr6-54457234-DEL-70
chr6-54534618-DEL-716
chr6-54539112-DEL-82
chr6-54565699-DEL-102
chr6-54566846-DEL-401
chr6-54594606-DEL-398
chr6-54603034-DEL-198
chr6-54604665-DEL-237
chr6-54639782-DEL-127
chr6-54644189-DEL-466
chr6-54647447-DEL-204
chr6-54702545-DEL-520
chr6-54761916-DEL-63
chr6-54809779-DEL-195
chr6-54844893-DEL-61
chr6-54861111-DEL-60
chr6-54940035-DEL-1419
chr6-54951293-DEL-56
chr6-54953947-DEL-676
chr6-54954703-DEL-104
chr6-54956865-DEL-54
chr6-54971741-DEL-63
chr6-54972238-DEL-391
chr6-55007762-DEL-468
chr6-55102300-DEL-111
chr6-55110807-DEL-75
chr6-55145898-DEL-96
chr6-55177151-DEL-206
chr6-55183028-DEL-930
chr6-55188627-DEL-54
chr6-55207129-DEL-54
chr6-55285559-DEL-399
chr6-55354403-DEL-70
chr6-55363055-DEL-72
chr6-55401327-DEL-119
chr6-55534257-DEL-86
chr6-55564929-DEL-64
chr6-55607668-DEL-69
chr6-55698434-DEL-68
chr6-55735724-DEL-1935
chr6-55763185-DEL-78
chr6-55797550-DEL-1201
chr6-55802202-DEL-59
chr6-55818045-DEL-59
chr6-55820970-DEL-414
chr6-55846180-DEL-722
chr6-55862556-DEL-250
chr6-55868963-DEL-140
chr6-55869434-DEL-1549
chr6-55903929-DEL-132
chr6-55957060-DEL-487
chr6-55964942-DEL-1452
chr6-55967203-DEL-470
chr6-56025360-DEL-235
chr6-56030863-DEL-2465
chr6-56048851-DEL-58
chr6-56093521-DEL-1578
chr6-56105662-DEL-486
chr6-56125058-DEL-1137
chr6-56130291-DEL-112

chr6-56155740-DEL-51
chr6-56179668-DEL-1392
chr6-56185719-DEL-317
chr6-56217509-DEL-8172
chr6-56256415-DEL-52
chr6-56276438-DEL-212
chr6-56335981-DEL-52
chr6-56368944-DEL-4136
chr6-56385172-DEL-1038
chr6-56419703-DEL-194
chr6-56461255-DEL-510
chr6-56495567-DEL-51
chr6-56501621-DEL-367
chr6-56511344-DEL-81
chr6-56523112-DEL-1217
chr6-56533420-DEL-1174
chr6-56554813-DEL-1216
chr6-56567821-DEL-201
chr6-56589071-DEL-53
chr6-56624163-DEL-221
chr6-56656574-DEL-70
chr6-56670648-DEL-406
chr6-56710138-DEL-212
chr6-56711884-DEL-57
chr6-56799441-DEL-54
chr6-56824376-DEL-745
chr6-56825048-DEL-74
chr6-56838930-DEL-3040
chr6-56857212-DEL-577
chr6-56917991-DEL-268
chr6-56921336-DEL-4158
chr6-56945233-DEL-58
chr6-57027387-DEL-2815
chr6-57029535-DEL-234
chr6-57063658-DEL-7325
chr6-57106920-DEL-247
chr6-57149672-DEL-1792
chr6-57162670-DEL-186
chr6-57192341-DEL-136
chr6-57300460-DEL-452
chr6-57303001-DEL-169
chr6-57320596-DEL-15700
chr6-57385696-DEL-119
chr6-57413931-DEL-390
chr6-57430207-DEL-1280
chr6-57432786-DEL-5215
chr6-57438029-DEL-1469
chr6-57454935-DEL-158
chr6-57466379-DEL-58
chr6-57469376-DEL-88
chr6-57478187-DEL-2784
chr6-57506630-DEL-62
chr6-57542208-DEL-92
chr6-57545088-DEL-79
chr6-57552973-DEL-1048
chr6-57559820-DEL-128
chr6-57578239-DEL-51
chr6-57606158-DEL-7823
chr6-57627848-DEL-859
chr6-57634258-DEL-456
chr6-57644374-DEL-6293
chr6-57652279-DEL-328
chr6-57668439-DEL-65
chr6-57677226-DEL-435
chr6-57698150-DEL-56
chr6-57735109-DEL-1540
chr6-57737448-DEL-80
chr6-57751382-DEL-71
chr6-57764400-DEL-52
chr6-57780220-DEL-217
chr6-57788992-DEL-114
chr6-57815199-DEL-739
chr6-57837746-DEL-1239
chr6-57844111-DEL-1047
chr6-57871962-DEL-102
chr6-57877936-DEL-550
chr6-57951169-DEL-202

chr6-57954025-DEL-89
chr6-57999287-DEL-614
chr6-58061408-DEL-6321
chr6-58061409-DEL-197
chr6-58080810-DEL-63
chr6-58116507-DEL-409
chr6-58124565-DEL-6624
chr6-58129318-DEL-141
chr6-58137187-DEL-631
chr6-58138218-DEL-116
chr6-58139608-DEL-290
chr6-58140710-DEL-58
chr6-58144802-DEL-353
chr6-58150714-DEL-204
chr6-58197267-DEL-101
chr6-58238918-DEL-116
chr6-58269674-DEL-6229
chr6-58300818-DEL-3259
chr6-58305766-DEL-142
chr6-58307351-DEL-3640
chr6-58335444-DEL-198
chr6-58353720-DEL-5772
chr6-58355959-DEL-1893
chr6-58361267-DEL-1891
chr6-58382873-DEL-145
chr6-58401575-DEL-75
chr6-58401771-DEL-6783
chr6-58408282-DEL-208
chr6-58425427-DEL-3456
chr6-58429692-DEL-237
chr6-58434092-DEL-52
chr6-58436163-DEL-5251
chr6-58441672-DEL-234
chr6-58456769-DEL-141
chr6-58466971-DEL-3355
chr6-58494645-DEL-990
chr6-58494859-DEL-493
chr6-58498981-DEL-7304
chr6-58501220-DEL-84
chr6-58505647-DEL-51
chr6-58510830-DEL-1978
chr6-58517853-DEL-3674
chr6-58521529-DEL-278
chr6-58522272-DEL-960
chr6-58532813-DEL-1901
chr6-58538516-DEL-6281
chr6-58547261-DEL-1368
chr6-58565989-DEL-156
chr6-58589402-DEL-197
chr6-58645025-DEL-142
chr6-58657871-DEL-62
chr6-58665096-DEL-725
chr6-58677839-DEL-328
chr6-58678390-DEL-2325
chr6-58681094-DEL-7614
chr6-58681920-DEL-90
chr6-58698002-DEL-339
chr6-58701263-DEL-3095
chr6-58712034-DEL-94
chr6-58734701-DEL-5346
chr6-58789953-DEL-153
chr6-58791228-DEL-184
chr6-58806877-DEL-58
chr6-58858152-DEL-113
chr6-58934922-DEL-905
chr6-58960026-DEL-161
chr6-58993399-DEL-280
chr6-59063115-DEL-1038
chr6-59071031-DEL-1967
chr6-59073257-DEL-338
chr6-59079951-DEL-88
chr6-59113311-DEL-602
chr6-59116375-DEL-1082
chr6-59118055-DEL-100
chr6-59171794-DEL-294
chr6-59177003-DEL-211
chr6-59186871-DEL-111

chr6-59202668-DEL-66
chr6-59202692-DEL-3924
chr6-59206637-DEL-1015
chr6-59215025-DEL-2376
chr6-59240639-DEL-1183
chr6-59336699-DEL-522
chr6-59354668-DEL-387
chr6-59361759-DEL-618
chr6-59436232-DEL-8298
chr6-59484830-DEL-315
chr6-59538157-DEL-652
chr6-59553873-DEL-68
chr6-59558762-DEL-2133
chr6-59559978-DEL-486
chr6-59561452-DEL-7084
chr6-59572378-DEL-3274
chr6-59587129-DEL-181
chr6-59607827-DEL-6332
chr6-59626324-DEL-180
chr6-59652167-DEL-6154
chr6-59665338-DEL-25237
chr6-59670149-DEL-55
chr6-59690987-DEL-56
chr6-59697644-DEL-51
chr6-59699253-DEL-55
chr6-59718919-DEL-56
chr6-59727006-DEL-15702
chr6-59769908-DEL-1257
chr6-59773288-DEL-74
chr6-59793457-DEL-1148
chr6-59802264-DEL-6017
chr6-59833490-DEL-322
chr6-59837104-DEL-383
chr6-59904859-DEL-84
chr6-59907547-DEL-2684
chr6-59907547-DEL-2694
chr6-59918775-DEL-63
chr6-59946799-DEL-60
chr6-59952683-DEL-461
chr6-59957362-DEL-1644
chr6-60010014-DEL-8341
chr6-60022494-DEL-332
chr6-60043262-DEL-113
chr6-60049292-DEL-6977
chr6-60051093-DEL-5003
chr6-60072010-DEL-6305
chr6-60126911-DEL-53
chr6-60134080-DEL-55
chr6-60139154-DEL-492
chr6-60139944-DEL-334
chr6-60141738-DEL-63
chr6-60142924-DEL-695
chr6-60157486-DEL-106
chr6-60161197-DEL-6813
chr6-60169052-DEL-382
chr6-60171119-DEL-65
chr6-60172712-DEL-270
chr6-60174138-DEL-1205
chr6-60177148-DEL-293
chr6-60188878-DEL-58
chr6-60193398-DEL-125
chr6-60208477-DEL-733
chr6-60215483-DEL-359
chr6-60216914-DEL-6111
chr6-60224312-DEL-2042
chr6-60229981-DEL-206
chr6-60231699-DEL-133
chr6-60236290-DEL-6483
chr6-60250451-DEL-84
chr6-60253407-DEL-198
chr6-60258896-DEL-130
chr6-60265030-DEL-6148
chr6-60268461-DEL-193
chr6-60273067-DEL-64
chr6-60276172-DEL-6213
chr6-60284495-DEL-214
chr6-60289109-DEL-7595

chr6-60289161-DEL-212
chr6-60290229-DEL-424
chr6-60367786-DEL-121
chr6-60375907-DEL-344
chr6-60401859-DEL-1486
chr6-60432454-DEL-110
chr6-60438410-DEL-199
chr6-60444528-DEL-477
chr6-60471030-DEL-54
chr6-60473623-DEL-1044
chr6-60499059-DEL-137
chr6-60500488-DEL-2106
chr6-60515529-DEL-12486
chr6-60535265-DEL-759
chr6-60542738-DEL-1134
chr6-60542744-DEL-889
chr6-60542831-DEL-120
chr6-60543670-DEL-441
chr6-60559238-DEL-3891
chr6-60594821-DEL-97
chr6-60596447-DEL-6371
chr6-60596679-DEL-198
chr6-60607139-DEL-76
chr6-60625408-DEL-66
chr6-60629906-DEL-51
chr6-60645359-DEL-807
chr6-60646197-DEL-12127
chr6-60666737-DEL-6445
chr6-60676408-DEL-69
chr6-60706690-DEL-67
chr6-60749906-DEL-495
chr6-60755024-DEL-816
chr6-60834665-DEL-90
chr6-60843585-DEL-279
chr6-60850759-DEL-61
chr6-60851478-DEL-5546
chr6-60856616-DEL-236
chr6-60862254-DEL-54
chr6-60866416-DEL-505
chr6-60867241-DEL-2634
chr6-60874669-DEL-713
chr6-60880457-DEL-1184
chr6-60880526-DEL-592
chr6-60884657-DEL-313
chr6-60894698-DEL-191
chr6-60914647-DEL-3535
chr6-60982537-DEL-824
chr6-61024896-DEL-1789
chr6-61031759-DEL-838
chr6-61065385-DEL-6271
chr6-61101245-DEL-91
chr6-61138376-DEL-51
chr6-61170322-DEL-826
chr6-61172774-DEL-1784
chr6-61174408-DEL-336
chr6-61238328-DEL-53
chr6-61313149-DEL-187
chr6-61317561-DEL-474
chr6-61336986-DEL-618
chr6-61339017-DEL-250
chr6-61369146-DEL-57
chr6-61410564-DEL-2773
chr6-61416461-DEL-909
chr6-61421515-DEL-1352
chr6-61438860-DEL-1091
chr6-61464694-DEL-559
chr6-61470554-DEL-54
chr6-61474980-DEL-759
chr6-61513936-DEL-67
chr6-61524032-DEL-1584
chr6-61528926-DEL-645
chr6-61578464-DEL-108
chr6-61609482-DEL-868
chr6-61615117-DEL-89
chr6-61622343-DEL-77
chr6-61622997-DEL-63
chr6-61688104-DEL-4686

chr6-61697414-DEL-151
chr6-61748534-DEL-225
chr6-61779156-DEL-98
chr6-61794162-DEL-94
chr6-61834065-DEL-191
chr6-61876494-DEL-662
chr6-61880857-DEL-84
chr6-61912509-DEL-237
chr6-61929668-DEL-818
chr6-61936325-DEL-133
chr6-61938873-DEL-66
chr6-61948160-DEL-3861
chr6-61961936-DEL-208
chr6-61965188-DEL-262
chr6-62010152-DEL-1442
chr6-62037780-DEL-357
chr6-62047567-DEL-752
chr6-62059884-DEL-283
chr6-62062781-DEL-52
chr6-62077435-DEL-1983
chr6-62098811-DEL-281
chr6-62128983-DEL-107
chr6-62147294-DEL-69
chr6-62151198-DEL-412
chr6-62152430-DEL-231
chr6-62215864-DEL-525
chr6-62232220-DEL-421
chr6-62235697-DEL-163
chr6-62265621-DEL-2827
chr6-62269563-DEL-129
chr6-62274096-DEL-53
chr6-62295455-DEL-63
chr6-62300690-DEL-276
chr6-62308934-DEL-4059
chr6-62314155-DEL-71
chr6-62325398-DEL-135
chr6-62327278-DEL-152
chr6-62333981-DEL-563
chr6-62347329-DEL-1684
chr6-62352191-DEL-1011
chr6-62355179-DEL-57
chr6-62381003-DEL-64
chr6-62391033-DEL-272
chr6-62431703-DEL-1022
chr6-62441159-DEL-693
chr6-62441934-DEL-728
chr6-62446227-DEL-343
chr6-62457433-DEL-392
chr6-62498580-DEL-420
chr6-62519793-DEL-305
chr6-62521859-DEL-323
chr6-62532445-DEL-891
chr6-62536465-DEL-6215
chr6-62544503-DEL-649
chr6-62577188-DEL-363
chr6-62587947-DEL-2017
chr6-62596589-DEL-209
chr6-62614021-DEL-1739
chr6-62641146-DEL-2291
chr6-62677675-DEL-112
chr6-62696692-DEL-591
chr6-62699956-DEL-3853
chr6-62707277-DEL-456
chr6-62718758-DEL-110
chr6-62723942-DEL-58
chr6-62751342-DEL-1828
chr6-62765522-DEL-70
chr6-62790219-DEL-368
chr6-62798354-DEL-6300
chr6-62826790-DEL-735
chr6-62848883-DEL-746
chr6-62854084-DEL-137
chr6-62864313-DEL-328
chr6-62865094-DEL-58
chr6-62893093-DEL-552
chr6-62917196-DEL-807
chr6-62952460-DEL-55

chr6-62965866-DEL-6434
chr6-62971999-DEL-212
chr6-62973155-DEL-77
chr6-62983573-DEL-4685
chr6-62991388-DEL-8616
chr6-63032172-DEL-6227
chr6-63042481-DEL-516
chr6-63046609-DEL-195
chr6-63052063-DEL-4288
chr6-63085342-DEL-62
chr6-63151700-DEL-2847
chr6-63167238-DEL-377
chr6-63175488-DEL-1247
chr6-63191635-DEL-168
chr6-63212765-DEL-1119
chr6-63225065-DEL-96
chr6-63259514-DEL-474
chr6-63328159-DEL-3987
chr6-63339202-DEL-52
chr6-63346468-DEL-534
chr6-63360299-DEL-711
chr6-63362524-DEL-375
chr6-63375432-DEL-1520
chr6-63382566-DEL-298
chr6-63437622-DEL-117
chr6-63465710-DEL-262
chr6-63466479-DEL-126
chr6-63475487-DEL-82
chr6-63482944-DEL-680
chr6-63532219-DEL-54
chr6-63613530-DEL-388
chr6-63627578-DEL-68
chr6-63635524-DEL-491
chr6-63672143-DEL-6223
chr6-63683280-DEL-394
chr6-63687656-DEL-90
chr6-63720718-DEL-267
chr6-63746346-DEL-1984
chr6-63755440-DEL-761
chr6-63758181-DEL-825
chr6-63785740-DEL-352
chr6-63794276-DEL-128
chr6-63809362-DEL-189
chr6-63843564-DEL-52
chr6-63892248-DEL-153
chr6-63906467-DEL-501
chr6-63919242-DEL-56
chr6-63953020-DEL-933
chr6-63965982-DEL-1452
chr6-64018343-DEL-498
chr6-64081525-DEL-751
chr6-64083942-DEL-555
chr6-64105498-DEL-159
chr6-64186320-DEL-279
chr6-64212557-DEL-142
chr6-64215227-DEL-58
chr6-64236739-DEL-116
chr6-64238808-DEL-400
chr6-64308716-DEL-7253
chr6-64308912-DEL-68
chr6-64393097-DEL-92
chr6-64397848-DEL-95
chr6-64438987-DEL-266
chr6-64451975-DEL-810
chr6-64505596-DEL-164
chr6-64530158-DEL-539
chr6-64534934-DEL-430
chr6-64571245-DEL-4835
chr6-64712939-DEL-194
chr6-64714948-DEL-641
chr6-64736984-DEL-1824
chr6-64745656-DEL-2438
chr6-64794245-DEL-386
chr6-64820259-DEL-93
chr6-64828672-DEL-150
chr6-64850055-DEL-60
chr6-64895876-DEL-795

chr6-64909952-DEL-460
chr6-64918850-DEL-76
chr6-64942795-DEL-54
chr6-64950957-DEL-76
chr6-64960275-DEL-52
chr6-64964150-DEL-62
chr6-64970190-DEL-89
chr6-64976443-DEL-494
chr6-64977874-DEL-162
chr6-64980671-DEL-102
chr6-64981857-DEL-212
chr6-64987547-DEL-121
chr6-64991207-DEL-334
chr6-64996200-DEL-185
chr6-65000693-DEL-76
chr6-65002542-DEL-214
chr6-65003257-DEL-185
chr6-65007187-DEL-52
chr6-65007547-DEL-636
chr6-65013647-DEL-201
chr6-65022246-DEL-207
chr6-65036402-DEL-2754
chr6-65047072-DEL-183
chr6-65077409-DEL-6231
chr6-65093218-DEL-1170
chr6-65098663-DEL-73
chr6-65138996-DEL-299
chr6-65142929-DEL-674
chr6-65147523-DEL-602
chr6-65159139-DEL-186
chr6-65164138-DEL-932
chr6-65167280-DEL-116
chr6-65179400-DEL-6490
chr6-65192309-DEL-6886
chr6-65223822-DEL-1250
chr6-65244480-DEL-440
chr6-65275740-DEL-6192
chr6-65329793-DEL-523
chr6-65361660-DEL-7379
chr6-65422355-DEL-61
chr6-65427128-DEL-66
chr6-65445815-DEL-88
chr6-65453507-DEL-1135
chr6-65455589-DEL-954
chr6-65488605-DEL-56
chr6-65493275-DEL-401
chr6-65534368-DEL-84
chr6-65552544-DEL-61
chr6-65580422-DEL-87
chr6-65586142-DEL-66
chr6-65589877-DEL-985
chr6-65677420-DEL-454
chr6-65696213-DEL-54
chr6-65699957-DEL-1571
chr6-65708200-DEL-83
chr6-65743581-DEL-192
chr6-65762474-DEL-555
chr6-65790213-DEL-138
chr6-65824841-DEL-330
chr6-65831335-DEL-86
chr6-65846489-DEL-98
chr6-65874324-DEL-117
chr6-65915795-DEL-100
chr6-65931104-DEL-484
chr6-65934091-DEL-78
chr6-65944908-DEL-357
chr6-65946909-DEL-82
chr6-65967358-DEL-86
chr6-65975121-DEL-176
chr6-66088736-DEL-400
chr6-66103615-DEL-197
chr6-66131505-DEL-1405
chr6-66141265-DEL-797
chr6-66148123-DEL-792
chr6-66162113-DEL-164
chr6-66189259-DEL-257
chr6-66194230-DEL-70

chr6-66227625-DEL-377
chr6-66265620-DEL-74
chr6-66270673-DEL-190
chr6-66288651-DEL-329
chr6-66289620-DEL-160
chr6-66292176-DEL-110
chr6-66294247-DEL-6301
chr6-66308356-DEL-198
chr6-66313517-DEL-3178
chr6-66327268-DEL-5427
chr6-66413619-DEL-431
chr6-66424753-DEL-1303
chr6-66446584-DEL-194
chr6-66499703-DEL-70
chr6-66506331-DEL-102
chr6-66581563-DEL-72
chr6-66591910-DEL-99
chr6-66608300-DEL-310
chr6-66628867-DEL-122
chr6-66647530-DEL-291
chr6-66647552-DEL-1041
chr6-66647921-DEL-1228
chr6-66648204-DEL-2487
chr6-66648618-DEL-514
chr6-66649205-DEL-1194
chr6-66720836-DEL-2158
chr6-66812104-DEL-545
chr6-66816481-DEL-58
chr6-66817662-DEL-52
chr6-66839659-DEL-77
chr6-66841714-DEL-178
chr6-66940699-DEL-253
chr6-67072244-DEL-66
chr6-67073134-DEL-53
chr6-67088909-DEL-291
chr6-67127507-DEL-172
chr6-67139876-DEL-78
chr6-67140041-DEL-1260
chr6-67152516-DEL-54
chr6-67158404-DEL-534
chr6-67159222-DEL-52
chr6-67164201-DEL-553
chr6-67175400-DEL-1801
chr6-67194374-DEL-3923
chr6-67225874-DEL-187
chr6-67308907-DEL-159
chr6-67311643-DEL-80
chr6-67317596-DEL-105
chr6-67340651-DEL-398
chr6-67351303-DEL-383
chr6-67363177-DEL-4593
chr6-67374255-DEL-1038
chr6-67386470-DEL-535
chr6-67425130-DEL-1070
chr6-67445931-DEL-71
chr6-67470570-DEL-346
chr6-67471614-DEL-212
chr6-67497096-DEL-2316
chr6-67516111-DEL-113
chr6-67536084-DEL-55
chr6-67552345-DEL-18646
chr6-67553295-DEL-7323
chr6-67559888-DEL-636
chr6-67562914-DEL-6636
chr6-67569255-DEL-214
chr6-67580507-DEL-2823
chr6-67587779-DEL-282
chr6-67608311-DEL-178
chr6-67611896-DEL-143
chr6-67612782-DEL-942
chr6-67618251-DEL-115
chr6-67658845-DEL-51
chr6-67662647-DEL-1308
chr6-67674038-DEL-59
chr6-67716282-DEL-591
chr6-67716983-DEL-2681
chr6-67728002-DEL-364

chr6-67809433-DEL-149
chr6-67812302-DEL-222
chr6-67823239-DEL-211
chr6-67827179-DEL-1720
chr6-67834062-DEL-57
chr6-67877362-DEL-319
chr6-67900627-DEL-226
chr6-67903542-DEL-55
chr6-67913381-DEL-5520
chr6-67961635-DEL-3019
chr6-67974820-DEL-1663
chr6-68048246-DEL-55
chr6-68072599-DEL-2924
chr6-68080430-DEL-159
chr6-68100783-DEL-9463
chr6-68105350-DEL-1512
chr6-68110851-DEL-119
chr6-68111685-DEL-653
chr6-68113439-DEL-6451
chr6-68144355-DEL-5535
chr6-68151147-DEL-1906
chr6-68153904-DEL-516
chr6-68156661-DEL-75
chr6-68161284-DEL-1168
chr6-68164692-DEL-7044
chr6-68171111-DEL-77
chr6-68180375-DEL-87
chr6-68183422-DEL-797
chr6-68188281-DEL-1888
chr6-68198836-DEL-6281
chr6-68204988-DEL-127
chr6-68207150-DEL-6181
chr6-68216264-DEL-25147
chr6-68219369-DEL-9901
chr6-68250224-DEL-8715
chr6-68260188-DEL-7198
chr6-68269152-DEL-124
chr6-68290544-DEL-2664
chr6-68306229-DEL-630
chr6-68318135-DEL-1653
chr6-68330319-DEL-165
chr6-68332719-DEL-6946
chr6-68358461-DEL-175
chr6-68359244-DEL-34191
chr6-68404100-DEL-323
chr6-68422051-DEL-13443
chr6-68504194-DEL-303
chr6-68549897-DEL-47480
chr6-68602576-DEL-202
chr6-68633471-DEL-687
chr6-68640382-DEL-1544
chr6-68643599-DEL-79
chr6-68658409-DEL-2890
chr6-68661605-DEL-69
chr6-68664230-DEL-518
chr6-68665774-DEL-112
chr6-68668383-DEL-66
chr6-68686449-DEL-8318
chr6-68687463-DEL-481
chr6-68698208-DEL-2969
chr6-68702316-DEL-179
chr6-68703473-DEL-121
chr6-68707461-DEL-309
chr6-68737868-DEL-239
chr6-68747093-DEL-1552
chr6-68759449-DEL-3854
chr6-68764074-DEL-55
chr6-68765634-DEL-135
chr6-68766709-DEL-3932
chr6-68774495-DEL-7010
chr6-68775840-DEL-3658
chr6-68792562-DEL-3717
chr6-68814132-DEL-959
chr6-68908487-DEL-52
chr6-68947416-DEL-6124
chr6-68958153-DEL-204
chr6-68994380-DEL-244

chr6-68994632-DEL-6432
chr6-69006505-DEL-6926
chr6-69012752-DEL-212
chr6-69051329-DEL-94
chr6-69136027-DEL-6317
chr6-69152993-DEL-56
chr6-69187749-DEL-205
chr6-69193990-DEL-102
chr6-69205103-DEL-410
chr6-69208660-DEL-9957
chr6-69221357-DEL-318
chr6-69232899-DEL-480
chr6-69233761-DEL-8041
chr6-69233903-DEL-1259
chr6-69238679-DEL-849
chr6-69243058-DEL-1716
chr6-69251900-DEL-61
chr6-69270191-DEL-7333
chr6-69293029-DEL-5321
chr6-69303299-DEL-81
chr6-69357562-DEL-1940
chr6-69366252-DEL-75
chr6-69392053-DEL-1725
chr6-69440927-DEL-6184
chr6-69452077-DEL-9564
chr6-69462958-DEL-10600
chr6-69464689-DEL-6443
chr6-69475349-DEL-170
chr6-69486081-DEL-555
chr6-69489072-DEL-516
chr6-69492928-DEL-225
chr6-69502489-DEL-51
chr6-69545221-DEL-172
chr6-69545228-DEL-172
chr6-69547309-DEL-207
chr6-69563589-DEL-59
chr6-69567012-DEL-6303
chr6-69603916-DEL-3545
chr6-69606815-DEL-1719
chr6-69616303-DEL-92
chr6-69617395-DEL-221
chr6-69618473-DEL-1114
chr6-69620325-DEL-95
chr6-69622253-DEL-56
chr6-69758029-DEL-74
chr6-69773929-DEL-6473
chr6-69815888-DEL-668
chr6-69848242-DEL-3270
chr6-69990870-DEL-6499
chr6-70023889-DEL-6679
chr6-70030759-DEL-126
chr6-70041642-DEL-224
chr6-70052096-DEL-7767
chr6-70065084-DEL-6313
chr6-70111848-DEL-5046
chr6-70138740-DEL-1330
chr6-70163435-DEL-6281
chr6-70235434-DEL-53
chr6-70246645-DEL-77
chr6-70247954-DEL-16400
chr6-70267459-DEL-52
chr6-70303509-DEL-3390
chr6-70367699-DEL-174
chr6-70405624-DEL-22960
chr6-70446540-DEL-1329
chr6-70458985-DEL-52
chr6-70460159-DEL-6740
chr6-70473513-DEL-1915
chr6-70474801-DEL-325
chr6-70478568-DEL-30623
chr6-70501164-DEL-6861
chr6-70509495-DEL-87
chr6-70511474-DEL-5449
chr6-70513691-DEL-142
chr6-70565827-DEL-670
chr6-70571839-DEL-294
chr6-70597310-DEL-6363

chr6-70619088-DEL-93
chr6-70653691-DEL-85
chr6-70664880-DEL-124
chr6-70665890-DEL-111
chr6-70668676-DEL-4502
chr6-70675647-DEL-324
chr6-70676216-DEL-1599
chr6-70676561-DEL-99
chr6-70677943-DEL-610
chr6-70726055-DEL-140
chr6-70738596-DEL-200
chr6-70754534-DEL-76
chr6-70757316-DEL-165
chr6-70760937-DEL-396
chr6-70773289-DEL-117
chr6-70776207-DEL-64
chr6-70779383-DEL-1215
chr6-70785580-DEL-72
chr6-70786815-DEL-78
chr6-70789280-DEL-152
chr6-70796674-DEL-648
chr6-70811956-DEL-58
chr6-70814561-DEL-190
chr6-70826170-DEL-177
chr6-70844261-DEL-160
chr6-70866850-DEL-62
chr6-70867444-DEL-351
chr6-70874388-DEL-69
chr6-70887839-DEL-58
chr6-70904254-DEL-750
chr6-70922439-DEL-64
chr6-70929632-DEL-74
chr6-70990947-DEL-52
chr6-71012370-DEL-3337
chr6-71020783-DEL-272
chr6-71022621-DEL-1494
chr6-71053117-DEL-51
chr6-71061296-DEL-151
chr6-71076265-DEL-181
chr6-71099701-DEL-123
chr6-71112764-DEL-92
chr6-71122988-DEL-3061
chr6-71134567-DEL-5146
chr6-71145680-DEL-82
chr6-71145902-DEL-238
chr6-71147217-DEL-306
chr6-71154311-DEL-85
chr6-71174388-DEL-59
chr6-71200338-DEL-334
chr6-71201121-DEL-60
chr6-71201521-DEL-506
chr6-71203188-DEL-203
chr6-71244210-DEL-87
chr6-71264363-DEL-5204
chr6-71284102-DEL-336
chr6-71287150-DEL-1887
chr6-71287157-DEL-1138
chr6-71288339-DEL-691
chr6-71308448-DEL-210
chr6-71318896-DEL-60
chr6-71332849-DEL-56
chr6-71347125-DEL-116
chr6-71377908-DEL-63
chr6-71380671-DEL-233
chr6-71408123-DEL-65
chr6-71412056-DEL-398
chr6-71440865-DEL-201
chr6-71444723-DEL-91
chr6-71467947-DEL-1016
chr6-71540329-DEL-70
chr6-71582302-DEL-210
chr6-71617970-DEL-60
chr6-71618366-DEL-753
chr6-71640052-DEL-146
chr6-71670503-DEL-159
chr6-71680304-DEL-134
chr6-71682913-DEL-67

chr6-71731151-DEL-72
chr6-71747502-DEL-52
chr6-71766736-DEL-2404
chr6-71774535-DEL-85
chr6-71776399-DEL-98
chr6-71925986-DEL-162
chr6-71977353-DEL-148
chr6-71992903-DEL-79
chr6-71993499-DEL-1241
chr6-71996095-DEL-409
chr6-71998663-DEL-1356
chr6-72007554-DEL-6560
chr6-72037722-DEL-852
chr6-72051180-DEL-120
chr6-72067458-DEL-313
chr6-72077227-DEL-55
chr6-72093826-DEL-62
chr6-72095976-DEL-719
chr6-72104066-DEL-363
chr6-72123001-DEL-266
chr6-72137778-DEL-167
chr6-72140950-DEL-52
chr6-72203992-DEL-138
chr6-72224880-DEL-84
chr6-72232039-DEL-61
chr6-72241204-DEL-166
chr6-72264518-DEL-63
chr6-72267058-DEL-61
chr6-72301969-DEL-179
chr6-72306344-DEL-68
chr6-72320034-DEL-273
chr6-72371191-DEL-425
chr6-72388735-DEL-1523
chr6-72428833-DEL-61
chr6-72447417-DEL-192
chr6-72474333-DEL-74
chr6-72514324-DEL-569
chr6-72541989-DEL-190
chr6-72598572-DEL-112
chr6-72605229-DEL-144
chr6-72609325-DEL-119
chr6-72621942-DEL-70
chr6-72623462-DEL-80
chr6-72675280-DEL-386
chr6-72703474-DEL-104
chr6-72718651-DEL-175
chr6-72779133-DEL-61
chr6-72805232-DEL-339
chr6-72828220-DEL-2171
chr6-72854159-DEL-553
chr6-72856749-DEL-1352
chr6-72894129-DEL-175
chr6-72919577-DEL-72
chr6-72968806-DEL-66
chr6-72989115-DEL-155
chr6-73030322-DEL-76
chr6-73052053-DEL-52
chr6-73057269-DEL-1888
chr6-73080666-DEL-104
chr6-73084015-DEL-208
chr6-73144859-DEL-100
chr6-73152711-DEL-73
chr6-73177588-DEL-1967
chr6-73179576-DEL-1213
chr6-73321376-DEL-2201
chr6-73321394-DEL-286
chr6-73323658-DEL-2321
chr6-73326098-DEL-2402
chr6-73329779-DEL-59
chr6-73359148-DEL-75
chr6-73366866-DEL-74
chr6-73369052-DEL-1883
chr6-73381063-DEL-2871
chr6-73432177-DEL-133
chr6-73465174-DEL-339
chr6-73486691-DEL-399
chr6-73487996-DEL-330

chr6-73497046-DEL-84
chr6-73574839-DEL-52
chr6-73592087-DEL-1086
chr6-73596202-DEL-594
chr6-73604472-DEL-873
chr6-73648318-DEL-1298
chr6-73663249-DEL-435
chr6-73671629-DEL-100
chr6-73686850-DEL-101
chr6-73713186-DEL-8302
chr6-73714218-DEL-6796
chr6-73714222-DEL-5973
chr6-73731358-DEL-56
chr6-73747492-DEL-698
chr6-73802317-DEL-192
chr6-73823811-DEL-205
chr6-73826087-DEL-88
chr6-73827604-DEL-1331
chr6-73905934-DEL-547
chr6-73906828-DEL-557
chr6-73930141-DEL-69
chr6-73933002-DEL-324
chr6-73958102-DEL-51
chr6-74005293-DEL-1651
chr6-74008256-DEL-65
chr6-74097673-DEL-58
chr6-74103793-DEL-2538
chr6-74113435-DEL-7049
chr6-74134353-DEL-6239
chr6-74149858-DEL-88
chr6-74168978-DEL-1473
chr6-74176855-DEL-313
chr6-74195545-DEL-78
chr6-74212951-DEL-200
chr6-74222755-DEL-783
chr6-74248093-DEL-125
chr6-74252733-DEL-1209
chr6-74354960-DEL-161
chr6-74360309-DEL-6239
chr6-74372013-DEL-74
chr6-74391022-DEL-569
chr6-74414669-DEL-87
chr6-74418281-DEL-756
chr6-74434404-DEL-161
chr6-74437226-DEL-943
chr6-74453787-DEL-198
chr6-74467276-DEL-63
chr6-74478384-DEL-204
chr6-74483872-DEL-791
chr6-74492589-DEL-6209
chr6-74502727-DEL-6455
chr6-74530329-DEL-223
chr6-74533398-DEL-62
chr6-74559200-DEL-81
chr6-74585457-DEL-183
chr6-74640749-DEL-1436
chr6-74647241-DEL-1863
chr6-74659094-DEL-1383
chr6-74676276-DEL-52
chr6-74729633-DEL-2080
chr6-74733195-DEL-2355
chr6-74736372-DEL-548
chr6-74757979-DEL-53
chr6-74765524-DEL-1399
chr6-74855400-DEL-54
chr6-74886418-DEL-173
chr6-74911289-DEL-115
chr6-74911308-DEL-6379
chr6-74924882-DEL-3483
chr6-74970958-DEL-80
chr6-74973397-DEL-6470
chr6-74973402-DEL-13387
chr6-74973537-DEL-782
chr6-74975333-DEL-3734
chr6-74979943-DEL-1799
chr6-74981804-DEL-4985
chr6-75001153-DEL-581

chr6-75030782-DEL-2795
chr6-75034241-DEL-219
chr6-75054803-DEL-6159
chr6-75063931-DEL-60
chr6-75075187-DEL-274
chr6-75077368-DEL-798
chr6-75079163-DEL-66
chr6-75089637-DEL-182
chr6-75098119-DEL-1263
chr6-75098119-DEL-749
chr6-75098701-DEL-1129
chr6-75174107-DEL-63
chr6-75180284-DEL-1196
chr6-75196636-DEL-691
chr6-75229228-DEL-688
chr6-75231512-DEL-389
chr6-75265008-DEL-63
chr6-75272992-DEL-2130
chr6-75293336-DEL-5561
chr6-75310301-DEL-1857
chr6-75312171-DEL-3529
chr6-75315919-DEL-98
chr6-75318548-DEL-876
chr6-75320092-DEL-6693
chr6-75381150-DEL-59
chr6-75383373-DEL-225
chr6-75392407-DEL-1053
chr6-75419946-DEL-109
chr6-75426624-DEL-2019
chr6-75430987-DEL-66
chr6-75453078-DEL-286
chr6-75454602-DEL-61
chr6-75476254-DEL-2402
chr6-75479340-DEL-492
chr6-75486913-DEL-216
chr6-75492149-DEL-910
chr6-75495245-DEL-55
chr6-75497393-DEL-231
chr6-75529678-DEL-1221
chr6-75585225-DEL-141
chr6-75593406-DEL-429
chr6-75594645-DEL-5811
chr6-75603883-DEL-792
chr6-75634365-DEL-171
chr6-75639752-DEL-2236
chr6-75658750-DEL-80
chr6-75659325-DEL-1641
chr6-75672061-DEL-160
chr6-75686782-DEL-4783
chr6-75695436-DEL-982
chr6-75780385-DEL-172
chr6-75820660-DEL-56
chr6-75828315-DEL-457
chr6-75856666-DEL-2483
chr6-75862828-DEL-209
chr6-75878392-DEL-1379
chr6-75879623-DEL-122
chr6-75902254-DEL-640
chr6-75903589-DEL-89
chr6-75936869-DEL-3154
chr6-75949119-DEL-1367
chr6-75953114-DEL-301
chr6-75983056-DEL-60
chr6-75989760-DEL-155
chr6-76059893-DEL-64
chr6-76080469-DEL-56
chr6-76086951-DEL-1424
chr6-76090543-DEL-116
chr6-76155719-DEL-59
chr6-76165846-DEL-722
chr6-76184752-DEL-52
chr6-76187940-DEL-748
chr6-76192584-DEL-4411
chr6-76261171-DEL-738
chr6-76293628-DEL-73
chr6-76296467-DEL-2502
chr6-76299015-DEL-819

chr6-76300029-DEL-198
chr6-76304945-DEL-130
chr6-76311778-DEL-62
chr6-76373748-DEL-939
chr6-76378833-DEL-362
chr6-76409531-DEL-228
chr6-76512452-DEL-6325
chr6-76569798-DEL-318
chr6-76582422-DEL-2030
chr6-76605927-DEL-1328
chr6-76622054-DEL-6183
chr6-76622064-DEL-625
chr6-76641497-DEL-57
chr6-76667811-DEL-56
chr6-76694194-DEL-52
chr6-76705165-DEL-6632
chr6-76718414-DEL-172
chr6-76742382-DEL-85
chr6-76775156-DEL-549
chr6-76777144-DEL-434
chr6-76820163-DEL-86
chr6-76824105-DEL-6998
chr6-76834715-DEL-58
chr6-76841274-DEL-61
chr6-76865986-DEL-52
chr6-76880580-DEL-260
chr6-76887704-DEL-52
chr6-76889092-DEL-88
chr6-76901470-DEL-223
chr6-76935368-DEL-111
chr6-76945380-DEL-83
chr6-76977722-DEL-59
chr6-76993327-DEL-108
chr6-77028169-DEL-64
chr6-77029118-DEL-1573
chr6-77033884-DEL-81
chr6-77048618-DEL-1889
chr6-77051366-DEL-481
chr6-77104821-DEL-65
chr6-77113655-DEL-96
chr6-77129833-DEL-279
chr6-77149978-DEL-93
chr6-77158528-DEL-253
chr6-77164931-DEL-100
chr6-77201847-DEL-66
chr6-77219034-DEL-61
chr6-77232591-DEL-144
chr6-77249759-DEL-609
chr6-77264008-DEL-104
chr6-77283023-DEL-52
chr6-77303156-DEL-2517
chr6-77329053-DEL-64
chr6-77379391-DEL-288
chr6-77434020-DEL-121
chr6-77457017-DEL-419
chr6-77501018-DEL-52
chr6-77539963-DEL-733
chr6-77562766-DEL-321
chr6-77567486-DEL-51
chr6-77573043-DEL-91
chr6-77632551-DEL-221
chr6-77638573-DEL-142
chr6-77683280-DEL-617
chr6-77705997-DEL-512
chr6-77731529-DEL-484
chr6-77834865-DEL-1015
chr6-77844656-DEL-167
chr6-77859889-DEL-622
chr6-77891829-DEL-55
chr6-77905982-DEL-53
chr6-77910840-DEL-1163
chr6-77933882-DEL-884
chr6-77934063-DEL-416
chr6-77950041-DEL-60
chr6-77959773-DEL-138
chr6-77980810-DEL-376
chr6-77983075-DEL-52

chr6-77996893-DEL-283
chr6-78013865-DEL-6223
chr6-78016834-DEL-1337
chr6-78070356-DEL-2828
chr6-78084251-DEL-575
chr6-78085666-DEL-61
chr6-78108465-DEL-346
chr6-78122083-DEL-1158
chr6-78143824-DEL-107
chr6-78147204-DEL-67
chr6-78167716-DEL-109
chr6-78174944-DEL-652
chr6-78185085-DEL-178
chr6-78186999-DEL-3140
chr6-78203788-DEL-394
chr6-78204749-DEL-60
chr6-78207290-DEL-193
chr6-78237037-DEL-210
chr6-78294387-DEL-64
chr6-78295174-DEL-276
chr6-78301874-DEL-1474
chr6-78358381-DEL-134
chr6-78363473-DEL-158
chr6-78364947-DEL-384
chr6-78364972-DEL-1098
chr6-78372560-DEL-211
chr6-78397180-DEL-2536
chr6-78422018-DEL-787
chr6-78436152-DEL-533
chr6-78447284-DEL-401
chr6-78477218-DEL-1524
chr6-78506074-DEL-1251
chr6-78509361-DEL-188
chr6-78545696-DEL-95
chr6-78559120-DEL-53
chr6-78568024-DEL-78
chr6-78583233-DEL-1869
chr6-78592323-DEL-88
chr6-78652316-DEL-53
chr6-78701537-DEL-57
chr6-78723189-DEL-212
chr6-78724415-DEL-343
chr6-78788947-DEL-61
chr6-78805240-DEL-617
chr6-78818117-DEL-142
chr6-78824366-DEL-109
chr6-78836694-DEL-403
chr6-78878249-DEL-327
chr6-78888344-DEL-103
chr6-78906237-DEL-56
chr6-78908224-DEL-606
chr6-78909090-DEL-845
chr6-78915628-DEL-8120
chr6-78930318-DEL-375
chr6-78977023-DEL-88
chr6-78979149-DEL-54
chr6-78999427-DEL-2955
chr6-79014890-DEL-3607
chr6-79059813-DEL-633
chr6-79064979-DEL-107
chr6-79080816-DEL-95
chr6-79082985-DEL-1016
chr6-79087615-DEL-935
chr6-79115947-DEL-443
chr6-79165958-DEL-71
chr6-79167260-DEL-203
chr6-79175012-DEL-60
chr6-79206454-DEL-182
chr6-79223837-DEL-52
chr6-79240016-DEL-2431
chr6-79270710-DEL-55
chr6-79274366-DEL-1335
chr6-79276726-DEL-54
chr6-79291744-DEL-157
chr6-79302912-DEL-68
chr6-79347650-DEL-61
chr6-79357068-DEL-1031

chr6-79363608-DEL-61
chr6-79372477-DEL-165
chr6-79374139-DEL-156
chr6-79412567-DEL-64
chr6-79427715-DEL-5931
chr6-79433647-DEL-121
chr6-79438123-DEL-84
chr6-79456021-DEL-61
chr6-79471660-DEL-2560
chr6-79491117-DEL-483
chr6-79506750-DEL-215
chr6-79513054-DEL-79
chr6-79514264-DEL-444
chr6-79521786-DEL-208
chr6-79531471-DEL-62
chr6-79553763-DEL-751
chr6-79554534-DEL-1020
chr6-79556226-DEL-217
chr6-79582910-DEL-189
chr6-79591375-DEL-179
chr6-79623410-DEL-64
chr6-79657348-DEL-1177
chr6-79660594-DEL-6060
chr6-79673472-DEL-52
chr6-79675748-DEL-183
chr6-79681380-DEL-534
chr6-79698215-DEL-509
chr6-79708341-DEL-52
chr6-79709967-DEL-294
chr6-79713912-DEL-117
chr6-79717354-DEL-186
chr6-79736735-DEL-740
chr6-79744365-DEL-127
chr6-79747976-DEL-114
chr6-79755510-DEL-684
chr6-79800746-DEL-2452
chr6-79829908-DEL-1112
chr6-79834851-DEL-76
chr6-79847541-DEL-455
chr6-79903158-DEL-77
chr6-79936712-DEL-9130
chr6-79941196-DEL-2833
chr6-79965501-DEL-91
chr6-80010012-DEL-73
chr6-80018616-DEL-502
chr6-80032262-DEL-104
chr6-80071146-DEL-66
chr6-80093220-DEL-1445
chr6-80104416-DEL-1877
chr6-80120679-DEL-136
chr6-80151241-DEL-52
chr6-80184949-DEL-482
chr6-80206204-DEL-1151
chr6-80218931-DEL-53
chr6-80255885-DEL-107
chr6-80363540-DEL-52
chr6-80371110-DEL-6129
chr6-80382031-DEL-74
chr6-80407798-DEL-505
chr6-80427635-DEL-89
chr6-80435709-DEL-642
chr6-80442544-DEL-693
chr6-80451692-DEL-51
chr6-80466329-DEL-63
chr6-80468447-DEL-701
chr6-80468542-DEL-1292
chr6-80469022-DEL-701
chr6-80469481-DEL-296
chr6-80499642-DEL-203
chr6-80501906-DEL-708
chr6-80515704-DEL-68
chr6-80519801-DEL-63
chr6-80526096-DEL-160
chr6-80539306-DEL-155
chr6-80557114-DEL-494
chr6-80586988-DEL-56
chr6-80593463-DEL-391

chr6-80595981-DEL-52
chr6-80606321-DEL-673
chr6-80625933-DEL-559
chr6-80636486-DEL-94
chr6-80649065-DEL-805
chr6-80666152-DEL-51
chr6-80669674-DEL-363
chr6-80672824-DEL-115
chr6-80755856-DEL-57
chr6-80820348-DEL-169
chr6-80838402-DEL-62
chr6-80850089-DEL-3971
chr6-80883375-DEL-261
chr6-80911258-DEL-411
chr6-80918046-DEL-72
chr6-80938815-DEL-61
chr6-80954050-DEL-206
chr6-80956403-DEL-63
chr6-80957429-DEL-54
chr6-80964118-DEL-57
chr6-80983133-DEL-119
chr6-80991858-DEL-2804
chr6-81007321-DEL-3779
chr6-81051704-DEL-52
chr6-81054800-DEL-79
chr6-81068462-DEL-343
chr6-81071003-DEL-440
chr6-81077603-DEL-53
chr6-81081169-DEL-116
chr6-81081884-DEL-152
chr6-81110154-DEL-64
chr6-81115034-DEL-1537
chr6-81116104-DEL-487
chr6-81173279-DEL-75
chr6-81195216-DEL-142
chr6-81221662-DEL-20007
chr6-81229112-DEL-124
chr6-81270794-DEL-156
chr6-81292100-DEL-82
chr6-81299892-DEL-208
chr6-81306619-DEL-71
chr6-81306940-DEL-1562
chr6-81327470-DEL-114
chr6-81343349-DEL-161
chr6-81366261-DEL-144
chr6-81367426-DEL-72
chr6-81372043-DEL-59
chr6-81376489-DEL-206
chr6-81382594-DEL-5024
chr6-81389318-DEL-110
chr6-81396968-DEL-141
chr6-81409514-DEL-130
chr6-81414749-DEL-206
chr6-81457578-DEL-204
chr6-81495821-DEL-1359
chr6-81497706-DEL-252
chr6-81505616-DEL-476
chr6-81528478-DEL-68
chr6-81620841-DEL-126
chr6-81626747-DEL-57
chr6-81658557-DEL-62
chr6-81679760-DEL-219
chr6-81684614-DEL-111
chr6-81697058-DEL-57
chr6-81730793-DEL-57
chr6-81755346-DEL-559
chr6-81817677-DEL-60
chr6-81869826-DEL-124
chr6-81889042-DEL-56
chr6-82007974-DEL-58
chr6-82012129-DEL-174
chr6-82024515-DEL-434
chr6-82025654-DEL-74
chr6-82039510-DEL-55
chr6-82040494-DEL-63
chr6-82043172-DEL-163
chr6-82058012-DEL-128

chr6-82082675-DEL-124
chr6-82100607-DEL-237
chr6-82111248-DEL-534
chr6-82124805-DEL-790
chr6-82138691-DEL-6539
chr6-82170258-DEL-109
chr6-82181297-DEL-103
chr6-82206641-DEL-219
chr6-82208742-DEL-399
chr6-82245477-DEL-345
chr6-82292194-DEL-174
chr6-82312047-DEL-172
chr6-82313692-DEL-57
chr6-82320802-DEL-1513
chr6-82320802-DEL-2159
chr6-82323043-DEL-1113
chr6-82324216-DEL-3772
chr6-82345127-DEL-112
chr6-82346747-DEL-438
chr6-82355760-DEL-140
chr6-82370967-DEL-3547
chr6-82392107-DEL-78
chr6-82460303-DEL-182
chr6-82467717-DEL-60
chr6-82471803-DEL-1014
chr6-82496089-DEL-56
chr6-82559998-DEL-118
chr6-82571785-DEL-57
chr6-82578270-DEL-978
chr6-82602960-DEL-212
chr6-82635837-DEL-723
chr6-82651604-DEL-110
chr6-82683209-DEL-112
chr6-82684567-DEL-190
chr6-82686208-DEL-204
chr6-82689972-DEL-222
chr6-82695997-DEL-683
chr6-82736023-DEL-602
chr6-82756893-DEL-63
chr6-82786104-DEL-106
chr6-82806088-DEL-146
chr6-82809133-DEL-72
chr6-82810292-DEL-60
chr6-82813041-DEL-54
chr6-82836630-DEL-542
chr6-82844864-DEL-358
chr6-82853104-DEL-116
chr6-82866299-DEL-69
chr6-82878657-DEL-83
chr6-82884442-DEL-564
chr6-82891727-DEL-290
chr6-82900275-DEL-77
chr6-82917105-DEL-137
chr6-82929593-DEL-55
chr6-82931797-DEL-60
chr6-82954371-DEL-1168
chr6-82959708-DEL-100
chr6-82962962-DEL-145
chr6-82984097-DEL-322
chr6-83023658-DEL-52
chr6-83082002-DEL-373
chr6-83132855-DEL-52
chr6-83153942-DEL-302
chr6-83190597-DEL-253
chr6-83192931-DEL-1414
chr6-83193516-DEL-110
chr6-83194543-DEL-1324
chr6-83194560-DEL-515
chr6-83195673-DEL-126
chr6-83204745-DEL-52
chr6-83209031-DEL-52
chr6-83216482-DEL-57
chr6-83224972-DEL-98
chr6-83248382-DEL-187
chr6-83258190-DEL-179
chr6-83263563-DEL-399
chr6-83310327-DEL-71

chr6-83320881-DEL-544
chr6-83330164-DEL-78
chr6-83365967-DEL-85
chr6-83371869-DEL-193
chr6-83378244-DEL-308
chr6-83382210-DEL-71
chr6-83468284-DEL-64
chr6-83471667-DEL-92
chr6-83514989-DEL-51
chr6-83550238-DEL-424
chr6-83612446-DEL-110
chr6-83672629-DEL-139
chr6-83700983-DEL-66
chr6-83716160-DEL-207
chr6-83736144-DEL-60
chr6-83813644-DEL-203
chr6-83816231-DEL-52
chr6-83820753-DEL-204
chr6-83831218-DEL-158
chr6-83841231-DEL-835
chr6-83857550-DEL-1747
chr6-83880823-DEL-1290
chr6-83926302-DEL-62
chr6-83935921-DEL-64
chr6-83937440-DEL-220
chr6-83980206-DEL-202
chr6-83990271-DEL-853
chr6-84013014-DEL-198
chr6-84056519-DEL-140
chr6-84059954-DEL-108
chr6-84067263-DEL-84
chr6-84104267-DEL-177
chr6-84112471-DEL-269
chr6-84115877-DEL-127
chr6-84122340-DEL-51
chr6-84140318-DEL-104
chr6-84150095-DEL-254
chr6-84166798-DEL-141
chr6-84173991-DEL-118
chr6-84191638-DEL-155
chr6-84219177-DEL-52
chr6-84229926-DEL-71
chr6-84234111-DEL-111
chr6-84250821-DEL-545
chr6-84303431-DEL-120
chr6-84352083-DEL-234
chr6-84415268-DEL-754
chr6-84415776-DEL-250
chr6-84418503-DEL-100
chr6-84502962-DEL-1347
chr6-84589304-DEL-404
chr6-84616628-DEL-77
chr6-84767822-DEL-74
chr6-84790934-DEL-132
chr6-84799940-DEL-1919
chr6-84803099-DEL-3416
chr6-84836473-DEL-196
chr6-84874290-DEL-164
chr6-84898220-DEL-997
chr6-84931773-DEL-136
chr6-84933054-DEL-53
chr6-85021328-DEL-263
chr6-85028022-DEL-3263
chr6-85097104-DEL-488
chr6-85119906-DEL-107
chr6-85135693-DEL-401
chr6-85140437-DEL-97
chr6-85145064-DEL-67
chr6-85213076-DEL-99
chr6-85273684-DEL-626
chr6-85286278-DEL-72
chr6-85335650-DEL-1019
chr6-85336262-DEL-213
chr6-85360437-DEL-174
chr6-85371835-DEL-259
chr6-85373590-DEL-187
chr6-85390780-DEL-87

chr6-85454692-DEL-66
chr6-85465928-DEL-131
chr6-85479718-DEL-56
chr6-85517358-DEL-104
chr6-85522959-DEL-217
chr6-85544902-DEL-1038
chr6-85546553-DEL-631
chr6-85556252-DEL-198
chr6-85578977-DEL-363
chr6-85632434-DEL-66
chr6-85668974-DEL-796
chr6-85692960-DEL-397
chr6-85694694-DEL-2904
chr6-85697663-DEL-496
chr6-85698203-DEL-2952
chr6-85701463-DEL-518
chr6-85702350-DEL-542
chr6-85711131-DEL-5840
chr6-85741302-DEL-89
chr6-85745501-DEL-403
chr6-85770399-DEL-796
chr6-85793077-DEL-55
chr6-85814652-DEL-64
chr6-85815027-DEL-4830
chr6-85817833-DEL-136
chr6-85820309-DEL-55
chr6-85832984-DEL-70
chr6-85918507-DEL-486
chr6-85920302-DEL-863
chr6-85933037-DEL-66
chr6-85942745-DEL-466
chr6-85953222-DEL-4938
chr6-85974389-DEL-5715
chr6-86009504-DEL-697
chr6-86026084-DEL-55
chr6-86067994-DEL-80
chr6-86108923-DEL-177
chr6-86117477-DEL-132
chr6-86140864-DEL-106
chr6-86170459-DEL-59
chr6-86174444-DEL-108
chr6-86206780-DEL-144
chr6-86216695-DEL-76
chr6-86236613-DEL-481
chr6-86253653-DEL-1743
chr6-86258259-DEL-496
chr6-86264032-DEL-218
chr6-86284544-DEL-2716
chr6-86296675-DEL-1013
chr6-86302038-DEL-263
chr6-86320230-DEL-388
chr6-86322228-DEL-51
chr6-86331583-DEL-56
chr6-86390963-DEL-179
chr6-86451395-DEL-189
chr6-86459290-DEL-195
chr6-86472334-DEL-183
chr6-86477370-DEL-939
chr6-86479571-DEL-194
chr6-86496196-DEL-174
chr6-86506559-DEL-159
chr6-86523455-DEL-180
chr6-86533086-DEL-210
chr6-86541964-DEL-117
chr6-86570871-DEL-64
chr6-86580444-DEL-234
chr6-86615988-DEL-55
chr6-86617570-DEL-58
chr6-86695184-DEL-52
chr6-86734025-DEL-130
chr6-86742374-DEL-206
chr6-86763285-DEL-209
chr6-86775086-DEL-154
chr6-86836591-DEL-76
chr6-86838547-DEL-84
chr6-86852974-DEL-60
chr6-86858835-DEL-58

chr6-86868310-DEL-845
chr6-86884888-DEL-93
chr6-86903950-DEL-114
chr6-86934387-DEL-1136
chr6-87016490-DEL-132
chr6-87022665-DEL-2628
chr6-87027633-DEL-175
chr6-87088492-DEL-99
chr6-87114839-DEL-154
chr6-87141936-DEL-73
chr6-87148565-DEL-1370
chr6-87202083-DEL-56
chr6-87206141-DEL-56
chr6-87211734-DEL-81
chr6-87257980-DEL-143
chr6-87280839-DEL-110
chr6-87304680-DEL-184
chr6-87317974-DEL-60
chr6-87320797-DEL-266
chr6-87374021-DEL-111
chr6-87387263-DEL-170
chr6-87391092-DEL-101
chr6-87391969-DEL-83
chr6-87440764-DEL-90
chr6-87443668-DEL-62
chr6-87446867-DEL-506
chr6-87476476-DEL-146
chr6-87484599-DEL-186
chr6-87520664-DEL-455
chr6-87521559-DEL-55
chr6-87535313-DEL-140
chr6-87575096-DEL-58
chr6-87577218-DEL-165
chr6-87607253-DEL-58
chr6-87629712-DEL-56
chr6-87652973-DEL-1252
chr6-87750963-DEL-93
chr6-87752259-DEL-172
chr6-87763079-DEL-100
chr6-87777781-DEL-126
chr6-87792248-DEL-239
chr6-87805127-DEL-348
chr6-87832132-DEL-196
chr6-87839297-DEL-335
chr6-87862719-DEL-126
chr6-87897340-DEL-66
chr6-87913890-DEL-259
chr6-87917708-DEL-2716
chr6-87927553-DEL-200
chr6-87928368-DEL-184
chr6-87929487-DEL-380
chr6-87974225-DEL-211
chr6-87990991-DEL-55
chr6-87992470-DEL-1992
chr6-87998841-DEL-3653
chr6-88012211-DEL-64
chr6-88022790-DEL-1666
chr6-88028508-DEL-188
chr6-88043564-DEL-53
chr6-88047444-DEL-216
chr6-88100546-DEL-65
chr6-88118423-DEL-1761
chr6-88122288-DEL-255
chr6-88146111-DEL-57
chr6-88156495-DEL-71
chr6-88156774-DEL-969
chr6-88157963-DEL-739
chr6-88187864-DEL-221
chr6-88188369-DEL-96
chr6-88195561-DEL-65
chr6-88218146-DEL-134
chr6-88246363-DEL-53
chr6-88360654-DEL-69
chr6-88369188-DEL-64
chr6-88403324-DEL-81
chr6-88405690-DEL-213
chr6-88407769-DEL-67

chr6-88412790-DEL-149
chr6-88423864-DEL-169
chr6-88471608-DEL-435
chr6-88471695-DEL-744
chr6-88472096-DEL-335
chr6-88488997-DEL-64
chr6-88512576-DEL-70
chr6-88535605-DEL-172
chr6-88573437-DEL-96
chr6-88609342-DEL-502
chr6-88622160-DEL-68
chr6-88630670-DEL-1583
chr6-88646679-DEL-1927
chr6-88646679-DEL-2550
chr6-88648677-DEL-3720
chr6-88649271-DEL-3106
chr6-88651106-DEL-1268
chr6-88668903-DEL-69
chr6-88673188-DEL-211
chr6-88725848-DEL-750
chr6-88740143-DEL-140
chr6-88753427-DEL-84
chr6-88775171-DEL-516
chr6-88832591-DEL-163
chr6-88839449-DEL-201
chr6-88845845-DEL-282
chr6-88875021-DEL-53
chr6-88891686-DEL-633
chr6-88904406-DEL-121
chr6-88962771-DEL-464
chr6-88981418-DEL-60
chr6-89004084-DEL-174
chr6-89018839-DEL-200
chr6-89031858-DEL-55
chr6-89056997-DEL-209
chr6-89063151-DEL-120
chr6-89103787-DEL-85
chr6-89104392-DEL-289
chr6-89157152-DEL-496
chr6-89165729-DEL-486
chr6-89167637-DEL-957
chr6-89174122-DEL-485
chr6-89183131-DEL-189
chr6-89198444-DEL-58
chr6-89206892-DEL-360
chr6-89256454-DEL-246
chr6-89385183-DEL-54
chr6-89475809-DEL-59
chr6-89499784-DEL-88
chr6-89579667-DEL-1656
chr6-89589926-DEL-84
chr6-89595760-DEL-1611
chr6-89630243-DEL-53
chr6-89635214-DEL-334
chr6-89663354-DEL-402
chr6-89687964-DEL-318
chr6-89700083-DEL-207
chr6-89708975-DEL-1569
chr6-89728573-DEL-66
chr6-89730856-DEL-418
chr6-89732027-DEL-8799
chr6-89748248-DEL-306
chr6-89755108-DEL-665
chr6-89759701-DEL-1148
chr6-89759741-DEL-6218
chr6-89800100-DEL-13310
chr6-89806038-DEL-1948
chr6-89812449-DEL-391
chr6-89842288-DEL-823
chr6-89855796-DEL-371
chr6-89880408-DEL-2303
chr6-89888684-DEL-6315
chr6-90020323-DEL-1228
chr6-90045591-DEL-170
chr6-90058111-DEL-155
chr6-90060153-DEL-1319
chr6-90066070-DEL-8290

chr6-90083382-DEL-58
chr6-90117210-DEL-1665
chr6-90148047-DEL-132
chr6-90188142-DEL-4868
chr6-90220511-DEL-68
chr6-90268183-DEL-820
chr6-90269529-DEL-3653
chr6-90303198-DEL-329
chr6-90342644-DEL-62
chr6-90395386-DEL-207
chr6-90412984-DEL-316
chr6-90478119-DEL-57
chr6-90492280-DEL-277
chr6-90493905-DEL-125
chr6-90494492-DEL-1160
chr6-90494789-DEL-356
chr6-90495446-DEL-332
chr6-90506325-DEL-71
chr6-90512435-DEL-54
chr6-90522747-DEL-88
chr6-90550676-DEL-504
chr6-90555939-DEL-62
chr6-90570777-DEL-58
chr6-90577391-DEL-112
chr6-90824200-DEL-65
chr6-90860763-DEL-56
chr6-90882533-DEL-1564
chr6-90942336-DEL-384
chr6-90985321-DEL-147
chr6-91006936-DEL-79
chr6-91009472-DEL-62
chr6-91021114-DEL-220
chr6-91040572-DEL-67
chr6-91041027-DEL-61
chr6-91118327-DEL-180
chr6-91152124-DEL-62
chr6-91189294-DEL-68
chr6-91250339-DEL-111
chr6-91256610-DEL-79
chr6-91257131-DEL-558
chr6-91257881-DEL-109
chr6-91339669-DEL-177
chr6-91349224-DEL-78
chr6-91355214-DEL-1016
chr6-91368522-DEL-68
chr6-91375915-DEL-65
chr6-91412140-DEL-118
chr6-91421948-DEL-1170
chr6-91433913-DEL-354
chr6-91439105-DEL-651
chr6-91473608-DEL-399
chr6-91543728-DEL-216
chr6-91556231-DEL-60
chr6-91564664-DEL-52
chr6-91619028-DEL-64
chr6-91643865-DEL-208
chr6-91748363-DEL-64
chr6-91803108-DEL-644
chr6-91803827-DEL-66
chr6-91807331-DEL-56
chr6-91833122-DEL-249
chr6-91862490-DEL-334
chr6-91873205-DEL-52
chr6-91886831-DEL-143
chr6-91890523-DEL-52
chr6-91930905-DEL-197
chr6-91935344-DEL-116
chr6-91954655-DEL-224
chr6-91976583-DEL-317
chr6-92006808-DEL-146
chr6-92024062-DEL-56
chr6-92026348-DEL-254
chr6-92034367-DEL-54
chr6-92097778-DEL-155
chr6-92105614-DEL-1114
chr6-92107030-DEL-750
chr6-92164711-DEL-130

chr6-92248100-DEL-62
chr6-92252693-DEL-1960
chr6-92337456-DEL-246
chr6-92346218-DEL-59
chr6-92369782-DEL-549
chr6-92414309-DEL-267
chr6-92417661-DEL-646
chr6-92426607-DEL-84
chr6-92434076-DEL-142
chr6-92477748-DEL-2699
chr6-92505656-DEL-761
chr6-92541880-DEL-1345
chr6-92573845-DEL-97
chr6-92610625-DEL-396
chr6-92742681-DEL-79
chr6-92751728-DEL-70
chr6-92767210-DEL-169
chr6-92803314-DEL-118
chr6-92832785-DEL-62
chr6-92853856-DEL-1365
chr6-92862711-DEL-227
chr6-92885342-DEL-66
chr6-92927501-DEL-70
chr6-92951432-DEL-97
chr6-92994667-DEL-57
chr6-93007045-DEL-59
chr6-93041012-DEL-188
chr6-93089829-DEL-110
chr6-93118377-DEL-97
chr6-93122880-DEL-1203
chr6-93172451-DEL-334
chr6-93183238-DEL-107
chr6-93219683-DEL-1106
chr6-93230101-DEL-58
chr6-93265221-DEL-60
chr6-93269791-DEL-58
chr6-93341080-DEL-296
chr6-93381555-DEL-4242
chr6-93424174-DEL-334
chr6-93436191-DEL-213
chr6-93466602-DEL-1425
chr6-93495830-DEL-77
chr6-93530500-DEL-356
chr6-93559759-DEL-179
chr6-93564482-DEL-1171
chr6-93580469-DEL-74
chr6-93582335-DEL-827
chr6-93614648-DEL-185
chr6-93621730-DEL-802
chr6-93718863-DEL-63
chr6-93729879-DEL-318
chr6-93742671-DEL-142
chr6-93755489-DEL-56
chr6-93759896-DEL-108
chr6-93775073-DEL-252
chr6-93795591-DEL-85
chr6-93801108-DEL-114
chr6-93856995-DEL-2674
chr6-93877051-DEL-67
chr6-93933364-DEL-290
chr6-93982067-DEL-896
chr6-94000196-DEL-87
chr6-94031808-DEL-96
chr6-94060209-DEL-1079
chr6-94098857-DEL-68
chr6-94109617-DEL-52
chr6-94149706-DEL-255
chr6-94165066-DEL-1143
chr6-94193223-DEL-61
chr6-94206933-DEL-80
chr6-94209704-DEL-146
chr6-94212036-DEL-76
chr6-94215656-DEL-113
chr6-94231108-DEL-63
chr6-94243768-DEL-64
chr6-94269206-DEL-665
chr6-94282748-DEL-202

chr6-94297931-DEL-57
chr6-94298569-DEL-79
chr6-94301069-DEL-305
chr6-94311961-DEL-175
chr6-94318043-DEL-206
chr6-94319156-DEL-492
chr6-94372799-DEL-169
chr6-94379826-DEL-62
chr6-94388051-DEL-796
chr6-94392415-DEL-186
chr6-94393025-DEL-1145
chr6-94464098-DEL-345
chr6-94466343-DEL-4719
chr6-94482868-DEL-625
chr6-94495334-DEL-92
chr6-94557462-DEL-96
chr6-94563846-DEL-106
chr6-94588164-DEL-176
chr6-94712708-DEL-66
chr6-94747787-DEL-94
chr6-94779624-DEL-178
chr6-94801733-DEL-285
chr6-94802849-DEL-431
chr6-94817287-DEL-69
chr6-94873861-DEL-408
chr6-94880486-DEL-364
chr6-94890762-DEL-52
chr6-94903263-DEL-442
chr6-94933274-DEL-54
chr6-94936649-DEL-66
chr6-94957958-DEL-74
chr6-94965727-DEL-1898
chr6-94972373-DEL-170
chr6-94993743-DEL-51
chr6-95006648-DEL-6410
chr6-95006742-DEL-895
chr6-95008091-DEL-4822
chr6-95013092-DEL-57
chr6-95030207-DEL-110
chr6-95032872-DEL-104
chr6-95034038-DEL-241
chr6-95038845-DEL-335
chr6-95087608-DEL-158
chr6-95100952-DEL-320
chr6-95109920-DEL-1344
chr6-95182812-DEL-251
chr6-95185264-DEL-182
chr6-95216591-DEL-214
chr6-95244050-DEL-71
chr6-95258089-DEL-54
chr6-95277457-DEL-60
chr6-95361644-DEL-60
chr6-95386324-DEL-134
chr6-95386727-DEL-6417
chr6-95443952-DEL-726
chr6-95473536-DEL-71
chr6-95475869-DEL-121
chr6-95491835-DEL-97
chr6-95519491-DEL-112
chr6-95587456-DEL-1194
chr6-95607488-DEL-56
chr6-95632267-DEL-70
chr6-95668153-DEL-52
chr6-95677635-DEL-52
chr6-95701448-DEL-123
chr6-95730656-DEL-1391
chr6-95765858-DEL-199
chr6-95817776-DEL-212
chr6-95819985-DEL-220
chr6-95829974-DEL-315
chr6-95837419-DEL-6340
chr6-95869979-DEL-51
chr6-95885004-DEL-71
chr6-95921455-DEL-227
chr6-95935475-DEL-68
chr6-95936371-DEL-346
chr6-95938525-DEL-62

chr6-95957687-DEL-214
chr6-95976125-DEL-64
chr6-96009825-DEL-522
chr6-96095025-DEL-107
chr6-96147884-DEL-70
chr6-96170387-DEL-1431
chr6-96208305-DEL-106
chr6-96283160-DEL-312
chr6-96296765-DEL-280
chr6-96320206-DEL-94
chr6-96329606-DEL-1141
chr6-96384831-DEL-440
chr6-96393189-DEL-145
chr6-96399998-DEL-66
chr6-96443321-DEL-53
chr6-96462753-DEL-68
chr6-96488565-DEL-74
chr6-96584661-DEL-1380
chr6-96593090-DEL-211
chr6-96593817-DEL-6419
chr6-96665064-DEL-1076
chr6-96692786-DEL-96
chr6-96696242-DEL-58
chr6-96752911-DEL-8157
chr6-96752984-DEL-1984
chr6-96754203-DEL-1983
chr6-96763900-DEL-152
chr6-96791407-DEL-656
chr6-96793133-DEL-353
chr6-96797660-DEL-2349
chr6-96815066-DEL-53
chr6-96830579-DEL-279
chr6-96846074-DEL-60
chr6-96869999-DEL-3193
chr6-96898875-DEL-6402
chr6-96900565-DEL-3778
chr6-96905281-DEL-310
chr6-96923922-DEL-173
chr6-96933111-DEL-1431
chr6-96987263-DEL-562
chr6-96987980-DEL-364
chr6-97058323-DEL-54
chr6-97063392-DEL-4315
chr6-97064700-DEL-3015
chr6-97067077-DEL-360
chr6-97081753-DEL-1959
chr6-97084487-DEL-1148
chr6-97094942-DEL-88
chr6-97097294-DEL-66
chr6-97098389-DEL-139
chr6-97139229-DEL-74
chr6-97144775-DEL-60
chr6-97194580-DEL-51
chr6-97204835-DEL-174
chr6-97210353-DEL-944
chr6-97217899-DEL-1924
chr6-97221850-DEL-679
chr6-97249101-DEL-54
chr6-97251842-DEL-83
chr6-97267058-DEL-1557
chr6-97297445-DEL-121
chr6-97314522-DEL-730
chr6-97320288-DEL-209
chr6-97362537-DEL-52
chr6-97387556-DEL-56
chr6-97393471-DEL-112
chr6-97491105-DEL-560
chr6-97496053-DEL-214
chr6-97513543-DEL-228
chr6-97516562-DEL-466
chr6-97520894-DEL-622
chr6-97535265-DEL-68
chr6-97545107-DEL-1133
chr6-97562130-DEL-853
chr6-97595246-DEL-63
chr6-97639753-DEL-935
chr6-97640708-DEL-1763

chr6-97642253-DEL-217
chr6-97684982-DEL-86
chr6-97690337-DEL-199
chr6-97703828-DEL-262
chr6-97711259-DEL-70
chr6-97777077-DEL-192
chr6-97784935-DEL-80
chr6-97828363-DEL-1074
chr6-97899405-DEL-171
chr6-97974770-DEL-448
chr6-98004756-DEL-157
chr6-98065471-DEL-281
chr6-98092352-DEL-183
chr6-98137881-DEL-2037
chr6-98140459-DEL-408
chr6-98159097-DEL-2728
chr6-98166551-DEL-55
chr6-98196688-DEL-61
chr6-98205257-DEL-68
chr6-98246300-DEL-84
chr6-98254023-DEL-2280
chr6-98302398-DEL-135
chr6-98304763-DEL-64
chr6-98309574-DEL-137
chr6-98327628-DEL-62
chr6-98333273-DEL-54
chr6-98364324-DEL-722
chr6-98366840-DEL-308
chr6-98371227-DEL-54
chr6-98451970-DEL-358
chr6-98521684-DEL-958
chr6-98524376-DEL-445
chr6-98616741-DEL-101
chr6-98618105-DEL-177
chr6-98620809-DEL-232
chr6-98710347-DEL-143
chr6-98714111-DEL-1030
chr6-98739481-DEL-64
chr6-98740147-DEL-1681
chr6-98782927-DEL-51
chr6-98787652-DEL-368
chr6-98813558-DEL-378
chr6-98814352-DEL-225
chr6-98858331-DEL-5980
chr6-98887363-DEL-2620
chr6-98902146-DEL-54
chr6-98942670-DEL-93
chr6-99014260-DEL-198
chr6-99018021-DEL-1107
chr6-99038850-DEL-84
chr6-99039807-DEL-112
chr6-99145326-DEL-174
chr6-99156351-DEL-68
chr6-99208125-DEL-216
chr6-99216265-DEL-130
chr6-99219113-DEL-157
chr6-99250506-DEL-54
chr6-99250545-DEL-791
chr6-99270847-DEL-224
chr6-99280744-DEL-224
chr6-99290248-DEL-348
chr6-99292155-DEL-111
chr6-99295262-DEL-213
chr6-99303920-DEL-170
chr6-99314046-DEL-100
chr6-99319972-DEL-142
chr6-99333551-DEL-374
chr6-99403896-DEL-67
chr6-99408534-DEL-245
chr6-99414923-DEL-155
chr6-99508600-DEL-66
chr6-99514555-DEL-81
chr6-99526061-DEL-58
chr6-99538941-DEL-129
chr6-99548810-DEL-65
chr6-99607240-DEL-80
chr6-99611765-DEL-87

chr6-99619008-DEL-64
chr6-99635123-DEL-446
chr6-99682486-DEL-71
chr6-99695530-DEL-262
chr6-99708110-DEL-75
chr6-99782865-DEL-168
chr6-99807772-DEL-277
chr6-99828175-DEL-87
chr6-99835560-DEL-106
chr6-99859943-DEL-310
chr6-99884098-DEL-1549
chr6-99907579-DEL-344
chr6-99925043-DEL-514
chr6-99930603-DEL-58
chr6-99938478-DEL-64
chr6-99966379-DEL-202
chr6-99986161-DEL-72
chr6-99993186-DEL-56
chr6-99999784-DEL-130
chr6-100001274-DEL-58
chr6-100012875-DEL-4934
chr6-100038027-DEL-64
chr6-100098946-DEL-210
chr6-100138037-DEL-193
chr6-100145904-DEL-181
chr6-100170319-DEL-337
chr6-100181973-DEL-131
chr6-100187198-DEL-59
chr6-100212016-DEL-253
chr6-100223783-DEL-427
chr6-100274902-DEL-60
chr6-100280360-DEL-129
chr6-100287870-DEL-66
chr6-100394378-DEL-64
chr6-100400960-DEL-126
chr6-100441474-DEL-431
chr6-100442391-DEL-96
chr6-100444753-DEL-113
chr6-100445183-DEL-102
chr6-100483661-DEL-58
chr6-100507544-DEL-68
chr6-100523570-DEL-200
chr6-100612107-DEL-116
chr6-100617080-DEL-88
chr6-100643246-DEL-112
chr6-100653546-DEL-256
chr6-100700064-DEL-74
chr6-100706476-DEL-55
chr6-100713731-DEL-3200
chr6-100721794-DEL-69
chr6-100724863-DEL-59
chr6-100725973-DEL-60
chr6-100734656-DEL-424
chr6-100746877-DEL-2582
chr6-100760786-DEL-1117
chr6-100783038-DEL-52
chr6-100821221-DEL-164
chr6-100864900-DEL-327
chr6-100868790-DEL-111
chr6-100871265-DEL-476
chr6-100895377-DEL-55
chr6-100956782-DEL-309
chr6-101027107-DEL-75
chr6-101027930-DEL-2137
chr6-101037949-DEL-1255
chr6-101050164-DEL-88
chr6-101052231-DEL-127
chr6-101069000-DEL-56
chr6-101076603-DEL-113
chr6-101135076-DEL-72
chr6-101155441-DEL-274
chr6-101156187-DEL-311
chr6-101230432-DEL-55
chr6-101274849-DEL-337
chr6-101278390-DEL-612
chr6-101301286-DEL-223
chr6-101352980-DEL-55

chr6-101415956-DEL-1164
chr6-101434995-DEL-63
chr6-101448134-DEL-71
chr6-101462332-DEL-501
chr6-101493329-DEL-546
chr6-101547455-DEL-72
chr6-101548065-DEL-6155
chr6-101562749-DEL-6948
chr6-101574444-DEL-3031
chr6-101585824-DEL-96
chr6-101595657-DEL-53
chr6-101689180-DEL-744
chr6-101714986-DEL-974
chr6-101716866-DEL-83
chr6-101717310-DEL-6199
chr6-101731816-DEL-506
chr6-101732314-DEL-6545
chr6-101732319-DEL-2511
chr6-101732560-DEL-215
chr6-101734866-DEL-2113
chr6-101736996-DEL-1864
chr6-101765741-DEL-203
chr6-101783829-DEL-52
chr6-101831759-DEL-108
chr6-101834229-DEL-99
chr6-101841475-DEL-354
chr6-101888345-DEL-54
chr6-101906705-DEL-437
chr6-101911379-DEL-96
chr6-101913109-DEL-57
chr6-101914251-DEL-216
chr6-101951956-DEL-575
chr6-101956781-DEL-127
chr6-101960788-DEL-75
chr6-101963324-DEL-2122
chr6-101987660-DEL-1004
chr6-101993915-DEL-593
chr6-101997984-DEL-106
chr6-102045908-DEL-508
chr6-102067972-DEL-1078
chr6-102077291-DEL-60
chr6-102168615-DEL-105
chr6-102191233-DEL-231
chr6-102196162-DEL-1878
chr6-102248074-DEL-1244
chr6-102259045-DEL-76
chr6-102261837-DEL-55
chr6-102330541-DEL-311
chr6-102346038-DEL-210
chr6-102396334-DEL-159
chr6-102397564-DEL-1041
chr6-102497271-DEL-140
chr6-102506045-DEL-64
chr6-102512019-DEL-106
chr6-102558667-DEL-465
chr6-102568400-DEL-90
chr6-102578739-DEL-1286
chr6-102586997-DEL-2995
chr6-102638475-DEL-1614
chr6-102641777-DEL-648
chr6-102643890-DEL-6599
chr6-102656149-DEL-310
chr6-102660919-DEL-533
chr6-102661181-DEL-6661
chr6-102661733-DEL-150
chr6-102686445-DEL-108
chr6-102701752-DEL-68
chr6-102708154-DEL-100
chr6-102710377-DEL-992
chr6-102754996-DEL-987
chr6-102755814-DEL-170
chr6-102758941-DEL-9575
chr6-102761725-DEL-3536
chr6-102827282-DEL-3077
chr6-102831384-DEL-58
chr6-102881622-DEL-54
chr6-102883734-DEL-476

chr6-102885622-DEL-54
chr6-102886530-DEL-54
chr6-102891202-DEL-141
chr6-102895936-DEL-11185
chr6-102934673-DEL-66
chr6-102972083-DEL-480
chr6-102977070-DEL-1392
chr6-102987327-DEL-58
chr6-103028949-DEL-524
chr6-103123057-DEL-104
chr6-103130348-DEL-1377
chr6-103132078-DEL-66
chr6-103188589-DEL-390
chr6-103217119-DEL-229
chr6-103262331-DEL-1214
chr6-103295378-DEL-363
chr6-103320077-DEL-95
chr6-103356343-DEL-68
chr6-103393367-DEL-76
chr6-103402300-DEL-62
chr6-103417520-DEL-69
chr6-103436499-DEL-588
chr6-103452490-DEL-2098
chr6-103454795-DEL-6224
chr6-103479485-DEL-268
chr6-103481857-DEL-439
chr6-103500451-DEL-62
chr6-103509499-DEL-1454
chr6-103558436-DEL-214
chr6-103564687-DEL-500
chr6-103574045-DEL-1160
chr6-103582860-DEL-490
chr6-103602205-DEL-257
chr6-103629222-DEL-1070
chr6-103630287-DEL-132
chr6-103680208-DEL-475
chr6-103725837-DEL-52
chr6-103727624-DEL-1226
chr6-103774643-DEL-62
chr6-103782745-DEL-73
chr6-103797479-DEL-779
chr6-103816809-DEL-66
chr6-103829541-DEL-183
chr6-103834356-DEL-90
chr6-103925863-DEL-1033
chr6-103947241-DEL-1258
chr6-103957241-DEL-95
chr6-103962689-DEL-75
chr6-103968103-DEL-63
chr6-103974617-DEL-85
chr6-103995417-DEL-659
chr6-104010087-DEL-514
chr6-104020003-DEL-65
chr6-104026759-DEL-59
chr6-104028437-DEL-94
chr6-104038533-DEL-245
chr6-104069757-DEL-1279
chr6-104083131-DEL-1030
chr6-104096170-DEL-181
chr6-104101007-DEL-56
chr6-104106539-DEL-72
chr6-104143027-DEL-3765
chr6-104150376-DEL-353
chr6-104152059-DEL-60
chr6-104158959-DEL-64
chr6-104210662-DEL-90
chr6-104217066-DEL-697
chr6-104228801-DEL-188
chr6-104256562-DEL-85
chr6-104281091-DEL-325
chr6-104297947-DEL-854
chr6-104353385-DEL-72
chr6-104362097-DEL-212
chr6-104377678-DEL-1423
chr6-104383987-DEL-262
chr6-104404562-DEL-57
chr6-104405548-DEL-51

chr6-104415403-DEL-469
chr6-104420115-DEL-75
chr6-104443755-DEL-945
chr6-104475283-DEL-110
chr6-104492361-DEL-3294
chr6-104510281-DEL-132
chr6-104535744-DEL-831
chr6-104610012-DEL-337
chr6-104650060-DEL-175
chr6-104654197-DEL-852
chr6-104696262-DEL-1758
chr6-104722159-DEL-913
chr6-104733760-DEL-109
chr6-104757512-DEL-75
chr6-104783862-DEL-74
chr6-104791591-DEL-142
chr6-104807620-DEL-371
chr6-104808831-DEL-188
chr6-104821315-DEL-184
chr6-104825720-DEL-1298
chr6-104828111-DEL-105
chr6-104839110-DEL-298
chr6-104872300-DEL-74
chr6-104875113-DEL-2536
chr6-104876215-DEL-90
chr6-104883496-DEL-1230
chr6-104889167-DEL-446
chr6-104895454-DEL-993
chr6-104911871-DEL-491
chr6-104930760-DEL-1294
chr6-104931101-DEL-2145
chr6-104932419-DEL-853
chr6-104937029-DEL-197
chr6-104937880-DEL-198
chr6-104938506-DEL-212
chr6-104939292-DEL-99
chr6-104957494-DEL-102
chr6-104958159-DEL-85
chr6-105048931-DEL-318
chr6-105058662-DEL-173
chr6-105084805-DEL-54
chr6-105085079-DEL-411
chr6-105087999-DEL-256
chr6-105093771-DEL-195
chr6-105102293-DEL-96
chr6-105114287-DEL-58
chr6-105117951-DEL-218
chr6-105125784-DEL-1165
chr6-105126655-DEL-252
chr6-105138295-DEL-54
chr6-105143775-DEL-216
chr6-105152882-DEL-573
chr6-105197902-DEL-1409
chr6-105205440-DEL-182
chr6-105207957-DEL-554
chr6-105244750-DEL-3531
chr6-105266221-DEL-945
chr6-105272191-DEL-252
chr6-105291474-DEL-6504
chr6-105312035-DEL-6198
chr6-105312444-DEL-1792
chr6-105323656-DEL-80
chr6-105333695-DEL-170
chr6-105351307-DEL-184
chr6-105376931-DEL-68
chr6-105385508-DEL-127
chr6-105398159-DEL-6364
chr6-105436730-DEL-425
chr6-105437858-DEL-79
chr6-105438238-DEL-7690
chr6-105448804-DEL-375
chr6-105455269-DEL-249
chr6-105457724-DEL-69
chr6-105473567-DEL-66
chr6-105492242-DEL-55
chr6-105493290-DEL-55
chr6-105499443-DEL-207

chr6-105506425-DEL-1698
chr6-105513059-DEL-54
chr6-105523128-DEL-174
chr6-105539654-DEL-114
chr6-105544552-DEL-277
chr6-105560839-DEL-66
chr6-105597107-DEL-4830
chr6-105601973-DEL-1512
chr6-105639962-DEL-411
chr6-105666347-DEL-166
chr6-105696166-DEL-142
chr6-105696535-DEL-1896
chr6-105709027-DEL-179
chr6-105761978-DEL-515
chr6-105772493-DEL-212
chr6-105778186-DEL-70
chr6-105822135-DEL-56
chr6-105827014-DEL-387
chr6-105837674-DEL-867
chr6-105842085-DEL-54
chr6-105853457-DEL-120
chr6-105861772-DEL-4744
chr6-105882781-DEL-51
chr6-105898030-DEL-1848
chr6-105926846-DEL-579
chr6-105974793-DEL-70
chr6-105984971-DEL-1259
chr6-106004963-DEL-51
chr6-106026471-DEL-108
chr6-106046395-DEL-56
chr6-106076107-DEL-422
chr6-106077942-DEL-1466
chr6-106080080-DEL-55
chr6-106084206-DEL-278
chr6-106096965-DEL-59
chr6-106115839-DEL-722
chr6-106156681-DEL-137
chr6-106177269-DEL-95
chr6-106179435-DEL-3744
chr6-106181531-DEL-137
chr6-106193126-DEL-52
chr6-106199795-DEL-225
chr6-106249632-DEL-100
chr6-106250389-DEL-54
chr6-106250849-DEL-620
chr6-106299794-DEL-64
chr6-106326574-DEL-58
chr6-106337272-DEL-97
chr6-106343407-DEL-1015
chr6-106343985-DEL-83
chr6-106414199-DEL-58
chr6-106501461-DEL-170
chr6-106502765-DEL-124
chr6-106530505-DEL-373
chr6-106530537-DEL-176
chr6-106531196-DEL-153
chr6-106531232-DEL-590
chr6-106531723-DEL-198
chr6-106591076-DEL-299
chr6-106625919-DEL-1191
chr6-106733708-DEL-63
chr6-106791827-DEL-149
chr6-106800453-DEL-72
chr6-106811044-DEL-1148
chr6-106818738-DEL-66
chr6-106849733-DEL-120
chr6-106857186-DEL-83
chr6-106894951-DEL-79
chr6-106903377-DEL-96
chr6-106974364-DEL-172
chr6-106982395-DEL-1167
chr6-107036672-DEL-3027
chr6-107073218-DEL-7272
chr6-107128772-DEL-264
chr6-107136786-DEL-137
chr6-107188448-DEL-51
chr6-107222483-DEL-4721

chr6-107223151-DEL-6887
chr6-107238529-DEL-827
chr6-107253831-DEL-587
chr6-107261195-DEL-6227
chr6-107271110-DEL-366
chr6-107271477-DEL-6625
chr6-107325409-DEL-69
chr6-107330338-DEL-1429
chr6-107331656-DEL-146
chr6-107354347-DEL-81
chr6-107378133-DEL-629
chr6-107389762-DEL-228
chr6-107391766-DEL-298
chr6-107433524-DEL-123
chr6-107443780-DEL-194
chr6-107454456-DEL-208
chr6-107457020-DEL-59
chr6-107478764-DEL-198
chr6-107529381-DEL-67
chr6-107550288-DEL-92
chr6-107560438-DEL-381
chr6-107636748-DEL-566
chr6-107647760-DEL-6890
chr6-107654170-DEL-212
chr6-107677393-DEL-74
chr6-107755267-DEL-504
chr6-107757039-DEL-75
chr6-107780190-DEL-52
chr6-107786704-DEL-81
chr6-107789211-DEL-414
chr6-107807419-DEL-6138
chr6-107833435-DEL-6249
chr6-107847018-DEL-310
chr6-107858627-DEL-6285
chr6-107868248-DEL-67
chr6-107879627-DEL-136
chr6-107880554-DEL-6221
chr6-107891089-DEL-6439
chr6-107916972-DEL-527
chr6-107917080-DEL-5065
chr6-107918945-DEL-120
chr6-107952262-DEL-754
chr6-107955506-DEL-182
chr6-107963965-DEL-6678
chr6-107974426-DEL-349
chr6-108000577-DEL-52
chr6-108007189-DEL-159
chr6-108007980-DEL-1969
chr6-108011391-DEL-2466
chr6-108070197-DEL-150
chr6-108093939-DEL-166
chr6-108095849-DEL-776
chr6-108125075-DEL-54
chr6-108148241-DEL-208
chr6-108181971-DEL-60
chr6-108234456-DEL-216
chr6-108237187-DEL-926
chr6-108313311-DEL-110
chr6-108329847-DEL-56
chr6-108331014-DEL-54
chr6-108441978-DEL-53
chr6-108462560-DEL-165
chr6-108485157-DEL-67
chr6-108500775-DEL-929
chr6-108511638-DEL-62
chr6-108578279-DEL-201
chr6-108594429-DEL-190
chr6-108598125-DEL-344
chr6-108624588-DEL-252
chr6-108629674-DEL-608
chr6-108648600-DEL-84
chr6-108650476-DEL-107
chr6-108651527-DEL-117
chr6-108692105-DEL-2214
chr6-108693850-DEL-57
chr6-108707522-DEL-88
chr6-108736621-DEL-248

chr6-108738939-DEL-456
chr6-108745334-DEL-119
chr6-108763634-DEL-384
chr6-108785811-DEL-61
chr6-108823956-DEL-56
chr6-108832035-DEL-58
chr6-108845698-DEL-72
chr6-108846775-DEL-56
chr6-108853423-DEL-57
chr6-108857954-DEL-54
chr6-108907810-DEL-3753
chr6-108949381-DEL-1790
chr6-108972896-DEL-277
chr6-109029932-DEL-1008
chr6-109032172-DEL-56
chr6-109043973-DEL-6556
chr6-109069397-DEL-6781
chr6-109084828-DEL-133
chr6-109142257-DEL-154
chr6-109181991-DEL-53
chr6-109238166-DEL-92
chr6-109240388-DEL-191
chr6-109243733-DEL-107
chr6-109324785-DEL-163
chr6-109334327-DEL-10540
chr6-109353155-DEL-1430
chr6-109371574-DEL-106
chr6-109379058-DEL-205
chr6-109382582-DEL-52
chr6-109390817-DEL-110
chr6-109398630-DEL-6388
chr6-109427829-DEL-1881
chr6-109469776-DEL-617
chr6-109494323-DEL-236
chr6-109581722-DEL-108
chr6-109604716-DEL-55
chr6-109608208-DEL-330
chr6-109612015-DEL-81
chr6-109639973-DEL-1111
chr6-109683245-DEL-7112
chr6-109742096-DEL-994
chr6-109748368-DEL-3042
chr6-109756808-DEL-52
chr6-109804283-DEL-125
chr6-109843111-DEL-52
chr6-109854756-DEL-614
chr6-109865360-DEL-52
chr6-109897522-DEL-347
chr6-109917617-DEL-971
chr6-110015277-DEL-52
chr6-110024811-DEL-260
chr6-110026689-DEL-2261
chr6-110033324-DEL-127
chr6-110043973-DEL-56
chr6-110069816-DEL-562
chr6-110078392-DEL-109
chr6-110079289-DEL-903
chr6-110084382-DEL-51
chr6-110086982-DEL-66
chr6-110151203-DEL-62
chr6-110178152-DEL-820
chr6-110189758-DEL-3683
chr6-110230947-DEL-432
chr6-110231704-DEL-309
chr6-110233147-DEL-590
chr6-110272511-DEL-87
chr6-110282275-DEL-1293
chr6-110294390-DEL-205
chr6-110320175-DEL-72
chr6-110330899-DEL-66
chr6-110333171-DEL-57
chr6-110360103-DEL-56
chr6-110374097-DEL-84
chr6-110377625-DEL-127
chr6-110403030-DEL-94
chr6-110421695-DEL-188
chr6-110490610-DEL-427

chr6-110501499-DEL-440
chr6-110513668-DEL-1587
chr6-110541186-DEL-74
chr6-110565191-DEL-278
chr6-110566108-DEL-270
chr6-110567313-DEL-1483
chr6-110572242-DEL-178
chr6-110591220-DEL-549
chr6-110593153-DEL-234
chr6-110603692-DEL-341
chr6-110609129-DEL-396
chr6-110659771-DEL-68
chr6-110660953-DEL-52
chr6-110664986-DEL-78
chr6-110665861-DEL-65
chr6-110678286-DEL-590
chr6-110717546-DEL-179
chr6-110729279-DEL-51
chr6-110730275-DEL-52
chr6-110747270-DEL-83
chr6-110761344-DEL-508
chr6-110785780-DEL-156
chr6-110796687-DEL-58
chr6-110812890-DEL-351
chr6-110817004-DEL-308
chr6-110817555-DEL-835
chr6-110821691-DEL-72
chr6-110829415-DEL-368
chr6-110850779-DEL-1424
chr6-110875938-DEL-51
chr6-110881637-DEL-1386
chr6-110886327-DEL-74
chr6-110934097-DEL-116
chr6-110963070-DEL-342
chr6-110988116-DEL-59
chr6-110989993-DEL-177
chr6-111032005-DEL-604
chr6-111047712-DEL-4923
chr6-111047803-DEL-195
chr6-111048044-DEL-2950
chr6-111051019-DEL-1614
chr6-111061147-DEL-3237
chr6-111095807-DEL-60
chr6-111109504-DEL-69
chr6-111116146-DEL-230
chr6-111130670-DEL-7272
chr6-111151619-DEL-817
chr6-111165855-DEL-373
chr6-111173021-DEL-1410
chr6-111225795-DEL-172
chr6-111226936-DEL-53
chr6-111237864-DEL-147
chr6-111239915-DEL-562
chr6-111273231-DEL-58
chr6-111304452-DEL-50
chr6-111328784-DEL-107
chr6-111359945-DEL-5317
chr6-111368313-DEL-6131
chr6-111382774-DEL-56
chr6-111392480-DEL-6415
chr6-111398736-DEL-346
chr6-111399376-DEL-52
chr6-111416503-DEL-239
chr6-111422374-DEL-1724
chr6-111459964-DEL-191
chr6-111482657-DEL-105
chr6-111530755-DEL-129
chr6-111531569-DEL-92
chr6-111586837-DEL-59
chr6-111631071-DEL-64
chr6-111643192-DEL-11859
chr6-111644872-DEL-8838
chr6-111667778-DEL-2450
chr6-111669670-DEL-575
chr6-111684104-DEL-167
chr6-111687342-DEL-4051
chr6-111699299-DEL-951

chr6-111736785-DEL-60
chr6-111752188-DEL-95
chr6-111754251-DEL-531
chr6-111765364-DEL-205
chr6-111765905-DEL-1030
chr6-111780230-DEL-342
chr6-111795245-DEL-81
chr6-111825452-DEL-84
chr6-111868407-DEL-85
chr6-111925456-DEL-128
chr6-111951107-DEL-3540
chr6-111960365-DEL-241
chr6-111969404-DEL-752
chr6-111971304-DEL-317
chr6-112022373-DEL-54
chr6-112058613-DEL-145
chr6-112067938-DEL-66
chr6-112086336-DEL-52
chr6-112087507-DEL-380
chr6-112111641-DEL-71
chr6-112118648-DEL-1186
chr6-112134934-DEL-263
chr6-112137858-DEL-120
chr6-112156508-DEL-6319
chr6-112163865-DEL-84
chr6-112171007-DEL-312
chr6-112181415-DEL-1385
chr6-112201486-DEL-2392
chr6-112201560-DEL-577
chr6-112202146-DEL-1754
chr6-112227939-DEL-558
chr6-112229222-DEL-674
chr6-112230155-DEL-1008
chr6-112230338-DEL-398
chr6-112236428-DEL-96
chr6-112315533-DEL-812
chr6-112335958-DEL-1564
chr6-112340360-DEL-78
chr6-112365506-DEL-208
chr6-112373234-DEL-250
chr6-112410839-DEL-64
chr6-112438025-DEL-106
chr6-112475020-DEL-62
chr6-112476197-DEL-256
chr6-112503953-DEL-496
chr6-112505000-DEL-104
chr6-112571565-DEL-54
chr6-112576025-DEL-85
chr6-112580046-DEL-299
chr6-112586798-DEL-170
chr6-112587216-DEL-460
chr6-112588154-DEL-166
chr6-112612725-DEL-60
chr6-112626566-DEL-72
chr6-112655646-DEL-438
chr6-112675191-DEL-116
chr6-112691231-DEL-76
chr6-112708011-DEL-93
chr6-112751845-DEL-116
chr6-112801829-DEL-161
chr6-112836513-DEL-172
chr6-112871269-DEL-100
chr6-112874840-DEL-112
chr6-112961360-DEL-82
chr6-112982822-DEL-186
chr6-112997841-DEL-65
chr6-113003872-DEL-156
chr6-113016481-DEL-130
chr6-113016741-DEL-514
chr6-113028739-DEL-247
chr6-113075025-DEL-66
chr6-113104904-DEL-67
chr6-113139400-DEL-59
chr6-113140396-DEL-65
chr6-113176358-DEL-1893
chr6-113178467-DEL-164
chr6-113200824-DEL-82

chr6-113207660-DEL-53
chr6-113223564-DEL-522
chr6-113226610-DEL-135
chr6-113235431-DEL-215
chr6-113244159-DEL-51
chr6-113267464-DEL-84
chr6-113309790-DEL-161
chr6-113322967-DEL-56
chr6-113327840-DEL-219
chr6-113331099-DEL-225
chr6-113331841-DEL-59
chr6-113350887-DEL-54
chr6-113378913-DEL-181
chr6-113381274-DEL-343
chr6-113412101-DEL-177
chr6-113450631-DEL-84
chr6-113481667-DEL-84
chr6-113482449-DEL-170
chr6-113503249-DEL-133
chr6-113507000-DEL-122
chr6-113537207-DEL-110
chr6-113580556-DEL-141
chr6-113582605-DEL-143
chr6-113619375-DEL-182
chr6-113651410-DEL-95
chr6-113664070-DEL-85
chr6-113694951-DEL-74
chr6-113713836-DEL-79
chr6-113769741-DEL-298
chr6-113784710-DEL-51
chr6-113805310-DEL-121
chr6-113904036-DEL-968
chr6-113927946-DEL-80
chr6-113962345-DEL-57
chr6-113980157-DEL-174
chr6-113987976-DEL-336
chr6-114008872-DEL-77
chr6-114025922-DEL-123
chr6-114113023-DEL-61
chr6-114152100-DEL-609
chr6-114232889-DEL-144
chr6-114245031-DEL-136
chr6-114275541-DEL-158
chr6-114303713-DEL-954
chr6-114306131-DEL-234
chr6-114317914-DEL-1360
chr6-114318670-DEL-60
chr6-114324525-DEL-140
chr6-114373935-DEL-293
chr6-114464385-DEL-177
chr6-114467407-DEL-4698
chr6-114469360-DEL-1678
chr6-114471088-DEL-100
chr6-114475707-DEL-88
chr6-114478189-DEL-70
chr6-114499684-DEL-57
chr6-114502236-DEL-2405
chr6-114536086-DEL-60
chr6-114545811-DEL-136
chr6-114548314-DEL-64
chr6-114550743-DEL-58
chr6-114553241-DEL-75
chr6-114567760-DEL-103
chr6-114575306-DEL-2506
chr6-114592591-DEL-63
chr6-114594933-DEL-115
chr6-114600020-DEL-52
chr6-114602425-DEL-81
chr6-114604808-DEL-118
chr6-114612371-DEL-56
chr6-114624995-DEL-199
chr6-114652335-DEL-61
chr6-114717010-DEL-3414
chr6-114742957-DEL-88
chr6-114767548-DEL-156
chr6-114801660-DEL-117
chr6-114802238-DEL-205

chr6-114803172-DEL-54
chr6-114812938-DEL-60
chr6-114822278-DEL-86
chr6-114830125-DEL-155
chr6-114848172-DEL-1435
chr6-114849443-DEL-345
chr6-114866913-DEL-57
chr6-114916735-DEL-150
chr6-114947300-DEL-54
chr6-114968856-DEL-52
chr6-114991759-DEL-201
chr6-115034089-DEL-52
chr6-115058896-DEL-1150
chr6-115065039-DEL-70
chr6-115066955-DEL-70
chr6-115083696-DEL-217
chr6-115090430-DEL-196
chr6-115117339-DEL-79
chr6-115147763-DEL-93
chr6-115179184-DEL-171
chr6-115179772-DEL-197
chr6-115203507-DEL-52
chr6-115205711-DEL-1586
chr6-115217575-DEL-97
chr6-115232966-DEL-80
chr6-115330858-DEL-595
chr6-115356265-DEL-70
chr6-115403936-DEL-704
chr6-115449363-DEL-282
chr6-115455810-DEL-400
chr6-115483393-DEL-55
chr6-115526614-DEL-55
chr6-115553983-DEL-62
chr6-115577183-DEL-163
chr6-115628958-DEL-1318
chr6-115646046-DEL-199
chr6-115656568-DEL-144
chr6-115661031-DEL-205
chr6-115666005-DEL-490
chr6-115683757-DEL-58
chr6-115686501-DEL-52
chr6-115703817-DEL-198
chr6-115714753-DEL-664
chr6-115723803-DEL-463
chr6-115743179-DEL-189
chr6-115750009-DEL-77
chr6-115756314-DEL-175
chr6-115796429-DEL-537
chr6-115845456-DEL-3527
chr6-115847139-DEL-4150
chr6-115849004-DEL-595
chr6-115897809-DEL-281
chr6-115899219-DEL-64
chr6-115928501-DEL-2738
chr6-115980691-DEL-145
chr6-115987598-DEL-53
chr6-116006800-DEL-202
chr6-116041669-DEL-198
chr6-116042806-DEL-182
chr6-116088559-DEL-2998
chr6-116096145-DEL-52
chr6-116105105-DEL-1170
chr6-116126733-DEL-1263
chr6-116128722-DEL-450
chr6-116141538-DEL-295
chr6-116145166-DEL-204
chr6-116166091-DEL-95
chr6-116205140-DEL-90
chr6-116213856-DEL-207
chr6-116215728-DEL-806
chr6-116219074-DEL-124
chr6-116223541-DEL-140
chr6-116227994-DEL-964
chr6-116228143-DEL-297
chr6-116228770-DEL-62
chr6-116240267-DEL-75
chr6-116245269-DEL-96

chr6-116266397-DEL-401
chr6-116273156-DEL-378
chr6-116275359-DEL-124
chr6-116283897-DEL-75
chr6-116294931-DEL-58
chr6-116303096-DEL-192
chr6-116322449-DEL-190
chr6-116323022-DEL-750
chr6-116407494-DEL-68
chr6-116429662-DEL-77
chr6-116459286-DEL-2275
chr6-116464587-DEL-203
chr6-116466288-DEL-57
chr6-116475202-DEL-164
chr6-116483185-DEL-816
chr6-116492089-DEL-68
chr6-116495950-DEL-72
chr6-116509564-DEL-147
chr6-116549512-DEL-3684
chr6-116562617-DEL-55
chr6-116579158-DEL-62
chr6-116600746-DEL-197
chr6-116633106-DEL-162
chr6-116633565-DEL-518
chr6-116652860-DEL-52
chr6-116657144-DEL-58
chr6-116704359-DEL-6174
chr6-116715249-DEL-56
chr6-116725319-DEL-510
chr6-116725387-DEL-1342
chr6-116734015-DEL-191
chr6-116774876-DEL-108
chr6-116813300-DEL-7630
chr6-116832976-DEL-144
chr6-116860189-DEL-428
chr6-116862378-DEL-56
chr6-116886724-DEL-198
chr6-116892855-DEL-55
chr6-116932359-DEL-340
chr6-116942210-DEL-149
chr6-116978591-DEL-71
chr6-117000228-DEL-57
chr6-117004516-DEL-99
chr6-117014365-DEL-190
chr6-117030262-DEL-228
chr6-117031431-DEL-1240
chr6-117039592-DEL-147
chr6-117041555-DEL-126
chr6-117042351-DEL-2213
chr6-117064421-DEL-411
chr6-117090167-DEL-53
chr6-117127729-DEL-53
chr6-117168128-DEL-93
chr6-117188503-DEL-2757
chr6-117189005-DEL-994
chr6-117195044-DEL-184
chr6-117198833-DEL-54
chr6-117214762-DEL-341
chr6-117235610-DEL-2671
chr6-117235612-DEL-761
chr6-117236488-DEL-296
chr6-117237869-DEL-1627
chr6-117238499-DEL-997
chr6-117270114-DEL-204
chr6-117281643-DEL-67
chr6-117338906-DEL-174
chr6-117348117-DEL-853
chr6-117364888-DEL-1017
chr6-117365996-DEL-418
chr6-117377156-DEL-214
chr6-117390690-DEL-74
chr6-117435669-DEL-53
chr6-117440416-DEL-90
chr6-117446347-DEL-214
chr6-117526949-DEL-860
chr6-117528759-DEL-223
chr6-117537465-DEL-52

chr6-117601624-DEL-163
chr6-117631864-DEL-56
chr6-117651119-DEL-234
chr6-117681372-DEL-53
chr6-117688812-DEL-162
chr6-117712294-DEL-84
chr6-117717744-DEL-57
chr6-117749282-DEL-116
chr6-117758850-DEL-101
chr6-117782430-DEL-784
chr6-117782635-DEL-673
chr6-117806619-DEL-92
chr6-117807121-DEL-164
chr6-117815663-DEL-1757
chr6-117821648-DEL-123
chr6-117836823-DEL-152
chr6-117841706-DEL-80
chr6-117851191-DEL-248
chr6-117872041-DEL-167
chr6-117881402-DEL-1221
chr6-117900078-DEL-207
chr6-117903506-DEL-329
chr6-117926810-DEL-1532
chr6-117937181-DEL-797
chr6-117958929-DEL-53
chr6-117983579-DEL-65
chr6-117989810-DEL-64
chr6-117996303-DEL-56
chr6-118013223-DEL-118
chr6-118015436-DEL-2576
chr6-118015929-DEL-549
chr6-118028386-DEL-1342
chr6-118037800-DEL-3204
chr6-118039348-DEL-106
chr6-118060108-DEL-55
chr6-118063799-DEL-57
chr6-118077185-DEL-161
chr6-118120051-DEL-80
chr6-118138390-DEL-74
chr6-118143640-DEL-152
chr6-118194784-DEL-74
chr6-118214168-DEL-202
chr6-118246812-DEL-247
chr6-118318437-DEL-82
chr6-118353648-DEL-606
chr6-118354579-DEL-59
chr6-118366761-DEL-360
chr6-118386393-DEL-202
chr6-118442560-DEL-67
chr6-118473531-DEL-412
chr6-118474326-DEL-698
chr6-118475240-DEL-1097
chr6-118489042-DEL-482
chr6-118618019-DEL-68
chr6-118646513-DEL-54
chr6-118649043-DEL-64
chr6-118662679-DEL-137
chr6-118686133-DEL-72
chr6-118689647-DEL-64
chr6-118736561-DEL-605
chr6-118762337-DEL-80
chr6-118770935-DEL-56
chr6-118779005-DEL-66
chr6-118787064-DEL-181
chr6-118857043-DEL-70
chr6-118940855-DEL-185
chr6-118947864-DEL-686
chr6-118969854-DEL-753
chr6-118999962-DEL-4027
chr6-119007745-DEL-56
chr6-119021770-DEL-124
chr6-119044869-DEL-57
chr6-119057713-DEL-1041
chr6-119064195-DEL-223
chr6-119070237-DEL-286
chr6-119079702-DEL-53
chr6-119089144-DEL-131

chr6-119112092-DEL-1614
chr6-119144859-DEL-52
chr6-119179094-DEL-199
chr6-119184526-DEL-3520
chr6-119207049-DEL-258
chr6-119209859-DEL-166
chr6-119220490-DEL-390
chr6-119226445-DEL-331
chr6-119234624-DEL-198
chr6-119241717-DEL-78
chr6-119278062-DEL-52
chr6-119309363-DEL-146
chr6-119313893-DEL-170
chr6-119370670-DEL-108
chr6-119380848-DEL-58
chr6-119393239-DEL-176
chr6-119431252-DEL-2339
chr6-119443044-DEL-202
chr6-119446304-DEL-63
chr6-119446999-DEL-885
chr6-119451633-DEL-288
chr6-119470242-DEL-132
chr6-119476209-DEL-78
chr6-119478425-DEL-84
chr6-119489071-DEL-98
chr6-119509779-DEL-151
chr6-119512126-DEL-161
chr6-119517196-DEL-82
chr6-119524674-DEL-215
chr6-119535426-DEL-58
chr6-119588782-DEL-67
chr6-119613343-DEL-198
chr6-119651896-DEL-90
chr6-119667141-DEL-129
chr6-119723803-DEL-53
chr6-119724518-DEL-408
chr6-119732485-DEL-73
chr6-119809475-DEL-141
chr6-119812590-DEL-76
chr6-119849660-DEL-3699
chr6-119858448-DEL-194
chr6-119865652-DEL-75
chr6-119872562-DEL-154
chr6-119898638-DEL-149
chr6-120011220-DEL-81
chr6-120025264-DEL-170
chr6-120035284-DEL-447
chr6-120065450-DEL-319
chr6-120124660-DEL-401
chr6-120137548-DEL-84
chr6-120189077-DEL-103
chr6-120216786-DEL-169
chr6-120245357-DEL-201
chr6-120248089-DEL-57
chr6-120254647-DEL-60
chr6-120293060-DEL-202
chr6-120293358-DEL-519
chr6-120303062-DEL-180
chr6-120315522-DEL-57
chr6-120325974-DEL-66
chr6-120336067-DEL-183
chr6-120342032-DEL-59
chr6-120360482-DEL-199
chr6-120400463-DEL-1109
chr6-120411518-DEL-199
chr6-120426928-DEL-180
chr6-120434667-DEL-199
chr6-120442913-DEL-73
chr6-120484985-DEL-180
chr6-120491055-DEL-61
chr6-120496434-DEL-662
chr6-120523586-DEL-201
chr6-120541349-DEL-113
chr6-120545311-DEL-99
chr6-120547484-DEL-1738
chr6-120548745-DEL-978
chr6-120551426-DEL-2040

chr6-120567818-DEL-88
chr6-120609302-DEL-390
chr6-120631231-DEL-398
chr6-120637878-DEL-1599
chr6-120697945-DEL-2494
chr6-120747363-DEL-189
chr6-120783076-DEL-206
chr6-120797897-DEL-68
chr6-120802028-DEL-1539
chr6-120817663-DEL-739
chr6-120823116-DEL-114
chr6-120896791-DEL-3203
chr6-120899945-DEL-324
chr6-120925609-DEL-64
chr6-121003754-DEL-137
chr6-121023416-DEL-320
chr6-121024104-DEL-294
chr6-121025112-DEL-242
chr6-121042079-DEL-99
chr6-121066275-DEL-438
chr6-121098081-DEL-195
chr6-121166257-DEL-210
chr6-121167456-DEL-337
chr6-121200944-DEL-578
chr6-121234328-DEL-132
chr6-121251994-DEL-62
chr6-121296364-DEL-145
chr6-121317557-DEL-796
chr6-121396221-DEL-437
chr6-121429218-DEL-401
chr6-121465277-DEL-165
chr6-121493549-DEL-1396
chr6-121503589-DEL-352
chr6-121530385-DEL-530
chr6-121547771-DEL-3657
chr6-121555757-DEL-401
chr6-121627024-DEL-3779
chr6-121627162-DEL-2939
chr6-121627163-DEL-1175
chr6-121628884-DEL-251
chr6-121629136-DEL-1764
chr6-121629219-DEL-882
chr6-121665503-DEL-733
chr6-121672981-DEL-624
chr6-121686637-DEL-1524
chr6-121705813-DEL-344
chr6-121732848-DEL-142
chr6-121739745-DEL-137
chr6-121915098-DEL-51
chr6-121915825-DEL-539
chr6-121955567-DEL-2249
chr6-121962809-DEL-2767
chr6-121967039-DEL-219
chr6-121974304-DEL-5388
chr6-121990419-DEL-6566
chr6-122007899-DEL-213
chr6-122107080-DEL-570
chr6-122167485-DEL-1179
chr6-122214478-DEL-153
chr6-122216019-DEL-357
chr6-122221004-DEL-1030
chr6-122229580-DEL-96
chr6-122230041-DEL-443
chr6-122230929-DEL-79
chr6-122241990-DEL-81
chr6-122249642-DEL-267
chr6-122258622-DEL-554
chr6-122277208-DEL-71
chr6-122279007-DEL-1133
chr6-122283094-DEL-488
chr6-122324310-DEL-160
chr6-122341010-DEL-74
chr6-122360763-DEL-541
chr6-122363766-DEL-715
chr6-122366860-DEL-280
chr6-122372028-DEL-72
chr6-122404583-DEL-1052

chr6-122407599-DEL-2010
chr6-122466593-DEL-114
chr6-122485129-DEL-56
chr6-122492591-DEL-181
chr6-122495715-DEL-61
chr6-122534596-DEL-241
chr6-122538550-DEL-94
chr6-122546773-DEL-824
chr6-122553365-DEL-115
chr6-122565825-DEL-613
chr6-122580260-DEL-198
chr6-122582736-DEL-108
chr6-122592708-DEL-341
chr6-122594071-DEL-1487
chr6-122603588-DEL-71
chr6-122608198-DEL-586
chr6-122617553-DEL-505
chr6-122641280-DEL-68
chr6-122646142-DEL-120
chr6-122657131-DEL-249
chr6-122672916-DEL-982
chr6-122674168-DEL-587
chr6-122680272-DEL-67
chr6-122694104-DEL-197
chr6-122697506-DEL-2638
chr6-122710290-DEL-234
chr6-122712923-DEL-165
chr6-122725476-DEL-386
chr6-122731287-DEL-387
chr6-122734427-DEL-235
chr6-122743887-DEL-201
chr6-122750430-DEL-83
chr6-122759658-DEL-1688
chr6-122760238-DEL-52
chr6-122767397-DEL-300
chr6-122768485-DEL-1888
chr6-122776341-DEL-745
chr6-122783206-DEL-100
chr6-122825992-DEL-64
chr6-122837154-DEL-599
chr6-122842985-DEL-404
chr6-122844430-DEL-115
chr6-122871472-DEL-193
chr6-122912421-DEL-1026
chr6-122958180-DEL-75
chr6-122960310-DEL-1488
chr6-122972773-DEL-612
chr6-123006207-DEL-211
chr6-123014036-DEL-5692
chr6-123024387-DEL-1025
chr6-123044622-DEL-78
chr6-123044833-DEL-1434
chr6-123056196-DEL-6228
chr6-123077272-DEL-2497
chr6-123087499-DEL-735
chr6-123088308-DEL-1939
chr6-123107053-DEL-8341
chr6-123115126-DEL-226
chr6-123155079-DEL-1315
chr6-123167819-DEL-5131
chr6-123190458-DEL-300
chr6-123226549-DEL-515
chr6-123234040-DEL-231
chr6-123255688-DEL-52
chr6-123277042-DEL-213
chr6-123333958-DEL-72
chr6-123336411-DEL-197
chr6-123336697-DEL-10910
chr6-123336703-DEL-4344
chr6-123343207-DEL-4263
chr6-123347694-DEL-450
chr6-123348271-DEL-140
chr6-123349406-DEL-1306
chr6-123364888-DEL-60
chr6-123377930-DEL-208
chr6-123432882-DEL-925
chr6-123456519-DEL-135

chr6-123456766-DEL-849
chr6-123472144-DEL-80
chr6-123475690-DEL-1592
chr6-123480730-DEL-681
chr6-123491945-DEL-126
chr6-123499274-DEL-228
chr6-123502392-DEL-5570
chr6-123514013-DEL-82
chr6-123531922-DEL-91
chr6-123570456-DEL-197
chr6-123590678-DEL-6107
chr6-123592914-DEL-1592
chr6-123604917-DEL-1557
chr6-123609474-DEL-109
chr6-123640961-DEL-342
chr6-123683819-DEL-820
chr6-123699486-DEL-396
chr6-123699923-DEL-2968
chr6-123725271-DEL-723
chr6-123726128-DEL-1584
chr6-123727991-DEL-1233
chr6-123731894-DEL-7751
chr6-123741392-DEL-4695
chr6-123783506-DEL-56
chr6-123790642-DEL-58
chr6-123861899-DEL-147
chr6-123876009-DEL-162
chr6-123882856-DEL-109
chr6-123907045-DEL-213
chr6-123908089-DEL-332
chr6-123911168-DEL-134
chr6-123946759-DEL-2165
chr6-123972150-DEL-70
chr6-124055137-DEL-187
chr6-124063617-DEL-574
chr6-124068606-DEL-53
chr6-124128571-DEL-6190
chr6-124135093-DEL-203
chr6-124137172-DEL-69
chr6-124164056-DEL-64
chr6-124175555-DEL-340
chr6-124188354-DEL-428
chr6-124188860-DEL-428
chr6-124230028-DEL-259
chr6-124245431-DEL-1308
chr6-124253246-DEL-73
chr6-124262261-DEL-634
chr6-124265535-DEL-5429
chr6-124274269-DEL-1447
chr6-124276959-DEL-148
chr6-124291334-DEL-418
chr6-124320183-DEL-130
chr6-124346920-DEL-73
chr6-124386767-DEL-61
chr6-124395175-DEL-57
chr6-124405227-DEL-66
chr6-124418976-DEL-1015
chr6-124419055-DEL-420
chr6-124420855-DEL-1297
chr6-124429533-DEL-176
chr6-124443097-DEL-449
chr6-124447634-DEL-142
chr6-124455838-DEL-337
chr6-124491594-DEL-572
chr6-124512439-DEL-149
chr6-124522801-DEL-464
chr6-124534197-DEL-1142
chr6-124542457-DEL-62
chr6-124542731-DEL-13197
chr6-124547498-DEL-132
chr6-124572959-DEL-2163
chr6-124591320-DEL-3543
chr6-124628283-DEL-165
chr6-124629579-DEL-204
chr6-124657169-DEL-54
chr6-124678812-DEL-170
chr6-124699789-DEL-76

chr6-124748196-DEL-600
chr6-124749851-DEL-138
chr6-124753255-DEL-117
chr6-124818913-DEL-80
chr6-124821333-DEL-71
chr6-124871336-DEL-1310
chr6-124902582-DEL-192
chr6-124904839-DEL-73
chr6-124916910-DEL-464
chr6-124961306-DEL-68
chr6-124962390-DEL-206
chr6-124963264-DEL-271
chr6-124966490-DEL-191
chr6-124985361-DEL-201
chr6-125017631-DEL-390
chr6-125029940-DEL-56
chr6-125049103-DEL-68
chr6-125080479-DEL-195
chr6-125082543-DEL-312
chr6-125129762-DEL-67
chr6-125134029-DEL-94
chr6-125157162-DEL-282
chr6-125181711-DEL-185
chr6-125187167-DEL-204
chr6-125219026-DEL-105
chr6-125225960-DEL-3149
chr6-125236622-DEL-197
chr6-125252915-DEL-80
chr6-125268922-DEL-144
chr6-125278116-DEL-289
chr6-125284635-DEL-83
chr6-125295548-DEL-53
chr6-125322077-DEL-63
chr6-125367264-DEL-183
chr6-125367400-DEL-1384
chr6-125367417-DEL-10569
chr6-125369645-DEL-10591
chr6-125374167-DEL-2582
chr6-125392932-DEL-1797
chr6-125397905-DEL-148
chr6-125399531-DEL-76
chr6-125401599-DEL-167
chr6-125404752-DEL-87
chr6-125405033-DEL-605
chr6-125406266-DEL-66
chr6-125413092-DEL-816
chr6-125416568-DEL-152
chr6-125433523-DEL-199
chr6-125443977-DEL-79
chr6-125445888-DEL-85
chr6-125461495-DEL-221
chr6-125467888-DEL-197
chr6-125513253-DEL-70
chr6-125518702-DEL-54
chr6-125527909-DEL-236
chr6-125558773-DEL-85
chr6-125576833-DEL-120
chr6-125596356-DEL-1541
chr6-125596597-DEL-64
chr6-125613575-DEL-114
chr6-125616499-DEL-404
chr6-125634352-DEL-339
chr6-125644675-DEL-54
chr6-125658204-DEL-120
chr6-125677056-DEL-400
chr6-125686608-DEL-622
chr6-125688751-DEL-72
chr6-125726337-DEL-74
chr6-125747579-DEL-75
chr6-125751198-DEL-91
chr6-125764778-DEL-1438
chr6-125781234-DEL-77
chr6-125820978-DEL-1846
chr6-125823757-DEL-109
chr6-125853928-DEL-170
chr6-125854361-DEL-225
chr6-125889148-DEL-82

chr6-125916121-DEL-78
chr6-125927414-DEL-58
chr6-125938649-DEL-400
chr6-125955380-DEL-153
chr6-125965403-DEL-319
chr6-126009232-DEL-1139
chr6-126057332-DEL-64
chr6-126104577-DEL-51
chr6-126130375-DEL-200
chr6-126164027-DEL-113
chr6-126166862-DEL-80
chr6-126205051-DEL-333
chr6-126211531-DEL-364
chr6-126213131-DEL-138
chr6-126216644-DEL-98
chr6-126234050-DEL-6198
chr6-126270890-DEL-60
chr6-126274650-DEL-157
chr6-126325068-DEL-90
chr6-126341359-DEL-68
chr6-126395514-DEL-1785
chr6-126403514-DEL-54
chr6-126408873-DEL-155
chr6-126411502-DEL-132
chr6-126412227-DEL-316
chr6-126427170-DEL-65
chr6-126440033-DEL-157
chr6-126459768-DEL-56
chr6-126472677-DEL-60
chr6-126475727-DEL-54
chr6-126476747-DEL-69
chr6-126479310-DEL-54
chr6-126493507-DEL-675
chr6-126516379-DEL-320
chr6-126550531-DEL-3503
chr6-126567465-DEL-98
chr6-126579359-DEL-74
chr6-126612946-DEL-56
chr6-126693398-DEL-183
chr6-126727708-DEL-1157
chr6-126734575-DEL-599
chr6-126737951-DEL-91
chr6-126759779-DEL-70
chr6-126761862-DEL-55
chr6-126784374-DEL-192
chr6-126790051-DEL-77
chr6-126796298-DEL-111
chr6-126802616-DEL-51
chr6-126807475-DEL-87
chr6-126808820-DEL-140
chr6-126817806-DEL-90
chr6-126819528-DEL-111
chr6-126826515-DEL-56
chr6-126831571-DEL-76
chr6-126875047-DEL-211
chr6-126936086-DEL-1860
chr6-126936152-DEL-806
chr6-126937115-DEL-248
chr6-126937450-DEL-434
chr6-126947681-DEL-204
chr6-126952674-DEL-201
chr6-126984304-DEL-864
chr6-126994406-DEL-53
chr6-126999801-DEL-161
chr6-127006885-DEL-86
chr6-127025103-DEL-176
chr6-127046080-DEL-142
chr6-127110596-DEL-64
chr6-127157343-DEL-54
chr6-127215414-DEL-64
chr6-127222517-DEL-74
chr6-127262170-DEL-318
chr6-127273071-DEL-72
chr6-127283265-DEL-97
chr6-127292625-DEL-63
chr6-127313264-DEL-84
chr6-127324597-DEL-92

chr6-127337138-DEL-57
chr6-127342962-DEL-62
chr6-127345850-DEL-765
chr6-127357174-DEL-175
chr6-127358907-DEL-204
chr6-127366109-DEL-89
chr6-127413442-DEL-722
chr6-127419320-DEL-206
chr6-127438249-DEL-203
chr6-127462235-DEL-58
chr6-127476408-DEL-172
chr6-127483727-DEL-72
chr6-127500131-DEL-450
chr6-127540428-DEL-58
chr6-127541576-DEL-57
chr6-127542415-DEL-104
chr6-127561692-DEL-201
chr6-127570464-DEL-161
chr6-127571511-DEL-165
chr6-127583080-DEL-169
chr6-127592609-DEL-940
chr6-127598119-DEL-70
chr6-127613336-DEL-70
chr6-127630418-DEL-67
chr6-127667745-DEL-62
chr6-127684973-DEL-91
chr6-127711742-DEL-84
chr6-127722129-DEL-500
chr6-127763888-DEL-108
chr6-127764878-DEL-84
chr6-127859483-DEL-92
chr6-127899513-DEL-101
chr6-127916827-DEL-475
chr6-127917588-DEL-497
chr6-127917903-DEL-616
chr6-127918619-DEL-75
chr6-127931446-DEL-115
chr6-127934721-DEL-70
chr6-127936630-DEL-472
chr6-127974263-DEL-57
chr6-127991668-DEL-95
chr6-127994653-DEL-463
chr6-127996682-DEL-79
chr6-128063144-DEL-140
chr6-128092595-DEL-237
chr6-128104856-DEL-297
chr6-128128535-DEL-145
chr6-128152209-DEL-58
chr6-128175024-DEL-58
chr6-128184727-DEL-204
chr6-128190506-DEL-902
chr6-128193230-DEL-300
chr6-128206680-DEL-108
chr6-128210426-DEL-293
chr6-128231215-DEL-85
chr6-128301429-DEL-154
chr6-128310011-DEL-574
chr6-128315717-DEL-86
chr6-128365195-DEL-237
chr6-128396101-DEL-206
chr6-128410334-DEL-90
chr6-128417912-DEL-273
chr6-128424893-DEL-82
chr6-128435702-DEL-514
chr6-128443942-DEL-209
chr6-128499459-DEL-74
chr6-128532559-DEL-76
chr6-128578742-DEL-2506
chr6-128589605-DEL-5033
chr6-128602494-DEL-85
chr6-128694978-DEL-395
chr6-128725703-DEL-184
chr6-128752537-DEL-878
chr6-128763661-DEL-113
chr6-128772558-DEL-64
chr6-128806013-DEL-71
chr6-128806803-DEL-2438

chr6-128829744-DEL-1230
chr6-128832017-DEL-115
chr6-128834947-DEL-275
chr6-128900640-DEL-3925
chr6-128905976-DEL-438
chr6-128911559-DEL-114
chr6-128922836-DEL-3954
chr6-128923688-DEL-883
chr6-129026450-DEL-106
chr6-129036590-DEL-5494
chr6-129051612-DEL-994
chr6-129203301-DEL-1698
chr6-129226939-DEL-304
chr6-129230710-DEL-60
chr6-129276004-DEL-709
chr6-129315329-DEL-97
chr6-129429898-DEL-68
chr6-129452006-DEL-332
chr6-129458030-DEL-86
chr6-129463668-DEL-65
chr6-129469265-DEL-157
chr6-129516066-DEL-474
chr6-129524402-DEL-74
chr6-129557564-DEL-883
chr6-129602664-DEL-94
chr6-129605819-DEL-870
chr6-129608269-DEL-3339
chr6-129615574-DEL-94
chr6-129629767-DEL-1239
chr6-129647293-DEL-2212
chr6-129669005-DEL-56
chr6-129678894-DEL-149
chr6-129706726-DEL-395
chr6-129738258-DEL-5881
chr6-129765281-DEL-6377
chr6-129771683-DEL-2384
chr6-129776026-DEL-205
chr6-129825484-DEL-8010
chr6-129833956-DEL-212
chr6-129854091-DEL-2858
chr6-129877836-DEL-50
chr6-129878488-DEL-632
chr6-129880055-DEL-181
chr6-129907518-DEL-5038
chr6-129923566-DEL-532
chr6-129924914-DEL-376
chr6-129926818-DEL-727
chr6-129936898-DEL-1020
chr6-129939770-DEL-3711
chr6-129945967-DEL-430
chr6-129947867-DEL-683
chr6-129957646-DEL-52
chr6-129959277-DEL-2191
chr6-129963165-DEL-68
chr6-129966313-DEL-216
chr6-129966589-DEL-3024
chr6-129984747-DEL-54
chr6-130038128-DEL-58
chr6-130041263-DEL-507
chr6-130042492-DEL-199
chr6-130048918-DEL-342
chr6-130050307-DEL-1041
chr6-130052317-DEL-1558
chr6-130056070-DEL-85
chr6-130057826-DEL-56
chr6-130064599-DEL-96
chr6-130067056-DEL-282
chr6-130072099-DEL-82
chr6-130081849-DEL-454
chr6-130082478-DEL-62
chr6-130083674-DEL-1553
chr6-130086242-DEL-430
chr6-130087944-DEL-73
chr6-130091208-DEL-754
chr6-130097250-DEL-96
chr6-130099221-DEL-119
chr6-130119739-DEL-61

chr6-130136653-DEL-4339
chr6-130138219-DEL-65
chr6-130140289-DEL-242
chr6-130145348-DEL-438
chr6-130149940-DEL-136
chr6-130151482-DEL-73
chr6-130154658-DEL-759
chr6-130157678-DEL-56
chr6-130160670-DEL-96
chr6-130163131-DEL-108
chr6-130181304-DEL-190
chr6-130182168-DEL-2338
chr6-130195389-DEL-56
chr6-130199359-DEL-310
chr6-130204209-DEL-55
chr6-130239225-DEL-3830
chr6-130247950-DEL-56
chr6-130251381-DEL-71
chr6-130255453-DEL-595
chr6-130263210-DEL-59
chr6-130282460-DEL-365
chr6-130298359-DEL-498
chr6-130299273-DEL-753
chr6-130317129-DEL-110
chr6-130333608-DEL-959
chr6-130335752-DEL-751
chr6-130336910-DEL-503
chr6-130339329-DEL-61
chr6-130372405-DEL-135
chr6-130401188-DEL-568
chr6-130414262-DEL-1808
chr6-130421152-DEL-314
chr6-130439359-DEL-3158
chr6-130460356-DEL-252
chr6-130460956-DEL-84
chr6-130464495-DEL-616
chr6-130471861-DEL-6183
chr6-130474399-DEL-288
chr6-130506277-DEL-1081
chr6-130508632-DEL-1193
chr6-130510188-DEL-150
chr6-130532399-DEL-165
chr6-130536915-DEL-89
chr6-130555730-DEL-126
chr6-130603758-DEL-274
chr6-130610145-DEL-13984
chr6-130614032-DEL-199
chr6-130633771-DEL-292
chr6-130644668-DEL-63
chr6-130660682-DEL-6386
chr6-130685348-DEL-375
chr6-130689490-DEL-329
chr6-130690336-DEL-193
chr6-130692707-DEL-243
chr6-130705835-DEL-168
chr6-130708231-DEL-456
chr6-130708713-DEL-2193
chr6-130726759-DEL-6205
chr6-130734651-DEL-275
chr6-130745922-DEL-51
chr6-130755246-DEL-76
chr6-130809503-DEL-874
chr6-130831275-DEL-413
chr6-130869355-DEL-3309
chr6-130882287-DEL-243
chr6-130887091-DEL-76
chr6-130895398-DEL-1602
chr6-130902396-DEL-7085
chr6-130966998-DEL-6258
chr6-130996016-DEL-72
chr6-131044781-DEL-394
chr6-131086051-DEL-6145
chr6-131132089-DEL-4816
chr6-131177809-DEL-4549
chr6-131204847-DEL-107
chr6-131255996-DEL-605
chr6-131273653-DEL-175

chr6-131278004-DEL-72
chr6-131311788-DEL-62
chr6-131324533-DEL-576
chr6-131401189-DEL-59
chr6-131403533-DEL-1933
chr6-131417016-DEL-7043
chr6-131455041-DEL-71
chr6-131460954-DEL-615
chr6-131461962-DEL-56
chr6-131484852-DEL-54
chr6-131494838-DEL-1658
chr6-131505590-DEL-440
chr6-131517174-DEL-330
chr6-131517923-DEL-1204
chr6-131559292-DEL-1216
chr6-131571636-DEL-6436
chr6-131578752-DEL-8872
chr6-131579293-DEL-13402
chr6-131613793-DEL-98
chr6-131613856-DEL-2167
chr6-131616061-DEL-1346
chr6-131617448-DEL-166
chr6-131617655-DEL-1344
chr6-131619365-DEL-531
chr6-131646747-DEL-210
chr6-131657665-DEL-58
chr6-131672836-DEL-100
chr6-131679360-DEL-6179
chr6-131687267-DEL-1310
chr6-131690217-DEL-6136
chr6-131719650-DEL-51
chr6-131730400-DEL-4774
chr6-131736596-DEL-3285
chr6-131739520-DEL-3441
chr6-131743417-DEL-219
chr6-131743873-DEL-496
chr6-131774647-DEL-208
chr6-131786738-DEL-211
chr6-131788641-DEL-886
chr6-131804086-DEL-67
chr6-131850972-DEL-1002
chr6-131873537-DEL-179
chr6-131875626-DEL-6069
chr6-131919773-DEL-4097
chr6-131931369-DEL-208
chr6-131976953-DEL-60
chr6-132032781-DEL-111
chr6-132091871-DEL-165
chr6-132118678-DEL-4630
chr6-132128305-DEL-289
chr6-132142627-DEL-6303
chr6-132182552-DEL-2977
chr6-132184708-DEL-57
chr6-132245934-DEL-5668
chr6-132250345-DEL-2569
chr6-132287290-DEL-2434
chr6-132318318-DEL-2607
chr6-132320679-DEL-3238
chr6-132344007-DEL-707
chr6-132346720-DEL-1443
chr6-132367349-DEL-3007
chr6-132392430-DEL-358
chr6-132440966-DEL-5556
chr6-132480186-DEL-354
chr6-132497993-DEL-56
chr6-132506138-DEL-95
chr6-132519359-DEL-82
chr6-132543215-DEL-360
chr6-132577153-DEL-123
chr6-132579969-DEL-56
chr6-132580785-DEL-2436
chr6-132612246-DEL-7821
chr6-132696727-DEL-179
chr6-132718191-DEL-3133
chr6-132723753-DEL-2264
chr6-132727324-DEL-1127
chr6-132727338-DEL-70

chr6-132759963-DEL-133
chr6-132781950-DEL-60
chr6-132783617-DEL-103
chr6-132805897-DEL-828
chr6-132818419-DEL-63
chr6-132828660-DEL-408
chr6-132849996-DEL-2326
chr6-132875985-DEL-106
chr6-132909612-DEL-1842
chr6-132917981-DEL-548
chr6-132932010-DEL-626
chr6-132932662-DEL-1802
chr6-132939436-DEL-1161
chr6-132940825-DEL-1341
chr6-132942174-DEL-4984
chr6-132949828-DEL-81
chr6-132952377-DEL-2527
chr6-132953284-DEL-506
chr6-132955881-DEL-861
chr6-132971236-DEL-765
chr6-132991550-DEL-103
chr6-133008267-DEL-2446
chr6-133043803-DEL-237
chr6-133045595-DEL-3913
chr6-133055073-DEL-2545
chr6-133061377-DEL-275
chr6-133066353-DEL-2233
chr6-133075105-DEL-384
chr6-133076372-DEL-646
chr6-133110344-DEL-74
chr6-133195547-DEL-207
chr6-133198670-DEL-2966
chr6-133206003-DEL-65
chr6-133221202-DEL-2427
chr6-133226725-DEL-60
chr6-133231906-DEL-70
chr6-133253065-DEL-1529
chr6-133254330-DEL-1421
chr6-133256328-DEL-72
chr6-133262261-DEL-70
chr6-133279371-DEL-124
chr6-133311195-DEL-416
chr6-133313073-DEL-60
chr6-133336807-DEL-55
chr6-133375818-DEL-10765
chr6-133420969-DEL-1692
chr6-133424211-DEL-12063
chr6-133444384-DEL-108
chr6-133446633-DEL-65
chr6-133475511-DEL-75
chr6-133517602-DEL-440
chr6-133524602-DEL-791
chr6-133530478-DEL-798
chr6-133543620-DEL-200
chr6-133545580-DEL-984
chr6-133547317-DEL-59
chr6-133553813-DEL-473
chr6-133564139-DEL-86
chr6-133625832-DEL-3506
chr6-133630920-DEL-108
chr6-133658193-DEL-1848
chr6-133664137-DEL-1165
chr6-133666346-DEL-5367
chr6-133713529-DEL-58
chr6-133722154-DEL-6514
chr6-133765397-DEL-4767
chr6-133809315-DEL-6668
chr6-133829549-DEL-63
chr6-133832503-DEL-78
chr6-133836705-DEL-764
chr6-133846456-DEL-1328
chr6-133848516-DEL-264
chr6-133852200-DEL-203
chr6-133869103-DEL-1684
chr6-133876682-DEL-433
chr6-133891455-DEL-351
chr6-133892240-DEL-1625

chr6-134067633-DEL-53
chr6-134084457-DEL-3793
chr6-134189705-DEL-2846
chr6-134209044-DEL-52
chr6-134218868-DEL-217
chr6-134291798-DEL-308
chr6-134301055-DEL-110
chr6-134301997-DEL-89
chr6-134313162-DEL-1057
chr6-134316375-DEL-398
chr6-134339322-DEL-1362
chr6-134359397-DEL-52
chr6-134363748-DEL-72
chr6-134367436-DEL-1275
chr6-134371900-DEL-76
chr6-134381342-DEL-57
chr6-134417865-DEL-729
chr6-134499520-DEL-1369
chr6-134514224-DEL-99
chr6-134547093-DEL-274
chr6-134561599-DEL-55
chr6-134566806-DEL-67
chr6-134573391-DEL-236
chr6-134575163-DEL-78
chr6-134597551-DEL-483
chr6-134614759-DEL-54
chr6-134616216-DEL-73
chr6-134627986-DEL-1684
chr6-134646009-DEL-51
chr6-134655889-DEL-98
chr6-134719794-DEL-198
chr6-134747965-DEL-610
chr6-134774511-DEL-313
chr6-134796729-DEL-66
chr6-134804579-DEL-119
chr6-134816508-DEL-400
chr6-134835595-DEL-67
chr6-134840808-DEL-195
chr6-134843623-DEL-206
chr6-134883099-DEL-216
chr6-134908730-DEL-180
chr6-134913127-DEL-82
chr6-134916933-DEL-720
chr6-134924752-DEL-69
chr6-134941610-DEL-182
chr6-134942646-DEL-164
chr6-134973835-DEL-78
chr6-134986566-DEL-64
chr6-135006564-DEL-567
chr6-135011081-DEL-79
chr6-135014278-DEL-605
chr6-135030416-DEL-68
chr6-135031294-DEL-233
chr6-135036034-DEL-183
chr6-135098152-DEL-122
chr6-135127690-DEL-55
chr6-135141466-DEL-207
chr6-135154817-DEL-59
chr6-135163529-DEL-100
chr6-135190159-DEL-1216
chr6-135198842-DEL-111
chr6-135208134-DEL-112
chr6-135245101-DEL-165
chr6-135292663-DEL-54
chr6-135350801-DEL-145
chr6-135376374-DEL-163
chr6-135425022-DEL-53
chr6-135429126-DEL-58
chr6-135485236-DEL-373
chr6-135503398-DEL-76
chr6-135521253-DEL-51
chr6-135584862-DEL-1228
chr6-135594739-DEL-2796
chr6-135652035-DEL-57
chr6-135684152-DEL-94
chr6-135727505-DEL-78
chr6-135759268-DEL-157

chr6-135764779-DEL-54
chr6-135803727-DEL-230
chr6-135818572-DEL-96
chr6-135831976-DEL-111
chr6-135841705-DEL-126
chr6-135853411-DEL-86
chr6-135872384-DEL-63
chr6-135881841-DEL-60
chr6-135927811-DEL-110
chr6-135955553-DEL-60
chr6-135962716-DEL-106
chr6-136007812-DEL-63
chr6-136029016-DEL-55
chr6-136030288-DEL-56
chr6-136041834-DEL-3063
chr6-136046568-DEL-182
chr6-136052737-DEL-104
chr6-136066090-DEL-323
chr6-136104260-DEL-53
chr6-136174525-DEL-1387
chr6-136202831-DEL-132
chr6-136210712-DEL-178
chr6-136214751-DEL-54
chr6-136237768-DEL-53
chr6-136266648-DEL-1889
chr6-136274084-DEL-149
chr6-136282584-DEL-156
chr6-136308467-DEL-54
chr6-136311371-DEL-68
chr6-136329538-DEL-88
chr6-136357984-DEL-276
chr6-136382877-DEL-80
chr6-136386724-DEL-259
chr6-136391385-DEL-821
chr6-136403680-DEL-69
chr6-136407284-DEL-282
chr6-136409632-DEL-132
chr6-136411134-DEL-152
chr6-136422539-DEL-128
chr6-136425407-DEL-199
chr6-136435596-DEL-190
chr6-136449002-DEL-52
chr6-136478665-DEL-55
chr6-136595243-DEL-125
chr6-136608948-DEL-91
chr6-136620862-DEL-232
chr6-136631456-DEL-66
chr6-136646874-DEL-97
chr6-136654196-DEL-96
chr6-136659471-DEL-71
chr6-136660946-DEL-62
chr6-136661818-DEL-681
chr6-136693494-DEL-52
chr6-136705243-DEL-59
chr6-136738233-DEL-165
chr6-136740345-DEL-56
chr6-136784498-DEL-73
chr6-136785952-DEL-86
chr6-136795324-DEL-282
chr6-136825089-DEL-197
chr6-136828260-DEL-601
chr6-136856873-DEL-60
chr6-136862107-DEL-56
chr6-136876881-DEL-53
chr6-136885498-DEL-93
chr6-136909454-DEL-54
chr6-136916232-DEL-118
chr6-136932923-DEL-56
chr6-136936826-DEL-135
chr6-136954815-DEL-209
chr6-136965196-DEL-54
chr6-137001944-DEL-71
chr6-137003053-DEL-60
chr6-137009831-DEL-215
chr6-137057054-DEL-104
chr6-137080538-DEL-1052
chr6-137089256-DEL-336

chr6-137110499-DEL-56
chr6-137117808-DEL-66
chr6-137181157-DEL-1722
chr6-137190762-DEL-52
chr6-137222422-DEL-100
chr6-137240939-DEL-748
chr6-137262685-DEL-110
chr6-137268274-DEL-70
chr6-137311787-DEL-63
chr6-137329067-DEL-980
chr6-137333506-DEL-309
chr6-137337254-DEL-494
chr6-137339599-DEL-87
chr6-137363928-DEL-133
chr6-137379804-DEL-1520
chr6-137383527-DEL-73
chr6-137398647-DEL-59
chr6-137408775-DEL-52
chr6-137451022-DEL-66
chr6-137488176-DEL-828
chr6-137502734-DEL-143
chr6-137506916-DEL-56
chr6-137535013-DEL-158
chr6-137536163-DEL-7758
chr6-137539315-DEL-95
chr6-137543335-DEL-110
chr6-137550014-DEL-486
chr6-137554821-DEL-60
chr6-137580051-DEL-111
chr6-137599126-DEL-52
chr6-137622946-DEL-83
chr6-137660806-DEL-78
chr6-137662608-DEL-1592
chr6-137669621-DEL-163
chr6-137697548-DEL-173
chr6-137700118-DEL-505
chr6-137707939-DEL-101
chr6-137780956-DEL-70
chr6-137803520-DEL-176
chr6-137804054-DEL-152
chr6-137820248-DEL-88
chr6-137823357-DEL-56
chr6-137836236-DEL-56
chr6-137857234-DEL-58
chr6-137858803-DEL-126
chr6-137880734-DEL-1790
chr6-137932005-DEL-572
chr6-137951507-DEL-135
chr6-137990376-DEL-605
chr6-138027601-DEL-70
chr6-138038927-DEL-501
chr6-138050878-DEL-140
chr6-138062529-DEL-671
chr6-138062559-DEL-208
chr6-138082061-DEL-87
chr6-138152968-DEL-1016
chr6-138160999-DEL-214
chr6-138177279-DEL-155
chr6-138179789-DEL-399
chr6-138199244-DEL-1488
chr6-138201089-DEL-76
chr6-138206997-DEL-75
chr6-138219551-DEL-94
chr6-138224837-DEL-2106
chr6-138231038-DEL-74
chr6-138236746-DEL-146
chr6-138267897-DEL-52
chr6-138269439-DEL-488
chr6-138269685-DEL-568
chr6-138303490-DEL-65
chr6-138383274-DEL-52
chr6-138407231-DEL-142
chr6-138416034-DEL-94
chr6-138419714-DEL-80
chr6-138471507-DEL-189
chr6-138501807-DEL-58
chr6-138546491-DEL-998

chr6-138554350-DEL-95
chr6-138581622-DEL-200
chr6-138618672-DEL-97
chr6-138664692-DEL-316
chr6-138669548-DEL-58
chr6-138693009-DEL-59
chr6-138695608-DEL-77
chr6-138739766-DEL-186
chr6-138744313-DEL-339
chr6-138796111-DEL-2308
chr6-138822308-DEL-79
chr6-138846269-DEL-216
chr6-138848175-DEL-129
chr6-138859799-DEL-77
chr6-138872186-DEL-109
chr6-138903715-DEL-111
chr6-138921592-DEL-62
chr6-138953358-DEL-220
chr6-138957617-DEL-102
chr6-138961708-DEL-183
chr6-138962131-DEL-314
chr6-139038982-DEL-130
chr6-139043813-DEL-117
chr6-139066447-DEL-116
chr6-139090784-DEL-58
chr6-139093085-DEL-533
chr6-139123753-DEL-1533
chr6-139137867-DEL-61
chr6-139166177-DEL-106
chr6-139176744-DEL-200
chr6-139215188-DEL-58
chr6-139226133-DEL-68
chr6-139239874-DEL-56
chr6-139268055-DEL-141
chr6-139268330-DEL-558
chr6-139291747-DEL-376
chr6-139300998-DEL-145
chr6-139301837-DEL-237
chr6-139335054-DEL-6191
chr6-139350236-DEL-6017
chr6-139361495-DEL-1690
chr6-139363797-DEL-344
chr6-139392686-DEL-151
chr6-139420040-DEL-359
chr6-139434254-DEL-62
chr6-139442203-DEL-85
chr6-139448194-DEL-1490
chr6-139467473-DEL-991
chr6-139472311-DEL-60
chr6-139503134-DEL-200
chr6-139506879-DEL-4116
chr6-139511164-DEL-298
chr6-139540131-DEL-3424
chr6-139545449-DEL-52
chr6-139558660-DEL-1587
chr6-139605178-DEL-169
chr6-139608977-DEL-623
chr6-139640169-DEL-129
chr6-139640501-DEL-104
chr6-139644323-DEL-76
chr6-139649988-DEL-288
chr6-139651245-DEL-8060
chr6-139676485-DEL-63
chr6-139711363-DEL-119
chr6-139726252-DEL-80
chr6-139782076-DEL-3396
chr6-139813007-DEL-236
chr6-139849455-DEL-1155
chr6-139885901-DEL-423
chr6-139885996-DEL-972
chr6-139886610-DEL-324
chr6-139897604-DEL-225
chr6-139909919-DEL-162
chr6-139952480-DEL-63
chr6-139981283-DEL-250
chr6-139996376-DEL-52
chr6-140002737-DEL-210

chr6-140073080-DEL-66
chr6-140086525-DEL-2322
chr6-140090120-DEL-128
chr6-140125286-DEL-112
chr6-140126821-DEL-125
chr6-140127341-DEL-110
chr6-140129461-DEL-75
chr6-140143451-DEL-132
chr6-140151659-DEL-1151
chr6-140152556-DEL-525
chr6-140160953-DEL-442
chr6-140180466-DEL-221
chr6-140186905-DEL-251
chr6-140202143-DEL-95
chr6-140221066-DEL-493
chr6-140232273-DEL-84
chr6-140274424-DEL-205
chr6-140286686-DEL-75
chr6-140321783-DEL-62
chr6-140324580-DEL-284
chr6-140334802-DEL-402
chr6-140337744-DEL-102
chr6-140357184-DEL-174
chr6-140421805-DEL-565
chr6-140479850-DEL-204
chr6-140487373-DEL-52
chr6-140540429-DEL-216
chr6-140553148-DEL-163
chr6-140554941-DEL-144
chr6-140562977-DEL-68
chr6-140578173-DEL-192
chr6-140578912-DEL-124
chr6-140589919-DEL-75
chr6-140590643-DEL-212
chr6-140604307-DEL-654
chr6-140604989-DEL-2463
chr6-140605753-DEL-94
chr6-140611092-DEL-94
chr6-140615833-DEL-306
chr6-140618772-DEL-92
chr6-140638501-DEL-184
chr6-140647531-DEL-79
chr6-140647677-DEL-502
chr6-140656826-DEL-91
chr6-140672001-DEL-57
chr6-140686737-DEL-7314
chr6-140698913-DEL-439
chr6-140707160-DEL-57
chr6-140743647-DEL-144
chr6-140754191-DEL-71
chr6-140764784-DEL-151
chr6-140792171-DEL-131
chr6-140805333-DEL-1293
chr6-140809035-DEL-4849
chr6-140814085-DEL-513
chr6-140815226-DEL-457
chr6-140848905-DEL-87
chr6-140857540-DEL-112
chr6-140882363-DEL-172
chr6-140910953-DEL-58
chr6-140918503-DEL-127
chr6-140944537-DEL-60
chr6-140956057-DEL-65
chr6-140970834-DEL-1248
chr6-140984920-DEL-2035
chr6-140993004-DEL-440
chr6-141019116-DEL-155
chr6-141019662-DEL-1436
chr6-141023068-DEL-293
chr6-141027789-DEL-163
chr6-141032525-DEL-880
chr6-141058180-DEL-78
chr6-141067221-DEL-2254
chr6-141148493-DEL-947
chr6-141152592-DEL-66
chr6-141204665-DEL-1260
chr6-141216450-DEL-83

chr6-141233868-DEL-716
chr6-141242541-DEL-51
chr6-141266465-DEL-96
chr6-141272606-DEL-95
chr6-141279909-DEL-431
chr6-141284091-DEL-76
chr6-141301381-DEL-10828
chr6-141312216-DEL-56
chr6-141330379-DEL-238
chr6-141333289-DEL-97
chr6-141340122-DEL-232
chr6-141344508-DEL-118
chr6-141379077-DEL-69
chr6-141438594-DEL-57
chr6-141479071-DEL-76
chr6-141484303-DEL-435
chr6-141499512-DEL-417
chr6-141537193-DEL-5156
chr6-141569448-DEL-401
chr6-141579136-DEL-56
chr6-141595710-DEL-58
chr6-141646317-DEL-4287
chr6-141685637-DEL-54
chr6-141715292-DEL-501
chr6-141730650-DEL-308
chr6-141738292-DEL-908
chr6-141748258-DEL-831
chr6-141782970-DEL-717
chr6-141788930-DEL-137
chr6-141791294-DEL-90
chr6-141806763-DEL-4106
chr6-141823767-DEL-66
chr6-141838930-DEL-909
chr6-141844975-DEL-2633
chr6-141908564-DEL-196
chr6-141925284-DEL-852
chr6-141947822-DEL-257
chr6-141979023-DEL-100
chr6-141995475-DEL-448
chr6-142012567-DEL-6712
chr6-142018938-DEL-204
chr6-142020174-DEL-191
chr6-142023541-DEL-6189
chr6-142070154-DEL-93
chr6-142076992-DEL-723
chr6-142103591-DEL-3927
chr6-142113721-DEL-70
chr6-142117897-DEL-101
chr6-142122343-DEL-101
chr6-142138509-DEL-1816
chr6-142171123-DEL-78
chr6-142173364-DEL-181
chr6-142176230-DEL-92
chr6-142194262-DEL-53
chr6-142195046-DEL-51
chr6-142215838-DEL-4276
chr6-142249711-DEL-664
chr6-142255491-DEL-55
chr6-142263008-DEL-58
chr6-142263856-DEL-336
chr6-142266145-DEL-269
chr6-142326307-DEL-428
chr6-142342541-DEL-352
chr6-142342567-DEL-3311
chr6-142367928-DEL-268
chr6-142388036-DEL-103
chr6-142389564-DEL-808
chr6-142446321-DEL-206
chr6-142452025-DEL-101
chr6-142468973-DEL-476
chr6-142471400-DEL-160
chr6-142472613-DEL-3831
chr6-142481083-DEL-400
chr6-142500857-DEL-898
chr6-142543237-DEL-106
chr6-142555780-DEL-1874
chr6-142565937-DEL-51

chr6-142572088-DEL-270
chr6-142600279-DEL-89
chr6-142616536-DEL-283
chr6-142627922-DEL-104
chr6-142629332-DEL-72
chr6-142644329-DEL-184
chr6-142653565-DEL-72
chr6-142657473-DEL-231
chr6-142676290-DEL-1661
chr6-142687779-DEL-80
chr6-142690770-DEL-228
chr6-142696378-DEL-233
chr6-142696398-DEL-499
chr6-142716589-DEL-66
chr6-142749716-DEL-60
chr6-142755007-DEL-99
chr6-142767576-DEL-51
chr6-142789829-DEL-52
chr6-142792325-DEL-66
chr6-142813944-DEL-124
chr6-142837659-DEL-223
chr6-142842623-DEL-60
chr6-142842775-DEL-6332
chr6-142858390-DEL-52
chr6-142869799-DEL-68
chr6-142883501-DEL-52
chr6-142886320-DEL-172
chr6-142899659-DEL-55
chr6-142903003-DEL-85
chr6-142916374-DEL-72
chr6-142924146-DEL-90
chr6-142924785-DEL-155
chr6-142929443-DEL-282
chr6-142930750-DEL-74
chr6-142959842-DEL-2196
chr6-142979603-DEL-104
chr6-142996811-DEL-56
chr6-143002927-DEL-216
chr6-143021325-DEL-200
chr6-143039278-DEL-215
chr6-143042279-DEL-102
chr6-143067670-DEL-6603
chr6-143082306-DEL-5149
chr6-143084504-DEL-151
chr6-143098664-DEL-134
chr6-143101504-DEL-67
chr6-143110524-DEL-882
chr6-143141809-DEL-490
chr6-143147726-DEL-202
chr6-143164360-DEL-69
chr6-143196258-DEL-51
chr6-143218734-DEL-1105
chr6-143260753-DEL-67
chr6-143275278-DEL-70
chr6-143294011-DEL-84
chr6-143299807-DEL-381
chr6-143330232-DEL-89
chr6-143334545-DEL-262
chr6-143343325-DEL-72
chr6-143381658-DEL-124
chr6-143399014-DEL-415
chr6-143410457-DEL-51
chr6-143436631-DEL-143
chr6-143439134-DEL-94
chr6-143443380-DEL-459
chr6-143465026-DEL-107
chr6-143465937-DEL-53
chr6-143491746-DEL-153
chr6-143524491-DEL-110
chr6-143556080-DEL-127
chr6-143589149-DEL-142
chr6-143592608-DEL-67
chr6-143620444-DEL-93
chr6-143633902-DEL-53
chr6-143634391-DEL-69
chr6-143666953-DEL-192
chr6-143675077-DEL-205

chr6-143695935-DEL-138
chr6-143714086-DEL-343
chr6-143728608-DEL-182
chr6-143741859-DEL-62
chr6-143748443-DEL-1957
chr6-143751052-DEL-618
chr6-143756569-DEL-961
chr6-143766279-DEL-80
chr6-143776897-DEL-60
chr6-143782619-DEL-122
chr6-143783905-DEL-351
chr6-143792010-DEL-503
chr6-143804783-DEL-58
chr6-143836447-DEL-288
chr6-143849271-DEL-117
chr6-143854454-DEL-86
chr6-143856818-DEL-192
chr6-143862933-DEL-358
chr6-143883951-DEL-167
chr6-143902036-DEL-227
chr6-143914440-DEL-59
chr6-143934663-DEL-66
chr6-143941487-DEL-214
chr6-143955195-DEL-182
chr6-143956387-DEL-81
chr6-143961952-DEL-181
chr6-143965520-DEL-75
chr6-144017750-DEL-68
chr6-144046755-DEL-62
chr6-144048162-DEL-52
chr6-144070978-DEL-119
chr6-144078720-DEL-117
chr6-144110412-DEL-65
chr6-144165280-DEL-1485
chr6-144177984-DEL-56
chr6-144194178-DEL-59
chr6-144273534-DEL-60
chr6-144306766-DEL-68
chr6-144316539-DEL-72
chr6-144326452-DEL-128
chr6-144345027-DEL-478
chr6-144402317-DEL-124
chr6-144427910-DEL-88
chr6-144441369-DEL-199
chr6-144465557-DEL-58
chr6-144476723-DEL-146
chr6-144481250-DEL-109
chr6-144494924-DEL-75
chr6-144518658-DEL-64
chr6-144528551-DEL-106
chr6-144534681-DEL-123
chr6-144537868-DEL-90
chr6-144540901-DEL-532
chr6-144558388-DEL-82
chr6-144562463-DEL-120
chr6-144570952-DEL-78
chr6-144575437-DEL-655
chr6-144605287-DEL-68
chr6-144613574-DEL-354
chr6-144614368-DEL-448
chr6-144614581-DEL-640
chr6-144615445-DEL-416
chr6-144702477-DEL-90
chr6-144712943-DEL-94
chr6-144735441-DEL-75
chr6-144737910-DEL-60
chr6-144745395-DEL-77
chr6-144762774-DEL-67
chr6-144771053-DEL-312
chr6-144781741-DEL-540
chr6-144784998-DEL-56
chr6-144785688-DEL-1191
chr6-144797033-DEL-578
chr6-144799237-DEL-431
chr6-144821426-DEL-64
chr6-144872547-DEL-285
chr6-144895433-DEL-502

chr6-144915135-DEL-231
chr6-144915681-DEL-76
chr6-144950249-DEL-87
chr6-144956921-DEL-54
chr6-144960647-DEL-55
chr6-144961329-DEL-60
chr6-144962169-DEL-109
chr6-144970698-DEL-2780
chr6-144991196-DEL-321
chr6-144998354-DEL-56
chr6-145017971-DEL-659
chr6-145025344-DEL-535
chr6-145026160-DEL-1086
chr6-145029860-DEL-52
chr6-145037531-DEL-115
chr6-145041468-DEL-199
chr6-145056353-DEL-178
chr6-145069245-DEL-52
chr6-145076511-DEL-54
chr6-145078798-DEL-51
chr6-145088117-DEL-62
chr6-145095559-DEL-54
chr6-145126989-DEL-375
chr6-145134690-DEL-678
chr6-145157446-DEL-59
chr6-145188734-DEL-1267
chr6-145200788-DEL-124
chr6-145226752-DEL-82
chr6-145268911-DEL-77
chr6-145275723-DEL-52
chr6-145294418-DEL-55
chr6-145324218-DEL-109
chr6-145385846-DEL-230
chr6-145397707-DEL-641
chr6-145404116-DEL-52
chr6-145420549-DEL-51
chr6-145439306-DEL-240
chr6-145450837-DEL-814
chr6-145454058-DEL-156
chr6-145457009-DEL-91
chr6-145460292-DEL-876
chr6-145510781-DEL-882
chr6-145512339-DEL-62
chr6-145515739-DEL-80
chr6-145541649-DEL-933
chr6-145554774-DEL-151
chr6-145577399-DEL-87
chr6-145585483-DEL-329
chr6-145618988-DEL-90
chr6-145627593-DEL-1016
chr6-145628655-DEL-418
chr6-145628895-DEL-636
chr6-145635619-DEL-504
chr6-145638220-DEL-71
chr6-145652971-DEL-62
chr6-145654241-DEL-64
chr6-145688151-DEL-53
chr6-145730189-DEL-1456
chr6-145730481-DEL-65
chr6-145730546-DEL-1511
chr6-145730721-DEL-612
chr6-145730761-DEL-1095
chr6-145730794-DEL-280
chr6-145732290-DEL-247
chr6-145740934-DEL-58
chr6-145749639-DEL-139
chr6-145758067-DEL-61
chr6-145783264-DEL-104
chr6-145785597-DEL-200
chr6-145789240-DEL-164
chr6-145829066-DEL-397
chr6-145841546-DEL-3069
chr6-145841885-DEL-953
chr6-145842424-DEL-6231
chr6-145844634-DEL-4031
chr6-145915551-DEL-1577
chr6-145965037-DEL-186

chr6-145965439-DEL-1114
chr6-145965700-DEL-66
chr6-145966222-DEL-354
chr6-145972358-DEL-63
chr6-146003417-DEL-317
chr6-146034446-DEL-83
chr6-146098604-DEL-399
chr6-146114064-DEL-107
chr6-146152707-DEL-483
chr6-146258322-DEL-142
chr6-146280359-DEL-110
chr6-146293080-DEL-86
chr6-146303383-DEL-73
chr6-146309716-DEL-336
chr6-146337743-DEL-54
chr6-146342033-DEL-70
chr6-146346613-DEL-105
chr6-146348777-DEL-3368
chr6-146370566-DEL-345
chr6-146410366-DEL-168
chr6-146411080-DEL-58
chr6-146415064-DEL-65
chr6-146424111-DEL-427
chr6-146434181-DEL-107
chr6-146440672-DEL-199
chr6-146441223-DEL-51
chr6-146509690-DEL-352
chr6-146520695-DEL-68
chr6-146541417-DEL-62
chr6-146545259-DEL-263
chr6-146567320-DEL-311
chr6-146579376-DEL-63
chr6-146608703-DEL-5116
chr6-146609406-DEL-75
chr6-146614141-DEL-4067
chr6-146616143-DEL-664
chr6-146682367-DEL-54
chr6-146684926-DEL-1254
chr6-146709776-DEL-357
chr6-146730869-DEL-69
chr6-146756795-DEL-132
chr6-146935538-DEL-203
chr6-146941883-DEL-76
chr6-146955029-DEL-142
chr6-146995219-DEL-52
chr6-147020466-DEL-80
chr6-147042896-DEL-168
chr6-147055273-DEL-174
chr6-147071891-DEL-334
chr6-147105917-DEL-61
chr6-147131567-DEL-1428
chr6-147138247-DEL-151
chr6-147142785-DEL-51
chr6-147171793-DEL-246
chr6-147189909-DEL-55
chr6-147221122-DEL-59
chr6-147241602-DEL-625
chr6-147243123-DEL-139
chr6-147297220-DEL-52
chr6-147298477-DEL-109
chr6-147308722-DEL-524
chr6-147330681-DEL-391
chr6-147332237-DEL-132
chr6-147410672-DEL-106
chr6-147427337-DEL-79
chr6-147439499-DEL-71
chr6-147444309-DEL-1517
chr6-147445971-DEL-53
chr6-147466832-DEL-60
chr6-147473131-DEL-98
chr6-147495340-DEL-1664
chr6-147518192-DEL-104
chr6-147539270-DEL-375
chr6-147567766-DEL-59
chr6-147569153-DEL-2134
chr6-147599658-DEL-61
chr6-147610098-DEL-72

chr6-147621380-DEL-152
chr6-147626527-DEL-84
chr6-147636866-DEL-63
chr6-147666377-DEL-54
chr6-147690367-DEL-65
chr6-147693035-DEL-167
chr6-147709107-DEL-56
chr6-147740517-DEL-57
chr6-147746538-DEL-186
chr6-147753278-DEL-109
chr6-147776737-DEL-81
chr6-147816818-DEL-744
chr6-147892402-DEL-80
chr6-147898973-DEL-968
chr6-147914716-DEL-416
chr6-147914909-DEL-1984
chr6-147916378-DEL-154
chr6-147957516-DEL-78
chr6-147963697-DEL-78
chr6-147967172-DEL-127
chr6-147976212-DEL-4285
chr6-147998120-DEL-204
chr6-148027474-DEL-5189
chr6-148037960-DEL-385
chr6-148098982-DEL-351
chr6-148124393-DEL-87
chr6-148149343-DEL-271
chr6-148195930-DEL-66
chr6-148214163-DEL-276
chr6-148324348-DEL-56
chr6-148361949-DEL-70
chr6-148408855-DEL-61
chr6-148434719-DEL-97
chr6-148446053-DEL-250
chr6-148469086-DEL-512
chr6-148482880-DEL-583
chr6-148506639-DEL-5871
chr6-148522110-DEL-84
chr6-148533564-DEL-1197
chr6-148536102-DEL-142
chr6-148536519-DEL-5323
chr6-148536762-DEL-233
chr6-148540379-DEL-68
chr6-148542961-DEL-90
chr6-148543029-DEL-2892
chr6-148543430-DEL-70
chr6-148549729-DEL-3124
chr6-148574086-DEL-829
chr6-148587926-DEL-561
chr6-148599596-DEL-195
chr6-148610303-DEL-146
chr6-148624201-DEL-76
chr6-148640269-DEL-80
chr6-148675540-DEL-72
chr6-148692056-DEL-109
chr6-148696396-DEL-99
chr6-148794528-DEL-293
chr6-148802348-DEL-320
chr6-148813669-DEL-439
chr6-148831212-DEL-187
chr6-148832563-DEL-276
chr6-148842481-DEL-154
chr6-148843842-DEL-187
chr6-148877481-DEL-6889
chr6-148916751-DEL-51
chr6-148922616-DEL-4639
chr6-148970331-DEL-149
chr6-148977984-DEL-202
chr6-148999701-DEL-204
chr6-149021835-DEL-199
chr6-149026002-DEL-648
chr6-149027857-DEL-52
chr6-149068644-DEL-544
chr6-149084441-DEL-89
chr6-149107628-DEL-52
chr6-149121293-DEL-159
chr6-149150039-DEL-764

chr6-149154687-DEL-362
chr6-149161270-DEL-752
chr6-149203122-DEL-1478
chr6-149206339-DEL-182
chr6-149214023-DEL-212
chr6-149215792-DEL-133
chr6-149226209-DEL-58
chr6-149244148-DEL-524
chr6-149259874-DEL-545
chr6-149302440-DEL-3141
chr6-149303734-DEL-360
chr6-149308583-DEL-202
chr6-149313660-DEL-69
chr6-149322545-DEL-185
chr6-149328139-DEL-3108
chr6-149328285-DEL-1537
chr6-149334462-DEL-80
chr6-149360825-DEL-434
chr6-149369430-DEL-268
chr6-149387463-DEL-78
chr6-149427204-DEL-958
chr6-149435669-DEL-1055
chr7-3052965-DEL-128
chr7-3148813-DEL-2019
chr7-3158231-DEL-140
chr7-3158490-DEL-3475
chr7-3196889-DEL-173
chr7-3216653-DEL-68
chr7-3223536-DEL-2340
chr7-3225761-DEL-98
chr7-3233873-DEL-207
chr7-3240450-DEL-2142
chr7-3250335-DEL-61
chr7-3251547-DEL-210
chr7-3256535-DEL-219
chr7-3264638-DEL-61
chr7-3270307-DEL-61
chr7-3294488-DEL-761
chr7-3301513-DEL-1107
chr7-3301948-DEL-140
chr7-3355730-DEL-52
chr7-3360224-DEL-146
chr7-3364953-DEL-804
chr7-3373422-DEL-54
chr7-3393285-DEL-395
chr7-3400692-DEL-148
chr7-3403657-DEL-54
chr7-3411214-DEL-530
chr7-3412391-DEL-58
chr7-3423380-DEL-71
chr7-3450222-DEL-202
chr7-3452361-DEL-175
chr7-3452886-DEL-56
chr7-3488395-DEL-204
chr7-3525965-DEL-400
chr7-3529606-DEL-326
chr7-3556970-DEL-104
chr7-3567195-DEL-1502
chr7-3567258-DEL-472
chr7-3567549-DEL-829
chr7-3576003-DEL-216
chr7-3608012-DEL-204
chr7-3610833-DEL-328
chr7-3611745-DEL-77
chr7-3630812-DEL-215
chr7-3652489-DEL-108
chr7-3667706-DEL-156
chr7-3711455-DEL-189
chr7-3719960-DEL-19722
chr7-3743547-DEL-205
chr7-3744929-DEL-331
chr7-3761827-DEL-402
chr7-3787537-DEL-1735
chr7-3792128-DEL-6738
chr7-3792558-DEL-214
chr7-3820398-DEL-86
chr7-3821882-DEL-1887

chr7-3831531-DEL-180
chr7-3850091-DEL-331
chr7-3851489-DEL-231
chr7-3875369-DEL-74
chr7-3885398-DEL-1735
chr7-3889988-DEL-6954
chr7-3890130-DEL-214
chr7-3904334-DEL-60
chr7-3935624-DEL-629
chr7-3975443-DEL-6291
chr7-4060581-DEL-4972
chr7-4067046-DEL-68
chr7-4081365-DEL-162
chr7-4107672-DEL-126
chr7-4112094-DEL-114
chr7-4134919-DEL-61
chr7-4153033-DEL-51
chr7-4170566-DEL-56
chr7-4187522-DEL-576
chr7-4188523-DEL-7599
chr7-4262881-DEL-214
chr7-4277632-DEL-150
chr7-4290001-DEL-250
chr7-4294133-DEL-204
chr7-4302649-DEL-955
chr7-4328392-DEL-322
chr7-4329992-DEL-339
chr7-4334388-DEL-68
chr7-4345048-DEL-90
chr7-4357810-DEL-340
chr7-4361587-DEL-162
chr7-4369437-DEL-444
chr7-4391082-DEL-5263
chr7-4398079-DEL-164
chr7-4420966-DEL-4943
chr7-4431322-DEL-122
chr7-4451804-DEL-58
chr7-4453601-DEL-66
chr7-4456529-DEL-177
chr7-4458142-DEL-53
chr7-4480173-DEL-87
chr7-4510923-DEL-74
chr7-4520468-DEL-211
chr7-4521643-DEL-80
chr7-4542215-DEL-85
chr7-4551904-DEL-141
chr7-4559124-DEL-110
chr7-4563571-DEL-129
chr7-4564751-DEL-310
chr7-4566822-DEL-82
chr7-4573660-DEL-120
chr7-4578044-DEL-5336
chr7-4584303-DEL-1512
chr7-4585961-DEL-6422
chr7-4622609-DEL-422
chr7-4625681-DEL-71
chr7-4628292-DEL-61
chr7-4652616-DEL-194
chr7-4653714-DEL-555
chr7-4675797-DEL-144
chr7-4678905-DEL-460
chr7-4684040-DEL-310
chr7-4699352-DEL-88
chr7-4714381-DEL-204
chr7-4717579-DEL-60
chr7-4735695-DEL-150
chr7-4737428-DEL-55
chr7-4753285-DEL-551
chr7-4758661-DEL-100
chr7-4766471-DEL-155
chr7-4770938-DEL-98
chr7-4799348-DEL-57
chr7-4809494-DEL-54
chr7-4816519-DEL-57
chr7-4854609-DEL-65
chr7-4859895-DEL-72
chr7-4864120-DEL-207

chr7-4864655-DEL-60
chr7-4866042-DEL-50
chr7-4878118-DEL-178
chr7-4881627-DEL-3080
chr7-4881634-DEL-99
chr7-4898537-DEL-56
chr7-4954113-DEL-57
chr7-4958159-DEL-89
chr7-4962442-DEL-204
chr7-4964562-DEL-505
chr7-4969909-DEL-83
chr7-4974369-DEL-90
chr7-5001273-DEL-59
chr7-5014130-DEL-54
chr7-5027923-DEL-120
chr7-5030290-DEL-58
chr7-5047640-DEL-308
chr7-5050700-DEL-538
chr7-5085843-DEL-90
chr7-5127025-DEL-97
chr7-5130851-DEL-1249
chr7-5178068-DEL-6228
chr7-5205604-DEL-6871
chr7-5205635-DEL-424
chr7-5214401-DEL-401
chr7-5223244-DEL-7993
chr7-5272018-DEL-7368
chr7-5284093-DEL-147
chr7-5295900-DEL-4587
chr7-5318844-DEL-75
chr7-5329408-DEL-106
chr7-5333148-DEL-70
chr7-5347519-DEL-99
chr7-5348586-DEL-85
chr7-5356924-DEL-87
chr7-5357312-DEL-305
chr7-5358013-DEL-613
chr7-5371143-DEL-7499
chr7-5371253-DEL-6383
chr7-5471823-DEL-258
chr7-5473411-DEL-56
chr7-5480124-DEL-52
chr7-5556324-DEL-6585
chr7-5564673-DEL-291
chr7-5588821-DEL-222
chr7-5594362-DEL-2458
chr7-5597578-DEL-200
chr7-5616100-DEL-1222
chr7-5618016-DEL-1683
chr7-5630071-DEL-102
chr7-5654224-DEL-68
chr7-5763638-DEL-2302
chr7-5826294-DEL-66
chr7-5850238-DEL-6791
chr7-5855802-DEL-84
chr7-5872457-DEL-3601
chr7-5877183-DEL-5107
chr7-5963836-DEL-66
chr7-5975714-DEL-72
chr7-5990495-DEL-3543
chr7-5994948-DEL-270
chr7-5999219-DEL-2391
chr7-6001549-DEL-161
chr7-6002658-DEL-4779
chr7-6003276-DEL-815
chr7-6012992-DEL-89
chr7-6024443-DEL-99
chr7-6024552-DEL-1802
chr7-6029245-DEL-183
chr7-6032178-DEL-126
chr7-6043877-DEL-375
chr7-6052986-DEL-6135
chr7-6060192-DEL-716
chr7-6077778-DEL-264
chr7-6077920-DEL-6639
chr7-6084241-DEL-214
chr7-6179604-DEL-72

chr7-6204840-DEL-315
chr7-6204920-DEL-53
chr7-6243937-DEL-85
chr7-6256868-DEL-194
chr7-6259241-DEL-96
chr7-6262997-DEL-51
chr7-6303978-DEL-108
chr7-6324364-DEL-51
chr7-6328879-DEL-97
chr7-6340172-DEL-205
chr7-6351604-DEL-381
chr7-6352221-DEL-108
chr7-6363253-DEL-79
chr7-6368854-DEL-88
chr7-6387999-DEL-658
chr7-6389376-DEL-181
chr7-6395652-DEL-59
chr7-6399961-DEL-84
chr7-6404002-DEL-1119
chr7-6408728-DEL-107
chr7-6410843-DEL-194
chr7-6415372-DEL-201
chr7-6477603-DEL-129
chr7-6480822-DEL-2040
chr7-6505527-DEL-681
chr7-6507043-DEL-347
chr7-6520478-DEL-56
chr7-6521000-DEL-60
chr7-6541579-DEL-194
chr7-6545097-DEL-1378
chr7-6552333-DEL-1287
chr7-6553037-DEL-349
chr7-6578641-DEL-7959
chr7-6586684-DEL-118
chr7-6593720-DEL-7295
chr7-6593772-DEL-206
chr7-6606141-DEL-66
chr7-6643443-DEL-3155
chr7-6737956-DEL-61
chr7-6755894-DEL-73
chr7-6812805-DEL-79
chr7-6841666-DEL-98
chr7-6851390-DEL-308
chr7-6869953-DEL-151
chr7-6874789-DEL-116
chr7-6897109-DEL-90
chr7-6907616-DEL-158
chr7-6931698-DEL-823
chr7-6933452-DEL-91
chr7-6951073-DEL-162
chr7-6960181-DEL-554
chr7-6974700-DEL-204
chr7-6977040-DEL-114
chr7-6984637-DEL-116
chr7-7011243-DEL-6648
chr7-7021834-DEL-143
chr7-7022639-DEL-1382
chr7-7024075-DEL-98
chr7-7024756-DEL-523
chr7-7045245-DEL-125
chr7-7072345-DEL-4079
chr7-7076462-DEL-6627
chr7-7092666-DEL-1083
chr7-7094988-DEL-17761
chr7-7106284-DEL-204
chr7-7115960-DEL-1303
chr7-7125475-DEL-698
chr7-7129770-DEL-132
chr7-7140544-DEL-377
chr7-7152329-DEL-566
chr7-7161536-DEL-6985
chr7-7167786-DEL-745
chr7-7179318-DEL-2253
chr7-7186799-DEL-141
chr7-7191781-DEL-124
chr7-7219402-DEL-6464
chr7-7238213-DEL-217

chr7-7263773-DEL-139
chr7-7271002-DEL-3102
chr7-7271005-DEL-2018
chr7-7273058-DEL-1043
chr7-7274395-DEL-51
chr7-7274785-DEL-4377
chr7-7275141-DEL-199
chr7-7276311-DEL-2784
chr7-7460192-DEL-2823
chr7-7476098-DEL-52
chr7-7562574-DEL-111
chr7-7626435-DEL-3877
chr7-7648298-DEL-2830
chr7-7651316-DEL-274
chr7-7680279-DEL-56
chr7-7752851-DEL-53
chr7-7783476-DEL-154
chr7-7804384-DEL-53
chr7-7808969-DEL-53
chr7-7893523-DEL-1944
chr7-7987993-DEL-60
chr7-8028081-DEL-2482
chr7-8035571-DEL-51
chr7-8048782-DEL-2481
chr7-8081215-DEL-74
chr7-8095154-DEL-149
chr7-8180444-DEL-2822
chr7-8273206-DEL-52
chr7-8287996-DEL-54
chr7-8293435-DEL-182
chr7-8307865-DEL-609
chr7-8356885-DEL-53
chr7-8360555-DEL-90
chr7-8387499-DEL-104
chr7-8407857-DEL-3719
chr7-8411410-DEL-9643
chr7-8495640-DEL-64
chr7-8583261-DEL-154
chr7-8599676-DEL-4949
chr7-8676865-DEL-65
chr7-8774995-DEL-273
chr7-8783041-DEL-6213
chr7-8846643-DEL-598
chr7-8895011-DEL-545
chr7-9016630-DEL-340
chr7-9078981-DEL-7121
chr7-9153817-DEL-53
chr7-9154815-DEL-7353
chr7-9167484-DEL-77
chr7-9195895-DEL-1953
chr7-9253747-DEL-5524
chr7-9320992-DEL-65
chr7-9455678-DEL-54
chr7-9521573-DEL-270
chr7-9540985-DEL-362
chr7-9640030-DEL-548
chr7-9652668-DEL-9372
chr7-9665667-DEL-2338
chr7-9690422-DEL-846
chr7-9761382-DEL-3581
chr7-9810723-DEL-1490
chr7-9888838-DEL-54
chr7-9891180-DEL-492
chr7-9925387-DEL-1176
chr7-9956280-DEL-668
chr7-9958840-DEL-468
chr7-9961174-DEL-6339
chr7-9977945-DEL-67
chr7-9983835-DEL-280
chr7-10015611-DEL-245
chr7-10025439-DEL-67
chr7-10038900-DEL-121
chr7-10041885-DEL-369
chr7-10048763-DEL-1430
chr7-10050703-DEL-536
chr7-10051979-DEL-1036
chr7-10053068-DEL-2010

chr7-10056404-DEL-3791
chr7-10071421-DEL-402
chr7-10075415-DEL-60
chr7-10091489-DEL-55
chr7-10107572-DEL-124
chr7-10107820-DEL-980
chr7-10109806-DEL-190
chr7-10111382-DEL-96
chr7-10114566-DEL-128
chr7-10130820-DEL-168
chr7-10166632-DEL-413
chr7-10193826-DEL-412
chr7-10205057-DEL-10105
chr7-10205758-DEL-235
chr7-10226787-DEL-197
chr7-10274775-DEL-471
chr7-10293024-DEL-3888
chr7-10310220-DEL-53
chr7-10326983-DEL-57
chr7-10338002-DEL-336
chr7-10353197-DEL-6271
chr7-10366631-DEL-53
chr7-10368462-DEL-1335
chr7-10385185-DEL-1887
chr7-10388275-DEL-1453
chr7-10390213-DEL-688
chr7-10410194-DEL-75
chr7-10465525-DEL-102
chr7-10534918-DEL-57
chr7-10558718-DEL-6413
chr7-10576686-DEL-3518
chr7-10586374-DEL-1559
chr7-10633539-DEL-556
chr7-10641892-DEL-189
chr7-10643255-DEL-7133
chr7-10690948-DEL-597
chr7-10735286-DEL-83
chr7-10755409-DEL-5979
chr7-10767211-DEL-444
chr7-10776849-DEL-5386
chr7-10782022-DEL-431
chr7-10788201-DEL-4373
chr7-10800158-DEL-1723
chr7-10837167-DEL-1204
chr7-10883338-DEL-2224
chr7-10900637-DEL-353
chr7-10903869-DEL-6444
chr7-10903925-DEL-253
chr7-10974351-DEL-97
chr7-10976586-DEL-55
chr7-10987009-DEL-1237
chr7-11020989-DEL-2570
chr7-11082994-DEL-284
chr7-11085737-DEL-250
chr7-11088529-DEL-11723
chr7-11108977-DEL-219
chr7-11127479-DEL-83
chr7-11168785-DEL-434
chr7-11178589-DEL-195
chr7-11180023-DEL-151
chr7-11182805-DEL-3561
chr7-11190906-DEL-198
chr7-11193277-DEL-1081
chr7-11195288-DEL-252
chr7-11195954-DEL-83
chr7-11240107-DEL-9311
chr7-11296413-DEL-150
chr7-11307453-DEL-9326
chr7-11329639-DEL-69
chr7-11334483-DEL-193
chr7-11335429-DEL-5538
chr7-11348462-DEL-537
chr7-11353731-DEL-922
chr7-11378460-DEL-416
chr7-11425618-DEL-5744
chr7-11444024-DEL-171
chr7-11453981-DEL-1083

chr7-11530554-DEL-416
chr7-11533899-DEL-53
chr7-11534746-DEL-564
chr7-11583134-DEL-75
chr7-11591446-DEL-1839
chr7-11595073-DEL-1952
chr7-11625033-DEL-3095
chr7-11627046-DEL-6397
chr7-11632915-DEL-1159
chr7-11643627-DEL-110
chr7-11646728-DEL-2466
chr7-11649227-DEL-5444
chr7-11663537-DEL-4040
chr7-11670190-DEL-98
chr7-11680071-DEL-145
chr7-11719476-DEL-689
chr7-11744803-DEL-176
chr7-11746994-DEL-177
chr7-11748825-DEL-88
chr7-11750009-DEL-100
chr7-11750715-DEL-549
chr7-11772245-DEL-1582
chr7-11784958-DEL-6235
chr7-11794918-DEL-769
chr7-11796297-DEL-2481
chr7-11799706-DEL-715
chr7-11803780-DEL-6201
chr7-11811739-DEL-128
chr7-11815003-DEL-5608
chr7-11832584-DEL-670
chr7-11883455-DEL-91
chr7-11887754-DEL-4931
chr7-11898086-DEL-3290
chr7-11905575-DEL-59
chr7-11921382-DEL-96
chr7-11935406-DEL-299
chr7-11964129-DEL-192
chr7-11970957-DEL-1697
chr7-11974409-DEL-61
chr7-12046139-DEL-220
chr7-12064890-DEL-3058
chr7-12066870-DEL-1719
chr7-12068819-DEL-19883
chr7-12103991-DEL-17871
chr7-12123278-DEL-1697
chr7-12128508-DEL-3564
chr7-12180135-DEL-52
chr7-12183495-DEL-761
chr7-12183680-DEL-56
chr7-12184454-DEL-2468
chr7-12190806-DEL-578
chr7-12234312-DEL-2734
chr7-12260448-DEL-6452
chr7-12260461-DEL-3646
chr7-12264075-DEL-2808
chr7-12270295-DEL-11863
chr7-12300317-DEL-6198
chr7-12322077-DEL-7551
chr7-12327401-DEL-1291
chr7-12373472-DEL-70
chr7-12377966-DEL-6210
chr7-12403356-DEL-65
chr7-12412694-DEL-522
chr7-12432133-DEL-3305
chr7-12449323-DEL-317
chr7-12450754-DEL-718
chr7-12456024-DEL-204
chr7-12465960-DEL-53
chr7-12472100-DEL-90
chr7-12497822-DEL-53
chr7-12508999-DEL-54
chr7-12524867-DEL-134
chr7-12549565-DEL-208
chr7-12557978-DEL-326
chr7-12559539-DEL-498
chr7-12590470-DEL-157
chr7-12608706-DEL-617

chr7-12612277-DEL-413
chr7-12654954-DEL-293
chr7-12665033-DEL-57
chr7-12667017-DEL-57
chr7-12671326-DEL-160
chr7-12683854-DEL-211
chr7-12687802-DEL-189
chr7-12689290-DEL-122
chr7-12749612-DEL-231
chr7-12793491-DEL-2961
chr7-12799146-DEL-69
chr7-12800374-DEL-1518
chr7-12813717-DEL-165
chr7-12822163-DEL-67
chr7-12824360-DEL-60
chr7-12840296-DEL-173
chr7-12846074-DEL-197
chr7-12881902-DEL-192
chr7-12902482-DEL-432
chr7-12920512-DEL-6401
chr7-12943088-DEL-322
chr7-12959614-DEL-6745
chr7-13043838-DEL-378
chr7-13046659-DEL-178
chr7-13068483-DEL-164
chr7-13095550-DEL-146
chr7-13112029-DEL-56
chr7-13142913-DEL-102
chr7-13148227-DEL-1431
chr7-13178187-DEL-6472
chr7-13190514-DEL-54
chr7-13209873-DEL-5151
chr7-13215012-DEL-6180
chr7-13223057-DEL-275
chr7-13234161-DEL-2587
chr7-13237236-DEL-6187
chr7-13276841-DEL-117
chr7-13279641-DEL-117
chr7-13296522-DEL-500
chr7-13312101-DEL-8405
chr7-13340946-DEL-54
chr7-13348092-DEL-6292
chr7-13383583-DEL-704
chr7-13420011-DEL-435
chr7-13424237-DEL-59
chr7-13449434-DEL-2414
chr7-13464544-DEL-80
chr7-13466955-DEL-86
chr7-13511599-DEL-51
chr7-13512359-DEL-118
chr7-13515696-DEL-75
chr7-13579628-DEL-52
chr7-13605729-DEL-132
chr7-13605903-DEL-4987
chr7-13620540-DEL-10863
chr7-13625984-DEL-114
chr7-13631167-DEL-215
chr7-13654324-DEL-90
chr7-13660007-DEL-73
chr7-13660354-DEL-330
chr7-13661277-DEL-114
chr7-13699112-DEL-1696
chr7-13700768-DEL-1659
chr7-13765455-DEL-107
chr7-13793670-DEL-6371
chr7-13811943-DEL-92
chr7-13874956-DEL-465
chr7-13909338-DEL-11259
chr7-13922427-DEL-2892
chr7-13928000-DEL-56
chr7-13972611-DEL-324
chr7-13998890-DEL-1030
chr7-14049457-DEL-56
chr7-14056191-DEL-205
chr7-14082641-DEL-201
chr7-14088610-DEL-4387
chr7-14104198-DEL-6186

chr7-14118246-DEL-6008
chr7-14137441-DEL-6751
chr7-14170958-DEL-292
chr7-14177504-DEL-56
chr7-14221204-DEL-1870
chr7-14241726-DEL-3984
chr7-14319314-DEL-364
chr7-14336573-DEL-3184
chr7-14349054-DEL-760
chr7-14356232-DEL-1356
chr7-14364809-DEL-52
chr7-14398719-DEL-220
chr7-14406633-DEL-196
chr7-14415940-DEL-946
chr7-14432992-DEL-88
chr7-14453223-DEL-239
chr7-14457388-DEL-53
chr7-14470717-DEL-9828
chr7-14494317-DEL-135
chr7-14594701-DEL-141
chr7-14595692-DEL-1557
chr7-14631604-DEL-140
chr7-14643019-DEL-5643
chr7-14685173-DEL-6554
chr7-14686481-DEL-6655
chr7-14707552-DEL-5193
chr7-14729655-DEL-138
chr7-14730638-DEL-5157
chr7-14731930-DEL-3601
chr7-14735761-DEL-440
chr7-14750350-DEL-6620
chr7-14777968-DEL-853
chr7-14855760-DEL-132
chr7-14858495-DEL-8619
chr7-15041724-DEL-84
chr7-15154308-DEL-63
chr7-15196084-DEL-7333
chr7-15196189-DEL-4131
chr7-15200740-DEL-1409
chr7-15202423-DEL-167
chr7-15223982-DEL-711
chr7-15230195-DEL-144
chr7-15231184-DEL-1557
chr7-15275244-DEL-3503
chr7-15329802-DEL-7079
chr7-15388056-DEL-6060
chr7-15413978-DEL-6148
chr7-15462140-DEL-520
chr7-15470020-DEL-2192
chr7-15516071-DEL-460
chr7-15571203-DEL-174
chr7-15571561-DEL-530
chr7-15572169-DEL-576
chr7-15575766-DEL-155
chr7-15596733-DEL-168
chr7-15598938-DEL-53
chr7-15616121-DEL-174
chr7-15622842-DEL-822
chr7-15641253-DEL-627
chr7-15649840-DEL-164
chr7-15652086-DEL-212
chr7-15653204-DEL-75
chr7-15663425-DEL-244
chr7-15699203-DEL-51
chr7-15732633-DEL-52
chr7-15742758-DEL-106
chr7-15745176-DEL-106
chr7-15763085-DEL-105
chr7-15763666-DEL-205
chr7-15818797-DEL-65
chr7-15856459-DEL-177
chr7-15910750-DEL-106
chr7-15913343-DEL-114
chr7-15915056-DEL-106
chr7-15931555-DEL-296
chr7-15953861-DEL-67
chr7-15959419-DEL-82

chr7-15984391-DEL-60
chr7-15986097-DEL-407
chr7-15987164-DEL-231
chr7-15991466-DEL-259
chr7-15997031-DEL-100
chr7-16013685-DEL-203
chr7-16026594-DEL-195
chr7-16041349-DEL-51
chr7-16065925-DEL-83
chr7-16074640-DEL-2471
chr7-16087103-DEL-2009
chr7-16105377-DEL-150
chr7-16114456-DEL-178
chr7-16117465-DEL-443
chr7-16132316-DEL-198
chr7-16176195-DEL-75
chr7-16216935-DEL-635
chr7-16237124-DEL-73
chr7-16258731-DEL-80
chr7-16354750-DEL-79
chr7-16360875-DEL-55
chr7-16366242-DEL-373
chr7-16389394-DEL-407
chr7-16393090-DEL-680
chr7-16463363-DEL-342
chr7-16464578-DEL-701
chr7-16466966-DEL-202
chr7-16479236-DEL-106
chr7-16479739-DEL-275
chr7-16486586-DEL-77
chr7-16490810-DEL-177
chr7-16493489-DEL-1205
chr7-16499562-DEL-98
chr7-16504667-DEL-234
chr7-16512093-DEL-130
chr7-16537787-DEL-81
chr7-16538675-DEL-1324
chr7-16576441-DEL-52
chr7-16585132-DEL-165
chr7-16598819-DEL-186
chr7-16654512-DEL-52
chr7-16666655-DEL-197
chr7-16669044-DEL-177
chr7-16699464-DEL-63
chr7-16712043-DEL-293
chr7-16735219-DEL-52
chr7-16774463-DEL-52
chr7-16780534-DEL-261
chr7-16783301-DEL-205
chr7-16786243-DEL-175
chr7-16791511-DEL-117
chr7-16798196-DEL-105
chr7-16819269-DEL-52
chr7-16831704-DEL-91
chr7-16834930-DEL-239
chr7-16843190-DEL-202
chr7-16843702-DEL-7002
chr7-16845892-DEL-6241
chr7-16855404-DEL-6300
chr7-16865365-DEL-2457
chr7-16874441-DEL-655
chr7-16886843-DEL-390
chr7-16903948-DEL-67
chr7-16916463-DEL-6602
chr7-16921207-DEL-1987
chr7-16923545-DEL-6305
chr7-16930176-DEL-66
chr7-16930728-DEL-82
chr7-16932720-DEL-374
chr7-16949384-DEL-347
chr7-16955678-DEL-721
chr7-16958088-DEL-72
chr7-16958571-DEL-63
chr7-16959228-DEL-5582
chr7-16975812-DEL-6786
chr7-16982643-DEL-93
chr7-16997559-DEL-7943

chr7-17027563-DEL-5725
chr7-17033516-DEL-2568
chr7-17044079-DEL-1467
chr7-17044225-DEL-527
chr7-17059271-DEL-278
chr7-17084647-DEL-190
chr7-17089547-DEL-1274
chr7-17095310-DEL-147
chr7-17110563-DEL-7202
chr7-17118203-DEL-1481
chr7-17120642-DEL-10041
chr7-17142689-DEL-2573
chr7-17153263-DEL-1467
chr7-17158193-DEL-1852
chr7-17173774-DEL-2153
chr7-17177239-DEL-2160
chr7-17178497-DEL-287
chr7-17188738-DEL-5183
chr7-17190331-DEL-2749
chr7-17201989-DEL-1595
chr7-17204264-DEL-473
chr7-17221415-DEL-180
chr7-17256636-DEL-73
chr7-17264368-DEL-5218
chr7-17265996-DEL-2749
chr7-17270703-DEL-516
chr7-17271759-DEL-5410
chr7-17272377-DEL-76
chr7-17302371-DEL-180
chr7-17315481-DEL-1882
chr7-17317466-DEL-69
chr7-17385267-DEL-836
chr7-17422565-DEL-6759
chr7-17432167-DEL-349
chr7-17433855-DEL-1549
chr7-17456089-DEL-197
chr7-17471561-DEL-77
chr7-17480714-DEL-102
chr7-17493952-DEL-321
chr7-17501523-DEL-2478
chr7-17547490-DEL-53
chr7-17554844-DEL-198
chr7-17578497-DEL-6351
chr7-17585077-DEL-1337
chr7-17587042-DEL-52
chr7-17588731-DEL-937
chr7-17590964-DEL-6308
chr7-17624643-DEL-104
chr7-17650668-DEL-69
chr7-17682515-DEL-668
chr7-17721188-DEL-1709
chr7-17727343-DEL-3070
chr7-17732934-DEL-573
chr7-17736622-DEL-3275
chr7-17749509-DEL-1493
chr7-17774882-DEL-444
chr7-17818232-DEL-444
chr7-17829738-DEL-4337
chr7-17836913-DEL-51
chr7-17841604-DEL-6606
chr7-17875166-DEL-53
chr7-17921880-DEL-6575
chr7-17928448-DEL-3723
chr7-17932733-DEL-3658
chr7-18004604-DEL-101
chr7-18012578-DEL-65
chr7-18015318-DEL-3478
chr7-18022346-DEL-3753
chr7-18031741-DEL-141
chr7-18036647-DEL-51
chr7-18059778-DEL-138
chr7-18061973-DEL-7814
chr7-18066715-DEL-58
chr7-18072358-DEL-3830
chr7-18112726-DEL-13242
chr7-18146711-DEL-425
chr7-18147543-DEL-7254

chr7-18186308-DEL-1356
chr7-18186331-DEL-934
chr7-18187159-DEL-506
chr7-18187359-DEL-303
chr7-18188798-DEL-220
chr7-18199264-DEL-57
chr7-18235464-DEL-4307
chr7-18246776-DEL-78
chr7-18273438-DEL-2816
chr7-18431475-DEL-7040
chr7-18440522-DEL-83
chr7-18448777-DEL-156
chr7-18449471-DEL-129
chr7-18467652-DEL-70
chr7-18479851-DEL-217
chr7-18484060-DEL-432
chr7-18551600-DEL-220
chr7-18564575-DEL-76
chr7-18575847-DEL-117
chr7-18584190-DEL-166
chr7-18584851-DEL-166
chr7-18591322-DEL-57
chr7-18603303-DEL-3209
chr7-18609729-DEL-61
chr7-18645623-DEL-158
chr7-18655821-DEL-76
chr7-18666603-DEL-52
chr7-18672127-DEL-53
chr7-18675126-DEL-53
chr7-18679799-DEL-265
chr7-18683266-DEL-52
chr7-18700573-DEL-53
chr7-18706165-DEL-54
chr7-18707322-DEL-328
chr7-18709079-DEL-129
chr7-18710867-DEL-54
chr7-18713625-DEL-52
chr7-18751071-DEL-323
chr7-18803347-DEL-1203
chr7-18849912-DEL-159
chr7-18896396-DEL-52
chr7-18952845-DEL-52
chr7-18984136-DEL-82
chr7-19009179-DEL-110
chr7-19021715-DEL-255
chr7-19029119-DEL-64
chr7-19034138-DEL-99
chr7-19062480-DEL-51
chr7-19063644-DEL-772
chr7-19067253-DEL-54
chr7-19107618-DEL-1316
chr7-19152188-DEL-51
chr7-19153796-DEL-185
chr7-19156122-DEL-1133
chr7-19169190-DEL-106
chr7-19170485-DEL-52
chr7-19171492-DEL-95
chr7-19188360-DEL-107
chr7-19213261-DEL-217
chr7-19216859-DEL-135
chr7-19219936-DEL-106
chr7-19227194-DEL-88
chr7-19246903-DEL-1156
chr7-19261620-DEL-899
chr7-19272090-DEL-777
chr7-19289216-DEL-246
chr7-19305052-DEL-54
chr7-19374257-DEL-169
chr7-19388249-DEL-149
chr7-19405839-DEL-1128
chr7-19405843-DEL-612
chr7-19417576-DEL-54
chr7-19421717-DEL-52
chr7-19421974-DEL-63
chr7-19429961-DEL-51
chr7-19461962-DEL-166
chr7-19464892-DEL-256

chr7-19494620-DEL-106
chr7-19512109-DEL-211
chr7-19533598-DEL-222
chr7-19534124-DEL-232
chr7-19549644-DEL-54
chr7-19582357-DEL-511
chr7-19603977-DEL-72
chr7-19617906-DEL-168
chr7-19618694-DEL-212
chr7-19624843-DEL-200
chr7-19629060-DEL-78
chr7-19629684-DEL-64
chr7-19642434-DEL-53
chr7-19656650-DEL-51
chr7-19662458-DEL-20100
chr7-19685445-DEL-59
chr7-19687265-DEL-53
chr7-19720770-DEL-77
chr7-19722201-DEL-550
chr7-19725294-DEL-70
chr7-19729455-DEL-189
chr7-19764797-DEL-835
chr7-19768470-DEL-1482
chr7-19794381-DEL-230
chr7-19808730-DEL-16857
chr7-19830863-DEL-389
chr7-19840182-DEL-5528
chr7-20179854-DEL-76
chr7-20264907-DEL-1394
chr7-20479526-DEL-52
chr7-20500864-DEL-395
chr7-20567119-DEL-204
chr7-20635763-DEL-3689
chr7-20640366-DEL-507
chr7-20804200-DEL-6797
chr7-20810560-DEL-212
chr7-20845842-DEL-8750
chr7-20849823-DEL-243
chr7-20850671-DEL-1599
chr7-20853231-DEL-204
chr7-20854163-DEL-9376
chr7-20898075-DEL-1598
chr7-20908017-DEL-553
chr7-20910453-DEL-6424
chr7-20925551-DEL-197
chr7-20927719-DEL-61
chr7-20931869-DEL-8789
chr7-20954400-DEL-5929
chr7-20974518-DEL-268
chr7-20994801-DEL-376
chr7-21024439-DEL-196
chr7-21040320-DEL-203
chr7-21051698-DEL-97
chr7-21086069-DEL-12849
chr7-21086100-DEL-84
chr7-21111473-DEL-79
chr7-21113524-DEL-81
chr7-21114456-DEL-1074
chr7-21146982-DEL-67
chr7-21156512-DEL-926
chr7-21167105-DEL-1177
chr7-21184101-DEL-158
chr7-21189125-DEL-703
chr7-21388193-DEL-8779
chr7-21410677-DEL-5927
chr7-21632131-DEL-52
chr7-21637432-DEL-703
chr7-21638231-DEL-1333
chr7-21657120-DEL-132
chr7-22117584-DEL-6406
chr7-22124635-DEL-1710
chr7-22139505-DEL-2963
chr7-22140980-DEL-76
chr7-22143850-DEL-6380
chr7-22365994-DEL-341
chr7-22368615-DEL-193
chr7-22463349-DEL-2112

chr7-22464953-DEL-6203
chr7-22471799-DEL-7062
chr7-22472245-DEL-5353
chr7-22475472-DEL-830
chr7-22480792-DEL-4763
chr7-22483384-DEL-2900
chr7-22486297-DEL-1516
chr7-22520094-DEL-9476
chr7-22524766-DEL-243
chr7-22816989-DEL-11950
chr7-22855846-DEL-6427
chr7-22858523-DEL-2218
chr7-22886941-DEL-110
chr7-22907856-DEL-362
chr7-22927938-DEL-194
chr7-22959115-DEL-1353
chr7-22978760-DEL-1177
chr7-22993274-DEL-56
chr7-22997610-DEL-84
chr7-23006921-DEL-8526
chr7-23023314-DEL-8775
chr7-23036833-DEL-900
chr7-23056432-DEL-1818
chr7-23073331-DEL-56
chr7-23082817-DEL-76
chr7-23087107-DEL-283
chr7-23150622-DEL-51
chr7-23192533-DEL-535
chr7-23234140-DEL-325
chr7-23285185-DEL-203
chr7-23286544-DEL-4472
chr7-23295020-DEL-3560
chr7-23298677-DEL-938
chr7-23304844-DEL-235
chr7-23313159-DEL-66
chr7-23321602-DEL-342
chr7-23355112-DEL-63
chr7-23387268-DEL-1475
chr7-23445054-DEL-5900
chr7-23452152-DEL-146
chr7-23491820-DEL-9060
chr7-23528083-DEL-2162
chr7-23537036-DEL-12054
chr7-23575963-DEL-4836
chr7-23583434-DEL-347
chr7-23618837-DEL-56
chr7-23687430-DEL-630
chr7-23701992-DEL-65
chr7-23709018-DEL-1263
chr7-23709574-DEL-23309
chr7-23709816-DEL-6017
chr7-23710917-DEL-6937
chr7-23725617-DEL-6226
chr7-23759733-DEL-5567
chr7-23772188-DEL-270
chr7-23776383-DEL-53
chr7-23810817-DEL-495
chr7-23824647-DEL-528
chr7-23830774-DEL-233
chr7-23885232-DEL-115
chr7-23996090-DEL-84
chr7-24025115-DEL-51
chr7-24043755-DEL-56
chr7-24061363-DEL-170
chr7-24067457-DEL-121
chr7-24071676-DEL-57
chr7-24074775-DEL-1394
chr7-24075567-DEL-159
chr7-24088603-DEL-102
chr7-24096050-DEL-438
chr7-24105581-DEL-189
chr7-24110910-DEL-216
chr7-24124593-DEL-320
chr7-24139024-DEL-187
chr7-24139346-DEL-550
chr7-24143915-DEL-158
chr7-24147320-DEL-104

chr7-24152494-DEL-176
chr7-24160655-DEL-79
chr7-24163122-DEL-54
chr7-24170513-DEL-53
chr7-24195046-DEL-53
chr7-24215615-DEL-585
chr7-24219956-DEL-131
chr7-24221799-DEL-370
chr7-24226951-DEL-239
chr7-24288584-DEL-52
chr7-24289478-DEL-54
chr7-24304194-DEL-208
chr7-24346402-DEL-54
chr7-24358648-DEL-54
chr7-24361752-DEL-59
chr7-24383109-DEL-407
chr7-24402247-DEL-173
chr7-24415010-DEL-162
chr7-24429685-DEL-404
chr7-24433699-DEL-196
chr7-24458904-DEL-81
chr7-24461770-DEL-54
chr7-24490100-DEL-301
chr7-24510269-DEL-226
chr7-24562736-DEL-645
chr7-24565726-DEL-263
chr7-24590899-DEL-64
chr7-24613608-DEL-688
chr7-24663135-DEL-66
chr7-24682072-DEL-51
chr7-24697719-DEL-174
chr7-24699113-DEL-183
chr7-24723106-DEL-76
chr7-24730100-DEL-1152
chr7-24740125-DEL-276
chr7-24795374-DEL-76
chr7-24810797-DEL-367
chr7-24824815-DEL-90
chr7-24836459-DEL-742
chr7-24839923-DEL-79
chr7-24843091-DEL-68
chr7-24857739-DEL-54
chr7-24865336-DEL-56
chr7-24890894-DEL-72
chr7-24895242-DEL-275
chr7-24897974-DEL-60
chr7-24898935-DEL-200
chr7-24913155-DEL-189
chr7-24916249-DEL-174
chr7-24958698-DEL-318
chr7-24958822-DEL-838
chr7-24959034-DEL-339
chr7-24965776-DEL-597
chr7-25004412-DEL-58
chr7-25010096-DEL-92
chr7-25019710-DEL-71
chr7-25031919-DEL-68
chr7-25032407-DEL-103
chr7-25035935-DEL-56
chr7-25050695-DEL-203
chr7-25052542-DEL-102
chr7-25061741-DEL-105
chr7-25095580-DEL-58
chr7-25104927-DEL-2109
chr7-25116204-DEL-146
chr7-25117704-DEL-128
chr7-25123286-DEL-195
chr7-25145717-DEL-80
chr7-25155623-DEL-98
chr7-25156399-DEL-2976
chr7-25188279-DEL-203
chr7-25218999-DEL-599
chr7-25226060-DEL-122
chr7-25230267-DEL-1596
chr7-25230391-DEL-5670
chr7-25267753-DEL-53
chr7-25269156-DEL-5186

chr7-25282041-DEL-65
chr7-25285721-DEL-4068
chr7-25310650-DEL-54
chr7-25311413-DEL-54
chr7-25312766-DEL-126
chr7-25317857-DEL-93
chr7-25347797-DEL-107
chr7-25352250-DEL-52
chr7-25371517-DEL-205
chr7-25379112-DEL-90
chr7-25391188-DEL-108
chr7-25398975-DEL-53
chr7-25406112-DEL-61
chr7-25434623-DEL-213
chr7-25474877-DEL-152
chr7-25476734-DEL-319
chr7-25484628-DEL-348
chr7-25495478-DEL-101
chr7-25506630-DEL-56
chr7-25512073-DEL-203
chr7-25513651-DEL-106
chr7-25519686-DEL-439
chr7-25525723-DEL-993
chr7-25560416-DEL-165
chr7-25565064-DEL-241
chr7-25584841-DEL-81
chr7-25588929-DEL-109
chr7-25591602-DEL-411
chr7-25599138-DEL-117
chr7-25603459-DEL-164
chr7-25621712-DEL-389
chr7-25622437-DEL-2411
chr7-25626089-DEL-213
chr7-25646703-DEL-620
chr7-25660623-DEL-347
chr7-25661860-DEL-112
chr7-25662656-DEL-849
chr7-25691690-DEL-349
chr7-25706153-DEL-134
chr7-25728319-DEL-2560
chr7-25730756-DEL-344
chr7-25743894-DEL-6490
chr7-25789072-DEL-274
chr7-25828425-DEL-3576
chr7-25850443-DEL-344
chr7-25885428-DEL-128
chr7-25901238-DEL-2261
chr7-25905280-DEL-1043
chr7-25908858-DEL-2566
chr7-25959575-DEL-53
chr7-25967640-DEL-1052
chr7-25983193-DEL-6620
chr7-26009491-DEL-159
chr7-26118153-DEL-98
chr7-26129959-DEL-505
chr7-26140491-DEL-100
chr7-26167282-DEL-181
chr7-26169820-DEL-1407
chr7-26188522-DEL-1501
chr7-26206154-DEL-146
chr7-26213357-DEL-59
chr7-26213888-DEL-615
chr7-26216310-DEL-271
chr7-26256527-DEL-55
chr7-26277607-DEL-282
chr7-26292041-DEL-5506
chr7-26322904-DEL-317
chr7-26340638-DEL-370
chr7-26351415-DEL-6403
chr7-26362846-DEL-59
chr7-26382889-DEL-106
chr7-26405365-DEL-5065
chr7-26446365-DEL-70
chr7-26457498-DEL-525
chr7-26464048-DEL-100
chr7-26479082-DEL-185
chr7-26537664-DEL-149

chr7-26567615-DEL-4475
chr7-26622242-DEL-6468
chr7-26630864-DEL-52
chr7-26677094-DEL-1525
chr7-26680887-DEL-66
chr7-26706364-DEL-5068
chr7-26710242-DEL-3851
chr7-26725721-DEL-96
chr7-26756679-DEL-53
chr7-26764439-DEL-2746
chr7-26771161-DEL-104
chr7-26788136-DEL-56
chr7-26794412-DEL-794
chr7-26796155-DEL-1609
chr7-26835279-DEL-156
chr7-26849401-DEL-204
chr7-26850537-DEL-55
chr7-26856493-DEL-55
chr7-26861996-DEL-69
chr7-26883070-DEL-213
chr7-26916980-DEL-138
chr7-26921948-DEL-53
chr7-26938780-DEL-165
chr7-26971214-DEL-261
chr7-26990290-DEL-122
chr7-26992311-DEL-181
chr7-27010336-DEL-54
chr7-27014534-DEL-807
chr7-27043745-DEL-86
chr7-27070155-DEL-59
chr7-27072936-DEL-212
chr7-27080500-DEL-2075
chr7-27089438-DEL-106
chr7-27099265-DEL-181
chr7-27159686-DEL-100
chr7-27170467-DEL-96
chr7-27184493-DEL-96
chr7-27194228-DEL-266
chr7-27194913-DEL-64
chr7-27205012-DEL-60
chr7-27208934-DEL-207
chr7-27210501-DEL-301
chr7-27229715-DEL-53
chr7-27245408-DEL-198
chr7-27256279-DEL-1179
chr7-27256456-DEL-567
chr7-27256921-DEL-603
chr7-27263090-DEL-286
chr7-27268488-DEL-113
chr7-27275965-DEL-52
chr7-27314652-DEL-405
chr7-27334701-DEL-90
chr7-27344310-DEL-204
chr7-27348454-DEL-220
chr7-27351929-DEL-64
chr7-27355492-DEL-90
chr7-27387146-DEL-120
chr7-27388584-DEL-88
chr7-27397276-DEL-279
chr7-27405243-DEL-305
chr7-27411265-DEL-208
chr7-27412149-DEL-7206
chr7-27418731-DEL-77
chr7-27451503-DEL-199
chr7-27468888-DEL-407
chr7-27475103-DEL-590
chr7-27498050-DEL-132
chr7-27504841-DEL-182
chr7-27508519-DEL-235
chr7-27523339-DEL-149
chr7-27526044-DEL-1688
chr7-27548804-DEL-208
chr7-27581797-DEL-2250
chr7-27594184-DEL-3414
chr7-27598939-DEL-644
chr7-27599555-DEL-1849
chr7-27630535-DEL-1075

chr7-27630799-DEL-93
chr7-27646079-DEL-205
chr7-27659731-DEL-336
chr7-27671968-DEL-197
chr7-27699935-DEL-1684
chr7-27709554-DEL-202
chr7-27759557-DEL-460
chr7-27782564-DEL-204
chr7-27795353-DEL-2208
chr7-27795381-DEL-2683
chr7-27795745-DEL-609
chr7-27796440-DEL-137
chr7-27797091-DEL-540
chr7-27797828-DEL-237
chr7-27849636-DEL-231
chr7-27859908-DEL-198
chr7-27864428-DEL-141
chr7-27901296-DEL-116
chr7-27901956-DEL-317
chr7-27915912-DEL-163
chr7-27928190-DEL-111
chr7-27943235-DEL-396
chr7-27944140-DEL-164
chr7-27944413-DEL-396
chr7-27944486-DEL-181
chr7-27946279-DEL-66
chr7-27946930-DEL-165
chr7-27948593-DEL-365
chr7-27963645-DEL-53
chr7-27974998-DEL-74
chr7-27995322-DEL-303
chr7-28012233-DEL-604
chr7-28027095-DEL-198
chr7-28032135-DEL-181
chr7-28033580-DEL-201
chr7-28034761-DEL-141
chr7-28041540-DEL-270
chr7-28042460-DEL-209
chr7-28060827-DEL-269
chr7-28087341-DEL-86
chr7-28096817-DEL-204
chr7-28125406-DEL-200
chr7-28128163-DEL-218
chr7-28141796-DEL-181
chr7-28143005-DEL-70
chr7-28162596-DEL-200
chr7-28178467-DEL-730
chr7-28204597-DEL-64
chr7-28231025-DEL-177
chr7-28236081-DEL-106
chr7-28249304-DEL-199
chr7-28252020-DEL-146
chr7-28267749-DEL-110
chr7-28279978-DEL-305
chr7-28321593-DEL-106
chr7-28338185-DEL-369
chr7-28340118-DEL-70
chr7-28349463-DEL-51
chr7-28355227-DEL-106
chr7-28366490-DEL-116
chr7-28375689-DEL-253
chr7-28380251-DEL-51
chr7-28381439-DEL-86
chr7-28382187-DEL-434
chr7-28402428-DEL-200
chr7-28405215-DEL-211
chr7-28408869-DEL-64
chr7-28419035-DEL-60
chr7-28444956-DEL-68
chr7-28447442-DEL-102
chr7-28451073-DEL-61
chr7-28490308-DEL-89
chr7-28493628-DEL-80
chr7-28506084-DEL-996
chr7-28506944-DEL-51
chr7-28522578-DEL-61
chr7-28529064-DEL-115

chr7-28542958-DEL-57
chr7-28555285-DEL-91
chr7-28574554-DEL-445
chr7-28583438-DEL-53
chr7-28621710-DEL-52
chr7-28665624-DEL-60
chr7-28714514-DEL-102
chr7-28722322-DEL-72
chr7-28727430-DEL-153
chr7-28729763-DEL-132
chr7-28733818-DEL-354
chr7-28734602-DEL-481
chr7-28735781-DEL-1685
chr7-28736518-DEL-73
chr7-28739632-DEL-65
chr7-28741979-DEL-137
chr7-28751172-DEL-86
chr7-28779872-DEL-325
chr7-28843826-DEL-52
chr7-28844869-DEL-214
chr7-28876988-DEL-201
chr7-28894143-DEL-233
chr7-28917730-DEL-186
chr7-28936685-DEL-179
chr7-28960762-DEL-148
chr7-28967158-DEL-991
chr7-28968397-DEL-413
chr7-28974024-DEL-674
chr7-29058953-DEL-120
chr7-29080426-DEL-55
chr7-29089062-DEL-138
chr7-29113586-DEL-57
chr7-29119483-DEL-63
chr7-29122224-DEL-487
chr7-29143095-DEL-102
chr7-29145079-DEL-53
chr7-29185873-DEL-51
chr7-29201970-DEL-106
chr7-29206849-DEL-204
chr7-29240976-DEL-96
chr7-29248591-DEL-407
chr7-29275175-DEL-83
chr7-29275736-DEL-185
chr7-29277403-DEL-111
chr7-29279463-DEL-186
chr7-29296576-DEL-331
chr7-29312666-DEL-73
chr7-29328139-DEL-519
chr7-29330498-DEL-6047
chr7-29351224-DEL-163
chr7-29360667-DEL-138
chr7-29369933-DEL-1190
chr7-29433870-DEL-1944
chr7-29436502-DEL-346
chr7-29439902-DEL-169
chr7-29445315-DEL-6128
chr7-29459445-DEL-386
chr7-29498548-DEL-220
chr7-29504569-DEL-6396
chr7-29516621-DEL-175
chr7-29519333-DEL-2139
chr7-29532473-DEL-2608
chr7-29536134-DEL-346
chr7-29538305-DEL-401
chr7-29564166-DEL-255
chr7-29567899-DEL-3035
chr7-29569121-DEL-881
chr7-29583063-DEL-1498
chr7-29584052-DEL-80
chr7-29596974-DEL-860
chr7-29618639-DEL-52
chr7-29630207-DEL-210
chr7-29661051-DEL-153
chr7-29661613-DEL-94
chr7-29663492-DEL-154
chr7-29668663-DEL-84
chr7-29673413-DEL-386

chr7-29673824-DEL-4331
chr7-29681303-DEL-56
chr7-29682163-DEL-135
chr7-29691513-DEL-148
chr7-29694759-DEL-127
chr7-29729035-DEL-204
chr7-29749132-DEL-238
chr7-29752249-DEL-981
chr7-29807506-DEL-109
chr7-29828326-DEL-199
chr7-29840035-DEL-63
chr7-29881756-DEL-66
chr7-29901542-DEL-580
chr7-29912766-DEL-625
chr7-29921547-DEL-588
chr7-29981851-DEL-229
chr7-29987886-DEL-1121
chr7-29992619-DEL-176
chr7-30000021-DEL-103
chr7-30003697-DEL-688
chr7-30029712-DEL-125
chr7-30084494-DEL-185
chr7-30150352-DEL-246
chr7-30168199-DEL-149
chr7-30170696-DEL-81
chr7-30187264-DEL-104
chr7-30191441-DEL-136
chr7-30195217-DEL-75
chr7-30216133-DEL-52
chr7-30230617-DEL-72
chr7-30341091-DEL-85
chr7-30354989-DEL-53
chr7-30373110-DEL-610
chr7-30383265-DEL-176
chr7-30407166-DEL-158
chr7-30421097-DEL-58
chr7-30423408-DEL-101
chr7-30443885-DEL-474
chr7-30462246-DEL-68
chr7-30496550-DEL-222
chr7-30505902-DEL-53
chr7-30510976-DEL-71
chr7-30530487-DEL-90
chr7-30531095-DEL-470
chr7-30532732-DEL-53
chr7-30541134-DEL-154
chr7-30544946-DEL-173
chr7-30571385-DEL-54
chr7-30581974-DEL-106
chr7-30587929-DEL-130
chr7-30604990-DEL-106
chr7-30615394-DEL-160
chr7-30622787-DEL-107
chr7-30625438-DEL-165
chr7-30626850-DEL-8119
chr7-30737861-DEL-106
chr7-30747147-DEL-59
chr7-30790546-DEL-53
chr7-30805383-DEL-98
chr7-30811473-DEL-52
chr7-30879898-DEL-97
chr7-30884483-DEL-1383
chr7-30888997-DEL-179
chr7-30888997-DEL-78908
chr7-30923938-DEL-6294
chr7-30936691-DEL-22051
chr7-30973132-DEL-59
chr7-31005848-DEL-4104
chr7-31012310-DEL-1416
chr7-31016501-DEL-7274
chr7-31030429-DEL-59
chr7-31040900-DEL-210
chr7-31076624-DEL-385
chr7-31078206-DEL-1095
chr7-31082295-DEL-1130
chr7-31089912-DEL-245
chr7-31154880-DEL-678

chr7-31157304-DEL-3240
chr7-31168177-DEL-204
chr7-31168746-DEL-1787
chr7-31183976-DEL-1155
chr7-31201582-DEL-139
chr7-31210641-DEL-54
chr7-31252419-DEL-6303
chr7-31252683-DEL-4274
chr7-31262004-DEL-60
chr7-31287536-DEL-1075
chr7-31287563-DEL-831
chr7-31291665-DEL-5264
chr7-31312320-DEL-329
chr7-31322124-DEL-57
chr7-31374829-DEL-2877
chr7-31386660-DEL-67
chr7-31392527-DEL-138
chr7-31419754-DEL-99
chr7-31442184-DEL-1944
chr7-31464185-DEL-53
chr7-31489508-DEL-87
chr7-31525102-DEL-2130
chr7-31545987-DEL-617
chr7-31555813-DEL-6772
chr7-31558891-DEL-4311
chr7-31561114-DEL-202
chr7-31568884-DEL-65
chr7-31592267-DEL-266
chr7-31631890-DEL-8835
chr7-31645537-DEL-54
chr7-31720319-DEL-2722
chr7-31722443-DEL-202
chr7-31745966-DEL-53
chr7-31787060-DEL-332
chr7-31842255-DEL-340
chr7-31854671-DEL-72
chr7-31855600-DEL-131
chr7-31872844-DEL-92
chr7-31875077-DEL-109
chr7-31884025-DEL-5582
chr7-31901426-DEL-258
chr7-31902532-DEL-1394
chr7-31923862-DEL-344
chr7-31926619-DEL-68
chr7-31943328-DEL-105
chr7-31961752-DEL-57
chr7-31969875-DEL-6757
chr7-31969890-DEL-212
chr7-31991437-DEL-56
chr7-32019563-DEL-3481
chr7-32029860-DEL-56
chr7-32038834-DEL-128
chr7-32056323-DEL-693
chr7-32059895-DEL-161
chr7-32062573-DEL-52
chr7-32062983-DEL-2970
chr7-32066774-DEL-53
chr7-32069892-DEL-4027
chr7-32095508-DEL-67
chr7-32101373-DEL-138
chr7-32214696-DEL-826
chr7-32286486-DEL-922
chr7-32288160-DEL-1096
chr7-32290222-DEL-397
chr7-32318021-DEL-266
chr7-32358369-DEL-62
chr7-32432869-DEL-52
chr7-32492495-DEL-89
chr7-32659764-DEL-52
chr7-32719424-DEL-72
chr7-32798971-DEL-196
chr7-32801043-DEL-255
chr7-32802138-DEL-1396
chr7-32805531-DEL-2564
chr7-32840483-DEL-109
chr7-32864141-DEL-258
chr7-32865248-DEL-1395

chr7-32886574-DEL-344
chr7-32890261-DEL-55
chr7-32916494-DEL-203
chr7-32929402-DEL-2394
chr7-32939626-DEL-3036
chr7-32972581-DEL-3479
chr7-33016178-DEL-2993
chr7-33020804-DEL-812
chr7-33052853-DEL-138
chr7-33078921-DEL-1879
chr7-33093748-DEL-1345
chr7-33095281-DEL-265
chr7-33135628-DEL-697
chr7-33138623-DEL-161
chr7-33140975-DEL-5479
chr7-33147678-DEL-56
chr7-33225856-DEL-494
chr7-33250465-DEL-3398
chr7-33267184-DEL-1595
chr7-33275729-DEL-482
chr7-33301968-DEL-401
chr7-33321562-DEL-447
chr7-33328670-DEL-1332
chr7-33336428-DEL-13191
chr7-33380892-DEL-1473
chr7-33459616-DEL-649
chr7-33461500-DEL-851
chr7-33495843-DEL-1298
chr7-33509332-DEL-610
chr7-33513701-DEL-5095
chr7-33529396-DEL-7462
chr7-33537499-DEL-74
chr7-33538807-DEL-6322
chr7-33579214-DEL-318
chr7-33585278-DEL-6221
chr7-33593336-DEL-423
chr7-33594041-DEL-2417
chr7-33605433-DEL-588
chr7-33667010-DEL-899
chr7-33668653-DEL-6881
chr7-33674338-DEL-498
chr7-33681867-DEL-332
chr7-33730645-DEL-7235
chr7-33730680-DEL-46087
chr7-33839293-DEL-188
chr7-33842484-DEL-52
chr7-33873054-DEL-277
chr7-33875060-DEL-454
chr7-33877730-DEL-334
chr7-33885361-DEL-175
chr7-33903705-DEL-72
chr7-33991541-DEL-87
chr7-33996527-DEL-2124
chr7-34046627-DEL-103
chr7-34062344-DEL-209
chr7-34066547-DEL-79
chr7-34071475-DEL-91
chr7-34085543-DEL-51
chr7-34128043-DEL-402
chr7-34142845-DEL-204
chr7-34174172-DEL-79
chr7-34194051-DEL-179
chr7-34222690-DEL-65
chr7-34265655-DEL-90
chr7-34282874-DEL-66
chr7-34304916-DEL-110
chr7-34308855-DEL-163
chr7-34311386-DEL-52
chr7-34319577-DEL-53
chr7-34322683-DEL-218
chr7-34324642-DEL-657
chr7-34344720-DEL-117
chr7-34359195-DEL-59
chr7-34368419-DEL-142
chr7-34377180-DEL-176
chr7-34385077-DEL-91
chr7-34391616-DEL-64

chr7-34412931-DEL-370
chr7-34415709-DEL-115
chr7-34424128-DEL-213
chr7-34428545-DEL-88
chr7-34439244-DEL-674
chr7-34515290-DEL-265
chr7-34586337-DEL-131
chr7-34592139-DEL-163
chr7-34606974-DEL-64
chr7-34676885-DEL-351
chr7-34703540-DEL-69
chr7-34803685-DEL-352
chr7-34805347-DEL-269
chr7-34809742-DEL-843
chr7-34824492-DEL-54
chr7-34832873-DEL-148
chr7-34859213-DEL-622
chr7-34906329-DEL-621
chr7-34929575-DEL-220
chr7-34944960-DEL-138
chr7-34950581-DEL-118
chr7-34956095-DEL-158
chr7-34967349-DEL-54
chr7-34970885-DEL-53
chr7-34978660-DEL-396
chr7-34979499-DEL-78
chr7-35018955-DEL-102
chr7-35026411-DEL-1345
chr7-35042703-DEL-71
chr7-35052415-DEL-206
chr7-35065771-DEL-74
chr7-35079466-DEL-64
chr7-35097808-DEL-80
chr7-35101087-DEL-205
chr7-35126781-DEL-198
chr7-35135400-DEL-93
chr7-35140911-DEL-939
chr7-35162369-DEL-463
chr7-35197786-DEL-65
chr7-35212874-DEL-200
chr7-35240442-DEL-399
chr7-35244767-DEL-195
chr7-35261402-DEL-335
chr7-35269703-DEL-99
chr7-35270125-DEL-185
chr7-35272026-DEL-238
chr7-35293867-DEL-104
chr7-35310395-DEL-103
chr7-35313571-DEL-216
chr7-35331306-DEL-160
chr7-35336774-DEL-155
chr7-35354742-DEL-67
chr7-35360469-DEL-277
chr7-35365806-DEL-1102
chr7-35405435-DEL-77
chr7-35416420-DEL-239
chr7-35448473-DEL-102
chr7-35461280-DEL-122
chr7-35464484-DEL-424
chr7-35481875-DEL-60
chr7-35495270-DEL-81
chr7-35549678-DEL-53
chr7-35562027-DEL-60
chr7-35605774-DEL-139
chr7-35607610-DEL-91
chr7-35637373-DEL-53
chr7-35639294-DEL-175
chr7-35661131-DEL-101
chr7-35668939-DEL-103
chr7-35698678-DEL-79
chr7-35701235-DEL-52
chr7-35709512-DEL-73
chr7-35717560-DEL-561
chr7-35803189-DEL-195
chr7-35806854-DEL-72
chr7-35840750-DEL-664
chr7-35861032-DEL-157

chr7-35861706-DEL-110
chr7-35872294-DEL-112
chr7-35893556-DEL-66
chr7-35932413-DEL-1166
chr7-35957722-DEL-180
chr7-35979806-DEL-60
chr7-36106188-DEL-52
chr7-36152726-DEL-186
chr7-36196383-DEL-53
chr7-36250841-DEL-84
chr7-36262827-DEL-983
chr7-36272055-DEL-801
chr7-36278676-DEL-62
chr7-36298081-DEL-286
chr7-36314316-DEL-52
chr7-36321607-DEL-55
chr7-36387086-DEL-58
chr7-36435093-DEL-53
chr7-36461569-DEL-198
chr7-36505093-DEL-59
chr7-36511855-DEL-107
chr7-36534522-DEL-102
chr7-36581377-DEL-54
chr7-36591178-DEL-289
chr7-36634130-DEL-426
chr7-36721642-DEL-343
chr7-36732108-DEL-80
chr7-36753982-DEL-68
chr7-36772083-DEL-131
chr7-36781594-DEL-1419
chr7-36814569-DEL-108
chr7-36831600-DEL-125
chr7-36866528-DEL-353
chr7-36867475-DEL-474
chr7-36947170-DEL-66
chr7-36952339-DEL-761
chr7-37093041-DEL-467
chr7-37107728-DEL-154
chr7-37114274-DEL-56
chr7-37155722-DEL-1439
chr7-37190767-DEL-213
chr7-37222932-DEL-136
chr7-37241081-DEL-119
chr7-37280999-DEL-64
chr7-37305882-DEL-58
chr7-37313572-DEL-616
chr7-37347511-DEL-54
chr7-37369336-DEL-738
chr7-37383609-DEL-57
chr7-37407125-DEL-51
chr7-37457071-DEL-56
chr7-37473770-DEL-58
chr7-37479962-DEL-98
chr7-37495142-DEL-75
chr7-37503856-DEL-70
chr7-37504613-DEL-64
chr7-37521475-DEL-65
chr7-37526238-DEL-134
chr7-37530415-DEL-51
chr7-37553013-DEL-52
chr7-37576359-DEL-66
chr7-37577349-DEL-198
chr7-37583224-DEL-109
chr7-37605453-DEL-414
chr7-37629686-DEL-53
chr7-37724816-DEL-950
chr7-37747693-DEL-78
chr7-37749010-DEL-119
chr7-37828351-DEL-124
chr7-37853022-DEL-2514
chr7-37856154-DEL-70
chr7-37860848-DEL-58
chr7-37872140-DEL-407
chr7-37876883-DEL-103
chr7-37879981-DEL-2447
chr7-37886343-DEL-616
chr7-37897927-DEL-1124

chr7-37936293-DEL-341
chr7-37948592-DEL-1624
chr7-37950700-DEL-163
chr7-37959456-DEL-51
chr7-37968828-DEL-200
chr7-37985587-DEL-101
chr7-37998237-DEL-3993
chr7-38000196-DEL-5545
chr7-38087645-DEL-585
chr7-38094246-DEL-1360
chr7-38109499-DEL-714
chr7-38120905-DEL-123
chr7-38155008-DEL-3581
chr7-38160169-DEL-708
chr7-38162406-DEL-663
chr7-38167300-DEL-494
chr7-38200582-DEL-1212
chr7-38217727-DEL-197
chr7-38224149-DEL-55
chr7-38224935-DEL-50
chr7-38235669-DEL-718
chr7-38237063-DEL-410
chr7-38240834-DEL-76
chr7-38248364-DEL-223
chr7-38256342-DEL-76
chr7-38281702-DEL-71
chr7-38297349-DEL-2824
chr7-38301912-DEL-56
chr7-38318735-DEL-694
chr7-38319546-DEL-5987
chr7-38327413-DEL-106
chr7-38386693-DEL-701
chr7-38388069-DEL-410
chr7-38398923-DEL-2046
chr7-38410144-DEL-1337
chr7-38410167-DEL-62
chr7-38417958-DEL-64
chr7-38428028-DEL-127
chr7-38430230-DEL-7902
chr7-38442363-DEL-495
chr7-38515769-DEL-355
chr7-38533127-DEL-73
chr7-38562794-DEL-100
chr7-38576397-DEL-664
chr7-38581288-DEL-495
chr7-38665237-DEL-59
chr7-38674774-DEL-152
chr7-38677902-DEL-451
chr7-38685624-DEL-60
chr7-38767189-DEL-495
chr7-38803117-DEL-53
chr7-38805007-DEL-84
chr7-38807560-DEL-524
chr7-38819307-DEL-142
chr7-38826160-DEL-68
chr7-38831198-DEL-72
chr7-38838723-DEL-409
chr7-38842391-DEL-104
chr7-38843171-DEL-121
chr7-38878626-DEL-610
chr7-38881584-DEL-666
chr7-38990853-DEL-51
chr7-39002889-DEL-1337
chr7-39029399-DEL-6471
chr7-39039735-DEL-495
chr7-39047360-DEL-62
chr7-39054656-DEL-605
chr7-39055722-DEL-87
chr7-39065139-DEL-141
chr7-39066784-DEL-134
chr7-39068397-DEL-3761
chr7-39072443-DEL-671
chr7-39080952-DEL-403
chr7-39089075-DEL-87
chr7-39108916-DEL-57
chr7-39129570-DEL-1085
chr7-39157982-DEL-6479

chr7-39250346-DEL-210
chr7-39279254-DEL-412
chr7-39299471-DEL-123
chr7-39300598-DEL-70
chr7-39309875-DEL-766
chr7-39317652-DEL-107
chr7-39329464-DEL-54
chr7-39329918-DEL-373
chr7-39331093-DEL-54
chr7-39343705-DEL-66
chr7-39397990-DEL-149
chr7-39405144-DEL-455
chr7-39565738-DEL-441
chr7-39604686-DEL-78
chr7-39628756-DEL-1138
chr7-39633691-DEL-5533
chr7-39643064-DEL-55
chr7-39665656-DEL-383
chr7-39693522-DEL-61
chr7-39698220-DEL-72
chr7-39699762-DEL-88
chr7-39710739-DEL-51
chr7-39732616-DEL-1983
chr7-39780742-DEL-2881
chr7-39781393-DEL-75
chr7-39838773-DEL-1319
chr7-39872460-DEL-79
chr7-39873161-DEL-195
chr7-39904620-DEL-94
chr7-39911030-DEL-381
chr7-39913617-DEL-59
chr7-39927269-DEL-62
chr7-39977402-DEL-417
chr7-40013302-DEL-129
chr7-40039221-DEL-54
chr7-40040998-DEL-153
chr7-40050043-DEL-351
chr7-40052511-DEL-1627
chr7-40086530-DEL-59
chr7-40167715-DEL-60
chr7-40213610-DEL-336
chr7-40214742-DEL-105
chr7-40253918-DEL-51
chr7-40275968-DEL-71
chr7-40279373-DEL-194
chr7-40282100-DEL-168
chr7-40317524-DEL-705
chr7-40352439-DEL-176
chr7-40364320-DEL-178
chr7-40370439-DEL-513
chr7-40391877-DEL-83
chr7-40410014-DEL-80
chr7-40448415-DEL-68
chr7-40498118-DEL-209
chr7-40503013-DEL-163
chr7-40513526-DEL-227
chr7-40567556-DEL-52
chr7-40597108-DEL-66
chr7-40633103-DEL-2142
chr7-40645184-DEL-73
chr7-40645590-DEL-2329
chr7-40710411-DEL-54
chr7-40712800-DEL-4046
chr7-40727154-DEL-161
chr7-40732793-DEL-4864
chr7-40739777-DEL-233
chr7-40744295-DEL-60
chr7-40758462-DEL-74
chr7-40761341-DEL-34798
chr7-40807914-DEL-80
chr7-40809936-DEL-104
chr7-40815169-DEL-9784
chr7-40829251-DEL-157
chr7-40830751-DEL-67
chr7-40839342-DEL-5608
chr7-40864200-DEL-130
chr7-40881835-DEL-78

chr7-40929359-DEL-77
chr7-40965795-DEL-146
chr7-40976730-DEL-798
chr7-40991014-DEL-193
chr7-41000661-DEL-71
chr7-41001244-DEL-235
chr7-41015751-DEL-3481
chr7-41051523-DEL-2615
chr7-41062062-DEL-8798
chr7-41078641-DEL-3810
chr7-41083309-DEL-1825
chr7-41091649-DEL-1185
chr7-41095699-DEL-119
chr7-41101303-DEL-89
chr7-41145752-DEL-122
chr7-41147700-DEL-52
chr7-41169715-DEL-58
chr7-41180664-DEL-8064
chr7-41195865-DEL-84
chr7-41200452-DEL-81
chr7-41251523-DEL-214
chr7-41269911-DEL-54
chr7-41281578-DEL-57
chr7-41286968-DEL-585
chr7-41291537-DEL-75
chr7-41313392-DEL-202
chr7-41331062-DEL-2430
chr7-41333542-DEL-7155
chr7-41347521-DEL-71
chr7-41367607-DEL-243
chr7-41388536-DEL-120
chr7-41393355-DEL-39392
chr7-41432530-DEL-839
chr7-41441097-DEL-188
chr7-41454988-DEL-2101
chr7-41457804-DEL-826
chr7-41465409-DEL-213
chr7-41474355-DEL-6010
chr7-41519546-DEL-203
chr7-41526029-DEL-252
chr7-41546448-DEL-73
chr7-41569846-DEL-5692
chr7-41605547-DEL-2360
chr7-41652882-DEL-1008
chr7-41670144-DEL-1302
chr7-41676894-DEL-6815
chr7-41683753-DEL-7960
chr7-41697702-DEL-201
chr7-41702226-DEL-215
chr7-41738054-DEL-157
chr7-41768831-DEL-1929
chr7-41774393-DEL-1303
chr7-41777802-DEL-6308
chr7-41786551-DEL-68
chr7-41875773-DEL-81
chr7-41877464-DEL-6214
chr7-41892124-DEL-876
chr7-41918507-DEL-676
chr7-42002995-DEL-80
chr7-42014271-DEL-6402
chr7-42022086-DEL-164
chr7-42024289-DEL-855
chr7-42024377-DEL-421
chr7-42025676-DEL-83
chr7-42029111-DEL-946
chr7-42038046-DEL-9963
chr7-42053395-DEL-650
chr7-42067641-DEL-197
chr7-42083912-DEL-2470
chr7-42089935-DEL-400
chr7-42122503-DEL-89
chr7-42127413-DEL-453
chr7-42128631-DEL-354
chr7-42132525-DEL-669
chr7-42141144-DEL-199
chr7-42174199-DEL-52
chr7-42175318-DEL-193

chr7-42183738-DEL-3607
chr7-42188965-DEL-80
chr7-42203060-DEL-193
chr7-42223724-DEL-438
chr7-42262041-DEL-252
chr7-42296089-DEL-189
chr7-42322412-DEL-400
chr7-42356176-DEL-118
chr7-42392673-DEL-183
chr7-42398017-DEL-67
chr7-42402560-DEL-67
chr7-42408838-DEL-547
chr7-42410039-DEL-495
chr7-42412190-DEL-857
chr7-42466166-DEL-520
chr7-42491383-DEL-2355
chr7-42494504-DEL-193
chr7-42533636-DEL-11583
chr7-42537168-DEL-10308
chr7-42552291-DEL-196
chr7-42593251-DEL-183
chr7-42606202-DEL-60
chr7-42617895-DEL-3399
chr7-42622071-DEL-8441
chr7-42632196-DEL-336
chr7-42706555-DEL-114
chr7-42711163-DEL-53
chr7-42712355-DEL-103
chr7-42725078-DEL-55
chr7-42739857-DEL-263
chr7-42742286-DEL-3134
chr7-42747999-DEL-388
chr7-42752658-DEL-73
chr7-42759285-DEL-190
chr7-42765489-DEL-37687
chr7-42796521-DEL-71
chr7-42818002-DEL-1478
chr7-42822763-DEL-837
chr7-42829189-DEL-498
chr7-42831078-DEL-6370
chr7-42845352-DEL-7397
chr7-42851636-DEL-51
chr7-42861491-DEL-440
chr7-42893345-DEL-62
chr7-42903574-DEL-78
chr7-42903906-DEL-1020
chr7-42924038-DEL-55
chr7-42932296-DEL-72
chr7-42937025-DEL-903
chr7-42939268-DEL-229
chr7-42991852-DEL-70
chr7-42997332-DEL-51
chr7-43018850-DEL-83
chr7-43021572-DEL-149
chr7-43046703-DEL-99
chr7-43056913-DEL-94
chr7-43077459-DEL-75
chr7-43079768-DEL-70
chr7-43083798-DEL-110
chr7-43104359-DEL-427
chr7-43141109-DEL-151
chr7-43145594-DEL-58
chr7-43149584-DEL-80
chr7-43274395-DEL-77
chr7-43284493-DEL-112
chr7-43304901-DEL-294
chr7-43320432-DEL-323
chr7-43326425-DEL-114
chr7-43405815-DEL-255
chr7-43413517-DEL-52
chr7-43417947-DEL-52
chr7-43444851-DEL-68
chr7-43484255-DEL-880
chr7-43518071-DEL-150
chr7-43522238-DEL-531
chr7-43523838-DEL-197
chr7-43529391-DEL-62

chr7-43549546-DEL-363
chr7-43549924-DEL-2534
chr7-43575350-DEL-57
chr7-43609916-DEL-72
chr7-43637156-DEL-1187
chr7-43698918-DEL-56
chr7-43735484-DEL-2997
chr7-43792975-DEL-6185
chr7-43802277-DEL-763
chr7-43803099-DEL-120
chr7-43843922-DEL-160
chr7-43846820-DEL-64
chr7-43853750-DEL-83
chr7-43854319-DEL-318
chr7-43881068-DEL-79
chr7-43994599-DEL-104
chr7-44011187-DEL-63
chr7-44014447-DEL-94
chr7-44046917-DEL-87
chr7-44047848-DEL-73
chr7-44062082-DEL-67
chr7-44086186-DEL-149
chr7-44097815-DEL-120
chr7-44100242-DEL-323
chr7-44104412-DEL-58
chr7-44131348-DEL-53
chr7-44166542-DEL-164
chr7-44194013-DEL-201
chr7-44204651-DEL-98
chr7-44207403-DEL-544
chr7-44246042-DEL-89
chr7-44270237-DEL-271
chr7-44308479-DEL-549
chr7-44315765-DEL-106
chr7-44330536-DEL-64
chr7-44333902-DEL-53
chr7-44350496-DEL-102
chr7-44362062-DEL-52
chr7-44379807-DEL-184
chr7-44387991-DEL-220
chr7-44419899-DEL-143
chr7-44421715-DEL-64
chr7-44431437-DEL-66
chr7-44435147-DEL-2446
chr7-44475950-DEL-69
chr7-44482960-DEL-176
chr7-44511396-DEL-80
chr7-44539527-DEL-176
chr7-44560949-DEL-56
chr7-44562051-DEL-60
chr7-44584814-DEL-74
chr7-44602988-DEL-55
chr7-44612743-DEL-194
chr7-44640055-DEL-74
chr7-44694718-DEL-145
chr7-44742215-DEL-96
chr7-44743997-DEL-70
chr7-44747884-DEL-52
chr7-44759180-DEL-336
chr7-44762373-DEL-63
chr7-44782589-DEL-60
chr7-44839772-DEL-55
chr7-44851018-DEL-937
chr7-44875683-DEL-107
chr7-44879590-DEL-285
chr7-44901802-DEL-228
chr7-44905064-DEL-300
chr7-44911436-DEL-58
chr7-44917464-DEL-674
chr7-45023913-DEL-174
chr7-45046631-DEL-3423
chr7-45050703-DEL-199
chr7-45089846-DEL-214
chr7-45091508-DEL-276
chr7-45129952-DEL-207
chr7-45178136-DEL-92
chr7-45193765-DEL-787

chr7-45194374-DEL-204
chr7-45201156-DEL-163
chr7-45204838-DEL-196
chr7-45230359-DEL-958
chr7-45233277-DEL-177
chr7-45261739-DEL-102
chr7-45262446-DEL-70
chr7-45265462-DEL-65
chr7-45275822-DEL-179
chr7-45283601-DEL-85
chr7-45295678-DEL-72
chr7-45316551-DEL-384
chr7-45318048-DEL-62
chr7-45335164-DEL-81
chr7-45346209-DEL-263
chr7-45382142-DEL-68
chr7-45403117-DEL-52
chr7-45409397-DEL-293
chr7-45413142-DEL-200
chr7-45491022-DEL-3257
chr7-45601027-DEL-132
chr7-45611293-DEL-60
chr7-45640212-DEL-71
chr7-45665541-DEL-120
chr7-45672569-DEL-63
chr7-45676649-DEL-58
chr7-45681286-DEL-53
chr7-45691374-DEL-340
chr7-45714463-DEL-108
chr7-45741030-DEL-90
chr7-45744256-DEL-1250
chr7-45764955-DEL-69
chr7-45779788-DEL-83
chr7-45784464-DEL-133
chr7-45812897-DEL-58
chr7-45824023-DEL-629
chr7-45829903-DEL-224
chr7-45837586-DEL-64
chr7-45838259-DEL-56
chr7-45881858-DEL-184
chr7-45882267-DEL-412
chr7-45885700-DEL-102
chr7-45942911-DEL-54
chr7-45944695-DEL-341
chr7-45946735-DEL-85
chr7-45979597-DEL-57
chr7-46001003-DEL-78
chr7-46036871-DEL-78
chr7-46105186-DEL-74
chr7-46110518-DEL-102
chr7-46118232-DEL-62
chr7-46121422-DEL-72
chr7-46138582-DEL-306
chr7-46155694-DEL-440
chr7-46169781-DEL-453
chr7-46176541-DEL-73
chr7-46212118-DEL-141
chr7-46264813-DEL-72
chr7-46269345-DEL-152
chr7-46273334-DEL-147
chr7-46276063-DEL-895
chr7-46298545-DEL-505
chr7-46310417-DEL-107
chr7-46330805-DEL-99
chr7-46363540-DEL-62
chr7-46374282-DEL-2627
chr7-46428021-DEL-339
chr7-46433138-DEL-87
chr7-46477939-DEL-208
chr7-46497753-DEL-101
chr7-46513278-DEL-106
chr7-46539698-DEL-161
chr7-46588553-DEL-110
chr7-46619082-DEL-1723
chr7-46622354-DEL-166
chr7-46632429-DEL-80
chr7-46651576-DEL-185

chr7-46673643-DEL-1030
chr7-46683394-DEL-878
chr7-46695547-DEL-122
chr7-46706776-DEL-62
chr7-46725225-DEL-529
chr7-46763055-DEL-52
chr7-46779493-DEL-153
chr7-46788480-DEL-61
chr7-46794069-DEL-336
chr7-46794674-DEL-51
chr7-46803949-DEL-5184
chr7-46806138-DEL-198
chr7-46822780-DEL-263
chr7-46824648-DEL-6165
chr7-46866250-DEL-51
chr7-46880091-DEL-163
chr7-46882715-DEL-520
chr7-46890776-DEL-2443
chr7-46890896-DEL-287
chr7-46906107-DEL-56
chr7-46908864-DEL-6619
chr7-46952061-DEL-83
chr7-46952738-DEL-2526
chr7-46956947-DEL-149
chr7-46965014-DEL-68
chr7-46974413-DEL-85
chr7-46982933-DEL-761
chr7-47057386-DEL-70
chr7-47063560-DEL-4185
chr7-47068994-DEL-202
chr7-47070701-DEL-374
chr7-47091774-DEL-346
chr7-47105428-DEL-131
chr7-47108181-DEL-362
chr7-47117678-DEL-13923
chr7-47128653-DEL-1983
chr7-47131206-DEL-3023
chr7-47152122-DEL-55
chr7-47171214-DEL-323
chr7-47178065-DEL-1333
chr7-47179758-DEL-474
chr7-47207144-DEL-738
chr7-47209407-DEL-59
chr7-47212557-DEL-314
chr7-47213736-DEL-338
chr7-47245590-DEL-80
chr7-47255240-DEL-856
chr7-47263634-DEL-5709
chr7-47284965-DEL-65
chr7-47307270-DEL-103
chr7-47314635-DEL-146
chr7-47317951-DEL-77
chr7-47324859-DEL-60
chr7-47347080-DEL-1438
chr7-47357022-DEL-51
chr7-47372893-DEL-61
chr7-47404044-DEL-59
chr7-47429675-DEL-407
chr7-47448309-DEL-56
chr7-47449123-DEL-716
chr7-47470164-DEL-627
chr7-47484592-DEL-1581
chr7-47550452-DEL-12796
chr7-47550902-DEL-1009
chr7-47551013-DEL-648
chr7-47563019-DEL-75
chr7-47581880-DEL-696
chr7-47584301-DEL-1618
chr7-47602500-DEL-844
chr7-47676532-DEL-199
chr7-47735569-DEL-744
chr7-47794418-DEL-7672
chr7-47861599-DEL-1999
chr7-47869222-DEL-190
chr7-47900342-DEL-6479
chr7-47908105-DEL-532
chr7-47911255-DEL-28823

chr7-47920736-DEL-58
chr7-47928232-DEL-53
chr7-47932399-DEL-499
chr7-47937068-DEL-225
chr7-47954034-DEL-212
chr7-47968418-DEL-9918
chr7-47999640-DEL-1210
chr7-48013678-DEL-2002
chr7-48030308-DEL-6349
chr7-48042180-DEL-83
chr7-48050030-DEL-837
chr7-48052651-DEL-7215
chr7-48052702-DEL-5883
chr7-48062077-DEL-6580
chr7-48062315-DEL-214
chr7-48131261-DEL-127
chr7-48133906-DEL-118
chr7-48137086-DEL-67
chr7-48141756-DEL-67
chr7-48160609-DEL-53
chr7-48182773-DEL-66
chr7-48255031-DEL-2989
chr7-48274350-DEL-66
chr7-48279720-DEL-96
chr7-48342048-DEL-979
chr7-48355356-DEL-79
chr7-48365094-DEL-959
chr7-48377529-DEL-437
chr7-48383619-DEL-832
chr7-48444226-DEL-202
chr7-48451861-DEL-54
chr7-48467546-DEL-164
chr7-48553435-DEL-568
chr7-48562376-DEL-278
chr7-48566238-DEL-312
chr7-48583854-DEL-294
chr7-48595777-DEL-456
chr7-48612880-DEL-67
chr7-48645988-DEL-238
chr7-48669873-DEL-62
chr7-48712710-DEL-82
chr7-48718371-DEL-100
chr7-48732872-DEL-279
chr7-48734056-DEL-66
chr7-48745650-DEL-494
chr7-48747280-DEL-121
chr7-48765180-DEL-165
chr7-48780354-DEL-58
chr7-48804267-DEL-431
chr7-48824636-DEL-105
chr7-48865388-DEL-129
chr7-48920389-DEL-57
chr7-48958644-DEL-59
chr7-48959788-DEL-108
chr7-49008203-DEL-75
chr7-49076501-DEL-70
chr7-49110176-DEL-898
chr7-49114802-DEL-52
chr7-49117176-DEL-92
chr7-49136249-DEL-113
chr7-49141945-DEL-158
chr7-49186818-DEL-1380
chr7-49205782-DEL-56
chr7-49206906-DEL-179
chr7-49215418-DEL-553
chr7-49231597-DEL-758
chr7-49246173-DEL-135
chr7-49273270-DEL-138
chr7-49298486-DEL-81
chr7-49330909-DEL-376
chr7-49360528-DEL-54
chr7-49377968-DEL-676
chr7-49392662-DEL-59
chr7-49481534-DEL-197
chr7-49521380-DEL-1728
chr7-49534985-DEL-52
chr7-49540335-DEL-57

chr7-49578846-DEL-96
chr7-49620610-DEL-58
chr7-49671679-DEL-52
chr7-49688920-DEL-102
chr7-49689725-DEL-59
chr7-49732960-DEL-1277
chr7-49752486-DEL-1257
chr7-49753847-DEL-829
chr7-49771873-DEL-6386
chr7-49776244-DEL-8484
chr7-49826569-DEL-232
chr7-49860206-DEL-50
chr7-49892922-DEL-946
chr7-49896430-DEL-57
chr7-49984497-DEL-134
chr7-50000287-DEL-973
chr7-50023705-DEL-86
chr7-50032339-DEL-140
chr7-50066762-DEL-55
chr7-50079810-DEL-445
chr7-50087133-DEL-100
chr7-50093670-DEL-152
chr7-50100538-DEL-81
chr7-50110649-DEL-266
chr7-50129372-DEL-75
chr7-50146634-DEL-954
chr7-50147269-DEL-405
chr7-50163505-DEL-215
chr7-50184260-DEL-1556
chr7-50195782-DEL-118
chr7-50224483-DEL-230
chr7-50228915-DEL-310
chr7-50232225-DEL-993
chr7-50246747-DEL-1240
chr7-50261656-DEL-60
chr7-50323352-DEL-371
chr7-50341412-DEL-153
chr7-50361970-DEL-128
chr7-50365154-DEL-54
chr7-50411751-DEL-698
chr7-50436991-DEL-513
chr7-50460755-DEL-169
chr7-50548931-DEL-66
chr7-50568931-DEL-70
chr7-50571784-DEL-54
chr7-50595605-DEL-79
chr7-50604357-DEL-79
chr7-50613103-DEL-79
chr7-50621856-DEL-79
chr7-50630609-DEL-80
chr7-50633169-DEL-10817
chr7-50639511-DEL-78
chr7-50646227-DEL-194
chr7-50651720-DEL-2363
chr7-50685568-DEL-52
chr7-50797293-DEL-3166
chr7-50806530-DEL-802
chr7-50807496-DEL-751
chr7-50809088-DEL-1430
chr7-50813101-DEL-6176
chr7-50827463-DEL-150
chr7-50854591-DEL-78
chr7-50889366-DEL-64
chr7-50921662-DEL-2577
chr7-50943655-DEL-903
chr7-50964792-DEL-1749
chr7-50976868-DEL-159
chr7-50977644-DEL-464
chr7-51018128-DEL-715
chr7-51028258-DEL-2153
chr7-51031615-DEL-293
chr7-51032660-DEL-502
chr7-51039185-DEL-67
chr7-51044350-DEL-205
chr7-51046096-DEL-347
chr7-51081239-DEL-163
chr7-51081847-DEL-2536

chr7-51198450-DEL-2525
chr7-51287531-DEL-529
chr7-51306490-DEL-52
chr7-51325127-DEL-539
chr7-51333001-DEL-52
chr7-51337637-DEL-254
chr7-51354180-DEL-68
chr7-51361483-DEL-426
chr7-51362319-DEL-242
chr7-51362627-DEL-1059
chr7-51400362-DEL-164
chr7-51441864-DEL-122
chr7-51449265-DEL-2426
chr7-51451904-DEL-10527
chr7-51470616-DEL-121
chr7-51558862-DEL-2179
chr7-51574584-DEL-68
chr7-51600788-DEL-139
chr7-51625607-DEL-259
chr7-51625875-DEL-4038
chr7-51653340-DEL-174
chr7-51718919-DEL-116
chr7-51723918-DEL-514
chr7-51738003-DEL-507
chr7-51757398-DEL-395
chr7-51766641-DEL-1269
chr7-51775354-DEL-1023
chr7-51820065-DEL-94
chr7-51852132-DEL-66
chr7-51865650-DEL-494
chr7-51884056-DEL-52
chr7-51887003-DEL-557
chr7-51905730-DEL-145
chr7-51917403-DEL-118
chr7-51925545-DEL-79
chr7-51925854-DEL-51
chr7-51932855-DEL-216
chr7-51933063-DEL-6216
chr7-51948167-DEL-54
chr7-51949064-DEL-99
chr7-51975255-DEL-91
chr7-51984933-DEL-1712
chr7-51991967-DEL-56
chr7-52004046-DEL-60
chr7-52044445-DEL-69
chr7-52052557-DEL-66
chr7-52061781-DEL-1432
chr7-52108440-DEL-699
chr7-52126740-DEL-117
chr7-52139877-DEL-86
chr7-52156026-DEL-307
chr7-52172368-DEL-838
chr7-52173711-DEL-58
chr7-52187135-DEL-52
chr7-52205196-DEL-55
chr7-52208951-DEL-337
chr7-52217735-DEL-271
chr7-52259919-DEL-158
chr7-52272446-DEL-166
chr7-52285228-DEL-66
chr7-52305318-DEL-140
chr7-52371496-DEL-76
chr7-52430884-DEL-78
chr7-52438373-DEL-6493
chr7-52450068-DEL-6457
chr7-52459647-DEL-922
chr7-52471841-DEL-60
chr7-52474881-DEL-206
chr7-52479524-DEL-6258
chr7-52486158-DEL-1326
chr7-52489104-DEL-74
chr7-52496597-DEL-79
chr7-52512712-DEL-708
chr7-52528469-DEL-102
chr7-52551904-DEL-204
chr7-52576806-DEL-2058
chr7-52589117-DEL-563

chr7-52593516-DEL-2300
chr7-52602741-DEL-51
chr7-52620698-DEL-319
chr7-52649713-DEL-312
chr7-52695459-DEL-258
chr7-52701777-DEL-66
chr7-52702967-DEL-121
chr7-52738651-DEL-76
chr7-52744535-DEL-977
chr7-52755397-DEL-52
chr7-52761829-DEL-400
chr7-52769143-DEL-214
chr7-52773038-DEL-81
chr7-52786680-DEL-6327
chr7-52786695-DEL-6873
chr7-52793007-DEL-4898
chr7-52806056-DEL-771
chr7-52812803-DEL-7389
chr7-52861740-DEL-64
chr7-52862802-DEL-949
chr7-52865091-DEL-387
chr7-52867710-DEL-2755
chr7-52921848-DEL-110
chr7-52951652-DEL-57
chr7-52952925-DEL-487
chr7-52955506-DEL-2570
chr7-52958527-DEL-10488
chr7-52963155-DEL-1487
chr7-52964686-DEL-1150
chr7-52965054-DEL-111
chr7-52965430-DEL-4166
chr7-52973410-DEL-625
chr7-53027834-DEL-6299
chr7-53034477-DEL-104
chr7-53035869-DEL-3224
chr7-53049072-DEL-3429
chr7-53066072-DEL-59
chr7-53068779-DEL-108
chr7-53087069-DEL-1258
chr7-53088500-DEL-1423
chr7-53097360-DEL-86
chr7-53103267-DEL-52
chr7-53122146-DEL-3446
chr7-53125765-DEL-744
chr7-53129261-DEL-309
chr7-53151833-DEL-1614
chr7-53165057-DEL-256
chr7-53172417-DEL-1610
chr7-53180474-DEL-1021
chr7-53185640-DEL-426
chr7-53188933-DEL-2200
chr7-53189964-DEL-159
chr7-53217651-DEL-599
chr7-53218030-DEL-495
chr7-53219265-DEL-192
chr7-53219870-DEL-211
chr7-53230519-DEL-856
chr7-53240510-DEL-896
chr7-53245153-DEL-92
chr7-53254477-DEL-1027
chr7-53284308-DEL-376
chr7-53287398-DEL-1421
chr7-53301188-DEL-93
chr7-53372846-DEL-66
chr7-53408850-DEL-876
chr7-53409839-DEL-430
chr7-53512133-DEL-645
chr7-53544783-DEL-336
chr7-53567815-DEL-758
chr7-53596962-DEL-123
chr7-53619592-DEL-52
chr7-53620200-DEL-231
chr7-53630648-DEL-1424
chr7-53638137-DEL-1285
chr7-53656337-DEL-899
chr7-53663477-DEL-374
chr7-53685289-DEL-78

chr7-53724171-DEL-507
chr7-53725006-DEL-1938
chr7-53730718-DEL-717
chr7-53733818-DEL-156
chr7-53744980-DEL-227
chr7-53757573-DEL-65
chr7-53762048-DEL-1173
chr7-53763288-DEL-836
chr7-53829601-DEL-1803
chr7-53853963-DEL-107
chr7-53869009-DEL-68
chr7-53894856-DEL-294
chr7-53958672-DEL-85
chr7-53978632-DEL-58
chr7-53996776-DEL-4238
chr7-54001050-DEL-1103
chr7-54009279-DEL-2267
chr7-54028421-DEL-72
chr7-54049124-DEL-62
chr7-54062426-DEL-1396
chr7-54071718-DEL-648
chr7-54074766-DEL-203
chr7-54095714-DEL-57
chr7-54130989-DEL-224
chr7-54133703-DEL-130
chr7-54136193-DEL-989
chr7-54137069-DEL-55
chr7-54159856-DEL-911
chr7-54163105-DEL-56
chr7-54164637-DEL-214
chr7-54173781-DEL-58
chr7-54192915-DEL-5628
chr7-54194207-DEL-121
chr7-54198231-DEL-206
chr7-54218023-DEL-150
chr7-54222024-DEL-2461
chr7-54239811-DEL-297
chr7-54243221-DEL-58
chr7-54254116-DEL-1112
chr7-54278935-DEL-213
chr7-54282391-DEL-384
chr7-54289895-DEL-300
chr7-54322585-DEL-5975
chr7-54328272-DEL-182
chr7-54329839-DEL-355
chr7-54338018-DEL-100
chr7-54369898-DEL-1702
chr7-54401909-DEL-582
chr7-54404760-DEL-1209
chr7-54475637-DEL-258
chr7-54490591-DEL-64
chr7-54572756-DEL-6859
chr7-54584662-DEL-50
chr7-54587018-DEL-410
chr7-54588755-DEL-6953
chr7-54599333-DEL-630
chr7-54608155-DEL-2118
chr7-54633959-DEL-76
chr7-54643661-DEL-1079
chr7-54670014-DEL-372
chr7-54732205-DEL-622
chr7-54735812-DEL-76
chr7-54736698-DEL-54
chr7-54794876-DEL-129
chr7-54805502-DEL-3155
chr7-54806576-DEL-582
chr7-54853000-DEL-98
chr7-54875975-DEL-1009
chr7-54879706-DEL-58
chr7-54881847-DEL-270
chr7-54904693-DEL-818
chr7-54907185-DEL-498
chr7-54924086-DEL-492
chr7-54940514-DEL-959
chr7-54998959-DEL-81
chr7-55013509-DEL-900
chr7-55020153-DEL-279

chr7-55021267-DEL-408
chr7-55034742-DEL-4934
chr7-55044048-DEL-1260
chr7-55053690-DEL-641
chr7-55071285-DEL-372
chr7-55111983-DEL-459
chr7-55113368-DEL-213
chr7-55136111-DEL-1496
chr7-55167889-DEL-606
chr7-55172290-DEL-221
chr7-55195050-DEL-243
chr7-55205264-DEL-70
chr7-55218809-DEL-313
chr7-55241510-DEL-82
chr7-55244478-DEL-354
chr7-55244841-DEL-1174
chr7-55248073-DEL-8451
chr7-55248294-DEL-428
chr7-55264819-DEL-415
chr7-55276134-DEL-6196
chr7-55302492-DEL-591
chr7-55303231-DEL-10695
chr7-55316920-DEL-202
chr7-55319838-DEL-134
chr7-55331078-DEL-2153
chr7-55336532-DEL-13755
chr7-55339132-DEL-6005
chr7-55347747-DEL-1333
chr7-55357514-DEL-171
chr7-55358136-DEL-13076
chr7-55364575-DEL-602
chr7-55406904-DEL-1594
chr7-55438081-DEL-52
chr7-55461000-DEL-283
chr7-55483651-DEL-1751
chr7-55510116-DEL-124
chr7-55526023-DEL-78
chr7-55539054-DEL-58
chr7-55553587-DEL-59
chr7-55584534-DEL-197
chr7-55613157-DEL-2740
chr7-55620901-DEL-174
chr7-55642974-DEL-223
chr7-55652250-DEL-172
chr7-55688078-DEL-60
chr7-55732019-DEL-193
chr7-55761013-DEL-811
chr7-55809927-DEL-152
chr7-55809950-DEL-2963
chr7-55810378-DEL-1976
chr7-55810638-DEL-1956
chr7-55812400-DEL-228
chr7-55875815-DEL-282
chr7-55888104-DEL-603
chr7-55947158-DEL-74
chr7-56004636-DEL-621
chr7-56070747-DEL-86
chr7-56087519-DEL-459
chr7-56088267-DEL-84
chr7-56089898-DEL-464
chr7-56091157-DEL-22229
chr7-56093038-DEL-70
chr7-56094111-DEL-59
chr7-56205148-DEL-341
chr7-56220016-DEL-361
chr7-56238544-DEL-87
chr7-56256091-DEL-62
chr7-56259565-DEL-7570
chr7-56295097-DEL-75
chr7-56328283-DEL-956
chr7-56329207-DEL-6179
chr7-56337689-DEL-76
chr7-56353399-DEL-269
chr7-56414193-DEL-63
chr7-56434259-DEL-724
chr7-56540454-DEL-254
chr7-56546603-DEL-56

chr7-56570105-DEL-9754
chr7-56579745-DEL-305
chr7-56668015-DEL-6254
chr7-56684898-DEL-311
chr7-56721733-DEL-105
chr7-56768217-DEL-708
chr7-56781913-DEL-247
chr7-56865634-DEL-66
chr7-56951820-DEL-2062
chr7-56954471-DEL-221
chr7-56963036-DEL-2391
chr7-56977294-DEL-55
chr7-56982957-DEL-801
chr7-57003303-DEL-183
chr7-57024946-DEL-250
chr7-57077992-DEL-518
chr7-57079438-DEL-81
chr7-57084481-DEL-54
chr7-57093295-DEL-1164
chr7-57121258-DEL-269
chr7-57130975-DEL-317
chr7-57178870-DEL-60
chr7-57190341-DEL-519
chr7-57204943-DEL-161
chr7-57215550-DEL-65
chr7-57237372-DEL-124
chr7-57244233-DEL-172
chr7-57281585-DEL-57
chr7-57291072-DEL-329
chr7-57292197-DEL-389
chr7-57342614-DEL-928
chr7-57359638-DEL-63
chr7-57445099-DEL-69
chr7-57485835-DEL-64
chr7-57486537-DEL-759
chr7-57502707-DEL-222
chr7-57577658-DEL-3165
chr7-57608111-DEL-52
chr7-57618447-DEL-371
chr7-57687767-DEL-6852
chr7-57707277-DEL-208
chr7-57720450-DEL-6497
chr7-57750244-DEL-7435
chr7-57767905-DEL-7240
chr7-57775434-DEL-166
chr7-57784029-DEL-184
chr7-57792674-DEL-147
chr7-57797976-DEL-774
chr7-57810739-DEL-6480
chr7-57821085-DEL-6556
chr7-57839993-DEL-212
chr7-57884833-DEL-6940
chr7-57913436-DEL-729
chr7-57920405-DEL-220
chr7-57921613-DEL-3013
chr7-57925496-DEL-6131
chr7-57932977-DEL-552
chr7-57954197-DEL-80
chr7-57965012-DEL-1938
chr7-57998274-DEL-281
chr7-58050228-DEL-76
chr7-58058399-DEL-529
chr7-58080766-DEL-664
chr7-58109118-DEL-1620
chr7-58133904-DEL-87
chr7-58146622-DEL-1893
chr7-58150447-DEL-284
chr7-58200291-DEL-624
chr7-58203844-DEL-124
chr7-58211204-DEL-130
chr7-58211970-DEL-589
chr7-58214299-DEL-124
chr7-58225491-DEL-420
chr7-58228367-DEL-904
chr7-58230464-DEL-1000
chr7-58238582-DEL-205
chr7-58245102-DEL-62

chr7-58251551-DEL-2048
chr7-58288841-DEL-1323
chr7-58298231-DEL-85
chr7-58301199-DEL-1995
chr7-58338551-DEL-164
chr7-58342446-DEL-51
chr7-58358190-DEL-129
chr7-58373889-DEL-58
chr7-58382998-DEL-1329
chr7-58396828-DEL-5236
chr7-58422202-DEL-218
chr7-58485480-DEL-53
chr7-58508060-DEL-101
chr7-58510141-DEL-57
chr7-58544329-DEL-6113
chr7-58588656-DEL-62
chr7-58651748-DEL-487
chr7-58653791-DEL-72
chr7-58668372-DEL-112
chr7-58681877-DEL-417
chr7-58710015-DEL-1683
chr7-58741953-DEL-4454
chr7-58746428-DEL-1985
chr7-58749402-DEL-106
chr7-58759636-DEL-1697
chr7-58781324-DEL-2631
chr7-58785449-DEL-2987
chr7-58786335-DEL-73
chr7-58792819-DEL-83
chr7-58809371-DEL-2098
chr7-58852853-DEL-79
chr7-58964520-DEL-77
chr7-58973970-DEL-73
chr7-58995965-DEL-54
chr7-58998748-DEL-548
chr7-59003433-DEL-1869
chr7-59013801-DEL-3740
chr7-59019555-DEL-1869
chr7-59021715-DEL-3737
chr7-59054472-DEL-1864
chr7-59055384-DEL-5605
chr7-59058626-DEL-3739
chr7-59063893-DEL-897
chr7-59071931-DEL-1868
chr7-59074090-DEL-1870
chr7-59079011-DEL-1871
chr7-59103782-DEL-7692
chr7-59106503-DEL-2078
chr7-59111360-DEL-1872
chr7-59119157-DEL-1869
chr7-59135183-DEL-1868
chr7-59146973-DEL-409
chr7-59148448-DEL-9157
chr7-59148602-DEL-7303
chr7-59220107-DEL-1871
chr7-59221513-DEL-1872
chr7-59266609-DEL-1326
chr7-59282460-DEL-802
chr7-59309561-DEL-735
chr7-59326933-DEL-2487
chr7-59328775-DEL-2545
chr7-59333714-DEL-2122
chr7-59335833-DEL-7086
chr7-59342917-DEL-425
chr7-59346687-DEL-50
chr7-59354351-DEL-2524
chr7-59364473-DEL-2529
chr7-59525246-DEL-327
chr7-59543048-DEL-2510
chr7-59555711-DEL-638
chr7-59557369-DEL-1195
chr7-59557636-DEL-6387
chr7-59558629-DEL-2711
chr7-59561383-DEL-4480
chr7-59572240-DEL-54
chr7-59590753-DEL-195
chr7-59591519-DEL-2768

chr7-59592282-DEL-891
chr7-59606125-DEL-178
chr7-59609275-DEL-3133
chr7-59612628-DEL-109
chr7-59614470-DEL-1272
chr7-59614734-DEL-849
chr7-59625761-DEL-76
chr7-59646121-DEL-682
chr7-59647576-DEL-71
chr7-59666023-DEL-516
chr7-59675292-DEL-91
chr7-59708557-DEL-167
chr7-59712945-DEL-941
chr7-59757681-DEL-68
chr7-59762023-DEL-899
chr7-59800533-DEL-52
chr7-59812937-DEL-212
chr7-59814628-DEL-89
chr7-59841820-DEL-11638
chr7-59851911-DEL-1090
chr7-59883818-DEL-583
chr7-59913495-DEL-273
chr7-59922280-DEL-81
chr7-59923542-DEL-6305
chr7-59950288-DEL-404
chr7-59959470-DEL-70
chr7-59966683-DEL-2059
chr7-59971842-DEL-120
chr7-59978825-DEL-891
chr7-59983123-DEL-884
chr7-59994486-DEL-6807
chr7-60003165-DEL-3705
chr7-60043299-DEL-140
chr7-60056337-DEL-57
chr7-60065105-DEL-6333
chr7-60077928-DEL-821
chr7-60081629-DEL-118
chr7-60085723-DEL-868
chr7-60089440-DEL-447
chr7-60100610-DEL-3700
chr7-60154162-DEL-3403
chr7-60154507-DEL-582
chr7-60161387-DEL-741
chr7-60163592-DEL-88
chr7-60167384-DEL-165
chr7-60168296-DEL-1383
chr7-60170767-DEL-100
chr7-60172909-DEL-2411
chr7-60180788-DEL-1064
chr7-60199020-DEL-312
chr7-60207901-DEL-659
chr7-60210769-DEL-118
chr7-60215289-DEL-3586
chr7-60251233-DEL-1179
chr7-60261946-DEL-644
chr7-60287186-DEL-65
chr7-60294112-DEL-861
chr7-60297677-DEL-2546
chr7-60301205-DEL-164
chr7-60302043-DEL-1389
chr7-60306661-DEL-1521
chr7-60340686-DEL-822
chr7-60345708-DEL-27850
chr7-60357582-DEL-289
chr7-60358873-DEL-1264
chr7-60372479-DEL-578
chr7-60378897-DEL-998
chr7-60388344-DEL-904
chr7-60388450-DEL-54
chr7-60397726-DEL-68
chr7-60398854-DEL-741
chr7-60402938-DEL-791
chr7-60403881-DEL-679
chr7-60405102-DEL-860
chr7-60412657-DEL-7204
chr7-60419323-DEL-79
chr7-60422612-DEL-145

chr7-60434214-DEL-70
chr7-60456059-DEL-121
chr7-60460143-DEL-873
chr7-60464406-DEL-543
chr7-60465992-DEL-438
chr7-60507874-DEL-1257
chr7-60517031-DEL-166
chr7-60527727-DEL-60
chr7-60540007-DEL-57
chr7-60564526-DEL-58
chr7-60596892-DEL-126
chr7-60597862-DEL-1326
chr7-60602987-DEL-54
chr7-60609494-DEL-190
chr7-60631810-DEL-69
chr7-60634352-DEL-76
chr7-60646438-DEL-204
chr7-60660987-DEL-60
chr7-60667232-DEL-86
chr7-60712290-DEL-1276
chr7-60718585-DEL-1450
chr7-60756504-DEL-1623
chr7-60763241-DEL-174
chr7-60765414-DEL-517
chr7-60775922-DEL-156
chr7-60792556-DEL-5020
chr7-60795602-DEL-133
chr7-60802974-DEL-100
chr7-60826324-DEL-627
chr7-60852586-DEL-56
chr7-60860026-DEL-1321
chr7-60862155-DEL-357
chr7-60864299-DEL-921
chr7-60866744-DEL-413
chr7-60881339-DEL-107
chr7-60884349-DEL-3875
chr7-60910103-DEL-665
chr7-60943940-DEL-184
chr7-60968556-DEL-7477
chr7-60973591-DEL-1307
chr7-60995901-DEL-26949
chr7-61006846-DEL-13359
chr7-61020251-DEL-2593
chr7-61026126-DEL-64
chr7-61027665-DEL-6884
chr7-61027734-DEL-95
chr7-61036313-DEL-85
chr7-61036349-DEL-4550
chr7-61055027-DEL-149
chr7-61077980-DEL-1786
chr7-61080651-DEL-1315
chr7-61082852-DEL-4695
chr7-61083833-DEL-7242
chr7-61097668-DEL-1958
chr7-61099845-DEL-3325
chr7-61108628-DEL-97
chr7-61126946-DEL-497
chr7-61204433-DEL-65
chr7-61219611-DEL-424
chr7-61227231-DEL-93
chr7-61249191-DEL-1096
chr7-61260353-DEL-188
chr7-61272929-DEL-288
chr7-61334396-DEL-52
chr7-61341962-DEL-242
chr7-61351182-DEL-99
chr7-61352396-DEL-377
chr7-61353886-DEL-357
chr7-61355170-DEL-217
chr7-61390693-DEL-1023
chr7-61392219-DEL-807
chr7-61452944-DEL-61
chr7-61488068-DEL-356
chr7-61489079-DEL-1907
chr7-61579459-DEL-61
chr7-61607269-DEL-1318
chr7-61657212-DEL-707

chr7-61660929-DEL-4991
chr7-61662507-DEL-2801
chr7-61707304-DEL-489
chr7-61714360-DEL-398
chr7-61733097-DEL-2244
chr7-61771367-DEL-1258
chr7-61787938-DEL-71
chr7-61807844-DEL-66
chr7-61844152-DEL-408
chr7-61853258-DEL-68
chr7-61862060-DEL-76
chr7-61911947-DEL-72
chr7-61966982-DEL-226
chr7-61975468-DEL-139
chr7-61987255-DEL-75
chr7-62019262-DEL-179
chr7-62033017-DEL-602
chr7-62093504-DEL-287
chr7-62101891-DEL-95
chr7-62118502-DEL-2298
chr7-62119108-DEL-2294
chr7-62125987-DEL-491
chr7-62168903-DEL-268
chr7-62170882-DEL-112
chr7-62208654-DEL-1775
chr7-62209156-DEL-1054
chr7-62232777-DEL-113
chr7-62237099-DEL-63
chr7-62251620-DEL-69
chr7-62264735-DEL-435
chr7-62268832-DEL-12604
chr7-62270013-DEL-5903
chr7-62279169-DEL-199
chr7-62301149-DEL-1875
chr7-62343914-DEL-5476
chr7-62378984-DEL-3888
chr7-62382797-DEL-226
chr7-62405029-DEL-6022
chr7-62420771-DEL-205
chr7-62441383-DEL-93
chr7-62444614-DEL-6398
chr7-62459105-DEL-145
chr7-62463832-DEL-203
chr7-62472137-DEL-81
chr7-62503468-DEL-6354
chr7-62511528-DEL-153
chr7-62559015-DEL-61
chr7-62562801-DEL-192
chr7-62563206-DEL-1252
chr7-62564732-DEL-194
chr7-62566129-DEL-185
chr7-62579694-DEL-59
chr7-62589225-DEL-359
chr7-62606871-DEL-210
chr7-62659499-DEL-106
chr7-62661766-DEL-111
chr7-62665048-DEL-87
chr7-62666646-DEL-1508
chr7-62673829-DEL-414
chr7-62686179-DEL-63
chr7-62717443-DEL-80
chr7-62718742-DEL-71
chr7-62737701-DEL-63
chr7-62738736-DEL-352
chr7-62786956-DEL-197
chr7-62811480-DEL-103
chr7-62854463-DEL-104
chr7-62893654-DEL-51
chr7-62922032-DEL-6350
chr7-62934808-DEL-6202
chr7-62979049-DEL-68
chr7-63018803-DEL-52
chr7-63044854-DEL-6370
chr7-63075661-DEL-1848
chr7-63122533-DEL-53
chr7-63135815-DEL-429
chr7-63143430-DEL-132

chr7-63151986-DEL-83
chr7-63166582-DEL-1741
chr7-63193742-DEL-196
chr7-63218682-DEL-52
chr7-63219289-DEL-1924
chr7-63227423-DEL-352
chr7-63247728-DEL-794
chr7-63252709-DEL-1585
chr7-63253957-DEL-68
chr7-63334937-DEL-54
chr7-63349917-DEL-697
chr7-63398743-DEL-244
chr7-63450222-DEL-562
chr7-63477721-DEL-179
chr7-63523054-DEL-53
chr7-63578412-DEL-111
chr7-63595655-DEL-599
chr7-63598260-DEL-98
chr7-63718421-DEL-100
chr7-63724265-DEL-84
chr7-63728029-DEL-63
chr7-63736307-DEL-543
chr7-63740075-DEL-1339
chr7-63741974-DEL-155
chr7-63765337-DEL-62
chr7-63765649-DEL-6327
chr7-63775770-DEL-61
chr7-63810891-DEL-52
chr7-63823076-DEL-78
chr7-63824698-DEL-56
chr7-63831977-DEL-615
chr7-63832010-DEL-54
chr7-63861654-DEL-103
chr7-63866292-DEL-105
chr7-63896905-DEL-72
chr7-63930911-DEL-443
chr7-63941749-DEL-57
chr7-63980886-DEL-168
chr7-63990718-DEL-63
chr7-64013884-DEL-102
chr7-64016141-DEL-110
chr7-64021694-DEL-93
chr7-64050607-DEL-76
chr7-64115595-DEL-30128
chr7-64115603-DEL-606
chr7-64115828-DEL-232
chr7-64116433-DEL-324
chr7-64117031-DEL-325
chr7-64117544-DEL-1804
chr7-64119289-DEL-277
chr7-64120039-DEL-138
chr7-64120783-DEL-1108
chr7-64121040-DEL-324
chr7-64122218-DEL-975
chr7-64122315-DEL-70
chr7-64122562-DEL-374
chr7-64123521-DEL-1944
chr7-64123610-DEL-186
chr7-64124380-DEL-601
chr7-64125014-DEL-93
chr7-64126245-DEL-183
chr7-64128005-DEL-93
chr7-64129508-DEL-322
chr7-64130390-DEL-92
chr7-64132630-DEL-137
chr7-64135770-DEL-185
chr7-64135802-DEL-556
chr7-64136611-DEL-323
chr7-64137527-DEL-137
chr7-64137872-DEL-978
chr7-64138068-DEL-233
chr7-64139365-DEL-972
chr7-64139991-DEL-139
chr7-64141261-DEL-139
chr7-64142385-DEL-89
chr7-64142971-DEL-186
chr7-64144134-DEL-93

chr7-64149701-DEL-134
chr7-64164779-DEL-119
chr7-64169638-DEL-76
chr7-64188861-DEL-82
chr7-64217556-DEL-58
chr7-64236705-DEL-794
chr7-64243640-DEL-144
chr7-64249754-DEL-6995
chr7-64259791-DEL-51
chr7-64261084-DEL-6417
chr7-64288022-DEL-67
chr7-64332359-DEL-72
chr7-64344340-DEL-66
chr7-64420745-DEL-78
chr7-64455307-DEL-283
chr7-64461982-DEL-440
chr7-64467951-DEL-807
chr7-64507815-DEL-63
chr7-64515546-DEL-1741
chr7-64575419-DEL-6329
chr7-64582619-DEL-66
chr7-64597734-DEL-52
chr7-64638645-DEL-92
chr7-64695897-DEL-54
chr7-64703220-DEL-150
chr7-64756420-DEL-4492
chr7-64780808-DEL-116
chr7-64796731-DEL-82
chr7-64802065-DEL-52
chr7-64805136-DEL-67
chr7-64810929-DEL-54
chr7-64835370-DEL-12878
chr7-64851150-DEL-139
chr7-64869849-DEL-943
chr7-64871652-DEL-184
chr7-64919063-DEL-53
chr7-64924600-DEL-2583
chr7-64927430-DEL-281
chr7-64937641-DEL-992
chr7-64941855-DEL-73
chr7-64993885-DEL-173
chr7-64999917-DEL-60
chr7-65053844-DEL-200
chr7-65069543-DEL-1518
chr7-65104630-DEL-201
chr7-65112020-DEL-279
chr7-65133225-DEL-790
chr7-65195083-DEL-53
chr7-65211008-DEL-193
chr7-65215375-DEL-131
chr7-65219312-DEL-117
chr7-65243638-DEL-56
chr7-65269705-DEL-53
chr7-65346095-DEL-518
chr7-65368898-DEL-7142
chr7-65405998-DEL-1469
chr7-65408865-DEL-748
chr7-65409860-DEL-168
chr7-65449445-DEL-1759
chr7-65464241-DEL-298
chr7-65492887-DEL-2109
chr7-65496020-DEL-204
chr7-65523619-DEL-108
chr7-65534852-DEL-274
chr7-65610307-DEL-66
chr7-65678296-DEL-64
chr7-65762006-DEL-213
chr7-65795532-DEL-135
chr7-65795646-DEL-133
chr7-65801504-DEL-4675
chr7-65876022-DEL-102
chr7-65880201-DEL-54
chr7-65881256-DEL-60
chr7-65941032-DEL-599
chr7-65950311-DEL-76
chr7-65962482-DEL-51
chr7-66000344-DEL-57

chr7-66011367-DEL-825
chr7-66028566-DEL-67
chr7-66029708-DEL-82
chr7-66160643-DEL-268
chr7-66167008-DEL-95
chr7-66169986-DEL-53
chr7-66198276-DEL-13213
chr7-66204333-DEL-102
chr7-66231748-DEL-104
chr7-66255054-DEL-69
chr7-66272393-DEL-58
chr7-66283720-DEL-104
chr7-66288455-DEL-66
chr7-66292664-DEL-132
chr7-66321808-DEL-64
chr7-66345375-DEL-97
chr7-66389681-DEL-64
chr7-66391559-DEL-672
chr7-66400787-DEL-80
chr7-66403462-DEL-178
chr7-66411884-DEL-57
chr7-66432802-DEL-82
chr7-66460949-DEL-63
chr7-66471069-DEL-108
chr7-66473605-DEL-82
chr7-66562133-DEL-153
chr7-66564933-DEL-565
chr7-66568100-DEL-246
chr7-66593155-DEL-99
chr7-66674069-DEL-1050
chr7-66695624-DEL-133
chr7-66706028-DEL-69
chr7-66744545-DEL-86
chr7-66766617-DEL-79
chr7-66772523-DEL-61
chr7-66779959-DEL-153
chr7-66839269-DEL-1374
chr7-66903985-DEL-723
chr7-66904461-DEL-116
chr7-66948033-DEL-6184
chr7-66973555-DEL-88
chr7-67010846-DEL-170
chr7-67042061-DEL-1736
chr7-67107622-DEL-52
chr7-67130645-DEL-256
chr7-67136154-DEL-171
chr7-67142067-DEL-62
chr7-67144771-DEL-102
chr7-67145615-DEL-51
chr7-67206668-DEL-95
chr7-67213316-DEL-217
chr7-67233222-DEL-53
chr7-67236592-DEL-115
chr7-67269421-DEL-90
chr7-67270842-DEL-113
chr7-67299123-DEL-115
chr7-67333520-DEL-81
chr7-67365369-DEL-2995
chr7-67370640-DEL-906
chr7-67394963-DEL-84
chr7-67407531-DEL-212
chr7-67408057-DEL-268
chr7-67413701-DEL-248
chr7-67436847-DEL-612
chr7-67464225-DEL-96
chr7-67473730-DEL-101
chr7-67508164-DEL-72
chr7-67512409-DEL-55
chr7-67517918-DEL-185
chr7-67518170-DEL-474
chr7-67528445-DEL-161
chr7-67537582-DEL-55
chr7-67538934-DEL-54
chr7-67552330-DEL-1067
chr7-67565089-DEL-391
chr7-67565606-DEL-1225
chr7-67612483-DEL-1070

chr7-67640772-DEL-199
chr7-67644311-DEL-366
chr7-67662931-DEL-172
chr7-67685662-DEL-89
chr7-67715328-DEL-69
chr7-67731501-DEL-150
chr7-67747751-DEL-85
chr7-67790806-DEL-127
chr7-67797424-DEL-99
chr7-67809012-DEL-123
chr7-67821446-DEL-316
chr7-67883509-DEL-55
chr7-67884436-DEL-212
chr7-67890588-DEL-108
chr7-67892613-DEL-236
chr7-67901955-DEL-190
chr7-67914845-DEL-55
chr7-67927667-DEL-423
chr7-67937026-DEL-74
chr7-67975055-DEL-469
chr7-67977140-DEL-51
chr7-67983268-DEL-102
chr7-67990644-DEL-181
chr7-67999551-DEL-127
chr7-68013667-DEL-56
chr7-68086372-DEL-99
chr7-68137920-DEL-582
chr7-68141234-DEL-67
chr7-68151309-DEL-70
chr7-68157932-DEL-105
chr7-68168078-DEL-561
chr7-68198525-DEL-314
chr7-68204292-DEL-64
chr7-68213297-DEL-572
chr7-68214854-DEL-70
chr7-68236683-DEL-740
chr7-68248884-DEL-112
chr7-68297719-DEL-287
chr7-68298213-DEL-506
chr7-68307422-DEL-133
chr7-68320997-DEL-77
chr7-68327306-DEL-72
chr7-68328682-DEL-328
chr7-68339330-DEL-925
chr7-68363659-DEL-1144
chr7-68363987-DEL-779
chr7-68380560-DEL-284
chr7-68381564-DEL-278
chr7-68383346-DEL-54
chr7-68529090-DEL-234
chr7-68561287-DEL-136
chr7-68593375-DEL-58
chr7-68600292-DEL-51
chr7-68607034-DEL-321
chr7-68620606-DEL-63
chr7-68626272-DEL-352
chr7-68675569-DEL-125
chr7-68688534-DEL-96
chr7-68716407-DEL-2480
chr7-68723673-DEL-54
chr7-68759451-DEL-206
chr7-68788587-DEL-84
chr7-68804890-DEL-1665
chr7-68840315-DEL-899
chr7-68846977-DEL-466
chr7-68860247-DEL-55
chr7-68913457-DEL-82
chr7-68941918-DEL-509
chr7-68959708-DEL-7686
chr7-68974289-DEL-1057
chr7-68980304-DEL-64
chr7-68981361-DEL-56
chr7-68998325-DEL-1175
chr7-69031332-DEL-51
chr7-69065590-DEL-1213
chr7-69097841-DEL-68
chr7-69103846-DEL-15481

chr7-69106044-DEL-1491
chr7-69112405-DEL-55
chr7-69137877-DEL-64
chr7-69176102-DEL-51
chr7-69183973-DEL-52
chr7-69212560-DEL-641
chr7-69213525-DEL-709
chr7-69246821-DEL-647
chr7-69272495-DEL-837
chr7-69283534-DEL-56
chr7-69325445-DEL-53
chr7-69337756-DEL-2094
chr7-69351097-DEL-80
chr7-69366454-DEL-64
chr7-69406350-DEL-56
chr7-69413990-DEL-965
chr7-69421823-DEL-1096
chr7-69480685-DEL-1660
chr7-69504432-DEL-90
chr7-69509846-DEL-586
chr7-69512674-DEL-109
chr7-69518829-DEL-201
chr7-69524412-DEL-1834
chr7-69536467-DEL-143
chr7-69548331-DEL-1967
chr7-69564069-DEL-81
chr7-69601789-DEL-132
chr7-69617550-DEL-948
chr7-69618595-DEL-394
chr7-69619037-DEL-1429
chr7-69654884-DEL-138
chr7-69667170-DEL-62
chr7-69672317-DEL-1243
chr7-69672444-DEL-154
chr7-69733551-DEL-1041
chr7-69775841-DEL-82
chr7-69792605-DEL-58
chr7-69850719-DEL-58
chr7-69855711-DEL-60
chr7-69862935-DEL-397
chr7-69864972-DEL-96
chr7-69879991-DEL-129
chr7-69900013-DEL-270
chr7-69945861-DEL-58
chr7-69951809-DEL-68
chr7-70025091-DEL-86
chr7-70088273-DEL-103
chr7-70117925-DEL-76
chr7-70124401-DEL-80
chr7-70160931-DEL-62
chr7-70167297-DEL-83
chr7-70172097-DEL-672
chr7-70182496-DEL-452
chr7-70206646-DEL-58
chr7-70261667-DEL-714
chr7-70265065-DEL-833
chr7-70277473-DEL-148
chr7-70286103-DEL-92
chr7-70353632-DEL-270
chr7-70423526-DEL-1738
chr7-70447530-DEL-53
chr7-70477034-DEL-164
chr7-70562251-DEL-1451
chr7-70643563-DEL-64
chr7-70661499-DEL-52
chr7-70678962-DEL-205
chr7-70716289-DEL-804
chr7-70790261-DEL-55
chr7-70841471-DEL-53
chr7-70845926-DEL-103
chr7-70903434-DEL-73
chr7-70925867-DEL-56
chr7-70928249-DEL-75
chr7-70936488-DEL-145
chr7-70940556-DEL-109
chr7-70982427-DEL-79
chr7-70984357-DEL-54

chr7-70986597-DEL-55
chr7-71009344-DEL-65
chr7-71057428-DEL-140
chr7-71087832-DEL-191
chr7-71096582-DEL-85
chr7-71104662-DEL-519
chr7-71118654-DEL-104
chr7-71206991-DEL-54
chr7-71222074-DEL-68
chr7-71223631-DEL-2987
chr7-71289762-DEL-67
chr7-71342726-DEL-2568
chr7-71375875-DEL-201
chr7-71407057-DEL-853
chr7-71409685-DEL-183
chr7-71417210-DEL-731
chr7-71439347-DEL-94
chr7-71462407-DEL-650
chr7-71530469-DEL-53
chr7-71581036-DEL-303
chr7-71616050-DEL-69
chr7-71708585-DEL-4348
chr7-71713015-DEL-56
chr7-71746739-DEL-129
chr7-71785641-DEL-58
chr7-71792939-DEL-87
chr7-71825421-DEL-1296
chr7-71900224-DEL-505
chr7-71911812-DEL-207
chr7-71967349-DEL-52
chr7-71967788-DEL-415
chr7-71989584-DEL-177
chr7-72059603-DEL-457
chr7-72063957-DEL-475
chr7-72065047-DEL-62
chr7-72067964-DEL-58
chr7-72073329-DEL-1296
chr7-72085956-DEL-925
chr7-72099860-DEL-584
chr7-72102547-DEL-9151
chr7-72113715-DEL-100
chr7-72116208-DEL-68
chr7-72122899-DEL-476
chr7-72147700-DEL-171
chr7-72161477-DEL-414
chr7-72177203-DEL-193
chr7-72194139-DEL-145
chr7-72200319-DEL-196
chr7-72264725-DEL-228
chr7-72268782-DEL-1041
chr7-72316403-DEL-145
chr7-72322494-DEL-2166
chr7-72322526-DEL-784
chr7-72323334-DEL-1295
chr7-72327892-DEL-1005
chr7-72331239-DEL-60
chr7-72334911-DEL-292
chr7-72394302-DEL-60
chr7-72403928-DEL-226
chr7-72408820-DEL-121
chr7-72421677-DEL-566
chr7-72429376-DEL-746
chr7-72437745-DEL-80
chr7-72442136-DEL-98
chr7-72463660-DEL-100
chr7-72466892-DEL-1481
chr7-72488637-DEL-179
chr7-72503188-DEL-266
chr7-72505921-DEL-55
chr7-72507242-DEL-64
chr7-72531013-DEL-2621
chr7-72533777-DEL-2732
chr7-72644155-DEL-65
chr7-72717346-DEL-1101
chr7-72726006-DEL-6207
chr7-72735989-DEL-75
chr7-72749149-DEL-137

chr7-72763968-DEL-269
chr7-72853360-DEL-70
chr7-72879347-DEL-153
chr7-72908484-DEL-1253
chr7-72915834-DEL-1209
chr7-72929524-DEL-52
chr7-72974463-DEL-59
chr7-72995671-DEL-193
chr7-73051926-DEL-259
chr7-73065670-DEL-209
chr7-73073781-DEL-52
chr7-73084149-DEL-102
chr7-73111409-DEL-440
chr7-73135544-DEL-358
chr7-73144217-DEL-3001
chr7-73145205-DEL-207
chr7-73198951-DEL-222
chr7-73224289-DEL-267
chr7-73232213-DEL-56
chr7-73234722-DEL-54
chr7-73242074-DEL-421
chr7-73281595-DEL-661
chr7-73282798-DEL-1417
chr7-73284145-DEL-104
chr7-73321945-DEL-6516
chr7-73384802-DEL-51
chr7-73411140-DEL-70
chr7-73413546-DEL-991
chr7-73433748-DEL-390
chr7-73512450-DEL-1507
chr7-73522987-DEL-924
chr7-73532840-DEL-96
chr7-73535368-DEL-308
chr7-73660395-DEL-124
chr7-73718719-DEL-102
chr7-73731136-DEL-6492
chr7-73746702-DEL-248
chr7-73753549-DEL-1083
chr7-73756180-DEL-70
chr7-73758654-DEL-60
chr7-73783866-DEL-62
chr7-73846873-DEL-63
chr7-73851004-DEL-11236
chr7-73859286-DEL-2586
chr7-73863771-DEL-56
chr7-73870050-DEL-182
chr7-73877699-DEL-62
chr7-73923961-DEL-363
chr7-73937519-DEL-235
chr7-73982583-DEL-1890
chr7-73993926-DEL-99
chr7-74000156-DEL-82
chr7-74000157-DEL-1135
chr7-74000494-DEL-942
chr7-74000572-DEL-282
chr7-74000944-DEL-480
chr7-74013900-DEL-126
chr7-74053798-DEL-184
chr7-74077870-DEL-181
chr7-74103261-DEL-742
chr7-74123644-DEL-70
chr7-74126618-DEL-654
chr7-74131279-DEL-60
chr7-74136134-DEL-74
chr7-74144222-DEL-222
chr7-74183189-DEL-67
chr7-74200038-DEL-61
chr7-74217505-DEL-233
chr7-74250591-DEL-120
chr7-74275934-DEL-72
chr7-74291860-DEL-64
chr7-74305979-DEL-103
chr7-74327353-DEL-154
chr7-74353901-DEL-67
chr7-74403710-DEL-318
chr7-74408182-DEL-467
chr7-74423026-DEL-1346

chr7-74457357-DEL-227
chr7-74494621-DEL-66
chr7-74501009-DEL-316
chr7-74569351-DEL-78
chr7-74575450-DEL-3654
chr7-74576877-DEL-517
chr7-74597321-DEL-1102
chr7-74644197-DEL-468
chr7-74646895-DEL-66
chr7-74666981-DEL-612
chr7-74670671-DEL-1134
chr7-74679088-DEL-267
chr7-74731313-DEL-1125
chr7-74752567-DEL-176
chr7-74793780-DEL-1196
chr7-74807693-DEL-3072
chr7-74810993-DEL-3351
chr7-74827505-DEL-226
chr7-74830243-DEL-9063
chr7-74874662-DEL-130
chr7-74897367-DEL-103
chr7-74902571-DEL-218
chr7-74915429-DEL-57
chr7-74931251-DEL-56
chr7-74933685-DEL-72
chr7-74956459-DEL-72
chr7-74959670-DEL-103
chr7-74965872-DEL-438
chr7-74972463-DEL-1191
chr7-74974030-DEL-82
chr7-74983243-DEL-72
chr7-75007143-DEL-146
chr7-75012946-DEL-223
chr7-75026923-DEL-53
chr7-75040088-DEL-107
chr7-75046754-DEL-74
chr7-75071127-DEL-1297
chr7-75087639-DEL-678
chr7-75093293-DEL-74
chr7-75136393-DEL-1628
chr7-75157070-DEL-7697
chr7-75190459-DEL-8790
chr7-75238302-DEL-1599
chr7-75239903-DEL-914
chr7-75240838-DEL-864
chr7-75241845-DEL-77
chr7-75276062-DEL-72
chr7-75308994-DEL-74
chr7-75325574-DEL-690
chr7-75341349-DEL-73
chr7-75377175-DEL-55
chr7-75390427-DEL-469
chr7-75405537-DEL-56
chr7-75410994-DEL-168
chr7-75414273-DEL-409
chr7-75433326-DEL-68
chr7-75439201-DEL-561
chr7-75439828-DEL-129
chr7-75473062-DEL-3293
chr7-75473070-DEL-770
chr7-75473900-DEL-672
chr7-75475084-DEL-771
chr7-75475915-DEL-190
chr7-75487113-DEL-3816
chr7-75534066-DEL-340
chr7-75582337-DEL-85
chr7-75594524-DEL-6466
chr7-75622886-DEL-99
chr7-75638971-DEL-7499
chr7-75645739-DEL-424
chr7-75654909-DEL-59
chr7-75684873-DEL-291
chr7-75686904-DEL-814
chr7-75715894-DEL-13406
chr7-75745347-DEL-295
chr7-75753936-DEL-2669
chr7-75788584-DEL-56

chr7-75796290-DEL-67
chr7-75813334-DEL-102
chr7-75816162-DEL-64
chr7-75828435-DEL-59
chr7-75841730-DEL-836
chr7-75858627-DEL-2352
chr7-75884152-DEL-1513
chr7-75890589-DEL-9295
chr7-75916155-DEL-2436
chr7-75931334-DEL-483
chr7-75934516-DEL-64
chr7-75957672-DEL-202
chr7-75968211-DEL-128
chr7-76005457-DEL-74
chr7-76130104-DEL-179
chr7-76133583-DEL-793
chr7-76140796-DEL-2241
chr7-76142130-DEL-919
chr7-76165812-DEL-250
chr7-76168264-DEL-344
chr7-76228596-DEL-255
chr7-76268327-DEL-89
chr7-76307752-DEL-192
chr7-76325647-DEL-1812
chr7-76343107-DEL-114
chr7-76359603-DEL-201
chr7-76386838-DEL-6344
chr7-76414871-DEL-96
chr7-76434740-DEL-110
chr7-76460561-DEL-96
chr7-76500536-DEL-110
chr7-76500708-DEL-134
chr7-76529032-DEL-101
chr7-76550898-DEL-188
chr7-76576539-DEL-938
chr7-76612715-DEL-443
chr7-76619084-DEL-16270
chr7-76622578-DEL-157
chr7-76653339-DEL-170
chr7-76666612-DEL-401
chr7-76674314-DEL-219
chr7-76698193-DEL-53
chr7-76701303-DEL-76
chr7-76712095-DEL-52
chr7-76715677-DEL-63
chr7-76770228-DEL-68
chr7-76792864-DEL-839
chr7-76795113-DEL-91
chr7-76800498-DEL-724
chr7-76804875-DEL-222
chr7-76844636-DEL-106
chr7-76879545-DEL-56
chr7-76904546-DEL-762
chr7-76921339-DEL-2482
chr7-76921932-DEL-8615
chr7-76924585-DEL-111
chr7-76927192-DEL-462
chr7-76957169-DEL-2284
chr7-76967709-DEL-1125
chr7-76971996-DEL-98
chr7-76984360-DEL-212
chr7-77003752-DEL-332
chr7-77034855-DEL-61
chr7-77037934-DEL-6281
chr7-77037946-DEL-3351
chr7-77043286-DEL-911
chr7-77048652-DEL-180
chr7-77055937-DEL-62
chr7-77064561-DEL-94
chr7-77071216-DEL-83
chr7-77079049-DEL-574
chr7-77101276-DEL-62
chr7-77107182-DEL-1204
chr7-77119522-DEL-6195
chr7-77127911-DEL-844
chr7-77129692-DEL-60
chr7-77137970-DEL-765

chr7-77151930-DEL-294
chr7-77182836-DEL-101
chr7-77239144-DEL-336
chr7-77248467-DEL-64
chr7-77255800-DEL-1064
chr7-77268230-DEL-52
chr7-77279861-DEL-453
chr7-77297859-DEL-440
chr7-77302109-DEL-2818
chr7-77450863-DEL-91
chr7-77490096-DEL-59
chr7-77496149-DEL-66
chr7-77565065-DEL-1364
chr7-77566705-DEL-1224
chr7-77591568-DEL-64
chr7-77598268-DEL-59
chr7-77602454-DEL-5994
chr7-77603229-DEL-6010
chr7-77609677-DEL-922
chr7-77627686-DEL-116
chr7-77680319-DEL-136
chr7-77682860-DEL-4181
chr7-77778960-DEL-57
chr7-77789713-DEL-62
chr7-77812772-DEL-1268
chr7-77828496-DEL-240
chr7-77832391-DEL-285
chr7-77851492-DEL-199
chr7-77863646-DEL-1888
chr7-77874664-DEL-1364
chr7-77882932-DEL-163
chr7-77889501-DEL-510
chr7-77940503-DEL-58
chr7-78041007-DEL-260
chr7-78065819-DEL-212
chr7-78083756-DEL-10480
chr7-78084839-DEL-779
chr7-78093591-DEL-182
chr7-78097071-DEL-52
chr7-78113035-DEL-288
chr7-78117577-DEL-52
chr7-78120233-DEL-138
chr7-78163069-DEL-93
chr7-78214972-DEL-121
chr7-78262282-DEL-325
chr7-78291915-DEL-84
chr7-78429538-DEL-161
chr7-78452329-DEL-66
chr7-78459102-DEL-1443
chr7-78461307-DEL-193
chr7-78465403-DEL-88
chr7-78466959-DEL-745
chr7-78480953-DEL-460
chr7-78481340-DEL-2076
chr7-78481389-DEL-738
chr7-78482129-DEL-1586
chr7-78482459-DEL-78
chr7-78483042-DEL-160
chr7-78499502-DEL-762
chr7-78500623-DEL-161
chr7-78541930-DEL-91
chr7-78577382-DEL-605
chr7-78581301-DEL-54
chr7-78610928-DEL-69
chr7-78645435-DEL-65
chr7-78654578-DEL-845
chr7-78655159-DEL-686
chr7-78673009-DEL-1384
chr7-78696307-DEL-82
chr7-78697617-DEL-51
chr7-78711864-DEL-730
chr7-78773825-DEL-310
chr7-78806399-DEL-79
chr7-78831703-DEL-1226
chr7-78841724-DEL-75
chr7-78851451-DEL-63
chr7-78965109-DEL-6705

chr7-79006762-DEL-739
chr7-79046827-DEL-119
chr7-79075453-DEL-225
chr7-79146226-DEL-72
chr7-79179029-DEL-126
chr7-79217948-DEL-190
chr7-79225827-DEL-60
chr7-79246228-DEL-80
chr7-79252739-DEL-76
chr7-79283046-DEL-52
chr7-79300286-DEL-68
chr7-79302361-DEL-156
chr7-79338151-DEL-219
chr7-79355093-DEL-452
chr7-79374427-DEL-54
chr7-79378112-DEL-72
chr7-79384142-DEL-52
chr7-79397037-DEL-203
chr7-79432707-DEL-218
chr7-79439780-DEL-746
chr7-79467708-DEL-132
chr7-79468657-DEL-375
chr7-79478123-DEL-2432
chr7-79478648-DEL-1296
chr7-79532904-DEL-308
chr7-79555516-DEL-222
chr7-79574613-DEL-155
chr7-79575144-DEL-269
chr7-79592968-DEL-2839
chr7-79595969-DEL-55
chr7-79598979-DEL-66
chr7-79602541-DEL-106
chr7-79610035-DEL-276
chr7-79622579-DEL-52
chr7-79627628-DEL-96
chr7-79645287-DEL-222
chr7-79660424-DEL-123
chr7-79666146-DEL-1288
chr7-79689324-DEL-52
chr7-79726603-DEL-55
chr7-79759825-DEL-106
chr7-79785359-DEL-83
chr7-79796326-DEL-160
chr7-79811358-DEL-888
chr7-79824801-DEL-179
chr7-79828627-DEL-442
chr7-79872239-DEL-63
chr7-79889673-DEL-116
chr7-79892872-DEL-206
chr7-79912448-DEL-316
chr7-80004458-DEL-183
chr7-80046059-DEL-198
chr7-80097842-DEL-90
chr7-80127617-DEL-1396
chr7-80148262-DEL-207
chr7-80188563-DEL-203
chr7-80195254-DEL-196
chr7-80196889-DEL-82
chr7-80200349-DEL-62
chr7-80251638-DEL-158
chr7-80257969-DEL-864
chr7-80283716-DEL-1995
chr7-80295994-DEL-96
chr7-80312812-DEL-63
chr7-80326471-DEL-267
chr7-80341643-DEL-446
chr7-80351930-DEL-132
chr7-80358171-DEL-1717
chr7-80358381-DEL-364
chr7-80359134-DEL-104
chr7-80419141-DEL-75
chr7-80438613-DEL-53
chr7-80443348-DEL-1166
chr7-80443493-DEL-397
chr7-80444057-DEL-152
chr7-80450925-DEL-218
chr7-80456578-DEL-209

chr7-80476370-DEL-85
chr7-80509270-DEL-282
chr7-80517740-DEL-524
chr7-80519234-DEL-95
chr7-80531368-DEL-1956
chr7-80552739-DEL-353
chr7-80582238-DEL-120
chr7-80600660-DEL-136
chr7-80609708-DEL-2851
chr7-80661010-DEL-1005
chr7-80685876-DEL-66
chr7-80689418-DEL-53
chr7-80704342-DEL-68
chr7-80723721-DEL-52
chr7-80732588-DEL-705
chr7-80735638-DEL-220
chr7-80737168-DEL-504
chr7-80737931-DEL-2173
chr7-80768781-DEL-68
chr7-80780308-DEL-92
chr7-80785553-DEL-60
chr7-80800836-DEL-131
chr7-80822135-DEL-277
chr7-80823457-DEL-604
chr7-80843259-DEL-207
chr7-80849422-DEL-55
chr7-80884256-DEL-136
chr7-80889101-DEL-68
chr7-80891376-DEL-200
chr7-80901523-DEL-151
chr7-80955046-DEL-446
chr7-80967276-DEL-110
chr7-80973371-DEL-243
chr7-80982157-DEL-85
chr7-81015113-DEL-276
chr7-81037539-DEL-6309
chr7-81038240-DEL-93
chr7-81055823-DEL-3825
chr7-81081601-DEL-207
chr7-81127916-DEL-75
chr7-81153169-DEL-615
chr7-81198188-DEL-80
chr7-81230225-DEL-649
chr7-81268979-DEL-457
chr7-81283299-DEL-711
chr7-81315117-DEL-107
chr7-81369165-DEL-119
chr7-81402513-DEL-54
chr7-81421600-DEL-214
chr7-81434910-DEL-62
chr7-81435472-DEL-83
chr7-81437214-DEL-215
chr7-81488148-DEL-204
chr7-81503797-DEL-91
chr7-81506279-DEL-64
chr7-81539807-DEL-68
chr7-81578611-DEL-58
chr7-81596873-DEL-390
chr7-81616815-DEL-147
chr7-81625996-DEL-54
chr7-81661303-DEL-54
chr7-81668508-DEL-693
chr7-81701925-DEL-104
chr7-81705079-DEL-53
chr7-81712581-DEL-57
chr7-81735689-DEL-1516
chr7-81739699-DEL-59
chr7-81743836-DEL-695
chr7-81759106-DEL-94
chr7-81772528-DEL-1635
chr7-81790514-DEL-84
chr7-81801382-DEL-508
chr7-81804030-DEL-282
chr7-81812472-DEL-544
chr7-81816514-DEL-4909
chr7-81834491-DEL-335
chr7-81843715-DEL-2842

chr7-81843870-DEL-128
chr7-81846730-DEL-608
chr7-81848472-DEL-79
chr7-81859343-DEL-57
chr7-81863103-DEL-212
chr7-81863440-DEL-424
chr7-81870555-DEL-180
chr7-81892266-DEL-987
chr7-81907272-DEL-151
chr7-81916484-DEL-58
chr7-81932477-DEL-167
chr7-81949671-DEL-242
chr7-81997398-DEL-69
chr7-82022613-DEL-65
chr7-82024231-DEL-151
chr7-82082836-DEL-444
chr7-82127970-DEL-307
chr7-82174662-DEL-70
chr7-82229869-DEL-68
chr7-82267830-DEL-175
chr7-82343027-DEL-1087
chr7-82377239-DEL-749
chr7-82378951-DEL-1342
chr7-82462882-DEL-230
chr7-82471510-DEL-95
chr7-82482231-DEL-115
chr7-82485642-DEL-51
chr7-82504213-DEL-53
chr7-82511499-DEL-254
chr7-82541742-DEL-1678
chr7-82565003-DEL-200
chr7-82614080-DEL-61
chr7-82645194-DEL-1892
chr7-82660527-DEL-266
chr7-82667697-DEL-82
chr7-82701394-DEL-109
chr7-82705054-DEL-82
chr7-82715665-DEL-2746
chr7-82731524-DEL-297
chr7-82750702-DEL-88
chr7-82755744-DEL-715
chr7-82774238-DEL-84
chr7-82783428-DEL-2919
chr7-82800192-DEL-57
chr7-82813711-DEL-64
chr7-82821167-DEL-90
chr7-82893293-DEL-76
chr7-82898244-DEL-56
chr7-82975413-DEL-131
chr7-82992836-DEL-2425
chr7-83002320-DEL-3095
chr7-83078119-DEL-195
chr7-83090633-DEL-123
chr7-83147302-DEL-173
chr7-83152517-DEL-1987
chr7-83228451-DEL-1668
chr7-83306970-DEL-58
chr7-83346095-DEL-399
chr7-83347974-DEL-52
chr7-83368602-DEL-105
chr7-83389986-DEL-57
chr7-83417580-DEL-62
chr7-83432532-DEL-6142
chr7-83441738-DEL-178
chr7-83454716-DEL-153
chr7-83474793-DEL-208
chr7-83536939-DEL-99
chr7-83547622-DEL-126
chr7-83553272-DEL-6014
chr7-83557841-DEL-76
chr7-83561711-DEL-159
chr7-83565554-DEL-227
chr7-83583358-DEL-66
chr7-83588112-DEL-1088
chr7-83623708-DEL-52
chr7-83666799-DEL-245
chr7-83679044-DEL-213

chr7-83684431-DEL-67
chr7-83696047-DEL-1555
chr7-83711561-DEL-76
chr7-83726320-DEL-97
chr7-83732999-DEL-144
chr7-83740367-DEL-58
chr7-83742130-DEL-432
chr7-83744059-DEL-3621
chr7-83751063-DEL-1591
chr7-83780404-DEL-95
chr7-83805418-DEL-66
chr7-83811255-DEL-284
chr7-83871056-DEL-56
chr7-83891756-DEL-201
chr7-83900965-DEL-3007
chr7-83943488-DEL-66
chr7-83955496-DEL-126
chr7-84014946-DEL-50
chr7-84015932-DEL-335
chr7-84038662-DEL-161
chr7-84040363-DEL-178
chr7-84062484-DEL-165
chr7-84114400-DEL-97
chr7-84127309-DEL-199
chr7-84133987-DEL-2642
chr7-84139016-DEL-58
chr7-84148579-DEL-215
chr7-84179622-DEL-58
chr7-84198753-DEL-1059
chr7-84228763-DEL-123
chr7-84242682-DEL-109
chr7-84243613-DEL-114
chr7-84316758-DEL-221
chr7-84398173-DEL-368
chr7-84434077-DEL-71
chr7-84457692-DEL-447
chr7-84461892-DEL-335
chr7-84476165-DEL-2522
chr7-84481108-DEL-329
chr7-84484311-DEL-5335
chr7-84513373-DEL-3002
chr7-84530149-DEL-771
chr7-84552806-DEL-3070
chr7-84584693-DEL-58
chr7-84603682-DEL-91
chr7-84606446-DEL-2206
chr7-84630900-DEL-197
chr7-84709089-DEL-1302
chr7-84752570-DEL-90
chr7-84765100-DEL-69
chr7-84770479-DEL-55
chr7-84813647-DEL-911
chr7-84847154-DEL-73
chr7-84849331-DEL-68
chr7-84850670-DEL-443
chr7-84856928-DEL-66
chr7-84888972-DEL-6066
chr7-84903290-DEL-144
chr7-84928308-DEL-6762
chr7-84935503-DEL-982
chr7-84980583-DEL-1189
chr7-84981871-DEL-831
chr7-84983023-DEL-489
chr7-84983827-DEL-1704
chr7-84994461-DEL-86
chr7-85007405-DEL-58
chr7-85023270-DEL-4534
chr7-85047004-DEL-86
chr7-85048567-DEL-67
chr7-85049450-DEL-2256
chr7-85052616-DEL-67
chr7-85065701-DEL-6001
chr7-85071906-DEL-1864
chr7-85078878-DEL-60
chr7-85086380-DEL-6216
chr7-85094684-DEL-1522
chr7-85102864-DEL-8679

chr7-85113694-DEL-964
chr7-85116883-DEL-7999
chr7-85116955-DEL-4529
chr7-85132723-DEL-57
chr7-85134810-DEL-3424
chr7-85138543-DEL-634
chr7-85143479-DEL-80
chr7-85148211-DEL-2789
chr7-85161226-DEL-5389
chr7-85174368-DEL-4627
chr7-85179395-DEL-1870
chr7-85183596-DEL-1330
chr7-85184618-DEL-653
chr7-85193997-DEL-6882
chr7-85247810-DEL-516
chr7-85335124-DEL-1511
chr7-85354048-DEL-634
chr7-85365554-DEL-8206
chr7-85374738-DEL-9025
chr7-85406334-DEL-12749
chr7-85430754-DEL-342
chr7-85443865-DEL-171
chr7-85446019-DEL-117
chr7-85449513-DEL-547
chr7-85451984-DEL-1684
chr7-85453319-DEL-717
chr7-85467469-DEL-329
chr7-85470538-DEL-79
chr7-85470892-DEL-2045
chr7-85491360-DEL-550
chr7-85564583-DEL-167
chr7-85575046-DEL-9119
chr7-85584844-DEL-144
chr7-85657672-DEL-1897
chr7-85667941-DEL-58
chr7-85743734-DEL-4263
chr7-85788025-DEL-4178
chr7-85792301-DEL-1179
chr7-85799825-DEL-93
chr7-85812656-DEL-597
chr7-85824582-DEL-399
chr7-85837603-DEL-69
chr7-85867935-DEL-2526
chr7-85886325-DEL-2232
chr7-85905937-DEL-657
chr7-85923050-DEL-312
chr7-85968281-DEL-166
chr7-86027964-DEL-862
chr7-86053416-DEL-771
chr7-86056817-DEL-1168
chr7-86063322-DEL-193
chr7-86085554-DEL-1998
chr7-86099506-DEL-60
chr7-86115662-DEL-252
chr7-86119245-DEL-52
chr7-86120508-DEL-2921
chr7-86125094-DEL-47023
chr7-86127522-DEL-73
chr7-86132305-DEL-363
chr7-86144786-DEL-51
chr7-86178034-DEL-357
chr7-86187461-DEL-958
chr7-86202205-DEL-55
chr7-86214019-DEL-7765
chr7-86264574-DEL-58
chr7-86276215-DEL-303
chr7-86345929-DEL-235
chr7-86352593-DEL-149
chr7-86352892-DEL-768
chr7-86355423-DEL-6209
chr7-86364773-DEL-64
chr7-86371010-DEL-917
chr7-86387356-DEL-70
chr7-86390069-DEL-69
chr7-86402475-DEL-638
chr7-86445661-DEL-195
chr7-86446475-DEL-104

chr7-86465568-DEL-175
chr7-86483274-DEL-377
chr7-86487409-DEL-101
chr7-86492265-DEL-1455
chr7-86492894-DEL-7320
chr7-86495030-DEL-6004
chr7-86502146-DEL-3085
chr7-86521971-DEL-2471
chr7-86527055-DEL-402
chr7-86547067-DEL-2551
chr7-86605162-DEL-78
chr7-86640535-DEL-85
chr7-86682604-DEL-2088
chr7-86694473-DEL-1492
chr7-86703783-DEL-1817
chr7-86711333-DEL-480
chr7-86786921-DEL-270
chr7-86804877-DEL-70
chr7-86806242-DEL-455
chr7-86808021-DEL-202
chr7-86809039-DEL-7188
chr7-86816487-DEL-390
chr7-86821371-DEL-82
chr7-86839492-DEL-333
chr7-86851926-DEL-412
chr7-86894086-DEL-82
chr7-86908307-DEL-446
chr7-86922435-DEL-76
chr7-86949440-DEL-765
chr7-87054816-DEL-58
chr7-87058062-DEL-110
chr7-87064452-DEL-6429
chr7-87075780-DEL-205
chr7-87080351-DEL-187
chr7-87084253-DEL-740
chr7-87092927-DEL-60
chr7-87172460-DEL-69
chr7-87176124-DEL-200
chr7-87179559-DEL-110
chr7-87225182-DEL-111
chr7-87273857-DEL-105
chr7-87288016-DEL-75
chr7-87344327-DEL-4336
chr7-87351177-DEL-13841
chr7-87360343-DEL-534
chr7-87371692-DEL-203
chr7-87390941-DEL-240
chr7-87393036-DEL-95
chr7-87400746-DEL-1286
chr7-87408599-DEL-162
chr7-87442454-DEL-188
chr7-87457807-DEL-58
chr7-87461952-DEL-649
chr7-87470596-DEL-2271
chr7-87473018-DEL-88
chr7-87501199-DEL-162
chr7-87505479-DEL-72
chr7-87520508-DEL-109
chr7-87579496-DEL-189
chr7-87621594-DEL-402
chr7-87650352-DEL-209
chr7-87677737-DEL-844
chr7-87702941-DEL-94
chr7-87715424-DEL-87
chr7-87730925-DEL-608
chr7-87762924-DEL-113
chr7-87793648-DEL-179
chr7-87814016-DEL-56
chr7-87816344-DEL-6789
chr7-87826657-DEL-235
chr7-87831915-DEL-51
chr7-87838031-DEL-59
chr7-87853859-DEL-70
chr7-87881600-DEL-178
chr7-87913715-DEL-58
chr7-87915741-DEL-2996
chr7-87915833-DEL-218

chr7-87916293-DEL-1350
chr7-87916652-DEL-1643
chr7-87916965-DEL-74
chr7-87918357-DEL-71
chr7-87949150-DEL-84
chr7-87954259-DEL-799
chr7-87970883-DEL-549
chr7-87974185-DEL-112
chr7-87983629-DEL-3707
chr7-87993403-DEL-1297
chr7-88008458-DEL-225
chr7-88014168-DEL-1197
chr7-88016760-DEL-87
chr7-88035522-DEL-160
chr7-88043513-DEL-54
chr7-88049283-DEL-100
chr7-88053512-DEL-1032
chr7-88059307-DEL-1300
chr7-88060645-DEL-1553
chr7-88066929-DEL-1447
chr7-88077750-DEL-986
chr7-88094970-DEL-422
chr7-88100229-DEL-746
chr7-88105579-DEL-55
chr7-88115233-DEL-82
chr7-88207697-DEL-662
chr7-88209182-DEL-1739
chr7-88219131-DEL-457
chr7-88220481-DEL-82
chr7-88226678-DEL-51
chr7-88273719-DEL-317
chr7-88287251-DEL-448
chr7-88373026-DEL-154
chr7-88414805-DEL-1924
chr7-88440099-DEL-153
chr7-88454806-DEL-1018
chr7-88463086-DEL-1575
chr7-88471510-DEL-271
chr7-88482385-DEL-64
chr7-88506388-DEL-1201
chr7-88513428-DEL-6208
chr7-88537445-DEL-219
chr7-88537743-DEL-3130
chr7-88540921-DEL-1535
chr7-88596655-DEL-55
chr7-88597107-DEL-200
chr7-88600911-DEL-454
chr7-88637190-DEL-6346
chr7-88648624-DEL-6576
chr7-88695732-DEL-98
chr7-88697122-DEL-594
chr7-88710505-DEL-1375
chr7-88721277-DEL-52
chr7-88735160-DEL-53
chr7-88743133-DEL-440
chr7-88747816-DEL-202
chr7-88793218-DEL-265
chr7-88837655-DEL-1254
chr7-88851877-DEL-186
chr7-88892933-DEL-51
chr7-88899853-DEL-328
chr7-88913228-DEL-105
chr7-88915701-DEL-187
chr7-88917075-DEL-55
chr7-88927324-DEL-287
chr7-88938786-DEL-374
chr7-88939229-DEL-553
chr7-88948763-DEL-98
chr7-88963545-DEL-157
chr7-88983995-DEL-219
chr7-89074210-DEL-58
chr7-89080401-DEL-60
chr7-89099467-DEL-205
chr7-89109871-DEL-62
chr7-89119377-DEL-194
chr7-89123991-DEL-128
chr7-89126549-DEL-68

chr7-89139163-DEL-67
chr7-89152391-DEL-175
chr7-89167807-DEL-120
chr7-89181931-DEL-55
chr7-89216113-DEL-56
chr7-89221280-DEL-54
chr7-89231400-DEL-6412
chr7-89254092-DEL-115
chr7-89279288-DEL-77
chr7-89324285-DEL-1108
chr7-89343341-DEL-1626
chr7-89421640-DEL-52
chr7-89437395-DEL-227
chr7-89437928-DEL-604
chr7-89444031-DEL-1573
chr7-89445461-DEL-61
chr7-89467654-DEL-55
chr7-89488287-DEL-473
chr7-89521596-DEL-2018
chr7-89524363-DEL-145
chr7-89529658-DEL-758
chr7-89547376-DEL-170
chr7-89578801-DEL-188
chr7-89592282-DEL-292
chr7-89600501-DEL-516
chr7-89609231-DEL-53
chr7-89613352-DEL-52
chr7-89614177-DEL-180
chr7-89622347-DEL-209
chr7-89631629-DEL-59
chr7-89657033-DEL-368
chr7-89664183-DEL-56
chr7-89667455-DEL-189
chr7-89677684-DEL-185
chr7-89730479-DEL-3109
chr7-89755892-DEL-56
chr7-89789779-DEL-113
chr7-89791721-DEL-1212
chr7-89802655-DEL-53
chr7-89817141-DEL-313
chr7-89846753-DEL-84
chr7-89878390-DEL-177
chr7-89884670-DEL-53
chr7-89904619-DEL-1903
chr7-89941139-DEL-3051
chr7-89945105-DEL-69
chr7-89974256-DEL-61
chr7-90023993-DEL-83
chr7-90034659-DEL-68
chr7-90037487-DEL-85
chr7-90052477-DEL-166
chr7-90054596-DEL-1347
chr7-90103685-DEL-403
chr7-90112465-DEL-54
chr7-90120937-DEL-289
chr7-90128433-DEL-67
chr7-90135274-DEL-127
chr7-90154312-DEL-54
chr7-90175598-DEL-9279
chr7-90192124-DEL-52
chr7-90195178-DEL-216
chr7-90245274-DEL-193
chr7-90267957-DEL-201
chr7-90271902-DEL-2258
chr7-90290205-DEL-54
chr7-90336059-DEL-377
chr7-90356334-DEL-301
chr7-90358934-DEL-62
chr7-90365999-DEL-6280
chr7-90384560-DEL-329
chr7-90391505-DEL-692
chr7-90409824-DEL-2112
chr7-90412416-DEL-2540
chr7-90427514-DEL-1151
chr7-90445272-DEL-1554
chr7-90454218-DEL-7264
chr7-90463216-DEL-126

chr7-90491227-DEL-809
chr7-90515161-DEL-1082
chr7-90549286-DEL-927
chr7-90571575-DEL-2356
chr7-90678017-DEL-84
chr7-90699713-DEL-162
chr7-90709547-DEL-210
chr7-90790633-DEL-801
chr7-90811834-DEL-4944
chr7-90857173-DEL-92
chr7-90860729-DEL-255
chr7-90862735-DEL-186
chr7-90878353-DEL-168
chr7-90885771-DEL-1378
chr7-90887598-DEL-379
chr7-90902646-DEL-140
chr7-90920304-DEL-318
chr7-90936343-DEL-915
chr7-90941301-DEL-291
chr7-90956604-DEL-291
chr7-91002559-DEL-118
chr7-91024498-DEL-245
chr7-91041915-DEL-164
chr7-91060330-DEL-60
chr7-91067341-DEL-128
chr7-91105964-DEL-276
chr7-91117927-DEL-55
chr7-91126607-DEL-90
chr7-91153690-DEL-60
chr7-91162719-DEL-810
chr7-91213443-DEL-77
chr7-91214381-DEL-937
chr7-91217339-DEL-6778
chr7-91277664-DEL-94
chr7-91280388-DEL-1114
chr7-91291710-DEL-242
chr7-91347025-DEL-68
chr7-91401299-DEL-52
chr7-91412763-DEL-199
chr7-91431554-DEL-538
chr7-91457818-DEL-1578
chr7-91459482-DEL-76
chr7-91459830-DEL-501
chr7-91469269-DEL-161
chr7-91511905-DEL-111
chr7-91523734-DEL-59
chr7-91596655-DEL-6875
chr7-91601992-DEL-819
chr7-91714277-DEL-128
chr7-91721303-DEL-201
chr7-91758094-DEL-60
chr7-91766221-DEL-76
chr7-91792526-DEL-458
chr7-91795841-DEL-701
chr7-91796960-DEL-132
chr7-91840611-DEL-3072
chr7-91843826-DEL-178
chr7-91850830-DEL-59
chr7-91931972-DEL-61
chr7-91956626-DEL-53
chr7-91969312-DEL-97
chr7-91983949-DEL-150
chr7-91985218-DEL-162
chr7-92027230-DEL-499
chr7-92046867-DEL-65
chr7-92059208-DEL-638
chr7-92116729-DEL-132
chr7-92117272-DEL-160
chr7-92139569-DEL-154
chr7-92157627-DEL-2227
chr7-92173424-DEL-721
chr7-92188664-DEL-848
chr7-92192260-DEL-156
chr7-92200645-DEL-66
chr7-92217832-DEL-303
chr7-92222766-DEL-828
chr7-92254670-DEL-897

chr7-92323015-DEL-222
chr7-92330239-DEL-53
chr7-92362241-DEL-63
chr7-92377662-DEL-54
chr7-92396440-DEL-165
chr7-92434162-DEL-64
chr7-92435679-DEL-9185
chr7-92464979-DEL-111
chr7-92483439-DEL-152
chr7-92602052-DEL-73
chr7-92640575-DEL-52
chr7-92642053-DEL-448
chr7-92663614-DEL-2761
chr7-92676242-DEL-1161
chr7-92685728-DEL-51
chr7-92705684-DEL-53
chr7-92718965-DEL-129
chr7-92735452-DEL-271
chr7-92737807-DEL-189
chr7-92758106-DEL-342
chr7-92787465-DEL-235
chr7-92805084-DEL-6393
chr7-92816146-DEL-6618
chr7-92816971-DEL-950
chr7-92828155-DEL-719
chr7-92830370-DEL-112
chr7-92836211-DEL-1064
chr7-92837836-DEL-6236
chr7-92847785-DEL-1930
chr7-92850660-DEL-431
chr7-92853406-DEL-482
chr7-92856070-DEL-2012
chr7-92893823-DEL-111
chr7-92904194-DEL-89
chr7-92917247-DEL-1063
chr7-92954821-DEL-89
chr7-92965502-DEL-218
chr7-92998957-DEL-161
chr7-93041371-DEL-133
chr7-93071504-DEL-126
chr7-93082114-DEL-4274
chr7-93120984-DEL-84
chr7-93128948-DEL-231
chr7-93132954-DEL-241
chr7-93140515-DEL-1307
chr7-93150062-DEL-564
chr7-93161867-DEL-115
chr7-93170581-DEL-6232
chr7-93223044-DEL-877
chr7-93235799-DEL-177
chr7-93251668-DEL-60
chr7-93265811-DEL-119
chr7-93303131-DEL-230
chr7-93347421-DEL-480
chr7-93348613-DEL-7922
chr7-93358339-DEL-54
chr7-93360617-DEL-6300
chr7-93408603-DEL-164
chr7-93440634-DEL-60
chr7-93486209-DEL-108
chr7-93495371-DEL-904
chr7-93497201-DEL-1170
chr7-93498805-DEL-328
chr7-93515150-DEL-741
chr7-93516038-DEL-264
chr7-93536598-DEL-79
chr7-93540478-DEL-144
chr7-93544241-DEL-159
chr7-93562193-DEL-492
chr7-93572254-DEL-199
chr7-93573005-DEL-1898
chr7-93616832-DEL-106
chr7-93617620-DEL-6211
chr7-93624860-DEL-400
chr7-93625932-DEL-1770
chr7-93630549-DEL-669
chr7-93636677-DEL-3464

chr7-93640362-DEL-3331
chr7-93650321-DEL-57
chr7-93662780-DEL-424
chr7-93682311-DEL-68
chr7-93683096-DEL-1782
chr7-93699799-DEL-513
chr7-93727917-DEL-51
chr7-93770053-DEL-206
chr7-93772960-DEL-108
chr7-93811165-DEL-299
chr7-93813310-DEL-52
chr7-93825279-DEL-1989
chr7-93846854-DEL-56
chr7-93857067-DEL-71
chr7-93890842-DEL-7116
chr7-93902352-DEL-60
chr7-93903774-DEL-847
chr7-93907748-DEL-60
chr7-93911579-DEL-261
chr7-93934159-DEL-789
chr7-93937110-DEL-159
chr7-93941548-DEL-268
chr7-94002703-DEL-781
chr7-94014301-DEL-1331
chr7-94023580-DEL-72
chr7-94037610-DEL-608
chr7-94038708-DEL-3670
chr7-94042857-DEL-1624
chr7-94048368-DEL-117
chr7-94076072-DEL-56
chr7-94156671-DEL-743
chr7-94171289-DEL-293
chr7-94179586-DEL-52
chr7-94225215-DEL-3807
chr7-94248115-DEL-204
chr7-94260154-DEL-180
chr7-94280390-DEL-3387
chr7-94293173-DEL-132
chr7-94329573-DEL-177
chr7-94340648-DEL-94
chr7-94345662-DEL-171
chr7-94358488-DEL-72
chr7-94359988-DEL-86
chr7-94438788-DEL-6140
chr7-94484742-DEL-1275
chr7-94504788-DEL-117
chr7-94516271-DEL-198
chr7-94551331-DEL-341
chr7-94570009-DEL-88
chr7-94582007-DEL-67
chr7-94587161-DEL-519
chr7-94587906-DEL-57
chr7-94588688-DEL-617
chr7-94640892-DEL-58
chr7-94646293-DEL-64
chr7-94650605-DEL-256
chr7-94651776-DEL-97
chr7-94662751-DEL-247
chr7-94669281-DEL-48215
chr7-94678986-DEL-102
chr7-94694578-DEL-652
chr7-94695331-DEL-918
chr7-94731087-DEL-53
chr7-94734190-DEL-82
chr7-94756142-DEL-6199
chr7-94776904-DEL-255
chr7-94785885-DEL-258
chr7-94799541-DEL-116
chr7-94801137-DEL-438
chr7-94810291-DEL-249
chr7-94814027-DEL-401
chr7-94826407-DEL-3156
chr7-94829829-DEL-444
chr7-94830443-DEL-170
chr7-94836651-DEL-6264
chr7-94849410-DEL-533
chr7-94920123-DEL-56238

chr7-94920401-DEL-142
chr7-94958842-DEL-1654
chr7-94982753-DEL-399
chr7-94987100-DEL-6018
chr7-94994309-DEL-203
chr7-95036243-DEL-82
chr7-95039098-DEL-5382
chr7-95078169-DEL-58
chr7-95127236-DEL-1296
chr7-95161024-DEL-140
chr7-95247497-DEL-110
chr7-95249906-DEL-110
chr7-95269553-DEL-56
chr7-95284147-DEL-751
chr7-95284987-DEL-66
chr7-95286878-DEL-346
chr7-95298991-DEL-251
chr7-95309505-DEL-60
chr7-95365194-DEL-57
chr7-95380107-DEL-70
chr7-95415268-DEL-194
chr7-95416899-DEL-159
chr7-95432504-DEL-110
chr7-95538154-DEL-90
chr7-95547674-DEL-720
chr7-95589196-DEL-58
chr7-95632100-DEL-372
chr7-95671457-DEL-283
chr7-95723621-DEL-297
chr7-95744659-DEL-68
chr7-95810605-DEL-57
chr7-95875252-DEL-60
chr7-95886019-DEL-55
chr7-95916924-DEL-112
chr7-95958997-DEL-83
chr7-96022320-DEL-144
chr7-96098707-DEL-54
chr7-96192606-DEL-1343
chr7-96200440-DEL-62
chr7-96220912-DEL-148
chr7-96242640-DEL-134
chr7-96252799-DEL-55
chr7-96279099-DEL-85
chr7-96279638-DEL-507
chr7-96286446-DEL-127
chr7-96302956-DEL-73
chr7-96324201-DEL-177
chr7-96334470-DEL-1265
chr7-96349640-DEL-61
chr7-96470500-DEL-210
chr7-96485561-DEL-53
chr7-96500981-DEL-90
chr7-96502510-DEL-104
chr7-96510271-DEL-235
chr7-96571770-DEL-54
chr7-96580252-DEL-200
chr7-96604051-DEL-54
chr7-96630944-DEL-53
chr7-96638761-DEL-180
chr7-96702886-DEL-230
chr7-96715932-DEL-180
chr7-96742190-DEL-346
chr7-96769832-DEL-53
chr7-96800946-DEL-592
chr7-96828417-DEL-96
chr7-96834491-DEL-60
chr7-96902851-DEL-2273
chr7-96909797-DEL-676
chr7-96921845-DEL-15573
chr7-96937923-DEL-2443
chr7-96939909-DEL-937
chr7-96982234-DEL-150
chr7-97012056-DEL-167
chr7-97023719-DEL-173
chr7-97044224-DEL-58
chr7-97080472-DEL-169
chr7-97081255-DEL-486

chr7-97082253-DEL-80
chr7-97088616-DEL-52
chr7-97092088-DEL-1406
chr7-97092208-DEL-628
chr7-97103556-DEL-70
chr7-97112214-DEL-936
chr7-97114533-DEL-148
chr7-97149758-DEL-62
chr7-97172027-DEL-68
chr7-97250524-DEL-53
chr7-97256627-DEL-67
chr7-97272842-DEL-61
chr7-97297497-DEL-74
chr7-97321975-DEL-77
chr7-97357890-DEL-276
chr7-97360453-DEL-86
chr7-97369872-DEL-57
chr7-97371687-DEL-84
chr7-97397707-DEL-131
chr7-97400012-DEL-201
chr7-97487575-DEL-138
chr7-97490918-DEL-180
chr7-97516107-DEL-204
chr7-97521949-DEL-658
chr7-97528863-DEL-209
chr7-97529944-DEL-374
chr7-97536753-DEL-76
chr7-97598213-DEL-51
chr7-97651369-DEL-1769
chr7-97653971-DEL-112
chr7-97680469-DEL-171
chr7-97709300-DEL-55
chr7-97804136-DEL-206
chr7-97806724-DEL-74
chr7-97829903-DEL-207
chr7-97831943-DEL-141
chr7-97836643-DEL-315
chr7-97858780-DEL-1491
chr7-97866770-DEL-56
chr7-97876239-DEL-58
chr7-97890237-DEL-567
chr7-97903211-DEL-97
chr7-97905732-DEL-749
chr7-97916582-DEL-53
chr7-97991339-DEL-65
chr7-97995770-DEL-54
chr7-98023257-DEL-163
chr7-98036233-DEL-138
chr7-98054631-DEL-53
chr7-98079873-DEL-71
chr7-98115169-DEL-461
chr7-98177206-DEL-243
chr7-98194378-DEL-280
chr7-98194921-DEL-1101
chr7-98220026-DEL-51
chr7-98384737-DEL-392
chr7-98398242-DEL-109
chr7-98403190-DEL-160
chr7-98430559-DEL-203
chr7-98451516-DEL-129
chr7-98483118-DEL-92
chr7-98504578-DEL-78
chr7-98569627-DEL-317
chr7-98570149-DEL-201
chr7-98609178-DEL-409
chr7-98610659-DEL-171
chr7-98666710-DEL-62
chr7-98708127-DEL-93
chr7-98739814-DEL-885
chr7-98741733-DEL-144
chr7-98750835-DEL-78
chr7-98760392-DEL-1492
chr7-98798579-DEL-1207
chr7-98804296-DEL-345
chr7-98932525-DEL-94
chr7-98946107-DEL-165
chr7-98946302-DEL-720

chr7-98946307-DEL-2903
chr7-98947139-DEL-54
chr7-98947443-DEL-247
chr7-98948215-DEL-360
chr7-98949017-DEL-194
chr7-99006822-DEL-2073
chr7-99017253-DEL-72
chr7-99019750-DEL-74
chr7-99041587-DEL-54
chr7-99072866-DEL-66
chr7-99080115-DEL-60
chr7-99093034-DEL-58
chr7-99101142-DEL-379
chr7-99139701-DEL-64
chr7-99140921-DEL-68
chr7-99142851-DEL-185
chr7-99144171-DEL-59
chr7-99180576-DEL-51
chr7-99212591-DEL-366
chr7-99292353-DEL-55
chr7-99377398-DEL-199
chr7-99405102-DEL-143
chr7-99434012-DEL-76
chr7-99473651-DEL-52
chr7-99484284-DEL-214
chr7-99493179-DEL-204
chr7-99514112-DEL-116
chr7-99553357-DEL-102
chr7-99583629-DEL-203
chr7-99591773-DEL-294
chr7-99594222-DEL-861
chr7-99595835-DEL-57
chr7-99609800-DEL-96
chr7-99677355-DEL-68
chr7-99719384-DEL-65
chr7-99737741-DEL-481
chr7-99774263-DEL-330
chr7-99780409-DEL-64
chr7-99801672-DEL-1974
chr7-99808107-DEL-91
chr7-99835953-DEL-54
chr7-99845354-DEL-62
chr7-99861401-DEL-52
chr7-99884865-DEL-137
chr7-99941295-DEL-364
chr7-99962134-DEL-90
chr7-99970795-DEL-4419
chr7-99988183-DEL-171
chr7-100040888-DEL-57
chr7-100084397-DEL-90
chr7-100096695-DEL-167
chr7-100113957-DEL-339
chr7-100131637-DEL-66
chr7-100174285-DEL-119
chr7-100175316-DEL-211
chr7-100183455-DEL-61
chr7-100193813-DEL-512
chr7-100196723-DEL-152
chr7-100244321-DEL-2501
chr7-100281918-DEL-232
chr7-100291389-DEL-211
chr7-100311996-DEL-159
chr7-100314871-DEL-52
chr7-100343247-DEL-1080
chr7-100359962-DEL-202
chr7-100375128-DEL-995
chr7-100379185-DEL-129
chr7-100394880-DEL-1688
chr7-100401023-DEL-219
chr7-100424093-DEL-56
chr7-100432346-DEL-74
chr7-100443251-DEL-238
chr7-100515108-DEL-76
chr7-100554993-DEL-138
chr7-100648507-DEL-1680
chr7-100680924-DEL-132
chr7-100681670-DEL-236

chr7-100683325-DEL-138
chr7-100685680-DEL-52
chr7-100729894-DEL-80
chr7-100731201-DEL-57
chr7-100761264-DEL-649
chr7-100776837-DEL-55
chr7-100803798-DEL-550
chr7-100871498-DEL-157
chr7-100893841-DEL-551
chr7-100899856-DEL-195
chr7-100903480-DEL-203
chr7-100904025-DEL-310
chr7-100917490-DEL-214
chr7-100959041-DEL-103
chr7-100963091-DEL-202
chr7-100964337-DEL-892
chr7-100965759-DEL-112
chr7-100977608-DEL-23017
chr7-101020775-DEL-132
chr7-101055932-DEL-118
chr7-101067453-DEL-184
chr7-101129112-DEL-92
chr7-101150638-DEL-69
chr7-101178553-DEL-77
chr7-101189884-DEL-58
chr7-101217351-DEL-2243
chr7-101241221-DEL-1093
chr7-101253193-DEL-398
chr7-101260095-DEL-55
chr7-101271022-DEL-202
chr7-101280216-DEL-61
chr7-101297941-DEL-51
chr7-101303575-DEL-218
chr7-101306130-DEL-86
chr7-101309895-DEL-64
chr7-101359723-DEL-67
chr7-101377992-DEL-531
chr7-101442360-DEL-67
chr7-101473984-DEL-182
chr7-101541673-DEL-309
chr7-101584231-DEL-61
chr7-101592096-DEL-2144
chr7-101614885-DEL-87
chr7-101627138-DEL-1368
chr7-101654047-DEL-148
chr7-101700322-DEL-90
chr7-101718639-DEL-306
chr7-101719346-DEL-397
chr7-101725783-DEL-209
chr7-101763867-DEL-216
chr7-101791852-DEL-200
chr7-101794146-DEL-204
chr7-101806903-DEL-100
chr7-101816912-DEL-80
chr7-101820784-DEL-195
chr7-101826705-DEL-207
chr7-101846392-DEL-510
chr7-101851229-DEL-202
chr7-101861806-DEL-991
chr7-101879415-DEL-79
chr7-101882493-DEL-90
chr7-101963046-DEL-144
chr7-101984082-DEL-69
chr7-102040189-DEL-586
chr7-102107111-DEL-125
chr7-102119284-DEL-3521
chr7-102122820-DEL-2241
chr7-102140411-DEL-51
chr7-102140779-DEL-59
chr7-102150283-DEL-487
chr7-102177966-DEL-207
chr7-102270742-DEL-2085
chr7-102271493-DEL-1967
chr7-102271554-DEL-6375
chr7-102275210-DEL-208
chr7-102292331-DEL-7047
chr7-102320768-DEL-1324

chr7-102381158-DEL-112
chr7-102433895-DEL-308
chr7-102443900-DEL-3647
chr7-102451560-DEL-1426
chr7-102462001-DEL-52
chr7-102525143-DEL-66
chr7-102599574-DEL-1500
chr7-102601087-DEL-1232
chr7-102628163-DEL-1302
chr7-102644885-DEL-2560
chr7-102648382-DEL-5442
chr7-102648523-DEL-5303
chr7-102664214-DEL-97
chr7-102674037-DEL-64
chr7-102715549-DEL-313
chr7-102746510-DEL-322
chr7-102750365-DEL-201
chr7-102754220-DEL-7077
chr7-102768203-DEL-839
chr7-102882975-DEL-58
chr7-102902126-DEL-2311
chr7-102902662-DEL-2436
chr7-102902752-DEL-1198
chr7-102908716-DEL-644
chr7-102938893-DEL-4551
chr7-102944254-DEL-114
chr7-103176258-DEL-19320
chr7-103177738-DEL-6153
chr7-103177750-DEL-5019
chr7-103182854-DEL-1124
chr7-103225711-DEL-5072
chr7-103245866-DEL-60
chr7-103247631-DEL-943
chr7-103335536-DEL-5237
chr7-103343202-DEL-208
chr7-103345035-DEL-5538
chr7-103357095-DEL-117
chr7-103442530-DEL-687
chr7-103461202-DEL-52
chr7-103462055-DEL-358
chr7-103464669-DEL-59
chr7-103465191-DEL-6212
chr7-103529538-DEL-70
chr7-103559870-DEL-123
chr7-103598824-DEL-58
chr7-103610091-DEL-6341
chr7-103630187-DEL-6148
chr7-103636512-DEL-339
chr7-103656437-DEL-88
chr7-103664231-DEL-1081
chr7-103821202-DEL-190
chr7-103832234-DEL-677
chr7-103879414-DEL-75
chr7-103900291-DEL-6483
chr7-103906731-DEL-197
chr7-103916043-DEL-7127
chr7-103927011-DEL-62
chr7-103930773-DEL-51
chr7-103934689-DEL-205
chr7-103958396-DEL-51
chr7-103959070-DEL-943
chr7-103962394-DEL-204
chr7-103967864-DEL-187
chr7-103968174-DEL-5121
chr7-103970869-DEL-577
chr7-103989169-DEL-55
chr7-104000960-DEL-76
chr7-104064856-DEL-3206
chr7-104127747-DEL-177
chr7-104138536-DEL-4088
chr7-104146980-DEL-6659
chr7-104158384-DEL-128
chr7-104162586-DEL-452
chr7-104194269-DEL-222
chr7-104228217-DEL-65
chr7-104273530-DEL-102
chr7-104276434-DEL-6157

chr7-104297468-DEL-137
chr7-104358271-DEL-12873
chr7-104415034-DEL-696
chr7-104428371-DEL-3244
chr7-104431104-DEL-68
chr7-104449482-DEL-4142
chr7-104461378-DEL-2986
chr7-104472064-DEL-76
chr7-104476366-DEL-6656
chr7-104569079-DEL-1332
chr7-104602968-DEL-120
chr7-104622475-DEL-1206
chr7-104691647-DEL-502
chr7-104698858-DEL-225
chr7-104702820-DEL-112
chr7-104730673-DEL-63
chr7-104737816-DEL-54
chr7-104743152-DEL-1523
chr7-104756295-DEL-523
chr7-104759153-DEL-1440
chr7-104760688-DEL-4385
chr7-104766315-DEL-191
chr7-104773093-DEL-861
chr7-104777733-DEL-247
chr7-104813349-DEL-3581
chr7-104838060-DEL-54
chr7-104850330-DEL-256
chr7-104965400-DEL-426
chr7-105012705-DEL-1071
chr7-105023599-DEL-53
chr7-105071232-DEL-280
chr7-105102771-DEL-198
chr7-105107363-DEL-238
chr7-105108843-DEL-171
chr7-105125158-DEL-91
chr7-105134946-DEL-132
chr7-105137439-DEL-79
chr7-105142144-DEL-463
chr7-105143318-DEL-70
chr7-105173994-DEL-1845
chr7-105182331-DEL-74
chr7-105184437-DEL-77
chr7-105186592-DEL-927
chr7-105199675-DEL-104
chr7-105221039-DEL-52
chr7-105247571-DEL-72
chr7-105291779-DEL-167
chr7-105488894-DEL-65
chr7-105493594-DEL-79
chr7-105703351-DEL-1316
chr7-105704978-DEL-70
chr7-105708674-DEL-80
chr7-105721956-DEL-100
chr7-105722697-DEL-215
chr7-105796798-DEL-58
chr7-105796896-DEL-1894
chr7-105810986-DEL-770
chr7-105852480-DEL-3410
chr7-105856329-DEL-310
chr7-105861499-DEL-13151
chr7-105903654-DEL-87
chr7-105906040-DEL-302
chr7-105909005-DEL-4793
chr7-105920648-DEL-546
chr7-105960828-DEL-5240
chr7-105977354-DEL-6240
chr7-106102611-DEL-185
chr7-106141476-DEL-62
chr7-106236931-DEL-568
chr7-106256542-DEL-7440
chr7-106266567-DEL-244
chr7-106463675-DEL-72
chr7-106500311-DEL-476
chr7-106680922-DEL-6662
chr7-106706729-DEL-1774
chr7-106709033-DEL-60
chr7-106711418-DEL-298

chr7-106712544-DEL-77
chr7-106744005-DEL-1294
chr7-106895596-DEL-178
chr7-106897755-DEL-226
chr7-106918654-DEL-528
chr7-106925683-DEL-218
chr7-106936811-DEL-151
chr7-106946342-DEL-104
chr7-106961153-DEL-1214
chr7-107007289-DEL-85
chr7-107032255-DEL-1812
chr7-107052819-DEL-289
chr7-107070455-DEL-201
chr7-107085057-DEL-74
chr7-107089154-DEL-90
chr7-107124175-DEL-106
chr7-107213006-DEL-99
chr7-107216248-DEL-87
chr7-107236244-DEL-83
chr7-107239230-DEL-217
chr7-107261881-DEL-137
chr7-107291368-DEL-157
chr7-107311518-DEL-2233
chr7-107340281-DEL-174
chr7-107361046-DEL-114
chr7-107362440-DEL-684
chr7-107377010-DEL-916
chr7-107399282-DEL-68
chr7-107407961-DEL-1655
chr7-107408219-DEL-201
chr7-107408970-DEL-88
chr7-107409510-DEL-126
chr7-107464925-DEL-108
chr7-107470311-DEL-6458
chr7-107511132-DEL-268
chr7-107550088-DEL-74
chr7-107555054-DEL-57
chr7-107574236-DEL-6420
chr7-107591115-DEL-15297
chr7-107606344-DEL-6468
chr7-107625761-DEL-212
chr7-107626130-DEL-1422
chr7-107626150-DEL-636
chr7-107642550-DEL-6639
chr7-107667753-DEL-85
chr7-107667953-DEL-536
chr7-107687926-DEL-108
chr7-107730853-DEL-56
chr7-107732131-DEL-4824
chr7-107756326-DEL-54
chr7-107762524-DEL-61
chr7-107777484-DEL-63
chr7-107782053-DEL-95
chr7-107792858-DEL-702
chr7-107795342-DEL-10942
chr7-107817190-DEL-57
chr7-107819184-DEL-2957
chr7-107824124-DEL-86
chr7-107824737-DEL-623
chr7-107826821-DEL-1432
chr7-107832268-DEL-3856
chr7-107834354-DEL-206
chr7-107863166-DEL-94
chr7-107920692-DEL-79
chr7-107946980-DEL-185
chr7-107976421-DEL-53
chr7-108039170-DEL-6231
chr7-108046864-DEL-215
chr7-108058106-DEL-54
chr7-108060232-DEL-959
chr7-108074167-DEL-440
chr7-108080001-DEL-677
chr7-108081958-DEL-281
chr7-108087858-DEL-6605
chr7-108098630-DEL-6990
chr7-108098691-DEL-401
chr7-108099007-DEL-400

chr7-108114683-DEL-283
chr7-108129400-DEL-6297
chr7-108143636-DEL-4499
chr7-108148415-DEL-7449
chr7-108159563-DEL-813
chr7-108176793-DEL-567
chr7-108194202-DEL-6251
chr7-108204365-DEL-323
chr7-108205112-DEL-1429
chr7-108221356-DEL-1141
chr7-108223304-DEL-116
chr7-108238457-DEL-84
chr7-108242182-DEL-2236
chr7-108253921-DEL-288
chr7-108258608-DEL-64
chr7-108260774-DEL-1972
chr7-108270257-DEL-731
chr7-108280811-DEL-1116
chr7-108293534-DEL-156
chr7-108337016-DEL-515
chr7-108347089-DEL-399
chr7-108376598-DEL-224
chr7-108398183-DEL-1956
chr7-108417747-DEL-6717
chr7-108420803-DEL-2022
chr7-108430687-DEL-2700
chr7-108432716-DEL-482
chr7-108433348-DEL-224
chr7-108438512-DEL-4401
chr7-108438653-DEL-3073
chr7-108451311-DEL-313
chr7-108453786-DEL-1172
chr7-108477020-DEL-100
chr7-108506660-DEL-244
chr7-108513973-DEL-72
chr7-108522396-DEL-122
chr7-108526594-DEL-56
chr7-108547685-DEL-203
chr7-108577165-DEL-1769
chr7-108623025-DEL-288
chr7-108652580-DEL-655
chr7-108653984-DEL-171
chr7-108668185-DEL-662
chr7-108739950-DEL-80
chr7-108782897-DEL-82
chr7-108830366-DEL-83
chr7-108851917-DEL-226
chr7-108915753-DEL-64
chr7-109029534-DEL-62
chr7-109046092-DEL-389
chr7-109090674-DEL-54
chr7-109098288-DEL-55
chr7-109100995-DEL-204
chr7-109112738-DEL-223
chr7-109122799-DEL-59
chr7-109184135-DEL-222
chr7-109187911-DEL-75
chr7-109196355-DEL-515
chr7-109247306-DEL-201
chr7-109264150-DEL-208
chr7-109276504-DEL-1147
chr7-109277363-DEL-191
chr7-109283590-DEL-734
chr7-109327697-DEL-158
chr7-109331961-DEL-798
chr7-109352553-DEL-95
chr7-109356875-DEL-92
chr7-109388775-DEL-2771
chr7-109444451-DEL-199
chr7-109445238-DEL-619
chr7-109445765-DEL-202
chr7-109465748-DEL-95
chr7-109496756-DEL-172
chr7-109503395-DEL-112
chr7-109519760-DEL-53
chr7-109530273-DEL-686
chr7-109569692-DEL-78

chr7-109584436-DEL-81
chr7-109589289-DEL-1085
chr7-109604745-DEL-70
chr7-109628406-DEL-160
chr7-109656807-DEL-86
chr7-109658606-DEL-470
chr7-109665530-DEL-749
chr7-109691430-DEL-113
chr7-109706546-DEL-99
chr7-109715509-DEL-183
chr7-109735100-DEL-206
chr7-109738022-DEL-112
chr7-109771455-DEL-190
chr7-109771705-DEL-1358
chr7-109773465-DEL-197
chr7-109780957-DEL-501
chr7-109783062-DEL-100
chr7-109793018-DEL-115
chr7-109793176-DEL-589
chr7-109816273-DEL-138
chr7-109850432-DEL-1220
chr7-109892843-DEL-493
chr7-109916673-DEL-119
chr7-109937476-DEL-55
chr7-109948518-DEL-76
chr7-109973845-DEL-528
chr7-109983610-DEL-88
chr7-110025424-DEL-87
chr7-110034579-DEL-746
chr7-110047796-DEL-303
chr7-110069013-DEL-209
chr7-110075676-DEL-841
chr7-110112513-DEL-71
chr7-110173485-DEL-424
chr7-110188597-DEL-54
chr7-110194867-DEL-203
chr7-110205196-DEL-107
chr7-110263828-DEL-68
chr7-110332757-DEL-1608
chr7-110336477-DEL-268
chr7-110398479-DEL-74
chr7-110405576-DEL-226
chr7-110464469-DEL-150
chr7-110464832-DEL-545
chr7-110472735-DEL-627
chr7-110511657-DEL-140
chr7-110517862-DEL-58
chr7-110538546-DEL-1110
chr7-110559246-DEL-578
chr7-110585645-DEL-1064
chr7-110608286-DEL-157
chr7-110619123-DEL-52
chr7-110620958-DEL-113
chr7-110630209-DEL-161
chr7-110649860-DEL-148
chr7-110652728-DEL-75
chr7-110712003-DEL-64
chr7-110714715-DEL-65
chr7-110732463-DEL-102
chr7-110765136-DEL-303
chr7-110820478-DEL-140
chr7-110822255-DEL-64
chr7-110828383-DEL-1120
chr7-110828396-DEL-583
chr7-110885308-DEL-3049
chr7-110898544-DEL-61
chr7-110916593-DEL-149
chr7-110930388-DEL-74
chr7-110938201-DEL-63
chr7-110957070-DEL-658
chr7-110987711-DEL-1002
chr7-111003149-DEL-143
chr7-111026952-DEL-1506
chr7-111041275-DEL-208
chr7-111056819-DEL-60
chr7-111127365-DEL-148
chr7-111132138-DEL-126

chr7-111136593-DEL-70
chr7-111190153-DEL-112
chr7-111261093-DEL-69
chr7-111265487-DEL-51
chr7-111281161-DEL-86
chr7-111284156-DEL-609
chr7-111325686-DEL-79
chr7-111332334-DEL-2388
chr7-111334649-DEL-331
chr7-111364586-DEL-234
chr7-111395284-DEL-62
chr7-111402888-DEL-120
chr7-111404206-DEL-64
chr7-111411507-DEL-119
chr7-111484210-DEL-811
chr7-111485244-DEL-1482
chr7-111496925-DEL-74
chr7-111499368-DEL-118
chr7-111543845-DEL-97
chr7-111582062-DEL-104
chr7-111597593-DEL-237
chr7-111619447-DEL-592
chr7-111747079-DEL-55
chr7-111747803-DEL-146
chr7-111749274-DEL-231
chr7-111774200-DEL-429
chr7-111816936-DEL-1861
chr7-111902337-DEL-138
chr7-111933947-DEL-82
chr7-111944767-DEL-71
chr7-111947654-DEL-54
chr7-111959174-DEL-842
chr7-111994832-DEL-236
chr7-112046923-DEL-68
chr7-112052230-DEL-146
chr7-112122296-DEL-676
chr7-112139161-DEL-385
chr7-112140032-DEL-91
chr7-112145111-DEL-310
chr7-112148750-DEL-161
chr7-112186976-DEL-469
chr7-112236316-DEL-270
chr7-112236645-DEL-150
chr7-112237546-DEL-120
chr7-112241576-DEL-57
chr7-112256543-DEL-362
chr7-112305762-DEL-160
chr7-112313555-DEL-68
chr7-112340546-DEL-78
chr7-112348486-DEL-96
chr7-112368192-DEL-318
chr7-112378688-DEL-92
chr7-112387750-DEL-18679
chr7-112489941-DEL-362
chr7-112535360-DEL-62
chr7-112566127-DEL-52
chr7-112590295-DEL-526
chr7-112619109-DEL-374
chr7-112645103-DEL-1280
chr7-112683401-DEL-58
chr7-112691300-DEL-148
chr7-112693668-DEL-76
chr7-112699227-DEL-56
chr7-112700061-DEL-1833
chr7-112715736-DEL-56
chr7-112773490-DEL-110
chr7-112834239-DEL-674
chr7-112859134-DEL-76
chr7-112867412-DEL-129
chr7-112875714-DEL-90
chr7-112888023-DEL-56
chr7-112890134-DEL-81
chr7-112906840-DEL-73
chr7-112917595-DEL-4176
chr7-112952372-DEL-70
chr7-112974579-DEL-94
chr7-112982260-DEL-56

chr7-112989984-DEL-216
chr7-113013282-DEL-53
chr7-113063697-DEL-51
chr7-113076386-DEL-187
chr7-113129059-DEL-55
chr7-113147726-DEL-269
chr7-113179857-DEL-1217
chr7-113185343-DEL-104
chr7-113209165-DEL-65
chr7-113214196-DEL-117
chr7-113226248-DEL-62
chr7-113266390-DEL-60
chr7-113290918-DEL-60
chr7-113292243-DEL-359
chr7-113297451-DEL-269
chr7-113340900-DEL-470
chr7-113363226-DEL-70
chr7-113375985-DEL-97
chr7-113376499-DEL-54
chr7-113377723-DEL-151
chr7-113408658-DEL-321
chr7-113436389-DEL-213
chr7-113449117-DEL-308
chr7-113512259-DEL-365
chr7-113514443-DEL-399
chr7-113659957-DEL-543
chr7-113672456-DEL-171
chr7-113698687-DEL-163
chr7-113711140-DEL-267
chr7-113740447-DEL-456
chr7-113766922-DEL-190
chr7-113789892-DEL-970
chr7-113804213-DEL-238
chr7-113822793-DEL-1836
chr7-113848002-DEL-53
chr7-113859020-DEL-210
chr7-113865506-DEL-119
chr7-113867083-DEL-1529
chr7-113874081-DEL-53
chr7-113879514-DEL-93
chr7-113882537-DEL-89
chr7-113901771-DEL-120
chr7-113905048-DEL-365
chr7-113905860-DEL-414
chr7-113922737-DEL-124
chr7-113925145-DEL-247
chr7-113937597-DEL-62
chr7-113947788-DEL-198
chr7-113982365-DEL-2942
chr7-114034097-DEL-157
chr7-114082235-DEL-80
chr7-114101490-DEL-87
chr7-114108838-DEL-95
chr7-114109661-DEL-966
chr7-114134121-DEL-79
chr7-114142327-DEL-262
chr7-114172507-DEL-399
chr7-114174900-DEL-80
chr7-114200155-DEL-55
chr7-114245765-DEL-916
chr7-114268548-DEL-1272
chr7-114271382-DEL-124
chr7-114273858-DEL-194
chr7-114300518-DEL-53
chr7-114332753-DEL-114
chr7-114334655-DEL-1346
chr7-114342342-DEL-286
chr7-114358274-DEL-80
chr7-114362735-DEL-394
chr7-114400280-DEL-115
chr7-114440612-DEL-732
chr7-114465436-DEL-83
chr7-114472250-DEL-102
chr7-114509385-DEL-88
chr7-114543781-DEL-55
chr7-114588829-DEL-140
chr7-114648791-DEL-81

chr7-114669337-DEL-57
chr7-114722497-DEL-1771
chr7-114750156-DEL-156
chr7-114754534-DEL-57
chr7-114780486-DEL-223
chr7-114794728-DEL-60
chr7-114827935-DEL-77
chr7-114844466-DEL-122
chr7-114845293-DEL-637
chr7-114855549-DEL-1678
chr7-114871655-DEL-64
chr7-114904034-DEL-70
chr7-114925226-DEL-2051
chr7-115017109-DEL-751
chr7-115048010-DEL-52
chr7-115094703-DEL-905
chr7-115102509-DEL-655
chr7-115146689-DEL-65
chr7-115161355-DEL-51
chr7-115199065-DEL-58
chr7-115205453-DEL-121
chr7-115274942-DEL-713
chr7-115278268-DEL-1185
chr7-115298946-DEL-100
chr7-115333729-DEL-80
chr7-115397796-DEL-60
chr7-115417456-DEL-410
chr7-115486369-DEL-69
chr7-115506761-DEL-230
chr7-115616185-DEL-93
chr7-115617446-DEL-228
chr7-115670044-DEL-1923
chr7-115688477-DEL-79
chr7-115807638-DEL-389
chr7-115816153-DEL-286
chr7-115819918-DEL-186
chr7-115829740-DEL-77
chr7-115890705-DEL-154
chr7-115914385-DEL-585
chr7-115915005-DEL-134
chr7-115915406-DEL-391
chr7-115921049-DEL-164
chr7-115925711-DEL-257
chr7-115935223-DEL-208
chr7-115942973-DEL-60
chr7-115969295-DEL-191
chr7-115969976-DEL-482
chr7-115981519-DEL-4855
chr7-116057672-DEL-88
chr7-116064042-DEL-72
chr7-116067233-DEL-108
chr7-116069878-DEL-765
chr7-116070705-DEL-829
chr7-116106270-DEL-170
chr7-116135470-DEL-363
chr7-116151191-DEL-1666
chr7-116178441-DEL-168
chr7-116187153-DEL-150
chr7-116193653-DEL-703
chr7-116255058-DEL-87
chr7-116333362-DEL-67
chr7-116350410-DEL-97
chr7-116368198-DEL-59
chr7-116397326-DEL-834
chr7-116438055-DEL-76
chr7-116476882-DEL-463
chr7-116528440-DEL-67
chr7-116554948-DEL-1602
chr7-116556824-DEL-139
chr7-116577822-DEL-934
chr7-116581708-DEL-171
chr7-116587423-DEL-58
chr7-116601147-DEL-324
chr7-116632193-DEL-328
chr7-116666841-DEL-908
chr7-116680894-DEL-220
chr7-116715231-DEL-394

chr7-116726989-DEL-530
chr7-116727586-DEL-1929
chr7-116780924-DEL-66
chr7-116863430-DEL-835
chr7-116883755-DEL-70
chr7-116905958-DEL-78
chr7-116943562-DEL-66
chr7-116995526-DEL-63
chr7-117030266-DEL-64
chr7-117033752-DEL-1543
chr7-117036441-DEL-72
chr7-117056558-DEL-139
chr7-117075853-DEL-52
chr7-117096077-DEL-74
chr7-117098167-DEL-54
chr7-117115982-DEL-52
chr7-117126111-DEL-96
chr7-117277875-DEL-181
chr7-117336735-DEL-67
chr7-117350300-DEL-994
chr7-117353845-DEL-52
chr7-117429134-DEL-56
chr7-117429508-DEL-1505
chr7-117434797-DEL-1527
chr7-117434838-DEL-1549
chr7-117436366-DEL-570
chr7-117451421-DEL-63
chr7-117476214-DEL-1144
chr7-117485736-DEL-185
chr7-117547762-DEL-848
chr7-117613264-DEL-64
chr7-117637204-DEL-59
chr7-117693033-DEL-191
chr7-117722427-DEL-64
chr7-117786309-DEL-422
chr7-117808851-DEL-186
chr7-117810893-DEL-72
chr7-117814028-DEL-169
chr7-117821440-DEL-67
chr7-117853171-DEL-61
chr7-117860630-DEL-77
chr7-117865583-DEL-119
chr7-117867385-DEL-6195
chr7-117910254-DEL-244
chr7-117914138-DEL-176
chr7-117924533-DEL-198
chr7-117944283-DEL-534
chr7-118051280-DEL-1299
chr7-118052625-DEL-206
chr7-118092821-DEL-178
chr7-118100810-DEL-361
chr7-118104709-DEL-64
chr7-118105245-DEL-56
chr7-118141554-DEL-93
chr7-118162191-DEL-102
chr7-118174768-DEL-130
chr7-118180087-DEL-75
chr7-118188535-DEL-1251
chr7-118195792-DEL-94
chr7-118202417-DEL-150
chr7-118218209-DEL-52
chr7-118230897-DEL-188
chr7-118242029-DEL-54
chr7-118259614-DEL-1872
chr7-118284769-DEL-50
chr7-118302085-DEL-195
chr7-118327565-DEL-66
chr7-118356834-DEL-70
chr7-118379553-DEL-73
chr7-118389364-DEL-64
chr7-118410074-DEL-123
chr7-118414915-DEL-602
chr7-118420198-DEL-172
chr7-118460384-DEL-797
chr7-118516645-DEL-727
chr7-118526222-DEL-99
chr7-118577452-DEL-985

chr7-118612281-DEL-66
chr7-118614234-DEL-100
chr7-118627017-DEL-81
chr7-118628236-DEL-770
chr7-118637757-DEL-72
chr7-118641980-DEL-1098
chr7-118666816-DEL-79
chr7-118681429-DEL-630
chr7-118710549-DEL-5212
chr7-118775560-DEL-698
chr7-118808449-DEL-98
chr7-118873652-DEL-253
chr7-118969756-DEL-109
chr7-119011584-DEL-53
chr7-119028131-DEL-106
chr7-119055926-DEL-60
chr7-119066716-DEL-105
chr7-119116514-DEL-72
chr7-119159794-DEL-57
chr7-119183017-DEL-1327
chr7-119256375-DEL-156
chr7-119276162-DEL-1452
chr7-119282891-DEL-114
chr7-119287638-DEL-155
chr7-119309215-DEL-65
chr7-119325076-DEL-52
chr7-119328331-DEL-293
chr7-119330659-DEL-3723
chr7-119341517-DEL-166
chr7-119358616-DEL-450
chr7-119362869-DEL-177
chr7-119369375-DEL-52
chr7-119408228-DEL-149
chr7-119450792-DEL-197
chr7-119478830-DEL-371
chr7-119500404-DEL-185
chr7-119509978-DEL-55
chr7-119552274-DEL-100
chr7-119553745-DEL-411
chr7-119576028-DEL-389
chr7-119582276-DEL-68
chr7-119583489-DEL-625
chr7-119593802-DEL-71
chr7-119621197-DEL-81
chr7-119622611-DEL-62
chr7-119624919-DEL-56
chr7-119626082-DEL-268
chr7-119736889-DEL-63
chr7-119739749-DEL-654
chr7-119759232-DEL-101
chr7-119801482-DEL-84
chr7-119821168-DEL-174
chr7-119833641-DEL-75
chr7-119841271-DEL-126
chr7-119855328-DEL-6830
chr7-119864138-DEL-61
chr7-119905167-DEL-1188
chr7-119945579-DEL-58
chr7-119951174-DEL-53
chr7-119958381-DEL-218
chr7-119965695-DEL-2257
chr7-119972760-DEL-68
chr7-119975609-DEL-87
chr7-119996980-DEL-271
chr7-119997263-DEL-1975
chr7-119997638-DEL-842
chr7-120016252-DEL-1094
chr7-120021476-DEL-256
chr7-120041728-DEL-6192
chr7-120061823-DEL-847
chr7-120062711-DEL-1901
chr7-120085130-DEL-114
chr7-120173446-DEL-397
chr7-120177411-DEL-160
chr7-120194054-DEL-1009
chr7-120198343-DEL-84
chr7-120207376-DEL-2137

chr7-120213364-DEL-64
chr7-120228518-DEL-214
chr7-120237317-DEL-53
chr7-120273681-DEL-62
chr7-120287918-DEL-53
chr7-120295994-DEL-51
chr7-120353788-DEL-69
chr7-120370577-DEL-52
chr7-120375663-DEL-59
chr7-120384418-DEL-66
chr7-120388141-DEL-120
chr7-120449193-DEL-110
chr7-120467093-DEL-79
chr7-120491727-DEL-147
chr7-120543251-DEL-56
chr7-120545521-DEL-238
chr7-120545546-DEL-782
chr7-120546343-DEL-85
chr7-120552587-DEL-101
chr7-120582296-DEL-65
chr7-120607657-DEL-160
chr7-120660424-DEL-154
chr7-120668634-DEL-204
chr7-120684049-DEL-72
chr7-120720369-DEL-82
chr7-120726090-DEL-271
chr7-120729952-DEL-59
chr7-120738039-DEL-554
chr7-120763617-DEL-339
chr7-120802947-DEL-1871
chr7-120828676-DEL-933
chr7-120835858-DEL-156
chr7-120841957-DEL-51
chr7-120844947-DEL-274
chr7-120861392-DEL-55
chr7-120929863-DEL-1005
chr7-120959576-DEL-73
chr7-121054805-DEL-62
chr7-121071841-DEL-196
chr7-121075049-DEL-184
chr7-121110612-DEL-81
chr7-121117162-DEL-376
chr7-121166957-DEL-93
chr7-121172875-DEL-248
chr7-121174142-DEL-89
chr7-121179149-DEL-90
chr7-121229344-DEL-103
chr7-121231412-DEL-70
chr7-121250319-DEL-74
chr7-121300768-DEL-304
chr7-121352204-DEL-90
chr7-121365910-DEL-304
chr7-121388683-DEL-225
chr7-121416358-DEL-66
chr7-121432881-DEL-248
chr7-121441844-DEL-115
chr7-121443224-DEL-68
chr7-121449209-DEL-106
chr7-121459670-DEL-112
chr7-121461779-DEL-72
chr7-121503732-DEL-248
chr7-121505099-DEL-205
chr7-121508916-DEL-165
chr7-121564219-DEL-69
chr7-121578009-DEL-72
chr7-121626146-DEL-55
chr7-121650827-DEL-1022
chr7-121664412-DEL-93
chr7-121714063-DEL-528
chr7-121718296-DEL-399
chr7-121722647-DEL-77
chr7-121822885-DEL-51
chr7-121856838-DEL-398
chr7-121903731-DEL-69
chr7-121930427-DEL-58
chr7-121994014-DEL-340
chr7-122001096-DEL-123

chr7-122008176-DEL-275
chr7-122019483-DEL-52
chr7-122057883-DEL-154
chr7-122073399-DEL-397
chr7-122119492-DEL-735
chr7-122140588-DEL-520
chr7-122143012-DEL-53
chr7-122149889-DEL-54
chr7-122268247-DEL-59
chr7-122331088-DEL-110
chr7-122343672-DEL-72
chr7-122367009-DEL-68
chr7-122385716-DEL-57
chr7-122425132-DEL-397
chr7-122432080-DEL-1436
chr7-122481585-DEL-840
chr7-122481595-DEL-60
chr7-122483245-DEL-56
chr7-122518176-DEL-60
chr7-122529248-DEL-404
chr7-122532809-DEL-150
chr7-122533687-DEL-207
chr7-122540112-DEL-80
chr7-122546996-DEL-55
chr7-122558697-DEL-5423
chr7-122565181-DEL-56
chr7-122566747-DEL-242
chr7-122610418-DEL-179
chr7-122615627-DEL-398
chr7-122619504-DEL-4586
chr7-122619987-DEL-211
chr7-122620053-DEL-2693
chr7-122620570-DEL-301
chr7-122625073-DEL-78
chr7-122657416-DEL-1217
chr7-122660079-DEL-65
chr7-122669754-DEL-7104
chr7-122731845-DEL-471
chr7-122758245-DEL-1565
chr7-122776598-DEL-74
chr7-122812847-DEL-219
chr7-122815486-DEL-207
chr7-122848693-DEL-108
chr7-122860426-DEL-166
chr7-122901233-DEL-128
chr7-122928675-DEL-74
chr7-122962551-DEL-99
chr7-122996962-DEL-53
chr7-123015031-DEL-448
chr7-123043585-DEL-82
chr7-123048427-DEL-175
chr7-123068713-DEL-164
chr7-123112110-DEL-330
chr7-123119233-DEL-59
chr7-123120626-DEL-88
chr7-123172226-DEL-54
chr7-123222283-DEL-4429
chr7-123249689-DEL-1408
chr7-123253913-DEL-708
chr7-123263507-DEL-615
chr7-123313945-DEL-2266
chr7-123317183-DEL-83
chr7-123334008-DEL-367
chr7-123363511-DEL-9867
chr7-123373426-DEL-145
chr7-123403702-DEL-540
chr7-123404411-DEL-865
chr7-123421355-DEL-87
chr7-123426810-DEL-51
chr7-123481948-DEL-75
chr7-123538228-DEL-321
chr7-123560341-DEL-54
chr7-123570971-DEL-346
chr7-123617325-DEL-53
chr7-123617881-DEL-169
chr7-123624387-DEL-142
chr7-123658752-DEL-120

chr7-123694633-DEL-173
chr7-123696708-DEL-131
chr7-123716043-DEL-60
chr7-123746785-DEL-355
chr7-123753050-DEL-2507
chr7-123763737-DEL-108
chr7-123814333-DEL-210
chr7-123820576-DEL-106
chr7-123830465-DEL-62
chr7-123843583-DEL-401
chr7-123867491-DEL-90
chr7-123869812-DEL-73
chr7-123881278-DEL-364
chr7-123882325-DEL-386
chr7-123901838-DEL-68
chr7-123912331-DEL-97
chr7-123916806-DEL-54
chr7-123929523-DEL-63
chr7-123936311-DEL-179
chr7-123969192-DEL-61
chr7-123974285-DEL-78
chr7-123991717-DEL-248
chr7-124027276-DEL-51
chr7-124079105-DEL-679
chr7-124115356-DEL-1580
chr7-124133731-DEL-1137
chr7-124142885-DEL-1368
chr7-124144480-DEL-188
chr7-124179196-DEL-92
chr7-124185968-DEL-872
chr7-124203005-DEL-193
chr7-124208421-DEL-96
chr7-124238074-DEL-718
chr7-124241377-DEL-163
chr7-124242885-DEL-788
chr7-124249107-DEL-56
chr7-124321298-DEL-2905
chr7-124493251-DEL-60
chr7-124537341-DEL-220
chr7-124544430-DEL-497
chr7-124552597-DEL-56
chr7-124576208-DEL-2214
chr7-124606918-DEL-1416
chr7-124634253-DEL-132
chr7-124656776-DEL-1425
chr7-124660303-DEL-196
chr7-124703782-DEL-540
chr7-124704137-DEL-795
chr7-124726781-DEL-95
chr7-124743708-DEL-72
chr7-124749990-DEL-329
chr7-124782399-DEL-254
chr7-124821947-DEL-64
chr7-124830457-DEL-101
chr7-124866875-DEL-148
chr7-124873515-DEL-52
chr7-124887459-DEL-554
chr7-124893128-DEL-93
chr7-124911678-DEL-62
chr7-124913278-DEL-3011
chr7-124916875-DEL-1858
chr7-124962010-DEL-59
chr7-124985399-DEL-407
chr7-124986660-DEL-52
chr7-124995530-DEL-109
chr7-125002085-DEL-58
chr7-125005492-DEL-61
chr7-125038170-DEL-120
chr7-125061698-DEL-595
chr7-125061778-DEL-626
chr7-125062849-DEL-415
chr7-125079649-DEL-238
chr7-125144870-DEL-911
chr7-125211280-DEL-229
chr7-125211877-DEL-852
chr7-125224804-DEL-166
chr7-125249165-DEL-91

chr7-125264523-DEL-80
chr7-125313055-DEL-160
chr7-125376156-DEL-884
chr7-125407184-DEL-857
chr7-125408867-DEL-399
chr7-125488560-DEL-76
chr7-125574939-DEL-142
chr7-125581869-DEL-176
chr7-125587545-DEL-65
chr7-125609309-DEL-59
chr7-125645845-DEL-60
chr7-125674000-DEL-56
chr7-125717217-DEL-75
chr7-125746706-DEL-454
chr7-125747266-DEL-266
chr7-125753528-DEL-66
chr7-125757849-DEL-76
chr7-125795506-DEL-69
chr7-125800547-DEL-220
chr7-125813371-DEL-62
chr7-125815630-DEL-62
chr7-125825449-DEL-133
chr7-125853703-DEL-52
chr7-125854767-DEL-516
chr7-125903848-DEL-179
chr7-125921159-DEL-145
chr7-125921594-DEL-71
chr7-125922585-DEL-187
chr7-125941922-DEL-321
chr7-125966145-DEL-84
chr7-125984940-DEL-147
chr7-126012092-DEL-61
chr7-126015688-DEL-92
chr7-126019295-DEL-51
chr7-126024440-DEL-218
chr7-126026044-DEL-181
chr7-126032387-DEL-88
chr7-126038761-DEL-145
chr7-126040045-DEL-181
chr7-126069850-DEL-54
chr7-126076338-DEL-281
chr7-126104757-DEL-270
chr7-126128627-DEL-182
chr7-126130856-DEL-1846
chr7-126239793-DEL-58
chr7-126247464-DEL-54
chr7-126253363-DEL-260
chr7-126258523-DEL-91
chr7-126313515-DEL-70
chr7-126314599-DEL-91
chr7-126316312-DEL-68
chr7-126321142-DEL-51
chr7-126331742-DEL-168
chr7-126340789-DEL-114
chr7-126343121-DEL-104
chr7-126389900-DEL-153
chr7-126410009-DEL-203
chr7-126432997-DEL-398
chr7-126443040-DEL-178
chr7-126462973-DEL-54
chr7-126493945-DEL-125
chr7-126499128-DEL-244
chr7-126524677-DEL-184
chr7-126526123-DEL-198
chr7-126554145-DEL-154
chr7-126659825-DEL-78
chr7-126684740-DEL-4648
chr7-126748156-DEL-72
chr7-126749122-DEL-341
chr7-126752304-DEL-203
chr7-126760772-DEL-198
chr7-126772356-DEL-191
chr7-126786451-DEL-183
chr7-126823500-DEL-87
chr7-126825856-DEL-84
chr7-126828969-DEL-498
chr7-126848094-DEL-428

chr7-126863588-DEL-104
chr7-126886138-DEL-213
chr7-126888278-DEL-51
chr7-126941445-DEL-61
chr7-127010938-DEL-75
chr7-127015011-DEL-2635
chr7-127032288-DEL-172
chr7-127033107-DEL-185
chr7-127038908-DEL-61
chr7-127052953-DEL-286
chr7-127057191-DEL-63
chr7-127107242-DEL-153
chr7-127118318-DEL-200
chr7-127153355-DEL-213
chr7-127230152-DEL-203
chr7-127232197-DEL-1882
chr7-127237207-DEL-204
chr7-127248629-DEL-76
chr7-127250696-DEL-217
chr7-127255119-DEL-206
chr7-127255795-DEL-178
chr7-127257572-DEL-544
chr7-127274124-DEL-228
chr7-127281454-DEL-578
chr7-127282436-DEL-106
chr7-127283527-DEL-71
chr7-127286848-DEL-202
chr7-127287455-DEL-68
chr7-127290025-DEL-84
chr7-127338831-DEL-203
chr7-127360922-DEL-400
chr7-127393672-DEL-205
chr7-127402518-DEL-51
chr7-127405160-DEL-142
chr7-127419199-DEL-285
chr7-127420388-DEL-59
chr7-127424832-DEL-207
chr7-127432756-DEL-170
chr7-127446806-DEL-173
chr7-127447215-DEL-321
chr7-127457101-DEL-75
chr7-127463442-DEL-90
chr7-127472574-DEL-233
chr7-127512689-DEL-205
chr7-127513018-DEL-411
chr7-127541829-DEL-76
chr7-127555244-DEL-1257
chr7-127596526-DEL-208
chr7-127612032-DEL-382
chr7-127619778-DEL-807
chr7-127642335-DEL-250
chr7-127670733-DEL-2185
chr7-127718198-DEL-76
chr7-127722538-DEL-2394
chr7-127731119-DEL-74
chr7-127785876-DEL-858
chr7-127786380-DEL-137
chr7-127787203-DEL-396
chr7-127798274-DEL-906
chr7-127813819-DEL-77
chr7-127818413-DEL-398
chr7-127898579-DEL-199
chr7-127925987-DEL-177
chr7-127928206-DEL-1488
chr7-127936324-DEL-76
chr7-127942127-DEL-702
chr7-127949580-DEL-121
chr7-127997819-DEL-204
chr7-128036624-DEL-65
chr7-128060600-DEL-65
chr7-128110487-DEL-54
chr7-128135743-DEL-101
chr7-128170638-DEL-202
chr7-128194151-DEL-146
chr7-128205880-DEL-201
chr7-128230468-DEL-57
chr7-128277953-DEL-72

chr7-128288060-DEL-57
chr7-128290214-DEL-55
chr7-128290844-DEL-116
chr7-128323794-DEL-113
chr7-128328837-DEL-718
chr7-128339510-DEL-125
chr7-128343476-DEL-148
chr7-128344969-DEL-51
chr7-128347393-DEL-94
chr7-128412423-DEL-52
chr7-128418630-DEL-93
chr7-128431207-DEL-1156
chr7-128432505-DEL-323
chr7-128471804-DEL-864
chr7-128483338-DEL-1543
chr7-128487844-DEL-59
chr7-128489838-DEL-1100
chr7-128561144-DEL-64
chr7-128574785-DEL-6018
chr7-128596489-DEL-69
chr7-128618745-DEL-356
chr7-128620664-DEL-4515
chr7-128624582-DEL-2765
chr7-128631705-DEL-3853
chr7-128657170-DEL-53
chr7-128664331-DEL-3470
chr7-128711861-DEL-107
chr7-128738811-DEL-59
chr7-128743446-DEL-224
chr7-128786101-DEL-226
chr7-128788148-DEL-52
chr7-128790774-DEL-1226
chr7-128806519-DEL-80
chr7-128890320-DEL-62
chr7-128926903-DEL-52
chr7-128940958-DEL-56
chr7-128969069-DEL-434
chr7-128972923-DEL-226
chr7-128975948-DEL-66
chr7-128992061-DEL-256
chr7-129002779-DEL-123
chr7-129029623-DEL-71
chr7-129040000-DEL-96
chr7-129050591-DEL-68
chr7-129052335-DEL-4194
chr7-129059024-DEL-60
chr7-129087311-DEL-124
chr7-129089901-DEL-220
chr7-129093423-DEL-3617
chr7-129101611-DEL-201
chr7-129122377-DEL-61
chr7-129153823-DEL-132
chr7-129157104-DEL-1037
chr7-129161146-DEL-9923
chr7-129183302-DEL-52
chr7-129192461-DEL-123
chr7-129197977-DEL-549
chr7-129281639-DEL-224
chr7-129286115-DEL-809
chr7-129308189-DEL-80
chr7-129319433-DEL-60
chr7-129321494-DEL-65
chr7-129362101-DEL-1649
chr7-129373347-DEL-153
chr7-129377747-DEL-86
chr7-129386182-DEL-68
chr7-129419182-DEL-64
chr7-129420433-DEL-114
chr7-129423794-DEL-69
chr7-129426362-DEL-56
chr7-129464218-DEL-77
chr7-129470736-DEL-2027
chr7-129471295-DEL-2042
chr7-129494594-DEL-128
chr7-129538126-DEL-65
chr7-129553579-DEL-60
chr7-129557723-DEL-86

chr7-129563354-DEL-258
chr7-129577692-DEL-68
chr7-129578737-DEL-143
chr7-129598695-DEL-83
chr7-129599864-DEL-669
chr7-129666947-DEL-110
chr7-129699724-DEL-731
chr7-129728173-DEL-95
chr7-129825159-DEL-739
chr7-129827409-DEL-92
chr7-129871960-DEL-53
chr7-129896989-DEL-267
chr7-129945232-DEL-10908
chr7-129959086-DEL-497
chr7-129997074-DEL-163
chr7-130003509-DEL-69
chr7-130015717-DEL-182
chr7-130029269-DEL-79
chr7-130052131-DEL-1483
chr7-130059165-DEL-673
chr7-130090074-DEL-444
chr7-130096469-DEL-4432
chr7-130128487-DEL-67
chr7-130132752-DEL-494
chr7-130147886-DEL-96
chr7-130160604-DEL-304
chr7-130193301-DEL-3090
chr7-130194458-DEL-3049
chr7-130286449-DEL-916
chr7-130286630-DEL-255
chr7-130287108-DEL-95
chr7-130288339-DEL-103
chr7-130372961-DEL-87
chr7-130374256-DEL-555
chr7-130501520-DEL-630
chr7-130517573-DEL-179
chr7-130594196-DEL-243
chr7-130624234-DEL-399
chr7-130630462-DEL-171
chr7-130731174-DEL-56
chr7-130733864-DEL-61
chr7-130762814-DEL-407
chr7-130763885-DEL-1504
chr7-130780996-DEL-362
chr7-130782982-DEL-104
chr7-130792707-DEL-151
chr7-130807693-DEL-84
chr7-130811041-DEL-51
chr7-130813551-DEL-54
chr7-130879727-DEL-1301
chr7-130886393-DEL-56
chr7-130915170-DEL-105
chr7-130915506-DEL-325
chr7-130930623-DEL-120
chr7-130934930-DEL-100
chr7-130954511-DEL-74
chr7-131005559-DEL-204
chr7-131018221-DEL-337
chr7-131028124-DEL-152
chr7-131032696-DEL-201
chr7-131089481-DEL-214
chr7-131105831-DEL-1412
chr7-131132583-DEL-294
chr7-131182906-DEL-96
chr7-131191006-DEL-60
chr7-131198934-DEL-109
chr7-131212317-DEL-141
chr7-131234398-DEL-71
chr7-131235468-DEL-96
chr7-131237797-DEL-366
chr7-131240180-DEL-581
chr7-131316137-DEL-176
chr7-131345552-DEL-201
chr7-131429510-DEL-368
chr7-131433766-DEL-655
chr7-131487590-DEL-65
chr7-131525764-DEL-73

chr7-131526851-DEL-239
chr7-131531833-DEL-66
chr7-131535870-DEL-436
chr7-131559500-DEL-216
chr7-131564634-DEL-68
chr7-131582687-DEL-1463
chr7-131640126-DEL-62
chr7-131656168-DEL-95
chr7-131663139-DEL-61
chr7-131698717-DEL-592
chr7-131726997-DEL-84
chr7-131742273-DEL-101
chr7-131765098-DEL-623
chr7-131816959-DEL-336
chr7-131826361-DEL-125
chr7-131830526-DEL-54
chr7-131844486-DEL-114
chr7-131894765-DEL-81
chr7-131903809-DEL-75
chr7-131921201-DEL-563
chr7-131948033-DEL-87
chr7-131963554-DEL-205
chr7-131968389-DEL-102
chr7-132003904-DEL-243
chr7-132053252-DEL-652
chr7-132113047-DEL-60
chr7-132116489-DEL-171
chr7-132121890-DEL-227
chr7-132127423-DEL-197
chr7-132136664-DEL-373
chr7-132160594-DEL-127
chr7-132221012-DEL-194
chr7-132222885-DEL-141
chr7-132251524-DEL-57
chr7-132254141-DEL-60
chr7-132256615-DEL-108
chr7-132261795-DEL-64
chr7-132268964-DEL-666
chr7-132279486-DEL-74
chr7-132315079-DEL-60
chr7-132318414-DEL-194
chr7-132326088-DEL-123
chr7-132336422-DEL-496
chr7-132347986-DEL-592
chr7-132377415-DEL-58
chr7-132435742-DEL-212
chr7-132482658-DEL-198
chr7-132514245-DEL-69
chr7-132515733-DEL-6203
chr7-132522603-DEL-69
chr7-132570861-DEL-480
chr7-132581635-DEL-143
chr7-132611769-DEL-61
chr7-132645745-DEL-110
chr7-132706015-DEL-52
chr7-132739874-DEL-624
chr7-132747992-DEL-510
chr7-132748570-DEL-221
chr7-132749785-DEL-190
chr7-132765890-DEL-63
chr7-132779510-DEL-100
chr7-132789266-DEL-305
chr7-132839978-DEL-64
chr7-132861819-DEL-332
chr7-132867612-DEL-108
chr7-132880270-DEL-55
chr7-132885323-DEL-545
chr7-132910819-DEL-70
chr7-132931573-DEL-206
chr7-132933714-DEL-724
chr7-132947672-DEL-100
chr7-132961380-DEL-329
chr7-132984471-DEL-196
chr7-133038007-DEL-52
chr7-133050571-DEL-160
chr7-133090345-DEL-1120
chr7-133090630-DEL-895

chr7-133122112-DEL-57
chr7-133125776-DEL-109
chr7-133180105-DEL-188
chr7-133189840-DEL-104
chr7-133204407-DEL-404
chr7-133223714-DEL-98
chr7-133271543-DEL-198
chr7-133277922-DEL-54
chr7-133287271-DEL-499
chr7-133305538-DEL-698
chr7-133306115-DEL-120
chr7-133312204-DEL-485
chr7-133328341-DEL-64
chr7-133329285-DEL-397
chr7-133340144-DEL-545
chr7-133366498-DEL-58
chr7-133385466-DEL-2847
chr7-133385970-DEL-2027
chr7-133386607-DEL-302
chr7-133387368-DEL-767
chr7-133388460-DEL-810
chr7-133412700-DEL-1836
chr7-133413137-DEL-418
chr7-133413258-DEL-810
chr7-133413334-DEL-1206
chr7-133413875-DEL-662
chr7-133416036-DEL-198
chr7-133449327-DEL-71
chr7-133465577-DEL-76
chr7-133466438-DEL-207
chr7-133467543-DEL-180
chr7-133515633-DEL-1361
chr7-133560182-DEL-226
chr7-133699782-DEL-68
chr7-133735884-DEL-75
chr7-133769157-DEL-59
chr7-133784109-DEL-151
chr7-133785746-DEL-248
chr7-133798479-DEL-136
chr7-133802541-DEL-60
chr7-133882368-DEL-193
chr7-133917777-DEL-187
chr7-133924891-DEL-102
chr7-133926159-DEL-108
chr7-133930910-DEL-6211
chr7-133952588-DEL-124
chr7-133954603-DEL-72
chr7-133996404-DEL-64
chr7-134001556-DEL-210
chr7-134008736-DEL-1249
chr7-134061355-DEL-188
chr7-134132324-DEL-562
chr7-134133458-DEL-72
chr7-134142783-DEL-2437
chr7-134155954-DEL-55
chr7-134231032-DEL-1009
chr7-134284646-DEL-98
chr7-134296421-DEL-142
chr7-134323553-DEL-660
chr7-134378433-DEL-82
chr7-134410159-DEL-599
chr7-134447033-DEL-122
chr7-134499818-DEL-133
chr7-134505939-DEL-106
chr7-134522789-DEL-198
chr7-134527293-DEL-58
chr7-134563068-DEL-76
chr7-134610260-DEL-138
chr7-134667732-DEL-52
chr7-134697071-DEL-288
chr7-134749443-DEL-195
chr7-134787082-DEL-165
chr7-134841500-DEL-333
chr7-134853940-DEL-434
chr7-134860182-DEL-332
chr7-134900320-DEL-112
chr7-134951817-DEL-54

chr7-135010593-DEL-532
chr7-135018080-DEL-166
chr7-135039797-DEL-394
chr7-135084952-DEL-1496
chr7-135106857-DEL-1014
chr7-135111709-DEL-85
chr7-135121988-DEL-118
chr7-135131859-DEL-82
chr7-135160679-DEL-161
chr7-135187871-DEL-51
chr7-135191255-DEL-74
chr7-135217151-DEL-52
chr7-135298938-DEL-1398
chr7-135299620-DEL-711
chr7-135324161-DEL-595
chr7-135379166-DEL-81
chr7-135391880-DEL-86
chr7-135401884-DEL-116
chr7-135421057-DEL-850
chr7-135482076-DEL-256
chr7-135512867-DEL-112
chr7-135524304-DEL-1073
chr7-135558582-DEL-790
chr7-135600820-DEL-215
chr7-135640566-DEL-120
chr7-135644165-DEL-82
chr7-135670020-DEL-63
chr7-135693121-DEL-131
chr7-135723101-DEL-73
chr7-135738876-DEL-955
chr7-135748550-DEL-159
chr7-135788161-DEL-74
chr7-135847446-DEL-56
chr7-135849381-DEL-71
chr7-135851583-DEL-60
chr7-135862875-DEL-624
chr7-135909829-DEL-3044
chr7-135986310-DEL-80
chr7-136112937-DEL-1096
chr7-136116945-DEL-84
chr7-136143013-DEL-53
chr7-136177854-DEL-72
chr7-136211225-DEL-141
chr7-136236049-DEL-53
chr7-136255007-DEL-64
chr7-136267945-DEL-1002
chr7-136333906-DEL-102
chr7-136349937-DEL-185
chr7-136362534-DEL-81
chr7-136378078-DEL-80
chr7-136409371-DEL-119
chr7-136409524-DEL-927
chr7-136412534-DEL-1846
chr7-136458197-DEL-55
chr7-136469426-DEL-2997
chr7-136469572-DEL-1739
chr7-136479985-DEL-51
chr7-136492508-DEL-707
chr7-136498149-DEL-377
chr7-136501734-DEL-1798
chr7-136619222-DEL-152
chr7-136655485-DEL-153
chr7-136695145-DEL-72
chr7-136716487-DEL-140
chr7-136717320-DEL-56
chr7-136737405-DEL-69
chr7-136740706-DEL-138
chr7-136817217-DEL-312
chr7-136834896-DEL-56
chr7-136834911-DEL-955
chr7-136835308-DEL-224
chr7-136873222-DEL-133
chr7-136877141-DEL-486
chr7-136933060-DEL-86
chr7-136936832-DEL-174
chr7-136940874-DEL-92
chr7-136946299-DEL-144

chr7-136964528-DEL-52
chr7-137009256-DEL-1627
chr7-137057942-DEL-132
chr7-137071617-DEL-162
chr7-137076415-DEL-565
chr7-137090654-DEL-155
chr7-137097286-DEL-58
chr7-137112665-DEL-88
chr7-137115104-DEL-107
chr7-137118497-DEL-221
chr7-137159676-DEL-601
chr7-137187043-DEL-463
chr7-137219627-DEL-79
chr7-137242547-DEL-54
chr7-137257822-DEL-54
chr7-137285953-DEL-55
chr7-137305923-DEL-224
chr7-137312303-DEL-186
chr7-137367035-DEL-87
chr7-137368243-DEL-93
chr7-137370914-DEL-58
chr7-137378390-DEL-108
chr7-137393454-DEL-2028
chr7-137399852-DEL-77
chr7-137400890-DEL-158
chr7-137457892-DEL-198
chr7-137560865-DEL-187
chr7-137573682-DEL-371
chr7-137602808-DEL-2225
chr7-137617573-DEL-52
chr7-137622207-DEL-579
chr7-137652073-DEL-193
chr7-137663450-DEL-805
chr7-137664852-DEL-92
chr7-137675543-DEL-149
chr7-137697953-DEL-61
chr7-137764348-DEL-224
chr7-137788868-DEL-57
chr7-137895426-DEL-128
chr7-137900387-DEL-112
chr7-137903177-DEL-80
chr7-137917904-DEL-1569
chr7-137931990-DEL-122
chr7-137973526-DEL-205
chr7-138004188-DEL-371
chr7-138038280-DEL-51
chr7-138061471-DEL-52
chr7-138110955-DEL-54
chr7-138119834-DEL-382
chr7-138138442-DEL-388
chr7-138164748-DEL-52
chr7-138170269-DEL-73
chr7-138191595-DEL-147
chr7-138237604-DEL-2896
chr7-138257256-DEL-72
chr7-138278821-DEL-66
chr7-138282819-DEL-263
chr7-138287392-DEL-195
chr7-138337685-DEL-68
chr7-138394530-DEL-299
chr7-138402878-DEL-184
chr7-138431703-DEL-408
chr7-138450822-DEL-170
chr7-138485384-DEL-52
chr7-138487303-DEL-741
chr7-138522395-DEL-88
chr7-138527991-DEL-156
chr7-138530659-DEL-177
chr7-138598959-DEL-365
chr7-138630761-DEL-86
chr7-138634186-DEL-137
chr7-138677180-DEL-52
chr7-138760767-DEL-144
chr7-138769515-DEL-81
chr7-138777623-DEL-81
chr7-138783303-DEL-745
chr7-138786076-DEL-131

chr7-138786990-DEL-74
chr7-138900519-DEL-407
chr7-138901646-DEL-177
chr7-138930394-DEL-75
chr7-139001123-DEL-400
chr7-139004429-DEL-2663
chr7-139027743-DEL-1098
chr7-139028309-DEL-74
chr7-139090898-DEL-63
chr7-139133382-DEL-58
chr7-139188582-DEL-67
chr7-139199451-DEL-87
chr7-139200463-DEL-185
chr7-139218718-DEL-63
chr7-139223445-DEL-123
chr7-139237731-DEL-56
chr7-139249107-DEL-275
chr7-139250540-DEL-1774
chr7-139256518-DEL-52
chr7-139256937-DEL-545
chr7-139278733-DEL-84
chr7-139315941-DEL-975
chr7-139332503-DEL-97
chr7-139365630-DEL-52
chr7-139432189-DEL-195
chr7-139442326-DEL-75
chr7-139495481-DEL-91
chr7-139505153-DEL-55
chr7-139541527-DEL-312
chr7-139548919-DEL-184
chr7-139580700-DEL-76
chr7-139592872-DEL-70
chr7-139594665-DEL-6584
chr7-139601249-DEL-5937
chr7-139622176-DEL-167
chr7-139672314-DEL-101
chr7-139674083-DEL-58
chr7-139722449-DEL-117
chr7-139735713-DEL-214
chr7-139736888-DEL-54
chr7-139799590-DEL-125
chr7-139812113-DEL-931
chr7-139814847-DEL-6239
chr7-139830301-DEL-446
chr7-139841367-DEL-6673
chr7-139848012-DEL-64
chr7-139880723-DEL-318
chr7-139881011-DEL-6437
chr7-139895647-DEL-419
chr7-139900912-DEL-67
chr7-139911935-DEL-158
chr7-139922699-DEL-87
chr7-139943302-DEL-58
chr7-139944050-DEL-379
chr7-139944820-DEL-471
chr7-139946836-DEL-56
chr7-139969672-DEL-2366
chr7-139984057-DEL-2534
chr7-140003637-DEL-56
chr7-140004177-DEL-2261
chr7-140021753-DEL-112
chr7-140068392-DEL-103
chr7-140096575-DEL-1624
chr7-140101041-DEL-236
chr7-140122801-DEL-221
chr7-140137707-DEL-1136
chr7-140160371-DEL-5666
chr7-140183860-DEL-578
chr7-140221792-DEL-417
chr7-140246211-DEL-491
chr7-140252168-DEL-749
chr7-140257189-DEL-1409
chr7-140314171-DEL-62
chr7-140331779-DEL-397
chr7-140332376-DEL-754
chr7-140378094-DEL-60
chr7-140432267-DEL-184

chr7-140446907-DEL-116
chr7-140453015-DEL-51
chr7-140481015-DEL-349
chr7-140482007-DEL-102
chr7-140498129-DEL-612
chr7-140530580-DEL-188
chr7-140542754-DEL-358
chr7-140553702-DEL-95
chr7-140575911-DEL-867
chr7-140576433-DEL-125
chr7-140580630-DEL-100
chr7-140620096-DEL-192
chr7-140624741-DEL-56
chr7-140635511-DEL-57
chr7-140644069-DEL-96
chr7-140648936-DEL-233
chr7-140656377-DEL-56
chr7-140663674-DEL-64
chr7-140665740-DEL-296
chr7-140681271-DEL-60
chr7-140690823-DEL-202
chr7-140702618-DEL-204
chr7-140725914-DEL-73
chr7-140729515-DEL-197
chr7-140739191-DEL-264
chr7-140756068-DEL-59
chr7-140756658-DEL-386
chr7-140757698-DEL-732
chr7-140760164-DEL-73
chr7-140763452-DEL-558
chr7-140857105-DEL-144
chr7-140862422-DEL-68
chr7-140895603-DEL-85
chr7-140897711-DEL-55
chr7-140965650-DEL-202
chr7-140979581-DEL-64
chr7-140983125-DEL-254
chr7-141028991-DEL-669
chr7-141031309-DEL-80
chr7-141032607-DEL-179
chr7-141083285-DEL-2138
chr7-141086143-DEL-114
chr7-141087431-DEL-142
chr7-141093175-DEL-116
chr7-141124978-DEL-205
chr7-141134800-DEL-95
chr7-141135261-DEL-198
chr7-141238877-DEL-304
chr7-141239134-DEL-252
chr7-141255327-DEL-201
chr7-141285237-DEL-56
chr7-141291775-DEL-720
chr7-141293230-DEL-186
chr7-141293775-DEL-81
chr7-141295010-DEL-129
chr7-141314425-DEL-104
chr7-141321814-DEL-81
chr7-141324202-DEL-139
chr7-141373332-DEL-84
chr7-141407513-DEL-408
chr7-141416046-DEL-941
chr7-141416164-DEL-54
chr7-141452655-DEL-210
chr7-141486544-DEL-79
chr7-141528467-DEL-2092
chr7-141559644-DEL-2462
chr7-141602799-DEL-99
chr7-141661144-DEL-107
chr7-141665987-DEL-62
chr7-141702723-DEL-51
chr7-141728770-DEL-492
chr7-141741581-DEL-52
chr7-141752925-DEL-53
chr7-141755588-DEL-288
chr7-141763153-DEL-132
chr7-141798951-DEL-182
chr7-141821313-DEL-1317

chr7-141834177-DEL-114
chr7-141847629-DEL-81
chr7-141920833-DEL-184
chr7-141978490-DEL-116
chr7-141990106-DEL-395
chr7-142007437-DEL-59
chr7-142009539-DEL-55
chr7-142073294-DEL-80
chr7-142083985-DEL-72
chr7-142114477-DEL-83
chr7-142150327-DEL-78
chr7-142159433-DEL-56
chr7-142227269-DEL-243
chr7-142252587-DEL-56
chr7-142271929-DEL-115
chr7-142310921-DEL-54
chr7-142322548-DEL-16671
chr7-142391679-DEL-7082
chr7-142421588-DEL-193
chr7-142491359-DEL-73
chr7-142498209-DEL-481
chr7-142543602-DEL-100
chr7-142579996-DEL-65
chr7-142597187-DEL-302
chr7-142666096-DEL-67
chr7-142826160-DEL-189
chr7-142834958-DEL-125
chr7-142880037-DEL-51
chr7-142888732-DEL-164
chr7-142985195-DEL-56
chr7-143009692-DEL-94
chr7-143040229-DEL-113
chr7-143177550-DEL-175
chr7-143181184-DEL-53
chr7-143187355-DEL-372
chr7-143189439-DEL-756
chr7-143190465-DEL-51
chr7-143216621-DEL-178
chr7-143253899-DEL-68
chr7-143267583-DEL-169
chr7-143279273-DEL-118
chr7-143315721-DEL-91
chr7-143340932-DEL-73
chr7-143341996-DEL-67
chr7-143355730-DEL-56
chr7-143403087-DEL-14192
chr7-143457155-DEL-61
chr7-143502334-DEL-66
chr7-143514290-DEL-103
chr7-143551848-DEL-92
chr7-143587958-DEL-815
chr7-143641316-DEL-209
chr7-143654640-DEL-62
chr7-143655287-DEL-55
chr7-143687761-DEL-285
chr7-143709169-DEL-53
chr7-143796858-DEL-357
chr7-143808600-DEL-168
chr7-143846613-DEL-57
chr7-143858316-DEL-59
chr7-143863293-DEL-191
chr7-143876161-DEL-170
chr7-143884422-DEL-108
chr7-143886236-DEL-1975
chr7-143893944-DEL-89
chr7-143925302-DEL-58
chr7-143942518-DEL-262
chr7-143984594-DEL-57
chr7-143987976-DEL-71
chr7-143988149-DEL-1132
chr7-144018218-DEL-216
chr7-144051667-DEL-66
chr7-144095257-DEL-59
chr7-144096331-DEL-223
chr7-144109106-DEL-53
chr7-144133038-DEL-78
chr7-144137572-DEL-72

chr7-144144194-DEL-59
chr7-144157912-DEL-53
chr7-144174360-DEL-60
chr7-144213482-DEL-80
chr7-144226079-DEL-91
chr7-144246305-DEL-103
chr7-144271420-DEL-63
chr7-144279253-DEL-2604
chr7-144281252-DEL-612
chr7-144284955-DEL-154
chr7-144331770-DEL-661
chr7-144385701-DEL-58
chr7-144406425-DEL-58
chr7-144434922-DEL-187
chr7-144454855-DEL-576
chr7-144466982-DEL-163
chr7-144513157-DEL-184
chr7-144575650-DEL-52
chr7-144597108-DEL-65
chr7-144606854-DEL-289
chr7-144715556-DEL-1181
chr7-144758633-DEL-200
chr7-144773987-DEL-59
chr7-144780120-DEL-255
chr7-144800007-DEL-1868
chr7-144829477-DEL-137
chr7-144834985-DEL-384
chr7-144841803-DEL-400
chr7-144849245-DEL-939
chr7-144851856-DEL-384
chr7-144873034-DEL-58
chr7-144893645-DEL-67
chr8-3067494-DEL-2046
chr8-3099374-DEL-81
chr8-3132050-DEL-33097
chr8-3165702-DEL-1007
chr8-3186289-DEL-899
chr8-3226480-DEL-245
chr8-3244345-DEL-96
chr8-3249569-DEL-192
chr8-3306459-DEL-69
chr8-3328926-DEL-60
chr8-3341376-DEL-174
chr8-3342101-DEL-217
chr8-3367647-DEL-141
chr8-3369169-DEL-439
chr8-3371636-DEL-711
chr8-3387891-DEL-86
chr8-3421627-DEL-58
chr8-3452553-DEL-100
chr8-3482277-DEL-318
chr8-3485057-DEL-171
chr8-3492317-DEL-154
chr8-3516201-DEL-333
chr8-3519151-DEL-63
chr8-3557009-DEL-70
chr8-3560104-DEL-171
chr8-3604188-DEL-103
chr8-3607754-DEL-202
chr8-3612597-DEL-265
chr8-3635661-DEL-52
chr8-3694973-DEL-60
chr8-3735992-DEL-99
chr8-3744424-DEL-220
chr8-3819160-DEL-3252
chr8-3822630-DEL-6488
chr8-3833179-DEL-748
chr8-3835457-DEL-15151
chr8-3843132-DEL-315
chr8-3844970-DEL-53
chr8-3845445-DEL-91
chr8-3846272-DEL-564
chr8-3848226-DEL-3302
chr8-3850979-DEL-870
chr8-3856700-DEL-2879
chr8-3868407-DEL-1186
chr8-3873176-DEL-17584

chr8-3906788-DEL-227
chr8-3949469-DEL-2049
chr8-3951931-DEL-7217
chr8-3960500-DEL-6433
chr8-4006486-DEL-59
chr8-4120642-DEL-857
chr8-4124638-DEL-498
chr8-4125348-DEL-6178
chr8-4137214-DEL-2709
chr8-4150120-DEL-686
chr8-4151092-DEL-1313
chr8-4157075-DEL-537
chr8-4235315-DEL-195
chr8-4314761-DEL-118
chr8-4344010-DEL-52
chr8-4378646-DEL-2026
chr8-4381462-DEL-3417
chr8-4395972-DEL-110
chr8-4419619-DEL-54
chr8-4422928-DEL-9174
chr8-4437100-DEL-165
chr8-4442386-DEL-6414
chr8-4472703-DEL-400
chr8-4476349-DEL-489
chr8-4490921-DEL-18300
chr8-4518363-DEL-52
chr8-4533127-DEL-198
chr8-4537482-DEL-84
chr8-4582388-DEL-199
chr8-4582940-DEL-630
chr8-4595818-DEL-6591
chr8-4606244-DEL-55
chr8-4617366-DEL-154
chr8-4662168-DEL-434
chr8-4671832-DEL-2101
chr8-4675533-DEL-69
chr8-4706647-DEL-107
chr8-4707066-DEL-117
chr8-4731079-DEL-6905
chr8-4737309-DEL-709
chr8-4768411-DEL-476
chr8-4771307-DEL-1131
chr8-4778884-DEL-129
chr8-4821257-DEL-3477
chr8-4824516-DEL-442
chr8-4902605-DEL-6334
chr8-4903703-DEL-4940
chr8-4909809-DEL-1163
chr8-4919743-DEL-1450
chr8-4929672-DEL-107
chr8-4936132-DEL-7088
chr8-4947319-DEL-233
chr8-4949156-DEL-213
chr8-4979031-DEL-1137
chr8-4981483-DEL-73
chr8-5078586-DEL-515
chr8-5079370-DEL-219
chr8-5084379-DEL-76
chr8-5093462-DEL-1115
chr8-5159469-DEL-67
chr8-5218730-DEL-100
chr8-5231963-DEL-55
chr8-5247943-DEL-6178
chr8-5267390-DEL-8306
chr8-5309705-DEL-1021
chr8-5366898-DEL-1603
chr8-5382589-DEL-199
chr8-5405161-DEL-675
chr8-5435064-DEL-303
chr8-5447609-DEL-80
chr8-5505644-DEL-75
chr8-5525755-DEL-828
chr8-5548976-DEL-1439
chr8-5552219-DEL-3675
chr8-5669514-DEL-61
chr8-5683441-DEL-2128
chr8-5727315-DEL-60

chr8-5749863-DEL-64
chr8-5757479-DEL-6090
chr8-5803157-DEL-60
chr8-5835401-DEL-1494
chr8-5884643-DEL-78
chr8-5885574-DEL-6238
chr8-5936781-DEL-129
chr8-5967592-DEL-400
chr8-5999215-DEL-67
chr8-6004330-DEL-1044
chr8-6005705-DEL-678
chr8-6006566-DEL-399
chr8-6011213-DEL-52
chr8-6038625-DEL-1732
chr8-6050428-DEL-653
chr8-6064250-DEL-204
chr8-6086104-DEL-109
chr8-6091993-DEL-58
chr8-6103399-DEL-374
chr8-6110116-DEL-312
chr8-6136668-DEL-73
chr8-6145754-DEL-554
chr8-6146657-DEL-204
chr8-6153981-DEL-54
chr8-6167741-DEL-64
chr8-6184554-DEL-234
chr8-6187112-DEL-152
chr8-6227303-DEL-75
chr8-6302297-DEL-222
chr8-6306921-DEL-655
chr8-6370713-DEL-60
chr8-6423462-DEL-58
chr8-6427792-DEL-1355
chr8-6486015-DEL-56
chr8-6487113-DEL-62
chr8-6499915-DEL-85
chr8-6508579-DEL-1665
chr8-6522360-DEL-56
chr8-6531569-DEL-159
chr8-6555732-DEL-52
chr8-6560789-DEL-234
chr8-6586761-DEL-466
chr8-6638475-DEL-447
chr8-6704385-DEL-286
chr8-6714550-DEL-967
chr8-6755986-DEL-151
chr8-6781256-DEL-8692
chr8-6796764-DEL-374
chr8-6846036-DEL-2950
chr8-6907617-DEL-76
chr8-6937390-DEL-2151
chr8-6947217-DEL-1071
chr8-6952765-DEL-237
chr8-6982866-DEL-80
chr8-7021186-DEL-154
chr8-7058387-DEL-177
chr8-7100208-DEL-108
chr8-7106293-DEL-1136
chr8-7120457-DEL-77
chr8-7129740-DEL-1971
chr8-7197855-DEL-1416
chr8-7215226-DEL-200
chr8-7216490-DEL-64
chr8-7242590-DEL-196
chr8-7244130-DEL-2844
chr8-7298683-DEL-5985
chr8-7331231-DEL-432
chr8-7346680-DEL-765
chr8-7383146-DEL-5968
chr8-7419207-DEL-2859
chr8-7428452-DEL-80
chr8-7492507-DEL-1338
chr8-7509791-DEL-204
chr8-7520198-DEL-400
chr8-7545282-DEL-54
chr8-7548991-DEL-156
chr8-7560884-DEL-55

chr8-7573449-DEL-70
chr8-7592214-DEL-6497
chr8-7602453-DEL-630
chr8-7615993-DEL-975
chr8-7621303-DEL-197
chr8-7682328-DEL-1308
chr8-7711855-DEL-52
chr8-7722097-DEL-254
chr8-7723624-DEL-247
chr8-7737552-DEL-56
chr8-7742794-DEL-6196
chr8-7797134-DEL-439
chr8-7799142-DEL-8450
chr8-7806360-DEL-917
chr8-7817655-DEL-81
chr8-7875527-DEL-285
chr8-7906315-DEL-328
chr8-7907976-DEL-62
chr8-8027546-DEL-56
chr8-8043330-DEL-54
chr8-8092929-DEL-95
chr8-8109575-DEL-133
chr8-8206950-DEL-2078
chr8-8214258-DEL-403
chr8-8251034-DEL-1151
chr8-8327759-DEL-112
chr8-8330379-DEL-1007
chr8-8346107-DEL-120
chr8-8386437-DEL-313
chr8-8403704-DEL-116
chr8-8423008-DEL-170
chr8-8444957-DEL-118
chr8-8461435-DEL-167
chr8-8483047-DEL-884
chr8-8497221-DEL-102
chr8-8509334-DEL-259
chr8-8536509-DEL-58
chr8-8544381-DEL-58
chr8-8545115-DEL-123
chr8-8550890-DEL-70
chr8-8562402-DEL-374
chr8-8592062-DEL-175
chr8-8629240-DEL-335
chr8-8664882-DEL-194
chr8-8723434-DEL-58
chr8-8723661-DEL-898
chr8-8746165-DEL-55
chr8-8793830-DEL-1831
chr8-8799337-DEL-128
chr8-8800171-DEL-393
chr8-8803752-DEL-99
chr8-8848107-DEL-1752
chr8-8848389-DEL-867
chr8-8848412-DEL-215
chr8-8849221-DEL-1088
chr8-8866566-DEL-2321
chr8-8875404-DEL-52
chr8-8924725-DEL-189
chr8-8928306-DEL-211
chr8-8932659-DEL-61
chr8-8938716-DEL-53
chr8-8972092-DEL-64
chr8-8983580-DEL-358
chr8-9004385-DEL-56
chr8-9008241-DEL-128
chr8-9032518-DEL-78
chr8-9038300-DEL-63
chr8-9048324-DEL-53
chr8-9063098-DEL-208
chr8-9080027-DEL-54
chr8-9145573-DEL-56
chr8-9148651-DEL-116
chr8-9157895-DEL-52
chr8-9214908-DEL-585
chr8-9229307-DEL-144
chr8-9263924-DEL-72
chr8-9308205-DEL-93

chr8-9313348-DEL-51
chr8-9322557-DEL-1292
chr8-9329566-DEL-145
chr8-9331327-DEL-50
chr8-9362622-DEL-90
chr8-9379277-DEL-54
chr8-9394238-DEL-176
chr8-9435794-DEL-114
chr8-9466411-DEL-115
chr8-9470489-DEL-335
chr8-9471603-DEL-102
chr8-9488400-DEL-51
chr8-9573422-DEL-203
chr8-9615200-DEL-518
chr8-9628332-DEL-300
chr8-9662967-DEL-344
chr8-9681633-DEL-66
chr8-9692481-DEL-97
chr8-9702088-DEL-60
chr8-9707798-DEL-1268
chr8-9711336-DEL-86
chr8-9741083-DEL-636
chr8-9741520-DEL-537
chr8-9744319-DEL-350
chr8-9755007-DEL-58
chr8-9782124-DEL-103
chr8-9795619-DEL-962
chr8-9799577-DEL-1619
chr8-9806830-DEL-189
chr8-9810689-DEL-54
chr8-9811099-DEL-629
chr8-9812595-DEL-69
chr8-9825173-DEL-72
chr8-9833054-DEL-3209
chr8-9870056-DEL-52
chr8-9880513-DEL-1201
chr8-9931231-DEL-81
chr8-9941438-DEL-60
chr8-9942836-DEL-141
chr8-9976274-DEL-66
chr8-10009544-DEL-109
chr8-10013004-DEL-205
chr8-10060885-DEL-252
chr8-10070142-DEL-452
chr8-10074270-DEL-204
chr8-10077875-DEL-56
chr8-10104308-DEL-99
chr8-10124604-DEL-205
chr8-10128294-DEL-72
chr8-10178111-DEL-148
chr8-10201657-DEL-219
chr8-10218830-DEL-122
chr8-10276207-DEL-60
chr8-10297292-DEL-95
chr8-10302455-DEL-1854
chr8-10303159-DEL-120
chr8-10316618-DEL-228
chr8-10335840-DEL-52
chr8-10337194-DEL-699
chr8-10342876-DEL-100
chr8-10355970-DEL-230
chr8-10382133-DEL-459
chr8-10402399-DEL-482
chr8-10414458-DEL-124
chr8-10424963-DEL-61
chr8-10447278-DEL-465
chr8-10457698-DEL-54
chr8-10547430-DEL-192
chr8-10585986-DEL-218
chr8-10614367-DEL-231
chr8-10620250-DEL-63
chr8-10633297-DEL-109
chr8-10637387-DEL-66
chr8-10649552-DEL-130
chr8-10668068-DEL-103
chr8-10704578-DEL-188
chr8-10710287-DEL-88

chr8-10717767-DEL-340
chr8-10722838-DEL-71
chr8-10726605-DEL-150
chr8-10732628-DEL-65
chr8-10741259-DEL-734
chr8-10754846-DEL-52
chr8-10755088-DEL-368
chr8-10775235-DEL-257
chr8-10830176-DEL-144
chr8-10847121-DEL-65
chr8-10865228-DEL-122
chr8-10910094-DEL-144
chr8-10923230-DEL-210
chr8-10935247-DEL-95
chr8-10987215-DEL-72
chr8-11013950-DEL-88
chr8-11017022-DEL-234
chr8-11019981-DEL-54
chr8-11030764-DEL-253
chr8-11074082-DEL-60
chr8-11117088-DEL-62
chr8-11171129-DEL-144
chr8-11175969-DEL-525
chr8-11187154-DEL-884
chr8-11195087-DEL-345
chr8-11220200-DEL-88
chr8-11247533-DEL-110
chr8-11370772-DEL-120
chr8-11387940-DEL-1100
chr8-11402292-DEL-72
chr8-11409147-DEL-9992
chr8-11448871-DEL-359
chr8-11482128-DEL-114
chr8-11515805-DEL-313
chr8-11537178-DEL-67
chr8-11538442-DEL-203
chr8-11551787-DEL-170
chr8-11577218-DEL-141
chr8-11617421-DEL-152
chr8-11626973-DEL-269
chr8-11639416-DEL-6546
chr8-11663476-DEL-160
chr8-11682146-DEL-61
chr8-11697618-DEL-144
chr8-11700971-DEL-178
chr8-11702884-DEL-189
chr8-11795306-DEL-53
chr8-11846364-DEL-152
chr8-11859974-DEL-65
chr8-11875667-DEL-71
chr8-11900712-DEL-252
chr8-11917470-DEL-52
chr8-11923032-DEL-107
chr8-11935331-DEL-336
chr8-11971770-DEL-69
chr8-11981235-DEL-6169
chr8-12005969-DEL-119
chr8-12006961-DEL-124
chr8-12019975-DEL-54
chr8-12026699-DEL-172
chr8-12035767-DEL-59
chr8-12053329-DEL-92
chr8-12054975-DEL-237
chr8-12062045-DEL-83
chr8-12068697-DEL-97
chr8-12070645-DEL-462
chr8-12088990-DEL-61
chr8-12092662-DEL-54
chr8-12114408-DEL-964
chr8-12167708-DEL-65
chr8-12178906-DEL-78
chr8-12180964-DEL-303
chr8-12181410-DEL-2578
chr8-12181661-DEL-114
chr8-12182060-DEL-1356
chr8-12182705-DEL-82
chr8-12183641-DEL-201

chr8-12195566-DEL-65
chr8-12238533-DEL-94
chr8-12253507-DEL-752
chr8-12315487-DEL-87
chr8-12324359-DEL-70
chr8-12324768-DEL-3342
chr8-12356728-DEL-156
chr8-12361677-DEL-198
chr8-12362295-DEL-58
chr8-12386038-DEL-589
chr8-12406799-DEL-1884
chr8-12483536-DEL-62
chr8-12506186-DEL-138
chr8-12511142-DEL-75
chr8-12549542-DEL-67
chr8-12550766-DEL-82
chr8-12569781-DEL-71
chr8-12582878-DEL-204
chr8-12609609-DEL-75
chr8-12615001-DEL-52
chr8-12633406-DEL-123
chr8-12636730-DEL-1074
chr8-12637142-DEL-300
chr8-12653253-DEL-417
chr8-12672148-DEL-81
chr8-12682877-DEL-57
chr8-12701820-DEL-91
chr8-12773737-DEL-231
chr8-12774025-DEL-1630
chr8-12816464-DEL-108
chr8-12828197-DEL-533
chr8-12837559-DEL-61
chr8-12884512-DEL-55
chr8-12898483-DEL-143
chr8-12986231-DEL-343
chr8-13019808-DEL-145
chr8-13049520-DEL-121
chr8-13073200-DEL-74
chr8-13082453-DEL-101
chr8-13100939-DEL-66
chr8-13106215-DEL-72
chr8-13117374-DEL-228
chr8-13119419-DEL-865
chr8-13119598-DEL-160
chr8-13120304-DEL-190
chr8-13125566-DEL-165
chr8-13178284-DEL-199
chr8-13181003-DEL-206
chr8-13198751-DEL-62
chr8-13211656-DEL-177
chr8-13244022-DEL-82
chr8-13260195-DEL-258
chr8-13289957-DEL-80
chr8-13296701-DEL-159
chr8-13300776-DEL-87
chr8-13325217-DEL-1086
chr8-13344163-DEL-98
chr8-13367403-DEL-380
chr8-13377651-DEL-389
chr8-13426616-DEL-266
chr8-13503595-DEL-76
chr8-13507419-DEL-54
chr8-13533923-DEL-103
chr8-13552048-DEL-239
chr8-13616647-DEL-71
chr8-13657307-DEL-247
chr8-13657348-DEL-535
chr8-13658521-DEL-84
chr8-13733185-DEL-579
chr8-13743116-DEL-93
chr8-13760404-DEL-5353
chr8-13769298-DEL-5044
chr8-13779324-DEL-52
chr8-13821962-DEL-134
chr8-13849993-DEL-175
chr8-13853520-DEL-1582
chr8-13863182-DEL-637

chr8-13863473-DEL-2054
chr8-13864074-DEL-1071
chr8-13946140-DEL-692
chr8-13953475-DEL-1892
chr8-13959607-DEL-88
chr8-13967845-DEL-1504
chr8-13967890-DEL-513
chr8-13967954-DEL-2187
chr8-13968613-DEL-1406
chr8-13970183-DEL-262
chr8-13993432-DEL-122
chr8-14007754-DEL-63
chr8-14010145-DEL-440
chr8-14020319-DEL-207
chr8-14040306-DEL-140
chr8-14044103-DEL-144
chr8-14057483-DEL-160
chr8-14057510-DEL-765
chr8-14088055-DEL-171
chr8-14088593-DEL-53
chr8-14094583-DEL-55
chr8-14112134-DEL-87
chr8-14118065-DEL-323
chr8-14154701-DEL-148
chr8-14205471-DEL-93
chr8-14207763-DEL-53
chr8-14268208-DEL-299
chr8-14308134-DEL-53
chr8-14311209-DEL-456
chr8-14317961-DEL-51
chr8-14344223-DEL-4848
chr8-14355400-DEL-107
chr8-14406231-DEL-330
chr8-14410805-DEL-80
chr8-14428394-DEL-68
chr8-14470880-DEL-2060
chr8-14484915-DEL-116
chr8-14500197-DEL-122
chr8-14529154-DEL-99
chr8-14627907-DEL-52
chr8-14633996-DEL-98
chr8-14636153-DEL-64
chr8-14655104-DEL-91
chr8-14667857-DEL-76
chr8-14722389-DEL-72
chr8-14733093-DEL-65
chr8-14754367-DEL-53
chr8-14773243-DEL-123
chr8-14779041-DEL-424
chr8-14783223-DEL-285
chr8-14786428-DEL-71
chr8-14817767-DEL-69
chr8-14834404-DEL-2308
chr8-14836767-DEL-3004
chr8-14853244-DEL-3862
chr8-14870920-DEL-110
chr8-14935274-DEL-64
chr8-14941568-DEL-89
chr8-14986851-DEL-135
chr8-15036233-DEL-64
chr8-15044874-DEL-71
chr8-15059978-DEL-1147
chr8-15089154-DEL-72
chr8-15091975-DEL-81
chr8-15094237-DEL-143
chr8-15121553-DEL-402
chr8-15152270-DEL-101
chr8-15192068-DEL-467
chr8-15198208-DEL-51
chr8-15245147-DEL-54
chr8-15245611-DEL-1756
chr8-15272799-DEL-79
chr8-15283911-DEL-217
chr8-15290615-DEL-67
chr8-15295952-DEL-159
chr8-15309437-DEL-529
chr8-15348730-DEL-102

chr8-15353712-DEL-131
chr8-15439091-DEL-168
chr8-15469927-DEL-62
chr8-15473585-DEL-198
chr8-15511036-DEL-115
chr8-15536172-DEL-60
chr8-15540546-DEL-52
chr8-15555641-DEL-94
chr8-15595252-DEL-1179
chr8-15622878-DEL-141
chr8-15626326-DEL-127
chr8-15629506-DEL-52
chr8-15640124-DEL-813
chr8-15640796-DEL-140
chr8-15664449-DEL-6201
chr8-15673729-DEL-506
chr8-15682306-DEL-64
chr8-15682743-DEL-244
chr8-15685618-DEL-6428
chr8-15693317-DEL-118
chr8-15728271-DEL-348
chr8-15729550-DEL-1411
chr8-15730996-DEL-490
chr8-15801143-DEL-13377
chr8-15814503-DEL-3594
chr8-15902405-DEL-52
chr8-15983144-DEL-203
chr8-16007221-DEL-166
chr8-16037262-DEL-100
chr8-16105470-DEL-95
chr8-16106089-DEL-57
chr8-16123458-DEL-212
chr8-16173091-DEL-95
chr8-16201894-DEL-311
chr8-16237882-DEL-58
chr8-16249334-DEL-58
chr8-16296850-DEL-1530
chr8-16298731-DEL-730
chr8-16310838-DEL-7166
chr8-16324702-DEL-1030
chr8-16336379-DEL-68
chr8-16394150-DEL-70
chr8-16430327-DEL-511
chr8-16455670-DEL-61741
chr8-16487438-DEL-7778
chr8-16541776-DEL-2597
chr8-16567278-DEL-930
chr8-16568995-DEL-2343
chr8-16593529-DEL-64
chr8-16685079-DEL-251
chr8-16706140-DEL-80
chr8-16717117-DEL-144
chr8-16737947-DEL-268
chr8-16753080-DEL-939
chr8-16786530-DEL-69
chr8-16803746-DEL-594
chr8-16827104-DEL-86
chr8-16833589-DEL-65
chr8-16884124-DEL-68
chr8-16940784-DEL-372
chr8-16979181-DEL-164
chr8-16982091-DEL-59
chr8-17049822-DEL-1770
chr8-17074110-DEL-55
chr8-17114906-DEL-121
chr8-17183363-DEL-86
chr8-17184258-DEL-811
chr8-17189144-DEL-252
chr8-17190359-DEL-200
chr8-17198268-DEL-153
chr8-17215446-DEL-81
chr8-17252364-DEL-66
chr8-17273321-DEL-107
chr8-17301220-DEL-316
chr8-17323392-DEL-80
chr8-17326652-DEL-1419
chr8-17332785-DEL-64

chr8-17408325-DEL-69
chr8-17415808-DEL-116
chr8-17421711-DEL-77
chr8-17425651-DEL-909
chr8-17491016-DEL-1699
chr8-17507528-DEL-243
chr8-17525852-DEL-829
chr8-17527328-DEL-71
chr8-17533091-DEL-212
chr8-17611333-DEL-79
chr8-17626268-DEL-490
chr8-17643619-DEL-119
chr8-17650518-DEL-51
chr8-17670835-DEL-68
chr8-17673251-DEL-2067
chr8-17677452-DEL-590
chr8-17725100-DEL-360
chr8-17769660-DEL-104
chr8-17817711-DEL-229
chr8-17821522-DEL-59
chr8-17844516-DEL-189
chr8-17849416-DEL-120
chr8-17917214-DEL-205
chr8-17980383-DEL-331
chr8-18024585-DEL-54
chr8-18095893-DEL-6403
chr8-18158001-DEL-434
chr8-18167612-DEL-78
chr8-18182910-DEL-539
chr8-18226688-DEL-156
chr8-18233010-DEL-913
chr8-18235883-DEL-477
chr8-18248222-DEL-6209
chr8-18270053-DEL-212
chr8-18278486-DEL-234
chr8-18282067-DEL-58
chr8-18282897-DEL-6290
chr8-18421060-DEL-129
chr8-18460453-DEL-119
chr8-18482135-DEL-6719
chr8-18488975-DEL-146
chr8-18551124-DEL-253
chr8-18589187-DEL-199
chr8-18602953-DEL-104
chr8-18617217-DEL-309
chr8-18618214-DEL-63
chr8-18619371-DEL-2560
chr8-18642982-DEL-202
chr8-18650312-DEL-544
chr8-18655049-DEL-396
chr8-18658660-DEL-272
chr8-18703606-DEL-412
chr8-18706423-DEL-209
chr8-18722446-DEL-55
chr8-18747577-DEL-142
chr8-18815665-DEL-100
chr8-18821607-DEL-831
chr8-18825430-DEL-69
chr8-18898672-DEL-323
chr8-18945228-DEL-258
chr8-18973968-DEL-59
chr8-19026252-DEL-80
chr8-19047114-DEL-528
chr8-19066520-DEL-1599
chr8-19103974-DEL-6300
chr8-19148740-DEL-3009
chr8-19155532-DEL-209
chr8-19156200-DEL-65
chr8-19179116-DEL-1057
chr8-19223420-DEL-75
chr8-19245788-DEL-543
chr8-19249014-DEL-56
chr8-19252592-DEL-1771
chr8-19261032-DEL-526
chr8-19315558-DEL-1398
chr8-19347445-DEL-145
chr8-19348414-DEL-440

chr8-19352641-DEL-597
chr8-19364755-DEL-198
chr8-19395549-DEL-4548
chr8-19402735-DEL-263
chr8-19404650-DEL-599
chr8-19419488-DEL-4640
chr8-19424715-DEL-4149
chr8-19444790-DEL-2881
chr8-19473553-DEL-390
chr8-19484370-DEL-2075
chr8-19488525-DEL-81
chr8-19490320-DEL-59
chr8-19500238-DEL-351
chr8-19510079-DEL-54
chr8-19521441-DEL-209
chr8-19542910-DEL-724
chr8-19545268-DEL-120
chr8-19549274-DEL-13205
chr8-19558832-DEL-3617
chr8-19561890-DEL-159
chr8-19562588-DEL-1166
chr8-19580257-DEL-470
chr8-19620453-DEL-56
chr8-19626889-DEL-2269
chr8-19646384-DEL-135
chr8-19663560-DEL-56
chr8-19679808-DEL-375
chr8-19809911-DEL-51
chr8-19810613-DEL-982
chr8-19813059-DEL-806
chr8-19817734-DEL-60
chr8-19833367-DEL-1533
chr8-20023747-DEL-963
chr8-20113061-DEL-982
chr8-20119644-DEL-7476
chr8-20131702-DEL-207
chr8-20174700-DEL-51
chr8-20507594-DEL-273
chr8-20508959-DEL-981
chr8-20574421-DEL-72
chr8-20578649-DEL-76
chr8-20582064-DEL-59
chr8-20619929-DEL-2526
chr8-20647024-DEL-312
chr8-20660978-DEL-154
chr8-20696075-DEL-470
chr8-20696927-DEL-634
chr8-20727873-DEL-582
chr8-21094764-DEL-3207
chr8-21103196-DEL-60
chr8-21161993-DEL-225
chr8-21189416-DEL-51
chr8-21200915-DEL-669
chr8-21203994-DEL-102
chr8-21205115-DEL-312
chr8-21205785-DEL-156
chr8-21206119-DEL-517
chr8-21206339-DEL-465
chr8-21217521-DEL-513
chr8-21221081-DEL-360
chr8-21225883-DEL-989
chr8-21236504-DEL-1748
chr8-21237532-DEL-360
chr8-21238714-DEL-313
chr8-21239441-DEL-312
chr8-21239556-DEL-835
chr8-21240811-DEL-204
chr8-21242520-DEL-153
chr8-21245365-DEL-156
chr8-21246210-DEL-207
chr8-21246340-DEL-156
chr8-21333179-DEL-622
chr8-21340955-DEL-17096
chr8-21341867-DEL-76
chr8-21348738-DEL-63
chr8-21350127-DEL-76
chr8-21358305-DEL-853

chr8-21393945-DEL-51
chr8-21399062-DEL-2016
chr8-21402983-DEL-2440
chr8-21460303-DEL-13051
chr8-21464304-DEL-963
chr8-21478272-DEL-3570
chr8-21480188-DEL-64
chr8-21502391-DEL-490
chr8-21505400-DEL-206
chr8-21557311-DEL-197
chr8-21560778-DEL-1834
chr8-21563652-DEL-394
chr8-21597041-DEL-6188
chr8-21616678-DEL-6407
chr8-21641262-DEL-1122
chr8-21643633-DEL-656
chr8-21646817-DEL-270
chr8-21658593-DEL-6233
chr8-21668664-DEL-83
chr8-21671529-DEL-1158
chr8-21679412-DEL-1370
chr8-21698836-DEL-6003
chr8-21700833-DEL-167
chr8-21713899-DEL-5652
chr8-21715304-DEL-71
chr8-21733393-DEL-645
chr8-21735062-DEL-7193
chr8-21748109-DEL-6468
chr8-21748114-DEL-374818
chr8-21768603-DEL-109
chr8-21775231-DEL-1378
chr8-21779358-DEL-997
chr8-21781913-DEL-261
chr8-21794185-DEL-856
chr8-21862505-DEL-2859
chr8-21867459-DEL-60
chr8-21875186-DEL-86
chr8-21887699-DEL-193
chr8-21922604-DEL-192
chr8-21924196-DEL-395
chr8-21942432-DEL-2822
chr8-21946087-DEL-80
chr8-21953312-DEL-55
chr8-21993255-DEL-2847
chr8-22005928-DEL-84
chr8-22034013-DEL-3837
chr8-22040015-DEL-113
chr8-22042850-DEL-2949
chr8-22062154-DEL-395
chr8-22071558-DEL-5626
chr8-22080389-DEL-2811
chr8-22084043-DEL-80
chr8-22086712-DEL-619
chr8-22092230-DEL-6131
chr8-22104349-DEL-749
chr8-22106741-DEL-889
chr8-22113762-DEL-831
chr8-22116521-DEL-6412
chr8-22136969-DEL-105
chr8-22150274-DEL-261
chr8-22160910-DEL-5488
chr8-22165708-DEL-80
chr8-22168504-DEL-5989
chr8-22183570-DEL-5584
chr8-22200689-DEL-541
chr8-22201277-DEL-6196
chr8-22226822-DEL-8017
chr8-22257781-DEL-72
chr8-22302453-DEL-255
chr8-22303651-DEL-167
chr8-22307612-DEL-110
chr8-22314646-DEL-171
chr8-22321718-DEL-74
chr8-22324330-DEL-4631
chr8-22379537-DEL-617
chr8-22414351-DEL-80
chr8-22427083-DEL-3577

chr8-22432154-DEL-85
chr8-22443581-DEL-76
chr8-22489853-DEL-67
chr8-22493443-DEL-161
chr8-22530473-DEL-92
chr8-22532930-DEL-59
chr8-22565868-DEL-76
chr8-22579051-DEL-66
chr8-22594046-DEL-962
chr8-22637911-DEL-86
chr8-22668043-DEL-59
chr8-22679664-DEL-200
chr8-22686482-DEL-112
chr8-22688416-DEL-68
chr8-22713781-DEL-215
chr8-22727953-DEL-59
chr8-22765326-DEL-131
chr8-22777092-DEL-400
chr8-22787366-DEL-53
chr8-22819848-DEL-839
chr8-22827939-DEL-165
chr8-22841081-DEL-71
chr8-22871029-DEL-56
chr8-22886242-DEL-660
chr8-22898831-DEL-192
chr8-22908823-DEL-858
chr8-22909431-DEL-123
chr8-22913631-DEL-209
chr8-22933713-DEL-180
chr8-22947162-DEL-111
chr8-22956873-DEL-1237
chr8-22971604-DEL-60
chr8-23002259-DEL-163
chr8-23015712-DEL-54
chr8-23065722-DEL-98
chr8-23093903-DEL-211
chr8-23099537-DEL-193
chr8-23102528-DEL-76
chr8-23107930-DEL-210
chr8-23112259-DEL-54
chr8-23128375-DEL-695
chr8-23139469-DEL-428
chr8-23174246-DEL-143
chr8-23179525-DEL-987
chr8-23184575-DEL-357
chr8-23257875-DEL-97
chr8-23265991-DEL-473
chr8-23291636-DEL-120
chr8-23305161-DEL-63
chr8-23309410-DEL-58
chr8-23310042-DEL-57
chr8-23311591-DEL-104
chr8-23326185-DEL-308
chr8-23380723-DEL-213
chr8-23393732-DEL-182
chr8-23460703-DEL-916
chr8-23462786-DEL-262
chr8-23526276-DEL-57
chr8-23556665-DEL-539
chr8-23574400-DEL-60
chr8-23630767-DEL-219
chr8-23675802-DEL-1243
chr8-23677972-DEL-269
chr8-23689683-DEL-112
chr8-23702525-DEL-142
chr8-23704574-DEL-69
chr8-23706082-DEL-83
chr8-23767905-DEL-126
chr8-23884984-DEL-125
chr8-23887068-DEL-65
chr8-23946229-DEL-60
chr8-24043491-DEL-75
chr8-24059762-DEL-134
chr8-24073791-DEL-158
chr8-24083051-DEL-401
chr8-24093804-DEL-54
chr8-24137686-DEL-154

chr8-24149976-DEL-348
chr8-24203648-DEL-76
chr8-24223653-DEL-94
chr8-24238552-DEL-143
chr8-24247500-DEL-119
chr8-24257297-DEL-92
chr8-24261342-DEL-96
chr8-24307904-DEL-152
chr8-24377143-DEL-301
chr8-24424818-DEL-112
chr8-24443333-DEL-80
chr8-24453437-DEL-64
chr8-24455424-DEL-1093
chr8-24474341-DEL-125
chr8-24496990-DEL-157
chr8-24518205-DEL-54
chr8-24523710-DEL-1343
chr8-24524963-DEL-69
chr8-24572787-DEL-105
chr8-24576891-DEL-57
chr8-24598076-DEL-313
chr8-24609809-DEL-69
chr8-24614998-DEL-60
chr8-24727002-DEL-204
chr8-24737581-DEL-265
chr8-24755405-DEL-124
chr8-24805805-DEL-1537
chr8-24836994-DEL-195
chr8-24849177-DEL-60
chr8-24853541-DEL-82
chr8-24858230-DEL-341
chr8-24874331-DEL-52
chr8-24882527-DEL-505
chr8-24912860-DEL-83
chr8-24916370-DEL-60
chr8-24944706-DEL-75
chr8-25018203-DEL-62
chr8-25018753-DEL-2879
chr8-25023426-DEL-115
chr8-25041199-DEL-76
chr8-25056630-DEL-61
chr8-25078632-DEL-75
chr8-25084351-DEL-72
chr8-25085009-DEL-62
chr8-25085841-DEL-71
chr8-25092907-DEL-96
chr8-25099269-DEL-80
chr8-25123870-DEL-154
chr8-25128275-DEL-52
chr8-25129230-DEL-80
chr8-25130093-DEL-1106
chr8-25130140-DEL-280
chr8-25130827-DEL-295
chr8-25144615-DEL-609
chr8-25147139-DEL-135
chr8-25171508-DEL-60
chr8-25172797-DEL-264
chr8-25197556-DEL-75
chr8-25206015-DEL-327
chr8-25213088-DEL-135
chr8-25217851-DEL-75
chr8-25235485-DEL-607
chr8-25241217-DEL-12928
chr8-25280521-DEL-417
chr8-25294932-DEL-277
chr8-25298378-DEL-260
chr8-25315053-DEL-78
chr8-25316583-DEL-173
chr8-25323244-DEL-415
chr8-25328159-DEL-61
chr8-25329316-DEL-52
chr8-25341005-DEL-74
chr8-25344303-DEL-53
chr8-25348492-DEL-1646
chr8-25355718-DEL-1541
chr8-25364604-DEL-78
chr8-25373225-DEL-218

chr8-25374817-DEL-360
chr8-25389457-DEL-865
chr8-25404573-DEL-244
chr8-25425904-DEL-59
chr8-25427489-DEL-60
chr8-25434703-DEL-996
chr8-25443305-DEL-164
chr8-25464379-DEL-87
chr8-25473038-DEL-2247
chr8-25483961-DEL-85
chr8-25502309-DEL-52
chr8-25508930-DEL-560
chr8-25521441-DEL-261
chr8-25540507-DEL-300
chr8-25557104-DEL-180
chr8-25566168-DEL-57
chr8-25580128-DEL-51
chr8-25584838-DEL-54
chr8-25589394-DEL-119
chr8-25597258-DEL-2270
chr8-25620891-DEL-5777
chr8-25637137-DEL-95
chr8-25656600-DEL-75
chr8-25701168-DEL-202
chr8-25725692-DEL-50
chr8-25732769-DEL-58
chr8-25749841-DEL-51
chr8-25754760-DEL-2170
chr8-25774883-DEL-74
chr8-25787593-DEL-345
chr8-25803533-DEL-58
chr8-25807946-DEL-51
chr8-25827535-DEL-1119
chr8-25836085-DEL-633
chr8-25854592-DEL-608
chr8-25860386-DEL-120
chr8-25867712-DEL-118
chr8-25871197-DEL-111
chr8-25873948-DEL-57
chr8-25877688-DEL-67
chr8-25889661-DEL-70
chr8-25917473-DEL-106
chr8-25921209-DEL-195
chr8-25924929-DEL-60
chr8-25936979-DEL-366
chr8-25944259-DEL-532
chr8-25966106-DEL-2246
chr8-25974244-DEL-73
chr8-25978301-DEL-101
chr8-25981082-DEL-113
chr8-25991473-DEL-59
chr8-25992190-DEL-316
chr8-25996476-DEL-79
chr8-26022817-DEL-63
chr8-26037462-DEL-192
chr8-26067769-DEL-98
chr8-26068936-DEL-52
chr8-26090736-DEL-324
chr8-26116838-DEL-248
chr8-26185085-DEL-2089
chr8-26187681-DEL-79
chr8-26230891-DEL-118
chr8-26251838-DEL-273
chr8-26325468-DEL-264
chr8-26344137-DEL-221
chr8-26355770-DEL-85
chr8-26356603-DEL-112
chr8-26357354-DEL-88
chr8-26358814-DEL-236
chr8-26377881-DEL-963
chr8-26403249-DEL-305
chr8-26459624-DEL-370
chr8-26483237-DEL-115
chr8-26492100-DEL-113
chr8-26506425-DEL-473
chr8-26539026-DEL-129
chr8-26539608-DEL-649

chr8-26543853-DEL-554
chr8-26634507-DEL-669
chr8-26663134-DEL-81
chr8-26664619-DEL-313
chr8-26670496-DEL-577
chr8-26683841-DEL-120
chr8-26693494-DEL-246
chr8-26705069-DEL-185
chr8-26713890-DEL-81
chr8-26715034-DEL-72
chr8-26786100-DEL-120
chr8-26849870-DEL-137
chr8-26872597-DEL-126
chr8-26895414-DEL-167
chr8-26898278-DEL-60
chr8-26912188-DEL-52
chr8-26933513-DEL-200
chr8-26943260-DEL-88
chr8-26957032-DEL-87
chr8-26989024-DEL-318
chr8-26997193-DEL-164
chr8-27052079-DEL-57
chr8-27137338-DEL-107
chr8-27145121-DEL-653
chr8-27172433-DEL-2858
chr8-27174767-DEL-507
chr8-27191856-DEL-68
chr8-27249080-DEL-108
chr8-27249686-DEL-113
chr8-27304233-DEL-153
chr8-27491000-DEL-78
chr8-27492413-DEL-57
chr8-27557887-DEL-258
chr8-27563455-DEL-110
chr8-27586603-DEL-120
chr8-27610100-DEL-138
chr8-27636887-DEL-235
chr8-27678137-DEL-176
chr8-27680572-DEL-1649
chr8-27682524-DEL-2384
chr8-27695237-DEL-190
chr8-27699331-DEL-382
chr8-27719719-DEL-626
chr8-27727642-DEL-593
chr8-27733274-DEL-703
chr8-27734924-DEL-252
chr8-27742293-DEL-456
chr8-27752902-DEL-72
chr8-27756501-DEL-56
chr8-27765805-DEL-62
chr8-27769128-DEL-88
chr8-27772357-DEL-4912
chr8-27779686-DEL-667
chr8-27793751-DEL-8817
chr8-27803857-DEL-19668
chr8-27803921-DEL-445
chr8-27805385-DEL-152
chr8-27830082-DEL-211
chr8-27852153-DEL-80
chr8-27867366-DEL-528
chr8-27870760-DEL-57
chr8-27876906-DEL-63
chr8-27906452-DEL-62
chr8-27926710-DEL-71
chr8-27934632-DEL-60
chr8-27948987-DEL-400
chr8-27950630-DEL-432
chr8-27966240-DEL-2805
chr8-27973729-DEL-903
chr8-27974606-DEL-6206
chr8-28005560-DEL-1047
chr8-28046323-DEL-196
chr8-28050789-DEL-6616
chr8-28051008-DEL-201
chr8-28063411-DEL-127
chr8-28074892-DEL-96
chr8-28093915-DEL-238

chr8-28095483-DEL-1084
chr8-28098674-DEL-59
chr8-28102066-DEL-85
chr8-28102977-DEL-1397
chr8-28105904-DEL-56
chr8-28116687-DEL-3786
chr8-28136351-DEL-1541
chr8-28137943-DEL-1498
chr8-28161294-DEL-353
chr8-28165140-DEL-103
chr8-28183170-DEL-394
chr8-28192855-DEL-1010
chr8-28198603-DEL-91
chr8-28212924-DEL-58
chr8-28227487-DEL-64
chr8-28280471-DEL-497
chr8-28286230-DEL-1955
chr8-28293026-DEL-6633
chr8-28425305-DEL-65
chr8-28459831-DEL-2861
chr8-28467042-DEL-69
chr8-28490662-DEL-959
chr8-28506319-DEL-6420
chr8-28516093-DEL-6396
chr8-28559850-DEL-99
chr8-28563316-DEL-92
chr8-28605516-DEL-65
chr8-28621736-DEL-58
chr8-28649273-DEL-166
chr8-28693224-DEL-89
chr8-28715757-DEL-714
chr8-28717048-DEL-587
chr8-28718696-DEL-89
chr8-28728873-DEL-280
chr8-28740183-DEL-1887
chr8-28743686-DEL-3936
chr8-28788117-DEL-151
chr8-28795402-DEL-1246
chr8-28853924-DEL-143
chr8-28860058-DEL-58
chr8-28871743-DEL-113
chr8-28921071-DEL-683
chr8-28930783-DEL-654
chr8-28936536-DEL-96
chr8-28945859-DEL-215
chr8-28969383-DEL-53
chr8-28985549-DEL-56
chr8-29013863-DEL-72
chr8-29017569-DEL-60
chr8-29027228-DEL-55
chr8-29038986-DEL-385
chr8-29062320-DEL-89
chr8-29090214-DEL-149
chr8-29097477-DEL-120
chr8-29114803-DEL-2294
chr8-29119112-DEL-129
chr8-29147213-DEL-126
chr8-29149884-DEL-153
chr8-29158699-DEL-249
chr8-29237668-DEL-136
chr8-29252220-DEL-61
chr8-29296802-DEL-937
chr8-29340438-DEL-119
chr8-29420251-DEL-173
chr8-29442519-DEL-394
chr8-29453571-DEL-233
chr8-29523643-DEL-108
chr8-29573483-DEL-168
chr8-29598891-DEL-1408
chr8-29609346-DEL-448
chr8-29614317-DEL-54
chr8-29617797-DEL-199
chr8-29698289-DEL-62
chr8-29720696-DEL-104
chr8-29776222-DEL-786
chr8-29804068-DEL-60
chr8-29834643-DEL-4180

chr8-29878884-DEL-56
chr8-29887696-DEL-2598
chr8-29914565-DEL-58
chr8-29956357-DEL-252
chr8-30006174-DEL-68
chr8-30057887-DEL-51
chr8-30065458-DEL-87
chr8-30098998-DEL-10176
chr8-30111896-DEL-975
chr8-30129345-DEL-1065
chr8-30150963-DEL-53
chr8-30156472-DEL-2726
chr8-30168017-DEL-821
chr8-30181086-DEL-51
chr8-30277221-DEL-63
chr8-30283299-DEL-3931
chr8-30301624-DEL-107
chr8-30312033-DEL-80
chr8-30324451-DEL-62
chr8-30377622-DEL-75
chr8-30393586-DEL-170
chr8-30399944-DEL-54
chr8-30402553-DEL-64
chr8-30404911-DEL-1062
chr8-30483671-DEL-64
chr8-30501225-DEL-310
chr8-30525903-DEL-6435
chr8-30533728-DEL-65
chr8-30555911-DEL-3688
chr8-30562270-DEL-106
chr8-30571849-DEL-121
chr8-30575513-DEL-803
chr8-30581954-DEL-59
chr8-30583580-DEL-180
chr8-30583631-DEL-7093
chr8-30583996-DEL-482
chr8-30599593-DEL-51
chr8-30612605-DEL-63
chr8-30612934-DEL-526
chr8-30614245-DEL-41039
chr8-30685678-DEL-103
chr8-30714723-DEL-95
chr8-30734574-DEL-79
chr8-30738724-DEL-367
chr8-30758155-DEL-517
chr8-30767772-DEL-67
chr8-30781442-DEL-86
chr8-30787564-DEL-2758
chr8-30791351-DEL-54
chr8-30870355-DEL-59
chr8-30873969-DEL-77
chr8-30905566-DEL-77
chr8-30915872-DEL-829
chr8-30929407-DEL-138
chr8-31014255-DEL-63
chr8-31021149-DEL-351
chr8-31039187-DEL-1087
chr8-31067268-DEL-179
chr8-31079946-DEL-1088
chr8-31105444-DEL-6269
chr8-31113127-DEL-154
chr8-31129365-DEL-372
chr8-31146848-DEL-59
chr8-31162584-DEL-174
chr8-31167351-DEL-166
chr8-31167936-DEL-958
chr8-31167945-DEL-223
chr8-31168741-DEL-111
chr8-31189308-DEL-157
chr8-31249541-DEL-68
chr8-31265309-DEL-52
chr8-31274793-DEL-114
chr8-31277186-DEL-1244
chr8-31309625-DEL-78
chr8-31330523-DEL-383
chr8-31365737-DEL-135
chr8-31391801-DEL-77

chr8-31399728-DEL-183
chr8-31432520-DEL-2641
chr8-31490813-DEL-56
chr8-31494475-DEL-53
chr8-31505761-DEL-1996
chr8-31537624-DEL-6716
chr8-31552808-DEL-196
chr8-31560979-DEL-100
chr8-31574361-DEL-143
chr8-31587675-DEL-73
chr8-31591817-DEL-966
chr8-31626272-DEL-200
chr8-31632242-DEL-115
chr8-31634859-DEL-60
chr8-31663037-DEL-2545
chr8-31682955-DEL-57
chr8-31683123-DEL-1145
chr8-31717092-DEL-1004
chr8-31719860-DEL-56
chr8-31774218-DEL-73
chr8-31787923-DEL-193
chr8-31801963-DEL-666
chr8-31815704-DEL-195
chr8-31827862-DEL-64
chr8-31834717-DEL-2837
chr8-31872613-DEL-8335
chr8-31872684-DEL-2270
chr8-31892745-DEL-1138
chr8-31900501-DEL-78
chr8-31910351-DEL-316
chr8-31943871-DEL-88
chr8-31958030-DEL-731
chr8-31962185-DEL-83
chr8-31965681-DEL-1271
chr8-31975838-DEL-108
chr8-31990833-DEL-232
chr8-32040111-DEL-62
chr8-32046436-DEL-2591
chr8-32092031-DEL-1350
chr8-32139688-DEL-52
chr8-32150002-DEL-67
chr8-32178109-DEL-1340
chr8-32184505-DEL-117
chr8-32186264-DEL-276
chr8-32197575-DEL-220
chr8-32235418-DEL-52
chr8-32242948-DEL-87
chr8-32243778-DEL-60
chr8-32248202-DEL-80
chr8-32257539-DEL-55
chr8-32278158-DEL-2033
chr8-32280304-DEL-531
chr8-32280588-DEL-729
chr8-32281906-DEL-330
chr8-32354574-DEL-64
chr8-32398785-DEL-78
chr8-32445203-DEL-60
chr8-32456285-DEL-65
chr8-32466347-DEL-53
chr8-32478634-DEL-52
chr8-32478660-DEL-1088
chr8-32530389-DEL-75
chr8-32569522-DEL-1120
chr8-32584698-DEL-66
chr8-32588371-DEL-77
chr8-32702644-DEL-52
chr8-32744448-DEL-1777
chr8-32744458-DEL-464
chr8-32744997-DEL-1228
chr8-32792351-DEL-209
chr8-32817085-DEL-90
chr8-32838662-DEL-133
chr8-32944352-DEL-54
chr8-32962563-DEL-52
chr8-32972950-DEL-1347
chr8-32988336-DEL-2126
chr8-32999334-DEL-192

chr8-33001899-DEL-140
chr8-33017095-DEL-94
chr8-33020760-DEL-304
chr8-33106955-DEL-59
chr8-33128055-DEL-270
chr8-33143683-DEL-191
chr8-33172193-DEL-82
chr8-33177517-DEL-99
chr8-33188685-DEL-375
chr8-33199339-DEL-80
chr8-33252906-DEL-72
chr8-33268547-DEL-100
chr8-33269574-DEL-64
chr8-33278475-DEL-249
chr8-33300591-DEL-54
chr8-33312243-DEL-301
chr8-33337132-DEL-51
chr8-33338262-DEL-52
chr8-33356688-DEL-155
chr8-33366357-DEL-277
chr8-33399503-DEL-52
chr8-33404945-DEL-88
chr8-33465746-DEL-242
chr8-33488459-DEL-79
chr8-33523022-DEL-62
chr8-33525587-DEL-177
chr8-33533619-DEL-101
chr8-33536585-DEL-319
chr8-33538172-DEL-74
chr8-33555031-DEL-74
chr8-33565478-DEL-85
chr8-33575049-DEL-108
chr8-33582996-DEL-64
chr8-33597024-DEL-53
chr8-33650920-DEL-71
chr8-33654818-DEL-54
chr8-33657854-DEL-6358
chr8-33665712-DEL-1746
chr8-33671768-DEL-775
chr8-33687084-DEL-59
chr8-33687546-DEL-435
chr8-33691625-DEL-1190
chr8-33692836-DEL-1214
chr8-33696389-DEL-853
chr8-33697338-DEL-3397
chr8-33701033-DEL-158
chr8-33713545-DEL-6637
chr8-33721334-DEL-1462
chr8-33739338-DEL-216
chr8-33754031-DEL-460
chr8-33771715-DEL-393
chr8-33783440-DEL-114
chr8-33829170-DEL-72
chr8-33829330-DEL-1244
chr8-33845284-DEL-415
chr8-33849513-DEL-336
chr8-33859862-DEL-4751
chr8-33894820-DEL-64
chr8-33910432-DEL-96
chr8-33919535-DEL-1637
chr8-33935008-DEL-1837
chr8-33965247-DEL-434
chr8-33971227-DEL-305
chr8-34004816-DEL-152
chr8-34008282-DEL-58
chr8-34022009-DEL-2722
chr8-34051457-DEL-720
chr8-34052254-DEL-114
chr8-34057198-DEL-1816
chr8-34058428-DEL-1066
chr8-34129692-DEL-118
chr8-34148466-DEL-172
chr8-34176019-DEL-653
chr8-34233008-DEL-716
chr8-34245986-DEL-210
chr8-34290434-DEL-114
chr8-34390248-DEL-3709

chr8-34393219-DEL-143
chr8-34401648-DEL-179
chr8-34439167-DEL-283
chr8-34449250-DEL-174
chr8-34479150-DEL-71
chr8-34485881-DEL-588
chr8-34494438-DEL-220
chr8-34510949-DEL-56
chr8-34546698-DEL-63
chr8-34554982-DEL-2037
chr8-34562461-DEL-242
chr8-34569675-DEL-130
chr8-34625732-DEL-197
chr8-34626387-DEL-75
chr8-34629985-DEL-213
chr8-34659514-DEL-53
chr8-34660878-DEL-53
chr8-34696586-DEL-131
chr8-34715586-DEL-59
chr8-34716725-DEL-135
chr8-34740800-DEL-566
chr8-34753189-DEL-590
chr8-34758418-DEL-145
chr8-34762891-DEL-492
chr8-34792408-DEL-422
chr8-34806593-DEL-60
chr8-34875048-DEL-133
chr8-34903369-DEL-51
chr8-34964182-DEL-905
chr8-34975953-DEL-233
chr8-34978605-DEL-1746
chr8-34979281-DEL-225
chr8-34999025-DEL-378
chr8-35007042-DEL-85
chr8-35034089-DEL-68
chr8-35051389-DEL-63
chr8-35091497-DEL-298
chr8-35132130-DEL-1460
chr8-35137491-DEL-79
chr8-35163890-DEL-57
chr8-35164544-DEL-1101
chr8-35169132-DEL-1190
chr8-35182829-DEL-383
chr8-35187621-DEL-60
chr8-35193877-DEL-53
chr8-35204204-DEL-54
chr8-35213286-DEL-734
chr8-35231612-DEL-55
chr8-35289463-DEL-200
chr8-35340869-DEL-64
chr8-35368349-DEL-54
chr8-35411266-DEL-402
chr8-35439328-DEL-67
chr8-35448377-DEL-146
chr8-35455288-DEL-4165
chr8-35478182-DEL-55
chr8-35493025-DEL-74
chr8-35525803-DEL-79
chr8-35544747-DEL-184
chr8-35549700-DEL-54
chr8-35570676-DEL-405
chr8-35629476-DEL-897
chr8-35635689-DEL-51
chr8-35674664-DEL-97
chr8-35686157-DEL-93
chr8-35704877-DEL-53
chr8-35733657-DEL-1550
chr8-35737565-DEL-52
chr8-35763120-DEL-67
chr8-35771179-DEL-95
chr8-35785153-DEL-87
chr8-35791231-DEL-149
chr8-35792203-DEL-160
chr8-35798217-DEL-68
chr8-35813576-DEL-51
chr8-35814969-DEL-66
chr8-35863521-DEL-209

chr8-35920818-DEL-62
chr8-35939939-DEL-1600
chr8-35990970-DEL-66
chr8-35992959-DEL-1052
chr8-35994088-DEL-80
chr8-36027481-DEL-101
chr8-36039941-DEL-864
chr8-36040006-DEL-105
chr8-36042902-DEL-130
chr8-36045877-DEL-270
chr8-36063079-DEL-74
chr8-36064889-DEL-65
chr8-36068778-DEL-300
chr8-36123281-DEL-124
chr8-36156805-DEL-260
chr8-36164496-DEL-126
chr8-36225848-DEL-239
chr8-36233845-DEL-137
chr8-36241279-DEL-78
chr8-36312450-DEL-70
chr8-36377283-DEL-98
chr8-36379066-DEL-351
chr8-36433167-DEL-96
chr8-36478926-DEL-1129
chr8-36488812-DEL-177
chr8-36490039-DEL-230
chr8-36492784-DEL-534
chr8-36525566-DEL-72
chr8-36528177-DEL-119
chr8-36534536-DEL-59
chr8-36546554-DEL-55
chr8-36636270-DEL-153
chr8-36690451-DEL-71
chr8-36700666-DEL-60
chr8-36712238-DEL-75
chr8-36732804-DEL-136
chr8-36752981-DEL-888
chr8-36775000-DEL-68
chr8-36778255-DEL-54
chr8-36788476-DEL-150
chr8-36794652-DEL-98
chr8-36802161-DEL-773
chr8-36815350-DEL-2665
chr8-36837483-DEL-936
chr8-36866998-DEL-85
chr8-36871652-DEL-495
chr8-36973536-DEL-1657
chr8-36984833-DEL-99
chr8-36989422-DEL-55
chr8-36999112-DEL-166
chr8-37021301-DEL-414
chr8-37043023-DEL-934
chr8-37062006-DEL-172
chr8-37064668-DEL-84
chr8-37189266-DEL-74
chr8-37191655-DEL-68
chr8-37205894-DEL-56
chr8-37224090-DEL-60
chr8-37227808-DEL-1814
chr8-37239151-DEL-221
chr8-37245711-DEL-52
chr8-37263314-DEL-66
chr8-37268003-DEL-1372
chr8-37289374-DEL-90
chr8-37345341-DEL-1209
chr8-37513850-DEL-2385
chr8-37550209-DEL-74
chr8-37560005-DEL-193
chr8-37579718-DEL-60
chr8-37583401-DEL-102
chr8-37586930-DEL-955
chr8-37588212-DEL-1641
chr8-37635760-DEL-68
chr8-37638104-DEL-95
chr8-37646570-DEL-323
chr8-37681754-DEL-58
chr8-37709584-DEL-2486

chr8-37735753-DEL-440
chr8-37739866-DEL-87
chr8-37761664-DEL-67
chr8-37769466-DEL-54
chr8-37770727-DEL-67
chr8-37771964-DEL-51
chr8-37773712-DEL-68
chr8-37794078-DEL-1271
chr8-37799547-DEL-1430
chr8-37807199-DEL-75
chr8-37840687-DEL-75
chr8-37852351-DEL-101
chr8-37873807-DEL-220
chr8-37895361-DEL-56
chr8-37911081-DEL-1419
chr8-37912870-DEL-1129
chr8-37937224-DEL-498
chr8-37948085-DEL-7883
chr8-37953662-DEL-1464
chr8-37954217-DEL-543
chr8-37979619-DEL-960
chr8-38066856-DEL-1905
chr8-38089719-DEL-2800
chr8-38091440-DEL-240
chr8-38094407-DEL-2777
chr8-38094415-DEL-6514
chr8-38099026-DEL-3200
chr8-38197161-DEL-1732
chr8-38264389-DEL-109
chr8-38267044-DEL-1289
chr8-38312573-DEL-6348
chr8-38342770-DEL-94
chr8-38361116-DEL-435
chr8-38367042-DEL-162
chr8-38383333-DEL-1236
chr8-38389906-DEL-123
chr8-38404667-DEL-110
chr8-38425089-DEL-118
chr8-38473237-DEL-108
chr8-38478714-DEL-901
chr8-38484882-DEL-971
chr8-38508351-DEL-55
chr8-38523267-DEL-71
chr8-38546249-DEL-418
chr8-38549615-DEL-60
chr8-38578607-DEL-1140
chr8-38600015-DEL-274
chr8-38614167-DEL-3229
chr8-38640093-DEL-1368
chr8-38660235-DEL-792
chr8-38674510-DEL-2099
chr8-38678207-DEL-2283
chr8-38709640-DEL-471
chr8-38727169-DEL-4283
chr8-38734863-DEL-2133
chr8-38735023-DEL-51
chr8-38749141-DEL-154
chr8-38753313-DEL-68
chr8-38779701-DEL-326
chr8-38785420-DEL-72
chr8-38832508-DEL-107
chr8-38875332-DEL-6397
chr8-38885415-DEL-940
chr8-38894670-DEL-107
chr8-38958197-DEL-312
chr8-39032468-DEL-67
chr8-39084557-DEL-358
chr8-39136154-DEL-3992
chr8-39148785-DEL-730
chr8-39156370-DEL-58
chr8-39166559-DEL-65
chr8-39179372-DEL-6145
chr8-39219274-DEL-2312
chr8-39263864-DEL-60
chr8-39266144-DEL-68
chr8-39273978-DEL-57
chr8-39275254-DEL-1412

chr8-39276864-DEL-179
chr8-39295017-DEL-6174
chr8-39307203-DEL-156
chr8-39319980-DEL-3140
chr8-39329192-DEL-442
chr8-39339340-DEL-176
chr8-39341420-DEL-144
chr8-39352160-DEL-66
chr8-39386872-DEL-536
chr8-39404049-DEL-64
chr8-39407346-DEL-2042
chr8-39419886-DEL-6400
chr8-39440106-DEL-319
chr8-39456109-DEL-472
chr8-39508633-DEL-55
chr8-39512368-DEL-238
chr8-39526529-DEL-54
chr8-39549463-DEL-1401
chr8-39567803-DEL-53
chr8-39599472-DEL-64
chr8-39600051-DEL-253
chr8-39600054-DEL-197
chr8-39600180-DEL-8847
chr8-39600476-DEL-125
chr8-39601117-DEL-6570
chr8-39601189-DEL-175
chr8-39601641-DEL-258
chr8-39602160-DEL-368
chr8-39602609-DEL-106
chr8-39602897-DEL-4353
chr8-39602989-DEL-4371
chr8-39603470-DEL-3312
chr8-39607464-DEL-253
chr8-39607986-DEL-101
chr8-39608485-DEL-77
chr8-39611427-DEL-129
chr8-39627182-DEL-2186
chr8-39635900-DEL-886
chr8-39639288-DEL-66
chr8-39644946-DEL-2108
chr8-39654971-DEL-862
chr8-39669819-DEL-569
chr8-39689469-DEL-181
chr8-39791952-DEL-190
chr8-39819960-DEL-274
chr8-39843103-DEL-64
chr8-39845880-DEL-6550
chr8-39847158-DEL-545
chr8-39892616-DEL-108
chr8-39922848-DEL-622
chr8-39961835-DEL-409
chr8-39970996-DEL-6227
chr8-39977750-DEL-53
chr8-39989804-DEL-92
chr8-40016783-DEL-126
chr8-40021081-DEL-56
chr8-40027737-DEL-160
chr8-40042969-DEL-735
chr8-40062187-DEL-377
chr8-40091825-DEL-74
chr8-40103554-DEL-288
chr8-40122244-DEL-562
chr8-40134782-DEL-58
chr8-40139301-DEL-194
chr8-40221301-DEL-411
chr8-40235857-DEL-62
chr8-40236746-DEL-195
chr8-40263828-DEL-100
chr8-40266798-DEL-353
chr8-40303122-DEL-77
chr8-40341330-DEL-138
chr8-40341473-DEL-162
chr8-40341883-DEL-1508
chr8-40343481-DEL-255
chr8-40347530-DEL-183
chr8-40384496-DEL-335
chr8-40462115-DEL-1553

chr8-40464512-DEL-669
chr8-40478892-DEL-189
chr8-40489233-DEL-2504
chr8-40557097-DEL-61
chr8-40592656-DEL-4880
chr8-40631849-DEL-121
chr8-40811253-DEL-118
chr8-40821559-DEL-197
chr8-40846179-DEL-73
chr8-40852521-DEL-58
chr8-40867342-DEL-57
chr8-40928064-DEL-168
chr8-40951253-DEL-201
chr8-40999685-DEL-89
chr8-41095986-DEL-194
chr8-41098163-DEL-443
chr8-41124536-DEL-542
chr8-41148359-DEL-2209
chr8-41153356-DEL-172
chr8-41163095-DEL-836
chr8-41165696-DEL-858
chr8-41181867-DEL-172
chr8-41183639-DEL-1671
chr8-41223678-DEL-2124
chr8-41256093-DEL-201
chr8-41263829-DEL-415
chr8-41290762-DEL-6881
chr8-41336400-DEL-1048
chr8-41385567-DEL-63
chr8-41406290-DEL-4343
chr8-41411513-DEL-2779
chr8-41531145-DEL-61
chr8-41624344-DEL-75
chr8-41647553-DEL-77
chr8-41674139-DEL-77
chr8-41691151-DEL-62
chr8-41713234-DEL-89
chr8-41772950-DEL-54
chr8-41780222-DEL-60
chr8-41783021-DEL-204
chr8-41791464-DEL-791
chr8-41822997-DEL-61
chr8-41883229-DEL-136
chr8-41894793-DEL-85
chr8-41903603-DEL-173
chr8-41914141-DEL-68
chr8-41918067-DEL-6034
chr8-41925697-DEL-154
chr8-41943162-DEL-152
chr8-41967470-DEL-186
chr8-41974333-DEL-177
chr8-41986679-DEL-63
chr8-41996313-DEL-370
chr8-41996403-DEL-217
chr8-42051938-DEL-927
chr8-42054476-DEL-84
chr8-42076650-DEL-69
chr8-42094659-DEL-183
chr8-42154098-DEL-432
chr8-42177245-DEL-6367
chr8-42261310-DEL-364
chr8-42356282-DEL-772
chr8-42384964-DEL-51
chr8-42388705-DEL-126
chr8-42389005-DEL-937
chr8-42394512-DEL-204
chr8-42433895-DEL-283
chr8-42446375-DEL-441
chr8-42452375-DEL-162
chr8-42510906-DEL-4204
chr8-42515256-DEL-2659
chr8-42528059-DEL-6174
chr8-42539453-DEL-77
chr8-42556991-DEL-58
chr8-42599380-DEL-696
chr8-42606681-DEL-86
chr8-42607361-DEL-63

chr8-42654926-DEL-1450
chr8-42692337-DEL-167
chr8-42695688-DEL-109
chr8-42761960-DEL-108
chr8-42786846-DEL-99
chr8-42788916-DEL-82
chr8-42792051-DEL-955
chr8-42811250-DEL-130
chr8-42811516-DEL-2271
chr8-42836228-DEL-80
chr8-42869495-DEL-2503
chr8-42872081-DEL-374
chr8-42876184-DEL-208
chr8-42905058-DEL-1302
chr8-42910908-DEL-71
chr8-42919451-DEL-470
chr8-42950593-DEL-66
chr8-42975269-DEL-84
chr8-42979455-DEL-59
chr8-42983416-DEL-122
chr8-43013469-DEL-5123
chr8-43022891-DEL-6350
chr8-43032051-DEL-6958
chr8-43087927-DEL-93
chr8-43099450-DEL-189
chr8-43104654-DEL-59
chr8-43113494-DEL-262
chr8-43130791-DEL-146
chr8-43164690-DEL-78
chr8-43168270-DEL-249
chr8-43199529-DEL-866
chr8-43229900-DEL-1071
chr8-43243501-DEL-81
chr8-43252368-DEL-1486
chr8-43254703-DEL-67
chr8-43291800-DEL-4516
chr8-43296618-DEL-415
chr8-43299416-DEL-173
chr8-43301804-DEL-6360
chr8-43310489-DEL-54
chr8-43311093-DEL-6378
chr8-43388411-DEL-64
chr8-43414723-DEL-60
chr8-43557110-DEL-56
chr8-43609824-DEL-104
chr8-43631416-DEL-766
chr8-43635453-DEL-103
chr8-43692360-DEL-726
chr8-43733592-DEL-59
chr8-43764503-DEL-122
chr8-43774550-DEL-78
chr8-43782082-DEL-57
chr8-43815617-DEL-66
chr8-43836705-DEL-467
chr8-43837564-DEL-67
chr8-43862339-DEL-92
chr8-43864756-DEL-68
chr8-43879031-DEL-400
chr8-43919105-DEL-1648
chr8-43939970-DEL-73
chr8-43959981-DEL-602
chr8-43972533-DEL-325
chr8-43989788-DEL-6418
chr8-44007197-DEL-538
chr8-44010371-DEL-668
chr8-44015432-DEL-175
chr8-44020972-DEL-234
chr8-44050468-DEL-6498
chr8-44072086-DEL-154
chr8-44073189-DEL-25798
chr8-44092599-DEL-6388
chr8-44103100-DEL-161
chr8-44109800-DEL-6167
chr8-44132399-DEL-12616
chr8-44159297-DEL-867
chr8-44161170-DEL-2380
chr8-44193748-DEL-4220

chr8-44198184-DEL-287
chr8-44206523-DEL-941
chr8-44222131-DEL-4045
chr8-44226422-DEL-1410
chr8-44263795-DEL-2427
chr8-44269895-DEL-7472
chr8-44270592-DEL-1996
chr8-44287164-DEL-112
chr8-44304803-DEL-86
chr8-44331762-DEL-144
chr8-44341717-DEL-56
chr8-44353794-DEL-367
chr8-44365604-DEL-56
chr8-44424710-DEL-6313
chr8-44426558-DEL-1098
chr8-44433443-DEL-58
chr8-44434730-DEL-228
chr8-44447778-DEL-112
chr8-44464180-DEL-885
chr8-44475972-DEL-66
chr8-44476796-DEL-57
chr8-44486069-DEL-297
chr8-44492300-DEL-90
chr8-44508405-DEL-1168
chr8-44580276-DEL-56
chr8-44589050-DEL-164
chr8-44601161-DEL-107
chr8-44603732-DEL-74
chr8-44621765-DEL-66
chr8-44641042-DEL-52
chr8-44685272-DEL-891
chr8-44686874-DEL-6923
chr8-44695965-DEL-6239
chr8-44711261-DEL-106
chr8-44728868-DEL-57
chr8-44745462-DEL-54
chr8-44816198-DEL-130
chr8-44853920-DEL-72
chr8-44859763-DEL-577
chr8-44932623-DEL-131
chr8-44985052-DEL-58
chr8-45008104-DEL-2697
chr8-45036142-DEL-2732
chr8-45038900-DEL-3503
chr8-45078820-DEL-58
chr8-45086574-DEL-347
chr8-45135950-DEL-361
chr8-45175599-DEL-336
chr8-45197659-DEL-97
chr8-45290962-DEL-117
chr8-45296251-DEL-2486
chr8-45305049-DEL-116
chr8-45305739-DEL-67
chr8-45309403-DEL-6007
chr8-45318396-DEL-54
chr8-45378809-DEL-5333
chr8-45470701-DEL-90
chr8-45490811-DEL-162
chr8-45518012-DEL-53
chr8-45521046-DEL-83
chr8-45534442-DEL-78
chr8-45553037-DEL-70
chr8-45557214-DEL-1111
chr8-45580500-DEL-1114
chr8-45620135-DEL-60
chr8-45625669-DEL-161
chr8-45667381-DEL-7215
chr8-45686377-DEL-671
chr8-45740330-DEL-184
chr8-45741053-DEL-56
chr8-45767339-DEL-774
chr8-45866595-DEL-214
chr8-45874248-DEL-55
chr8-45875037-DEL-6435
chr8-45883898-DEL-55
chr8-45884267-DEL-1399
chr8-45895948-DEL-4051

chr8-45916050-DEL-1196
chr8-45918309-DEL-163
chr8-45924705-DEL-543
chr8-45927709-DEL-538
chr8-45965821-DEL-252
chr8-45978020-DEL-736
chr8-45996899-DEL-73
chr8-46004342-DEL-74
chr8-46046193-DEL-64
chr8-46084237-DEL-112
chr8-46101741-DEL-65
chr8-46107539-DEL-194
chr8-46155716-DEL-65
chr8-46202612-DEL-107
chr8-46238517-DEL-257
chr8-46288814-DEL-233
chr8-46299760-DEL-107
chr8-46309278-DEL-130
chr8-46323902-DEL-61
chr8-46365320-DEL-122
chr8-46374639-DEL-73
chr8-46384183-DEL-395
chr8-46401020-DEL-1285
chr8-46419471-DEL-359
chr8-46441539-DEL-103
chr8-46461707-DEL-166
chr8-46474586-DEL-109
chr8-46476381-DEL-53
chr8-46480020-DEL-63
chr8-46488310-DEL-156
chr8-46505729-DEL-88
chr8-46514049-DEL-171
chr8-46514924-DEL-76
chr8-46528588-DEL-1240
chr8-46530120-DEL-216
chr8-46566736-DEL-367
chr8-46674095-DEL-94
chr8-46674716-DEL-628
chr8-46688778-DEL-56
chr8-46705681-DEL-2270
chr8-46714037-DEL-244
chr8-46721150-DEL-243
chr8-46797027-DEL-111
chr8-46804185-DEL-72
chr8-46809866-DEL-3181
chr8-46831459-DEL-61
chr8-46852094-DEL-245
chr8-46871988-DEL-114
chr8-46876912-DEL-165
chr8-46907485-DEL-60
chr8-46934646-DEL-106
chr8-46969730-DEL-106
chr8-47047234-DEL-388
chr8-47050253-DEL-74
chr8-47125556-DEL-137
chr8-47149092-DEL-68
chr8-47151867-DEL-195
chr8-47157658-DEL-206
chr8-47161531-DEL-112
chr8-47167356-DEL-90
chr8-47181649-DEL-1558
chr8-47183682-DEL-93
chr8-47200277-DEL-110
chr8-47218929-DEL-58
chr8-47221419-DEL-63
chr8-47232940-DEL-136
chr8-47274939-DEL-606
chr8-47335593-DEL-132
chr8-47365241-DEL-57
chr8-47379489-DEL-144
chr8-47438318-DEL-620
chr8-47442372-DEL-58
chr8-47456830-DEL-722
chr8-47457580-DEL-66
chr8-47457713-DEL-244
chr8-47459309-DEL-109
chr8-47499923-DEL-147

chr8-47500690-DEL-172
chr8-47503962-DEL-52
chr8-47536931-DEL-5539
chr8-47618216-DEL-186
chr8-47618321-DEL-2210
chr8-47626636-DEL-869
chr8-47654013-DEL-202
chr8-47657647-DEL-163
chr8-47707186-DEL-134
chr8-47727712-DEL-597
chr8-47749682-DEL-88
chr8-47751342-DEL-79
chr8-47763367-DEL-76
chr8-47785017-DEL-327
chr8-47816813-DEL-119
chr8-47821516-DEL-284
chr8-47826070-DEL-173
chr8-47828692-DEL-60
chr8-47847753-DEL-127
chr8-47876604-DEL-115
chr8-47889182-DEL-153
chr8-47919215-DEL-77
chr8-47925890-DEL-77
chr8-47944307-DEL-84
chr8-47953010-DEL-71
chr8-47963380-DEL-452
chr8-48055826-DEL-51
chr8-48063365-DEL-178
chr8-48084607-DEL-204
chr8-48085259-DEL-594
chr8-48086132-DEL-1007
chr8-48087209-DEL-419
chr8-48087952-DEL-208
chr8-48100077-DEL-96
chr8-48115109-DEL-208
chr8-48117179-DEL-51
chr8-48167405-DEL-51
chr8-48182616-DEL-59
chr8-48202345-DEL-62
chr8-48211267-DEL-927
chr8-48232299-DEL-209
chr8-48244715-DEL-95
chr8-48256342-DEL-91
chr8-48258313-DEL-1085
chr8-48261323-DEL-1652
chr8-48288988-DEL-966
chr8-48289570-DEL-147
chr8-48295305-DEL-172
chr8-48303503-DEL-54
chr8-48326203-DEL-280
chr8-48331711-DEL-60
chr8-48332972-DEL-52
chr8-48340547-DEL-149
chr8-48347958-DEL-54
chr8-48362521-DEL-93
chr8-48374227-DEL-302
chr8-48394482-DEL-53
chr8-48396983-DEL-1832
chr8-48407732-DEL-312
chr8-48408411-DEL-56
chr8-48426202-DEL-193
chr8-48429411-DEL-238
chr8-48446547-DEL-573
chr8-48487196-DEL-196
chr8-48495572-DEL-222
chr8-48531911-DEL-503
chr8-48541781-DEL-94
chr8-48564216-DEL-75
chr8-48565362-DEL-56
chr8-48576881-DEL-504
chr8-48582007-DEL-1597
chr8-48598525-DEL-167
chr8-48610038-DEL-52
chr8-48621766-DEL-75
chr8-48622007-DEL-344
chr8-48627978-DEL-855
chr8-48629079-DEL-91

chr8-48631533-DEL-53
chr8-48639677-DEL-197
chr8-48695400-DEL-58
chr8-48699932-DEL-969
chr8-48700803-DEL-237
chr8-48784810-DEL-84
chr8-48812095-DEL-1501
chr8-48842854-DEL-132
chr8-48869098-DEL-109
chr8-48894616-DEL-93
chr8-48896842-DEL-92
chr8-48931403-DEL-482
chr8-48935853-DEL-151
chr8-48948498-DEL-381
chr8-48992609-DEL-179
chr8-49018553-DEL-656
chr8-49025444-DEL-86
chr8-49036302-DEL-81
chr8-49054976-DEL-403
chr8-49059631-DEL-81
chr8-49155872-DEL-166
chr8-49178073-DEL-59
chr8-49203862-DEL-53
chr8-49230574-DEL-76
chr8-49285679-DEL-70
chr8-49291992-DEL-84
chr8-49302102-DEL-54
chr8-49322486-DEL-93
chr8-49429992-DEL-146
chr8-49486188-DEL-63
chr8-49487756-DEL-87
chr8-49488329-DEL-56
chr8-49548266-DEL-78
chr8-49576717-DEL-236
chr8-49645876-DEL-64
chr8-49652454-DEL-52
chr8-49686187-DEL-55
chr8-49727954-DEL-70
chr8-49730942-DEL-658
chr8-49821053-DEL-253
chr8-49822364-DEL-156
chr8-49823005-DEL-184
chr8-49831415-DEL-88
chr8-49869701-DEL-54
chr8-49902408-DEL-162
chr8-49924567-DEL-210
chr8-49955053-DEL-273
chr8-49976833-DEL-396
chr8-50011885-DEL-1982
chr8-50068749-DEL-610
chr8-50163763-DEL-103
chr8-50180203-DEL-224
chr8-50190123-DEL-296
chr8-50198926-DEL-479
chr8-50256409-DEL-56
chr8-50268825-DEL-2876
chr8-50296447-DEL-1256
chr8-50299300-DEL-78
chr8-50303737-DEL-1760
chr8-50312489-DEL-12143
chr8-50314863-DEL-55
chr8-50337564-DEL-127
chr8-50398659-DEL-142
chr8-50420675-DEL-265
chr8-50457363-DEL-103
chr8-50460671-DEL-100
chr8-50470857-DEL-67
chr8-50477459-DEL-163
chr8-50520838-DEL-88
chr8-50547035-DEL-335
chr8-50547904-DEL-62
chr8-50558456-DEL-863
chr8-50566856-DEL-369
chr8-50578994-DEL-72
chr8-50586799-DEL-86
chr8-50624107-DEL-59
chr8-50635687-DEL-152

chr8-50640187-DEL-58
chr8-50641454-DEL-52
chr8-50764245-DEL-203
chr8-50772920-DEL-203
chr8-50780822-DEL-264
chr8-50802927-DEL-2507
chr8-50861528-DEL-567
chr8-50892172-DEL-236
chr8-50898435-DEL-490
chr8-50910766-DEL-435
chr8-50915134-DEL-594
chr8-50921682-DEL-109
chr8-50926915-DEL-82
chr8-50927671-DEL-162
chr8-50946303-DEL-202
chr8-50958601-DEL-431
chr8-50972842-DEL-88
chr8-50992009-DEL-320
chr8-51030595-DEL-173
chr8-51039522-DEL-79
chr8-51043490-DEL-307
chr8-51075665-DEL-74
chr8-51122545-DEL-1254
chr8-51247143-DEL-66
chr8-51255094-DEL-70
chr8-51266827-DEL-435
chr8-51296029-DEL-261
chr8-51300860-DEL-1108
chr8-51312705-DEL-880
chr8-51334317-DEL-379
chr8-51384513-DEL-224
chr8-51385781-DEL-6211
chr8-51406159-DEL-688
chr8-51416047-DEL-284
chr8-51428275-DEL-86
chr8-51455516-DEL-1334
chr8-51462923-DEL-64
chr8-51545926-DEL-174
chr8-51582796-DEL-929
chr8-51585424-DEL-331
chr8-51587102-DEL-855
chr8-51608758-DEL-51
chr8-51615965-DEL-525
chr8-51621895-DEL-54
chr8-51643455-DEL-571
chr8-51671834-DEL-55
chr8-51679757-DEL-150
chr8-51682792-DEL-1037
chr8-51753729-DEL-366
chr8-51763178-DEL-371
chr8-51775908-DEL-13667
chr8-51780331-DEL-149
chr8-51782929-DEL-52
chr8-51798876-DEL-303
chr8-51799676-DEL-212
chr8-51800260-DEL-212
chr8-51837365-DEL-77
chr8-51870724-DEL-840
chr8-51929693-DEL-181
chr8-51941524-DEL-424
chr8-51943880-DEL-53
chr8-51948206-DEL-79
chr8-51950782-DEL-83
chr8-51958179-DEL-1179
chr8-51975369-DEL-54
chr8-52015860-DEL-58
chr8-52032236-DEL-6319
chr8-52061509-DEL-1949
chr8-52073750-DEL-421
chr8-52115338-DEL-1200
chr8-52152900-DEL-2537
chr8-52161794-DEL-58
chr8-52204541-DEL-271
chr8-52253180-DEL-115
chr8-52272184-DEL-56
chr8-52310861-DEL-57
chr8-52318960-DEL-317

chr8-52326827-DEL-106
chr8-52331558-DEL-884
chr8-52371078-DEL-98
chr8-52390585-DEL-72
chr8-52410336-DEL-190
chr8-52445926-DEL-1492
chr8-52461936-DEL-2773
chr8-52469212-DEL-2182
chr8-52514492-DEL-201
chr8-52522506-DEL-2058
chr8-52550385-DEL-2662
chr8-52555998-DEL-196
chr8-52557039-DEL-1124
chr8-52582751-DEL-62
chr8-52611142-DEL-60
chr8-52618228-DEL-441
chr8-52622085-DEL-5112
chr8-52637264-DEL-1474
chr8-52643447-DEL-1565
chr8-52645076-DEL-1990
chr8-52661045-DEL-53
chr8-52670585-DEL-427
chr8-52708699-DEL-185
chr8-52737978-DEL-6177
chr8-52747670-DEL-74
chr8-52757433-DEL-1484
chr8-52757696-DEL-424
chr8-52761079-DEL-213
chr8-52791031-DEL-228
chr8-52796306-DEL-413
chr8-52799584-DEL-78
chr8-52802944-DEL-52
chr8-52805330-DEL-548
chr8-52820611-DEL-54
chr8-52835389-DEL-65
chr8-52843719-DEL-82
chr8-52853850-DEL-621
chr8-52872871-DEL-5157
chr8-52959286-DEL-1706
chr8-53018858-DEL-144
chr8-53035854-DEL-17150
chr8-53065235-DEL-2436
chr8-53072791-DEL-79
chr8-53132464-DEL-2285
chr8-53154575-DEL-103
chr8-53210202-DEL-2554
chr8-53210415-DEL-2954
chr8-53216562-DEL-60
chr8-53249299-DEL-204
chr8-53259334-DEL-1197
chr8-53266914-DEL-300
chr8-53275803-DEL-957
chr8-53281207-DEL-1419
chr8-53289195-DEL-1030
chr8-53315335-DEL-296
chr8-53321292-DEL-170
chr8-53324599-DEL-242
chr8-53379749-DEL-738
chr8-53386133-DEL-52
chr8-53392005-DEL-254
chr8-53409480-DEL-6402
chr8-53409542-DEL-13605
chr8-53424845-DEL-72
chr8-53436432-DEL-69
chr8-53439043-DEL-6040
chr8-53524779-DEL-553
chr8-53525897-DEL-506
chr8-53559073-DEL-289
chr8-53563416-DEL-78
chr8-53584418-DEL-6221
chr8-53661705-DEL-1127
chr8-53664023-DEL-184
chr8-53668484-DEL-523
chr8-53732898-DEL-150
chr8-53744491-DEL-398
chr8-53803231-DEL-55
chr8-53888160-DEL-79

chr8-53889142-DEL-142
chr8-53930860-DEL-76
chr8-53937147-DEL-53
chr8-53967520-DEL-55
chr8-53984972-DEL-72
chr8-53985985-DEL-627
chr8-54003131-DEL-379
chr8-54011617-DEL-1324
chr8-54018797-DEL-136
chr8-54043130-DEL-52
chr8-54045944-DEL-843
chr8-54046192-DEL-56
chr8-54049785-DEL-62
chr8-54050942-DEL-336
chr8-54054600-DEL-191
chr8-54079162-DEL-655
chr8-54125717-DEL-65
chr8-54207385-DEL-451
chr8-54210376-DEL-992
chr8-54229831-DEL-109
chr8-54238844-DEL-736
chr8-54239741-DEL-137
chr8-54249270-DEL-1459
chr8-54278732-DEL-182
chr8-54299801-DEL-312
chr8-54307751-DEL-1933
chr8-54309593-DEL-305
chr8-54376019-DEL-1124
chr8-54391663-DEL-148
chr8-54425259-DEL-6247
chr8-54451235-DEL-248
chr8-54466677-DEL-205
chr8-54486541-DEL-230
chr8-54494369-DEL-486
chr8-54499563-DEL-1036
chr8-54512713-DEL-135
chr8-54516365-DEL-178
chr8-54535614-DEL-100
chr8-54536578-DEL-827
chr8-54559690-DEL-53
chr8-54583478-DEL-119
chr8-54594301-DEL-114
chr8-54604452-DEL-873
chr8-54619088-DEL-53
chr8-54631931-DEL-136
chr8-54666567-DEL-987
chr8-54690587-DEL-264
chr8-54694975-DEL-7135
chr8-54700341-DEL-200
chr8-54715615-DEL-74
chr8-54728219-DEL-52
chr8-54729569-DEL-3617
chr8-54749930-DEL-4711
chr8-54751088-DEL-1848
chr8-54757660-DEL-197
chr8-54766953-DEL-413
chr8-54781271-DEL-807
chr8-54805330-DEL-73
chr8-54866295-DEL-220
chr8-54879316-DEL-247
chr8-54912558-DEL-1856
chr8-54932044-DEL-70
chr8-54989459-DEL-656
chr8-55049390-DEL-120
chr8-55069054-DEL-318
chr8-55069782-DEL-6344
chr8-55070090-DEL-117
chr8-55071595-DEL-5282
chr8-55077116-DEL-1357
chr8-55080377-DEL-348
chr8-55081755-DEL-195
chr8-55090354-DEL-210
chr8-55160124-DEL-206
chr8-55188673-DEL-62
chr8-55207057-DEL-608
chr8-55238512-DEL-1918
chr8-55242214-DEL-63

chr8-55312271-DEL-85
chr8-55331911-DEL-62
chr8-55345932-DEL-451
chr8-55350391-DEL-196
chr8-55371821-DEL-90
chr8-55387673-DEL-5895
chr8-55420145-DEL-58
chr8-55443672-DEL-66
chr8-55445914-DEL-73
chr8-55481270-DEL-185
chr8-55490139-DEL-102
chr8-55509063-DEL-384
chr8-55537060-DEL-77
chr8-55838270-DEL-1867
chr8-55846014-DEL-2994
chr8-55849425-DEL-484
chr8-55901187-DEL-7033
chr8-55947152-DEL-52
chr8-55974013-DEL-57
chr8-55984892-DEL-68
chr8-55985319-DEL-130
chr8-55998184-DEL-64
chr8-56011636-DEL-73
chr8-56016973-DEL-4539
chr8-56031628-DEL-6056
chr8-56073905-DEL-264
chr8-56080006-DEL-70
chr8-56092761-DEL-1712
chr8-56123316-DEL-52
chr8-56134115-DEL-6037
chr8-56193688-DEL-100
chr8-56196135-DEL-6774
chr8-56259174-DEL-6229
chr8-56277093-DEL-65
chr8-56290438-DEL-12442
chr8-56326954-DEL-85
chr8-56345685-DEL-2615
chr8-56361441-DEL-1808
chr8-56372670-DEL-349
chr8-56400552-DEL-645
chr8-56434656-DEL-152
chr8-56449520-DEL-187
chr8-56530820-DEL-276
chr8-56534407-DEL-1367
chr8-56555580-DEL-6164
chr8-56570469-DEL-189
chr8-56676411-DEL-151
chr8-56713967-DEL-145
chr8-56719317-DEL-224
chr8-56724375-DEL-62
chr8-56754461-DEL-112
chr8-56762416-DEL-56
chr8-56781836-DEL-64
chr8-56790633-DEL-1317
chr8-56802637-DEL-8814
chr8-56838378-DEL-464
chr8-56850607-DEL-3974
chr8-56900029-DEL-258
chr8-56939031-DEL-75
chr8-56956056-DEL-472
chr8-56959245-DEL-491
chr8-56986115-DEL-65
chr8-56994125-DEL-248
chr8-57048188-DEL-210
chr8-57079561-DEL-51
chr8-57131490-DEL-64
chr8-57150308-DEL-244
chr8-57165074-DEL-1573
chr8-57178709-DEL-324
chr8-57190930-DEL-462
chr8-57196171-DEL-204
chr8-57213205-DEL-88
chr8-57246376-DEL-9876
chr8-57259068-DEL-5632
chr8-57294030-DEL-1016
chr8-57295063-DEL-314
chr8-57335074-DEL-70

chr8-57341224-DEL-132
chr8-57364616-DEL-1361
chr8-57379070-DEL-57
chr8-57395965-DEL-12915
chr8-57425729-DEL-60
chr8-57452612-DEL-128
chr8-57473164-DEL-6191
chr8-57479916-DEL-1086
chr8-57504192-DEL-103
chr8-57551422-DEL-179
chr8-57559472-DEL-84
chr8-57561823-DEL-502
chr8-57571049-DEL-192
chr8-57600479-DEL-141
chr8-57604883-DEL-138
chr8-57611116-DEL-872
chr8-57623844-DEL-6445
chr8-57633723-DEL-1193
chr8-57635626-DEL-6941
chr8-57648260-DEL-2745
chr8-57665061-DEL-101
chr8-57702920-DEL-686
chr8-57728733-DEL-115
chr8-57730457-DEL-84
chr8-57737781-DEL-182
chr8-57739472-DEL-63
chr8-57797562-DEL-275
chr8-57900096-DEL-110
chr8-57907082-DEL-1116
chr8-57916699-DEL-86
chr8-57928522-DEL-123
chr8-57941995-DEL-113
chr8-57957765-DEL-71
chr8-57969559-DEL-210
chr8-57973994-DEL-186
chr8-58083168-DEL-82
chr8-58116797-DEL-94
chr8-58162022-DEL-56
chr8-58180541-DEL-108
chr8-58222604-DEL-985
chr8-58246385-DEL-184
chr8-58261576-DEL-120
chr8-58267631-DEL-205
chr8-58304842-DEL-70
chr8-58403785-DEL-86
chr8-58484699-DEL-62
chr8-58527278-DEL-56
chr8-58528609-DEL-1769
chr8-58536165-DEL-123
chr8-58539510-DEL-175
chr8-58548605-DEL-208
chr8-58560975-DEL-383
chr8-58596545-DEL-200
chr8-58606872-DEL-6470
chr8-58617053-DEL-6860
chr8-58637176-DEL-233
chr8-58656942-DEL-73
chr8-58671732-DEL-79
chr8-58711950-DEL-66
chr8-58794785-DEL-62
chr8-58828350-DEL-63
chr8-58839187-DEL-155
chr8-58854687-DEL-442
chr8-58856032-DEL-5601
chr8-58867834-DEL-213
chr8-58874589-DEL-62
chr8-58876667-DEL-84
chr8-58877138-DEL-858
chr8-58884948-DEL-10424
chr8-58908713-DEL-216
chr8-58927823-DEL-1486
chr8-58951448-DEL-347
chr8-59034170-DEL-1751
chr8-59069545-DEL-1974
chr8-59077449-DEL-290
chr8-59102868-DEL-682
chr8-59130602-DEL-148

chr8-59149022-DEL-260
chr8-59212169-DEL-52
chr8-59221211-DEL-177
chr8-59236962-DEL-1053
chr8-59273952-DEL-86
chr8-59280569-DEL-139
chr8-59320352-DEL-53
chr8-59356912-DEL-233
chr8-59367611-DEL-60
chr8-59390739-DEL-181
chr8-59392541-DEL-1774
chr8-59415931-DEL-394
chr8-59436853-DEL-74
chr8-59449348-DEL-561
chr8-59452496-DEL-540
chr8-59458575-DEL-130
chr8-59499508-DEL-271
chr8-59508156-DEL-261
chr8-59520910-DEL-853
chr8-59522136-DEL-52
chr8-59542094-DEL-1916
chr8-59550858-DEL-63
chr8-59566877-DEL-79
chr8-59573658-DEL-77
chr8-59605701-DEL-901
chr8-59624591-DEL-58
chr8-59629724-DEL-70
chr8-59679946-DEL-455
chr8-59684494-DEL-1227
chr8-59696919-DEL-56
chr8-59698819-DEL-119
chr8-59735502-DEL-61
chr8-59737806-DEL-136
chr8-59798491-DEL-175
chr8-59801360-DEL-59
chr8-59810097-DEL-6131
chr8-59831338-DEL-5629
chr8-59840001-DEL-1102
chr8-59864357-DEL-356
chr8-59874850-DEL-514
chr8-59921302-DEL-76
chr8-59940742-DEL-132
chr8-60021102-DEL-4938
chr8-60031734-DEL-1334
chr8-60032813-DEL-522
chr8-60181085-DEL-102
chr8-60207405-DEL-164
chr8-60219373-DEL-297
chr8-60227103-DEL-87
chr8-60261374-DEL-168
chr8-60273841-DEL-62
chr8-60286446-DEL-89
chr8-60309593-DEL-52
chr8-60334070-DEL-203
chr8-60341504-DEL-6019
chr8-60367816-DEL-68
chr8-60381214-DEL-51
chr8-60381374-DEL-437
chr8-60384458-DEL-150
chr8-60388810-DEL-362
chr8-60390885-DEL-111
chr8-60410362-DEL-117
chr8-60411045-DEL-291
chr8-60464079-DEL-198
chr8-60470161-DEL-3828
chr8-60518205-DEL-1225
chr8-60541187-DEL-246
chr8-60547298-DEL-54
chr8-60554975-DEL-7318
chr8-60598348-DEL-6299
chr8-60598481-DEL-200
chr8-60608733-DEL-445
chr8-60614561-DEL-59
chr8-60631961-DEL-534
chr8-60717803-DEL-573
chr8-60760273-DEL-838
chr8-60797509-DEL-71

chr8-60810798-DEL-6017
chr8-60826781-DEL-59
chr8-60862566-DEL-2520
chr8-60873894-DEL-198
chr8-60879404-DEL-353
chr8-60887296-DEL-197
chr8-60917300-DEL-554
chr8-60920220-DEL-365
chr8-60922295-DEL-54
chr8-60934051-DEL-60
chr8-60937101-DEL-104
chr8-60940353-DEL-885
chr8-60961821-DEL-136
chr8-60966231-DEL-59
chr8-60970486-DEL-56
chr8-61010846-DEL-253
chr8-61028697-DEL-1787
chr8-61034349-DEL-1431
chr8-61040099-DEL-5239
chr8-61088367-DEL-1048
chr8-61095005-DEL-73
chr8-61100880-DEL-396
chr8-61137545-DEL-9383
chr8-61179571-DEL-139
chr8-61233006-DEL-94
chr8-61253601-DEL-153
chr8-61262232-DEL-269
chr8-61338792-DEL-239
chr8-61361441-DEL-70
chr8-61368183-DEL-69
chr8-61401517-DEL-484
chr8-61406305-DEL-71
chr8-61410141-DEL-240
chr8-61410711-DEL-126
chr8-61474898-DEL-72
chr8-61530666-DEL-207
chr8-61531830-DEL-437
chr8-61538964-DEL-94
chr8-61544236-DEL-56
chr8-61558976-DEL-369
chr8-61578800-DEL-128
chr8-61591784-DEL-634
chr8-61602389-DEL-52
chr8-61619450-DEL-56
chr8-61638706-DEL-2170
chr8-61651586-DEL-348
chr8-61657616-DEL-737
chr8-61658380-DEL-57
chr8-61666988-DEL-72
chr8-61669488-DEL-56
chr8-61683499-DEL-228
chr8-61720493-DEL-56
chr8-61752687-DEL-219
chr8-61766519-DEL-2541
chr8-61771463-DEL-104
chr8-61781314-DEL-136
chr8-61854254-DEL-87
chr8-61941380-DEL-721
chr8-61944914-DEL-425
chr8-61994095-DEL-238
chr8-61997433-DEL-1418
chr8-62015147-DEL-176
chr8-62020718-DEL-608
chr8-62054613-DEL-325
chr8-62112249-DEL-108
chr8-62143805-DEL-152
chr8-62145048-DEL-80
chr8-62158394-DEL-60
chr8-62186526-DEL-305
chr8-62191008-DEL-4555
chr8-62205277-DEL-53
chr8-62283415-DEL-146
chr8-62289962-DEL-190
chr8-62310955-DEL-200
chr8-62312263-DEL-706
chr8-62365641-DEL-102
chr8-62371589-DEL-53

chr8-62384678-DEL-64
chr8-62386872-DEL-743
chr8-62409001-DEL-112
chr8-62421226-DEL-102
chr8-62457180-DEL-74
chr8-62459522-DEL-125
chr8-62463874-DEL-685
chr8-62510386-DEL-58
chr8-62566512-DEL-3404
chr8-62605805-DEL-62
chr8-62637418-DEL-100
chr8-62638086-DEL-54
chr8-62667943-DEL-366
chr8-62681218-DEL-52
chr8-62727336-DEL-68
chr8-62730292-DEL-198
chr8-62756297-DEL-58
chr8-62779406-DEL-5196
chr8-62786891-DEL-377
chr8-62796890-DEL-2238
chr8-62839329-DEL-199
chr8-62841302-DEL-3698
chr8-62873329-DEL-1313
chr8-62880238-DEL-193
chr8-62911998-DEL-456
chr8-62928915-DEL-156
chr8-62938524-DEL-1345
chr8-62946849-DEL-83
chr8-62989860-DEL-68
chr8-62999825-DEL-378
chr8-63009617-DEL-446
chr8-63026498-DEL-1050
chr8-63034200-DEL-448
chr8-63088487-DEL-77
chr8-63110828-DEL-233
chr8-63115039-DEL-477
chr8-63140611-DEL-150
chr8-63150113-DEL-60
chr8-63192913-DEL-80
chr8-63272505-DEL-5339
chr8-63284950-DEL-54
chr8-63297835-DEL-112
chr8-63312453-DEL-640
chr8-63337599-DEL-6314
chr8-63341515-DEL-7021
chr8-63346492-DEL-1338
chr8-63350316-DEL-428
chr8-63389959-DEL-195
chr8-63429618-DEL-205
chr8-63451544-DEL-352
chr8-63452653-DEL-1094
chr8-63468559-DEL-3427
chr8-63482872-DEL-54
chr8-63485238-DEL-183
chr8-63489822-DEL-184
chr8-63500489-DEL-6277
chr8-63511498-DEL-527
chr8-63584724-DEL-52
chr8-63636227-DEL-864
chr8-63638357-DEL-64
chr8-63663676-DEL-52
chr8-63707143-DEL-102
chr8-63734632-DEL-109
chr8-63741048-DEL-76
chr8-63744810-DEL-172
chr8-63750484-DEL-1852
chr8-63751012-DEL-8067
chr8-63762075-DEL-17314
chr8-63770914-DEL-8475
chr8-63786596-DEL-789
chr8-63833856-DEL-82
chr8-63834217-DEL-254
chr8-63840318-DEL-6152
chr8-63916325-DEL-257
chr8-63926818-DEL-6349
chr8-63933646-DEL-2687
chr8-63965471-DEL-68

chr8-63989185-DEL-60
chr8-63990823-DEL-941
chr8-64002625-DEL-125
chr8-64011818-DEL-63
chr8-64079611-DEL-3751
chr8-64092197-DEL-165
chr8-64148736-DEL-51
chr8-64167969-DEL-494
chr8-64168642-DEL-1321
chr8-64219888-DEL-419
chr8-64249386-DEL-2296
chr8-64253440-DEL-84
chr8-64268707-DEL-6151
chr8-64296935-DEL-505
chr8-64302848-DEL-6800
chr8-64336580-DEL-614
chr8-64337053-DEL-1635
chr8-64337322-DEL-237
chr8-64337654-DEL-1034
chr8-64364708-DEL-4062
chr8-64385349-DEL-3291
chr8-64390768-DEL-447
chr8-64427923-DEL-3016
chr8-64474432-DEL-60
chr8-64503314-DEL-254
chr8-64538200-DEL-95
chr8-64565412-DEL-77
chr8-64580555-DEL-73
chr8-64582693-DEL-1337
chr8-64613815-DEL-252
chr8-64625491-DEL-396
chr8-64636203-DEL-3754
chr8-64661379-DEL-177
chr8-64704090-DEL-1518
chr8-64705868-DEL-487
chr8-64709078-DEL-123
chr8-64715425-DEL-70
chr8-64721249-DEL-392
chr8-64722088-DEL-517
chr8-64753906-DEL-1099
chr8-64756227-DEL-54
chr8-64797178-DEL-263
chr8-64812957-DEL-1452
chr8-64831861-DEL-407
chr8-64858063-DEL-51
chr8-64862560-DEL-72
chr8-64888327-DEL-69
chr8-64898863-DEL-424
chr8-64899443-DEL-212
chr8-64899802-DEL-212
chr8-64906400-DEL-1782
chr8-64909927-DEL-51
chr8-64942058-DEL-68
chr8-64950230-DEL-115
chr8-64951831-DEL-54
chr8-64966049-DEL-89
chr8-64987739-DEL-377
chr8-65025641-DEL-516
chr8-65026219-DEL-516
chr8-65026948-DEL-517
chr8-65027624-DEL-56
chr8-65062917-DEL-65
chr8-65064833-DEL-829
chr8-65069183-DEL-63
chr8-65084614-DEL-718
chr8-65097711-DEL-2104
chr8-65109284-DEL-131
chr8-65181699-DEL-155
chr8-65182389-DEL-71
chr8-65234375-DEL-66
chr8-65274887-DEL-306
chr8-65298781-DEL-88
chr8-65330046-DEL-397
chr8-65330847-DEL-63
chr8-65338666-DEL-327
chr8-65357557-DEL-1275
chr8-65362517-DEL-15963

chr8-65386763-DEL-308
chr8-65389097-DEL-199
chr8-65391468-DEL-55
chr8-65403291-DEL-1258
chr8-65422687-DEL-260
chr8-65468214-DEL-470
chr8-65469623-DEL-182
chr8-65479787-DEL-171
chr8-65497800-DEL-54
chr8-65505032-DEL-146
chr8-65513208-DEL-56
chr8-65522668-DEL-206
chr8-65562003-DEL-101
chr8-65566976-DEL-2414
chr8-65576867-DEL-1543
chr8-65599696-DEL-222
chr8-65605053-DEL-109
chr8-65609840-DEL-195
chr8-65637789-DEL-5209
chr8-65662263-DEL-66
chr8-65674749-DEL-182
chr8-65683058-DEL-963
chr8-65685817-DEL-2492
chr8-65717518-DEL-51
chr8-65741624-DEL-72
chr8-65746865-DEL-13722
chr8-65747935-DEL-68
chr8-65748291-DEL-10174
chr8-65760803-DEL-8367
chr8-65763229-DEL-288
chr8-65764333-DEL-96
chr8-65765024-DEL-1746
chr8-65769172-DEL-6873
chr8-65771018-DEL-282
chr8-65771996-DEL-4614
chr8-65794643-DEL-3099
chr8-65818997-DEL-61
chr8-65833263-DEL-2110
chr8-65846810-DEL-192
chr8-65850175-DEL-176
chr8-65851786-DEL-56
chr8-65856024-DEL-6855
chr8-65882380-DEL-127
chr8-65887775-DEL-6672
chr8-65898172-DEL-75
chr8-65910018-DEL-3018
chr8-65925654-DEL-556
chr8-65932599-DEL-674
chr8-65955916-DEL-68
chr8-65971751-DEL-3317
chr8-65986495-DEL-4211
chr8-66023430-DEL-199
chr8-66030813-DEL-6203
chr8-66043418-DEL-9032
chr8-66118885-DEL-62
chr8-66119339-DEL-152
chr8-66164933-DEL-288
chr8-66166473-DEL-88
chr8-66177989-DEL-150
chr8-66182892-DEL-752
chr8-66194986-DEL-60
chr8-66198575-DEL-58
chr8-66202459-DEL-6031
chr8-66239839-DEL-1054
chr8-66256401-DEL-1582
chr8-66269627-DEL-123
chr8-66278450-DEL-1593
chr8-66302849-DEL-68
chr8-66308073-DEL-113
chr8-66314107-DEL-197
chr8-66329402-DEL-111
chr8-66343724-DEL-148
chr8-66349220-DEL-84
chr8-66365190-DEL-255
chr8-66369000-DEL-611
chr8-66371368-DEL-68
chr8-66397913-DEL-66

chr8-66408004-DEL-58
chr8-66413962-DEL-90
chr8-66414725-DEL-908
chr8-66450611-DEL-4492
chr8-66462877-DEL-72
chr8-66481580-DEL-130
chr8-66501597-DEL-1002
chr8-66507174-DEL-103
chr8-66519692-DEL-400
chr8-66528146-DEL-56
chr8-66532791-DEL-755
chr8-66557187-DEL-986
chr8-66559484-DEL-222
chr8-66560556-DEL-642
chr8-66585057-DEL-602
chr8-66609375-DEL-499
chr8-66617061-DEL-542
chr8-66617795-DEL-462
chr8-66644000-DEL-92
chr8-66647239-DEL-56
chr8-66669389-DEL-65
chr8-66674096-DEL-735
chr8-66707720-DEL-944
chr8-66710028-DEL-3749
chr8-66733142-DEL-142
chr8-66749659-DEL-6060
chr8-66761068-DEL-58
chr8-66765251-DEL-105
chr8-66770973-DEL-1665
chr8-66803802-DEL-968
chr8-66833020-DEL-52
chr8-66850850-DEL-82
chr8-66853757-DEL-167
chr8-66857074-DEL-265
chr8-66869296-DEL-758
chr8-66909424-DEL-226
chr8-66940114-DEL-135
chr8-66945672-DEL-184
chr8-66957092-DEL-88
chr8-67003238-DEL-94
chr8-67015329-DEL-144
chr8-67029526-DEL-2616
chr8-67063327-DEL-63
chr8-67083898-DEL-84
chr8-67117545-DEL-326
chr8-67139175-DEL-122
chr8-67196714-DEL-334
chr8-67231127-DEL-97
chr8-67267423-DEL-158
chr8-67297802-DEL-640
chr8-67313378-DEL-213
chr8-67349287-DEL-66
chr8-67396811-DEL-587
chr8-67426588-DEL-1149
chr8-67430887-DEL-387
chr8-67431621-DEL-92
chr8-67457813-DEL-6412
chr8-67468568-DEL-99
chr8-67541939-DEL-3284
chr8-67558930-DEL-263
chr8-67571320-DEL-10778
chr8-67582702-DEL-8685
chr8-67585817-DEL-7303
chr8-67589328-DEL-1101
chr8-67590296-DEL-230
chr8-67595520-DEL-51
chr8-67608984-DEL-52
chr8-67612767-DEL-142
chr8-67624839-DEL-1509
chr8-67670313-DEL-80
chr8-67678632-DEL-2035
chr8-67711748-DEL-549
chr8-67716089-DEL-310
chr8-67739885-DEL-609
chr8-67751472-DEL-685
chr8-67759050-DEL-249
chr8-67822419-DEL-6497

chr8-67849950-DEL-1047
chr8-67851815-DEL-152
chr8-67860880-DEL-6276
chr8-67884422-DEL-55
chr8-67905000-DEL-827
chr8-67974401-DEL-213
chr8-68036467-DEL-51
chr8-68081814-DEL-511
chr8-68086525-DEL-438
chr8-68124435-DEL-125
chr8-68130218-DEL-329
chr8-68131177-DEL-175
chr8-68140687-DEL-80
chr8-68194798-DEL-560
chr8-68228727-DEL-6449
chr8-68238262-DEL-137
chr8-68240867-DEL-64
chr8-68245357-DEL-7507
chr8-68315906-DEL-528
chr8-68328980-DEL-149
chr8-68345941-DEL-52
chr8-68383856-DEL-75
chr8-68392757-DEL-110
chr8-68410619-DEL-83
chr8-68438245-DEL-880
chr8-68456357-DEL-859
chr8-68471618-DEL-74
chr8-68492927-DEL-79
chr8-68494332-DEL-61
chr8-68521612-DEL-54
chr8-68523128-DEL-58
chr8-68531316-DEL-255
chr8-68544545-DEL-330
chr8-68619282-DEL-108
chr8-68652033-DEL-115
chr8-68662217-DEL-68
chr8-68681967-DEL-64
chr8-68744645-DEL-569
chr8-68805572-DEL-1383
chr8-68822507-DEL-66
chr8-68832160-DEL-151
chr8-68846207-DEL-71
chr8-68849377-DEL-742
chr8-68916425-DEL-2702
chr8-68920024-DEL-1189
chr8-68989564-DEL-52
chr8-68991243-DEL-784
chr8-69047007-DEL-82
chr8-69048399-DEL-268
chr8-69068297-DEL-68
chr8-69087982-DEL-1560
chr8-69088199-DEL-60
chr8-69092873-DEL-450
chr8-69097945-DEL-310
chr8-69115202-DEL-133
chr8-69156673-DEL-51
chr8-69213460-DEL-131
chr8-69243403-DEL-410
chr8-69244652-DEL-85
chr8-69300559-DEL-320
chr8-69336804-DEL-105
chr8-69358095-DEL-83
chr8-69374099-DEL-637
chr8-69374839-DEL-1141
chr8-69380764-DEL-172
chr8-69382946-DEL-53
chr8-69404288-DEL-1017
chr8-69451074-DEL-231
chr8-69456317-DEL-199
chr8-69460272-DEL-64
chr8-69461214-DEL-218
chr8-69484592-DEL-490
chr8-69514530-DEL-75
chr8-69544005-DEL-110
chr8-69551907-DEL-907
chr8-69553024-DEL-203
chr8-69601274-DEL-51

chr8-69649665-DEL-1413
chr8-69667213-DEL-198
chr8-69684072-DEL-4193
chr8-69692885-DEL-1494
chr8-69694409-DEL-785
chr8-69710075-DEL-147
chr8-69720007-DEL-239
chr8-69801194-DEL-904
chr8-69807112-DEL-537
chr8-69814395-DEL-246
chr8-69815049-DEL-820
chr8-69831854-DEL-56
chr8-69833247-DEL-58
chr8-69868415-DEL-2262
chr8-69870818-DEL-1750
chr8-69888539-DEL-1356
chr8-69899356-DEL-64
chr8-69907761-DEL-7835
chr8-69912662-DEL-2103
chr8-69937943-DEL-358
chr8-69991125-DEL-175
chr8-69996776-DEL-318
chr8-70001367-DEL-329
chr8-70007890-DEL-5984
chr8-70021814-DEL-57
chr8-70030407-DEL-127
chr8-70058443-DEL-393
chr8-70069883-DEL-183
chr8-70072012-DEL-59
chr8-70088236-DEL-871
chr8-70096098-DEL-2880
chr8-70121908-DEL-6135
chr8-70165214-DEL-323
chr8-70170403-DEL-486
chr8-70183740-DEL-163
chr8-70198762-DEL-470
chr8-70235396-DEL-6293
chr8-70243261-DEL-103
chr8-70250147-DEL-116
chr8-70309146-DEL-369
chr8-70319538-DEL-259
chr8-70331606-DEL-194
chr8-70351703-DEL-88
chr8-70357261-DEL-78
chr8-70369711-DEL-284
chr8-70398002-DEL-80
chr8-70411418-DEL-197
chr8-70412292-DEL-54
chr8-70416590-DEL-111
chr8-70437290-DEL-64
chr8-70441807-DEL-180
chr8-70459045-DEL-89
chr8-70463074-DEL-831
chr8-70513766-DEL-201
chr8-70526022-DEL-300
chr8-70527459-DEL-52
chr8-70562096-DEL-204
chr8-70632140-DEL-457
chr8-70683228-DEL-227
chr8-70753437-DEL-158
chr8-70807731-DEL-203
chr8-70812525-DEL-57
chr8-70817859-DEL-203
chr8-70843222-DEL-56
chr8-70883672-DEL-54
chr8-70907804-DEL-57
chr8-70940312-DEL-98
chr8-70963348-DEL-304
chr8-70986169-DEL-60
chr8-70989665-DEL-97
chr8-71007773-DEL-146
chr8-71020341-DEL-2301
chr8-71028806-DEL-68
chr8-71032038-DEL-244
chr8-71036042-DEL-72
chr8-71071753-DEL-69
chr8-71072760-DEL-225

chr8-71080042-DEL-408
chr8-71094160-DEL-194
chr8-71114993-DEL-2546
chr8-71119191-DEL-195
chr8-71121981-DEL-99
chr8-71171549-DEL-842
chr8-71237541-DEL-220
chr8-71269227-DEL-102
chr8-71277034-DEL-211
chr8-71300085-DEL-52
chr8-71313830-DEL-114
chr8-71362504-DEL-830
chr8-71375376-DEL-56
chr8-71413643-DEL-52
chr8-71418873-DEL-314
chr8-71419702-DEL-152
chr8-71421757-DEL-531
chr8-71422817-DEL-304
chr8-71423441-DEL-912
chr8-71424607-DEL-114
chr8-71426031-DEL-76
chr8-71426877-DEL-304
chr8-71428560-DEL-304
chr8-71428674-DEL-304
chr8-71429309-DEL-301
chr8-71431062-DEL-145
chr8-71431801-DEL-76
chr8-71432475-DEL-226
chr8-71434083-DEL-379
chr8-71434741-DEL-190
chr8-71435872-DEL-334
chr8-71436187-DEL-785
chr8-71437482-DEL-258
chr8-71445903-DEL-238
chr8-71446970-DEL-269
chr8-71448773-DEL-331
chr8-71450336-DEL-76
chr8-71453556-DEL-114
chr8-71454638-DEL-1129
chr8-71456665-DEL-190
chr8-71458493-DEL-76
chr8-71459214-DEL-255
chr8-71461616-DEL-243
chr8-71473343-DEL-101
chr8-71490356-DEL-203
chr8-71505527-DEL-2037
chr8-71510424-DEL-1543
chr8-71589535-DEL-56
chr8-71625925-DEL-95
chr8-71631096-DEL-1910
chr8-71633717-DEL-1980
chr8-71640110-DEL-613
chr8-71641911-DEL-145
chr8-71650696-DEL-3965
chr8-71684230-DEL-84
chr8-71696006-DEL-2939
chr8-71710487-DEL-664
chr8-71761601-DEL-81
chr8-71816769-DEL-206
chr8-71827641-DEL-345
chr8-71870865-DEL-1112
chr8-71873460-DEL-4863
chr8-71882579-DEL-135
chr8-71883132-DEL-3457
chr8-71891105-DEL-140
chr8-71899480-DEL-333
chr8-71912658-DEL-74
chr8-71920408-DEL-200
chr8-71956058-DEL-328
chr8-71986585-DEL-152
chr8-71990434-DEL-410
chr8-72003965-DEL-71
chr8-72011611-DEL-85
chr8-72027670-DEL-1097
chr8-72029121-DEL-101
chr8-72035531-DEL-98
chr8-72053924-DEL-646

chr8-72074165-DEL-151
chr8-72093105-DEL-396
chr8-72158316-DEL-210
chr8-72202876-DEL-64
chr8-72209726-DEL-305
chr8-72239133-DEL-5625
chr8-72245640-DEL-4468
chr8-72258686-DEL-463
chr8-72260444-DEL-606
chr8-72570278-DEL-205
chr8-72576559-DEL-100
chr8-72579796-DEL-51
chr8-72620412-DEL-411
chr8-72625563-DEL-1713
chr8-72675445-DEL-776
chr8-72687693-DEL-168
chr8-72710193-DEL-1139
chr8-72792481-DEL-297
chr8-72810508-DEL-80
chr8-72822180-DEL-259
chr8-72870305-DEL-1529
chr8-72894657-DEL-68
chr8-72916823-DEL-153
chr8-72932899-DEL-196
chr8-72945661-DEL-174
chr8-72963200-DEL-84
chr8-72967301-DEL-82
chr8-72970263-DEL-54
chr8-72979357-DEL-76
chr8-72980888-DEL-53
chr8-72982067-DEL-4728
chr8-72986867-DEL-124
chr8-72987823-DEL-73
chr8-72990229-DEL-53
chr8-72996929-DEL-782
chr8-73010117-DEL-330
chr8-73017663-DEL-100
chr8-73018503-DEL-391
chr8-73086085-DEL-82
chr8-73092445-DEL-891
chr8-73126449-DEL-205
chr8-73143516-DEL-53
chr8-73164464-DEL-531
chr8-73187453-DEL-1021
chr8-73189381-DEL-167
chr8-73210974-DEL-73
chr8-73226193-DEL-210
chr8-73346910-DEL-388
chr8-73347736-DEL-59
chr8-73350287-DEL-123
chr8-73356319-DEL-60
chr8-73360645-DEL-60
chr8-73361063-DEL-971
chr8-73363766-DEL-65
chr8-73365169-DEL-100
chr8-73367166-DEL-59
chr8-73368060-DEL-60
chr8-73368565-DEL-211
chr8-73369497-DEL-118
chr8-73370337-DEL-678
chr8-73391262-DEL-78
chr8-73410655-DEL-2985
chr8-73469990-DEL-495
chr8-73479018-DEL-566
chr8-73506774-DEL-495
chr8-73521288-DEL-1146
chr8-73527236-DEL-84
chr8-73580361-DEL-1324
chr8-73581770-DEL-314
chr8-73582285-DEL-2680
chr8-73590057-DEL-8275
chr8-73614082-DEL-62
chr8-73634124-DEL-3724
chr8-73652428-DEL-5825
chr8-73653762-DEL-119
chr8-73665922-DEL-248
chr8-73667936-DEL-201

chr8-73670747-DEL-345
chr8-73676107-DEL-54
chr8-73693393-DEL-9935
chr8-73703602-DEL-133
chr8-73704186-DEL-5190
chr8-73706811-DEL-1937
chr8-73741302-DEL-262
chr8-73748530-DEL-56
chr8-73752772-DEL-844
chr8-73761692-DEL-110
chr8-73764371-DEL-897
chr8-73772367-DEL-2542
chr8-73793137-DEL-54
chr8-73816219-DEL-247
chr8-73826103-DEL-1925
chr8-73873490-DEL-191
chr8-73925810-DEL-65
chr8-73931841-DEL-3292
chr8-73935738-DEL-879
chr8-73941278-DEL-68
chr8-73960955-DEL-116
chr8-73969253-DEL-54
chr8-74009195-DEL-2627
chr8-74036505-DEL-94
chr8-74089789-DEL-135
chr8-74090969-DEL-10847
chr8-74118460-DEL-6278
chr8-74128987-DEL-116
chr8-74140593-DEL-52
chr8-74148783-DEL-429
chr8-74153287-DEL-73
chr8-74156285-DEL-102
chr8-74165623-DEL-76
chr8-74194425-DEL-399
chr8-74195858-DEL-3372
chr8-74196249-DEL-59
chr8-74210714-DEL-98
chr8-74218489-DEL-6484
chr8-74256746-DEL-59
chr8-74264304-DEL-6414
chr8-74276368-DEL-399
chr8-74330424-DEL-207
chr8-74332433-DEL-1340
chr8-74345608-DEL-307
chr8-74366558-DEL-54
chr8-74370031-DEL-159
chr8-74382147-DEL-418
chr8-74422606-DEL-6168
chr8-74514365-DEL-1755
chr8-74519805-DEL-4618
chr8-74529968-DEL-53
chr8-74535340-DEL-426
chr8-74540188-DEL-676
chr8-74561839-DEL-376
chr8-74570388-DEL-400
chr8-74573473-DEL-54
chr8-74576208-DEL-72
chr8-74593732-DEL-56
chr8-74606221-DEL-69
chr8-74758929-DEL-751
chr8-74769981-DEL-7434
chr8-74804824-DEL-58
chr8-74863975-DEL-120
chr8-74881074-DEL-73
chr8-74893456-DEL-1407
chr8-74897784-DEL-616
chr8-74913770-DEL-1674
chr8-74915867-DEL-10595
chr8-74953035-DEL-829
chr8-74959073-DEL-118
chr8-75004744-DEL-109
chr8-75052998-DEL-351
chr8-75076297-DEL-7090
chr8-75086693-DEL-2971
chr8-75112779-DEL-272
chr8-75160659-DEL-156
chr8-75177105-DEL-355

chr8-75191596-DEL-775
chr8-75205877-DEL-2733
chr8-75225354-DEL-167
chr8-75262730-DEL-3763
chr8-75267684-DEL-6314
chr8-75292030-DEL-2259
chr8-75296294-DEL-88
chr8-75334273-DEL-285
chr8-75335276-DEL-224
chr8-75335862-DEL-515
chr8-75366590-DEL-102
chr8-75370148-DEL-3303
chr8-75373467-DEL-2939
chr8-75409147-DEL-110
chr8-75431905-DEL-206
chr8-75433724-DEL-3714
chr8-75444358-DEL-207
chr8-75466810-DEL-247
chr8-75495008-DEL-57
chr8-75538510-DEL-84
chr8-75542025-DEL-1345
chr8-75557528-DEL-82
chr8-75584371-DEL-114
chr8-75610433-DEL-588
chr8-75629059-DEL-52
chr8-75633354-DEL-208
chr8-75649133-DEL-363
chr8-75658643-DEL-102
chr8-75665720-DEL-51
chr8-75672199-DEL-202
chr8-75684697-DEL-66
chr8-75701816-DEL-250
chr8-75729626-DEL-536
chr8-75731862-DEL-319
chr8-75732928-DEL-94
chr8-75735937-DEL-206
chr8-75736348-DEL-2059
chr8-75807924-DEL-777
chr8-75813648-DEL-56
chr8-75858167-DEL-280
chr8-75865822-DEL-200
chr8-75872729-DEL-201
chr8-75876371-DEL-98
chr8-75912420-DEL-116
chr8-75916102-DEL-75
chr8-75922459-DEL-103
chr8-75958094-DEL-1492
chr8-75960471-DEL-80
chr8-75961357-DEL-164
chr8-75969935-DEL-852
chr8-75987857-DEL-2239
chr8-75993204-DEL-119
chr8-76000934-DEL-89
chr8-76009107-DEL-358
chr8-76009139-DEL-520
chr8-76010293-DEL-203
chr8-76011192-DEL-111
chr8-76020049-DEL-67
chr8-76027638-DEL-6213
chr8-76069183-DEL-399
chr8-76083772-DEL-117886
chr8-76116563-DEL-225
chr8-76117443-DEL-153
chr8-76125119-DEL-1005
chr8-76160635-DEL-81
chr8-76165681-DEL-107
chr8-76167607-DEL-93
chr8-76179920-DEL-2996
chr8-76214809-DEL-2007
chr8-76219533-DEL-61
chr8-76229601-DEL-56
chr8-76244416-DEL-1987
chr8-76249061-DEL-882
chr8-76249083-DEL-1024
chr8-76249971-DEL-292
chr8-76253398-DEL-981
chr8-76254641-DEL-354

chr8-76255826-DEL-793
chr8-76270041-DEL-51
chr8-76281278-DEL-68
chr8-76285625-DEL-204
chr8-76288272-DEL-96
chr8-76289289-DEL-369
chr8-76293835-DEL-6759
chr8-76336646-DEL-219
chr8-76382355-DEL-365
chr8-76396463-DEL-52
chr8-76403457-DEL-58
chr8-76409813-DEL-240
chr8-76426922-DEL-490
chr8-76436929-DEL-591
chr8-76445114-DEL-123
chr8-76449467-DEL-279
chr8-76452069-DEL-91
chr8-76458469-DEL-1783
chr8-76517554-DEL-49973
chr8-76566617-DEL-78
chr8-76569150-DEL-779
chr8-76572525-DEL-5404
chr8-76608402-DEL-2903
chr8-76630621-DEL-199
chr8-76672071-DEL-721
chr8-76676263-DEL-273
chr8-76685006-DEL-824
chr8-76691896-DEL-108
chr8-76694112-DEL-2891
chr8-76702570-DEL-593
chr8-76704132-DEL-306
chr8-76736730-DEL-64
chr8-76784890-DEL-81
chr8-76790098-DEL-577
chr8-76799279-DEL-55
chr8-76821608-DEL-9720
chr8-76844526-DEL-158
chr8-76891230-DEL-1322
chr8-76898858-DEL-8220
chr8-76914597-DEL-65
chr8-76966531-DEL-54
chr8-76971338-DEL-96
chr8-76984500-DEL-94
chr8-76985263-DEL-243
chr8-76995481-DEL-464
chr8-76996852-DEL-258
chr8-77008745-DEL-52
chr8-77030757-DEL-2018
chr8-77043259-DEL-3338
chr8-77060410-DEL-6219
chr8-77060482-DEL-199
chr8-77071358-DEL-166
chr8-77126387-DEL-881
chr8-77138435-DEL-499
chr8-77138970-DEL-7300
chr8-77151101-DEL-64
chr8-77161067-DEL-268
chr8-77163117-DEL-472
chr8-77177207-DEL-52
chr8-77186811-DEL-178
chr8-77187593-DEL-73
chr8-77188458-DEL-6754
chr8-77194786-DEL-214
chr8-77201971-DEL-53
chr8-77206824-DEL-80
chr8-77213461-DEL-105
chr8-77220003-DEL-56
chr8-77220740-DEL-54
chr8-77233732-DEL-100
chr8-77234388-DEL-73
chr8-77241689-DEL-96
chr8-77263446-DEL-163
chr8-77276911-DEL-1404
chr8-77308342-DEL-542
chr8-77346254-DEL-288
chr8-77376709-DEL-1703
chr8-77391841-DEL-1865

chr8-77401217-DEL-1529
chr8-77401625-DEL-759
chr8-77402684-DEL-60
chr8-77423790-DEL-51
chr8-77436219-DEL-174
chr8-77482197-DEL-114
chr8-77486955-DEL-67
chr8-77497220-DEL-530
chr8-77504258-DEL-142
chr8-77528496-DEL-1475
chr8-77576659-DEL-3125
chr8-77590578-DEL-348
chr8-77602421-DEL-66
chr8-77608018-DEL-120
chr8-77641202-DEL-102
chr8-77661898-DEL-99
chr8-77670203-DEL-940
chr8-77755199-DEL-81
chr8-77777511-DEL-297
chr8-77845861-DEL-85
chr8-77889545-DEL-182
chr8-77904503-DEL-2198
chr8-77916681-DEL-154
chr8-77948050-DEL-216
chr8-77948846-DEL-88
chr8-77958143-DEL-107
chr8-77959791-DEL-2587
chr8-77987649-DEL-68
chr8-77995117-DEL-164
chr8-78046913-DEL-117
chr8-78152105-DEL-54
chr8-78175476-DEL-520
chr8-78197732-DEL-439
chr8-78238271-DEL-172
chr8-78269094-DEL-198
chr8-78278741-DEL-192
chr8-78287844-DEL-255
chr8-78291198-DEL-66
chr8-78301819-DEL-403
chr8-78314374-DEL-1782
chr8-78384342-DEL-176
chr8-78394914-DEL-149
chr8-78414407-DEL-534
chr8-78427140-DEL-808
chr8-78432622-DEL-1158
chr8-78433368-DEL-57
chr8-78437790-DEL-90
chr8-78465989-DEL-228
chr8-78475251-DEL-80
chr8-78496137-DEL-259
chr8-78500378-DEL-70
chr8-78534765-DEL-729
chr8-78548320-DEL-61
chr8-78565412-DEL-100
chr8-78567474-DEL-324
chr8-78567581-DEL-359
chr8-78575312-DEL-56
chr8-78580751-DEL-85
chr8-78591287-DEL-4202
chr8-78618117-DEL-208
chr8-78623052-DEL-184
chr8-78625486-DEL-52
chr8-78627459-DEL-2904
chr8-78633733-DEL-6932
chr8-78643606-DEL-1517
chr8-78652339-DEL-1583
chr8-78652435-DEL-60
chr8-78654266-DEL-148
chr8-78704288-DEL-137
chr8-78743777-DEL-54
chr8-78763593-DEL-462
chr8-78769965-DEL-357
chr8-78818487-DEL-53
chr8-78844605-DEL-52
chr8-78913540-DEL-62
chr8-78957747-DEL-385
chr8-79006696-DEL-76

chr8-79013865-DEL-65
chr8-79023365-DEL-76
chr8-79043703-DEL-327
chr8-79066636-DEL-251
chr8-79079143-DEL-138
chr8-79104725-DEL-1680
chr8-79111709-DEL-311
chr8-79141486-DEL-195
chr8-79200799-DEL-574
chr8-79212238-DEL-346
chr8-79278856-DEL-231
chr8-79318359-DEL-187
chr8-79318784-DEL-317
chr8-79320794-DEL-68
chr8-79328748-DEL-257
chr8-79329303-DEL-176
chr8-79339750-DEL-264
chr8-79350811-DEL-64
chr8-79360277-DEL-256
chr8-79365444-DEL-576
chr8-79397142-DEL-184
chr8-79406498-DEL-56
chr8-79418866-DEL-76
chr8-79425707-DEL-255
chr8-79436193-DEL-180
chr8-79436371-DEL-5355
chr8-79442534-DEL-65
chr8-79449653-DEL-60
chr8-79470402-DEL-144
chr8-79479532-DEL-140
chr8-79483477-DEL-63
chr8-79498788-DEL-55
chr8-79502711-DEL-448
chr8-79515098-DEL-87
chr8-79520686-DEL-337
chr8-79521561-DEL-110
chr8-79546805-DEL-62
chr8-79553525-DEL-302
chr8-79576548-DEL-266
chr8-79600855-DEL-236
chr8-79634362-DEL-706
chr8-79666070-DEL-59
chr8-79669197-DEL-2058
chr8-79690219-DEL-104
chr8-79690677-DEL-179
chr8-79703590-DEL-173
chr8-79711399-DEL-228
chr8-79712417-DEL-1226
chr8-79724983-DEL-351
chr8-79750537-DEL-131
chr8-79761952-DEL-180
chr8-79813794-DEL-76
chr8-79855329-DEL-94
chr8-79866650-DEL-537
chr8-79870826-DEL-72
chr8-79874160-DEL-301
chr8-79886760-DEL-67
chr8-79910049-DEL-182
chr8-79935174-DEL-55
chr8-79940548-DEL-208
chr8-79952587-DEL-86
chr8-79979220-DEL-142
chr8-79988888-DEL-115
chr8-80027592-DEL-160
chr8-80030951-DEL-123
chr8-80049056-DEL-70
chr8-80139778-DEL-194
chr8-80170188-DEL-56
chr8-80174609-DEL-75
chr8-80183886-DEL-159
chr8-80235006-DEL-131
chr8-80259431-DEL-95
chr8-80262198-DEL-124
chr8-80285070-DEL-120
chr8-80298771-DEL-3327
chr8-80313543-DEL-132
chr8-80314378-DEL-1385

chr8-80318329-DEL-939
chr8-80341252-DEL-99
chr8-80348310-DEL-163
chr8-80353997-DEL-865
chr8-80414656-DEL-88
chr8-80418683-DEL-488
chr8-80494391-DEL-198
chr8-80500935-DEL-367
chr8-80518872-DEL-71
chr8-80577829-DEL-145
chr8-80601605-DEL-74
chr8-80626490-DEL-330
chr8-80646019-DEL-900
chr8-80646859-DEL-237
chr8-80672948-DEL-102
chr8-80676073-DEL-59
chr8-80688870-DEL-68
chr8-80718234-DEL-64
chr8-80745210-DEL-54
chr8-80752685-DEL-55
chr8-80763054-DEL-68
chr8-80765031-DEL-167
chr8-80773928-DEL-180
chr8-80870479-DEL-566
chr8-80914968-DEL-60
chr8-80945857-DEL-3920
chr8-80986589-DEL-74
chr8-81004842-DEL-62
chr8-81082792-DEL-329
chr8-81086539-DEL-116
chr8-81141824-DEL-60
chr8-81210077-DEL-68
chr8-81213815-DEL-59
chr8-81238050-DEL-64
chr8-81242281-DEL-195
chr8-81243124-DEL-56
chr8-81307008-DEL-560
chr8-81311421-DEL-290
chr8-81312379-DEL-57
chr8-81328414-DEL-55
chr8-81331849-DEL-193
chr8-81345569-DEL-1552
chr8-81351733-DEL-505
chr8-81359353-DEL-130
chr8-81363492-DEL-8228
chr8-81393411-DEL-556
chr8-81394326-DEL-213
chr8-81417998-DEL-72
chr8-81424361-DEL-499
chr8-81451226-DEL-52
chr8-81556260-DEL-71
chr8-81580599-DEL-106
chr8-81584115-DEL-173
chr8-81626519-DEL-100
chr8-81630610-DEL-664
chr8-81635612-DEL-136
chr8-81659212-DEL-2557
chr8-81671061-DEL-66
chr8-81684089-DEL-170
chr8-81698826-DEL-307
chr8-81705158-DEL-509
chr8-81724186-DEL-193
chr8-81728834-DEL-984
chr8-81729961-DEL-1390
chr8-81744921-DEL-208
chr8-81774051-DEL-67
chr8-81782778-DEL-57
chr8-81787883-DEL-73
chr8-81811603-DEL-367
chr8-81811649-DEL-65
chr8-81823129-DEL-71
chr8-81852507-DEL-4603
chr8-81877536-DEL-1243
chr8-81896233-DEL-582
chr8-81911791-DEL-162
chr8-81944534-DEL-186
chr8-81975410-DEL-119

chr8-81976749-DEL-55
chr8-82002742-DEL-74
chr8-82049575-DEL-643
chr8-82053527-DEL-208
chr8-82059755-DEL-52
chr8-82098025-DEL-59
chr8-82101507-DEL-233
chr8-82109106-DEL-93
chr8-82111719-DEL-192
chr8-82180418-DEL-143
chr8-82227593-DEL-970
chr8-82234544-DEL-85
chr8-82252236-DEL-983
chr8-82291623-DEL-202
chr8-82304382-DEL-66
chr8-82313789-DEL-173
chr8-82325194-DEL-73
chr8-82326856-DEL-76
chr8-82328970-DEL-284
chr8-82334352-DEL-68
chr8-82380327-DEL-189
chr8-82382521-DEL-144
chr8-82428122-DEL-82
chr8-82429804-DEL-1804
chr8-82447189-DEL-198
chr8-82447938-DEL-6434
chr8-82453125-DEL-1200
chr8-82453688-DEL-637
chr8-82459526-DEL-251
chr8-82484789-DEL-805
chr8-82517740-DEL-67
chr8-82597922-DEL-99
chr8-82625050-DEL-94
chr8-82646976-DEL-485
chr8-82661545-DEL-300
chr8-82663491-DEL-229
chr8-82741035-DEL-2658
chr8-82769987-DEL-84
chr8-82843706-DEL-262
chr8-82858855-DEL-69
chr8-82871915-DEL-70
chr8-82945552-DEL-239
chr8-82977767-DEL-80
chr8-83007469-DEL-166
chr8-83020392-DEL-813
chr8-83079109-DEL-860
chr8-83097805-DEL-648
chr8-83122288-DEL-2312
chr8-83140615-DEL-568
chr8-83161943-DEL-2995
chr8-83165021-DEL-552
chr8-83197836-DEL-739
chr8-83199420-DEL-159
chr8-83201539-DEL-968
chr8-83208098-DEL-52
chr8-83209791-DEL-526
chr8-83223935-DEL-54
chr8-83252625-DEL-1258
chr8-83258060-DEL-95
chr8-83259740-DEL-86
chr8-83272141-DEL-143
chr8-83297303-DEL-520
chr8-83319310-DEL-5086
chr8-83346965-DEL-56
chr8-83359321-DEL-6366
chr8-83371264-DEL-6573
chr8-83380707-DEL-63
chr8-83389469-DEL-642
chr8-83402339-DEL-1821
chr8-83445706-DEL-205
chr8-83529599-DEL-6195
chr8-83542309-DEL-54
chr8-83543078-DEL-169
chr8-83545315-DEL-6654
chr8-83566730-DEL-56
chr8-83577849-DEL-166
chr8-83588395-DEL-73

chr8-83593474-DEL-60
chr8-83655112-DEL-117
chr8-83667455-DEL-561
chr8-83676404-DEL-61
chr8-83681090-DEL-898
chr8-83745398-DEL-6338
chr8-83746928-DEL-6335
chr8-83756837-DEL-93
chr8-83797282-DEL-2636
chr8-83802504-DEL-135
chr8-83849106-DEL-5210
chr8-83850423-DEL-1955
chr8-83854977-DEL-332
chr8-83857708-DEL-57
chr8-83878342-DEL-315
chr8-83882521-DEL-90
chr8-83897300-DEL-234
chr8-83899883-DEL-58
chr8-83915431-DEL-209
chr8-83920396-DEL-57
chr8-83922913-DEL-166
chr8-83925143-DEL-85
chr8-83951329-DEL-72
chr8-83955078-DEL-1709
chr8-83955169-DEL-2340
chr8-83956892-DEL-879
chr8-83956894-DEL-1266
chr8-83957827-DEL-332
chr8-83989064-DEL-524
chr8-84005813-DEL-73
chr8-84014954-DEL-57
chr8-84022380-DEL-554
chr8-84053381-DEL-121
chr8-84120333-DEL-168
chr8-84141273-DEL-4516
chr8-84149042-DEL-525
chr8-84157764-DEL-53
chr8-84174036-DEL-52
chr8-84177182-DEL-275
chr8-84189706-DEL-140
chr8-84233155-DEL-3556
chr8-84253669-DEL-1279
chr8-84290796-DEL-98
chr8-84306265-DEL-91
chr8-84328233-DEL-74
chr8-84364736-DEL-128
chr8-84366649-DEL-270
chr8-84418030-DEL-1111
chr8-84418235-DEL-184
chr8-84422099-DEL-56
chr8-84434108-DEL-91
chr8-84435198-DEL-213
chr8-84437555-DEL-199
chr8-84470416-DEL-207
chr8-84481526-DEL-197
chr8-84483363-DEL-192
chr8-84489771-DEL-590
chr8-84542975-DEL-65
chr8-84544368-DEL-113
chr8-84545012-DEL-1228
chr8-84548554-DEL-132
chr8-84555849-DEL-77
chr8-84578352-DEL-181
chr8-84585867-DEL-92
chr8-84589847-DEL-243
chr8-84592966-DEL-140
chr8-84611351-DEL-95
chr8-84631246-DEL-52
chr8-84676715-DEL-246
chr8-84678392-DEL-227
chr8-84712115-DEL-207
chr8-84751345-DEL-62
chr8-84837328-DEL-2383
chr8-84847284-DEL-57
chr8-84850218-DEL-318
chr8-84884296-DEL-51
chr8-84885319-DEL-165

chr8-84903634-DEL-54
chr8-84904424-DEL-168
chr8-84912489-DEL-66
chr8-84957707-DEL-79
chr8-84969912-DEL-202
chr8-85002968-DEL-126
chr8-85022338-DEL-68
chr8-85038109-DEL-206
chr8-85044169-DEL-52
chr8-85057516-DEL-61
chr8-85112133-DEL-63
chr8-85126422-DEL-68
chr8-85148727-DEL-178
chr8-85159752-DEL-103
chr8-85195257-DEL-245
chr8-85214600-DEL-77
chr8-85219455-DEL-164
chr8-85222176-DEL-102
chr8-85239712-DEL-104
chr8-85254928-DEL-765
chr8-85280309-DEL-98
chr8-85297219-DEL-51
chr8-85329676-DEL-68
chr8-85335743-DEL-145
chr8-85339832-DEL-52
chr8-85346626-DEL-84
chr8-85367763-DEL-206
chr8-85375149-DEL-1801
chr8-85378372-DEL-836
chr8-85379360-DEL-958
chr8-85380341-DEL-483
chr8-85418413-DEL-212
chr8-85421727-DEL-51
chr8-85466457-DEL-351
chr8-85543771-DEL-60
chr8-85544774-DEL-603
chr8-85555110-DEL-2305
chr8-85555507-DEL-417
chr8-85576596-DEL-202
chr8-85578430-DEL-1882
chr8-85632785-DEL-221
chr8-85652865-DEL-84
chr8-85666739-DEL-206
chr8-85669454-DEL-158
chr8-85688512-DEL-385
chr8-85736754-DEL-198
chr8-85761978-DEL-102
chr8-85766229-DEL-131
chr8-85786356-DEL-174
chr8-85797230-DEL-88
chr8-85804704-DEL-206
chr8-85840520-DEL-68
chr8-85840981-DEL-94
chr8-85852478-DEL-6392
chr8-85862657-DEL-77
chr8-85863697-DEL-1468
chr8-85863834-DEL-5014
chr8-85865458-DEL-2993
chr8-85881962-DEL-54
chr8-85885733-DEL-7480
chr8-85888820-DEL-216
chr8-85890457-DEL-205
chr8-85900745-DEL-208
chr8-85971214-DEL-6158
chr8-86007801-DEL-51
chr8-86018522-DEL-120
chr8-86078997-DEL-87
chr8-86105300-DEL-652
chr8-86117165-DEL-99
chr8-86149071-DEL-208
chr8-86162753-DEL-59
chr8-86170070-DEL-71
chr8-86171168-DEL-80
chr8-86179586-DEL-206
chr8-86186792-DEL-405
chr8-86199244-DEL-417
chr8-86207507-DEL-628

chr8-86226244-DEL-296
chr8-86256771-DEL-75
chr8-86271705-DEL-57
chr8-86289448-DEL-135
chr8-86331849-DEL-114
chr8-86375557-DEL-709
chr8-86382911-DEL-1651
chr8-86446221-DEL-360
chr8-86454818-DEL-204
chr8-86469159-DEL-73
chr8-86504245-DEL-67
chr8-86507297-DEL-87
chr8-86514739-DEL-143
chr8-86523623-DEL-2111
chr8-86526929-DEL-57
chr8-86578184-DEL-252
chr8-86603811-DEL-79
chr8-86608463-DEL-54
chr8-86613776-DEL-86
chr8-86633236-DEL-168
chr8-86633453-DEL-1447
chr8-86634883-DEL-1586
chr8-86656907-DEL-101
chr8-86675184-DEL-2223
chr8-86686209-DEL-144
chr8-86687971-DEL-109
chr8-86704648-DEL-1992
chr8-86706331-DEL-735
chr8-86706349-DEL-1243
chr8-86707103-DEL-869
chr8-86707843-DEL-1786
chr8-86709169-DEL-484
chr8-86715869-DEL-4940
chr8-86730416-DEL-53
chr8-86772789-DEL-141
chr8-86773819-DEL-176
chr8-86777229-DEL-2742
chr8-86783724-DEL-95
chr8-86793358-DEL-9407
chr8-86795030-DEL-132
chr8-86818522-DEL-933
chr8-86834439-DEL-200
chr8-86845380-DEL-462
chr8-86856046-DEL-1345
chr8-86856656-DEL-3927
chr8-86882484-DEL-69
chr8-86886787-DEL-191
chr8-86896760-DEL-52
chr8-86909892-DEL-1286
chr8-86958637-DEL-140
chr8-86963474-DEL-62
chr8-86989885-DEL-63
chr8-87040241-DEL-438
chr8-87044243-DEL-378
chr8-87064155-DEL-116
chr8-87068429-DEL-179
chr8-87105260-DEL-212
chr8-87126500-DEL-52
chr8-87130838-DEL-404
chr8-87149268-DEL-210
chr8-87161712-DEL-174
chr8-87172032-DEL-51
chr8-87189200-DEL-688
chr8-87198171-DEL-243
chr8-87234133-DEL-51
chr8-87245101-DEL-373
chr8-87251787-DEL-1030
chr8-87291522-DEL-241
chr8-87313668-DEL-60
chr8-87377425-DEL-407
chr8-87416566-DEL-88
chr8-87419851-DEL-119
chr8-87428845-DEL-2653
chr8-87431956-DEL-91
chr8-87507410-DEL-52
chr8-87531963-DEL-75
chr8-87536627-DEL-1785

chr8-87538491-DEL-1558
chr8-87572324-DEL-115
chr8-87585627-DEL-88
chr8-87605026-DEL-3754
chr8-87607013-DEL-169
chr8-87619658-DEL-98
chr8-87646283-DEL-203
chr8-87668663-DEL-72
chr8-87689481-DEL-162
chr8-87726318-DEL-120
chr8-87750840-DEL-110
chr8-87777472-DEL-128
chr8-87797168-DEL-74
chr8-87805600-DEL-63
chr8-87806659-DEL-1522
chr8-87836424-DEL-144
chr8-87918206-DEL-58
chr8-87943013-DEL-316
chr8-87959220-DEL-708
chr8-87972564-DEL-1420
chr8-88004242-DEL-174
chr8-88029280-DEL-264
chr8-88035819-DEL-1479
chr8-88071155-DEL-61
chr8-88247224-DEL-134
chr8-88275107-DEL-1421
chr8-88322384-DEL-1447
chr8-88487867-DEL-204
chr8-88523427-DEL-63
chr8-88526418-DEL-63
chr8-88552565-DEL-81
chr8-88575300-DEL-69
chr8-88592709-DEL-1496
chr8-88612450-DEL-114
chr8-88646527-DEL-198
chr8-88715364-DEL-67
chr8-88762133-DEL-56
chr8-88781456-DEL-219
chr8-88785889-DEL-109
chr8-88796780-DEL-58
chr8-88837303-DEL-205
chr8-88870097-DEL-165
chr8-88885442-DEL-57
chr8-88886568-DEL-85
chr8-88890125-DEL-385
chr8-88908517-DEL-176
chr8-88911289-DEL-425
chr8-88915699-DEL-177
chr8-88923262-DEL-57
chr8-88946194-DEL-104
chr8-88958196-DEL-54
chr8-88962411-DEL-223
chr8-88971662-DEL-283
chr8-88995489-DEL-138
chr8-89030993-DEL-52
chr8-89031972-DEL-104
chr8-89047505-DEL-60
chr8-89069828-DEL-195
chr8-89080536-DEL-490
chr8-89082448-DEL-188
chr8-89097994-DEL-194
chr8-89130713-DEL-1131
chr8-89131897-DEL-498
chr8-89136211-DEL-51
chr8-89139341-DEL-54
chr8-89173196-DEL-60
chr8-89201996-DEL-934
chr8-89205125-DEL-72
chr8-89223230-DEL-64
chr8-89234973-DEL-358
chr8-89326931-DEL-202
chr8-89333519-DEL-78
chr8-89354776-DEL-55
chr8-89413999-DEL-104
chr8-89525911-DEL-54
chr8-89543820-DEL-100
chr8-89643115-DEL-56

chr8-89677687-DEL-206
chr8-89688946-DEL-353
chr8-89711879-DEL-52
chr8-89723111-DEL-167
chr8-89725289-DEL-496
chr8-89731521-DEL-181
chr8-89803657-DEL-1349
chr8-89815590-DEL-52
chr8-89817822-DEL-106
chr8-89817974-DEL-614
chr8-89823457-DEL-1502
chr8-89824979-DEL-1566
chr8-89842401-DEL-604
chr8-89843127-DEL-2169
chr8-89853120-DEL-56
chr8-89854293-DEL-78
chr8-89860171-DEL-64
chr8-89875597-DEL-127
chr8-89885682-DEL-56
chr8-89916835-DEL-57
chr8-89940656-DEL-118
chr8-89957130-DEL-995
chr8-89959610-DEL-5760
chr8-89962218-DEL-93
chr8-90069343-DEL-220
chr8-90097017-DEL-65
chr8-90131635-DEL-648
chr8-90269124-DEL-70
chr8-90307222-DEL-195
chr8-90331732-DEL-66
chr8-90351579-DEL-59
chr8-90363457-DEL-988
chr8-90397081-DEL-285
chr8-90462255-DEL-99
chr8-90481761-DEL-59
chr8-90552628-DEL-1137
chr8-90593882-DEL-78
chr8-90604530-DEL-997
chr8-90659840-DEL-1213
chr8-90661586-DEL-2239
chr8-90691337-DEL-65
chr8-90693818-DEL-71
chr8-90707559-DEL-63
chr8-90726476-DEL-52
chr8-90752091-DEL-70
chr8-90783512-DEL-502
chr8-90801165-DEL-71
chr8-90812900-DEL-53
chr8-90814908-DEL-75
chr8-90823314-DEL-53
chr8-90853663-DEL-52
chr8-90884845-DEL-66
chr8-90887874-DEL-141
chr8-90895062-DEL-126
chr8-90904979-DEL-152
chr8-90939051-DEL-119
chr8-90942126-DEL-6838
chr8-90951320-DEL-2731
chr8-91032026-DEL-232
chr8-91055937-DEL-51
chr8-91070737-DEL-160
chr8-91182258-DEL-56
chr8-91205845-DEL-131
chr8-91210720-DEL-78
chr8-91212153-DEL-176
chr8-91213674-DEL-90
chr8-91221204-DEL-54
chr8-91267811-DEL-53
chr8-91273147-DEL-57
chr8-91291485-DEL-140
chr8-91299087-DEL-1248
chr8-91300094-DEL-386
chr8-91407012-DEL-70
chr8-91428198-DEL-56
chr8-91432859-DEL-236
chr8-91442319-DEL-125
chr8-91497609-DEL-51

chr8-91498231-DEL-332
chr8-91499058-DEL-4448
chr8-91499444-DEL-101
chr8-91504998-DEL-253
chr8-91507528-DEL-100
chr8-91509302-DEL-155
chr8-91515882-DEL-121
chr8-91519807-DEL-56
chr8-91523312-DEL-60
chr8-91551601-DEL-281
chr8-91562462-DEL-200
chr8-91563532-DEL-120
chr8-91572869-DEL-362
chr8-91584828-DEL-104
chr8-91587115-DEL-70
chr8-91606014-DEL-65
chr8-91691982-DEL-199
chr8-91733822-DEL-60
chr8-91747275-DEL-848
chr8-91762012-DEL-199
chr8-91784423-DEL-54
chr8-91795217-DEL-52
chr8-91820575-DEL-123
chr8-91831370-DEL-57
chr8-91876230-DEL-202
chr8-91895896-DEL-105
chr8-91908356-DEL-66
chr8-91909881-DEL-66
chr8-91963411-DEL-65
chr8-92002108-DEL-66
chr8-92048307-DEL-56
chr8-92059791-DEL-97
chr8-92090446-DEL-52
chr8-92111897-DEL-2125
chr8-92140844-DEL-417
chr8-92149979-DEL-64
chr8-92154337-DEL-112
chr8-92183379-DEL-75
chr8-92186065-DEL-58
chr8-92197415-DEL-283
chr8-92203325-DEL-117
chr8-92224742-DEL-253
chr8-92233109-DEL-59
chr8-92253495-DEL-53
chr8-92281143-DEL-56
chr8-92282217-DEL-2162
chr8-92290874-DEL-212
chr8-92331062-DEL-56
chr8-92395540-DEL-103
chr8-92400315-DEL-60
chr8-92536572-DEL-266
chr8-92538917-DEL-65
chr8-92550409-DEL-52
chr8-92556203-DEL-58
chr8-92645535-DEL-176
chr8-92648473-DEL-52
chr8-92671815-DEL-54
chr8-92682448-DEL-108
chr8-92749699-DEL-52
chr8-92794784-DEL-52
chr8-92801369-DEL-61
chr8-92816838-DEL-201
chr8-92853491-DEL-81
chr8-92857309-DEL-173
chr8-92889861-DEL-3381
chr8-92896195-DEL-73
chr8-92901915-DEL-62
chr8-92924340-DEL-65
chr8-92942590-DEL-247
chr8-92956504-DEL-193
chr8-92993760-DEL-75
chr8-93019686-DEL-81
chr8-93034547-DEL-265
chr8-93036028-DEL-109
chr8-93048162-DEL-549
chr8-93102280-DEL-57
chr8-93105892-DEL-53

chr8-93112139-DEL-55
chr8-93120915-DEL-68
chr8-93130564-DEL-59
chr8-93150794-DEL-82
chr8-93151268-DEL-527
chr8-93197623-DEL-120
chr8-93221586-DEL-138
chr8-93240284-DEL-1911
chr8-93273179-DEL-106
chr8-93306092-DEL-532
chr8-93307259-DEL-62
chr8-93320782-DEL-2781
chr8-93337954-DEL-107
chr8-93346866-DEL-113
chr8-93414616-DEL-87
chr8-93422995-DEL-309
chr8-93431095-DEL-62
chr8-93452215-DEL-169
chr8-93455850-DEL-79
chr8-93459693-DEL-172
chr8-93476925-DEL-57
chr8-93531406-DEL-82
chr8-93533688-DEL-210
chr8-93651966-DEL-52
chr8-93682287-DEL-135
chr8-93746047-DEL-55
chr8-93767596-DEL-114
chr8-93770439-DEL-199
chr8-93846412-DEL-76
chr8-93860542-DEL-54
chr8-93912057-DEL-154
chr8-93962036-DEL-701
chr8-93982149-DEL-553
chr8-93987953-DEL-76
chr8-94025479-DEL-214
chr8-94036226-DEL-262
chr8-94040862-DEL-101
chr8-94091413-DEL-982
chr8-94113065-DEL-2807
chr8-94155850-DEL-854
chr8-94163191-DEL-444
chr8-94176312-DEL-3531
chr8-94179913-DEL-3380
chr8-94193781-DEL-3044
chr8-94205949-DEL-6570
chr8-94212282-DEL-530
chr8-94222309-DEL-239
chr8-94248078-DEL-293
chr8-94253574-DEL-56
chr8-94266980-DEL-514
chr8-94293538-DEL-173
chr8-94310140-DEL-212
chr8-94334909-DEL-3094
chr8-94379694-DEL-63
chr8-94415908-DEL-94
chr8-94423358-DEL-189
chr8-94460103-DEL-400
chr8-94484472-DEL-207
chr8-94547436-DEL-53
chr8-94591240-DEL-1145
chr8-94607642-DEL-60
chr8-94640705-DEL-416
chr8-94713627-DEL-325
chr8-94746635-DEL-545
chr8-94748331-DEL-69
chr8-94772189-DEL-77
chr8-94783846-DEL-97
chr8-94848452-DEL-81
chr8-94857997-DEL-92
chr8-94883474-DEL-91
chr8-94894839-DEL-1432
chr8-94896298-DEL-1513
chr8-94902071-DEL-259
chr8-94904666-DEL-84
chr8-94921473-DEL-147
chr8-94927046-DEL-1088
chr8-94929104-DEL-1541

chr8-94943703-DEL-230
chr8-94960188-DEL-90
chr8-95006006-DEL-58
chr8-95023242-DEL-62
chr8-95077895-DEL-104
chr8-95093388-DEL-72
chr8-95133946-DEL-93
chr8-95143250-DEL-51
chr8-95164883-DEL-158
chr8-95175411-DEL-51
chr8-95180070-DEL-113
chr8-95210779-DEL-225
chr8-95229316-DEL-251
chr8-95230621-DEL-200
chr8-95241871-DEL-105
chr8-95247593-DEL-387
chr8-95331622-DEL-170
chr8-95351023-DEL-152
chr8-95370419-DEL-2503
chr8-95374246-DEL-252
chr8-95388658-DEL-276
chr8-95414762-DEL-131
chr8-95417363-DEL-68
chr8-95425032-DEL-105
chr8-95495136-DEL-56
chr8-95517751-DEL-206
chr8-95522016-DEL-254
chr8-95525784-DEL-605
chr8-95534634-DEL-122
chr8-95556457-DEL-437
chr8-95616880-DEL-59
chr8-95653404-DEL-209
chr8-95687313-DEL-171
chr8-95748082-DEL-685
chr8-95750461-DEL-83
chr8-95753404-DEL-171
chr8-95760339-DEL-842
chr8-95789486-DEL-104
chr8-95805065-DEL-153
chr8-95865648-DEL-64
chr8-95885997-DEL-164
chr8-95894043-DEL-85
chr8-95917768-DEL-340
chr8-95943554-DEL-414
chr8-95977239-DEL-317
chr8-95998833-DEL-103
chr8-95999801-DEL-96
chr8-96048278-DEL-59
chr8-96062048-DEL-140
chr8-96077347-DEL-204
chr8-96158010-DEL-304
chr8-96189988-DEL-62
chr8-96221566-DEL-105
chr8-96222361-DEL-94
chr8-96261343-DEL-338
chr8-96264889-DEL-83
chr8-96274871-DEL-286
chr8-96291480-DEL-246
chr8-96332814-DEL-476
chr8-96334779-DEL-90
chr8-96367683-DEL-180
chr8-96372281-DEL-1018
chr8-96377291-DEL-174
chr8-96392634-DEL-181
chr8-96444191-DEL-60
chr8-96453382-DEL-209
chr8-96464441-DEL-815
chr8-96508464-DEL-1286
chr8-96508980-DEL-77
chr8-96518090-DEL-212
chr8-96521062-DEL-2928
chr8-96536614-DEL-72
chr8-96541434-DEL-193
chr8-96541435-DEL-513
chr8-96557173-DEL-763
chr8-96560875-DEL-413
chr8-96573990-DEL-413

chr8-96576689-DEL-83
chr8-96581364-DEL-189
chr8-96581756-DEL-2061
chr8-96584942-DEL-401
chr8-96611371-DEL-353
chr8-96618779-DEL-53
chr8-96646261-DEL-136
chr8-96652602-DEL-177
chr8-96685756-DEL-448
chr8-96810407-DEL-224
chr8-96811256-DEL-1967
chr8-96815874-DEL-728
chr8-96821742-DEL-127
chr8-96851964-DEL-60
chr8-96860217-DEL-539
chr8-96909120-DEL-514
chr8-96927235-DEL-413
chr8-96967486-DEL-1068
chr8-96972008-DEL-126
chr8-96984796-DEL-507
chr8-96986939-DEL-5279
chr8-97012147-DEL-1085
chr8-97019464-DEL-62
chr8-97059876-DEL-6449
chr8-97094846-DEL-100
chr8-97127830-DEL-204
chr8-97143143-DEL-6243
chr8-97182122-DEL-104
chr8-97265638-DEL-58
chr8-97327437-DEL-896
chr8-97354088-DEL-109
chr8-97363155-DEL-69
chr8-97364213-DEL-851
chr8-97370602-DEL-1840
chr8-97385570-DEL-174
chr8-97388411-DEL-6578
chr8-97388978-DEL-263
chr8-97389784-DEL-292
chr8-97392785-DEL-85
chr8-97393318-DEL-1007
chr8-97436344-DEL-106
chr8-97456326-DEL-1454
chr8-97474554-DEL-135
chr8-97486087-DEL-210
chr8-97517174-DEL-1586
chr8-97542806-DEL-647
chr8-97559640-DEL-55
chr8-97560630-DEL-221
chr8-97566631-DEL-54
chr8-97591960-DEL-76
chr8-97596634-DEL-600
chr8-97602537-DEL-399
chr8-97610611-DEL-496
chr8-97619884-DEL-885
chr8-97631704-DEL-521
chr8-97632194-DEL-313
chr8-97692306-DEL-177
chr8-97695373-DEL-208
chr8-97705691-DEL-70
chr8-97714806-DEL-139
chr8-97733492-DEL-1066
chr8-97757859-DEL-72
chr8-97761125-DEL-470
chr8-97779721-DEL-54
chr8-97782561-DEL-136
chr8-97792228-DEL-2289
chr8-97797567-DEL-392
chr8-97802924-DEL-2222
chr8-97804884-DEL-487
chr8-97826317-DEL-71
chr8-97834021-DEL-235
chr8-97845536-DEL-411
chr8-97846692-DEL-767
chr8-97854902-DEL-141
chr8-97904379-DEL-899
chr8-97905274-DEL-369
chr8-97986585-DEL-414

chr8-98044757-DEL-180
chr8-98070380-DEL-401
chr8-98081578-DEL-140
chr8-98087531-DEL-79
chr8-98093125-DEL-468
chr8-98100951-DEL-68
chr8-98178498-DEL-1616
chr8-98193680-DEL-118
chr8-98209723-DEL-1336
chr8-98229006-DEL-51
chr8-98239365-DEL-135
chr8-98246751-DEL-1482
chr8-98288846-DEL-69
chr8-98297652-DEL-56
chr8-98317206-DEL-1403
chr8-98318455-DEL-155
chr8-98330716-DEL-389
chr8-98354879-DEL-56
chr8-98388931-DEL-1085
chr8-98391020-DEL-58
chr8-98431499-DEL-53
chr8-98440982-DEL-101
chr8-98467680-DEL-153
chr8-98485552-DEL-202
chr8-98510014-DEL-67
chr8-98522244-DEL-329
chr8-98522281-DEL-6621
chr8-98535814-DEL-553
chr8-98542952-DEL-496
chr8-98548833-DEL-790
chr8-98576167-DEL-60
chr8-98579205-DEL-1969
chr8-98601778-DEL-270
chr8-98613569-DEL-1106
chr8-98617457-DEL-1149
chr8-98632828-DEL-1102
chr8-98659756-DEL-375
chr8-98725177-DEL-652
chr8-98747082-DEL-60
chr8-98755315-DEL-6223
chr8-98781721-DEL-790
chr8-98794048-DEL-400
chr8-98837366-DEL-5306
chr8-98918569-DEL-198
chr8-98919772-DEL-52
chr8-98925643-DEL-533
chr8-98942827-DEL-52
chr8-98947252-DEL-349
chr8-98950665-DEL-786
chr8-98963450-DEL-980
chr8-98983972-DEL-177
chr8-98989446-DEL-190
chr8-99017638-DEL-57
chr8-99029671-DEL-3321
chr8-99127362-DEL-171
chr8-99139204-DEL-85
chr8-99164229-DEL-129
chr8-99186959-DEL-119
chr8-99245081-DEL-68
chr8-99258221-DEL-261
chr8-99270187-DEL-105
chr8-99277584-DEL-1575
chr8-99283434-DEL-2518
chr8-99288615-DEL-1320
chr8-99301052-DEL-150
chr8-99316056-DEL-99
chr8-99334564-DEL-67
chr8-99372170-DEL-65
chr8-99375351-DEL-133
chr8-99378571-DEL-462
chr8-99386235-DEL-63
chr8-99392843-DEL-676
chr8-99401837-DEL-520
chr8-99418536-DEL-52
chr8-99434907-DEL-10413
chr8-99484208-DEL-364
chr8-99485834-DEL-59

chr8-99511120-DEL-57
chr8-99544773-DEL-58
chr8-99556720-DEL-77
chr8-99561096-DEL-134
chr8-99564293-DEL-8543
chr8-99574981-DEL-66
chr8-99607125-DEL-94
chr8-99628244-DEL-418
chr8-99663515-DEL-62
chr8-99681456-DEL-225
chr8-99700897-DEL-161
chr8-99707142-DEL-509
chr8-99718716-DEL-61
chr8-99745015-DEL-569
chr8-99864294-DEL-389
chr8-99925727-DEL-160
chr8-99939980-DEL-174
chr8-99951910-DEL-83
chr8-99979805-DEL-222
chr8-99998567-DEL-140
chr8-100017743-DEL-112
chr8-100020467-DEL-55
chr8-100022297-DEL-1544
chr8-100077946-DEL-82
chr8-100097660-DEL-51
chr8-100103745-DEL-1244
chr8-100108790-DEL-59
chr8-100123610-DEL-79
chr8-100180384-DEL-497
chr8-100193631-DEL-212
chr8-100208691-DEL-208
chr8-100239454-DEL-134
chr8-100242093-DEL-2343
chr8-100255263-DEL-210
chr8-100273018-DEL-647
chr8-100273946-DEL-1334
chr8-100282820-DEL-752
chr8-100337098-DEL-5224
chr8-100340568-DEL-366
chr8-100359515-DEL-70
chr8-100360241-DEL-75
chr8-100373678-DEL-77
chr8-100441141-DEL-54
chr8-100442662-DEL-2333
chr8-100448371-DEL-314
chr8-100486357-DEL-719
chr8-100490171-DEL-77
chr8-100530503-DEL-624
chr8-100531477-DEL-178
chr8-100549951-DEL-95
chr8-100586681-DEL-6025
chr8-100586691-DEL-907
chr8-100593060-DEL-231
chr8-100665925-DEL-240
chr8-100678526-DEL-2280
chr8-100678693-DEL-826
chr8-100688244-DEL-394
chr8-100701052-DEL-62
chr8-100748898-DEL-2719
chr8-100800890-DEL-1207
chr8-100823377-DEL-1788
chr8-100858560-DEL-51
chr8-100870299-DEL-97
chr8-100887746-DEL-2167
chr8-100897628-DEL-189
chr8-100918307-DEL-294
chr8-100931632-DEL-193
chr8-100986848-DEL-101
chr8-100991932-DEL-78
chr8-100996539-DEL-528
chr8-101067876-DEL-799
chr8-101101642-DEL-4925
chr8-101114995-DEL-1406
chr8-101144439-DEL-436
chr8-101160060-DEL-121
chr8-101160467-DEL-1389
chr8-101161874-DEL-998

chr8-101207344-DEL-123
chr8-101224649-DEL-897
chr8-101238810-DEL-1390
chr8-101250718-DEL-680
chr8-101260972-DEL-72
chr8-101276930-DEL-2067
chr8-101331715-DEL-2254
chr8-101409649-DEL-1320
chr8-101415274-DEL-1893
chr8-101419486-DEL-202
chr8-101419884-DEL-1303
chr8-101421171-DEL-58
chr8-101428827-DEL-55
chr8-101438131-DEL-56
chr8-101464739-DEL-176
chr8-101466902-DEL-52
chr8-101497374-DEL-73
chr8-101535425-DEL-463
chr8-101547248-DEL-456
chr8-101549956-DEL-3382
chr8-101552412-DEL-261
chr8-101594479-DEL-63
chr8-101606030-DEL-3526
chr8-101616360-DEL-688
chr8-101625053-DEL-53
chr8-101636738-DEL-7220
chr8-101659177-DEL-410
chr8-101664267-DEL-551
chr8-101679164-DEL-938
chr8-101685137-DEL-64
chr8-101688198-DEL-6040
chr8-101726791-DEL-71
chr8-101733603-DEL-401
chr8-101772797-DEL-2392
chr8-101800976-DEL-56
chr8-101812972-DEL-258
chr8-101823220-DEL-75
chr8-101846681-DEL-363
chr8-101869476-DEL-1557
chr8-102008221-DEL-1840
chr8-102012112-DEL-223
chr8-102021725-DEL-77
chr8-102023053-DEL-11474
chr8-102026258-DEL-867
chr8-102027227-DEL-227
chr8-102049566-DEL-1796
chr8-102096283-DEL-886
chr8-102101211-DEL-6370
chr8-102141888-DEL-358
chr8-102169734-DEL-306
chr8-102182684-DEL-172
chr8-102198907-DEL-63
chr8-102213602-DEL-60
chr8-102232215-DEL-236
chr8-102236742-DEL-490
chr8-102250127-DEL-596
chr8-102272686-DEL-605
chr8-102314065-DEL-199
chr8-102326320-DEL-4101
chr8-102337038-DEL-2597
chr8-102360680-DEL-165
chr8-102401585-DEL-54
chr8-102425941-DEL-82
chr8-102457305-DEL-4220
chr8-102469954-DEL-840
chr8-102471026-DEL-361
chr8-102515154-DEL-2024
chr8-102536224-DEL-1209
chr8-102543483-DEL-1897
chr8-102571922-DEL-91
chr8-102626797-DEL-1734
chr8-102659158-DEL-472
chr8-102668682-DEL-239
chr8-102685713-DEL-192
chr8-102689698-DEL-247
chr8-102698870-DEL-126
chr8-102701795-DEL-9728

chr8-102723254-DEL-692
chr8-102778633-DEL-1175
chr8-102821671-DEL-1302
chr8-102838895-DEL-140
chr8-102851749-DEL-10723
chr8-102878002-DEL-71
chr8-102887242-DEL-54
chr8-102887600-DEL-5008
chr8-102908622-DEL-60
chr8-102927569-DEL-285
chr8-102985072-DEL-128
chr8-102999255-DEL-101
chr8-103015806-DEL-174
chr8-103037417-DEL-76
chr8-103043377-DEL-200
chr8-103081450-DEL-89
chr8-103110782-DEL-1167
chr8-103125285-DEL-219
chr8-103177488-DEL-55
chr8-103213781-DEL-56
chr8-103231681-DEL-1405
chr8-103231750-DEL-118
chr8-103254770-DEL-127
chr8-103288639-DEL-343
chr8-103311385-DEL-694
chr8-103399088-DEL-69
chr8-103423176-DEL-1106
chr8-103427701-DEL-66
chr8-103454699-DEL-66
chr8-103534739-DEL-101
chr8-103563139-DEL-123
chr8-103586328-DEL-51
chr8-103614066-DEL-1530
chr8-103614067-DEL-3766
chr8-103615619-DEL-3259
chr8-103618945-DEL-1182
chr8-103622371-DEL-52
chr8-103626738-DEL-196
chr8-103632818-DEL-445
chr8-103643468-DEL-803
chr8-103695092-DEL-57
chr8-103781639-DEL-470
chr8-103785998-DEL-703
chr8-103792140-DEL-107
chr8-103822784-DEL-68
chr8-103824070-DEL-51
chr8-103828016-DEL-95
chr8-103839874-DEL-61
chr8-103841338-DEL-54
chr8-103894596-DEL-140
chr8-103894887-DEL-749
chr8-103975793-DEL-453
chr8-103978310-DEL-3134
chr8-104023999-DEL-56
chr8-104025841-DEL-78
chr8-104198646-DEL-7714
chr8-104240586-DEL-73
chr8-104294434-DEL-206
chr8-104327237-DEL-845
chr8-104339280-DEL-70
chr8-104341888-DEL-60
chr8-104373186-DEL-61
chr8-104383160-DEL-54
chr8-104408028-DEL-72
chr8-104418353-DEL-84
chr8-104444762-DEL-88
chr8-104454813-DEL-1047
chr8-104506902-DEL-51
chr8-104527953-DEL-111
chr8-104569187-DEL-154
chr8-104581342-DEL-90
chr8-104615071-DEL-134
chr8-104657528-DEL-247
chr8-104670457-DEL-298
chr8-104705172-DEL-51
chr8-104747913-DEL-157
chr8-104756710-DEL-72

chr8-104787608-DEL-75
chr8-104793376-DEL-442
chr8-104801740-DEL-107
chr8-104819820-DEL-72
chr8-104863188-DEL-55
chr8-104873965-DEL-858
chr8-104886989-DEL-105
chr8-104894878-DEL-503
chr8-104935962-DEL-97
chr8-104936234-DEL-1153
chr8-104938847-DEL-91
chr8-104985049-DEL-358
chr8-104991739-DEL-400
chr8-105003597-DEL-3843
chr8-105061726-DEL-278
chr8-105133261-DEL-129
chr8-105205389-DEL-84
chr8-105220338-DEL-292
chr8-105242281-DEL-52
chr8-105245131-DEL-132
chr8-105248802-DEL-182
chr8-105250726-DEL-160
chr8-105263410-DEL-107
chr8-105291374-DEL-59
chr8-105299209-DEL-52
chr8-105309144-DEL-204
chr8-105339061-DEL-144
chr8-105352221-DEL-68
chr8-105353633-DEL-226
chr8-105393284-DEL-200
chr8-105444567-DEL-126
chr8-105463258-DEL-68
chr8-105474495-DEL-4237
chr8-105497571-DEL-4416
chr8-105525545-DEL-181
chr8-105534279-DEL-688
chr8-105584309-DEL-6856
chr8-105601112-DEL-944
chr8-105602045-DEL-62
chr8-105603888-DEL-6648
chr8-105620138-DEL-6299
chr8-105636396-DEL-116
chr8-105678988-DEL-73
chr8-105762355-DEL-208
chr8-105775011-DEL-116
chr8-105795943-DEL-877
chr8-105797451-DEL-923
chr8-105808836-DEL-219
chr8-105812736-DEL-64
chr8-105827925-DEL-598
chr8-105830075-DEL-54
chr8-105836992-DEL-65
chr8-105846087-DEL-78
chr8-105859284-DEL-138
chr8-105938967-DEL-448
chr8-105950495-DEL-112
chr8-105970797-DEL-145
chr8-105971666-DEL-207
chr8-105984748-DEL-77
chr8-106028229-DEL-517
chr8-106047125-DEL-225
chr8-106070791-DEL-54
chr8-106137439-DEL-332
chr8-106154654-DEL-203
chr8-106179901-DEL-76
chr8-106183061-DEL-111
chr8-106215688-DEL-132
chr8-106261942-DEL-387
chr8-106274452-DEL-614
chr8-106311869-DEL-995
chr8-106354758-DEL-196
chr8-106401800-DEL-203
chr8-106464217-DEL-70
chr8-106478128-DEL-312
chr8-106479592-DEL-78
chr8-106491024-DEL-79
chr8-106515571-DEL-60

chr8-106521678-DEL-1284
chr8-106529170-DEL-206
chr8-106532879-DEL-207
chr8-106543045-DEL-183
chr8-106543806-DEL-236
chr8-106550815-DEL-167
chr8-106558355-DEL-1532
chr8-106561372-DEL-204
chr8-106577892-DEL-90
chr8-106591470-DEL-55
chr8-106613219-DEL-113
chr8-106622825-DEL-184
chr8-106640832-DEL-432
chr8-106645029-DEL-232
chr8-106651336-DEL-266
chr8-106655678-DEL-63
chr8-106664849-DEL-80
chr8-106682878-DEL-114
chr8-106697569-DEL-205
chr8-106703296-DEL-206
chr8-106738390-DEL-2887
chr8-106746802-DEL-208
chr8-106752387-DEL-87
chr8-106765265-DEL-69
chr8-106769286-DEL-71
chr8-106774340-DEL-206
chr8-106795640-DEL-97
chr8-106812503-DEL-71
chr8-106842957-DEL-211
chr8-106846970-DEL-460
chr8-106874018-DEL-408
chr8-106901074-DEL-205
chr8-106915096-DEL-293
chr8-106917095-DEL-227
chr8-106920504-DEL-196
chr8-106929513-DEL-245
chr8-106947429-DEL-144
chr8-106971899-DEL-209
chr8-106974660-DEL-181
chr8-106987959-DEL-70
chr8-107004721-DEL-92
chr8-107011995-DEL-138
chr8-107035447-DEL-168
chr8-107092023-DEL-84
chr8-107098685-DEL-58
chr8-107110943-DEL-87
chr8-107121926-DEL-539
chr8-107137310-DEL-147
chr8-107137949-DEL-164
chr8-107143790-DEL-98
chr8-107150043-DEL-54
chr8-107163646-DEL-193
chr8-107168856-DEL-159
chr8-107183528-DEL-106
chr8-107197447-DEL-64
chr8-107225864-DEL-147
chr8-107234187-DEL-92
chr8-107242897-DEL-170
chr8-107284367-DEL-190
chr8-107299553-DEL-751
chr8-107328920-DEL-73
chr8-107350665-DEL-53
chr8-107370201-DEL-111
chr8-107433963-DEL-191
chr8-107442344-DEL-219
chr8-107448437-DEL-236
chr8-107455525-DEL-466
chr8-107467689-DEL-62
chr8-107473232-DEL-402
chr8-107479061-DEL-343
chr8-107492907-DEL-277
chr8-107511335-DEL-146
chr8-107526225-DEL-97
chr8-107540155-DEL-136
chr8-107574505-DEL-81
chr8-107630510-DEL-60
chr8-107636108-DEL-1296

chr8-107636316-DEL-78
chr8-107640632-DEL-106
chr8-107647541-DEL-51
chr8-107660430-DEL-138
chr8-107677012-DEL-91
chr8-107677808-DEL-167
chr8-107688199-DEL-551
chr8-107692950-DEL-206
chr8-107722977-DEL-118
chr8-107728416-DEL-63
chr8-107730395-DEL-123
chr8-107735226-DEL-125
chr8-107740088-DEL-2794
chr8-107741307-DEL-143
chr8-107748118-DEL-860
chr8-107749929-DEL-695
chr8-107760450-DEL-211
chr8-107763693-DEL-1169
chr8-107781204-DEL-362
chr8-107793912-DEL-334
chr8-107793929-DEL-317
chr8-107815169-DEL-58
chr8-107815578-DEL-196
chr8-107835855-DEL-95
chr8-107855361-DEL-87
chr8-107880973-DEL-205
chr8-107883205-DEL-104
chr8-107950464-DEL-51
chr8-107957878-DEL-56
chr8-107973322-DEL-275
chr8-108002581-DEL-58
chr8-108003120-DEL-55
chr8-108015031-DEL-84
chr8-108045359-DEL-179
chr8-108079998-DEL-172
chr8-108141663-DEL-111
chr8-108162086-DEL-345
chr8-108180809-DEL-343
chr8-108207576-DEL-65
chr8-108227976-DEL-365
chr8-108232342-DEL-78
chr8-108298954-DEL-81
chr8-108410787-DEL-64
chr8-108437549-DEL-106
chr8-108449713-DEL-74
chr8-108451020-DEL-185
chr8-108472025-DEL-93
chr8-108495296-DEL-170
chr8-108531623-DEL-65
chr8-108565385-DEL-1096
chr8-108585961-DEL-53
chr8-108586429-DEL-164
chr8-108600869-DEL-79
chr8-108602795-DEL-151
chr8-108604804-DEL-177
chr8-108607320-DEL-52
chr8-108693107-DEL-58
chr8-108752492-DEL-356
chr8-108804482-DEL-248
chr8-108810209-DEL-65
chr8-108837175-DEL-51
chr8-108868698-DEL-55
chr8-108929838-DEL-83
chr8-108984960-DEL-87
chr8-108992556-DEL-447
chr8-109024489-DEL-56
chr8-109032994-DEL-89
chr8-109048282-DEL-1817
chr8-109115814-DEL-115
chr8-109138904-DEL-78
chr8-109194930-DEL-51
chr8-109216583-DEL-69
chr8-109225193-DEL-176
chr8-109251178-DEL-94
chr8-109313463-DEL-70
chr8-109350163-DEL-88
chr8-109361388-DEL-1769

chr8-109387839-DEL-66
chr8-109465650-DEL-197
chr8-109536234-DEL-90
chr8-109583059-DEL-54
chr8-109619642-DEL-431
chr8-109633826-DEL-68
chr8-109642610-DEL-274
chr8-109659171-DEL-53
chr8-109697099-DEL-64
chr8-109702469-DEL-817
chr8-109712252-DEL-692
chr8-109752294-DEL-316
chr8-109764751-DEL-77
chr8-109837016-DEL-62
chr8-109846595-DEL-72
chr8-109892375-DEL-206
chr8-109924648-DEL-513
chr8-109985386-DEL-54
chr8-110001609-DEL-52
chr8-110018984-DEL-2646
chr8-110078194-DEL-79
chr8-110109728-DEL-90
chr8-110117724-DEL-577
chr8-110157734-DEL-62
chr8-110167332-DEL-51
chr8-110172999-DEL-2599
chr8-110181083-DEL-716
chr8-110251514-DEL-78
chr8-110270147-DEL-419
chr8-110298575-DEL-204
chr8-110305010-DEL-58
chr8-110324396-DEL-162
chr8-110343757-DEL-127
chr8-110349920-DEL-155
chr8-110389499-DEL-166
chr8-110390116-DEL-111
chr8-110406448-DEL-128
chr8-110446167-DEL-222
chr8-110466594-DEL-6240
chr8-110494613-DEL-107
chr8-110533437-DEL-128
chr8-110543484-DEL-194
chr8-110561648-DEL-69
chr8-110564450-DEL-78
chr8-110612752-DEL-156
chr8-110621687-DEL-278
chr8-110626421-DEL-142
chr8-110706386-DEL-985
chr8-110740116-DEL-289
chr8-110745601-DEL-93
chr8-110752208-DEL-237
chr8-110820175-DEL-1797
chr8-110821534-DEL-442
chr8-110833228-DEL-94
chr8-110842542-DEL-75
chr8-110843252-DEL-103
chr8-110858440-DEL-146
chr8-110859801-DEL-449
chr8-110863116-DEL-133
chr8-110878279-DEL-174
chr8-110890675-DEL-436
chr8-110918933-DEL-919
chr8-110934299-DEL-60
chr8-110956266-DEL-62
chr8-110989973-DEL-71
chr8-110999148-DEL-123
chr8-111040907-DEL-178
chr8-111045712-DEL-2684
chr8-111079651-DEL-96
chr8-111083450-DEL-72
chr8-111084655-DEL-61
chr8-111112931-DEL-973
chr8-111142091-DEL-79
chr8-111152146-DEL-1717
chr8-111160554-DEL-280
chr8-111182770-DEL-187
chr8-111226983-DEL-140

chr8-111248140-DEL-107
chr8-111248935-DEL-536
chr8-111279367-DEL-237
chr8-111297866-DEL-328
chr8-111349138-DEL-223
chr8-111365803-DEL-168
chr8-111383832-DEL-228
chr8-111394405-DEL-139
chr8-111411833-DEL-394
chr8-111448478-DEL-117
chr8-111460223-DEL-62
chr8-111471882-DEL-218
chr8-111480514-DEL-253
chr8-111496135-DEL-69
chr8-111499876-DEL-356
chr8-111505140-DEL-1186
chr8-111505892-DEL-106
chr8-111565897-DEL-210
chr8-111581022-DEL-288
chr8-111585341-DEL-91
chr8-111587226-DEL-259
chr8-111600043-DEL-103
chr8-111623127-DEL-192
chr8-111632484-DEL-259
chr8-111643024-DEL-684
chr8-111679953-DEL-205
chr8-111746287-DEL-210
chr8-111748673-DEL-126
chr8-111749310-DEL-208
chr8-111751961-DEL-790
chr8-111753444-DEL-199
chr8-111758643-DEL-88
chr8-111766978-DEL-80
chr8-111814450-DEL-179
chr8-111817609-DEL-181
chr8-111818967-DEL-424
chr8-111824157-DEL-107
chr8-111836828-DEL-183
chr8-111860129-DEL-198
chr8-111868379-DEL-190
chr8-111893879-DEL-135
chr8-111934011-DEL-205
chr8-111993025-DEL-204
chr8-112012401-DEL-51
chr8-112051739-DEL-370
chr8-112105947-DEL-1579
chr8-112109808-DEL-54
chr8-112110131-DEL-366
chr8-112126045-DEL-394
chr8-112134099-DEL-162
chr8-112191530-DEL-72
chr8-112224980-DEL-70
chr8-112233077-DEL-85
chr8-112252333-DEL-192
chr8-112261237-DEL-689
chr8-112271398-DEL-76
chr8-112283453-DEL-112
chr8-112313146-DEL-884
chr8-112328562-DEL-77
chr8-112336348-DEL-68
chr8-112340645-DEL-1319
chr8-112341416-DEL-142
chr8-112362773-DEL-2341
chr8-112362861-DEL-54
chr8-112364576-DEL-225
chr8-112365153-DEL-1092
chr8-112370910-DEL-59
chr8-112379828-DEL-95
chr8-112397979-DEL-60
chr8-112425187-DEL-201
chr8-112448401-DEL-150
chr8-112476212-DEL-499
chr8-112482477-DEL-149
chr8-112499102-DEL-257
chr8-112523634-DEL-192
chr8-112524851-DEL-56
chr8-112552974-DEL-60

chr8-112606349-DEL-443
chr8-112624713-DEL-382
chr8-112661275-DEL-170
chr8-112664956-DEL-221
chr8-112673193-DEL-339
chr8-112675200-DEL-139
chr8-112682452-DEL-173
chr8-112696429-DEL-168
chr8-112699645-DEL-969
chr8-112704374-DEL-65
chr8-112717193-DEL-183
chr8-112728641-DEL-75
chr8-112762153-DEL-476
chr8-112766240-DEL-118
chr8-112779950-DEL-764
chr8-112783535-DEL-68
chr8-112799128-DEL-1531
chr8-112818851-DEL-378
chr8-112842954-DEL-91
chr8-112849077-DEL-118
chr8-112865144-DEL-67
chr8-112867053-DEL-61
chr8-112872390-DEL-571
chr8-112884835-DEL-101
chr8-112909691-DEL-62
chr8-112973113-DEL-324
chr8-112975831-DEL-57
chr8-112998491-DEL-87
chr8-113004365-DEL-104
chr8-113015867-DEL-206
chr8-113123635-DEL-80
chr8-113132028-DEL-194
chr8-113134330-DEL-51
chr8-113174856-DEL-468
chr8-113176580-DEL-122
chr8-113274027-DEL-116
chr8-113348755-DEL-1183
chr8-113397124-DEL-75
chr8-113407500-DEL-81
chr8-113409677-DEL-170
chr8-113411101-DEL-931
chr8-113458714-DEL-284
chr8-113469213-DEL-63
chr8-113483401-DEL-202
chr8-113511029-DEL-64
chr8-113532715-DEL-64
chr8-113545432-DEL-91
chr8-113564209-DEL-64
chr8-113612815-DEL-55
chr8-113661519-DEL-94
chr8-113663896-DEL-95
chr8-113724516-DEL-66
chr8-113728234-DEL-536
chr8-113734959-DEL-93
chr8-113781945-DEL-1681
chr8-113783635-DEL-1794
chr8-113835573-DEL-111
chr8-113852847-DEL-191
chr8-113887370-DEL-211
chr8-113897861-DEL-78
chr8-113958811-DEL-75
chr8-113991939-DEL-337
chr8-114000719-DEL-66
chr8-114018667-DEL-63
chr8-114060945-DEL-1265
chr8-114083854-DEL-56
chr8-114093079-DEL-1203
chr8-114170023-DEL-70
chr8-114186928-DEL-839
chr8-114216860-DEL-390
chr8-114263855-DEL-926
chr8-114274182-DEL-69
chr8-114349727-DEL-59
chr8-114360616-DEL-118
chr8-114420600-DEL-60
chr8-114421958-DEL-54
chr8-114426895-DEL-1592

chr8-114453401-DEL-91
chr8-114461630-DEL-116
chr8-114508787-DEL-83
chr8-114512624-DEL-260
chr8-114521703-DEL-1196
chr8-114534429-DEL-320
chr8-114560361-DEL-68
chr8-114586291-DEL-291
chr8-114631757-DEL-818
chr8-114658993-DEL-113
chr8-114665799-DEL-52
chr8-114718267-DEL-62
chr8-114756773-DEL-59
chr8-114785254-DEL-114
chr8-114787305-DEL-176
chr8-114848970-DEL-136
chr8-114871035-DEL-178
chr8-114876317-DEL-763
chr8-114897959-DEL-484
chr8-114898774-DEL-55
chr8-114911167-DEL-143
chr8-114914611-DEL-56
chr8-114926143-DEL-118
chr8-114931429-DEL-187
chr8-114934789-DEL-62
chr8-114974326-DEL-70
chr8-114986278-DEL-99
chr8-115000754-DEL-168
chr8-115015635-DEL-61
chr8-115029362-DEL-55
chr8-115069791-DEL-116
chr8-115073190-DEL-179
chr8-115149822-DEL-110
chr8-115150867-DEL-420
chr8-115163421-DEL-62
chr8-115183950-DEL-126
chr8-115214705-DEL-372
chr8-115217206-DEL-1989
chr8-115220413-DEL-54
chr8-115265926-DEL-72
chr8-115273934-DEL-131
chr8-115298242-DEL-56
chr8-115312136-DEL-85
chr8-115316485-DEL-99
chr8-115326218-DEL-972
chr8-115347251-DEL-1023
chr8-115358285-DEL-109
chr8-115391428-DEL-255
chr8-115489613-DEL-98
chr8-115500095-DEL-54
chr8-115523835-DEL-478
chr8-115542226-DEL-58
chr8-115546730-DEL-81
chr8-115564260-DEL-58
chr8-115583189-DEL-740
chr8-115592074-DEL-106
chr8-115596918-DEL-81
chr8-115608351-DEL-246
chr8-115618150-DEL-361
chr8-115620210-DEL-160
chr8-115628588-DEL-63
chr8-115644361-DEL-52
chr8-115649837-DEL-252
chr8-115661168-DEL-758
chr8-115692832-DEL-225
chr8-115694058-DEL-165
chr8-115697407-DEL-146
chr8-115774138-DEL-82
chr8-115836859-DEL-81
chr8-115838354-DEL-54
chr8-115841849-DEL-75
chr8-115850376-DEL-65
chr8-115874935-DEL-59
chr8-115879194-DEL-182
chr8-115881767-DEL-160
chr8-115883901-DEL-93
chr8-115909150-DEL-138

chr8-115928098-DEL-52
chr8-115934152-DEL-542
chr8-116024050-DEL-92
chr8-116024783-DEL-87
chr8-116027957-DEL-196
chr8-116039041-DEL-62
chr8-116050191-DEL-139
chr8-116065795-DEL-214
chr8-116098151-DEL-77
chr8-116104542-DEL-176
chr8-116113279-DEL-230
chr8-116151228-DEL-954
chr8-116190030-DEL-60
chr8-116274323-DEL-625
chr8-116280328-DEL-198
chr8-116283625-DEL-54
chr8-116284637-DEL-267
chr8-116304293-DEL-69
chr8-116306795-DEL-345
chr8-116315645-DEL-120
chr8-116320524-DEL-1246
chr8-116364671-DEL-182
chr8-116371588-DEL-133
chr8-116380547-DEL-419
chr8-116395622-DEL-72
chr8-116519223-DEL-147
chr8-116535148-DEL-546
chr8-116554529-DEL-240
chr8-116561267-DEL-60
chr8-116563316-DEL-171
chr8-116597777-DEL-66
chr8-116689322-DEL-54
chr8-116728369-DEL-968
chr8-116733249-DEL-65
chr8-116733882-DEL-250
chr8-116784013-DEL-497
chr8-116844097-DEL-468
chr8-116854027-DEL-64
chr8-116864397-DEL-776
chr8-116865002-DEL-95
chr8-116899738-DEL-54
chr8-116953588-DEL-74
chr8-116966909-DEL-100
chr8-116969454-DEL-51
chr8-116973491-DEL-78
chr8-116976535-DEL-236
chr8-116999147-DEL-204
chr8-117005803-DEL-60
chr8-117007354-DEL-119
chr8-117048335-DEL-62
chr8-117055354-DEL-72
chr8-117062291-DEL-579
chr8-117071996-DEL-338
chr8-117076472-DEL-982
chr8-117093196-DEL-59
chr8-117134593-DEL-199
chr8-117152414-DEL-93
chr8-117158012-DEL-51
chr8-117191605-DEL-148
chr8-117218561-DEL-6186
chr8-117247050-DEL-1037
chr8-117250177-DEL-69
chr8-117253489-DEL-98
chr8-117258437-DEL-228
chr8-117261745-DEL-305
chr8-117270995-DEL-58
chr8-117277559-DEL-305
chr8-117278582-DEL-98
chr8-117286128-DEL-1846
chr8-117286186-DEL-2753
chr8-117286262-DEL-334
chr8-117287102-DEL-402
chr8-117287479-DEL-370
chr8-117288050-DEL-553
chr8-117288571-DEL-370
chr8-117289841-DEL-73
chr8-117299780-DEL-68

chr8-117302257-DEL-58
chr8-117313703-DEL-591
chr8-117334657-DEL-408
chr8-117343610-DEL-418
chr8-117352796-DEL-312
chr8-117359067-DEL-60
chr8-117366519-DEL-450
chr8-117404198-DEL-152
chr8-117404358-DEL-4754
chr8-117410390-DEL-51
chr8-117430717-DEL-896
chr8-117430895-DEL-1006
chr8-117431165-DEL-68
chr8-117431545-DEL-173
chr8-117438065-DEL-56
chr8-117463419-DEL-69
chr8-117468313-DEL-110
chr8-117468617-DEL-359
chr8-117478664-DEL-1679
chr8-117524053-DEL-81
chr8-117526010-DEL-115
chr8-117554125-DEL-60
chr8-117583188-DEL-763
chr8-117592074-DEL-72
chr8-117621564-DEL-100
chr8-117623345-DEL-361
chr8-117624088-DEL-1223
chr8-117629117-DEL-1049
chr8-117640647-DEL-152
chr8-117711665-DEL-404
chr8-117717101-DEL-170
chr8-117720847-DEL-540
chr8-117727581-DEL-77
chr8-117747891-DEL-2043
chr8-117755732-DEL-244
chr8-117775146-DEL-383
chr8-117789501-DEL-148
chr8-117800968-DEL-69
chr8-117810189-DEL-89
chr8-117814805-DEL-96
chr8-117821464-DEL-70
chr8-117828294-DEL-398
chr8-117841825-DEL-54
chr8-117844961-DEL-56
chr8-117855167-DEL-64
chr8-117945853-DEL-135
chr8-117946812-DEL-183
chr8-117949111-DEL-1396
chr8-117998230-DEL-77
chr8-118002000-DEL-72
chr8-118037548-DEL-195
chr8-118077469-DEL-182
chr8-118125355-DEL-56
chr8-118128447-DEL-234
chr8-118139649-DEL-64
chr8-118156289-DEL-139
chr8-118179206-DEL-376
chr8-118184626-DEL-115
chr8-118217366-DEL-51
chr8-118218865-DEL-472
chr8-118235946-DEL-52
chr8-118242462-DEL-802
chr8-118249584-DEL-146
chr8-118262923-DEL-255
chr8-118264063-DEL-67
chr8-118275794-DEL-101
chr8-118306735-DEL-211
chr8-118340260-DEL-246
chr8-118381179-DEL-52
chr8-118418703-DEL-74
chr8-118452223-DEL-137
chr8-118463926-DEL-202
chr8-118471103-DEL-176
chr8-118484255-DEL-79
chr8-118503304-DEL-73
chr8-118505801-DEL-1232
chr8-118534853-DEL-52

chr8-118535878-DEL-98
chr8-118542949-DEL-268
chr8-118553152-DEL-608
chr8-118561419-DEL-85
chr8-118564874-DEL-147
chr8-118591529-DEL-132
chr8-118637514-DEL-60
chr8-118680541-DEL-58
chr8-118694503-DEL-79
chr8-118696584-DEL-116
chr8-118726934-DEL-558
chr8-118739661-DEL-100
chr8-118756687-DEL-92
chr8-118782136-DEL-60
chr8-118797664-DEL-756
chr8-118843514-DEL-206
chr8-118857047-DEL-53
chr8-118875772-DEL-79
chr8-118882446-DEL-52
chr8-118942323-DEL-57
chr8-119053081-DEL-947
chr8-119076180-DEL-128
chr8-119163213-DEL-234
chr8-119170656-DEL-80
chr8-119182894-DEL-96
chr8-119202211-DEL-111
chr8-119256548-DEL-132
chr8-119281743-DEL-72
chr8-119306274-DEL-88
chr8-119315522-DEL-53
chr8-119338978-DEL-72
chr8-119343337-DEL-1095
chr8-119351229-DEL-58
chr8-119366324-DEL-56
chr8-119375234-DEL-55
chr8-119395319-DEL-88
chr8-119399615-DEL-63
chr8-119404247-DEL-104
chr8-119409583-DEL-105
chr8-119417556-DEL-74
chr8-119418619-DEL-210
chr8-119424519-DEL-63
chr8-119533956-DEL-57
chr8-119534669-DEL-51
chr8-119571541-DEL-170
chr8-119576389-DEL-58
chr8-119599829-DEL-64
chr8-119657660-DEL-68
chr8-119663587-DEL-835
chr8-119714534-DEL-577
chr8-119724476-DEL-1042
chr8-119724482-DEL-70
chr8-119734444-DEL-73
chr8-119740736-DEL-782
chr8-119743992-DEL-75
chr8-119744875-DEL-58
chr8-119758051-DEL-178
chr8-119787329-DEL-65
chr8-119816137-DEL-372
chr8-119828943-DEL-345
chr8-119842002-DEL-197
chr8-119858419-DEL-74
chr8-119886977-DEL-103
chr8-119893605-DEL-770
chr8-119922658-DEL-380
chr8-119946956-DEL-66
chr8-119987589-DEL-379
chr8-120002240-DEL-421
chr8-120004740-DEL-61
chr8-120042504-DEL-125
chr8-120106918-DEL-62
chr8-120131457-DEL-466
chr8-120142436-DEL-72
chr8-120143273-DEL-68
chr8-120143877-DEL-108
chr8-120153931-DEL-198
chr8-120190315-DEL-76

chr8-120195180-DEL-340
chr8-120205075-DEL-51
chr8-120221764-DEL-77
chr8-120234273-DEL-175
chr8-120237850-DEL-60
chr8-120241129-DEL-87
chr8-120266643-DEL-331
chr8-120305078-DEL-218
chr8-120311712-DEL-1589
chr8-120328830-DEL-554
chr8-120334925-DEL-66
chr8-120367218-DEL-209
chr8-120373817-DEL-51
chr8-120379226-DEL-224
chr8-120398776-DEL-152
chr8-120419194-DEL-315
chr8-120462367-DEL-584
chr8-120466928-DEL-136
chr8-120513430-DEL-52
chr8-120513774-DEL-58
chr8-120560558-DEL-77
chr8-120578246-DEL-63
chr8-120613580-DEL-163
chr8-120624599-DEL-155
chr8-120635251-DEL-269
chr8-120645207-DEL-120
chr8-120688027-DEL-112
chr8-120694688-DEL-65
chr8-120796023-DEL-204
chr8-120799599-DEL-58
chr8-120804210-DEL-171
chr8-120806912-DEL-106
chr8-120814990-DEL-857
chr8-120815354-DEL-208
chr8-120834954-DEL-191
chr8-120859684-DEL-228
chr8-120866689-DEL-81
chr8-120909734-DEL-260
chr8-120911553-DEL-54
chr8-120940362-DEL-177
chr8-120947271-DEL-62
chr8-120996961-DEL-58
chr8-121032029-DEL-170
chr8-121121690-DEL-108
chr8-121156273-DEL-210
chr8-121173054-DEL-110
chr8-121180655-DEL-73
chr8-121184036-DEL-190
chr8-121191004-DEL-58
chr8-121224838-DEL-121
chr8-121228700-DEL-278
chr8-121233351-DEL-161
chr8-121237112-DEL-58
chr8-121238852-DEL-247
chr8-121245179-DEL-56
chr8-121249505-DEL-52
chr8-121271173-DEL-60
chr8-121314303-DEL-179
chr8-121321733-DEL-262
chr8-121375691-DEL-104
chr8-121401250-DEL-451
chr8-121415485-DEL-64
chr8-121416412-DEL-129
chr8-121417381-DEL-2145
chr8-121439872-DEL-641
chr8-121447050-DEL-69
chr8-121476511-DEL-104
chr8-121491849-DEL-93
chr8-121498364-DEL-71
chr8-121541665-DEL-372
chr8-121556770-DEL-118
chr8-121562240-DEL-84
chr8-121600090-DEL-56
chr8-121601471-DEL-85
chr8-121633105-DEL-93
chr8-121671317-DEL-125
chr8-121688558-DEL-271

chr8-121721917-DEL-693
chr8-121734589-DEL-256
chr8-121739232-DEL-59
chr8-121751691-DEL-72
chr8-121798777-DEL-156
chr8-121831851-DEL-203
chr8-121836493-DEL-747
chr8-121871025-DEL-74
chr8-121893398-DEL-76
chr8-121900087-DEL-56
chr8-121912177-DEL-245
chr8-121931823-DEL-1935
chr8-121971826-DEL-140
chr8-121976363-DEL-123
chr8-121979518-DEL-98
chr8-121994415-DEL-84
chr8-122012381-DEL-108
chr8-122044786-DEL-662
chr8-122053957-DEL-61
chr8-122065851-DEL-192
chr8-122111071-DEL-90
chr8-122113174-DEL-122
chr8-122156646-DEL-53
chr8-122165644-DEL-96
chr8-122169191-DEL-102
chr8-122169606-DEL-164
chr8-122173991-DEL-54
chr8-122174235-DEL-1226
chr8-122179672-DEL-377
chr8-122192885-DEL-139
chr8-122199776-DEL-201
chr8-122200304-DEL-159
chr8-122202879-DEL-52
chr8-122205461-DEL-60
chr8-122209928-DEL-367
chr8-122217965-DEL-166
chr8-122243213-DEL-88
chr8-122263503-DEL-328
chr8-122287608-DEL-67
chr8-122297676-DEL-52
chr8-122339589-DEL-324
chr8-122340466-DEL-136
chr8-122361794-DEL-357
chr8-122365642-DEL-153
chr8-122367490-DEL-143
chr8-122368460-DEL-63
chr8-122406499-DEL-758
chr8-122416270-DEL-167
chr8-122420402-DEL-54
chr8-122434807-DEL-604
chr8-122438440-DEL-232
chr8-122473282-DEL-126
chr8-122493638-DEL-1005
chr8-122503585-DEL-54
chr8-122510911-DEL-79
chr8-122531133-DEL-87
chr8-122541902-DEL-1544
chr8-122548820-DEL-5608
chr8-122549017-DEL-2552
chr8-122549021-DEL-4029
chr8-122549067-DEL-1020
chr8-122549379-DEL-2142
chr8-122549381-DEL-1275
chr8-122549482-DEL-918
chr8-122550807-DEL-714
chr8-122551828-DEL-204
chr8-122552287-DEL-1225
chr8-122552343-DEL-126
chr8-122552859-DEL-102
chr8-122553262-DEL-601
chr8-122553816-DEL-397
chr8-122596264-DEL-93
chr8-122597543-DEL-94
chr8-122644040-DEL-216
chr8-122658584-DEL-245
chr8-122699100-DEL-1769
chr8-122706892-DEL-154

chr8-122738413-DEL-368
chr8-122834996-DEL-387
chr8-122866440-DEL-195
chr8-122894720-DEL-752
chr8-122914538-DEL-111
chr8-122939142-DEL-69
chr8-122940572-DEL-77
chr8-122983935-DEL-119
chr8-123022476-DEL-440
chr8-123038819-DEL-68
chr8-123049392-DEL-233
chr8-123059375-DEL-90
chr8-123071326-DEL-237
chr8-123084381-DEL-60
chr8-123207310-DEL-204
chr8-123232564-DEL-55
chr8-123248249-DEL-179
chr8-123292154-DEL-229
chr8-123295235-DEL-346
chr8-123344654-DEL-119
chr8-123387220-DEL-242
chr8-123518996-DEL-59
chr8-123526809-DEL-521
chr8-123530484-DEL-92
chr8-123535405-DEL-1121
chr8-123535406-DEL-675
chr8-123544782-DEL-53
chr8-123549296-DEL-396
chr8-123550433-DEL-208
chr8-123593293-DEL-55
chr8-123595800-DEL-65
chr8-123598082-DEL-171
chr8-123643111-DEL-76
chr8-123665835-DEL-167
chr8-123684861-DEL-103
chr8-123692619-DEL-175
chr8-123694939-DEL-63
chr8-123703229-DEL-352
chr8-123707453-DEL-242
chr8-123710718-DEL-91
chr8-123739482-DEL-345
chr8-123756017-DEL-384
chr8-123762205-DEL-77
chr8-123763607-DEL-184
chr8-123764731-DEL-179
chr8-123773406-DEL-91
chr8-123780929-DEL-54
chr8-123782580-DEL-85
chr8-123788029-DEL-144
chr8-123801196-DEL-185
chr8-123825721-DEL-118
chr8-123827416-DEL-81
chr8-123855574-DEL-54
chr8-123872385-DEL-102
chr8-123950924-DEL-172
chr8-123987738-DEL-412
chr8-123988856-DEL-201
chr8-124004135-DEL-220
chr8-124021292-DEL-171
chr8-124046760-DEL-81
chr8-124049650-DEL-53
chr8-124069513-DEL-652
chr8-124123859-DEL-186
chr8-124132448-DEL-72
chr8-124142255-DEL-207
chr8-124146284-DEL-163
chr8-124150964-DEL-384
chr8-124165905-DEL-475
chr8-124168248-DEL-209
chr8-124171436-DEL-54
chr8-124173854-DEL-68
chr8-124178704-DEL-191
chr8-124209179-DEL-95
chr8-124250678-DEL-60
chr8-124252113-DEL-118
chr8-124258004-DEL-688
chr8-124265047-DEL-127

chr8-124266728-DEL-143
chr8-124268463-DEL-115
chr8-124270142-DEL-131
chr8-124271821-DEL-137
chr8-124273510-DEL-143
chr8-124275206-DEL-141
chr8-124276902-DEL-143
chr8-124278611-DEL-137
chr8-124280332-DEL-123
chr8-124282006-DEL-143
chr8-124329532-DEL-115
chr8-124331206-DEL-135
chr8-124332897-DEL-123
chr8-124334555-DEL-143
chr8-124336283-DEL-129
chr8-124337962-DEL-143
chr8-124339659-DEL-145
chr8-124341383-DEL-115
chr8-124343053-DEL-137
chr8-124344776-DEL-141
chr8-124346484-DEL-139
chr8-124348204-DEL-123
chr8-124350166-DEL-914
chr8-124358455-DEL-193
chr8-124359944-DEL-53
chr8-124390727-DEL-88
chr8-124392926-DEL-54
chr8-124413178-DEL-143
chr8-124424745-DEL-172
chr8-124446148-DEL-664
chr8-124497143-DEL-181
chr8-124508754-DEL-222
chr8-124521033-DEL-2111
chr8-124526614-DEL-62
chr8-124527557-DEL-830
chr8-124529966-DEL-63
chr8-124562617-DEL-434
chr8-124579385-DEL-199
chr8-124580523-DEL-486
chr8-124588256-DEL-224
chr8-124595463-DEL-54
chr8-124610367-DEL-74
chr8-124658443-DEL-73
chr8-124672464-DEL-156
chr8-124714640-DEL-185
chr8-124721087-DEL-312
chr8-124730678-DEL-979
chr8-124818523-DEL-76
chr8-124827011-DEL-119
chr8-124847020-DEL-5030
chr8-124854049-DEL-6436
chr8-124861241-DEL-65
chr8-124893955-DEL-149
chr8-124904343-DEL-64
chr8-124931270-DEL-643
chr8-124952257-DEL-52
chr8-125011562-DEL-140
chr8-125019453-DEL-57
chr8-125038321-DEL-227
chr8-125057017-DEL-154
chr8-125091022-DEL-96
chr8-125099932-DEL-225
chr8-125103462-DEL-251
chr8-125104076-DEL-218
chr8-125128509-DEL-509
chr8-125140725-DEL-90
chr8-125148572-DEL-52
chr8-125149955-DEL-76
chr8-125151958-DEL-247
chr8-125198325-DEL-153
chr8-125221912-DEL-75
chr8-125224469-DEL-112
chr8-125225106-DEL-129
chr8-125240802-DEL-1986
chr8-125246880-DEL-308
chr8-125262070-DEL-70
chr8-125288684-DEL-76

chr8-125319250-DEL-55
chr8-125328702-DEL-332
chr8-125352217-DEL-296
chr8-125385041-DEL-52
chr8-125385690-DEL-200
chr8-125393400-DEL-52
chr8-125401423-DEL-244
chr8-125443660-DEL-52
chr8-125444412-DEL-720
chr8-125444780-DEL-137
chr8-125461195-DEL-487
chr8-125462321-DEL-59
chr8-125495743-DEL-10664
chr8-125533563-DEL-227
chr8-125544629-DEL-97
chr8-125551007-DEL-208
chr8-125569629-DEL-208
chr8-125579444-DEL-78
chr8-125593771-DEL-156
chr8-125603160-DEL-87
chr8-125711636-DEL-79
chr8-125747567-DEL-172
chr8-125747959-DEL-360
chr8-125749237-DEL-302
chr8-125766536-DEL-299
chr8-125783146-DEL-923
chr8-125786811-DEL-74
chr8-125797974-DEL-746
chr8-125802294-DEL-353
chr8-125804579-DEL-98
chr8-125817728-DEL-103
chr8-125819048-DEL-507
chr8-125826263-DEL-487
chr8-125828398-DEL-72
chr8-125831431-DEL-132
chr8-125846892-DEL-206
chr8-125849391-DEL-118
chr8-125854845-DEL-225
chr8-125870550-DEL-247
chr8-125888963-DEL-176
chr8-125919109-DEL-157
chr8-125937380-DEL-133
chr8-125972870-DEL-74
chr8-126059497-DEL-60
chr8-126088984-DEL-52
chr8-126098765-DEL-229
chr8-126105143-DEL-284
chr8-126110468-DEL-121
chr8-126124867-DEL-137
chr8-126138220-DEL-379
chr8-126144579-DEL-64
chr8-126146456-DEL-1519
chr8-126147292-DEL-222
chr8-126156807-DEL-120
chr8-126158319-DEL-53
chr8-126204562-DEL-141
chr8-126213146-DEL-373
chr8-126228128-DEL-217
chr8-126232942-DEL-61
chr8-126264315-DEL-465
chr8-126264415-DEL-164
chr8-126310095-DEL-345
chr8-126318404-DEL-84
chr8-126326484-DEL-85
chr8-126349087-DEL-112
chr8-126377758-DEL-358
chr8-126382104-DEL-145
chr8-126395861-DEL-127
chr8-126406765-DEL-245
chr8-126413850-DEL-591
chr8-126447712-DEL-81
chr8-126507088-DEL-58
chr8-126512935-DEL-147
chr8-126531594-DEL-340
chr8-126532679-DEL-384
chr8-126546230-DEL-56
chr8-126548496-DEL-53

chr8-126558125-DEL-84
chr8-126561695-DEL-256
chr8-126569848-DEL-56
chr8-126596836-DEL-77
chr8-126628350-DEL-62
chr8-126631721-DEL-186
chr8-126654512-DEL-88
chr8-126665863-DEL-76
chr8-126667817-DEL-55
chr8-126691055-DEL-80
chr8-126691745-DEL-110
chr8-126731376-DEL-270
chr8-126742077-DEL-60
chr8-126747377-DEL-94
chr8-126763961-DEL-610
chr8-126765423-DEL-55
chr8-126770144-DEL-56
chr8-126771807-DEL-77
chr8-126809270-DEL-58
chr8-126810549-DEL-386
chr8-126820935-DEL-811
chr8-126822549-DEL-63
chr8-126825819-DEL-15588
chr8-126845908-DEL-55
chr8-126889051-DEL-70
chr8-126925098-DEL-72
chr8-126931682-DEL-83
chr8-126934766-DEL-65
chr8-126943840-DEL-401
chr8-126950518-DEL-6392
chr8-126995615-DEL-98
chr8-127011423-DEL-94
chr8-127060472-DEL-1154
chr8-127087412-DEL-100
chr8-127089785-DEL-94
chr8-127100018-DEL-90
chr8-127103418-DEL-130
chr8-127105107-DEL-76
chr8-127112013-DEL-67
chr8-127186534-DEL-797
chr8-127192214-DEL-496
chr8-127193427-DEL-264
chr8-127217420-DEL-52
chr8-127229623-DEL-111
chr8-127242399-DEL-138
chr8-127279758-DEL-89
chr8-127319027-DEL-60
chr8-127332865-DEL-56
chr8-127333078-DEL-366
chr8-127334674-DEL-70
chr8-127385707-DEL-96
chr8-127521766-DEL-177
chr8-127527406-DEL-55
chr8-127550646-DEL-118
chr8-127583229-DEL-146
chr8-127596852-DEL-84
chr8-127617727-DEL-596
chr8-127631127-DEL-130
chr8-127677012-DEL-60
chr8-127686813-DEL-59
chr8-127707359-DEL-73
chr8-127723497-DEL-199
chr8-127733090-DEL-287
chr8-127764195-DEL-5209
chr8-127767501-DEL-87
chr8-127769090-DEL-1623
chr8-127772401-DEL-1418
chr8-127808990-DEL-56
chr8-127859890-DEL-80
chr8-127885720-DEL-298
chr8-127897015-DEL-123
chr8-127972979-DEL-64
chr8-127982810-DEL-55
chr8-127983969-DEL-264
chr8-127988218-DEL-58
chr8-128007283-DEL-358
chr8-128044734-DEL-59

chr8-128054450-DEL-134
chr8-128090246-DEL-101
chr8-128094110-DEL-91
chr8-128174961-DEL-211
chr8-128193094-DEL-201
chr8-128229521-DEL-68
chr8-128240997-DEL-641
chr8-128242964-DEL-55
chr8-128263828-DEL-262
chr8-128342910-DEL-80
chr8-128345935-DEL-60
chr8-128356772-DEL-53
chr8-128384184-DEL-60
chr8-128386134-DEL-146
chr8-128389997-DEL-978
chr8-128455918-DEL-222
chr8-128477108-DEL-60
chr8-128571601-DEL-886
chr8-128590145-DEL-428
chr8-128590797-DEL-89
chr8-128649329-DEL-80
chr8-128661090-DEL-72
chr8-128682746-DEL-97
chr8-128735241-DEL-79
chr8-128776659-DEL-567
chr8-128868421-DEL-1393
chr8-128887853-DEL-135
chr8-128893845-DEL-699
chr8-128894766-DEL-314
chr8-128903204-DEL-62
chr8-128986279-DEL-314
chr8-128996621-DEL-1181
chr8-128997434-DEL-131
chr8-129021483-DEL-640
chr8-129045412-DEL-70
chr8-129069303-DEL-375
chr8-129106126-DEL-264
chr8-129173534-DEL-84
chr8-129175307-DEL-150
chr8-129183583-DEL-56
chr8-129250316-DEL-57
chr8-129254559-DEL-66
chr8-129263898-DEL-52
chr8-129272771-DEL-92
chr8-129294917-DEL-1775
chr8-129296931-DEL-195
chr8-129304166-DEL-419
chr8-129339075-DEL-770
chr8-129341039-DEL-1052
chr8-129366283-DEL-52
chr8-129407752-DEL-120
chr8-129425929-DEL-1364
chr8-129465735-DEL-3223
chr8-129494485-DEL-358
chr8-129498411-DEL-2866
chr8-129502986-DEL-3390
chr8-129503684-DEL-4228
chr8-129504226-DEL-2254
chr8-129548974-DEL-241
chr8-129555997-DEL-254
chr8-129564668-DEL-163
chr8-129571212-DEL-7005
chr8-129581582-DEL-147
chr8-129587118-DEL-883
chr8-129588037-DEL-520
chr8-129597630-DEL-1131
chr8-129609888-DEL-61
chr8-129616023-DEL-55
chr8-129643544-DEL-73
chr8-129667882-DEL-332
chr8-129675349-DEL-6359
chr8-129682378-DEL-55
chr8-129684364-DEL-73
chr8-129687188-DEL-107
chr8-129722652-DEL-438
chr8-129748731-DEL-1292
chr8-129756558-DEL-6313

chr8-129772121-DEL-54
chr8-129784422-DEL-66
chr8-129808526-DEL-1222
chr8-129823955-DEL-5504
chr8-129841358-DEL-932
chr8-129842462-DEL-369
chr8-129842933-DEL-730
chr8-129844570-DEL-6239
chr8-129862192-DEL-269
chr8-129889235-DEL-835
chr8-129974127-DEL-354
chr8-129979403-DEL-1450
chr9-3011600-DEL-176
chr9-3051854-DEL-302
chr9-3057371-DEL-848
chr9-3059987-DEL-93
chr9-3061047-DEL-89
chr9-3064678-DEL-62
chr9-3065561-DEL-420
chr9-3070137-DEL-1531
chr9-3074109-DEL-58
chr9-3075704-DEL-139
chr9-3095649-DEL-601
chr9-3096675-DEL-230
chr9-3108730-DEL-223
chr9-3113709-DEL-67
chr9-3141498-DEL-2782
chr9-3145732-DEL-6269
chr9-3163195-DEL-14157
chr9-3174989-DEL-134
chr9-3215333-DEL-503
chr9-3239289-DEL-185
chr9-3249896-DEL-207
chr9-3310549-DEL-193
chr9-3316577-DEL-914
chr9-3317550-DEL-204
chr9-3324950-DEL-884
chr9-3331806-DEL-498
chr9-3339426-DEL-158
chr9-3342119-DEL-198
chr9-3349837-DEL-242
chr9-3354524-DEL-688
chr9-3392144-DEL-966
chr9-3397266-DEL-53
chr9-3436000-DEL-1627
chr9-3452459-DEL-1909
chr9-3476405-DEL-61
chr9-3483290-DEL-95
chr9-3490425-DEL-3881
chr9-3533691-DEL-135
chr9-3584803-DEL-159
chr9-3628654-DEL-71
chr9-3642449-DEL-7336
chr9-3674691-DEL-6378
chr9-3743528-DEL-6241
chr9-3751558-DEL-68
chr9-3802006-DEL-6192
chr9-3817554-DEL-622
chr9-3823236-DEL-83
chr9-3824925-DEL-400
chr9-3827856-DEL-208
chr9-3833236-DEL-57
chr9-3836424-DEL-6347
chr9-3880993-DEL-7559
chr9-3908600-DEL-5508
chr9-3939450-DEL-8139
chr9-3939462-DEL-8139
chr9-3995654-DEL-1459
chr9-4050294-DEL-112
chr9-4076120-DEL-101
chr9-4090400-DEL-1868
chr9-4094805-DEL-285
chr9-4094873-DEL-2178
chr9-4100632-DEL-1054
chr9-4101790-DEL-1578
chr9-4116461-DEL-167
chr9-4131384-DEL-53

chr9-4202396-DEL-467
chr9-4238240-DEL-7491
chr9-4240873-DEL-4391
chr9-4249826-DEL-203
chr9-4256300-DEL-57
chr9-4272665-DEL-366
chr9-4275933-DEL-611
chr9-4293840-DEL-902
chr9-4303558-DEL-109
chr9-4310050-DEL-71
chr9-4353175-DEL-57
chr9-4387055-DEL-5712
chr9-4401202-DEL-293
chr9-4401597-DEL-2548
chr9-4404855-DEL-1364
chr9-4407327-DEL-267
chr9-4420604-DEL-228
chr9-4509184-DEL-2530
chr9-4512748-DEL-207
chr9-4548527-DEL-243
chr9-4565717-DEL-57
chr9-4578162-DEL-480
chr9-4592052-DEL-99
chr9-4614800-DEL-205
chr9-4650664-DEL-990
chr9-4652974-DEL-1649
chr9-4654637-DEL-1733
chr9-4668245-DEL-151
chr9-4705720-DEL-75
chr9-4731460-DEL-4486
chr9-4739927-DEL-136
chr9-4771748-DEL-212
chr9-4777638-DEL-62
chr9-4814676-DEL-62
chr9-4842725-DEL-13174
chr9-4846700-DEL-8845
chr9-4914900-DEL-2040
chr9-4938533-DEL-195
chr9-5017231-DEL-293
chr9-5019649-DEL-91
chr9-5033627-DEL-146
chr9-5045858-DEL-92
chr9-5053832-DEL-181
chr9-5055765-DEL-395
chr9-5057192-DEL-937
chr9-5062672-DEL-5570
chr9-5073446-DEL-3137
chr9-5076106-DEL-64
chr9-5077159-DEL-99
chr9-5088849-DEL-114
chr9-5090353-DEL-75
chr9-5098696-DEL-76791
chr9-5132527-DEL-114
chr9-5153531-DEL-745
chr9-5187500-DEL-54
chr9-5193980-DEL-1639
chr9-5201104-DEL-1338
chr9-5223430-DEL-4341
chr9-5227527-DEL-628
chr9-5233143-DEL-102
chr9-5246909-DEL-79
chr9-5259063-DEL-4832
chr9-5274034-DEL-2153
chr9-5275876-DEL-2318
chr9-5292786-DEL-222
chr9-5309184-DEL-222
chr9-5323985-DEL-76
chr9-5354201-DEL-171
chr9-5366245-DEL-931
chr9-5377053-DEL-106
chr9-5380605-DEL-227
chr9-5419514-DEL-67
chr9-5421292-DEL-490
chr9-5425933-DEL-345
chr9-5467686-DEL-468
chr9-5490873-DEL-161
chr9-5491635-DEL-89

chr9-5529197-DEL-239
chr9-5530708-DEL-601
chr9-5548470-DEL-7006
chr9-5560010-DEL-51
chr9-5576213-DEL-288
chr9-5578076-DEL-538
chr9-5591940-DEL-2158
chr9-5649756-DEL-307
chr9-5657394-DEL-121
chr9-5658732-DEL-5507
chr9-5682837-DEL-56
chr9-5690899-DEL-1187
chr9-5692526-DEL-830
chr9-5692806-DEL-3164
chr9-5730551-DEL-151
chr9-5755653-DEL-731
chr9-5764218-DEL-169
chr9-5792023-DEL-52
chr9-5852316-DEL-65
chr9-5861427-DEL-2223
chr9-5895760-DEL-290
chr9-5903687-DEL-3803
chr9-5905102-DEL-64
chr9-5905551-DEL-165
chr9-5911704-DEL-153
chr9-5926969-DEL-3759
chr9-5930071-DEL-191
chr9-5955062-DEL-73
chr9-5956172-DEL-434
chr9-5976253-DEL-2152
chr9-5990478-DEL-363
chr9-6042953-DEL-3076
chr9-6060027-DEL-69
chr9-6067064-DEL-11522
chr9-6083591-DEL-121
chr9-6088830-DEL-312
chr9-6121149-DEL-632
chr9-6142986-DEL-74
chr9-6146433-DEL-2448
chr9-6160444-DEL-797
chr9-6219134-DEL-472
chr9-6241231-DEL-694
chr9-6251424-DEL-62
chr9-6272014-DEL-628
chr9-6273985-DEL-148
chr9-6324356-DEL-60
chr9-6393033-DEL-581
chr9-6419587-DEL-398
chr9-6439606-DEL-1929
chr9-6442487-DEL-4116
chr9-6449048-DEL-803
chr9-6452896-DEL-565
chr9-6459845-DEL-806
chr9-6471461-DEL-80
chr9-6513742-DEL-4570
chr9-6529931-DEL-83
chr9-6536807-DEL-114
chr9-6542985-DEL-251
chr9-6551304-DEL-91
chr9-6583101-DEL-3164
chr9-6589519-DEL-822
chr9-6665538-DEL-194
chr9-6675736-DEL-281
chr9-6678833-DEL-383
chr9-6680483-DEL-96
chr9-6692103-DEL-11766
chr9-6694760-DEL-8101
chr9-6710175-DEL-1261
chr9-6715899-DEL-1387
chr9-6734079-DEL-108
chr9-6756132-DEL-572
chr9-6761091-DEL-60724
chr9-6762890-DEL-326
chr9-6791284-DEL-650
chr9-6799804-DEL-213
chr9-6801224-DEL-57
chr9-6802713-DEL-60

chr9-6817903-DEL-165
chr9-6820395-DEL-122
chr9-6865905-DEL-2836
chr9-6866102-DEL-1903
chr9-6868019-DEL-721
chr9-6956777-DEL-2125
chr9-6959706-DEL-117
chr9-6960098-DEL-312
chr9-6962254-DEL-545
chr9-6967949-DEL-8605
chr9-7010656-DEL-178
chr9-7018590-DEL-393
chr9-7027905-DEL-641
chr9-7036004-DEL-635
chr9-7038351-DEL-96
chr9-7045601-DEL-89
chr9-7088852-DEL-1890
chr9-7091471-DEL-6178
chr9-7108577-DEL-810
chr9-7116080-DEL-241
chr9-7162947-DEL-103
chr9-7198291-DEL-2752
chr9-7231476-DEL-2895
chr9-7245703-DEL-865
chr9-7246612-DEL-612
chr9-7285000-DEL-718
chr9-7303969-DEL-94
chr9-7305096-DEL-521
chr9-7308416-DEL-82
chr9-7394430-DEL-487
chr9-7402108-DEL-149
chr9-7407033-DEL-8832
chr9-7477618-DEL-5970
chr9-7489255-DEL-247
chr9-7492125-DEL-526
chr9-7494411-DEL-5418
chr9-7569060-DEL-807
chr9-7576126-DEL-179
chr9-7641483-DEL-289
chr9-7657745-DEL-119
chr9-7680026-DEL-324
chr9-7687029-DEL-83
chr9-7716493-DEL-82
chr9-7737681-DEL-70
chr9-7751673-DEL-675
chr9-7783622-DEL-116
chr9-7798238-DEL-92
chr9-7857906-DEL-214
chr9-7871719-DEL-518
chr9-7925389-DEL-54
chr9-7939564-DEL-174
chr9-7951074-DEL-144
chr9-7958990-DEL-202
chr9-7988755-DEL-165
chr9-7991805-DEL-213
chr9-7993837-DEL-71
chr9-7997609-DEL-52
chr9-8029342-DEL-1293
chr9-8029574-DEL-367
chr9-8030304-DEL-53
chr9-8044246-DEL-135
chr9-8057234-DEL-217
chr9-8065136-DEL-1999
chr9-8081265-DEL-563
chr9-8097662-DEL-189
chr9-8111701-DEL-339
chr9-8117254-DEL-378
chr9-8237572-DEL-248
chr9-8243182-DEL-950
chr9-8287923-DEL-11573
chr9-8300465-DEL-63
chr9-8302476-DEL-929
chr9-8311245-DEL-67
chr9-8314210-DEL-6304
chr9-8327799-DEL-327
chr9-8354888-DEL-6222
chr9-8366365-DEL-18402

chr9-8371548-DEL-3836
chr9-8377427-DEL-6786
chr9-8442673-DEL-149
chr9-8463085-DEL-82
chr9-8477506-DEL-6256
chr9-8506553-DEL-841
chr9-8508035-DEL-401
chr9-8537206-DEL-845
chr9-8566407-DEL-276
chr9-8575190-DEL-273
chr9-8617022-DEL-885
chr9-8646759-DEL-58
chr9-8650476-DEL-57
chr9-8652623-DEL-131
chr9-8659859-DEL-7725
chr9-8667695-DEL-2243
chr9-8669964-DEL-999
chr9-8711820-DEL-60
chr9-8743779-DEL-1185
chr9-8745286-DEL-504
chr9-8770680-DEL-2558
chr9-8837428-DEL-81
chr9-8888094-DEL-700
chr9-8895843-DEL-336
chr9-8913729-DEL-84
chr9-8914691-DEL-52
chr9-8959081-DEL-90
chr9-8976664-DEL-232
chr9-9010664-DEL-73
chr9-9071148-DEL-471
chr9-9071880-DEL-208
chr9-9078856-DEL-611
chr9-9085174-DEL-1179
chr9-9114646-DEL-561
chr9-9123459-DEL-64
chr9-9152709-DEL-708
chr9-9153002-DEL-215
chr9-9178640-DEL-64
chr9-9192442-DEL-56
chr9-9281733-DEL-3155
chr9-9299648-DEL-66
chr9-9310753-DEL-656
chr9-9319750-DEL-1653
chr9-9325880-DEL-6057
chr9-9331396-DEL-87
chr9-9333407-DEL-1459
chr9-9348128-DEL-74
chr9-9361023-DEL-401
chr9-9372584-DEL-61
chr9-9397362-DEL-6223
chr9-9453693-DEL-82
chr9-9466163-DEL-355
chr9-9468261-DEL-6253
chr9-9479771-DEL-6336
chr9-9487723-DEL-213
chr9-9594161-DEL-57
chr9-9595771-DEL-212
chr9-9627095-DEL-65
chr9-9631331-DEL-499
chr9-9741391-DEL-77
chr9-9762460-DEL-612
chr9-9815615-DEL-880
chr9-9846591-DEL-197
chr9-9880262-DEL-294
chr9-9905547-DEL-117
chr9-9906669-DEL-645
chr9-9940893-DEL-3423
chr9-9941387-DEL-189
chr9-9941496-DEL-1242
chr9-9955918-DEL-1269
chr9-10029580-DEL-54
chr9-10050586-DEL-282
chr9-10061357-DEL-267
chr9-10084894-DEL-117
chr9-10149374-DEL-123
chr9-10153017-DEL-597
chr9-10248641-DEL-236

chr9-10263234-DEL-189
chr9-10377848-DEL-90
chr9-10415235-DEL-184
chr9-10432550-DEL-1166
chr9-10436714-DEL-60
chr9-10438656-DEL-198
chr9-10446183-DEL-518
chr9-10474682-DEL-75
chr9-10479418-DEL-1115
chr9-10509210-DEL-76
chr9-10525028-DEL-193
chr9-10545506-DEL-52
chr9-10565027-DEL-70
chr9-10566507-DEL-53
chr9-10579973-DEL-116
chr9-10592147-DEL-441
chr9-10624956-DEL-275
chr9-10639445-DEL-109
chr9-10651940-DEL-55
chr9-10652449-DEL-67
chr9-10681016-DEL-69
chr9-10697743-DEL-1090
chr9-10747666-DEL-643
chr9-10779865-DEL-183
chr9-10791954-DEL-1912
chr9-10796858-DEL-70
chr9-10832849-DEL-270
chr9-10869927-DEL-305
chr9-10917154-DEL-1645
chr9-10923044-DEL-1831
chr9-10933437-DEL-70
chr9-10934555-DEL-436
chr9-10940577-DEL-492
chr9-10941700-DEL-2210
chr9-10952917-DEL-70
chr9-10961763-DEL-1453
chr9-10978321-DEL-189
chr9-11021354-DEL-88
chr9-11028795-DEL-60
chr9-11065168-DEL-831
chr9-11106846-DEL-557
chr9-11107839-DEL-5997
chr9-11140630-DEL-218
chr9-11209426-DEL-767
chr9-11211105-DEL-1510
chr9-11212988-DEL-192
chr9-11292163-DEL-933
chr9-11297173-DEL-207
chr9-11333846-DEL-840
chr9-11346212-DEL-838
chr9-11380236-DEL-112
chr9-11399079-DEL-1230
chr9-11408783-DEL-63
chr9-11419025-DEL-400
chr9-11439031-DEL-827
chr9-11471273-DEL-61
chr9-11485453-DEL-5176
chr9-11488930-DEL-275
chr9-11508859-DEL-1248
chr9-11514500-DEL-563
chr9-11521908-DEL-93
chr9-11528971-DEL-1283
chr9-11545280-DEL-149
chr9-11560328-DEL-72
chr9-11596110-DEL-93
chr9-11605302-DEL-224
chr9-11632505-DEL-1950
chr9-11632508-DEL-1507
chr9-11640828-DEL-104
chr9-11678628-DEL-138
chr9-11699517-DEL-64
chr9-11699712-DEL-650
chr9-11706904-DEL-652
chr9-11745685-DEL-1408
chr9-11751627-DEL-52
chr9-11766308-DEL-3041
chr9-11768374-DEL-88

chr9-11771165-DEL-1309
chr9-11781700-DEL-550
chr9-11811907-DEL-14652
chr9-11839080-DEL-6445
chr9-11876200-DEL-102
chr9-11925203-DEL-529
chr9-11979734-DEL-77
chr9-11985074-DEL-74
chr9-11992264-DEL-1225
chr9-12021301-DEL-5305
chr9-12127434-DEL-16602
chr9-12163408-DEL-2837
chr9-12165160-DEL-126
chr9-12172935-DEL-6081
chr9-12184104-DEL-53
chr9-12197974-DEL-70
chr9-12205976-DEL-53
chr9-12216099-DEL-382
chr9-12218996-DEL-107
chr9-12221641-DEL-3557
chr9-12242733-DEL-289
chr9-12257114-DEL-434
chr9-12257757-DEL-1051
chr9-12265180-DEL-823
chr9-12266198-DEL-65
chr9-12266716-DEL-70
chr9-12281702-DEL-5893
chr9-12300032-DEL-56
chr9-12325200-DEL-896
chr9-12327453-DEL-191
chr9-12341521-DEL-53
chr9-12346784-DEL-66
chr9-12350609-DEL-329
chr9-12360906-DEL-1611
chr9-12364311-DEL-3042
chr9-12409145-DEL-6415
chr9-12416053-DEL-1397
chr9-12441715-DEL-438
chr9-12462225-DEL-66
chr9-12476925-DEL-51
chr9-12502491-DEL-100
chr9-12563914-DEL-1716
chr9-12565769-DEL-2239
chr9-12567671-DEL-321
chr9-12569897-DEL-4085
chr9-12594814-DEL-169
chr9-12635898-DEL-88
chr9-12638526-DEL-2149
chr9-12673545-DEL-2801
chr9-12687437-DEL-545
chr9-12689474-DEL-85
chr9-12696752-DEL-4309
chr9-12738387-DEL-266
chr9-12768265-DEL-5630
chr9-12780030-DEL-66
chr9-12794588-DEL-116
chr9-12797324-DEL-153
chr9-12799494-DEL-775
chr9-12801647-DEL-90
chr9-12802396-DEL-6218
chr9-12823345-DEL-551
chr9-12897722-DEL-444
chr9-12906029-DEL-8303
chr9-12911495-DEL-265
chr9-12933160-DEL-222
chr9-12936289-DEL-2408
chr9-12941366-DEL-558
chr9-12943355-DEL-67
chr9-12989487-DEL-98
chr9-12997905-DEL-6529
chr9-13004203-DEL-203
chr9-13013229-DEL-87
chr9-13036270-DEL-9340
chr9-13039375-DEL-3106
chr9-13065605-DEL-61
chr9-13079610-DEL-190
chr9-13101717-DEL-91

chr9-13105744-DEL-343
chr9-13117934-DEL-442
chr9-13138654-DEL-439
chr9-13173518-DEL-145
chr9-13209485-DEL-4674
chr9-13275819-DEL-4895
chr9-13287021-DEL-60
chr9-13309968-DEL-98
chr9-13325536-DEL-452
chr9-13354572-DEL-144
chr9-13355641-DEL-149
chr9-13357942-DEL-61
chr9-13415064-DEL-299
chr9-13431464-DEL-185
chr9-13445718-DEL-468
chr9-13466400-DEL-411
chr9-13480903-DEL-390
chr9-13490639-DEL-77
chr9-13491983-DEL-79
chr9-13501019-DEL-848
chr9-13526315-DEL-77
chr9-13538951-DEL-56
chr9-13580612-DEL-435
chr9-13583506-DEL-206
chr9-13602821-DEL-1439
chr9-13650546-DEL-75
chr9-13655238-DEL-381
chr9-13657804-DEL-214
chr9-13684279-DEL-946
chr9-13691443-DEL-201
chr9-13776903-DEL-116
chr9-13811689-DEL-1091
chr9-13817255-DEL-122
chr9-13866508-DEL-218
chr9-13868588-DEL-80
chr9-13906077-DEL-464
chr9-13907038-DEL-407
chr9-13934222-DEL-54
chr9-13937455-DEL-60
chr9-13974348-DEL-397
chr9-13989944-DEL-293
chr9-13998120-DEL-96
chr9-14030749-DEL-120
chr9-14060360-DEL-187
chr9-14082799-DEL-89
chr9-14090430-DEL-356
chr9-14103269-DEL-109
chr9-14117225-DEL-219
chr9-14120739-DEL-81
chr9-14143703-DEL-400
chr9-14144713-DEL-84
chr9-14151802-DEL-156
chr9-14163418-DEL-232
chr9-14204069-DEL-352
chr9-14209189-DEL-79
chr9-14259261-DEL-71
chr9-14264197-DEL-87
chr9-14324498-DEL-2297
chr9-14337574-DEL-2987
chr9-14352801-DEL-1463
chr9-14436728-DEL-203
chr9-14454710-DEL-51
chr9-14480583-DEL-60
chr9-14492674-DEL-53
chr9-14500879-DEL-55
chr9-14506102-DEL-201
chr9-14511689-DEL-494
chr9-14544785-DEL-55
chr9-14578424-DEL-59
chr9-14605894-DEL-54
chr9-14610913-DEL-56
chr9-14611926-DEL-210
chr9-14700296-DEL-964
chr9-14749366-DEL-209
chr9-14828442-DEL-100
chr9-14829103-DEL-86
chr9-14833753-DEL-229

chr9-14836979-DEL-698
chr9-14839411-DEL-5039
chr9-14851465-DEL-520
chr9-14874731-DEL-198
chr9-14913978-DEL-68
chr9-14916730-DEL-93
chr9-14924986-DEL-205
chr9-14928899-DEL-96
chr9-14934474-DEL-60
chr9-14941538-DEL-58
chr9-14946730-DEL-360
chr9-14955075-DEL-136
chr9-14959169-DEL-204
chr9-14987698-DEL-52
chr9-14993421-DEL-67
chr9-15006565-DEL-142
chr9-15008160-DEL-141
chr9-15009716-DEL-269
chr9-15016149-DEL-62
chr9-15039492-DEL-54
chr9-15045549-DEL-2103
chr9-15076575-DEL-165
chr9-15112406-DEL-404
chr9-15113411-DEL-442
chr9-15116567-DEL-162
chr9-15118881-DEL-60
chr9-15124112-DEL-172
chr9-15149913-DEL-68
chr9-15163056-DEL-336
chr9-15184196-DEL-53
chr9-15188414-DEL-104
chr9-15201151-DEL-554
chr9-15211884-DEL-146
chr9-15222188-DEL-259
chr9-15227343-DEL-163
chr9-15254274-DEL-153
chr9-15303306-DEL-191
chr9-15316037-DEL-1906
chr9-15351749-DEL-4798
chr9-15398452-DEL-186
chr9-15401879-DEL-152
chr9-15413421-DEL-66
chr9-15430209-DEL-87
chr9-15479815-DEL-2453
chr9-15495046-DEL-55
chr9-15508575-DEL-510
chr9-15524969-DEL-98
chr9-15551022-DEL-441
chr9-15591521-DEL-148
chr9-15597734-DEL-248
chr9-15611226-DEL-351
chr9-15642510-DEL-207
chr9-15653785-DEL-85
chr9-15687122-DEL-242
chr9-15710158-DEL-460
chr9-15711160-DEL-88
chr9-15739201-DEL-1808
chr9-15773536-DEL-57
chr9-15812947-DEL-797
chr9-15817088-DEL-254
chr9-15839905-DEL-155
chr9-15852781-DEL-197
chr9-15868081-DEL-90
chr9-15902229-DEL-245
chr9-15924002-DEL-649
chr9-15928059-DEL-1308
chr9-15994534-DEL-1156
chr9-16035901-DEL-108
chr9-16045896-DEL-63
chr9-16108681-DEL-108
chr9-16132624-DEL-54
chr9-16173770-DEL-144
chr9-16226862-DEL-171
chr9-16233596-DEL-779
chr9-16245126-DEL-654
chr9-16267830-DEL-69
chr9-16303803-DEL-61

chr9-16307212-DEL-60
chr9-16344282-DEL-143
chr9-16374308-DEL-522
chr9-16376318-DEL-2062
chr9-16398025-DEL-247
chr9-16405854-DEL-513
chr9-16406787-DEL-672
chr9-16415646-DEL-199
chr9-16416989-DEL-265
chr9-16434075-DEL-702
chr9-16460401-DEL-284
chr9-16518334-DEL-214
chr9-16540797-DEL-977
chr9-16543886-DEL-354
chr9-16569238-DEL-51
chr9-16579205-DEL-291
chr9-16580769-DEL-92
chr9-16582361-DEL-157
chr9-16606082-DEL-566
chr9-16608894-DEL-53
chr9-16616213-DEL-1496
chr9-16627252-DEL-5174
chr9-16655081-DEL-502
chr9-16656834-DEL-576
chr9-16679988-DEL-3088
chr9-16689217-DEL-23390
chr9-16701863-DEL-2298
chr9-16705392-DEL-433
chr9-16723355-DEL-734
chr9-16730050-DEL-805
chr9-16765390-DEL-7283
chr9-16791465-DEL-68
chr9-16796100-DEL-64
chr9-16849885-DEL-65
chr9-16859026-DEL-2835
chr9-16882796-DEL-1921
chr9-16910215-DEL-566
chr9-16916418-DEL-210
chr9-16919005-DEL-1692
chr9-16927437-DEL-136
chr9-16945071-DEL-823
chr9-16970075-DEL-90
chr9-16982062-DEL-63
chr9-16985130-DEL-106
chr9-17000552-DEL-62
chr9-17001421-DEL-200
chr9-17012843-DEL-1638
chr9-17028753-DEL-1250
chr9-17076200-DEL-126
chr9-17086182-DEL-56
chr9-17149935-DEL-876
chr9-17153402-DEL-599
chr9-17171718-DEL-2929
chr9-17193297-DEL-84
chr9-17206147-DEL-272
chr9-17222983-DEL-5985
chr9-17233194-DEL-2779
chr9-17263590-DEL-1845
chr9-17290856-DEL-801
chr9-17292100-DEL-442
chr9-17293168-DEL-1963
chr9-17336403-DEL-445
chr9-17372288-DEL-5658
chr9-17419116-DEL-135
chr9-17434434-DEL-468
chr9-17492720-DEL-819
chr9-17494634-DEL-1648
chr9-17495793-DEL-6079
chr9-17535758-DEL-168
chr9-17537655-DEL-2852
chr9-17553161-DEL-223
chr9-17555158-DEL-1363
chr9-17556468-DEL-223
chr9-17570240-DEL-849
chr9-17653799-DEL-257
chr9-17688459-DEL-124
chr9-17705513-DEL-982

chr9-17734899-DEL-157
chr9-17738478-DEL-2441
chr9-17745880-DEL-54
chr9-17798031-DEL-84
chr9-17837815-DEL-84
chr9-17839559-DEL-350
chr9-17845581-DEL-235
chr9-17847229-DEL-934
chr9-17854831-DEL-1210
chr9-17864549-DEL-709
chr9-17876832-DEL-226
chr9-17887042-DEL-4624
chr9-17892459-DEL-59
chr9-17903869-DEL-1666
chr9-17942374-DEL-83
chr9-17952563-DEL-52
chr9-17959987-DEL-499
chr9-18002093-DEL-757
chr9-18006963-DEL-2729
chr9-18010189-DEL-1375
chr9-18028281-DEL-13383
chr9-18043099-DEL-1385
chr9-18057608-DEL-706
chr9-18083841-DEL-54
chr9-18091481-DEL-1970
chr9-18094619-DEL-11860
chr9-18113284-DEL-885
chr9-18120828-DEL-339
chr9-18208122-DEL-2112
chr9-18219091-DEL-51
chr9-18223848-DEL-74
chr9-18254001-DEL-253
chr9-18266611-DEL-55
chr9-18282149-DEL-65
chr9-18308095-DEL-217
chr9-18323860-DEL-60
chr9-18325028-DEL-2660
chr9-18366007-DEL-82
chr9-18400192-DEL-6236
chr9-18412956-DEL-6321
chr9-18424839-DEL-84
chr9-18494168-DEL-57
chr9-18520679-DEL-165
chr9-18533893-DEL-80
chr9-18538170-DEL-663
chr9-18539421-DEL-165
chr9-18579572-DEL-84
chr9-18641035-DEL-79
chr9-18689216-DEL-70
chr9-18702290-DEL-6205
chr9-18736140-DEL-705
chr9-18745047-DEL-462
chr9-18751909-DEL-265
chr9-18757244-DEL-343
chr9-18767162-DEL-115
chr9-18789553-DEL-269
chr9-18794405-DEL-201
chr9-18805048-DEL-2759
chr9-18805056-DEL-6215
chr9-18807831-DEL-3441
chr9-18827729-DEL-461
chr9-18852123-DEL-1216
chr9-18854047-DEL-171
chr9-18901359-DEL-6025
chr9-18917500-DEL-302
chr9-18926784-DEL-3554
chr9-18932495-DEL-95
chr9-18963729-DEL-139
chr9-18966940-DEL-6237
chr9-18982096-DEL-52
chr9-18995799-DEL-1964
chr9-19004710-DEL-4616
chr9-19015207-DEL-1337
chr9-19059845-DEL-538
chr9-19061403-DEL-1023
chr9-19071078-DEL-6766
chr9-19102619-DEL-744

chr9-19103533-DEL-93
chr9-19117524-DEL-5504
chr9-19129535-DEL-13569
chr9-19152538-DEL-147
chr9-19166054-DEL-575
chr9-19174690-DEL-1484
chr9-19190886-DEL-66
chr9-19207174-DEL-628
chr9-19213913-DEL-50
chr9-19216374-DEL-70
chr9-19231602-DEL-6208
chr9-19253807-DEL-58
chr9-19259677-DEL-1036
chr9-19265387-DEL-201
chr9-19291094-DEL-7826
chr9-19296622-DEL-3975
chr9-19299960-DEL-231
chr9-19309134-DEL-61
chr9-19310476-DEL-67
chr9-19311747-DEL-715
chr9-19311762-DEL-197
chr9-19414849-DEL-199
chr9-19426098-DEL-386
chr9-19429023-DEL-1369
chr9-19437488-DEL-3538
chr9-19454226-DEL-58
chr9-19464004-DEL-1977
chr9-19505655-DEL-7296
chr9-19513706-DEL-6306
chr9-19523393-DEL-203
chr9-19544423-DEL-6066
chr9-19576895-DEL-959
chr9-19577739-DEL-351
chr9-19578447-DEL-5941
chr9-19603674-DEL-56
chr9-19603730-DEL-505
chr9-19614792-DEL-633
chr9-19627681-DEL-57
chr9-19637260-DEL-3064
chr9-19686906-DEL-60
chr9-19716806-DEL-203
chr9-19720745-DEL-740
chr9-19776248-DEL-4099
chr9-19780942-DEL-621
chr9-19796239-DEL-571
chr9-19804649-DEL-88
chr9-19839098-DEL-523
chr9-19866634-DEL-12558
chr9-19867561-DEL-456
chr9-19868207-DEL-8697
chr9-19889035-DEL-5912
chr9-19890402-DEL-10326
chr9-19903085-DEL-2187
chr9-19906783-DEL-155
chr9-19909909-DEL-300
chr9-19910798-DEL-721
chr9-19915434-DEL-652
chr9-19921063-DEL-210
chr9-19947813-DEL-281
chr9-19972067-DEL-11060
chr9-20003157-DEL-51
chr9-20008726-DEL-266
chr9-20029448-DEL-3132
chr9-20052633-DEL-1645
chr9-20077135-DEL-119
chr9-20126447-DEL-6898
chr9-20133294-DEL-7090
chr9-20146801-DEL-194
chr9-20156142-DEL-492
chr9-20186229-DEL-55
chr9-20265916-DEL-6483
chr9-20277605-DEL-961
chr9-20279053-DEL-6403
chr9-20305547-DEL-620
chr9-20354115-DEL-417
chr9-20360244-DEL-463
chr9-20365080-DEL-996

chr9-20372234-DEL-64
chr9-20466583-DEL-579
chr9-20467580-DEL-183
chr9-20481374-DEL-123
chr9-20481514-DEL-5390
chr9-20492121-DEL-54
chr9-20493805-DEL-66
chr9-20525568-DEL-819
chr9-20526269-DEL-121
chr9-20567091-DEL-103
chr9-20585237-DEL-192
chr9-20597712-DEL-88
chr9-20604720-DEL-77
chr9-20614479-DEL-506
chr9-20628946-DEL-251
chr9-20671894-DEL-70
chr9-20675129-DEL-80
chr9-20680559-DEL-380
chr9-20688613-DEL-152
chr9-20724816-DEL-70
chr9-20741710-DEL-239
chr9-20743507-DEL-86
chr9-20744824-DEL-62
chr9-20749770-DEL-52
chr9-20752484-DEL-156
chr9-20768107-DEL-51
chr9-20787603-DEL-56
chr9-20790635-DEL-97
chr9-20792508-DEL-1528
chr9-20794097-DEL-1538
chr9-20798522-DEL-185
chr9-20806875-DEL-51
chr9-20810826-DEL-106
chr9-20812218-DEL-291
chr9-20817696-DEL-237
chr9-20830575-DEL-61
chr9-20867564-DEL-168
chr9-20907223-DEL-209
chr9-20909616-DEL-130
chr9-20916631-DEL-142
chr9-20924255-DEL-99
chr9-20929245-DEL-246
chr9-20952523-DEL-82
chr9-20959569-DEL-494
chr9-20960647-DEL-53
chr9-20965799-DEL-1610
chr9-21014385-DEL-55
chr9-21079001-DEL-63
chr9-21107660-DEL-75
chr9-21129014-DEL-74
chr9-21143593-DEL-72
chr9-21166728-DEL-566
chr9-21175351-DEL-263
chr9-21194359-DEL-239
chr9-21204958-DEL-531
chr9-21225134-DEL-197
chr9-21227575-DEL-108
chr9-21240816-DEL-247
chr9-21273383-DEL-122
chr9-21285163-DEL-199
chr9-21293335-DEL-181
chr9-21302129-DEL-201
chr9-21323901-DEL-180
chr9-21324890-DEL-205
chr9-21330173-DEL-197
chr9-21332465-DEL-161
chr9-21339816-DEL-143
chr9-21344296-DEL-60
chr9-21362637-DEL-52
chr9-21369207-DEL-64
chr9-21387216-DEL-142
chr9-21415431-DEL-52
chr9-21427727-DEL-57
chr9-21444615-DEL-984
chr9-21446556-DEL-131
chr9-21452643-DEL-193
chr9-21470923-DEL-203

chr9-21510154-DEL-166
chr9-21520984-DEL-207
chr9-21540175-DEL-1215
chr9-21541864-DEL-51
chr9-21593314-DEL-199
chr9-21609742-DEL-211
chr9-21628215-DEL-688
chr9-21631136-DEL-107
chr9-21649953-DEL-98
chr9-21652137-DEL-405
chr9-21677253-DEL-53
chr9-21694227-DEL-193
chr9-21694818-DEL-798
chr9-21705008-DEL-90
chr9-21719434-DEL-69
chr9-21720396-DEL-192
chr9-21726587-DEL-140
chr9-21748775-DEL-52
chr9-21768141-DEL-198
chr9-21773754-DEL-439
chr9-21806408-DEL-96
chr9-21807221-DEL-192
chr9-21833037-DEL-210
chr9-21856919-DEL-85
chr9-21878009-DEL-907
chr9-21878686-DEL-1872
chr9-21880601-DEL-1089
chr9-21932588-DEL-927
chr9-21965318-DEL-60
chr9-21971824-DEL-72
chr9-22005459-DEL-74
chr9-22009635-DEL-104
chr9-22035620-DEL-172
chr9-22044830-DEL-94
chr9-22055530-DEL-291
chr9-22055912-DEL-1970
chr9-22059065-DEL-363
chr9-22059804-DEL-199
chr9-22087515-DEL-184
chr9-22094650-DEL-69
chr9-22096827-DEL-730
chr9-22101002-DEL-1175
chr9-22101074-DEL-132
chr9-22101151-DEL-2921
chr9-22101865-DEL-388
chr9-22102380-DEL-120
chr9-22103206-DEL-614
chr9-22103709-DEL-191
chr9-22111677-DEL-168
chr9-22119849-DEL-148
chr9-22123655-DEL-186
chr9-22133669-DEL-212
chr9-22138547-DEL-546
chr9-22140117-DEL-194
chr9-22165505-DEL-90
chr9-22172299-DEL-52
chr9-22201212-DEL-94
chr9-22210148-DEL-140
chr9-22211362-DEL-281
chr9-22227159-DEL-243
chr9-22231248-DEL-1523
chr9-22253719-DEL-463
chr9-22275068-DEL-141
chr9-22276791-DEL-58
chr9-22303924-DEL-125
chr9-22307862-DEL-1213
chr9-22331002-DEL-82
chr9-22339192-DEL-162
chr9-22368070-DEL-125
chr9-22395664-DEL-123
chr9-22411154-DEL-444
chr9-22464662-DEL-84
chr9-22496107-DEL-336
chr9-22507403-DEL-88
chr9-22521963-DEL-578
chr9-22525863-DEL-138
chr9-22527597-DEL-2509

chr9-22547093-DEL-308
chr9-22588001-DEL-336
chr9-22622022-DEL-114
chr9-22622871-DEL-917
chr9-22624572-DEL-141
chr9-22669356-DEL-120
chr9-22699957-DEL-480
chr9-22707503-DEL-128
chr9-22721123-DEL-1408
chr9-22742745-DEL-70
chr9-22749237-DEL-99
chr9-22751961-DEL-1761
chr9-22751966-DEL-307
chr9-22753411-DEL-314
chr9-22753536-DEL-3479
chr9-22780461-DEL-194
chr9-22787212-DEL-108
chr9-22793800-DEL-3074
chr9-22822720-DEL-231
chr9-22828945-DEL-1035
chr9-22859705-DEL-2910
chr9-22866769-DEL-1419
chr9-22869013-DEL-52
chr9-22891636-DEL-1767
chr9-22935673-DEL-52
chr9-22991533-DEL-88
chr9-23020562-DEL-115
chr9-23028495-DEL-819
chr9-23046707-DEL-70
chr9-23063713-DEL-91
chr9-23070568-DEL-401
chr9-23155949-DEL-69
chr9-23197965-DEL-6454
chr9-23220685-DEL-64
chr9-23239071-DEL-148
chr9-23274165-DEL-62
chr9-23388355-DEL-84
chr9-23433640-DEL-538
chr9-23447125-DEL-63
chr9-23459249-DEL-390
chr9-23529102-DEL-337
chr9-23584730-DEL-148
chr9-23586951-DEL-66
chr9-23589625-DEL-6524
chr9-23595903-DEL-201
chr9-23643043-DEL-80
chr9-23657193-DEL-3162
chr9-23694641-DEL-54
chr9-23731195-DEL-70
chr9-23744437-DEL-383
chr9-23751687-DEL-204
chr9-23771812-DEL-335
chr9-23789434-DEL-64
chr9-23840192-DEL-212
chr9-23905889-DEL-237
chr9-23912149-DEL-64
chr9-23928091-DEL-593
chr9-24015113-DEL-182
chr9-24041608-DEL-630
chr9-24046895-DEL-56
chr9-24059411-DEL-214
chr9-24068160-DEL-158
chr9-24113998-DEL-393
chr9-24146160-DEL-852
chr9-24151787-DEL-260
chr9-24169169-DEL-207
chr9-24188585-DEL-51
chr9-24232043-DEL-156
chr9-24233506-DEL-6526
chr9-24240545-DEL-211
chr9-24299604-DEL-166
chr9-24398282-DEL-121
chr9-24432157-DEL-244
chr9-24444693-DEL-187
chr9-24451105-DEL-70
chr9-24500703-DEL-113
chr9-24506197-DEL-108

chr9-24513090-DEL-5740
chr9-24551627-DEL-73
chr9-24619751-DEL-99
chr9-24641558-DEL-904
chr9-24707683-DEL-207
chr9-24727009-DEL-155
chr9-24748045-DEL-72
chr9-24776341-DEL-438
chr9-24835178-DEL-1519
chr9-24843997-DEL-3180
chr9-24860021-DEL-77
chr9-24863903-DEL-897
chr9-24869684-DEL-62
chr9-24879681-DEL-279
chr9-24931449-DEL-53
chr9-24939806-DEL-215
chr9-24940369-DEL-118
chr9-24956550-DEL-112
chr9-25002041-DEL-84
chr9-25012402-DEL-411
chr9-25021283-DEL-134
chr9-25030530-DEL-1999
chr9-25040355-DEL-383
chr9-25053672-DEL-87
chr9-25129767-DEL-331
chr9-25166067-DEL-63
chr9-25252088-DEL-1251
chr9-25264227-DEL-82
chr9-25276029-DEL-611
chr9-25338786-DEL-158
chr9-25368209-DEL-60
chr9-25408761-DEL-172
chr9-25414013-DEL-330
chr9-25436662-DEL-100
chr9-25448704-DEL-438
chr9-25473305-DEL-125
chr9-25491990-DEL-127
chr9-25530907-DEL-75
chr9-25536112-DEL-220
chr9-25544609-DEL-71
chr9-25547398-DEL-255
chr9-25549460-DEL-6232
chr9-25575287-DEL-882
chr9-25581536-DEL-630
chr9-25582218-DEL-1375
chr9-25591284-DEL-199
chr9-25599761-DEL-598
chr9-25641673-DEL-52
chr9-25654611-DEL-609
chr9-25702407-DEL-356
chr9-25703472-DEL-1514
chr9-25710255-DEL-253
chr9-25723659-DEL-94
chr9-25724032-DEL-492
chr9-25753964-DEL-6080
chr9-25765706-DEL-130
chr9-25792087-DEL-137
chr9-25833740-DEL-69
chr9-25839245-DEL-6220
chr9-25850564-DEL-206
chr9-25853924-DEL-64
chr9-25877995-DEL-5978
chr9-25925580-DEL-52
chr9-25935473-DEL-3592
chr9-25985935-DEL-222
chr9-25990508-DEL-285
chr9-25993196-DEL-1488
chr9-26038365-DEL-202
chr9-26038813-DEL-1203
chr9-26048418-DEL-77
chr9-26081717-DEL-892
chr9-26103140-DEL-1046
chr9-26120385-DEL-1361
chr9-26126729-DEL-2129
chr9-26128922-DEL-655
chr9-26135067-DEL-53
chr9-26169330-DEL-741

chr9-26173711-DEL-55
chr9-26175690-DEL-105
chr9-26191368-DEL-964
chr9-26226532-DEL-58
chr9-26274342-DEL-764
chr9-26299890-DEL-4714
chr9-26356311-DEL-395
chr9-26359318-DEL-52
chr9-26376421-DEL-345
chr9-26378832-DEL-80
chr9-26404575-DEL-80
chr9-26417934-DEL-6249
chr9-26432109-DEL-97
chr9-26459387-DEL-95
chr9-26497924-DEL-1234
chr9-26585971-DEL-857
chr9-26607467-DEL-1531
chr9-26628814-DEL-114
chr9-26644188-DEL-56
chr9-26647870-DEL-503
chr9-26668659-DEL-56
chr9-26696285-DEL-52
chr9-26704702-DEL-60
chr9-26709001-DEL-74
chr9-26734083-DEL-90
chr9-26734339-DEL-69
chr9-26747064-DEL-139
chr9-26756009-DEL-401
chr9-26757898-DEL-147
chr9-26761687-DEL-238
chr9-26803688-DEL-164
chr9-26858809-DEL-87
chr9-26904483-DEL-51
chr9-26975860-DEL-186
chr9-26989984-DEL-84
chr9-27043744-DEL-556
chr9-27122985-DEL-56
chr9-27124470-DEL-72
chr9-27129541-DEL-65
chr9-27175560-DEL-80
chr9-27227297-DEL-600
chr9-27281172-DEL-162
chr9-27299623-DEL-67
chr9-27371926-DEL-209
chr9-27380217-DEL-52
chr9-27381762-DEL-704
chr9-27384285-DEL-123
chr9-27425172-DEL-52
chr9-27458467-DEL-334
chr9-27468933-DEL-103
chr9-27499541-DEL-68
chr9-27514084-DEL-69
chr9-27578242-DEL-1629
chr9-27590098-DEL-6155
chr9-27607727-DEL-202
chr9-27624325-DEL-180
chr9-27626880-DEL-577
chr9-27686478-DEL-866
chr9-27690599-DEL-105
chr9-27735021-DEL-75
chr9-27806319-DEL-115
chr9-27808730-DEL-105
chr9-27814820-DEL-56
chr9-27821579-DEL-53
chr9-27923416-DEL-55
chr9-27933678-DEL-60
chr9-27933988-DEL-703
chr9-27998523-DEL-88
chr9-28003303-DEL-100
chr9-28085549-DEL-57
chr9-28106384-DEL-656
chr9-28126907-DEL-144
chr9-28150554-DEL-137
chr9-28152267-DEL-186
chr9-28175690-DEL-457
chr9-28244408-DEL-2428
chr9-28268347-DEL-242

chr9-28331172-DEL-117
chr9-28397623-DEL-403
chr9-28410315-DEL-57
chr9-28419236-DEL-1829
chr9-28438586-DEL-111
chr9-28440965-DEL-324
chr9-28476779-DEL-56
chr9-28482306-DEL-1182
chr9-28493642-DEL-60
chr9-28605146-DEL-75
chr9-28606092-DEL-66
chr9-28682212-DEL-52
chr9-28691836-DEL-72
chr9-28710587-DEL-515
chr9-28743931-DEL-59
chr9-28758177-DEL-78
chr9-28768767-DEL-833
chr9-28775081-DEL-230
chr9-28807956-DEL-72
chr9-28813629-DEL-68
chr9-28820957-DEL-245
chr9-28851654-DEL-238
chr9-28900635-DEL-131
chr9-28949472-DEL-326
chr9-28983649-DEL-78
chr9-28992730-DEL-130
chr9-29003317-DEL-75
chr9-29021622-DEL-169
chr9-29070710-DEL-2599
chr9-29080304-DEL-112
chr9-29106221-DEL-1814
chr9-29114289-DEL-75
chr9-29124996-DEL-75
chr9-29220140-DEL-764
chr9-29225080-DEL-54
chr9-29234065-DEL-2320
chr9-29331283-DEL-59
chr9-29353803-DEL-98
chr9-29363807-DEL-496
chr9-29364981-DEL-52
chr9-29377617-DEL-172
chr9-29395413-DEL-189
chr9-29396228-DEL-128
chr9-29417129-DEL-296
chr9-29425141-DEL-399
chr9-29451083-DEL-154
chr9-29455007-DEL-155
chr9-29459402-DEL-51
chr9-29488385-DEL-195
chr9-29511590-DEL-360
chr9-29617641-DEL-321
chr9-29780991-DEL-240
chr9-29869670-DEL-315
chr9-29879774-DEL-56
chr9-29885593-DEL-87
chr9-29887373-DEL-213
chr9-29900143-DEL-57
chr9-29944920-DEL-194
chr9-29962478-DEL-72
chr9-29980561-DEL-460
chr9-29995208-DEL-94
chr9-30004559-DEL-66
chr9-30010625-DEL-56
chr9-30022723-DEL-182
chr9-30081057-DEL-99
chr9-30132737-DEL-400
chr9-30149314-DEL-130
chr9-30157414-DEL-60
chr9-30176145-DEL-56
chr9-30203485-DEL-56
chr9-30222275-DEL-246
chr9-30224832-DEL-55
chr9-30270173-DEL-2477
chr9-30324817-DEL-59
chr9-30327011-DEL-397
chr9-30350309-DEL-78
chr9-30354884-DEL-69

chr9-30399809-DEL-203
chr9-30421067-DEL-64
chr9-30437420-DEL-2889
chr9-30453616-DEL-608
chr9-30470019-DEL-2003
chr9-30473653-DEL-224
chr9-30508359-DEL-548
chr9-30533006-DEL-603
chr9-30566579-DEL-345
chr9-30572643-DEL-120
chr9-30589036-DEL-106
chr9-30601705-DEL-5071
chr9-30607179-DEL-264
chr9-30623086-DEL-271
chr9-30696796-DEL-210
chr9-30716889-DEL-568
chr9-30772401-DEL-225
chr9-30825092-DEL-87
chr9-30844756-DEL-90
chr9-30936509-DEL-171
chr9-30954962-DEL-192
chr9-30966347-DEL-124
chr9-30975694-DEL-70
chr9-31034017-DEL-122
chr9-31035304-DEL-348
chr9-31076810-DEL-199
chr9-31101770-DEL-64
chr9-31105117-DEL-1936
chr9-31131235-DEL-87
chr9-31142508-DEL-197
chr9-31144561-DEL-858
chr9-31149550-DEL-58
chr9-31151253-DEL-370
chr9-31159287-DEL-76
chr9-31178594-DEL-60
chr9-31194138-DEL-313
chr9-31220688-DEL-1377
chr9-31248895-DEL-56
chr9-31261556-DEL-80
chr9-31270641-DEL-200
chr9-31273368-DEL-170
chr9-31281233-DEL-189
chr9-31295543-DEL-201
chr9-31296104-DEL-106
chr9-31324777-DEL-92
chr9-31354177-DEL-253
chr9-31355090-DEL-211
chr9-31379411-DEL-51
chr9-31392637-DEL-73
chr9-31395885-DEL-213
chr9-31399575-DEL-141
chr9-31409139-DEL-60
chr9-31428582-DEL-253
chr9-31451221-DEL-145
chr9-31491873-DEL-237
chr9-31505916-DEL-105
chr9-31564067-DEL-64
chr9-31582905-DEL-104
chr9-31596391-DEL-61
chr9-31614061-DEL-139
chr9-31618879-DEL-416
chr9-31679258-DEL-114
chr9-31682775-DEL-541
chr9-31683730-DEL-103
chr9-31722304-DEL-55
chr9-31740231-DEL-1000
chr9-31742838-DEL-167
chr9-31745193-DEL-127
chr9-31759580-DEL-288
chr9-31776625-DEL-507
chr9-31797403-DEL-52
chr9-31811494-DEL-149
chr9-31833671-DEL-439
chr9-31846564-DEL-1639
chr9-31861112-DEL-174
chr9-31862547-DEL-56
chr9-31878236-DEL-823

chr9-31912341-DEL-70
chr9-31948369-DEL-138
chr9-32026464-DEL-64
chr9-32032728-DEL-5649
chr9-32041934-DEL-250
chr9-32042556-DEL-186
chr9-32075302-DEL-140
chr9-32114472-DEL-786
chr9-32145342-DEL-70
chr9-32163237-DEL-930
chr9-32164687-DEL-60
chr9-32185605-DEL-96
chr9-32210236-DEL-55
chr9-32211497-DEL-216
chr9-32212593-DEL-153
chr9-32218580-DEL-51
chr9-32218954-DEL-1537
chr9-32219548-DEL-300
chr9-32219847-DEL-307
chr9-32241144-DEL-427
chr9-32264498-DEL-93
chr9-32265545-DEL-1121
chr9-32265546-DEL-955
chr9-32265597-DEL-214
chr9-32266804-DEL-460
chr9-32273725-DEL-2178
chr9-32275990-DEL-706
chr9-32286280-DEL-70
chr9-32370001-DEL-51
chr9-32379403-DEL-52
chr9-32388911-DEL-205
chr9-32427048-DEL-194
chr9-32447383-DEL-50
chr9-32470623-DEL-116
chr9-32486057-DEL-97
chr9-32519013-DEL-284
chr9-32522614-DEL-62
chr9-32560804-DEL-1388
chr9-32595025-DEL-246
chr9-32596139-DEL-59
chr9-32680487-DEL-1486
chr9-32702902-DEL-89
chr9-32709755-DEL-399
chr9-32710328-DEL-1090
chr9-32720028-DEL-3325
chr9-32740495-DEL-3036
chr9-32741082-DEL-77
chr9-32749085-DEL-60
chr9-32761040-DEL-192
chr9-32779880-DEL-57
chr9-32790054-DEL-1116
chr9-32835061-DEL-192
chr9-32877796-DEL-129
chr9-32950952-DEL-1001
chr9-32952114-DEL-208
chr9-32978416-DEL-76
chr9-33005842-DEL-873
chr9-33037739-DEL-201
chr9-33054898-DEL-6019
chr9-33064318-DEL-67
chr9-33130766-DEL-81
chr9-33199398-DEL-60
chr9-33249253-DEL-1521
chr9-33302960-DEL-12585
chr9-33309580-DEL-4685
chr9-33319892-DEL-99
chr9-33353103-DEL-57
chr9-33389476-DEL-1101
chr9-33405914-DEL-109
chr9-33424609-DEL-195
chr9-33438237-DEL-53
chr9-33451602-DEL-185
chr9-33469326-DEL-100
chr9-33516788-DEL-129
chr9-33517411-DEL-8470
chr9-33526357-DEL-60
chr9-33546988-DEL-101

chr9-33548070-DEL-115
chr9-33558177-DEL-114
chr9-33566666-DEL-102
chr9-33567991-DEL-565
chr9-33570809-DEL-506
chr9-33601132-DEL-228
chr9-33610139-DEL-234
chr9-33613240-DEL-66
chr9-33628213-DEL-55
chr9-33634941-DEL-70
chr9-33666620-DEL-1929
chr9-33727951-DEL-68
chr9-33751279-DEL-786
chr9-33781581-DEL-1180
chr9-33828405-DEL-116
chr9-33829736-DEL-306
chr9-33945230-DEL-447
chr9-33950698-DEL-1441
chr9-33956382-DEL-168
chr9-33967615-DEL-54
chr9-34010800-DEL-487
chr9-34035586-DEL-533
chr9-34038122-DEL-228
chr9-34077053-DEL-864
chr9-34086053-DEL-59
chr9-34199587-DEL-66
chr9-34247061-DEL-74
chr9-34275237-DEL-210
chr9-34314784-DEL-574
chr9-34320951-DEL-60
chr9-34389937-DEL-60
chr9-34402959-DEL-56
chr9-34423980-DEL-52
chr9-34453656-DEL-967
chr9-34482056-DEL-60
chr9-34538144-DEL-75
chr9-34550504-DEL-1921
chr9-34551161-DEL-1553
chr9-34552916-DEL-190
chr9-34557422-DEL-336
chr9-34635901-DEL-399
chr9-34697289-DEL-416
chr9-34774562-DEL-2196
chr9-34781816-DEL-156
chr9-34797144-DEL-151
chr9-34807461-DEL-612
chr9-34827752-DEL-52
chr9-34830978-DEL-64
chr9-34844720-DEL-174
chr9-34870596-DEL-80
chr9-34941442-DEL-69
chr9-34969665-DEL-69
chr9-34982422-DEL-116
chr9-34984843-DEL-211
chr9-34990127-DEL-103
chr9-34997367-DEL-97
chr9-35018123-DEL-399
chr9-35023713-DEL-147
chr9-35061482-DEL-111
chr9-35065880-DEL-207
chr9-35107719-DEL-466
chr9-35149115-DEL-108
chr9-35152676-DEL-2159
chr9-35164017-DEL-84
chr9-35281953-DEL-77
chr9-35356560-DEL-91
chr9-35362684-DEL-132
chr9-35364278-DEL-330
chr9-35410699-DEL-129
chr9-35462865-DEL-54
chr9-35496868-DEL-407
chr9-35514522-DEL-157
chr9-35522079-DEL-945
chr9-35535070-DEL-6471
chr9-35545151-DEL-99
chr9-35545936-DEL-96
chr9-35554066-DEL-184

chr9-35563066-DEL-547
chr9-35613841-DEL-2571
chr9-35618329-DEL-94
chr9-35650204-DEL-88
chr9-35743406-DEL-126
chr9-35784212-DEL-2318
chr9-35797188-DEL-856
chr9-35797966-DEL-212
chr9-35844203-DEL-246
chr9-35857813-DEL-6000
chr9-35863609-DEL-884
chr9-35864987-DEL-495
chr9-35882856-DEL-587
chr9-35917198-DEL-600
chr9-35932195-DEL-1981
chr9-35977748-DEL-10864
chr9-35977991-DEL-4744
chr9-35990009-DEL-65
chr9-35998679-DEL-9156
chr9-35999351-DEL-7479
chr9-36033656-DEL-135
chr9-36053709-DEL-4372
chr9-36070035-DEL-2077
chr9-36117637-DEL-1038
chr9-36136637-DEL-17632
chr9-36162397-DEL-138
chr9-36207254-DEL-984
chr9-36212696-DEL-64
chr9-36230245-DEL-284
chr9-36239947-DEL-535
chr9-36258402-DEL-4163
chr9-36287553-DEL-6491
chr9-36294603-DEL-2210
chr9-36302417-DEL-708
chr9-36324259-DEL-869
chr9-36366819-DEL-75
chr9-36371061-DEL-3581
chr9-36391417-DEL-1833
chr9-36403446-DEL-204
chr9-36420688-DEL-825
chr9-36422714-DEL-218
chr9-36466278-DEL-1527
chr9-36473586-DEL-6756
chr9-36490198-DEL-79
chr9-36531679-DEL-2834
chr9-36573681-DEL-599
chr9-36582210-DEL-6307
chr9-36593690-DEL-108
chr9-36619179-DEL-233
chr9-36662318-DEL-79
chr9-36666991-DEL-905
chr9-36711942-DEL-1096
chr9-36715994-DEL-2654
chr9-36732326-DEL-204
chr9-36738741-DEL-83
chr9-36798848-DEL-72
chr9-36822017-DEL-65
chr9-36843372-DEL-174
chr9-36864372-DEL-63
chr9-36917805-DEL-1369
chr9-36940192-DEL-52
chr9-36990396-DEL-757
chr9-36991382-DEL-1665
chr9-37028593-DEL-256
chr9-37032691-DEL-56
chr9-37066066-DEL-52
chr9-37125517-DEL-53
chr9-37156327-DEL-982
chr9-37183318-DEL-1752
chr9-37223087-DEL-441
chr9-37228623-DEL-207
chr9-37234609-DEL-1157
chr9-37323553-DEL-240
chr9-37337738-DEL-281
chr9-37353062-DEL-7175
chr9-37360179-DEL-203
chr9-37389470-DEL-87

chr9-37391005-DEL-55
chr9-37397364-DEL-306
chr9-37403265-DEL-88
chr9-37468978-DEL-62
chr9-37491852-DEL-57
chr9-37495456-DEL-60
chr9-37499677-DEL-271
chr9-37507147-DEL-70
chr9-37533630-DEL-222
chr9-37539815-DEL-70
chr9-37596021-DEL-96
chr9-37602943-DEL-163
chr9-37622274-DEL-60
chr9-37634292-DEL-106
chr9-37656954-DEL-56
chr9-37670430-DEL-154
chr9-37673166-DEL-5407
chr9-37713190-DEL-58
chr9-37733086-DEL-53
chr9-37764760-DEL-1291
chr9-37841602-DEL-79
chr9-37862211-DEL-749
chr9-37871485-DEL-261
chr9-37934304-DEL-87
chr9-37938138-DEL-79
chr9-37943715-DEL-299
chr9-37955210-DEL-134
chr9-37956922-DEL-351
chr9-37962349-DEL-77
chr9-37981242-DEL-328
chr9-38000888-DEL-184
chr9-38006490-DEL-66
chr9-38028909-DEL-213
chr9-38029279-DEL-1644
chr9-38059693-DEL-2412
chr9-38066938-DEL-238
chr9-38076303-DEL-215
chr9-38090876-DEL-16045
chr9-38111447-DEL-1449
chr9-38144576-DEL-266
chr9-38151004-DEL-10015
chr9-38152034-DEL-423
chr9-38161566-DEL-864
chr9-38191582-DEL-465
chr9-38207328-DEL-1451
chr9-38228247-DEL-77
chr9-38346610-DEL-132
chr9-38356399-DEL-60
chr9-38390371-DEL-134
chr9-38403467-DEL-54
chr9-38472461-DEL-201
chr9-38474880-DEL-7473
chr9-38501801-DEL-297
chr9-38502666-DEL-224
chr9-38522630-DEL-113
chr9-38529001-DEL-12619
chr9-38529338-DEL-221
chr9-38531867-DEL-1056
chr9-38543121-DEL-11910
chr9-38546161-DEL-108
chr9-38547177-DEL-67
chr9-38578832-DEL-94
chr9-38596405-DEL-220
chr9-38612264-DEL-84
chr9-38619628-DEL-69
chr9-38632487-DEL-1013
chr9-38656858-DEL-71
chr9-38693608-DEL-116
chr9-38697597-DEL-1319
chr9-38699852-DEL-88
chr9-38702400-DEL-276
chr9-38719306-DEL-912
chr9-38721064-DEL-65
chr9-38723591-DEL-1064
chr9-38779396-DEL-492
chr9-38790319-DEL-1049
chr9-38790489-DEL-260

chr9-38797559-DEL-34729
chr9-38800927-DEL-801
chr9-38802281-DEL-364
chr9-38806019-DEL-512
chr9-38807839-DEL-208
chr9-38826132-DEL-1439
chr9-38829218-DEL-99
chr9-38861980-DEL-150
chr9-38867750-DEL-668
chr9-38878307-DEL-3385
chr9-38904990-DEL-95
chr9-38961576-DEL-2008
chr9-38973577-DEL-54
chr9-38985675-DEL-1292
chr9-38996691-DEL-6740
chr9-39044989-DEL-1387
chr9-39046837-DEL-2358
chr9-39072792-DEL-2409
chr9-39091242-DEL-114
chr9-39098325-DEL-3199
chr9-39105807-DEL-2002
chr9-39107633-DEL-2364
chr9-39115285-DEL-407
chr9-39130991-DEL-77
chr9-39137364-DEL-1154
chr9-39149429-DEL-85
chr9-39167044-DEL-744
chr9-39175787-DEL-135
chr9-39214600-DEL-387
chr9-39239383-DEL-253
chr9-39257001-DEL-149
chr9-39257896-DEL-1664
chr9-39259960-DEL-11078
chr9-39259998-DEL-4731
chr9-39261691-DEL-232
chr9-39262014-DEL-882
chr9-39264865-DEL-6173
chr9-39277282-DEL-75
chr9-39281426-DEL-262
chr9-39310911-DEL-6421
chr9-39325026-DEL-3386
chr9-39341215-DEL-104
chr9-39345381-DEL-55
chr9-39345763-DEL-1026
chr9-39365656-DEL-476
chr9-39388696-DEL-1181
chr9-39425438-DEL-1060
chr9-39427354-DEL-175
chr9-39431533-DEL-585
chr9-39435646-DEL-138
chr9-39436724-DEL-6417
chr9-39448419-DEL-54
chr9-39457166-DEL-105
chr9-39467976-DEL-51
chr9-39491017-DEL-925
chr9-39492528-DEL-1313
chr9-39494076-DEL-958
chr9-39495101-DEL-275
chr9-39496520-DEL-306
chr9-39543332-DEL-344
chr9-39553115-DEL-1833
chr9-39561532-DEL-89
chr9-39576259-DEL-569
chr9-39602403-DEL-301
chr9-39615025-DEL-60
chr9-39662623-DEL-327
chr9-39671793-DEL-4947
chr9-39692703-DEL-52
chr9-39705420-DEL-159
chr9-39713999-DEL-215
chr9-39723539-DEL-3210
chr9-39740131-DEL-1075
chr9-39750487-DEL-94
chr9-39756404-DEL-266
chr9-39758549-DEL-340
chr9-39787089-DEL-10788
chr9-39823206-DEL-198

chr9-39840746-DEL-76
chr9-39850436-DEL-84
chr9-39859136-DEL-60
chr9-39864525-DEL-132
chr9-39876003-DEL-996
chr9-39896897-DEL-6198
chr9-39929476-DEL-53
chr9-39934824-DEL-87
chr9-39952066-DEL-6193
chr9-39965630-DEL-1102
chr9-39969673-DEL-940
chr9-39998021-DEL-247
chr9-39998948-DEL-222
chr9-40008817-DEL-439
chr9-40014989-DEL-159
chr9-40015692-DEL-3041
chr9-40019876-DEL-136
chr9-40032991-DEL-133
chr9-40069698-DEL-137
chr9-40071009-DEL-72
chr9-40080201-DEL-5312
chr9-40094208-DEL-232
chr9-40095013-DEL-500
chr9-40148259-DEL-52
chr9-40150676-DEL-202
chr9-40161439-DEL-58
chr9-40207518-DEL-71
chr9-40219791-DEL-335
chr9-40278906-DEL-92
chr9-40308063-DEL-55
chr9-40330082-DEL-233
chr9-40342007-DEL-57
chr9-40363174-DEL-100
chr9-40395159-DEL-54
chr9-40478840-DEL-68
chr9-40501517-DEL-2321
chr9-40546528-DEL-612
chr9-40549043-DEL-63
chr9-40557987-DEL-239
chr9-40579590-DEL-60
chr9-40630726-DEL-87
chr9-40648184-DEL-132
chr9-40664247-DEL-167
chr9-40666600-DEL-133
chr9-40680238-DEL-1263
chr9-40687089-DEL-191
chr9-40688138-DEL-52
chr9-40691083-DEL-1055
chr9-40698648-DEL-138
chr9-40702084-DEL-173
chr9-40708545-DEL-66
chr9-40731775-DEL-96
chr9-40735885-DEL-220
chr9-40737575-DEL-280
chr9-40756149-DEL-197
chr9-40826838-DEL-89
chr9-40847677-DEL-123
chr9-40853681-DEL-74
chr9-40854070-DEL-267
chr9-40879273-DEL-126
chr9-40885725-DEL-488
chr9-40898913-DEL-96
chr9-40901634-DEL-320
chr9-40930799-DEL-60
chr9-40969967-DEL-88
chr9-41007415-DEL-53
chr9-41015148-DEL-968
chr9-41074954-DEL-432
chr9-41104876-DEL-61
chr9-41106855-DEL-68
chr9-41125163-DEL-51
chr9-41128815-DEL-431
chr9-41141903-DEL-72
chr9-41154764-DEL-60
chr9-41167651-DEL-74
chr9-41230042-DEL-98
chr9-41291616-DEL-74

chr9-41310083-DEL-202
chr9-41313013-DEL-107
chr9-41317006-DEL-239
chr9-41326907-DEL-78
chr9-41327571-DEL-894
chr9-41335144-DEL-767
chr9-41426923-DEL-211
chr9-41463548-DEL-174
chr9-41480199-DEL-279
chr9-41513179-DEL-79
chr9-41540185-DEL-312
chr9-41560294-DEL-203
chr9-41564857-DEL-548
chr9-41602946-DEL-400
chr9-41680966-DEL-66
chr9-41682073-DEL-56
chr9-41682499-DEL-498
chr9-41748790-DEL-68
chr9-41753508-DEL-70
chr9-41754366-DEL-174
chr9-41756398-DEL-127
chr9-41758406-DEL-83
chr9-41791039-DEL-150
chr9-41829870-DEL-56
chr9-41860173-DEL-172
chr9-41910573-DEL-64
chr9-41923690-DEL-85
chr9-41936144-DEL-174
chr9-41951767-DEL-86
chr9-41961363-DEL-1050
chr9-41963085-DEL-615
chr9-42025424-DEL-115
chr9-42043716-DEL-76
chr9-42081661-DEL-98
chr9-42090070-DEL-74
chr9-42111219-DEL-91
chr9-42116173-DEL-59
chr9-42227067-DEL-62
chr9-42228055-DEL-2127
chr9-42313701-DEL-270
chr9-42319447-DEL-62
chr9-42349612-DEL-167
chr9-42384062-DEL-83
chr9-42409052-DEL-145
chr9-42436815-DEL-72
chr9-42452567-DEL-122
chr9-42457223-DEL-89
chr9-42505469-DEL-816
chr9-42530634-DEL-221
chr9-42541250-DEL-184
chr9-42555629-DEL-316
chr9-42575525-DEL-57
chr9-42600309-DEL-532
chr9-42606567-DEL-64
chr9-42610771-DEL-82
chr9-42646116-DEL-58
chr9-42706611-DEL-183
chr9-42727211-DEL-64
chr9-42747801-DEL-375
chr9-42748609-DEL-72
chr9-42766323-DEL-519
chr9-42784917-DEL-74
chr9-42795931-DEL-66
chr9-42804174-DEL-96
chr9-42830576-DEL-124
chr9-42858948-DEL-74
chr9-42859707-DEL-61
chr9-42863436-DEL-75
chr9-42871638-DEL-66
chr9-42876075-DEL-1664
chr9-42907394-DEL-547
chr9-42908355-DEL-93
chr9-42912526-DEL-183
chr9-42921810-DEL-114
chr9-42929999-DEL-979
chr9-42930052-DEL-411
chr9-42943576-DEL-65

chr9-42956371-DEL-740
chr9-42973791-DEL-96
chr9-43033183-DEL-227
chr9-43047674-DEL-149
chr9-43090394-DEL-57
chr9-43092905-DEL-51
chr9-43125743-DEL-93
chr9-43161832-DEL-111
chr9-43164412-DEL-68
chr9-43169267-DEL-57
chr9-43200629-DEL-116
chr9-43219121-DEL-178
chr9-43221333-DEL-103
chr9-43270996-DEL-110
chr9-43281403-DEL-116
chr9-43326468-DEL-1455
chr9-43332696-DEL-51
chr9-43337517-DEL-71
chr9-43338655-DEL-186
chr9-43356032-DEL-123
chr9-43357518-DEL-230
chr9-43361318-DEL-392
chr9-43378027-DEL-152
chr9-43399404-DEL-69
chr9-43409864-DEL-51
chr9-43438139-DEL-82
chr9-43470363-DEL-1224
chr9-43492721-DEL-147
chr9-43518532-DEL-97
chr9-43534592-DEL-56
chr9-43542113-DEL-403
chr9-43542805-DEL-60
chr9-43556539-DEL-195
chr9-43557739-DEL-1318
chr9-43588300-DEL-201
chr9-43689540-DEL-991
chr9-43713804-DEL-66
chr9-43725545-DEL-82
chr9-43731860-DEL-149
chr9-43780234-DEL-103
chr9-43788452-DEL-55
chr9-43837295-DEL-52
chr9-43839640-DEL-295
chr9-43920345-DEL-216
chr9-43930532-DEL-76
chr9-43931775-DEL-336
chr9-43936785-DEL-808
chr9-43965030-DEL-178
chr9-44026509-DEL-229
chr9-44033241-DEL-193
chr9-44055627-DEL-52
chr9-44067116-DEL-92
chr9-44085364-DEL-178
chr9-44112911-DEL-209
chr9-44208083-DEL-204
chr9-44211361-DEL-339
chr9-44219580-DEL-179
chr9-44249621-DEL-188
chr9-44262174-DEL-52
chr9-44273509-DEL-199
chr9-44288576-DEL-375
chr9-44320189-DEL-64
chr9-44353597-DEL-195
chr9-44463370-DEL-61
chr9-44466103-DEL-86
chr9-44468911-DEL-60
chr9-44471255-DEL-65
chr9-44472559-DEL-215
chr9-44474755-DEL-548
chr9-44477806-DEL-241
chr9-44479482-DEL-72
chr9-44500708-DEL-217
chr9-44501319-DEL-170
chr9-44537602-DEL-59
chr9-44554853-DEL-337
chr9-44572305-DEL-175
chr9-44642888-DEL-72

chr9-44649718-DEL-205
chr9-44682430-DEL-202
chr9-44693578-DEL-140
chr9-44772591-DEL-51
chr9-44775178-DEL-86
chr9-44886868-DEL-91
chr9-44920095-DEL-54
chr9-44926800-DEL-88
chr9-44932570-DEL-68
chr9-44937678-DEL-51
chr9-44944673-DEL-277
chr9-44946631-DEL-56
chr9-45012861-DEL-299
chr9-45016532-DEL-68
chr9-45022058-DEL-56
chr9-45034798-DEL-63
chr9-45040034-DEL-868
chr9-45047925-DEL-109
chr9-45071873-DEL-538
chr9-45072745-DEL-1219
chr9-45129872-DEL-52
chr9-45141305-DEL-341
chr9-45168012-DEL-158
chr9-45207903-DEL-636
chr9-45235655-DEL-86
chr9-45274641-DEL-711
chr9-45298596-DEL-221
chr9-45339825-DEL-115
chr9-45356216-DEL-60
chr9-45373790-DEL-1451
chr9-45374629-DEL-1286
chr9-45385930-DEL-99
chr9-45430216-DEL-578
chr9-45500912-DEL-221
chr9-45504161-DEL-251
chr9-45512599-DEL-51
chr9-45513448-DEL-117
chr9-45535656-DEL-67
chr9-45561950-DEL-83
chr9-45567349-DEL-83
chr9-45573677-DEL-340
chr9-45581603-DEL-83
chr9-45583622-DEL-91
chr9-45599949-DEL-70
chr9-45602354-DEL-66
chr9-45636952-DEL-200
chr9-45639032-DEL-167
chr9-45641163-DEL-275
chr9-45644509-DEL-168
chr9-45648085-DEL-376
chr9-45657324-DEL-68
chr9-45668940-DEL-1016
chr9-45701210-DEL-200
chr9-45781008-DEL-162
chr9-45810310-DEL-180
chr9-45860482-DEL-69
chr9-45893267-DEL-92
chr9-45911677-DEL-983
chr9-45922117-DEL-112
chr9-45931903-DEL-55
chr9-45935098-DEL-460
chr9-45936438-DEL-411
chr9-45959638-DEL-85
chr9-45968031-DEL-100
chr9-45971479-DEL-1197
chr9-45988769-DEL-828
chr9-46044197-DEL-410
chr9-46044403-DEL-2246
chr9-46044829-DEL-174
chr9-46045461-DEL-102
chr9-46046159-DEL-134
chr9-46047094-DEL-2522
chr9-46048203-DEL-323
chr9-46048714-DEL-514
chr9-46078024-DEL-392
chr9-46103763-DEL-123
chr9-46112094-DEL-174

chr9-46166948-DEL-198
chr9-46196620-DEL-498
chr9-46200517-DEL-200
chr9-46208483-DEL-152
chr9-46242340-DEL-423
chr9-46250805-DEL-1029
chr9-46251019-DEL-629
chr9-46251521-DEL-393
chr9-46273628-DEL-98
chr9-46278733-DEL-203
chr9-46284223-DEL-458
chr9-46325836-DEL-136
chr9-46398259-DEL-68
chr9-46419213-DEL-51
chr9-46434282-DEL-81
chr9-46458930-DEL-63
chr9-46486659-DEL-368
chr9-46563218-DEL-213
chr9-46563218-DEL-341
chr9-46597694-DEL-52
chr9-46598251-DEL-60
chr9-46636138-DEL-61
chr9-46684955-DEL-67
chr9-46712311-DEL-2564
chr9-46715487-DEL-1109
chr9-46722342-DEL-51
chr9-46749837-DEL-1126
chr9-46767968-DEL-1025
chr9-46780051-DEL-61
chr9-46832117-DEL-60
chr9-46852941-DEL-86
chr9-46857741-DEL-65
chr9-46912743-DEL-112
chr9-46940517-DEL-206
chr9-46946443-DEL-89
chr9-46981483-DEL-65
chr9-47014406-DEL-68
chr9-47027896-DEL-205
chr9-47079980-DEL-80
chr9-47096206-DEL-102
chr9-47108141-DEL-407
chr9-47108763-DEL-437
chr9-47113753-DEL-840
chr9-47125449-DEL-392
chr9-47128824-DEL-54
chr9-47141967-DEL-307
chr9-47153253-DEL-81
chr9-47239241-DEL-55
chr9-47253827-DEL-58
chr9-47290489-DEL-65
chr9-47319065-DEL-72
chr9-47331356-DEL-403
chr9-47404575-DEL-1472
chr9-47407559-DEL-142
chr9-47480344-DEL-130
chr9-47490045-DEL-149
chr9-47543839-DEL-211
chr9-47578423-DEL-52
chr9-47581118-DEL-54
chr9-47753862-DEL-104
chr9-47833539-DEL-142
chr9-47851060-DEL-378
chr9-47852866-DEL-61
chr9-47869919-DEL-893
chr9-47900997-DEL-3252
chr9-47908694-DEL-52
chr9-47925378-DEL-81
chr9-47930148-DEL-54
chr9-47931027-DEL-56
chr9-47950571-DEL-52
chr9-47964881-DEL-192
chr9-47984744-DEL-400
chr9-47999106-DEL-1174
chr9-48027710-DEL-55
chr9-48136337-DEL-66
chr9-48159520-DEL-62
chr9-48226547-DEL-140

chr9-48278839-DEL-67
chr9-48283002-DEL-210
chr9-48289877-DEL-2456
chr9-48302042-DEL-238
chr9-48368630-DEL-52
chr9-48383246-DEL-103
chr9-48399538-DEL-167
chr9-48412817-DEL-105
chr9-48426757-DEL-94
chr9-48437463-DEL-118
chr9-48439215-DEL-234
chr9-48486031-DEL-431
chr9-48489742-DEL-424
chr9-48515441-DEL-102
chr9-48521892-DEL-52
chr9-48531065-DEL-62
chr9-48545865-DEL-400
chr9-48575684-DEL-52
chr9-48591203-DEL-203
chr9-48594472-DEL-1012
chr9-48638090-DEL-60
chr9-48685807-DEL-52
chr9-48740580-DEL-60
chr9-48803438-DEL-97
chr9-48825021-DEL-54
chr9-48912165-DEL-218
chr9-48919123-DEL-75
chr9-48948590-DEL-51
chr9-48973883-DEL-66
chr9-48983700-DEL-215
chr9-49002412-DEL-205
chr9-49002950-DEL-114
chr9-49012021-DEL-63
chr9-49034114-DEL-55
chr9-49054619-DEL-71
chr9-49063132-DEL-92
chr9-49075860-DEL-358
chr9-49130343-DEL-67
chr9-49143803-DEL-106
chr9-49187208-DEL-69
chr9-49204560-DEL-178
chr9-49228188-DEL-151
chr9-49236898-DEL-6581
chr9-49243474-DEL-808
chr9-49272690-DEL-569
chr9-49328822-DEL-225
chr9-49348206-DEL-637
chr9-49424872-DEL-103
chr9-49434686-DEL-69
chr9-49436887-DEL-99
chr9-49465351-DEL-330
chr9-49516276-DEL-178
chr9-49524626-DEL-77
chr9-49540848-DEL-900
chr9-49628278-DEL-74
chr9-49674187-DEL-123
chr9-49690123-DEL-315
chr9-49723203-DEL-1590
chr9-49726671-DEL-69
chr9-49731697-DEL-1043
chr9-49731729-DEL-218
chr9-49732189-DEL-551
chr9-49737352-DEL-584
chr9-49751321-DEL-71
chr9-49766809-DEL-60
chr9-49769694-DEL-161
chr9-49811224-DEL-305
chr9-49827762-DEL-281
chr9-49831150-DEL-150
chr9-49840785-DEL-63
chr9-49848430-DEL-66
chr9-49856232-DEL-54
chr9-49858740-DEL-134
chr9-49889403-DEL-355
chr9-49916298-DEL-204
chr9-49986651-DEL-2466
chr9-49988233-DEL-899

chr9-50046941-DEL-86
chr9-50093935-DEL-108
chr9-50096351-DEL-417
chr9-50187167-DEL-1180
chr9-50216666-DEL-95
chr9-50235159-DEL-53
chr9-50241156-DEL-76
chr9-50246834-DEL-66
chr9-50262498-DEL-105
chr9-50269629-DEL-114
chr9-50331580-DEL-51
chr9-50350091-DEL-226
chr9-50370818-DEL-67
chr9-50376598-DEL-804
chr9-50377424-DEL-806
chr9-50421121-DEL-3854
chr9-50481924-DEL-60
chr9-50497970-DEL-74
chr9-50498628-DEL-2250
chr9-50505078-DEL-227
chr9-50517130-DEL-58
chr9-50519499-DEL-65
chr9-50551897-DEL-51
chr9-50586430-DEL-55
chr9-50597261-DEL-55
chr9-50631899-DEL-151
chr9-50718109-DEL-203
chr9-50727401-DEL-171
chr9-50736418-DEL-211
chr9-50744219-DEL-251
chr9-50813082-DEL-106
chr9-50834969-DEL-58
chr9-50845539-DEL-247
chr9-50888090-DEL-513
chr9-50956805-DEL-233
chr9-50957169-DEL-576
chr9-50966923-DEL-195
chr9-50970917-DEL-68
chr9-51010569-DEL-77
chr9-51011624-DEL-128
chr9-51022845-DEL-266
chr9-51039454-DEL-64
chr9-51047491-DEL-51
chr9-51073336-DEL-101
chr9-51137598-DEL-57
chr9-51172330-DEL-52
chr9-51179806-DEL-66
chr9-51252585-DEL-108
chr9-51276035-DEL-734
chr9-51285340-DEL-120
chr9-51337186-DEL-575
chr9-51340217-DEL-87
chr9-51362186-DEL-143
chr9-51376033-DEL-343
chr9-51404064-DEL-60
chr9-51407290-DEL-1579
chr9-51420981-DEL-2162
chr9-51429526-DEL-76
chr9-51446017-DEL-306
chr9-51457578-DEL-457
chr9-51535656-DEL-429
chr9-51547778-DEL-58
chr9-51568072-DEL-57
chr9-51586352-DEL-62
chr9-51594507-DEL-121
chr9-51612025-DEL-528
chr9-51623800-DEL-58
chr9-51640817-DEL-605
chr9-51650487-DEL-56
chr9-51658579-DEL-102
chr9-51681078-DEL-684
chr9-51681114-DEL-180
chr9-51681221-DEL-1007
chr9-51681347-DEL-683
chr9-51815152-DEL-185
chr9-51838340-DEL-116
chr9-51847341-DEL-587

chr9-51849195-DEL-52
chr9-51876986-DEL-3242
chr9-51893502-DEL-106
chr9-51910732-DEL-1086
chr9-51929632-DEL-486
chr9-51946787-DEL-659
chr9-51994922-DEL-160
chr9-52007813-DEL-213
chr9-52033304-DEL-191
chr9-52036928-DEL-361
chr9-52069084-DEL-64
chr9-52088957-DEL-55
chr9-52149552-DEL-59
chr9-52197884-DEL-114
chr9-52216018-DEL-449
chr9-52265988-DEL-104
chr9-52290976-DEL-1145
chr9-52292805-DEL-745
chr9-52293694-DEL-481
chr9-52294282-DEL-2355
chr9-52296369-DEL-212
chr9-52379110-DEL-166
chr9-52389093-DEL-878
chr9-52413264-DEL-89
chr9-52468524-DEL-215
chr9-52469498-DEL-149
chr9-52503154-DEL-101
chr9-52603835-DEL-3469
chr9-52614705-DEL-54
chr9-52660150-DEL-131
chr9-52661886-DEL-733
chr9-52684884-DEL-103
chr9-52710609-DEL-100
chr9-52736277-DEL-546
chr9-52765280-DEL-92
chr9-52790309-DEL-7896
chr9-52806286-DEL-15776
chr9-52818211-DEL-3852
chr9-52831801-DEL-368
chr9-52853500-DEL-80
chr9-52866809-DEL-60
chr9-52866883-DEL-1275
chr9-52899581-DEL-922
chr9-52938938-DEL-55
chr9-52974302-DEL-127
chr9-52974624-DEL-920
chr9-52997597-DEL-171
chr9-53001043-DEL-1936
chr9-53032006-DEL-55
chr9-53049854-DEL-105
chr9-53097869-DEL-354
chr9-53140840-DEL-2213
chr9-53152393-DEL-71
chr9-53205949-DEL-335
chr9-53221498-DEL-167
chr9-53231546-DEL-59
chr9-53237831-DEL-355
chr9-53243616-DEL-341
chr9-53277241-DEL-176
chr9-53294978-DEL-173
chr9-53305450-DEL-55
chr9-53305735-DEL-1284
chr9-53322279-DEL-431
chr9-53364324-DEL-201
chr9-53366149-DEL-159
chr9-53420507-DEL-613
chr9-53442170-DEL-220
chr9-53466781-DEL-227
chr9-53469982-DEL-52
chr9-53470614-DEL-200
chr9-53507254-DEL-299
chr9-53525068-DEL-503
chr9-53550482-DEL-78
chr9-53557095-DEL-70
chr9-53560840-DEL-855
chr9-53564677-DEL-784
chr9-53581745-DEL-168

chr9-53581844-DEL-1212
chr9-53629292-DEL-249
chr9-53631987-DEL-265
chr9-53666763-DEL-2037
chr9-53687110-DEL-299
chr9-53710310-DEL-104
chr9-53721054-DEL-447
chr9-53757406-DEL-52
chr9-53773108-DEL-88
chr9-53801570-DEL-466
chr9-53865070-DEL-399
chr9-53894564-DEL-408
chr9-53918642-DEL-66
chr9-53932287-DEL-52
chr9-53976978-DEL-715
chr9-54023097-DEL-236
chr9-54032342-DEL-70
chr9-54049498-DEL-569
chr9-54057644-DEL-956
chr9-54106418-DEL-2482
chr9-54179703-DEL-519
chr9-54245302-DEL-56
chr9-54325541-DEL-52
chr9-54341943-DEL-87
chr9-54373433-DEL-202
chr9-54468762-DEL-337
chr9-54486546-DEL-291
chr9-54596594-DEL-92
chr9-54673337-DEL-59
chr9-54695731-DEL-172
chr9-54705043-DEL-190
chr9-54717961-DEL-104
chr9-54740265-DEL-1136
chr9-54829233-DEL-192
chr9-54844112-DEL-92
chr9-54862399-DEL-60
chr9-54863905-DEL-105
chr9-54884930-DEL-130
chr9-54895404-DEL-143
chr9-54935044-DEL-207
chr9-54947140-DEL-66
chr9-54950621-DEL-140
chr9-55000696-DEL-116
chr9-55005639-DEL-109
chr9-55058566-DEL-68
chr9-55061690-DEL-52
chr9-55078110-DEL-609
chr9-55132105-DEL-56
chr9-55142860-DEL-74
chr9-55151039-DEL-223
chr9-55175230-DEL-534
chr9-55207566-DEL-302
chr9-55219463-DEL-61
chr9-55220554-DEL-70
chr9-55234132-DEL-100
chr9-55280953-DEL-183
chr9-55282183-DEL-267
chr9-55283722-DEL-274
chr9-55309071-DEL-78
chr9-55362357-DEL-114
chr9-55398044-DEL-158
chr9-55403363-DEL-986
chr9-55425725-DEL-1693
chr9-55432923-DEL-91
chr9-55469609-DEL-344
chr9-55479124-DEL-72
chr9-55480412-DEL-1337
chr9-55536573-DEL-68
chr9-55541301-DEL-8644
chr9-55545235-DEL-211
chr9-55548194-DEL-1751
chr9-55571970-DEL-1913
chr9-55616346-DEL-968
chr9-55616946-DEL-318
chr9-55644509-DEL-119
chr9-55662449-DEL-83
chr9-55670892-DEL-677

chr9-55675683-DEL-7622
chr9-55683538-DEL-208
chr9-55690633-DEL-63
chr9-55700245-DEL-497
chr9-55725413-DEL-5161
chr9-55730928-DEL-343
chr9-55735367-DEL-7865
chr9-55742735-DEL-875
chr9-55747024-DEL-449
chr9-55754205-DEL-130
chr9-55756747-DEL-661
chr9-55759293-DEL-3446
chr9-55782400-DEL-51
chr9-55786909-DEL-1275
chr9-55831183-DEL-136
chr9-55846488-DEL-1126
chr9-55881650-DEL-744
chr9-55891145-DEL-941
chr9-55902184-DEL-521
chr9-55911907-DEL-478
chr9-55914126-DEL-70
chr9-55918891-DEL-1955
chr9-55984567-DEL-341
chr9-56091172-DEL-64
chr9-56091880-DEL-2123
chr9-56096852-DEL-120
chr9-56158902-DEL-74
chr9-56261649-DEL-725
chr9-56306648-DEL-815
chr9-56335219-DEL-76
chr9-56344491-DEL-96
chr9-56423066-DEL-281
chr9-56432266-DEL-65
chr9-56434397-DEL-237
chr9-56442472-DEL-75
chr9-56445804-DEL-275
chr9-56452416-DEL-73
chr9-56465561-DEL-84
chr9-56472186-DEL-561
chr9-56642822-DEL-56
chr9-56661091-DEL-52
chr9-56676121-DEL-133
chr9-56716345-DEL-70
chr9-56768697-DEL-202
chr9-56800624-DEL-591
chr9-56828226-DEL-186
chr9-56857538-DEL-234
chr9-56863620-DEL-212
chr9-56879857-DEL-99
chr9-56921556-DEL-428
chr9-56948489-DEL-77
chr9-57005820-DEL-197
chr9-57028238-DEL-562
chr9-57029615-DEL-140
chr9-57059984-DEL-206
chr9-57060936-DEL-411
chr9-57066440-DEL-293
chr9-57095593-DEL-135
chr9-57107902-DEL-134
chr9-57120701-DEL-179
chr9-57124594-DEL-88
chr9-57158359-DEL-60
chr9-57218114-DEL-169
chr9-57222983-DEL-428
chr9-57229671-DEL-66
chr9-57230526-DEL-345
chr9-57238915-DEL-86
chr9-57241958-DEL-1835
chr9-57296558-DEL-162
chr9-57299775-DEL-111
chr9-57368423-DEL-256
chr9-57391188-DEL-719
chr9-57471749-DEL-198
chr9-57480221-DEL-216
chr9-57482418-DEL-418
chr9-57502890-DEL-177
chr9-57557029-DEL-225

chr9-57569267-DEL-155
chr9-57569799-DEL-244
chr9-57577611-DEL-68
chr9-57594100-DEL-8618
chr9-57645502-DEL-313
chr9-57652381-DEL-133
chr9-57677267-DEL-65
chr9-57680524-DEL-59
chr9-57688620-DEL-1628
chr9-57695402-DEL-122
chr9-57734554-DEL-401
chr9-57735310-DEL-73
chr9-57749112-DEL-128
chr9-57751799-DEL-2102
chr9-57754866-DEL-530
chr9-57760104-DEL-213
chr9-57762311-DEL-52
chr9-57790551-DEL-129
chr9-57792999-DEL-1255
chr9-57862550-DEL-126
chr9-57873470-DEL-89
chr9-57907094-DEL-146
chr9-57914935-DEL-210
chr9-57921172-DEL-51
chr9-57929665-DEL-436
chr9-57947666-DEL-135
chr9-57985486-DEL-281
chr9-58032320-DEL-86
chr9-58068214-DEL-451
chr9-58085540-DEL-54
chr9-58091190-DEL-178
chr9-58115835-DEL-201
chr9-58122097-DEL-58
chr9-58145594-DEL-95
chr9-58148048-DEL-60
chr9-58175938-DEL-76
chr9-58178672-DEL-210
chr9-58188416-DEL-239
chr9-58193227-DEL-197
chr9-58207200-DEL-60
chr9-58213558-DEL-856
chr9-58261437-DEL-1292
chr9-58292751-DEL-82
chr9-58298267-DEL-149
chr9-58356486-DEL-204
chr9-58379489-DEL-127
chr9-58414859-DEL-83
chr9-58418270-DEL-165
chr9-58446639-DEL-71
chr9-58457487-DEL-170
chr9-58473517-DEL-2125
chr9-58483437-DEL-1513
chr9-58497571-DEL-204
chr9-58498694-DEL-1528
chr9-58518134-DEL-66
chr9-58583813-DEL-120
chr9-58625350-DEL-73
chr9-58634772-DEL-77
chr9-58776775-DEL-161
chr9-58803050-DEL-743
chr9-58804953-DEL-108
chr9-58887176-DEL-69
chr9-58906871-DEL-56
chr9-58973203-DEL-140
chr9-59045466-DEL-56
chr9-59051968-DEL-55
chr9-59052551-DEL-90
chr9-59086434-DEL-170
chr9-59086876-DEL-849
chr9-59114515-DEL-586
chr9-59132847-DEL-4050
chr9-59158764-DEL-628
chr9-59189171-DEL-60
chr9-59189543-DEL-242
chr9-59191964-DEL-64
chr9-59205246-DEL-240
chr9-59213469-DEL-164

chr9-59227790-DEL-786
chr9-59234161-DEL-206
chr9-59264988-DEL-1926
chr9-59289310-DEL-397
chr9-59294487-DEL-357
chr9-59318669-DEL-98
chr9-59323413-DEL-70
chr9-59332259-DEL-194
chr9-59357023-DEL-256
chr9-59417941-DEL-290
chr9-59419953-DEL-54
chr9-59438646-DEL-54
chr9-59439615-DEL-218
chr9-59445977-DEL-199
chr9-59474966-DEL-794
chr9-59500667-DEL-198
chr9-59506409-DEL-59
chr9-59506930-DEL-63
chr9-59512529-DEL-197
chr9-59528693-DEL-209
chr9-59536955-DEL-387
chr9-59625324-DEL-140
chr9-59630200-DEL-72
chr9-59636471-DEL-52
chr9-59638341-DEL-102
chr9-59650500-DEL-72
chr9-59724613-DEL-148
chr9-59772536-DEL-178
chr9-59831371-DEL-59
chr9-59843303-DEL-532
chr9-59849754-DEL-52
chr9-59860348-DEL-130
chr9-59870259-DEL-2453
chr9-59883006-DEL-187
chr9-59885772-DEL-202
chr9-59911336-DEL-71
chr9-59936727-DEL-58
chr9-59941360-DEL-79
chr9-59949590-DEL-415
chr9-59962866-DEL-83
chr9-59984654-DEL-63
chr9-59990242-DEL-554
chr9-60010579-DEL-84
chr9-60042921-DEL-2466
chr9-60096990-DEL-116
chr9-60109091-DEL-67
chr9-60126763-DEL-74
chr9-60136939-DEL-338
chr9-60144431-DEL-91
chr9-60198791-DEL-830
chr9-60199573-DEL-52
chr9-60238562-DEL-85
chr9-60248629-DEL-674
chr9-60273292-DEL-83
chr9-60306608-DEL-56
chr9-60354485-DEL-108
chr9-60370557-DEL-175
chr9-60386783-DEL-52
chr9-60436207-DEL-250
chr9-60446352-DEL-1024
chr9-60446359-DEL-957
chr9-60447354-DEL-92
chr9-60462711-DEL-64
chr9-60469502-DEL-71
chr9-60497281-DEL-1172
chr9-60512832-DEL-527
chr9-60528031-DEL-97
chr9-60572758-DEL-301
chr9-60604152-DEL-351
chr9-60646888-DEL-63
chr9-60666509-DEL-107
chr9-60692422-DEL-4387
chr9-60697371-DEL-66
chr9-60704311-DEL-56
chr9-60783112-DEL-206
chr9-60785068-DEL-81
chr9-60802003-DEL-95

chr9-60825473-DEL-119
chr9-60867906-DEL-67
chr9-60884051-DEL-60
chr9-60907886-DEL-54
chr9-60928136-DEL-344
chr9-60931338-DEL-91
chr9-60935130-DEL-274
chr9-60985583-DEL-215
chr9-61027589-DEL-167
chr9-61035488-DEL-72
chr9-61044882-DEL-128
chr9-61047862-DEL-487
chr9-61054915-DEL-1982
chr9-61085732-DEL-144
chr9-61089646-DEL-53
chr9-61100834-DEL-168
chr9-61110022-DEL-72
chr9-61158529-DEL-51
chr9-61161445-DEL-534
chr9-61161601-DEL-416
chr9-61267976-DEL-121
chr9-61294855-DEL-56
chr9-61313337-DEL-58
chr9-61313818-DEL-668
chr9-61321387-DEL-131
chr9-61326891-DEL-273
chr9-61332081-DEL-118
chr9-61342239-DEL-59
chr9-61356805-DEL-61
chr9-61382392-DEL-60
chr9-61383247-DEL-178
chr9-61385820-DEL-90
chr9-61388432-DEL-56
chr9-61418170-DEL-54
chr9-61421046-DEL-1772
chr9-61474834-DEL-70
chr9-61481704-DEL-79
chr9-61498333-DEL-56
chr9-61514226-DEL-83
chr9-61515325-DEL-78
chr9-61516980-DEL-64
chr9-61595813-DEL-82
chr9-61641885-DEL-85
chr9-61675125-DEL-80
chr9-61725092-DEL-132
chr9-61776840-DEL-505
chr9-61815549-DEL-981
chr9-61850513-DEL-289
chr9-61854631-DEL-122
chr9-61861058-DEL-448
chr9-61864134-DEL-400
chr9-61869502-DEL-74
chr9-61884788-DEL-170
chr9-61903647-DEL-210
chr9-61912763-DEL-96
chr9-61918458-DEL-479
chr9-61919006-DEL-1730
chr9-61935549-DEL-375
chr9-61945438-DEL-71
chr9-61995326-DEL-310
chr9-62038658-DEL-1765
chr9-62074775-DEL-101
chr9-62078084-DEL-82
chr9-62089905-DEL-1254
chr9-62110155-DEL-67
chr9-62117242-DEL-56
chr9-62124781-DEL-194
chr9-62140689-DEL-171
chr9-62141187-DEL-523
chr9-62172921-DEL-71
chr9-62182201-DEL-160
chr9-62203492-DEL-80
chr9-62222561-DEL-80
chr9-62270877-DEL-1406
chr9-62278561-DEL-124
chr9-62306861-DEL-54
chr9-62326934-DEL-102

chr9-62337750-DEL-55
chr9-62354754-DEL-78
chr9-62383887-DEL-64
chr9-62389174-DEL-140
chr9-62395090-DEL-120
chr9-62428225-DEL-60
chr9-62435503-DEL-70
chr9-62453158-DEL-143
chr9-62469530-DEL-189
chr9-62485197-DEL-85
chr9-62486075-DEL-5445
chr9-62515861-DEL-60
chr9-62541902-DEL-401
chr9-62543952-DEL-103
chr9-62548124-DEL-140
chr9-62596513-DEL-97
chr9-62600353-DEL-96
chr9-62610781-DEL-144
chr9-62612868-DEL-107
chr9-62618535-DEL-429
chr9-62667388-DEL-52
chr9-62676269-DEL-64
chr9-62697880-DEL-214
chr9-62721483-DEL-254
chr9-62725211-DEL-59
chr9-62747240-DEL-469
chr9-62749503-DEL-228
chr9-62750862-DEL-737
chr9-62759809-DEL-66
chr9-62780215-DEL-79
chr9-62781860-DEL-60
chr9-62794887-DEL-86
chr9-62827108-DEL-203
chr9-62868208-DEL-71
chr9-62893182-DEL-317
chr9-62896637-DEL-159
chr9-62897466-DEL-312
chr9-62921637-DEL-62
chr9-62933547-DEL-159
chr9-62975601-DEL-197
chr9-62984807-DEL-64
chr9-62989361-DEL-66
chr9-63018440-DEL-52
chr9-63023881-DEL-2459
chr9-63115837-DEL-588
chr9-63241815-DEL-265
chr9-63260641-DEL-568
chr9-63265683-DEL-274
chr9-63277048-DEL-1250
chr9-63316069-DEL-277
chr9-63318665-DEL-106
chr9-63323773-DEL-317
chr9-63329434-DEL-1626
chr9-63337421-DEL-378
chr9-63345416-DEL-56
chr9-63363015-DEL-190
chr9-63418957-DEL-228
chr9-63460294-DEL-377
chr9-63472098-DEL-60
chr9-63477894-DEL-61
chr9-63498424-DEL-388
chr9-63511800-DEL-61
chr9-63539114-DEL-297
chr9-63544103-DEL-1590
chr9-63560220-DEL-376
chr9-63583502-DEL-120
chr9-63591113-DEL-105
chr9-63612587-DEL-184
chr9-63639219-DEL-243
chr9-63646855-DEL-54
chr9-63683029-DEL-58
chr9-63708706-DEL-316
chr9-63766660-DEL-70
chr9-63775132-DEL-331
chr9-63799439-DEL-70
chr9-63804137-DEL-62
chr9-63853150-DEL-309

chr9-63853789-DEL-1453
chr9-63878688-DEL-200
chr9-63882968-DEL-69
chr9-63987080-DEL-218
chr9-64009350-DEL-100
chr9-64011267-DEL-228
chr9-64013646-DEL-205
chr9-64017334-DEL-225
chr9-64029808-DEL-721
chr9-64030817-DEL-304
chr9-64031378-DEL-389
chr9-64035810-DEL-175
chr9-64046323-DEL-328
chr9-64061105-DEL-142
chr9-64061773-DEL-198
chr9-64090516-DEL-162
chr9-64131507-DEL-181
chr9-64150467-DEL-56
chr9-64164903-DEL-1499
chr9-64166121-DEL-63
chr9-64169660-DEL-1611
chr9-64193175-DEL-349
chr9-64202113-DEL-210
chr9-64204798-DEL-170
chr9-64212713-DEL-172
chr9-64235875-DEL-308
chr9-64238613-DEL-209
chr9-64286839-DEL-185
chr9-64358788-DEL-112
chr9-64388454-DEL-64
chr9-64388782-DEL-360
chr9-64403275-DEL-58
chr9-64404721-DEL-132
chr9-64427587-DEL-63
chr9-64464105-DEL-67
chr9-64471715-DEL-80
chr9-64549216-DEL-52
chr9-64579229-DEL-113
chr9-64594027-DEL-52
chr9-64617783-DEL-220
chr9-64655550-DEL-74
chr9-64657064-DEL-352
chr9-64727005-DEL-314
chr9-64741050-DEL-271
chr9-64757317-DEL-105
chr9-64764763-DEL-86
chr9-64765180-DEL-188
chr9-64767666-DEL-310
chr9-64789886-DEL-177
chr9-64807285-DEL-53
chr9-64821543-DEL-60
chr9-64854712-DEL-66
chr9-64862682-DEL-235
chr9-64871999-DEL-938
chr9-64891799-DEL-88
chr9-64914575-DEL-70
chr9-64967878-DEL-188
chr9-64993890-DEL-60
chr9-64995228-DEL-1451
chr9-64995465-DEL-97
chr9-65018628-DEL-75
chr9-65026741-DEL-200
chr9-65094949-DEL-400
chr9-65101755-DEL-68
chr9-65109932-DEL-54
chr9-65114948-DEL-175
chr9-65121276-DEL-56
chr9-65131528-DEL-64
chr9-65135468-DEL-90
chr9-65141437-DEL-216
chr9-65144420-DEL-320
chr9-65171500-DEL-94
chr9-65197114-DEL-57
chr9-65220566-DEL-183
chr9-65233805-DEL-190
chr9-65239815-DEL-208
chr9-65252630-DEL-473

chr9-65255839-DEL-551
chr9-65283170-DEL-214
chr9-65291911-DEL-95
chr9-65293377-DEL-100
chr9-65302197-DEL-66
chr9-65324270-DEL-51
chr9-65331313-DEL-109
chr9-65340954-DEL-56
chr9-65367194-DEL-68
chr9-65373214-DEL-681
chr9-65389993-DEL-124
chr9-65395549-DEL-55
chr9-65425250-DEL-53
chr9-65452304-DEL-52
chr9-65462880-DEL-214
chr9-65503568-DEL-201
chr9-65508520-DEL-197
chr9-65535725-DEL-51
chr9-65548735-DEL-58
chr9-65576486-DEL-656
chr9-65577168-DEL-586
chr9-65578185-DEL-207
chr9-65605160-DEL-169
chr9-65628410-DEL-60
chr9-65678222-DEL-71
chr9-65687416-DEL-247
chr9-65718922-DEL-54
chr9-65728356-DEL-76
chr9-65749058-DEL-54
chr9-65763212-DEL-195
chr9-65764269-DEL-317
chr9-65775073-DEL-982
chr9-65775305-DEL-236
chr9-65775418-DEL-894
chr9-65776143-DEL-1312
chr9-65776209-DEL-805
chr9-65776719-DEL-201
chr9-65777468-DEL-168
chr9-65788974-DEL-186
chr9-65794040-DEL-183
chr9-65800966-DEL-1570
chr9-65802980-DEL-203
chr9-65825424-DEL-265
chr9-65841901-DEL-251
chr9-65907636-DEL-250
chr9-65964409-DEL-136
chr9-65982999-DEL-76
chr9-66011623-DEL-51
chr9-66019931-DEL-78
chr9-66029101-DEL-56
chr9-66049205-DEL-119
chr9-66060363-DEL-219
chr9-66084745-DEL-84
chr9-66121785-DEL-57
chr9-66166499-DEL-219
chr9-66166839-DEL-769
chr9-66168612-DEL-404
chr9-66191827-DEL-52
chr9-66198440-DEL-147
chr9-66222016-DEL-323
chr9-66229848-DEL-82
chr9-66242375-DEL-87
chr9-66254072-DEL-58
chr9-66263776-DEL-164
chr9-66299550-DEL-84
chr9-66303139-DEL-135
chr9-66351696-DEL-77
chr9-66354458-DEL-128
chr9-66365027-DEL-68
chr9-66370542-DEL-981
chr9-66387129-DEL-344
chr9-66428042-DEL-3518
chr9-66439721-DEL-225
chr9-66458586-DEL-97
chr9-66462694-DEL-240
chr9-66531237-DEL-137
chr9-66538876-DEL-66

chr9-66556123-DEL-116
chr9-66566157-DEL-58
chr9-66624625-DEL-52
chr9-66657370-DEL-272
chr9-66696628-DEL-190
chr9-66711992-DEL-1518
chr9-66717215-DEL-1527
chr9-66727058-DEL-175
chr9-66730223-DEL-2622
chr9-66784274-DEL-58
chr9-66785050-DEL-189
chr9-66835496-DEL-74
chr9-66851981-DEL-707
chr9-66915401-DEL-63
chr9-66934428-DEL-83
chr9-66938659-DEL-52
chr9-66944429-DEL-323
chr9-66965555-DEL-58
chr9-66982997-DEL-1519
chr9-67009074-DEL-52
chr9-67047919-DEL-778
chr9-67079991-DEL-55
chr9-67152449-DEL-125
chr9-67186879-DEL-1461
chr9-67229637-DEL-292
chr9-67232256-DEL-131
chr9-67262915-DEL-56
chr9-67264478-DEL-1581
chr9-67310906-DEL-111
chr9-67314865-DEL-127
chr9-67347478-DEL-439
chr9-67347574-DEL-58
chr9-67348005-DEL-729
chr9-67348663-DEL-161
chr9-67373929-DEL-51
chr9-67387525-DEL-115
chr9-67393315-DEL-65
chr9-67399455-DEL-149
chr9-67400197-DEL-889
chr9-67405408-DEL-234
chr9-67421310-DEL-436
chr9-67425690-DEL-62
chr9-67433387-DEL-78
chr9-67459291-DEL-286
chr9-67460985-DEL-1460
chr9-67479016-DEL-145
chr9-67526059-DEL-139
chr9-67559894-DEL-2258
chr9-67569135-DEL-136
chr9-67586145-DEL-72
chr9-67594630-DEL-57
chr9-67601972-DEL-128
chr9-67677679-DEL-521
chr9-67695783-DEL-1860
chr9-67700405-DEL-224
chr9-67713009-DEL-338
chr9-67714027-DEL-59
chr9-67726232-DEL-280
chr9-67773982-DEL-62
chr9-67787958-DEL-87
chr9-67838028-DEL-56
chr9-67904103-DEL-99
chr9-67966818-DEL-204
chr9-68073181-DEL-98
chr9-68106252-DEL-197
chr9-68107215-DEL-589
chr9-68112060-DEL-1158
chr9-68129667-DEL-1250
chr9-68158713-DEL-58
chr9-68178182-DEL-62
chr9-68183447-DEL-56
chr9-68191944-DEL-56
chr9-68216131-DEL-102
chr9-68229956-DEL-1865
chr9-68267330-DEL-84
chr9-68280873-DEL-100
chr9-68292833-DEL-66

chr9-68339341-DEL-62
chr9-68379868-DEL-82
chr9-68434408-DEL-6197
chr9-68461452-DEL-56
chr9-68489931-DEL-410
chr9-68490864-DEL-269
chr9-68508958-DEL-55
chr9-68535119-DEL-70
chr9-68566179-DEL-896
chr9-68587914-DEL-53
chr9-68619464-DEL-167
chr9-68665470-DEL-109
chr9-68697787-DEL-146
chr9-68720284-DEL-138
chr9-68721875-DEL-73
chr9-68793853-DEL-76
chr9-68805126-DEL-84
chr9-68877183-DEL-57
chr9-68904297-DEL-72
chr9-68923800-DEL-70
chr9-68929262-DEL-58
chr9-68982634-DEL-83
chr9-69007451-DEL-135
chr9-69016774-DEL-74
chr9-69055329-DEL-414
chr9-69136431-DEL-105
chr9-69148464-DEL-65
chr9-69214823-DEL-70
chr9-69238531-DEL-65
chr9-69264930-DEL-93
chr9-69299867-DEL-111
chr9-69430228-DEL-313
chr9-69440128-DEL-638
chr9-69448278-DEL-110
chr9-69450804-DEL-137
chr9-69452711-DEL-170
chr9-69466244-DEL-2437
chr9-69499506-DEL-76
chr9-69503112-DEL-76
chr9-69514176-DEL-329
chr9-69542398-DEL-74
chr9-69560829-DEL-435
chr9-69639445-DEL-168
chr9-69717741-DEL-114
chr9-69767550-DEL-52
chr9-69870220-DEL-1037
chr9-69892223-DEL-119
chr9-69894270-DEL-332
chr9-69937800-DEL-53
chr9-69972353-DEL-135
chr9-69984435-DEL-98
chr9-69998209-DEL-2213
chr9-70006695-DEL-332
chr9-70009309-DEL-180
chr9-70021999-DEL-1051
chr9-70113785-DEL-52
chr9-70183357-DEL-52
chr9-70247993-DEL-57
chr9-70265625-DEL-56
chr9-70288403-DEL-59
chr9-70291524-DEL-432
chr9-70314876-DEL-61
chr9-70341719-DEL-214
chr9-70370392-DEL-6206
chr9-70378622-DEL-101
chr9-70384315-DEL-114
chr9-70397583-DEL-105
chr9-70397879-DEL-3417
chr9-70417994-DEL-180
chr9-70461895-DEL-310
chr9-70501411-DEL-62
chr9-70530440-DEL-199
chr9-70537818-DEL-92
chr9-70549593-DEL-53
chr9-70573245-DEL-160
chr9-70741346-DEL-73
chr9-70742513-DEL-54

chr9-70750468-DEL-1281
chr9-70757940-DEL-53
chr9-70759817-DEL-1951
chr9-70775126-DEL-966
chr9-70830346-DEL-1830
chr9-70835798-DEL-85
chr9-70895740-DEL-553
chr9-70924403-DEL-60
chr9-70928524-DEL-383
chr9-70933583-DEL-100
chr9-70985649-DEL-58
chr9-71009053-DEL-477
chr9-71012449-DEL-69
chr9-71055567-DEL-88
chr9-71059449-DEL-276
chr9-71067847-DEL-184
chr9-71111715-DEL-85
chr9-71187302-DEL-215
chr9-71193824-DEL-388
chr9-71231131-DEL-213
chr9-71235894-DEL-216
chr9-71240014-DEL-73
chr9-71252815-DEL-88
chr9-71257829-DEL-123
chr9-71291129-DEL-78
chr9-71325234-DEL-66
chr9-71371472-DEL-871
chr9-71373908-DEL-531
chr9-71375906-DEL-59
chr9-71395996-DEL-432
chr9-71398690-DEL-192
chr9-71404897-DEL-89
chr9-71411057-DEL-55
chr9-71419140-DEL-99
chr9-71423857-DEL-120
chr9-71460549-DEL-66
chr9-71467012-DEL-514
chr9-71586497-DEL-61
chr9-71589522-DEL-238
chr9-71593720-DEL-55
chr9-71607358-DEL-174
chr9-71625784-DEL-53
chr9-71675139-DEL-310
chr9-71688010-DEL-651
chr9-71712345-DEL-85
chr9-71714381-DEL-93
chr9-71725334-DEL-76
chr9-71726625-DEL-200
chr9-71728238-DEL-806
chr9-71737925-DEL-125
chr9-71758900-DEL-1519
chr9-71761688-DEL-1735
chr9-71772561-DEL-1124
chr9-71827504-DEL-75
chr9-71904083-DEL-127
chr9-71940505-DEL-58
chr9-72018168-DEL-51
chr9-72024161-DEL-53
chr9-72034242-DEL-76
chr9-72086624-DEL-140
chr9-72100145-DEL-212
chr9-72108759-DEL-75
chr9-72244698-DEL-381
chr9-72290459-DEL-56
chr9-72303156-DEL-57
chr9-72306162-DEL-68
chr9-72324825-DEL-241
chr9-72336711-DEL-196
chr9-72361721-DEL-63
chr9-72366457-DEL-151
chr9-72418474-DEL-175
chr9-72432410-DEL-549
chr9-72505622-DEL-67
chr9-72558090-DEL-201
chr9-72590315-DEL-200
chr9-72591103-DEL-217
chr9-72604729-DEL-228

chr9-72619673-DEL-182
chr9-72641060-DEL-76
chr9-72690249-DEL-4344
chr9-72694783-DEL-199
chr9-72697012-DEL-80
chr9-72701531-DEL-201
chr9-72705002-DEL-195
chr9-72712501-DEL-72
chr9-72728728-DEL-207
chr9-72731565-DEL-188
chr9-72746160-DEL-1699
chr9-72757876-DEL-188
chr9-72766689-DEL-935
chr9-72861323-DEL-193
chr9-72866639-DEL-114
chr9-72872015-DEL-113
chr9-72898683-DEL-162
chr9-72900144-DEL-716
chr9-72953748-DEL-221
chr9-72956863-DEL-55
chr9-72963608-DEL-56
chr9-72963918-DEL-1309
chr9-72966936-DEL-79
chr9-72968448-DEL-198
chr9-72969678-DEL-574
chr9-72970438-DEL-142
chr9-72973132-DEL-308
chr9-73025803-DEL-53
chr9-73048319-DEL-71
chr9-73064272-DEL-211
chr9-73067060-DEL-69
chr9-73096811-DEL-59
chr9-73117185-DEL-239
chr9-73121870-DEL-949
chr9-73151266-DEL-112
chr9-73157181-DEL-118
chr9-73180379-DEL-77
chr9-73194967-DEL-2253
chr9-73200153-DEL-168
chr9-73243206-DEL-56
chr9-73255576-DEL-192
chr9-73256196-DEL-3737
chr9-73276256-DEL-1010
chr9-73382833-DEL-128
chr9-73422350-DEL-105
chr9-73502857-DEL-62
chr9-73543891-DEL-108
chr9-73557375-DEL-480
chr9-73561115-DEL-153
chr9-73615289-DEL-90
chr9-73677232-DEL-52
chr9-73695653-DEL-106
chr9-73715777-DEL-355
chr9-73729205-DEL-93
chr9-73731956-DEL-208
chr9-73753004-DEL-205
chr9-73761273-DEL-99
chr9-73778782-DEL-139
chr9-73825134-DEL-98
chr9-73836297-DEL-86
chr9-73881004-DEL-160
chr9-73882149-DEL-152
chr9-73926832-DEL-81
chr9-73953472-DEL-155
chr9-73957273-DEL-82
chr9-73960434-DEL-295
chr9-73963465-DEL-367
chr9-73977656-DEL-131
chr9-73996514-DEL-3378
chr9-74030931-DEL-56
chr9-74116893-DEL-72
chr9-74121409-DEL-1122
chr9-74137632-DEL-1320
chr9-74167306-DEL-156
chr9-74224899-DEL-80
chr9-74286374-DEL-148
chr9-74385675-DEL-235

chr9-74391608-DEL-74
chr9-74402529-DEL-1049
chr9-74424956-DEL-1867
chr9-74487211-DEL-119
chr9-74495700-DEL-53
chr9-74500894-DEL-279
chr9-74506456-DEL-56
chr9-74508074-DEL-273
chr9-74525619-DEL-156
chr9-74585311-DEL-393
chr9-74611970-DEL-59
chr9-74617373-DEL-60
chr9-74619523-DEL-55
chr9-74624331-DEL-260
chr9-74667191-DEL-420
chr9-74690444-DEL-1653
chr9-74694212-DEL-1040
chr9-74731783-DEL-74
chr9-74741228-DEL-424
chr9-74743717-DEL-1040
chr9-74755256-DEL-78
chr9-74790848-DEL-192
chr9-74858084-DEL-116
chr9-74863696-DEL-936
chr9-74906867-DEL-140
chr9-74947208-DEL-283
chr9-74995721-DEL-201
chr9-75021776-DEL-88
chr9-75039642-DEL-173
chr9-75099995-DEL-770
chr9-75107903-DEL-86
chr9-75175214-DEL-116
chr9-75183392-DEL-76
chr9-75216484-DEL-64
chr9-75220509-DEL-124
chr9-75242612-DEL-202
chr9-75245779-DEL-209
chr9-75276837-DEL-437
chr9-75310760-DEL-57
chr9-75327853-DEL-52
chr9-75331525-DEL-172
chr9-75353313-DEL-199
chr9-75355785-DEL-164
chr9-75363850-DEL-58
chr9-75381787-DEL-279
chr9-75395794-DEL-311
chr9-75413070-DEL-555
chr9-75428426-DEL-165
chr9-75434516-DEL-171
chr9-75440497-DEL-377
chr9-75468460-DEL-74
chr9-75485369-DEL-52
chr9-75521608-DEL-156
chr9-75566787-DEL-177
chr9-75575501-DEL-155
chr9-75579096-DEL-59
chr9-75598360-DEL-1002
chr9-75616286-DEL-6904
chr9-75628020-DEL-6328
chr9-75637623-DEL-5612
chr9-75649583-DEL-1472
chr9-75656092-DEL-59
chr9-75660139-DEL-767
chr9-75705384-DEL-123
chr9-75740422-DEL-82
chr9-75774893-DEL-631
chr9-75787990-DEL-117
chr9-75797956-DEL-52
chr9-75798684-DEL-226
chr9-75801872-DEL-79
chr9-75822966-DEL-272
chr9-75850736-DEL-51
chr9-75902964-DEL-71
chr9-75970028-DEL-58
chr9-75990609-DEL-202
chr9-75996545-DEL-74
chr9-76018693-DEL-116

chr9-76019491-DEL-390
chr9-76030472-DEL-54
chr9-76035312-DEL-314
chr9-76052067-DEL-58
chr9-76054342-DEL-2311
chr9-76074451-DEL-90
chr9-76103308-DEL-108
chr9-76168271-DEL-1824
chr9-76198126-DEL-86
chr9-76199908-DEL-937
chr9-76200704-DEL-141
chr9-76213889-DEL-110
chr9-76221239-DEL-80
chr9-76233877-DEL-470
chr9-76237084-DEL-68
chr9-76240280-DEL-1132
chr9-76241457-DEL-52
chr9-76257631-DEL-747
chr9-76393144-DEL-559
chr9-76407795-DEL-120
chr9-76430828-DEL-419
chr9-76440343-DEL-399
chr9-76450004-DEL-148
chr9-76506426-DEL-179
chr9-76522846-DEL-1437
chr9-76525835-DEL-236
chr9-76529271-DEL-241
chr9-76652360-DEL-539
chr9-76711326-DEL-1974
chr9-76724813-DEL-67
chr9-76728110-DEL-321
chr9-76785205-DEL-580
chr9-76793687-DEL-531
chr9-76810722-DEL-797
chr9-76814309-DEL-62
chr9-76830354-DEL-195
chr9-76832547-DEL-875
chr9-76860361-DEL-91
chr9-76862683-DEL-132
chr9-76867158-DEL-66
chr9-76875611-DEL-66
chr9-76985034-DEL-847
chr9-76987307-DEL-2680
chr9-76991217-DEL-197
chr9-76993025-DEL-79
chr9-77001167-DEL-96
chr9-77035163-DEL-213
chr9-77059493-DEL-7464
chr9-77059701-DEL-130
chr9-77099665-DEL-1652
chr9-77117613-DEL-509
chr9-77130809-DEL-930
chr9-77144242-DEL-78
chr9-77145387-DEL-140
chr9-77147628-DEL-538
chr9-77182645-DEL-1006
chr9-77221047-DEL-598
chr9-77287101-DEL-149
chr9-77302656-DEL-108
chr9-77320394-DEL-128
chr9-77321527-DEL-150
chr9-77367825-DEL-76
chr9-77370107-DEL-52
chr9-77378967-DEL-60
chr9-77404889-DEL-876
chr9-77431358-DEL-1336
chr9-77438457-DEL-178
chr9-77439031-DEL-169
chr9-77439513-DEL-80
chr9-77456131-DEL-81
chr9-77483419-DEL-668
chr9-77484138-DEL-241
chr9-77484664-DEL-554
chr9-77489438-DEL-141
chr9-77514420-DEL-122
chr9-77526421-DEL-766
chr9-77529808-DEL-56

chr9-77537485-DEL-60
chr9-77573779-DEL-263
chr9-77582113-DEL-6212
chr9-77592013-DEL-63
chr9-77628213-DEL-58
chr9-77643120-DEL-514
chr9-77649030-DEL-66
chr9-77702302-DEL-363
chr9-77706609-DEL-405
chr9-77758067-DEL-60
chr9-77767972-DEL-17551
chr9-77815120-DEL-74
chr9-77816054-DEL-69
chr9-77835735-DEL-69
chr9-77840283-DEL-294
chr9-77880256-DEL-2130
chr9-77898575-DEL-51
chr9-77909700-DEL-411
chr9-77925646-DEL-140
chr9-77930141-DEL-200
chr9-77947702-DEL-221
chr9-77952049-DEL-338
chr9-77964584-DEL-66
chr9-77985418-DEL-233
chr9-77989497-DEL-439
chr9-78071440-DEL-107
chr9-78082321-DEL-202
chr9-78160032-DEL-144
chr9-78197400-DEL-73
chr9-78197895-DEL-959
chr9-78199816-DEL-583
chr9-78203031-DEL-67
chr9-78212885-DEL-128
chr9-78217974-DEL-164
chr9-78226782-DEL-90
chr9-78258739-DEL-161
chr9-78283131-DEL-56
chr9-78291928-DEL-201
chr9-78312386-DEL-156
chr9-78323457-DEL-175
chr9-78324197-DEL-321
chr9-78353305-DEL-123
chr9-78362173-DEL-233
chr9-78378356-DEL-200
chr9-78379161-DEL-96
chr9-78382993-DEL-189
chr9-78393588-DEL-767
chr9-78397653-DEL-206
chr9-78399529-DEL-57
chr9-78400878-DEL-857
chr9-78404561-DEL-225
chr9-78440400-DEL-79
chr9-78451371-DEL-406
chr9-78452516-DEL-275
chr9-78460943-DEL-194
chr9-78469680-DEL-7508
chr9-78482303-DEL-267
chr9-78485024-DEL-85
chr9-78501577-DEL-168
chr9-78510193-DEL-394
chr9-78540320-DEL-576
chr9-78547156-DEL-150
chr9-78595228-DEL-62
chr9-78599528-DEL-148
chr9-78607914-DEL-92
chr9-78652058-DEL-1600
chr9-78655529-DEL-324
chr9-78668450-DEL-494
chr9-78671313-DEL-3396
chr9-78675812-DEL-63
chr9-78719401-DEL-942
chr9-78766547-DEL-427
chr9-78798765-DEL-1228
chr9-78817982-DEL-853
chr9-78822979-DEL-1714
chr9-78826367-DEL-63
chr9-78834973-DEL-1137

chr9-78837632-DEL-2195
chr9-78854542-DEL-1695
chr9-78855739-DEL-260
chr9-78878541-DEL-314
chr9-78891969-DEL-78
chr9-78955833-DEL-1498
chr9-78973610-DEL-2376
chr9-78999636-DEL-125
chr9-79098444-DEL-108
chr9-79099379-DEL-269
chr9-79106331-DEL-613
chr9-79136003-DEL-157
chr9-79141684-DEL-56
chr9-79169474-DEL-714
chr9-79186377-DEL-817
chr9-79187919-DEL-929
chr9-79191829-DEL-158
chr9-79198515-DEL-102
chr9-79220499-DEL-397
chr9-79221679-DEL-158
chr9-79227737-DEL-167
chr9-79244774-DEL-57
chr9-79255107-DEL-2010
chr9-79258315-DEL-1185
chr9-79258316-DEL-71
chr9-79258356-DEL-915
chr9-79291556-DEL-81
chr9-79342068-DEL-114
chr9-79354768-DEL-674
chr9-79370094-DEL-408
chr9-79388500-DEL-71
chr9-79394204-DEL-4870
chr9-79432637-DEL-442
chr9-79439831-DEL-1043
chr9-79446130-DEL-5118
chr9-79446131-DEL-5626
chr9-79446389-DEL-2576
chr9-79448639-DEL-2560
chr9-79455177-DEL-587
chr9-79474091-DEL-825
chr9-79525768-DEL-125
chr9-79542993-DEL-990
chr9-79631614-DEL-60
chr9-79658857-DEL-606
chr9-79688237-DEL-164
chr9-79690263-DEL-255
chr9-79696590-DEL-94
chr9-79754777-DEL-338
chr9-79773836-DEL-52
chr9-79785981-DEL-55
chr9-79806815-DEL-10504
chr9-79831067-DEL-306
chr9-79840470-DEL-334
chr9-79847856-DEL-57
chr9-79926299-DEL-65
chr9-79954442-DEL-75
chr9-79964081-DEL-91
chr9-80011645-DEL-302
chr9-80026499-DEL-829
chr9-80054796-DEL-183
chr9-80056636-DEL-237
chr9-80058111-DEL-1669
chr9-80061660-DEL-156
chr9-80068309-DEL-64
chr9-80069302-DEL-58
chr9-80098939-DEL-350
chr9-80129092-DEL-277
chr9-80150019-DEL-206
chr9-80170503-DEL-62
chr9-80187435-DEL-95
chr9-80225676-DEL-457
chr9-80229158-DEL-406
chr9-80262206-DEL-68
chr9-80340904-DEL-57
chr9-80345219-DEL-54
chr9-80431662-DEL-116
chr9-80462141-DEL-66

chr9-80463618-DEL-172
chr9-80488058-DEL-755
chr9-80491592-DEL-168
chr9-80586590-DEL-77
chr9-80591217-DEL-969
chr9-80598121-DEL-801
chr9-80611321-DEL-2599
chr9-80616243-DEL-75
chr9-80663576-DEL-70
chr9-80684364-DEL-2925
chr9-80718999-DEL-54
chr9-80721259-DEL-78
chr9-80742303-DEL-1236
chr9-80744788-DEL-942
chr9-80804991-DEL-438
chr9-80852798-DEL-3318
chr9-80906289-DEL-58
chr9-80944860-DEL-7632
chr9-80946312-DEL-261
chr9-80946592-DEL-2806
chr9-80949440-DEL-1446
chr9-80954373-DEL-909
chr9-80982983-DEL-598
chr9-80983820-DEL-338
chr9-80988980-DEL-66
chr9-80995750-DEL-1702
chr9-81031325-DEL-220
chr9-81048594-DEL-114
chr9-81053169-DEL-826
chr9-81075598-DEL-1670
chr9-81078951-DEL-1340
chr9-81088808-DEL-206
chr9-81110580-DEL-340
chr9-81112217-DEL-1470
chr9-81161785-DEL-6197
chr9-81197906-DEL-88
chr9-81199909-DEL-4648
chr9-81226462-DEL-117
chr9-81228312-DEL-1161
chr9-81248827-DEL-401
chr9-81254197-DEL-2614
chr9-81267643-DEL-215
chr9-81282333-DEL-845
chr9-81299603-DEL-153
chr9-81304202-DEL-325
chr9-81304714-DEL-298
chr9-81312316-DEL-5739
chr9-81314690-DEL-1935
chr9-81321049-DEL-2187
chr9-81325536-DEL-739
chr9-81326378-DEL-2109
chr9-81328511-DEL-2620
chr9-81336923-DEL-82
chr9-81345437-DEL-2495
chr9-81357473-DEL-126
chr9-81362171-DEL-74
chr9-81376430-DEL-1654
chr9-81408318-DEL-405
chr9-81415030-DEL-1788
chr9-81449837-DEL-537
chr9-81517000-DEL-56
chr9-81548972-DEL-713
chr9-81559632-DEL-1109
chr9-81595651-DEL-391
chr9-81600732-DEL-1183
chr9-81609770-DEL-2779
chr9-81678327-DEL-52
chr9-81681948-DEL-316
chr9-81701704-DEL-877
chr9-81739135-DEL-70
chr9-81752409-DEL-116
chr9-81780390-DEL-616
chr9-81806475-DEL-505
chr9-81855921-DEL-1136
chr9-81886096-DEL-136
chr9-81900491-DEL-194
chr9-81934283-DEL-815

chr9-81945564-DEL-331
chr9-81954645-DEL-81
chr9-81962786-DEL-74
chr9-82009712-DEL-6271
chr9-82021868-DEL-242
chr9-82042666-DEL-54
chr9-82045925-DEL-2560
chr9-82049857-DEL-1355
chr9-82054118-DEL-1168
chr9-82055979-DEL-2451
chr9-82085991-DEL-66
chr9-82103046-DEL-1037
chr9-82105258-DEL-141
chr9-82173657-DEL-3452
chr9-82198511-DEL-50
chr9-82285761-DEL-60
chr9-82322319-DEL-558
chr9-82324884-DEL-6234
chr9-82337995-DEL-255
chr9-82407260-DEL-220
chr9-82408814-DEL-82
chr9-82413498-DEL-635
chr9-82429691-DEL-72
chr9-82466182-DEL-64
chr9-82467602-DEL-94
chr9-82468329-DEL-6481
chr9-82501631-DEL-1454
chr9-82501878-DEL-51
chr9-82521135-DEL-443
chr9-82538310-DEL-59
chr9-82539515-DEL-445
chr9-82560763-DEL-1461
chr9-82563212-DEL-138
chr9-82595676-DEL-384
chr9-82605415-DEL-93
chr9-82612315-DEL-4261
chr9-82616643-DEL-117
chr9-82620415-DEL-1644
chr9-82646868-DEL-54
chr9-82667837-DEL-192
chr9-82692855-DEL-58
chr9-82878719-DEL-115
chr9-82902866-DEL-1255
chr9-82909691-DEL-104
chr9-82933104-DEL-693
chr9-82937322-DEL-204
chr9-82963600-DEL-52
chr9-82972456-DEL-329
chr9-83020449-DEL-976
chr9-83038473-DEL-62
chr9-83046887-DEL-6126
chr9-83054498-DEL-5616
chr9-83088262-DEL-1324
chr9-83090254-DEL-109
chr9-83092971-DEL-972
chr9-83097287-DEL-76
chr9-83111232-DEL-60
chr9-83112126-DEL-103
chr9-83170212-DEL-977
chr9-83306256-DEL-633
chr9-83318076-DEL-116
chr9-83375735-DEL-166
chr9-83390816-DEL-966
chr9-83403681-DEL-239
chr9-83407387-DEL-59
chr9-83418143-DEL-87
chr9-83466992-DEL-377
chr9-83487054-DEL-61
chr9-83497760-DEL-170
chr9-83503487-DEL-56
chr9-83558267-DEL-4250
chr9-83563625-DEL-340
chr9-83578958-DEL-183
chr9-83617758-DEL-2393
chr9-83642931-DEL-52
chr9-83647920-DEL-76
chr9-83665972-DEL-52

chr9-83718347-DEL-156
chr9-83755354-DEL-1782
chr9-83789380-DEL-2045
chr9-83801863-DEL-191
chr9-83821736-DEL-59
chr9-83835867-DEL-1464
chr9-83857874-DEL-1055
chr9-83883517-DEL-192
chr9-83956460-DEL-399
chr9-83960795-DEL-95
chr9-84098212-DEL-310
chr9-84098896-DEL-10152
chr9-84123811-DEL-94
chr9-84128476-DEL-57
chr9-84153556-DEL-68
chr9-84159596-DEL-88
chr9-84187132-DEL-542
chr9-84239153-DEL-1339
chr9-84320253-DEL-51
chr9-84324799-DEL-1884
chr9-84324938-DEL-117
chr9-84371234-DEL-6140
chr9-84385231-DEL-179
chr9-84389285-DEL-2960
chr9-84394789-DEL-476
chr9-84429673-DEL-268
chr9-84432411-DEL-57
chr9-84495961-DEL-598
chr9-84505730-DEL-132
chr9-84506484-DEL-1168
chr9-84514617-DEL-51
chr9-84516966-DEL-103
chr9-84534029-DEL-285
chr9-84581364-DEL-383
chr9-84605725-DEL-89
chr9-84628548-DEL-1419
chr9-84672652-DEL-60
chr9-84674807-DEL-150
chr9-84694494-DEL-424
chr9-84695651-DEL-642
chr9-84720959-DEL-283
chr9-84747651-DEL-6204
chr9-84753560-DEL-187
chr9-84754830-DEL-207
chr9-84765975-DEL-6938
chr9-84780554-DEL-99
chr9-84805142-DEL-59
chr9-84822566-DEL-70
chr9-84835228-DEL-1871
chr9-84856963-DEL-985
chr9-84905561-DEL-639
chr9-84939986-DEL-1691
chr9-84943885-DEL-4539
chr9-84981034-DEL-1891
chr9-85011277-DEL-399
chr9-85016024-DEL-160
chr9-85025745-DEL-292
chr9-85034895-DEL-366
chr9-85041955-DEL-172
chr9-85085265-DEL-121
chr9-85097986-DEL-645
chr9-85117670-DEL-80
chr9-85126892-DEL-52
chr9-85127576-DEL-199
chr9-85152400-DEL-339
chr9-85193465-DEL-85
chr9-85213131-DEL-58
chr9-85245076-DEL-96
chr9-85248895-DEL-54
chr9-85253628-DEL-51
chr9-85288403-DEL-175
chr9-85317113-DEL-507
chr9-85384478-DEL-265
chr9-85408848-DEL-65
chr9-85412832-DEL-57
chr9-85432024-DEL-59
chr9-85436568-DEL-3964

chr9-85508198-DEL-6752
chr9-85508600-DEL-214
chr9-85643613-DEL-167
chr9-85654104-DEL-69
chr9-85662508-DEL-119
chr9-85671193-DEL-3500
chr9-85691047-DEL-115
chr9-85749995-DEL-727
chr9-85769728-DEL-83
chr9-85790087-DEL-64
chr9-85855158-DEL-9943
chr9-85865248-DEL-821
chr9-85879080-DEL-109
chr9-85880983-DEL-921
chr9-85934461-DEL-52
chr9-85992616-DEL-506
chr9-86011360-DEL-52
chr9-86035429-DEL-645
chr9-86073034-DEL-1010
chr9-86083049-DEL-60
chr9-86094903-DEL-1300
chr9-86104585-DEL-404
chr9-86109033-DEL-343
chr9-86135899-DEL-66
chr9-86171549-DEL-328
chr9-86223811-DEL-81
chr9-86266306-DEL-299
chr9-86275125-DEL-164
chr9-86292953-DEL-102
chr9-86352592-DEL-5040
chr9-86410882-DEL-2119
chr9-86449348-DEL-201
chr9-86458999-DEL-5194
chr9-86467818-DEL-400
chr9-86479096-DEL-260
chr9-86507705-DEL-414
chr9-86511603-DEL-636
chr9-86582172-DEL-197
chr9-86606372-DEL-660
chr9-86613605-DEL-53
chr9-86651124-DEL-241
chr9-86660760-DEL-452
chr9-86670201-DEL-219
chr9-86693863-DEL-52
chr9-86796474-DEL-855
chr9-86801678-DEL-75
chr9-86816881-DEL-115
chr9-86845459-DEL-333
chr9-86950955-DEL-194
chr9-86958321-DEL-186
chr9-86977922-DEL-125
chr9-86995773-DEL-500
chr9-87009070-DEL-2111
chr9-87017330-DEL-110
chr9-87029282-DEL-62
chr9-87034859-DEL-169
chr9-87051888-DEL-100
chr9-87052584-DEL-101
chr9-87052958-DEL-995
chr9-87058562-DEL-59
chr9-87072455-DEL-75
chr9-87093409-DEL-164
chr9-87139207-DEL-464
chr9-87140642-DEL-54
chr9-87145612-DEL-60
chr9-87173110-DEL-52
chr9-87176954-DEL-1311
chr9-87188831-DEL-595
chr9-87203078-DEL-94
chr9-87252266-DEL-262
chr9-87264127-DEL-76
chr9-87305733-DEL-280
chr9-87342032-DEL-3153
chr9-87342769-DEL-64
chr9-87369646-DEL-13342
chr9-87433443-DEL-177
chr9-87437631-DEL-495

chr9-87440982-DEL-218
chr9-87458701-DEL-6450
chr9-87486757-DEL-832
chr9-87492150-DEL-789
chr9-87498493-DEL-112
chr9-87503692-DEL-264
chr9-87504875-DEL-298
chr9-87513169-DEL-752
chr9-87526775-DEL-205
chr9-87578592-DEL-940
chr9-87635431-DEL-100
chr9-87667510-DEL-90
chr9-87673021-DEL-6143
chr9-87676264-DEL-2900
chr9-87679357-DEL-1493
chr9-87697194-DEL-6477
chr9-87708128-DEL-111
chr9-87714764-DEL-5459
chr9-87720529-DEL-1967
chr9-87720894-DEL-415
chr9-87751445-DEL-359
chr9-87777264-DEL-213
chr9-87809007-DEL-131
chr9-87816499-DEL-507
chr9-87825324-DEL-60
chr9-87861002-DEL-831
chr9-87870635-DEL-390
chr9-87871178-DEL-488
chr9-87880918-DEL-54
chr9-87883928-DEL-348
chr9-87898648-DEL-57
chr9-87925893-DEL-88
chr9-87939729-DEL-61
chr9-87958881-DEL-989
chr9-87969023-DEL-99
chr9-87993241-DEL-62
chr9-88028635-DEL-1627
chr9-88037557-DEL-295
chr9-88043151-DEL-1429
chr9-88070761-DEL-1254
chr9-88088918-DEL-957
chr9-88106448-DEL-166
chr9-88124093-DEL-2426
chr9-88148662-DEL-96
chr9-88156489-DEL-88
chr9-88184086-DEL-66
chr9-88194233-DEL-54
chr9-88231612-DEL-314
chr9-88237325-DEL-72
chr9-88269925-DEL-341
chr9-88287634-DEL-56
chr9-88292082-DEL-355
chr9-88315168-DEL-171
chr9-88372255-DEL-83
chr9-88383135-DEL-85
chr9-88384930-DEL-99
chr9-88388231-DEL-429
chr9-88389892-DEL-459
chr9-88390396-DEL-510
chr9-88390910-DEL-559
chr9-88409129-DEL-378
chr9-88413065-DEL-130
chr9-88458218-DEL-465
chr9-88474227-DEL-37936
chr9-88533994-DEL-209
chr9-88548099-DEL-6611
chr9-88555501-DEL-204
chr9-88615007-DEL-62
chr9-88620840-DEL-218
chr9-88692740-DEL-6413
chr9-88762919-DEL-1766
chr9-88771950-DEL-2981
chr9-88780848-DEL-217
chr9-88805182-DEL-7121
chr9-88808955-DEL-67
chr9-88856039-DEL-175
chr9-88890060-DEL-60

chr9-88895239-DEL-9473
chr9-88911362-DEL-61
chr9-88945596-DEL-80
chr9-88949835-DEL-7324
chr9-88950068-DEL-118
chr9-88961715-DEL-678
chr9-88969046-DEL-185
chr9-88986957-DEL-6202
chr9-89001482-DEL-57
chr9-89010090-DEL-2574
chr9-89010642-DEL-2572
chr9-89022903-DEL-217
chr9-89070342-DEL-413
chr9-89132234-DEL-6474
chr9-89176058-DEL-140
chr9-89202036-DEL-125
chr9-89221780-DEL-14715
chr9-89243801-DEL-417
chr9-89244828-DEL-76
chr9-89274433-DEL-82
chr9-89279821-DEL-51
chr9-89296943-DEL-60
chr9-89331435-DEL-90
chr9-89336084-DEL-55
chr9-89351668-DEL-644
chr9-89353549-DEL-60
chr9-89366057-DEL-67
chr9-89388128-DEL-62
chr9-89480623-DEL-72
chr9-89499753-DEL-218
chr9-89511573-DEL-85
chr9-89515518-DEL-129
chr9-89539291-DEL-58
chr9-89541363-DEL-825
chr9-89593516-DEL-1626
chr9-89595245-DEL-262
chr9-89624598-DEL-221
chr9-89625029-DEL-157
chr9-89646766-DEL-230
chr9-89693152-DEL-840
chr9-89739556-DEL-72
chr9-89748381-DEL-71
chr9-89763304-DEL-120
chr9-89780144-DEL-193
chr9-89782952-DEL-54
chr9-89840207-DEL-64
chr9-89844776-DEL-170
chr9-89861825-DEL-89
chr9-89955268-DEL-188
chr9-89964573-DEL-200
chr9-89965464-DEL-380
chr9-89975057-DEL-166
chr9-89989966-DEL-122
chr9-89999077-DEL-156
chr9-90013440-DEL-120
chr9-90051623-DEL-52
chr9-90098388-DEL-57
chr9-90100012-DEL-201
chr9-90139126-DEL-129
chr9-90167302-DEL-1040
chr9-90218001-DEL-312
chr9-90224490-DEL-423
chr9-90225233-DEL-6204
chr9-90238807-DEL-2469
chr9-90241362-DEL-426
chr9-90247868-DEL-1522
chr9-90260251-DEL-198
chr9-90272135-DEL-6126
chr9-90278512-DEL-221
chr9-90287727-DEL-211
chr9-90290606-DEL-58
chr9-90305137-DEL-735
chr9-90309461-DEL-52
chr9-90322785-DEL-442
chr9-90332746-DEL-102
chr9-90337493-DEL-1937
chr9-90373862-DEL-888

chr9-90381082-DEL-1360
chr9-90383956-DEL-1936
chr9-90433861-DEL-946
chr9-90442924-DEL-121
chr9-90444072-DEL-2304
chr9-90481783-DEL-107
chr9-90511431-DEL-1449
chr9-90528188-DEL-6329
chr9-90548373-DEL-64
chr9-90561637-DEL-58
chr9-90591765-DEL-565
chr9-90594799-DEL-56
chr9-90633834-DEL-1043
chr9-90638102-DEL-460
chr9-90663344-DEL-226
chr9-90665045-DEL-2102
chr9-90667212-DEL-310
chr9-90681447-DEL-94
chr9-90698363-DEL-368
chr9-90700527-DEL-472
chr9-90731569-DEL-188
chr9-90735786-DEL-13387
chr9-90753373-DEL-73
chr9-90756697-DEL-3378
chr9-90762239-DEL-467
chr9-90783902-DEL-78
chr9-90787585-DEL-462
chr9-90802076-DEL-209
chr9-90829960-DEL-92
chr9-90833890-DEL-419
chr9-90839532-DEL-203
chr9-90861507-DEL-76
chr9-90863440-DEL-336
chr9-90873107-DEL-67
chr9-90963879-DEL-504
chr9-90966568-DEL-56
chr9-91051005-DEL-52
chr9-91056263-DEL-126
chr9-91120936-DEL-5224
chr9-91215355-DEL-404
chr9-91216713-DEL-69
chr9-91222885-DEL-330
chr9-91227360-DEL-103
chr9-91421553-DEL-53
chr9-91476599-DEL-62
chr9-91487965-DEL-97
chr9-91500831-DEL-6957
chr9-91516198-DEL-293
chr9-91516476-DEL-6436
chr9-91522530-DEL-212
chr9-91525364-DEL-5778
chr9-91537503-DEL-297
chr9-91580972-DEL-362
chr9-91597310-DEL-100
chr9-91611998-DEL-57
chr9-91624408-DEL-177
chr9-91626710-DEL-66
chr9-91646662-DEL-80
chr9-91674973-DEL-1486
chr9-91748111-DEL-143
chr9-91753914-DEL-619
chr9-91768638-DEL-2242
chr9-91772885-DEL-323
chr9-91776449-DEL-444
chr9-91778372-DEL-77
chr9-91798774-DEL-742
chr9-91806934-DEL-2240
chr9-91834323-DEL-59
chr9-91836850-DEL-1972
chr9-91887577-DEL-92
chr9-91899143-DEL-55
chr9-91931620-DEL-52
chr9-91949740-DEL-190
chr9-91952747-DEL-6190
chr9-91970483-DEL-51
chr9-91973745-DEL-1473
chr9-91974090-DEL-13120

chr9-91996806-DEL-3729
chr9-92031321-DEL-322
chr9-92037430-DEL-58
chr9-92055207-DEL-67
chr9-92062051-DEL-807
chr9-92094954-DEL-1101
chr9-92105806-DEL-117
chr9-92120830-DEL-66
chr9-92134015-DEL-918
chr9-92143838-DEL-89
chr9-92172209-DEL-54
chr9-92203240-DEL-401
chr9-92215129-DEL-551
chr9-92225097-DEL-89
chr9-92227320-DEL-469
chr9-92228320-DEL-119
chr9-92238758-DEL-57
chr9-92245954-DEL-89
chr9-92246529-DEL-1075
chr9-92249342-DEL-213
chr9-92274384-DEL-3078
chr9-92304527-DEL-6503
chr9-92312596-DEL-64
chr9-92355875-DEL-718
chr9-92365211-DEL-314
chr9-92369440-DEL-53
chr9-92392136-DEL-313
chr9-92393302-DEL-255
chr9-92417055-DEL-633
chr9-92435740-DEL-220
chr9-92457583-DEL-91
chr9-92463965-DEL-898
chr9-92513285-DEL-763
chr9-92518121-DEL-114
chr9-92528432-DEL-61
chr9-92529429-DEL-442
chr9-92529902-DEL-1830
chr9-92531814-DEL-4741
chr9-92557354-DEL-292
chr9-92559252-DEL-90
chr9-92562627-DEL-61
chr9-92576099-DEL-271
chr9-92635642-DEL-52
chr9-92704200-DEL-84
chr9-92706101-DEL-1733
chr9-92730310-DEL-6203
chr9-92740501-DEL-602
chr9-92740697-DEL-403
chr9-92741428-DEL-1553
chr9-92741788-DEL-529
chr9-92741958-DEL-665
chr9-92773501-DEL-1024
chr9-92784220-DEL-1673
chr9-92810727-DEL-482
chr9-92905130-DEL-74
chr9-92911034-DEL-6365
chr9-92934424-DEL-52
chr9-92935683-DEL-119
chr9-92939284-DEL-53
chr9-92942408-DEL-397
chr9-92955494-DEL-253
chr9-92996568-DEL-544
chr9-93019553-DEL-6130
chr9-93025563-DEL-64
chr9-93037322-DEL-68
chr9-93043063-DEL-122
chr9-93095824-DEL-464
chr9-93120203-DEL-677
chr9-93124133-DEL-785
chr9-93133343-DEL-145
chr9-93230588-DEL-134
chr9-93241013-DEL-138
chr9-93242871-DEL-6183
chr9-93251868-DEL-165
chr9-93255237-DEL-80
chr9-93260236-DEL-6590
chr9-93260944-DEL-212

chr9-93273894-DEL-10048
chr9-93289067-DEL-353
chr9-93296033-DEL-2231
chr9-93351111-DEL-179
chr9-93353538-DEL-56
chr9-93377408-DEL-817
chr9-93380064-DEL-73
chr9-93451794-DEL-242
chr9-93451797-DEL-481
chr9-93455026-DEL-1633
chr9-93478834-DEL-327
chr9-93495972-DEL-6062
chr9-93502749-DEL-529
chr9-93514720-DEL-1143
chr9-93520475-DEL-60
chr9-93526262-DEL-7284
chr9-93544391-DEL-6172
chr9-93560939-DEL-918
chr9-93588939-DEL-1041
chr9-93590856-DEL-77
chr9-93596637-DEL-77
chr9-93606230-DEL-125
chr9-93618497-DEL-279
chr9-93631676-DEL-123
chr9-93652466-DEL-95
chr9-93655906-DEL-131
chr9-93685512-DEL-53
chr9-93707736-DEL-76
chr9-93724512-DEL-84
chr9-93762979-DEL-60
chr9-93766605-DEL-109
chr9-93789360-DEL-412
chr9-93793011-DEL-86
chr9-93799070-DEL-540
chr9-93835788-DEL-405
chr9-93855236-DEL-105
chr9-93878448-DEL-1564
chr9-93879573-DEL-390
chr9-93898428-DEL-77
chr9-93908782-DEL-54
chr9-93947717-DEL-70
chr9-93977033-DEL-1302
chr9-93979997-DEL-55
chr9-94065668-DEL-352
chr9-94073846-DEL-357
chr9-94091394-DEL-58
chr9-94098409-DEL-271
chr9-94111712-DEL-73
chr9-94175757-DEL-446
chr9-94191475-DEL-5797
chr9-94216227-DEL-103
chr9-94221097-DEL-315
chr9-94226807-DEL-507
chr9-94237212-DEL-306
chr9-94286973-DEL-58
chr9-94346003-DEL-74
chr9-94395055-DEL-1466
chr9-94448710-DEL-98
chr9-94472651-DEL-284
chr9-94478020-DEL-59
chr9-94478541-DEL-223
chr9-94491634-DEL-102
chr9-94500774-DEL-94
chr9-94533812-DEL-221
chr9-94537253-DEL-52
chr9-94538211-DEL-127
chr9-94538838-DEL-56
chr9-94552920-DEL-79
chr9-94576234-DEL-116
chr9-94590240-DEL-83
chr9-94606193-DEL-1144
chr9-94620623-DEL-71
chr9-94646632-DEL-116
chr9-94673962-DEL-998
chr9-94687836-DEL-368
chr9-94711449-DEL-173
chr9-94785054-DEL-298

chr9-94903232-DEL-1427
chr9-94903317-DEL-672
chr9-94904065-DEL-569
chr9-94904665-DEL-180
chr9-94929922-DEL-457
chr9-94973981-DEL-315
chr9-95053627-DEL-714
chr9-95127814-DEL-6192
chr9-95150224-DEL-182
chr9-95151386-DEL-200
chr9-95238522-DEL-1010
chr9-95296257-DEL-151
chr9-95360232-DEL-495
chr9-95381878-DEL-471
chr9-95407972-DEL-55
chr9-95416226-DEL-62
chr9-95417105-DEL-1033
chr9-95450541-DEL-265
chr9-95479067-DEL-108
chr9-95491301-DEL-579
chr9-95493106-DEL-1531
chr9-95505784-DEL-66
chr9-95507668-DEL-129
chr9-95558657-DEL-138
chr9-95569456-DEL-54
chr9-95657085-DEL-88
chr9-95659417-DEL-105
chr9-95669443-DEL-53
chr9-95685620-DEL-88
chr9-95690603-DEL-72
chr9-95722131-DEL-98
chr9-95730458-DEL-58
chr9-95763578-DEL-198
chr9-95800681-DEL-1222
chr9-95812653-DEL-203
chr9-95896450-DEL-214
chr9-95906646-DEL-499
chr9-95909538-DEL-105
chr9-95922441-DEL-1841
chr9-95929993-DEL-160
chr9-95949887-DEL-62
chr9-95959803-DEL-181
chr9-95962366-DEL-58
chr9-95963219-DEL-83
chr9-95969357-DEL-56
chr9-95975600-DEL-72
chr9-96020702-DEL-272
chr9-96037884-DEL-579
chr9-96038955-DEL-200
chr9-96045808-DEL-380
chr9-96070586-DEL-92
chr9-96085431-DEL-63
chr9-96103067-DEL-62
chr9-96122115-DEL-138
chr9-96145422-DEL-2227
chr9-96147761-DEL-62
chr9-96153748-DEL-220
chr9-96156935-DEL-381
chr9-96166610-DEL-80
chr9-96178597-DEL-187
chr9-96183626-DEL-51
chr9-96207664-DEL-62
chr9-96254493-DEL-1030
chr9-96256030-DEL-237
chr9-96260127-DEL-2106
chr9-96273196-DEL-123
chr9-96292221-DEL-1777
chr9-96300370-DEL-104
chr9-96357384-DEL-934
chr9-96364114-DEL-274
chr9-96367727-DEL-53
chr9-96370457-DEL-303
chr9-96390768-DEL-232
chr9-96398883-DEL-69
chr9-96428038-DEL-154
chr9-96461101-DEL-152
chr9-96496041-DEL-54

chr9-96508819-DEL-78
chr9-96517134-DEL-66
chr9-96538619-DEL-62
chr9-96559740-DEL-213
chr9-96648794-DEL-131
chr9-96675204-DEL-81
chr9-96675776-DEL-54
chr9-96705316-DEL-59
chr9-96742620-DEL-269
chr9-96798965-DEL-741
chr9-96818329-DEL-214
chr9-96828283-DEL-55
chr9-96882578-DEL-111
chr9-96884514-DEL-56
chr9-96916396-DEL-118
chr9-96918642-DEL-52
chr9-96919797-DEL-58
chr9-96931467-DEL-1006
chr9-96940211-DEL-329
chr9-96951810-DEL-264
chr9-96952318-DEL-208
chr9-96965791-DEL-263
chr9-96968940-DEL-623
chr9-96972738-DEL-88
chr9-96993777-DEL-1111
chr9-96999826-DEL-288
chr9-97018939-DEL-429
chr9-97032489-DEL-385
chr9-97058972-DEL-120
chr9-97076471-DEL-712
chr9-97089589-DEL-911
chr9-97127281-DEL-155
chr9-97170586-DEL-2799
chr9-97208958-DEL-12628
chr9-97209014-DEL-9787
chr9-97212670-DEL-6579
chr9-97219250-DEL-2402
chr9-97225603-DEL-61
chr9-97227782-DEL-872
chr9-97232123-DEL-117
chr9-97252716-DEL-92
chr9-97262745-DEL-77
chr9-97298607-DEL-68
chr9-97308634-DEL-54
chr9-97331089-DEL-120
chr9-97412051-DEL-69
chr9-97502252-DEL-61
chr9-97546046-DEL-64
chr9-97578412-DEL-70
chr9-97583091-DEL-203
chr9-97630167-DEL-1062
chr9-97671779-DEL-68
chr9-97674105-DEL-292
chr9-97785567-DEL-120
chr9-97808789-DEL-398
chr9-97838145-DEL-61
chr9-97839970-DEL-70
chr9-97853439-DEL-1067
chr9-97869443-DEL-3016
chr9-97880493-DEL-1270
chr9-97887106-DEL-80
chr9-97890215-DEL-1433
chr9-97891715-DEL-242
chr9-97903349-DEL-53
chr9-97922941-DEL-1445
chr9-97928192-DEL-64
chr9-97990124-DEL-52
chr9-98002917-DEL-205
chr9-98038636-DEL-119
chr9-98068313-DEL-64
chr9-98090597-DEL-742
chr9-98132987-DEL-426
chr9-98191272-DEL-134
chr9-98201327-DEL-1578
chr9-98253157-DEL-176
chr9-98294645-DEL-64
chr9-98295408-DEL-743

chr9-98300257-DEL-243
chr9-98359304-DEL-85
chr9-98390861-DEL-88
chr9-98399218-DEL-147
chr9-98408363-DEL-101
chr9-98477713-DEL-58
chr9-98505112-DEL-147
chr9-98535584-DEL-51
chr9-98559640-DEL-5839
chr9-98570012-DEL-2465
chr9-98614795-DEL-707
chr9-98626545-DEL-120
chr9-98636712-DEL-52
chr9-98648453-DEL-5292
chr9-98715785-DEL-811
chr9-98716371-DEL-99
chr9-98726751-DEL-64
chr9-98730830-DEL-209
chr9-98749280-DEL-94
chr9-98750785-DEL-55
chr9-98775914-DEL-15116
chr9-98784007-DEL-95
chr9-98792698-DEL-1460
chr9-98828476-DEL-60
chr9-98843039-DEL-67
chr9-98860523-DEL-236
chr9-98929150-DEL-58
chr9-98960950-DEL-226
chr9-98966083-DEL-136
chr9-99052643-DEL-200
chr9-99063153-DEL-76
chr9-99075016-DEL-159
chr9-99128443-DEL-2761
chr9-99202521-DEL-206
chr9-99243533-DEL-2119
chr9-99244945-DEL-233
chr9-99248968-DEL-51
chr9-99258165-DEL-103
chr9-99262438-DEL-1324
chr9-99264293-DEL-66
chr9-99274647-DEL-88
chr9-99294577-DEL-56
chr9-99296273-DEL-118
chr9-99326959-DEL-602
chr9-99327645-DEL-255
chr9-99337847-DEL-60
chr9-99340164-DEL-58
chr9-99354620-DEL-173
chr9-99358582-DEL-81
chr9-99382362-DEL-60
chr9-99427414-DEL-2281
chr9-99427703-DEL-70
chr9-99447745-DEL-56
chr9-99448343-DEL-168
chr9-99506828-DEL-162
chr9-99509853-DEL-242
chr9-99530692-DEL-89
chr9-99558895-DEL-1915
chr9-99569323-DEL-201
chr9-99586231-DEL-100
chr9-99634268-DEL-60
chr9-99636005-DEL-87
chr9-99648320-DEL-567
chr9-99666192-DEL-401
chr9-99684114-DEL-68
chr9-99699712-DEL-874
chr9-99703362-DEL-713
chr9-99785333-DEL-6488
chr9-99913468-DEL-130
chr9-99921648-DEL-284
chr9-99938248-DEL-517
chr9-99953498-DEL-361
chr9-99988157-DEL-268
chr9-100005496-DEL-76
chr9-100016922-DEL-58
chr9-100028203-DEL-235
chr9-100033213-DEL-1712

chr9-100074708-DEL-63
chr9-100126006-DEL-71
chr9-100175504-DEL-59
chr9-100201600-DEL-795
chr9-100211116-DEL-191
chr9-100212682-DEL-53
chr9-100243367-DEL-596
chr9-100268061-DEL-77
chr9-100277082-DEL-172
chr9-100281712-DEL-78
chr9-100305013-DEL-258
chr9-100305640-DEL-89
chr9-100309511-DEL-490
chr9-100324574-DEL-294
chr9-100345468-DEL-87
chr9-100346586-DEL-186
chr9-100360580-DEL-1688
chr9-100436416-DEL-783
chr9-100484522-DEL-335
chr9-100486940-DEL-72
chr9-100497295-DEL-250
chr9-100506219-DEL-2388
chr9-100506227-DEL-139
chr9-100510187-DEL-105
chr9-100514883-DEL-184
chr9-100577260-DEL-3519
chr9-100577296-DEL-575
chr9-100579613-DEL-1127
chr9-100585629-DEL-120
chr9-100590041-DEL-191
chr9-100591798-DEL-63
chr9-100600046-DEL-92
chr9-100654516-DEL-1707
chr9-100655545-DEL-932
chr9-100690918-DEL-319
chr9-100705165-DEL-185
chr9-100709202-DEL-600
chr9-100720911-DEL-2688
chr9-100740644-DEL-1632
chr9-100754086-DEL-143
chr9-100783334-DEL-2306
chr9-100793110-DEL-54
chr9-100815659-DEL-6142
chr9-100825826-DEL-54
chr9-100862776-DEL-400
chr9-100868408-DEL-224
chr9-100880958-DEL-193
chr9-100911327-DEL-340
chr9-100922778-DEL-163
chr9-100924206-DEL-108
chr9-100926416-DEL-206
chr9-100929205-DEL-163
chr9-100930961-DEL-52
chr9-100939669-DEL-502
chr9-100940667-DEL-86
chr9-100967644-DEL-1012
chr9-100967682-DEL-850
chr9-100968445-DEL-145
chr9-100970093-DEL-73
chr9-100974842-DEL-182
chr9-101005467-DEL-6160
chr9-101014304-DEL-55
chr9-101016444-DEL-103
chr9-101070920-DEL-54
chr9-101085469-DEL-326
chr9-101106758-DEL-495
chr9-101111074-DEL-656
chr9-101116411-DEL-67
chr9-101197587-DEL-60
chr9-101210435-DEL-60
chr9-101237838-DEL-295
chr9-101264459-DEL-400
chr9-101265792-DEL-3096
chr9-101287356-DEL-52
chr9-101367924-DEL-110
chr9-101370313-DEL-114
chr9-101391961-DEL-55

chr9-101415496-DEL-291
chr9-101425724-DEL-179
chr9-101438120-DEL-51
chr9-101457454-DEL-66
chr9-101468096-DEL-60
chr9-101471916-DEL-79
chr9-101510266-DEL-69
chr9-101520168-DEL-88
chr9-101522239-DEL-429
chr9-101547982-DEL-63
chr9-101549870-DEL-60
chr9-101598664-DEL-63
chr9-101606470-DEL-92
chr9-101616011-DEL-111
chr9-101655412-DEL-685
chr9-101659105-DEL-6160
chr9-101742670-DEL-138
chr9-101776431-DEL-144
chr9-101798512-DEL-103
chr9-101803316-DEL-66
chr9-101867563-DEL-142
chr9-101895123-DEL-74
chr9-101929319-DEL-124
chr9-101951799-DEL-69
chr9-101983444-DEL-279
chr9-102020513-DEL-106
chr9-102055248-DEL-74
chr9-102096568-DEL-863
chr9-102108815-DEL-671
chr9-102111047-DEL-227
chr9-102121639-DEL-703
chr9-102129348-DEL-151
chr9-102182473-DEL-105
chr9-102187001-DEL-91
chr9-102190853-DEL-816
chr9-102219637-DEL-966
chr9-102264663-DEL-54
chr9-102265795-DEL-103
chr9-102308891-DEL-234
chr9-102343973-DEL-165
chr9-102356983-DEL-161
chr9-102381970-DEL-66
chr9-102416421-DEL-67
chr9-102466706-DEL-203
chr9-102488126-DEL-172
chr9-102495638-DEL-110
chr9-102531356-DEL-60
chr9-102542198-DEL-56
chr9-102574368-DEL-335
chr9-102578730-DEL-52
chr9-102653874-DEL-78
chr9-102660983-DEL-427
chr9-102677005-DEL-209
chr9-102702437-DEL-65
chr9-102722660-DEL-3589
chr9-102732480-DEL-389
chr9-102732985-DEL-52
chr9-102751004-DEL-387
chr9-102792273-DEL-3546
chr9-102795440-DEL-85
chr9-102808012-DEL-77
chr9-102812535-DEL-327
chr9-102832088-DEL-3536
chr9-102839444-DEL-128
chr9-102839830-DEL-86
chr9-102867812-DEL-130
chr9-102873981-DEL-93
chr9-102931090-DEL-117
chr9-103003036-DEL-64
chr9-103008009-DEL-51
chr9-103010267-DEL-144
chr9-103022035-DEL-57
chr9-103042285-DEL-1123
chr9-103068452-DEL-238
chr9-103082903-DEL-650
chr9-103097734-DEL-60
chr9-103133570-DEL-65

chr9-103145751-DEL-328
chr9-103147371-DEL-68
chr9-103173415-DEL-2976
chr9-103173426-DEL-1728
chr9-103173545-DEL-4175
chr9-103173555-DEL-336
chr9-103173578-DEL-912
chr9-103173841-DEL-1152
chr9-103173922-DEL-936
chr9-103174153-DEL-1392
chr9-103174730-DEL-192
chr9-103175235-DEL-288
chr9-103175324-DEL-96
chr9-103175671-DEL-288
chr9-103175897-DEL-1919
chr9-103176566-DEL-336
chr9-103176674-DEL-1104
chr9-103176935-DEL-720
chr9-103198913-DEL-147
chr9-103227573-DEL-51
chr9-103250221-DEL-141
chr9-103265294-DEL-1682
chr9-103267685-DEL-550
chr9-103297843-DEL-89
chr9-103333583-DEL-612
chr9-103339569-DEL-102
chr9-103344008-DEL-101
chr9-103348599-DEL-1500
chr9-103370336-DEL-1311
chr9-103427304-DEL-106
chr9-103481512-DEL-498
chr9-103520740-DEL-66
chr9-103550895-DEL-76
chr9-103623033-DEL-931
chr9-103659266-DEL-54
chr9-103698506-DEL-152
chr9-103711229-DEL-120
chr9-103716441-DEL-1086
chr9-103717694-DEL-115
chr9-103727529-DEL-129
chr9-103764763-DEL-67
chr9-103767906-DEL-749
chr9-103774492-DEL-193
chr9-103793958-DEL-52
chr9-103798475-DEL-255
chr9-103805957-DEL-1130
chr9-103820142-DEL-56
chr9-103833887-DEL-60
chr9-103837493-DEL-540
chr9-103888090-DEL-54
chr9-103905942-DEL-75
chr9-103935844-DEL-400
chr9-103936436-DEL-164
chr9-104011662-DEL-153
chr9-104020772-DEL-134
chr9-104022161-DEL-119
chr9-104025501-DEL-171
chr9-104052538-DEL-72
chr9-104092293-DEL-142
chr9-104106659-DEL-52
chr9-104200981-DEL-53
chr9-104203944-DEL-51
chr9-104213790-DEL-87
chr9-104227347-DEL-438
chr9-104229681-DEL-53
chr9-104256023-DEL-903
chr9-104280641-DEL-62
chr9-104282388-DEL-541
chr9-104311499-DEL-182
chr9-104327067-DEL-861
chr9-104338372-DEL-69
chr9-104357211-DEL-214
chr9-104365476-DEL-54
chr9-104401270-DEL-220
chr9-104401665-DEL-1150
chr9-104425797-DEL-72
chr9-104452347-DEL-63

chr9-104455950-DEL-85
chr9-104457025-DEL-59
chr9-104480754-DEL-139
chr9-104488945-DEL-76
chr9-104498967-DEL-195
chr9-104500157-DEL-71
chr9-104502437-DEL-304
chr9-104508992-DEL-256
chr9-104533430-DEL-52
chr9-104537832-DEL-58
chr9-104552788-DEL-411
chr9-104568935-DEL-2943
chr9-104575360-DEL-55
chr9-104578640-DEL-655
chr9-104597830-DEL-54
chr9-104598628-DEL-165
chr9-104619575-DEL-53
chr9-104623178-DEL-71
chr9-104639166-DEL-476
chr9-104661221-DEL-119
chr9-104713605-DEL-65
chr9-104718966-DEL-498
chr9-104719963-DEL-399
chr9-104721156-DEL-102
chr9-104725291-DEL-1374
chr9-104727446-DEL-928
chr9-104733886-DEL-86
chr9-104757514-DEL-518
chr9-104770945-DEL-192
chr9-104783714-DEL-6171
chr9-104800176-DEL-149
chr9-104802120-DEL-168
chr9-104803373-DEL-1858
chr9-104804487-DEL-77
chr9-104863117-DEL-196
chr9-104885090-DEL-58
chr9-104890154-DEL-64
chr9-104895176-DEL-51
chr9-104913257-DEL-1222
chr9-104923473-DEL-77
chr9-104929473-DEL-68
chr9-104974348-DEL-161
chr9-104974879-DEL-744
chr9-104985956-DEL-134
chr9-104987498-DEL-1437
chr9-105002291-DEL-71
chr9-105024862-DEL-6205
chr9-105036279-DEL-129
chr9-105052490-DEL-53
chr9-105053471-DEL-309
chr9-105059440-DEL-235
chr9-105097733-DEL-515
chr9-105126089-DEL-317
chr9-105194824-DEL-93
chr9-105198867-DEL-63
chr9-105230508-DEL-85
chr9-105246003-DEL-1123
chr9-105248354-DEL-2720
chr9-105252390-DEL-254
chr9-105332977-DEL-414
chr9-105348902-DEL-508
chr9-105359744-DEL-3597
chr9-105363244-DEL-74
chr9-105376307-DEL-322
chr9-105392221-DEL-51
chr9-105399794-DEL-72
chr9-105428604-DEL-187
chr9-105437222-DEL-780
chr9-105454065-DEL-68
chr9-105463035-DEL-108
chr9-105463437-DEL-661
chr9-105479169-DEL-77
chr9-105500625-DEL-8364
chr9-105509203-DEL-830
chr9-105567448-DEL-205
chr9-105583391-DEL-112
chr9-105616519-DEL-231

chr9-105628668-DEL-63
chr9-105647786-DEL-126
chr9-105660611-DEL-100
chr9-105665325-DEL-52
chr9-105668336-DEL-107
chr9-105679998-DEL-71
chr9-105685303-DEL-614
chr9-105690377-DEL-59
chr9-105702118-DEL-56
chr9-105706381-DEL-54
chr9-105721778-DEL-54
chr9-105735295-DEL-1925
chr9-105793749-DEL-878
chr9-105813829-DEL-84
chr9-105950628-DEL-1287
chr9-105956229-DEL-184
chr9-105970372-DEL-206
chr9-105989070-DEL-330
chr9-105990884-DEL-58
chr9-106015882-DEL-60
chr9-106020223-DEL-106
chr9-106041007-DEL-158
chr9-106052113-DEL-60
chr9-106109613-DEL-1114
chr9-106116147-DEL-82
chr9-106119814-DEL-72
chr9-106121177-DEL-233
chr9-106135263-DEL-439
chr9-106141332-DEL-189
chr9-106164121-DEL-62
chr9-106293346-DEL-505
chr9-106303724-DEL-196
chr9-106343737-DEL-197
chr9-106365740-DEL-217
chr9-106368516-DEL-827
chr9-106413138-DEL-70
chr9-106431183-DEL-3348
chr9-106436173-DEL-110
chr9-106436864-DEL-64
chr9-106478634-DEL-82
chr9-106489186-DEL-116
chr9-106514044-DEL-62
chr9-106537166-DEL-398
chr9-106582000-DEL-78
chr9-106594816-DEL-52
chr9-106598364-DEL-80
chr9-106641974-DEL-1959
chr9-106650543-DEL-83
chr9-106651985-DEL-284
chr9-106676380-DEL-1376
chr9-106679810-DEL-1064
chr9-106807839-DEL-299
chr9-106822792-DEL-142
chr9-106827197-DEL-226
chr9-106853350-DEL-84
chr9-106858819-DEL-71
chr9-106898515-DEL-503
chr9-106910431-DEL-58
chr9-106914814-DEL-382
chr9-106930875-DEL-64
chr9-106932918-DEL-374
chr9-106934244-DEL-273
chr9-106946849-DEL-330
chr9-106975765-DEL-368
chr9-107005168-DEL-6598
chr9-107075364-DEL-208
chr9-107121391-DEL-827
chr9-107129655-DEL-52
chr9-107145758-DEL-71
chr9-107152524-DEL-89
chr9-107164427-DEL-350
chr9-107182061-DEL-107
chr9-107183777-DEL-270
chr9-107193529-DEL-207
chr9-107288917-DEL-132
chr9-107319575-DEL-94
chr9-107500206-DEL-205

chr9-107518747-DEL-56
chr9-107524506-DEL-199
chr9-107601770-DEL-60
chr9-107609257-DEL-54
chr9-107638143-DEL-1018
chr9-107642164-DEL-121
chr9-107658793-DEL-385
chr9-107663452-DEL-201
chr9-107694882-DEL-63
chr9-107755464-DEL-196
chr9-107773009-DEL-80
chr9-107791073-DEL-178
chr9-107799894-DEL-58
chr9-107850209-DEL-180
chr9-107865070-DEL-54
chr9-107882648-DEL-1772
chr9-107889206-DEL-72
chr9-107911676-DEL-160
chr9-107977470-DEL-295
chr9-108001398-DEL-150
chr9-108002370-DEL-115
chr9-108010081-DEL-210
chr9-108011383-DEL-486
chr9-108013320-DEL-218
chr9-108035307-DEL-1448
chr9-108044518-DEL-320
chr9-108059874-DEL-62
chr9-108075251-DEL-197
chr9-108104070-DEL-266
chr9-108117943-DEL-1166
chr9-108118226-DEL-803
chr9-108118949-DEL-288
chr9-108130371-DEL-243
chr9-108159239-DEL-213
chr9-108169668-DEL-310
chr9-108205708-DEL-1820
chr9-108220875-DEL-2970
chr9-108234557-DEL-81
chr9-108241686-DEL-81
chr9-108246658-DEL-67
chr9-108248519-DEL-52
chr9-108308637-DEL-170
chr9-108381289-DEL-190
chr9-108420393-DEL-183
chr9-108424132-DEL-82
chr9-108446419-DEL-73
chr9-108469898-DEL-212
chr9-108478265-DEL-541
chr9-108524966-DEL-172
chr9-108576200-DEL-295
chr9-108579417-DEL-1239
chr9-108579940-DEL-195
chr9-108588951-DEL-180
chr9-108597564-DEL-161
chr9-108601092-DEL-519
chr9-108614784-DEL-64
chr9-108644794-DEL-53
chr9-108655206-DEL-133
chr9-108740273-DEL-59
chr9-108748706-DEL-894
chr9-108749561-DEL-69
chr9-108761269-DEL-640
chr9-108776305-DEL-58
chr9-108794731-DEL-94
chr9-108831058-DEL-461
chr9-108848252-DEL-1649
chr9-108849320-DEL-1110
chr9-108850013-DEL-216
chr9-108876108-DEL-913
chr9-108958629-DEL-710
chr9-108960711-DEL-252
chr9-108988064-DEL-550
chr9-109027845-DEL-1365
chr9-109053953-DEL-902
chr9-109066783-DEL-973
chr9-109096294-DEL-1484
chr9-109098630-DEL-56

chr9-109109750-DEL-232
chr9-109130867-DEL-1874
chr9-109138915-DEL-848
chr9-109158823-DEL-112
chr9-109182708-DEL-109
chr9-109189976-DEL-6661
chr9-109198039-DEL-65
chr9-109226737-DEL-1243
chr9-109233553-DEL-4053
chr9-109239530-DEL-498
chr9-109243030-DEL-392
chr9-109274280-DEL-85
chr9-109297424-DEL-2891
chr9-109317941-DEL-134
chr9-109331488-DEL-105
chr9-109338378-DEL-7938
chr9-109359634-DEL-212
chr9-109372946-DEL-369
chr9-109373853-DEL-799
chr9-109413078-DEL-526
chr9-109473241-DEL-196
chr9-109535259-DEL-1807
chr9-109603872-DEL-54
chr9-109610328-DEL-111
chr9-109612188-DEL-2373
chr9-109616526-DEL-62
chr9-109643674-DEL-600
chr9-109644282-DEL-168
chr9-109672273-DEL-688
chr9-109710695-DEL-103
chr9-109725250-DEL-52
chr9-109755255-DEL-64
chr9-109767186-DEL-109
chr9-109776654-DEL-81
chr9-109783297-DEL-119
chr9-109806090-DEL-123
chr9-109813880-DEL-193
chr9-109837167-DEL-273
chr9-109846000-DEL-190
chr9-109851693-DEL-111
chr9-109859456-DEL-184
chr9-109888362-DEL-60
chr9-109937152-DEL-73
chr9-109939508-DEL-162
chr9-109953276-DEL-202
chr9-109975545-DEL-518
chr9-110004612-DEL-54
chr9-110026153-DEL-71
chr9-110048158-DEL-58
chr9-110063203-DEL-62
chr9-110064626-DEL-347
chr9-110069622-DEL-169
chr9-110098971-DEL-2672
chr9-110099701-DEL-302
chr9-110102011-DEL-1458
chr9-110103307-DEL-52
chr9-110107498-DEL-91
chr9-110120207-DEL-285
chr9-110147527-DEL-776
chr9-110154409-DEL-60
chr9-110160211-DEL-203
chr9-110166524-DEL-148
chr9-110183712-DEL-167
chr9-110193018-DEL-59
chr9-110194290-DEL-309
chr9-110239941-DEL-96
chr9-110243294-DEL-80
chr9-110272835-DEL-121
chr9-110273163-DEL-595
chr9-110274811-DEL-284
chr9-110279296-DEL-144
chr9-110299852-DEL-522
chr9-110315519-DEL-81
chr9-110317426-DEL-59
chr9-110327796-DEL-146
chr9-110331475-DEL-3511
chr9-110334680-DEL-306

chr9-110347832-DEL-61
chr9-110355703-DEL-164
chr9-110365617-DEL-59
chr9-110400513-DEL-377
chr9-110409229-DEL-89
chr9-110427455-DEL-253
chr9-110430828-DEL-208
chr9-110486992-DEL-76
chr9-110489286-DEL-929
chr9-110503215-DEL-288
chr9-110509595-DEL-202
chr9-110600112-DEL-78
chr9-110601169-DEL-60
chr9-110618150-DEL-63
chr9-110642711-DEL-213
chr9-110649855-DEL-132
chr9-110657071-DEL-102
chr9-110676752-DEL-1170
chr9-110716128-DEL-60
chr9-110732285-DEL-806
chr9-110733128-DEL-1384
chr9-110734906-DEL-809
chr9-110736152-DEL-153
chr9-110736986-DEL-253
chr9-110761104-DEL-417
chr9-110791851-DEL-131
chr9-110808332-DEL-94
chr9-110815773-DEL-185
chr9-110840085-DEL-127
chr9-110850614-DEL-80
chr9-110870245-DEL-58
chr9-110871914-DEL-52
chr9-110887665-DEL-325
chr9-110895784-DEL-434
chr9-110952016-DEL-64
chr9-110959988-DEL-1083
chr9-111005668-DEL-183
chr9-111031914-DEL-2063
chr9-111037878-DEL-126
chr9-111082841-DEL-332
chr9-111088460-DEL-74
chr9-111151237-DEL-54
chr9-111171526-DEL-59
chr9-111200216-DEL-232
chr9-111245357-DEL-55
chr9-111274317-DEL-195
chr9-111324096-DEL-130
chr9-111343505-DEL-218
chr9-111348205-DEL-387
chr9-111350778-DEL-1255
chr9-111382304-DEL-66
chr9-111394081-DEL-1464
chr9-111409247-DEL-784
chr9-111415921-DEL-791
chr9-111426449-DEL-1009
chr9-111436458-DEL-286
chr9-111445929-DEL-60
chr9-111545549-DEL-1314
chr9-111556475-DEL-129
chr9-111580959-DEL-295
chr9-111632043-DEL-58
chr9-111690609-DEL-161
chr9-111696869-DEL-72
chr9-111719068-DEL-339
chr9-111722109-DEL-82
chr9-111735219-DEL-1836
chr9-111773459-DEL-94
chr9-111803779-DEL-86
chr9-111811304-DEL-298
chr9-111812045-DEL-60
chr9-111818972-DEL-98
chr9-111840827-DEL-57
chr9-111875998-DEL-58
chr9-111877806-DEL-56
chr9-111880086-DEL-568
chr9-111884664-DEL-55
chr9-111890814-DEL-487

chr9-111897548-DEL-56
chr9-111898356-DEL-53
chr9-111916092-DEL-216
chr9-111932904-DEL-132
chr9-111942296-DEL-400
chr9-111951060-DEL-385
chr9-111967465-DEL-83
chr9-112013828-DEL-149
chr9-112088947-DEL-353
chr9-112129718-DEL-708
chr9-112134413-DEL-248
chr9-112152612-DEL-92
chr9-112166293-DEL-320
chr9-112170828-DEL-53
chr9-112183403-DEL-1038
chr9-112215132-DEL-178
chr9-112228772-DEL-68
chr9-112245132-DEL-2836
chr9-112248009-DEL-963
chr9-112298233-DEL-58
chr9-112355986-DEL-119
chr9-112358493-DEL-344
chr9-112388270-DEL-58
chr9-112408356-DEL-77
chr9-112412110-DEL-64
chr9-112418744-DEL-64
chr9-112430546-DEL-64
chr9-112431822-DEL-82
chr9-112464321-DEL-492
chr9-112486082-DEL-64
chr9-112498372-DEL-75
chr9-112499485-DEL-51
chr9-112539355-DEL-134
chr9-112558230-DEL-98
chr9-112571533-DEL-400
chr9-112609334-DEL-70
chr9-112636061-DEL-121
chr9-112676586-DEL-54
chr9-112683612-DEL-120
chr9-112752740-DEL-85
chr9-112801857-DEL-1473
chr9-112810419-DEL-530
chr9-112813416-DEL-6414
chr9-112829512-DEL-55
chr9-112834548-DEL-58
chr9-112876323-DEL-55
chr9-112906200-DEL-136
chr9-112934043-DEL-228
chr9-112951943-DEL-64
chr9-112969937-DEL-118
chr9-113045796-DEL-64
chr9-113055535-DEL-2206
chr9-113107930-DEL-69
chr9-113117376-DEL-108
chr9-113131459-DEL-56
chr9-113147435-DEL-996
chr9-113151887-DEL-57
chr9-113172765-DEL-87
chr9-113177751-DEL-1021
chr9-113179483-DEL-52
chr9-113187987-DEL-133
chr9-113229748-DEL-154
chr9-113278426-DEL-77
chr9-113322378-DEL-72
chr9-113327912-DEL-58
chr9-113345848-DEL-174
chr9-113354426-DEL-1142
chr9-113358385-DEL-208
chr9-113422439-DEL-2851
chr9-113432841-DEL-182
chr9-113437360-DEL-857
chr9-113438606-DEL-76
chr9-113456531-DEL-96
chr9-113457019-DEL-645
chr9-113472799-DEL-193
chr9-113473624-DEL-99
chr9-113475847-DEL-57

chr9-113512163-DEL-555
chr9-113516214-DEL-56
chr9-113545791-DEL-54
chr9-113552403-DEL-60
chr9-113597127-DEL-52
chr9-113608150-DEL-137
chr9-113615472-DEL-55
chr9-113616988-DEL-934
chr9-113618418-DEL-390
chr9-113654006-DEL-60
chr9-113657481-DEL-189
chr9-113668738-DEL-586
chr9-113682806-DEL-137
chr9-113699295-DEL-269
chr9-113723566-DEL-128
chr9-113751998-DEL-384
chr9-113818013-DEL-57
chr9-113826308-DEL-176
chr9-113838496-DEL-82
chr9-113872939-DEL-55
chr9-113884015-DEL-98
chr9-113941069-DEL-190
chr9-113970221-DEL-6326
chr9-113978460-DEL-60
chr9-114002532-DEL-912
chr9-114003155-DEL-417
chr9-114003595-DEL-5549
chr9-114004328-DEL-798
chr9-114004335-DEL-209
chr9-114005021-DEL-266
chr9-114007528-DEL-371
chr9-114044532-DEL-53
chr9-114060412-DEL-197
chr9-114064528-DEL-1953
chr9-114079407-DEL-228
chr9-114079513-DEL-95
chr9-114081563-DEL-133
chr9-114084995-DEL-3260
chr9-114088778-DEL-1176
chr9-114104193-DEL-1363
chr9-114108736-DEL-60
chr9-114110761-DEL-201
chr9-114112500-DEL-6495
chr9-114184787-DEL-173
chr9-114192756-DEL-655
chr9-114210909-DEL-215
chr9-114213132-DEL-166
chr9-114251549-DEL-137
chr9-114254022-DEL-59
chr9-114254265-DEL-2395
chr9-114256705-DEL-601
chr9-114267544-DEL-87
chr9-114288935-DEL-2272
chr9-114294083-DEL-120
chr9-114298453-DEL-212
chr9-114305150-DEL-155
chr9-114314686-DEL-273
chr9-114323075-DEL-353
chr9-114350634-DEL-137
chr9-114358363-DEL-239
chr9-114362321-DEL-210
chr9-114365245-DEL-203
chr9-114366929-DEL-130
chr9-114379681-DEL-56
chr9-114387184-DEL-192
chr9-114388012-DEL-59
chr9-114395304-DEL-72
chr9-114396500-DEL-267
chr9-114403034-DEL-85
chr9-114421309-DEL-597
chr9-114433083-DEL-228
chr9-114456094-DEL-52
chr9-114522426-DEL-387
chr9-114523105-DEL-194
chr9-114544344-DEL-59
chr9-114556290-DEL-521
chr9-114564888-DEL-285

chr9-114567649-DEL-141
chr9-114575206-DEL-129
chr9-114580755-DEL-63
chr9-114582803-DEL-57
chr9-114585049-DEL-54
chr9-114592909-DEL-66
chr9-114594702-DEL-87
chr9-114649076-DEL-180
chr9-114650496-DEL-60
chr9-114656748-DEL-60
chr9-114661174-DEL-464
chr9-114663885-DEL-59
chr9-114683307-DEL-103
chr9-114689286-DEL-52
chr9-114712116-DEL-433
chr9-114725992-DEL-1011
chr9-114751271-DEL-451
chr9-114753945-DEL-83
chr9-114777665-DEL-58
chr9-114779622-DEL-76
chr9-114785184-DEL-206
chr9-114799191-DEL-4501
chr9-114805528-DEL-82
chr9-114816962-DEL-289
chr9-114822529-DEL-184
chr9-114841289-DEL-126
chr9-114855968-DEL-1071
chr9-114903069-DEL-114
chr9-114917407-DEL-240
chr9-114924193-DEL-115
chr9-114942153-DEL-58
chr9-114963881-DEL-536
chr9-114967711-DEL-118
chr9-114982643-DEL-115
chr9-114989855-DEL-216
chr9-114993973-DEL-127
chr9-115003297-DEL-80
chr9-115016534-DEL-56
chr9-115031478-DEL-199
chr9-115034617-DEL-330
chr9-115045443-DEL-111
chr9-115052566-DEL-60
chr9-115089391-DEL-194
chr9-115098791-DEL-74
chr9-115102540-DEL-65
chr9-115127260-DEL-113
chr9-115161817-DEL-56
chr9-115176647-DEL-104
chr9-115179902-DEL-143
chr9-115214551-DEL-71
chr9-115222747-DEL-64
chr9-115225469-DEL-94
chr9-115229253-DEL-69
chr9-115240086-DEL-228
chr9-115242882-DEL-61
chr9-115264870-DEL-68
chr9-115266706-DEL-969
chr9-115312519-DEL-137
chr9-115325743-DEL-583
chr9-115326023-DEL-1327
chr9-115326542-DEL-1016
chr9-115326569-DEL-821
chr9-115367909-DEL-68
chr9-115370285-DEL-630
chr9-115429672-DEL-152
chr9-115436947-DEL-151
chr9-115443997-DEL-87
chr9-115464805-DEL-499
chr9-115523918-DEL-1074
chr9-115549955-DEL-191
chr9-115562689-DEL-68
chr9-115572718-DEL-505
chr9-115600480-DEL-449
chr9-115616099-DEL-71
chr9-115646907-DEL-197
chr9-115649308-DEL-1268
chr9-115656642-DEL-346

chr9-115659525-DEL-443
chr9-115672300-DEL-594
chr9-115693354-DEL-86
chr9-115693497-DEL-719
chr9-115698240-DEL-8569
chr9-115704049-DEL-372
chr9-115774468-DEL-87
chr9-115796478-DEL-54
chr9-115834664-DEL-377
chr9-115842437-DEL-81
chr9-115871389-DEL-214
chr9-115882469-DEL-58
chr9-115944124-DEL-186
chr9-115989771-DEL-1148
chr9-116005409-DEL-64
chr9-116015901-DEL-87
chr9-116029500-DEL-386
chr9-116045417-DEL-275
chr9-116059377-DEL-242
chr9-116086425-DEL-329
chr9-116140940-DEL-66
chr9-116172909-DEL-70
chr9-116191740-DEL-188
chr9-116196649-DEL-75
chr9-116200187-DEL-114
chr9-116217049-DEL-96
chr9-116222086-DEL-137
chr9-116227902-DEL-60
chr9-116235575-DEL-315
chr9-116238713-DEL-52
chr9-116239711-DEL-1368
chr9-116271017-DEL-143
chr9-116275029-DEL-198
chr9-116287108-DEL-74
chr9-116325043-DEL-86
chr9-116377058-DEL-202
chr9-116380642-DEL-272
chr9-116390652-DEL-4428
chr9-116416413-DEL-54
chr9-116423523-DEL-63
chr9-116445352-DEL-64
chr9-116451543-DEL-254
chr9-116467087-DEL-61
chr9-116471388-DEL-86
chr9-116484853-DEL-432
chr9-116492734-DEL-520
chr9-116554534-DEL-52
chr9-116560180-DEL-64
chr9-116575522-DEL-62
chr9-116582668-DEL-161
chr9-116625966-DEL-435
chr9-116627679-DEL-54
chr9-116661340-DEL-162
chr9-116666681-DEL-180
chr9-116666845-DEL-528
chr9-116702474-DEL-60
chr9-116703018-DEL-440
chr9-116760260-DEL-82
chr9-116795452-DEL-66
chr9-116805093-DEL-56
chr9-116817279-DEL-208
chr9-116827710-DEL-122
chr9-116875447-DEL-88
chr9-116879404-DEL-64
chr9-116898967-DEL-1967
chr9-116926666-DEL-58
chr9-116958289-DEL-223
chr9-116958817-DEL-96
chr9-116987536-DEL-52
chr9-117017153-DEL-69
chr9-117023413-DEL-66
chr9-117034829-DEL-76
chr9-117045944-DEL-56
chr9-117058030-DEL-60
chr9-117064126-DEL-132
chr9-117068736-DEL-293
chr9-117075616-DEL-97

chr9-117086931-DEL-87
chr9-117088221-DEL-66
chr9-117115219-DEL-326
chr9-117132812-DEL-56
chr9-117137098-DEL-70
chr9-117144306-DEL-239
chr9-117173141-DEL-488
chr9-117173995-DEL-265
chr9-117211824-DEL-78
chr9-117243026-DEL-469
chr9-117257092-DEL-213
chr9-117281971-DEL-173
chr9-117285154-DEL-1540
chr9-117295894-DEL-122
chr9-117304816-DEL-73
chr9-117347259-DEL-91
chr9-117350196-DEL-56
chr9-117368861-DEL-53
chr9-117382665-DEL-51
chr9-117392252-DEL-78
chr9-117416353-DEL-165
chr9-117421598-DEL-86
chr9-117432066-DEL-62
chr9-117435186-DEL-521
chr9-117449485-DEL-92
chr9-117454680-DEL-66
chr9-117457727-DEL-164
chr9-117461495-DEL-140
chr9-117465566-DEL-60
chr9-117471577-DEL-63
chr9-117476375-DEL-69
chr9-117488188-DEL-84
chr9-117488780-DEL-930
chr9-117504712-DEL-1028
chr9-117520923-DEL-88
chr9-117534639-DEL-128
chr9-117552806-DEL-80
chr9-117559468-DEL-402
chr9-117560827-DEL-118
chr9-117575786-DEL-60
chr9-117602899-DEL-1041
chr9-117619462-DEL-69
chr9-117626953-DEL-92
chr9-117629983-DEL-122
chr9-117651127-DEL-56
chr9-117659251-DEL-70
chr9-117668097-DEL-105
chr9-117687172-DEL-142
chr9-117694661-DEL-50
chr9-117745111-DEL-124
chr9-117754021-DEL-96
chr9-117783975-DEL-326
chr9-117816612-DEL-9353
chr9-117828447-DEL-53
chr9-117850849-DEL-1763
chr9-117871066-DEL-177
chr9-117887145-DEL-62
chr9-117916971-DEL-2040
chr9-117916988-DEL-332
chr9-117917036-DEL-198
chr9-117917115-DEL-4854
chr9-117918133-DEL-1942
chr9-117918548-DEL-217
chr9-117918807-DEL-2478
chr9-117919665-DEL-2376
chr9-117919888-DEL-1346
chr9-117920759-DEL-926
chr9-117921428-DEL-339
chr9-117940863-DEL-370
chr9-117942566-DEL-74
chr9-117943059-DEL-100
chr9-117945049-DEL-194
chr9-117963376-DEL-915
chr9-117975509-DEL-398
chr9-117999536-DEL-100
chr9-118019536-DEL-62
chr9-118030265-DEL-484

chr9-118047273-DEL-550
chr9-118061924-DEL-300
chr9-118069510-DEL-51
chr9-118093028-DEL-736
chr9-118127983-DEL-194
chr9-118151913-DEL-79
chr9-118171979-DEL-142
chr9-118181877-DEL-78
chr9-118191604-DEL-145
chr9-118219718-DEL-77
chr9-118227836-DEL-76
chr9-118264252-DEL-342
chr9-118271557-DEL-64
chr9-118279588-DEL-60
chr9-118283114-DEL-51
chr9-118286743-DEL-152
chr9-118319496-DEL-172
chr9-118319913-DEL-400
chr9-118331372-DEL-861
chr9-118334179-DEL-77
chr9-118378317-DEL-200
chr9-118382703-DEL-70
chr9-118383272-DEL-74
chr9-118417348-DEL-63
chr9-118423268-DEL-325
chr9-118430185-DEL-59
chr9-118434016-DEL-95
chr9-118449399-DEL-112
chr9-118469427-DEL-689
chr9-118485004-DEL-56
chr9-118493698-DEL-3127
chr9-118516447-DEL-96
chr9-118521067-DEL-144
chr9-118552364-DEL-67
chr9-118554805-DEL-1949
chr9-118584516-DEL-451
chr9-118627034-DEL-222
chr9-118649781-DEL-285
chr9-118659463-DEL-55
chr9-118666479-DEL-686
chr9-118680502-DEL-122
chr9-118774763-DEL-1321
chr9-118775101-DEL-52
chr9-118777560-DEL-98
chr9-118784667-DEL-68
chr9-118820100-DEL-68
chr9-118822211-DEL-82
chr9-118828074-DEL-309
chr9-118835978-DEL-70
chr9-118876747-DEL-79
chr9-118879148-DEL-71
chr9-118889111-DEL-218
chr9-118952803-DEL-96
chr9-119025676-DEL-56
chr9-119029322-DEL-582
chr9-119034329-DEL-741
chr9-119037530-DEL-86
chr9-119038906-DEL-1849
chr9-119042514-DEL-94
chr9-119050587-DEL-230
chr9-119111213-DEL-59
chr9-119125611-DEL-2844
chr9-119128514-DEL-1918
chr9-119156406-DEL-197
chr9-119190623-DEL-63
chr9-119195116-DEL-196
chr9-119200732-DEL-97
chr9-119202983-DEL-68
chr9-119203546-DEL-2009
chr9-119215787-DEL-87
chr9-119247498-DEL-114
chr9-119250584-DEL-51
chr9-119301434-DEL-72
chr9-119305617-DEL-380
chr9-119329777-DEL-90
chr9-119338999-DEL-72
chr9-119358425-DEL-415

chr9-119400037-DEL-60
chr9-119431551-DEL-123
chr9-119442308-DEL-80
chr9-119466651-DEL-160
chr9-119477715-DEL-96
chr9-119492227-DEL-65
chr9-119504077-DEL-88
chr9-119531894-DEL-304
chr9-119532401-DEL-88
chr9-119585645-DEL-53
chr9-119607857-DEL-90
chr9-119627025-DEL-70
chr9-119628999-DEL-59
chr9-119656968-DEL-169
chr9-119690270-DEL-14505
chr9-119712128-DEL-58
chr9-119715503-DEL-391
chr9-119742546-DEL-103
chr9-119748986-DEL-51
chr9-119757613-DEL-67
chr9-119759754-DEL-1257
chr9-119777812-DEL-1004
chr9-119805230-DEL-91
chr9-119816574-DEL-292
chr9-119819085-DEL-786
chr9-119825432-DEL-125
chr9-119835018-DEL-93
chr9-119897636-DEL-64
chr9-119914164-DEL-239
chr9-119924892-DEL-164
chr9-119943596-DEL-163
chr9-119964134-DEL-148
chr9-119977574-DEL-389
chr9-119987863-DEL-70
chr9-120056145-DEL-52
chr9-120058550-DEL-52
chr9-120063260-DEL-56
chr9-120074080-DEL-786
chr9-120116622-DEL-336
chr9-120184864-DEL-250
chr9-120200662-DEL-153
chr9-120214986-DEL-440
chr9-120236231-DEL-86
chr9-120236666-DEL-172
chr9-120266185-DEL-67
chr9-120275935-DEL-70
chr9-120283333-DEL-54
chr9-120287326-DEL-53
chr9-120288723-DEL-228
chr9-120305370-DEL-86
chr9-120310814-DEL-56
chr9-120365394-DEL-99
chr9-120378505-DEL-95
chr9-120404635-DEL-196
chr9-120416597-DEL-239
chr9-120452402-DEL-879
chr9-120460211-DEL-98
chr9-120477158-DEL-102
chr9-120484257-DEL-523
chr9-120517265-DEL-793
chr9-120517916-DEL-59
chr9-120549518-DEL-67
chr9-120554669-DEL-179
chr9-120557252-DEL-358
chr9-120581110-DEL-181
chr9-120600837-DEL-52
chr9-120611606-DEL-2578
chr9-120621906-DEL-447
chr9-120627040-DEL-120
chr9-120628670-DEL-1862
chr9-120645655-DEL-88
chr9-120646118-DEL-128
chr9-120675641-DEL-181
chr9-120689654-DEL-126
chr9-120715273-DEL-88
chr9-120727583-DEL-110
chr9-120732714-DEL-134

chr9-120811334-DEL-216
chr9-120821008-DEL-68
chr9-120824564-DEL-76
chr9-120864806-DEL-66
chr9-120869855-DEL-1678
chr9-120870082-DEL-1091
chr9-120870994-DEL-251
chr9-120918247-DEL-351
chr9-120933242-DEL-562
chr9-120944877-DEL-80
chr9-120948752-DEL-84
chr9-120949717-DEL-61
chr9-120961288-DEL-177
chr9-120964421-DEL-80
chr9-120965386-DEL-120
chr9-120976596-DEL-654
chr9-120980331-DEL-112
chr9-120992685-DEL-72
chr9-120999604-DEL-60
chr9-121014037-DEL-60
chr9-121017588-DEL-183
chr9-121019892-DEL-52
chr9-121041110-DEL-307
chr9-121049286-DEL-328
chr9-121049305-DEL-11144
chr9-121049833-DEL-18044
chr9-121050291-DEL-87
chr9-121050779-DEL-112
chr9-121050827-DEL-1285
chr9-121051407-DEL-83
chr9-121051736-DEL-417
chr9-121052636-DEL-286
chr9-121053396-DEL-139
chr9-121054185-DEL-255
chr9-121054774-DEL-174
chr9-121055609-DEL-162
chr9-121056186-DEL-174
chr9-121057577-DEL-248
chr9-121058124-DEL-1335
chr9-121058546-DEL-62
chr9-121059137-DEL-255
chr9-121059589-DEL-433
chr9-121060438-DEL-161
chr9-121060550-DEL-1236
chr9-121060884-DEL-1904
chr9-121060910-DEL-329
chr9-121061375-DEL-105
chr9-121061963-DEL-752
chr9-121062526-DEL-479
chr9-121062773-DEL-3123
chr9-121062816-DEL-5155
chr9-121062927-DEL-299
chr9-121064590-DEL-379
chr9-121065836-DEL-174
chr9-121066333-DEL-1348
chr9-121066446-DEL-323
chr9-121066507-DEL-143
chr9-121067761-DEL-211
chr9-121068286-DEL-151
chr9-121097699-DEL-722
chr9-121098801-DEL-1036
chr9-121098835-DEL-983
chr9-121099783-DEL-53
chr9-121122279-DEL-78
chr9-121124322-DEL-75
chr9-121136753-DEL-384
chr9-121149979-DEL-135
chr9-121168129-DEL-187
chr9-121192264-DEL-52
chr9-121194310-DEL-62
chr9-121198146-DEL-216
chr9-121207955-DEL-72
chr9-121210017-DEL-99
chr9-121217491-DEL-61
chr9-121218183-DEL-170
chr9-121256970-DEL-51
chr9-121291933-DEL-174

chr9-121332957-DEL-1416
chr9-121351096-DEL-55
chr9-121385642-DEL-175
chr9-121388604-DEL-102
chr9-121433552-DEL-528
chr9-121436156-DEL-118
chr9-121437230-DEL-392
chr9-121464342-DEL-82
chr9-121466383-DEL-78
chr9-121485370-DEL-75
chr9-121491037-DEL-213
chr9-121526776-DEL-225
chr9-121530220-DEL-172
chr9-121530745-DEL-808
chr9-121562028-DEL-58
chr9-121593656-DEL-100
chr9-121602579-DEL-80
chr9-121691141-DEL-58
chr9-121720998-DEL-197
chr9-121733991-DEL-587
chr9-121760309-DEL-867
chr9-121770598-DEL-68
chr9-121782714-DEL-616
chr9-121789741-DEL-400
chr9-121791589-DEL-61
chr9-121797743-DEL-880
chr9-121815981-DEL-54
chr9-121818039-DEL-54
chr9-121838546-DEL-96
chr9-121841412-DEL-80
chr9-121854004-DEL-276
chr9-121855708-DEL-86
chr9-121909251-DEL-226
chr9-121927438-DEL-224
chr9-121929143-DEL-160
chr9-121934696-DEL-131
chr9-121958789-DEL-307
chr9-121990466-DEL-1702
chr9-121990712-DEL-368
chr9-122001219-DEL-1706
chr9-122016531-DEL-1967
chr9-122027332-DEL-679
chr9-122030291-DEL-169
chr9-122044645-DEL-316
chr9-122053794-DEL-51
chr9-122074893-DEL-190
chr9-122086931-DEL-429
chr9-122092466-DEL-60
chr9-122169160-DEL-112
chr9-122176212-DEL-287
chr9-122176934-DEL-63
chr9-122204145-DEL-156
chr9-122219538-DEL-136
chr9-122278758-DEL-1681
chr9-122288272-DEL-1220
chr9-122304293-DEL-88
chr9-122318501-DEL-56
chr9-122320097-DEL-384
chr9-122335987-DEL-182
chr9-122343160-DEL-64
chr9-122356452-DEL-256
chr9-122362414-DEL-138
chr9-122371614-DEL-102
chr9-122372053-DEL-100
chr9-122373058-DEL-54
chr9-122374308-DEL-102
chr9-122403143-DEL-66
chr9-122435007-DEL-130
chr9-122435772-DEL-94
chr9-122470257-DEL-93
chr9-122477515-DEL-594
chr9-122501380-DEL-1133
chr9-122517124-DEL-56
chr9-122546494-DEL-2409
chr9-122590461-DEL-1249
chr9-122590591-DEL-758
chr9-122591135-DEL-61

chr9-122609218-DEL-93
chr9-122617289-DEL-56
chr9-122640408-DEL-246
chr9-122676888-DEL-548
chr9-122684167-DEL-545
chr9-122692531-DEL-302
chr9-122717049-DEL-90
chr9-122733351-DEL-2054
chr9-122733353-DEL-91
chr9-122733644-DEL-325
chr9-122734372-DEL-351
chr9-122735011-DEL-52
chr9-122736272-DEL-60
chr9-122795855-DEL-86
chr9-122821814-DEL-60
chr9-122858636-DEL-190
chr9-122859786-DEL-366
chr9-122868257-DEL-69
chr9-122878223-DEL-215
chr9-122893922-DEL-63
chr9-122943692-DEL-167
chr9-122951883-DEL-54
chr9-122979286-DEL-167
chr9-122997214-DEL-83
chr9-123010547-DEL-201
chr9-123031317-DEL-54
chr9-123040689-DEL-62
chr9-123053456-DEL-53
chr9-123073865-DEL-55
chr9-123077768-DEL-63
chr9-123080116-DEL-142
chr9-123093288-DEL-68
chr9-123114122-DEL-128
chr9-123137740-DEL-80
chr9-123180079-DEL-82
chr9-123182784-DEL-482
chr9-123198394-DEL-841
chr9-123200278-DEL-81
chr9-123224987-DEL-82
chr9-123243419-DEL-426
chr9-123275415-DEL-772
chr9-123276283-DEL-242
chr9-123279577-DEL-270
chr9-123292336-DEL-205
chr9-123401305-DEL-198
chr9-123565495-DEL-1699
chr9-123568853-DEL-80
chr9-123570571-DEL-183
chr9-123586399-DEL-192
chr9-123601632-DEL-60
chr9-123602677-DEL-51
chr9-123620976-DEL-112
chr9-123624993-DEL-72
chr9-123656064-DEL-62
chr9-123686378-DEL-773
chr9-123695327-DEL-1714
chr9-123702208-DEL-61
chr9-123702650-DEL-18548
chr9-123721435-DEL-489
chr9-123729737-DEL-1821
chr9-123733733-DEL-2219
chr9-123733865-DEL-238
chr9-123744562-DEL-78
chr9-123751427-DEL-436
chr9-123760040-DEL-57
chr9-123786882-DEL-2378
chr9-123792199-DEL-2143
chr9-123792255-DEL-69
chr9-123803340-DEL-210
chr9-123842081-DEL-52
chr9-123857683-DEL-196
chr9-123897888-DEL-56
chr9-123917018-DEL-930
chr9-123932427-DEL-568
chr9-124017003-DEL-61
chr9-124024399-DEL-126
chr9-124037120-DEL-2444

chr9-124110823-DEL-6204
chr9-124129209-DEL-64
chr9-124198900-DEL-69
chr9-124213486-DEL-979
chr9-124215413-DEL-15994
chrX-3055926-DEL-620
chrX-3246315-DEL-57
chrX-3431979-DEL-395
chrX-3477367-DEL-53
chrX-3737424-DEL-540
chrX-3744717-DEL-1627
chrX-3770274-DEL-64
chrX-3835754-DEL-3273
chrX-3866334-DEL-828
chrX-3935234-DEL-76
chrX-3953523-DEL-5612
chrX-4035562-DEL-78
chrX-4038177-DEL-79
chrX-4040229-DEL-1163
chrX-4041522-DEL-395
chrX-4043798-DEL-1596
chrX-4047681-DEL-53
chrX-4106847-DEL-52
chrX-4258505-DEL-4002
chrX-4347220-DEL-3230
chrX-4361846-DEL-5249
chrX-4395506-DEL-1907
chrX-4406309-DEL-4199
chrX-4500213-DEL-288
chrX-4502214-DEL-60
chrX-4516510-DEL-5248
chrX-4539388-DEL-58
chrX-4543624-DEL-2874
chrX-4722648-DEL-58
chrX-4828808-DEL-55
chrX-5013995-DEL-558
chrX-5040460-DEL-395
chrX-5090337-DEL-468
chrX-5155438-DEL-288
chrX-5207797-DEL-85
chrX-5235409-DEL-12187
chrX-5278062-DEL-5507
chrX-5361316-DEL-988
chrX-5362658-DEL-79
chrX-5386949-DEL-2139
chrX-5455493-DEL-332
chrX-5461054-DEL-7561
chrX-5473378-DEL-6466
chrX-5534268-DEL-3868
chrX-5546252-DEL-6376
chrX-5546572-DEL-199
chrX-5553583-DEL-480
chrX-5609570-DEL-4558
chrX-5710975-DEL-139
chrX-5755921-DEL-1899
chrX-5826922-DEL-939
chrX-5851728-DEL-215
chrX-5913903-DEL-125
chrX-5923415-DEL-120
chrX-5931316-DEL-419
chrX-5950823-DEL-56
chrX-5966893-DEL-388
chrX-5967545-DEL-703
chrX-5968645-DEL-99
chrX-5977963-DEL-1288
chrX-6034366-DEL-59
chrX-6091627-DEL-989
chrX-6095640-DEL-56
chrX-6109516-DEL-131
chrX-6137852-DEL-740
chrX-6200764-DEL-3345
chrX-6214063-DEL-451
chrX-6247676-DEL-311
chrX-6258132-DEL-288
chrX-6266708-DEL-53
chrX-6276712-DEL-105
chrX-6285783-DEL-2854

chrX-6297301-DEL-2741
chrX-6307378-DEL-108
chrX-6316151-DEL-111
chrX-6330869-DEL-66
chrX-6381081-DEL-3339
chrX-6404634-DEL-203
chrX-6409660-DEL-86
chrX-6423066-DEL-78
chrX-6447331-DEL-893
chrX-6501485-DEL-74
chrX-6541590-DEL-328
chrX-6616204-DEL-790
chrX-6631771-DEL-1068
chrX-6633554-DEL-1257
chrX-6640291-DEL-88
chrX-6690092-DEL-59
chrX-6698155-DEL-300
chrX-6766990-DEL-171
chrX-6822202-DEL-66
chrX-6836308-DEL-59
chrX-6850300-DEL-164
chrX-6856918-DEL-1640
chrX-6877278-DEL-491
chrX-6878940-DEL-539
chrX-6880178-DEL-92
chrX-6881743-DEL-102
chrX-6951180-DEL-261
chrX-6953804-DEL-475
chrX-7010243-DEL-62
chrX-7018932-DEL-539
chrX-7049108-DEL-195
chrX-7066269-DEL-52
chrX-7083669-DEL-218
chrX-7084464-DEL-74
chrX-7098163-DEL-1794
chrX-7100208-DEL-289
chrX-7120671-DEL-215
chrX-7209261-DEL-168
chrX-7265493-DEL-769
chrX-7284325-DEL-72
chrX-7293459-DEL-972
chrX-7314446-DEL-269
chrX-7325255-DEL-2282
chrX-7327592-DEL-211
chrX-7353120-DEL-163
chrX-7417682-DEL-828
chrX-7418778-DEL-58
chrX-7456786-DEL-71
chrX-7518727-DEL-78
chrX-7528616-DEL-62
chrX-7612035-DEL-55
chrX-7620847-DEL-400
chrX-7654730-DEL-84
chrX-7682182-DEL-497
chrX-7698813-DEL-64
chrX-7715881-DEL-181
chrX-7771562-DEL-175
chrX-7783968-DEL-241
chrX-7807028-DEL-80
chrX-7815415-DEL-107
chrX-7815864-DEL-212
chrX-7821122-DEL-202
chrX-7857180-DEL-184
chrX-7873788-DEL-266
chrX-7891140-DEL-149
chrX-7901321-DEL-72
chrX-7912373-DEL-72
chrX-7920957-DEL-76
chrX-7965133-DEL-96
chrX-7969180-DEL-59
chrX-7991320-DEL-108
chrX-8038495-DEL-288
chrX-8076807-DEL-1259
chrX-8087483-DEL-815
chrX-8099932-DEL-470
chrX-8112343-DEL-161
chrX-8121495-DEL-199

chrX-8169680-DEL-570
chrX-8278114-DEL-702
chrX-8322876-DEL-5886
chrX-8338252-DEL-27915
chrX-8378544-DEL-718
chrX-8416170-DEL-1655
chrX-8424074-DEL-180
chrX-8425746-DEL-112
chrX-8490469-DEL-510
chrX-8493467-DEL-72
chrX-8500844-DEL-647
chrX-8502795-DEL-1149
chrX-8503155-DEL-82
chrX-8522233-DEL-6249
chrX-8557228-DEL-1340
chrX-8631481-DEL-4826
chrX-8691397-DEL-1662
chrX-8703174-DEL-183
chrX-8738414-DEL-72
chrX-8742725-DEL-320
chrX-8743915-DEL-1034
chrX-8778172-DEL-1504
chrX-8807070-DEL-338
chrX-8830120-DEL-72
chrX-8843424-DEL-139
chrX-8858503-DEL-278
chrX-8922280-DEL-3036
chrX-8939040-DEL-3036
chrX-8945022-DEL-144
chrX-8975649-DEL-406
chrX-8990927-DEL-152
chrX-9022883-DEL-1078
chrX-9084507-DEL-59
chrX-9110600-DEL-325
chrX-9114678-DEL-71
chrX-9161086-DEL-922
chrX-9161187-DEL-1126
chrX-9161771-DEL-408
chrX-9163202-DEL-52
chrX-9211017-DEL-368
chrX-9222573-DEL-9576
chrX-9233411-DEL-4014
chrX-9248824-DEL-2499
chrX-9251334-DEL-7803
chrX-9251807-DEL-3324
chrX-9255613-DEL-6358
chrX-9259642-DEL-2329
chrX-9274408-DEL-54
chrX-9276466-DEL-107
chrX-9311725-DEL-99
chrX-9387764-DEL-3472
chrX-9419765-DEL-471
chrX-9440896-DEL-107
chrX-9445539-DEL-1035
chrX-9446831-DEL-548
chrX-9447545-DEL-53
chrX-9448249-DEL-322
chrX-9449109-DEL-628
chrX-9450194-DEL-53
chrX-9454077-DEL-54
chrX-9471568-DEL-268
chrX-9473180-DEL-422
chrX-9474455-DEL-107
chrX-9502147-DEL-54
chrX-9502955-DEL-140
chrX-9505391-DEL-2131
chrX-9512877-DEL-106
chrX-9515984-DEL-270
chrX-9524648-DEL-57
chrX-9542557-DEL-308
chrX-9572365-DEL-1909
chrX-9577429-DEL-2422
chrX-9584651-DEL-6409
chrX-9592655-DEL-109
chrX-9604493-DEL-268
chrX-9677271-DEL-80
chrX-9682802-DEL-200

chrX-9754324-DEL-82
chrX-9756017-DEL-6425
chrX-9765355-DEL-1985
chrX-9780717-DEL-116
chrX-9785368-DEL-6884
chrX-9812081-DEL-185
chrX-9824511-DEL-52
chrX-9830652-DEL-82
chrX-9832171-DEL-6490
chrX-9849731-DEL-838
chrX-9911884-DEL-151
chrX-9926245-DEL-67
chrX-9950620-DEL-1522
chrX-10026344-DEL-286
chrX-10029340-DEL-78
chrX-10062552-DEL-66
chrX-10077104-DEL-84
chrX-10097434-DEL-63
chrX-10116110-DEL-68
chrX-10129638-DEL-64
chrX-10132043-DEL-168
chrX-10148072-DEL-68
chrX-10168908-DEL-211
chrX-10200309-DEL-165
chrX-10225805-DEL-3870
chrX-10338899-DEL-1929
chrX-10340664-DEL-182
chrX-10443349-DEL-167
chrX-10516818-DEL-62
chrX-10523490-DEL-1566
chrX-10526711-DEL-2488
chrX-10560605-DEL-66
chrX-10565546-DEL-55
chrX-10579005-DEL-91
chrX-10615472-DEL-53
chrX-10661425-DEL-2835
chrX-10666395-DEL-540
chrX-10719936-DEL-51
chrX-10729047-DEL-218
chrX-10816371-DEL-6024
chrX-10826849-DEL-188
chrX-10831550-DEL-424
chrX-10843988-DEL-94
chrX-10876502-DEL-414
chrX-10883041-DEL-85
chrX-10896912-DEL-97
chrX-10911451-DEL-64
chrX-10939651-DEL-204
chrX-10953712-DEL-192
chrX-10958316-DEL-360
chrX-10979296-DEL-168
chrX-11028460-DEL-63
chrX-11042510-DEL-453
chrX-11066613-DEL-59
chrX-11068874-DEL-61
chrX-11168001-DEL-6846
chrX-11168420-DEL-154
chrX-11193757-DEL-155
chrX-11240767-DEL-70
chrX-11276172-DEL-72
chrX-11319455-DEL-51
chrX-11329004-DEL-68
chrX-11339427-DEL-102
chrX-11395921-DEL-66
chrX-11403106-DEL-70
chrX-11422173-DEL-51
chrX-11424008-DEL-90
chrX-11428813-DEL-93
chrX-11448176-DEL-981
chrX-11452279-DEL-53
chrX-11522385-DEL-540
chrX-11571149-DEL-131
chrX-11598591-DEL-82
chrX-11682846-DEL-199
chrX-11700144-DEL-225
chrX-11704672-DEL-82
chrX-11707686-DEL-255

chrX-11708255-DEL-140
chrX-11735771-DEL-93
chrX-11802485-DEL-242
chrX-11827434-DEL-66
chrX-11893470-DEL-58
chrX-11967643-DEL-238
chrX-12005432-DEL-52
chrX-12009975-DEL-155
chrX-12035250-DEL-62
chrX-12038458-DEL-65
chrX-12077146-DEL-90
chrX-12083810-DEL-156
chrX-12108675-DEL-2817
chrX-12131777-DEL-1528
chrX-12132574-DEL-180
chrX-12160186-DEL-52
chrX-12187259-DEL-205
chrX-12187925-DEL-90
chrX-12192919-DEL-72
chrX-12237211-DEL-770
chrX-12261013-DEL-2004
chrX-12272150-DEL-957
chrX-12307107-DEL-1447
chrX-12317578-DEL-358
chrX-12354540-DEL-155
chrX-12359214-DEL-62
chrX-12402057-DEL-470
chrX-12416664-DEL-64
chrX-12430577-DEL-473
chrX-12448176-DEL-66
chrX-12449324-DEL-3145
chrX-12463509-DEL-3189
chrX-12465042-DEL-205
chrX-12467436-DEL-281
chrX-12500867-DEL-170
chrX-12501583-DEL-236
chrX-12502123-DEL-1293
chrX-12509434-DEL-190
chrX-12530415-DEL-68
chrX-12530932-DEL-2234
chrX-12587546-DEL-65
chrX-12595018-DEL-200
chrX-12599225-DEL-67
chrX-12618342-DEL-188
chrX-12650028-DEL-91
chrX-12650777-DEL-172
chrX-12680937-DEL-60
chrX-12685471-DEL-85
chrX-12695226-DEL-73
chrX-12709465-DEL-647
chrX-12717874-DEL-52
chrX-12752747-DEL-65
chrX-12754917-DEL-52
chrX-12761070-DEL-56
chrX-12814203-DEL-67
chrX-12840252-DEL-132
chrX-12872285-DEL-591
chrX-12971751-DEL-68
chrX-12980970-DEL-127
chrX-13018844-DEL-138
chrX-13033857-DEL-143
chrX-13080817-DEL-159
chrX-13085474-DEL-108
chrX-13086631-DEL-181
chrX-13088149-DEL-66
chrX-13098477-DEL-212
chrX-13103458-DEL-52
chrX-13117239-DEL-124
chrX-13127335-DEL-134
chrX-13135888-DEL-373
chrX-13153121-DEL-74
chrX-13160562-DEL-297
chrX-13206061-DEL-53
chrX-13228131-DEL-116
chrX-13231846-DEL-1424
chrX-13252836-DEL-61
chrX-13253941-DEL-355

chrX-13260432-DEL-1440
chrX-13270823-DEL-198
chrX-13292293-DEL-199
chrX-13306999-DEL-167
chrX-13320904-DEL-139
chrX-13323896-DEL-53
chrX-13343252-DEL-66
chrX-13343918-DEL-124
chrX-13365583-DEL-58
chrX-13377082-DEL-58
chrX-13438822-DEL-158
chrX-13456829-DEL-85
chrX-13476917-DEL-184
chrX-13500165-DEL-102
chrX-13529412-DEL-97
chrX-13556539-DEL-79
chrX-13628980-DEL-196
chrX-13658622-DEL-90
chrX-13674565-DEL-130
chrX-13732595-DEL-88
chrX-13791463-DEL-78
chrX-13797765-DEL-1042
chrX-13810401-DEL-1217
chrX-13887650-DEL-197
chrX-13911531-DEL-496
chrX-14027953-DEL-130
chrX-14079513-DEL-771
chrX-14106209-DEL-62
chrX-14111099-DEL-66
chrX-14157287-DEL-1498
chrX-14176631-DEL-517
chrX-14184964-DEL-288
chrX-14189293-DEL-442
chrX-14232463-DEL-53
chrX-14248039-DEL-3231
chrX-14276857-DEL-6643
chrX-14285406-DEL-63
chrX-14292466-DEL-99
chrX-14324666-DEL-1693
chrX-14338779-DEL-401
chrX-14340111-DEL-2380
chrX-14380086-DEL-603
chrX-14426198-DEL-6273
chrX-14436500-DEL-8176
chrX-14436695-DEL-424
chrX-14437787-DEL-636
chrX-14451256-DEL-6728
chrX-14481962-DEL-268
chrX-14486764-DEL-115
chrX-14539195-DEL-90
chrX-14541226-DEL-1745
chrX-14558616-DEL-128
chrX-14582021-DEL-468
chrX-14671691-DEL-3592
chrX-14683741-DEL-204
chrX-14685803-DEL-5110
chrX-14714726-DEL-387
chrX-14737377-DEL-55
chrX-14773711-DEL-53
chrX-14816824-DEL-835
chrX-14818028-DEL-2616
chrX-14832159-DEL-3229
chrX-14842094-DEL-66
chrX-14846191-DEL-76
chrX-14873729-DEL-1493
chrX-14988669-DEL-1790
chrX-15001109-DEL-493
chrX-15004030-DEL-54
chrX-15021877-DEL-55
chrX-15043241-DEL-905
chrX-15085512-DEL-121
chrX-15153211-DEL-71
chrX-15155371-DEL-94
chrX-15210565-DEL-1404
chrX-15221153-DEL-134
chrX-15230542-DEL-64
chrX-15235961-DEL-58

chrX-15236807-DEL-1556
chrX-15291548-DEL-218
chrX-15353128-DEL-6354
chrX-15380341-DEL-324
chrX-15411174-DEL-249
chrX-15441768-DEL-119
chrX-15491455-DEL-205
chrX-15500361-DEL-86
chrX-15503647-DEL-4232
chrX-15551538-DEL-1231
chrX-15553207-DEL-2186
chrX-15555462-DEL-455
chrX-15556038-DEL-4357
chrX-15558293-DEL-3868
chrX-15597350-DEL-62
chrX-15606893-DEL-6212
chrX-15620147-DEL-637
chrX-15635608-DEL-96
chrX-15644411-DEL-709
chrX-15719303-DEL-538
chrX-15743235-DEL-2064
chrX-15746242-DEL-73
chrX-15764416-DEL-108
chrX-15799384-DEL-6403
chrX-15812621-DEL-211
chrX-15828847-DEL-7448
chrX-15837575-DEL-60
chrX-15855254-DEL-191
chrX-15903835-DEL-2443
chrX-15929488-DEL-560
chrX-15941924-DEL-6314
chrX-15951591-DEL-873
chrX-15954127-DEL-498
chrX-15958777-DEL-483
chrX-15973743-DEL-9629
chrX-15978321-DEL-888
chrX-15981375-DEL-394
chrX-15990441-DEL-642
chrX-16037644-DEL-205
chrX-16061575-DEL-58
chrX-16076606-DEL-6201
chrX-16084892-DEL-712
chrX-16105378-DEL-762
chrX-16105972-DEL-1224
chrX-16156876-DEL-99
chrX-16157898-DEL-268
chrX-16165036-DEL-277
chrX-16168774-DEL-2259
chrX-16172250-DEL-204
chrX-16186899-DEL-1591
chrX-16190286-DEL-2023
chrX-16196673-DEL-578
chrX-16197475-DEL-54
chrX-16205557-DEL-5502
chrX-16218348-DEL-112
chrX-16237267-DEL-528
chrX-16262262-DEL-268
chrX-16262780-DEL-6602
chrX-16286034-DEL-94
chrX-16289822-DEL-1037
chrX-16303980-DEL-51
chrX-16314716-DEL-425
chrX-16326290-DEL-93
chrX-16337391-DEL-6315
chrX-16347997-DEL-854
chrX-16355546-DEL-6213
chrX-16362215-DEL-2956
chrX-16376557-DEL-91
chrX-16404002-DEL-985
chrX-16416356-DEL-559
chrX-16417141-DEL-175
chrX-16421637-DEL-119
chrX-16435617-DEL-136
chrX-16439228-DEL-6667
chrX-16444846-DEL-781
chrX-16495087-DEL-566
chrX-16615839-DEL-383

chrX-16619724-DEL-705
chrX-16624998-DEL-519
chrX-16638642-DEL-167
chrX-16664420-DEL-1166
chrX-16670452-DEL-89
chrX-16710556-DEL-150
chrX-16724965-DEL-53
chrX-16765944-DEL-409
chrX-16773539-DEL-56
chrX-16859544-DEL-389
chrX-16894078-DEL-68
chrX-16913649-DEL-65
chrX-16934415-DEL-56
chrX-16975714-DEL-336
chrX-16992945-DEL-234
chrX-17007514-DEL-56
chrX-17058363-DEL-339
chrX-17104378-DEL-197
chrX-17154949-DEL-368
chrX-17161662-DEL-527
chrX-17228264-DEL-197
chrX-17373176-DEL-725
chrX-17376331-DEL-9337
chrX-17390076-DEL-486
chrX-17394049-DEL-77
chrX-17413089-DEL-83
chrX-17415631-DEL-388
chrX-17416234-DEL-61
chrX-17418130-DEL-65
chrX-17443805-DEL-3056
chrX-17474126-DEL-304
chrX-17495302-DEL-346
chrX-17507710-DEL-12726
chrX-17533362-DEL-982
chrX-17539987-DEL-163
chrX-17560279-DEL-64
chrX-17568562-DEL-226
chrX-17608935-DEL-3363
chrX-17631030-DEL-60
chrX-17708337-DEL-60
chrX-17721361-DEL-424
chrX-17722699-DEL-56
chrX-17732958-DEL-73
chrX-17743849-DEL-871
chrX-17749971-DEL-105
chrX-17764044-DEL-66
chrX-17773196-DEL-198
chrX-17782283-DEL-105
chrX-17796590-DEL-480
chrX-17804469-DEL-206
chrX-17822310-DEL-3886
chrX-17838032-DEL-1650
chrX-17842487-DEL-2734
chrX-17875801-DEL-105
chrX-17891112-DEL-6215
chrX-17902019-DEL-85
chrX-17933952-DEL-6585
chrX-17957849-DEL-1563
chrX-17980388-DEL-292
chrX-18019654-DEL-51
chrX-18035549-DEL-2211
chrX-18068563-DEL-222
chrX-18069406-DEL-709
chrX-18097804-DEL-57
chrX-18138694-DEL-359
chrX-18154255-DEL-14032
chrX-18155082-DEL-12873
chrX-18159561-DEL-6439
chrX-18168787-DEL-6440
chrX-18180300-DEL-58
chrX-18180750-DEL-59
chrX-18187354-DEL-1715
chrX-18190400-DEL-71
chrX-18194221-DEL-91
chrX-18201308-DEL-5571
chrX-18224508-DEL-230
chrX-18252478-DEL-1522

chrX-18338473-DEL-1827
chrX-18394443-DEL-169
chrX-18404662-DEL-7269
chrX-18404838-DEL-212
chrX-18421219-DEL-6225
chrX-18428364-DEL-54
chrX-18430688-DEL-104
chrX-18445828-DEL-165
chrX-18450263-DEL-6592
chrX-18459168-DEL-71
chrX-18463166-DEL-1698
chrX-18467562-DEL-58
chrX-18499360-DEL-58
chrX-18706080-DEL-223
chrX-18784933-DEL-605
chrX-18794664-DEL-530
chrX-18812849-DEL-1608
chrX-18831321-DEL-324
chrX-18832955-DEL-133
chrX-18837044-DEL-78
chrX-18854794-DEL-71
chrX-18857680-DEL-264
chrX-18927887-DEL-268
chrX-18947512-DEL-140
chrX-18968128-DEL-2057
chrX-18981673-DEL-68
chrX-18989330-DEL-6222
chrX-19017610-DEL-230
chrX-19043814-DEL-191
chrX-19064154-DEL-71
chrX-19109307-DEL-125
chrX-19134428-DEL-2980
chrX-19141123-DEL-813
chrX-19144564-DEL-57
chrX-19209539-DEL-67
chrX-19216270-DEL-263
chrX-19221444-DEL-309
chrX-19258216-DEL-462
chrX-19274758-DEL-181
chrX-19328448-DEL-193
chrX-19380774-DEL-1030
chrX-19382089-DEL-193
chrX-19393157-DEL-330
chrX-19396025-DEL-120
chrX-19434875-DEL-336
chrX-19439191-DEL-1078
chrX-19451866-DEL-507
chrX-19487387-DEL-62
chrX-19490604-DEL-136
chrX-19501033-DEL-390
chrX-19503714-DEL-1382
chrX-19505306-DEL-2976
chrX-19509141-DEL-4644
chrX-19517516-DEL-6506
chrX-19524942-DEL-116
chrX-19525742-DEL-804
chrX-19632745-DEL-299
chrX-19645161-DEL-1908
chrX-19647517-DEL-2973
chrX-19744739-DEL-613
chrX-19747468-DEL-57
chrX-19763139-DEL-69
chrX-19792028-DEL-3392
chrX-19798447-DEL-3072
chrX-19826268-DEL-1878
chrX-19833238-DEL-1997
chrX-19857229-DEL-157
chrX-19906954-DEL-80
chrX-19908510-DEL-425
chrX-19921738-DEL-470
chrX-19945720-DEL-1228
chrX-19983358-DEL-3180
chrX-20044605-DEL-1220
chrX-20078843-DEL-92
chrX-20080784-DEL-134
chrX-20104069-DEL-270
chrX-20107264-DEL-856

chrX-20127822-DEL-1512
chrX-20145801-DEL-74
chrX-20182659-DEL-68
chrX-20185319-DEL-612
chrX-20198066-DEL-144
chrX-20207263-DEL-162
chrX-20210801-DEL-109
chrX-20240196-DEL-104
chrX-20304158-DEL-176
chrX-20316143-DEL-102
chrX-20333539-DEL-346
chrX-20346775-DEL-1713
chrX-20399674-DEL-121
chrX-20446184-DEL-3943
chrX-20460511-DEL-6019
chrX-20479197-DEL-751
chrX-20521694-DEL-52
chrX-20526059-DEL-676
chrX-20605160-DEL-60
chrX-20611672-DEL-139
chrX-20618049-DEL-57
chrX-20620163-DEL-202
chrX-20621365-DEL-74
chrX-20628603-DEL-6420
chrX-20645414-DEL-1804
chrX-20652698-DEL-2774
chrX-20674824-DEL-367
chrX-20696277-DEL-371
chrX-20757232-DEL-51
chrX-20769119-DEL-196
chrX-20777281-DEL-53
chrX-20865400-DEL-6964
chrX-20881479-DEL-51
chrX-20909364-DEL-52
chrX-20975329-DEL-407
chrX-21052631-DEL-235
chrX-21054486-DEL-120
chrX-21062078-DEL-7525
chrX-21069821-DEL-2799
chrX-21073945-DEL-111
chrX-21075101-DEL-60
chrX-21101244-DEL-217
chrX-21102359-DEL-9947
chrX-21112033-DEL-214
chrX-21120857-DEL-7153
chrX-21127382-DEL-412
chrX-21147489-DEL-233
chrX-21150294-DEL-488
chrX-21157174-DEL-94
chrX-21198951-DEL-1646
chrX-21204693-DEL-186
chrX-21243673-DEL-79
chrX-21265037-DEL-303
chrX-21310586-DEL-69
chrX-21322120-DEL-104
chrX-21327106-DEL-1247
chrX-21393244-DEL-308
chrX-21404502-DEL-72
chrX-21485125-DEL-5453
chrX-21544304-DEL-137
chrX-21546790-DEL-122
chrX-21548481-DEL-69
chrX-21566816-DEL-675
chrX-21673496-DEL-1203
chrX-21675923-DEL-662
chrX-21677862-DEL-189
chrX-21712352-DEL-5675
chrX-21712442-DEL-7312
chrX-21718104-DEL-1650
chrX-21738340-DEL-117
chrX-21745886-DEL-218
chrX-21756326-DEL-206
chrX-21793057-DEL-831
chrX-21816838-DEL-71
chrX-21864149-DEL-358
chrX-21887231-DEL-9109
chrX-21887839-DEL-6783

chrX-21894248-DEL-208
chrX-21896651-DEL-154
chrX-21900528-DEL-9153
chrX-21953738-DEL-55
chrX-21978507-DEL-62
chrX-21996208-DEL-6819
chrX-22016865-DEL-916
chrX-22054861-DEL-2527
chrX-22081381-DEL-129
chrX-22111755-DEL-1915
chrX-22144533-DEL-94
chrX-22158447-DEL-312
chrX-22165302-DEL-62
chrX-22170423-DEL-403
chrX-22174139-DEL-175
chrX-22180480-DEL-805
chrX-22186006-DEL-1309
chrX-22189994-DEL-992
chrX-22193900-DEL-353
chrX-22213481-DEL-621
chrX-22228083-DEL-61
chrX-22234385-DEL-66
chrX-22241804-DEL-178
chrX-22243802-DEL-73
chrX-22263090-DEL-211
chrX-22288072-DEL-60
chrX-22299347-DEL-1655
chrX-22301754-DEL-1166
chrX-22330373-DEL-353
chrX-22339843-DEL-87
chrX-22365362-DEL-10691
chrX-22366538-DEL-7015
chrX-22378984-DEL-6439
chrX-22385602-DEL-63
chrX-22386404-DEL-538
chrX-22394241-DEL-6464
chrX-22425265-DEL-70
chrX-22430166-DEL-7178
chrX-22439867-DEL-6476
chrX-22460118-DEL-635
chrX-22468018-DEL-437
chrX-22468718-DEL-2774
chrX-22471305-DEL-187
chrX-22476652-DEL-57
chrX-22480186-DEL-54
chrX-22490522-DEL-131
chrX-22492439-DEL-194
chrX-22498836-DEL-175
chrX-22509059-DEL-60
chrX-22561906-DEL-93
chrX-22578249-DEL-60
chrX-22612539-DEL-58
chrX-22619291-DEL-337
chrX-22624927-DEL-571
chrX-22649857-DEL-114
chrX-22658804-DEL-60
chrX-22662800-DEL-392
chrX-22713319-DEL-64
chrX-22725710-DEL-640
chrX-22739330-DEL-529
chrX-22740324-DEL-9332
chrX-22799617-DEL-2224
chrX-22809396-DEL-1089
chrX-22855670-DEL-2771
chrX-22904033-DEL-460
chrX-22935544-DEL-1951
chrX-22953014-DEL-1246
chrX-23008349-DEL-6189
chrX-23023922-DEL-153
chrX-23046210-DEL-56
chrX-23167287-DEL-66
chrX-23221106-DEL-578
chrX-23255673-DEL-1413
chrX-23259716-DEL-107
chrX-23320268-DEL-486
chrX-23327601-DEL-791
chrX-23368018-DEL-56

chrX-23387330-DEL-5194
chrX-23392653-DEL-1178
chrX-23393729-DEL-386
chrX-23401037-DEL-7168
chrX-23413825-DEL-66
chrX-23444004-DEL-53
chrX-23448828-DEL-14838
chrX-23470381-DEL-170
chrX-23480109-DEL-90
chrX-23518966-DEL-2499
chrX-23519499-DEL-100
chrX-23574695-DEL-183
chrX-23612502-DEL-94
chrX-23686528-DEL-1739
chrX-23701714-DEL-230
chrX-23817595-DEL-71
chrX-23835880-DEL-56
chrX-23880772-DEL-2542
chrX-23900391-DEL-401
chrX-23944253-DEL-8756
chrX-23954543-DEL-189
chrX-23964617-DEL-97
chrX-23993322-DEL-807
chrX-24018143-DEL-191
chrX-24024917-DEL-400
chrX-24050982-DEL-807
chrX-24067728-DEL-678
chrX-24078249-DEL-60
chrX-24081180-DEL-110
chrX-24105652-DEL-627
chrX-24112484-DEL-331
chrX-24115217-DEL-7506
chrX-24163740-DEL-82
chrX-24349867-DEL-3300
chrX-24385960-DEL-8505
chrX-24396422-DEL-2710
chrX-24444553-DEL-5585
chrX-24460221-DEL-2201
chrX-24483433-DEL-4292
chrX-24539102-DEL-672
chrX-24568668-DEL-65
chrX-24570127-DEL-189
chrX-24663818-DEL-1585
chrX-24689720-DEL-2852
chrX-24690289-DEL-73
chrX-24697537-DEL-3647
chrX-25345958-DEL-2774
chrX-25557463-DEL-1724
chrX-25656599-DEL-165
chrX-25705319-DEL-8773
chrX-25801294-DEL-56
chrX-25861014-DEL-874
chrX-25862533-DEL-3039
chrX-25868498-DEL-17845
chrX-25888033-DEL-6500
chrX-25955954-DEL-56
chrX-25961554-DEL-90
chrX-26113029-DEL-112
chrX-26252003-DEL-484
chrX-26255380-DEL-535
chrX-26339822-DEL-51
chrX-26364977-DEL-3038
chrX-26443111-DEL-165
chrX-26443827-DEL-1277
chrX-26512203-DEL-1679
chrX-26534935-DEL-199
chrX-26545225-DEL-215
chrX-26758781-DEL-73
chrX-26808265-DEL-2113
chrX-26835607-DEL-193
chrX-26836290-DEL-1891
chrX-26844127-DEL-2730
chrX-26856385-DEL-54
chrX-28785817-DEL-66
chrX-29296718-DEL-417
chrX-29570298-DEL-865
chrX-29868221-DEL-208

chrX-29916885-DEL-18921
chrX-29953460-DEL-5376
chrX-30018451-DEL-2790
chrX-30801908-DEL-6443
chrX-30839216-DEL-64
chrX-30954978-DEL-5999
chrX-31196107-DEL-127
chrX-31249539-DEL-542
chrX-31309275-DEL-1046
chrX-31346821-DEL-1191
chrX-31355287-DEL-102
chrX-31362044-DEL-551
chrX-31393029-DEL-10095
chrX-31412210-DEL-6113
chrX-31420566-DEL-1202
chrX-31490047-DEL-7505
chrX-31509189-DEL-58
chrX-31597281-DEL-78
chrX-31619267-DEL-6570
chrX-31622803-DEL-8230
chrX-31635082-DEL-102
chrX-31701321-DEL-280
chrX-31786083-DEL-97
chrX-31789939-DEL-393
chrX-31886479-DEL-172
chrX-31987239-DEL-200
chrX-32014221-DEL-157
chrX-32018405-DEL-403
chrX-32050736-DEL-395
chrX-32082356-DEL-6932
chrX-32129120-DEL-6600
chrX-32217973-DEL-2957
chrX-32225654-DEL-78
chrX-32348968-DEL-167
chrX-32646878-DEL-5259
chrX-32658465-DEL-66
chrX-32835868-DEL-58
chrX-32848148-DEL-81
chrX-32852848-DEL-1162
chrX-32854140-DEL-396
chrX-32942981-DEL-3669
chrX-32950665-DEL-3019
chrX-33263774-DEL-377
chrX-33278789-DEL-5393
chrX-33307753-DEL-288
chrX-33368500-DEL-335
chrX-33378523-DEL-4759
chrX-33379079-DEL-4203
chrX-33693730-DEL-8852
chrX-33712403-DEL-378
chrX-33824894-DEL-556
chrX-33886735-DEL-82
chrX-33947028-DEL-118
chrX-34476859-DEL-4368
chrX-34494073-DEL-3518
chrX-34585881-DEL-403
chrX-34761745-DEL-10812
chrX-34780002-DEL-519
chrX-34808332-DEL-4851
chrX-34818860-DEL-5533
chrX-34820915-DEL-1482
chrX-34852378-DEL-109
chrX-34912230-DEL-55
chrX-34913453-DEL-2550
chrX-34929993-DEL-78
chrX-34956106-DEL-74
chrX-34957830-DEL-5981
chrX-34983978-DEL-3680
chrX-35028068-DEL-327
chrX-35043543-DEL-472
chrX-35047313-DEL-472
chrX-35097347-DEL-875
chrX-35100208-DEL-1342
chrX-35112514-DEL-194
chrX-35122606-DEL-196
chrX-35131872-DEL-154
chrX-35135337-DEL-444

chrX-35140220-DEL-746
chrX-35142393-DEL-72
chrX-35172950-DEL-398
chrX-35187742-DEL-381
chrX-35198170-DEL-80
chrX-35236264-DEL-285
chrX-35291376-DEL-80
chrX-35317690-DEL-142
chrX-35320818-DEL-80
chrX-35339381-DEL-68
chrX-35349728-DEL-89
chrX-35381116-DEL-66
chrX-35389104-DEL-71
chrX-35398415-DEL-187
chrX-35423355-DEL-163
chrX-35427444-DEL-56
chrX-35469450-DEL-202
chrX-35479335-DEL-310
chrX-35494914-DEL-168
chrX-35561638-DEL-800
chrX-35564422-DEL-2304
chrX-35659581-DEL-52
chrX-35684979-DEL-201
chrX-35690784-DEL-117
chrX-35693442-DEL-58
chrX-35704923-DEL-140
chrX-35723739-DEL-6599
chrX-35749732-DEL-397
chrX-35763290-DEL-124
chrX-35786660-DEL-228
chrX-35803334-DEL-2009
chrX-35806201-DEL-228
chrX-35818667-DEL-88
chrX-35823017-DEL-667
chrX-35828401-DEL-138
chrX-35831562-DEL-139
chrX-35858526-DEL-61
chrX-35897718-DEL-324
chrX-35924101-DEL-56
chrX-35969193-DEL-5305
chrX-35977616-DEL-158
chrX-35981901-DEL-400
chrX-35986425-DEL-1207
chrX-35998269-DEL-291
chrX-36008040-DEL-194
chrX-36013668-DEL-215
chrX-36043064-DEL-286
chrX-36083668-DEL-60
chrX-36110538-DEL-123
chrX-36167556-DEL-66
chrX-36189453-DEL-114
chrX-36194865-DEL-1037
chrX-36198472-DEL-199
chrX-36201632-DEL-704
chrX-36208671-DEL-64
chrX-36216256-DEL-105
chrX-36223538-DEL-118
chrX-36282251-DEL-441
chrX-36284381-DEL-72
chrX-36298939-DEL-208
chrX-36299500-DEL-160
chrX-36318218-DEL-63
chrX-36320347-DEL-310
chrX-36328450-DEL-1413
chrX-36330057-DEL-73
chrX-36338115-DEL-207
chrX-36344791-DEL-938
chrX-36351042-DEL-218
chrX-36377496-DEL-56
chrX-36394158-DEL-179
chrX-36396263-DEL-230
chrX-36399338-DEL-189
chrX-36416042-DEL-347
chrX-36418619-DEL-213
chrX-36433175-DEL-52
chrX-36437979-DEL-373
chrX-36440378-DEL-106

chrX-36446723-DEL-200
chrX-36458317-DEL-315
chrX-36463239-DEL-188
chrX-36492585-DEL-115
chrX-36529047-DEL-56
chrX-36610699-DEL-54
chrX-36622753-DEL-424
chrX-36667845-DEL-5263
chrX-36680951-DEL-580
chrX-36682209-DEL-434
chrX-36713634-DEL-6594
chrX-36722647-DEL-89
chrX-36735646-DEL-54
chrX-36745191-DEL-6202
chrX-36760133-DEL-104
chrX-36769281-DEL-126
chrX-36807006-DEL-846
chrX-36808752-DEL-76
chrX-36815407-DEL-5152
chrX-36846235-DEL-744
chrX-36846293-DEL-1426
chrX-36846752-DEL-867
chrX-36880012-DEL-54
chrX-36887625-DEL-74
chrX-36998601-DEL-212
chrX-37006923-DEL-345
chrX-37010914-DEL-408
chrX-37017743-DEL-64
chrX-37018912-DEL-421
chrX-37051426-DEL-232
chrX-37052392-DEL-554
chrX-37056792-DEL-206
chrX-37057848-DEL-578
chrX-37059695-DEL-587
chrX-37068270-DEL-53
chrX-37088770-DEL-2115
chrX-37123071-DEL-574
chrX-37142711-DEL-410
chrX-37152766-DEL-344
chrX-37184728-DEL-61
chrX-37197575-DEL-613
chrX-37207695-DEL-121
chrX-37211296-DEL-56
chrX-37231143-DEL-54
chrX-37268956-DEL-529
chrX-37271582-DEL-637
chrX-37292818-DEL-117
chrX-37311299-DEL-77
chrX-37325925-DEL-51
chrX-37328287-DEL-94
chrX-37350252-DEL-327
chrX-37379270-DEL-282
chrX-37398055-DEL-127
chrX-37419284-DEL-110
chrX-37437692-DEL-193
chrX-37467783-DEL-379
chrX-37470699-DEL-358
chrX-37516750-DEL-172
chrX-37522460-DEL-120
chrX-37522692-DEL-1291
chrX-37556203-DEL-137
chrX-37564399-DEL-1549
chrX-37574711-DEL-77
chrX-37624221-DEL-201
chrX-37657491-DEL-109
chrX-37658467-DEL-194
chrX-37728363-DEL-255
chrX-37749339-DEL-751
chrX-37750579-DEL-6389
chrX-37761702-DEL-63
chrX-37764796-DEL-86
chrX-37773853-DEL-1330
chrX-37784911-DEL-173
chrX-37801825-DEL-5503
chrX-37844117-DEL-4346
chrX-37870861-DEL-9936
chrX-37898800-DEL-331

chrX-37914332-DEL-401
chrX-37975971-DEL-7658
chrX-37998583-DEL-85
chrX-38008212-DEL-572
chrX-38052617-DEL-565
chrX-38070815-DEL-76
chrX-38074776-DEL-584
chrX-38085590-DEL-6282
chrX-38092782-DEL-79
chrX-38149084-DEL-68
chrX-38180333-DEL-911
chrX-38182035-DEL-1854
chrX-38221003-DEL-18375
chrX-38227205-DEL-1894
chrX-38245180-DEL-6434
chrX-38257720-DEL-165
chrX-38291723-DEL-1109
chrX-38294322-DEL-6697
chrX-38324906-DEL-759
chrX-38338027-DEL-129
chrX-38342812-DEL-5161
chrX-38350193-DEL-248
chrX-38357071-DEL-158
chrX-38387319-DEL-2385
chrX-38396635-DEL-70
chrX-38403503-DEL-10822
chrX-38433646-DEL-1115
chrX-38444803-DEL-177
chrX-38479814-DEL-86
chrX-38527950-DEL-6220
chrX-38544548-DEL-60
chrX-38582955-DEL-375
chrX-38585141-DEL-2790
chrX-38609182-DEL-404
chrX-38634945-DEL-1214
chrX-38678401-DEL-1214
chrX-38707061-DEL-903
chrX-38715961-DEL-3289
chrX-38717291-DEL-354
chrX-38721994-DEL-58
chrX-38756187-DEL-212
chrX-38756717-DEL-242
chrX-38769549-DEL-3239
chrX-38781390-DEL-2986
chrX-38798538-DEL-187
chrX-38806780-DEL-1800
chrX-38827426-DEL-580
chrX-38833830-DEL-5097
chrX-38834629-DEL-222
chrX-38838493-DEL-988
chrX-38881917-DEL-878
chrX-38883571-DEL-336
chrX-38894342-DEL-6469
chrX-38895880-DEL-97
chrX-38901186-DEL-1612
chrX-38904873-DEL-76
chrX-38918432-DEL-168
chrX-38935189-DEL-1110
chrX-38968764-DEL-138
chrX-38988885-DEL-117
chrX-38993654-DEL-102
chrX-39000580-DEL-394
chrX-39014195-DEL-324
chrX-39036704-DEL-61
chrX-39054694-DEL-148
chrX-39067836-DEL-1570
chrX-39095906-DEL-131
chrX-39117125-DEL-282
chrX-39164971-DEL-323
chrX-39176887-DEL-614
chrX-39186068-DEL-358
chrX-39187639-DEL-2309
chrX-39202631-DEL-6263
chrX-39214922-DEL-295
chrX-39247139-DEL-85
chrX-39263471-DEL-60
chrX-39340799-DEL-579

chrX-39346703-DEL-1801
chrX-39367920-DEL-58
chrX-39371085-DEL-333
chrX-39388476-DEL-2526
chrX-39412985-DEL-831
chrX-39453116-DEL-8737
chrX-39468834-DEL-681
chrX-39493503-DEL-1057
chrX-39504122-DEL-4931
chrX-39513276-DEL-455
chrX-39515102-DEL-358
chrX-39565165-DEL-6218
chrX-39574723-DEL-493
chrX-39589615-DEL-190
chrX-39628377-DEL-19536
chrX-39673384-DEL-207
chrX-39680755-DEL-3335
chrX-39692007-DEL-882
chrX-39694932-DEL-2653
chrX-39694937-DEL-1073
chrX-39696066-DEL-1519
chrX-39722380-DEL-6596
chrX-39738556-DEL-86
chrX-39751161-DEL-124
chrX-39758279-DEL-681
chrX-39781070-DEL-203
chrX-39783107-DEL-423
chrX-39836671-DEL-62
chrX-39858498-DEL-114
chrX-39867792-DEL-558
chrX-39878241-DEL-105
chrX-39929746-DEL-53
chrX-39953017-DEL-56
chrX-39953671-DEL-6516
chrX-39999875-DEL-1410
chrX-40027021-DEL-98
chrX-40030524-DEL-661
chrX-40046568-DEL-562
chrX-40073254-DEL-403
chrX-40082878-DEL-6034
chrX-40096143-DEL-7032
chrX-40128003-DEL-56
chrX-40143291-DEL-4690
chrX-40183214-DEL-6325
chrX-40201013-DEL-107
chrX-40222649-DEL-64
chrX-40237431-DEL-202
chrX-40320901-DEL-424
chrX-40350624-DEL-392
chrX-40381760-DEL-353
chrX-40417608-DEL-1283
chrX-40422590-DEL-157
chrX-40422908-DEL-747
chrX-40448499-DEL-151
chrX-40473604-DEL-60
chrX-40505325-DEL-97
chrX-40529667-DEL-73
chrX-40571338-DEL-424
chrX-40652890-DEL-117
chrX-40665336-DEL-71
chrX-40706756-DEL-81
chrX-40719015-DEL-52
chrX-40728078-DEL-903
chrX-40742459-DEL-374
chrX-40896844-DEL-216
chrX-40909486-DEL-602
chrX-40919862-DEL-827
chrX-40930922-DEL-161
chrX-40949808-DEL-97
chrX-40974578-DEL-96
chrX-40985787-DEL-81
chrX-41008379-DEL-221
chrX-41023077-DEL-1192
chrX-41076099-DEL-205
chrX-41089950-DEL-446
chrX-41092615-DEL-456
chrX-41127382-DEL-1300

chrX-41162703-DEL-1214
chrX-41179448-DEL-1714
chrX-41225956-DEL-461
chrX-41227091-DEL-201
chrX-41248009-DEL-65
chrX-41249973-DEL-782
chrX-41287289-DEL-51
chrX-41313342-DEL-84
chrX-41410683-DEL-74
chrX-41418649-DEL-602
chrX-41443922-DEL-316
chrX-41477964-DEL-474
chrX-41488627-DEL-76
chrX-41499282-DEL-2956
chrX-41503349-DEL-3052
chrX-41534219-DEL-141
chrX-41537055-DEL-1172
chrX-41787228-DEL-89
chrX-41810599-DEL-131
chrX-41925902-DEL-80
chrX-41947360-DEL-358
chrX-41967666-DEL-7570
chrX-41992615-DEL-778
chrX-42080729-DEL-480
chrX-42097020-DEL-623
chrX-42109861-DEL-6487
chrX-42120197-DEL-545
chrX-42149244-DEL-557
chrX-42189369-DEL-630
chrX-42196968-DEL-890
chrX-42258031-DEL-517
chrX-42265003-DEL-64
chrX-42297655-DEL-6879
chrX-42297784-DEL-424
chrX-42314731-DEL-6395
chrX-42415536-DEL-149
chrX-42446401-DEL-1158
chrX-42450548-DEL-1557
chrX-42492204-DEL-583
chrX-42508317-DEL-60
chrX-42552220-DEL-989
chrX-42556617-DEL-188
chrX-42562854-DEL-481
chrX-42563722-DEL-1292
chrX-42598634-DEL-788
chrX-42657283-DEL-204
chrX-42666748-DEL-311
chrX-42695026-DEL-58
chrX-42703533-DEL-1862
chrX-42714083-DEL-5096
chrX-42740845-DEL-58
chrX-42742617-DEL-8712
chrX-42755121-DEL-423
chrX-42774944-DEL-56
chrX-42786818-DEL-66
chrX-42844526-DEL-145
chrX-42847764-DEL-232
chrX-42848557-DEL-1446
chrX-42894369-DEL-1245
chrX-42916295-DEL-2719
chrX-42966094-DEL-6237
chrX-42996624-DEL-63
chrX-43010861-DEL-396
chrX-43015857-DEL-81
chrX-43016843-DEL-58
chrX-43028586-DEL-114
chrX-43096891-DEL-11171
chrX-43113679-DEL-476
chrX-43125357-DEL-310
chrX-43132082-DEL-9403
chrX-43156640-DEL-4894
chrX-43164251-DEL-216
chrX-43166275-DEL-2385
chrX-43175872-DEL-923
chrX-43177015-DEL-75
chrX-43190653-DEL-54
chrX-43192982-DEL-5964

chrX-43199558-DEL-958
chrX-43203536-DEL-6227
chrX-43248884-DEL-1731
chrX-43251461-DEL-121
chrX-43296238-DEL-817
chrX-43323184-DEL-9512
chrX-43333203-DEL-54
chrX-43336481-DEL-3147
chrX-43351387-DEL-7075
chrX-43367632-DEL-268
chrX-43372380-DEL-4932
chrX-43373328-DEL-7749
chrX-43379869-DEL-2600
chrX-43416321-DEL-3068
chrX-43463975-DEL-364
chrX-43540381-DEL-269
chrX-43549652-DEL-647
chrX-43560360-DEL-64
chrX-43565033-DEL-3352
chrX-43595262-DEL-53
chrX-43646096-DEL-56
chrX-43700235-DEL-1418
chrX-43707121-DEL-87
chrX-43744319-DEL-108
chrX-43744793-DEL-6338
chrX-43776976-DEL-397
chrX-43796903-DEL-1094
chrX-43806756-DEL-6587
chrX-43821295-DEL-6394
chrX-43854236-DEL-2221
chrX-43860575-DEL-3450
chrX-43874632-DEL-193
chrX-43901453-DEL-83
chrX-43918916-DEL-544
chrX-43937078-DEL-54
chrX-43937870-DEL-2456
chrX-43960542-DEL-494
chrX-43966505-DEL-60
chrX-43968037-DEL-107
chrX-43981556-DEL-907
chrX-43996111-DEL-452
chrX-44010595-DEL-95
chrX-44040916-DEL-118
chrX-44054257-DEL-774
chrX-44080677-DEL-267
chrX-44117124-DEL-124
chrX-44135840-DEL-7162
chrX-44144089-DEL-890
chrX-44161546-DEL-1504
chrX-44190517-DEL-6612
chrX-44190741-DEL-5014
chrX-44197322-DEL-571
chrX-44231269-DEL-1790
chrX-44283405-DEL-4504
chrX-44291458-DEL-312
chrX-44302848-DEL-210
chrX-44314339-DEL-659
chrX-44315118-DEL-938
chrX-44318167-DEL-6255
chrX-44324766-DEL-8427
chrX-44341292-DEL-8618
chrX-44393992-DEL-4251
chrX-44400510-DEL-77
chrX-44482397-DEL-66
chrX-44530885-DEL-65
chrX-44563520-DEL-1063
chrX-44565233-DEL-207
chrX-44571216-DEL-67
chrX-44604170-DEL-266
chrX-44609737-DEL-56
chrX-44610945-DEL-53
chrX-44613476-DEL-543
chrX-44621268-DEL-5961
chrX-44633302-DEL-922
chrX-44639255-DEL-1310
chrX-44650322-DEL-265
chrX-44661197-DEL-5315

chrX-44677240-DEL-2216
chrX-44701293-DEL-96
chrX-44704336-DEL-58
chrX-44710495-DEL-6625
chrX-44726991-DEL-66
chrX-44747960-DEL-217
chrX-44749372-DEL-2554
chrX-44749660-DEL-679
chrX-44750020-DEL-1713
chrX-44773733-DEL-1935
chrX-44844934-DEL-160
chrX-44856815-DEL-400
chrX-44913307-DEL-1939
chrX-44935992-DEL-4858
chrX-45002935-DEL-275
chrX-45010262-DEL-1770
chrX-45021299-DEL-97
chrX-45047394-DEL-3235
chrX-45052678-DEL-7498
chrX-45069173-DEL-56
chrX-45093720-DEL-194
chrX-45094335-DEL-1532
chrX-45120086-DEL-2326
chrX-45157717-DEL-1289
chrX-45173052-DEL-65
chrX-45292159-DEL-388
chrX-45297435-DEL-2330
chrX-45304086-DEL-529
chrX-45311957-DEL-68
chrX-45344654-DEL-158
chrX-45346304-DEL-53
chrX-45438649-DEL-1140
chrX-45459817-DEL-3886
chrX-45492460-DEL-1892
chrX-45501993-DEL-169
chrX-45520537-DEL-356
chrX-45566200-DEL-896
chrX-45581165-DEL-1610
chrX-45596617-DEL-699
chrX-45601762-DEL-59
chrX-45612638-DEL-4925
chrX-45664638-DEL-187
chrX-45689236-DEL-336
chrX-45703374-DEL-1587
chrX-45703387-DEL-53
chrX-45715504-DEL-198
chrX-45731980-DEL-597
chrX-45779376-DEL-480
chrX-45787476-DEL-74
chrX-45804578-DEL-2104
chrX-45811600-DEL-543
chrX-45847419-DEL-94
chrX-45858213-DEL-661
chrX-45889597-DEL-4061
chrX-45905119-DEL-6231
chrX-45913314-DEL-107
chrX-45915170-DEL-342
chrX-45936076-DEL-328
chrX-45955191-DEL-1614
chrX-46011600-DEL-1232
chrX-46033566-DEL-399
chrX-46066041-DEL-6684
chrX-46079328-DEL-6033
chrX-46112656-DEL-83
chrX-46120009-DEL-52
chrX-46152906-DEL-1605
chrX-46210836-DEL-559
chrX-46268683-DEL-51
chrX-46370807-DEL-909
chrX-46432447-DEL-198
chrX-46469733-DEL-181
chrX-46472774-DEL-436
chrX-46473549-DEL-55
chrX-46495192-DEL-166
chrX-46507023-DEL-7500
chrX-46523755-DEL-56
chrX-46609282-DEL-832

chrX-46626605-DEL-1528
chrX-46669600-DEL-60
chrX-46747782-DEL-396
chrX-46760780-DEL-693
chrX-46761729-DEL-1160
chrX-46774312-DEL-452
chrX-46809957-DEL-1095
chrX-46811856-DEL-553
chrX-46825299-DEL-148
chrX-46896149-DEL-125
chrX-46900597-DEL-52
chrX-46904638-DEL-59
chrX-46925337-DEL-599
chrX-46950951-DEL-876
chrX-46995906-DEL-288
chrX-46998859-DEL-995
chrX-47007856-DEL-72
chrX-47023072-DEL-73
chrX-47037659-DEL-330
chrX-47056868-DEL-2295
chrX-47068253-DEL-202
chrX-47074932-DEL-215
chrX-47084010-DEL-174
chrX-47089353-DEL-398
chrX-47120135-DEL-202
chrX-47128537-DEL-318
chrX-47144130-DEL-274
chrX-47150478-DEL-52
chrX-47167046-DEL-198
chrX-47173974-DEL-427
chrX-47188423-DEL-414
chrX-47256481-DEL-340
chrX-47263813-DEL-528
chrX-47301310-DEL-136
chrX-47304911-DEL-89
chrX-47311230-DEL-284
chrX-47313855-DEL-90
chrX-47365472-DEL-205
chrX-47390963-DEL-69
chrX-47438464-DEL-634
chrX-47500586-DEL-105
chrX-47560993-DEL-207
chrX-47581894-DEL-230
chrX-47589819-DEL-104
chrX-47596325-DEL-210
chrX-47619400-DEL-202
chrX-47668540-DEL-51
chrX-47694647-DEL-60
chrX-47707450-DEL-87
chrX-47720170-DEL-368
chrX-47734418-DEL-74
chrX-47738255-DEL-54
chrX-47805899-DEL-2149
chrX-47827075-DEL-64
chrX-47827546-DEL-6138
chrX-47835458-DEL-670
chrX-47844501-DEL-6665
chrX-47855453-DEL-4382
chrX-47871095-DEL-182
chrX-47935345-DEL-6238
chrX-47982194-DEL-143
chrX-47984985-DEL-56
chrX-48007389-DEL-1716
chrX-48033103-DEL-294
chrX-48038097-DEL-2459
chrX-48042163-DEL-657
chrX-48044894-DEL-469
chrX-48064250-DEL-450
chrX-48095068-DEL-2191
chrX-48105342-DEL-65
chrX-48109017-DEL-60
chrX-48130899-DEL-81
chrX-48149548-DEL-137
chrX-48260050-DEL-71
chrX-48261713-DEL-528
chrX-48286632-DEL-3883
chrX-48366537-DEL-266

chrX-48382142-DEL-242
chrX-48408259-DEL-887
chrX-48409382-DEL-172
chrX-48440701-DEL-212
chrX-48509600-DEL-3106
chrX-48526894-DEL-1010
chrX-48611871-DEL-64
chrX-48618085-DEL-1745
chrX-48649849-DEL-7820
chrX-48695119-DEL-9946
chrX-48730331-DEL-797
chrX-48736952-DEL-435
chrX-48749243-DEL-1190
chrX-48752676-DEL-550
chrX-48753785-DEL-4564
chrX-48765439-DEL-194
chrX-48774326-DEL-1423
chrX-48780922-DEL-58
chrX-48783022-DEL-201
chrX-48795351-DEL-223
chrX-48796227-DEL-628
chrX-48803468-DEL-6977
chrX-48828331-DEL-2589
chrX-48894479-DEL-860
chrX-48895721-DEL-127
chrX-48911187-DEL-107
chrX-48919378-DEL-399
chrX-48976241-DEL-56
chrX-48983869-DEL-1820
chrX-48985765-DEL-1580
chrX-48986224-DEL-5742
chrX-48997253-DEL-203
chrX-49020574-DEL-81
chrX-49032850-DEL-845
chrX-49033787-DEL-52
chrX-49068635-DEL-226
chrX-49086936-DEL-7001
chrX-49127805-DEL-660
chrX-49143343-DEL-390
chrX-49150786-DEL-123
chrX-49164014-DEL-393
chrX-49185020-DEL-52
chrX-49237254-DEL-3955
chrX-49246862-DEL-7774
chrX-49288966-DEL-596
chrX-49292592-DEL-168
chrX-49307207-DEL-353
chrX-49334653-DEL-1915
chrX-49336721-DEL-107
chrX-49337829-DEL-4476
chrX-49350380-DEL-2286
chrX-49353942-DEL-731
chrX-49395120-DEL-2408
chrX-49396041-DEL-1185
chrX-49430557-DEL-6409
chrX-49443443-DEL-6121
chrX-49455204-DEL-436
chrX-49467910-DEL-337
chrX-49491201-DEL-411
chrX-49492068-DEL-201
chrX-49494677-DEL-166
chrX-49502273-DEL-5175
chrX-49524701-DEL-237
chrX-49557858-DEL-358
chrX-49562944-DEL-133
chrX-49591307-DEL-53
chrX-49600959-DEL-6493
chrX-49622438-DEL-1447
chrX-49627010-DEL-2331
chrX-49640564-DEL-252
chrX-49654118-DEL-319
chrX-49752067-DEL-52
chrX-49770485-DEL-881
chrX-49771424-DEL-255
chrX-49772098-DEL-61
chrX-49776050-DEL-1467
chrX-49795892-DEL-55

chrX-49816861-DEL-255
chrX-49825360-DEL-101
chrX-49855625-DEL-6551
chrX-49898281-DEL-383
chrX-49911211-DEL-152
chrX-49924916-DEL-102
chrX-49926790-DEL-144
chrX-50007403-DEL-138
chrX-50058220-DEL-72
chrX-50085473-DEL-1153
chrX-50098301-DEL-119
chrX-50139085-DEL-72
chrX-50153221-DEL-272
chrX-50160769-DEL-399
chrX-50163636-DEL-92
chrX-50184906-DEL-1247
chrX-50194298-DEL-2343
chrX-50269638-DEL-210
chrX-50284281-DEL-90
chrX-50309350-DEL-79
chrX-50344977-DEL-6202
chrX-50367007-DEL-63
chrX-50414824-DEL-68
chrX-50415936-DEL-296
chrX-50417636-DEL-763
chrX-50427246-DEL-112
chrX-50438981-DEL-57
chrX-50439958-DEL-6227
chrX-50481818-DEL-54
chrX-50497686-DEL-256
chrX-50637915-DEL-130
chrX-50646904-DEL-97
chrX-50651330-DEL-97
chrX-50684490-DEL-164
chrX-50743205-DEL-72
chrX-50790034-DEL-57
chrX-50805680-DEL-89
chrX-50830078-DEL-70
chrX-50831027-DEL-1657
chrX-50855519-DEL-645
chrX-50872801-DEL-1270
chrX-50878730-DEL-53
chrX-50898738-DEL-345
chrX-50900972-DEL-551
chrX-50940052-DEL-90
chrX-50975478-DEL-171
chrX-50987643-DEL-54
chrX-51000759-DEL-966
chrX-51081857-DEL-1067
chrX-51098184-DEL-403
chrX-51130505-DEL-323
chrX-51133851-DEL-1050
chrX-51169128-DEL-136
chrX-51198091-DEL-70
chrX-51235224-DEL-173
chrX-51239123-DEL-295
chrX-51284016-DEL-400
chrX-51320727-DEL-82
chrX-51329659-DEL-55
chrX-51340968-DEL-391
chrX-51344412-DEL-594
chrX-51384721-DEL-197
chrX-51389170-DEL-57
chrX-51415826-DEL-55
chrX-51416277-DEL-197
chrX-51433801-DEL-83
chrX-51455680-DEL-89
chrX-51470742-DEL-52
chrX-51481271-DEL-98
chrX-51482768-DEL-72
chrX-51564835-DEL-54
chrX-51597254-DEL-75
chrX-51612375-DEL-296
chrX-51738945-DEL-279
chrX-51775251-DEL-384
chrX-51779307-DEL-199
chrX-51813551-DEL-964

chrX-51816624-DEL-310
chrX-51822287-DEL-82
chrX-51826929-DEL-200
chrX-51837178-DEL-61
chrX-51840996-DEL-54
chrX-51846492-DEL-407
chrX-51858657-DEL-58
chrX-51861992-DEL-405
chrX-51886789-DEL-71
chrX-51906029-DEL-58
chrX-51994223-DEL-87
chrX-52037671-DEL-131
chrX-52050194-DEL-98
chrX-52082254-DEL-443
chrX-52113776-DEL-61
chrX-52114214-DEL-440
chrX-52125264-DEL-72
chrX-52129481-DEL-80
chrX-52131731-DEL-1688
chrX-52225612-DEL-55
chrX-52249220-DEL-327
chrX-52278594-DEL-170
chrX-52310137-DEL-267
chrX-52384300-DEL-55
chrX-52496161-DEL-354
chrX-52511664-DEL-408
chrX-52552067-DEL-108
chrX-52552480-DEL-672
chrX-52565968-DEL-115
chrX-52577261-DEL-1238
chrX-52589169-DEL-146
chrX-52590427-DEL-60
chrX-52606449-DEL-58
chrX-52607001-DEL-1094
chrX-52727501-DEL-178
chrX-52752036-DEL-716
chrX-52758572-DEL-81
chrX-52776593-DEL-72
chrX-52804603-DEL-6360
chrX-52828914-DEL-1943
chrX-52897989-DEL-54
chrX-52942084-DEL-1715
chrX-52979531-DEL-4393
chrX-53023900-DEL-204
chrX-53057512-DEL-555
chrX-53091608-DEL-326
chrX-53107407-DEL-6772
chrX-53120963-DEL-3831
chrX-53134248-DEL-63
chrX-53229162-DEL-5436
chrX-53251297-DEL-558
chrX-53255620-DEL-239
chrX-53424404-DEL-1006
chrX-53460552-DEL-4393
chrX-53606444-DEL-1006
chrX-53610114-DEL-56
chrX-53613624-DEL-3830
chrX-53621193-DEL-1010
chrX-53697463-DEL-1007
chrX-53712207-DEL-1010
chrX-53818135-DEL-75
chrX-53819120-DEL-134
chrX-53883596-DEL-2629
chrX-53883618-DEL-661
chrX-53896066-DEL-280
chrX-53911929-DEL-1199
chrX-53992597-DEL-331
chrX-54046902-DEL-1006
chrX-54251072-DEL-142
chrX-54346909-DEL-4372
chrX-54353156-DEL-750
chrX-54407936-DEL-2017
chrX-54434484-DEL-493
chrX-54465452-DEL-1121
chrX-54469296-DEL-434
chrX-54470875-DEL-56
chrX-54472855-DEL-2149

chrX-54507448-DEL-586
chrX-54510380-DEL-62
chrX-54516891-DEL-64
chrX-54532974-DEL-1292
chrX-54539339-DEL-105
chrX-54605408-DEL-239
chrX-54646296-DEL-6787
chrX-54706174-DEL-858
chrX-54726335-DEL-10595
chrX-54754893-DEL-279
chrX-54765139-DEL-111
chrX-54856678-DEL-3695
chrX-54908180-DEL-692
chrX-54929874-DEL-6211
chrX-54942694-DEL-56
chrX-54967187-DEL-1752
chrX-54989150-DEL-402
chrX-55002286-DEL-56
chrX-55005778-DEL-161
chrX-55022309-DEL-124
chrX-55025060-DEL-1356
chrX-55026462-DEL-1015
chrX-55073002-DEL-403
chrX-55082563-DEL-915
chrX-55194061-DEL-124
chrX-55215393-DEL-428
chrX-55232516-DEL-153
chrX-55257276-DEL-390
chrX-55264964-DEL-153
chrX-55276989-DEL-68
chrX-55281412-DEL-2023
chrX-55284243-DEL-1943
chrX-55333200-DEL-204
chrX-55413952-DEL-96
chrX-55426554-DEL-189
chrX-55476075-DEL-1215
chrX-55570818-DEL-148
chrX-55580042-DEL-51
chrX-55603172-DEL-175
chrX-55614797-DEL-226
chrX-55725875-DEL-112
chrX-55726838-DEL-80
chrX-55783896-DEL-202
chrX-55814034-DEL-840
chrX-55859196-DEL-763
chrX-55882480-DEL-197
chrX-55900961-DEL-52
chrX-55941840-DEL-587
chrX-55985088-DEL-51
chrX-56020251-DEL-1130
chrX-56041381-DEL-393
chrX-56043525-DEL-198
chrX-56044533-DEL-394
chrX-56119625-DEL-280
chrX-56188568-DEL-149
chrX-56252831-DEL-53
chrX-56316506-DEL-68
chrX-56327222-DEL-119
chrX-56385909-DEL-389
chrX-56393636-DEL-57
chrX-56441751-DEL-89
chrX-56449770-DEL-232
chrX-56471601-DEL-1095
chrX-56571032-DEL-1935
chrX-56598760-DEL-511
chrX-56649918-DEL-424
chrX-56667634-DEL-703
chrX-56681924-DEL-439
chrX-56787303-DEL-64
chrX-56836026-DEL-198
chrX-56886268-DEL-52
chrX-56895137-DEL-767
chrX-56936301-DEL-226
chrX-56990643-DEL-202
chrX-57001938-DEL-136
chrX-57038984-DEL-455
chrX-57136340-DEL-56

chrX-57145328-DEL-73
chrX-57150070-DEL-62
chrX-57254212-DEL-6678
chrX-57274337-DEL-533
chrX-57317160-DEL-67
chrX-57323394-DEL-54
chrX-57359817-DEL-58
chrX-57383119-DEL-7729
chrX-57394179-DEL-609
chrX-57442253-DEL-6189
chrX-57479492-DEL-420
chrX-57533055-DEL-799
chrX-57548545-DEL-228
chrX-57553429-DEL-3732
chrX-57574098-DEL-692
chrX-57583869-DEL-219
chrX-57590679-DEL-683
chrX-57595859-DEL-1635
chrX-57599363-DEL-187
chrX-57627892-DEL-1021
chrX-57687951-DEL-1271
chrX-57700871-DEL-169
chrX-57703786-DEL-1358
chrX-57714653-DEL-1742
chrX-57725662-DEL-1347
chrX-57726667-DEL-531
chrX-57735000-DEL-1159
chrX-57737064-DEL-513
chrX-57741327-DEL-60
chrX-57790532-DEL-360
chrX-57817028-DEL-2374
chrX-57827976-DEL-18983
chrX-57865103-DEL-63
chrX-57881945-DEL-1050
chrX-57899916-DEL-96
chrX-57941749-DEL-1879
chrX-57997925-DEL-677
chrX-58036440-DEL-408
chrX-58102496-DEL-2102
chrX-58106941-DEL-61
chrX-58215582-DEL-217
chrX-58219262-DEL-54
chrX-58340461-DEL-997
chrX-58353433-DEL-280
chrX-58381063-DEL-89
chrX-58433981-DEL-56
chrX-58472203-DEL-902
chrX-58483850-DEL-424
chrX-58490035-DEL-3468
chrX-58516112-DEL-77
chrX-58567878-DEL-1811
chrX-58632022-DEL-1363
chrX-58632813-DEL-681
chrX-58696545-DEL-52
chrX-58698595-DEL-58
chrX-58712351-DEL-2596
chrX-58715305-DEL-6589
chrX-58721471-DEL-208
chrX-58724041-DEL-348
chrX-58740042-DEL-99
chrX-58785496-DEL-1637
chrX-58817036-DEL-2546
chrX-58822797-DEL-481
chrX-58843278-DEL-160
chrX-58925489-DEL-118
chrX-58935481-DEL-76
chrX-58963486-DEL-1292
chrX-58971430-DEL-400
chrX-58974148-DEL-6935
chrX-58981651-DEL-62
chrX-58989855-DEL-287
chrX-59009187-DEL-2075
chrX-59010805-DEL-929
chrX-59023771-DEL-1823
chrX-59025358-DEL-272
chrX-59036433-DEL-520
chrX-59111372-DEL-71

chrX-59144965-DEL-217
chrX-59171762-DEL-458
chrX-59179344-DEL-226
chrX-59215997-DEL-73
chrX-59218293-DEL-100
chrX-59223744-DEL-61
chrX-59235285-DEL-120
chrX-59238855-DEL-107
chrX-59261332-DEL-575
chrX-59271608-DEL-278
chrX-59380722-DEL-4467
chrX-59454206-DEL-247
chrX-59486200-DEL-2918
chrX-59491976-DEL-57
chrX-59510619-DEL-690
chrX-59517860-DEL-1630
chrX-59550486-DEL-225
chrX-59559888-DEL-78
chrX-59621275-DEL-161
chrX-59629006-DEL-4588
chrX-59638360-DEL-676
chrX-59641389-DEL-52
chrX-59700008-DEL-106
chrX-59704653-DEL-493
chrX-59720846-DEL-83
chrX-59759821-DEL-1367
chrX-59815184-DEL-158
chrX-59829079-DEL-168
chrX-59854542-DEL-64
chrX-59923306-DEL-214
chrX-59955026-DEL-203
chrX-59972450-DEL-52
chrX-59995550-DEL-6288
chrX-60065248-DEL-162
chrX-60068124-DEL-1260
chrX-60083423-DEL-51
chrX-60090775-DEL-116
chrX-60103056-DEL-2542
chrX-60128446-DEL-1499
chrX-60228238-DEL-216
chrX-60261886-DEL-53
chrX-60282355-DEL-58
chrX-60309892-DEL-87
chrX-60312255-DEL-1407
chrX-60357198-DEL-486
chrX-60372103-DEL-6303
chrX-60381672-DEL-8901
chrX-60383038-DEL-6376
chrX-60405046-DEL-128
chrX-60411425-DEL-1171
chrX-60457274-DEL-55
chrX-60490596-DEL-2190
chrX-60490680-DEL-831
chrX-60491540-DEL-1028
chrX-60492157-DEL-1176
chrX-60492735-DEL-152
chrX-60497619-DEL-1842
chrX-60567293-DEL-412
chrX-60569264-DEL-1083
chrX-60575845-DEL-456
chrX-60579532-DEL-204
chrX-60581581-DEL-500
chrX-60584845-DEL-2051
chrX-60596986-DEL-6186
chrX-60613023-DEL-210
chrX-60626947-DEL-589
chrX-60638404-DEL-85
chrX-60721891-DEL-61
chrX-60723580-DEL-80
chrX-60745150-DEL-1726
chrX-60759244-DEL-66
chrX-60828947-DEL-1668
chrX-60876027-DEL-544
chrX-60919725-DEL-336
chrX-60924327-DEL-63
chrX-60952388-DEL-83
chrX-60970073-DEL-59

chrX-61003668-DEL-347
chrX-61006040-DEL-8444
chrX-61013379-DEL-418
chrX-61017768-DEL-887
chrX-61044960-DEL-9220
chrX-61086040-DEL-79
chrX-61090178-DEL-54
chrX-61094538-DEL-1534
chrX-61097488-DEL-275
chrX-61108233-DEL-498
chrX-61109129-DEL-7023
chrX-61150406-DEL-9103
chrX-61160134-DEL-3051
chrX-61174470-DEL-252
chrX-61186829-DEL-79
chrX-61188857-DEL-204
chrX-61196799-DEL-299
chrX-61228748-DEL-320
chrX-61238996-DEL-373
chrX-61270201-DEL-306
chrX-61275795-DEL-174
chrX-61279924-DEL-308
chrX-61281487-DEL-2240
chrX-61283741-DEL-124
chrX-61293593-DEL-290
chrX-61301426-DEL-65
chrX-61368023-DEL-53
chrX-61387202-DEL-3600
chrX-61409200-DEL-130
chrX-61471505-DEL-93
chrX-61485576-DEL-52
chrX-61498283-DEL-60
chrX-61505802-DEL-1008
chrX-61507426-DEL-2549
chrX-61539702-DEL-414
chrX-61547186-DEL-2282
chrX-61565363-DEL-746
chrX-61591805-DEL-5231
chrX-61591961-DEL-775
chrX-61594062-DEL-4480
chrX-61595442-DEL-745
chrX-61596985-DEL-748
chrX-61597491-DEL-746
chrX-61612813-DEL-1177
chrX-61626296-DEL-71
chrX-61635975-DEL-54
chrX-61643763-DEL-685
chrX-61660631-DEL-6409
chrX-61669630-DEL-6948
chrX-61687539-DEL-1651
chrX-61688886-DEL-542
chrX-61699714-DEL-624
chrX-61722064-DEL-80
chrX-61732479-DEL-469
chrX-61735644-DEL-6237
chrX-61752831-DEL-638
chrX-61756354-DEL-400
chrX-61765687-DEL-67
chrX-61829367-DEL-120
chrX-61837136-DEL-14388
chrX-61859789-DEL-424
chrX-61871842-DEL-68
chrX-61872395-DEL-113
chrX-61896754-DEL-67
chrX-61898085-DEL-351
chrX-61898983-DEL-1066
chrX-61928394-DEL-6793
chrX-61935668-DEL-1092
chrX-61937889-DEL-6212
chrX-61945421-DEL-270
chrX-61947623-DEL-6547
chrX-61964625-DEL-56
chrX-61968194-DEL-805
chrX-61971134-DEL-11520
chrX-61974862-DEL-7411
chrX-61974876-DEL-1040
chrX-61983697-DEL-76

chrX-61988474-DEL-144
chrX-61989927-DEL-7439
chrX-61991537-DEL-2994
chrX-62013467-DEL-6251
chrX-62031902-DEL-1573
chrX-62059402-DEL-1337
chrX-62063830-DEL-60
chrX-62066665-DEL-437
chrX-62080557-DEL-920
chrX-62084514-DEL-2276
chrX-62085910-DEL-2249
chrX-62092070-DEL-52
chrX-62106905-DEL-134
chrX-62120305-DEL-59
chrX-62125919-DEL-52
chrX-62137316-DEL-62
chrX-62139486-DEL-230
chrX-62187439-DEL-52
chrX-62209551-DEL-2419
chrX-62269618-DEL-1069
chrX-62294475-DEL-54
chrX-62297964-DEL-2512
chrX-62306242-DEL-361
chrX-62307074-DEL-368
chrX-62307951-DEL-69
chrX-62314262-DEL-2033
chrX-62320379-DEL-1387
chrX-62361516-DEL-1142
chrX-62405323-DEL-3600
chrX-62408998-DEL-59
chrX-62415064-DEL-232
chrX-62426081-DEL-85
chrX-62437871-DEL-62
chrX-62451091-DEL-1155
chrX-62454136-DEL-194
chrX-62459057-DEL-130
chrX-62464674-DEL-519
chrX-62508823-DEL-456
chrX-62514800-DEL-2665
chrX-62518044-DEL-4626
chrX-62553171-DEL-208
chrX-62580959-DEL-386
chrX-62625429-DEL-82
chrX-62660320-DEL-408
chrX-62663816-DEL-121
chrX-62673668-DEL-208
chrX-62676993-DEL-582
chrX-62678129-DEL-6482
chrX-62678243-DEL-214
chrX-62713545-DEL-633
chrX-62778066-DEL-600
chrX-62787400-DEL-777
chrX-62804700-DEL-3076
chrX-62853427-DEL-3615
chrX-62874186-DEL-56
chrX-62876705-DEL-225
chrX-62879413-DEL-1370
chrX-62891945-DEL-71
chrX-62895970-DEL-71
chrX-62896520-DEL-6212
chrX-62922278-DEL-13660
chrX-62930475-DEL-6160
chrX-62959827-DEL-91
chrX-62969533-DEL-6227
chrX-62977341-DEL-203
chrX-63012061-DEL-728
chrX-63045570-DEL-228
chrX-63047168-DEL-428
chrX-63062488-DEL-107
chrX-63073579-DEL-7055
chrX-63080202-DEL-195
chrX-63080539-DEL-6618
chrX-63115625-DEL-170
chrX-63116174-DEL-718
chrX-63125678-DEL-3407
chrX-63129416-DEL-72
chrX-63135586-DEL-74

chrX-63140641-DEL-61
chrX-63143366-DEL-473
chrX-63202344-DEL-56
chrX-63245045-DEL-188
chrX-63297212-DEL-116
chrX-63390440-DEL-2655
chrX-63400062-DEL-5061
chrX-63423674-DEL-70
chrX-63424092-DEL-830
chrX-63456275-DEL-949
chrX-63486249-DEL-384
chrX-63499497-DEL-173
chrX-63513658-DEL-66
chrX-63540175-DEL-2344
chrX-63582874-DEL-6207
chrX-63640534-DEL-67
chrX-63666948-DEL-11288
chrX-63675927-DEL-206
chrX-63679337-DEL-220
chrX-63682469-DEL-76
chrX-63684794-DEL-77
chrX-63703068-DEL-634
chrX-63720041-DEL-3036
chrX-63723247-DEL-1801
chrX-63729358-DEL-1808
chrX-63751701-DEL-919
chrX-63789672-DEL-69
chrX-63802535-DEL-5594
chrX-63818554-DEL-795
chrX-63823422-DEL-81
chrX-63861844-DEL-104
chrX-63863325-DEL-942
chrX-63885155-DEL-837
chrX-63903862-DEL-264
chrX-63908610-DEL-202
chrX-63925589-DEL-76
chrX-63962437-DEL-52
chrX-63970630-DEL-125
chrX-63972966-DEL-1012
chrX-63996473-DEL-256
chrX-63997074-DEL-52
chrX-64004317-DEL-57
chrX-64029003-DEL-64
chrX-64074784-DEL-283
chrX-64099767-DEL-400
chrX-64115879-DEL-72
chrX-64117379-DEL-1758
chrX-64130189-DEL-9442
chrX-64130966-DEL-6326
chrX-64142124-DEL-66729
chrX-64154803-DEL-2574
chrX-64155713-DEL-378
chrX-64178300-DEL-1753
chrX-64183734-DEL-4530
chrX-64204939-DEL-1717
chrX-64208840-DEL-2820
chrX-64215820-DEL-1547
chrX-64220170-DEL-380
chrX-64221748-DEL-107
chrX-64286212-DEL-54
chrX-64315127-DEL-330
chrX-64337090-DEL-316
chrX-64339723-DEL-3782
chrX-64344204-DEL-2100
chrX-64357900-DEL-717
chrX-64416267-DEL-193
chrX-64446942-DEL-354
chrX-64459871-DEL-1081
chrX-64459916-DEL-52
chrX-64492890-DEL-1623
chrX-64500081-DEL-365
chrX-64508676-DEL-57
chrX-64512183-DEL-60
chrX-64532957-DEL-1802
chrX-64549170-DEL-5241
chrX-64555095-DEL-614
chrX-64556820-DEL-439

chrX-64568215-DEL-451
chrX-64577156-DEL-767
chrX-64627766-DEL-53
chrX-64681851-DEL-64
chrX-64703109-DEL-461
chrX-64709047-DEL-125
chrX-64712848-DEL-2663
chrX-64715531-DEL-2305
chrX-64734834-DEL-1000
chrX-64739840-DEL-121
chrX-64747243-DEL-596
chrX-64793626-DEL-1015
chrX-64816260-DEL-181
chrX-64822033-DEL-1049
chrX-64824050-DEL-62
chrX-64843398-DEL-128
chrX-64845286-DEL-79
chrX-64847673-DEL-960
chrX-64860363-DEL-569
chrX-64862650-DEL-208
chrX-64870078-DEL-1972
chrX-64894140-DEL-197
chrX-64929457-DEL-6056
chrX-64940683-DEL-3000
chrX-64982803-DEL-64
chrX-65010837-DEL-198
chrX-65018857-DEL-208
chrX-65026242-DEL-944
chrX-65030892-DEL-12633
chrX-65030963-DEL-214
chrX-65069043-DEL-170
chrX-65081420-DEL-52
chrX-65085384-DEL-84
chrX-65099210-DEL-51
chrX-65154636-DEL-492
chrX-65167936-DEL-5545
chrX-65172782-DEL-117
chrX-65188490-DEL-416
chrX-65233365-DEL-57
chrX-65273265-DEL-7135
chrX-65299751-DEL-899
chrX-65321509-DEL-62
chrX-65340814-DEL-764
chrX-65350853-DEL-958
chrX-65361813-DEL-648
chrX-65382484-DEL-352
chrX-65413148-DEL-563
chrX-65413813-DEL-629
chrX-65421733-DEL-1887
chrX-65424744-DEL-71
chrX-65444584-DEL-1596
chrX-65464946-DEL-303
chrX-65487273-DEL-825
chrX-65492408-DEL-497
chrX-65537123-DEL-8436
chrX-65575940-DEL-52
chrX-65577483-DEL-245
chrX-65594510-DEL-331
chrX-65615098-DEL-2158
chrX-65629009-DEL-106
chrX-65678931-DEL-706
chrX-65710899-DEL-1000
chrX-65733200-DEL-175
chrX-65769833-DEL-1152
chrX-65842042-DEL-666
chrX-65859158-DEL-1884
chrX-65870436-DEL-3252
chrX-65877249-DEL-3267
chrX-65886360-DEL-621
chrX-65891384-DEL-1100
chrX-65900509-DEL-1561
chrX-65916268-DEL-58
chrX-65942393-DEL-1186
chrX-65961903-DEL-1890
chrX-65964512-DEL-152
chrX-65968136-DEL-62
chrX-66008199-DEL-5110

chrX-66009463-DEL-106
chrX-66011933-DEL-148
chrX-66014327-DEL-1197
chrX-66020892-DEL-1874
chrX-66027186-DEL-66
chrX-66039005-DEL-473
chrX-66047323-DEL-333
chrX-66056864-DEL-7206
chrX-66067154-DEL-3183
chrX-66075354-DEL-5974
chrX-66182388-DEL-705
chrX-66185794-DEL-63
chrX-66187188-DEL-415
chrX-66202323-DEL-682
chrX-66216030-DEL-695
chrX-66235107-DEL-401
chrX-66249285-DEL-1624
chrX-66258552-DEL-502
chrX-66280182-DEL-186
chrX-66288885-DEL-100
chrX-66306091-DEL-656
chrX-66306762-DEL-597
chrX-66307458-DEL-3294
chrX-66310783-DEL-719
chrX-66311528-DEL-1261
chrX-66339409-DEL-802
chrX-66364108-DEL-234
chrX-66381593-DEL-67
chrX-66386229-DEL-6588
chrX-66406568-DEL-1027
chrX-66432254-DEL-229
chrX-66483439-DEL-224
chrX-66485838-DEL-356
chrX-66486777-DEL-71
chrX-66488089-DEL-345
chrX-66489372-DEL-4273
chrX-66505741-DEL-790
chrX-66514737-DEL-214
chrX-66517745-DEL-533
chrX-66528828-DEL-84
chrX-66534920-DEL-84
chrX-66535222-DEL-6397
chrX-66544949-DEL-278
chrX-66549663-DEL-2481
chrX-66577656-DEL-56
chrX-66582982-DEL-9348
chrX-66591907-DEL-200
chrX-66595989-DEL-263
chrX-66596967-DEL-194
chrX-66612885-DEL-1362
chrX-66661064-DEL-581
chrX-66673680-DEL-960
chrX-66675121-DEL-874
chrX-66715610-DEL-6361
chrX-66756802-DEL-672
chrX-66762221-DEL-1043
chrX-66827878-DEL-1465
chrX-66829760-DEL-193
chrX-66862157-DEL-451
chrX-66875063-DEL-177
chrX-66879476-DEL-118
chrX-66879884-DEL-251
chrX-66889192-DEL-59
chrX-66949194-DEL-6175
chrX-66997741-DEL-2213
chrX-67001793-DEL-4532
chrX-67011917-DEL-792
chrX-67034655-DEL-52
chrX-67064082-DEL-72
chrX-67106752-DEL-1596
chrX-67139291-DEL-6348
chrX-67152778-DEL-1600
chrX-67250437-DEL-571
chrX-67265125-DEL-885
chrX-67307485-DEL-6280
chrX-67325399-DEL-2408
chrX-67351194-DEL-2344

chrX-67401138-DEL-270
chrX-67418975-DEL-1322
chrX-67448284-DEL-3674
chrX-67470365-DEL-1052
chrX-67470679-DEL-6819
chrX-67490168-DEL-994
chrX-67513127-DEL-3783
chrX-67516924-DEL-5550
chrX-67566735-DEL-170
chrX-67599446-DEL-1727
chrX-67642227-DEL-720
chrX-67687678-DEL-221
chrX-67692487-DEL-873
chrX-67693428-DEL-157
chrX-67702512-DEL-197
chrX-67715449-DEL-944
chrX-67731113-DEL-692
chrX-67732709-DEL-170
chrX-67735466-DEL-647
chrX-67736601-DEL-60
chrX-67764783-DEL-1588
chrX-67785367-DEL-94
chrX-67791038-DEL-4731
chrX-67803154-DEL-1785
chrX-67809944-DEL-978
chrX-67874295-DEL-162
chrX-67888561-DEL-468
chrX-67901270-DEL-247
chrX-67902939-DEL-653
chrX-67911840-DEL-3612
chrX-67937059-DEL-56
chrX-67945967-DEL-800
chrX-67967823-DEL-364
chrX-67972675-DEL-78
chrX-67981343-DEL-141
chrX-68009617-DEL-112
chrX-68011008-DEL-602
chrX-68044685-DEL-5009
chrX-68067853-DEL-211
chrX-68100666-DEL-3291
chrX-68115732-DEL-497
chrX-68149940-DEL-87
chrX-68163652-DEL-191
chrX-68173549-DEL-422
chrX-68174648-DEL-728
chrX-68178039-DEL-61
chrX-68186509-DEL-935
chrX-68188528-DEL-2602
chrX-68194437-DEL-80
chrX-68230976-DEL-386
chrX-68289865-DEL-6660
chrX-68302623-DEL-402
chrX-68324869-DEL-512
chrX-68334512-DEL-52
chrX-68336164-DEL-400
chrX-68338367-DEL-2348
chrX-68348008-DEL-69
chrX-68386644-DEL-1916
chrX-68389426-DEL-56
chrX-68440931-DEL-74
chrX-68449120-DEL-1794
chrX-68533851-DEL-1443
chrX-68550391-DEL-224
chrX-68556369-DEL-68
chrX-68571118-DEL-135
chrX-68590126-DEL-1490
chrX-68600454-DEL-1326
chrX-68608835-DEL-106
chrX-68685758-DEL-184
chrX-68712246-DEL-190
chrX-68715206-DEL-1825
chrX-68720906-DEL-349
chrX-68734406-DEL-211
chrX-68746093-DEL-1199
chrX-68761486-DEL-94
chrX-68762810-DEL-3979
chrX-68819531-DEL-347

chrX-68839251-DEL-196
chrX-68888113-DEL-223
chrX-68961484-DEL-114
chrX-68981213-DEL-424
chrX-69026851-DEL-122
chrX-69039446-DEL-73
chrX-69045854-DEL-64
chrX-69062502-DEL-414
chrX-69082306-DEL-67
chrX-69098899-DEL-6644
chrX-69109692-DEL-6628
chrX-69124402-DEL-969
chrX-69125353-DEL-62
chrX-69126728-DEL-254
chrX-69150078-DEL-1587
chrX-69171973-DEL-102
chrX-69188130-DEL-806
chrX-69230490-DEL-4895
chrX-69287606-DEL-853
chrX-69290263-DEL-13492
chrX-69307251-DEL-11642
chrX-69317354-DEL-176
chrX-69326057-DEL-54
chrX-69335939-DEL-1393
chrX-69418042-DEL-91
chrX-69424148-DEL-1397
chrX-69439273-DEL-1347
chrX-69528015-DEL-10762
chrX-69544379-DEL-205
chrX-69553035-DEL-426
chrX-69668533-DEL-572
chrX-69704200-DEL-6277
chrX-69710162-DEL-64
chrX-69736117-DEL-2204
chrX-69762907-DEL-2734
chrX-69763512-DEL-778
chrX-69763597-DEL-1873
chrX-69879241-DEL-178
chrX-69885993-DEL-60
chrX-69903897-DEL-6532
chrX-69924079-DEL-119
chrX-69931557-DEL-852
chrX-69946733-DEL-373
chrX-70014430-DEL-71
chrX-70036679-DEL-467
chrX-70044569-DEL-60
chrX-70056760-DEL-153
chrX-70079026-DEL-154
chrX-70120252-DEL-72
chrX-70148538-DEL-52
chrX-70274574-DEL-71
chrX-70282442-DEL-217
chrX-70374832-DEL-126
chrX-70435423-DEL-100
chrX-70467779-DEL-98
chrX-70505940-DEL-71
chrX-70524595-DEL-150
chrX-70528559-DEL-400
chrX-70552524-DEL-179
chrX-70566911-DEL-110
chrX-70574482-DEL-57
chrX-70637860-DEL-67
chrX-70666581-DEL-125
chrX-70689028-DEL-97
chrX-70756306-DEL-60
chrX-70771835-DEL-427
chrX-70774013-DEL-256
chrX-70774888-DEL-64
chrX-70776600-DEL-606
chrX-70779548-DEL-287
chrX-70782387-DEL-382
chrX-70802954-DEL-1052
chrX-70804755-DEL-166
chrX-70835099-DEL-467
chrX-70849781-DEL-354
chrX-70884989-DEL-544
chrX-70918106-DEL-58

chrX-70939031-DEL-1887
chrX-70944588-DEL-267
chrX-70951556-DEL-218
chrX-70962252-DEL-709
chrX-70963005-DEL-299
chrX-70990150-DEL-87
chrX-71071406-DEL-1401
chrX-71140881-DEL-259
chrX-71166327-DEL-8128
chrX-71194093-DEL-1128
chrX-71201099-DEL-95
chrX-71222520-DEL-2903
chrX-71226166-DEL-513
chrX-71235044-DEL-1464
chrX-71241831-DEL-108
chrX-71275500-DEL-11541
chrX-71319738-DEL-220
chrX-71321288-DEL-88
chrX-71332529-DEL-558
chrX-71336201-DEL-221
chrX-71387571-DEL-58
chrX-71388276-DEL-420
chrX-71398711-DEL-2901
chrX-71419696-DEL-414
chrX-71422789-DEL-419
chrX-71433306-DEL-371
chrX-71442929-DEL-6394
chrX-71448943-DEL-199
chrX-71498142-DEL-478
chrX-71512627-DEL-75
chrX-71546938-DEL-200
chrX-71563024-DEL-66
chrX-71571086-DEL-2451
chrX-71599060-DEL-643
chrX-71621749-DEL-636
chrX-71634059-DEL-611
chrX-71668935-DEL-587
chrX-71720184-DEL-71
chrX-71765869-DEL-425
chrX-71798700-DEL-982
chrX-71827154-DEL-65
chrX-71843604-DEL-1324
chrX-71865419-DEL-239
chrX-71895009-DEL-68
chrX-71922466-DEL-190
chrX-71975780-DEL-6275
chrX-72039847-DEL-143
chrX-72084999-DEL-1750
chrX-72116232-DEL-81
chrX-72116844-DEL-526
chrX-72121141-DEL-177
chrX-72128208-DEL-78
chrX-72148356-DEL-55
chrX-72161363-DEL-140
chrX-72173965-DEL-444
chrX-72179295-DEL-444
chrX-72192189-DEL-140
chrX-72220572-DEL-130
chrX-72311690-DEL-7117
chrX-72375198-DEL-753
chrX-72381721-DEL-51
chrX-72382845-DEL-580
chrX-72403299-DEL-202
chrX-72426148-DEL-181
chrX-72446573-DEL-52
chrX-72448126-DEL-2305
chrX-72498769-DEL-54
chrX-72506868-DEL-73
chrX-72514151-DEL-154
chrX-72577571-DEL-52
chrX-72611697-DEL-56
chrX-72641562-DEL-428
chrX-72666281-DEL-345
chrX-72746568-DEL-121
chrX-72771002-DEL-77
chrX-72777152-DEL-76
chrX-72783122-DEL-410

chrX-72787928-DEL-51
chrX-72847697-DEL-500
chrX-72854563-DEL-223
chrX-72873809-DEL-394
chrX-72875303-DEL-1047
chrX-72877630-DEL-204
chrX-72942743-DEL-170
chrX-72975008-DEL-210
chrX-73016207-DEL-182
chrX-73024283-DEL-142
chrX-73041119-DEL-726
chrX-73056135-DEL-202
chrX-73096452-DEL-110
chrX-73107755-DEL-60
chrX-73111452-DEL-238
chrX-73114623-DEL-122
chrX-73214227-DEL-1330
chrX-73226748-DEL-2266
chrX-73228866-DEL-166
chrX-73250644-DEL-360
chrX-73304491-DEL-695
chrX-73370308-DEL-56
chrX-73404988-DEL-58
chrX-73425987-DEL-1336
chrX-73464997-DEL-205
chrX-73481473-DEL-710
chrX-73497621-DEL-14897
chrX-73522255-DEL-163
chrX-73541899-DEL-79
chrX-73555964-DEL-3784
chrX-73567590-DEL-203
chrX-73596691-DEL-3612
chrX-73611400-DEL-496
chrX-73658185-DEL-8154
chrX-73665424-DEL-212
chrX-73667465-DEL-73
chrX-73676519-DEL-214
chrX-73706670-DEL-55
chrX-73719039-DEL-66
chrX-73730733-DEL-3709
chrX-73732929-DEL-250
chrX-73752510-DEL-2252
chrX-73779306-DEL-59
chrX-73795670-DEL-4232
chrX-73799643-DEL-7593
chrX-73814190-DEL-230
chrX-73907107-DEL-16829
chrX-73934879-DEL-58
chrX-73939997-DEL-901
chrX-73945944-DEL-105
chrX-73951922-DEL-3113
chrX-73971226-DEL-74
chrX-73989531-DEL-195
chrX-74081415-DEL-373
chrX-74108253-DEL-152
chrX-74116281-DEL-608
chrX-74147458-DEL-388
chrX-74179257-DEL-62
chrX-74190332-DEL-80
chrX-74199504-DEL-117
chrX-74219621-DEL-188
chrX-74276452-DEL-2905
chrX-74317317-DEL-202
chrX-74332535-DEL-468
chrX-74400788-DEL-5590
chrX-74441377-DEL-1012
chrX-74474613-DEL-405
chrX-74485281-DEL-211
chrX-74487639-DEL-114
chrX-74579042-DEL-171
chrX-74588454-DEL-1588
chrX-74600138-DEL-410
chrX-74622944-DEL-499
chrX-74662878-DEL-2461
chrX-74663632-DEL-66
chrX-74691566-DEL-66
chrX-74692799-DEL-474

chrX-74693457-DEL-717
chrX-74694375-DEL-57
chrX-74695724-DEL-879
chrX-74698689-DEL-59
chrX-74705589-DEL-2447
chrX-74707327-DEL-431
chrX-74708107-DEL-59
chrX-74725326-DEL-977
chrX-74730216-DEL-1298
chrX-74734195-DEL-197
chrX-74772264-DEL-894
chrX-74773844-DEL-69
chrX-74773993-DEL-2845
chrX-74777892-DEL-158
chrX-74778738-DEL-67
chrX-74782900-DEL-968
chrX-74802066-DEL-968
chrX-74809777-DEL-159
chrX-74840165-DEL-79
chrX-74898327-DEL-289
chrX-74925095-DEL-303
chrX-74980297-DEL-9081
chrX-74982570-DEL-3129
chrX-74989554-DEL-302
chrX-74991183-DEL-2019
chrX-74994728-DEL-9467
chrX-75046645-DEL-54
chrX-75060304-DEL-354
chrX-75068194-DEL-581
chrX-75090041-DEL-2015
chrX-75094755-DEL-1075
chrX-75099236-DEL-77
chrX-75101723-DEL-2095
chrX-75146038-DEL-55
chrX-75198316-DEL-1323
chrX-75249934-DEL-14189
chrX-75269222-DEL-63
chrX-75270027-DEL-4554
chrX-75306828-DEL-91
chrX-75339762-DEL-769
chrX-75366339-DEL-2030
chrX-75397588-DEL-4363
chrX-75403530-DEL-11135
chrX-75485859-DEL-640
chrX-75491274-DEL-141
chrX-75507726-DEL-6398
chrX-75517031-DEL-51
chrX-75529249-DEL-1012
chrX-75590223-DEL-6182
chrX-75605209-DEL-58
chrX-75649558-DEL-1246
chrX-75661487-DEL-64
chrX-75714263-DEL-645
chrX-75720868-DEL-62
chrX-75745544-DEL-5666
chrX-75773779-DEL-203
chrX-75785053-DEL-2770
chrX-75810887-DEL-509
chrX-75835676-DEL-513
chrX-75842815-DEL-497
chrX-75846221-DEL-184
chrX-75869153-DEL-169
chrX-75895754-DEL-198
chrX-75920516-DEL-122
chrX-75959407-DEL-52
chrX-76010061-DEL-60
chrX-76031392-DEL-2842
chrX-76054756-DEL-56
chrX-76065373-DEL-63
chrX-76128829-DEL-54
chrX-76133332-DEL-81
chrX-76173307-DEL-434
chrX-76182966-DEL-2219
chrX-76189421-DEL-774
chrX-76215969-DEL-723
chrX-76216764-DEL-12224
chrX-76219025-DEL-350

chrX-76219086-DEL-5482
chrX-76286034-DEL-199
chrX-76297366-DEL-279
chrX-76301085-DEL-7960
chrX-76301101-DEL-5535
chrX-76301414-DEL-14664
chrX-76301576-DEL-212
chrX-76308262-DEL-6351
chrX-76309214-DEL-6865
chrX-76423819-DEL-91
chrX-76430321-DEL-64
chrX-76589397-DEL-1213
chrX-76717249-DEL-62
chrX-76722242-DEL-55
chrX-76729579-DEL-800
chrX-76762695-DEL-222
chrX-76783813-DEL-595
chrX-76798450-DEL-180
chrX-76906042-DEL-87
chrX-76938390-DEL-6233
chrX-76939053-DEL-5595
chrX-76944790-DEL-214
chrX-76989736-DEL-1638
chrX-77000991-DEL-158
chrX-77002716-DEL-17516
chrX-77057025-DEL-2210
chrX-77068897-DEL-6227
chrX-77085815-DEL-6531
chrX-77116950-DEL-65
chrX-77117643-DEL-2004
chrX-77120032-DEL-6052
chrX-77163471-DEL-1461
chrX-77167467-DEL-352
chrX-77217176-DEL-4768
chrX-77243085-DEL-113
chrX-77248154-DEL-4231
chrX-77249469-DEL-6472
chrX-77257208-DEL-52
chrX-77270270-DEL-202
chrX-77332639-DEL-258
chrX-77376642-DEL-8037
chrX-77423544-DEL-1126
chrX-77423803-DEL-858
chrX-77436746-DEL-74
chrX-77441528-DEL-123
chrX-77454096-DEL-740
chrX-77467075-DEL-1830
chrX-77484844-DEL-52
chrX-77515229-DEL-227
chrX-77520926-DEL-2507
chrX-77537567-DEL-221
chrX-77547309-DEL-381
chrX-77556097-DEL-761
chrX-77612560-DEL-955
chrX-77620888-DEL-51
chrX-77621189-DEL-444
chrX-77693474-DEL-54
chrX-77694673-DEL-5952
chrX-77700667-DEL-11738
chrX-77727463-DEL-2945
chrX-77747901-DEL-72
chrX-77752394-DEL-7119
chrX-77794398-DEL-5545
chrX-77844094-DEL-1472
chrX-77881370-DEL-3717
chrX-77886623-DEL-51
chrX-77902050-DEL-53
chrX-77911523-DEL-6076
chrX-77918074-DEL-152
chrX-77931161-DEL-8885
chrX-77990186-DEL-498
chrX-77991013-DEL-4907
chrX-77991335-DEL-522
chrX-78089516-DEL-60
chrX-78102378-DEL-1026
chrX-78124829-DEL-898
chrX-78163251-DEL-141

chrX-78178522-DEL-2010
chrX-78200776-DEL-60
chrX-78204268-DEL-207
chrX-78236937-DEL-4853
chrX-78285647-DEL-557
chrX-78294942-DEL-317
chrX-78311917-DEL-63
chrX-78330715-DEL-6518
chrX-78343897-DEL-79
chrX-78352413-DEL-722
chrX-78360029-DEL-85
chrX-78427406-DEL-1691
chrX-78431152-DEL-506
chrX-78442213-DEL-1230
chrX-78448330-DEL-1719
chrX-78460870-DEL-452
chrX-78464977-DEL-3918
chrX-78503042-DEL-1785
chrX-78551891-DEL-58
chrX-78555349-DEL-511
chrX-78579600-DEL-8457
chrX-78598950-DEL-671
chrX-78605102-DEL-3432
chrX-78653891-DEL-6214
chrX-78676458-DEL-171
chrX-78685697-DEL-226
chrX-78691813-DEL-1218
chrX-78719235-DEL-401
chrX-78738350-DEL-2840
chrX-78747243-DEL-229
chrX-78750878-DEL-527
chrX-78751451-DEL-1274
chrX-78753123-DEL-1065
chrX-78761370-DEL-214
chrX-78798325-DEL-1441
chrX-78806647-DEL-168
chrX-78823868-DEL-191
chrX-78826354-DEL-188
chrX-78829732-DEL-206
chrX-78861412-DEL-8808
chrX-78869859-DEL-441
chrX-78918998-DEL-498
chrX-78925331-DEL-214
chrX-78925976-DEL-100
chrX-78960782-DEL-296
chrX-78965180-DEL-6231
chrX-78984120-DEL-284
chrX-79009603-DEL-1895
chrX-79022405-DEL-96
chrX-79044312-DEL-1327
chrX-79066398-DEL-1342
chrX-79071115-DEL-671
chrX-79077705-DEL-70
chrX-79127198-DEL-72
chrX-79131303-DEL-6213
chrX-79133400-DEL-2364
chrX-79226989-DEL-885
chrX-79229586-DEL-397
chrX-79272589-DEL-114
chrX-79288582-DEL-365
chrX-79334458-DEL-388
chrX-79357492-DEL-695
chrX-79360569-DEL-601
chrX-79379309-DEL-2690
chrX-79400660-DEL-198
chrX-79403755-DEL-94
chrX-79405876-DEL-2354
chrX-79409377-DEL-6421
chrX-79422369-DEL-6195
chrX-79430383-DEL-2471
chrX-79436411-DEL-2103
chrX-79449380-DEL-198
chrX-79458123-DEL-4595
chrX-79485537-DEL-64
chrX-79487162-DEL-1436
chrX-79489817-DEL-1838
chrX-79538465-DEL-414

chrX-79542717-DEL-62
chrX-79546573-DEL-6392
chrX-79553550-DEL-1178
chrX-79556678-DEL-3568
chrX-79575138-DEL-68
chrX-79584760-DEL-6149
chrX-79597648-DEL-9614
chrX-79601316-DEL-5168
chrX-79684359-DEL-88
chrX-79699226-DEL-316
chrX-79702917-DEL-238
chrX-79731561-DEL-390
chrX-79734149-DEL-1722
chrX-79738531-DEL-1894
chrX-79779748-DEL-54
chrX-79780635-DEL-4797
chrX-79786782-DEL-396
chrX-79787469-DEL-11174
chrX-79790656-DEL-316
chrX-79792405-DEL-6660
chrX-79830969-DEL-72
chrX-79854368-DEL-373
chrX-79867959-DEL-6154
chrX-79909667-DEL-636
chrX-79914460-DEL-64
chrX-79933445-DEL-6449
chrX-79946149-DEL-106
chrX-79991369-DEL-256
chrX-80021650-DEL-9269
chrX-80022365-DEL-39369
chrX-80023328-DEL-2441
chrX-80031062-DEL-59
chrX-80069449-DEL-4881
chrX-80122230-DEL-67
chrX-80154314-DEL-2717
chrX-80182230-DEL-170
chrX-80194487-DEL-6366
chrX-80194562-DEL-200
chrX-80224679-DEL-6516
chrX-80231641-DEL-51
chrX-80238664-DEL-64
chrX-80302475-DEL-59
chrX-80326140-DEL-615
chrX-80347196-DEL-753
chrX-80368574-DEL-75
chrX-80372587-DEL-55
chrX-80375354-DEL-122
chrX-80402129-DEL-185
chrX-80421347-DEL-103
chrX-80427120-DEL-74
chrX-80427185-DEL-1459
chrX-80430590-DEL-4886
chrX-80559747-DEL-4649
chrX-80568261-DEL-500
chrX-80583148-DEL-243
chrX-80584556-DEL-183
chrX-80618580-DEL-400
chrX-80621525-DEL-109
chrX-80622319-DEL-1089
chrX-80684420-DEL-335
chrX-80720430-DEL-6798
chrX-80727270-DEL-1327
chrX-80762369-DEL-1862
chrX-80765740-DEL-56
chrX-80770124-DEL-480
chrX-80782299-DEL-20958
chrX-80787550-DEL-1249
chrX-80800595-DEL-10631
chrX-80834002-DEL-78
chrX-80875988-DEL-851
chrX-80878337-DEL-2330
chrX-80881606-DEL-1293
chrX-80918471-DEL-125
chrX-80920284-DEL-96
chrX-80964312-DEL-1228
chrX-80966038-DEL-2937
chrX-81000998-DEL-527

chrX-81006635-DEL-2308
chrX-81011678-DEL-94
chrX-81033212-DEL-249
chrX-81035013-DEL-633
chrX-81044712-DEL-659
chrX-81062652-DEL-710
chrX-81128363-DEL-5976
chrX-81143313-DEL-207
chrX-81254440-DEL-5448
chrX-81255957-DEL-315
chrX-81257002-DEL-2945
chrX-81268357-DEL-219
chrX-81276083-DEL-6645
chrX-81293469-DEL-669
chrX-81322776-DEL-1579
chrX-81378051-DEL-84
chrX-81409072-DEL-619
chrX-81420578-DEL-1836
chrX-81427637-DEL-1392
chrX-81468939-DEL-121
chrX-81478725-DEL-1317
chrX-81512899-DEL-543
chrX-81516593-DEL-331
chrX-81519366-DEL-51
chrX-81558849-DEL-65
chrX-81575869-DEL-6296
chrX-81587687-DEL-105
chrX-81610067-DEL-1903
chrX-81632101-DEL-2941
chrX-81637107-DEL-501
chrX-81642848-DEL-53
chrX-81653354-DEL-62
chrX-81674593-DEL-391
chrX-81690548-DEL-72
chrX-81694958-DEL-11719
chrX-81721293-DEL-57
chrX-81743536-DEL-68
chrX-81744601-DEL-181
chrX-81802491-DEL-2169
chrX-81835426-DEL-1923
chrX-81897770-DEL-52
chrX-81905207-DEL-134
chrX-81921809-DEL-235
chrX-81922991-DEL-2025
chrX-81929953-DEL-2725
chrX-81975366-DEL-142
chrX-81985905-DEL-221
chrX-82030751-DEL-51
chrX-82039072-DEL-98
chrX-82059482-DEL-54
chrX-82110312-DEL-104
chrX-82129758-DEL-272
chrX-82169277-DEL-60
chrX-82179414-DEL-284
chrX-82193383-DEL-123
chrX-82204877-DEL-59
chrX-82212704-DEL-58
chrX-82220708-DEL-52
chrX-82244527-DEL-194
chrX-82272241-DEL-652
chrX-82306892-DEL-98
chrX-82310269-DEL-578
chrX-82346404-DEL-295
chrX-82350862-DEL-4440
chrX-82357701-DEL-173
chrX-82367717-DEL-732
chrX-82370663-DEL-3617
chrX-82384576-DEL-1450
chrX-82397021-DEL-96
chrX-82405942-DEL-4917
chrX-82427468-DEL-493
chrX-82458409-DEL-56
chrX-82498738-DEL-472
chrX-82514160-DEL-110
chrX-82522221-DEL-82
chrX-82527892-DEL-235
chrX-82528871-DEL-375

chrX-82542542-DEL-10212
chrX-82546889-DEL-10236
chrX-82551429-DEL-3417
chrX-82569794-DEL-6565
chrX-82579034-DEL-71
chrX-82616309-DEL-70
chrX-82628808-DEL-839
chrX-82632187-DEL-78
chrX-82633468-DEL-674
chrX-82643469-DEL-120
chrX-82667598-DEL-1772
chrX-82668183-DEL-74
chrX-82711859-DEL-55
chrX-82736503-DEL-352
chrX-82740064-DEL-227
chrX-82767735-DEL-323
chrX-82769591-DEL-68
chrX-82778095-DEL-978
chrX-82823881-DEL-246
chrX-82847475-DEL-118
chrX-82860498-DEL-576
chrX-82892211-DEL-2282
chrX-82900945-DEL-185
chrX-82907744-DEL-199
chrX-82911563-DEL-905
chrX-82938323-DEL-211
chrX-82944059-DEL-171
chrX-82968761-DEL-93
chrX-83002292-DEL-328
chrX-83015739-DEL-744
chrX-83027696-DEL-772
chrX-83067596-DEL-131
chrX-83148104-DEL-58
chrX-83182293-DEL-1559
chrX-83222769-DEL-401
chrX-83232376-DEL-485
chrX-83264557-DEL-54
chrX-83278736-DEL-79
chrX-83300742-DEL-83
chrX-83309250-DEL-314
chrX-83323322-DEL-2027
chrX-83334327-DEL-6222
chrX-83366332-DEL-82
chrX-83379415-DEL-86
chrX-83422497-DEL-1187
chrX-83450182-DEL-1029
chrX-83460265-DEL-668
chrX-83511070-DEL-1276
chrX-83520550-DEL-1468
chrX-83523936-DEL-334
chrX-83529428-DEL-1762
chrX-83560643-DEL-88
chrX-83595990-DEL-425
chrX-83602520-DEL-2695
chrX-83612730-DEL-81
chrX-83619509-DEL-60
chrX-83620324-DEL-780
chrX-83675088-DEL-169
chrX-83723553-DEL-927
chrX-83806661-DEL-102
chrX-83858646-DEL-73
chrX-83881094-DEL-109
chrX-83907052-DEL-74
chrX-83940260-DEL-78
chrX-83944670-DEL-52
chrX-83946353-DEL-80
chrX-83977614-DEL-52
chrX-83988934-DEL-124
chrX-83992344-DEL-303
chrX-84056532-DEL-488
chrX-84110887-DEL-114
chrX-84138240-DEL-397
chrX-84140691-DEL-66
chrX-84142986-DEL-51
chrX-84163931-DEL-151
chrX-84172419-DEL-336
chrX-84207217-DEL-123

chrX-84208726-DEL-558
chrX-84217956-DEL-56
chrX-84220810-DEL-59
chrX-84255329-DEL-55
chrX-84379621-DEL-71
chrX-84394210-DEL-176
chrX-84396631-DEL-2077
chrX-84408845-DEL-3307
chrX-84417050-DEL-1448
chrX-84495467-DEL-497
chrX-84531676-DEL-3233
chrX-84534923-DEL-1121
chrX-84548753-DEL-6425
chrX-84573576-DEL-1231
chrX-84605459-DEL-65
chrX-84684609-DEL-246
chrX-84710867-DEL-199
chrX-84727267-DEL-153
chrX-84798969-DEL-151
chrX-84821438-DEL-115
chrX-84897444-DEL-66
chrX-84900351-DEL-372
chrX-84935028-DEL-60
chrX-84944459-DEL-262
chrX-84952380-DEL-432
chrX-84952809-DEL-983
chrX-84963007-DEL-68
chrX-84969493-DEL-267
chrX-84971007-DEL-92
chrX-84989421-DEL-59
chrX-85014590-DEL-297
chrX-85024194-DEL-112
chrX-85142358-DEL-7171
chrX-85161191-DEL-54
chrX-85164446-DEL-3632
chrX-85167181-DEL-58
chrX-85171191-DEL-796
chrX-85194783-DEL-277
chrX-85196814-DEL-58
chrX-85217145-DEL-601
chrX-85218331-DEL-66
chrX-85259818-DEL-122
chrX-85290033-DEL-107
chrX-85294549-DEL-309
chrX-85301036-DEL-3405
chrX-85331830-DEL-6919
chrX-85366324-DEL-90
chrX-85387012-DEL-114
chrX-85397712-DEL-62
chrX-85426915-DEL-61
chrX-85429711-DEL-62
chrX-85503646-DEL-62
chrX-85519378-DEL-411
chrX-85537827-DEL-1233
chrX-85547706-DEL-54
chrX-85558431-DEL-2464
chrX-85562574-DEL-138
chrX-85578844-DEL-1251
chrX-85617482-DEL-134
chrX-85641302-DEL-538
chrX-85658035-DEL-404
chrX-85691626-DEL-162
chrX-85707540-DEL-1811
chrX-85736341-DEL-152
chrX-85741197-DEL-5770
chrX-85758253-DEL-73
chrX-85759646-DEL-123
chrX-85836491-DEL-2636
chrX-85855063-DEL-56
chrX-85864779-DEL-8013
chrX-85904515-DEL-376
chrX-85935565-DEL-167
chrX-85983966-DEL-1257
chrX-86000529-DEL-830
chrX-86012118-DEL-71
chrX-86027059-DEL-99
chrX-86032234-DEL-1372

chrX-86036075-DEL-54
chrX-86073096-DEL-116
chrX-86107036-DEL-56
chrX-86113887-DEL-65
chrX-86133959-DEL-487
chrX-86162077-DEL-111
chrX-86184800-DEL-524
chrX-86247441-DEL-66
chrX-86259972-DEL-51
chrX-86263024-DEL-647
chrX-86287043-DEL-75
chrX-86292528-DEL-52
chrX-86343438-DEL-57
chrX-86386377-DEL-140
chrX-86392572-DEL-239
chrX-86403589-DEL-66
chrX-86404347-DEL-85
chrX-86427762-DEL-63
chrX-86459258-DEL-54
chrX-86477395-DEL-124
chrX-86561592-DEL-449
chrX-86580840-DEL-88
chrX-86603449-DEL-520
chrX-86608278-DEL-1255
chrX-86609550-DEL-338
chrX-86618051-DEL-477
chrX-86622098-DEL-914
chrX-86634253-DEL-1058
chrX-86638826-DEL-928
chrX-86650788-DEL-58
chrX-86672152-DEL-2735
chrX-86680089-DEL-689
chrX-86681542-DEL-656
chrX-86683726-DEL-513
chrX-86713504-DEL-941
chrX-86726385-DEL-76
chrX-86750157-DEL-64
chrX-86752762-DEL-55
chrX-86761433-DEL-408
chrX-86771013-DEL-513
chrX-86814176-DEL-779
chrX-86824491-DEL-804
chrX-86827326-DEL-230
chrX-86882374-DEL-424
chrX-86981943-DEL-320
chrX-87001635-DEL-80
chrX-87029470-DEL-1739
chrX-87066185-DEL-63
chrX-87074783-DEL-94
chrX-87131182-DEL-1018
chrX-87182638-DEL-501
chrX-87194598-DEL-98
chrX-87236683-DEL-59
chrX-87253065-DEL-1349
chrX-87263317-DEL-107
chrX-87267344-DEL-1722
chrX-87273721-DEL-6205
chrX-87285981-DEL-218
chrX-87318256-DEL-332
chrX-87337568-DEL-651
chrX-87390375-DEL-6417
chrX-87412010-DEL-69
chrX-87431964-DEL-195
chrX-87473366-DEL-68
chrX-87475870-DEL-1008
chrX-87478217-DEL-2441
chrX-87516390-DEL-101
chrX-87544237-DEL-1315
chrX-87587089-DEL-400
chrX-87592584-DEL-1085
chrX-87600093-DEL-56
chrX-87605265-DEL-5852
chrX-87616285-DEL-138
chrX-87621200-DEL-2462
chrX-87630597-DEL-108
chrX-87636137-DEL-75
chrX-87681635-DEL-124

chrX-87691866-DEL-1557
chrX-87696821-DEL-124
chrX-87721301-DEL-783
chrX-87724584-DEL-51
chrX-87725787-DEL-1636
chrX-87758530-DEL-186
chrX-87763061-DEL-1180
chrX-87783024-DEL-844
chrX-87790010-DEL-205
chrX-87801069-DEL-56
chrX-87801854-DEL-248
chrX-87807707-DEL-1396
chrX-87831400-DEL-1285
chrX-87872954-DEL-2633
chrX-87894226-DEL-248
chrX-87942205-DEL-57
chrX-87967006-DEL-6556
chrX-87984117-DEL-58
chrX-88002788-DEL-5021
chrX-88014543-DEL-78
chrX-88032399-DEL-1896
chrX-88048974-DEL-426
chrX-88051276-DEL-12481
chrX-88081038-DEL-670
chrX-88105907-DEL-5472
chrX-88114439-DEL-6903
chrX-88163834-DEL-80
chrX-88188473-DEL-114
chrX-88189922-DEL-163
chrX-88195624-DEL-2043
chrX-88218965-DEL-57
chrX-88220463-DEL-217
chrX-88237634-DEL-66
chrX-88247230-DEL-234
chrX-88256965-DEL-637
chrX-88260857-DEL-443
chrX-88284536-DEL-201
chrX-88369784-DEL-4308
chrX-88389435-DEL-59
chrX-88427295-DEL-285
chrX-88433973-DEL-103
chrX-88442831-DEL-219
chrX-88452356-DEL-259
chrX-88458326-DEL-72
chrX-88474703-DEL-102
chrX-88495060-DEL-225
chrX-88501154-DEL-776
chrX-88510853-DEL-2096
chrX-88512970-DEL-1275
chrX-88524277-DEL-56
chrX-88530051-DEL-13067
chrX-88531681-DEL-6823
chrX-88558807-DEL-331
chrX-88582661-DEL-1207
chrX-88584819-DEL-80
chrX-88587646-DEL-1196
chrX-88597756-DEL-394
chrX-88658358-DEL-2841
chrX-88661992-DEL-548
chrX-88674334-DEL-5926
chrX-88685688-DEL-8512
chrX-88698175-DEL-203
chrX-88708938-DEL-968
chrX-88761129-DEL-467
chrX-88801260-DEL-7451
chrX-88820862-DEL-160
chrX-88839902-DEL-1785
chrX-88847290-DEL-698
chrX-88874800-DEL-96
chrX-88886951-DEL-309
chrX-88890765-DEL-70
chrX-88891692-DEL-3728
chrX-88908680-DEL-129
chrX-88925895-DEL-1604
chrX-88928532-DEL-401
chrX-88929479-DEL-1934
chrX-88950090-DEL-1391

chrX-88958275-DEL-2030
chrX-88973055-DEL-401
chrX-88990327-DEL-391
chrX-89027539-DEL-242
chrX-89057450-DEL-62
chrX-89076850-DEL-192
chrX-89089438-DEL-56
chrX-89102764-DEL-62
chrX-89118670-DEL-731
chrX-89127643-DEL-1014
chrX-89183421-DEL-2261
chrX-89211462-DEL-384
chrX-89222538-DEL-1311
chrX-89242055-DEL-61
chrX-89277894-DEL-1680
chrX-89304547-DEL-102
chrX-89328780-DEL-862
chrX-89341074-DEL-86
chrX-89351323-DEL-6175
chrX-89369905-DEL-400
chrX-89373917-DEL-211
chrX-89385014-DEL-62
chrX-89399757-DEL-2493
chrX-89429846-DEL-786
chrX-89444297-DEL-2381
chrX-89480955-DEL-6351
chrX-89498543-DEL-53
chrX-89521904-DEL-180
chrX-89610438-DEL-1356
chrX-89619039-DEL-340
chrX-89627514-DEL-1339
chrX-89630652-DEL-71
chrX-89636112-DEL-5062
chrX-89642925-DEL-999
chrX-89659818-DEL-56
chrX-89688369-DEL-816
chrX-89737835-DEL-727
chrX-89747069-DEL-180
chrX-89748132-DEL-13942
chrX-89762635-DEL-1231
chrX-89818813-DEL-848
chrX-89827512-DEL-103
chrX-89833434-DEL-408
chrX-89835164-DEL-72
chrX-89858064-DEL-95
chrX-89897530-DEL-6185
chrX-89897610-DEL-198
chrX-89903716-DEL-14997
chrX-89904561-DEL-193
chrX-89905535-DEL-65
chrX-89916176-DEL-483
chrX-89956735-DEL-60
chrX-89965096-DEL-1499
chrX-90034775-DEL-201
chrX-90057202-DEL-433
chrX-90083055-DEL-52
chrX-90090764-DEL-72
chrX-90127822-DEL-60
chrX-90161613-DEL-58306
chrX-90164752-DEL-10155
chrX-90227209-DEL-54
chrX-90277944-DEL-672
chrX-90318931-DEL-236
chrX-90323130-DEL-1965
chrX-90330822-DEL-65
chrX-90337612-DEL-3368
chrX-90363696-DEL-350
chrX-90367439-DEL-4697
chrX-90369863-DEL-80
chrX-90369986-DEL-122
chrX-90377598-DEL-1314
chrX-90378007-DEL-2033
chrX-90385404-DEL-517
chrX-90390132-DEL-282
chrX-90394842-DEL-55
chrX-90546315-DEL-2105
chrX-90572979-DEL-5821

chrX-90591517-DEL-76
chrX-90740276-DEL-326
chrX-90763065-DEL-486
chrX-90787532-DEL-102
chrX-90787693-DEL-5787
chrX-90845232-DEL-2096
chrX-90875873-DEL-205
chrX-90878360-DEL-9724
chrX-90888922-DEL-7105
chrX-90904639-DEL-564
chrX-90917039-DEL-6398
chrX-90926068-DEL-475
chrX-90952920-DEL-67
chrX-90976595-DEL-414
chrX-90979956-DEL-385
chrX-91050828-DEL-198
chrX-91057036-DEL-188
chrX-91062821-DEL-1222
chrX-91063068-DEL-120
chrX-91064697-DEL-3043
chrX-91064923-DEL-184
chrX-91066943-DEL-228
chrX-91069618-DEL-204
chrX-91097640-DEL-856
chrX-91104371-DEL-64
chrX-91105731-DEL-10595
chrX-91126659-DEL-58
chrX-91142125-DEL-1109
chrX-91147299-DEL-150
chrX-91150078-DEL-223
chrX-91150808-DEL-742
chrX-91151621-DEL-1002
chrX-91168473-DEL-263
chrX-91172140-DEL-62
chrX-91175209-DEL-139
chrX-91209863-DEL-190
chrX-91214002-DEL-703
chrX-91225466-DEL-304
chrX-91229738-DEL-61
chrX-91239713-DEL-71
chrX-91250345-DEL-62
chrX-91285681-DEL-1228
chrX-91307172-DEL-13983
chrX-91335753-DEL-141
chrX-91347062-DEL-88
chrX-91347968-DEL-644
chrX-91375576-DEL-76
chrX-91396203-DEL-150
chrX-91399812-DEL-431
chrX-91411373-DEL-254
chrX-91427592-DEL-1297
chrX-91445051-DEL-80
chrX-91485295-DEL-120
chrX-91506811-DEL-1271
chrX-91542336-DEL-51
chrX-91567649-DEL-206
chrX-91587889-DEL-58
chrX-91615415-DEL-228
chrX-91658659-DEL-214
chrX-91683509-DEL-244
chrX-91693898-DEL-85
chrX-91756082-DEL-214
chrX-91811502-DEL-425
chrX-91844242-DEL-4348
chrX-91848609-DEL-1319
chrX-91863027-DEL-416
chrX-91864679-DEL-655
chrX-91894805-DEL-205
chrX-92004539-DEL-78
chrX-92004752-DEL-6488
chrX-92011239-DEL-767
chrX-92035431-DEL-2612
chrX-92036834-DEL-59
chrX-92043962-DEL-856
chrX-92065015-DEL-6270
chrX-92087894-DEL-274
chrX-92089536-DEL-68

chrX-92090114-DEL-584
chrX-92101672-DEL-486
chrX-92103243-DEL-2553
chrX-92125739-DEL-1995
chrX-92135912-DEL-610
chrX-92143513-DEL-2104
chrX-92205310-DEL-29514
chrX-92220230-DEL-2133
chrX-92231839-DEL-2976
chrX-92303670-DEL-277
chrX-92304290-DEL-78
chrX-92324497-DEL-54
chrX-92371168-DEL-58
chrX-92387037-DEL-173
chrX-92392857-DEL-174
chrX-92434467-DEL-58
chrX-92444278-DEL-306
chrX-92500350-DEL-80
chrX-92518450-DEL-368
chrX-92574707-DEL-52
chrX-92607689-DEL-120
chrX-92614216-DEL-772
chrX-92615729-DEL-266
chrX-92666657-DEL-427
chrX-92703593-DEL-806
chrX-92705308-DEL-199
chrX-92756636-DEL-197
chrX-92821792-DEL-52
chrX-92839553-DEL-172
chrX-92912551-DEL-107
chrX-92922723-DEL-116
chrX-92925726-DEL-400
chrX-92940476-DEL-54
chrX-92965762-DEL-210
chrX-93030853-DEL-148
chrX-93048656-DEL-95
chrX-93058880-DEL-112
chrX-93083489-DEL-174
chrX-93084390-DEL-170
chrX-93100755-DEL-1739
chrX-93113508-DEL-187
chrX-93136530-DEL-63
chrX-93173027-DEL-276
chrX-93203096-DEL-810
chrX-93212028-DEL-414
chrX-93212052-DEL-382
chrX-93212649-DEL-244
chrX-93227709-DEL-205
chrX-93231635-DEL-359
chrX-93248799-DEL-213
chrX-93266489-DEL-164
chrX-93269804-DEL-264
chrX-93288320-DEL-66
chrX-93289435-DEL-371
chrX-93304869-DEL-56
chrX-93329479-DEL-1103
chrX-93367759-DEL-774
chrX-93381093-DEL-52
chrX-93448900-DEL-1089
chrX-93472400-DEL-70
chrX-93478596-DEL-75
chrX-93502126-DEL-73
chrX-93513985-DEL-60
chrX-93514084-DEL-1486
chrX-93537889-DEL-1219
chrX-93542060-DEL-185
chrX-93542963-DEL-293
chrX-93553952-DEL-956
chrX-93557884-DEL-686
chrX-93562608-DEL-135
chrX-93567954-DEL-326
chrX-93571527-DEL-533
chrX-93669183-DEL-1002
chrX-93670457-DEL-222
chrX-93777649-DEL-56
chrX-93790546-DEL-536
chrX-93796073-DEL-55

chrX-93844034-DEL-2124
chrX-93866643-DEL-1967
chrX-93871161-DEL-3898
chrX-93886636-DEL-473
chrX-93888743-DEL-8541
chrX-93916059-DEL-530
chrX-93919364-DEL-650
chrX-93922188-DEL-1307
chrX-93922455-DEL-703
chrX-93961597-DEL-50001
chrX-94016659-DEL-2061
chrX-94018545-DEL-1372
chrX-94023249-DEL-695
chrX-94027542-DEL-76
chrX-94111185-DEL-223
chrX-94112662-DEL-204
chrX-94151641-DEL-87
chrX-94168506-DEL-396
chrX-94169221-DEL-282
chrX-94195689-DEL-72
chrX-94217904-DEL-897
chrX-94245541-DEL-75
chrX-94251973-DEL-404
chrX-94273659-DEL-611
chrX-94309386-DEL-8133
chrX-94319156-DEL-6141
chrX-94328932-DEL-6411
chrX-94346407-DEL-1175
chrX-94365498-DEL-1917
chrX-94377115-DEL-6171
chrX-94492401-DEL-106
chrX-94517099-DEL-209
chrX-94553165-DEL-58
chrX-94561897-DEL-60
chrX-94568156-DEL-300
chrX-94570069-DEL-55
chrX-94606536-DEL-894
chrX-94610779-DEL-53
chrX-94610923-DEL-6034
chrX-94616481-DEL-276
chrX-94631491-DEL-315
chrX-94641122-DEL-60
chrX-94696861-DEL-192
chrX-94714777-DEL-80
chrX-94733211-DEL-1150
chrX-94760709-DEL-500
chrX-94771799-DEL-395
chrX-94774959-DEL-94
chrX-94779928-DEL-802
chrX-94787349-DEL-6818
chrX-94787481-DEL-208
chrX-94794378-DEL-198
chrX-94795144-DEL-83
chrX-94797090-DEL-6366
chrX-94812096-DEL-1724
chrX-94832887-DEL-99
chrX-94883876-DEL-2501
chrX-94896657-DEL-118
chrX-94923122-DEL-56
chrX-94989591-DEL-416
chrX-94995190-DEL-189
chrX-94997303-DEL-85
chrX-94999737-DEL-477
chrX-95044828-DEL-108
chrX-95044835-DEL-1770
chrX-95074938-DEL-709
chrX-95081037-DEL-78
chrX-95082419-DEL-62
chrX-95099513-DEL-1324
chrX-95103305-DEL-1870
chrX-95105422-DEL-1458
chrX-95112164-DEL-368
chrX-95114466-DEL-96
chrX-95124589-DEL-76
chrX-95126391-DEL-224
chrX-95192411-DEL-403
chrX-95219202-DEL-121

chrX-95228180-DEL-975
chrX-95231666-DEL-310
chrX-95235099-DEL-1279
chrX-95247882-DEL-202
chrX-95289299-DEL-682
chrX-95290088-DEL-187
chrX-95296732-DEL-191
chrX-95297849-DEL-870
chrX-95307827-DEL-473
chrX-95310076-DEL-237
chrX-95317982-DEL-416
chrX-95338497-DEL-3059
chrX-95353621-DEL-1505
chrX-95380174-DEL-54
chrX-95386359-DEL-1328
chrX-95399605-DEL-294
chrX-95422878-DEL-6605
chrX-95436782-DEL-11732
chrX-95436782-DEL-6241
chrX-95445665-DEL-8361
chrX-95451156-DEL-2870
chrX-95463404-DEL-52
chrX-95466995-DEL-1354
chrX-95481222-DEL-5456
chrX-95507620-DEL-155
chrX-95508891-DEL-282
chrX-95565798-DEL-774
chrX-95602462-DEL-1176
chrX-95649041-DEL-163
chrX-95672542-DEL-2905
chrX-95688551-DEL-1529
chrX-95821017-DEL-66
chrX-95845573-DEL-2943
chrX-95854038-DEL-9839
chrX-95871435-DEL-6476
chrX-95886989-DEL-133
chrX-95905680-DEL-9775
chrX-95925064-DEL-316
chrX-95927083-DEL-826
chrX-95936549-DEL-176
chrX-95943894-DEL-325
chrX-95946973-DEL-66
chrX-95948975-DEL-89
chrX-95963689-DEL-2319
chrX-95988680-DEL-5042
chrX-95997871-DEL-71
chrX-96042570-DEL-221
chrX-96049259-DEL-1727
chrX-96120375-DEL-209
chrX-96153695-DEL-106
chrX-96191228-DEL-510
chrX-96192671-DEL-160
chrX-96214665-DEL-237
chrX-96245401-DEL-276
chrX-96250053-DEL-113
chrX-96266724-DEL-1617
chrX-96291783-DEL-1510
chrX-96298410-DEL-6234
chrX-96333070-DEL-474
chrX-96358727-DEL-110
chrX-96360592-DEL-52
chrX-96383107-DEL-1367
chrX-96394336-DEL-75
chrX-96435158-DEL-2355
chrX-96458551-DEL-774
chrX-96505546-DEL-8199
chrX-96517799-DEL-7623
chrX-96528004-DEL-214
chrX-96535300-DEL-199
chrX-96614379-DEL-281
chrX-96662002-DEL-1632
chrX-96664642-DEL-59
chrX-96710221-DEL-56
chrX-96771592-DEL-211
chrX-96789461-DEL-483
chrX-96812194-DEL-1505
chrX-96820213-DEL-102

chrX-96830385-DEL-420
chrX-96868101-DEL-837
chrX-96876934-DEL-8303
chrX-96880520-DEL-1031
chrX-96903030-DEL-52
chrX-96909856-DEL-9038
chrX-96921301-DEL-8812
chrX-96945798-DEL-676
chrX-96948196-DEL-469
chrX-96950768-DEL-9511
chrX-96951504-DEL-6553
chrX-97003166-DEL-57
chrX-97103631-DEL-780
chrX-97113501-DEL-320
chrX-97120883-DEL-399
chrX-97129652-DEL-2144
chrX-97138681-DEL-120
chrX-97149402-DEL-5398
chrX-97150158-DEL-54
chrX-97153493-DEL-123
chrX-97154801-DEL-115
chrX-97183019-DEL-2876
chrX-97211506-DEL-86
chrX-97222418-DEL-576
chrX-97239767-DEL-137
chrX-97254900-DEL-203
chrX-97276959-DEL-529
chrX-97294124-DEL-496
chrX-97317608-DEL-145
chrX-97406175-DEL-3718
chrX-97421744-DEL-336
chrX-97437457-DEL-68
chrX-97454822-DEL-937
chrX-97482318-DEL-6890
chrX-97527367-DEL-314
chrX-97530002-DEL-589
chrX-97534555-DEL-447
chrX-97540745-DEL-6243
chrX-97569058-DEL-84
chrX-97575167-DEL-160
chrX-97620633-DEL-523
chrX-97622854-DEL-7957
chrX-97627084-DEL-6271
chrX-97635441-DEL-4462
chrX-97649328-DEL-6225
chrX-97659667-DEL-76
chrX-97715244-DEL-350
chrX-97749616-DEL-465
chrX-97771932-DEL-588
chrX-97784172-DEL-1792
chrX-97786156-DEL-4869
chrX-97791697-DEL-13453
chrX-97854098-DEL-7051
chrX-97863559-DEL-1821
chrX-97868147-DEL-103
chrX-97895813-DEL-56
chrX-97939843-DEL-77
chrX-97951186-DEL-161
chrX-97975518-DEL-108
chrX-97976587-DEL-52
chrX-98006754-DEL-52
chrX-98042109-DEL-277
chrX-98050955-DEL-462
chrX-98136939-DEL-78
chrX-98158708-DEL-589
chrX-98245816-DEL-52
chrX-98311571-DEL-198
chrX-98328804-DEL-103
chrX-98372606-DEL-1999
chrX-98387056-DEL-62
chrX-98460450-DEL-129
chrX-98513564-DEL-62
chrX-98524729-DEL-51
chrX-98629774-DEL-117
chrX-98644262-DEL-128
chrX-98679798-DEL-1065
chrX-98690452-DEL-66

chrX-98709790-DEL-55
chrX-98749899-DEL-196
chrX-98755932-DEL-980
chrX-98760999-DEL-77
chrX-98770632-DEL-128
chrX-98803000-DEL-1864
chrX-98933679-DEL-64
chrX-98953463-DEL-728
chrX-98960178-DEL-515
chrX-99029670-DEL-322
chrX-99050944-DEL-208
chrX-99101221-DEL-643
chrX-99105239-DEL-55
chrX-99151957-DEL-624
chrX-99175648-DEL-4413
chrX-99191434-DEL-4156
chrX-99233050-DEL-99
chrX-99239598-DEL-2321
chrX-99240207-DEL-979
chrX-99385619-DEL-2191
chrX-99393314-DEL-237
chrX-99462189-DEL-190
chrX-99548354-DEL-72
chrX-99549756-DEL-51
chrX-99570135-DEL-68
chrX-99580694-DEL-119
chrX-99659578-DEL-51
chrX-99684470-DEL-54
chrX-99690975-DEL-66
chrX-99736024-DEL-204
chrX-99741764-DEL-1971
chrX-99815875-DEL-52
chrX-99856125-DEL-118
chrX-99862298-DEL-181
chrX-99865275-DEL-74
chrX-99927578-DEL-69
chrX-99999277-DEL-95
chrX-100046907-DEL-1740
chrX-100048696-DEL-1022
chrX-100053573-DEL-800
chrX-100072052-DEL-1604
chrX-100104543-DEL-100
chrX-100152977-DEL-70
chrX-100213100-DEL-191
chrX-100216991-DEL-56
chrX-100233207-DEL-179
chrX-100246002-DEL-883
chrX-100438631-DEL-64
chrX-100467804-DEL-174
chrX-100478564-DEL-165
chrX-100531729-DEL-71
chrX-100538911-DEL-2740
chrX-100540073-DEL-1545
chrX-100547935-DEL-177
chrX-100553346-DEL-227
chrX-100565401-DEL-206
chrX-100579039-DEL-242
chrX-100620476-DEL-490
chrX-100631235-DEL-206
chrX-100638267-DEL-225
chrX-100664463-DEL-86
chrX-100702831-DEL-150
chrX-100714187-DEL-214
chrX-100723986-DEL-66
chrX-100730812-DEL-216
chrX-100739839-DEL-327
chrX-100753702-DEL-6215
chrX-100762093-DEL-84
chrX-100768886-DEL-806
chrX-100817262-DEL-357
chrX-100868015-DEL-893
chrX-100896509-DEL-58
chrX-100951748-DEL-195
chrX-100971904-DEL-67
chrX-101059116-DEL-496
chrX-101083240-DEL-213
chrX-101133899-DEL-51

chrX-101144766-DEL-71
chrX-101167419-DEL-79
chrX-101195422-DEL-204
chrX-101198864-DEL-398
chrX-101208367-DEL-191
chrX-101208958-DEL-509
chrX-101212196-DEL-865
chrX-101220301-DEL-67
chrX-101239185-DEL-60
chrX-101248925-DEL-61
chrX-101254790-DEL-446
chrX-101258503-DEL-184
chrX-101259793-DEL-167
chrX-101262344-DEL-82
chrX-101264573-DEL-75
chrX-101269240-DEL-182
chrX-101270728-DEL-534
chrX-101339017-DEL-66
chrX-101340459-DEL-106
chrX-101374500-DEL-97
chrX-101434282-DEL-54
chrX-101453984-DEL-204
chrX-101499068-DEL-78
chrX-101556014-DEL-360
chrX-101572161-DEL-1149
chrX-101647230-DEL-194
chrX-101649926-DEL-90
chrX-101655910-DEL-345
chrX-101677817-DEL-329
chrX-101728372-DEL-422
chrX-101744860-DEL-72
chrX-101759219-DEL-2894
chrX-101776708-DEL-558
chrX-101787561-DEL-233
chrX-101812093-DEL-2849
chrX-101839185-DEL-60
chrX-101875904-DEL-82
chrX-101905990-DEL-498
chrX-101906578-DEL-912
chrX-101939721-DEL-107
chrX-101955373-DEL-78
chrX-101994658-DEL-1160
chrX-102006920-DEL-11675
chrX-102028268-DEL-111
chrX-102105980-DEL-196
chrX-102121317-DEL-279
chrX-102133271-DEL-61
chrX-102142770-DEL-60
chrX-102156357-DEL-2778
chrX-102163806-DEL-6461
chrX-102190753-DEL-59
chrX-102191203-DEL-1191
chrX-102203159-DEL-72
chrX-102211999-DEL-2130
chrX-102219698-DEL-341
chrX-102241637-DEL-70
chrX-102250836-DEL-231
chrX-102265016-DEL-80
chrX-102267178-DEL-336
chrX-102282536-DEL-2183
chrX-102285434-DEL-201
chrX-102288001-DEL-200
chrX-102288893-DEL-505
chrX-102295906-DEL-111
chrX-102307106-DEL-759
chrX-102310735-DEL-122
chrX-102318198-DEL-311
chrX-102354174-DEL-94
chrX-102375436-DEL-57
chrX-102407831-DEL-54
chrX-102426678-DEL-97
chrX-102431054-DEL-204
chrX-102438713-DEL-104
chrX-102453752-DEL-198
chrX-102462692-DEL-626
chrX-102490677-DEL-58
chrX-102491751-DEL-67

chrX-102492198-DEL-138
chrX-102492911-DEL-136
chrX-102498803-DEL-935
chrX-102498804-DEL-221
chrX-102499307-DEL-204
chrX-102501609-DEL-122
chrX-102502766-DEL-875
chrX-102510983-DEL-148
chrX-102537089-DEL-52
chrX-102552687-DEL-51
chrX-102587861-DEL-63
chrX-102588854-DEL-53
chrX-102589485-DEL-410
chrX-102629912-DEL-77
chrX-102641033-DEL-52
chrX-102655912-DEL-801
chrX-102656884-DEL-210
chrX-102669484-DEL-100
chrX-102677180-DEL-95
chrX-102708980-DEL-57
chrX-102722294-DEL-201
chrX-102730854-DEL-510
chrX-102793463-DEL-120
chrX-102878376-DEL-288
chrX-102879007-DEL-208
chrX-102893681-DEL-132
chrX-102902091-DEL-1299
chrX-102909063-DEL-54
chrX-102955701-DEL-459
chrX-103012957-DEL-239
chrX-103019794-DEL-186
chrX-103032917-DEL-577
chrX-103059869-DEL-60
chrX-103142754-DEL-193
chrX-103146623-DEL-147
chrX-103162494-DEL-107
chrX-103190478-DEL-111
chrX-103199414-DEL-1163
chrX-103228450-DEL-249
chrX-103256138-DEL-1804
chrX-103296318-DEL-468
chrX-103335601-DEL-56
chrX-103354831-DEL-982
chrX-103377628-DEL-2491
chrX-103387110-DEL-201
chrX-103396866-DEL-408
chrX-103527690-DEL-54
chrX-103538098-DEL-69
chrX-103578079-DEL-102
chrX-103637622-DEL-54
chrX-103666239-DEL-6402
chrX-103682548-DEL-6423
chrX-103761223-DEL-419
chrX-103801525-DEL-808
chrX-103819416-DEL-13429
chrX-103832988-DEL-359
chrX-103877697-DEL-62
chrX-103879040-DEL-870
chrX-103914361-DEL-8401
chrX-103919039-DEL-963
chrX-103922150-DEL-118
chrX-103925793-DEL-68
chrX-103927536-DEL-206
chrX-103948818-DEL-297
chrX-103964343-DEL-174
chrX-103995790-DEL-54
chrX-104010326-DEL-56
chrX-104033392-DEL-398
chrX-104036054-DEL-498
chrX-104037811-DEL-53
chrX-104063858-DEL-6167
chrX-104073477-DEL-61
chrX-104098403-DEL-65
chrX-104103355-DEL-844
chrX-104133398-DEL-104
chrX-104135820-DEL-126
chrX-104139113-DEL-193

chrX-104153927-DEL-101
chrX-104187819-DEL-938
chrX-104194691-DEL-205
chrX-104226437-DEL-62
chrX-104236320-DEL-637
chrX-104257607-DEL-70
chrX-104310903-DEL-206
chrX-104316350-DEL-112
chrX-104330542-DEL-673
chrX-104367354-DEL-1085
chrX-104386341-DEL-70
chrX-104403615-DEL-56
chrX-104429853-DEL-231
chrX-104467820-DEL-53
chrX-104495364-DEL-2589
chrX-104534332-DEL-3545
chrX-104564031-DEL-152
chrX-104565919-DEL-98
chrX-104566781-DEL-232
chrX-104567063-DEL-4439
chrX-104668396-DEL-72
chrX-104671941-DEL-361
chrX-104694781-DEL-994
chrX-104701543-DEL-52
chrX-104717793-DEL-97
chrX-104759225-DEL-54
chrX-104854923-DEL-170
chrX-104904981-DEL-118
chrX-104931641-DEL-196
chrX-104942594-DEL-207
chrX-104961858-DEL-169
chrX-104966806-DEL-151
chrX-104982563-DEL-210
chrX-105007081-DEL-53
chrX-105028194-DEL-132
chrX-105033968-DEL-63
chrX-105039945-DEL-182
chrX-105048252-DEL-199
chrX-105061007-DEL-126
chrX-105079159-DEL-136
chrX-105084833-DEL-163
chrX-105148109-DEL-69
chrX-105151086-DEL-643
chrX-105204556-DEL-2166
chrX-105247847-DEL-396
chrX-105274441-DEL-209
chrX-105281594-DEL-119
chrX-105312307-DEL-98
chrX-105320532-DEL-400
chrX-105335829-DEL-181
chrX-105379085-DEL-66
chrX-105433115-DEL-8616
chrX-105444119-DEL-189
chrX-105446161-DEL-4059
chrX-105464528-DEL-295
chrX-105471214-DEL-75
chrX-105494260-DEL-83
chrX-105496990-DEL-76
chrX-105562336-DEL-2210
chrX-105572781-DEL-536
chrX-105587877-DEL-52
chrX-105597723-DEL-3675
chrX-105603969-DEL-6085
chrX-105648185-DEL-207
chrX-105650343-DEL-968
chrX-105656940-DEL-791
chrX-105662841-DEL-1383
chrX-105664158-DEL-197
chrX-105683940-DEL-80
chrX-105723774-DEL-580
chrX-105725063-DEL-7506
chrX-105744990-DEL-197
chrX-105747904-DEL-225
chrX-105763299-DEL-360
chrX-105772231-DEL-7385
chrX-105818509-DEL-420
chrX-105845172-DEL-121

chrX-105865852-DEL-579
chrX-105867326-DEL-207
chrX-105921994-DEL-290
chrX-105932470-DEL-1520
chrX-105934661-DEL-222
chrX-105980195-DEL-53
chrX-105995036-DEL-140
chrX-106027526-DEL-99
chrX-106081970-DEL-113
chrX-106116588-DEL-772
chrX-106120425-DEL-1221
chrX-106150442-DEL-2727
chrX-106163635-DEL-270
chrX-106164339-DEL-164
chrX-106172674-DEL-323
chrX-106198378-DEL-57
chrX-106227381-DEL-2159
chrX-106232661-DEL-1156
chrX-106263568-DEL-131
chrX-106306360-DEL-256
chrX-106316405-DEL-1357
chrX-106392582-DEL-2030
chrX-106482063-DEL-978
chrX-106503562-DEL-97
chrX-106515517-DEL-52
chrX-106518472-DEL-84
chrX-106523740-DEL-439
chrX-106579882-DEL-439
chrX-106591282-DEL-7453
chrX-106610086-DEL-440
chrX-106630604-DEL-392
chrX-106641831-DEL-3635
chrX-106645613-DEL-424
chrX-106690575-DEL-1890
chrX-106788053-DEL-110
chrX-106790566-DEL-82
chrX-106794530-DEL-161
chrX-106857722-DEL-86
chrX-106875375-DEL-629
chrX-106928570-DEL-1071
chrX-106952233-DEL-100
chrX-106956820-DEL-53
chrX-106958920-DEL-72
chrX-106963808-DEL-1571
chrX-106972743-DEL-182
chrX-106972966-DEL-4219
chrX-107089950-DEL-348
chrX-107157926-DEL-1004
chrX-107159511-DEL-6259
chrX-107193955-DEL-87
chrX-107201708-DEL-177
chrX-107241914-DEL-234
chrX-107260075-DEL-6175
chrX-107296484-DEL-61
chrX-107300239-DEL-5382
chrX-107329980-DEL-59
chrX-107343576-DEL-200
chrX-107372780-DEL-1158
chrX-107374745-DEL-455
chrX-107401249-DEL-147
chrX-107433438-DEL-472
chrX-107454421-DEL-153
chrX-107498593-DEL-1799
chrX-107548685-DEL-74
chrX-107597711-DEL-237
chrX-107600671-DEL-127
chrX-107618004-DEL-6573
chrX-107668993-DEL-310
chrX-107685144-DEL-79
chrX-107704228-DEL-70
chrX-107710420-DEL-1059
chrX-107731117-DEL-3422
chrX-107759065-DEL-951
chrX-107762417-DEL-2006
chrX-107788666-DEL-53
chrX-107791609-DEL-153
chrX-107815280-DEL-104

chrX-107841801-DEL-73
chrX-107867138-DEL-57
chrX-107956011-DEL-477
chrX-107965947-DEL-506
chrX-108010091-DEL-579
chrX-108030481-DEL-132
chrX-108032939-DEL-169
chrX-108044111-DEL-63
chrX-108052617-DEL-340
chrX-108055072-DEL-60
chrX-108081238-DEL-5991
chrX-108255754-DEL-1115
chrX-108279694-DEL-386
chrX-108287598-DEL-54
chrX-108304791-DEL-154
chrX-108308732-DEL-6403
chrX-108316106-DEL-418
chrX-108320047-DEL-1700
chrX-108320903-DEL-10162
chrX-108323635-DEL-5918
chrX-108323939-DEL-173
chrX-108349767-DEL-394
chrX-108355253-DEL-139
chrX-108366704-DEL-512
chrX-108402078-DEL-283
chrX-108405754-DEL-3232
chrX-108479176-DEL-1606
chrX-108497816-DEL-72
chrX-108501151-DEL-437
chrX-108517580-DEL-6149
chrX-108532956-DEL-334
chrX-108559285-DEL-2227
chrX-108563383-DEL-834
chrX-108565120-DEL-125
chrX-108565867-DEL-70
chrX-108597127-DEL-449
chrX-108620002-DEL-340
chrX-108639864-DEL-156
chrX-108658428-DEL-779
chrX-108685270-DEL-3581
chrX-108702926-DEL-206
chrX-108732573-DEL-1466
chrX-108735186-DEL-4754
chrX-108764395-DEL-56
chrX-108785723-DEL-967
chrX-108792663-DEL-936
chrX-108810676-DEL-6279
chrX-108832409-DEL-52
chrX-108868485-DEL-6581
chrX-108874853-DEL-215
chrX-108874861-DEL-896
chrX-108884332-DEL-172
chrX-108900205-DEL-194
chrX-108914550-DEL-6179
chrX-108920733-DEL-416
chrX-108976306-DEL-56
chrX-108987419-DEL-120
chrX-109016150-DEL-139
chrX-109076073-DEL-203
chrX-109085786-DEL-6246
chrX-109093617-DEL-2164
chrX-109098630-DEL-51
chrX-109108509-DEL-129
chrX-109126229-DEL-2705
chrX-109141303-DEL-57
chrX-109146953-DEL-124
chrX-109172221-DEL-554
chrX-109210519-DEL-425
chrX-109211731-DEL-276
chrX-109215555-DEL-2435
chrX-109236177-DEL-87
chrX-109236389-DEL-1293
chrX-109259471-DEL-99
chrX-109317745-DEL-54
chrX-109454744-DEL-60
chrX-109469288-DEL-82
chrX-109476680-DEL-6369

chrX-109489069-DEL-6231
chrX-109495415-DEL-1056
chrX-109502729-DEL-325
chrX-109546202-DEL-9983
chrX-109547134-DEL-9046
chrX-109549037-DEL-2402
chrX-109586526-DEL-1263
chrX-109600680-DEL-433
chrX-109608025-DEL-232
chrX-109629998-DEL-405
chrX-109647174-DEL-1030
chrX-109664892-DEL-384
chrX-109675991-DEL-55
chrX-109688227-DEL-401
chrX-109706833-DEL-54
chrX-109807894-DEL-64
chrX-109884762-DEL-1677
chrX-109904631-DEL-38889
chrX-109911760-DEL-6202
chrX-109932796-DEL-381
chrX-109941219-DEL-390
chrX-109982351-DEL-1208
chrX-109983911-DEL-424
chrX-109995595-DEL-111
chrX-110021071-DEL-452
chrX-110048097-DEL-3349
chrX-110058135-DEL-77
chrX-110063582-DEL-214
chrX-110105471-DEL-1982
chrX-110133441-DEL-194
chrX-110168404-DEL-215
chrX-110174239-DEL-5201
chrX-110183401-DEL-1152
chrX-110220722-DEL-298
chrX-110227083-DEL-72
chrX-110231607-DEL-168
chrX-110241415-DEL-1019
chrX-110243979-DEL-395
chrX-110267147-DEL-688
chrX-110271379-DEL-559
chrX-110352882-DEL-145
chrX-110393693-DEL-150
chrX-110409926-DEL-57
chrX-110416086-DEL-291
chrX-110444279-DEL-76
chrX-110477943-DEL-56
chrX-110511343-DEL-134
chrX-110554045-DEL-367
chrX-110588541-DEL-801
chrX-110592529-DEL-234
chrX-110600093-DEL-385
chrX-110607707-DEL-53
chrX-110619055-DEL-2118
chrX-110645529-DEL-267
chrX-110673915-DEL-88
chrX-110690196-DEL-78
chrX-110727962-DEL-5449
chrX-110736044-DEL-717
chrX-110760196-DEL-54
chrX-110779889-DEL-818
chrX-110836465-DEL-1643
chrX-110943017-DEL-979
chrX-110944202-DEL-110
chrX-110969792-DEL-184
chrX-110970392-DEL-213
chrX-110971566-DEL-450
chrX-110984959-DEL-82
chrX-110987635-DEL-1951
chrX-111020297-DEL-303
chrX-111067872-DEL-1748
chrX-111070975-DEL-311
chrX-111144995-DEL-108
chrX-111162133-DEL-71
chrX-111174919-DEL-632
chrX-111194935-DEL-464
chrX-111245707-DEL-3941
chrX-111301519-DEL-762

chrX-111302556-DEL-119
chrX-111303301-DEL-78
chrX-111314033-DEL-300
chrX-111322181-DEL-3423
chrX-111336407-DEL-8905
chrX-111352447-DEL-1941
chrX-111354744-DEL-1174
chrX-111389618-DEL-6697
chrX-111445396-DEL-66
chrX-111454338-DEL-1424
chrX-111530984-DEL-370
chrX-111562440-DEL-227
chrX-111599625-DEL-52
chrX-111637140-DEL-6372
chrX-111661155-DEL-179
chrX-111689020-DEL-54
chrX-111741830-DEL-579
chrX-111764329-DEL-1179
chrX-111790029-DEL-435
chrX-111805872-DEL-83
chrX-111833559-DEL-396
chrX-111836704-DEL-1824
chrX-111911672-DEL-56
chrX-111942259-DEL-851
chrX-111943132-DEL-1420
chrX-111963150-DEL-53
chrX-112026436-DEL-13294
chrX-112031691-DEL-6094
chrX-112144764-DEL-159
chrX-112149135-DEL-586
chrX-112161699-DEL-623
chrX-112170300-DEL-53
chrX-112220948-DEL-61
chrX-112245151-DEL-600
chrX-112257754-DEL-569
chrX-112269071-DEL-60
chrX-112274292-DEL-60
chrX-112346875-DEL-1548
chrX-112349249-DEL-911
chrX-112372922-DEL-206
chrX-112381690-DEL-6071
chrX-112410609-DEL-73
chrX-112461513-DEL-533
chrX-112473081-DEL-2253
chrX-112502864-DEL-57
chrX-112507184-DEL-80
chrX-112513103-DEL-6360
chrX-112525978-DEL-367
chrX-112535125-DEL-599
chrX-112551043-DEL-1150
chrX-112584728-DEL-141
chrX-112631214-DEL-421
chrX-112671824-DEL-796
chrX-112679940-DEL-765
chrX-112699167-DEL-178
chrX-112713719-DEL-1517
chrX-112746046-DEL-1697
chrX-112768727-DEL-60
chrX-112780738-DEL-343
chrX-112784804-DEL-6835
chrX-112802060-DEL-515
chrX-112805104-DEL-5027
chrX-112852371-DEL-124
chrX-112854696-DEL-426
chrX-112867101-DEL-1646
chrX-112868807-DEL-820
chrX-112874685-DEL-8183
chrX-112902591-DEL-916
chrX-112903287-DEL-213
chrX-112927668-DEL-240
chrX-112940727-DEL-4677
chrX-112944793-DEL-150
chrX-112952512-DEL-78
chrX-112957358-DEL-3342
chrX-112966572-DEL-116
chrX-112992684-DEL-2994
chrX-113021417-DEL-139

chrX-113030491-DEL-64
chrX-113037089-DEL-8835
chrX-113037973-DEL-76
chrX-113048295-DEL-72
chrX-113049886-DEL-6468
chrX-113060587-DEL-340
chrX-113096782-DEL-5415
chrX-113125672-DEL-77
chrX-113143201-DEL-136
chrX-113145815-DEL-313
chrX-113150830-DEL-88
chrX-113170369-DEL-7355
chrX-113176863-DEL-211
chrX-113196010-DEL-1138
chrX-113220414-DEL-390
chrX-113230188-DEL-2179
chrX-113232958-DEL-66
chrX-113273716-DEL-195
chrX-113285042-DEL-358
chrX-113293462-DEL-51
chrX-113298751-DEL-252
chrX-113301478-DEL-1001
chrX-113302663-DEL-793
chrX-113303460-DEL-283
chrX-113304056-DEL-5273
chrX-113320171-DEL-11541
chrX-113338989-DEL-589
chrX-113371003-DEL-138
chrX-113371438-DEL-1898
chrX-113389716-DEL-910
chrX-113422148-DEL-151
chrX-113424681-DEL-52
chrX-113427588-DEL-105
chrX-113429231-DEL-96
chrX-113473920-DEL-82
chrX-113474085-DEL-930
chrX-113475621-DEL-791
chrX-113478091-DEL-894
chrX-113500819-DEL-1451
chrX-113515736-DEL-6158
chrX-113526950-DEL-54
chrX-113541214-DEL-118
chrX-113542778-DEL-511
chrX-113562785-DEL-180
chrX-113613430-DEL-6263
chrX-113624303-DEL-71
chrX-113639007-DEL-483
chrX-113640512-DEL-1558
chrX-113659794-DEL-14722
chrX-113659845-DEL-6893
chrX-113660168-DEL-159
chrX-113666066-DEL-8329
chrX-113672584-DEL-1639
chrX-113677401-DEL-301
chrX-113680035-DEL-70
chrX-113683672-DEL-55
chrX-113684560-DEL-6210
chrX-113693632-DEL-185
chrX-113704557-DEL-168
chrX-113720981-DEL-157
chrX-113771707-DEL-77
chrX-113774207-DEL-2339
chrX-113847703-DEL-189
chrX-113891930-DEL-500
chrX-113900754-DEL-6823
chrX-113909459-DEL-64
chrX-113911134-DEL-6330
chrX-113941456-DEL-472
chrX-113973866-DEL-1172
chrX-114001445-DEL-66
chrX-114016983-DEL-118
chrX-114022566-DEL-78
chrX-114022906-DEL-7745
chrX-114061138-DEL-131
chrX-114065778-DEL-6191
chrX-114072117-DEL-2489
chrX-114074724-DEL-311

chrX-114101506-DEL-954
chrX-114103802-DEL-72
chrX-114124110-DEL-2514
chrX-114150927-DEL-489
chrX-114160057-DEL-2855
chrX-114186131-DEL-140
chrX-114201760-DEL-62
chrX-114225652-DEL-84
chrX-114304411-DEL-64
chrX-114318307-DEL-3250
chrX-114337763-DEL-441
chrX-114340032-DEL-6154
chrX-114377657-DEL-1426
chrX-114391182-DEL-102
chrX-114400569-DEL-62
chrX-114434476-DEL-77
chrX-114442841-DEL-105
chrX-114467937-DEL-10637
chrX-114484034-DEL-7298
chrX-114491367-DEL-1272
chrX-114500585-DEL-1193
chrX-114507042-DEL-908
chrX-114517647-DEL-52
chrX-114548047-DEL-723
chrX-114570203-DEL-6422
chrX-114642320-DEL-118
chrX-114648755-DEL-7145
chrX-114655762-DEL-138
chrX-114658286-DEL-5773
chrX-114668210-DEL-66
chrX-114669755-DEL-178
chrX-114711636-DEL-58
chrX-114716860-DEL-453
chrX-114722131-DEL-781
chrX-114733911-DEL-884
chrX-114782291-DEL-630
chrX-114785376-DEL-1456
chrX-114799157-DEL-1866
chrX-114824862-DEL-301
chrX-114830538-DEL-93
chrX-114836630-DEL-196
chrX-114855408-DEL-56
chrX-114890643-DEL-127
chrX-114927922-DEL-2075
chrX-114937158-DEL-2052
chrX-114946020-DEL-1484
chrX-114950881-DEL-1002
chrX-114966410-DEL-56
chrX-114972339-DEL-476
chrX-114984149-DEL-52
chrX-114988825-DEL-59
chrX-115006366-DEL-77
chrX-115012414-DEL-105
chrX-115018894-DEL-591
chrX-115020622-DEL-71
chrX-115029542-DEL-533
chrX-115048753-DEL-98
chrX-115064099-DEL-55
chrX-115071027-DEL-84
chrX-115088137-DEL-690
chrX-115107062-DEL-846
chrX-115113651-DEL-1466
chrX-115114031-DEL-7224
chrX-115134800-DEL-6307
chrX-115175879-DEL-1232
chrX-115196803-DEL-269
chrX-115198266-DEL-477
chrX-115213227-DEL-205
chrX-115236078-DEL-2449
chrX-115239112-DEL-472
chrX-115247794-DEL-116
chrX-115250653-DEL-763
chrX-115288655-DEL-282
chrX-115332052-DEL-1828
chrX-115360367-DEL-64
chrX-115373266-DEL-55
chrX-115382773-DEL-515

chrX-115406679-DEL-957
chrX-115419374-DEL-2782
chrX-115426155-DEL-278
chrX-115469930-DEL-671
chrX-115489267-DEL-89
chrX-115520365-DEL-1840
chrX-115528399-DEL-51
chrX-115558507-DEL-124
chrX-115569970-DEL-201
chrX-115583853-DEL-489
chrX-115638306-DEL-632
chrX-115647091-DEL-162
chrX-115658555-DEL-101
chrX-115662208-DEL-292
chrX-115695864-DEL-262
chrX-115700214-DEL-1439
chrX-115768016-DEL-76
chrX-115806345-DEL-58
chrX-115837336-DEL-219
chrX-115848560-DEL-210
chrX-115858508-DEL-278
chrX-115859620-DEL-52
chrX-115872408-DEL-1760
chrX-115881861-DEL-1285
chrX-115920050-DEL-191
chrX-115948110-DEL-54
chrX-115990685-DEL-149
chrX-116011333-DEL-1754
chrX-116028797-DEL-6263
chrX-116040594-DEL-2438
chrX-116040607-DEL-3735
chrX-116043047-DEL-1295
chrX-116052022-DEL-3979
chrX-116063484-DEL-1826
chrX-116066399-DEL-145
chrX-116078436-DEL-1697
chrX-116110543-DEL-57
chrX-116111966-DEL-70
chrX-116112824-DEL-1167
chrX-116142018-DEL-2216
chrX-116151445-DEL-603
chrX-116171681-DEL-305
chrX-116178166-DEL-655
chrX-116181789-DEL-53
chrX-116215206-DEL-999
chrX-116219370-DEL-60
chrX-116261837-DEL-84
chrX-116265224-DEL-82
chrX-116303455-DEL-84
chrX-116357612-DEL-6633
chrX-116366263-DEL-85
chrX-116379985-DEL-56
chrX-116390196-DEL-586
chrX-116394681-DEL-899
chrX-116415194-DEL-625
chrX-116419587-DEL-123
chrX-116431236-DEL-1677
chrX-116442278-DEL-1191
chrX-116458982-DEL-3192
chrX-116475314-DEL-3912
chrX-116476939-DEL-100
chrX-116479088-DEL-2076
chrX-116494003-DEL-831
chrX-116498917-DEL-65
chrX-116526506-DEL-589
chrX-116527789-DEL-482
chrX-116534487-DEL-2686
chrX-116546187-DEL-499
chrX-116558256-DEL-57
chrX-116558353-DEL-2570
chrX-116566682-DEL-84
chrX-116584780-DEL-269
chrX-116586093-DEL-326
chrX-116591398-DEL-539
chrX-116603175-DEL-893
chrX-116604834-DEL-2163
chrX-116628361-DEL-2302

chrX-116630426-DEL-56
chrX-116633948-DEL-357
chrX-116667909-DEL-658
chrX-116679217-DEL-85
chrX-116702457-DEL-632
chrX-116708428-DEL-53
chrX-116728283-DEL-442
chrX-116730143-DEL-279
chrX-116739726-DEL-274
chrX-116742601-DEL-67
chrX-116764503-DEL-1241
chrX-116791663-DEL-74
chrX-116793770-DEL-684
chrX-116815988-DEL-511
chrX-116816899-DEL-983
chrX-116849965-DEL-157
chrX-116887082-DEL-193
chrX-116946649-DEL-1716
chrX-116976327-DEL-2038
chrX-116978996-DEL-2341
chrX-117011092-DEL-1365
chrX-117011212-DEL-1774
chrX-117021301-DEL-59
chrX-117031983-DEL-394
chrX-117034998-DEL-371
chrX-117071793-DEL-400
chrX-117089494-DEL-285
chrX-117097792-DEL-1665
chrX-117106770-DEL-72
chrX-117149820-DEL-128
chrX-117150948-DEL-532
chrX-117160889-DEL-194
chrX-117163046-DEL-59
chrX-117168627-DEL-119
chrX-117179416-DEL-71
chrX-117181796-DEL-2151
chrX-117182894-DEL-984
chrX-117190923-DEL-449
chrX-117192787-DEL-1548
chrX-117193133-DEL-106
chrX-117199072-DEL-4615
chrX-117205151-DEL-384
chrX-117216319-DEL-111
chrX-117232620-DEL-737
chrX-117234197-DEL-364
chrX-117238978-DEL-2719
chrX-117241310-DEL-603
chrX-117242668-DEL-1545
chrX-117286958-DEL-1490
chrX-117300076-DEL-6421
chrX-117305904-DEL-159
chrX-117308118-DEL-125
chrX-117343815-DEL-2550
chrX-117353758-DEL-225
chrX-117363612-DEL-54
chrX-117365939-DEL-407
chrX-117370120-DEL-206
chrX-117371609-DEL-2639
chrX-117396585-DEL-1161
chrX-117399213-DEL-631
chrX-117426395-DEL-1261
chrX-117436217-DEL-1518
chrX-117446121-DEL-2352
chrX-117448292-DEL-386
chrX-117463118-DEL-791
chrX-117468920-DEL-634
chrX-117515495-DEL-280
chrX-117551319-DEL-400
chrX-117555442-DEL-58
chrX-117569684-DEL-6190
chrX-117610406-DEL-6764
chrX-117630746-DEL-253
chrX-117686483-DEL-84
chrX-117697455-DEL-680
chrX-117708937-DEL-158
chrX-117744345-DEL-408
chrX-117755846-DEL-114

chrX-117773297-DEL-63014
chrX-117776647-DEL-1454
chrX-117790654-DEL-6215
chrX-117797637-DEL-79
chrX-117835391-DEL-429
chrX-117837391-DEL-873
chrX-117875243-DEL-474
chrX-117902896-DEL-140
chrX-117919330-DEL-5522
chrX-117935151-DEL-118
chrX-117973862-DEL-130
chrX-118055090-DEL-66
chrX-118064449-DEL-100
chrX-118090005-DEL-401
chrX-118094341-DEL-508
chrX-118116346-DEL-111
chrX-118122138-DEL-101
chrX-118130334-DEL-435
chrX-118134003-DEL-7952
chrX-118171300-DEL-3247
chrX-118179290-DEL-104
chrX-118195399-DEL-842
chrX-118222828-DEL-618
chrX-118264082-DEL-64
chrX-118362705-DEL-100
chrX-118373109-DEL-1050
chrX-118395153-DEL-2257
chrX-118421290-DEL-806
chrX-118478249-DEL-8958
chrX-118483164-DEL-5352
chrX-118484812-DEL-848
chrX-118489287-DEL-70
chrX-118492550-DEL-1050
chrX-118495391-DEL-5630
chrX-118507421-DEL-368
chrX-118556680-DEL-68
chrX-118570474-DEL-555
chrX-118644923-DEL-235
chrX-118658098-DEL-1577
chrX-118667093-DEL-958
chrX-118689656-DEL-6098
chrX-118698706-DEL-72
chrX-118711988-DEL-134
chrX-118736310-DEL-285
chrX-118780436-DEL-1624
chrX-118783526-DEL-147
chrX-118798974-DEL-1763
chrX-118803788-DEL-166
chrX-118806295-DEL-69
chrX-118812896-DEL-51
chrX-118814316-DEL-485
chrX-118852586-DEL-6568
chrX-118859725-DEL-53
chrX-118864486-DEL-63
chrX-118865378-DEL-1978
chrX-118886591-DEL-74
chrX-118894905-DEL-742
chrX-118896218-DEL-1437
chrX-118906105-DEL-789
chrX-118918407-DEL-59
chrX-118920390-DEL-4868
chrX-118937535-DEL-1771
chrX-118945103-DEL-9059
chrX-118955023-DEL-3781
chrX-118965436-DEL-536
chrX-118986749-DEL-83
chrX-119041977-DEL-70
chrX-119079258-DEL-202
chrX-119080869-DEL-1351
chrX-119082333-DEL-1420
chrX-119134126-DEL-354
chrX-119233558-DEL-1792
chrX-119248395-DEL-1539
chrX-119260475-DEL-6239
chrX-119267489-DEL-594
chrX-119325953-DEL-57
chrX-119342297-DEL-340

chrX-119380051-DEL-7409
chrX-119409324-DEL-213
chrX-119424775-DEL-98
chrX-119452698-DEL-935
chrX-119471134-DEL-574
chrX-119479040-DEL-436
chrX-119493813-DEL-163
chrX-119495948-DEL-60
chrX-119513619-DEL-122
chrX-119556860-DEL-78
chrX-119630710-DEL-56
chrX-119660431-DEL-6195
chrX-119689796-DEL-80
chrX-119724999-DEL-3922
chrX-119763335-DEL-58
chrX-119771085-DEL-279
chrX-119772614-DEL-1833
chrX-119794371-DEL-2570
chrX-119796955-DEL-5076
chrX-119811113-DEL-724
chrX-119822510-DEL-141
chrX-119877712-DEL-4311
chrX-119879558-DEL-3224
chrX-119909545-DEL-967
chrX-119979274-DEL-1016
chrX-119992076-DEL-260
chrX-120012568-DEL-1028
chrX-120035759-DEL-59
chrX-120059533-DEL-12413
chrX-120092768-DEL-70
chrX-120109668-DEL-109
chrX-120114550-DEL-800
chrX-120160196-DEL-241
chrX-120193355-DEL-7086
chrX-120209726-DEL-229
chrX-120240600-DEL-109
chrX-120243626-DEL-2650
chrX-120257016-DEL-190
chrX-120258739-DEL-358
chrX-120267483-DEL-2171
chrX-120271143-DEL-1253
chrX-120275725-DEL-7508
chrX-120283574-DEL-6264
chrX-120290542-DEL-202
chrX-120290941-DEL-8283
chrX-120299138-DEL-553
chrX-120309956-DEL-151
chrX-120340230-DEL-106
chrX-120362318-DEL-4111
chrX-120367243-DEL-1321
chrX-120384226-DEL-400
chrX-120385918-DEL-2289
chrX-120400883-DEL-944
chrX-120404088-DEL-358
chrX-120419382-DEL-6390
chrX-120472860-DEL-76
chrX-120484072-DEL-84
chrX-120528305-DEL-4113
chrX-120545541-DEL-893
chrX-120588540-DEL-282
chrX-120601143-DEL-53
chrX-120609859-DEL-507
chrX-120610394-DEL-2252
chrX-120636106-DEL-52
chrX-120639811-DEL-904
chrX-120671743-DEL-56
chrX-120679845-DEL-972
chrX-120690296-DEL-600
chrX-120721830-DEL-57
chrX-120739754-DEL-2098
chrX-120766611-DEL-796
chrX-120793545-DEL-198
chrX-120809197-DEL-2490
chrX-120817465-DEL-342
chrX-120824623-DEL-493
chrX-120854213-DEL-1725
chrX-120872398-DEL-7319

chrX-120911667-DEL-6160
chrX-120920264-DEL-727
chrX-120925781-DEL-119
chrX-120949376-DEL-877
chrX-120977152-DEL-408
chrX-120996348-DEL-4441
chrX-121010514-DEL-2272
chrX-121034055-DEL-66
chrX-121084214-DEL-748
chrX-121129330-DEL-58
chrX-121150069-DEL-355
chrX-121180796-DEL-540
chrX-121195195-DEL-13048
chrX-121208972-DEL-507
chrX-121218621-DEL-3242
chrX-121234324-DEL-6370
chrX-121249508-DEL-926
chrX-121284684-DEL-110
chrX-121295635-DEL-926
chrX-121296302-DEL-1644
chrX-121299723-DEL-1479
chrX-121314918-DEL-277
chrX-121327700-DEL-5840
chrX-121337460-DEL-1226
chrX-121366803-DEL-53
chrX-121400195-DEL-115
chrX-121402849-DEL-1022
chrX-121420954-DEL-54
chrX-121440136-DEL-112
chrX-121442905-DEL-3303
chrX-121457970-DEL-1809
chrX-121470359-DEL-58
chrX-121476281-DEL-2611
chrX-121499980-DEL-208
chrX-121500913-DEL-80
chrX-121538178-DEL-771
chrX-121547037-DEL-1891
chrX-121592290-DEL-7501
chrX-121599916-DEL-1912
chrX-121628066-DEL-1979
chrX-121679566-DEL-69
chrX-121685393-DEL-1951
chrX-121716532-DEL-97
chrX-121772089-DEL-6222
chrX-121785563-DEL-85
chrX-121846991-DEL-5098
chrX-121888632-DEL-6356
chrX-121901048-DEL-4568
chrX-121916208-DEL-100
chrX-121928821-DEL-381
chrX-121943736-DEL-196
chrX-121945804-DEL-583
chrX-121949149-DEL-685
chrX-121967453-DEL-79
chrX-121983103-DEL-52
chrX-121993673-DEL-1254
chrX-121994064-DEL-1258
chrX-122020980-DEL-6154
chrX-122058752-DEL-5152
chrX-122092681-DEL-77
chrX-122139340-DEL-1149
chrX-122141291-DEL-1149
chrX-122157672-DEL-132
chrX-122163042-DEL-7108
chrX-122175988-DEL-63
chrX-122177984-DEL-142
chrX-122179337-DEL-77
chrX-122400457-DEL-3319
chrX-122407395-DEL-99
chrX-122410702-DEL-2241
chrX-122468550-DEL-1171
chrX-122470720-DEL-7078
chrX-122506060-DEL-56
chrX-122521882-DEL-853
chrX-122547361-DEL-72
chrX-122608714-DEL-753
chrX-122615626-DEL-175

chrX-122621539-DEL-3601
chrX-122622829-DEL-300
chrX-122626738-DEL-63
chrX-122630097-DEL-68
chrX-122688427-DEL-2212
chrX-122791894-DEL-651
chrX-122834737-DEL-176
chrX-122839041-DEL-371
chrX-122839563-DEL-1262
chrX-122843357-DEL-3292
chrX-122850228-DEL-145
chrX-122853573-DEL-2241
chrX-122868920-DEL-2138
chrX-122869450-DEL-3068
chrX-122906079-DEL-4019
chrX-122945421-DEL-96
chrX-122988513-DEL-75
chrX-123020166-DEL-1353
chrX-123028003-DEL-187
chrX-123042921-DEL-95
chrX-123052354-DEL-18623
chrX-123056322-DEL-1197
chrX-123073229-DEL-14474
chrX-123080287-DEL-496
chrX-123110556-DEL-1137
chrX-123130676-DEL-2404
chrX-123135192-DEL-52
chrX-123139295-DEL-120
chrX-123140932-DEL-537
chrX-123148221-DEL-58
chrX-123149436-DEL-2731
chrX-123149590-DEL-3008
chrX-123165650-DEL-498
chrX-123165655-DEL-3971
chrX-123189763-DEL-5434
chrX-123264197-DEL-75
chrX-123273712-DEL-240
chrX-123280176-DEL-3325
chrX-123390939-DEL-1595
chrX-123399446-DEL-3337
chrX-123404487-DEL-751
chrX-123520348-DEL-1595
chrX-123528855-DEL-3337
chrX-123534722-DEL-669
chrX-124041972-DEL-1595
chrX-124050481-DEL-3338
chrX-124174362-DEL-4346
chrX-124184767-DEL-64
chrX-124253927-DEL-2240
chrX-124263111-DEL-3324
chrX-124268524-DEL-53
chrX-124270379-DEL-62
chrX-124373864-DEL-1595
chrX-124382373-DEL-3337
chrX-124384904-DEL-72
chrX-124466962-DEL-2240
chrX-124476150-DEL-3325
chrX-124483419-DEL-62
chrX-124491580-DEL-381
chrX-124593740-DEL-1595
chrX-124602248-DEL-3337
chrX-124776591-DEL-1595
chrX-124785099-DEL-3336
chrX-124809035-DEL-1149
chrX-124811328-DEL-1149
chrX-124905877-DEL-1595
chrX-124916904-DEL-71
chrX-124930273-DEL-762
chrX-124959518-DEL-4628
chrX-124973446-DEL-2296
chrX-124975834-DEL-7475
chrX-125008592-DEL-52
chrX-125021370-DEL-3125
chrX-125041169-DEL-112
chrX-125042726-DEL-6448
chrX-125050712-DEL-140
chrX-125076448-DEL-1490

chrX-125094677-DEL-213
chrX-125153732-DEL-451
chrX-125155228-DEL-143
chrX-125157292-DEL-59
chrX-125160242-DEL-17414
chrX-125170297-DEL-2507
chrX-125177322-DEL-196
chrX-125189005-DEL-2623
chrX-125209672-DEL-62
chrX-125228087-DEL-6388
chrX-125242227-DEL-235
chrX-125254884-DEL-8331
chrX-125256291-DEL-6650
chrX-125282582-DEL-6433
chrX-125304587-DEL-88
chrX-125309083-DEL-1473
chrX-125310488-DEL-7048
chrX-125316700-DEL-416
chrX-125343859-DEL-797
chrX-125350845-DEL-1675
chrX-125414364-DEL-783
chrX-125444997-DEL-6478
chrX-125492455-DEL-1699
chrX-125495443-DEL-5496
chrX-125512382-DEL-7194
chrX-125530282-DEL-182
chrX-125583812-DEL-1671
chrX-125656693-DEL-972
chrX-125662662-DEL-348
chrX-125665459-DEL-295
chrX-125679669-DEL-1057
chrX-125681405-DEL-1393
chrX-125689996-DEL-806
chrX-125690881-DEL-334
chrX-125693198-DEL-190
chrX-125693832-DEL-1433
chrX-125705990-DEL-363
chrX-125726221-DEL-2027
chrX-125749299-DEL-1154
chrX-125760840-DEL-7766
chrX-125800293-DEL-165
chrX-125808951-DEL-827
chrX-125852329-DEL-59
chrX-125905181-DEL-52
chrX-125908817-DEL-7294
chrX-125918077-DEL-214
chrX-125925712-DEL-8866
chrX-125926251-DEL-8302
chrX-125950905-DEL-1691
chrX-125972324-DEL-163
chrX-125978044-DEL-2715
chrX-125993919-DEL-92
chrX-126031658-DEL-201
chrX-126046066-DEL-6167
chrX-126072665-DEL-7090
chrX-126090935-DEL-877
chrX-126107412-DEL-273
chrX-126115690-DEL-126
chrX-126116714-DEL-769
chrX-126139349-DEL-54
chrX-126183696-DEL-3409
chrX-126191230-DEL-141
chrX-126194285-DEL-459
chrX-126194978-DEL-91
chrX-126201847-DEL-717
chrX-126205366-DEL-54
chrX-126212175-DEL-192
chrX-126238172-DEL-3131
chrX-126251898-DEL-274
chrX-126258256-DEL-9916
chrX-126273155-DEL-441
chrX-126305504-DEL-1244
chrX-126322015-DEL-218
chrX-126335145-DEL-197
chrX-126359071-DEL-1010
chrX-126360469-DEL-1716
chrX-126377370-DEL-54

chrX-126378325-DEL-804
chrX-126395057-DEL-135
chrX-126397578-DEL-835
chrX-126418103-DEL-404
chrX-126419899-DEL-239
chrX-126428814-DEL-965
chrX-126435472-DEL-6122
chrX-126442107-DEL-59
chrX-126443927-DEL-199
chrX-126464727-DEL-504
chrX-126596779-DEL-1893
chrX-126627677-DEL-222
chrX-126655468-DEL-1830
chrX-126675235-DEL-255
chrX-126679333-DEL-52
chrX-126686271-DEL-2116
chrX-126695286-DEL-64
chrX-126700550-DEL-1465
chrX-126720610-DEL-67
chrX-126737121-DEL-1137
chrX-126755870-DEL-677
chrX-126759373-DEL-2632
chrX-126798916-DEL-68
chrX-126836860-DEL-196
chrX-126838243-DEL-101
chrX-126858115-DEL-833
chrX-126858982-DEL-5717
chrX-126886557-DEL-779
chrX-126897374-DEL-1260
chrX-126911900-DEL-5521
chrX-126924894-DEL-211
chrX-126929445-DEL-200
chrX-126941264-DEL-282
chrX-126951160-DEL-80
chrX-126953209-DEL-179
chrX-126956556-DEL-315
chrX-127013322-DEL-1172
chrX-127017357-DEL-60
chrX-127018244-DEL-153
chrX-127019324-DEL-3691
chrX-127050949-DEL-787
chrX-127061314-DEL-88
chrX-127069444-DEL-118
chrX-127075191-DEL-78
chrX-127083319-DEL-83
chrX-127111025-DEL-70
chrX-127118230-DEL-725
chrX-127123972-DEL-5388
chrX-127154354-DEL-82
chrX-127177449-DEL-2792
chrX-127185998-DEL-103
chrX-127205628-DEL-281
chrX-127238702-DEL-165
chrX-127272763-DEL-275
chrX-127287952-DEL-1412
chrX-127295914-DEL-27986
chrX-127335679-DEL-278
chrX-127347126-DEL-381
chrX-127375950-DEL-1001
chrX-127377391-DEL-202
chrX-127389069-DEL-1112
chrX-127396316-DEL-303
chrX-127398099-DEL-1381
chrX-127405002-DEL-5191
chrX-127415239-DEL-957
chrX-127422253-DEL-53
chrX-127422826-DEL-206
chrX-127423319-DEL-492
chrX-127481173-DEL-59
chrX-127561417-DEL-192
chrX-127563740-DEL-296
chrX-127566294-DEL-60
chrX-127583804-DEL-688
chrX-127595058-DEL-89
chrX-127608683-DEL-6807
chrX-127672760-DEL-1558
chrX-127735460-DEL-445

chrX-127759143-DEL-53
chrX-127778383-DEL-6331
chrX-127790923-DEL-1342
chrX-127792400-DEL-5957
chrX-127797745-DEL-1927
chrX-127805657-DEL-6360
chrX-127816472-DEL-64
chrX-127833686-DEL-1123
chrX-127847179-DEL-286
chrX-127860508-DEL-6170
chrX-127875478-DEL-64
chrX-127894951-DEL-214
chrX-127918546-DEL-118
chrX-127984468-DEL-3339
chrX-127997989-DEL-162
chrX-128010866-DEL-221
chrX-128040792-DEL-766
chrX-128041629-DEL-1957
chrX-128058314-DEL-60
chrX-128115517-DEL-399
chrX-128118133-DEL-1176
chrX-128147313-DEL-266
chrX-128176584-DEL-6181
chrX-128192981-DEL-1287
chrX-128214482-DEL-628
chrX-128218529-DEL-170
chrX-128218961-DEL-3317
chrX-128273915-DEL-1076
chrX-128329376-DEL-19632
chrX-128331081-DEL-3895
chrX-128335354-DEL-147
chrX-128366764-DEL-53
chrX-128401283-DEL-80
chrX-128424465-DEL-593
chrX-128427340-DEL-176
chrX-128493640-DEL-56
chrX-128511547-DEL-1800
chrX-128526135-DEL-110
chrX-128613450-DEL-213
chrX-128616049-DEL-406
chrX-128665223-DEL-211
chrX-128671997-DEL-66
chrX-128706843-DEL-6215
chrX-128777646-DEL-71
chrX-128791617-DEL-212
chrX-128825975-DEL-53
chrX-128853865-DEL-259
chrX-128865127-DEL-247
chrX-128912442-DEL-4319
chrX-128921279-DEL-278
chrX-128966749-DEL-1486
chrX-128968336-DEL-1100
chrX-128974700-DEL-777
chrX-128975694-DEL-78
chrX-129014123-DEL-117
chrX-129015256-DEL-93
chrX-129021613-DEL-107
chrX-129035033-DEL-62
chrX-129063993-DEL-125
chrX-129065062-DEL-2249
chrX-129068016-DEL-203
chrX-129136782-DEL-85
chrX-129154147-DEL-166
chrX-129156092-DEL-66
chrX-129192827-DEL-951
chrX-129217031-DEL-94
chrX-129225326-DEL-172
chrX-129283057-DEL-79
chrX-129295370-DEL-332
chrX-129355966-DEL-401
chrX-129377792-DEL-881
chrX-129385735-DEL-189
chrX-129546954-DEL-361
chrX-129568866-DEL-62
chrX-129589354-DEL-671
chrX-129608758-DEL-527
chrX-129627074-DEL-79

chrX-129681161-DEL-5166
chrX-129693965-DEL-4893
chrX-129698901-DEL-217
chrX-129751071-DEL-6183
chrX-129786455-DEL-402
chrX-129816265-DEL-414
chrX-129836895-DEL-308
chrX-129854324-DEL-3012
chrX-129894636-DEL-92
chrX-129899118-DEL-87
chrX-129927513-DEL-76
chrX-129938194-DEL-57
chrX-129962515-DEL-66
chrX-130013764-DEL-67
chrX-130054249-DEL-1866
chrX-130057977-DEL-1096
chrX-130065829-DEL-408
chrX-130073649-DEL-81
chrX-130076366-DEL-86
chrX-130093783-DEL-79
chrX-130107283-DEL-67
chrX-130120175-DEL-1133
chrX-130140906-DEL-223
chrX-130144428-DEL-177
chrX-130145988-DEL-327
chrX-130185625-DEL-140
chrX-130278095-DEL-57
chrX-130296633-DEL-2412
chrX-130317274-DEL-88
chrX-130387142-DEL-396
chrX-130429625-DEL-1889
chrX-130440685-DEL-60
chrX-130457046-DEL-714
chrX-130490884-DEL-127
chrX-130529349-DEL-194
chrX-130550312-DEL-403
chrX-130564735-DEL-864
chrX-130584660-DEL-65
chrX-130600823-DEL-177
chrX-130604697-DEL-496
chrX-130614795-DEL-155
chrX-130616983-DEL-87
chrX-130653489-DEL-3080
chrX-130661086-DEL-204
chrX-130673022-DEL-56
chrX-130696146-DEL-69
chrX-130705531-DEL-1692
chrX-130739886-DEL-349
chrX-130740540-DEL-54
chrX-130777771-DEL-124
chrX-130781344-DEL-208
chrX-130806914-DEL-58
chrX-130828902-DEL-687
chrX-130862584-DEL-121
chrX-130874482-DEL-72
chrX-130878470-DEL-6538
chrX-130878689-DEL-214
chrX-130904572-DEL-260
chrX-130939241-DEL-1173
chrX-130955813-DEL-311
chrX-130984704-DEL-215
chrX-131032789-DEL-845
chrX-131096517-DEL-70
chrX-131100344-DEL-731
chrX-131130286-DEL-60
chrX-131204109-DEL-979
chrX-131205194-DEL-70
chrX-131210925-DEL-147
chrX-131231062-DEL-814
chrX-131266441-DEL-108
chrX-131318660-DEL-159
chrX-131334525-DEL-310
chrX-131449712-DEL-191
chrX-131471451-DEL-375
chrX-131478071-DEL-7752
chrX-131478187-DEL-6391
chrX-131486031-DEL-5022

chrX-131487429-DEL-2116
chrX-131490416-DEL-3010
chrX-131513502-DEL-302
chrX-131525507-DEL-69
chrX-131529356-DEL-993
chrX-131532346-DEL-1054
chrX-131557985-DEL-54
chrX-131572499-DEL-1533
chrX-131580493-DEL-320
chrX-131592495-DEL-77
chrX-131609248-DEL-90
chrX-131626425-DEL-920
chrX-131645955-DEL-6885
chrX-131666716-DEL-146
chrX-131676961-DEL-84
chrX-131692458-DEL-6395
chrX-131700150-DEL-58
chrX-131707096-DEL-148
chrX-131711122-DEL-366
chrX-131714504-DEL-77
chrX-131739592-DEL-149
chrX-131747279-DEL-477
chrX-131807807-DEL-1639
chrX-131832889-DEL-199
chrX-131893118-DEL-1293
chrX-131901033-DEL-4982
chrX-131907518-DEL-450
chrX-131910353-DEL-425
chrX-131918956-DEL-370
chrX-131923784-DEL-174
chrX-131954079-DEL-250
chrX-131964758-DEL-2499
chrX-131974085-DEL-72
chrX-131994091-DEL-330
chrX-131998263-DEL-52
chrX-132030139-DEL-153
chrX-132051857-DEL-536
chrX-132068710-DEL-2228
chrX-132076543-DEL-1511
chrX-132123188-DEL-759
chrX-132124008-DEL-1273
chrX-132125338-DEL-360
chrX-132137249-DEL-62
chrX-132178355-DEL-251
chrX-132190058-DEL-124
chrX-132196648-DEL-74
chrX-132207703-DEL-54
chrX-132218589-DEL-65
chrX-132278954-DEL-98
chrX-132292891-DEL-493
chrX-132304842-DEL-54
chrX-132335254-DEL-217
chrX-132337405-DEL-2477
chrX-132358061-DEL-1330
chrX-132363297-DEL-93
chrX-132371688-DEL-1909
chrX-132377692-DEL-445
chrX-132383710-DEL-1671
chrX-132385984-DEL-87
chrX-132422575-DEL-286
chrX-132438887-DEL-88
chrX-132442763-DEL-151
chrX-132448664-DEL-124
chrX-132475655-DEL-120
chrX-132507091-DEL-171
chrX-132522475-DEL-58
chrX-132601015-DEL-292
chrX-132662024-DEL-55
chrX-132684557-DEL-58
chrX-132704665-DEL-3532
chrX-132735591-DEL-66
chrX-132756407-DEL-111
chrX-132799339-DEL-99
chrX-132803997-DEL-2496
chrX-132804218-DEL-215
chrX-132875310-DEL-537
chrX-132904912-DEL-244

chrX-132909127-DEL-497
chrX-132923928-DEL-56
chrX-132936688-DEL-55
chrX-132944515-DEL-609
chrX-133015557-DEL-105
chrX-133030844-DEL-132
chrX-133044663-DEL-173
chrX-133047046-DEL-77
chrX-133050855-DEL-144
chrX-133068629-DEL-144
chrX-133095664-DEL-112
chrX-133120498-DEL-248
chrX-133122797-DEL-56
chrX-133139929-DEL-219
chrX-133150815-DEL-1676
chrX-133156573-DEL-58
chrX-133173525-DEL-107
chrX-133193690-DEL-60
chrX-133230770-DEL-137
chrX-133237712-DEL-131
chrX-133241726-DEL-55
chrX-133277907-DEL-51
chrX-133312696-DEL-2044
chrX-133332091-DEL-68
chrX-133333642-DEL-177
chrX-133392638-DEL-65
chrX-133412184-DEL-61
chrX-133426808-DEL-175
chrX-133427922-DEL-174
chrX-133429479-DEL-62
chrX-133453518-DEL-98
chrX-133475929-DEL-201
chrX-133479567-DEL-76
chrX-133480818-DEL-63
chrX-133495412-DEL-64
chrX-133498705-DEL-639
chrX-133530860-DEL-54
chrX-133534944-DEL-188
chrX-133559903-DEL-83
chrX-133574201-DEL-537
chrX-133579315-DEL-89
chrX-133665251-DEL-206
chrX-133671422-DEL-322
chrX-133735307-DEL-74
chrX-133792652-DEL-134
chrX-133804540-DEL-54
chrX-133832407-DEL-1309
chrX-133854449-DEL-844
chrX-133865865-DEL-54
chrX-133938070-DEL-819
chrX-133941327-DEL-12932
chrX-133941388-DEL-15763
chrX-133942902-DEL-931
chrX-133943009-DEL-285
chrX-133944062-DEL-76
chrX-133944821-DEL-456
chrX-133945419-DEL-456
chrX-133946916-DEL-57
chrX-133947911-DEL-285
chrX-133949107-DEL-133
chrX-133951952-DEL-323
chrX-133953427-DEL-835
chrX-133954172-DEL-3059
chrX-133954821-DEL-247
chrX-133955061-DEL-76
chrX-133955673-DEL-192
chrX-133972506-DEL-64
chrX-133974781-DEL-52
chrX-133987554-DEL-160
chrX-133996542-DEL-174
chrX-134000217-DEL-273
chrX-134001115-DEL-117
chrX-134001994-DEL-54
chrX-134014269-DEL-215
chrX-134056766-DEL-56
chrX-134064668-DEL-148
chrX-134075653-DEL-426

chrX-134083344-DEL-62
chrX-134084716-DEL-71
chrX-134085608-DEL-10188
chrX-134089183-DEL-96
chrX-134092083-DEL-344
chrX-134094498-DEL-85
chrX-134106337-DEL-124
chrX-134112597-DEL-2118
chrX-134148329-DEL-148
chrX-134153775-DEL-58
chrX-134154600-DEL-1432
chrX-134157439-DEL-1295
chrX-134161216-DEL-138
chrX-134183118-DEL-266
chrX-134186665-DEL-168
chrX-134193801-DEL-69
chrX-134232267-DEL-11450
chrX-134234459-DEL-685
chrX-134259860-DEL-430
chrX-134312849-DEL-104
chrX-134320815-DEL-86
chrX-134338292-DEL-50
chrX-134355417-DEL-7795
chrX-134374977-DEL-66
chrX-134381467-DEL-172
chrX-134384503-DEL-87
chrX-134390308-DEL-67
chrX-134468563-DEL-75
chrX-134482926-DEL-69
chrX-134510761-DEL-322
chrX-134554975-DEL-311
chrX-134586569-DEL-59
chrX-134587041-DEL-143
chrX-134595281-DEL-582
chrX-134597575-DEL-107
chrX-134617931-DEL-59
chrX-134630821-DEL-205
chrX-134639117-DEL-60
chrX-134672723-DEL-1486
chrX-134684670-DEL-75
chrX-134689382-DEL-88
chrX-134735477-DEL-197
chrX-134751414-DEL-1197
chrX-134810125-DEL-735
chrX-134838008-DEL-328
chrX-134859803-DEL-235
chrX-134870289-DEL-418
chrX-134887935-DEL-142
chrX-134889184-DEL-57
chrX-134901882-DEL-66
chrX-134929369-DEL-464
chrX-134930230-DEL-143
chrX-134942998-DEL-155
chrX-134956190-DEL-1157
chrX-135043569-DEL-85
chrX-135046319-DEL-529
chrX-135060952-DEL-363
chrX-135081218-DEL-79
chrX-135117623-DEL-54
chrX-135128370-DEL-92
chrX-135160271-DEL-198
chrX-135208103-DEL-58
chrX-135304887-DEL-890
chrX-135312596-DEL-506
chrX-135319101-DEL-632
chrX-135338149-DEL-66
chrX-135366476-DEL-79
chrX-135377350-DEL-9574
chrX-135397870-DEL-6787
chrX-135397884-DEL-5752
chrX-135405268-DEL-159
chrX-135413480-DEL-125
chrX-135417148-DEL-1281
chrX-135423931-DEL-601
chrX-135428476-DEL-716
chrX-135434231-DEL-54
chrX-135436787-DEL-164

chrX-135474114-DEL-293
chrX-135502818-DEL-1477
chrX-135505513-DEL-214
chrX-135537991-DEL-989
chrX-135544435-DEL-760
chrX-135550928-DEL-2857
chrX-135572900-DEL-64
chrX-135573629-DEL-938
chrX-135581462-DEL-72
chrX-135585177-DEL-304
chrX-135628771-DEL-163
chrX-135629813-DEL-478
chrX-135713774-DEL-502
chrX-135726685-DEL-52
chrX-135734444-DEL-177
chrX-135738438-DEL-66
chrX-135806651-DEL-744
chrX-135817403-DEL-399
chrX-135827161-DEL-155
chrX-135834535-DEL-964
chrX-135847080-DEL-70
chrX-135856411-DEL-108
chrX-135882653-DEL-52
chrX-135955695-DEL-69
chrX-135962710-DEL-477
chrX-135964724-DEL-106
chrX-135990628-DEL-53
chrX-136007586-DEL-2704
chrX-136010841-DEL-1403
chrX-136029057-DEL-3116
chrX-136033346-DEL-1371
chrX-136034354-DEL-766
chrX-136066898-DEL-816
chrX-136069043-DEL-93
chrX-136074944-DEL-212
chrX-136111131-DEL-62
chrX-136132778-DEL-449
chrX-136137526-DEL-54
chrX-136140308-DEL-57
chrX-136170277-DEL-450
chrX-136173322-DEL-65
chrX-136206286-DEL-1446
chrX-136216047-DEL-197
chrX-136285644-DEL-3550
chrX-136293628-DEL-1612
chrX-136312673-DEL-8202
chrX-136327834-DEL-212
chrX-136352112-DEL-524
chrX-136388137-DEL-2552
chrX-136396198-DEL-60
chrX-136397446-DEL-195
chrX-136409656-DEL-72
chrX-136430447-DEL-225
chrX-136432788-DEL-267
chrX-136436932-DEL-204
chrX-136506672-DEL-544
chrX-136523020-DEL-145
chrX-136526352-DEL-212
chrX-136609168-DEL-5311
chrX-136617942-DEL-134
chrX-136632174-DEL-412
chrX-136632970-DEL-1759
chrX-136645626-DEL-150
chrX-136753125-DEL-104
chrX-136754081-DEL-358
chrX-136807071-DEL-907
chrX-136816667-DEL-718
chrX-136823531-DEL-116
chrX-136839146-DEL-115
chrX-136845921-DEL-317
chrX-136856475-DEL-202
chrX-136888851-DEL-200
chrX-136889852-DEL-1020
chrX-136898841-DEL-2112
chrX-136990521-DEL-94
chrX-136992781-DEL-58
chrX-137012206-DEL-520

chrX-137027132-DEL-6213
chrX-137031058-DEL-79
chrX-137038431-DEL-70
chrX-137118697-DEL-373
chrX-137119237-DEL-186
chrX-137120094-DEL-351
chrX-137140595-DEL-94
chrX-137200259-DEL-708
chrX-137204752-DEL-88
chrX-137213154-DEL-162
chrX-137214158-DEL-71
chrX-137250014-DEL-423
chrX-137291320-DEL-863
chrX-137319399-DEL-85
chrX-137336837-DEL-1255
chrX-137353132-DEL-144
chrX-137417609-DEL-97
chrX-137441147-DEL-511
chrX-137465112-DEL-1087
chrX-137491566-DEL-2669
chrX-137500240-DEL-1583
chrX-137518947-DEL-196
chrX-137570293-DEL-1248
chrX-137605429-DEL-3777
chrX-137680001-DEL-79
chrX-137691998-DEL-895
chrX-137721274-DEL-81
chrX-137764978-DEL-62
chrX-137823383-DEL-88
chrX-137854436-DEL-63
chrX-137863076-DEL-64
chrX-137962525-DEL-77
chrX-138001670-DEL-114
chrX-138002350-DEL-387
chrX-138002883-DEL-51
chrX-138087748-DEL-1653
chrX-138101090-DEL-866
chrX-138124415-DEL-67
chrX-138127653-DEL-124
chrX-138158398-DEL-204
chrX-138160040-DEL-52
chrX-138195847-DEL-52
chrX-138203642-DEL-1724
chrX-138217441-DEL-56
chrX-138248054-DEL-170
chrX-138365493-DEL-768
chrX-138402141-DEL-1490
chrX-138515353-DEL-62
chrX-138517691-DEL-365
chrX-138541871-DEL-137
chrX-138548220-DEL-66
chrX-138558467-DEL-184
chrX-138614336-DEL-921
chrX-138619082-DEL-310
chrX-138625214-DEL-61
chrX-138665687-DEL-51
chrX-138682741-DEL-115
chrX-138727053-DEL-112
chrX-138746000-DEL-2241
chrX-138747651-DEL-55
chrX-138840446-DEL-80
chrX-138856418-DEL-108
chrX-138865094-DEL-55
chrX-138874533-DEL-1017
chrX-138901996-DEL-516
chrX-138933616-DEL-245
chrX-138939373-DEL-98
chrX-138980943-DEL-862
chrX-139025594-DEL-136
chrX-139061080-DEL-592
chrX-139075446-DEL-54
chrX-139128927-DEL-157
chrX-139173451-DEL-108
chrX-139247559-DEL-79
chrX-139301912-DEL-117
chrX-139308832-DEL-97
chrX-139433543-DEL-88

chrX-139536819-DEL-215
chrX-139539091-DEL-55
chrX-139551154-DEL-80
chrX-139558076-DEL-80
chrX-139595517-DEL-128
chrX-139614044-DEL-129
chrX-139638154-DEL-68
chrX-139655343-DEL-180
chrX-139672598-DEL-1051
chrX-139689449-DEL-548
chrX-139701381-DEL-187
chrX-139756091-DEL-54
chrX-139772567-DEL-86
chrX-139781984-DEL-1301
chrX-139785480-DEL-432
chrX-139898396-DEL-118
chrX-139916706-DEL-115
chrX-139953379-DEL-60
chrX-139993947-DEL-159
chrX-140022972-DEL-502
chrX-140029072-DEL-408
chrX-140030085-DEL-1023
chrX-140073128-DEL-399
chrX-140082888-DEL-751
chrX-140149862-DEL-2866
chrX-140214538-DEL-1840
chrX-140217957-DEL-575
chrX-140282579-DEL-504
chrX-140300941-DEL-144
chrX-140319731-DEL-1014
chrX-140320845-DEL-132
chrX-140322130-DEL-668
chrX-140350822-DEL-428
chrX-140357233-DEL-81
chrX-140397840-DEL-58
chrX-140404750-DEL-322
chrX-140417698-DEL-598
chrX-140459463-DEL-181
chrX-140460298-DEL-807
chrX-140471378-DEL-141
chrX-140472740-DEL-541
chrX-140474799-DEL-76
chrX-140557736-DEL-112
chrX-140574454-DEL-348
chrX-140598252-DEL-98
chrX-140611980-DEL-65
chrX-140614414-DEL-6116
chrX-140623690-DEL-58
chrX-140642466-DEL-72
chrX-140682512-DEL-5391
chrX-140682743-DEL-3004
chrX-140701630-DEL-2828
chrX-140723042-DEL-2486
chrX-140766435-DEL-236
chrX-140780160-DEL-515
chrX-140845275-DEL-810
chrX-140884750-DEL-353
chrX-140896839-DEL-63
chrX-140927114-DEL-4604
chrX-140927397-DEL-6416
chrX-140927535-DEL-201
chrX-140970809-DEL-52
chrX-141013980-DEL-112
chrX-141060723-DEL-804
chrX-141110467-DEL-183
chrX-141114653-DEL-62
chrX-141118351-DEL-105
chrX-141143790-DEL-58
chrX-141158441-DEL-85
chrX-141169411-DEL-128
chrX-141196176-DEL-3425
chrX-141217393-DEL-399
chrX-141281259-DEL-60
chrX-141293802-DEL-214
chrX-141295875-DEL-83
chrX-141305026-DEL-3018
chrX-141308617-DEL-107

chrX-141324944-DEL-52
chrX-141331083-DEL-1537
chrX-141346596-DEL-127
chrX-141353486-DEL-325
chrX-141371223-DEL-572
chrX-141375543-DEL-93
chrX-141522505-DEL-57
chrX-141583600-DEL-197
chrX-141584214-DEL-461
chrX-141627750-DEL-618
chrX-141721674-DEL-232
chrX-141760579-DEL-320
chrX-141792164-DEL-381
chrX-141840806-DEL-160
chrX-141855887-DEL-70
chrX-141911763-DEL-481
chrX-141913658-DEL-578
chrX-141925512-DEL-218
chrX-141938764-DEL-1332
chrX-141940104-DEL-705
chrX-141950157-DEL-52
chrX-141953024-DEL-11090
chrX-141983722-DEL-160
chrX-141992399-DEL-170
chrX-142003851-DEL-60
chrX-142006596-DEL-79
chrX-142037415-DEL-1231
chrX-142135693-DEL-139
chrX-142140521-DEL-62
chrX-142198720-DEL-1306
chrX-142209081-DEL-66
chrX-142252739-DEL-5700
chrX-142270973-DEL-200
chrX-142302425-DEL-58
chrX-142338864-DEL-64
chrX-142427402-DEL-93
chrX-142452658-DEL-291
chrX-142534032-DEL-3193
chrX-142541250-DEL-216
chrX-142587901-DEL-70
chrX-142646979-DEL-100
chrX-142700443-DEL-287
chrX-142735628-DEL-338
chrX-142739274-DEL-7922
chrX-142755588-DEL-6408
chrX-142774038-DEL-1749
chrX-142779858-DEL-496
chrX-142807828-DEL-6617
chrX-142826476-DEL-11948
chrX-142882420-DEL-491
chrX-142909866-DEL-2401
chrX-142911742-DEL-358
chrX-142951670-DEL-1888
chrX-142992626-DEL-500
chrX-143014961-DEL-72
chrX-143024419-DEL-174
chrX-143026541-DEL-206
chrX-143027235-DEL-58
chrX-143028476-DEL-55
chrX-143041131-DEL-1948
chrX-143046958-DEL-566
chrX-143048029-DEL-72
chrX-143067867-DEL-52
chrX-143080568-DEL-512
chrX-143123984-DEL-402
chrX-143143567-DEL-125
chrX-143145812-DEL-194
chrX-143175063-DEL-70
chrX-143201554-DEL-116
chrX-143223408-DEL-61
chrX-143235124-DEL-763
chrX-143260263-DEL-96
chrX-143260820-DEL-2146
chrX-143271716-DEL-211
chrX-143284615-DEL-96
chrX-143295822-DEL-101
chrX-143321811-DEL-1129

chrX-143333992-DEL-735
chrX-143339184-DEL-176
chrX-143345909-DEL-301
chrX-143365588-DEL-77
chrX-143369435-DEL-2470
chrX-143390936-DEL-1368
chrX-143416976-DEL-2384
chrX-143505190-DEL-167
chrX-143521705-DEL-67
chrX-143524294-DEL-552
chrX-143563002-DEL-680
chrX-143565539-DEL-203
chrX-143585736-DEL-85
chrX-143604386-DEL-207
chrX-143677190-DEL-714
chrX-143686499-DEL-140
chrX-143689982-DEL-203
chrX-143691300-DEL-113
chrX-143724276-DEL-64
chrX-143727017-DEL-401
chrX-143729681-DEL-289
chrX-143737154-DEL-3239
chrX-143744145-DEL-60
chrX-143763597-DEL-56
chrX-143772688-DEL-60
chrX-143791923-DEL-53
chrX-143963202-DEL-493
chrX-143981442-DEL-55
chrX-143986014-DEL-3392
chrX-144076018-DEL-659
chrX-144096513-DEL-8602
chrX-144182911-DEL-717
chrX-144219689-DEL-65
chrX-144227524-DEL-80
chrX-144365152-DEL-59
chrX-144374865-DEL-196
chrX-144380533-DEL-82
chrX-144390899-DEL-1109
chrX-144407688-DEL-996
chrX-144429375-DEL-200
chrX-144444164-DEL-3136
chrX-144458350-DEL-157
chrX-144458718-DEL-1305
chrX-144478614-DEL-52
chrX-144547919-DEL-88
chrX-144576502-DEL-1019
chrX-144580707-DEL-486
chrX-144583498-DEL-64
chrX-144624860-DEL-885
chrX-144634741-DEL-189
chrX-144651626-DEL-2710
chrX-144671369-DEL-749
chrX-144708637-DEL-62
chrX-144721461-DEL-690
chrX-144728399-DEL-172
chrX-144741094-DEL-202
chrX-144753339-DEL-677
chrX-144775952-DEL-1275
chrX-144797290-DEL-313
chrX-144799566-DEL-600
chrX-144817155-DEL-7168
chrX-144831455-DEL-281
chrX-144841015-DEL-52
chrX-144848989-DEL-1319
chrX-144886806-DEL-132
chrX-144899985-DEL-400
chrX-144906506-DEL-68
chrX-144911704-DEL-323
chrX-144914644-DEL-52
chrX-144931546-DEL-197
chrX-144940098-DEL-6186
chrX-144958822-DEL-1272
chrX-144973694-DEL-119
chrX-145052101-DEL-615
chrX-145088047-DEL-91
chrX-145098396-DEL-214
chrX-145130344-DEL-228

chrX-145207119-DEL-1497
chrX-145223782-DEL-64
chrX-145253369-DEL-291
chrX-145269615-DEL-72
chrX-145365474-DEL-4792
chrX-145439062-DEL-1110
chrX-145444964-DEL-199
chrX-145464140-DEL-1171
chrX-145493363-DEL-91
chrX-145587549-DEL-89
chrX-145588782-DEL-638
chrX-145611514-DEL-81
chrX-145620924-DEL-377
chrX-145641860-DEL-340
chrX-145642441-DEL-191
chrX-145678346-DEL-2098
chrX-145682417-DEL-12614
chrX-145707706-DEL-84
chrX-145713744-DEL-556
chrX-145720136-DEL-169
chrX-145735370-DEL-510
chrX-145752270-DEL-267
chrX-145765447-DEL-162
chrX-145781785-DEL-84
chrX-145806365-DEL-66
chrX-145833812-DEL-67
chrX-145861225-DEL-224
chrX-145891140-DEL-68
chrX-145925510-DEL-63
chrX-145932525-DEL-2733
chrX-145987833-DEL-8238
chrX-146013078-DEL-2122
chrX-146040409-DEL-189
chrX-146067285-DEL-10152
chrX-146067977-DEL-5495
chrX-146068543-DEL-7842
chrX-146071181-DEL-6241
chrX-146075919-DEL-4541
chrX-146077952-DEL-11973
chrX-146084204-DEL-6361
chrX-146133110-DEL-103
chrX-146142385-DEL-87
chrX-146178342-DEL-83
chrX-146208149-DEL-677
chrX-146209044-DEL-140
chrX-146245299-DEL-1125
chrX-146273212-DEL-1561
chrX-146317621-DEL-684
chrX-146339369-DEL-52
chrX-146350622-DEL-86
chrX-146360233-DEL-1160
chrX-146403927-DEL-1690
chrX-146414454-DEL-872
chrX-146422105-DEL-7410
chrX-146429735-DEL-332
chrX-146432838-DEL-6217
chrX-146497720-DEL-55
chrX-146549537-DEL-4265
chrX-146559089-DEL-53
chrX-146565756-DEL-1904
chrX-146595021-DEL-72
chrX-146677380-DEL-635
chrX-146685558-DEL-743
chrX-146694003-DEL-6206
chrX-146829735-DEL-60
chrX-146830285-DEL-4317
chrX-146881803-DEL-89
chrX-146904248-DEL-57
chrX-146957819-DEL-4735
chrX-146986204-DEL-72
chrX-146999606-DEL-62
chrX-147014561-DEL-68
chrX-147294932-DEL-72
chrX-147365643-DEL-51
chrX-147387418-DEL-4191
chrX-147390124-DEL-57
chrX-147435777-DEL-217

chrX-147437661-DEL-6068
chrX-147580678-DEL-299
chrX-147587643-DEL-6401
chrX-147628616-DEL-9802
chrX-147634774-DEL-1535
chrX-147640191-DEL-69
chrX-147659787-DEL-4823
chrX-147765686-DEL-89
chrX-147818587-DEL-848
chrX-147970334-DEL-579
chrX-148089749-DEL-52
chrX-148199251-DEL-650
chrX-148213119-DEL-561
chrX-148357812-DEL-2047
chrX-148360045-DEL-565
chrX-148361982-DEL-217
chrX-148363861-DEL-10833
chrX-148365435-DEL-13868
chrX-148433843-DEL-4290
chrX-148484628-DEL-463
chrX-148485522-DEL-2500
chrX-148513337-DEL-1982
chrX-148522046-DEL-52
chrX-148538262-DEL-1239
chrX-148607894-DEL-74
chrX-148608461-DEL-123
chrX-148611300-DEL-64
chrX-148613084-DEL-2643
chrX-148677355-DEL-344
chrX-148681971-DEL-105
chrX-148734686-DEL-3062
chrX-148786379-DEL-583
chrX-148849838-DEL-6209
chrX-148879382-DEL-57
chrX-148927606-DEL-12669
chrX-148972146-DEL-1060
chrX-149010134-DEL-186
chrX-149015407-DEL-4754
chrX-149020524-DEL-212
chrX-149042394-DEL-144
chrX-149095401-DEL-86
chrX-149106394-DEL-3780
chrX-149176386-DEL-58
chrX-149176837-DEL-403
chrX-149197752-DEL-52
chrX-149213821-DEL-604
chrX-149287069-DEL-1196
chrX-149365913-DEL-276
chrX-149369172-DEL-170
chrX-149496673-DEL-6393
chrX-149526591-DEL-203
chrX-149557931-DEL-164
chrX-149602262-DEL-1890
chrX-149605484-DEL-99
chrX-149659783-DEL-571
chrX-149727345-DEL-64
chrX-149764457-DEL-179
chrX-149767482-DEL-178
chrX-149777541-DEL-176
chrX-149781938-DEL-351
chrX-149788262-DEL-511
chrX-149790696-DEL-163
chrX-149795975-DEL-397
chrX-149816759-DEL-195
chrX-149825804-DEL-204
chrX-149847842-DEL-665
chrX-149858169-DEL-202
chrX-149894096-DEL-1325
chrX-149925073-DEL-80
chrX-149934286-DEL-136
chrX-149937287-DEL-112
chrX-149966138-DEL-92
chrX-149975679-DEL-212
chrX-149978554-DEL-181
chrX-149988650-DEL-109
chrX-150042743-DEL-320
chrX-150074818-DEL-145

chrX-150111982-DEL-438
chrX-150113128-DEL-2848
chrX-150150651-DEL-85
chrX-150183191-DEL-249
chrX-150242859-DEL-1498
chrX-150271639-DEL-1517
chrX-150295631-DEL-492
chrX-150358184-DEL-600
chrX-150383905-DEL-293
chrX-150387253-DEL-158
chrX-150392857-DEL-66
chrX-150423430-DEL-60
chrX-150429719-DEL-56
chrX-150446181-DEL-1896
chrX-150487062-DEL-736
chrX-150501914-DEL-276
chrX-150519476-DEL-1937
chrX-150521340-DEL-1766
chrX-150525833-DEL-214
chrX-150534199-DEL-199
chrX-150534872-DEL-98
chrX-150576688-DEL-224
chrX-150615770-DEL-315
chrX-150628222-DEL-59
chrX-150709402-DEL-715
chrX-150710216-DEL-60
chrX-150714210-DEL-162
chrX-150726566-DEL-1418
chrX-150728341-DEL-1695
chrX-150735039-DEL-6174
chrX-150741935-DEL-208
chrX-150750058-DEL-58
chrX-150756775-DEL-422
chrX-150761363-DEL-405
chrX-150769547-DEL-218
chrX-150911582-DEL-166
chrX-150966953-DEL-381
chrX-150973471-DEL-79
chrX-150994408-DEL-150
chrX-151015176-DEL-68
chrX-151062278-DEL-318
chrX-151064293-DEL-206
chrX-151074351-DEL-369
chrX-151096740-DEL-835
chrX-151118108-DEL-109
chrX-151132223-DEL-533
chrX-151155604-DEL-61
chrX-151164282-DEL-58
chrX-151173519-DEL-252
chrX-151174326-DEL-197
chrX-151236364-DEL-12407
chrX-151237012-DEL-1637
chrX-151304383-DEL-468
chrX-151332809-DEL-523
chrX-151370143-DEL-811
chrX-151371513-DEL-994
chrX-151372827-DEL-2505
chrX-151379356-DEL-372
chrX-151423902-DEL-173
chrX-151433437-DEL-516
chrX-151461051-DEL-69
chrX-151496558-DEL-217
chrX-151508904-DEL-228
chrX-151572186-DEL-53
chrX-151621627-DEL-4810
chrX-151649373-DEL-325
chrX-151660199-DEL-1652
chrX-151665309-DEL-95
chrX-151692460-DEL-729
chrX-151697963-DEL-122
chrX-151700925-DEL-74
chrX-151703328-DEL-324
chrX-151726814-DEL-440
chrX-151727315-DEL-117
chrX-151734835-DEL-9238
chrX-151867673-DEL-147
chrX-151880278-DEL-219

chrX-151881354-DEL-126
chrX-151894550-DEL-195
chrX-151897517-DEL-84
chrX-151958189-DEL-472
chrX-151967890-DEL-102
chrX-151974719-DEL-161
chrX-152022483-DEL-110
chrX-152116813-DEL-63
chrX-152136213-DEL-86
chrX-152220564-DEL-52
chrX-152228393-DEL-2526
chrX-152277145-DEL-156
chrX-152310561-DEL-58
chrX-152320445-DEL-870
chrX-152332642-DEL-95
chrX-152352061-DEL-6472
chrX-152367752-DEL-1114
chrX-152373731-DEL-200
chrX-152394519-DEL-350
chrX-152414950-DEL-970
chrX-152415719-DEL-204
chrX-152421629-DEL-140
chrX-152422848-DEL-109
chrX-152438151-DEL-95
chrX-152445637-DEL-76
chrX-152550050-DEL-2200
chrX-152569649-DEL-227
chrX-152581239-DEL-498
chrX-152624528-DEL-2888
chrX-152655757-DEL-129
chrX-152664599-DEL-731
chrX-152675440-DEL-74
chrX-152701805-DEL-235
chrX-152719062-DEL-54
chrX-152724578-DEL-2135
chrX-152752627-DEL-2168
chrX-152870337-DEL-7153
chrX-152885721-DEL-750
chrX-152887365-DEL-166
chrX-152893886-DEL-6082
chrX-152900971-DEL-686
chrX-152909016-DEL-276
chrX-152916326-DEL-198
chrX-152928786-DEL-201
chrX-152930713-DEL-276
chrX-152933146-DEL-52
chrX-152953666-DEL-6358
chrX-152973785-DEL-4183
chrX-152977166-DEL-1596
chrX-152995476-DEL-383
chrX-153022582-DEL-969
chrX-153023901-DEL-6209
chrX-153033089-DEL-620
chrX-153037453-DEL-1733
chrX-153063940-DEL-3020
chrX-153097145-DEL-64
chrX-153102266-DEL-61
chrX-153107774-DEL-2282
chrX-153109611-DEL-81
chrX-153133233-DEL-470
chrX-153151102-DEL-92
chrX-153175429-DEL-348
chrX-153185256-DEL-1942
chrX-153187315-DEL-261
chrX-153189369-DEL-254
chrX-153199395-DEL-1890
chrX-153203226-DEL-2702
chrX-153212177-DEL-461
chrX-153215180-DEL-1892
chrX-153222318-DEL-5299
chrX-153255445-DEL-56
chrX-153303256-DEL-51
chrX-153305944-DEL-1832
chrX-153313090-DEL-1339
chrX-153325073-DEL-252
chrX-153330065-DEL-154
chrX-153339891-DEL-5747

chrX-153341738-DEL-889
chrX-153341947-DEL-2857
chrX-153343421-DEL-1728
chrX-153394812-DEL-1718
chrX-153468200-DEL-3205
chrX-153502460-DEL-4663
chrX-153536982-DEL-390
chrX-153539779-DEL-77
chrX-153542382-DEL-2085
chrX-153543501-DEL-16332
chrX-153662461-DEL-339
chrX-153726702-DEL-363
chrX-153735251-DEL-820
chrX-153751120-DEL-6812
chrX-153776778-DEL-104
chrX-153825059-DEL-199
chrX-153860106-DEL-1516
chrX-153861383-DEL-620
chrX-153891964-DEL-413
chrX-153906201-DEL-89
chrX-153914532-DEL-701
chrX-153980057-DEL-169
chrX-154010879-DEL-212
chrX-154014664-DEL-7976
chrX-154015953-DEL-73
chrX-154034055-DEL-198
chrX-154088969-DEL-112
chrX-154092101-DEL-98
chrX-154096946-DEL-277
chrX-154114737-DEL-87
chrX-154132460-DEL-159
chrX-154133348-DEL-198
chrX-154146965-DEL-74
chrX-154182200-DEL-194
chrX-154230179-DEL-1157
chrX-154278641-DEL-138
chrX-154279388-DEL-4961
chrX-154286601-DEL-95
chrX-154290202-DEL-377
chrX-154412655-DEL-932
chrX-154436559-DEL-87
chrX-154462428-DEL-1486
chrX-154487379-DEL-4053
chrX-154529161-DEL-67
chrX-154529611-DEL-128
chrX-154530795-DEL-1376
chrX-154539226-DEL-84
chrX-154547263-DEL-243
chrX-154549068-DEL-143
chrX-154558318-DEL-1481
chrX-154596997-DEL-68
chrX-154662538-DEL-283
chrX-154664930-DEL-56
chrX-154678030-DEL-2712
chrX-154694027-DEL-1061
chrX-154733438-DEL-1643
chrX-154760001-DEL-516
chrX-154771075-DEL-59
chrX-154781249-DEL-357
chrX-154790212-DEL-129
chrX-154874249-DEL-67
chrX-154919836-DEL-51
chrX-154948572-DEL-526
chrX-154957173-DEL-1390
chrX-154993872-DEL-57
chrX-155036269-DEL-454
chrX-155115560-DEL-154
chrX-155126146-DEL-59
chrX-155130322-DEL-448
chrX-155132593-DEL-1271
chrX-155142905-DEL-73
chrX-155144634-DEL-3807
chrX-155169884-DEL-56
chrX-155175205-DEL-584
chrX-155226840-DEL-98
chrX-155252797-DEL-987
chrX-155267404-DEL-178

chrX-155285896-DEL-4396
chrX-155322243-DEL-236
chrX-155431129-DEL-524
chrX-155458811-DEL-1609
chrX-155465905-DEL-2401
chrX-155478719-DEL-200
chrX-155484974-DEL-686
chrX-155511347-DEL-6430
chrX-155580182-DEL-58
chrX-155582641-DEL-222
chrX-155599635-DEL-148
chrX-155622326-DEL-55
chrX-155651666-DEL-6196
chrX-155701271-DEL-203
chrX-155722591-DEL-53
chrX-155731755-DEL-347
chrX-155769899-DEL-90
chrX-155780572-DEL-486
chrX-155860655-DEL-581
chrX-155881837-DEL-1255
chrX-155909210-DEL-4085
chrX-155974582-DEL-411
chrX-155975828-DEL-371
chrX-156012144-DEL-2119
chrX-156020395-DEL-531
chrX-156030854-DEL-2711
chrX-156110426-DEL-54
chrX-156156376-DEL-256
chrX-156177680-DEL-140
chrX-156178592-DEL-242
chrX-156195486-DEL-422
chrX-156221751-DEL-196
chrX-156239420-DEL-160
chrX-156314719-DEL-51
chrX-156348168-DEL-87
chrX-156352750-DEL-258
chrX-156489712-DEL-60
chrX-156494773-DEL-52
chrX-156510310-DEL-85
chrX-156515972-DEL-66
chrX-156528191-DEL-1191
chrX-156576480-DEL-87
chrX-156651802-DEL-60
chrX-156682066-DEL-95
chrX-156711872-DEL-52
chrX-156736557-DEL-100
chrX-156765251-DEL-620
chrX-156780732-DEL-220
chrX-156804627-DEL-53
chrX-156881618-DEL-764
chrX-156902619-DEL-263
chrX-156923715-DEL-76
chrX-156958387-DEL-1328
chrX-156960709-DEL-53
chrX-156970171-DEL-358
chrX-156974030-DEL-352
chrX-156975253-DEL-307
chrX-157021459-DEL-74
chrX-157033343-DEL-104
chrX-157055074-DEL-1008
chrX-157069683-DEL-116
chrX-157113935-DEL-4770
chrX-157119325-DEL-158
chrX-157154959-DEL-56
chrX-157155880-DEL-52
chrX-157172308-DEL-132
chrX-157240614-DEL-3513
chrX-157277525-DEL-2324
chrX-157306823-DEL-1794
chrX-157338917-DEL-3582
chrX-157339498-DEL-639
chrX-157343476-DEL-182
chrX-157346998-DEL-1626
chrX-157363887-DEL-471
chrX-157418901-DEL-1193
chrX-157468451-DEL-6016
chrX-157478742-DEL-54

chrX-157535775-DEL-55
chrX-157562996-DEL-402
chrX-157572238-DEL-249
chrX-157578497-DEL-300
chrX-157597530-DEL-204
chrX-157625037-DEL-52
chrX-157660919-DEL-720
chrX-157663116-DEL-137
chrX-157684304-DEL-6245
chrX-157714275-DEL-432
chrX-157731995-DEL-97
chrX-157754437-DEL-61
chrX-157762200-DEL-2409
chrX-157790112-DEL-642
chrX-157877049-DEL-60
chrX-157884895-DEL-72
chrX-157886673-DEL-498
chrX-157903110-DEL-2150
chrX-157956921-DEL-981
chrX-157971888-DEL-363
chrX-158030045-DEL-567
chrX-158104845-DEL-1752
chrX-158132960-DEL-64
chrX-158135862-DEL-110
chrX-158139277-DEL-1295
chrX-158174029-DEL-112
chrX-158174984-DEL-127
chrX-158175564-DEL-334
chrX-158179583-DEL-269
chrX-158181174-DEL-6191
chrX-158189746-DEL-202
chrX-158223613-DEL-56
chrX-158250861-DEL-86
chrX-158259488-DEL-77
chrX-158268816-DEL-167
chrX-158269750-DEL-179
chrX-158270533-DEL-499
chrX-158284061-DEL-91
chrX-158305169-DEL-507
chrX-158318818-DEL-201
chrX-158322009-DEL-93
chrX-158332196-DEL-62
chrX-158353799-DEL-56
chrX-158393835-DEL-264
chrX-158439523-DEL-198
chrX-158440836-DEL-58
chrX-158556010-DEL-1289
chrX-158571314-DEL-400
chrX-158723543-DEL-85
chrX-158764084-DEL-387
chrX-158821533-DEL-66
chrX-158841284-DEL-213
chrX-158860614-DEL-237
chrX-158886804-DEL-103
chrX-158892571-DEL-163
chrX-158914911-DEL-95
chrX-158953418-DEL-333
chrX-158953423-DEL-2222
chrX-158991173-DEL-84
chrX-159004755-DEL-533
chrX-159050490-DEL-1557
chrX-159100464-DEL-156
chrX-159111775-DEL-238
chrX-159116859-DEL-1797
chrX-159146399-DEL-147
chrX-159219841-DEL-58
chrX-159259378-DEL-95
chrX-159282965-DEL-485
chrX-159492307-DEL-916
chrX-159522432-DEL-309
chrX-159565611-DEL-78
chrX-159571259-DEL-107
chrX-159579335-DEL-53
chrX-159600744-DEL-128
chrX-159610338-DEL-62
chrX-159636342-DEL-345
chrX-159645735-DEL-173

chrX-159661077-DEL-80
chrX-159663677-DEL-115
chrX-159692339-DEL-1191
chrX-159693051-DEL-100
chrX-159705710-DEL-78
chrX-159709407-DEL-500
chrX-159726367-DEL-71
chrX-159823057-DEL-479
chrX-159825451-DEL-258
chrX-159828010-DEL-60
chrX-159839298-DEL-2498
chrX-159841827-DEL-3907
chrX-159895723-DEL-1070
chrX-159928638-DEL-140
chrX-159986800-DEL-2027
chrX-160005045-DEL-83
chrX-160007410-DEL-1416
chrX-160008788-DEL-5523
chrX-160009255-DEL-100
chrX-160009500-DEL-2118
chrX-160061000-DEL-1992
chrX-160069929-DEL-58
chrX-160132157-DEL-71
chrX-160170812-DEL-3453
chrX-160174491-DEL-1010
chrX-160184152-DEL-2326
chrX-160273478-DEL-96
chrX-160329971-DEL-212
chrX-160331242-DEL-2294
chrX-160467179-DEL-61
chrX-160472232-DEL-86
chrX-160489146-DEL-6675
chrX-160492061-DEL-87
chrX-160507499-DEL-53
chrX-160543739-DEL-339
chrX-160557207-DEL-1437
chrX-160743353-DEL-52
chrX-160765581-DEL-332
chrX-160792307-DEL-89
chrX-160843284-DEL-210
chrX-160850936-DEL-189
chrX-160954431-DEL-544
chrX-160986453-DEL-955
chrX-161067914-DEL-56
chrX-161129907-DEL-66
chrX-161137774-DEL-175
chrX-161140803-DEL-929
chrX-161163451-DEL-72
chrX-161192900-DEL-68
chrX-161198223-DEL-64
chrX-161205394-DEL-2023
chrX-161258731-DEL-546
chrX-161266039-DEL-65
chrX-161349790-DEL-533
chrX-161350849-DEL-158
chrX-161380467-DEL-851
chrX-161385249-DEL-100
chrX-161389672-DEL-539
chrX-161397250-DEL-412
chrX-161416037-DEL-401
chrX-161421181-DEL-150
chrX-161462039-DEL-54
chrX-161465650-DEL-354
chrX-161484889-DEL-1871
chrX-161490638-DEL-89
chrX-161503738-DEL-187
chrX-161508097-DEL-176
chrX-161508437-DEL-869
chrX-161509043-DEL-237
chrX-161528754-DEL-589
chrX-161548437-DEL-204
chrX-161578806-DEL-1275
chrX-161582883-DEL-179
chrX-161586049-DEL-986
chrX-161587423-DEL-391
chrX-161589517-DEL-347
chrX-161606499-DEL-217

chrX-161616414-DEL-68
chrX-161622790-DEL-4177
chrX-161628388-DEL-225
chrX-161657954-DEL-202
chrX-161671544-DEL-72
chrX-161741774-DEL-91
chrX-161742468-DEL-144
chrX-161790671-DEL-66
chrX-161838420-DEL-100
chrX-161858379-DEL-1484
chrX-161891697-DEL-899
chrX-161902650-DEL-664
chrX-161907687-DEL-479
chrX-161914076-DEL-68
chrX-161932413-DEL-55
chrX-161934302-DEL-1175
chrX-161973494-DEL-1579
chrX-162021034-DEL-117
chrX-162122790-DEL-180
chrX-162135473-DEL-90
chrX-162137828-DEL-505
chrX-162186395-DEL-66
chrX-162202286-DEL-52
chrX-162204726-DEL-54
chrX-162260989-DEL-920
chrX-162280690-DEL-902
chrX-162301243-DEL-445
chrX-162340727-DEL-7326
chrX-162348789-DEL-5657
chrX-162369996-DEL-189
chrX-162377792-DEL-79
chrX-162427923-DEL-61
chrX-162457549-DEL-1320
chrX-162476820-DEL-122
chrX-162517831-DEL-79
chrX-162522852-DEL-177
chrX-162542198-DEL-259
chrX-162587983-DEL-670
chrX-162599007-DEL-51
chrX-162655160-DEL-3472
chrX-162670580-DEL-1553
chrX-162756342-DEL-805
chrX-162789056-DEL-659
chrX-162807711-DEL-146
chrX-162839784-DEL-60
chrX-162845249-DEL-57
chrX-162850707-DEL-174
chrX-162889390-DEL-109
chrX-162896006-DEL-527
chrX-162904785-DEL-72
chrX-162921631-DEL-421
chrX-162984817-DEL-196
chrX-163007923-DEL-1442
chrX-163020196-DEL-60
chrX-163081337-DEL-840
chrX-163108175-DEL-2006
chrX-163132460-DEL-275
chrX-163241339-DEL-116
chrX-163246416-DEL-532
chrX-163251044-DEL-61
chrX-163261709-DEL-1437
chrX-163272309-DEL-2020
chrX-163275417-DEL-796
chrX-163322069-DEL-118
chrX-163343281-DEL-486
chrX-163354857-DEL-58
chrX-163378369-DEL-606
chrX-163398015-DEL-212
chrX-163415290-DEL-958
chrX-163431874-DEL-84
chrX-163481753-DEL-73
chrX-163487472-DEL-53
chrX-163533376-DEL-6223
chrX-163574801-DEL-2487
chrX-163652155-DEL-56
chrX-163679652-DEL-976
chrX-163686684-DEL-96

chrX-163692426-DEL-55
chrX-163696799-DEL-445
chrX-163700945-DEL-54
chrX-163706470-DEL-501
chrX-163738556-DEL-614
chrX-163803234-DEL-124
chrX-163804596-DEL-241
chrX-163819837-DEL-51
chrX-163829414-DEL-679
chrX-163832353-DEL-916
chrX-163835832-DEL-71
chrX-163846568-DEL-781
chrX-163927885-DEL-51
chrX-163966056-DEL-73
chrX-163977599-DEL-52
chrX-163984492-DEL-737
chrX-164000011-DEL-61
chrX-164024196-DEL-91
chrX-164053726-DEL-93
chrX-164089376-DEL-68
chrX-164096296-DEL-65
chrX-164124100-DEL-439
chrX-164159346-DEL-459
chrX-164162882-DEL-52
chrX-164193429-DEL-6626
chrX-164193722-DEL-201
chrX-164203466-DEL-70
chrX-164218429-DEL-346
chrX-164240036-DEL-134
chrX-164257101-DEL-6283
chrX-164299042-DEL-911
chrX-164342272-DEL-2061
chrX-164425724-DEL-593
chrX-164427000-DEL-5715
chrX-164484178-DEL-6485
chrX-164484356-DEL-214
chrX-164518872-DEL-6215
chrX-164535638-DEL-2423
chrX-164573672-DEL-6225
chrX-164589260-DEL-15171
chrX-164589824-DEL-8752
chrX-164589826-DEL-8187
chrX-164613537-DEL-6748
chrX-164623392-DEL-78
chrX-164678532-DEL-87
chrX-164685916-DEL-236
chrX-164692228-DEL-821
chrX-164743147-DEL-271
chrX-164759017-DEL-96
chrX-164769254-DEL-4175
chrX-164772552-DEL-4758
chrX-164773629-DEL-941
chrX-164776971-DEL-437
chrX-164780092-DEL-359
chrX-164792351-DEL-51
chrX-164838857-DEL-239
chrX-164845922-DEL-6275
chrX-164874411-DEL-1603
chrX-164879414-DEL-401
chrX-164909982-DEL-2596
chrX-164921328-DEL-246
chrX-164934531-DEL-441
chrX-164951156-DEL-166
chrX-165130400-DEL-51
chrX-165194500-DEL-106
chrX-165350695-DEL-130
chrX-165351953-DEL-157
chrX-165402971-DEL-393
chrX-165417457-DEL-406
chrX-165452686-DEL-69
chrX-165453224-DEL-89
chrX-165506647-DEL-199
chrX-165545697-DEL-52
chrX-165624359-DEL-64
chrX-165656972-DEL-56
chrX-165669722-DEL-54
chrX-165680809-DEL-54

chrX-165720045-DEL-1761
chrX-165742108-DEL-1826
chrX-165756646-DEL-72
chrX-165769388-DEL-6284
chrX-165782459-DEL-63
chrX-165784924-DEL-80
chrX-165785544-DEL-196
chrX-165802346-DEL-208
chrX-165831659-DEL-61
chrX-165842852-DEL-553
chrX-165896166-DEL-76
chrX-165925867-DEL-84
chrX-165935266-DEL-191
chrX-165976443-DEL-84
chrX-166029075-DEL-6183
chrX-166039618-DEL-909
chrX-166040816-DEL-126
chrX-166041263-DEL-534
chrX-166097284-DEL-59
chrX-166108494-DEL-707
chrX-166120064-DEL-407
chrX-166127071-DEL-1706
chrX-166184956-DEL-211
chrX-166264166-DEL-2080
chrX-166310447-DEL-430
chrX-166319909-DEL-179
chrX-166338603-DEL-157
chrX-166381425-DEL-121
chrX-166438894-DEL-1736
chrX-166444086-DEL-714
chrX-166521744-DEL-62
chrX-166542480-DEL-485
chrX-166607724-DEL-924
chrX-166634467-DEL-1271
chrX-166660315-DEL-52
chrX-166704685-DEL-56
chrX-166709529-DEL-54
chrX-166726213-DEL-72
chrX-166740547-DEL-825
chrX-166769888-DEL-455
chrX-166772898-DEL-71
chrX-166781312-DEL-54
chrX-166782653-DEL-65
chrX-166786916-DEL-60
chrX-166798583-DEL-228
chrX-166802781-DEL-108
chrX-166805891-DEL-658
chrX-166809166-DEL-771
chrX-166826173-DEL-338
chrX-166860327-DEL-81
chrX-166883112-DEL-60
chrX-166891960-DEL-308
chrX-166936252-DEL-904
chrX-166967671-DEL-91
chrX-166975170-DEL-714
chrX-167074134-DEL-54
chrX-167100162-DEL-89
chrX-167135410-DEL-3202
chrX-167171289-DEL-57
chrX-167176702-DEL-58
chrX-167187946-DEL-2399
chrX-167192412-DEL-135
chrX-167253189-DEL-421
chrX-167270718-DEL-598
chrX-167271363-DEL-1220
chrX-167273622-DEL-286
chrX-167340780-DEL-1002
chrX-167343011-DEL-463
chrX-167416720-DEL-497
chrX-167501938-DEL-723
chrX-167502709-DEL-1174
chrX-167508144-DEL-80
chrX-167540936-DEL-81
chrX-167583116-DEL-205
chrX-167589722-DEL-584
chrX-167652647-DEL-60
chrX-167779431-DEL-64

chrX-167852789-DEL-56
chrX-167884321-DEL-97
chrX-168023983-DEL-563
chrX-168058685-DEL-110
chrX-168061702-DEL-79
chrX-168088266-DEL-170
chrX-168091329-DEL-65
chrX-168106661-DEL-171
chrX-168151321-DEL-51
chrX-168153682-DEL-186
chrX-168161463-DEL-281
chrX-168189938-DEL-1036
chrX-168190024-DEL-59
chrX-168194939-DEL-212
chrX-168207704-DEL-186
chrX-168224158-DEL-536
chrX-168281324-DEL-438
chrX-168298565-DEL-2474
chrX-168310699-DEL-1177
chrX-168316882-DEL-1031
chrX-168320109-DEL-306
chrX-168340223-DEL-2202
chrX-168343988-DEL-53
chrX-168355113-DEL-1527
chrX-168366139-DEL-54
chrX-168425382-DEL-60
chrX-168442868-DEL-64
chrX-168490275-DEL-90
chrX-168505179-DEL-59
chrX-168514795-DEL-103
chrX-168532926-DEL-247
chrX-168554412-DEL-607
chrX-168573825-DEL-89
chrX-168646510-DEL-2103
chrX-168652083-DEL-75
chrX-168706411-DEL-3408
chrX-168714148-DEL-34774
chrX-168726886-DEL-324
chrX-168741733-DEL-204
chrX-168753216-DEL-1821
chrX-168757982-DEL-70
chrX-168762246-DEL-252
chrX-168769162-DEL-310
chrX-168769829-DEL-323
chrX-168770916-DEL-117
chrX-168771329-DEL-300
chrX-168772519-DEL-123
chrX-168773972-DEL-62
chrX-168775074-DEL-966
chrX-168776463-DEL-2515
chrX-168776739-DEL-93
chrX-168777817-DEL-590
chrX-168778729-DEL-12462
chrX-168779948-DEL-493
chrX-168781095-DEL-149
chrX-168782034-DEL-217
chrX-168782356-DEL-207
chrX-168782709-DEL-249
chrX-168783817-DEL-123
chrX-168784172-DEL-252
chrX-168784700-DEL-62
chrX-168785567-DEL-86
chrX-168786326-DEL-124
chrX-168787379-DEL-184
chrX-168788052-DEL-174
chrX-168788539-DEL-1233
chrX-168791621-DEL-80
chrX-168792615-DEL-86
chrX-168794464-DEL-1353
chrX-168794473-DEL-5749
chrX-168794498-DEL-2474
chrX-168794533-DEL-2196
chrX-168794695-DEL-249
chrX-168794956-DEL-474
chrX-168795336-DEL-294
chrX-168795983-DEL-335
chrX-168796348-DEL-493

chrX-168797228-DEL-315
chrX-168798226-DEL-540
chrX-168798310-DEL-90
chrX-168798359-DEL-1282
chrX-168799042-DEL-630
chrX-168799097-DEL-944
chrX-168799943-DEL-135
chrX-168801354-DEL-95
chrX-168803157-DEL-697
chrX-168804080-DEL-51
chrX-168804187-DEL-131
chrX-168909728-DEL-74
chrX-168961087-DEL-150
chrX-169012537-DEL-91
chrX-169068908-DEL-341
chrX-169069916-DEL-213
chrX-169071192-DEL-281
chrX-169072179-DEL-144
chrX-169072410-DEL-68
chrX-169074331-DEL-434
chrX-169074332-DEL-1962
chrX-169074959-DEL-450
chrX-169075171-DEL-1130
chrX-169077147-DEL-217
chrX-169081929-DEL-741
chrX-169085845-DEL-90
chrX-169087941-DEL-79
chrX-169089479-DEL-62
chrX-169091605-DEL-119
chrX-169092693-DEL-105
chrX-169093748-DEL-80
chrX-169095320-DEL-168
chrX-169100866-DEL-58
chrX-169102914-DEL-176
chrX-169106938-DEL-76
chrX-169108050-DEL-87
chrX-169112239-DEL-131
chrX-169112717-DEL-122
chrX-169114188-DEL-1977
chrX-169114523-DEL-226
chrX-169115691-DEL-121
chrX-169117636-DEL-188
chrX-169118128-DEL-791
chrX-169118352-DEL-53
chrX-169121750-DEL-87
chrX-169177245-DEL-60
chrX-169178270-DEL-175
chrX-169180441-DEL-65
chrX-169181039-DEL-88
chrX-169183309-DEL-149
chrX-169184071-DEL-60
chrX-169187564-DEL-677
chrX-169190483-DEL-121
chrX-169259901-DEL-123
chrX-169261494-DEL-305
chrX-169262914-DEL-244
chrX-169269009-DEL-73
chrX-169281609-DEL-242
chrX-169300247-DEL-322
chrX-169304489-DEL-62
chrX-169307512-DEL-150
chrX-169312856-DEL-93
chrX-169313397-DEL-119

**Identified_By_JAX**
chr1-3146996-DEL-112
chr1-3147029-DEL-609
chr1-3173596-DEL-74
chr1-3348094-DEL-550
chr1-3525277-DEL-793
chr1-3645377-DEL-1504
chr1-3689877-DEL-2894
chr1-3817458-DEL-3877
chr1-3835532-DEL-132
chr1-3922796-DEL-212
chr1-3948029-DEL-2575
chr1-4014069-DEL-85
chr1-4022102-DEL-1903
chr1-4053616-DEL-5095
chr1-4326652-DEL-339
chr1-4326848-DEL-54
chr1-4326869-DEL-581
chr1-4327621-DEL-666
chr1-4328400-DEL-461
chr1-4450951-DEL-51
chr1-4509900-DEL-5169
chr1-4574684-DEL-69
chr1-4579449-DEL-97
chr1-4579507-DEL-308
chr1-4627201-DEL-10188
chr1-4746007-DEL-65
chr1-4756721-DEL-226
chr1-4782874-DEL-53
chr1-4799291-DEL-67
chr1-4825510-DEL-136
chr1-4828676-DEL-87
chr1-4835830-DEL-133
chr1-4881175-DEL-192
chr1-4886966-DEL-2594
chr1-4905100-DEL-592
chr1-4905167-DEL-181
chr1-4932001-DEL-165
chr1-4932751-DEL-1992
chr1-4936729-DEL-288
chr1-4948925-DEL-61
chr1-4949409-DEL-1562
chr1-4984338-DEL-246
chr1-5001464-DEL-59
chr1-5041511-DEL-1696
chr1-5142274-DEL-667
chr1-5147026-DEL-5395
chr1-5179654-DEL-4946
chr1-5185089-DEL-50
chr1-5346244-DEL-50
chr1-5346983-DEL-114
chr1-5493937-DEL-7061
chr1-5531609-DEL-599
chr1-5634772-DEL-2807
chr1-5905155-DEL-71
chr1-5991226-DEL-836
chr1-6073270-DEL-90
chr1-6215202-DEL-6243
chr1-6221144-DEL-199
chr1-6423430-DEL-999
chr1-6426422-DEL-789
chr1-6444126-DEL-86
chr1-6450492-DEL-2137
chr1-6454024-DEL-969
chr1-6495868-DEL-1329
chr1-6514116-DEL-1358
chr1-6519264-DEL-1880
chr1-6722364-DEL-78
chr1-6733081-DEL-1491
chr1-6811668-DEL-210
chr1-6987234-DEL-201
chr1-7016601-DEL-4182
chr1-7024773-DEL-625
chr1-7058966-DEL-186
chr1-7058972-DEL-60
chr1-7109713-DEL-253
chr1-7189183-DEL-242

chr1-7195099-DEL-5099
chr1-7418585-DEL-87
chr1-7429474-DEL-52
chr1-7429738-DEL-200
chr1-7464152-DEL-566
chr1-7475132-DEL-91
chr1-7531990-DEL-216
chr1-7566853-DEL-134
chr1-7567933-DEL-144
chr1-7724265-DEL-669
chr1-7745720-DEL-78
chr1-7746188-DEL-72
chr1-7800561-DEL-56
chr1-7826715-DEL-50
chr1-7827147-DEL-3235
chr1-7841104-DEL-54
chr1-7976204-DEL-1275
chr1-7982134-DEL-4483
chr1-7992966-DEL-16943
chr1-8009959-DEL-292
chr1-8010550-DEL-122
chr1-8161391-DEL-58
chr1-8163291-DEL-126
chr1-8163369-DEL-51
chr1-8167811-DEL-65
chr1-8323277-DEL-104
chr1-8392770-DEL-70
chr1-8422983-DEL-71
chr1-8441774-DEL-91
chr1-8493091-DEL-60
chr1-8494934-DEL-67
chr1-8526663-DEL-54
chr1-8698652-DEL-1709
chr1-8705210-DEL-6400
chr1-8758723-DEL-416
chr1-8833561-DEL-123
chr1-8857569-DEL-229
chr1-8886255-DEL-75
chr1-8910763-DEL-69
chr1-8918416-DEL-62
chr1-8928877-DEL-207
chr1-8929837-DEL-53
chr1-9006042-DEL-50
chr1-9039687-DEL-57
chr1-9163387-DEL-189
chr1-9398420-DEL-73
chr1-9398612-DEL-189
chr1-9417205-DEL-423
chr1-9496277-DEL-5110
chr1-9524419-DEL-78
chr1-9565087-DEL-566
chr1-9589367-DEL-108
chr1-9600802-DEL-200
chr1-9661735-DEL-113
chr1-9675994-DEL-56
chr1-9684002-DEL-162
chr1-9707107-DEL-261
chr1-9775548-DEL-533
chr1-9834348-DEL-136
chr1-9852209-DEL-72
chr1-9854632-DEL-163
chr1-9873758-DEL-62
chr1-9897484-DEL-115
chr1-9945661-DEL-54
chr1-9961085-DEL-164
chr1-10054135-DEL-949
chr1-10054573-DEL-474
chr1-10074887-DEL-208
chr1-10076908-DEL-54
chr1-10170925-DEL-322
chr1-10171297-DEL-227
chr1-10171343-DEL-643
chr1-10260924-DEL-1330
chr1-10291399-DEL-699
chr1-10361150-DEL-51
chr1-10452824-DEL-201
chr1-10506749-DEL-702

chr1-10548392-DEL-1276
chr1-10687896-DEL-124
chr1-10754217-DEL-1203
chr1-10829040-DEL-449
chr1-10829790-DEL-56
chr1-10927554-DEL-471
chr1-10933604-DEL-93
chr1-10938252-DEL-76
chr1-11085443-DEL-445
chr1-11178199-DEL-56
chr1-11199953-DEL-1182
chr1-11262242-DEL-61
chr1-11264681-DEL-196
chr1-11264965-DEL-98
chr1-11296986-DEL-3667
chr1-11309290-DEL-330
chr1-11422269-DEL-3402
chr1-11465386-DEL-76
chr1-11469010-DEL-148
chr1-11552074-DEL-265
chr1-11561450-DEL-50
chr1-11589755-DEL-3178
chr1-11591226-DEL-50
chr1-11614345-DEL-124
chr1-11677935-DEL-203
chr1-11680269-DEL-238
chr1-11680584-DEL-439
chr1-11711697-DEL-720
chr1-11725064-DEL-50
chr1-11725898-DEL-55
chr1-11762954-DEL-86
chr1-11870621-DEL-235
chr1-11875575-DEL-13334
chr1-12032654-DEL-130
chr1-12092997-DEL-53
chr1-12195945-DEL-218
chr1-12246978-DEL-1355
chr1-12268617-DEL-418
chr1-12379289-DEL-81
chr1-12436271-DEL-50
chr1-12459251-DEL-50
chr1-12714999-DEL-50
chr1-12727985-DEL-208
chr1-12829713-DEL-50
chr1-12844489-DEL-156
chr1-12910717-DEL-117
chr1-12911216-DEL-187
chr1-13048337-DEL-56
chr1-13066522-DEL-58
chr1-13079961-DEL-62
chr1-13102783-DEL-100
chr1-13103218-DEL-67
chr1-13121886-DEL-217
chr1-13133108-DEL-94
chr1-13152920-DEL-702
chr1-13169446-DEL-210
chr1-13215494-DEL-144
chr1-13239996-DEL-78
chr1-13357843-DEL-213
chr1-13388456-DEL-216
chr1-13397877-DEL-74
chr1-13407868-DEL-305
chr1-13407972-DEL-123
chr1-13409193-DEL-285
chr1-13409406-DEL-72
chr1-13438319-DEL-122
chr1-13569179-DEL-165
chr1-13569452-DEL-204
chr1-13584213-DEL-117
chr1-13772046-DEL-404
chr1-13808918-DEL-150
chr1-13809292-DEL-134
chr1-13866906-DEL-50
chr1-13882802-DEL-7995
chr1-13976072-DEL-62
chr1-14010777-DEL-662
chr1-14024793-DEL-1222

chr1-14026196-DEL-338
chr1-14075653-DEL-75
chr1-14267267-DEL-504
chr1-14267668-DEL-115
chr1-14275519-DEL-71
chr1-14309535-DEL-264
chr1-14316020-DEL-52
chr1-14408493-DEL-86
chr1-14421389-DEL-93
chr1-14428710-DEL-60
chr1-14430655-DEL-2009
chr1-14434605-DEL-328
chr1-14462871-DEL-247
chr1-14509177-DEL-204
chr1-14554192-DEL-112
chr1-14643316-DEL-1118
chr1-14666324-DEL-92
chr1-14668441-DEL-80
chr1-14715464-DEL-107
chr1-14715771-DEL-63
chr1-14771164-DEL-77
chr1-14963023-DEL-79
chr1-15043213-DEL-1409
chr1-15197757-DEL-151
chr1-15211901-DEL-87
chr1-15243276-DEL-1010
chr1-15396130-DEL-758
chr1-15404478-DEL-54
chr1-15469768-DEL-136
chr1-15492050-DEL-288
chr1-15492895-DEL-162
chr1-15531322-DEL-2760
chr1-15562300-DEL-676
chr1-15562319-DEL-1701
chr1-15578777-DEL-156
chr1-15665338-DEL-56
chr1-15694037-DEL-72
chr1-15699821-DEL-1682
chr1-15856320-DEL-187
chr1-16107209-DEL-636
chr1-16130933-DEL-78
chr1-16139846-DEL-3637
chr1-16143246-DEL-1520
chr1-16222602-DEL-50
chr1-16413085-DEL-112
chr1-16573942-DEL-200
chr1-16576571-DEL-50
chr1-16662924-DEL-579
chr1-16663269-DEL-177
chr1-16663671-DEL-84
chr1-16820950-DEL-83
chr1-16830564-DEL-145
chr1-16844858-DEL-2366
chr1-16854023-DEL-78
chr1-16855101-DEL-2689
chr1-16859899-DEL-100
chr1-16916041-DEL-155
chr1-17051920-DEL-881
chr1-17078513-DEL-83
chr1-17087205-DEL-76
chr1-17088411-DEL-204
chr1-17148580-DEL-249
chr1-17222696-DEL-264
chr1-17222926-DEL-51
chr1-17358769-DEL-6271
chr1-17426656-DEL-151
chr1-17466429-DEL-86
chr1-17468251-DEL-621
chr1-17530489-DEL-542
chr1-17594608-DEL-2924
chr1-17696112-DEL-87
chr1-17865384-DEL-116
chr1-17866110-DEL-224
chr1-18049207-DEL-86
chr1-18053396-DEL-3398
chr1-18128768-DEL-280
chr1-18148606-DEL-428

chr1-18160059-DEL-141
chr1-18257554-DEL-55
chr1-18400108-DEL-51
chr1-18513679-DEL-3189
chr1-18627054-DEL-103
chr1-18752068-DEL-336
chr1-18864202-DEL-52
chr1-18871853-DEL-50
chr1-18907513-DEL-64
chr1-18937415-DEL-817
chr1-18962878-DEL-626
chr1-19097903-DEL-184
chr1-19217737-DEL-202
chr1-19448928-DEL-151
chr1-19501527-DEL-144
chr1-19589244-DEL-1382
chr1-19745642-DEL-1031
chr1-19817442-DEL-552
chr1-19818043-DEL-929
chr1-19956394-DEL-1695
chr1-20069295-DEL-408
chr1-20112178-DEL-512
chr1-20145229-DEL-55
chr1-20181936-DEL-149
chr1-20216360-DEL-537
chr1-20216944-DEL-60
chr1-20221398-DEL-406
chr1-20257119-DEL-55
chr1-20272613-DEL-117
chr1-20383640-DEL-295
chr1-20383990-DEL-164
chr1-20406620-DEL-158
chr1-20561395-DEL-1930
chr1-20570654-DEL-70
chr1-20570719-DEL-280
chr1-20579566-DEL-192
chr1-20582507-DEL-70
chr1-20615030-DEL-631
chr1-20652957-DEL-350
chr1-20654689-DEL-173
chr1-20691279-DEL-233
chr1-20713201-DEL-963
chr1-20772574-DEL-1890
chr1-20789173-DEL-59
chr1-20922957-DEL-374
chr1-20938935-DEL-50
chr1-20941692-DEL-77
chr1-20996512-DEL-111
chr1-21107882-DEL-54
chr1-21125648-DEL-52
chr1-21163692-DEL-194
chr1-21211793-DEL-73
chr1-21214859-DEL-56
chr1-21221055-DEL-54
chr1-21271977-DEL-212
chr1-21280452-DEL-72
chr1-21280548-DEL-504
chr1-21416241-DEL-50
chr1-21441108-DEL-174
chr1-21496297-DEL-668
chr1-21596777-DEL-79
chr1-21642169-DEL-2784
chr1-21750973-DEL-2450
chr1-21789212-DEL-54
chr1-21809489-DEL-50
chr1-21843664-DEL-72
chr1-21904367-DEL-63
chr1-22042766-DEL-50
chr1-22169478-DEL-54
chr1-22172478-DEL-126
chr1-22269986-DEL-618
chr1-22301094-DEL-6161
chr1-22328040-DEL-72
chr1-22369185-DEL-139
chr1-22369841-DEL-81
chr1-22397816-DEL-348
chr1-22430928-DEL-146

chr1-22452565-DEL-1034
chr1-22630381-DEL-489
chr1-22706103-DEL-1909
chr1-22878127-DEL-80
chr1-23100021-DEL-308
chr1-23100219-DEL-66
chr1-23138236-DEL-74
chr1-23161514-DEL-58
chr1-23184887-DEL-65
chr1-23220133-DEL-785
chr1-23348127-DEL-94
chr1-23402561-DEL-77
chr1-23572480-DEL-212
chr1-23572955-DEL-64
chr1-23573163-DEL-76
chr1-23595278-DEL-833
chr1-23721642-DEL-110
chr1-23749211-DEL-232
chr1-23760920-DEL-290
chr1-23940085-DEL-4256
chr1-24220256-DEL-58
chr1-24220401-DEL-200
chr1-24307550-DEL-219
chr1-24326464-DEL-1597
chr1-24365966-DEL-65
chr1-24368473-DEL-364
chr1-24416419-DEL-126
chr1-24515095-DEL-70
chr1-24644664-DEL-50
chr1-24670186-DEL-67
chr1-24889603-DEL-77
chr1-25040379-DEL-1466
chr1-25261058-DEL-56
chr1-25510772-DEL-50
chr1-25588035-DEL-319
chr1-25638082-DEL-108
chr1-25758871-DEL-479
chr1-25969463-DEL-2545
chr1-25972201-DEL-59
chr1-25972318-DEL-114
chr1-26033967-DEL-92
chr1-26045250-DEL-2207
chr1-26048157-DEL-2212
chr1-26063968-DEL-50
chr1-26065122-DEL-88
chr1-26072736-DEL-211
chr1-26090468-DEL-3765
chr1-26139327-DEL-72
chr1-26186303-DEL-537
chr1-26205940-DEL-1248
chr1-26229397-DEL-694
chr1-26285607-DEL-54
chr1-26285607-DEL-156
chr1-26301036-DEL-4187
chr1-26336747-DEL-116
chr1-26621585-DEL-52
chr1-26669462-DEL-53
chr1-26681247-DEL-75
chr1-26684193-DEL-68
chr1-26797092-DEL-57
chr1-26812525-DEL-58
chr1-26916900-DEL-54
chr1-26938308-DEL-85
chr1-27145970-DEL-77
chr1-27233455-DEL-416
chr1-27233546-DEL-207
chr1-27299987-DEL-50
chr1-27471800-DEL-99
chr1-27539335-DEL-1504
chr1-27725256-DEL-82
chr1-27850636-DEL-883
chr1-27935063-DEL-52
chr1-27949338-DEL-1530
chr1-28150700-DEL-3883
chr1-28179583-DEL-50
chr1-28210039-DEL-4857
chr1-28241229-DEL-50

chr1-28282580-DEL-84
chr1-28300598-DEL-1331
chr1-28319414-DEL-62
chr1-28326825-DEL-424
chr1-28359770-DEL-61
chr1-28444401-DEL-52
chr1-28452604-DEL-1416
chr1-28470816-DEL-246
chr1-28489921-DEL-198
chr1-28528244-DEL-50
chr1-28618235-DEL-256
chr1-28699836-DEL-498
chr1-28705141-DEL-62
chr1-28779467-DEL-1700
chr1-28789791-DEL-132
chr1-28822077-DEL-3387
chr1-29062296-DEL-78
chr1-29062334-DEL-339
chr1-29202571-DEL-336
chr1-29259443-DEL-153
chr1-29280534-DEL-93
chr1-29343439-DEL-50
chr1-29351375-DEL-50
chr1-29513622-DEL-136
chr1-29513830-DEL-198
chr1-29640120-DEL-58
chr1-29643617-DEL-9067
chr1-29892660-DEL-169
chr1-29947291-DEL-945
chr1-30017683-DEL-50
chr1-30098533-DEL-116
chr1-30150698-DEL-305
chr1-30265408-DEL-58
chr1-30304722-DEL-1492
chr1-30410179-DEL-416
chr1-30753447-DEL-83
chr1-30814406-DEL-372
chr1-31003799-DEL-83
chr1-31094047-DEL-134
chr1-31166847-DEL-402
chr1-31206087-DEL-143
chr1-31248988-DEL-65
chr1-31303658-DEL-7554
chr1-31637451-DEL-69
chr1-31690018-DEL-77
chr1-31721131-DEL-200
chr1-31721497-DEL-77
chr1-31872774-DEL-128
chr1-31917857-DEL-52
chr1-31936427-DEL-54
chr1-31963498-DEL-771
chr1-31998656-DEL-4472
chr1-32024593-DEL-4418
chr1-32161546-DEL-50
chr1-32270243-DEL-350
chr1-32370647-DEL-93
chr1-32432573-DEL-367
chr1-32434463-DEL-730
chr1-32444633-DEL-1478
chr1-32449570-DEL-584
chr1-32452248-DEL-906
chr1-32456993-DEL-86
chr1-32583570-DEL-89
chr1-32657871-DEL-80
chr1-32734341-DEL-55
chr1-32979201-DEL-907
chr1-33077132-DEL-98
chr1-33122259-DEL-100
chr1-33130835-DEL-110
chr1-33139465-DEL-221
chr1-33146225-DEL-62
chr1-33279421-DEL-99
chr1-33365676-DEL-243
chr1-33447222-DEL-409
chr1-33518022-DEL-53
chr1-33525825-DEL-82
chr1-33561300-DEL-147

chr1-33606225-DEL-2106
chr1-33606437-DEL-234
chr1-33607726-DEL-78
chr1-33609111-DEL-858
chr1-33639343-DEL-261
chr1-33639566-DEL-53
chr1-33639755-DEL-939
chr1-33720797-DEL-82
chr1-33870712-DEL-99
chr1-33933829-DEL-330
chr1-33933889-DEL-60
chr1-33972799-DEL-54
chr1-33973012-DEL-53
chr1-33977667-DEL-54
chr1-33998501-DEL-100
chr1-34023436-DEL-106
chr1-34060031-DEL-50
chr1-34138777-DEL-73
chr1-34146547-DEL-734
chr1-34183078-DEL-186
chr1-34183091-DEL-387
chr1-34183472-DEL-218
chr1-34183511-DEL-84
chr1-34250480-DEL-268
chr1-34362441-DEL-246
chr1-34416240-DEL-104
chr1-34423558-DEL-264
chr1-34442927-DEL-251
chr1-34494660-DEL-110
chr1-34521029-DEL-66
chr1-34559431-DEL-266
chr1-34695678-DEL-307
chr1-34756500-DEL-79
chr1-34919370-DEL-195
chr1-34965083-DEL-63
chr1-34990554-DEL-96
chr1-35136083-DEL-50
chr1-35208080-DEL-288
chr1-35208223-DEL-96
chr1-35215299-DEL-3759
chr1-35244369-DEL-5201
chr1-35302910-DEL-124
chr1-35309687-DEL-63
chr1-35314117-DEL-1681
chr1-35400411-DEL-68
chr1-35410568-DEL-64
chr1-35421887-DEL-1437
chr1-35433794-DEL-164
chr1-35439425-DEL-50
chr1-35695744-DEL-244
chr1-35701205-DEL-54
chr1-35701497-DEL-80
chr1-35736065-DEL-172
chr1-35847492-DEL-60
chr1-35853102-DEL-50
chr1-35957347-DEL-345
chr1-36002953-DEL-62
chr1-36030666-DEL-169
chr1-36079929-DEL-55
chr1-36088315-DEL-201
chr1-36088676-DEL-324
chr1-36095537-DEL-65
chr1-36136457-DEL-315
chr1-36136487-DEL-93
chr1-36163401-DEL-116
chr1-36296140-DEL-52
chr1-36623528-DEL-391
chr1-36649401-DEL-147
chr1-36649401-DEL-637
chr1-36701006-DEL-100
chr1-36752657-DEL-79
chr1-36765918-DEL-334
chr1-36772569-DEL-54
chr1-36975095-DEL-425
chr1-37012440-DEL-77
chr1-37063484-DEL-156
chr1-37098055-DEL-140

chr1-37205597-DEL-470
chr1-37347791-DEL-56
chr1-37410390-DEL-58
chr1-37430703-DEL-64
chr1-37430767-DEL-144
chr1-37473818-DEL-407
chr1-37473930-DEL-58
chr1-37474019-DEL-186
chr1-37664121-DEL-63
chr1-37857021-DEL-52
chr1-37859070-DEL-3871
chr1-37902627-DEL-89
chr1-38018394-DEL-1477
chr1-38111215-DEL-263
chr1-38413450-DEL-76
chr1-38417800-DEL-137
chr1-38418123-DEL-158
chr1-38425410-DEL-229
chr1-38425416-DEL-638
chr1-38463706-DEL-145
chr1-38469087-DEL-297
chr1-38469625-DEL-63
chr1-38535212-DEL-130
chr1-38580034-DEL-153
chr1-38810896-DEL-50
chr1-38822524-DEL-54
chr1-38847971-DEL-594
chr1-38849268-DEL-99
chr1-38926605-DEL-50
chr1-39027497-DEL-165
chr1-39072670-DEL-739
chr1-39082960-DEL-722
chr1-39261750-DEL-217
chr1-39261929-DEL-86
chr1-39342280-DEL-70
chr1-39367957-DEL-71
chr1-39411998-DEL-395
chr1-39570834-DEL-420
chr1-39683430-DEL-69
chr1-39715071-DEL-184
chr1-39724389-DEL-87
chr1-39778907-DEL-86
chr1-39780285-DEL-54
chr1-39795553-DEL-161
chr1-39795852-DEL-183
chr1-39866309-DEL-168
chr1-39866317-DEL-81
chr1-39873060-DEL-124
chr1-39903252-DEL-127
chr1-39910792-DEL-56
chr1-39975137-DEL-117
chr1-39975137-DEL-297
chr1-39996843-DEL-62
chr1-40072258-DEL-50
chr1-40107545-DEL-67
chr1-40174710-DEL-128
chr1-40198490-DEL-111
chr1-40202274-DEL-69
chr1-40337705-DEL-146
chr1-40383636-DEL-95
chr1-40413283-DEL-50
chr1-40434025-DEL-142
chr1-40463675-DEL-50
chr1-40521434-DEL-105
chr1-40558164-DEL-101
chr1-40558448-DEL-60
chr1-40593369-DEL-547
chr1-40727577-DEL-52
chr1-40762301-DEL-533
chr1-40764013-DEL-283
chr1-40888110-DEL-163
chr1-40888129-DEL-72
chr1-40921311-DEL-92
chr1-40921628-DEL-232
chr1-40967166-DEL-57
chr1-40992079-DEL-104
chr1-41189223-DEL-50

chr1-41256754-DEL-897
chr1-41348779-DEL-67
chr1-41404346-DEL-437
chr1-41410340-DEL-830
chr1-41420301-DEL-50
chr1-41491995-DEL-210
chr1-41544930-DEL-51
chr1-41794965-DEL-386
chr1-41874249-DEL-97
chr1-41895230-DEL-120
chr1-41987609-DEL-1319
chr1-42033472-DEL-88
chr1-42285539-DEL-174
chr1-42290247-DEL-60
chr1-42315206-DEL-86
chr1-42419936-DEL-2454
chr1-42546128-DEL-60
chr1-42671882-DEL-178
chr1-42860441-DEL-50
chr1-42885069-DEL-65
chr1-43157986-DEL-537
chr1-43167956-DEL-85
chr1-43226540-DEL-104
chr1-43236862-DEL-175
chr1-43240026-DEL-378
chr1-43262688-DEL-50
chr1-43460106-DEL-72
chr1-43679876-DEL-1957
chr1-43817161-DEL-305
chr1-43836953-DEL-59
chr1-43922163-DEL-111
chr1-44025251-DEL-95
chr1-44191943-DEL-1326
chr1-44193763-DEL-208
chr1-44193771-DEL-56
chr1-44194003-DEL-88
chr1-44236241-DEL-76
chr1-44261559-DEL-5643
chr1-44272181-DEL-1066
chr1-44310611-DEL-2209
chr1-44357931-DEL-65
chr1-44377686-DEL-50
chr1-44479289-DEL-56
chr1-44500276-DEL-63
chr1-44501845-DEL-50
chr1-44612700-DEL-50
chr1-44707025-DEL-1691
chr1-44782164-DEL-50
chr1-44790818-DEL-334
chr1-44827871-DEL-50
chr1-44840452-DEL-2570
chr1-44898603-DEL-2138
chr1-45088698-DEL-637
chr1-45174295-DEL-94
chr1-45190651-DEL-58
chr1-45214983-DEL-298
chr1-45296246-DEL-55
chr1-45437447-DEL-931
chr1-45524184-DEL-56
chr1-45548939-DEL-142
chr1-45756494-DEL-1054
chr1-45876169-DEL-83
chr1-45915469-DEL-127
chr1-46066559-DEL-50
chr1-46130556-DEL-3583
chr1-46352551-DEL-400
chr1-46413778-DEL-56
chr1-46425481-DEL-1112
chr1-46480506-DEL-104
chr1-46524135-DEL-6627
chr1-46531418-DEL-164
chr1-46575497-DEL-56
chr1-46658781-DEL-4863
chr1-46667232-DEL-50
chr1-46747042-DEL-201
chr1-46822553-DEL-54
chr1-46954930-DEL-60

chr1-47061383-DEL-540
chr1-47072019-DEL-70
chr1-47157579-DEL-50
chr1-47274825-DEL-100
chr1-47283423-DEL-4277
chr1-47315305-DEL-183
chr1-47444729-DEL-1541
chr1-47479849-DEL-137
chr1-47711197-DEL-56
chr1-47723010-DEL-67
chr1-47792391-DEL-66
chr1-47826441-DEL-234
chr1-47826763-DEL-57
chr1-47906713-DEL-155
chr1-47912723-DEL-89
chr1-47922330-DEL-5203
chr1-48008830-DEL-78
chr1-48038741-DEL-66
chr1-48054594-DEL-772
chr1-48067142-DEL-88
chr1-48174540-DEL-60
chr1-48252325-DEL-51
chr1-48294241-DEL-97
chr1-48377051-DEL-1134
chr1-48569470-DEL-367
chr1-48596884-DEL-52
chr1-48710130-DEL-2912
chr1-48714081-DEL-52
chr1-48763592-DEL-52
chr1-48805143-DEL-275
chr1-49169141-DEL-379
chr1-49169686-DEL-201
chr1-49169689-DEL-710
chr1-49210909-DEL-51
chr1-49263228-DEL-60
chr1-49274509-DEL-69
chr1-49276149-DEL-51
chr1-49277828-DEL-58
chr1-49352597-DEL-64
chr1-49586507-DEL-64
chr1-49612828-DEL-293
chr1-49650021-DEL-128
chr1-49654136-DEL-78
chr1-49774771-DEL-78
chr1-49837876-DEL-1934
chr1-49889773-DEL-55
chr1-49936921-DEL-206
chr1-49938259-DEL-174
chr1-49945166-DEL-464
chr1-50024941-DEL-217
chr1-50033176-DEL-2629
chr1-50254742-DEL-3859
chr1-50303320-DEL-50
chr1-50406177-DEL-448
chr1-50501090-DEL-265
chr1-50569028-DEL-167
chr1-50715945-DEL-1887
chr1-50810979-DEL-65
chr1-50822687-DEL-50
chr1-50943531-DEL-101
chr1-51058338-DEL-191
chr1-51074112-DEL-51
chr1-51151239-DEL-144
chr1-51161848-DEL-52
chr1-51269159-DEL-354
chr1-51307047-DEL-51
chr1-51322492-DEL-66
chr1-51337755-DEL-77
chr1-51358822-DEL-69
chr1-51511574-DEL-539
chr1-51593573-DEL-2151
chr1-51649234-DEL-58
chr1-51722401-DEL-514
chr1-51849743-DEL-2217
chr1-51856744-DEL-78
chr1-52040666-DEL-56
chr1-52085582-DEL-649

chr1-52085708-DEL-50
chr1-52168350-DEL-197
chr1-52343422-DEL-72
chr1-52439087-DEL-6939
chr1-52569974-DEL-2152
chr1-52661062-DEL-1494
chr1-52794818-DEL-120
chr1-52810032-DEL-79
chr1-52810087-DEL-171
chr1-52810337-DEL-386
chr1-52810873-DEL-329
chr1-52908987-DEL-64
chr1-52909211-DEL-128
chr1-52926285-DEL-102
chr1-52926463-DEL-163
chr1-52978669-DEL-212
chr1-53142465-DEL-76
chr1-53195018-DEL-112
chr1-53206278-DEL-57
chr1-53218244-DEL-302
chr1-53219065-DEL-50
chr1-53219337-DEL-62
chr1-53234784-DEL-310
chr1-53274703-DEL-2802
chr1-53450173-DEL-54
chr1-53457785-DEL-60
chr1-53596648-DEL-416
chr1-53691415-DEL-3363
chr1-53771009-DEL-82
chr1-53898546-DEL-97
chr1-53898712-DEL-556
chr1-53983273-DEL-4104
chr1-54210813-DEL-1365
chr1-54247057-DEL-208
chr1-54247342-DEL-199
chr1-54333639-DEL-368
chr1-54588982-DEL-617
chr1-54621855-DEL-54
chr1-54650051-DEL-74
chr1-54650166-DEL-62
chr1-54698992-DEL-881
chr1-54702030-DEL-320
chr1-54748409-DEL-481
chr1-54793606-DEL-63
chr1-54859872-DEL-140
chr1-54859872-DEL-616
chr1-54873635-DEL-60
chr1-54921735-DEL-72
chr1-54996547-DEL-52
chr1-55048339-DEL-589
chr1-55071365-DEL-55
chr1-55136555-DEL-272
chr1-55174035-DEL-69
chr1-55208830-DEL-59
chr1-55274864-DEL-81
chr1-55552003-DEL-80
chr1-55665741-DEL-72
chr1-55701202-DEL-172
chr1-55799404-DEL-1337
chr1-55810205-DEL-18441
chr1-55850364-DEL-1069
chr1-56120653-DEL-112
chr1-56158953-DEL-1140
chr1-56195561-DEL-130
chr1-56280681-DEL-221
chr1-56473879-DEL-55
chr1-56668283-DEL-384
chr1-56721056-DEL-669
chr1-56799640-DEL-76
chr1-56841154-DEL-216
chr1-56841319-DEL-88
chr1-56931457-DEL-61
chr1-57061542-DEL-114
chr1-57061869-DEL-171
chr1-57172483-DEL-405
chr1-57172814-DEL-412
chr1-57310376-DEL-65

chr1-57332054-DEL-944
chr1-57495616-DEL-2551
chr1-57574062-DEL-298
chr1-57581718-DEL-70
chr1-57605840-DEL-201
chr1-57605850-DEL-435
chr1-57746440-DEL-58
chr1-57765900-DEL-100
chr1-57890749-DEL-76
chr1-57894049-DEL-172
chr1-57961327-DEL-300
chr1-58151208-DEL-58
chr1-58213946-DEL-341
chr1-58254720-DEL-53
chr1-58411072-DEL-62
chr1-58411320-DEL-126
chr1-58599598-DEL-203
chr1-58680129-DEL-9730
chr1-58747464-DEL-1039
chr1-58810204-DEL-342
chr1-58934419-DEL-88
chr1-58938164-DEL-51
chr1-58998380-DEL-208
chr1-59150671-DEL-407
chr1-59161660-DEL-125
chr1-59171994-DEL-317
chr1-59178636-DEL-170
chr1-59178670-DEL-68
chr1-59249589-DEL-73
chr1-59249688-DEL-467
chr1-59344265-DEL-248
chr1-59363369-DEL-867
chr1-59391650-DEL-90
chr1-59439775-DEL-630
chr1-59439908-DEL-270
chr1-59447470-DEL-128
chr1-59465237-DEL-122
chr1-59502377-DEL-555
chr1-59639652-DEL-77
chr1-59640655-DEL-98
chr1-59702319-DEL-51
chr1-59818608-DEL-58
chr1-60043142-DEL-62
chr1-60140038-DEL-198
chr1-60197037-DEL-50
chr1-60340051-DEL-50
chr1-60340392-DEL-208
chr1-60577632-DEL-83
chr1-60619713-DEL-67
chr1-60733728-DEL-138
chr1-60740318-DEL-74
chr1-60746638-DEL-654
chr1-60965922-DEL-369
chr1-60974186-DEL-50
chr1-61100495-DEL-67
chr1-61102525-DEL-66
chr1-61219300-DEL-90
chr1-61387269-DEL-53
chr1-61465599-DEL-50
chr1-61537121-DEL-424
chr1-61591112-DEL-60
chr1-61605864-DEL-74
chr1-61786324-DEL-50
chr1-61812667-DEL-56
chr1-61838079-DEL-98
chr1-61863681-DEL-52
chr1-61927593-DEL-1718
chr1-61986333-DEL-125
chr1-62151025-DEL-742
chr1-62175652-DEL-63
chr1-62211785-DEL-54
chr1-62234050-DEL-208
chr1-62334807-DEL-212
chr1-62580494-DEL-53
chr1-62640626-DEL-170
chr1-62643819-DEL-138
chr1-62847054-DEL-113

chr1-63010815-DEL-494
chr1-63010828-DEL-234
chr1-63259719-DEL-52
chr1-63298966-DEL-74
chr1-63775186-DEL-796
chr1-63775437-DEL-1189
chr1-63813016-DEL-456
chr1-63838205-DEL-3750
chr1-64208930-DEL-122
chr1-64493623-DEL-129
chr1-64564436-DEL-54
chr1-64719599-DEL-50
chr1-64799102-DEL-50
chr1-64799843-DEL-236
chr1-64837587-DEL-2035
chr1-64867659-DEL-63
chr1-64887543-DEL-82
chr1-64957438-DEL-50
chr1-65036052-DEL-158
chr1-65066597-DEL-68
chr1-65137195-DEL-60
chr1-65153237-DEL-1616
chr1-65155129-DEL-132
chr1-65161816-DEL-109
chr1-65424347-DEL-94
chr1-65424347-DEL-190
chr1-65515555-DEL-56
chr1-65534757-DEL-50
chr1-65715180-DEL-58
chr1-65755202-DEL-4593
chr1-65861746-DEL-313
chr1-65925513-DEL-76
chr1-65979026-DEL-50
chr1-65987269-DEL-1068
chr1-65988280-DEL-1067
chr1-66056763-DEL-7248
chr1-66115041-DEL-50
chr1-66535897-DEL-156
chr1-66580993-DEL-4009
chr1-66796678-DEL-139
chr1-66796996-DEL-68
chr1-66922680-DEL-1628
chr1-66924388-DEL-55
chr1-66967266-DEL-57
chr1-66967508-DEL-223
chr1-67200794-DEL-482
chr1-67467751-DEL-57
chr1-67506739-DEL-66
chr1-67653321-DEL-106
chr1-67750727-DEL-50
chr1-67920201-DEL-50
chr1-68151137-DEL-1386
chr1-68152984-DEL-322
chr1-68279524-DEL-54
chr1-68292187-DEL-91
chr1-68302898-DEL-50
chr1-68393036-DEL-50
chr1-68529329-DEL-153
chr1-68550596-DEL-83
chr1-68637610-DEL-214
chr1-68666118-DEL-1450
chr1-68771511-DEL-50
chr1-68805474-DEL-64
chr1-68806992-DEL-153
chr1-69185184-DEL-313
chr1-69310452-DEL-50
chr1-69627251-DEL-56
chr1-69862399-DEL-81
chr1-69997423-DEL-50
chr1-70003764-DEL-4775
chr1-70089246-DEL-106
chr1-70115041-DEL-176
chr1-70244342-DEL-597
chr1-70294241-DEL-245
chr1-70616721-DEL-125
chr1-70640351-DEL-161
chr1-70644033-DEL-196

chr1-70718744-DEL-294
chr1-70878996-DEL-56
chr1-70913379-DEL-1545
chr1-70963934-DEL-240
chr1-71328471-DEL-84
chr1-71444657-DEL-2012
chr1-71508202-DEL-96
chr1-71528458-DEL-83
chr1-71799953-DEL-110
chr1-71960543-DEL-64
chr1-71972322-DEL-1479
chr1-71983549-DEL-60
chr1-71985388-DEL-56
chr1-72012006-DEL-60
chr1-72267676-DEL-50
chr1-72279357-DEL-67
chr1-72335180-DEL-252
chr1-72420069-DEL-232
chr1-72540077-DEL-80
chr1-72592382-DEL-296
chr1-72760524-DEL-53
chr1-72760654-DEL-109
chr1-72764041-DEL-77
chr1-72781666-DEL-75
chr1-72798117-DEL-81
chr1-72811303-DEL-189
chr1-72814496-DEL-63
chr1-72834004-DEL-1897
chr1-72841846-DEL-77
chr1-72945814-DEL-113
chr1-72963564-DEL-2701
chr1-72970976-DEL-107
chr1-72980094-DEL-52
chr1-73021090-DEL-64
chr1-73022477-DEL-3571
chr1-73049348-DEL-53
chr1-73343969-DEL-462
chr1-73505025-DEL-92
chr1-73539485-DEL-58
chr1-73576216-DEL-54
chr1-73594817-DEL-258
chr1-73605172-DEL-106
chr1-73679564-DEL-53
chr1-73738392-DEL-56
chr1-73743269-DEL-52
chr1-73872462-DEL-53
chr1-73872698-DEL-53
chr1-73931592-DEL-107
chr1-73974302-DEL-113
chr1-74103684-DEL-163
chr1-74140441-DEL-201
chr1-74213330-DEL-202
chr1-74223711-DEL-99
chr1-74311484-DEL-290
chr1-74353284-DEL-332
chr1-74359784-DEL-73
chr1-74365377-DEL-794
chr1-74444933-DEL-68
chr1-74454451-DEL-69
chr1-74501062-DEL-181
chr1-74550471-DEL-464
chr1-74592811-DEL-59
chr1-74749843-DEL-370
chr1-74751226-DEL-203
chr1-74800675-DEL-820
chr1-74801917-DEL-54
chr1-74852373-DEL-218
chr1-74952911-DEL-66
chr1-75015491-DEL-83
chr1-75043966-DEL-159
chr1-75086638-DEL-574
chr1-75413747-DEL-103
chr1-75434127-DEL-65
chr1-75437399-DEL-81
chr1-75437756-DEL-554
chr1-75437968-DEL-254
chr1-75439058-DEL-157

chr1-75618032-DEL-58
chr1-75683998-DEL-50
chr1-75684449-DEL-3019
chr1-75728554-DEL-50
chr1-75831596-DEL-111
chr1-75934962-DEL-60
chr1-75959417-DEL-53
chr1-76208718-DEL-50
chr1-76324756-DEL-50
chr1-76364775-DEL-50
chr1-76453393-DEL-70
chr1-76548457-DEL-67
chr1-76774230-DEL-117
chr1-76780186-DEL-228
chr1-76780258-DEL-62
chr1-76935897-DEL-155
chr1-76940473-DEL-119
chr1-76991174-DEL-381
chr1-77001423-DEL-91
chr1-77018440-DEL-53
chr1-77052499-DEL-210
chr1-77083567-DEL-60
chr1-77122321-DEL-57
chr1-77147314-DEL-150
chr1-77339799-DEL-438
chr1-77445007-DEL-134
chr1-77551141-DEL-548
chr1-77553202-DEL-105
chr1-77582558-DEL-76
chr1-77583889-DEL-63
chr1-77682702-DEL-329
chr1-77773284-DEL-160
chr1-77878456-DEL-51
chr1-77887421-DEL-50
chr1-78051745-DEL-60
chr1-78155831-DEL-50
chr1-78281211-DEL-110
chr1-78374897-DEL-53
chr1-78601104-DEL-74
chr1-78611541-DEL-182
chr1-78611843-DEL-74
chr1-78616225-DEL-68
chr1-78616414-DEL-271
chr1-78668270-DEL-124
chr1-78673149-DEL-360
chr1-78673271-DEL-120
chr1-78673509-DEL-60
chr1-78734029-DEL-2782
chr1-78763064-DEL-844
chr1-78765609-DEL-108
chr1-78791781-DEL-50
chr1-78836323-DEL-89
chr1-78854355-DEL-156
chr1-78889096-DEL-110
chr1-78969331-DEL-52
chr1-79014158-DEL-5000
chr1-79076211-DEL-208
chr1-79078845-DEL-107
chr1-79113301-DEL-3421
chr1-79129034-DEL-129
chr1-79159296-DEL-2172
chr1-79304349-DEL-50
chr1-79353734-DEL-146
chr1-79441957-DEL-78
chr1-79445161-DEL-88
chr1-79593268-DEL-52
chr1-79774032-DEL-50
chr1-79813877-DEL-72
chr1-79889181-DEL-371
chr1-79924174-DEL-170
chr1-79964039-DEL-375
chr1-79979511-DEL-6411
chr1-79981491-DEL-896
chr1-80182717-DEL-50
chr1-80325220-DEL-1419
chr1-80376318-DEL-135
chr1-80477859-DEL-50

chr1-80595285-DEL-51
chr1-80609023-DEL-50
chr1-80683074-DEL-107
chr1-80692903-DEL-132
chr1-80744430-DEL-215
chr1-80744523-DEL-101
chr1-80744653-DEL-337
chr1-80744862-DEL-127
chr1-80751957-DEL-60
chr1-80806446-DEL-56
chr1-80866090-DEL-61
chr1-80875308-DEL-50
chr1-81014982-DEL-121
chr1-81117223-DEL-62
chr1-81421692-DEL-194
chr1-81597641-DEL-54
chr1-81733416-DEL-60
chr1-81982427-DEL-54
chr1-82064079-DEL-844
chr1-82455003-DEL-657
chr1-82455063-DEL-100
chr1-82455107-DEL-284
chr1-82455281-DEL-126
chr1-82521562-DEL-118
chr1-82542674-DEL-110
chr1-82572538-DEL-288
chr1-82591967-DEL-52
chr1-82625300-DEL-81
chr1-82848783-DEL-272
chr1-82924184-DEL-54
chr1-82933541-DEL-63
chr1-82978380-DEL-126
chr1-82978550-DEL-70
chr1-83008707-DEL-147
chr1-83020951-DEL-65
chr1-83081236-DEL-159
chr1-83148680-DEL-54
chr1-83207502-DEL-5672
chr1-83261269-DEL-61
chr1-83360201-DEL-193
chr1-83502620-DEL-67
chr1-83502766-DEL-78
chr1-83704905-DEL-62
chr1-83767442-DEL-55
chr1-83907296-DEL-55
chr1-83931472-DEL-96
chr1-84009181-DEL-81
chr1-84210765-DEL-148
chr1-84240118-DEL-53
chr1-84352873-DEL-1044
chr1-84375676-DEL-103
chr1-84393235-DEL-1513
chr1-84416408-DEL-60
chr1-84663276-DEL-2256
chr1-85002187-DEL-206
chr1-85031241-DEL-1082
chr1-85052061-DEL-302
chr1-85087294-DEL-4619
chr1-85088688-DEL-52
chr1-85102882-DEL-1269
chr1-85183798-DEL-121
chr1-85244435-DEL-1242
chr1-85470186-DEL-279
chr1-85478200-DEL-468
chr1-85487523-DEL-1737
chr1-85511776-DEL-486
chr1-85531719-DEL-53
chr1-85544133-DEL-128
chr1-85581694-DEL-54
chr1-85861675-DEL-71
chr1-85889172-DEL-59
chr1-85903834-DEL-58
chr1-85909907-DEL-481
chr1-85932156-DEL-68
chr1-85939366-DEL-4791
chr1-86034775-DEL-60
chr1-86069453-DEL-189

chr1-86193864-DEL-54
chr1-86205810-DEL-94
chr1-86388528-DEL-56
chr1-86401886-DEL-291
chr1-86524952-DEL-60
chr1-86678327-DEL-68
chr1-86717116-DEL-72
chr1-86831203-DEL-106
chr1-87058070-DEL-228
chr1-87319606-DEL-50
chr1-87422989-DEL-50
chr1-87581134-DEL-108
chr1-87750071-DEL-319
chr1-88036436-DEL-120
chr1-88126650-DEL-628
chr1-88150065-DEL-723
chr1-88161960-DEL-69
chr1-88226661-DEL-173
chr1-88249049-DEL-224
chr1-88328154-DEL-74
chr1-88379919-DEL-60
chr1-88413235-DEL-2144
chr1-88430917-DEL-129
chr1-88430917-DEL-282
chr1-88517196-DEL-56
chr1-88523744-DEL-90
chr1-88545710-DEL-78
chr1-88644170-DEL-119
chr1-88671573-DEL-127
chr1-88750302-DEL-200
chr1-88750308-DEL-94
chr1-88987058-DEL-60
chr1-89288010-DEL-305
chr1-89339125-DEL-150
chr1-89339167-DEL-65
chr1-89429475-DEL-1259
chr1-89430251-DEL-65
chr1-89953861-DEL-54
chr1-89974532-DEL-254
chr1-90008808-DEL-80
chr1-90113775-DEL-87
chr1-90115627-DEL-50
chr1-90309871-DEL-127
chr1-90314422-DEL-50
chr1-90327982-DEL-2694
chr1-90373195-DEL-2023
chr1-90373277-DEL-212
chr1-90373991-DEL-882
chr1-90374552-DEL-1725
chr1-90483468-DEL-69
chr1-90663728-DEL-52
chr1-90684250-DEL-50
chr1-90703524-DEL-63
chr1-90760049-DEL-52
chr1-90791753-DEL-64
chr1-90797753-DEL-72
chr1-90833055-DEL-102
chr1-90841264-DEL-144
chr1-90841584-DEL-56
chr1-90877751-DEL-264
chr1-90877913-DEL-51
chr1-90912776-DEL-2132
chr1-91349834-DEL-423
chr1-91350113-DEL-162
chr1-91419634-DEL-112
chr1-91576497-DEL-109
chr1-91610764-DEL-66
chr1-91709714-DEL-110
chr1-91732560-DEL-156
chr1-92191770-DEL-56
chr1-92258627-DEL-56
chr1-92303745-DEL-212
chr1-92313585-DEL-105
chr1-92427444-DEL-2118
chr1-93047090-DEL-55
chr1-93057520-DEL-128
chr1-93057809-DEL-62

chr1-93307995-DEL-54
chr1-93620037-DEL-51
chr1-93699148-DEL-123
chr1-93784628-DEL-415
chr1-93824494-DEL-53
chr1-93849252-DEL-963
chr1-93872053-DEL-352
chr1-93909088-DEL-51
chr1-93974180-DEL-52
chr1-93984241-DEL-1497
chr1-94174212-DEL-263
chr1-94733041-DEL-728
chr1-94763550-DEL-1021
chr1-94843360-DEL-350
chr1-94946586-DEL-140
chr1-95010434-DEL-50
chr1-95017397-DEL-275
chr1-95187969-DEL-96
chr1-95725140-DEL-55
chr1-95756335-DEL-72
chr1-95785738-DEL-54
chr1-95849749-DEL-58
chr1-95886631-DEL-3158
chr1-95901868-DEL-425
chr1-95901916-DEL-69
chr1-95901994-DEL-180
chr1-95902267-DEL-255
chr1-95920379-DEL-539
chr1-96066853-DEL-347
chr1-96270149-DEL-121
chr1-96359526-DEL-108
chr1-96379113-DEL-1775
chr1-96380986-DEL-427
chr1-96403115-DEL-1005
chr1-96479349-DEL-726
chr1-96519561-DEL-89
chr1-96570993-DEL-688
chr1-96593451-DEL-6044
chr1-96661386-DEL-1676
chr1-96667244-DEL-55
chr1-96681544-DEL-2230
chr1-96885969-DEL-74
chr1-97064251-DEL-163114
chr1-97714746-DEL-63
chr1-97937626-DEL-71
chr1-97938336-DEL-55
chr1-98003757-DEL-77
chr1-98078367-DEL-54
chr1-98153588-DEL-72
chr1-98170633-DEL-727
chr1-98174038-DEL-56
chr1-98174226-DEL-107
chr1-98278738-DEL-66
chr1-98297734-DEL-2802
chr1-98313261-DEL-214
chr1-98395097-DEL-115
chr1-98442411-DEL-62
chr1-98475779-DEL-68
chr1-98475824-DEL-139
chr1-98550700-DEL-174
chr1-98724129-DEL-3181
chr1-98728210-DEL-601
chr1-98801106-DEL-2007
chr1-98918305-DEL-122
chr1-98936803-DEL-1163
chr1-98955968-DEL-50
chr1-99002890-DEL-1307
chr1-99031585-DEL-185
chr1-99062825-DEL-140
chr1-99063191-DEL-213
chr1-99173768-DEL-85
chr1-99269347-DEL-266
chr1-99442979-DEL-50
chr1-99549021-DEL-59
chr1-99568454-DEL-913
chr1-99761883-DEL-50
chr1-99789097-DEL-81

chr1-99811447-DEL-100
chr1-99824819-DEL-814
chr1-99824954-DEL-345
chr1-99837358-DEL-1982
chr1-99880683-DEL-379
chr1-99903482-DEL-1468
chr1-99919682-DEL-95
chr1-100027404-DEL-229
chr1-100126759-DEL-1781
chr1-100284701-DEL-114
chr1-100338228-DEL-189
chr1-100848632-DEL-545
chr1-100862820-DEL-400
chr1-100863568-DEL-1304
chr1-100963416-DEL-114
chr1-100976448-DEL-158
chr1-101017173-DEL-4156
chr1-101030805-DEL-2604
chr1-101072323-DEL-132
chr1-101189627-DEL-68
chr1-101381433-DEL-53
chr1-101555566-DEL-61
chr1-101614360-DEL-50
chr1-101627541-DEL-72
chr1-101924690-DEL-83
chr1-101928728-DEL-50
chr1-101962558-DEL-474
chr1-101969198-DEL-760
chr1-102013885-DEL-6764
chr1-102030497-DEL-53
chr1-102159367-DEL-152
chr1-102182926-DEL-448
chr1-102183607-DEL-201
chr1-102183898-DEL-402
chr1-102197055-DEL-66
chr1-102258128-DEL-3339
chr1-102265254-DEL-606
chr1-102522867-DEL-12513
chr1-102525819-DEL-4074
chr1-102675808-DEL-3712
chr1-102808771-DEL-67
chr1-102863557-DEL-1822
chr1-102902001-DEL-199
chr1-102917655-DEL-52
chr1-102922026-DEL-6165
chr1-102959390-DEL-6758
chr1-102978676-DEL-66
chr1-103012442-DEL-52
chr1-103030397-DEL-60
chr1-103066824-DEL-152
chr1-103093958-DEL-589
chr1-103120447-DEL-52
chr1-103210858-DEL-6674
chr1-103218956-DEL-2958
chr1-103229845-DEL-60
chr1-103328746-DEL-54
chr1-103405529-DEL-53
chr1-103411003-DEL-63
chr1-103451986-DEL-165
chr1-103498229-DEL-205
chr1-103506159-DEL-50
chr1-103561738-DEL-57
chr1-103612226-DEL-242
chr1-103621706-DEL-264
chr1-103651615-DEL-54
chr1-103651974-DEL-194
chr1-103677555-DEL-164
chr1-103692001-DEL-597
chr1-103713400-DEL-57
chr1-103734706-DEL-53
chr1-103944170-DEL-675
chr1-104086811-DEL-147
chr1-104247194-DEL-106
chr1-104299286-DEL-1017
chr1-104462465-DEL-52
chr1-104485788-DEL-160
chr1-104583171-DEL-75

chr1-104828222-DEL-941
chr1-104910639-DEL-82
chr1-104970762-DEL-4758
chr1-105244429-DEL-138
chr1-105244487-DEL-60
chr1-105310194-DEL-130
chr1-105422544-DEL-3653
chr1-105603689-DEL-55
chr1-105604102-DEL-170
chr1-105837279-DEL-227
chr1-105862369-DEL-1160
chr1-105977909-DEL-109
chr1-105985233-DEL-77
chr1-106052508-DEL-62
chr1-106053853-DEL-409
chr1-106057157-DEL-114
chr1-106139394-DEL-57
chr1-106139499-DEL-585
chr1-106146966-DEL-147
chr1-106155387-DEL-235
chr1-106310271-DEL-256
chr1-106397174-DEL-300
chr1-106400706-DEL-880
chr1-106532973-DEL-60
chr1-106549004-DEL-1240
chr1-106757624-DEL-592
chr1-106774784-DEL-60
chr1-106832262-DEL-50
chr1-106940783-DEL-6118
chr1-106947608-DEL-173
chr1-107077523-DEL-857
chr1-107184770-DEL-575
chr1-107189366-DEL-1632
chr1-107266278-DEL-218
chr1-107266380-DEL-50
chr1-107331884-DEL-50
chr1-107367579-DEL-163
chr1-107368197-DEL-914
chr1-107402919-DEL-122
chr1-107498874-DEL-50
chr1-107588839-DEL-58
chr1-107668013-DEL-51
chr1-107668174-DEL-255
chr1-107668428-DEL-93
chr1-107696728-DEL-52
chr1-107702341-DEL-64
chr1-107952806-DEL-50
chr1-108036752-DEL-210
chr1-108224202-DEL-87
chr1-108326047-DEL-4443
chr1-108334012-DEL-1556
chr1-108377321-DEL-256
chr1-108534879-DEL-267
chr1-108640628-DEL-63
chr1-108705652-DEL-57
chr1-108705883-DEL-244
chr1-108816689-DEL-51
chr1-108876113-DEL-81
chr1-109003094-DEL-80
chr1-109014284-DEL-58
chr1-109018060-DEL-50
chr1-109142024-DEL-88
chr1-109183936-DEL-1591
chr1-109241743-DEL-55
chr1-109267446-DEL-76
chr1-109301009-DEL-889
chr1-109304355-DEL-474
chr1-109503862-DEL-56
chr1-109633214-DEL-4740
chr1-109653341-DEL-5546
chr1-109720692-DEL-2558
chr1-109843911-DEL-54
chr1-109965432-DEL-3346
chr1-110002784-DEL-116
chr1-110034511-DEL-80
chr1-110137386-DEL-302
chr1-110401241-DEL-69

chr1-110440234-DEL-1635
chr1-110546841-DEL-5808
chr1-110663873-DEL-1608
chr1-110759393-DEL-1223
chr1-110805675-DEL-3053
chr1-110863762-DEL-327
chr1-110964893-DEL-116
chr1-111025480-DEL-272
chr1-111050802-DEL-990
chr1-111051691-DEL-52
chr1-111065848-DEL-8485
chr1-111249953-DEL-50
chr1-111357310-DEL-6438
chr1-111358724-DEL-204
chr1-111360677-DEL-1217
chr1-111363790-DEL-101
chr1-111471473-DEL-858
chr1-111516514-DEL-54
chr1-111536340-DEL-50
chr1-111578903-DEL-116
chr1-111646260-DEL-418
chr1-111765556-DEL-50
chr1-111779211-DEL-54
chr1-111824764-DEL-101
chr1-111991850-DEL-52
chr1-112020040-DEL-401
chr1-112160885-DEL-4239
chr1-112181061-DEL-126
chr1-112237548-DEL-59
chr1-112345835-DEL-140
chr1-112359900-DEL-70
chr1-112366027-DEL-6466
chr1-112434346-DEL-228
chr1-112546086-DEL-2258
chr1-112579889-DEL-6351
chr1-112588082-DEL-56
chr1-112618033-DEL-89
chr1-112623122-DEL-1338
chr1-112630628-DEL-152
chr1-112648700-DEL-503
chr1-112799342-DEL-92
chr1-112952205-DEL-155
chr1-113065202-DEL-258
chr1-113154927-DEL-60
chr1-113164939-DEL-738
chr1-113168687-DEL-3478
chr1-113280710-DEL-186
chr1-113293970-DEL-1955
chr1-113374343-DEL-60
chr1-113557682-DEL-153
chr1-113618404-DEL-60
chr1-113648696-DEL-1627
chr1-113690963-DEL-184
chr1-113710666-DEL-702
chr1-113852198-DEL-2597
chr1-113859866-DEL-216
chr1-113930979-DEL-1374
chr1-114024812-DEL-1054
chr1-114029455-DEL-4571
chr1-114101898-DEL-1765
chr1-114117646-DEL-2131
chr1-114243522-DEL-53
chr1-114275389-DEL-56
chr1-114360511-DEL-81
chr1-114653451-DEL-83
chr1-114676665-DEL-50
chr1-114706407-DEL-50
chr1-114718014-DEL-869
chr1-114997959-DEL-65
chr1-115027997-DEL-201
chr1-115223415-DEL-72
chr1-115252849-DEL-64
chr1-115258614-DEL-1301
chr1-115260023-DEL-89
chr1-115272295-DEL-50
chr1-115427900-DEL-60
chr1-115453276-DEL-74

chr1-115580974-DEL-66
chr1-115632838-DEL-54
chr1-115682261-DEL-110
chr1-115718008-DEL-50
chr1-115764956-DEL-2362
chr1-115934672-DEL-83
chr1-116009120-DEL-52
chr1-116031572-DEL-1453
chr1-116247145-DEL-6936
chr1-116333142-DEL-50
chr1-116531100-DEL-66
chr1-116556022-DEL-50
chr1-116670961-DEL-2671
chr1-116726478-DEL-480
chr1-116746487-DEL-54
chr1-116917975-DEL-5005
chr1-116935712-DEL-130
chr1-117001275-DEL-857
chr1-117121396-DEL-50
chr1-117209068-DEL-101
chr1-117213151-DEL-279
chr1-117216301-DEL-626
chr1-117309832-DEL-170
chr1-117384446-DEL-122
chr1-117468994-DEL-113
chr1-117490146-DEL-928
chr1-117538455-DEL-64
chr1-117553325-DEL-1092
chr1-117554417-DEL-168
chr1-117629605-DEL-230882
chr1-117674960-DEL-2267
chr1-117765435-DEL-686
chr1-117772350-DEL-4418
chr1-117832382-DEL-65
chr1-117868978-DEL-85
chr1-117914119-DEL-84
chr1-118024751-DEL-208
chr1-118032166-DEL-76
chr1-118080618-DEL-75
chr1-118126606-DEL-84
chr1-118127819-DEL-252
chr1-118175751-DEL-952
chr1-118176829-DEL-394
chr1-118204303-DEL-4266
chr1-118269559-DEL-57
chr1-118299143-DEL-562
chr1-118425018-DEL-156
chr1-118486852-DEL-148
chr1-118668584-DEL-138
chr1-118680013-DEL-873
chr1-118698461-DEL-50
chr1-118789814-DEL-50
chr1-118813711-DEL-63
chr1-119029769-DEL-198
chr1-119038273-DEL-52
chr1-119045821-DEL-123
chr1-119046319-DEL-284
chr1-119329751-DEL-98
chr1-119416514-DEL-346
chr1-119858263-DEL-93
chr1-119889392-DEL-50
chr1-119917944-DEL-120
chr1-119917947-DEL-519
chr1-119970141-DEL-388
chr1-119974759-DEL-76
chr1-120021993-DEL-226
chr1-120024180-DEL-518
chr1-120084205-DEL-107
chr1-120117546-DEL-75
chr1-120288595-DEL-196
chr1-120306361-DEL-390
chr1-120345196-DEL-65
chr1-120360939-DEL-131
chr1-120371285-DEL-98
chr1-120387265-DEL-58
chr1-120388977-DEL-538
chr1-120413224-DEL-100

chr1-120423585-DEL-50
chr1-120561599-DEL-659
chr1-120586127-DEL-695
chr1-120673511-DEL-52
chr1-120702642-DEL-9025
chr1-120855126-DEL-106
chr1-120865659-DEL-279
chr1-120886968-DEL-304
chr1-120921299-DEL-1024
chr1-120921735-DEL-212
chr1-120939353-DEL-88
chr1-120939599-DEL-74
chr1-120939951-DEL-104
chr1-120953185-DEL-52
chr1-121042061-DEL-151
chr1-121056081-DEL-967
chr1-121093779-DEL-87
chr1-121114884-DEL-51
chr1-121286517-DEL-344
chr1-121287177-DEL-84
chr1-121297341-DEL-136
chr1-121331796-DEL-219
chr1-121331962-DEL-78
chr1-121382962-DEL-127
chr1-121603975-DEL-51
chr1-121725276-DEL-58
chr1-121813870-DEL-554
chr1-121851349-DEL-50
chr1-121945392-DEL-1689
chr1-122254739-DEL-68
chr1-122259925-DEL-80
chr1-122273465-DEL-97
chr1-122273671-DEL-106
chr1-122535482-DEL-2082
chr1-122554686-DEL-65
chr1-122579197-DEL-55
chr1-122582297-DEL-1841
chr1-122587194-DEL-94
chr1-122598004-DEL-51
chr1-122615444-DEL-2036
chr1-122635742-DEL-59
chr1-122688563-DEL-72
chr1-122700421-DEL-416
chr1-122750328-DEL-4103
chr1-122848769-DEL-4459
chr1-123186442-DEL-50
chr1-123194770-DEL-72
chr1-123344732-DEL-64
chr1-123415339-DEL-221
chr1-123586443-DEL-100
chr1-123612384-DEL-50
chr1-123695476-DEL-52
chr1-123724773-DEL-8599
chr1-123740726-DEL-84
chr1-123748156-DEL-180
chr1-123806320-DEL-349
chr1-123890746-DEL-52
chr1-123920312-DEL-58
chr1-124083133-DEL-160
chr1-124152976-DEL-421
chr1-124198178-DEL-1163
chr1-124272248-DEL-52
chr1-124334817-DEL-62
chr1-124424684-DEL-68
chr1-124495827-DEL-50
chr1-124533365-DEL-50
chr1-124585983-DEL-59
chr1-124586308-DEL-182
chr1-124670269-DEL-64
chr1-124700945-DEL-68
chr1-124826626-DEL-350
chr1-125153326-DEL-1431
chr1-125316774-DEL-90
chr1-125317935-DEL-83
chr1-125442183-DEL-72
chr1-125501279-DEL-974
chr1-125586517-DEL-118

chr1-126077624-DEL-50
chr1-126101352-DEL-81
chr1-126112592-DEL-1160
chr1-126156765-DEL-126
chr1-126185668-DEL-450
chr1-126203274-DEL-354
chr1-126218357-DEL-208
chr1-126224044-DEL-50
chr1-126250637-DEL-75
chr1-126250810-DEL-168
chr1-126413109-DEL-434
chr1-126451428-DEL-135
chr1-126470680-DEL-100
chr1-126539245-DEL-348
chr1-126539327-DEL-108
chr1-126674401-DEL-120
chr1-126877041-DEL-61
chr1-126953449-DEL-1189
chr1-127134838-DEL-450
chr1-127341005-DEL-55
chr1-127354730-DEL-128
chr1-127422354-DEL-60
chr1-127498894-DEL-85
chr1-127526668-DEL-3093
chr1-127618446-DEL-55
chr1-127723810-DEL-87
chr1-127792586-DEL-76
chr1-127856412-DEL-123
chr1-128009975-DEL-2712
chr1-128012810-DEL-2101
chr1-128117654-DEL-60
chr1-128243612-DEL-150
chr1-128244670-DEL-201
chr1-128244872-DEL-401
chr1-128393764-DEL-88
chr1-128488452-DEL-104
chr1-128540034-DEL-290
chr1-128540047-DEL-94
chr1-128540292-DEL-136
chr1-128637120-DEL-76
chr1-128674216-DEL-120
chr1-128716648-DEL-50
chr1-128900030-DEL-126
chr1-128900528-DEL-178
chr1-128945362-DEL-3164
chr1-128965203-DEL-130
chr1-129027854-DEL-91
chr1-129071333-DEL-53
chr1-129112247-DEL-1489
chr1-129184085-DEL-201
chr1-129239173-DEL-192
chr1-129267363-DEL-207
chr1-129428399-DEL-50
chr1-129713239-DEL-183
chr1-129742528-DEL-1482
chr1-129900232-DEL-57
chr1-129939429-DEL-84
chr1-129939710-DEL-141
chr1-129947332-DEL-183
chr1-129947425-DEL-90
chr1-130039387-DEL-201
chr1-130101986-DEL-50
chr1-130288295-DEL-106
chr1-130288461-DEL-52
chr1-130288719-DEL-144
chr1-130376952-DEL-51
chr1-130673829-DEL-50
chr1-130693104-DEL-164
chr1-130823677-DEL-54
chr1-130837470-DEL-51
chr1-130837470-DEL-701
chr1-130838164-DEL-200
chr1-130878183-DEL-90
chr1-131106617-DEL-228
chr1-131153598-DEL-183
chr1-131208988-DEL-261
chr1-131299607-DEL-3702

chr1-131325614-DEL-56
chr1-131470848-DEL-80
chr1-131622160-DEL-76
chr1-131637713-DEL-176
chr1-131640699-DEL-67
chr1-131661807-DEL-125
chr1-131662273-DEL-138
chr1-131749574-DEL-58
chr1-131749921-DEL-204
chr1-132015479-DEL-188
chr1-132015534-DEL-78
chr1-132112649-DEL-280
chr1-132141687-DEL-66
chr1-132193449-DEL-439
chr1-132237931-DEL-97
chr1-132238045-DEL-83
chr1-132277000-DEL-115
chr1-132321064-DEL-145
chr1-132330427-DEL-74
chr1-132338857-DEL-50
chr1-132349205-DEL-58
chr1-132353809-DEL-55
chr1-132401259-DEL-1295
chr1-132445505-DEL-331
chr1-132445723-DEL-109
chr1-132649864-DEL-162
chr1-132650008-DEL-72
chr1-132793780-DEL-108
chr1-132893022-DEL-126
chr1-132903342-DEL-50
chr1-132905954-DEL-58
chr1-132950829-DEL-52
chr1-132953458-DEL-779
chr1-132954314-DEL-72
chr1-133087612-DEL-55
chr1-133107388-DEL-99
chr1-133371663-DEL-54
chr1-133377731-DEL-51
chr1-133437658-DEL-59
chr1-133505617-DEL-66
chr1-133511339-DEL-58
chr1-133522537-DEL-1336
chr1-133542676-DEL-135
chr1-133821991-DEL-50
chr1-134247517-DEL-164
chr1-134369919-DEL-70
chr1-134377546-DEL-50
chr1-134524478-DEL-56
chr1-134595550-DEL-64
chr1-134605866-DEL-279
chr1-134802311-DEL-67
chr1-134802678-DEL-69
chr1-135001653-DEL-50
chr1-135027779-DEL-50
chr1-135215017-DEL-916
chr1-135217402-DEL-505
chr1-135243453-DEL-75
chr1-135249199-DEL-134
chr1-135635438-DEL-97
chr1-135681062-DEL-97
chr1-135701447-DEL-69
chr1-135725486-DEL-107
chr1-135843858-DEL-50
chr1-135989929-DEL-83
chr1-136018069-DEL-68
chr1-136018813-DEL-54
chr1-136048164-DEL-55
chr1-136264442-DEL-2022
chr1-136264960-DEL-814
chr1-136266203-DEL-261
chr1-136304001-DEL-104
chr1-136304355-DEL-205
chr1-136310062-DEL-91
chr1-136315530-DEL-679
chr1-136378126-DEL-1950
chr1-136411673-DEL-324
chr1-136411871-DEL-58

chr1-136413105-DEL-88
chr1-136448673-DEL-83
chr1-136456691-DEL-50
chr1-136575253-DEL-332
chr1-136590433-DEL-162
chr1-136630348-DEL-178
chr1-136675607-DEL-209
chr1-136683461-DEL-216
chr1-136688531-DEL-289
chr1-136688565-DEL-85
chr1-136786587-DEL-63
chr1-136795596-DEL-70
chr1-136799750-DEL-144
chr1-136904853-DEL-52
chr1-136939673-DEL-597
chr1-137056050-DEL-50
chr1-137251430-DEL-92
chr1-137251669-DEL-90
chr1-137339739-DEL-95
chr1-137425248-DEL-660
chr1-137454607-DEL-6247
chr1-137505062-DEL-709
chr1-137724242-DEL-190
chr1-137806072-DEL-60
chr1-137902690-DEL-104
chr1-137932337-DEL-72
chr1-137954072-DEL-344
chr1-138278364-DEL-352
chr1-138370248-DEL-2481
chr1-138421799-DEL-68
chr1-138526735-DEL-60
chr1-138611193-DEL-56
chr1-138617742-DEL-68
chr1-138628399-DEL-277
chr1-138630257-DEL-171
chr1-138712783-DEL-50
chr1-138999757-DEL-1558
chr1-139072376-DEL-384
chr1-139086179-DEL-90
chr1-139212977-DEL-294
chr1-139213518-DEL-274
chr1-139300386-DEL-323
chr1-139413260-DEL-58
chr1-139568437-DEL-50
chr1-139824191-DEL-56
chr1-140108340-DEL-75
chr1-140135092-DEL-54
chr1-140158283-DEL-54
chr1-140213656-DEL-732
chr1-140331699-DEL-64
chr1-140599743-DEL-121
chr1-140602976-DEL-138
chr1-140825487-DEL-58
chr1-140868921-DEL-120
chr1-140887382-DEL-145
chr1-140981809-DEL-310
chr1-140988885-DEL-454
chr1-141086350-DEL-127
chr1-141114358-DEL-13670
chr1-141167211-DEL-56
chr1-141254725-DEL-62
chr1-141403128-DEL-109
chr1-141432751-DEL-83
chr1-141467062-DEL-4336
chr1-141698389-DEL-538
chr1-141884155-DEL-60
chr1-141941093-DEL-205
chr1-141946410-DEL-78
chr1-142061575-DEL-57
chr1-142061731-DEL-100
chr1-142110026-DEL-60
chr1-142218165-DEL-596
chr1-142326163-DEL-2410
chr1-142334233-DEL-249
chr1-142493747-DEL-930
chr1-142539949-DEL-3716
chr1-142596522-DEL-541

chr1-142756733-DEL-434
chr1-142757232-DEL-377
chr1-142969309-DEL-443
chr1-143083113-DEL-763
chr1-143175417-DEL-4768
chr1-143284949-DEL-912
chr1-143439262-DEL-58
chr1-143441564-DEL-153
chr1-143468328-DEL-66
chr1-143582121-DEL-51
chr1-143637795-DEL-778
chr1-143638609-DEL-62
chr1-143709687-DEL-54
chr1-143722921-DEL-2481
chr1-143747291-DEL-266
chr1-143747549-DEL-92
chr1-143860994-DEL-92
chr1-143911582-DEL-286
chr1-144128493-DEL-2784
chr1-144288063-DEL-2639
chr1-144486299-DEL-938
chr1-144659204-DEL-3857
chr1-144676893-DEL-79
chr1-144677085-DEL-233
chr1-144753758-DEL-100
chr1-144967045-DEL-51
chr1-145593686-DEL-555
chr1-145668174-DEL-64
chr1-145668181-DEL-168
chr1-145856649-DEL-110
chr1-145900086-DEL-199
chr1-145981118-DEL-442
chr1-146071487-DEL-8827
chr1-146191036-DEL-513
chr1-146257937-DEL-2825
chr1-146296040-DEL-1900
chr1-146300542-DEL-804
chr1-146301376-DEL-665
chr1-146328693-DEL-140
chr1-146328718-DEL-52
chr1-146498791-DEL-50
chr1-146520730-DEL-50
chr1-146701803-DEL-271
chr1-146701898-DEL-66
chr1-146918060-DEL-1524
chr1-147014916-DEL-168
chr1-147200940-DEL-1076
chr1-147273728-DEL-458
chr1-147341054-DEL-53
chr1-147341556-DEL-1032
chr1-147426047-DEL-52
chr1-147428753-DEL-640
chr1-147434209-DEL-229
chr1-147603827-DEL-1692
chr1-147656400-DEL-3054
chr1-147708985-DEL-2677
chr1-147957536-DEL-208
chr1-147959525-DEL-50
chr1-148043990-DEL-54
chr1-148277449-DEL-76
chr1-148280879-DEL-888
chr1-148397545-DEL-424
chr1-148598559-DEL-660
chr1-148744528-DEL-76
chr1-148753669-DEL-6388
chr1-148813467-DEL-54
chr1-148835890-DEL-56
chr1-148866603-DEL-611
chr1-148880073-DEL-280
chr1-149174105-DEL-3033
chr1-149294683-DEL-1706
chr1-149487973-DEL-182
chr1-149651563-DEL-50
chr1-149795896-DEL-595
chr1-149853133-DEL-170
chr1-149867168-DEL-1124
chr1-149886992-DEL-74

chr1-149910938-DEL-1477
chr1-149921415-DEL-143
chr1-150252957-DEL-2803
chr1-150330660-DEL-96
chr1-150419348-DEL-62
chr1-150487657-DEL-50
chr1-150502363-DEL-66
chr1-150621777-DEL-354
chr1-150621785-DEL-118
chr1-150621996-DEL-685
chr1-150622277-DEL-317
chr1-150855087-DEL-50
chr1-150959739-DEL-299
chr1-150992723-DEL-170
chr1-151058276-DEL-15838
chr1-151069537-DEL-11742
chr1-151128448-DEL-69
chr1-151158922-DEL-480
chr1-151251895-DEL-2677
chr1-151450479-DEL-116
chr1-151463331-DEL-90
chr1-151463340-DEL-350
chr1-151618053-DEL-492
chr1-151640798-DEL-115
chr1-151676138-DEL-243
chr1-151676497-DEL-639
chr1-151677277-DEL-92
chr1-151677733-DEL-414
chr1-151677900-DEL-138
chr1-151678401-DEL-289
chr1-151716206-DEL-182
chr1-151735153-DEL-114
chr1-151752463-DEL-613
chr1-151812260-DEL-52
chr1-151918643-DEL-79
chr1-152014304-DEL-51
chr1-152118105-DEL-214
chr1-152139319-DEL-120
chr1-152143511-DEL-74
chr1-152185245-DEL-63
chr1-152242602-DEL-330
chr1-152280240-DEL-52
chr1-152509086-DEL-560
chr1-152518082-DEL-68
chr1-152529981-DEL-197
chr1-152603008-DEL-179
chr1-152611020-DEL-76
chr1-152639098-DEL-50
chr1-152694649-DEL-54
chr1-152953056-DEL-6934
chr1-153052071-DEL-79
chr1-153063354-DEL-74
chr1-153177883-DEL-130
chr1-153263264-DEL-9697
chr1-153277252-DEL-90
chr1-153277947-DEL-221
chr1-153281007-DEL-2679
chr1-153339806-DEL-63
chr1-153369564-DEL-310
chr1-153519074-DEL-213
chr1-153544509-DEL-52
chr1-153828053-DEL-224
chr1-153897359-DEL-138
chr1-153918196-DEL-69
chr1-153971673-DEL-219
chr1-154046942-DEL-53
chr1-154065385-DEL-83
chr1-154082763-DEL-1396
chr1-154365286-DEL-98
chr1-154431047-DEL-66
chr1-154637363-DEL-61
chr1-154673939-DEL-105
chr1-154734526-DEL-94
chr1-154734526-DEL-210
chr1-154735004-DEL-182
chr1-154757409-DEL-260
chr1-154757558-DEL-65

chr1-154795228-DEL-100
chr1-154864238-DEL-147
chr1-154895771-DEL-3537
chr1-154901258-DEL-63
chr1-154951325-DEL-50
chr1-154978290-DEL-96
chr1-154991009-DEL-362
chr1-155028761-DEL-135
chr1-155052692-DEL-52
chr1-155052826-DEL-79
chr1-155080139-DEL-64
chr1-155126924-DEL-100
chr1-155180077-DEL-55
chr1-155276858-DEL-172
chr1-155356034-DEL-112
chr1-155372419-DEL-209
chr1-155372545-DEL-104
chr1-155427205-DEL-61
chr1-155427332-DEL-68
chr1-155473904-DEL-109
chr1-155553490-DEL-700
chr1-155605002-DEL-165
chr1-155630381-DEL-72
chr1-155813068-DEL-59
chr1-155871635-DEL-215
chr1-155953529-DEL-444
chr1-156217908-DEL-64
chr1-156409029-DEL-97
chr1-156432026-DEL-98
chr1-156504262-DEL-109
chr1-156510257-DEL-109
chr1-156521426-DEL-89
chr1-156521709-DEL-295
chr1-156567416-DEL-83
chr1-156567594-DEL-914
chr1-156614609-DEL-85
chr1-156773945-DEL-148
chr1-156879817-DEL-52
chr1-156940753-DEL-70
chr1-156954700-DEL-669
chr1-157251334-DEL-70
chr1-157366455-DEL-128
chr1-157386559-DEL-168
chr1-157428118-DEL-112
chr1-157440429-DEL-64
chr1-157552984-DEL-64
chr1-157804794-DEL-294
chr1-157805082-DEL-612
chr1-157805387-DEL-54
chr1-157805437-DEL-126
chr1-157920075-DEL-58
chr1-158074945-DEL-325
chr1-158089410-DEL-160
chr1-158102884-DEL-174
chr1-158365074-DEL-52
chr1-158482782-DEL-120
chr1-158482835-DEL-52
chr1-158483102-DEL-274
chr1-158839295-DEL-55
chr1-158867217-DEL-50
chr1-159038151-DEL-54
chr1-159196559-DEL-50
chr1-159411949-DEL-1178
chr1-159620685-DEL-149
chr1-159620907-DEL-152
chr1-159620999-DEL-60
chr1-159744362-DEL-80
chr1-159773781-DEL-701
chr1-160249176-DEL-208
chr1-160300284-DEL-54
chr1-160335248-DEL-60
chr1-160556983-DEL-96
chr1-160573342-DEL-54
chr1-160634502-DEL-312
chr1-160818449-DEL-63
chr1-160866242-DEL-356
chr1-160924845-DEL-66

chr1-161086422-DEL-61
chr1-161144365-DEL-4160
chr1-161151997-DEL-93
chr1-161226692-DEL-156
chr1-161242877-DEL-50
chr1-161348713-DEL-173
chr1-161426782-DEL-56
chr1-161826079-DEL-127
chr1-161974511-DEL-55
chr1-162163282-DEL-60
chr1-162242253-DEL-213
chr1-162242433-DEL-50
chr1-162356562-DEL-50
chr1-162491398-DEL-128
chr1-162664554-DEL-85
chr1-162799624-DEL-56
chr1-162864209-DEL-68
chr1-163210308-DEL-72
chr1-163284117-DEL-158
chr1-163284526-DEL-3819
chr1-163365899-DEL-51
chr1-163508253-DEL-90
chr1-163623840-DEL-85
chr1-163787636-DEL-55
chr1-163815003-DEL-89
chr1-163847951-DEL-1452
chr1-163871577-DEL-132
chr1-163930860-DEL-2042
chr1-163935753-DEL-86
chr1-164143556-DEL-50
chr1-164152533-DEL-101
chr1-164156375-DEL-393
chr1-164156377-DEL-109
chr1-164158140-DEL-1293
chr1-164163318-DEL-50
chr1-164186537-DEL-501
chr1-164306141-DEL-337
chr1-164324838-DEL-1439
chr1-164363161-DEL-600
chr1-164469018-DEL-226
chr1-164535697-DEL-142
chr1-164596262-DEL-84
chr1-164841749-DEL-75
chr1-164853578-DEL-199
chr1-164861575-DEL-445
chr1-164862698-DEL-52
chr1-164953690-DEL-1109
chr1-164968326-DEL-54
chr1-165067885-DEL-181
chr1-165156468-DEL-77
chr1-165180921-DEL-166
chr1-165507651-DEL-3908
chr1-165507853-DEL-114
chr1-165586518-DEL-670
chr1-165595210-DEL-366
chr1-165899750-DEL-50
chr1-166039396-DEL-7089
chr1-166179581-DEL-7956
chr1-166211871-DEL-202
chr1-166223066-DEL-65
chr1-166477386-DEL-58
chr1-166638310-DEL-51
chr1-166707430-DEL-50
chr1-166903744-DEL-64
chr1-167047809-DEL-142
chr1-167108270-DEL-57
chr1-167147631-DEL-144
chr1-167165974-DEL-373
chr1-167172943-DEL-173
chr1-167604370-DEL-84
chr1-167696742-DEL-66
chr1-167791399-DEL-75
chr1-167882117-DEL-50
chr1-168152739-DEL-66
chr1-168161737-DEL-111
chr1-168231515-DEL-78
chr1-168329632-DEL-106

chr1-168370055-DEL-50
chr1-168390330-DEL-87
chr1-168393998-DEL-57
chr1-168430207-DEL-143
chr1-168636664-DEL-608
chr1-168638252-DEL-166
chr1-168638662-DEL-64
chr1-168740026-DEL-314
chr1-168772712-DEL-50
chr1-168824439-DEL-98
chr1-168845399-DEL-50
chr1-168941227-DEL-50
chr1-169025134-DEL-66
chr1-169098078-DEL-58
chr1-169173354-DEL-56
chr1-169173515-DEL-163
chr1-169333895-DEL-81
chr1-169341247-DEL-53
chr1-169345177-DEL-337
chr1-169345336-DEL-152
chr1-169345548-DEL-263
chr1-169475591-DEL-70
chr1-169536843-DEL-89
chr1-169734456-DEL-108
chr1-169902643-DEL-71
chr1-169969127-DEL-167
chr1-170103867-DEL-69
chr1-170254568-DEL-50
chr1-170281534-DEL-60
chr1-170311177-DEL-248
chr1-170356464-DEL-108
chr1-170448388-DEL-52
chr1-170526330-DEL-50
chr1-170652836-DEL-71
chr1-170674773-DEL-151
chr1-170747545-DEL-66
chr1-170786979-DEL-53
chr1-170815474-DEL-158
chr1-170815875-DEL-216
chr1-170816031-DEL-72
chr1-170873730-DEL-434
chr1-170894380-DEL-54
chr1-170895391-DEL-78
chr1-171028007-DEL-50
chr1-171036665-DEL-149
chr1-171174140-DEL-50
chr1-171260803-DEL-131
chr1-171284096-DEL-78
chr1-171285662-DEL-224
chr1-171299405-DEL-136
chr1-171302722-DEL-101
chr1-171359716-DEL-73
chr1-171369214-DEL-329
chr1-171418477-DEL-459
chr1-171588141-DEL-164
chr1-171642555-DEL-547
chr1-171805509-DEL-95
chr1-171805746-DEL-150
chr1-171890484-DEL-131
chr1-171928309-DEL-64
chr1-172134405-DEL-56
chr1-172423062-DEL-52
chr1-172601842-DEL-75
chr1-172636515-DEL-83
chr1-172639331-DEL-76
chr1-172640823-DEL-54
chr1-172733415-DEL-433
chr1-172822137-DEL-200
chr1-172822165-DEL-65
chr1-172847647-DEL-50
chr1-173009898-DEL-50
chr1-173079051-DEL-59
chr1-173299275-DEL-282
chr1-173326881-DEL-200
chr1-173366866-DEL-7408
chr1-173436345-DEL-419
chr1-173539911-DEL-65

chr1-173660691-DEL-214
chr1-173745675-DEL-2400
chr1-173750531-DEL-52
chr1-173800108-DEL-468
chr1-173837505-DEL-147
chr1-173837514-DEL-435
chr1-173837828-DEL-126
chr1-173865016-DEL-52
chr1-173867873-DEL-50
chr1-173954988-DEL-2438
chr1-174015453-DEL-1436
chr1-174153316-DEL-3462
chr1-174246700-DEL-100
chr1-174302544-DEL-63
chr1-174318535-DEL-56
chr1-174417920-DEL-142
chr1-174418075-DEL-50
chr1-174447955-DEL-50
chr1-174551250-DEL-300
chr1-174551527-DEL-92
chr1-174715908-DEL-75
chr1-174848820-DEL-54
chr1-175149746-DEL-52
chr1-175220668-DEL-59
chr1-175628969-DEL-77
chr1-175785307-DEL-148
chr1-175825110-DEL-50
chr1-175875244-DEL-1714
chr1-175960659-DEL-103
chr1-176085411-DEL-60
chr1-176185629-DEL-1531
chr1-176301494-DEL-125
chr1-176381763-DEL-212
chr1-176406357-DEL-2717
chr1-176411123-DEL-3790
chr1-176480731-DEL-148
chr1-176480816-DEL-58
chr1-176549645-DEL-331
chr1-176549829-DEL-130
chr1-176646993-DEL-159
chr1-176718941-DEL-93
chr1-176802530-DEL-93
chr1-176868175-DEL-171
chr1-177206344-DEL-73
chr1-177377214-DEL-1809
chr1-177456408-DEL-195
chr1-177495531-DEL-72
chr1-177534727-DEL-1971
chr1-177549916-DEL-69
chr1-177567867-DEL-125
chr1-177807628-DEL-190
chr1-177813956-DEL-336
chr1-177824173-DEL-63
chr1-177838121-DEL-214
chr1-177905034-DEL-14047
chr1-178344797-DEL-73
chr1-178349308-DEL-52
chr1-178587779-DEL-50
chr1-178669597-DEL-51
chr1-178899847-DEL-81
chr1-178990650-DEL-322
chr1-178990915-DEL-97
chr1-178991203-DEL-536
chr1-179007495-DEL-54
chr1-179430058-DEL-140
chr1-179430911-DEL-100
chr1-179498994-DEL-140
chr1-179499195-DEL-157
chr1-179542969-DEL-80
chr1-179619062-DEL-137
chr1-179665569-DEL-68
chr1-179667294-DEL-189
chr1-179730178-DEL-422
chr1-179730827-DEL-790
chr1-179731072-DEL-1203
chr1-179732098-DEL-1144
chr1-179794275-DEL-328

chr1-179794402-DEL-133
chr1-179891771-DEL-96
chr1-180069673-DEL-225
chr1-180070213-DEL-1924
chr1-180070597-DEL-672
chr1-180071021-DEL-124
chr1-180071948-DEL-50
chr1-180239309-DEL-56
chr1-180239518-DEL-187
chr1-180315546-DEL-190
chr1-180371095-DEL-926
chr1-180373274-DEL-62
chr1-180415063-DEL-139
chr1-180417362-DEL-1075
chr1-180439643-DEL-4414
chr1-180499569-DEL-66
chr1-180506468-DEL-232
chr1-180543314-DEL-52
chr1-180577688-DEL-56
chr1-180593777-DEL-82
chr1-180594054-DEL-63
chr1-180614152-DEL-50
chr1-180757802-DEL-202
chr1-180836739-DEL-164
chr1-180847440-DEL-82
chr1-180907661-DEL-55
chr1-181018374-DEL-409
chr1-181360149-DEL-54
chr1-181391344-DEL-495
chr1-181391535-DEL-82
chr1-181395852-DEL-185
chr1-181457770-DEL-100
chr1-181502059-DEL-100
chr1-181570552-DEL-200
chr1-181571326-DEL-160
chr1-181576120-DEL-50
chr1-181579155-DEL-120
chr1-181683025-DEL-54
chr1-181890769-DEL-93
chr1-181978781-DEL-90
chr1-181994573-DEL-51
chr1-181994761-DEL-173
chr1-182014000-DEL-119
chr1-182113273-DEL-94
chr1-182332695-DEL-110
chr1-182456928-DEL-126
chr1-182457309-DEL-54
chr1-182705026-DEL-291
chr1-182844605-DEL-64
chr1-182884960-DEL-553
chr1-182887462-DEL-373
chr1-182943383-DEL-78
chr1-183040232-DEL-710
chr1-183124141-DEL-148
chr1-183172388-DEL-81
chr1-183227531-DEL-784
chr1-183257366-DEL-104
chr1-183308618-DEL-50
chr1-183316530-DEL-72
chr1-183329963-DEL-108
chr1-183330216-DEL-358
chr1-183349454-DEL-570
chr1-183350220-DEL-75
chr1-183360765-DEL-176
chr1-183391803-DEL-69
chr1-183392010-DEL-107
chr1-183559717-DEL-50
chr1-183694537-DEL-50
chr1-183888422-DEL-141
chr1-184077399-DEL-58
chr1-184140446-DEL-63
chr1-184225528-DEL-56
chr1-184291147-DEL-65
chr1-184434922-DEL-620
chr1-184473359-DEL-98
chr1-184523598-DEL-79
chr1-184546420-DEL-650

chr1-184620913-DEL-118
chr1-184621350-DEL-53
chr1-184688616-DEL-393
chr1-184688739-DEL-118
chr1-184763508-DEL-94
chr1-185051292-DEL-50
chr1-185190421-DEL-182
chr1-185190683-DEL-159
chr1-185405898-DEL-122
chr1-185406163-DEL-144
chr1-185462753-DEL-77
chr1-185482544-DEL-72
chr1-185511763-DEL-67
chr1-185691813-DEL-92
chr1-185720800-DEL-99
chr1-185749545-DEL-335
chr1-185750032-DEL-118
chr1-185827158-DEL-52
chr1-186092195-DEL-62
chr1-186180587-DEL-276
chr1-186441943-DEL-612
chr1-186442460-DEL-369
chr1-186462111-DEL-3216
chr1-186462489-DEL-62
chr1-186546162-DEL-76
chr1-186557869-DEL-66
chr1-186587798-DEL-669
chr1-186620201-DEL-50
chr1-186725660-DEL-1133
chr1-186903660-DEL-368
chr1-186919645-DEL-50
chr1-186982886-DEL-1458
chr1-187071282-DEL-236
chr1-187071559-DEL-166
chr1-187087004-DEL-682
chr1-187121206-DEL-53
chr1-187124934-DEL-848
chr1-187267010-DEL-125
chr1-187348731-DEL-111
chr1-187448846-DEL-75
chr1-187547794-DEL-98
chr1-187552932-DEL-90
chr1-187625749-DEL-121
chr1-187805640-DEL-167
chr1-187957563-DEL-1594
chr1-188056924-DEL-68
chr1-188057486-DEL-72
chr1-188057858-DEL-101
chr1-188058044-DEL-61
chr1-188058100-DEL-399
chr1-188070453-DEL-3110
chr1-188242375-DEL-83
chr1-188242662-DEL-184
chr1-188385993-DEL-172
chr1-188386081-DEL-73
chr1-188414330-DEL-132
chr1-188425208-DEL-257
chr1-188463449-DEL-271
chr1-188463556-DEL-50
chr1-188522323-DEL-76
chr1-188526384-DEL-96
chr1-188539077-DEL-154
chr1-188601492-DEL-52
chr1-188601820-DEL-162
chr1-188622992-DEL-72
chr1-188623228-DEL-108
chr1-188625839-DEL-71
chr1-188628210-DEL-180
chr1-188665090-DEL-50
chr1-188705546-DEL-106
chr1-188762857-DEL-110
chr1-188861546-DEL-92
chr1-188926522-DEL-120
chr1-188927171-DEL-120
chr1-188975626-DEL-147
chr1-189032886-DEL-128
chr1-189033226-DEL-256

chr1-189233108-DEL-108
chr1-189269866-DEL-76
chr1-189270079-DEL-170
chr1-189306680-DEL-57
chr1-189339890-DEL-168
chr1-189356422-DEL-50
chr1-189425037-DEL-59
chr1-189432962-DEL-68
chr1-189466306-DEL-216
chr1-189500914-DEL-50
chr1-189521667-DEL-62
chr1-189612151-DEL-54
chr1-189659726-DEL-73
chr1-189712704-DEL-354
chr1-189716739-DEL-55
chr1-189716890-DEL-69
chr1-189738368-DEL-136
chr1-189748772-DEL-76
chr1-189773832-DEL-51
chr1-189776741-DEL-71
chr1-189802459-DEL-90
chr1-189884717-DEL-130
chr1-189935288-DEL-148
chr1-189944178-DEL-464
chr1-189950178-DEL-50
chr1-190024512-DEL-837
chr1-190032936-DEL-1474
chr1-190035930-DEL-175
chr1-190275575-DEL-112
chr1-190406803-DEL-423
chr1-190488224-DEL-676
chr1-190488724-DEL-68
chr1-190513004-DEL-1059
chr1-190522600-DEL-208
chr1-190583193-DEL-408
chr1-190583750-DEL-351
chr1-190603065-DEL-260
chr1-190613384-DEL-50
chr1-190649689-DEL-166
chr1-190679290-DEL-132
chr1-190733314-DEL-55
chr1-190745841-DEL-222
chr1-190783816-DEL-65
chr1-191101869-DEL-291
chr1-191155252-DEL-61
chr1-191205341-DEL-178
chr1-191287494-DEL-586
chr1-191379219-DEL-60
chr1-191490808-DEL-94
chr1-191491045-DEL-153
chr1-191586256-DEL-52
chr1-191600431-DEL-432
chr1-191884664-DEL-117
chr1-191896665-DEL-97
chr1-191900914-DEL-53
chr1-191912990-DEL-240
chr1-191937239-DEL-392
chr1-192142759-DEL-58
chr1-192147598-DEL-6609
chr1-192230046-DEL-180
chr1-192265737-DEL-64
chr1-192311825-DEL-3380
chr1-192342324-DEL-96
chr1-192344805-DEL-755
chr1-192352081-DEL-434
chr1-192352854-DEL-252
chr1-192398257-DEL-58
chr1-192398714-DEL-50
chr1-192601115-DEL-124
chr1-192734100-DEL-417
chr1-192809501-DEL-250
chr1-193028071-DEL-79
chr1-193057157-DEL-130
chr1-193180253-DEL-91
chr1-193183400-DEL-134
chr1-193452957-DEL-3498
chr1-193459885-DEL-50

chr1-193757578-DEL-641
chr1-193825943-DEL-1197
chr1-193885466-DEL-144
chr1-194431930-DEL-492
chr1-194738319-DEL-416
chr1-194843968-DEL-77
chr1-194870842-DEL-594
chr1-194895218-DEL-3874
chr1-194964245-DEL-408
chr10-3062791-DEL-3204
chr10-3320128-DEL-51
chr10-3360828-DEL-84
chr10-3456305-DEL-69
chr10-3482279-DEL-52
chr10-3482391-DEL-86
chr10-3746342-DEL-120
chr10-3787195-DEL-54
chr10-3824458-DEL-178
chr10-3841380-DEL-74
chr10-3857187-DEL-506
chr10-3876528-DEL-176
chr10-3878704-DEL-207
chr10-3889541-DEL-53
chr10-3912139-DEL-63
chr10-3936798-DEL-1814
chr10-3950871-DEL-619
chr10-3960709-DEL-433
chr10-4005504-DEL-54
chr10-4012124-DEL-2937
chr10-4087429-DEL-184
chr10-4108416-DEL-68
chr10-4191341-DEL-545
chr10-4195400-DEL-419
chr10-4195671-DEL-148
chr10-4208671-DEL-170
chr10-4249336-DEL-1690
chr10-4264818-DEL-309
chr10-4268647-DEL-102
chr10-4295096-DEL-699
chr10-4324996-DEL-66
chr10-4336080-DEL-51
chr10-4429234-DEL-128
chr10-4430507-DEL-874
chr10-4453142-DEL-61
chr10-4457362-DEL-110
chr10-4477105-DEL-143
chr10-4488168-DEL-59
chr10-4680290-DEL-256
chr10-4753150-DEL-62
chr10-4753514-DEL-212
chr10-4833867-DEL-382
chr10-4861272-DEL-210
chr10-4861292-DEL-424
chr10-4861858-DEL-212
chr10-4956525-DEL-105
chr10-5030930-DEL-2374
chr10-5031514-DEL-535
chr10-5040322-DEL-212
chr10-5107920-DEL-59
chr10-5108067-DEL-265
chr10-5163084-DEL-53
chr10-5207573-DEL-142
chr10-5207651-DEL-64
chr10-5211203-DEL-82
chr10-5269395-DEL-205
chr10-5305309-DEL-62
chr10-5374852-DEL-50
chr10-5387390-DEL-61
chr10-5405234-DEL-54
chr10-5450079-DEL-84
chr10-5518268-DEL-68
chr10-5603557-DEL-74
chr10-5664845-DEL-109
chr10-5761394-DEL-248
chr10-5825133-DEL-913
chr10-5833206-DEL-698
chr10-5847790-DEL-228

chr10-5858252-DEL-77
chr10-5865702-DEL-121
chr10-5891653-DEL-232
chr10-5906519-DEL-50
chr10-5929837-DEL-177
chr10-5931394-DEL-13487
chr10-6081520-DEL-54
chr10-6095544-DEL-246
chr10-6095568-DEL-75
chr10-6137981-DEL-54
chr10-6171565-DEL-54
chr10-6263142-DEL-1024
chr10-6354971-DEL-2674
chr10-6429388-DEL-599
chr10-6464835-DEL-99
chr10-6480929-DEL-787
chr10-6580558-DEL-330
chr10-6698996-DEL-106
chr10-6699405-DEL-53
chr10-6846215-DEL-58
chr10-6875576-DEL-74
chr10-6998971-DEL-190
chr10-6998989-DEL-65
chr10-7097017-DEL-2002
chr10-7099466-DEL-84
chr10-7099590-DEL-1335
chr10-7118206-DEL-238
chr10-7140277-DEL-65
chr10-7166963-DEL-53
chr10-7182577-DEL-108
chr10-7509463-DEL-2760
chr10-7608817-DEL-760
chr10-7778843-DEL-140
chr10-7783802-DEL-64
chr10-7874103-DEL-58
chr10-7912610-DEL-131
chr10-7938886-DEL-106
chr10-7988115-DEL-96
chr10-8093861-DEL-60
chr10-8168917-DEL-469
chr10-8269518-DEL-1035
chr10-8290625-DEL-82
chr10-8329854-DEL-518
chr10-8378304-DEL-104
chr10-8389051-DEL-534
chr10-8456129-DEL-50
chr10-8486458-DEL-617
chr10-8554298-DEL-99
chr10-8571476-DEL-98
chr10-8571560-DEL-480
chr10-8585022-DEL-68
chr10-8593641-DEL-402
chr10-8594108-DEL-163
chr10-8600191-DEL-60
chr10-8602258-DEL-213
chr10-8639522-DEL-590
chr10-8705150-DEL-129
chr10-8793090-DEL-50
chr10-8831049-DEL-52
chr10-8838008-DEL-86
chr10-8893653-DEL-70
chr10-8893941-DEL-181
chr10-9023958-DEL-158
chr10-9032102-DEL-66
chr10-9073245-DEL-1762
chr10-9300632-DEL-364
chr10-9315587-DEL-59
chr10-9421866-DEL-103
chr10-9457091-DEL-76
chr10-9496108-DEL-337
chr10-9496362-DEL-60
chr10-9545830-DEL-50
chr10-9582964-DEL-2799
chr10-9586961-DEL-120
chr10-9587334-DEL-110
chr10-9729210-DEL-936
chr10-9739651-DEL-481

chr10-9751380-DEL-2338
chr10-9758127-DEL-112
chr10-9819890-DEL-52
chr10-9856492-DEL-50
chr10-10004958-DEL-277
chr10-10008754-DEL-51
chr10-10210993-DEL-86
chr10-10354378-DEL-1491
chr10-10503524-DEL-63
chr10-10568680-DEL-130
chr10-10808563-DEL-52
chr10-10842910-DEL-79
chr10-10848010-DEL-54
chr10-10865983-DEL-156
chr10-10899449-DEL-62
chr10-10905983-DEL-56
chr10-10921189-DEL-61
chr10-10931394-DEL-102
chr10-10999310-DEL-381
chr10-11193985-DEL-74
chr10-11218369-DEL-103
chr10-11253956-DEL-93
chr10-11263853-DEL-139
chr10-11276855-DEL-5556
chr10-11309829-DEL-474
chr10-11399518-DEL-1430
chr10-11441140-DEL-96
chr10-11448885-DEL-3804
chr10-11511567-DEL-639
chr10-11512213-DEL-107
chr10-11568189-DEL-10218
chr10-11751910-DEL-107
chr10-11825270-DEL-513
chr10-11835335-DEL-106
chr10-11891818-DEL-117
chr10-11892294-DEL-134
chr10-12272811-DEL-57
chr10-12283752-DEL-60
chr10-12330575-DEL-81
chr10-12366452-DEL-761
chr10-12368190-DEL-50
chr10-12519651-DEL-50
chr10-12523517-DEL-85
chr10-12529440-DEL-72
chr10-12564794-DEL-153
chr10-12590341-DEL-130
chr10-12896732-DEL-148
chr10-12998052-DEL-50
chr10-13006444-DEL-116
chr10-13107981-DEL-132
chr10-13304186-DEL-52
chr10-13318569-DEL-56
chr10-13366142-DEL-1144
chr10-13510766-DEL-95
chr10-13554802-DEL-80
chr10-13660512-DEL-212
chr10-13660779-DEL-548
chr10-13660855-DEL-218
chr10-13747809-DEL-51
chr10-13772999-DEL-3517
chr10-13777505-DEL-444
chr10-14102076-DEL-51
chr10-14121587-DEL-7182
chr10-14241635-DEL-1136
chr10-14271560-DEL-157
chr10-14382947-DEL-60
chr10-14413494-DEL-158
chr10-14450393-DEL-106
chr10-14453206-DEL-50
chr10-14490400-DEL-80
chr10-14521718-DEL-73
chr10-14538568-DEL-1483
chr10-14679468-DEL-118
chr10-14686327-DEL-97
chr10-14752543-DEL-124
chr10-14869826-DEL-154
chr10-14917154-DEL-56

chr10-14963266-DEL-52
chr10-15000246-DEL-2716
chr10-15012633-DEL-73
chr10-15012978-DEL-57
chr10-15021443-DEL-144
chr10-15067453-DEL-2462
chr10-15067453-DEL-4926
chr10-15127279-DEL-60
chr10-15183952-DEL-94
chr10-15236043-DEL-325
chr10-15248916-DEL-1251
chr10-15330182-DEL-50
chr10-15448556-DEL-811
chr10-15450536-DEL-4938
chr10-15510672-DEL-212
chr10-15557906-DEL-1886
chr10-15594888-DEL-3553
chr10-15821476-DEL-115
chr10-15821835-DEL-120
chr10-15839871-DEL-2884
chr10-15929371-DEL-188
chr10-16006248-DEL-5938
chr10-16010691-DEL-178
chr10-16068916-DEL-146
chr10-16130424-DEL-56
chr10-16152462-DEL-54
chr10-16253990-DEL-50
chr10-16446964-DEL-853
chr10-16511467-DEL-1060
chr10-16568989-DEL-2368
chr10-16676034-DEL-284
chr10-16738354-DEL-101
chr10-16829604-DEL-50
chr10-16854317-DEL-109
chr10-16969324-DEL-52
chr10-17249235-DEL-89
chr10-17256025-DEL-50
chr10-17337644-DEL-2071
chr10-17510875-DEL-50
chr10-18003389-DEL-100
chr10-18008281-DEL-671
chr10-18021906-DEL-267
chr10-18031234-DEL-615
chr10-18182630-DEL-50
chr10-18198883-DEL-86
chr10-18500692-DEL-52
chr10-18513964-DEL-78
chr10-18549147-DEL-58
chr10-18581319-DEL-507
chr10-18589198-DEL-83
chr10-18688778-DEL-102
chr10-18700749-DEL-79
chr10-18714080-DEL-64
chr10-18820748-DEL-73
chr10-18968341-DEL-141
chr10-18968595-DEL-90
chr10-18985274-DEL-1705
chr10-19008149-DEL-5388
chr10-19036168-DEL-142
chr10-19082776-DEL-210
chr10-19147068-DEL-1487
chr10-19151502-DEL-90
chr10-19151723-DEL-197
chr10-19303469-DEL-625
chr10-19303752-DEL-77
chr10-19383752-DEL-56
chr10-19407564-DEL-57
chr10-19430372-DEL-2849
chr10-19456175-DEL-60
chr10-19456218-DEL-205
chr10-19473870-DEL-587
chr10-19559441-DEL-243
chr10-19583650-DEL-93
chr10-19614064-DEL-84
chr10-19640276-DEL-1271
chr10-19642973-DEL-1685
chr10-19699346-DEL-113

chr10-19744591-DEL-101
chr10-19748204-DEL-214
chr10-19749622-DEL-671
chr10-19795854-DEL-58
chr10-19854863-DEL-2620
chr10-19864204-DEL-166
chr10-19973710-DEL-66
chr10-19993783-DEL-54
chr10-20013132-DEL-170
chr10-20140751-DEL-71
chr10-20247276-DEL-1990
chr10-20265229-DEL-377
chr10-20357412-DEL-56
chr10-20392771-DEL-50
chr10-20411300-DEL-59
chr10-20464945-DEL-214
chr10-20499268-DEL-258
chr10-20511504-DEL-319
chr10-20573005-DEL-126
chr10-20628971-DEL-55
chr10-20642263-DEL-50
chr10-20660386-DEL-60
chr10-20713444-DEL-82
chr10-20721104-DEL-51
chr10-20751227-DEL-423
chr10-20791811-DEL-804
chr10-20873227-DEL-60
chr10-20927442-DEL-86
chr10-20936278-DEL-185
chr10-20952734-DEL-62
chr10-21119743-DEL-56
chr10-21303327-DEL-68
chr10-21343147-DEL-190
chr10-21371765-DEL-194
chr10-21373902-DEL-52
chr10-21376300-DEL-60
chr10-21444116-DEL-63
chr10-21526374-DEL-1018
chr10-21526558-DEL-80
chr10-21571854-DEL-59
chr10-21641563-DEL-71
chr10-21696713-DEL-63
chr10-21739551-DEL-104
chr10-21751248-DEL-158
chr10-21751879-DEL-53
chr10-21754397-DEL-56
chr10-21762648-DEL-50
chr10-21772045-DEL-2018
chr10-21801259-DEL-142
chr10-21801914-DEL-55
chr10-21827236-DEL-101
chr10-21835750-DEL-747
chr10-21839616-DEL-91
chr10-22154210-DEL-66
chr10-22155535-DEL-329
chr10-22164956-DEL-1487
chr10-22173820-DEL-1401
chr10-22252151-DEL-50
chr10-22284278-DEL-81
chr10-22324205-DEL-72
chr10-22346331-DEL-923
chr10-22441839-DEL-61
chr10-22465399-DEL-891
chr10-22479498-DEL-64
chr10-22487404-DEL-107
chr10-22589386-DEL-55
chr10-22620692-DEL-106
chr10-22620898-DEL-130
chr10-22621324-DEL-238
chr10-22668235-DEL-985
chr10-22728622-DEL-57
chr10-22760025-DEL-365
chr10-22760321-DEL-168
chr10-22772608-DEL-666
chr10-22780682-DEL-57
chr10-22811309-DEL-332
chr10-22876002-DEL-138

chr10-22882384-DEL-1796
chr10-23154288-DEL-157
chr10-23218863-DEL-255
chr10-23219103-DEL-51
chr10-23232942-DEL-1046
chr10-23483756-DEL-100
chr10-23545069-DEL-1126
chr10-23573846-DEL-174
chr10-23574050-DEL-702
chr10-23639793-DEL-149
chr10-23667172-DEL-55
chr10-23669545-DEL-62
chr10-23684256-DEL-560
chr10-23698133-DEL-88
chr10-23772669-DEL-64
chr10-23794918-DEL-213
chr10-23803931-DEL-100
chr10-23804296-DEL-233
chr10-23920763-DEL-167
chr10-23922553-DEL-60
chr10-24046812-DEL-50
chr10-24097837-DEL-91
chr10-24140586-DEL-4664
chr10-24151202-DEL-50
chr10-24230138-DEL-882
chr10-24245585-DEL-228
chr10-24343890-DEL-1077
chr10-24406689-DEL-63
chr10-24413734-DEL-51
chr10-24417406-DEL-129
chr10-24476383-DEL-608
chr10-24476586-DEL-117
chr10-24477007-DEL-99
chr10-24530602-DEL-86
chr10-24547109-DEL-59
chr10-24581965-DEL-135
chr10-24689397-DEL-138
chr10-24690233-DEL-170
chr10-24714114-DEL-126
chr10-24755220-DEL-238
chr10-24795026-DEL-132
chr10-24795173-DEL-265
chr10-24799929-DEL-50
chr10-24826478-DEL-60
chr10-24868521-DEL-1767
chr10-24884614-DEL-1484
chr10-24906333-DEL-1057
chr10-24935257-DEL-55
chr10-24968191-DEL-174
chr10-24968452-DEL-232
chr10-24973811-DEL-53
chr10-25063283-DEL-404
chr10-25090301-DEL-1448
chr10-25093012-DEL-694
chr10-25140361-DEL-100
chr10-25157629-DEL-100
chr10-25158219-DEL-50
chr10-25213154-DEL-207
chr10-25334600-DEL-565
chr10-25334807-DEL-189
chr10-25349845-DEL-736
chr10-25351090-DEL-664
chr10-25351720-DEL-98
chr10-25436798-DEL-117
chr10-25444553-DEL-707
chr10-25604030-DEL-595
chr10-25689982-DEL-220
chr10-25715722-DEL-64
chr10-25737430-DEL-440
chr10-25782487-DEL-50
chr10-25803307-DEL-50
chr10-25905368-DEL-178
chr10-26035093-DEL-4363
chr10-26245115-DEL-50
chr10-26441503-DEL-50
chr10-26552741-DEL-1526
chr10-26557290-DEL-162

chr10-26559975-DEL-4244
chr10-26565353-DEL-102
chr10-26614200-DEL-51
chr10-26636893-DEL-52
chr10-26642737-DEL-66
chr10-26642909-DEL-114
chr10-26647652-DEL-54
chr10-26683883-DEL-63
chr10-26823748-DEL-257
chr10-26841461-DEL-64
chr10-27054510-DEL-86
chr10-27208210-DEL-85
chr10-27338039-DEL-2210
chr10-27451370-DEL-73
chr10-27520675-DEL-2146
chr10-27560947-DEL-50
chr10-27619851-DEL-1074
chr10-27644554-DEL-60
chr10-27865872-DEL-298
chr10-27898979-DEL-3201
chr10-27914430-DEL-67
chr10-27970732-DEL-124
chr10-27970780-DEL-52
chr10-28114983-DEL-92
chr10-28123605-DEL-3601
chr10-28147110-DEL-50
chr10-28292270-DEL-60
chr10-28335127-DEL-70
chr10-28412570-DEL-2844
chr10-28484901-DEL-50
chr10-28488112-DEL-52
chr10-28496310-DEL-102
chr10-28525847-DEL-76
chr10-28700975-DEL-143
chr10-28953031-DEL-6032
chr10-29069569-DEL-52
chr10-29160730-DEL-120
chr10-29292593-DEL-56
chr10-29316918-DEL-201
chr10-29413664-DEL-54
chr10-29574414-DEL-2296
chr10-29574729-DEL-648
chr10-29640910-DEL-261
chr10-29848256-DEL-137
chr10-30092013-DEL-424
chr10-30115633-DEL-776
chr10-30256347-DEL-63
chr10-30382670-DEL-61
chr10-30466336-DEL-434
chr10-30466624-DEL-124
chr10-30544805-DEL-150
chr10-30554740-DEL-142
chr10-30700570-DEL-52
chr10-30764189-DEL-1029
chr10-30765488-DEL-345
chr10-31078732-DEL-144
chr10-31091876-DEL-60
chr10-31275675-DEL-213
chr10-31316132-DEL-2041
chr10-31328479-DEL-929
chr10-31533037-DEL-314
chr10-31595930-DEL-155
chr10-31624645-DEL-62
chr10-31684434-DEL-596
chr10-31686081-DEL-50
chr10-31691732-DEL-50
chr10-31691811-DEL-6253
chr10-31754321-DEL-1183
chr10-31823669-DEL-51
chr10-31931203-DEL-72
chr10-32118240-DEL-219
chr10-32139722-DEL-881
chr10-32187475-DEL-8323
chr10-32229805-DEL-802
chr10-32231317-DEL-589
chr10-32299598-DEL-232
chr10-32299711-DEL-58

chr10-32363924-DEL-90
chr10-32518958-DEL-2130
chr10-32606416-DEL-103
chr10-32635254-DEL-102
chr10-32659762-DEL-517
chr10-32674678-DEL-415
chr10-32733311-DEL-232
chr10-32777455-DEL-68
chr10-32794418-DEL-6872
chr10-32841603-DEL-353
chr10-32882873-DEL-5935
chr10-32893144-DEL-92
chr10-32901399-DEL-61
chr10-32928868-DEL-57
chr10-33065134-DEL-169
chr10-33073903-DEL-52
chr10-33081444-DEL-2427
chr10-33115910-DEL-213
chr10-33230627-DEL-3324
chr10-33250200-DEL-50
chr10-33275485-DEL-53
chr10-33333175-DEL-289
chr10-33454986-DEL-1110
chr10-33474899-DEL-54
chr10-33608908-DEL-1899
chr10-33687144-DEL-69
chr10-33711550-DEL-3591
chr10-33722440-DEL-76
chr10-33854392-DEL-139
chr10-33948437-DEL-478
chr10-33995857-DEL-261
chr10-34146824-DEL-169
chr10-34165943-DEL-1113
chr10-34230063-DEL-284
chr10-34253919-DEL-207
chr10-34344257-DEL-190
chr10-34348505-DEL-138
chr10-34443835-DEL-964
chr10-34544980-DEL-56
chr10-34600908-DEL-68
chr10-34612838-DEL-116
chr10-34634590-DEL-88
chr10-34634979-DEL-61
chr10-34742511-DEL-222
chr10-34820803-DEL-82
chr10-34822116-DEL-98
chr10-34837000-DEL-116
chr10-34966239-DEL-112
chr10-35066722-DEL-3781
chr10-35076750-DEL-95
chr10-35224096-DEL-54
chr10-35241805-DEL-462
chr10-35242482-DEL-323
chr10-35339992-DEL-56
chr10-35532098-DEL-63
chr10-35553095-DEL-208
chr10-35606247-DEL-67
chr10-35623422-DEL-850
chr10-35652036-DEL-130
chr10-35842071-DEL-4385
chr10-35848427-DEL-66
chr10-35899035-DEL-3937
chr10-36027338-DEL-52
chr10-36178402-DEL-125
chr10-36553139-DEL-321
chr10-36786486-DEL-53
chr10-36786638-DEL-265
chr10-36878422-DEL-81
chr10-36913548-DEL-6461
chr10-36969085-DEL-61
chr10-37049698-DEL-58
chr10-37143748-DEL-305
chr10-37297647-DEL-122
chr10-37577737-DEL-3022
chr10-37750316-DEL-50
chr10-37751600-DEL-50
chr10-37780501-DEL-62

chr10-37915218-DEL-52
chr10-38156070-DEL-57
chr10-38175795-DEL-436
chr10-38262816-DEL-205
chr10-38266876-DEL-104
chr10-38307904-DEL-1391
chr10-38319531-DEL-55
chr10-38376624-DEL-8225
chr10-38521279-DEL-2107
chr10-38536794-DEL-773
chr10-38574902-DEL-51
chr10-38733546-DEL-71
chr10-38917787-DEL-363
chr10-38983190-DEL-84
chr10-39002051-DEL-4201
chr10-39015804-DEL-135
chr10-39035241-DEL-68
chr10-39036532-DEL-54
chr10-39036675-DEL-84
chr10-39078902-DEL-56
chr10-39308045-DEL-494
chr10-39351728-DEL-84
chr10-39474759-DEL-105
chr10-39495130-DEL-233
chr10-39541601-DEL-104
chr10-39675249-DEL-263
chr10-39687236-DEL-107
chr10-39751547-DEL-450
chr10-39795041-DEL-279
chr10-39795871-DEL-248
chr10-39829208-DEL-62
chr10-39847385-DEL-680
chr10-39867666-DEL-164
chr10-39892781-DEL-353
chr10-39918438-DEL-111
chr10-39960969-DEL-52
chr10-39991014-DEL-52
chr10-40004179-DEL-52
chr10-40030721-DEL-477
chr10-40030898-DEL-53
chr10-40031640-DEL-263
chr10-40031708-DEL-64
chr10-40065413-DEL-50
chr10-40078352-DEL-1684
chr10-40151131-DEL-50
chr10-40167333-DEL-196
chr10-40328623-DEL-90
chr10-40328698-DEL-299
chr10-40328892-DEL-147
chr10-40341352-DEL-126
chr10-40361750-DEL-64
chr10-40374111-DEL-58
chr10-40384366-DEL-54
chr10-40423393-DEL-52
chr10-40485276-DEL-1509
chr10-40522534-DEL-512
chr10-40526956-DEL-557
chr10-40558781-DEL-62
chr10-40705697-DEL-151
chr10-40706047-DEL-186
chr10-40793973-DEL-408
chr10-40815182-DEL-76
chr10-40894365-DEL-92
chr10-40959602-DEL-75
chr10-40971115-DEL-207
chr10-40971235-DEL-80
chr10-41039860-DEL-233
chr10-41068074-DEL-1656
chr10-41071088-DEL-78
chr10-41110766-DEL-395
chr10-41185984-DEL-5686
chr10-41192454-DEL-68
chr10-41339112-DEL-202
chr10-41436030-DEL-314
chr10-41522767-DEL-70
chr10-41667069-DEL-77
chr10-41700827-DEL-272

chr10-41775576-DEL-235
chr10-41775582-DEL-94
chr10-41995959-DEL-138
chr10-42013295-DEL-304
chr10-42027336-DEL-2044
chr10-42051132-DEL-105
chr10-42055470-DEL-271
chr10-42191044-DEL-407
chr10-42200057-DEL-50
chr10-42211743-DEL-279
chr10-42277536-DEL-780
chr10-42277997-DEL-720
chr10-42278424-DEL-210
chr10-42278536-DEL-990
chr10-42279242-DEL-270
chr10-42322810-DEL-50
chr10-42413020-DEL-2232
chr10-42415619-DEL-88
chr10-42586265-DEL-127
chr10-42728623-DEL-260
chr10-42728668-DEL-63
chr10-42728855-DEL-102
chr10-42768111-DEL-66
chr10-42800462-DEL-50
chr10-42931218-DEL-223
chr10-42931395-DEL-110
chr10-42933422-DEL-100
chr10-42939574-DEL-74
chr10-42978862-DEL-98
chr10-42991477-DEL-50
chr10-43079686-DEL-50
chr10-43143502-DEL-2921
chr10-43157101-DEL-99
chr10-43176072-DEL-90
chr10-43183450-DEL-112
chr10-43307206-DEL-55
chr10-43326627-DEL-100
chr10-43380602-DEL-84
chr10-43429053-DEL-463
chr10-43430428-DEL-1301
chr10-43486078-DEL-73
chr10-43494308-DEL-130
chr10-43554586-DEL-51
chr10-43576079-DEL-238
chr10-43632624-DEL-58
chr10-43814423-DEL-116
chr10-43847256-DEL-78
chr10-43851941-DEL-78
chr10-43857859-DEL-357
chr10-43861548-DEL-51
chr10-43868117-DEL-70
chr10-43890761-DEL-316
chr10-43917951-DEL-306
chr10-44007406-DEL-228
chr10-44007463-DEL-87
chr10-44267139-DEL-52
chr10-44274311-DEL-183
chr10-44610731-DEL-214
chr10-44658665-DEL-114
chr10-44703496-DEL-62
chr10-44710857-DEL-74
chr10-44743019-DEL-101
chr10-44803717-DEL-126
chr10-44884181-DEL-125
chr10-44893272-DEL-66
chr10-44992009-DEL-230
chr10-44992567-DEL-99
chr10-45011803-DEL-153
chr10-45122960-DEL-51
chr10-45164177-DEL-60
chr10-45167830-DEL-76
chr10-45282599-DEL-82
chr10-45299671-DEL-2564
chr10-45302451-DEL-60
chr10-45428957-DEL-112
chr10-45488700-DEL-115
chr10-45501058-DEL-211

chr10-45502124-DEL-679
chr10-45503183-DEL-87
chr10-45688306-DEL-50
chr10-45757837-DEL-156
chr10-45809673-DEL-78
chr10-45966615-DEL-54
chr10-46094333-DEL-54
chr10-46118224-DEL-70
chr10-46248144-DEL-94
chr10-46305553-DEL-58
chr10-46307952-DEL-1404
chr10-46344373-DEL-3633
chr10-46394430-DEL-86
chr10-46462855-DEL-208
chr10-46493755-DEL-126
chr10-46608963-DEL-67
chr10-46710372-DEL-973
chr10-46730850-DEL-290
chr10-46762523-DEL-2564
chr10-46772985-DEL-57
chr10-46794960-DEL-1453
chr10-46905390-DEL-138
chr10-47043388-DEL-114
chr10-47043955-DEL-225
chr10-47162863-DEL-225
chr10-47251754-DEL-1337
chr10-47302503-DEL-53
chr10-47325168-DEL-162
chr10-47403444-DEL-982
chr10-47406642-DEL-103
chr10-47436220-DEL-243
chr10-47460956-DEL-64
chr10-47493651-DEL-10889
chr10-47527858-DEL-1100
chr10-47579427-DEL-117
chr10-47586494-DEL-4052
chr10-47646957-DEL-81
chr10-47704033-DEL-620
chr10-47792449-DEL-567
chr10-47908296-DEL-2792
chr10-47948440-DEL-187
chr10-48010681-DEL-102
chr10-48032733-DEL-3479
chr10-48125126-DEL-61
chr10-48323048-DEL-1521
chr10-48383086-DEL-67
chr10-48384057-DEL-683
chr10-48410202-DEL-1484
chr10-48430044-DEL-131
chr10-48480447-DEL-113
chr10-48480458-DEL-56
chr10-48531366-DEL-424
chr10-48533536-DEL-193
chr10-48608063-DEL-217
chr10-48608111-DEL-88
chr10-48608374-DEL-52
chr10-48650857-DEL-209
chr10-48650858-DEL-102
chr10-48651239-DEL-231
chr10-48740063-DEL-666
chr10-48855464-DEL-432
chr10-48863146-DEL-134
chr10-48903325-DEL-186
chr10-48996170-DEL-110
chr10-49112766-DEL-50
chr10-49112892-DEL-755
chr10-49260900-DEL-77
chr10-49261062-DEL-110
chr10-49261356-DEL-78
chr10-49298737-DEL-1664
chr10-49443710-DEL-83
chr10-49456326-DEL-327
chr10-49461481-DEL-543
chr10-49614249-DEL-149
chr10-49629211-DEL-161
chr10-49724900-DEL-364
chr10-49729795-DEL-53

chr10-49877211-DEL-50
chr10-49934448-DEL-78
chr10-50197141-DEL-82
chr10-50197383-DEL-259
chr10-50305390-DEL-315
chr10-50309263-DEL-54
chr10-50367393-DEL-4157
chr10-50431019-DEL-66
chr10-50812254-DEL-60
chr10-50890262-DEL-302
chr10-51178111-DEL-1332
chr10-51217018-DEL-550
chr10-51248815-DEL-314
chr10-51352432-DEL-10125
chr10-51422833-DEL-1291
chr10-51424893-DEL-87
chr10-51426376-DEL-50
chr10-51474798-DEL-50
chr10-51566205-DEL-72
chr10-51620142-DEL-2011
chr10-51678084-DEL-50
chr10-51700824-DEL-252
chr10-51713092-DEL-94
chr10-51879636-DEL-201
chr10-51903313-DEL-50
chr10-51945064-DEL-1958
chr10-51963224-DEL-493
chr10-51992554-DEL-420
chr10-51992694-DEL-94
chr10-52081650-DEL-8173
chr10-52149832-DEL-89
chr10-52154031-DEL-61
chr10-52161712-DEL-50
chr10-52171117-DEL-163
chr10-52171337-DEL-56
chr10-52179314-DEL-56
chr10-52198758-DEL-949
chr10-52202374-DEL-3999
chr10-52245400-DEL-140
chr10-52397379-DEL-201
chr10-52430791-DEL-200
chr10-52511248-DEL-203
chr10-52522329-DEL-80
chr10-52563427-DEL-1302
chr10-52618321-DEL-113
chr10-52678540-DEL-61
chr10-52705122-DEL-126
chr10-52746895-DEL-65
chr10-52772955-DEL-91
chr10-52812156-DEL-60
chr10-52812214-DEL-158
chr10-52893120-DEL-54
chr10-52944072-DEL-63
chr10-53047536-DEL-52
chr10-53196483-DEL-53
chr10-53310244-DEL-52
chr10-53318435-DEL-614
chr10-53431965-DEL-2466
chr10-53440268-DEL-58
chr10-53457151-DEL-76
chr10-53518900-DEL-54
chr10-53529487-DEL-476
chr10-53651208-DEL-96
chr10-53659500-DEL-618
chr10-53719559-DEL-99
chr10-53803074-DEL-156
chr10-53803139-DEL-64
chr10-53860984-DEL-56
chr10-53863750-DEL-54
chr10-53871019-DEL-104
chr10-53945385-DEL-87
chr10-53986803-DEL-60
chr10-54040272-DEL-50
chr10-54067317-DEL-1788
chr10-54192797-DEL-66
chr10-54208427-DEL-50
chr10-54219007-DEL-95

chr10-54277221-DEL-54
chr10-54288737-DEL-485
chr10-54336647-DEL-1159
chr10-54375755-DEL-106
chr10-54592179-DEL-50
chr10-54609979-DEL-54
chr10-54750901-DEL-60
chr10-54788875-DEL-97
chr10-54791628-DEL-50
chr10-55065534-DEL-2125
chr10-55121247-DEL-2144
chr10-55192182-DEL-236
chr10-55331992-DEL-56
chr10-55364956-DEL-69
chr10-55380217-DEL-212
chr10-55390102-DEL-121
chr10-55514063-DEL-60
chr10-55603935-DEL-212
chr10-55621034-DEL-201
chr10-55847377-DEL-62
chr10-55862317-DEL-60
chr10-55977136-DEL-812
chr10-56014201-DEL-6224
chr10-56042466-DEL-148
chr10-56079199-DEL-50
chr10-56092607-DEL-52
chr10-56108205-DEL-57
chr10-56126091-DEL-474
chr10-56226694-DEL-56
chr10-56429860-DEL-878
chr10-56434942-DEL-87
chr10-56618106-DEL-537
chr10-56647875-DEL-56
chr10-56709022-DEL-170
chr10-56713668-DEL-56
chr10-56714911-DEL-66
chr10-56770159-DEL-3198
chr10-56886810-DEL-148
chr10-56925781-DEL-680
chr10-57027739-DEL-2028
chr10-57137850-DEL-267
chr10-57137853-DEL-63
chr10-57138168-DEL-68
chr10-57138395-DEL-105
chr10-57172394-DEL-736
chr10-57243612-DEL-524
chr10-57280297-DEL-152
chr10-57314707-DEL-456
chr10-57316232-DEL-185
chr10-57326648-DEL-6142
chr10-57370044-DEL-187
chr10-57370650-DEL-247
chr10-57422114-DEL-161
chr10-57436128-DEL-103
chr10-57589405-DEL-66
chr10-58023537-DEL-3681
chr10-58065164-DEL-57
chr10-58117452-DEL-151
chr10-58145266-DEL-3563
chr10-58161858-DEL-593
chr10-58182184-DEL-2448
chr10-58399761-DEL-79
chr10-58406539-DEL-116
chr10-58420231-DEL-65
chr10-58420370-DEL-225
chr10-58448299-DEL-237
chr10-58498267-DEL-604
chr10-58500216-DEL-83
chr10-58512365-DEL-118
chr10-58544097-DEL-65
chr10-58604477-DEL-3572
chr10-58675758-DEL-125
chr10-58760823-DEL-144
chr10-58760919-DEL-528
chr10-58761399-DEL-96
chr10-58766679-DEL-100
chr10-58853585-DEL-78

chr10-58869724-DEL-603
chr10-58957079-DEL-60
chr10-59093991-DEL-79
chr10-59181407-DEL-60
chr10-59314121-DEL-96
chr10-59322276-DEL-62
chr10-59332547-DEL-186
chr10-59334960-DEL-108
chr10-59358624-DEL-114
chr10-59442428-DEL-267
chr10-59486257-DEL-294
chr10-59503014-DEL-1331
chr10-59521692-DEL-125
chr10-59521848-DEL-151
chr10-59531377-DEL-3252
chr10-59703569-DEL-943
chr10-59766494-DEL-1591
chr10-59768133-DEL-56
chr10-59799282-DEL-50
chr10-59844586-DEL-74
chr10-59901211-DEL-76
chr10-59917611-DEL-122
chr10-59947929-DEL-55
chr10-59947930-DEL-117
chr10-60020479-DEL-70
chr10-60088069-DEL-76
chr10-60088337-DEL-56
chr10-60254347-DEL-59
chr10-60304497-DEL-54
chr10-60311430-DEL-70
chr10-60537656-DEL-93
chr10-60689699-DEL-560
chr10-60845144-DEL-112
chr10-60878322-DEL-498
chr10-60906123-DEL-50
chr10-60918353-DEL-52
chr10-60919524-DEL-105
chr10-60919693-DEL-231
chr10-61052866-DEL-60
chr10-61063748-DEL-52
chr10-61095154-DEL-54
chr10-61095374-DEL-96
chr10-61107326-DEL-51
chr10-61113440-DEL-99
chr10-61239416-DEL-65
chr10-61249986-DEL-92
chr10-61259592-DEL-53
chr10-61295334-DEL-177
chr10-61295621-DEL-279
chr10-61327429-DEL-50
chr10-61415595-DEL-142
chr10-61539840-DEL-230
chr10-61647179-DEL-98
chr10-61796735-DEL-82
chr10-61804389-DEL-192
chr10-62042441-DEL-134
chr10-62042912-DEL-143
chr10-62147955-DEL-166
chr10-62164571-DEL-362
chr10-62220322-DEL-55
chr10-62220457-DEL-292
chr10-62329343-DEL-92
chr10-62390484-DEL-244
chr10-62409623-DEL-84
chr10-62414904-DEL-178
chr10-62611191-DEL-50
chr10-62739480-DEL-56
chr10-62745923-DEL-1941
chr10-62760648-DEL-216
chr10-62770649-DEL-60
chr10-62772102-DEL-50
chr10-62804155-DEL-62
chr10-62804285-DEL-215
chr10-62868034-DEL-50
chr10-62873285-DEL-52
chr10-62929492-DEL-60
chr10-63027726-DEL-79

chr10-63221440-DEL-128
chr10-63221655-DEL-203
chr10-63372456-DEL-66
chr10-63404043-DEL-191
chr10-63409541-DEL-157
chr10-63413403-DEL-59
chr10-63547557-DEL-398
chr10-63891141-DEL-192
chr10-63891159-DEL-84
chr10-64120422-DEL-51
chr10-64185152-DEL-74
chr10-64239803-DEL-69
chr10-64290716-DEL-155
chr10-64497538-DEL-60
chr10-64573338-DEL-62
chr10-64612681-DEL-8864
chr10-64649810-DEL-96
chr10-64650070-DEL-112
chr10-64888542-DEL-50
chr10-64959894-DEL-352
chr10-64960189-DEL-84
chr10-64963516-DEL-50
chr10-64975972-DEL-64
chr10-65135046-DEL-56
chr10-65353901-DEL-97
chr10-65418859-DEL-50
chr10-65571923-DEL-70
chr10-65601524-DEL-438
chr10-65894332-DEL-54
chr10-65918359-DEL-780
chr10-65972415-DEL-190
chr10-66108256-DEL-169
chr10-66325703-DEL-70
chr10-66329740-DEL-90
chr10-66329923-DEL-169
chr10-66330024-DEL-68
chr10-66347122-DEL-258
chr10-66348001-DEL-74
chr10-66389187-DEL-61
chr10-66411540-DEL-64
chr10-66592527-DEL-73
chr10-66607147-DEL-187
chr10-66665577-DEL-240
chr10-66723788-DEL-108
chr10-66806538-DEL-1253
chr10-66844374-DEL-65
chr10-67095580-DEL-347
chr10-67099106-DEL-557
chr10-67099233-DEL-174
chr10-67099648-DEL-75
chr10-67099698-DEL-310
chr10-67122913-DEL-50
chr10-67230484-DEL-76
chr10-67264805-DEL-50
chr10-67284788-DEL-2818
chr10-67479964-DEL-119
chr10-67540224-DEL-54
chr10-67644398-DEL-283
chr10-67644694-DEL-226
chr10-67645135-DEL-179
chr10-68060987-DEL-300
chr10-68515068-DEL-62
chr10-68585141-DEL-58
chr10-68597260-DEL-2140
chr10-68597353-DEL-4281
chr10-68668785-DEL-72
chr10-68697508-DEL-168
chr10-68716994-DEL-64
chr10-68723144-DEL-50
chr10-68763683-DEL-86
chr10-68778072-DEL-100
chr10-68840070-DEL-475
chr10-68853015-DEL-63
chr10-68855638-DEL-75
chr10-68903107-DEL-50
chr10-68903129-DEL-134
chr10-68938356-DEL-373

chr10-68938786-DEL-56
chr10-68948437-DEL-188
chr10-69039825-DEL-192
chr10-69060714-DEL-56
chr10-69166276-DEL-58
chr10-69233761-DEL-77
chr10-69268015-DEL-54
chr10-69293044-DEL-694
chr10-69313235-DEL-52
chr10-69395838-DEL-110
chr10-69400863-DEL-123
chr10-69411722-DEL-56
chr10-69450308-DEL-134
chr10-69450408-DEL-52
chr10-69505901-DEL-74
chr10-69505948-DEL-152
chr10-69508964-DEL-108
chr10-69554265-DEL-51
chr10-69570953-DEL-132
chr10-69571158-DEL-82
chr10-69583247-DEL-71
chr10-69590652-DEL-76
chr10-69601985-DEL-67
chr10-69731331-DEL-50
chr10-69743812-DEL-96
chr10-69785299-DEL-51
chr10-69952782-DEL-56
chr10-70107571-DEL-64
chr10-70136085-DEL-497
chr10-70147553-DEL-301
chr10-70147675-DEL-93
chr10-70147966-DEL-93
chr10-70187422-DEL-194
chr10-70249457-DEL-106
chr10-70261219-DEL-104
chr10-70336772-DEL-50
chr10-70605051-DEL-167
chr10-70605315-DEL-52
chr10-70688639-DEL-322
chr10-70732911-DEL-98
chr10-70733130-DEL-104
chr10-70768533-DEL-465
chr10-70837403-DEL-58
chr10-70858741-DEL-67
chr10-70918447-DEL-84
chr10-71033055-DEL-50
chr10-71076639-DEL-181
chr10-71103707-DEL-143
chr10-71105007-DEL-89
chr10-71151283-DEL-3684
chr10-71238859-DEL-1937
chr10-71259470-DEL-55
chr10-71266534-DEL-54
chr10-71328630-DEL-60
chr10-71525761-DEL-50
chr10-71554118-DEL-161
chr10-71554521-DEL-80
chr10-71579239-DEL-60
chr10-71607482-DEL-74
chr10-71710643-DEL-163
chr10-71721126-DEL-56
chr10-71738367-DEL-3128
chr10-71762573-DEL-466
chr10-71762706-DEL-91
chr10-71812762-DEL-76
chr10-71953617-DEL-84
chr10-72010897-DEL-198
chr10-72104428-DEL-102
chr10-72128733-DEL-58
chr10-72133014-DEL-73
chr10-72244362-DEL-52
chr10-72327114-DEL-361
chr10-72327115-DEL-161
chr10-72328737-DEL-214
chr10-72364957-DEL-95
chr10-72716610-DEL-56
chr10-72785321-DEL-140

chr10-72928408-DEL-54
chr10-72941486-DEL-109
chr10-72989466-DEL-60
chr10-73178517-DEL-85
chr10-73195431-DEL-52
chr10-73273016-DEL-56
chr10-73420243-DEL-56
chr10-73509367-DEL-2787
chr10-73607924-DEL-50
chr10-73624585-DEL-1126
chr10-73649374-DEL-1055
chr10-73663485-DEL-60
chr10-73668665-DEL-71
chr10-73762305-DEL-53
chr10-73831725-DEL-72
chr10-73834438-DEL-90
chr10-73901028-DEL-95
chr10-73944553-DEL-62
chr10-73950093-DEL-2506
chr10-74035754-DEL-101
chr10-74053619-DEL-52
chr10-74155226-DEL-247
chr10-74307048-DEL-76
chr10-74354708-DEL-50
chr10-74486727-DEL-50
chr10-74575117-DEL-7314
chr10-74586501-DEL-269
chr10-74590109-DEL-1519
chr10-74607116-DEL-2225
chr10-74703985-DEL-143
chr10-74704013-DEL-52
chr10-74728316-DEL-275
chr10-74728441-DEL-125
chr10-74756102-DEL-79
chr10-74880680-DEL-52
chr10-75008125-DEL-50
chr10-75176466-DEL-121
chr10-75255164-DEL-112
chr10-75386635-DEL-52
chr10-75543648-DEL-72
chr10-75559705-DEL-275
chr10-75559789-DEL-55
chr10-75627096-DEL-116
chr10-75627319-DEL-144
chr10-75645362-DEL-58
chr10-75699966-DEL-6463
chr10-75893367-DEL-51
chr10-75907773-DEL-278
chr10-75908703-DEL-405
chr10-75952284-DEL-68
chr10-76023071-DEL-144
chr10-76082162-DEL-291
chr10-76142112-DEL-78
chr10-76146249-DEL-659
chr10-76146403-DEL-139
chr10-76146887-DEL-134
chr10-76147125-DEL-451
chr10-76147872-DEL-637
chr10-76148011-DEL-52
chr10-76160865-DEL-1398
chr10-76166564-DEL-420
chr10-76169273-DEL-56
chr10-76194621-DEL-52
chr10-76401638-DEL-248
chr10-76455371-DEL-52
chr10-76467431-DEL-84
chr10-76467627-DEL-55
chr10-76523853-DEL-85
chr10-76845527-DEL-156
chr10-76920817-DEL-54
chr10-76953184-DEL-94
chr10-76985479-DEL-130
chr10-76992157-DEL-56
chr10-77066805-DEL-76
chr10-77090026-DEL-129
chr10-77153357-DEL-52
chr10-77164137-DEL-129

chr10-77164483-DEL-66
chr10-77213077-DEL-270
chr10-77444843-DEL-79
chr10-77606300-DEL-11217
chr10-77612240-DEL-5598
chr10-77782896-DEL-167
chr10-77785689-DEL-737
chr10-77794462-DEL-1131
chr10-77967922-DEL-56
chr10-77974283-DEL-76
chr10-77986138-DEL-235
chr10-78181135-DEL-53
chr10-78214735-DEL-1459
chr10-78216392-DEL-202
chr10-78230086-DEL-64
chr10-78290407-DEL-146
chr10-78313844-DEL-52
chr10-78342995-DEL-234
chr10-78438430-DEL-70
chr10-78440851-DEL-51
chr10-78503640-DEL-54
chr10-78567313-DEL-4314
chr10-78608948-DEL-77
chr10-78660298-DEL-460
chr10-79043497-DEL-4890
chr10-79085737-DEL-199
chr10-79137255-DEL-2502
chr10-79424370-DEL-1037
chr10-79425483-DEL-77
chr10-79677439-DEL-53
chr10-79795519-DEL-91
chr10-79924426-DEL-201
chr10-80049269-DEL-156
chr10-80049325-DEL-429
chr10-80117768-DEL-56
chr10-80300141-DEL-148
chr10-80324028-DEL-78
chr10-80340488-DEL-50
chr10-80485949-DEL-50
chr10-80519993-DEL-198
chr10-80658727-DEL-54
chr10-80696151-DEL-211
chr10-80728386-DEL-201
chr10-80728395-DEL-96
chr10-81005326-DEL-88
chr10-81042986-DEL-60
chr10-81343129-DEL-102
chr10-81493573-DEL-145
chr10-81573221-DEL-199
chr10-81642032-DEL-84
chr10-82085680-DEL-2832
chr10-82246429-DEL-73
chr10-82247291-DEL-6361
chr10-82353191-DEL-1493
chr10-82481475-DEL-61
chr10-82625029-DEL-5674
chr10-82639003-DEL-71
chr10-82641591-DEL-382
chr10-82647980-DEL-58
chr10-82723638-DEL-55
chr10-82734669-DEL-160
chr10-82759763-DEL-90
chr10-82884555-DEL-142
chr10-82973678-DEL-445
chr10-83042725-DEL-1226
chr10-83075022-DEL-66
chr10-83081850-DEL-62
chr10-83098495-DEL-80
chr10-83144364-DEL-1066
chr10-83739669-DEL-69
chr10-83740071-DEL-61
chr10-83824430-DEL-950
chr10-83958993-DEL-86
chr10-83959278-DEL-113
chr10-84020428-DEL-83
chr10-84038158-DEL-59
chr10-84067695-DEL-50

chr10-84342390-DEL-50
chr10-84358170-DEL-120
chr10-84462758-DEL-1080
chr10-84470221-DEL-108
chr10-84491116-DEL-378
chr10-84491201-DEL-131
chr10-84499952-DEL-178
chr10-84602808-DEL-351
chr10-84644166-DEL-62
chr10-84719533-DEL-50
chr10-84779960-DEL-94
chr10-85082556-DEL-56
chr10-85117221-DEL-290
chr10-85228942-DEL-83
chr10-85251263-DEL-4069
chr10-85357194-DEL-156
chr10-85729633-DEL-1264
chr10-85730936-DEL-275
chr10-85761664-DEL-78
chr10-85791856-DEL-396
chr10-85932708-DEL-393
chr10-85961226-DEL-83
chr10-85967707-DEL-69
chr10-85991475-DEL-1456
chr10-86041218-DEL-89
chr10-86051573-DEL-54
chr10-86286046-DEL-57
chr10-86407567-DEL-1284
chr10-86547580-DEL-58
chr10-86623847-DEL-140
chr10-86676156-DEL-57
chr10-86681266-DEL-84
chr10-86715916-DEL-50
chr10-86728567-DEL-56
chr10-86764897-DEL-50
chr10-86796758-DEL-317
chr10-86802708-DEL-111
chr10-86860658-DEL-114
chr10-86978253-DEL-3159
chr10-87037920-DEL-59
chr10-87117518-DEL-138
chr10-87117653-DEL-63
chr10-87273097-DEL-4745
chr10-87306790-DEL-50
chr10-87307393-DEL-56
chr10-87418169-DEL-181
chr10-87418440-DEL-172
chr10-87537189-DEL-98
chr10-87537457-DEL-191
chr10-87600385-DEL-51
chr10-87752461-DEL-88
chr10-87846200-DEL-560
chr10-87866772-DEL-157
chr10-88030156-DEL-64
chr10-88066715-DEL-50
chr10-88167850-DEL-78
chr10-88183745-DEL-53
chr10-88287737-DEL-107
chr10-88327891-DEL-50
chr10-88384484-DEL-85
chr10-88463154-DEL-102
chr10-88494193-DEL-54
chr10-88494370-DEL-157
chr10-88520073-DEL-51
chr10-88715838-DEL-92
chr10-88769750-DEL-6577
chr10-88783114-DEL-224
chr10-88794643-DEL-1302
chr10-88843436-DEL-481
chr10-89069356-DEL-695
chr10-89105707-DEL-130
chr10-89191070-DEL-236
chr10-89448167-DEL-64
chr10-89610345-DEL-50
chr10-89649140-DEL-680
chr10-89660548-DEL-487
chr10-89661004-DEL-360

chr10-89661184-DEL-63
chr10-89789889-DEL-9150
chr10-89867094-DEL-191
chr10-89872867-DEL-107
chr10-89971605-DEL-113
chr10-89996136-DEL-463
chr10-90023169-DEL-172
chr10-90039726-DEL-124
chr10-90092965-DEL-500
chr10-90093315-DEL-100
chr10-90095260-DEL-138
chr10-90322692-DEL-50
chr10-90497955-DEL-1933
chr10-90722163-DEL-50
chr10-90786543-DEL-50
chr10-91005762-DEL-101
chr10-91053435-DEL-144
chr10-91110590-DEL-217
chr10-91200910-DEL-69
chr10-91239454-DEL-464
chr10-91379889-DEL-166
chr10-91650322-DEL-2652
chr10-91683014-DEL-102
chr10-91741421-DEL-50
chr10-91809663-DEL-97
chr10-91997910-DEL-50
chr10-92143434-DEL-74
chr10-92361248-DEL-65
chr10-92390969-DEL-50
chr10-92418761-DEL-52
chr10-92430281-DEL-55
chr10-92480997-DEL-51
chr10-92576168-DEL-158
chr10-92861064-DEL-87
chr10-92879441-DEL-66
chr10-92934945-DEL-83
chr10-92977487-DEL-404
chr10-92985838-DEL-72
chr10-92990098-DEL-113
chr10-93034247-DEL-185
chr10-93037776-DEL-144
chr10-93058644-DEL-174
chr10-93058828-DEL-688
chr10-93158542-DEL-171
chr10-93158867-DEL-185
chr10-93221709-DEL-124
chr10-93263898-DEL-157
chr10-93264013-DEL-57
chr10-93269318-DEL-125
chr10-93353273-DEL-93
chr10-93455498-DEL-156
chr10-93461171-DEL-143
chr10-93500035-DEL-66
chr10-93562691-DEL-690
chr10-93610790-DEL-435
chr10-93616690-DEL-64
chr10-93642822-DEL-202
chr10-93687989-DEL-205
chr10-93765715-DEL-146
chr10-93844493-DEL-3729
chr10-94053589-DEL-115
chr10-94079435-DEL-201
chr10-94109018-DEL-131
chr10-94122575-DEL-100
chr10-94124348-DEL-52
chr10-94248764-DEL-56
chr10-94399717-DEL-84
chr10-94460781-DEL-135
chr10-94705250-DEL-114
chr10-94766170-DEL-98
chr10-94792459-DEL-100
chr10-94831800-DEL-102
chr10-94865819-DEL-68
chr10-94900811-DEL-259
chr10-94921023-DEL-69
chr10-95001308-DEL-2058
chr10-95015251-DEL-58

chr10-95036941-DEL-69
chr10-95098643-DEL-125
chr10-95122238-DEL-115
chr10-95137414-DEL-94
chr10-95141515-DEL-230
chr10-95254331-DEL-855
chr10-95264802-DEL-198
chr10-95347641-DEL-75
chr10-95395014-DEL-55
chr10-95396620-DEL-1308
chr10-95420048-DEL-1049
chr10-95533574-DEL-609
chr10-95536153-DEL-78
chr10-95587390-DEL-70
chr10-95607935-DEL-72
chr10-95648344-DEL-64
chr10-95648568-DEL-468
chr10-95702735-DEL-155
chr10-96015505-DEL-67
chr10-96072195-DEL-114
chr10-96362060-DEL-63
chr10-96376578-DEL-88
chr10-96411717-DEL-255
chr10-96420744-DEL-70
chr10-96485468-DEL-160
chr10-96545043-DEL-78
chr10-96619714-DEL-97
chr10-96665485-DEL-364
chr10-96727378-DEL-67
chr10-96799274-DEL-62
chr10-96835110-DEL-199
chr10-96861521-DEL-2543
chr10-97167898-DEL-140
chr10-97241845-DEL-66
chr10-97381163-DEL-116
chr10-97461706-DEL-385
chr10-97631680-DEL-253
chr10-98011337-DEL-802
chr10-98054461-DEL-317
chr10-98123628-DEL-163
chr10-98133853-DEL-125
chr10-98145714-DEL-1044
chr10-98173308-DEL-201
chr10-98247332-DEL-182
chr10-98247377-DEL-56
chr10-98254146-DEL-61
chr10-98254670-DEL-50
chr10-98332961-DEL-1360
chr10-98355770-DEL-83
chr10-98368546-DEL-56
chr10-98441039-DEL-1291
chr10-98489127-DEL-88
chr10-98612584-DEL-450
chr10-98649273-DEL-1467
chr10-98711593-DEL-89
chr10-98892123-DEL-58
chr10-98892873-DEL-1477
chr10-98950655-DEL-570
chr10-98965882-DEL-50
chr10-99006527-DEL-108
chr10-99006572-DEL-418
chr10-99041370-DEL-325
chr10-99164367-DEL-83
chr10-99198755-DEL-56
chr10-99241407-DEL-60
chr10-99253882-DEL-251
chr10-99301592-DEL-1657
chr10-99378884-DEL-309
chr10-99379258-DEL-71
chr10-99590341-DEL-57
chr10-99590789-DEL-206
chr10-99737983-DEL-313
chr10-99749902-DEL-114
chr10-99755443-DEL-150
chr10-99790397-DEL-145
chr10-99892923-DEL-56
chr10-100040203-DEL-242

chr10-100051719-DEL-1828
chr10-100139430-DEL-57
chr10-100139622-DEL-1871
chr10-100140496-DEL-251
chr10-100141328-DEL-219
chr10-100356008-DEL-484
chr10-100468605-DEL-55
chr10-100666971-DEL-2150
chr10-100680217-DEL-460
chr10-100680709-DEL-1352
chr10-100684798-DEL-61
chr10-100765313-DEL-52
chr10-100782978-DEL-50
chr10-100841316-DEL-71
chr10-100860360-DEL-56
chr10-100888378-DEL-90
chr10-101090860-DEL-863
chr10-101115597-DEL-64
chr10-101141004-DEL-325
chr10-101212926-DEL-52
chr10-101223529-DEL-1280
chr10-101314859-DEL-1117
chr10-101444305-DEL-81
chr10-101519089-DEL-113
chr10-101519390-DEL-452
chr10-101529156-DEL-50
chr10-101665362-DEL-58
chr10-101689307-DEL-201
chr10-101723078-DEL-50
chr10-101772336-DEL-946
chr10-101792956-DEL-977
chr10-101812828-DEL-829
chr10-101905211-DEL-55
chr10-101930771-DEL-672
chr10-101973150-DEL-74
chr10-101997300-DEL-77
chr10-102006975-DEL-143
chr10-102019611-DEL-252
chr10-102023001-DEL-67
chr10-102179995-DEL-1775
chr10-102186974-DEL-157
chr10-102312311-DEL-74
chr10-102321807-DEL-70
chr10-102347172-DEL-53
chr10-102472782-DEL-55
chr10-102566953-DEL-77
chr10-102684207-DEL-83
chr10-102721458-DEL-360
chr10-102721628-DEL-68
chr10-102729519-DEL-130
chr10-102754229-DEL-97
chr10-103028226-DEL-50
chr10-103069000-DEL-192
chr10-103098552-DEL-8710
chr10-103133795-DEL-377
chr10-103163452-DEL-50
chr10-103309586-DEL-1950
chr10-103324738-DEL-52
chr10-103325655-DEL-624
chr10-103458046-DEL-87
chr10-103477141-DEL-1616
chr10-103734843-DEL-203
chr10-103740095-DEL-3121
chr10-103928095-DEL-466
chr10-103994524-DEL-42553
chr10-104324838-DEL-246
chr10-104338764-DEL-50
chr10-104360695-DEL-838
chr10-104370173-DEL-52
chr10-104760698-DEL-1276
chr10-104951239-DEL-699
chr10-104975339-DEL-895
chr10-104979010-DEL-59
chr10-104979103-DEL-62
chr10-105056406-DEL-152
chr10-105256357-DEL-63
chr10-105286581-DEL-50

chr10-105324262-DEL-87
chr10-105351870-DEL-58
chr10-105357271-DEL-83
chr10-105437553-DEL-481
chr10-105448831-DEL-71
chr10-105449053-DEL-504
chr10-105493018-DEL-52
chr10-105554556-DEL-51
chr10-105602707-DEL-445
chr10-105636002-DEL-64
chr10-105642791-DEL-50
chr10-105702316-DEL-75
chr10-105738663-DEL-80
chr10-105766639-DEL-52
chr10-105782438-DEL-224
chr10-105811848-DEL-217
chr10-105876149-DEL-153
chr10-105877968-DEL-218
chr10-105915559-DEL-109
chr10-106108351-DEL-51
chr10-106263314-DEL-140
chr10-106280668-DEL-138
chr10-106284430-DEL-221
chr10-106360965-DEL-56
chr10-106393708-DEL-65
chr10-106430213-DEL-123
chr10-106496140-DEL-2763
chr10-106560554-DEL-51
chr10-106838411-DEL-143
chr10-106921731-DEL-78
chr10-106965486-DEL-7323
chr10-107007606-DEL-301
chr10-107101989-DEL-584
chr10-107214448-DEL-297
chr10-107273796-DEL-134
chr10-107295617-DEL-54
chr10-107382702-DEL-50
chr10-107382704-DEL-156
chr10-107439685-DEL-83
chr10-107478194-DEL-85
chr10-107497009-DEL-79
chr10-107497156-DEL-429
chr10-107497455-DEL-86
chr10-107589311-DEL-63
chr10-107657102-DEL-52
chr10-107731866-DEL-81
chr10-107732066-DEL-56
chr10-107878651-DEL-325
chr10-107923041-DEL-120
chr10-107923887-DEL-69
chr10-107948492-DEL-116
chr10-107990375-DEL-282
chr10-107990591-DEL-90
chr10-107996447-DEL-78
chr10-108036367-DEL-144
chr10-108094601-DEL-151
chr10-108124268-DEL-55
chr10-108183703-DEL-118
chr10-108196196-DEL-78
chr10-108266966-DEL-213
chr10-108274045-DEL-70
chr10-108342422-DEL-997
chr10-108363812-DEL-239
chr10-108369594-DEL-60
chr10-108465402-DEL-110
chr10-108465759-DEL-89
chr10-108554798-DEL-54
chr10-108591239-DEL-101
chr10-108647640-DEL-50
chr10-108704327-DEL-1570
chr10-108725888-DEL-665
chr10-108803456-DEL-60
chr10-108831963-DEL-1493
chr10-108902998-DEL-1739
chr10-108994508-DEL-95
chr10-109186364-DEL-2172
chr10-109252066-DEL-2213

chr10-109289364-DEL-137
chr10-109450517-DEL-6147
chr10-109496669-DEL-52
chr10-109498057-DEL-56
chr10-109626219-DEL-1047
chr10-109686338-DEL-326
chr10-109793377-DEL-74
chr10-109800650-DEL-75
chr10-109959002-DEL-171
chr10-109959252-DEL-92
chr10-109962315-DEL-76
chr10-109977634-DEL-97
chr10-109986051-DEL-1002
chr10-110068711-DEL-62
chr10-110109769-DEL-90
chr10-110109896-DEL-240
chr10-110224260-DEL-71
chr10-110328128-DEL-50
chr10-110339860-DEL-12629
chr10-110552594-DEL-52
chr10-110654183-DEL-88
chr10-110742800-DEL-113
chr10-110747046-DEL-55
chr10-110855415-DEL-51
chr10-110871487-DEL-195
chr10-110918076-DEL-1617
chr10-110927979-DEL-996
chr10-110940559-DEL-80
chr10-111026636-DEL-352
chr10-111039858-DEL-52
chr10-111053605-DEL-132
chr10-111068459-DEL-69
chr10-111161902-DEL-159
chr10-111170943-DEL-76
chr10-111219087-DEL-99
chr10-111287339-DEL-212
chr10-111287634-DEL-847
chr10-111371937-DEL-90
chr10-111377877-DEL-60
chr10-111408162-DEL-1981
chr10-111457793-DEL-160
chr10-111562482-DEL-59
chr10-111582681-DEL-1150
chr10-111584420-DEL-56
chr10-111633724-DEL-66
chr10-111676659-DEL-74
chr10-111688187-DEL-280
chr10-111762890-DEL-236
chr10-111783430-DEL-82
chr10-111839911-DEL-56
chr10-111941399-DEL-59
chr10-111974559-DEL-321
chr10-112003142-DEL-7668
chr10-112010811-DEL-58
chr10-112159107-DEL-7857
chr10-112159240-DEL-220
chr10-112260998-DEL-211
chr10-112467151-DEL-50
chr10-112486160-DEL-51
chr10-112510559-DEL-54
chr10-112535097-DEL-639
chr10-112657794-DEL-456
chr10-112957454-DEL-72
chr10-113041120-DEL-543
chr10-113074917-DEL-102
chr10-113075032-DEL-538
chr10-113161791-DEL-84
chr10-113285113-DEL-261
chr10-113304698-DEL-53
chr10-113476315-DEL-1255
chr10-113476516-DEL-101
chr10-113477237-DEL-251
chr10-113747882-DEL-150
chr10-113750037-DEL-107
chr10-113751929-DEL-202
chr10-113752633-DEL-55
chr10-113837498-DEL-52

chr10-113949754-DEL-68
chr10-114096434-DEL-50
chr10-114114388-DEL-898
chr10-114122903-DEL-61
chr10-114152707-DEL-666
chr10-114198310-DEL-50
chr10-114263320-DEL-52
chr10-114263668-DEL-98
chr10-114286924-DEL-51
chr10-114287256-DEL-59
chr10-114371397-DEL-424
chr10-114451086-DEL-89
chr10-114484664-DEL-63
chr10-114558363-DEL-51
chr10-114683953-DEL-60
chr10-114723565-DEL-50
chr10-114867135-DEL-1173
chr10-115006876-DEL-58
chr10-115027922-DEL-2907
chr10-115076139-DEL-186
chr10-115088897-DEL-484
chr10-115119301-DEL-52
chr10-115241972-DEL-50
chr10-115271341-DEL-95
chr10-115271565-DEL-176
chr10-115271938-DEL-61
chr10-115319074-DEL-145
chr10-115320027-DEL-88
chr10-115408901-DEL-274
chr10-115438301-DEL-64
chr10-115466004-DEL-60
chr10-115559611-DEL-1156
chr10-115575666-DEL-50
chr10-115731113-DEL-200
chr10-115756516-DEL-122
chr10-115824956-DEL-354
chr10-115834857-DEL-292
chr10-115834926-DEL-130
chr10-115893600-DEL-108
chr10-115943655-DEL-62
chr10-116031715-DEL-120
chr10-116075951-DEL-225
chr10-116148026-DEL-208
chr10-116253087-DEL-57
chr10-116268135-DEL-89
chr10-116337364-DEL-58
chr10-116337372-DEL-179
chr10-116394952-DEL-167
chr10-116432267-DEL-108
chr10-116432429-DEL-392
chr10-116503373-DEL-88
chr10-116662244-DEL-51
chr10-116682046-DEL-84
chr10-116743196-DEL-2045
chr10-116748348-DEL-50
chr10-116759900-DEL-234
chr10-116760010-DEL-60
chr10-116846218-DEL-59
chr10-116853232-DEL-570
chr10-116876635-DEL-228
chr10-116894359-DEL-108
chr10-116894603-DEL-156
chr10-117010164-DEL-102
chr10-117026457-DEL-52
chr10-117049450-DEL-54
chr10-117165029-DEL-72
chr10-117201487-DEL-157
chr10-117227425-DEL-273
chr10-117235482-DEL-72
chr10-117316858-DEL-146
chr10-117370478-DEL-348
chr10-117418727-DEL-92
chr10-117450675-DEL-50
chr10-117456012-DEL-76
chr10-117472511-DEL-442
chr10-117486959-DEL-129
chr10-117513211-DEL-153

chr10-117529224-DEL-216
chr10-117554542-DEL-84
chr10-117559281-DEL-1018
chr10-117561118-DEL-71
chr10-117561881-DEL-61
chr10-117609675-DEL-114
chr10-117625403-DEL-178
chr10-117810058-DEL-60
chr10-117815206-DEL-60
chr10-117894176-DEL-424
chr10-117894176-DEL-888
chr10-117924447-DEL-833
chr10-117989150-DEL-102
chr10-118035621-DEL-50
chr10-118074185-DEL-3179
chr10-118126704-DEL-321
chr10-118162808-DEL-21710
chr10-118340514-DEL-180
chr10-118349199-DEL-234
chr10-118380972-DEL-136
chr10-118381276-DEL-158
chr10-118381293-DEL-58
chr10-118385412-DEL-107
chr10-118430946-DEL-107
chr10-118461225-DEL-310
chr10-118487009-DEL-119
chr10-118489353-DEL-75
chr10-118526170-DEL-5135
chr10-118544937-DEL-282
chr10-118563698-DEL-50
chr10-118592284-DEL-50
chr10-118744497-DEL-68
chr10-118788079-DEL-186
chr10-118796953-DEL-65
chr10-118886711-DEL-51
chr10-118888022-DEL-62
chr10-118902155-DEL-1215
chr10-118906059-DEL-130
chr10-118913942-DEL-203
chr10-118930581-DEL-62
chr10-118974007-DEL-60
chr10-119213198-DEL-343
chr10-119338857-DEL-78
chr10-119516948-DEL-97
chr10-119620583-DEL-684
chr10-119675699-DEL-58
chr10-119695731-DEL-414
chr10-119729401-DEL-588
chr10-119729634-DEL-1148
chr10-119749020-DEL-50
chr10-119789231-DEL-129
chr10-119856764-DEL-68
chr10-119884760-DEL-337
chr10-119918071-DEL-138
chr10-119918152-DEL-54
chr10-119979728-DEL-169
chr10-120058665-DEL-85
chr10-120058762-DEL-62
chr10-120061042-DEL-50
chr10-120114369-DEL-188
chr10-120114906-DEL-71
chr10-120202174-DEL-120
chr10-120245997-DEL-232
chr10-120246085-DEL-51
chr10-120255642-DEL-74
chr10-120257658-DEL-94
chr10-120282788-DEL-66
chr10-120336484-DEL-72
chr10-120347583-DEL-84
chr10-120437897-DEL-96
chr10-120438158-DEL-484
chr10-120520449-DEL-121
chr10-120597015-DEL-70
chr10-120748703-DEL-128
chr10-120758231-DEL-50
chr10-120780417-DEL-191
chr10-120790144-DEL-62

chr10-120793242-DEL-108
chr10-120803794-DEL-168
chr10-120811075-DEL-1120
chr10-120812372-DEL-58
chr10-120856008-DEL-92
chr10-120906106-DEL-142
chr10-120910149-DEL-67
chr10-120935186-DEL-50
chr10-120964382-DEL-51
chr10-121005689-DEL-56
chr10-121032394-DEL-50
chr10-121038055-DEL-2469
chr10-121048972-DEL-54
chr10-121129013-DEL-246
chr10-121205537-DEL-807
chr10-121237506-DEL-108
chr10-121259577-DEL-50
chr10-121274959-DEL-1822
chr10-121300131-DEL-63
chr10-121314060-DEL-3218
chr10-121321241-DEL-248
chr10-121412595-DEL-60
chr10-121412706-DEL-210
chr10-121412967-DEL-180
chr10-121415218-DEL-232
chr10-121415612-DEL-462
chr10-121483849-DEL-188
chr10-121548847-DEL-1332
chr10-121725524-DEL-290
chr10-121760176-DEL-110
chr10-121789511-DEL-121
chr10-121825283-DEL-64
chr10-121825540-DEL-115
chr10-122146049-DEL-50
chr10-122215331-DEL-101
chr10-122274692-DEL-51
chr10-122295324-DEL-194
chr10-122300915-DEL-1720
chr10-122324920-DEL-371
chr10-122329468-DEL-74
chr10-122404077-DEL-52
chr10-122598769-DEL-96
chr10-122622233-DEL-332
chr10-122743015-DEL-77
chr10-122750657-DEL-231
chr10-122760233-DEL-50
chr10-122882992-DEL-480
chr10-122883674-DEL-162
chr10-122918450-DEL-90
chr10-123022250-DEL-83
chr10-123319943-DEL-217
chr10-123320489-DEL-217
chr10-123320994-DEL-75
chr10-123351071-DEL-86
chr10-123368493-DEL-77
chr10-123385131-DEL-56
chr10-123551720-DEL-54
chr10-123606903-DEL-78
chr10-123654580-DEL-57
chr10-123715477-DEL-2647
chr10-123815306-DEL-1408
chr10-123941229-DEL-98
chr10-123944839-DEL-58
chr10-123950690-DEL-55
chr10-124060994-DEL-76
chr10-124214263-DEL-72
chr10-124228121-DEL-577
chr10-124276304-DEL-159
chr10-124304144-DEL-162
chr10-124304649-DEL-54
chr10-124321063-DEL-52
chr10-124373945-DEL-478
chr10-124452693-DEL-232
chr10-124688998-DEL-66
chr10-124825096-DEL-142
chr10-124825342-DEL-165
chr10-124825370-DEL-77

chr10-124828751-DEL-59
chr10-124846277-DEL-62
chr10-124859716-DEL-50
chr10-124971138-DEL-70
chr10-125045061-DEL-69
chr10-125236385-DEL-66
chr10-125236595-DEL-249
chr10-125323496-DEL-102
chr10-125359840-DEL-70
chr10-125388917-DEL-76
chr10-125415833-DEL-70
chr10-125416037-DEL-298
chr10-125471646-DEL-507
chr10-125498819-DEL-60
chr10-125528026-DEL-204
chr10-125528083-DEL-51
chr10-125550714-DEL-6290
chr10-125580332-DEL-52
chr10-125609846-DEL-103
chr10-125690871-DEL-213
chr10-125690935-DEL-96
chr10-125714363-DEL-211
chr10-125728691-DEL-69
chr10-125740100-DEL-56
chr10-125843572-DEL-63
chr10-125913911-DEL-59
chr10-125913973-DEL-169
chr10-125914353-DEL-253
chr10-125914516-DEL-81
chr10-125928130-DEL-96
chr10-125928367-DEL-88
chr10-125944166-DEL-50
chr10-125975829-DEL-155
chr10-125983302-DEL-83
chr10-125989357-DEL-92
chr10-126051300-DEL-146
chr10-126074188-DEL-71
chr10-126148268-DEL-52
chr10-126181367-DEL-478
chr10-126182016-DEL-81
chr10-126193375-DEL-50
chr10-126293557-DEL-51
chr10-126380453-DEL-51
chr10-126389326-DEL-74
chr10-126430736-DEL-76
chr10-126497203-DEL-72
chr10-126517130-DEL-64
chr10-126552888-DEL-610
chr10-126580101-DEL-540
chr10-126661065-DEL-289
chr10-126730522-DEL-177
chr10-126788041-DEL-51
chr10-126794051-DEL-199
chr10-126984678-DEL-87
chr10-127077297-DEL-354
chr10-127145223-DEL-105
chr10-127228278-DEL-72
chr10-127228521-DEL-206
chr10-127493675-DEL-106
chr10-127676207-DEL-946
chr10-127680746-DEL-86
chr10-127767985-DEL-71
chr10-127768153-DEL-180
chr10-127783266-DEL-1631
chr10-127815764-DEL-84
chr10-127833194-DEL-82
chr10-127854498-DEL-219
chr10-127991941-DEL-166
chr10-128002533-DEL-825
chr10-128024897-DEL-226
chr10-128024964-DEL-66
chr10-128033846-DEL-55
chr10-128111248-DEL-383
chr10-128126534-DEL-72
chr10-128168420-DEL-258
chr10-128313923-DEL-65
chr10-128374876-DEL-202

chr10-128375233-DEL-814
chr10-128444134-DEL-92
chr10-128445524-DEL-178
chr10-128445859-DEL-193
chr10-128449902-DEL-114
chr10-128454052-DEL-312
chr10-128491840-DEL-188
chr10-128618914-DEL-116
chr10-128798420-DEL-96
chr10-128838596-DEL-63
chr10-128838691-DEL-164
chr10-128862595-DEL-76
chr10-128886868-DEL-8603
chr10-128949865-DEL-438
chr10-129115149-DEL-1826
chr10-129176642-DEL-214
chr10-129321328-DEL-65
chr10-129618300-DEL-52
chr10-129654823-DEL-2767
chr10-129703003-DEL-110
chr10-129705275-DEL-1645
chr10-129801353-DEL-8867
chr10-129809449-DEL-8876
chr10-129863410-DEL-58
chr10-129949640-DEL-69
chr10-129949833-DEL-207
chr10-129990895-DEL-50
chr10-130056944-DEL-94
chr10-130298971-DEL-2978
chr10-130299580-DEL-1485
chr11-3064274-DEL-123
chr11-3149372-DEL-1192
chr11-3206137-DEL-372
chr11-3407608-DEL-4238
chr11-3526852-DEL-386
chr11-3527331-DEL-86
chr11-3549620-DEL-278
chr11-3552193-DEL-91
chr11-3552442-DEL-365
chr11-3737287-DEL-105
chr11-3738230-DEL-218
chr11-3774501-DEL-600
chr11-3802590-DEL-416
chr11-3929648-DEL-152
chr11-3958125-DEL-180
chr11-3994812-DEL-1001
chr11-4021871-DEL-467
chr11-4051022-DEL-134
chr11-4054144-DEL-68
chr11-4054481-DEL-130
chr11-4175435-DEL-282
chr11-4175682-DEL-60
chr11-4510426-DEL-101
chr11-4522863-DEL-1058
chr11-4580540-DEL-209
chr11-4585615-DEL-193
chr11-4645821-DEL-186
chr11-4648429-DEL-78
chr11-4648696-DEL-134
chr11-4803472-DEL-405
chr11-4857353-DEL-116
chr11-4929741-DEL-266
chr11-5004703-DEL-58
chr11-5096670-DEL-226
chr11-5097036-DEL-74
chr11-5136182-DEL-120
chr11-5150372-DEL-72
chr11-5153348-DEL-54
chr11-5249146-DEL-314
chr11-5405131-DEL-55
chr11-5416649-DEL-300
chr11-5421044-DEL-356
chr11-5437378-DEL-50
chr11-5633899-DEL-54
chr11-5700835-DEL-54
chr11-5727595-DEL-140
chr11-5765520-DEL-59

chr11-5774341-DEL-380
chr11-5841603-DEL-50
chr11-5877468-DEL-101
chr11-5962517-DEL-248
chr11-6037369-DEL-50
chr11-6300931-DEL-55
chr11-6300969-DEL-220
chr11-6301387-DEL-64
chr11-6329514-DEL-63
chr11-6363085-DEL-355
chr11-6363284-DEL-82
chr11-6397244-DEL-110
chr11-6427537-DEL-197
chr11-6481478-DEL-4420
chr11-6487094-DEL-201
chr11-6524417-DEL-398
chr11-6599292-DEL-54
chr11-6599675-DEL-94
chr11-6733582-DEL-53
chr11-6733594-DEL-212
chr11-6737968-DEL-416
chr11-6888428-DEL-193
chr11-6985316-DEL-57
chr11-7056304-DEL-112
chr11-7281834-DEL-208
chr11-7349555-DEL-2358
chr11-7463695-DEL-50
chr11-7685413-DEL-56
chr11-7691904-DEL-1448
chr11-7738182-DEL-635
chr11-7791883-DEL-3118
chr11-7983737-DEL-3436
chr11-8000322-DEL-62
chr11-8064931-DEL-68
chr11-8186436-DEL-197
chr11-8444596-DEL-267
chr11-8485179-DEL-58
chr11-8501542-DEL-196
chr11-8515197-DEL-454
chr11-8519942-DEL-64
chr11-8558421-DEL-105
chr11-8694790-DEL-1995
chr11-8698598-DEL-493
chr11-8808074-DEL-54
chr11-8826498-DEL-300
chr11-8826840-DEL-238
chr11-8882569-DEL-143
chr11-8898418-DEL-195
chr11-8905262-DEL-208
chr11-8977222-DEL-72
chr11-9019645-DEL-1339
chr11-9139769-DEL-102
chr11-9195314-DEL-222
chr11-9207184-DEL-95
chr11-9238824-DEL-69
chr11-9250185-DEL-98
chr11-9314981-DEL-201
chr11-9340721-DEL-442
chr11-9447159-DEL-2114
chr11-9550145-DEL-104
chr11-9614778-DEL-132
chr11-9632597-DEL-72
chr11-9694660-DEL-6276
chr11-9716660-DEL-2834
chr11-9769233-DEL-1273
chr11-9825058-DEL-182
chr11-9843305-DEL-68
chr11-9890514-DEL-2374
chr11-9903638-DEL-77
chr11-9991695-DEL-181
chr11-10086347-DEL-162
chr11-10146709-DEL-156
chr11-10147071-DEL-585
chr11-10226368-DEL-56
chr11-10363302-DEL-50
chr11-10521425-DEL-100
chr11-10547324-DEL-82

chr11-10574072-DEL-4942
chr11-10638381-DEL-146
chr11-10661110-DEL-50
chr11-10678972-DEL-121
chr11-10683136-DEL-62
chr11-10683322-DEL-401
chr11-10752417-DEL-60
chr11-10796517-DEL-92
chr11-10851345-DEL-64
chr11-10899769-DEL-98
chr11-10942502-DEL-612
chr11-11033266-DEL-60
chr11-11049155-DEL-63
chr11-11143170-DEL-372
chr11-11190288-DEL-89
chr11-11258331-DEL-83
chr11-11280449-DEL-108
chr11-11285178-DEL-90
chr11-11356086-DEL-246
chr11-11386751-DEL-1309
chr11-11411738-DEL-82
chr11-11451171-DEL-145
chr11-11513184-DEL-194
chr11-11661231-DEL-51
chr11-11687915-DEL-80
chr11-11868219-DEL-65
chr11-12355773-DEL-79
chr11-12386687-DEL-182
chr11-12449394-DEL-60
chr11-12534165-DEL-176
chr11-12627344-DEL-85
chr11-12683902-DEL-50
chr11-12778933-DEL-1530
chr11-12979128-DEL-671
chr11-13021461-DEL-904
chr11-13178375-DEL-3887
chr11-13195494-DEL-272
chr11-13235498-DEL-159
chr11-13329088-DEL-6659
chr11-13365873-DEL-93
chr11-13378114-DEL-286
chr11-13460316-DEL-56
chr11-13641828-DEL-51
chr11-13701694-DEL-61
chr11-13799299-DEL-55
chr11-13956848-DEL-72
chr11-14030950-DEL-163
chr11-14077952-DEL-214
chr11-14161612-DEL-5435
chr11-14215014-DEL-81
chr11-14243270-DEL-53
chr11-14445914-DEL-134
chr11-14519055-DEL-2515
chr11-14556504-DEL-380
chr11-14583022-DEL-50
chr11-14764138-DEL-65
chr11-15043340-DEL-3131
chr11-15048853-DEL-286
chr11-15061129-DEL-51
chr11-15143946-DEL-228
chr11-15201580-DEL-51
chr11-15213990-DEL-3618
chr11-15240127-DEL-14000
chr11-15344726-DEL-50
chr11-15350944-DEL-609
chr11-15370536-DEL-386
chr11-15379324-DEL-306
chr11-15526163-DEL-78
chr11-15557831-DEL-506
chr11-15563659-DEL-1249
chr11-15628519-DEL-94
chr11-15696738-DEL-404
chr11-15888762-DEL-1343
chr11-15920913-DEL-50
chr11-15946564-DEL-70
chr11-15975123-DEL-844
chr11-16052636-DEL-88

chr11-16121427-DEL-118
chr11-16146301-DEL-141
chr11-16149372-DEL-1311
chr11-16165592-DEL-113
chr11-16402860-DEL-5503
chr11-16492278-DEL-53
chr11-16508615-DEL-68
chr11-16608122-DEL-124
chr11-16829157-DEL-848
chr11-16991451-DEL-127
chr11-16993296-DEL-145
chr11-17070427-DEL-1228
chr11-17097592-DEL-50
chr11-17102596-DEL-373
chr11-17103053-DEL-197
chr11-17158435-DEL-175
chr11-17164687-DEL-68
chr11-17166837-DEL-207
chr11-17251921-DEL-2010
chr11-17269822-DEL-50
chr11-17346721-DEL-83
chr11-17415554-DEL-936
chr11-17415664-DEL-132
chr11-17416229-DEL-141
chr11-17416700-DEL-99
chr11-17483576-DEL-2063
chr11-17880193-DEL-203
chr11-17971007-DEL-488
chr11-18001901-DEL-1014
chr11-18124691-DEL-50
chr11-18156134-DEL-437
chr11-18207397-DEL-112
chr11-18207578-DEL-116
chr11-18282055-DEL-240
chr11-18282199-DEL-96
chr11-18376236-DEL-1279
chr11-18409541-DEL-445
chr11-18471058-DEL-67
chr11-18660380-DEL-112
chr11-18950288-DEL-61
chr11-19085476-DEL-50
chr11-19172634-DEL-54
chr11-19217944-DEL-967
chr11-19351479-DEL-124
chr11-19442267-DEL-105
chr11-19475666-DEL-102
chr11-19522252-DEL-146
chr11-19745972-DEL-178
chr11-19754052-DEL-1040
chr11-19756674-DEL-1403
chr11-19777914-DEL-52
chr11-19918221-DEL-246
chr11-19982150-DEL-107
chr11-20011336-DEL-281
chr11-20099656-DEL-126
chr11-20131830-DEL-62
chr11-20133928-DEL-80
chr11-20206063-DEL-1486
chr11-20260451-DEL-640
chr11-20260960-DEL-80
chr11-20358754-DEL-217
chr11-20376798-DEL-1217
chr11-20397706-DEL-81
chr11-20424146-DEL-85
chr11-20435041-DEL-61
chr11-20491051-DEL-50
chr11-20532267-DEL-96
chr11-20532287-DEL-480
chr11-20532720-DEL-61
chr11-20624582-DEL-120
chr11-20804939-DEL-98
chr11-20805125-DEL-82
chr11-20857376-DEL-50
chr11-20882319-DEL-60
chr11-20882493-DEL-129
chr11-20883150-DEL-70
chr11-20886541-DEL-58

chr11-20886576-DEL-194
chr11-20983599-DEL-56
chr11-21006700-DEL-331
chr11-21126412-DEL-112
chr11-21148528-DEL-55
chr11-21156052-DEL-7895
chr11-21179343-DEL-1869
chr11-21285494-DEL-127
chr11-21405697-DEL-150
chr11-21463513-DEL-132
chr11-21467817-DEL-257
chr11-21487228-DEL-93
chr11-21543966-DEL-1099
chr11-21679581-DEL-50
chr11-21715630-DEL-55
chr11-21803816-DEL-848
chr11-21888530-DEL-60
chr11-22152316-DEL-1136
chr11-22231281-DEL-117
chr11-22276351-DEL-62
chr11-22457124-DEL-253
chr11-22481179-DEL-775
chr11-22493442-DEL-3804
chr11-22545705-DEL-104
chr11-22565468-DEL-110
chr11-22589251-DEL-60
chr11-22615667-DEL-2490
chr11-22655793-DEL-54
chr11-22688258-DEL-92
chr11-22711339-DEL-3228
chr11-22791855-DEL-182
chr11-22947952-DEL-56
chr11-22989602-DEL-1520
chr11-22992436-DEL-100
chr11-23023559-DEL-148
chr11-23048151-DEL-180
chr11-23063105-DEL-200
chr11-23094812-DEL-50
chr11-23117095-DEL-54
chr11-23299973-DEL-462
chr11-23329368-DEL-102
chr11-23491277-DEL-1072
chr11-23491553-DEL-532
chr11-23491673-DEL-172
chr11-23536928-DEL-52
chr11-23560239-DEL-54
chr11-23584246-DEL-56
chr11-23585393-DEL-116
chr11-23759560-DEL-88
chr11-23767043-DEL-457
chr11-23808396-DEL-454
chr11-23837283-DEL-63
chr11-23860205-DEL-207
chr11-23870411-DEL-198
chr11-23963963-DEL-118
chr11-24053976-DEL-51
chr11-24124580-DEL-135
chr11-24125059-DEL-140
chr11-24219752-DEL-208
chr11-24360975-DEL-407
chr11-24576904-DEL-761
chr11-24842931-DEL-51
chr11-25321410-DEL-5717
chr11-25433881-DEL-90
chr11-25599040-DEL-90
chr11-25608365-DEL-50
chr11-25630748-DEL-50
chr11-25778180-DEL-52
chr11-25819747-DEL-50
chr11-26133811-DEL-215
chr11-26288841-DEL-1416
chr11-26291534-DEL-50
chr11-26301938-DEL-50
chr11-26589158-DEL-284
chr11-26959320-DEL-3421
chr11-27034966-DEL-1402
chr11-27144604-DEL-53

chr11-27246066-DEL-147
chr11-27246435-DEL-2934
chr11-27393950-DEL-75
chr11-27395817-DEL-185
chr11-27410866-DEL-3169
chr11-27459266-DEL-50
chr11-27507185-DEL-99
chr11-27818006-DEL-38622
chr11-27861584-DEL-9280
chr11-27951310-DEL-50
chr11-28006179-DEL-50
chr11-28014606-DEL-1073
chr11-28029751-DEL-66
chr11-28074942-DEL-562
chr11-28185293-DEL-5499
chr11-28211210-DEL-94
chr11-28281956-DEL-52
chr11-28322910-DEL-52
chr11-28424489-DEL-1203
chr11-28777854-DEL-1598
chr11-28913085-DEL-1878
chr11-29154129-DEL-50
chr11-29187245-DEL-90
chr11-29216772-DEL-124
chr11-29246190-DEL-64
chr11-29470855-DEL-4524
chr11-29661588-DEL-106
chr11-29703332-DEL-52
chr11-29762417-DEL-1037
chr11-29767172-DEL-59
chr11-29826225-DEL-127
chr11-29900943-DEL-68
chr11-29901331-DEL-63
chr11-29911150-DEL-199
chr11-29948738-DEL-85
chr11-30243287-DEL-110
chr11-30243308-DEL-52
chr11-30273687-DEL-179
chr11-30332062-DEL-221
chr11-30416919-DEL-1707
chr11-30460944-DEL-3356
chr11-30464380-DEL-215
chr11-30508557-DEL-56
chr11-30508772-DEL-132
chr11-30509839-DEL-52
chr11-30552090-DEL-5426
chr11-30559002-DEL-61
chr11-30598176-DEL-139
chr11-30664118-DEL-50
chr11-30696195-DEL-109
chr11-30812799-DEL-1620
chr11-30831749-DEL-131
chr11-30883402-DEL-64
chr11-30985810-DEL-214
chr11-31198922-DEL-5697
chr11-31236230-DEL-227
chr11-31236273-DEL-104
chr11-31238003-DEL-184
chr11-31238431-DEL-118
chr11-31293753-DEL-1876
chr11-31358823-DEL-289
chr11-31359151-DEL-90
chr11-31396322-DEL-55
chr11-31410209-DEL-675
chr11-31431854-DEL-326
chr11-31567631-DEL-52
chr11-31596597-DEL-50
chr11-31621052-DEL-175
chr11-31661193-DEL-105
chr11-31673899-DEL-50
chr11-31678006-DEL-111
chr11-31678012-DEL-251
chr11-31678413-DEL-114
chr11-31795311-DEL-60
chr11-31872629-DEL-92
chr11-31901520-DEL-78
chr11-31901752-DEL-196

chr11-31908542-DEL-50
chr11-31950848-DEL-152
chr11-32013333-DEL-50
chr11-32069027-DEL-4766
chr11-32074774-DEL-180
chr11-32115176-DEL-62
chr11-32142355-DEL-61
chr11-32209649-DEL-105
chr11-32209926-DEL-105
chr11-32223590-DEL-170
chr11-32308840-DEL-52
chr11-32442017-DEL-50
chr11-32529392-DEL-53
chr11-32657098-DEL-51
chr11-32675790-DEL-102
chr11-32758035-DEL-61
chr11-32770878-DEL-62
chr11-32771194-DEL-77
chr11-32792926-DEL-86
chr11-32869672-DEL-207
chr11-32914813-DEL-74
chr11-33364782-DEL-54
chr11-33397161-DEL-98
chr11-33397198-DEL-291
chr11-33420507-DEL-99
chr11-33511712-DEL-192
chr11-33535355-DEL-50
chr11-33552719-DEL-342
chr11-33612791-DEL-51
chr11-33645342-DEL-52
chr11-33660264-DEL-260
chr11-33660733-DEL-104
chr11-33678141-DEL-388
chr11-33776100-DEL-202
chr11-33809246-DEL-59
chr11-33959219-DEL-156
chr11-33979940-DEL-466
chr11-34036747-DEL-60
chr11-34098947-DEL-84
chr11-34124458-DEL-129
chr11-34164058-DEL-63
chr11-34170382-DEL-102
chr11-34173135-DEL-373
chr11-34173202-DEL-141
chr11-34270657-DEL-190
chr11-34303718-DEL-62
chr11-34365171-DEL-77
chr11-34518412-DEL-73
chr11-34520728-DEL-242
chr11-34525333-DEL-396
chr11-34555121-DEL-240
chr11-34557685-DEL-71
chr11-34562616-DEL-71
chr11-34581177-DEL-137
chr11-34657224-DEL-52
chr11-34658041-DEL-268
chr11-34887539-DEL-104
chr11-34901079-DEL-114
chr11-35036283-DEL-307
chr11-35037033-DEL-155
chr11-35039393-DEL-160
chr11-35039464-DEL-58
chr11-35054942-DEL-434
chr11-35138741-DEL-57
chr11-35175562-DEL-50
chr11-35202227-DEL-175
chr11-35270214-DEL-50
chr11-35316794-DEL-54
chr11-35379411-DEL-251
chr11-35408639-DEL-283
chr11-35624161-DEL-50
chr11-35637081-DEL-87
chr11-35651121-DEL-50
chr11-35654253-DEL-1740
chr11-35654695-DEL-105
chr11-35655064-DEL-616
chr11-35656238-DEL-69

chr11-35693135-DEL-1326
chr11-35834697-DEL-264
chr11-35885784-DEL-50
chr11-35907832-DEL-52
chr11-35907949-DEL-312
chr11-35966617-DEL-116
chr11-35993334-DEL-54
chr11-35993483-DEL-247
chr11-36003853-DEL-2529
chr11-36088841-DEL-163
chr11-36174644-DEL-61
chr11-36174654-DEL-196
chr11-36320615-DEL-122
chr11-36389276-DEL-168
chr11-36413262-DEL-148
chr11-36413281-DEL-50
chr11-36500996-DEL-50
chr11-36682496-DEL-52
chr11-36826585-DEL-74
chr11-37003487-DEL-216
chr11-37003835-DEL-170
chr11-37417891-DEL-65
chr11-37451285-DEL-86
chr11-37477106-DEL-372
chr11-37613022-DEL-51
chr11-37643522-DEL-424
chr11-37728623-DEL-2275
chr11-37897038-DEL-52
chr11-37897289-DEL-64
chr11-38032301-DEL-50
chr11-38060256-DEL-50
chr11-38508619-DEL-114
chr11-38596414-DEL-833
chr11-38821566-DEL-54
chr11-38925739-DEL-614
chr11-38947967-DEL-50
chr11-38981613-DEL-50
chr11-39060399-DEL-670
chr11-39450297-DEL-59
chr11-39477212-DEL-63
chr11-39478210-DEL-56
chr11-39512110-DEL-2084
chr11-39549664-DEL-186
chr11-39551595-DEL-70
chr11-39709968-DEL-1018
chr11-40083930-DEL-154
chr11-40182617-DEL-54
chr11-40234548-DEL-1505
chr11-40239510-DEL-415
chr11-40239718-DEL-415
chr11-40393912-DEL-52
chr11-40435622-DEL-411
chr11-40549922-DEL-522
chr11-40688483-DEL-1286
chr11-40800690-DEL-201
chr11-40831505-DEL-60
chr11-40983426-DEL-264
chr11-41022394-DEL-2919
chr11-41267780-DEL-55
chr11-41338505-DEL-72
chr11-41713665-DEL-103
chr11-41832168-DEL-90
chr11-42011858-DEL-52
chr11-42121370-DEL-109
chr11-42156278-DEL-589
chr11-42217552-DEL-64
chr11-42229113-DEL-2258
chr11-42238055-DEL-62
chr11-42263472-DEL-1637
chr11-42276597-DEL-70
chr11-42280545-DEL-440
chr11-42577238-DEL-1758
chr11-42598098-DEL-1709
chr11-42610057-DEL-5135
chr11-42787130-DEL-54
chr11-42806469-DEL-202
chr11-42806767-DEL-104

chr11-42936237-DEL-62
chr11-42936404-DEL-57
chr11-42936417-DEL-126
chr11-43017974-DEL-92
chr11-43153923-DEL-78
chr11-43250971-DEL-57
chr11-43294325-DEL-52
chr11-43345802-DEL-7280
chr11-43393030-DEL-1516
chr11-43393559-DEL-56
chr11-43398347-DEL-66
chr11-43398560-DEL-197
chr11-43497135-DEL-56
chr11-43680325-DEL-152
chr11-43707613-DEL-85
chr11-43922632-DEL-50
chr11-43942440-DEL-50
chr11-44183567-DEL-228
chr11-44184088-DEL-183
chr11-44236516-DEL-118
chr11-44275135-DEL-2268
chr11-44331011-DEL-238
chr11-44338525-DEL-1272
chr11-44386423-DEL-120
chr11-44386463-DEL-259
chr11-44391066-DEL-95
chr11-44727007-DEL-564
chr11-44731473-DEL-64
chr11-44820164-DEL-1561
chr11-45061848-DEL-50
chr11-45243366-DEL-50
chr11-45522442-DEL-74
chr11-45524851-DEL-256
chr11-45565575-DEL-116
chr11-45614446-DEL-155
chr11-45755574-DEL-1266
chr11-45831007-DEL-239
chr11-45869594-DEL-69
chr11-45905047-DEL-201
chr11-45958926-DEL-81
chr11-45959236-DEL-158
chr11-46100276-DEL-117
chr11-46207257-DEL-60
chr11-46406991-DEL-61
chr11-46407781-DEL-192
chr11-46407797-DEL-81
chr11-46438020-DEL-196
chr11-46464230-DEL-114
chr11-46480349-DEL-248
chr11-46480510-DEL-53
chr11-46531546-DEL-63
chr11-46586938-DEL-95
chr11-46587220-DEL-170
chr11-46620816-DEL-70
chr11-46624821-DEL-104
chr11-46695723-DEL-82
chr11-46695836-DEL-675
chr11-46801742-DEL-198
chr11-46846645-DEL-142
chr11-46846735-DEL-60
chr11-46848254-DEL-563
chr11-46884181-DEL-78
chr11-47100681-DEL-57
chr11-47465189-DEL-56
chr11-47531733-DEL-343
chr11-47585246-DEL-52
chr11-47595581-DEL-387
chr11-47650619-DEL-179
chr11-47779068-DEL-262
chr11-47828262-DEL-1133
chr11-47902588-DEL-50
chr11-48143353-DEL-52
chr11-48235520-DEL-1573
chr11-48286234-DEL-56
chr11-48342734-DEL-214
chr11-48370522-DEL-146
chr11-48399418-DEL-617

chr11-48606206-DEL-83
chr11-48624164-DEL-1117
chr11-48966848-DEL-52
chr11-48978439-DEL-50
chr11-48989399-DEL-217
chr11-49031636-DEL-180
chr11-49037729-DEL-52
chr11-49118427-DEL-334
chr11-49118841-DEL-222
chr11-49119245-DEL-117
chr11-49119554-DEL-60
chr11-49119896-DEL-160
chr11-49208156-DEL-188
chr11-49441364-DEL-125
chr11-49622594-DEL-237
chr11-49631865-DEL-84
chr11-49735598-DEL-88
chr11-49855764-DEL-85
chr11-49869823-DEL-175
chr11-49921933-DEL-84
chr11-50023845-DEL-942
chr11-50058227-DEL-183
chr11-50158807-DEL-55
chr11-50228684-DEL-1136
chr11-50303429-DEL-541
chr11-50310095-DEL-83
chr11-50310317-DEL-54
chr11-50339284-DEL-50
chr11-50457627-DEL-50
chr11-50634122-DEL-58
chr11-50669710-DEL-84
chr11-50671976-DEL-55
chr11-50766798-DEL-344
chr11-50766909-DEL-155
chr11-50824747-DEL-197
chr11-50903636-DEL-396
chr11-50907534-DEL-28432
chr11-51078037-DEL-50
chr11-51240808-DEL-528
chr11-51261978-DEL-100
chr11-51271530-DEL-124
chr11-51428564-DEL-121
chr11-51456198-DEL-149
chr11-51465794-DEL-123
chr11-51470245-DEL-50
chr11-51487885-DEL-780
chr11-51521124-DEL-113
chr11-51684088-DEL-94
chr11-51832742-DEL-72
chr11-51832742-DEL-162
chr11-51921071-DEL-800
chr11-51922337-DEL-523
chr11-52064380-DEL-54
chr11-52095646-DEL-611
chr11-52096127-DEL-101
chr11-52179522-DEL-159
chr11-52179827-DEL-127
chr11-52186462-DEL-1210
chr11-52207931-DEL-738
chr11-52225222-DEL-275
chr11-52281993-DEL-429
chr11-52315821-DEL-188
chr11-52319772-DEL-378
chr11-52444342-DEL-1864
chr11-52516043-DEL-57
chr11-52548122-DEL-59
chr11-52572342-DEL-3150
chr11-52591610-DEL-50
chr11-52791508-DEL-74
chr11-52792954-DEL-65
chr11-52872422-DEL-187
chr11-52902834-DEL-60
chr11-53066015-DEL-364
chr11-53116434-DEL-195
chr11-53403977-DEL-50
chr11-53428920-DEL-496
chr11-53429467-DEL-182

chr11-53444929-DEL-6354
chr11-53817630-DEL-1892
chr11-53834960-DEL-62
chr11-53875601-DEL-73
chr11-53907618-DEL-90
chr11-53975359-DEL-82
chr11-54199390-DEL-55
chr11-54261182-DEL-52
chr11-54322510-DEL-58
chr11-54375740-DEL-71
chr11-54573350-DEL-733
chr11-54577501-DEL-140
chr11-54721175-DEL-142
chr11-54735247-DEL-302
chr11-54883770-DEL-50
chr11-54905042-DEL-50
chr11-54929838-DEL-142
chr11-54929870-DEL-55
chr11-54936987-DEL-52
chr11-55021680-DEL-91
chr11-55029064-DEL-182
chr11-55029700-DEL-186
chr11-55045817-DEL-646
chr11-55064432-DEL-291
chr11-55356729-DEL-70
chr11-55432037-DEL-50
chr11-55521274-DEL-50
chr11-55644090-DEL-261
chr11-55793830-DEL-95
chr11-55927978-DEL-79
chr11-55963821-DEL-56
chr11-56003669-DEL-56
chr11-56012050-DEL-52
chr11-56095220-DEL-424
chr11-56113402-DEL-62
chr11-56125065-DEL-158
chr11-56149981-DEL-229
chr11-56177456-DEL-86
chr11-56184603-DEL-3613
chr11-56265911-DEL-1963
chr11-56440473-DEL-92
chr11-56480478-DEL-2060
chr11-56587569-DEL-55
chr11-56645562-DEL-70
chr11-56734735-DEL-71
chr11-56744008-DEL-50
chr11-56789893-DEL-52
chr11-56861321-DEL-50
chr11-56922910-DEL-75
chr11-57004776-DEL-106
chr11-57009981-DEL-50
chr11-57014559-DEL-1047
chr11-57028462-DEL-58
chr11-57032372-DEL-300
chr11-57227504-DEL-66
chr11-57242830-DEL-508
chr11-57397219-DEL-94
chr11-57524644-DEL-60
chr11-57564653-DEL-120
chr11-57799889-DEL-50
chr11-57817949-DEL-172
chr11-57829471-DEL-88
chr11-57837696-DEL-68
chr11-57837953-DEL-207
chr11-57892440-DEL-50
chr11-57979346-DEL-50
chr11-57997462-DEL-108
chr11-57997685-DEL-106
chr11-58143597-DEL-71
chr11-58174911-DEL-78
chr11-58246869-DEL-51
chr11-58272271-DEL-184
chr11-58281978-DEL-144
chr11-58303088-DEL-71
chr11-58333509-DEL-108
chr11-58334787-DEL-80
chr11-58352924-DEL-60

chr11-58367237-DEL-50
chr11-58463227-DEL-506
chr11-58603356-DEL-3200
chr11-58642247-DEL-74
chr11-58650828-DEL-62
chr11-58654820-DEL-628
chr11-58803494-DEL-206
chr11-58826084-DEL-190
chr11-58894748-DEL-1091
chr11-58895669-DEL-169
chr11-58896249-DEL-210
chr11-58896249-DEL-456
chr11-58896529-DEL-120
chr11-58986987-DEL-125
chr11-58991983-DEL-56
chr11-59253971-DEL-82
chr11-59336898-DEL-85
chr11-59467144-DEL-141
chr11-59561630-DEL-136
chr11-59684387-DEL-88
chr11-59687234-DEL-194
chr11-59844946-DEL-1836
chr11-60130278-DEL-98
chr11-60138156-DEL-120
chr11-60201022-DEL-50
chr11-60215627-DEL-78
chr11-60216441-DEL-732
chr11-60341989-DEL-421
chr11-60380657-DEL-198
chr11-60406558-DEL-58
chr11-60455706-DEL-421
chr11-60514923-DEL-91
chr11-60517128-DEL-828
chr11-60518344-DEL-52
chr11-60519720-DEL-206
chr11-60520167-DEL-322
chr11-60530557-DEL-839
chr11-60565801-DEL-131
chr11-60571187-DEL-746
chr11-60580334-DEL-2025
chr11-60580409-DEL-55
chr11-60757342-DEL-50
chr11-60855502-DEL-82
chr11-60887713-DEL-378
chr11-60949868-DEL-85
chr11-60990373-DEL-570
chr11-60990977-DEL-70
chr11-61019264-DEL-52
chr11-61101567-DEL-50
chr11-61101573-DEL-105
chr11-61124897-DEL-50
chr11-61127134-DEL-1073
chr11-61151760-DEL-188
chr11-61166940-DEL-170
chr11-61209153-DEL-162
chr11-61232446-DEL-80
chr11-61262079-DEL-82
chr11-61299293-DEL-54
chr11-61348680-DEL-91
chr11-61518015-DEL-50
chr11-61545176-DEL-55
chr11-61548674-DEL-118
chr11-61665230-DEL-57
chr11-61710945-DEL-66
chr11-61713359-DEL-206
chr11-61754464-DEL-82
chr11-61869310-DEL-91
chr11-61908668-DEL-403
chr11-61908797-DEL-62
chr11-62151034-DEL-140
chr11-62376474-DEL-51
chr11-62377298-DEL-88
chr11-62405714-DEL-232
chr11-62405714-DEL-609
chr11-62437197-DEL-214
chr11-62459804-DEL-431
chr11-62460262-DEL-75

chr11-62639862-DEL-2505
chr11-62654156-DEL-136
chr11-62757071-DEL-125
chr11-62813224-DEL-50
chr11-62851414-DEL-63
chr11-62851542-DEL-89
chr11-62851669-DEL-97
chr11-62902885-DEL-131
chr11-62906163-DEL-83
chr11-63119815-DEL-620
chr11-63162953-DEL-1573
chr11-63176239-DEL-53
chr11-63224897-DEL-145
chr11-63297609-DEL-118
chr11-63309767-DEL-168
chr11-63310377-DEL-171
chr11-63310831-DEL-109
chr11-63352237-DEL-64
chr11-63352454-DEL-179
chr11-63499063-DEL-99
chr11-63514905-DEL-53
chr11-63521638-DEL-846
chr11-63521685-DEL-376
chr11-63557047-DEL-139
chr11-63566610-DEL-57
chr11-63659393-DEL-677
chr11-63711143-DEL-477
chr11-63722881-DEL-50
chr11-63742694-DEL-445
chr11-63743506-DEL-1470
chr11-63746887-DEL-1511
chr11-63757653-DEL-81
chr11-64138967-DEL-68
chr11-64184921-DEL-533
chr11-64279373-DEL-56
chr11-64305092-DEL-864
chr11-64458639-DEL-80
chr11-64506730-DEL-50
chr11-64510851-DEL-144
chr11-64586654-DEL-122
chr11-64604306-DEL-96
chr11-64759646-DEL-142
chr11-64779734-DEL-152
chr11-64800081-DEL-56
chr11-65094276-DEL-68
chr11-65166959-DEL-2835
chr11-65324712-DEL-52
chr11-65425326-DEL-64
chr11-65502989-DEL-59
chr11-65518477-DEL-2038
chr11-65618862-DEL-71
chr11-65682177-DEL-255
chr11-65782264-DEL-88
chr11-65787990-DEL-6322
chr11-65934013-DEL-112
chr11-65958655-DEL-96
chr11-65989521-DEL-145
chr11-66129037-DEL-192
chr11-66312000-DEL-66
chr11-66329071-DEL-142
chr11-66329109-DEL-372
chr11-66431941-DEL-131
chr11-66511495-DEL-108
chr11-66668252-DEL-138
chr11-66672160-DEL-59
chr11-66699128-DEL-64
chr11-66711152-DEL-175
chr11-66725134-DEL-50
chr11-66734730-DEL-85
chr11-66902628-DEL-88
chr11-66982342-DEL-117
chr11-67012694-DEL-202
chr11-67012997-DEL-70
chr11-67052724-DEL-401
chr11-67103261-DEL-62
chr11-67414249-DEL-125
chr11-67414281-DEL-58

chr11-67437166-DEL-79
chr11-67463390-DEL-105
chr11-67506850-DEL-80
chr11-67508657-DEL-54
chr11-67554016-DEL-64
chr11-67658775-DEL-119
chr11-67817860-DEL-176
chr11-67840617-DEL-214
chr11-67982300-DEL-170
chr11-68044087-DEL-224
chr11-68048601-DEL-928
chr11-68097236-DEL-504
chr11-68112569-DEL-122
chr11-68175779-DEL-93
chr11-68367377-DEL-101
chr11-68464348-DEL-178
chr11-68464529-DEL-64
chr11-68473858-DEL-158
chr11-68614234-DEL-155
chr11-68624296-DEL-88
chr11-68687498-DEL-148
chr11-68687599-DEL-333
chr11-68801090-DEL-66
chr11-68801571-DEL-144
chr11-68816517-DEL-52
chr11-68816528-DEL-135
chr11-68876109-DEL-211
chr11-69136113-DEL-52
chr11-69207103-DEL-65
chr11-69227151-DEL-274
chr11-69227535-DEL-55
chr11-69229490-DEL-216
chr11-69376605-DEL-63
chr11-69378828-DEL-175
chr11-69424599-DEL-50
chr11-69441768-DEL-394
chr11-69441795-DEL-79
chr11-69466317-DEL-101
chr11-69615212-DEL-1102
chr11-69615492-DEL-372
chr11-69615506-DEL-124
chr11-69974298-DEL-72
chr11-69980332-DEL-57
chr11-70026874-DEL-107
chr11-70165536-DEL-369
chr11-70165969-DEL-735
chr11-70198892-DEL-269
chr11-70252135-DEL-6757
chr11-70261643-DEL-58
chr11-70261667-DEL-252
chr11-70261825-DEL-94
chr11-70284082-DEL-116
chr11-70319849-DEL-410
chr11-70527028-DEL-282
chr11-70627544-DEL-127
chr11-70867656-DEL-92
chr11-70915247-DEL-50
chr11-70948644-DEL-234
chr11-70956541-DEL-3510
chr11-70988143-DEL-71
chr11-71068566-DEL-2465
chr11-71073765-DEL-72
chr11-71074038-DEL-149
chr11-71078169-DEL-158
chr11-71164723-DEL-201
chr11-71167934-DEL-66
chr11-71210893-DEL-480
chr11-71211412-DEL-873
chr11-71215690-DEL-3968
chr11-71498372-DEL-129
chr11-71550712-DEL-63
chr11-71557670-DEL-429
chr11-71557787-DEL-195
chr11-71618961-DEL-50
chr11-71623958-DEL-55
chr11-71712704-DEL-66
chr11-71763236-DEL-62

chr11-71885053-DEL-71
chr11-71931523-DEL-50
chr11-71993393-DEL-217
chr11-71993775-DEL-113
chr11-72073329-DEL-88
chr11-72073543-DEL-326
chr11-72189403-DEL-1482
chr11-72202807-DEL-248
chr11-72312847-DEL-144
chr11-72430513-DEL-50
chr11-72452795-DEL-81
chr11-72583919-DEL-55
chr11-72740533-DEL-125
chr11-72740778-DEL-95
chr11-73030676-DEL-174
chr11-73036698-DEL-91
chr11-73052803-DEL-56
chr11-73091527-DEL-55
chr11-73100050-DEL-60
chr11-73206077-DEL-428
chr11-73283726-DEL-59
chr11-73322999-DEL-12535
chr11-73495966-DEL-52
chr11-73496867-DEL-198
chr11-73675337-DEL-4399
chr11-73709199-DEL-388
chr11-73746286-DEL-266
chr11-73888506-DEL-65
chr11-74073084-DEL-76
chr11-74162940-DEL-150
chr11-74197007-DEL-4998
chr11-74429008-DEL-127
chr11-74486532-DEL-266
chr11-74503088-DEL-52
chr11-74842281-DEL-118
chr11-74978097-DEL-680
chr11-75078127-DEL-640
chr11-75108185-DEL-50
chr11-75113377-DEL-270
chr11-75179505-DEL-180
chr11-75274003-DEL-254
chr11-75409292-DEL-55
chr11-75614335-DEL-222
chr11-75700196-DEL-366
chr11-75710898-DEL-347
chr11-75830853-DEL-145
chr11-75869635-DEL-52
chr11-75872645-DEL-300
chr11-75929126-DEL-50
chr11-76047572-DEL-136
chr11-76094436-DEL-71
chr11-76161551-DEL-253
chr11-76407528-DEL-61
chr11-76520978-DEL-53
chr11-76543283-DEL-403
chr11-76579782-DEL-60
chr11-76590082-DEL-911
chr11-76607631-DEL-105
chr11-76611582-DEL-50
chr11-76901694-DEL-110
chr11-76929917-DEL-70
chr11-77003056-DEL-95
chr11-77221038-DEL-78
chr11-77540420-DEL-58
chr11-77540771-DEL-197
chr11-77568038-DEL-72
chr11-77570060-DEL-356
chr11-77580144-DEL-50
chr11-77638458-DEL-314
chr11-77666169-DEL-52
chr11-77694637-DEL-76
chr11-77740400-DEL-59
chr11-77740687-DEL-50
chr11-77809679-DEL-225
chr11-77934163-DEL-143
chr11-77934406-DEL-173
chr11-77934688-DEL-168

chr11-78002359-DEL-74
chr11-78075618-DEL-79
chr11-78129466-DEL-311
chr11-78130264-DEL-222
chr11-78187064-DEL-197
chr11-78351935-DEL-277
chr11-78448917-DEL-52
chr11-78467081-DEL-737
chr11-78483101-DEL-1004
chr11-78548759-DEL-244
chr11-78549128-DEL-272
chr11-78570817-DEL-82
chr11-78739047-DEL-50
chr11-78782136-DEL-219
chr11-78873386-DEL-352
chr11-78930792-DEL-174
chr11-78958623-DEL-51
chr11-79045548-DEL-126
chr11-79070777-DEL-50
chr11-79099042-DEL-57
chr11-79171684-DEL-58
chr11-79204268-DEL-164
chr11-79243767-DEL-207
chr11-79243986-DEL-51
chr11-79350993-DEL-165
chr11-79351057-DEL-55
chr11-79476004-DEL-155
chr11-79623176-DEL-161
chr11-79623229-DEL-54
chr11-79639684-DEL-81
chr11-79682135-DEL-478
chr11-79703289-DEL-112
chr11-79709980-DEL-131
chr11-79839593-DEL-50
chr11-79859740-DEL-577
chr11-79918834-DEL-221
chr11-79931328-DEL-50
chr11-79983168-DEL-1219
chr11-80093584-DEL-311
chr11-80112034-DEL-108
chr11-80240171-DEL-145
chr11-80315982-DEL-111
chr11-80357057-DEL-453
chr11-80376176-DEL-93
chr11-80376305-DEL-97
chr11-80417580-DEL-1144
chr11-80448825-DEL-473
chr11-80454822-DEL-383
chr11-80458812-DEL-198
chr11-80476536-DEL-87
chr11-80574526-DEL-263
chr11-80574660-DEL-129
chr11-80806818-DEL-70
chr11-80836224-DEL-52
chr11-81073135-DEL-110
chr11-81163579-DEL-51
chr11-81280572-DEL-54
chr11-81339135-DEL-174
chr11-81362737-DEL-56
chr11-81458589-DEL-108
chr11-81520184-DEL-120
chr11-81620957-DEL-1086
chr11-81666648-DEL-52
chr11-81747444-DEL-255
chr11-81767520-DEL-66
chr11-81793586-DEL-77
chr11-81801385-DEL-57
chr11-82045992-DEL-69
chr11-82185321-DEL-52
chr11-82286249-DEL-50
chr11-82364560-DEL-391
chr11-82548859-DEL-78
chr11-82548885-DEL-571
chr11-82620343-DEL-63
chr11-82658372-DEL-158
chr11-82658739-DEL-221
chr11-82821027-DEL-84

chr11-82823992-DEL-153
chr11-82891447-DEL-58
chr11-82892852-DEL-97
chr11-82895748-DEL-101
chr11-82950392-DEL-1542
chr11-82974312-DEL-950
chr11-83446046-DEL-340
chr11-83446511-DEL-282
chr11-83450038-DEL-372
chr11-83470953-DEL-79
chr11-83586360-DEL-60
chr11-83708316-DEL-50
chr11-83720213-DEL-62
chr11-83730775-DEL-60
chr11-83808337-DEL-60
chr11-84005246-DEL-55
chr11-84103124-DEL-357
chr11-84339709-DEL-134
chr11-84399844-DEL-156
chr11-84402595-DEL-193
chr11-84459844-DEL-54
chr11-84525727-DEL-56
chr11-84784287-DEL-328
chr11-84784463-DEL-83
chr11-84829964-DEL-60
chr11-84833359-DEL-58
chr11-85116589-DEL-1818
chr11-85160598-DEL-50
chr11-85330305-DEL-2437
chr11-85395205-DEL-267
chr11-85395483-DEL-252
chr11-85395590-DEL-54
chr11-85480873-DEL-81
chr11-85493072-DEL-89
chr11-85565062-DEL-94
chr11-85567318-DEL-200
chr11-85598852-DEL-54
chr11-85607638-DEL-184
chr11-85822947-DEL-71
chr11-85843017-DEL-58
chr11-86068676-DEL-416
chr11-86344922-DEL-52
chr11-86368602-DEL-65
chr11-86444535-DEL-50
chr11-86455554-DEL-51
chr11-86468910-DEL-99
chr11-86521910-DEL-217
chr11-86531643-DEL-285
chr11-86603820-DEL-50
chr11-86665619-DEL-65
chr11-86665835-DEL-210
chr11-86671347-DEL-169
chr11-86690645-DEL-61
chr11-86690808-DEL-202
chr11-86728893-DEL-54
chr11-87060130-DEL-1111
chr11-87153550-DEL-217
chr11-87343538-DEL-204
chr11-87354919-DEL-474
chr11-87544910-DEL-60
chr11-87648850-DEL-50
chr11-87773389-DEL-50
chr11-87780953-DEL-50
chr11-87788802-DEL-127
chr11-87791384-DEL-99
chr11-87863067-DEL-73
chr11-87945975-DEL-272
chr11-88015654-DEL-57
chr11-88040524-DEL-198
chr11-88081177-DEL-50
chr11-88081718-DEL-124
chr11-88120216-DEL-82
chr11-88207600-DEL-62
chr11-88583599-DEL-173
chr11-88641123-DEL-111
chr11-88670832-DEL-69
chr11-88756495-DEL-1395

chr11-88801415-DEL-50
chr11-88833178-DEL-149
chr11-88838615-DEL-57
chr11-88846966-DEL-54
chr11-88860440-DEL-139
chr11-88860585-DEL-69
chr11-88876727-DEL-56
chr11-88940115-DEL-297
chr11-88970752-DEL-66
chr11-89016076-DEL-183
chr11-89113458-DEL-50
chr11-89260091-DEL-51
chr11-89272391-DEL-52
chr11-89305150-DEL-58
chr11-89523337-DEL-86
chr11-89591787-DEL-107
chr11-89652602-DEL-836
chr11-89690061-DEL-716
chr11-89721385-DEL-929
chr11-89726215-DEL-57
chr11-89754939-DEL-52
chr11-89793460-DEL-64
chr11-89800947-DEL-214
chr11-90052665-DEL-270
chr11-90074514-DEL-219
chr11-90146185-DEL-127
chr11-90170898-DEL-142
chr11-90192804-DEL-175
chr11-90297661-DEL-60
chr11-90316363-DEL-337
chr11-90398452-DEL-2822
chr11-90467933-DEL-68
chr11-90645722-DEL-88
chr11-90680625-DEL-50
chr11-90704484-DEL-253
chr11-90704938-DEL-473
chr11-90802724-DEL-2677
chr11-90869434-DEL-215
chr11-90996484-DEL-4608
chr11-91059453-DEL-103
chr11-91082536-DEL-56
chr11-91157853-DEL-609
chr11-91190822-DEL-92
chr11-91233247-DEL-5345
chr11-91238118-DEL-93
chr11-91264299-DEL-50
chr11-91270495-DEL-50
chr11-91362417-DEL-73
chr11-91391346-DEL-112
chr11-91436556-DEL-50
chr11-91468886-DEL-94
chr11-91579827-DEL-80
chr11-91695134-DEL-252
chr11-91753969-DEL-170
chr11-91754177-DEL-278
chr11-91754262-DEL-55
chr11-91781081-DEL-857
chr11-91806087-DEL-264
chr11-91861403-DEL-159
chr11-91915892-DEL-62
chr11-91916067-DEL-108
chr11-91920251-DEL-511
chr11-91938584-DEL-57
chr11-92150503-DEL-2656
chr11-92153437-DEL-1427
chr11-92396311-DEL-70
chr11-92577292-DEL-72
chr11-92868860-DEL-53
chr11-92882581-DEL-181
chr11-92896025-DEL-50
chr11-92914034-DEL-243
chr11-92939854-DEL-50
chr11-93100222-DEL-50
chr11-93135629-DEL-57
chr11-93202196-DEL-64
chr11-93395515-DEL-102
chr11-93467147-DEL-1440

chr11-93517973-DEL-111
chr11-93545038-DEL-52
chr11-93576351-DEL-53
chr11-93579441-DEL-56
chr11-93593299-DEL-56
chr11-93614018-DEL-177
chr11-93780655-DEL-148
chr11-93892081-DEL-361
chr11-93892616-DEL-79
chr11-93892892-DEL-283
chr11-94000614-DEL-134
chr11-94008509-DEL-273
chr11-94062059-DEL-74
chr11-94077005-DEL-50
chr11-94077704-DEL-50
chr11-94115589-DEL-359
chr11-94131521-DEL-52
chr11-94132011-DEL-130
chr11-94260434-DEL-66
chr11-94418927-DEL-254
chr11-94446644-DEL-50
chr11-94482756-DEL-159
chr11-94485440-DEL-727
chr11-94491207-DEL-50
chr11-94508780-DEL-1566
chr11-94539614-DEL-107
chr11-94683492-DEL-15559
chr11-94692537-DEL-15600
chr11-94805378-DEL-50
chr11-94850634-DEL-50
chr11-95097878-DEL-138
chr11-95098086-DEL-140
chr11-95209461-DEL-141
chr11-95357064-DEL-262
chr11-95357960-DEL-69
chr11-95427937-DEL-175
chr11-95667348-DEL-137
chr11-95689817-DEL-729
chr11-95690084-DEL-262
chr11-95690484-DEL-55
chr11-95767531-DEL-97
chr11-95799391-DEL-1347
chr11-95803314-DEL-65
chr11-95803460-DEL-66
chr11-95828218-DEL-50
chr11-96113331-DEL-75
chr11-96113426-DEL-211
chr11-96359981-DEL-74
chr11-96365550-DEL-148
chr11-96382413-DEL-776
chr11-96563192-DEL-50
chr11-96758864-DEL-91
chr11-96968736-DEL-209
chr11-97225162-DEL-77
chr11-97230348-DEL-73
chr11-97512251-DEL-170
chr11-97529956-DEL-250
chr11-97530528-DEL-234
chr11-97530955-DEL-126
chr11-97531182-DEL-132
chr11-97626909-DEL-50
chr11-97642514-DEL-99
chr11-97735397-DEL-63
chr11-97736218-DEL-151
chr11-97771736-DEL-219
chr11-97840608-DEL-72
chr11-97873012-DEL-402
chr11-97873158-DEL-67
chr11-98010276-DEL-90
chr11-98011361-DEL-339
chr11-98088320-DEL-66
chr11-98295102-DEL-129
chr11-98399199-DEL-108
chr11-98399501-DEL-75
chr11-98495481-DEL-177
chr11-98495814-DEL-52
chr11-98560170-DEL-50

chr11-98684481-DEL-101
chr11-98721515-DEL-176
chr11-98910519-DEL-250
chr11-98911666-DEL-188
chr11-98949521-DEL-54
chr11-99021924-DEL-76
chr11-99145801-DEL-71
chr11-99149176-DEL-69
chr11-99178118-DEL-120
chr11-99256645-DEL-58
chr11-99256775-DEL-50
chr11-99298616-DEL-152
chr11-99298728-DEL-420
chr11-99298972-DEL-60
chr11-99375728-DEL-50
chr11-99424134-DEL-62
chr11-99663375-DEL-75
chr11-99680084-DEL-58
chr11-99687348-DEL-231
chr11-99687393-DEL-72
chr11-99690303-DEL-101
chr11-99690458-DEL-125
chr11-99694761-DEL-90
chr11-99695784-DEL-56
chr11-99695958-DEL-161
chr11-99696034-DEL-56
chr11-99716234-DEL-847
chr11-99741823-DEL-57
chr11-99779974-DEL-409
chr11-99903030-DEL-52
chr11-99973558-DEL-56
chr11-100019574-DEL-196
chr11-100126115-DEL-72
chr11-100174194-DEL-117
chr11-100206144-DEL-58
chr11-100237806-DEL-138
chr11-100258124-DEL-1220
chr11-100371455-DEL-50
chr11-100507670-DEL-57
chr11-100539841-DEL-84
chr11-100573765-DEL-50
chr11-100785685-DEL-108
chr11-100787618-DEL-57
chr11-100820930-DEL-56
chr11-100835440-DEL-52
chr11-100876145-DEL-107
chr11-100880363-DEL-70
chr11-100896158-DEL-102
chr11-101015531-DEL-62
chr11-101093055-DEL-81
chr11-101099717-DEL-74
chr11-101245131-DEL-565
chr11-101257749-DEL-149
chr11-101300123-DEL-60
chr11-101312300-DEL-115
chr11-101351883-DEL-154
chr11-101431860-DEL-185
chr11-101516957-DEL-173
chr11-101537089-DEL-1936
chr11-101710780-DEL-50
chr11-101738817-DEL-50
chr11-101759747-DEL-221
chr11-101904400-DEL-96
chr11-101937776-DEL-78
chr11-102000777-DEL-110
chr11-102005623-DEL-63
chr11-102025919-DEL-2827
chr11-102026859-DEL-102
chr11-102314566-DEL-78
chr11-102374894-DEL-93
chr11-102523993-DEL-1908
chr11-102763876-DEL-421
chr11-102763998-DEL-84
chr11-102764460-DEL-87
chr11-102781404-DEL-358
chr11-102782042-DEL-112
chr11-102904489-DEL-644

chr11-102938743-DEL-458
chr11-102973726-DEL-166
chr11-103205497-DEL-136
chr11-103311621-DEL-310
chr11-103401507-DEL-164
chr11-103418411-DEL-58
chr11-103444489-DEL-1167
chr11-103476757-DEL-61
chr11-103562669-DEL-74
chr11-103651674-DEL-864
chr11-103675036-DEL-61
chr11-103734653-DEL-60
chr11-103753571-DEL-82
chr11-103813562-DEL-59
chr11-103818715-DEL-497
chr11-103881180-DEL-57
chr11-103924838-DEL-52
chr11-104041434-DEL-53
chr11-104148944-DEL-57
chr11-104167521-DEL-192
chr11-104167875-DEL-613
chr11-104216471-DEL-51
chr11-104252280-DEL-50
chr11-104291850-DEL-1891
chr11-104373895-DEL-116
chr11-104378089-DEL-76
chr11-104386849-DEL-858
chr11-104402743-DEL-50
chr11-104596863-DEL-129
chr11-104621283-DEL-108
chr11-104638695-DEL-383
chr11-104639031-DEL-84
chr11-104705512-DEL-184
chr11-104705803-DEL-156
chr11-104757452-DEL-89
chr11-104769407-DEL-53
chr11-104935917-DEL-214
chr11-104952045-DEL-52
chr11-105006679-DEL-408
chr11-105011701-DEL-1968
chr11-105044032-DEL-54
chr11-105052658-DEL-1159
chr11-105060130-DEL-190
chr11-105150226-DEL-180
chr11-105162347-DEL-141
chr11-105163238-DEL-55
chr11-105446200-DEL-116
chr11-105549377-DEL-51
chr11-105567155-DEL-62
chr11-105742963-DEL-525
chr11-105763358-DEL-1129
chr11-105844829-DEL-647
chr11-105846033-DEL-97
chr11-105917129-DEL-66
chr11-105961159-DEL-88
chr11-106007840-DEL-50
chr11-106029481-DEL-77
chr11-106364487-DEL-50
chr11-106469080-DEL-374
chr11-106475011-DEL-420
chr11-106555646-DEL-245
chr11-106606569-DEL-52
chr11-106609719-DEL-166
chr11-106616375-DEL-593
chr11-106616897-DEL-167
chr11-106646261-DEL-56
chr11-106646267-DEL-125
chr11-106667577-DEL-79
chr11-106706134-DEL-62
chr11-106834675-DEL-6139
chr11-106835747-DEL-2644
chr11-106846043-DEL-60
chr11-106851822-DEL-213
chr11-106894216-DEL-1418
chr11-106939282-DEL-731
chr11-106939365-DEL-2154
chr11-106941275-DEL-158

chr11-106950534-DEL-112
chr11-107040214-DEL-63
chr11-107049097-DEL-203
chr11-107089640-DEL-60
chr11-107123487-DEL-217
chr11-107167140-DEL-62
chr11-107292407-DEL-262
chr11-107312914-DEL-54
chr11-107318074-DEL-355
chr11-107407126-DEL-204
chr11-107418241-DEL-57
chr11-107487049-DEL-138
chr11-107487059-DEL-68
chr11-107502133-DEL-66
chr11-107637182-DEL-51
chr11-107641438-DEL-165
chr11-107672135-DEL-56
chr11-107675462-DEL-472
chr11-107682472-DEL-68
chr11-107716586-DEL-860
chr11-107749761-DEL-52
chr11-107749983-DEL-242
chr11-107754173-DEL-51
chr11-107770274-DEL-52
chr11-107808360-DEL-57
chr11-107820004-DEL-50
chr11-107845464-DEL-80
chr11-108153211-DEL-88
chr11-108351186-DEL-56
chr11-108416933-DEL-300
chr11-108416933-DEL-620
chr11-108539533-DEL-50
chr11-108796621-DEL-872
chr11-108917259-DEL-51
chr11-108919296-DEL-160
chr11-108987104-DEL-240
chr11-109005230-DEL-52
chr11-109010042-DEL-1695
chr11-109031721-DEL-180
chr11-109033644-DEL-54
chr11-109042548-DEL-88
chr11-109055854-DEL-680
chr11-109055869-DEL-76
chr11-109056416-DEL-73
chr11-109056660-DEL-56
chr11-109056711-DEL-330
chr11-109061321-DEL-156
chr11-109111264-DEL-441
chr11-109127555-DEL-86
chr11-109162086-DEL-932
chr11-109162882-DEL-74
chr11-109163137-DEL-441
chr11-109179516-DEL-94
chr11-109238023-DEL-50
chr11-109322408-DEL-58
chr11-109351848-DEL-54
chr11-109406697-DEL-372
chr11-109428651-DEL-228
chr11-109428787-DEL-92
chr11-109455999-DEL-141
chr11-109456350-DEL-198
chr11-109517326-DEL-1389
chr11-109526227-DEL-74
chr11-109581727-DEL-96
chr11-109669338-DEL-51
chr11-109687148-DEL-225
chr11-109689876-DEL-2330
chr11-109706243-DEL-71
chr11-109714977-DEL-491
chr11-109730951-DEL-76
chr11-109760621-DEL-111
chr11-109804454-DEL-51
chr11-109844110-DEL-56
chr11-109864915-DEL-60
chr11-109878674-DEL-341
chr11-109910463-DEL-4821
chr11-109923268-DEL-294

chr11-110086586-DEL-63
chr11-110230292-DEL-59
chr11-110276161-DEL-151
chr11-110277628-DEL-139
chr11-110277846-DEL-57
chr11-110358401-DEL-1083
chr11-110358519-DEL-279
chr11-110358860-DEL-372
chr11-110359141-DEL-62
chr11-110432326-DEL-50
chr11-110432438-DEL-90
chr11-110432664-DEL-115
chr11-110450153-DEL-64
chr11-110450365-DEL-581
chr11-110474833-DEL-364
chr11-110516058-DEL-100
chr11-110573114-DEL-50
chr11-110615929-DEL-80
chr11-110640418-DEL-199
chr11-110715693-DEL-98
chr11-110723274-DEL-61
chr11-110759111-DEL-71
chr11-110792999-DEL-58
chr11-110837757-DEL-61
chr11-110874891-DEL-1199
chr11-110897218-DEL-196
chr11-110999328-DEL-71
chr11-111002218-DEL-50
chr11-111003741-DEL-50
chr11-111018574-DEL-51
chr11-111090634-DEL-86
chr11-111179027-DEL-767
chr11-111273464-DEL-57
chr11-111287830-DEL-339
chr11-111331827-DEL-78
chr11-111357776-DEL-1033
chr11-111361476-DEL-88
chr11-111373048-DEL-245
chr11-111438314-DEL-53
chr11-111613162-DEL-971
chr11-111620312-DEL-136
chr11-111639793-DEL-50
chr11-111670987-DEL-576
chr11-111691259-DEL-117
chr11-111746634-DEL-114
chr11-111772300-DEL-187
chr11-111772303-DEL-64
chr11-111811795-DEL-72
chr11-111938061-DEL-1213
chr11-111955023-DEL-64
chr11-111989879-DEL-1118
chr11-112226805-DEL-50
chr11-112230118-DEL-77
chr11-112420678-DEL-72
chr11-112452550-DEL-51
chr11-112571636-DEL-50
chr11-112585175-DEL-77
chr11-112606493-DEL-50
chr11-112620688-DEL-104
chr11-112625675-DEL-88
chr11-112625679-DEL-178
chr11-112625986-DEL-127
chr11-112769160-DEL-54
chr11-112777509-DEL-60
chr11-112802811-DEL-56
chr11-112838588-DEL-69
chr11-112957858-DEL-56
chr11-112957858-DEL-119
chr11-113062159-DEL-53
chr11-113139409-DEL-60
chr11-113192075-DEL-518
chr11-113222831-DEL-50
chr11-113265216-DEL-59
chr11-113312945-DEL-62
chr11-113375132-DEL-82
chr11-113400890-DEL-184
chr11-113403089-DEL-77

chr11-113434807-DEL-84
chr11-113695057-DEL-52
chr11-113736029-DEL-152
chr11-113771027-DEL-93
chr11-113867474-DEL-57
chr11-113900993-DEL-230
chr11-114005466-DEL-98
chr11-114021925-DEL-86
chr11-114039758-DEL-50
chr11-114136257-DEL-97
chr11-114159675-DEL-386
chr11-114186917-DEL-286
chr11-114224853-DEL-52
chr11-114239794-DEL-102
chr11-114250062-DEL-55
chr11-114262770-DEL-118
chr11-114272017-DEL-224
chr11-114272232-DEL-50
chr11-114280256-DEL-63
chr11-114303150-DEL-66
chr11-114404307-DEL-51
chr11-114436666-DEL-50
chr11-114541648-DEL-175
chr11-114553480-DEL-58
chr11-114565168-DEL-248
chr11-114627697-DEL-56
chr11-114701522-DEL-50
chr11-114729663-DEL-54
chr11-114729851-DEL-135
chr11-114752211-DEL-57
chr11-114898244-DEL-4285
chr11-114908208-DEL-326
chr11-114909764-DEL-57
chr11-114918591-DEL-79
chr11-114918915-DEL-326
chr11-114926447-DEL-13642
chr11-114929348-DEL-79
chr11-114935499-DEL-66
chr11-115023988-DEL-88
chr11-115192797-DEL-55
chr11-115258958-DEL-57
chr11-115276708-DEL-52
chr11-115368465-DEL-560
chr11-115377243-DEL-208
chr11-115610778-DEL-303
chr11-115616159-DEL-63
chr11-115616385-DEL-193
chr11-115619218-DEL-302
chr11-115619750-DEL-110
chr11-115625899-DEL-101
chr11-115640779-DEL-50
chr11-115662249-DEL-92
chr11-115662575-DEL-2188
chr11-115681282-DEL-220
chr11-115834090-DEL-71
chr11-115849160-DEL-408
chr11-115849391-DEL-51
chr11-115930637-DEL-199
chr11-115934472-DEL-133
chr11-115965402-DEL-59
chr11-115983470-DEL-125
chr11-116014948-DEL-80
chr11-116028443-DEL-67
chr11-116031474-DEL-660
chr11-116032442-DEL-386
chr11-116059988-DEL-2550
chr11-116060290-DEL-68
chr11-116167146-DEL-88
chr11-116177904-DEL-82
chr11-116277971-DEL-156
chr11-116304919-DEL-55
chr11-116330443-DEL-58
chr11-116347657-DEL-60
chr11-116386859-DEL-429
chr11-116406808-DEL-91
chr11-116416404-DEL-139
chr11-116416572-DEL-60

chr11-116502667-DEL-50
chr11-116598832-DEL-84
chr11-116630045-DEL-26669
chr11-116661407-DEL-54
chr11-116774716-DEL-204
chr11-116779142-DEL-110
chr11-116800451-DEL-60
chr11-116800932-DEL-90
chr11-116862146-DEL-50
chr11-116862228-DEL-147
chr11-116910047-DEL-50
chr11-116943766-DEL-138
chr11-116943806-DEL-66
chr11-116958437-DEL-207
chr11-116977901-DEL-54
chr11-116987055-DEL-152
chr11-117087955-DEL-72
chr11-117198397-DEL-104
chr11-117259611-DEL-106
chr11-117274977-DEL-50
chr11-117342490-DEL-162
chr11-117342839-DEL-60
chr11-117343045-DEL-90
chr11-117354486-DEL-96
chr11-117385202-DEL-52
chr11-117390634-DEL-5136
chr11-117458340-DEL-98
chr11-117521653-DEL-206
chr11-117526763-DEL-102
chr11-117743648-DEL-86
chr11-117794523-DEL-153
chr11-117809770-DEL-50
chr11-117838353-DEL-143
chr11-118045981-DEL-572
chr11-118050605-DEL-37602
chr11-118082050-DEL-240
chr11-118247467-DEL-112
chr11-118247611-DEL-50
chr11-118286351-DEL-120
chr11-118332539-DEL-370
chr11-118510820-DEL-72
chr11-118659237-DEL-53
chr11-118675841-DEL-124
chr11-118692013-DEL-66
chr11-118692146-DEL-462
chr11-118692441-DEL-132
chr11-118713834-DEL-134
chr11-118764227-DEL-417
chr11-118960243-DEL-60
chr11-118977856-DEL-89
chr11-118988905-DEL-52
chr11-119003451-DEL-96
chr11-119003761-DEL-181
chr11-119085269-DEL-58
chr11-119206705-DEL-158
chr11-119244555-DEL-105
chr11-119251430-DEL-169
chr11-119264125-DEL-63
chr11-119269063-DEL-347
chr11-119282460-DEL-50
chr11-119319650-DEL-115
chr11-119350139-DEL-50
chr11-119359308-DEL-65
chr11-119396600-DEL-78
chr11-119473068-DEL-349
chr11-119669502-DEL-186
chr11-119670116-DEL-404
chr11-119704404-DEL-65
chr11-119750381-DEL-152
chr11-119886220-DEL-50
chr11-120044871-DEL-69
chr11-120059600-DEL-59
chr11-120095424-DEL-207
chr11-120140054-DEL-866
chr11-120219367-DEL-61
chr11-120232911-DEL-433
chr11-120391910-DEL-799

chr11-120397532-DEL-180
chr11-120404670-DEL-85
chr11-120415118-DEL-60
chr11-120522283-DEL-50
chr11-120633976-DEL-480
chr11-120634433-DEL-173
chr11-120634508-DEL-71
chr11-120665740-DEL-497
chr11-120672022-DEL-133
chr11-120987033-DEL-162
chr11-121043359-DEL-144
chr11-121072820-DEL-77
chr11-121076125-DEL-52
chr11-121268301-DEL-50
chr11-121768617-DEL-50
chr11-121772276-DEL-60
chr11-121820774-DEL-102
chr12-3146550-DEL-54
chr12-3246585-DEL-11714
chr12-3325788-DEL-50
chr12-3370862-DEL-86
chr12-3405988-DEL-70
chr12-3407309-DEL-280
chr12-3446108-DEL-575
chr12-3446142-DEL-105
chr12-3446593-DEL-106
chr12-3518650-DEL-52
chr12-3567462-DEL-140
chr12-3629653-DEL-1588
chr12-3639632-DEL-1049
chr12-3862089-DEL-136
chr12-3896992-DEL-250
chr12-3897094-DEL-80
chr12-3903318-DEL-52
chr12-3975178-DEL-58
chr12-4107967-DEL-55
chr12-4113022-DEL-52
chr12-4113148-DEL-139
chr12-4139403-DEL-122
chr12-4163425-DEL-670
chr12-4168235-DEL-80
chr12-4202525-DEL-282
chr12-4212383-DEL-152
chr12-4222338-DEL-336
chr12-4237866-DEL-275
chr12-4269564-DEL-1792
chr12-4271019-DEL-217
chr12-4326421-DEL-107
chr12-4342175-DEL-760
chr12-4408917-DEL-56
chr12-4443988-DEL-58
chr12-4480809-DEL-496
chr12-4486930-DEL-480
chr12-4487392-DEL-384
chr12-4516692-DEL-340
chr12-4562163-DEL-90
chr12-4677639-DEL-51
chr12-4744910-DEL-186
chr12-4829235-DEL-194
chr12-4830035-DEL-93
chr12-4845283-DEL-130
chr12-4865508-DEL-200
chr12-4875592-DEL-102
chr12-4927640-DEL-126
chr12-4957594-DEL-91
chr12-5012274-DEL-274
chr12-5076372-DEL-61
chr12-5100556-DEL-743
chr12-5101851-DEL-3374
chr12-5107924-DEL-1239
chr12-5109936-DEL-61
chr12-5111202-DEL-311
chr12-5174933-DEL-63
chr12-5375404-DEL-56
chr12-5413393-DEL-65
chr12-5556125-DEL-352
chr12-5556431-DEL-334

chr12-5946978-DEL-238
chr12-5960713-DEL-73
chr12-6066113-DEL-1148
chr12-6226233-DEL-51
chr12-6246961-DEL-70
chr12-6273593-DEL-856
chr12-6329203-DEL-74
chr12-6329368-DEL-322
chr12-6370341-DEL-64
chr12-6556005-DEL-341
chr12-6588285-DEL-154
chr12-6593100-DEL-4430
chr12-6625326-DEL-53
chr12-6685497-DEL-846
chr12-6692414-DEL-779
chr12-6736765-DEL-191
chr12-6749969-DEL-160
chr12-6750354-DEL-115
chr12-6924175-DEL-74
chr12-7010357-DEL-86
chr12-7044967-DEL-144
chr12-7193088-DEL-8575
chr12-7383393-DEL-197
chr12-7482851-DEL-1273
chr12-7568395-DEL-70
chr12-7691560-DEL-1511
chr12-7715035-DEL-168
chr12-7808603-DEL-53
chr12-7940360-DEL-150
chr12-7987931-DEL-1669
chr12-8228438-DEL-416
chr12-8418345-DEL-62
chr12-8449499-DEL-116
chr12-8483427-DEL-147
chr12-8483650-DEL-64
chr12-8492996-DEL-52
chr12-8509512-DEL-54
chr12-8539802-DEL-1712
chr12-8539917-DEL-127
chr12-8540551-DEL-474
chr12-8667228-DEL-143
chr12-8810236-DEL-175
chr12-8865182-DEL-77
chr12-9013970-DEL-260
chr12-9014313-DEL-111
chr12-9014584-DEL-1323
chr12-9018032-DEL-532
chr12-9054741-DEL-112
chr12-9071131-DEL-172
chr12-9095474-DEL-84
chr12-9366772-DEL-162
chr12-9457502-DEL-2654
chr12-9475312-DEL-221
chr12-9475955-DEL-1206
chr12-9493729-DEL-1297
chr12-9494171-DEL-220
chr12-9494660-DEL-200
chr12-9636369-DEL-54
chr12-9695383-DEL-50
chr12-9769176-DEL-269
chr12-9868219-DEL-54
chr12-9999864-DEL-1246
chr12-10001613-DEL-104
chr12-10168138-DEL-53
chr12-10216656-DEL-76
chr12-10216840-DEL-87
chr12-10906183-DEL-72
chr12-10953686-DEL-63
chr12-10955303-DEL-72
chr12-10986964-DEL-54
chr12-11011639-DEL-1428
chr12-11100810-DEL-63
chr12-11116236-DEL-56
chr12-11218661-DEL-69
chr12-11231779-DEL-245
chr12-11513151-DEL-141
chr12-11667752-DEL-100

chr12-11694834-DEL-1386
chr12-11844451-DEL-310
chr12-11900677-DEL-201
chr12-11918258-DEL-105
chr12-11930702-DEL-54
chr12-12061770-DEL-99
chr12-12237456-DEL-216
chr12-12339452-DEL-65
chr12-12380675-DEL-2473
chr12-12400885-DEL-130
chr12-12420619-DEL-326
chr12-12420873-DEL-290
chr12-12431350-DEL-50
chr12-12455393-DEL-68
chr12-12584465-DEL-55
chr12-12844764-DEL-347
chr12-12848394-DEL-84
chr12-12879507-DEL-73
chr12-12905844-DEL-494
chr12-12922123-DEL-154
chr12-12961974-DEL-408
chr12-13112490-DEL-1486
chr12-13263443-DEL-280
chr12-13309674-DEL-356
chr12-13362075-DEL-2787
chr12-13504651-DEL-2019
chr12-13562345-DEL-557
chr12-13651901-DEL-226
chr12-13652082-DEL-50
chr12-13653085-DEL-81
chr12-13866030-DEL-56
chr12-13866450-DEL-116
chr12-14016506-DEL-310
chr12-14047463-DEL-636
chr12-14155378-DEL-50
chr12-14232512-DEL-52
chr12-14328920-DEL-71
chr12-14387374-DEL-76
chr12-14418900-DEL-80
chr12-14419080-DEL-68
chr12-14419580-DEL-64
chr12-14709480-DEL-548
chr12-14841022-DEL-416
chr12-14965216-DEL-57
chr12-15046013-DEL-143
chr12-15100680-DEL-109
chr12-15117117-DEL-372
chr12-15624937-DEL-50
chr12-15743024-DEL-119
chr12-15862064-DEL-138
chr12-15875239-DEL-922
chr12-15892692-DEL-56
chr12-15939497-DEL-74
chr12-15953472-DEL-60
chr12-16041018-DEL-120
chr12-16065446-DEL-127
chr12-16216381-DEL-1264
chr12-16366345-DEL-1458
chr12-16388167-DEL-74
chr12-16430996-DEL-154
chr12-16571482-DEL-50
chr12-16654660-DEL-501
chr12-16682609-DEL-62
chr12-16688497-DEL-210
chr12-17133681-DEL-152
chr12-17134075-DEL-114
chr12-17243836-DEL-295
chr12-17244060-DEL-96
chr12-17440352-DEL-966
chr12-17446782-DEL-56
chr12-17487865-DEL-136
chr12-17488262-DEL-72
chr12-17490035-DEL-50
chr12-17493979-DEL-50
chr12-17519034-DEL-117
chr12-17553495-DEL-174
chr12-17669053-DEL-62

chr12-17896175-DEL-74
chr12-17979292-DEL-75
chr12-18385730-DEL-391
chr12-18388762-DEL-58
chr12-18396817-DEL-460
chr12-18407952-DEL-90
chr12-18408313-DEL-82
chr12-18409615-DEL-70
chr12-18902800-DEL-565
chr12-18906352-DEL-210
chr12-18908607-DEL-57
chr12-18913510-DEL-61
chr12-18978513-DEL-62
chr12-18983888-DEL-66
chr12-18989311-DEL-435
chr12-19011888-DEL-178
chr12-19048396-DEL-108
chr12-19056666-DEL-99
chr12-19247337-DEL-352
chr12-19484072-DEL-82
chr12-19908945-DEL-826
chr12-19929400-DEL-573
chr12-20376534-DEL-60
chr12-20464388-DEL-322
chr12-20482503-DEL-185
chr12-20683629-DEL-314
chr12-20854379-DEL-329
chr12-20949444-DEL-386
chr12-21066057-DEL-84
chr12-21125262-DEL-237
chr12-21208525-DEL-179
chr12-21300031-DEL-72
chr12-21402816-DEL-55
chr12-21452299-DEL-304
chr12-21503267-DEL-191
chr12-21536605-DEL-70
chr12-21675094-DEL-63
chr12-21943755-DEL-630
chr12-22043468-DEL-336
chr12-22075896-DEL-173
chr12-22449850-DEL-315
chr12-22567208-DEL-65
chr12-22668411-DEL-287
chr12-22682741-DEL-52
chr12-22686707-DEL-2508
chr12-22723725-DEL-3036
chr12-22774543-DEL-438
chr12-23114669-DEL-52
chr12-23164008-DEL-236
chr12-23174833-DEL-127
chr12-23247576-DEL-64
chr12-23247730-DEL-100
chr12-23379957-DEL-206
chr12-23455658-DEL-50
chr12-23478100-DEL-318
chr12-23709909-DEL-195
chr12-23714458-DEL-1574
chr12-23731368-DEL-572
chr12-23734937-DEL-318
chr12-24331870-DEL-110
chr12-24395667-DEL-2732
chr12-24612080-DEL-71
chr12-24726667-DEL-472
chr12-24809758-DEL-51
chr12-24930641-DEL-200
chr12-24938952-DEL-111
chr12-24995241-DEL-50
chr12-25100277-DEL-62
chr12-25191218-DEL-71
chr12-25262460-DEL-119
chr12-25297600-DEL-54
chr12-25322051-DEL-162
chr12-25335113-DEL-3814
chr12-25463448-DEL-193
chr12-25463540-DEL-56
chr12-25463706-DEL-101
chr12-25641497-DEL-3792

chr12-25686117-DEL-1482
chr12-25850863-DEL-201
chr12-26085032-DEL-208
chr12-26099649-DEL-1417
chr12-26106506-DEL-54
chr12-26235921-DEL-1626
chr12-26281070-DEL-93
chr12-26338350-DEL-1497
chr12-26346273-DEL-56
chr12-26397050-DEL-73
chr12-26412140-DEL-78
chr12-26504454-DEL-50
chr12-26642661-DEL-153
chr12-26893851-DEL-100
chr12-26893978-DEL-223
chr12-26918721-DEL-495
chr12-27133045-DEL-52
chr12-27291471-DEL-1338
chr12-27292020-DEL-2627
chr12-27312376-DEL-52
chr12-27558307-DEL-2464
chr12-27568550-DEL-120
chr12-27568602-DEL-55
chr12-27661891-DEL-180
chr12-27719059-DEL-90
chr12-27728717-DEL-180
chr12-27740427-DEL-192
chr12-27740586-DEL-523
chr12-27740645-DEL-204
chr12-27740663-DEL-60
chr12-27761788-DEL-57
chr12-27765351-DEL-141
chr12-27769081-DEL-948
chr12-27797905-DEL-1196
chr12-27819309-DEL-54
chr12-28036871-DEL-91
chr12-28139360-DEL-68
chr12-28204864-DEL-78
chr12-28288896-DEL-84
chr12-28411329-DEL-101
chr12-28446156-DEL-186
chr12-28462909-DEL-52
chr12-28530132-DEL-246
chr12-28625313-DEL-62
chr12-28697503-DEL-1049
chr12-28794143-DEL-54
chr12-28794461-DEL-206
chr12-28804596-DEL-453
chr12-28864658-DEL-53
chr12-28910675-DEL-105
chr12-28910958-DEL-108
chr12-28912271-DEL-293
chr12-28937027-DEL-330
chr12-29097850-DEL-394
chr12-29098064-DEL-503
chr12-29196272-DEL-200
chr12-29196632-DEL-62
chr12-29214532-DEL-110
chr12-29316095-DEL-66
chr12-29977853-DEL-52
chr12-30104404-DEL-56
chr12-30240588-DEL-2108
chr12-30262367-DEL-263
chr12-30405316-DEL-264
chr12-30469865-DEL-100
chr12-30628736-DEL-233
chr12-30741340-DEL-165
chr12-30741429-DEL-69
chr12-30781187-DEL-188
chr12-30864834-DEL-1114
chr12-30973156-DEL-324
chr12-31020790-DEL-116
chr12-31052020-DEL-177
chr12-31124481-DEL-141
chr12-31128376-DEL-85
chr12-31134723-DEL-117
chr12-31196921-DEL-57

chr12-31260187-DEL-50
chr12-31272070-DEL-176
chr12-31320080-DEL-219
chr12-31443685-DEL-90
chr12-31560955-DEL-194
chr12-31603862-DEL-51
chr12-31655923-DEL-455
chr12-31677184-DEL-116
chr12-31685966-DEL-178
chr12-31768623-DEL-95
chr12-31779447-DEL-518
chr12-31820038-DEL-190
chr12-31820480-DEL-193
chr12-31875956-DEL-74
chr12-31898952-DEL-116
chr12-31921830-DEL-60
chr12-31924934-DEL-93
chr12-31971449-DEL-60
chr12-32062845-DEL-56
chr12-32159094-DEL-4061
chr12-32200806-DEL-58
chr12-32271913-DEL-56
chr12-32275657-DEL-50
chr12-32291132-DEL-378
chr12-32301241-DEL-119
chr12-32315363-DEL-1970
chr12-32351542-DEL-196
chr12-32460253-DEL-957
chr12-32467847-DEL-970
chr12-32468264-DEL-424
chr12-32558670-DEL-9200
chr12-32635955-DEL-432
chr12-32636081-DEL-96
chr12-32636771-DEL-192
chr12-32720885-DEL-79
chr12-32898195-DEL-88
chr12-32952328-DEL-3767
chr12-32974026-DEL-565
chr12-33056694-DEL-89
chr12-33066418-DEL-50
chr12-33223457-DEL-74
chr12-33253082-DEL-50
chr12-33273734-DEL-124
chr12-33344717-DEL-81
chr12-33383409-DEL-52
chr12-33457744-DEL-2480
chr12-33458871-DEL-230
chr12-33832942-DEL-80
chr12-33866053-DEL-54
chr12-33873797-DEL-55
chr12-33966719-DEL-64
chr12-34004356-DEL-76
chr12-34310144-DEL-176
chr12-34402861-DEL-56
chr12-34417668-DEL-226
chr12-34417933-DEL-96
chr12-34506396-DEL-52
chr12-34519861-DEL-50
chr12-34584267-DEL-110
chr12-34636311-DEL-74
chr12-34729871-DEL-94
chr12-34898887-DEL-60
chr12-34900659-DEL-84
chr12-34948679-DEL-114
chr12-34968477-DEL-67
chr12-35004501-DEL-124
chr12-35046654-DEL-341
chr12-35074891-DEL-354
chr12-35075006-DEL-159
chr12-35127699-DEL-224
chr12-35471471-DEL-68
chr12-35529668-DEL-1156
chr12-35850917-DEL-13586
chr12-35937345-DEL-128
chr12-35937817-DEL-56
chr12-36011366-DEL-59
chr12-36155313-DEL-1628

chr12-36156722-DEL-74
chr12-36228353-DEL-192
chr12-36264125-DEL-121
chr12-36438646-DEL-84
chr12-36500162-DEL-104
chr12-36514441-DEL-340
chr12-36582134-DEL-185
chr12-36649637-DEL-60
chr12-36729105-DEL-1867
chr12-36877867-DEL-136
chr12-36877891-DEL-56
chr12-37303409-DEL-60
chr12-37611675-DEL-123
chr12-37707100-DEL-60
chr12-37740843-DEL-77
chr12-37808553-DEL-2253
chr12-37860054-DEL-60
chr12-37884782-DEL-52
chr12-37943348-DEL-75
chr12-37995240-DEL-3644
chr12-38124748-DEL-1462
chr12-38391440-DEL-52
chr12-38502883-DEL-50
chr12-38541226-DEL-638
chr12-38541243-DEL-90
chr12-38630692-DEL-102
chr12-38879404-DEL-60
chr12-38946260-DEL-670
chr12-39057956-DEL-74
chr12-39143566-DEL-533
chr12-39183145-DEL-54
chr12-39210216-DEL-64
chr12-39289015-DEL-3113
chr12-39473690-DEL-83
chr12-39765810-DEL-62
chr12-39768130-DEL-464
chr12-39808701-DEL-50
chr12-39809509-DEL-55
chr12-40000853-DEL-50
chr12-40029577-DEL-75
chr12-40035367-DEL-358
chr12-40131043-DEL-67
chr12-40256846-DEL-57
chr12-40358576-DEL-59
chr12-40380749-DEL-2938
chr12-40652389-DEL-72
chr12-40652534-DEL-224
chr12-40682079-DEL-1824
chr12-40698537-DEL-290
chr12-40989638-DEL-109
chr12-41000068-DEL-50
chr12-41054120-DEL-1890
chr12-41212817-DEL-741
chr12-41267558-DEL-102
chr12-41301605-DEL-52
chr12-41347379-DEL-8781
chr12-41347740-DEL-3670
chr12-41435508-DEL-350
chr12-41579075-DEL-330
chr12-41617204-DEL-2881
chr12-41630908-DEL-62
chr12-41737960-DEL-54
chr12-41856830-DEL-338
chr12-41951750-DEL-321
chr12-41982626-DEL-740
chr12-42097365-DEL-127
chr12-42328519-DEL-159
chr12-42361105-DEL-1372
chr12-42368179-DEL-120
chr12-42410745-DEL-57
chr12-42410879-DEL-322
chr12-42440397-DEL-54
chr12-42614266-DEL-60
chr12-42926431-DEL-77
chr12-43127505-DEL-2268
chr12-43153479-DEL-26534
chr12-43258039-DEL-256

chr12-43262600-DEL-2099
chr12-43267047-DEL-1399
chr12-43354873-DEL-69
chr12-43401651-DEL-74
chr12-43459417-DEL-4174
chr12-43469576-DEL-1981
chr12-43730761-DEL-268
chr12-43927825-DEL-50
chr12-43950291-DEL-230
chr12-43952532-DEL-84
chr12-44063006-DEL-58
chr12-44067742-DEL-84
chr12-44087879-DEL-59
chr12-44189506-DEL-103
chr12-44298589-DEL-3987
chr12-44432534-DEL-62
chr12-44796541-DEL-58
chr12-44952248-DEL-57
chr12-45010830-DEL-113
chr12-45137835-DEL-58
chr12-45157055-DEL-914
chr12-45228033-DEL-1073
chr12-45251770-DEL-4247
chr12-45299676-DEL-66
chr12-45333664-DEL-77
chr12-45366518-DEL-207
chr12-45383537-DEL-2061
chr12-45449205-DEL-185
chr12-45449205-DEL-407
chr12-45459406-DEL-120
chr12-45463841-DEL-2332
chr12-45472430-DEL-533
chr12-45473028-DEL-77
chr12-45594527-DEL-139
chr12-45678318-DEL-6196
chr12-45890951-DEL-1083
chr12-45893143-DEL-57
chr12-46057656-DEL-1759
chr12-46088845-DEL-1115
chr12-46213430-DEL-3347
chr12-46236209-DEL-177
chr12-46344102-DEL-1103
chr12-46399626-DEL-737
chr12-46495895-DEL-62
chr12-46509021-DEL-614
chr12-46605771-DEL-2050
chr12-46726620-DEL-50
chr12-46822732-DEL-457
chr12-46829650-DEL-65
chr12-46888861-DEL-50
chr12-46921247-DEL-61
chr12-46926038-DEL-615
chr12-46927378-DEL-175
chr12-47043751-DEL-66
chr12-47043895-DEL-426
chr12-47074120-DEL-2466
chr12-47088836-DEL-498
chr12-47349871-DEL-1027
chr12-47407128-DEL-199
chr12-47443822-DEL-136
chr12-47542342-DEL-58
chr12-47628148-DEL-1706
chr12-47715213-DEL-1969
chr12-47741137-DEL-54
chr12-47854172-DEL-325
chr12-47930160-DEL-288
chr12-48040940-DEL-198
chr12-48060895-DEL-51
chr12-48113052-DEL-120
chr12-48221874-DEL-169
chr12-48286283-DEL-50
chr12-48348915-DEL-181
chr12-48404520-DEL-1091
chr12-48502457-DEL-79
chr12-48516043-DEL-520
chr12-48603363-DEL-1621
chr12-48604960-DEL-334

chr12-48857849-DEL-67
chr12-48945479-DEL-140
chr12-49068269-DEL-115
chr12-49293875-DEL-1278
chr12-49343644-DEL-52
chr12-49735882-DEL-50
chr12-50022149-DEL-70
chr12-50449478-DEL-6491
chr12-50616336-DEL-551
chr12-50620181-DEL-110
chr12-50899671-DEL-59
chr12-50928539-DEL-144
chr12-51084617-DEL-54
chr12-51097774-DEL-100
chr12-51304474-DEL-70
chr12-51369307-DEL-67
chr12-51447789-DEL-239
chr12-51520888-DEL-186
chr12-51582875-DEL-658
chr12-51590929-DEL-102
chr12-51621656-DEL-98
chr12-51633883-DEL-55
chr12-51799890-DEL-143
chr12-51868552-DEL-336
chr12-51924926-DEL-531
chr12-51938785-DEL-50
chr12-51949158-DEL-581
chr12-52072035-DEL-54
chr12-52105214-DEL-255
chr12-52438389-DEL-51
chr12-52455325-DEL-461
chr12-52625896-DEL-50
chr12-52717579-DEL-50
chr12-52774799-DEL-151
chr12-52814627-DEL-156
chr12-52849232-DEL-432
chr12-52870109-DEL-51
chr12-52873507-DEL-56
chr12-52901165-DEL-84
chr12-52953936-DEL-184
chr12-52973608-DEL-5374
chr12-52973856-DEL-311
chr12-52979166-DEL-82
chr12-53153374-DEL-64
chr12-53178722-DEL-70
chr12-53224659-DEL-264
chr12-53352050-DEL-77
chr12-53395702-DEL-501
chr12-53395998-DEL-242
chr12-53396313-DEL-54
chr12-53471919-DEL-132
chr12-53520413-DEL-584
chr12-53674926-DEL-1005
chr12-53754046-DEL-444
chr12-53789098-DEL-322
chr12-53820598-DEL-131
chr12-53847695-DEL-200
chr12-53985331-DEL-1020
chr12-54202078-DEL-86
chr12-54305267-DEL-154
chr12-54306182-DEL-60
chr12-54364461-DEL-56
chr12-54420207-DEL-66
chr12-54438074-DEL-2978
chr12-54466726-DEL-57
chr12-54558447-DEL-404
chr12-54589448-DEL-169
chr12-54607205-DEL-186
chr12-54621238-DEL-686
chr12-54687443-DEL-60
chr12-54743801-DEL-197
chr12-54766690-DEL-52
chr12-54858079-DEL-75
chr12-54916298-DEL-68
chr12-54946431-DEL-53
chr12-54999412-DEL-78
chr12-55074609-DEL-206

chr12-55074988-DEL-296
chr12-55110166-DEL-60
chr12-55385500-DEL-227
chr12-55603298-DEL-165
chr12-55603501-DEL-55
chr12-55792318-DEL-139
chr12-55810626-DEL-66
chr12-55836361-DEL-77
chr12-55872251-DEL-128
chr12-55886557-DEL-55
chr12-55924917-DEL-66
chr12-55955885-DEL-184
chr12-55976229-DEL-405
chr12-55976413-DEL-64
chr12-56098974-DEL-64
chr12-56145185-DEL-50
chr12-56146396-DEL-53
chr12-56211445-DEL-3357
chr12-56227402-DEL-2838
chr12-56248919-DEL-91
chr12-56250813-DEL-67
chr12-56319882-DEL-56
chr12-56327700-DEL-106
chr12-56395830-DEL-1164
chr12-56425112-DEL-143
chr12-56456207-DEL-89
chr12-56514158-DEL-55
chr12-56663186-DEL-141
chr12-56754216-DEL-106
chr12-56931103-DEL-106
chr12-56957888-DEL-2817
chr12-56998099-DEL-50
chr12-57064944-DEL-148
chr12-57199347-DEL-72
chr12-57260626-DEL-70
chr12-57261070-DEL-59
chr12-57351479-DEL-116
chr12-57436139-DEL-157
chr12-57436333-DEL-249
chr12-57485709-DEL-51
chr12-57798865-DEL-530
chr12-57924804-DEL-90
chr12-58028086-DEL-76
chr12-58167649-DEL-409
chr12-58191589-DEL-62
chr12-58273259-DEL-199
chr12-58293766-DEL-657
chr12-58557380-DEL-50
chr12-58703631-DEL-51
chr12-58944601-DEL-50
chr12-58971903-DEL-177
chr12-59028218-DEL-54
chr12-59081759-DEL-2755
chr12-59145917-DEL-65
chr12-59160208-DEL-1122
chr12-59167015-DEL-65
chr12-59184843-DEL-324
chr12-59246941-DEL-174
chr12-59310546-DEL-70
chr12-59523799-DEL-137
chr12-59690187-DEL-307
chr12-59800857-DEL-559
chr12-59801189-DEL-731
chr12-59921523-DEL-1665
chr12-59994196-DEL-198
chr12-60000784-DEL-10064
chr12-60074762-DEL-176
chr12-60082882-DEL-197
chr12-60082904-DEL-73
chr12-60118673-DEL-128
chr12-60192622-DEL-52
chr12-60209352-DEL-54
chr12-60247462-DEL-124
chr12-60381197-DEL-72
chr12-60422707-DEL-90
chr12-60902875-DEL-60
chr12-60902915-DEL-159

chr12-61016113-DEL-72
chr12-61104347-DEL-103
chr12-61117785-DEL-3040
chr12-61232820-DEL-93
chr12-61285365-DEL-119
chr12-61367601-DEL-59
chr12-61588308-DEL-127
chr12-61601309-DEL-903
chr12-61614978-DEL-56
chr12-61615277-DEL-2738
chr12-61703301-DEL-303
chr12-61735349-DEL-52
chr12-61763266-DEL-192
chr12-61855289-DEL-50
chr12-61934129-DEL-59
chr12-62057888-DEL-50
chr12-62085315-DEL-50
chr12-62133990-DEL-512
chr12-62149687-DEL-1669
chr12-62154875-DEL-201
chr12-62198369-DEL-3641
chr12-62245693-DEL-194
chr12-62260103-DEL-106
chr12-62346033-DEL-265
chr12-62347147-DEL-74
chr12-62471501-DEL-2566
chr12-62527306-DEL-229
chr12-62621065-DEL-4540
chr12-62704980-DEL-94
chr12-62712875-DEL-52
chr12-62769487-DEL-155
chr12-62828135-DEL-171
chr12-62931946-DEL-149
chr12-62973115-DEL-2631
chr12-63070824-DEL-401
chr12-63127444-DEL-50
chr12-63247071-DEL-825
chr12-63260997-DEL-642
chr12-63341117-DEL-2834
chr12-63360043-DEL-581
chr12-63366721-DEL-2495
chr12-63516557-DEL-50
chr12-63532650-DEL-401
chr12-63534868-DEL-72
chr12-63535068-DEL-732
chr12-63620531-DEL-4214
chr12-63696171-DEL-91
chr12-63711456-DEL-1170
chr12-63711607-DEL-51
chr12-63733582-DEL-946
chr12-63761576-DEL-368
chr12-64023779-DEL-153
chr12-64088133-DEL-75
chr12-64178768-DEL-1446
chr12-64217065-DEL-51
chr12-64240594-DEL-1368
chr12-64267466-DEL-59
chr12-64370314-DEL-181
chr12-64447033-DEL-82
chr12-64447224-DEL-53
chr12-64447520-DEL-1464
chr12-64646616-DEL-60
chr12-64699258-DEL-85
chr12-64699430-DEL-363
chr12-64710948-DEL-3185
chr12-64714244-DEL-897
chr12-64784268-DEL-4037
chr12-64796424-DEL-72
chr12-64840899-DEL-50
chr12-65138200-DEL-220
chr12-65170157-DEL-96
chr12-65242658-DEL-208
chr12-65254462-DEL-118
chr12-65290830-DEL-78
chr12-65349100-DEL-61
chr12-65387665-DEL-55
chr12-65434334-DEL-50

chr12-65461771-DEL-169
chr12-65545631-DEL-169
chr12-65570714-DEL-549
chr12-65691936-DEL-2613
chr12-65887870-DEL-540
chr12-65924648-DEL-108
chr12-65937741-DEL-172
chr12-66017737-DEL-55
chr12-66218598-DEL-53
chr12-66229293-DEL-58
chr12-66416392-DEL-50
chr12-66447167-DEL-565
chr12-66531283-DEL-56
chr12-66715545-DEL-54
chr12-66798712-DEL-942
chr12-67084560-DEL-185
chr12-67441835-DEL-50
chr12-67442023-DEL-80
chr12-67570940-DEL-177
chr12-67571326-DEL-75
chr12-67796358-DEL-189
chr12-67796685-DEL-204
chr12-67809593-DEL-442
chr12-67881730-DEL-416
chr12-67962235-DEL-66
chr12-68113130-DEL-1338
chr12-68335493-DEL-1321
chr12-68360622-DEL-141
chr12-68440399-DEL-56
chr12-68454366-DEL-567
chr12-68580809-DEL-156
chr12-68590178-DEL-252
chr12-68832595-DEL-76
chr12-68883918-DEL-126
chr12-69074780-DEL-73
chr12-69100991-DEL-234
chr12-69123953-DEL-77
chr12-69193764-DEL-1172
chr12-69242300-DEL-171
chr12-69322941-DEL-588
chr12-69372125-DEL-1570
chr12-69406482-DEL-162
chr12-69433005-DEL-92
chr12-69448372-DEL-51
chr12-69488074-DEL-362
chr12-69488107-DEL-159
chr12-69490634-DEL-73
chr12-69506241-DEL-235
chr12-69506582-DEL-64
chr12-69535652-DEL-84
chr12-69556265-DEL-234
chr12-69556284-DEL-693
chr12-69564789-DEL-50
chr12-69655206-DEL-190
chr12-69714985-DEL-2523
chr12-69792737-DEL-66
chr12-69801580-DEL-51
chr12-69830970-DEL-100
chr12-69842316-DEL-5013
chr12-69920948-DEL-64
chr12-70002588-DEL-370
chr12-70036752-DEL-200
chr12-70075379-DEL-159
chr12-70112651-DEL-514
chr12-70193951-DEL-98
chr12-70315569-DEL-64
chr12-70376629-DEL-2134
chr12-70516938-DEL-84
chr12-70547626-DEL-56
chr12-70600384-DEL-1649
chr12-70697033-DEL-66
chr12-70730355-DEL-232
chr12-70730367-DEL-68
chr12-70878774-DEL-174
chr12-70929819-DEL-64
chr12-71135294-DEL-5731
chr12-71136264-DEL-133

chr12-71152964-DEL-218
chr12-71184893-DEL-51
chr12-71240412-DEL-234
chr12-71312565-DEL-179
chr12-71404902-DEL-1185
chr12-71492639-DEL-145
chr12-71515228-DEL-196
chr12-71663599-DEL-222
chr12-71681306-DEL-51
chr12-71698508-DEL-64
chr12-71710375-DEL-296
chr12-71809290-DEL-105
chr12-71809459-DEL-84
chr12-72021160-DEL-58
chr12-72075541-DEL-75
chr12-72269343-DEL-88
chr12-72287710-DEL-177
chr12-72364907-DEL-58
chr12-72414949-DEL-91
chr12-72466599-DEL-50
chr12-72672349-DEL-128
chr12-72681571-DEL-1422
chr12-72683228-DEL-134
chr12-72720630-DEL-284
chr12-72723328-DEL-50
chr12-72790186-DEL-296
chr12-72791869-DEL-57
chr12-72898445-DEL-153
chr12-72937299-DEL-84
chr12-72937327-DEL-276
chr12-72964364-DEL-120
chr12-73034410-DEL-54
chr12-73057618-DEL-73
chr12-73110913-DEL-64
chr12-73118834-DEL-61
chr12-73161680-DEL-98
chr12-73177759-DEL-104
chr12-73186373-DEL-156
chr12-73284080-DEL-91
chr12-73395175-DEL-152
chr12-73460835-DEL-66
chr12-73472981-DEL-72
chr12-73472989-DEL-196
chr12-73486203-DEL-62
chr12-73541482-DEL-75
chr12-73574972-DEL-160
chr12-73603174-DEL-96
chr12-73627930-DEL-50
chr12-73665331-DEL-71
chr12-73741849-DEL-523
chr12-73756373-DEL-55
chr12-74058777-DEL-72
chr12-74059040-DEL-52
chr12-74266949-DEL-768
chr12-74431549-DEL-59
chr12-74930480-DEL-68
chr12-74930715-DEL-263
chr12-75093193-DEL-86
chr12-75165088-DEL-123
chr12-75226654-DEL-2058
chr12-75341075-DEL-50
chr12-75384196-DEL-76
chr12-75451818-DEL-165
chr12-75454319-DEL-100
chr12-75502605-DEL-196
chr12-75660910-DEL-54
chr12-75700169-DEL-51
chr12-75759312-DEL-121
chr12-75875641-DEL-127
chr12-75895964-DEL-465
chr12-76086450-DEL-134
chr12-76203387-DEL-245
chr12-76203447-DEL-54
chr12-76207445-DEL-93
chr12-76207702-DEL-93
chr12-76226693-DEL-474
chr12-76226793-DEL-148

chr12-76236350-DEL-618
chr12-76236632-DEL-204
chr12-76258813-DEL-100
chr12-76375824-DEL-51
chr12-76376197-DEL-72
chr12-76468394-DEL-69
chr12-76508999-DEL-142
chr12-76615671-DEL-241
chr12-76683580-DEL-180
chr12-76732599-DEL-50
chr12-76824261-DEL-200
chr12-76827997-DEL-144
chr12-76904751-DEL-52
chr12-76968343-DEL-82
chr12-76977316-DEL-424
chr12-77034251-DEL-55
chr12-77048108-DEL-55
chr12-77143598-DEL-54
chr12-77182603-DEL-206
chr12-77228216-DEL-274
chr12-77340782-DEL-226
chr12-77371662-DEL-419
chr12-77379615-DEL-199
chr12-77485852-DEL-63
chr12-77485852-DEL-306
chr12-77486047-DEL-152
chr12-77554543-DEL-115
chr12-77606345-DEL-50
chr12-77649058-DEL-154
chr12-77674301-DEL-76
chr12-77709490-DEL-68
chr12-77734234-DEL-56
chr12-77798705-DEL-174
chr12-77798785-DEL-76
chr12-77844161-DEL-52
chr12-77895678-DEL-68
chr12-77929740-DEL-2191
chr12-78039470-DEL-62
chr12-78123367-DEL-54
chr12-78276384-DEL-50
chr12-78453180-DEL-5214
chr12-78483893-DEL-50
chr12-78617486-DEL-50
chr12-78620639-DEL-54
chr12-78707495-DEL-227
chr12-78709989-DEL-230
chr12-78891172-DEL-201
chr12-78949618-DEL-166
chr12-78987117-DEL-591
chr12-78987989-DEL-60
chr12-78987989-DEL-170
chr12-79272068-DEL-57
chr12-79272235-DEL-190
chr12-79300157-DEL-60
chr12-79306108-DEL-302
chr12-79306361-DEL-58
chr12-79503822-DEL-50
chr12-79547508-DEL-50
chr12-79608450-DEL-7398
chr12-79646739-DEL-88
chr12-79662011-DEL-78
chr12-79671933-DEL-50
chr12-79686407-DEL-276
chr12-79724255-DEL-504
chr12-79985770-DEL-84
chr12-80006059-DEL-64
chr12-80010781-DEL-64
chr12-80039060-DEL-440
chr12-80039069-DEL-156
chr12-80043384-DEL-50
chr12-80200170-DEL-156
chr12-80202314-DEL-50
chr12-80287460-DEL-98
chr12-80427268-DEL-76
chr12-80509542-DEL-97
chr12-80586134-DEL-51
chr12-80644666-DEL-96

chr12-80666055-DEL-3363
chr12-80764393-DEL-125
chr12-80835701-DEL-398
chr12-80881849-DEL-1136
chr12-80919659-DEL-146
chr12-80930333-DEL-58
chr12-80958237-DEL-5311
chr12-81038383-DEL-3354
chr12-81057338-DEL-210
chr12-81118661-DEL-120
chr12-81328615-DEL-1417
chr12-81753558-DEL-86
chr12-81814251-DEL-74
chr12-81860399-DEL-160
chr12-81882157-DEL-95
chr12-81882673-DEL-132
chr12-81885705-DEL-50
chr12-81985049-DEL-102
chr12-82032683-DEL-60
chr12-82074874-DEL-13279
chr12-82102099-DEL-8495
chr12-82170583-DEL-1505
chr12-82174107-DEL-72
chr12-82210447-DEL-426
chr12-82257934-DEL-862
chr12-82328163-DEL-1795
chr12-82328624-DEL-237
chr12-82364695-DEL-984
chr12-82406209-DEL-161
chr12-82437703-DEL-405
chr12-82439188-DEL-105
chr12-82524292-DEL-60
chr12-82654516-DEL-90
chr12-82686418-DEL-212
chr12-82744701-DEL-67
chr12-82812493-DEL-62
chr12-82817948-DEL-235
chr12-82898904-DEL-173
chr12-82974869-DEL-161
chr12-83017855-DEL-58
chr12-83041597-DEL-55
chr12-83063010-DEL-89
chr12-83071148-DEL-78
chr12-83083096-DEL-73
chr12-83153681-DEL-50
chr12-83209931-DEL-54
chr12-83211570-DEL-196
chr12-83242663-DEL-260
chr12-83243062-DEL-124
chr12-83631869-DEL-288
chr12-83648967-DEL-50
chr12-83801226-DEL-1213
chr12-83807151-DEL-759
chr12-84012249-DEL-132
chr12-84012489-DEL-51
chr12-84053662-DEL-56
chr12-84107275-DEL-165
chr12-84107371-DEL-75
chr12-84133334-DEL-87
chr12-84158571-DEL-930
chr12-84169306-DEL-444
chr12-84316735-DEL-629
chr12-84317004-DEL-60
chr12-84317210-DEL-151
chr12-84319668-DEL-254
chr12-84329538-DEL-348
chr12-84441633-DEL-62
chr12-84450926-DEL-211
chr12-84556457-DEL-106
chr12-84592318-DEL-187
chr12-84631504-DEL-107
chr12-84664798-DEL-76
chr12-84671200-DEL-50
chr12-84776437-DEL-162
chr12-84776580-DEL-72
chr12-84857261-DEL-62
chr12-84885796-DEL-92

chr12-84929988-DEL-879
chr12-84988924-DEL-412
chr12-85128036-DEL-50
chr12-85178915-DEL-109
chr12-85188949-DEL-86
chr12-85210014-DEL-84
chr12-85230269-DEL-55
chr12-85263487-DEL-338
chr12-85386561-DEL-56
chr12-85398787-DEL-52
chr12-85565882-DEL-82
chr12-85684734-DEL-62
chr12-85791303-DEL-52
chr12-85797004-DEL-88
chr12-85914714-DEL-130
chr12-85951893-DEL-50
chr12-85977238-DEL-327
chr12-85984566-DEL-77
chr12-86204509-DEL-67
chr12-86418544-DEL-50
chr12-86562231-DEL-241
chr12-86572205-DEL-56
chr12-86572205-DEL-196
chr12-86579827-DEL-65
chr12-86598760-DEL-201
chr12-86638884-DEL-80
chr12-86684371-DEL-59
chr12-86716799-DEL-54
chr12-86812132-DEL-142
chr12-86819324-DEL-58
chr12-86874183-DEL-50
chr12-86892965-DEL-207
chr12-86949189-DEL-391
chr12-86991377-DEL-55
chr12-87026204-DEL-209
chr12-87083461-DEL-71
chr12-87097460-DEL-341
chr12-87142758-DEL-50
chr12-87263983-DEL-134
chr12-87337694-DEL-55
chr12-87560858-DEL-534
chr12-87569156-DEL-102
chr12-87603931-DEL-71
chr12-87832219-DEL-78
chr12-87921454-DEL-56
chr12-87922123-DEL-84
chr12-88016674-DEL-54
chr12-88080826-DEL-193
chr12-88081711-DEL-2590
chr12-88218533-DEL-213
chr12-88234149-DEL-52
chr12-88236076-DEL-757
chr12-88249776-DEL-182
chr12-88261517-DEL-238
chr12-88336353-DEL-51
chr12-88349249-DEL-492
chr12-88489949-DEL-132
chr12-88496402-DEL-442
chr12-88509949-DEL-64
chr12-88612926-DEL-983
chr12-88706150-DEL-77
chr12-89004105-DEL-500
chr12-89067976-DEL-116
chr12-89133960-DEL-211
chr12-89168528-DEL-68
chr12-89676868-DEL-62
chr12-89676997-DEL-90
chr12-89913504-DEL-2087
chr12-90205967-DEL-60
chr12-90377215-DEL-50
chr12-90422352-DEL-58
chr12-90422739-DEL-133
chr12-90495444-DEL-59
chr12-90545851-DEL-217
chr12-90719635-DEL-52
chr12-90763651-DEL-71
chr12-90936967-DEL-154

chr12-91058407-DEL-906
chr12-91084179-DEL-4155
chr12-91099345-DEL-86
chr12-91201957-DEL-1020
chr12-91202092-DEL-60
chr12-91239911-DEL-114
chr12-91251952-DEL-69
chr12-91252056-DEL-100
chr12-91259827-DEL-100
chr12-91298192-DEL-54
chr12-91441080-DEL-424
chr12-91575978-DEL-60
chr12-91684016-DEL-60
chr12-91686834-DEL-64
chr12-91806051-DEL-60
chr12-91931707-DEL-91
chr12-91942681-DEL-57
chr12-92024791-DEL-1161
chr12-92101010-DEL-951
chr12-92114455-DEL-401
chr12-92165110-DEL-66
chr12-92206310-DEL-85
chr12-92310353-DEL-64
chr12-92389593-DEL-662
chr12-92390590-DEL-145
chr12-92433485-DEL-50
chr12-92463894-DEL-436
chr12-92591004-DEL-1400
chr12-92656817-DEL-50
chr12-92687350-DEL-104
chr12-92687378-DEL-260
chr12-92691995-DEL-1854
chr12-92727845-DEL-1437
chr12-92805204-DEL-6174
chr12-93030926-DEL-5036
chr12-93052113-DEL-72
chr12-93150241-DEL-1082
chr12-93187574-DEL-323
chr12-93199010-DEL-4470
chr12-93352068-DEL-6196
chr12-93433637-DEL-68
chr12-93462580-DEL-152
chr12-93462623-DEL-54
chr12-93496812-DEL-1163
chr12-93543898-DEL-66
chr12-93713068-DEL-120
chr12-93873194-DEL-176
chr12-93887381-DEL-208
chr12-94074010-DEL-185
chr12-94243300-DEL-50
chr12-94245211-DEL-199
chr12-94245517-DEL-111
chr12-94245670-DEL-179
chr12-94289504-DEL-121
chr12-94448763-DEL-136
chr12-94628528-DEL-1121
chr12-94638421-DEL-52
chr12-94655448-DEL-314
chr12-94655794-DEL-60
chr12-94706750-DEL-1370
chr12-94710209-DEL-2909
chr12-94743015-DEL-64
chr12-94782943-DEL-144
chr12-94822441-DEL-62
chr12-94849049-DEL-52
chr12-94875193-DEL-51
chr12-94924214-DEL-61
chr12-94956441-DEL-4362
chr12-95006176-DEL-118
chr12-95026748-DEL-65
chr12-95053894-DEL-137
chr12-95124962-DEL-547
chr12-95145159-DEL-1541
chr12-95170256-DEL-146
chr12-95223458-DEL-236
chr12-95248328-DEL-703
chr12-95276664-DEL-104

chr12-95294316-DEL-238
chr12-95397268-DEL-201
chr12-95458816-DEL-78
chr12-95495633-DEL-2870
chr12-95500511-DEL-100
chr12-95525121-DEL-50
chr12-95540292-DEL-222
chr12-95580774-DEL-1108
chr12-95646350-DEL-57
chr12-95792391-DEL-2529
chr12-95832991-DEL-1259
chr12-95917833-DEL-58
chr12-95973152-DEL-201
chr12-96001412-DEL-64
chr12-96040278-DEL-192
chr12-96083438-DEL-108
chr12-96145827-DEL-1569
chr12-96149090-DEL-52
chr12-96159052-DEL-74
chr12-96412187-DEL-1552
chr12-96477753-DEL-424
chr12-96500649-DEL-121
chr12-96508036-DEL-243
chr12-96556056-DEL-62
chr12-96635853-DEL-50
chr12-96787495-DEL-115
chr12-96801979-DEL-66
chr12-96842745-DEL-50
chr12-96906199-DEL-555
chr12-96952585-DEL-183
chr12-96952595-DEL-65
chr12-97007136-DEL-3325
chr12-97199995-DEL-72
chr12-97279724-DEL-79
chr12-97318815-DEL-2200
chr12-97321480-DEL-161
chr12-97407509-DEL-73
chr12-97407640-DEL-60
chr12-97463462-DEL-2059
chr12-97630168-DEL-50
chr12-97653757-DEL-181
chr12-97721234-DEL-50
chr12-97845991-DEL-104
chr12-97907816-DEL-562
chr12-98048196-DEL-69
chr12-98204966-DEL-2970
chr12-98270529-DEL-108
chr12-98379082-DEL-221
chr12-98471166-DEL-55
chr12-98772000-DEL-1833
chr12-98877261-DEL-51
chr12-98916405-DEL-280
chr12-98936938-DEL-275
chr12-98965344-DEL-51
chr12-99072678-DEL-73
chr12-99092414-DEL-70
chr12-99134489-DEL-1556
chr12-99169728-DEL-565
chr12-99174099-DEL-54
chr12-99200963-DEL-101
chr12-99221310-DEL-204
chr12-99221704-DEL-219
chr12-99376224-DEL-171
chr12-99441433-DEL-278
chr12-99471105-DEL-144
chr12-99471369-DEL-3160
chr12-99471811-DEL-144
chr12-99473185-DEL-1970
chr12-99474222-DEL-766
chr12-99608272-DEL-50
chr12-99671651-DEL-156
chr12-99806405-DEL-175
chr12-99806695-DEL-69
chr12-99927948-DEL-65
chr12-99994174-DEL-1776
chr12-100049528-DEL-242
chr12-100187679-DEL-65

chr12-100208468-DEL-62
chr12-100208558-DEL-171
chr12-100241922-DEL-57
chr12-100309112-DEL-51
chr12-100343821-DEL-207
chr12-100352849-DEL-143
chr12-100355428-DEL-52
chr12-100448441-DEL-185
chr12-100454183-DEL-520
chr12-100457200-DEL-957
chr12-100567314-DEL-147
chr12-100655739-DEL-63
chr12-100694551-DEL-56
chr12-100694764-DEL-124
chr12-100758497-DEL-53
chr12-100758594-DEL-64
chr12-100784742-DEL-382
chr12-100822083-DEL-78
chr12-100861062-DEL-50
chr12-100933770-DEL-50
chr12-101505337-DEL-1000
chr12-101571183-DEL-50
chr12-101702384-DEL-78
chr12-101709000-DEL-95
chr12-101772447-DEL-50
chr12-101806166-DEL-262
chr12-101806483-DEL-52
chr12-101806716-DEL-118
chr12-101807011-DEL-243
chr12-101807110-DEL-1033
chr12-101807827-DEL-312
chr12-101843307-DEL-198
chr12-101920336-DEL-423
chr12-101927723-DEL-3717
chr12-102021467-DEL-76
chr12-102037426-DEL-9069
chr12-102089833-DEL-1705
chr12-102101122-DEL-109
chr12-102184650-DEL-150
chr12-102279984-DEL-79
chr12-102310697-DEL-238
chr12-102310750-DEL-112
chr12-102343697-DEL-131
chr12-102343792-DEL-56
chr12-102357513-DEL-77
chr12-102367114-DEL-74
chr12-102413100-DEL-640
chr12-102416174-DEL-52
chr12-102431850-DEL-159
chr12-102451231-DEL-69
chr12-102461153-DEL-52
chr12-102461384-DEL-132
chr12-102481616-DEL-140
chr12-102496597-DEL-1507
chr12-102595959-DEL-680
chr12-102625430-DEL-55
chr12-102626090-DEL-58
chr12-102626381-DEL-335
chr12-103069191-DEL-259
chr12-103141243-DEL-96
chr12-103174985-DEL-102
chr12-103175471-DEL-192
chr12-103179428-DEL-174
chr12-103230701-DEL-88
chr12-103260846-DEL-50
chr12-103263440-DEL-56
chr12-103352735-DEL-58
chr12-103487494-DEL-89
chr12-103505925-DEL-64
chr12-103521304-DEL-204
chr12-103575829-DEL-60
chr12-103793468-DEL-236
chr12-103858168-DEL-60
chr12-103898837-DEL-3288
chr12-104019682-DEL-272
chr12-104019836-DEL-118
chr12-104028829-DEL-145

chr12-104035293-DEL-61
chr12-104077933-DEL-61
chr12-104086360-DEL-66
chr12-104335691-DEL-52
chr12-104371368-DEL-90
chr12-104385160-DEL-1732
chr12-104408596-DEL-290
chr12-104434416-DEL-60
chr12-104478199-DEL-78
chr12-104478219-DEL-172
chr12-104478536-DEL-237
chr12-104555219-DEL-60
chr12-104611153-DEL-144
chr12-104770483-DEL-96
chr12-104799401-DEL-104
chr12-104849713-DEL-268
chr12-104854224-DEL-64
chr12-104877428-DEL-84
chr12-104909879-DEL-134
chr12-104914411-DEL-96
chr12-104989373-DEL-248
chr12-104989858-DEL-259
chr12-104990213-DEL-51
chr12-105001634-DEL-182
chr12-105005828-DEL-54
chr12-105107828-DEL-189
chr12-105148103-DEL-106
chr12-105148567-DEL-437
chr12-105299695-DEL-52
chr12-105365657-DEL-52
chr12-105428696-DEL-60
chr12-105469854-DEL-1764
chr12-105489087-DEL-969
chr12-105490747-DEL-78
chr12-105510930-DEL-102
chr12-105621809-DEL-52
chr12-105668674-DEL-368
chr12-105749755-DEL-179
chr12-105876766-DEL-168
chr12-105876840-DEL-336
chr12-105913590-DEL-176
chr12-105914440-DEL-104
chr12-105919983-DEL-110
chr12-105920066-DEL-294
chr12-105920261-DEL-126
chr12-106072333-DEL-56
chr12-106078010-DEL-142
chr12-106108569-DEL-97
chr12-106154239-DEL-117
chr12-106173038-DEL-71
chr12-106202143-DEL-88
chr12-106204180-DEL-81
chr12-106217594-DEL-102
chr12-106304501-DEL-55
chr12-106333937-DEL-115
chr12-106374933-DEL-114
chr12-106375211-DEL-230
chr12-106375312-DEL-671
chr12-106375880-DEL-145
chr12-106571582-DEL-148
chr12-106571882-DEL-168
chr12-106597831-DEL-53
chr12-106643763-DEL-216
chr12-106982550-DEL-57
chr12-107042226-DEL-64
chr12-107077508-DEL-318
chr12-107078063-DEL-75
chr12-107174103-DEL-202
chr12-107185116-DEL-60
chr12-107187525-DEL-50
chr12-107302759-DEL-591
chr12-107415184-DEL-85
chr12-107422435-DEL-64
chr12-107456366-DEL-50
chr12-107460650-DEL-280
chr12-107460887-DEL-326
chr12-107461030-DEL-128

chr12-107507352-DEL-62
chr12-107522533-DEL-426
chr12-107522804-DEL-426
chr12-107563972-DEL-55
chr12-107649295-DEL-1327
chr12-107808913-DEL-112
chr12-107830754-DEL-677
chr12-107990053-DEL-132
chr12-107995280-DEL-84
chr12-108013009-DEL-88
chr12-108037217-DEL-50
chr12-108051627-DEL-293
chr12-108051720-DEL-76
chr12-108059353-DEL-164
chr12-108180558-DEL-187
chr12-108248841-DEL-71
chr12-108254096-DEL-189
chr12-108290777-DEL-94
chr12-108291564-DEL-61
chr12-108315662-DEL-93
chr12-108324794-DEL-300
chr12-108324916-DEL-98
chr12-108422128-DEL-2390
chr12-108804431-DEL-50
chr12-108804537-DEL-156
chr12-108946279-DEL-62
chr12-108968368-DEL-167
chr12-109043196-DEL-288
chr12-109086060-DEL-428
chr12-109249990-DEL-50
chr12-109389511-DEL-69
chr12-109396784-DEL-129
chr12-109396807-DEL-54
chr12-109445427-DEL-215
chr12-109597053-DEL-93
chr12-109769393-DEL-493
chr12-109769488-DEL-87
chr12-109769886-DEL-145
chr12-109836435-DEL-63
chr12-109931897-DEL-54
chr12-109977473-DEL-1051
chr12-109979451-DEL-58
chr12-109979824-DEL-185
chr12-109980824-DEL-80
chr12-110007988-DEL-2005
chr12-110019567-DEL-95
chr12-110040988-DEL-54
chr12-110057747-DEL-102
chr12-110067045-DEL-50
chr12-110115486-DEL-154
chr12-110116815-DEL-168
chr12-110129078-DEL-57
chr12-110205112-DEL-105
chr12-110334896-DEL-1535
chr12-110335723-DEL-65
chr12-110336881-DEL-614
chr12-110468050-DEL-50
chr12-110676094-DEL-56
chr12-110686243-DEL-201
chr12-110700963-DEL-64
chr12-110865813-DEL-50
chr12-111035794-DEL-314
chr12-111035897-DEL-76
chr12-111108746-DEL-210
chr12-111144753-DEL-174
chr12-111144840-DEL-77
chr12-111247280-DEL-50
chr12-111256576-DEL-58
chr12-111477683-DEL-76
chr12-111495745-DEL-95
chr12-111696572-DEL-700
chr12-111704657-DEL-164
chr12-111708541-DEL-72
chr12-111762691-DEL-890
chr12-111816399-DEL-646
chr12-111834204-DEL-440
chr12-111834839-DEL-160

chr12-111895482-DEL-110
chr12-111976424-DEL-210
chr12-111976574-DEL-60
chr12-112173110-DEL-98
chr12-112263697-DEL-629
chr12-112374178-DEL-80
chr12-112376204-DEL-50
chr12-112380914-DEL-150
chr12-112402844-DEL-114
chr12-112411509-DEL-618
chr12-112452950-DEL-390
chr12-112513600-DEL-194
chr12-112527297-DEL-110
chr12-112572945-DEL-137
chr12-112575966-DEL-96
chr12-112666033-DEL-68
chr12-112679455-DEL-220
chr12-112807874-DEL-60
chr12-112826624-DEL-86
chr12-112860042-DEL-76
chr12-112922742-DEL-107
chr12-112980335-DEL-53
chr12-112986360-DEL-92
chr12-113012059-DEL-282
chr12-113079425-DEL-129
chr12-113188961-DEL-132
chr12-113189032-DEL-53
chr12-113202692-DEL-50
chr12-113211499-DEL-2931
chr12-113255973-DEL-144
chr12-113256811-DEL-51
chr12-113257214-DEL-155
chr12-113257428-DEL-182
chr12-113274244-DEL-245
chr12-113274799-DEL-293
chr12-113282294-DEL-69
chr12-113297879-DEL-50
chr12-113411661-DEL-145
chr12-113576449-DEL-119
chr12-113618543-DEL-98
chr12-113619722-DEL-86
chr12-113653303-DEL-2567
chr12-113661222-DEL-137
chr12-113701818-DEL-56
chr12-113701818-DEL-136
chr12-113707953-DEL-1478
chr12-113730478-DEL-26315
chr12-113736595-DEL-81
chr12-113852170-DEL-498
chr12-113902390-DEL-55
chr12-113904630-DEL-80
chr12-113907019-DEL-170
chr12-113910439-DEL-130
chr12-113912731-DEL-2108
chr12-113913321-DEL-229
chr12-113966815-DEL-240
chr12-113971866-DEL-235
chr12-113972088-DEL-82
chr12-113977974-DEL-198
chr12-113979272-DEL-1088
chr12-114015331-DEL-55
chr12-114122616-DEL-11626
chr12-114204947-DEL-208
chr12-114233669-DEL-56
chr12-114259020-DEL-2772
chr12-114350859-DEL-65
chr12-114375985-DEL-498
chr12-114378589-DEL-946
chr12-114379910-DEL-311
chr12-114410785-DEL-533
chr12-114420839-DEL-114
chr12-114453887-DEL-127
chr12-114464304-DEL-50
chr12-114488480-DEL-154
chr12-114488809-DEL-75
chr12-114505250-DEL-127
chr12-114553764-DEL-51

chr12-114555648-DEL-127
chr12-114580591-DEL-50
chr12-114584048-DEL-137
chr12-114587160-DEL-204
chr12-114611028-DEL-114
chr12-114611238-DEL-362
chr12-114614345-DEL-3158
chr12-114629094-DEL-4833
chr12-114641868-DEL-60
chr12-114662263-DEL-189
chr12-114673205-DEL-857
chr12-114732752-DEL-950
chr12-114752462-DEL-189
chr12-114783406-DEL-328
chr12-114803228-DEL-69
chr12-114806989-DEL-59
chr12-114808286-DEL-36239
chr12-114844446-DEL-853
chr12-114873594-DEL-1357
chr12-114911581-DEL-654
chr12-114912877-DEL-55
chr12-114913617-DEL-211
chr12-114951967-DEL-654
chr12-114989640-DEL-1108
chr12-114995702-DEL-407
chr12-115012107-DEL-329
chr12-115153515-DEL-1203
chr12-115160733-DEL-77
chr12-115217413-DEL-93
chr12-115432269-DEL-131
chr12-115504003-DEL-323
chr12-115514754-DEL-95
chr12-115516554-DEL-52
chr12-115519035-DEL-823
chr12-115530690-DEL-1657
chr12-115557564-DEL-50
chr12-115568875-DEL-3140
chr12-115572086-DEL-53
chr12-115598586-DEL-230
chr12-115598747-DEL-84
chr12-115606081-DEL-129
chr12-115613372-DEL-203
chr12-115660374-DEL-62
chr12-115758139-DEL-1314
chr12-115767119-DEL-173
chr12-115801001-DEL-54
chr12-115811519-DEL-13986
chr12-115812658-DEL-159
chr12-115812659-DEL-74
chr12-115820854-DEL-5606
chr12-115828664-DEL-74
chr12-115855499-DEL-58
chr12-115900015-DEL-194
chr12-115962363-DEL-1318
chr12-115998091-DEL-154
chr12-116025486-DEL-1024
chr12-116070002-DEL-1328
chr12-116185768-DEL-60
chr12-116212066-DEL-802
chr12-116212345-DEL-166
chr12-116213533-DEL-1454
chr12-116214692-DEL-295
chr12-116215323-DEL-322
chr12-116215328-DEL-77
chr12-116239053-DEL-52
chr12-116262341-DEL-53
chr12-116283139-DEL-52
chr12-116345783-DEL-58
chr12-116535699-DEL-294
chr12-116540392-DEL-300
chr12-116543276-DEL-50
chr12-116607467-DEL-81
chr12-116616179-DEL-53
chr12-116628900-DEL-52
chr12-116642069-DEL-54
chr12-116652999-DEL-8897
chr12-116662964-DEL-57

chr12-116663334-DEL-6164
chr12-116902597-DEL-84
chr12-116944947-DEL-108
chr12-116970804-DEL-149
chr12-116970944-DEL-343
chr12-116971546-DEL-116
chr12-117090062-DEL-50
chr12-117183082-DEL-74
chr12-117202970-DEL-147
chr12-117358439-DEL-660
chr12-117359570-DEL-1515
chr12-117363746-DEL-50
chr12-117406662-DEL-84
chr12-117409264-DEL-81
chr12-117578137-DEL-170
chr12-117579653-DEL-138
chr12-117664538-DEL-361
chr12-117699433-DEL-110
chr12-117812905-DEL-175
chr12-117825663-DEL-384
chr12-117840533-DEL-50
chr12-118049816-DEL-260
chr12-118317118-DEL-5659
chr12-118396921-DEL-66
chr12-118513806-DEL-64
chr12-118678755-DEL-72
chr12-118679892-DEL-114
chr12-118757888-DEL-3171
chr12-118878612-DEL-50
chr12-119323907-DEL-497
chr12-119420032-DEL-213
chr12-119516210-DEL-60
chr12-119734507-DEL-51
chr12-119819082-DEL-3512
chr12-119830926-DEL-309
chr12-119831067-DEL-72
chr12-119952945-DEL-4200
chr13-3200664-DEL-51
chr13-3398385-DEL-3598
chr13-3402771-DEL-4627
chr13-3499406-DEL-3223
chr13-3524748-DEL-1456
chr13-3845590-DEL-50
chr13-3882966-DEL-56
chr13-3901431-DEL-69
chr13-3901808-DEL-107
chr13-3903965-DEL-351
chr13-3912724-DEL-50
chr13-4014336-DEL-104
chr13-4014409-DEL-557
chr13-4014775-DEL-178
chr13-4197787-DEL-97
chr13-4469687-DEL-733
chr13-4655899-DEL-138
chr13-4656369-DEL-50
chr13-4656394-DEL-398
chr13-4877679-DEL-2454
chr13-4951743-DEL-70
chr13-4985137-DEL-56
chr13-5132509-DEL-501
chr13-5431121-DEL-3117
chr13-5459633-DEL-4377
chr13-5685388-DEL-74
chr13-5856296-DEL-75
chr13-5976884-DEL-65
chr13-5982275-DEL-1745
chr13-6043346-DEL-80
chr13-6095541-DEL-56
chr13-6213897-DEL-100
chr13-6299299-DEL-422
chr13-6361152-DEL-69
chr13-6365447-DEL-319
chr13-6502035-DEL-85
chr13-6502196-DEL-214
chr13-6527987-DEL-63
chr13-6552483-DEL-50
chr13-6656277-DEL-6120

chr13-6798138-DEL-52
chr13-7154149-DEL-230
chr13-7195927-DEL-52
chr13-7331166-DEL-57
chr13-7442203-DEL-80
chr13-7651788-DEL-3603
chr13-7779392-DEL-1288
chr13-8025986-DEL-2996
chr13-8360103-DEL-176
chr13-8533694-DEL-79
chr13-8681189-DEL-50
chr13-8731385-DEL-62
chr13-8785695-DEL-54
chr13-8805628-DEL-285
chr13-8819922-DEL-227
chr13-8820320-DEL-81
chr13-8924839-DEL-196
chr13-8991291-DEL-114
chr13-8992991-DEL-60
chr13-9059381-DEL-203
chr13-9082632-DEL-186
chr13-9309708-DEL-62
chr13-9447790-DEL-1270
chr13-9524370-DEL-53
chr13-9707806-DEL-52
chr13-9767992-DEL-54
chr13-9933599-DEL-766
chr13-9934218-DEL-108
chr13-9934617-DEL-67
chr13-9939750-DEL-159
chr13-9972757-DEL-312
chr13-9973093-DEL-501
chr13-10038792-DEL-56
chr13-10049488-DEL-70
chr13-10094994-DEL-58
chr13-10129057-DEL-111
chr13-10175867-DEL-52
chr13-10176077-DEL-58
chr13-10252766-DEL-105
chr13-10271987-DEL-50
chr13-10413529-DEL-54
chr13-10474136-DEL-204
chr13-10499934-DEL-110
chr13-10558744-DEL-128
chr13-10566019-DEL-1487
chr13-10628674-DEL-2650
chr13-10642510-DEL-52
chr13-10670340-DEL-61
chr13-10700890-DEL-76
chr13-10865744-DEL-54
chr13-10880331-DEL-1573
chr13-10960095-DEL-79
chr13-11042516-DEL-71
chr13-11067977-DEL-85
chr13-11142085-DEL-2381
chr13-11153361-DEL-1297
chr13-11210303-DEL-198
chr13-11410699-DEL-56
chr13-11549421-DEL-55
chr13-11722478-DEL-69
chr13-11789035-DEL-1487
chr13-11896579-DEL-165
chr13-11904886-DEL-76
chr13-11936920-DEL-183
chr13-12007150-DEL-625
chr13-12007818-DEL-665
chr13-12081505-DEL-312
chr13-12152353-DEL-962
chr13-12184890-DEL-193
chr13-12202352-DEL-212
chr13-12241910-DEL-110
chr13-12443785-DEL-142
chr13-12480085-DEL-62
chr13-12576198-DEL-78
chr13-12611338-DEL-512
chr13-12620773-DEL-210
chr13-12749191-DEL-1234

chr13-12778780-DEL-79
chr13-13151951-DEL-2903
chr13-13157172-DEL-998
chr13-13258140-DEL-294
chr13-13449726-DEL-648
chr13-13516166-DEL-55
chr13-13522155-DEL-115
chr13-13522405-DEL-51
chr13-13581548-DEL-75
chr13-13696256-DEL-863
chr13-13702198-DEL-234
chr13-13702361-DEL-108
chr13-13702653-DEL-150
chr13-13843947-DEL-63
chr13-13940363-DEL-104
chr13-14035220-DEL-293
chr13-14108292-DEL-264
chr13-14253270-DEL-2033
chr13-14272110-DEL-55
chr13-14280852-DEL-51
chr13-14315532-DEL-603
chr13-14389489-DEL-2246
chr13-14466333-DEL-547
chr13-14522456-DEL-158
chr13-14533465-DEL-50
chr13-14670362-DEL-323
chr13-14787238-DEL-79
chr13-14986523-DEL-84
chr13-15047517-DEL-3561
chr13-15089426-DEL-60
chr13-15174110-DEL-60
chr13-15188496-DEL-114
chr13-15213918-DEL-57
chr13-15486060-DEL-1004
chr13-15685083-DEL-84
chr13-15741552-DEL-52
chr13-15936455-DEL-61
chr13-15973503-DEL-50
chr13-16005994-DEL-5696
chr13-16471364-DEL-58
chr13-16707514-DEL-377
chr13-16908925-DEL-76
chr13-17110945-DEL-117
chr13-17125497-DEL-333
chr13-17171022-DEL-66
chr13-17317488-DEL-964
chr13-17452561-DEL-682
chr13-17688125-DEL-608
chr13-17923336-DEL-124
chr13-18258096-DEL-491
chr13-18294602-DEL-52
chr13-18385945-DEL-58
chr13-18397086-DEL-90
chr13-18562632-DEL-50
chr13-18613220-DEL-50
chr13-18679849-DEL-1194
chr13-18791623-DEL-50
chr13-18953711-DEL-98
chr13-18953711-DEL-294
chr13-18960641-DEL-53
chr13-18960853-DEL-106
chr13-18961095-DEL-51
chr13-19272504-DEL-80
chr13-19328634-DEL-84
chr13-19347000-DEL-14191
chr13-19449785-DEL-158
chr13-19599997-DEL-130
chr13-19619519-DEL-60
chr13-19645531-DEL-519
chr13-19756384-DEL-56
chr13-19757669-DEL-281
chr13-19759675-DEL-126
chr13-20103163-DEL-566
chr13-20104151-DEL-151
chr13-20173203-DEL-64
chr13-20214078-DEL-118
chr13-20329017-DEL-224

chr13-20434891-DEL-114
chr13-20492592-DEL-100
chr13-20531759-DEL-56
chr13-20539722-DEL-207
chr13-20616179-DEL-85
chr13-20692506-DEL-2801
chr13-20752263-DEL-52
chr13-20968868-DEL-112
chr13-21198600-DEL-54
chr13-21206081-DEL-123
chr13-21206429-DEL-101
chr13-21210388-DEL-1145
chr13-21415205-DEL-86
chr13-21462894-DEL-84
chr13-21492504-DEL-2040
chr13-21564360-DEL-83
chr13-21615543-DEL-89
chr13-21843618-DEL-125
chr13-21930354-DEL-76
chr13-22026809-DEL-69
chr13-22164710-DEL-50
chr13-22217279-DEL-1137
chr13-22280358-DEL-127
chr13-22341299-DEL-6339
chr13-22514342-DEL-50
chr13-22527768-DEL-167
chr13-22527906-DEL-69
chr13-22666238-DEL-151
chr13-22826777-DEL-10085
chr13-22857716-DEL-1914
chr13-22868210-DEL-7751
chr13-22877213-DEL-186
chr13-22891765-DEL-1331
chr13-23083857-DEL-41475
chr13-23126132-DEL-436
chr13-23136952-DEL-1801
chr13-23273751-DEL-1688
chr13-23389928-DEL-2411
chr13-23444812-DEL-66
chr13-23557480-DEL-50
chr13-23589384-DEL-2151
chr13-23624682-DEL-1252
chr13-23835905-DEL-78
chr13-23925622-DEL-118
chr13-24022290-DEL-135
chr13-24063437-DEL-54
chr13-24070834-DEL-50
chr13-24096144-DEL-194
chr13-24292067-DEL-72
chr13-24300449-DEL-51
chr13-24340121-DEL-84
chr13-24357413-DEL-105
chr13-24365223-DEL-632
chr13-24366700-DEL-57
chr13-24582092-DEL-4739
chr13-24595140-DEL-150
chr13-24714825-DEL-12300
chr13-24766703-DEL-1391
chr13-24851203-DEL-1027
chr13-24865827-DEL-154
chr13-25011464-DEL-135
chr13-25013485-DEL-111
chr13-25118158-DEL-71
chr13-25213157-DEL-50
chr13-25273493-DEL-65
chr13-25274089-DEL-59
chr13-25330529-DEL-84
chr13-25369229-DEL-50
chr13-25376791-DEL-430
chr13-25376824-DEL-86
chr13-25485713-DEL-50
chr13-25486391-DEL-52
chr13-25513875-DEL-81
chr13-25530811-DEL-52
chr13-25745524-DEL-252
chr13-25865765-DEL-57
chr13-25865798-DEL-126

chr13-25883448-DEL-81
chr13-25916400-DEL-122
chr13-25916408-DEL-54
chr13-25942229-DEL-104
chr13-25986676-DEL-569
chr13-26056375-DEL-289
chr13-26121319-DEL-2619
chr13-26517385-DEL-65
chr13-26518139-DEL-202
chr13-26705847-DEL-6294
chr13-26733710-DEL-56
chr13-26798901-DEL-1752
chr13-26844942-DEL-386
chr13-27072772-DEL-107
chr13-27072775-DEL-299
chr13-27073070-DEL-99
chr13-27093700-DEL-210
chr13-27183461-DEL-65
chr13-27268316-DEL-5029
chr13-27289135-DEL-25326
chr13-27326343-DEL-1541
chr13-27377813-DEL-68
chr13-27377817-DEL-146
chr13-27377941-DEL-56
chr13-27381590-DEL-108
chr13-27412223-DEL-1211
chr13-27507636-DEL-417
chr13-27603292-DEL-1459
chr13-27604920-DEL-860
chr13-27609973-DEL-822
chr13-27629821-DEL-63
chr13-27646542-DEL-1108
chr13-27675283-DEL-94
chr13-27725679-DEL-66
chr13-27732155-DEL-57
chr13-27736218-DEL-2563
chr13-27930886-DEL-62
chr13-27931206-DEL-145
chr13-28046577-DEL-930
chr13-28080940-DEL-3132
chr13-28109846-DEL-82
chr13-28426888-DEL-64
chr13-28659792-DEL-208
chr13-28815770-DEL-58
chr13-28861597-DEL-220
chr13-28861767-DEL-67
chr13-28963279-DEL-53
chr13-29056853-DEL-79
chr13-29132068-DEL-146
chr13-29177182-DEL-57
chr13-29256020-DEL-212
chr13-29484463-DEL-88
chr13-29502297-DEL-2088
chr13-29546765-DEL-54
chr13-29799713-DEL-142
chr13-29812944-DEL-147
chr13-29851174-DEL-80
chr13-29873325-DEL-953
chr13-29873892-DEL-455
chr13-29934179-DEL-58
chr13-29934640-DEL-64
chr13-30033394-DEL-214
chr13-30118701-DEL-200
chr13-30312345-DEL-2262
chr13-30314543-DEL-230
chr13-30319123-DEL-52
chr13-30327889-DEL-67
chr13-30328242-DEL-182
chr13-30405087-DEL-50
chr13-30434090-DEL-238
chr13-30434112-DEL-118
chr13-30570625-DEL-50
chr13-30881564-DEL-161
chr13-30883117-DEL-266
chr13-30983651-DEL-54
chr13-31019869-DEL-162
chr13-31104687-DEL-50

chr13-31182443-DEL-72
chr13-31294758-DEL-766
chr13-31400004-DEL-575
chr13-31400234-DEL-125
chr13-31512100-DEL-208
chr13-31517626-DEL-362
chr13-31788557-DEL-52
chr13-31820147-DEL-64
chr13-31876261-DEL-50
chr13-31965363-DEL-82
chr13-32620150-DEL-60
chr13-32688537-DEL-1705
chr13-32846313-DEL-454
chr13-33040551-DEL-132
chr13-33272062-DEL-850
chr13-33273080-DEL-200
chr13-33342112-DEL-440
chr13-33713678-DEL-323
chr13-33840461-DEL-3592
chr13-34252094-DEL-107
chr13-34533733-DEL-50
chr13-34603531-DEL-354
chr13-34821828-DEL-242
chr13-34821848-DEL-88
chr13-35036852-DEL-100
chr13-35167289-DEL-82
chr13-35222420-DEL-231
chr13-35626511-DEL-300
chr13-35814451-DEL-201
chr13-35883612-DEL-188
chr13-35970063-DEL-619
chr13-36069642-DEL-208
chr13-36289259-DEL-129
chr13-36390132-DEL-90
chr13-36564611-DEL-205
chr13-36581863-DEL-50
chr13-36659341-DEL-80
chr13-36787295-DEL-59
chr13-36839579-DEL-81
chr13-36841558-DEL-76
chr13-36944659-DEL-112
chr13-36978109-DEL-3193
chr13-37106844-DEL-2440
chr13-37147564-DEL-912
chr13-37162513-DEL-222
chr13-37209412-DEL-56
chr13-37281092-DEL-377
chr13-37331150-DEL-60
chr13-37433027-DEL-202
chr13-37444123-DEL-1116
chr13-37465447-DEL-58
chr13-37559298-DEL-51
chr13-37661853-DEL-50
chr13-37697390-DEL-61
chr13-37874393-DEL-75
chr13-37959687-DEL-238
chr13-37994039-DEL-692
chr13-38075816-DEL-369
chr13-38111022-DEL-66
chr13-38188158-DEL-208
chr13-38188218-DEL-918
chr13-38188700-DEL-84
chr13-38223738-DEL-274
chr13-38345354-DEL-95
chr13-38350545-DEL-150
chr13-38425533-DEL-50
chr13-38457656-DEL-254
chr13-38499269-DEL-67
chr13-38522099-DEL-483
chr13-38554138-DEL-96
chr13-38555194-DEL-1785
chr13-38608650-DEL-92
chr13-38635777-DEL-655
chr13-38654675-DEL-897
chr13-38660375-DEL-52
chr13-38826481-DEL-216
chr13-38826579-DEL-729

chr13-38969859-DEL-81
chr13-39048486-DEL-68
chr13-39052711-DEL-65
chr13-39056665-DEL-719
chr13-39060187-DEL-688
chr13-39218556-DEL-57
chr13-39262586-DEL-50
chr13-39287766-DEL-63
chr13-39334554-DEL-156
chr13-39386465-DEL-688
chr13-39388272-DEL-227
chr13-39486451-DEL-52
chr13-39541823-DEL-61
chr13-39571188-DEL-543
chr13-39580197-DEL-269
chr13-39580554-DEL-234
chr13-39583915-DEL-56
chr13-39591500-DEL-72
chr13-39622673-DEL-162
chr13-39644308-DEL-103
chr13-39659629-DEL-70
chr13-39712841-DEL-234
chr13-39723624-DEL-58
chr13-39757861-DEL-62
chr13-39893046-DEL-67
chr13-39995553-DEL-84
chr13-40068188-DEL-50
chr13-40083870-DEL-338
chr13-40093043-DEL-6713
chr13-40427729-DEL-63
chr13-40551192-DEL-57
chr13-40652480-DEL-1148
chr13-40655954-DEL-51
chr13-40701701-DEL-77
chr13-40717241-DEL-108
chr13-40763980-DEL-400
chr13-40823904-DEL-200
chr13-40849338-DEL-58
chr13-40852143-DEL-286
chr13-40985026-DEL-6201
chr13-41003789-DEL-206
chr13-41051556-DEL-126
chr13-41114661-DEL-3093
chr13-41476905-DEL-72
chr13-41571416-DEL-208
chr13-41924931-DEL-60
chr13-41965347-DEL-228
chr13-41967208-DEL-52
chr13-42039974-DEL-131
chr13-42060432-DEL-68
chr13-42160385-DEL-60
chr13-42175462-DEL-173
chr13-42177418-DEL-120
chr13-42216418-DEL-355
chr13-42216695-DEL-78
chr13-42229628-DEL-181
chr13-42248875-DEL-68
chr13-42349667-DEL-63
chr13-42383821-DEL-275
chr13-42401954-DEL-116
chr13-42421442-DEL-54
chr13-42468981-DEL-56
chr13-42507158-DEL-95
chr13-42517517-DEL-53
chr13-42543846-DEL-91
chr13-42627012-DEL-178
chr13-42706756-DEL-188
chr13-42745394-DEL-67
chr13-42773021-DEL-66
chr13-42910146-DEL-50
chr13-42960485-DEL-1661
chr13-42973388-DEL-145
chr13-43181644-DEL-135
chr13-43193778-DEL-107
chr13-43283285-DEL-64
chr13-43295541-DEL-77
chr13-43321020-DEL-62

chr13-43501170-DEL-742
chr13-43535697-DEL-67
chr13-43753548-DEL-50
chr13-43813717-DEL-90
chr13-44429966-DEL-71
chr13-44515147-DEL-68
chr13-44600468-DEL-67
chr13-44629142-DEL-324
chr13-44681852-DEL-140
chr13-44737270-DEL-50
chr13-44746928-DEL-222
chr13-44801971-DEL-62
chr13-44936052-DEL-120
chr13-45013347-DEL-2736
chr13-45286057-DEL-50
chr13-45737840-DEL-104
chr13-45800612-DEL-566
chr13-45801214-DEL-671
chr13-45916542-DEL-122
chr13-45943511-DEL-250
chr13-45992455-DEL-84
chr13-46031611-DEL-177
chr13-46295344-DEL-60
chr13-46350422-DEL-70
chr13-46470026-DEL-1894
chr13-46680694-DEL-50
chr13-46729541-DEL-201
chr13-46757356-DEL-224
chr13-46818967-DEL-274
chr13-46956787-DEL-111
chr13-46990616-DEL-56
chr13-46996239-DEL-150
chr13-47142669-DEL-620
chr13-47216677-DEL-175
chr13-47384154-DEL-157
chr13-47630960-DEL-1038
chr13-47862149-DEL-50
chr13-47987937-DEL-430
chr13-48041583-DEL-153
chr13-48302874-DEL-468
chr13-48327695-DEL-55
chr13-48428940-DEL-56
chr13-48587553-DEL-82
chr13-48602695-DEL-140
chr13-48710849-DEL-50
chr13-48994580-DEL-253
chr13-49060614-DEL-162
chr13-49079287-DEL-98
chr13-49183635-DEL-120
chr13-49207685-DEL-102
chr13-49208139-DEL-221
chr13-49524136-DEL-133
chr13-49551231-DEL-259
chr13-49552921-DEL-634
chr13-49553149-DEL-211
chr13-49651572-DEL-53
chr13-49949800-DEL-50
chr13-49976708-DEL-78
chr13-50009270-DEL-70
chr13-50009485-DEL-340
chr13-50127569-DEL-109
chr13-50127928-DEL-341
chr13-50206453-DEL-58
chr13-50646362-DEL-56
chr13-50744323-DEL-2140
chr13-50778729-DEL-54
chr13-50830932-DEL-116
chr13-50940498-DEL-4302
chr13-50946908-DEL-262
chr13-51117150-DEL-199
chr13-51157261-DEL-131
chr13-51219437-DEL-677
chr13-51315485-DEL-116
chr13-51366558-DEL-63
chr13-51447553-DEL-107
chr13-51457975-DEL-302
chr13-51458438-DEL-113

chr13-51712946-DEL-56
chr13-51788359-DEL-4568
chr13-51818213-DEL-168
chr13-51871304-DEL-405
chr13-51871385-DEL-189
chr13-51983533-DEL-204
chr13-52028767-DEL-51
chr13-52167094-DEL-92
chr13-52193031-DEL-52
chr13-52202654-DEL-1714
chr13-52319974-DEL-1241
chr13-52390877-DEL-184
chr13-52436895-DEL-107
chr13-52449982-DEL-105
chr13-52474065-DEL-496
chr13-52498266-DEL-57
chr13-52507349-DEL-265
chr13-52591548-DEL-1380
chr13-52649143-DEL-52
chr13-52668958-DEL-180
chr13-52914153-DEL-56
chr13-52914210-DEL-119
chr13-53213492-DEL-156
chr13-53213561-DEL-331
chr13-53267585-DEL-58
chr13-53344254-DEL-76
chr13-53344394-DEL-198
chr13-53361051-DEL-77
chr13-53404037-DEL-86
chr13-53404498-DEL-56
chr13-53422881-DEL-54
chr13-53423942-DEL-54
chr13-53458774-DEL-54
chr13-53546091-DEL-64
chr13-53546214-DEL-64
chr13-53550527-DEL-52
chr13-53637899-DEL-334
chr13-53638021-DEL-144
chr13-53660272-DEL-85
chr13-53712466-DEL-66
chr13-53760226-DEL-50
chr13-53816475-DEL-70
chr13-54173765-DEL-813
chr13-54174060-DEL-336
chr13-54277071-DEL-64
chr13-54335797-DEL-56
chr13-54349552-DEL-132
chr13-54398167-DEL-50
chr13-54600556-DEL-1321
chr13-54602194-DEL-74
chr13-54755743-DEL-111
chr13-54957006-DEL-52
chr13-54960872-DEL-56
chr13-54978630-DEL-102
chr13-55040872-DEL-64
chr13-55117754-DEL-53
chr13-55119594-DEL-117
chr13-55178375-DEL-164
chr13-55189833-DEL-50
chr13-55233548-DEL-652
chr13-55237639-DEL-97
chr13-55240042-DEL-124
chr13-55246047-DEL-60
chr13-55345068-DEL-173
chr13-55345291-DEL-1558
chr13-55479621-DEL-185
chr13-55488768-DEL-2309
chr13-55515708-DEL-55
chr13-55641366-DEL-115
chr13-55753829-DEL-73
chr13-55760386-DEL-1855
chr13-55834486-DEL-376
chr13-55843699-DEL-63
chr13-55861886-DEL-206
chr13-55912554-DEL-154
chr13-55925049-DEL-1063
chr13-56217406-DEL-78

chr13-56232101-DEL-2037
chr13-56310419-DEL-70
chr13-56365360-DEL-132
chr13-56389861-DEL-52
chr13-56415269-DEL-55
chr13-56565037-DEL-139
chr13-56594625-DEL-50
chr13-56620248-DEL-54
chr13-56634437-DEL-394
chr13-56672665-DEL-50
chr13-56799556-DEL-61
chr13-56801300-DEL-237
chr13-56818828-DEL-98
chr13-56892484-DEL-626
chr13-56893297-DEL-203
chr13-56931267-DEL-55
chr13-56948941-DEL-771
chr13-56949020-DEL-50
chr13-57135114-DEL-66
chr13-57152172-DEL-115
chr13-57166236-DEL-72
chr13-57172867-DEL-50
chr13-57229990-DEL-99
chr13-57262780-DEL-63
chr13-57275002-DEL-328
chr13-57329862-DEL-137
chr13-57463228-DEL-58
chr13-57467995-DEL-106
chr13-57524862-DEL-155
chr13-57557961-DEL-80
chr13-57563985-DEL-108
chr13-57593143-DEL-70
chr13-57633565-DEL-54
chr13-57696521-DEL-61
chr13-57780062-DEL-641
chr13-57806439-DEL-475
chr13-57932838-DEL-249
chr13-57967805-DEL-119
chr13-57982396-DEL-68
chr13-57987282-DEL-53
chr13-58007549-DEL-56
chr13-58047949-DEL-99
chr13-58058069-DEL-330
chr13-58119983-DEL-70
chr13-58187031-DEL-281
chr13-58188973-DEL-1102
chr13-58191889-DEL-106
chr13-58264901-DEL-1517
chr13-58494261-DEL-410
chr13-58494419-DEL-94
chr13-58496296-DEL-1329
chr13-58496877-DEL-411
chr13-58657418-DEL-51
chr13-58680843-DEL-580
chr13-58683371-DEL-59
chr13-58714984-DEL-52
chr13-58726624-DEL-64
chr13-58727046-DEL-64
chr13-58790105-DEL-461
chr13-58790159-DEL-197
chr13-58801994-DEL-320
chr13-58859679-DEL-72
chr13-58859967-DEL-90
chr13-58884267-DEL-90
chr13-58892475-DEL-66
chr13-58922080-DEL-148
chr13-58971693-DEL-1821
chr13-59060316-DEL-100
chr13-59060475-DEL-202
chr13-59168317-DEL-219
chr13-59180016-DEL-68
chr13-59192213-DEL-561
chr13-59202502-DEL-113
chr13-59286747-DEL-756
chr13-59355392-DEL-53
chr13-59357094-DEL-69
chr13-59400547-DEL-125

chr13-59576346-DEL-158
chr13-59625787-DEL-157
chr13-59660693-DEL-208
chr13-59673626-DEL-126
chr13-59673643-DEL-336
chr13-59709371-DEL-350
chr13-59806887-DEL-100
chr13-59886038-DEL-173
chr13-59998587-DEL-150
chr13-60047108-DEL-67
chr13-60102611-DEL-300
chr13-60102872-DEL-55
chr13-60115101-DEL-111
chr13-60150756-DEL-2861
chr13-60222349-DEL-54
chr13-60222350-DEL-427
chr13-60224682-DEL-57
chr13-60307600-DEL-56
chr13-60307768-DEL-330
chr13-60317704-DEL-122
chr13-60338348-DEL-70
chr13-60386042-DEL-602
chr13-60405633-DEL-58
chr13-60489012-DEL-72
chr13-60590603-DEL-210
chr13-60710052-DEL-54
chr13-60710679-DEL-63
chr13-60778491-DEL-50
chr13-60778702-DEL-50
chr13-60828944-DEL-667
chr13-60828967-DEL-261
chr13-60829460-DEL-145
chr13-60898704-DEL-64
chr13-60922120-DEL-78
chr13-60936394-DEL-96
chr13-60943955-DEL-114
chr13-60981649-DEL-1201
chr13-61031286-DEL-3688
chr13-61057578-DEL-427
chr13-61090186-DEL-168
chr13-61113915-DEL-129
chr13-61117955-DEL-22860
chr13-61135088-DEL-176
chr13-61230276-DEL-5579
chr13-61305853-DEL-195
chr13-61320968-DEL-98
chr13-61611238-DEL-7020
chr13-61638721-DEL-128
chr13-61682603-DEL-80
chr13-61738413-DEL-105
chr13-61756915-DEL-199
chr13-61812725-DEL-52
chr13-61858801-DEL-331
chr13-61893358-DEL-84
chr13-62137322-DEL-385
chr13-62259852-DEL-350
chr13-62262080-DEL-358
chr13-62268313-DEL-97
chr13-62319873-DEL-83
chr13-62361612-DEL-212
chr13-62385818-DEL-87
chr13-62517405-DEL-336
chr13-62567024-DEL-848
chr13-62683783-DEL-408
chr13-62710936-DEL-89
chr13-62711506-DEL-188
chr13-62848499-DEL-84
chr13-62884466-DEL-50
chr13-62977406-DEL-1498
chr13-63023708-DEL-97
chr13-63077488-DEL-94
chr13-63174882-DEL-86
chr13-63191097-DEL-199
chr13-63191386-DEL-301
chr13-63353932-DEL-1244
chr13-63512010-DEL-68
chr13-63527043-DEL-190

chr13-63527165-DEL-760
chr13-63527932-DEL-190
chr13-63725278-DEL-188
chr13-63844502-DEL-195
chr13-63844606-DEL-62
chr13-63844953-DEL-113
chr13-63897359-DEL-50
chr13-63901969-DEL-2092
chr13-63909988-DEL-384
chr13-63977156-DEL-50
chr13-64046249-DEL-57
chr13-64154228-DEL-90
chr13-64192300-DEL-203
chr13-64192345-DEL-79
chr13-64192575-DEL-52
chr13-64194673-DEL-120
chr13-64236074-DEL-50
chr13-64242031-DEL-120
chr13-64371248-DEL-52
chr13-64375796-DEL-52
chr13-64402228-DEL-60
chr13-64420159-DEL-145
chr13-64442635-DEL-512
chr13-64452167-DEL-119
chr13-64469689-DEL-670
chr13-64476449-DEL-135
chr13-64493555-DEL-50
chr13-64497799-DEL-72
chr13-64550715-DEL-195
chr13-64644353-DEL-50
chr13-64694661-DEL-786
chr13-64710783-DEL-152
chr13-64724935-DEL-51
chr13-64738825-DEL-65
chr13-64801040-DEL-740
chr13-64807930-DEL-1007
chr13-64829388-DEL-1149
chr13-64838462-DEL-68
chr13-64868898-DEL-50
chr13-64937354-DEL-56
chr13-65087633-DEL-205
chr13-65162289-DEL-82
chr13-65209927-DEL-187
chr13-65360174-DEL-335
chr13-65360221-DEL-152
chr13-65483038-DEL-99
chr13-65486153-DEL-420
chr13-65486298-DEL-168
chr13-65486576-DEL-252
chr13-65486744-DEL-84
chr13-65494292-DEL-142
chr13-65513761-DEL-855
chr13-65522915-DEL-3435
chr13-65524370-DEL-50
chr13-65644968-DEL-94
chr13-65694750-DEL-504
chr13-65811185-DEL-222
chr13-65894296-DEL-127
chr13-65894352-DEL-60
chr13-65907297-DEL-104
chr13-66055529-DEL-85
chr13-66078577-DEL-64
chr13-66246444-DEL-514
chr13-66311465-DEL-168
chr13-66426702-DEL-93
chr13-66471855-DEL-179
chr13-66472588-DEL-135
chr13-66473797-DEL-610
chr13-66628138-DEL-318
chr13-66676695-DEL-503
chr13-66789019-DEL-667
chr13-66789345-DEL-90
chr13-66940880-DEL-84
chr13-66943590-DEL-211
chr13-67000043-DEL-840
chr13-67001042-DEL-381
chr13-67037855-DEL-3985

chr13-67115500-DEL-176
chr13-67115541-DEL-731
chr13-67201659-DEL-2023
chr13-67297030-DEL-499
chr13-67402123-DEL-74
chr13-67502433-DEL-66
chr13-67524012-DEL-24006
chr13-67579949-DEL-119
chr13-67583641-DEL-628
chr13-67601582-DEL-1102
chr13-67613383-DEL-681
chr13-67646758-DEL-134
chr13-67725553-DEL-66
chr13-67835007-DEL-50
chr13-67863771-DEL-204
chr13-67946014-DEL-52
chr13-67988787-DEL-1678
chr13-68009161-DEL-530
chr13-68013681-DEL-1887
chr13-68075302-DEL-114
chr13-68100026-DEL-78
chr13-68212694-DEL-1047
chr13-68343576-DEL-59
chr13-68452809-DEL-66
chr13-68455571-DEL-408
chr13-68491644-DEL-86
chr13-68495155-DEL-5567
chr13-68524299-DEL-136
chr13-68524606-DEL-136
chr13-68613772-DEL-315
chr13-68614161-DEL-181
chr13-68742146-DEL-85
chr13-68771572-DEL-144
chr13-68866592-DEL-330
chr13-68866816-DEL-106
chr13-68874496-DEL-353
chr13-68883346-DEL-64
chr13-68883470-DEL-67
chr13-68969963-DEL-58
chr13-69190198-DEL-56
chr13-69266090-DEL-122
chr13-69302784-DEL-581
chr13-69335918-DEL-60
chr13-69388991-DEL-105
chr13-69407358-DEL-58
chr13-69418749-DEL-182
chr13-69485079-DEL-225
chr13-69522295-DEL-86
chr13-69579886-DEL-144
chr13-69580006-DEL-66
chr13-69598208-DEL-55
chr13-69602818-DEL-53
chr13-69656619-DEL-124
chr13-69949660-DEL-52
chr13-69979133-DEL-84
chr13-70010830-DEL-90
chr13-70051842-DEL-256
chr13-70052408-DEL-315
chr13-70057354-DEL-64
chr13-70057625-DEL-88
chr13-70154503-DEL-87
chr13-70180270-DEL-236
chr13-70253800-DEL-189
chr13-70548760-DEL-2058
chr13-70566689-DEL-79
chr13-70627281-DEL-180
chr13-70641414-DEL-76
chr13-70678084-DEL-116
chr13-70870695-DEL-1294
chr13-70895142-DEL-168
chr13-70898062-DEL-1142
chr13-70924924-DEL-91
chr13-70934741-DEL-50
chr13-70980066-DEL-61
chr13-71042414-DEL-107
chr13-71042704-DEL-270
chr13-71162138-DEL-58

chr13-71166403-DEL-239
chr13-71166837-DEL-442
chr13-71529557-DEL-427
chr13-71529788-DEL-213
chr13-71554646-DEL-76
chr13-71567563-DEL-80
chr13-71960092-DEL-57
chr13-72058180-DEL-103
chr13-72093293-DEL-983
chr13-72093858-DEL-192
chr13-72094102-DEL-440
chr13-72154157-DEL-1979
chr13-72228928-DEL-51
chr13-72459867-DEL-84
chr13-72460111-DEL-190
chr13-72462345-DEL-66
chr13-72474348-DEL-71
chr13-72474368-DEL-146
chr13-72499556-DEL-95
chr13-72543053-DEL-50
chr13-72601403-DEL-3267
chr13-72636011-DEL-286
chr13-72636101-DEL-1108
chr13-72636650-DEL-232
chr13-72637177-DEL-325
chr13-72637467-DEL-53
chr13-72683007-DEL-81
chr13-72796239-DEL-505
chr13-72836561-DEL-102
chr13-72907811-DEL-126
chr13-72942582-DEL-122
chr13-72971264-DEL-62
chr13-73063657-DEL-106
chr13-73117584-DEL-51
chr13-73154330-DEL-83
chr13-73210343-DEL-50
chr13-73250263-DEL-51
chr13-73369441-DEL-147
chr13-73426602-DEL-59
chr13-73529936-DEL-254
chr13-73570810-DEL-56
chr13-73636285-DEL-51
chr13-73668835-DEL-310
chr13-73749040-DEL-160
chr13-73784057-DEL-175
chr13-73870060-DEL-63
chr13-74032217-DEL-233
chr13-74056230-DEL-1396
chr13-74195546-DEL-81
chr13-74379228-DEL-1513
chr13-74434923-DEL-154
chr13-74503256-DEL-144
chr13-74677147-DEL-877
chr13-74678834-DEL-734
chr13-74915143-DEL-50
chr13-74948952-DEL-1085
chr13-74971418-DEL-78
chr13-75000690-DEL-73
chr13-75000742-DEL-193
chr13-75199383-DEL-5097
chr13-75314747-DEL-2124
chr13-75547718-DEL-386
chr13-75700616-DEL-312
chr13-75728102-DEL-50
chr13-75851645-DEL-388
chr13-75931717-DEL-561
chr13-76017135-DEL-60
chr13-76103529-DEL-72
chr13-76165833-DEL-464
chr13-76215951-DEL-75
chr13-76262763-DEL-68
chr13-76346421-DEL-52
chr13-76487315-DEL-489
chr13-76507694-DEL-867
chr13-76524578-DEL-129
chr13-76524717-DEL-52
chr13-76525199-DEL-52

chr13-76622235-DEL-4270
chr13-76699761-DEL-99
chr13-76988814-DEL-233
chr13-77080169-DEL-50
chr13-77257809-DEL-150
chr13-77454041-DEL-1082
chr13-77569989-DEL-1527
chr13-77587030-DEL-135
chr13-77668549-DEL-54
chr13-77685949-DEL-212
chr13-77871693-DEL-77
chr13-77872393-DEL-353
chr13-77872785-DEL-343
chr13-77885286-DEL-126
chr13-77952731-DEL-89
chr13-78068684-DEL-270
chr13-78112427-DEL-1074
chr13-78422723-DEL-73
chr13-78506817-DEL-398
chr13-78824255-DEL-420
chr13-78961322-DEL-56
chr13-79060066-DEL-459
chr13-79132012-DEL-102
chr13-79159605-DEL-50
chr13-79161974-DEL-1460
chr13-79221941-DEL-100
chr13-79257578-DEL-213
chr13-79396242-DEL-105
chr13-79447047-DEL-52
chr13-79478971-DEL-213
chr13-79590421-DEL-144
chr13-79607842-DEL-1848
chr13-79620812-DEL-2574
chr13-79834118-DEL-54
chr13-79928909-DEL-498
chr13-79940518-DEL-90
chr13-80172561-DEL-212
chr13-80238450-DEL-59
chr13-80341550-DEL-56
chr13-80411074-DEL-68
chr13-80430034-DEL-185
chr13-80541471-DEL-186
chr13-80675644-DEL-149
chr13-80684864-DEL-1488
chr13-80976889-DEL-89
chr13-81146465-DEL-51
chr13-81151161-DEL-1185
chr13-81165177-DEL-130
chr13-81457631-DEL-343
chr13-81457843-DEL-401
chr13-81494714-DEL-59
chr13-81563077-DEL-58
chr13-81621762-DEL-393
chr13-81700779-DEL-2351
chr13-81837845-DEL-69
chr13-81849818-DEL-740
chr13-81896085-DEL-51
chr13-82168403-DEL-1586
chr13-82241386-DEL-51
chr13-82333245-DEL-394
chr13-82390493-DEL-292
chr13-82483931-DEL-213
chr13-82483990-DEL-433
chr13-82671145-DEL-6272
chr13-82704118-DEL-799
chr13-82870506-DEL-4095
chr13-83051044-DEL-261
chr13-83190296-DEL-172
chr13-83208166-DEL-3304
chr13-83245554-DEL-98
chr13-83299310-DEL-296
chr13-83356851-DEL-70
chr13-83405293-DEL-2310
chr13-83409177-DEL-181
chr13-83459939-DEL-104
chr13-83487751-DEL-124
chr13-83785472-DEL-916

chr13-83948912-DEL-80
chr13-83984984-DEL-50
chr13-83989343-DEL-133
chr13-84009755-DEL-64
chr13-84085107-DEL-342
chr13-84085483-DEL-96
chr13-84144336-DEL-133
chr13-84184605-DEL-60
chr13-84435788-DEL-204
chr13-84467922-DEL-50
chr13-84483469-DEL-87
chr13-84531649-DEL-4766
chr13-84550595-DEL-312
chr13-84575901-DEL-211
chr13-85137640-DEL-82
chr13-85466721-DEL-50
chr13-85472311-DEL-624
chr13-85502962-DEL-82
chr13-85517842-DEL-504
chr13-85538603-DEL-89
chr13-85538945-DEL-54
chr13-85609340-DEL-287
chr13-85620005-DEL-90
chr13-85630063-DEL-81
chr13-85964509-DEL-55
chr13-85994357-DEL-3573
chr13-86008314-DEL-282
chr13-86038077-DEL-122
chr13-86162690-DEL-131
chr13-86269826-DEL-6193
chr13-86299496-DEL-199
chr13-86304365-DEL-106
chr13-86319097-DEL-66
chr13-86412365-DEL-194
chr13-86504523-DEL-562
chr13-86516312-DEL-52
chr13-86610833-DEL-156
chr13-86624840-DEL-818
chr13-86631019-DEL-55
chr13-86661560-DEL-1243
chr13-86685110-DEL-938
chr13-86709695-DEL-134
chr13-86709935-DEL-56
chr13-86824087-DEL-2142
chr13-86833922-DEL-4834
chr13-86937430-DEL-56
chr13-86957500-DEL-2081
chr13-86970921-DEL-2760
chr13-87051409-DEL-3461
chr13-87209625-DEL-646
chr13-87289645-DEL-3669
chr13-87346576-DEL-689
chr13-87350668-DEL-50
chr13-87354529-DEL-159
chr13-87354592-DEL-50
chr13-87401069-DEL-598
chr13-87401277-DEL-81
chr13-87432011-DEL-70
chr13-87692326-DEL-84
chr13-87768141-DEL-420
chr13-87921950-DEL-124
chr13-87930940-DEL-86
chr13-88012070-DEL-93
chr13-88026005-DEL-128
chr13-88026801-DEL-155
chr13-88046900-DEL-3712
chr13-88095146-DEL-79
chr13-88105770-DEL-67
chr13-88215001-DEL-50
chr13-88220317-DEL-102
chr13-88321127-DEL-50
chr13-88471787-DEL-112
chr13-88500202-DEL-3573
chr13-88934398-DEL-64
chr13-89203337-DEL-50
chr13-89323892-DEL-219
chr13-89404331-DEL-1401

chr13-89493358-DEL-374
chr13-89502184-DEL-68
chr13-89719841-DEL-800
chr13-89724356-DEL-2228
chr13-89734887-DEL-764
chr13-89814775-DEL-165
chr13-89814978-DEL-159
chr13-89952405-DEL-115
chr13-89982743-DEL-880
chr13-90154557-DEL-68
chr13-90269175-DEL-71
chr13-90348802-DEL-100
chr13-90404587-DEL-60
chr13-90416259-DEL-858
chr13-90451156-DEL-84
chr13-90785655-DEL-60
chr13-90953371-DEL-163
chr13-90983393-DEL-72
chr13-91005209-DEL-345
chr13-91005591-DEL-200
chr13-91165070-DEL-74
chr13-91172239-DEL-72
chr13-91205692-DEL-50
chr13-91229708-DEL-105
chr13-91232631-DEL-88
chr13-91232681-DEL-812
chr13-91322344-DEL-454
chr13-91322999-DEL-281
chr13-91363540-DEL-2475
chr13-91496726-DEL-54
chr13-91506364-DEL-72
chr13-91552426-DEL-145
chr13-91786730-DEL-255
chr13-91786797-DEL-51
chr13-91928475-DEL-114
chr13-91956105-DEL-200
chr13-92051658-DEL-68
chr13-92248419-DEL-54
chr13-92287896-DEL-216
chr13-92304016-DEL-444
chr13-92328480-DEL-144
chr13-92347128-DEL-225
chr13-92353568-DEL-52
chr13-92366277-DEL-62
chr13-92366408-DEL-254
chr13-92366545-DEL-63
chr13-92501222-DEL-252
chr13-92508245-DEL-6840
chr13-92543470-DEL-66
chr13-92625497-DEL-1138
chr13-92626323-DEL-132
chr13-92635224-DEL-53
chr13-92752730-DEL-68
chr13-92782953-DEL-286
chr13-92783021-DEL-82
chr13-92790146-DEL-161
chr13-92790472-DEL-520
chr13-92790679-DEL-724
chr13-92791348-DEL-90
chr13-92939963-DEL-199
chr13-92968334-DEL-50
chr13-93134623-DEL-107
chr13-93134775-DEL-50
chr13-93159451-DEL-171
chr13-93218905-DEL-50
chr13-93277928-DEL-50
chr13-93362091-DEL-444
chr13-93490800-DEL-528
chr13-93508133-DEL-143
chr13-93576549-DEL-82
chr13-93605633-DEL-67
chr13-93684644-DEL-203
chr13-93711297-DEL-122
chr13-93751459-DEL-192
chr13-93765186-DEL-774
chr13-93772505-DEL-56
chr13-93777990-DEL-69

chr13-93795140-DEL-2326
chr13-93818296-DEL-368
chr13-93893094-DEL-205
chr13-93978082-DEL-149
chr13-93978138-DEL-68
chr13-93978351-DEL-709
chr13-93978559-DEL-228
chr13-93979761-DEL-91
chr13-94099698-DEL-375
chr13-94217061-DEL-1252
chr13-94232564-DEL-99
chr13-94312315-DEL-77
chr13-94312446-DEL-145
chr13-94358997-DEL-60
chr13-94387174-DEL-55
chr13-94417306-DEL-99
chr13-94568034-DEL-68
chr13-94650545-DEL-515
chr13-94717826-DEL-314
chr13-94764632-DEL-242
chr13-94834796-DEL-84
chr13-95030467-DEL-70
chr13-95115754-DEL-95
chr13-95240688-DEL-151
chr13-95306452-DEL-236
chr13-95545182-DEL-84
chr13-95606487-DEL-82
chr13-95623032-DEL-130
chr13-95635590-DEL-72
chr13-95646880-DEL-217
chr13-95685312-DEL-67
chr13-95715189-DEL-50
chr13-95790997-DEL-69
chr13-95887034-DEL-217
chr13-95949005-DEL-103
chr13-95962121-DEL-98
chr13-96077918-DEL-58
chr13-96232190-DEL-60
chr13-96257811-DEL-176
chr13-96283392-DEL-3204
chr13-96284721-DEL-385
chr13-96304281-DEL-52
chr13-96340168-DEL-287
chr13-96351929-DEL-50
chr13-96506482-DEL-58
chr13-96624986-DEL-96
chr13-96712604-DEL-853
chr13-96750654-DEL-156
chr13-96839876-DEL-130
chr13-96841851-DEL-720
chr13-96851897-DEL-59
chr13-96941934-DEL-50
chr13-97025815-DEL-3153
chr13-97059978-DEL-231
chr13-97097869-DEL-92
chr13-97155775-DEL-959
chr13-97156944-DEL-1614
chr13-97256820-DEL-707
chr13-97298726-DEL-228
chr13-97445032-DEL-58
chr13-97505609-DEL-225
chr13-97521252-DEL-53
chr13-97562782-DEL-69
chr13-97631914-DEL-72
chr13-97714877-DEL-640
chr13-97714887-DEL-86
chr13-97829048-DEL-50
chr13-97878823-DEL-50
chr13-97948923-DEL-105
chr13-98044321-DEL-64
chr13-98107887-DEL-88
chr13-98145949-DEL-58
chr13-98170848-DEL-238
chr13-98231299-DEL-976
chr13-98299853-DEL-210
chr13-98300302-DEL-147
chr13-98317579-DEL-1840

chr13-98328500-DEL-127
chr13-98349022-DEL-79
chr13-98459314-DEL-173
chr13-98469492-DEL-74
chr13-98481532-DEL-50
chr13-98512442-DEL-52
chr13-98538388-DEL-1274
chr13-98562194-DEL-336
chr13-98562283-DEL-132
chr13-98804723-DEL-160
chr13-98821789-DEL-136
chr13-98822009-DEL-152
chr13-98822432-DEL-50
chr13-98840673-DEL-65
chr13-98874476-DEL-256
chr13-98887514-DEL-405
chr13-98930638-DEL-313
chr13-98953430-DEL-107
chr13-98962436-DEL-302
chr13-98972576-DEL-350
chr13-99009308-DEL-1611
chr13-99069831-DEL-119
chr13-99070007-DEL-313
chr13-99133365-DEL-466
chr13-99166021-DEL-267
chr13-99166343-DEL-69
chr13-99221132-DEL-216
chr13-99221726-DEL-192
chr13-99221736-DEL-566
chr13-99267332-DEL-7127
chr13-99293691-DEL-91
chr13-99337855-DEL-120
chr13-99384575-DEL-57
chr13-99421625-DEL-173
chr13-99459161-DEL-382
chr13-99459218-DEL-190
chr13-99485755-DEL-56
chr13-99496765-DEL-190
chr13-99497108-DEL-72
chr13-99525355-DEL-226
chr13-99588268-DEL-160
chr13-99696303-DEL-3386
chr13-99703840-DEL-234
chr13-99707488-DEL-58
chr13-99888980-DEL-69
chr13-99912443-DEL-50
chr13-99939419-DEL-312
chr13-99995205-DEL-293
chr13-100018330-DEL-80
chr13-100038159-DEL-988
chr13-100131976-DEL-224
chr13-100330822-DEL-119
chr13-100361059-DEL-94
chr13-100370418-DEL-208
chr13-100370892-DEL-208
chr13-100434263-DEL-128
chr13-100461788-DEL-275
chr13-100462789-DEL-824
chr13-100536126-DEL-93
chr13-100537383-DEL-55
chr13-100540175-DEL-276
chr13-100556163-DEL-558
chr13-100632609-DEL-73
chr13-100662105-DEL-201
chr13-100678588-DEL-56
chr13-100745332-DEL-332
chr13-100745756-DEL-91
chr13-100833439-DEL-50
chr13-100843248-DEL-1470
chr13-100884977-DEL-50
chr13-100938846-DEL-71
chr13-101010885-DEL-89
chr13-101025076-DEL-399
chr13-101081822-DEL-420
chr13-101133654-DEL-104
chr13-101191957-DEL-398
chr13-101264497-DEL-50

chr13-101495644-DEL-62
chr13-101531610-DEL-114
chr13-101531878-DEL-61
chr13-101532276-DEL-90
chr13-101582761-DEL-171
chr13-101719493-DEL-1488
chr13-101811371-DEL-74
chr13-101840378-DEL-831
chr13-101865593-DEL-274
chr13-102203112-DEL-50
chr13-102222155-DEL-72
chr13-102493510-DEL-158
chr13-102493826-DEL-77
chr13-102578869-DEL-623
chr13-102584748-DEL-70
chr13-102866362-DEL-160
chr13-102882712-DEL-763
chr13-103019460-DEL-51
chr13-103114315-DEL-86
chr13-103114919-DEL-83
chr13-103199132-DEL-50
chr13-103253883-DEL-63
chr13-103254090-DEL-434
chr13-103412508-DEL-348
chr13-103416744-DEL-56
chr13-103457005-DEL-439
chr13-103457134-DEL-97
chr13-103457347-DEL-97
chr13-103527019-DEL-471
chr13-103537305-DEL-201
chr13-103677229-DEL-77
chr13-103697397-DEL-1082
chr13-103718682-DEL-214
chr13-103759245-DEL-96
chr13-103831498-DEL-52
chr13-103845385-DEL-54
chr13-104066447-DEL-7345
chr13-104218279-DEL-676
chr13-104230023-DEL-80
chr13-104298048-DEL-72
chr13-104358862-DEL-144
chr13-104358880-DEL-60
chr13-104487272-DEL-90
chr13-104592829-DEL-66
chr13-104638765-DEL-175
chr13-104641766-DEL-54
chr13-104797197-DEL-58
chr13-104797399-DEL-201
chr13-104809924-DEL-86
chr13-105253911-DEL-55
chr13-105255316-DEL-342
chr13-105316508-DEL-53
chr13-105447133-DEL-128
chr13-105603452-DEL-50
chr13-105652581-DEL-326
chr13-105701585-DEL-62
chr13-105707473-DEL-54
chr13-105732910-DEL-430
chr13-105939109-DEL-1496
chr13-105995756-DEL-554
chr13-106011526-DEL-394
chr13-106113186-DEL-59
chr13-106145125-DEL-50
chr13-106274887-DEL-2311
chr13-106408298-DEL-1861
chr13-106544312-DEL-50
chr13-106641053-DEL-78
chr13-106800113-DEL-64
chr13-106803449-DEL-112
chr13-106937044-DEL-50
chr13-106942036-DEL-150
chr13-106989632-DEL-50
chr13-106996046-DEL-52
chr13-107160612-DEL-128
chr13-107176203-DEL-538
chr13-107249078-DEL-380
chr13-107255909-DEL-242

chr13-107256123-DEL-350
chr13-107302181-DEL-222
chr13-107321267-DEL-56
chr13-107340952-DEL-87
chr13-107404859-DEL-1546
chr13-107405437-DEL-90
chr13-107622874-DEL-308
chr13-107623019-DEL-152
chr13-107623103-DEL-72
chr13-107668386-DEL-377
chr13-107686323-DEL-59
chr13-107689955-DEL-126
chr13-107750499-DEL-567
chr13-107975669-DEL-600
chr13-107976251-DEL-150
chr13-107994790-DEL-78
chr13-108203295-DEL-344
chr13-108203317-DEL-114
chr13-108203547-DEL-138
chr13-108319770-DEL-54
chr13-108393472-DEL-50
chr13-108465072-DEL-711
chr13-108546374-DEL-51
chr13-108812638-DEL-54
chr13-108891191-DEL-68
chr13-108911765-DEL-50
chr13-108983527-DEL-50
chr13-109006732-DEL-56
chr13-109018816-DEL-93
chr13-109018924-DEL-245
chr13-109116849-DEL-1211
chr13-109167020-DEL-1361
chr13-109185732-DEL-81
chr13-109240772-DEL-3009
chr13-109411378-DEL-93
chr13-109544469-DEL-122
chr13-109650721-DEL-66
chr13-109684051-DEL-55
chr13-109694196-DEL-73
chr13-109694247-DEL-190
chr13-109694350-DEL-85
chr13-109707563-DEL-56
chr13-109737829-DEL-770
chr13-109763556-DEL-10125
chr13-110019009-DEL-50
chr13-110076632-DEL-619
chr13-110171206-DEL-83
chr13-110245953-DEL-287
chr13-110267445-DEL-67
chr13-110287576-DEL-188
chr13-110352746-DEL-254
chr13-110400591-DEL-152
chr13-110446214-DEL-214
chr13-110593080-DEL-729
chr13-110860327-DEL-162
chr13-110860424-DEL-64
chr13-110880533-DEL-170
chr13-110880562-DEL-83
chr13-110897005-DEL-1603
chr13-111117573-DEL-50
chr13-111193086-DEL-60
chr13-111193118-DEL-176
chr13-111366692-DEL-50
chr13-111458401-DEL-116
chr13-111479067-DEL-110
chr13-111522630-DEL-142
chr13-111526956-DEL-207
chr13-111608193-DEL-218
chr13-111688720-DEL-95
chr13-111708467-DEL-576
chr13-111708582-DEL-222
chr13-111708861-DEL-206
chr13-111733711-DEL-73
chr13-111911048-DEL-66
chr13-111911372-DEL-66
chr13-111965075-DEL-82
chr13-111965251-DEL-72

chr13-112026291-DEL-498
chr13-112027202-DEL-82
chr13-112099117-DEL-186
chr13-112167161-DEL-54
chr13-112243144-DEL-767
chr13-112247461-DEL-1348
chr13-112290903-DEL-816
chr13-112377486-DEL-56
chr13-112392079-DEL-52
chr13-112457275-DEL-50
chr13-112465236-DEL-236
chr13-112465280-DEL-72
chr13-112516412-DEL-438
chr13-112558784-DEL-353
chr13-112558826-DEL-124
chr13-112567082-DEL-1736
chr13-112567169-DEL-310
chr13-112568594-DEL-62
chr13-112587154-DEL-70
chr13-112588913-DEL-176
chr13-112614531-DEL-68
chr13-112676383-DEL-107
chr13-112690455-DEL-110
chr13-112709130-DEL-251
chr13-112711884-DEL-76
chr13-112773004-DEL-77
chr13-112810289-DEL-120
chr13-112826873-DEL-114
chr13-112854312-DEL-51
chr13-112866389-DEL-341
chr13-112880962-DEL-62
chr13-112980652-DEL-661
chr13-113160339-DEL-61
chr13-113255420-DEL-73
chr13-113300279-DEL-505
chr13-113322311-DEL-205
chr13-113322618-DEL-61
chr13-113519538-DEL-538
chr13-113532513-DEL-98
chr13-113534451-DEL-63
chr13-113669135-DEL-145
chr13-113711945-DEL-50
chr13-113809683-DEL-242
chr13-113829266-DEL-160
chr13-113847643-DEL-52
chr13-113875285-DEL-72
chr13-113880533-DEL-50
chr13-113998121-DEL-84
chr13-114077240-DEL-180
chr13-114210117-DEL-311
chr13-114494788-DEL-66
chr13-114514875-DEL-88
chr13-114618982-DEL-147
chr13-114652718-DEL-203
chr13-114718482-DEL-58
chr13-114918332-DEL-321
chr13-114938099-DEL-100
chr13-114940622-DEL-251
chr13-114940902-DEL-459
chr13-114968366-DEL-53
chr13-114981519-DEL-72
chr13-114990022-DEL-60
chr13-115078312-DEL-1959
chr13-115199103-DEL-52
chr13-115356844-DEL-82
chr13-115361443-DEL-78
chr13-115381391-DEL-51
chr13-115382322-DEL-89
chr13-115442135-DEL-56
chr13-115590007-DEL-3375
chr13-115627145-DEL-96
chr13-115638960-DEL-738
chr13-115640807-DEL-200
chr13-115749803-DEL-2322
chr13-115758860-DEL-52
chr13-115804197-DEL-151
chr13-115891063-DEL-1418

chr13-115930681-DEL-62
chr13-116016062-DEL-164
chr13-116229034-DEL-2963
chr13-116318029-DEL-354
chr13-116364903-DEL-146
chr13-116365085-DEL-117
chr13-116408764-DEL-143
chr13-116466542-DEL-60
chr13-116521521-DEL-69
chr13-116546189-DEL-102
chr13-116647893-DEL-1130
chr13-116818918-DEL-59
chr13-117027477-DEL-96
chr13-117139153-DEL-311
chr13-117145931-DEL-113
chr13-117185729-DEL-420
chr13-117191472-DEL-50
chr13-117288900-DEL-119
chr13-117454010-DEL-78
chr13-117577585-DEL-4251
chr13-117640804-DEL-2818
chr13-117667820-DEL-50
chr13-117887743-DEL-50
chr13-117960959-DEL-52
chr13-118020367-DEL-91
chr13-118102407-DEL-161
chr13-118216279-DEL-50
chr13-118245227-DEL-155
chr13-118285247-DEL-3794
chr13-118472940-DEL-78
chr13-118782076-DEL-64
chr13-118835418-DEL-89
chr13-118844235-DEL-74
chr13-119050863-DEL-60
chr13-119095595-DEL-126
chr13-119128628-DEL-4985
chr13-119141632-DEL-70
chr13-119160254-DEL-96
chr13-119185008-DEL-92
chr13-119259369-DEL-413
chr13-119540155-DEL-351
chr13-119564715-DEL-151
chr13-119566217-DEL-61
chr13-119624161-DEL-188
chr13-119628999-DEL-439015
chr13-120072885-DEL-1469
chr13-120075954-DEL-444
chr13-120078505-DEL-997
chr13-120079332-DEL-164
chr13-120202641-DEL-52
chr13-120209457-DEL-1094
chr13-120268340-DEL-50
chr13-120347608-DEL-72
chr13-120382801-DEL-57
chr13-120470077-DEL-240
chr13-120499058-DEL-1509
chr13-120645407-DEL-405
chr14-3155991-DEL-86
chr14-3407183-DEL-792
chr14-3572781-DEL-68
chr14-3621641-DEL-50
chr14-3765766-DEL-2314
chr14-4248691-DEL-131
chr14-4436452-DEL-72
chr14-4537449-DEL-2128
chr14-4544397-DEL-244
chr14-4545006-DEL-195
chr14-4545255-DEL-58
chr14-4609889-DEL-51
chr14-4976293-DEL-257
chr14-5113075-DEL-230
chr14-5163322-DEL-985
chr14-5219587-DEL-71
chr14-5247071-DEL-50
chr14-5257756-DEL-50
chr14-5259814-DEL-1204
chr14-5288763-DEL-111

chr14-5384289-DEL-267
chr14-5408061-DEL-301
chr14-5426404-DEL-90
chr14-5438605-DEL-51645
chr14-5443586-DEL-423
chr14-5605649-DEL-54
chr14-5623867-DEL-204
chr14-5624003-DEL-72
chr14-5893293-DEL-80
chr14-6010029-DEL-84
chr14-6010088-DEL-280
chr14-6066795-DEL-52
chr14-6124103-DEL-93
chr14-6312098-DEL-1292
chr14-6320924-DEL-5082
chr14-6379587-DEL-129
chr14-6379653-DEL-63
chr14-6595991-DEL-272
chr14-6775705-DEL-400
chr14-6782566-DEL-159
chr14-6786489-DEL-1179
chr14-6805388-DEL-50
chr14-6925289-DEL-453
chr14-7119964-DEL-272
chr14-7267256-DEL-58
chr14-7387698-DEL-72
chr14-7521432-DEL-91
chr14-7533474-DEL-1094
chr14-7675086-DEL-61
chr14-7677376-DEL-296
chr14-7677594-DEL-112
chr14-7783807-DEL-50
chr14-7788852-DEL-156
chr14-7795891-DEL-62
chr14-7951987-DEL-52
chr14-8008442-DEL-50
chr14-8419174-DEL-1097
chr14-8532720-DEL-4424
chr14-8575859-DEL-55
chr14-8670231-DEL-855
chr14-8866682-DEL-572
chr14-8986430-DEL-1382
chr14-9160324-DEL-371
chr14-9185943-DEL-54
chr14-9195390-DEL-1125
chr14-9210459-DEL-235
chr14-9238640-DEL-68
chr14-9268523-DEL-783
chr14-9325313-DEL-75
chr14-9325766-DEL-1460
chr14-9328902-DEL-112
chr14-9329473-DEL-75
chr14-9438818-DEL-1030
chr14-9527253-DEL-63
chr14-9535754-DEL-73
chr14-9579436-DEL-52
chr14-9582692-DEL-66
chr14-9675858-DEL-147
chr14-9721937-DEL-138
chr14-9873709-DEL-112
chr14-9875572-DEL-89
chr14-9901057-DEL-60
chr14-10025942-DEL-52
chr14-10067847-DEL-862
chr14-10087886-DEL-80
chr14-10154996-DEL-73
chr14-10331227-DEL-117
chr14-10514490-DEL-55
chr14-10514785-DEL-61
chr14-10531187-DEL-64
chr14-10642655-DEL-110
chr14-10643072-DEL-456
chr14-10810712-DEL-286
chr14-10918657-DEL-143
chr14-10989340-DEL-936
chr14-10995158-DEL-50
chr14-11069605-DEL-89

chr14-11073771-DEL-88
chr14-11114972-DEL-253
chr14-11199105-DEL-1802
chr14-11201294-DEL-191
chr14-11201345-DEL-84
chr14-11212602-DEL-51
chr14-11227007-DEL-54
chr14-11419872-DEL-2196
chr14-11420196-DEL-108
chr14-11422583-DEL-468
chr14-11431848-DEL-128
chr14-11492455-DEL-144
chr14-11584551-DEL-466
chr14-11649956-DEL-57
chr14-11672585-DEL-96
chr14-11802697-DEL-50
chr14-11844806-DEL-58
chr14-11965416-DEL-3349
chr14-12115954-DEL-116
chr14-12197333-DEL-2118
chr14-12258401-DEL-640
chr14-12382983-DEL-214
chr14-12524916-DEL-50
chr14-12779118-DEL-51
chr14-12783977-DEL-205
chr14-12915008-DEL-50
chr14-12986066-DEL-50
chr14-12986268-DEL-62
chr14-13043082-DEL-1567
chr14-13079681-DEL-224
chr14-13216190-DEL-56
chr14-13321024-DEL-52
chr14-13384603-DEL-686
chr14-13466988-DEL-148
chr14-13507114-DEL-82
chr14-13649692-DEL-56
chr14-13804785-DEL-118
chr14-13923219-DEL-94
chr14-13947393-DEL-166
chr14-13973386-DEL-50
chr14-13982019-DEL-1248
chr14-13997543-DEL-93
chr14-14005934-DEL-66
chr14-14044917-DEL-82
chr14-14109873-DEL-678
chr14-14256704-DEL-296
chr14-14261437-DEL-54
chr14-14293753-DEL-177
chr14-14395694-DEL-6334
chr14-14452150-DEL-165
chr14-14452249-DEL-66
chr14-14480819-DEL-51
chr14-14571811-DEL-160
chr14-14685141-DEL-94
chr14-14706963-DEL-50
chr14-14808531-DEL-108
chr14-15197135-DEL-216
chr14-15266659-DEL-231
chr14-15468950-DEL-202
chr14-16218771-DEL-115
chr14-18249317-DEL-56
chr14-18941294-DEL-953
chr14-19264400-DEL-96
chr14-19265654-DEL-122
chr14-19638053-DEL-50
chr14-19783304-DEL-59
chr14-19819304-DEL-58
chr14-19840591-DEL-60
chr14-19840857-DEL-60
chr14-20015386-DEL-68
chr14-20031585-DEL-356
chr14-20050754-DEL-155
chr14-20083078-DEL-68
chr14-20167801-DEL-252
chr14-20168194-DEL-102
chr14-20237979-DEL-268
chr14-20270488-DEL-66

chr14-20304936-DEL-362
chr14-20306266-DEL-324
chr14-20306497-DEL-111
chr14-20342822-DEL-114
chr14-20402997-DEL-127
chr14-20469003-DEL-488
chr14-20469530-DEL-54
chr14-20469994-DEL-269
chr14-20470372-DEL-235
chr14-20525038-DEL-80
chr14-20525149-DEL-371
chr14-20525478-DEL-51
chr14-20819915-DEL-50
chr14-20872543-DEL-1321
chr14-20946682-DEL-634
chr14-21017184-DEL-54
chr14-21073068-DEL-863
chr14-21250440-DEL-99
chr14-21284652-DEL-50
chr14-21330018-DEL-71
chr14-21359148-DEL-61
chr14-21487007-DEL-77
chr14-21733505-DEL-82
chr14-21776332-DEL-182
chr14-21781637-DEL-92
chr14-21806937-DEL-388
chr14-21887044-DEL-140
chr14-21961962-DEL-54
chr14-21971452-DEL-573
chr14-22059752-DEL-50
chr14-22103517-DEL-124
chr14-22130792-DEL-66
chr14-22264967-DEL-147
chr14-22282674-DEL-178
chr14-22346633-DEL-54
chr14-22398902-DEL-114
chr14-22674828-DEL-78
chr14-22805315-DEL-180
chr14-22883216-DEL-2388
chr14-22949465-DEL-725
chr14-22985104-DEL-662
chr14-22988386-DEL-50
chr14-23065457-DEL-51
chr14-23083904-DEL-213
chr14-23146944-DEL-220
chr14-23459246-DEL-132
chr14-23460394-DEL-214
chr14-23585014-DEL-3965
chr14-23604101-DEL-59
chr14-23667884-DEL-77
chr14-23668046-DEL-140
chr14-23813513-DEL-2069
chr14-23900921-DEL-155
chr14-24104977-DEL-54
chr14-24121551-DEL-88
chr14-24171901-DEL-2904
chr14-24391105-DEL-69
chr14-24593797-DEL-56
chr14-24906510-DEL-84
chr14-24930045-DEL-74
chr14-24948772-DEL-50
chr14-24954358-DEL-284
chr14-25038078-DEL-60
chr14-25144543-DEL-255
chr14-25144699-DEL-101
chr14-25212992-DEL-52
chr14-25330741-DEL-555
chr14-25391149-DEL-50
chr14-25467590-DEL-122
chr14-25556981-DEL-56
chr14-25675473-DEL-382
chr14-25675601-DEL-72
chr14-25704658-DEL-204
chr14-25754819-DEL-76
chr14-25792344-DEL-54
chr14-25796829-DEL-50
chr14-25815325-DEL-1053

chr14-25827894-DEL-208
chr14-25861599-DEL-54
chr14-25912086-DEL-1731
chr14-25914602-DEL-164
chr14-25915210-DEL-391
chr14-25915267-DEL-129
chr14-25915686-DEL-184
chr14-25922275-DEL-71
chr14-25990653-DEL-220
chr14-25998396-DEL-62
chr14-26071415-DEL-4147
chr14-26168298-DEL-87
chr14-26290019-DEL-155
chr14-26461212-DEL-135
chr14-26462413-DEL-81
chr14-26467385-DEL-58
chr14-26467520-DEL-52
chr14-26468216-DEL-81
chr14-26476249-DEL-108
chr14-26573831-DEL-78
chr14-26634697-DEL-112
chr14-26758111-DEL-925
chr14-26785209-DEL-59
chr14-26843131-DEL-784
chr14-26961167-DEL-124
chr14-26967870-DEL-101
chr14-26970515-DEL-142
chr14-27250432-DEL-60
chr14-27250509-DEL-126
chr14-27256367-DEL-1719
chr14-27284342-DEL-14085
chr14-27371314-DEL-59
chr14-27415320-DEL-327
chr14-27484663-DEL-81
chr14-27535316-DEL-60
chr14-27554508-DEL-67
chr14-27587744-DEL-6107
chr14-27721344-DEL-298
chr14-27975937-DEL-152
chr14-28019436-DEL-52
chr14-28112810-DEL-918
chr14-28358643-DEL-88
chr14-28359438-DEL-84
chr14-28415874-DEL-50
chr14-28464036-DEL-55
chr14-28617963-DEL-329
chr14-28618063-DEL-149
chr14-28619680-DEL-87
chr14-28622335-DEL-62
chr14-28634724-DEL-52
chr14-28639966-DEL-5701
chr14-28722604-DEL-151
chr14-28765996-DEL-1171
chr14-29013551-DEL-53
chr14-29204665-DEL-52
chr14-29295482-DEL-208
chr14-29459512-DEL-62
chr14-29474373-DEL-50
chr14-29482140-DEL-105
chr14-29745592-DEL-124
chr14-29830769-DEL-1275
chr14-30187063-DEL-810
chr14-30187222-DEL-143
chr14-30433525-DEL-92
chr14-30545162-DEL-100
chr14-30648672-DEL-50
chr14-30662315-DEL-55
chr14-30670913-DEL-680
chr14-30684430-DEL-240
chr14-30691405-DEL-122
chr14-31237374-DEL-71
chr14-31302993-DEL-305
chr14-31303397-DEL-294
chr14-31304935-DEL-50
chr14-31596311-DEL-357
chr14-31613024-DEL-54
chr14-31639357-DEL-528

chr14-31702566-DEL-291
chr14-31725659-DEL-62
chr14-31729678-DEL-1827
chr14-31744599-DEL-80
chr14-32050657-DEL-76
chr14-32051674-DEL-125
chr14-32073365-DEL-234
chr14-32080143-DEL-307
chr14-32084328-DEL-1080
chr14-32096224-DEL-86
chr14-32131062-DEL-96
chr14-32171945-DEL-575
chr14-32205761-DEL-635
chr14-32232417-DEL-315
chr14-32243619-DEL-362
chr14-32280840-DEL-117
chr14-32334697-DEL-1018
chr14-32379703-DEL-202
chr14-32497542-DEL-57
chr14-32497647-DEL-168
chr14-32577092-DEL-50
chr14-32751278-DEL-1012
chr14-32856714-DEL-887
chr14-32892864-DEL-92
chr14-32902787-DEL-146
chr14-32908633-DEL-1849
chr14-32919763-DEL-61
chr14-32939820-DEL-64
chr14-32963446-DEL-927
chr14-32965701-DEL-116
chr14-33009519-DEL-51
chr14-33014996-DEL-156
chr14-33150257-DEL-763
chr14-33174497-DEL-52
chr14-33505185-DEL-68
chr14-33661917-DEL-104
chr14-33673153-DEL-77
chr14-33734353-DEL-1308
chr14-33786615-DEL-120
chr14-33876376-DEL-54
chr14-33970259-DEL-56
chr14-34063338-DEL-51
chr14-34068761-DEL-219
chr14-34268339-DEL-55
chr14-34301779-DEL-123
chr14-34328322-DEL-488
chr14-34334104-DEL-50
chr14-34421044-DEL-160
chr14-34466204-DEL-52
chr14-34502745-DEL-495
chr14-34516968-DEL-50
chr14-34531320-DEL-54
chr14-34634202-DEL-5213
chr14-34689567-DEL-72
chr14-34689795-DEL-120
chr14-34709358-DEL-254
chr14-34726042-DEL-50
chr14-34854882-DEL-628
chr14-34888121-DEL-57
chr14-34894569-DEL-1713
chr14-34955988-DEL-88
chr14-35204322-DEL-94
chr14-35279969-DEL-56
chr14-35388305-DEL-125
chr14-35390109-DEL-261
chr14-35390271-DEL-87
chr14-35392315-DEL-1347
chr14-35415008-DEL-54
chr14-35543511-DEL-101
chr14-35571879-DEL-161
chr14-35624182-DEL-219
chr14-35686913-DEL-1123
chr14-35704933-DEL-59
chr14-35736466-DEL-56
chr14-35740903-DEL-6322
chr14-35852609-DEL-62
chr14-36019490-DEL-182

chr14-36628405-DEL-140
chr14-36669258-DEL-382
chr14-36694282-DEL-184
chr14-36696498-DEL-233
chr14-36719467-DEL-54
chr14-36730273-DEL-65
chr14-36777699-DEL-63
chr14-36777817-DEL-83
chr14-36900132-DEL-160
chr14-36982876-DEL-214
chr14-37268262-DEL-16918
chr14-37334639-DEL-2111
chr14-37352777-DEL-1247
chr14-37615000-DEL-197
chr14-37773102-DEL-591
chr14-38002446-DEL-4219
chr14-38006849-DEL-1322
chr14-38017777-DEL-54
chr14-38208697-DEL-65
chr14-38250682-DEL-112
chr14-38270958-DEL-4354
chr14-38282308-DEL-3272
chr14-38429879-DEL-50
chr14-38539612-DEL-1908
chr14-38606013-DEL-73
chr14-38621755-DEL-50
chr14-38621788-DEL-141
chr14-38621873-DEL-56
chr14-38769727-DEL-58
chr14-38961734-DEL-84
chr14-39132257-DEL-60
chr14-39157394-DEL-170
chr14-39377852-DEL-411
chr14-39498395-DEL-56
chr14-39513112-DEL-50
chr14-39671050-DEL-50
chr14-39720778-DEL-51
chr14-39759406-DEL-114
chr14-39955239-DEL-129
chr14-40133038-DEL-50
chr14-40149058-DEL-2485
chr14-40269208-DEL-212
chr14-40282756-DEL-158
chr14-40334264-DEL-1103
chr14-40385213-DEL-66
chr14-40446540-DEL-4228
chr14-40541260-DEL-148
chr14-40543607-DEL-808
chr14-40593800-DEL-53
chr14-40682740-DEL-967
chr14-40683083-DEL-165
chr14-40683542-DEL-165
chr14-40754135-DEL-76
chr14-40806791-DEL-363
chr14-40858690-DEL-50
chr14-40866802-DEL-99
chr14-41356124-DEL-912
chr14-41430660-DEL-1271
chr14-42002912-DEL-338
chr14-42847260-DEL-52
chr14-42853622-DEL-483
chr14-42987778-DEL-64
chr14-43133958-DEL-56
chr14-43150621-DEL-201
chr14-43198706-DEL-1454
chr14-43212652-DEL-830
chr14-43812776-DEL-56
chr14-44229889-DEL-60
chr14-45163809-DEL-137
chr14-45164910-DEL-1490
chr14-45167510-DEL-895
chr14-45172069-DEL-815
chr14-45249941-DEL-60
chr14-45289188-DEL-364
chr14-45331343-DEL-75
chr14-45604468-DEL-64
chr14-45630594-DEL-76

chr14-45636944-DEL-50
chr14-45679403-DEL-377
chr14-45733292-DEL-424
chr14-45781850-DEL-52
chr14-45805361-DEL-83
chr14-45806653-DEL-637
chr14-45926290-DEL-281
chr14-45926529-DEL-72
chr14-46007299-DEL-114
chr14-46194835-DEL-80
chr14-46513715-DEL-145
chr14-46587779-DEL-582
chr14-46694696-DEL-52
chr14-46818219-DEL-52
chr14-46921978-DEL-463
chr14-46928298-DEL-108
chr14-46951660-DEL-60
chr14-46976563-DEL-195
chr14-46976582-DEL-76
chr14-46976742-DEL-200
chr14-46976848-DEL-91
chr14-47223349-DEL-67
chr14-47294606-DEL-78
chr14-47299019-DEL-132
chr14-47369600-DEL-417
chr14-47369694-DEL-115
chr14-47375840-DEL-50
chr14-47381466-DEL-57
chr14-47493947-DEL-202
chr14-47494516-DEL-62
chr14-47705670-DEL-102
chr14-47705874-DEL-201
chr14-47909380-DEL-73
chr14-47935617-DEL-1128
chr14-48109962-DEL-739
chr14-48110835-DEL-158
chr14-48167992-DEL-64
chr14-48183388-DEL-66
chr14-48336322-DEL-50
chr14-48452683-DEL-1089
chr14-48642029-DEL-67
chr14-48713009-DEL-50
chr14-48713359-DEL-50
chr14-49007324-DEL-105
chr14-49079486-DEL-84
chr14-49079904-DEL-55
chr14-49121727-DEL-84
chr14-49222588-DEL-106
chr14-49273963-DEL-86
chr14-49353595-DEL-50
chr14-49456592-DEL-213
chr14-49764305-DEL-72
chr14-49901391-DEL-76
chr14-49901560-DEL-64
chr14-50098401-DEL-68
chr14-50122122-DEL-214
chr14-50162052-DEL-195
chr14-50194802-DEL-72
chr14-50194808-DEL-202
chr14-50326363-DEL-64
chr14-50392151-DEL-60
chr14-50703441-DEL-90
chr14-50708006-DEL-108
chr14-50708036-DEL-341
chr14-50708233-DEL-122
chr14-50838042-DEL-4508
chr14-50843894-DEL-3192
chr14-50989882-DEL-50
chr14-51063882-DEL-63
chr14-51190411-DEL-13263
chr14-51199166-DEL-973
chr14-51211505-DEL-3144
chr14-51364477-DEL-50
chr14-51370554-DEL-54
chr14-51448278-DEL-2656
chr14-51534038-DEL-832
chr14-51674850-DEL-110

chr14-51675013-DEL-61
chr14-51849712-DEL-8332
chr14-51921230-DEL-94
chr14-51972379-DEL-65
chr14-52051683-DEL-1890
chr14-52137827-DEL-210
chr14-52332874-DEL-67
chr14-52333722-DEL-61
chr14-52498004-DEL-60
chr14-52518598-DEL-272
chr14-52573305-DEL-153
chr14-52573490-DEL-678
chr14-52602686-DEL-3056
chr14-52617754-DEL-210
chr14-52621327-DEL-287
chr14-52655052-DEL-637
chr14-52717203-DEL-598
chr14-52717523-DEL-92
chr14-52747861-DEL-2152
chr14-52881314-DEL-208
chr14-52881418-DEL-60
chr14-52910474-DEL-6551
chr14-52957606-DEL-109
chr14-53193127-DEL-75
chr14-53194100-DEL-50
chr14-53271001-DEL-489
chr14-53480109-DEL-50
chr14-53514595-DEL-56
chr14-53528468-DEL-2382
chr14-53556531-DEL-888
chr14-53575221-DEL-497
chr14-53750543-DEL-511
chr14-53751535-DEL-482
chr14-53801090-DEL-8381
chr14-53822961-DEL-90
chr14-53838598-DEL-52
chr14-53871580-DEL-115
chr14-54539040-DEL-50
chr14-54567215-DEL-88
chr14-54580463-DEL-153
chr14-54653495-DEL-56
chr14-54687200-DEL-68
chr14-54694349-DEL-177
chr14-54803003-DEL-108
chr14-54803293-DEL-209
chr14-54928702-DEL-1271
chr14-54946185-DEL-82
chr14-55073188-DEL-375
chr14-55137238-DEL-64
chr14-55209279-DEL-63
chr14-55209441-DEL-244
chr14-55214860-DEL-90
chr14-55215273-DEL-165
chr14-55404664-DEL-228
chr14-55404696-DEL-106
chr14-55699023-DEL-109
chr14-55849561-DEL-50
chr14-55925434-DEL-152
chr14-56074018-DEL-55
chr14-56074241-DEL-201
chr14-56178550-DEL-50
chr14-56381742-DEL-211
chr14-56555467-DEL-71
chr14-56564889-DEL-66
chr14-56797862-DEL-288
chr14-56804412-DEL-115
chr14-56971359-DEL-108
chr14-57319585-DEL-499
chr14-57625922-DEL-58
chr14-57659015-DEL-199
chr14-57742637-DEL-50
chr14-57797022-DEL-392
chr14-57966009-DEL-666
chr14-58077261-DEL-330
chr14-58117517-DEL-204
chr14-58157933-DEL-204
chr14-58184143-DEL-380

chr14-58230925-DEL-240
chr14-58265514-DEL-288
chr14-58267067-DEL-760
chr14-58348393-DEL-88
chr14-58562237-DEL-204
chr14-58570882-DEL-50
chr14-58587448-DEL-88
chr14-58850677-DEL-662
chr14-59058689-DEL-51
chr14-59079150-DEL-64
chr14-59415545-DEL-59
chr14-59517247-DEL-44173
chr14-59889764-DEL-132
chr14-59905685-DEL-50
chr14-59917200-DEL-184
chr14-59926521-DEL-72
chr14-60229230-DEL-192
chr14-60229336-DEL-2232
chr14-60278172-DEL-941
chr14-60674726-DEL-885
chr14-60735055-DEL-135
chr14-61232154-DEL-76
chr14-61481462-DEL-181
chr14-61499400-DEL-1160
chr14-61550679-DEL-1358
chr14-61582996-DEL-245
chr14-61708604-DEL-291
chr14-61723446-DEL-251
chr14-61741675-DEL-81
chr14-61802426-DEL-194
chr14-61817908-DEL-6460
chr14-61843460-DEL-180
chr14-61855549-DEL-137
chr14-61994863-DEL-52
chr14-62109594-DEL-54
chr14-62234921-DEL-90
chr14-62272571-DEL-92
chr14-62398512-DEL-208
chr14-62398968-DEL-64
chr14-62419314-DEL-209
chr14-62465910-DEL-78
chr14-62515564-DEL-56
chr14-62560079-DEL-50
chr14-62574337-DEL-84
chr14-62617902-DEL-57
chr14-62639584-DEL-616
chr14-62664269-DEL-51
chr14-62664368-DEL-51
chr14-62724975-DEL-128
chr14-62752025-DEL-50
chr14-62804327-DEL-51
chr14-63048532-DEL-216
chr14-63261522-DEL-858
chr14-63376710-DEL-256
chr14-63420276-DEL-52
chr14-63517754-DEL-192
chr14-63517754-DEL-390
chr14-63571537-DEL-80
chr14-63663770-DEL-64
chr14-63740887-DEL-60
chr14-63919800-DEL-104
chr14-64252317-DEL-72
chr14-64344376-DEL-234
chr14-64441972-DEL-54
chr14-64524157-DEL-64
chr14-64643516-DEL-50
chr14-64710467-DEL-70
chr14-64882928-DEL-169
chr14-65011447-DEL-60
chr14-65057877-DEL-69
chr14-65175443-DEL-1289
chr14-65212992-DEL-168
chr14-65223892-DEL-1110
chr14-65224772-DEL-629
chr14-65273453-DEL-108
chr14-65282190-DEL-50
chr14-65344531-DEL-214

chr14-65626864-DEL-50
chr14-65633209-DEL-50
chr14-65778545-DEL-195
chr14-65831807-DEL-133
chr14-65835943-DEL-86
chr14-65862431-DEL-97
chr14-65862712-DEL-96
chr14-65863423-DEL-146
chr14-65863723-DEL-239
chr14-65867762-DEL-98
chr14-65926306-DEL-63
chr14-65935644-DEL-57
chr14-65947209-DEL-98
chr14-66023443-DEL-319
chr14-66033514-DEL-1438
chr14-66123149-DEL-382
chr14-66144207-DEL-175
chr14-66318792-DEL-316
chr14-66472656-DEL-54
chr14-66644742-DEL-59
chr14-66746939-DEL-64
chr14-66750302-DEL-60
chr14-66804534-DEL-59
chr14-66941576-DEL-337
chr14-67000249-DEL-120
chr14-67003770-DEL-80
chr14-67013114-DEL-284
chr14-67144688-DEL-62
chr14-67157696-DEL-50
chr14-67209364-DEL-136
chr14-67383435-DEL-69
chr14-67399733-DEL-182
chr14-67441243-DEL-150
chr14-67478902-DEL-54
chr14-67527129-DEL-50
chr14-67636546-DEL-74
chr14-67676391-DEL-109
chr14-67752570-DEL-88
chr14-67763516-DEL-52
chr14-67999372-DEL-323
chr14-68111143-DEL-60
chr14-68114004-DEL-75
chr14-68259170-DEL-67
chr14-68297370-DEL-83
chr14-68404231-DEL-87
chr14-68404590-DEL-155
chr14-68470690-DEL-53
chr14-68470921-DEL-223
chr14-68546910-DEL-1410
chr14-68547809-DEL-317
chr14-68572396-DEL-61
chr14-68713317-DEL-2003
chr14-68713491-DEL-804
chr14-68760488-DEL-134
chr14-68881926-DEL-78
chr14-68975630-DEL-229
chr14-68975646-DEL-68
chr14-69107584-DEL-2614
chr14-69115642-DEL-461
chr14-69127603-DEL-80
chr14-69146104-DEL-195
chr14-69199881-DEL-68
chr14-69403587-DEL-1026
chr14-69556709-DEL-71
chr14-69565925-DEL-56
chr14-69598510-DEL-192
chr14-69753701-DEL-248
chr14-69784175-DEL-56
chr14-69799176-DEL-189
chr14-69816760-DEL-224
chr14-69830906-DEL-201
chr14-69834147-DEL-66
chr14-69925502-DEL-310
chr14-69962078-DEL-2464
chr14-69995997-DEL-68
chr14-70079573-DEL-54
chr14-70102297-DEL-78

chr14-70166916-DEL-190
chr14-70167208-DEL-65
chr14-70473675-DEL-87
chr14-70502859-DEL-270
chr14-70805956-DEL-298
chr14-70963980-DEL-54
chr14-70964212-DEL-267
chr14-70998705-DEL-186
chr14-71264459-DEL-664
chr14-71328538-DEL-353
chr14-71485935-DEL-57
chr14-71712333-DEL-82
chr14-71712515-DEL-77
chr14-72092893-DEL-56
chr14-72140746-DEL-50
chr14-72274971-DEL-574
chr14-72393373-DEL-130
chr14-72393454-DEL-63
chr14-72515843-DEL-50
chr14-72527190-DEL-92
chr14-72589763-DEL-150
chr14-72686580-DEL-171
chr14-72706331-DEL-76
chr14-72757725-DEL-6353
chr14-72874498-DEL-54
chr14-72919922-DEL-300
chr14-72981202-DEL-56
chr14-73165126-DEL-61
chr14-73250844-DEL-63
chr14-73321831-DEL-346
chr14-73321914-DEL-58
chr14-73322319-DEL-178
chr14-73386013-DEL-224
chr14-73392771-DEL-189
chr14-73566331-DEL-52
chr14-73642472-DEL-266
chr14-73642548-DEL-76
chr14-73670131-DEL-76
chr14-73684322-DEL-68
chr14-73691564-DEL-72
chr14-73709490-DEL-69
chr14-73719884-DEL-79
chr14-73729767-DEL-876
chr14-73877776-DEL-84
chr14-73936654-DEL-83
chr14-73997022-DEL-78
chr14-74202398-DEL-75
chr14-74292761-DEL-207
chr14-74360112-DEL-315
chr14-74590199-DEL-66
chr14-74642352-DEL-407
chr14-74728116-DEL-360
chr14-74785612-DEL-1492
chr14-75085321-DEL-51
chr14-75101297-DEL-68
chr14-75211433-DEL-50
chr14-75246625-DEL-102
chr14-75263649-DEL-2581
chr14-75519474-DEL-341
chr14-75761248-DEL-51
chr14-75824272-DEL-189
chr14-75906612-DEL-71
chr14-75948234-DEL-353
chr14-76038663-DEL-87
chr14-76041592-DEL-76
chr14-76042381-DEL-112
chr14-76060451-DEL-1025
chr14-76372911-DEL-60
chr14-76377010-DEL-671
chr14-76587010-DEL-120
chr14-76587036-DEL-58
chr14-76726473-DEL-129
chr14-77194067-DEL-56
chr14-77285857-DEL-3350
chr14-77314475-DEL-207
chr14-77348198-DEL-104
chr14-77391361-DEL-96

chr14-77749693-DEL-100
chr14-77937522-DEL-318
chr14-78017788-DEL-213
chr14-78028903-DEL-52
chr14-78049984-DEL-217
chr14-78383530-DEL-50
chr14-78494560-DEL-368
chr14-78673868-DEL-51
chr14-78681161-DEL-63
chr14-78712080-DEL-222
chr14-78852194-DEL-57
chr14-78882336-DEL-52
chr14-78944371-DEL-105
chr14-78994766-DEL-50
chr14-79065824-DEL-548
chr14-79222422-DEL-809
chr14-79223705-DEL-133
chr14-79254265-DEL-1129
chr14-79365643-DEL-50
chr14-79428596-DEL-1446
chr14-79455050-DEL-57
chr14-79601907-DEL-561
chr14-79713701-DEL-240
chr14-79821235-DEL-75
chr14-79900926-DEL-1236
chr14-80334677-DEL-134
chr14-80800937-DEL-127
chr14-80924128-DEL-110
chr14-81037840-DEL-828
chr14-81353845-DEL-60
chr14-81414935-DEL-3657
chr14-81639372-DEL-122
chr14-81701947-DEL-50
chr14-81721814-DEL-50
chr14-81784943-DEL-1534
chr14-81964508-DEL-52
chr14-81968927-DEL-62
chr14-81995207-DEL-123
chr14-81995257-DEL-56
chr14-82259974-DEL-56
chr14-82321429-DEL-636
chr14-83510080-DEL-50
chr14-83615144-DEL-164
chr14-84429911-DEL-1911
chr14-84543094-DEL-50
chr14-84605256-DEL-418
chr14-85397205-DEL-66
chr14-85534504-DEL-186
chr14-85676331-DEL-122
chr14-85888570-DEL-50
chr14-86241333-DEL-191
chr14-86300129-DEL-50
chr14-86383343-DEL-77
chr14-86386994-DEL-50
chr14-86497350-DEL-87
chr14-86759603-DEL-288
chr14-86919095-DEL-91
chr14-87082139-DEL-269
chr14-87336674-DEL-50
chr14-87402294-DEL-60
chr14-87465803-DEL-58
chr14-87487136-DEL-108
chr14-87524405-DEL-162
chr14-87556403-DEL-157
chr14-87565878-DEL-50
chr14-87692516-DEL-56
chr14-87700262-DEL-216
chr14-87700617-DEL-130
chr14-87715663-DEL-90
chr14-87715669-DEL-333
chr14-87715818-DEL-55
chr14-87834587-DEL-50
chr14-87890947-DEL-140
chr14-87891351-DEL-323
chr14-87891377-DEL-157
chr14-88087078-DEL-50
chr14-88218665-DEL-50

chr14-88381663-DEL-50
chr14-88387182-DEL-51
chr14-88401172-DEL-4805
chr14-88406733-DEL-6183
chr14-88509218-DEL-137
chr14-88667588-DEL-367
chr14-88683554-DEL-110
chr14-88774359-DEL-62
chr14-88785953-DEL-920
chr14-88810920-DEL-5068
chr14-89062023-DEL-52
chr14-89097985-DEL-145
chr14-89131883-DEL-257
chr14-89215026-DEL-53
chr14-89278796-DEL-7515
chr14-89443012-DEL-54
chr14-89511121-DEL-84
chr14-89551998-DEL-80
chr14-89917311-DEL-779
chr14-89923115-DEL-412
chr14-89923492-DEL-126
chr14-89947076-DEL-241
chr14-90059108-DEL-50
chr14-90346883-DEL-50
chr14-90467928-DEL-52
chr14-90569868-DEL-291
chr14-90611851-DEL-148
chr14-90696139-DEL-141
chr14-90737379-DEL-437
chr14-90737764-DEL-52
chr14-90850834-DEL-60
chr14-90949918-DEL-65
chr14-91237207-DEL-50
chr14-91280408-DEL-708
chr14-91294247-DEL-913
chr14-91326143-DEL-50
chr14-91565273-DEL-58
chr14-91652294-DEL-50
chr14-91765061-DEL-50
chr14-91935560-DEL-1085
chr14-91992161-DEL-6221
chr14-92019055-DEL-279
chr14-92044040-DEL-402
chr14-92069495-DEL-206
chr14-92190248-DEL-466
chr14-92312323-DEL-67
chr14-92327003-DEL-601
chr14-92628848-DEL-171
chr14-92664889-DEL-5491
chr14-92773249-DEL-7589
chr14-92876626-DEL-100
chr14-93058436-DEL-50
chr14-93359214-DEL-58
chr14-93410242-DEL-118
chr14-93529190-DEL-60
chr14-93592542-DEL-6817
chr14-93647518-DEL-50
chr14-93686553-DEL-242
chr14-93950302-DEL-165
chr14-94193647-DEL-51
chr14-94194490-DEL-68
chr14-94271959-DEL-77
chr14-94287346-DEL-259
chr14-94409746-DEL-2435
chr14-94587069-DEL-216
chr14-94740577-DEL-52
chr14-94761296-DEL-51
chr14-94818926-DEL-50
chr14-94914318-DEL-164
chr14-94924627-DEL-52
chr14-94924770-DEL-236
chr14-94941365-DEL-50
chr14-95066924-DEL-115
chr14-95245594-DEL-340
chr14-95245832-DEL-1821
chr14-95246314-DEL-327
chr14-95246715-DEL-102

chr14-95247403-DEL-309
chr14-95355764-DEL-142
chr14-95368890-DEL-2076
chr14-95408884-DEL-56
chr14-95493131-DEL-104
chr14-95495384-DEL-498
chr14-95513707-DEL-50
chr14-95523298-DEL-457
chr14-95542487-DEL-78
chr14-95600221-DEL-5994
chr14-95721930-DEL-181
chr14-95822495-DEL-71
chr14-95970892-DEL-694
chr14-96198355-DEL-50
chr14-96244385-DEL-98
chr14-96321061-DEL-125
chr14-96321384-DEL-66
chr14-96460688-DEL-4171
chr14-96472711-DEL-51
chr14-96553063-DEL-50
chr14-96588886-DEL-107
chr14-96589121-DEL-82
chr14-96616907-DEL-50
chr14-96737129-DEL-50
chr14-96742435-DEL-52
chr14-96825912-DEL-50
chr14-96870732-DEL-82
chr14-96870846-DEL-440
chr14-96882797-DEL-2150
chr14-96923085-DEL-50
chr14-97100163-DEL-128
chr14-97123499-DEL-50
chr14-97197901-DEL-1234
chr14-97444527-DEL-52
chr14-97444901-DEL-124
chr14-97736238-DEL-160
chr14-97811280-DEL-347
chr14-97855879-DEL-76
chr14-98028982-DEL-50
chr14-98048209-DEL-54
chr14-98562351-DEL-76
chr14-98916242-DEL-70
chr14-98990451-DEL-63
chr14-99008378-DEL-92
chr14-99307927-DEL-154
chr14-99308084-DEL-326
chr14-99423550-DEL-238
chr14-99509372-DEL-50
chr14-99523498-DEL-60
chr14-99525686-DEL-91
chr14-99590007-DEL-50
chr14-99702124-DEL-350
chr14-99728626-DEL-66
chr14-99820980-DEL-307
chr14-99821172-DEL-118
chr14-99935441-DEL-72
chr14-100031690-DEL-71
chr14-100171354-DEL-109
chr14-100239323-DEL-58
chr14-100282437-DEL-50
chr14-100360431-DEL-313
chr14-100400875-DEL-1182
chr14-100503617-DEL-84
chr14-100503847-DEL-78
chr14-100535882-DEL-91
chr14-100543668-DEL-580
chr14-100643315-DEL-50
chr14-100706425-DEL-77
chr14-100806978-DEL-218
chr14-100918507-DEL-1091
chr14-100959293-DEL-160
chr14-100974838-DEL-208
chr14-100986310-DEL-184
chr14-101042423-DEL-2211
chr14-101148535-DEL-60
chr14-101262680-DEL-50
chr14-101452421-DEL-72

chr14-101478466-DEL-534
chr14-101576730-DEL-2151
chr14-101670671-DEL-1779
chr14-101758174-DEL-58
chr14-101925524-DEL-120
chr14-101941206-DEL-543
chr14-102327393-DEL-50
chr14-102390617-DEL-942
chr14-102438744-DEL-2160
chr14-102459165-DEL-2127
chr14-102965329-DEL-66
chr14-103106757-DEL-3792
chr14-103290503-DEL-189
chr14-103300132-DEL-198
chr14-103452000-DEL-124
chr14-103696349-DEL-110
chr14-103734161-DEL-50
chr14-103801595-DEL-150
chr14-103857599-DEL-586
chr14-103871556-DEL-87
chr14-103914656-DEL-692
chr14-103945322-DEL-74
chr14-104006397-DEL-56
chr14-104221143-DEL-75
chr14-104309035-DEL-212
chr14-104709388-DEL-50
chr14-104843933-DEL-97
chr14-104920548-DEL-72
chr14-104941302-DEL-428
chr14-105089623-DEL-57
chr14-105277161-DEL-796
chr14-105279458-DEL-50
chr14-105303568-DEL-572
chr14-105303695-DEL-224
chr14-105526382-DEL-1051
chr14-105526384-DEL-107
chr14-105527007-DEL-211
chr14-105536254-DEL-57
chr14-105593379-DEL-58
chr14-105714260-DEL-69
chr14-106031761-DEL-123
chr14-106067406-DEL-169
chr14-106349482-DEL-5677
chr14-106401209-DEL-58
chr14-106401441-DEL-54
chr14-106417906-DEL-72
chr14-106500472-DEL-129
chr14-106587808-DEL-2436
chr14-106642487-DEL-1354
chr14-106685149-DEL-21218
chr14-106734757-DEL-84
chr14-107167022-DEL-448
chr14-107464482-DEL-55
chr14-107477305-DEL-640
chr14-107481872-DEL-224
chr14-107611295-DEL-185
chr14-107646902-DEL-50
chr14-107647125-DEL-120
chr14-107673227-DEL-149
chr14-107796540-DEL-71
chr14-108095932-DEL-51
chr14-108571124-DEL-80
chr14-108646324-DEL-187
chr14-108671735-DEL-625
chr14-108885029-DEL-2609
chr14-108891766-DEL-1276
chr14-109017511-DEL-84
chr14-109098017-DEL-66
chr14-109336498-DEL-672
chr14-109336654-DEL-190
chr14-109381458-DEL-52
chr14-109559150-DEL-149
chr14-109581839-DEL-1697
chr14-109612169-DEL-76
chr14-109624049-DEL-100
chr14-109796407-DEL-7015
chr14-109827448-DEL-556

chr14-109851193-DEL-390
chr14-109925280-DEL-50
chr14-109997070-DEL-462
chr14-110116843-DEL-50
chr14-110313367-DEL-125
chr14-110426702-DEL-820
chr14-110502412-DEL-170
chr14-110513174-DEL-548
chr14-110531727-DEL-52
chr14-110557438-DEL-90
chr14-110810223-DEL-565
chr14-110863242-DEL-52
chr14-110888622-DEL-68
chr14-110921755-DEL-60
chr14-111132950-DEL-304
chr14-111134169-DEL-71
chr14-111233078-DEL-76
chr14-111269489-DEL-828
chr14-111504281-DEL-70
chr14-111582440-DEL-64
chr14-111646323-DEL-50
chr14-111649399-DEL-183
chr14-111649570-DEL-70
chr14-111826910-DEL-55
chr14-111830774-DEL-75
chr14-111981453-DEL-50
chr14-112066055-DEL-125
chr14-112180362-DEL-50
chr14-112207417-DEL-66
chr14-112211519-DEL-105
chr14-112250529-DEL-124
chr14-112296964-DEL-1095
chr14-112427670-DEL-59
chr14-112440935-DEL-67
chr14-112626911-DEL-113
chr14-112674000-DEL-1694
chr14-112679487-DEL-52
chr14-112704791-DEL-53
chr14-112828744-DEL-1572
chr14-113094418-DEL-89
chr14-113101623-DEL-53
chr14-113466673-DEL-559
chr14-113499176-DEL-158
chr14-113515976-DEL-183
chr14-113610388-DEL-194
chr14-113624202-DEL-2183
chr14-113662481-DEL-144
chr14-113699708-DEL-1081
chr14-113740875-DEL-174
chr14-113742004-DEL-351
chr14-113787745-DEL-52
chr14-113819672-DEL-4830
chr14-113915973-DEL-2540
chr14-114206696-DEL-580
chr14-114225861-DEL-64
chr14-114227957-DEL-2394
chr14-114374288-DEL-104
chr14-114375712-DEL-50
chr14-114425317-DEL-3156
chr14-114530771-DEL-108
chr14-114537533-DEL-88
chr14-114673054-DEL-100
chr14-114714216-DEL-3629
chr14-114727664-DEL-245
chr14-114908111-DEL-1423
chr14-115027054-DEL-51
chr14-115064394-DEL-1355
chr14-115096281-DEL-377
chr14-115304661-DEL-60
chr14-115354033-DEL-208
chr14-115491213-DEL-50
chr14-115586411-DEL-386
chr14-115642810-DEL-120
chr14-115728687-DEL-62
chr14-116156920-DEL-213
chr14-116234258-DEL-186
chr14-116261461-DEL-3504

chr14-116378267-DEL-50
chr14-116470473-DEL-412
chr14-116531215-DEL-189
chr14-116738319-DEL-94
chr14-116742646-DEL-106
chr14-116840629-DEL-125
chr14-116976233-DEL-164
chr14-116976343-DEL-54
chr14-116982690-DEL-132
chr14-117022081-DEL-50
chr14-117034172-DEL-100
chr14-117049779-DEL-97
chr14-117207969-DEL-58
chr14-117397018-DEL-384
chr14-117480549-DEL-279
chr14-117480863-DEL-306
chr14-117506681-DEL-64
chr14-117520319-DEL-349
chr14-117587772-DEL-189
chr14-117613064-DEL-206
chr14-117613358-DEL-67
chr14-117623632-DEL-521
chr14-117664779-DEL-655
chr14-117678795-DEL-70
chr14-117692454-DEL-250
chr14-117705993-DEL-1113
chr14-117751985-DEL-152
chr14-117842539-DEL-268
chr14-117866332-DEL-87
chr14-117957974-DEL-353
chr14-117958223-DEL-58
chr14-118044911-DEL-68
chr14-118069664-DEL-56
chr14-118169842-DEL-93
chr14-118233710-DEL-405
chr14-118233991-DEL-148
chr14-118234306-DEL-69
chr14-118236772-DEL-288
chr14-118237143-DEL-399
chr14-118237428-DEL-178
chr14-118297614-DEL-56
chr14-118313339-DEL-720
chr14-118337226-DEL-56
chr14-118343466-DEL-50
chr14-118393232-DEL-338
chr14-118401188-DEL-50
chr14-118607458-DEL-50
chr14-118707719-DEL-110
chr14-118730822-DEL-51
chr14-118767618-DEL-169
chr14-118784613-DEL-482
chr14-118785607-DEL-173
chr14-118809166-DEL-76
chr14-118878179-DEL-56
chr14-118878297-DEL-140
chr14-118878402-DEL-56
chr14-118906904-DEL-55
chr14-118958228-DEL-56
chr14-119040649-DEL-62
chr14-119049029-DEL-478
chr14-119050032-DEL-201
chr14-119055255-DEL-392
chr14-119055556-DEL-56
chr14-119065538-DEL-824
chr14-119100040-DEL-78
chr14-119191053-DEL-122
chr14-119219792-DEL-60
chr14-119219881-DEL-63
chr14-119220124-DEL-50
chr14-119265418-DEL-63
chr14-119271107-DEL-562
chr14-119271170-DEL-185
chr14-119271257-DEL-89
chr14-119271445-DEL-228
chr14-119271776-DEL-185
chr14-119272764-DEL-230
chr14-119282713-DEL-422

chr14-119318304-DEL-175
chr14-119323788-DEL-73
chr14-119360732-DEL-286
chr14-119367328-DEL-138
chr14-119382409-DEL-52
chr14-119384576-DEL-481
chr14-119394111-DEL-68
chr14-119419441-DEL-72
chr14-119441839-DEL-60
chr14-119461893-DEL-202
chr14-119493197-DEL-782
chr14-119551330-DEL-117
chr14-119596638-DEL-98
chr14-119644325-DEL-50
chr14-119669022-DEL-244
chr14-119737248-DEL-132
chr14-119737340-DEL-62
chr14-119738317-DEL-70
chr14-119746786-DEL-1131
chr14-119761354-DEL-247
chr14-119764014-DEL-119
chr14-119827363-DEL-160
chr14-119847473-DEL-51
chr14-119847853-DEL-1392
chr14-119911595-DEL-54
chr14-119948732-DEL-90
chr14-120011586-DEL-52
chr14-120064932-DEL-337
chr14-120143311-DEL-52
chr14-120156225-DEL-1056
chr14-120211605-DEL-308
chr14-120245814-DEL-181
chr14-120248969-DEL-216
chr14-120269808-DEL-309
chr14-120288143-DEL-107
chr14-120303601-DEL-453
chr14-120416825-DEL-400
chr14-120541415-DEL-55
chr14-120587708-DEL-227
chr14-120677459-DEL-59
chr14-120677488-DEL-224
chr14-120700010-DEL-92
chr14-120740067-DEL-453
chr14-120797972-DEL-50
chr14-120803085-DEL-63
chr14-120808248-DEL-158
chr14-120911822-DEL-92
chr14-120913866-DEL-324
chr14-120920549-DEL-166
chr14-121038241-DEL-1893
chr14-121041607-DEL-3648
chr14-121066257-DEL-284
chr14-121066329-DEL-820
chr14-121076172-DEL-140
chr14-121101495-DEL-202
chr14-121193378-DEL-1407
chr14-121197056-DEL-225
chr14-121225752-DEL-63
chr14-121240766-DEL-90
chr14-121253269-DEL-50
chr14-121256490-DEL-80
chr14-121269946-DEL-63
chr14-121294234-DEL-62
chr14-121322173-DEL-245
chr14-121322220-DEL-98
chr14-121427753-DEL-267
chr14-121506588-DEL-78
chr14-121511786-DEL-282
chr14-121702928-DEL-226
chr14-121720701-DEL-68
chr14-121720701-DEL-465
chr14-121721065-DEL-104
chr14-121721112-DEL-397
chr14-121783037-DEL-271
chr14-121807250-DEL-51
chr14-121972190-DEL-61
chr14-121972381-DEL-202

chr14-122010517-DEL-232
chr14-122023111-DEL-1342
chr14-122080036-DEL-167
chr14-122134878-DEL-122
chr14-122205689-DEL-80
chr14-122288992-DEL-171
chr14-122404726-DEL-935
chr14-122433000-DEL-114
chr14-122475499-DEL-210
chr14-122475793-DEL-246
chr14-122475919-DEL-120
chr14-122597947-DEL-122
chr14-122832687-DEL-378
chr14-122842131-DEL-60
chr14-122901865-DEL-185
chr14-122902838-DEL-60
chr14-122939916-DEL-64
chr14-123339221-DEL-208
chr14-123556622-DEL-1066
chr14-123628566-DEL-174
chr14-123702713-DEL-688
chr14-123792959-DEL-70
chr14-123871380-DEL-92
chr14-123991310-DEL-52
chr14-124030347-DEL-208
chr14-124074681-DEL-58
chr14-124262592-DEL-129
chr14-124301794-DEL-1175
chr14-124317115-DEL-1602
chr14-124359270-DEL-50
chr14-124425158-DEL-783
chr14-124604692-DEL-106
chr14-124854433-DEL-1440
chr15-3248395-DEL-390
chr15-3442271-DEL-50
chr15-3442626-DEL-60
chr15-3571412-DEL-65
chr15-3612302-DEL-50
chr15-3974371-DEL-1750
chr15-4142438-DEL-230
chr15-4232139-DEL-50
chr15-4274761-DEL-63
chr15-4340260-DEL-205
chr15-4340328-DEL-78
chr15-4476194-DEL-51
chr15-4493072-DEL-150
chr15-4823650-DEL-108
chr15-4860601-DEL-76
chr15-4894692-DEL-199
chr15-4902448-DEL-237
chr15-4905669-DEL-52
chr15-5120343-DEL-55
chr15-5162515-DEL-81
chr15-5190346-DEL-747
chr15-5191402-DEL-58
chr15-5311928-DEL-59
chr15-5360289-DEL-423
chr15-5605888-DEL-60
chr15-5633838-DEL-63
chr15-5673542-DEL-68
chr15-5673686-DEL-172
chr15-5673714-DEL-408
chr15-5776060-DEL-68
chr15-6013749-DEL-1215
chr15-6033819-DEL-670
chr15-6083588-DEL-879
chr15-6330269-DEL-141
chr15-6381269-DEL-234
chr15-6381479-DEL-235
chr15-6426766-DEL-52
chr15-6545433-DEL-196
chr15-6548442-DEL-152
chr15-6557157-DEL-1272
chr15-6558722-DEL-271
chr15-6559189-DEL-769
chr15-6577710-DEL-87
chr15-6608398-DEL-54

chr15-6615952-DEL-91
chr15-6626977-DEL-120
chr15-6637757-DEL-264
chr15-6640489-DEL-435
chr15-6902361-DEL-238
chr15-6929085-DEL-58
chr15-6936099-DEL-52
chr15-6987426-DEL-193
chr15-7022559-DEL-50
chr15-7182340-DEL-139
chr15-7336292-DEL-136
chr15-7453088-DEL-205
chr15-7453255-DEL-68
chr15-7599111-DEL-1168
chr15-7602998-DEL-56
chr15-7809032-DEL-265
chr15-7817367-DEL-220
chr15-7927348-DEL-260
chr15-7927376-DEL-691
chr15-7932597-DEL-330
chr15-7947993-DEL-161
chr15-8021378-DEL-128
chr15-8021798-DEL-64
chr15-8028393-DEL-85
chr15-8096859-DEL-520
chr15-8116388-DEL-105
chr15-8149686-DEL-159
chr15-8257471-DEL-92
chr15-8510036-DEL-160
chr15-8523901-DEL-101
chr15-8547261-DEL-2664
chr15-8548358-DEL-190
chr15-8550749-DEL-219
chr15-8551125-DEL-52
chr15-8561640-DEL-62
chr15-8561745-DEL-66
chr15-8586512-DEL-551
chr15-8597100-DEL-72
chr15-8610410-DEL-141
chr15-8630233-DEL-66
chr15-8677703-DEL-62
chr15-8708371-DEL-104
chr15-8708572-DEL-180
chr15-8742754-DEL-64
chr15-8761078-DEL-246
chr15-8780856-DEL-54
chr15-8834751-DEL-206
chr15-8843855-DEL-208
chr15-8876354-DEL-94
chr15-8928098-DEL-314
chr15-8932128-DEL-53
chr15-9019009-DEL-51
chr15-9035724-DEL-224
chr15-9035826-DEL-625
chr15-9045868-DEL-2891
chr15-9047939-DEL-2554
chr15-9247359-DEL-631
chr15-9314188-DEL-7211
chr15-9422581-DEL-84
chr15-9824106-DEL-52
chr15-9886128-DEL-56
chr15-9888102-DEL-557
chr15-9992426-DEL-66
chr15-10228063-DEL-52
chr15-10466033-DEL-264
chr15-10533621-DEL-70
chr15-10552610-DEL-50
chr15-10579943-DEL-252
chr15-10590348-DEL-903
chr15-10614698-DEL-74
chr15-10658086-DEL-84
chr15-10998888-DEL-66
chr15-11004368-DEL-418
chr15-11060573-DEL-128
chr15-11063655-DEL-78
chr15-11095139-DEL-69
chr15-11189416-DEL-159

chr15-11212463-DEL-81
chr15-11329986-DEL-459
chr15-11330062-DEL-204
chr15-11480013-DEL-238
chr15-11480626-DEL-81
chr15-11535916-DEL-145
chr15-11611147-DEL-149
chr15-11892645-DEL-144
chr15-12033686-DEL-537
chr15-12044118-DEL-3496
chr15-12065713-DEL-70
chr15-12081758-DEL-231
chr15-12095946-DEL-66
chr15-12098469-DEL-402
chr15-12099030-DEL-94
chr15-12099463-DEL-312
chr15-12105709-DEL-53
chr15-12200747-DEL-80
chr15-12249899-DEL-111
chr15-12256613-DEL-205
chr15-12570078-DEL-61
chr15-12570256-DEL-65
chr15-12572873-DEL-63
chr15-12607711-DEL-352
chr15-12607775-DEL-96
chr15-12632156-DEL-134
chr15-12646902-DEL-164
chr15-12741453-DEL-284
chr15-12784192-DEL-61
chr15-12795423-DEL-169
chr15-12872898-DEL-111
chr15-12873289-DEL-74
chr15-13003113-DEL-458
chr15-13003184-DEL-952
chr15-13046946-DEL-124
chr15-13046946-DEL-310
chr15-13047383-DEL-62
chr15-13100323-DEL-185
chr15-13226564-DEL-228
chr15-13237696-DEL-156
chr15-13383505-DEL-66
chr15-13542521-DEL-60
chr15-13586304-DEL-51
chr15-13704399-DEL-52
chr15-13770588-DEL-284
chr15-13778020-DEL-50
chr15-13798227-DEL-1149
chr15-13798988-DEL-429
chr15-13841462-DEL-901
chr15-14176600-DEL-174
chr15-14234030-DEL-1185
chr15-14308549-DEL-114
chr15-14394353-DEL-65
chr15-14480556-DEL-4099
chr15-14524326-DEL-52
chr15-14524753-DEL-251
chr15-14985269-DEL-154
chr15-14985706-DEL-101
chr15-15002278-DEL-58
chr15-15051494-DEL-8196
chr15-15479018-DEL-534
chr15-15541643-DEL-83
chr15-15712141-DEL-109
chr15-15776784-DEL-58
chr15-15828886-DEL-52
chr15-15947305-DEL-76
chr15-16068777-DEL-531
chr15-16082226-DEL-2709
chr15-16088686-DEL-197
chr15-16124762-DEL-4512
chr15-16166846-DEL-78
chr15-16321778-DEL-51
chr15-16815127-DEL-88
chr15-16939792-DEL-1268
chr15-17253128-DEL-227
chr15-17267491-DEL-212
chr15-17276526-DEL-1485

chr15-17284601-DEL-50
chr15-17599883-DEL-62
chr15-17661501-DEL-64
chr15-17701170-DEL-528
chr15-17702167-DEL-84
chr15-17722032-DEL-499
chr15-17771974-DEL-336
chr15-17795085-DEL-184
chr15-17819661-DEL-140
chr15-17858861-DEL-77
chr15-17877801-DEL-1145
chr15-17988832-DEL-559
chr15-18030662-DEL-1376
chr15-18058710-DEL-50
chr15-18174986-DEL-571
chr15-18177080-DEL-1312
chr15-18310055-DEL-2103
chr15-18312347-DEL-305
chr15-18359634-DEL-417
chr15-18385657-DEL-355
chr15-18411308-DEL-52
chr15-18411467-DEL-103
chr15-18441878-DEL-104
chr15-18450556-DEL-351
chr15-18537358-DEL-51
chr15-18551198-DEL-50
chr15-18640931-DEL-112
chr15-18861345-DEL-586
chr15-18875997-DEL-809
chr15-18954753-DEL-50
chr15-18982255-DEL-293
chr15-18999010-DEL-60
chr15-18999178-DEL-160
chr15-19020254-DEL-192
chr15-19135155-DEL-1947
chr15-19221682-DEL-58
chr15-19234188-DEL-103
chr15-19294784-DEL-92
chr15-19338375-DEL-62
chr15-19355169-DEL-62
chr15-19376030-DEL-80
chr15-19378166-DEL-783
chr15-19386297-DEL-90
chr15-19448051-DEL-6687
chr15-19483989-DEL-889
chr15-19710029-DEL-68
chr15-19718325-DEL-188
chr15-19774391-DEL-144
chr15-19801159-DEL-96
chr15-19951281-DEL-3627
chr15-20370613-DEL-2575
chr15-20377581-DEL-50
chr15-20468980-DEL-3825
chr15-20496999-DEL-119
chr15-20669909-DEL-191
chr15-20909675-DEL-285
chr15-20913436-DEL-533
chr15-20915122-DEL-5958
chr15-20966820-DEL-72
chr15-21028564-DEL-203
chr15-21035831-DEL-199
chr15-21101990-DEL-264
chr15-21454612-DEL-81
chr15-21861864-DEL-707
chr15-21864576-DEL-12153
chr15-21875976-DEL-212
chr15-21876335-DEL-212
chr15-21899411-DEL-51
chr15-21918122-DEL-6909
chr15-21992081-DEL-54
chr15-22211515-DEL-1012
chr15-22246059-DEL-68
chr15-22410872-DEL-129
chr15-22411149-DEL-65
chr15-22411161-DEL-131
chr15-22629301-DEL-68
chr15-22660731-DEL-1283

chr15-22799428-DEL-66
chr15-22799718-DEL-770
chr15-22930845-DEL-570
chr15-22947942-DEL-184
chr15-23114792-DEL-104
chr15-23263659-DEL-289
chr15-23297357-DEL-212
chr15-23343056-DEL-5312
chr15-23384408-DEL-245
chr15-23461045-DEL-428
chr15-23461611-DEL-90
chr15-23462409-DEL-6293
chr15-23651278-DEL-62
chr15-23695033-DEL-58
chr15-23719109-DEL-3416
chr15-23741996-DEL-3491
chr15-23776492-DEL-204
chr15-23861133-DEL-57
chr15-23913284-DEL-830
chr15-24269124-DEL-60
chr15-24289333-DEL-734
chr15-24356890-DEL-69
chr15-24470131-DEL-58
chr15-24582667-DEL-329
chr15-24583306-DEL-556
chr15-24889592-DEL-125
chr15-24945655-DEL-399
chr15-24949333-DEL-283
chr15-25127863-DEL-77
chr15-25199790-DEL-66
chr15-25255613-DEL-74
chr15-25506402-DEL-53
chr15-25551319-DEL-81
chr15-25553431-DEL-151
chr15-25553691-DEL-101
chr15-25560344-DEL-71
chr15-25570301-DEL-60
chr15-25591352-DEL-66
chr15-25651813-DEL-1023
chr15-25873987-DEL-588
chr15-25954356-DEL-80
chr15-25959022-DEL-117
chr15-26029425-DEL-124
chr15-26050975-DEL-50
chr15-26111358-DEL-109
chr15-26163520-DEL-60
chr15-26219397-DEL-71
chr15-26219601-DEL-55
chr15-26257064-DEL-969
chr15-26258109-DEL-56
chr15-26282546-DEL-1561
chr15-26304138-DEL-89
chr15-26397683-DEL-458
chr15-26428244-DEL-137
chr15-26441233-DEL-593
chr15-26554148-DEL-753
chr15-26623362-DEL-92
chr15-26665145-DEL-157
chr15-26672699-DEL-280
chr15-26681316-DEL-212
chr15-26709484-DEL-240
chr15-26918190-DEL-686
chr15-26918571-DEL-305
chr15-26941462-DEL-56
chr15-27112676-DEL-59
chr15-27115311-DEL-63
chr15-27119082-DEL-4226
chr15-27206187-DEL-66
chr15-27214447-DEL-76
chr15-27269608-DEL-59
chr15-27304602-DEL-63
chr15-27347215-DEL-393
chr15-27347420-DEL-81
chr15-27397262-DEL-205
chr15-27459802-DEL-338
chr15-27599881-DEL-2519
chr15-27663157-DEL-72

chr15-27682386-DEL-118
chr15-27687825-DEL-76
chr15-27753623-DEL-84
chr15-27854013-DEL-50
chr15-27914383-DEL-190
chr15-27935603-DEL-84
chr15-27956908-DEL-55
chr15-27969591-DEL-66
chr15-27971153-DEL-70
chr15-27971286-DEL-315
chr15-27971318-DEL-1288
chr15-27971716-DEL-413
chr15-27972290-DEL-302
chr15-27972734-DEL-51
chr15-28010820-DEL-76
chr15-28078143-DEL-664
chr15-28083158-DEL-91
chr15-28117453-DEL-1194
chr15-28118750-DEL-606
chr15-28134688-DEL-80
chr15-28134713-DEL-165
chr15-28139628-DEL-63
chr15-28190609-DEL-57
chr15-28205673-DEL-630
chr15-28317117-DEL-51
chr15-28565605-DEL-12668
chr15-28614264-DEL-78
chr15-28767703-DEL-83
chr15-28909818-DEL-210
chr15-28941015-DEL-2170
chr15-28959162-DEL-61
chr15-28966808-DEL-281
chr15-29021311-DEL-57
chr15-29058448-DEL-133
chr15-29150116-DEL-2847
chr15-29162266-DEL-265
chr15-29257764-DEL-269
chr15-29284548-DEL-7118
chr15-29331360-DEL-491
chr15-29353184-DEL-194
chr15-29569155-DEL-3297
chr15-29605824-DEL-63
chr15-29626307-DEL-596
chr15-29629305-DEL-76
chr15-29648330-DEL-6250
chr15-29656429-DEL-204
chr15-29663740-DEL-277
chr15-29819747-DEL-95
chr15-29897020-DEL-72
chr15-29957246-DEL-77
chr15-29969930-DEL-2078
chr15-30041165-DEL-290
chr15-30114215-DEL-118
chr15-30115136-DEL-552
chr15-30433048-DEL-53
chr15-30539870-DEL-123
chr15-30540537-DEL-50
chr15-30547091-DEL-52
chr15-30730487-DEL-74
chr15-30898878-DEL-163
chr15-30903995-DEL-50
chr15-31041939-DEL-85
chr15-31110576-DEL-50
chr15-31175903-DEL-80
chr15-31175972-DEL-558
chr15-31229988-DEL-367
chr15-31249691-DEL-204
chr15-31249945-DEL-93
chr15-31337995-DEL-50
chr15-31516648-DEL-180
chr15-31652252-DEL-161
chr15-31652252-DEL-529
chr15-31690275-DEL-70
chr15-31696905-DEL-6611
chr15-31714389-DEL-312
chr15-31714485-DEL-102
chr15-31736571-DEL-63

chr15-31834791-DEL-445
chr15-31906208-DEL-60
chr15-32077199-DEL-142
chr15-32228975-DEL-96
chr15-32229074-DEL-297
chr15-32229259-DEL-94
chr15-32287020-DEL-492
chr15-32296313-DEL-58
chr15-32357386-DEL-719
chr15-32454158-DEL-54
chr15-32709977-DEL-108
chr15-32879973-DEL-206
chr15-33094730-DEL-2588
chr15-33216936-DEL-3742
chr15-33238072-DEL-50
chr15-33307991-DEL-52
chr15-33532887-DEL-50
chr15-33541725-DEL-55
chr15-33704179-DEL-2119
chr15-33762526-DEL-51
chr15-33814258-DEL-3011
chr15-33865870-DEL-87
chr15-33905549-DEL-116
chr15-33995707-DEL-183
chr15-34020519-DEL-417
chr15-34051370-DEL-158
chr15-34069176-DEL-80
chr15-34073530-DEL-2470
chr15-34118501-DEL-50
chr15-34147868-DEL-104
chr15-34149507-DEL-672
chr15-34213169-DEL-61
chr15-34241654-DEL-53
chr15-34374864-DEL-57
chr15-34445371-DEL-1038
chr15-34455217-DEL-113
chr15-34505609-DEL-127
chr15-34630441-DEL-240
chr15-34862504-DEL-52
chr15-34900622-DEL-50
chr15-35345933-DEL-1919
chr15-35614862-DEL-144
chr15-35685438-DEL-572
chr15-35686009-DEL-619
chr15-35758776-DEL-96
chr15-35908605-DEL-598
chr15-35908891-DEL-936
chr15-35909733-DEL-162
chr15-36001993-DEL-104
chr15-36002399-DEL-84
chr15-36028047-DEL-232
chr15-36124120-DEL-114
chr15-36202439-DEL-588
chr15-36202496-DEL-251
chr15-36203275-DEL-126
chr15-36214581-DEL-78
chr15-36214593-DEL-252
chr15-36324201-DEL-95
chr15-36464656-DEL-1215
chr15-36472500-DEL-210
chr15-36522416-DEL-51
chr15-36531421-DEL-195
chr15-36531846-DEL-66
chr15-36531868-DEL-135
chr15-36553015-DEL-50
chr15-36575349-DEL-85
chr15-36575356-DEL-1248
chr15-36603649-DEL-107
chr15-36642412-DEL-77
chr15-36643700-DEL-165
chr15-36663826-DEL-65
chr15-36749276-DEL-67
chr15-36866893-DEL-298
chr15-36882823-DEL-192
chr15-36909012-DEL-88
chr15-36909012-DEL-286
chr15-36929803-DEL-91

chr15-36930921-DEL-56
chr15-36976597-DEL-357
chr15-36977265-DEL-121
chr15-37005853-DEL-123
chr15-37021062-DEL-67
chr15-37021631-DEL-206
chr15-37021896-DEL-257
chr15-37059380-DEL-164
chr15-37108105-DEL-547
chr15-37124922-DEL-220
chr15-37125515-DEL-64
chr15-37156126-DEL-104
chr15-37173100-DEL-215
chr15-37175530-DEL-1140
chr15-37175624-DEL-134
chr15-37175923-DEL-68
chr15-37176153-DEL-202
chr15-37176470-DEL-235
chr15-37203524-DEL-200
chr15-37226415-DEL-72
chr15-37272667-DEL-104
chr15-37518300-DEL-75
chr15-37541865-DEL-61
chr15-37541921-DEL-219
chr15-37541922-DEL-945
chr15-37542609-DEL-192
chr15-37668736-DEL-105
chr15-37764634-DEL-105
chr15-37950889-DEL-86
chr15-37988032-DEL-239
chr15-38120374-DEL-56
chr15-38186616-DEL-92
chr15-38206868-DEL-102
chr15-38251329-DEL-80
chr15-38273255-DEL-2958
chr15-38305533-DEL-604
chr15-38305585-DEL-107
chr15-38308984-DEL-54
chr15-38315855-DEL-217
chr15-38522362-DEL-198
chr15-38541598-DEL-339
chr15-38695351-DEL-82
chr15-38701321-DEL-246
chr15-38734751-DEL-98
chr15-38802764-DEL-219
chr15-38808399-DEL-439
chr15-38838045-DEL-265
chr15-39057357-DEL-68
chr15-39102650-DEL-62
chr15-39102763-DEL-216
chr15-39205136-DEL-151
chr15-39285595-DEL-93
chr15-39337666-DEL-55
chr15-39387221-DEL-54
chr15-39506035-DEL-61
chr15-39605291-DEL-72
chr15-39613154-DEL-161
chr15-39613410-DEL-92
chr15-39692445-DEL-166
chr15-39727384-DEL-84
chr15-39787186-DEL-1436
chr15-39828284-DEL-60
chr15-39860605-DEL-200
chr15-39860674-DEL-74
chr15-39879374-DEL-72
chr15-39889043-DEL-96
chr15-39949491-DEL-50
chr15-40166463-DEL-184
chr15-40212843-DEL-85
chr15-40269162-DEL-78
chr15-40301199-DEL-316
chr15-40301543-DEL-464
chr15-40455033-DEL-181
chr15-40910642-DEL-204
chr15-41021516-DEL-141
chr15-41021594-DEL-69
chr15-41071465-DEL-260

chr15-41107711-DEL-6501
chr15-41109360-DEL-2638
chr15-41114261-DEL-306
chr15-41266831-DEL-50
chr15-41499297-DEL-141
chr15-41671813-DEL-60
chr15-41775749-DEL-117
chr15-41775906-DEL-82
chr15-41781019-DEL-275
chr15-41877432-DEL-506
chr15-41979728-DEL-50
chr15-42016309-DEL-229
chr15-42214136-DEL-209
chr15-42214136-DEL-433
chr15-42214297-DEL-3968
chr15-42352274-DEL-339
chr15-42361397-DEL-240
chr15-42402408-DEL-124
chr15-42437711-DEL-50
chr15-42503982-DEL-52
chr15-42515142-DEL-246
chr15-42822219-DEL-116
chr15-42895950-DEL-62
chr15-42902842-DEL-52
chr15-42932785-DEL-78
chr15-43053085-DEL-584
chr15-43070261-DEL-457
chr15-43365258-DEL-962
chr15-43651187-DEL-2637
chr15-43665504-DEL-129
chr15-43792638-DEL-138
chr15-43800728-DEL-68
chr15-43839891-DEL-50
chr15-43988890-DEL-204
chr15-43989636-DEL-53
chr15-44147464-DEL-186
chr15-44179486-DEL-66
chr15-44303777-DEL-52
chr15-44411853-DEL-2220
chr15-44662376-DEL-74
chr15-44676889-DEL-84
chr15-44725403-DEL-2529
chr15-44832975-DEL-461
chr15-44988511-DEL-60
chr15-45116643-DEL-5098
chr15-45167929-DEL-3032
chr15-45203253-DEL-192
chr15-45379819-DEL-1674
chr15-45470819-DEL-67
chr15-45470989-DEL-229
chr15-45488860-DEL-53
chr15-45648888-DEL-645
chr15-45799278-DEL-312
chr15-46053998-DEL-1588
chr15-46095092-DEL-299
chr15-46112898-DEL-1644
chr15-46189454-DEL-486
chr15-46346720-DEL-50
chr15-46398616-DEL-50
chr15-46410088-DEL-5869
chr15-46569707-DEL-5224
chr15-46609226-DEL-58
chr15-46907121-DEL-303
chr15-46963380-DEL-599
chr15-46995054-DEL-2654
chr15-47004887-DEL-126
chr15-47197926-DEL-1115
chr15-47254203-DEL-263
chr15-47255843-DEL-2724
chr15-47283389-DEL-298
chr15-47339357-DEL-56
chr15-47360597-DEL-261
chr15-47397347-DEL-629
chr15-47465722-DEL-878
chr15-47468261-DEL-160
chr15-47555157-DEL-417
chr15-47570780-DEL-316

chr15-47606689-DEL-141
chr15-47656987-DEL-245
chr15-47688709-DEL-370
chr15-47691076-DEL-58
chr15-47691211-DEL-1140
chr15-47698098-DEL-253
chr15-47735017-DEL-100
chr15-48036737-DEL-7578
chr15-48117926-DEL-498
chr15-48289553-DEL-3366
chr15-48325106-DEL-56
chr15-48397154-DEL-56
chr15-48429725-DEL-50
chr15-48435966-DEL-227
chr15-48445693-DEL-58
chr15-48471216-DEL-66
chr15-48473932-DEL-79
chr15-48494013-DEL-50
chr15-48582458-DEL-513
chr15-48600924-DEL-1837
chr15-48668422-DEL-1211
chr15-48703806-DEL-50
chr15-48708547-DEL-78
chr15-48803988-DEL-183
chr15-48804021-DEL-74
chr15-48804384-DEL-64
chr15-48826764-DEL-239
chr15-48946442-DEL-3843
chr15-49016034-DEL-2374
chr15-49145137-DEL-58
chr15-49168331-DEL-195
chr15-49191582-DEL-66
chr15-49277879-DEL-204
chr15-49285207-DEL-60
chr15-49311903-DEL-79
chr15-49382589-DEL-819
chr15-49493038-DEL-113
chr15-49505509-DEL-247
chr15-49514950-DEL-77
chr15-49520903-DEL-687
chr15-49570174-DEL-610
chr15-49577569-DEL-68
chr15-49624619-DEL-53
chr15-49748003-DEL-50
chr15-49948485-DEL-144
chr15-49960801-DEL-51
chr15-49980349-DEL-56
chr15-50090483-DEL-115
chr15-50107045-DEL-69
chr15-50127327-DEL-50
chr15-50592469-DEL-66
chr15-50720953-DEL-57
chr15-50877464-DEL-79
chr15-50893012-DEL-82
chr15-51061648-DEL-50
chr15-51114129-DEL-94
chr15-51258355-DEL-54
chr15-51317439-DEL-248
chr15-51347113-DEL-138
chr15-51431983-DEL-56
chr15-51467398-DEL-72
chr15-51517826-DEL-52
chr15-51646294-DEL-61
chr15-51710917-DEL-90
chr15-51721043-DEL-80
chr15-51727675-DEL-167
chr15-51772949-DEL-235
chr15-51891037-DEL-214
chr15-51969218-DEL-72
chr15-51982261-DEL-99
chr15-51997554-DEL-116
chr15-52134828-DEL-201
chr15-52155685-DEL-430
chr15-52302790-DEL-730
chr15-52395371-DEL-1795
chr15-52419400-DEL-76
chr15-52421730-DEL-56

chr15-52490782-DEL-167
chr15-52538425-DEL-212
chr15-52776361-DEL-58
chr15-52782270-DEL-206
chr15-52787506-DEL-601
chr15-52874722-DEL-96
chr15-52979604-DEL-83
chr15-53118987-DEL-100
chr15-53169458-DEL-52
chr15-53190204-DEL-64
chr15-53190514-DEL-110
chr15-53248083-DEL-78
chr15-53539028-DEL-90
chr15-53569177-DEL-50
chr15-53670784-DEL-54
chr15-53676702-DEL-107
chr15-53736892-DEL-50
chr15-53790053-DEL-1655
chr15-53840126-DEL-1483
chr15-53917567-DEL-276
chr15-54079630-DEL-1985
chr15-54089537-DEL-86
chr15-54201021-DEL-252
chr15-54316319-DEL-135
chr15-54351694-DEL-260
chr15-54351907-DEL-72
chr15-54383001-DEL-50
chr15-54446236-DEL-132
chr15-54452044-DEL-151
chr15-54778111-DEL-56
chr15-54794004-DEL-132
chr15-54812537-DEL-295
chr15-54884676-DEL-133
chr15-54884698-DEL-343
chr15-54884918-DEL-77
chr15-55022523-DEL-2198
chr15-55092070-DEL-186
chr15-55142298-DEL-627
chr15-55157277-DEL-1120
chr15-55278546-DEL-52
chr15-55463412-DEL-70
chr15-55477123-DEL-266
chr15-55639656-DEL-293
chr15-55722983-DEL-311
chr15-55722985-DEL-104
chr15-55743995-DEL-77
chr15-55818761-DEL-52
chr15-55832589-DEL-54
chr15-55842756-DEL-88
chr15-55846096-DEL-52
chr15-56085528-DEL-74
chr15-56118692-DEL-5843
chr15-56166701-DEL-58
chr15-56319822-DEL-230
chr15-56481944-DEL-58
chr15-56538966-DEL-230
chr15-56549800-DEL-126
chr15-56550033-DEL-66
chr15-56712945-DEL-82
chr15-56730093-DEL-62
chr15-56731975-DEL-780
chr15-56749075-DEL-212
chr15-56982912-DEL-74
chr15-57001658-DEL-64
chr15-57042158-DEL-154
chr15-57119269-DEL-108
chr15-57142702-DEL-214
chr15-57172642-DEL-5289
chr15-57256612-DEL-810
chr15-57342745-DEL-1861
chr15-57352816-DEL-1902
chr15-57508376-DEL-145
chr15-57554991-DEL-50
chr15-57607208-DEL-156
chr15-57647881-DEL-102
chr15-57795217-DEL-628
chr15-57803501-DEL-168

chr15-57820642-DEL-69
chr15-57860721-DEL-1050
chr15-57860963-DEL-160
chr15-57861139-DEL-68
chr15-57861591-DEL-368
chr15-57861887-DEL-112
chr15-57994296-DEL-825
chr15-58003291-DEL-1069
chr15-58102832-DEL-1890
chr15-58111045-DEL-96
chr15-58115148-DEL-50
chr15-58258813-DEL-88
chr15-58258965-DEL-179
chr15-58309729-DEL-1792
chr15-58340739-DEL-86
chr15-58383469-DEL-53
chr15-58386093-DEL-224
chr15-58394254-DEL-258
chr15-58422169-DEL-66
chr15-58434859-DEL-72
chr15-58475072-DEL-90
chr15-58527782-DEL-62
chr15-58632406-DEL-50
chr15-58781502-DEL-3829
chr15-58861730-DEL-50
chr15-59155439-DEL-1049
chr15-59219385-DEL-420
chr15-59234533-DEL-148
chr15-59303481-DEL-2857
chr15-59303867-DEL-908
chr15-59305241-DEL-1481
chr15-59366956-DEL-55
chr15-59419992-DEL-74
chr15-59547698-DEL-51
chr15-59548820-DEL-195
chr15-59684896-DEL-87
chr15-59760577-DEL-50
chr15-59807739-DEL-58
chr15-59940408-DEL-1138
chr15-59988063-DEL-1972
chr15-59997701-DEL-52
chr15-60129314-DEL-72
chr15-60134397-DEL-84
chr15-60368234-DEL-50
chr15-60478809-DEL-100
chr15-60517543-DEL-242
chr15-60552989-DEL-234
chr15-60555261-DEL-842
chr15-60576696-DEL-78
chr15-60582857-DEL-1239
chr15-60743008-DEL-489
chr15-61114565-DEL-50
chr15-61151002-DEL-7499
chr15-61206477-DEL-99
chr15-61257206-DEL-70
chr15-61352566-DEL-54
chr15-61358620-DEL-530
chr15-61365387-DEL-50
chr15-61408502-DEL-107
chr15-61415785-DEL-76
chr15-61461297-DEL-317
chr15-61531997-DEL-110
chr15-61567603-DEL-65
chr15-61633319-DEL-50
chr15-61650551-DEL-50
chr15-61890579-DEL-54
chr15-61899780-DEL-63
chr15-62098135-DEL-2448
chr15-62284340-DEL-76
chr15-62314723-DEL-472
chr15-62314975-DEL-177
chr15-62315726-DEL-50
chr15-62478493-DEL-2374
chr15-62734857-DEL-439
chr15-62735246-DEL-2043
chr15-62916701-DEL-58
chr15-63210298-DEL-61

chr15-63281389-DEL-56
chr15-63281787-DEL-188
chr15-63281849-DEL-71
chr15-63315643-DEL-94
chr15-63385739-DEL-1738
chr15-63444540-DEL-260
chr15-63444833-DEL-222
chr15-63458568-DEL-174
chr15-63469951-DEL-1429
chr15-63544825-DEL-92
chr15-63668699-DEL-1423
chr15-63719163-DEL-67117
chr15-63811960-DEL-126
chr15-63812368-DEL-213
chr15-63964162-DEL-161
chr15-63978469-DEL-98
chr15-64040978-DEL-60
chr15-64129040-DEL-61
chr15-64147457-DEL-122
chr15-64438711-DEL-52
chr15-64534674-DEL-90
chr15-64551448-DEL-3640
chr15-64630522-DEL-612
chr15-64630592-DEL-200
chr15-64702731-DEL-50
chr15-64813122-DEL-117
chr15-64825834-DEL-93
chr15-64933422-DEL-94
chr15-65083156-DEL-52
chr15-65083165-DEL-106
chr15-65146803-DEL-760
chr15-65210407-DEL-73
chr15-65235364-DEL-630
chr15-65277722-DEL-209
chr15-65334766-DEL-267
chr15-65390879-DEL-116
chr15-65400435-DEL-1213
chr15-65455233-DEL-94
chr15-65535032-DEL-132
chr15-65561706-DEL-60
chr15-65896740-DEL-187
chr15-66042617-DEL-78
chr15-66248082-DEL-432
chr15-66318896-DEL-54
chr15-66490216-DEL-3549
chr15-66524500-DEL-209
chr15-66680056-DEL-158
chr15-66992391-DEL-53
chr15-67127166-DEL-84
chr15-67204263-DEL-53
chr15-67388867-DEL-974
chr15-67464162-DEL-292
chr15-67507582-DEL-5751
chr15-67644839-DEL-54
chr15-67839073-DEL-399
chr15-68008373-DEL-859
chr15-68008562-DEL-389
chr15-68028545-DEL-730
chr15-68083992-DEL-87
chr15-68115795-DEL-150
chr15-68419877-DEL-58
chr15-68505312-DEL-409
chr15-68602112-DEL-1093
chr15-68641711-DEL-656
chr15-68643585-DEL-1623
chr15-68693450-DEL-90
chr15-68830388-DEL-60
chr15-69028868-DEL-129
chr15-69034560-DEL-1486
chr15-69054076-DEL-50
chr15-69200313-DEL-179
chr15-69201197-DEL-189
chr15-69279815-DEL-3900
chr15-69310575-DEL-2455
chr15-69339438-DEL-804
chr15-69412952-DEL-564
chr15-69481869-DEL-4204

chr15-69553117-DEL-54
chr15-69559101-DEL-50
chr15-69700498-DEL-325
chr15-69848829-DEL-66
chr15-69954835-DEL-214
chr15-70132626-DEL-85
chr15-70138936-DEL-52
chr15-70226837-DEL-120
chr15-70303702-DEL-269
chr15-70303799-DEL-81
chr15-70304026-DEL-104
chr15-70309624-DEL-77
chr15-70397842-DEL-56
chr15-70415476-DEL-59
chr15-70484644-DEL-164
chr15-70518908-DEL-76
chr15-70519079-DEL-1497
chr15-70524379-DEL-75
chr15-70548342-DEL-315
chr15-70608644-DEL-1348
chr15-70738801-DEL-836
chr15-70743904-DEL-85
chr15-70745019-DEL-78
chr15-70810166-DEL-66
chr15-70819121-DEL-74
chr15-70842712-DEL-56
chr15-70843558-DEL-80
chr15-70986707-DEL-50
chr15-71083034-DEL-75
chr15-71135552-DEL-132
chr15-71221647-DEL-1143
chr15-71253143-DEL-74
chr15-71282384-DEL-3108
chr15-71367779-DEL-54
chr15-71367865-DEL-143
chr15-71419313-DEL-50
chr15-71488334-DEL-63
chr15-71500757-DEL-76
chr15-71587388-DEL-183
chr15-71608805-DEL-470
chr15-71609189-DEL-471
chr15-71634142-DEL-794
chr15-71643879-DEL-15537
chr15-71828213-DEL-80
chr15-71979782-DEL-82
chr15-71992585-DEL-50
chr15-72066558-DEL-50
chr15-72106734-DEL-59
chr15-72120686-DEL-1696
chr15-72120686-DEL-3393
chr15-72394505-DEL-306
chr15-72395299-DEL-101
chr15-72395765-DEL-305
chr15-72457035-DEL-780
chr15-72655233-DEL-55
chr15-72849318-DEL-66
chr15-72868350-DEL-384
chr15-72869099-DEL-768
chr15-72869654-DEL-224
chr15-72869983-DEL-84
chr15-73139671-DEL-100
chr15-73217667-DEL-138
chr15-73290184-DEL-610
chr15-73373226-DEL-179
chr15-73403842-DEL-130
chr15-73646680-DEL-104
chr15-73713020-DEL-102
chr15-73777689-DEL-50
chr15-74046940-DEL-212
chr15-74059435-DEL-59
chr15-74059614-DEL-120
chr15-74072661-DEL-1744
chr15-74159664-DEL-135
chr15-74187509-DEL-62
chr15-74187694-DEL-67
chr15-74211223-DEL-1414
chr15-74232575-DEL-7167

chr15-74424297-DEL-365
chr15-74441908-DEL-50
chr15-74585883-DEL-171
chr15-74825717-DEL-70
chr15-74877882-DEL-75
chr15-74932750-DEL-1426
chr15-75036401-DEL-11182
chr15-75251304-DEL-139
chr15-75267374-DEL-980
chr15-75355785-DEL-48903
chr15-75450653-DEL-488
chr15-75450996-DEL-88
chr15-75461989-DEL-1240
chr15-75753234-DEL-152
chr15-75898189-DEL-642
chr15-75898240-DEL-257
chr15-75898497-DEL-254
chr15-75974423-DEL-7725
chr15-76044068-DEL-51
chr15-76094791-DEL-64
chr15-76457683-DEL-51
chr15-76465357-DEL-124
chr15-76693956-DEL-64
chr15-76697326-DEL-76
chr15-76745845-DEL-180
chr15-76762948-DEL-164
chr15-76790845-DEL-126
chr15-76813813-DEL-1346
chr15-76823921-DEL-104
chr15-76857385-DEL-522
chr15-76877662-DEL-74
chr15-76885535-DEL-892
chr15-76923471-DEL-119
chr15-77030796-DEL-279
chr15-77163584-DEL-158
chr15-77200764-DEL-50
chr15-77212882-DEL-50
chr15-77256265-DEL-76
chr15-77256912-DEL-211
chr15-77269940-DEL-1330
chr15-77274796-DEL-666
chr15-77314846-DEL-54
chr15-77350401-DEL-165
chr15-77382645-DEL-6437
chr15-77556980-DEL-650
chr15-77681006-DEL-92
chr15-77738083-DEL-483
chr15-78055214-DEL-153
chr15-78097820-DEL-52
chr15-78179828-DEL-50
chr15-78358791-DEL-148
chr15-78397982-DEL-51
chr15-78477826-DEL-56
chr15-78534109-DEL-54
chr15-78620456-DEL-362
chr15-78623379-DEL-1586
chr15-78684114-DEL-63
chr15-78818316-DEL-188
chr15-78819378-DEL-153
chr15-78826019-DEL-87
chr15-78987109-DEL-70
chr15-79009917-DEL-125
chr15-79186612-DEL-124
chr15-79202414-DEL-78
chr15-79210309-DEL-50
chr15-79222952-DEL-90
chr15-79300495-DEL-89
chr15-79606231-DEL-372
chr15-79614556-DEL-681
chr15-79614706-DEL-149
chr15-79631173-DEL-63
chr15-79661552-DEL-6932
chr15-79667056-DEL-131
chr15-79667402-DEL-97
chr15-79696820-DEL-234
chr15-79725320-DEL-51
chr15-79771015-DEL-64

chr15-79790029-DEL-71
chr15-79793498-DEL-242
chr15-79796188-DEL-68
chr15-79826613-DEL-58
chr15-79871719-DEL-56
chr15-79918244-DEL-147
chr15-79930704-DEL-67
chr15-79930768-DEL-183
chr15-79981909-DEL-51
chr15-80093458-DEL-50
chr15-80124173-DEL-766
chr15-80124273-DEL-285
chr15-80125003-DEL-1504
chr15-80125973-DEL-534
chr15-80164292-DEL-56
chr15-80197808-DEL-108
chr15-80217870-DEL-297
chr15-80237415-DEL-58
chr15-80496219-DEL-107
chr15-80512506-DEL-54
chr15-80636016-DEL-160
chr15-80696244-DEL-150
chr15-80804328-DEL-162
chr15-80805640-DEL-163
chr15-80835526-DEL-399
chr15-80842154-DEL-511
chr15-80845282-DEL-202
chr15-80849121-DEL-640
chr15-80868123-DEL-288
chr15-80875602-DEL-191
chr15-80994474-DEL-53
chr15-81060225-DEL-99
chr15-81150258-DEL-118
chr15-81194110-DEL-273
chr15-81194336-DEL-68
chr15-81369608-DEL-61
chr15-81369978-DEL-181
chr15-81431333-DEL-331
chr15-81438238-DEL-64
chr15-81598646-DEL-537
chr15-81654170-DEL-179
chr15-81807197-DEL-96
chr15-81828957-DEL-1176
chr15-81902210-DEL-269
chr15-81904526-DEL-176
chr15-81937604-DEL-4108
chr15-81941304-DEL-185
chr15-81941575-DEL-358
chr15-81958165-DEL-89
chr15-81974384-DEL-138
chr15-82002851-DEL-54
chr15-82010204-DEL-307
chr15-82101961-DEL-66
chr15-82246212-DEL-155
chr15-82442019-DEL-2214
chr15-82509196-DEL-113
chr15-82763249-DEL-237
chr15-82904715-DEL-92
chr15-82956798-DEL-585
chr15-83048376-DEL-63
chr15-83048603-DEL-327
chr15-83068543-DEL-57
chr15-83141056-DEL-870
chr15-83142378-DEL-395
chr15-83175616-DEL-166
chr15-83185717-DEL-1122
chr15-83194107-DEL-59
chr15-83207464-DEL-58
chr15-83314789-DEL-85
chr15-83368963-DEL-120
chr15-83369122-DEL-52
chr15-83388343-DEL-64
chr15-83388551-DEL-64
chr15-83506531-DEL-61
chr15-83628944-DEL-766
chr15-83629365-DEL-286
chr15-83709882-DEL-50

chr15-83710754-DEL-242
chr15-83711106-DEL-154
chr15-83723146-DEL-60
chr15-83805019-DEL-74
chr15-83875333-DEL-65
chr15-83875470-DEL-50
chr15-83884873-DEL-66
chr15-83905470-DEL-66
chr15-83914999-DEL-175
chr15-83915280-DEL-195
chr15-83920111-DEL-431
chr15-83922706-DEL-99
chr15-83922742-DEL-365
chr15-83923331-DEL-1302
chr15-83994922-DEL-126
chr15-84031914-DEL-1647
chr15-84106305-DEL-81
chr15-84159374-DEL-69
chr15-84243736-DEL-162
chr15-84246654-DEL-299
chr15-84302264-DEL-77
chr15-84361733-DEL-236
chr15-84409109-DEL-132
chr15-84419378-DEL-2237
chr15-84426386-DEL-562
chr15-84518393-DEL-50
chr15-84519517-DEL-110
chr15-84519683-DEL-74
chr15-84519879-DEL-248
chr15-84534363-DEL-68
chr15-84599879-DEL-55
chr15-84601307-DEL-51
chr15-84601832-DEL-54
chr15-84666359-DEL-219
chr15-84677605-DEL-210
chr15-84750450-DEL-1347
chr15-84767098-DEL-113
chr15-84794622-DEL-203
chr15-84854818-DEL-980
chr15-84854838-DEL-380
chr15-84855318-DEL-420
chr15-84888799-DEL-51
chr15-84907222-DEL-129
chr15-84912466-DEL-63
chr15-84991807-DEL-96
chr15-85069604-DEL-54
chr15-85208102-DEL-165
chr15-85212988-DEL-66
chr15-85241684-DEL-52
chr15-85338523-DEL-83
chr15-85365339-DEL-72
chr15-85400031-DEL-377
chr15-85447660-DEL-56
chr15-85475006-DEL-208
chr15-85520239-DEL-66
chr15-85651562-DEL-135
chr15-85669493-DEL-379
chr15-85754411-DEL-86
chr15-85758211-DEL-238
chr15-85762642-DEL-59
chr15-85800445-DEL-254
chr15-85800472-DEL-102
chr15-85807647-DEL-53
chr15-85937350-DEL-128
chr15-86093504-DEL-2897
chr15-86100060-DEL-95
chr15-86111220-DEL-1126
chr15-86129669-DEL-620
chr15-86129811-DEL-163
chr15-86209516-DEL-154
chr15-86209708-DEL-754
chr15-86210403-DEL-152
chr15-86287266-DEL-609
chr15-86406074-DEL-495
chr15-86549418-DEL-85
chr15-86589556-DEL-208
chr15-86613186-DEL-68

chr15-86637657-DEL-300
chr15-86702939-DEL-91
chr15-86769332-DEL-475
chr15-86770537-DEL-84
chr15-86789398-DEL-464
chr15-86798383-DEL-218
chr15-86925537-DEL-90
chr15-86977761-DEL-82
chr15-87011861-DEL-2819
chr15-87067115-DEL-52
chr15-87067184-DEL-176
chr15-87105126-DEL-109
chr15-87120727-DEL-230
chr15-87264483-DEL-652
chr15-87368980-DEL-101
chr15-87385653-DEL-145
chr15-87385699-DEL-61
chr15-87445292-DEL-70
chr15-87447104-DEL-56
chr15-87454470-DEL-54
chr15-87466072-DEL-51
chr15-87548983-DEL-144
chr15-87575436-DEL-119
chr15-87678448-DEL-72
chr15-87845421-DEL-430
chr15-87845423-DEL-1405
chr15-87845430-DEL-90
chr15-87845935-DEL-71
chr15-87856515-DEL-73
chr15-87856663-DEL-84
chr15-87995379-DEL-246
chr15-87999330-DEL-452
chr15-88071230-DEL-96
chr15-88104246-DEL-308
chr15-88142386-DEL-58
chr15-88216473-DEL-334
chr15-88239412-DEL-90
chr15-88240785-DEL-75
chr15-88308839-DEL-103
chr15-88308865-DEL-1259
chr15-88309184-DEL-505
chr15-88309888-DEL-109
chr15-88409688-DEL-164
chr15-88409769-DEL-62
chr15-88410059-DEL-98
chr15-88447500-DEL-78
chr15-88514871-DEL-114
chr15-88679119-DEL-1181
chr15-88681706-DEL-273
chr15-88681728-DEL-56
chr15-88682087-DEL-140
chr15-88698205-DEL-207
chr15-88711037-DEL-80
chr15-88711182-DEL-89
chr15-88711539-DEL-59
chr15-88722812-DEL-75
chr15-88724839-DEL-56
chr15-88791601-DEL-56
chr15-88828652-DEL-193
chr15-88859462-DEL-420
chr15-88878655-DEL-140
chr15-89017542-DEL-192
chr15-89018411-DEL-64
chr15-89103066-DEL-673
chr15-89103873-DEL-233
chr15-89118170-DEL-82
chr15-89130917-DEL-1556
chr15-89131326-DEL-480
chr15-89133274-DEL-1620
chr15-89215258-DEL-1183
chr15-89321544-DEL-829
chr15-89364677-DEL-50
chr15-89424200-DEL-57
chr15-89600080-DEL-86
chr15-89600250-DEL-55
chr15-89720456-DEL-220
chr15-89800362-DEL-56

chr15-89801822-DEL-2012
chr15-89842050-DEL-199
chr15-89895544-DEL-85
chr15-89900469-DEL-166
chr15-89936229-DEL-1419
chr15-90022733-DEL-101
chr15-90054847-DEL-130
chr15-90100515-DEL-151
chr15-90167945-DEL-58
chr15-90261978-DEL-176
chr15-90262943-DEL-1222
chr15-90269998-DEL-2936
chr15-90543237-DEL-247
chr15-90616300-DEL-116
chr15-90677189-DEL-55
chr15-90677503-DEL-5422
chr15-90733426-DEL-1431
chr15-90760063-DEL-239
chr15-90773671-DEL-575
chr15-90808000-DEL-1755
chr15-90815085-DEL-95
chr15-90827285-DEL-284
chr15-90869274-DEL-50
chr15-90990484-DEL-221
chr15-91088124-DEL-145
chr15-91118265-DEL-98
chr15-91176660-DEL-84
chr15-91185170-DEL-1412
chr15-91226613-DEL-63
chr15-91275102-DEL-69
chr15-91308490-DEL-97
chr15-91320206-DEL-446
chr15-91356406-DEL-52
chr15-91394246-DEL-207
chr15-91407942-DEL-177
chr15-91407995-DEL-77
chr15-91451803-DEL-64
chr15-91525671-DEL-69
chr15-91570684-DEL-244
chr15-91670508-DEL-336
chr15-91682419-DEL-87
chr15-91889136-DEL-1040
chr15-91894782-DEL-84
chr15-91912223-DEL-7959
chr15-91941123-DEL-224
chr15-91941148-DEL-94
chr15-92082054-DEL-297
chr15-92200343-DEL-106
chr15-92205396-DEL-108
chr15-92217410-DEL-165
chr15-92337715-DEL-78
chr15-92511592-DEL-4530
chr15-92529043-DEL-194
chr15-92659682-DEL-61
chr15-92665167-DEL-510
chr15-92665316-DEL-95
chr15-92680083-DEL-55
chr15-92680949-DEL-66
chr15-92683975-DEL-122
chr15-92705231-DEL-58
chr15-92705384-DEL-53
chr15-92721942-DEL-205
chr15-92808439-DEL-80
chr15-92978949-DEL-55
chr15-93005055-DEL-2481
chr15-93083846-DEL-809
chr15-93127921-DEL-60
chr15-93274481-DEL-62
chr15-93301346-DEL-195
chr15-93310484-DEL-329
chr15-93329229-DEL-287
chr15-93365870-DEL-953
chr15-93407569-DEL-100
chr15-93407823-DEL-66
chr15-93449206-DEL-50
chr15-93569865-DEL-50
chr15-93582852-DEL-54

chr15-93583156-DEL-58
chr15-93615061-DEL-76
chr15-93650227-DEL-116
chr15-93762532-DEL-94
chr15-93829118-DEL-99
chr15-93845185-DEL-315
chr15-93899908-DEL-69
chr15-93910769-DEL-166
chr15-93975902-DEL-185
chr15-93986369-DEL-56
chr15-94126844-DEL-172
chr15-94261173-DEL-486
chr15-94282168-DEL-670
chr15-94282586-DEL-63
chr15-94343229-DEL-99
chr15-94343509-DEL-215
chr15-94343798-DEL-100
chr15-94349961-DEL-70
chr15-94373788-DEL-54
chr15-94388125-DEL-58
chr15-94388261-DEL-222
chr15-94450795-DEL-59
chr15-94453282-DEL-60
chr15-94491176-DEL-97
chr15-94525359-DEL-238
chr15-94583463-DEL-68
chr15-94622484-DEL-487
chr15-94633447-DEL-56
chr15-94720705-DEL-54
chr15-94767783-DEL-60
chr15-94798564-DEL-70
chr15-94928753-DEL-333
chr15-94962973-DEL-50
chr15-94969103-DEL-189
chr15-95022699-DEL-51
chr15-95098327-DEL-53
chr15-95159364-DEL-439
chr15-95166290-DEL-284
chr15-95181527-DEL-62
chr15-95189544-DEL-104
chr15-95273318-DEL-249
chr15-95337244-DEL-205
chr15-95367419-DEL-106
chr15-95367481-DEL-582
chr15-95420315-DEL-203
chr15-95459566-DEL-562
chr15-95464564-DEL-67
chr15-95523487-DEL-60
chr15-95575106-DEL-60
chr15-95577135-DEL-184
chr15-95605815-DEL-90
chr15-95621450-DEL-50
chr15-95745678-DEL-50
chr15-95750531-DEL-300
chr15-95968504-DEL-643
chr15-95979557-DEL-54
chr15-95981856-DEL-112
chr15-95982044-DEL-80
chr15-96001325-DEL-55
chr15-96046939-DEL-104
chr15-96047204-DEL-58
chr15-96126527-DEL-474
chr15-96274670-DEL-54
chr15-96411629-DEL-1738
chr15-96503461-DEL-54
chr15-96547289-DEL-56
chr15-96553960-DEL-89
chr15-96721683-DEL-131
chr15-96766529-DEL-80
chr15-96832450-DEL-156
chr15-96832527-DEL-338
chr15-96833125-DEL-52
chr15-96840926-DEL-214
chr15-96841101-DEL-54
chr15-96922333-DEL-78
chr15-96984181-DEL-66
chr15-97003364-DEL-65

chr15-97022112-DEL-249
chr15-97030839-DEL-226
chr15-97047618-DEL-110
chr15-97172065-DEL-76
chr15-97186822-DEL-50
chr15-97385406-DEL-176
chr15-97425010-DEL-54
chr15-97464514-DEL-463
chr15-97482357-DEL-50
chr15-97560277-DEL-270
chr15-97560656-DEL-232
chr15-97560766-DEL-86
chr15-97622213-DEL-175
chr15-97701268-DEL-54
chr15-97762730-DEL-172
chr15-97810829-DEL-58
chr15-97818461-DEL-95
chr15-97826923-DEL-55
chr15-97856508-DEL-625
chr15-97856542-DEL-242
chr15-97857001-DEL-138
chr15-97865419-DEL-216
chr15-97920163-DEL-196
chr15-97920273-DEL-86
chr15-98017685-DEL-171
chr15-98095592-DEL-148
chr15-98147381-DEL-62
chr15-98207423-DEL-312
chr15-98252378-DEL-60
chr15-98266704-DEL-60
chr15-98391013-DEL-62
chr15-98396478-DEL-269
chr15-98399273-DEL-70
chr15-98399394-DEL-56
chr15-98401517-DEL-113
chr15-98482448-DEL-105
chr15-98546952-DEL-108
chr15-98592236-DEL-63
chr15-98603490-DEL-56
chr15-98636409-DEL-70
chr15-98680789-DEL-66
chr15-98683867-DEL-677
chr15-98739443-DEL-55
chr15-98772742-DEL-69
chr15-98790522-DEL-117
chr15-98835649-DEL-92
chr15-98835674-DEL-341
chr15-98901467-DEL-130
chr15-98904243-DEL-193
chr15-98932571-DEL-2316
chr15-98945045-DEL-487
chr15-98959740-DEL-119
chr15-98971717-DEL-203
chr15-99005625-DEL-189
chr15-99005926-DEL-355
chr15-99027331-DEL-295
chr15-99037206-DEL-60
chr15-99067465-DEL-64
chr15-99087919-DEL-4389
chr15-99094376-DEL-70
chr15-99129283-DEL-87
chr15-99166652-DEL-56
chr15-99278578-DEL-430
chr15-99345853-DEL-84
chr15-99450086-DEL-211
chr15-99450767-DEL-809
chr15-99606026-DEL-148
chr15-99663756-DEL-208
chr15-99719853-DEL-241
chr15-99731495-DEL-56
chr15-99762386-DEL-52
chr15-99809699-DEL-62
chr15-99895815-DEL-87
chr15-99925656-DEL-381
chr15-100098328-DEL-497
chr15-100165315-DEL-110
chr15-100204977-DEL-103

chr15-100219526-DEL-126
chr15-100226181-DEL-19343
chr15-100254267-DEL-297
chr15-100268277-DEL-60
chr15-100314687-DEL-100
chr15-100429075-DEL-316
chr15-100429276-DEL-317
chr15-100470495-DEL-315
chr15-100482221-DEL-169
chr15-100483062-DEL-231
chr15-100495842-DEL-168
chr15-100540591-DEL-84
chr15-100544996-DEL-1323
chr15-100553591-DEL-137
chr15-100666965-DEL-135
chr15-100747215-DEL-435
chr15-100765115-DEL-156
chr15-100765398-DEL-100
chr15-100802496-DEL-91
chr15-100862898-DEL-88
chr15-101021652-DEL-96
chr15-101200984-DEL-140
chr15-101244821-DEL-60
chr15-101367394-DEL-56
chr15-101572277-DEL-79
chr15-101572356-DEL-152
chr15-101581841-DEL-767
chr15-101582859-DEL-603
chr15-101639334-DEL-80
chr15-101688117-DEL-181
chr15-101728367-DEL-78
chr15-101856202-DEL-132
chr15-101864750-DEL-152
chr15-101864946-DEL-64
chr15-101910373-DEL-54
chr15-102079639-DEL-578
chr15-102108344-DEL-62
chr15-102288585-DEL-116
chr15-102351858-DEL-1527
chr15-102363766-DEL-195
chr15-102405900-DEL-54
chr15-102535837-DEL-50
chr15-102558175-DEL-50
chr15-102590059-DEL-79
chr15-102640100-DEL-238
chr15-102665867-DEL-207
chr15-102766254-DEL-502
chr15-103050581-DEL-691
chr15-103219690-DEL-191
chr15-103281373-DEL-51
chr15-103478776-DEL-1507
chr15-103512915-DEL-1465
chr15-103551593-DEL-218
chr15-103778724-DEL-1274
chr15-103963716-DEL-205
chr16-3055863-DEL-720
chr16-3056161-DEL-230
chr16-3068019-DEL-2261
chr16-3117649-DEL-202
chr16-3169653-DEL-66
chr16-3174530-DEL-4387
chr16-3197971-DEL-97
chr16-3315071-DEL-521
chr16-3361299-DEL-7880
chr16-3466300-DEL-229
chr16-3480213-DEL-60
chr16-3551017-DEL-86
chr16-3669462-DEL-82
chr16-3784670-DEL-68
chr16-4041335-DEL-51
chr16-4157218-DEL-120
chr16-4182298-DEL-116
chr16-4193374-DEL-165
chr16-4210377-DEL-98
chr16-4294908-DEL-210
chr16-4362040-DEL-120
chr16-4603528-DEL-147

chr16-4614778-DEL-54
chr16-4638528-DEL-186
chr16-4651068-DEL-120
chr16-4753301-DEL-54
chr16-4912609-DEL-145
chr16-4955422-DEL-116
chr16-5026101-DEL-51
chr16-5029216-DEL-83
chr16-5064174-DEL-122
chr16-5064602-DEL-62
chr16-5170254-DEL-117
chr16-5200811-DEL-50
chr16-5221339-DEL-116
chr16-5221816-DEL-52
chr16-5302339-DEL-298
chr16-5308603-DEL-86
chr16-5327536-DEL-60
chr16-5346394-DEL-616
chr16-5386215-DEL-174
chr16-5437769-DEL-539
chr16-5536118-DEL-72
chr16-5544233-DEL-90
chr16-5588601-DEL-51
chr16-5588739-DEL-93
chr16-5601226-DEL-56
chr16-5601547-DEL-65
chr16-5601560-DEL-161
chr16-5634595-DEL-52
chr16-5637076-DEL-50
chr16-5637990-DEL-50
chr16-5665129-DEL-52
chr16-5826913-DEL-80
chr16-5863602-DEL-50
chr16-5922334-DEL-65
chr16-5933564-DEL-22303
chr16-5988541-DEL-53
chr16-6057806-DEL-2274
chr16-6204855-DEL-1852
chr16-6264465-DEL-108
chr16-6448663-DEL-52
chr16-6753681-DEL-60
chr16-6844666-DEL-728
chr16-6872193-DEL-96
chr16-6919121-DEL-74
chr16-7005592-DEL-50
chr16-7066555-DEL-232
chr16-7096250-DEL-164
chr16-7344608-DEL-51
chr16-7364383-DEL-252
chr16-7471855-DEL-70
chr16-7491689-DEL-56
chr16-7500526-DEL-85
chr16-7543338-DEL-601
chr16-7630105-DEL-2724
chr16-7755581-DEL-58
chr16-7755620-DEL-203
chr16-7756343-DEL-58
chr16-7795838-DEL-138
chr16-7798834-DEL-210
chr16-7799312-DEL-76
chr16-7835311-DEL-148
chr16-7849000-DEL-50
chr16-7901949-DEL-177
chr16-7902275-DEL-89
chr16-7919235-DEL-1173
chr16-7929062-DEL-85
chr16-7979730-DEL-102
chr16-7991977-DEL-149
chr16-8111531-DEL-163
chr16-8171386-DEL-60
chr16-8182292-DEL-50
chr16-8210287-DEL-84
chr16-8305366-DEL-139
chr16-8310417-DEL-105
chr16-8334195-DEL-72
chr16-8337405-DEL-69
chr16-8345242-DEL-52

chr16-8350047-DEL-110
chr16-8419714-DEL-74
chr16-8443077-DEL-192
chr16-8575771-DEL-231
chr16-8587634-DEL-68
chr16-8588038-DEL-122
chr16-8616785-DEL-222
chr16-8717103-DEL-161
chr16-8717567-DEL-98
chr16-8722765-DEL-59
chr16-8726006-DEL-168
chr16-8734322-DEL-56
chr16-8919462-DEL-986
chr16-9041536-DEL-206
chr16-9042102-DEL-304
chr16-9172862-DEL-481
chr16-9257081-DEL-56
chr16-9283443-DEL-528
chr16-9362338-DEL-228
chr16-9366303-DEL-72
chr16-9401317-DEL-336
chr16-9456254-DEL-54
chr16-9540870-DEL-221
chr16-9540884-DEL-445
chr16-9574194-DEL-51
chr16-9638587-DEL-99
chr16-9649877-DEL-50
chr16-9712564-DEL-68
chr16-9737278-DEL-120
chr16-9838498-DEL-119
chr16-9904808-DEL-117
chr16-10132454-DEL-365
chr16-10193493-DEL-347
chr16-10205629-DEL-62
chr16-10214863-DEL-50
chr16-10381891-DEL-477
chr16-10458719-DEL-60
chr16-10717852-DEL-67
chr16-10744479-DEL-69
chr16-10761829-DEL-52
chr16-10792660-DEL-184
chr16-10793005-DEL-118
chr16-10813165-DEL-99
chr16-10835040-DEL-136
chr16-10929336-DEL-80
chr16-11009334-DEL-846
chr16-11080576-DEL-200
chr16-11091860-DEL-54
chr16-11179016-DEL-393
chr16-11179151-DEL-97
chr16-11191819-DEL-156
chr16-11241337-DEL-285
chr16-11284048-DEL-139
chr16-11404875-DEL-244
chr16-11446959-DEL-64
chr16-11452431-DEL-957
chr16-11474657-DEL-190
chr16-11495715-DEL-273
chr16-11504448-DEL-990
chr16-11504919-DEL-88
chr16-11505551-DEL-268
chr16-11518775-DEL-909
chr16-11737109-DEL-53
chr16-11742129-DEL-104
chr16-11757142-DEL-500
chr16-11760696-DEL-86
chr16-11780428-DEL-74
chr16-11784901-DEL-161
chr16-11854062-DEL-55
chr16-12016194-DEL-54
chr16-12293032-DEL-55
chr16-12356070-DEL-139
chr16-12461329-DEL-55
chr16-12461481-DEL-224
chr16-12516543-DEL-167
chr16-12516625-DEL-50
chr16-12553026-DEL-136

chr16-12579623-DEL-94
chr16-12669851-DEL-93
chr16-12836511-DEL-250
chr16-12836559-DEL-77
chr16-12836918-DEL-115
chr16-12846119-DEL-119
chr16-12846341-DEL-69
chr16-13020245-DEL-669
chr16-13028845-DEL-340
chr16-13117085-DEL-183
chr16-13299452-DEL-2893
chr16-13313568-DEL-66
chr16-13363420-DEL-466
chr16-13363691-DEL-119
chr16-13404908-DEL-51
chr16-13440767-DEL-2077
chr16-13451380-DEL-331
chr16-13475808-DEL-92
chr16-13510712-DEL-112
chr16-13513172-DEL-510
chr16-13539511-DEL-174
chr16-13568750-DEL-69
chr16-13569088-DEL-64
chr16-13573454-DEL-174
chr16-13602103-DEL-137
chr16-13642844-DEL-51
chr16-13642850-DEL-255
chr16-13670081-DEL-2083
chr16-13995819-DEL-60
chr16-14010960-DEL-54
chr16-14066853-DEL-77
chr16-14076108-DEL-71
chr16-14259201-DEL-1090
chr16-14259284-DEL-140
chr16-14259778-DEL-186
chr16-14260122-DEL-75
chr16-14320733-DEL-52
chr16-14320856-DEL-65
chr16-14683278-DEL-1473
chr16-14876671-DEL-2116
chr16-15044616-DEL-4108
chr16-15051003-DEL-1683
chr16-15079766-DEL-871
chr16-15164754-DEL-69
chr16-15399419-DEL-2756
chr16-15459854-DEL-357
chr16-15483464-DEL-85
chr16-15490740-DEL-50
chr16-15536074-DEL-122
chr16-15578352-DEL-369
chr16-15588024-DEL-61
chr16-15716956-DEL-60
chr16-15717135-DEL-150
chr16-15762265-DEL-1269
chr16-15811794-DEL-3993
chr16-15896178-DEL-50
chr16-15940842-DEL-1521
chr16-15995373-DEL-3613
chr16-16220497-DEL-2747
chr16-16526050-DEL-380
chr16-16581715-DEL-206
chr16-16654748-DEL-5554
chr16-16765135-DEL-100
chr16-16771956-DEL-1800
chr16-16816783-DEL-211
chr16-16964409-DEL-117
chr16-17010140-DEL-64
chr16-17068985-DEL-60
chr16-17087330-DEL-88
chr16-17105959-DEL-58
chr16-17360035-DEL-50
chr16-17422616-DEL-72
chr16-17528882-DEL-245
chr16-17551651-DEL-94
chr16-17551901-DEL-329
chr16-17605939-DEL-52
chr16-17696416-DEL-72

chr16-17758409-DEL-954
chr16-17758802-DEL-422
chr16-17765160-DEL-420
chr16-17810011-DEL-3586
chr16-17868143-DEL-112
chr16-17940259-DEL-57
chr16-18113368-DEL-205
chr16-18131105-DEL-182
chr16-18150519-DEL-728
chr16-18180858-DEL-767
chr16-18184811-DEL-2497
chr16-18252511-DEL-371
chr16-18343554-DEL-454
chr16-18411127-DEL-95
chr16-18500017-DEL-977
chr16-18510627-DEL-61
chr16-18556378-DEL-145
chr16-18577740-DEL-72
chr16-18659210-DEL-211
chr16-18659617-DEL-285
chr16-18665612-DEL-170
chr16-18679852-DEL-1490
chr16-18775149-DEL-234
chr16-18830004-DEL-3145
chr16-18898484-DEL-750
chr16-19216652-DEL-2365
chr16-19244760-DEL-95
chr16-19311022-DEL-90
chr16-19340948-DEL-58
chr16-19467001-DEL-4122
chr16-19555437-DEL-3670
chr16-19610543-DEL-69
chr16-19711381-DEL-96
chr16-19758519-DEL-6401
chr16-19768555-DEL-204
chr16-19843593-DEL-197
chr16-19888781-DEL-52
chr16-19945094-DEL-90
chr16-19948337-DEL-126
chr16-19948392-DEL-464
chr16-20084622-DEL-1440
chr16-20086070-DEL-159
chr16-20086807-DEL-144
chr16-20181925-DEL-451
chr16-20182518-DEL-105
chr16-20208761-DEL-149
chr16-20242900-DEL-51
chr16-20258303-DEL-154
chr16-20341259-DEL-65
chr16-20585042-DEL-50
chr16-20731978-DEL-65
chr16-20755851-DEL-51
chr16-20776086-DEL-120
chr16-20899025-DEL-168
chr16-20956281-DEL-176
chr16-20964539-DEL-176
chr16-20985298-DEL-60
chr16-21049891-DEL-66
chr16-21094041-DEL-84
chr16-21160850-DEL-261
chr16-21187507-DEL-185
chr16-21209387-DEL-54
chr16-21270509-DEL-118
chr16-21300673-DEL-84
chr16-21303472-DEL-2990
chr16-21319237-DEL-143
chr16-21370236-DEL-50
chr16-21417857-DEL-2570
chr16-21420423-DEL-479
chr16-21436656-DEL-61
chr16-21477041-DEL-474
chr16-21497450-DEL-144
chr16-21497707-DEL-120
chr16-21538041-DEL-101
chr16-21545089-DEL-274
chr16-21615594-DEL-50
chr16-21678646-DEL-55

chr16-21681428-DEL-985
chr16-21946430-DEL-204
chr16-21954699-DEL-222
chr16-22035127-DEL-71
chr16-22296191-DEL-446
chr16-22296463-DEL-140
chr16-22330443-DEL-99
chr16-22348053-DEL-385
chr16-22360635-DEL-96
chr16-22381726-DEL-50
chr16-22521758-DEL-300
chr16-22551716-DEL-2118
chr16-22612585-DEL-54
chr16-22649661-DEL-102
chr16-22808700-DEL-246
chr16-22808813-DEL-122
chr16-23038697-DEL-50
chr16-23081475-DEL-74
chr16-23100727-DEL-166
chr16-23232587-DEL-69
chr16-23473423-DEL-207
chr16-23483570-DEL-54
chr16-23545950-DEL-56
chr16-23547880-DEL-1088
chr16-23621715-DEL-62
chr16-23860800-DEL-84
chr16-23934778-DEL-1196
chr16-23937567-DEL-14395
chr16-23972378-DEL-78
chr16-24075116-DEL-183
chr16-24276743-DEL-75
chr16-24364152-DEL-50
chr16-24424381-DEL-57
chr16-24442141-DEL-162
chr16-24693997-DEL-184
chr16-24855277-DEL-315
chr16-25045049-DEL-60
chr16-25229886-DEL-108
chr16-25238982-DEL-180
chr16-25239082-DEL-68
chr16-25271425-DEL-124
chr16-25569110-DEL-50
chr16-25650912-DEL-121
chr16-25884175-DEL-143
chr16-26012475-DEL-1163
chr16-26014043-DEL-544
chr16-26049201-DEL-65
chr16-26088301-DEL-1492
chr16-26092881-DEL-84
chr16-26136091-DEL-6370
chr16-26202093-DEL-123
chr16-26249443-DEL-83
chr16-26267315-DEL-50
chr16-26387121-DEL-54
chr16-26573615-DEL-58
chr16-26627189-DEL-6460
chr16-26685851-DEL-212
chr16-26717520-DEL-81
chr16-26726060-DEL-2579
chr16-26739605-DEL-50
chr16-26748455-DEL-103
chr16-26812210-DEL-305
chr16-26871706-DEL-50
chr16-26927016-DEL-2066
chr16-26985976-DEL-180
chr16-27013319-DEL-103
chr16-27035145-DEL-2146
chr16-27048533-DEL-62
chr16-27065736-DEL-552
chr16-27137554-DEL-2218
chr16-27149853-DEL-135
chr16-27158766-DEL-57
chr16-27242601-DEL-206
chr16-27427013-DEL-201
chr16-27431761-DEL-64
chr16-27441703-DEL-56
chr16-27520958-DEL-206

chr16-27521232-DEL-60
chr16-27587725-DEL-54
chr16-27822420-DEL-1991
chr16-27853468-DEL-50
chr16-27893354-DEL-56
chr16-27957358-DEL-154
chr16-27965491-DEL-317
chr16-28043015-DEL-83
chr16-28075442-DEL-96
chr16-28163984-DEL-70
chr16-28249901-DEL-50
chr16-28280893-DEL-192
chr16-28344515-DEL-116
chr16-28408183-DEL-1565
chr16-28497799-DEL-52
chr16-28771404-DEL-91
chr16-29122516-DEL-233
chr16-29156529-DEL-407
chr16-29236845-DEL-62
chr16-29281569-DEL-51
chr16-29291764-DEL-55
chr16-29321380-DEL-50
chr16-29334735-DEL-198
chr16-29334980-DEL-149
chr16-29401544-DEL-438
chr16-29401606-DEL-94
chr16-29401672-DEL-1097
chr16-29402270-DEL-312
chr16-29402393-DEL-63
chr16-29403155-DEL-256
chr16-29412137-DEL-171
chr16-29506057-DEL-64
chr16-29510948-DEL-68
chr16-29624323-DEL-105
chr16-29637443-DEL-70
chr16-29691091-DEL-50
chr16-29696909-DEL-243
chr16-29731614-DEL-115
chr16-29760652-DEL-54
chr16-29851690-DEL-68
chr16-29973963-DEL-57
chr16-30043548-DEL-83
chr16-30051581-DEL-68
chr16-30066172-DEL-306
chr16-30192353-DEL-133
chr16-30295600-DEL-68
chr16-30366409-DEL-234
chr16-30366423-DEL-114
chr16-30462760-DEL-84
chr16-30468921-DEL-129
chr16-30536291-DEL-304
chr16-30747678-DEL-426
chr16-30759385-DEL-72
chr16-30891762-DEL-62
chr16-30969226-DEL-117
chr16-31012211-DEL-360
chr16-31053471-DEL-172
chr16-31267750-DEL-54
chr16-31293468-DEL-216
chr16-31383299-DEL-78
chr16-31442766-DEL-260
chr16-31463000-DEL-114
chr16-31463193-DEL-304
chr16-31532122-DEL-50
chr16-31659114-DEL-138
chr16-31730667-DEL-1240
chr16-31801198-DEL-174
chr16-31804991-DEL-51
chr16-31950388-DEL-390
chr16-31977205-DEL-190
chr16-32089323-DEL-106
chr16-32137556-DEL-166
chr16-32309007-DEL-783
chr16-32417500-DEL-124
chr16-32429678-DEL-80
chr16-32459726-DEL-161
chr16-32881763-DEL-211

chr16-32907977-DEL-132
chr16-32907988-DEL-52
chr16-32942664-DEL-50
chr16-32979346-DEL-50
chr16-33038130-DEL-99
chr16-33094116-DEL-51
chr16-33227179-DEL-109
chr16-33359328-DEL-69
chr16-33414368-DEL-8298
chr16-33541433-DEL-85
chr16-33685338-DEL-69
chr16-33804112-DEL-60
chr16-33804241-DEL-178
chr16-33827077-DEL-819
chr16-33895687-DEL-118
chr16-33953958-DEL-68
chr16-34066859-DEL-233
chr16-34075656-DEL-75
chr16-34153387-DEL-61
chr16-34153389-DEL-144
chr16-34177060-DEL-76
chr16-34206909-DEL-128
chr16-34229725-DEL-676
chr16-34419617-DEL-56
chr16-34463546-DEL-102
chr16-34472508-DEL-50
chr16-34581229-DEL-77
chr16-34819121-DEL-226
chr16-34911447-DEL-150
chr16-34911556-DEL-50
chr16-35018244-DEL-76
chr16-35348298-DEL-484
chr16-35385991-DEL-64
chr16-35393613-DEL-360
chr16-35464673-DEL-284
chr16-35472475-DEL-90
chr16-35481194-DEL-72
chr16-35521071-DEL-378
chr16-35521088-DEL-84
chr16-35882644-DEL-207
chr16-35882728-DEL-69
chr16-35932774-DEL-50
chr16-35960777-DEL-87
chr16-35981723-DEL-214
chr16-35981931-DEL-80
chr16-36035926-DEL-316
chr16-36048798-DEL-1118
chr16-36078225-DEL-27915
chr16-36143815-DEL-74
chr16-36336889-DEL-259
chr16-36342222-DEL-84
chr16-36435958-DEL-52
chr16-36458490-DEL-1532
chr16-36479998-DEL-106
chr16-36533802-DEL-299
chr16-36564390-DEL-351
chr16-36638813-DEL-564
chr16-36639697-DEL-59
chr16-36876479-DEL-73
chr16-37043677-DEL-132
chr16-37065285-DEL-60
chr16-37189098-DEL-50
chr16-37397458-DEL-60
chr16-37432839-DEL-60
chr16-37523135-DEL-87
chr16-37561049-DEL-62
chr16-37602259-DEL-213
chr16-37633220-DEL-240
chr16-37685856-DEL-73
chr16-37686079-DEL-88
chr16-37862375-DEL-55
chr16-37902283-DEL-77
chr16-38046208-DEL-350
chr16-38063348-DEL-68
chr16-38102483-DEL-70
chr16-38173745-DEL-193
chr16-38320323-DEL-203

chr16-38320549-DEL-82
chr16-38324902-DEL-60
chr16-38640307-DEL-119
chr16-38651549-DEL-5209
chr16-38667168-DEL-56
chr16-38868177-DEL-60
chr16-38904332-DEL-57
chr16-39149479-DEL-83
chr16-39234809-DEL-323
chr16-39561091-DEL-198
chr16-39822823-DEL-3819
chr16-39853537-DEL-50
chr16-40008094-DEL-60
chr16-40068464-DEL-600
chr16-40070228-DEL-77
chr16-40070399-DEL-60
chr16-40140178-DEL-114
chr16-40192262-DEL-649
chr16-40518274-DEL-86
chr16-40676604-DEL-68
chr16-41038092-DEL-104
chr16-41102147-DEL-56
chr16-41292575-DEL-202
chr16-41367731-DEL-2961
chr16-41668732-DEL-92
chr16-41711861-DEL-183
chr16-41858963-DEL-50
chr16-41972656-DEL-67
chr16-42087585-DEL-108
chr16-42384788-DEL-258
chr16-42588217-DEL-54
chr16-42881163-DEL-142
chr16-43289106-DEL-88
chr16-43320195-DEL-50
chr16-43504687-DEL-112
chr16-43532692-DEL-72
chr16-43596231-DEL-399
chr16-43596279-DEL-198
chr16-43598916-DEL-1491
chr16-43733070-DEL-1224
chr16-43734579-DEL-439
chr16-44072557-DEL-457
chr16-44123996-DEL-55
chr16-44140954-DEL-127
chr16-44216569-DEL-630
chr16-44219416-DEL-494
chr16-44295027-DEL-122
chr16-44346402-DEL-52
chr16-44407532-DEL-85
chr16-44495335-DEL-263
chr16-44495688-DEL-707
chr16-44523885-DEL-50
chr16-44626901-DEL-66
chr16-44774937-DEL-50
chr16-44887715-DEL-54
chr16-44995810-DEL-76
chr16-45122505-DEL-52
chr16-45142396-DEL-101
chr16-45175974-DEL-140
chr16-45349665-DEL-131
chr16-45397391-DEL-373
chr16-45488289-DEL-155
chr16-45519063-DEL-55
chr16-45528323-DEL-107
chr16-45730381-DEL-376
chr16-45764860-DEL-56
chr16-45836404-DEL-2088
chr16-45882087-DEL-50
chr16-45910343-DEL-640
chr16-45923832-DEL-2407
chr16-45964198-DEL-52
chr16-45979016-DEL-145
chr16-46354294-DEL-64
chr16-46354444-DEL-330
chr16-46354638-DEL-56
chr16-46376279-DEL-99
chr16-46378445-DEL-52

chr16-46440961-DEL-65
chr16-46463892-DEL-424
chr16-46538229-DEL-377
chr16-46735565-DEL-111
chr16-46841021-DEL-63
chr16-46937504-DEL-63
chr16-47037568-DEL-212
chr16-47310294-DEL-1116
chr16-47461009-DEL-117
chr16-47468407-DEL-66
chr16-47468629-DEL-216
chr16-47508642-DEL-62
chr16-47548341-DEL-229
chr16-47568471-DEL-1225
chr16-47579384-DEL-229
chr16-47640554-DEL-12465
chr16-47876489-DEL-78
chr16-47945457-DEL-533
chr16-47946051-DEL-63
chr16-47950666-DEL-183
chr16-48068227-DEL-274
chr16-48070634-DEL-398
chr16-48070978-DEL-60
chr16-48118430-DEL-1193
chr16-48173934-DEL-108
chr16-48282465-DEL-463
chr16-48305118-DEL-982
chr16-48358961-DEL-9354
chr16-48400325-DEL-438
chr16-48409090-DEL-96
chr16-48409463-DEL-249
chr16-48509466-DEL-4619
chr16-48518665-DEL-56
chr16-48900829-DEL-56
chr16-48940649-DEL-117
chr16-48968823-DEL-1491
chr16-49117723-DEL-50
chr16-49381090-DEL-121
chr16-49439939-DEL-103
chr16-49454405-DEL-694
chr16-49508625-DEL-113
chr16-49513473-DEL-78
chr16-49939084-DEL-126
chr16-49939367-DEL-108
chr16-50116083-DEL-468
chr16-50188066-DEL-82
chr16-50192906-DEL-50
chr16-50404707-DEL-50
chr16-50421750-DEL-51
chr16-50458615-DEL-80
chr16-50472842-DEL-67
chr16-50515857-DEL-249
chr16-50688808-DEL-82
chr16-50743930-DEL-314
chr16-50793067-DEL-87
chr16-50851088-DEL-85
chr16-50863048-DEL-1149
chr16-50881858-DEL-1485
chr16-51005283-DEL-1532
chr16-51053377-DEL-597
chr16-51463273-DEL-99
chr16-51475051-DEL-58
chr16-51475191-DEL-1179
chr16-51476428-DEL-66
chr16-51476859-DEL-64
chr16-51883119-DEL-390
chr16-51904077-DEL-110
chr16-51981094-DEL-116
chr16-52267985-DEL-52
chr16-52372806-DEL-67
chr16-52393390-DEL-2012
chr16-52478852-DEL-56
chr16-52501189-DEL-362
chr16-52572943-DEL-8395
chr16-52575910-DEL-106
chr16-52622809-DEL-791
chr16-52634871-DEL-57

chr16-52711693-DEL-148
chr16-52900967-DEL-91
chr16-52939560-DEL-455
chr16-53204757-DEL-61
chr16-53250547-DEL-72
chr16-53250977-DEL-1680
chr16-53253876-DEL-485
chr16-53432292-DEL-95
chr16-53443561-DEL-52
chr16-53503316-DEL-2926
chr16-53664338-DEL-765
chr16-53725207-DEL-1527
chr16-53782067-DEL-569
chr16-53783665-DEL-4916
chr16-53905022-DEL-206
chr16-53982960-DEL-359
chr16-54145536-DEL-64
chr16-54156613-DEL-92
chr16-54156785-DEL-204
chr16-54264667-DEL-178
chr16-54272931-DEL-50
chr16-54335782-DEL-376
chr16-54688147-DEL-212
chr16-54751017-DEL-2297
chr16-54847179-DEL-1912
chr16-54861032-DEL-60
chr16-54861468-DEL-138
chr16-54887784-DEL-308
chr16-54951489-DEL-1766
chr16-55151128-DEL-108
chr16-55151428-DEL-234
chr16-55337232-DEL-153
chr16-55526384-DEL-71
chr16-55571617-DEL-203
chr16-55672862-DEL-68
chr16-55674654-DEL-354
chr16-55674701-DEL-70
chr16-55674937-DEL-70
chr16-55675180-DEL-340
chr16-55676953-DEL-84
chr16-55750302-DEL-145
chr16-55773127-DEL-60
chr16-55837826-DEL-55
chr16-55910106-DEL-51
chr16-55916835-DEL-125
chr16-55979662-DEL-60
chr16-55985087-DEL-560
chr16-56029546-DEL-270
chr16-56277159-DEL-594
chr16-56277190-DEL-119
chr16-56277442-DEL-113
chr16-56628960-DEL-67
chr16-56796280-DEL-55
chr16-56993918-DEL-161
chr16-57031039-DEL-50
chr16-57082911-DEL-154
chr16-57082977-DEL-60
chr16-57163856-DEL-177
chr16-57299993-DEL-126
chr16-57300178-DEL-55
chr16-57387322-DEL-164
chr16-57387462-DEL-79
chr16-57387687-DEL-162
chr16-57387691-DEL-55
chr16-57722406-DEL-60
chr16-57730274-DEL-50
chr16-57818472-DEL-210
chr16-57818635-DEL-92
chr16-57946167-DEL-50
chr16-58085138-DEL-738
chr16-58116613-DEL-637
chr16-58188395-DEL-7826
chr16-58284059-DEL-106
chr16-58392115-DEL-1493
chr16-58421905-DEL-55
chr16-58441978-DEL-204
chr16-58461502-DEL-1003

chr16-58518456-DEL-120
chr16-58610995-DEL-156
chr16-58797694-DEL-6687
chr16-58964577-DEL-240
chr16-59032255-DEL-53178
chr16-59059919-DEL-64
chr16-59156343-DEL-199
chr16-59316437-DEL-5038
chr16-59370218-DEL-52
chr16-59429293-DEL-55
chr16-59429521-DEL-106
chr16-59477451-DEL-56
chr16-59512707-DEL-76
chr16-59757986-DEL-50
chr16-59997628-DEL-50
chr16-60010303-DEL-1311
chr16-60012678-DEL-51
chr16-60141533-DEL-53
chr16-60156054-DEL-52
chr16-60339211-DEL-63
chr16-60391360-DEL-574
chr16-60484022-DEL-64
chr16-60595470-DEL-1650
chr16-60618721-DEL-96
chr16-60644051-DEL-123
chr16-60673081-DEL-705
chr16-60762712-DEL-58
chr16-60868768-DEL-50
chr16-60954498-DEL-104
chr16-61190125-DEL-204
chr16-61290702-DEL-92
chr16-61343549-DEL-1656
chr16-61345652-DEL-50
chr16-61389340-DEL-54
chr16-61621295-DEL-294
chr16-61621540-DEL-79
chr16-61645710-DEL-506
chr16-61768603-DEL-200
chr16-61794474-DEL-109
chr16-61867088-DEL-56
chr16-61917400-DEL-3951
chr16-62054024-DEL-148
chr16-62056670-DEL-279
chr16-62143938-DEL-897
chr16-62294957-DEL-85
chr16-62390736-DEL-212
chr16-62484680-DEL-267
chr16-62484883-DEL-64
chr16-62535468-DEL-129
chr16-62535751-DEL-86
chr16-62635370-DEL-50
chr16-62653495-DEL-176
chr16-62653818-DEL-318
chr16-62779759-DEL-50
chr16-62789178-DEL-1217
chr16-62840526-DEL-54
chr16-62915744-DEL-50
chr16-63061558-DEL-117
chr16-63198788-DEL-1146
chr16-63218796-DEL-65
chr16-63280734-DEL-1593
chr16-63280781-DEL-87
chr16-63280786-DEL-222
chr16-63282512-DEL-176
chr16-63283274-DEL-80
chr16-63653344-DEL-1627
chr16-63653861-DEL-753
chr16-63655525-DEL-552
chr16-63777841-DEL-85
chr16-63951410-DEL-60
chr16-63951414-DEL-125
chr16-63998291-DEL-117
chr16-64015003-DEL-5273
chr16-64057404-DEL-1264
chr16-64104409-DEL-401
chr16-64200578-DEL-82
chr16-64218645-DEL-2077

chr16-64268246-DEL-52
chr16-64270632-DEL-212
chr16-64270644-DEL-5246
chr16-64311993-DEL-94
chr16-64365467-DEL-431
chr16-64506692-DEL-74
chr16-64730934-DEL-53
chr16-64754762-DEL-422
chr16-64784804-DEL-316
chr16-64784972-DEL-122
chr16-64785217-DEL-144
chr16-64800165-DEL-197
chr16-64854866-DEL-376
chr16-64855568-DEL-106
chr16-65087839-DEL-1549
chr16-65150101-DEL-63
chr16-65331202-DEL-156
chr16-65456756-DEL-51
chr16-65494065-DEL-624
chr16-65505088-DEL-117
chr16-65604852-DEL-50
chr16-65631367-DEL-50
chr16-65706741-DEL-72
chr16-65799686-DEL-837
chr16-65854062-DEL-4367
chr16-65897899-DEL-50
chr16-65986952-DEL-77
chr16-66115158-DEL-56
chr16-66250675-DEL-60
chr16-66372218-DEL-442
chr16-66372242-DEL-126
chr16-66372512-DEL-208
chr16-66406633-DEL-51
chr16-66682255-DEL-58
chr16-66745955-DEL-58
chr16-67194359-DEL-267
chr16-67205538-DEL-84
chr16-67205756-DEL-92
chr16-67225719-DEL-116
chr16-67361917-DEL-757
chr16-67448445-DEL-58
chr16-67450128-DEL-50
chr16-67457136-DEL-73
chr16-67579772-DEL-542
chr16-67638890-DEL-3041
chr16-67801831-DEL-60
chr16-67820446-DEL-2928
chr16-67829431-DEL-1601
chr16-67834902-DEL-64
chr16-67871351-DEL-50
chr16-67897197-DEL-206
chr16-67904731-DEL-338
chr16-67912936-DEL-464
chr16-68010772-DEL-76
chr16-68099537-DEL-538
chr16-68100248-DEL-65
chr16-68124979-DEL-60
chr16-68125249-DEL-76
chr16-68160869-DEL-54
chr16-68209418-DEL-50
chr16-68214038-DEL-101
chr16-68326230-DEL-157
chr16-68424107-DEL-54
chr16-68493442-DEL-71
chr16-68620465-DEL-1129
chr16-68666685-DEL-80
chr16-68783455-DEL-50
chr16-68854789-DEL-72
chr16-69005681-DEL-212
chr16-69254344-DEL-74
chr16-69265375-DEL-70
chr16-69269763-DEL-50
chr16-69292320-DEL-751
chr16-69328401-DEL-1015
chr16-69377330-DEL-51
chr16-69399177-DEL-52
chr16-69416287-DEL-568

chr16-69502993-DEL-214
chr16-69728314-DEL-304
chr16-69872545-DEL-718
chr16-69941719-DEL-84
chr16-69941770-DEL-1218
chr16-70012878-DEL-50
chr16-70020802-DEL-80
chr16-70098022-DEL-56
chr16-70118256-DEL-104
chr16-70276960-DEL-194
chr16-70277222-DEL-88
chr16-70464284-DEL-208
chr16-70585529-DEL-157
chr16-70669474-DEL-180
chr16-70669526-DEL-74
chr16-70669874-DEL-240
chr16-70681026-DEL-3745
chr16-70687856-DEL-119
chr16-71042057-DEL-208
chr16-71080495-DEL-70
chr16-71154886-DEL-2270
chr16-71278748-DEL-416
chr16-71315937-DEL-420
chr16-71332573-DEL-2295
chr16-71348491-DEL-2019
chr16-71462983-DEL-277
chr16-71503167-DEL-2659
chr16-71541931-DEL-77
chr16-71554969-DEL-467
chr16-71616812-DEL-132
chr16-71668688-DEL-264
chr16-71691537-DEL-88
chr16-71760304-DEL-55
chr16-71773068-DEL-694
chr16-71782300-DEL-310
chr16-71804287-DEL-223
chr16-71837926-DEL-56
chr16-71903552-DEL-91
chr16-71916832-DEL-59
chr16-72234516-DEL-144
chr16-72277450-DEL-152
chr16-72313324-DEL-51
chr16-72356366-DEL-66
chr16-72366605-DEL-77
chr16-72376240-DEL-265
chr16-72450071-DEL-63
chr16-72481602-DEL-70
chr16-72585852-DEL-330
chr16-72642311-DEL-64
chr16-72767547-DEL-120
chr16-72767578-DEL-56
chr16-72856439-DEL-54
chr16-72875338-DEL-56
chr16-72944977-DEL-55
chr16-73049345-DEL-194
chr16-73055982-DEL-7213
chr16-73161789-DEL-99
chr16-73186175-DEL-1133
chr16-73338612-DEL-50
chr16-73517843-DEL-72
chr16-73630148-DEL-59
chr16-73645194-DEL-92
chr16-73665533-DEL-97
chr16-73971260-DEL-328
chr16-74096669-DEL-202
chr16-74336780-DEL-120
chr16-74471630-DEL-128
chr16-74557192-DEL-56
chr16-74557341-DEL-102
chr16-74720317-DEL-68
chr16-74756493-DEL-56
chr16-74809915-DEL-73
chr16-74986351-DEL-123
chr16-75023450-DEL-90
chr16-75030790-DEL-336
chr16-75043610-DEL-524
chr16-75070743-DEL-66

chr16-75085954-DEL-67
chr16-75088711-DEL-96
chr16-75102033-DEL-78
chr16-75102493-DEL-1560
chr16-75104301-DEL-247
chr16-75105730-DEL-62
chr16-75111172-DEL-74
chr16-75126100-DEL-3192
chr16-75134969-DEL-64
chr16-75416739-DEL-2361
chr16-75465751-DEL-786
chr16-75643923-DEL-218
chr16-75692291-DEL-1025
chr16-75702307-DEL-80
chr16-75719860-DEL-55
chr16-75842064-DEL-144
chr16-75853899-DEL-3758
chr16-75887179-DEL-100
chr16-75900826-DEL-50
chr16-76064610-DEL-180
chr16-76084014-DEL-72
chr16-76276938-DEL-405
chr16-76277049-DEL-54
chr16-76302489-DEL-91
chr16-76302645-DEL-230
chr16-76302938-DEL-71
chr16-76323010-DEL-943
chr16-76324946-DEL-1360
chr16-76341474-DEL-177
chr16-76357820-DEL-604
chr16-76404647-DEL-92
chr16-76459736-DEL-80
chr16-76459957-DEL-149
chr16-76717087-DEL-2230
chr16-76736975-DEL-62
chr16-76790200-DEL-53
chr16-76979494-DEL-53
chr16-77014578-DEL-375
chr16-77140954-DEL-50
chr16-77674929-DEL-52
chr16-77751405-DEL-2435
chr16-77777022-DEL-554
chr16-77782235-DEL-192
chr16-77782329-DEL-56
chr16-77788539-DEL-204
chr16-77874551-DEL-466
chr16-77875104-DEL-54
chr16-77922918-DEL-157
chr16-77969256-DEL-2471
chr16-77972177-DEL-1528
chr16-77998065-DEL-126
chr16-78014194-DEL-55
chr16-78049446-DEL-474
chr16-78049723-DEL-136
chr16-78067841-DEL-96
chr16-78075478-DEL-68
chr16-78089153-DEL-64
chr16-78233599-DEL-327
chr16-78233696-DEL-152
chr16-78247831-DEL-161
chr16-78390713-DEL-60
chr16-78495601-DEL-50
chr16-78546539-DEL-50
chr16-78613335-DEL-625
chr16-78708539-DEL-799
chr16-78816948-DEL-112
chr16-78866601-DEL-267
chr16-78893784-DEL-158
chr16-78913221-DEL-2974
chr16-78942767-DEL-210
chr16-78952321-DEL-138
chr16-79207306-DEL-93
chr16-79264515-DEL-535
chr16-79333439-DEL-123
chr16-79333832-DEL-171
chr16-79334151-DEL-247
chr16-79334306-DEL-100

chr16-79336455-DEL-50
chr16-79485625-DEL-55
chr16-79593589-DEL-53
chr16-79643074-DEL-75
chr16-79832992-DEL-55
chr16-80051960-DEL-55
chr16-80065192-DEL-2285
chr16-80288353-DEL-221
chr16-80474328-DEL-201
chr16-80660454-DEL-100
chr16-80669572-DEL-1110
chr16-80681916-DEL-130
chr16-80733134-DEL-1323
chr16-80744615-DEL-52
chr16-80839255-DEL-2914
chr16-80958456-DEL-50
chr16-81066232-DEL-69
chr16-81155370-DEL-264
chr16-81169219-DEL-1187
chr16-81171148-DEL-2234
chr16-81177070-DEL-4146
chr16-81218269-DEL-149
chr16-81224997-DEL-2957
chr16-81254591-DEL-71
chr16-81462061-DEL-62
chr16-81606024-DEL-67
chr16-81621942-DEL-9653
chr16-81650934-DEL-524
chr16-81696384-DEL-84
chr16-81706404-DEL-582
chr16-81799503-DEL-184
chr16-81866127-DEL-2891
chr16-81898090-DEL-1667
chr16-82002477-DEL-143
chr16-82070587-DEL-1592
chr16-82082987-DEL-263
chr16-82304697-DEL-212
chr16-82395706-DEL-292
chr16-82521834-DEL-2395
chr16-82588033-DEL-65
chr16-82601069-DEL-179
chr16-82715311-DEL-204
chr16-82742615-DEL-78
chr16-82848295-DEL-113
chr16-82855145-DEL-68
chr16-83093411-DEL-85
chr16-83150632-DEL-1639
chr16-83338036-DEL-137
chr16-83480704-DEL-50
chr16-83502793-DEL-69
chr16-83520860-DEL-60
chr16-83553088-DEL-66
chr16-83614052-DEL-1619
chr16-83697515-DEL-256
chr16-83763366-DEL-78
chr16-83859308-DEL-1295
chr16-84034543-DEL-50
chr16-84161977-DEL-297
chr16-84162202-DEL-69
chr16-84193420-DEL-138
chr16-84203003-DEL-1106
chr16-84236630-DEL-201
chr16-84242922-DEL-1660
chr16-84339374-DEL-68
chr16-84361212-DEL-176
chr16-84369726-DEL-87
chr16-84370355-DEL-197
chr16-84417713-DEL-76
chr16-84464014-DEL-385
chr16-84464689-DEL-54
chr16-84537293-DEL-50
chr16-84541043-DEL-51
chr16-84565507-DEL-98
chr16-84565826-DEL-230
chr16-84659814-DEL-153
chr16-84694418-DEL-51
chr16-84738988-DEL-212

chr16-84771996-DEL-54
chr16-84779539-DEL-1155
chr16-84877097-DEL-69
chr16-84909751-DEL-86
chr16-84935362-DEL-50
chr16-84958321-DEL-247
chr16-84958656-DEL-72
chr16-84958902-DEL-66
chr16-85117428-DEL-200
chr16-85208357-DEL-1168
chr16-85232831-DEL-124
chr16-85246540-DEL-2548
chr16-85267917-DEL-140
chr16-85422829-DEL-107
chr16-85437160-DEL-333
chr16-85491614-DEL-57
chr16-85497528-DEL-57
chr16-85509427-DEL-298
chr16-85578966-DEL-1463
chr16-85586154-DEL-64
chr16-85616038-DEL-82
chr16-85620108-DEL-103
chr16-85620216-DEL-1334
chr16-85785300-DEL-949
chr16-85797592-DEL-182
chr16-86027138-DEL-6718
chr16-86027952-DEL-1551
chr16-86100360-DEL-581
chr16-86190945-DEL-52
chr16-86670875-DEL-210
chr16-86814545-DEL-50
chr16-86962150-DEL-400
chr16-87090206-DEL-93
chr16-87106074-DEL-82
chr16-87158222-DEL-392
chr16-87169333-DEL-1715
chr16-87183005-DEL-139
chr16-87183068-DEL-479
chr16-87253608-DEL-337
chr16-87305511-DEL-206
chr16-87305518-DEL-711
chr16-87305919-DEL-250
chr16-87305968-DEL-55
chr16-87306329-DEL-320
chr16-87306985-DEL-130
chr16-87357247-DEL-1128
chr16-87389348-DEL-63
chr16-87389418-DEL-165
chr16-87397040-DEL-242
chr16-87397430-DEL-279
chr16-87445961-DEL-72
chr16-87446299-DEL-75
chr16-87539219-DEL-55
chr16-87747984-DEL-58
chr16-87778265-DEL-69
chr16-87802125-DEL-420
chr16-88029422-DEL-60
chr16-88043283-DEL-60
chr16-88189156-DEL-604
chr16-88236338-DEL-188
chr16-88236454-DEL-92
chr16-88242187-DEL-52
chr16-88267291-DEL-76
chr16-88316665-DEL-54
chr16-88316844-DEL-586
chr16-88321512-DEL-1640
chr16-88362893-DEL-111
chr16-88522326-DEL-50
chr16-88530560-DEL-109
chr16-88723833-DEL-660
chr16-88806214-DEL-120
chr16-88853675-DEL-2038
chr16-88880288-DEL-63
chr16-88950855-DEL-74
chr16-88981035-DEL-3034
chr16-89017455-DEL-5007
chr16-89108850-DEL-164

chr16-89159158-DEL-301
chr16-89168169-DEL-1274
chr16-89207547-DEL-295
chr16-89224630-DEL-57
chr16-89388499-DEL-57
chr16-89433476-DEL-214
chr16-89472756-DEL-50
chr16-89500782-DEL-70
chr16-89528535-DEL-52
chr16-89538165-DEL-54
chr16-89557938-DEL-57
chr16-89658982-DEL-257
chr16-89675536-DEL-200
chr16-89702389-DEL-292
chr16-89748401-DEL-380
chr16-89760141-DEL-550
chr16-89767468-DEL-106
chr16-89921105-DEL-1912
chr16-89930133-DEL-123
chr16-89956091-DEL-112
chr16-89974752-DEL-207
chr16-89979085-DEL-239
chr16-89985722-DEL-545
chr16-90069794-DEL-64
chr16-90090070-DEL-244
chr16-90093737-DEL-79
chr16-90098082-DEL-183
chr16-90152594-DEL-237
chr16-90163220-DEL-142
chr16-90225065-DEL-52
chr16-90245687-DEL-332
chr16-90369057-DEL-3812
chr16-90425793-DEL-62
chr16-90576900-DEL-65
chr16-90603533-DEL-95
chr16-90603745-DEL-380
chr16-90604159-DEL-95
chr16-90691788-DEL-128
chr16-90699422-DEL-66
chr16-90737655-DEL-112
chr16-90873018-DEL-56
chr16-90884074-DEL-981
chr16-90908473-DEL-451
chr16-90928369-DEL-101
chr16-90963486-DEL-65
chr16-91036859-DEL-110
chr16-91036970-DEL-52
chr16-91061946-DEL-166
chr16-91062286-DEL-235
chr16-91146693-DEL-71
chr16-91214737-DEL-105
chr16-91218107-DEL-114
chr16-91235578-DEL-60
chr16-91251673-DEL-931
chr16-91331466-DEL-158
chr16-91355848-DEL-220
chr16-91364170-DEL-232
chr16-91543913-DEL-180
chr16-91546845-DEL-429
chr16-91595870-DEL-84
chr16-91621372-DEL-73
chr16-91749952-DEL-132
chr16-91782499-DEL-372
chr16-91850920-DEL-230
chr16-91851264-DEL-117
chr16-91890709-DEL-427
chr16-91934421-DEL-134
chr16-91961278-DEL-57
chr16-92042404-DEL-98
chr16-92045634-DEL-153
chr16-92045685-DEL-51
chr16-92051653-DEL-952
chr16-92052077-DEL-422
chr16-92052342-DEL-579
chr16-92053030-DEL-212
chr16-92120524-DEL-254
chr16-92142641-DEL-60

chr16-92144506-DEL-256
chr16-92144887-DEL-96
chr16-92255453-DEL-50
chr16-92310625-DEL-75
chr16-92318483-DEL-64
chr16-92324587-DEL-50
chr16-92358723-DEL-88
chr16-92358959-DEL-134
chr16-92366964-DEL-62
chr16-92511293-DEL-72
chr16-92554994-DEL-73
chr16-92571151-DEL-234
chr16-92662645-DEL-185
chr16-92663328-DEL-109
chr16-92675807-DEL-172
chr16-92676094-DEL-100
chr16-92735956-DEL-111
chr16-92804398-DEL-51
chr16-92837429-DEL-58
chr16-92890427-DEL-50
chr16-92914647-DEL-128
chr16-92933249-DEL-321
chr16-93088419-DEL-70
chr16-93106095-DEL-72
chr16-93186652-DEL-72
chr16-93243990-DEL-50
chr16-93261048-DEL-368
chr16-93275209-DEL-88
chr16-93276757-DEL-215
chr16-93307337-DEL-343
chr16-93320863-DEL-52
chr16-93322803-DEL-194
chr16-93376656-DEL-146
chr16-93376885-DEL-164
chr16-93434429-DEL-63
chr16-93443682-DEL-89
chr16-93451608-DEL-404
chr16-93510325-DEL-483
chr16-93523624-DEL-370
chr16-93569095-DEL-306
chr16-93569527-DEL-132
chr16-93569577-DEL-54
chr16-93575125-DEL-200
chr16-93592198-DEL-132
chr16-93710172-DEL-120
chr16-93724306-DEL-139
chr16-93742961-DEL-598
chr16-93743671-DEL-104
chr16-93789306-DEL-3017
chr16-93791575-DEL-1560
chr16-93792292-DEL-1938
chr16-93796686-DEL-734
chr16-93811617-DEL-122
chr16-93842995-DEL-75
chr16-93862063-DEL-436
chr16-93916637-DEL-103
chr16-94072525-DEL-367
chr16-94100700-DEL-111
chr16-94125277-DEL-55
chr16-94134642-DEL-210
chr16-94134642-DEL-455
chr16-94152330-DEL-1459
chr16-94430869-DEL-52
chr16-94470009-DEL-1405
chr16-94499251-DEL-319
chr16-94499686-DEL-145
chr16-94500056-DEL-87
chr16-94542220-DEL-110
chr16-94542421-DEL-77
chr16-94542616-DEL-110
chr16-94576290-DEL-93
chr16-94671121-DEL-94
chr16-94681398-DEL-50
chr16-94682399-DEL-50
chr16-94724711-DEL-420
chr16-94760341-DEL-305
chr16-94849945-DEL-205

chr16-94863774-DEL-110
chr16-94891320-DEL-84
chr16-94891531-DEL-148
chr16-94891775-DEL-132
chr16-94897985-DEL-151
chr16-94910477-DEL-84
chr16-94932002-DEL-55
chr16-94983903-DEL-86
chr16-95055895-DEL-301
chr16-95056675-DEL-60
chr16-95111537-DEL-50
chr16-95156827-DEL-80
chr16-95192996-DEL-50
chr16-95287969-DEL-70
chr16-95525270-DEL-71
chr16-95543812-DEL-55
chr16-95575408-DEL-152
chr16-95618098-DEL-58
chr16-95694763-DEL-84
chr16-95732055-DEL-64
chr16-95806784-DEL-218
chr16-95813925-DEL-59
chr16-95942772-DEL-54
chr16-95963921-DEL-94
chr16-96005245-DEL-152
chr16-96005571-DEL-332
chr16-96008013-DEL-70
chr16-96026563-DEL-208
chr16-96079503-DEL-77
chr16-96106769-DEL-52
chr16-96148165-DEL-77
chr16-96153727-DEL-58
chr16-96177103-DEL-85
chr16-96177351-DEL-136
chr16-96192884-DEL-84
chr16-96193125-DEL-1229
chr16-96221833-DEL-555
chr16-96524437-DEL-134
chr16-96524814-DEL-80
chr16-96598090-DEL-61
chr16-96733164-DEL-146
chr16-96734397-DEL-498
chr16-96757690-DEL-140
chr16-96757827-DEL-112
chr16-97441664-DEL-64
chr16-97461280-DEL-644
chr16-97736196-DEL-62
chr17-3053020-DEL-57
chr17-3122469-DEL-2626
chr17-3239343-DEL-143
chr17-3382613-DEL-50
chr17-3399239-DEL-65
chr17-3434659-DEL-78
chr17-3435046-DEL-162
chr17-3499638-DEL-304
chr17-3499760-DEL-134
chr17-3506490-DEL-64
chr17-3540392-DEL-82
chr17-3544768-DEL-124
chr17-3622149-DEL-1303
chr17-3623411-DEL-198
chr17-3623445-DEL-73
chr17-3643004-DEL-96
chr17-3655247-DEL-72
chr17-3735207-DEL-195
chr17-3824888-DEL-90
chr17-3954976-DEL-60
chr17-4013280-DEL-328
chr17-4290451-DEL-1465
chr17-4427149-DEL-284
chr17-4427809-DEL-130
chr17-4456478-DEL-109
chr17-4493094-DEL-57
chr17-4557095-DEL-56
chr17-4569691-DEL-127
chr17-4575754-DEL-50
chr17-4587286-DEL-238

chr17-4662373-DEL-2592
chr17-4798125-DEL-1377
chr17-4844813-DEL-78
chr17-4846082-DEL-78
chr17-4861419-DEL-4979
chr17-4861520-DEL-430
chr17-4865550-DEL-424
chr17-4865808-DEL-114
chr17-4987358-DEL-1147
chr17-5035935-DEL-114
chr17-5061824-DEL-106
chr17-5069794-DEL-180
chr17-5070064-DEL-92
chr17-5085811-DEL-95
chr17-5088749-DEL-1224
chr17-5143654-DEL-92
chr17-5196392-DEL-538
chr17-5196419-DEL-1827
chr17-5197478-DEL-1553
chr17-5198787-DEL-243
chr17-5208840-DEL-230
chr17-5208840-DEL-736
chr17-5217125-DEL-51
chr17-5235377-DEL-50
chr17-5264021-DEL-54
chr17-5291312-DEL-99
chr17-5291660-DEL-264
chr17-5303566-DEL-134
chr17-5322958-DEL-74
chr17-5343399-DEL-338
chr17-5406253-DEL-504
chr17-5406335-DEL-112
chr17-5494590-DEL-133
chr17-5494624-DEL-271
chr17-5494975-DEL-195
chr17-5495257-DEL-98
chr17-5508783-DEL-56
chr17-5521487-DEL-50
chr17-5633642-DEL-52
chr17-5643224-DEL-124
chr17-5669935-DEL-1060
chr17-5672265-DEL-66
chr17-5717405-DEL-124
chr17-5732674-DEL-81
chr17-5753416-DEL-145
chr17-5755659-DEL-1565
chr17-5768385-DEL-264
chr17-5785763-DEL-1883
chr17-5791624-DEL-254
chr17-5795944-DEL-130
chr17-5799434-DEL-114
chr17-5807448-DEL-61
chr17-5812264-DEL-208
chr17-5818720-DEL-535
chr17-5823522-DEL-5721
chr17-5850442-DEL-63
chr17-5861557-DEL-184
chr17-5885938-DEL-83
chr17-5901403-DEL-102
chr17-5914346-DEL-837
chr17-5914498-DEL-1716
chr17-5914605-DEL-398
chr17-5914852-DEL-75
chr17-5915367-DEL-538
chr17-5924716-DEL-56
chr17-5930611-DEL-3662
chr17-5933574-DEL-1657
chr17-5938417-DEL-189
chr17-5942352-DEL-77
chr17-5960225-DEL-274
chr17-5999791-DEL-82
chr17-6011910-DEL-145
chr17-6064692-DEL-78
chr17-6259544-DEL-112
chr17-6319114-DEL-300
chr17-6445262-DEL-266
chr17-6764850-DEL-88

chr17-6795583-DEL-258
chr17-6796216-DEL-194
chr17-6901052-DEL-65
chr17-6923773-DEL-106
chr17-6930337-DEL-9430
chr17-6950327-DEL-1719
chr17-6957207-DEL-53
chr17-6960374-DEL-1368
chr17-7105915-DEL-4849
chr17-7202197-DEL-2423
chr17-7224804-DEL-249
chr17-7239319-DEL-139
chr17-7251905-DEL-134
chr17-7257799-DEL-72
chr17-7356868-DEL-100
chr17-7356868-DEL-225
chr17-7358726-DEL-62
chr17-7620967-DEL-51
chr17-7742247-DEL-239
chr17-7818732-DEL-95
chr17-7823256-DEL-167
chr17-7865061-DEL-432
chr17-7919092-DEL-325
chr17-7919139-DEL-68
chr17-7919370-DEL-492
chr17-7935648-DEL-169
chr17-7962383-DEL-258
chr17-7985858-DEL-226
chr17-7993136-DEL-255
chr17-7997752-DEL-134
chr17-8003886-DEL-89
chr17-8003886-DEL-223
chr17-8004384-DEL-54
chr17-8033741-DEL-109
chr17-8126385-DEL-295
chr17-8126526-DEL-67
chr17-8127633-DEL-569
chr17-8128623-DEL-312
chr17-8128633-DEL-123
chr17-8130563-DEL-506
chr17-8130699-DEL-137
chr17-8174987-DEL-225
chr17-8175449-DEL-50
chr17-8175462-DEL-350
chr17-8360899-DEL-138
chr17-8429661-DEL-50
chr17-8565707-DEL-165
chr17-8565903-DEL-421
chr17-8566293-DEL-106
chr17-8583244-DEL-122
chr17-8604803-DEL-50
chr17-8763043-DEL-73
chr17-8847589-DEL-132
chr17-8955735-DEL-144
chr17-9044242-DEL-105
chr17-9061434-DEL-150
chr17-9071910-DEL-68
chr17-9398601-DEL-309
chr17-9452134-DEL-70
chr17-9457251-DEL-90
chr17-9457466-DEL-432
chr17-9615791-DEL-50
chr17-9622447-DEL-54
chr17-9806571-DEL-1492
chr17-9895106-DEL-50
chr17-9898831-DEL-629
chr17-9899224-DEL-243
chr17-9963674-DEL-218
chr17-10074323-DEL-70
chr17-10214938-DEL-99
chr17-10363732-DEL-127
chr17-10542880-DEL-75
chr17-10628471-DEL-62
chr17-10663845-DEL-986
chr17-10684968-DEL-87
chr17-10809703-DEL-135
chr17-10866629-DEL-70

chr17-11012796-DEL-141
chr17-11063026-DEL-53
chr17-11099460-DEL-1565
chr17-11268175-DEL-79
chr17-11288003-DEL-154
chr17-11329769-DEL-140
chr17-11355361-DEL-1612
chr17-11382558-DEL-703
chr17-11386257-DEL-86
chr17-11451431-DEL-134
chr17-11513772-DEL-4145
chr17-11544039-DEL-66
chr17-11546972-DEL-275
chr17-11547112-DEL-100
chr17-11547743-DEL-300
chr17-11574080-DEL-61
chr17-11574188-DEL-160
chr17-11603797-DEL-206
chr17-11610653-DEL-1363
chr17-11620096-DEL-60
chr17-11817945-DEL-57
chr17-11825430-DEL-50
chr17-11878371-DEL-64
chr17-11904632-DEL-60
chr17-12004212-DEL-50
chr17-12046787-DEL-2981
chr17-12235885-DEL-624
chr17-12442303-DEL-3202
chr17-12443336-DEL-93
chr17-12453604-DEL-70
chr17-12464478-DEL-935
chr17-12465204-DEL-75
chr17-12465212-DEL-334
chr17-12465471-DEL-75
chr17-12502576-DEL-4491
chr17-12602372-DEL-63
chr17-12666241-DEL-271
chr17-12667975-DEL-1978
chr17-12690562-DEL-185
chr17-12730946-DEL-90
chr17-12776290-DEL-204
chr17-12808692-DEL-57
chr17-12823449-DEL-204
chr17-12854962-DEL-1116
chr17-12854975-DEL-63
chr17-12855364-DEL-322
chr17-12855818-DEL-191
chr17-13100912-DEL-50
chr17-13116993-DEL-51
chr17-13117230-DEL-393
chr17-13117289-DEL-50
chr17-13151364-DEL-97
chr17-13195663-DEL-170
chr17-13240924-DEL-53
chr17-13241138-DEL-58
chr17-13241296-DEL-95
chr17-13285529-DEL-100
chr17-13289514-DEL-103
chr17-13396144-DEL-56
chr17-13580548-DEL-993
chr17-13680747-DEL-56
chr17-13760479-DEL-50
chr17-13899181-DEL-336
chr17-14074888-DEL-379
chr17-14180658-DEL-401
chr17-14205384-DEL-60
chr17-14215477-DEL-58256
chr17-14219251-DEL-342
chr17-14413965-DEL-50
chr17-14445302-DEL-1131
chr17-14449651-DEL-87
chr17-14471381-DEL-183
chr17-14523907-DEL-80
chr17-14570442-DEL-60
chr17-14665606-DEL-166
chr17-14665650-DEL-64
chr17-14674365-DEL-102

chr17-14674754-DEL-248
chr17-14692371-DEL-54
chr17-14897463-DEL-74
chr17-14928255-DEL-52
chr17-14943804-DEL-50
chr17-15023575-DEL-1629
chr17-15181071-DEL-20463
chr17-15181071-DEL-62992
chr17-15263794-DEL-11449
chr17-15706302-DEL-3184
chr17-15706537-DEL-1101
chr17-15707891-DEL-197
chr17-15708976-DEL-70
chr17-15709602-DEL-52
chr17-15776928-DEL-108
chr17-15806400-DEL-206
chr17-15806564-DEL-50
chr17-15943022-DEL-114
chr17-16023285-DEL-83
chr17-16073596-DEL-1554
chr17-16237432-DEL-258
chr17-16364662-DEL-90
chr17-16562196-DEL-141
chr17-16664718-DEL-51
chr17-16727754-DEL-578
chr17-16845398-DEL-78
chr17-16948953-DEL-1379
chr17-17248244-DEL-76
chr17-17308913-DEL-336
chr17-17309521-DEL-85
chr17-17365079-DEL-156
chr17-17365960-DEL-504
chr17-17366841-DEL-342
chr17-17376411-DEL-8300
chr17-17390298-DEL-306
chr17-17515118-DEL-2373
chr17-17725589-DEL-376
chr17-17725812-DEL-611
chr17-17774853-DEL-212
chr17-17796652-DEL-987
chr17-17966786-DEL-1873
chr17-17974487-DEL-50
chr17-18101219-DEL-50
chr17-18155745-DEL-82
chr17-18223199-DEL-54
chr17-18310277-DEL-5491
chr17-18416676-DEL-130
chr17-18552365-DEL-60
chr17-18657909-DEL-73
chr17-19325842-DEL-1992
chr17-19357276-DEL-56
chr17-19638880-DEL-67
chr17-19663698-DEL-53
chr17-19691721-DEL-103
chr17-19768502-DEL-2125
chr17-19812953-DEL-7264
chr17-19942495-DEL-27708
chr17-20034982-DEL-328
chr17-20044157-DEL-510
chr17-20111033-DEL-4477
chr17-20203249-DEL-74
chr17-20264052-DEL-240
chr17-20271273-DEL-1349
chr17-20508647-DEL-340
chr17-20539041-DEL-58
chr17-20541732-DEL-335
chr17-20548021-DEL-400
chr17-20556871-DEL-1227
chr17-20559848-DEL-631
chr17-20562203-DEL-2173
chr17-20603760-DEL-488
chr17-20632238-DEL-370
chr17-20676914-DEL-60
chr17-20717991-DEL-72
chr17-20726461-DEL-76
chr17-20767166-DEL-301
chr17-20822195-DEL-50

chr17-20842644-DEL-51
chr17-20880726-DEL-88
chr17-20885321-DEL-54
chr17-20933648-DEL-599
chr17-20967846-DEL-8127
chr17-21047644-DEL-73
chr17-21116924-DEL-85
chr17-21183134-DEL-2297
chr17-21220289-DEL-102
chr17-21393905-DEL-23168
chr17-21621089-DEL-91
chr17-21829318-DEL-421
chr17-21836685-DEL-1788
chr17-21913143-DEL-1634
chr17-21948819-DEL-222
chr17-22268284-DEL-55
chr17-22272188-DEL-781
chr17-22369275-DEL-446
chr17-22436527-DEL-116
chr17-22581391-DEL-625
chr17-22775610-DEL-2012
chr17-22804434-DEL-5604
chr17-22822791-DEL-402
chr17-22846573-DEL-102
chr17-22921742-DEL-816
chr17-22924567-DEL-1164
chr17-22948094-DEL-84
chr17-22952931-DEL-67
chr17-23124999-DEL-4352
chr17-23142895-DEL-12720
chr17-23201783-DEL-148033
chr17-23208051-DEL-68619
chr17-23354262-DEL-279
chr17-23415106-DEL-66
chr17-23445012-DEL-860
chr17-23470483-DEL-8561
chr17-23682274-DEL-198
chr17-23790729-DEL-206
chr17-23813066-DEL-391
chr17-23844240-DEL-958
chr17-23920479-DEL-328
chr17-24005562-DEL-165
chr17-24125153-DEL-51
chr17-24128203-DEL-316
chr17-24168644-DEL-784
chr17-24208073-DEL-86
chr17-24250752-DEL-53
chr17-24252503-DEL-203
chr17-24253766-DEL-58
chr17-24261167-DEL-58
chr17-24315023-DEL-253
chr17-24315746-DEL-59
chr17-24384916-DEL-110
chr17-24399026-DEL-126
chr17-24451650-DEL-53
chr17-24474452-DEL-80
chr17-24474595-DEL-680
chr17-24474680-DEL-201
chr17-24475072-DEL-200
chr17-24475633-DEL-680
chr17-24476036-DEL-200
chr17-24487153-DEL-331
chr17-24678718-DEL-1491
chr17-24681293-DEL-297
chr17-24743608-DEL-96
chr17-24758699-DEL-75
chr17-24759019-DEL-216
chr17-25253779-DEL-52
chr17-25265542-DEL-451
chr17-25280157-DEL-50
chr17-25316466-DEL-50
chr17-25316921-DEL-373
chr17-25414259-DEL-85
chr17-25544914-DEL-105
chr17-25583571-DEL-52
chr17-25670767-DEL-60
chr17-25761252-DEL-199

chr17-25764228-DEL-56
chr17-25779278-DEL-327
chr17-25779386-DEL-74
chr17-25819394-DEL-167
chr17-25870563-DEL-195
chr17-25870860-DEL-300
chr17-25877144-DEL-538
chr17-25878043-DEL-95
chr17-25893180-DEL-76
chr17-25919096-DEL-253
chr17-26131742-DEL-185
chr17-26147100-DEL-102
chr17-26151314-DEL-436
chr17-26151640-DEL-55
chr17-26193998-DEL-50
chr17-26266990-DEL-5270
chr17-26284532-DEL-192
chr17-26293780-DEL-96
chr17-26301410-DEL-50
chr17-26305655-DEL-331
chr17-26326144-DEL-1503
chr17-26433819-DEL-100
chr17-26510524-DEL-6867
chr17-26527069-DEL-68
chr17-26538973-DEL-90
chr17-26541730-DEL-249
chr17-26541898-DEL-108
chr17-26543169-DEL-86
chr17-26549442-DEL-206
chr17-26549718-DEL-273
chr17-26567542-DEL-69
chr17-26665795-DEL-55
chr17-26691237-DEL-216
chr17-26767063-DEL-123
chr17-26767327-DEL-61
chr17-26767360-DEL-221
chr17-26784758-DEL-164
chr17-26784764-DEL-73
chr17-26852033-DEL-87
chr17-26983486-DEL-276
chr17-26983751-DEL-138
chr17-27077367-DEL-140
chr17-27077636-DEL-50
chr17-27080292-DEL-52
chr17-27311084-DEL-60
chr17-27311511-DEL-96
chr17-27365337-DEL-50
chr17-27384946-DEL-170
chr17-27385184-DEL-68
chr17-27402284-DEL-244
chr17-27402288-DEL-61
chr17-27402396-DEL-81
chr17-27748568-DEL-203
chr17-27751955-DEL-440
chr17-27755081-DEL-276
chr17-27759802-DEL-60
chr17-27818381-DEL-342
chr17-27834226-DEL-64
chr17-27892460-DEL-153
chr17-27918579-DEL-66
chr17-27918796-DEL-65
chr17-27918975-DEL-99
chr17-27948247-DEL-102
chr17-27963066-DEL-52
chr17-28028481-DEL-363
chr17-28180884-DEL-1445
chr17-28371501-DEL-69
chr17-28511899-DEL-50
chr17-28577752-DEL-146
chr17-28577782-DEL-54
chr17-28589721-DEL-210
chr17-28673363-DEL-56
chr17-28693626-DEL-97
chr17-28828056-DEL-351
chr17-28834002-DEL-191
chr17-29001139-DEL-1044
chr17-29067470-DEL-318

chr17-29199768-DEL-254
chr17-29220360-DEL-149
chr17-29273872-DEL-63
chr17-29296669-DEL-97
chr17-29301696-DEL-172
chr17-29320025-DEL-1371
chr17-29320066-DEL-5486
chr17-29320240-DEL-67
chr17-29322513-DEL-2162
chr17-29322943-DEL-3988
chr17-29351802-DEL-152
chr17-29417144-DEL-676
chr17-29417241-DEL-262
chr17-29505338-DEL-82
chr17-29511968-DEL-189
chr17-29553363-DEL-59
chr17-29553511-DEL-181
chr17-29627700-DEL-68
chr17-29704075-DEL-78
chr17-29735174-DEL-72
chr17-30081740-DEL-279
chr17-30160896-DEL-96
chr17-30189493-DEL-54
chr17-30302080-DEL-435
chr17-30308100-DEL-74
chr17-30359331-DEL-622
chr17-30476175-DEL-238
chr17-30520963-DEL-182
chr17-30521101-DEL-720
chr17-30521653-DEL-120
chr17-30593310-DEL-743
chr17-30624785-DEL-64
chr17-30712241-DEL-113
chr17-30730942-DEL-110
chr17-30843814-DEL-2521
chr17-30873988-DEL-511
chr17-30886101-DEL-53
chr17-30904466-DEL-108
chr17-30935147-DEL-66
chr17-30982100-DEL-56
chr17-31005572-DEL-96
chr17-31011986-DEL-338
chr17-31017689-DEL-144
chr17-31084469-DEL-50
chr17-31115420-DEL-101
chr17-31115860-DEL-101
chr17-31136903-DEL-67
chr17-31137480-DEL-171
chr17-31211851-DEL-67
chr17-31212037-DEL-55
chr17-31214830-DEL-64
chr17-31430530-DEL-105
chr17-31510467-DEL-99
chr17-31510467-DEL-231
chr17-31578314-DEL-60
chr17-31659566-DEL-58
chr17-31682456-DEL-83
chr17-31696811-DEL-1658
chr17-31698499-DEL-80
chr17-31756747-DEL-58
chr17-31756910-DEL-73
chr17-31775120-DEL-50
chr17-31920582-DEL-306
chr17-32041770-DEL-119
chr17-32085661-DEL-96
chr17-32119955-DEL-66
chr17-32236850-DEL-73
chr17-32430290-DEL-426
chr17-32518180-DEL-1317
chr17-32624850-DEL-1090
chr17-32626961-DEL-179
chr17-32640647-DEL-1370
chr17-32641033-DEL-146
chr17-32646572-DEL-81
chr17-32649025-DEL-92
chr17-32649189-DEL-340
chr17-32676320-DEL-82

chr17-32774664-DEL-24598
chr17-32931155-DEL-50
chr17-33047432-DEL-336
chr17-33059082-DEL-60
chr17-33090124-DEL-217
chr17-33090781-DEL-77
chr17-33117625-DEL-326
chr17-33126074-DEL-186
chr17-33145180-DEL-50
chr17-33239460-DEL-72
chr17-33241549-DEL-102
chr17-33376948-DEL-3021
chr17-33670232-DEL-52
chr17-33719219-DEL-96
chr17-33760784-DEL-124
chr17-33822728-DEL-127
chr17-33824179-DEL-82
chr17-34009450-DEL-504
chr17-34009975-DEL-84
chr17-34014875-DEL-121
chr17-34182758-DEL-95
chr17-34183279-DEL-448
chr17-34199620-DEL-181
chr17-34209720-DEL-54
chr17-34286662-DEL-126
chr17-34421054-DEL-60
chr17-34421220-DEL-103
chr17-34421422-DEL-226
chr17-34472917-DEL-154
chr17-34484836-DEL-856
chr17-34492307-DEL-90
chr17-34499672-DEL-371
chr17-34500176-DEL-54
chr17-34549469-DEL-99
chr17-34572569-DEL-224
chr17-34607060-DEL-144
chr17-34681750-DEL-1493
chr17-34698294-DEL-65
chr17-34717140-DEL-223
chr17-34717920-DEL-332
chr17-34717943-DEL-158
chr17-34732296-DEL-112
chr17-34741402-DEL-4342
chr17-34858065-DEL-50
chr17-34957575-DEL-134
chr17-34964666-DEL-79504
chr17-34987543-DEL-176
chr17-35061116-DEL-60
chr17-35253255-DEL-192
chr17-35289566-DEL-50
chr17-35370278-DEL-313
chr17-35489407-DEL-137
chr17-35507442-DEL-80
chr17-35534499-DEL-572
chr17-35641624-DEL-80
chr17-35648815-DEL-326
chr17-35653162-DEL-122750
chr17-35779006-DEL-180
chr17-35799684-DEL-435
chr17-35800174-DEL-140
chr17-35806035-DEL-144
chr17-35806806-DEL-5206
chr17-35842671-DEL-50
chr17-35851393-DEL-58
chr17-35884972-DEL-80
chr17-35930752-DEL-450
chr17-35933035-DEL-972
chr17-35973397-DEL-152
chr17-36014510-DEL-52
chr17-36020830-DEL-196
chr17-36034408-DEL-90
chr17-36129763-DEL-214
chr17-36187321-DEL-456
chr17-36195608-DEL-383
chr17-36202252-DEL-612
chr17-36295609-DEL-50
chr17-36445388-DEL-66

chr17-36450108-DEL-56
chr17-36450290-DEL-260
chr17-36450473-DEL-68
chr17-36464864-DEL-1579
chr17-36504495-DEL-1369
chr17-36506994-DEL-163
chr17-36524472-DEL-83
chr17-36525243-DEL-68
chr17-36544367-DEL-347
chr17-36730524-DEL-689
chr17-36875327-DEL-50
chr17-36883262-DEL-200
chr17-36965964-DEL-29984
chr17-37024036-DEL-2916
chr17-37092487-DEL-617
chr17-37196872-DEL-72
chr17-37336760-DEL-1326
chr17-37345173-DEL-74
chr17-37497273-DEL-62
chr17-37614137-DEL-68
chr17-37622278-DEL-75
chr17-37692903-DEL-879
chr17-38533443-DEL-60
chr17-38686318-DEL-84
chr17-38881729-DEL-1196
chr17-38883984-DEL-66
chr17-38886374-DEL-64
chr17-38895475-DEL-84
chr17-39116412-DEL-393
chr17-39211772-DEL-6514
chr17-39839138-DEL-132
chr17-39929785-DEL-80
chr17-39947616-DEL-429
chr17-39959071-DEL-21086
chr17-39974940-DEL-21001
chr17-40017350-DEL-95
chr17-40079455-DEL-214
chr17-40144414-DEL-120
chr17-40351293-DEL-54
chr17-40426432-DEL-2167
chr17-40630792-DEL-187
chr17-40852055-DEL-427
chr17-40852266-DEL-644
chr17-40866458-DEL-72
chr17-41205535-DEL-206
chr17-41363596-DEL-409
chr17-41420793-DEL-50
chr17-41431405-DEL-84
chr17-41445642-DEL-72
chr17-41445946-DEL-56
chr17-41695569-DEL-398
chr17-42016446-DEL-56
chr17-42174292-DEL-80
chr17-42224177-DEL-1283
chr17-42241591-DEL-957
chr17-42258297-DEL-636
chr17-42267358-DEL-218
chr17-42379728-DEL-4083
chr17-42437666-DEL-595
chr17-42543731-DEL-66
chr17-42551493-DEL-238
chr17-42594269-DEL-62
chr17-42689991-DEL-669
chr17-42856957-DEL-58
chr17-42869884-DEL-1126
chr17-42889266-DEL-850
chr17-42902068-DEL-631
chr17-42915174-DEL-140
chr17-42984804-DEL-494
chr17-43010219-DEL-50
chr17-43013310-DEL-50
chr17-43102865-DEL-243
chr17-43161516-DEL-95
chr17-43210578-DEL-50
chr17-43216715-DEL-90
chr17-43243609-DEL-57
chr17-43243984-DEL-120

chr17-43265264-DEL-216
chr17-43270822-DEL-56
chr17-43278189-DEL-102
chr17-43297618-DEL-57
chr17-43301375-DEL-51
chr17-43361107-DEL-212
chr17-43361249-DEL-78
chr17-43362389-DEL-285
chr17-43405972-DEL-110
chr17-43409499-DEL-842
chr17-43503878-DEL-127
chr17-43596570-DEL-50
chr17-43608390-DEL-50
chr17-43614920-DEL-425
chr17-43615076-DEL-58
chr17-43617046-DEL-285
chr17-43617123-DEL-55
chr17-43626497-DEL-206
chr17-43628725-DEL-64
chr17-43629878-DEL-260
chr17-43801328-DEL-515
chr17-43872198-DEL-179
chr17-43925756-DEL-51
chr17-44054715-DEL-50
chr17-44162100-DEL-1638
chr17-44197973-DEL-54
chr17-44213597-DEL-50
chr17-44295959-DEL-110
chr17-44342960-DEL-50
chr17-44359268-DEL-244
chr17-44419650-DEL-6250
chr17-44431978-DEL-548
chr17-44500251-DEL-50
chr17-44510491-DEL-54
chr17-44542871-DEL-1544
chr17-44711302-DEL-114
chr17-44771690-DEL-60
chr17-44805979-DEL-65
chr17-44864950-DEL-55
chr17-44913045-DEL-273
chr17-44913413-DEL-168
chr17-45163782-DEL-3385
chr17-45257801-DEL-60
chr17-45290303-DEL-50
chr17-45475593-DEL-142
chr17-45475631-DEL-51
chr17-45482419-DEL-50
chr17-45539119-DEL-116
chr17-45592249-DEL-91
chr17-45661607-DEL-132
chr17-45769932-DEL-65
chr17-45790957-DEL-264
chr17-45794180-DEL-50
chr17-45797610-DEL-63
chr17-45831027-DEL-50
chr17-45908554-DEL-82
chr17-46225984-DEL-383
chr17-46251908-DEL-50
chr17-46294294-DEL-90
chr17-46322020-DEL-80
chr17-46406213-DEL-56
chr17-46411657-DEL-54
chr17-46432079-DEL-560
chr17-46444937-DEL-115
chr17-46448210-DEL-121
chr17-46545732-DEL-305
chr17-46546376-DEL-93
chr17-46603801-DEL-54
chr17-46607275-DEL-62
chr17-46622288-DEL-752
chr17-46641361-DEL-123
chr17-46646171-DEL-54
chr17-46744668-DEL-66
chr17-46781985-DEL-934
chr17-46787110-DEL-67
chr17-46821557-DEL-50
chr17-46892469-DEL-51

chr17-46966114-DEL-82
chr17-47077985-DEL-238
chr17-47078104-DEL-68
chr17-47095427-DEL-168
chr17-47105883-DEL-73
chr17-47146553-DEL-57
chr17-47146699-DEL-57
chr17-47210781-DEL-66
chr17-47241586-DEL-60
chr17-47261336-DEL-1109
chr17-47302487-DEL-82
chr17-47345481-DEL-62
chr17-47428108-DEL-1028
chr17-47429629-DEL-1467
chr17-47508265-DEL-74
chr17-47518120-DEL-63
chr17-47518345-DEL-87
chr17-47527271-DEL-60
chr17-47581046-DEL-172
chr17-47680839-DEL-624
chr17-47733744-DEL-50
chr17-47740891-DEL-203
chr17-47814247-DEL-422
chr17-47820674-DEL-150
chr17-47820764-DEL-60
chr17-47893160-DEL-63
chr17-47901288-DEL-54
chr17-47953346-DEL-56
chr17-48067188-DEL-54
chr17-48134837-DEL-81
chr17-48206133-DEL-66
chr17-48224625-DEL-148
chr17-48316473-DEL-156
chr17-48316545-DEL-77
chr17-48394359-DEL-231
chr17-48422501-DEL-102
chr17-48467220-DEL-621
chr17-48475218-DEL-317
chr17-48520709-DEL-55
chr17-48523009-DEL-304
chr17-48526081-DEL-108
chr17-48528347-DEL-52
chr17-48528960-DEL-6543
chr17-48529067-DEL-213
chr17-48710955-DEL-54
chr17-48747876-DEL-125
chr17-48786596-DEL-236
chr17-48792545-DEL-64
chr17-48817222-DEL-160
chr17-48831282-DEL-123
chr17-48952430-DEL-50
chr17-49037166-DEL-506
chr17-49037302-DEL-66
chr17-49044431-DEL-181
chr17-49163411-DEL-86
chr17-49216258-DEL-58
chr17-49218701-DEL-144
chr17-49268408-DEL-116
chr17-49298075-DEL-52
chr17-49319798-DEL-198
chr17-49320203-DEL-234
chr17-49428106-DEL-1137
chr17-49452078-DEL-64
chr17-49455686-DEL-217
chr17-49456322-DEL-188
chr17-49504478-DEL-126
chr17-49504664-DEL-56
chr17-49507091-DEL-96
chr17-49512030-DEL-50
chr17-49522019-DEL-50
chr17-49531464-DEL-72
chr17-49570882-DEL-50
chr17-49587172-DEL-331
chr17-49614957-DEL-82
chr17-49615252-DEL-203
chr17-50031793-DEL-291
chr17-50035867-DEL-171

chr17-50166846-DEL-117
chr17-50224249-DEL-4071
chr17-50232234-DEL-207
chr17-50232935-DEL-223
chr17-50233615-DEL-60
chr17-50233959-DEL-373
chr17-50248057-DEL-86
chr17-50368859-DEL-85
chr17-50408054-DEL-66
chr17-50422863-DEL-80
chr17-50423448-DEL-256
chr17-50539084-DEL-145
chr17-50569406-DEL-50
chr17-50652597-DEL-64
chr17-50652605-DEL-186
chr17-50653011-DEL-214
chr17-50706640-DEL-132
chr17-50791391-DEL-138
chr17-50791461-DEL-64
chr17-50800938-DEL-56
chr17-50836242-DEL-318
chr17-50938312-DEL-126
chr17-50939091-DEL-175
chr17-50960979-DEL-161
chr17-50966517-DEL-56
chr17-51079911-DEL-258
chr17-51241353-DEL-51
chr17-51266700-DEL-485
chr17-51345250-DEL-4470
chr17-51428807-DEL-1940
chr17-51620996-DEL-325
chr17-51621306-DEL-65
chr17-51621534-DEL-66
chr17-51874790-DEL-104
chr17-52021577-DEL-147
chr17-52102825-DEL-85
chr17-52158872-DEL-50
chr17-52206694-DEL-54
chr17-52376718-DEL-85
chr17-52457867-DEL-50
chr17-52554315-DEL-287
chr17-52633322-DEL-2385
chr17-52809931-DEL-237
chr17-52835365-DEL-78
chr17-53094056-DEL-52
chr17-53282411-DEL-184
chr17-53324650-DEL-53
chr17-53530500-DEL-998
chr17-53646039-DEL-1336
chr17-53909200-DEL-5695
chr17-53936663-DEL-69
chr17-54042144-DEL-859
chr17-54078819-DEL-60
chr17-54092369-DEL-933
chr17-54103662-DEL-187
chr17-54167314-DEL-66
chr17-54204111-DEL-665
chr17-54204479-DEL-668
chr17-54318655-DEL-178
chr17-54419842-DEL-416
chr17-54540537-DEL-3302
chr17-54626383-DEL-68
chr17-54648444-DEL-140
chr17-54661633-DEL-215
chr17-54730210-DEL-68
chr17-54750419-DEL-728
chr17-54752056-DEL-50
chr17-54816420-DEL-156
chr17-54829117-DEL-116
chr17-54832623-DEL-90
chr17-54906438-DEL-679
chr17-54950774-DEL-122
chr17-54960596-DEL-72
chr17-54975259-DEL-76
chr17-54984919-DEL-58
chr17-55000168-DEL-1292
chr17-55028000-DEL-3824

chr17-55066578-DEL-2912
chr17-55080969-DEL-426
chr17-55088596-DEL-2085
chr17-55108914-DEL-67
chr17-55147820-DEL-153
chr17-55286250-DEL-56
chr17-55343897-DEL-97
chr17-55362783-DEL-50
chr17-55490732-DEL-2476
chr17-55606800-DEL-6212
chr17-55625507-DEL-140
chr17-55628676-DEL-981
chr17-55640681-DEL-64
chr17-55640859-DEL-156
chr17-55641081-DEL-54
chr17-55659071-DEL-658
chr17-55676509-DEL-796
chr17-55732513-DEL-1113
chr17-55733652-DEL-199
chr17-55778148-DEL-57
chr17-55983217-DEL-4144
chr17-56036977-DEL-3147
chr17-56083165-DEL-1255
chr17-56412297-DEL-497
chr17-56413013-DEL-198
chr17-56413632-DEL-396
chr17-56435858-DEL-143
chr17-56506941-DEL-50
chr17-56690049-DEL-105
chr17-56690734-DEL-188
chr17-56690787-DEL-90
chr17-56717465-DEL-93
chr17-56729702-DEL-446
chr17-56729897-DEL-90
chr17-56808679-DEL-258
chr17-56937003-DEL-858
chr17-57025866-DEL-146
chr17-57114802-DEL-53
chr17-57124118-DEL-298
chr17-57148573-DEL-93
chr17-57232739-DEL-52
chr17-57263494-DEL-114
chr17-57336257-DEL-52
chr17-57392164-DEL-64
chr17-57400136-DEL-927
chr17-57409580-DEL-65
chr17-57425619-DEL-98
chr17-57462808-DEL-924
chr17-57462836-DEL-56
chr17-57491250-DEL-90
chr17-57509569-DEL-54
chr17-57569114-DEL-50
chr17-57602832-DEL-100
chr17-57616547-DEL-114
chr17-57909644-DEL-85
chr17-57983373-DEL-58
chr17-58019554-DEL-829
chr17-58020722-DEL-1249
chr17-58033708-DEL-55
chr17-58101198-DEL-145
chr17-58110816-DEL-474
chr17-58134479-DEL-235
chr17-58227263-DEL-109
chr17-58237493-DEL-268
chr17-58259924-DEL-84
chr17-58322134-DEL-224
chr17-58416357-DEL-1813
chr17-58434265-DEL-1687
chr17-58444453-DEL-431
chr17-58478094-DEL-6558
chr17-58490343-DEL-52
chr17-58609410-DEL-1818
chr17-58732155-DEL-76
chr17-58732273-DEL-159
chr17-58787967-DEL-91
chr17-58874435-DEL-108
chr17-58970087-DEL-3241

chr17-59012790-DEL-56
chr17-59089457-DEL-54
chr17-59093589-DEL-105
chr17-59139025-DEL-95
chr17-59168033-DEL-53
chr17-59174783-DEL-50
chr17-59316399-DEL-50
chr17-59352110-DEL-423
chr17-59369744-DEL-53
chr17-59380670-DEL-56
chr17-59409815-DEL-1770
chr17-59439515-DEL-284
chr17-59702292-DEL-136
chr17-59702349-DEL-367
chr17-59774289-DEL-54
chr17-59810508-DEL-2094
chr17-59866341-DEL-61
chr17-59977931-DEL-1048
chr17-59984455-DEL-427
chr17-59988552-DEL-74
chr17-60242785-DEL-50
chr17-60264662-DEL-64
chr17-60266702-DEL-172
chr17-60297323-DEL-468
chr17-60370565-DEL-238
chr17-60387966-DEL-54
chr17-60388427-DEL-391
chr17-60393321-DEL-300
chr17-60414782-DEL-94
chr17-60501449-DEL-87
chr17-60516341-DEL-63
chr17-60518761-DEL-620
chr17-60657739-DEL-93
chr17-60710276-DEL-82
chr17-60715223-DEL-272
chr17-60783652-DEL-1963
chr17-60813134-DEL-877
chr17-60842891-DEL-2576
chr17-60843513-DEL-60
chr17-60848246-DEL-231
chr17-60858605-DEL-534
chr17-61038900-DEL-172
chr17-61164196-DEL-94
chr17-61185357-DEL-50
chr17-61254653-DEL-279
chr17-61303817-DEL-1701
chr17-61586149-DEL-492
chr17-61626412-DEL-50
chr17-61635104-DEL-6004
chr17-61668863-DEL-148
chr17-61712493-DEL-473
chr17-61796872-DEL-63
chr17-61799923-DEL-848
chr17-61961536-DEL-207
chr17-62212677-DEL-144
chr17-62221754-DEL-314
chr17-62279270-DEL-79
chr17-62519766-DEL-53
chr17-62533611-DEL-58
chr17-62533815-DEL-84
chr17-62614351-DEL-130
chr17-62621373-DEL-66
chr17-62622016-DEL-1005
chr17-62650063-DEL-1170
chr17-62695419-DEL-1973
chr17-62707643-DEL-1493
chr17-62870463-DEL-230
chr17-62975050-DEL-203
chr17-62998675-DEL-54
chr17-63220973-DEL-997
chr17-63357990-DEL-87
chr17-63428855-DEL-78
chr17-63574182-DEL-121
chr17-63598647-DEL-364
chr17-63598681-DEL-104
chr17-63703314-DEL-144
chr17-63732774-DEL-1013

chr17-63867410-DEL-308
chr17-63909766-DEL-50
chr17-63948979-DEL-1056
chr17-63956870-DEL-172
chr17-63999577-DEL-324
chr17-64003471-DEL-50
chr17-64034103-DEL-277
chr17-64182127-DEL-69
chr17-64184699-DEL-181
chr17-64219449-DEL-189
chr17-64243570-DEL-54
chr17-64256411-DEL-64
chr17-64396604-DEL-50
chr17-64448892-DEL-285
chr17-64478337-DEL-54
chr17-64540377-DEL-497
chr17-64628561-DEL-58
chr17-64696146-DEL-873
chr17-64707455-DEL-81
chr17-64735049-DEL-902
chr17-64759569-DEL-365
chr17-64765066-DEL-76
chr17-64780035-DEL-60
chr17-64780231-DEL-172
chr17-64872057-DEL-116
chr17-64894733-DEL-165
chr17-64895109-DEL-172
chr17-64946577-DEL-244
chr17-64955830-DEL-352
chr17-65098053-DEL-50
chr17-65181805-DEL-81
chr17-65227488-DEL-81
chr17-65306224-DEL-83
chr17-65493777-DEL-51
chr17-65588465-DEL-165
chr17-65621921-DEL-3480
chr17-65767421-DEL-225
chr17-65868992-DEL-218
chr17-65869212-DEL-256
chr17-65872197-DEL-946
chr17-65886478-DEL-112
chr17-65963790-DEL-116
chr17-65988332-DEL-50
chr17-66013892-DEL-52
chr17-66015113-DEL-94
chr17-66036174-DEL-52
chr17-66042675-DEL-66
chr17-66069674-DEL-50
chr17-66154128-DEL-424
chr17-66228129-DEL-51
chr17-66235938-DEL-76
chr17-66250302-DEL-158
chr17-66336631-DEL-943
chr17-66410546-DEL-125
chr17-66493907-DEL-91
chr17-66537520-DEL-123
chr17-66537864-DEL-102
chr17-66538150-DEL-57
chr17-66571154-DEL-79
chr17-66671420-DEL-56
chr17-66679259-DEL-185
chr17-66761072-DEL-61
chr17-66761193-DEL-118
chr17-66767437-DEL-191
chr17-66799785-DEL-144
chr17-66842250-DEL-278
chr17-66842872-DEL-205
chr17-66852803-DEL-966
chr17-66921811-DEL-122
chr17-66921985-DEL-343
chr17-66922265-DEL-118
chr17-66953687-DEL-165
chr17-66953921-DEL-62
chr17-66954755-DEL-50
chr17-67039587-DEL-808
chr17-67152858-DEL-546
chr17-67281322-DEL-68

chr17-67347195-DEL-58
chr17-67368150-DEL-64
chr17-67431881-DEL-210
chr17-67443104-DEL-59
chr17-67537562-DEL-492
chr17-67538081-DEL-88
chr17-67557228-DEL-96
chr17-67564250-DEL-58
chr17-67564411-DEL-338
chr17-67587399-DEL-1016
chr17-67663950-DEL-125
chr17-67726694-DEL-132
chr17-67726703-DEL-290
chr17-67747317-DEL-94
chr17-67787569-DEL-86
chr17-67822852-DEL-64
chr17-67907141-DEL-90
chr17-68148396-DEL-65
chr17-68148529-DEL-137
chr17-68152882-DEL-74
chr17-68167345-DEL-137
chr17-68212375-DEL-66
chr17-68278776-DEL-114
chr17-68284742-DEL-442
chr17-68299355-DEL-162
chr17-68300646-DEL-274
chr17-68550912-DEL-224
chr17-68585602-DEL-60
chr17-68591275-DEL-53
chr17-68615624-DEL-636
chr17-68627302-DEL-78
chr17-68857268-DEL-106
chr17-68884274-DEL-72
chr17-69012097-DEL-59
chr17-69012468-DEL-147
chr17-69012826-DEL-146
chr17-69067081-DEL-715
chr17-69118239-DEL-67
chr17-69162039-DEL-90
chr17-69208254-DEL-56
chr17-69248755-DEL-6742
chr17-69331447-DEL-69
chr17-69450846-DEL-180
chr17-69451038-DEL-70
chr17-69486025-DEL-159
chr17-69486116-DEL-54
chr17-69571658-DEL-1118
chr17-69604048-DEL-56
chr17-69605261-DEL-525
chr17-69605676-DEL-159
chr17-69606006-DEL-56
chr17-69740442-DEL-64
chr17-69798577-DEL-585
chr17-69850589-DEL-69
chr17-69884539-DEL-58
chr17-69913734-DEL-76
chr17-70033593-DEL-121
chr17-70033887-DEL-90
chr17-70152834-DEL-58
chr17-70186798-DEL-365
chr17-70257000-DEL-91
chr17-70373204-DEL-581
chr17-70449582-DEL-102
chr17-70450058-DEL-124
chr17-70570972-DEL-54
chr17-70710206-DEL-523
chr17-70737050-DEL-1279
chr17-70807006-DEL-90
chr17-70836221-DEL-60
chr17-70950417-DEL-282
chr17-70981741-DEL-949
chr17-70984390-DEL-1951
chr17-71070797-DEL-183
chr17-71071033-DEL-392
chr17-71110799-DEL-494
chr17-71111375-DEL-584
chr17-71117739-DEL-56

chr17-71129794-DEL-102
chr17-71144845-DEL-162
chr17-71148863-DEL-69
chr17-71207741-DEL-7412
chr17-71226585-DEL-130
chr17-71266435-DEL-112
chr17-71335635-DEL-110
chr17-71335864-DEL-148
chr17-71335865-DEL-50
chr17-71355146-DEL-3493
chr17-71365081-DEL-2690
chr17-71570020-DEL-50
chr17-71574005-DEL-52
chr17-71588787-DEL-56
chr17-71601237-DEL-184
chr17-71745127-DEL-205
chr17-71745982-DEL-144
chr17-71746212-DEL-76
chr17-71746683-DEL-95
chr17-71846188-DEL-100
chr17-71936143-DEL-51
chr17-71971848-DEL-92
chr17-72038996-DEL-81
chr17-72189828-DEL-60
chr17-72190152-DEL-942
chr17-72220627-DEL-451
chr17-72220874-DEL-204
chr17-72253293-DEL-52
chr17-72266533-DEL-56
chr17-72379515-DEL-80
chr17-72386264-DEL-132
chr17-72469028-DEL-50
chr17-72496415-DEL-90
chr17-72499557-DEL-56
chr17-72520795-DEL-1600
chr17-72527565-DEL-73
chr17-72625342-DEL-1003
chr17-72628858-DEL-53
chr17-72808399-DEL-51
chr17-72986513-DEL-117
chr17-72999759-DEL-52
chr17-73012033-DEL-50
chr17-73111274-DEL-351
chr17-73126018-DEL-189
chr17-73128328-DEL-58
chr17-73187692-DEL-90
chr17-73206137-DEL-219
chr17-73263325-DEL-56
chr17-73265209-DEL-106
chr17-73380876-DEL-54
chr17-73387159-DEL-68
chr17-73422766-DEL-223
chr17-73652376-DEL-174
chr17-73652749-DEL-127
chr17-73719918-DEL-137
chr17-73720006-DEL-66
chr17-73772727-DEL-184
chr17-73815502-DEL-95
chr17-73819739-DEL-52
chr17-73892448-DEL-63
chr17-73892453-DEL-210
chr17-73986435-DEL-322
chr17-74049243-DEL-52
chr17-74055583-DEL-50
chr17-74092043-DEL-146
chr17-74172773-DEL-106
chr17-74299990-DEL-89
chr17-74337000-DEL-54
chr17-74337148-DEL-131
chr17-74340198-DEL-51
chr17-74376214-DEL-76
chr17-74384209-DEL-67
chr17-74466380-DEL-50
chr17-74485782-DEL-1167
chr17-74489643-DEL-105
chr17-74498644-DEL-80
chr17-74586509-DEL-225

chr17-74608584-DEL-72
chr17-74625873-DEL-221
chr17-74636260-DEL-722
chr17-74784077-DEL-731
chr17-74816922-DEL-5465
chr17-74847851-DEL-1762
chr17-74858975-DEL-1348
chr17-74912448-DEL-50
chr17-75227545-DEL-111
chr17-75227554-DEL-224
chr17-75269152-DEL-62
chr17-75317107-DEL-50
chr17-75317128-DEL-167
chr17-75317387-DEL-117
chr17-75317504-DEL-50
chr17-75374873-DEL-161
chr17-75430838-DEL-63
chr17-75753781-DEL-116
chr17-75753898-DEL-52
chr17-75806473-DEL-56
chr17-75825551-DEL-50
chr17-75903675-DEL-3198
chr17-75945755-DEL-50
chr17-75971174-DEL-53
chr17-76017131-DEL-54
chr17-76055070-DEL-265
chr17-76125193-DEL-55
chr17-76127201-DEL-58
chr17-76267602-DEL-58
chr17-76292081-DEL-84
chr17-76340355-DEL-61
chr17-76384861-DEL-1241
chr17-76519813-DEL-1302
chr17-76553027-DEL-139
chr17-76720266-DEL-57
chr17-76755390-DEL-1220
chr17-76789476-DEL-52
chr17-76789574-DEL-57
chr17-76853167-DEL-395
chr17-76871922-DEL-1873
chr17-76948987-DEL-167
chr17-76963587-DEL-1075
chr17-77012504-DEL-52
chr17-77055334-DEL-358
chr17-77332989-DEL-50
chr17-77376650-DEL-64
chr17-77396636-DEL-56
chr17-77407279-DEL-70
chr17-77407434-DEL-102
chr17-77453237-DEL-50
chr17-77458507-DEL-1739
chr17-77565250-DEL-56
chr17-77647406-DEL-55
chr17-77650343-DEL-62
chr17-77742505-DEL-54
chr17-77747015-DEL-55
chr17-77816270-DEL-291
chr17-77940873-DEL-50
chr17-77959976-DEL-3090
chr17-78039504-DEL-224
chr17-78045513-DEL-338
chr17-78045962-DEL-105
chr17-78138472-DEL-1155
chr17-78163242-DEL-63
chr17-78293201-DEL-1033
chr17-78352595-DEL-1318
chr17-78368311-DEL-54
chr17-78394806-DEL-138
chr17-78394814-DEL-58
chr17-78403250-DEL-548
chr17-78408360-DEL-76
chr17-78769713-DEL-78
chr17-78875943-DEL-68
chr17-78879566-DEL-299
chr17-78907966-DEL-56
chr17-78929840-DEL-59
chr17-78968986-DEL-68

chr17-79079229-DEL-68
chr17-79083194-DEL-70
chr17-79153607-DEL-209
chr17-79153615-DEL-81
chr17-79159631-DEL-2377
chr17-79184139-DEL-139
chr17-79199064-DEL-63
chr17-79211233-DEL-98
chr17-79359339-DEL-54
chr17-79366930-DEL-54
chr17-79394826-DEL-677
chr17-79408461-DEL-126
chr17-79413960-DEL-1527
chr17-79481682-DEL-220
chr17-79481883-DEL-66
chr17-79515996-DEL-179
chr17-79516394-DEL-262
chr17-79516445-DEL-77
chr17-79524274-DEL-1872
chr17-79546070-DEL-76
chr17-79578311-DEL-1100
chr17-79628623-DEL-116
chr17-79697855-DEL-95
chr17-79816160-DEL-108
chr17-79879592-DEL-167
chr17-80050193-DEL-98
chr17-80100781-DEL-3172
chr17-80105745-DEL-60
chr17-80185656-DEL-142
chr17-80186326-DEL-120
chr17-80258630-DEL-62
chr17-80274166-DEL-50
chr17-80280240-DEL-138
chr17-80315138-DEL-56
chr17-80458285-DEL-201
chr17-80548550-DEL-52
chr17-80639265-DEL-54
chr17-80645868-DEL-92
chr17-80647917-DEL-77
chr17-80650787-DEL-66
chr17-80775929-DEL-89
chr17-80775992-DEL-188
chr17-80827661-DEL-128
chr17-80906371-DEL-216
chr17-80951129-DEL-96
chr17-80981612-DEL-239
chr17-81004426-DEL-55
chr17-81044512-DEL-56
chr17-81052043-DEL-52
chr17-81072979-DEL-603
chr17-81075062-DEL-110
chr17-81112492-DEL-150
chr17-81161392-DEL-7080
chr17-81262114-DEL-100
chr17-81262114-DEL-224
chr17-81262536-DEL-284
chr17-81317589-DEL-53
chr17-81338127-DEL-222
chr17-81382885-DEL-58
chr17-81581670-DEL-50
chr17-81591756-DEL-230
chr17-81601037-DEL-412
chr17-81764647-DEL-50
chr17-81819842-DEL-139
chr17-81904414-DEL-1015
chr17-81917053-DEL-304
chr17-81972671-DEL-52
chr17-81972755-DEL-258
chr17-82074111-DEL-69
chr17-82184829-DEL-716
chr17-82191299-DEL-142
chr17-82191577-DEL-527
chr17-82192053-DEL-54
chr17-82198236-DEL-803
chr17-82199150-DEL-1959
chr17-82228628-DEL-52
chr17-82241202-DEL-296

chr17-82640623-DEL-1065
chr17-82786059-DEL-147
chr17-82859865-DEL-92
chr17-83024490-DEL-941
chr17-83081512-DEL-4040
chr17-83118550-DEL-57
chr17-83118581-DEL-174
chr17-83118810-DEL-273
chr17-83118933-DEL-123
chr17-83142728-DEL-175
chr17-83187816-DEL-1583
chr17-83264502-DEL-138
chr17-83350443-DEL-72
chr17-83350445-DEL-184
chr17-83370075-DEL-522
chr17-83370358-DEL-129
chr17-83370860-DEL-115
chr17-83412054-DEL-84
chr17-83415270-DEL-153
chr17-83475377-DEL-2340
chr17-83475438-DEL-122
chr17-83475794-DEL-199
chr17-83476171-DEL-561
chr17-83476426-DEL-182
chr17-83477102-DEL-669
chr17-83601665-DEL-58
chr17-83607371-DEL-376
chr17-83607403-DEL-62
chr17-83634506-DEL-60
chr17-83651069-DEL-74
chr17-83655183-DEL-54
chr17-83664940-DEL-661
chr17-83696567-DEL-62
chr17-83710142-DEL-571
chr17-83710368-DEL-233
chr17-83728728-DEL-104
chr17-83898499-DEL-249
chr17-83899371-DEL-62
chr17-83901039-DEL-79
chr17-83920152-DEL-218
chr17-83929942-DEL-175
chr17-84005622-DEL-108
chr17-84028668-DEL-368
chr17-84076427-DEL-61
chr17-84163205-DEL-70
chr17-84434621-DEL-88
chr17-84520368-DEL-52
chr17-84542265-DEL-401
chr17-84542428-DEL-104
chr17-84569060-DEL-57
chr17-84593161-DEL-146
chr17-84593359-DEL-56
chr17-84608384-DEL-147
chr17-84608660-DEL-116
chr17-84695160-DEL-606
chr17-84710228-DEL-172
chr17-84766713-DEL-2589
chr17-84766827-DEL-52
chr17-84813750-DEL-138
chr17-84814048-DEL-94
chr17-84846556-DEL-138
chr17-84854971-DEL-78
chr17-84872135-DEL-419
chr17-84895399-DEL-187
chr17-84921292-DEL-196
chr17-84921601-DEL-300
chr17-84944568-DEL-56
chr17-84946016-DEL-192
chr17-84974526-DEL-402
chr17-85077148-DEL-149
chr17-85093899-DEL-80
chr17-85100066-DEL-137
chr17-85109349-DEL-200
chr17-85149369-DEL-110
chr17-85165460-DEL-76
chr17-85206986-DEL-54
chr17-85227019-DEL-135

chr17-85308278-DEL-912
chr17-85308393-DEL-227
chr17-85315315-DEL-99
chr17-85347014-DEL-78
chr17-85363007-DEL-69
chr17-85374309-DEL-58
chr17-85436520-DEL-58
chr17-85463271-DEL-62
chr17-85521632-DEL-62
chr17-85545990-DEL-109
chr17-85547962-DEL-59
chr17-85667723-DEL-54
chr17-85672525-DEL-50
chr17-85677168-DEL-74
chr17-85692437-DEL-76
chr17-85703477-DEL-50
chr17-85785192-DEL-151
chr17-85812516-DEL-66
chr17-85945885-DEL-249
chr17-86004606-DEL-62
chr17-86019905-DEL-120
chr17-86020131-DEL-381
chr17-86095888-DEL-50
chr17-86211361-DEL-88
chr17-86531327-DEL-200
chr17-86597609-DEL-110
chr17-86664575-DEL-500
chr17-86732201-DEL-91
chr17-86734911-DEL-62
chr17-86735240-DEL-1119
chr17-86750346-DEL-124
chr17-86784837-DEL-93
chr17-86823602-DEL-61
chr17-86876199-DEL-495
chr17-86929613-DEL-62
chr17-87220272-DEL-53
chr17-87278129-DEL-168
chr17-87288154-DEL-508
chr17-87319802-DEL-1155
chr17-87320783-DEL-165
chr17-87355500-DEL-81
chr17-87359403-DEL-139
chr17-87379634-DEL-52
chr17-87410669-DEL-97
chr17-87516161-DEL-646
chr17-87516166-DEL-1760
chr17-87517154-DEL-232
chr17-87557623-DEL-122
chr17-87605096-DEL-104
chr17-87608731-DEL-73
chr17-87626595-DEL-57
chr17-87702282-DEL-156
chr17-87723115-DEL-212
chr17-87723121-DEL-63
chr17-87729013-DEL-174
chr17-87888646-DEL-52
chr17-87889234-DEL-81
chr17-87941089-DEL-265
chr17-87987599-DEL-99
chr17-87987806-DEL-165
chr17-88020648-DEL-54
chr17-88020725-DEL-113
chr17-88020797-DEL-341
chr17-88021324-DEL-341
chr17-88024101-DEL-50
chr17-88042463-DEL-119
chr17-88056456-DEL-5026
chr17-88062013-DEL-72
chr17-88069809-DEL-236
chr17-88069931-DEL-62
chr17-88081281-DEL-50
chr17-88086823-DEL-111
chr17-88087058-DEL-71
chr17-88120423-DEL-237
chr17-88120754-DEL-84
chr17-88143673-DEL-110
chr17-88143677-DEL-234

chr17-88143982-DEL-60
chr17-88153662-DEL-405
chr17-88154067-DEL-57
chr17-88162375-DEL-5393
chr17-88175819-DEL-1520
chr17-88227043-DEL-484
chr17-88309738-DEL-118
chr17-88377318-DEL-50
chr17-88411057-DEL-221
chr17-88535617-DEL-58
chr17-88638794-DEL-69
chr17-88664079-DEL-173
chr17-88820353-DEL-67
chr17-88820584-DEL-256
chr17-88831957-DEL-144
chr17-88839722-DEL-58
chr17-88873368-DEL-285
chr17-88909974-DEL-52
chr17-88950735-DEL-254
chr17-88951070-DEL-350
chr17-88984663-DEL-133
chr17-89017118-DEL-504
chr17-89024774-DEL-307
chr17-89025769-DEL-78
chr17-89082626-DEL-234
chr17-89127618-DEL-200
chr17-89153543-DEL-109
chr17-89181352-DEL-54
chr17-89271669-DEL-105
chr17-89308366-DEL-93
chr17-89498158-DEL-111
chr17-89583898-DEL-156
chr17-89655899-DEL-125
chr17-89755440-DEL-408
chr17-89770533-DEL-58
chr17-90060142-DEL-142
chr17-90268404-DEL-88
chr17-90292049-DEL-333
chr17-90313806-DEL-96
chr17-90338340-DEL-53
chr17-90377987-DEL-50
chr17-90447938-DEL-50
chr17-90480248-DEL-171
chr17-90564410-DEL-58
chr17-90699495-DEL-132
chr17-90751996-DEL-60
chr17-90752316-DEL-60
chr17-90899962-DEL-60
chr17-91230523-DEL-270
chr17-91339656-DEL-480
chr17-91341051-DEL-63
chr17-91379319-DEL-50
chr17-91389192-DEL-176
chr17-91521961-DEL-1764
chr17-91527315-DEL-200
chr17-91527315-DEL-632
chr17-91568731-DEL-55
chr17-91793881-DEL-3306
chr17-91834371-DEL-125
chr17-91885755-DEL-493
chr17-91921775-DEL-53
chr17-91928318-DEL-651
chr17-92254805-DEL-50
chr17-92376355-DEL-64
chr17-92377346-DEL-1094
chr17-92378396-DEL-70
chr17-92408213-DEL-309
chr17-92509386-DEL-3348
chr17-92563469-DEL-529
chr17-92600455-DEL-1461
chr17-92651944-DEL-3608
chr17-92655872-DEL-678
chr17-92665479-DEL-110
chr17-92693694-DEL-76
chr17-92731185-DEL-1853
chr17-92760225-DEL-3695
chr17-92793199-DEL-2131

chr17-92910135-DEL-91
chr17-93020929-DEL-3234
chr17-93140702-DEL-102
chr17-93164417-DEL-64
chr17-93191485-DEL-55
chr17-93225932-DEL-84
chr17-93286400-DEL-57
chr17-93505087-DEL-54
chr17-93800436-DEL-5663
chr17-93823225-DEL-64
chr17-94007597-DEL-1048
chr17-94337601-DEL-55
chr17-94464553-DEL-141
chr17-94488372-DEL-50
chr17-94500508-DEL-198
chr17-94535101-DEL-2994
chr17-94657021-DEL-853
chr17-94693903-DEL-57
chr17-94745180-DEL-63
chr17-94762966-DEL-54
chr17-94763074-DEL-180
chr17-94763262-DEL-412
chr17-94763482-DEL-192
chr17-94998787-DEL-386
chr17-95052183-DEL-53
chr17-95067083-DEL-25045
chr17-95100806-DEL-115
chr17-95144985-DEL-54
chr18-3004431-DEL-300
chr18-3120213-DEL-88
chr18-3355660-DEL-60
chr18-3394773-DEL-180
chr18-3799213-DEL-89
chr18-3799379-DEL-52
chr18-3880093-DEL-1492
chr18-3883615-DEL-106
chr18-3908867-DEL-65
chr18-3960283-DEL-223
chr18-3962443-DEL-170
chr18-3990900-DEL-50
chr18-4065255-DEL-122
chr18-4147292-DEL-78
chr18-4152444-DEL-60
chr18-4177187-DEL-77
chr18-4177866-DEL-62
chr18-4255189-DEL-61
chr18-4687052-DEL-76
chr18-4692909-DEL-1565
chr18-4714912-DEL-152
chr18-4728496-DEL-313
chr18-4734619-DEL-96
chr18-4984525-DEL-528
chr18-5027053-DEL-140
chr18-5062515-DEL-202
chr18-5065392-DEL-50
chr18-5109635-DEL-3895
chr18-5144101-DEL-241
chr18-5144258-DEL-74
chr18-5144438-DEL-453
chr18-5187650-DEL-259
chr18-5187768-DEL-812
chr18-5325605-DEL-76
chr18-5531334-DEL-1474
chr18-5748053-DEL-93
chr18-5751439-DEL-212
chr18-5777653-DEL-132
chr18-5788585-DEL-1288
chr18-5788797-DEL-292
chr18-5789190-DEL-275
chr18-5789836-DEL-543
chr18-5790334-DEL-60
chr18-5947748-DEL-51
chr18-5962323-DEL-67
chr18-6081550-DEL-297
chr18-6081940-DEL-558
chr18-6151487-DEL-50
chr18-6157024-DEL-1218

chr18-6157080-DEL-185
chr18-6157907-DEL-120
chr18-6158556-DEL-57
chr18-6410177-DEL-266
chr18-6437527-DEL-50
chr18-6641295-DEL-116
chr18-6643785-DEL-263
chr18-6765816-DEL-86
chr18-6796372-DEL-7146
chr18-6979161-DEL-65
chr18-7070106-DEL-3894
chr18-7093114-DEL-472
chr18-7113483-DEL-489
chr18-7113592-DEL-133
chr18-7113915-DEL-144
chr18-7174408-DEL-145
chr18-7401671-DEL-120
chr18-7432000-DEL-50
chr18-7437221-DEL-64
chr18-7484400-DEL-52
chr18-7549537-DEL-275
chr18-7556502-DEL-255
chr18-7608194-DEL-61
chr18-7608948-DEL-750
chr18-7611968-DEL-51
chr18-7628524-DEL-211
chr18-7754298-DEL-440
chr18-7755154-DEL-60
chr18-7790343-DEL-62
chr18-7823973-DEL-282
chr18-7858987-DEL-2200
chr18-8223162-DEL-2469
chr18-8233293-DEL-268
chr18-8283838-DEL-74
chr18-8626741-DEL-157
chr18-8627019-DEL-268
chr18-8683840-DEL-1138
chr18-8685086-DEL-140
chr18-8707453-DEL-56
chr18-8722792-DEL-4393
chr18-8981782-DEL-819
chr18-8986407-DEL-97
chr18-9035830-DEL-82
chr18-9042600-DEL-111
chr18-9042797-DEL-50
chr18-9098950-DEL-72
chr18-9122372-DEL-57
chr18-9487319-DEL-1785
chr18-9489352-DEL-147
chr18-9678385-DEL-356
chr18-9678796-DEL-141
chr18-9683039-DEL-3678
chr18-9686498-DEL-2404
chr18-9733030-DEL-1530
chr18-9733416-DEL-646
chr18-9789804-DEL-123
chr18-9836672-DEL-112
chr18-9916835-DEL-1340
chr18-10195559-DEL-50
chr18-10265792-DEL-64
chr18-10428947-DEL-98
chr18-10431134-DEL-72
chr18-10431459-DEL-236
chr18-10546066-DEL-70
chr18-10548459-DEL-164
chr18-10687268-DEL-7170
chr18-10698512-DEL-123
chr18-10734859-DEL-1448
chr18-10823081-DEL-399
chr18-10888788-DEL-160
chr18-10889025-DEL-197
chr18-10934667-DEL-176
chr18-10947712-DEL-76
chr18-10954994-DEL-50
chr18-10971051-DEL-56
chr18-11010768-DEL-352
chr18-11062336-DEL-50

chr18-11228924-DEL-90
chr18-11275696-DEL-2171
chr18-11303146-DEL-295
chr18-11303376-DEL-55
chr18-11411050-DEL-174
chr18-11423540-DEL-3662
chr18-11555211-DEL-62
chr18-11567784-DEL-124
chr18-11611910-DEL-50
chr18-11622468-DEL-100
chr18-11679447-DEL-73
chr18-11738818-DEL-191
chr18-12025898-DEL-248
chr18-12031779-DEL-355
chr18-12056946-DEL-58
chr18-12056946-DEL-118
chr18-12138850-DEL-204
chr18-12145550-DEL-186
chr18-12145794-DEL-262
chr18-12150267-DEL-710
chr18-12163309-DEL-125
chr18-12166511-DEL-220
chr18-12166702-DEL-97
chr18-12166903-DEL-681
chr18-12167129-DEL-105
chr18-12224744-DEL-4512
chr18-12305664-DEL-416
chr18-12307174-DEL-54
chr18-12336593-DEL-132
chr18-12498893-DEL-532
chr18-12534948-DEL-50
chr18-12544178-DEL-140
chr18-12585342-DEL-134
chr18-12585821-DEL-136
chr18-12606499-DEL-2253
chr18-12683057-DEL-185
chr18-12752935-DEL-872
chr18-12753167-DEL-1284
chr18-12754090-DEL-488
chr18-12834670-DEL-92
chr18-12853187-DEL-280
chr18-12853357-DEL-80
chr18-12853595-DEL-120
chr18-12856907-DEL-503
chr18-12857275-DEL-54
chr18-12907867-DEL-51
chr18-13034772-DEL-1179
chr18-13080398-DEL-62
chr18-13083650-DEL-50
chr18-13107154-DEL-214
chr18-13175867-DEL-765
chr18-13257335-DEL-88
chr18-13382470-DEL-71
chr18-13383819-DEL-142
chr18-13429638-DEL-840
chr18-13431066-DEL-150
chr18-13431215-DEL-700
chr18-13810157-DEL-165
chr18-14054879-DEL-58
chr18-14055047-DEL-66
chr18-14073092-DEL-152
chr18-14160212-DEL-56
chr18-14215614-DEL-50
chr18-14392540-DEL-59
chr18-14436271-DEL-140
chr18-14448509-DEL-53
chr18-14497718-DEL-65
chr18-14540846-DEL-51
chr18-14593762-DEL-1746
chr18-14724719-DEL-2275
chr18-14785986-DEL-60
chr18-14793734-DEL-72
chr18-14899196-DEL-1520
chr18-15046819-DEL-382
chr18-15046881-DEL-137
chr18-15051081-DEL-311
chr18-15051331-DEL-140

chr18-15153684-DEL-63
chr18-15246813-DEL-146
chr18-15247337-DEL-90
chr18-15247337-DEL-303
chr18-15310161-DEL-165
chr18-15329711-DEL-171
chr18-15333149-DEL-550
chr18-15338138-DEL-60
chr18-15338488-DEL-62
chr18-15355904-DEL-78
chr18-15355990-DEL-160
chr18-15356112-DEL-54
chr18-15441321-DEL-93
chr18-15488707-DEL-734
chr18-15491835-DEL-60
chr18-15567357-DEL-286
chr18-15567617-DEL-104
chr18-15574195-DEL-104
chr18-15995856-DEL-68
chr18-16021028-DEL-2048
chr18-16134858-DEL-150
chr18-16229223-DEL-54
chr18-16293691-DEL-73
chr18-16294059-DEL-2433
chr18-16377837-DEL-356
chr18-16484659-DEL-1007
chr18-16490287-DEL-62
chr18-16517759-DEL-50
chr18-16555994-DEL-1314
chr18-16584001-DEL-50
chr18-16585797-DEL-62
chr18-16636346-DEL-407
chr18-16716406-DEL-51
chr18-16733152-DEL-1242
chr18-16792165-DEL-722
chr18-17000711-DEL-1347
chr18-17041949-DEL-1816
chr18-17054405-DEL-1538
chr18-17104657-DEL-974
chr18-17404118-DEL-3799
chr18-17508559-DEL-1823
chr18-17583754-DEL-94
chr18-17619901-DEL-57
chr18-17641177-DEL-90
chr18-17641729-DEL-150
chr18-17668799-DEL-68
chr18-17804465-DEL-1467
chr18-17908158-DEL-4280
chr18-17927131-DEL-181
chr18-17960067-DEL-206
chr18-18024327-DEL-2259
chr18-18242406-DEL-60
chr18-18408950-DEL-673
chr18-18422847-DEL-1258
chr18-18526314-DEL-86
chr18-18747000-DEL-395
chr18-18749786-DEL-2314
chr18-18778016-DEL-1676
chr18-18834156-DEL-519
chr18-18924096-DEL-209
chr18-18991488-DEL-1478
chr18-18998603-DEL-54
chr18-19070158-DEL-425
chr18-19093486-DEL-351
chr18-19095061-DEL-2671
chr18-19108084-DEL-10191
chr18-19119879-DEL-7667
chr18-19154009-DEL-964
chr18-19187929-DEL-541
chr18-19188560-DEL-66
chr18-19230301-DEL-152
chr18-19301357-DEL-1054
chr18-19358239-DEL-1436
chr18-19367252-DEL-64
chr18-19394058-DEL-205
chr18-19510744-DEL-5674
chr18-19628989-DEL-123

chr18-19666203-DEL-2386
chr18-19745526-DEL-208
chr18-19974699-DEL-398
chr18-20034309-DEL-87
chr18-20081407-DEL-56
chr18-20129992-DEL-82
chr18-20130177-DEL-178
chr18-20130407-DEL-150
chr18-20418989-DEL-97
chr18-20463151-DEL-415
chr18-20621740-DEL-549
chr18-20659402-DEL-72
chr18-20720961-DEL-130
chr18-20722813-DEL-106
chr18-20754603-DEL-241
chr18-20756289-DEL-989
chr18-20812279-DEL-245
chr18-20812558-DEL-60
chr18-20880435-DEL-86
chr18-20899625-DEL-868
chr18-21011530-DEL-127
chr18-21045796-DEL-100
chr18-21045827-DEL-229
chr18-21108509-DEL-100
chr18-21108796-DEL-198
chr18-21129918-DEL-50
chr18-21171409-DEL-67
chr18-21182733-DEL-177
chr18-21194619-DEL-5809
chr18-21303904-DEL-85
chr18-21359222-DEL-184
chr18-21439356-DEL-150
chr18-21455241-DEL-107
chr18-21561216-DEL-50
chr18-21766914-DEL-53
chr18-21918826-DEL-58
chr18-21962782-DEL-55
chr18-22000068-DEL-61
chr18-22320978-DEL-102
chr18-22393623-DEL-50
chr18-22850848-DEL-52
chr18-22906784-DEL-925
chr18-23081990-DEL-82
chr18-23104742-DEL-179
chr18-23146852-DEL-62
chr18-23237048-DEL-356
chr18-23265570-DEL-67
chr18-23265595-DEL-173
chr18-23371769-DEL-2938
chr18-23382426-DEL-1487
chr18-23411962-DEL-50
chr18-23427169-DEL-147
chr18-23451571-DEL-144
chr18-23516564-DEL-90
chr18-23537917-DEL-3280
chr18-23607134-DEL-338
chr18-23629078-DEL-59
chr18-23693590-DEL-292
chr18-23714491-DEL-50
chr18-23907959-DEL-79
chr18-23919304-DEL-60
chr18-24055431-DEL-103
chr18-24298234-DEL-2133
chr18-24392214-DEL-111
chr18-24434326-DEL-50
chr18-24439088-DEL-64
chr18-24496326-DEL-51
chr18-24497669-DEL-88
chr18-24533677-DEL-64
chr18-24563039-DEL-60
chr18-24587879-DEL-118
chr18-24588229-DEL-253
chr18-24614245-DEL-83
chr18-24615521-DEL-78
chr18-24804795-DEL-290
chr18-24805431-DEL-135
chr18-24806229-DEL-146

chr18-24809970-DEL-308
chr18-24815448-DEL-64
chr18-24881643-DEL-68
chr18-24976862-DEL-2571
chr18-25053347-DEL-135
chr18-25204469-DEL-61
chr18-25392321-DEL-255
chr18-25407477-DEL-1675
chr18-25488932-DEL-87
chr18-25715090-DEL-103
chr18-25715556-DEL-123
chr18-25759887-DEL-82
chr18-25895132-DEL-54
chr18-25922538-DEL-128
chr18-26058553-DEL-2245
chr18-26143528-DEL-64
chr18-26147295-DEL-56
chr18-26189424-DEL-180
chr18-26194731-DEL-65
chr18-26207141-DEL-255
chr18-26417976-DEL-973
chr18-26419044-DEL-476
chr18-26453401-DEL-485
chr18-26567213-DEL-50
chr18-26748821-DEL-72
chr18-26954720-DEL-50
chr18-26998177-DEL-100
chr18-27153406-DEL-52
chr18-27188443-DEL-202
chr18-27272253-DEL-70
chr18-27273477-DEL-741
chr18-27280167-DEL-381
chr18-27326943-DEL-108
chr18-27327196-DEL-81
chr18-27356919-DEL-829
chr18-27488564-DEL-62
chr18-27519707-DEL-61
chr18-27519934-DEL-137
chr18-27565319-DEL-1024
chr18-27640122-DEL-774
chr18-27710467-DEL-332
chr18-27717107-DEL-82
chr18-27733739-DEL-3172
chr18-27829882-DEL-2095
chr18-27849509-DEL-144
chr18-27849713-DEL-64
chr18-27918387-DEL-917
chr18-28011680-DEL-552
chr18-28071641-DEL-278
chr18-28122358-DEL-50
chr18-28192067-DEL-2977
chr18-28209517-DEL-54
chr18-28217936-DEL-88
chr18-28291322-DEL-322
chr18-28299294-DEL-1493
chr18-28343104-DEL-327
chr18-28463231-DEL-50
chr18-28510570-DEL-184
chr18-28533050-DEL-210
chr18-28582631-DEL-54
chr18-28593354-DEL-201
chr18-28705570-DEL-490
chr18-28821628-DEL-86
chr18-29035476-DEL-56
chr18-29176325-DEL-532
chr18-29176351-DEL-68
chr18-29176627-DEL-136
chr18-29201215-DEL-195
chr18-29201843-DEL-7574
chr18-29346213-DEL-65
chr18-29346530-DEL-50
chr18-29355814-DEL-80
chr18-29648998-DEL-52
chr18-29732511-DEL-84
chr18-29928084-DEL-50
chr18-30000849-DEL-65
chr18-30095971-DEL-50

chr18-30139947-DEL-52
chr18-30140923-DEL-118
chr18-30182552-DEL-160
chr18-30331929-DEL-1142
chr18-30492234-DEL-87
chr18-30562083-DEL-56
chr18-30657666-DEL-51
chr18-30722078-DEL-145
chr18-30803494-DEL-189
chr18-30809706-DEL-1626
chr18-30852899-DEL-115
chr18-30900996-DEL-1200
chr18-30982226-DEL-62
chr18-31024601-DEL-308
chr18-31038450-DEL-68
chr18-31038880-DEL-219
chr18-31052271-DEL-70
chr18-31067405-DEL-112
chr18-31192203-DEL-56
chr18-31246458-DEL-54
chr18-31560589-DEL-117
chr18-31560664-DEL-244
chr18-31587386-DEL-98
chr18-31614292-DEL-100
chr18-31626418-DEL-76
chr18-31651440-DEL-1160
chr18-31681250-DEL-50
chr18-31827395-DEL-930
chr18-31828321-DEL-60
chr18-31835755-DEL-94
chr18-31840996-DEL-714
chr18-31952471-DEL-55
chr18-31966177-DEL-83
chr18-32053664-DEL-58
chr18-32144647-DEL-92
chr18-32261115-DEL-106
chr18-32317472-DEL-746
chr18-32317688-DEL-127
chr18-32318118-DEL-100
chr18-32327188-DEL-66
chr18-32354016-DEL-351
chr18-32572643-DEL-63
chr18-32654546-DEL-636
chr18-32686172-DEL-218
chr18-32696087-DEL-328
chr18-32750411-DEL-113
chr18-32903462-DEL-11835
chr18-32911904-DEL-11827
chr18-32936011-DEL-90
chr18-32983712-DEL-208
chr18-33275506-DEL-940
chr18-33404178-DEL-57
chr18-33463310-DEL-428
chr18-33561275-DEL-645
chr18-33648263-DEL-155
chr18-33654888-DEL-68
chr18-33654916-DEL-198
chr18-33694729-DEL-180
chr18-33719367-DEL-1397
chr18-33815957-DEL-1926
chr18-33924326-DEL-315
chr18-33934840-DEL-50
chr18-33943658-DEL-147
chr18-33953859-DEL-83
chr18-33956151-DEL-50
chr18-33973707-DEL-1610
chr18-33985436-DEL-85
chr18-33991084-DEL-544
chr18-33991161-DEL-204
chr18-34017604-DEL-94
chr18-34025465-DEL-354
chr18-34043462-DEL-94
chr18-34087492-DEL-70
chr18-34177147-DEL-251
chr18-34195472-DEL-181
chr18-34205564-DEL-54
chr18-34206442-DEL-229

chr18-34322408-DEL-590
chr18-34373369-DEL-71
chr18-34457022-DEL-117
chr18-34479612-DEL-102
chr18-34652714-DEL-57
chr18-34703851-DEL-102
chr18-34754703-DEL-52
chr18-34777370-DEL-172
chr18-34842722-DEL-89
chr18-34923336-DEL-131
chr18-34954311-DEL-68
chr18-35026052-DEL-83
chr18-35134539-DEL-194
chr18-35183919-DEL-156
chr18-35388708-DEL-195
chr18-35416318-DEL-72
chr18-35797063-DEL-63
chr18-35827560-DEL-1431
chr18-36004168-DEL-50
chr18-36073820-DEL-139
chr18-36132633-DEL-408
chr18-36132973-DEL-136
chr18-36183660-DEL-63
chr18-36183977-DEL-76
chr18-36359438-DEL-179
chr18-36382657-DEL-210
chr18-36382956-DEL-272
chr18-36656507-DEL-50
chr18-36658200-DEL-50
chr18-36681669-DEL-69
chr18-36830318-DEL-66
chr18-36855756-DEL-80
chr18-36950372-DEL-1440
chr18-37030873-DEL-2622
chr18-37049196-DEL-205
chr18-37162615-DEL-77
chr18-37180279-DEL-53
chr18-37217634-DEL-3551
chr18-37377603-DEL-195
chr18-37377892-DEL-8536
chr18-37387047-DEL-4801
chr18-37411883-DEL-89
chr18-37419296-DEL-66
chr18-37753492-DEL-54
chr18-37765866-DEL-400
chr18-37879248-DEL-201
chr18-37916024-DEL-141
chr18-37931028-DEL-56
chr18-38293914-DEL-50
chr18-38351441-DEL-53
chr18-38423487-DEL-217
chr18-38439473-DEL-66
chr18-38457795-DEL-268
chr18-38554770-DEL-101
chr18-38645356-DEL-76
chr18-38687460-DEL-1914
chr18-38756703-DEL-1992
chr18-38791672-DEL-62
chr18-38793008-DEL-86
chr18-38793169-DEL-53
chr18-38794917-DEL-160
chr18-38883965-DEL-72
chr18-38906802-DEL-93
chr18-39068942-DEL-424
chr18-39251457-DEL-150
chr18-39273090-DEL-50
chr18-39311889-DEL-574
chr18-39312422-DEL-164
chr18-39349451-DEL-67
chr18-39438598-DEL-65
chr18-39519624-DEL-140
chr18-39632864-DEL-54
chr18-39823806-DEL-54
chr18-39969807-DEL-63
chr18-40066037-DEL-2758
chr18-40085910-DEL-1118
chr18-40133046-DEL-6141

chr18-40253309-DEL-1011
chr18-40409771-DEL-53
chr18-40557389-DEL-1460
chr18-40772554-DEL-66
chr18-40801738-DEL-56
chr18-40804571-DEL-4883
chr18-40950004-DEL-82
chr18-40970314-DEL-60
chr18-40984887-DEL-50
chr18-41000295-DEL-179
chr18-41078049-DEL-4594
chr18-41288864-DEL-2932
chr18-41343827-DEL-77
chr18-41347109-DEL-200
chr18-41589450-DEL-60
chr18-41892703-DEL-214
chr18-41976539-DEL-324
chr18-41976613-DEL-85
chr18-41976933-DEL-322
chr18-42030302-DEL-392
chr18-42197601-DEL-57
chr18-42231059-DEL-52
chr18-42231197-DEL-171
chr18-42270585-DEL-54
chr18-42302760-DEL-54
chr18-42332760-DEL-58
chr18-42341678-DEL-57
chr18-42356782-DEL-225
chr18-42517533-DEL-70
chr18-42649911-DEL-501
chr18-42714148-DEL-156
chr18-42744113-DEL-51
chr18-42786417-DEL-225
chr18-42901852-DEL-66
chr18-43055188-DEL-58
chr18-43060082-DEL-88
chr18-43189201-DEL-76
chr18-43305197-DEL-60
chr18-43325746-DEL-268
chr18-43360862-DEL-56
chr18-43388048-DEL-63
chr18-43763989-DEL-90
chr18-43789896-DEL-98
chr18-43789914-DEL-202
chr18-43790052-DEL-64
chr18-43825077-DEL-344
chr18-43825232-DEL-144
chr18-43968885-DEL-332
chr18-43974564-DEL-67
chr18-44127650-DEL-53
chr18-44239135-DEL-73
chr18-44331181-DEL-2873
chr18-44361048-DEL-1760
chr18-44539110-DEL-59
chr18-44559134-DEL-111
chr18-44681340-DEL-57
chr18-44681377-DEL-228
chr18-44681542-DEL-63
chr18-44880583-DEL-1003
chr18-44913402-DEL-146
chr18-44970901-DEL-1060
chr18-45245174-DEL-168
chr18-45333240-DEL-138
chr18-45480371-DEL-251
chr18-45480426-DEL-54
chr18-45616793-DEL-76
chr18-45819895-DEL-173
chr18-45963800-DEL-454
chr18-46130916-DEL-1448
chr18-46132890-DEL-54
chr18-46173697-DEL-77
chr18-46288489-DEL-491
chr18-46294283-DEL-63
chr18-46340626-DEL-142
chr18-46340697-DEL-54
chr18-46417222-DEL-483
chr18-46450253-DEL-6280

chr18-46513147-DEL-100
chr18-46513395-DEL-243
chr18-46519168-DEL-260
chr18-46519314-DEL-126
chr18-46522940-DEL-1298
chr18-46675406-DEL-471
chr18-46684816-DEL-234
chr18-46685182-DEL-138
chr18-46718538-DEL-50
chr18-46839431-DEL-57
chr18-46924510-DEL-1489
chr18-47011889-DEL-64
chr18-47130881-DEL-361
chr18-47490628-DEL-50
chr18-47744909-DEL-79
chr18-47760995-DEL-90
chr18-47763927-DEL-66
chr18-47806567-DEL-203
chr18-48280322-DEL-460
chr18-48415169-DEL-81
chr18-48613719-DEL-286
chr18-48699470-DEL-191
chr18-48711290-DEL-55
chr18-48715533-DEL-133
chr18-48727660-DEL-1630
chr18-48777627-DEL-130
chr18-48831333-DEL-50
chr18-48899973-DEL-108
chr18-48900045-DEL-306
chr18-48950258-DEL-378
chr18-48950752-DEL-668
chr18-48951268-DEL-208
chr18-49057109-DEL-340
chr18-49127248-DEL-50
chr18-49148236-DEL-871
chr18-49222369-DEL-103
chr18-49434752-DEL-86
chr18-49520835-DEL-2474
chr18-49579494-DEL-227
chr18-49579582-DEL-58
chr18-49619099-DEL-124
chr18-49741871-DEL-136
chr18-49747609-DEL-201
chr18-49802393-DEL-6144
chr18-49867496-DEL-71
chr18-49869842-DEL-69
chr18-49879872-DEL-64
chr18-49895312-DEL-528
chr18-49901644-DEL-54
chr18-49926507-DEL-106
chr18-50119574-DEL-64
chr18-50165277-DEL-58
chr18-50235323-DEL-322
chr18-50235726-DEL-401
chr18-50253201-DEL-348
chr18-50253878-DEL-166
chr18-50254384-DEL-1350
chr18-50271277-DEL-124
chr18-50354366-DEL-140
chr18-50354458-DEL-428
chr18-50477705-DEL-21710
chr18-50492628-DEL-428
chr18-50493772-DEL-58
chr18-50674517-DEL-56
chr18-50994285-DEL-198
chr18-51041804-DEL-4477
chr18-51103111-DEL-18904
chr18-51425336-DEL-79
chr18-51595100-DEL-59
chr18-51595100-DEL-149
chr18-51741816-DEL-63
chr18-51790518-DEL-4706
chr18-51841745-DEL-71
chr18-51872403-DEL-60
chr18-51993877-DEL-150
chr18-52005267-DEL-66
chr18-52073364-DEL-212

chr18-52119357-DEL-50
chr18-52204945-DEL-187
chr18-52221108-DEL-1186
chr18-52222730-DEL-334
chr18-52354901-DEL-110
chr18-52373376-DEL-392
chr18-52490884-DEL-160
chr18-52508622-DEL-88
chr18-52508903-DEL-70
chr18-52510624-DEL-3561
chr18-52557104-DEL-294
chr18-52557482-DEL-1160
chr18-52574536-DEL-194
chr18-52587849-DEL-157
chr18-52803107-DEL-307
chr18-52889949-DEL-140
chr18-52979564-DEL-389
chr18-53009906-DEL-212
chr18-53017159-DEL-52
chr18-53034053-DEL-82
chr18-53114273-DEL-52
chr18-53162255-DEL-93
chr18-53195237-DEL-846
chr18-53195464-DEL-420
chr18-53195884-DEL-288
chr18-53217821-DEL-237
chr18-53217904-DEL-97
chr18-53226819-DEL-102
chr18-53298431-DEL-780
chr18-53301356-DEL-60
chr18-53307963-DEL-64
chr18-53315731-DEL-61
chr18-53347093-DEL-780
chr18-53363067-DEL-72
chr18-53363067-DEL-240
chr18-53436762-DEL-54
chr18-53437027-DEL-218
chr18-53437567-DEL-162
chr18-53488100-DEL-240
chr18-53488655-DEL-240
chr18-53636297-DEL-2132
chr18-53636312-DEL-453
chr18-53636914-DEL-351
chr18-53637377-DEL-1052
chr18-53637633-DEL-253
chr18-53638197-DEL-148
chr18-53638921-DEL-74
chr18-53724809-DEL-50
chr18-53727669-DEL-50
chr18-53833030-DEL-2960
chr18-54093315-DEL-252
chr18-54093809-DEL-108
chr18-54112317-DEL-54
chr18-54156914-DEL-201
chr18-54193629-DEL-168
chr18-54230952-DEL-397
chr18-54231438-DEL-125
chr18-54687132-DEL-55
chr18-54728136-DEL-60
chr18-54774046-DEL-232
chr18-54858662-DEL-75
chr18-54913353-DEL-146
chr18-55169980-DEL-60
chr18-55270940-DEL-90
chr18-55305204-DEL-75
chr18-55367009-DEL-50
chr18-55439503-DEL-153
chr18-55439503-DEL-321
chr18-55504590-DEL-152
chr18-55504644-DEL-68
chr18-55548363-DEL-84
chr18-55688074-DEL-50
chr18-55741235-DEL-64
chr18-55775011-DEL-50
chr18-55792880-DEL-175
chr18-55841226-DEL-82
chr18-55870369-DEL-715

chr18-56065890-DEL-150
chr18-56078895-DEL-82
chr18-56103669-DEL-137
chr18-56294348-DEL-70
chr18-56302297-DEL-169
chr18-56324220-DEL-1424
chr18-56698366-DEL-110
chr18-56737639-DEL-203
chr18-56750521-DEL-50
chr18-56815534-DEL-135
chr18-56817298-DEL-309
chr18-56890089-DEL-90
chr18-56897363-DEL-129
chr18-57006738-DEL-435
chr18-57018325-DEL-55
chr18-57126819-DEL-368
chr18-57130828-DEL-1422
chr18-57184042-DEL-64
chr18-57201419-DEL-100
chr18-57251425-DEL-1149
chr18-57263393-DEL-116
chr18-57298445-DEL-819
chr18-57377381-DEL-66
chr18-57381578-DEL-78
chr18-57432681-DEL-1724
chr18-57439286-DEL-50
chr18-57463893-DEL-80
chr18-57552721-DEL-57
chr18-57580862-DEL-358
chr18-57640025-DEL-67
chr18-57703303-DEL-83
chr18-57723006-DEL-130
chr18-57817933-DEL-245
chr18-57818371-DEL-84
chr18-57876602-DEL-177
chr18-57916186-DEL-187
chr18-57999704-DEL-134
chr18-58037553-DEL-1265
chr18-58102091-DEL-122
chr18-58123866-DEL-948
chr18-58161855-DEL-110
chr18-58184647-DEL-51
chr18-58227166-DEL-54
chr18-58398423-DEL-258
chr18-58398490-DEL-58
chr18-58398789-DEL-192
chr18-58398946-DEL-75
chr18-58634822-DEL-120
chr18-58635030-DEL-128
chr18-58643456-DEL-82
chr18-58650473-DEL-215
chr18-58748572-DEL-168
chr18-58808765-DEL-395
chr18-58927578-DEL-216
chr18-58977061-DEL-50
chr18-59048333-DEL-96
chr18-59110074-DEL-1102
chr18-59114894-DEL-80
chr18-59149909-DEL-121
chr18-59150126-DEL-100
chr18-59443412-DEL-100
chr18-59468854-DEL-193
chr18-59519588-DEL-58
chr18-59577820-DEL-94
chr18-59688394-DEL-67
chr18-59764085-DEL-246
chr18-59777811-DEL-108
chr18-59827582-DEL-54
chr18-60004993-DEL-68
chr18-60086980-DEL-212
chr18-60259420-DEL-50
chr18-60402265-DEL-184
chr18-60482147-DEL-602
chr18-60594098-DEL-183
chr18-60745491-DEL-140
chr18-60893682-DEL-81
chr18-60919726-DEL-60

chr18-60922968-DEL-1299
chr18-60923815-DEL-610
chr18-60926364-DEL-377
chr18-61001802-DEL-337
chr18-61029001-DEL-216
chr18-61029030-DEL-462
chr18-61060262-DEL-132
chr18-61100598-DEL-54
chr18-61267249-DEL-112
chr18-61395089-DEL-77
chr18-61415926-DEL-80
chr18-61453693-DEL-63
chr18-61486388-DEL-50
chr18-61489689-DEL-120
chr18-61492163-DEL-740
chr18-61552047-DEL-53
chr18-61634609-DEL-231
chr18-61643845-DEL-122
chr18-61666499-DEL-178
chr18-61667066-DEL-283
chr18-61746014-DEL-228
chr18-61844029-DEL-78
chr18-61902320-DEL-100
chr18-61950104-DEL-50
chr18-62107953-DEL-60
chr18-62162253-DEL-73
chr18-62301355-DEL-123
chr18-62301430-DEL-51
chr18-62488083-DEL-244
chr18-62527296-DEL-405
chr18-62810071-DEL-50
chr18-62840280-DEL-64
chr18-62840407-DEL-567
chr18-62896779-DEL-104
chr18-63025950-DEL-489
chr18-63092753-DEL-168
chr18-63240753-DEL-172
chr18-63286785-DEL-150
chr18-63382526-DEL-1785
chr18-63436134-DEL-261
chr18-63492317-DEL-78
chr18-63512290-DEL-324
chr18-63533424-DEL-50
chr18-63660231-DEL-243
chr18-63667764-DEL-1862
chr18-63673688-DEL-416
chr18-63845361-DEL-90
chr18-63845546-DEL-180
chr18-64066009-DEL-54
chr18-64099805-DEL-200
chr18-64234354-DEL-103
chr18-64308543-DEL-62
chr18-64314536-DEL-52
chr18-64419281-DEL-134
chr18-64517062-DEL-135
chr18-64585389-DEL-50
chr18-64607347-DEL-234
chr18-64620145-DEL-50
chr18-64629358-DEL-317
chr18-64656559-DEL-1013
chr18-64701795-DEL-62
chr18-64701877-DEL-92
chr18-64703288-DEL-159
chr18-64722214-DEL-220
chr18-64722494-DEL-75
chr18-64876093-DEL-163
chr18-64877503-DEL-50
chr18-64901614-DEL-109
chr18-64901664-DEL-235
chr18-64901738-DEL-53
chr18-64943386-DEL-67
chr18-64959768-DEL-1535
chr18-64960016-DEL-50
chr18-64965376-DEL-60
chr18-64981442-DEL-67
chr18-65138157-DEL-65
chr18-65147921-DEL-60

chr18-65271440-DEL-82
chr18-65271440-DEL-184
chr18-65271455-DEL-372
chr18-65271741-DEL-86
chr18-65360772-DEL-54
chr18-65514230-DEL-322
chr18-65556893-DEL-1552
chr18-65623123-DEL-241
chr18-65681047-DEL-883
chr18-65750166-DEL-246
chr18-65829253-DEL-855
chr18-65859919-DEL-51
chr18-66011884-DEL-54
chr18-66087430-DEL-1193
chr18-66119311-DEL-270
chr18-66163387-DEL-109
chr18-66220249-DEL-62
chr18-66224316-DEL-196
chr18-66315397-DEL-103
chr18-66324412-DEL-87
chr18-66345467-DEL-50
chr18-66349021-DEL-190
chr18-66394931-DEL-1050
chr18-66515349-DEL-77
chr18-66516107-DEL-90
chr18-66540643-DEL-103
chr18-66616739-DEL-58
chr18-66651327-DEL-78
chr18-66656183-DEL-54
chr18-66747591-DEL-50
chr18-66773890-DEL-75
chr18-66815254-DEL-50
chr18-66892127-DEL-410
chr18-66892271-DEL-146
chr18-66893424-DEL-54
chr18-66980279-DEL-62
chr18-67014311-DEL-1592
chr18-67243153-DEL-3035
chr18-67262932-DEL-373
chr18-67280790-DEL-156
chr18-67285114-DEL-206
chr18-67337253-DEL-51
chr18-67373240-DEL-65
chr18-67392078-DEL-64
chr18-67419630-DEL-768
chr18-67486351-DEL-57
chr18-67486428-DEL-54
chr18-67489295-DEL-77
chr18-67492368-DEL-71
chr18-67514110-DEL-617
chr18-67522492-DEL-74
chr18-67527919-DEL-60
chr18-67527923-DEL-170
chr18-67603021-DEL-1938
chr18-67666668-DEL-949
chr18-67684430-DEL-226
chr18-67747468-DEL-75
chr18-67757445-DEL-53
chr18-67761080-DEL-503
chr18-67790978-DEL-383
chr18-67893287-DEL-69
chr18-67969877-DEL-252
chr18-68022160-DEL-201
chr18-68205133-DEL-155
chr18-68249817-DEL-68
chr18-68297285-DEL-74
chr18-68384932-DEL-54
chr18-68405963-DEL-169
chr18-68494706-DEL-60
chr18-68520108-DEL-208
chr18-68554979-DEL-50
chr18-68568593-DEL-84
chr18-68605519-DEL-203
chr18-68607490-DEL-206
chr18-68652027-DEL-276
chr18-68679006-DEL-59
chr18-68730846-DEL-670

chr18-68758391-DEL-92
chr18-68904354-DEL-467
chr18-68911525-DEL-3970
chr18-68945492-DEL-841
chr18-69058185-DEL-52
chr18-69071658-DEL-282
chr18-69071783-DEL-130
chr18-69075083-DEL-148
chr18-69163821-DEL-113
chr18-69170193-DEL-1713
chr18-69242547-DEL-68
chr18-69242597-DEL-187
chr18-69242772-DEL-85
chr18-69271394-DEL-107
chr18-69324173-DEL-68
chr18-69362691-DEL-255
chr18-69363049-DEL-629
chr18-69373898-DEL-60
chr18-69377713-DEL-69
chr18-69395189-DEL-1350
chr18-69409267-DEL-219
chr18-69590825-DEL-85
chr18-69742455-DEL-50
chr18-69748107-DEL-1254
chr18-69826481-DEL-75
chr18-69855276-DEL-92
chr18-69944027-DEL-407
chr18-69944096-DEL-66
chr18-69944331-DEL-520
chr18-69974799-DEL-74
chr18-70035852-DEL-151
chr18-70119020-DEL-55
chr18-70193712-DEL-67
chr18-70195917-DEL-276
chr18-70329452-DEL-1758
chr18-70383098-DEL-149
chr18-70457074-DEL-2746
chr18-70499109-DEL-1129
chr18-70525926-DEL-52
chr18-70539813-DEL-189
chr18-70539860-DEL-75
chr18-70545344-DEL-71
chr18-70588301-DEL-56
chr18-70594291-DEL-53
chr18-70614828-DEL-98
chr18-70617660-DEL-1027
chr18-70756618-DEL-126
chr18-70892332-DEL-830
chr18-70968822-DEL-67
chr18-71061269-DEL-208
chr18-71186368-DEL-50
chr18-71191735-DEL-122
chr18-71225569-DEL-80
chr18-71238172-DEL-235
chr18-71424803-DEL-252
chr18-71556704-DEL-207
chr18-71594155-DEL-569
chr18-71909424-DEL-86
chr18-72024061-DEL-90
chr18-72040239-DEL-62
chr18-72178397-DEL-50
chr18-72352556-DEL-57
chr18-72452484-DEL-58
chr18-72482033-DEL-62
chr18-72522717-DEL-110
chr18-72680664-DEL-76
chr18-72710249-DEL-424
chr18-72724167-DEL-161
chr18-72735112-DEL-943
chr18-72763460-DEL-63
chr18-72781331-DEL-1575
chr18-72787476-DEL-4534
chr18-72905876-DEL-1081
chr18-72979047-DEL-256
chr18-73137117-DEL-75
chr18-73188914-DEL-121
chr18-73364174-DEL-5384

chr18-73421977-DEL-320
chr18-73422251-DEL-114
chr18-73429044-DEL-163
chr18-73444742-DEL-70
chr18-73500285-DEL-80
chr18-73556649-DEL-690
chr18-73556649-DEL-1800
chr18-73557745-DEL-360
chr18-73573108-DEL-65
chr18-73574082-DEL-56
chr18-73574194-DEL-157
chr18-73580781-DEL-459
chr18-73582188-DEL-124
chr18-73785555-DEL-84
chr18-73822808-DEL-1114
chr18-73822874-DEL-128
chr18-73822903-DEL-458
chr18-73826419-DEL-62
chr18-73905643-DEL-50
chr18-73905776-DEL-50
chr18-73960353-DEL-56
chr18-73962188-DEL-140
chr18-74008156-DEL-93
chr18-74126061-DEL-84
chr18-74149246-DEL-72
chr18-74165133-DEL-51
chr18-74205999-DEL-92
chr18-74327640-DEL-51
chr18-74597038-DEL-85
chr18-74637545-DEL-60
chr18-74686066-DEL-1642
chr18-74746377-DEL-669
chr18-74773160-DEL-60
chr18-74871747-DEL-96
chr18-74896083-DEL-507
chr18-74938556-DEL-58
chr18-74994607-DEL-104
chr18-75013718-DEL-184
chr18-75013753-DEL-699
chr18-75013890-DEL-72
chr18-75014108-DEL-142
chr18-75014437-DEL-120
chr18-75211226-DEL-166
chr18-75218775-DEL-88
chr18-75219499-DEL-252
chr18-75240236-DEL-96
chr18-75275725-DEL-75
chr18-75275914-DEL-57
chr18-75276047-DEL-57
chr18-75401358-DEL-153
chr18-75415613-DEL-63
chr18-75448720-DEL-58
chr18-75558540-DEL-54
chr18-75569078-DEL-100
chr18-75613431-DEL-124
chr18-75742376-DEL-63
chr18-75748452-DEL-381
chr18-75852585-DEL-704
chr18-75877460-DEL-50
chr18-76083196-DEL-71
chr18-76143551-DEL-157
chr18-76174443-DEL-153
chr18-76174710-DEL-55
chr18-76174786-DEL-126
chr18-76239700-DEL-82
chr18-76239962-DEL-134
chr18-76299099-DEL-108
chr18-76307503-DEL-54
chr18-76359584-DEL-153
chr18-76377105-DEL-129
chr18-76472963-DEL-195
chr18-76548359-DEL-834
chr18-76593231-DEL-102
chr18-76627873-DEL-78
chr18-76655502-DEL-592
chr18-76655864-DEL-626
chr18-76662689-DEL-104

chr18-76663677-DEL-490
chr18-76685359-DEL-228
chr18-76689537-DEL-466
chr18-76721750-DEL-110
chr18-76724501-DEL-537
chr18-76725075-DEL-945
chr18-76741556-DEL-111
chr18-76768046-DEL-98
chr18-76778063-DEL-60
chr18-76858261-DEL-52
chr18-76928805-DEL-213
chr18-77044792-DEL-678
chr18-77131800-DEL-130
chr18-77156981-DEL-120
chr18-77183348-DEL-400
chr18-77277229-DEL-344
chr18-77277402-DEL-130
chr18-77324197-DEL-317
chr18-77348777-DEL-75
chr18-77385570-DEL-88
chr18-77507415-DEL-131
chr18-77526301-DEL-105
chr18-77545320-DEL-135
chr18-77585169-DEL-235
chr18-77609506-DEL-72
chr18-77618616-DEL-245
chr18-77678483-DEL-66
chr18-77709788-DEL-54
chr18-77824553-DEL-112
chr18-77828010-DEL-76
chr18-77839742-DEL-206
chr18-77841344-DEL-190
chr18-77859047-DEL-152
chr18-77875106-DEL-94
chr18-77878662-DEL-438
chr18-77932096-DEL-58
chr18-77939507-DEL-168
chr18-77971905-DEL-74
chr18-77972008-DEL-111
chr18-78008594-DEL-54
chr18-78037283-DEL-181
chr18-78043185-DEL-221
chr18-78059716-DEL-345
chr18-78095708-DEL-95
chr18-78171297-DEL-51
chr18-78201388-DEL-53
chr18-78211887-DEL-306
chr18-78275088-DEL-60
chr18-78314224-DEL-1789
chr18-78413438-DEL-185
chr18-78641445-DEL-174
chr18-78687910-DEL-1377
chr18-78739643-DEL-636
chr18-78791703-DEL-584
chr18-78795972-DEL-60
chr18-79020749-DEL-69
chr18-79028513-DEL-168
chr18-79071607-DEL-94
chr18-79238643-DEL-1014
chr18-79391142-DEL-68
chr18-79457307-DEL-64
chr18-79484751-DEL-472
chr18-79572203-DEL-59
chr18-79740327-DEL-268
chr18-79743144-DEL-55
chr18-79768008-DEL-208
chr18-79853283-DEL-3849
chr18-79970686-DEL-84
chr18-79970929-DEL-109
chr18-79973253-DEL-71
chr18-80055116-DEL-13458
chr18-80058766-DEL-6514
chr18-80204441-DEL-95
chr18-80259570-DEL-319
chr18-80259735-DEL-62
chr18-80307793-DEL-205
chr18-80344423-DEL-66

chr18-80362815-DEL-994
chr18-80428471-DEL-121
chr18-80548217-DEL-55
chr18-80763641-DEL-122
chr18-80763775-DEL-58
chr18-80953919-DEL-55
chr18-80958364-DEL-1810
chr18-80997403-DEL-55
chr18-81072398-DEL-81
chr18-81098394-DEL-108
chr18-81139422-DEL-338
chr18-81139708-DEL-468
chr18-81274164-DEL-81
chr18-81310386-DEL-58
chr18-81330361-DEL-50
chr18-81447353-DEL-252
chr18-81541737-DEL-51
chr18-81597233-DEL-851
chr18-81643706-DEL-332
chr18-81662979-DEL-2885
chr18-81696438-DEL-59
chr18-81707885-DEL-56
chr18-81744483-DEL-54
chr18-81755223-DEL-363
chr18-81756074-DEL-506
chr18-81756084-DEL-221
chr18-81817307-DEL-50
chr18-81838305-DEL-136
chr18-81838749-DEL-204
chr18-81916042-DEL-56
chr18-81916152-DEL-90
chr18-81916522-DEL-60
chr18-82036143-DEL-77
chr18-82040415-DEL-103
chr18-82207255-DEL-120
chr18-82258102-DEL-104
chr18-82280650-DEL-70
chr18-82296972-DEL-425
chr18-82320751-DEL-50
chr18-82321106-DEL-71
chr18-82437899-DEL-75
chr18-82548108-DEL-53
chr18-82650009-DEL-288
chr18-82726574-DEL-74
chr18-82737966-DEL-75
chr18-82764891-DEL-58
chr18-82776832-DEL-57
chr18-82796989-DEL-51
chr18-82893391-DEL-64
chr18-82942204-DEL-96
chr18-82948734-DEL-50
chr18-82970920-DEL-139
chr18-83002046-DEL-124
chr18-83003792-DEL-54
chr18-83036748-DEL-96
chr18-83064080-DEL-276
chr18-83074954-DEL-54
chr18-83075115-DEL-146
chr18-83102990-DEL-178
chr18-83104014-DEL-177
chr18-83105468-DEL-112
chr18-83120832-DEL-104
chr18-83121129-DEL-119
chr18-83156433-DEL-78
chr18-83156454-DEL-162
chr18-83156767-DEL-158
chr18-83157152-DEL-60
chr18-83274231-DEL-127
chr18-83290509-DEL-57
chr18-83344110-DEL-52
chr18-83376863-DEL-87
chr18-83388290-DEL-109
chr18-83434736-DEL-282
chr18-83451814-DEL-74
chr18-83455330-DEL-54
chr18-83511409-DEL-60
chr18-83559318-DEL-66

chr18-83559556-DEL-214
chr18-83586483-DEL-144
chr18-83716811-DEL-381
chr18-83803931-DEL-88
chr18-83855133-DEL-99
chr18-83905355-DEL-68
chr18-84000670-DEL-81
chr18-84128969-DEL-1098
chr18-84160565-DEL-834
chr18-84175919-DEL-640
chr18-84230997-DEL-96
chr18-84408007-DEL-180
chr18-84521626-DEL-467
chr18-84581850-DEL-1449
chr18-84581881-DEL-690
chr18-84582922-DEL-498
chr18-84661063-DEL-102
chr18-84661453-DEL-67
chr18-84672264-DEL-146
chr18-84672492-DEL-290
chr18-84672519-DEL-132
chr18-84689106-DEL-798
chr18-84690510-DEL-72
chr18-84690648-DEL-95
chr18-84690891-DEL-72
chr18-84752309-DEL-51
chr18-84848940-DEL-2658
chr18-84848978-DEL-666
chr18-84849467-DEL-115
chr18-84873204-DEL-394
chr18-84881246-DEL-60
chr18-85019956-DEL-6265
chr18-85027103-DEL-83
chr18-85027310-DEL-280
chr18-85088149-DEL-183
chr18-85266622-DEL-84
chr18-85404732-DEL-50
chr18-85524809-DEL-56
chr18-85533314-DEL-126
chr18-85625056-DEL-85
chr18-85635392-DEL-3990
chr18-85780272-DEL-150
chr18-85808645-DEL-1061
chr18-85845566-DEL-50
chr18-86273350-DEL-51
chr18-86389816-DEL-368
chr18-86631135-DEL-1736
chr18-86749345-DEL-251
chr18-86951006-DEL-134
chr18-86995123-DEL-53
chr18-87023685-DEL-64
chr18-87023874-DEL-3754
chr18-87133714-DEL-422
chr18-87213780-DEL-67
chr18-87386728-DEL-1057
chr18-87406777-DEL-50
chr18-87497009-DEL-452
chr18-87733689-DEL-50
chr18-87781793-DEL-96
chr18-87858759-DEL-60
chr18-87927126-DEL-161
chr18-88079197-DEL-2657
chr18-88123749-DEL-234
chr18-88145714-DEL-60
chr18-88281435-DEL-811
chr18-88380742-DEL-54
chr18-88585948-DEL-5822
chr18-88591422-DEL-398
chr18-89052585-DEL-232
chr18-89387021-DEL-129
chr18-89538532-DEL-1218
chr18-89784208-DEL-83
chr18-89878130-DEL-6530
chr18-89885065-DEL-1963
chr18-89899881-DEL-150
chr18-89935662-DEL-207
chr18-90302207-DEL-2081

chr18-90391958-DEL-424
chr18-90423178-DEL-616
chr18-90488373-DEL-1852
chr18-90571229-DEL-7104
chr19-3133057-DEL-3989
chr19-3377379-DEL-404
chr19-3470357-DEL-278
chr19-3491608-DEL-88
chr19-3491713-DEL-390
chr19-3491802-DEL-150
chr19-3492012-DEL-59
chr19-3645912-DEL-80
chr19-3657254-DEL-98
chr19-3684519-DEL-158
chr19-3722638-DEL-50
chr19-3786949-DEL-158
chr19-3788849-DEL-620
chr19-3788879-DEL-177
chr19-3877135-DEL-144
chr19-3907332-DEL-59
chr19-4017585-DEL-50
chr19-4026865-DEL-68
chr19-4072677-DEL-92
chr19-4074060-DEL-162
chr19-4074398-DEL-108
chr19-4074454-DEL-51
chr19-4079678-DEL-52
chr19-4268550-DEL-50
chr19-4329006-DEL-167
chr19-4478843-DEL-50
chr19-4512870-DEL-86
chr19-4521531-DEL-108
chr19-4636883-DEL-70
chr19-4684874-DEL-50
chr19-4759403-DEL-260
chr19-4759525-DEL-58
chr19-4868265-DEL-1888
chr19-4868403-DEL-628
chr19-4878045-DEL-250
chr19-5213166-DEL-536
chr19-5246255-DEL-56
chr19-5255050-DEL-481
chr19-5291472-DEL-245
chr19-5291509-DEL-97
chr19-5306805-DEL-118
chr19-5366796-DEL-2998
chr19-5492996-DEL-50
chr19-5591756-DEL-52
chr19-5643195-DEL-143
chr19-5650629-DEL-114
chr19-5675754-DEL-52
chr19-5681665-DEL-54
chr19-5720937-DEL-52
chr19-5728137-DEL-217
chr19-5736867-DEL-168
chr19-5814909-DEL-50
chr19-5822471-DEL-1241
chr19-5863384-DEL-54
chr19-5864095-DEL-216
chr19-5872113-DEL-75
chr19-5873761-DEL-95
chr19-5880831-DEL-344
chr19-5904075-DEL-440
chr19-6054177-DEL-241
chr19-6054602-DEL-109
chr19-6054653-DEL-54
chr19-6055094-DEL-159
chr19-6092083-DEL-479
chr19-6160696-DEL-495
chr19-6182881-DEL-828
chr19-6225314-DEL-500
chr19-6240214-DEL-158
chr19-6277348-DEL-120
chr19-6298464-DEL-657
chr19-6377855-DEL-175
chr19-6379341-DEL-138
chr19-6445047-DEL-71

chr19-6543953-DEL-153
chr19-6558733-DEL-306
chr19-6608655-DEL-100
chr19-6608852-DEL-411
chr19-6609139-DEL-103
chr19-6624720-DEL-126
chr19-6628761-DEL-108
chr19-6679685-DEL-1179
chr19-6681335-DEL-223
chr19-6703909-DEL-52
chr19-6717658-DEL-126
chr19-6739507-DEL-50645
chr19-6988276-DEL-68
chr19-6994392-DEL-168
chr19-7090390-DEL-103
chr19-7092959-DEL-116
chr19-7100870-DEL-151
chr19-7101368-DEL-456
chr19-7101450-DEL-211
chr19-7167856-DEL-51
chr19-7196529-DEL-986
chr19-7289323-DEL-104
chr19-7329773-DEL-95
chr19-7338886-DEL-485
chr19-7354571-DEL-90
chr19-7361252-DEL-62
chr19-7378363-DEL-224
chr19-7379404-DEL-744
chr19-7491700-DEL-2071
chr19-7547781-DEL-50
chr19-7547959-DEL-115
chr19-7615126-DEL-96
chr19-7694128-DEL-1508
chr19-7704539-DEL-722
chr19-7730794-DEL-176
chr19-7930834-DEL-151
chr19-7937403-DEL-80
chr19-7974627-DEL-3389
chr19-8060191-DEL-327
chr19-8120414-DEL-56
chr19-8182651-DEL-463
chr19-8190762-DEL-1393
chr19-8221012-DEL-6804
chr19-8470146-DEL-11904
chr19-8500237-DEL-2904
chr19-8704490-DEL-207
chr19-8704702-DEL-209
chr19-8842592-DEL-219
chr19-8860716-DEL-255
chr19-8927011-DEL-492
chr19-8927156-DEL-175
chr19-8927478-DEL-147
chr19-9129298-DEL-1364
chr19-9585413-DEL-84
chr19-9630543-DEL-1833
chr19-9633809-DEL-124
chr19-9731235-DEL-190
chr19-9805892-DEL-8693
chr19-9945170-DEL-524
chr19-9964605-DEL-102
chr19-10028946-DEL-100
chr19-10189433-DEL-50
chr19-10252064-DEL-468
chr19-10301380-DEL-84
chr19-10445896-DEL-169
chr19-10448772-DEL-222
chr19-10449091-DEL-222
chr19-10450222-DEL-81
chr19-10485767-DEL-167
chr19-10486974-DEL-133
chr19-10681920-DEL-108
chr19-10692381-DEL-62
chr19-10694901-DEL-180
chr19-10728372-DEL-109
chr19-10728393-DEL-54
chr19-10863445-DEL-711
chr19-10926167-DEL-120

chr19-10959389-DEL-68
chr19-10976475-DEL-64
chr19-11050791-DEL-164
chr19-11061827-DEL-8042
chr19-11086449-DEL-197
chr19-11100139-DEL-2675
chr19-11108758-DEL-112
chr19-11165271-DEL-78
chr19-11239209-DEL-64
chr19-11280848-DEL-105
chr19-11284904-DEL-50
chr19-11370331-DEL-72
chr19-11396364-DEL-40160
chr19-11410042-DEL-88
chr19-11429666-DEL-86
chr19-11527346-DEL-50
chr19-11708623-DEL-3519
chr19-11761051-DEL-69
chr19-11871326-DEL-156
chr19-11966942-DEL-279
chr19-11994364-DEL-663
chr19-12015412-DEL-62
chr19-12033186-DEL-14935
chr19-12135466-DEL-1138
chr19-12153104-DEL-62878
chr19-12250236-DEL-386
chr19-12318735-DEL-78
chr19-12530105-DEL-107
chr19-12615025-DEL-84
chr19-12821025-DEL-85
chr19-12829168-DEL-104
chr19-12845810-DEL-9009
chr19-13080905-DEL-88
chr19-13081136-DEL-61
chr19-13131807-DEL-90
chr19-13221190-DEL-1887
chr19-13231413-DEL-169
chr19-13365857-DEL-55
chr19-13494663-DEL-10229
chr19-13501093-DEL-220
chr19-13601726-DEL-140
chr19-13678854-DEL-50
chr19-13715444-DEL-86
chr19-13977112-DEL-201
chr19-14076519-DEL-189
chr19-14139293-DEL-52
chr19-14141456-DEL-54
chr19-14193116-DEL-403
chr19-14264473-DEL-235
chr19-14264770-DEL-336
chr19-14343501-DEL-170
chr19-14354006-DEL-185
chr19-14384358-DEL-52
chr19-14536335-DEL-56
chr19-14536491-DEL-170
chr19-14595419-DEL-56
chr19-14844192-DEL-224
chr19-14897464-DEL-50
chr19-15129857-DEL-58
chr19-15225174-DEL-142
chr19-15243368-DEL-277
chr19-15425401-DEL-236
chr19-15461110-DEL-50
chr19-15603197-DEL-82
chr19-15776678-DEL-536
chr19-15953020-DEL-152
chr19-15956733-DEL-54
chr19-15979842-DEL-82
chr19-16029435-DEL-50
chr19-16042909-DEL-50
chr19-16115558-DEL-60
chr19-16122975-DEL-58
chr19-16172240-DEL-57
chr19-16204645-DEL-191
chr19-16424477-DEL-341
chr19-16482704-DEL-61
chr19-16566263-DEL-696

chr19-16640013-DEL-68
chr19-16691903-DEL-68
chr19-16735265-DEL-318
chr19-16755007-DEL-159
chr19-16755182-DEL-60
chr19-16778627-DEL-56
chr19-16795617-DEL-175
chr19-16795682-DEL-60
chr19-17016309-DEL-105
chr19-17016467-DEL-57
chr19-17022346-DEL-557
chr19-17023376-DEL-389
chr19-17110928-DEL-106
chr19-17116525-DEL-153
chr19-17123060-DEL-397
chr19-17222634-DEL-150
chr19-17254611-DEL-58
chr19-17254894-DEL-228
chr19-17335467-DEL-5175
chr19-17354835-DEL-185
chr19-17407063-DEL-199
chr19-17407129-DEL-97
chr19-17407392-DEL-53
chr19-17502305-DEL-52
chr19-17503837-DEL-60
chr19-17618246-DEL-143
chr19-17645817-DEL-119
chr19-17646072-DEL-67
chr19-17693496-DEL-58
chr19-17701586-DEL-61
chr19-17714397-DEL-50
chr19-17840098-DEL-66
chr19-17957362-DEL-50
chr19-17985989-DEL-53
chr19-18018826-DEL-284
chr19-18069680-DEL-131
chr19-18069861-DEL-63
chr19-18191378-DEL-88
chr19-18227634-DEL-158
chr19-18257810-DEL-79
chr19-18362987-DEL-138
chr19-18576790-DEL-57
chr19-18678300-DEL-101
chr19-18723050-DEL-98
chr19-18771198-DEL-398
chr19-18815929-DEL-103
chr19-19147583-DEL-84
chr19-19438353-DEL-78
chr19-19440497-DEL-1075
chr19-19447280-DEL-102
chr19-19575926-DEL-52
chr19-19727616-DEL-136
chr19-19806063-DEL-1780
chr19-19884023-DEL-51
chr19-19938588-DEL-1212
chr19-20003707-DEL-102
chr19-20045084-DEL-99
chr19-20058028-DEL-1419
chr19-20181405-DEL-594
chr19-20400338-DEL-284
chr19-20479093-DEL-53
chr19-20492721-DEL-52
chr19-20563093-DEL-164
chr19-20708394-DEL-182
chr19-20853796-DEL-68
chr19-20866843-DEL-1667
chr19-20897059-DEL-183
chr19-20903336-DEL-50
chr19-20975794-DEL-353
chr19-21079337-DEL-110
chr19-21154438-DEL-576
chr19-21179716-DEL-52
chr19-21187015-DEL-76
chr19-21195705-DEL-212
chr19-21199935-DEL-1123
chr19-21352523-DEL-52
chr19-21428899-DEL-1294

chr19-21481189-DEL-72
chr19-21496172-DEL-176
chr19-21531879-DEL-102
chr19-21605745-DEL-50
chr19-21605776-DEL-201
chr19-21761645-DEL-50
chr19-21762856-DEL-723
chr19-21810158-DEL-73
chr19-21886371-DEL-74
chr19-21889056-DEL-80
chr19-21897793-DEL-14076
chr19-21933541-DEL-634
chr19-21990619-DEL-72
chr19-22017367-DEL-178
chr19-22055041-DEL-50
chr19-22067776-DEL-268
chr19-22113436-DEL-57
chr19-22427604-DEL-149
chr19-22501336-DEL-179
chr19-22565210-DEL-695
chr19-22565767-DEL-110
chr19-22630066-DEL-50
chr19-22669168-DEL-52
chr19-22720274-DEL-81
chr19-22831369-DEL-63
chr19-22864592-DEL-108
chr19-23011084-DEL-50
chr19-23076034-DEL-180
chr19-23282263-DEL-3879
chr19-23389574-DEL-135
chr19-23548837-DEL-134
chr19-23549054-DEL-129
chr19-23595031-DEL-2971
chr19-23702312-DEL-64
chr19-23816502-DEL-55
chr19-23929619-DEL-75
chr19-23959121-DEL-244
chr19-23962331-DEL-249
chr19-23962478-DEL-102
chr19-23969784-DEL-50
chr19-23984017-DEL-84
chr19-24123052-DEL-64
chr19-24133715-DEL-50
chr19-24140198-DEL-50
chr19-24147043-DEL-518
chr19-24199478-DEL-90
chr19-24274039-DEL-252
chr19-24300125-DEL-126
chr19-24326996-DEL-139
chr19-24327889-DEL-192
chr19-24353027-DEL-157
chr19-24479395-DEL-158
chr19-24551092-DEL-50
chr19-24564443-DEL-57
chr19-24602422-DEL-74
chr19-24746071-DEL-1240
chr19-24747042-DEL-390
chr19-24796205-DEL-556
chr19-24823535-DEL-128
chr19-24835397-DEL-79
chr19-24889595-DEL-6305
chr19-25034771-DEL-234
chr19-25095142-DEL-1232
chr19-25102155-DEL-112
chr19-25202684-DEL-84
chr19-25228344-DEL-592
chr19-25251161-DEL-194
chr19-25273903-DEL-75
chr19-25275586-DEL-52
chr19-25287095-DEL-136
chr19-25310724-DEL-100
chr19-25348351-DEL-325
chr19-25348479-DEL-1053
chr19-25349032-DEL-372
chr19-25349440-DEL-160
chr19-25429904-DEL-392
chr19-25434846-DEL-282

chr19-25442335-DEL-314
chr19-25448244-DEL-168
chr19-25449164-DEL-311
chr19-25449250-DEL-101
chr19-25490810-DEL-242
chr19-25554644-DEL-79
chr19-25572625-DEL-152
chr19-25614010-DEL-247
chr19-25614175-DEL-52
chr19-25688032-DEL-453
chr19-25814711-DEL-74
chr19-25872125-DEL-66
chr19-25873535-DEL-461
chr19-25963218-DEL-158
chr19-26035649-DEL-288
chr19-26041121-DEL-51
chr19-26041272-DEL-156
chr19-26136142-DEL-141
chr19-26273188-DEL-52
chr19-26369102-DEL-98
chr19-26514102-DEL-304
chr19-26533462-DEL-55
chr19-26533677-DEL-1266
chr19-26553187-DEL-880
chr19-26649068-DEL-54
chr19-26658815-DEL-50
chr19-26661563-DEL-62
chr19-26798833-DEL-504
chr19-26923160-DEL-403
chr19-27058490-DEL-459
chr19-27069896-DEL-660
chr19-27172647-DEL-56
chr19-27326910-DEL-50
chr19-27516582-DEL-51
chr19-27539281-DEL-77
chr19-27612035-DEL-141
chr19-27645024-DEL-54
chr19-27658306-DEL-85
chr19-27680659-DEL-229
chr19-27728782-DEL-424
chr19-27747064-DEL-255
chr19-27833419-DEL-50
chr19-28038617-DEL-64
chr19-28085647-DEL-64
chr19-28101105-DEL-1108
chr19-28219324-DEL-525
chr19-28378117-DEL-60
chr19-28503762-DEL-275
chr19-28503940-DEL-124
chr19-28504002-DEL-322
chr19-28504437-DEL-91
chr19-28667940-DEL-55
chr19-28713492-DEL-135
chr19-28938285-DEL-158
chr19-29074022-DEL-1275
chr19-29080434-DEL-58
chr19-29114837-DEL-124
chr19-29383102-DEL-75
chr19-29428474-DEL-245
chr19-29455397-DEL-68
chr19-29491577-DEL-310
chr19-29504134-DEL-56
chr19-29603555-DEL-311
chr19-29617816-DEL-202
chr19-29640104-DEL-502
chr19-29670112-DEL-2486
chr19-29674174-DEL-169
chr19-29829683-DEL-399
chr19-29830750-DEL-50
chr19-29961158-DEL-243
chr19-30125659-DEL-53
chr19-30144003-DEL-77
chr19-30187587-DEL-1109
chr19-30202944-DEL-209
chr19-30205800-DEL-66
chr19-30213067-DEL-423
chr19-30220155-DEL-56

chr19-30228489-DEL-311
chr19-30228682-DEL-128
chr19-30238769-DEL-166
chr19-30365535-DEL-1488
chr19-30375357-DEL-112
chr19-30401792-DEL-60
chr19-30401809-DEL-134
chr19-30455546-DEL-224
chr19-30455659-DEL-111
chr19-30478281-DEL-1084
chr19-30490181-DEL-56
chr19-30677867-DEL-174
chr19-30804246-DEL-68
chr19-30896020-DEL-330
chr19-30959024-DEL-50
chr19-31015195-DEL-337
chr19-31107105-DEL-78
chr19-31111195-DEL-57
chr19-31115615-DEL-534
chr19-31116318-DEL-57
chr19-31441564-DEL-50
chr19-31462028-DEL-126
chr19-31470027-DEL-92
chr19-31665438-DEL-54
chr19-31843804-DEL-55
chr19-31862048-DEL-2344
chr19-31869611-DEL-72
chr19-31921959-DEL-131
chr19-31955978-DEL-78
chr19-31976675-DEL-194
chr19-31979259-DEL-63
chr19-32057658-DEL-360
chr19-32109365-DEL-50
chr19-32109636-DEL-50
chr19-32123713-DEL-196
chr19-32278742-DEL-122
chr19-32284793-DEL-153
chr19-32368880-DEL-110
chr19-32479596-DEL-356
chr19-32505474-DEL-86
chr19-32699273-DEL-138
chr19-32723909-DEL-61
chr19-32728373-DEL-132
chr19-32744187-DEL-56
chr19-32852548-DEL-62
chr19-32982833-DEL-68
chr19-33053192-DEL-54
chr19-33055252-DEL-704
chr19-33394162-DEL-95
chr19-33399139-DEL-401
chr19-33434080-DEL-312
chr19-33457908-DEL-8978
chr19-33473267-DEL-50
chr19-33543979-DEL-982
chr19-33560245-DEL-50
chr19-33629082-DEL-160429
chr19-33649666-DEL-1786
chr19-33729443-DEL-5322
chr19-33786305-DEL-6357
chr19-33796177-DEL-219
chr19-33839635-DEL-66
chr19-33840077-DEL-5932
chr19-33847698-DEL-137
chr19-33866239-DEL-59
chr19-33866560-DEL-50
chr19-33928908-DEL-3175
chr19-33998031-DEL-634
chr19-34002796-DEL-6266
chr19-34082848-DEL-199
chr19-34369605-DEL-236
chr19-34426823-DEL-50
chr19-34463592-DEL-5910
chr19-34464806-DEL-1339
chr19-34475292-DEL-156
chr19-34569195-DEL-24309
chr19-34588504-DEL-5738
chr19-34662225-DEL-51

chr19-34762399-DEL-453
chr19-34956073-DEL-130
chr19-34956321-DEL-54
chr19-34956906-DEL-126
chr19-34957111-DEL-143
chr19-34993810-DEL-50
chr19-35003735-DEL-59
chr19-35006602-DEL-60
chr19-35073199-DEL-94
chr19-35131058-DEL-52
chr19-35132839-DEL-122
chr19-35133002-DEL-138
chr19-35244973-DEL-50
chr19-35430308-DEL-53
chr19-35447983-DEL-102
chr19-35508954-DEL-977
chr19-35562158-DEL-6783
chr19-35577012-DEL-93
chr19-35590565-DEL-127
chr19-35644004-DEL-204
chr19-35862686-DEL-69
chr19-35866296-DEL-763
chr19-35866605-DEL-185
chr19-36047795-DEL-2519
chr19-36050350-DEL-373
chr19-36082491-DEL-129
chr19-36135374-DEL-50
chr19-36136096-DEL-3083
chr19-36187177-DEL-60
chr19-36362845-DEL-97
chr19-36490269-DEL-117
chr19-36502861-DEL-115
chr19-36568762-DEL-50
chr19-36605655-DEL-80
chr19-36651910-DEL-84
chr19-36671196-DEL-148
chr19-36685010-DEL-82
chr19-36797812-DEL-52
chr19-36950780-DEL-52
chr19-36952823-DEL-1343
chr19-37012686-DEL-78
chr19-37276179-DEL-1396
chr19-37319348-DEL-217
chr19-37396151-DEL-50
chr19-37432916-DEL-637
chr19-37433164-DEL-278
chr19-37439745-DEL-159
chr19-37465940-DEL-340
chr19-37710871-DEL-249
chr19-37721914-DEL-293
chr19-37792254-DEL-128
chr19-37796730-DEL-53
chr19-37804175-DEL-74
chr19-37981885-DEL-138
chr19-38010528-DEL-53
chr19-38017607-DEL-858
chr19-38196093-DEL-54
chr19-38196453-DEL-53
chr19-38299206-DEL-51
chr19-38306524-DEL-173
chr19-38326305-DEL-54
chr19-38332239-DEL-207
chr19-38379926-DEL-99
chr19-38410914-DEL-833
chr19-38411782-DEL-98
chr19-38412056-DEL-90
chr19-38429259-DEL-2062
chr19-38445340-DEL-59
chr19-38463385-DEL-119
chr19-38478413-DEL-5282
chr19-38635136-DEL-50
chr19-38780788-DEL-50
chr19-38794269-DEL-64
chr19-38926380-DEL-106
chr19-38966138-DEL-1092
chr19-38985297-DEL-63
chr19-39034338-DEL-2535

chr19-39040877-DEL-2542
chr19-39342776-DEL-1981
chr19-39486604-DEL-74
chr19-39560354-DEL-481
chr19-39636665-DEL-54
chr19-39728629-DEL-213
chr19-39735765-DEL-58
chr19-39761074-DEL-75524
chr19-39810221-DEL-628
chr19-39823936-DEL-1484
chr19-39898099-DEL-2223
chr19-39986429-DEL-50
chr19-40123070-DEL-114
chr19-40175389-DEL-52
chr19-40215591-DEL-113
chr19-40294707-DEL-72
chr19-40317168-DEL-491
chr19-40373131-DEL-134
chr19-40472414-DEL-51
chr19-40558383-DEL-78
chr19-40694293-DEL-2270
chr19-40725828-DEL-62
chr19-40761703-DEL-452
chr19-40804801-DEL-156
chr19-40830622-DEL-1021
chr19-40843898-DEL-177
chr19-40845492-DEL-295
chr19-40879221-DEL-53
chr19-40932980-DEL-72
chr19-40988371-DEL-50
chr19-41054135-DEL-124
chr19-41082339-DEL-96
chr19-41083035-DEL-129
chr19-41092636-DEL-131
chr19-41115798-DEL-100
chr19-41119402-DEL-1484
chr19-41174210-DEL-128
chr19-41257495-DEL-54
chr19-41280021-DEL-689
chr19-41329932-DEL-80
chr19-41363329-DEL-169
chr19-41436948-DEL-427
chr19-41480861-DEL-50
chr19-41621750-DEL-107
chr19-41632181-DEL-129
chr19-41633630-DEL-69
chr19-41749319-DEL-114
chr19-41757716-DEL-132
chr19-41837667-DEL-153
chr19-41861535-DEL-212
chr19-41861572-DEL-70
chr19-41892602-DEL-220
chr19-42000759-DEL-69
chr19-42000939-DEL-207
chr19-42077528-DEL-122
chr19-42094694-DEL-1340
chr19-42253715-DEL-102
chr19-42284995-DEL-50
chr19-42341603-DEL-53
chr19-42480893-DEL-51
chr19-42545663-DEL-571
chr19-42657533-DEL-54
chr19-42687754-DEL-84
chr19-42687754-DEL-189
chr19-42703773-DEL-162
chr19-42704158-DEL-139
chr19-42761266-DEL-50
chr19-42838376-DEL-182
chr19-43018216-DEL-123
chr19-43072807-DEL-163
chr19-43210560-DEL-1839
chr19-43413067-DEL-892
chr19-43481796-DEL-70
chr19-43511527-DEL-283
chr19-43525142-DEL-687
chr19-43529932-DEL-164
chr19-43537860-DEL-135

chr19-43620460-DEL-89
chr19-43622651-DEL-141
chr19-43673722-DEL-54
chr19-43682974-DEL-200
chr19-43682977-DEL-50
chr19-43729982-DEL-94
chr19-43785479-DEL-60
chr19-43925992-DEL-98
chr19-44008960-DEL-114
chr19-44114378-DEL-84
chr19-44139340-DEL-214
chr19-44204402-DEL-122
chr19-44225960-DEL-50
chr19-44257512-DEL-102
chr19-44305800-DEL-82
chr19-44306147-DEL-177
chr19-44310754-DEL-361
chr19-44319694-DEL-158
chr19-44479829-DEL-225
chr19-44480505-DEL-66
chr19-44518310-DEL-50
chr19-44633652-DEL-53
chr19-44639662-DEL-50
chr19-44648796-DEL-93
chr19-44915667-DEL-52
chr19-45006275-DEL-63
chr19-45117897-DEL-116
chr19-45119020-DEL-224
chr19-45180996-DEL-53
chr19-45181898-DEL-100
chr19-45182070-DEL-140
chr19-45193966-DEL-69
chr19-45248394-DEL-201
chr19-45248634-DEL-191
chr19-45261893-DEL-272
chr19-45312439-DEL-218
chr19-45312776-DEL-110
chr19-45544754-DEL-193
chr19-45696325-DEL-154
chr19-45696675-DEL-136
chr19-45703150-DEL-116
chr19-45913065-DEL-50
chr19-45944625-DEL-128
chr19-45973595-DEL-76
chr19-45974264-DEL-200
chr19-46000234-DEL-59
chr19-46011563-DEL-1040
chr19-46075238-DEL-182
chr19-46077877-DEL-85
chr19-46078981-DEL-52
chr19-46096768-DEL-182
chr19-46101454-DEL-93
chr19-46217555-DEL-157
chr19-46282207-DEL-56
chr19-46330591-DEL-245
chr19-46330890-DEL-203
chr19-46452623-DEL-68
chr19-46481486-DEL-203
chr19-46527482-DEL-200
chr19-46587048-DEL-104
chr19-46594172-DEL-76
chr19-46599913-DEL-105
chr19-46615046-DEL-319
chr19-46685898-DEL-129
chr19-46688320-DEL-51
chr19-46714102-DEL-61
chr19-46738676-DEL-124
chr19-46739601-DEL-162
chr19-46739996-DEL-243
chr19-46816317-DEL-65
chr19-46831123-DEL-147
chr19-46980158-DEL-50
chr19-46989416-DEL-111
chr19-46995116-DEL-114
chr19-47047006-DEL-508
chr19-47074040-DEL-261
chr19-47090293-DEL-94

chr19-47130682-DEL-50
chr19-47135735-DEL-63
chr19-47237473-DEL-171
chr19-47283271-DEL-70
chr19-47296478-DEL-225
chr19-47511327-DEL-128
chr19-47622412-DEL-365
chr19-47622962-DEL-246
chr19-47679474-DEL-57
chr19-47736741-DEL-214
chr19-47742861-DEL-318
chr19-47742889-DEL-140
chr19-47744725-DEL-50
chr19-47793328-DEL-214
chr19-47810380-DEL-110
chr19-47814915-DEL-104
chr19-47815165-DEL-78
chr19-47822962-DEL-54
chr19-47901152-DEL-66
chr19-47945339-DEL-78
chr19-47945550-DEL-51
chr19-47960793-DEL-67
chr19-47968444-DEL-55
chr19-48000552-DEL-50
chr19-48039962-DEL-58
chr19-48042133-DEL-168
chr19-48042359-DEL-228
chr19-48043030-DEL-115
chr19-48053688-DEL-188
chr19-48053792-DEL-87
chr19-48102521-DEL-274
chr19-48165993-DEL-271
chr19-48166007-DEL-132
chr19-48166216-DEL-321
chr19-48166360-DEL-58
chr19-48189832-DEL-110
chr19-48256627-DEL-100
chr19-48390826-DEL-210
chr19-48397234-DEL-287
chr19-48397340-DEL-113
chr19-48420800-DEL-106
chr19-48458162-DEL-50
chr19-48537432-DEL-98
chr19-48545734-DEL-56
chr19-48546104-DEL-133
chr19-48802795-DEL-174
chr19-48877180-DEL-55
chr19-48998258-DEL-65
chr19-49090328-DEL-1374
chr19-49121217-DEL-790
chr19-49178713-DEL-124
chr19-49215277-DEL-52
chr19-49367717-DEL-52
chr19-49384019-DEL-74
chr19-49434427-DEL-141
chr19-49621360-DEL-102
chr19-49656355-DEL-66
chr19-49656491-DEL-182
chr19-49683935-DEL-95
chr19-49718295-DEL-61
chr19-49918822-DEL-53
chr19-49918941-DEL-54
chr19-49939062-DEL-2329
chr19-49973583-DEL-288
chr19-50016746-DEL-150
chr19-50172864-DEL-53
chr19-50175627-DEL-54
chr19-50251886-DEL-104
chr19-50301994-DEL-595
chr19-50312398-DEL-104
chr19-50776200-DEL-60
chr19-50826821-DEL-207
chr19-50890610-DEL-194
chr19-50890865-DEL-56
chr19-51148790-DEL-1974
chr19-51153950-DEL-130
chr19-51161846-DEL-1711

chr19-51236790-DEL-2632
chr19-51288895-DEL-195
chr19-51301551-DEL-50
chr19-51409675-DEL-68
chr19-51423661-DEL-117
chr19-51497142-DEL-144
chr19-51497194-DEL-57
chr19-51674796-DEL-56
chr19-51706230-DEL-1062
chr19-51940410-DEL-3076
chr19-51943711-DEL-212
chr19-51950525-DEL-231
chr19-51975144-DEL-806
chr19-52039633-DEL-168
chr19-52053488-DEL-69
chr19-52097895-DEL-205
chr19-52142691-DEL-216
chr19-52207419-DEL-150
chr19-52233062-DEL-81
chr19-52381004-DEL-76
chr19-52399040-DEL-50
chr19-52399086-DEL-136
chr19-52401191-DEL-118
chr19-52449103-DEL-139
chr19-52449369-DEL-143
chr19-52449458-DEL-64
chr19-52458370-DEL-207
chr19-52524018-DEL-188
chr19-52534060-DEL-556
chr19-52552488-DEL-50
chr19-52739669-DEL-69
chr19-52823310-DEL-182
chr19-52844368-DEL-52
chr19-52862665-DEL-211
chr19-52865336-DEL-150
chr19-52904053-DEL-77
chr19-53084680-DEL-128
chr19-53092331-DEL-62
chr19-53105130-DEL-424
chr19-53150582-DEL-56
chr19-53219041-DEL-50
chr19-53336123-DEL-618
chr19-53337214-DEL-158
chr19-53337747-DEL-180
chr19-53397363-DEL-128
chr19-53485947-DEL-68
chr19-53511646-DEL-955
chr19-53511920-DEL-2409
chr19-53513345-DEL-981
chr19-53514512-DEL-420
chr19-53528228-DEL-257
chr19-53528329-DEL-62
chr19-53564895-DEL-60
chr19-53587787-DEL-150
chr19-53644054-DEL-50
chr19-53712350-DEL-209
chr19-53763749-DEL-523
chr19-53763817-DEL-1438
chr19-53765261-DEL-449
chr19-53826670-DEL-76
chr19-53856272-DEL-90
chr19-53927259-DEL-56
chr19-53996747-DEL-60
chr19-53997048-DEL-67
chr19-54075071-DEL-76
chr19-54114148-DEL-62
chr19-54136128-DEL-255
chr19-54137050-DEL-400
chr19-54139097-DEL-901
chr19-54139347-DEL-308
chr19-54140436-DEL-110
chr19-54143862-DEL-72
chr19-54150557-DEL-4681
chr19-54170261-DEL-54
chr19-54185170-DEL-364
chr19-54195657-DEL-88
chr19-54197503-DEL-123

chr19-54234739-DEL-97
chr19-54238985-DEL-56
chr19-54371165-DEL-77
chr19-54477032-DEL-100
chr19-54477269-DEL-229
chr19-54490910-DEL-50
chr19-54689880-DEL-1818
chr19-54706793-DEL-63
chr19-54759960-DEL-134
chr19-54799139-DEL-2305
chr19-54844339-DEL-69
chr19-54943989-DEL-66
chr19-54967864-DEL-64
chr19-55029427-DEL-50
chr19-55040269-DEL-80
chr19-55051460-DEL-173
chr19-55051646-DEL-74
chr19-55099872-DEL-56
chr19-55118778-DEL-140
chr19-55131520-DEL-155
chr19-55137313-DEL-72
chr19-55153418-DEL-170
chr19-55153661-DEL-116
chr19-55185866-DEL-68
chr19-55227177-DEL-212
chr19-55230581-DEL-131
chr19-55292534-DEL-112
chr19-55334955-DEL-401
chr19-55356920-DEL-504
chr19-55357041-DEL-196
chr19-55445975-DEL-87
chr19-55503382-DEL-238
chr19-55503807-DEL-363
chr19-55504001-DEL-125
chr19-55674193-DEL-54
chr19-55684393-DEL-147
chr19-55684776-DEL-64
chr19-55712453-DEL-114
chr19-55719800-DEL-68
chr19-55758879-DEL-192
chr19-55831329-DEL-138
chr19-56026381-DEL-55
chr19-56029890-DEL-63
chr19-56058459-DEL-76
chr19-56131629-DEL-235
chr19-56184169-DEL-58
chr19-56264543-DEL-52
chr19-56311030-DEL-336
chr19-56311167-DEL-56
chr19-56311771-DEL-56
chr19-56344698-DEL-106
chr19-56371718-DEL-51
chr19-56409202-DEL-51
chr19-56451952-DEL-62
chr19-56477361-DEL-356
chr19-56477705-DEL-223
chr19-56630704-DEL-90
chr19-56678919-DEL-60
chr19-56697931-DEL-206
chr19-56721657-DEL-50
chr19-56746822-DEL-242
chr19-56785421-DEL-366
chr19-56785490-DEL-160
chr19-56883083-DEL-135
chr19-56921370-DEL-82
chr19-56921478-DEL-246
chr19-56956482-DEL-102
chr19-56960338-DEL-192
chr19-56960989-DEL-66
chr19-56967568-DEL-198
chr19-57084861-DEL-52
chr19-57102129-DEL-54
chr19-57164471-DEL-100
chr19-57179405-DEL-51
chr19-57181651-DEL-129
chr19-57234139-DEL-50
chr19-57235576-DEL-51

chr19-57236880-DEL-61
chr19-57239047-DEL-50
chr19-57334020-DEL-584
chr19-57335968-DEL-134
chr19-57336233-DEL-54
chr19-57436984-DEL-529
chr19-57437359-DEL-151
chr19-57437661-DEL-59
chr19-57437796-DEL-205
chr19-57439608-DEL-97
chr19-57509668-DEL-54
chr19-57570826-DEL-89
chr19-57613066-DEL-80
chr19-57613464-DEL-120
chr19-57650385-DEL-106
chr19-57659321-DEL-60
chr19-57659609-DEL-440
chr19-57660001-DEL-80
chr19-57660222-DEL-440
chr19-57660321-DEL-120
chr19-57697503-DEL-68
chr19-57697833-DEL-169
chr19-57703734-DEL-157
chr19-57703830-DEL-61
chr19-57747635-DEL-140
chr19-57855396-DEL-384
chr19-57925995-DEL-99
chr19-57971305-DEL-58
chr19-57982832-DEL-150
chr19-58131991-DEL-50
chr19-58132896-DEL-211
chr19-58138085-DEL-60
chr19-58220746-DEL-132
chr19-58220943-DEL-100
chr19-58231661-DEL-54
chr19-58271883-DEL-451
chr19-58280208-DEL-105
chr19-58280459-DEL-269
chr19-58306904-DEL-136
chr19-58320937-DEL-119
chr19-58334957-DEL-294
chr19-58477854-DEL-50
chr19-58558157-DEL-82
chr19-58562125-DEL-153
chr19-58674044-DEL-90
chr19-58760291-DEL-144
chr19-58821114-DEL-63
chr19-58955625-DEL-204
chr19-58955761-DEL-68
chr19-59033609-DEL-52
chr19-59188677-DEL-84
chr19-59188939-DEL-63
chr19-59188955-DEL-325
chr19-59201506-DEL-792
chr19-59367979-DEL-177
chr19-59459160-DEL-50
chr19-59519529-DEL-66
chr19-59519685-DEL-294
chr19-59541115-DEL-58
chr19-59541248-DEL-79
chr19-59654439-DEL-64
chr19-59749240-DEL-78
chr19-59873479-DEL-55
chr19-59873526-DEL-136
chr19-59951329-DEL-59
chr19-60117207-DEL-52
chr19-60283390-DEL-52
chr19-60336750-DEL-208
chr19-60398961-DEL-112
chr19-60712892-DEL-70
chr19-60789453-DEL-52
chr19-60841516-DEL-134
chr19-60892519-DEL-1060
chr19-60913200-DEL-62
chr2-3138721-DEL-62
chr2-3265328-DEL-50
chr2-3277195-DEL-64

chr2-3352006-DEL-78
chr2-3529891-DEL-336
chr2-3530811-DEL-206
chr2-3531896-DEL-777
chr2-3577123-DEL-54
chr2-3615790-DEL-187
chr2-3674689-DEL-66
chr2-3734766-DEL-150
chr2-3786010-DEL-322
chr2-3793599-DEL-128
chr2-3835544-DEL-4446
chr2-3851900-DEL-261
chr2-3921421-DEL-136
chr2-3921448-DEL-67
chr2-3965859-DEL-3398
chr2-3977421-DEL-72
chr2-4138698-DEL-50
chr2-4141397-DEL-138
chr2-4149042-DEL-140
chr2-4170867-DEL-52
chr2-4219683-DEL-1322
chr2-4224663-DEL-132
chr2-4224915-DEL-54
chr2-4259134-DEL-83
chr2-4323465-DEL-191
chr2-4458789-DEL-75
chr2-4527137-DEL-254
chr2-4569281-DEL-136
chr2-4576479-DEL-50
chr2-4602837-DEL-69
chr2-4619210-DEL-83
chr2-4623516-DEL-163
chr2-4623615-DEL-64
chr2-4623997-DEL-196
chr2-4624329-DEL-97
chr2-4658202-DEL-88
chr2-4658240-DEL-220
chr2-4670674-DEL-112
chr2-4670898-DEL-252
chr2-4749491-DEL-624
chr2-4815068-DEL-69
chr2-4829960-DEL-678
chr2-4847295-DEL-152
chr2-4850235-DEL-588
chr2-4877724-DEL-104
chr2-4957903-DEL-431
chr2-4982018-DEL-393
chr2-4997042-DEL-106
chr2-5014519-DEL-561
chr2-5030286-DEL-343
chr2-5060236-DEL-78
chr2-5088362-DEL-155
chr2-5124314-DEL-62
chr2-5163318-DEL-50
chr2-5178734-DEL-693
chr2-5193612-DEL-185
chr2-5231212-DEL-100
chr2-5248747-DEL-237
chr2-5272760-DEL-421
chr2-5273595-DEL-4814
chr2-5301401-DEL-68
chr2-5323509-DEL-491
chr2-5363675-DEL-123
chr2-5380886-DEL-372
chr2-5380931-DEL-52
chr2-5381207-DEL-304
chr2-5385439-DEL-1376
chr2-5388373-DEL-71
chr2-5412664-DEL-55
chr2-5458429-DEL-208
chr2-5458437-DEL-90
chr2-5479690-DEL-452
chr2-5481170-DEL-55
chr2-5488135-DEL-52
chr2-5495860-DEL-68
chr2-5546911-DEL-529
chr2-5550706-DEL-1094

chr2-5606173-DEL-57
chr2-5640732-DEL-412
chr2-5672517-DEL-75
chr2-5700699-DEL-947
chr2-5738876-DEL-86
chr2-5739084-DEL-86
chr2-5809996-DEL-907
chr2-5844266-DEL-218
chr2-5846355-DEL-158
chr2-5921641-DEL-207
chr2-5930368-DEL-1563
chr2-5976023-DEL-199
chr2-5979592-DEL-181
chr2-5990508-DEL-2244
chr2-6004460-DEL-201
chr2-6063310-DEL-557
chr2-6066800-DEL-65
chr2-6083403-DEL-808
chr2-6095061-DEL-196
chr2-6108338-DEL-96
chr2-6184482-DEL-261
chr2-6201875-DEL-50
chr2-6207628-DEL-2327
chr2-6267095-DEL-206
chr2-6276981-DEL-50
chr2-6305222-DEL-101
chr2-6307925-DEL-64
chr2-6317887-DEL-680
chr2-6332145-DEL-1005
chr2-6349457-DEL-148
chr2-6349590-DEL-383
chr2-6349814-DEL-159
chr2-6360156-DEL-66
chr2-6370133-DEL-50
chr2-6370322-DEL-96
chr2-6422188-DEL-105
chr2-6494094-DEL-66
chr2-6494320-DEL-79
chr2-6503863-DEL-54
chr2-6588158-DEL-89
chr2-6642634-DEL-77
chr2-6661039-DEL-125
chr2-6679141-DEL-60
chr2-6679353-DEL-76
chr2-6686538-DEL-52
chr2-6713280-DEL-72
chr2-6771477-DEL-56
chr2-6777870-DEL-3922
chr2-6914329-DEL-915
chr2-7090290-DEL-63
chr2-7096601-DEL-68
chr2-7137452-DEL-931
chr2-7138500-DEL-726
chr2-7149139-DEL-205
chr2-7208505-DEL-245
chr2-7293174-DEL-524
chr2-7303180-DEL-60
chr2-7350784-DEL-135
chr2-7391447-DEL-90
chr2-7517208-DEL-50
chr2-7539359-DEL-50
chr2-7555305-DEL-220
chr2-7605168-DEL-66
chr2-7770798-DEL-53
chr2-7786806-DEL-329
chr2-7799782-DEL-226
chr2-7841771-DEL-50
chr2-7971540-DEL-50
chr2-8144062-DEL-62
chr2-8167813-DEL-50
chr2-8194293-DEL-848
chr2-8222160-DEL-272
chr2-8414414-DEL-275
chr2-8423567-DEL-521
chr2-8488020-DEL-144
chr2-8490566-DEL-2121
chr2-8563845-DEL-62

chr2-8570440-DEL-1835
chr2-8571324-DEL-83
chr2-8576123-DEL-114
chr2-8695076-DEL-56
chr2-8700053-DEL-82
chr2-8937024-DEL-286
chr2-8937814-DEL-52
chr2-9075342-DEL-369
chr2-9084582-DEL-1475
chr2-9201181-DEL-153
chr2-9215009-DEL-89
chr2-9308733-DEL-62
chr2-9464238-DEL-96
chr2-9519006-DEL-208
chr2-9532706-DEL-462
chr2-9538138-DEL-112
chr2-9593849-DEL-50
chr2-9597814-DEL-68
chr2-9665791-DEL-54
chr2-9665791-DEL-180
chr2-9778552-DEL-106
chr2-9779992-DEL-89
chr2-9780684-DEL-120
chr2-9780833-DEL-121
chr2-9850315-DEL-307
chr2-9868755-DEL-109
chr2-9896287-DEL-233
chr2-9909680-DEL-184
chr2-9997507-DEL-80
chr2-10035161-DEL-106
chr2-10118625-DEL-178
chr2-10118960-DEL-98
chr2-10160360-DEL-114
chr2-10162768-DEL-84
chr2-10286790-DEL-166
chr2-10405007-DEL-305
chr2-10405429-DEL-52
chr2-10467003-DEL-146
chr2-10500200-DEL-58
chr2-10509751-DEL-176
chr2-10558815-DEL-624
chr2-10564141-DEL-64
chr2-10623397-DEL-54
chr2-10683866-DEL-2364
chr2-10765505-DEL-101
chr2-10765905-DEL-80
chr2-10796679-DEL-424
chr2-10814793-DEL-108
chr2-10925249-DEL-78
chr2-11059566-DEL-52
chr2-11067019-DEL-259
chr2-11165178-DEL-68
chr2-11165972-DEL-51
chr2-11222472-DEL-50
chr2-11391594-DEL-92
chr2-11391924-DEL-260
chr2-11500871-DEL-63
chr2-11509909-DEL-80
chr2-11515678-DEL-1396
chr2-11564665-DEL-254
chr2-11598504-DEL-160
chr2-11632270-DEL-63
chr2-11644143-DEL-250
chr2-11663544-DEL-62
chr2-11666516-DEL-72
chr2-11749557-DEL-152
chr2-11791325-DEL-195
chr2-11791389-DEL-56
chr2-11873447-DEL-77
chr2-11910222-DEL-356
chr2-11910682-DEL-632
chr2-11945972-DEL-4171
chr2-11992206-DEL-1062
chr2-12053511-DEL-142
chr2-12073990-DEL-174
chr2-12074364-DEL-76
chr2-12127514-DEL-65

chr2-12235217-DEL-52
chr2-12269095-DEL-78
chr2-12312753-DEL-50
chr2-12392292-DEL-65
chr2-12411326-DEL-53
chr2-12411330-DEL-113
chr2-12478161-DEL-63
chr2-12612994-DEL-64
chr2-12681486-DEL-832
chr2-12758657-DEL-1604
chr2-12940737-DEL-236
chr2-13063731-DEL-94
chr2-13170030-DEL-69
chr2-13348482-DEL-251
chr2-13394434-DEL-50
chr2-13546826-DEL-52
chr2-13663268-DEL-211
chr2-13769293-DEL-65
chr2-13808671-DEL-285
chr2-13808837-DEL-132
chr2-13827212-DEL-1180
chr2-13844833-DEL-174
chr2-13856587-DEL-60
chr2-13979228-DEL-201
chr2-14001901-DEL-60
chr2-14002024-DEL-140
chr2-14059818-DEL-72
chr2-14104708-DEL-853
chr2-14163739-DEL-1768
chr2-14200358-DEL-81
chr2-14205348-DEL-98
chr2-14270991-DEL-56
chr2-14459591-DEL-188
chr2-14480054-DEL-60
chr2-14517137-DEL-1183
chr2-15031230-DEL-78
chr2-15101903-DEL-70
chr2-15376120-DEL-126
chr2-15426205-DEL-50
chr2-15653409-DEL-65
chr2-15659645-DEL-99
chr2-15661865-DEL-522
chr2-15704274-DEL-424
chr2-15830166-DEL-109
chr2-15938933-DEL-157
chr2-16218276-DEL-75
chr2-16520647-DEL-56
chr2-16520849-DEL-90
chr2-16595513-DEL-72
chr2-16752838-DEL-64
chr2-17100963-DEL-84
chr2-17102888-DEL-304
chr2-17172252-DEL-141
chr2-17209297-DEL-72
chr2-17344692-DEL-149
chr2-17485975-DEL-67
chr2-17491134-DEL-6181
chr2-17559623-DEL-400
chr2-17559734-DEL-100
chr2-17671246-DEL-80
chr2-17764999-DEL-2721
chr2-17799417-DEL-52
chr2-17893327-DEL-61
chr2-17893562-DEL-187
chr2-17912171-DEL-85
chr2-18109613-DEL-79
chr2-18109780-DEL-57
chr2-18174102-DEL-120
chr2-18230814-DEL-344
chr2-18259567-DEL-1761
chr2-18291174-DEL-5141
chr2-18399093-DEL-762
chr2-18467117-DEL-73
chr2-18501047-DEL-54
chr2-18544210-DEL-87
chr2-18579445-DEL-96
chr2-18584900-DEL-50

chr2-18595443-DEL-1248
chr2-18644771-DEL-63
chr2-18644928-DEL-165
chr2-18651541-DEL-199
chr2-18783535-DEL-157
chr2-18809660-DEL-648
chr2-18809689-DEL-120
chr2-18810097-DEL-72
chr2-19053724-DEL-52
chr2-19140137-DEL-63
chr2-19224630-DEL-50
chr2-19261725-DEL-57
chr2-19267152-DEL-176
chr2-19371527-DEL-55
chr2-19378531-DEL-304
chr2-19562779-DEL-65
chr2-19618301-DEL-56
chr2-19698958-DEL-110
chr2-19736721-DEL-133
chr2-19798586-DEL-971
chr2-19830723-DEL-1890
chr2-19843883-DEL-1263
chr2-19855016-DEL-152
chr2-19855280-DEL-61
chr2-19873203-DEL-236
chr2-19873572-DEL-207
chr2-19913345-DEL-50
chr2-19965588-DEL-1775
chr2-20034256-DEL-56
chr2-20050697-DEL-867
chr2-20102316-DEL-120
chr2-20181304-DEL-4112
chr2-20193604-DEL-542
chr2-20347308-DEL-52
chr2-20428880-DEL-1565
chr2-20464997-DEL-154
chr2-20480071-DEL-68
chr2-20540877-DEL-186
chr2-20582518-DEL-150
chr2-20629528-DEL-315
chr2-20672248-DEL-210
chr2-20686486-DEL-4620
chr2-20692085-DEL-179
chr2-20725436-DEL-80
chr2-20893190-DEL-424
chr2-20895146-DEL-393
chr2-20980092-DEL-52
chr2-20991593-DEL-112
chr2-20997001-DEL-174
chr2-21022250-DEL-52
chr2-21043792-DEL-300
chr2-21129246-DEL-56
chr2-21236438-DEL-70
chr2-21335194-DEL-473
chr2-21342383-DEL-50
chr2-21380838-DEL-80
chr2-21551148-DEL-355
chr2-21557142-DEL-98
chr2-21574516-DEL-61
chr2-21574691-DEL-62
chr2-21607760-DEL-201
chr2-21671537-DEL-92
chr2-21693308-DEL-3087
chr2-21720254-DEL-3557
chr2-21737222-DEL-3652
chr2-21760046-DEL-95
chr2-21849803-DEL-969
chr2-21875972-DEL-3403
chr2-22094480-DEL-52
chr2-22115315-DEL-1494
chr2-22279423-DEL-399
chr2-22285641-DEL-99
chr2-22412913-DEL-1162
chr2-22432946-DEL-656
chr2-22536925-DEL-51
chr2-22546395-DEL-85
chr2-22588850-DEL-348

chr2-22604907-DEL-60
chr2-22631863-DEL-50
chr2-22726245-DEL-3856
chr2-22883034-DEL-75
chr2-22917661-DEL-72
chr2-22950279-DEL-2725
chr2-23207134-DEL-82
chr2-23325111-DEL-75
chr2-23386372-DEL-67
chr2-23392035-DEL-850
chr2-23470783-DEL-155
chr2-23499924-DEL-2318
chr2-23576148-DEL-59
chr2-23583660-DEL-1912
chr2-23628017-DEL-1013
chr2-23684338-DEL-4761
chr2-23731045-DEL-1038
chr2-23742428-DEL-66
chr2-23859477-DEL-50
chr2-24190328-DEL-582
chr2-24192289-DEL-103
chr2-24395774-DEL-50
chr2-24669443-DEL-104
chr2-24708633-DEL-64
chr2-24708736-DEL-130
chr2-24746156-DEL-53
chr2-24746284-DEL-157
chr2-24761131-DEL-568
chr2-24762652-DEL-216
chr2-24822523-DEL-221
chr2-24937253-DEL-140
chr2-25048635-DEL-422
chr2-25049007-DEL-116
chr2-25051029-DEL-404
chr2-25148490-DEL-77
chr2-25191030-DEL-114
chr2-25191560-DEL-54
chr2-25257897-DEL-105
chr2-25258216-DEL-65
chr2-25342269-DEL-90
chr2-25379407-DEL-56
chr2-25405467-DEL-210
chr2-25443545-DEL-211
chr2-25499929-DEL-308
chr2-25500216-DEL-69
chr2-25522183-DEL-65
chr2-25630886-DEL-670
chr2-25631229-DEL-150
chr2-25643818-DEL-57
chr2-25836213-DEL-62
chr2-25986754-DEL-190
chr2-26159027-DEL-53
chr2-26176888-DEL-215
chr2-26381589-DEL-69
chr2-26501754-DEL-173
chr2-26552746-DEL-74
chr2-26588480-DEL-6798
chr2-26624966-DEL-82
chr2-26725158-DEL-114
chr2-26743946-DEL-204
chr2-26825752-DEL-57
chr2-26878032-DEL-75
chr2-26931879-DEL-50
chr2-27007658-DEL-128
chr2-27019404-DEL-748
chr2-27091691-DEL-260
chr2-27196014-DEL-516
chr2-27207973-DEL-52
chr2-27273084-DEL-585
chr2-27370988-DEL-64
chr2-27376312-DEL-227
chr2-27475964-DEL-93
chr2-27476348-DEL-73
chr2-27559672-DEL-80
chr2-27707485-DEL-78
chr2-27783146-DEL-100
chr2-27783916-DEL-82

chr2-27828893-DEL-107
chr2-28074546-DEL-67
chr2-28257913-DEL-52
chr2-28262010-DEL-72
chr2-28283731-DEL-144
chr2-28283731-DEL-384
chr2-28348889-DEL-348
chr2-28349787-DEL-70
chr2-28490897-DEL-380
chr2-28492814-DEL-492
chr2-28492943-DEL-222
chr2-28498666-DEL-52
chr2-29024932-DEL-89
chr2-29417934-DEL-60
chr2-29458851-DEL-51
chr2-29486105-DEL-67
chr2-29528643-DEL-119
chr2-29598500-DEL-57
chr2-29644891-DEL-252
chr2-29648036-DEL-93
chr2-29648411-DEL-149
chr2-29648466-DEL-73
chr2-29666496-DEL-195
chr2-29696363-DEL-63
chr2-29778569-DEL-288
chr2-29798257-DEL-280
chr2-29836680-DEL-87
chr2-29941398-DEL-94
chr2-29955356-DEL-51
chr2-30165875-DEL-96
chr2-30345035-DEL-325
chr2-30382472-DEL-120
chr2-30382827-DEL-147
chr2-30383191-DEL-66
chr2-30383963-DEL-406
chr2-30384069-DEL-187
chr2-30384419-DEL-96
chr2-30433286-DEL-50
chr2-30523955-DEL-55
chr2-30525901-DEL-130
chr2-30600428-DEL-56
chr2-30655173-DEL-132
chr2-30813461-DEL-60
chr2-30887583-DEL-262
chr2-30924779-DEL-232
chr2-30988506-DEL-190
chr2-31117658-DEL-107
chr2-31217483-DEL-187
chr2-31280267-DEL-88
chr2-31337933-DEL-165
chr2-31338120-DEL-236
chr2-31338294-DEL-109
chr2-31352700-DEL-79
chr2-31610667-DEL-1743
chr2-31648228-DEL-128
chr2-31684164-DEL-121
chr2-31694961-DEL-649
chr2-31743387-DEL-281
chr2-31767777-DEL-165
chr2-32022753-DEL-141
chr2-32075684-DEL-55
chr2-32299902-DEL-170
chr2-32363364-DEL-3988
chr2-32371886-DEL-212
chr2-32394316-DEL-104
chr2-32751351-DEL-50
chr2-32817863-DEL-51
chr2-32841859-DEL-308
chr2-32842217-DEL-218
chr2-32854549-DEL-68
chr2-32861334-DEL-68
chr2-32898872-DEL-288
chr2-32951399-DEL-100
chr2-33099013-DEL-2458
chr2-33177987-DEL-75
chr2-33189130-DEL-51
chr2-33189388-DEL-559

chr2-33205055-DEL-77
chr2-33303604-DEL-70
chr2-33473127-DEL-51
chr2-33511163-DEL-320
chr2-33511281-DEL-80
chr2-33600126-DEL-50
chr2-33631839-DEL-119
chr2-33635252-DEL-188
chr2-33722916-DEL-271
chr2-33735149-DEL-68
chr2-33874669-DEL-1389
chr2-33885191-DEL-774
chr2-33886398-DEL-763
chr2-33887267-DEL-2233
chr2-33903887-DEL-206
chr2-33940565-DEL-572
chr2-33941367-DEL-88
chr2-34476747-DEL-1230
chr2-34654449-DEL-397
chr2-34751426-DEL-108
chr2-34830785-DEL-56
chr2-34855861-DEL-357
chr2-34878395-DEL-114
chr2-35012228-DEL-166
chr2-35258050-DEL-127
chr2-35258225-DEL-84
chr2-35304941-DEL-324
chr2-35331073-DEL-51
chr2-35430571-DEL-81
chr2-35433943-DEL-272
chr2-35615427-DEL-107
chr2-35666231-DEL-68
chr2-35681995-DEL-764
chr2-35692104-DEL-316
chr2-35698537-DEL-76
chr2-35807309-DEL-143
chr2-35810234-DEL-153
chr2-35829528-DEL-84
chr2-35861132-DEL-486
chr2-35887392-DEL-2111
chr2-35997240-DEL-68
chr2-36021807-DEL-61
chr2-36096036-DEL-174
chr2-36100743-DEL-54
chr2-36269645-DEL-985
chr2-36277384-DEL-85
chr2-36286352-DEL-805
chr2-36304361-DEL-117
chr2-36354241-DEL-586
chr2-36375341-DEL-724
chr2-36544981-DEL-216
chr2-36585003-DEL-86
chr2-36618264-DEL-1492
chr2-36637175-DEL-229
chr2-36708970-DEL-83
chr2-36789893-DEL-2561
chr2-36891856-DEL-56
chr2-36988135-DEL-1890
chr2-37112164-DEL-54
chr2-37144104-DEL-2017
chr2-37223409-DEL-2835
chr2-37298489-DEL-56
chr2-37501910-DEL-53
chr2-37611639-DEL-2617
chr2-37628462-DEL-139
chr2-37761443-DEL-50
chr2-38015334-DEL-149
chr2-38015634-DEL-80
chr2-38222583-DEL-224
chr2-38397771-DEL-59
chr2-38566726-DEL-1049
chr2-38572887-DEL-173
chr2-38647906-DEL-63
chr2-38711282-DEL-55
chr2-38746598-DEL-323
chr2-38749523-DEL-891
chr2-38759679-DEL-85

chr2-38793817-DEL-108
chr2-38842356-DEL-54
chr2-38865927-DEL-156
chr2-38934453-DEL-1827
chr2-39110402-DEL-70
chr2-39166632-DEL-1140
chr2-39182023-DEL-87
chr2-39218011-DEL-72
chr2-39222953-DEL-58
chr2-39276452-DEL-2934
chr2-39400341-DEL-50
chr2-39486376-DEL-142
chr2-39509925-DEL-78
chr2-39696923-DEL-58
chr2-39708594-DEL-480
chr2-39746609-DEL-78
chr2-39746776-DEL-150
chr2-39985326-DEL-2533
chr2-40113717-DEL-1772
chr2-40122821-DEL-71
chr2-40189365-DEL-348
chr2-40225655-DEL-54
chr2-40448601-DEL-1006
chr2-40523695-DEL-235
chr2-40524273-DEL-78
chr2-40524297-DEL-209
chr2-40524588-DEL-387
chr2-40534562-DEL-101
chr2-40546426-DEL-163
chr2-40596615-DEL-263
chr2-40627770-DEL-51
chr2-40741912-DEL-229
chr2-41341556-DEL-71
chr2-41382990-DEL-949
chr2-41573492-DEL-50
chr2-41584217-DEL-4298
chr2-41590772-DEL-149
chr2-41633145-DEL-96
chr2-41664478-DEL-581
chr2-41924155-DEL-174
chr2-41942791-DEL-821
chr2-41948520-DEL-122
chr2-42015948-DEL-50
chr2-42177745-DEL-60
chr2-42244686-DEL-112
chr2-42263068-DEL-50
chr2-42350576-DEL-541
chr2-42369086-DEL-63
chr2-42369218-DEL-136
chr2-42563314-DEL-710
chr2-42613067-DEL-50
chr2-42757503-DEL-241
chr2-42834021-DEL-281
chr2-42862211-DEL-996
chr2-43013154-DEL-225
chr2-43128941-DEL-537
chr2-43428610-DEL-50
chr2-43567180-DEL-51
chr2-43641784-DEL-148
chr2-43669194-DEL-75
chr2-43779221-DEL-64
chr2-43853760-DEL-50
chr2-43935211-DEL-77
chr2-44027936-DEL-1177
chr2-44280455-DEL-248
chr2-44342889-DEL-80
chr2-44530695-DEL-54
chr2-44654421-DEL-8831
chr2-44935623-DEL-152
chr2-45064865-DEL-74
chr2-45080888-DEL-197
chr2-45081291-DEL-250
chr2-45132179-DEL-86
chr2-45185132-DEL-137
chr2-45241821-DEL-507
chr2-45243493-DEL-122
chr2-45310396-DEL-52

chr2-45358243-DEL-94
chr2-45431104-DEL-13290
chr2-45451219-DEL-204
chr2-45488544-DEL-217
chr2-45784231-DEL-54
chr2-45816108-DEL-50
chr2-45820715-DEL-70
chr2-46056477-DEL-97
chr2-46171911-DEL-337
chr2-46171968-DEL-68
chr2-46233252-DEL-4491
chr2-46280590-DEL-56
chr2-46314299-DEL-50
chr2-46527543-DEL-128
chr2-46570360-DEL-209
chr2-46574589-DEL-450
chr2-46661195-DEL-260
chr2-46724424-DEL-7032
chr2-46843203-DEL-93
chr2-46843351-DEL-223
chr2-47018846-DEL-212
chr2-47474122-DEL-50
chr2-47657625-DEL-248
chr2-47687539-DEL-563
chr2-47924615-DEL-57
chr2-47996305-DEL-81
chr2-48023790-DEL-50
chr2-48261705-DEL-180
chr2-48450895-DEL-406
chr2-48451047-DEL-163
chr2-48478877-DEL-51
chr2-48796601-DEL-640
chr2-48880757-DEL-114
chr2-48946808-DEL-1147
chr2-48949692-DEL-3702
chr2-49077060-DEL-869
chr2-49137706-DEL-104
chr2-49137862-DEL-300
chr2-49138426-DEL-80
chr2-49176937-DEL-290
chr2-49193566-DEL-145
chr2-49263358-DEL-102
chr2-49296480-DEL-79
chr2-49357753-DEL-139
chr2-49461877-DEL-70
chr2-49479636-DEL-66
chr2-49498030-DEL-702
chr2-49514451-DEL-84
chr2-49516888-DEL-64
chr2-49572432-DEL-149
chr2-49654889-DEL-50
chr2-49674978-DEL-170
chr2-49729324-DEL-54
chr2-49759695-DEL-107
chr2-49926205-DEL-103
chr2-49984400-DEL-768
chr2-50046815-DEL-58
chr2-50057129-DEL-340
chr2-50153210-DEL-1014
chr2-50204288-DEL-1018
chr2-50280093-DEL-499
chr2-50358075-DEL-64
chr2-50455910-DEL-68
chr2-50594070-DEL-78
chr2-50595620-DEL-130
chr2-50823602-DEL-155
chr2-50866928-DEL-62
chr2-50867019-DEL-155
chr2-50976789-DEL-91
chr2-50978491-DEL-195
chr2-51042978-DEL-60
chr2-51092727-DEL-132
chr2-51290036-DEL-145
chr2-51293938-DEL-222
chr2-51444716-DEL-87
chr2-51468974-DEL-63
chr2-51488934-DEL-381

chr2-51515607-DEL-783
chr2-51516670-DEL-68
chr2-51532927-DEL-50
chr2-51550730-DEL-170
chr2-51681687-DEL-55
chr2-51836800-DEL-1155
chr2-51851972-DEL-54
chr2-51901722-DEL-110
chr2-51923525-DEL-81
chr2-52305964-DEL-69
chr2-52441423-DEL-77
chr2-52549307-DEL-461
chr2-52587609-DEL-1170
chr2-52588259-DEL-270
chr2-52607492-DEL-236
chr2-52608178-DEL-152
chr2-52628101-DEL-53
chr2-52679050-DEL-52
chr2-52713723-DEL-338
chr2-52727564-DEL-56
chr2-52742161-DEL-100
chr2-52784242-DEL-359
chr2-52930004-DEL-144
chr2-52940819-DEL-81
chr2-53056364-DEL-61
chr2-53086438-DEL-81
chr2-53101786-DEL-446
chr2-53181486-DEL-1850
chr2-53261949-DEL-112
chr2-53366952-DEL-82
chr2-53417601-DEL-356
chr2-53488383-DEL-57
chr2-53488709-DEL-2265
chr2-53556382-DEL-325
chr2-53935554-DEL-51
chr2-53938789-DEL-86
chr2-54017884-DEL-56
chr2-54040978-DEL-50
chr2-54051586-DEL-64
chr2-54303447-DEL-50
chr2-54349543-DEL-378
chr2-54361147-DEL-1734
chr2-54497882-DEL-902
chr2-54625881-DEL-1748
chr2-54660822-DEL-336
chr2-54681937-DEL-426
chr2-54806641-DEL-6929
chr2-54834309-DEL-68
chr2-54842047-DEL-2499
chr2-55083346-DEL-52
chr2-55142317-DEL-54
chr2-55162460-DEL-1357
chr2-55163921-DEL-1383
chr2-55413246-DEL-470
chr2-55585169-DEL-1351
chr2-55595260-DEL-272
chr2-55605278-DEL-51
chr2-55892363-DEL-1152
chr2-55893230-DEL-4972
chr2-56059292-DEL-61
chr2-56305278-DEL-131
chr2-56338594-DEL-64
chr2-56371411-DEL-1554
chr2-56376511-DEL-132
chr2-56423179-DEL-50
chr2-56587655-DEL-112
chr2-56689277-DEL-424
chr2-56689764-DEL-1060
chr2-56690805-DEL-212
chr2-56700373-DEL-53
chr2-56923439-DEL-2336
chr2-56961934-DEL-203
chr2-56964751-DEL-1251
chr2-57091242-DEL-316
chr2-57102451-DEL-107
chr2-57227046-DEL-124
chr2-57345724-DEL-597

chr2-57502510-DEL-1039
chr2-57577216-DEL-51
chr2-57777418-DEL-73
chr2-57845095-DEL-266
chr2-58008574-DEL-1523
chr2-58010416-DEL-1898
chr2-58101152-DEL-2937
chr2-58111738-DEL-1077
chr2-58114641-DEL-2583
chr2-58261295-DEL-50
chr2-58277180-DEL-78
chr2-58289595-DEL-52
chr2-58303065-DEL-1103
chr2-58316066-DEL-216
chr2-58327722-DEL-56
chr2-58334611-DEL-52
chr2-58537760-DEL-222
chr2-58588144-DEL-85
chr2-58651894-DEL-295
chr2-58774763-DEL-1645
chr2-58777506-DEL-416
chr2-58849289-DEL-186
chr2-58918402-DEL-297
chr2-58947266-DEL-1911
chr2-58947283-DEL-352
chr2-58948193-DEL-460
chr2-58949517-DEL-145
chr2-59115989-DEL-64
chr2-59227368-DEL-373
chr2-59230645-DEL-142
chr2-59498049-DEL-952
chr2-59535025-DEL-63
chr2-59554182-DEL-223
chr2-59554312-DEL-78
chr2-59560828-DEL-117
chr2-59569879-DEL-52
chr2-59620478-DEL-127
chr2-59652236-DEL-141
chr2-59700354-DEL-250
chr2-59700527-DEL-100
chr2-59759319-DEL-664
chr2-59760149-DEL-94
chr2-59834352-DEL-204
chr2-59871246-DEL-131
chr2-59924255-DEL-173
chr2-59940124-DEL-290
chr2-59965964-DEL-247
chr2-59985160-DEL-52
chr2-59995387-DEL-148
chr2-60100747-DEL-396
chr2-60153149-DEL-74
chr2-60219951-DEL-106
chr2-60403984-DEL-216
chr2-60457415-DEL-134
chr2-60462218-DEL-806
chr2-60505783-DEL-58
chr2-60508521-DEL-1031
chr2-60679773-DEL-53
chr2-60921469-DEL-455
chr2-60921594-DEL-175
chr2-60925325-DEL-422
chr2-60925815-DEL-577
chr2-60947545-DEL-76
chr2-60949755-DEL-54
chr2-61008019-DEL-78
chr2-61047694-DEL-165
chr2-61076834-DEL-141
chr2-61079286-DEL-80
chr2-61122412-DEL-80
chr2-61249659-DEL-420
chr2-61253342-DEL-50
chr2-61403425-DEL-423
chr2-61507307-DEL-81
chr2-62027580-DEL-1750
chr2-62304458-DEL-60
chr2-62394831-DEL-110
chr2-62506341-DEL-55

chr2-62534134-DEL-69
chr2-62580872-DEL-1462
chr2-62615989-DEL-52
chr2-62915070-DEL-161
chr2-63038333-DEL-80
chr2-63270694-DEL-261
chr2-63289209-DEL-54
chr2-63296994-DEL-106
chr2-63440798-DEL-1069
chr2-63955298-DEL-108
chr2-63958535-DEL-403
chr2-63993991-DEL-62
chr2-64039977-DEL-445
chr2-64041067-DEL-2365
chr2-64173535-DEL-67
chr2-64198683-DEL-140
chr2-64526227-DEL-57
chr2-64638800-DEL-56
chr2-64772144-DEL-158
chr2-64772196-DEL-76
chr2-64829506-DEL-138
chr2-65080620-DEL-50
chr2-65095753-DEL-186
chr2-65150703-DEL-132
chr2-65173385-DEL-1193
chr2-65303724-DEL-102
chr2-65496680-DEL-52
chr2-65625651-DEL-999
chr2-65625681-DEL-108
chr2-65626752-DEL-144
chr2-65632333-DEL-56
chr2-65870645-DEL-870
chr2-65876438-DEL-551
chr2-65886219-DEL-6233
chr2-66019618-DEL-66
chr2-66117542-DEL-58
chr2-66121358-DEL-153
chr2-66212515-DEL-52
chr2-66359041-DEL-431
chr2-66421560-DEL-55
chr2-66422130-DEL-496
chr2-66452688-DEL-736
chr2-66454286-DEL-1686
chr2-66482428-DEL-1821
chr2-66585225-DEL-60
chr2-66623217-DEL-50
chr2-66698374-DEL-1466
chr2-66728552-DEL-53
chr2-66775561-DEL-98
chr2-66804536-DEL-96
chr2-66920446-DEL-205
chr2-66974210-DEL-217
chr2-67017494-DEL-140
chr2-67120945-DEL-58
chr2-67127657-DEL-68
chr2-67134905-DEL-227
chr2-67136925-DEL-55
chr2-67149760-DEL-199
chr2-67158239-DEL-77
chr2-67172727-DEL-100
chr2-67172884-DEL-61
chr2-67210191-DEL-276
chr2-67213411-DEL-50
chr2-67352479-DEL-87
chr2-67357446-DEL-445
chr2-67367228-DEL-1305
chr2-67489649-DEL-1089
chr2-67602895-DEL-64
chr2-67652016-DEL-78
chr2-67653848-DEL-62
chr2-67704383-DEL-57
chr2-67763848-DEL-2395
chr2-67837325-DEL-257
chr2-67898780-DEL-66
chr2-67912353-DEL-216
chr2-67915377-DEL-72
chr2-68032015-DEL-50

chr2-68065529-DEL-350
chr2-68070455-DEL-200
chr2-68112303-DEL-53
chr2-68172929-DEL-59
chr2-68252679-DEL-851
chr2-68252703-DEL-111
chr2-68276295-DEL-97
chr2-68322805-DEL-53
chr2-68486944-DEL-209
chr2-68504946-DEL-205
chr2-68694001-DEL-94
chr2-68788083-DEL-50
chr2-68844775-DEL-168
chr2-68847018-DEL-60
chr2-68851062-DEL-92
chr2-68854260-DEL-57
chr2-68870431-DEL-66
chr2-68904256-DEL-95
chr2-68932149-DEL-50
chr2-68943464-DEL-104
chr2-68956971-DEL-686
chr2-68999939-DEL-50
chr2-69177593-DEL-1413
chr2-69187336-DEL-54
chr2-69204535-DEL-80
chr2-69221251-DEL-55
chr2-69222181-DEL-261
chr2-69392097-DEL-192
chr2-69409920-DEL-82
chr2-69435182-DEL-59
chr2-69559393-DEL-53
chr2-69718485-DEL-373
chr2-69718531-DEL-151
chr2-69760539-DEL-829
chr2-69890777-DEL-38929
chr2-69932127-DEL-2304
chr2-70106863-DEL-57
chr2-70243226-DEL-147
chr2-70295314-DEL-51
chr2-70295665-DEL-118
chr2-70357403-DEL-130
chr2-70365533-DEL-52
chr2-70409257-DEL-68
chr2-70467945-DEL-117
chr2-70522607-DEL-81
chr2-70540221-DEL-50
chr2-70684625-DEL-50
chr2-70721701-DEL-65
chr2-70721773-DEL-2029
chr2-70728471-DEL-57
chr2-70794052-DEL-289
chr2-70804037-DEL-231
chr2-70840969-DEL-161
chr2-70965290-DEL-69
chr2-71040313-DEL-99
chr2-71116713-DEL-169
chr2-71138890-DEL-81
chr2-71180936-DEL-122
chr2-71181196-DEL-105
chr2-71207126-DEL-777
chr2-71414000-DEL-189
chr2-71440212-DEL-100
chr2-71443127-DEL-50
chr2-71460916-DEL-159
chr2-71466798-DEL-50
chr2-71516233-DEL-244
chr2-71524559-DEL-94
chr2-71567573-DEL-287
chr2-71607724-DEL-54
chr2-71688863-DEL-66
chr2-71736657-DEL-394
chr2-71789826-DEL-219
chr2-71789872-DEL-51
chr2-71825733-DEL-436
chr2-71839057-DEL-859
chr2-71975452-DEL-51
chr2-72063169-DEL-1167

chr2-72315679-DEL-52
chr2-72366327-DEL-67
chr2-72549476-DEL-1605
chr2-73056455-DEL-68
chr2-73056479-DEL-269
chr2-73056680-DEL-100
chr2-73285759-DEL-135
chr2-73376954-DEL-54
chr2-73456760-DEL-92
chr2-73480875-DEL-53
chr2-73615023-DEL-64
chr2-73824212-DEL-56
chr2-73828443-DEL-88
chr2-73838183-DEL-130
chr2-73967362-DEL-204
chr2-74081604-DEL-569
chr2-74081609-DEL-50
chr2-74115324-DEL-51
chr2-74458138-DEL-54
chr2-74786217-DEL-115
chr2-75020291-DEL-200
chr2-75041727-DEL-712
chr2-75209055-DEL-258
chr2-75263898-DEL-60
chr2-75285925-DEL-119
chr2-75296598-DEL-50
chr2-75427123-DEL-66
chr2-75578380-DEL-156
chr2-75578658-DEL-260
chr2-75619785-DEL-50
chr2-75634272-DEL-146
chr2-75651971-DEL-170
chr2-75717978-DEL-164
chr2-75799289-DEL-86
chr2-75802370-DEL-119
chr2-75911942-DEL-114
chr2-76367915-DEL-78
chr2-76368104-DEL-309
chr2-76368241-DEL-104
chr2-76482865-DEL-1056
chr2-76487409-DEL-108
chr2-76735565-DEL-50
chr2-76824392-DEL-213
chr2-76832943-DEL-81
chr2-76988986-DEL-338
chr2-77033162-DEL-188
chr2-77199261-DEL-51
chr2-77369323-DEL-65
chr2-77544619-DEL-56
chr2-77581935-DEL-185
chr2-77582217-DEL-89
chr2-77656489-DEL-64
chr2-77685801-DEL-1495
chr2-77716846-DEL-408
chr2-77717353-DEL-60
chr2-77718703-DEL-64
chr2-77817110-DEL-52
chr2-77986388-DEL-2114
chr2-77992477-DEL-266
chr2-78041737-DEL-132
chr2-78250112-DEL-253
chr2-78259299-DEL-52
chr2-78312089-DEL-729
chr2-78481580-DEL-55
chr2-78582274-DEL-51
chr2-78620406-DEL-81
chr2-78640113-DEL-66
chr2-78658450-DEL-1824
chr2-78734859-DEL-138
chr2-78821378-DEL-3286
chr2-78936009-DEL-177
chr2-78938961-DEL-222
chr2-78950326-DEL-59
chr2-78985716-DEL-87
chr2-79001028-DEL-59
chr2-79010625-DEL-306
chr2-79030780-DEL-92

chr2-79116819-DEL-111
chr2-79422408-DEL-50
chr2-79426812-DEL-84
chr2-79674137-DEL-690
chr2-79730936-DEL-92
chr2-79767312-DEL-423
chr2-79803541-DEL-60
chr2-80004208-DEL-50
chr2-80019529-DEL-50
chr2-80135778-DEL-1050
chr2-80281858-DEL-2593
chr2-80284999-DEL-60
chr2-80287187-DEL-133
chr2-80413535-DEL-751
chr2-80433543-DEL-82
chr2-80546513-DEL-63
chr2-80640421-DEL-1439
chr2-80707188-DEL-52
chr2-80883371-DEL-196
chr2-80963632-DEL-374
chr2-81040638-DEL-1714
chr2-81287572-DEL-106
chr2-81353896-DEL-2006
chr2-81356055-DEL-62
chr2-81356394-DEL-136
chr2-81416849-DEL-131
chr2-81541440-DEL-212
chr2-81558237-DEL-126
chr2-81581749-DEL-66
chr2-81603309-DEL-2776
chr2-81606558-DEL-189
chr2-81628112-DEL-50
chr2-81704278-DEL-50
chr2-81758966-DEL-50
chr2-81806957-DEL-1299
chr2-81919827-DEL-102
chr2-81949906-DEL-52
chr2-81958521-DEL-1500
chr2-82070556-DEL-192
chr2-82177437-DEL-59
chr2-82261297-DEL-208
chr2-82678652-DEL-1649
chr2-82747292-DEL-2736
chr2-82871738-DEL-199
chr2-83091830-DEL-750
chr2-83102644-DEL-50
chr2-83154477-DEL-4150
chr2-83177088-DEL-128
chr2-83179187-DEL-55
chr2-83268973-DEL-204
chr2-83281645-DEL-56
chr2-83286572-DEL-851
chr2-83337866-DEL-161
chr2-83533088-DEL-204
chr2-83543154-DEL-1386
chr2-83647907-DEL-55
chr2-83648188-DEL-55
chr2-83717273-DEL-6295
chr2-83783931-DEL-15389
chr2-83784117-DEL-34860
chr2-83868050-DEL-337
chr2-83908035-DEL-1608
chr2-84024522-DEL-53
chr2-84024614-DEL-136
chr2-84137856-DEL-276
chr2-84164761-DEL-635
chr2-84203068-DEL-233
chr2-84292382-DEL-60
chr2-84352011-DEL-106
chr2-84384769-DEL-209
chr2-84436652-DEL-291
chr2-84547031-DEL-331
chr2-84725211-DEL-121
chr2-84751089-DEL-121
chr2-85250633-DEL-50
chr2-85406139-DEL-50
chr2-85509993-DEL-52

chr2-85618697-DEL-103
chr2-85724914-DEL-1494
chr2-85786647-DEL-98
chr2-85884194-DEL-28874
chr2-85959976-DEL-78
chr2-86077156-DEL-53
chr2-86117100-DEL-1041
chr2-86133015-DEL-2726
chr2-86161564-DEL-70
chr2-86216218-DEL-11601
chr2-86217008-DEL-28492
chr2-86315544-DEL-235
chr2-86453907-DEL-138
chr2-86557733-DEL-364
chr2-86727347-DEL-50
chr2-86793296-DEL-125
chr2-86805116-DEL-4152
chr2-86850821-DEL-114
chr2-86852978-DEL-423
chr2-86995464-DEL-1957
chr2-87001982-DEL-1254
chr2-87003425-DEL-50
chr2-87021229-DEL-50
chr2-87021456-DEL-468
chr2-87049128-DEL-140
chr2-87052076-DEL-22324
chr2-87086930-DEL-522
chr2-87112001-DEL-1515
chr2-87134682-DEL-193
chr2-87184469-DEL-57
chr2-87204790-DEL-937
chr2-87221973-DEL-368
chr2-87292760-DEL-913
chr2-87293803-DEL-914
chr2-87301790-DEL-214
chr2-87364160-DEL-72
chr2-87741172-DEL-659
chr2-87778375-DEL-329
chr2-87787256-DEL-326
chr2-87788030-DEL-431
chr2-87894846-DEL-64
chr2-87914039-DEL-1167
chr2-87960783-DEL-137
chr2-87966079-DEL-584
chr2-88131500-DEL-5945
chr2-88360627-DEL-78
chr2-88415848-DEL-60
chr2-88443797-DEL-4124
chr2-88469968-DEL-236
chr2-88494076-DEL-7001
chr2-88606017-DEL-5386
chr2-88892081-DEL-107
chr2-89042242-DEL-32056
chr2-89067447-DEL-8026
chr2-89142245-DEL-345
chr2-89216247-DEL-70
chr2-89298364-DEL-6535
chr2-89335111-DEL-309
chr2-89426719-DEL-115
chr2-89504029-DEL-56
chr2-89766285-DEL-3049
chr2-90111966-DEL-371
chr2-90128601-DEL-60
chr2-90166917-DEL-77411
chr2-90206088-DEL-5357
chr2-90233249-DEL-2171
chr2-90402116-DEL-172
chr2-90440769-DEL-73
chr2-90452337-DEL-296
chr2-90452753-DEL-477
chr2-90456857-DEL-52
chr2-90813650-DEL-126
chr2-91021747-DEL-85
chr2-91070571-DEL-200
chr2-91212168-DEL-56
chr2-91224052-DEL-162
chr2-91332545-DEL-82

chr2-91462141-DEL-94
chr2-91478766-DEL-84
chr2-91524154-DEL-60
chr2-91600623-DEL-90
chr2-91647791-DEL-53
chr2-91667069-DEL-66
chr2-91883980-DEL-55
chr2-91901103-DEL-56
chr2-91913884-DEL-74
chr2-91923979-DEL-96
chr2-91972777-DEL-94
chr2-91996704-DEL-145
chr2-92449381-DEL-63
chr2-92491106-DEL-423
chr2-92491372-DEL-147
chr2-92568982-DEL-187
chr2-92584832-DEL-50
chr2-92622340-DEL-3542
chr2-92872618-DEL-67
chr2-93052343-DEL-192
chr2-93135585-DEL-56
chr2-93191799-DEL-69
chr2-93200391-DEL-55
chr2-93314659-DEL-164
chr2-93332805-DEL-92
chr2-93377808-DEL-91
chr2-93390542-DEL-101
chr2-93505180-DEL-55
chr2-93579253-DEL-64
chr2-93598629-DEL-50
chr2-93603503-DEL-56
chr2-93645032-DEL-72
chr2-93650705-DEL-180
chr2-93703478-DEL-87
chr2-93726944-DEL-167
chr2-93770074-DEL-271
chr2-93770395-DEL-106
chr2-93952721-DEL-376
chr2-93971322-DEL-195
chr2-94017084-DEL-270
chr2-94179114-DEL-216
chr2-94219119-DEL-628
chr2-94270008-DEL-50
chr2-94292412-DEL-202
chr2-94395401-DEL-1132
chr2-94627924-DEL-50
chr2-94675129-DEL-66
chr2-94680167-DEL-5757
chr2-94923446-DEL-102
chr2-95018600-DEL-76
chr2-95081996-DEL-3622
chr2-95117240-DEL-3411
chr2-95177093-DEL-167
chr2-95298612-DEL-1806
chr2-95316857-DEL-121
chr2-95389657-DEL-1312
chr2-95391089-DEL-740
chr2-95408707-DEL-52
chr2-95409491-DEL-51
chr2-95441975-DEL-3469
chr2-95467023-DEL-309
chr2-95551467-DEL-50
chr2-95688802-DEL-51
chr2-95752340-DEL-52
chr2-96042073-DEL-50
chr2-96106859-DEL-64
chr2-96107049-DEL-195
chr2-96457628-DEL-303
chr2-96554777-DEL-54
chr2-96611078-DEL-84
chr2-96617095-DEL-2786
chr2-96661343-DEL-232
chr2-96693009-DEL-97
chr2-96754860-DEL-50
chr2-96760848-DEL-471
chr2-96809546-DEL-1576
chr2-96826976-DEL-52

chr2-97268201-DEL-62
chr2-97369782-DEL-78
chr2-97607279-DEL-439
chr2-97640371-DEL-54
chr2-97721873-DEL-129
chr2-97722096-DEL-194
chr2-97746862-DEL-87
chr2-97750405-DEL-50
chr2-97750650-DEL-183
chr2-97764153-DEL-1894
chr2-97830970-DEL-97
chr2-97848634-DEL-50
chr2-97866060-DEL-563
chr2-97928380-DEL-732
chr2-97939982-DEL-64
chr2-97940230-DEL-104
chr2-97970933-DEL-117
chr2-98066413-DEL-265
chr2-98084179-DEL-130
chr2-98090807-DEL-99
chr2-98315104-DEL-1309
chr2-98317127-DEL-88
chr2-98429491-DEL-55
chr2-98449808-DEL-344
chr2-98462646-DEL-208
chr2-98489778-DEL-1547
chr2-98501110-DEL-767
chr2-98554908-DEL-1153
chr2-98591188-DEL-120
chr2-98636693-DEL-669
chr2-98668967-DEL-369
chr2-98746963-DEL-2473
chr2-98815612-DEL-2169
chr2-98899895-DEL-53
chr2-98910024-DEL-115
chr2-98997336-DEL-440
chr2-99001787-DEL-120
chr2-99102975-DEL-52
chr2-99178458-DEL-56
chr2-99208974-DEL-290
chr2-99277442-DEL-88
chr2-99307334-DEL-938
chr2-99312676-DEL-59
chr2-99339212-DEL-53
chr2-99399964-DEL-1571
chr2-99401950-DEL-216
chr2-99457461-DEL-368
chr2-99549071-DEL-50
chr2-99733437-DEL-6827
chr2-99784939-DEL-168
chr2-99785950-DEL-131
chr2-99822364-DEL-8751
chr2-99831163-DEL-59
chr2-99867739-DEL-1329
chr2-99997728-DEL-77
chr2-100000858-DEL-104
chr2-100009220-DEL-72
chr2-100702278-DEL-53
chr2-100714383-DEL-1002
chr2-100755996-DEL-109
chr2-100852620-DEL-106
chr2-100853701-DEL-233
chr2-101046782-DEL-72
chr2-101354219-DEL-55
chr2-101354494-DEL-348
chr2-101366143-DEL-55
chr2-101399276-DEL-294
chr2-101487997-DEL-126
chr2-101503332-DEL-52
chr2-101560623-DEL-68
chr2-101780162-DEL-120
chr2-101789034-DEL-51
chr2-101810385-DEL-137
chr2-101888800-DEL-181
chr2-101888976-DEL-50
chr2-101897002-DEL-760
chr2-101897077-DEL-200

chr2-102198977-DEL-382
chr2-102199014-DEL-127
chr2-102495396-DEL-88
chr2-102866750-DEL-425
chr2-102929636-DEL-76
chr2-103145034-DEL-55
chr2-103317236-DEL-142
chr2-103359589-DEL-204
chr2-103429030-DEL-546
chr2-103682211-DEL-5661
chr2-103709317-DEL-203
chr2-103736160-DEL-62
chr2-103794069-DEL-578
chr2-103795230-DEL-163
chr2-103798276-DEL-124
chr2-103828106-DEL-82
chr2-103850080-DEL-149
chr2-103877037-DEL-310
chr2-103877647-DEL-208
chr2-103877864-DEL-102
chr2-103913699-DEL-136
chr2-103915530-DEL-980
chr2-104016806-DEL-190
chr2-104095081-DEL-54
chr2-104167271-DEL-67
chr2-104189118-DEL-236
chr2-104194259-DEL-113
chr2-104315955-DEL-88
chr2-104328836-DEL-111
chr2-104359990-DEL-56
chr2-104704728-DEL-291
chr2-104803246-DEL-52
chr2-104889873-DEL-200
chr2-104922372-DEL-50
chr2-105005001-DEL-480
chr2-105005061-DEL-160
chr2-105042651-DEL-54
chr2-105043836-DEL-168
chr2-105088829-DEL-234
chr2-105210900-DEL-155
chr2-105251001-DEL-120
chr2-105298407-DEL-270
chr2-105303096-DEL-50
chr2-105326777-DEL-51
chr2-105377477-DEL-54
chr2-105377546-DEL-432
chr2-105394855-DEL-162
chr2-105440270-DEL-137
chr2-105454566-DEL-244
chr2-105475325-DEL-118
chr2-105475949-DEL-172
chr2-105476010-DEL-84
chr2-105599382-DEL-60
chr2-105717863-DEL-259
chr2-105794757-DEL-95
chr2-105971692-DEL-725
chr2-105987825-DEL-50
chr2-106270835-DEL-122
chr2-106271213-DEL-72
chr2-106312767-DEL-1011
chr2-106352245-DEL-164
chr2-106397108-DEL-2605
chr2-106510969-DEL-54
chr2-106672292-DEL-125
chr2-106749885-DEL-116
chr2-106956206-DEL-69
chr2-107034566-DEL-157
chr2-107242291-DEL-51
chr2-107280841-DEL-66
chr2-107420689-DEL-59
chr2-107729028-DEL-749
chr2-108031330-DEL-4561
chr2-108038470-DEL-1187
chr2-108080794-DEL-66
chr2-108091354-DEL-1464
chr2-108117220-DEL-272
chr2-108208420-DEL-54

chr2-108300163-DEL-654
chr2-108769762-DEL-76
chr2-109217769-DEL-186
chr2-109294090-DEL-1182
chr2-109296452-DEL-1075
chr2-109324659-DEL-72
chr2-109410207-DEL-531
chr2-109410693-DEL-62
chr2-109753015-DEL-73
chr2-109802876-DEL-568
chr2-109882615-DEL-51
chr2-110004601-DEL-223
chr2-110005211-DEL-126
chr2-110023825-DEL-2108
chr2-110326995-DEL-6857
chr2-110396702-DEL-106
chr2-110655876-DEL-216
chr2-110681251-DEL-93
chr2-110811654-DEL-89
chr2-110822537-DEL-120
chr2-110839040-DEL-5714
chr2-110840630-DEL-1752
chr2-110845696-DEL-342
chr2-110953077-DEL-214
chr2-111119455-DEL-1173
chr2-111242028-DEL-1086
chr2-111416521-DEL-3871
chr2-111423749-DEL-5886
chr2-111479246-DEL-24487
chr2-111628192-DEL-2882
chr2-111752661-DEL-2238
chr2-111799854-DEL-192
chr2-111933191-DEL-514
chr2-111990918-DEL-268
chr2-112097859-DEL-144
chr2-112163003-DEL-52
chr2-112227422-DEL-51
chr2-112417067-DEL-122
chr2-112434250-DEL-183
chr2-112548237-DEL-85
chr2-112582855-DEL-118
chr2-112583097-DEL-198
chr2-112583498-DEL-76
chr2-112699931-DEL-579
chr2-112910683-DEL-114
chr2-112971007-DEL-133
chr2-113035576-DEL-98
chr2-113115677-DEL-112
chr2-113153085-DEL-198
chr2-113307497-DEL-50
chr2-113313754-DEL-70
chr2-113568223-DEL-131
chr2-113591331-DEL-70
chr2-113597620-DEL-197
chr2-113613749-DEL-52
chr2-113630062-DEL-76
chr2-113662633-DEL-188
chr2-113786412-DEL-1913
chr2-113791990-DEL-50
chr2-114415557-DEL-1387
chr2-114542824-DEL-82
chr2-114663263-DEL-1459
chr2-114811273-DEL-50
chr2-114850012-DEL-316
chr2-114866212-DEL-189
chr2-114866219-DEL-65
chr2-115088439-DEL-243
chr2-115219312-DEL-76
chr2-115304763-DEL-334
chr2-115304834-DEL-120
chr2-115321701-DEL-54
chr2-115624034-DEL-827
chr2-116091995-DEL-80
chr2-116211541-DEL-89
chr2-116224043-DEL-1965
chr2-116244753-DEL-56
chr2-116365854-DEL-124

chr2-116376961-DEL-662
chr2-116386951-DEL-606
chr2-116388036-DEL-145
chr2-116524933-DEL-101
chr2-116594631-DEL-262
chr2-116726208-DEL-50
chr2-116732479-DEL-56
chr2-116743514-DEL-54
chr2-116752065-DEL-60
chr2-116904733-DEL-70
chr2-116905259-DEL-112
chr2-116948142-DEL-328
chr2-116948360-DEL-381
chr2-116948486-DEL-133
chr2-116948561-DEL-58
chr2-116973152-DEL-76
chr2-117063390-DEL-1596
chr2-117063619-DEL-304
chr2-117064189-DEL-684
chr2-117218329-DEL-52
chr2-117278656-DEL-72
chr2-117278768-DEL-106
chr2-117296036-DEL-69
chr2-117377659-DEL-21712
chr2-117508678-DEL-117
chr2-117613657-DEL-2546
chr2-117729646-DEL-91
chr2-117752933-DEL-147
chr2-117776799-DEL-50
chr2-117791838-DEL-61
chr2-117808894-DEL-145
chr2-117838966-DEL-80
chr2-118042543-DEL-4980
chr2-118065207-DEL-491
chr2-118132399-DEL-1080
chr2-118132697-DEL-127
chr2-118135132-DEL-80
chr2-118135285-DEL-60
chr2-118160701-DEL-188
chr2-118173658-DEL-51
chr2-118394548-DEL-674
chr2-118465179-DEL-126
chr2-118529053-DEL-321
chr2-118550842-DEL-1735
chr2-118632078-DEL-136
chr2-118643063-DEL-130
chr2-118666844-DEL-251
chr2-118677509-DEL-256
chr2-118685991-DEL-2561
chr2-118767589-DEL-70
chr2-118784557-DEL-556
chr2-118785945-DEL-50
chr2-118824013-DEL-128
chr2-118923445-DEL-94
chr2-119092289-DEL-93
chr2-119226763-DEL-684
chr2-119234500-DEL-67
chr2-119287930-DEL-100
chr2-119315663-DEL-181
chr2-119316280-DEL-205
chr2-119316680-DEL-74
chr2-119419745-DEL-73
chr2-119535328-DEL-476
chr2-119599171-DEL-307
chr2-119667159-DEL-55
chr2-119710875-DEL-374
chr2-119711149-DEL-66
chr2-119736011-DEL-127
chr2-119736567-DEL-80
chr2-119859563-DEL-144
chr2-120078497-DEL-59
chr2-120078805-DEL-110
chr2-120102028-DEL-74
chr2-120104602-DEL-180
chr2-120206213-DEL-50
chr2-120268952-DEL-54
chr2-120320851-DEL-66

chr2-120320953-DEL-114
chr2-120368041-DEL-211
chr2-120453004-DEL-153
chr2-120474931-DEL-126
chr2-120481606-DEL-778
chr2-120500474-DEL-290
chr2-120511700-DEL-50
chr2-120914362-DEL-145
chr2-120976070-DEL-215
chr2-121109741-DEL-210
chr2-121175269-DEL-55
chr2-121318749-DEL-56
chr2-121351242-DEL-302
chr2-121352825-DEL-165
chr2-121410867-DEL-61
chr2-121553760-DEL-76
chr2-121625945-DEL-130
chr2-121647573-DEL-96
chr2-121650954-DEL-56
chr2-121753032-DEL-50
chr2-121910027-DEL-276
chr2-121936935-DEL-135
chr2-122155715-DEL-149
chr2-122216137-DEL-96
chr2-122216939-DEL-1260
chr2-122221098-DEL-58
chr2-122237054-DEL-243
chr2-122248326-DEL-112
chr2-122248438-DEL-54
chr2-122310141-DEL-126
chr2-122593177-DEL-171
chr2-122640925-DEL-360
chr2-122661601-DEL-76
chr2-122932669-DEL-130
chr2-123028111-DEL-99
chr2-123223871-DEL-50
chr2-123355884-DEL-5588
chr2-123421223-DEL-138
chr2-123429589-DEL-51
chr2-123576529-DEL-56
chr2-123599710-DEL-50
chr2-123619028-DEL-202
chr2-123697296-DEL-53
chr2-123774312-DEL-415
chr2-123892046-DEL-1532
chr2-123965438-DEL-50
chr2-124012244-DEL-178
chr2-124192003-DEL-3279
chr2-124415987-DEL-331
chr2-124643034-DEL-2401
chr2-124699228-DEL-602
chr2-124699616-DEL-271
chr2-124741058-DEL-180
chr2-124900542-DEL-65
chr2-124900555-DEL-133
chr2-124902165-DEL-64
chr2-124975287-DEL-122
chr2-125081998-DEL-625
chr2-125171949-DEL-64
chr2-125176889-DEL-72
chr2-125445982-DEL-50
chr2-125608441-DEL-510
chr2-125608739-DEL-204
chr2-125659793-DEL-60
chr2-125659799-DEL-252
chr2-125740780-DEL-285
chr2-125760225-DEL-195
chr2-125845533-DEL-58
chr2-125870764-DEL-199
chr2-125871149-DEL-58
chr2-126072785-DEL-54
chr2-126287389-DEL-1482
chr2-126340522-DEL-50
chr2-126348605-DEL-50
chr2-126568292-DEL-52
chr2-126586358-DEL-166
chr2-126586442-DEL-82

chr2-126621293-DEL-50
chr2-126746444-DEL-50
chr2-126753589-DEL-474
chr2-126859260-DEL-707
chr2-126881788-DEL-420
chr2-126939784-DEL-154
chr2-126977910-DEL-52
chr2-127012633-DEL-56
chr2-127134996-DEL-76
chr2-127144666-DEL-336
chr2-127144914-DEL-112
chr2-127145489-DEL-637
chr2-127170029-DEL-64
chr2-127217468-DEL-52
chr2-127317923-DEL-90
chr2-127318065-DEL-95
chr2-127319628-DEL-377
chr2-127505992-DEL-100
chr2-127528892-DEL-66
chr2-127737320-DEL-69
chr2-127737326-DEL-575
chr2-127738389-DEL-90
chr2-128094917-DEL-54
chr2-128148427-DEL-53
chr2-128150199-DEL-52
chr2-128164954-DEL-220
chr2-128251591-DEL-84
chr2-128263434-DEL-60
chr2-128572631-DEL-68
chr2-128631790-DEL-74
chr2-128720152-DEL-76
chr2-128738756-DEL-98
chr2-128923702-DEL-100
chr2-128962779-DEL-150
chr2-128962841-DEL-50
chr2-128996734-DEL-68
chr2-129005276-DEL-109
chr2-129075445-DEL-92
chr2-129080019-DEL-69
chr2-129095626-DEL-50
chr2-129106610-DEL-62
chr2-129106610-DEL-496
chr2-129130038-DEL-50
chr2-129212872-DEL-70
chr2-129241796-DEL-50
chr2-129296328-DEL-214
chr2-129662049-DEL-50
chr2-129782923-DEL-119
chr2-129938925-DEL-88
chr2-130050385-DEL-372
chr2-130050391-DEL-91
chr2-130050953-DEL-165
chr2-130051451-DEL-53
chr2-130079403-DEL-51
chr2-130171063-DEL-145
chr2-130181026-DEL-401
chr2-130352116-DEL-146
chr2-130411103-DEL-3058
chr2-130530598-DEL-63
chr2-130610737-DEL-54
chr2-130622695-DEL-50
chr2-130716268-DEL-50
chr2-130716530-DEL-453
chr2-130730377-DEL-623
chr2-130744831-DEL-64
chr2-131127307-DEL-320
chr2-131136671-DEL-384
chr2-131146716-DEL-442
chr2-131163079-DEL-154
chr2-131209815-DEL-74
chr2-131216355-DEL-52
chr2-131236639-DEL-156
chr2-131236960-DEL-60
chr2-131316116-DEL-81
chr2-131400704-DEL-100
chr2-131434058-DEL-1850
chr2-131473035-DEL-52

chr2-131706290-DEL-52
chr2-131804526-DEL-80
chr2-131817020-DEL-180
chr2-131840813-DEL-444
chr2-131840895-DEL-54
chr2-131857211-DEL-140
chr2-131910198-DEL-227
chr2-131910239-DEL-98
chr2-131917774-DEL-164
chr2-131918023-DEL-121
chr2-131948338-DEL-7536
chr2-131959733-DEL-107
chr2-132010816-DEL-1972
chr2-132015388-DEL-61
chr2-132155617-DEL-130
chr2-132185202-DEL-139
chr2-132192561-DEL-77
chr2-132244264-DEL-14237
chr2-132268052-DEL-174
chr2-132294118-DEL-4385
chr2-132301568-DEL-4652
chr2-132317107-DEL-80
chr2-132365311-DEL-51
chr2-132373539-DEL-349
chr2-132399200-DEL-6732
chr2-132489456-DEL-211
chr2-132553185-DEL-111
chr2-132553414-DEL-315
chr2-132553659-DEL-70
chr2-132569801-DEL-261
chr2-132581240-DEL-60
chr2-132621158-DEL-102
chr2-132643925-DEL-116
chr2-132681841-DEL-63
chr2-132727521-DEL-5977
chr2-132774273-DEL-110
chr2-132917497-DEL-303
chr2-132954395-DEL-138
chr2-132966562-DEL-56
chr2-132966636-DEL-140
chr2-132984692-DEL-5065
chr2-133075802-DEL-234
chr2-133076502-DEL-70
chr2-133415767-DEL-86
chr2-133423043-DEL-164
chr2-133606535-DEL-128
chr2-133687501-DEL-232
chr2-133687530-DEL-78
chr2-133815999-DEL-72
chr2-134034700-DEL-268
chr2-134156920-DEL-1298
chr2-134317422-DEL-65
chr2-134579722-DEL-75
chr2-134579889-DEL-304
chr2-134632266-DEL-316
chr2-134679623-DEL-68
chr2-134837470-DEL-50
chr2-134936990-DEL-50
chr2-135008987-DEL-792
chr2-135043474-DEL-172
chr2-135216214-DEL-52
chr2-135216280-DEL-186
chr2-135252061-DEL-50
chr2-135332117-DEL-121
chr2-135365658-DEL-50
chr2-135387187-DEL-51
chr2-135519971-DEL-831
chr2-135528185-DEL-250
chr2-135551541-DEL-58
chr2-135696425-DEL-149
chr2-135769486-DEL-81
chr2-135801010-DEL-101
chr2-135975313-DEL-2983
chr2-136037072-DEL-91
chr2-136060325-DEL-158
chr2-136096968-DEL-131
chr2-136250966-DEL-156

chr2-136678973-DEL-687
chr2-136705365-DEL-267
chr2-136740965-DEL-69
chr2-136936672-DEL-100
chr2-136955164-DEL-61
chr2-137245756-DEL-1551
chr2-137524211-DEL-502
chr2-137577265-DEL-98
chr2-137890840-DEL-724
chr2-137963528-DEL-80
chr2-137974152-DEL-175
chr2-138085373-DEL-94
chr2-138335985-DEL-231
chr2-138636489-DEL-328
chr2-138640668-DEL-63
chr2-138664773-DEL-225
chr2-138755977-DEL-78
chr2-138802332-DEL-448
chr2-139237233-DEL-113
chr2-139435607-DEL-1894
chr2-139438456-DEL-90
chr2-139578644-DEL-56
chr2-139623839-DEL-60
chr2-139640395-DEL-108
chr2-139645218-DEL-58
chr2-139718790-DEL-1169
chr2-139732934-DEL-84
chr2-139831687-DEL-266
chr2-139903745-DEL-81
chr2-139917730-DEL-50
chr2-139918566-DEL-88
chr2-139958133-DEL-98
chr2-140043784-DEL-50
chr2-140313129-DEL-138
chr2-140313142-DEL-61
chr2-140360625-DEL-1711
chr2-140382244-DEL-214
chr2-140448152-DEL-2377
chr2-140516559-DEL-120
chr2-140567779-DEL-124
chr2-140745713-DEL-379
chr2-140914721-DEL-86
chr2-140940668-DEL-142
chr2-140991879-DEL-93
chr2-141050163-DEL-172
chr2-141213945-DEL-93
chr2-141237614-DEL-51
chr2-141452765-DEL-78
chr2-141481809-DEL-113
chr2-141517600-DEL-65
chr2-141805682-DEL-70
chr2-142008653-DEL-51
chr2-142030115-DEL-141
chr2-142068526-DEL-93
chr2-142104862-DEL-57
chr2-142159976-DEL-237
chr2-142191099-DEL-91
chr2-142255955-DEL-253
chr2-142391210-DEL-132
chr2-142394663-DEL-118
chr2-142419200-DEL-133
chr2-142603369-DEL-98
chr2-142643736-DEL-62
chr2-142708387-DEL-504
chr2-142708950-DEL-58
chr2-143136384-DEL-333
chr2-143204550-DEL-231
chr2-143208965-DEL-50
chr2-143274395-DEL-4358
chr2-143339867-DEL-633
chr2-143491690-DEL-98
chr2-143570767-DEL-2276
chr2-143588444-DEL-66
chr2-143611760-DEL-3360
chr2-143671779-DEL-66
chr2-143674476-DEL-155
chr2-143675762-DEL-334

chr2-143702101-DEL-64
chr2-143715350-DEL-131
chr2-143715833-DEL-861
chr2-143717425-DEL-70
chr2-143830534-DEL-54
chr2-143899642-DEL-87
chr2-143923600-DEL-53
chr2-143931816-DEL-56
chr2-143931990-DEL-136
chr2-143946523-DEL-72
chr2-144063829-DEL-423
chr2-144084302-DEL-54
chr2-144177463-DEL-722
chr2-144189769-DEL-55
chr2-144304572-DEL-76
chr2-144349580-DEL-72
chr2-144404911-DEL-148
chr2-144441350-DEL-90
chr2-144529145-DEL-145
chr2-144674425-DEL-58
chr2-144929557-DEL-132
chr2-145393407-DEL-471
chr2-145642876-DEL-72
chr2-145738065-DEL-62
chr2-145844380-DEL-199
chr2-145948597-DEL-52
chr2-146371555-DEL-267
chr2-146445978-DEL-2432
chr2-146573546-DEL-1198
chr2-146631847-DEL-462
chr2-146765673-DEL-80
chr2-146801983-DEL-246
chr2-146830459-DEL-81
chr2-146863227-DEL-1419
chr2-147014357-DEL-52
chr2-147018039-DEL-165
chr2-147018512-DEL-94
chr2-147037870-DEL-168
chr2-147037955-DEL-75
chr2-147052715-DEL-68
chr2-147067442-DEL-243
chr2-147067760-DEL-78
chr2-147067836-DEL-158
chr2-147225123-DEL-77
chr2-147239726-DEL-138
chr2-147272582-DEL-50
chr2-147280972-DEL-1643
chr2-147322704-DEL-815
chr2-147504079-DEL-1151
chr2-147521144-DEL-51
chr2-148091437-DEL-447
chr2-148147745-DEL-50
chr2-148157926-DEL-50
chr2-148302739-DEL-212
chr2-148932954-DEL-98
chr2-148941471-DEL-144
chr2-148982330-DEL-163
chr2-148982397-DEL-60
chr2-149014296-DEL-422
chr2-149080804-DEL-383
chr2-149207025-DEL-54
chr2-149207351-DEL-69
chr2-149319606-DEL-51
chr2-149394370-DEL-117
chr2-149404361-DEL-50
chr2-149406119-DEL-733
chr2-149464967-DEL-1464
chr2-149636035-DEL-52
chr2-149949509-DEL-644
chr2-150096698-DEL-252
chr2-150109672-DEL-252
chr2-150110092-DEL-1176
chr2-150215540-DEL-385
chr2-150246014-DEL-76
chr2-150337059-DEL-81
chr2-150663683-DEL-447
chr2-150663738-DEL-74

chr2-150741603-DEL-270
chr2-150764583-DEL-50
chr2-150767554-DEL-139
chr2-150833295-DEL-690
chr2-150864917-DEL-52
chr2-151047180-DEL-3371
chr2-151175752-DEL-6891
chr2-151295213-DEL-1129
chr2-151458065-DEL-180
chr2-151552005-DEL-50
chr2-151633366-DEL-52
chr2-151667440-DEL-84
chr2-151758381-DEL-648
chr2-151792944-DEL-1434
chr2-151856187-DEL-1394
chr2-151864546-DEL-764
chr2-151868095-DEL-1185
chr2-151960831-DEL-88
chr2-152036320-DEL-1096
chr2-152163845-DEL-71
chr2-152195442-DEL-1124
chr2-152204246-DEL-274
chr2-152231103-DEL-141
chr2-152231195-DEL-430
chr2-152235151-DEL-70
chr2-152292044-DEL-63
chr2-152293337-DEL-55
chr2-152534730-DEL-62
chr2-152806160-DEL-203
chr2-152886533-DEL-95
chr2-152921750-DEL-233
chr2-152936085-DEL-2292
chr2-152936104-DEL-936
chr2-152938275-DEL-552
chr2-152938706-DEL-2104
chr2-153015584-DEL-51
chr2-153105271-DEL-62
chr2-153123988-DEL-234
chr2-153201360-DEL-203
chr2-153231906-DEL-4726
chr2-153268554-DEL-205
chr2-153268665-DEL-83
chr2-153281541-DEL-146
chr2-153400932-DEL-151
chr2-153657126-DEL-126
chr2-153699329-DEL-69
chr2-153699572-DEL-69
chr2-153715527-DEL-689
chr2-153715716-DEL-253
chr2-153716005-DEL-254
chr2-153725592-DEL-204
chr2-153782246-DEL-120
chr2-153782470-DEL-64
chr2-153784009-DEL-198
chr2-153796894-DEL-100
chr2-153826318-DEL-52
chr2-153910547-DEL-53
chr2-154092450-DEL-58
chr2-154109236-DEL-49379
chr2-154257474-DEL-233
chr2-154275033-DEL-66
chr2-154512716-DEL-1349
chr2-154644444-DEL-1540
chr2-154753055-DEL-50
chr2-154764874-DEL-235
chr2-154768619-DEL-198
chr2-154769472-DEL-68
chr2-154769819-DEL-189
chr2-154881041-DEL-54
chr2-154918962-DEL-211
chr2-154961490-DEL-230
chr2-155063946-DEL-120
chr2-155167636-DEL-90
chr2-155240203-DEL-134
chr2-155561035-DEL-127
chr2-155561327-DEL-52
chr2-155568066-DEL-50

chr2-155620852-DEL-133
chr2-155627320-DEL-61
chr2-155679703-DEL-64
chr2-155743514-DEL-133
chr2-155762056-DEL-177
chr2-155868440-DEL-82
chr2-155897485-DEL-119
chr2-155897690-DEL-84
chr2-155900139-DEL-68
chr2-155949971-DEL-63
chr2-155954802-DEL-58
chr2-155978487-DEL-1927
chr2-156072887-DEL-224
chr2-156072911-DEL-88
chr2-156116779-DEL-559
chr2-156116826-DEL-199
chr2-156117118-DEL-181
chr2-156160944-DEL-50
chr2-156179093-DEL-84
chr2-156208295-DEL-58
chr2-156208477-DEL-68
chr2-156325986-DEL-510
chr2-156432351-DEL-51
chr2-156507102-DEL-50
chr2-156524141-DEL-54
chr2-156543784-DEL-173
chr2-156551972-DEL-186
chr2-156618497-DEL-160
chr2-156618697-DEL-112
chr2-156646579-DEL-74
chr2-156659645-DEL-63
chr2-156702768-DEL-310
chr2-156703016-DEL-147
chr2-156722858-DEL-268
chr2-156876757-DEL-53
chr2-156927626-DEL-62
chr2-156947340-DEL-50
chr2-156951584-DEL-208
chr2-157007506-DEL-319
chr2-157079527-DEL-102
chr2-157079694-DEL-51
chr2-157079857-DEL-87
chr2-157084809-DEL-53
chr2-157108024-DEL-1987
chr2-157116675-DEL-59
chr2-157250619-DEL-189
chr2-157287029-DEL-126
chr2-157287127-DEL-326
chr2-157363330-DEL-114
chr2-157427514-DEL-5277
chr2-157462400-DEL-4465
chr2-157787347-DEL-349
chr2-157800839-DEL-130
chr2-157802466-DEL-119
chr2-157881267-DEL-60
chr2-157892446-DEL-98
chr2-157892498-DEL-350
chr2-157968330-DEL-184
chr2-157979015-DEL-50
chr2-157990580-DEL-72
chr2-158112906-DEL-52
chr2-158242655-DEL-78
chr2-158396594-DEL-497
chr2-158397121-DEL-144
chr2-158536037-DEL-250
chr2-158574580-DEL-50
chr2-158579203-DEL-64
chr2-158589290-DEL-84
chr2-158591993-DEL-59
chr2-158593590-DEL-59
chr2-158681131-DEL-308
chr2-158723441-DEL-144
chr2-158736238-DEL-568
chr2-158736389-DEL-226
chr2-158975855-DEL-262
chr2-158991203-DEL-50
chr2-159064635-DEL-237

chr2-159064644-DEL-51
chr2-159066627-DEL-1857
chr2-159267340-DEL-76
chr2-159344292-DEL-302
chr2-159447736-DEL-88
chr2-159475155-DEL-92
chr2-159475376-DEL-57
chr2-159478237-DEL-208
chr2-159514940-DEL-50
chr2-159589777-DEL-51
chr2-159713888-DEL-1177
chr2-159788258-DEL-110
chr2-159885104-DEL-58
chr2-159977216-DEL-80
chr2-160284163-DEL-112
chr2-160312206-DEL-58
chr2-160312300-DEL-58
chr2-160329347-DEL-102
chr2-160408873-DEL-187
chr2-160512896-DEL-59
chr2-160559456-DEL-194
chr2-160563240-DEL-96
chr2-160717674-DEL-2037
chr2-160761702-DEL-150
chr2-160799465-DEL-519
chr2-161014943-DEL-53
chr2-161015962-DEL-54
chr2-161015980-DEL-144
chr2-161033189-DEL-50
chr2-161110498-DEL-124
chr2-161161876-DEL-281
chr2-161222524-DEL-58
chr2-161299305-DEL-469
chr2-161403923-DEL-151
chr2-161404018-DEL-56
chr2-161404430-DEL-114
chr2-161524061-DEL-50
chr2-161639490-DEL-102
chr2-161639823-DEL-66
chr2-161640990-DEL-102
chr2-161641156-DEL-299
chr2-161775248-DEL-73
chr2-161823275-DEL-67
chr2-161823421-DEL-96
chr2-161851654-DEL-1518
chr2-161914248-DEL-54
chr2-161929682-DEL-80
chr2-161953989-DEL-262
chr2-161988058-DEL-204
chr2-162200129-DEL-323
chr2-162626241-DEL-1370
chr2-162769549-DEL-104
chr2-162797442-DEL-234
chr2-162803875-DEL-55
chr2-162804927-DEL-64
chr2-162879742-DEL-782
chr2-162880052-DEL-390
chr2-163023574-DEL-50
chr2-163165522-DEL-150
chr2-163484566-DEL-593
chr2-163543469-DEL-106
chr2-163543487-DEL-530
chr2-163646675-DEL-123
chr2-163790527-DEL-58
chr2-163811944-DEL-131
chr2-163839453-DEL-3143
chr2-163849135-DEL-94
chr2-163849224-DEL-329
chr2-163849453-DEL-94
chr2-163915528-DEL-185
chr2-163929980-DEL-50
chr2-164183188-DEL-103
chr2-164204812-DEL-52
chr2-164239372-DEL-414
chr2-164239482-DEL-144
chr2-164239937-DEL-162
chr2-164349525-DEL-224

chr2-164353879-DEL-64
chr2-164362251-DEL-159
chr2-164376278-DEL-88
chr2-164378243-DEL-116
chr2-164475724-DEL-68
chr2-164505991-DEL-198
chr2-164535062-DEL-447
chr2-164535109-DEL-120
chr2-164535388-DEL-213
chr2-164546910-DEL-82
chr2-164571408-DEL-98
chr2-164575811-DEL-170
chr2-164616366-DEL-505
chr2-164616671-DEL-166
chr2-164671679-DEL-435
chr2-164684766-DEL-167
chr2-164853555-DEL-938
chr2-164871599-DEL-551
chr2-165016056-DEL-64
chr2-165016114-DEL-140
chr2-165198909-DEL-124
chr2-165224957-DEL-96
chr2-165272949-DEL-120
chr2-165353057-DEL-109
chr2-165353463-DEL-89
chr2-165408469-DEL-114
chr2-165456851-DEL-144
chr2-165457307-DEL-118
chr2-165457675-DEL-119
chr2-165457837-DEL-250
chr2-165457943-DEL-113
chr2-165461413-DEL-59
chr2-165537794-DEL-69
chr2-165543340-DEL-80
chr2-165543889-DEL-60
chr2-165638155-DEL-68
chr2-165644017-DEL-161
chr2-165644160-DEL-72
chr2-165645821-DEL-98
chr2-165749997-DEL-107
chr2-165757758-DEL-217
chr2-165784437-DEL-132
chr2-165792500-DEL-73
chr2-165804767-DEL-60
chr2-165868267-DEL-70
chr2-165926741-DEL-135
chr2-166003080-DEL-132
chr2-166006697-DEL-62
chr2-166082319-DEL-56
chr2-166127648-DEL-244
chr2-166231561-DEL-252
chr2-166332148-DEL-54
chr2-166446131-DEL-248
chr2-166456735-DEL-206
chr2-166522826-DEL-1185
chr2-166575624-DEL-160
chr2-166602301-DEL-66
chr2-166630524-DEL-159
chr2-166672847-DEL-600
chr2-166673099-DEL-80
chr2-166712401-DEL-65
chr2-166712441-DEL-132
chr2-166725493-DEL-648
chr2-166725557-DEL-284
chr2-166725906-DEL-51
chr2-166725977-DEL-158
chr2-166760606-DEL-50
chr2-166762003-DEL-54
chr2-166809920-DEL-92
chr2-166810118-DEL-191
chr2-166887215-DEL-324
chr2-166887593-DEL-54
chr2-166916699-DEL-627
chr2-166927349-DEL-152
chr2-166927709-DEL-56
chr2-167020795-DEL-100
chr2-167020959-DEL-204

chr2-167058785-DEL-64
chr2-167102817-DEL-72
chr2-167124416-DEL-7596
chr2-167239749-DEL-95
chr2-167362710-DEL-4264
chr2-167432962-DEL-110
chr2-167514002-DEL-76
chr2-167592332-DEL-176
chr2-167592824-DEL-487
chr2-167593160-DEL-98
chr2-167627959-DEL-404
chr2-167677378-DEL-60
chr2-167712421-DEL-61
chr2-167719089-DEL-76
chr2-167726353-DEL-56
chr2-167734120-DEL-75
chr2-167812298-DEL-681
chr2-167832717-DEL-163
chr2-167840598-DEL-50
chr2-167851310-DEL-92
chr2-167857134-DEL-380
chr2-168061211-DEL-240
chr2-168079296-DEL-93
chr2-168107369-DEL-88
chr2-168154032-DEL-52
chr2-168155496-DEL-312
chr2-168203699-DEL-166
chr2-168203939-DEL-116
chr2-168204073-DEL-54
chr2-168233060-DEL-71
chr2-168252502-DEL-62
chr2-168337888-DEL-268
chr2-168356722-DEL-68
chr2-168356879-DEL-164
chr2-168382388-DEL-71
chr2-168461103-DEL-231
chr2-168487142-DEL-150
chr2-168512875-DEL-74
chr2-168519349-DEL-2007
chr2-168521282-DEL-74
chr2-168526590-DEL-53
chr2-168535651-DEL-59
chr2-168545112-DEL-157
chr2-168557244-DEL-143
chr2-168589250-DEL-112
chr2-168589260-DEL-822
chr2-168629639-DEL-748
chr2-168638536-DEL-80
chr2-168668993-DEL-177
chr2-168763266-DEL-81
chr2-168816181-DEL-5603
chr2-168837463-DEL-91
chr2-168840158-DEL-61
chr2-168841696-DEL-242
chr2-168854304-DEL-147
chr2-168854414-DEL-69
chr2-168910672-DEL-51
chr2-168944662-DEL-50
chr2-168946598-DEL-60
chr2-168947502-DEL-267
chr2-168957823-DEL-77
chr2-168990775-DEL-64
chr2-169012389-DEL-66
chr2-169017871-DEL-54
chr2-169041508-DEL-550
chr2-169079664-DEL-101
chr2-169133853-DEL-50
chr2-169218324-DEL-51
chr2-169257340-DEL-51
chr2-169263532-DEL-60
chr2-169319883-DEL-379
chr2-169320046-DEL-102
chr2-169327217-DEL-336
chr2-169354717-DEL-54
chr2-169354856-DEL-92
chr2-169405733-DEL-72
chr2-169423926-DEL-83

chr2-169429446-DEL-610
chr2-169429897-DEL-88
chr2-169435092-DEL-2237
chr2-169459617-DEL-900
chr2-169485706-DEL-96
chr2-169490734-DEL-392
chr2-169514143-DEL-643
chr2-169544277-DEL-58
chr2-169550866-DEL-50
chr2-169598701-DEL-134
chr2-169646825-DEL-50
chr2-169677628-DEL-114
chr2-169699082-DEL-80
chr2-169749498-DEL-132
chr2-169786667-DEL-336
chr2-169786769-DEL-80
chr2-169822041-DEL-176
chr2-169822276-DEL-88
chr2-169868327-DEL-101
chr2-169880147-DEL-88
chr2-169905281-DEL-171
chr2-170053251-DEL-120
chr2-170085246-DEL-108
chr2-170162639-DEL-264
chr2-170162994-DEL-259
chr2-170171010-DEL-65
chr2-170202761-DEL-71
chr2-170203229-DEL-80
chr2-170215869-DEL-50
chr2-170251017-DEL-55
chr2-170253341-DEL-55
chr2-170259394-DEL-841
chr2-170275554-DEL-3159
chr2-170311759-DEL-65
chr2-170315718-DEL-58
chr2-170347172-DEL-71
chr2-170361048-DEL-203
chr2-170365560-DEL-126
chr2-170390684-DEL-64
chr2-170424027-DEL-76
chr2-170452241-DEL-68
chr2-170468513-DEL-51
chr2-170506656-DEL-59
chr2-170507190-DEL-64
chr2-170523553-DEL-184
chr2-170524027-DEL-174
chr2-170556134-DEL-51
chr2-170556205-DEL-103
chr2-170571282-DEL-216
chr2-170601047-DEL-81
chr2-170722498-DEL-699
chr2-170753331-DEL-64
chr2-170819485-DEL-86
chr2-170839602-DEL-453
chr2-170839674-DEL-222
chr2-170846989-DEL-62
chr2-170876690-DEL-298
chr2-170913897-DEL-329
chr2-170962173-DEL-225
chr2-171005386-DEL-267
chr2-171052520-DEL-460
chr2-171091065-DEL-2106
chr2-171091218-DEL-648
chr2-171093920-DEL-90
chr2-171109579-DEL-68
chr2-171132978-DEL-190
chr2-171133086-DEL-750
chr2-171159305-DEL-132
chr2-171170918-DEL-1277
chr2-171182001-DEL-157
chr2-171220545-DEL-309
chr2-171232459-DEL-357
chr2-171232502-DEL-56
chr2-171232811-DEL-151
chr2-171232967-DEL-53
chr2-171252779-DEL-139
chr2-171253821-DEL-348

chr2-171253995-DEL-87
chr2-171315580-DEL-58
chr2-171338295-DEL-354
chr2-171347383-DEL-858
chr2-171382421-DEL-211
chr2-171384660-DEL-192
chr2-171400399-DEL-160
chr2-171426481-DEL-96
chr2-171467256-DEL-726
chr2-171474719-DEL-60
chr2-171550316-DEL-58
chr2-171645246-DEL-98
chr2-171656911-DEL-59
chr2-171667985-DEL-1374
chr2-171674884-DEL-128
chr2-171704886-DEL-420
chr2-171705609-DEL-203
chr2-171729827-DEL-506
chr2-171734700-DEL-450
chr2-171741181-DEL-120
chr2-171788003-DEL-507
chr2-171788364-DEL-78
chr2-171811374-DEL-99
chr2-171836488-DEL-337
chr2-171847592-DEL-54
chr2-171856920-DEL-77
chr2-171871545-DEL-87
chr2-171925285-DEL-509
chr2-171927216-DEL-120
chr2-171940149-DEL-3069
chr2-171945275-DEL-64
chr2-171982915-DEL-127
chr2-171983118-DEL-56
chr2-172000251-DEL-342
chr2-172021122-DEL-202
chr2-172043581-DEL-319
chr2-172046291-DEL-365
chr2-172046293-DEL-761
chr2-172059954-DEL-249
chr2-172066328-DEL-180
chr2-172097504-DEL-196
chr2-172121255-DEL-1151
chr2-172122627-DEL-288
chr2-172151756-DEL-581
chr2-172169665-DEL-96
chr2-172175592-DEL-109
chr2-172325270-DEL-98
chr2-172346124-DEL-50
chr2-172408634-DEL-50
chr2-172436901-DEL-248
chr2-172437265-DEL-222
chr2-172466705-DEL-78
chr2-172501117-DEL-51
chr2-172518964-DEL-132
chr2-172556134-DEL-96
chr2-172683642-DEL-170
chr2-172755432-DEL-58
chr2-172806672-DEL-112
chr2-172806803-DEL-96
chr2-172810013-DEL-50
chr2-172898680-DEL-64
chr2-172920465-DEL-108
chr2-172920537-DEL-50
chr2-172941699-DEL-67
chr2-172941881-DEL-228
chr2-172947087-DEL-50
chr2-172969954-DEL-201
chr2-172988079-DEL-147
chr2-173004708-DEL-216
chr2-173042593-DEL-1336
chr2-173051415-DEL-120
chr2-173154317-DEL-140
chr2-173242409-DEL-105
chr2-173243635-DEL-1640
chr2-173257962-DEL-66
chr2-173348890-DEL-52
chr2-173372266-DEL-81

chr2-173384071-DEL-300
chr2-173469250-DEL-108
chr2-173477539-DEL-223
chr2-173509853-DEL-65
chr2-173549367-DEL-291
chr2-173549569-DEL-86
chr2-173594035-DEL-677
chr2-173594636-DEL-67
chr2-173603630-DEL-93
chr2-173611736-DEL-80
chr2-173730532-DEL-1753
chr2-173747763-DEL-1694
chr2-173809695-DEL-116
chr2-174131485-DEL-80
chr2-174202139-DEL-476
chr2-174202168-DEL-204
chr2-174240974-DEL-106
chr2-174244516-DEL-107
chr2-174300706-DEL-78
chr2-174365872-DEL-56
chr2-174387080-DEL-68
chr2-174387218-DEL-1540
chr2-174425653-DEL-134
chr2-174459219-DEL-62
chr2-174525272-DEL-50
chr2-174573047-DEL-96
chr2-174855321-DEL-196
chr2-175010801-DEL-252
chr2-175011105-DEL-84
chr2-176547999-DEL-362
chr2-177200827-DEL-84
chr2-177261784-DEL-84
chr2-177411585-DEL-84
chr2-177594264-DEL-1530
chr2-177598804-DEL-816
chr2-177769222-DEL-116
chr2-177827344-DEL-54
chr2-177838290-DEL-134
chr2-177844200-DEL-240
chr2-177870872-DEL-1233
chr2-177879583-DEL-80
chr2-177940673-DEL-101
chr2-177941147-DEL-129
chr2-178042002-DEL-52
chr2-178104174-DEL-126
chr2-178141055-DEL-351
chr2-178145378-DEL-50
chr2-178170014-DEL-290
chr2-178170753-DEL-232
chr2-178184427-DEL-63
chr2-178235995-DEL-388
chr2-178281231-DEL-96
chr2-178317461-DEL-495
chr2-178317597-DEL-62
chr2-178341027-DEL-137
chr2-178365938-DEL-68
chr2-178371619-DEL-855
chr2-178404742-DEL-402
chr2-178422649-DEL-481
chr2-178422933-DEL-148
chr2-178428069-DEL-213
chr2-178438087-DEL-76
chr2-178439293-DEL-68
chr2-178458714-DEL-55
chr2-178539960-DEL-52
chr2-178582416-DEL-50
chr2-178644339-DEL-1628
chr2-178688150-DEL-120
chr2-178783335-DEL-132
chr2-178783440-DEL-363
chr2-178783803-DEL-66
chr2-178785505-DEL-52
chr2-178837601-DEL-108
chr2-178860141-DEL-50
chr2-178951282-DEL-363
chr2-178968265-DEL-1648
chr2-179033905-DEL-65

chr2-179077483-DEL-60
chr2-179122229-DEL-92
chr2-179152070-DEL-72
chr2-179155521-DEL-127
chr2-179196170-DEL-82
chr2-179296919-DEL-150
chr2-179358225-DEL-338
chr2-179380291-DEL-165
chr2-179380744-DEL-123
chr2-179387470-DEL-87
chr2-179387689-DEL-136
chr2-179400377-DEL-60
chr2-179486406-DEL-100
chr2-179532261-DEL-85
chr2-179561101-DEL-324
chr2-179612041-DEL-105
chr2-179706354-DEL-88
chr2-179706415-DEL-256
chr2-179706807-DEL-2078
chr2-179706979-DEL-726
chr2-179708257-DEL-632
chr2-179895135-DEL-406
chr2-179910893-DEL-66
chr2-179936963-DEL-450
chr2-180036377-DEL-63
chr2-180080879-DEL-50
chr2-180229549-DEL-60
chr2-180229635-DEL-236
chr2-180237006-DEL-108
chr2-180391792-DEL-264
chr2-180392603-DEL-264
chr2-180423433-DEL-75
chr2-180445889-DEL-59
chr2-180790172-DEL-82
chr2-180986673-DEL-158
chr2-181069703-DEL-465
chr2-181070000-DEL-72
chr2-181121121-DEL-158
chr2-181235132-DEL-50
chr2-181392379-DEL-563
chr2-181420627-DEL-197
chr2-181443098-DEL-51
chr2-181513009-DEL-59
chr3-3075496-DEL-63
chr3-3271123-DEL-1182
chr3-3413779-DEL-201
chr3-3924408-DEL-1758
chr3-3928967-DEL-199
chr3-4050648-DEL-56
chr3-4069730-DEL-81
chr3-4091237-DEL-2783
chr3-4175031-DEL-126
chr3-4362972-DEL-405
chr3-4368094-DEL-68
chr3-4370041-DEL-1479
chr3-4421875-DEL-198
chr3-4451474-DEL-7145
chr3-4547124-DEL-131
chr3-4547218-DEL-60
chr3-4588602-DEL-289
chr3-4588675-DEL-70
chr3-4740037-DEL-67
chr3-4897194-DEL-93
chr3-4938734-DEL-580
chr3-4955140-DEL-51
chr3-5180515-DEL-383
chr3-5327007-DEL-65
chr3-5525243-DEL-50
chr3-5842791-DEL-954
chr3-5877439-DEL-85
chr3-6046047-DEL-54
chr3-6145343-DEL-2607
chr3-6235580-DEL-89
chr3-6439544-DEL-66
chr3-6559367-DEL-57
chr3-6578510-DEL-203
chr3-6587901-DEL-52

chr3-6618375-DEL-2317
chr3-6621009-DEL-3256
chr3-6842612-DEL-137
chr3-6863535-DEL-2135
chr3-7147941-DEL-6477
chr3-7224350-DEL-70
chr3-7236425-DEL-133
chr3-7253498-DEL-352
chr3-7315373-DEL-55
chr3-7420406-DEL-200
chr3-7453123-DEL-50
chr3-7545983-DEL-169
chr3-7573366-DEL-132
chr3-7657072-DEL-262
chr3-7692436-DEL-53
chr3-7896001-DEL-2144
chr3-7946046-DEL-52
chr3-8032382-DEL-121
chr3-8144550-DEL-1472
chr3-8151116-DEL-1740
chr3-8164665-DEL-74
chr3-8207759-DEL-50
chr3-8257883-DEL-150
chr3-8258188-DEL-311
chr3-8278443-DEL-494
chr3-8291079-DEL-79
chr3-8308825-DEL-53
chr3-8320224-DEL-52
chr3-8332967-DEL-66
chr3-8530774-DEL-56
chr3-8552887-DEL-282
chr3-8565063-DEL-58
chr3-8598265-DEL-71
chr3-8598367-DEL-120
chr3-8612071-DEL-50
chr3-8684708-DEL-1681
chr3-8766714-DEL-64
chr3-8828825-DEL-258
chr3-8835849-DEL-107
chr3-8841181-DEL-64
chr3-8858928-DEL-462
chr3-8858996-DEL-96
chr3-8859184-DEL-112
chr3-8897166-DEL-416
chr3-8931831-DEL-61
chr3-8946866-DEL-96
chr3-8980782-DEL-321
chr3-8986837-DEL-57
chr3-9006209-DEL-335
chr3-9006847-DEL-112
chr3-9017817-DEL-97
chr3-9050233-DEL-113
chr3-9050643-DEL-68
chr3-9122801-DEL-675
chr3-9135061-DEL-62
chr3-9135454-DEL-70
chr3-9237124-DEL-265
chr3-9237510-DEL-533
chr3-9237597-DEL-69
chr3-9274772-DEL-612
chr3-9322175-DEL-89
chr3-9325435-DEL-790
chr3-9358254-DEL-185
chr3-9393014-DEL-56
chr3-9481023-DEL-55
chr3-9516020-DEL-144
chr3-9516020-DEL-618
chr3-9516497-DEL-98
chr3-9516621-DEL-270
chr3-9517043-DEL-243
chr3-9519459-DEL-53
chr3-9566847-DEL-480
chr3-9592902-DEL-150
chr3-9592908-DEL-72
chr3-9593139-DEL-354
chr3-9597687-DEL-584
chr3-9629229-DEL-272

chr3-9636109-DEL-63
chr3-9651224-DEL-90
chr3-9667514-DEL-50
chr3-9667569-DEL-393
chr3-9683682-DEL-78
chr3-9683933-DEL-119
chr3-9701037-DEL-81
chr3-9701199-DEL-285
chr3-9707087-DEL-356
chr3-9748289-DEL-144
chr3-9842582-DEL-240
chr3-9867854-DEL-630
chr3-9932677-DEL-198
chr3-9932923-DEL-145
chr3-9973674-DEL-468
chr3-9984394-DEL-335
chr3-9990314-DEL-88
chr3-9992963-DEL-54
chr3-10014740-DEL-86
chr3-10099400-DEL-190
chr3-10123747-DEL-116
chr3-10215148-DEL-800
chr3-10263113-DEL-50
chr3-10341297-DEL-122
chr3-10341320-DEL-246
chr3-10389480-DEL-64
chr3-10402615-DEL-81
chr3-10412977-DEL-1235
chr3-10419632-DEL-89
chr3-10438330-DEL-56
chr3-10546997-DEL-6395
chr3-10593848-DEL-3221
chr3-10777826-DEL-94
chr3-10778190-DEL-1577
chr3-11173675-DEL-58
chr3-11215016-DEL-50
chr3-11240198-DEL-656
chr3-11310543-DEL-129
chr3-11471674-DEL-54
chr3-11579594-DEL-64
chr3-11956463-DEL-59
chr3-12182030-DEL-2034
chr3-12221175-DEL-74
chr3-12257038-DEL-147
chr3-12387802-DEL-55
chr3-12394016-DEL-486
chr3-12460776-DEL-54
chr3-12488475-DEL-4700
chr3-12565558-DEL-54
chr3-12638724-DEL-6193
chr3-12684783-DEL-201
chr3-12728608-DEL-100
chr3-12741964-DEL-58
chr3-12789608-DEL-89
chr3-12801396-DEL-423
chr3-12848871-DEL-65
chr3-12924890-DEL-54
chr3-12928175-DEL-3936
chr3-13008495-DEL-2093
chr3-13036886-DEL-1615
chr3-13043237-DEL-1444
chr3-13069588-DEL-66
chr3-13112042-DEL-70
chr3-13173651-DEL-2914
chr3-13199544-DEL-120
chr3-13269973-DEL-51
chr3-13321526-DEL-597
chr3-13324779-DEL-116
chr3-13383133-DEL-230
chr3-13492173-DEL-77
chr3-13586562-DEL-182
chr3-13598714-DEL-51
chr3-13629991-DEL-1484
chr3-13837254-DEL-62
chr3-14005600-DEL-997
chr3-14007601-DEL-1224
chr3-14047018-DEL-175

chr3-14065240-DEL-2562
chr3-14146081-DEL-50
chr3-14414892-DEL-2611
chr3-14418341-DEL-211
chr3-14455441-DEL-212
chr3-14457290-DEL-62
chr3-14529130-DEL-3678
chr3-14544156-DEL-68
chr3-14607852-DEL-103
chr3-14636519-DEL-50
chr3-14665498-DEL-126
chr3-14699552-DEL-110
chr3-14753108-DEL-413
chr3-14849644-DEL-202
chr3-14849647-DEL-80
chr3-14850010-DEL-50
chr3-14940495-DEL-109
chr3-14945947-DEL-70
chr3-14977180-DEL-858
chr3-15045857-DEL-6800
chr3-15172022-DEL-433
chr3-15180832-DEL-252
chr3-15182178-DEL-84
chr3-15184387-DEL-75
chr3-15201328-DEL-170
chr3-15209815-DEL-82
chr3-15294376-DEL-1016
chr3-15296347-DEL-60
chr3-15424230-DEL-430
chr3-15479281-DEL-537
chr3-15483416-DEL-56
chr3-15606373-DEL-64
chr3-15649545-DEL-619
chr3-15663785-DEL-58
chr3-15829450-DEL-59
chr3-15848855-DEL-51
chr3-15990300-DEL-74
chr3-15991829-DEL-1279
chr3-16018246-DEL-31265
chr3-16046299-DEL-61
chr3-16049315-DEL-327
chr3-16093118-DEL-117
chr3-16093163-DEL-57
chr3-16106283-DEL-117
chr3-16119661-DEL-54
chr3-16124098-DEL-70
chr3-16139308-DEL-56
chr3-16247076-DEL-1367
chr3-16251335-DEL-61
chr3-16352561-DEL-6552
chr3-16424881-DEL-50
chr3-16441026-DEL-895
chr3-16462896-DEL-63
chr3-16506157-DEL-52
chr3-16554421-DEL-104
chr3-16612170-DEL-420
chr3-16697527-DEL-1648
chr3-16932329-DEL-84
chr3-16962570-DEL-105
chr3-17048544-DEL-643
chr3-17075277-DEL-1190
chr3-17186486-DEL-334
chr3-17357508-DEL-116
chr3-17754939-DEL-4596
chr3-17773019-DEL-571
chr3-17788665-DEL-411
chr3-17820193-DEL-426
chr3-17981753-DEL-127
chr3-18075919-DEL-120
chr3-18189890-DEL-50
chr3-18221609-DEL-61
chr3-18450289-DEL-182
chr3-18468050-DEL-294
chr3-18558727-DEL-768
chr3-18598809-DEL-99
chr3-18610607-DEL-208
chr3-18638841-DEL-199

chr3-18709359-DEL-109
chr3-18820128-DEL-140
chr3-19020647-DEL-1437
chr3-19133989-DEL-54
chr3-19149071-DEL-399
chr3-19200129-DEL-768
chr3-19201098-DEL-470
chr3-19206517-DEL-281
chr3-19261611-DEL-238
chr3-19292890-DEL-64
chr3-19389827-DEL-392
chr3-19671599-DEL-1637
chr3-19736684-DEL-241
chr3-19737163-DEL-390
chr3-19772587-DEL-155
chr3-19867816-DEL-7472
chr3-19971922-DEL-4156
chr3-20027729-DEL-146
chr3-20048601-DEL-693
chr3-20078548-DEL-172
chr3-20154682-DEL-974
chr3-20240624-DEL-68
chr3-20292284-DEL-52
chr3-20363693-DEL-249
chr3-20390115-DEL-259
chr3-20577462-DEL-201
chr3-20782348-DEL-992
chr3-20784951-DEL-53
chr3-20934061-DEL-372
chr3-21038067-DEL-285
chr3-21056461-DEL-2805
chr3-21081572-DEL-60
chr3-21188798-DEL-86
chr3-21192914-DEL-1392
chr3-21220300-DEL-2741
chr3-21335049-DEL-2505
chr3-21339905-DEL-50
chr3-21449737-DEL-175
chr3-21500280-DEL-161
chr3-21589700-DEL-63
chr3-21827927-DEL-129
chr3-21836781-DEL-1412
chr3-21893372-DEL-544
chr3-22019238-DEL-50
chr3-22165456-DEL-136
chr3-22385493-DEL-62
chr3-22484050-DEL-1253
chr3-22523627-DEL-242
chr3-22566480-DEL-2453
chr3-22627199-DEL-561
chr3-22805542-DEL-1351
chr3-22865333-DEL-50
chr3-22973980-DEL-305
chr3-22986966-DEL-214
chr3-23108973-DEL-54
chr3-23310077-DEL-355
chr3-23332129-DEL-231
chr3-23393199-DEL-58
chr3-23475178-DEL-55
chr3-23675298-DEL-109
chr3-23701353-DEL-1041
chr3-23759908-DEL-57
chr3-23788058-DEL-948
chr3-23837435-DEL-96
chr3-23965158-DEL-52
chr3-24060129-DEL-56
chr3-24161319-DEL-50
chr3-24410603-DEL-78
chr3-24508164-DEL-94
chr3-24600985-DEL-323
chr3-24601332-DEL-1862
chr3-24711207-DEL-4326
chr3-24758722-DEL-62
chr3-24807989-DEL-1089
chr3-24833210-DEL-58
chr3-24835284-DEL-172
chr3-24874054-DEL-89

chr3-24874665-DEL-132
chr3-24883763-DEL-349
chr3-24993667-DEL-52
chr3-24999047-DEL-52
chr3-25247673-DEL-51
chr3-25330211-DEL-2530
chr3-25382706-DEL-632
chr3-25386749-DEL-60
chr3-25622968-DEL-69
chr3-25630860-DEL-342
chr3-25660531-DEL-84
chr3-25692782-DEL-1068
chr3-25728740-DEL-50
chr3-25751963-DEL-1137
chr3-25823314-DEL-54
chr3-25989618-DEL-313
chr3-26099939-DEL-124
chr3-26156392-DEL-53
chr3-26306271-DEL-84
chr3-26508710-DEL-66
chr3-26514526-DEL-61
chr3-26543144-DEL-68
chr3-26547945-DEL-1250
chr3-26563634-DEL-50
chr3-26655772-DEL-52
chr3-26656006-DEL-103
chr3-26665965-DEL-815
chr3-26668738-DEL-2718
chr3-26683835-DEL-434
chr3-26685381-DEL-262
chr3-26685498-DEL-60
chr3-26685682-DEL-78
chr3-26774562-DEL-53
chr3-26779063-DEL-62
chr3-26823321-DEL-50
chr3-26912329-DEL-6569
chr3-26947311-DEL-4730
chr3-26971917-DEL-367
chr3-26998646-DEL-199
chr3-27045340-DEL-2750
chr3-27051518-DEL-959
chr3-27052731-DEL-333
chr3-27077344-DEL-76
chr3-27130230-DEL-630
chr3-27149466-DEL-459
chr3-27212908-DEL-128
chr3-27214231-DEL-60
chr3-27230934-DEL-60
chr3-27270663-DEL-135
chr3-27358597-DEL-1461
chr3-27396558-DEL-132
chr3-27494929-DEL-175
chr3-27651269-DEL-64
chr3-27715320-DEL-68
chr3-27795142-DEL-456
chr3-27850499-DEL-95
chr3-27954214-DEL-56
chr3-27955118-DEL-55
chr3-28002210-DEL-273
chr3-28065527-DEL-102
chr3-28111291-DEL-51
chr3-28287692-DEL-77
chr3-28342353-DEL-68
chr3-28395417-DEL-424
chr3-28428733-DEL-74
chr3-28449334-DEL-61
chr3-28535853-DEL-65
chr3-28562357-DEL-326
chr3-28758318-DEL-68
chr3-28758318-DEL-236
chr3-28785212-DEL-125
chr3-28791653-DEL-2424
chr3-28950510-DEL-78
chr3-28963106-DEL-63
chr3-28976067-DEL-56
chr3-29174303-DEL-79
chr3-29288518-DEL-393

chr3-29288629-DEL-86
chr3-29329365-DEL-144
chr3-29497550-DEL-1002
chr3-29886380-DEL-327
chr3-29897709-DEL-135
chr3-29897709-DEL-459
chr3-29975293-DEL-57
chr3-29993438-DEL-174
chr3-30044502-DEL-50
chr3-30178692-DEL-60
chr3-30239602-DEL-84
chr3-30286345-DEL-81
chr3-30613144-DEL-56
chr3-30614884-DEL-50
chr3-30633123-DEL-462
chr3-30645068-DEL-94
chr3-30757367-DEL-1092
chr3-30787225-DEL-597
chr3-30826991-DEL-832
chr3-30920559-DEL-51
chr3-30994859-DEL-113
chr3-31124431-DEL-400
chr3-31164024-DEL-50
chr3-31183543-DEL-315
chr3-31186425-DEL-103
chr3-31265899-DEL-78
chr3-31385287-DEL-3356
chr3-31419621-DEL-50
chr3-31507177-DEL-71
chr3-31544877-DEL-514
chr3-31619355-DEL-370
chr3-31632827-DEL-76
chr3-31720244-DEL-136
chr3-31763089-DEL-480
chr3-31891024-DEL-163
chr3-32000440-DEL-865
chr3-32104018-DEL-318
chr3-32159433-DEL-60
chr3-32185982-DEL-166
chr3-32192388-DEL-575
chr3-32214867-DEL-53
chr3-32312770-DEL-280
chr3-32319185-DEL-84
chr3-32537273-DEL-54
chr3-32544028-DEL-84
chr3-32551974-DEL-162
chr3-32630151-DEL-187
chr3-32630409-DEL-91
chr3-32718256-DEL-721
chr3-32767761-DEL-782
chr3-32783528-DEL-58
chr3-32799863-DEL-164
chr3-32896862-DEL-68
chr3-32937263-DEL-66
chr3-32945289-DEL-54
chr3-33163526-DEL-145
chr3-33242615-DEL-68
chr3-33272932-DEL-66
chr3-33336729-DEL-3254
chr3-33411696-DEL-120
chr3-33498422-DEL-879
chr3-33637134-DEL-56
chr3-33670969-DEL-126
chr3-33974216-DEL-54
chr3-33989763-DEL-95
chr3-33992427-DEL-202
chr3-34054258-DEL-53
chr3-34188029-DEL-97
chr3-34247784-DEL-184
chr3-34287459-DEL-130
chr3-34317452-DEL-86
chr3-34479145-DEL-96
chr3-34508343-DEL-105
chr3-34678639-DEL-89
chr3-34836874-DEL-230
chr3-34857522-DEL-300
chr3-34857989-DEL-118

chr3-34859936-DEL-82
chr3-34895995-DEL-152
chr3-34896230-DEL-132
chr3-34921044-DEL-52
chr3-34948770-DEL-88
chr3-35002204-DEL-74
chr3-35014415-DEL-114
chr3-35033359-DEL-242
chr3-35053707-DEL-50
chr3-35275572-DEL-251
chr3-35338695-DEL-8447
chr3-35456028-DEL-141
chr3-35533460-DEL-216
chr3-35533718-DEL-110
chr3-35533893-DEL-194
chr3-35719448-DEL-936
chr3-35777012-DEL-137
chr3-35778558-DEL-96
chr3-35827062-DEL-4114
chr3-35950543-DEL-57
chr3-35950602-DEL-230
chr3-36056315-DEL-60
chr3-36058328-DEL-52
chr3-36097339-DEL-150
chr3-36097540-DEL-261
chr3-36246055-DEL-512
chr3-36277052-DEL-54
chr3-36278667-DEL-56
chr3-36298063-DEL-108
chr3-36350482-DEL-54
chr3-36382441-DEL-92
chr3-36499753-DEL-174
chr3-36653289-DEL-50
chr3-36686510-DEL-65
chr3-36707608-DEL-66
chr3-36816505-DEL-62
chr3-36816534-DEL-161
chr3-36907375-DEL-208
chr3-37225369-DEL-62
chr3-37416092-DEL-50
chr3-37709529-DEL-97
chr3-37771163-DEL-50
chr3-37974634-DEL-174
chr3-38020604-DEL-128
chr3-38237560-DEL-54
chr3-38245294-DEL-147
chr3-38245527-DEL-69
chr3-38264278-DEL-7414
chr3-38271232-DEL-218
chr3-38312840-DEL-85
chr3-38344320-DEL-50
chr3-38369273-DEL-79
chr3-38392792-DEL-79
chr3-38392837-DEL-269
chr3-38393039-DEL-95
chr3-38579661-DEL-72
chr3-38701524-DEL-77
chr3-38701656-DEL-69
chr3-38715578-DEL-128
chr3-38733394-DEL-50
chr3-38828402-DEL-94
chr3-38832550-DEL-1473
chr3-38873413-DEL-84
chr3-38920158-DEL-1571
chr3-38934108-DEL-462
chr3-38972310-DEL-69
chr3-38980546-DEL-83
chr3-38980655-DEL-188
chr3-38980796-DEL-67
chr3-39052823-DEL-50
chr3-39079158-DEL-81
chr3-39079294-DEL-777
chr3-39123997-DEL-114
chr3-39191347-DEL-53
chr3-39320782-DEL-50
chr3-39540653-DEL-1257
chr3-39623507-DEL-160

chr3-39623717-DEL-54
chr3-39635403-DEL-99
chr3-39717502-DEL-1213
chr3-39737248-DEL-91
chr3-39757111-DEL-128
chr3-40010798-DEL-311
chr3-40013501-DEL-137
chr3-40062510-DEL-54
chr3-40167217-DEL-102
chr3-40233509-DEL-50
chr3-40270585-DEL-1821
chr3-40292269-DEL-58
chr3-40303225-DEL-61
chr3-40356525-DEL-278
chr3-40365766-DEL-104
chr3-40494962-DEL-58
chr3-40506446-DEL-156
chr3-40506468-DEL-72
chr3-40527909-DEL-96
chr3-40589118-DEL-120
chr3-40604423-DEL-316
chr3-40604460-DEL-888
chr3-40605644-DEL-144
chr3-40627411-DEL-102
chr3-40663998-DEL-5321
chr3-40893341-DEL-62
chr3-40981125-DEL-54
chr3-41184503-DEL-52
chr3-41247943-DEL-56
chr3-41248059-DEL-85
chr3-41336833-DEL-60
chr3-41399569-DEL-60
chr3-41404474-DEL-58
chr3-41537525-DEL-50
chr3-41583227-DEL-72
chr3-41635542-DEL-50
chr3-41662376-DEL-55
chr3-41802106-DEL-134
chr3-41871969-DEL-50
chr3-42119592-DEL-1052
chr3-42181360-DEL-182
chr3-42238307-DEL-88
chr3-42497368-DEL-190
chr3-42567477-DEL-124
chr3-42571353-DEL-71
chr3-42575223-DEL-375
chr3-42581940-DEL-344
chr3-42582277-DEL-74
chr3-42593086-DEL-1491
chr3-42656678-DEL-939
chr3-42687865-DEL-147
chr3-42866779-DEL-294
chr3-42868248-DEL-56
chr3-42902606-DEL-97
chr3-43492817-DEL-60
chr3-43574939-DEL-897
chr3-43646078-DEL-50
chr3-43734534-DEL-173
chr3-43855993-DEL-2790
chr3-43888808-DEL-56
chr3-44476159-DEL-62
chr3-44602649-DEL-3629
chr3-44649020-DEL-314
chr3-44871139-DEL-170
chr3-44901497-DEL-64
chr3-44945531-DEL-469
chr3-44946062-DEL-348
chr3-44948234-DEL-52
chr3-44963980-DEL-132
chr3-45028800-DEL-1476
chr3-45044042-DEL-61
chr3-45103570-DEL-95
chr3-45115271-DEL-201
chr3-45167204-DEL-52
chr3-45233445-DEL-55
chr3-45238663-DEL-57
chr3-45263070-DEL-50

chr3-45452291-DEL-50
chr3-45502391-DEL-92
chr3-45540098-DEL-201
chr3-45565093-DEL-50
chr3-45644597-DEL-50
chr3-45754674-DEL-130
chr3-45772183-DEL-3404
chr3-45813186-DEL-489
chr3-46007296-DEL-52
chr3-46154204-DEL-1988
chr3-46193346-DEL-5614
chr3-46235624-DEL-59
chr3-46711313-DEL-64
chr3-46781773-DEL-368
chr3-46811177-DEL-90
chr3-46820785-DEL-55
chr3-46825511-DEL-52
chr3-46960085-DEL-50
chr3-47006851-DEL-2782
chr3-47067747-DEL-63
chr3-47316853-DEL-755
chr3-47326404-DEL-6224
chr3-47371244-DEL-2936
chr3-47458770-DEL-147
chr3-47546258-DEL-200
chr3-47554824-DEL-91
chr3-47585087-DEL-69
chr3-47633072-DEL-1700
chr3-47700769-DEL-60
chr3-47700864-DEL-210
chr3-47719133-DEL-201
chr3-47920576-DEL-795
chr3-47999590-DEL-74
chr3-48058441-DEL-50
chr3-48068108-DEL-238
chr3-48106687-DEL-126
chr3-48129225-DEL-57
chr3-48129320-DEL-192
chr3-48129566-DEL-255
chr3-48135574-DEL-104
chr3-48236183-DEL-119
chr3-48307616-DEL-127
chr3-48317607-DEL-72
chr3-48388360-DEL-203
chr3-48523972-DEL-140
chr3-48593823-DEL-67
chr3-48672163-DEL-370
chr3-48684471-DEL-163
chr3-48796202-DEL-102
chr3-48804157-DEL-58
chr3-48951267-DEL-53
chr3-48951472-DEL-54
chr3-48955623-DEL-5824
chr3-49016989-DEL-164
chr3-49185195-DEL-55
chr3-49201275-DEL-50
chr3-49209710-DEL-65
chr3-49261563-DEL-50
chr3-49261599-DEL-419
chr3-49262283-DEL-70
chr3-49266515-DEL-2063
chr3-49342442-DEL-141
chr3-49359949-DEL-124
chr3-49418751-DEL-208
chr3-49430840-DEL-615
chr3-49484758-DEL-70
chr3-49496777-DEL-60
chr3-49562801-DEL-117
chr3-49563018-DEL-63
chr3-49639693-DEL-52
chr3-49681727-DEL-7109
chr3-49888957-DEL-67
chr3-50006122-DEL-113
chr3-50015604-DEL-143
chr3-50088083-DEL-1119
chr3-50114553-DEL-444
chr3-50194444-DEL-61

chr3-50210487-DEL-99
chr3-50210837-DEL-56
chr3-50234735-DEL-56
chr3-50469436-DEL-50
chr3-50616954-DEL-639
chr3-50617230-DEL-191
chr3-50696856-DEL-56
chr3-50696947-DEL-97
chr3-50720581-DEL-745
chr3-50794698-DEL-515
chr3-50797461-DEL-62
chr3-50821252-DEL-60
chr3-50894537-DEL-102
chr3-51079398-DEL-61
chr3-51120920-DEL-214
chr3-51179836-DEL-72
chr3-51180068-DEL-178
chr3-51206099-DEL-135
chr3-51206332-DEL-60
chr3-51210556-DEL-53
chr3-51267361-DEL-2176
chr3-51388270-DEL-53
chr3-51394535-DEL-105
chr3-51403643-DEL-147
chr3-51431529-DEL-68
chr3-51480312-DEL-50
chr3-51614855-DEL-328
chr3-51614884-DEL-55
chr3-51673406-DEL-842
chr3-51691189-DEL-336
chr3-51915880-DEL-98
chr3-51916100-DEL-125
chr3-51916109-DEL-318
chr3-51976177-DEL-52
chr3-52085825-DEL-87
chr3-52169695-DEL-54
chr3-52192959-DEL-54
chr3-52235399-DEL-688
chr3-52236576-DEL-301
chr3-52261251-DEL-157
chr3-52314435-DEL-75
chr3-52356886-DEL-202
chr3-52365072-DEL-120
chr3-52393439-DEL-65
chr3-52451174-DEL-3794
chr3-52490049-DEL-330
chr3-52574094-DEL-67
chr3-52666272-DEL-1227
chr3-52830030-DEL-476
chr3-52834712-DEL-93
chr3-52932405-DEL-105
chr3-53006071-DEL-166
chr3-53141024-DEL-72
chr3-53181755-DEL-70
chr3-53192262-DEL-101
chr3-53284047-DEL-104
chr3-53296038-DEL-1571
chr3-53343141-DEL-61
chr3-53350633-DEL-58
chr3-53391361-DEL-57
chr3-53400795-DEL-179
chr3-53406343-DEL-196
chr3-53409574-DEL-174
chr3-53419090-DEL-118
chr3-53422346-DEL-71
chr3-53439289-DEL-174
chr3-53457659-DEL-167
chr3-53590848-DEL-216
chr3-53591096-DEL-102
chr3-53653671-DEL-316
chr3-53785453-DEL-60
chr3-53829769-DEL-238
chr3-53859693-DEL-68
chr3-54036103-DEL-214
chr3-54067064-DEL-62
chr3-54079050-DEL-54
chr3-54183344-DEL-342

chr3-54327275-DEL-191
chr3-54390697-DEL-404
chr3-54395826-DEL-54
chr3-54469676-DEL-256
chr3-54479932-DEL-90
chr3-54586055-DEL-399
chr3-54601394-DEL-221
chr3-54610790-DEL-409
chr3-54837705-DEL-52
chr3-54904101-DEL-675
chr3-54934349-DEL-84
chr3-55111274-DEL-191
chr3-55118097-DEL-1105
chr3-55248788-DEL-50
chr3-55251357-DEL-69
chr3-55251500-DEL-223
chr3-55275066-DEL-75
chr3-55276519-DEL-54
chr3-55323659-DEL-176
chr3-55347787-DEL-80
chr3-55365709-DEL-50
chr3-55506652-DEL-66
chr3-55521305-DEL-54
chr3-55568696-DEL-98
chr3-55757132-DEL-141
chr3-55844682-DEL-61
chr3-55908914-DEL-940
chr3-55971522-DEL-563
chr3-56116718-DEL-213
chr3-56140089-DEL-123
chr3-56140193-DEL-57
chr3-56169035-DEL-69
chr3-56224314-DEL-69
chr3-56276554-DEL-176
chr3-56281819-DEL-1507
chr3-56404996-DEL-56
chr3-56530848-DEL-236
chr3-56607322-DEL-76
chr3-56638813-DEL-188
chr3-56707382-DEL-462
chr3-56719119-DEL-1105
chr3-56721178-DEL-52
chr3-56747113-DEL-180
chr3-56797785-DEL-100
chr3-56823209-DEL-13623
chr3-56823211-DEL-5870
chr3-56859442-DEL-828
chr3-56932601-DEL-68
chr3-57162400-DEL-2906
chr3-57315735-DEL-80
chr3-57315999-DEL-74
chr3-57531713-DEL-53
chr3-57535510-DEL-194
chr3-57535834-DEL-114
chr3-57563361-DEL-912
chr3-57682865-DEL-52
chr3-57764545-DEL-11124
chr3-57799071-DEL-294
chr3-57805362-DEL-52
chr3-57834026-DEL-1560
chr3-57904461-DEL-488
chr3-57913129-DEL-64
chr3-57973433-DEL-97
chr3-57973635-DEL-1892
chr3-57975996-DEL-252
chr3-57982043-DEL-78
chr3-58027811-DEL-126
chr3-58160697-DEL-154
chr3-58261119-DEL-80
chr3-58381909-DEL-406
chr3-58382156-DEL-111
chr3-58440407-DEL-328
chr3-58441964-DEL-198
chr3-58443205-DEL-187
chr3-58507754-DEL-2273
chr3-58553223-DEL-68
chr3-58573266-DEL-54

chr3-58771182-DEL-50
chr3-58841323-DEL-50
chr3-58878587-DEL-113
chr3-58903486-DEL-58
chr3-58903662-DEL-58
chr3-59016729-DEL-670
chr3-59026651-DEL-177
chr3-59153785-DEL-54
chr3-59176694-DEL-2607
chr3-59330403-DEL-1364
chr3-59536295-DEL-1871
chr3-59662766-DEL-222
chr3-59736326-DEL-57
chr3-60008944-DEL-67
chr3-60024453-DEL-184
chr3-60025431-DEL-65
chr3-60025470-DEL-979
chr3-60053531-DEL-492
chr3-60054706-DEL-858
chr3-60078208-DEL-121
chr3-60221149-DEL-1666
chr3-60335708-DEL-60
chr3-60469584-DEL-176
chr3-60593341-DEL-934
chr3-60615781-DEL-54
chr3-60620715-DEL-99
chr3-60625369-DEL-61
chr3-60625601-DEL-83
chr3-60641700-DEL-58
chr3-60736957-DEL-52
chr3-61114195-DEL-573
chr3-61241082-DEL-1630
chr3-61329691-DEL-54
chr3-61419774-DEL-1062
chr3-61527265-DEL-1068
chr3-61558079-DEL-435
chr3-61588728-DEL-64
chr3-61738083-DEL-50
chr3-61841990-DEL-2119
chr3-61852011-DEL-165
chr3-61856917-DEL-265
chr3-61945700-DEL-201
chr3-61980085-DEL-2377
chr3-62066150-DEL-66
chr3-62066586-DEL-219
chr3-62168874-DEL-336
chr3-62186177-DEL-61
chr3-62649687-DEL-212
chr3-62649955-DEL-212
chr3-62786531-DEL-429
chr3-62821531-DEL-632
chr3-62945558-DEL-170
chr3-62978575-DEL-302
chr3-63257156-DEL-53
chr3-63262023-DEL-100
chr3-63396054-DEL-667
chr3-63419714-DEL-215
chr3-63420700-DEL-68
chr3-63478965-DEL-130
chr3-63563092-DEL-116
chr3-63573702-DEL-98
chr3-63580898-DEL-569
chr3-63583216-DEL-212
chr3-63654011-DEL-2877
chr3-64079369-DEL-115
chr3-64143083-DEL-58
chr3-64158003-DEL-53
chr3-64409667-DEL-237
chr3-64412146-DEL-420
chr3-64610938-DEL-540
chr3-64926931-DEL-68
chr3-64936985-DEL-179
chr3-64937196-DEL-50
chr3-64964673-DEL-68
chr3-64967894-DEL-50
chr3-64987607-DEL-136
chr3-65046260-DEL-298

chr3-65046624-DEL-1903
chr3-65168305-DEL-126
chr3-65206134-DEL-58
chr3-65368737-DEL-53
chr3-65374196-DEL-94
chr3-65374198-DEL-196
chr3-65594541-DEL-5761
chr3-65676774-DEL-201
chr3-65676908-DEL-67
chr3-65677192-DEL-67
chr3-65682044-DEL-50
chr3-65703023-DEL-84
chr3-65772989-DEL-298
chr3-65786038-DEL-76
chr3-65838167-DEL-809
chr3-65907307-DEL-107
chr3-65949224-DEL-165
chr3-65949387-DEL-56
chr3-65978831-DEL-5824
chr3-66019901-DEL-101
chr3-66067486-DEL-256
chr3-66067721-DEL-50
chr3-66166859-DEL-264
chr3-66176276-DEL-6260
chr3-66244079-DEL-75
chr3-66274098-DEL-144
chr3-66392941-DEL-74
chr3-66426554-DEL-52
chr3-66471470-DEL-55
chr3-66577704-DEL-206
chr3-66601099-DEL-86
chr3-66601263-DEL-300
chr3-66630288-DEL-336
chr3-66690437-DEL-679
chr3-66722315-DEL-72
chr3-66742249-DEL-80
chr3-67081525-DEL-53
chr3-67170051-DEL-135
chr3-67170969-DEL-272
chr3-67206303-DEL-59
chr3-67272822-DEL-58
chr3-67382250-DEL-59
chr3-67389728-DEL-427
chr3-67412154-DEL-77
chr3-67412407-DEL-152
chr3-67498397-DEL-60
chr3-67512960-DEL-53
chr3-67544739-DEL-93
chr3-67601628-DEL-139
chr3-67752860-DEL-75
chr3-67753581-DEL-1088
chr3-67769958-DEL-2679
chr3-67790845-DEL-50
chr3-67822422-DEL-52
chr3-67848403-DEL-677
chr3-67956437-DEL-60
chr3-68247209-DEL-814
chr3-68271167-DEL-748
chr3-68274375-DEL-82
chr3-68304736-DEL-91
chr3-68391258-DEL-98
chr3-68409929-DEL-69
chr3-68423556-DEL-109
chr3-68425810-DEL-125
chr3-68476749-DEL-358
chr3-68587405-DEL-2371
chr3-68706822-DEL-88
chr3-68942329-DEL-169
chr3-69060682-DEL-177
chr3-69060867-DEL-56
chr3-69061210-DEL-396
chr3-69069008-DEL-50
chr3-69081594-DEL-50
chr3-69205498-DEL-1634
chr3-69230733-DEL-72
chr3-69251633-DEL-356
chr3-69251650-DEL-129

chr3-69252330-DEL-461
chr3-69252674-DEL-97
chr3-69252913-DEL-542
chr3-69255088-DEL-411
chr3-69262254-DEL-76
chr3-69279780-DEL-667
chr3-69299370-DEL-135
chr3-69320287-DEL-84
chr3-69385182-DEL-51
chr3-69406865-DEL-52
chr3-69412576-DEL-52
chr3-69427467-DEL-140
chr3-69522239-DEL-352
chr3-69532383-DEL-58
chr3-69675572-DEL-180
chr3-69679041-DEL-884
chr3-69857773-DEL-207
chr3-69922478-DEL-94
chr3-69922692-DEL-60
chr3-70108528-DEL-54
chr3-70551614-DEL-121
chr3-70575988-DEL-55
chr3-70622738-DEL-79
chr3-70656314-DEL-50
chr3-70704167-DEL-64
chr3-70874576-DEL-79
chr3-70894521-DEL-160
chr3-70900373-DEL-696
chr3-70904366-DEL-55
chr3-70932553-DEL-231
chr3-70993496-DEL-1244
chr3-70999623-DEL-52
chr3-71192762-DEL-57
chr3-71218306-DEL-72
chr3-71240580-DEL-50
chr3-71270703-DEL-666
chr3-71477918-DEL-135
chr3-71661978-DEL-202
chr3-71719189-DEL-51
chr3-71777295-DEL-69
chr3-71806338-DEL-3439
chr3-71906467-DEL-52
chr3-71927636-DEL-59
chr3-72126677-DEL-102
chr3-72146174-DEL-262
chr3-72167030-DEL-104
chr3-72272707-DEL-70
chr3-72339429-DEL-420
chr3-72518371-DEL-50
chr3-72718440-DEL-101
chr3-72809063-DEL-62
chr3-73053632-DEL-53
chr3-73414146-DEL-1494
chr3-73762849-DEL-7684
chr3-73826097-DEL-1653
chr3-73858087-DEL-209
chr3-73905619-DEL-1170
chr3-73908469-DEL-66
chr3-73909620-DEL-1606
chr3-73959326-DEL-102
chr3-73975088-DEL-70
chr3-74004866-DEL-484
chr3-74008592-DEL-605
chr3-74052331-DEL-678
chr3-74138674-DEL-427
chr3-74214041-DEL-52
chr3-74224029-DEL-77
chr3-74236652-DEL-50
chr3-74290337-DEL-433
chr3-74312173-DEL-84
chr3-74382710-DEL-14595
chr3-74403662-DEL-215
chr3-74415485-DEL-382
chr3-74440887-DEL-7274
chr3-74441568-DEL-2050
chr3-74474434-DEL-243
chr3-74477800-DEL-739

chr3-74534183-DEL-202
chr3-74562965-DEL-146
chr3-74588657-DEL-205
chr3-74656830-DEL-59
chr3-74668711-DEL-53
chr3-74682912-DEL-58
chr3-74704192-DEL-50
chr3-74721251-DEL-446
chr3-74890425-DEL-5311
chr3-74906373-DEL-50
chr3-75044737-DEL-62
chr3-75083817-DEL-55
chr3-75133559-DEL-66
chr3-75170855-DEL-56
chr3-75190461-DEL-1434
chr3-75240172-DEL-51
chr3-75335315-DEL-173
chr3-75417125-DEL-55
chr3-75438796-DEL-51
chr3-75519667-DEL-96
chr3-75591969-DEL-341
chr3-75609780-DEL-55
chr3-75695893-DEL-694
chr3-75824194-DEL-590
chr3-75834703-DEL-52
chr3-75854020-DEL-145
chr3-75870522-DEL-53
chr3-75888400-DEL-61
chr3-75888650-DEL-298
chr3-75891922-DEL-60
chr3-75979713-DEL-54
chr3-76042816-DEL-155
chr3-76105307-DEL-52
chr3-76105448-DEL-157
chr3-76127701-DEL-64
chr3-76307733-DEL-70
chr3-76431494-DEL-58
chr3-76541613-DEL-255
chr3-76655069-DEL-60
chr3-76673492-DEL-66
chr3-76730611-DEL-76
chr3-76943795-DEL-54
chr3-76966804-DEL-92
chr3-77014151-DEL-204
chr3-77107539-DEL-113
chr3-77198938-DEL-5154
chr3-77551338-DEL-1177
chr3-77572737-DEL-50
chr3-77574080-DEL-8265
chr3-77633076-DEL-65
chr3-77798103-DEL-50
chr3-77834159-DEL-2111
chr3-77844208-DEL-69
chr3-77872106-DEL-73
chr3-77874903-DEL-2989
chr3-77939318-DEL-64
chr3-77982414-DEL-3711
chr3-78042619-DEL-50
chr3-78112936-DEL-1141
chr3-78169299-DEL-339
chr3-78187973-DEL-57
chr3-78188023-DEL-191
chr3-78207638-DEL-74
chr3-78226659-DEL-471
chr3-78245789-DEL-199
chr3-78246395-DEL-398
chr3-78293254-DEL-53
chr3-78302899-DEL-2259
chr3-78304520-DEL-65
chr3-78319364-DEL-102
chr3-78558083-DEL-69
chr3-78703136-DEL-50
chr3-78782232-DEL-61
chr3-78817187-DEL-55
chr3-78853280-DEL-686
chr3-78853289-DEL-210
chr3-78853780-DEL-186

chr3-78853851-DEL-84
chr3-78878318-DEL-93
chr3-78909289-DEL-3445
chr3-78926179-DEL-80
chr3-78983308-DEL-50
chr3-79134140-DEL-52
chr3-79143852-DEL-310
chr3-79150003-DEL-194
chr3-79150098-DEL-82
chr3-79151721-DEL-50
chr3-79212879-DEL-155
chr3-79304018-DEL-78
chr3-79304370-DEL-190
chr3-79332580-DEL-90
chr3-79334635-DEL-208
chr3-79416378-DEL-176
chr3-79429789-DEL-186
chr3-79491609-DEL-65
chr3-79541632-DEL-212
chr3-79671484-DEL-6700
chr3-79726285-DEL-250
chr3-79751080-DEL-176
chr3-79778788-DEL-314
chr3-79778967-DEL-126
chr3-79803663-DEL-230
chr3-79906587-DEL-228
chr3-79919329-DEL-2269
chr3-79919440-DEL-877
chr3-80223007-DEL-66
chr3-80233796-DEL-72
chr3-80246279-DEL-90
chr3-80338917-DEL-330
chr3-80347035-DEL-1704
chr3-80360038-DEL-204
chr3-80409709-DEL-1177
chr3-80641034-DEL-52
chr3-80741153-DEL-105
chr3-80756216-DEL-383
chr3-80774085-DEL-320
chr3-80774091-DEL-120
chr3-80813496-DEL-58
chr3-80816442-DEL-55
chr3-80834251-DEL-2130
chr3-80902621-DEL-780
chr3-81072306-DEL-52
chr3-81096958-DEL-211
chr3-81099202-DEL-50
chr3-81109187-DEL-244
chr3-81143371-DEL-131
chr3-81145748-DEL-4502
chr3-81317539-DEL-1458
chr3-81340510-DEL-84
chr3-81482857-DEL-874
chr3-81489098-DEL-345
chr3-81502424-DEL-1090
chr3-81525595-DEL-4737
chr3-81561825-DEL-60
chr3-81598649-DEL-1839
chr3-81631991-DEL-51
chr3-81698918-DEL-58
chr3-81730470-DEL-228
chr3-81750239-DEL-158
chr3-81794406-DEL-50
chr3-81878259-DEL-114
chr3-81902372-DEL-54
chr3-81942437-DEL-132
chr3-81993410-DEL-1001
chr3-82092630-DEL-50
chr3-82737987-DEL-301
chr3-82838002-DEL-100
chr3-82861932-DEL-138
chr3-82971208-DEL-709
chr3-82971308-DEL-330
chr3-83060238-DEL-50
chr3-83129096-DEL-167
chr3-83427677-DEL-1244
chr3-83479342-DEL-78

chr3-83499655-DEL-80
chr3-83499791-DEL-82
chr3-83578261-DEL-74
chr3-83626526-DEL-60
chr3-83724616-DEL-50
chr3-83753027-DEL-2466
chr3-83753839-DEL-113
chr3-83811269-DEL-119
chr3-83815135-DEL-122
chr3-83840804-DEL-87
chr3-83862988-DEL-272
chr3-83924639-DEL-172
chr3-83925084-DEL-70
chr3-84110917-DEL-184
chr3-84230582-DEL-62
chr3-84496201-DEL-91
chr3-84526984-DEL-1321
chr3-84528342-DEL-93
chr3-84532578-DEL-52
chr3-84563577-DEL-287
chr3-84570927-DEL-67
chr3-84571014-DEL-154
chr3-84619499-DEL-51
chr3-84751897-DEL-304
chr3-84752263-DEL-306
chr3-84756967-DEL-2484
chr3-84915953-DEL-50
chr3-84978438-DEL-80
chr3-85037187-DEL-79
chr3-85037224-DEL-287
chr3-85038089-DEL-119
chr3-85127033-DEL-62
chr3-85185515-DEL-370
chr3-85346499-DEL-417
chr3-85408308-DEL-151
chr3-85675766-DEL-56
chr3-85763613-DEL-66
chr3-85871132-DEL-87
chr3-85964899-DEL-64
chr3-85975775-DEL-249
chr3-86081497-DEL-547
chr3-86085559-DEL-209
chr3-86128617-DEL-198
chr3-86309216-DEL-164
chr3-86544235-DEL-124
chr3-86723017-DEL-55
chr3-86732964-DEL-115
chr3-86732994-DEL-401
chr3-86750294-DEL-190
chr3-86976386-DEL-3633
chr3-87085346-DEL-164
chr3-87129047-DEL-56
chr3-87210513-DEL-1060
chr3-87214138-DEL-221
chr3-87319938-DEL-216
chr3-87387445-DEL-50
chr3-87472358-DEL-60
chr3-87580135-DEL-2440
chr3-87667876-DEL-242
chr3-87727720-DEL-53
chr3-87733163-DEL-360
chr3-87804645-DEL-412
chr3-87919172-DEL-74
chr3-87947570-DEL-54
chr3-88003642-DEL-63
chr3-88111458-DEL-59
chr3-88177289-DEL-616
chr3-88274796-DEL-14888
chr3-88281564-DEL-5715
chr3-88287205-DEL-5830
chr3-88311574-DEL-59
chr3-88369861-DEL-260
chr3-88372365-DEL-50
chr3-88386217-DEL-234
chr3-88488980-DEL-546
chr3-88504127-DEL-62
chr3-88686031-DEL-62151

chr3-88735237-DEL-50
chr3-88846616-DEL-292
chr3-89018932-DEL-330
chr3-89106701-DEL-91
chr3-89106703-DEL-414
chr3-89108786-DEL-89
chr3-89277037-DEL-114
chr3-89373414-DEL-181
chr3-89445782-DEL-314
chr3-89452878-DEL-347
chr3-89569473-DEL-184
chr3-89635400-DEL-1130
chr3-89639014-DEL-301
chr3-89655827-DEL-190
chr3-89656042-DEL-76
chr3-89673433-DEL-90
chr3-89674495-DEL-100
chr3-89743666-DEL-164
chr3-89743876-DEL-85
chr3-89745696-DEL-58
chr3-89775216-DEL-248
chr3-89789497-DEL-54
chr3-89810001-DEL-77
chr3-89827152-DEL-53
chr3-89837052-DEL-262
chr3-89837065-DEL-84
chr3-89849431-DEL-84
chr3-89862236-DEL-90
chr3-90207490-DEL-95
chr3-90275314-DEL-321
chr3-90275515-DEL-632
chr3-90288322-DEL-52
chr3-90313258-DEL-56
chr3-90574559-DEL-162
chr3-90574696-DEL-51
chr3-91047887-DEL-91
chr3-91185547-DEL-5395
chr3-91315478-DEL-2129
chr3-91322562-DEL-466
chr3-91364174-DEL-1274
chr3-91398738-DEL-481
chr3-91399928-DEL-2524
chr3-91407860-DEL-51
chr3-91444986-DEL-90
chr3-91446451-DEL-2190
chr3-91597609-DEL-154290
chr3-91897052-DEL-1466
chr3-91975599-DEL-50
chr3-92053171-DEL-60
chr3-92086894-DEL-364
chr3-92147500-DEL-758
chr3-92187301-DEL-17870
chr3-92379546-DEL-4046
chr3-92385466-DEL-203
chr3-92397820-DEL-144
chr3-92434144-DEL-1457
chr3-92455130-DEL-257
chr3-92633332-DEL-114
chr3-92694158-DEL-256
chr3-92848790-DEL-112
chr3-92966747-DEL-2945
chr3-92993723-DEL-198
chr3-93000620-DEL-1150
chr3-93118981-DEL-3996
chr3-93231329-DEL-510
chr3-93232011-DEL-531
chr3-93232996-DEL-531
chr3-93285110-DEL-50
chr3-93303764-DEL-72
chr3-93304688-DEL-72
chr3-93317497-DEL-130
chr3-93353003-DEL-60
chr3-93432681-DEL-92
chr3-93555632-DEL-477
chr3-93580719-DEL-51
chr3-93682256-DEL-324
chr3-93683821-DEL-338

chr3-93736570-DEL-60
chr3-93748957-DEL-51
chr3-93942007-DEL-64
chr3-93991889-DEL-187
chr3-94088015-DEL-152
chr3-94315839-DEL-114
chr3-94350409-DEL-57
chr3-94415111-DEL-755
chr3-94516040-DEL-79
chr3-94673623-DEL-687
chr3-94710296-DEL-53
chr3-94874894-DEL-320
chr3-94969863-DEL-81
chr3-94998750-DEL-59
chr3-95027709-DEL-225
chr3-95264949-DEL-51
chr3-95303345-DEL-5473
chr3-95310895-DEL-355
chr3-95311104-DEL-50
chr3-95323901-DEL-513
chr3-95330642-DEL-145
chr3-95336615-DEL-119
chr3-95369071-DEL-232
chr3-95378890-DEL-586
chr3-95431877-DEL-58
chr3-95555068-DEL-125
chr3-95624069-DEL-50
chr3-95772987-DEL-112
chr3-95785018-DEL-50
chr3-95818861-DEL-119
chr3-95842930-DEL-206
chr3-95900554-DEL-59
chr3-95923993-DEL-60
chr3-95933372-DEL-360
chr3-96007977-DEL-160
chr3-96194760-DEL-117
chr3-96229032-DEL-52
chr3-96259720-DEL-51
chr3-96288997-DEL-51
chr3-96349720-DEL-270
chr3-96642405-DEL-91
chr3-96658876-DEL-50
chr3-96742627-DEL-88
chr3-96748674-DEL-987
chr3-96748684-DEL-2706
chr3-96750221-DEL-1043
chr3-96917517-DEL-175
chr3-97063223-DEL-233
chr3-97167963-DEL-192
chr3-97168170-DEL-288
chr3-97190248-DEL-67
chr3-97222681-DEL-564
chr3-97402460-DEL-60
chr3-97444562-DEL-547
chr3-97455849-DEL-56
chr3-97465967-DEL-116
chr3-97486331-DEL-71
chr3-97539194-DEL-76
chr3-97553300-DEL-225
chr3-97596490-DEL-51
chr3-97739929-DEL-87
chr3-97844188-DEL-64
chr3-97896885-DEL-87
chr3-97896894-DEL-541
chr3-97897160-DEL-94
chr3-97975988-DEL-124
chr3-98062935-DEL-3342
chr3-98121630-DEL-63
chr3-98122085-DEL-69
chr3-98165411-DEL-82
chr3-98165563-DEL-205
chr3-98165849-DEL-82
chr3-98180789-DEL-50
chr3-98268587-DEL-372
chr3-98271081-DEL-188
chr3-98339616-DEL-308
chr3-98563631-DEL-72

chr3-98851558-DEL-245
chr3-99001895-DEL-208
chr3-99318815-DEL-66
chr3-99475902-DEL-56
chr3-99729391-DEL-63
chr3-100130409-DEL-54
chr3-100131214-DEL-54
chr3-100212796-DEL-207
chr3-100244148-DEL-721
chr3-100333022-DEL-50
chr3-100447374-DEL-71
chr3-100458127-DEL-60
chr3-100458127-DEL-240
chr3-100618761-DEL-810
chr3-100648398-DEL-69
chr3-100661077-DEL-838
chr3-100717269-DEL-50
chr3-100750346-DEL-416
chr3-100803451-DEL-52
chr3-100888209-DEL-65
chr3-100931817-DEL-67
chr3-101005873-DEL-60
chr3-101073980-DEL-903
chr3-101133830-DEL-56
chr3-101358674-DEL-139
chr3-101376833-DEL-133
chr3-101396218-DEL-62
chr3-101440373-DEL-50
chr3-101442719-DEL-66
chr3-101566062-DEL-133
chr3-101576646-DEL-50
chr3-101646542-DEL-111
chr3-101656970-DEL-56
chr3-101666506-DEL-134
chr3-101729937-DEL-50
chr3-101755986-DEL-142
chr3-101815218-DEL-70
chr3-101889810-DEL-132
chr3-101925841-DEL-105
chr3-101935570-DEL-841
chr3-101951688-DEL-153
chr3-101978637-DEL-70
chr3-102083015-DEL-50
chr3-102108334-DEL-52
chr3-102162878-DEL-80
chr3-102318573-DEL-372
chr3-102445732-DEL-300
chr3-102462922-DEL-134
chr3-102488163-DEL-50
chr3-102501989-DEL-132
chr3-102501995-DEL-440
chr3-102573722-DEL-91
chr3-102608093-DEL-136
chr3-102837384-DEL-80
chr3-102850801-DEL-56
chr3-102861510-DEL-80
chr3-102867079-DEL-120
chr3-102919151-DEL-1433
chr3-103047967-DEL-134
chr3-103048847-DEL-50
chr3-103105877-DEL-195
chr3-103284491-DEL-184
chr3-103302583-DEL-224
chr3-103302827-DEL-448
chr3-103668292-DEL-319
chr3-103668293-DEL-133
chr3-103668520-DEL-135
chr3-103693155-DEL-99
chr3-103756200-DEL-54
chr3-103863029-DEL-50
chr3-103863239-DEL-100
chr3-104213047-DEL-51
chr3-104254351-DEL-50
chr3-104356592-DEL-3855
chr3-104454678-DEL-52
chr3-104487420-DEL-2002
chr3-104598671-DEL-74

chr3-104618681-DEL-190
chr3-104705879-DEL-144
chr3-104749374-DEL-76
chr3-104935705-DEL-1696
chr3-104937619-DEL-112
chr3-104985736-DEL-185
chr3-105110129-DEL-87
chr3-105168986-DEL-79
chr3-105247371-DEL-2353
chr3-105330363-DEL-58
chr3-105348353-DEL-852
chr3-105449605-DEL-230
chr3-105466001-DEL-71
chr3-105810901-DEL-52
chr3-106022848-DEL-367
chr3-106023126-DEL-157
chr3-106581874-DEL-502
chr3-106623347-DEL-50
chr3-106844834-DEL-66
chr3-106987650-DEL-55
chr3-107041434-DEL-50
chr3-107137185-DEL-50
chr3-107180638-DEL-128
chr3-107302705-DEL-3438
chr3-107356026-DEL-62
chr3-107385688-DEL-350
chr3-107386290-DEL-812
chr3-107387431-DEL-169
chr3-107387621-DEL-56
chr3-107458319-DEL-240
chr3-107823396-DEL-52
chr3-107901648-DEL-51
chr3-107908467-DEL-1955
chr3-107943196-DEL-140
chr3-107945185-DEL-144
chr3-107996528-DEL-334
chr3-108187786-DEL-137
chr3-108316524-DEL-210
chr3-108414714-DEL-909
chr3-108419027-DEL-68
chr3-108435367-DEL-577
chr3-108456442-DEL-50
chr3-108619730-DEL-64
chr3-108712050-DEL-52
chr3-108714958-DEL-59
chr3-108715124-DEL-101
chr3-108724060-DEL-178
chr3-108744253-DEL-50
chr3-108774666-DEL-145
chr3-108975207-DEL-394
chr3-108992021-DEL-54
chr3-109011641-DEL-78
chr3-109126988-DEL-62
chr3-109224780-DEL-60
chr3-109237175-DEL-249
chr3-109256490-DEL-102
chr3-109284501-DEL-75
chr3-109288271-DEL-88
chr3-109302091-DEL-116
chr3-109912795-DEL-56
chr3-109914768-DEL-1205
chr3-109984134-DEL-720
chr3-109999490-DEL-56
chr3-110173075-DEL-71
chr3-110233570-DEL-214
chr3-110256285-DEL-105
chr3-110343082-DEL-330
chr3-110374410-DEL-52
chr3-110377876-DEL-60
chr3-110386206-DEL-144
chr3-110755596-DEL-715
chr3-111090971-DEL-3262
chr3-111369387-DEL-261
chr3-111553353-DEL-91
chr3-111646585-DEL-22385
chr3-111649617-DEL-847
chr3-111667928-DEL-621

chr3-111678439-DEL-273
chr3-111701318-DEL-4937
chr3-111713293-DEL-100
chr3-111790852-DEL-58
chr3-111801795-DEL-92
chr3-112348478-DEL-1187
chr3-112397219-DEL-164
chr3-112418067-DEL-189
chr3-112479592-DEL-54
chr3-112650382-DEL-1835
chr3-112840939-DEL-193
chr3-112880159-DEL-74
chr3-112944115-DEL-50
chr3-113020872-DEL-268
chr3-113148965-DEL-98
chr3-113194226-DEL-127314
chr3-113461044-DEL-10456
chr3-113580316-DEL-51
chr3-113656899-DEL-659
chr3-113751589-DEL-148
chr3-113837156-DEL-95
chr3-113908720-DEL-50
chr3-114004316-DEL-56
chr3-114040440-DEL-146
chr3-114256277-DEL-60
chr3-114463292-DEL-6206
chr3-114658414-DEL-81
chr3-114786879-DEL-51
chr3-114969186-DEL-1374
chr3-115185135-DEL-2229
chr3-115249342-DEL-50
chr3-115259059-DEL-64
chr3-115259221-DEL-131
chr3-115373110-DEL-86
chr3-115475187-DEL-357
chr3-115583107-DEL-84
chr3-115583128-DEL-196
chr3-115640476-DEL-180
chr3-115640724-DEL-291
chr3-115669721-DEL-95
chr3-115880160-DEL-60
chr3-115943991-DEL-197
chr3-115944289-DEL-377
chr3-115952732-DEL-189
chr3-116065248-DEL-50
chr3-116094980-DEL-57
chr3-116106760-DEL-54
chr3-116115350-DEL-50
chr3-116115375-DEL-187
chr3-116123315-DEL-65
chr3-116123367-DEL-131
chr3-116266282-DEL-264
chr3-116266300-DEL-120
chr3-116279086-DEL-58
chr3-116315965-DEL-93
chr3-116469928-DEL-82
chr3-116486132-DEL-126
chr3-116488334-DEL-1766
chr3-116560565-DEL-473
chr3-116562155-DEL-114
chr3-116591283-DEL-224
chr3-116618039-DEL-56
chr3-116636643-DEL-493
chr3-116636928-DEL-203
chr3-116637182-DEL-116
chr3-116662639-DEL-304
chr3-116662925-DEL-145
chr3-116684977-DEL-189
chr3-116767470-DEL-83
chr3-116833635-DEL-50
chr3-116852977-DEL-493
chr3-116912483-DEL-98
chr3-116927507-DEL-57
chr3-116935853-DEL-50
chr3-116974572-DEL-56
chr3-117047514-DEL-2656
chr3-117143345-DEL-387

chr3-117208329-DEL-83
chr3-117266256-DEL-59
chr3-117415926-DEL-214
chr3-117416234-DEL-138
chr3-117513177-DEL-213
chr3-117537541-DEL-52
chr3-117537905-DEL-108
chr3-117591626-DEL-108
chr3-117591680-DEL-418
chr3-117591916-DEL-190
chr3-117644381-DEL-72
chr3-117651389-DEL-50
chr3-117810506-DEL-244
chr3-117847740-DEL-778
chr3-117900526-DEL-1876
chr3-117948534-DEL-84
chr3-118010612-DEL-102
chr3-118020368-DEL-126
chr3-118047271-DEL-1621
chr3-118281630-DEL-53
chr3-118446559-DEL-1048
chr3-118501430-DEL-66
chr3-118525380-DEL-52
chr3-118635130-DEL-64
chr3-118680833-DEL-56
chr3-118740138-DEL-798
chr3-118760175-DEL-75
chr3-118785365-DEL-50
chr3-118820110-DEL-168
chr3-118924058-DEL-282
chr3-119058916-DEL-56
chr3-119110742-DEL-82
chr3-119177409-DEL-58
chr3-119312652-DEL-420
chr3-119415247-DEL-181
chr3-119420820-DEL-363
chr3-119425405-DEL-77
chr3-119474929-DEL-72
chr3-119532405-DEL-81
chr3-119810935-DEL-64
chr3-119954086-DEL-66
chr3-120045137-DEL-129
chr3-120047392-DEL-687
chr3-120063132-DEL-51
chr3-120170714-DEL-127
chr3-120177858-DEL-52
chr3-120178926-DEL-449
chr3-120220725-DEL-74
chr3-120429422-DEL-242
chr3-120429777-DEL-50
chr3-120591958-DEL-50
chr3-120608185-DEL-228
chr3-120651064-DEL-50
chr3-120661074-DEL-100
chr3-120681937-DEL-54
chr3-120745328-DEL-82
chr3-120752735-DEL-720
chr3-120787075-DEL-52
chr3-120801216-DEL-50
chr3-120840802-DEL-54
chr3-120954942-DEL-86
chr3-120976074-DEL-99
chr3-120976262-DEL-109
chr3-121093229-DEL-366
chr3-121247033-DEL-311
chr3-121384807-DEL-180
chr3-121536574-DEL-702
chr3-121564235-DEL-459
chr3-121567044-DEL-138
chr3-121607257-DEL-112
chr3-121650864-DEL-111
chr3-121676104-DEL-636
chr3-121676219-DEL-159
chr3-121684512-DEL-114
chr3-121684670-DEL-168
chr3-121715077-DEL-104
chr3-121717932-DEL-52

chr3-121718270-DEL-117
chr3-121720333-DEL-86
chr3-121756223-DEL-174
chr3-121814493-DEL-121
chr3-121816971-DEL-174
chr3-121940070-DEL-53
chr3-121976771-DEL-50
chr3-121987179-DEL-303
chr3-122021414-DEL-11526
chr3-122100190-DEL-497
chr3-122103720-DEL-156
chr3-122104834-DEL-194
chr3-122122363-DEL-140
chr3-122209574-DEL-212
chr3-122241839-DEL-64
chr3-122307136-DEL-348
chr3-122404993-DEL-160
chr3-122432454-DEL-53
chr3-122439806-DEL-52
chr3-122439972-DEL-68
chr3-122598847-DEL-539
chr3-122619118-DEL-90
chr3-122684419-DEL-182
chr3-122785801-DEL-50
chr3-122856940-DEL-286
chr3-122857060-DEL-88
chr3-122879408-DEL-175
chr3-122917302-DEL-68
chr3-122964099-DEL-2122
chr3-123166875-DEL-425
chr3-123175243-DEL-845
chr3-123180214-DEL-258
chr3-123199654-DEL-50
chr3-123304785-DEL-215
chr3-123377563-DEL-326
chr3-123377609-DEL-102
chr3-123401495-DEL-50
chr3-123405866-DEL-240
chr3-123470440-DEL-1286
chr3-123471063-DEL-1391
chr3-123599819-DEL-58
chr3-123670477-DEL-54
chr3-123791285-DEL-52
chr3-123791321-DEL-145
chr3-123887248-DEL-324
chr3-124004654-DEL-53
chr3-124053015-DEL-101
chr3-124087420-DEL-102
chr3-124162048-DEL-349
chr3-124197674-DEL-52
chr3-124251588-DEL-82
chr3-124284295-DEL-50
chr3-124324813-DEL-167
chr3-124470238-DEL-54
chr3-124509445-DEL-60
chr3-124525203-DEL-63
chr3-124620481-DEL-1175
chr3-124622149-DEL-3875
chr3-124762334-DEL-128
chr3-124801198-DEL-55
chr3-124834929-DEL-52
chr3-124951370-DEL-50
chr3-125093382-DEL-132
chr3-125093462-DEL-64
chr3-125211711-DEL-50
chr3-125217529-DEL-1484
chr3-125248652-DEL-66
chr3-125313722-DEL-407
chr3-125346461-DEL-1361
chr3-125364138-DEL-53
chr3-125369117-DEL-72
chr3-125388896-DEL-536
chr3-125437540-DEL-862
chr3-125453653-DEL-72
chr3-125494768-DEL-52
chr3-125848846-DEL-58
chr3-125921028-DEL-4093

chr3-125974557-DEL-72
chr3-125998637-DEL-223
chr3-126091849-DEL-70
chr3-126451287-DEL-59
chr3-126485383-DEL-167
chr3-126513987-DEL-52
chr3-126514270-DEL-232
chr3-126517867-DEL-129
chr3-126550291-DEL-64
chr3-126649792-DEL-8374
chr3-126820645-DEL-344
chr3-126877997-DEL-56
chr3-126935707-DEL-1944
chr3-126939409-DEL-1242
chr3-127009733-DEL-4378
chr3-127028854-DEL-677
chr3-127098371-DEL-74
chr3-127118583-DEL-110
chr3-127151571-DEL-54
chr3-127154018-DEL-72
chr3-127233284-DEL-193
chr3-127258888-DEL-52
chr3-127264665-DEL-58
chr3-127274821-DEL-54
chr3-127459879-DEL-50
chr3-127515605-DEL-289
chr3-127557098-DEL-51
chr3-127595907-DEL-50
chr3-127608046-DEL-63
chr3-127621332-DEL-329
chr3-127712654-DEL-102
chr3-127716164-DEL-50
chr3-127736065-DEL-222
chr3-127741050-DEL-90
chr3-127764016-DEL-242
chr3-127764075-DEL-68
chr3-127813505-DEL-160
chr3-127813609-DEL-58
chr3-127814621-DEL-281
chr3-127815092-DEL-68
chr3-127868302-DEL-88
chr3-127949376-DEL-50
chr3-127997166-DEL-1443
chr3-128385155-DEL-50
chr3-128529922-DEL-241
chr3-128537579-DEL-96
chr3-128566523-DEL-330
chr3-128566881-DEL-95
chr3-128733253-DEL-481
chr3-128776296-DEL-72
chr3-128818675-DEL-7096
chr3-128851058-DEL-2125
chr3-128903937-DEL-64
chr3-128987621-DEL-67
chr3-129030645-DEL-137
chr3-129030851-DEL-51
chr3-129118717-DEL-56
chr3-129118863-DEL-222
chr3-129146831-DEL-130
chr3-129151067-DEL-50
chr3-129184394-DEL-55
chr3-129219182-DEL-1217
chr3-129230049-DEL-750
chr3-129247663-DEL-152
chr3-129289047-DEL-62
chr3-129302040-DEL-53
chr3-129302059-DEL-142
chr3-129402611-DEL-1509
chr3-129477602-DEL-230
chr3-129700309-DEL-53
chr3-129735955-DEL-60
chr3-129877120-DEL-737
chr3-130170342-DEL-82
chr3-130235495-DEL-776
chr3-130558799-DEL-1291
chr3-130657577-DEL-145
chr3-130693047-DEL-50

chr3-130724498-DEL-132
chr3-130732836-DEL-157
chr3-130817310-DEL-50
chr3-130855819-DEL-197
chr3-131069448-DEL-210
chr3-131082808-DEL-159
chr3-131093872-DEL-61
chr3-131111260-DEL-237
chr3-131117513-DEL-159
chr3-131117918-DEL-80
chr3-131155375-DEL-50
chr3-131219719-DEL-966
chr3-131296840-DEL-1536
chr3-131416187-DEL-5888
chr3-131530805-DEL-501
chr3-131531249-DEL-171
chr3-131531687-DEL-791
chr3-131531746-DEL-59
chr3-131532403-DEL-69
chr3-131551535-DEL-52
chr3-131596107-DEL-172
chr3-131631606-DEL-54
chr3-131631696-DEL-70
chr3-131735707-DEL-321
chr3-131744667-DEL-52
chr3-131826167-DEL-256
chr3-131846220-DEL-74
chr3-131916634-DEL-50
chr3-132045424-DEL-118
chr3-132096986-DEL-54
chr3-132231198-DEL-50
chr3-132376172-DEL-70
chr3-132441467-DEL-80
chr3-132490152-DEL-108
chr3-132504768-DEL-794
chr3-132599637-DEL-54
chr3-132645230-DEL-1009
chr3-132670321-DEL-78
chr3-132695346-DEL-4616
chr3-132853183-DEL-54
chr3-132854318-DEL-116
chr3-132896074-DEL-356
chr3-133012789-DEL-167
chr3-133057195-DEL-1544
chr3-133057295-DEL-524
chr3-133058393-DEL-658
chr3-133084304-DEL-162
chr3-133091857-DEL-50
chr3-133106922-DEL-1893
chr3-133195509-DEL-50
chr3-133271581-DEL-4590
chr3-133384750-DEL-60
chr3-133436547-DEL-51
chr3-133436825-DEL-70
chr3-133488021-DEL-981
chr3-133497657-DEL-71
chr3-133643896-DEL-669
chr3-133645891-DEL-124
chr3-133763167-DEL-101
chr3-133809612-DEL-674
chr3-133812682-DEL-120
chr3-133847747-DEL-60
chr3-134019024-DEL-75
chr3-134031553-DEL-164
chr3-134034630-DEL-68
chr3-134220548-DEL-62
chr3-134404558-DEL-55
chr3-134422747-DEL-142
chr3-134604699-DEL-258
chr3-134632258-DEL-447
chr3-134644047-DEL-367
chr3-134645697-DEL-4095
chr3-134661960-DEL-5142
chr3-134820064-DEL-413
chr3-134900551-DEL-4175
chr3-134980903-DEL-66
chr3-134991657-DEL-2302

chr3-135005162-DEL-50
chr3-135048618-DEL-150
chr3-135048694-DEL-74
chr3-135048888-DEL-76
chr3-135282914-DEL-546
chr3-135283226-DEL-250
chr3-135305363-DEL-54
chr3-135386307-DEL-83
chr3-135475310-DEL-5610
chr3-135564720-DEL-99
chr3-135660402-DEL-50
chr3-135683839-DEL-2896
chr3-135716856-DEL-3970
chr3-135753797-DEL-148
chr3-135811931-DEL-1915
chr3-135954722-DEL-66
chr3-136005458-DEL-239
chr3-136101891-DEL-242
chr3-136233912-DEL-68
chr3-136240096-DEL-498
chr3-136294513-DEL-305
chr3-136383111-DEL-87
chr3-136508879-DEL-102
chr3-136536206-DEL-143
chr3-136878598-DEL-50
chr3-137016815-DEL-108
chr3-137024217-DEL-1362
chr3-137137072-DEL-50
chr3-137277510-DEL-96
chr3-137277874-DEL-159
chr3-137316442-DEL-50
chr3-137390986-DEL-1864
chr3-137451455-DEL-120
chr3-137517749-DEL-147
chr3-137619817-DEL-189
chr3-137620218-DEL-265
chr3-137734172-DEL-54
chr3-138026259-DEL-63
chr3-138050533-DEL-50
chr3-138168777-DEL-507
chr3-138191042-DEL-53
chr3-138199280-DEL-171
chr3-138272811-DEL-56
chr3-138306924-DEL-1483
chr3-138437855-DEL-136
chr3-138511512-DEL-124
chr3-138547491-DEL-195
chr3-138612378-DEL-130
chr3-138733790-DEL-50
chr3-138751800-DEL-495
chr3-138759950-DEL-96
chr3-138983722-DEL-1481
chr3-139049869-DEL-177
chr3-139504276-DEL-50
chr3-139518299-DEL-525
chr3-139521109-DEL-466
chr3-139622224-DEL-242
chr3-139651418-DEL-58
chr3-139893719-DEL-190
chr3-140318871-DEL-147
chr3-140395451-DEL-1569
chr3-140440822-DEL-50
chr3-140449470-DEL-64
chr3-140470619-DEL-9039
chr3-140476979-DEL-416
chr3-140598716-DEL-50
chr3-140604164-DEL-86
chr3-140785305-DEL-5957
chr3-140831339-DEL-1434
chr3-141000715-DEL-967
chr3-141022137-DEL-61
chr3-141131288-DEL-50
chr3-141200856-DEL-56
chr3-141249328-DEL-195
chr3-141249771-DEL-165
chr3-141340160-DEL-55
chr3-141389212-DEL-60

chr3-141389308-DEL-174
chr3-141596877-DEL-67
chr3-141596957-DEL-119
chr3-141628160-DEL-56
chr3-142003502-DEL-188
chr3-142003601-DEL-81
chr3-142034889-DEL-69
chr3-142109348-DEL-56
chr3-142138090-DEL-591
chr3-142163666-DEL-138
chr3-142245200-DEL-56
chr3-142316031-DEL-15542
chr3-142365287-DEL-280
chr3-142373057-DEL-13084
chr3-142389163-DEL-156
chr3-142402448-DEL-66
chr3-142479237-DEL-104
chr3-142653332-DEL-93
chr3-142731055-DEL-6271
chr3-142804404-DEL-189
chr3-142894421-DEL-260
chr3-143044796-DEL-80
chr3-143136586-DEL-1541
chr3-143375797-DEL-56
chr3-143432475-DEL-177
chr3-143432714-DEL-75
chr3-143440984-DEL-180
chr3-143450536-DEL-50
chr3-143472199-DEL-774
chr3-143587863-DEL-57
chr3-143626109-DEL-52
chr3-143640387-DEL-108
chr3-143670093-DEL-313
chr3-143756886-DEL-66
chr3-143783139-DEL-122
chr3-143814957-DEL-76
chr3-143877729-DEL-126
chr3-143949418-DEL-129
chr3-143955401-DEL-813
chr3-143955622-DEL-103
chr3-143956329-DEL-131
chr3-144037614-DEL-90
chr3-144108809-DEL-80
chr3-144118173-DEL-55
chr3-144157572-DEL-290
chr3-144174201-DEL-82
chr3-144194739-DEL-408
chr3-144232760-DEL-58
chr3-144252840-DEL-478
chr3-144253080-DEL-537
chr3-144264578-DEL-77
chr3-144299815-DEL-153
chr3-144338843-DEL-63
chr3-144338870-DEL-132
chr3-144340242-DEL-57
chr3-144345070-DEL-62
chr3-144392324-DEL-82
chr3-144400380-DEL-80
chr3-144415808-DEL-61
chr3-144415808-DEL-187
chr3-144419256-DEL-56
chr3-144426351-DEL-53
chr3-144437938-DEL-1863
chr3-144443961-DEL-6378
chr3-144444282-DEL-234
chr3-144450922-DEL-528
chr3-144451143-DEL-66
chr3-144509751-DEL-99
chr3-144509760-DEL-726
chr3-144509956-DEL-99
chr3-144510281-DEL-429
chr3-144528604-DEL-54
chr3-144528604-DEL-378
chr3-144534319-DEL-1059
chr3-144570363-DEL-64
chr3-144621526-DEL-412
chr3-144623884-DEL-601

chr3-144868837-DEL-272
chr3-144871286-DEL-79
chr3-144889144-DEL-255
chr3-144924396-DEL-591
chr3-144934785-DEL-54
chr3-145057912-DEL-468
chr3-145058042-DEL-124
chr3-145146469-DEL-124
chr3-145368595-DEL-1385
chr3-145405360-DEL-132
chr3-145431848-DEL-90
chr3-145513236-DEL-50
chr3-145549730-DEL-90
chr3-145556410-DEL-87
chr3-145566152-DEL-234
chr3-145664995-DEL-51
chr3-145682255-DEL-1544
chr3-145718440-DEL-78
chr3-145757952-DEL-88
chr3-145758270-DEL-175
chr3-145767736-DEL-104
chr3-145829863-DEL-155
chr3-145866814-DEL-68
chr3-145896698-DEL-86
chr3-145960796-DEL-158
chr3-145983746-DEL-53
chr3-146025459-DEL-231
chr3-146103146-DEL-260
chr3-146138900-DEL-141
chr3-146146522-DEL-63
chr3-146146800-DEL-208
chr3-146182643-DEL-175
chr3-146183192-DEL-1007
chr3-146183406-DEL-368
chr3-146183774-DEL-262
chr3-146196042-DEL-68
chr3-146265787-DEL-84
chr3-146377287-DEL-780
chr3-146401620-DEL-117
chr3-146442163-DEL-466
chr3-146442297-DEL-115
chr3-146442821-DEL-147
chr3-146608707-DEL-110
chr3-146614154-DEL-60
chr3-146631733-DEL-75
chr3-146638553-DEL-481
chr3-146653769-DEL-51
chr3-146694145-DEL-775
chr3-146850606-DEL-164
chr3-146853909-DEL-50
chr3-146855899-DEL-463
chr3-146957279-DEL-50
chr3-147097290-DEL-56
chr3-147140363-DEL-66
chr3-147154216-DEL-133
chr3-147224601-DEL-58
chr3-147225037-DEL-54
chr3-147245793-DEL-95
chr3-147274026-DEL-51
chr3-147373025-DEL-758
chr3-147457060-DEL-86
chr3-147520516-DEL-582
chr3-147527299-DEL-125
chr3-147576567-DEL-207
chr3-147580661-DEL-577
chr3-147768660-DEL-355
chr3-147851121-DEL-88
chr3-147851440-DEL-7060
chr3-147866097-DEL-55
chr3-147925067-DEL-104
chr3-147928998-DEL-62
chr3-148019128-DEL-50
chr3-148055874-DEL-173
chr3-148057163-DEL-50
chr3-148137027-DEL-156
chr3-148146108-DEL-53
chr3-148171594-DEL-120

chr3-148190717-DEL-87
chr3-148191988-DEL-271
chr3-148291132-DEL-129
chr3-148332110-DEL-1880
chr3-148345403-DEL-159
chr3-148365679-DEL-1083
chr3-148391803-DEL-50
chr3-148504272-DEL-75
chr3-148514174-DEL-52
chr3-148556159-DEL-177
chr3-148610916-DEL-236
chr3-148756533-DEL-62
chr3-148764281-DEL-376
chr3-148764455-DEL-96
chr3-148764807-DEL-150
chr3-148827219-DEL-1023
chr3-148888454-DEL-80
chr3-148937969-DEL-69
chr3-149007465-DEL-254
chr3-149033017-DEL-72
chr3-149079107-DEL-50
chr3-149143373-DEL-167
chr3-149191997-DEL-50
chr3-149232323-DEL-246
chr3-149252936-DEL-82
chr3-149285520-DEL-458
chr3-149419698-DEL-288
chr3-149419986-DEL-64
chr3-149443287-DEL-92
chr3-149450493-DEL-92
chr3-149556916-DEL-503
chr3-149585342-DEL-53
chr3-149825926-DEL-81
chr3-149836504-DEL-194
chr3-149846887-DEL-202
chr3-149887339-DEL-308
chr3-149971415-DEL-50
chr3-150010999-DEL-68
chr3-150031063-DEL-54
chr3-150154874-DEL-66
chr3-150216649-DEL-63
chr3-150216793-DEL-397
chr3-150217247-DEL-471
chr3-150242074-DEL-166
chr3-150263154-DEL-75
chr3-150263361-DEL-148
chr3-150299608-DEL-50
chr3-150322557-DEL-143
chr3-150433414-DEL-581
chr3-150438501-DEL-1070
chr3-150608470-DEL-858
chr3-150614326-DEL-280
chr3-150644479-DEL-111
chr3-150670101-DEL-887
chr3-150679573-DEL-419
chr3-150701042-DEL-56
chr3-150761017-DEL-179
chr3-150869302-DEL-50
chr3-150900633-DEL-2328
chr3-150974761-DEL-67
chr3-150993474-DEL-53
chr3-151042604-DEL-221
chr3-151147368-DEL-70
chr3-151155903-DEL-60
chr3-151239521-DEL-151
chr3-151270100-DEL-56
chr3-151406246-DEL-290
chr3-151492278-DEL-50
chr3-151508993-DEL-96
chr3-151516726-DEL-196
chr3-151520224-DEL-71
chr3-151520551-DEL-63
chr3-151529327-DEL-104
chr3-151603404-DEL-58
chr3-151618092-DEL-56
chr3-151717635-DEL-6661
chr3-151758548-DEL-944

chr3-151779668-DEL-280
chr3-151814091-DEL-60
chr3-151826377-DEL-76
chr3-151834690-DEL-155
chr3-151881433-DEL-58
chr3-151937377-DEL-74
chr3-151991064-DEL-190
chr3-152018887-DEL-185
chr3-152019348-DEL-871
chr3-152028193-DEL-52
chr3-152075451-DEL-80
chr3-152143851-DEL-195
chr3-152188820-DEL-155
chr3-152191757-DEL-213
chr3-152191757-DEL-573
chr3-152196741-DEL-132
chr3-152208268-DEL-506
chr3-152235783-DEL-689
chr3-152250237-DEL-149
chr3-152252040-DEL-239
chr3-152261829-DEL-90
chr3-152268700-DEL-321
chr3-152295009-DEL-396
chr3-152295216-DEL-88
chr3-152314293-DEL-52
chr3-152355457-DEL-237
chr3-152355720-DEL-190
chr3-152388713-DEL-644
chr3-152422889-DEL-93
chr3-152427403-DEL-111
chr3-152439065-DEL-58
chr3-152447696-DEL-225
chr3-152461386-DEL-547
chr3-152484832-DEL-98
chr3-152490211-DEL-138
chr3-152491867-DEL-660
chr3-152524924-DEL-61
chr3-152556882-DEL-105
chr3-152568061-DEL-89
chr3-152569296-DEL-53
chr3-152621512-DEL-126
chr3-152670484-DEL-281
chr3-152709783-DEL-118
chr3-152738910-DEL-52
chr3-152748211-DEL-68
chr3-152765163-DEL-98
chr3-152773871-DEL-66
chr3-152774340-DEL-126
chr3-152796914-DEL-1328
chr3-152813848-DEL-430
chr3-152909408-DEL-105
chr3-152998777-DEL-58
chr3-153017830-DEL-99
chr3-153036591-DEL-988
chr3-153038937-DEL-512
chr3-153038983-DEL-72
chr3-153116665-DEL-190
chr3-153119694-DEL-74
chr3-153139874-DEL-62
chr3-153158691-DEL-1119
chr3-153162238-DEL-66
chr3-153229864-DEL-60
chr3-153246810-DEL-91
chr3-153249027-DEL-222
chr3-153317450-DEL-60
chr3-153330093-DEL-459
chr3-153332817-DEL-88
chr3-153356676-DEL-56
chr3-153391371-DEL-398
chr3-153403270-DEL-355
chr3-153410602-DEL-168
chr3-153419476-DEL-52
chr3-153439190-DEL-64
chr3-153439316-DEL-144
chr3-153458869-DEL-1248
chr3-153465618-DEL-60
chr3-153475791-DEL-90

chr3-153494232-DEL-338
chr3-153499878-DEL-233
chr3-153536439-DEL-202
chr3-153605401-DEL-106
chr3-153636621-DEL-200
chr3-153656689-DEL-87
chr3-153677449-DEL-57
chr3-153685659-DEL-199
chr3-153685777-DEL-80
chr3-153713239-DEL-284
chr3-153721758-DEL-214
chr3-153772050-DEL-86
chr3-153772538-DEL-515
chr3-153777634-DEL-223
chr3-153777841-DEL-279
chr3-153779882-DEL-233
chr3-153826144-DEL-70
chr3-153826333-DEL-128
chr3-153836953-DEL-76
chr3-153986893-DEL-96
chr3-154021454-DEL-231
chr3-154054657-DEL-361
chr3-154069650-DEL-66
chr3-154105338-DEL-91
chr3-154120529-DEL-576
chr3-154121356-DEL-657
chr3-154121456-DEL-153
chr3-154121915-DEL-99
chr3-154128337-DEL-1093
chr3-154160755-DEL-135
chr3-154178432-DEL-576
chr3-154178772-DEL-512
chr3-154246937-DEL-774
chr3-154280665-DEL-54
chr3-154283840-DEL-444
chr3-154287384-DEL-212
chr3-154314680-DEL-318
chr3-154359931-DEL-50
chr3-154365028-DEL-59
chr3-154381874-DEL-54
chr3-154435258-DEL-74
chr3-154552490-DEL-132
chr3-154579303-DEL-368
chr3-154590369-DEL-166
chr3-154607527-DEL-3436
chr3-154652465-DEL-60
chr3-154677671-DEL-7426
chr3-154688755-DEL-553
chr3-154852178-DEL-6047
chr3-154947349-DEL-52
chr3-154981746-DEL-613
chr3-155098741-DEL-104
chr3-155219127-DEL-1127
chr3-155308160-DEL-95
chr3-155413744-DEL-57
chr3-155530979-DEL-80
chr3-155633102-DEL-1217
chr3-155716839-DEL-50
chr3-155813796-DEL-304
chr3-155896811-DEL-108
chr3-155902426-DEL-162
chr3-155914211-DEL-767
chr3-155941012-DEL-52
chr3-155950092-DEL-291
chr3-156121239-DEL-60
chr3-156142039-DEL-588
chr3-156407469-DEL-72
chr3-156414822-DEL-50
chr3-156514387-DEL-432
chr3-156540202-DEL-467
chr3-156641547-DEL-55
chr3-156711728-DEL-105
chr3-156798829-DEL-86
chr3-156939515-DEL-89
chr3-157056182-DEL-69
chr3-157081962-DEL-737
chr3-157146651-DEL-1243

chr3-157247085-DEL-120
chr3-157321509-DEL-56
chr3-157338551-DEL-131
chr3-157385118-DEL-82
chr3-157482948-DEL-73
chr3-157506505-DEL-132
chr3-157506506-DEL-65
chr3-157511724-DEL-2820
chr3-157533699-DEL-189
chr3-157645900-DEL-2207
chr3-157952535-DEL-89
chr3-158083122-DEL-56
chr3-158155959-DEL-108
chr3-158195659-DEL-50
chr3-158381638-DEL-2472
chr3-158473039-DEL-173
chr3-158486175-DEL-1369
chr3-158494214-DEL-50
chr3-158503847-DEL-1250
chr3-158517945-DEL-66
chr3-158796741-DEL-50
chr3-158898265-DEL-564
chr3-159000784-DEL-5460
chr3-159032671-DEL-50
chr3-159116326-DEL-1889
chr3-159200635-DEL-63
chr3-159244850-DEL-357
chr3-159306456-DEL-369
chr3-159306898-DEL-52
chr3-159484575-DEL-355
chr3-159484845-DEL-85
chr3-159537527-DEL-80
chr4-3055654-DEL-53
chr4-3060170-DEL-832
chr4-3061346-DEL-159
chr4-3061828-DEL-334
chr4-3064251-DEL-972
chr4-3065531-DEL-501
chr4-3082597-DEL-230
chr4-3083386-DEL-63
chr4-3085178-DEL-146
chr4-3164623-DEL-66
chr4-3175480-DEL-260
chr4-3178683-DEL-154
chr4-3218240-DEL-65
chr4-3228541-DEL-68
chr4-3230121-DEL-1253
chr4-3232928-DEL-143
chr4-3240066-DEL-315
chr4-3269766-DEL-152
chr4-3300281-DEL-80
chr4-3382831-DEL-2068
chr4-3543961-DEL-2287
chr4-3613603-DEL-490
chr4-3613826-DEL-104
chr4-3613963-DEL-350
chr4-3657399-DEL-121
chr4-3661557-DEL-178
chr4-3661749-DEL-76
chr4-3697485-DEL-60
chr4-3778940-DEL-1172
chr4-3791865-DEL-61
chr4-3806554-DEL-3078
chr4-4023918-DEL-68
chr4-4072890-DEL-95
chr4-4120268-DEL-2627
chr4-4149257-DEL-162
chr4-4334242-DEL-305
chr4-4446190-DEL-2422
chr4-4469978-DEL-192
chr4-4480960-DEL-186
chr4-4481613-DEL-187
chr4-4482574-DEL-2475
chr4-4506920-DEL-50
chr4-4557528-DEL-90
chr4-4632702-DEL-182
chr4-4640622-DEL-1799

chr4-4642693-DEL-1091
chr4-4686803-DEL-98
chr4-4715084-DEL-72
chr4-4724577-DEL-136
chr4-4807665-DEL-60
chr4-4809065-DEL-56
chr4-4976793-DEL-603
chr4-4987964-DEL-50
chr4-5077062-DEL-70
chr4-5119998-DEL-54
chr4-5135920-DEL-109
chr4-5162625-DEL-68
chr4-5443050-DEL-65
chr4-5492678-DEL-56
chr4-5635269-DEL-65
chr4-5718351-DEL-872
chr4-5748510-DEL-105
chr4-5784031-DEL-50
chr4-5968298-DEL-1288
chr4-5977314-DEL-88
chr4-6051368-DEL-501
chr4-6264724-DEL-55
chr4-6301216-DEL-54
chr4-6431563-DEL-92
chr4-6431588-DEL-433
chr4-6482027-DEL-64
chr4-6685464-DEL-81
chr4-6705057-DEL-63
chr4-6784326-DEL-2114
chr4-6858102-DEL-1314
chr4-7007395-DEL-319
chr4-7033403-DEL-216
chr4-7234527-DEL-160
chr4-7294727-DEL-10296
chr4-7422701-DEL-50
chr4-7451419-DEL-511
chr4-7459904-DEL-50
chr4-7461658-DEL-4162
chr4-7502481-DEL-964
chr4-7523306-DEL-2114
chr4-7616435-DEL-216
chr4-7659635-DEL-2490
chr4-7781632-DEL-553
chr4-7813867-DEL-1203
chr4-7838737-DEL-1095
chr4-7868648-DEL-50
chr4-8081084-DEL-72
chr4-8104394-DEL-118
chr4-8117029-DEL-7448
chr4-8133479-DEL-3376
chr4-8152084-DEL-196
chr4-8157364-DEL-169
chr4-8168749-DEL-444
chr4-8178132-DEL-109
chr4-8235600-DEL-303
chr4-8316501-DEL-78
chr4-8366126-DEL-334
chr4-8367784-DEL-54
chr4-8380363-DEL-139
chr4-8512978-DEL-1493
chr4-8590607-DEL-76
chr4-8608190-DEL-134
chr4-8622439-DEL-4305
chr4-8730843-DEL-444
chr4-8731293-DEL-52
chr4-8740143-DEL-84
chr4-8758658-DEL-239
chr4-8759059-DEL-91
chr4-8780444-DEL-98
chr4-8817232-DEL-100
chr4-8878553-DEL-55
chr4-8894786-DEL-53
chr4-9021676-DEL-235
chr4-9084505-DEL-68
chr4-9086564-DEL-146
chr4-9167538-DEL-236
chr4-9182982-DEL-480

chr4-9206664-DEL-51
chr4-9206741-DEL-459
chr4-9305981-DEL-74
chr4-9309298-DEL-630
chr4-9309322-DEL-175
chr4-9309815-DEL-70
chr4-9432522-DEL-337
chr4-9442128-DEL-84
chr4-9547128-DEL-152
chr4-9620559-DEL-128
chr4-9742673-DEL-118
chr4-9748932-DEL-103
chr4-9749007-DEL-51
chr4-9769658-DEL-822
chr4-9827022-DEL-608
chr4-9827673-DEL-108
chr4-9895345-DEL-385
chr4-9958769-DEL-1024
chr4-10046039-DEL-733
chr4-10078023-DEL-880
chr4-10111023-DEL-208
chr4-10199732-DEL-50
chr4-10207732-DEL-126
chr4-10277804-DEL-78
chr4-10371047-DEL-1889
chr4-10461190-DEL-280
chr4-10576263-DEL-50
chr4-10582738-DEL-1130
chr4-10694150-DEL-325
chr4-10734806-DEL-348
chr4-10737696-DEL-1920
chr4-10819780-DEL-75
chr4-10819912-DEL-468
chr4-10900593-DEL-247
chr4-10905375-DEL-1370
chr4-10906991-DEL-209
chr4-10907287-DEL-126
chr4-10907397-DEL-214
chr4-10939232-DEL-315
chr4-10949999-DEL-206
chr4-10972379-DEL-88
chr4-11027912-DEL-113
chr4-11045948-DEL-216
chr4-11053936-DEL-1530
chr4-11054741-DEL-405
chr4-11065364-DEL-342
chr4-11065413-DEL-148
chr4-11106689-DEL-816
chr4-11174676-DEL-58
chr4-11220115-DEL-185
chr4-11238828-DEL-302
chr4-11341923-DEL-175
chr4-11397152-DEL-93
chr4-11441504-DEL-1515
chr4-11458331-DEL-1180
chr4-11464108-DEL-410
chr4-11532005-DEL-381
chr4-11545814-DEL-73
chr4-11566317-DEL-242
chr4-11571066-DEL-323
chr4-11571313-DEL-112
chr4-11648670-DEL-878
chr4-11665438-DEL-62
chr4-11718762-DEL-317
chr4-11718795-DEL-153
chr4-11735081-DEL-301
chr4-11767918-DEL-768
chr4-11776099-DEL-340
chr4-11819995-DEL-64
chr4-11884575-DEL-931
chr4-11948763-DEL-160
chr4-11987260-DEL-58
chr4-12024994-DEL-67
chr4-12037435-DEL-291
chr4-12037580-DEL-68
chr4-12048686-DEL-372
chr4-12139629-DEL-69

chr4-12152094-DEL-312
chr4-12152352-DEL-78
chr4-12155020-DEL-122
chr4-12174772-DEL-2018
chr4-12357690-DEL-62
chr4-12358026-DEL-452
chr4-12358357-DEL-96
chr4-12395932-DEL-105
chr4-12445667-DEL-3850
chr4-12484603-DEL-676
chr4-12551210-DEL-230
chr4-12627331-DEL-50
chr4-12696694-DEL-50
chr4-12701293-DEL-132
chr4-12701377-DEL-52
chr4-12715723-DEL-64
chr4-12807400-DEL-135
chr4-12934389-DEL-64
chr4-13282399-DEL-54
chr4-13295869-DEL-210
chr4-13717141-DEL-52
chr4-13939474-DEL-64
chr4-13973702-DEL-230
chr4-14078244-DEL-50
chr4-14201236-DEL-313
chr4-14238377-DEL-122
chr4-14312003-DEL-1230
chr4-14561249-DEL-50
chr4-14728774-DEL-50
chr4-14744764-DEL-51
chr4-14753388-DEL-52
chr4-14753719-DEL-63
chr4-14786152-DEL-261
chr4-14791092-DEL-2017
chr4-14866151-DEL-196
chr4-15095587-DEL-533
chr4-15118752-DEL-276
chr4-15120045-DEL-378
chr4-15128303-DEL-54
chr4-15320468-DEL-576
chr4-15376767-DEL-60
chr4-15596662-DEL-2837
chr4-15629868-DEL-540
chr4-15725240-DEL-50
chr4-15862046-DEL-476
chr4-15883118-DEL-59
chr4-15952256-DEL-70
chr4-15954239-DEL-114
chr4-15954485-DEL-66
chr4-16021867-DEL-325
chr4-16125443-DEL-855
chr4-16147148-DEL-2939
chr4-16246939-DEL-2387
chr4-16425228-DEL-170
chr4-16485038-DEL-63
chr4-16507863-DEL-72
chr4-16634888-DEL-1779
chr4-17065891-DEL-5199
chr4-17072174-DEL-418
chr4-17377381-DEL-51
chr4-17466195-DEL-1023
chr4-17707151-DEL-78
chr4-17711681-DEL-5193
chr4-17720591-DEL-884
chr4-17768110-DEL-53
chr4-17933029-DEL-1475
chr4-17960992-DEL-137
chr4-18230952-DEL-66
chr4-18280317-DEL-110
chr4-18333982-DEL-13527
chr4-18347867-DEL-1167
chr4-18528772-DEL-893
chr4-18530638-DEL-52
chr4-18535894-DEL-326
chr4-18536440-DEL-291
chr4-18578500-DEL-66
chr4-18664389-DEL-56

chr4-18726162-DEL-677
chr4-18775384-DEL-165
chr4-18805535-DEL-84
chr4-18828564-DEL-58
chr4-18848514-DEL-297
chr4-18848684-DEL-54
chr4-18864056-DEL-72
chr4-18873721-DEL-362
chr4-18882482-DEL-80
chr4-18896584-DEL-66
chr4-19023208-DEL-2065
chr4-19028003-DEL-82
chr4-19030845-DEL-2801
chr4-19043323-DEL-53
chr4-19051937-DEL-63
chr4-19067553-DEL-1944
chr4-19083828-DEL-63
chr4-19123864-DEL-795
chr4-19134388-DEL-327
chr4-19148947-DEL-8073
chr4-19177467-DEL-552
chr4-19229381-DEL-1763
chr4-19274300-DEL-4956
chr4-19277771-DEL-66
chr4-19279327-DEL-106
chr4-19289642-DEL-55
chr4-19307329-DEL-50
chr4-19337137-DEL-565
chr4-19349693-DEL-303
chr4-19362220-DEL-50
chr4-19376046-DEL-80
chr4-19478436-DEL-115
chr4-19490491-DEL-54
chr4-19666111-DEL-1847
chr4-19688172-DEL-360
chr4-19729698-DEL-114
chr4-19734346-DEL-52
chr4-19755794-DEL-86
chr4-19787027-DEL-605
chr4-19917995-DEL-88
chr4-19994342-DEL-76
chr4-20001606-DEL-1251
chr4-20003034-DEL-89
chr4-20013933-DEL-58
chr4-20086041-DEL-515
chr4-20125715-DEL-211
chr4-20141773-DEL-176
chr4-20196590-DEL-56
chr4-20417799-DEL-50
chr4-20575349-DEL-498
chr4-20671346-DEL-62
chr4-20757036-DEL-526
chr4-20964327-DEL-874
chr4-21003017-DEL-53
chr4-21050397-DEL-53
chr4-21050974-DEL-182
chr4-21126770-DEL-8595
chr4-21153524-DEL-3968
chr4-21287187-DEL-325
chr4-21362983-DEL-155
chr4-21363196-DEL-406
chr4-21367754-DEL-64
chr4-21408294-DEL-84
chr4-21408805-DEL-184
chr4-21513308-DEL-2776
chr4-21579065-DEL-7820
chr4-21586975-DEL-54
chr4-21588825-DEL-1054
chr4-21592970-DEL-89
chr4-21624664-DEL-292
chr4-21630237-DEL-362
chr4-21634310-DEL-1162
chr4-21665046-DEL-92
chr4-21743681-DEL-113
chr4-21829102-DEL-84
chr4-21925670-DEL-1427
chr4-21937859-DEL-2069

chr4-21964079-DEL-88
chr4-21992223-DEL-135
chr4-22026494-DEL-696
chr4-22026791-DEL-96
chr4-22076688-DEL-720
chr4-22078261-DEL-68
chr4-22340183-DEL-57
chr4-22346785-DEL-50
chr4-22347176-DEL-79
chr4-22472703-DEL-60
chr4-22503156-DEL-104
chr4-22519756-DEL-493
chr4-22556267-DEL-139
chr4-22641021-DEL-94
chr4-22707515-DEL-52
chr4-22889339-DEL-122
chr4-22908644-DEL-774
chr4-22957015-DEL-50
chr4-23067968-DEL-135
chr4-23151508-DEL-93
chr4-23236087-DEL-1462
chr4-23288619-DEL-2762
chr4-23372905-DEL-78
chr4-23392090-DEL-79
chr4-23542677-DEL-93
chr4-23566264-DEL-100
chr4-23582855-DEL-63
chr4-23694799-DEL-119
chr4-23837493-DEL-1251
chr4-23883521-DEL-881
chr4-23884622-DEL-399
chr4-23935984-DEL-108
chr4-23974585-DEL-133
chr4-23981268-DEL-102
chr4-24014667-DEL-182
chr4-24023746-DEL-2727
chr4-24103068-DEL-57
chr4-24193816-DEL-212
chr4-24194488-DEL-212
chr4-24204286-DEL-553
chr4-24258247-DEL-621
chr4-24354970-DEL-69
chr4-24366063-DEL-2224
chr4-24397237-DEL-192
chr4-24410224-DEL-55
chr4-24488529-DEL-77
chr4-24493354-DEL-543
chr4-24548464-DEL-92
chr4-24613671-DEL-51
chr4-24624871-DEL-93
chr4-24666635-DEL-82
chr4-24751630-DEL-65
chr4-24898824-DEL-64
chr4-25009537-DEL-90
chr4-25009871-DEL-83
chr4-25030723-DEL-252
chr4-25084883-DEL-697
chr4-25230536-DEL-690
chr4-25280161-DEL-51
chr4-25306483-DEL-228
chr4-25333373-DEL-66
chr4-25375277-DEL-329
chr4-25438693-DEL-50
chr4-25512924-DEL-1699
chr4-25750125-DEL-94
chr4-25756677-DEL-50
chr4-25801268-DEL-205
chr4-25801282-DEL-80
chr4-25958476-DEL-4441
chr4-26078041-DEL-59
chr4-26128765-DEL-709
chr4-26129165-DEL-1054
chr4-26136361-DEL-165
chr4-26195445-DEL-427
chr4-26287271-DEL-66
chr4-26410865-DEL-1062
chr4-26444321-DEL-458

chr4-26508816-DEL-1247
chr4-26601147-DEL-218
chr4-26624433-DEL-386
chr4-26671830-DEL-586
chr4-26675041-DEL-2103
chr4-26689218-DEL-608
chr4-26695279-DEL-92
chr4-26712163-DEL-231
chr4-26712454-DEL-199
chr4-26718402-DEL-117
chr4-26776623-DEL-50
chr4-26875791-DEL-1145
chr4-26954398-DEL-2426
chr4-27029825-DEL-6104
chr4-27087759-DEL-52
chr4-27096323-DEL-2058
chr4-27104719-DEL-590
chr4-27155822-DEL-92
chr4-27283797-DEL-61
chr4-27299529-DEL-50
chr4-27405807-DEL-757
chr4-27423574-DEL-120
chr4-27436760-DEL-80
chr4-27551888-DEL-750
chr4-27791217-DEL-75
chr4-27861177-DEL-243
chr4-27877198-DEL-998
chr4-27996920-DEL-2076
chr4-28033121-DEL-5266
chr4-28061786-DEL-1281
chr4-28163699-DEL-105
chr4-28198313-DEL-134
chr4-28207512-DEL-116
chr4-28208341-DEL-135
chr4-28282634-DEL-88
chr4-28285065-DEL-272
chr4-28395266-DEL-50
chr4-28446583-DEL-215
chr4-28485626-DEL-60
chr4-28493942-DEL-397
chr4-28567163-DEL-381
chr4-28590715-DEL-232
chr4-28592641-DEL-367
chr4-28765670-DEL-2926
chr4-28878668-DEL-70
chr4-28888510-DEL-84
chr4-28954286-DEL-52
chr4-28995140-DEL-103
chr4-28995470-DEL-213
chr4-29041835-DEL-245
chr4-29095883-DEL-156
chr4-29102329-DEL-62
chr4-29166684-DEL-50
chr4-29233704-DEL-1494
chr4-29336937-DEL-759
chr4-29405182-DEL-577
chr4-29516996-DEL-52
chr4-29588926-DEL-64
chr4-29626146-DEL-52
chr4-29751793-DEL-199
chr4-29765049-DEL-50
chr4-29831156-DEL-105
chr4-29834719-DEL-76
chr4-29906091-DEL-61
chr4-29913515-DEL-850
chr4-29952044-DEL-56
chr4-30114272-DEL-56
chr4-30170080-DEL-204
chr4-30296216-DEL-255
chr4-30333849-DEL-583
chr4-30419713-DEL-245
chr4-30479370-DEL-224
chr4-30506088-DEL-1083
chr4-30532985-DEL-5402
chr4-30580132-DEL-1582
chr4-30631640-DEL-2372
chr4-30637665-DEL-114

chr4-30657376-DEL-186
chr4-30746590-DEL-75
chr4-30772540-DEL-70
chr4-30886722-DEL-1285
chr4-30937885-DEL-66
chr4-30982832-DEL-79
chr4-31124960-DEL-3696
chr4-31199533-DEL-50
chr4-31230107-DEL-172
chr4-31254681-DEL-849
chr4-31303560-DEL-165
chr4-31347316-DEL-1218
chr4-31414911-DEL-2374
chr4-31470322-DEL-1592
chr4-31483339-DEL-58
chr4-31527783-DEL-208
chr4-31567525-DEL-198
chr4-31597321-DEL-2663
chr4-31599307-DEL-75
chr4-31629699-DEL-53
chr4-31692935-DEL-60
chr4-31772196-DEL-180
chr4-31828175-DEL-169
chr4-31831323-DEL-169
chr4-31878324-DEL-56
chr4-31986986-DEL-60
chr4-32049677-DEL-50
chr4-32082643-DEL-50
chr4-32088631-DEL-839
chr4-32179406-DEL-158
chr4-32185502-DEL-251
chr4-32185624-DEL-63
chr4-32232236-DEL-680
chr4-32255666-DEL-54
chr4-32294803-DEL-189
chr4-32314974-DEL-77
chr4-32341518-DEL-172
chr4-32395470-DEL-368
chr4-32527179-DEL-135
chr4-32527179-DEL-276
chr4-32590758-DEL-326
chr4-32606147-DEL-71
chr4-32658662-DEL-556
chr4-32744247-DEL-81
chr4-32777449-DEL-67
chr4-32811908-DEL-54
chr4-32825922-DEL-62
chr4-32841715-DEL-254
chr4-32865263-DEL-180
chr4-32902920-DEL-609
chr4-32931638-DEL-287
chr4-32934506-DEL-200
chr4-32945890-DEL-56
chr4-32946137-DEL-231
chr4-32946483-DEL-60
chr4-32969417-DEL-506
chr4-33038940-DEL-75
chr4-33073444-DEL-343
chr4-33077287-DEL-57
chr4-33097185-DEL-82
chr4-33097497-DEL-61
chr4-33199579-DEL-398
chr4-33213525-DEL-50
chr4-33286230-DEL-54
chr4-33349373-DEL-75
chr4-33374515-DEL-488
chr4-33375024-DEL-365
chr4-33540817-DEL-50
chr4-33551759-DEL-85
chr4-33551986-DEL-56
chr4-33648603-DEL-232
chr4-33674958-DEL-102
chr4-33729573-DEL-339
chr4-33753883-DEL-64
chr4-33789946-DEL-178
chr4-33802351-DEL-71
chr4-34228578-DEL-171

chr4-34242530-DEL-153
chr4-34250882-DEL-1009
chr4-34394123-DEL-103
chr4-34465220-DEL-139
chr4-34531125-DEL-55
chr4-34586071-DEL-96
chr4-34649204-DEL-60
chr4-34649547-DEL-136
chr4-34736999-DEL-170
chr4-34856885-DEL-116
chr4-34875864-DEL-207
chr4-34876210-DEL-239
chr4-34938431-DEL-50
chr4-34997170-DEL-190
chr4-35062714-DEL-150
chr4-35132632-DEL-341
chr4-35176459-DEL-80
chr4-35542165-DEL-920
chr4-35657458-DEL-3541
chr4-35662804-DEL-221
chr4-35820483-DEL-80
chr4-35839882-DEL-120
chr4-35848166-DEL-73
chr4-35866810-DEL-416
chr4-36046267-DEL-60
chr4-36133877-DEL-1286
chr4-36184133-DEL-681
chr4-36214654-DEL-79
chr4-36214767-DEL-463
chr4-36392414-DEL-90
chr4-36535655-DEL-60
chr4-36606396-DEL-4999
chr4-36620558-DEL-59
chr4-36833842-DEL-216
chr4-36859840-DEL-714
chr4-36863971-DEL-56
chr4-37027747-DEL-50
chr4-37058707-DEL-94
chr4-37166675-DEL-369
chr4-37172581-DEL-89
chr4-37176175-DEL-488
chr4-37251259-DEL-327
chr4-37315515-DEL-350
chr4-37458213-DEL-50
chr4-37589968-DEL-2356
chr4-37647564-DEL-50
chr4-37805616-DEL-3438
chr4-37842877-DEL-2385
chr4-37846013-DEL-152
chr4-37940964-DEL-998
chr4-38023021-DEL-1494
chr4-38086949-DEL-1303
chr4-38265197-DEL-50
chr4-38270715-DEL-444
chr4-38278031-DEL-52
chr4-38525505-DEL-61
chr4-38597153-DEL-696
chr4-38597775-DEL-222
chr4-38605641-DEL-2084
chr4-38703279-DEL-99
chr4-38705612-DEL-153
chr4-38715616-DEL-71
chr4-38725451-DEL-1825
chr4-38729780-DEL-2015
chr4-38801781-DEL-50
chr4-38884017-DEL-1015
chr4-38963947-DEL-1155
chr4-38966041-DEL-1253
chr4-38985648-DEL-266
chr4-38985679-DEL-95
chr4-38986138-DEL-252
chr4-39031681-DEL-66
chr4-39101927-DEL-53
chr4-39153777-DEL-141
chr4-39222218-DEL-679
chr4-39304992-DEL-349
chr4-39340354-DEL-67

chr4-39347635-DEL-6310
chr4-39442431-DEL-152
chr4-39448064-DEL-430
chr4-39535774-DEL-644
chr4-39548176-DEL-126
chr4-39556512-DEL-1423
chr4-39593758-DEL-108
chr4-39629210-DEL-183
chr4-39688431-DEL-266
chr4-39828427-DEL-57
chr4-39832118-DEL-2155
chr4-39840791-DEL-283
chr4-39914132-DEL-70
chr4-40091755-DEL-58
chr4-40164356-DEL-54
chr4-40169568-DEL-153
chr4-40312986-DEL-271
chr4-40332604-DEL-92
chr4-40356315-DEL-280
chr4-40356495-DEL-100
chr4-40356822-DEL-168
chr4-40385019-DEL-180
chr4-40385134-DEL-54
chr4-40414576-DEL-18014
chr4-40499738-DEL-67
chr4-40502998-DEL-1093
chr4-40637884-DEL-138
chr4-40650418-DEL-51
chr4-40677572-DEL-52
chr4-40677842-DEL-199
chr4-40685497-DEL-441
chr4-40744452-DEL-225
chr4-40766524-DEL-153
chr4-40799208-DEL-98
chr4-40933873-DEL-69
chr4-40935703-DEL-57
chr4-41058752-DEL-50
chr4-41082375-DEL-442
chr4-41246083-DEL-92
chr4-41323571-DEL-202
chr4-41326301-DEL-113
chr4-41376395-DEL-70
chr4-41437904-DEL-451
chr4-41482443-DEL-370
chr4-41496018-DEL-98
chr4-41588622-DEL-188
chr4-41592871-DEL-588
chr4-41593019-DEL-210
chr4-41611396-DEL-139
chr4-41666146-DEL-57
chr4-41743670-DEL-178
chr4-42205782-DEL-58
chr4-42490009-DEL-51
chr4-42790903-DEL-194
chr4-42806858-DEL-804
chr4-42838858-DEL-52
chr4-42914778-DEL-67
chr4-42928036-DEL-50
chr4-42932771-DEL-111
chr4-42936577-DEL-130
chr4-42991338-DEL-1643
chr4-43046499-DEL-111
chr4-43052504-DEL-73
chr4-43080065-DEL-260
chr4-43083079-DEL-2205
chr4-43113833-DEL-67
chr4-43131724-DEL-144
chr4-43211690-DEL-91
chr4-43387879-DEL-542
chr4-43388708-DEL-72
chr4-43433623-DEL-970
chr4-43434680-DEL-601
chr4-43436356-DEL-2191
chr4-43471829-DEL-290
chr4-43481536-DEL-72
chr4-43599404-DEL-126
chr4-43691423-DEL-50

chr4-43708287-DEL-436
chr4-43714497-DEL-104
chr4-43802789-DEL-54
chr4-43804691-DEL-77
chr4-43871295-DEL-63
chr4-43880922-DEL-96
chr4-44063287-DEL-1489
chr4-44329145-DEL-59
chr4-44393836-DEL-112
chr4-44678579-DEL-60
chr4-44738040-DEL-58
chr4-45063917-DEL-80
chr4-45076187-DEL-480
chr4-45081233-DEL-53
chr4-45139817-DEL-99
chr4-45232046-DEL-63
chr4-45232245-DEL-63
chr4-45240954-DEL-63
chr4-45289588-DEL-55
chr4-45311967-DEL-530
chr4-45312108-DEL-118
chr4-45313165-DEL-742
chr4-45365364-DEL-50
chr4-45417857-DEL-93
chr4-45585032-DEL-50
chr4-45594347-DEL-80
chr4-45652138-DEL-54
chr4-45818383-DEL-115
chr4-45926588-DEL-137
chr4-45959540-DEL-70
chr4-45969474-DEL-50
chr4-46029214-DEL-96
chr4-46029262-DEL-262
chr4-46216029-DEL-58
chr4-46219259-DEL-150
chr4-46289324-DEL-172
chr4-46289456-DEL-56
chr4-46428487-DEL-136
chr4-46439206-DEL-1562
chr4-46481882-DEL-108
chr4-46481907-DEL-531
chr4-46482481-DEL-203
chr4-46519531-DEL-52
chr4-46544621-DEL-62
chr4-46557881-DEL-50
chr4-46640651-DEL-480
chr4-46705593-DEL-117
chr4-46723035-DEL-73
chr4-46733187-DEL-58
chr4-46754916-DEL-791
chr4-46773090-DEL-255
chr4-46773379-DEL-288
chr4-46778055-DEL-61
chr4-46861634-DEL-86
chr4-46871299-DEL-50
chr4-47165015-DEL-749
chr4-47180306-DEL-82
chr4-47242544-DEL-91
chr4-47285316-DEL-51
chr4-47313379-DEL-154
chr4-47390845-DEL-122
chr4-47423011-DEL-54
chr4-47430893-DEL-1446
chr4-47905082-DEL-252
chr4-47969059-DEL-733
chr4-47992383-DEL-104
chr4-48035832-DEL-399
chr4-48035910-DEL-60
chr4-48221443-DEL-50
chr4-48521834-DEL-119
chr4-48619084-DEL-50
chr4-48655758-DEL-50
chr4-48686001-DEL-53
chr4-48686147-DEL-66
chr4-48844782-DEL-866
chr4-48850160-DEL-5520
chr4-48972177-DEL-120

chr4-48974731-DEL-2864
chr4-49144400-DEL-56
chr4-49170836-DEL-237
chr4-49191255-DEL-58
chr4-49228669-DEL-3336
chr4-49344820-DEL-71
chr4-49413169-DEL-486
chr4-49423199-DEL-3285
chr4-49466656-DEL-636
chr4-49557384-DEL-70
chr4-49569571-DEL-1156
chr4-49731505-DEL-832
chr4-49813097-DEL-265
chr4-50369575-DEL-248
chr4-50441522-DEL-99
chr4-50856204-DEL-662
chr4-51186402-DEL-219
chr4-51283179-DEL-56
chr4-51385162-DEL-6487
chr4-51389001-DEL-1523
chr4-51489775-DEL-99
chr4-51619307-DEL-206
chr4-51752061-DEL-2714
chr4-51831357-DEL-208
chr4-51860755-DEL-111
chr4-51867587-DEL-2782
chr4-51956901-DEL-4640
chr4-52086131-DEL-636
chr4-52086767-DEL-636
chr4-52087108-DEL-212
chr4-52291745-DEL-924
chr4-52330353-DEL-68
chr4-52544060-DEL-50
chr4-52600973-DEL-69
chr4-52658086-DEL-222
chr4-52768739-DEL-103
chr4-52821216-DEL-379
chr4-52860192-DEL-60
chr4-52933338-DEL-88
chr4-52969589-DEL-54
chr4-52969920-DEL-131
chr4-53058186-DEL-424
chr4-53098104-DEL-247
chr4-53182332-DEL-64
chr4-53201431-DEL-186
chr4-53214927-DEL-271
chr4-53364200-DEL-60
chr4-53375558-DEL-60
chr4-53457476-DEL-124
chr4-53485625-DEL-50
chr4-53601848-DEL-1285
chr4-53668445-DEL-212
chr4-53772176-DEL-2128
chr4-53810958-DEL-6243
chr4-53933979-DEL-57
chr4-53936746-DEL-86
chr4-54035096-DEL-50
chr4-54072273-DEL-56
chr4-54101232-DEL-72
chr4-54114423-DEL-104
chr4-54166731-DEL-88
chr4-54191388-DEL-726
chr4-54249260-DEL-1826
chr4-54364303-DEL-60
chr4-54440996-DEL-339
chr4-54626283-DEL-129
chr4-54767922-DEL-55
chr4-54927759-DEL-120
chr4-54928049-DEL-259
chr4-54933674-DEL-1120
chr4-55107068-DEL-71
chr4-55159716-DEL-54
chr4-55171633-DEL-50
chr4-55268252-DEL-100
chr4-55430957-DEL-52
chr4-55459028-DEL-52
chr4-55688925-DEL-78

chr4-55787996-DEL-355
chr4-55788398-DEL-58
chr4-55792775-DEL-72
chr4-55801127-DEL-54
chr4-55829260-DEL-1146
chr4-55891967-DEL-55
chr4-55999510-DEL-79
chr4-56054572-DEL-56
chr4-56124341-DEL-50
chr4-56173513-DEL-71
chr4-56201650-DEL-78
chr4-56293717-DEL-90
chr4-56356286-DEL-482
chr4-56428889-DEL-118
chr4-56447778-DEL-51
chr4-56579953-DEL-74
chr4-56736066-DEL-110
chr4-56737509-DEL-85
chr4-56855493-DEL-82
chr4-56919477-DEL-189
chr4-56933869-DEL-50
chr4-56970582-DEL-162
chr4-57124167-DEL-68
chr4-57199230-DEL-50
chr4-57248119-DEL-65
chr4-57340491-DEL-77
chr4-57384212-DEL-106
chr4-57410646-DEL-78
chr4-57541733-DEL-56
chr4-57570365-DEL-8298
chr4-57605198-DEL-208
chr4-57623555-DEL-238
chr4-57623793-DEL-90
chr4-57767510-DEL-82
chr4-57791743-DEL-78
chr4-57811475-DEL-103
chr4-57870121-DEL-210
chr4-57964703-DEL-393
chr4-58026093-DEL-114
chr4-58058894-DEL-550
chr4-58082093-DEL-83
chr4-58239783-DEL-88
chr4-58322767-DEL-109
chr4-58412855-DEL-412
chr4-58421867-DEL-247
chr4-58430633-DEL-53
chr4-58490518-DEL-51
chr4-58579855-DEL-110
chr4-58600058-DEL-135
chr4-58654790-DEL-350
chr4-58676319-DEL-622
chr4-58717269-DEL-136
chr4-58778900-DEL-50
chr4-58866849-DEL-164
chr4-58871119-DEL-172
chr4-58951574-DEL-172
chr4-59000897-DEL-838
chr4-59020766-DEL-153
chr4-59064222-DEL-54
chr4-59133836-DEL-74
chr4-59232122-DEL-57
chr4-59300760-DEL-214
chr4-59303147-DEL-4432
chr4-59406206-DEL-82
chr4-59516663-DEL-486
chr4-59516768-DEL-176
chr4-59517079-DEL-173
chr4-59713973-DEL-3355
chr4-59740627-DEL-372
chr4-59972167-DEL-279
chr4-60071226-DEL-175
chr4-60181952-DEL-50
chr4-60698188-DEL-175
chr4-61549218-DEL-209
chr4-61554083-DEL-100
chr4-61660903-DEL-1318
chr4-61738551-DEL-94

chr4-61921500-DEL-3177
chr4-62171539-DEL-54
chr4-62298735-DEL-84
chr4-62377802-DEL-294
chr4-62631766-DEL-2060
chr4-62757864-DEL-56
chr4-63177657-DEL-98
chr4-63250620-DEL-147
chr4-63507291-DEL-73
chr4-63520123-DEL-182
chr4-63520526-DEL-116
chr4-63538507-DEL-78
chr4-63549724-DEL-50
chr4-63581820-DEL-57
chr4-63582052-DEL-569
chr4-63582145-DEL-114
chr4-63624298-DEL-55
chr4-63627713-DEL-149
chr4-64021836-DEL-52
chr4-64074741-DEL-570
chr4-64131974-DEL-991
chr4-64189450-DEL-50
chr4-64268923-DEL-126
chr4-64305575-DEL-124
chr4-64447986-DEL-140
chr4-64507303-DEL-322
chr4-64522835-DEL-50
chr4-64714719-DEL-1079
chr4-64800773-DEL-60
chr4-64805609-DEL-710
chr4-64944156-DEL-176
chr4-65059662-DEL-161
chr4-65257990-DEL-103
chr4-65258070-DEL-423
chr4-65258381-DEL-85
chr4-65517853-DEL-185
chr4-65578044-DEL-50
chr4-65693039-DEL-89
chr4-65726185-DEL-623
chr4-65840992-DEL-63
chr4-65873235-DEL-3487
chr4-66350655-DEL-64
chr4-66407504-DEL-4857
chr4-66457942-DEL-50
chr4-66506253-DEL-50
chr4-66820608-DEL-5089
chr4-66877959-DEL-64
chr4-66890906-DEL-617
chr4-67058410-DEL-66
chr4-67203954-DEL-52
chr4-67274371-DEL-138
chr4-67607469-DEL-187
chr4-67664481-DEL-78
chr4-67733981-DEL-853
chr4-67746206-DEL-50
chr4-67834247-DEL-212
chr4-67873963-DEL-72
chr4-68023261-DEL-115
chr4-68029566-DEL-53
chr4-68143502-DEL-214
chr4-68252071-DEL-66
chr4-68314456-DEL-51
chr4-68344583-DEL-64
chr4-68497440-DEL-68
chr4-68711551-DEL-490
chr4-68746051-DEL-60
chr4-68825551-DEL-62
chr4-68869685-DEL-107
chr4-68932780-DEL-420
chr4-69095733-DEL-60
chr4-69146638-DEL-62
chr4-69195850-DEL-128
chr4-69277960-DEL-112
chr4-69373115-DEL-50
chr4-69573402-DEL-77
chr4-69586939-DEL-653
chr4-69841801-DEL-51

chr4-69986736-DEL-402
chr4-70252086-DEL-168
chr4-70513222-DEL-287
chr4-70531400-DEL-1037
chr4-70556783-DEL-58
chr4-70569823-DEL-1886
chr4-70571899-DEL-1247
chr4-70598089-DEL-332
chr4-70663122-DEL-83
chr4-70770197-DEL-209
chr4-70770490-DEL-64
chr4-70877065-DEL-145
chr4-70910028-DEL-17146
chr4-70998908-DEL-1258
chr4-71012451-DEL-60
chr4-71281432-DEL-112
chr4-71383094-DEL-172
chr4-71588132-DEL-52
chr4-71733219-DEL-4192
chr4-71761068-DEL-7109
chr4-71927323-DEL-1886
chr4-71944880-DEL-126
chr4-72429046-DEL-306
chr4-72831613-DEL-50
chr4-73062588-DEL-66
chr4-73181359-DEL-132
chr4-73188689-DEL-416
chr4-73283662-DEL-94
chr4-73328439-DEL-84
chr4-73551558-DEL-60557
chr4-73631154-DEL-199
chr4-73816823-DEL-36464
chr4-73964337-DEL-54
chr4-73993045-DEL-6516
chr4-74043893-DEL-161
chr4-74057725-DEL-256
chr4-74139787-DEL-184
chr4-74263920-DEL-612
chr4-74284456-DEL-64
chr4-74302738-DEL-50
chr4-74325393-DEL-50
chr4-74362456-DEL-50
chr4-74391414-DEL-62
chr4-74406152-DEL-1995
chr4-74426773-DEL-56
chr4-74547491-DEL-2121
chr4-74564187-DEL-9412
chr4-74733082-DEL-50
chr4-74745292-DEL-9155
chr4-75018840-DEL-2397
chr4-75147391-DEL-265
chr4-75164817-DEL-1116
chr4-75290024-DEL-618
chr4-75290771-DEL-719
chr4-75362906-DEL-60
chr4-75505988-DEL-605
chr4-76024164-DEL-78
chr4-76132140-DEL-208
chr4-76325538-DEL-94
chr4-76390731-DEL-52
chr4-76471770-DEL-51
chr4-76482065-DEL-50
chr4-76521835-DEL-53
chr4-76676185-DEL-1761
chr4-76745856-DEL-68
chr4-76885531-DEL-52
chr4-76912781-DEL-436
chr4-76931466-DEL-222
chr4-77054855-DEL-1222
chr4-77060907-DEL-2109
chr4-77084361-DEL-262
chr4-77187286-DEL-169
chr4-77218820-DEL-1091
chr4-77220143-DEL-883
chr4-77281622-DEL-84
chr4-77295438-DEL-787
chr4-77411334-DEL-96

chr4-77411465-DEL-130
chr4-77662789-DEL-51
chr4-77668790-DEL-2891
chr4-77733191-DEL-224
chr4-77769075-DEL-58
chr4-77886070-DEL-1882
chr4-77919753-DEL-50
chr4-78006862-DEL-237
chr4-78068709-DEL-50
chr4-78146475-DEL-59
chr4-78241250-DEL-139
chr4-78258713-DEL-51
chr4-78274157-DEL-511
chr4-78376065-DEL-73
chr4-78378465-DEL-154
chr4-78473776-DEL-1159
chr4-78506922-DEL-73
chr4-78537820-DEL-109
chr4-78589816-DEL-90
chr4-78654876-DEL-2162
chr4-78778746-DEL-50
chr4-78821559-DEL-917
chr4-78848633-DEL-1109
chr4-78890347-DEL-1089
chr4-79030352-DEL-66
chr4-79237552-DEL-103
chr4-79572967-DEL-50
chr4-79577012-DEL-147
chr4-79577159-DEL-306
chr4-79727767-DEL-122
chr4-79744585-DEL-112
chr4-79909018-DEL-50
chr4-80040926-DEL-76
chr4-80060866-DEL-50
chr4-80102501-DEL-66
chr4-80263783-DEL-68
chr4-80525510-DEL-92
chr4-80602914-DEL-2236
chr4-80605202-DEL-1241
chr4-80711387-DEL-475
chr4-80822632-DEL-1470
chr4-80869214-DEL-53
chr4-81055443-DEL-187
chr4-81055446-DEL-78
chr4-81067564-DEL-419
chr4-81074697-DEL-461
chr4-81247973-DEL-840
chr4-81258348-DEL-1298
chr4-81261746-DEL-3972
chr4-81314748-DEL-201
chr4-81314770-DEL-69
chr4-81318321-DEL-116
chr4-81380068-DEL-823
chr4-81389276-DEL-61
chr4-81403430-DEL-1178
chr4-81444600-DEL-70
chr4-81505625-DEL-146
chr4-81717222-DEL-50
chr4-81733813-DEL-123
chr4-81734677-DEL-88
chr4-81734691-DEL-234
chr4-81794992-DEL-4558
chr4-81814939-DEL-87
chr4-81833929-DEL-54
chr4-81859053-DEL-198
chr4-81869541-DEL-400
chr4-81905270-DEL-57
chr4-81928745-DEL-2449
chr4-81931380-DEL-6135
chr4-81932625-DEL-2430
chr4-82022720-DEL-1133
chr4-82104286-DEL-61
chr4-82272801-DEL-84
chr4-82448213-DEL-50
chr4-82476344-DEL-50
chr4-82571810-DEL-389
chr4-82577622-DEL-58

chr4-82579035-DEL-120
chr4-82579178-DEL-829
chr4-82635864-DEL-50
chr4-82866045-DEL-60
chr4-82951724-DEL-62
chr4-82956082-DEL-2241
chr4-82984547-DEL-208
chr4-82993203-DEL-828
chr4-82994312-DEL-1581
chr4-83132256-DEL-50
chr4-83178615-DEL-50
chr4-83310111-DEL-60
chr4-83319117-DEL-842
chr4-83345574-DEL-117
chr4-83423078-DEL-106
chr4-83506769-DEL-122
chr4-83517342-DEL-50
chr4-83541492-DEL-144
chr4-83619638-DEL-101
chr4-83660025-DEL-90
chr4-83709014-DEL-86
chr4-83759993-DEL-944
chr4-83813175-DEL-57
chr4-84043744-DEL-56
chr4-84143796-DEL-708
chr4-84403751-DEL-286
chr4-84428405-DEL-61
chr4-84657604-DEL-183
chr4-84657734-DEL-69
chr4-84717957-DEL-64
chr4-84782222-DEL-223
chr4-84805318-DEL-500
chr4-84890279-DEL-71
chr4-85140215-DEL-279
chr4-85140524-DEL-705
chr4-85170923-DEL-78
chr4-85201655-DEL-65
chr4-85228581-DEL-64
chr4-85245611-DEL-430
chr4-85333570-DEL-50
chr4-85348849-DEL-395
chr4-85429473-DEL-188
chr4-85456941-DEL-311
chr4-85459032-DEL-86
chr4-85460526-DEL-2556
chr4-85672950-DEL-166
chr4-85677505-DEL-121
chr4-85724780-DEL-137
chr4-85957070-DEL-54
chr4-85993961-DEL-50
chr4-86082021-DEL-1785
chr4-86088164-DEL-538
chr4-86116881-DEL-270
chr4-86191003-DEL-479
chr4-86228281-DEL-71
chr4-86354767-DEL-868
chr4-86361214-DEL-52
chr4-86361526-DEL-321
chr4-86376589-DEL-2980
chr4-86387346-DEL-7429
chr4-86596581-DEL-203
chr4-86596896-DEL-130
chr4-86599292-DEL-66
chr4-86638910-DEL-61
chr4-86651908-DEL-721
chr4-86651913-DEL-62
chr4-86715341-DEL-76
chr4-86748894-DEL-1674
chr4-86778914-DEL-54
chr4-86808300-DEL-742
chr4-86823687-DEL-74
chr4-86844281-DEL-71
chr4-86877306-DEL-119
chr4-86921454-DEL-130
chr4-87101812-DEL-191
chr4-87159536-DEL-121
chr4-87267911-DEL-66

chr4-87284110-DEL-60
chr4-87285117-DEL-222
chr4-87407369-DEL-59
chr4-87532594-DEL-6413
chr4-87564329-DEL-84
chr4-87779709-DEL-54
chr4-87794351-DEL-53
chr4-87808662-DEL-233
chr4-87808842-DEL-113
chr4-87840458-DEL-50
chr4-87916100-DEL-54
chr4-88165902-DEL-330
chr4-88280340-DEL-3546
chr4-88558679-DEL-53
chr4-88560667-DEL-56
chr4-88563966-DEL-252
chr4-88577757-DEL-111
chr4-88616302-DEL-119
chr4-88616721-DEL-76
chr4-88673092-DEL-36682
chr4-88733990-DEL-50
chr4-88754863-DEL-213
chr4-88772555-DEL-1168
chr4-88779927-DEL-778
chr4-88802276-DEL-66
chr4-88822774-DEL-86
chr4-88871203-DEL-53
chr4-88876363-DEL-170
chr4-88876478-DEL-59
chr4-88882920-DEL-108
chr4-88883186-DEL-50
chr4-88892092-DEL-76
chr4-88911571-DEL-61
chr4-88958653-DEL-159
chr4-89103436-DEL-7103
chr4-89125374-DEL-1230
chr4-89156216-DEL-54
chr4-89182935-DEL-137
chr4-89203188-DEL-69
chr4-89203297-DEL-184
chr4-89203596-DEL-127
chr4-89269120-DEL-146
chr4-89305754-DEL-107
chr4-89376529-DEL-74
chr4-89434923-DEL-52
chr4-89463096-DEL-90
chr4-89463330-DEL-57
chr4-89627431-DEL-232
chr4-89678070-DEL-13984
chr4-89762894-DEL-86
chr4-89765826-DEL-3207
chr4-89786083-DEL-72
chr4-89968766-DEL-79
chr4-90006421-DEL-1265
chr4-90109267-DEL-60
chr4-90321893-DEL-93
chr4-90715713-DEL-64
chr4-90846996-DEL-60
chr4-90867670-DEL-424
chr4-90897333-DEL-244
chr4-91051889-DEL-65
chr4-91073264-DEL-1950
chr4-91133517-DEL-57
chr4-91247671-DEL-192
chr4-91280019-DEL-347
chr4-91327830-DEL-232
chr4-91835768-DEL-1648
chr4-91860663-DEL-7091
chr4-92067695-DEL-2163
chr4-92241833-DEL-85
chr4-92290171-DEL-51
chr4-92385655-DEL-1283
chr4-92498954-DEL-462
chr4-92516640-DEL-1767
chr4-93068169-DEL-70
chr4-93068195-DEL-154
chr4-93086312-DEL-50

chr4-93122801-DEL-69
chr4-93135406-DEL-50
chr4-93298147-DEL-1128
chr4-93535083-DEL-66
chr4-93635066-DEL-1888
chr4-93723961-DEL-60
chr4-93733532-DEL-62
chr4-93886083-DEL-51
chr4-94245886-DEL-1472
chr4-94338511-DEL-56
chr4-94370878-DEL-566
chr4-94559533-DEL-1533
chr4-94612975-DEL-56
chr4-94764886-DEL-1218
chr4-94993890-DEL-82
chr4-95047796-DEL-80
chr4-95099164-DEL-51
chr4-95187315-DEL-68
chr4-95314678-DEL-52
chr4-95384908-DEL-622
chr4-95561946-DEL-86
chr4-95706656-DEL-81
chr4-95706658-DEL-402
chr4-95706980-DEL-192
chr4-95776865-DEL-1636
chr4-95841293-DEL-81
chr4-95992461-DEL-1487
chr4-96009956-DEL-81
chr4-96010143-DEL-69
chr4-96122274-DEL-52
chr4-96137320-DEL-55
chr4-96179543-DEL-1567
chr4-96238666-DEL-54
chr4-96548871-DEL-54
chr4-96905708-DEL-54
chr4-96962108-DEL-2130
chr4-97299396-DEL-747
chr4-97351249-DEL-813
chr4-97392434-DEL-117
chr4-97680898-DEL-58
chr4-97847140-DEL-50
chr4-97926404-DEL-124
chr4-97926499-DEL-328
chr4-97977158-DEL-128
chr4-97979396-DEL-50
chr4-98041762-DEL-60
chr4-98067994-DEL-53
chr4-98108046-DEL-54
chr4-98130979-DEL-78
chr4-98197082-DEL-4540
chr4-98223001-DEL-100
chr4-98230274-DEL-152
chr4-98230274-DEL-342
chr4-98232623-DEL-512
chr4-98244395-DEL-82
chr4-98294722-DEL-167
chr4-98303579-DEL-219
chr4-98472912-DEL-76
chr4-98513223-DEL-90
chr4-98559598-DEL-52
chr4-98597237-DEL-61
chr4-98603737-DEL-497
chr4-98603758-DEL-229
chr4-98714579-DEL-56
chr4-98752728-DEL-73
chr4-98754175-DEL-50
chr4-98762932-DEL-267
chr4-98810823-DEL-82
chr4-98845351-DEL-195
chr4-98866565-DEL-127
chr4-98867048-DEL-977
chr4-98937796-DEL-76
chr4-98986209-DEL-85
chr4-99028403-DEL-133
chr4-99048211-DEL-126
chr4-99089240-DEL-247
chr4-99089415-DEL-52

chr4-99136684-DEL-210
chr4-99142816-DEL-57
chr4-99142837-DEL-228
chr4-99389394-DEL-60
chr4-99415584-DEL-67
chr4-99420662-DEL-96
chr4-99420662-DEL-270
chr4-99444198-DEL-251
chr4-99452489-DEL-63
chr4-99467778-DEL-254
chr4-99507858-DEL-132
chr4-99518874-DEL-116
chr4-99520575-DEL-80
chr4-99559285-DEL-181
chr4-99591841-DEL-584
chr4-99592218-DEL-110
chr4-99694612-DEL-4196
chr4-99733939-DEL-438
chr4-99892617-DEL-141
chr4-99968225-DEL-56
chr4-99980583-DEL-54
chr4-100010553-DEL-229
chr4-100038212-DEL-51
chr4-100038522-DEL-64
chr4-100052002-DEL-50
chr4-100052187-DEL-169
chr4-100096502-DEL-58
chr4-100167451-DEL-67
chr4-100170986-DEL-375
chr4-100284707-DEL-123
chr4-100313415-DEL-5255
chr4-100337377-DEL-118
chr4-100395641-DEL-220
chr4-100395746-DEL-54
chr4-100481493-DEL-364
chr4-100481630-DEL-156
chr4-100526331-DEL-81
chr4-100553156-DEL-1528
chr4-100573244-DEL-52
chr4-100643107-DEL-55
chr4-100752953-DEL-1447
chr4-100797221-DEL-118
chr4-100887025-DEL-50
chr4-100889179-DEL-99
chr4-100901586-DEL-144
chr4-100922010-DEL-1519
chr4-100931878-DEL-171
chr4-100933024-DEL-103
chr4-101105767-DEL-280
chr4-101129312-DEL-96
chr4-101129408-DEL-288
chr4-101170345-DEL-1364
chr4-101180595-DEL-62
chr4-101189296-DEL-82
chr4-101325707-DEL-1609
chr4-101338578-DEL-66
chr4-101402092-DEL-65
chr4-101416839-DEL-251
chr4-101453502-DEL-112
chr4-101453726-DEL-56
chr4-101453737-DEL-168
chr4-101457982-DEL-652
chr4-101677289-DEL-197
chr4-101697304-DEL-135
chr4-102033008-DEL-53
chr4-102054060-DEL-112
chr4-102054265-DEL-56
chr4-102138865-DEL-103
chr4-102236571-DEL-54
chr4-102237951-DEL-135
chr4-102241295-DEL-55
chr4-102323331-DEL-96
chr4-102351179-DEL-845
chr4-102522145-DEL-120
chr4-102578556-DEL-50
chr4-102590943-DEL-63
chr4-102687401-DEL-924

chr4-102722627-DEL-70
chr4-102817567-DEL-600
chr4-102876032-DEL-6556
chr4-102885149-DEL-1499
chr4-102915778-DEL-66
chr4-102945645-DEL-391
chr4-103030048-DEL-50
chr4-103060001-DEL-245
chr4-103128371-DEL-733
chr4-103180842-DEL-560
chr4-103230915-DEL-6341
chr4-103368728-DEL-70
chr4-103495692-DEL-50
chr4-103613785-DEL-57
chr4-104193101-DEL-56
chr4-104239683-DEL-204
chr4-104311018-DEL-405
chr4-104529019-DEL-62
chr4-104617265-DEL-115
chr4-104712232-DEL-166
chr4-104730988-DEL-50
chr4-104737898-DEL-154
chr4-104783474-DEL-382
chr4-104838199-DEL-76
chr4-104861913-DEL-54
chr4-104879667-DEL-533
chr4-104884149-DEL-57
chr4-104894533-DEL-105
chr4-104939925-DEL-57
chr4-104944756-DEL-127
chr4-104951238-DEL-61
chr4-105200851-DEL-59
chr4-105200851-DEL-236
chr4-105532204-DEL-3498
chr4-105535732-DEL-240
chr4-105583297-DEL-84
chr4-105592680-DEL-424
chr4-105703715-DEL-70
chr4-105910376-DEL-50
chr4-105989466-DEL-82
chr4-106034464-DEL-201
chr4-106108600-DEL-50
chr4-106115751-DEL-66
chr4-106235936-DEL-60
chr4-106317290-DEL-91
chr4-106322561-DEL-72
chr4-106458825-DEL-58
chr4-106538577-DEL-106
chr4-106542299-DEL-1558
chr4-106547057-DEL-117
chr4-106598615-DEL-50
chr4-106616026-DEL-193
chr4-106625412-DEL-108
chr4-106682965-DEL-86
chr4-106692947-DEL-138
chr4-106692987-DEL-58
chr4-106701921-DEL-584
chr4-106733706-DEL-158
chr4-106735792-DEL-143
chr4-106735988-DEL-457
chr4-106966167-DEL-276
chr4-107062928-DEL-55
chr4-107085046-DEL-3958
chr4-107153040-DEL-160
chr4-107158431-DEL-68
chr4-107215127-DEL-337
chr4-107262044-DEL-135
chr4-107325560-DEL-105
chr4-107403472-DEL-50
chr4-107456067-DEL-972
chr4-107487761-DEL-573
chr4-107511182-DEL-50
chr4-107538192-DEL-1633
chr4-107546137-DEL-60
chr4-107682606-DEL-2690
chr4-107824832-DEL-65
chr4-107942813-DEL-54

chr4-107960265-DEL-1652
chr4-108057669-DEL-50
chr4-108126920-DEL-132
chr4-108181200-DEL-84
chr4-108280873-DEL-168
chr4-108307677-DEL-75
chr4-108351244-DEL-50
chr4-108411761-DEL-343
chr4-108411785-DEL-83
chr4-108435227-DEL-148
chr4-108435452-DEL-366
chr4-108518791-DEL-50
chr4-108650514-DEL-182
chr4-108744633-DEL-64
chr4-108803424-DEL-126
chr4-108808604-DEL-117
chr4-108824449-DEL-51
chr4-108861229-DEL-63
chr4-109017039-DEL-66
chr4-109157267-DEL-800
chr4-109260481-DEL-416
chr4-109274186-DEL-679
chr4-109298297-DEL-94
chr4-109395471-DEL-6245
chr4-109544993-DEL-93
chr4-109675260-DEL-60
chr4-109858393-DEL-102
chr4-109967400-DEL-60
chr4-110000761-DEL-136
chr4-110075165-DEL-1069
chr4-110155279-DEL-152
chr4-110240939-DEL-62
chr4-110393701-DEL-127
chr4-110408841-DEL-205
chr4-110659357-DEL-74
chr4-110787485-DEL-1692
chr4-110816861-DEL-200
chr4-110950317-DEL-1909
chr4-110959438-DEL-1890
chr4-111050653-DEL-381
chr4-111080362-DEL-90
chr4-111316345-DEL-54
chr4-111318981-DEL-85
chr4-111455672-DEL-53
chr4-111545882-DEL-50
chr4-111649855-DEL-1100
chr4-111847966-DEL-1544
chr4-112556359-DEL-114
chr4-112639144-DEL-1427
chr4-112649319-DEL-124
chr4-113488935-DEL-263
chr4-113501561-DEL-58
chr4-113517236-DEL-300
chr4-113518257-DEL-116
chr4-113526160-DEL-91
chr4-113530910-DEL-95
chr4-113546699-DEL-52
chr4-113568927-DEL-481
chr4-113580302-DEL-132
chr4-113583186-DEL-2735
chr4-113627649-DEL-68
chr4-113776018-DEL-4047
chr4-113798653-DEL-50
chr4-113831330-DEL-104
chr4-113899188-DEL-1100
chr4-113912016-DEL-204
chr4-114002156-DEL-141
chr4-114021486-DEL-50
chr4-114205704-DEL-339
chr4-114331577-DEL-623
chr4-114593541-DEL-140
chr4-114778390-DEL-92
chr4-114813046-DEL-179
chr4-114965352-DEL-56
chr4-115046176-DEL-1493
chr4-115113402-DEL-1757
chr4-115125982-DEL-1191

chr4-115167104-DEL-1981
chr4-115193443-DEL-106514
chr4-115282314-DEL-6194
chr4-115308744-DEL-2633
chr4-115507516-DEL-60
chr4-115523654-DEL-4417
chr4-115538346-DEL-2259
chr4-115617582-DEL-50
chr4-115705869-DEL-708
chr4-115902116-DEL-480
chr4-115902216-DEL-992
chr4-116005620-DEL-158
chr4-116025141-DEL-391
chr4-116093955-DEL-197
chr4-116155364-DEL-96
chr4-116155477-DEL-384
chr4-116283957-DEL-103
chr4-116373793-DEL-3241
chr4-116375389-DEL-169
chr4-116394908-DEL-50
chr4-116476953-DEL-153
chr4-116523052-DEL-1371
chr4-116539272-DEL-72
chr4-116559496-DEL-102
chr4-116570835-DEL-64
chr4-116653268-DEL-110
chr4-116867124-DEL-143
chr4-117256571-DEL-64
chr4-117281959-DEL-122
chr4-117305624-DEL-68
chr4-117366184-DEL-60
chr4-117366783-DEL-70
chr4-117401335-DEL-53
chr4-117564367-DEL-462
chr4-117565049-DEL-84
chr4-117565259-DEL-84
chr4-117654480-DEL-163
chr4-117764346-DEL-189
chr4-117829430-DEL-56
chr4-117829559-DEL-125
chr4-117870007-DEL-53
chr4-117932098-DEL-214
chr4-117932427-DEL-53
chr4-117959696-DEL-74
chr4-118055737-DEL-62
chr4-118110136-DEL-124
chr4-118157631-DEL-54
chr4-118228810-DEL-140
chr4-118260877-DEL-93
chr4-118261180-DEL-364
chr4-118261726-DEL-120
chr4-118279639-DEL-190
chr4-118308707-DEL-72
chr4-118404564-DEL-1369
chr4-118412967-DEL-719
chr4-118412999-DEL-319
chr4-118444140-DEL-112
chr4-118479582-DEL-186
chr4-118500037-DEL-335
chr4-118540820-DEL-5545
chr4-118652410-DEL-196
chr4-118674438-DEL-50
chr4-118690503-DEL-66
chr4-118730707-DEL-531
chr4-118743328-DEL-69
chr4-118759111-DEL-56
chr4-118799119-DEL-1529
chr4-118832583-DEL-250
chr4-119068339-DEL-55
chr4-119077338-DEL-178
chr4-119077481-DEL-88
chr4-119213535-DEL-158
chr4-119226241-DEL-171
chr4-119246816-DEL-174
chr4-119252505-DEL-202
chr4-119343403-DEL-262
chr4-119350345-DEL-177

chr4-119390659-DEL-111
chr4-119405191-DEL-362
chr4-119571014-DEL-58
chr4-119698569-DEL-386
chr4-119748122-DEL-74
chr4-119858886-DEL-59
chr4-119886935-DEL-82
chr4-119898125-DEL-131
chr4-120004248-DEL-154
chr4-120077503-DEL-254
chr4-120077755-DEL-76
chr4-120113625-DEL-56
chr4-120257827-DEL-937
chr4-120388426-DEL-127
chr4-120502406-DEL-232
chr4-120502435-DEL-105
chr4-120534503-DEL-340
chr4-120721731-DEL-261
chr4-120756924-DEL-104
chr4-120804596-DEL-78
chr4-120837095-DEL-135
chr4-120977314-DEL-177
chr4-121077241-DEL-141
chr4-121192988-DEL-95
chr4-121292023-DEL-72
chr4-121324873-DEL-198
chr4-121374141-DEL-64
chr4-121562736-DEL-2295
chr4-121571761-DEL-1154
chr4-121608819-DEL-50
chr4-121832356-DEL-55
chr4-122014750-DEL-109
chr4-122407794-DEL-55
chr4-122444381-DEL-121
chr4-122478283-DEL-3760
chr4-122483507-DEL-688
chr4-122495355-DEL-130
chr4-122508540-DEL-237
chr4-122644627-DEL-64
chr4-122689102-DEL-1783
chr4-122721297-DEL-60
chr4-122818791-DEL-70
chr4-122818948-DEL-133
chr4-122819243-DEL-70
chr4-122902385-DEL-58
chr4-122914901-DEL-307
chr4-122972617-DEL-144
chr4-123032392-DEL-60
chr4-123163360-DEL-942
chr4-123208995-DEL-158
chr4-123233462-DEL-69
chr4-123233479-DEL-166
chr4-123233699-DEL-68
chr4-123267526-DEL-149
chr4-123339262-DEL-52
chr4-123434496-DEL-290
chr4-123548140-DEL-153
chr4-123596964-DEL-6851
chr4-123603845-DEL-322
chr4-123614020-DEL-126
chr4-123632813-DEL-50
chr4-123647301-DEL-360
chr4-123649427-DEL-224
chr4-123660274-DEL-315
chr4-123660631-DEL-66
chr4-123756410-DEL-164
chr4-123914296-DEL-85
chr4-123969245-DEL-60
chr4-124005437-DEL-58
chr4-124017923-DEL-50
chr4-124046239-DEL-109
chr4-124049077-DEL-137
chr4-124049106-DEL-276
chr4-124049536-DEL-84
chr4-124049554-DEL-253
chr4-124169963-DEL-88
chr4-124217238-DEL-93

chr4-124230344-DEL-51
chr4-124264110-DEL-56
chr4-124354485-DEL-286
chr4-124354607-DEL-132
chr4-124443193-DEL-60
chr4-124460796-DEL-375
chr4-124719226-DEL-52
chr4-124795063-DEL-124
chr4-124802462-DEL-78
chr4-124885509-DEL-50
chr4-124919782-DEL-145
chr4-124929761-DEL-464
chr4-124930089-DEL-60
chr4-125041009-DEL-99
chr4-125075879-DEL-67
chr4-125103715-DEL-196
chr4-125193108-DEL-50
chr4-125228346-DEL-630
chr4-125283619-DEL-162
chr4-125299052-DEL-69
chr4-125511308-DEL-50
chr4-125565431-DEL-118
chr4-125587296-DEL-125
chr4-125831921-DEL-229
chr4-125862861-DEL-72
chr4-125870661-DEL-76
chr4-125889109-DEL-83
chr4-125989818-DEL-206
chr4-126049614-DEL-70
chr4-126083570-DEL-248
chr4-126168382-DEL-253
chr4-126193394-DEL-172
chr4-126308075-DEL-176
chr4-126308099-DEL-400
chr4-126308407-DEL-152
chr4-126345736-DEL-57
chr4-126369945-DEL-1140
chr4-126407040-DEL-299
chr4-126407061-DEL-143
chr4-126414480-DEL-94
chr4-126482746-DEL-84
chr4-126567646-DEL-120
chr4-126657071-DEL-61
chr4-126698588-DEL-50
chr4-126720841-DEL-104
chr4-126872235-DEL-69
chr4-127026600-DEL-50
chr4-127052230-DEL-192
chr4-127102370-DEL-101
chr4-127271556-DEL-66
chr4-127375453-DEL-96
chr4-127431940-DEL-84
chr4-127475566-DEL-90
chr4-127597605-DEL-52
chr4-127963170-DEL-64
chr4-128002499-DEL-408
chr4-128002521-DEL-85
chr4-128060852-DEL-115
chr4-128065873-DEL-70
chr4-128066056-DEL-79
chr4-128086120-DEL-79
chr4-128115361-DEL-237
chr4-128132542-DEL-54
chr4-128159909-DEL-680
chr4-128204425-DEL-62
chr4-128251685-DEL-52
chr4-128279759-DEL-314
chr4-128279842-DEL-149
chr4-128348862-DEL-53
chr4-128524010-DEL-120
chr4-128758973-DEL-63
chr4-128759417-DEL-185
chr4-128799613-DEL-90
chr4-128846828-DEL-50
chr4-128892168-DEL-89
chr4-128922967-DEL-1031
chr4-128923009-DEL-384

chr4-128923743-DEL-393
chr4-128985305-DEL-256
chr4-128993230-DEL-203
chr4-129040902-DEL-219
chr4-129092877-DEL-53
chr4-129170193-DEL-78
chr4-129414464-DEL-59
chr4-129422939-DEL-205
chr4-129423169-DEL-91
chr4-129423429-DEL-155
chr4-129436945-DEL-1034
chr4-129454792-DEL-196
chr4-129454898-DEL-88
chr4-129496644-DEL-2000
chr4-129561722-DEL-195
chr4-129584359-DEL-52
chr4-129694274-DEL-58
chr4-129805087-DEL-66
chr4-129810457-DEL-50
chr4-129810599-DEL-140
chr4-129851427-DEL-354
chr4-129896800-DEL-152
chr4-129926467-DEL-72
chr4-129928815-DEL-917
chr4-129965215-DEL-226
chr4-129965266-DEL-58
chr4-130017203-DEL-50
chr4-130017500-DEL-186
chr4-130044492-DEL-61
chr4-130085825-DEL-50
chr4-130173870-DEL-52
chr4-130215224-DEL-126
chr4-130285675-DEL-359
chr4-130285677-DEL-83
chr4-130348429-DEL-316
chr4-130369063-DEL-78
chr4-130511749-DEL-446
chr4-130521156-DEL-53
chr4-130537732-DEL-72
chr4-130577377-DEL-256
chr4-130613979-DEL-131
chr4-130616282-DEL-148
chr4-130616652-DEL-182
chr4-130618382-DEL-86
chr4-130821503-DEL-96
chr4-130826379-DEL-120
chr4-130859288-DEL-171
chr4-130859569-DEL-50
chr4-130889980-DEL-50
chr4-130911886-DEL-92
chr4-130944141-DEL-54
chr4-130949422-DEL-7119
chr4-131074767-DEL-120
chr4-131075316-DEL-54
chr4-131109973-DEL-130
chr4-131111692-DEL-93
chr4-131201669-DEL-50
chr4-131273505-DEL-78
chr4-131531980-DEL-196
chr4-131532302-DEL-65
chr4-131660677-DEL-202
chr4-131857317-DEL-66
chr4-131902912-DEL-60
chr4-132144305-DEL-1627
chr4-132165720-DEL-289
chr4-132166167-DEL-59
chr4-132177663-DEL-201
chr4-132275865-DEL-118
chr4-132339688-DEL-100
chr4-132424411-DEL-70
chr4-132441921-DEL-222
chr4-132486054-DEL-602
chr4-132487224-DEL-166
chr4-132669511-DEL-332
chr4-132681549-DEL-81
chr4-132687431-DEL-78
chr4-132962509-DEL-57

chr4-133011191-DEL-188
chr4-133128672-DEL-50
chr4-133146324-DEL-120
chr4-133382627-DEL-207
chr4-133464321-DEL-52
chr4-133490129-DEL-111
chr4-133503419-DEL-50
chr4-133510134-DEL-52
chr4-133563269-DEL-204
chr4-133563323-DEL-74
chr4-133585994-DEL-178
chr4-133616833-DEL-142
chr4-133663971-DEL-206
chr4-133670957-DEL-189
chr4-133763737-DEL-207
chr4-133764014-DEL-54
chr4-133828722-DEL-166
chr4-133916452-DEL-1146
chr4-133953389-DEL-53
chr4-133953421-DEL-125
chr4-134047958-DEL-52
chr4-134107834-DEL-471
chr4-134141482-DEL-60
chr4-134225968-DEL-132
chr4-134247497-DEL-84
chr4-134312127-DEL-249
chr4-134312393-DEL-50
chr4-134312430-DEL-119
chr4-134314297-DEL-56
chr4-134350844-DEL-67
chr4-134437980-DEL-77
chr4-134556417-DEL-56
chr4-134571737-DEL-156
chr4-134660975-DEL-158
chr4-134665206-DEL-650
chr4-134679142-DEL-76
chr4-134679864-DEL-52
chr4-134697915-DEL-137
chr4-134700366-DEL-315
chr4-134720679-DEL-91
chr4-134932362-DEL-50
chr4-135006311-DEL-188
chr4-135038893-DEL-159
chr4-135060749-DEL-77
chr4-135062524-DEL-89
chr4-135078247-DEL-50
chr4-135134426-DEL-284
chr4-135157653-DEL-126
chr4-135226620-DEL-93
chr4-135239899-DEL-56
chr4-135250326-DEL-110
chr4-135406071-DEL-378
chr4-135511533-DEL-188
chr4-135526675-DEL-51
chr4-135613986-DEL-200
chr4-135707425-DEL-149
chr4-135725421-DEL-1462
chr4-135791567-DEL-107
chr4-135806320-DEL-1451
chr4-135892823-DEL-65
chr4-135990076-DEL-62
chr4-136027393-DEL-195
chr4-136126074-DEL-134
chr4-136159744-DEL-50
chr4-136194139-DEL-128
chr4-136197200-DEL-61
chr4-136357715-DEL-192
chr4-136537073-DEL-180
chr4-136546627-DEL-62
chr4-136596080-DEL-206
chr4-136598651-DEL-129
chr4-136750601-DEL-50
chr4-136774588-DEL-245
chr4-136780738-DEL-60
chr4-136843748-DEL-56
chr4-136875336-DEL-50
chr4-136985781-DEL-107

chr4-136988048-DEL-50
chr4-137071883-DEL-81
chr4-137090658-DEL-401
chr4-137101993-DEL-561
chr4-137116812-DEL-1195
chr4-137118380-DEL-321
chr4-137138106-DEL-56
chr4-137138635-DEL-50
chr4-137142687-DEL-195
chr4-137142775-DEL-67
chr4-137145502-DEL-887
chr4-137153838-DEL-176
chr4-137218558-DEL-417
chr4-137422096-DEL-279
chr4-137621815-DEL-210
chr4-137622150-DEL-201
chr4-137661116-DEL-145
chr4-137661444-DEL-294
chr4-137697362-DEL-8470
chr4-137759724-DEL-50
chr4-137771151-DEL-336
chr4-137784726-DEL-146
chr4-137789287-DEL-173
chr4-137911471-DEL-50
chr4-138029694-DEL-66
chr4-138140843-DEL-276
chr4-138208727-DEL-62
chr4-138230357-DEL-482
chr4-138269233-DEL-85
chr4-138309577-DEL-54
chr4-138388396-DEL-234
chr4-138389986-DEL-108
chr4-138445558-DEL-99
chr4-138516134-DEL-50
chr4-138535528-DEL-50
chr4-138570533-DEL-202
chr4-138587995-DEL-220
chr4-138595406-DEL-66
chr4-138616758-DEL-111
chr4-138624503-DEL-70
chr4-138748633-DEL-60
chr4-138752386-DEL-142
chr4-138752386-DEL-1476
chr4-138753061-DEL-440
chr4-138753598-DEL-230
chr4-138755075-DEL-93
chr4-138865109-DEL-448
chr4-138937222-DEL-233
chr4-139046310-DEL-63
chr4-139046483-DEL-134
chr4-139063799-DEL-710
chr4-139066313-DEL-119
chr4-139123170-DEL-160
chr4-139379505-DEL-52
chr4-139448467-DEL-52
chr4-139456157-DEL-323
chr4-139502078-DEL-130
chr4-139606210-DEL-230
chr4-139606286-DEL-60
chr4-139606388-DEL-182
chr4-139606553-DEL-73
chr4-139638379-DEL-198
chr4-139707873-DEL-54
chr4-139860354-DEL-140
chr4-140044317-DEL-318
chr4-140045767-DEL-54
chr4-140120338-DEL-176
chr4-140144882-DEL-185
chr4-140217169-DEL-84
chr4-140217446-DEL-99
chr4-140272878-DEL-80
chr4-140273257-DEL-200
chr4-140381484-DEL-56
chr4-140459340-DEL-180
chr4-140613038-DEL-69
chr4-140624430-DEL-54
chr4-140650516-DEL-105

chr4-140650523-DEL-266
chr4-140650699-DEL-67
chr4-140736630-DEL-279
chr4-140813688-DEL-209
chr4-141081989-DEL-336
chr4-141082442-DEL-213
chr4-141142742-DEL-50
chr4-141142963-DEL-121
chr4-141146149-DEL-56
chr4-141167270-DEL-748
chr4-141169787-DEL-95
chr4-141232320-DEL-56
chr4-141232488-DEL-199
chr4-141366789-DEL-90
chr4-141388040-DEL-74
chr4-141393617-DEL-67
chr4-141395165-DEL-82
chr4-141416309-DEL-55
chr4-141416688-DEL-152
chr4-141477841-DEL-171
chr4-141477916-DEL-81
chr4-141500698-DEL-232
chr4-141535230-DEL-1619
chr4-141563337-DEL-86
chr4-141726377-DEL-92
chr4-141726455-DEL-291
chr4-141821366-DEL-53
chr4-141925398-DEL-384
chr4-141967923-DEL-79
chr4-141970860-DEL-64
chr4-141970944-DEL-108
chr4-141973169-DEL-80
chr4-141999843-DEL-212
chr4-142000069-DEL-412
chr4-142000364-DEL-165
chr4-142157202-DEL-90
chr4-142362366-DEL-60
chr4-142447383-DEL-86
chr4-142493781-DEL-96
chr4-142525116-DEL-149
chr4-142567910-DEL-92
chr4-142577721-DEL-132
chr4-142584929-DEL-4228
chr4-142596321-DEL-72
chr4-142596665-DEL-206
chr4-142810064-DEL-94
chr4-142948067-DEL-130
chr4-142961197-DEL-57
chr4-143044234-DEL-95
chr4-143057731-DEL-98
chr4-143113927-DEL-116
chr4-143185921-DEL-11848
chr4-143336285-DEL-72
chr4-143336447-DEL-115
chr4-143440060-DEL-729
chr4-143442002-DEL-120
chr4-143463809-DEL-358
chr4-143465204-DEL-112
chr4-143472363-DEL-145
chr4-143480804-DEL-2921
chr4-143491744-DEL-56
chr4-143628071-DEL-139
chr4-143777648-DEL-83588
chr4-143907293-DEL-473
chr4-143983303-DEL-171
chr4-143987371-DEL-562
chr4-144034649-DEL-155
chr4-144061097-DEL-60
chr4-144078942-DEL-150
chr4-144264981-DEL-54
chr4-144426958-DEL-50
chr4-144483364-DEL-88
chr4-144519110-DEL-167
chr4-144561562-DEL-884
chr4-144562093-DEL-1115
chr4-144562185-DEL-182
chr4-144622561-DEL-66

chr4-144650888-DEL-80
chr4-144686740-DEL-177
chr4-144689996-DEL-50
chr4-144733805-DEL-214
chr4-144880676-DEL-54
chr4-145016725-DEL-416
chr4-145017011-DEL-182
chr4-145020623-DEL-107
chr4-145075571-DEL-1291
chr4-145108412-DEL-50
chr4-145141996-DEL-195
chr4-145159222-DEL-155
chr4-145176245-DEL-84
chr4-145237790-DEL-220
chr4-145240654-DEL-123
chr4-145249867-DEL-51
chr4-145303391-DEL-503
chr4-145306342-DEL-72
chr4-145310749-DEL-95
chr4-145332736-DEL-440
chr4-145349663-DEL-756
chr4-145351517-DEL-174
chr4-145369537-DEL-196
chr4-145506787-DEL-94
chr4-145533170-DEL-742
chr4-146057932-DEL-50
chr4-146077282-DEL-327
chr4-146082142-DEL-502
chr4-146092940-DEL-219
chr4-146135768-DEL-52
chr4-146199410-DEL-274
chr4-146204927-DEL-1395
chr4-146208030-DEL-79
chr4-146254069-DEL-84
chr4-146310684-DEL-438
chr4-146464713-DEL-73
chr4-146492653-DEL-97
chr4-146559533-DEL-159
chr4-146565159-DEL-212
chr4-146614572-DEL-70
chr4-147005777-DEL-52
chr4-147006015-DEL-117
chr4-147074815-DEL-169
chr4-147182315-DEL-50
chr4-147190522-DEL-227
chr4-147409009-DEL-159
chr4-147413152-DEL-204
chr4-147436130-DEL-52
chr4-147497247-DEL-142
chr4-147498818-DEL-61
chr4-147583974-DEL-116
chr4-147597124-DEL-671
chr4-147599875-DEL-183
chr4-147601668-DEL-222
chr4-147632040-DEL-550
chr4-147663727-DEL-486
chr4-147769334-DEL-276
chr4-147839303-DEL-336
chr4-147876800-DEL-196
chr4-147877589-DEL-68
chr4-147893877-DEL-171
chr4-147926429-DEL-96
chr4-147928716-DEL-51
chr4-147938635-DEL-64
chr4-147943631-DEL-171
chr4-147943850-DEL-62
chr4-148038975-DEL-227
chr4-148039354-DEL-52
chr4-148049242-DEL-136
chr4-148049311-DEL-61
chr4-148087879-DEL-62
chr4-148087894-DEL-186
chr4-148208018-DEL-50
chr4-148364463-DEL-88
chr4-148417428-DEL-60
chr4-148455484-DEL-141
chr4-148471792-DEL-88

chr4-148487382-DEL-76
chr4-148506863-DEL-62
chr4-148544956-DEL-56
chr4-148545165-DEL-183
chr4-148804706-DEL-104
chr4-148836473-DEL-51
chr4-149094976-DEL-100
chr4-149179854-DEL-93
chr4-149196363-DEL-66
chr4-149196457-DEL-355
chr4-149312511-DEL-230
chr4-149434752-DEL-84
chr4-149464896-DEL-96
chr4-149532627-DEL-89
chr4-149532918-DEL-181
chr4-149575321-DEL-216
chr4-149581610-DEL-116
chr4-149613094-DEL-182
chr4-149778668-DEL-130
chr4-149850682-DEL-83
chr4-149850873-DEL-89
chr4-149853317-DEL-717
chr4-149890876-DEL-123
chr4-149902667-DEL-266
chr4-149920794-DEL-689
chr4-149945315-DEL-124
chr4-149984596-DEL-117
chr4-150002189-DEL-53
chr4-150061007-DEL-88
chr4-150101971-DEL-75
chr4-150108455-DEL-151
chr4-150131387-DEL-75
chr4-150232114-DEL-52
chr4-150246442-DEL-82
chr4-150358276-DEL-214
chr4-150409174-DEL-82
chr4-150503355-DEL-203
chr4-150562927-DEL-53
chr4-150598255-DEL-170
chr4-150598643-DEL-148
chr4-150598853-DEL-586
chr4-150599899-DEL-96
chr4-150725720-DEL-63
chr4-150730480-DEL-190
chr4-150731286-DEL-560
chr4-150752963-DEL-351
chr4-150763370-DEL-119
chr4-150768757-DEL-50
chr4-150778795-DEL-64
chr4-150814123-DEL-97
chr4-150816536-DEL-92
chr4-150848502-DEL-261
chr4-150848718-DEL-58
chr4-150864298-DEL-58
chr4-150926017-DEL-144
chr4-151072315-DEL-463
chr4-151072777-DEL-52
chr4-151256250-DEL-68
chr4-151256263-DEL-204
chr4-151315821-DEL-609
chr4-151316886-DEL-80
chr4-151317598-DEL-1304
chr4-151335836-DEL-64
chr4-151336012-DEL-108
chr4-151345063-DEL-490
chr4-151393307-DEL-344
chr4-151410050-DEL-90
chr4-151433875-DEL-58
chr4-151493531-DEL-51
chr4-151501588-DEL-67
chr4-151515281-DEL-68
chr4-151592255-DEL-106
chr4-151608621-DEL-355
chr4-151608805-DEL-171
chr4-151616532-DEL-176
chr4-151616532-DEL-396
chr4-151706038-DEL-178

chr4-151706207-DEL-483
chr4-151711635-DEL-674
chr4-151714140-DEL-171
chr4-151721650-DEL-72
chr4-151737840-DEL-126
chr4-151815539-DEL-58
chr4-151815849-DEL-88
chr4-151829415-DEL-87
chr4-151888473-DEL-134
chr4-151895796-DEL-59
chr4-151975586-DEL-66
chr4-152006826-DEL-58
chr4-152011975-DEL-222
chr4-152184340-DEL-77
chr4-152288603-DEL-174
chr4-152314933-DEL-76
chr4-152318822-DEL-63
chr4-152331209-DEL-2008
chr4-152454073-DEL-279
chr4-152470704-DEL-77
chr4-152528364-DEL-70
chr4-152548682-DEL-110
chr4-152609555-DEL-114
chr4-152651547-DEL-111
chr4-152703018-DEL-125
chr4-152703062-DEL-55
chr4-152703348-DEL-111
chr4-152703632-DEL-126
chr4-152717592-DEL-52
chr4-152737437-DEL-261
chr4-152737777-DEL-542
chr4-152738159-DEL-160
chr4-152744972-DEL-727
chr4-152898379-DEL-235
chr4-152898527-DEL-78
chr4-152953195-DEL-56
chr4-152978174-DEL-448
chr4-153000093-DEL-74
chr4-153075987-DEL-240
chr4-153076037-DEL-60
chr4-153076417-DEL-342
chr4-153097252-DEL-187
chr4-153097336-DEL-51
chr4-153157841-DEL-57
chr4-153161291-DEL-63
chr4-153172107-DEL-292
chr4-153172651-DEL-56
chr4-153182060-DEL-154
chr4-153220806-DEL-61
chr4-153224738-DEL-342
chr4-153232278-DEL-894
chr4-153237500-DEL-110
chr4-153242536-DEL-51
chr4-153279629-DEL-471
chr4-153288100-DEL-526
chr4-153299164-DEL-114
chr4-153339026-DEL-138
chr4-153344366-DEL-50
chr4-153351888-DEL-50
chr4-153357406-DEL-78
chr4-153422872-DEL-82
chr4-153474508-DEL-221
chr4-153481748-DEL-369
chr4-153593411-DEL-50
chr4-153593436-DEL-200
chr4-153652393-DEL-307
chr4-153758616-DEL-63
chr4-153769855-DEL-222
chr4-153777740-DEL-51
chr4-153892138-DEL-63
chr4-153892192-DEL-253
chr4-153892487-DEL-126
chr4-153973109-DEL-1248
chr4-154052424-DEL-152
chr4-154066592-DEL-85
chr4-154070472-DEL-64
chr4-154084750-DEL-1158

chr4-154085890-DEL-80
chr4-154087340-DEL-90
chr4-154089675-DEL-59
chr4-154116597-DEL-64
chr4-154124665-DEL-83
chr4-154134496-DEL-200
chr4-154134608-DEL-56
chr4-154134850-DEL-56
chr4-154136096-DEL-1201
chr4-154147788-DEL-102
chr4-154160016-DEL-122
chr4-154204532-DEL-86
chr4-154211159-DEL-87
chr4-154233574-DEL-290
chr4-154233812-DEL-125
chr4-154237735-DEL-287
chr4-154267674-DEL-50
chr4-154295098-DEL-133
chr4-154345882-DEL-97
chr4-154462443-DEL-73
chr4-154571291-DEL-94
chr4-154602219-DEL-51
chr4-154602231-DEL-167
chr4-154619970-DEL-53
chr4-154757308-DEL-60
chr4-154764903-DEL-612
chr4-154840720-DEL-68
chr4-154861272-DEL-76
chr4-154922309-DEL-189
chr4-154922567-DEL-108
chr4-154946858-DEL-51
chr4-154957138-DEL-300
chr4-154962620-DEL-1159
chr4-154996726-DEL-72
chr4-155009277-DEL-300
chr4-155019740-DEL-98
chr4-155031367-DEL-236
chr4-155041206-DEL-58
chr4-155297083-DEL-160
chr4-155313520-DEL-54
chr4-155344014-DEL-111
chr4-155359631-DEL-54
chr4-155363948-DEL-250
chr4-155413121-DEL-203
chr4-155434165-DEL-1003
chr4-155447905-DEL-395
chr4-155476787-DEL-88
chr4-155529885-DEL-117
chr4-155529972-DEL-51
chr4-155533675-DEL-519
chr4-155539050-DEL-90
chr4-155540850-DEL-1743
chr4-155563238-DEL-234
chr4-155690072-DEL-733
chr4-155837862-DEL-103
chr4-155880176-DEL-77
chr4-155880183-DEL-1270
chr4-155923003-DEL-101
chr4-155962514-DEL-137
chr4-156013446-DEL-1773
chr4-156013985-DEL-172
chr4-156060042-DEL-1321
chr4-156069152-DEL-336
chr4-156081032-DEL-394
chr4-156085891-DEL-52
chr4-156085928-DEL-512
chr4-156086151-DEL-96
chr4-156095353-DEL-50
chr4-156121342-DEL-171
chr4-156121506-DEL-70
chr4-156121932-DEL-200
chr4-156125974-DEL-63
chr4-156221590-DEL-494
chr4-156287521-DEL-66
chr4-156297787-DEL-1687
chr4-156320951-DEL-299
chr4-156373069-DEL-408

chr4-156494823-DEL-71
chr4-156564599-DEL-333
chr4-156566960-DEL-169
chr5-3071454-DEL-1303
chr5-3485821-DEL-1463
chr5-3485893-DEL-229
chr5-3487269-DEL-58
chr5-3559359-DEL-74
chr5-3572227-DEL-198
chr5-3660358-DEL-810
chr5-3660700-DEL-945
chr5-4012472-DEL-120
chr5-4012735-DEL-60
chr5-4013026-DEL-62
chr5-4102566-DEL-64
chr5-4102907-DEL-389
chr5-4116725-DEL-216
chr5-4222762-DEL-3077
chr5-4244447-DEL-90
chr5-4292091-DEL-50
chr5-4402428-DEL-1475
chr5-4542014-DEL-116
chr5-4588348-DEL-100
chr5-4704023-DEL-57
chr5-4705570-DEL-52
chr5-4963568-DEL-99
chr5-5000596-DEL-95
chr5-5056138-DEL-1934
chr5-5070062-DEL-401
chr5-5136255-DEL-185
chr5-5179438-DEL-64
chr5-5223757-DEL-92
chr5-5278227-DEL-151
chr5-5278314-DEL-510
chr5-5402296-DEL-763
chr5-5406982-DEL-50
chr5-5503476-DEL-50
chr5-5537913-DEL-86
chr5-5540989-DEL-58
chr5-5762356-DEL-50
chr5-5861684-DEL-87
chr5-5867945-DEL-2782
chr5-5904153-DEL-8650
chr5-6104793-DEL-153
chr5-6148819-DEL-1759
chr5-6187158-DEL-50
chr5-6187423-DEL-5325
chr5-6321966-DEL-3742
chr5-6333589-DEL-122
chr5-6521823-DEL-204
chr5-6561458-DEL-1621
chr5-6665356-DEL-203
chr5-6857762-DEL-50
chr5-7039398-DEL-114
chr5-7103927-DEL-54
chr5-7162549-DEL-392
chr5-7319238-DEL-79
chr5-7319392-DEL-125
chr5-7395606-DEL-50
chr5-7516002-DEL-2213
chr5-7589493-DEL-50
chr5-7623922-DEL-2861
chr5-7645885-DEL-50
chr5-7673156-DEL-50
chr5-7703575-DEL-50
chr5-7805776-DEL-320
chr5-8003577-DEL-57
chr5-8013415-DEL-1946
chr5-8115717-DEL-64
chr5-8117350-DEL-51
chr5-8146808-DEL-545
chr5-8161199-DEL-220
chr5-8421717-DEL-61
chr5-8538931-DEL-220
chr5-8592730-DEL-78
chr5-8599966-DEL-64
chr5-8605212-DEL-504

chr5-8620305-DEL-65
chr5-8794186-DEL-825
chr5-8839976-DEL-57
chr5-8870454-DEL-244
chr5-8994496-DEL-465
chr5-8994545-DEL-95
chr5-8994879-DEL-150
chr5-9074691-DEL-2038
chr5-9083557-DEL-198
chr5-9086514-DEL-177
chr5-9089038-DEL-174
chr5-9097249-DEL-182
chr5-9139170-DEL-116
chr5-9238506-DEL-61
chr5-9312906-DEL-95
chr5-9340747-DEL-2526
chr5-9355369-DEL-3049
chr5-9431145-DEL-214
chr5-9431573-DEL-62
chr5-9502684-DEL-476
chr5-9626052-DEL-78
chr5-9798886-DEL-100
chr5-9809019-DEL-108
chr5-9813132-DEL-64
chr5-9861045-DEL-52
chr5-9871145-DEL-620
chr5-9918766-DEL-160
chr5-9967259-DEL-50
chr5-10054314-DEL-1459
chr5-10103452-DEL-65
chr5-10154701-DEL-1231
chr5-10201960-DEL-106
chr5-10255490-DEL-115
chr5-10260913-DEL-5138
chr5-10275600-DEL-906
chr5-10277045-DEL-224
chr5-10296030-DEL-812
chr5-10464323-DEL-4107
chr5-10464328-DEL-91
chr5-10660620-DEL-67
chr5-10716846-DEL-567
chr5-10717742-DEL-122
chr5-10882450-DEL-2520
chr5-10931193-DEL-722
chr5-11955120-DEL-150
chr5-12170056-DEL-52
chr5-12181381-DEL-1643
chr5-12194767-DEL-528
chr5-12314586-DEL-495
chr5-12481674-DEL-50
chr5-12531339-DEL-510
chr5-12531642-DEL-177
chr5-12585668-DEL-430
chr5-12679326-DEL-2912
chr5-12719067-DEL-1624
chr5-12833215-DEL-67
chr5-12900008-DEL-1926
chr5-12943918-DEL-70
chr5-13006640-DEL-1357
chr5-13038113-DEL-99
chr5-13055127-DEL-77
chr5-13092201-DEL-1274
chr5-13100352-DEL-117
chr5-13114916-DEL-579
chr5-13127995-DEL-50
chr5-13128008-DEL-206
chr5-13140072-DEL-300
chr5-13149457-DEL-5237
chr5-13305951-DEL-60
chr5-13364974-DEL-3086
chr5-13425921-DEL-153
chr5-13455971-DEL-234
chr5-13513248-DEL-107
chr5-13715799-DEL-50
chr5-13813452-DEL-82
chr5-14000246-DEL-72
chr5-14015299-DEL-50

chr5-14100654-DEL-98
chr5-14316577-DEL-62
chr5-14320799-DEL-1111
chr5-14342932-DEL-329
chr5-14378276-DEL-205
chr5-14395297-DEL-51
chr5-14506301-DEL-2205
chr5-14643973-DEL-50
chr5-14833879-DEL-60
chr5-15061454-DEL-398
chr5-15722235-DEL-261
chr5-15772877-DEL-234
chr5-15852746-DEL-881
chr5-15854408-DEL-335
chr5-15893314-DEL-71
chr5-15940073-DEL-50
chr5-15951530-DEL-103
chr5-16016981-DEL-1338
chr5-16030960-DEL-4435
chr5-16064838-DEL-102
chr5-16133698-DEL-55
chr5-16571407-DEL-657
chr5-16571939-DEL-50
chr5-16572284-DEL-88
chr5-16582105-DEL-644
chr5-16596441-DEL-71
chr5-16698743-DEL-54
chr5-16698878-DEL-112
chr5-16754260-DEL-183
chr5-16916470-DEL-5714
chr5-17220231-DEL-162
chr5-17220294-DEL-66
chr5-17388445-DEL-72
chr5-17450016-DEL-424
chr5-17451738-DEL-424
chr5-17486290-DEL-58
chr5-17517847-DEL-5872
chr5-17696180-DEL-53
chr5-17838377-DEL-147
chr5-17946346-DEL-104
chr5-18017863-DEL-102
chr5-18027603-DEL-50
chr5-18030791-DEL-54
chr5-18194803-DEL-134
chr5-18195020-DEL-66
chr5-18226626-DEL-139
chr5-18263317-DEL-63
chr5-18391558-DEL-70
chr5-18507968-DEL-1434
chr5-18522301-DEL-120
chr5-18664779-DEL-63
chr5-18857410-DEL-453
chr5-18857926-DEL-63
chr5-18968296-DEL-132
chr5-19141465-DEL-227
chr5-19142760-DEL-94
chr5-19189312-DEL-807
chr5-19277699-DEL-971
chr5-19472614-DEL-53
chr5-19497102-DEL-1662
chr5-19562491-DEL-624
chr5-19607307-DEL-165
chr5-19629382-DEL-3187
chr5-19708419-DEL-5830
chr5-19828032-DEL-378
chr5-19839615-DEL-63
chr5-19842366-DEL-52
chr5-19920717-DEL-57
chr5-19981492-DEL-71
chr5-20064825-DEL-50
chr5-20088593-DEL-2413
chr5-20108292-DEL-342
chr5-20108467-DEL-97
chr5-20135775-DEL-50
chr5-20326198-DEL-6417
chr5-20463214-DEL-69
chr5-20542339-DEL-1547

chr5-20566876-DEL-795
chr5-20649490-DEL-55
chr5-20693805-DEL-56
chr5-20877435-DEL-598
chr5-20911847-DEL-475
chr5-20912363-DEL-66
chr5-20954188-DEL-112
chr5-20991735-DEL-240
chr5-21002151-DEL-594
chr5-21070158-DEL-2087
chr5-21103743-DEL-3534
chr5-21182194-DEL-195
chr5-21183507-DEL-928
chr5-21183981-DEL-66
chr5-21198623-DEL-342
chr5-21204799-DEL-185
chr5-21277309-DEL-230
chr5-21277680-DEL-337
chr5-21346133-DEL-163
chr5-21373613-DEL-50
chr5-21396378-DEL-51
chr5-21471661-DEL-132
chr5-21473058-DEL-653
chr5-21527934-DEL-139
chr5-21555105-DEL-548
chr5-21561004-DEL-329
chr5-21585227-DEL-88
chr5-21585454-DEL-66
chr5-21592904-DEL-58
chr5-21627188-DEL-130
chr5-21674048-DEL-50
chr5-21696960-DEL-50
chr5-21805302-DEL-176
chr5-21936156-DEL-86
chr5-22011579-DEL-295
chr5-22024899-DEL-84
chr5-22092578-DEL-99
chr5-22111182-DEL-264
chr5-22111738-DEL-340
chr5-22112410-DEL-172
chr5-22112492-DEL-84
chr5-22113824-DEL-749
chr5-22133429-DEL-96
chr5-22228839-DEL-102
chr5-22230078-DEL-54
chr5-22387348-DEL-84
chr5-22649536-DEL-72
chr5-22652138-DEL-946
chr5-22696973-DEL-4037
chr5-22699724-DEL-851
chr5-22702135-DEL-154
chr5-22718296-DEL-1570
chr5-22738504-DEL-129
chr5-22741073-DEL-286
chr5-22859687-DEL-2058
chr5-22925093-DEL-78
chr5-22994212-DEL-385
chr5-23200838-DEL-749
chr5-23205516-DEL-211
chr5-23205681-DEL-66
chr5-23226827-DEL-50
chr5-23246092-DEL-54
chr5-23374115-DEL-217
chr5-23464477-DEL-89
chr5-23470014-DEL-3892
chr5-23482155-DEL-250
chr5-23483155-DEL-187
chr5-23589207-DEL-93
chr5-23589354-DEL-620
chr5-23655645-DEL-2220
chr5-23692759-DEL-57
chr5-23717346-DEL-52
chr5-23721519-DEL-90
chr5-23754341-DEL-76
chr5-23810059-DEL-50
chr5-23931404-DEL-3699
chr5-23972613-DEL-224

chr5-23987376-DEL-50
chr5-24026118-DEL-165
chr5-24042432-DEL-567
chr5-24050587-DEL-64
chr5-24097991-DEL-190
chr5-24157836-DEL-77
chr5-24161461-DEL-150
chr5-24161553-DEL-60
chr5-24219485-DEL-76
chr5-24250783-DEL-187
chr5-24341203-DEL-205
chr5-24397376-DEL-116
chr5-24410184-DEL-56
chr5-24410675-DEL-120
chr5-24428500-DEL-234
chr5-24430198-DEL-65
chr5-24447093-DEL-50
chr5-24463600-DEL-155
chr5-24475261-DEL-54
chr5-24515155-DEL-114
chr5-24515348-DEL-96
chr5-24819272-DEL-114
chr5-24819484-DEL-114
chr5-24832106-DEL-60
chr5-24846568-DEL-219
chr5-24921074-DEL-60
chr5-25027710-DEL-58
chr5-25070546-DEL-133
chr5-25070776-DEL-251
chr5-25070916-DEL-111
chr5-25073450-DEL-50
chr5-25073576-DEL-162
chr5-25110790-DEL-118
chr5-25185574-DEL-72
chr5-25185744-DEL-165
chr5-25267031-DEL-66
chr5-25288246-DEL-376
chr5-25288494-DEL-407
chr5-25317440-DEL-123
chr5-25335511-DEL-609
chr5-25401334-DEL-71
chr5-25401335-DEL-367
chr5-25671507-DEL-79
chr5-25767909-DEL-50
chr5-25824183-DEL-243
chr5-25854541-DEL-1703
chr5-25909011-DEL-93
chr5-25944951-DEL-52
chr5-25947374-DEL-50
chr5-25970789-DEL-173
chr5-25970873-DEL-75
chr5-26020792-DEL-660
chr5-26021122-DEL-1361
chr5-26038401-DEL-146
chr5-26050202-DEL-59
chr5-26077224-DEL-54
chr5-26080423-DEL-107
chr5-26081123-DEL-104
chr5-26112980-DEL-90
chr5-26114452-DEL-75
chr5-26129298-DEL-95
chr5-26265386-DEL-698
chr5-26390156-DEL-407
chr5-26390784-DEL-243
chr5-26436789-DEL-287
chr5-26529057-DEL-778
chr5-26747141-DEL-214
chr5-26747608-DEL-246
chr5-27242796-DEL-63
chr5-27623147-DEL-86
chr5-27632703-DEL-459
chr5-27675605-DEL-85
chr5-27675827-DEL-69
chr5-27731003-DEL-200
chr5-27804106-DEL-50
chr5-27853625-DEL-123
chr5-27962563-DEL-548

chr5-27962948-DEL-142
chr5-28054114-DEL-56
chr5-28070977-DEL-5982
chr5-28077532-DEL-55
chr5-28077818-DEL-55
chr5-28080196-DEL-64
chr5-28141219-DEL-677
chr5-28190191-DEL-5586
chr5-28241411-DEL-101
chr5-28251185-DEL-1526
chr5-28303326-DEL-1087
chr5-28304386-DEL-159
chr5-28359881-DEL-136
chr5-28461343-DEL-67
chr5-28533989-DEL-56
chr5-28935777-DEL-52
chr5-29075315-DEL-300
chr5-29132943-DEL-423
chr5-29161563-DEL-748
chr5-29415832-DEL-77
chr5-29531420-DEL-52
chr5-29909571-DEL-364
chr5-30120722-DEL-281
chr5-30146257-DEL-224
chr5-30147791-DEL-211
chr5-30188500-DEL-88
chr5-30192042-DEL-546
chr5-30246004-DEL-185
chr5-30414866-DEL-200
chr5-30509433-DEL-65
chr5-30509585-DEL-133
chr5-30530541-DEL-58
chr5-30698230-DEL-411
chr5-30739889-DEL-230
chr5-30739892-DEL-106
chr5-30856203-DEL-50
chr5-31113787-DEL-794
chr5-31115006-DEL-52
chr5-31125138-DEL-92
chr5-31169497-DEL-240
chr5-31302189-DEL-173
chr5-31368810-DEL-126
chr5-31371075-DEL-53
chr5-31504442-DEL-87
chr5-31515480-DEL-50
chr5-31704808-DEL-250
chr5-31732339-DEL-75
chr5-31778610-DEL-69
chr5-31846552-DEL-88
chr5-32105171-DEL-369
chr5-32105212-DEL-943
chr5-32105472-DEL-451
chr5-32105783-DEL-575
chr5-32173198-DEL-135
chr5-32639110-DEL-64
chr5-32693989-DEL-198
chr5-32694608-DEL-62
chr5-32698526-DEL-367
chr5-32704138-DEL-230
chr5-32708851-DEL-455
chr5-32872210-DEL-557
chr5-32872545-DEL-193
chr5-33071850-DEL-59
chr5-33078202-DEL-202
chr5-33078445-DEL-116
chr5-33198284-DEL-61
chr5-33203961-DEL-54
chr5-33215666-DEL-153
chr5-33275761-DEL-238
chr5-33294460-DEL-50
chr5-33772463-DEL-194
chr5-33774138-DEL-146
chr5-33942974-DEL-1481
chr5-34103526-DEL-61
chr5-34178200-DEL-2602
chr5-34195531-DEL-56
chr5-34301590-DEL-217

chr5-34742085-DEL-549
chr5-34805278-DEL-118
chr5-34912065-DEL-66
chr5-34924781-DEL-52
chr5-34940594-DEL-105
chr5-35102650-DEL-100
chr5-35193727-DEL-230
chr5-35285149-DEL-117
chr5-35408361-DEL-81
chr5-35440139-DEL-70
chr5-35473402-DEL-252
chr5-35495808-DEL-293
chr5-35677876-DEL-141
chr5-35678074-DEL-285
chr5-35758405-DEL-50
chr5-35881918-DEL-1041
chr5-35930254-DEL-68
chr5-36047912-DEL-54
chr5-36048029-DEL-71
chr5-36074819-DEL-224
chr5-36116349-DEL-423
chr5-36117009-DEL-150
chr5-36171154-DEL-2132
chr5-36193902-DEL-86
chr5-36199959-DEL-88
chr5-36278722-DEL-50
chr5-36298544-DEL-570
chr5-36340864-DEL-120
chr5-36340864-DEL-268
chr5-36341063-DEL-74
chr5-36429542-DEL-219
chr5-36438438-DEL-226
chr5-36608648-DEL-50
chr5-36624943-DEL-119
chr5-36706522-DEL-108
chr5-36753144-DEL-78
chr5-36788128-DEL-30000
chr5-36864988-DEL-70
chr5-36865421-DEL-525
chr5-36912513-DEL-195
chr5-36925384-DEL-52
chr5-36925692-DEL-62
chr5-36937344-DEL-137
chr5-37108821-DEL-66
chr5-37167275-DEL-245
chr5-37167315-DEL-82
chr5-37176949-DEL-890
chr5-37316319-DEL-50
chr5-37333229-DEL-301
chr5-37339090-DEL-78
chr5-37357847-DEL-101
chr5-37396461-DEL-54
chr5-37421952-DEL-132
chr5-37422223-DEL-209
chr5-37445551-DEL-131
chr5-37586345-DEL-55
chr5-37650912-DEL-87
chr5-37969178-DEL-90
chr5-37979002-DEL-90
chr5-37992860-DEL-143
chr5-38180298-DEL-408
chr5-38180507-DEL-168
chr5-38326641-DEL-81
chr5-38327368-DEL-121
chr5-38497769-DEL-50
chr5-38540945-DEL-601
chr5-38575430-DEL-50
chr5-38643948-DEL-311
chr5-38814170-DEL-159
chr5-38894111-DEL-128
chr5-38909378-DEL-116
chr5-39105889-DEL-103
chr5-39224191-DEL-249
chr5-39224507-DEL-309
chr5-39224992-DEL-50
chr5-39276441-DEL-122
chr5-39339708-DEL-52

chr5-39427889-DEL-276
chr5-39524939-DEL-53
chr5-39527356-DEL-2582
chr5-39577366-DEL-92
chr5-39659122-DEL-2371
chr5-39749132-DEL-80
chr5-40112305-DEL-70
chr5-40115247-DEL-662
chr5-40198236-DEL-316
chr5-40202217-DEL-404
chr5-40480398-DEL-50
chr5-40704976-DEL-644
chr5-40718651-DEL-158
chr5-40743923-DEL-90
chr5-40880381-DEL-50
chr5-41087164-DEL-52
chr5-41198408-DEL-92
chr5-41203537-DEL-68
chr5-41476275-DEL-1875
chr5-41544656-DEL-1866
chr5-41645677-DEL-8295
chr5-41731670-DEL-267
chr5-41731702-DEL-75
chr5-41731841-DEL-96
chr5-41755443-DEL-50
chr5-41932157-DEL-66
chr5-42001950-DEL-128
chr5-42004988-DEL-73
chr5-42017580-DEL-352
chr5-42117614-DEL-94
chr5-42117845-DEL-61
chr5-42129615-DEL-74
chr5-42129627-DEL-158
chr5-42145590-DEL-54
chr5-42162554-DEL-5289
chr5-42345784-DEL-212
chr5-42479032-DEL-130
chr5-42479143-DEL-50
chr5-42706576-DEL-70
chr5-42819566-DEL-2052
chr5-42912212-DEL-1270
chr5-42949783-DEL-518
chr5-42983169-DEL-13349
chr5-43218938-DEL-240
chr5-43277296-DEL-96
chr5-43331047-DEL-361
chr5-43410463-DEL-1374
chr5-43461993-DEL-63
chr5-43464952-DEL-274
chr5-43470304-DEL-94
chr5-43476398-DEL-198
chr5-43603613-DEL-72
chr5-43718428-DEL-240
chr5-43718742-DEL-138
chr5-43718826-DEL-54
chr5-43781754-DEL-150
chr5-43820349-DEL-51
chr5-44110167-DEL-2421
chr5-44115968-DEL-1036
chr5-44222593-DEL-299
chr5-44443494-DEL-216
chr5-44470063-DEL-2999
chr5-44518608-DEL-190
chr5-44764981-DEL-163
chr5-44765945-DEL-51
chr5-44840690-DEL-144
chr5-44902082-DEL-84
chr5-44914705-DEL-1602
chr5-44920039-DEL-133
chr5-45025334-DEL-520
chr5-45025880-DEL-805
chr5-45087784-DEL-204
chr5-45130193-DEL-75
chr5-45219649-DEL-70
chr5-45243028-DEL-1496
chr5-45392643-DEL-52
chr5-45609849-DEL-152

chr5-45657323-DEL-150
chr5-45782287-DEL-50
chr5-45841980-DEL-166
chr5-45869198-DEL-50
chr5-46022333-DEL-50
chr5-46047219-DEL-60
chr5-46152435-DEL-792
chr5-46157366-DEL-672
chr5-46249284-DEL-56
chr5-46438146-DEL-543
chr5-46515092-DEL-51
chr5-46575528-DEL-50
chr5-46701809-DEL-96
chr5-46703592-DEL-690
chr5-46727221-DEL-60
chr5-46759243-DEL-126
chr5-46817533-DEL-115
chr5-46887384-DEL-1063
chr5-46918855-DEL-6239
chr5-46925319-DEL-188
chr5-47044037-DEL-55
chr5-47176811-DEL-2254
chr5-47235389-DEL-52
chr5-47511806-DEL-60
chr5-47594799-DEL-89
chr5-47765348-DEL-171
chr5-47765361-DEL-64
chr5-47800705-DEL-53
chr5-47845486-DEL-60
chr5-47915435-DEL-83
chr5-47915502-DEL-270
chr5-47974691-DEL-191
chr5-48111868-DEL-1544
chr5-48156498-DEL-360
chr5-48310905-DEL-54
chr5-48384235-DEL-204
chr5-48866058-DEL-1897
chr5-49152322-DEL-50
chr5-49261580-DEL-167
chr5-49348508-DEL-73
chr5-49474435-DEL-299
chr5-49523953-DEL-206
chr5-49678403-DEL-76
chr5-49680743-DEL-56
chr5-49745096-DEL-113
chr5-49950145-DEL-59
chr5-50026766-DEL-592
chr5-50077084-DEL-68
chr5-50096138-DEL-54
chr5-50109379-DEL-2342
chr5-50361056-DEL-1866
chr5-50476006-DEL-424
chr5-50501910-DEL-51
chr5-50552474-DEL-265
chr5-50582789-DEL-128
chr5-50678352-DEL-1296
chr5-50840619-DEL-270
chr5-50963813-DEL-888
chr5-51276016-DEL-66
chr5-51306470-DEL-54
chr5-51316982-DEL-92
chr5-51358896-DEL-537
chr5-51451385-DEL-86
chr5-51545671-DEL-50
chr5-51806684-DEL-226
chr5-51851689-DEL-60
chr5-51872840-DEL-138
chr5-51879741-DEL-125
chr5-51879746-DEL-373
chr5-51950807-DEL-50
chr5-52249085-DEL-54
chr5-52507334-DEL-132
chr5-52578903-DEL-54
chr5-52657564-DEL-68
chr5-52668125-DEL-77
chr5-52709016-DEL-118
chr5-52735766-DEL-50

chr5-52753421-DEL-50
chr5-52875722-DEL-50
chr5-52931907-DEL-1931
chr5-53075861-DEL-453
chr5-53106544-DEL-58
chr5-53171595-DEL-158
chr5-53191884-DEL-738
chr5-53191895-DEL-240
chr5-53192495-DEL-288
chr5-53195198-DEL-74
chr5-53251736-DEL-51
chr5-53398172-DEL-66
chr5-53459789-DEL-216
chr5-53465482-DEL-5969
chr5-53542354-DEL-52
chr5-53735871-DEL-148
chr5-53740096-DEL-122
chr5-53740522-DEL-250
chr5-53825151-DEL-50
chr5-53833257-DEL-160
chr5-53931939-DEL-83
chr5-53974620-DEL-60
chr5-54096286-DEL-102
chr5-54490820-DEL-253
chr5-54493333-DEL-60
chr5-54695278-DEL-121
chr5-54695560-DEL-195
chr5-54744656-DEL-54
chr5-54945708-DEL-235
chr5-54967926-DEL-90
chr5-55130713-DEL-1384
chr5-55174419-DEL-114
chr5-55207628-DEL-117
chr5-55563528-DEL-196
chr5-55721022-DEL-64
chr5-55747599-DEL-268
chr5-55768746-DEL-1493
chr5-55780494-DEL-52
chr5-55808429-DEL-175
chr5-55934755-DEL-298
chr5-55935615-DEL-136
chr5-56010130-DEL-78
chr5-56023605-DEL-69
chr5-56085833-DEL-50
chr5-56126329-DEL-50
chr5-56196574-DEL-52
chr5-56298225-DEL-62
chr5-56595935-DEL-90
chr5-56723586-DEL-50
chr5-56754783-DEL-195
chr5-56790551-DEL-84
chr5-57085476-DEL-1072
chr5-57137667-DEL-186
chr5-57214914-DEL-1110
chr5-57596684-DEL-50
chr5-57916677-DEL-244
chr5-57917049-DEL-210
chr5-58235911-DEL-527
chr5-58353610-DEL-125
chr5-58363396-DEL-50
chr5-58391771-DEL-3096
chr5-58477889-DEL-143
chr5-58737364-DEL-115
chr5-58810339-DEL-1332
chr5-58933378-DEL-594
chr5-58945844-DEL-72
chr5-58949201-DEL-76
chr5-58950334-DEL-104
chr5-58984766-DEL-2658
chr5-58985453-DEL-66
chr5-59024820-DEL-1174
chr5-59055825-DEL-64
chr5-59109123-DEL-198
chr5-59114597-DEL-559
chr5-59208712-DEL-175
chr5-59242636-DEL-1429
chr5-59338578-DEL-58

chr5-59504092-DEL-1214
chr5-59505586-DEL-369
chr5-59531297-DEL-945
chr5-59610625-DEL-2077
chr5-59615979-DEL-106
chr5-59641709-DEL-200
chr5-59664001-DEL-574
chr5-59684995-DEL-1416
chr5-59719238-DEL-967
chr5-59741237-DEL-243
chr5-59812685-DEL-70
chr5-59819199-DEL-56
chr5-59862958-DEL-357
chr5-59905581-DEL-84
chr5-59930736-DEL-64
chr5-59976146-DEL-60
chr5-60022463-DEL-1967
chr5-60075691-DEL-1075
chr5-60106288-DEL-7464
chr5-60123816-DEL-66
chr5-60164551-DEL-513
chr5-60173850-DEL-60
chr5-60235438-DEL-244
chr5-60398337-DEL-56
chr5-60415744-DEL-99
chr5-60425369-DEL-399
chr5-60446135-DEL-161
chr5-60446837-DEL-2514
chr5-60462460-DEL-599
chr5-60517590-DEL-64
chr5-60529691-DEL-154
chr5-60585381-DEL-50
chr5-60670818-DEL-1247
chr5-60698415-DEL-89
chr5-60904607-DEL-50
chr5-60935841-DEL-122
chr5-60937229-DEL-526
chr5-60940398-DEL-1013
chr5-60995359-DEL-68
chr5-61008756-DEL-50
chr5-61022950-DEL-90
chr5-61197374-DEL-605
chr5-61207291-DEL-188
chr5-61217330-DEL-1084
chr5-61257441-DEL-2792
chr5-61328441-DEL-1271
chr5-61389667-DEL-214
chr5-61415566-DEL-50
chr5-61450558-DEL-212
chr5-61677524-DEL-353
chr5-61678258-DEL-1114
chr5-61682751-DEL-221
chr5-61717704-DEL-278
chr5-61809399-DEL-666
chr5-61864464-DEL-1606
chr5-61933299-DEL-26949
chr5-61994610-DEL-101
chr5-62015353-DEL-800
chr5-62035220-DEL-2471
chr5-62186850-DEL-5605
chr5-62324944-DEL-90
chr5-62644061-DEL-15103
chr5-62681574-DEL-58
chr5-62755427-DEL-64
chr5-62755558-DEL-420
chr5-62756043-DEL-130
chr5-62909237-DEL-1430
chr5-62929996-DEL-2785
chr5-63009735-DEL-50
chr5-63016460-DEL-697
chr5-63111230-DEL-1166
chr5-63134667-DEL-3780
chr5-63369285-DEL-75
chr5-63402338-DEL-1516
chr5-63443321-DEL-666
chr5-63505930-DEL-50
chr5-63527798-DEL-238

chr5-63570146-DEL-142
chr5-63663023-DEL-1111
chr5-63725225-DEL-74
chr5-63733142-DEL-570
chr5-63840313-DEL-92
chr5-63900627-DEL-167
chr5-63955639-DEL-68
chr5-64004627-DEL-278
chr5-64110671-DEL-140
chr5-64157979-DEL-83
chr5-64183673-DEL-178
chr5-64301373-DEL-629
chr5-64385874-DEL-674
chr5-64416479-DEL-96
chr5-64480377-DEL-75
chr5-64551341-DEL-60
chr5-64576398-DEL-62
chr5-64607132-DEL-172
chr5-64761875-DEL-76
chr5-64794584-DEL-76
chr5-64893675-DEL-50
chr5-64917340-DEL-379
chr5-64930395-DEL-53
chr5-64936093-DEL-53
chr5-64937239-DEL-52
chr5-65039147-DEL-222
chr5-65042040-DEL-53
chr5-65114826-DEL-176
chr5-65201599-DEL-192
chr5-65272432-DEL-80
chr5-65286604-DEL-54
chr5-65389255-DEL-255
chr5-65389512-DEL-298
chr5-65424383-DEL-55
chr5-65467555-DEL-472
chr5-65604750-DEL-91
chr5-65782438-DEL-134
chr5-65832085-DEL-1623
chr5-65911226-DEL-60
chr5-65946207-DEL-66
chr5-65946402-DEL-56
chr5-66008921-DEL-103
chr5-66034122-DEL-2157
chr5-66036141-DEL-50
chr5-66074881-DEL-79
chr5-66086234-DEL-82
chr5-66107042-DEL-86
chr5-66141343-DEL-151
chr5-66146086-DEL-154
chr5-66151642-DEL-50
chr5-66181065-DEL-74
chr5-66360064-DEL-240
chr5-66395836-DEL-150
chr5-66445883-DEL-695
chr5-66488214-DEL-1916
chr5-66623813-DEL-58
chr5-66684024-DEL-65
chr5-66686593-DEL-54
chr5-66761586-DEL-135
chr5-66786601-DEL-76
chr5-66787631-DEL-660
chr5-66800544-DEL-114
chr5-66831263-DEL-61
chr5-66854141-DEL-120
chr5-66854244-DEL-50
chr5-66896735-DEL-56
chr5-66985191-DEL-187
chr5-67019943-DEL-60
chr5-67043684-DEL-58
chr5-67046750-DEL-503
chr5-67065968-DEL-76
chr5-67083897-DEL-50
chr5-67178761-DEL-80
chr5-67295229-DEL-90
chr5-67312936-DEL-140
chr5-67407040-DEL-62
chr5-67472842-DEL-113

chr5-67533568-DEL-3337
chr5-67610504-DEL-64
chr5-67672195-DEL-87
chr5-67674394-DEL-80
chr5-67674519-DEL-174
chr5-67719543-DEL-60
chr5-67726738-DEL-50
chr5-67756896-DEL-50
chr5-67949147-DEL-426
chr5-68015908-DEL-456
chr5-68031696-DEL-55
chr5-68097525-DEL-1275
chr5-68134920-DEL-878
chr5-68237693-DEL-1905
chr5-68256260-DEL-624
chr5-68263727-DEL-75
chr5-68350309-DEL-51
chr5-68362907-DEL-503
chr5-68378642-DEL-208
chr5-68395128-DEL-65
chr5-68446446-DEL-162
chr5-68560955-DEL-66
chr5-68575309-DEL-52
chr5-68626209-DEL-589
chr5-68666987-DEL-183
chr5-68672746-DEL-158
chr5-68685066-DEL-50
chr5-68688101-DEL-807
chr5-68703865-DEL-90
chr5-68708898-DEL-2018
chr5-68721784-DEL-60
chr5-68745543-DEL-208
chr5-68758748-DEL-50
chr5-68785679-DEL-610
chr5-68825062-DEL-1789
chr5-68828356-DEL-572
chr5-68835650-DEL-56
chr5-68848900-DEL-1816
chr5-68953525-DEL-75
chr5-68982876-DEL-194
chr5-69006709-DEL-1305
chr5-69084417-DEL-65
chr5-69092324-DEL-65
chr5-69124163-DEL-62
chr5-69173089-DEL-82
chr5-69360337-DEL-146
chr5-69363866-DEL-128
chr5-69397641-DEL-465
chr5-69399623-DEL-73
chr5-69399995-DEL-979
chr5-69473681-DEL-474
chr5-69548241-DEL-1517
chr5-69571856-DEL-99
chr5-69688941-DEL-428
chr5-70023310-DEL-76
chr5-70049289-DEL-1056
chr5-70094242-DEL-675
chr5-70106877-DEL-64
chr5-70169178-DEL-52
chr5-70202879-DEL-6011
chr5-70209257-DEL-111
chr5-70212502-DEL-97
chr5-70253187-DEL-2238
chr5-70330704-DEL-544
chr5-70344365-DEL-73
chr5-70393958-DEL-3102
chr5-70400077-DEL-1778
chr5-70432253-DEL-66
chr5-70498891-DEL-1558
chr5-70553303-DEL-2552
chr5-70622803-DEL-152
chr5-70625436-DEL-1054
chr5-70691267-DEL-1431
chr5-70743621-DEL-108
chr5-70924475-DEL-1081
chr5-70962321-DEL-313
chr5-70986633-DEL-54

chr5-71069747-DEL-336
chr5-71112084-DEL-62
chr5-71141064-DEL-489
chr5-71195099-DEL-52
chr5-71307113-DEL-50
chr5-71309266-DEL-52
chr5-71491732-DEL-51
chr5-71496424-DEL-137
chr5-71749065-DEL-114
chr5-72215375-DEL-145
chr5-72215489-DEL-69
chr5-72215719-DEL-78
chr5-72219724-DEL-62
chr5-72309827-DEL-62
chr5-72335989-DEL-68
chr5-72424560-DEL-61
chr5-72428475-DEL-500
chr5-72429155-DEL-222
chr5-72468539-DEL-724
chr5-72545577-DEL-86
chr5-72555692-DEL-1033
chr5-72633163-DEL-52
chr5-72666760-DEL-87
chr5-72741922-DEL-95
chr5-72742216-DEL-230
chr5-72742236-DEL-90
chr5-72750985-DEL-50
chr5-72814625-DEL-259
chr5-72841053-DEL-418
chr5-72843590-DEL-244
chr5-72994979-DEL-509
chr5-73049287-DEL-405
chr5-73050543-DEL-60
chr5-73063127-DEL-622
chr5-73102030-DEL-2137
chr5-73146758-DEL-152
chr5-73158603-DEL-337
chr5-73527509-DEL-56
chr5-73556501-DEL-69
chr5-73560067-DEL-51
chr5-73594939-DEL-51
chr5-73601676-DEL-3579
chr5-73605879-DEL-129
chr5-73611108-DEL-50
chr5-73613152-DEL-50
chr5-73720254-DEL-532
chr5-73758443-DEL-50
chr5-73840438-DEL-75
chr5-73856394-DEL-282
chr5-73856722-DEL-118
chr5-73856756-DEL-54
chr5-73918470-DEL-225
chr5-73961338-DEL-70
chr5-73964272-DEL-66
chr5-74035312-DEL-158
chr5-74076092-DEL-997
chr5-74091687-DEL-359
chr5-74137493-DEL-164
chr5-74138112-DEL-199
chr5-74138994-DEL-226
chr5-74155643-DEL-112
chr5-74156058-DEL-383
chr5-74204384-DEL-58
chr5-74274182-DEL-75
chr5-74301080-DEL-134
chr5-74389702-DEL-1201
chr5-74432460-DEL-108
chr5-74468947-DEL-324
chr5-74510616-DEL-112
chr5-74510843-DEL-389
chr5-74511096-DEL-84
chr5-74600052-DEL-207
chr5-74646807-DEL-1388
chr5-74666877-DEL-125
chr5-74670136-DEL-66
chr5-74688696-DEL-51
chr5-74731886-DEL-50

chr5-74775874-DEL-119
chr5-74880224-DEL-93
chr5-74884746-DEL-174
chr5-74942972-DEL-50
chr5-74951321-DEL-90
chr5-74979074-DEL-70
chr5-75059250-DEL-50
chr5-75148498-DEL-2699
chr5-75148975-DEL-81
chr5-75158419-DEL-135
chr5-75271503-DEL-88
chr5-75281479-DEL-52
chr5-75388608-DEL-64
chr5-75457139-DEL-443
chr5-75488412-DEL-105
chr5-75495690-DEL-59
chr5-75540857-DEL-200
chr5-75585948-DEL-400
chr5-75608218-DEL-87
chr5-75612863-DEL-307
chr5-75612902-DEL-1080
chr5-75613347-DEL-466
chr5-75622587-DEL-548
chr5-75622896-DEL-108
chr5-75714223-DEL-51
chr5-75728241-DEL-107
chr5-75748182-DEL-56
chr5-75821097-DEL-204
chr5-75842554-DEL-120
chr5-75890102-DEL-194
chr5-75990140-DEL-177
chr5-76022122-DEL-52
chr5-76037791-DEL-353
chr5-76127840-DEL-50
chr5-76190240-DEL-87
chr5-76217327-DEL-54
chr5-76242958-DEL-81
chr5-76254602-DEL-88
chr5-76267239-DEL-51
chr5-76376077-DEL-105
chr5-76431750-DEL-885
chr5-76483242-DEL-71
chr5-76499052-DEL-59
chr5-76560175-DEL-1158
chr5-76573320-DEL-813
chr5-76574317-DEL-621
chr5-76742092-DEL-67
chr5-76865178-DEL-83
chr5-76904640-DEL-237
chr5-76904678-DEL-69
chr5-76905794-DEL-1484
chr5-76905971-DEL-444
chr5-76906287-DEL-86
chr5-76909724-DEL-128
chr5-76912304-DEL-917
chr5-76924302-DEL-1244
chr5-76949502-DEL-569
chr5-76950521-DEL-54
chr5-77000074-DEL-56
chr5-77149819-DEL-63
chr5-77152610-DEL-89
chr5-77201280-DEL-134
chr5-77291342-DEL-134
chr5-77300239-DEL-1135
chr5-77313660-DEL-51
chr5-77355725-DEL-272
chr5-77355735-DEL-80
chr5-77376497-DEL-53
chr5-77376681-DEL-206
chr5-77407863-DEL-97
chr5-77448079-DEL-257
chr5-77453909-DEL-86
chr5-77460397-DEL-157
chr5-77497639-DEL-67
chr5-77518685-DEL-882
chr5-77522019-DEL-90
chr5-77557474-DEL-6418

chr5-77559760-DEL-788
chr5-77618599-DEL-126
chr5-77618900-DEL-302
chr5-77704603-DEL-405
chr5-77753894-DEL-60
chr5-77753921-DEL-180
chr5-77782518-DEL-172
chr5-77823129-DEL-246
chr5-77832145-DEL-158
chr5-77852541-DEL-62
chr5-77876973-DEL-385
chr5-77880356-DEL-99
chr5-77982583-DEL-99
chr5-77988729-DEL-50
chr5-78008760-DEL-19245
chr5-78011338-DEL-1467
chr5-78021176-DEL-585
chr5-78031285-DEL-1031
chr5-78309124-DEL-519
chr5-78309363-DEL-246
chr5-78309367-DEL-96
chr5-78419093-DEL-60
chr5-78509234-DEL-50
chr5-78525336-DEL-1935
chr5-78545373-DEL-68
chr5-78548055-DEL-403
chr5-78626811-DEL-149
chr5-78676337-DEL-84
chr5-78761462-DEL-67
chr5-78779248-DEL-56
chr5-78967507-DEL-5381
chr5-78975066-DEL-515
chr5-79173371-DEL-3432
chr5-79238939-DEL-51
chr5-79239784-DEL-198
chr5-79295783-DEL-59
chr5-79302410-DEL-96
chr5-79385775-DEL-98
chr5-79552977-DEL-54
chr5-79573252-DEL-100
chr5-79593363-DEL-245
chr5-79597885-DEL-90
chr5-79687833-DEL-2411
chr5-79728501-DEL-2967
chr5-79744404-DEL-136
chr5-79745926-DEL-4746
chr5-79821409-DEL-3413
chr5-79839141-DEL-117
chr5-79883396-DEL-67
chr5-79959929-DEL-656
chr5-79963808-DEL-6953
chr5-80075326-DEL-72
chr5-80120822-DEL-108
chr5-80124740-DEL-227
chr5-80202562-DEL-1062
chr5-80246022-DEL-54
chr5-80270206-DEL-50
chr5-80335047-DEL-4120
chr5-80347759-DEL-618
chr5-80407320-DEL-3406
chr5-80430669-DEL-52
chr5-80537909-DEL-2165
chr5-80579369-DEL-50
chr5-80591404-DEL-221
chr5-80664753-DEL-63
chr5-80803468-DEL-1889
chr5-80843000-DEL-59
chr5-80918643-DEL-142
chr5-80928372-DEL-2548
chr5-81074740-DEL-2251
chr5-81103500-DEL-61
chr5-81119292-DEL-79
chr5-81186188-DEL-52
chr5-81225840-DEL-457
chr5-81229182-DEL-658
chr5-81258964-DEL-124
chr5-81587598-DEL-123

chr5-81785338-DEL-60
chr5-81876563-DEL-186
chr5-81954144-DEL-364
chr5-82238483-DEL-837
chr5-82449143-DEL-6620
chr5-82503126-DEL-355
chr5-82605763-DEL-389
chr5-82626916-DEL-279
chr5-82740870-DEL-131
chr5-82801552-DEL-201
chr5-82824282-DEL-8775
chr5-82872025-DEL-1597
chr5-82912005-DEL-203
chr5-82924078-DEL-872
chr5-82925605-DEL-62
chr5-82972694-DEL-61
chr5-83074958-DEL-1582
chr5-83089830-DEL-144
chr5-83128710-DEL-64
chr5-83129131-DEL-82
chr5-83204589-DEL-60
chr5-83238138-DEL-68
chr5-83344648-DEL-146
chr5-83358360-DEL-50
chr5-83465231-DEL-5992
chr5-83482126-DEL-430
chr5-83537232-DEL-212
chr5-83619337-DEL-1187
chr5-83649298-DEL-182
chr5-83708545-DEL-150
chr5-83718620-DEL-168
chr5-83912814-DEL-50
chr5-83975758-DEL-1801
chr5-83977932-DEL-52
chr5-84011683-DEL-56
chr5-84111081-DEL-3879
chr5-84115577-DEL-1275
chr5-84127411-DEL-50
chr5-84189029-DEL-106
chr5-84236910-DEL-1125
chr5-84251169-DEL-137
chr5-84285666-DEL-56
chr5-84525678-DEL-424
chr5-84595984-DEL-2307
chr5-84666748-DEL-94
chr5-84666992-DEL-71
chr5-84714612-DEL-79
chr5-84733841-DEL-55
chr5-84794352-DEL-453
chr5-84813012-DEL-155
chr5-84851531-DEL-161
chr5-84901637-DEL-6514
chr5-84919273-DEL-86
chr5-85130078-DEL-50
chr5-85211369-DEL-1045
chr5-85257765-DEL-1294
chr5-85330831-DEL-214
chr5-85332806-DEL-52
chr5-85507351-DEL-964
chr5-85539932-DEL-1811
chr5-85546959-DEL-2987
chr5-85637671-DEL-60
chr5-85645905-DEL-64
chr5-85813578-DEL-66
chr5-85889610-DEL-380
chr5-85970368-DEL-6053
chr5-86074820-DEL-1836
chr5-86076760-DEL-356
chr5-86101695-DEL-74
chr5-86115490-DEL-451
chr5-86237240-DEL-1341
chr5-86291125-DEL-175
chr5-86330486-DEL-54
chr5-86348519-DEL-6569
chr5-86443529-DEL-188
chr5-86447126-DEL-51
chr5-86460122-DEL-62

chr5-86574264-DEL-254
chr5-86694832-DEL-1012
chr5-86865561-DEL-115
chr5-86898344-DEL-50
chr5-86950429-DEL-129
chr5-86987993-DEL-223
chr5-87093295-DEL-1666
chr5-87120204-DEL-109
chr5-87259864-DEL-857
chr5-87342410-DEL-50
chr5-87366450-DEL-455
chr5-87463440-DEL-52
chr5-87475658-DEL-90
chr5-87565267-DEL-247
chr5-87677666-DEL-62
chr5-87788678-DEL-1000
chr5-87792164-DEL-8395
chr5-87852174-DEL-272
chr5-87852390-DEL-105
chr5-87892636-DEL-78
chr5-87944963-DEL-2477
chr5-87947898-DEL-252
chr5-88043690-DEL-50
chr5-88328573-DEL-2508
chr5-88333158-DEL-99
chr5-88345710-DEL-2122
chr5-88364320-DEL-52
chr5-88476058-DEL-78
chr5-88507493-DEL-2816
chr5-88512253-DEL-209
chr5-88513714-DEL-1034
chr5-88544143-DEL-78
chr5-88552998-DEL-75
chr5-88585116-DEL-154
chr5-88597100-DEL-739
chr5-88624317-DEL-374
chr5-88649936-DEL-66
chr5-88678308-DEL-90
chr5-88690163-DEL-66
chr5-88696484-DEL-2842
chr5-88759547-DEL-212
chr5-88768287-DEL-117
chr5-88770582-DEL-1787
chr5-88772477-DEL-575
chr5-88818720-DEL-143
chr5-88926527-DEL-156
chr5-88948667-DEL-117
chr5-88964067-DEL-183
chr5-89009409-DEL-56
chr5-89009711-DEL-81
chr5-89018464-DEL-1699
chr5-89030874-DEL-50
chr5-89070491-DEL-52
chr5-89126226-DEL-894
chr5-89156143-DEL-80
chr5-89156452-DEL-52
chr5-89272733-DEL-62
chr5-89393007-DEL-66
chr5-89530289-DEL-792
chr5-89616736-DEL-64
chr5-89660704-DEL-2449
chr5-89767755-DEL-1638
chr5-89768012-DEL-168
chr5-89769523-DEL-126
chr5-89794672-DEL-55
chr5-89811147-DEL-238
chr5-89903587-DEL-272
chr5-89994886-DEL-1034
chr5-90051477-DEL-50
chr5-90058158-DEL-222
chr5-90115621-DEL-71
chr5-90124459-DEL-57
chr5-90133630-DEL-105
chr5-90268987-DEL-128
chr5-90302501-DEL-1234
chr5-90317356-DEL-6249
chr5-90344126-DEL-50

chr5-90352406-DEL-50
chr5-90407206-DEL-59
chr5-90437995-DEL-289
chr5-90440387-DEL-94
chr5-90468834-DEL-1739
chr5-90479121-DEL-57
chr5-90533801-DEL-537
chr5-90549390-DEL-50
chr5-90601767-DEL-83
chr5-90657390-DEL-197
chr5-90664875-DEL-100
chr5-90668517-DEL-133
chr5-90668828-DEL-72
chr5-90846121-DEL-929
chr5-90888209-DEL-61
chr5-90983711-DEL-1401
chr5-91000885-DEL-61
chr5-91029873-DEL-121
chr5-91086183-DEL-130
chr5-91106922-DEL-138
chr5-91106962-DEL-50
chr5-91255030-DEL-72
chr5-91310995-DEL-797
chr5-91486921-DEL-1686
chr5-91495699-DEL-55
chr5-91550074-DEL-50
chr5-91582658-DEL-90
chr5-91611333-DEL-176
chr5-91629507-DEL-130
chr5-91786440-DEL-310
chr5-91787919-DEL-609
chr5-91827963-DEL-336
chr5-91835835-DEL-128
chr5-91893779-DEL-79
chr5-91922261-DEL-136
chr5-91942725-DEL-52
chr5-92003420-DEL-246
chr5-92040415-DEL-132
chr5-92043930-DEL-198
chr5-92080054-DEL-215
chr5-92164602-DEL-64
chr5-92171797-DEL-106
chr5-92175396-DEL-69
chr5-92257088-DEL-467
chr5-92259191-DEL-116
chr5-92261695-DEL-88
chr5-92384969-DEL-55
chr5-92408260-DEL-57
chr5-92510656-DEL-64
chr5-92560812-DEL-221
chr5-92618308-DEL-199
chr5-92660892-DEL-58
chr5-92730523-DEL-154
chr5-92817528-DEL-1810
chr5-92902695-DEL-3822
chr5-92966217-DEL-250
chr5-93011915-DEL-84
chr5-93119141-DEL-100
chr5-93203878-DEL-182
chr5-93204858-DEL-73
chr5-93234520-DEL-79
chr5-93250567-DEL-629
chr5-93291977-DEL-243
chr5-93343769-DEL-50
chr5-93425930-DEL-122
chr5-95667409-DEL-221
chr5-95668876-DEL-143
chr5-95969449-DEL-201
chr5-96004667-DEL-201
chr5-96009486-DEL-1287
chr5-96031581-DEL-82
chr5-96142631-DEL-1196
chr5-96204423-DEL-6826
chr5-96235797-DEL-66
chr5-96244260-DEL-77
chr5-96257270-DEL-81
chr5-96332190-DEL-80

chr5-96400121-DEL-100
chr5-96403018-DEL-64
chr5-96433837-DEL-4554
chr5-96461188-DEL-65
chr5-96500279-DEL-91
chr5-96509952-DEL-266
chr5-96540900-DEL-782
chr5-96540998-DEL-123
chr5-96541555-DEL-116
chr5-96549968-DEL-128
chr5-96589268-DEL-50
chr5-96594877-DEL-87
chr5-96760555-DEL-57
chr5-96831721-DEL-401
chr5-96889501-DEL-59
chr5-96987792-DEL-118
chr5-97011465-DEL-148
chr5-97063425-DEL-50
chr5-97168988-DEL-120
chr5-97277245-DEL-89
chr5-97319575-DEL-70
chr5-97360172-DEL-50
chr5-97394800-DEL-687
chr5-97412056-DEL-210
chr5-97428400-DEL-61
chr5-97446351-DEL-151
chr5-97486473-DEL-54
chr5-97504058-DEL-99
chr5-97563349-DEL-194
chr5-97711142-DEL-94
chr5-97763322-DEL-498
chr5-97767088-DEL-848
chr5-97806037-DEL-111
chr5-97807353-DEL-898
chr5-97915080-DEL-266
chr5-97923085-DEL-63
chr5-97931800-DEL-563
chr5-98006149-DEL-50
chr5-98038062-DEL-143
chr5-98038935-DEL-60
chr5-98065455-DEL-195
chr5-98099473-DEL-50
chr5-98132491-DEL-53
chr5-98208137-DEL-525
chr5-98208515-DEL-147
chr5-98242711-DEL-650
chr5-98276319-DEL-58
chr5-98277232-DEL-68
chr5-98309177-DEL-982
chr5-98322650-DEL-199
chr5-98322977-DEL-95
chr5-98348728-DEL-50
chr5-98399216-DEL-52
chr5-98406975-DEL-81
chr5-98443893-DEL-139
chr5-98494999-DEL-59
chr5-98598372-DEL-252
chr5-98675513-DEL-6792
chr5-98684960-DEL-972
chr5-98715495-DEL-51
chr5-98781099-DEL-54
chr5-98830897-DEL-50
chr5-98857087-DEL-98
chr5-98961050-DEL-361
chr5-98978345-DEL-368
chr5-98994832-DEL-253
chr5-99154983-DEL-305
chr5-99242886-DEL-1764
chr5-99312471-DEL-92
chr5-99363274-DEL-60
chr5-99429428-DEL-118
chr5-99477931-DEL-78
chr5-99506763-DEL-102
chr5-99655493-DEL-140
chr5-99676796-DEL-247
chr5-99708431-DEL-52
chr5-99714854-DEL-672

chr5-99732670-DEL-248
chr5-99741416-DEL-1070
chr5-99788254-DEL-167
chr5-99791352-DEL-1048
chr5-99800320-DEL-210
chr5-99806993-DEL-52
chr5-99908308-DEL-60
chr5-99952756-DEL-88
chr5-99978893-DEL-944
chr5-99990116-DEL-167
chr5-99990219-DEL-59
chr5-100037459-DEL-86
chr5-100040348-DEL-76
chr5-100045922-DEL-100
chr5-100065938-DEL-50
chr5-100086465-DEL-50
chr5-100106087-DEL-50
chr5-100134329-DEL-7166
chr5-100145335-DEL-3336
chr5-100145831-DEL-1363
chr5-100165183-DEL-369
chr5-100167800-DEL-406
chr5-100171039-DEL-944
chr5-100197018-DEL-77
chr5-100252936-DEL-132
chr5-100278722-DEL-866
chr5-100314235-DEL-61
chr5-100330417-DEL-1853
chr5-100355519-DEL-283
chr5-100355624-DEL-671
chr5-100356188-DEL-119
chr5-100425018-DEL-56
chr5-100452686-DEL-56
chr5-100476693-DEL-163
chr5-100490079-DEL-59
chr5-100522598-DEL-75
chr5-100679509-DEL-73
chr5-100686272-DEL-328
chr5-100686719-DEL-79
chr5-100700318-DEL-229
chr5-100702109-DEL-57
chr5-100703227-DEL-50
chr5-100908130-DEL-72
chr5-100909141-DEL-116
chr5-100975389-DEL-74
chr5-101059232-DEL-74
chr5-101089904-DEL-128
chr5-101171332-DEL-220
chr5-101183123-DEL-192
chr5-101297570-DEL-733
chr5-101306348-DEL-86
chr5-101307944-DEL-181
chr5-101317285-DEL-276
chr5-101394860-DEL-58
chr5-101403333-DEL-507
chr5-101408441-DEL-90
chr5-101470185-DEL-67
chr5-101484636-DEL-92
chr5-101509942-DEL-274
chr5-101564929-DEL-100
chr5-101565139-DEL-102
chr5-101566558-DEL-503
chr5-101572094-DEL-407
chr5-101572672-DEL-286
chr5-101582006-DEL-62
chr5-101606741-DEL-252
chr5-101694273-DEL-68
chr5-101736450-DEL-72
chr5-101749694-DEL-62
chr5-101770690-DEL-1050
chr5-101781969-DEL-54
chr5-101782235-DEL-699
chr5-101782349-DEL-108
chr5-101782596-DEL-135
chr5-101783194-DEL-810
chr5-101858833-DEL-124
chr5-101895760-DEL-52

chr5-101922375-DEL-192
chr5-101948619-DEL-182
chr5-101954587-DEL-117
chr5-101961216-DEL-105
chr5-102038833-DEL-120
chr5-102064772-DEL-398
chr5-102064927-DEL-70
chr5-102087428-DEL-183
chr5-102112059-DEL-880
chr5-102126538-DEL-429
chr5-102132991-DEL-248
chr5-102170414-DEL-60
chr5-102349445-DEL-55
chr5-102356106-DEL-788
chr5-102382273-DEL-54
chr5-102394781-DEL-72
chr5-102404520-DEL-215
chr5-102406925-DEL-88
chr5-102421765-DEL-60
chr5-102487661-DEL-129
chr5-102515440-DEL-66
chr5-102570181-DEL-121
chr5-102591694-DEL-1374
chr5-102602279-DEL-50
chr5-102685133-DEL-58
chr5-102695408-DEL-154
chr5-102722794-DEL-138
chr5-102747406-DEL-1432
chr5-102782832-DEL-72
chr5-102800275-DEL-68
chr5-102800633-DEL-160
chr5-102838843-DEL-53
chr5-102839203-DEL-224
chr5-102869774-DEL-89
chr5-102901363-DEL-52
chr5-103021267-DEL-64
chr5-103076501-DEL-109
chr5-103099429-DEL-96
chr5-103115678-DEL-119
chr5-103116093-DEL-177
chr5-103130345-DEL-50
chr5-103174915-DEL-237
chr5-103191017-DEL-91
chr5-103244815-DEL-52
chr5-103405933-DEL-186
chr5-103441407-DEL-69
chr5-103458660-DEL-64
chr5-103459146-DEL-240
chr5-103498550-DEL-163
chr5-103502484-DEL-235
chr5-103614646-DEL-150
chr5-103619919-DEL-208
chr5-103625436-DEL-51
chr5-103655697-DEL-1448
chr5-103692664-DEL-99
chr5-103704976-DEL-58
chr5-103723945-DEL-126
chr5-103724098-DEL-363
chr5-103752775-DEL-51
chr5-103874399-DEL-82
chr5-103874743-DEL-222
chr5-103978771-DEL-131
chr5-104006700-DEL-319
chr5-104040403-DEL-150
chr5-104045270-DEL-594
chr5-104049405-DEL-130
chr5-104050922-DEL-1538
chr5-104079704-DEL-50
chr5-104084909-DEL-297
chr5-104090620-DEL-131
chr5-104098767-DEL-53
chr5-104165394-DEL-162
chr5-104199953-DEL-1134
chr5-104213523-DEL-89
chr5-104242100-DEL-50
chr5-104250722-DEL-97
chr5-104272437-DEL-1110

chr5-104277310-DEL-539
chr5-104475783-DEL-87
chr5-104478827-DEL-1167
chr5-104488548-DEL-144
chr5-104570630-DEL-71
chr5-104576846-DEL-66
chr5-104641005-DEL-54
chr5-104786238-DEL-252
chr5-104786537-DEL-406
chr5-104897888-DEL-252
chr5-104901125-DEL-56
chr5-104916595-DEL-96
chr5-104952470-DEL-347
chr5-104962672-DEL-84
chr5-105101305-DEL-398
chr5-105115160-DEL-66
chr5-105216488-DEL-3169
chr5-105382687-DEL-170
chr5-105387521-DEL-51
chr5-105409273-DEL-66
chr5-105409574-DEL-55
chr5-105437693-DEL-52
chr5-105686942-DEL-184
chr5-105687064-DEL-60
chr5-105691516-DEL-86
chr5-105709244-DEL-55
chr5-105711557-DEL-104
chr5-105773084-DEL-96
chr5-105786403-DEL-133
chr5-105919305-DEL-958
chr5-105947174-DEL-111
chr5-105947222-DEL-261
chr5-105947417-DEL-66
chr5-105977811-DEL-52
chr5-105986246-DEL-304
chr5-105987007-DEL-80
chr5-106005822-DEL-608
chr5-106005966-DEL-168
chr5-106012700-DEL-425
chr5-106020960-DEL-55
chr5-106067721-DEL-223
chr5-106074598-DEL-51
chr5-106077432-DEL-480
chr5-106077794-DEL-61
chr5-106096178-DEL-85
chr5-106107279-DEL-1069
chr5-106116763-DEL-86
chr5-106137796-DEL-64
chr5-106171111-DEL-214
chr5-106173963-DEL-205
chr5-106174260-DEL-114
chr5-106236483-DEL-100
chr5-106276059-DEL-114
chr5-106276244-DEL-189
chr5-106300863-DEL-197
chr5-106301093-DEL-687
chr5-106303525-DEL-1298
chr5-106304995-DEL-673
chr5-106324642-DEL-54
chr5-106379078-DEL-52
chr5-106483971-DEL-1872
chr5-106485564-DEL-120
chr5-106551374-DEL-476
chr5-106555623-DEL-55
chr5-106761180-DEL-70
chr5-106816571-DEL-371
chr5-106816620-DEL-125
chr5-106956307-DEL-124
chr5-106985235-DEL-76
chr5-107044414-DEL-98
chr5-107069501-DEL-492
chr5-107090490-DEL-1728
chr5-107097535-DEL-254
chr5-107125632-DEL-53
chr5-107135914-DEL-288
chr5-107369046-DEL-54
chr5-107369151-DEL-354

chr5-107473061-DEL-298
chr5-107517010-DEL-91
chr5-107519571-DEL-156
chr5-107519639-DEL-76
chr5-107534371-DEL-54
chr5-107602611-DEL-114
chr5-107740009-DEL-58
chr5-107762672-DEL-57
chr5-107778720-DEL-50
chr5-107778876-DEL-255
chr5-107779683-DEL-1192
chr5-107817321-DEL-66
chr5-107832107-DEL-255
chr5-107839452-DEL-53
chr5-108029706-DEL-52
chr5-108070965-DEL-763
chr5-108211902-DEL-64
chr5-108242388-DEL-206
chr5-108371730-DEL-57
chr5-108420590-DEL-1043
chr5-108541717-DEL-59
chr5-108545656-DEL-51
chr5-108830113-DEL-140
chr5-108859913-DEL-112
chr5-108909009-DEL-1750
chr5-108962547-DEL-2782
chr5-108967432-DEL-1625
chr5-108970511-DEL-600
chr5-108979180-DEL-66
chr5-109096560-DEL-74
chr5-109263524-DEL-158
chr5-109265650-DEL-878
chr5-109430024-DEL-2925
chr5-109614158-DEL-5843
chr5-109640819-DEL-17006
chr5-109656067-DEL-51
chr5-109677098-DEL-60
chr5-109689752-DEL-56
chr5-109738369-DEL-51
chr5-109801838-DEL-237
chr5-109969624-DEL-55
chr5-109985204-DEL-1118
chr5-110047514-DEL-407
chr5-110115813-DEL-194
chr5-110121937-DEL-6355
chr5-110133532-DEL-384
chr5-110134001-DEL-100
chr5-110134920-DEL-202
chr5-110200051-DEL-171
chr5-110240058-DEL-253
chr5-110244851-DEL-1691
chr5-110257512-DEL-2910
chr5-110257555-DEL-220
chr5-110297272-DEL-113
chr5-110320319-DEL-116
chr5-110348725-DEL-52
chr5-110456203-DEL-413
chr5-110499570-DEL-1009
chr5-110553040-DEL-599
chr5-110553786-DEL-157
chr5-110613462-DEL-223
chr5-110625025-DEL-3155
chr5-110715204-DEL-69
chr5-110800581-DEL-56
chr5-110898616-DEL-104
chr5-110951119-DEL-176
chr5-111106449-DEL-72
chr5-111110094-DEL-621
chr5-111110540-DEL-676
chr5-111110715-DEL-1464
chr5-111111786-DEL-1081
chr5-111111853-DEL-110
chr5-111112341-DEL-297
chr5-111143981-DEL-148
chr5-111277754-DEL-913
chr5-111282779-DEL-93
chr5-111297430-DEL-50

chr5-111331503-DEL-166
chr5-111430359-DEL-69
chr5-111540250-DEL-54
chr5-111588915-DEL-122
chr5-111598312-DEL-88
chr5-111617859-DEL-204
chr5-111656203-DEL-78
chr5-111687680-DEL-50
chr5-111693598-DEL-50
chr5-111706762-DEL-399
chr5-111707358-DEL-119
chr5-111723871-DEL-170
chr5-111723927-DEL-56
chr5-111792132-DEL-56
chr5-111822353-DEL-75
chr5-111874182-DEL-118
chr5-111900811-DEL-131
chr5-111905573-DEL-204
chr5-111924599-DEL-120
chr5-111924605-DEL-54
chr5-111924777-DEL-54
chr5-111934403-DEL-296
chr5-111945828-DEL-150
chr5-111960147-DEL-132
chr5-112145376-DEL-62
chr5-112145517-DEL-64
chr5-112267701-DEL-118
chr5-112305666-DEL-95
chr5-112325931-DEL-90
chr5-112417489-DEL-50
chr5-112487014-DEL-62
chr5-112519382-DEL-57
chr5-112625506-DEL-52
chr5-112646407-DEL-207
chr5-112688946-DEL-60
chr5-112710482-DEL-198
chr5-112751173-DEL-82
chr5-112805329-DEL-397
chr5-112814465-DEL-50
chr5-112824211-DEL-123
chr5-112873490-DEL-74
chr5-112889107-DEL-189
chr5-112919137-DEL-186
chr5-112937082-DEL-57
chr5-112948946-DEL-710
chr5-112966413-DEL-95
chr5-113036893-DEL-924
chr5-113038405-DEL-50
chr5-113079187-DEL-50
chr5-113114896-DEL-53
chr5-113186684-DEL-78
chr5-113199285-DEL-97
chr5-113325568-DEL-89
chr5-113344344-DEL-1969
chr5-113359297-DEL-126
chr5-113390234-DEL-59
chr5-113431463-DEL-60
chr5-113548439-DEL-120
chr5-113655579-DEL-120
chr5-113812033-DEL-88
chr5-113829565-DEL-1057
chr5-113830763-DEL-58
chr5-113846722-DEL-225
chr5-113860078-DEL-73
chr5-113918188-DEL-66
chr5-113945110-DEL-65
chr5-114013380-DEL-124
chr5-114061874-DEL-215
chr5-114062324-DEL-228
chr5-114062529-DEL-68
chr5-114137242-DEL-50
chr5-114177432-DEL-115
chr5-114233937-DEL-335
chr5-114234369-DEL-52
chr5-114234468-DEL-647
chr5-114258732-DEL-84
chr5-114324911-DEL-138

chr5-114470849-DEL-434
chr5-114479214-DEL-168
chr5-114479395-DEL-57
chr5-114534943-DEL-84
chr5-114846514-DEL-78
chr5-114884089-DEL-482
chr5-114906017-DEL-68
chr5-114906085-DEL-137
chr5-114911757-DEL-69
chr5-114953398-DEL-10408
chr5-115027981-DEL-5204
chr5-115082472-DEL-75
chr5-115095359-DEL-1413
chr5-115098941-DEL-90
chr5-115143448-DEL-51
chr5-115187710-DEL-71
chr5-115248987-DEL-279
chr5-115257927-DEL-215
chr5-115259818-DEL-102
chr5-115259947-DEL-50
chr5-115292076-DEL-1042
chr5-115345858-DEL-58
chr5-115365211-DEL-62
chr5-115457032-DEL-122
chr5-115515277-DEL-2658
chr5-115525636-DEL-55
chr5-115526046-DEL-144
chr5-115626372-DEL-675
chr5-115898925-DEL-2577
chr5-115947502-DEL-378
chr5-115969837-DEL-82
chr5-116028840-DEL-140
chr5-116036977-DEL-60
chr5-116055487-DEL-52
chr5-116136669-DEL-78
chr5-116136696-DEL-248
chr5-116136955-DEL-74
chr5-116193377-DEL-184
chr5-116229408-DEL-120
chr5-116281541-DEL-51
chr5-116329298-DEL-86
chr5-116343429-DEL-72
chr5-116378727-DEL-55
chr5-116379625-DEL-115
chr5-116392919-DEL-100
chr5-116423274-DEL-54
chr5-116467076-DEL-60
chr5-116481676-DEL-276
chr5-116611054-DEL-94
chr5-116776822-DEL-58
chr5-116878406-DEL-61
chr5-117166189-DEL-53
chr5-117236960-DEL-87
chr5-117369591-DEL-76
chr5-117584486-DEL-187
chr5-117658468-DEL-109
chr5-117685395-DEL-211
chr5-117738544-DEL-56
chr5-117802127-DEL-91
chr5-117811278-DEL-66
chr5-117854038-DEL-50
chr5-118002621-DEL-77
chr5-118038110-DEL-160
chr5-118038158-DEL-54
chr5-118145152-DEL-178
chr5-118145155-DEL-57
chr5-118159592-DEL-196
chr5-118162055-DEL-206
chr5-118189901-DEL-67
chr5-118321403-DEL-112
chr5-118321527-DEL-52
chr5-118340328-DEL-106
chr5-118452572-DEL-60
chr5-118456819-DEL-52
chr5-118569841-DEL-69
chr5-118570130-DEL-56
chr5-118648174-DEL-53

chr5-118709179-DEL-59
chr5-118886063-DEL-60
chr5-118932563-DEL-123
chr5-118991458-DEL-121
chr5-119019661-DEL-205
chr5-119064776-DEL-153
chr5-119092775-DEL-62
chr5-119121771-DEL-56
chr5-119265438-DEL-286
chr5-119383893-DEL-64
chr5-119408491-DEL-226
chr5-119408500-DEL-54
chr5-119472382-DEL-159
chr5-119481821-DEL-82
chr5-119489239-DEL-274
chr5-119489659-DEL-147
chr5-119489771-DEL-56
chr5-119506472-DEL-114
chr5-119614936-DEL-150
chr5-119633793-DEL-66
chr5-119748211-DEL-86
chr5-119856198-DEL-329
chr5-119858443-DEL-52
chr5-119904043-DEL-162
chr5-119918384-DEL-109
chr5-120032859-DEL-60
chr5-120074435-DEL-133
chr5-120236987-DEL-156
chr5-120336773-DEL-101
chr5-120389085-DEL-728
chr5-120441925-DEL-104
chr5-120457023-DEL-80
chr5-120482885-DEL-228
chr5-120545608-DEL-51
chr5-120564228-DEL-62
chr5-120612872-DEL-59
chr5-120699688-DEL-75
chr5-120783836-DEL-60
chr5-120804027-DEL-72
chr5-120883397-DEL-120
chr5-120884067-DEL-50
chr5-120996137-DEL-52
chr5-121023199-DEL-179
chr5-121044252-DEL-60
chr5-121045908-DEL-83
chr5-121046048-DEL-106
chr5-121094184-DEL-89
chr5-121173681-DEL-285
chr5-121193538-DEL-50
chr5-121225425-DEL-52
chr5-121248132-DEL-60
chr5-121262819-DEL-255
chr5-121265073-DEL-126
chr5-121361729-DEL-50
chr5-121470480-DEL-124
chr5-121693413-DEL-4073
chr5-122171787-DEL-54
chr5-122229224-DEL-883
chr5-122237724-DEL-64
chr5-122264157-DEL-122
chr5-122264231-DEL-54
chr5-122323048-DEL-78
chr5-122364980-DEL-208
chr5-122659627-DEL-693
chr5-122659821-DEL-296
chr5-122785883-DEL-473
chr5-122787221-DEL-90
chr5-122919328-DEL-1745
chr5-122968300-DEL-218
chr5-123022337-DEL-583
chr5-123064905-DEL-53
chr5-123174187-DEL-55
chr5-123199357-DEL-226
chr5-123210095-DEL-853
chr5-123251363-DEL-57
chr5-123321871-DEL-83
chr5-123384673-DEL-492

chr5-123385031-DEL-696
chr5-123458702-DEL-129
chr5-123657165-DEL-97
chr5-123706825-DEL-310
chr5-123758408-DEL-202
chr5-123810604-DEL-111
chr5-123853488-DEL-189
chr5-123993173-DEL-62
chr5-123993436-DEL-432
chr5-124041544-DEL-69
chr5-124054718-DEL-75
chr5-124072796-DEL-62
chr5-124125939-DEL-106
chr5-124191772-DEL-212
chr5-124208826-DEL-452
chr5-124473166-DEL-86
chr5-124473517-DEL-187
chr5-124701148-DEL-1749
chr5-124786079-DEL-1116
chr5-124786871-DEL-144
chr5-124787159-DEL-252
chr5-124788802-DEL-225
chr5-124855697-DEL-118
chr5-125049680-DEL-1136
chr5-125071805-DEL-55
chr5-125243386-DEL-338
chr5-125320017-DEL-54
chr5-125326971-DEL-470
chr5-125337359-DEL-1162
chr5-125474787-DEL-773
chr5-125478149-DEL-108
chr5-125528451-DEL-52
chr5-125573561-DEL-791
chr5-125582748-DEL-168
chr5-125583626-DEL-1410
chr5-125608207-DEL-52
chr5-125638386-DEL-131
chr5-125672774-DEL-95
chr5-125672970-DEL-62
chr5-125817065-DEL-80
chr5-125818820-DEL-68
chr5-125825420-DEL-51
chr5-126047402-DEL-1010
chr5-126058088-DEL-56
chr5-126183148-DEL-59
chr5-126184278-DEL-133
chr5-126295176-DEL-77
chr5-126364777-DEL-91
chr5-126393352-DEL-112
chr5-126393462-DEL-226
chr5-126550294-DEL-218
chr5-126606760-DEL-164
chr5-126684476-DEL-74
chr5-126736209-DEL-75
chr5-126820422-DEL-50
chr5-126903824-DEL-628
chr5-126977249-DEL-240
chr5-127019056-DEL-52
chr5-127146632-DEL-52
chr5-127244120-DEL-131
chr5-127262737-DEL-94
chr5-127588425-DEL-82
chr5-127705282-DEL-99
chr5-127732164-DEL-503
chr5-127792372-DEL-298
chr5-127881893-DEL-762
chr5-127893120-DEL-75
chr5-128002660-DEL-58
chr5-128007566-DEL-248
chr5-128079886-DEL-51
chr5-128189911-DEL-53
chr5-128199435-DEL-65
chr5-128392654-DEL-224
chr5-128450378-DEL-166
chr5-128597373-DEL-63
chr5-128640946-DEL-50
chr5-128798875-DEL-6329

chr5-128805260-DEL-242
chr5-128899698-DEL-53
chr5-128920767-DEL-67
chr5-128951654-DEL-222
chr5-129000544-DEL-114
chr5-129025029-DEL-159
chr5-129034072-DEL-169
chr5-129049871-DEL-330
chr5-129052251-DEL-105
chr5-129092077-DEL-59
chr5-129181471-DEL-96
chr5-129212720-DEL-108
chr5-129234664-DEL-110
chr5-129265112-DEL-56
chr5-129287224-DEL-375
chr5-129287979-DEL-64
chr5-129305382-DEL-52
chr5-129331020-DEL-265
chr5-129331234-DEL-352
chr5-129331660-DEL-127
chr5-129402178-DEL-204
chr5-129402352-DEL-50
chr5-129463784-DEL-398
chr5-129468999-DEL-50
chr5-129480268-DEL-1263
chr5-129691168-DEL-422
chr5-129691634-DEL-227
chr5-129692115-DEL-210
chr5-129711968-DEL-201
chr5-129787260-DEL-50
chr5-129871864-DEL-250
chr5-129873436-DEL-122
chr5-129894181-DEL-70
chr5-129894277-DEL-177
chr5-129903426-DEL-252
chr5-130152100-DEL-51
chr5-130231850-DEL-61
chr5-130331371-DEL-52
chr5-130337383-DEL-107
chr5-130360935-DEL-58
chr5-130410852-DEL-87
chr5-130436704-DEL-193
chr5-130473281-DEL-50
chr5-130498489-DEL-70
chr5-130502290-DEL-50
chr5-130508275-DEL-53
chr5-130537228-DEL-52
chr5-130558276-DEL-209
chr5-130558350-DEL-53
chr5-130560599-DEL-57
chr5-130586253-DEL-55
chr5-130698032-DEL-53
chr5-130781936-DEL-82
chr5-130793384-DEL-118
chr5-130829977-DEL-173
chr5-130950117-DEL-152
chr5-130981991-DEL-53
chr5-130999900-DEL-208
chr5-131024814-DEL-53
chr5-131078173-DEL-171
chr5-131092509-DEL-114
chr5-131128794-DEL-53
chr5-131158397-DEL-215
chr5-131290746-DEL-64
chr5-131339324-DEL-53
chr5-131457914-DEL-53
chr5-131526452-DEL-159
chr5-131536199-DEL-51
chr5-131574794-DEL-107
chr5-131652084-DEL-206
chr5-131661700-DEL-470
chr5-131661830-DEL-1504
chr5-131661843-DEL-188
chr5-131662230-DEL-329
chr5-131667764-DEL-104
chr5-131677378-DEL-106
chr5-131742561-DEL-228

chr5-131826798-DEL-60
chr5-131928037-DEL-125
chr5-131996318-DEL-78
chr5-132262843-DEL-9490
chr5-132396198-DEL-50
chr5-132440368-DEL-102
chr5-132489626-DEL-54
chr5-132619823-DEL-106
chr5-132620061-DEL-57
chr5-132624600-DEL-106
chr5-132694026-DEL-60
chr5-132735170-DEL-53
chr5-132766000-DEL-212
chr5-132777597-DEL-53
chr5-132780087-DEL-106
chr5-132780321-DEL-131
chr5-132783226-DEL-221
chr5-132795346-DEL-53
chr5-132807781-DEL-64
chr5-132854132-DEL-50
chr5-132855560-DEL-53
chr5-132863877-DEL-158
chr5-132876212-DEL-567
chr5-132876490-DEL-61
chr5-132905414-DEL-51
chr5-132938470-DEL-80
chr5-133014047-DEL-53
chr5-133021164-DEL-74
chr5-133028542-DEL-688
chr5-133144373-DEL-183
chr5-133156813-DEL-60
chr5-133246159-DEL-57
chr5-133253799-DEL-70
chr5-133452416-DEL-50
chr5-133489590-DEL-7051
chr5-133515613-DEL-121
chr5-133524082-DEL-89
chr5-133524089-DEL-705
chr5-133524467-DEL-220
chr5-133538199-DEL-83
chr5-133603208-DEL-58
chr5-133606049-DEL-53
chr5-133688573-DEL-264
chr5-133715512-DEL-316
chr5-133755493-DEL-91
chr5-133755651-DEL-758
chr5-133756913-DEL-915
chr5-133764259-DEL-62
chr5-133858058-DEL-1032
chr5-133977729-DEL-180
chr5-133988289-DEL-69
chr5-133988488-DEL-360
chr5-134009826-DEL-85
chr5-134027879-DEL-218
chr5-134030382-DEL-55
chr5-134045675-DEL-680
chr5-134064978-DEL-324
chr5-134072154-DEL-266
chr5-134132907-DEL-265
chr5-134154272-DEL-269
chr5-134166678-DEL-106
chr5-134188910-DEL-201
chr5-134257931-DEL-97
chr5-134326263-DEL-183
chr5-134356425-DEL-83
chr5-134364930-DEL-226
chr5-134372325-DEL-1064
chr5-134376453-DEL-153
chr5-134403943-DEL-106
chr5-134407705-DEL-256
chr5-134417737-DEL-53
chr5-134461500-DEL-53
chr5-134529453-DEL-105
chr5-134543381-DEL-174
chr5-134556234-DEL-517
chr5-134556642-DEL-109
chr5-134590201-DEL-482

chr5-134622567-DEL-53
chr5-134623176-DEL-96
chr5-134639631-DEL-157
chr5-134639737-DEL-51
chr5-134677827-DEL-53
chr5-134678337-DEL-70
chr5-134684374-DEL-312
chr5-134802266-DEL-157
chr5-134825534-DEL-318
chr5-134836272-DEL-216
chr5-134837187-DEL-96
chr5-134843613-DEL-106
chr5-134935112-DEL-142
chr5-134968235-DEL-796
chr5-134988639-DEL-65
chr5-134991299-DEL-50
chr5-135032032-DEL-551
chr5-135032676-DEL-494
chr5-135056461-DEL-53
chr5-135058904-DEL-106
chr5-135072136-DEL-77
chr5-135097503-DEL-145
chr5-135109960-DEL-53
chr5-135206907-DEL-55
chr5-135286218-DEL-52
chr5-135296191-DEL-148
chr5-135296611-DEL-53
chr5-135306159-DEL-426
chr5-135316515-DEL-53
chr5-135318554-DEL-220
chr5-135388160-DEL-53
chr5-135388429-DEL-53
chr5-135434107-DEL-62
chr5-135441770-DEL-211
chr5-135464307-DEL-160
chr5-135475247-DEL-53
chr5-135480699-DEL-245
chr5-135480703-DEL-53
chr5-135497778-DEL-53
chr5-135498278-DEL-370
chr5-135512744-DEL-52
chr5-135523017-DEL-52
chr5-135540939-DEL-120
chr5-135556227-DEL-53
chr5-135589780-DEL-265
chr5-135606601-DEL-212
chr5-135621646-DEL-203
chr5-135622974-DEL-72
chr5-135623196-DEL-381
chr5-135716294-DEL-368
chr5-135750074-DEL-63
chr5-135783004-DEL-105
chr5-135783165-DEL-306
chr5-135827922-DEL-119
chr5-135866440-DEL-195
chr5-135866688-DEL-186
chr5-135867138-DEL-62
chr5-135998555-DEL-256
chr5-136077160-DEL-72
chr5-136081912-DEL-106
chr5-136095417-DEL-164
chr5-136098248-DEL-106
chr5-136109298-DEL-53
chr5-136180038-DEL-106
chr5-136232829-DEL-105
chr5-136284054-DEL-53
chr5-136287349-DEL-53
chr5-136455194-DEL-52
chr5-136486229-DEL-60
chr5-136495995-DEL-218
chr5-136520243-DEL-88
chr5-136546049-DEL-60
chr5-136576572-DEL-52
chr5-136675284-DEL-53
chr5-136678872-DEL-53
chr5-136689641-DEL-53
chr5-136692124-DEL-85

chr5-136731891-DEL-212
chr5-136756283-DEL-169
chr5-136756647-DEL-89
chr5-136782159-DEL-53
chr5-136787229-DEL-416
chr5-136797328-DEL-159
chr5-136846762-DEL-166
chr5-136855715-DEL-159
chr5-136855993-DEL-580
chr5-136861244-DEL-94
chr5-136861256-DEL-253
chr5-136895462-DEL-121
chr5-136904392-DEL-54
chr5-136922966-DEL-54
chr5-136931061-DEL-52
chr5-136957455-DEL-186
chr5-136966937-DEL-52
chr5-136978605-DEL-315
chr5-136987091-DEL-53
chr5-137045371-DEL-107
chr5-137047439-DEL-162
chr5-137087736-DEL-53
chr5-137087945-DEL-106
chr5-137088654-DEL-182
chr5-137115511-DEL-123
chr5-137115920-DEL-325
chr5-137117601-DEL-156
chr5-137121483-DEL-51
chr5-137152946-DEL-377
chr5-137171950-DEL-51
chr5-137178731-DEL-72
chr5-137179284-DEL-252
chr5-137191768-DEL-50
chr5-137194892-DEL-319
chr5-137217436-DEL-900
chr5-137224209-DEL-90
chr5-137255268-DEL-2709
chr5-137259563-DEL-144
chr5-137320270-DEL-254
chr5-137322496-DEL-196
chr5-137343318-DEL-524
chr5-137395320-DEL-106
chr5-137452940-DEL-212
chr5-137462596-DEL-237
chr5-137479137-DEL-50
chr5-137592005-DEL-50
chr5-137595004-DEL-62
chr5-137604085-DEL-184
chr5-137615492-DEL-100
chr5-137663223-DEL-53
chr5-137666227-DEL-212
chr5-137674174-DEL-285
chr5-137686518-DEL-140
chr5-137734865-DEL-52
chr5-137753034-DEL-391
chr5-137810803-DEL-3057
chr5-137825221-DEL-188
chr5-137836268-DEL-203
chr5-137845748-DEL-15566
chr5-137855403-DEL-13710
chr5-137882658-DEL-55
chr5-137967477-DEL-94
chr5-138019977-DEL-15971
chr5-138118983-DEL-2040
chr5-138147410-DEL-1040
chr5-138157464-DEL-929
chr5-138174645-DEL-50
chr5-138178720-DEL-50
chr5-138238944-DEL-50
chr5-138279578-DEL-50
chr5-138462288-DEL-824
chr5-138463136-DEL-90
chr5-138608414-DEL-222
chr5-138613698-DEL-88
chr5-138655640-DEL-66
chr5-138702166-DEL-54
chr5-138759313-DEL-53

chr5-138785777-DEL-192
chr5-138812555-DEL-87
chr5-138823417-DEL-90
chr5-138878469-DEL-52
chr5-138919096-DEL-75
chr5-138921239-DEL-98
chr5-138941048-DEL-51
chr5-138954283-DEL-78
chr5-139023409-DEL-104
chr5-139026845-DEL-120
chr5-139047826-DEL-159
chr5-139054290-DEL-81
chr5-139058103-DEL-53
chr5-139068889-DEL-107
chr5-139090016-DEL-167
chr5-139090311-DEL-136
chr5-139112895-DEL-213
chr5-139174699-DEL-207
chr5-139175058-DEL-52
chr5-139175483-DEL-111
chr5-139179969-DEL-120
chr5-139232951-DEL-106
chr5-139261773-DEL-141
chr5-139292071-DEL-53
chr5-139437959-DEL-56
chr5-139439803-DEL-78
chr5-139508801-DEL-263
chr5-139512092-DEL-52
chr5-139565265-DEL-154
chr5-139593263-DEL-420
chr5-139594233-DEL-70
chr5-139596299-DEL-60
chr5-139597566-DEL-207
chr5-139630186-DEL-61
chr5-139637650-DEL-1244
chr5-139641863-DEL-1449
chr5-139656497-DEL-52
chr5-139700623-DEL-207
chr5-139722753-DEL-279
chr5-139758863-DEL-106
chr5-139763762-DEL-50
chr5-139802223-DEL-202
chr5-139802553-DEL-1431
chr5-139828402-DEL-141
chr5-139838413-DEL-105
chr5-139840474-DEL-50
chr5-139884165-DEL-53
chr5-139922881-DEL-373
chr5-139923334-DEL-52
chr5-139952781-DEL-52
chr5-139997510-DEL-106
chr5-139997846-DEL-158
chr5-140030565-DEL-64
chr5-140052132-DEL-162
chr5-140063246-DEL-103
chr5-140071435-DEL-55
chr5-140259473-DEL-69
chr5-140285433-DEL-462
chr5-140312368-DEL-55
chr5-140312843-DEL-226
chr5-140354023-DEL-159
chr5-140369824-DEL-56
chr5-140376032-DEL-67
chr5-140377043-DEL-141
chr5-140393560-DEL-598
chr5-140393623-DEL-80
chr5-140419044-DEL-206
chr5-140424497-DEL-905
chr5-140435020-DEL-67
chr5-140469872-DEL-264
chr5-140480396-DEL-53
chr5-140482102-DEL-103
chr5-140486926-DEL-105
chr5-140498076-DEL-211
chr5-140504048-DEL-53
chr5-140515569-DEL-5132
chr5-140540846-DEL-118

chr5-140541188-DEL-57
chr5-140554912-DEL-54
chr5-140694891-DEL-53
chr5-140695191-DEL-203
chr5-140725167-DEL-157
chr5-140790848-DEL-74
chr5-140806016-DEL-81
chr5-140820097-DEL-84
chr5-140868786-DEL-262
chr5-140868810-DEL-105
chr5-140880144-DEL-91
chr5-140880897-DEL-208
chr5-140887753-DEL-54
chr5-140901841-DEL-154
chr5-140953127-DEL-85
chr5-140953628-DEL-86
chr5-141001364-DEL-54
chr5-141073890-DEL-108
chr5-141118113-DEL-60
chr5-141131698-DEL-1128
chr5-141140643-DEL-76
chr5-141158925-DEL-138
chr5-141170483-DEL-157
chr5-141174285-DEL-62
chr5-141175370-DEL-70
chr5-141214619-DEL-51
chr5-141352875-DEL-82
chr5-141381413-DEL-106
chr5-141516708-DEL-3611
chr5-141554157-DEL-62
chr5-141574233-DEL-55
chr5-141576068-DEL-63
chr5-141633032-DEL-1291
chr5-141660791-DEL-66
chr5-141709308-DEL-51
chr5-141751987-DEL-160
chr5-141968052-DEL-143
chr5-141977800-DEL-145
chr5-141991339-DEL-96
chr5-142010132-DEL-876
chr5-142065993-DEL-136
chr5-142066081-DEL-56
chr5-142069162-DEL-159
chr5-142081288-DEL-52
chr5-142110565-DEL-60
chr5-142137705-DEL-52
chr5-142154446-DEL-53
chr5-142164729-DEL-62
chr5-142174134-DEL-106
chr5-142193552-DEL-54
chr5-142202614-DEL-50
chr5-142229671-DEL-210
chr5-142229830-DEL-51
chr5-142244948-DEL-53
chr5-142287956-DEL-166
chr5-142314784-DEL-222
chr5-142324796-DEL-113
chr5-142324807-DEL-252
chr5-142367877-DEL-121
chr5-142376060-DEL-75
chr5-142414205-DEL-76
chr5-142428899-DEL-130
chr5-142456727-DEL-264
chr5-142469792-DEL-435
chr5-142469823-DEL-111
chr5-142470280-DEL-52
chr5-142570733-DEL-53
chr5-142598947-DEL-56
chr5-142666902-DEL-94
chr5-142684652-DEL-275
chr5-142690695-DEL-211
chr5-142693127-DEL-96
chr5-142706347-DEL-366
chr5-142708744-DEL-171
chr5-142727580-DEL-52
chr5-142767759-DEL-51
chr5-142794727-DEL-265

chr5-142813847-DEL-53
chr5-142815335-DEL-172
chr5-142825406-DEL-283
chr5-142831039-DEL-313
chr5-142844854-DEL-52
chr5-142868940-DEL-106
chr5-142909891-DEL-107
chr5-142914004-DEL-152
chr5-142937112-DEL-51
chr5-142951261-DEL-53
chr5-142965722-DEL-200
chr5-142969894-DEL-52
chr5-143056244-DEL-1276
chr5-143119590-DEL-549
chr5-143120163-DEL-53
chr5-143137908-DEL-438
chr5-143177154-DEL-84
chr5-143189538-DEL-60
chr5-143193975-DEL-275
chr5-143257023-DEL-84
chr5-143283432-DEL-124
chr5-143290970-DEL-65
chr5-143306895-DEL-121
chr5-143332953-DEL-422
chr5-143364573-DEL-159
chr5-143368895-DEL-57
chr5-143384639-DEL-99
chr5-143399773-DEL-594
chr5-143422067-DEL-732
chr5-143462358-DEL-182
chr5-143526182-DEL-50
chr5-143589598-DEL-262
chr5-143589799-DEL-262
chr5-143591645-DEL-62
chr5-143600950-DEL-160
chr5-143793936-DEL-194
chr5-143807302-DEL-247
chr5-143807912-DEL-172
chr5-143808071-DEL-50
chr5-143830185-DEL-84
chr5-143833181-DEL-50
chr5-143868682-DEL-205
chr5-143876533-DEL-2084
chr5-143966306-DEL-158
chr5-143967585-DEL-330
chr5-143975127-DEL-50
chr5-143992437-DEL-165
chr5-144022200-DEL-56
chr5-144088242-DEL-79
chr5-144104366-DEL-299
chr5-144108920-DEL-50
chr5-144117268-DEL-78
chr5-144145979-DEL-153
chr5-144145988-DEL-54
chr5-144167975-DEL-52
chr5-144182698-DEL-105
chr5-144182795-DEL-52
chr5-144214710-DEL-138
chr5-144237639-DEL-152
chr5-144243201-DEL-452
chr5-144276870-DEL-237
chr5-144303426-DEL-157
chr5-144312861-DEL-112
chr5-144315745-DEL-60
chr5-144336123-DEL-159
chr5-144340867-DEL-52
chr5-144351032-DEL-198
chr5-144351336-DEL-414
chr5-144351528-DEL-204
chr5-144363380-DEL-50
chr5-144367304-DEL-202
chr5-144376929-DEL-97
chr5-144387171-DEL-135
chr5-144396219-DEL-52
chr5-144409403-DEL-54
chr5-144415039-DEL-116
chr5-144431605-DEL-75

chr5-144437580-DEL-64
chr5-144485749-DEL-55
chr5-144528752-DEL-154
chr5-144553283-DEL-66
chr5-144570198-DEL-74
chr5-144570261-DEL-452
chr5-144574565-DEL-683
chr5-144636181-DEL-52
chr5-144636306-DEL-156
chr5-144639513-DEL-69
chr5-144716907-DEL-54
chr5-144757538-DEL-50
chr5-144793877-DEL-52
chr5-144793877-DEL-156
chr5-144798368-DEL-4363
chr5-144924539-DEL-89
chr5-144937245-DEL-234
chr5-144968126-DEL-78
chr5-144996486-DEL-50
chr5-145001447-DEL-68
chr5-145012251-DEL-209
chr5-145012290-DEL-52
chr5-145034421-DEL-94
chr5-145134829-DEL-52
chr5-145140056-DEL-187
chr5-145184837-DEL-65
chr5-145289649-DEL-4288
chr5-145352656-DEL-64
chr5-145405018-DEL-58
chr5-145459719-DEL-205
chr5-145463690-DEL-138
chr5-145498271-DEL-104
chr5-145502460-DEL-822
chr5-145502806-DEL-360
chr5-145504479-DEL-366
chr5-145539529-DEL-75
chr5-145608595-DEL-2863
chr5-145615179-DEL-579
chr5-145716718-DEL-603
chr5-145920429-DEL-60
chr5-146024879-DEL-362
chr5-146028396-DEL-78
chr5-146042592-DEL-55
chr5-146043969-DEL-352
chr5-146143040-DEL-425
chr5-146264760-DEL-348
chr5-146282660-DEL-478
chr5-146282806-DEL-124
chr5-146314326-DEL-92
chr5-146343857-DEL-672
chr5-146359326-DEL-341
chr5-146400479-DEL-80
chr5-146400761-DEL-200
chr5-146401025-DEL-80
chr5-146457393-DEL-30111
chr5-146469913-DEL-191
chr5-146488455-DEL-190
chr5-146511014-DEL-2030
chr5-146530149-DEL-66
chr5-146662733-DEL-67
chr5-146675337-DEL-78
chr5-146721307-DEL-83
chr5-146733665-DEL-1339
chr5-146762464-DEL-63
chr5-146792692-DEL-70
chr5-146840835-DEL-50
chr5-146845891-DEL-54
chr5-146849907-DEL-196
chr5-146855615-DEL-54
chr5-146859196-DEL-191
chr5-146865368-DEL-50
chr5-146918721-DEL-79
chr5-146937074-DEL-955
chr5-146982399-DEL-72
chr5-147023354-DEL-50
chr5-147028724-DEL-560
chr5-147049467-DEL-399

chr5-147049493-DEL-163
chr5-147049843-DEL-77
chr5-147059381-DEL-54
chr5-147074719-DEL-52
chr5-147099370-DEL-118
chr5-147179135-DEL-52
chr5-147296902-DEL-305
chr5-147340626-DEL-369
chr5-147342328-DEL-144
chr5-147358047-DEL-116
chr5-147358480-DEL-145
chr5-147397065-DEL-50
chr5-147471252-DEL-730
chr5-147542397-DEL-56
chr5-147650956-DEL-104
chr5-147686689-DEL-54
chr5-147689150-DEL-150
chr5-147714147-DEL-114
chr5-147744775-DEL-3074
chr5-147768932-DEL-64
chr5-147777703-DEL-350
chr5-147795127-DEL-250
chr5-147815771-DEL-55
chr5-147833050-DEL-60
chr5-147865302-DEL-78
chr5-147896225-DEL-310
chr5-147900766-DEL-1656
chr5-147910473-DEL-432
chr5-147911915-DEL-127
chr5-147964353-DEL-58
chr5-147983790-DEL-54
chr5-148002613-DEL-296
chr5-148011732-DEL-92
chr5-148036697-DEL-120
chr5-148060783-DEL-97
chr5-148105821-DEL-50
chr5-148182863-DEL-400
chr5-148214530-DEL-80
chr5-148260174-DEL-2159
chr5-148327665-DEL-64
chr5-148333271-DEL-68
chr5-148432407-DEL-62
chr5-148548998-DEL-1118
chr5-148593071-DEL-73
chr5-148593427-DEL-67
chr5-148597148-DEL-224
chr5-148608479-DEL-68
chr5-148651684-DEL-54
chr5-148672451-DEL-408
chr5-148672902-DEL-320
chr5-148716122-DEL-92
chr5-148730394-DEL-256
chr5-148740915-DEL-50
chr5-148818052-DEL-106
chr5-148843972-DEL-114
chr5-148850844-DEL-153
chr5-148857054-DEL-360
chr5-148857092-DEL-100
chr5-148975059-DEL-148
chr5-149019230-DEL-168
chr5-149019483-DEL-174
chr5-149020113-DEL-50
chr5-149034322-DEL-121
chr5-149108484-DEL-100
chr5-149108564-DEL-120
chr5-149210978-DEL-50
chr5-149282595-DEL-270
chr5-149282752-DEL-113
chr5-149306664-DEL-140
chr5-149318995-DEL-50
chr5-149410924-DEL-62
chr5-149442881-DEL-106
chr5-149465235-DEL-52
chr5-149467086-DEL-84
chr5-149469955-DEL-66
chr5-149469967-DEL-268
chr5-149474366-DEL-225

chr5-149498373-DEL-60
chr5-149622197-DEL-69
chr5-149634474-DEL-50
chr5-149659655-DEL-120
chr5-149662497-DEL-137
chr5-149705127-DEL-51
chr5-149818236-DEL-105
chr5-149837149-DEL-101
chr5-149862539-DEL-59
chr5-149867552-DEL-240
chr5-149904462-DEL-198
chr5-149906027-DEL-156
chr5-150055378-DEL-140
chr5-150100319-DEL-293
chr5-150132774-DEL-63
chr5-150177414-DEL-195
chr5-150250486-DEL-128
chr5-150273795-DEL-864
chr5-150335016-DEL-928
chr5-150335851-DEL-272
chr5-150367563-DEL-948
chr5-150629434-DEL-348
chr5-150744317-DEL-128
chr5-150746243-DEL-61
chr5-150769571-DEL-54
chr5-150775477-DEL-88
chr5-150821362-DEL-90
chr5-150826698-DEL-114
chr5-150848508-DEL-50
chr5-150848665-DEL-139
chr5-150868344-DEL-172
chr5-150920290-DEL-61
chr5-150923003-DEL-71
chr5-150971422-DEL-418
chr5-151044146-DEL-83
chr5-151058307-DEL-335
chr5-151088835-DEL-94
chr5-151189262-DEL-54
chr5-151223089-DEL-1387
chr5-151316760-DEL-52
chr5-151331283-DEL-1458
chr5-151556001-DEL-80
chr5-151638540-DEL-91
chr6-3155419-DEL-55
chr6-3192347-DEL-96
chr6-3320477-DEL-299
chr6-3392534-DEL-38930
chr6-3419734-DEL-1247
chr6-3444838-DEL-156
chr6-3455830-DEL-50
chr6-3461394-DEL-252
chr6-3558019-DEL-55
chr6-3617866-DEL-127
chr6-3849408-DEL-136
chr6-3853442-DEL-968
chr6-4156381-DEL-52
chr6-4242767-DEL-154
chr6-4296597-DEL-66
chr6-4321903-DEL-201
chr6-4387120-DEL-144
chr6-4496787-DEL-319
chr6-4576664-DEL-73
chr6-4583313-DEL-54
chr6-4606452-DEL-50
chr6-4608787-DEL-634
chr6-4923735-DEL-505
chr6-4992906-DEL-182
chr6-5055503-DEL-50
chr6-5079656-DEL-128
chr6-5107148-DEL-110
chr6-5149810-DEL-599
chr6-5196642-DEL-88
chr6-5254011-DEL-70
chr6-5262839-DEL-61
chr6-5340565-DEL-50
chr6-5343410-DEL-50
chr6-5356848-DEL-273

chr6-5577818-DEL-243
chr6-5617495-DEL-56
chr6-5638574-DEL-50
chr6-5688609-DEL-108
chr6-5720328-DEL-145
chr6-5875421-DEL-50
chr6-5914843-DEL-78
chr6-5958409-DEL-270
chr6-6170845-DEL-162
chr6-6171007-DEL-189
chr6-6244510-DEL-200
chr6-6274714-DEL-657
chr6-6279350-DEL-862
chr6-6282067-DEL-132
chr6-6282193-DEL-270
chr6-6283712-DEL-640
chr6-6284768-DEL-288
chr6-6423689-DEL-2397
chr6-6469224-DEL-82
chr6-6490011-DEL-201
chr6-6645731-DEL-155
chr6-6645991-DEL-168
chr6-6646336-DEL-280
chr6-6646851-DEL-90
chr6-6788964-DEL-56
chr6-6907147-DEL-120
chr6-7231323-DEL-554
chr6-7283068-DEL-4910
chr6-7405905-DEL-134
chr6-7444000-DEL-57
chr6-7462802-DEL-528
chr6-7567890-DEL-78
chr6-7614109-DEL-399
chr6-7628952-DEL-74
chr6-7658670-DEL-74
chr6-7667742-DEL-91
chr6-7793540-DEL-52
chr6-7795275-DEL-2080
chr6-7809650-DEL-50
chr6-7813776-DEL-96
chr6-7883187-DEL-72
chr6-7916870-DEL-67
chr6-7958376-DEL-76
chr6-7960832-DEL-64
chr6-7962895-DEL-124
chr6-8099573-DEL-622
chr6-8099581-DEL-260
chr6-8138296-DEL-50
chr6-8165448-DEL-159
chr6-8177206-DEL-114
chr6-8177218-DEL-259
chr6-8414368-DEL-176
chr6-8451993-DEL-60
chr6-8607596-DEL-75
chr6-8621617-DEL-1373
chr6-8623128-DEL-50
chr6-8728818-DEL-581
chr6-8779827-DEL-329
chr6-8821527-DEL-50
chr6-8873982-DEL-226
chr6-9179778-DEL-120
chr6-9222111-DEL-51
chr6-9490758-DEL-1888
chr6-9557658-DEL-64
chr6-9780540-DEL-907
chr6-9974733-DEL-55
chr6-9984509-DEL-1735
chr6-10083547-DEL-52
chr6-10229019-DEL-158
chr6-10251658-DEL-83
chr6-10251873-DEL-128
chr6-10337977-DEL-74
chr6-10478560-DEL-57
chr6-10814857-DEL-75
chr6-11189010-DEL-1488
chr6-11341045-DEL-111
chr6-11415104-DEL-1806

chr6-11462466-DEL-64
chr6-11483204-DEL-60
chr6-11608822-DEL-58
chr6-11813552-DEL-69
chr6-11858525-DEL-26217
chr6-11874447-DEL-22013
chr6-12066431-DEL-1975
chr6-12097562-DEL-438
chr6-12139462-DEL-167
chr6-12233432-DEL-1882
chr6-12250797-DEL-71
chr6-12425897-DEL-6839
chr6-12459921-DEL-272
chr6-12497704-DEL-64
chr6-12604609-DEL-58
chr6-12662473-DEL-84
chr6-12717024-DEL-55
chr6-12782581-DEL-50
chr6-12784472-DEL-409
chr6-13014436-DEL-63
chr6-13417465-DEL-62
chr6-13561948-DEL-76
chr6-13600471-DEL-190
chr6-13693971-DEL-76
chr6-13853591-DEL-744
chr6-13899369-DEL-3593
chr6-13929086-DEL-50
chr6-13949217-DEL-352
chr6-14142982-DEL-78
chr6-14174062-DEL-50
chr6-14244840-DEL-2153
chr6-14264713-DEL-75
chr6-14282533-DEL-54
chr6-14413761-DEL-1146
chr6-14420162-DEL-68
chr6-14566566-DEL-1206
chr6-14603282-DEL-2847
chr6-14612015-DEL-100
chr6-14612470-DEL-109
chr6-14639953-DEL-116
chr6-14840575-DEL-1031
chr6-14927220-DEL-312
chr6-15078230-DEL-196
chr6-15174637-DEL-327
chr6-15279493-DEL-93
chr6-15365851-DEL-94
chr6-15380459-DEL-54
chr6-15472349-DEL-91
chr6-15495567-DEL-56
chr6-15511806-DEL-223
chr6-15512116-DEL-507
chr6-15527334-DEL-266
chr6-15618047-DEL-75
chr6-15835036-DEL-60
chr6-15991463-DEL-52
chr6-16112515-DEL-66
chr6-16135262-DEL-51
chr6-16179849-DEL-199
chr6-16203492-DEL-2521
chr6-16209992-DEL-364
chr6-16219204-DEL-261
chr6-16229588-DEL-114
chr6-16322850-DEL-59
chr6-16340775-DEL-58
chr6-16454873-DEL-385
chr6-16488839-DEL-678
chr6-16512496-DEL-62
chr6-16652827-DEL-5390
chr6-16909040-DEL-50
chr6-17004232-DEL-407
chr6-17004272-DEL-203
chr6-17032653-DEL-127
chr6-17123153-DEL-69
chr6-17174658-DEL-1003
chr6-17322915-DEL-54
chr6-17326332-DEL-69
chr6-17335441-DEL-1745

chr6-17371210-DEL-93
chr6-17389867-DEL-60
chr6-17492811-DEL-65
chr6-17526505-DEL-341
chr6-17564652-DEL-154
chr6-17681763-DEL-217
chr6-17717507-DEL-50
chr6-17719197-DEL-415
chr6-17727752-DEL-69
chr6-17902626-DEL-56
chr6-17935904-DEL-702
chr6-17936204-DEL-324
chr6-17936903-DEL-108
chr6-17937034-DEL-54
chr6-17999823-DEL-298
chr6-18000033-DEL-78
chr6-18052914-DEL-1366
chr6-18094197-DEL-52
chr6-18537684-DEL-50
chr6-18638475-DEL-1242
chr6-18675424-DEL-476
chr6-18675981-DEL-1612
chr6-18684618-DEL-50
chr6-18697679-DEL-229
chr6-18755129-DEL-133
chr6-18804873-DEL-159
chr6-19099738-DEL-50
chr6-19103852-DEL-52
chr6-19204454-DEL-90
chr6-19341004-DEL-66
chr6-19478061-DEL-971
chr6-19479613-DEL-666
chr6-19516261-DEL-147
chr6-19766526-DEL-50
chr6-19784965-DEL-107
chr6-19784976-DEL-330
chr6-19827291-DEL-63
chr6-19827499-DEL-190
chr6-19845338-DEL-469
chr6-19884604-DEL-85
chr6-20021611-DEL-558
chr6-20044625-DEL-58
chr6-20045988-DEL-2646
chr6-20103136-DEL-57
chr6-20119055-DEL-765
chr6-20295535-DEL-199
chr6-20321678-DEL-2465
chr6-20372812-DEL-114
chr6-20509089-DEL-60
chr6-20528247-DEL-199
chr6-20570071-DEL-1097
chr6-20572967-DEL-50
chr6-20600690-DEL-1317
chr6-20775306-DEL-313
chr6-20922409-DEL-3420
chr6-20950818-DEL-1082
chr6-20954578-DEL-53
chr6-21041043-DEL-195
chr6-21099226-DEL-74
chr6-21132718-DEL-172
chr6-21140236-DEL-134
chr6-21140339-DEL-58
chr6-21194301-DEL-125
chr6-21209029-DEL-158
chr6-21380119-DEL-67
chr6-21539676-DEL-264
chr6-21539859-DEL-118
chr6-21766520-DEL-391
chr6-21766675-DEL-176
chr6-21860342-DEL-73
chr6-21881014-DEL-1975
chr6-21888083-DEL-102
chr6-21951913-DEL-50
chr6-22065051-DEL-123
chr6-22119765-DEL-260
chr6-22125154-DEL-6668
chr6-22228381-DEL-690

chr6-22240441-DEL-1519
chr6-22260624-DEL-56
chr6-22382020-DEL-60
chr6-22386015-DEL-186
chr6-22393904-DEL-843
chr6-22399751-DEL-1539
chr6-22427198-DEL-302
chr6-22448270-DEL-59
chr6-22466857-DEL-60
chr6-22478402-DEL-1746
chr6-22492759-DEL-50
chr6-22513874-DEL-96
chr6-22528939-DEL-85
chr6-22551545-DEL-363
chr6-22551555-DEL-750
chr6-22648220-DEL-1386
chr6-22750666-DEL-33071
chr6-22840272-DEL-77
chr6-22871373-DEL-60
chr6-22924417-DEL-54
chr6-22953629-DEL-50
chr6-23079535-DEL-110
chr6-23097750-DEL-126
chr6-23121789-DEL-98
chr6-23123972-DEL-229
chr6-23156792-DEL-289
chr6-23178596-DEL-64
chr6-23364890-DEL-212
chr6-23378624-DEL-97
chr6-23432902-DEL-128
chr6-23459942-DEL-180
chr6-23505043-DEL-81
chr6-23525159-DEL-212
chr6-23541046-DEL-60
chr6-23586815-DEL-135
chr6-23604823-DEL-185
chr6-23605181-DEL-6211
chr6-24039141-DEL-65
chr6-24084197-DEL-943
chr6-24118831-DEL-50
chr6-24161290-DEL-56
chr6-24161509-DEL-197
chr6-24356976-DEL-866
chr6-24470919-DEL-50
chr6-24491956-DEL-212
chr6-24608562-DEL-52
chr6-24644718-DEL-80
chr6-24732421-DEL-89
chr6-24755118-DEL-268
chr6-24851887-DEL-97
chr6-24885280-DEL-903
chr6-24907106-DEL-104
chr6-24956912-DEL-50
chr6-24956993-DEL-137
chr6-25036695-DEL-67
chr6-25079224-DEL-51
chr6-25362983-DEL-64
chr6-25394561-DEL-195
chr6-25436794-DEL-258
chr6-25602681-DEL-525
chr6-25652830-DEL-90
chr6-25711361-DEL-217
chr6-25711412-DEL-74
chr6-25745779-DEL-52
chr6-25758353-DEL-273
chr6-25837174-DEL-58
chr6-26003681-DEL-463
chr6-26040393-DEL-2367
chr6-26062562-DEL-188
chr6-26111992-DEL-55
chr6-26364397-DEL-2066
chr6-26528153-DEL-59
chr6-26564218-DEL-71
chr6-26765131-DEL-52
chr6-26766125-DEL-50
chr6-26822207-DEL-226
chr6-26859867-DEL-389

chr6-26868981-DEL-1839
chr6-26962115-DEL-2800
chr6-27075926-DEL-64
chr6-27140136-DEL-2514
chr6-27280843-DEL-311
chr6-27306183-DEL-1466
chr6-27422833-DEL-171
chr6-27515117-DEL-142
chr6-27767751-DEL-152
chr6-27922795-DEL-72
chr6-28083597-DEL-52
chr6-28138591-DEL-51
chr6-28347098-DEL-131
chr6-28436867-DEL-51
chr6-28476752-DEL-76
chr6-28531654-DEL-184
chr6-28701541-DEL-50
chr6-28990028-DEL-54
chr6-29030666-DEL-172
chr6-29086791-DEL-55
chr6-29233497-DEL-152
chr6-29272125-DEL-181
chr6-29283973-DEL-96
chr6-29373997-DEL-50
chr6-29479909-DEL-66
chr6-29487671-DEL-235
chr6-29519590-DEL-58
chr6-29748813-DEL-173
chr6-29749141-DEL-50
chr6-29879618-DEL-50
chr6-29991059-DEL-128
chr6-29991129-DEL-270
chr6-30011648-DEL-167
chr6-30011964-DEL-140
chr6-30012175-DEL-210
chr6-30019064-DEL-52
chr6-30019630-DEL-182
chr6-30038491-DEL-101
chr6-30038730-DEL-207
chr6-30061644-DEL-72
chr6-30108259-DEL-57
chr6-30157738-DEL-568
chr6-30158334-DEL-104
chr6-30461930-DEL-88
chr6-30486425-DEL-124
chr6-30522636-DEL-74
chr6-30535879-DEL-124
chr6-30535953-DEL-55
chr6-30591519-DEL-91
chr6-30852008-DEL-2506
chr6-30951135-DEL-55
chr6-30953555-DEL-65
chr6-30973373-DEL-4059
chr6-31012550-DEL-1426
chr6-31022154-DEL-675
chr6-31034187-DEL-836
chr6-31056775-DEL-161
chr6-31155756-DEL-57
chr6-31326783-DEL-114
chr6-31507587-DEL-699
chr6-31558419-DEL-150
chr6-31603993-DEL-55
chr6-31610829-DEL-75
chr6-31628787-DEL-125
chr6-31650656-DEL-147
chr6-31653548-DEL-57
chr6-31680937-DEL-58
chr6-31776305-DEL-5218
chr6-31808025-DEL-127
chr6-31835610-DEL-732
chr6-31843333-DEL-106
chr6-31862424-DEL-130
chr6-32012347-DEL-1486
chr6-32055788-DEL-3743
chr6-32071138-DEL-212
chr6-32091852-DEL-775
chr6-32106003-DEL-1202

chr6-32243917-DEL-108
chr6-32497600-DEL-78
chr6-32688276-DEL-54
chr6-32772409-DEL-60
chr6-32989272-DEL-131
chr6-33050513-DEL-179
chr6-33061635-DEL-65
chr6-33061850-DEL-248
chr6-33096159-DEL-54
chr6-33151544-DEL-2402
chr6-33179279-DEL-3513
chr6-33247781-DEL-96
chr6-33267855-DEL-1774
chr6-33393129-DEL-130
chr6-33455831-DEL-50
chr6-33486628-DEL-50
chr6-33595664-DEL-111
chr6-33858568-DEL-60
chr6-33920123-DEL-160
chr6-33995008-DEL-55
chr6-34090116-DEL-204
chr6-34175758-DEL-60
chr6-34296488-DEL-67
chr6-34351825-DEL-508
chr6-34603411-DEL-54
chr6-34829333-DEL-131
chr6-34898383-DEL-62
chr6-35260652-DEL-188
chr6-35263653-DEL-323
chr6-35268605-DEL-162
chr6-35346742-DEL-105
chr6-35396186-DEL-82
chr6-35427711-DEL-545
chr6-35478064-DEL-73
chr6-35511892-DEL-202
chr6-35661478-DEL-2582
chr6-35790180-DEL-120
chr6-36034736-DEL-1003
chr6-36055785-DEL-56
chr6-36118558-DEL-4626
chr6-36132873-DEL-2265
chr6-36212591-DEL-376
chr6-36607721-DEL-107
chr6-36957192-DEL-1911
chr6-37058622-DEL-660
chr6-37065777-DEL-191
chr6-37065925-DEL-56
chr6-37152906-DEL-70
chr6-37204004-DEL-208
chr6-37238555-DEL-50
chr6-37258157-DEL-378
chr6-37390094-DEL-54
chr6-37552919-DEL-4395
chr6-37555767-DEL-1964
chr6-37558739-DEL-499
chr6-37559309-DEL-680
chr6-37575426-DEL-122
chr6-37656215-DEL-133
chr6-37664110-DEL-941
chr6-37674308-DEL-1015
chr6-37774086-DEL-64
chr6-38016257-DEL-59
chr6-38150867-DEL-72
chr6-38167211-DEL-206
chr6-38340263-DEL-65
chr6-38345028-DEL-68
chr6-38350367-DEL-245
chr6-38357098-DEL-53
chr6-38521030-DEL-221
chr6-38551757-DEL-1174
chr6-38552968-DEL-96
chr6-38655478-DEL-87
chr6-38691842-DEL-50
chr6-38904562-DEL-57
chr6-39282371-DEL-148
chr6-39287221-DEL-144
chr6-39292977-DEL-56

chr6-39314600-DEL-54
chr6-39317662-DEL-678
chr6-39337395-DEL-511
chr6-39370134-DEL-68
chr6-39429427-DEL-411
chr6-39431242-DEL-218
chr6-39458459-DEL-213
chr6-39518408-DEL-88
chr6-39674380-DEL-410
chr6-39837102-DEL-51
chr6-39932700-DEL-50
chr6-39999026-DEL-140
chr6-40203440-DEL-57
chr6-40305340-DEL-78
chr6-40494964-DEL-210
chr6-40500223-DEL-962
chr6-40568805-DEL-66
chr6-40577196-DEL-1129
chr6-40596847-DEL-449
chr6-40609819-DEL-59
chr6-40657598-DEL-26297
chr6-40695266-DEL-1506
chr6-40699539-DEL-201
chr6-40848839-DEL-90
chr6-40929993-DEL-53
chr6-41173084-DEL-161
chr6-41190574-DEL-50
chr6-41259220-DEL-197
chr6-41373068-DEL-66542
chr6-41407234-DEL-2041
chr6-41450774-DEL-787
chr6-41507243-DEL-53
chr6-41649811-DEL-66
chr6-41741853-DEL-78
chr6-41900384-DEL-64
chr6-42072044-DEL-1710
chr6-42146792-DEL-2048
chr6-42158011-DEL-52
chr6-42398981-DEL-72
chr6-42486824-DEL-61
chr6-42581206-DEL-60
chr6-42696668-DEL-1167
chr6-42711170-DEL-89
chr6-42808563-DEL-140
chr6-42817169-DEL-95
chr6-42876011-DEL-98
chr6-42978305-DEL-200
chr6-42983357-DEL-775
chr6-42984385-DEL-2192
chr6-42986130-DEL-68
chr6-42990526-DEL-86
chr6-42998817-DEL-87
chr6-43162465-DEL-50
chr6-43463432-DEL-353
chr6-43475013-DEL-2759
chr6-43529058-DEL-1280
chr6-43605033-DEL-1944
chr6-43694163-DEL-100
chr6-43894042-DEL-210
chr6-43949822-DEL-170
chr6-44240038-DEL-128
chr6-44241591-DEL-2944
chr6-44246134-DEL-2266
chr6-44389394-DEL-185
chr6-44452477-DEL-57
chr6-44596978-DEL-458
chr6-44601206-DEL-51
chr6-44762424-DEL-416
chr6-44845440-DEL-132
chr6-44921262-DEL-102
chr6-44974361-DEL-6855
chr6-45041192-DEL-270
chr6-45210722-DEL-56
chr6-45387537-DEL-1050
chr6-45404041-DEL-357
chr6-45483663-DEL-302
chr6-45495557-DEL-97

chr6-45588835-DEL-56
chr6-45616222-DEL-1104
chr6-45784623-DEL-127
chr6-46061473-DEL-50
chr6-46204504-DEL-560
chr6-46228106-DEL-56
chr6-46258346-DEL-66
chr6-46276561-DEL-799
chr6-46305980-DEL-6720
chr6-46317217-DEL-500
chr6-46796932-DEL-2258
chr6-46808670-DEL-145
chr6-46871716-DEL-865
chr6-46915301-DEL-155
chr6-46949601-DEL-152
chr6-47083986-DEL-104
chr6-47106582-DEL-37862
chr6-47136529-DEL-280
chr6-47137294-DEL-171
chr6-47137586-DEL-113
chr6-47158973-DEL-51
chr6-47178823-DEL-54
chr6-47258014-DEL-159
chr6-47262952-DEL-605
chr6-47329668-DEL-55
chr6-47628966-DEL-93403
chr6-47760006-DEL-313
chr6-47808140-DEL-174
chr6-47889048-DEL-2482
chr6-47914719-DEL-382
chr6-47951045-DEL-221
chr6-48009944-DEL-98
chr6-48200624-DEL-59
chr6-48222479-DEL-586
chr6-48266277-DEL-2713
chr6-48281689-DEL-14037
chr6-48351527-DEL-7067
chr6-48398785-DEL-90
chr6-48628997-DEL-61
chr6-48731616-DEL-2887
chr6-48731753-DEL-559
chr6-48936069-DEL-96
chr6-48944637-DEL-113
chr6-48946554-DEL-341
chr6-48946592-DEL-155
chr6-49004243-DEL-55
chr6-49107264-DEL-396
chr6-49107661-DEL-50
chr6-49134919-DEL-370
chr6-49173825-DEL-52
chr6-49225329-DEL-254
chr6-49260708-DEL-89
chr6-49261028-DEL-171
chr6-49302077-DEL-131
chr6-49315033-DEL-259
chr6-49322655-DEL-1293
chr6-49361448-DEL-177
chr6-49385200-DEL-136
chr6-49456938-DEL-828
chr6-49484449-DEL-324
chr6-49486174-DEL-230
chr6-49502094-DEL-74
chr6-49566653-DEL-280
chr6-49566888-DEL-287
chr6-49684528-DEL-244
chr6-49684906-DEL-53
chr6-49862816-DEL-1595
chr6-50042379-DEL-199
chr6-50042819-DEL-155
chr6-50046691-DEL-106
chr6-50136171-DEL-57
chr6-50144460-DEL-55
chr6-50173864-DEL-273
chr6-50431162-DEL-172
chr6-50435097-DEL-6257
chr6-50488938-DEL-2106
chr6-50499243-DEL-90

chr6-50584809-DEL-61
chr6-50612665-DEL-620
chr6-50615930-DEL-255
chr6-50626890-DEL-94
chr6-50709233-DEL-60
chr6-51069028-DEL-150
chr6-51084432-DEL-50
chr6-51216461-DEL-57
chr6-51239138-DEL-5203
chr6-51350973-DEL-132
chr6-51395045-DEL-229
chr6-51395141-DEL-88
chr6-51419956-DEL-54
chr6-51480259-DEL-223
chr6-51480540-DEL-691
chr6-51481388-DEL-441
chr6-51491168-DEL-53
chr6-51536412-DEL-52
chr6-51540609-DEL-140
chr6-51627090-DEL-88
chr6-51627381-DEL-581
chr6-51638060-DEL-50
chr6-51690500-DEL-308
chr6-51725493-DEL-95
chr6-51770560-DEL-76
chr6-51798266-DEL-223
chr6-51918411-DEL-455
chr6-51941675-DEL-50
chr6-51963215-DEL-78
chr6-52057055-DEL-320
chr6-52057240-DEL-57
chr6-52057464-DEL-129
chr6-52057649-DEL-110
chr6-52066040-DEL-1590
chr6-52074742-DEL-124
chr6-52258521-DEL-1947
chr6-52304283-DEL-55
chr6-52325403-DEL-2625
chr6-52328384-DEL-322
chr6-52337292-DEL-127
chr6-52348630-DEL-215
chr6-52613494-DEL-3005
chr6-52757437-DEL-85
chr6-52801123-DEL-81
chr6-52922025-DEL-81
chr6-52922203-DEL-50
chr6-52922284-DEL-56
chr6-52922626-DEL-223
chr6-52939360-DEL-269
chr6-53026426-DEL-258
chr6-53128782-DEL-147
chr6-53485619-DEL-50
chr6-53536173-DEL-232
chr6-53708551-DEL-54
chr6-53749854-DEL-224
chr6-53910773-DEL-83
chr6-54138978-DEL-50
chr6-54143456-DEL-85
chr6-54145796-DEL-214
chr6-54166035-DEL-250
chr6-54190360-DEL-130
chr6-54240025-DEL-144
chr6-54265354-DEL-231
chr6-54369853-DEL-86
chr6-54409432-DEL-54
chr6-54433304-DEL-408
chr6-54464000-DEL-60
chr6-54565577-DEL-51
chr6-54565899-DEL-85
chr6-54639738-DEL-86
chr6-54639891-DEL-113
chr6-54648961-DEL-327
chr6-54673371-DEL-499
chr6-54701075-DEL-110
chr6-54822348-DEL-236
chr6-54859025-DEL-54
chr6-54953908-DEL-494

chr6-54954623-DEL-182
chr6-54957265-DEL-111
chr6-55007810-DEL-52
chr6-55008008-DEL-466
chr6-55034128-DEL-226
chr6-55068466-DEL-59
chr6-55131355-DEL-50
chr6-55192240-DEL-98
chr6-55205871-DEL-64
chr6-55220011-DEL-57
chr6-55327315-DEL-50
chr6-55347456-DEL-188
chr6-55354298-DEL-68
chr6-55554841-DEL-51
chr6-55903969-DEL-65
chr6-55965294-DEL-92
chr6-56065273-DEL-160
chr6-56213617-DEL-51
chr6-56276491-DEL-214
chr6-56303330-DEL-116
chr6-56488311-DEL-101
chr6-56568847-DEL-156
chr6-56685154-DEL-1071
chr6-56813939-DEL-152
chr6-56824300-DEL-722
chr6-57029600-DEL-234
chr6-57207189-DEL-115
chr6-57351695-DEL-214
chr6-57400817-DEL-413
chr6-57454955-DEL-196
chr6-57458351-DEL-206
chr6-57469828-DEL-321
chr6-57559873-DEL-111
chr6-57668370-DEL-108
chr6-57673428-DEL-50
chr6-57733372-DEL-1710
chr6-57751286-DEL-72
chr6-58038201-DEL-2455
chr6-58129209-DEL-142
chr6-58137646-DEL-58
chr6-58138331-DEL-116
chr6-58140864-DEL-56
chr6-58150628-DEL-204
chr6-58197453-DEL-80
chr6-58494642-DEL-412
chr6-58505311-DEL-56
chr6-58510848-DEL-11539
chr6-58518225-DEL-5431
chr6-58560162-DEL-54
chr6-58574089-DEL-56
chr6-58700192-DEL-61
chr6-58701319-DEL-3038
chr6-59293479-DEL-68
chr6-59775621-DEL-2131
chr6-59790154-DEL-98
chr6-59799091-DEL-121
chr6-59808291-DEL-2288
chr6-59819436-DEL-182
chr6-59826246-DEL-109
chr6-59828364-DEL-59
chr6-59855656-DEL-56
chr6-59874430-DEL-69
chr6-59876952-DEL-838
chr6-59895624-DEL-1412
chr6-59904511-DEL-84
chr6-59905176-DEL-50
chr6-59907561-DEL-2776
chr6-59914006-DEL-666
chr6-59945643-DEL-126
chr6-59966064-DEL-2045
chr6-59978034-DEL-559
chr6-59994279-DEL-861
chr6-59996276-DEL-2928
chr6-60047978-DEL-149
chr6-60085150-DEL-84
chr6-60103539-DEL-132
chr6-60111222-DEL-196

chr6-60114742-DEL-1069
chr6-60140341-DEL-57
chr6-60206246-DEL-81
chr6-60426074-DEL-113
chr6-60441194-DEL-7090
chr6-60448680-DEL-122
chr6-60498352-DEL-203
chr6-60535275-DEL-612
chr6-60542771-DEL-1340
chr6-60594726-DEL-65
chr6-60596554-DEL-199
chr6-60625767-DEL-104
chr6-60750049-DEL-496
chr6-60815606-DEL-149
chr6-60832625-DEL-541
chr6-60856501-DEL-64
chr6-60856585-DEL-204
chr6-60856788-DEL-64
chr6-60867241-DEL-2583
chr6-60880458-DEL-589
chr6-60900078-DEL-176
chr6-60999044-DEL-80
chr6-61098561-DEL-54
chr6-61320209-DEL-661
chr6-61321950-DEL-52
chr6-61622150-DEL-215
chr6-61697348-DEL-82
chr6-61718065-DEL-551
chr6-61722751-DEL-3008
chr6-61731575-DEL-53
chr6-61779109-DEL-87
chr6-61917743-DEL-60
chr6-61993087-DEL-317
chr6-62033310-DEL-51
chr6-62038075-DEL-350
chr6-62091039-DEL-913
chr6-62101751-DEL-63
chr6-62128981-DEL-56
chr6-62129090-DEL-71
chr6-62171678-DEL-61
chr6-62203112-DEL-60
chr6-62265683-DEL-2703
chr6-62292111-DEL-52
chr6-62335361-DEL-77
chr6-62421856-DEL-52
chr6-62457844-DEL-101
chr6-62473519-DEL-181
chr6-62544560-DEL-215
chr6-62545226-DEL-373
chr6-62583504-DEL-1579
chr6-62623954-DEL-50
chr6-62952511-DEL-52
chr6-62971808-DEL-199
chr6-62972238-DEL-57
chr6-63040397-DEL-1246
chr6-63207640-DEL-1186
chr6-63240804-DEL-285
chr6-63346241-DEL-535
chr6-63418660-DEL-474
chr6-63437614-DEL-243
chr6-63503367-DEL-321
chr6-63506994-DEL-56
chr6-63513425-DEL-568
chr6-63530678-DEL-51
chr6-63555328-DEL-894
chr6-63720582-DEL-129
chr6-63720582-DEL-267
chr6-63886595-DEL-57
chr6-63886633-DEL-209
chr6-63892217-DEL-125
chr6-64018587-DEL-53
chr6-64308723-DEL-6813
chr6-64398039-DEL-59
chr6-64508004-DEL-401
chr6-64820106-DEL-122
chr6-64820385-DEL-337
chr6-64820689-DEL-66

chr6-64849798-DEL-171
chr6-64895932-DEL-760
chr6-64897194-DEL-50
chr6-64950904-DEL-52
chr6-64977879-DEL-274
chr6-64978300-DEL-400
chr6-64979967-DEL-158
chr6-65234944-DEL-1606
chr6-65244766-DEL-84
chr6-65269489-DEL-52
chr6-65269750-DEL-242
chr6-65361510-DEL-7381
chr6-65656005-DEL-51
chr6-65824861-DEL-55
chr6-65825182-DEL-165
chr6-65874215-DEL-546
chr6-65874641-DEL-78
chr6-66140003-DEL-483
chr6-66143094-DEL-2417
chr6-66150873-DEL-724
chr6-66282828-DEL-349
chr6-66608299-DEL-117
chr6-66608709-DEL-504
chr6-66632073-DEL-207
chr6-66939773-DEL-2717
chr6-66960237-DEL-100
chr6-66960588-DEL-100
chr6-67139938-DEL-1364
chr6-67184382-DEL-222
chr6-67366269-DEL-242
chr6-67516053-DEL-171
chr6-67658654-DEL-53
chr6-67692469-DEL-256
chr6-67709259-DEL-711
chr6-67710971-DEL-70
chr6-67728038-DEL-424
chr6-67778820-DEL-258
chr6-67866795-DEL-1984
chr6-67871189-DEL-58
chr6-67875228-DEL-1099
chr6-67888197-DEL-50
chr6-67902139-DEL-50
chr6-68001808-DEL-137
chr6-68019498-DEL-19717
chr6-68040236-DEL-54
chr6-68047258-DEL-5549
chr6-68080513-DEL-60
chr6-68254937-DEL-1518
chr6-68256580-DEL-407
chr6-68269211-DEL-112
chr6-68270233-DEL-2147
chr6-68275283-DEL-2031
chr6-68291042-DEL-665
chr6-68293136-DEL-654
chr6-68299144-DEL-1770
chr6-68308473-DEL-82
chr6-68312130-DEL-267
chr6-68317259-DEL-119
chr6-68317746-DEL-120
chr6-68318345-DEL-101
chr6-68326106-DEL-2893
chr6-68367428-DEL-371
chr6-68393024-DEL-220
chr6-68404392-DEL-111
chr6-68404404-DEL-314
chr6-68404591-DEL-130
chr6-68412227-DEL-56
chr6-68451668-DEL-50
chr6-68495971-DEL-3963
chr6-68518519-DEL-79
chr6-68544691-DEL-2260
chr6-68723333-DEL-276
chr6-68723864-DEL-711
chr6-68733934-DEL-3939
chr6-68737856-DEL-79
chr6-68738190-DEL-401
chr6-68744542-DEL-244

chr6-68958369-DEL-58
chr6-69006335-DEL-161283
chr6-69024330-DEL-3842
chr6-69194070-DEL-84
chr6-69360930-DEL-392
chr6-69485703-DEL-926
chr6-69493309-DEL-225
chr6-69676608-DEL-848
chr6-69757399-DEL-48308
chr6-69759957-DEL-8589
chr6-69807051-DEL-61
chr6-69816202-DEL-69
chr6-69838960-DEL-68
chr6-70041681-DEL-284
chr6-70085716-DEL-2415
chr6-70198549-DEL-174
chr6-70215574-DEL-3034
chr6-70220091-DEL-90
chr6-70243706-DEL-453
chr6-70285141-DEL-6168
chr6-70392281-DEL-1250
chr6-70395269-DEL-234
chr6-70585577-DEL-79
chr6-70592665-DEL-410
chr6-70619006-DEL-94
chr6-70676529-DEL-80
chr6-70682052-DEL-2553
chr6-70802351-DEL-159
chr6-70866142-DEL-69
chr6-70867605-DEL-193
chr6-70867914-DEL-130
chr6-70952399-DEL-54
chr6-70971073-DEL-66
chr6-71145969-DEL-902
chr6-71147078-DEL-92
chr6-71147277-DEL-498
chr6-71287181-DEL-1828
chr6-71639965-DEL-276
chr6-71639997-DEL-94
chr6-71731237-DEL-56
chr6-71887258-DEL-57
chr6-71999859-DEL-58
chr6-72013482-DEL-64
chr6-72013753-DEL-64
chr6-72067463-DEL-57
chr6-72067613-DEL-249
chr6-72122949-DEL-238
chr6-72150313-DEL-50
chr6-72150539-DEL-54
chr6-72264461-DEL-64
chr6-72270171-DEL-84
chr6-72283193-DEL-134
chr6-72320035-DEL-156
chr6-72364260-DEL-94
chr6-72447480-DEL-168
chr6-72482351-DEL-56
chr6-72525730-DEL-50
chr6-72541611-DEL-945
chr6-72542000-DEL-107
chr6-72589715-DEL-214
chr6-72621998-DEL-60
chr6-72703079-DEL-53
chr6-72804610-DEL-72
chr6-72879566-DEL-34330
chr6-72894044-DEL-97
chr6-72947348-DEL-52
chr6-72950789-DEL-277
chr6-73057597-DEL-1488
chr6-73080794-DEL-83
chr6-73099570-DEL-136
chr6-73099648-DEL-58
chr6-73177595-DEL-3183
chr6-73279504-DEL-105
chr6-73279594-DEL-51
chr6-73321376-DEL-489
chr6-73359210-DEL-59
chr6-73379483-DEL-210

chr6-73492936-DEL-542
chr6-73497005-DEL-210
chr6-73660187-DEL-156
chr6-73662234-DEL-1687
chr6-73714714-DEL-6628
chr6-73729602-DEL-1722
chr6-73859725-DEL-179
chr6-73883301-DEL-129
chr6-73889029-DEL-50
chr6-73930337-DEL-67
chr6-74168770-DEL-1473
chr6-74214549-DEL-1132
chr6-74222844-DEL-781
chr6-74319611-DEL-77
chr6-74340596-DEL-214
chr6-74401609-DEL-50
chr6-74440726-DEL-335
chr6-74454197-DEL-153
chr6-74656638-DEL-4488
chr6-74915305-DEL-50
chr6-74946474-DEL-51
chr6-74970857-DEL-77
chr6-74973526-DEL-1393
chr6-75030789-DEL-2354
chr6-75098118-DEL-1664
chr6-75109203-DEL-73
chr6-75289458-DEL-50
chr6-75315913-DEL-188
chr6-75581008-DEL-1268
chr6-75634524-DEL-335
chr6-75742530-DEL-1756
chr6-75754432-DEL-778
chr6-75759129-DEL-818
chr6-75870209-DEL-164
chr6-75949105-DEL-1325
chr6-75983423-DEL-265
chr6-76080293-DEL-672
chr6-76160290-DEL-1162
chr6-76236812-DEL-56
chr6-76296469-DEL-3800
chr6-76311778-DEL-128
chr6-76342702-DEL-102
chr6-76760777-DEL-50
chr6-76888152-DEL-152
chr6-76928247-DEL-77
chr6-76935355-DEL-70
chr6-76963691-DEL-62
chr6-77089319-DEL-52
chr6-77219091-DEL-60
chr6-77379328-DEL-242
chr6-77422553-DEL-90
chr6-77433942-DEL-110
chr6-77708840-DEL-135
chr6-77774482-DEL-54
chr6-77781164-DEL-50
chr6-77888720-DEL-121
chr6-77934122-DEL-520
chr6-77981115-DEL-222
chr6-78198166-DEL-458
chr6-78299931-DEL-1775
chr6-78412706-DEL-625
chr6-78759967-DEL-2580
chr6-78812604-DEL-128
chr6-78821477-DEL-356
chr6-78848410-DEL-427
chr6-79014619-DEL-1644
chr6-79270448-DEL-72
chr6-79284032-DEL-6233
chr6-79295067-DEL-2973
chr6-79437917-DEL-142
chr6-79438003-DEL-56
chr6-80009722-DEL-50
chr6-80342778-DEL-160
chr6-80468950-DEL-958
chr6-80525953-DEL-629
chr6-80526029-DEL-117
chr6-80724684-DEL-2175

chr6-80755995-DEL-82
chr6-80817202-DEL-50
chr6-80956422-DEL-109
chr6-80957561-DEL-62
chr6-81007628-DEL-3275
chr6-81012243-DEL-75
chr6-81070986-DEL-962
chr6-81073456-DEL-100
chr6-81081885-DEL-50
chr6-81092536-DEL-62
chr6-81223257-DEL-50
chr6-81267981-DEL-180
chr6-81344067-DEL-50
chr6-81391894-DEL-106
chr6-81479950-DEL-200
chr6-81658481-DEL-54
chr6-81792781-DEL-121
chr6-81889596-DEL-54
chr6-81958956-DEL-297
chr6-81967040-DEL-919
chr6-82064772-DEL-124
chr6-82082668-DEL-70
chr6-82084332-DEL-66
chr6-82100786-DEL-128
chr6-82101205-DEL-110
chr6-82111325-DEL-458
chr6-82245187-DEL-1460
chr6-82345128-DEL-168
chr6-82346540-DEL-345
chr6-82347022-DEL-162
chr6-82663079-DEL-58
chr6-82663734-DEL-1763
chr6-82664407-DEL-798
chr6-82685944-DEL-136
chr6-82686345-DEL-68
chr6-82695754-DEL-649
chr6-82805766-DEL-377
chr6-82806078-DEL-390
chr6-82844793-DEL-437
chr6-82931332-DEL-218
chr6-83002553-DEL-110
chr6-83002687-DEL-54
chr6-83023477-DEL-72
chr6-83104141-DEL-202
chr6-83192931-DEL-94
chr6-83193152-DEL-1033
chr6-83193204-DEL-291
chr6-83194510-DEL-608
chr6-83195458-DEL-348
chr6-83195565-DEL-126
chr6-83258097-DEL-54
chr6-83258255-DEL-115
chr6-83307407-DEL-983
chr6-83471725-DEL-97
chr6-83498109-DEL-183
chr6-83508278-DEL-366
chr6-83560522-DEL-50
chr6-83625517-DEL-59
chr6-83656243-DEL-199
chr6-83672606-DEL-84
chr6-83736338-DEL-70
chr6-83821164-DEL-51
chr6-84112453-DEL-216
chr6-84166866-DEL-96
chr6-84250822-DEL-419
chr6-84251328-DEL-105
chr6-84415216-DEL-568
chr6-84415927-DEL-114
chr6-84745727-DEL-1188
chr6-85021429-DEL-182
chr6-85065313-DEL-374
chr6-85647457-DEL-54044
chr6-85671097-DEL-114
chr6-85686923-DEL-1392
chr6-85727372-DEL-47383
chr6-85817728-DEL-177
chr6-85916963-DEL-490

chr6-85933085-DEL-82
chr6-85979866-DEL-51
chr6-86107331-DEL-91
chr6-86164100-DEL-613
chr6-86206914-DEL-142
chr6-86223968-DEL-51
chr6-86422054-DEL-88
chr6-86482084-DEL-72
chr6-86490906-DEL-220
chr6-86570945-DEL-90
chr6-86593804-DEL-1105
chr6-86617654-DEL-66
chr6-86760841-DEL-202
chr6-86887286-DEL-50
chr6-87027479-DEL-166
chr6-87151227-DEL-81
chr6-87211547-DEL-81
chr6-87211799-DEL-162
chr6-87211888-DEL-54
chr6-87270778-DEL-1052
chr6-87307075-DEL-90
chr6-87374081-DEL-148
chr6-87387203-DEL-66
chr6-87498861-DEL-70
chr6-87520655-DEL-721
chr6-87521494-DEL-51
chr6-87577315-DEL-82
chr6-87639053-DEL-50
chr6-87752203-DEL-258
chr6-88010929-DEL-90
chr6-88040252-DEL-137
chr6-88050439-DEL-114
chr6-88051382-DEL-244
chr6-88187865-DEL-100
chr6-88188312-DEL-192
chr6-88471607-DEL-375
chr6-88472375-DEL-144
chr6-88498673-DEL-119
chr6-88531876-DEL-52
chr6-88554863-DEL-108
chr6-88646684-DEL-5656
chr6-88712096-DEL-53
chr6-88722909-DEL-136
chr6-88962702-DEL-464
chr6-88974428-DEL-905
chr6-88997332-DEL-1746
chr6-89040482-DEL-3258
chr6-89055886-DEL-63
chr6-89104361-DEL-370
chr6-89104501-DEL-175
chr6-89106400-DEL-52
chr6-89256361-DEL-226
chr6-89479694-DEL-69
chr6-89635116-DEL-62
chr6-89688219-DEL-64
chr6-89812612-DEL-114
chr6-89855749-DEL-180
chr6-89995968-DEL-50
chr6-90550672-DEL-52
chr6-91041051-DEL-426
chr6-91100766-DEL-316
chr6-91100768-DEL-76
chr6-91234208-DEL-54
chr6-91617199-DEL-53
chr6-91617748-DEL-167
chr6-91812604-DEL-64
chr6-92006962-DEL-79
chr6-92171653-DEL-72
chr6-92247817-DEL-228
chr6-92248189-DEL-66
chr6-92251661-DEL-80
chr6-92382209-DEL-95
chr6-92383113-DEL-234
chr6-92900910-DEL-50
chr6-93019652-DEL-59
chr6-93119697-DEL-132
chr6-93172589-DEL-1084

chr6-93183160-DEL-76
chr6-93189158-DEL-50
chr6-93207981-DEL-141
chr6-93269748-DEL-111
chr6-93426231-DEL-842
chr6-93489695-DEL-50
chr6-93502654-DEL-50
chr6-93759965-DEL-162
chr6-93775091-DEL-180
chr6-93775271-DEL-72
chr6-93933405-DEL-222
chr6-94193160-DEL-580
chr6-94205688-DEL-1547
chr6-94297718-DEL-271
chr6-94305024-DEL-2588
chr6-94318091-DEL-135
chr6-94318341-DEL-51
chr6-94379905-DEL-54
chr6-94460906-DEL-59
chr6-94712825-DEL-99
chr6-94779698-DEL-141
chr6-94801734-DEL-54
chr6-94801823-DEL-285
chr6-94832885-DEL-51
chr6-94873877-DEL-316
chr6-94965696-DEL-1869
chr6-95032869-DEL-56
chr6-95038077-DEL-52
chr6-95074273-DEL-61
chr6-95087549-DEL-218
chr6-95087559-DEL-104
chr6-95126412-DEL-50
chr6-95185457-DEL-216
chr6-95238966-DEL-54
chr6-95357042-DEL-1266
chr6-95443653-DEL-84
chr6-95519665-DEL-72
chr6-95607538-DEL-77
chr6-95667708-DEL-55
chr6-95935364-DEL-67
chr6-95975757-DEL-438
chr6-96147859-DEL-144
chr6-96155902-DEL-50
chr6-96284237-DEL-223
chr6-96553872-DEL-50
chr6-96648883-DEL-54
chr6-96665221-DEL-279
chr6-96696101-DEL-196
chr6-96696193-DEL-50
chr6-96735925-DEL-296
chr6-96870069-DEL-3111
chr6-96873447-DEL-51
chr6-96898888-DEL-5443
chr6-96905341-DEL-251
chr6-96987206-DEL-498
chr6-96987488-DEL-120
chr6-96987981-DEL-416
chr6-97064701-DEL-2406
chr6-97067223-DEL-495
chr6-97094871-DEL-90
chr6-97141166-DEL-1812
chr6-97198417-DEL-168
chr6-97314634-DEL-618
chr6-97429173-DEL-60
chr6-97448783-DEL-79
chr6-97472129-DEL-52
chr6-97501336-DEL-66
chr6-97684935-DEL-54
chr6-97703528-DEL-328
chr6-97703660-DEL-134
chr6-97703955-DEL-144
chr6-97711195-DEL-111
chr6-97747002-DEL-50
chr6-98198682-DEL-50
chr6-98260682-DEL-57
chr6-98474550-DEL-190
chr6-98710298-DEL-130

chr6-98857883-DEL-4745
chr6-99017599-DEL-56
chr6-99203206-DEL-62
chr6-99259998-DEL-50
chr6-99333504-DEL-330
chr6-99387249-DEL-112
chr6-99574338-DEL-100
chr6-99782746-DEL-144
chr6-99807970-DEL-94
chr6-99828051-DEL-110
chr6-99859906-DEL-294
chr6-100280421-DEL-66
chr6-100439093-DEL-885
chr6-100441243-DEL-479
chr6-100444248-DEL-309
chr6-100444705-DEL-164
chr6-100444817-DEL-938
chr6-100445183-DEL-380
chr6-100556219-DEL-51
chr6-100643188-DEL-87
chr6-100899814-DEL-151
chr6-100905239-DEL-109
chr6-100934937-DEL-56
chr6-101050288-DEL-128
chr6-101076545-DEL-113
chr6-101301333-DEL-100
chr6-101301336-DEL-240
chr6-101478271-DEL-83
chr6-101576810-DEL-463
chr6-101809210-DEL-50
chr6-101888053-DEL-642
chr6-101963491-DEL-2102
chr6-102024915-DEL-60
chr6-102196479-DEL-1477
chr6-102207418-DEL-461
chr6-102253269-DEL-52
chr6-102558666-DEL-346
chr6-102559180-DEL-118
chr6-102613501-DEL-1590
chr6-102640175-DEL-1490
chr6-102761441-DEL-3536
chr6-102827464-DEL-1539
chr6-102858583-DEL-1137
chr6-102934544-DEL-268
chr6-103113220-DEL-151
chr6-103120050-DEL-55
chr6-103125199-DEL-1085
chr6-103340606-DEL-70
chr6-103344761-DEL-70
chr6-103467173-DEL-244
chr6-103479647-DEL-82
chr6-103558555-DEL-214
chr6-103564702-DEL-76
chr6-103630155-DEL-157
chr6-103630425-DEL-74
chr6-103651680-DEL-53
chr6-103764405-DEL-59
chr6-103834859-DEL-806
chr6-104096287-DEL-66
chr6-104100919-DEL-55
chr6-104221381-DEL-50
chr6-104232791-DEL-96
chr6-104297713-DEL-157
chr6-104298189-DEL-366
chr6-104353335-DEL-70
chr6-104404957-DEL-54
chr6-104622693-DEL-60
chr6-104707584-DEL-56
chr6-104958215-DEL-50
chr6-105291770-DEL-6702
chr6-105334030-DEL-63
chr6-105376892-DEL-122
chr6-105777946-DEL-56
chr6-105853515-DEL-63
chr6-106530573-DEL-220
chr6-106531120-DEL-1166
chr6-106531579-DEL-462

chr6-107408391-DEL-154
chr6-107454682-DEL-60
chr6-107606258-DEL-421
chr6-107756956-DEL-95
chr6-107858654-DEL-6234
chr6-107883336-DEL-219
chr6-107951933-DEL-968
chr6-107955426-DEL-230
chr6-107955549-DEL-88
chr6-108008074-DEL-1969
chr6-108061315-DEL-78
chr6-108069982-DEL-100
chr6-108070248-DEL-100
chr6-108124989-DEL-74
chr6-108139967-DEL-158
chr6-108182030-DEL-59
chr6-108233759-DEL-50
chr6-108234444-DEL-62
chr6-108287699-DEL-134
chr6-108331744-DEL-52
chr6-108462547-DEL-239
chr6-108693782-DEL-54
chr6-108739047-DEL-349
chr6-108749636-DEL-56
chr6-108763670-DEL-271
chr6-108824048-DEL-168
chr6-108907895-DEL-3898
chr6-108959381-DEL-368
chr6-109030941-DEL-22779
chr6-109181940-DEL-54
chr6-109229743-DEL-64
chr6-109324755-DEL-68
chr6-109324910-DEL-80
chr6-109354410-DEL-343
chr6-109354648-DEL-327
chr6-109591993-DEL-876
chr6-109695115-DEL-71
chr6-109700608-DEL-83
chr6-109756000-DEL-117
chr6-109884525-DEL-1915
chr6-109895715-DEL-179
chr6-109965994-DEL-50
chr6-109981008-DEL-8372
chr6-110033385-DEL-52
chr6-110037084-DEL-53
chr6-110230947-DEL-1065
chr6-110233193-DEL-539
chr6-110294599-DEL-119
chr6-110330767-DEL-50
chr6-110420623-DEL-64
chr6-110476435-DEL-448
chr6-110541252-DEL-80
chr6-110761368-DEL-425
chr6-110779795-DEL-92
chr6-110785787-DEL-588
chr6-110796629-DEL-53
chr6-110796646-DEL-128
chr6-110875850-DEL-51
chr6-110934028-DEL-111
chr6-111018733-DEL-439
chr6-111095749-DEL-119
chr6-111155932-DEL-78
chr6-111296803-DEL-50
chr6-111304493-DEL-82
chr6-111392497-DEL-6585
chr6-111434963-DEL-159
chr6-111490992-DEL-56
chr6-111642797-DEL-8853
chr6-111663111-DEL-62
chr6-111752135-DEL-76
chr6-111780231-DEL-128
chr6-111780668-DEL-342
chr6-111780713-DEL-102
chr6-111819997-DEL-130
chr6-111960490-DEL-241
chr6-111979620-DEL-51
chr6-112075797-DEL-50

chr6-112126056-DEL-52
chr6-112137855-DEL-171
chr6-112201619-DEL-374
chr6-112201619-DEL-1908
chr6-112201905-DEL-154
chr6-112202433-DEL-198
chr6-112203165-DEL-164
chr6-112227938-DEL-3202
chr6-112340515-DEL-66
chr6-112504938-DEL-105
chr6-112507917-DEL-567
chr6-112587747-DEL-50
chr6-112653044-DEL-54
chr6-112655657-DEL-316
chr6-112655701-DEL-97
chr6-112675104-DEL-114
chr6-112836437-DEL-186
chr6-112836806-DEL-111
chr6-112837087-DEL-118
chr6-112941294-DEL-83
chr6-112982887-DEL-162
chr6-113104931-DEL-134
chr6-113140213-DEL-194
chr6-113140222-DEL-64
chr6-113167902-DEL-50
chr6-113187239-DEL-117
chr6-113201203-DEL-106
chr6-113329202-DEL-56
chr6-113333441-DEL-73
chr6-113433833-DEL-132
chr6-113580611-DEL-256
chr6-113898354-DEL-58
chr6-113921672-DEL-50
chr6-113928009-DEL-79
chr6-114001175-DEL-53
chr6-114145123-DEL-107
chr6-114214429-DEL-363
chr6-114232837-DEL-65
chr6-114254270-DEL-304
chr6-114317924-DEL-1611
chr6-114329074-DEL-150
chr6-114492411-DEL-62
chr6-114592487-DEL-74
chr6-114599965-DEL-96
chr6-114604893-DEL-58
chr6-114612265-DEL-88
chr6-114886457-DEL-208
chr6-114977023-DEL-236
chr6-115065128-DEL-228
chr6-115068965-DEL-278
chr6-115263360-DEL-74
chr6-115526637-DEL-83
chr6-115604746-DEL-64
chr6-115749854-DEL-80
chr6-115862812-DEL-57
chr6-115897810-DEL-1289
chr6-115897861-DEL-628
chr6-115898030-DEL-65
chr6-116166033-DEL-64
chr6-116166033-DEL-144
chr6-116215706-DEL-918
chr6-116227990-DEL-911
chr6-116228144-DEL-77
chr6-116228657-DEL-174
chr6-116238067-DEL-50
chr6-116275360-DEL-62
chr6-116323053-DEL-575
chr6-116429653-DEL-171
chr6-116633314-DEL-174
chr6-116633736-DEL-326
chr6-116740738-DEL-121
chr6-116790168-DEL-73
chr6-116832977-DEL-54
chr6-116833244-DEL-144
chr6-116860404-DEL-214
chr6-116885410-DEL-2642
chr6-116942377-DEL-50

chr6-116978894-DEL-50
chr6-116995510-DEL-54
chr6-117014366-DEL-51
chr6-117014663-DEL-190
chr6-117030355-DEL-116
chr6-117235615-DEL-3880
chr6-117334459-DEL-176
chr6-117338975-DEL-78
chr6-117364964-DEL-1095
chr6-117377047-DEL-214
chr6-117592581-DEL-122
chr6-117650920-DEL-246
chr6-117704057-DEL-260
chr6-117705475-DEL-289
chr6-117706722-DEL-445
chr6-117718009-DEL-229
chr6-117749623-DEL-100
chr6-117758859-DEL-202
chr6-117769862-DEL-1706
chr6-117770913-DEL-353
chr6-117782485-DEL-756
chr6-117782497-DEL-308
chr6-117815765-DEL-202
chr6-117841631-DEL-209
chr6-117855440-DEL-178
chr6-117959834-DEL-575
chr6-118039265-DEL-152
chr6-118354655-DEL-67
chr6-118366770-DEL-432
chr6-118408132-DEL-50
chr6-118458826-DEL-898
chr6-118473485-DEL-1543
chr6-118553246-DEL-456
chr6-118598214-DEL-130
chr6-118648947-DEL-64
chr6-118833564-DEL-53
chr6-118947016-DEL-60
chr6-118991726-DEL-189
chr6-119021829-DEL-95
chr6-119063274-DEL-213
chr6-119064178-DEL-162
chr6-119076264-DEL-108
chr6-119134287-DEL-668
chr6-119207036-DEL-170
chr6-119209934-DEL-83
chr6-119234641-DEL-112
chr6-119283418-DEL-67
chr6-119309428-DEL-197
chr6-119370490-DEL-108
chr6-119431185-DEL-2318
chr6-119451562-DEL-288
chr6-119451654-DEL-128
chr6-119476132-DEL-62
chr6-119478569-DEL-76
chr6-119489013-DEL-157
chr6-119620439-DEL-56
chr6-119794922-DEL-2517
chr6-119865372-DEL-685
chr6-120028503-DEL-52
chr6-120030734-DEL-102
chr6-120053147-DEL-938
chr6-120132343-DEL-88
chr6-120161904-DEL-64
chr6-120517490-DEL-50
chr6-120547480-DEL-2243
chr6-120572530-DEL-65
chr6-120631549-DEL-188
chr6-120765356-DEL-762
chr6-120801882-DEL-440
chr6-120805976-DEL-60
chr6-120876455-DEL-155
chr6-120959986-DEL-211
chr6-121025012-DEL-272
chr6-121104992-DEL-103
chr6-121157995-DEL-984
chr6-121164741-DEL-98
chr6-121175411-DEL-50

chr6-121176423-DEL-78
chr6-121234332-DEL-52
chr6-121341889-DEL-189
chr6-121465376-DEL-56
chr6-121483315-DEL-93
chr6-121503947-DEL-199
chr6-121580653-DEL-260
chr6-121627025-DEL-3989
chr6-121629220-DEL-1386
chr6-122027818-DEL-72
chr6-122129723-DEL-44313
chr6-122190272-DEL-11688
chr6-122218952-DEL-449
chr6-122249841-DEL-69
chr6-122324446-DEL-78
chr6-122332943-DEL-126
chr6-122363849-DEL-715
chr6-122380012-DEL-53
chr6-122492839-DEL-66
chr6-122580252-DEL-89
chr6-122580527-DEL-198
chr6-122592638-DEL-357
chr6-122617604-DEL-506
chr6-122657029-DEL-54
chr6-122657145-DEL-294
chr6-122767340-DEL-390
chr6-122767398-DEL-180
chr6-122792153-DEL-50
chr6-122875022-DEL-50
chr6-123024904-DEL-161
chr6-123044870-DEL-1462
chr6-123084911-DEL-2588
chr6-123201848-DEL-2061
chr6-123206063-DEL-52
chr6-123226456-DEL-451
chr6-123226500-DEL-215
chr6-123226910-DEL-145
chr6-123289527-DEL-58
chr6-123364650-DEL-243
chr6-123493276-DEL-208
chr6-123551464-DEL-64
chr6-123676746-DEL-32349
chr6-123696205-DEL-125
chr6-123861816-DEL-184
chr6-123861968-DEL-51
chr6-123876256-DEL-162
chr6-123933856-DEL-192
chr6-124099093-DEL-1071
chr6-124129688-DEL-2337
chr6-124163991-DEL-73
chr6-124186122-DEL-1327
chr6-124320120-DEL-81
chr6-124320221-DEL-209
chr6-124320626-DEL-179
chr6-124337324-DEL-71
chr6-124419343-DEL-268
chr6-124429653-DEL-218
chr6-124447672-DEL-71
chr6-124473760-DEL-56
chr6-124487730-DEL-630
chr6-124512574-DEL-54
chr6-124554256-DEL-3825
chr6-124554735-DEL-205
chr6-124555198-DEL-123
chr6-124616207-DEL-56
chr6-124749932-DEL-50
chr6-124804866-DEL-65
chr6-124904803-DEL-128
chr6-124917230-DEL-53
chr6-124993805-DEL-92
chr6-125007643-DEL-51
chr6-125082503-DEL-389
chr6-125082779-DEL-116
chr6-125088532-DEL-233
chr6-125311199-DEL-54
chr6-125370059-DEL-7927
chr6-125380150-DEL-147

chr6-125393674-DEL-1136
chr6-125405013-DEL-542
chr6-125409234-DEL-281
chr6-125416604-DEL-50
chr6-125419752-DEL-80
chr6-125451365-DEL-55
chr6-125523000-DEL-70
chr6-125558840-DEL-67
chr6-125588185-DEL-81
chr6-125596524-DEL-159
chr6-125616627-DEL-124
chr6-125658212-DEL-66
chr6-125709283-DEL-125
chr6-125756356-DEL-1087
chr6-125853929-DEL-60
chr6-125867444-DEL-54
chr6-125955230-DEL-280
chr6-125965232-DEL-60
chr6-125965332-DEL-340
chr6-126057274-DEL-126
chr6-126408953-DEL-96
chr6-126440114-DEL-66
chr6-126454194-DEL-52
chr6-126505763-DEL-54
chr6-126516299-DEL-400
chr6-126550610-DEL-3424
chr6-126690592-DEL-111
chr6-126722193-DEL-138
chr6-126796204-DEL-108
chr6-126936124-DEL-527
chr6-127283080-DEL-151
chr6-127324516-DEL-74
chr6-127349335-DEL-53
chr6-127358962-DEL-158
chr6-127362751-DEL-77
chr6-127398215-DEL-54
chr6-127419437-DEL-152
chr6-127508664-DEL-70
chr6-127524789-DEL-70
chr6-127596312-DEL-180
chr6-127859455-DEL-60
chr6-127900969-DEL-72
chr6-127917590-DEL-725
chr6-127918733-DEL-105
chr6-127922535-DEL-240
chr6-127934781-DEL-94
chr6-127973038-DEL-128
chr6-128003103-DEL-234
chr6-128063340-DEL-139
chr6-128210450-DEL-86
chr6-128210566-DEL-152
chr6-128417714-DEL-819
chr6-128417911-DEL-210
chr6-128418431-DEL-64
chr6-128424930-DEL-122
chr6-128602468-DEL-56
chr6-128606047-DEL-161
chr6-128624290-DEL-94
chr6-128642068-DEL-1094
chr6-128695317-DEL-328
chr6-128702078-DEL-52
chr6-128725851-DEL-170
chr6-128763660-DEL-51
chr6-128772895-DEL-73
chr6-128824283-DEL-1422
chr6-128845124-DEL-80
chr6-128854715-DEL-144
chr6-128855783-DEL-2008
chr6-128881018-DEL-498
chr6-128902056-DEL-9981
chr6-128904907-DEL-1688
chr6-128982585-DEL-69
chr6-129338956-DEL-65
chr6-129393871-DEL-65
chr6-129582855-DEL-1405
chr6-129606741-DEL-51
chr6-129763552-DEL-31292

chr6-129790721-DEL-50
chr6-129859455-DEL-113
chr6-129963497-DEL-72
chr6-130012900-DEL-71
chr6-130028900-DEL-17026
chr6-130032567-DEL-57
chr6-130037578-DEL-52
chr6-130061878-DEL-56
chr6-130066417-DEL-97906
chr6-130072197-DEL-64
chr6-130073726-DEL-2619
chr6-130091951-DEL-81
chr6-130093986-DEL-50
chr6-130118804-DEL-227
chr6-130150008-DEL-68
chr6-130163550-DEL-135
chr6-130235401-DEL-738
chr6-130273979-DEL-310
chr6-130282469-DEL-63
chr6-130296567-DEL-199
chr6-130316339-DEL-616
chr6-130504754-DEL-85
chr6-130532449-DEL-99
chr6-130555165-DEL-420
chr6-130555832-DEL-84
chr6-130556403-DEL-84
chr6-130607134-DEL-3128
chr6-130614117-DEL-199
chr6-130644344-DEL-769
chr6-130646871-DEL-1258
chr6-130653284-DEL-168
chr6-130673000-DEL-51
chr6-130681457-DEL-57
chr6-130688064-DEL-156
chr6-130710623-DEL-274
chr6-130764013-DEL-84
chr6-130769344-DEL-183
chr6-130772376-DEL-243
chr6-130790616-DEL-661
chr6-130809568-DEL-809
chr6-130887023-DEL-87
chr6-130897483-DEL-534
chr6-130956282-DEL-55
chr6-130981828-DEL-319
chr6-130996222-DEL-202
chr6-131021006-DEL-2878
chr6-131129825-DEL-4778
chr6-131132226-DEL-4815
chr6-131241453-DEL-120
chr6-131294787-DEL-50
chr6-131324532-DEL-82
chr6-131324714-DEL-374
chr6-131325152-DEL-120
chr6-131462009-DEL-60
chr6-131494831-DEL-57
chr6-131509671-DEL-523
chr6-131613389-DEL-527
chr6-131743684-DEL-200
chr6-131931286-DEL-116
chr6-131931518-DEL-60
chr6-132032425-DEL-172
chr6-132085598-DEL-595
chr6-132091909-DEL-111
chr6-132092336-DEL-126
chr6-132101963-DEL-1782
chr6-132119307-DEL-249541
chr6-132169880-DEL-3289
chr6-132170371-DEL-322
chr6-132183639-DEL-104795
chr6-132184433-DEL-330
chr6-132184771-DEL-54
chr6-132185127-DEL-57
chr6-132213383-DEL-234
chr6-132237586-DEL-52
chr6-132295516-DEL-1010
chr6-132318312-DEL-5563
chr6-132346352-DEL-1447

chr6-132416023-DEL-164
chr6-132453113-DEL-52
chr6-132531803-DEL-351
chr6-132531807-DEL-111
chr6-132577020-DEL-84
chr6-132577267-DEL-165
chr6-132579875-DEL-59
chr6-132696685-DEL-64
chr6-132875910-DEL-61
chr6-132900550-DEL-50
chr6-132953420-DEL-66
chr6-133012508-DEL-1132
chr6-133029461-DEL-103
chr6-133157882-DEL-160
chr6-133175616-DEL-188
chr6-133250603-DEL-830
chr6-133344606-DEL-1545
chr6-133459362-DEL-71
chr6-133462499-DEL-2368
chr6-133470794-DEL-2863
chr6-133739549-DEL-92
chr6-133767099-DEL-307
chr6-133845865-DEL-57
chr6-133848520-DEL-147
chr6-133848722-DEL-264
chr6-133853311-DEL-2816
chr6-133869082-DEL-1578
chr6-133870682-DEL-104
chr6-133914195-DEL-93
chr6-133951557-DEL-72
chr6-133956456-DEL-126
chr6-133965524-DEL-58
chr6-134048203-DEL-54
chr6-134056054-DEL-334
chr6-134276665-DEL-422
chr6-134280584-DEL-50
chr6-134284058-DEL-58
chr6-134371935-DEL-114
chr6-134644123-DEL-264
chr6-134652887-DEL-220
chr6-134700525-DEL-58
chr6-134804847-DEL-154
chr6-134924182-DEL-60
chr6-134935171-DEL-560
chr6-134946903-DEL-52
chr6-134946935-DEL-257
chr6-134947356-DEL-69
chr6-134947717-DEL-84
chr6-134981618-DEL-1258
chr6-135036043-DEL-320
chr6-135098045-DEL-123
chr6-135106175-DEL-65
chr6-135127604-DEL-120
chr6-135141407-DEL-188
chr6-135149784-DEL-755
chr6-135157196-DEL-56
chr6-135182075-DEL-706
chr6-135184401-DEL-557
chr6-135190086-DEL-286
chr6-135244993-DEL-180
chr6-135245211-DEL-56
chr6-135292518-DEL-57
chr6-135503442-DEL-87
chr6-135656234-DEL-162
chr6-135827543-DEL-84
chr6-135962729-DEL-168
chr6-135962806-DEL-76
chr6-135962932-DEL-421
chr6-135980205-DEL-67
chr6-136026400-DEL-64
chr6-136099899-DEL-69
chr6-136118234-DEL-54
chr6-136202836-DEL-77
chr6-136210736-DEL-71
chr6-136210962-DEL-95
chr6-136255438-DEL-60
chr6-136286360-DEL-637

chr6-136308182-DEL-143
chr6-136308195-DEL-67
chr6-136315961-DEL-421
chr6-136329581-DEL-100
chr6-136382682-DEL-98
chr6-136383152-DEL-685
chr6-136383225-DEL-109
chr6-136386807-DEL-103
chr6-136391335-DEL-820
chr6-136403610-DEL-65
chr6-136407273-DEL-130
chr6-136407356-DEL-51
chr6-136410005-DEL-215
chr6-136456847-DEL-275
chr6-136560551-DEL-348
chr6-136560795-DEL-364
chr6-136595353-DEL-101
chr6-136635902-DEL-165
chr6-136646962-DEL-60
chr6-136661815-DEL-470
chr6-136771378-DEL-109
chr6-136799292-DEL-208
chr6-136909508-DEL-62
chr6-136944755-DEL-84
chr6-136968967-DEL-1175
chr6-136976563-DEL-937
chr6-137095144-DEL-57
chr6-137101626-DEL-53
chr6-137163400-DEL-62
chr6-137222479-DEL-96
chr6-137262768-DEL-69
chr6-137268097-DEL-153
chr6-137268099-DEL-55
chr6-137268361-DEL-62
chr6-137317656-DEL-62
chr6-137393585-DEL-100
chr6-137444186-DEL-66
chr6-137575659-DEL-79
chr6-137580120-DEL-108
chr6-137632859-DEL-52
chr6-137669685-DEL-100
chr6-137695310-DEL-270
chr6-137697765-DEL-323
chr6-137699957-DEL-692
chr6-137708024-DEL-63
chr6-137709248-DEL-979
chr6-137719307-DEL-60
chr6-137804124-DEL-151
chr6-137823201-DEL-55
chr6-137836267-DEL-76
chr6-137868220-DEL-132
chr6-137946629-DEL-614
chr6-137951561-DEL-52
chr6-137985325-DEL-1219
chr6-138027456-DEL-108
chr6-138039105-DEL-129
chr6-138039284-DEL-56
chr6-138062560-DEL-72
chr6-138197618-DEL-3304
chr6-138201252-DEL-232
chr6-138269449-DEL-229
chr6-138270338-DEL-117
chr6-138324501-DEL-1008
chr6-138333567-DEL-50
chr6-138407218-DEL-72
chr6-138419537-DEL-52
chr6-138432354-DEL-332
chr6-138693069-DEL-72
chr6-138739973-DEL-50
chr6-138789111-DEL-890
chr6-138803652-DEL-50
chr6-138820213-DEL-1293
chr6-138846351-DEL-167
chr6-138950714-DEL-50
chr6-138957699-DEL-103
chr6-138962140-DEL-770
chr6-139039003-DEL-65

chr6-139173985-DEL-59
chr6-139203761-DEL-89
chr6-139392728-DEL-83
chr6-139589134-DEL-72
chr6-139676642-DEL-60
chr6-139696646-DEL-58
chr6-139711267-DEL-52
chr6-139787575-DEL-92
chr6-139788073-DEL-56
chr6-139885886-DEL-947
chr6-139885971-DEL-73
chr6-139952519-DEL-131
chr6-139952546-DEL-63
chr6-139981093-DEL-149
chr6-139981334-DEL-175
chr6-140129401-DEL-56
chr6-140143452-DEL-220
chr6-140180461-DEL-370
chr6-140186941-DEL-183
chr6-140237332-DEL-50
chr6-140325465-DEL-66
chr6-140335053-DEL-62
chr6-140357185-DEL-54
chr6-140377696-DEL-471
chr6-140446318-DEL-54
chr6-140563058-DEL-60
chr6-140563914-DEL-459
chr6-140564630-DEL-720
chr6-140604327-DEL-3124
chr6-140605541-DEL-106
chr6-140605825-DEL-77
chr6-140615870-DEL-144
chr6-140638601-DEL-66
chr6-140672008-DEL-252
chr6-140697496-DEL-116
chr6-140702782-DEL-208
chr6-140743601-DEL-241
chr6-140743700-DEL-92
chr6-140764986-DEL-52
chr6-140780535-DEL-208
chr6-140792059-DEL-142
chr6-140848037-DEL-58
chr6-140857456-DEL-96
chr6-140882356-DEL-60
chr6-140882551-DEL-120
chr6-140909994-DEL-50
chr6-140910390-DEL-66
chr6-140944906-DEL-50
chr6-141058262-DEL-50
chr6-141082000-DEL-50
chr6-141234044-DEL-510
chr6-141234244-DEL-222
chr6-141333423-DEL-87
chr6-141338419-DEL-50
chr6-141538651-DEL-78
chr6-141715256-DEL-326
chr6-141715291-DEL-141
chr6-141715572-DEL-63
chr6-141715770-DEL-172
chr6-141748532-DEL-511
chr6-141759822-DEL-293
chr6-141788923-DEL-82
chr6-141804859-DEL-1436
chr6-141876813-DEL-62
chr6-141946780-DEL-229
chr6-142018684-DEL-74
chr6-142122439-DEL-50
chr6-142147905-DEL-672
chr6-142266164-DEL-214
chr6-142326308-DEL-316
chr6-142342520-DEL-692
chr6-142342520-DEL-3256
chr6-142342754-DEL-1260
chr6-142345391-DEL-571
chr6-142369817-DEL-58
chr6-142422464-DEL-78
chr6-142422768-DEL-131

chr6-142520057-DEL-66
chr6-142520383-DEL-82
chr6-142543145-DEL-96
chr6-142555044-DEL-50
chr6-142565634-DEL-50
chr6-142568918-DEL-83
chr6-142570359-DEL-130
chr6-142586019-DEL-216
chr6-142650500-DEL-82
chr6-142657525-DEL-95
chr6-142696321-DEL-563
chr6-142749627-DEL-61
chr6-142784031-DEL-85
chr6-142885469-DEL-578
chr6-142899710-DEL-57
chr6-142902637-DEL-121
chr6-142946129-DEL-216
chr6-142953602-DEL-69
chr6-142979656-DEL-64
chr6-143002928-DEL-72
chr6-143021433-DEL-58
chr6-143141713-DEL-506
chr6-143215187-DEL-541
chr6-143334597-DEL-260
chr6-143343168-DEL-194
chr6-143399279-DEL-75
chr6-143408667-DEL-56
chr6-143436659-DEL-194
chr6-143465107-DEL-76
chr6-143519984-DEL-104
chr6-143637342-DEL-219
chr6-143696037-DEL-67
chr6-143728675-DEL-116
chr6-143782699-DEL-94
chr6-143791980-DEL-86
chr6-143792187-DEL-205
chr6-143836484-DEL-180
chr6-143839873-DEL-527
chr6-143849227-DEL-57
chr6-143849423-DEL-117
chr6-143883971-DEL-98
chr6-143956564-DEL-59
chr6-144110473-DEL-65
chr6-144213914-DEL-67
chr6-144269712-DEL-163
chr6-144402192-DEL-91
chr6-144471085-DEL-170
chr6-144481189-DEL-59
chr6-144540644-DEL-457
chr6-144613300-DEL-96
chr6-144613728-DEL-256
chr6-144796513-DEL-89
chr6-144799273-DEL-240
chr6-144821354-DEL-79
chr6-144821780-DEL-60
chr6-144930645-DEL-72
chr6-144932208-DEL-57
chr6-144958977-DEL-1026
chr6-144991493-DEL-50
chr6-145041092-DEL-200
chr6-145041331-DEL-199
chr6-145041500-DEL-80
chr6-145056409-DEL-164
chr6-145078633-DEL-51
chr6-145088171-DEL-50
chr6-145216375-DEL-117
chr6-145220384-DEL-123
chr6-145268883-DEL-50
chr6-145370226-DEL-88
chr6-145385749-DEL-373
chr6-145388060-DEL-100
chr6-145453598-DEL-702
chr6-145515676-DEL-72
chr6-145599166-DEL-66
chr6-145627578-DEL-2176
chr6-145710110-DEL-135
chr6-145711635-DEL-1247

chr6-145715178-DEL-147
chr6-145715430-DEL-63
chr6-145729503-DEL-115
chr6-145729536-DEL-778
chr6-145729750-DEL-1806
chr6-145730140-DEL-131
chr6-145730845-DEL-133
chr6-145731974-DEL-629
chr6-145732421-DEL-132
chr6-145749651-DEL-64
chr6-145753293-DEL-50
chr6-145785701-DEL-72
chr6-146099541-DEL-54
chr6-146152707-DEL-74
chr6-146152898-DEL-130
chr6-146152902-DEL-629
chr6-146276674-DEL-101
chr6-146280307-DEL-114
chr6-146341317-DEL-71
chr6-146370606-DEL-402
chr6-146370740-DEL-172
chr6-146424020-DEL-427
chr6-146541448-DEL-93
chr6-146567369-DEL-313
chr6-146614135-DEL-4149
chr6-146709897-DEL-237
chr6-146720680-DEL-270
chr6-146730546-DEL-51
chr6-146763017-DEL-58
chr6-146941797-DEL-69
chr6-147020386-DEL-240
chr6-147060208-DEL-54
chr6-147138099-DEL-381
chr6-147138315-DEL-148
chr6-147243032-DEL-135
chr6-147297394-DEL-54
chr6-147346687-DEL-92
chr6-147371246-DEL-69
chr6-147378776-DEL-284
chr6-147410447-DEL-180
chr6-147431080-DEL-52
chr6-147452848-DEL-1900
chr6-147578939-DEL-54
chr6-147589975-DEL-50
chr6-147609670-DEL-71
chr6-147621459-DEL-220
chr6-147621621-DEL-64
chr6-147669537-DEL-790
chr6-147708899-DEL-737
chr6-147816672-DEL-250
chr6-147817039-DEL-378
chr6-147914688-DEL-768
chr6-147914740-DEL-2112
chr6-147916186-DEL-96
chr6-147916378-DEL-448
chr6-148360289-DEL-54
chr6-148543884-DEL-60
chr6-148597898-DEL-54
chr6-148607160-DEL-1732
chr6-148610276-DEL-116
chr6-148624305-DEL-114
chr6-148640376-DEL-58
chr6-149265582-DEL-58
chr7-3211565-DEL-112
chr7-3219087-DEL-64
chr7-3240601-DEL-185
chr7-3241628-DEL-658
chr7-3260161-DEL-82
chr7-3281096-DEL-79
chr7-3292152-DEL-180
chr7-3301683-DEL-893
chr7-3327704-DEL-129
chr7-3393286-DEL-535
chr7-3418686-DEL-51
chr7-3424173-DEL-118
chr7-3465540-DEL-164
chr7-3478638-DEL-51

chr7-3557067-DEL-108
chr7-3566486-DEL-1939
chr7-3567511-DEL-602
chr7-3576004-DEL-270
chr7-3578100-DEL-372
chr7-3586947-DEL-52
chr7-3587070-DEL-51
chr7-3684338-DEL-240
chr7-3736747-DEL-56
chr7-3738713-DEL-158229
chr7-3746389-DEL-174
chr7-3846084-DEL-53064
chr7-3928414-DEL-75
chr7-4072770-DEL-60
chr7-4094349-DEL-82
chr7-4107607-DEL-234
chr7-4107747-DEL-108
chr7-4112113-DEL-53
chr7-4290013-DEL-174
chr7-4327261-DEL-1571
chr7-4391235-DEL-5143
chr7-4431131-DEL-343
chr7-4458089-DEL-53
chr7-4511010-DEL-130
chr7-4551890-DEL-92
chr7-4563662-DEL-94
chr7-4564124-DEL-100
chr7-4652333-DEL-228
chr7-4653517-DEL-536
chr7-4684207-DEL-155
chr7-4699378-DEL-178
chr7-4758518-DEL-652
chr7-4758622-DEL-86
chr7-4803169-DEL-113
chr7-4852594-DEL-73
chr7-4872011-DEL-80
chr7-4887903-DEL-243
chr7-4958362-DEL-50
chr7-4961297-DEL-283
chr7-4976782-DEL-206
chr7-5016701-DEL-74
chr7-5043075-DEL-933
chr7-5050693-DEL-463
chr7-5051049-DEL-116
chr7-5332755-DEL-111
chr7-5356985-DEL-120
chr7-5357361-DEL-64
chr7-5518762-DEL-164
chr7-5556305-DEL-3778
chr7-5600270-DEL-158
chr7-5663351-DEL-3384
chr7-5712405-DEL-1083
chr7-5719344-DEL-6433
chr7-5748218-DEL-67
chr7-5763143-DEL-3384
chr7-5773803-DEL-2432
chr7-5784076-DEL-16161
chr7-5793324-DEL-95
chr7-5833584-DEL-63
chr7-5873198-DEL-196
chr7-5960913-DEL-10610
chr7-6041611-DEL-68
chr7-6042089-DEL-2767
chr7-6077427-DEL-7343
chr7-6204563-DEL-529
chr7-6240395-DEL-292
chr7-6257287-DEL-140
chr7-6259167-DEL-161
chr7-6262976-DEL-110
chr7-6301494-DEL-81
chr7-6303951-DEL-180
chr7-6312345-DEL-62
chr7-6324178-DEL-82
chr7-6351692-DEL-277
chr7-6368807-DEL-188
chr7-6389462-DEL-101
chr7-6436351-DEL-410

chr7-6541579-DEL-105
chr7-6552398-DEL-1223
chr7-6828316-DEL-190
chr7-6851479-DEL-76
chr7-6870004-DEL-166
chr7-6874830-DEL-318
chr7-6874903-DEL-121
chr7-7106228-DEL-379
chr7-7191779-DEL-50
chr7-7192074-DEL-108
chr7-7427041-DEL-76
chr7-7539798-DEL-388
chr7-7947848-DEL-1562
chr7-8293446-DEL-255
chr7-9816642-DEL-50
chr7-10048766-DEL-6399
chr7-10125993-DEL-196
chr7-10126232-DEL-95
chr7-10205625-DEL-411
chr7-10295466-DEL-8430
chr7-10388274-DEL-2629
chr7-10407564-DEL-945
chr7-10408906-DEL-1190
chr7-10461948-DEL-1393
chr7-10478178-DEL-58
chr7-10515687-DEL-1635
chr7-10533676-DEL-336
chr7-10535652-DEL-219
chr7-10860286-DEL-162
chr7-10875806-DEL-101
chr7-10876898-DEL-557
chr7-10891822-DEL-392
chr7-11054669-DEL-317
chr7-11082657-DEL-74
chr7-11123110-DEL-4915
chr7-11378713-DEL-208
chr7-11438376-DEL-692
chr7-11670098-DEL-96
chr7-11680040-DEL-299
chr7-11697366-DEL-72
chr7-11719274-DEL-692
chr7-11744954-DEL-65
chr7-11824249-DEL-871
chr7-11832485-DEL-520
chr7-11872370-DEL-105
chr7-11879382-DEL-1989
chr7-11883394-DEL-74
chr7-11921340-DEL-85
chr7-11974351-DEL-62
chr7-12399747-DEL-1888
chr7-12472105-DEL-148
chr7-12549289-DEL-99
chr7-12653274-DEL-107
chr7-12653863-DEL-723
chr7-12654879-DEL-282
chr7-12671336-DEL-86
chr7-12749430-DEL-437
chr7-12881879-DEL-128
chr7-12920357-DEL-6987
chr7-12958323-DEL-5590
chr7-12984631-DEL-355
chr7-13110105-DEL-53
chr7-13142899-DEL-52
chr7-13146987-DEL-6322
chr7-13214580-DEL-440
chr7-13233751-DEL-3118
chr7-13296469-DEL-501
chr7-13605755-DEL-80
chr7-13654400-DEL-54
chr7-13700112-DEL-1661
chr7-13713036-DEL-52
chr7-13763424-DEL-105
chr7-13765282-DEL-94
chr7-13812032-DEL-105
chr7-13901768-DEL-16081
chr7-13904394-DEL-192
chr7-14109511-DEL-33813

chr7-14220863-DEL-2539
chr7-14223716-DEL-2684
chr7-14241881-DEL-3985
chr7-14356442-DEL-83
chr7-14432638-DEL-196
chr7-14432944-DEL-76
chr7-14481168-DEL-219
chr7-14556798-DEL-2418
chr7-14557038-DEL-78
chr7-14609694-DEL-2959
chr7-14691401-DEL-204
chr7-14701382-DEL-505
chr7-15047061-DEL-60
chr7-15054201-DEL-50
chr7-15540030-DEL-5889
chr7-15571398-DEL-537
chr7-15572556-DEL-216
chr7-15651999-DEL-159
chr7-15700135-DEL-54
chr7-15704990-DEL-209
chr7-15712315-DEL-53
chr7-15732474-DEL-418
chr7-15742758-DEL-159
chr7-15745255-DEL-53
chr7-15763089-DEL-53
chr7-15915136-DEL-107
chr7-15924252-DEL-78
chr7-15931528-DEL-420
chr7-15931595-DEL-140
chr7-15932109-DEL-72
chr7-15955288-DEL-99
chr7-15986080-DEL-372
chr7-16076411-DEL-274
chr7-16076492-DEL-104
chr7-16180113-DEL-50
chr7-16193476-DEL-53
chr7-16353566-DEL-826
chr7-16460597-DEL-73
chr7-16470087-DEL-64
chr7-16479220-DEL-187
chr7-16479320-DEL-54
chr7-16484123-DEL-54
chr7-16488483-DEL-83
chr7-16587728-DEL-57
chr7-16632573-DEL-54
chr7-16668217-DEL-344
chr7-16668892-DEL-584
chr7-16701366-DEL-1986
chr7-16710704-DEL-210
chr7-16721202-DEL-57
chr7-16729710-DEL-120
chr7-16780616-DEL-315
chr7-16783374-DEL-206
chr7-16798079-DEL-106
chr7-16835928-DEL-530
chr7-16836130-DEL-52
chr7-16865137-DEL-2441
chr7-17187401-DEL-72
chr7-17264462-DEL-4428
chr7-17296739-DEL-54
chr7-17480672-DEL-64
chr7-17565748-DEL-2774
chr7-17927204-DEL-883
chr7-18059724-DEL-151
chr7-18066552-DEL-57
chr7-18066675-DEL-121
chr7-18093761-DEL-6347
chr7-18103054-DEL-130083
chr7-18105937-DEL-921
chr7-18112026-DEL-487
chr7-18113066-DEL-17118
chr7-18123420-DEL-19445
chr7-18147097-DEL-66
chr7-18162863-DEL-71
chr7-18169172-DEL-83
chr7-18187755-DEL-229
chr7-18234165-DEL-222

chr7-18246862-DEL-62
chr7-18329289-DEL-199
chr7-18329655-DEL-40690
chr7-18575903-DEL-78
chr7-18591452-DEL-63
chr7-18645675-DEL-52
chr7-18645675-DEL-157
chr7-18671721-DEL-159
chr7-18671929-DEL-52
chr7-18679642-DEL-158
chr7-18731020-DEL-53
chr7-18904929-DEL-371
chr7-18938990-DEL-121
chr7-19032917-DEL-233
chr7-19085251-DEL-53
chr7-19132369-DEL-52
chr7-19144077-DEL-134
chr7-19152049-DEL-53
chr7-19169244-DEL-53
chr7-19170478-DEL-103
chr7-19179727-DEL-262
chr7-19181443-DEL-57
chr7-19214540-DEL-53
chr7-19220002-DEL-54
chr7-19279382-DEL-182
chr7-19406094-DEL-65
chr7-19406094-DEL-275
chr7-19406820-DEL-144
chr7-19421927-DEL-53
chr7-19422435-DEL-61
chr7-19494689-DEL-54
chr7-19512162-DEL-159
chr7-19573540-DEL-457
chr7-19599386-DEL-107
chr7-19601576-DEL-53
chr7-19618849-DEL-53
chr7-19624851-DEL-57
chr7-19624992-DEL-200
chr7-19626270-DEL-173
chr7-19640029-DEL-281
chr7-19642783-DEL-106
chr7-19661836-DEL-56
chr7-19861067-DEL-118
chr7-19863830-DEL-370
chr7-19877562-DEL-356
chr7-20640308-DEL-507
chr7-20682390-DEL-1124
chr7-20687108-DEL-360
chr7-22528434-DEL-204
chr7-22823650-DEL-4422
chr7-22957180-DEL-78
chr7-23023170-DEL-9220
chr7-23360081-DEL-98
chr7-23400590-DEL-149
chr7-23518929-DEL-7003
chr7-23558991-DEL-1813
chr7-23589704-DEL-121
chr7-23609909-DEL-50
chr7-23652815-DEL-483
chr7-23772378-DEL-266
chr7-23772611-DEL-58
chr7-23776289-DEL-103
chr7-23830115-DEL-1254
chr7-23831940-DEL-51
chr7-23885179-DEL-128
chr7-23909464-DEL-50
chr7-23913614-DEL-107
chr7-23984304-DEL-87
chr7-24043698-DEL-52
chr7-24074984-DEL-1183
chr7-24077323-DEL-50
chr7-24088535-DEL-170
chr7-24096078-DEL-70
chr7-24108099-DEL-202
chr7-24143984-DEL-53
chr7-24160579-DEL-106
chr7-24170665-DEL-106

chr7-24195127-DEL-53
chr7-24315544-DEL-53
chr7-24433321-DEL-72
chr7-24433321-DEL-432
chr7-24435010-DEL-1265
chr7-24458700-DEL-226
chr7-24510316-DEL-140
chr7-24551291-DEL-51
chr7-24562655-DEL-710
chr7-24613266-DEL-685
chr7-24667123-DEL-83
chr7-24697482-DEL-215
chr7-24697559-DEL-66
chr7-24697792-DEL-102
chr7-24750606-DEL-54
chr7-24810871-DEL-313
chr7-24822798-DEL-1224
chr7-24824068-DEL-415
chr7-24824544-DEL-1044
chr7-24824726-DEL-180
chr7-24825653-DEL-933
chr7-24836147-DEL-519
chr7-24857564-DEL-204
chr7-24958678-DEL-664
chr7-24959123-DEL-430
chr7-25010131-DEL-153
chr7-25052464-DEL-119
chr7-25269289-DEL-5187
chr7-25311254-DEL-214
chr7-25311333-DEL-53
chr7-25476708-DEL-266
chr7-25484645-DEL-174
chr7-25484720-DEL-79
chr7-25513426-DEL-83
chr7-25599103-DEL-231
chr7-25660685-DEL-188
chr7-25660824-DEL-230
chr7-25705823-DEL-379
chr7-25738340-DEL-450
chr7-25802115-DEL-164
chr7-25855693-DEL-1488
chr7-26136259-DEL-54
chr7-26277606-DEL-92
chr7-26292048-DEL-5178
chr7-26340388-DEL-309
chr7-26340868-DEL-383
chr7-26351421-DEL-5905
chr7-26434492-DEL-50
chr7-26464317-DEL-100
chr7-26524406-DEL-54
chr7-26550462-DEL-112313
chr7-26567627-DEL-96
chr7-26709079-DEL-5009
chr7-26780699-DEL-184
chr7-26835174-DEL-106
chr7-26849731-DEL-62
chr7-26939051-DEL-55
chr7-26971215-DEL-80
chr7-26971377-DEL-261
chr7-26990344-DEL-72
chr7-27158027-DEL-52
chr7-27170519-DEL-88
chr7-27194321-DEL-53
chr7-27256352-DEL-540
chr7-27256577-DEL-1073
chr7-27262998-DEL-492
chr7-27263404-DEL-82
chr7-27293736-DEL-58
chr7-27348655-DEL-424
chr7-27348911-DEL-189
chr7-27352019-DEL-58
chr7-27382985-DEL-53
chr7-27382985-DEL-212
chr7-27387207-DEL-60
chr7-27397221-DEL-279
chr7-27405683-DEL-58
chr7-27418666-DEL-77

chr7-27458002-DEL-511
chr7-27461929-DEL-256
chr7-27468930-DEL-61
chr7-27478688-DEL-704
chr7-27508518-DEL-92
chr7-27594462-DEL-2967
chr7-27598609-DEL-108
chr7-27599109-DEL-391
chr7-27795524-DEL-68
chr7-27796102-DEL-1358
chr7-27796136-DEL-235
chr7-27796852-DEL-812
chr7-27849630-DEL-180
chr7-27888185-DEL-50
chr7-27901929-DEL-371
chr7-27924540-DEL-1485
chr7-27943028-DEL-66
chr7-27945611-DEL-66
chr7-27946280-DEL-132
chr7-27981978-DEL-55
chr7-27982778-DEL-109
chr7-28042489-DEL-260
chr7-28061075-DEL-52
chr7-28142939-DEL-71
chr7-28177786-DEL-159
chr7-28220335-DEL-118
chr7-28231201-DEL-164
chr7-28253208-DEL-158
chr7-28312126-DEL-320
chr7-28321745-DEL-54
chr7-28323241-DEL-273
chr7-28338471-DEL-51
chr7-28367704-DEL-53
chr7-28368650-DEL-1445
chr7-28380268-DEL-90
chr7-28382244-DEL-398
chr7-28405254-DEL-316
chr7-28427820-DEL-1680
chr7-28452576-DEL-54
chr7-28506785-DEL-99
chr7-28527146-DEL-1766
chr7-28576328-DEL-55
chr7-28631779-DEL-50
chr7-28679509-DEL-200
chr7-28703335-DEL-53
chr7-28727377-DEL-75
chr7-28736438-DEL-142
chr7-28741904-DEL-137
chr7-28763073-DEL-52
chr7-28763142-DEL-159
chr7-28765391-DEL-159
chr7-28800748-DEL-53
chr7-28843863-DEL-156
chr7-28843908-DEL-53
chr7-29058960-DEL-66
chr7-29088946-DEL-87
chr7-29089017-DEL-202
chr7-29113341-DEL-163
chr7-29119258-DEL-92
chr7-29143117-DEL-204
chr7-29145237-DEL-106
chr7-29273003-DEL-62
chr7-29360879-DEL-54
chr7-29509858-DEL-457
chr7-29516533-DEL-220
chr7-29535991-DEL-389
chr7-29564205-DEL-103
chr7-29630207-DEL-71
chr7-29630375-DEL-210
chr7-29656061-DEL-1623
chr7-29674695-DEL-170
chr7-29675592-DEL-2338
chr7-29729064-DEL-134
chr7-29888803-DEL-211
chr7-29901464-DEL-268
chr7-29992639-DEL-107
chr7-30003632-DEL-793

chr7-30079971-DEL-699
chr7-30097393-DEL-52
chr7-30109967-DEL-243
chr7-30150326-DEL-82
chr7-30150761-DEL-120
chr7-30175323-DEL-53
chr7-30203844-DEL-159
chr7-30204122-DEL-106
chr7-30207224-DEL-107
chr7-30230673-DEL-66
chr7-30238507-DEL-58
chr7-30410589-DEL-50
chr7-30467242-DEL-106
chr7-30472691-DEL-53
chr7-30494737-DEL-50
chr7-30500327-DEL-1462
chr7-30530518-DEL-4076
chr7-30531142-DEL-235
chr7-30571318-DEL-53
chr7-30582006-DEL-211
chr7-30604991-DEL-212
chr7-30605046-DEL-105
chr7-30737831-DEL-54
chr7-30805438-DEL-106
chr7-30822902-DEL-159
chr7-30865398-DEL-1566
chr7-30879963-DEL-88
chr7-31066651-DEL-2825
chr7-31560978-DEL-64
chr7-31722251-DEL-413
chr7-31760022-DEL-89
chr7-31809797-DEL-64
chr7-32962359-DEL-1746
chr7-33052905-DEL-276
chr7-33093744-DEL-1689
chr7-33095538-DEL-265
chr7-33115354-DEL-7050
chr7-33593953-DEL-1848
chr7-33698681-DEL-214
chr7-34156674-DEL-82
chr7-34173939-DEL-159
chr7-34173969-DEL-53
chr7-34193900-DEL-282
chr7-34268693-DEL-51
chr7-34305085-DEL-55
chr7-34322824-DEL-130
chr7-34377254-DEL-167
chr7-34412904-DEL-53
chr7-34413064-DEL-318
chr7-34415639-DEL-139
chr7-34424182-DEL-54
chr7-34427652-DEL-55
chr7-34439119-DEL-209
chr7-34439370-DEL-465
chr7-34515186-DEL-265
chr7-34652799-DEL-77
chr7-34659308-DEL-1286
chr7-34803495-DEL-462
chr7-34859214-DEL-268
chr7-34909255-DEL-70
chr7-34941636-DEL-127
chr7-34944837-DEL-196
chr7-34944961-DEL-84
chr7-34950754-DEL-112
chr7-34950814-DEL-50
chr7-34956196-DEL-212
chr7-34989704-DEL-90
chr7-35101088-DEL-126
chr7-35101408-DEL-205
chr7-35212795-DEL-181
chr7-35258752-DEL-2857
chr7-35272093-DEL-206
chr7-35329801-DEL-68
chr7-35331403-DEL-159
chr7-35360470-DEL-160
chr7-35360693-DEL-87
chr7-35416293-DEL-142

chr7-35464615-DEL-372
chr7-35464660-DEL-53
chr7-35605781-DEL-88
chr7-35646722-DEL-50
chr7-35668999-DEL-62
chr7-35724256-DEL-422
chr7-35872257-DEL-174
chr7-35893557-DEL-180
chr7-35979675-DEL-159
chr7-35991393-DEL-64
chr7-36124607-DEL-78
chr7-36154712-DEL-159
chr7-36208428-DEL-52
chr7-36242607-DEL-55
chr7-36272158-DEL-698
chr7-36275595-DEL-52
chr7-36278607-DEL-51
chr7-36362541-DEL-50
chr7-36369136-DEL-52
chr7-36410848-DEL-63
chr7-36733146-DEL-86
chr7-36772111-DEL-72
chr7-36814562-DEL-50
chr7-36866569-DEL-424
chr7-36866623-DEL-72
chr7-36866642-DEL-163
chr7-36878686-DEL-58
chr7-36906908-DEL-84
chr7-36910201-DEL-120
chr7-37052921-DEL-104
chr7-37107729-DEL-50
chr7-37108102-DEL-56
chr7-37108179-DEL-152
chr7-37127519-DEL-268
chr7-37195180-DEL-50
chr7-37221378-DEL-112
chr7-37241057-DEL-75
chr7-37281052-DEL-68
chr7-37383564-DEL-109
chr7-37479960-DEL-162
chr7-37494945-DEL-186
chr7-37495203-DEL-60
chr7-37504644-DEL-110
chr7-37504723-DEL-52
chr7-37530397-DEL-148
chr7-37576026-DEL-66
chr7-37576490-DEL-892
chr7-37577315-DEL-265
chr7-37747887-DEL-80
chr7-37819660-DEL-63
chr7-37819871-DEL-151
chr7-37822237-DEL-50
chr7-37823323-DEL-50
chr7-37860728-DEL-60
chr7-37974780-DEL-199
chr7-37985243-DEL-70
chr7-37998236-DEL-1136
chr7-38057377-DEL-56
chr7-38057671-DEL-123
chr7-38109444-DEL-715
chr7-38121290-DEL-175
chr7-38152383-DEL-147
chr7-38205466-DEL-60
chr7-38209057-DEL-106
chr7-38217693-DEL-126
chr7-38217698-DEL-262
chr7-38819401-DEL-272
chr7-38877465-DEL-1917
chr7-38915134-DEL-815
chr7-39081169-DEL-201
chr7-39250435-DEL-90
chr7-39278772-DEL-120
chr7-39285932-DEL-66
chr7-39300511-DEL-72
chr7-39303954-DEL-1830
chr7-39305894-DEL-113
chr7-39310103-DEL-2093

chr7-39317535-DEL-52
chr7-39343706-DEL-118
chr7-39343762-DEL-58
chr7-39397889-DEL-150
chr7-39413008-DEL-50
chr7-39413123-DEL-50
chr7-39483234-DEL-70
chr7-39484186-DEL-56
chr7-39710680-DEL-112
chr7-39859864-DEL-50
chr7-39872411-DEL-70
chr7-39873081-DEL-126
chr7-39936985-DEL-55
chr7-39997430-DEL-50
chr7-40028492-DEL-92
chr7-40050239-DEL-195
chr7-40088288-DEL-347
chr7-40397178-DEL-50
chr7-40645600-DEL-109
chr7-40809989-DEL-64
chr7-40821403-DEL-208
chr7-40847853-DEL-102
chr7-40881784-DEL-90
chr7-40885648-DEL-195
chr7-40929429-DEL-71
chr7-41033210-DEL-52
chr7-41179230-DEL-5851
chr7-41209274-DEL-50
chr7-41383815-DEL-80
chr7-41414023-DEL-1202
chr7-41465348-DEL-214
chr7-41795549-DEL-50
chr7-41968904-DEL-11814
chr7-42188902-DEL-79
chr7-42215960-DEL-2534
chr7-42361361-DEL-445
chr7-42502764-DEL-53
chr7-42648258-DEL-117
chr7-42652790-DEL-241
chr7-42654766-DEL-295
chr7-42772869-DEL-740
chr7-42819393-DEL-10188
chr7-42829432-DEL-53
chr7-42893451-DEL-70
chr7-42937026-DEL-728
chr7-43007129-DEL-50
chr7-43056552-DEL-74
chr7-43056952-DEL-108
chr7-43249026-DEL-295
chr7-43272462-DEL-359
chr7-43387047-DEL-289
chr7-43405834-DEL-128
chr7-43413443-DEL-65
chr7-43529488-DEL-63
chr7-43609833-DEL-96
chr7-43814242-DEL-641
chr7-43838413-DEL-4368
chr7-44047706-DEL-161
chr7-44047785-DEL-74
chr7-44052138-DEL-52
chr7-44132522-DEL-60
chr7-44195023-DEL-456
chr7-44229382-DEL-212
chr7-44235653-DEL-53
chr7-44270444-DEL-75
chr7-44309634-DEL-53
chr7-44349578-DEL-64
chr7-44387992-DEL-513
chr7-44482961-DEL-88
chr7-44511262-DEL-120
chr7-44612995-DEL-134
chr7-44714684-DEL-52
chr7-44904983-DEL-300
chr7-44943212-DEL-50
chr7-44975078-DEL-53
chr7-45046813-DEL-3406
chr7-45119503-DEL-200

chr7-45127472-DEL-53
chr7-45194375-DEL-86
chr7-45194692-DEL-203
chr7-45212108-DEL-50
chr7-45232805-DEL-50
chr7-45317998-DEL-60
chr7-45397158-DEL-52
chr7-45405704-DEL-175
chr7-45405958-DEL-124
chr7-45553038-DEL-120
chr7-45574154-DEL-54
chr7-45601015-DEL-88
chr7-45611081-DEL-561
chr7-45614489-DEL-263
chr7-45640431-DEL-71
chr7-45698734-DEL-60
chr7-45708230-DEL-50
chr7-45764886-DEL-92
chr7-45784525-DEL-253
chr7-45800584-DEL-58
chr7-45885753-DEL-78
chr7-46152135-DEL-58
chr7-46196649-DEL-393
chr7-46276211-DEL-748
chr7-46374359-DEL-2540
chr7-46632022-DEL-207
chr7-46659544-DEL-243
chr7-46659983-DEL-1014
chr7-46676916-DEL-50
chr7-46779035-DEL-520
chr7-46794609-DEL-117
chr7-46805996-DEL-408
chr7-46806324-DEL-138
chr7-46871607-DEL-1831
chr7-46890842-DEL-275
chr7-46906053-DEL-56
chr7-47171078-DEL-324
chr7-47209470-DEL-3992
chr7-47215091-DEL-46063
chr7-47264317-DEL-325
chr7-47307333-DEL-56
chr7-47314630-DEL-98
chr7-47327389-DEL-197
chr7-47338592-DEL-2624
chr7-47345812-DEL-3304
chr7-47365869-DEL-2313
chr7-47372979-DEL-65
chr7-47376256-DEL-2199
chr7-47398002-DEL-51
chr7-47400397-DEL-2337
chr7-47421490-DEL-531
chr7-47436317-DEL-1449
chr7-47561718-DEL-117
chr7-47562051-DEL-196
chr7-47590660-DEL-100
chr7-47619648-DEL-161
chr7-47621591-DEL-77
chr7-47623958-DEL-50
chr7-47643676-DEL-294
chr7-47668026-DEL-309
chr7-47676509-DEL-140
chr7-47692744-DEL-191
chr7-47712296-DEL-50
chr7-47735559-DEL-693
chr7-48133806-DEL-75
chr7-48226444-DEL-50
chr7-48353986-DEL-1633
chr7-48595750-DEL-538
chr7-48628154-DEL-60
chr7-48635001-DEL-56
chr7-48718425-DEL-94
chr7-48926834-DEL-59
chr7-48957004-DEL-162
chr7-49035616-DEL-68
chr7-49067313-DEL-102
chr7-49304068-DEL-58
chr7-49488088-DEL-3117

chr7-49540314-DEL-323
chr7-49688818-DEL-99
chr7-49860114-DEL-94
chr7-49892981-DEL-878
chr7-50017209-DEL-2477
chr7-50027097-DEL-108
chr7-50066907-DEL-50
chr7-50087048-DEL-66
chr7-50225165-DEL-106
chr7-50261571-DEL-58
chr7-50285227-DEL-244
chr7-50302345-DEL-50
chr7-50303729-DEL-113
chr7-50319724-DEL-64
chr7-50341392-DEL-208
chr7-50341413-DEL-102
chr7-50510434-DEL-71
chr7-50568847-DEL-60
chr7-50640016-DEL-2230
chr7-50642098-DEL-1889
chr7-50684486-DEL-149
chr7-50719081-DEL-95
chr7-50827281-DEL-57
chr7-50827388-DEL-173
chr7-50928910-DEL-874
chr7-50932372-DEL-50
chr7-50976604-DEL-87
chr7-50976719-DEL-319
chr7-51045897-DEL-279
chr7-51441856-DEL-70
chr7-51451691-DEL-590
chr7-51718798-DEL-71
chr7-51719026-DEL-89
chr7-51852028-DEL-60
chr7-51884107-DEL-68
chr7-51925227-DEL-117
chr7-51996436-DEL-1042
chr7-52030998-DEL-391
chr7-52052506-DEL-70
chr7-52126799-DEL-97
chr7-52146054-DEL-76
chr7-52308816-DEL-144
chr7-52322968-DEL-127
chr7-52324767-DEL-53
chr7-52371552-DEL-60
chr7-52372657-DEL-53
chr7-52376363-DEL-386
chr7-52528176-DEL-570
chr7-52528521-DEL-91
chr7-52576807-DEL-2340
chr7-52628049-DEL-290
chr7-52922061-DEL-122
chr7-53068776-DEL-159
chr7-53217494-DEL-1335
chr7-53219238-DEL-1773
chr7-53240514-DEL-523
chr7-53372632-DEL-66
chr7-53395136-DEL-616
chr7-53419746-DEL-6229
chr7-53598443-DEL-4170
chr7-53606023-DEL-65
chr7-53630699-DEL-1372
chr7-53699271-DEL-63
chr7-53712820-DEL-50
chr7-53836850-DEL-858
chr7-53863069-DEL-84
chr7-53996808-DEL-5346
chr7-54071943-DEL-56
chr7-54119518-DEL-381
chr7-54119966-DEL-1970
chr7-54159739-DEL-51
chr7-54437855-DEL-14949
chr7-54475607-DEL-214
chr7-54735605-DEL-1048
chr7-54735926-DEL-212
chr7-54748824-DEL-50
chr7-54857528-DEL-181

chr7-54929155-DEL-50
chr7-55071207-DEL-332
chr7-55135788-DEL-1897
chr7-55159398-DEL-85
chr7-55241477-DEL-56
chr7-55244595-DEL-233
chr7-55248667-DEL-214
chr7-55276846-DEL-3507
chr7-55551868-DEL-56
chr7-55620892-DEL-376
chr7-55687994-DEL-58
chr7-55809930-DEL-1900
chr7-55835092-DEL-87
chr7-55935322-DEL-58
chr7-56064069-DEL-50
chr7-56157561-DEL-1538
chr7-56209574-DEL-3246
chr7-56433125-DEL-65
chr7-56434274-DEL-1474
chr7-56528128-DEL-2481
chr7-56549169-DEL-54
chr7-56570124-DEL-9079
chr7-56639627-DEL-1783
chr7-56668054-DEL-6268
chr7-57005448-DEL-53
chr7-57244267-DEL-86
chr7-57310171-DEL-78
chr7-57389982-DEL-50
chr7-57502720-DEL-614
chr7-57618499-DEL-375
chr7-57618957-DEL-615
chr7-57628672-DEL-1371
chr7-57741096-DEL-164
chr7-58120506-DEL-52
chr7-58157854-DEL-2674
chr7-58175949-DEL-4469
chr7-58185165-DEL-103
chr7-58185515-DEL-2203
chr7-58194200-DEL-54
chr7-58214216-DEL-104
chr7-58216617-DEL-51
chr7-58238433-DEL-206
chr7-58485494-DEL-100
chr7-58645881-DEL-139
chr7-58668453-DEL-52
chr7-58785730-DEL-2494
chr7-58819996-DEL-60
chr7-58843508-DEL-50
chr7-58964658-DEL-54
chr7-58964897-DEL-63
chr7-59003091-DEL-2916
chr7-59579554-DEL-807
chr7-59612543-DEL-110
chr7-59614470-DEL-850
chr7-59614856-DEL-1316
chr7-59712978-DEL-877
chr7-59932110-DEL-2946
chr7-60132552-DEL-4549
chr7-60142369-DEL-809
chr7-60147061-DEL-3545
chr7-60201386-DEL-236
chr7-60379479-DEL-350
chr7-60434999-DEL-899
chr7-60471064-DEL-120
chr7-60471138-DEL-50
chr7-60516453-DEL-55
chr7-60516789-DEL-183
chr7-60574791-DEL-153
chr7-60708586-DEL-52
chr7-60763308-DEL-161
chr7-60807190-DEL-233
chr7-60912040-DEL-52
chr7-60973433-DEL-1307
chr7-61027641-DEL-52
chr7-61028055-DEL-64
chr7-61036450-DEL-4451
chr7-61083622-DEL-7269

chr7-61098577-DEL-62
chr7-61256071-DEL-4544
chr7-61260422-DEL-119
chr7-61273105-DEL-112
chr7-61487988-DEL-357
chr7-61540400-DEL-51
chr7-61566377-DEL-682
chr7-61661396-DEL-5384
chr7-61687395-DEL-60
chr7-61695109-DEL-51
chr7-61748918-DEL-735
chr7-62039458-DEL-54
chr7-62118038-DEL-2302
chr7-62232733-DEL-272
chr7-62270008-DEL-6399
chr7-62382826-DEL-156
chr7-62420850-DEL-63
chr7-62503744-DEL-199
chr7-62557375-DEL-50
chr7-62579585-DEL-91
chr7-62705903-DEL-56
chr7-62706017-DEL-52
chr7-62718400-DEL-729
chr7-62811393-DEL-103
chr7-62978550-DEL-74
chr7-63035736-DEL-60
chr7-63179268-DEL-60
chr7-63262438-DEL-51
chr7-63390837-DEL-62
chr7-63463286-DEL-81
chr7-63664809-DEL-56
chr7-63717993-DEL-124
chr7-63719116-DEL-247
chr7-63724470-DEL-63
chr7-63734904-DEL-314
chr7-63821663-DEL-241
chr7-63861989-DEL-159
chr7-63866385-DEL-70
chr7-63949821-DEL-339
chr7-64000107-DEL-61
chr7-64013885-DEL-51
chr7-64050559-DEL-52
chr7-64115495-DEL-15920
chr7-64115829-DEL-700
chr7-64149693-DEL-257
chr7-64149760-DEL-126
chr7-64164858-DEL-80
chr7-64169621-DEL-334
chr7-64188821-DEL-162
chr7-64305667-DEL-75
chr7-64332397-DEL-97
chr7-64582555-DEL-130
chr7-64777023-DEL-86
chr7-64780859-DEL-64
chr7-64923749-DEL-56
chr7-64925089-DEL-2164
chr7-65102190-DEL-1177
chr7-65186355-DEL-56
chr7-65187685-DEL-56
chr7-65189225-DEL-81
chr7-65330931-DEL-51
chr7-65369146-DEL-205
chr7-65400553-DEL-54
chr7-65527521-DEL-52
chr7-65544230-DEL-112
chr7-65576328-DEL-162
chr7-65743884-DEL-2945
chr7-65795108-DEL-980
chr7-65795508-DEL-100
chr7-65796187-DEL-267
chr7-65807451-DEL-50
chr7-65941019-DEL-559
chr7-65969083-DEL-96
chr7-66000477-DEL-76
chr7-66011487-DEL-560
chr7-66024051-DEL-58
chr7-66029701-DEL-156

chr7-66035941-DEL-50
chr7-66160550-DEL-297
chr7-66204380-DEL-106
chr7-66311752-DEL-50
chr7-66389607-DEL-96
chr7-66432747-DEL-163
chr7-66513446-DEL-475
chr7-66795249-DEL-56
chr7-66866412-DEL-1347
chr7-66904122-DEL-746
chr7-67233225-DEL-106
chr7-67236540-DEL-106
chr7-67394919-DEL-68
chr7-67413216-DEL-59
chr7-67565089-DEL-1746
chr7-67658177-DEL-101
chr7-67699928-DEL-58
chr7-67715239-DEL-63
chr7-67715263-DEL-144
chr7-67731446-DEL-50
chr7-67731721-DEL-100
chr7-67821472-DEL-85
chr7-67866708-DEL-671
chr7-67893688-DEL-50
chr7-67901957-DEL-930
chr7-68086288-DEL-204
chr7-68086429-DEL-51
chr7-68116275-DEL-82
chr7-68157984-DEL-53
chr7-68162381-DEL-50
chr7-68236684-DEL-843
chr7-68237261-DEL-264
chr7-68285821-DEL-67
chr7-68296141-DEL-51
chr7-68363458-DEL-1377
chr7-68463832-DEL-314
chr7-68496227-DEL-56
chr7-68707153-DEL-203
chr7-68762628-DEL-984
chr7-68788273-DEL-50
chr7-68808396-DEL-95
chr7-68905331-DEL-55
chr7-68913366-DEL-95
chr7-68970451-DEL-647
chr7-68984764-DEL-52
chr7-69074418-DEL-95
chr7-69280976-DEL-606
chr7-69288843-DEL-176
chr7-69862932-DEL-336
chr7-70167258-DEL-63
chr7-70286076-DEL-68
chr7-70516852-DEL-52
chr7-70634576-DEL-3145
chr7-70678873-DEL-170
chr7-70807154-DEL-52
chr7-71096627-DEL-124
chr7-71510198-DEL-1491
chr7-71692925-DEL-1945
chr7-71792097-DEL-54
chr7-71967623-DEL-416
chr7-72105333-DEL-273
chr7-72147679-DEL-257
chr7-72147938-DEL-297
chr7-72252375-DEL-60
chr7-72258496-DEL-118
chr7-72264460-DEL-54
chr7-72316276-DEL-146
chr7-72322529-DEL-2100
chr7-72327800-DEL-1005
chr7-72338054-DEL-2132
chr7-72403523-DEL-520
chr7-72403545-DEL-180
chr7-72403983-DEL-172
chr7-72428201-DEL-147
chr7-72488569-DEL-135
chr7-72532319-DEL-1702
chr7-72564574-DEL-4354

chr7-72649948-DEL-53
chr7-72686311-DEL-70
chr7-72886626-DEL-75
chr7-72913444-DEL-131
chr7-72917179-DEL-1544
chr7-72974543-DEL-307
chr7-72995755-DEL-50
chr7-72995974-DEL-193
chr7-73063937-DEL-50
chr7-73224315-DEL-148
chr7-73251355-DEL-50
chr7-73296183-DEL-234
chr7-73322188-DEL-6071
chr7-73392368-DEL-50
chr7-73411111-DEL-140
chr7-73480211-DEL-366
chr7-73562804-DEL-190
chr7-73718660-DEL-170
chr7-73756109-DEL-68
chr7-73783761-DEL-54
chr7-73810014-DEL-81
chr7-73863625-DEL-72
chr7-73863625-DEL-156
chr7-73878071-DEL-54
chr7-73898910-DEL-78
chr7-73942348-DEL-205
chr7-74000005-DEL-88
chr7-74000297-DEL-141
chr7-74000440-DEL-853
chr7-74001237-DEL-188
chr7-74053782-DEL-246
chr7-74080065-DEL-328
chr7-74080502-DEL-307
chr7-74173430-DEL-65
chr7-74327300-DEL-155
chr7-74457360-DEL-72
chr7-74550516-DEL-216
chr7-74587216-DEL-50
chr7-74679085-DEL-211
chr7-74679152-DEL-63
chr7-74717051-DEL-170
chr7-74752668-DEL-116
chr7-74793822-DEL-60
chr7-74794415-DEL-74
chr7-74810987-DEL-3756
chr7-74814779-DEL-2859
chr7-74827523-DEL-175
chr7-74874744-DEL-54
chr7-74902457-DEL-222
chr7-74902745-DEL-288
chr7-74922477-DEL-70
chr7-74956408-DEL-152
chr7-75088007-DEL-541
chr7-75139432-DEL-88
chr7-75257409-DEL-50
chr7-75429853-DEL-54
chr7-75473063-DEL-3049
chr7-75476243-DEL-112
chr7-75496665-DEL-61
chr7-75570852-DEL-50
chr7-75645113-DEL-424
chr7-75645736-DEL-212
chr7-75736092-DEL-54
chr7-75745348-DEL-221
chr7-75754002-DEL-1882
chr7-75890762-DEL-9122
chr7-76460532-DEL-60
chr7-76468504-DEL-50
chr7-76500243-DEL-553
chr7-76500491-DEL-211
chr7-76501017-DEL-134
chr7-76537549-DEL-106
chr7-76674823-DEL-50
chr7-76701314-DEL-120
chr7-76884359-DEL-64
chr7-77101492-DEL-480
chr7-77225554-DEL-180

chr7-77255801-DEL-479
chr7-77256316-DEL-546
chr7-77279789-DEL-67
chr7-77380536-DEL-62
chr7-77400902-DEL-972
chr7-77416576-DEL-1134
chr7-77495475-DEL-111
chr7-77496332-DEL-66
chr7-77828524-DEL-99
chr7-77828715-DEL-51
chr7-78151051-DEL-50
chr7-78159127-DEL-69
chr7-78185917-DEL-62
chr7-78264738-DEL-66
chr7-78452064-DEL-189
chr7-78480890-DEL-1894
chr7-78481808-DEL-566
chr7-78481846-DEL-120
chr7-78482916-DEL-538
chr7-78483670-DEL-88
chr7-78654551-DEL-959
chr7-78655068-DEL-671
chr7-78655225-DEL-274
chr7-78655722-DEL-114
chr7-78697645-DEL-86
chr7-78773824-DEL-144
chr7-78774050-DEL-376
chr7-78774308-DEL-166
chr7-79246312-DEL-57
chr7-79439833-DEL-696
chr7-79467733-DEL-162
chr7-79478537-DEL-1080
chr7-79555592-DEL-147
chr7-79558474-DEL-195
chr7-79574186-DEL-120
chr7-79613010-DEL-1116
chr7-79622522-DEL-50
chr7-79627580-DEL-50
chr7-79645192-DEL-312
chr7-79824815-DEL-114
chr7-79871517-DEL-64
chr7-79921351-DEL-155
chr7-79922879-DEL-454
chr7-80097618-DEL-315
chr7-80097708-DEL-90
chr7-80197248-DEL-82
chr7-80200344-DEL-164
chr7-80313258-DEL-146
chr7-80358237-DEL-208
chr7-80358302-DEL-520
chr7-80374287-DEL-60
chr7-80443142-DEL-1034
chr7-80443708-DEL-482
chr7-80456835-DEL-52
chr7-80528579-DEL-61
chr7-80587037-DEL-55
chr7-80768562-DEL-66
chr7-80779817-DEL-187
chr7-80779818-DEL-621
chr7-80780183-DEL-62
chr7-80822161-DEL-80
chr7-80823126-DEL-1209
chr7-80884342-DEL-136
chr7-80967185-DEL-74
chr7-81211562-DEL-86
chr7-81239662-DEL-193
chr7-81287506-DEL-201
chr7-81315172-DEL-65
chr7-81592136-DEL-110
chr7-81661149-DEL-106
chr7-81803993-DEL-206
chr7-81804260-DEL-70
chr7-81816655-DEL-4907
chr7-81859317-DEL-112
chr7-81932442-DEL-54
chr7-81971100-DEL-540
chr7-82001739-DEL-55

chr7-82069854-DEL-50
chr7-82174678-DEL-210
chr7-82213831-DEL-97
chr7-82331120-DEL-130
chr7-82357565-DEL-132
chr7-82504051-DEL-124
chr7-82701195-DEL-209
chr7-82701401-DEL-50
chr7-82705206-DEL-52
chr7-82814299-DEL-51
chr7-83120985-DEL-50
chr7-83203112-DEL-408
chr7-83228935-DEL-963
chr7-83238432-DEL-62
chr7-83346080-DEL-222
chr7-83346496-DEL-109
chr7-83398997-DEL-1036
chr7-83417596-DEL-139
chr7-83462616-DEL-54
chr7-83474772-DEL-312
chr7-83666567-DEL-79
chr7-83742130-DEL-324
chr7-83963830-DEL-639
chr7-84014827-DEL-97
chr7-84038685-DEL-189
chr7-84040365-DEL-77
chr7-84040561-DEL-178
chr7-84062485-DEL-66
chr7-84148824-DEL-215
chr7-84237613-DEL-99
chr7-84273994-DEL-59
chr7-84285537-DEL-1163
chr7-84372446-DEL-149
chr7-84409152-DEL-53
chr7-84420015-DEL-115
chr7-84420153-DEL-163
chr7-84596689-DEL-159
chr7-84670083-DEL-1628
chr7-84827065-DEL-36009
chr7-85005056-DEL-65
chr7-85023151-DEL-4535
chr7-85083181-DEL-249
chr7-85138020-DEL-214
chr7-85183608-DEL-1654
chr7-85248203-DEL-52
chr7-85302983-DEL-1046
chr7-85508154-DEL-111
chr7-85513803-DEL-1917
chr7-85584401-DEL-73
chr7-85607592-DEL-152
chr7-85613450-DEL-144
chr7-85743879-DEL-4258
chr7-85836683-DEL-90
chr7-85897027-DEL-100
chr7-85998856-DEL-5432
chr7-86052364-DEL-235
chr7-86054483-DEL-411
chr7-86055398-DEL-929
chr7-86081252-DEL-340
chr7-86181545-DEL-1309
chr7-86212685-DEL-23475
chr7-86394317-DEL-56
chr7-86446535-DEL-64
chr7-86544646-DEL-1375
chr7-86547680-DEL-54
chr7-86580040-DEL-50
chr7-86652948-DEL-50
chr7-86657186-DEL-3286
chr7-86672098-DEL-54
chr7-86694088-DEL-1492
chr7-86729402-DEL-5933
chr7-86796155-DEL-50
chr7-86837365-DEL-141
chr7-86837459-DEL-50
chr7-86850774-DEL-193
chr7-86865170-DEL-1702
chr7-86985717-DEL-125

chr7-87013304-DEL-776
chr7-87053482-DEL-482
chr7-87056896-DEL-125
chr7-87058421-DEL-154
chr7-87107424-DEL-54
chr7-87206360-DEL-195
chr7-87207940-DEL-367
chr7-87211024-DEL-50
chr7-87270423-DEL-451
chr7-87350005-DEL-58
chr7-87457687-DEL-85
chr7-87551995-DEL-424
chr7-87762906-DEL-52
chr7-87827170-DEL-64
chr7-87915797-DEL-363
chr7-87915797-DEL-2594
chr7-87915833-DEL-108
chr7-88027800-DEL-299
chr7-88042465-DEL-50
chr7-88043469-DEL-113
chr7-88059307-DEL-1213
chr7-88060550-DEL-1648
chr7-88123781-DEL-54
chr7-88158872-DEL-52
chr7-88226729-DEL-51
chr7-88326752-DEL-1563
chr7-88372991-DEL-87
chr7-88440106-DEL-83
chr7-88537327-DEL-5108
chr7-88639134-DEL-1405
chr7-88793466-DEL-58
chr7-88860596-DEL-1345
chr7-88915713-DEL-94
chr7-89021850-DEL-50
chr7-89371781-DEL-114
chr7-89379165-DEL-446
chr7-89400155-DEL-2249
chr7-89553514-DEL-50
chr7-89593043-DEL-400
chr7-89653033-DEL-63
chr7-89662117-DEL-100
chr7-89664699-DEL-61
chr7-89732027-DEL-55
chr7-89876634-DEL-60
chr7-89878424-DEL-85
chr7-89932472-DEL-68
chr7-90010172-DEL-176
chr7-90111392-DEL-84
chr7-90112283-DEL-4248
chr7-90179683-DEL-2940
chr7-90191984-DEL-103
chr7-90225734-DEL-77
chr7-90245244-DEL-135
chr7-90267803-DEL-201
chr7-90274163-DEL-297
chr7-90356057-DEL-71
chr7-90453130-DEL-1499
chr7-90657650-DEL-153
chr7-90878322-DEL-84
chr7-90882973-DEL-122
chr7-91002673-DEL-76
chr7-91106906-DEL-51
chr7-91153544-DEL-294
chr7-91212348-DEL-52
chr7-91291677-DEL-51
chr7-91291800-DEL-377
chr7-91292201-DEL-54
chr7-91327726-DEL-54
chr7-91412807-DEL-101
chr7-91457872-DEL-1520
chr7-91469299-DEL-77
chr7-91601915-DEL-821
chr7-91714330-DEL-98
chr7-91758580-DEL-195
chr7-91792574-DEL-286
chr7-91796833-DEL-156
chr7-91983961-DEL-60

chr7-92136394-DEL-58
chr7-92210938-DEL-116
chr7-92281304-DEL-329
chr7-92289637-DEL-64
chr7-92330091-DEL-63
chr7-92363982-DEL-103
chr7-92434100-DEL-61
chr7-92468164-DEL-72
chr7-92483440-DEL-57
chr7-92536311-DEL-70
chr7-92705547-DEL-55
chr7-92737755-DEL-182
chr7-92787437-DEL-188
chr7-92931465-DEL-251
chr7-92954745-DEL-170
chr7-92954772-DEL-56
chr7-92968976-DEL-3332
chr7-93292793-DEL-399
chr7-93299717-DEL-50
chr7-93401534-DEL-96
chr7-93524718-DEL-99
chr7-93580662-DEL-2086
chr7-93606946-DEL-120
chr7-93616961-DEL-51
chr7-93649412-DEL-640
chr7-93681475-DEL-860
chr7-93697453-DEL-62
chr7-93846801-DEL-52
chr7-93857111-DEL-104
chr7-94048323-DEL-79
chr7-94076033-DEL-155
chr7-94150551-DEL-2012
chr7-94179528-DEL-92
chr7-94260226-DEL-84
chr7-94260304-DEL-61
chr7-94293053-DEL-173
chr7-94293074-DEL-79
chr7-94439712-DEL-5524
chr7-94504852-DEL-115
chr7-94782969-DEL-55
chr7-94826409-DEL-4200
chr7-94920272-DEL-162
chr7-94920488-DEL-180
chr7-94920543-DEL-57
chr7-95127223-DEL-1230
chr7-95249833-DEL-66
chr7-95284148-DEL-668
chr7-95284920-DEL-62
chr7-95287210-DEL-50
chr7-95674656-DEL-56
chr7-95810486-DEL-76
chr7-95840701-DEL-2219
chr7-96181127-DEL-50
chr7-96242725-DEL-130
chr7-96286496-DEL-75
chr7-96286864-DEL-90
chr7-96407389-DEL-71
chr7-96470462-DEL-342
chr7-96510372-DEL-177
chr7-96570097-DEL-82
chr7-96580314-DEL-139
chr7-96633087-DEL-69
chr7-96715717-DEL-610
chr7-96982235-DEL-75
chr7-97050858-DEL-50
chr7-97074978-DEL-50
chr7-97092208-DEL-1279
chr7-97108232-DEL-88
chr7-97180862-DEL-292
chr7-97413141-DEL-50
chr7-97445610-DEL-128
chr7-97453316-DEL-442
chr7-97583881-DEL-53
chr7-97653918-DEL-166
chr7-97654028-DEL-54
chr7-97804290-DEL-56
chr7-97806890-DEL-61

chr7-97830113-DEL-56
chr7-97836579-DEL-118
chr7-97836812-DEL-315
chr7-97858997-DEL-1491
chr7-97866705-DEL-86
chr7-97968252-DEL-143
chr7-98177205-DEL-151
chr7-98262029-DEL-65
chr7-98336335-DEL-52
chr7-98403145-DEL-128
chr7-98516144-DEL-50
chr7-98613026-DEL-70
chr7-98708084-DEL-62
chr7-98750897-DEL-56
chr7-98804295-DEL-190
chr7-98804531-DEL-58
chr7-98889491-DEL-51
chr7-98945810-DEL-415
chr7-98945810-DEL-1604
chr7-98945865-DEL-110
chr7-98946246-DEL-2985
chr7-98947414-DEL-1492
chr7-98947909-DEL-110
chr7-98980580-DEL-170
chr7-98980688-DEL-70
chr7-99166403-DEL-50
chr7-99405212-DEL-93
chr7-99609705-DEL-64
chr7-99637521-DEL-74
chr7-99707876-DEL-128
chr7-99798680-DEL-491
chr7-99819900-DEL-84
chr7-99821214-DEL-4177
chr7-99957450-DEL-78
chr7-100033221-DEL-258
chr7-100131554-DEL-58
chr7-100175393-DEL-135
chr7-100301860-DEL-306
chr7-100311939-DEL-150
chr7-100323422-DEL-2358
chr7-100348646-DEL-56
chr7-100388156-DEL-388
chr7-100397006-DEL-145
chr7-100431131-DEL-565
chr7-100603661-DEL-169
chr7-100675516-DEL-50
chr7-100965562-DEL-54
chr7-101020329-DEL-200
chr7-101067413-DEL-237
chr7-101179795-DEL-54
chr7-101204996-DEL-25359
chr7-101217730-DEL-2251
chr7-101306066-DEL-87
chr7-101591937-DEL-2144
chr7-101627189-DEL-1052
chr7-101791909-DEL-113
chr7-101911861-DEL-50
chr7-101935990-DEL-53
chr7-101963101-DEL-83
chr7-101985333-DEL-434
chr7-102003131-DEL-175
chr7-102042319-DEL-52
chr7-102068664-DEL-194
chr7-102088457-DEL-1510
chr7-102141144-DEL-60
chr7-102237673-DEL-52
chr7-102381159-DEL-55
chr7-102383895-DEL-64
chr7-102409266-DEL-242
chr7-102430005-DEL-64
chr7-102715545-DEL-529
chr7-102866437-DEL-50
chr7-102902198-DEL-2229
chr7-102938898-DEL-4162
chr7-103027743-DEL-543
chr7-103106555-DEL-96
chr7-103115098-DEL-124

chr7-103212445-DEL-1991
chr7-103225879-DEL-4545
chr7-103242485-DEL-4869
chr7-103380625-DEL-50
chr7-103529713-DEL-54
chr7-103559987-DEL-69
chr7-103595360-DEL-1389
chr7-103682947-DEL-212
chr7-103683539-DEL-211
chr7-103700503-DEL-50
chr7-103704830-DEL-54
chr7-103705420-DEL-1256
chr7-103710311-DEL-76
chr7-103737726-DEL-204
chr7-103793070-DEL-50
chr7-103821222-DEL-63
chr7-103821514-DEL-190
chr7-103857749-DEL-170
chr7-103857769-DEL-60
chr7-103879355-DEL-77
chr7-103898509-DEL-81828
chr7-103899353-DEL-914
chr7-103913668-DEL-50
chr7-103988997-DEL-58
chr7-104000883-DEL-88
chr7-104020835-DEL-183
chr7-104065044-DEL-2713
chr7-104077203-DEL-57
chr7-104126516-DEL-214
chr7-104197972-DEL-78
chr7-104297354-DEL-247
chr7-104370168-DEL-642
chr7-104445910-DEL-8248
chr7-104554719-DEL-1104
chr7-104734854-DEL-35741
chr7-104860789-DEL-272
chr7-104992514-DEL-988
chr7-105107486-DEL-127
chr7-105121714-DEL-50
chr7-105142148-DEL-411
chr7-105142387-DEL-51
chr7-105247654-DEL-54
chr7-105497196-DEL-420
chr7-105850465-DEL-570
chr7-106009998-DEL-1464
chr7-106034550-DEL-3860
chr7-106102612-DEL-83
chr7-106102890-DEL-190
chr7-106102903-DEL-81
chr7-106131530-DEL-2623
chr7-106141415-DEL-50
chr7-106236934-DEL-469
chr7-106237071-DEL-168
chr7-106255335-DEL-90
chr7-106270819-DEL-82
chr7-106429175-DEL-171
chr7-106574206-DEL-50
chr7-106599642-DEL-157
chr7-106680233-DEL-6662
chr7-106704514-DEL-22529
chr7-106711604-DEL-69
chr7-106885523-DEL-116
chr7-106936812-DEL-86
chr7-106937150-DEL-151
chr7-106946218-DEL-184
chr7-106946242-DEL-84
chr7-106946511-DEL-121
chr7-107082865-DEL-92
chr7-107174780-DEL-57
chr7-107381635-DEL-76
chr7-107407905-DEL-1247
chr7-107407932-DEL-239
chr7-107408506-DEL-62
chr7-107408925-DEL-65
chr7-107409092-DEL-525
chr7-107466475-DEL-1774
chr7-107567310-DEL-22968

chr7-107601484-DEL-51
chr7-107612498-DEL-214
chr7-107689013-DEL-2026
chr7-107896388-DEL-322
chr7-107936596-DEL-54
chr7-107963602-DEL-75
chr7-108388039-DEL-408
chr7-108458239-DEL-262
chr7-108492578-DEL-52
chr7-108547557-DEL-85
chr7-108622854-DEL-517
chr7-108830329-DEL-66
chr7-108857702-DEL-68
chr7-109046088-DEL-109
chr7-109046266-DEL-389
chr7-109098132-DEL-66
chr7-109187806-DEL-153
chr7-109251008-DEL-50
chr7-109283807-DEL-377
chr7-109334596-DEL-54
chr7-109444526-DEL-144
chr7-109445006-DEL-78
chr7-109445206-DEL-672
chr7-109445868-DEL-100
chr7-109465851-DEL-66
chr7-109551957-DEL-99
chr7-109552602-DEL-67
chr7-109589229-DEL-1086
chr7-109658875-DEL-56
chr7-109691594-DEL-234
chr7-109892786-DEL-634
chr7-109937395-DEL-55
chr7-109948405-DEL-253
chr7-109948630-DEL-78
chr7-110043898-DEL-910
chr7-110511615-DEL-98
chr7-110523884-DEL-50
chr7-110617809-DEL-1257
chr7-110620917-DEL-78
chr7-110700434-DEL-61
chr7-110732344-DEL-221
chr7-110786376-DEL-52
chr7-110786507-DEL-50
chr7-110820551-DEL-52
chr7-110820681-DEL-108
chr7-110828426-DEL-1033
chr7-110847002-DEL-224
chr7-110956964-DEL-69
chr7-110957177-DEL-524
chr7-110982620-DEL-942
chr7-110983643-DEL-2508
chr7-111056749-DEL-60
chr7-111127365-DEL-84
chr7-111136514-DEL-77
chr7-111334973-DEL-160
chr7-111402894-DEL-60
chr7-111734060-DEL-241
chr7-111734229-DEL-73
chr7-111789799-DEL-51
chr7-111876395-DEL-234
chr7-111973150-DEL-104
chr7-112099499-DEL-68
chr7-112139984-DEL-61
chr7-112193835-DEL-58
chr7-112236505-DEL-420
chr7-112535014-DEL-67
chr7-112693705-DEL-116
chr7-112700599-DEL-98
chr7-112714403-DEL-243
chr7-112772024-DEL-184
chr7-112990019-DEL-136
chr7-113193144-DEL-51
chr7-113214212-DEL-52
chr7-113232581-DEL-100
chr7-113250117-DEL-55
chr7-113363361-DEL-50
chr7-113376546-DEL-96

chr7-113682287-DEL-50
chr7-113697265-DEL-141
chr7-113746410-DEL-448
chr7-113746803-DEL-96
chr7-113780063-DEL-294
chr7-113937598-DEL-124
chr7-113949048-DEL-1457
chr7-114176864-DEL-53
chr7-114271478-DEL-58
chr7-114342481-DEL-145
chr7-114362750-DEL-335
chr7-114472353-DEL-73
chr7-114591196-DEL-50
chr7-114648739-DEL-67
chr7-114856620-DEL-50
chr7-115101376-DEL-511
chr7-115102524-DEL-596
chr7-115333651-DEL-81
chr7-115417029-DEL-411
chr7-115714183-DEL-154
chr7-115773234-DEL-292
chr7-115785139-DEL-314
chr7-115807702-DEL-206
chr7-115807956-DEL-73
chr7-115914473-DEL-1577
chr7-116001792-DEL-2785
chr7-116063033-DEL-50
chr7-116241198-DEL-478
chr7-116505815-DEL-96
chr7-116601128-DEL-267
chr7-116601146-DEL-126
chr7-116601528-DEL-69
chr7-116601887-DEL-98
chr7-116623276-DEL-65
chr7-116675765-DEL-64
chr7-116680656-DEL-230
chr7-116750857-DEL-78
chr7-116921526-DEL-64
chr7-117156421-DEL-50
chr7-117336692-DEL-58
chr7-117353884-DEL-67
chr7-117434822-DEL-2098
chr7-117485900-DEL-278
chr7-117665848-DEL-50
chr7-117667951-DEL-50
chr7-117693005-DEL-109
chr7-117786241-DEL-428
chr7-117786690-DEL-340
chr7-117809100-DEL-74
chr7-117851447-DEL-50
chr7-117914334-DEL-102
chr7-117924595-DEL-113
chr7-117947941-DEL-57
chr7-118068909-DEL-68
chr7-118117613-DEL-69
chr7-118141308-DEL-169
chr7-118277422-DEL-170
chr7-118277752-DEL-50
chr7-118353401-DEL-57
chr7-118357211-DEL-90
chr7-118402727-DEL-120
chr7-118412473-DEL-136
chr7-118452779-DEL-1863
chr7-118512532-DEL-138
chr7-118586914-DEL-63
chr7-118969831-DEL-74
chr7-119011695-DEL-50
chr7-119094881-DEL-106
chr7-119358669-DEL-385
chr7-119561890-DEL-87
chr7-119608199-DEL-112
chr7-119626066-DEL-65
chr7-119626241-DEL-129
chr7-119739537-DEL-136
chr7-119739912-DEL-269
chr7-119833567-DEL-114
chr7-119864078-DEL-61

chr7-119958373-DEL-94
chr7-120061822-DEL-2791
chr7-120088581-DEL-222
chr7-120144850-DEL-68
chr7-120171202-DEL-79
chr7-120190943-DEL-51
chr7-120198287-DEL-89
chr7-120207382-DEL-2040
chr7-120209502-DEL-103
chr7-120213469-DEL-66
chr7-120245573-DEL-1584
chr7-120302003-DEL-56
chr7-120388462-DEL-243
chr7-120458770-DEL-228
chr7-120545581-DEL-272
chr7-120545586-DEL-832
chr7-120552613-DEL-158
chr7-120582356-DEL-62
chr7-120600892-DEL-155
chr7-120607586-DEL-252
chr7-120607626-DEL-100
chr7-120679695-DEL-144
chr7-120696003-DEL-824
chr7-120720364-DEL-140
chr7-120726308-DEL-81
chr7-120844932-DEL-136
chr7-120896189-DEL-403
chr7-120902267-DEL-366
chr7-121124945-DEL-110
chr7-121155477-DEL-85
chr7-121166745-DEL-73
chr7-121172936-DEL-244
chr7-121179191-DEL-126
chr7-121179200-DEL-62
chr7-121201235-DEL-57
chr7-121229252-DEL-58
chr7-121271775-DEL-487
chr7-121366151-DEL-80
chr7-121416292-DEL-66
chr7-121435777-DEL-79
chr7-121503670-DEL-249
chr7-121503733-DEL-124
chr7-121528097-DEL-76
chr7-121564272-DEL-51
chr7-121564677-DEL-92
chr7-121651484-DEL-150
chr7-121696377-DEL-75
chr7-121718126-DEL-570
chr7-121725267-DEL-51
chr7-121782797-DEL-151
chr7-121854347-DEL-50
chr7-121856943-DEL-285
chr7-121902231-DEL-52
chr7-122140835-DEL-832
chr7-122284185-DEL-54
chr7-122481578-DEL-780
chr7-122481608-DEL-132
chr7-122482022-DEL-76
chr7-122482069-DEL-204
chr7-122482522-DEL-72
chr7-122541978-DEL-193
chr7-122566931-DEL-257
chr7-122620262-DEL-2514
chr7-122815985-DEL-74
chr7-122901182-DEL-152
chr7-123015350-DEL-63
chr7-123018805-DEL-222
chr7-123043458-DEL-127
chr7-123092419-DEL-210
chr7-123119134-DEL-99
chr7-123431427-DEL-50
chr7-123533347-DEL-80
chr7-123617916-DEL-101
chr7-123694715-DEL-133
chr7-123753709-DEL-54
chr7-123763709-DEL-80
chr7-123938394-DEL-782

chr7-124552256-DEL-64
chr7-124552759-DEL-65
chr7-124556000-DEL-123
chr7-124557201-DEL-186
chr7-124596727-DEL-52
chr7-124703767-DEL-423
chr7-124759824-DEL-1093
chr7-124880112-DEL-159
chr7-125002019-DEL-60
chr7-125031939-DEL-1889
chr7-125038113-DEL-84
chr7-125061778-DEL-1013
chr7-125117775-DEL-50
chr7-125357843-DEL-50
chr7-125488382-DEL-51
chr7-125575014-DEL-92
chr7-125581861-DEL-128
chr7-125610312-DEL-77
chr7-125674005-DEL-108
chr7-125696396-DEL-50
chr7-125757809-DEL-220
chr7-125856639-DEL-52
chr7-125918388-DEL-211
chr7-125920762-DEL-179
chr7-125921489-DEL-75
chr7-125941947-DEL-402
chr7-125966155-DEL-164
chr7-126032398-DEL-130
chr7-126138515-DEL-143
chr7-126240860-DEL-1300
chr7-126247397-DEL-78
chr7-126499129-DEL-164
chr7-126688782-DEL-52
chr7-126848082-DEL-125
chr7-126848447-DEL-76
chr7-126883957-DEL-84
chr7-126885157-DEL-517
chr7-126886139-DEL-65
chr7-126886526-DEL-214
chr7-126919550-DEL-843
chr7-126984370-DEL-169
chr7-127042458-DEL-70
chr7-127052962-DEL-205
chr7-127269947-DEL-73
chr7-127319648-DEL-106
chr7-127320119-DEL-526
chr7-127358285-DEL-358
chr7-127360850-DEL-109
chr7-127360879-DEL-255
chr7-127361316-DEL-139
chr7-127419471-DEL-213
chr7-127432596-DEL-510
chr7-127446807-DEL-338
chr7-127518404-DEL-57
chr7-127541643-DEL-201
chr7-127668582-DEL-50
chr7-127718335-DEL-50
chr7-127718640-DEL-64
chr7-127718849-DEL-50
chr7-127787117-DEL-482
chr7-127798299-DEL-548
chr7-127842874-DEL-76
chr7-127868675-DEL-92
chr7-127882518-DEL-138
chr7-127912803-DEL-72
chr7-127954299-DEL-51
chr7-127997453-DEL-50
chr7-128222930-DEL-50
chr7-128298367-DEL-61
chr7-128399573-DEL-52
chr7-128407366-DEL-821
chr7-128489639-DEL-1463
chr7-128526402-DEL-83
chr7-128561169-DEL-141
chr7-128657198-DEL-168
chr7-128657213-DEL-65
chr7-128664471-DEL-3030

chr7-128794222-DEL-160
chr7-128806447-DEL-81
chr7-128950475-DEL-750
chr7-128970658-DEL-56
chr7-128976023-DEL-70
chr7-129002695-DEL-162
chr7-129050631-DEL-112
chr7-129059025-DEL-120
chr7-129153718-DEL-51
chr7-129153731-DEL-228
chr7-129153937-DEL-93
chr7-129319326-DEL-54
chr7-129332968-DEL-98
chr7-129386454-DEL-60
chr7-129469932-DEL-2045
chr7-129471409-DEL-1899
chr7-129533131-DEL-305
chr7-129553469-DEL-90
chr7-129557843-DEL-58
chr7-129559496-DEL-52
chr7-129666914-DEL-75
chr7-129677049-DEL-57
chr7-129717903-DEL-289
chr7-129739155-DEL-551
chr7-129896462-DEL-56
chr7-129917484-DEL-217
chr7-129979081-DEL-50
chr7-129979457-DEL-144
chr7-130003275-DEL-63
chr7-130003658-DEL-90
chr7-130015593-DEL-92
chr7-130133039-DEL-55
chr7-130163614-DEL-52
chr7-130286417-DEL-161
chr7-130340699-DEL-157
chr7-130456386-DEL-84
chr7-130615459-DEL-54
chr7-130733718-DEL-197
chr7-130771708-DEL-781
chr7-130781002-DEL-560
chr7-130808806-DEL-52
chr7-130839106-DEL-61
chr7-130858403-DEL-75
chr7-130878547-DEL-158
chr7-130914820-DEL-50
chr7-130915541-DEL-258
chr7-131005560-DEL-68
chr7-131018224-DEL-57
chr7-131018371-DEL-338
chr7-131059887-DEL-76
chr7-131119367-DEL-82
chr7-131173758-DEL-1532
chr7-131212333-DEL-63
chr7-131235551-DEL-84
chr7-131235597-DEL-240
chr7-131240169-DEL-1161
chr7-131465131-DEL-74
chr7-131500909-DEL-50
chr7-131525701-DEL-100
chr7-131559501-DEL-144
chr7-131570673-DEL-66
chr7-131731597-DEL-899
chr7-131738665-DEL-53
chr7-131817180-DEL-196
chr7-131826559-DEL-117
chr7-131830553-DEL-108
chr7-132058450-DEL-105
chr7-132116578-DEL-249
chr7-132223094-DEL-153
chr7-132256621-DEL-162
chr7-132256670-DEL-54
chr7-132347358-DEL-51
chr7-132482659-DEL-132
chr7-132642195-DEL-126
chr7-132705956-DEL-79
chr7-132748028-DEL-448
chr7-132847267-DEL-50

chr7-132855224-DEL-102
chr7-132872320-DEL-2274
chr7-132884526-DEL-81
chr7-132935656-DEL-55
chr7-132947618-DEL-117
chr7-132961109-DEL-205
chr7-133050572-DEL-52
chr7-133090399-DEL-1012
chr7-133091520-DEL-54
chr7-133204341-DEL-287
chr7-133305513-DEL-565
chr7-133312209-DEL-609
chr7-133385465-DEL-1524
chr7-133413137-DEL-544
chr7-133413277-DEL-1242
chr7-133413402-DEL-104
chr7-133413765-DEL-288
chr7-133465392-DEL-50
chr7-133493513-DEL-58
chr7-133547610-DEL-105
chr7-133560261-DEL-150
chr7-133608775-DEL-74
chr7-133686287-DEL-375
chr7-133785323-DEL-433
chr7-133785555-DEL-124
chr7-133802457-DEL-94
chr7-133834879-DEL-52
chr7-133882344-DEL-54
chr7-133882437-DEL-115
chr7-134127686-DEL-94
chr7-134239753-DEL-161
chr7-134243017-DEL-123
chr7-134274629-DEL-50
chr7-134284722-DEL-150
chr7-134298746-DEL-60
chr7-134323553-DEL-580
chr7-134446852-DEL-248
chr7-134610245-DEL-60
chr7-134648585-DEL-67
chr7-134649845-DEL-275
chr7-134684497-DEL-2296
chr7-134717843-DEL-52
chr7-134749455-DEL-123
chr7-134787083-DEL-243
chr7-134787118-DEL-112
chr7-135131641-DEL-72
chr7-135131641-DEL-172
chr7-135191112-DEL-75
chr7-135246111-DEL-84
chr7-135401892-DEL-58
chr7-135600873-DEL-163
chr7-135602508-DEL-50
chr7-135613534-DEL-1324
chr7-135640520-DEL-100
chr7-135701934-DEL-2099
chr7-135704263-DEL-287
chr7-135748488-DEL-269
chr7-135755093-DEL-61
chr7-135848951-DEL-50
chr7-136029289-DEL-1078
chr7-136326970-DEL-59
chr7-136469499-DEL-2775
chr7-136498242-DEL-266
chr7-136568308-DEL-648
chr7-136716429-DEL-140
chr7-136737395-DEL-141
chr7-136740951-DEL-138
chr7-136784126-DEL-68
chr7-136817130-DEL-208
chr7-136834778-DEL-956
chr7-136835459-DEL-224
chr7-136933061-DEL-136
chr7-136933180-DEL-62
chr7-136940967-DEL-50
chr7-136964429-DEL-80
chr7-137000259-DEL-55
chr7-137071557-DEL-163

chr7-137115337-DEL-105
chr7-137118361-DEL-340
chr7-137148208-DEL-58
chr7-137159627-DEL-596
chr7-137177775-DEL-76
chr7-137242079-DEL-51
chr7-137282472-DEL-58
chr7-137312339-DEL-93
chr7-137368297-DEL-61
chr7-137374192-DEL-268
chr7-137427382-DEL-849
chr7-137457989-DEL-165
chr7-137561401-DEL-51
chr7-137663444-DEL-54
chr7-137663540-DEL-696
chr7-137664821-DEL-60
chr7-137683628-DEL-86
chr7-137782843-DEL-123
chr7-137879272-DEL-208
chr7-137900416-DEL-136
chr7-137932110-DEL-77
chr7-137973575-DEL-220
chr7-137973928-DEL-216
chr7-137973960-DEL-65
chr7-138129863-DEL-101
chr7-138191480-DEL-196
chr7-138211791-DEL-694
chr7-138217522-DEL-50
chr7-138278362-DEL-85
chr7-138282736-DEL-132
chr7-138402880-DEL-115
chr7-138416910-DEL-66
chr7-138431595-DEL-408
chr7-138431772-DEL-68
chr7-138518841-DEL-53
chr7-138530734-DEL-333
chr7-138587380-DEL-50
chr7-138596294-DEL-50
chr7-138619806-DEL-50
chr7-138649549-DEL-51
chr7-138661449-DEL-58
chr7-138760531-DEL-325
chr7-138769283-DEL-466
chr7-138769395-DEL-57
chr7-138769833-DEL-105
chr7-139027812-DEL-444
chr7-139028200-DEL-110
chr7-139028521-DEL-289
chr7-139132973-DEL-51
chr7-139200429-DEL-97
chr7-139237528-DEL-56
chr7-139239421-DEL-675
chr7-139290542-DEL-221
chr7-139332586-DEL-64
chr7-139431694-DEL-78
chr7-139432124-DEL-105
chr7-139443350-DEL-50
chr7-139445592-DEL-124
chr7-139473459-DEL-77
chr7-139555067-DEL-53
chr7-139711682-DEL-50
chr7-139728924-DEL-65
chr7-139735714-DEL-88
chr7-139735875-DEL-104
chr7-139841374-DEL-6502
chr7-139943360-DEL-65
chr7-140092493-DEL-197
chr7-140233852-DEL-3108
chr7-140331684-DEL-78
chr7-140331891-DEL-284
chr7-140559299-DEL-50
chr7-140620134-DEL-102
chr7-140648937-DEL-66
chr7-140649134-DEL-234
chr7-140703684-DEL-141
chr7-140977639-DEL-110
chr7-140995797-DEL-60

chr7-141083105-DEL-2136
chr7-141218342-DEL-819
chr7-141239048-DEL-295
chr7-141292319-DEL-78
chr7-141321480-DEL-219
chr7-141321765-DEL-226
chr7-141328321-DEL-52
chr7-141362415-DEL-206
chr7-141527358-DEL-2090
chr7-141698003-DEL-78
chr7-141728857-DEL-300
chr7-141729152-DEL-66
chr7-141729356-DEL-66
chr7-141743378-DEL-90
chr7-141755267-DEL-210
chr7-141755690-DEL-78
chr7-141755912-DEL-66
chr7-141755930-DEL-222
chr7-141872657-DEL-56
chr7-142019075-DEL-176
chr7-142271867-DEL-178
chr7-142275132-DEL-52
chr7-142498300-DEL-388
chr7-143040231-DEL-58
chr7-143042000-DEL-56
chr7-143189353-DEL-962
chr7-143190241-DEL-52
chr7-143190446-DEL-218
chr7-143201046-DEL-51
chr7-143216749-DEL-299
chr7-143514372-DEL-97
chr7-143586941-DEL-50
chr7-143687867-DEL-262
chr7-143808754-DEL-114
chr7-143818315-DEL-61
chr7-143846648-DEL-88
chr7-143858241-DEL-67
chr7-143880840-DEL-64
chr7-143942381-DEL-200
chr7-143942601-DEL-444
chr7-144346211-DEL-52
chr7-144406456-DEL-84
chr7-144423961-DEL-50
chr7-144670227-DEL-128
chr8-3244538-DEL-96
chr8-3249673-DEL-60
chr8-3362998-DEL-73
chr8-3492342-DEL-51
chr8-3735959-DEL-66
chr8-3742073-DEL-296
chr8-3856707-DEL-2453
chr8-3992889-DEL-52
chr8-4141192-DEL-223
chr8-4239725-DEL-139
chr8-4262036-DEL-62
chr8-4262452-DEL-91
chr8-4395964-DEL-66
chr8-4417380-DEL-1948
chr8-4707027-DEL-60
chr8-4707154-DEL-53
chr8-4733801-DEL-4360
chr8-4790029-DEL-52
chr8-4803685-DEL-412
chr8-4807751-DEL-3777
chr8-5079323-DEL-279
chr8-5079710-DEL-64
chr8-5084460-DEL-97
chr8-5135231-DEL-166
chr8-5150302-DEL-64
chr8-5382323-DEL-57
chr8-5382629-DEL-54
chr8-5447579-DEL-158
chr8-5669569-DEL-56
chr8-5818299-DEL-97
chr8-5835239-DEL-1494
chr8-5884045-DEL-3528
chr8-5936501-DEL-57

chr8-5937009-DEL-210
chr8-6118492-DEL-74
chr8-6187195-DEL-166
chr8-6316968-DEL-69
chr8-6531702-DEL-201
chr8-6608944-DEL-498
chr8-6659826-DEL-119
chr8-6705761-DEL-63
chr8-7058268-DEL-132
chr8-7138740-DEL-606
chr8-7216425-DEL-72
chr8-7299168-DEL-59
chr8-7331397-DEL-87
chr8-7331418-DEL-210
chr8-7602241-DEL-584
chr8-7679530-DEL-68
chr8-7737193-DEL-54
chr8-7942136-DEL-56
chr8-8181312-DEL-54
chr8-8295078-DEL-50
chr8-8336598-DEL-1455
chr8-8360066-DEL-54
chr8-8509284-DEL-100
chr8-8592069-DEL-98
chr8-8649232-DEL-79
chr8-8789085-DEL-52
chr8-8794690-DEL-216
chr8-8848010-DEL-2349
chr8-8848116-DEL-868
chr8-8849130-DEL-182
chr8-8857836-DEL-155
chr8-8872228-DEL-63
chr8-8955896-DEL-51
chr8-9062989-DEL-307
chr8-9063117-DEL-92
chr8-9104214-DEL-67
chr8-9331268-DEL-130
chr8-9435834-DEL-146
chr8-9532068-DEL-52
chr8-9594300-DEL-99
chr8-9632519-DEL-128
chr8-9666321-DEL-52
chr8-9741063-DEL-321
chr8-9741302-DEL-118
chr8-9741888-DEL-105
chr8-9741978-DEL-51
chr8-9778530-DEL-77
chr8-9806526-DEL-81
chr8-9806891-DEL-58
chr8-9812864-DEL-69
chr8-9850366-DEL-56
chr8-9880747-DEL-966
chr8-9901985-DEL-76
chr8-9958412-DEL-68
chr8-10074324-DEL-151
chr8-10178209-DEL-111
chr8-10240004-DEL-115
chr8-10240429-DEL-54
chr8-10250027-DEL-68
chr8-10302457-DEL-1361
chr8-10353769-DEL-55
chr8-10457853-DEL-141
chr8-10483262-DEL-93
chr8-10486191-DEL-59
chr8-10633350-DEL-270
chr8-10649447-DEL-236
chr8-10726419-DEL-375
chr8-10732706-DEL-82
chr8-10755432-DEL-81
chr8-10775299-DEL-89
chr8-10775574-DEL-279
chr8-10847163-DEL-98
chr8-10900492-DEL-50
chr8-10910153-DEL-72
chr8-10923239-DEL-294
chr8-10935611-DEL-57
chr8-10968105-DEL-225

chr8-11183742-DEL-83
chr8-11200932-DEL-95
chr8-11220150-DEL-170
chr8-11370673-DEL-126
chr8-11449500-DEL-77
chr8-11577168-DEL-131
chr8-11581385-DEL-1153
chr8-11717130-DEL-56
chr8-11724660-DEL-210
chr8-11935527-DEL-54
chr8-11969694-DEL-2182
chr8-11971702-DEL-70
chr8-12017289-DEL-130
chr8-12017506-DEL-129
chr8-12026786-DEL-86
chr8-12054754-DEL-192
chr8-12092582-DEL-80
chr8-12106678-DEL-52
chr8-12178602-DEL-273
chr8-12180928-DEL-72
chr8-12181102-DEL-2326
chr8-12181609-DEL-390
chr8-12312123-DEL-50
chr8-12315334-DEL-116
chr8-12405931-DEL-113
chr8-12550286-DEL-60
chr8-12636668-DEL-900
chr8-12637212-DEL-375
chr8-12744047-DEL-51
chr8-12766642-DEL-60
chr8-12773851-DEL-86
chr8-12828075-DEL-492
chr8-13117240-DEL-317
chr8-13119430-DEL-183
chr8-13119459-DEL-1208
chr8-13125637-DEL-147
chr8-13178285-DEL-92
chr8-13178649-DEL-199
chr8-13254061-DEL-68
chr8-13300690-DEL-51
chr8-13435969-DEL-52
chr8-13503602-DEL-122
chr8-13653486-DEL-199
chr8-13657004-DEL-63
chr8-13657595-DEL-284
chr8-13658479-DEL-63
chr8-13660284-DEL-57
chr8-13733256-DEL-462
chr8-13746942-DEL-72
chr8-13862849-DEL-2248
chr8-13896149-DEL-4687
chr8-13953481-DEL-1492
chr8-13968131-DEL-1875
chr8-13970245-DEL-161
chr8-13990678-DEL-50
chr8-13993459-DEL-68
chr8-14010034-DEL-595
chr8-14017114-DEL-94
chr8-14057700-DEL-591
chr8-14058832-DEL-50
chr8-14088144-DEL-91
chr8-14428395-DEL-136
chr8-14428681-DEL-118
chr8-14500236-DEL-139
chr8-14557724-DEL-195
chr8-14627955-DEL-91
chr8-14654803-DEL-57
chr8-14779036-DEL-245
chr8-14779489-DEL-181
chr8-14840490-DEL-2312
chr8-14853377-DEL-3419
chr8-14932563-DEL-58
chr8-15045039-DEL-134
chr8-15064969-DEL-50
chr8-15272709-DEL-79
chr8-15273143-DEL-57
chr8-15284099-DEL-145

chr8-15295885-DEL-160
chr8-15309543-DEL-502
chr8-15310757-DEL-81
chr8-15322977-DEL-66
chr8-15391691-DEL-51
chr8-15512050-DEL-129
chr8-15622998-DEL-142
chr8-15622999-DEL-64
chr8-15626396-DEL-95
chr8-15673623-DEL-116
chr8-15705817-DEL-1200
chr8-15728156-DEL-116
chr8-15728199-DEL-344
chr8-15729576-DEL-1907
chr8-15784124-DEL-56
chr8-15819281-DEL-2976
chr8-15947214-DEL-81
chr8-15991609-DEL-78
chr8-16028836-DEL-94
chr8-16037328-DEL-62
chr8-16054033-DEL-61
chr8-16127606-DEL-367
chr8-16249359-DEL-86
chr8-16542101-DEL-2256
chr8-16640015-DEL-99
chr8-16650694-DEL-419
chr8-16717102-DEL-64
chr8-16827031-DEL-82
chr8-16940823-DEL-318
chr8-16979226-DEL-54
chr8-17093115-DEL-62
chr8-17113873-DEL-79
chr8-17114901-DEL-55
chr8-17115009-DEL-62
chr8-17172244-DEL-68
chr8-17186682-DEL-66
chr8-17200335-DEL-203
chr8-17236763-DEL-53
chr8-17404911-DEL-50
chr8-17415748-DEL-2970
chr8-17421600-DEL-346
chr8-17670903-DEL-54
chr8-17713162-DEL-426
chr8-17769753-DEL-56
chr8-17844610-DEL-195
chr8-17844750-DEL-73
chr8-17849534-DEL-180
chr8-17917276-DEL-221
chr8-18024479-DEL-102
chr8-18840509-DEL-54
chr8-18887900-DEL-83
chr8-19003098-DEL-86
chr8-19020015-DEL-67
chr8-19248910-DEL-64
chr8-19259510-DEL-52
chr8-19280385-DEL-203
chr8-19299016-DEL-1463
chr8-19364691-DEL-198
chr8-19510518-DEL-50
chr8-19561957-DEL-159
chr8-19572515-DEL-9841
chr8-19625776-DEL-141
chr8-19641430-DEL-2128
chr8-19644149-DEL-1346
chr8-19682864-DEL-1055
chr8-19689632-DEL-6489
chr8-20621035-DEL-1445
chr8-20672141-DEL-204
chr8-21052431-DEL-468
chr8-21065419-DEL-156
chr8-21074675-DEL-56
chr8-21129240-DEL-210
chr8-21188756-DEL-102
chr8-21348078-DEL-8260
chr8-21533768-DEL-68
chr8-21593355-DEL-540
chr8-21622805-DEL-199

chr8-21670301-DEL-68
chr8-21724481-DEL-619
chr8-22006453-DEL-6487
chr8-22302092-DEL-617
chr8-22341925-DEL-2371
chr8-22530415-DEL-52
chr8-22530916-DEL-86
chr8-22542940-DEL-300
chr8-22568434-DEL-117
chr8-22576378-DEL-561
chr8-22596876-DEL-52
chr8-22619050-DEL-50
chr8-22688099-DEL-175
chr8-22688673-DEL-54
chr8-22713542-DEL-58
chr8-22713717-DEL-93
chr8-22896911-DEL-2152
chr8-22908875-DEL-736
chr8-22908876-DEL-245
chr8-22933804-DEL-131
chr8-22976318-DEL-50
chr8-23002349-DEL-111
chr8-23051663-DEL-50
chr8-23093929-DEL-121
chr8-23107931-DEL-110
chr8-23108263-DEL-210
chr8-23174371-DEL-72
chr8-23310074-DEL-86
chr8-23494842-DEL-60
chr8-23574443-DEL-100
chr8-23654308-DEL-50
chr8-23825315-DEL-3755
chr8-23869343-DEL-52
chr8-23885049-DEL-96
chr8-23887207-DEL-69
chr8-23930125-DEL-79
chr8-24045279-DEL-351
chr8-24073666-DEL-102
chr8-24073854-DEL-96
chr8-24247425-DEL-102
chr8-24286392-DEL-701
chr8-24304419-DEL-146
chr8-24350835-DEL-143
chr8-24421809-DEL-50
chr8-24443411-DEL-52
chr8-24474463-DEL-79
chr8-24496599-DEL-56
chr8-24614916-DEL-73
chr8-24756897-DEL-342
chr8-24765133-DEL-151
chr8-24765635-DEL-50
chr8-24837097-DEL-339
chr8-24884584-DEL-213
chr8-24912834-DEL-52
chr8-24970162-DEL-142
chr8-25092974-DEL-68
chr8-25099182-DEL-52
chr8-25130058-DEL-1078
chr8-25130393-DEL-196
chr8-25183348-DEL-50
chr8-25213143-DEL-81
chr8-25232269-DEL-50
chr8-25245503-DEL-1338
chr8-25298412-DEL-155
chr8-25315147-DEL-75
chr8-25482719-DEL-51
chr8-25508899-DEL-627
chr8-25556731-DEL-154
chr8-25557120-DEL-55
chr8-25580125-DEL-113
chr8-25702906-DEL-172
chr8-25725604-DEL-76
chr8-25728939-DEL-86
chr8-25732980-DEL-59
chr8-25756334-DEL-136
chr8-25756804-DEL-192
chr8-25787629-DEL-173

chr8-25845096-DEL-74
chr8-25854831-DEL-57
chr8-25860652-DEL-56
chr8-25871155-DEL-96
chr8-25917494-DEL-50
chr8-25962044-DEL-64
chr8-25986253-DEL-227
chr8-25992099-DEL-163
chr8-25992169-DEL-80
chr8-25992225-DEL-342
chr8-25992422-DEL-118
chr8-26097232-DEL-51
chr8-26147401-DEL-80
chr8-26274459-DEL-63
chr8-26435856-DEL-54
chr8-26459568-DEL-371
chr8-26476177-DEL-50
chr8-26506667-DEL-102
chr8-26506899-DEL-54
chr8-26540425-DEL-133
chr8-26663000-DEL-110
chr8-26664623-DEL-112
chr8-26713276-DEL-727
chr8-26796411-DEL-66
chr8-26808532-DEL-68
chr8-26825840-DEL-60
chr8-26933510-DEL-70
chr8-26997329-DEL-156
chr8-27011525-DEL-50
chr8-27266821-DEL-1107
chr8-27270883-DEL-2377
chr8-27304218-DEL-111
chr8-27336298-DEL-50
chr8-27368578-DEL-85
chr8-27480171-DEL-66
chr8-27490873-DEL-126
chr8-27557983-DEL-163
chr8-27563482-DEL-210
chr8-27564173-DEL-434
chr8-27636035-DEL-226
chr8-27668669-DEL-84
chr8-27682113-DEL-50
chr8-27734920-DEL-929
chr8-27765728-DEL-63
chr8-27765912-DEL-127
chr8-27768277-DEL-47879
chr8-27815544-DEL-884
chr8-28046329-DEL-59
chr8-28050958-DEL-402
chr8-28063429-DEL-75
chr8-28093814-DEL-348
chr8-28093824-DEL-118
chr8-28136384-DEL-3053
chr8-28150812-DEL-164
chr8-28192802-DEL-107
chr8-28244317-DEL-50
chr8-28439799-DEL-843
chr8-28621685-DEL-82
chr8-28663715-DEL-274
chr8-28715946-DEL-1737
chr8-28717842-DEL-3886
chr8-28753958-DEL-55
chr8-28795461-DEL-1247
chr8-28853832-DEL-68
chr8-28853998-DEL-70
chr8-29013805-DEL-51
chr8-29014278-DEL-58
chr8-29027095-DEL-87
chr8-29090153-DEL-106
chr8-29090632-DEL-149
chr8-29350269-DEL-50
chr8-29368969-DEL-483
chr8-29442514-DEL-313
chr8-29442875-DEL-67
chr8-29453571-DEL-285
chr8-29583308-DEL-125
chr8-29647568-DEL-50

chr8-29720768-DEL-115
chr8-29834857-DEL-3719
chr8-29914616-DEL-78
chr8-29993181-DEL-58
chr8-30081949-DEL-368
chr8-30236939-DEL-72
chr8-30255706-DEL-54
chr8-30312435-DEL-52
chr8-30393107-DEL-1529
chr8-30421632-DEL-123
chr8-30430889-DEL-208
chr8-30481761-DEL-51
chr8-30562057-DEL-70
chr8-30562330-DEL-60
chr8-30583817-DEL-6910
chr8-30642594-DEL-57
chr8-30646594-DEL-325
chr8-30647924-DEL-66
chr8-30677466-DEL-76
chr8-30785136-DEL-2809
chr8-31083170-DEL-50
chr8-31084564-DEL-187
chr8-31167860-DEL-140
chr8-31168238-DEL-77
chr8-31168418-DEL-71
chr8-31415298-DEL-110
chr8-31428744-DEL-150
chr8-31456892-DEL-60
chr8-31560907-DEL-159
chr8-31587628-DEL-54
chr8-31616808-DEL-50
chr8-31717838-DEL-2804
chr8-31827756-DEL-72
chr8-31828035-DEL-67
chr8-31831717-DEL-52
chr8-31900564-DEL-80
chr8-31910322-DEL-232
chr8-31942450-DEL-295
chr8-31943826-DEL-55
chr8-31962189-DEL-132
chr8-31990834-DEL-163
chr8-31991256-DEL-232
chr8-32084995-DEL-70
chr8-32171524-DEL-208
chr8-32183466-DEL-417
chr8-32241020-DEL-154
chr8-32248137-DEL-65
chr8-32294373-DEL-63
chr8-32354522-DEL-62
chr8-32522669-DEL-124
chr8-32584234-DEL-64
chr8-32744467-DEL-1714
chr8-32792352-DEL-121
chr8-32792577-DEL-206
chr8-32944323-DEL-144
chr8-32957850-DEL-77
chr8-33020658-DEL-398
chr8-33020673-DEL-50
chr8-33104052-DEL-265
chr8-33254842-DEL-74
chr8-33269672-DEL-58
chr8-33277072-DEL-61
chr8-33343643-DEL-207
chr8-33404914-DEL-58
chr8-33407748-DEL-64
chr8-33408193-DEL-1736
chr8-33413358-DEL-227
chr8-33456012-DEL-905
chr8-33465747-DEL-120
chr8-33465956-DEL-242
chr8-33523024-DEL-297
chr8-33690073-DEL-57
chr8-33753901-DEL-343
chr8-33894754-DEL-65
chr8-33949974-DEL-6278
chr8-33983129-DEL-93
chr8-34057150-DEL-1798

chr8-34188970-DEL-195
chr8-34210709-DEL-142
chr8-34245689-DEL-52
chr8-34343041-DEL-56
chr8-34393220-DEL-85
chr8-34393359-DEL-143
chr8-34459633-DEL-187
chr8-34494508-DEL-221
chr8-34500101-DEL-198
chr8-34548462-DEL-50
chr8-34569644-DEL-65
chr8-34628554-DEL-125
chr8-34629289-DEL-524
chr8-34644832-DEL-80
chr8-34660915-DEL-187
chr8-34745163-DEL-92
chr8-34758543-DEL-54
chr8-34794755-DEL-1503
chr8-34834006-DEL-52
chr8-34850586-DEL-56
chr8-34957041-DEL-51
chr8-34978892-DEL-244
chr8-34979624-DEL-122
chr8-35013112-DEL-178
chr8-35013624-DEL-60
chr8-35037197-DEL-75
chr8-35108839-DEL-53
chr8-35183773-DEL-755
chr8-35224432-DEL-50
chr8-35250382-DEL-56
chr8-35393843-DEL-70
chr8-35419719-DEL-3351
chr8-35448342-DEL-59
chr8-35452949-DEL-91
chr8-35455288-DEL-4107
chr8-35482562-DEL-50
chr8-35525759-DEL-56
chr8-35623301-DEL-207
chr8-35629475-DEL-795
chr8-35703312-DEL-50
chr8-35785049-DEL-98
chr8-35792200-DEL-91
chr8-35792379-DEL-160
chr8-35815069-DEL-66
chr8-35846689-DEL-61
chr8-36027533-DEL-58
chr8-36039878-DEL-170
chr8-36040107-DEL-80
chr8-36062819-DEL-83
chr8-36123222-DEL-115
chr8-36201320-DEL-1004
chr8-36461987-DEL-1261
chr8-36465709-DEL-97
chr8-36606125-DEL-275
chr8-36606126-DEL-60
chr8-36690394-DEL-72
chr8-36837414-DEL-940
chr8-36866120-DEL-288
chr8-36989337-DEL-113
chr8-37026049-DEL-56
chr8-37168694-DEL-64
chr8-37189203-DEL-105
chr8-37189286-DEL-240
chr8-37200392-DEL-6765
chr8-37325802-DEL-56
chr8-37559729-DEL-143
chr8-37559906-DEL-69
chr8-37563086-DEL-10542
chr8-37604406-DEL-51
chr8-37635904-DEL-58
chr8-37646543-DEL-65
chr8-37646643-DEL-253
chr8-37873386-DEL-743
chr8-37873756-DEL-220
chr8-37907284-DEL-52
chr8-37954076-DEL-1132
chr8-38094624-DEL-213

chr8-38138073-DEL-625
chr8-38261863-DEL-71
chr8-38305962-DEL-743
chr8-38342757-DEL-56
chr8-38364348-DEL-123
chr8-38373308-DEL-154
chr8-38443964-DEL-1974
chr8-38473237-DEL-50
chr8-38473333-DEL-65
chr8-38508378-DEL-117
chr8-38809637-DEL-50
chr8-38925676-DEL-549
chr8-39010425-DEL-50
chr8-39040666-DEL-50
chr8-39075391-DEL-1776
chr8-39219091-DEL-2654
chr8-39319900-DEL-3113
chr8-39338154-DEL-2654
chr8-39404091-DEL-84
chr8-39457727-DEL-743
chr8-39512863-DEL-82
chr8-39545504-DEL-2116
chr8-39547715-DEL-2471
chr8-39600286-DEL-72
chr8-39603392-DEL-3527
chr8-39669834-DEL-498
chr8-39681292-DEL-713
chr8-39774417-DEL-66
chr8-40053191-DEL-305
chr8-40075370-DEL-52
chr8-40114131-DEL-1059
chr8-40182922-DEL-208
chr8-40390824-DEL-367
chr8-40502411-DEL-1093
chr8-40555086-DEL-50
chr8-40592650-DEL-5317
chr8-40631826-DEL-64
chr8-40635676-DEL-95
chr8-40636119-DEL-488
chr8-40821486-DEL-198
chr8-40869392-DEL-82
chr8-41256193-DEL-52
chr8-41256197-DEL-169
chr8-41503507-DEL-50
chr8-41674267-DEL-145
chr8-41822763-DEL-472
chr8-41887639-DEL-617
chr8-41901503-DEL-136
chr8-41903696-DEL-161
chr8-41996313-DEL-63
chr8-41996532-DEL-382
chr8-42076477-DEL-293
chr8-42384820-DEL-53
chr8-42408244-DEL-52
chr8-42596306-DEL-52
chr8-42692250-DEL-90
chr8-42692444-DEL-56
chr8-42762013-DEL-72
chr8-42871979-DEL-157
chr8-42872069-DEL-56
chr8-42872237-DEL-370
chr8-42904467-DEL-1878
chr8-42911001-DEL-51
chr8-43043465-DEL-475
chr8-43087863-DEL-97
chr8-43130874-DEL-148
chr8-43132990-DEL-50
chr8-43148909-DEL-591
chr8-43299422-DEL-876
chr8-43358310-DEL-50
chr8-43506316-DEL-50
chr8-43519229-DEL-86
chr8-43550672-DEL-98
chr8-43553859-DEL-191
chr8-43666286-DEL-132
chr8-43786615-DEL-153
chr8-43815618-DEL-120

chr8-43837597-DEL-159
chr8-43976874-DEL-48473
chr8-44055009-DEL-780
chr8-44073191-DEL-26194
chr8-44097307-DEL-1065
chr8-44103126-DEL-14687
chr8-44172517-DEL-52
chr8-44194583-DEL-81
chr8-44202572-DEL-191
chr8-44224430-DEL-125
chr8-44287225-DEL-63
chr8-44621385-DEL-60
chr8-44666216-DEL-50
chr8-44666771-DEL-50
chr8-44853863-DEL-66
chr8-44859566-DEL-578
chr8-45164444-DEL-64
chr8-45175123-DEL-196
chr8-45175563-DEL-214
chr8-45175918-DEL-202
chr8-45244000-DEL-349
chr8-45369332-DEL-50
chr8-45553179-DEL-59
chr8-45610746-DEL-1610
chr8-45666519-DEL-1935
chr8-45671547-DEL-138
chr8-45706600-DEL-76
chr8-45716499-DEL-158
chr8-45858343-DEL-52
chr8-45883695-DEL-76
chr8-45895945-DEL-3936
chr8-46289035-DEL-214
chr8-46303212-DEL-140
chr8-46384148-DEL-263
chr8-46450145-DEL-92
chr8-46472745-DEL-60
chr8-46488382-DEL-78
chr8-46513566-DEL-912
chr8-46514544-DEL-646
chr8-46714112-DEL-64
chr8-46757533-DEL-60
chr8-46809915-DEL-2845
chr8-46824434-DEL-94
chr8-46852006-DEL-106
chr8-46934185-DEL-568
chr8-46935652-DEL-132
chr8-46981374-DEL-71
chr8-46984229-DEL-60
chr8-47097301-DEL-152
chr8-47098555-DEL-99
chr8-47110008-DEL-13853
chr8-47148968-DEL-92
chr8-47181530-DEL-1558
chr8-47300783-DEL-60
chr8-47438382-DEL-445
chr8-47456214-DEL-344
chr8-47456454-DEL-127
chr8-47456590-DEL-508
chr8-47456862-DEL-62
chr8-47457150-DEL-341
chr8-47457436-DEL-93
chr8-47458060-DEL-244
chr8-47500214-DEL-82
chr8-47500877-DEL-69
chr8-47536902-DEL-5481
chr8-47568111-DEL-1268
chr8-47570839-DEL-255
chr8-47582519-DEL-72
chr8-47783598-DEL-112
chr8-47821395-DEL-366
chr8-47886166-DEL-86
chr8-47889174-DEL-52
chr8-47890000-DEL-50
chr8-48099965-DEL-120
chr8-48289066-DEL-308
chr8-48304338-DEL-50
chr8-48313758-DEL-53

chr8-48374294-DEL-125
chr8-48482691-DEL-74
chr8-48495516-DEL-238
chr8-48512446-DEL-429
chr8-48526679-DEL-111
chr8-48559905-DEL-53
chr8-48621549-DEL-62
chr8-48621671-DEL-58
chr8-48695049-DEL-120
chr8-48698106-DEL-51
chr8-48700015-DEL-1154
chr8-48722664-DEL-50
chr8-48784811-DEL-140
chr8-48842917-DEL-156
chr8-48876377-DEL-50
chr8-49361500-DEL-51
chr8-49371270-DEL-170
chr8-49418559-DEL-68
chr8-49427938-DEL-64
chr8-49430368-DEL-57
chr8-49540854-DEL-844
chr8-49548457-DEL-108
chr8-49725766-DEL-318
chr8-49904424-DEL-247
chr8-50067517-DEL-50
chr8-50068699-DEL-493
chr8-50107378-DEL-114
chr8-50166064-DEL-160
chr8-50190149-DEL-58
chr8-50190371-DEL-76
chr8-50222379-DEL-54
chr8-50420774-DEL-209
chr8-50490693-DEL-57
chr8-50578948-DEL-59
chr8-50767747-DEL-240
chr8-50803078-DEL-194
chr8-50856239-DEL-252
chr8-51039456-DEL-88
chr8-51075603-DEL-77
chr8-51108536-DEL-1551
chr8-51180713-DEL-76
chr8-51263672-DEL-62
chr8-51360246-DEL-131
chr8-51384486-DEL-56
chr8-51414249-DEL-63
chr8-51736633-DEL-955
chr8-51786805-DEL-50
chr8-51799878-DEL-212
chr8-51839489-DEL-81
chr8-52272247-DEL-50
chr8-52366252-DEL-2054
chr8-52427239-DEL-119
chr8-52550407-DEL-2259
chr8-52570174-DEL-113
chr8-52590223-DEL-50
chr8-52618456-DEL-80
chr8-52684492-DEL-135
chr8-52757418-DEL-212
chr8-52820522-DEL-145
chr8-52905077-DEL-50
chr8-53074416-DEL-160
chr8-53095702-DEL-70
chr8-53153757-DEL-60
chr8-53244903-DEL-59
chr8-53248244-DEL-135
chr8-53256262-DEL-51
chr8-53260116-DEL-6566
chr8-53281554-DEL-68
chr8-53511856-DEL-1015
chr8-53587625-DEL-404
chr8-53663463-DEL-1422
chr8-53901887-DEL-160
chr8-53967503-DEL-124
chr8-54054597-DEL-142
chr8-54115584-DEL-77
chr8-54197776-DEL-2885
chr8-54238874-DEL-100

chr8-54294283-DEL-2616
chr8-54309723-DEL-156
chr8-54425029-DEL-1486
chr8-54486534-DEL-185
chr8-54594447-DEL-66
chr8-54830939-DEL-1036
chr8-55050269-DEL-50
chr8-55069019-DEL-199
chr8-55069272-DEL-100
chr8-55070233-DEL-117
chr8-55072264-DEL-5284
chr8-55088967-DEL-61
chr8-55104655-DEL-399
chr8-55105081-DEL-1253
chr8-55231833-DEL-50
chr8-55371728-DEL-67
chr8-55397513-DEL-54
chr8-55490203-DEL-123
chr8-55881747-DEL-311
chr8-55893939-DEL-54
chr8-55958478-DEL-3348
chr8-56108576-DEL-50
chr8-56123035-DEL-54
chr8-56387862-DEL-62
chr8-56403319-DEL-564
chr8-56413944-DEL-58
chr8-56461028-DEL-413
chr8-56555163-DEL-58
chr8-56606364-DEL-2835
chr8-56620744-DEL-571
chr8-56685973-DEL-66
chr8-56781768-DEL-99
chr8-57076948-DEL-176
chr8-57130852-DEL-284
chr8-57151117-DEL-51
chr8-57298711-DEL-52
chr8-57353853-DEL-97
chr8-57452528-DEL-111
chr8-57559143-DEL-225
chr8-57648139-DEL-479
chr8-57685575-DEL-87
chr8-57698777-DEL-51
chr8-57844911-DEL-1225
chr8-58218037-DEL-123
chr8-58275903-DEL-97
chr8-58464553-DEL-4730
chr8-58486305-DEL-3446
chr8-58509395-DEL-80
chr8-58532780-DEL-96
chr8-58563755-DEL-68
chr8-58839276-DEL-66
chr8-58842144-DEL-424
chr8-58896504-DEL-64
chr8-58946535-DEL-66
chr8-58971141-DEL-355
chr8-59017544-DEL-58
chr8-59415879-DEL-291
chr8-59499620-DEL-280
chr8-59592604-DEL-50
chr8-59624706-DEL-59
chr8-59653306-DEL-497
chr8-59690280-DEL-56
chr8-59802075-DEL-80
chr8-59808871-DEL-679
chr8-59920937-DEL-3356
chr8-60309622-DEL-74
chr8-60370222-DEL-257
chr8-60376765-DEL-147
chr8-60387231-DEL-983
chr8-60410223-DEL-61
chr8-60463995-DEL-88
chr8-60463995-DEL-308
chr8-60551801-DEL-50
chr8-60552221-DEL-693
chr8-60598428-DEL-200
chr8-60600505-DEL-4104
chr8-60614683-DEL-81

chr8-60684292-DEL-698
chr8-60711847-DEL-451
chr8-60826806-DEL-120
chr8-60862546-DEL-2470
chr8-60873442-DEL-160
chr8-60923693-DEL-276
chr8-60925978-DEL-214
chr8-60961823-DEL-64
chr8-61010690-DEL-204
chr8-61094259-DEL-81
chr8-61094868-DEL-118
chr8-61262295-DEL-130
chr8-61553914-DEL-2048
chr8-61619419-DEL-160
chr8-61657673-DEL-258
chr8-61658014-DEL-610
chr8-61658655-DEL-54
chr8-61662711-DEL-50
chr8-61683385-DEL-87
chr8-61683551-DEL-192
chr8-61771330-DEL-265
chr8-61781265-DEL-204
chr8-61781401-DEL-68
chr8-62286419-DEL-908
chr8-62289963-DEL-95
chr8-62290163-DEL-191
chr8-62365373-DEL-204
chr8-62541279-DEL-703
chr8-62601262-DEL-311
chr8-62603228-DEL-1029
chr8-62657885-DEL-104
chr8-62707188-DEL-53
chr8-62730270-DEL-352
chr8-62784930-DEL-5171
chr8-62796968-DEL-2241
chr8-62880248-DEL-239
chr8-63396015-DEL-1477
chr8-63451559-DEL-1633
chr8-63462920-DEL-57
chr8-63468280-DEL-3850
chr8-63684232-DEL-1787
chr8-63719213-DEL-79
chr8-63740901-DEL-67
chr8-63750898-DEL-8669
chr8-64092259-DEL-104
chr8-64281725-DEL-1313
chr8-64336636-DEL-2052
chr8-64365091-DEL-4057
chr8-64385435-DEL-3205
chr8-64580652-DEL-82
chr8-64704119-DEL-2196
chr8-64706572-DEL-4668
chr8-64720345-DEL-50
chr8-64741102-DEL-1831
chr8-64883345-DEL-212
chr8-64884602-DEL-60
chr8-65027677-DEL-57
chr8-65053926-DEL-76
chr8-65062917-DEL-121
chr8-65096230-DEL-69
chr8-65275050-DEL-196
chr8-65336817-DEL-50
chr8-65386665-DEL-287
chr8-65391074-DEL-50
chr8-65463916-DEL-50
chr8-65469656-DEL-92
chr8-65568386-DEL-87
chr8-65576922-DEL-1544
chr8-65578402-DEL-252
chr8-65609795-DEL-160
chr8-65609824-DEL-74
chr8-65674564-DEL-52
chr8-65741556-DEL-74
chr8-65819118-DEL-90
chr8-65875425-DEL-58
chr8-65879045-DEL-50
chr8-66176801-DEL-312

chr8-66269590-DEL-85
chr8-66270782-DEL-943
chr8-66280198-DEL-65
chr8-66343827-DEL-218
chr8-66410790-DEL-2015
chr8-66507225-DEL-75
chr8-66512909-DEL-53
chr8-66513328-DEL-53
chr8-66517796-DEL-4062
chr8-66532792-DEL-652
chr8-66535134-DEL-51
chr8-66535142-DEL-636
chr8-66535706-DEL-72
chr8-66602074-DEL-3507
chr8-66634040-DEL-78
chr8-66710072-DEL-3756
chr8-66767314-DEL-399
chr8-66906723-DEL-683
chr8-66909449-DEL-154
chr8-66945657-DEL-104
chr8-67083854-DEL-72
chr8-67097737-DEL-1464
chr8-67139207-DEL-57
chr8-67267485-DEL-133
chr8-67313147-DEL-84
chr8-67313147-DEL-252
chr8-67406455-DEL-208
chr8-67739920-DEL-3384
chr8-67821038-DEL-7874
chr8-67840364-DEL-430
chr8-67851909-DEL-77
chr8-67974449-DEL-79
chr8-68036526-DEL-81
chr8-68078357-DEL-362
chr8-68124544-DEL-50
chr8-68126381-DEL-139
chr8-68134526-DEL-846
chr8-68392739-DEL-51
chr8-68449123-DEL-54
chr8-68490481-DEL-52
chr8-68623812-DEL-4712
chr8-68629262-DEL-387
chr8-68631109-DEL-87
chr8-68668400-DEL-149
chr8-68681966-DEL-160
chr8-69068457-DEL-51
chr8-69144356-DEL-70
chr8-69358186-DEL-53
chr8-69374098-DEL-1876
chr8-69380582-DEL-80
chr8-69380883-DEL-54
chr8-69397579-DEL-132
chr8-69544381-DEL-110
chr8-69720008-DEL-77
chr8-69720174-DEL-239
chr8-69720242-DEL-99
chr8-69720624-DEL-741
chr8-69899822-DEL-69409
chr8-69978804-DEL-58
chr8-70022472-DEL-4970
chr8-70181221-DEL-145
chr8-70382876-DEL-114
chr8-70434960-DEL-80
chr8-70464713-DEL-132
chr8-70670731-DEL-202
chr8-70881616-DEL-52
chr8-70881750-DEL-268
chr8-70887669-DEL-143
chr8-70897272-DEL-134
chr8-70926278-DEL-560
chr8-70940367-DEL-82
chr8-70985965-DEL-60
chr8-71138188-DEL-50
chr8-71401093-DEL-52
chr8-71433996-DEL-73
chr8-71447024-DEL-113
chr8-71454268-DEL-1457

chr8-71473288-DEL-94
chr8-71511073-DEL-13256
chr8-71641791-DEL-324
chr8-71748752-DEL-59
chr8-71862173-DEL-54
chr8-71882598-DEL-80
chr8-71945756-DEL-339
chr8-71954018-DEL-407
chr8-71956058-DEL-248
chr8-71956546-DEL-163
chr8-71982164-DEL-54
chr8-72004020-DEL-52
chr8-72035588-DEL-74
chr8-72053860-DEL-644
chr8-72062991-DEL-68
chr8-72158317-DEL-110
chr8-72158688-DEL-209
chr8-72202973-DEL-54
chr8-72258659-DEL-103
chr8-72259387-DEL-50
chr8-72718590-DEL-57
chr8-72809562-DEL-56
chr8-72821346-DEL-114
chr8-72963094-DEL-50
chr8-73045717-DEL-159
chr8-73187101-DEL-90
chr8-73187864-DEL-205
chr8-73346886-DEL-149
chr8-73347196-DEL-240
chr8-73359884-DEL-8657
chr8-73359905-DEL-60
chr8-73587299-DEL-58
chr8-73642362-DEL-344
chr8-73653692-DEL-60
chr8-73665970-DEL-170
chr8-73683922-DEL-432
chr8-74112918-DEL-218
chr8-74153060-DEL-68
chr8-74198907-DEL-2279
chr8-74210642-DEL-97
chr8-74224473-DEL-199
chr8-74456589-DEL-128
chr8-74456604-DEL-304
chr8-74497809-DEL-1279
chr8-74593570-DEL-78
chr8-74615027-DEL-4108
chr8-74685087-DEL-1263
chr8-74806461-DEL-51
chr8-74944949-DEL-83
chr8-75049662-DEL-3226
chr8-75146816-DEL-64
chr8-75334271-DEL-101
chr8-75334522-DEL-315
chr8-75335277-DEL-1437
chr8-75409044-DEL-81
chr8-75409313-DEL-132
chr8-75431795-DEL-204
chr8-75431853-DEL-612
chr8-75610319-DEL-392
chr8-75649106-DEL-314
chr8-75793617-DEL-191
chr8-75866048-DEL-78
chr8-75866309-DEL-81
chr8-75882117-DEL-50
chr8-75900174-DEL-161
chr8-75912348-DEL-117
chr8-75922381-DEL-90
chr8-76009106-DEL-64
chr8-76009315-DEL-355
chr8-76009369-DEL-164
chr8-76116389-DEL-1475
chr8-76231767-DEL-84
chr8-76243282-DEL-194
chr8-76243860-DEL-2079
chr8-76249071-DEL-1177
chr8-76426881-DEL-108
chr8-76426970-DEL-393

chr8-76512787-DEL-52
chr8-76517021-DEL-4408
chr8-76564873-DEL-98
chr8-76630552-DEL-198
chr8-76658674-DEL-62
chr8-76689219-DEL-115
chr8-76691964-DEL-68
chr8-76844479-DEL-138
chr8-77008678-DEL-78
chr8-77126397-DEL-1381
chr8-77194931-DEL-214
chr8-77233787-DEL-84
chr8-77238479-DEL-62
chr8-77341388-DEL-77
chr8-77346191-DEL-188
chr8-77346311-DEL-68
chr8-77346491-DEL-52
chr8-77401218-DEL-1453
chr8-77423852-DEL-73
chr8-77608019-DEL-60
chr8-77791125-DEL-126
chr8-77791161-DEL-60
chr8-77947760-DEL-217
chr8-77995226-DEL-81
chr8-77999752-DEL-55
chr8-78113408-DEL-56
chr8-78118300-DEL-2911
chr8-78148562-DEL-813
chr8-78152014-DEL-81
chr8-78175399-DEL-442
chr8-78190938-DEL-65
chr8-78231344-DEL-2131
chr8-78231347-DEL-68
chr8-78246558-DEL-54
chr8-78276777-DEL-1998
chr8-78277266-DEL-68
chr8-78378710-DEL-60
chr8-78395200-DEL-90
chr8-78414388-DEL-68
chr8-78414669-DEL-284
chr8-78414713-DEL-98
chr8-78437708-DEL-73
chr8-78514791-DEL-70
chr8-78547475-DEL-446
chr8-78547663-DEL-111
chr8-78547720-DEL-54
chr8-78547868-DEL-212
chr8-78548027-DEL-53
chr8-78548150-DEL-194
chr8-78548484-DEL-57
chr8-78550853-DEL-163
chr8-78567551-DEL-576
chr8-78567580-DEL-126
chr8-78567994-DEL-216
chr8-78568138-DEL-72
chr8-78588431-DEL-171
chr8-78747495-DEL-213
chr8-78769981-DEL-3343
chr8-78793178-DEL-107
chr8-78886138-DEL-52
chr8-78977769-DEL-246
chr8-79024316-DEL-60
chr8-79079144-DEL-72
chr8-79196127-DEL-53
chr8-79200720-DEL-55
chr8-79212539-DEL-62
chr8-79279123-DEL-90
chr8-79318342-DEL-763
chr8-79319185-DEL-185
chr8-79319482-DEL-95
chr8-79328757-DEL-137
chr8-79339529-DEL-57
chr8-79339839-DEL-176
chr8-79365583-DEL-198
chr8-79424555-DEL-52
chr8-79438002-DEL-64
chr8-79470388-DEL-362

chr8-79498724-DEL-51
chr8-79520738-DEL-71
chr8-79521295-DEL-265
chr8-79521713-DEL-71
chr8-79553539-DEL-251
chr8-79610854-DEL-50
chr8-79611190-DEL-57
chr8-79618798-DEL-89
chr8-79640161-DEL-60
chr8-79711424-DEL-78
chr8-79725102-DEL-81
chr8-79761845-DEL-228
chr8-79855236-DEL-96
chr8-79979248-DEL-54
chr8-79988919-DEL-138
chr8-80262189-DEL-51
chr8-80285032-DEL-80
chr8-80322914-DEL-84
chr8-80330678-DEL-204
chr8-80348479-DEL-55
chr8-80543142-DEL-134
chr8-80543219-DEL-50
chr8-80646790-DEL-657
chr8-80647045-DEL-210
chr8-80688561-DEL-104
chr8-80740004-DEL-59
chr8-80762025-DEL-1695
chr8-80765032-DEL-90
chr8-80765276-DEL-167
chr8-80943733-DEL-50
chr8-81002140-DEL-349
chr8-81242231-DEL-193
chr8-81243301-DEL-62
chr8-81349900-DEL-1379
chr8-81383393-DEL-777
chr8-81417928-DEL-220
chr8-81424548-DEL-53
chr8-81470318-DEL-68
chr8-81515975-DEL-70
chr8-81584187-DEL-50
chr8-81584191-DEL-115
chr8-81602595-DEL-50
chr8-81630530-DEL-50
chr8-81641768-DEL-51
chr8-81646993-DEL-887
chr8-81671711-DEL-50
chr8-81811376-DEL-56
chr8-81811737-DEL-309
chr8-81811878-DEL-65
chr8-81811915-DEL-137
chr8-81822324-DEL-51
chr8-81975333-DEL-209
chr8-81975339-DEL-99
chr8-81976839-DEL-100
chr8-82049577-DEL-266
chr8-82049892-DEL-365
chr8-82109022-DEL-96
chr8-82118765-DEL-50
chr8-82180419-DEL-68
chr8-82234535-DEL-340
chr8-82277778-DEL-1666
chr8-82329071-DEL-239
chr8-82329227-DEL-56
chr8-82472647-DEL-57
chr8-82556221-DEL-123
chr8-82556280-DEL-50
chr8-82697132-DEL-203
chr8-82734354-DEL-129
chr8-82757193-DEL-88
chr8-82758696-DEL-60
chr8-82782841-DEL-2373
chr8-82843704-DEL-115
chr8-82843979-DEL-62
chr8-82945732-DEL-83
chr8-83013576-DEL-75
chr8-83259680-DEL-78
chr8-83400614-DEL-62

chr8-83441869-DEL-3552
chr8-83453580-DEL-587
chr8-83593910-DEL-56
chr8-83744657-DEL-6339
chr8-83848227-DEL-1958
chr8-83945959-DEL-50
chr8-83955175-DEL-801
chr8-83956382-DEL-331
chr8-83957003-DEL-490
chr8-83957891-DEL-269
chr8-84005981-DEL-53
chr8-84093830-DEL-151
chr8-84133111-DEL-60
chr8-84157711-DEL-106
chr8-84189587-DEL-141
chr8-84233154-DEL-3351
chr8-84290734-DEL-64
chr8-84329322-DEL-358
chr8-84329877-DEL-167
chr8-84351421-DEL-894
chr8-84361053-DEL-51
chr8-84364549-DEL-54
chr8-84364689-DEL-218
chr8-84483394-DEL-132
chr8-84548597-DEL-140
chr8-84548673-DEL-56
chr8-84589881-DEL-51
chr8-84590076-DEL-243
chr8-84593027-DEL-76
chr8-84632360-DEL-54
chr8-84676870-DEL-56
chr8-84709775-DEL-104
chr8-84840029-DEL-266
chr8-84845888-DEL-50
chr8-84847133-DEL-84
chr8-84847170-DEL-172
chr8-84919980-DEL-123
chr8-84923249-DEL-273
chr8-84972353-DEL-50
chr8-85036260-DEL-120
chr8-85097346-DEL-142
chr8-85119524-DEL-75
chr8-85161397-DEL-99
chr8-85172248-DEL-286
chr8-85173044-DEL-2260
chr8-85178747-DEL-7413
chr8-85219486-DEL-61
chr8-85219691-DEL-75
chr8-85379373-DEL-1645
chr8-85653353-DEL-50
chr8-85669496-DEL-88
chr8-85726234-DEL-118
chr8-85790084-DEL-249
chr8-85841033-DEL-86
chr8-86128351-DEL-50
chr8-86186849-DEL-330
chr8-86224309-DEL-419
chr8-86226243-DEL-188
chr8-86334850-DEL-50
chr8-86517159-DEL-229
chr8-86630392-DEL-72
chr8-86675399-DEL-2009
chr8-86704647-DEL-4989
chr8-86777049-DEL-371
chr8-86783697-DEL-62
chr8-86805276-DEL-800
chr8-86845377-DEL-185
chr8-86853638-DEL-58
chr8-86864810-DEL-748
chr8-87022877-DEL-1467
chr8-87024247-DEL-155
chr8-87044244-DEL-189
chr8-87161607-DEL-94
chr8-87198326-DEL-135
chr8-87300724-DEL-56
chr8-87309064-DEL-289
chr8-87313743-DEL-54

chr8-87390033-DEL-485
chr8-87419902-DEL-63
chr8-87503281-DEL-1476
chr8-87508801-DEL-74
chr8-87646272-DEL-161
chr8-87665005-DEL-50
chr8-87731545-DEL-50
chr8-87818666-DEL-60
chr8-87909994-DEL-1907
chr8-87948956-DEL-526
chr8-88052744-DEL-78
chr8-88202579-DEL-1364
chr8-88487135-DEL-51
chr8-88552515-DEL-51
chr8-88573585-DEL-261
chr8-88597632-DEL-52
chr8-88626165-DEL-81
chr8-88705218-DEL-90
chr8-88781329-DEL-93
chr8-88781594-DEL-138
chr8-88785809-DEL-110
chr8-88908518-DEL-99
chr8-88908756-DEL-177
chr8-88911374-DEL-425
chr8-88915286-DEL-296
chr8-88915524-DEL-441
chr8-88938932-DEL-104
chr8-88988287-DEL-53
chr8-89166344-DEL-50
chr8-89303671-DEL-126
chr8-89333364-DEL-52
chr8-89343446-DEL-54
chr8-89373728-DEL-197
chr8-89418515-DEL-50
chr8-89525856-DEL-55
chr8-89543369-DEL-145
chr8-89543731-DEL-66
chr8-89693634-DEL-552
chr8-89742806-DEL-50
chr8-89815650-DEL-204
chr8-89842241-DEL-771
chr8-89885816-DEL-100
chr8-89962133-DEL-89
chr8-89972677-DEL-126
chr8-90210993-DEL-76
chr8-90211191-DEL-52
chr8-90307164-DEL-189
chr8-90462317-DEL-115
chr8-90481701-DEL-60
chr8-90593792-DEL-96
chr8-90604397-DEL-989
chr8-90832748-DEL-50
chr8-90875996-DEL-54
chr8-90884764-DEL-90
chr8-90895053-DEL-69
chr8-91011736-DEL-53
chr8-91070805-DEL-141
chr8-91206009-DEL-113
chr8-91210683-DEL-63
chr8-91213644-DEL-63
chr8-91291471-DEL-90
chr8-91359953-DEL-169
chr8-91406700-DEL-190
chr8-91407092-DEL-50
chr8-91428420-DEL-52
chr8-91432860-DEL-155
chr8-91433106-DEL-69
chr8-91448300-DEL-52
chr8-91461314-DEL-56
chr8-91498230-DEL-1279
chr8-91517479-DEL-54
chr8-91547364-DEL-72
chr8-91587183-DEL-52
chr8-91733896-DEL-106
chr8-91818254-DEL-1772
chr8-91820422-DEL-84
chr8-91820450-DEL-249

chr8-91831255-DEL-58
chr8-91895795-DEL-104
chr8-91909911-DEL-106
chr8-91910210-DEL-52
chr8-92002161-DEL-52
chr8-92060237-DEL-153
chr8-92060295-DEL-60
chr8-92063063-DEL-125
chr8-92140653-DEL-438
chr8-92140902-DEL-186
chr8-92149907-DEL-268
chr8-92183344-DEL-50
chr8-92193711-DEL-54
chr8-92197356-DEL-295
chr8-92203290-DEL-50
chr8-92203389-DEL-478
chr8-92210317-DEL-137
chr8-92290799-DEL-182
chr8-92347800-DEL-175
chr8-92452171-DEL-810
chr8-92500828-DEL-66
chr8-92505483-DEL-68
chr8-92536630-DEL-208
chr8-92583499-DEL-56
chr8-92647293-DEL-213
chr8-92902002-DEL-188
chr8-92911557-DEL-74
chr8-92942643-DEL-191
chr8-92987931-DEL-64
chr8-93097405-DEL-157
chr8-93120829-DEL-84
chr8-93197417-DEL-60
chr8-93197580-DEL-60
chr8-93221576-DEL-201
chr8-93315020-DEL-498
chr8-93346936-DEL-88
chr8-93393674-DEL-921
chr8-93414594-DEL-54
chr8-93414704-DEL-261
chr8-93455795-DEL-80
chr8-93493059-DEL-50
chr8-93624804-DEL-339
chr8-93640187-DEL-97
chr8-93648872-DEL-219
chr8-93730801-DEL-51
chr8-94003454-DEL-1190
chr8-94036452-DEL-234
chr8-94079249-DEL-52
chr8-94102291-DEL-126
chr8-94163577-DEL-99
chr8-94176331-DEL-6908
chr8-94178366-DEL-1436
chr8-94205941-DEL-6272
chr8-94222413-DEL-84
chr8-94248902-DEL-73
chr8-94253329-DEL-687
chr8-94260421-DEL-58
chr8-94309684-DEL-494
chr8-94379695-DEL-126
chr8-94412542-DEL-77
chr8-94423201-DEL-183
chr8-94457570-DEL-221
chr8-94496241-DEL-52
chr8-94685672-DEL-303
chr8-94713821-DEL-100
chr8-94724829-DEL-50
chr8-94746716-DEL-545
chr8-94748240-DEL-138
chr8-94890545-DEL-82
chr8-94900980-DEL-58
chr8-94924034-DEL-59
chr8-94933845-DEL-60
chr8-94943801-DEL-133
chr8-95077961-DEL-78
chr8-95133810-DEL-72
chr8-95229200-DEL-139
chr8-95229452-DEL-116

chr8-95230737-DEL-120
chr8-95243342-DEL-52
chr8-95249921-DEL-922
chr8-95268567-DEL-54
chr8-95495068-DEL-108
chr8-95532749-DEL-111
chr8-95733055-DEL-54
chr8-95956647-DEL-460
chr8-96118607-DEL-220
chr8-96133872-DEL-58
chr8-96149939-DEL-101
chr8-96157993-DEL-256
chr8-96244027-DEL-484
chr8-96291591-DEL-123
chr8-96312386-DEL-50
chr8-96400715-DEL-61
chr8-96412489-DEL-314
chr8-96466043-DEL-100
chr8-96541316-DEL-791
chr8-96652552-DEL-348
chr8-96652601-DEL-109
chr8-96652742-DEL-146
chr8-96679252-DEL-56
chr8-96685724-DEL-84
chr8-96685837-DEL-370
chr8-96750390-DEL-58
chr8-96812255-DEL-2914
chr8-96821795-DEL-58
chr8-96877328-DEL-84
chr8-96972086-DEL-134
chr8-97045230-DEL-58
chr8-97385619-DEL-119
chr8-97385839-DEL-67
chr8-97396117-DEL-53
chr8-97485952-DEL-119
chr8-97486023-DEL-272
chr8-97486199-DEL-99
chr8-97649572-DEL-57
chr8-97649722-DEL-231
chr8-97692070-DEL-208
chr8-97692368-DEL-121
chr8-97714839-DEL-73
chr8-97732414-DEL-64
chr8-97824803-DEL-243
chr8-98044673-DEL-100
chr8-98044758-DEL-260
chr8-98069160-DEL-79
chr8-98087139-DEL-97
chr8-98188862-DEL-50
chr8-98209704-DEL-1294
chr8-98474154-DEL-574
chr8-98485624-DEL-819
chr8-98522616-DEL-6289
chr8-98565791-DEL-82
chr8-98635916-DEL-50
chr8-98660431-DEL-100
chr8-98918609-DEL-126
chr8-98963559-DEL-981
chr8-99013614-DEL-1604
chr8-99018732-DEL-62
chr8-99050211-DEL-50
chr8-99066301-DEL-2760
chr8-99132313-DEL-52
chr8-99164143-DEL-90
chr8-99186935-DEL-72
chr8-99245012-DEL-67
chr8-99301134-DEL-90
chr8-99375403-DEL-69
chr8-99406852-DEL-1936
chr8-99441541-DEL-94
chr8-99492121-DEL-1082
chr8-99495522-DEL-58
chr8-99628037-DEL-76
chr8-99634603-DEL-171
chr8-99681531-DEL-65
chr8-99713366-DEL-75
chr8-99808772-DEL-56

chr8-99925755-DEL-90
chr8-99986034-DEL-1209
chr8-99998568-DEL-60
chr8-100022225-DEL-68
chr8-100034346-DEL-58
chr8-100165849-DEL-60
chr8-100213620-DEL-940
chr8-100224299-DEL-56
chr8-100255351-DEL-120
chr8-100261021-DEL-529
chr8-100360374-DEL-57
chr8-100422338-DEL-3388
chr8-100436491-DEL-316
chr8-100470111-DEL-52
chr8-100492437-DEL-1679
chr8-100536197-DEL-2694
chr8-100574935-DEL-102
chr8-100586671-DEL-6427
chr8-100630906-DEL-50
chr8-100677443-DEL-3202
chr8-100700858-DEL-112
chr8-100749224-DEL-2239
chr8-100772818-DEL-98
chr8-100858283-DEL-200
chr8-100858747-DEL-204
chr8-100880049-DEL-1118
chr8-100918397-DEL-96
chr8-100986823-DEL-60
chr8-101054225-DEL-489
chr8-101066320-DEL-72
chr8-101103401-DEL-3765
chr8-101108660-DEL-52
chr8-101250721-DEL-602
chr8-101318958-DEL-186
chr8-101322455-DEL-51
chr8-101466960-DEL-78
chr8-101552294-DEL-362
chr8-101629963-DEL-50
chr8-101655422-DEL-52
chr8-102075266-DEL-60
chr8-102190287-DEL-54
chr8-102213267-DEL-146
chr8-102258510-DEL-50
chr8-102273877-DEL-2338
chr8-102366035-DEL-57
chr8-102366563-DEL-448
chr8-102475409-DEL-50
chr8-102521814-DEL-131
chr8-102603099-DEL-59
chr8-102667736-DEL-61
chr8-102965325-DEL-281
chr8-102985060-DEL-72
chr8-103074007-DEL-285
chr8-103081334-DEL-164
chr8-103086464-DEL-138
chr8-103194912-DEL-684
chr8-103231751-DEL-57
chr8-103262591-DEL-67
chr8-103288723-DEL-212
chr8-103310068-DEL-58
chr8-103500387-DEL-53
chr8-103624486-DEL-69
chr8-103663008-DEL-58
chr8-103758201-DEL-52
chr8-103841270-DEL-59
chr8-103944332-DEL-54
chr8-104024035-DEL-74
chr8-104158065-DEL-52
chr8-104240458-DEL-108
chr8-104646611-DEL-1451
chr8-104738281-DEL-2152
chr8-104886564-DEL-70
chr8-104886936-DEL-107
chr8-104895077-DEL-132
chr8-104938876-DEL-135
chr8-105061761-DEL-168
chr8-105094219-DEL-123

chr8-105101359-DEL-97
chr8-105299013-DEL-247
chr8-105352282-DEL-81
chr8-105382881-DEL-53
chr8-105492674-DEL-343
chr8-105575925-DEL-12079
chr8-105584309-DEL-26227
chr8-105859371-DEL-51
chr8-105887017-DEL-54
chr8-105889598-DEL-157
chr8-105938763-DEL-768
chr8-105960287-DEL-662
chr8-105971909-DEL-126
chr8-106010943-DEL-54
chr8-106047576-DEL-405
chr8-106137440-DEL-168
chr8-106137444-DEL-386
chr8-106153210-DEL-75
chr8-106154846-DEL-350
chr8-106262044-DEL-68
chr8-106275867-DEL-252
chr8-106515848-DEL-176
chr8-106519421-DEL-50
chr8-106543936-DEL-101
chr8-106551758-DEL-83
chr8-106752323-DEL-87
chr8-106774381-DEL-108
chr8-106795598-DEL-296
chr8-106917890-DEL-153
chr8-106918084-DEL-64
chr8-106920716-DEL-56
chr8-107035481-DEL-98
chr8-107144817-DEL-51
chr8-107280688-DEL-83
chr8-107287734-DEL-113
chr8-107343381-DEL-517
chr8-107448328-DEL-244
chr8-107493294-DEL-52
chr8-107511280-DEL-147
chr8-107534841-DEL-51
chr8-107540122-DEL-102
chr8-107540138-DEL-442
chr8-107636104-DEL-750
chr8-107636275-DEL-156
chr8-107637012-DEL-118
chr8-107640634-DEL-190
chr8-107730356-DEL-82
chr8-107747901-DEL-70
chr8-107793716-DEL-339
chr8-107815988-DEL-58
chr8-107855286-DEL-79
chr8-107868622-DEL-199
chr8-107895711-DEL-359
chr8-107939892-DEL-50
chr8-108106545-DEL-1043
chr8-108144265-DEL-137
chr8-108180914-DEL-203
chr8-108181095-DEL-50
chr8-108235878-DEL-50
chr8-108381013-DEL-170
chr8-108381096-DEL-68
chr8-108437499-DEL-58
chr8-108472035-DEL-159
chr8-108500416-DEL-172
chr8-108602820-DEL-205
chr8-108607413-DEL-120
chr8-108651602-DEL-98
chr8-108667854-DEL-124
chr8-108773768-DEL-884
chr8-108895344-DEL-50
chr8-108912754-DEL-62
chr8-109116266-DEL-93
chr8-109225126-DEL-135
chr8-109350106-DEL-52
chr8-109361383-DEL-1348
chr8-109387749-DEL-130
chr8-109414735-DEL-201

chr8-109467121-DEL-698
chr8-109561391-DEL-208
chr8-109568608-DEL-185
chr8-109665774-DEL-392
chr8-109886805-DEL-101
chr8-110039201-DEL-159
chr8-110251462-DEL-100
chr8-110349865-DEL-78
chr8-110390178-DEL-223
chr8-110446168-DEL-148
chr8-110498260-DEL-60
chr8-110621853-DEL-90
chr8-110814585-DEL-120
chr8-110842580-DEL-95
chr8-110858303-DEL-104
chr8-110862924-DEL-264
chr8-110878385-DEL-92
chr8-110920824-DEL-52
chr8-110972737-DEL-595
chr8-110999103-DEL-202
chr8-111012677-DEL-437
chr8-111084501-DEL-124
chr8-111160394-DEL-304
chr8-111214767-DEL-108
chr8-111280190-DEL-50
chr8-111383842-DEL-57
chr8-111394328-DEL-140
chr8-111412085-DEL-50
chr8-111449952-DEL-58
chr8-111471896-DEL-105
chr8-111489527-DEL-104
chr8-111623180-DEL-88
chr8-111633941-DEL-1435
chr8-111748596-DEL-122
chr8-111934028-DEL-143
chr8-111993002-DEL-1453
chr8-112028057-DEL-80
chr8-112040568-DEL-56
chr8-112048092-DEL-52
chr8-112125956-DEL-1775
chr8-112134148-DEL-190
chr8-112168803-DEL-1308
chr8-112170555-DEL-86
chr8-112176714-DEL-50
chr8-112178106-DEL-295
chr8-112182271-DEL-70
chr8-112250861-DEL-54
chr8-112280329-DEL-51
chr8-112283332-DEL-262
chr8-112283400-DEL-78
chr8-112340647-DEL-1701
chr8-112362737-DEL-3443
chr8-112364561-DEL-164
chr8-112448349-DEL-275
chr8-112562843-DEL-191
chr8-112570549-DEL-79
chr8-112606356-DEL-487
chr8-112620660-DEL-559
chr8-112630056-DEL-67
chr8-112630635-DEL-71
chr8-112665122-DEL-55
chr8-112675199-DEL-74
chr8-112682512-DEL-52
chr8-112712556-DEL-209
chr8-112766264-DEL-166
chr8-112779297-DEL-56
chr8-112779866-DEL-52
chr8-112779881-DEL-553
chr8-112783476-DEL-112
chr8-112842931-DEL-57
chr8-112849517-DEL-106
chr8-112865064-DEL-81
chr8-112956143-DEL-98
chr8-112973172-DEL-334
chr8-113032599-DEL-52
chr8-113123578-DEL-68
chr8-113258824-DEL-259

chr8-113259252-DEL-63
chr8-113274068-DEL-62
chr8-113391146-DEL-51
chr8-113413891-DEL-81
chr8-113447854-DEL-50
chr8-113640246-DEL-55
chr8-113776769-DEL-56
chr8-113781945-DEL-3477
chr8-113836073-DEL-80
chr8-113852965-DEL-76
chr8-113860486-DEL-56
chr8-113965916-DEL-112
chr8-114035550-DEL-52
chr8-114154876-DEL-200
chr8-114169963-DEL-119
chr8-114178532-DEL-60
chr8-114183710-DEL-964
chr8-114200304-DEL-88
chr8-114216864-DEL-333
chr8-114256207-DEL-128
chr8-114260346-DEL-130
chr8-114349761-DEL-80
chr8-114461524-DEL-232
chr8-114461777-DEL-66
chr8-114527312-DEL-292
chr8-114563383-DEL-55
chr8-114584861-DEL-78
chr8-114756527-DEL-63
chr8-114756712-DEL-215
chr8-114757046-DEL-126
chr8-114773225-DEL-76
chr8-114849059-DEL-58
chr8-114870958-DEL-220
chr8-114911284-DEL-51
chr8-114914404-DEL-238
chr8-114914520-DEL-98
chr8-114926297-DEL-102
chr8-115000617-DEL-57
chr8-115015584-DEL-64
chr8-115183936-DEL-88
chr8-115214400-DEL-193
chr8-115214724-DEL-120
chr8-115214916-DEL-162
chr8-115214928-DEL-64
chr8-115232442-DEL-52
chr8-115390713-DEL-1862
chr8-115391029-DEL-176
chr8-115465811-DEL-104
chr8-115488828-DEL-56
chr8-115489549-DEL-116
chr8-115558248-DEL-60
chr8-115608217-DEL-646
chr8-115649964-DEL-160
chr8-115694168-DEL-196
chr8-115697462-DEL-52
chr8-115796720-DEL-69
chr8-115909211-DEL-78
chr8-116036002-DEL-78
chr8-116050180-DEL-70
chr8-116104489-DEL-170
chr8-116108692-DEL-51
chr8-116113459-DEL-113
chr8-116134918-DEL-60
chr8-116189920-DEL-57
chr8-116274338-DEL-478
chr8-116280329-DEL-66
chr8-116315382-DEL-111
chr8-116315714-DEL-52
chr8-116360264-DEL-50
chr8-116371644-DEL-83
chr8-116371785-DEL-58
chr8-116569239-DEL-59
chr8-116604483-DEL-787
chr8-116715778-DEL-118
chr8-116733963-DEL-169
chr8-116749060-DEL-92
chr8-116862464-DEL-50

chr8-116999153-DEL-146
chr8-117005714-DEL-99
chr8-117062147-DEL-527
chr8-117080358-DEL-50
chr8-117093065-DEL-91
chr8-117139746-DEL-183
chr8-117209395-DEL-102
chr8-117233945-DEL-121
chr8-117250094-DEL-159
chr8-117261357-DEL-547
chr8-117261507-DEL-262
chr8-117278054-DEL-74
chr8-117278667-DEL-123
chr8-117286136-DEL-2266
chr8-117286674-DEL-485
chr8-117286874-DEL-235
chr8-117287428-DEL-168
chr8-117304873-DEL-275
chr8-117333122-DEL-268
chr8-117334412-DEL-441
chr8-117343858-DEL-96
chr8-117352817-DEL-356
chr8-117392389-DEL-50
chr8-117430748-DEL-377
chr8-117430781-DEL-68
chr8-117431409-DEL-442
chr8-117534287-DEL-52
chr8-117539311-DEL-56
chr8-117554126-DEL-518
chr8-117608453-DEL-68
chr8-117621374-DEL-309
chr8-117621441-DEL-110
chr8-117720808-DEL-525
chr8-117738427-DEL-103
chr8-117775119-DEL-59
chr8-117958892-DEL-70
chr8-118076935-DEL-199
chr8-118249697-DEL-94
chr8-118315418-DEL-54
chr8-118366132-DEL-192
chr8-118425442-DEL-50
chr8-118591598-DEL-58
chr8-118677855-DEL-59
chr8-118727032-DEL-460
chr8-118727171-DEL-60
chr8-118810741-DEL-985
chr8-119037872-DEL-50
chr8-119079450-DEL-82
chr8-119439576-DEL-50
chr8-119482035-DEL-84
chr8-119533893-DEL-72
chr8-119537478-DEL-50
chr8-119613024-DEL-59
chr8-119714516-DEL-149
chr8-119714707-DEL-421
chr8-119744998-DEL-66
chr8-119887083-DEL-110
chr8-119980587-DEL-108
chr8-120042505-DEL-71
chr8-120042868-DEL-125
chr8-120043245-DEL-362
chr8-120132044-DEL-50
chr8-120143792-DEL-164
chr8-120153899-DEL-129
chr8-120154139-DEL-131
chr8-120235807-DEL-65
chr8-120236059-DEL-413
chr8-120288066-DEL-332
chr8-120328598-DEL-545
chr8-120379224-DEL-120
chr8-120398922-DEL-119
chr8-120411601-DEL-460
chr8-120419196-DEL-408
chr8-120419513-DEL-63
chr8-120466998-DEL-96
chr8-120514056-DEL-59
chr8-120560489-DEL-52

chr8-120560628-DEL-102
chr8-120563398-DEL-55
chr8-120573017-DEL-1127
chr8-120645186-DEL-200
chr8-120815713-DEL-63
chr8-120859696-DEL-177
chr8-120940499-DEL-117
chr8-121121691-DEL-54
chr8-121142083-DEL-99
chr8-121228574-DEL-108
chr8-121228698-DEL-406
chr8-121244143-DEL-60
chr8-121504880-DEL-612
chr8-121734418-DEL-64
chr8-121971827-DEL-84
chr8-121972665-DEL-92
chr8-121994571-DEL-84
chr8-122045020-DEL-59
chr8-122065852-DEL-72
chr8-122193385-DEL-92
chr8-122257034-DEL-60
chr8-122263561-DEL-199
chr8-122264628-DEL-1022
chr8-122339679-DEL-250
chr8-122353329-DEL-84
chr8-122367460-DEL-99
chr8-122406503-DEL-672
chr8-122438407-DEL-172
chr8-122548868-DEL-4080
chr8-122596172-DEL-94
chr8-122605323-DEL-50
chr8-122643987-DEL-396
chr8-122644149-DEL-108
chr8-122704359-DEL-525
chr8-122706659-DEL-124
chr8-122939078-DEL-66
chr8-122940553-DEL-192
chr8-122940656-DEL-89
chr8-122953638-DEL-216
chr8-123038781-DEL-136
chr8-123059147-DEL-229
chr8-123207101-DEL-225
chr8-123334105-DEL-169
chr8-123465314-DEL-111
chr8-123526820-DEL-638
chr8-123535493-DEL-460
chr8-123535897-DEL-630
chr8-123684821-DEL-156
chr8-123692603-DEL-50
chr8-123710746-DEL-121
chr8-123788016-DEL-64
chr8-123788386-DEL-96
chr8-124004136-DEL-440
chr8-124042610-DEL-90
chr8-124051754-DEL-84
chr8-124178682-DEL-331
chr8-124258074-DEL-689
chr8-124266638-DEL-234
chr8-124268367-DEL-212
chr8-124271729-DEL-230
chr8-124273445-DEL-226
chr8-124275123-DEL-224
chr8-124285340-DEL-250
chr8-124288755-DEL-246
chr8-124303946-DEL-210
chr8-124327723-DEL-234
chr8-124331111-DEL-230
chr8-124332810-DEL-210
chr8-124462899-DEL-50
chr8-124482391-DEL-135
chr8-124508497-DEL-56
chr8-124508513-DEL-165
chr8-124508832-DEL-166
chr8-124510979-DEL-50
chr8-124521321-DEL-68
chr8-124610532-DEL-72
chr8-124687899-DEL-87

chr8-124706486-DEL-50
chr8-124810619-DEL-50
chr8-124886066-DEL-99
chr8-124893849-DEL-145
chr8-124894720-DEL-190
chr8-124904449-DEL-64
chr8-125011689-DEL-58
chr8-125025102-DEL-97
chr8-125027840-DEL-56
chr8-125056852-DEL-67
chr8-125057072-DEL-87
chr8-125095520-DEL-50
chr8-125117041-DEL-209
chr8-125150024-DEL-158
chr8-125288769-DEL-107
chr8-125299130-DEL-51
chr8-125328749-DEL-350
chr8-125328835-DEL-144
chr8-125413971-DEL-74
chr8-125413975-DEL-826
chr8-125417815-DEL-64
chr8-125444413-DEL-137
chr8-125444875-DEL-86
chr8-125444875-DEL-258
chr8-125569630-DEL-59
chr8-125569848-DEL-208
chr8-125569931-DEL-103
chr8-125672979-DEL-120
chr8-125749326-DEL-256
chr8-125768949-DEL-56
chr8-125782915-DEL-1165
chr8-125782933-DEL-193
chr8-125786735-DEL-114
chr8-125819408-DEL-50
chr8-125831402-DEL-88
chr8-125843892-DEL-809
chr8-125846428-DEL-539
chr8-125846492-DEL-167
chr8-125846873-DEL-164
chr8-125877238-DEL-62
chr8-125888907-DEL-261
chr8-125918134-DEL-208
chr8-125998675-DEL-72
chr8-126003498-DEL-76
chr8-126102410-DEL-50
chr8-126110469-DEL-52
chr8-126110801-DEL-52
chr8-126158624-DEL-52
chr8-126204498-DEL-579
chr8-126264367-DEL-306
chr8-126264442-DEL-111
chr8-126264817-DEL-87
chr8-126265991-DEL-59
chr8-126348911-DEL-281
chr8-126377759-DEL-56
chr8-126377865-DEL-358
chr8-126382142-DEL-62
chr8-126382187-DEL-206
chr8-126413829-DEL-471
chr8-126507860-DEL-60
chr8-126531496-DEL-875
chr8-126531804-DEL-1259
chr8-126537461-DEL-50
chr8-126628293-DEL-72
chr8-126633252-DEL-50
chr8-126640674-DEL-130
chr8-126667775-DEL-108
chr8-126669803-DEL-62
chr8-126710018-DEL-747
chr8-126731377-DEL-210
chr8-126741054-DEL-50
chr8-126741943-DEL-90
chr8-126747671-DEL-94
chr8-126750477-DEL-59
chr8-126802465-DEL-55
chr8-126805934-DEL-58
chr8-126810319-DEL-448

chr8-126821298-DEL-51
chr8-126845874-DEL-110
chr8-126889147-DEL-56
chr8-126924717-DEL-56
chr8-126925157-DEL-52
chr8-126926974-DEL-1965
chr8-127011542-DEL-64
chr8-127087124-DEL-1355
chr8-127192145-DEL-802
chr8-127193136-DEL-77
chr8-127193584-DEL-72
chr8-127214783-DEL-59
chr8-127279825-DEL-90
chr8-127563566-DEL-70
chr8-127583266-DEL-300
chr8-127587261-DEL-52
chr8-127617975-DEL-120
chr8-127693066-DEL-163
chr8-127705326-DEL-52
chr8-127733028-DEL-272
chr8-127764195-DEL-9622
chr8-127885603-DEL-78
chr8-127885758-DEL-58
chr8-127885951-DEL-133
chr8-127983944-DEL-198
chr8-128054449-DEL-52
chr8-128054730-DEL-112
chr8-128093928-DEL-184
chr8-128107506-DEL-671
chr8-128175033-DEL-210
chr8-128352763-DEL-52
chr8-128386091-DEL-108
chr8-128455938-DEL-51
chr8-128477050-DEL-61
chr8-128583971-DEL-268
chr8-128590148-DEL-6140
chr8-128888055-DEL-134
chr8-128903001-DEL-267
chr8-128904118-DEL-68
chr8-128913202-DEL-61
chr8-128997398-DEL-57
chr8-128997494-DEL-72
chr8-129263822-DEL-78
chr8-129407841-DEL-66
chr8-129430725-DEL-65
chr8-129505945-DEL-1968
chr8-129554882-DEL-57
chr8-129757732-DEL-81
chr8-129841344-DEL-2314
chr8-129862193-DEL-126
chr8-129862424-DEL-269
chr9-3047204-DEL-244
chr9-3051194-DEL-62
chr9-3051548-DEL-150
chr9-3125436-DEL-875
chr9-3194443-DEL-70
chr9-3310725-DEL-1560
chr9-3471899-DEL-940
chr9-3481957-DEL-5572
chr9-3485421-DEL-520
chr9-3553834-DEL-1188
chr9-3573021-DEL-78
chr9-3722571-DEL-1020
chr9-3781259-DEL-5888
chr9-3939695-DEL-8139
chr9-3985746-DEL-85
chr9-4009204-DEL-979
chr9-4010518-DEL-260
chr9-4011874-DEL-813
chr9-4053876-DEL-519
chr9-4094787-DEL-2179
chr9-4100635-DEL-2887
chr9-4131196-DEL-128
chr9-4306453-DEL-217
chr9-4447459-DEL-61
chr9-4448878-DEL-576
chr9-4565606-DEL-200

chr9-4565657-DEL-91
chr9-4572990-DEL-80
chr9-4581998-DEL-663
chr9-4587699-DEL-288
chr9-4659488-DEL-238
chr9-4735477-DEL-1279
chr9-4739897-DEL-200
chr9-4915300-DEL-1641
chr9-5046064-DEL-124
chr9-5075028-DEL-1910
chr9-5131356-DEL-1506
chr9-5219529-DEL-1213
chr9-5376989-DEL-114
chr9-5491615-DEL-53
chr9-5559410-DEL-55
chr9-5566186-DEL-1817
chr9-5645140-DEL-147
chr9-5657467-DEL-52
chr9-5750711-DEL-50
chr9-5755416-DEL-270
chr9-5807665-DEL-1958
chr9-5829910-DEL-65
chr9-5860570-DEL-295
chr9-5942691-DEL-83
chr9-5980341-DEL-59
chr9-6075696-DEL-10851
chr9-6087139-DEL-8429
chr9-6126076-DEL-765
chr9-6194815-DEL-58
chr9-6257493-DEL-183
chr9-6339632-DEL-105
chr9-6413881-DEL-3048
chr9-6452923-DEL-614
chr9-6584291-DEL-1921
chr9-6665579-DEL-111
chr9-6708337-DEL-84
chr9-6778790-DEL-225
chr9-6802529-DEL-108
chr9-6822635-DEL-2611
chr9-6865790-DEL-177
chr9-6866118-DEL-2623
chr9-6894656-DEL-310
chr9-6916850-DEL-276
chr9-7108633-DEL-1146
chr9-7333677-DEL-92
chr9-7461813-DEL-159
chr9-7462904-DEL-62
chr9-7553084-DEL-290
chr9-7563697-DEL-59
chr9-7840304-DEL-64
chr9-7857744-DEL-129
chr9-8029807-DEL-423
chr9-8030359-DEL-321
chr9-8037652-DEL-258
chr9-8044299-DEL-102
chr9-8327845-DEL-196
chr9-8462178-DEL-58
chr9-8547165-DEL-52
chr9-8575199-DEL-78
chr9-8641281-DEL-50
chr9-9070854-DEL-1344
chr9-9123557-DEL-106
chr9-9152573-DEL-839
chr9-9192404-DEL-112
chr9-9262932-DEL-60
chr9-9336723-DEL-562
chr9-9359491-DEL-178
chr9-9545478-DEL-50
chr9-9594271-DEL-104
chr9-9619815-DEL-2077
chr9-9871589-DEL-55
chr9-9941384-DEL-1355
chr9-10069756-DEL-76
chr9-10085004-DEL-72
chr9-10248442-DEL-52
chr9-10270232-DEL-90
chr9-10280332-DEL-54

chr9-10351101-DEL-50
chr9-10446188-DEL-202
chr9-10472354-DEL-132
chr9-10474634-DEL-51
chr9-10509158-DEL-67
chr9-10525042-DEL-128
chr9-10525107-DEL-386
chr9-10525365-DEL-128
chr9-10564959-DEL-139
chr9-10699427-DEL-72
chr9-11121956-DEL-915
chr9-11242776-DEL-390
chr9-11456183-DEL-2162
chr9-11473510-DEL-5167
chr9-11508607-DEL-1247
chr9-11591592-DEL-632
chr9-11605449-DEL-78
chr9-11632509-DEL-1056
chr9-11678548-DEL-76
chr9-11849706-DEL-105
chr9-11938103-DEL-915
chr9-11987653-DEL-680
chr9-11994928-DEL-54
chr9-12021296-DEL-5786
chr9-12273582-DEL-3765
chr9-12415565-DEL-60
chr9-12416107-DEL-901
chr9-12475570-DEL-64
chr9-12563914-DEL-1412
chr9-12565443-DEL-2689
chr9-12768261-DEL-5988
chr9-12909656-DEL-1143
chr9-12911931-DEL-113
chr9-13007517-DEL-85
chr9-13039410-DEL-3126
chr9-13068770-DEL-330
chr9-13079471-DEL-131
chr9-13182128-DEL-110
chr9-13234891-DEL-220
chr9-13325011-DEL-138
chr9-13325701-DEL-293
chr9-13351304-DEL-1044
chr9-13415061-DEL-179
chr9-13445622-DEL-216
chr9-13501018-DEL-110
chr9-13501151-DEL-729
chr9-13580327-DEL-752
chr9-13906108-DEL-1166
chr9-14030796-DEL-155
chr9-14094912-DEL-9792
chr9-14149105-DEL-3724
chr9-14174700-DEL-2662
chr9-14222994-DEL-50
chr9-14249063-DEL-117
chr9-14277893-DEL-330
chr9-14351310-DEL-57
chr9-14356274-DEL-376
chr9-14487895-DEL-132
chr9-14511999-DEL-93
chr9-14559671-DEL-58
chr9-14612069-DEL-105
chr9-14828958-DEL-72
chr9-14833805-DEL-161
chr9-14928768-DEL-168
chr9-14941194-DEL-170
chr9-14946671-DEL-380
chr9-14955081-DEL-76
chr9-14995190-DEL-50
chr9-15009703-DEL-182
chr9-15009993-DEL-66
chr9-15015943-DEL-557
chr9-15039571-DEL-90
chr9-15068865-DEL-68
chr9-15092594-DEL-60
chr9-15117782-DEL-50
chr9-15119927-DEL-50
chr9-15163003-DEL-150

chr9-15163221-DEL-184
chr9-15198326-DEL-52
chr9-15227214-DEL-330
chr9-15227215-DEL-164
chr9-15310626-DEL-1237
chr9-15320469-DEL-549
chr9-15355933-DEL-140
chr9-15383942-DEL-64
chr9-15398441-DEL-53
chr9-15398453-DEL-292
chr9-15414460-DEL-53
chr9-15426378-DEL-108
chr9-15502686-DEL-69
chr9-15519209-DEL-919
chr9-15591633-DEL-74
chr9-15642409-DEL-245
chr9-15695024-DEL-73
chr9-15711086-DEL-89
chr9-15733046-DEL-1174
chr9-15739117-DEL-1808
chr9-15773480-DEL-58
chr9-15857117-DEL-77
chr9-15868124-DEL-98
chr9-15902323-DEL-141
chr9-15971551-DEL-286
chr9-16034281-DEL-50
chr9-16107915-DEL-50
chr9-16233625-DEL-672
chr9-16240933-DEL-56
chr9-16264059-DEL-72
chr9-16281373-DEL-243
chr9-16328997-DEL-123
chr9-16416987-DEL-190
chr9-16530646-DEL-72
chr9-16569280-DEL-100
chr9-16712211-DEL-115
chr9-16761669-DEL-56
chr9-16784720-DEL-63
chr9-16787204-DEL-479
chr9-16801108-DEL-52
chr9-16831248-DEL-81
chr9-16858672-DEL-50
chr9-16859021-DEL-3004
chr9-16948424-DEL-2962
chr9-17065921-DEL-707
chr9-17076139-DEL-126
chr9-17106007-DEL-1273
chr9-17135687-DEL-150
chr9-17137622-DEL-50
chr9-17148022-DEL-1702
chr9-17224475-DEL-4481
chr9-17231890-DEL-55
chr9-17290355-DEL-4800
chr9-17372509-DEL-5428
chr9-17414143-DEL-90
chr9-17493055-DEL-69
chr9-17524582-DEL-137
chr9-17526377-DEL-50
chr9-17554923-DEL-1598
chr9-17554943-DEL-237
chr9-17555246-DEL-255
chr9-17562385-DEL-50
chr9-17599121-DEL-853
chr9-17601668-DEL-8914
chr9-17602056-DEL-483
chr9-17619086-DEL-1071
chr9-17688530-DEL-80
chr9-17689327-DEL-9155
chr9-17691179-DEL-3454
chr9-17698882-DEL-988
chr9-17839558-DEL-258
chr9-17913541-DEL-54
chr9-17942324-DEL-82
chr9-18069622-DEL-2172
chr9-18089188-DEL-443
chr9-18120241-DEL-75
chr9-18323813-DEL-52

chr9-18402020-DEL-332
chr9-18424920-DEL-58
chr9-18425526-DEL-3716
chr9-18591461-DEL-71
chr9-18607320-DEL-235
chr9-18701867-DEL-7161
chr9-18716949-DEL-2870
chr9-18796106-DEL-58
chr9-18805110-DEL-6164
chr9-18927155-DEL-3076
chr9-18940841-DEL-410
chr9-19103143-DEL-844
chr9-19259533-DEL-1036
chr9-19278894-DEL-23265
chr9-19311763-DEL-92
chr9-19311951-DEL-610
chr9-19314228-DEL-1647
chr9-19478211-DEL-1148
chr9-19480059-DEL-1191
chr9-19532925-DEL-244
chr9-19544435-DEL-6211
chr9-19661122-DEL-52
chr9-20146603-DEL-194
chr9-20235311-DEL-1620
chr9-20299337-DEL-6572
chr9-20340117-DEL-76
chr9-20340411-DEL-52
chr9-20361859-DEL-185
chr9-20372574-DEL-134
chr9-20393739-DEL-456
chr9-20444113-DEL-149
chr9-20492221-DEL-123
chr9-20496843-DEL-666
chr9-20594877-DEL-75
chr9-20604689-DEL-57
chr9-20680618-DEL-220
chr9-20681198-DEL-60
chr9-20733054-DEL-50
chr9-20785381-DEL-50
chr9-20792483-DEL-3075
chr9-20909575-DEL-228
chr9-20929242-DEL-51
chr9-20959906-DEL-190
chr9-21030524-DEL-69
chr9-21107740-DEL-99
chr9-21128954-DEL-64
chr9-21175392-DEL-198
chr9-21204887-DEL-608
chr9-21245339-DEL-75
chr9-21273308-DEL-130
chr9-21330005-DEL-82
chr9-21330012-DEL-318
chr9-21330337-DEL-85
chr9-21400401-DEL-53
chr9-21445386-DEL-74
chr9-21539990-DEL-196
chr9-21629768-DEL-291
chr9-21652113-DEL-368
chr9-21676750-DEL-198
chr9-21697346-DEL-395
chr9-21704766-DEL-92
chr9-21814657-DEL-976
chr9-22044897-DEL-94
chr9-22055593-DEL-96
chr9-22055818-DEL-205
chr9-22102848-DEL-140
chr9-22103937-DEL-122
chr9-22133856-DEL-212
chr9-22211202-DEL-367
chr9-22231232-DEL-203
chr9-22231579-DEL-1192
chr9-22303701-DEL-57
chr9-22303712-DEL-180
chr9-22304023-DEL-206
chr9-22474441-DEL-56
chr9-22496033-DEL-420
chr9-22496108-DEL-126

chr9-22507468-DEL-80
chr9-22522192-DEL-134
chr9-22547041-DEL-132
chr9-22622880-DEL-264
chr9-22622925-DEL-696
chr9-22623406-DEL-258
chr9-22669321-DEL-80
chr9-22694023-DEL-246
chr9-22717908-DEL-998
chr9-22727991-DEL-93
chr9-22752031-DEL-1697
chr9-22780652-DEL-56
chr9-22935679-DEL-293
chr9-23020611-DEL-117
chr9-23066690-DEL-51
chr9-23232909-DEL-175
chr9-23264560-DEL-51
chr9-23388253-DEL-70
chr9-23459128-DEL-584
chr9-23462394-DEL-113
chr9-23595648-DEL-402
chr9-23731241-DEL-129
chr9-23800891-DEL-2077
chr9-23887811-DEL-2432
chr9-23890291-DEL-935
chr9-23912029-DEL-104
chr9-23961870-DEL-325
chr9-23962122-DEL-78
chr9-24059101-DEL-214
chr9-24067009-DEL-964
chr9-24169824-DEL-137
chr9-24211187-DEL-52
chr9-24489840-DEL-50
chr9-24493754-DEL-56
chr9-24619828-DEL-67
chr9-24833089-DEL-50
chr9-24843952-DEL-3319
chr9-24853564-DEL-140
chr9-24939548-DEL-1796
chr9-24961388-DEL-132
chr9-24962346-DEL-593
chr9-25070551-DEL-64
chr9-25167667-DEL-131
chr9-25266472-DEL-64
chr9-25324339-DEL-302
chr9-25387615-DEL-57
chr9-25408688-DEL-86
chr9-25436533-DEL-160
chr9-25491863-DEL-246
chr9-25492066-DEL-106
chr9-25557395-DEL-1216
chr9-25669499-DEL-214
chr9-25710314-DEL-145
chr9-25723758-DEL-52
chr9-25857949-DEL-53
chr9-25938704-DEL-651
chr9-26135240-DEL-98
chr9-26191367-DEL-1249
chr9-26432032-DEL-67
chr9-26461964-DEL-72
chr9-26509948-DEL-296
chr9-26704515-DEL-179
chr9-26857318-DEL-73
chr9-26975927-DEL-124
chr9-27236768-DEL-268
chr9-27637276-DEL-126
chr9-27766727-DEL-1661
chr9-27808695-DEL-69
chr9-28003164-DEL-54
chr9-28246608-DEL-147
chr9-28356978-DEL-50
chr9-28642152-DEL-175
chr9-28691837-DEL-144
chr9-28910061-DEL-50
chr9-29005615-DEL-50
chr9-29071462-DEL-533
chr9-29106389-DEL-1638

chr9-29248766-DEL-50
chr9-29353824-DEL-150
chr9-29395180-DEL-338
chr9-29395185-DEL-124
chr9-29395775-DEL-105
chr9-29395914-DEL-273
chr9-29396515-DEL-74
chr9-29435005-DEL-12089
chr9-29544033-DEL-50
chr9-29591986-DEL-51
chr9-29617510-DEL-216
chr9-29617588-DEL-463
chr9-29617941-DEL-104
chr9-29869613-DEL-63
chr9-29869613-DEL-420
chr9-29900054-DEL-73
chr9-30081868-DEL-50
chr9-30178607-DEL-52
chr9-30222181-DEL-359
chr9-30246694-DEL-74
chr9-30421003-DEL-134
chr9-30567770-DEL-53
chr9-30572655-DEL-54
chr9-30572867-DEL-80
chr9-30664153-DEL-95
chr9-30736449-DEL-118
chr9-30801779-DEL-54
chr9-30849384-DEL-72
chr9-30849567-DEL-50
chr9-30871202-DEL-615
chr9-30871451-DEL-140
chr9-31101714-DEL-96
chr9-31142498-DEL-78
chr9-31176392-DEL-729
chr9-31194380-DEL-476
chr9-31200197-DEL-50
chr9-31208599-DEL-50
chr9-31261733-DEL-110
chr9-31310964-DEL-141
chr9-31336547-DEL-96
chr9-31412541-DEL-64
chr9-31413885-DEL-58
chr9-31428649-DEL-284
chr9-31451222-DEL-61
chr9-31451331-DEL-233
chr9-31451514-DEL-74
chr9-31491851-DEL-136
chr9-31491950-DEL-303
chr9-31499403-DEL-50
chr9-31582789-DEL-134
chr9-31679258-DEL-60
chr9-31682754-DEL-322
chr9-31683176-DEL-342
chr9-31683726-DEL-230
chr9-31683883-DEL-73
chr9-31910923-DEL-185
chr9-32041747-DEL-1057
chr9-32041838-DEL-63
chr9-32042742-DEL-62
chr9-32075280-DEL-68
chr9-32218932-DEL-1357
chr9-32219412-DEL-525
chr9-32219982-DEL-292
chr9-32220162-DEL-142
chr9-32241198-DEL-60
chr9-32241271-DEL-142
chr9-32264449-DEL-155
chr9-32267085-DEL-740
chr9-32421674-DEL-50
chr9-32485972-DEL-189
chr9-32500540-DEL-53
chr9-32595004-DEL-161
chr9-32595425-DEL-246
chr9-32775365-DEL-1895
chr9-32788156-DEL-3856
chr9-32834930-DEL-110
chr9-32886118-DEL-945

chr9-32963058-DEL-50
chr9-32969624-DEL-122
chr9-33156226-DEL-369
chr9-33237200-DEL-448
chr9-33309273-DEL-4685
chr9-33319911-DEL-161
chr9-33415786-DEL-163
chr9-33555297-DEL-357
chr9-33601127-DEL-76
chr9-33601156-DEL-171
chr9-33814764-DEL-116
chr9-33829655-DEL-237
chr9-33829815-DEL-111
chr9-33837989-DEL-1553
chr9-33851878-DEL-50
chr9-33931990-DEL-521
chr9-33950123-DEL-188
chr9-34205487-DEL-118
chr9-34476547-DEL-50
chr9-34565224-DEL-188
chr9-34797005-DEL-124
chr9-34807864-DEL-84
chr9-34843496-DEL-549
chr9-35147806-DEL-50
chr9-35325950-DEL-54
chr9-35338363-DEL-50
chr9-35385589-DEL-74
chr9-35403675-DEL-153
chr9-35404517-DEL-496
chr9-35410669-DEL-102
chr9-35545764-DEL-755
chr9-35613712-DEL-52
chr9-35643459-DEL-981
chr9-35692191-DEL-149
chr9-35770306-DEL-84
chr9-35833330-DEL-207
chr9-35857680-DEL-9233
chr9-35863612-DEL-1227
chr9-35865325-DEL-495
chr9-35882983-DEL-56
chr9-35978854-DEL-4744
chr9-36162208-DEL-603
chr9-36162296-DEL-177
chr9-36162408-DEL-65
chr9-36212827-DEL-108
chr9-36392819-DEL-58
chr9-36534117-DEL-1520
chr9-36593665-DEL-50
chr9-36666843-DEL-1192
chr9-36681101-DEL-54
chr9-36712978-DEL-65
chr9-36958382-DEL-56
chr9-37156439-DEL-869
chr9-37491846-DEL-116
chr9-37499752-DEL-226
chr9-37539385-DEL-106
chr9-37651230-DEL-54
chr9-37758926-DEL-75
chr9-37786698-DEL-66
chr9-37841354-DEL-257
chr9-37841650-DEL-105
chr9-37891421-DEL-54
chr9-37944611-DEL-4042
chr9-38001191-DEL-184
chr9-38066939-DEL-563
chr9-38067611-DEL-162
chr9-38109880-DEL-126
chr9-38237187-DEL-83
chr9-38262055-DEL-91
chr9-38308182-DEL-92
chr9-38382723-DEL-72
chr9-38390331-DEL-107
chr9-38463015-DEL-214
chr9-38500607-DEL-145
chr9-38611962-DEL-70
chr9-38699917-DEL-72
chr9-38731842-DEL-150

chr9-38734510-DEL-564
chr9-38736319-DEL-522
chr9-38779777-DEL-386
chr9-38879488-DEL-1140
chr9-38973523-DEL-115
chr9-39072561-DEL-2409
chr9-39183614-DEL-1102
chr9-39345927-DEL-838
chr9-39457129-DEL-162
chr9-39646065-DEL-75
chr9-39750460-DEL-59
chr9-39756947-DEL-52
chr9-39776915-DEL-62
chr9-39850225-DEL-392
chr9-39934774-DEL-52
chr9-39966016-DEL-98
chr9-40027227-DEL-269
chr9-40076329-DEL-127
chr9-40080085-DEL-121
chr9-40247804-DEL-74
chr9-40532574-DEL-130
chr9-40557886-DEL-116
chr9-40630784-DEL-55
chr9-40648313-DEL-127
chr9-40684298-DEL-83
chr9-40702184-DEL-180
chr9-40731702-DEL-105
chr9-40826840-DEL-170
chr9-40836457-DEL-51
chr9-40847724-DEL-71
chr9-40853965-DEL-119
chr9-40930831-DEL-116
chr9-41096544-DEL-221
chr9-41097526-DEL-55
chr9-41112620-DEL-175
chr9-41125101-DEL-56
chr9-41159085-DEL-50
chr9-41199958-DEL-51
chr9-41208077-DEL-50
chr9-41310096-DEL-104
chr9-41317007-DEL-186
chr9-41317302-DEL-239
chr9-41427245-DEL-64
chr9-41458351-DEL-868
chr9-41479694-DEL-50
chr9-41618979-DEL-165
chr9-41682012-DEL-52
chr9-41730564-DEL-50
chr9-41753309-DEL-104
chr9-41753997-DEL-550
chr9-41756556-DEL-81
chr9-41787868-DEL-50
chr9-41848265-DEL-122
chr9-41890967-DEL-50
chr9-41951715-DEL-87
chr9-42025467-DEL-63
chr9-42111148-DEL-91
chr9-42115941-DEL-59
chr9-42116015-DEL-63
chr9-42349658-DEL-210
chr9-42400080-DEL-50
chr9-42408980-DEL-194
chr9-42409099-DEL-68
chr9-42555447-DEL-272
chr9-42784933-DEL-148
chr9-42921747-DEL-115
chr9-42930061-DEL-194
chr9-42930444-DEL-216
chr9-43058675-DEL-78
chr9-43096978-DEL-74
chr9-43332559-DEL-114
chr9-43369847-DEL-58
chr9-43588331-DEL-62
chr9-43588534-DEL-192
chr9-43780218-DEL-54
chr9-43794252-DEL-60
chr9-43839528-DEL-76

chr9-43854324-DEL-54
chr9-43930486-DEL-66
chr9-43931874-DEL-288
chr9-43989271-DEL-575
chr9-44025673-DEL-50
chr9-44100342-DEL-77
chr9-44344024-DEL-53
chr9-44368136-DEL-64
chr9-44370419-DEL-50
chr9-44415989-DEL-54
chr9-44450009-DEL-68
chr9-44481035-DEL-66
chr9-44566027-DEL-430
chr9-44775195-DEL-174
chr9-44814019-DEL-238
chr9-44932509-DEL-92
chr9-44998568-DEL-209
chr9-45133988-DEL-111
chr9-45304364-DEL-69
chr9-45385900-DEL-50
chr9-45641160-DEL-84
chr9-45687686-DEL-97
chr9-45701211-DEL-53
chr9-45701448-DEL-200
chr9-45893175-DEL-93
chr9-45971470-DEL-1388
chr9-46044265-DEL-444
chr9-46048800-DEL-238
chr9-46103340-DEL-52
chr9-46104192-DEL-123
chr9-46158140-DEL-103
chr9-46158454-DEL-56
chr9-46189942-DEL-70
chr9-46242433-DEL-142
chr9-46250873-DEL-963
chr9-46251251-DEL-242
chr9-46268976-DEL-120
chr9-46273563-DEL-104
chr9-46278792-DEL-102
chr9-46284165-DEL-324
chr9-46325892-DEL-176
chr9-46330561-DEL-72
chr9-46382288-DEL-72
chr9-46408098-DEL-52
chr9-46419220-DEL-95
chr9-46457593-DEL-50
chr9-46563234-DEL-147
chr9-46563292-DEL-455
chr9-46606912-DEL-1240
chr9-46613363-DEL-56
chr9-46912732-DEL-54
chr9-47142024-DEL-125
chr9-47209810-DEL-147
chr9-47210388-DEL-400
chr9-47480400-DEL-110
chr9-47588100-DEL-250
chr9-47608470-DEL-50
chr9-47753837-DEL-72
chr9-47753917-DEL-212
chr9-47901017-DEL-3374
chr9-48121164-DEL-54
chr9-48168003-DEL-62
chr9-48226562-DEL-476
chr9-48275447-DEL-81
chr9-48278901-DEL-159
chr9-48282931-DEL-54
chr9-48282949-DEL-337
chr9-48283149-DEL-137
chr9-48289881-DEL-2748
chr9-48301874-DEL-224
chr9-48322196-DEL-160
chr9-48486132-DEL-401
chr9-48522104-DEL-145
chr9-48591183-DEL-169
chr9-48803516-DEL-74
chr9-48865138-DEL-190
chr9-49145797-DEL-56

chr9-49206030-DEL-183
chr9-49237041-DEL-208
chr9-49465287-DEL-328
chr9-49682434-DEL-82
chr9-49731732-DEL-527
chr9-49732590-DEL-151
chr9-49766810-DEL-180
chr9-49769777-DEL-79
chr9-49856178-DEL-50
chr9-49863558-DEL-150
chr9-49916545-DEL-96
chr9-50093606-DEL-125
chr9-50262584-DEL-100
chr9-50309217-DEL-88
chr9-50331562-DEL-103
chr9-50376592-DEL-751
chr9-50718097-DEL-68
chr9-50813126-DEL-124
chr9-50834810-DEL-142
chr9-50835074-DEL-93
chr9-50838194-DEL-3501
chr9-51039352-DEL-64
chr9-51285400-DEL-56
chr9-51285504-DEL-64
chr9-51535275-DEL-363
chr9-51654326-DEL-104
chr9-51658538-DEL-54
chr9-51681113-DEL-756
chr9-51681689-DEL-360
chr9-51798017-DEL-61
chr9-51815107-DEL-92
chr9-51815246-DEL-139
chr9-51849393-DEL-942
chr9-51893637-DEL-55
chr9-51897862-DEL-639
chr9-51946248-DEL-70
chr9-51946597-DEL-285
chr9-51982465-DEL-172
chr9-51982981-DEL-62
chr9-52100523-DEL-1546
chr9-52119089-DEL-471
chr9-52119656-DEL-361
chr9-52197460-DEL-461
chr9-52215966-DEL-319
chr9-52216335-DEL-133
chr9-52265989-DEL-51
chr9-52418728-DEL-52
chr9-52503432-DEL-169
chr9-52614734-DEL-121
chr9-52660212-DEL-52
chr9-52684927-DEL-51
chr9-52711015-DEL-191
chr9-52801333-DEL-11927
chr9-52969333-DEL-646
chr9-52997566-DEL-143
chr9-53106427-DEL-7423
chr9-53237935-DEL-163
chr9-53305839-DEL-1286
chr9-53380653-DEL-78
chr9-53442159-DEL-53
chr9-53442238-DEL-171
chr9-53499221-DEL-68
chr9-53507251-DEL-118
chr9-53561122-DEL-254
chr9-53629356-DEL-182
chr9-53664604-DEL-2565
chr9-53687082-DEL-200
chr9-53687332-DEL-90
chr9-53776535-DEL-1549
chr9-53918533-DEL-214
chr9-54022252-DEL-50
chr9-54023406-DEL-288
chr9-54235086-DEL-121
chr9-54310060-DEL-50
chr9-54390649-DEL-657
chr9-54395246-DEL-400
chr9-54455058-DEL-150

chr9-54464741-DEL-60
chr9-54648701-DEL-140
chr9-54650790-DEL-69
chr9-54664814-DEL-126
chr9-54822240-DEL-152
chr9-54950672-DEL-121
chr9-54963522-DEL-54
chr9-55175190-DEL-319
chr9-55207642-DEL-153
chr9-55284143-DEL-62
chr9-55616345-DEL-909
chr9-55833980-DEL-128
chr9-55913892-DEL-98
chr9-56360098-DEL-58
chr9-56433959-DEL-188
chr9-56452329-DEL-60
chr9-56465696-DEL-84
chr9-56511287-DEL-216
chr9-56553083-DEL-54
chr9-56661344-DEL-57
chr9-56888603-DEL-134
chr9-56915123-DEL-93
chr9-57095518-DEL-108
chr9-57291268-DEL-58
chr9-57356311-DEL-65
chr9-57390231-DEL-812
chr9-57503175-DEL-52
chr9-57722469-DEL-85
chr9-57725302-DEL-51
chr9-57790567-DEL-67
chr9-57792995-DEL-850
chr9-57929779-DEL-469
chr9-57947628-DEL-78
chr9-57959360-DEL-146
chr9-58085483-DEL-53
chr9-58209149-DEL-191
chr9-58446606-DEL-52
chr9-58506527-DEL-50
chr9-58583882-DEL-85
chr9-58669014-DEL-52
chr9-58803033-DEL-686
chr9-58803051-DEL-286
chr9-59086545-DEL-60
chr9-59114496-DEL-465
chr9-59114985-DEL-116
chr9-59189694-DEL-68
chr9-59420266-DEL-127
chr9-59577149-DEL-189
chr9-59625142-DEL-98
chr9-59625142-DEL-294
chr9-59724505-DEL-91
chr9-60423597-DEL-50
chr9-60446410-DEL-990
chr9-60447747-DEL-52
chr9-60488986-DEL-50
chr9-60494549-DEL-50
chr9-60525485-DEL-239
chr9-60571795-DEL-131
chr9-60572732-DEL-57
chr9-60697308-DEL-64
chr9-60769447-DEL-120
chr9-60783233-DEL-76
chr9-60785204-DEL-50
chr9-60851832-DEL-50
chr9-60874282-DEL-51
chr9-61035187-DEL-148
chr9-61060501-DEL-72
chr9-61089583-DEL-461
chr9-61134196-DEL-536
chr9-61161392-DEL-580
chr9-61261621-DEL-551
chr9-61361971-DEL-61
chr9-61386024-DEL-56
chr9-61638751-DEL-390
chr9-61651976-DEL-103
chr9-61666249-DEL-222
chr9-61675501-DEL-137

chr9-61711921-DEL-568
chr9-61850422-DEL-582
chr9-61908583-DEL-114
chr9-62053648-DEL-65
chr9-62161921-DEL-60
chr9-62217762-DEL-61
chr9-62237294-DEL-499
chr9-62259976-DEL-63
chr9-62301855-DEL-56
chr9-62356767-DEL-2933
chr9-62419812-DEL-216
chr9-62453090-DEL-70
chr9-62453272-DEL-52
chr9-62512802-DEL-4906
chr9-62542152-DEL-401
chr9-62596490-DEL-62
chr9-62782014-DEL-151
chr9-62933302-DEL-173
chr9-63077834-DEL-50
chr9-63154091-DEL-50
chr9-63265663-DEL-430
chr9-63265741-DEL-117
chr9-63265898-DEL-77
chr9-63337556-DEL-65
chr9-63411647-DEL-523
chr9-63460243-DEL-378
chr9-63498244-DEL-384
chr9-63639370-DEL-82
chr9-63683060-DEL-87
chr9-63721597-DEL-64
chr9-63766731-DEL-58
chr9-63853146-DEL-2075
chr9-63933591-DEL-187
chr9-63971349-DEL-253
chr9-64013628-DEL-171
chr9-64029813-DEL-1529
chr9-64185434-DEL-166
chr9-64193263-DEL-257
chr9-64235946-DEL-110
chr9-64236192-DEL-86
chr9-64259715-DEL-50
chr9-64358328-DEL-525
chr9-64404673-DEL-66
chr9-64579079-DEL-170
chr9-64650591-DEL-72
chr9-64676213-DEL-189
chr9-64740990-DEL-117
chr9-64741221-DEL-96
chr9-64749067-DEL-50
chr9-64757323-DEL-51
chr9-64789887-DEL-68
chr9-64790075-DEL-173
chr9-64995489-DEL-147
chr9-65220725-DEL-179
chr9-65239947-DEL-208
chr9-65244680-DEL-261
chr9-65255835-DEL-420
chr9-65255848-DEL-80
chr9-65302330-DEL-120
chr9-65323707-DEL-93
chr9-65395333-DEL-55
chr9-65577246-DEL-182
chr9-65776176-DEL-1143
chr9-65776499-DEL-66
chr9-65776701-DEL-134
chr9-65776779-DEL-338
chr9-65954765-DEL-55
chr9-66084252-DEL-50
chr9-66144347-DEL-87
chr9-66166501-DEL-371
chr9-66167201-DEL-377
chr9-66226828-DEL-63
chr9-66242003-DEL-191
chr9-66242312-DEL-198
chr9-66556124-DEL-62
chr9-66556236-DEL-116
chr9-66581500-DEL-50

chr9-66596171-DEL-84
chr9-66784938-DEL-190
chr9-66785073-DEL-57
chr9-66940509-DEL-50
chr9-66965382-DEL-156
chr9-66970108-DEL-192
chr9-66970228-DEL-72
chr9-67044860-DEL-1635
chr9-67075358-DEL-54
chr9-67342680-DEL-3532
chr9-67346148-DEL-56
chr9-67347422-DEL-470
chr9-67348036-DEL-414
chr9-67348067-DEL-177
chr9-67348766-DEL-96
chr9-67405501-DEL-217
chr9-67459157-DEL-523
chr9-67459505-DEL-94
chr9-67490501-DEL-134
chr9-67569084-DEL-87
chr9-67583325-DEL-70
chr9-67678259-DEL-52
chr9-67700406-DEL-152
chr9-67915037-DEL-787
chr9-68061427-DEL-212
chr9-68061482-DEL-105
chr9-68162436-DEL-66
chr9-68216071-DEL-61
chr9-68360199-DEL-3513
chr9-68401252-DEL-6298
chr9-68554747-DEL-87
chr9-68578772-DEL-85
chr9-68665466-DEL-50
chr9-68720252-DEL-117
chr9-68923541-DEL-70
chr9-68923746-DEL-148
chr9-68929379-DEL-50
chr9-68981897-DEL-910
chr9-69007167-DEL-65
chr9-69010984-DEL-68
chr9-69027438-DEL-108
chr9-69148375-DEL-75
chr9-69148450-DEL-161
chr9-69262692-DEL-584
chr9-69299845-DEL-56
chr9-69347582-DEL-67
chr9-69488589-DEL-71
chr9-69506762-DEL-134
chr9-69514232-DEL-92
chr9-69542448-DEL-75
chr9-69639386-DEL-74
chr9-69639392-DEL-256
chr9-69767578-DEL-130
chr9-69767659-DEL-63
chr9-69800369-DEL-2277
chr9-69846505-DEL-59
chr9-69988764-DEL-50
chr9-70009353-DEL-104
chr9-70091303-DEL-175
chr9-70098897-DEL-2821
chr9-70163308-DEL-175
chr9-70174231-DEL-96
chr9-70179067-DEL-938
chr9-70244425-DEL-500
chr9-70530263-DEL-452
chr9-70543518-DEL-112
chr9-70553604-DEL-155
chr9-70650684-DEL-50
chr9-70660473-DEL-54
chr9-70742553-DEL-87
chr9-70751677-DEL-70
chr9-70895679-DEL-199
chr9-70895746-DEL-429
chr9-70896029-DEL-912
chr9-70960378-DEL-52
chr9-71202277-DEL-72
chr9-71205430-DEL-73

chr9-71230991-DEL-1265
chr9-71273528-DEL-160
chr9-71352522-DEL-51
chr9-71365518-DEL-68
chr9-71379645-DEL-56
chr9-71396034-DEL-366
chr9-71500777-DEL-61
chr9-71510412-DEL-89
chr9-71514938-DEL-524
chr9-71589659-DEL-102
chr9-71658226-DEL-60
chr9-71707756-DEL-153
chr9-71707769-DEL-70
chr9-71713566-DEL-69
chr9-72024016-DEL-287
chr9-72086549-DEL-148
chr9-72170605-DEL-53
chr9-72174634-DEL-5188
chr9-72199616-DEL-218
chr9-72290395-DEL-56
chr9-72380743-DEL-127
chr9-72425808-DEL-231
chr9-72535634-DEL-214
chr9-72553743-DEL-82
chr9-72575240-DEL-202
chr9-72662485-DEL-116
chr9-72673603-DEL-182
chr9-72730188-DEL-50
chr9-72759505-DEL-85
chr9-72842324-DEL-179
chr9-72849211-DEL-62
chr9-72966811-DEL-83
chr9-72970065-DEL-184
chr9-72974245-DEL-68
chr9-73064414-DEL-324
chr9-73151263-DEL-52
chr9-73189657-DEL-50
chr9-73195132-DEL-1614
chr9-73264658-DEL-1627
chr9-73455332-DEL-72
chr9-73543846-DEL-162
chr9-73561128-DEL-101
chr9-73605064-DEL-63
chr9-73715846-DEL-261
chr9-73715856-DEL-74
chr9-73732036-DEL-208
chr9-73761183-DEL-108
chr9-73778821-DEL-177
chr9-73832309-DEL-64
chr9-73953525-DEL-70
chr9-73963528-DEL-68
chr9-73977778-DEL-85
chr9-74066444-DEL-50
chr9-74153336-DEL-1014
chr9-74402735-DEL-109
chr9-74402952-DEL-164
chr9-74487106-DEL-162
chr9-74487268-DEL-65
chr9-74488703-DEL-50
chr9-74495766-DEL-70
chr9-74507930-DEL-283
chr9-74598143-DEL-75
chr9-74611917-DEL-59
chr9-74617065-DEL-348
chr9-74617173-DEL-129
chr9-74624378-DEL-138
chr9-74627704-DEL-67
chr9-74720206-DEL-73
chr9-74741087-DEL-424
chr9-75107838-DEL-86
chr9-75220504-DEL-76
chr9-75363691-DEL-317
chr9-75428479-DEL-60
chr9-75434384-DEL-172
chr9-75483090-DEL-50
chr9-75740362-DEL-145
chr9-75774974-DEL-632

chr9-75798444-DEL-335
chr9-76018685-DEL-68
chr9-76084935-DEL-52
chr9-76199887-DEL-798
chr9-76200795-DEL-141
chr9-76407763-DEL-100
chr9-76449952-DEL-158
chr9-76529278-DEL-133
chr9-76529583-DEL-241
chr9-76589759-DEL-228
chr9-76652264-DEL-541
chr9-76652681-DEL-533
chr9-76744455-DEL-938
chr9-76785175-DEL-537
chr9-76942271-DEL-86
chr9-77035253-DEL-123
chr9-77035411-DEL-90
chr9-77059621-DEL-130
chr9-77092598-DEL-2386
chr9-77144283-DEL-114
chr9-77145380-DEL-89
chr9-77205756-DEL-202
chr9-77205762-DEL-67
chr9-77347307-DEL-173
chr9-77367912-DEL-142
chr9-77439163-DEL-774
chr9-77439216-DEL-200
chr9-77439482-DEL-320
chr9-77439962-DEL-174
chr9-77462321-DEL-53
chr9-77479119-DEL-1633
chr9-77489503-DEL-140
chr9-77515415-DEL-662
chr9-77529883-DEL-54
chr9-77573344-DEL-152
chr9-77591695-DEL-64
chr9-77591696-DEL-320
chr9-77591952-DEL-64
chr9-77628236-DEL-132
chr9-77643289-DEL-502
chr9-77724947-DEL-145
chr9-77840347-DEL-147
chr9-77925688-DEL-62
chr9-77925718-DEL-146
chr9-77940635-DEL-72
chr9-77985504-DEL-50
chr9-78136649-DEL-4586
chr9-78164602-DEL-73
chr9-78181490-DEL-16302
chr9-78213089-DEL-62
chr9-78287826-DEL-88
chr9-78352604-DEL-82
chr9-78394036-DEL-541
chr9-78418395-DEL-181
chr9-78455239-DEL-50
chr9-78501583-DEL-103
chr9-78547027-DEL-242
chr9-78595229-DEL-180
chr9-78698811-DEL-50
chr9-78711473-DEL-1053
chr9-78774863-DEL-2401
chr9-78973340-DEL-2759
chr9-79045341-DEL-58
chr9-79075986-DEL-56
chr9-79213898-DEL-52
chr9-79287485-DEL-50
chr9-79310156-DEL-204
chr9-79342016-DEL-119
chr9-79346463-DEL-59
chr9-79393988-DEL-4867
chr9-79445818-DEL-3137
chr9-79468592-DEL-61
chr9-79532412-DEL-54
chr9-79545010-DEL-50
chr9-79631450-DEL-197
chr9-79631484-DEL-94
chr9-79860574-DEL-66

chr9-80117797-DEL-66
chr9-80129060-DEL-222
chr9-80170445-DEL-63
chr9-80431660-DEL-62
chr9-80467623-DEL-1790
chr9-80586550-DEL-66
chr9-80616313-DEL-54
chr9-80718921-DEL-208
chr9-80810318-DEL-50
chr9-80825766-DEL-3209
chr9-80859756-DEL-50
chr9-80862238-DEL-87
chr9-80988927-DEL-132
chr9-80992964-DEL-65
chr9-81007900-DEL-67
chr9-81031335-DEL-132
chr9-81048860-DEL-64
chr9-81267529-DEL-819
chr9-81272878-DEL-5977
chr9-81282786-DEL-96
chr9-81314745-DEL-16398
chr9-81336844-DEL-88
chr9-81453991-DEL-58
chr9-81559577-DEL-1165
chr9-81595421-DEL-140
chr9-81605793-DEL-1174
chr9-81678269-DEL-56
chr9-81893837-DEL-50
chr9-81900516-DEL-52
chr9-82022569-DEL-3298
chr9-82069509-DEL-190
chr9-82102997-DEL-999
chr9-82146825-DEL-58
chr9-82198585-DEL-56
chr9-82276460-DEL-62
chr9-82562905-DEL-486
chr9-82605479-DEL-90
chr9-82616546-DEL-59
chr9-82687249-DEL-74
chr9-82692667-DEL-484
chr9-82878781-DEL-133
chr9-82901696-DEL-2367
chr9-82972444-DEL-156
chr9-83093067-DEL-880
chr9-83294799-DEL-75
chr9-83407335-DEL-60
chr9-83466980-DEL-330
chr9-83487303-DEL-70
chr9-83523273-DEL-273
chr9-83589070-DEL-146
chr9-83642940-DEL-158
chr9-83877600-DEL-2605
chr9-83906652-DEL-393
chr9-83921500-DEL-80
chr9-83929698-DEL-384
chr9-84010909-DEL-66
chr9-84055121-DEL-2544
chr9-84254481-DEL-212
chr9-84498168-DEL-1185
chr9-84505787-DEL-50
chr9-84723991-DEL-50
chr9-84726281-DEL-172
chr9-85085210-DEL-138
chr9-85140044-DEL-2783
chr9-85148264-DEL-641
chr9-85253697-DEL-60
chr9-85317166-DEL-456
chr9-85390125-DEL-62
chr9-85393432-DEL-165
chr9-85445647-DEL-66
chr9-85508235-DEL-428
chr9-85530232-DEL-56
chr9-85547275-DEL-1870
chr9-85635324-DEL-2293
chr9-85637740-DEL-72
chr9-85638300-DEL-151
chr9-85716717-DEL-805

chr9-85778100-DEL-77
chr9-85846924-DEL-203
chr9-85851837-DEL-115
chr9-85934643-DEL-77
chr9-86154289-DEL-339
chr9-86207369-DEL-203
chr9-86223849-DEL-162
chr9-86303842-DEL-70
chr9-86313188-DEL-67
chr9-86386901-DEL-54
chr9-86400510-DEL-122
chr9-86410811-DEL-2120
chr9-86470579-DEL-72
chr9-86479323-DEL-293
chr9-86516380-DEL-56
chr9-86524562-DEL-360
chr9-86723822-DEL-181
chr9-86950988-DEL-350
chr9-87029150-DEL-228
chr9-87029196-DEL-58
chr9-87035308-DEL-190
chr9-87087444-DEL-629
chr9-87157077-DEL-681
chr9-87238902-DEL-63
chr9-87246151-DEL-53
chr9-87294350-DEL-109
chr9-87294656-DEL-75
chr9-87305850-DEL-56
chr9-87355817-DEL-1779
chr9-87498111-DEL-60
chr9-87679735-DEL-1493
chr9-87825271-DEL-57
chr9-87897976-DEL-122
chr9-87898161-DEL-571
chr9-87918464-DEL-50
chr9-87992337-DEL-287
chr9-88026170-DEL-129
chr9-88028642-DEL-4799
chr9-88043733-DEL-149
chr9-88070857-DEL-1060
chr9-88106533-DEL-166
chr9-88148650-DEL-148
chr9-88388551-DEL-99
chr9-88475473-DEL-676
chr9-88484544-DEL-345
chr9-88513305-DEL-255
chr9-88534123-DEL-145
chr9-88591136-DEL-744
chr9-88669218-DEL-2301
chr9-88676742-DEL-231
chr9-88676766-DEL-115
chr9-88727387-DEL-52
chr9-88856040-DEL-105
chr9-88895912-DEL-1284
chr9-88997431-DEL-54
chr9-89017678-DEL-307
chr9-89017966-DEL-76
chr9-89023158-DEL-1206
chr9-89145614-DEL-1253
chr9-89161964-DEL-13106
chr9-89220719-DEL-54
chr9-89251478-DEL-81
chr9-89257078-DEL-332
chr9-89289316-DEL-2876
chr9-89372209-DEL-60
chr9-89385475-DEL-769
chr9-89432918-DEL-90
chr9-89681609-DEL-1368
chr9-89748304-DEL-97
chr9-89762134-DEL-116
chr9-89763333-DEL-148
chr9-89844271-DEL-986
chr9-89844279-DEL-161
chr9-89844651-DEL-311
chr9-89960838-DEL-347
chr9-90013264-DEL-183
chr9-90013304-DEL-53

chr9-90030609-DEL-1379
chr9-90033568-DEL-101
chr9-90058785-DEL-84
chr9-90139179-DEL-482
chr9-90164825-DEL-70
chr9-90230784-DEL-649
chr9-90247887-DEL-52
chr9-90260251-DEL-396
chr9-90288246-DEL-70
chr9-90324555-DEL-60
chr9-90332797-DEL-52
chr9-90395333-DEL-133
chr9-90422532-DEL-467
chr9-90423830-DEL-2496
chr9-90431211-DEL-54
chr9-90441885-DEL-100
chr9-90502779-DEL-54
chr9-90547523-DEL-343
chr9-90783250-DEL-88
chr9-90963753-DEL-924
chr9-90964094-DEL-64
chr9-90966569-DEL-112
chr9-91150260-DEL-1593
chr9-91301634-DEL-56
chr9-91336991-DEL-50
chr9-91342503-DEL-136
chr9-91381804-DEL-52
chr9-91399557-DEL-50
chr9-91488028-DEL-120
chr9-91537802-DEL-120
chr9-91675112-DEL-1487
chr9-91757331-DEL-56
chr9-91803278-DEL-66
chr9-92105298-DEL-169
chr9-92105337-DEL-68
chr9-92105557-DEL-344
chr9-92138462-DEL-208
chr9-92138642-DEL-64
chr9-92215207-DEL-473
chr9-92313783-DEL-332
chr9-92314020-DEL-367
chr9-92344952-DEL-61
chr9-92457426-DEL-122
chr9-92523280-DEL-52
chr9-92558996-DEL-89
chr9-92566539-DEL-754
chr9-92576165-DEL-221
chr9-92611612-DEL-52
chr9-92645303-DEL-50
chr9-92773620-DEL-1024
chr9-93241096-DEL-56
chr9-93263758-DEL-692
chr9-93264815-DEL-1119
chr9-93451794-DEL-976
chr9-93521848-DEL-50
chr9-93574396-DEL-79
chr9-93604549-DEL-60
chr9-93652340-DEL-219
chr9-93776597-DEL-67
chr9-93855171-DEL-101
chr9-93878449-DEL-1169
chr9-93879679-DEL-288
chr9-93982389-DEL-83
chr9-93997589-DEL-55
chr9-94153271-DEL-167
chr9-94196994-DEL-201
chr9-94245273-DEL-320
chr9-94329438-DEL-291
chr9-94394978-DEL-1465
chr9-94463287-DEL-72
chr9-94466728-DEL-50
chr9-94478501-DEL-159
chr9-94534085-DEL-153
chr9-94576235-DEL-174
chr9-94590251-DEL-137
chr9-94719691-DEL-80
chr9-94761895-DEL-50

chr9-94769259-DEL-132
chr9-94777782-DEL-52
chr9-94780860-DEL-357
chr9-94872909-DEL-78
chr9-94886948-DEL-1703
chr9-94903239-DEL-1602
chr9-94962498-DEL-52
chr9-94991697-DEL-71
chr9-95022381-DEL-51
chr9-95054746-DEL-64
chr9-95151260-DEL-201
chr9-95237737-DEL-76
chr9-95281251-DEL-54
chr9-95313208-DEL-342
chr9-95316309-DEL-50
chr9-95450542-DEL-212
chr9-95505552-DEL-506
chr9-95536113-DEL-87
chr9-95558557-DEL-276
chr9-95648438-DEL-215
chr9-95659492-DEL-70
chr9-95669381-DEL-103
chr9-95708431-DEL-280
chr9-95906892-DEL-53
chr9-95959807-DEL-81
chr9-95960080-DEL-181
chr9-96147523-DEL-249
chr9-96260203-DEL-151
chr9-96273325-DEL-54
chr9-96292368-DEL-714
chr9-96300312-DEL-54
chr9-96338213-DEL-50
chr9-96370519-DEL-177
chr9-96398987-DEL-447
chr9-96452035-DEL-256
chr9-96461079-DEL-102
chr9-96675015-DEL-84
chr9-96752104-DEL-83
chr9-96755887-DEL-72
chr9-96756962-DEL-50
chr9-96807921-DEL-70
chr9-96818485-DEL-66
chr9-96882640-DEL-128
chr9-96886373-DEL-1213
chr9-96965659-DEL-132
chr9-96965723-DEL-330
chr9-96999816-DEL-195
chr9-97000105-DEL-96
chr9-97034769-DEL-682
chr9-97058695-DEL-174
chr9-97059249-DEL-66
chr9-97102675-DEL-67
chr9-97154756-DEL-52
chr9-97244252-DEL-111
chr9-97375324-DEL-114
chr9-97583092-DEL-58
chr9-97781792-DEL-62
chr9-97835632-DEL-263
chr9-97981247-DEL-72
chr9-98057744-DEL-344
chr9-98260757-DEL-70
chr9-98264068-DEL-52
chr9-98390862-DEL-176
chr9-98405237-DEL-60
chr9-98405988-DEL-52
chr9-98408351-DEL-52
chr9-98408557-DEL-65
chr9-98503841-DEL-90
chr9-98559653-DEL-5950
chr9-98598653-DEL-749
chr9-98615224-DEL-257
chr9-98648471-DEL-5201
chr9-98671379-DEL-176
chr9-98737749-DEL-108
chr9-98770694-DEL-1796
chr9-98828388-DEL-72
chr9-98953950-DEL-146

chr9-98965970-DEL-50
chr9-98966006-DEL-127
chr9-99157045-DEL-59
chr9-99376330-DEL-84
chr9-99382311-DEL-72
chr9-99427802-DEL-140
chr9-99568039-DEL-464
chr9-99578863-DEL-60
chr9-99586216-DEL-195
chr9-99586302-DEL-68
chr9-99601282-DEL-54
chr9-99684044-DEL-70
chr9-99692754-DEL-50
chr9-99703442-DEL-634
chr9-99724149-DEL-122
chr9-99921649-DEL-109
chr9-99921809-DEL-284
chr9-99964959-DEL-54
chr9-99988183-DEL-203
chr9-100000101-DEL-109
chr9-100005595-DEL-74
chr9-100100946-DEL-525
chr9-100244036-DEL-157
chr9-100258734-DEL-72
chr9-100324569-DEL-232
chr9-100324831-DEL-58
chr9-100345659-DEL-71
chr9-100383707-DEL-63
chr9-100486907-DEL-52
chr9-100685428-DEL-7128
chr9-100788990-DEL-100
chr9-100842539-DEL-1755
chr9-100844661-DEL-849
chr9-100939739-DEL-88
chr9-100939781-DEL-550
chr9-100940223-DEL-82
chr9-100940429-DEL-412
chr9-100940584-DEL-176
chr9-100967718-DEL-780
chr9-100998779-DEL-62
chr9-101103395-DEL-1405
chr9-101197616-DEL-104
chr9-101287216-DEL-104
chr9-101287256-DEL-266
chr9-101295991-DEL-50
chr9-101325590-DEL-102
chr9-101367175-DEL-76
chr9-101370456-DEL-62
chr9-101429299-DEL-50
chr9-101457199-DEL-68
chr9-101457713-DEL-60
chr9-101510223-DEL-58
chr9-102177262-DEL-50
chr9-102264463-DEL-594
chr9-102264490-DEL-108
chr9-102309122-DEL-70
chr9-102453273-DEL-50
chr9-102523114-DEL-52
chr9-102523497-DEL-62
chr9-102694022-DEL-117
chr9-102732466-DEL-182
chr9-102732544-DEL-390
chr9-102795318-DEL-118
chr9-102839957-DEL-86
chr9-102899432-DEL-51
chr9-102931181-DEL-117
chr9-102980286-DEL-52
chr9-103002978-DEL-65
chr9-103163248-DEL-83
chr9-103172944-DEL-190
chr9-103173781-DEL-3840
chr9-103173794-DEL-1344
chr9-103174178-DEL-624
chr9-103177058-DEL-720
chr9-103201747-DEL-50
chr9-103258663-DEL-899
chr9-103268372-DEL-52

chr9-103333644-DEL-452
chr9-103374678-DEL-54
chr9-103374732-DEL-376
chr9-103521793-DEL-56
chr9-103529543-DEL-50
chr9-103557272-DEL-52
chr9-103724927-DEL-58
chr9-103888414-DEL-51
chr9-103903579-DEL-50
chr9-103924188-DEL-67
chr9-104078565-DEL-72
chr9-104092350-DEL-66
chr9-104229354-DEL-74
chr9-104232490-DEL-50
chr9-104243222-DEL-52
chr9-104382288-DEL-163
chr9-104399156-DEL-57
chr9-104401453-DEL-214
chr9-104441484-DEL-96
chr9-104509710-DEL-52
chr9-104509993-DEL-130
chr9-104733867-DEL-54
chr9-104802099-DEL-123
chr9-104803323-DEL-1842
chr9-104807451-DEL-56
chr9-104833051-DEL-52
chr9-104842928-DEL-768
chr9-104858530-DEL-2472
chr9-104863117-DEL-312
chr9-104938368-DEL-94
chr9-105053549-DEL-301
chr9-105053668-DEL-113
chr9-105084820-DEL-59
chr9-105085192-DEL-87
chr9-105137103-DEL-56
chr9-105158013-DEL-840
chr9-105211673-DEL-208
chr9-105237524-DEL-50
chr9-105318961-DEL-68
chr9-105332859-DEL-414
chr9-105346024-DEL-91
chr9-105357037-DEL-50
chr9-105366004-DEL-52
chr9-105416826-DEL-121
chr9-105442526-DEL-75
chr9-105462827-DEL-620
chr9-105702256-DEL-60
chr9-105860469-DEL-77
chr9-105871253-DEL-1273
chr9-105989071-DEL-263
chr9-105989501-DEL-56
chr9-106141267-DEL-193
chr9-106149155-DEL-260
chr9-106293454-DEL-97
chr9-106380054-DEL-162
chr9-106435494-DEL-90
chr9-106513963-DEL-86
chr9-106748540-DEL-172
chr9-106853107-DEL-446
chr9-106975617-DEL-170
chr9-107121463-DEL-700
chr9-107183780-DEL-88
chr9-107250582-DEL-69
chr9-107630509-DEL-56
chr9-107760847-DEL-58
chr9-107952777-DEL-51
chr9-107953259-DEL-90
chr9-108010286-DEL-1130
chr9-108011052-DEL-134
chr9-108011510-DEL-544
chr9-108044782-DEL-60
chr9-108104197-DEL-259
chr9-108118065-DEL-965
chr9-108118128-DEL-60
chr9-108118208-DEL-450
chr9-108118553-DEL-124
chr9-108119230-DEL-81

chr9-108142942-DEL-239
chr9-108153516-DEL-50
chr9-108169857-DEL-91
chr9-108220876-DEL-2895
chr9-108223804-DEL-218
chr9-108308671-DEL-50
chr9-108422626-DEL-85
chr9-108478614-DEL-53
chr9-108483082-DEL-50
chr9-108572766-DEL-359
chr9-108580025-DEL-145
chr9-108740364-DEL-55
chr9-108740796-DEL-91
chr9-108826091-DEL-473
chr9-108848251-DEL-2103
chr9-108848262-DEL-64
chr9-108848496-DEL-527
chr9-108849807-DEL-616
chr9-109027321-DEL-428
chr9-109086411-DEL-215
chr9-109109982-DEL-107859
chr9-109200290-DEL-96
chr9-109372998-DEL-1654
chr9-109427091-DEL-50
chr9-109663272-DEL-58
chr9-109672368-DEL-466
chr9-109693610-DEL-153
chr9-109782719-DEL-66
chr9-110001019-DEL-136
chr9-110092013-DEL-590
chr9-110099787-DEL-123
chr9-110142111-DEL-129
chr9-110146786-DEL-185
chr9-110147539-DEL-668
chr9-110273414-DEL-545
chr9-110273521-DEL-80
chr9-110274274-DEL-1310
chr9-110274412-DEL-239
chr9-110274733-DEL-376
chr9-110274993-DEL-125
chr9-110279369-DEL-72
chr9-110534168-DEL-409
chr9-110550615-DEL-50
chr9-110676754-DEL-1269
chr9-110725775-DEL-50
chr9-110732310-DEL-3995
chr9-110765654-DEL-244
chr9-110808180-DEL-192
chr9-110815512-DEL-186
chr9-110934393-DEL-52
chr9-111005465-DEL-114
chr9-111082685-DEL-82
chr9-111082783-DEL-582
chr9-111082971-DEL-166
chr9-111083447-DEL-84
chr9-111324059-DEL-228
chr9-111343564-DEL-124
chr9-111415922-DEL-726
chr9-111471140-DEL-1017
chr9-111683081-DEL-270
chr9-111690334-DEL-134
chr9-111890881-DEL-555
chr9-111890947-DEL-66
chr9-111915798-DEL-55
chr9-111915917-DEL-295
chr9-111935139-DEL-91
chr9-112013822-DEL-73
chr9-112015450-DEL-55
chr9-112228706-DEL-102
chr9-112245138-DEL-3801
chr9-112267402-DEL-64
chr9-112274614-DEL-196
chr9-112304730-DEL-165
chr9-112416990-DEL-82
chr9-112430139-DEL-464
chr9-112492049-DEL-320
chr9-112517637-DEL-50

chr9-112636648-DEL-52
chr9-112684538-DEL-541
chr9-112828015-DEL-1313
chr9-112927924-DEL-52
chr9-112933683-DEL-76
chr9-112934044-DEL-62
chr9-112934165-DEL-228
chr9-112969947-DEL-192
chr9-112972404-DEL-1309
chr9-113022217-DEL-572
chr9-113057195-DEL-75
chr9-113107384-DEL-60
chr9-113133890-DEL-377
chr9-113242392-DEL-84
chr9-113312032-DEL-56
chr9-113322426-DEL-82
chr9-113327540-DEL-96
chr9-113380520-DEL-52
chr9-113392470-DEL-54
chr9-113545858-DEL-76
chr9-113723579-DEL-66
chr9-113742163-DEL-52
chr9-113752287-DEL-138
chr9-113826391-DEL-307
chr9-113883835-DEL-215
chr9-113940934-DEL-193
chr9-113969926-DEL-7995
chr9-113998460-DEL-174
chr9-114044167-DEL-264
chr9-114079355-DEL-380
chr9-114125323-DEL-393
chr9-114241501-DEL-173
chr9-114251442-DEL-134
chr9-114314907-DEL-64
chr9-114347736-DEL-187
chr9-114362535-DEL-210
chr9-114395188-DEL-236
chr9-114395297-DEL-1735
chr9-114399115-DEL-53
chr9-114401393-DEL-2301
chr9-114402959-DEL-79
chr9-114481069-DEL-566
chr9-114564953-DEL-221
chr9-114575099-DEL-130
chr9-114584967-DEL-70
chr9-114662444-DEL-54
chr9-114685667-DEL-717
chr9-114782106-DEL-960
chr9-114816961-DEL-145
chr9-114822582-DEL-188
chr9-114822806-DEL-280
chr9-114849836-DEL-56
chr9-114855708-DEL-1041
chr9-114891095-DEL-340
chr9-114903183-DEL-139
chr9-114919977-DEL-266
chr9-114920402-DEL-58
chr9-114971365-DEL-119
chr9-114990192-DEL-148
chr9-114993911-DEL-236
chr9-115045393-DEL-109
chr9-115075638-DEL-70
chr9-115168489-DEL-103
chr9-115181108-DEL-70
chr9-115202456-DEL-1533
chr9-115206674-DEL-59
chr9-115222622-DEL-58
chr9-115225658-DEL-62
chr9-115229115-DEL-58
chr9-115229121-DEL-380
chr9-115229257-DEL-189
chr9-115240313-DEL-194
chr9-115326491-DEL-860
chr9-115382538-DEL-50
chr9-115407260-DEL-70
chr9-115425125-DEL-57
chr9-115429674-DEL-88

chr9-115451347-DEL-50
chr9-115489483-DEL-186
chr9-115549936-DEL-51
chr9-115550220-DEL-191
chr9-115600690-DEL-199
chr9-115600735-DEL-99
chr9-115659589-DEL-50
chr9-115665263-DEL-101
chr9-115665537-DEL-64
chr9-115665658-DEL-226
chr9-115680983-DEL-102
chr9-115693529-DEL-348
chr9-115693947-DEL-270
chr9-115834590-DEL-378
chr9-115841500-DEL-764
chr9-115842730-DEL-81
chr9-115857835-DEL-66
chr9-115871363-DEL-106
chr9-115965518-DEL-202
chr9-116043998-DEL-245
chr9-116075818-DEL-819
chr9-116131288-DEL-85
chr9-116222225-DEL-81
chr9-116227143-DEL-50
chr9-116232129-DEL-503
chr9-116303498-DEL-65
chr9-116324769-DEL-102
chr9-116324891-DEL-297
chr9-116445298-DEL-52
chr9-116447708-DEL-68
chr9-116535804-DEL-50
chr9-116554482-DEL-53
chr9-116639913-DEL-1085
chr9-116645071-DEL-140
chr9-116661451-DEL-176
chr9-116666553-DEL-88
chr9-116666679-DEL-278
chr9-116667166-DEL-208
chr9-116669467-DEL-96
chr9-116687620-DEL-50
chr9-116862259-DEL-63
chr9-116901342-DEL-83
chr9-116913662-DEL-50
chr9-116959000-DEL-60
chr9-117029971-DEL-125
chr9-117064199-DEL-74
chr9-117065998-DEL-1928
chr9-117074351-DEL-112
chr9-117115296-DEL-250
chr9-117136999-DEL-123
chr9-117173068-DEL-244
chr9-117220307-DEL-261
chr9-117295834-DEL-78
chr9-117388102-DEL-113
chr9-117416286-DEL-205
chr9-117416563-DEL-57
chr9-117466935-DEL-135
chr9-117471587-DEL-148
chr9-117619374-DEL-69
chr9-117629766-DEL-106
chr9-117630903-DEL-1265
chr9-117650809-DEL-89
chr9-117679137-DEL-50
chr9-117687255-DEL-114
chr9-117694676-DEL-98
chr9-117698419-DEL-52
chr9-117789403-DEL-50
chr9-117917187-DEL-2042
chr9-117917383-DEL-4420
chr9-117940850-DEL-74
chr9-117941062-DEL-331
chr9-117941284-DEL-109
chr9-118021982-DEL-52
chr9-118062075-DEL-50
chr9-118128118-DEL-68
chr9-118151686-DEL-153
chr9-118286779-DEL-58

chr9-118286801-DEL-140
chr9-118382697-DEL-322
chr9-118423290-DEL-130
chr9-118444250-DEL-65
chr9-118449511-DEL-113
chr9-118459938-DEL-54
chr9-118498049-DEL-242
chr9-118503963-DEL-54
chr9-118532759-DEL-182
chr9-118533077-DEL-279
chr9-118541242-DEL-51
chr9-118632149-DEL-306
chr9-118790279-DEL-65
chr9-118836096-DEL-64
chr9-118856504-DEL-72
chr9-118867131-DEL-91
chr9-119050619-DEL-355
chr9-119103542-DEL-114
chr9-119175878-DEL-64
chr9-119187558-DEL-142
chr9-119194982-DEL-196
chr9-119215780-DEL-170
chr9-119305801-DEL-88
chr9-119338821-DEL-108
chr9-119339010-DEL-239
chr9-119431377-DEL-373
chr9-119431428-DEL-125
chr9-119466749-DEL-138
chr9-119639498-DEL-72
chr9-119639847-DEL-210
chr9-119715480-DEL-230
chr9-119819296-DEL-826
chr9-119827460-DEL-130
chr9-119878189-DEL-61
chr9-119885272-DEL-90
chr9-119914202-DEL-70
chr9-119914351-DEL-150
chr9-119952035-DEL-50
chr9-119964119-DEL-274
chr9-119971279-DEL-104
chr9-120002014-DEL-60
chr9-120055093-DEL-53
chr9-120076664-DEL-248
chr9-120195332-DEL-54
chr9-120196548-DEL-66
chr9-120236654-DEL-112
chr9-120236880-DEL-56
chr9-120249742-DEL-50
chr9-120288806-DEL-204
chr9-120310743-DEL-54
chr9-120378314-DEL-287
chr9-120414575-DEL-68
chr9-120416604-DEL-137
chr9-120417633-DEL-96
chr9-120517190-DEL-794
chr9-120543548-DEL-702
chr9-120557269-DEL-432
chr9-120557275-DEL-57
chr9-120581629-DEL-100
chr9-120671621-DEL-82
chr9-120715069-DEL-118
chr9-120727465-DEL-57
chr9-120811275-DEL-189
chr9-120839123-DEL-116
chr9-120839130-DEL-56
chr9-120850655-DEL-64
chr9-120869856-DEL-1090
chr9-120918131-DEL-429
chr9-120922880-DEL-511
chr9-120937322-DEL-507
chr9-120948713-DEL-55
chr9-120957206-DEL-104
chr9-120960076-DEL-817
chr9-120961348-DEL-98
chr9-120964515-DEL-68
chr9-120977152-DEL-104
chr9-120981926-DEL-52

chr9-121014022-DEL-210
chr9-121019990-DEL-96
chr9-121037540-DEL-58
chr9-121045952-DEL-222
chr9-121049179-DEL-19276
chr9-121049219-DEL-8113
chr9-121068240-DEL-87
chr9-121071765-DEL-122
chr9-121075672-DEL-59
chr9-121080798-DEL-62
chr9-121097700-DEL-430
chr9-121097933-DEL-849
chr9-121097952-DEL-137
chr9-121098204-DEL-345
chr9-121098460-DEL-293
chr9-121098668-DEL-544
chr9-121098958-DEL-943
chr9-121108515-DEL-79
chr9-121130899-DEL-208
chr9-121136800-DEL-309
chr9-121189047-DEL-63
chr9-121208007-DEL-58
chr9-121261760-DEL-558
chr9-121263622-DEL-116
chr9-121270045-DEL-80
chr9-121314533-DEL-84
chr9-121351685-DEL-60
chr9-121388633-DEL-136
chr9-121436067-DEL-91
chr9-121473422-DEL-388
chr9-121509923-DEL-249
chr9-121516825-DEL-69
chr9-121526750-DEL-135
chr9-121530720-DEL-363
chr9-121530870-DEL-735
chr9-121552212-DEL-78
chr9-121602523-DEL-53
chr9-121603782-DEL-114
chr9-121627536-DEL-60
chr9-121691082-DEL-62
chr9-121702175-DEL-52
chr9-121702774-DEL-84
chr9-121752098-DEL-87
chr9-121758756-DEL-161
chr9-121760484-DEL-525
chr9-121797686-DEL-560
chr9-121820871-DEL-178
chr9-121841580-DEL-108
chr9-121854166-DEL-69
chr9-121927357-DEL-236
chr9-121933491-DEL-50
chr9-121934529-DEL-66
chr9-121990583-DEL-1381
chr9-121990621-DEL-138
chr9-122040157-DEL-70
chr9-122204146-DEL-82
chr9-122343113-DEL-148
chr9-122369772-DEL-206
chr9-122370200-DEL-3096
chr9-122394124-DEL-59
chr9-122435015-DEL-65
chr9-122506076-DEL-53
chr9-122590417-DEL-513
chr9-122591615-DEL-59
chr9-122733593-DEL-624
chr9-122737238-DEL-305
chr9-122737776-DEL-57
chr9-122858816-DEL-96
chr9-122858928-DEL-81
chr9-122943686-DEL-91
chr9-122997146-DEL-147
chr9-123005880-DEL-57
chr9-123080157-DEL-88
chr9-123113944-DEL-184
chr9-123114123-DEL-64
chr9-123176032-DEL-50
chr9-123180023-DEL-456

chr9-123183018-DEL-192
chr9-123243420-DEL-340
chr9-123299841-DEL-74
chr9-123620853-DEL-220
chr9-123629081-DEL-91
chr9-123701971-DEL-123
chr9-123735417-DEL-502
chr9-123751129-DEL-57
chr9-123796074-DEL-58
chr9-123819739-DEL-76
chr9-123917191-DEL-975
chr9-123932450-DEL-482
chr9-124074008-DEL-54
chr9-124129374-DEL-64
chrX-4465129-DEL-161
chrX-4547712-DEL-572
chrX-5546504-DEL-200
chrX-5550460-DEL-18359
chrX-5625346-DEL-54
chrX-5851771-DEL-129
chrX-5937565-DEL-50
chrX-6147057-DEL-431
chrX-6247584-DEL-312
chrX-6307381-DEL-54
chrX-6316052-DEL-210
chrX-6360526-DEL-115
chrX-6404635-DEL-102
chrX-6541611-DEL-172
chrX-6541901-DEL-193
chrX-6806729-DEL-117
chrX-6850317-DEL-54
chrX-6879769-DEL-57
chrX-6880331-DEL-135
chrX-6937736-DEL-169
chrX-7048995-DEL-84
chrX-7049165-DEL-164
chrX-7254446-DEL-86
chrX-7681944-DEL-999
chrX-7771658-DEL-161
chrX-7775230-DEL-185
chrX-7873658-DEL-247
chrX-7912816-DEL-1592
chrX-7991381-DEL-104
chrX-8038495-DEL-231
chrX-8205959-DEL-371
chrX-8245075-DEL-118
chrX-8402491-DEL-682
chrX-8416172-DEL-192
chrX-8416941-DEL-1465
chrX-8441474-DEL-189
chrX-8499594-DEL-1892
chrX-8502072-DEL-4443
chrX-8635321-DEL-77
chrX-8691391-DEL-7806
chrX-8849720-DEL-114
chrX-8858244-DEL-599
chrX-8858301-DEL-251
chrX-8858640-DEL-131
chrX-8894060-DEL-92
chrX-8897271-DEL-610
chrX-8990890-DEL-116
chrX-9010144-DEL-60
chrX-9161116-DEL-1092
chrX-9161230-DEL-417
chrX-9211034-DEL-52
chrX-9268119-DEL-162
chrX-9275046-DEL-141
chrX-9440745-DEL-54
chrX-9447168-DEL-738
chrX-9447918-DEL-55
chrX-9448098-DEL-54
chrX-9504327-DEL-54
chrX-9512488-DEL-54
chrX-9668466-DEL-602
chrX-9909948-DEL-50
chrX-10077220-DEL-82
chrX-10407764-DEL-107

chrX-10427175-DEL-50
chrX-10433565-DEL-50
chrX-10495911-DEL-180
chrX-10578934-DEL-94
chrX-10731189-DEL-51
chrX-10789741-DEL-77
chrX-10855378-DEL-237
chrX-10935226-DEL-2910
chrX-10979442-DEL-85
chrX-11007574-DEL-379
chrX-11149713-DEL-54
chrX-11164619-DEL-1760
chrX-11194105-DEL-297
chrX-11274987-DEL-64
chrX-11329650-DEL-54
chrX-11340641-DEL-51
chrX-11584745-DEL-62
chrX-11598439-DEL-124
chrX-11662118-DEL-50
chrX-11705493-DEL-84
chrX-11802380-DEL-294
chrX-11898905-DEL-136
chrX-12077052-DEL-192
chrX-12132575-DEL-90
chrX-12280503-DEL-54
chrX-12296767-DEL-61
chrX-12430375-DEL-1657
chrX-12650944-DEL-196
chrX-12856013-DEL-50
chrX-13033496-DEL-5229
chrX-13040253-DEL-20191
chrX-13041722-DEL-10095
chrX-13067889-DEL-132
chrX-13087812-DEL-266
chrX-13117277-DEL-64
chrX-13253956-DEL-76
chrX-13254250-DEL-94
chrX-13260424-DEL-1391
chrX-13307133-DEL-55
chrX-13321148-DEL-336
chrX-13378493-DEL-52
chrX-13438943-DEL-52
chrX-13785506-DEL-783
chrX-13918328-DEL-99
chrX-14205730-DEL-125
chrX-14314792-DEL-59
chrX-14373211-DEL-50
chrX-14562321-DEL-1467
chrX-14822180-DEL-17089
chrX-14873723-DEL-1893
chrX-15001020-DEL-494
chrX-15112736-DEL-708
chrX-15113486-DEL-823
chrX-15221199-DEL-78
chrX-15287650-DEL-53
chrX-15551552-DEL-8832
chrX-15597021-DEL-484
chrX-15999654-DEL-82
chrX-16105365-DEL-990
chrX-16269041-DEL-201
chrX-16619779-DEL-572
chrX-16975590-DEL-337
chrX-17058341-DEL-55
chrX-17058444-DEL-290
chrX-17249535-DEL-4483
chrX-17352921-DEL-56
chrX-17361152-DEL-447
chrX-17362152-DEL-4536
chrX-17413359-DEL-78
chrX-17415722-DEL-886
chrX-17487829-DEL-320
chrX-17614074-DEL-6205
chrX-17842590-DEL-2329
chrX-17960199-DEL-50
chrX-18098133-DEL-56
chrX-18137426-DEL-81
chrX-18181208-DEL-59

chrX-18201514-DEL-5367
chrX-18404797-DEL-1060
chrX-18491580-DEL-214
chrX-19371001-DEL-50
chrX-19396080-DEL-123
chrX-19396114-DEL-58
chrX-20198067-DEL-66
chrX-20447251-DEL-360
chrX-20526116-DEL-66
chrX-20605136-DEL-116
chrX-20611294-DEL-396
chrX-20621413-DEL-136
chrX-20622353-DEL-652
chrX-20628739-DEL-201
chrX-20668336-DEL-195
chrX-20756802-DEL-490
chrX-20809084-DEL-748
chrX-21123978-DEL-1629
chrX-21127210-DEL-208
chrX-21322082-DEL-63
chrX-21711507-DEL-1645
chrX-21896584-DEL-155
chrX-22175021-DEL-50
chrX-22855774-DEL-2756
chrX-23033721-DEL-1490
chrX-23339372-DEL-2320
chrX-23444087-DEL-54
chrX-23480508-DEL-87
chrX-23519427-DEL-73
chrX-23557212-DEL-50
chrX-23574817-DEL-122
chrX-24018307-DEL-64
chrX-24562341-DEL-200
chrX-24567639-DEL-1340
chrX-24593835-DEL-115
chrX-25569647-DEL-195
chrX-25756092-DEL-163
chrX-26063703-DEL-446
chrX-30914133-DEL-1899
chrX-30919580-DEL-211
chrX-31410447-DEL-6907
chrX-31422261-DEL-1822
chrX-32014231-DEL-300
chrX-33724389-DEL-88
chrX-34958901-DEL-1109
chrX-34960058-DEL-1319
chrX-35133165-DEL-50
chrX-35160018-DEL-2174
chrX-35172977-DEL-325
chrX-35224121-DEL-50
chrX-35320661-DEL-118
chrX-35417390-DEL-138
chrX-35494782-DEL-116
chrX-35542507-DEL-50
chrX-35961145-DEL-1031
chrX-36043065-DEL-64
chrX-36394138-DEL-120
chrX-36438001-DEL-124
chrX-36438717-DEL-80
chrX-36440191-DEL-82
chrX-36492684-DEL-55
chrX-36537623-DEL-439
chrX-36760074-DEL-62
chrX-36765906-DEL-84
chrX-36830264-DEL-127
chrX-36832819-DEL-105
chrX-36833569-DEL-1094
chrX-36846156-DEL-1265
chrX-36855816-DEL-54
chrX-36862438-DEL-60
chrX-36862859-DEL-480
chrX-37010921-DEL-288
chrX-37051357-DEL-194
chrX-37052694-DEL-868
chrX-37058782-DEL-259
chrX-37268905-DEL-581
chrX-37292906-DEL-57

chrX-37350253-DEL-229
chrX-37352614-DEL-876
chrX-37522517-DEL-132
chrX-37569557-DEL-202
chrX-37574635-DEL-77
chrX-37892214-DEL-4211
chrX-37898874-DEL-261
chrX-37928051-DEL-117
chrX-38036151-DEL-229
chrX-38245185-DEL-5936
chrX-38388693-DEL-307
chrX-38674272-DEL-1101
chrX-38781619-DEL-2442
chrX-38976325-DEL-1628
chrX-38986076-DEL-890
chrX-39105237-DEL-56
chrX-39214970-DEL-187
chrX-39215203-DEL-149
chrX-39234196-DEL-50
chrX-39239368-DEL-66
chrX-39379932-DEL-57
chrX-39393341-DEL-11218
chrX-39858457-DEL-98
chrX-40082404-DEL-6395
chrX-40222535-DEL-50
chrX-40270652-DEL-50
chrX-40833351-DEL-214
chrX-40930692-DEL-166
chrX-41250267-DEL-175
chrX-41285209-DEL-564
chrX-41330278-DEL-68
chrX-41418648-DEL-517
chrX-41420265-DEL-210
chrX-41421453-DEL-53
chrX-41499287-DEL-2478
chrX-41504608-DEL-103
chrX-41575420-DEL-76
chrX-41763226-DEL-58
chrX-41810647-DEL-78
chrX-41925972-DEL-54
chrX-42012093-DEL-704
chrX-42013130-DEL-2839
chrX-42026102-DEL-54
chrX-42297845-DEL-212
chrX-42508292-DEL-175
chrX-42552641-DEL-673
chrX-42598022-DEL-70
chrX-42610449-DEL-54
chrX-43433916-DEL-50
chrX-43439997-DEL-50
chrX-43546265-DEL-50
chrX-43821350-DEL-1652
chrX-43953066-DEL-3379
chrX-43956861-DEL-4440
chrX-43958727-DEL-8491
chrX-43968027-DEL-55
chrX-44100234-DEL-212
chrX-44650271-DEL-89
chrX-44934928-DEL-5462
chrX-45010274-DEL-1425
chrX-45215085-DEL-58
chrX-45359707-DEL-1001
chrX-45492465-DEL-1492
chrX-45502453-DEL-70
chrX-45664721-DEL-87
chrX-45787547-DEL-66
chrX-45846240-DEL-63
chrX-45876629-DEL-936
chrX-45891477-DEL-1410
chrX-45936077-DEL-213
chrX-45936451-DEL-328
chrX-46071065-DEL-826
chrX-46152791-DEL-2004
chrX-46705032-DEL-104
chrX-46825325-DEL-91
chrX-46995923-DEL-191
chrX-47004373-DEL-234

chrX-47037660-DEL-60
chrX-47074163-DEL-50
chrX-47120203-DEL-54
chrX-47263854-DEL-340
chrX-47263858-DEL-60
chrX-47350665-DEL-52
chrX-47581759-DEL-215
chrX-47681680-DEL-255
chrX-47779843-DEL-1237
chrX-47984851-DEL-163
chrX-48130777-DEL-68
chrX-48130831-DEL-147
chrX-48149601-DEL-102
chrX-48373932-DEL-50
chrX-48576309-DEL-52
chrX-48650719-DEL-1519
chrX-48677904-DEL-58
chrX-48738721-DEL-105
chrX-48911106-DEL-54
chrX-49004860-DEL-50
chrX-49086870-DEL-7069
chrX-49169708-DEL-50
chrX-49266090-DEL-140
chrX-49351039-DEL-1484
chrX-49356553-DEL-963
chrX-49395980-DEL-1185
chrX-49672475-DEL-13606
chrX-49816861-DEL-116
chrX-49993083-DEL-152
chrX-50942937-DEL-9076
chrX-51098122-DEL-393
chrX-51169159-DEL-50
chrX-51822256-DEL-148
chrX-51846407-DEL-427
chrX-51851129-DEL-54
chrX-51938419-DEL-165
chrX-52037804-DEL-83
chrX-52158266-DEL-156
chrX-52449063-DEL-60
chrX-52485800-DEL-110
chrX-52587583-DEL-10867
chrX-52590126-DEL-423
chrX-52758822-DEL-196
chrX-52897588-DEL-664
chrX-54369530-DEL-115
chrX-54505778-DEL-611
chrX-54510208-DEL-258
chrX-54519351-DEL-599
chrX-54533019-DEL-1304
chrX-54761342-DEL-91224
chrX-54940555-DEL-30037
chrX-54976513-DEL-325
chrX-55025079-DEL-2372
chrX-55215522-DEL-151
chrX-55257257-DEL-133
chrX-55447920-DEL-51
chrX-55622168-DEL-56
chrX-56059633-DEL-1695
chrX-56305203-DEL-55
chrX-56440726-DEL-1202
chrX-56551773-DEL-706
chrX-56603677-DEL-277
chrX-56640472-DEL-194
chrX-56703152-DEL-64
chrX-56704068-DEL-254
chrX-56835954-DEL-165
chrX-56836350-DEL-133
chrX-56927981-DEL-93
chrX-56928343-DEL-77
chrX-57038503-DEL-265
chrX-57044167-DEL-52
chrX-57054221-DEL-53
chrX-57229181-DEL-52
chrX-57335486-DEL-2061
chrX-57370175-DEL-63
chrX-57461219-DEL-51
chrX-57542795-DEL-53

chrX-57816918-DEL-2782
chrX-58088242-DEL-1097
chrX-58103496-DEL-274
chrX-58106590-DEL-97
chrX-58108147-DEL-79
chrX-58128025-DEL-60
chrX-58236903-DEL-190
chrX-58644967-DEL-895
chrX-58649136-DEL-103
chrX-58717795-DEL-669
chrX-58903240-DEL-54
chrX-58910314-DEL-1270
chrX-58935537-DEL-67
chrX-58960205-DEL-82
chrX-58974199-DEL-225
chrX-59009199-DEL-2528
chrX-59235445-DEL-193
chrX-59352320-DEL-50
chrX-59559998-DEL-68
chrX-59700007-DEL-54
chrX-59815197-DEL-76
chrX-59922998-DEL-55
chrX-59960944-DEL-62
chrX-60065323-DEL-194
chrX-60129425-DEL-2482
chrX-60228042-DEL-141
chrX-60266471-DEL-208
chrX-60383065-DEL-2856
chrX-60510663-DEL-702
chrX-60612869-DEL-211
chrX-60638326-DEL-74
chrX-60849472-DEL-52
chrX-61094131-DEL-1941
chrX-61255280-DEL-50
chrX-61273156-DEL-56
chrX-61382254-DEL-923
chrX-61396445-DEL-371
chrX-61474614-DEL-89
chrX-61561904-DEL-3220
chrX-61564728-DEL-748
chrX-61571703-DEL-26427
chrX-61595643-DEL-745
chrX-61635199-DEL-67
chrX-61649417-DEL-312
chrX-61686864-DEL-66
chrX-61689939-DEL-391
chrX-61695241-DEL-5892
chrX-61899815-DEL-1159
chrX-61980172-DEL-1607
chrX-62002974-DEL-410
chrX-62277465-DEL-521
chrX-62314277-DEL-1604
chrX-62371534-DEL-1437
chrX-62839170-DEL-82
chrX-62876764-DEL-477
chrX-62925621-DEL-8658
chrX-63023120-DEL-1217
chrX-63074863-DEL-63
chrX-63080107-DEL-199
chrX-63080358-DEL-201
chrX-63091950-DEL-70
chrX-63140285-DEL-51
chrX-63179034-DEL-50
chrX-63344879-DEL-1154
chrX-63486723-DEL-198
chrX-63597133-DEL-60
chrX-63676134-DEL-206
chrX-63767907-DEL-4192
chrX-63861859-DEL-52
chrX-64225064-DEL-477
chrX-64271820-DEL-3271
chrX-65188504-DEL-208
chrX-65286814-DEL-50
chrX-65319578-DEL-60
chrX-65539422-DEL-1053
chrX-65615175-DEL-3047
chrX-65886493-DEL-621

chrX-65966428-DEL-52
chrX-66077490-DEL-2235
chrX-66149968-DEL-873
chrX-66187125-DEL-416
chrX-66288795-DEL-100
chrX-66306753-DEL-4473
chrX-66386488-DEL-400
chrX-66432609-DEL-110
chrX-66592004-DEL-215
chrX-66831110-DEL-4576
chrX-67004036-DEL-61
chrX-67470857-DEL-203
chrX-67484405-DEL-533
chrX-67513134-DEL-9338
chrX-67527057-DEL-63
chrX-67528775-DEL-55
chrX-67692559-DEL-1115
chrX-67736602-DEL-112
chrX-67901369-DEL-2196
chrX-67972589-DEL-98
chrX-68139499-DEL-54
chrX-68290031-DEL-6494
chrX-68556254-DEL-57
chrX-68556304-DEL-103
chrX-68666773-DEL-53
chrX-68991943-DEL-2638
chrX-69361170-DEL-60
chrX-69528144-DEL-10776
chrX-69668748-DEL-132
chrX-70056784-DEL-90
chrX-70524512-DEL-94
chrX-70566840-DEL-88
chrX-70570787-DEL-62
chrX-70577529-DEL-697
chrX-70666523-DEL-175
chrX-70666543-DEL-75
chrX-70677429-DEL-60
chrX-70688877-DEL-90
chrX-70689084-DEL-102
chrX-70708821-DEL-63
chrX-70774754-DEL-96
chrX-70872541-DEL-193
chrX-70962269-DEL-1035
chrX-70962538-DEL-50
chrX-71145840-DEL-1084
chrX-71166113-DEL-239
chrX-71233602-DEL-1478
chrX-71268541-DEL-367
chrX-71507953-DEL-816
chrX-71805047-DEL-1017
chrX-71895062-DEL-64
chrX-72039127-DEL-57
chrX-72128151-DEL-82
chrX-72129097-DEL-60
chrX-72147204-DEL-1507
chrX-72306384-DEL-50
chrX-72499882-DEL-102
chrX-72783123-DEL-201
chrX-72783383-DEL-410
chrX-72869865-DEL-60
chrX-72927510-DEL-52
chrX-73025133-DEL-2408
chrX-73177335-DEL-1374
chrX-73617803-DEL-50
chrX-73786868-DEL-158
chrX-73871910-DEL-289
chrX-73939556-DEL-849
chrX-73940438-DEL-954
chrX-74004037-DEL-65
chrX-74238881-DEL-3513
chrX-74270876-DEL-50
chrX-74403057-DEL-1363
chrX-74597359-DEL-2828
chrX-74663358-DEL-476
chrX-74674226-DEL-2497
chrX-74705107-DEL-717
chrX-74705140-DEL-2438

chrX-74777953-DEL-258
chrX-74861983-DEL-50
chrX-74982767-DEL-3130
chrX-75414754-DEL-50
chrX-75509356-DEL-4757
chrX-75516943-DEL-71
chrX-75632755-DEL-4961
chrX-75642396-DEL-144
chrX-75700954-DEL-8523
chrX-75920228-DEL-84
chrX-76031926-DEL-379
chrX-76688120-DEL-998
chrX-76909744-DEL-77
chrX-77145086-DEL-3661
chrX-77376552-DEL-15584
chrX-77423585-DEL-860
chrX-77441547-DEL-222
chrX-77466781-DEL-1830
chrX-77531236-DEL-50
chrX-77706821-DEL-6653
chrX-77749900-DEL-51
chrX-77844314-DEL-1472
chrX-77990722-DEL-4907
chrX-78186476-DEL-912
chrX-78494738-DEL-66
chrX-78504251-DEL-59
chrX-78565139-DEL-16669
chrX-78600530-DEL-56
chrX-78757216-DEL-618
chrX-78835073-DEL-1307
chrX-79009730-DEL-1494
chrX-79227007-DEL-3028
chrX-79290204-DEL-60
chrX-79545464-DEL-52
chrX-79696454-DEL-54
chrX-79738921-DEL-2901
chrX-79845324-DEL-57
chrX-79946127-DEL-54
chrX-80144207-DEL-58
chrX-80724453-DEL-1374
chrX-80762632-DEL-1462
chrX-80765452-DEL-431
chrX-80765694-DEL-113
chrX-81282039-DEL-214
chrX-81315898-DEL-70
chrX-81336807-DEL-162
chrX-81364912-DEL-160
chrX-81467091-DEL-332
chrX-81606864-DEL-2392
chrX-81660016-DEL-1809
chrX-81704879-DEL-1079
chrX-82053430-DEL-50
chrX-82129772-DEL-177
chrX-82221576-DEL-58
chrX-82312144-DEL-60
chrX-82518613-DEL-50
chrX-82527940-DEL-141
chrX-82528297-DEL-187
chrX-82542607-DEL-10248
chrX-82584290-DEL-52
chrX-82633723-DEL-64
chrX-82890035-DEL-159
chrX-82891531-DEL-303
chrX-82891959-DEL-52
chrX-82932208-DEL-914
chrX-82938369-DEL-424
chrX-82944008-DEL-171
chrX-82944484-DEL-124
chrX-82990506-DEL-374
chrX-82993455-DEL-60
chrX-83002226-DEL-342
chrX-83041511-DEL-2456
chrX-83433052-DEL-208
chrX-83447078-DEL-50
chrX-83616039-DEL-60
chrX-83881080-DEL-63
chrX-83988943-DEL-52

chrX-84289956-DEL-50
chrX-84554682-DEL-199
chrX-84648568-DEL-50
chrX-84684800-DEL-200
chrX-84693185-DEL-50
chrX-84710954-DEL-199
chrX-84939096-DEL-609
chrX-84952096-DEL-939
chrX-85014579-DEL-423
chrX-85098677-DEL-66
chrX-85167234-DEL-55
chrX-85217189-DEL-517
chrX-85294700-DEL-87
chrX-85736258-DEL-80
chrX-85755588-DEL-3243
chrX-86012064-DEL-106
chrX-86122071-DEL-1720
chrX-86243339-DEL-206
chrX-86378959-DEL-1350
chrX-86528222-DEL-3943
chrX-86540965-DEL-52
chrX-86608280-DEL-1601
chrX-86656704-DEL-59
chrX-86681525-DEL-1109
chrX-86713592-DEL-942
chrX-86827406-DEL-127
chrX-87014578-DEL-52
chrX-87518363-DEL-1491
chrX-87520024-DEL-5478
chrX-87805235-DEL-3868
chrX-87831053-DEL-2679
chrX-87922333-DEL-624
chrX-88220620-DEL-68
chrX-88253065-DEL-2075
chrX-88255086-DEL-559
chrX-88761236-DEL-359
chrX-89089464-DEL-106
chrX-89183513-DEL-2264
chrX-89241656-DEL-60
chrX-89369044-DEL-50
chrX-89419967-DEL-54
chrX-89695135-DEL-703
chrX-89742588-DEL-70
chrX-90056207-DEL-1424
chrX-90121804-DEL-2704
chrX-90192908-DEL-132
chrX-90369434-DEL-57
chrX-90370109-DEL-165
chrX-90370834-DEL-757
chrX-90385403-DEL-200
chrX-90385692-DEL-834
chrX-90398367-DEL-110
chrX-90459463-DEL-68
chrX-90645884-DEL-52
chrX-90680133-DEL-113
chrX-90962476-DEL-864
chrX-90997388-DEL-1929
chrX-91000175-DEL-1866
chrX-91005401-DEL-1878
chrX-91010689-DEL-1865
chrX-91014610-DEL-1217
chrX-91035427-DEL-9750
chrX-91062961-DEL-57
chrX-91065318-DEL-57
chrX-91070218-DEL-55
chrX-91071616-DEL-88
chrX-91072858-DEL-92
chrX-91128552-DEL-51
chrX-91129117-DEL-831
chrX-91239461-DEL-50
chrX-91250416-DEL-105
chrX-91376237-DEL-52
chrX-91683512-DEL-60
chrX-92083261-DEL-52
chrX-92252493-DEL-56
chrX-92323492-DEL-54
chrX-92475359-DEL-55

chrX-92846686-DEL-58
chrX-92847355-DEL-69
chrX-92952823-DEL-54
chrX-92965917-DEL-56
chrX-93009647-DEL-52
chrX-93173082-DEL-222
chrX-93211976-DEL-321
chrX-93212447-DEL-53
chrX-93212719-DEL-175
chrX-93562652-DEL-79
chrX-93875869-DEL-263
chrX-93887199-DEL-807
chrX-94028832-DEL-443
chrX-94258939-DEL-437
chrX-94365374-DEL-1485
chrX-94517217-DEL-50
chrX-94606711-DEL-566
chrX-94714894-DEL-60
chrX-94922993-DEL-108
chrX-95024819-DEL-595
chrX-95124105-DEL-58
chrX-95426883-DEL-13846
chrX-95436800-DEL-745
chrX-95507620-DEL-90
chrX-95843887-DEL-228
chrX-96268378-DEL-837
chrX-96285231-DEL-204
chrX-96346197-DEL-1612
chrX-96349505-DEL-11441
chrX-96358782-DEL-68
chrX-96382860-DEL-1615
chrX-96884197-DEL-931
chrX-96903064-DEL-634
chrX-97120869-DEL-51
chrX-97272136-DEL-620
chrX-97317517-DEL-99
chrX-97527563-DEL-60
chrX-97649261-DEL-5097
chrX-97786220-DEL-4817
chrX-97920528-DEL-69
chrX-98020186-DEL-1062
chrX-98079582-DEL-1691
chrX-98102048-DEL-4438
chrX-98399987-DEL-50
chrX-98620322-DEL-50
chrX-98631456-DEL-371
chrX-98749900-DEL-117
chrX-98750318-DEL-196
chrX-99109891-DEL-50
chrX-99335630-DEL-650
chrX-99804945-DEL-1816
chrX-100046908-DEL-2760
chrX-100053284-DEL-121
chrX-100157999-DEL-61
chrX-100158028-DEL-161
chrX-100213475-DEL-989
chrX-100289697-DEL-1904
chrX-100538969-DEL-2654
chrX-100651689-DEL-50
chrX-100664371-DEL-76
chrX-100675500-DEL-54
chrX-100739759-DEL-409
chrX-100739890-DEL-162
chrX-101082789-DEL-788
chrX-101343592-DEL-487
chrX-101556040-DEL-300
chrX-101556045-DEL-750
chrX-101626175-DEL-2737
chrX-101785581-DEL-18851
chrX-101787230-DEL-68
chrX-101838992-DEL-64
chrX-101875765-DEL-75
chrX-101905872-DEL-691
chrX-101906025-DEL-50
chrX-101906812-DEL-645
chrX-101907353-DEL-125
chrX-101911038-DEL-559

chrX-101942668-DEL-50
chrX-101976456-DEL-6133
chrX-102028263-DEL-50
chrX-102106071-DEL-56
chrX-102121441-DEL-135
chrX-102121518-DEL-58
chrX-102222476-DEL-122
chrX-102295521-DEL-166
chrX-102295908-DEL-51
chrX-102310700-DEL-231
chrX-102453807-DEL-839
chrX-102492566-DEL-305
chrX-102498806-DEL-84
chrX-102499137-DEL-578
chrX-102657080-DEL-57
chrX-102708099-DEL-53
chrX-102708906-DEL-247
chrX-102792449-DEL-50
chrX-102793427-DEL-60
chrX-102909125-DEL-69
chrX-103009398-DEL-472
chrX-103036067-DEL-50
chrX-103040527-DEL-50
chrX-103106379-DEL-1687
chrX-103248661-DEL-51
chrX-103312297-DEL-2375
chrX-103701831-DEL-50
chrX-103839544-DEL-14831
chrX-103878392-DEL-5767
chrX-103918824-DEL-964
chrX-103922242-DEL-322
chrX-104088278-DEL-183
chrX-104242342-DEL-258
chrX-104579888-DEL-52
chrX-104601103-DEL-66
chrX-104785430-DEL-3690
chrX-104882411-DEL-62
chrX-105008815-DEL-178
chrX-105110908-DEL-334
chrX-105187790-DEL-70
chrX-105281723-DEL-57
chrX-105284584-DEL-445
chrX-105312275-DEL-74
chrX-105379198-DEL-72
chrX-105433836-DEL-2363
chrX-105437798-DEL-8834
chrX-105464612-DEL-141
chrX-105783819-DEL-50
chrX-105932341-DEL-1518
chrX-106027579-DEL-72
chrX-106592867-DEL-2040
chrX-106686657-DEL-1790
chrX-106690583-DEL-1490
chrX-107597709-DEL-50
chrX-107597882-DEL-51
chrX-107600654-DEL-90
chrX-107616639-DEL-7867
chrX-107788419-DEL-276
chrX-107790524-DEL-151
chrX-107790535-DEL-73
chrX-108349717-DEL-197
chrX-108477889-DEL-50
chrX-108920732-DEL-285
chrX-109075891-DEL-402
chrX-109172029-DEL-204
chrX-109548828-DEL-2402
chrX-109586432-DEL-1357
chrX-109715664-DEL-1616
chrX-109875064-DEL-184
chrX-110048194-DEL-2974
chrX-110271274-DEL-560
chrX-110697854-DEL-234
chrX-111070734-DEL-310
chrX-111194582-DEL-1177
chrX-111337193-DEL-75
chrX-111928334-DEL-54
chrX-112230869-DEL-54

chrX-112269015-DEL-75
chrX-112350157-DEL-50
chrX-112513616-DEL-1078
chrX-112858769-DEL-755
chrX-112867126-DEL-2477
chrX-112874459-DEL-8183
chrX-113143074-DEL-142
chrX-113415664-DEL-1661
chrX-113609824-DEL-59
chrX-113683582-DEL-60
chrX-113838653-DEL-50
chrX-113876596-DEL-52
chrX-114061142-DEL-72
chrX-114227428-DEL-179
chrX-114320389-DEL-80
chrX-114570302-DEL-214
chrX-114589259-DEL-1413
chrX-114598692-DEL-910
chrX-114638823-DEL-54
chrX-114713081-DEL-2796
chrX-115382855-DEL-55
chrX-115419381-DEL-2421
chrX-115567566-DEL-3139
chrX-115695818-DEL-353
chrX-116181546-DEL-498
chrX-116419536-DEL-139
chrX-117011209-DEL-1363
chrX-117072446-DEL-3586
chrX-117173234-DEL-1526
chrX-117199924-DEL-882
chrX-117207215-DEL-56
chrX-117515574-DEL-263
chrX-117546543-DEL-5719
chrX-117621791-DEL-3030
chrX-117708842-DEL-60
chrX-117708902-DEL-132
chrX-117928628-DEL-2171
chrX-117970216-DEL-98
chrX-117999077-DEL-1240
chrX-118116502-DEL-63
chrX-118263975-DEL-129
chrX-119206144-DEL-50
chrX-119239229-DEL-50
chrX-119424612-DEL-97
chrX-119441275-DEL-50
chrX-119492103-DEL-60
chrX-119493845-DEL-50
chrX-119544154-DEL-82
chrX-119743785-DEL-1571
chrX-119794378-DEL-7647
chrX-119822520-DEL-530
chrX-120977359-DEL-207
chrX-120981395-DEL-54
chrX-120995584-DEL-186
chrX-121046148-DEL-73
chrX-121296124-DEL-1644
chrX-121634429-DEL-212
chrX-121638651-DEL-605
chrX-121916261-DEL-65
chrX-121928741-DEL-382
chrX-121946186-DEL-567
chrX-121965136-DEL-51
chrX-121993204-DEL-1259
chrX-122171783-DEL-299
chrX-122405357-DEL-63
chrX-122860340-DEL-2469
chrX-122954222-DEL-2029
chrX-123125404-DEL-142
chrX-123129056-DEL-2417
chrX-123250221-DEL-263
chrX-123278501-DEL-63
chrX-124924058-DEL-88
chrX-124930165-DEL-381
chrX-125288690-DEL-199
chrX-125317054-DEL-208
chrX-125464574-DEL-246
chrX-125665115-DEL-263

chrX-125993993-DEL-55
chrX-126082734-DEL-120
chrX-126139391-DEL-75
chrX-126191255-DEL-83
chrX-126257649-DEL-50
chrX-126537507-DEL-50
chrX-126871256-DEL-50
chrX-127336115-DEL-3104
chrX-127388995-DEL-1124
chrX-127575906-DEL-52
chrX-127579044-DEL-55
chrX-127626499-DEL-50
chrX-127886147-DEL-72
chrX-127918648-DEL-66
chrX-128058363-DEL-72
chrX-128330830-DEL-3896
chrX-128511587-DEL-1706
chrX-128582978-DEL-60
chrX-128617824-DEL-50
chrX-128702284-DEL-50
chrX-129098507-DEL-1326
chrX-129203957-DEL-52
chrX-129225039-DEL-118
chrX-129225165-DEL-280
chrX-129240070-DEL-78
chrX-129617280-DEL-64
chrX-130054255-DEL-1479
chrX-130317341-DEL-60
chrX-130387153-DEL-102
chrX-130399156-DEL-1415
chrX-130423259-DEL-5627
chrX-130476115-DEL-2514
chrX-130490589-DEL-541
chrX-130490758-DEL-96
chrX-130516007-DEL-1666
chrX-130584536-DEL-589
chrX-130715283-DEL-1722
chrX-130878562-DEL-214
chrX-130895949-DEL-1132
chrX-131438579-DEL-1078
chrX-131653408-DEL-52
chrX-131714386-DEL-594
chrX-131877420-DEL-50
chrX-132189990-DEL-257
chrX-132247775-DEL-374
chrX-132608880-DEL-91
chrX-132692335-DEL-54
chrX-132904904-DEL-413
chrX-132909518-DEL-54
chrX-133193735-DEL-165
chrX-133221463-DEL-102
chrX-133277547-DEL-50
chrX-133426670-DEL-520
chrX-133640463-DEL-53
chrX-133671560-DEL-160
chrX-133792754-DEL-123
chrX-133799888-DEL-87
chrX-133821283-DEL-833
chrX-133987515-DEL-120
chrX-133987916-DEL-80
chrX-133996483-DEL-115
chrX-133996658-DEL-58
chrX-134207051-DEL-67
chrX-134259861-DEL-344
chrX-134459175-DEL-172
chrX-134462226-DEL-53
chrX-134468518-DEL-54
chrX-134586120-DEL-84
chrX-134587189-DEL-111
chrX-134769402-DEL-70
chrX-134807374-DEL-52
chrX-134901695-DEL-70
chrX-134901809-DEL-206
chrX-135043580-DEL-175
chrX-135081343-DEL-98
chrX-135171362-DEL-54
chrX-135413384-DEL-56

chrX-135429570-DEL-553
chrX-135436849-DEL-120
chrX-135465598-DEL-784
chrX-135505424-DEL-179
chrX-135533713-DEL-785
chrX-135726616-DEL-80
chrX-135734407-DEL-93
chrX-135738525-DEL-75
chrX-135847028-DEL-83
chrX-136009474-DEL-2771
chrX-136022486-DEL-52
chrX-136057229-DEL-1010
chrX-136066964-DEL-57
chrX-136070197-DEL-212
chrX-136128627-DEL-1171
chrX-136132742-DEL-63
chrX-136140587-DEL-68
chrX-136476165-DEL-108
chrX-136816689-DEL-504
chrX-136941745-DEL-590
chrX-136983826-DEL-56
chrX-137088066-DEL-102
chrX-137141900-DEL-51
chrX-137448065-DEL-1077
chrX-137455647-DEL-328
chrX-137477758-DEL-60
chrX-138110516-DEL-55
chrX-138168938-DEL-50
chrX-138420973-DEL-70
chrX-138665490-DEL-93
chrX-138832291-DEL-171
chrX-139050200-DEL-55
chrX-139245115-DEL-72
chrX-139451394-DEL-54
chrX-139536612-DEL-777
chrX-139537037-DEL-196
chrX-139756031-DEL-156
chrX-139785559-DEL-293
chrX-139791024-DEL-53
chrX-139808294-DEL-38922
chrX-140029131-DEL-331
chrX-140030034-DEL-1639
chrX-140300955-DEL-66
chrX-140335840-DEL-3351
chrX-140350941-DEL-214
chrX-140574448-DEL-107
chrX-140574689-DEL-114
chrX-140970680-DEL-93
chrX-141240446-DEL-1823
chrX-141304987-DEL-3098
chrX-141462714-DEL-71
chrX-141632508-DEL-50
chrX-141983731-DEL-91
chrX-142232925-DEL-54
chrX-142302369-DEL-60
chrX-142302409-DEL-136
chrX-142369406-DEL-70
chrX-142452612-DEL-243
chrX-142544404-DEL-218
chrX-142636816-DEL-50
chrX-142807981-DEL-6609
chrX-142845308-DEL-4898
chrX-142992878-DEL-502
chrX-143047960-DEL-72
chrX-143088547-DEL-4115
chrX-143124206-DEL-50
chrX-143133694-DEL-497
chrX-143501352-DEL-1186
chrX-143521740-DEL-84
chrX-143578019-DEL-4834
chrX-143604270-DEL-208
chrX-143772694-DEL-132
chrX-143772738-DEL-65
chrX-143802296-DEL-65
chrX-144310577-DEL-51
chrX-144389538-DEL-130
chrX-144407691-DEL-3398

chrX-144547885-DEL-210
chrX-144643615-DEL-5437
chrX-144668377-DEL-69
chrX-144720395-DEL-1273
chrX-144776054-DEL-1197
chrX-144931665-DEL-196
chrX-145041480-DEL-78
chrX-145098298-DEL-214
chrX-145364081-DEL-2488
chrX-145405744-DEL-50
chrX-145578954-DEL-881
chrX-145620738-DEL-869
chrX-145679147-DEL-7296
chrX-145707643-DEL-168
chrX-145707762-DEL-60
chrX-145735351-DEL-54
chrX-145735430-DEL-459
chrX-145765509-DEL-105
chrX-145781849-DEL-76
chrX-145896547-DEL-2333
chrX-146037486-DEL-1051
chrX-146040410-DEL-84
chrX-146130737-DEL-56
chrX-146549543-DEL-14026
chrX-146734964-DEL-457
chrX-148688718-DEL-435
chrX-148875652-DEL-324
chrX-148881058-DEL-1964
chrX-148881087-DEL-100
chrX-149155880-DEL-1911
chrX-149526440-DEL-204
chrX-149602525-DEL-1490
chrX-149934370-DEL-56
chrX-150835396-DEL-53
chrX-150973415-DEL-80
chrX-151095794-DEL-638
chrX-151115244-DEL-50
chrX-151136129-DEL-52
chrX-151173728-DEL-156
chrX-151615276-DEL-1900
chrX-151645331-DEL-1404
chrX-151651542-DEL-771
chrX-151661200-DEL-124
chrX-151727299-DEL-55
chrX-151867647-DEL-216
chrX-151867767-DEL-105
chrX-151880745-DEL-72
chrX-151881407-DEL-123
chrX-151881437-DEL-59
chrX-152022244-DEL-128
chrX-152039902-DEL-50
chrX-152310510-DEL-82
chrX-152421657-DEL-256
chrX-152421809-DEL-116
chrX-152581486-DEL-53
chrX-152788594-DEL-21185
chrX-152810595-DEL-53546
chrX-152841181-DEL-8129
chrX-152887458-DEL-52
chrX-152890279-DEL-21222
chrX-152908373-DEL-21683
chrX-152930283-DEL-263
chrX-152956723-DEL-70249
chrX-153107651-DEL-2653
chrX-153203105-DEL-3233
chrX-153291143-DEL-68
chrX-153305491-DEL-53
chrX-153341668-DEL-3500
chrX-153582998-DEL-2873
chrX-153687180-DEL-24398
chrX-153762624-DEL-67271
chrX-153791090-DEL-5313
chrX-153793107-DEL-10567
chrX-153805888-DEL-395
chrX-153806410-DEL-1315
chrX-153806788-DEL-420
chrX-153809991-DEL-61141

chrX-153858353-DEL-3147
chrX-154088876-DEL-112
chrX-154242569-DEL-212
chrX-154532081-DEL-63
chrX-154773326-DEL-141
chrX-155196586-DEL-50
chrX-155436156-DEL-74
chrX-155605715-DEL-2143
chrX-155757904-DEL-2373
chrX-155769837-DEL-130
chrX-155910597-DEL-80
chrX-156177647-DEL-65
chrX-156239141-DEL-346
chrX-156443882-DEL-54
chrX-156576392-DEL-168
chrX-156576396-DEL-60
chrX-156576600-DEL-64
chrX-156692522-DEL-50
chrX-156755937-DEL-1124
chrX-157172309-DEL-72
chrX-157452264-DEL-825
chrX-157487865-DEL-50
chrX-157572291-DEL-185
chrX-157682658-DEL-8250
chrX-157902967-DEL-2523
chrX-157918544-DEL-1593
chrX-158007255-DEL-153
chrX-158136091-DEL-110
chrX-158175646-DEL-298
chrX-158299717-DEL-502
chrX-158840724-DEL-92
chrX-158840961-DEL-170
chrX-158841258-DEL-315
chrX-158841436-DEL-137
chrX-158908360-DEL-303
chrX-158965449-DEL-51
chrX-159171864-DEL-51
chrX-159241370-DEL-102
chrX-159692340-DEL-1376
chrX-159823084-DEL-2560
chrX-159893943-DEL-89
chrX-159928566-DEL-152
chrX-160157411-DEL-117
chrX-160175287-DEL-52
chrX-160223031-DEL-50
chrX-160489127-DEL-6628
chrX-160578764-DEL-205
chrX-160610948-DEL-50
chrX-160863520-DEL-817
chrX-161129779-DEL-84
chrX-161398419-DEL-50
chrX-161421216-DEL-196
chrX-161437848-DEL-50
chrX-161599513-DEL-750
chrX-161671490-DEL-72
chrX-161671532-DEL-172
chrX-161673219-DEL-323
chrX-161914004-DEL-112
chrX-162138109-DEL-398
chrX-162202310-DEL-88
chrX-162377872-DEL-78
chrX-162427827-DEL-120
chrX-162442418-DEL-844
chrX-162738305-DEL-936
chrX-162844163-DEL-51
chrX-162929618-DEL-791
chrX-162984837-DEL-124
chrX-163246712-DEL-153
chrX-163320036-DEL-65
chrX-163803571-DEL-222
chrX-163819728-DEL-228
chrX-163950737-DEL-50
chrX-163966155-DEL-72
chrX-164052826-DEL-157
chrX-164193488-DEL-402
chrX-164257039-DEL-6368
chrX-164387017-DEL-52

chrX-164432276-DEL-63
chrX-164454946-DEL-64
chrX-164484222-DEL-214
chrX-164630773-DEL-52
chrX-164792304-DEL-193
chrX-164921449-DEL-234
chrX-164959997-DEL-58
chrX-164960091-DEL-221
chrX-165100230-DEL-741
chrX-165351874-DEL-77
chrX-165545767-DEL-57
chrX-165553732-DEL-187
chrX-165696113-DEL-94
chrX-165799199-DEL-55
chrX-165802381-DEL-156
chrX-165880810-DEL-61
chrX-165881573-DEL-54
chrX-165976355-DEL-90
chrX-166381495-DEL-52
chrX-166802706-DEL-108
chrX-167192492-DEL-82
chrX-167198418-DEL-50
chrX-167632645-DEL-60
chrX-168119198-DEL-100
chrX-168207736-DEL-68
chrX-168344240-DEL-96
chrX-168356893-DEL-768
chrX-168604794-DEL-60
chrX-168769167-DEL-217
chrX-168770859-DEL-246
chrX-168771096-DEL-110
chrX-168772585-DEL-94
chrX-168774191-DEL-150
chrX-168775380-DEL-151
chrX-168776312-DEL-448
chrX-168776708-DEL-8706
chrX-168781664-DEL-587
chrX-168781694-DEL-61
chrX-169071276-DEL-272
chrX-169071280-DEL-125
chrX-169072130-DEL-101
chrX-169072580-DEL-144
chrX-169072615-DEL-342
chrX-169074960-DEL-249
chrX-169077304-DEL-148
chrX-169080517-DEL-102
chrX-169081288-DEL-156
chrX-169088410-DEL-60
chrX-169093831-DEL-155
chrX-169095098-DEL-62
chrX-169097494-DEL-187
chrX-169099166-DEL-175
chrX-169106623-DEL-365
chrX-169107880-DEL-63
chrX-169110387-DEL-59
chrX-169112198-DEL-309
chrX-169118428-DEL-65
chrX-169118709-DEL-57
chrX-169120265-DEL-147244
chrX-169290481-DEL-154
chrX-169295323-DEL-65

| Identified_By_Us | Identified_By_JAX |
|---|---|
| chr1-3094511-INS-51 | chr1-3094511-INS-51 |
| chr1-3097335-INS-3649 | chr1-3097326-INS-3643 |
| chr1-3112403-INS-79 | chr1-3112403-INS-79 |
| chr1-3119164-INS-68 | chr1-3119164-INS-72 |
| chr1-3120383-INS-154 | chr1-3120398-INS-151 |
| chr1-3121617-INS-122 | chr1-3121617-INS-115 |
| chr1-3145507-INS-1414 | chr1-3145508-INS-1415 |
| chr1-3154150-INS-71 | chr1-3154150-INS-72 |
| chr1-3182711-INS-957 | chr1-3182711-INS-957 |
| chr1-3190610-INS-4249 | chr1-3190611-INS-4250 |
| chr1-3217060-INS-382 | chr1-3217060-INS-382 |
| chr1-3254096-INS-1900 | chr1-3254097-INS-1900 |
| chr1-3293696-INS-6815 | chr1-3293697-INS-6821 |
| chr1-3318996-INS-379 | chr1-3318996-INS-379 |
| chr1-3338996-INS-66 | chr1-3338994-INS-70 |
| chr1-3387471-INS-807 | chr1-3387471-INS-806 |
| chr1-3414867-INS-213 | chr1-3414867-INS-213 |
| chr1-3418719-INS-6645 | chr1-3418719-INS-6645 |
| chr1-3420673-INS-1338 | chr1-3420674-INS-1339 |
| chr1-3442769-INS-6452 | chr1-3442770-INS-6452 |
| chr1-3460439-INS-52 | chr1-3460439-INS-50 |
| chr1-3481651-INS-52 | chr1-3481651-INS-54 |
| chr1-3482935-INS-59 | chr1-3482935-INS-59 |
| chr1-3493163-INS-56 | chr1-3493163-INS-62 |
| chr1-3582679-INS-1888 | chr1-3582680-INS-1888 |
| chr1-3609129-INS-328 | chr1-3609129-INS-328 |
| chr1-3614795-INS-6407 | chr1-3614796-INS-6407 |
| chr1-3646827-INS-92 | chr1-3646827-INS-107 |
| chr1-3655363-INS-58 | chr1-3655372-INS-66 |
| chr1-3673454-INS-90 | chr1-3673454-INS-90 |
| chr1-3743350-INS-58 | chr1-3743348-INS-100 |
| chr1-3751560-INS-354 | chr1-3751560-INS-355 |
| chr1-3756578-INS-62 | chr1-3756578-INS-64 |
| chr1-3766593-INS-427 | chr1-3766593-INS-427 |
| chr1-3771457-INS-1479 | chr1-3771458-INS-1479 |
| chr1-3774707-INS-77 | chr1-3774707-INS-77 |
| chr1-3776024-INS-479 | chr1-3776024-INS-479 |
| chr1-3779167-INS-2004 | chr1-3779168-INS-1998 |
| chr1-3786876-INS-401 | chr1-3786876-INS-401 |
| chr1-3814827-INS-420 | chr1-3814827-INS-420 |
| chr1-3825820-INS-307 | chr1-3825817-INS-307 |
| chr1-3830625-INS-431 | chr1-3830625-INS-459 |
| chr1-3831816-INS-167 | chr1-3831817-INS-169 |
| chr1-3857993-INS-423 | chr1-3857993-INS-423 |
| chr1-3864271-INS-2888 | chr1-3864272-INS-2888 |
| chr1-3890444-INS-228 | chr1-3890444-INS-229 |
| chr1-3964647-INS-167 | chr1-3964647-INS-165 |
| chr1-3983706-INS-216 | chr1-3983723-INS-204 |
| chr1-3986474-INS-66 | chr1-3986477-INS-56 |
| chr1-3994921-INS-99 | chr1-3994921-INS-90 |
| chr1-4021361-INS-221 | chr1-4021361-INS-221 |
| chr1-4031167-INS-527 | chr1-4031167-INS-527 |
| chr1-4047357-INS-60 | chr1-4047357-INS-60 |
| chr1-4058066-INS-8672 | chr1-4058067-INS-8673 |
| chr1-4065987-INS-198 | chr1-4065987-INS-198 |
| chr1-4071098-INS-52 | chr1-4071098-INS-52 |
| chr1-4079303-INS-317 | chr1-4079298-INS-318 |
| chr1-4084771-INS-83 | chr1-4084771-INS-83 |
| chr1-4114494-INS-524 | chr1-4114494-INS-524 |
| chr1-4115829-INS-984 | chr1-4115829-INS-980 |
| chr1-4124707-INS-172 | chr1-4124703-INS-176 |
| chr1-4159116-INS-7150 | chr1-4159117-INS-7150 |
| chr1-4188151-INS-664 | chr1-4188151-INS-664 |
| chr1-4214777-INS-55 | chr1-4214754-INS-58 |
| chr1-4245293-INS-61 | chr1-4245272-INS-62 |
| chr1-4312310-INS-1893 | chr1-4312311-INS-1893 |
| chr1-4319733-INS-257 | chr1-4319736-INS-244 |
| chr1-4394082-INS-6463 | chr1-4394083-INS-6463 |
| chr1-4413670-INS-59 | chr1-4413670-INS-94 |
| chr1-4421459-INS-80 | chr1-4421459-INS-80 |
| chr1-4437588-INS-6195 | chr1-4437589-INS-6195 |
| chr1-4455194-INS-1099 | chr1-4455195-INS-1100 |
| chr1-4457578-INS-696 | chr1-4457578-INS-696 |
| chr1-4488773-INS-51 | chr1-4488773-INS-58 |
| chr1-4512058-INS-6403 | chr1-4512059-INS-6403 |
| chr1-4550983-INS-126 | chr1-4550983-INS-128 |

| | |
|---|---|
| chr1-4580736-INS-131 | chr1-4580736-INS-138 |
| chr1-4602871-INS-169 | chr1-4602871-INS-169 |
| chr1-4621081-INS-498 | chr1-4621081-INS-498 |
| chr1-4664818-INS-6243 | chr1-4664819-INS-6243 |
| chr1-4685456-INS-814 | chr1-4685456-INS-814 |
| chr1-4706488-INS-253 | chr1-4706488-INS-253 |
| chr1-4718423-INS-142 | chr1-4718423-INS-142 |
| chr1-4721851-INS-137 | chr1-4721851-INS-137 |
| chr1-4733111-INS-359 | chr1-4733112-INS-359 |
| chr1-4746373-INS-64 | chr1-4746375-INS-70 |
| chr1-4751713-INS-1348 | chr1-4751714-INS-1348 |
| chr1-4757107-INS-506 | chr1-4757107-INS-506 |
| chr1-4768363-INS-142 | chr1-4768363-INS-142 |
| chr1-4774324-INS-6370 | chr1-4774325-INS-6370 |
| chr1-4776168-INS-708 | chr1-4776168-INS-708 |
| chr1-4781147-INS-246 | chr1-4781147-INS-246 |
| chr1-4796259-INS-58 | chr1-4796259-INS-103 |
| chr1-4835887-INS-97 | chr1-4835887-INS-87 |
| chr1-4842792-INS-206 | chr1-4842792-INS-206 |
| chr1-4846259-INS-213 | chr1-4846259-INS-214 |
| chr1-4861078-INS-222 | chr1-4861078-INS-222 |
| chr1-4868984-INS-185 | chr1-4868984-INS-187 |
| chr1-4871806-INS-201 | chr1-4871806-INS-201 |
| chr1-4886519-INS-176 | chr1-4886519-INS-176 |
| chr1-4888674-INS-215 | chr1-4888674-INS-215 |
| chr1-4900004-INS-194 | chr1-4900004-INS-197 |
| chr1-4904000-INS-216 | chr1-4904001-INS-216 |
| chr1-4907943-INS-199 | chr1-4907943-INS-199 |
| chr1-4917901-INS-200 | chr1-4917901-INS-200 |
| chr1-4922693-INS-204 | chr1-4922693-INS-202 |
| chr1-4924022-INS-110 | chr1-4924022-INS-102 |
| chr1-4956223-INS-70 | chr1-4956223-INS-64 |
| chr1-5026960-INS-713 | chr1-5026960-INS-712 |
| chr1-5047161-INS-102 | chr1-5047161-INS-92 |
| chr1-5048952-INS-1187 | chr1-5048953-INS-1185 |
| chr1-5049747-INS-1361 | chr1-5049748-INS-1362 |
| chr1-5066770-INS-2277 | chr1-5066771-INS-2278 |
| chr1-5071570-INS-5226 | chr1-5071571-INS-5226 |
| chr1-5073864-INS-841 | chr1-5073864-INS-840 |
| chr1-5077096-INS-199 | chr1-5077096-INS-199 |
| chr1-5102185-INS-2848 | chr1-5102186-INS-2848 |
| chr1-5106080-INS-1131 | chr1-5106081-INS-1131 |
| chr1-5123127-INS-6380 | chr1-5123128-INS-6381 |
| chr1-5137059-INS-326 | chr1-5137059-INS-326 |
| chr1-5143522-INS-56 | chr1-5143523-INS-52 |
| chr1-5163683-INS-62 | chr1-5163683-INS-62 |
| chr1-5184468-INS-57 | chr1-5184468-INS-57 |
| chr1-5206270-INS-610 | chr1-5206270-INS-609 |
| chr1-5227403-INS-1102 | chr1-5227404-INS-1103 |
| chr1-5268618-INS-7099 | chr1-5268619-INS-7099 |
| chr1-5294719-INS-52 | chr1-5294750-INS-50 |
| chr1-5308052-INS-80 | chr1-5308067-INS-96 |
| chr1-5311896-INS-475 | chr1-5311896-INS-473 |
| chr1-5313049-INS-4438 | chr1-5313050-INS-4438 |
| chr1-5330203-INS-8419 | chr1-5330204-INS-8419 |
| chr1-5347417-INS-76 | chr1-5347417-INS-76 |
| chr1-5352354-INS-78 | chr1-5352354-INS-78 |
| chr1-5555589-INS-575 | chr1-5555589-INS-575 |
| chr1-5586707-INS-64 | chr1-5586707-INS-64 |
| chr1-5590870-INS-178 | chr1-5590870-INS-172 |
| chr1-5595984-INS-5312 | chr1-5595985-INS-5312 |
| chr1-5649115-INS-436 | chr1-5649115-INS-436 |
| chr1-5687949-INS-76 | chr1-5687966-INS-72 |
| chr1-5765632-INS-792 | chr1-5765632-INS-792 |
| chr1-5799625-INS-6732 | chr1-5799626-INS-6732 |
| chr1-5817416-INS-81 | chr1-5817416-INS-81 |
| chr1-5831396-INS-111 | chr1-5831396-INS-111 |
| chr1-5854180-INS-703 | chr1-5854180-INS-704 |
| chr1-5903725-INS-335 | chr1-5903725-INS-333 |
| chr1-5935593-INS-56 | chr1-5935593-INS-58 |
| chr1-5946121-INS-380 | chr1-5946122-INS-366 |
| chr1-5959903-INS-6407 | chr1-5959904-INS-6409 |
| chr1-6005410-INS-1313 | chr1-6005411-INS-1315 |
| chr1-6012047-INS-59 | chr1-6012006-INS-50 |
| chr1-6031309-INS-55 | chr1-6031309-INS-70 |
| chr1-6045092-INS-1557 | chr1-6045093-INS-1557 |
| chr1-6075564-INS-748 | chr1-6075564-INS-748 |

| | |
|---|---|
| chr1-6085880-INS-328 | chr1-6085880-INS-328 |
| chr1-6104908-INS-519 | chr1-6104908-INS-519 |
| chr1-6111647-INS-6036 | chr1-6111648-INS-6036 |
| chr1-6131599-INS-85 | chr1-6131599-INS-85 |
| chr1-6156492-INS-61 | chr1-6156492-INS-73 |
| chr1-6162737-INS-343 | chr1-6162737-INS-357 |
| chr1-6169192-INS-218 | chr1-6169192-INS-218 |
| chr1-6188347-INS-219 | chr1-6188347-INS-220 |
| chr1-6205638-INS-164 | chr1-6205638-INS-164 |
| chr1-6213942-INS-88 | chr1-6213942-INS-88 |
| chr1-6224597-INS-1344 | chr1-6224598-INS-1345 |
| chr1-6229530-INS-1892 | chr1-6229531-INS-1892 |
| chr1-6267156-INS-94 | chr1-6267156-INS-98 |
| chr1-6268351-INS-157 | chr1-6268351-INS-157 |
| chr1-6277391-INS-189 | chr1-6277391-INS-189 |
| chr1-6305990-INS-188 | chr1-6305990-INS-192 |
| chr1-6384342-INS-4456 | chr1-6384343-INS-4456 |
| chr1-6464450-INS-5407 | chr1-6464451-INS-5407 |
| chr1-6466091-INS-68 | chr1-6466091-INS-71 |
| chr1-6482352-INS-416 | chr1-6482352-INS-416 |
| chr1-6491355-INS-66 | chr1-6491355-INS-62 |
| chr1-6494335-INS-112 | chr1-6494335-INS-112 |
| chr1-6495874-INS-151 | chr1-6495874-INS-151 |
| chr1-6526497-INS-68 | chr1-6526497-INS-68 |
| chr1-6532221-INS-57 | chr1-6532221-INS-57 |
| chr1-6540041-INS-4897 | chr1-6540042-INS-4897 |
| chr1-6553312-INS-6244 | chr1-6553313-INS-6244 |
| chr1-6623341-INS-252 | chr1-6623341-INS-252 |
| chr1-6634154-INS-1145 | chr1-6634155-INS-1146 |
| chr1-6639113-INS-204 | chr1-6639148-INS-203 |
| chr1-6687456-INS-341 | chr1-6687456-INS-342 |
| chr1-6701051-INS-537 | chr1-6701051-INS-537 |
| chr1-6708873-INS-56 | chr1-6708873-INS-54 |
| chr1-6716472-INS-525 | chr1-6716472-INS-525 |
| chr1-6751753-INS-343 | chr1-6751753-INS-343 |
| chr1-6774420-INS-351 | chr1-6774420-INS-351 |
| chr1-6776527-INS-1302 | chr1-6776528-INS-1303 |
| chr1-6793736-INS-98 | chr1-6793734-INS-100 |
| chr1-6848093-INS-218 | chr1-6848093-INS-218 |
| chr1-6856507-INS-6300 | chr1-6856508-INS-6300 |
| chr1-6868294-INS-88 | chr1-6868295-INS-92 |
| chr1-6871580-INS-64 | chr1-6871580-INS-100 |
| chr1-6874301-INS-54 | chr1-6874301-INS-54 |
| chr1-6881364-INS-1886 | chr1-6881365-INS-1885 |
| chr1-6904939-INS-104 | chr1-6904939-INS-103 |
| chr1-6926589-INS-54 | chr1-6926619-INS-58 |
| chr1-6926619-INS-940 | chr1-6926619-INS-940 |
| chr1-6947005-INS-59 | chr1-6947009-INS-94 |
| chr1-6951139-INS-230 | chr1-6951139-INS-230 |
| chr1-6953633-INS-1658 | chr1-6953632-INS-1650 |
| chr1-6955838-INS-2371 | chr1-6955839-INS-2375 |
| chr1-7003997-INS-84 | chr1-7003997-INS-84 |
| chr1-7009206-INS-572 | chr1-7009206-INS-572 |
| chr1-7013831-INS-165 | chr1-7013831-INS-165 |
| chr1-7040668-INS-84 | chr1-7040668-INS-80 |
| chr1-7053079-INS-6075 | chr1-7053080-INS-6075 |
| chr1-7053999-INS-211 | chr1-7053999-INS-211 |
| chr1-7077776-INS-111 | chr1-7077776-INS-115 |
| chr1-7080427-INS-81 | chr1-7080427-INS-81 |
| chr1-7082372-INS-475 | chr1-7082372-INS-475 |
| chr1-7087742-INS-56 | chr1-7087742-INS-56 |
| chr1-7093605-INS-135 | chr1-7093605-INS-129 |
| chr1-7106685-INS-192 | chr1-7106685-INS-192 |
| chr1-7129219-INS-5265 | chr1-7129220-INS-5265 |
| chr1-7138111-INS-366 | chr1-7138110-INS-339 |
| chr1-7142427-INS-204 | chr1-7142427-INS-204 |
| chr1-7230120-INS-6888 | chr1-7230121-INS-6889 |
| chr1-7232813-INS-159 | chr1-7232813-INS-159 |
| chr1-7282805-INS-170 | chr1-7282805-INS-169 |
| chr1-7283576-INS-2376 | chr1-7283577-INS-2376 |
| chr1-7313346-INS-348 | chr1-7313346-INS-348 |
| chr1-7315957-INS-1893 | chr1-7315958-INS-1893 |
| chr1-7358120-INS-214 | chr1-7358120-INS-214 |
| chr1-7361976-INS-4815 | chr1-7361977-INS-4815 |
| chr1-7375618-INS-6975 | chr1-7375619-INS-6975 |
| chr1-7384765-INS-7327 | chr1-7384766-INS-7327 |
| chr1-7391722-INS-116 | chr1-7391722-INS-73 |

| | |
|---|---|
| chr1-7496954-INS-62 | chr1-7496954-INS-62 |
| chr1-7535857-INS-321 | chr1-7535857-INS-328 |
| chr1-7568235-INS-119 | chr1-7568235-INS-119 |
| chr1-7573133-INS-565 | chr1-7573133-INS-565 |
| chr1-7584942-INS-75 | chr1-7584938-INS-91 |
| chr1-7600064-INS-473 | chr1-7600064-INS-473 |
| chr1-7608418-INS-1256 | chr1-7608419-INS-1256 |
| chr1-7619623-INS-435 | chr1-7619623-INS-435 |
| chr1-7620677-INS-2959 | chr1-7620678-INS-2959 |
| chr1-7620995-INS-691 | chr1-7620995-INS-691 |
| chr1-7658161-INS-2888 | chr1-7658162-INS-2888 |
| chr1-7660917-INS-7291 | chr1-7660918-INS-7291 |
| chr1-7699904-INS-506 | chr1-7699904-INS-506 |
| chr1-7718559-INS-217 | chr1-7718559-INS-217 |
| chr1-7719713-INS-51 | chr1-7719713-INS-51 |
| chr1-7723649-INS-1867 | chr1-7723650-INS-1867 |
| chr1-7726162-INS-402 | chr1-7726162-INS-402 |
| chr1-7753824-INS-55 | chr1-7753824-INS-55 |
| chr1-7776337-INS-3107 | chr1-7776338-INS-3107 |
| chr1-7798136-INS-328 | chr1-7798136-INS-328 |
| chr1-7803778-INS-226 | chr1-7803778-INS-226 |
| chr1-7826603-INS-128 | chr1-7826603-INS-128 |
| chr1-7838917-INS-2296 | chr1-7838918-INS-2296 |
| chr1-7845112-INS-144 | chr1-7845122-INS-164 |
| chr1-7853010-INS-300 | chr1-7853010-INS-300 |
| chr1-7925497-INS-6775 | chr1-7925498-INS-6776 |
| chr1-7973565-INS-392 | chr1-7973565-INS-392 |
| chr1-7986101-INS-152 | chr1-7986117-INS-165 |
| chr1-8041061-INS-73 | chr1-8041061-INS-81 |
| chr1-8126714-INS-6540 | chr1-8126715-INS-6540 |
| chr1-8135236-INS-6357 | chr1-8135237-INS-6358 |
| chr1-8156595-INS-7274 | chr1-8156596-INS-7274 |
| chr1-8156695-INS-3476 | chr1-8156696-INS-3476 |
| chr1-8171335-INS-51 | chr1-8171335-INS-53 |
| chr1-8179260-INS-543 | chr1-8179260-INS-543 |
| chr1-8193018-INS-62 | chr1-8193018-INS-62 |
| chr1-8204873-INS-130 | chr1-8204873-INS-130 |
| chr1-8207037-INS-85 | chr1-8207042-INS-80 |
| chr1-8225038-INS-68 | chr1-8225038-INS-80 |
| chr1-8237994-INS-53 | chr1-8237994-INS-57 |
| chr1-8250655-INS-73 | chr1-8250655-INS-73 |
| chr1-8262262-INS-7256 | chr1-8262263-INS-7256 |
| chr1-8272786-INS-2965 | chr1-8272787-INS-2965 |
| chr1-8342645-INS-7210 | chr1-8342646-INS-7210 |
| chr1-8355077-INS-74 | chr1-8355077-INS-78 |
| chr1-8387792-INS-138 | chr1-8387792-INS-137 |
| chr1-8404661-INS-6368 | chr1-8404662-INS-6368 |
| chr1-8413755-INS-200 | chr1-8413755-INS-200 |
| chr1-8425934-INS-4787 | chr1-8425935-INS-4787 |
| chr1-8437219-INS-58 | chr1-8437219-INS-52 |
| chr1-8503735-INS-60 | chr1-8503735-INS-58 |
| chr1-8532733-INS-6514 | chr1-8532734-INS-6515 |
| chr1-8571096-INS-56 | chr1-8571096-INS-53 |
| chr1-8571440-INS-446 | chr1-8571440-INS-446 |
| chr1-8625128-INS-198 | chr1-8625128-INS-198 |
| chr1-8628085-INS-55 | chr1-8628085-INS-55 |
| chr1-8628354-INS-6282 | chr1-8628355-INS-6284 |
| chr1-8638576-INS-176 | chr1-8638576-INS-176 |
| chr1-8656859-INS-99 | chr1-8656859-INS-99 |
| chr1-8661172-INS-6300 | chr1-8661173-INS-6300 |
| chr1-8668151-INS-132 | chr1-8668145-INS-135 |
| chr1-8674002-INS-620 | chr1-8674002-INS-622 |
| chr1-8721915-INS-3918 | chr1-8721916-INS-3917 |
| chr1-8722460-INS-256 | chr1-8722460-INS-256 |
| chr1-8723761-INS-124 | chr1-8723753-INS-125 |
| chr1-8734128-INS-68 | chr1-8734128-INS-68 |
| chr1-8752868-INS-158 | chr1-8752868-INS-158 |
| chr1-8813401-INS-52 | chr1-8813401-INS-56 |
| chr1-8862541-INS-6565 | chr1-8862542-INS-6565 |
| chr1-8889307-INS-60 | chr1-8889307-INS-64 |
| chr1-8890177-INS-6548 | chr1-8890178-INS-6548 |
| chr1-8892103-INS-60 | chr1-8892103-INS-60 |
| chr1-8901958-INS-204 | chr1-8901958-INS-204 |
| chr1-8910474-INS-166 | chr1-8910474-INS-166 |
| chr1-8915982-INS-8198 | chr1-8915982-INS-8198 |
| chr1-8920752-INS-219 | chr1-8920752-INS-219 |
| chr1-8941197-INS-731 | chr1-8941197-INS-727 |

| | |
|---|---|
| chr1-8968588-INS-4440 | chr1-8968589-INS-4440 |
| chr1-8997855-INS-66 | chr1-8997856-INS-66 |
| chr1-9016459-INS-70 | chr1-9016459-INS-70 |
| chr1-9022487-INS-4460 | chr1-9022488-INS-4460 |
| chr1-9027170-INS-252 | chr1-9027170-INS-252 |
| chr1-9033717-INS-852 | chr1-9033718-INS-852 |
| chr1-9040145-INS-583 | chr1-9040145-INS-583 |
| chr1-9043560-INS-195 | chr1-9043560-INS-195 |
| chr1-9081152-INS-214 | chr1-9081152-INS-214 |
| chr1-9082407-INS-51 | chr1-9082407-INS-51 |
| chr1-9155980-INS-52 | chr1-9155980-INS-54 |
| chr1-9167945-INS-325 | chr1-9167945-INS-325 |
| chr1-9191231-INS-201 | chr1-9191231-INS-201 |
| chr1-9208991-INS-7112 | chr1-9208992-INS-7112 |
| chr1-9210871-INS-6986 | chr1-9210872-INS-6986 |
| chr1-9225706-INS-774 | chr1-9225706-INS-774 |
| chr1-9233827-INS-6742 | chr1-9233828-INS-6742 |
| chr1-9281886-INS-6409 | chr1-9281887-INS-6409 |
| chr1-9282418-INS-116 | chr1-9282418-INS-116 |
| chr1-9284170-INS-106 | chr1-9284170-INS-106 |
| chr1-9290973-INS-4250 | chr1-9290974-INS-4250 |
| chr1-9292298-INS-202 | chr1-9292298-INS-202 |
| chr1-9295530-INS-68 | chr1-9295530-INS-76 |
| chr1-9329930-INS-56 | chr1-9329930-INS-56 |
| chr1-9350070-INS-59 | chr1-9350070-INS-55 |
| chr1-9352038-INS-82 | chr1-9352038-INS-82 |
| chr1-9355005-INS-629 | chr1-9355005-INS-628 |
| chr1-9377769-INS-151 | chr1-9377769-INS-155 |
| chr1-9406396-INS-6845 | chr1-9406397-INS-6848 |
| chr1-9417158-INS-6485 | chr1-9417159-INS-6485 |
| chr1-9452032-INS-808 | chr1-9452032-INS-808 |
| chr1-9473294-INS-220 | chr1-9473294-INS-204 |
| chr1-9479742-INS-85 | chr1-9479742-INS-85 |
| chr1-9484069-INS-7028 | chr1-9484070-INS-7029 |
| chr1-9487288-INS-65 | chr1-9487288-INS-65 |
| chr1-9509119-INS-192 | chr1-9509119-INS-192 |
| chr1-9528322-INS-192 | chr1-9528322-INS-187 |
| chr1-9567046-INS-300 | chr1-9567046-INS-300 |
| chr1-9578054-INS-371 | chr1-9578054-INS-373 |
| chr1-9621038-INS-370 | chr1-9621038-INS-370 |
| chr1-9651290-INS-5209 | chr1-9651291-INS-5209 |
| chr1-9668539-INS-2566 | chr1-9668540-INS-2566 |
| chr1-9668872-INS-987 | chr1-9668872-INS-987 |
| chr1-9684093-INS-136 | chr1-9684093-INS-140 |
| chr1-9694435-INS-147 | chr1-9694435-INS-145 |
| chr1-9776006-INS-268 | chr1-9776006-INS-249 |
| chr1-9782257-INS-196 | chr1-9782257-INS-196 |
| chr1-9799969-INS-846 | chr1-9799969-INS-846 |
| chr1-9802798-INS-194 | chr1-9802798-INS-196 |
| chr1-9813870-INS-224 | chr1-9813870-INS-225 |
| chr1-9823662-INS-183 | chr1-9823662-INS-183 |
| chr1-9825562-INS-62 | chr1-9825562-INS-62 |
| chr1-9840253-INS-287 | chr1-9840253-INS-288 |
| chr1-9853662-INS-74 | chr1-9853660-INS-66 |
| chr1-9860334-INS-7364 | chr1-9860335-INS-7364 |
| chr1-9863057-INS-213 | chr1-9863057-INS-213 |
| chr1-9869058-INS-76 | chr1-9869095-INS-76 |
| chr1-9879869-INS-136 | chr1-9879869-INS-137 |
| chr1-9944600-INS-1266 | chr1-9944601-INS-1266 |
| chr1-9950048-INS-810 | chr1-9950048-INS-844 |
| chr1-9955252-INS-424 | chr1-9955256-INS-427 |
| chr1-9957008-INS-201 | chr1-9957008-INS-201 |
| chr1-9964157-INS-88 | chr1-9964157-INS-88 |
| chr1-9965130-INS-770 | chr1-9965130-INS-770 |
| chr1-9971430-INS-287 | chr1-9971430-INS-287 |
| chr1-9983935-INS-198 | chr1-9983935-INS-198 |
| chr1-9987587-INS-7103 | chr1-9987588-INS-7103 |
| chr1-10000117-INS-207 | chr1-10000117-INS-207 |
| chr1-10000888-INS-862 | chr1-10000888-INS-862 |
| chr1-10006675-INS-58 | chr1-10006675-INS-60 |
| chr1-10025887-INS-57 | chr1-10025887-INS-57 |
| chr1-10029347-INS-6467 | chr1-10029348-INS-6467 |
| chr1-10033327-INS-199 | chr1-10033327-INS-199 |
| chr1-10043723-INS-86 | chr1-10043723-INS-86 |
| chr1-10050609-INS-74 | chr1-10050582-INS-64 |
| chr1-10087387-INS-204 | chr1-10087387-INS-206 |
| chr1-10122475-INS-62 | chr1-10122475-INS-62 |

| | |
|---|---|
| chr1-10310325-INS-108 | chr1-10310325-INS-108 |
| chr1-10311149-INS-60 | chr1-10311150-INS-60 |
| chr1-10314939-INS-2952 | chr1-10314940-INS-2952 |
| chr1-10327445-INS-518 | chr1-10327445-INS-518 |
| chr1-10329729-INS-1036 | chr1-10329755-INS-1036 |
| chr1-10349465-INS-249 | chr1-10349465-INS-249 |
| chr1-10356718-INS-81 | chr1-10356718-INS-81 |
| chr1-10365347-INS-6262 | chr1-10365348-INS-6262 |
| chr1-10376321-INS-60 | chr1-10376321-INS-60 |
| chr1-10378935-INS-1127 | chr1-10378942-INS-1121 |
| chr1-10421700-INS-102 | chr1-10421700-INS-96 |
| chr1-10494254-INS-2857 | chr1-10494254-INS-2857 |
| chr1-10494255-INS-188 | chr1-10494255-INS-188 |
| chr1-10498051-INS-400 | chr1-10498051-INS-400 |
| chr1-10506324-INS-239 | chr1-10506324-INS-239 |
| chr1-10527807-INS-56 | chr1-10527807-INS-56 |
| chr1-10534084-INS-60 | chr1-10534084-INS-60 |
| chr1-10551348-INS-70 | chr1-10551348-INS-70 |
| chr1-10557448-INS-161 | chr1-10557448-INS-159 |
| chr1-10646446-INS-83 | chr1-10646416-INS-90 |
| chr1-10654999-INS-2187 | chr1-10655000-INS-2189 |
| chr1-10656351-INS-80 | chr1-10656351-INS-80 |
| chr1-10680022-INS-332 | chr1-10680016-INS-330 |
| chr1-10696580-INS-7197 | chr1-10696581-INS-7198 |
| chr1-10699007-INS-61 | chr1-10699007-INS-61 |
| chr1-10716200-INS-433 | chr1-10716200-INS-433 |
| chr1-10724720-INS-54 | chr1-10724720-INS-56 |
| chr1-10729951-INS-6376 | chr1-10729952-INS-6376 |
| chr1-10737767-INS-94 | chr1-10737767-INS-88 |
| chr1-10742596-INS-52 | chr1-10742596-INS-50 |
| chr1-10743876-INS-1756 | chr1-10743877-INS-1756 |
| chr1-10763846-INS-2495 | chr1-10763847-INS-2495 |
| chr1-10785682-INS-6514 | chr1-10785683-INS-6515 |
| chr1-10830849-INS-70 | chr1-10830849-INS-70 |
| chr1-10844415-INS-68 | chr1-10844415-INS-68 |
| chr1-10845395-INS-5655 | chr1-10845396-INS-5655 |
| chr1-10847931-INS-266 | chr1-10847936-INS-274 |
| chr1-10863112-INS-201 | chr1-10863112-INS-201 |
| chr1-10872535-INS-998 | chr1-10872535-INS-998 |
| chr1-10908892-INS-5487 | chr1-10908893-INS-5487 |
| chr1-10927407-INS-288 | chr1-10927407-INS-296 |
| chr1-10930070-INS-1698 | chr1-10930071-INS-1698 |
| chr1-10931368-INS-615 | chr1-10931368-INS-620 |
| chr1-10938846-INS-123 | chr1-10938846-INS-135 |
| chr1-10972328-INS-421 | chr1-10972328-INS-421 |
| chr1-11000487-INS-70 | chr1-11000482-INS-70 |
| chr1-11031064-INS-102 | chr1-11031064-INS-102 |
| chr1-11035873-INS-180 | chr1-11035874-INS-180 |
| chr1-11038206-INS-98 | chr1-11038206-INS-102 |
| chr1-11046355-INS-53 | chr1-11046355-INS-53 |
| chr1-11056598-INS-74 | chr1-11056590-INS-100 |
| chr1-11080401-INS-171 | chr1-11080401-INS-176 |
| chr1-11082954-INS-52 | chr1-11082954-INS-52 |
| chr1-11086423-INS-139 | chr1-11086433-INS-102 |
| chr1-11093937-INS-89 | chr1-11093937-INS-93 |
| chr1-11108625-INS-2739 | chr1-11108626-INS-2739 |
| chr1-11118788-INS-2370 | chr1-11118789-INS-2370 |
| chr1-11130907-INS-64 | chr1-11130907-INS-64 |
| chr1-11143403-INS-1305 | chr1-11143404-INS-1305 |
| chr1-11160200-INS-69 | chr1-11160200-INS-69 |
| chr1-11164319-INS-53 | chr1-11164319-INS-53 |
| chr1-11168108-INS-215 | chr1-11168108-INS-215 |
| chr1-11171039-INS-116 | chr1-11171039-INS-118 |
| chr1-11185198-INS-83 | chr1-11185198-INS-83 |
| chr1-11194892-INS-52 | chr1-11194844-INS-68 |
| chr1-11231770-INS-51 | chr1-11231778-INS-52 |
| chr1-11243439-INS-194 | chr1-11243439-INS-194 |
| chr1-11245233-INS-162 | chr1-11245233-INS-162 |
| chr1-11280509-INS-747 | chr1-11280509-INS-752 |
| chr1-11283924-INS-463 | chr1-11283924-INS-480 |
| chr1-11286641-INS-558 | chr1-11286641-INS-559 |
| chr1-11290289-INS-62 | chr1-11290289-INS-66 |
| chr1-11294962-INS-61 | chr1-11294962-INS-61 |
| chr1-11297130-INS-85 | chr1-11297130-INS-85 |
| chr1-11306504-INS-1915 | chr1-11306505-INS-1916 |
| chr1-11351556-INS-596 | chr1-11351556-INS-598 |
| chr1-11367531-INS-6891 | chr1-11367532-INS-6892 |

| | |
|---|---|
| chr1-11374194-INS-6404 | chr1-11374206-INS-6391 |
| chr1-11375777-INS-61 | chr1-11375777-INS-61 |
| chr1-11381843-INS-1417 | chr1-11381844-INS-1417 |
| chr1-11384110-INS-65 | chr1-11384110-INS-65 |
| chr1-11390895-INS-96 | chr1-11390933-INS-66 |
| chr1-11393624-INS-390 | chr1-11393624-INS-390 |
| chr1-11409315-INS-96 | chr1-11409311-INS-108 |
| chr1-11413417-INS-190 | chr1-11413417-INS-186 |
| chr1-11441549-INS-75 | chr1-11441549-INS-75 |
| chr1-11445608-INS-2068 | chr1-11445609-INS-2067 |
| chr1-11452684-INS-227 | chr1-11452684-INS-227 |
| chr1-11471211-INS-109 | chr1-11471211-INS-109 |
| chr1-11472512-INS-69 | chr1-11472512-INS-79 |
| chr1-11504415-INS-563 | chr1-11504415-INS-568 |
| chr1-11531576-INS-255 | chr1-11531576-INS-254 |
| chr1-11560955-INS-3185 | chr1-11560956-INS-3185 |
| chr1-11578264-INS-392 | chr1-11578264-INS-392 |
| chr1-11581053-INS-52 | chr1-11581053-INS-56 |
| chr1-11590814-INS-204 | chr1-11590814-INS-205 |
| chr1-11599049-INS-538 | chr1-11599049-INS-538 |
| chr1-11601625-INS-2868 | chr1-11601626-INS-2868 |
| chr1-11611937-INS-116 | chr1-11611937-INS-116 |
| chr1-11632766-INS-98 | chr1-11632764-INS-144 |
| chr1-11670519-INS-412 | chr1-11670519-INS-411 |
| chr1-11711853-INS-58 | chr1-11711853-INS-58 |
| chr1-11716942-INS-54 | chr1-11716942-INS-54 |
| chr1-11726395-INS-63 | chr1-11726395-INS-63 |
| chr1-11755654-INS-734 | chr1-11755654-INS-728 |
| chr1-11768643-INS-270 | chr1-11768643-INS-270 |
| chr1-11811633-INS-72 | chr1-11811633-INS-70 |
| chr1-11831080-INS-62 | chr1-11831080-INS-68 |
| chr1-11837559-INS-295 | chr1-11837559-INS-295 |
| chr1-11873040-INS-228 | chr1-11873040-INS-228 |
| chr1-11874091-INS-90 | chr1-11874094-INS-78 |
| chr1-11898262-INS-502 | chr1-11898262-INS-502 |
| chr1-11910022-INS-72 | chr1-11910020-INS-77 |
| chr1-11916821-INS-2145 | chr1-11916822-INS-2159 |
| chr1-11924008-INS-197 | chr1-11924008-INS-197 |
| chr1-12026442-INS-424 | chr1-12026442-INS-424 |
| chr1-12034473-INS-204 | chr1-12034473-INS-204 |
| chr1-12037169-INS-56 | chr1-12037133-INS-50 |
| chr1-12052457-INS-5528 | chr1-12052458-INS-5528 |
| chr1-12083323-INS-77 | chr1-12083323-INS-77 |
| chr1-12106119-INS-75 | chr1-12106128-INS-54 |
| chr1-12119797-INS-6791 | chr1-12119798-INS-6792 |
| chr1-12228194-INS-6416 | chr1-12228195-INS-6416 |
| chr1-12232867-INS-1772 | chr1-12232868-INS-1776 |
| chr1-12240884-INS-160 | chr1-12240885-INS-167 |
| chr1-12270726-INS-415 | chr1-12270726-INS-415 |
| chr1-12286891-INS-401 | chr1-12286891-INS-401 |
| chr1-12288147-INS-194 | chr1-12288147-INS-196 |
| chr1-12300817-INS-132 | chr1-12300818-INS-135 |
| chr1-12367475-INS-203 | chr1-12367475-INS-203 |
| chr1-12396832-INS-62 | chr1-12396832-INS-58 |
| chr1-12414700-INS-681 | chr1-12414700-INS-681 |
| chr1-12440259-INS-116 | chr1-12440259-INS-116 |
| chr1-12458403-INS-116 | chr1-12458387-INS-129 |
| chr1-12466176-INS-163 | chr1-12466176-INS-163 |
| chr1-12493041-INS-101 | chr1-12493041-INS-101 |
| chr1-12517716-INS-6232 | chr1-12517717-INS-6232 |
| chr1-12534731-INS-6482 | chr1-12534732-INS-6482 |
| chr1-12573282-INS-58 | chr1-12573282-INS-58 |
| chr1-12640568-INS-542 | chr1-12640568-INS-542 |
| chr1-12646926-INS-7163 | chr1-12646927-INS-7163 |
| chr1-12657777-INS-7172 | chr1-12657778-INS-7172 |
| chr1-12661586-INS-5513 | chr1-12661587-INS-5513 |
| chr1-12670138-INS-218 | chr1-12670138-INS-219 |
| chr1-12683930-INS-84 | chr1-12683930-INS-86 |
| chr1-12696012-INS-148 | chr1-12696012-INS-128 |
| chr1-12700856-INS-208 | chr1-12700864-INS-208 |
| chr1-12702871-INS-298 | chr1-12702871-INS-296 |
| chr1-12703934-INS-733 | chr1-12703935-INS-752 |
| chr1-12710254-INS-75 | chr1-12710254-INS-76 |
| chr1-12713960-INS-211 | chr1-12713959-INS-224 |
| chr1-12747525-INS-144 | chr1-12747525-INS-144 |
| chr1-12760327-INS-64 | chr1-12760327-INS-64 |
| chr1-12788001-INS-59 | chr1-12788001-INS-56 |

| | |
|---|---|
| chr1-12789570-INS-173 | chr1-12789570-INS-175 |
| chr1-12802247-INS-52 | chr1-12802247-INS-58 |
| chr1-12811594-INS-68 | chr1-12811604-INS-51 |
| chr1-12820364-INS-201 | chr1-12820364-INS-202 |
| chr1-12834023-INS-207 | chr1-12834023-INS-207 |
| chr1-12844718-INS-51 | chr1-12844719-INS-81 |
| chr1-12866362-INS-203 | chr1-12866362-INS-203 |
| chr1-12888135-INS-219 | chr1-12888135-INS-219 |
| chr1-12912395-INS-65 | chr1-12912395-INS-65 |
| chr1-12920764-INS-223 | chr1-12920764-INS-222 |
| chr1-12933001-INS-99 | chr1-12933001-INS-99 |
| chr1-12948048-INS-126 | chr1-12948048-INS-126 |
| chr1-12954049-INS-193 | chr1-12954049-INS-196 |
| chr1-12960851-INS-216 | chr1-12960851-INS-216 |
| chr1-12962193-INS-104 | chr1-12962193-INS-105 |
| chr1-12971929-INS-6468 | chr1-12971930-INS-6468 |
| chr1-12981061-INS-214 | chr1-12981061-INS-213 |
| chr1-13000232-INS-610 | chr1-13000232-INS-610 |
| chr1-13006716-INS-345 | chr1-13006716-INS-345 |
| chr1-13009098-INS-120 | chr1-13009098-INS-120 |
| chr1-13018849-INS-209 | chr1-13018849-INS-209 |
| chr1-13032784-INS-172 | chr1-13032784-INS-172 |
| chr1-13046594-INS-187 | chr1-13046594-INS-188 |
| chr1-13053041-INS-96 | chr1-13053041-INS-100 |
| chr1-13056608-INS-303 | chr1-13056608-INS-303 |
| chr1-13058535-INS-74 | chr1-13058535-INS-74 |
| chr1-13079972-INS-1985 | chr1-13079973-INS-2000 |
| chr1-13095945-INS-660 | chr1-13095945-INS-662 |
| chr1-13098408-INS-237 | chr1-13098408-INS-237 |
| chr1-13102901-INS-167 | chr1-13102901-INS-169 |
| chr1-13106858-INS-69 | chr1-13106858-INS-69 |
| chr1-13116120-INS-6466 | chr1-13116121-INS-6466 |
| chr1-13117675-INS-205 | chr1-13117675-INS-205 |
| chr1-13133980-INS-699 | chr1-13133980-INS-698 |
| chr1-13134726-INS-75 | chr1-13134694-INS-60 |
| chr1-13147620-INS-106 | chr1-13147620-INS-106 |
| chr1-13156871-INS-161 | chr1-13156872-INS-174 |
| chr1-13173119-INS-178 | chr1-13173119-INS-178 |
| chr1-13175478-INS-75 | chr1-13175478-INS-75 |
| chr1-13177222-INS-54 | chr1-13177222-INS-54 |
| chr1-13180318-INS-212 | chr1-13180318-INS-212 |
| chr1-13182910-INS-175 | chr1-13182910-INS-175 |
| chr1-13187103-INS-1491 | chr1-13187104-INS-1491 |
| chr1-13198023-INS-82 | chr1-13198049-INS-92 |
| chr1-13201251-INS-64 | chr1-13201251-INS-64 |
| chr1-13202974-INS-71 | chr1-13202972-INS-71 |
| chr1-13208435-INS-318 | chr1-13208438-INS-311 |
| chr1-13220682-INS-214 | chr1-13220682-INS-217 |
| chr1-13226952-INS-405 | chr1-13226952-INS-405 |
| chr1-13245963-INS-54 | chr1-13245963-INS-54 |
| chr1-13310323-INS-741 | chr1-13310323-INS-741 |
| chr1-13325378-INS-226 | chr1-13325378-INS-226 |
| chr1-13328503-INS-438 | chr1-13328503-INS-438 |
| chr1-13338903-INS-424 | chr1-13338903-INS-424 |
| chr1-13356464-INS-2610 | chr1-13356465-INS-2612 |
| chr1-13368799-INS-164 | chr1-13368799-INS-166 |
| chr1-13404098-INS-52 | chr1-13404098-INS-54 |
| chr1-13431835-INS-52 | chr1-13431835-INS-52 |
| chr1-13454834-INS-6415 | chr1-13454835-INS-6415 |
| chr1-13455402-INS-180 | chr1-13455408-INS-196 |
| chr1-13484162-INS-52 | chr1-13484162-INS-52 |
| chr1-13499979-INS-68 | chr1-13499979-INS-72 |
| chr1-13512114-INS-75 | chr1-13512114-INS-75 |
| chr1-13515993-INS-200 | chr1-13515993-INS-200 |
| chr1-13520981-INS-198 | chr1-13520981-INS-198 |
| chr1-13524541-INS-124 | chr1-13524515-INS-126 |
| chr1-13536900-INS-52 | chr1-13536901-INS-51 |
| chr1-13569585-INS-52 | chr1-13569585-INS-52 |
| chr1-13602710-INS-323 | chr1-13602710-INS-321 |
| chr1-13629882-INS-1035 | chr1-13629883-INS-1035 |
| chr1-13655475-INS-52 | chr1-13655475-INS-54 |
| chr1-13663477-INS-6442 | chr1-13663478-INS-6442 |
| chr1-13691251-INS-58 | chr1-13691251-INS-58 |
| chr1-13692873-INS-56 | chr1-13692873-INS-52 |
| chr1-13698345-INS-214 | chr1-13698345-INS-214 |
| chr1-13700838-INS-295 | chr1-13700838-INS-295 |
| chr1-13703487-INS-181 | chr1-13703486-INS-180 |

| | |
|---|---|
| chr1-13717114-INS-60 | chr1-13717114-INS-60 |
| chr1-13727340-INS-196 | chr1-13727340-INS-196 |
| chr1-13733800-INS-53 | chr1-13733800-INS-53 |
| chr1-13767444-INS-65 | chr1-13767444-INS-71 |
| chr1-13783768-INS-129 | chr1-13783768-INS-129 |
| chr1-13788445-INS-57 | chr1-13788448-INS-57 |
| chr1-13799863-INS-57 | chr1-13799864-INS-56 |
| chr1-13812325-INS-114 | chr1-13812325-INS-114 |
| chr1-13814317-INS-342 | chr1-13814317-INS-342 |
| chr1-13831204-INS-6696 | chr1-13831205-INS-6697 |
| chr1-13871604-INS-52 | chr1-13871604-INS-50 |
| chr1-13894870-INS-3204 | chr1-13894871-INS-3204 |
| chr1-13901448-INS-646 | chr1-13901448-INS-646 |
| chr1-13904020-INS-1330 | chr1-13904021-INS-1330 |
| chr1-13915226-INS-571 | chr1-13915226-INS-573 |
| chr1-13920255-INS-57 | chr1-13920255-INS-57 |
| chr1-13953011-INS-3665 | chr1-13953012-INS-3689 |
| chr1-13978354-INS-546 | chr1-13978354-INS-556 |
| chr1-13991463-INS-62 | chr1-13991463-INS-60 |
| chr1-13993801-INS-62 | chr1-13993801-INS-62 |
| chr1-14010800-INS-60 | chr1-14010800-INS-60 |
| chr1-14029990-INS-53 | chr1-14029989-INS-70 |
| chr1-14035892-INS-57 | chr1-14035892-INS-57 |
| chr1-14044906-INS-1888 | chr1-14044907-INS-1888 |
| chr1-14062939-INS-54 | chr1-14062939-INS-66 |
| chr1-14090974-INS-761 | chr1-14090974-INS-761 |
| chr1-14136203-INS-84 | chr1-14136203-INS-84 |
| chr1-14151079-INS-706 | chr1-14151079-INS-707 |
| chr1-14178015-INS-86 | chr1-14178065-INS-56 |
| chr1-14219268-INS-54 | chr1-14219278-INS-77 |
| chr1-14227700-INS-400 | chr1-14227700-INS-400 |
| chr1-14235724-INS-83 | chr1-14235724-INS-60 |
| chr1-14255459-INS-60 | chr1-14255459-INS-56 |
| chr1-14285820-INS-91 | chr1-14285820-INS-91 |
| chr1-14292893-INS-374 | chr1-14292893-INS-374 |
| chr1-14298262-INS-89 | chr1-14298240-INS-100 |
| chr1-14317145-INS-194 | chr1-14317145-INS-197 |
| chr1-14358962-INS-61 | chr1-14358962-INS-61 |
| chr1-14396968-INS-60 | chr1-14396968-INS-60 |
| chr1-14452671-INS-6897 | chr1-14452672-INS-6897 |
| chr1-14467462-INS-52 | chr1-14467462-INS-52 |
| chr1-14506627-INS-237 | chr1-14506627-INS-237 |
| chr1-14509206-INS-59 | chr1-14509206-INS-56 |
| chr1-14525076-INS-135 | chr1-14525076-INS-135 |
| chr1-14542292-INS-92 | chr1-14542292-INS-92 |
| chr1-14555028-INS-96 | chr1-14555028-INS-96 |
| chr1-14656009-INS-5192 | chr1-14656010-INS-5192 |
| chr1-14659574-INS-1681 | chr1-14659572-INS-1671 |
| chr1-14672314-INS-95 | chr1-14672310-INS-107 |
| chr1-14679337-INS-1960 | chr1-14679338-INS-1964 |
| chr1-14680960-INS-3402 | chr1-14680961-INS-3402 |
| chr1-14683638-INS-128 | chr1-14683638-INS-128 |
| chr1-14688603-INS-1997 | chr1-14688604-INS-1997 |
| chr1-14714230-INS-1096 | chr1-14714231-INS-1098 |
| chr1-14714236-INS-530 | chr1-14714236-INS-530 |
| chr1-14715436-INS-55 | chr1-14715436-INS-55 |
| chr1-14728164-INS-325 | chr1-14728164-INS-325 |
| chr1-14729297-INS-50 | chr1-14729297-INS-52 |
| chr1-14729604-INS-1283 | chr1-14729605-INS-1284 |
| chr1-14734425-INS-387 | chr1-14734425-INS-387 |
| chr1-14741060-INS-205 | chr1-14741060-INS-205 |
| chr1-14745238-INS-79 | chr1-14745238-INS-78 |
| chr1-14767669-INS-432 | chr1-14767669-INS-432 |
| chr1-14770857-INS-665 | chr1-14770857-INS-664 |
| chr1-14833852-INS-393 | chr1-14833850-INS-397 |
| chr1-14838703-INS-74 | chr1-14838703-INS-72 |
| chr1-14842690-INS-65 | chr1-14842688-INS-110 |
| chr1-14847933-INS-342 | chr1-14847933-INS-342 |
| chr1-14877276-INS-8877 | chr1-14877277-INS-8876 |
| chr1-14881807-INS-54 | chr1-14881807-INS-54 |
| chr1-14887697-INS-80 | chr1-14887697-INS-56 |
| chr1-14897781-INS-150 | chr1-14897776-INS-174 |
| chr1-14903296-INS-1286 | chr1-14903297-INS-1287 |
| chr1-14915845-INS-1418 | chr1-14915846-INS-1418 |
| chr1-14918823-INS-354 | chr1-14918823-INS-354 |
| chr1-14938410-INS-100 | chr1-14938418-INS-103 |
| chr1-14939159-INS-438 | chr1-14939159-INS-432 |

| | |
|---|---|
| chr1-14951181-INS-1581 | chr1-14951182-INS-1580 |
| chr1-14953036-INS-389 | chr1-14953036-INS-385 |
| chr1-14972419-INS-316 | chr1-14972419-INS-315 |
| chr1-15009298-INS-789 | chr1-15009298-INS-789 |
| chr1-15019904-INS-58 | chr1-15019888-INS-52 |
| chr1-15071953-INS-1246 | chr1-15071954-INS-1246 |
| chr1-15075888-INS-217 | chr1-15075888-INS-217 |
| chr1-15105174-INS-439 | chr1-15105174-INS-439 |
| chr1-15129108-INS-63 | chr1-15129108-INS-63 |
| chr1-15145568-INS-96 | chr1-15145571-INS-99 |
| chr1-15164315-INS-80 | chr1-15164315-INS-80 |
| chr1-15204591-INS-985 | chr1-15204591-INS-985 |
| chr1-15205572-INS-60 | chr1-15205568-INS-81 |
| chr1-15206924-INS-187 | chr1-15206924-INS-188 |
| chr1-15217327-INS-72 | chr1-15217327-INS-72 |
| chr1-15236021-INS-81 | chr1-15236018-INS-88 |
| chr1-15270986-INS-397 | chr1-15270986-INS-397 |
| chr1-15314923-INS-1280 | chr1-15314924-INS-1280 |
| chr1-15321973-INS-325 | chr1-15322017-INS-310 |
| chr1-15347319-INS-65 | chr1-15347319-INS-65 |
| chr1-15418844-INS-529 | chr1-15418844-INS-529 |
| chr1-15440677-INS-156 | chr1-15440677-INS-156 |
| chr1-15451340-INS-442 | chr1-15451340-INS-420 |
| chr1-15455913-INS-54 | chr1-15455922-INS-56 |
| chr1-15468834-INS-795 | chr1-15468835-INS-759 |
| chr1-15474351-INS-53 | chr1-15474351-INS-53 |
| chr1-15539769-INS-364 | chr1-15539769-INS-364 |
| chr1-15542287-INS-419 | chr1-15542327-INS-375 |
| chr1-15552883-INS-288 | chr1-15552883-INS-288 |
| chr1-15567822-INS-204 | chr1-15567822-INS-204 |
| chr1-15577240-INS-146 | chr1-15577240-INS-148 |
| chr1-15600024-INS-192 | chr1-15600024-INS-192 |
| chr1-15630154-INS-223 | chr1-15630154-INS-223 |
| chr1-15631568-INS-570 | chr1-15631568-INS-569 |
| chr1-15637423-INS-54 | chr1-15637415-INS-70 |
| chr1-15643092-INS-448 | chr1-15643092-INS-448 |
| chr1-15661035-INS-68 | chr1-15661033-INS-56 |
| chr1-15667434-INS-204 | chr1-15667434-INS-204 |
| chr1-15676887-INS-6225 | chr1-15676873-INS-6224 |
| chr1-15683319-INS-178 | chr1-15683319-INS-179 |
| chr1-15690130-INS-142 | chr1-15690130-INS-142 |
| chr1-15691421-INS-297 | chr1-15691421-INS-274 |
| chr1-15693278-INS-82 | chr1-15693278-INS-86 |
| chr1-15694937-INS-60 | chr1-15694937-INS-106 |
| chr1-15715654-INS-90 | chr1-15715654-INS-135 |
| chr1-15747784-INS-6642 | chr1-15747785-INS-6642 |
| chr1-15759933-INS-6887 | chr1-15759934-INS-6889 |
| chr1-15765315-INS-443 | chr1-15765315-INS-443 |
| chr1-15766804-INS-184 | chr1-15766804-INS-184 |
| chr1-15769030-INS-2801 | chr1-15769031-INS-2801 |
| chr1-15776543-INS-114 | chr1-15776537-INS-132 |
| chr1-15842174-INS-230 | chr1-15842174-INS-230 |
| chr1-15850204-INS-72 | chr1-15850204-INS-72 |
| chr1-15853329-INS-216 | chr1-15853329-INS-216 |
| chr1-15863724-INS-56 | chr1-15863724-INS-56 |
| chr1-15903402-INS-69 | chr1-15903402-INS-69 |
| chr1-15911323-INS-86 | chr1-15911323-INS-79 |
| chr1-15930416-INS-78 | chr1-15930416-INS-74 |
| chr1-15945655-INS-52 | chr1-15945655-INS-58 |
| chr1-15982705-INS-60 | chr1-15982705-INS-72 |
| chr1-16005408-INS-91 | chr1-16005404-INS-88 |
| chr1-16026325-INS-201 | chr1-16026325-INS-201 |
| chr1-16033800-INS-221 | chr1-16033800-INS-221 |
| chr1-16034744-INS-63 | chr1-16034744-INS-60 |
| chr1-16045211-INS-194 | chr1-16045211-INS-194 |
| chr1-16048090-INS-105 | chr1-16048081-INS-141 |
| chr1-16095501-INS-65 | chr1-16095501-INS-50 |
| chr1-16099584-INS-52 | chr1-16099584-INS-52 |
| chr1-16111838-INS-79 | chr1-16111838-INS-79 |
| chr1-16114329-INS-505 | chr1-16114329-INS-505 |
| chr1-16130934-INS-609 | chr1-16130936-INS-562 |
| chr1-16133653-INS-184 | chr1-16133654-INS-185 |
| chr1-16140343-INS-399 | chr1-16140343-INS-399 |
| chr1-16144381-INS-100 | chr1-16144381-INS-100 |
| chr1-16151971-INS-236 | chr1-16151971-INS-237 |
| chr1-16158653-INS-836 | chr1-16158644-INS-839 |
| chr1-16164806-INS-5595 | chr1-16164807-INS-5596 |

| | |
|---|---|
| chr1-16203422-INS-138 | chr1-16203395-INS-173 |
| chr1-16211965-INS-7483 | chr1-16211966-INS-7483 |
| chr1-16213972-INS-724 | chr1-16213972-INS-729 |
| chr1-16223791-INS-196 | chr1-16223791-INS-196 |
| chr1-16224465-INS-75 | chr1-16224465-INS-75 |
| chr1-16245216-INS-316 | chr1-16245229-INS-307 |
| chr1-16266027-INS-1016 | chr1-16266028-INS-1017 |
| chr1-16275740-INS-6714 | chr1-16275740-INS-6714 |
| chr1-16324541-INS-106 | chr1-16324543-INS-90 |
| chr1-16387497-INS-54 | chr1-16387497-INS-80 |
| chr1-16390923-INS-64 | chr1-16390923-INS-64 |
| chr1-16447349-INS-116 | chr1-16447352-INS-120 |
| chr1-16489939-INS-6147 | chr1-16489940-INS-6148 |
| chr1-16502504-INS-209 | chr1-16502504-INS-209 |
| chr1-16513640-INS-61 | chr1-16513652-INS-80 |
| chr1-16537592-INS-5391 | chr1-16537593-INS-5391 |
| chr1-16542954-INS-54 | chr1-16542954-INS-54 |
| chr1-16559157-INS-748 | chr1-16559157-INS-755 |
| chr1-16564645-INS-519 | chr1-16564645-INS-519 |
| chr1-16573822-INS-132 | chr1-16573822-INS-133 |
| chr1-16606826-INS-291 | chr1-16606826-INS-282 |
| chr1-16610956-INS-503 | chr1-16610956-INS-503 |
| chr1-16618672-INS-96 | chr1-16618672-INS-96 |
| chr1-16621250-INS-58 | chr1-16621233-INS-66 |
| chr1-16624729-INS-55 | chr1-16624720-INS-55 |
| chr1-16640141-INS-62 | chr1-16640141-INS-66 |
| chr1-16663061-INS-1922 | chr1-16663062-INS-1922 |
| chr1-16716922-INS-452 | chr1-16716922-INS-452 |
| chr1-16747739-INS-184 | chr1-16747739-INS-179 |
| chr1-16754216-INS-211 | chr1-16754214-INS-210 |
| chr1-16755615-INS-51 | chr1-16755609-INS-51 |
| chr1-16774474-INS-435 | chr1-16774474-INS-435 |
| chr1-16780169-INS-95 | chr1-16780169-INS-95 |
| chr1-16795457-INS-245 | chr1-16795457-INS-252 |
| chr1-16821270-INS-86 | chr1-16821270-INS-92 |
| chr1-16825544-INS-73 | chr1-16825544-INS-77 |
| chr1-16827872-INS-294 | chr1-16827872-INS-294 |
| chr1-16830514-INS-156 | chr1-16830515-INS-185 |
| chr1-16854258-INS-273 | chr1-16854258-INS-277 |
| chr1-16884590-INS-81 | chr1-16884590-INS-99 |
| chr1-16912318-INS-197 | chr1-16912318-INS-197 |
| chr1-16924873-INS-65 | chr1-16924894-INS-53 |
| chr1-16933644-INS-437 | chr1-16933644-INS-437 |
| chr1-16974953-INS-56 | chr1-16974953-INS-60 |
| chr1-16998525-INS-252 | chr1-16998525-INS-252 |
| chr1-17035859-INS-73 | chr1-17035845-INS-86 |
| chr1-17040180-INS-55 | chr1-17040180-INS-55 |
| chr1-17087447-INS-105 | chr1-17087474-INS-73 |
| chr1-17178484-INS-147 | chr1-17178484-INS-147 |
| chr1-17195451-INS-405 | chr1-17195451-INS-405 |
| chr1-17260900-INS-309 | chr1-17260900-INS-309 |
| chr1-17262846-INS-209 | chr1-17262846-INS-209 |
| chr1-17263806-INS-159 | chr1-17263806-INS-159 |
| chr1-17305474-INS-450 | chr1-17305474-INS-452 |
| chr1-17312014-INS-5509 | chr1-17312015-INS-5509 |
| chr1-17340457-INS-83 | chr1-17340454-INS-86 |
| chr1-17350977-INS-53 | chr1-17350977-INS-53 |
| chr1-17354957-INS-434 | chr1-17354957-INS-434 |
| chr1-17357557-INS-114 | chr1-17357557-INS-114 |
| chr1-17392951-INS-631 | chr1-17392962-INS-639 |
| chr1-17410302-INS-102 | chr1-17410302-INS-102 |
| chr1-17413424-INS-458 | chr1-17413424-INS-458 |
| chr1-17471272-INS-51 | chr1-17471272-INS-51 |
| chr1-17481335-INS-84 | chr1-17481335-INS-84 |
| chr1-17485117-INS-6274 | chr1-17485118-INS-6259 |
| chr1-17507625-INS-6223 | chr1-17507626-INS-6223 |
| chr1-17512873-INS-256 | chr1-17512873-INS-256 |
| chr1-17526987-INS-90 | chr1-17526987-INS-92 |
| chr1-17528227-INS-418 | chr1-17528227-INS-420 |
| chr1-17555992-INS-548 | chr1-17555992-INS-549 |
| chr1-17584181-INS-154 | chr1-17584163-INS-129 |
| chr1-17655124-INS-6441 | chr1-17655125-INS-6441 |
| chr1-17683177-INS-360 | chr1-17683177-INS-360 |
| chr1-17687969-INS-259 | chr1-17687969-INS-260 |
| chr1-17689580-INS-757 | chr1-17689564-INS-745 |
| chr1-17710994-INS-125 | chr1-17710994-INS-125 |
| chr1-17726401-INS-659 | chr1-17726401-INS-659 |

| | |
|---|---|
| chr1-17743142-INS-57 | chr1-17743142-INS-57 |
| chr1-17746034-INS-194 | chr1-17746034-INS-194 |
| chr1-17807760-INS-98 | chr1-17807761-INS-68 |
| chr1-17827814-INS-54 | chr1-17827814-INS-56 |
| chr1-17833677-INS-52 | chr1-17833676-INS-52 |
| chr1-17842129-INS-76 | chr1-17842129-INS-76 |
| chr1-17870780-INS-2809 | chr1-17870781-INS-2810 |
| chr1-17873055-INS-54 | chr1-17873055-INS-52 |
| chr1-17874261-INS-6366 | chr1-17874262-INS-6367 |
| chr1-17877734-INS-182 | chr1-17877734-INS-182 |
| chr1-17881886-INS-911 | chr1-17881886-INS-911 |
| chr1-17885357-INS-52 | chr1-17885357-INS-52 |
| chr1-17894731-INS-7336 | chr1-17894732-INS-7336 |
| chr1-17898144-INS-322 | chr1-17898147-INS-302 |
| chr1-17909527-INS-108 | chr1-17909528-INS-106 |
| chr1-17918362-INS-227 | chr1-17918362-INS-201 |
| chr1-17947311-INS-78 | chr1-17947311-INS-78 |
| chr1-17954246-INS-711 | chr1-17954246-INS-711 |
| chr1-17994414-INS-60 | chr1-17994414-INS-51 |
| chr1-18011593-INS-695 | chr1-18011593-INS-694 |
| chr1-18023526-INS-199 | chr1-18023526-INS-199 |
| chr1-18064332-INS-1374 | chr1-18064333-INS-1374 |
| chr1-18070899-INS-96 | chr1-18070899-INS-72 |
| chr1-18079919-INS-300 | chr1-18079919-INS-300 |
| chr1-18082115-INS-58 | chr1-18082096-INS-58 |
| chr1-18083521-INS-151 | chr1-18083521-INS-133 |
| chr1-18096834-INS-83 | chr1-18096834-INS-83 |
| chr1-18101823-INS-72 | chr1-18101823-INS-72 |
| chr1-18125739-INS-3488 | chr1-18125740-INS-3473 |
| chr1-18125740-INS-530 | chr1-18125740-INS-530 |
| chr1-18137244-INS-119 | chr1-18137244-INS-119 |
| chr1-18142950-INS-526 | chr1-18142950-INS-526 |
| chr1-18184867-INS-197 | chr1-18184847-INS-205 |
| chr1-18184977-INS-665 | chr1-18184975-INS-667 |
| chr1-18193810-INS-631 | chr1-18193810-INS-631 |
| chr1-18200130-INS-123 | chr1-18200130-INS-95 |
| chr1-18232562-INS-6413 | chr1-18232563-INS-6417 |
| chr1-18241489-INS-572 | chr1-18241489-INS-571 |
| chr1-18243390-INS-6976 | chr1-18243391-INS-6978 |
| chr1-18289950-INS-5364 | chr1-18289951-INS-5365 |
| chr1-18297269-INS-159 | chr1-18297274-INS-171 |
| chr1-18308341-INS-565 | chr1-18308341-INS-559 |
| chr1-18316957-INS-221 | chr1-18316957-INS-222 |
| chr1-18339540-INS-5199 | chr1-18339541-INS-5199 |
| chr1-18395416-INS-6674 | chr1-18395417-INS-6675 |
| chr1-18449039-INS-55 | chr1-18449039-INS-58 |
| chr1-18472416-INS-58 | chr1-18472416-INS-58 |
| chr1-18482722-INS-55 | chr1-18482722-INS-60 |
| chr1-18506570-INS-149 | chr1-18506572-INS-149 |
| chr1-18506668-INS-631 | chr1-18506668-INS-638 |
| chr1-18509363-INS-57 | chr1-18509363-INS-63 |
| chr1-18517159-INS-58 | chr1-18517157-INS-59 |
| chr1-18535597-INS-207 | chr1-18535597-INS-210 |
| chr1-18624982-INS-94 | chr1-18624982-INS-95 |
| chr1-18625591-INS-80 | chr1-18625573-INS-62 |
| chr1-18669094-INS-102 | chr1-18669094-INS-102 |
| chr1-18684258-INS-94 | chr1-18684258-INS-96 |
| chr1-18703246-INS-102 | chr1-18703246-INS-144 |
| chr1-18708246-INS-364 | chr1-18708246-INS-364 |
| chr1-18716951-INS-1071 | chr1-18716952-INS-1071 |
| chr1-18720670-INS-53 | chr1-18720670-INS-53 |
| chr1-18739354-INS-213 | chr1-18739354-INS-214 |
| chr1-18750493-INS-77 | chr1-18750493-INS-72 |
| chr1-18790870-INS-1177 | chr1-18790847-INS-1176 |
| chr1-18805290-INS-91 | chr1-18805290-INS-77 |
| chr1-18820705-INS-58 | chr1-18820679-INS-54 |
| chr1-18824938-INS-174 | chr1-18824938-INS-174 |
| chr1-18849349-INS-5257 | chr1-18849350-INS-5257 |
| chr1-18894174-INS-56 | chr1-18894174-INS-56 |
| chr1-18895214-INS-6423 | chr1-18895215-INS-6423 |
| chr1-18897873-INS-78 | chr1-18897843-INS-111 |
| chr1-18902108-INS-55 | chr1-18902108-INS-55 |
| chr1-18923484-INS-6925 | chr1-18923485-INS-6925 |
| chr1-18927049-INS-562 | chr1-18927049-INS-562 |
| chr1-18929342-INS-64 | chr1-18929342-INS-50 |
| chr1-18939864-INS-57 | chr1-18939864-INS-60 |
| chr1-18954521-INS-252 | chr1-18954521-INS-252 |

| | |
|---|---|
| chr1-18976226-INS-194 | chr1-18976226-INS-207 |
| chr1-18982753-INS-242 | chr1-18982753-INS-241 |
| chr1-18997950-INS-208 | chr1-18997950-INS-208 |
| chr1-19002957-INS-3110 | chr1-19002958-INS-3110 |
| chr1-19005723-INS-465 | chr1-19005723-INS-467 |
| chr1-19038346-INS-56 | chr1-19038346-INS-56 |
| chr1-19050613-INS-272 | chr1-19050613-INS-273 |
| chr1-19051426-INS-6466 | chr1-19051427-INS-6466 |
| chr1-19076650-INS-222 | chr1-19076650-INS-226 |
| chr1-19091865-INS-6368 | chr1-19091866-INS-6368 |
| chr1-19125034-INS-8354 | chr1-19125035-INS-8355 |
| chr1-19128205-INS-56 | chr1-19128205-INS-60 |
| chr1-19145283-INS-650 | chr1-19145283-INS-650 |
| chr1-19151592-INS-553 | chr1-19151592-INS-553 |
| chr1-19155888-INS-7085 | chr1-19155889-INS-7085 |
| chr1-19204575-INS-64 | chr1-19204569-INS-50 |
| chr1-19210276-INS-139 | chr1-19210276-INS-139 |
| chr1-19214824-INS-232 | chr1-19214824-INS-233 |
| chr1-19235535-INS-103 | chr1-19235535-INS-148 |
| chr1-19249536-INS-83 | chr1-19249536-INS-63 |
| chr1-19268320-INS-233 | chr1-19268320-INS-265 |
| chr1-19270455-INS-681 | chr1-19270455-INS-682 |
| chr1-19273046-INS-6915 | chr1-19273047-INS-6930 |
| chr1-19280840-INS-89 | chr1-19280836-INS-50 |
| chr1-19303795-INS-58 | chr1-19303793-INS-60 |
| chr1-19311951-INS-201 | chr1-19311951-INS-185 |
| chr1-19328531-INS-277 | chr1-19328529-INS-309 |
| chr1-19340996-INS-111 | chr1-19340989-INS-93 |
| chr1-19351213-INS-7353 | chr1-19351214-INS-7354 |
| chr1-19364041-INS-70 | chr1-19364017-INS-109 |
| chr1-19364984-INS-204 | chr1-19364984-INS-203 |
| chr1-19419567-INS-7120 | chr1-19419568-INS-7120 |
| chr1-19426167-INS-1718 | chr1-19426168-INS-1718 |
| chr1-19486792-INS-3639 | chr1-19486793-INS-3639 |
| chr1-19491055-INS-714 | chr1-19491055-INS-714 |
| chr1-19496092-INS-1794 | chr1-19496093-INS-1794 |
| chr1-19511030-INS-201 | chr1-19511030-INS-201 |
| chr1-19522181-INS-54 | chr1-19522181-INS-54 |
| chr1-19545106-INS-6223 | chr1-19545107-INS-6224 |
| chr1-19652621-INS-53 | chr1-19652621-INS-61 |
| chr1-19667077-INS-342 | chr1-19667077-INS-342 |
| chr1-19669613-INS-80 | chr1-19669612-INS-69 |
| chr1-19672477-INS-449 | chr1-19672477-INS-455 |
| chr1-19690235-INS-105 | chr1-19690241-INS-121 |
| chr1-19728206-INS-51 | chr1-19728206-INS-51 |
| chr1-19746711-INS-56 | chr1-19746711-INS-56 |
| chr1-19747241-INS-1008 | chr1-19747242-INS-1008 |
| chr1-19791080-INS-801 | chr1-19791080-INS-801 |
| chr1-19813744-INS-57 | chr1-19813744-INS-97 |
| chr1-19821475-INS-342 | chr1-19821475-INS-342 |
| chr1-19835071-INS-2526 | chr1-19835072-INS-2526 |
| chr1-19853228-INS-72 | chr1-19853246-INS-86 |
| chr1-19868686-INS-517 | chr1-19868686-INS-517 |
| chr1-19888586-INS-912 | chr1-19888586-INS-912 |
| chr1-19985923-INS-103 | chr1-19985923-INS-97 |
| chr1-20001157-INS-144 | chr1-20001155-INS-146 |
| chr1-20019044-INS-60 | chr1-20019044-INS-84 |
| chr1-20033314-INS-711 | chr1-20033314-INS-711 |
| chr1-20043004-INS-201 | chr1-20043004-INS-201 |
| chr1-20052930-INS-4302 | chr1-20052931-INS-4302 |
| chr1-20055727-INS-84 | chr1-20055727-INS-84 |
| chr1-20112391-INS-3338 | chr1-20112392-INS-3338 |
| chr1-20121625-INS-126 | chr1-20121625-INS-126 |
| chr1-20135600-INS-106 | chr1-20135600-INS-105 |
| chr1-20141482-INS-94 | chr1-20141482-INS-94 |
| chr1-20185953-INS-1322 | chr1-20185954-INS-1319 |
| chr1-20216926-INS-56 | chr1-20216919-INS-106 |
| chr1-20232461-INS-5733 | chr1-20232462-INS-5733 |
| chr1-20242555-INS-52 | chr1-20242555-INS-56 |
| chr1-20270265-INS-181 | chr1-20270265-INS-181 |
| chr1-20306611-INS-58 | chr1-20306580-INS-71 |
| chr1-20409071-INS-70 | chr1-20409071-INS-70 |
| chr1-20419844-INS-366 | chr1-20419845-INS-321 |
| chr1-20476375-INS-7582 | chr1-20476376-INS-7582 |
| chr1-20535596-INS-71 | chr1-20535596-INS-71 |
| chr1-20542180-INS-279 | chr1-20542180-INS-279 |
| chr1-20545449-INS-74 | chr1-20545449-INS-92 |

| | |
|---|---|
| chr1-20579878-INS-283 | chr1-20579878-INS-280 |
| chr1-20588520-INS-459 | chr1-20588527-INS-469 |
| chr1-20609653-INS-355 | chr1-20609653-INS-360 |
| chr1-20612314-INS-6457 | chr1-20612315-INS-6457 |
| chr1-20618272-INS-54 | chr1-20618272-INS-54 |
| chr1-20619426-INS-74 | chr1-20619426-INS-74 |
| chr1-20640190-INS-72 | chr1-20640190-INS-72 |
| chr1-20641517-INS-57 | chr1-20641517-INS-59 |
| chr1-20651793-INS-193 | chr1-20651793-INS-192 |
| chr1-20683986-INS-144 | chr1-20683986-INS-144 |
| chr1-20694471-INS-114 | chr1-20694438-INS-135 |
| chr1-20728084-INS-84 | chr1-20728084-INS-86 |
| chr1-20731387-INS-87 | chr1-20731387-INS-87 |
| chr1-20732890-INS-7976 | chr1-20732891-INS-7976 |
| chr1-20735394-INS-114 | chr1-20735380-INS-116 |
| chr1-20750782-INS-553 | chr1-20750782-INS-554 |
| chr1-20765364-INS-3862 | chr1-20765365-INS-3862 |
| chr1-20767452-INS-338 | chr1-20767452-INS-338 |
| chr1-20809061-INS-330 | chr1-20809061-INS-330 |
| chr1-20812574-INS-100 | chr1-20812572-INS-145 |
| chr1-20816005-INS-237 | chr1-20816005-INS-237 |
| chr1-20843250-INS-790 | chr1-20843232-INS-816 |
| chr1-20844282-INS-204 | chr1-20844282-INS-205 |
| chr1-20877580-INS-83 | chr1-20877584-INS-84 |
| chr1-20881428-INS-130 | chr1-20881390-INS-130 |
| chr1-20901777-INS-69 | chr1-20901777-INS-67 |
| chr1-20906699-INS-292 | chr1-20906699-INS-296 |
| chr1-20911919-INS-124 | chr1-20911919-INS-125 |
| chr1-20917535-INS-76 | chr1-20917535-INS-73 |
| chr1-20926661-INS-64 | chr1-20926665-INS-76 |
| chr1-20952406-INS-713 | chr1-20952407-INS-713 |
| chr1-20955494-INS-66 | chr1-20955494-INS-66 |
| chr1-20957891-INS-104 | chr1-20957891-INS-108 |
| chr1-20964097-INS-84 | chr1-20964097-INS-84 |
| chr1-20980312-INS-208 | chr1-20980308-INS-204 |
| chr1-21010665-INS-69 | chr1-21010638-INS-72 |
| chr1-21041852-INS-208 | chr1-21041852-INS-208 |
| chr1-21046230-INS-243 | chr1-21046230-INS-243 |
| chr1-21058604-INS-123 | chr1-21058604-INS-123 |
| chr1-21092254-INS-1893 | chr1-21092255-INS-1893 |
| chr1-21099324-INS-570 | chr1-21099324-INS-571 |
| chr1-21100782-INS-166 | chr1-21100782-INS-201 |
| chr1-21151597-INS-3033 | chr1-21151598-INS-3033 |
| chr1-21173453-INS-418 | chr1-21173453-INS-416 |
| chr1-21194218-INS-128 | chr1-21194218-INS-123 |
| chr1-21197975-INS-60 | chr1-21197975-INS-65 |
| chr1-21216642-INS-63 | chr1-21216642-INS-93 |
| chr1-21218695-INS-102 | chr1-21218695-INS-102 |
| chr1-21244664-INS-52 | chr1-21244664-INS-52 |
| chr1-21282467-INS-179 | chr1-21282467-INS-179 |
| chr1-21308309-INS-64 | chr1-21308308-INS-70 |
| chr1-21320733-INS-212 | chr1-21320734-INS-213 |
| chr1-21324921-INS-302 | chr1-21324928-INS-303 |
| chr1-21326987-INS-358 | chr1-21326987-INS-358 |
| chr1-21343824-INS-72 | chr1-21343824-INS-72 |
| chr1-21349559-INS-216 | chr1-21349559-INS-216 |
| chr1-21355569-INS-562 | chr1-21355569-INS-563 |
| chr1-21358149-INS-61 | chr1-21358149-INS-61 |
| chr1-21367104-INS-102 | chr1-21367104-INS-104 |
| chr1-21380631-INS-178 | chr1-21380631-INS-178 |
| chr1-21384173-INS-404 | chr1-21384173-INS-415 |
| chr1-21400382-INS-5369 | chr1-21400383-INS-5331 |
| chr1-21409764-INS-660 | chr1-21409764-INS-660 |
| chr1-21427484-INS-799 | chr1-21427484-INS-799 |
| chr1-21431874-INS-6524 | chr1-21431875-INS-6524 |
| chr1-21434873-INS-4495 | chr1-21434874-INS-4496 |
| chr1-21473467-INS-148 | chr1-21473467-INS-148 |
| chr1-21476540-INS-217 | chr1-21476540-INS-217 |
| chr1-21488165-INS-90 | chr1-21488166-INS-92 |
| chr1-21497525-INS-1429 | chr1-21497526-INS-1447 |
| chr1-21497828-INS-118 | chr1-21497828-INS-118 |
| chr1-21500237-INS-185 | chr1-21500264-INS-136 |
| chr1-21501862-INS-52 | chr1-21501862-INS-52 |
| chr1-21504392-INS-88 | chr1-21504392-INS-88 |
| chr1-21512021-INS-200 | chr1-21512021-INS-201 |
| chr1-21524243-INS-441 | chr1-21524243-INS-441 |
| chr1-21545388-INS-220 | chr1-21545388-INS-200 |

| | |
|---|---|
| chr1-21559850-INS-72 | chr1-21559861-INS-88 |
| chr1-21562728-INS-60 | chr1-21562728-INS-60 |
| chr1-21574644-INS-58 | chr1-21574635-INS-54 |
| chr1-21597070-INS-172 | chr1-21597070-INS-171 |
| chr1-21611062-INS-7117 | chr1-21611063-INS-7117 |
| chr1-21631177-INS-1407 | chr1-21631178-INS-1407 |
| chr1-21672156-INS-6350 | chr1-21672157-INS-6351 |
| chr1-21697422-INS-6789 | chr1-21697423-INS-6789 |
| chr1-21703120-INS-958 | chr1-21703120-INS-958 |
| chr1-21724060-INS-372 | chr1-21724060-INS-372 |
| chr1-21731972-INS-51 | chr1-21731980-INS-73 |
| chr1-21740326-INS-578 | chr1-21740326-INS-593 |
| chr1-21745058-INS-974 | chr1-21745058-INS-974 |
| chr1-21754069-INS-356 | chr1-21754069-INS-356 |
| chr1-21755316-INS-490 | chr1-21755316-INS-491 |
| chr1-21761023-INS-147 | chr1-21761023-INS-149 |
| chr1-21768746-INS-6468 | chr1-21768747-INS-6468 |
| chr1-21792911-INS-68 | chr1-21792911-INS-72 |
| chr1-21796932-INS-51 | chr1-21796932-INS-53 |
| chr1-21825784-INS-77 | chr1-21825781-INS-88 |
| chr1-21851650-INS-377 | chr1-21851650-INS-375 |
| chr1-21865796-INS-1012 | chr1-21865797-INS-1016 |
| chr1-21868498-INS-146 | chr1-21868498-INS-146 |
| chr1-21883719-INS-147 | chr1-21883719-INS-147 |
| chr1-21885400-INS-1457 | chr1-21885401-INS-1456 |
| chr1-21897053-INS-1901 | chr1-21897054-INS-1902 |
| chr1-21903508-INS-1218 | chr1-21903509-INS-1218 |
| chr1-21922340-INS-397 | chr1-21922340-INS-397 |
| chr1-21937051-INS-96 | chr1-21937051-INS-68 |
| chr1-21964461-INS-110 | chr1-21964461-INS-110 |
| chr1-21966040-INS-398 | chr1-21966040-INS-398 |
| chr1-22010160-INS-72 | chr1-22010160-INS-73 |
| chr1-22014647-INS-104 | chr1-22014647-INS-104 |
| chr1-22040107-INS-52 | chr1-22040107-INS-54 |
| chr1-22056302-INS-606 | chr1-22056302-INS-606 |
| chr1-22078578-INS-339 | chr1-22078575-INS-344 |
| chr1-22099066-INS-152 | chr1-22099066-INS-157 |
| chr1-22133421-INS-76 | chr1-22133421-INS-78 |
| chr1-22134313-INS-450 | chr1-22134309-INS-472 |
| chr1-22151608-INS-213 | chr1-22151608-INS-215 |
| chr1-22157577-INS-106 | chr1-22157577-INS-106 |
| chr1-22172474-INS-266 | chr1-22172474-INS-265 |
| chr1-22174727-INS-96 | chr1-22174727-INS-72 |
| chr1-22177047-INS-65 | chr1-22177047-INS-80 |
| chr1-22214351-INS-599 | chr1-22214344-INS-606 |
| chr1-22216509-INS-358 | chr1-22216509-INS-358 |
| chr1-22254600-INS-52 | chr1-22254601-INS-60 |
| chr1-22266730-INS-70 | chr1-22266730-INS-70 |
| chr1-22276176-INS-70 | chr1-22276175-INS-111 |
| chr1-22318103-INS-2157 | chr1-22318104-INS-2157 |
| chr1-22328340-INS-72 | chr1-22328377-INS-68 |
| chr1-22331212-INS-342 | chr1-22331214-INS-330 |
| chr1-22360965-INS-202 | chr1-22360965-INS-202 |
| chr1-22369014-INS-55 | chr1-22369014-INS-55 |
| chr1-22371749-INS-335 | chr1-22371749-INS-334 |
| chr1-22377790-INS-60 | chr1-22377784-INS-52 |
| chr1-22384493-INS-89 | chr1-22384493-INS-89 |
| chr1-22404730-INS-6401 | chr1-22404731-INS-6401 |
| chr1-22424367-INS-56 | chr1-22424351-INS-96 |
| chr1-22425460-INS-573 | chr1-22425460-INS-573 |
| chr1-22436194-INS-875 | chr1-22436194-INS-875 |
| chr1-22489556-INS-62 | chr1-22489574-INS-68 |
| chr1-22506002-INS-6705 | chr1-22506003-INS-6704 |
| chr1-22531865-INS-66 | chr1-22531865-INS-75 |
| chr1-22553076-INS-497 | chr1-22553076-INS-497 |
| chr1-22591044-INS-66 | chr1-22591044-INS-66 |
| chr1-22604865-INS-63 | chr1-22604865-INS-53 |
| chr1-22613758-INS-125 | chr1-22613773-INS-141 |
| chr1-22622772-INS-62 | chr1-22622773-INS-52 |
| chr1-22643356-INS-886 | chr1-22643356-INS-886 |
| chr1-22661702-INS-98 | chr1-22661702-INS-114 |
| chr1-22671999-INS-3562 | chr1-22672000-INS-3563 |
| chr1-22675549-INS-52 | chr1-22675549-INS-52 |
| chr1-22686828-INS-74 | chr1-22686828-INS-72 |
| chr1-22741122-INS-107 | chr1-22741122-INS-82 |
| chr1-22750028-INS-593 | chr1-22750028-INS-593 |
| chr1-22754502-INS-249 | chr1-22754501-INS-252 |

| | |
|---|---|
| chr1-22783662-INS-1383 | chr1-22783663-INS-1388 |
| chr1-22821577-INS-2952 | chr1-22821578-INS-2952 |
| chr1-22822304-INS-212 | chr1-22822305-INS-212 |
| chr1-22839112-INS-87 | chr1-22839112-INS-87 |
| chr1-22849544-INS-137 | chr1-22849544-INS-141 |
| chr1-22866667-INS-80 | chr1-22866667-INS-67 |
| chr1-22914940-INS-58 | chr1-22914940-INS-54 |
| chr1-22926904-INS-70 | chr1-22926954-INS-65 |
| chr1-22962826-INS-208 | chr1-22962826-INS-208 |
| chr1-22970030-INS-775 | chr1-22970030-INS-775 |
| chr1-22972679-INS-864 | chr1-22972676-INS-865 |
| chr1-22975129-INS-894 | chr1-22975129-INS-895 |
| chr1-22978557-INS-59 | chr1-22978557-INS-59 |
| chr1-23025811-INS-55 | chr1-23025811-INS-55 |
| chr1-23028942-INS-52 | chr1-23028942-INS-50 |
| chr1-23036782-INS-7747 | chr1-23036783-INS-7748 |
| chr1-23053928-INS-74 | chr1-23053929-INS-68 |
| chr1-23088493-INS-85 | chr1-23088493-INS-85 |
| chr1-23100110-INS-176 | chr1-23100110-INS-176 |
| chr1-23120685-INS-57 | chr1-23120685-INS-60 |
| chr1-23130483-INS-112 | chr1-23130503-INS-84 |
| chr1-23184052-INS-197 | chr1-23184052-INS-194 |
| chr1-23190586-INS-92 | chr1-23190586-INS-92 |
| chr1-23234527-INS-59 | chr1-23234518-INS-71 |
| chr1-23282930-INS-211 | chr1-23282930-INS-207 |
| chr1-23311590-INS-3318 | chr1-23311591-INS-3321 |
| chr1-23357094-INS-60 | chr1-23357094-INS-60 |
| chr1-23382935-INS-389 | chr1-23382935-INS-389 |
| chr1-23391783-INS-61 | chr1-23391783-INS-65 |
| chr1-23402563-INS-91 | chr1-23402563-INS-95 |
| chr1-23420296-INS-51 | chr1-23420296-INS-51 |
| chr1-23425268-INS-402 | chr1-23425268-INS-402 |
| chr1-23434147-INS-3138 | chr1-23434148-INS-3138 |
| chr1-23448158-INS-76 | chr1-23448158-INS-76 |
| chr1-23466405-INS-126 | chr1-23466405-INS-126 |
| chr1-23467486-INS-88 | chr1-23467486-INS-88 |
| chr1-23487406-INS-286 | chr1-23487404-INS-294 |
| chr1-23488430-INS-575 | chr1-23488430-INS-576 |
| chr1-23498672-INS-206 | chr1-23498672-INS-206 |
| chr1-23525591-INS-50 | chr1-23525549-INS-74 |
| chr1-23535211-INS-120 | chr1-23535211-INS-125 |
| chr1-23569445-INS-824 | chr1-23569445-INS-824 |
| chr1-23573653-INS-326 | chr1-23573653-INS-326 |
| chr1-23594126-INS-390 | chr1-23594126-INS-390 |
| chr1-23603290-INS-600 | chr1-23603290-INS-601 |
| chr1-23614707-INS-94 | chr1-23614660-INS-99 |
| chr1-23624655-INS-73 | chr1-23624649-INS-75 |
| chr1-23633753-INS-148 | chr1-23633753-INS-111 |
| chr1-23639650-INS-889 | chr1-23639650-INS-889 |
| chr1-23640539-INS-567 | chr1-23640539-INS-568 |
| chr1-23653405-INS-102 | chr1-23653405-INS-102 |
| chr1-23679821-INS-110 | chr1-23679824-INS-113 |
| chr1-23700814-INS-167 | chr1-23700814-INS-167 |
| chr1-23702575-INS-542 | chr1-23702575-INS-542 |
| chr1-23711178-INS-85 | chr1-23711180-INS-85 |
| chr1-23721768-INS-52 | chr1-23721769-INS-56 |
| chr1-23724141-INS-169 | chr1-23724141-INS-169 |
| chr1-23751885-INS-759 | chr1-23751885-INS-759 |
| chr1-23755386-INS-114 | chr1-23755386-INS-114 |
| chr1-23758743-INS-534 | chr1-23758743-INS-534 |
| chr1-23803187-INS-56 | chr1-23803187-INS-56 |
| chr1-23828875-INS-54 | chr1-23828875-INS-54 |
| chr1-23898038-INS-713 | chr1-23898038-INS-715 |
| chr1-23913331-INS-169 | chr1-23913331-INS-165 |
| chr1-23945268-INS-208 | chr1-23945268-INS-208 |
| chr1-23955418-INS-60 | chr1-23955419-INS-70 |
| chr1-23971028-INS-206 | chr1-23971028-INS-203 |
| chr1-23972156-INS-67 | chr1-23972152-INS-62 |
| chr1-23988151-INS-194 | chr1-23988151-INS-194 |
| chr1-23999644-INS-218 | chr1-23999644-INS-218 |
| chr1-24003555-INS-539 | chr1-24003555-INS-539 |
| chr1-24014996-INS-203 | chr1-24014996-INS-204 |
| chr1-24016394-INS-60 | chr1-24016394-INS-60 |
| chr1-24033926-INS-464 | chr1-24033926-INS-452 |
| chr1-24048688-INS-217 | chr1-24048688-INS-217 |
| chr1-24064660-INS-1886 | chr1-24064661-INS-1886 |
| chr1-24107301-INS-95 | chr1-24107314-INS-93 |

| | |
|---|---|
| chr1-24108795-INS-56 | chr1-24108809-INS-68 |
| chr1-24143084-INS-7235 | chr1-24143085-INS-7235 |
| chr1-24152142-INS-325 | chr1-24152142-INS-325 |
| chr1-24152850-INS-52 | chr1-24152850-INS-54 |
| chr1-24163567-INS-90 | chr1-24163573-INS-89 |
| chr1-24174070-INS-233 | chr1-24174070-INS-233 |
| chr1-24189769-INS-57 | chr1-24189769-INS-59 |
| chr1-24191707-INS-2487 | chr1-24191708-INS-2487 |
| chr1-24198039-INS-5005 | chr1-24198040-INS-5005 |
| chr1-24229061-INS-77 | chr1-24229061-INS-77 |
| chr1-24231778-INS-248 | chr1-24231778-INS-252 |
| chr1-24305183-INS-70 | chr1-24305185-INS-106 |
| chr1-24327192-INS-67 | chr1-24327192-INS-63 |
| chr1-24334351-INS-199 | chr1-24334351-INS-199 |
| chr1-24353719-INS-188 | chr1-24353719-INS-186 |
| chr1-24373424-INS-342 | chr1-24373424-INS-342 |
| chr1-24386825-INS-90 | chr1-24386826-INS-66 |
| chr1-24425054-INS-6806 | chr1-24425055-INS-6806 |
| chr1-24439182-INS-600 | chr1-24439182-INS-600 |
| chr1-24441144-INS-65 | chr1-24441144-INS-66 |
| chr1-24450849-INS-1326 | chr1-24450850-INS-1328 |
| chr1-24476836-INS-60 | chr1-24476836-INS-58 |
| chr1-24478641-INS-73 | chr1-24478637-INS-121 |
| chr1-24494489-INS-79 | chr1-24494489-INS-78 |
| chr1-24504519-INS-348 | chr1-24504519-INS-349 |
| chr1-24515099-INS-146 | chr1-24515099-INS-146 |
| chr1-24535518-INS-208 | chr1-24535518-INS-208 |
| chr1-24544131-INS-62 | chr1-24544131-INS-62 |
| chr1-24544957-INS-54 | chr1-24544957-INS-54 |
| chr1-24550267-INS-173 | chr1-24550266-INS-174 |
| chr1-24575523-INS-464 | chr1-24575507-INS-464 |
| chr1-24610088-INS-54 | chr1-24610100-INS-52 |
| chr1-24613517-INS-52 | chr1-24613517-INS-50 |
| chr1-24628403-INS-332 | chr1-24628403-INS-333 |
| chr1-24637190-INS-3326 | chr1-24637191-INS-3326 |
| chr1-24639915-INS-71 | chr1-24639915-INS-71 |
| chr1-24686064-INS-192 | chr1-24686064-INS-180 |
| chr1-24686890-INS-406 | chr1-24686890-INS-406 |
| chr1-24706368-INS-69 | chr1-24706368-INS-69 |
| chr1-24713968-INS-315 | chr1-24713968-INS-299 |
| chr1-24738563-INS-70 | chr1-24738599-INS-70 |
| chr1-24778186-INS-1268 | chr1-24778187-INS-1268 |
| chr1-24782587-INS-76 | chr1-24782587-INS-74 |
| chr1-24806085-INS-59 | chr1-24806085-INS-59 |
| chr1-24813808-INS-87 | chr1-24813804-INS-85 |
| chr1-24816722-INS-56 | chr1-24816722-INS-60 |
| chr1-24823938-INS-52 | chr1-24823938-INS-52 |
| chr1-24826156-INS-398 | chr1-24826156-INS-398 |
| chr1-24849871-INS-884 | chr1-24849871-INS-888 |
| chr1-24871879-INS-68 | chr1-24871883-INS-66 |
| chr1-24874599-INS-117 | chr1-24874599-INS-117 |
| chr1-24890010-INS-868 | chr1-24890010-INS-869 |
| chr1-24892167-INS-69 | chr1-24892168-INS-64 |
| chr1-24906531-INS-103 | chr1-24906531-INS-66 |
| chr1-24915368-INS-62 | chr1-24915372-INS-64 |
| chr1-24936404-INS-7933 | chr1-24936405-INS-7934 |
| chr1-24949310-INS-79 | chr1-24949320-INS-92 |
| chr1-24954860-INS-78 | chr1-24954860-INS-78 |
| chr1-24981250-INS-88 | chr1-24981230-INS-118 |
| chr1-24989806-INS-76 | chr1-24989806-INS-76 |
| chr1-24992185-INS-54 | chr1-24992211-INS-55 |
| chr1-25019219-INS-2325 | chr1-25019220-INS-2325 |
| chr1-25019231-INS-309 | chr1-25019231-INS-310 |
| chr1-25044504-INS-83 | chr1-25044504-INS-77 |
| chr1-25064815-INS-66 | chr1-25064820-INS-69 |
| chr1-25090866-INS-51 | chr1-25090866-INS-51 |
| chr1-25157802-INS-146 | chr1-25157791-INS-122 |
| chr1-25159222-INS-60 | chr1-25159222-INS-60 |
| chr1-25186599-INS-79 | chr1-25186599-INS-79 |
| chr1-25207491-INS-183 | chr1-25207491-INS-184 |
| chr1-25210767-INS-66 | chr1-25210740-INS-66 |
| chr1-25238809-INS-69 | chr1-25238810-INS-69 |
| chr1-25262573-INS-56 | chr1-25262573-INS-52 |
| chr1-25316111-INS-6952 | chr1-25316112-INS-6953 |
| chr1-25330667-INS-56 | chr1-25330669-INS-61 |
| chr1-25332248-INS-174 | chr1-25332248-INS-174 |
| chr1-25339605-INS-3631 | chr1-25339606-INS-3631 |

| | |
|---|---|
| chr1-25342363-INS-171 | chr1-25342363-INS-151 |
| chr1-25353047-INS-1482 | chr1-25353048-INS-1482 |
| chr1-25380033-INS-52 | chr1-25380033-INS-52 |
| chr1-25439924-INS-58 | chr1-25439924-INS-62 |
| chr1-25442306-INS-214 | chr1-25442306-INS-214 |
| chr1-25463403-INS-195 | chr1-25463418-INS-186 |
| chr1-25475207-INS-51 | chr1-25475207-INS-58 |
| chr1-25553932-INS-1892 | chr1-25553933-INS-1892 |
| chr1-25553933-INS-400 | chr1-25553933-INS-400 |
| chr1-25567370-INS-169 | chr1-25567370-INS-169 |
| chr1-25571223-INS-215 | chr1-25571223-INS-216 |
| chr1-25576867-INS-67 | chr1-25576867-INS-75 |
| chr1-25612604-INS-6232 | chr1-25612605-INS-6232 |
| chr1-25648092-INS-344 | chr1-25648092-INS-343 |
| chr1-25687943-INS-158 | chr1-25687943-INS-158 |
| chr1-25718574-INS-3124 | chr1-25718575-INS-3124 |
| chr1-25720595-INS-52 | chr1-25720595-INS-52 |
| chr1-25756651-INS-60 | chr1-25756651-INS-56 |
| chr1-25766476-INS-69 | chr1-25766501-INS-68 |
| chr1-25797298-INS-991 | chr1-25797298-INS-989 |
| chr1-25824014-INS-52 | chr1-25824014-INS-50 |
| chr1-25847923-INS-122 | chr1-25847917-INS-128 |
| chr1-25856597-INS-53 | chr1-25856597-INS-59 |
| chr1-25902588-INS-199 | chr1-25902588-INS-201 |
| chr1-25927204-INS-381 | chr1-25927204-INS-381 |
| chr1-25964929-INS-100 | chr1-25964929-INS-100 |
| chr1-25979141-INS-53 | chr1-25979141-INS-53 |
| chr1-25980218-INS-97 | chr1-25980218-INS-97 |
| chr1-26031811-INS-203 | chr1-26031811-INS-203 |
| chr1-26035517-INS-710 | chr1-26035517-INS-710 |
| chr1-26065170-INS-55 | chr1-26065170-INS-55 |
| chr1-26065348-INS-586 | chr1-26065348-INS-586 |
| chr1-26075475-INS-129 | chr1-26075475-INS-129 |
| chr1-26114818-INS-6794 | chr1-26114819-INS-6794 |
| chr1-26119207-INS-50 | chr1-26119226-INS-51 |
| chr1-26130476-INS-733 | chr1-26130476-INS-738 |
| chr1-26134059-INS-191 | chr1-26134056-INS-196 |
| chr1-26149784-INS-2760 | chr1-26149785-INS-2760 |
| chr1-26177009-INS-360 | chr1-26177009-INS-360 |
| chr1-26183291-INS-2681 | chr1-26183292-INS-2683 |
| chr1-26229427-INS-315 | chr1-26229424-INS-312 |
| chr1-26241629-INS-64 | chr1-26241615-INS-65 |
| chr1-26245421-INS-58 | chr1-26245421-INS-62 |
| chr1-26265741-INS-59 | chr1-26265741-INS-59 |
| chr1-26272510-INS-52 | chr1-26272510-INS-56 |
| chr1-26277618-INS-6231 | chr1-26277619-INS-6232 |
| chr1-26280926-INS-621 | chr1-26280926-INS-624 |
| chr1-26304269-INS-98 | chr1-26304269-INS-106 |
| chr1-26318681-INS-6452 | chr1-26318682-INS-6452 |
| chr1-26325011-INS-51 | chr1-26325011-INS-51 |
| chr1-26365055-INS-504 | chr1-26365055-INS-504 |
| chr1-26432362-INS-111 | chr1-26432362-INS-117 |
| chr1-26443488-INS-336 | chr1-26443488-INS-336 |
| chr1-26465071-INS-52 | chr1-26465056-INS-50 |
| chr1-26501899-INS-225 | chr1-26501899-INS-225 |
| chr1-26557113-INS-115 | chr1-26557113-INS-115 |
| chr1-26578113-INS-184 | chr1-26578113-INS-185 |
| chr1-26604925-INS-1491 | chr1-26604945-INS-1491 |
| chr1-26631637-INS-204 | chr1-26631637-INS-204 |
| chr1-26634144-INS-68 | chr1-26634144-INS-56 |
| chr1-26647865-INS-115 | chr1-26647865-INS-87 |
| chr1-26656968-INS-284 | chr1-26656968-INS-284 |
| chr1-26658074-INS-538 | chr1-26658074-INS-538 |
| chr1-26667293-INS-103 | chr1-26667293-INS-103 |
| chr1-26669483-INS-55 | chr1-26669484-INS-55 |
| chr1-26685840-INS-189 | chr1-26685840-INS-189 |
| chr1-26718034-INS-257 | chr1-26718034-INS-257 |
| chr1-26743166-INS-80 | chr1-26743166-INS-80 |
| chr1-26773368-INS-137 | chr1-26773368-INS-133 |
| chr1-26788450-INS-187 | chr1-26788462-INS-215 |
| chr1-26810708-INS-285 | chr1-26810708-INS-285 |
| chr1-26844100-INS-213 | chr1-26844100-INS-213 |
| chr1-26857051-INS-6895 | chr1-26857052-INS-6895 |
| chr1-26891306-INS-843 | chr1-26891306-INS-843 |
| chr1-26894424-INS-1165 | chr1-26894425-INS-1165 |
| chr1-26896958-INS-193 | chr1-26896958-INS-194 |
| chr1-26904969-INS-54 | chr1-26904969-INS-60 |

| | |
|---|---|
| chr1-26929630-INS-60 | chr1-26929630-INS-59 |
| chr1-26938336-INS-53 | chr1-26938336-INS-60 |
| chr1-26945809-INS-995 | chr1-26945839-INS-1014 |
| chr1-26949887-INS-54 | chr1-26949887-INS-54 |
| chr1-27008778-INS-6636 | chr1-27008779-INS-6636 |
| chr1-27009526-INS-1122 | chr1-27009527-INS-1122 |
| chr1-27029359-INS-188 | chr1-27029321-INS-181 |
| chr1-27050333-INS-384 | chr1-27050333-INS-384 |
| chr1-27058962-INS-94 | chr1-27058962-INS-94 |
| chr1-27064309-INS-894 | chr1-27064309-INS-894 |
| chr1-27071624-INS-159 | chr1-27071624-INS-159 |
| chr1-27078412-INS-102 | chr1-27078412-INS-94 |
| chr1-27095917-INS-57 | chr1-27095918-INS-53 |
| chr1-27122489-INS-629 | chr1-27122490-INS-632 |
| chr1-27126717-INS-62 | chr1-27126717-INS-58 |
| chr1-27145120-INS-331 | chr1-27145120-INS-328 |
| chr1-27158512-INS-1802 | chr1-27158513-INS-1801 |
| chr1-27181446-INS-744 | chr1-27181446-INS-744 |
| chr1-27244635-INS-343 | chr1-27244635-INS-343 |
| chr1-27260940-INS-81 | chr1-27260941-INS-60 |
| chr1-27266916-INS-54 | chr1-27266916-INS-54 |
| chr1-27270288-INS-60 | chr1-27270282-INS-72 |
| chr1-27291312-INS-345 | chr1-27291312-INS-344 |
| chr1-27302221-INS-329 | chr1-27302221-INS-329 |
| chr1-27337151-INS-81 | chr1-27337151-INS-80 |
| chr1-27339688-INS-3667 | chr1-27339689-INS-3667 |
| chr1-27342024-INS-11683 | chr1-27342025-INS-11709 |
| chr1-27353737-INS-72 | chr1-27353737-INS-72 |
| chr1-27368540-INS-481 | chr1-27368540-INS-482 |
| chr1-27418268-INS-82 | chr1-27418268-INS-59 |
| chr1-27436394-INS-56 | chr1-27436394-INS-58 |
| chr1-27439074-INS-54 | chr1-27439074-INS-52 |
| chr1-27450822-INS-345 | chr1-27450822-INS-345 |
| chr1-27537401-INS-64 | chr1-27537420-INS-65 |
| chr1-27563069-INS-2358 | chr1-27563070-INS-2356 |
| chr1-27645970-INS-80 | chr1-27645970-INS-94 |
| chr1-27660855-INS-7165 | chr1-27660856-INS-7165 |
| chr1-27680955-INS-7136 | chr1-27680956-INS-7136 |
| chr1-27734114-INS-60 | chr1-27734106-INS-60 |
| chr1-27749094-INS-1604 | chr1-27749095-INS-1604 |
| chr1-27759142-INS-130 | chr1-27759142-INS-130 |
| chr1-27847899-INS-52 | chr1-27847899-INS-52 |
| chr1-27907057-INS-1276 | chr1-27907058-INS-1265 |
| chr1-27914159-INS-486 | chr1-27914159-INS-488 |
| chr1-27947040-INS-158 | chr1-27947040-INS-155 |
| chr1-27967260-INS-6596 | chr1-27967261-INS-6597 |
| chr1-27974504-INS-6191 | chr1-27974499-INS-6184 |
| chr1-27979953-INS-1061 | chr1-27979954-INS-1061 |
| chr1-27982132-INS-51 | chr1-27982132-INS-51 |
| chr1-28012446-INS-177 | chr1-28012446-INS-177 |
| chr1-28030491-INS-1893 | chr1-28030492-INS-1893 |
| chr1-28053352-INS-3115 | chr1-28053353-INS-3114 |
| chr1-28058235-INS-6485 | chr1-28058236-INS-6484 |
| chr1-28060657-INS-4545 | chr1-28060658-INS-4545 |
| chr1-28079454-INS-859 | chr1-28079452-INS-871 |
| chr1-28084486-INS-561 | chr1-28084487-INS-560 |
| chr1-28109960-INS-66 | chr1-28109960-INS-68 |
| chr1-28158711-INS-210 | chr1-28158711-INS-210 |
| chr1-28249112-INS-169 | chr1-28249112-INS-179 |
| chr1-28252402-INS-87 | chr1-28252402-INS-81 |
| chr1-28259214-INS-396 | chr1-28259214-INS-396 |
| chr1-28314567-INS-6449 | chr1-28314568-INS-6449 |
| chr1-28364560-INS-84 | chr1-28364560-INS-78 |
| chr1-28388083-INS-99 | chr1-28388084-INS-102 |
| chr1-28454550-INS-6193 | chr1-28454551-INS-6193 |
| chr1-28454702-INS-59 | chr1-28454702-INS-61 |
| chr1-28528984-INS-1335 | chr1-28528986-INS-1360 |
| chr1-28539733-INS-6076 | chr1-28539734-INS-6079 |
| chr1-28576722-INS-161 | chr1-28576722-INS-161 |
| chr1-28588871-INS-894 | chr1-28588871-INS-889 |
| chr1-28615713-INS-324 | chr1-28615713-INS-324 |
| chr1-28667747-INS-60 | chr1-28667745-INS-62 |
| chr1-28668822-INS-107 | chr1-28668822-INS-107 |
| chr1-28677682-INS-105 | chr1-28677682-INS-99 |
| chr1-28698801-INS-66 | chr1-28698801-INS-66 |
| chr1-28713346-INS-1888 | chr1-28713347-INS-1888 |
| chr1-28743871-INS-1366 | chr1-28743879-INS-1354 |

| | |
|---|---|
| chr1-28793145-INS-93 | chr1-28793145-INS-93 |
| chr1-28798243-INS-352 | chr1-28798243-INS-376 |
| chr1-28830992-INS-7128 | chr1-28830993-INS-7128 |
| chr1-28864183-INS-53 | chr1-28864194-INS-66 |
| chr1-28865689-INS-95 | chr1-28865689-INS-108 |
| chr1-28871805-INS-7551 | chr1-28871806-INS-7553 |
| chr1-28918293-INS-54 | chr1-28918293-INS-52 |
| chr1-28918391-INS-6278 | chr1-28918392-INS-6275 |
| chr1-28946653-INS-68 | chr1-28946653-INS-68 |
| chr1-28976762-INS-1534 | chr1-28976763-INS-1536 |
| chr1-29003498-INS-869 | chr1-29003498-INS-827 |
| chr1-29005690-INS-1473 | chr1-29005691-INS-1473 |
| chr1-29014434-INS-56 | chr1-29014434-INS-60 |
| chr1-29021216-INS-1065 | chr1-29021217-INS-1064 |
| chr1-29029994-INS-1747 | chr1-29029995-INS-1747 |
| chr1-29062843-INS-128 | chr1-29062843-INS-135 |
| chr1-29071560-INS-119 | chr1-29071560-INS-96 |
| chr1-29073717-INS-58 | chr1-29073717-INS-58 |
| chr1-29078049-INS-7152 | chr1-29078050-INS-7152 |
| chr1-29088187-INS-159 | chr1-29088186-INS-150 |
| chr1-29097851-INS-441 | chr1-29097851-INS-441 |
| chr1-29114757-INS-80 | chr1-29114757-INS-80 |
| chr1-29119621-INS-76 | chr1-29119621-INS-76 |
| chr1-29168868-INS-5792 | chr1-29168869-INS-5792 |
| chr1-29183896-INS-272 | chr1-29183896-INS-272 |
| chr1-29184806-INS-501 | chr1-29184806-INS-501 |
| chr1-29188874-INS-72 | chr1-29188866-INS-96 |
| chr1-29219345-INS-6468 | chr1-29219346-INS-6468 |
| chr1-29226485-INS-270 | chr1-29226485-INS-270 |
| chr1-29259963-INS-80 | chr1-29259963-INS-68 |
| chr1-29260003-INS-4918 | chr1-29260004-INS-4918 |
| chr1-29264338-INS-62 | chr1-29264340-INS-54 |
| chr1-29269731-INS-57 | chr1-29269731-INS-61 |
| chr1-29279609-INS-569 | chr1-29279609-INS-570 |
| chr1-29285141-INS-63 | chr1-29285141-INS-67 |
| chr1-29286205-INS-97 | chr1-29286205-INS-97 |
| chr1-29289753-INS-51 | chr1-29289753-INS-58 |
| chr1-29357558-INS-158 | chr1-29357556-INS-150 |
| chr1-29378470-INS-102 | chr1-29378470-INS-100 |
| chr1-29418979-INS-926 | chr1-29418979-INS-943 |
| chr1-29438921-INS-102 | chr1-29438921-INS-120 |
| chr1-29455620-INS-61 | chr1-29455622-INS-62 |
| chr1-29490175-INS-1742 | chr1-29490176-INS-1718 |
| chr1-29547890-INS-995 | chr1-29547890-INS-996 |
| chr1-29586828-INS-280 | chr1-29586828-INS-281 |
| chr1-29590747-INS-2075 | chr1-29590748-INS-2074 |
| chr1-29615453-INS-1244 | chr1-29615454-INS-1243 |
| chr1-29636565-INS-53 | chr1-29636565-INS-53 |
| chr1-29639857-INS-607 | chr1-29639857-INS-606 |
| chr1-29677869-INS-159 | chr1-29677869-INS-151 |
| chr1-29689521-INS-274 | chr1-29689521-INS-278 |
| chr1-29719645-INS-79 | chr1-29719645-INS-79 |
| chr1-29737971-INS-1890 | chr1-29737972-INS-1890 |
| chr1-29765748-INS-4593 | chr1-29765749-INS-4594 |
| chr1-29802887-INS-6914 | chr1-29802888-INS-6914 |
| chr1-29851314-INS-6483 | chr1-29851315-INS-6483 |
| chr1-29897164-INS-92 | chr1-29897164-INS-74 |
| chr1-29909520-INS-7140 | chr1-29909521-INS-7140 |
| chr1-29948773-INS-55 | chr1-29948773-INS-73 |
| chr1-29950710-INS-312 | chr1-29950710-INS-312 |
| chr1-29954493-INS-970 | chr1-29954493-INS-970 |
| chr1-29963444-INS-956 | chr1-29963444-INS-978 |
| chr1-29972724-INS-625 | chr1-29972739-INS-647 |
| chr1-30003228-INS-220 | chr1-30003228-INS-221 |
| chr1-30013021-INS-56 | chr1-30013021-INS-50 |
| chr1-30020943-INS-62 | chr1-30020943-INS-62 |
| chr1-30022496-INS-52 | chr1-30022496-INS-52 |
| chr1-30058551-INS-835 | chr1-30058551-INS-839 |
| chr1-30072694-INS-3020 | chr1-30072695-INS-3000 |
| chr1-30105071-INS-190 | chr1-30105071-INS-190 |
| chr1-30124989-INS-128 | chr1-30124989-INS-128 |
| chr1-30129740-INS-6656 | chr1-30129741-INS-6656 |
| chr1-30145736-INS-6297 | chr1-30145737-INS-6297 |
| chr1-30153607-INS-92 | chr1-30153598-INS-113 |
| chr1-30170709-INS-136 | chr1-30170709-INS-88 |
| chr1-30195345-INS-2562 | chr1-30195346-INS-2562 |
| chr1-30213093-INS-62 | chr1-30213093-INS-54 |

| | |
|---|---|
| chr1-30225966-INS-84 | chr1-30225924-INS-102 |
| chr1-30248602-INS-247 | chr1-30248602-INS-243 |
| chr1-30250327-INS-58 | chr1-30250327-INS-58 |
| chr1-30253618-INS-83 | chr1-30253618-INS-83 |
| chr1-30339435-INS-295 | chr1-30339435-INS-295 |
| chr1-30362345-INS-80 | chr1-30362345-INS-65 |
| chr1-30368283-INS-94 | chr1-30368288-INS-66 |
| chr1-30413591-INS-5387 | chr1-30413592-INS-5387 |
| chr1-30443333-INS-54 | chr1-30443293-INS-68 |
| chr1-30447228-INS-827 | chr1-30447228-INS-829 |
| chr1-30452521-INS-8743 | chr1-30452522-INS-8743 |
| chr1-30458450-INS-7218 | chr1-30458451-INS-7219 |
| chr1-30495351-INS-1720 | chr1-30495352-INS-1720 |
| chr1-30522806-INS-1268 | chr1-30522807-INS-1269 |
| chr1-30536417-INS-1819 | chr1-30536425-INS-1816 |
| chr1-30554712-INS-6302 | chr1-30554713-INS-6288 |
| chr1-30557127-INS-81 | chr1-30557127-INS-81 |
| chr1-30668745-INS-70 | chr1-30668745-INS-70 |
| chr1-30723120-INS-397 | chr1-30723120-INS-397 |
| chr1-30737443-INS-72 | chr1-30737443-INS-50 |
| chr1-30750380-INS-283 | chr1-30750365-INS-289 |
| chr1-30764181-INS-8327 | chr1-30764182-INS-8326 |
| chr1-30769861-INS-214 | chr1-30769861-INS-214 |
| chr1-30814139-INS-109 | chr1-30814139-INS-96 |
| chr1-30818173-INS-75 | chr1-30818173-INS-55 |
| chr1-30819972-INS-113 | chr1-30819972-INS-113 |
| chr1-30836674-INS-70 | chr1-30836674-INS-70 |
| chr1-30899433-INS-186 | chr1-30899433-INS-187 |
| chr1-30920443-INS-53 | chr1-30920443-INS-53 |
| chr1-30926513-INS-127 | chr1-30926516-INS-92 |
| chr1-30927425-INS-531 | chr1-30927425-INS-531 |
| chr1-30928597-INS-234 | chr1-30928597-INS-221 |
| chr1-30951122-INS-169 | chr1-30951122-INS-161 |
| chr1-30955792-INS-7212 | chr1-30955793-INS-7212 |
| chr1-30966883-INS-56 | chr1-30966883-INS-56 |
| chr1-30969985-INS-156 | chr1-30969985-INS-156 |
| chr1-31003119-INS-139 | chr1-31003131-INS-139 |
| chr1-31003805-INS-71 | chr1-31003805-INS-71 |
| chr1-31093367-INS-117 | chr1-31093365-INS-78 |
| chr1-31102481-INS-6619 | chr1-31102482-INS-6621 |
| chr1-31113563-INS-81 | chr1-31113563-INS-54 |
| chr1-31140279-INS-2452 | chr1-31140280-INS-2453 |
| chr1-31161857-INS-187 | chr1-31161857-INS-187 |
| chr1-31171787-INS-384 | chr1-31171787-INS-383 |
| chr1-31178128-INS-52 | chr1-31178128-INS-52 |
| chr1-31180469-INS-53 | chr1-31180469-INS-53 |
| chr1-31190674-INS-6483 | chr1-31190669-INS-6473 |
| chr1-31192908-INS-773 | chr1-31192908-INS-774 |
| chr1-31228835-INS-66 | chr1-31228835-INS-66 |
| chr1-31237164-INS-109 | chr1-31237164-INS-108 |
| chr1-31323031-INS-4174 | chr1-31323032-INS-4174 |
| chr1-31470855-INS-66 | chr1-31470855-INS-66 |
| chr1-31625686-INS-7173 | chr1-31625687-INS-7173 |
| chr1-31674384-INS-70 | chr1-31674384-INS-70 |
| chr1-31675596-INS-168 | chr1-31675642-INS-128 |
| chr1-31690926-INS-508 | chr1-31690926-INS-512 |
| chr1-31708806-INS-830 | chr1-31708812-INS-831 |
| chr1-31718216-INS-56 | chr1-31718216-INS-56 |
| chr1-31718856-INS-421 | chr1-31718856-INS-421 |
| chr1-31720008-INS-238 | chr1-31720008-INS-238 |
| chr1-31781452-INS-872 | chr1-31781452-INS-876 |
| chr1-31793703-INS-7064 | chr1-31793704-INS-7064 |
| chr1-31905671-INS-213 | chr1-31905671-INS-213 |
| chr1-32017674-INS-81 | chr1-32017674-INS-81 |
| chr1-32019952-INS-76 | chr1-32019952-INS-80 |
| chr1-32058629-INS-540 | chr1-32058629-INS-540 |
| chr1-32117989-INS-74 | chr1-32117989-INS-80 |
| chr1-32139908-INS-622 | chr1-32139908-INS-622 |
| chr1-32142734-INS-1650 | chr1-32142735-INS-1650 |
| chr1-32152448-INS-57 | chr1-32152448-INS-57 |
| chr1-32157208-INS-63 | chr1-32157208-INS-63 |
| chr1-32250386-INS-884 | chr1-32250390-INS-900 |
| chr1-32270705-INS-72 | chr1-32270705-INS-72 |
| chr1-32279381-INS-7155 | chr1-32279382-INS-7156 |
| chr1-32312604-INS-373 | chr1-32312604-INS-374 |
| chr1-32342221-INS-501 | chr1-32342221-INS-503 |
| chr1-32398060-INS-112 | chr1-32398060-INS-104 |

| | |
|---|---|
| chr1-32418697-INS-88 | chr1-32418693-INS-86 |
| chr1-32456456-INS-472 | chr1-32456456-INS-472 |
| chr1-32485053-INS-90 | chr1-32485049-INS-94 |
| chr1-32509233-INS-6429 | chr1-32509234-INS-6430 |
| chr1-32513290-INS-864 | chr1-32513289-INS-891 |
| chr1-32516495-INS-56 | chr1-32516495-INS-74 |
| chr1-32533186-INS-6737 | chr1-32533187-INS-6736 |
| chr1-32547555-INS-64 | chr1-32547555-INS-66 |
| chr1-32581889-INS-6344 | chr1-32581890-INS-6319 |
| chr1-32593682-INS-6668 | chr1-32593683-INS-6666 |
| chr1-32620189-INS-83 | chr1-32620189-INS-83 |
| chr1-32627725-INS-7390 | chr1-32627726-INS-7385 |
| chr1-32645697-INS-54 | chr1-32645705-INS-50 |
| chr1-32647193-INS-3967 | chr1-32647194-INS-3968 |
| chr1-32653745-INS-219 | chr1-32653745-INS-219 |
| chr1-32665630-INS-773 | chr1-32665630-INS-773 |
| chr1-32685674-INS-148 | chr1-32685674-INS-151 |
| chr1-32712842-INS-59 | chr1-32712842-INS-59 |
| chr1-32725234-INS-76 | chr1-32725234-INS-78 |
| chr1-32740968-INS-2211 | chr1-32740969-INS-2211 |
| chr1-32747327-INS-84 | chr1-32747332-INS-72 |
| chr1-32767314-INS-114 | chr1-32767314-INS-123 |
| chr1-32767890-INS-1495 | chr1-32767891-INS-1495 |
| chr1-32770903-INS-75 | chr1-32770903-INS-75 |
| chr1-32782214-INS-52 | chr1-32782214-INS-80 |
| chr1-32786676-INS-1559 | chr1-32786677-INS-1559 |
| chr1-32788414-INS-104 | chr1-32788417-INS-95 |
| chr1-32836570-INS-450 | chr1-32836570-INS-450 |
| chr1-32872055-INS-133 | chr1-32872055-INS-97 |
| chr1-32878164-INS-107 | chr1-32878164-INS-147 |
| chr1-32878777-INS-55 | chr1-32878777-INS-55 |
| chr1-32887333-INS-72 | chr1-32887333-INS-72 |
| chr1-32904246-INS-68 | chr1-32904246-INS-64 |
| chr1-32907122-INS-357 | chr1-32907122-INS-364 |
| chr1-32913833-INS-684 | chr1-32913833-INS-684 |
| chr1-32922920-INS-628 | chr1-32922920-INS-624 |
| chr1-32933946-INS-497 | chr1-32933946-INS-497 |
| chr1-32942659-INS-72 | chr1-32942659-INS-50 |
| chr1-32944620-INS-963 | chr1-32944620-INS-963 |
| chr1-32951272-INS-4505 | chr1-32951273-INS-4506 |
| chr1-32961011-INS-114 | chr1-32961011-INS-90 |
| chr1-32968239-INS-64 | chr1-32968239-INS-52 |
| chr1-32978446-INS-261 | chr1-32978446-INS-261 |
| chr1-32987671-INS-119 | chr1-32987672-INS-74 |
| chr1-32997971-INS-6290 | chr1-32997972-INS-6290 |
| chr1-33002053-INS-80 | chr1-33002053-INS-80 |
| chr1-33013965-INS-419 | chr1-33013965-INS-418 |
| chr1-33046179-INS-6413 | chr1-33046180-INS-6413 |
| chr1-33051671-INS-308 | chr1-33051671-INS-308 |
| chr1-33079473-INS-100 | chr1-33079473-INS-92 |
| chr1-33093006-INS-172 | chr1-33092997-INS-180 |
| chr1-33113656-INS-291 | chr1-33113629-INS-296 |
| chr1-33129281-INS-77 | chr1-33129277-INS-72 |
| chr1-33144975-INS-65 | chr1-33144975-INS-67 |
| chr1-33149420-INS-54 | chr1-33149458-INS-60 |
| chr1-33180416-INS-234 | chr1-33180416-INS-236 |
| chr1-33188872-INS-60 | chr1-33188872-INS-88 |
| chr1-33189274-INS-2587 | chr1-33189275-INS-2587 |
| chr1-33198283-INS-117 | chr1-33198283-INS-81 |
| chr1-33209620-INS-2597 | chr1-33209621-INS-2597 |
| chr1-33214618-INS-66 | chr1-33214624-INS-56 |
| chr1-33216324-INS-60 | chr1-33216324-INS-75 |
| chr1-33217392-INS-3122 | chr1-33217393-INS-3122 |
| chr1-33221774-INS-205 | chr1-33221774-INS-205 |
| chr1-33225392-INS-6799 | chr1-33225393-INS-6800 |
| chr1-33233117-INS-6227 | chr1-33233118-INS-6226 |
| chr1-33241921-INS-6582 | chr1-33241922-INS-6583 |
| chr1-33247908-INS-798 | chr1-33247908-INS-798 |
| chr1-33255274-INS-82 | chr1-33255274-INS-82 |
| chr1-33257585-INS-586 | chr1-33257585-INS-587 |
| chr1-33285977-INS-7270 | chr1-33285978-INS-7271 |
| chr1-33323966-INS-506 | chr1-33323966-INS-510 |
| chr1-33330258-INS-51 | chr1-33330258-INS-50 |
| chr1-33331022-INS-51 | chr1-33331013-INS-52 |
| chr1-33369134-INS-7112 | chr1-33369135-INS-7114 |
| chr1-33376537-INS-68 | chr1-33376537-INS-61 |
| chr1-33385822-INS-6834 | chr1-33385823-INS-6830 |

| | |
|---|---|
| chr1-33395177-INS-2974 | chr1-33395178-INS-2974 |
| chr1-33403057-INS-56 | chr1-33403057-INS-56 |
| chr1-33406783-INS-454 | chr1-33406783-INS-454 |
| chr1-33417413-INS-303 | chr1-33417413-INS-304 |
| chr1-33422536-INS-132 | chr1-33422527-INS-133 |
| chr1-33429313-INS-203 | chr1-33429313-INS-177 |
| chr1-33454408-INS-56 | chr1-33454409-INS-56 |
| chr1-33475895-INS-223 | chr1-33475895-INS-223 |
| chr1-33487339-INS-80 | chr1-33487330-INS-52 |
| chr1-33492004-INS-445 | chr1-33492004-INS-445 |
| chr1-33493000-INS-58 | chr1-33493000-INS-58 |
| chr1-33500148-INS-68 | chr1-33500148-INS-67 |
| chr1-33503285-INS-66 | chr1-33503285-INS-78 |
| chr1-33504104-INS-572 | chr1-33504104-INS-572 |
| chr1-33510191-INS-60 | chr1-33510191-INS-60 |
| chr1-33518552-INS-830 | chr1-33518552-INS-830 |
| chr1-33523051-INS-64 | chr1-33523065-INS-84 |
| chr1-33531888-INS-139 | chr1-33531888-INS-139 |
| chr1-33545372-INS-58 | chr1-33545372-INS-58 |
| chr1-33557934-INS-431 | chr1-33557934-INS-427 |
| chr1-33577172-INS-582 | chr1-33577172-INS-580 |
| chr1-33580426-INS-290 | chr1-33580430-INS-319 |
| chr1-33594097-INS-6820 | chr1-33594098-INS-6820 |
| chr1-33638146-INS-69 | chr1-33638146-INS-73 |
| chr1-33640210-INS-63 | chr1-33640210-INS-63 |
| chr1-33641256-INS-1021 | chr1-33641257-INS-1021 |
| chr1-33655992-INS-52 | chr1-33655992-INS-52 |
| chr1-33670133-INS-70 | chr1-33670133-INS-70 |
| chr1-33685545-INS-59 | chr1-33685542-INS-66 |
| chr1-33696314-INS-6057 | chr1-33696315-INS-6057 |
| chr1-33721695-INS-59 | chr1-33721715-INS-63 |
| chr1-33737738-INS-86 | chr1-33737734-INS-102 |
| chr1-33761110-INS-437 | chr1-33761110-INS-437 |
| chr1-33770088-INS-58 | chr1-33770088-INS-58 |
| chr1-33790079-INS-143 | chr1-33790079-INS-176 |
| chr1-33792537-INS-62 | chr1-33792537-INS-52 |
| chr1-33822866-INS-87 | chr1-33822866-INS-87 |
| chr1-33850485-INS-65 | chr1-33850485-INS-60 |
| chr1-33887969-INS-54 | chr1-33887969-INS-55 |
| chr1-33889785-INS-310 | chr1-33889786-INS-264 |
| chr1-33895173-INS-202 | chr1-33895173-INS-202 |
| chr1-33901307-INS-74 | chr1-33901337-INS-66 |
| chr1-33905911-INS-554 | chr1-33905907-INS-557 |
| chr1-33917821-INS-576 | chr1-33917821-INS-545 |
| chr1-33922733-INS-212 | chr1-33922733-INS-212 |
| chr1-33927000-INS-59 | chr1-33927000-INS-60 |
| chr1-33943793-INS-52 | chr1-33943794-INS-56 |
| chr1-33961540-INS-202 | chr1-33961540-INS-203 |
| chr1-33965064-INS-209 | chr1-33965064-INS-209 |
| chr1-33977642-INS-64 | chr1-33977646-INS-60 |
| chr1-34009255-INS-206 | chr1-34009255-INS-206 |
| chr1-34010113-INS-504 | chr1-34010113-INS-505 |
| chr1-34016010-INS-2678 | chr1-34016011-INS-2681 |
| chr1-34022614-INS-74 | chr1-34022614-INS-74 |
| chr1-34089533-INS-55 | chr1-34089533-INS-64 |
| chr1-34131788-INS-210 | chr1-34131788-INS-210 |
| chr1-34135642-INS-54 | chr1-34135642-INS-56 |
| chr1-34156998-INS-113 | chr1-34156998-INS-113 |
| chr1-34166232-INS-64 | chr1-34166232-INS-72 |
| chr1-34174632-INS-70 | chr1-34174672-INS-74 |
| chr1-34182089-INS-166 | chr1-34182090-INS-166 |
| chr1-34195568-INS-88 | chr1-34195584-INS-90 |
| chr1-34209178-INS-110 | chr1-34209178-INS-110 |
| chr1-34250800-INS-67 | chr1-34250800-INS-67 |
| chr1-34266152-INS-141 | chr1-34266152-INS-141 |
| chr1-34331537-INS-60 | chr1-34331547-INS-70 |
| chr1-34333071-INS-369 | chr1-34333071-INS-369 |
| chr1-34350232-INS-151 | chr1-34350232-INS-151 |
| chr1-34359272-INS-100 | chr1-34359272-INS-100 |
| chr1-34383721-INS-207 | chr1-34383721-INS-207 |
| chr1-34385784-INS-64 | chr1-34385784-INS-64 |
| chr1-34393346-INS-72 | chr1-34393346-INS-68 |
| chr1-34407404-INS-263 | chr1-34407404-INS-263 |
| chr1-34426006-INS-497 | chr1-34426006-INS-497 |
| chr1-34428382-INS-238 | chr1-34428402-INS-232 |
| chr1-34429674-INS-64 | chr1-34429649-INS-101 |
| chr1-34439236-INS-6690 | chr1-34439237-INS-6690 |

| | |
|---|---|
| chr1-34454134-INS-3134 | chr1-34454135-INS-3177 |
| chr1-34467431-INS-4204 | chr1-34467432-INS-4204 |
| chr1-34472799-INS-69 | chr1-34472799-INS-69 |
| chr1-34482359-INS-142 | chr1-34482359-INS-142 |
| chr1-34484167-INS-59 | chr1-34484167-INS-60 |
| chr1-34493577-INS-215 | chr1-34493577-INS-215 |
| chr1-34497999-INS-55 | chr1-34497999-INS-55 |
| chr1-34504417-INS-64 | chr1-34504417-INS-64 |
| chr1-34509571-INS-60 | chr1-34509569-INS-85 |
| chr1-34513923-INS-71 | chr1-34513923-INS-71 |
| chr1-34516113-INS-198 | chr1-34516133-INS-171 |
| chr1-34519289-INS-59 | chr1-34519289-INS-59 |
| chr1-34520294-INS-6412 | chr1-34520295-INS-6412 |
| chr1-34521865-INS-211 | chr1-34521865-INS-197 |
| chr1-34546160-INS-783 | chr1-34546160-INS-783 |
| chr1-34551826-INS-392 | chr1-34551826-INS-418 |
| chr1-34558902-INS-204 | chr1-34558902-INS-204 |
| chr1-34559664-INS-69 | chr1-34559664-INS-69 |
| chr1-34561033-INS-63 | chr1-34561033-INS-63 |
| chr1-34564486-INS-212 | chr1-34564486-INS-213 |
| chr1-34565845-INS-209 | chr1-34565845-INS-210 |
| chr1-34577969-INS-751 | chr1-34577969-INS-754 |
| chr1-34584174-INS-513 | chr1-34584174-INS-513 |
| chr1-34585953-INS-69 | chr1-34585953-INS-68 |
| chr1-34598018-INS-77 | chr1-34598018-INS-83 |
| chr1-34604298-INS-432 | chr1-34604298-INS-400 |
| chr1-34606134-INS-208 | chr1-34606134-INS-204 |
| chr1-34612604-INS-433 | chr1-34612604-INS-433 |
| chr1-34616573-INS-462 | chr1-34616574-INS-462 |
| chr1-34617142-INS-109 | chr1-34617101-INS-104 |
| chr1-34622449-INS-100 | chr1-34622449-INS-100 |
| chr1-34633210-INS-356 | chr1-34633210-INS-356 |
| chr1-34641563-INS-172 | chr1-34641563-INS-172 |
| chr1-34656850-INS-60 | chr1-34656848-INS-61 |
| chr1-34659521-INS-66 | chr1-34659521-INS-66 |
| chr1-34663241-INS-175 | chr1-34663242-INS-175 |
| chr1-34666607-INS-138 | chr1-34666607-INS-144 |
| chr1-34746611-INS-88 | chr1-34746611-INS-88 |
| chr1-34781336-INS-118 | chr1-34781366-INS-94 |
| chr1-34782830-INS-80 | chr1-34782830-INS-80 |
| chr1-34800833-INS-61 | chr1-34800833-INS-61 |
| chr1-34812609-INS-1889 | chr1-34812610-INS-1889 |
| chr1-34818565-INS-61 | chr1-34818565-INS-65 |
| chr1-34833381-INS-61 | chr1-34833381-INS-61 |
| chr1-34837162-INS-94 | chr1-34837154-INS-96 |
| chr1-34867132-INS-53 | chr1-34867132-INS-53 |
| chr1-34882288-INS-68 | chr1-34882288-INS-68 |
| chr1-34901942-INS-51 | chr1-34901942-INS-51 |
| chr1-34903050-INS-1355 | chr1-34903051-INS-1348 |
| chr1-34922636-INS-53 | chr1-34922648-INS-85 |
| chr1-34931597-INS-5380 | chr1-34931598-INS-5380 |
| chr1-34937968-INS-4329 | chr1-34937969-INS-4329 |
| chr1-34942107-INS-1282 | chr1-34942108-INS-1282 |
| chr1-34960017-INS-56 | chr1-34960017-INS-56 |
| chr1-34965184-INS-66 | chr1-34965166-INS-73 |
| chr1-34973836-INS-52 | chr1-34973825-INS-56 |
| chr1-34981199-INS-89 | chr1-34981199-INS-65 |
| chr1-34983522-INS-60 | chr1-34983522-INS-100 |
| chr1-34995498-INS-3113 | chr1-34995499-INS-3113 |
| chr1-35009045-INS-52 | chr1-35009045-INS-52 |
| chr1-35027653-INS-6759 | chr1-35027654-INS-6759 |
| chr1-35063669-INS-462 | chr1-35063669-INS-462 |
| chr1-35075952-INS-54 | chr1-35075953-INS-52 |
| chr1-35077730-INS-9231 | chr1-35077731-INS-9233 |
| chr1-35100129-INS-69 | chr1-35100129-INS-61 |
| chr1-35100510-INS-342 | chr1-35100510-INS-342 |
| chr1-35103431-INS-575 | chr1-35103431-INS-575 |
| chr1-35108976-INS-321 | chr1-35108976-INS-321 |
| chr1-35117472-INS-110 | chr1-35117472-INS-133 |
| chr1-35126365-INS-389 | chr1-35126365-INS-389 |
| chr1-35127614-INS-54 | chr1-35127614-INS-74 |
| chr1-35136014-INS-400 | chr1-35136014-INS-400 |
| chr1-35167084-INS-372 | chr1-35167084-INS-379 |
| chr1-35186795-INS-75 | chr1-35186771-INS-78 |
| chr1-35192245-INS-52 | chr1-35192245-INS-54 |
| chr1-35215694-INS-232 | chr1-35215694-INS-256 |
| chr1-35226678-INS-82 | chr1-35226674-INS-50 |

| | |
|---|---|
| chr1-35229502-INS-281 | chr1-35229502-INS-281 |
| chr1-35229502-INS-900 | chr1-35229502-INS-902 |
| chr1-35271287-INS-76 | chr1-35271287-INS-76 |
| chr1-35298034-INS-160 | chr1-35298034-INS-162 |
| chr1-35302828-INS-315 | chr1-35302829-INS-315 |
| chr1-35316836-INS-688 | chr1-35316836-INS-688 |
| chr1-35339709-INS-81 | chr1-35339709-INS-84 |
| chr1-35342470-INS-60 | chr1-35342470-INS-63 |
| chr1-35354401-INS-150 | chr1-35354401-INS-150 |
| chr1-35366927-INS-116 | chr1-35366927-INS-108 |
| chr1-35389338-INS-155 | chr1-35389338-INS-155 |
| chr1-35400437-INS-84 | chr1-35400437-INS-84 |
| chr1-35403000-INS-687 | chr1-35403000-INS-687 |
| chr1-35419024-INS-77 | chr1-35418996-INS-103 |
| chr1-35446915-INS-60 | chr1-35446915-INS-60 |
| chr1-35469103-INS-64 | chr1-35469103-INS-66 |
| chr1-35476578-INS-64 | chr1-35476578-INS-64 |
| chr1-35476920-INS-4868 | chr1-35476921-INS-4868 |
| chr1-35558536-INS-56 | chr1-35558536-INS-58 |
| chr1-35563474-INS-52 | chr1-35563474-INS-52 |
| chr1-35567961-INS-386 | chr1-35567961-INS-389 |
| chr1-35570886-INS-61 | chr1-35570900-INS-57 |
| chr1-35592972-INS-77 | chr1-35592972-INS-77 |
| chr1-35607844-INS-390 | chr1-35607844-INS-390 |
| chr1-35611689-INS-73 | chr1-35611689-INS-73 |
| chr1-35624648-INS-1210 | chr1-35624649-INS-1210 |
| chr1-35628571-INS-137 | chr1-35628571-INS-137 |
| chr1-35680390-INS-72 | chr1-35680390-INS-72 |
| chr1-35691838-INS-194 | chr1-35691838-INS-178 |
| chr1-35699355-INS-125 | chr1-35699355-INS-120 |
| chr1-35704632-INS-69 | chr1-35704633-INS-57 |
| chr1-35709719-INS-615 | chr1-35709719-INS-603 |
| chr1-35716023-INS-65 | chr1-35716023-INS-65 |
| chr1-35718150-INS-65 | chr1-35718150-INS-65 |
| chr1-35729472-INS-575 | chr1-35729472-INS-581 |
| chr1-35736027-INS-74 | chr1-35736028-INS-64 |
| chr1-35742443-INS-94 | chr1-35742471-INS-56 |
| chr1-35756580-INS-81 | chr1-35756580-INS-81 |
| chr1-35781806-INS-62 | chr1-35781806-INS-62 |
| chr1-35789645-INS-108 | chr1-35789645-INS-112 |
| chr1-35793803-INS-184 | chr1-35793803-INS-184 |
| chr1-35816351-INS-85 | chr1-35816351-INS-85 |
| chr1-35844390-INS-1164 | chr1-35844391-INS-1164 |
| chr1-35847975-INS-69 | chr1-35847947-INS-60 |
| chr1-35861094-INS-209 | chr1-35861094-INS-212 |
| chr1-35870950-INS-69 | chr1-35870963-INS-83 |
| chr1-35882681-INS-7676 | chr1-35882682-INS-7676 |
| chr1-35882682-INS-575 | chr1-35882682-INS-575 |
| chr1-35883492-INS-138 | chr1-35883483-INS-114 |
| chr1-35898151-INS-6320 | chr1-35898152-INS-6320 |
| chr1-35904922-INS-126 | chr1-35904922-INS-126 |
| chr1-35917053-INS-801 | chr1-35917057-INS-803 |
| chr1-35926227-INS-67 | chr1-35926227-INS-59 |
| chr1-35948799-INS-750 | chr1-35948799-INS-750 |
| chr1-35953524-INS-129 | chr1-35953524-INS-129 |
| chr1-35976871-INS-56 | chr1-35976871-INS-50 |
| chr1-35979272-INS-52 | chr1-35979272-INS-54 |
| chr1-35988062-INS-107 | chr1-35988062-INS-107 |
| chr1-35997617-INS-420 | chr1-35997612-INS-417 |
| chr1-36001084-INS-64 | chr1-36001084-INS-64 |
| chr1-36007471-INS-70 | chr1-36007471-INS-70 |
| chr1-36010076-INS-69 | chr1-36010077-INS-81 |
| chr1-36013947-INS-470 | chr1-36013947-INS-490 |
| chr1-36033892-INS-894 | chr1-36033892-INS-894 |
| chr1-36035566-INS-96 | chr1-36035566-INS-96 |
| chr1-36037067-INS-54 | chr1-36037067-INS-54 |
| chr1-36037376-INS-528 | chr1-36037376-INS-529 |
| chr1-36039509-INS-186 | chr1-36039509-INS-174 |
| chr1-36044420-INS-166 | chr1-36044420-INS-166 |
| chr1-36061527-INS-318 | chr1-36061520-INS-336 |
| chr1-36082931-INS-206 | chr1-36082931-INS-206 |
| chr1-36083546-INS-1893 | chr1-36083547-INS-1893 |
| chr1-36085699-INS-245 | chr1-36085699-INS-245 |
| chr1-36096850-INS-264 | chr1-36096850-INS-264 |
| chr1-36098993-INS-57 | chr1-36098993-INS-57 |
| chr1-36101238-INS-52 | chr1-36101238-INS-52 |
| chr1-36156374-INS-7736 | chr1-36156375-INS-7737 |

| | |
|---|---|
| chr1-36181589-INS-96 | chr1-36181589-INS-96 |
| chr1-36188975-INS-74 | chr1-36188977-INS-56 |
| chr1-36198885-INS-94 | chr1-36198885-INS-94 |
| chr1-36203129-INS-98 | chr1-36203129-INS-98 |
| chr1-36205684-INS-172 | chr1-36205684-INS-172 |
| chr1-36219893-INS-4107 | chr1-36219894-INS-4106 |
| chr1-36227135-INS-923 | chr1-36227152-INS-931 |
| chr1-36245230-INS-215 | chr1-36245230-INS-215 |
| chr1-36247784-INS-436 | chr1-36247784-INS-436 |
| chr1-36255117-INS-195 | chr1-36255117-INS-195 |
| chr1-36260029-INS-73 | chr1-36260029-INS-73 |
| chr1-36262065-INS-56 | chr1-36262069-INS-54 |
| chr1-36263788-INS-347 | chr1-36263788-INS-347 |
| chr1-36271376-INS-928 | chr1-36271376-INS-928 |
| chr1-36285980-INS-106 | chr1-36285980-INS-106 |
| chr1-36286368-INS-1332 | chr1-36286369-INS-1332 |
| chr1-36297514-INS-218 | chr1-36297514-INS-218 |
| chr1-36299318-INS-51 | chr1-36299334-INS-53 |
| chr1-36302337-INS-124 | chr1-36302358-INS-79 |
| chr1-36306584-INS-204 | chr1-36306584-INS-204 |
| chr1-36314355-INS-107 | chr1-36314355-INS-107 |
| chr1-36316172-INS-66 | chr1-36316172-INS-66 |
| chr1-36324655-INS-99 | chr1-36324655-INS-91 |
| chr1-36326946-INS-891 | chr1-36326946-INS-891 |
| chr1-36344385-INS-84 | chr1-36344385-INS-86 |
| chr1-36354084-INS-8976 | chr1-36354085-INS-8976 |
| chr1-36400903-INS-221 | chr1-36400903-INS-221 |
| chr1-36431841-INS-6442 | chr1-36431842-INS-6442 |
| chr1-36477257-INS-206 | chr1-36477257-INS-205 |
| chr1-36489082-INS-212 | chr1-36489082-INS-212 |
| chr1-36491852-INS-109 | chr1-36491852-INS-109 |
| chr1-36533774-INS-6452 | chr1-36533775-INS-6452 |
| chr1-36534397-INS-291 | chr1-36534397-INS-291 |
| chr1-36541384-INS-85 | chr1-36541384-INS-63 |
| chr1-36549692-INS-1528 | chr1-36549697-INS-1572 |
| chr1-36556984-INS-207 | chr1-36556984-INS-206 |
| chr1-36585320-INS-56 | chr1-36585320-INS-56 |
| chr1-36633223-INS-202 | chr1-36633224-INS-202 |
| chr1-36664099-INS-128 | chr1-36664099-INS-128 |
| chr1-36682380-INS-236 | chr1-36682380-INS-236 |
| chr1-36706971-INS-72 | chr1-36706985-INS-70 |
| chr1-36735320-INS-199 | chr1-36735320-INS-200 |
| chr1-36737033-INS-107 | chr1-36737033-INS-107 |
| chr1-36755457-INS-216 | chr1-36755457-INS-216 |
| chr1-36768060-INS-105 | chr1-36768060-INS-105 |
| chr1-36772036-INS-183 | chr1-36772049-INS-144 |
| chr1-36794731-INS-54 | chr1-36794731-INS-54 |
| chr1-36827325-INS-216 | chr1-36827325-INS-217 |
| chr1-36836798-INS-60 | chr1-36836798-INS-64 |
| chr1-36855353-INS-498 | chr1-36855353-INS-498 |
| chr1-36861592-INS-52 | chr1-36861592-INS-52 |
| chr1-36888583-INS-55 | chr1-36888583-INS-55 |
| chr1-36899358-INS-106 | chr1-36899337-INS-119 |
| chr1-36925845-INS-54 | chr1-36925846-INS-58 |
| chr1-36963783-INS-127 | chr1-36963783-INS-127 |
| chr1-36984004-INS-179 | chr1-36984004-INS-179 |
| chr1-36989243-INS-165 | chr1-36989243-INS-165 |
| chr1-37013627-INS-160 | chr1-37013627-INS-172 |
| chr1-37021168-INS-64 | chr1-37021169-INS-78 |
| chr1-37030328-INS-178 | chr1-37030328-INS-178 |
| chr1-37038744-INS-201 | chr1-37038744-INS-199 |
| chr1-37044989-INS-198 | chr1-37044989-INS-198 |
| chr1-37051567-INS-200 | chr1-37051567-INS-200 |
| chr1-37071479-INS-52 | chr1-37071515-INS-52 |
| chr1-37086159-INS-219 | chr1-37086159-INS-197 |
| chr1-37088505-INS-394 | chr1-37088505-INS-394 |
| chr1-37118199-INS-194 | chr1-37118199-INS-194 |
| chr1-37119410-INS-105 | chr1-37119428-INS-108 |
| chr1-37131423-INS-6440 | chr1-37131424-INS-6440 |
| chr1-37145259-INS-143 | chr1-37145259-INS-143 |
| chr1-37153695-INS-109 | chr1-37153695-INS-126 |
| chr1-37157601-INS-68 | chr1-37157561-INS-73 |
| chr1-37202826-INS-737 | chr1-37202826-INS-737 |
| chr1-37215671-INS-390 | chr1-37215671-INS-390 |
| chr1-37232970-INS-329 | chr1-37232970-INS-329 |
| chr1-37235846-INS-57 | chr1-37235847-INS-57 |
| chr1-37243485-INS-159 | chr1-37243486-INS-159 |

| | |
|---|---|
| chr1-37253255-INS-56 | chr1-37253255-INS-56 |
| chr1-37253659-INS-5508 | chr1-37253660-INS-5492 |
| chr1-37267015-INS-106 | chr1-37267015-INS-108 |
| chr1-37277764-INS-639 | chr1-37277765-INS-636 |
| chr1-37289373-INS-6442 | chr1-37289374-INS-6443 |
| chr1-37291649-INS-409 | chr1-37291649-INS-409 |
| chr1-37293458-INS-9249 | chr1-37293459-INS-9249 |
| chr1-37345263-INS-213 | chr1-37345263-INS-213 |
| chr1-37402504-INS-128 | chr1-37402504-INS-128 |
| chr1-37499023-INS-1709 | chr1-37499024-INS-1718 |
| chr1-37522555-INS-199 | chr1-37522555-INS-201 |
| chr1-37560329-INS-748 | chr1-37560329-INS-750 |
| chr1-37585893-INS-1882 | chr1-37585894-INS-1882 |
| chr1-37623858-INS-99 | chr1-37623818-INS-118 |
| chr1-37626807-INS-97 | chr1-37626820-INS-107 |
| chr1-37636183-INS-52 | chr1-37636183-INS-54 |
| chr1-37681653-INS-129 | chr1-37681653-INS-129 |
| chr1-37682619-INS-450 | chr1-37682619-INS-450 |
| chr1-37684420-INS-207 | chr1-37684420-INS-206 |
| chr1-37694614-INS-165 | chr1-37694614-INS-166 |
| chr1-37702173-INS-203 | chr1-37702173-INS-201 |
| chr1-37734161-INS-1761 | chr1-37734162-INS-1762 |
| chr1-37746989-INS-69 | chr1-37746989-INS-69 |
| chr1-37782645-INS-60 | chr1-37782645-INS-54 |
| chr1-37924637-INS-239 | chr1-37924637-INS-239 |
| chr1-37933287-INS-190 | chr1-37933287-INS-190 |
| chr1-37934855-INS-419 | chr1-37934855-INS-419 |
| chr1-37944571-INS-90 | chr1-37944562-INS-55 |
| chr1-37969110-INS-61 | chr1-37969110-INS-96 |
| chr1-38011962-INS-111 | chr1-38011962-INS-111 |
| chr1-38013400-INS-2725 | chr1-38013401-INS-2725 |
| chr1-38014363-INS-88 | chr1-38014363-INS-88 |
| chr1-38033791-INS-148 | chr1-38033791-INS-148 |
| chr1-38064208-INS-2000 | chr1-38064209-INS-2002 |
| chr1-38199529-INS-80 | chr1-38199529-INS-80 |
| chr1-38209762-INS-6653 | chr1-38209763-INS-6654 |
| chr1-38227724-INS-354 | chr1-38227718-INS-335 |
| chr1-38248044-INS-159 | chr1-38248044-INS-181 |
| chr1-38260084-INS-148 | chr1-38260084-INS-192 |
| chr1-38261882-INS-6442 | chr1-38261883-INS-6442 |
| chr1-38266561-INS-185 | chr1-38266561-INS-185 |
| chr1-38273273-INS-102 | chr1-38273273-INS-102 |
| chr1-38283042-INS-56 | chr1-38283042-INS-56 |
| chr1-38303648-INS-88 | chr1-38303648-INS-88 |
| chr1-38308356-INS-215 | chr1-38308356-INS-215 |
| chr1-38314436-INS-1892 | chr1-38314437-INS-1892 |
| chr1-38321479-INS-4578 | chr1-38321480-INS-4578 |
| chr1-38326932-INS-106 | chr1-38326932-INS-151 |
| chr1-38335446-INS-52 | chr1-38335446-INS-54 |
| chr1-38351548-INS-58 | chr1-38351548-INS-58 |
| chr1-38352388-INS-217 | chr1-38352388-INS-218 |
| chr1-38354331-INS-117 | chr1-38354331-INS-117 |
| chr1-38362019-INS-209 | chr1-38362019-INS-213 |
| chr1-38365464-INS-1045 | chr1-38365465-INS-1045 |
| chr1-38381340-INS-64 | chr1-38381340-INS-64 |
| chr1-38415461-INS-54 | chr1-38415479-INS-59 |
| chr1-38420384-INS-57 | chr1-38420383-INS-75 |
| chr1-38439507-INS-70 | chr1-38439507-INS-87 |
| chr1-38458876-INS-219 | chr1-38458876-INS-220 |
| chr1-38461532-INS-308 | chr1-38461532-INS-308 |
| chr1-38502430-INS-399 | chr1-38502430-INS-399 |
| chr1-38548645-INS-6818 | chr1-38548646-INS-6820 |
| chr1-38552868-INS-52 | chr1-38552868-INS-52 |
| chr1-38556684-INS-220 | chr1-38556684-INS-220 |
| chr1-38621193-INS-67 | chr1-38621193-INS-67 |
| chr1-38629386-INS-72 | chr1-38629386-INS-60 |
| chr1-38653313-INS-138 | chr1-38653313-INS-93 |
| chr1-38669730-INS-142 | chr1-38669730-INS-142 |
| chr1-38680141-INS-685 | chr1-38680141-INS-685 |
| chr1-38712454-INS-54 | chr1-38712446-INS-52 |
| chr1-38731430-INS-55 | chr1-38731430-INS-55 |
| chr1-38740174-INS-76 | chr1-38740174-INS-74 |
| chr1-38742048-INS-655 | chr1-38742054-INS-663 |
| chr1-38743187-INS-400 | chr1-38743187-INS-400 |
| chr1-38772646-INS-75 | chr1-38772646-INS-92 |
| chr1-38772965-INS-271 | chr1-38772979-INS-270 |
| chr1-38786159-INS-405 | chr1-38786159-INS-405 |

| | |
|---|---|
| chr1-38805186-INS-61 | chr1-38805188-INS-62 |
| chr1-38811632-INS-102 | chr1-38811632-INS-70 |
| chr1-38819225-INS-445 | chr1-38819225-INS-445 |
| chr1-38855295-INS-62 | chr1-38855295-INS-62 |
| chr1-38863731-INS-802 | chr1-38863734-INS-794 |
| chr1-38870078-INS-77 | chr1-38870078-INS-77 |
| chr1-38878463-INS-439 | chr1-38878463-INS-437 |
| chr1-38882048-INS-86 | chr1-38882046-INS-92 |
| chr1-38884273-INS-206 | chr1-38884273-INS-205 |
| chr1-38887114-INS-68 | chr1-38887114-INS-68 |
| chr1-38897275-INS-56 | chr1-38897275-INS-56 |
| chr1-38921414-INS-58 | chr1-38921414-INS-58 |
| chr1-38929037-INS-6744 | chr1-38929038-INS-6744 |
| chr1-38938169-INS-64 | chr1-38938169-INS-58 |
| chr1-38941580-INS-112 | chr1-38941580-INS-113 |
| chr1-38949057-INS-73 | chr1-38949057-INS-78 |
| chr1-38954728-INS-2647 | chr1-38954729-INS-2673 |
| chr1-38959773-INS-491 | chr1-38959773-INS-491 |
| chr1-38960828-INS-399 | chr1-38960828-INS-399 |
| chr1-38968645-INS-68 | chr1-38968643-INS-54 |
| chr1-38970230-INS-1888 | chr1-38970231-INS-1888 |
| chr1-38976554-INS-88 | chr1-38976554-INS-88 |
| chr1-38996115-INS-963 | chr1-38996115-INS-963 |
| chr1-39020830-INS-60 | chr1-39020830-INS-60 |
| chr1-39036340-INS-70 | chr1-39036340-INS-54 |
| chr1-39047138-INS-199 | chr1-39047138-INS-207 |
| chr1-39052885-INS-448 | chr1-39052885-INS-448 |
| chr1-39058191-INS-60 | chr1-39058218-INS-60 |
| chr1-39063103-INS-54 | chr1-39063103-INS-54 |
| chr1-39064017-INS-118 | chr1-39064017-INS-118 |
| chr1-39068686-INS-100 | chr1-39068686-INS-100 |
| chr1-39091978-INS-149 | chr1-39091977-INS-194 |
| chr1-39115546-INS-492 | chr1-39115546-INS-492 |
| chr1-39128425-INS-4078 | chr1-39128426-INS-4078 |
| chr1-39143431-INS-203 | chr1-39143431-INS-203 |
| chr1-39156526-INS-60 | chr1-39156526-INS-50 |
| chr1-39159615-INS-52 | chr1-39159592-INS-56 |
| chr1-39178251-INS-493 | chr1-39178251-INS-495 |
| chr1-39180014-INS-57 | chr1-39180014-INS-51 |
| chr1-39185288-INS-138 | chr1-39185288-INS-138 |
| chr1-39197407-INS-121 | chr1-39197407-INS-125 |
| chr1-39284035-INS-173 | chr1-39284035-INS-173 |
| chr1-39286410-INS-76 | chr1-39286410-INS-76 |
| chr1-39308527-INS-2732 | chr1-39308528-INS-2724 |
| chr1-39310462-INS-100 | chr1-39310462-INS-64 |
| chr1-39313248-INS-83 | chr1-39313238-INS-72 |
| chr1-39316047-INS-160 | chr1-39316047-INS-161 |
| chr1-39329934-INS-80 | chr1-39329934-INS-116 |
| chr1-39354138-INS-213 | chr1-39354138-INS-213 |
| chr1-39372639-INS-7248 | chr1-39372640-INS-7247 |
| chr1-39396630-INS-5067 | chr1-39396631-INS-5067 |
| chr1-39413664-INS-261 | chr1-39413664-INS-263 |
| chr1-39426214-INS-58 | chr1-39426214-INS-54 |
| chr1-39427452-INS-606 | chr1-39427455-INS-606 |
| chr1-39440442-INS-6403 | chr1-39440443-INS-6403 |
| chr1-39444734-INS-318 | chr1-39444734-INS-318 |
| chr1-39447277-INS-129 | chr1-39447277-INS-129 |
| chr1-39456530-INS-218 | chr1-39456530-INS-218 |
| chr1-39466718-INS-60 | chr1-39466718-INS-60 |
| chr1-39477287-INS-114 | chr1-39477261-INS-114 |
| chr1-39483900-INS-74 | chr1-39483872-INS-74 |
| chr1-39485278-INS-102 | chr1-39485278-INS-102 |
| chr1-39522012-INS-6392 | chr1-39522013-INS-6392 |
| chr1-39525299-INS-200 | chr1-39525299-INS-201 |
| chr1-39528064-INS-86 | chr1-39528064-INS-86 |
| chr1-39529878-INS-95 | chr1-39529878-INS-72 |
| chr1-39536402-INS-200 | chr1-39536411-INS-197 |
| chr1-39547070-INS-120 | chr1-39547070-INS-120 |
| chr1-39551198-INS-245 | chr1-39551198-INS-245 |
| chr1-39562032-INS-54 | chr1-39562033-INS-66 |
| chr1-39564838-INS-61 | chr1-39564830-INS-60 |
| chr1-39571069-INS-150 | chr1-39571064-INS-172 |
| chr1-39619529-INS-118 | chr1-39619524-INS-128 |
| chr1-39620603-INS-59 | chr1-39620603-INS-51 |
| chr1-39623458-INS-56 | chr1-39623458-INS-56 |
| chr1-39629059-INS-890 | chr1-39629057-INS-896 |
| chr1-39633733-INS-118 | chr1-39633733-INS-122 |

| | |
|---|---|
| chr1-39634965-INS-493 | chr1-39634965-INS-493 |
| chr1-39641486-INS-62 | chr1-39641486-INS-62 |
| chr1-39709299-INS-52 | chr1-39709299-INS-52 |
| chr1-39710112-INS-5549 | chr1-39710113-INS-5549 |
| chr1-39710294-INS-619 | chr1-39710294-INS-620 |
| chr1-39725057-INS-6430 | chr1-39725058-INS-6432 |
| chr1-39731518-INS-54 | chr1-39731519-INS-68 |
| chr1-39739586-INS-126 | chr1-39739589-INS-95 |
| chr1-39740222-INS-226 | chr1-39740222-INS-228 |
| chr1-39765429-INS-1890 | chr1-39765430-INS-1890 |
| chr1-39767122-INS-217 | chr1-39767122-INS-217 |
| chr1-39772055-INS-776 | chr1-39772055-INS-777 |
| chr1-39774415-INS-68 | chr1-39774411-INS-60 |
| chr1-39778777-INS-7149 | chr1-39778778-INS-7149 |
| chr1-39784954-INS-64 | chr1-39784954-INS-54 |
| chr1-39791864-INS-84 | chr1-39791864-INS-60 |
| chr1-39804746-INS-401 | chr1-39804746-INS-401 |
| chr1-39806407-INS-441 | chr1-39806407-INS-441 |
| chr1-39825821-INS-581 | chr1-39825821-INS-586 |
| chr1-39869825-INS-51 | chr1-39869825-INS-51 |
| chr1-39873060-INS-62 | chr1-39873060-INS-62 |
| chr1-39891782-INS-143 | chr1-39891782-INS-143 |
| chr1-39892932-INS-61 | chr1-39892932-INS-61 |
| chr1-39903195-INS-67 | chr1-39903195-INS-99 |
| chr1-39918059-INS-62 | chr1-39918059-INS-62 |
| chr1-39968684-INS-53 | chr1-39968684-INS-54 |
| chr1-39975146-INS-62 | chr1-39975146-INS-63 |
| chr1-39979757-INS-96 | chr1-39979757-INS-108 |
| chr1-39991851-INS-83 | chr1-39991853-INS-83 |
| chr1-40032234-INS-64 | chr1-40032234-INS-64 |
| chr1-40044229-INS-56 | chr1-40044229-INS-56 |
| chr1-40052681-INS-70 | chr1-40052681-INS-66 |
| chr1-40055288-INS-337 | chr1-40055283-INS-336 |
| chr1-40057951-INS-215 | chr1-40057951-INS-215 |
| chr1-40059335-INS-1310 | chr1-40059336-INS-1311 |
| chr1-40078699-INS-51 | chr1-40078699-INS-51 |
| chr1-40083909-INS-451 | chr1-40083909-INS-451 |
| chr1-40090139-INS-487 | chr1-40090139-INS-487 |
| chr1-40094243-INS-78 | chr1-40094243-INS-78 |
| chr1-40101531-INS-132 | chr1-40101531-INS-162 |
| chr1-40108083-INS-496 | chr1-40108083-INS-503 |
| chr1-40154026-INS-55 | chr1-40154026-INS-55 |
| chr1-40171798-INS-92 | chr1-40171804-INS-123 |
| chr1-40177333-INS-322 | chr1-40177333-INS-322 |
| chr1-40191882-INS-70 | chr1-40191882-INS-80 |
| chr1-40223637-INS-101 | chr1-40223637-INS-101 |
| chr1-40256218-INS-6377 | chr1-40256219-INS-6377 |
| chr1-40277126-INS-143 | chr1-40277126-INS-143 |
| chr1-40284186-INS-228 | chr1-40284182-INS-222 |
| chr1-40311843-INS-54 | chr1-40311843-INS-54 |
| chr1-40324634-INS-54 | chr1-40324634-INS-98 |
| chr1-40326573-INS-238 | chr1-40326573-INS-205 |
| chr1-40328380-INS-66 | chr1-40328380-INS-66 |
| chr1-40338435-INS-652 | chr1-40338435-INS-653 |
| chr1-40343849-INS-84 | chr1-40343849-INS-70 |
| chr1-40360823-INS-64 | chr1-40360823-INS-64 |
| chr1-40368681-INS-76 | chr1-40368669-INS-80 |
| chr1-40413977-INS-92 | chr1-40413977-INS-92 |
| chr1-40417363-INS-642 | chr1-40417363-INS-647 |
| chr1-40427197-INS-2513 | chr1-40427198-INS-2513 |
| chr1-40445831-INS-163 | chr1-40445831-INS-120 |
| chr1-40454379-INS-94 | chr1-40454379-INS-94 |
| chr1-40514761-INS-83 | chr1-40514727-INS-72 |
| chr1-40528072-INS-124 | chr1-40528072-INS-122 |
| chr1-40531780-INS-320 | chr1-40531780-INS-276 |
| chr1-40536239-INS-96 | chr1-40536239-INS-96 |
| chr1-40596083-INS-96 | chr1-40596083-INS-96 |
| chr1-40661785-INS-91 | chr1-40661785-INS-91 |
| chr1-40676636-INS-6733 | chr1-40676637-INS-6733 |
| chr1-40716035-INS-74 | chr1-40716037-INS-84 |
| chr1-40724098-INS-123 | chr1-40724098-INS-123 |
| chr1-40746888-INS-57 | chr1-40746888-INS-53 |
| chr1-40767581-INS-75 | chr1-40767558-INS-76 |
| chr1-40779488-INS-348 | chr1-40779513-INS-372 |
| chr1-40784398-INS-460 | chr1-40784398-INS-460 |
| chr1-40821390-INS-132 | chr1-40821390-INS-146 |
| chr1-40851381-INS-209 | chr1-40851393-INS-198 |

| | |
|---|---|
| chr1-40882950-INS-51 | chr1-40882951-INS-50 |
| chr1-40907956-INS-579 | chr1-40907956-INS-569 |
| chr1-40915115-INS-146 | chr1-40915103-INS-150 |
| chr1-40918136-INS-77 | chr1-40918136-INS-77 |
| chr1-40919203-INS-373 | chr1-40919203-INS-374 |
| chr1-40931753-INS-69 | chr1-40931753-INS-71 |
| chr1-40966098-INS-453 | chr1-40966098-INS-451 |
| chr1-40969930-INS-896 | chr1-40969930-INS-894 |
| chr1-40991399-INS-776 | chr1-40991413-INS-799 |
| chr1-41044002-INS-186 | chr1-41044002-INS-158 |
| chr1-41136378-INS-485 | chr1-41136378-INS-485 |
| chr1-41146894-INS-65 | chr1-41146894-INS-65 |
| chr1-41148207-INS-64 | chr1-41148207-INS-64 |
| chr1-41159077-INS-116 | chr1-41159077-INS-116 |
| chr1-41194301-INS-83 | chr1-41194301-INS-63 |
| chr1-41223111-INS-84 | chr1-41223111-INS-84 |
| chr1-41233174-INS-6631 | chr1-41233175-INS-6631 |
| chr1-41237665-INS-836 | chr1-41237665-INS-836 |
| chr1-41246126-INS-670 | chr1-41246129-INS-643 |
| chr1-41256279-INS-1897 | chr1-41256280-INS-1897 |
| chr1-41293910-INS-68 | chr1-41293910-INS-68 |
| chr1-41295738-INS-344 | chr1-41295738-INS-344 |
| chr1-41303208-INS-122 | chr1-41303206-INS-82 |
| chr1-41315803-INS-83 | chr1-41315803-INS-74 |
| chr1-41327472-INS-54 | chr1-41327472-INS-73 |
| chr1-41350336-INS-95 | chr1-41350336-INS-95 |
| chr1-41352169-INS-4982 | chr1-41352170-INS-4983 |
| chr1-41352641-INS-155 | chr1-41352641-INS-179 |
| chr1-41367466-INS-257 | chr1-41367466-INS-257 |
| chr1-41400058-INS-7297 | chr1-41400059-INS-7297 |
| chr1-41404804-INS-88 | chr1-41404804-INS-87 |
| chr1-41413598-INS-55 | chr1-41413598-INS-55 |
| chr1-41418134-INS-52 | chr1-41418134-INS-50 |
| chr1-41435135-INS-123 | chr1-41435182-INS-119 |
| chr1-41455937-INS-116 | chr1-41455937-INS-116 |
| chr1-41477604-INS-54 | chr1-41477604-INS-54 |
| chr1-41491141-INS-193 | chr1-41491141-INS-198 |
| chr1-41491141-INS-756 | chr1-41491141-INS-777 |
| chr1-41515946-INS-2316 | chr1-41515947-INS-2317 |
| chr1-41560679-INS-3183 | chr1-41560680-INS-3183 |
| chr1-41573359-INS-52 | chr1-41573359-INS-52 |
| chr1-41577833-INS-353 | chr1-41577821-INS-361 |
| chr1-41603548-INS-81 | chr1-41603548-INS-81 |
| chr1-41612515-INS-84 | chr1-41612504-INS-80 |
| chr1-41614834-INS-86 | chr1-41614834-INS-92 |
| chr1-41615880-INS-72 | chr1-41615880-INS-75 |
| chr1-41623490-INS-267 | chr1-41623490-INS-267 |
| chr1-41625693-INS-1741 | chr1-41625694-INS-1741 |
| chr1-41628802-INS-56 | chr1-41628821-INS-59 |
| chr1-41639640-INS-869 | chr1-41639640-INS-868 |
| chr1-41648577-INS-109 | chr1-41648561-INS-115 |
| chr1-41658318-INS-6644 | chr1-41658319-INS-6644 |
| chr1-41682724-INS-438 | chr1-41682724-INS-438 |
| chr1-41717328-INS-53 | chr1-41717328-INS-54 |
| chr1-41771271-INS-75 | chr1-41771271-INS-75 |
| chr1-41772413-INS-6440 | chr1-41772414-INS-6440 |
| chr1-41773582-INS-255 | chr1-41773582-INS-262 |
| chr1-41774308-INS-6241 | chr1-41774309-INS-6241 |
| chr1-41795325-INS-71 | chr1-41795325-INS-53 |
| chr1-41828684-INS-6488 | chr1-41828685-INS-6488 |
| chr1-41851981-INS-60 | chr1-41851981-INS-60 |
| chr1-41869315-INS-98 | chr1-41869315-INS-98 |
| chr1-41891887-INS-587 | chr1-41891887-INS-578 |
| chr1-41893000-INS-110 | chr1-41893002-INS-68 |
| chr1-41911241-INS-4603 | chr1-41911242-INS-4604 |
| chr1-41973013-INS-74 | chr1-41973013-INS-74 |
| chr1-42002510-INS-578 | chr1-42002510-INS-582 |
| chr1-42114313-INS-73 | chr1-42114352-INS-69 |
| chr1-42128286-INS-5176 | chr1-42128287-INS-5176 |
| chr1-42146378-INS-52 | chr1-42146378-INS-54 |
| chr1-42151867-INS-180 | chr1-42151867-INS-180 |
| chr1-42221132-INS-1310 | chr1-42221133-INS-1309 |
| chr1-42233397-INS-314 | chr1-42233397-INS-316 |
| chr1-42242471-INS-3304 | chr1-42242472-INS-3304 |
| chr1-42250651-INS-63 | chr1-42250651-INS-63 |
| chr1-42261754-INS-4090 | chr1-42261755-INS-4090 |
| chr1-42323387-INS-56 | chr1-42323387-INS-54 |

| | |
|---|---|
| chr1-42403831-INS-75 | chr1-42403831-INS-55 |
| chr1-42417103-INS-110 | chr1-42417113-INS-68 |
| chr1-42489413-INS-716 | chr1-42489413-INS-714 |
| chr1-42498746-INS-85 | chr1-42498753-INS-51 |
| chr1-42570858-INS-78 | chr1-42570858-INS-78 |
| chr1-42596033-INS-126 | chr1-42596033-INS-125 |
| chr1-42602206-INS-98 | chr1-42602206-INS-90 |
| chr1-42616833-INS-92 | chr1-42616836-INS-54 |
| chr1-42626308-INS-197 | chr1-42626308-INS-197 |
| chr1-42637894-INS-56 | chr1-42637894-INS-58 |
| chr1-42649702-INS-86 | chr1-42649702-INS-72 |
| chr1-42660332-INS-2426 | chr1-42660333-INS-2426 |
| chr1-42667981-INS-85 | chr1-42667981-INS-86 |
| chr1-42684397-INS-2999 | chr1-42684398-INS-2999 |
| chr1-42699150-INS-99 | chr1-42699149-INS-81 |
| chr1-42759298-INS-1891 | chr1-42759299-INS-1891 |
| chr1-42768878-INS-872 | chr1-42768878-INS-867 |
| chr1-42774309-INS-55 | chr1-42774309-INS-55 |
| chr1-42776528-INS-57 | chr1-42776542-INS-50 |
| chr1-42787440-INS-576 | chr1-42787440-INS-576 |
| chr1-42798747-INS-163 | chr1-42798747-INS-154 |
| chr1-42804395-INS-63 | chr1-42804395-INS-63 |
| chr1-42807541-INS-1415 | chr1-42807542-INS-1415 |
| chr1-42831817-INS-51 | chr1-42831817-INS-57 |
| chr1-42843086-INS-55 | chr1-42843086-INS-56 |
| chr1-42850344-INS-54 | chr1-42850344-INS-54 |
| chr1-42863041-INS-70 | chr1-42863041-INS-76 |
| chr1-42890942-INS-53 | chr1-42890987-INS-54 |
| chr1-42983757-INS-76 | chr1-42983757-INS-76 |
| chr1-43006998-INS-197 | chr1-43006998-INS-197 |
| chr1-43025144-INS-842 | chr1-43025144-INS-841 |
| chr1-43040291-INS-1045 | chr1-43040292-INS-1045 |
| chr1-43047752-INS-1919 | chr1-43047753-INS-1919 |
| chr1-43061369-INS-54 | chr1-43061370-INS-52 |
| chr1-43098661-INS-189 | chr1-43098661-INS-182 |
| chr1-43104945-INS-147 | chr1-43104908-INS-157 |
| chr1-43108882-INS-74 | chr1-43108882-INS-66 |
| chr1-43113561-INS-572 | chr1-43113561-INS-565 |
| chr1-43119893-INS-205 | chr1-43119893-INS-209 |
| chr1-43121836-INS-54 | chr1-43121834-INS-100 |
| chr1-43126021-INS-64 | chr1-43126034-INS-66 |
| chr1-43140231-INS-121 | chr1-43140231-INS-121 |
| chr1-43158293-INS-264 | chr1-43158293-INS-264 |
| chr1-43162204-INS-135 | chr1-43162204-INS-135 |
| chr1-43172391-INS-114 | chr1-43172391-INS-114 |
| chr1-43177862-INS-977 | chr1-43177862-INS-977 |
| chr1-43210185-INS-480 | chr1-43210193-INS-486 |
| chr1-43234142-INS-161 | chr1-43234150-INS-163 |
| chr1-43238224-INS-96 | chr1-43238224-INS-96 |
| chr1-43239974-INS-65 | chr1-43239968-INS-65 |
| chr1-43244171-INS-155 | chr1-43244171-INS-150 |
| chr1-43256161-INS-460 | chr1-43256161-INS-460 |
| chr1-43262667-INS-64 | chr1-43262667-INS-64 |
| chr1-43271759-INS-56 | chr1-43271745-INS-64 |
| chr1-43272831-INS-63 | chr1-43272831-INS-63 |
| chr1-43273835-INS-78 | chr1-43273844-INS-81 |
| chr1-43278075-INS-1253 | chr1-43278075-INS-1254 |
| chr1-43307981-INS-7082 | chr1-43307982-INS-7082 |
| chr1-43315080-INS-207 | chr1-43315080-INS-207 |
| chr1-43328527-INS-4858 | chr1-43328528-INS-4858 |
| chr1-43331633-INS-6158 | chr1-43331634-INS-6158 |
| chr1-43340852-INS-200 | chr1-43340852-INS-178 |
| chr1-43378610-INS-200 | chr1-43378610-INS-200 |
| chr1-43428370-INS-196 | chr1-43428370-INS-190 |
| chr1-43442536-INS-102 | chr1-43442536-INS-100 |
| chr1-43447335-INS-180 | chr1-43447355-INS-210 |
| chr1-43449689-INS-438 | chr1-43449689-INS-438 |
| chr1-43498764-INS-56 | chr1-43498764-INS-56 |
| chr1-43501213-INS-664 | chr1-43501213-INS-648 |
| chr1-43515645-INS-953 | chr1-43515645-INS-953 |
| chr1-43523977-INS-58 | chr1-43523977-INS-58 |
| chr1-43527579-INS-174 | chr1-43527579-INS-174 |
| chr1-43530692-INS-88 | chr1-43530690-INS-90 |
| chr1-43539251-INS-908 | chr1-43539251-INS-908 |
| chr1-43559227-INS-54 | chr1-43559227-INS-66 |
| chr1-43571512-INS-206 | chr1-43571512-INS-206 |
| chr1-43595693-INS-64 | chr1-43595693-INS-66 |

| | |
|---|---|
| chr1-43628240-INS-829 | chr1-43628240-INS-830 |
| chr1-43657738-INS-6807 | chr1-43657738-INS-6806 |
| chr1-43693375-INS-66 | chr1-43693375-INS-66 |
| chr1-43696093-INS-175 | chr1-43696097-INS-170 |
| chr1-43709567-INS-8446 | chr1-43709568-INS-8446 |
| chr1-43711959-INS-60 | chr1-43711966-INS-56 |
| chr1-43717448-INS-211 | chr1-43717448-INS-211 |
| chr1-43739142-INS-6567 | chr1-43739143-INS-6567 |
| chr1-43750598-INS-163 | chr1-43750598-INS-163 |
| chr1-43780204-INS-73 | chr1-43780208-INS-73 |
| chr1-43827376-INS-54 | chr1-43827376-INS-60 |
| chr1-43852349-INS-612 | chr1-43852349-INS-612 |
| chr1-43872804-INS-6992 | chr1-43872805-INS-6992 |
| chr1-43901650-INS-4061 | chr1-43901651-INS-4061 |
| chr1-43914161-INS-57 | chr1-43914161-INS-77 |
| chr1-43943630-INS-64 | chr1-43943630-INS-64 |
| chr1-43952932-INS-885 | chr1-43952932-INS-851 |
| chr1-43971802-INS-781 | chr1-43971804-INS-783 |
| chr1-43998183-INS-61 | chr1-43998183-INS-61 |
| chr1-44129948-INS-6674 | chr1-44129949-INS-6674 |
| chr1-44183845-INS-71 | chr1-44183845-INS-80 |
| chr1-44189537-INS-211 | chr1-44189537-INS-211 |
| chr1-44190760-INS-201 | chr1-44190760-INS-202 |
| chr1-44202509-INS-528 | chr1-44202509-INS-528 |
| chr1-44214959-INS-80 | chr1-44214960-INS-66 |
| chr1-44218843-INS-113 | chr1-44218843-INS-115 |
| chr1-44224240-INS-58 | chr1-44224240-INS-58 |
| chr1-44226234-INS-155 | chr1-44226235-INS-115 |
| chr1-44232575-INS-79 | chr1-44232575-INS-67 |
| chr1-44243137-INS-680 | chr1-44243137-INS-680 |
| chr1-44244716-INS-104 | chr1-44244716-INS-86 |
| chr1-44246567-INS-89 | chr1-44246567-INS-89 |
| chr1-44252090-INS-180 | chr1-44252090-INS-179 |
| chr1-44254464-INS-131 | chr1-44254466-INS-127 |
| chr1-44265211-INS-6223 | chr1-44265212-INS-6223 |
| chr1-44275481-INS-55 | chr1-44275482-INS-57 |
| chr1-44284240-INS-201 | chr1-44284240-INS-201 |
| chr1-44292188-INS-77 | chr1-44292171-INS-103 |
| chr1-44299502-INS-55 | chr1-44299502-INS-56 |
| chr1-44300712-INS-76 | chr1-44300712-INS-76 |
| chr1-44329906-INS-2661 | chr1-44329907-INS-2663 |
| chr1-44358820-INS-980 | chr1-44358820-INS-980 |
| chr1-44369217-INS-378 | chr1-44369218-INS-373 |
| chr1-44429402-INS-82 | chr1-44429402-INS-82 |
| chr1-44436657-INS-78 | chr1-44436657-INS-86 |
| chr1-44440698-INS-279 | chr1-44440698-INS-279 |
| chr1-44443150-INS-64 | chr1-44443150-INS-64 |
| chr1-44448071-INS-62 | chr1-44448071-INS-62 |
| chr1-44459561-INS-7613 | chr1-44459562-INS-7614 |
| chr1-44473894-INS-6312 | chr1-44473895-INS-6297 |
| chr1-44479210-INS-56 | chr1-44479201-INS-65 |
| chr1-44501076-INS-6791 | chr1-44501077-INS-6791 |
| chr1-44513592-INS-440 | chr1-44513592-INS-440 |
| chr1-44550798-INS-499 | chr1-44550798-INS-522 |
| chr1-44555574-INS-5353 | chr1-44555575-INS-5353 |
| chr1-44593612-INS-58 | chr1-44593612-INS-58 |
| chr1-44612680-INS-100 | chr1-44612672-INS-108 |
| chr1-44673067-INS-6407 | chr1-44673068-INS-6409 |
| chr1-44684549-INS-64 | chr1-44684549-INS-64 |
| chr1-44688024-INS-57 | chr1-44688024-INS-57 |
| chr1-44701261-INS-3083 | chr1-44701262-INS-3083 |
| chr1-44701262-INS-363 | chr1-44701262-INS-363 |
| chr1-44723796-INS-51 | chr1-44723790-INS-56 |
| chr1-44730738-INS-268 | chr1-44730738-INS-268 |
| chr1-44754567-INS-60 | chr1-44754567-INS-105 |
| chr1-44777914-INS-426 | chr1-44777914-INS-427 |
| chr1-44781553-INS-80 | chr1-44781553-INS-80 |
| chr1-44814945-INS-83 | chr1-44814945-INS-83 |
| chr1-44823055-INS-108 | chr1-44823055-INS-108 |
| chr1-44826028-INS-93 | chr1-44826060-INS-56 |
| chr1-44855945-INS-399 | chr1-44855945-INS-399 |
| chr1-44857955-INS-2630 | chr1-44857956-INS-2630 |
| chr1-44866770-INS-80 | chr1-44866759-INS-55 |
| chr1-44882292-INS-64 | chr1-44882292-INS-64 |
| chr1-44984908-INS-207 | chr1-44984908-INS-207 |
| chr1-44989513-INS-483 | chr1-44989513-INS-483 |
| chr1-44990443-INS-75 | chr1-44990443-INS-75 |

| | |
|---|---|
| chr1-45023975-INS-77 | chr1-45023939-INS-66 |
| chr1-45033154-INS-2281 | chr1-45033155-INS-2279 |
| chr1-45049518-INS-7124 | chr1-45049519-INS-7124 |
| chr1-45068669-INS-590 | chr1-45068669-INS-590 |
| chr1-45072264-INS-199 | chr1-45072264-INS-199 |
| chr1-45086627-INS-211 | chr1-45086627-INS-211 |
| chr1-45149133-INS-730 | chr1-45149133-INS-730 |
| chr1-45167081-INS-469 | chr1-45167081-INS-469 |
| chr1-45169019-INS-53 | chr1-45169019-INS-53 |
| chr1-45182850-INS-169 | chr1-45182851-INS-124 |
| chr1-45197306-INS-98 | chr1-45197306-INS-98 |
| chr1-45205124-INS-208 | chr1-45205124-INS-208 |
| chr1-45217032-INS-69 | chr1-45217032-INS-69 |
| chr1-45232192-INS-110 | chr1-45232200-INS-120 |
| chr1-45250811-INS-196 | chr1-45250811-INS-196 |
| chr1-45253136-INS-56 | chr1-45253136-INS-56 |
| chr1-45258581-INS-110 | chr1-45258581-INS-110 |
| chr1-45312796-INS-279 | chr1-45312796-INS-261 |
| chr1-45322941-INS-58 | chr1-45322940-INS-56 |
| chr1-45334611-INS-196 | chr1-45334641-INS-170 |
| chr1-45341634-INS-61 | chr1-45341644-INS-50 |
| chr1-45352621-INS-65 | chr1-45352628-INS-68 |
| chr1-45369369-INS-61 | chr1-45369376-INS-50 |
| chr1-45381456-INS-213 | chr1-45381456-INS-213 |
| chr1-45412413-INS-3176 | chr1-45412414-INS-3176 |
| chr1-45524168-INS-325 | chr1-45524169-INS-289 |
| chr1-45539771-INS-54 | chr1-45539771-INS-54 |
| chr1-45544428-INS-54 | chr1-45544428-INS-54 |
| chr1-45552660-INS-58 | chr1-45552660-INS-52 |
| chr1-45556678-INS-494 | chr1-45556684-INS-495 |
| chr1-45564856-INS-214 | chr1-45564856-INS-214 |
| chr1-45578900-INS-533 | chr1-45578900-INS-533 |
| chr1-45586115-INS-89 | chr1-45586115-INS-52 |
| chr1-45589556-INS-203 | chr1-45589556-INS-192 |
| chr1-45606308-INS-52 | chr1-45606308-INS-54 |
| chr1-45621845-INS-1810 | chr1-45621846-INS-1811 |
| chr1-45632027-INS-209 | chr1-45632027-INS-209 |
| chr1-45647539-INS-224 | chr1-45647539-INS-226 |
| chr1-45656755-INS-3198 | chr1-45656756-INS-3198 |
| chr1-45679537-INS-136 | chr1-45679583-INS-132 |
| chr1-45680309-INS-2710 | chr1-45680310-INS-2710 |
| chr1-45686445-INS-6653 | chr1-45686446-INS-6653 |
| chr1-45687445-INS-1107 | chr1-45687446-INS-1133 |
| chr1-45693113-INS-7256 | chr1-45693114-INS-7256 |
| chr1-45693491-INS-214 | chr1-45693491-INS-214 |
| chr1-45706605-INS-496 | chr1-45706605-INS-496 |
| chr1-45710730-INS-2125 | chr1-45710731-INS-2125 |
| chr1-45723414-INS-52 | chr1-45723412-INS-52 |
| chr1-45732518-INS-84 | chr1-45732518-INS-84 |
| chr1-45758189-INS-119 | chr1-45758189-INS-135 |
| chr1-45854566-INS-90 | chr1-45854566-INS-90 |
| chr1-45879268-INS-212 | chr1-45879268-INS-212 |
| chr1-45901899-INS-203 | chr1-45901899-INS-203 |
| chr1-45933519-INS-88 | chr1-45933519-INS-88 |
| chr1-45937710-INS-185 | chr1-45937710-INS-199 |
| chr1-45957697-INS-67 | chr1-45957697-INS-67 |
| chr1-45975633-INS-689 | chr1-45975654-INS-655 |
| chr1-45989473-INS-961 | chr1-45989473-INS-961 |
| chr1-46042003-INS-66 | chr1-46042003-INS-66 |
| chr1-46057534-INS-430 | chr1-46057534-INS-434 |
| chr1-46099682-INS-6228 | chr1-46099683-INS-6228 |
| chr1-46128875-INS-67 | chr1-46128856-INS-50 |
| chr1-46155258-INS-57 | chr1-46155258-INS-57 |
| chr1-46179723-INS-6476 | chr1-46179724-INS-6476 |
| chr1-46193227-INS-3056 | chr1-46193228-INS-3056 |
| chr1-46198207-INS-50 | chr1-46198207-INS-54 |
| chr1-46229513-INS-203 | chr1-46229513-INS-203 |
| chr1-46236477-INS-54 | chr1-46236495-INS-60 |
| chr1-46242503-INS-54 | chr1-46242503-INS-65 |
| chr1-46290289-INS-255 | chr1-46290289-INS-255 |
| chr1-46309032-INS-6868 | chr1-46309033-INS-6867 |
| chr1-46347659-INS-6607 | chr1-46347660-INS-6608 |
| chr1-46351591-INS-4508 | chr1-46351592-INS-4508 |
| chr1-46387602-INS-5238 | chr1-46387603-INS-5238 |
| chr1-46404006-INS-72 | chr1-46404027-INS-51 |
| chr1-46404431-INS-400 | chr1-46404431-INS-401 |
| chr1-46422416-INS-204 | chr1-46422416-INS-204 |

| | |
|---|---|
| chr1-46424716-INS-297 | chr1-46424716-INS-297 |
| chr1-46433392-INS-174 | chr1-46433392-INS-159 |
| chr1-46466493-INS-156 | chr1-46466493-INS-157 |
| chr1-46542198-INS-69 | chr1-46542198-INS-69 |
| chr1-46570256-INS-181 | chr1-46570256-INS-181 |
| chr1-46575497-INS-58 | chr1-46575497-INS-58 |
| chr1-46584093-INS-60 | chr1-46584093-INS-60 |
| chr1-46622552-INS-212 | chr1-46622552-INS-212 |
| chr1-46674161-INS-1490 | chr1-46674158-INS-1496 |
| chr1-46682468-INS-580 | chr1-46682468-INS-580 |
| chr1-46758970-INS-212 | chr1-46758970-INS-212 |
| chr1-46784020-INS-7354 | chr1-46784021-INS-7354 |
| chr1-46813939-INS-328 | chr1-46813939-INS-329 |
| chr1-46815992-INS-790 | chr1-46815992-INS-791 |
| chr1-46822234-INS-54 | chr1-46822234-INS-56 |
| chr1-46831610-INS-204 | chr1-46831610-INS-204 |
| chr1-46843408-INS-518 | chr1-46843408-INS-518 |
| chr1-46855216-INS-72 | chr1-46855216-INS-72 |
| chr1-46860684-INS-388 | chr1-46860684-INS-401 |
| chr1-46872886-INS-180 | chr1-46872866-INS-162 |
| chr1-46908512-INS-204 | chr1-46908512-INS-204 |
| chr1-46914592-INS-255 | chr1-46914592-INS-255 |
| chr1-46942293-INS-60 | chr1-46942293-INS-60 |
| chr1-46952596-INS-1203 | chr1-46952570-INS-1196 |
| chr1-46954430-INS-57 | chr1-46954443-INS-53 |
| chr1-46971038-INS-135 | chr1-46971038-INS-135 |
| chr1-46980584-INS-223 | chr1-46980584-INS-223 |
| chr1-46990111-INS-66 | chr1-46990111-INS-52 |
| chr1-47000362-INS-663 | chr1-47000362-INS-664 |
| chr1-47009698-INS-90 | chr1-47009693-INS-78 |
| chr1-47048080-INS-1281 | chr1-47048081-INS-1281 |
| chr1-47072272-INS-7116 | chr1-47072273-INS-7118 |
| chr1-47072533-INS-76 | chr1-47072533-INS-76 |
| chr1-47085109-INS-640 | chr1-47085109-INS-640 |
| chr1-47103885-INS-6497 | chr1-47103886-INS-6497 |
| chr1-47135005-INS-119 | chr1-47135005-INS-119 |
| chr1-47159414-INS-100 | chr1-47159419-INS-68 |
| chr1-47246411-INS-487 | chr1-47246411-INS-487 |
| chr1-47290003-INS-407 | chr1-47290003-INS-407 |
| chr1-47293520-INS-6987 | chr1-47293521-INS-6986 |
| chr1-47368941-INS-515 | chr1-47368941-INS-515 |
| chr1-47384487-INS-134 | chr1-47384487-INS-134 |
| chr1-47420869-INS-70 | chr1-47420869-INS-72 |
| chr1-47425329-INS-61 | chr1-47425329-INS-63 |
| chr1-47431398-INS-84 | chr1-47431398-INS-74 |
| chr1-47434239-INS-137 | chr1-47434240-INS-90 |
| chr1-47438142-INS-498 | chr1-47438142-INS-498 |
| chr1-47536095-INS-4573 | chr1-47536096-INS-4573 |
| chr1-47540931-INS-70 | chr1-47540925-INS-78 |
| chr1-47556997-INS-224 | chr1-47556997-INS-224 |
| chr1-47564695-INS-1914 | chr1-47564696-INS-1915 |
| chr1-47609319-INS-1186 | chr1-47609320-INS-1186 |
| chr1-47626851-INS-424 | chr1-47626851-INS-424 |
| chr1-47633578-INS-64 | chr1-47633578-INS-96 |
| chr1-47701569-INS-211 | chr1-47701569-INS-211 |
| chr1-47735356-INS-185 | chr1-47735356-INS-184 |
| chr1-47747186-INS-282 | chr1-47747184-INS-286 |
| chr1-47768790-INS-64 | chr1-47768790-INS-58 |
| chr1-47769246-INS-917 | chr1-47769246-INS-922 |
| chr1-47805121-INS-725 | chr1-47805121-INS-724 |
| chr1-47816093-INS-113 | chr1-47816093-INS-122 |
| chr1-47826378-INS-156 | chr1-47826378-INS-156 |
| chr1-47848800-INS-6669 | chr1-47848801-INS-6670 |
| chr1-47868431-INS-6359 | chr1-47868432-INS-6359 |
| chr1-47869609-INS-724 | chr1-47869609-INS-724 |
| chr1-47875063-INS-144 | chr1-47875063-INS-141 |
| chr1-47883715-INS-678 | chr1-47883715-INS-681 |
| chr1-47940777-INS-52 | chr1-47940790-INS-65 |
| chr1-47945903-INS-586 | chr1-47945903-INS-586 |
| chr1-47981079-INS-1892 | chr1-47981080-INS-1892 |
| chr1-48037618-INS-52 | chr1-48037618-INS-56 |
| chr1-48041226-INS-181 | chr1-48041226-INS-181 |
| chr1-48059545-INS-6410 | chr1-48059546-INS-6410 |
| chr1-48071114-INS-196 | chr1-48071114-INS-181 |
| chr1-48083011-INS-193 | chr1-48083000-INS-199 |
| chr1-48093313-INS-3231 | chr1-48093314-INS-3231 |
| chr1-48120451-INS-121 | chr1-48120451-INS-122 |

| | |
|---|---|
| chr1-48144583-INS-1692 | chr1-48144567-INS-1701 |
| chr1-48186345-INS-64 | chr1-48186348-INS-55 |
| chr1-48188510-INS-220 | chr1-48188510-INS-220 |
| chr1-48194047-INS-83 | chr1-48194031-INS-72 |
| chr1-48199548-INS-81 | chr1-48199548-INS-57 |
| chr1-48211887-INS-70 | chr1-48211871-INS-102 |
| chr1-48230319-INS-819 | chr1-48230319-INS-819 |
| chr1-48284856-INS-128 | chr1-48284856-INS-127 |
| chr1-48331356-INS-198 | chr1-48331356-INS-198 |
| chr1-48359102-INS-7055 | chr1-48359103-INS-7055 |
| chr1-48447521-INS-54 | chr1-48447521-INS-56 |
| chr1-48492392-INS-408 | chr1-48492392-INS-408 |
| chr1-48550402-INS-527 | chr1-48550402-INS-527 |
| chr1-48568355-INS-655 | chr1-48568355-INS-655 |
| chr1-48578167-INS-737 | chr1-48578167-INS-730 |
| chr1-48580724-INS-70 | chr1-48580724-INS-74 |
| chr1-48594455-INS-199 | chr1-48594455-INS-199 |
| chr1-48615392-INS-390 | chr1-48615392-INS-390 |
| chr1-48639480-INS-55 | chr1-48639480-INS-55 |
| chr1-48642517-INS-2485 | chr1-48642518-INS-2485 |
| chr1-48692961-INS-1284 | chr1-48692962-INS-1284 |
| chr1-48769748-INS-106 | chr1-48769748-INS-68 |
| chr1-48774254-INS-930 | chr1-48774254-INS-913 |
| chr1-48775794-INS-744 | chr1-48775794-INS-744 |
| chr1-48785203-INS-1055 | chr1-48785204-INS-1055 |
| chr1-48789031-INS-247 | chr1-48789031-INS-247 |
| chr1-48798485-INS-1291 | chr1-48798486-INS-1302 |
| chr1-48802355-INS-260 | chr1-48802355-INS-263 |
| chr1-48830707-INS-175 | chr1-48830705-INS-175 |
| chr1-48866587-INS-6444 | chr1-48866588-INS-6445 |
| chr1-48901162-INS-4087 | chr1-48901163-INS-4087 |
| chr1-48917193-INS-53 | chr1-48917193-INS-53 |
| chr1-48924708-INS-52 | chr1-48924707-INS-63 |
| chr1-48937687-INS-1321 | chr1-48937688-INS-1321 |
| chr1-48996183-INS-3247 | chr1-48996184-INS-3247 |
| chr1-49024690-INS-688 | chr1-49024690-INS-683 |
| chr1-49033329-INS-462 | chr1-49033329-INS-462 |
| chr1-49070785-INS-7165 | chr1-49070786-INS-7165 |
| chr1-49077421-INS-93 | chr1-49077463-INS-51 |
| chr1-49088426-INS-53 | chr1-49088426-INS-53 |
| chr1-49098488-INS-89 | chr1-49098488-INS-95 |
| chr1-49103371-INS-310 | chr1-49103365-INS-318 |
| chr1-49116585-INS-133 | chr1-49116585-INS-133 |
| chr1-49140864-INS-7577 | chr1-49140865-INS-7580 |
| chr1-49162990-INS-158 | chr1-49162990-INS-158 |
| chr1-49169167-INS-1127 | chr1-49169170-INS-1121 |
| chr1-49285741-INS-55 | chr1-49285741-INS-84 |
| chr1-49308013-INS-1751 | chr1-49308014-INS-1751 |
| chr1-49329582-INS-432 | chr1-49329582-INS-432 |
| chr1-49331629-INS-60 | chr1-49331621-INS-52 |
| chr1-49398372-INS-189 | chr1-49398371-INS-188 |
| chr1-49400094-INS-6395 | chr1-49400095-INS-6388 |
| chr1-49410367-INS-7167 | chr1-49410368-INS-7167 |
| chr1-49447343-INS-52 | chr1-49447339-INS-58 |
| chr1-49447394-INS-152 | chr1-49447390-INS-136 |
| chr1-49448710-INS-59 | chr1-49448710-INS-80 |
| chr1-49484887-INS-520 | chr1-49484887-INS-520 |
| chr1-49533304-INS-867 | chr1-49533304-INS-856 |
| chr1-49544440-INS-502 | chr1-49544440-INS-498 |
| chr1-49569130-INS-2034 | chr1-49569131-INS-2037 |
| chr1-49578555-INS-92 | chr1-49578555-INS-92 |
| chr1-49582889-INS-183 | chr1-49582889-INS-183 |
| chr1-49615442-INS-767 | chr1-49615442-INS-767 |
| chr1-49619443-INS-1049 | chr1-49619444-INS-1049 |
| chr1-49701437-INS-62 | chr1-49701420-INS-92 |
| chr1-49713484-INS-424 | chr1-49713480-INS-423 |
| chr1-49726186-INS-68 | chr1-49726186-INS-64 |
| chr1-49792368-INS-203 | chr1-49792368-INS-200 |
| chr1-49806765-INS-62 | chr1-49806764-INS-60 |
| chr1-49811571-INS-835 | chr1-49811571-INS-845 |
| chr1-49843835-INS-186 | chr1-49843835-INS-176 |
| chr1-49852833-INS-10534 | chr1-49852834-INS-10535 |
| chr1-49853022-INS-54 | chr1-49853046-INS-50 |
| chr1-49867653-INS-3790 | chr1-49867654-INS-3790 |
| chr1-49876662-INS-693 | chr1-49876662-INS-693 |
| chr1-49877691-INS-52 | chr1-49877691-INS-56 |
| chr1-49918328-INS-60 | chr1-49918328-INS-64 |

| | |
|---|---|
| chr1-49928996-INS-562 | chr1-49928997-INS-559 |
| chr1-49937160-INS-169 | chr1-49937161-INS-169 |
| chr1-49967988-INS-511 | chr1-49967988-INS-511 |
| chr1-49985201-INS-64 | chr1-49985211-INS-60 |
| chr1-49987440-INS-193 | chr1-49987440-INS-193 |
| chr1-50016933-INS-66 | chr1-50016933-INS-66 |
| chr1-50039298-INS-6693 | chr1-50039299-INS-6694 |
| chr1-50059770-INS-438 | chr1-50059770-INS-438 |
| chr1-50088967-INS-54 | chr1-50088967-INS-54 |
| chr1-50090635-INS-2766 | chr1-50090636-INS-2774 |
| chr1-50092975-INS-361 | chr1-50092975-INS-361 |
| chr1-50093355-INS-6440 | chr1-50093356-INS-6440 |
| chr1-50102442-INS-138 | chr1-50102442-INS-144 |
| chr1-50121011-INS-82 | chr1-50120962-INS-84 |
| chr1-50125475-INS-339 | chr1-50125475-INS-338 |
| chr1-50130464-INS-582 | chr1-50130464-INS-583 |
| chr1-50133951-INS-523 | chr1-50133951-INS-523 |
| chr1-50148894-INS-84 | chr1-50148894-INS-84 |
| chr1-50165153-INS-209 | chr1-50165153-INS-209 |
| chr1-50173178-INS-72 | chr1-50173178-INS-68 |
| chr1-50175374-INS-480 | chr1-50175374-INS-484 |
| chr1-50183575-INS-54 | chr1-50183576-INS-52 |
| chr1-50226153-INS-366 | chr1-50226153-INS-365 |
| chr1-50249005-INS-120 | chr1-50249007-INS-120 |
| chr1-50252458-INS-201 | chr1-50252458-INS-201 |
| chr1-50261623-INS-961 | chr1-50261623-INS-961 |
| chr1-50263148-INS-1156 | chr1-50263149-INS-1153 |
| chr1-50266515-INS-64 | chr1-50266515-INS-60 |
| chr1-50283412-INS-494 | chr1-50283412-INS-494 |
| chr1-50300070-INS-83 | chr1-50300090-INS-95 |
| chr1-50307115-INS-322 | chr1-50307115-INS-322 |
| chr1-50417932-INS-845 | chr1-50417932-INS-845 |
| chr1-50436525-INS-153 | chr1-50436525-INS-153 |
| chr1-50463728-INS-6487 | chr1-50463729-INS-6487 |
| chr1-50468965-INS-3164 | chr1-50468966-INS-3165 |
| chr1-50470365-INS-423 | chr1-50470365-INS-423 |
| chr1-50509389-INS-52 | chr1-50509389-INS-52 |
| chr1-50525252-INS-54 | chr1-50525252-INS-54 |
| chr1-50537053-INS-2566 | chr1-50537054-INS-2566 |
| chr1-50545083-INS-108 | chr1-50545083-INS-104 |
| chr1-50549734-INS-696 | chr1-50549734-INS-696 |
| chr1-50589750-INS-482 | chr1-50589750-INS-482 |
| chr1-50600815-INS-53 | chr1-50600815-INS-53 |
| chr1-50643747-INS-54 | chr1-50643748-INS-79 |
| chr1-50644967-INS-65 | chr1-50644967-INS-86 |
| chr1-50658334-INS-70 | chr1-50658334-INS-70 |
| chr1-50661042-INS-6387 | chr1-50661043-INS-6387 |
| chr1-50676207-INS-5546 | chr1-50676208-INS-5546 |
| chr1-50716484-INS-740 | chr1-50716484-INS-740 |
| chr1-50737940-INS-180 | chr1-50737940-INS-181 |
| chr1-50777851-INS-1453 | chr1-50777852-INS-1453 |
| chr1-50779492-INS-52 | chr1-50779492-INS-52 |
| chr1-50782896-INS-54 | chr1-50782896-INS-54 |
| chr1-50787770-INS-2481 | chr1-50787771-INS-2480 |
| chr1-50793362-INS-6287 | chr1-50793363-INS-6287 |
| chr1-50794166-INS-357 | chr1-50794166-INS-357 |
| chr1-50800236-INS-453 | chr1-50800236-INS-453 |
| chr1-50815880-INS-2411 | chr1-50815881-INS-2411 |
| chr1-50835223-INS-98 | chr1-50835223-INS-98 |
| chr1-50844735-INS-160 | chr1-50844735-INS-160 |
| chr1-50845309-INS-7341 | chr1-50845310-INS-7343 |
| chr1-50865519-INS-4665 | chr1-50865520-INS-4665 |
| chr1-50868491-INS-445 | chr1-50868491-INS-448 |
| chr1-50871210-INS-58 | chr1-50871213-INS-67 |
| chr1-50890676-INS-64 | chr1-50890676-INS-64 |
| chr1-50891527-INS-206 | chr1-50891527-INS-206 |
| chr1-50943522-INS-54 | chr1-50943522-INS-57 |
| chr1-50959742-INS-431 | chr1-50959742-INS-434 |
| chr1-50991842-INS-141 | chr1-50991842-INS-141 |
| chr1-51027602-INS-6129 | chr1-51027603-INS-6129 |
| chr1-51028960-INS-90 | chr1-51028960-INS-90 |
| chr1-51089450-INS-6259 | chr1-51089446-INS-6268 |
| chr1-51114246-INS-1614 | chr1-51114247-INS-1614 |
| chr1-51168311-INS-56 | chr1-51168320-INS-66 |
| chr1-51183019-INS-1948 | chr1-51183020-INS-1948 |
| chr1-51201366-INS-115 | chr1-51201366-INS-115 |
| chr1-51208343-INS-873 | chr1-51208343-INS-873 |

| | |
|---|---|
| chr1-51215632-INS-114 | chr1-51215658-INS-80 |
| chr1-51223468-INS-96 | chr1-51223442-INS-126 |
| chr1-51239678-INS-471 | chr1-51239678-INS-472 |
| chr1-51247511-INS-2529 | chr1-51247512-INS-2571 |
| chr1-51267009-INS-57 | chr1-51267009-INS-60 |
| chr1-51305514-INS-670 | chr1-51305514-INS-672 |
| chr1-51339241-INS-52 | chr1-51339241-INS-58 |
| chr1-51371993-INS-90 | chr1-51371993-INS-92 |
| chr1-51379456-INS-55 | chr1-51379436-INS-55 |
| chr1-51380237-INS-2826 | chr1-51380238-INS-2826 |
| chr1-51387553-INS-783 | chr1-51387553-INS-773 |
| chr1-51394278-INS-367 | chr1-51394278-INS-367 |
| chr1-51411362-INS-59 | chr1-51411379-INS-87 |
| chr1-51435511-INS-213 | chr1-51435511-INS-192 |
| chr1-51458950-INS-560 | chr1-51458950-INS-560 |
| chr1-51515011-INS-197 | chr1-51515003-INS-202 |
| chr1-51545053-INS-384 | chr1-51545053-INS-386 |
| chr1-51557143-INS-877 | chr1-51557142-INS-877 |
| chr1-51560157-INS-199 | chr1-51560157-INS-200 |
| chr1-51593207-INS-918 | chr1-51593207-INS-918 |
| chr1-51641638-INS-76 | chr1-51641638-INS-58 |
| chr1-51664112-INS-881 | chr1-51664112-INS-881 |
| chr1-51664220-INS-7156 | chr1-51664221-INS-7156 |
| chr1-51668085-INS-70 | chr1-51668087-INS-100 |
| chr1-51691009-INS-85 | chr1-51691009-INS-85 |
| chr1-51702521-INS-100 | chr1-51702521-INS-99 |
| chr1-51711282-INS-210 | chr1-51711282-INS-212 |
| chr1-51713335-INS-958 | chr1-51713335-INS-958 |
| chr1-51725985-INS-5433 | chr1-51725986-INS-5433 |
| chr1-51737914-INS-486 | chr1-51737914-INS-486 |
| chr1-51747689-INS-3863 | chr1-51747690-INS-3863 |
| chr1-51747690-INS-430 | chr1-51747690-INS-430 |
| chr1-51762981-INS-80 | chr1-51762981-INS-80 |
| chr1-51764398-INS-571 | chr1-51764398-INS-571 |
| chr1-51765109-INS-62 | chr1-51765109-INS-62 |
| chr1-51772063-INS-403 | chr1-51772063-INS-394 |
| chr1-51784378-INS-1420 | chr1-51784379-INS-1432 |
| chr1-51786715-INS-352 | chr1-51786715-INS-352 |
| chr1-51822244-INS-109 | chr1-51822244-INS-91 |
| chr1-51829530-INS-166 | chr1-51829530-INS-166 |
| chr1-51835786-INS-75 | chr1-51835786-INS-75 |
| chr1-51840458-INS-122 | chr1-51840458-INS-122 |
| chr1-51842549-INS-172 | chr1-51842591-INS-152 |
| chr1-51884549-INS-203 | chr1-51884549-INS-203 |
| chr1-51967171-INS-651 | chr1-51967171-INS-651 |
| chr1-51978012-INS-51 | chr1-51978012-INS-55 |
| chr1-51979822-INS-74 | chr1-51979822-INS-74 |
| chr1-51988217-INS-56 | chr1-51988217-INS-56 |
| chr1-51988984-INS-84 | chr1-51988985-INS-84 |
| chr1-52031060-INS-61 | chr1-52031060-INS-61 |
| chr1-52042357-INS-66 | chr1-52042357-INS-66 |
| chr1-52056345-INS-71 | chr1-52056345-INS-71 |
| chr1-52058197-INS-401 | chr1-52058197-INS-401 |
| chr1-52121098-INS-2050 | chr1-52121099-INS-2052 |
| chr1-52124860-INS-51 | chr1-52124861-INS-55 |
| chr1-52129111-INS-66 | chr1-52129136-INS-78 |
| chr1-52138510-INS-487 | chr1-52138510-INS-487 |
| chr1-52201319-INS-580 | chr1-52201319-INS-580 |
| chr1-52295771-INS-204 | chr1-52295771-INS-204 |
| chr1-52318941-INS-84 | chr1-52318926-INS-84 |
| chr1-52320373-INS-4145 | chr1-52320374-INS-4145 |
| chr1-52331179-INS-63 | chr1-52331179-INS-63 |
| chr1-52335248-INS-128 | chr1-52335254-INS-131 |
| chr1-52348146-INS-52 | chr1-52348146-INS-92 |
| chr1-52384543-INS-60 | chr1-52384543-INS-60 |
| chr1-52396699-INS-328 | chr1-52396699-INS-328 |
| chr1-52469933-INS-52 | chr1-52469917-INS-50 |
| chr1-52477032-INS-6613 | chr1-52477033-INS-6613 |
| chr1-52505345-INS-80 | chr1-52505345-INS-80 |
| chr1-52522383-INS-341 | chr1-52522381-INS-341 |
| chr1-52529008-INS-66 | chr1-52529008-INS-66 |
| chr1-52545742-INS-215 | chr1-52545742-INS-217 |
| chr1-52556126-INS-184 | chr1-52556126-INS-184 |
| chr1-52585479-INS-324 | chr1-52585479-INS-324 |
| chr1-52596693-INS-521 | chr1-52596693-INS-523 |
| chr1-52606335-INS-53 | chr1-52606335-INS-53 |
| chr1-52610507-INS-61 | chr1-52610507-INS-65 |

| | |
|---|---|
| chr1-52618735-INS-5189 | chr1-52618736-INS-5189 |
| chr1-52624497-INS-6634 | chr1-52624498-INS-6635 |
| chr1-52651210-INS-4890 | chr1-52651211-INS-4890 |
| chr1-52665799-INS-2950 | chr1-52665800-INS-2950 |
| chr1-52694183-INS-1466 | chr1-52694184-INS-1466 |
| chr1-52704923-INS-209 | chr1-52704923-INS-203 |
| chr1-52707330-INS-55 | chr1-52707330-INS-55 |
| chr1-52708703-INS-170 | chr1-52708703-INS-173 |
| chr1-52717408-INS-50 | chr1-52717408-INS-54 |
| chr1-52722199-INS-157 | chr1-52722199-INS-157 |
| chr1-52725400-INS-117 | chr1-52725400-INS-117 |
| chr1-52753577-INS-7079 | chr1-52753578-INS-7079 |
| chr1-52755845-INS-182 | chr1-52755845-INS-184 |
| chr1-52790047-INS-60 | chr1-52790047-INS-65 |
| chr1-52808616-INS-71 | chr1-52808616-INS-85 |
| chr1-52814456-INS-103 | chr1-52814462-INS-107 |
| chr1-52836509-INS-171 | chr1-52836509-INS-148 |
| chr1-52842692-INS-402 | chr1-52842692-INS-402 |
| chr1-52853384-INS-102 | chr1-52853384-INS-102 |
| chr1-52871478-INS-189 | chr1-52871478-INS-190 |
| chr1-52886451-INS-76 | chr1-52886451-INS-76 |
| chr1-52889973-INS-123 | chr1-52889973-INS-81 |
| chr1-52894689-INS-88 | chr1-52894689-INS-88 |
| chr1-52896040-INS-214 | chr1-52896040-INS-213 |
| chr1-52909948-INS-213 | chr1-52909948-INS-213 |
| chr1-52921604-INS-115 | chr1-52921604-INS-115 |
| chr1-52922800-INS-927 | chr1-52922800-INS-927 |
| chr1-52923521-INS-335 | chr1-52923521-INS-333 |
| chr1-52934602-INS-732 | chr1-52934602-INS-751 |
| chr1-52943412-INS-442 | chr1-52943412-INS-442 |
| chr1-52953652-INS-811 | chr1-52953652-INS-811 |
| chr1-52973423-INS-211 | chr1-52973423-INS-210 |
| chr1-53005490-INS-212 | chr1-53005490-INS-212 |
| chr1-53031825-INS-332 | chr1-53031825-INS-331 |
| chr1-53055062-INS-576 | chr1-53055062-INS-576 |
| chr1-53057257-INS-70 | chr1-53057257-INS-70 |
| chr1-53067263-INS-944 | chr1-53067263-INS-944 |
| chr1-53078909-INS-65 | chr1-53078913-INS-58 |
| chr1-53091000-INS-130 | chr1-53091000-INS-130 |
| chr1-53092475-INS-431 | chr1-53092475-INS-431 |
| chr1-53108485-INS-7760 | chr1-53108486-INS-7759 |
| chr1-53109945-INS-481 | chr1-53109945-INS-483 |
| chr1-53142154-INS-99 | chr1-53142143-INS-108 |
| chr1-53177967-INS-1622 | chr1-53177968-INS-1622 |
| chr1-53191058-INS-1377 | chr1-53191059-INS-1377 |
| chr1-53195099-INS-63 | chr1-53195126-INS-60 |
| chr1-53224731-INS-960 | chr1-53224731-INS-960 |
| chr1-53235017-INS-186 | chr1-53235017-INS-186 |
| chr1-53271076-INS-198 | chr1-53271076-INS-176 |
| chr1-53303472-INS-418 | chr1-53303472-INS-425 |
| chr1-53309906-INS-197 | chr1-53309906-INS-197 |
| chr1-53338360-INS-1890 | chr1-53338361-INS-1890 |
| chr1-53342678-INS-229 | chr1-53342678-INS-225 |
| chr1-53367206-INS-219 | chr1-53367206-INS-218 |
| chr1-53376443-INS-8267 | chr1-53376444-INS-8267 |
| chr1-53396354-INS-110 | chr1-53396354-INS-110 |
| chr1-53409702-INS-538 | chr1-53409702-INS-538 |
| chr1-53412826-INS-506 | chr1-53412826-INS-506 |
| chr1-53422050-INS-143 | chr1-53422096-INS-99 |
| chr1-53443051-INS-56 | chr1-53443051-INS-76 |
| chr1-53446241-INS-351 | chr1-53446241-INS-338 |
| chr1-53448724-INS-51 | chr1-53448724-INS-51 |
| chr1-53456876-INS-349 | chr1-53456876-INS-349 |
| chr1-53459482-INS-100 | chr1-53459483-INS-98 |
| chr1-53468177-INS-158 | chr1-53468179-INS-158 |
| chr1-53471646-INS-211 | chr1-53471646-INS-212 |
| chr1-53491853-INS-57 | chr1-53491853-INS-57 |
| chr1-53499700-INS-183 | chr1-53499700-INS-184 |
| chr1-53579285-INS-3147 | chr1-53579286-INS-3147 |
| chr1-53611697-INS-120 | chr1-53611650-INS-144 |
| chr1-53647530-INS-185 | chr1-53647527-INS-185 |
| chr1-53652820-INS-51 | chr1-53652820-INS-51 |
| chr1-53690758-INS-457 | chr1-53690758-INS-459 |
| chr1-53714043-INS-6847 | chr1-53714044-INS-6848 |
| chr1-53728248-INS-64 | chr1-53728256-INS-60 |
| chr1-53757084-INS-7478 | chr1-53757085-INS-7478 |
| chr1-53761275-INS-3731 | chr1-53761276-INS-3731 |

| | |
|---|---|
| chr1-53783998-INS-407 | chr1-53783998-INS-406 |
| chr1-53795206-INS-210 | chr1-53795206-INS-206 |
| chr1-53800760-INS-67 | chr1-53800760-INS-67 |
| chr1-53808326-INS-450 | chr1-53808336-INS-417 |
| chr1-53812403-INS-194 | chr1-53812403-INS-194 |
| chr1-53817276-INS-556 | chr1-53817276-INS-556 |
| chr1-53819029-INS-351 | chr1-53819029-INS-351 |
| chr1-53841473-INS-1545 | chr1-53841474-INS-1545 |
| chr1-53907332-INS-200 | chr1-53907332-INS-200 |
| chr1-53912073-INS-199 | chr1-53912073-INS-199 |
| chr1-53920635-INS-208 | chr1-53920635-INS-208 |
| chr1-53932476-INS-74 | chr1-53932476-INS-50 |
| chr1-53952627-INS-197 | chr1-53952618-INS-228 |
| chr1-53956388-INS-1431 | chr1-53956389-INS-1431 |
| chr1-53979383-INS-1919 | chr1-53979384-INS-1919 |
| chr1-53986228-INS-170 | chr1-53986228-INS-171 |
| chr1-54050038-INS-6310 | chr1-54050039-INS-6314 |
| chr1-54057384-INS-227 | chr1-54057384-INS-253 |
| chr1-54057798-INS-364 | chr1-54057798-INS-364 |
| chr1-54112710-INS-68 | chr1-54112710-INS-68 |
| chr1-54115308-INS-91 | chr1-54115312-INS-58 |
| chr1-54132871-INS-10086 | chr1-54132872-INS-10087 |
| chr1-54144903-INS-2963 | chr1-54144904-INS-2963 |
| chr1-54213298-INS-68 | chr1-54213299-INS-50 |
| chr1-54219339-INS-403 | chr1-54219339-INS-439 |
| chr1-54238992-INS-5549 | chr1-54238993-INS-5549 |
| chr1-54254236-INS-104 | chr1-54254236-INS-104 |
| chr1-54280213-INS-1203 | chr1-54280208-INS-1185 |
| chr1-54301015-INS-223 | chr1-54301015-INS-199 |
| chr1-54348838-INS-204 | chr1-54348838-INS-204 |
| chr1-54395691-INS-54 | chr1-54395691-INS-64 |
| chr1-54412491-INS-90 | chr1-54412491-INS-90 |
| chr1-54460096-INS-199 | chr1-54460096-INS-199 |
| chr1-54480309-INS-92 | chr1-54480307-INS-92 |
| chr1-54500169-INS-628 | chr1-54500169-INS-642 |
| chr1-54527662-INS-224 | chr1-54527662-INS-224 |
| chr1-54537820-INS-148 | chr1-54537820-INS-151 |
| chr1-54544922-INS-364 | chr1-54544922-INS-364 |
| chr1-54557509-INS-208 | chr1-54557509-INS-209 |
| chr1-54585994-INS-203 | chr1-54585994-INS-199 |
| chr1-54621164-INS-91 | chr1-54621164-INS-91 |
| chr1-54638161-INS-636 | chr1-54638161-INS-635 |
| chr1-54652804-INS-55 | chr1-54652804-INS-55 |
| chr1-54656800-INS-103 | chr1-54656800-INS-103 |
| chr1-54659702-INS-203 | chr1-54659702-INS-203 |
| chr1-54696687-INS-181 | chr1-54696687-INS-182 |
| chr1-54703314-INS-177 | chr1-54703312-INS-177 |
| chr1-54729436-INS-7086 | chr1-54729437-INS-7086 |
| chr1-54753747-INS-180 | chr1-54753747-INS-180 |
| chr1-54755695-INS-62 | chr1-54755675-INS-96 |
| chr1-54762740-INS-919 | chr1-54762740-INS-919 |
| chr1-54769882-INS-267 | chr1-54769885-INS-266 |
| chr1-54785910-INS-58 | chr1-54785910-INS-62 |
| chr1-54787761-INS-62 | chr1-54787761-INS-62 |
| chr1-54792865-INS-58 | chr1-54792865-INS-58 |
| chr1-54795427-INS-100 | chr1-54795427-INS-100 |
| chr1-54810280-INS-497 | chr1-54810280-INS-497 |
| chr1-54812429-INS-225 | chr1-54812429-INS-226 |
| chr1-54813504-INS-120 | chr1-54813504-INS-120 |
| chr1-54831393-INS-120 | chr1-54831393-INS-121 |
| chr1-54848343-INS-6927 | chr1-54848344-INS-6928 |
| chr1-54849391-INS-53 | chr1-54849391-INS-53 |
| chr1-54853732-INS-316 | chr1-54853732-INS-320 |
| chr1-54855934-INS-2360 | chr1-54855935-INS-2361 |
| chr1-54857789-INS-54 | chr1-54857789-INS-75 |
| chr1-54873635-INS-90 | chr1-54873635-INS-90 |
| chr1-54876757-INS-78 | chr1-54876758-INS-78 |
| chr1-54881993-INS-1086 | chr1-54881994-INS-1086 |
| chr1-54902123-INS-54 | chr1-54902124-INS-62 |
| chr1-54905139-INS-6377 | chr1-54905140-INS-6379 |
| chr1-54906078-INS-92 | chr1-54906078-INS-92 |
| chr1-54922367-INS-429 | chr1-54922367-INS-429 |
| chr1-54943449-INS-174 | chr1-54943449-INS-174 |
| chr1-54973413-INS-196 | chr1-54973413-INS-196 |
| chr1-54975235-INS-643 | chr1-54975235-INS-643 |
| chr1-54980111-INS-60 | chr1-54980111-INS-60 |
| chr1-54986059-INS-7115 | chr1-54986060-INS-7118 |

| | |
|---|---|
| chr1-54990644-INS-51 | chr1-54990644-INS-51 |
| chr1-55005787-INS-95 | chr1-55005787-INS-95 |
| chr1-55033284-INS-176 | chr1-55033284-INS-176 |
| chr1-55038459-INS-53 | chr1-55038426-INS-61 |
| chr1-55050178-INS-168 | chr1-55050172-INS-138 |
| chr1-55051689-INS-58 | chr1-55051689-INS-58 |
| chr1-55057475-INS-706 | chr1-55057475-INS-706 |
| chr1-55061909-INS-212 | chr1-55061909-INS-216 |
| chr1-55064290-INS-405 | chr1-55064290-INS-398 |
| chr1-55072291-INS-51 | chr1-55072291-INS-51 |
| chr1-55080581-INS-144 | chr1-55080581-INS-144 |
| chr1-55085224-INS-4087 | chr1-55085225-INS-4087 |
| chr1-55085446-INS-199 | chr1-55085446-INS-199 |
| chr1-55108322-INS-174 | chr1-55108322-INS-174 |
| chr1-55130856-INS-210 | chr1-55130856-INS-210 |
| chr1-55140602-INS-682 | chr1-55140602-INS-694 |
| chr1-55150124-INS-64 | chr1-55150125-INS-80 |
| chr1-55158528-INS-209 | chr1-55158528-INS-209 |
| chr1-55164367-INS-185 | chr1-55164367-INS-185 |
| chr1-55195940-INS-62 | chr1-55195940-INS-66 |
| chr1-55198484-INS-116 | chr1-55198484-INS-118 |
| chr1-55199008-INS-4392 | chr1-55199009-INS-4392 |
| chr1-55216913-INS-102 | chr1-55216911-INS-67 |
| chr1-55221697-INS-434 | chr1-55221697-INS-434 |
| chr1-55228079-INS-75 | chr1-55228059-INS-66 |
| chr1-55248346-INS-98 | chr1-55248346-INS-98 |
| chr1-55273511-INS-76 | chr1-55273511-INS-76 |
| chr1-55274308-INS-246 | chr1-55274308-INS-231 |
| chr1-55324092-INS-60 | chr1-55324092-INS-60 |
| chr1-55336418-INS-204 | chr1-55336418-INS-202 |
| chr1-55351719-INS-209 | chr1-55351719-INS-209 |
| chr1-55372150-INS-3038 | chr1-55372151-INS-3038 |
| chr1-55386417-INS-7126 | chr1-55386418-INS-7126 |
| chr1-55389534-INS-66 | chr1-55389534-INS-66 |
| chr1-55429950-INS-1164 | chr1-55429951-INS-1165 |
| chr1-55438787-INS-817 | chr1-55438787-INS-830 |
| chr1-55443332-INS-88 | chr1-55443332-INS-88 |
| chr1-55451960-INS-1088 | chr1-55451961-INS-1089 |
| chr1-55458488-INS-282 | chr1-55458488-INS-282 |
| chr1-55458626-INS-1213 | chr1-55458635-INS-1209 |
| chr1-55478719-INS-168 | chr1-55478719-INS-205 |
| chr1-55489810-INS-52 | chr1-55489811-INS-50 |
| chr1-55495725-INS-56 | chr1-55495729-INS-56 |
| chr1-55524171-INS-68 | chr1-55524171-INS-68 |
| chr1-55549390-INS-2391 | chr1-55549391-INS-2391 |
| chr1-55553964-INS-55 | chr1-55553964-INS-61 |
| chr1-55558029-INS-620 | chr1-55558029-INS-620 |
| chr1-55576912-INS-7119 | chr1-55576913-INS-7119 |
| chr1-55579010-INS-496 | chr1-55579010-INS-496 |
| chr1-55583565-INS-81 | chr1-55583560-INS-51 |
| chr1-55605863-INS-54 | chr1-55605863-INS-54 |
| chr1-55610813-INS-54 | chr1-55610813-INS-54 |
| chr1-55677221-INS-233 | chr1-55677229-INS-223 |
| chr1-55682733-INS-80 | chr1-55682733-INS-80 |
| chr1-55716122-INS-5505 | chr1-55716123-INS-5505 |
| chr1-55726071-INS-264 | chr1-55726071-INS-266 |
| chr1-55731401-INS-62 | chr1-55731401-INS-62 |
| chr1-55742944-INS-123 | chr1-55742944-INS-123 |
| chr1-55746532-INS-57 | chr1-55746532-INS-62 |
| chr1-55757140-INS-151 | chr1-55757140-INS-171 |
| chr1-55764390-INS-498 | chr1-55764390-INS-498 |
| chr1-55766967-INS-55 | chr1-55766967-INS-55 |
| chr1-55770014-INS-498 | chr1-55770014-INS-498 |
| chr1-55772402-INS-60 | chr1-55772402-INS-60 |
| chr1-55776448-INS-56 | chr1-55776445-INS-87 |
| chr1-55781629-INS-5167 | chr1-55781630-INS-5167 |
| chr1-55784095-INS-959 | chr1-55784095-INS-959 |
| chr1-55794345-INS-394 | chr1-55794345-INS-394 |
| chr1-55859449-INS-72 | chr1-55859449-INS-68 |
| chr1-55897979-INS-269 | chr1-55897979-INS-269 |
| chr1-55913960-INS-76 | chr1-55913960-INS-76 |
| chr1-56005832-INS-717 | chr1-56005832-INS-717 |
| chr1-56047828-INS-56 | chr1-56047826-INS-62 |
| chr1-56059839-INS-59 | chr1-56059849-INS-83 |
| chr1-56079145-INS-52 | chr1-56079145-INS-52 |
| chr1-56142535-INS-162 | chr1-56142535-INS-168 |
| chr1-56143638-INS-132 | chr1-56143639-INS-128 |

| | |
|---|---|
| chr1-56207199-INS-51 | chr1-56207199-INS-51 |
| chr1-56224054-INS-146 | chr1-56224054-INS-146 |
| chr1-56228293-INS-199 | chr1-56228293-INS-199 |
| chr1-56229981-INS-1678 | chr1-56229982-INS-1680 |
| chr1-56252145-INS-52 | chr1-56252145-INS-58 |
| chr1-56268738-INS-2852 | chr1-56268739-INS-2853 |
| chr1-56269941-INS-90 | chr1-56269946-INS-55 |
| chr1-56308439-INS-287 | chr1-56308439-INS-295 |
| chr1-56312025-INS-76 | chr1-56312025-INS-74 |
| chr1-56327561-INS-74 | chr1-56327561-INS-78 |
| chr1-56330826-INS-144 | chr1-56330826-INS-148 |
| chr1-56340317-INS-444 | chr1-56340317-INS-444 |
| chr1-56345434-INS-87 | chr1-56345434-INS-87 |
| chr1-56348309-INS-6454 | chr1-56348310-INS-6454 |
| chr1-56356668-INS-200 | chr1-56356668-INS-200 |
| chr1-56360166-INS-144 | chr1-56360169-INS-138 |
| chr1-56381590-INS-1608 | chr1-56381591-INS-1608 |
| chr1-56419567-INS-212 | chr1-56419567-INS-212 |
| chr1-56445577-INS-54 | chr1-56445604-INS-56 |
| chr1-56455063-INS-6421 | chr1-56455064-INS-6421 |
| chr1-56457515-INS-858 | chr1-56457515-INS-858 |
| chr1-56459702-INS-60 | chr1-56459702-INS-60 |
| chr1-56503983-INS-636 | chr1-56503983-INS-636 |
| chr1-56510804-INS-443 | chr1-56510804-INS-450 |
| chr1-56576002-INS-2966 | chr1-56576003-INS-2966 |
| chr1-56577917-INS-208 | chr1-56577917-INS-210 |
| chr1-56588627-INS-56 | chr1-56588627-INS-68 |
| chr1-56594369-INS-6842 | chr1-56594370-INS-6842 |
| chr1-56610109-INS-6501 | chr1-56610110-INS-6496 |
| chr1-56620386-INS-99 | chr1-56620386-INS-104 |
| chr1-56629241-INS-78 | chr1-56629241-INS-112 |
| chr1-56635186-INS-212 | chr1-56635186-INS-213 |
| chr1-56639362-INS-155 | chr1-56639362-INS-155 |
| chr1-56652698-INS-56 | chr1-56652698-INS-56 |
| chr1-56671160-INS-432 | chr1-56671160-INS-431 |
| chr1-56679389-INS-102 | chr1-56679389-INS-101 |
| chr1-56682558-INS-376 | chr1-56682558-INS-376 |
| chr1-56712109-INS-72 | chr1-56712105-INS-79 |
| chr1-56723253-INS-532 | chr1-56723253-INS-532 |
| chr1-56732248-INS-933 | chr1-56732238-INS-942 |
| chr1-56735024-INS-52 | chr1-56735024-INS-88 |
| chr1-56799622-INS-120 | chr1-56799626-INS-94 |
| chr1-56829779-INS-400 | chr1-56829779-INS-400 |
| chr1-56868424-INS-400 | chr1-56868424-INS-400 |
| chr1-56876609-INS-111 | chr1-56876609-INS-111 |
| chr1-56888533-INS-58 | chr1-56888533-INS-58 |
| chr1-56914423-INS-62 | chr1-56914379-INS-63 |
| chr1-56931485-INS-56 | chr1-56931468-INS-102 |
| chr1-56959250-INS-60 | chr1-56959243-INS-72 |
| chr1-56992770-INS-119 | chr1-56992770-INS-85 |
| chr1-57046437-INS-64 | chr1-57046425-INS-56 |
| chr1-57060676-INS-1247 | chr1-57060677-INS-1241 |
| chr1-57088084-INS-754 | chr1-57088084-INS-755 |
| chr1-57123127-INS-456 | chr1-57123127-INS-456 |
| chr1-57151450-INS-80 | chr1-57151450-INS-116 |
| chr1-57154900-INS-282 | chr1-57154900-INS-282 |
| chr1-57156663-INS-202 | chr1-57156663-INS-201 |
| chr1-57172846-INS-6445 | chr1-57172847-INS-6443 |
| chr1-57174783-INS-51 | chr1-57174783-INS-51 |
| chr1-57194702-INS-206 | chr1-57194702-INS-206 |
| chr1-57210225-INS-56 | chr1-57210198-INS-64 |
| chr1-57228888-INS-74 | chr1-57228888-INS-74 |
| chr1-57258860-INS-80 | chr1-57258860-INS-66 |
| chr1-57327704-INS-7136 | chr1-57327705-INS-7136 |
| chr1-57366256-INS-371 | chr1-57366267-INS-380 |
| chr1-57373601-INS-247 | chr1-57373567-INS-257 |
| chr1-57375314-INS-80 | chr1-57375327-INS-74 |
| chr1-57377231-INS-120 | chr1-57377231-INS-120 |
| chr1-57390926-INS-68 | chr1-57390926-INS-68 |
| chr1-57427006-INS-217 | chr1-57427006-INS-218 |
| chr1-57473325-INS-75 | chr1-57473325-INS-64 |
| chr1-57484547-INS-74 | chr1-57484547-INS-74 |
| chr1-57486767-INS-85 | chr1-57486767-INS-90 |
| chr1-57492423-INS-195 | chr1-57492423-INS-195 |
| chr1-57497842-INS-171 | chr1-57497842-INS-171 |
| chr1-57499602-INS-98 | chr1-57499602-INS-98 |
| chr1-57526167-INS-125 | chr1-57526167-INS-75 |

| | |
|---|---|
| chr1-57537925-INS-100 | chr1-57537924-INS-53 |
| chr1-57545431-INS-197 | chr1-57545431-INS-197 |
| chr1-57547705-INS-62 | chr1-57547705-INS-62 |
| chr1-57559718-INS-162 | chr1-57559718-INS-162 |
| chr1-57579933-INS-54 | chr1-57579933-INS-54 |
| chr1-57591311-INS-55 | chr1-57591311-INS-59 |
| chr1-57631683-INS-64 | chr1-57631683-INS-64 |
| chr1-57635919-INS-241 | chr1-57635919-INS-241 |
| chr1-57670024-INS-66 | chr1-57670024-INS-72 |
| chr1-57743787-INS-2905 | chr1-57743788-INS-2905 |
| chr1-57770382-INS-1893 | chr1-57770383-INS-1893 |
| chr1-57828100-INS-75 | chr1-57828100-INS-76 |
| chr1-57874055-INS-6665 | chr1-57874056-INS-6662 |
| chr1-57893378-INS-84 | chr1-57893378-INS-84 |
| chr1-57936963-INS-195 | chr1-57936963-INS-195 |
| chr1-57968884-INS-209 | chr1-57968879-INS-226 |
| chr1-57998082-INS-91 | chr1-57998082-INS-91 |
| chr1-58026983-INS-1546 | chr1-58026984-INS-1546 |
| chr1-58039935-INS-91 | chr1-58039935-INS-91 |
| chr1-58049466-INS-196 | chr1-58049466-INS-195 |
| chr1-58059658-INS-74 | chr1-58059658-INS-80 |
| chr1-58065851-INS-60 | chr1-58065851-INS-60 |
| chr1-58067586-INS-735 | chr1-58067586-INS-735 |
| chr1-58083874-INS-150 | chr1-58083874-INS-169 |
| chr1-58093157-INS-61 | chr1-58093157-INS-61 |
| chr1-58122592-INS-64 | chr1-58122592-INS-59 |
| chr1-58141271-INS-84 | chr1-58141271-INS-92 |
| chr1-58144379-INS-62 | chr1-58144379-INS-62 |
| chr1-58156163-INS-274 | chr1-58156163-INS-271 |
| chr1-58187212-INS-95 | chr1-58187212-INS-95 |
| chr1-58196153-INS-101 | chr1-58196153-INS-103 |
| chr1-58203307-INS-840 | chr1-58203294-INS-828 |
| chr1-58203819-INS-105 | chr1-58203817-INS-92 |
| chr1-58228969-INS-124 | chr1-58228970-INS-164 |
| chr1-58254720-INS-58 | chr1-58254720-INS-58 |
| chr1-58264631-INS-103 | chr1-58264632-INS-72 |
| chr1-58268874-INS-269 | chr1-58268874-INS-263 |
| chr1-58281140-INS-198 | chr1-58281140-INS-198 |
| chr1-58306984-INS-6910 | chr1-58306985-INS-6918 |
| chr1-58310105-INS-206 | chr1-58310106-INS-177 |
| chr1-58311414-INS-4479 | chr1-58311415-INS-4479 |
| chr1-58336128-INS-54 | chr1-58336129-INS-54 |
| chr1-58361392-INS-118 | chr1-58361393-INS-118 |
| chr1-58362246-INS-63 | chr1-58362240-INS-54 |
| chr1-58375403-INS-408 | chr1-58375403-INS-408 |
| chr1-58406420-INS-60 | chr1-58406435-INS-60 |
| chr1-58429423-INS-212 | chr1-58429423-INS-212 |
| chr1-58431478-INS-56 | chr1-58431479-INS-53 |
| chr1-58451694-INS-215 | chr1-58451694-INS-216 |
| chr1-58452659-INS-230 | chr1-58452659-INS-230 |
| chr1-58456631-INS-330 | chr1-58456631-INS-378 |
| chr1-58507988-INS-488 | chr1-58507988-INS-490 |
| chr1-58511169-INS-199 | chr1-58511169-INS-199 |
| chr1-58512315-INS-470 | chr1-58512315-INS-469 |
| chr1-58537347-INS-2263 | chr1-58537348-INS-2272 |
| chr1-58548272-INS-5201 | chr1-58548273-INS-5198 |
| chr1-58553182-INS-211 | chr1-58553182-INS-212 |
| chr1-58566540-INS-229 | chr1-58566540-INS-230 |
| chr1-58574744-INS-203 | chr1-58574744-INS-204 |
| chr1-58579598-INS-207 | chr1-58579598-INS-207 |
| chr1-58609315-INS-74 | chr1-58609315-INS-74 |
| chr1-58666641-INS-7178 | chr1-58666642-INS-7178 |
| chr1-58675446-INS-2667 | chr1-58675447-INS-2667 |
| chr1-58724387-INS-132 | chr1-58724387-INS-132 |
| chr1-58736038-INS-85 | chr1-58736037-INS-112 |
| chr1-58736479-INS-5437 | chr1-58736480-INS-5437 |
| chr1-58738655-INS-148 | chr1-58738655-INS-150 |
| chr1-58757060-INS-130 | chr1-58757060-INS-130 |
| chr1-58758676-INS-82 | chr1-58758676-INS-82 |
| chr1-58763813-INS-86 | chr1-58763813-INS-86 |
| chr1-58775689-INS-193 | chr1-58775689-INS-217 |
| chr1-58778272-INS-98 | chr1-58778272-INS-78 |
| chr1-58803031-INS-6909 | chr1-58803033-INS-6900 |
| chr1-58819282-INS-202 | chr1-58819282-INS-203 |
| chr1-58845747-INS-55 | chr1-58845747-INS-55 |
| chr1-58866157-INS-57 | chr1-58866157-INS-57 |
| chr1-58900146-INS-55 | chr1-58900146-INS-55 |

| | |
|---|---|
| chr1-58909934-INS-201 | chr1-58909934-INS-201 |
| chr1-58919538-INS-486 | chr1-58919538-INS-486 |
| chr1-58953298-INS-890 | chr1-58953298-INS-870 |
| chr1-58955721-INS-56 | chr1-58955722-INS-57 |
| chr1-58998441-INS-69 | chr1-58998441-INS-69 |
| chr1-59040855-INS-79 | chr1-59040848-INS-101 |
| chr1-59049147-INS-76 | chr1-59049147-INS-52 |
| chr1-59066490-INS-170 | chr1-59066490-INS-170 |
| chr1-59120329-INS-238 | chr1-59120330-INS-238 |
| chr1-59122646-INS-83 | chr1-59122646-INS-106 |
| chr1-59139401-INS-565 | chr1-59139401-INS-565 |
| chr1-59164048-INS-132 | chr1-59164048-INS-99 |
| chr1-59167487-INS-64 | chr1-59167533-INS-63 |
| chr1-59169790-INS-73 | chr1-59169790-INS-73 |
| chr1-59178738-INS-68 | chr1-59178738-INS-68 |
| chr1-59217189-INS-60 | chr1-59217189-INS-60 |
| chr1-59248762-INS-203 | chr1-59248762-INS-202 |
| chr1-59262395-INS-129 | chr1-59262395-INS-131 |
| chr1-59302128-INS-169 | chr1-59302128-INS-169 |
| chr1-59304116-INS-94 | chr1-59304105-INS-86 |
| chr1-59318637-INS-118 | chr1-59318637-INS-106 |
| chr1-59321451-INS-86 | chr1-59321451-INS-86 |
| chr1-59333795-INS-115 | chr1-59333795-INS-115 |
| chr1-59338446-INS-92 | chr1-59338446-INS-92 |
| chr1-59343335-INS-213 | chr1-59343335-INS-213 |
| chr1-59344408-INS-116 | chr1-59344433-INS-108 |
| chr1-59367945-INS-54 | chr1-59367947-INS-54 |
| chr1-59379109-INS-100 | chr1-59379158-INS-98 |
| chr1-59391679-INS-74 | chr1-59391679-INS-64 |
| chr1-59393209-INS-259 | chr1-59393209-INS-260 |
| chr1-59399027-INS-198 | chr1-59399027-INS-199 |
| chr1-59423425-INS-204 | chr1-59423425-INS-204 |
| chr1-59427246-INS-5511 | chr1-59427247-INS-5511 |
| chr1-59437141-INS-131 | chr1-59437165-INS-105 |
| chr1-59442961-INS-66 | chr1-59442961-INS-87 |
| chr1-59445100-INS-53 | chr1-59445100-INS-53 |
| chr1-59470605-INS-198 | chr1-59470605-INS-201 |
| chr1-59484773-INS-73 | chr1-59484781-INS-52 |
| chr1-59486756-INS-101 | chr1-59486756-INS-80 |
| chr1-59489289-INS-235 | chr1-59489289-INS-235 |
| chr1-59518771-INS-120 | chr1-59518771-INS-120 |
| chr1-59548620-INS-58 | chr1-59548624-INS-74 |
| chr1-59580082-INS-992 | chr1-59580082-INS-994 |
| chr1-59593818-INS-56 | chr1-59593831-INS-60 |
| chr1-59607444-INS-111 | chr1-59607444-INS-111 |
| chr1-59635481-INS-601 | chr1-59635481-INS-609 |
| chr1-59635489-INS-66 | chr1-59635489-INS-72 |
| chr1-59642850-INS-78 | chr1-59642850-INS-80 |
| chr1-59647978-INS-52 | chr1-59647979-INS-52 |
| chr1-59649619-INS-58 | chr1-59649619-INS-54 |
| chr1-59659987-INS-204 | chr1-59659987-INS-204 |
| chr1-59662003-INS-56 | chr1-59662004-INS-56 |
| chr1-59664667-INS-322 | chr1-59664667-INS-342 |
| chr1-59668923-INS-211 | chr1-59668923-INS-211 |
| chr1-59679497-INS-236 | chr1-59679497-INS-236 |
| chr1-59685139-INS-208 | chr1-59685139-INS-209 |
| chr1-59698522-INS-202 | chr1-59698522-INS-202 |
| chr1-59705406-INS-80 | chr1-59705406-INS-80 |
| chr1-59735043-INS-214 | chr1-59735043-INS-213 |
| chr1-59737389-INS-214 | chr1-59737389-INS-214 |
| chr1-59742562-INS-206 | chr1-59742562-INS-206 |
| chr1-59752225-INS-199 | chr1-59752225-INS-198 |
| chr1-59755468-INS-220 | chr1-59755468-INS-220 |
| chr1-59761329-INS-216 | chr1-59761329-INS-216 |
| chr1-59762813-INS-182 | chr1-59762813-INS-170 |
| chr1-59763051-INS-691 | chr1-59763051-INS-691 |
| chr1-59766444-INS-182 | chr1-59766425-INS-201 |
| chr1-59770636-INS-195 | chr1-59770636-INS-195 |
| chr1-59789296-INS-208 | chr1-59789299-INS-206 |
| chr1-59792403-INS-202 | chr1-59792403-INS-202 |
| chr1-59795027-INS-142 | chr1-59795027-INS-142 |
| chr1-59798927-INS-265 | chr1-59798927-INS-270 |
| chr1-59814526-INS-211 | chr1-59814526-INS-211 |
| chr1-59822136-INS-95 | chr1-59822129-INS-86 |
| chr1-59825224-INS-51 | chr1-59825222-INS-55 |
| chr1-59839207-INS-54 | chr1-59839223-INS-58 |
| chr1-59926842-INS-5545 | chr1-59926843-INS-5545 |

| | |
|---|---|
| chr1-59937037-INS-128 | chr1-59937056-INS-152 |
| chr1-59964813-INS-255 | chr1-59964813-INS-258 |
| chr1-59968487-INS-2128 | chr1-59968488-INS-2128 |
| chr1-59969542-INS-56 | chr1-59969542-INS-51 |
| chr1-60014430-INS-60 | chr1-60014456-INS-59 |
| chr1-60034557-INS-96 | chr1-60034557-INS-96 |
| chr1-60043402-INS-66 | chr1-60043414-INS-76 |
| chr1-60048068-INS-60 | chr1-60048068-INS-60 |
| chr1-60052444-INS-735 | chr1-60052444-INS-735 |
| chr1-60054340-INS-330 | chr1-60054340-INS-330 |
| chr1-60060828-INS-55 | chr1-60060828-INS-55 |
| chr1-60062653-INS-74 | chr1-60062653-INS-78 |
| chr1-60071709-INS-213 | chr1-60071709-INS-213 |
| chr1-60079816-INS-210 | chr1-60079816-INS-210 |
| chr1-60090028-INS-197 | chr1-60090028-INS-199 |
| chr1-60109960-INS-66 | chr1-60109960-INS-66 |
| chr1-60111845-INS-552 | chr1-60111845-INS-552 |
| chr1-60126487-INS-98 | chr1-60126487-INS-98 |
| chr1-60143706-INS-68 | chr1-60143706-INS-72 |
| chr1-60161091-INS-204 | chr1-60161091-INS-204 |
| chr1-60163245-INS-220 | chr1-60163245-INS-220 |
| chr1-60178054-INS-113 | chr1-60178054-INS-87 |
| chr1-60188898-INS-96 | chr1-60188898-INS-96 |
| chr1-60200940-INS-189 | chr1-60200940-INS-193 |
| chr1-60211814-INS-88 | chr1-60211810-INS-53 |
| chr1-60217919-INS-62 | chr1-60217919-INS-52 |
| chr1-60236255-INS-363 | chr1-60236255-INS-363 |
| chr1-60260706-INS-1122 | chr1-60260707-INS-1123 |
| chr1-60333392-INS-238 | chr1-60333392-INS-239 |
| chr1-60339823-INS-208 | chr1-60339823-INS-208 |
| chr1-60342862-INS-403 | chr1-60342862-INS-403 |
| chr1-60348894-INS-59 | chr1-60348894-INS-59 |
| chr1-60353267-INS-1290 | chr1-60353268-INS-1289 |
| chr1-60380475-INS-426 | chr1-60380475-INS-426 |
| chr1-60402973-INS-52 | chr1-60402973-INS-54 |
| chr1-60406815-INS-168 | chr1-60406815-INS-170 |
| chr1-60413849-INS-779 | chr1-60413849-INS-768 |
| chr1-60423007-INS-381 | chr1-60423007-INS-381 |
| chr1-60425956-INS-634 | chr1-60425956-INS-634 |
| chr1-60507727-INS-136 | chr1-60507727-INS-146 |
| chr1-60583705-INS-6664 | chr1-60583706-INS-6665 |
| chr1-60586307-INS-1559 | chr1-60586308-INS-1549 |
| chr1-60613032-INS-84 | chr1-60613032-INS-114 |
| chr1-60614524-INS-216 | chr1-60614524-INS-188 |
| chr1-60640456-INS-98 | chr1-60640456-INS-86 |
| chr1-60652836-INS-248 | chr1-60652836-INS-248 |
| chr1-60658295-INS-180 | chr1-60658295-INS-180 |
| chr1-60664305-INS-57 | chr1-60664305-INS-61 |
| chr1-60675831-INS-184 | chr1-60675831-INS-184 |
| chr1-60692893-INS-210 | chr1-60692893-INS-210 |
| chr1-60707960-INS-226 | chr1-60707981-INS-233 |
| chr1-60727239-INS-201 | chr1-60727212-INS-201 |
| chr1-60742802-INS-73 | chr1-60742802-INS-97 |
| chr1-60747444-INS-92 | chr1-60747444-INS-96 |
| chr1-60748287-INS-2283 | chr1-60748288-INS-2283 |
| chr1-60782123-INS-81 | chr1-60782123-INS-102 |
| chr1-60850049-INS-1449 | chr1-60850050-INS-1453 |
| chr1-60854751-INS-7093 | chr1-60854752-INS-7094 |
| chr1-60871193-INS-6467 | chr1-60871194-INS-6467 |
| chr1-60873883-INS-121 | chr1-60873883-INS-120 |
| chr1-60884423-INS-149 | chr1-60884423-INS-149 |
| chr1-60934153-INS-50 | chr1-60934153-INS-50 |
| chr1-60942755-INS-507 | chr1-60942755-INS-507 |
| chr1-60958807-INS-7025 | chr1-60958808-INS-7025 |
| chr1-60968044-INS-222 | chr1-60968044-INS-220 |
| chr1-60989272-INS-115 | chr1-60989265-INS-118 |
| chr1-60991638-INS-422 | chr1-60991638-INS-422 |
| chr1-60992713-INS-83 | chr1-60992735-INS-57 |
| chr1-61032238-INS-70 | chr1-61032238-INS-70 |
| chr1-61040990-INS-134 | chr1-61040990-INS-130 |
| chr1-61043463-INS-401 | chr1-61043463-INS-400 |
| chr1-61057064-INS-66 | chr1-61057064-INS-66 |
| chr1-61085644-INS-396 | chr1-61085644-INS-396 |
| chr1-61100511-INS-92 | chr1-61100490-INS-93 |
| chr1-61118332-INS-2830 | chr1-61118333-INS-2803 |
| chr1-61122735-INS-565 | chr1-61122735-INS-566 |
| chr1-61132555-INS-60 | chr1-61132555-INS-60 |

| | |
|---|---|
| chr1-61133476-INS-1022 | chr1-61133477-INS-1009 |
| chr1-61156871-INS-104 | chr1-61156871-INS-106 |
| chr1-61161688-INS-1130 | chr1-61161689-INS-1130 |
| chr1-61197407-INS-104 | chr1-61197406-INS-60 |
| chr1-61206367-INS-576 | chr1-61206367-INS-576 |
| chr1-61219488-INS-56 | chr1-61219486-INS-58 |
| chr1-61224571-INS-189 | chr1-61224571-INS-189 |
| chr1-61224775-INS-553 | chr1-61224770-INS-568 |
| chr1-61258544-INS-401 | chr1-61258544-INS-400 |
| chr1-61293232-INS-6808 | chr1-61293233-INS-6808 |
| chr1-61294389-INS-601 | chr1-61294390-INS-603 |
| chr1-61298719-INS-68 | chr1-61298715-INS-50 |
| chr1-61327577-INS-1295 | chr1-61327578-INS-1298 |
| chr1-61334345-INS-62 | chr1-61334345-INS-92 |
| chr1-61354042-INS-421 | chr1-61354042-INS-420 |
| chr1-61388697-INS-54 | chr1-61388697-INS-56 |
| chr1-61409189-INS-62 | chr1-61409189-INS-62 |
| chr1-61432232-INS-145 | chr1-61432231-INS-158 |
| chr1-61438051-INS-149 | chr1-61438051-INS-146 |
| chr1-61449043-INS-1890 | chr1-61449044-INS-1890 |
| chr1-61450766-INS-188 | chr1-61450766-INS-188 |
| chr1-61458587-INS-715 | chr1-61458587-INS-716 |
| chr1-61538125-INS-212 | chr1-61538125-INS-212 |
| chr1-61588163-INS-344 | chr1-61588163-INS-344 |
| chr1-61638170-INS-6928 | chr1-61638171-INS-6928 |
| chr1-61640616-INS-1998 | chr1-61640617-INS-1999 |
| chr1-61652930-INS-90 | chr1-61652930-INS-90 |
| chr1-61657632-INS-62 | chr1-61657632-INS-62 |
| chr1-61701389-INS-84 | chr1-61701400-INS-61 |
| chr1-61702070-INS-334 | chr1-61702070-INS-334 |
| chr1-61737063-INS-99 | chr1-61737063-INS-99 |
| chr1-61747741-INS-60 | chr1-61747741-INS-72 |
| chr1-61772599-INS-6349 | chr1-61772600-INS-6349 |
| chr1-61791129-INS-56 | chr1-61791129-INS-58 |
| chr1-61817762-INS-95 | chr1-61817762-INS-95 |
| chr1-61820497-INS-6113 | chr1-61820498-INS-6114 |
| chr1-61822278-INS-78 | chr1-61822278-INS-78 |
| chr1-61830729-INS-116 | chr1-61830729-INS-116 |
| chr1-61884876-INS-198 | chr1-61884876-INS-218 |
| chr1-61909820-INS-4778 | chr1-61909821-INS-4778 |
| chr1-61949408-INS-104 | chr1-61949408-INS-104 |
| chr1-61971177-INS-80 | chr1-61971177-INS-90 |
| chr1-62010177-INS-608 | chr1-62010177-INS-610 |
| chr1-62012141-INS-60 | chr1-62012141-INS-60 |
| chr1-62052330-INS-2707 | chr1-62052331-INS-2707 |
| chr1-62131850-INS-56 | chr1-62131841-INS-52 |
| chr1-62155948-INS-53 | chr1-62155948-INS-53 |
| chr1-62164655-INS-108 | chr1-62164655-INS-108 |
| chr1-62178162-INS-200 | chr1-62178132-INS-206 |
| chr1-62204018-INS-217 | chr1-62204018-INS-218 |
| chr1-62207234-INS-104 | chr1-62207234-INS-104 |
| chr1-62234007-INS-209 | chr1-62234007-INS-201 |
| chr1-62241569-INS-56 | chr1-62241569-INS-58 |
| chr1-62280198-INS-62 | chr1-62280199-INS-67 |
| chr1-62283147-INS-53 | chr1-62283147-INS-53 |
| chr1-62284447-INS-100 | chr1-62284447-INS-100 |
| chr1-62291558-INS-83 | chr1-62291558-INS-83 |
| chr1-62329783-INS-151 | chr1-62329783-INS-155 |
| chr1-62359094-INS-124 | chr1-62359094-INS-128 |
| chr1-62408433-INS-542 | chr1-62408433-INS-542 |
| chr1-62414205-INS-52 | chr1-62414205-INS-54 |
| chr1-62450461-INS-174 | chr1-62450462-INS-183 |
| chr1-62470499-INS-72 | chr1-62470499-INS-72 |
| chr1-62529253-INS-60 | chr1-62529269-INS-50 |
| chr1-62530137-INS-5838 | chr1-62530138-INS-5844 |
| chr1-62537178-INS-164 | chr1-62537178-INS-124 |
| chr1-62591969-INS-505 | chr1-62591969-INS-505 |
| chr1-62622784-INS-218 | chr1-62622784-INS-215 |
| chr1-62628094-INS-59 | chr1-62628094-INS-59 |
| chr1-62640696-INS-76 | chr1-62640696-INS-76 |
| chr1-62641974-INS-64 | chr1-62641974-INS-64 |
| chr1-62673737-INS-412 | chr1-62673737-INS-412 |
| chr1-62700933-INS-236 | chr1-62700933-INS-240 |
| chr1-62706682-INS-56 | chr1-62706690-INS-52 |
| chr1-62728651-INS-183 | chr1-62728651-INS-183 |
| chr1-62729719-INS-215 | chr1-62729719-INS-215 |
| chr1-62750464-INS-82 | chr1-62750464-INS-77 |

| | |
|---|---|
| chr1-62766194-INS-216 | chr1-62766194-INS-216 |
| chr1-62772693-INS-73 | chr1-62772693-INS-70 |
| chr1-62788940-INS-7138 | chr1-62788941-INS-7138 |
| chr1-62873312-INS-6017 | chr1-62873313-INS-6022 |
| chr1-62878618-INS-62 | chr1-62878618-INS-66 |
| chr1-62881767-INS-181 | chr1-62881767-INS-182 |
| chr1-62894165-INS-116 | chr1-62894122-INS-110 |
| chr1-62899442-INS-90 | chr1-62899442-INS-92 |
| chr1-62901900-INS-1886 | chr1-62901901-INS-1886 |
| chr1-62924957-INS-80 | chr1-62924929-INS-80 |
| chr1-62928582-INS-6473 | chr1-62928599-INS-6466 |
| chr1-62967963-INS-72 | chr1-62967961-INS-94 |
| chr1-62991459-INS-72 | chr1-62991459-INS-86 |
| chr1-63013946-INS-380 | chr1-63013941-INS-395 |
| chr1-63015188-INS-84 | chr1-63015188-INS-84 |
| chr1-63025474-INS-118 | chr1-63025474-INS-116 |
| chr1-63028102-INS-205 | chr1-63028102-INS-210 |
| chr1-63030232-INS-823 | chr1-63030232-INS-824 |
| chr1-63032853-INS-52 | chr1-63032853-INS-101 |
| chr1-63036556-INS-359 | chr1-63036556-INS-359 |
| chr1-63052956-INS-5219 | chr1-63052957-INS-5219 |
| chr1-63076394-INS-6413 | chr1-63076395-INS-6413 |
| chr1-63083419-INS-52 | chr1-63083419-INS-60 |
| chr1-63085672-INS-90 | chr1-63085672-INS-90 |
| chr1-63100793-INS-218 | chr1-63100788-INS-220 |
| chr1-63104508-INS-213 | chr1-63104508-INS-215 |
| chr1-63129185-INS-124 | chr1-63129185-INS-127 |
| chr1-63136254-INS-143 | chr1-63136254-INS-151 |
| chr1-63151536-INS-64 | chr1-63151536-INS-66 |
| chr1-63164088-INS-749 | chr1-63164088-INS-749 |
| chr1-63165082-INS-116 | chr1-63165082-INS-88 |
| chr1-63187125-INS-262 | chr1-63187126-INS-247 |
| chr1-63188275-INS-173 | chr1-63188275-INS-176 |
| chr1-63213574-INS-175 | chr1-63213574-INS-175 |
| chr1-63218917-INS-203 | chr1-63218917-INS-203 |
| chr1-63227408-INS-198 | chr1-63227408-INS-198 |
| chr1-63236827-INS-54 | chr1-63236841-INS-90 |
| chr1-63251544-INS-76 | chr1-63251543-INS-74 |
| chr1-63263714-INS-324 | chr1-63263687-INS-343 |
| chr1-63322047-INS-62 | chr1-63322047-INS-62 |
| chr1-63328316-INS-255 | chr1-63328316-INS-255 |
| chr1-63366729-INS-972 | chr1-63366729-INS-971 |
| chr1-63417571-INS-178 | chr1-63417570-INS-171 |
| chr1-63432613-INS-5420 | chr1-63432614-INS-5420 |
| chr1-63444267-INS-208 | chr1-63444267-INS-212 |
| chr1-63445332-INS-106 | chr1-63445335-INS-107 |
| chr1-63451803-INS-56 | chr1-63451802-INS-91 |
| chr1-63463154-INS-907 | chr1-63463154-INS-907 |
| chr1-63472874-INS-112 | chr1-63472874-INS-112 |
| chr1-63476956-INS-79 | chr1-63476956-INS-79 |
| chr1-63480930-INS-434 | chr1-63480930-INS-434 |
| chr1-63518431-INS-4404 | chr1-63518432-INS-4404 |
| chr1-63534949-INS-70 | chr1-63534971-INS-55 |
| chr1-63556369-INS-74 | chr1-63556370-INS-74 |
| chr1-63566280-INS-132 | chr1-63566280-INS-176 |
| chr1-63585330-INS-113 | chr1-63585330-INS-113 |
| chr1-63610652-INS-56 | chr1-63610654-INS-70 |
| chr1-63616620-INS-216 | chr1-63616620-INS-217 |
| chr1-63668958-INS-3305 | chr1-63668959-INS-3305 |
| chr1-63679542-INS-74 | chr1-63679520-INS-70 |
| chr1-63684302-INS-144 | chr1-63684285-INS-126 |
| chr1-63708291-INS-58 | chr1-63708283-INS-68 |
| chr1-63726796-INS-116 | chr1-63726797-INS-102 |
| chr1-63730785-INS-580 | chr1-63730785-INS-580 |
| chr1-63746593-INS-6944 | chr1-63746594-INS-6944 |
| chr1-63759245-INS-56 | chr1-63759245-INS-56 |
| chr1-63772586-INS-77 | chr1-63772586-INS-77 |
| chr1-63827263-INS-76 | chr1-63827262-INS-54 |
| chr1-63855696-INS-83 | chr1-63855677-INS-78 |
| chr1-63860338-INS-53 | chr1-63860351-INS-50 |
| chr1-63862272-INS-64 | chr1-63862246-INS-108 |
| chr1-63911673-INS-410 | chr1-63911673-INS-411 |
| chr1-63963300-INS-161 | chr1-63963300-INS-161 |
| chr1-63979880-INS-52 | chr1-63979887-INS-53 |
| chr1-63984712-INS-70 | chr1-63984719-INS-98 |
| chr1-63999712-INS-56 | chr1-63999712-INS-51 |
| chr1-64005959-INS-941 | chr1-64005959-INS-941 |

| | |
|---|---|
| chr1-64010050-INS-80 | chr1-64010050-INS-79 |
| chr1-64011580-INS-2583 | chr1-64011581-INS-2583 |
| chr1-64023037-INS-76 | chr1-64023037-INS-76 |
| chr1-64049058-INS-325 | chr1-64049059-INS-325 |
| chr1-64049156-INS-6380 | chr1-64049157-INS-6381 |
| chr1-64052901-INS-123 | chr1-64052900-INS-123 |
| chr1-64122388-INS-78 | chr1-64122388-INS-78 |
| chr1-64132083-INS-69 | chr1-64132083-INS-71 |
| chr1-64141617-INS-56 | chr1-64141617-INS-56 |
| chr1-64145888-INS-1122 | chr1-64145889-INS-1122 |
| chr1-64173204-INS-60 | chr1-64173204-INS-60 |
| chr1-64198192-INS-296 | chr1-64198192-INS-297 |
| chr1-64213331-INS-499 | chr1-64213331-INS-498 |
| chr1-64237326-INS-5550 | chr1-64237327-INS-5550 |
| chr1-64246322-INS-52 | chr1-64246322-INS-54 |
| chr1-64253424-INS-54 | chr1-64253424-INS-54 |
| chr1-64257403-INS-129 | chr1-64257403-INS-129 |
| chr1-64284408-INS-384 | chr1-64284408-INS-384 |
| chr1-64359585-INS-74 | chr1-64359585-INS-74 |
| chr1-64380109-INS-106 | chr1-64380109-INS-107 |
| chr1-64387357-INS-206 | chr1-64387357-INS-208 |
| chr1-64392143-INS-66 | chr1-64392143-INS-62 |
| chr1-64402464-INS-118 | chr1-64402464-INS-126 |
| chr1-64447053-INS-88 | chr1-64447053-INS-100 |
| chr1-64460745-INS-59 | chr1-64460745-INS-63 |
| chr1-64480579-INS-128 | chr1-64480579-INS-79 |
| chr1-64507361-INS-103 | chr1-64507361-INS-112 |
| chr1-64513991-INS-588 | chr1-64513991-INS-588 |
| chr1-64533894-INS-55 | chr1-64533894-INS-55 |
| chr1-64563527-INS-73 | chr1-64563527-INS-71 |
| chr1-64585287-INS-120 | chr1-64585285-INS-124 |
| chr1-64618907-INS-438 | chr1-64618907-INS-437 |
| chr1-64626689-INS-183 | chr1-64626689-INS-183 |
| chr1-64667346-INS-58 | chr1-64667302-INS-65 |
| chr1-64673799-INS-85 | chr1-64673799-INS-85 |
| chr1-64679339-INS-142 | chr1-64679339-INS-142 |
| chr1-64694879-INS-194 | chr1-64694879-INS-196 |
| chr1-64710558-INS-72 | chr1-64710558-INS-72 |
| chr1-64803691-INS-58 | chr1-64803691-INS-60 |
| chr1-64805264-INS-326 | chr1-64805264-INS-326 |
| chr1-64819815-INS-59 | chr1-64819815-INS-59 |
| chr1-64845283-INS-177 | chr1-64845283-INS-152 |
| chr1-64861238-INS-167 | chr1-64861238-INS-167 |
| chr1-64902524-INS-401 | chr1-64902524-INS-401 |
| chr1-64994355-INS-194 | chr1-64994355-INS-194 |
| chr1-65010563-INS-92 | chr1-65010563-INS-92 |
| chr1-65053060-INS-54 | chr1-65053060-INS-56 |
| chr1-65060460-INS-61 | chr1-65060454-INS-88 |
| chr1-65075164-INS-213 | chr1-65075164-INS-213 |
| chr1-65082221-INS-119 | chr1-65082221-INS-96 |
| chr1-65088091-INS-55 | chr1-65088091-INS-55 |
| chr1-65103357-INS-87 | chr1-65103357-INS-54 |
| chr1-65110053-INS-194 | chr1-65110053-INS-200 |
| chr1-65111594-INS-200 | chr1-65111594-INS-200 |
| chr1-65123796-INS-58 | chr1-65123796-INS-58 |
| chr1-65138865-INS-206 | chr1-65138865-INS-206 |
| chr1-65147851-INS-256 | chr1-65147852-INS-256 |
| chr1-65148440-INS-52 | chr1-65148440-INS-52 |
| chr1-65155651-INS-87 | chr1-65155652-INS-87 |
| chr1-65163565-INS-319 | chr1-65163565-INS-320 |
| chr1-65167813-INS-109 | chr1-65167813-INS-109 |
| chr1-65175410-INS-205 | chr1-65175410-INS-205 |
| chr1-65182526-INS-69 | chr1-65182526-INS-67 |
| chr1-65187182-INS-346 | chr1-65187185-INS-347 |
| chr1-65196515-INS-87 | chr1-65196515-INS-107 |
| chr1-65290876-INS-129 | chr1-65290861-INS-101 |
| chr1-65292285-INS-197 | chr1-65292285-INS-197 |
| chr1-65323015-INS-139 | chr1-65323015-INS-172 |
| chr1-65344644-INS-102 | chr1-65344634-INS-96 |
| chr1-65375255-INS-60 | chr1-65375255-INS-60 |
| chr1-65391298-INS-2881 | chr1-65391299-INS-2881 |
| chr1-65397669-INS-180 | chr1-65397669-INS-180 |
| chr1-65405246-INS-56 | chr1-65405198-INS-68 |
| chr1-65410906-INS-622 | chr1-65410906-INS-622 |
| chr1-65486574-INS-532 | chr1-65486574-INS-532 |
| chr1-65498024-INS-6199 | chr1-65498025-INS-6199 |
| chr1-65501125-INS-957 | chr1-65501125-INS-957 |

| | |
|---|---|
| chr1-65517853-INS-54 | chr1-65517853-INS-54 |
| chr1-65531855-INS-441 | chr1-65531856-INS-441 |
| chr1-65534780-INS-192 | chr1-65534791-INS-214 |
| chr1-65541455-INS-6273 | chr1-65541456-INS-6273 |
| chr1-65542329-INS-318 | chr1-65542330-INS-318 |
| chr1-65545770-INS-397 | chr1-65545770-INS-397 |
| chr1-65551625-INS-75 | chr1-65551625-INS-80 |
| chr1-65616074-INS-206 | chr1-65616074-INS-208 |
| chr1-65639633-INS-6572 | chr1-65639634-INS-6572 |
| chr1-65697308-INS-7293 | chr1-65697309-INS-7293 |
| chr1-65710341-INS-55 | chr1-65710341-INS-55 |
| chr1-65720528-INS-639 | chr1-65720528-INS-639 |
| chr1-65723954-INS-250 | chr1-65723954-INS-250 |
| chr1-65737127-INS-52 | chr1-65737127-INS-52 |
| chr1-65769974-INS-100 | chr1-65769974-INS-102 |
| chr1-65777435-INS-384 | chr1-65777435-INS-384 |
| chr1-65781570-INS-905 | chr1-65781570-INS-905 |
| chr1-65795889-INS-4664 | chr1-65795890-INS-4664 |
| chr1-65804512-INS-68 | chr1-65804512-INS-68 |
| chr1-65807778-INS-8172 | chr1-65807778-INS-8175 |
| chr1-65819874-INS-67 | chr1-65819874-INS-67 |
| chr1-65828002-INS-206 | chr1-65828002-INS-206 |
| chr1-65828933-INS-2899 | chr1-65828934-INS-2895 |
| chr1-65832069-INS-3328 | chr1-65832070-INS-3328 |
| chr1-65861398-INS-237 | chr1-65861398-INS-237 |
| chr1-65878898-INS-565 | chr1-65878898-INS-565 |
| chr1-65893199-INS-59 | chr1-65893199-INS-59 |
| chr1-65894871-INS-81 | chr1-65894887-INS-51 |
| chr1-65902912-INS-60 | chr1-65902911-INS-56 |
| chr1-65923435-INS-82 | chr1-65923435-INS-90 |
| chr1-65947761-INS-122 | chr1-65947761-INS-124 |
| chr1-65968537-INS-57 | chr1-65968537-INS-53 |
| chr1-65992375-INS-60 | chr1-65992371-INS-62 |
| chr1-66103991-INS-351 | chr1-66103991-INS-351 |
| chr1-66105607-INS-901 | chr1-66105607-INS-901 |
| chr1-66138294-INS-81 | chr1-66138294-INS-81 |
| chr1-66139701-INS-1655 | chr1-66139702-INS-1655 |
| chr1-66148600-INS-159 | chr1-66148600-INS-152 |
| chr1-66175313-INS-401 | chr1-66175310-INS-400 |
| chr1-66184677-INS-61 | chr1-66184677-INS-63 |
| chr1-66189384-INS-58 | chr1-66189384-INS-58 |
| chr1-66190530-INS-1240 | chr1-66190531-INS-1240 |
| chr1-66196213-INS-53 | chr1-66196216-INS-57 |
| chr1-66207666-INS-131 | chr1-66207666-INS-131 |
| chr1-66230369-INS-142 | chr1-66230377-INS-173 |
| chr1-66256017-INS-292 | chr1-66256017-INS-292 |
| chr1-66282324-INS-117 | chr1-66282324-INS-117 |
| chr1-66291824-INS-70 | chr1-66291824-INS-70 |
| chr1-66355969-INS-60 | chr1-66355962-INS-104 |
| chr1-66359209-INS-54 | chr1-66359209-INS-54 |
| chr1-66379797-INS-51 | chr1-66379752-INS-51 |
| chr1-66446512-INS-130 | chr1-66446512-INS-130 |
| chr1-66474753-INS-70 | chr1-66474749-INS-72 |
| chr1-66497601-INS-6489 | chr1-66497602-INS-6489 |
| chr1-66511883-INS-58 | chr1-66511883-INS-62 |
| chr1-66532811-INS-305 | chr1-66532811-INS-306 |
| chr1-66592940-INS-144 | chr1-66592940-INS-144 |
| chr1-66623621-INS-3715 | chr1-66623622-INS-3715 |
| chr1-66649624-INS-173 | chr1-66649624-INS-173 |
| chr1-66658297-INS-6407 | chr1-66658298-INS-6407 |
| chr1-66673746-INS-235 | chr1-66673746-INS-235 |
| chr1-66694659-INS-54 | chr1-66694659-INS-54 |
| chr1-66755378-INS-76 | chr1-66755378-INS-76 |
| chr1-66791799-INS-76 | chr1-66791775-INS-63 |
| chr1-66795807-INS-104 | chr1-66795808-INS-104 |
| chr1-66804245-INS-54 | chr1-66804245-INS-50 |
| chr1-66807771-INS-3885 | chr1-66807772-INS-3885 |
| chr1-66832368-INS-455 | chr1-66832368-INS-493 |
| chr1-66851522-INS-214 | chr1-66851522-INS-215 |
| chr1-66867612-INS-60 | chr1-66867612-INS-64 |
| chr1-66876451-INS-439 | chr1-66876451-INS-439 |
| chr1-66878388-INS-52 | chr1-66878382-INS-53 |
| chr1-66881342-INS-69 | chr1-66881342-INS-69 |
| chr1-66891697-INS-401 | chr1-66891697-INS-401 |
| chr1-66894072-INS-186 | chr1-66894072-INS-186 |
| chr1-66896150-INS-276 | chr1-66896150-INS-276 |
| chr1-66914198-INS-194 | chr1-66914198-INS-197 |

| | |
|---|---|
| chr1-66920858-INS-308 | chr1-66920856-INS-310 |
| chr1-66924466-INS-493 | chr1-66924467-INS-485 |
| chr1-66938281-INS-251 | chr1-66938281-INS-253 |
| chr1-66942063-INS-166 | chr1-66942063-INS-166 |
| chr1-66944145-INS-57 | chr1-66944146-INS-55 |
| chr1-66961757-INS-64 | chr1-66961757-INS-66 |
| chr1-66982262-INS-172 | chr1-66982262-INS-144 |
| chr1-66984909-INS-224 | chr1-66984909-INS-224 |
| chr1-66999457-INS-481 | chr1-66999457-INS-481 |
| chr1-67030404-INS-209 | chr1-67030404-INS-209 |
| chr1-67041321-INS-60 | chr1-67041321-INS-64 |
| chr1-67047900-INS-1863 | chr1-67047901-INS-1863 |
| chr1-67086885-INS-688 | chr1-67086885-INS-690 |
| chr1-67112220-INS-216 | chr1-67112221-INS-216 |
| chr1-67129907-INS-189 | chr1-67129907-INS-215 |
| chr1-67137634-INS-87 | chr1-67137634-INS-87 |
| chr1-67156605-INS-206 | chr1-67156605-INS-207 |
| chr1-67174059-INS-51 | chr1-67174059-INS-51 |
| chr1-67193598-INS-51 | chr1-67193598-INS-63 |
| chr1-67213877-INS-489 | chr1-67213877-INS-489 |
| chr1-67241029-INS-7075 | chr1-67241030-INS-7075 |
| chr1-67253947-INS-84 | chr1-67253947-INS-84 |
| chr1-67302936-INS-64 | chr1-67302936-INS-61 |
| chr1-67334952-INS-354 | chr1-67334952-INS-354 |
| chr1-67343929-INS-80 | chr1-67343929-INS-80 |
| chr1-67361937-INS-6492 | chr1-67361938-INS-6492 |
| chr1-67384078-INS-60 | chr1-67384078-INS-57 |
| chr1-67427181-INS-6715 | chr1-67427182-INS-6718 |
| chr1-67432620-INS-67 | chr1-67432620-INS-67 |
| chr1-67459621-INS-163 | chr1-67459605-INS-170 |
| chr1-67460712-INS-101 | chr1-67460712-INS-101 |
| chr1-67464232-INS-394 | chr1-67464232-INS-394 |
| chr1-67465072-INS-127 | chr1-67465068-INS-133 |
| chr1-67470480-INS-60 | chr1-67470480-INS-60 |
| chr1-67478042-INS-57 | chr1-67478042-INS-69 |
| chr1-67499596-INS-352 | chr1-67499596-INS-352 |
| chr1-67506764-INS-66 | chr1-67506764-INS-66 |
| chr1-67520289-INS-3786 | chr1-67520290-INS-3786 |
| chr1-67528407-INS-1157 | chr1-67528416-INS-1147 |
| chr1-67531341-INS-70 | chr1-67531359-INS-68 |
| chr1-67535558-INS-211 | chr1-67535558-INS-211 |
| chr1-67551598-INS-138 | chr1-67551598-INS-138 |
| chr1-67567242-INS-462 | chr1-67567242-INS-462 |
| chr1-67599416-INS-104 | chr1-67599416-INS-104 |
| chr1-67636111-INS-54 | chr1-67636134-INS-65 |
| chr1-67645830-INS-458 | chr1-67645830-INS-458 |
| chr1-67647392-INS-109 | chr1-67647392-INS-125 |
| chr1-67656340-INS-60 | chr1-67656340-INS-60 |
| chr1-67664913-INS-198 | chr1-67664913-INS-198 |
| chr1-67668735-INS-323 | chr1-67668735-INS-323 |
| chr1-67702511-INS-427 | chr1-67702511-INS-427 |
| chr1-67708851-INS-188 | chr1-67708851-INS-188 |
| chr1-67743018-INS-6676 | chr1-67743019-INS-6678 |
| chr1-67751500-INS-1529 | chr1-67751501-INS-1531 |
| chr1-67804347-INS-1789 | chr1-67804348-INS-1789 |
| chr1-67836650-INS-1462 | chr1-67836651-INS-1463 |
| chr1-67839011-INS-70 | chr1-67839011-INS-72 |
| chr1-67850074-INS-58 | chr1-67850074-INS-56 |
| chr1-67859821-INS-662 | chr1-67859821-INS-662 |
| chr1-67867610-INS-333 | chr1-67867610-INS-334 |
| chr1-67872787-INS-81 | chr1-67872783-INS-79 |
| chr1-67882288-INS-63 | chr1-67882289-INS-69 |
| chr1-67900127-INS-3309 | chr1-67900128-INS-3309 |
| chr1-67923056-INS-488 | chr1-67923056-INS-488 |
| chr1-67967328-INS-171 | chr1-67967328-INS-171 |
| chr1-68011816-INS-503 | chr1-68011816-INS-503 |
| chr1-68025944-INS-7166 | chr1-68025945-INS-7166 |
| chr1-68053901-INS-4657 | chr1-68053902-INS-4658 |
| chr1-68069485-INS-246 | chr1-68069485-INS-246 |
| chr1-68091252-INS-52 | chr1-68091253-INS-72 |
| chr1-68097135-INS-849 | chr1-68097135-INS-849 |
| chr1-68104362-INS-450 | chr1-68104372-INS-455 |
| chr1-68109467-INS-82 | chr1-68109467-INS-82 |
| chr1-68134053-INS-553 | chr1-68134053-INS-554 |
| chr1-68155584-INS-220 | chr1-68155584-INS-210 |
| chr1-68177016-INS-94 | chr1-68177016-INS-78 |
| chr1-68200876-INS-6803 | chr1-68200877-INS-6803 |

| | |
|---|---|
| chr1-68211583-INS-107 | chr1-68211583-INS-107 |
| chr1-68234146-INS-55 | chr1-68234146-INS-70 |
| chr1-68242011-INS-56 | chr1-68242003-INS-53 |
| chr1-68248660-INS-893 | chr1-68248660-INS-893 |
| chr1-68314831-INS-94 | chr1-68314831-INS-94 |
| chr1-68319754-INS-583 | chr1-68319754-INS-583 |
| chr1-68326347-INS-400 | chr1-68326347-INS-400 |
| chr1-68331918-INS-52 | chr1-68331918-INS-50 |
| chr1-68364962-INS-65 | chr1-68364962-INS-63 |
| chr1-68371708-INS-2023 | chr1-68371709-INS-2023 |
| chr1-68410140-INS-407 | chr1-68410136-INS-410 |
| chr1-68415418-INS-75 | chr1-68415418-INS-75 |
| chr1-68431393-INS-578 | chr1-68431393-INS-578 |
| chr1-68455203-INS-197 | chr1-68455203-INS-197 |
| chr1-68458213-INS-336 | chr1-68458215-INS-334 |
| chr1-68517667-INS-115 | chr1-68517667-INS-115 |
| chr1-68678541-INS-76 | chr1-68678541-INS-76 |
| chr1-68693568-INS-688 | chr1-68693568-INS-688 |
| chr1-68696740-INS-937 | chr1-68696740-INS-937 |
| chr1-68708194-INS-182 | chr1-68708194-INS-183 |
| chr1-68711732-INS-898 | chr1-68711732-INS-896 |
| chr1-68714958-INS-402 | chr1-68714958-INS-402 |
| chr1-68721167-INS-72 | chr1-68721197-INS-80 |
| chr1-68754658-INS-326 | chr1-68754658-INS-326 |
| chr1-68767719-INS-217 | chr1-68767719-INS-217 |
| chr1-68780045-INS-142 | chr1-68780045-INS-142 |
| chr1-68787355-INS-59 | chr1-68787368-INS-55 |
| chr1-68797257-INS-1061 | chr1-68797258-INS-1061 |
| chr1-68813893-INS-60 | chr1-68813859-INS-64 |
| chr1-68832839-INS-98 | chr1-68832839-INS-98 |
| chr1-68848040-INS-464 | chr1-68848040-INS-464 |
| chr1-68849621-INS-70 | chr1-68849618-INS-69 |
| chr1-68855522-INS-315 | chr1-68855522-INS-315 |
| chr1-68933076-INS-544 | chr1-68933076-INS-544 |
| chr1-68961672-INS-77 | chr1-68961658-INS-74 |
| chr1-68963437-INS-1496 | chr1-68963438-INS-1534 |
| chr1-68972757-INS-96 | chr1-68972755-INS-98 |
| chr1-68990625-INS-55 | chr1-68990625-INS-57 |
| chr1-68992127-INS-86 | chr1-68992127-INS-52 |
| chr1-68993390-INS-87 | chr1-68993390-INS-87 |
| chr1-69005593-INS-93 | chr1-69005574-INS-119 |
| chr1-69028217-INS-206 | chr1-69028217-INS-208 |
| chr1-69042288-INS-74 | chr1-69042288-INS-68 |
| chr1-69065439-INS-54 | chr1-69065439-INS-62 |
| chr1-69131168-INS-534 | chr1-69131168-INS-534 |
| chr1-69152667-INS-53 | chr1-69152656-INS-57 |
| chr1-69197576-INS-402 | chr1-69197576-INS-402 |
| chr1-69217480-INS-353 | chr1-69217480-INS-353 |
| chr1-69241542-INS-85 | chr1-69241542-INS-85 |
| chr1-69247206-INS-546 | chr1-69247206-INS-546 |
| chr1-69280997-INS-874 | chr1-69280997-INS-873 |
| chr1-69332769-INS-1254 | chr1-69332770-INS-1254 |
| chr1-69372936-INS-1647 | chr1-69372937-INS-1649 |
| chr1-69377239-INS-178 | chr1-69377239-INS-178 |
| chr1-69393142-INS-64 | chr1-69393142-INS-62 |
| chr1-69414977-INS-94 | chr1-69414987-INS-75 |
| chr1-69419706-INS-61 | chr1-69419706-INS-56 |
| chr1-69424187-INS-69 | chr1-69424195-INS-65 |
| chr1-69426491-INS-1890 | chr1-69426492-INS-1890 |
| chr1-69450365-INS-112 | chr1-69450365-INS-116 |
| chr1-69470514-INS-386 | chr1-69470514-INS-386 |
| chr1-69484027-INS-88 | chr1-69484027-INS-92 |
| chr1-69495901-INS-54 | chr1-69495901-INS-54 |
| chr1-69529971-INS-70 | chr1-69529971-INS-86 |
| chr1-69594856-INS-205 | chr1-69594856-INS-197 |
| chr1-69606203-INS-145 | chr1-69606199-INS-113 |
| chr1-69657670-INS-1021 | chr1-69657671-INS-1022 |
| chr1-69680738-INS-68 | chr1-69680738-INS-66 |
| chr1-69746339-INS-124 | chr1-69746339-INS-87 |
| chr1-69747071-INS-3129 | chr1-69747072-INS-3128 |
| chr1-69783992-INS-1197 | chr1-69783993-INS-1195 |
| chr1-69806637-INS-52 | chr1-69806637-INS-52 |
| chr1-69815424-INS-6611 | chr1-69815425-INS-6635 |
| chr1-69854847-INS-53 | chr1-69854847-INS-52 |
| chr1-69863752-INS-68 | chr1-69863752-INS-72 |
| chr1-69870897-INS-78 | chr1-69870897-INS-78 |
| chr1-69911152-INS-183 | chr1-69911152-INS-192 |

| | |
|---|---|
| chr1-69916489-INS-6670 | chr1-69916490-INS-6670 |
| chr1-69933804-INS-54 | chr1-69933804-INS-56 |
| chr1-69956444-INS-64 | chr1-69956444-INS-59 |
| chr1-69990454-INS-75 | chr1-69990461-INS-66 |
| chr1-70024266-INS-1020 | chr1-70024267-INS-998 |
| chr1-70040741-INS-552 | chr1-70040741-INS-550 |
| chr1-70104522-INS-2369 | chr1-70104523-INS-2369 |
| chr1-70108968-INS-79 | chr1-70108968-INS-61 |
| chr1-70114271-INS-211 | chr1-70114271-INS-233 |
| chr1-70128477-INS-56 | chr1-70128475-INS-64 |
| chr1-70133735-INS-98 | chr1-70133735-INS-98 |
| chr1-70144342-INS-500 | chr1-70144346-INS-500 |
| chr1-70179352-INS-1295 | chr1-70179353-INS-1295 |
| chr1-70199753-INS-69 | chr1-70199753-INS-69 |
| chr1-70203155-INS-75 | chr1-70203168-INS-82 |
| chr1-70209923-INS-6183 | chr1-70209924-INS-6183 |
| chr1-70218027-INS-58 | chr1-70218027-INS-58 |
| chr1-70238847-INS-52 | chr1-70238847-INS-54 |
| chr1-70244811-INS-288 | chr1-70244811-INS-288 |
| chr1-70285468-INS-62 | chr1-70285468-INS-62 |
| chr1-70288401-INS-187 | chr1-70288401-INS-187 |
| chr1-70291878-INS-69 | chr1-70291875-INS-68 |
| chr1-70338481-INS-1526 | chr1-70338482-INS-1523 |
| chr1-70341984-INS-51 | chr1-70341984-INS-51 |
| chr1-70382199-INS-4510 | chr1-70382200-INS-4510 |
| chr1-70393479-INS-8856 | chr1-70393480-INS-8856 |
| chr1-70393599-INS-2492 | chr1-70393600-INS-2492 |
| chr1-70407404-INS-1025 | chr1-70407405-INS-1025 |
| chr1-70414067-INS-597 | chr1-70414064-INS-594 |
| chr1-70455710-INS-65 | chr1-70455710-INS-65 |
| chr1-70464716-INS-1279 | chr1-70464717-INS-1283 |
| chr1-70475643-INS-66 | chr1-70475643-INS-56 |
| chr1-70508884-INS-307 | chr1-70508884-INS-307 |
| chr1-70617721-INS-55 | chr1-70617735-INS-90 |
| chr1-70624102-INS-697 | chr1-70624102-INS-698 |
| chr1-70636794-INS-87 | chr1-70636794-INS-88 |
| chr1-70646058-INS-793 | chr1-70646058-INS-793 |
| chr1-70661871-INS-97 | chr1-70661871-INS-97 |
| chr1-70679985-INS-208 | chr1-70679985-INS-208 |
| chr1-70747492-INS-1163 | chr1-70747493-INS-1157 |
| chr1-70781073-INS-141 | chr1-70781061-INS-116 |
| chr1-70800536-INS-56 | chr1-70800549-INS-68 |
| chr1-70801856-INS-103 | chr1-70801856-INS-91 |
| chr1-70834590-INS-508 | chr1-70834590-INS-508 |
| chr1-70894563-INS-6442 | chr1-70894564-INS-6442 |
| chr1-70949901-INS-374 | chr1-70949901-INS-343 |
| chr1-70954883-INS-52 | chr1-70954883-INS-52 |
| chr1-70969811-INS-497 | chr1-70969811-INS-514 |
| chr1-70988514-INS-427 | chr1-70988514-INS-461 |
| chr1-71008764-INS-231 | chr1-71008764-INS-231 |
| chr1-71010897-INS-825 | chr1-71010896-INS-844 |
| chr1-71065955-INS-58 | chr1-71065955-INS-58 |
| chr1-71070107-INS-855 | chr1-71070107-INS-854 |
| chr1-71072087-INS-85 | chr1-71072065-INS-63 |
| chr1-71075856-INS-72 | chr1-71075856-INS-72 |
| chr1-71080351-INS-325 | chr1-71080351-INS-325 |
| chr1-71089809-INS-1891 | chr1-71089810-INS-1891 |
| chr1-71155185-INS-7165 | chr1-71155186-INS-7165 |
| chr1-71165034-INS-2575 | chr1-71165035-INS-2575 |
| chr1-71192653-INS-78 | chr1-71192653-INS-78 |
| chr1-71205146-INS-101 | chr1-71205118-INS-110 |
| chr1-71212503-INS-1073 | chr1-71212504-INS-1073 |
| chr1-71232691-INS-6571 | chr1-71232692-INS-6571 |
| chr1-71262938-INS-2734 | chr1-71262954-INS-2736 |
| chr1-71269195-INS-106 | chr1-71269197-INS-101 |
| chr1-71274989-INS-82 | chr1-71274989-INS-74 |
| chr1-71299842-INS-76 | chr1-71299837-INS-120 |
| chr1-71302445-INS-327 | chr1-71302445-INS-327 |
| chr1-71328544-INS-54 | chr1-71328544-INS-82 |
| chr1-71329895-INS-57 | chr1-71329897-INS-57 |
| chr1-71361323-INS-88 | chr1-71361323-INS-92 |
| chr1-71398222-INS-624 | chr1-71398222-INS-624 |
| chr1-71431328-INS-112 | chr1-71431328-INS-116 |
| chr1-71446651-INS-52 | chr1-71446651-INS-52 |
| chr1-71459611-INS-52 | chr1-71459611-INS-52 |
| chr1-71461411-INS-424 | chr1-71461411-INS-424 |
| chr1-71492141-INS-86 | chr1-71492141-INS-56 |

| | |
|---|---|
| chr1-71503243-INS-414 | chr1-71503243-INS-416 |
| chr1-71510369-INS-57 | chr1-71510369-INS-93 |
| chr1-71517128-INS-6605 | chr1-71517129-INS-6605 |
| chr1-71535720-INS-79 | chr1-71535720-INS-79 |
| chr1-71539059-INS-338 | chr1-71539059-INS-338 |
| chr1-71580384-INS-1144 | chr1-71580384-INS-1140 |
| chr1-71581497-INS-2447 | chr1-71581498-INS-2447 |
| chr1-71599272-INS-170 | chr1-71599272-INS-170 |
| chr1-71600639-INS-163 | chr1-71600639-INS-163 |
| chr1-71645717-INS-56 | chr1-71645717-INS-58 |
| chr1-71646914-INS-122 | chr1-71646928-INS-140 |
| chr1-71670360-INS-161 | chr1-71670360-INS-161 |
| chr1-71686414-INS-196 | chr1-71686414-INS-200 |
| chr1-71698113-INS-145 | chr1-71698113-INS-137 |
| chr1-71703836-INS-66 | chr1-71703836-INS-66 |
| chr1-71708608-INS-197 | chr1-71708608-INS-201 |
| chr1-71711293-INS-217 | chr1-71711293-INS-217 |
| chr1-71739425-INS-109 | chr1-71739425-INS-112 |
| chr1-71748831-INS-4117 | chr1-71748832-INS-4117 |
| chr1-71756844-INS-93 | chr1-71756844-INS-93 |
| chr1-71766048-INS-1227 | chr1-71766049-INS-1228 |
| chr1-71771369-INS-92 | chr1-71771369-INS-52 |
| chr1-71777715-INS-78 | chr1-71777715-INS-103 |
| chr1-71781456-INS-518 | chr1-71781456-INS-494 |
| chr1-71798278-INS-204 | chr1-71798278-INS-204 |
| chr1-71798833-INS-235 | chr1-71798833-INS-236 |
| chr1-71808794-INS-53 | chr1-71808794-INS-55 |
| chr1-71813278-INS-152 | chr1-71813278-INS-152 |
| chr1-71829649-INS-195 | chr1-71829649-INS-195 |
| chr1-71869818-INS-309 | chr1-71869818-INS-309 |
| chr1-71906292-INS-62 | chr1-71906292-INS-62 |
| chr1-71923857-INS-82 | chr1-71923839-INS-63 |
| chr1-71960528-INS-147 | chr1-71960528-INS-120 |
| chr1-71979191-INS-53 | chr1-71979191-INS-51 |
| chr1-71987386-INS-283 | chr1-71987386-INS-293 |
| chr1-71999505-INS-306 | chr1-71999505-INS-312 |
| chr1-72016563-INS-364 | chr1-72016563-INS-363 |
| chr1-72022376-INS-76 | chr1-72022376-INS-80 |
| chr1-72026071-INS-126 | chr1-72026051-INS-102 |
| chr1-72058676-INS-548 | chr1-72058676-INS-539 |
| chr1-72061070-INS-4447 | chr1-72061071-INS-4447 |
| chr1-72069380-INS-345 | chr1-72069380-INS-340 |
| chr1-72074623-INS-443 | chr1-72074623-INS-443 |
| chr1-72088393-INS-65 | chr1-72088370-INS-65 |
| chr1-72093660-INS-2573 | chr1-72093661-INS-2573 |
| chr1-72137890-INS-81 | chr1-72137890-INS-90 |
| chr1-72141783-INS-3941 | chr1-72141774-INS-3935 |
| chr1-72143577-INS-1892 | chr1-72143578-INS-1892 |
| chr1-72152407-INS-844 | chr1-72152407-INS-843 |
| chr1-72159196-INS-401 | chr1-72159196-INS-401 |
| chr1-72175274-INS-251 | chr1-72175274-INS-250 |
| chr1-72189465-INS-66 | chr1-72189465-INS-66 |
| chr1-72193253-INS-72 | chr1-72193253-INS-72 |
| chr1-72200478-INS-76 | chr1-72200478-INS-80 |
| chr1-72204058-INS-237 | chr1-72204059-INS-237 |
| chr1-72213964-INS-92 | chr1-72213964-INS-124 |
| chr1-72215983-INS-62 | chr1-72215983-INS-66 |
| chr1-72243671-INS-1060 | chr1-72243672-INS-1060 |
| chr1-72245404-INS-195 | chr1-72245404-INS-186 |
| chr1-72249089-INS-203 | chr1-72249089-INS-203 |
| chr1-72253031-INS-6442 | chr1-72253032-INS-6442 |
| chr1-72256971-INS-223 | chr1-72256972-INS-208 |
| chr1-72260225-INS-1888 | chr1-72260226-INS-1888 |
| chr1-72264374-INS-56 | chr1-72264374-INS-56 |
| chr1-72279268-INS-143 | chr1-72279317-INS-106 |
| chr1-72281332-INS-331 | chr1-72281333-INS-317 |
| chr1-72284496-INS-215 | chr1-72284496-INS-215 |
| chr1-72287270-INS-212 | chr1-72287270-INS-212 |
| chr1-72298292-INS-116 | chr1-72298292-INS-117 |
| chr1-72302908-INS-205 | chr1-72302908-INS-209 |
| chr1-72305164-INS-73 | chr1-72305164-INS-73 |
| chr1-72307768-INS-174 | chr1-72307768-INS-174 |
| chr1-72309149-INS-195 | chr1-72309149-INS-195 |
| chr1-72335315-INS-66 | chr1-72335315-INS-66 |
| chr1-72362221-INS-229 | chr1-72362221-INS-229 |
| chr1-72372314-INS-1894 | chr1-72372315-INS-1894 |
| chr1-72374595-INS-216 | chr1-72374595-INS-216 |

| | |
|---|---|
| chr1-72405206-INS-62 | chr1-72405206-INS-62 |
| chr1-72494927-INS-199 | chr1-72494927-INS-199 |
| chr1-72508479-INS-141 | chr1-72508479-INS-141 |
| chr1-72514055-INS-58 | chr1-72514055-INS-56 |
| chr1-72587510-INS-201 | chr1-72587510-INS-201 |
| chr1-72588723-INS-972 | chr1-72588724-INS-974 |
| chr1-72591162-INS-123 | chr1-72591162-INS-123 |
| chr1-72597139-INS-571 | chr1-72597139-INS-608 |
| chr1-72601635-INS-70 | chr1-72601659-INS-52 |
| chr1-72631532-INS-76 | chr1-72631532-INS-76 |
| chr1-72641167-INS-57 | chr1-72641169-INS-57 |
| chr1-72644719-INS-287 | chr1-72644719-INS-287 |
| chr1-72683355-INS-403 | chr1-72683355-INS-403 |
| chr1-72683907-INS-7129 | chr1-72683908-INS-7129 |
| chr1-72696664-INS-61 | chr1-72696664-INS-61 |
| chr1-72702921-INS-977 | chr1-72702921-INS-977 |
| chr1-72716321-INS-52 | chr1-72716321-INS-64 |
| chr1-72754060-INS-101 | chr1-72754060-INS-101 |
| chr1-72762577-INS-94 | chr1-72762577-INS-96 |
| chr1-72773893-INS-98 | chr1-72773920-INS-64 |
| chr1-72782189-INS-203 | chr1-72782189-INS-203 |
| chr1-72837285-INS-55 | chr1-72837285-INS-55 |
| chr1-72841923-INS-56 | chr1-72841923-INS-52 |
| chr1-72868332-INS-52 | chr1-72868332-INS-54 |
| chr1-72877563-INS-195 | chr1-72877563-INS-195 |
| chr1-72932992-INS-123 | chr1-72932992-INS-87 |
| chr1-72951030-INS-60 | chr1-72951031-INS-68 |
| chr1-72958557-INS-391 | chr1-72958557-INS-390 |
| chr1-72960087-INS-52 | chr1-72960087-INS-56 |
| chr1-72965321-INS-117 | chr1-72965321-INS-81 |
| chr1-72980153-INS-60 | chr1-72980146-INS-68 |
| chr1-72988922-INS-68 | chr1-72988922-INS-96 |
| chr1-72998902-INS-169 | chr1-72998902-INS-179 |
| chr1-73008590-INS-52 | chr1-73008592-INS-54 |
| chr1-73012719-INS-52 | chr1-73012717-INS-56 |
| chr1-73021335-INS-6387 | chr1-73021336-INS-6387 |
| chr1-73025687-INS-116 | chr1-73025687-INS-116 |
| chr1-73065054-INS-66 | chr1-73065054-INS-66 |
| chr1-73074386-INS-66 | chr1-73074386-INS-50 |
| chr1-73085806-INS-62 | chr1-73085807-INS-98 |
| chr1-73090213-INS-387 | chr1-73090213-INS-387 |
| chr1-73123343-INS-72 | chr1-73123343-INS-68 |
| chr1-73142285-INS-6382 | chr1-73142286-INS-6382 |
| chr1-73144040-INS-53 | chr1-73144041-INS-59 |
| chr1-73167163-INS-831 | chr1-73167166-INS-831 |
| chr1-73179345-INS-65 | chr1-73179345-INS-60 |
| chr1-73217407-INS-7200 | chr1-73217408-INS-7200 |
| chr1-73232155-INS-137 | chr1-73232155-INS-133 |
| chr1-73243750-INS-104 | chr1-73243750-INS-107 |
| chr1-73254795-INS-461 | chr1-73254795-INS-461 |
| chr1-73277748-INS-730 | chr1-73277748-INS-725 |
| chr1-73294597-INS-532 | chr1-73294597-INS-532 |
| chr1-73315441-INS-572 | chr1-73315441-INS-572 |
| chr1-73355606-INS-64 | chr1-73355606-INS-66 |
| chr1-73358024-INS-1422 | chr1-73358025-INS-1422 |
| chr1-73399358-INS-99 | chr1-73399358-INS-102 |
| chr1-73415593-INS-72 | chr1-73415593-INS-74 |
| chr1-73517574-INS-1103 | chr1-73517575-INS-1103 |
| chr1-73524044-INS-128 | chr1-73524044-INS-128 |
| chr1-73543976-INS-60 | chr1-73543947-INS-64 |
| chr1-73558385-INS-7105 | chr1-73558386-INS-7105 |
| chr1-73575485-INS-51 | chr1-73575485-INS-51 |
| chr1-73594457-INS-208 | chr1-73594457-INS-208 |
| chr1-73605157-INS-72 | chr1-73605157-INS-74 |
| chr1-73622179-INS-214 | chr1-73622179-INS-214 |
| chr1-73654315-INS-413 | chr1-73654315-INS-418 |
| chr1-73684051-INS-56 | chr1-73684051-INS-64 |
| chr1-73685328-INS-110 | chr1-73685328-INS-106 |
| chr1-73697150-INS-52 | chr1-73697150-INS-52 |
| chr1-73711993-INS-85 | chr1-73711993-INS-91 |
| chr1-73738666-INS-68 | chr1-73738666-INS-56 |
| chr1-73743036-INS-139 | chr1-73743036-INS-189 |
| chr1-73823427-INS-84 | chr1-73823427-INS-90 |
| chr1-73839869-INS-511 | chr1-73839870-INS-552 |
| chr1-73849593-INS-52 | chr1-73849593-INS-58 |
| chr1-73870562-INS-56 | chr1-73870562-INS-56 |
| chr1-73876062-INS-55 | chr1-73876062-INS-55 |

| | |
|---|---|
| chr1-73890147-INS-256 | chr1-73890147-INS-256 |
| chr1-73954420-INS-130 | chr1-73954420-INS-94 |
| chr1-74009071-INS-67 | chr1-74009069-INS-53 |
| chr1-74013057-INS-203 | chr1-74013057-INS-203 |
| chr1-74031404-INS-553 | chr1-74031404-INS-553 |
| chr1-74041981-INS-76 | chr1-74041981-INS-114 |
| chr1-74109709-INS-212 | chr1-74109709-INS-211 |
| chr1-74115126-INS-401 | chr1-74115126-INS-401 |
| chr1-74133009-INS-54 | chr1-74133009-INS-58 |
| chr1-74155154-INS-5514 | chr1-74155155-INS-5514 |
| chr1-74163004-INS-5386 | chr1-74163005-INS-5386 |
| chr1-74193965-INS-84 | chr1-74193965-INS-88 |
| chr1-74222959-INS-117 | chr1-74222964-INS-120 |
| chr1-74249617-INS-390 | chr1-74249617-INS-390 |
| chr1-74250546-INS-157 | chr1-74250546-INS-157 |
| chr1-74259644-INS-105 | chr1-74259644-INS-107 |
| chr1-74269987-INS-355 | chr1-74269987-INS-355 |
| chr1-74272762-INS-201 | chr1-74272762-INS-205 |
| chr1-74282186-INS-55 | chr1-74282186-INS-55 |
| chr1-74294853-INS-80 | chr1-74294851-INS-81 |
| chr1-74305623-INS-62 | chr1-74305623-INS-86 |
| chr1-74308586-INS-181 | chr1-74308586-INS-181 |
| chr1-74350549-INS-201 | chr1-74350549-INS-215 |
| chr1-74366740-INS-270 | chr1-74366744-INS-288 |
| chr1-74380197-INS-66 | chr1-74380197-INS-66 |
| chr1-74383225-INS-229 | chr1-74383225-INS-229 |
| chr1-74393682-INS-107 | chr1-74393682-INS-110 |
| chr1-74405375-INS-196 | chr1-74405375-INS-202 |
| chr1-74407114-INS-203 | chr1-74407114-INS-203 |
| chr1-74445080-INS-56 | chr1-74445080-INS-76 |
| chr1-74463222-INS-198 | chr1-74463222-INS-198 |
| chr1-74468757-INS-64 | chr1-74468757-INS-64 |
| chr1-74491451-INS-54 | chr1-74491451-INS-54 |
| chr1-74512908-INS-55 | chr1-74512907-INS-55 |
| chr1-74526889-INS-209 | chr1-74526889-INS-209 |
| chr1-74532649-INS-209 | chr1-74532649-INS-210 |
| chr1-74536319-INS-180 | chr1-74536319-INS-180 |
| chr1-74572204-INS-212 | chr1-74572204-INS-213 |
| chr1-74592604-INS-89 | chr1-74592605-INS-54 |
| chr1-74600226-INS-52 | chr1-74600226-INS-52 |
| chr1-74619383-INS-262 | chr1-74619383-INS-260 |
| chr1-74635673-INS-177 | chr1-74635673-INS-167 |
| chr1-74756160-INS-2340 | chr1-74756160-INS-2338 |
| chr1-74804584-INS-56 | chr1-74804584-INS-58 |
| chr1-74853662-INS-72 | chr1-74853662-INS-66 |
| chr1-74860279-INS-216 | chr1-74860279-INS-216 |
| chr1-74888515-INS-70 | chr1-74888515-INS-69 |
| chr1-74910649-INS-202 | chr1-74910649-INS-203 |
| chr1-74937610-INS-51 | chr1-74937608-INS-68 |
| chr1-75015554-INS-85 | chr1-75015554-INS-85 |
| chr1-75078233-INS-219 | chr1-75078233-INS-219 |
| chr1-75085038-INS-333 | chr1-75085038-INS-333 |
| chr1-75113836-INS-163 | chr1-75113836-INS-171 |
| chr1-75135456-INS-173 | chr1-75135456-INS-173 |
| chr1-75141728-INS-69 | chr1-75141747-INS-94 |
| chr1-75144706-INS-90 | chr1-75144731-INS-77 |
| chr1-75171594-INS-52 | chr1-75171594-INS-52 |
| chr1-75199428-INS-392 | chr1-75199428-INS-395 |
| chr1-75212479-INS-206 | chr1-75212479-INS-206 |
| chr1-75214977-INS-247 | chr1-75214977-INS-247 |
| chr1-75256597-INS-54 | chr1-75256597-INS-66 |
| chr1-75265751-INS-270 | chr1-75265751-INS-269 |
| chr1-75273356-INS-171 | chr1-75273357-INS-172 |
| chr1-75278778-INS-56 | chr1-75278778-INS-60 |
| chr1-75308625-INS-82 | chr1-75308625-INS-82 |
| chr1-75355403-INS-212 | chr1-75355403-INS-210 |
| chr1-75373322-INS-52 | chr1-75373322-INS-58 |
| chr1-75410052-INS-6442 | chr1-75410053-INS-6442 |
| chr1-75494672-INS-2870 | chr1-75494673-INS-2839 |
| chr1-75545800-INS-84 | chr1-75545800-INS-78 |
| chr1-75550140-INS-326 | chr1-75550140-INS-326 |
| chr1-75560435-INS-96 | chr1-75560435-INS-96 |
| chr1-75568183-INS-60 | chr1-75568183-INS-60 |
| chr1-75588064-INS-80 | chr1-75588060-INS-80 |
| chr1-75588162-INS-6257 | chr1-75588153-INS-6238 |
| chr1-75590598-INS-54 | chr1-75590598-INS-56 |
| chr1-75602891-INS-51 | chr1-75602891-INS-51 |

| | |
|---|---|
| chr1-75604461-INS-563 | chr1-75604461-INS-563 |
| chr1-75610878-INS-128 | chr1-75610878-INS-128 |
| chr1-75616383-INS-66 | chr1-75616383-INS-66 |
| chr1-75619295-INS-315 | chr1-75619295-INS-315 |
| chr1-75621062-INS-80 | chr1-75621082-INS-64 |
| chr1-75680018-INS-506 | chr1-75680018-INS-506 |
| chr1-75702689-INS-163 | chr1-75702685-INS-165 |
| chr1-75704357-INS-92 | chr1-75704357-INS-92 |
| chr1-75714459-INS-379 | chr1-75714459-INS-379 |
| chr1-75716485-INS-683 | chr1-75716485-INS-681 |
| chr1-75717023-INS-575 | chr1-75717023-INS-574 |
| chr1-75727724-INS-54 | chr1-75727735-INS-50 |
| chr1-75743909-INS-313 | chr1-75743909-INS-313 |
| chr1-75793905-INS-116 | chr1-75793905-INS-116 |
| chr1-75798079-INS-51 | chr1-75798079-INS-51 |
| chr1-75826006-INS-6480 | chr1-75826007-INS-6481 |
| chr1-75847897-INS-68 | chr1-75847897-INS-68 |
| chr1-75906143-INS-103 | chr1-75906165-INS-125 |
| chr1-75920210-INS-5550 | chr1-75920211-INS-5550 |
| chr1-75925523-INS-51 | chr1-75925532-INS-57 |
| chr1-75956197-INS-400 | chr1-75956197-INS-400 |
| chr1-75959370-INS-87 | chr1-75959405-INS-82 |
| chr1-75975902-INS-52 | chr1-75975902-INS-54 |
| chr1-75990739-INS-478 | chr1-75990739-INS-478 |
| chr1-76014549-INS-442 | chr1-76014549-INS-437 |
| chr1-76143690-INS-85 | chr1-76143690-INS-85 |
| chr1-76161002-INS-6856 | chr1-76161003-INS-6855 |
| chr1-76169322-INS-80 | chr1-76169322-INS-80 |
| chr1-76178428-INS-64 | chr1-76178428-INS-68 |
| chr1-76219174-INS-58 | chr1-76219174-INS-56 |
| chr1-76221676-INS-1770 | chr1-76221677-INS-1769 |
| chr1-76255559-INS-66 | chr1-76255559-INS-66 |
| chr1-76255909-INS-272 | chr1-76255924-INS-268 |
| chr1-76320756-INS-401 | chr1-76320756-INS-401 |
| chr1-76374844-INS-76 | chr1-76374844-INS-64 |
| chr1-76378072-INS-147 | chr1-76378072-INS-147 |
| chr1-76452748-INS-341 | chr1-76452748-INS-341 |
| chr1-76455920-INS-63 | chr1-76455920-INS-57 |
| chr1-76457648-INS-57 | chr1-76457649-INS-61 |
| chr1-76482622-INS-622 | chr1-76482622-INS-622 |
| chr1-76484010-INS-1371 | chr1-76484011-INS-1371 |
| chr1-76487833-INS-127 | chr1-76487840-INS-114 |
| chr1-76493812-INS-7433 | chr1-76493813-INS-7433 |
| chr1-76523913-INS-65 | chr1-76523913-INS-68 |
| chr1-76582846-INS-116 | chr1-76582842-INS-132 |
| chr1-76585303-INS-9593 | chr1-76585304-INS-9593 |
| chr1-76669508-INS-6903 | chr1-76669509-INS-6903 |
| chr1-76676770-INS-536 | chr1-76676770-INS-536 |
| chr1-76693617-INS-2850 | chr1-76693618-INS-2851 |
| chr1-76696844-INS-58 | chr1-76696844-INS-60 |
| chr1-76698725-INS-223 | chr1-76698725-INS-223 |
| chr1-76704531-INS-130 | chr1-76704531-INS-130 |
| chr1-76723075-INS-83 | chr1-76723075-INS-89 |
| chr1-76733276-INS-228 | chr1-76733248-INS-225 |
| chr1-76737986-INS-89 | chr1-76737992-INS-89 |
| chr1-76773819-INS-74 | chr1-76773819-INS-68 |
| chr1-76798095-INS-4453 | chr1-76798096-INS-4453 |
| chr1-76851399-INS-174 | chr1-76851399-INS-170 |
| chr1-76856430-INS-82 | chr1-76856430-INS-67 |
| chr1-76857518-INS-913 | chr1-76857518-INS-913 |
| chr1-76870788-INS-6230 | chr1-76870789-INS-6231 |
| chr1-76911028-INS-422 | chr1-76911028-INS-386 |
| chr1-76918588-INS-206 | chr1-76918588-INS-206 |
| chr1-76925445-INS-54 | chr1-76925445-INS-60 |
| chr1-76932447-INS-1943 | chr1-76932448-INS-1938 |
| chr1-76942428-INS-88 | chr1-76942428-INS-50 |
| chr1-76955573-INS-82 | chr1-76955573-INS-88 |
| chr1-76967686-INS-120 | chr1-76967686-INS-120 |
| chr1-77001451-INS-79 | chr1-77001451-INS-80 |
| chr1-77004089-INS-615 | chr1-77004089-INS-615 |
| chr1-77012935-INS-801 | chr1-77012935-INS-801 |
| chr1-77021514-INS-64 | chr1-77021515-INS-75 |
| chr1-77047391-INS-105 | chr1-77047393-INS-140 |
| chr1-77111434-INS-64 | chr1-77111434-INS-64 |
| chr1-77111974-INS-628 | chr1-77111974-INS-628 |
| chr1-77116358-INS-59 | chr1-77116358-INS-59 |
| chr1-77116937-INS-172 | chr1-77116937-INS-176 |

| | |
|---|---|
| chr1-77129901-INS-894 | chr1-77129901-INS-893 |
| chr1-77147086-INS-76 | chr1-77147092-INS-56 |
| chr1-77195420-INS-103 | chr1-77195420-INS-103 |
| chr1-77207100-INS-5275 | chr1-77207101-INS-5275 |
| chr1-77210609-INS-145 | chr1-77210634-INS-145 |
| chr1-77216976-INS-74 | chr1-77216954-INS-54 |
| chr1-77230808-INS-81 | chr1-77230811-INS-75 |
| chr1-77300690-INS-152 | chr1-77300690-INS-152 |
| chr1-77307265-INS-64 | chr1-77307265-INS-72 |
| chr1-77321424-INS-2991 | chr1-77321425-INS-2990 |
| chr1-77337915-INS-91 | chr1-77337915-INS-98 |
| chr1-77343250-INS-54 | chr1-77343247-INS-56 |
| chr1-77372414-INS-52 | chr1-77372415-INS-56 |
| chr1-77419149-INS-202 | chr1-77419149-INS-201 |
| chr1-77428840-INS-117 | chr1-77428840-INS-117 |
| chr1-77434880-INS-53 | chr1-77434880-INS-55 |
| chr1-77441526-INS-217 | chr1-77441526-INS-215 |
| chr1-77462908-INS-72 | chr1-77462908-INS-56 |
| chr1-77472896-INS-208 | chr1-77472896-INS-208 |
| chr1-77474721-INS-347 | chr1-77474721-INS-349 |
| chr1-77475749-INS-51 | chr1-77475699-INS-100 |
| chr1-77476006-INS-388 | chr1-77476006-INS-389 |
| chr1-77496146-INS-207 | chr1-77496146-INS-207 |
| chr1-77508416-INS-69 | chr1-77508416-INS-69 |
| chr1-77524284-INS-97 | chr1-77524284-INS-102 |
| chr1-77536211-INS-4561 | chr1-77536212-INS-4561 |
| chr1-77543054-INS-112 | chr1-77543058-INS-117 |
| chr1-77555835-INS-126 | chr1-77555835-INS-117 |
| chr1-77567504-INS-1793 | chr1-77567505-INS-1793 |
| chr1-77597676-INS-56 | chr1-77597676-INS-56 |
| chr1-77608244-INS-355 | chr1-77608244-INS-355 |
| chr1-77619772-INS-53 | chr1-77619772-INS-63 |
| chr1-77626929-INS-280 | chr1-77626929-INS-280 |
| chr1-77661874-INS-6728 | chr1-77661874-INS-6729 |
| chr1-77675650-INS-1587 | chr1-77675651-INS-1587 |
| chr1-77675904-INS-321 | chr1-77675904-INS-321 |
| chr1-77681385-INS-56 | chr1-77681385-INS-60 |
| chr1-77700841-INS-399 | chr1-77700841-INS-384 |
| chr1-77762734-INS-6842 | chr1-77762735-INS-6842 |
| chr1-77771635-INS-88 | chr1-77771635-INS-60 |
| chr1-77776830-INS-87 | chr1-77776855-INS-62 |
| chr1-77835401-INS-1301 | chr1-77835420-INS-1276 |
| chr1-77857804-INS-82 | chr1-77857804-INS-82 |
| chr1-77859168-INS-153 | chr1-77859168-INS-153 |
| chr1-77937380-INS-68 | chr1-77937352-INS-82 |
| chr1-77938286-INS-2780 | chr1-77938287-INS-2780 |
| chr1-78001226-INS-206 | chr1-78001226-INS-206 |
| chr1-78057395-INS-6849 | chr1-78057396-INS-6849 |
| chr1-78096478-INS-80 | chr1-78096478-INS-56 |
| chr1-78104773-INS-55 | chr1-78104773-INS-55 |
| chr1-78136615-INS-114 | chr1-78136615-INS-113 |
| chr1-78167946-INS-54 | chr1-78167946-INS-54 |
| chr1-78199193-INS-92 | chr1-78199193-INS-92 |
| chr1-78208301-INS-764 | chr1-78208307-INS-714 |
| chr1-78212132-INS-363 | chr1-78212132-INS-363 |
| chr1-78248345-INS-194 | chr1-78248334-INS-186 |
| chr1-78250582-INS-51 | chr1-78250554-INS-52 |
| chr1-78298459-INS-62 | chr1-78298459-INS-62 |
| chr1-78334595-INS-6838 | chr1-78334609-INS-6829 |
| chr1-78343096-INS-447 | chr1-78343096-INS-447 |
| chr1-78351405-INS-138 | chr1-78351405-INS-134 |
| chr1-78356337-INS-558 | chr1-78356337-INS-561 |
| chr1-78412853-INS-496 | chr1-78412853-INS-496 |
| chr1-78418082-INS-75 | chr1-78418084-INS-61 |
| chr1-78433826-INS-123 | chr1-78433826-INS-123 |
| chr1-78458188-INS-529 | chr1-78458188-INS-530 |
| chr1-78459580-INS-427 | chr1-78459580-INS-427 |
| chr1-78488881-INS-247 | chr1-78488919-INS-199 |
| chr1-78497508-INS-3148 | chr1-78497509-INS-3147 |
| chr1-78515256-INS-6443 | chr1-78515257-INS-6443 |
| chr1-78536573-INS-238 | chr1-78536573-INS-244 |
| chr1-78543114-INS-5734 | chr1-78543115-INS-5734 |
| chr1-78554272-INS-203 | chr1-78554272-INS-203 |
| chr1-78577806-INS-95 | chr1-78577806-INS-95 |
| chr1-78580131-INS-197 | chr1-78580121-INS-197 |
| chr1-78646556-INS-100 | chr1-78646556-INS-100 |
| chr1-78663779-INS-65 | chr1-78663779-INS-70 |

| | |
|---|---|
| chr1-78674990-INS-52 | chr1-78674990-INS-62 |
| chr1-78700314-INS-90 | chr1-78700314-INS-90 |
| chr1-78715992-INS-218 | chr1-78715993-INS-218 |
| chr1-78719133-INS-59 | chr1-78719133-INS-59 |
| chr1-78719479-INS-1886 | chr1-78719480-INS-1886 |
| chr1-78750391-INS-5774 | chr1-78750392-INS-5774 |
| chr1-78765733-INS-199 | chr1-78765718-INS-240 |
| chr1-78784167-INS-509 | chr1-78784167-INS-510 |
| chr1-78791902-INS-661 | chr1-78791902-INS-660 |
| chr1-78801225-INS-309 | chr1-78801225-INS-308 |
| chr1-78803387-INS-1301 | chr1-78803388-INS-1301 |
| chr1-78824053-INS-89 | chr1-78824059-INS-64 |
| chr1-78825799-INS-4379 | chr1-78825800-INS-4379 |
| chr1-78839244-INS-54 | chr1-78839283-INS-52 |
| chr1-78848296-INS-173 | chr1-78848296-INS-173 |
| chr1-78850486-INS-6248 | chr1-78850487-INS-6248 |
| chr1-78855063-INS-210 | chr1-78855063-INS-195 |
| chr1-78866037-INS-72 | chr1-78866035-INS-51 |
| chr1-78896078-INS-6393 | chr1-78896079-INS-6394 |
| chr1-78898633-INS-158 | chr1-78898633-INS-158 |
| chr1-78921902-INS-184 | chr1-78921902-INS-184 |
| chr1-78927576-INS-816 | chr1-78927576-INS-820 |
| chr1-78946703-INS-6298 | chr1-78946704-INS-6298 |
| chr1-78957682-INS-678 | chr1-78957682-INS-678 |
| chr1-78967004-INS-5463 | chr1-78967004-INS-5464 |
| chr1-78968482-INS-58 | chr1-78968465-INS-50 |
| chr1-78980797-INS-50 | chr1-78980797-INS-50 |
| chr1-78984014-INS-7016 | chr1-78984015-INS-7018 |
| chr1-79027346-INS-489 | chr1-79027346-INS-489 |
| chr1-79030487-INS-95 | chr1-79030487-INS-95 |
| chr1-79042643-INS-68 | chr1-79042643-INS-60 |
| chr1-79045599-INS-567 | chr1-79045599-INS-568 |
| chr1-79071821-INS-812 | chr1-79071821-INS-812 |
| chr1-79129037-INS-1466 | chr1-79129038-INS-1463 |
| chr1-79138744-INS-6215 | chr1-79138745-INS-6215 |
| chr1-79178944-INS-219 | chr1-79178944-INS-219 |
| chr1-79195939-INS-52 | chr1-79195939-INS-52 |
| chr1-79210759-INS-6213 | chr1-79210760-INS-6213 |
| chr1-79223388-INS-622 | chr1-79223388-INS-622 |
| chr1-79244956-INS-70 | chr1-79244956-INS-72 |
| chr1-79251522-INS-7211 | chr1-79251523-INS-7211 |
| chr1-79253371-INS-3591 | chr1-79253372-INS-3592 |
| chr1-79253775-INS-69 | chr1-79253775-INS-69 |
| chr1-79263509-INS-7124 | chr1-79263510-INS-7124 |
| chr1-79273077-INS-164 | chr1-79273077-INS-164 |
| chr1-79274099-INS-1295 | chr1-79274100-INS-1295 |
| chr1-79274910-INS-6839 | chr1-79274911-INS-6840 |
| chr1-79279375-INS-3389 | chr1-79279376-INS-3389 |
| chr1-79279376-INS-553 | chr1-79279376-INS-553 |
| chr1-79294377-INS-78 | chr1-79294390-INS-86 |
| chr1-79298230-INS-52 | chr1-79298230-INS-52 |
| chr1-79300207-INS-7524 | chr1-79300208-INS-7523 |
| chr1-79304572-INS-157 | chr1-79304572-INS-160 |
| chr1-79321210-INS-6019 | chr1-79321211-INS-6019 |
| chr1-79322732-INS-334 | chr1-79322732-INS-333 |
| chr1-79323813-INS-69 | chr1-79323813-INS-69 |
| chr1-79326904-INS-65 | chr1-79326904-INS-65 |
| chr1-79331500-INS-954 | chr1-79331500-INS-954 |
| chr1-79339839-INS-84 | chr1-79339836-INS-84 |
| chr1-79354774-INS-675 | chr1-79354774-INS-675 |
| chr1-79392498-INS-50 | chr1-79392498-INS-64 |
| chr1-79396814-INS-238 | chr1-79396805-INS-216 |
| chr1-79409474-INS-709 | chr1-79409474-INS-710 |
| chr1-79410500-INS-509 | chr1-79410499-INS-508 |
| chr1-79411070-INS-96 | chr1-79411070-INS-84 |
| chr1-79420832-INS-57 | chr1-79420832-INS-57 |
| chr1-79443478-INS-471 | chr1-79443478-INS-471 |
| chr1-79448594-INS-405 | chr1-79448594-INS-405 |
| chr1-79450771-INS-76 | chr1-79450771-INS-76 |
| chr1-79460914-INS-71 | chr1-79460914-INS-67 |
| chr1-79480231-INS-6983 | chr1-79480232-INS-6983 |
| chr1-79491409-INS-964 | chr1-79491409-INS-964 |
| chr1-79499574-INS-940 | chr1-79499574-INS-940 |
| chr1-79503222-INS-217 | chr1-79503222-INS-218 |
| chr1-79505996-INS-6431 | chr1-79505997-INS-6432 |
| chr1-79516721-INS-1327 | chr1-79516721-INS-1325 |
| chr1-79519129-INS-2078 | chr1-79519130-INS-2080 |

| | |
|---|---|
| chr1-79541477-INS-940 | chr1-79541477-INS-940 |
| chr1-79543905-INS-162 | chr1-79543879-INS-171 |
| chr1-79545730-INS-361 | chr1-79545730-INS-361 |
| chr1-79604792-INS-66 | chr1-79604792-INS-66 |
| chr1-79606720-INS-156 | chr1-79606719-INS-164 |
| chr1-79618953-INS-56 | chr1-79618953-INS-56 |
| chr1-79620375-INS-841 | chr1-79620375-INS-838 |
| chr1-79623207-INS-169 | chr1-79623207-INS-168 |
| chr1-79645476-INS-62 | chr1-79645476-INS-66 |
| chr1-79652296-INS-7111 | chr1-79652297-INS-7111 |
| chr1-79654978-INS-200 | chr1-79654978-INS-200 |
| chr1-79655263-INS-1772 | chr1-79655264-INS-1773 |
| chr1-79663908-INS-534 | chr1-79663908-INS-534 |
| chr1-79683526-INS-93 | chr1-79683526-INS-96 |
| chr1-79738033-INS-175 | chr1-79738033-INS-176 |
| chr1-79767975-INS-895 | chr1-79767975-INS-901 |
| chr1-79777969-INS-54 | chr1-79777970-INS-68 |
| chr1-79778093-INS-2934 | chr1-79778094-INS-2934 |
| chr1-79779351-INS-177 | chr1-79779360-INS-167 |
| chr1-79796215-INS-1541 | chr1-79796216-INS-1541 |
| chr1-79817071-INS-292 | chr1-79817071-INS-292 |
| chr1-79841116-INS-423 | chr1-79841116-INS-424 |
| chr1-79845315-INS-196 | chr1-79845315-INS-232 |
| chr1-79849184-INS-415 | chr1-79849184-INS-415 |
| chr1-79849956-INS-4662 | chr1-79849957-INS-4662 |
| chr1-79861321-INS-76 | chr1-79861279-INS-99 |
| chr1-79874951-INS-70 | chr1-79874951-INS-56 |
| chr1-79877128-INS-58 | chr1-79877128-INS-60 |
| chr1-79886128-INS-172 | chr1-79886128-INS-172 |
| chr1-79926430-INS-353 | chr1-79926430-INS-353 |
| chr1-79937621-INS-264 | chr1-79937621-INS-262 |
| chr1-79971384-INS-255 | chr1-79971384-INS-255 |
| chr1-79972182-INS-6286 | chr1-79972183-INS-6286 |
| chr1-79976257-INS-58 | chr1-79976257-INS-58 |
| chr1-80012366-INS-186 | chr1-80012366-INS-186 |
| chr1-80025653-INS-590 | chr1-80025653-INS-592 |
| chr1-80073017-INS-6604 | chr1-80073018-INS-6605 |
| chr1-80093328-INS-88 | chr1-80093328-INS-92 |
| chr1-80096921-INS-90 | chr1-80096921-INS-74 |
| chr1-80110602-INS-55 | chr1-80110602-INS-59 |
| chr1-80128110-INS-2041 | chr1-80128111-INS-2078 |
| chr1-80131098-INS-754 | chr1-80131098-INS-754 |
| chr1-80136820-INS-216 | chr1-80136820-INS-219 |
| chr1-80139107-INS-239 | chr1-80139107-INS-239 |
| chr1-80156864-INS-203 | chr1-80156864-INS-205 |
| chr1-80172090-INS-114 | chr1-80172090-INS-116 |
| chr1-80178420-INS-208 | chr1-80178424-INS-207 |
| chr1-80212803-INS-208 | chr1-80212803-INS-208 |
| chr1-80256306-INS-2912 | chr1-80256307-INS-2912 |
| chr1-80290079-INS-229 | chr1-80290079-INS-219 |
| chr1-80303699-INS-220 | chr1-80303699-INS-176 |
| chr1-80305536-INS-204 | chr1-80305536-INS-204 |
| chr1-80326661-INS-104 | chr1-80326661-INS-92 |
| chr1-80359124-INS-87 | chr1-80359124-INS-85 |
| chr1-80380123-INS-177 | chr1-80380123-INS-178 |
| chr1-80389856-INS-77 | chr1-80389856-INS-81 |
| chr1-80392868-INS-213 | chr1-80392868-INS-213 |
| chr1-80393375-INS-156 | chr1-80393366-INS-156 |
| chr1-80401328-INS-52 | chr1-80401326-INS-84 |
| chr1-80402287-INS-333 | chr1-80402287-INS-334 |
| chr1-80410225-INS-1544 | chr1-80410226-INS-1544 |
| chr1-80414577-INS-118 | chr1-80414577-INS-118 |
| chr1-80421097-INS-5499 | chr1-80421098-INS-5499 |
| chr1-80445070-INS-6402 | chr1-80445057-INS-6391 |
| chr1-80452592-INS-7476 | chr1-80452593-INS-7477 |
| chr1-80463451-INS-53 | chr1-80463469-INS-65 |
| chr1-80465624-INS-102 | chr1-80465612-INS-91 |
| chr1-80510694-INS-53 | chr1-80510694-INS-53 |
| chr1-80527269-INS-362 | chr1-80527269-INS-362 |
| chr1-80538360-INS-792 | chr1-80538360-INS-794 |
| chr1-80540670-INS-55 | chr1-80540670-INS-57 |
| chr1-80548388-INS-79 | chr1-80548388-INS-60 |
| chr1-80550672-INS-696 | chr1-80550672-INS-698 |
| chr1-80565726-INS-72 | chr1-80565726-INS-72 |
| chr1-80582467-INS-6257 | chr1-80582468-INS-6259 |
| chr1-80588893-INS-53 | chr1-80588893-INS-53 |
| chr1-80594956-INS-6761 | chr1-80594957-INS-6761 |

| | |
|---|---|
| chr1-80605416-INS-202 | chr1-80605456-INS-208 |
| chr1-80627969-INS-1753 | chr1-80627970-INS-1753 |
| chr1-80633977-INS-74 | chr1-80633977-INS-80 |
| chr1-80646381-INS-1291 | chr1-80646382-INS-1291 |
| chr1-80694947-INS-145 | chr1-80694948-INS-96 |
| chr1-80695909-INS-56 | chr1-80695909-INS-72 |
| chr1-80708963-INS-637 | chr1-80708963-INS-637 |
| chr1-80714440-INS-321 | chr1-80714440-INS-322 |
| chr1-80717748-INS-63 | chr1-80717743-INS-60 |
| chr1-80743075-INS-6845 | chr1-80743076-INS-6845 |
| chr1-80749882-INS-206 | chr1-80749882-INS-208 |
| chr1-80754010-INS-58 | chr1-80753983-INS-62 |
| chr1-80769794-INS-96 | chr1-80769792-INS-60 |
| chr1-80776773-INS-93 | chr1-80776773-INS-69 |
| chr1-80819983-INS-69 | chr1-80819983-INS-72 |
| chr1-80843151-INS-452 | chr1-80843151-INS-453 |
| chr1-80877625-INS-730 | chr1-80877625-INS-730 |
| chr1-80888319-INS-56 | chr1-80888319-INS-52 |
| chr1-80889276-INS-6193 | chr1-80889277-INS-6195 |
| chr1-80900423-INS-7975 | chr1-80900424-INS-7975 |
| chr1-80923793-INS-76 | chr1-80923793-INS-76 |
| chr1-80924720-INS-4483 | chr1-80924721-INS-4483 |
| chr1-80985011-INS-682 | chr1-80985011-INS-683 |
| chr1-80986058-INS-2682 | chr1-80986059-INS-2683 |
| chr1-81045134-INS-877 | chr1-81045134-INS-877 |
| chr1-81071714-INS-6487 | chr1-81071715-INS-6488 |
| chr1-81116469-INS-1629 | chr1-81116470-INS-1629 |
| chr1-81119810-INS-7787 | chr1-81119811-INS-7787 |
| chr1-81181406-INS-425 | chr1-81181407-INS-427 |
| chr1-81184393-INS-53 | chr1-81184393-INS-57 |
| chr1-81254926-INS-6589 | chr1-81254927-INS-6591 |
| chr1-81257389-INS-54 | chr1-81257389-INS-50 |
| chr1-81298721-INS-83 | chr1-81298721-INS-88 |
| chr1-81358848-INS-63 | chr1-81358848-INS-83 |
| chr1-81392998-INS-669 | chr1-81392998-INS-669 |
| chr1-81397872-INS-214 | chr1-81397872-INS-214 |
| chr1-81410147-INS-73 | chr1-81410147-INS-77 |
| chr1-81425479-INS-371 | chr1-81425479-INS-370 |
| chr1-81447724-INS-343 | chr1-81447724-INS-343 |
| chr1-81449834-INS-132 | chr1-81449834-INS-124 |
| chr1-81472041-INS-2227 | chr1-81472042-INS-2227 |
| chr1-81484503-INS-2748 | chr1-81484504-INS-2755 |
| chr1-81492595-INS-3195 | chr1-81492596-INS-3195 |
| chr1-81500058-INS-974 | chr1-81500058-INS-974 |
| chr1-81506598-INS-178 | chr1-81506598-INS-177 |
| chr1-81556575-INS-96 | chr1-81556573-INS-80 |
| chr1-81589373-INS-120 | chr1-81589373-INS-95 |
| chr1-81593682-INS-2163 | chr1-81593683-INS-2163 |
| chr1-81647082-INS-70 | chr1-81647082-INS-70 |
| chr1-81670247-INS-467 | chr1-81670247-INS-467 |
| chr1-81699174-INS-202 | chr1-81699174-INS-202 |
| chr1-81700827-INS-233 | chr1-81700784-INS-245 |
| chr1-81731282-INS-5543 | chr1-81731283-INS-5527 |
| chr1-81735106-INS-79 | chr1-81735107-INS-77 |
| chr1-81742876-INS-52 | chr1-81742876-INS-50 |
| chr1-81765108-INS-80 | chr1-81765108-INS-80 |
| chr1-81804542-INS-555 | chr1-81804543-INS-555 |
| chr1-81806305-INS-342 | chr1-81806305-INS-342 |
| chr1-81817561-INS-6694 | chr1-81817562-INS-6695 |
| chr1-81881154-INS-118 | chr1-81881154-INS-109 |
| chr1-81890764-INS-182 | chr1-81890764-INS-187 |
| chr1-81897747-INS-120 | chr1-81897747-INS-128 |
| chr1-81903778-INS-60 | chr1-81903774-INS-60 |
| chr1-81919188-INS-87 | chr1-81919188-INS-79 |
| chr1-81922300-INS-69 | chr1-81922330-INS-67 |
| chr1-81943037-INS-849 | chr1-81943036-INS-849 |
| chr1-81946665-INS-61 | chr1-81946665-INS-56 |
| chr1-81981913-INS-6298 | chr1-81981914-INS-6298 |
| chr1-81989606-INS-6601 | chr1-81989607-INS-6601 |
| chr1-81999120-INS-72 | chr1-81999116-INS-52 |
| chr1-82014833-INS-2260 | chr1-82014834-INS-2260 |
| chr1-82042213-INS-5086 | chr1-82042214-INS-5086 |
| chr1-82102461-INS-2007 | chr1-82102462-INS-2010 |
| chr1-82109430-INS-66 | chr1-82109438-INS-60 |
| chr1-82116431-INS-570 | chr1-82116431-INS-569 |
| chr1-82129251-INS-162 | chr1-82129252-INS-162 |
| chr1-82147862-INS-77 | chr1-82147862-INS-77 |

| | |
|---|---|
| chr1-82199824-INS-811 | chr1-82199824-INS-811 |
| chr1-82200440-INS-6577 | chr1-82200441-INS-6577 |
| chr1-82230105-INS-69 | chr1-82230105-INS-69 |
| chr1-82284731-INS-135 | chr1-82284740-INS-119 |
| chr1-82286797-INS-66 | chr1-82286796-INS-97 |
| chr1-82338986-INS-1361 | chr1-82338987-INS-1394 |
| chr1-82346427-INS-64 | chr1-82346432-INS-58 |
| chr1-82360107-INS-62 | chr1-82360107-INS-64 |
| chr1-82382155-INS-207 | chr1-82382155-INS-206 |
| chr1-82424838-INS-131 | chr1-82424838-INS-131 |
| chr1-82430647-INS-66 | chr1-82430647-INS-57 |
| chr1-82439580-INS-3084 | chr1-82439581-INS-3084 |
| chr1-82443983-INS-60 | chr1-82443983-INS-60 |
| chr1-82449971-INS-146 | chr1-82449971-INS-146 |
| chr1-82462956-INS-529 | chr1-82462956-INS-529 |
| chr1-82468150-INS-206 | chr1-82468150-INS-206 |
| chr1-82517145-INS-586 | chr1-82517145-INS-591 |
| chr1-82537731-INS-90 | chr1-82537731-INS-87 |
| chr1-82545340-INS-60 | chr1-82545340-INS-52 |
| chr1-82546541-INS-498 | chr1-82546541-INS-498 |
| chr1-82588597-INS-54 | chr1-82588643-INS-52 |
| chr1-82617312-INS-81 | chr1-82617288-INS-97 |
| chr1-82630868-INS-142 | chr1-82630868-INS-155 |
| chr1-82637451-INS-51 | chr1-82637473-INS-58 |
| chr1-82653607-INS-6532 | chr1-82653608-INS-6532 |
| chr1-82667928-INS-110 | chr1-82667928-INS-110 |
| chr1-82671669-INS-72 | chr1-82671671-INS-76 |
| chr1-82682163-INS-2468 | chr1-82682164-INS-2465 |
| chr1-82702451-INS-1388 | chr1-82702452-INS-1389 |
| chr1-82735761-INS-62 | chr1-82735761-INS-50 |
| chr1-82742580-INS-643 | chr1-82742577-INS-646 |
| chr1-82746694-INS-199 | chr1-82746694-INS-198 |
| chr1-82746858-INS-1293 | chr1-82746859-INS-1292 |
| chr1-82758248-INS-55 | chr1-82758248-INS-59 |
| chr1-82770190-INS-845 | chr1-82770190-INS-845 |
| chr1-82781691-INS-245 | chr1-82781691-INS-245 |
| chr1-82782925-INS-83 | chr1-82782925-INS-83 |
| chr1-82791770-INS-60 | chr1-82791770-INS-60 |
| chr1-82806394-INS-251 | chr1-82806394-INS-278 |
| chr1-82831265-INS-70 | chr1-82831265-INS-70 |
| chr1-82846488-INS-136 | chr1-82846487-INS-108 |
| chr1-82862843-INS-86 | chr1-82862836-INS-112 |
| chr1-82865405-INS-51 | chr1-82865405-INS-51 |
| chr1-82872141-INS-78 | chr1-82872141-INS-82 |
| chr1-82892455-INS-599 | chr1-82892455-INS-595 |
| chr1-82923179-INS-189 | chr1-82923179-INS-155 |
| chr1-82938140-INS-153 | chr1-82938140-INS-153 |
| chr1-82954265-INS-72 | chr1-82954265-INS-72 |
| chr1-82955304-INS-65 | chr1-82955304-INS-91 |
| chr1-83015089-INS-126 | chr1-83015089-INS-125 |
| chr1-83057675-INS-74 | chr1-83057675-INS-74 |
| chr1-83064707-INS-498 | chr1-83064707-INS-498 |
| chr1-83084825-INS-52 | chr1-83084825-INS-52 |
| chr1-83100094-INS-99 | chr1-83100094-INS-98 |
| chr1-83101624-INS-74 | chr1-83101647-INS-74 |
| chr1-83104659-INS-211 | chr1-83104659-INS-211 |
| chr1-83106923-INS-6302 | chr1-83106924-INS-6302 |
| chr1-83148755-INS-64 | chr1-83148755-INS-64 |
| chr1-83156950-INS-59 | chr1-83156950-INS-59 |
| chr1-83170048-INS-54 | chr1-83170048-INS-60 |
| chr1-83199388-INS-52 | chr1-83199388-INS-54 |
| chr1-83223722-INS-79 | chr1-83223742-INS-79 |
| chr1-83225511-INS-74 | chr1-83225511-INS-66 |
| chr1-83233045-INS-117 | chr1-83233045-INS-125 |
| chr1-83235756-INS-60 | chr1-83235752-INS-68 |
| chr1-83239000-INS-122 | chr1-83239000-INS-122 |
| chr1-83261421-INS-86 | chr1-83261421-INS-89 |
| chr1-83270832-INS-209 | chr1-83270832-INS-209 |
| chr1-83290487-INS-68 | chr1-83290487-INS-52 |
| chr1-83298548-INS-76 | chr1-83298550-INS-56 |
| chr1-83315627-INS-536 | chr1-83315627-INS-536 |
| chr1-83360277-INS-400 | chr1-83360277-INS-430 |
| chr1-83399154-INS-492 | chr1-83399154-INS-492 |
| chr1-83410017-INS-386 | chr1-83410018-INS-385 |
| chr1-83414478-INS-457 | chr1-83414478-INS-457 |
| chr1-83415874-INS-6713 | chr1-83415875-INS-6713 |
| chr1-83424522-INS-64 | chr1-83424522-INS-68 |

| | |
|---|---|
| chr1-83495153-INS-213 | chr1-83495125-INS-214 |
| chr1-83502143-INS-390 | chr1-83502143-INS-390 |
| chr1-83519073-INS-1015 | chr1-83519074-INS-1012 |
| chr1-83531511-INS-451 | chr1-83531511-INS-452 |
| chr1-83549994-INS-62 | chr1-83549984-INS-60 |
| chr1-83573062-INS-100 | chr1-83573062-INS-146 |
| chr1-83588639-INS-565 | chr1-83588639-INS-565 |
| chr1-83608391-INS-99 | chr1-83608391-INS-91 |
| chr1-83620385-INS-55 | chr1-83620385-INS-58 |
| chr1-83622800-INS-57 | chr1-83622800-INS-57 |
| chr1-83630694-INS-6285 | chr1-83630695-INS-6283 |
| chr1-83648318-INS-67 | chr1-83648318-INS-67 |
| chr1-83649612-INS-206 | chr1-83649612-INS-206 |
| chr1-83650872-INS-201 | chr1-83650872-INS-201 |
| chr1-83681782-INS-101 | chr1-83681782-INS-101 |
| chr1-83685282-INS-476 | chr1-83685282-INS-476 |
| chr1-83704246-INS-334 | chr1-83704246-INS-334 |
| chr1-83706005-INS-82 | chr1-83706005-INS-82 |
| chr1-83714503-INS-66 | chr1-83714507-INS-57 |
| chr1-83715900-INS-58 | chr1-83715875-INS-60 |
| chr1-83718386-INS-384 | chr1-83718386-INS-384 |
| chr1-83747040-INS-64 | chr1-83747040-INS-68 |
| chr1-83758457-INS-1235 | chr1-83758458-INS-1235 |
| chr1-83761501-INS-215 | chr1-83761501-INS-215 |
| chr1-83773610-INS-53 | chr1-83773610-INS-53 |
| chr1-83789225-INS-1476 | chr1-83789226-INS-1477 |
| chr1-83792257-INS-103 | chr1-83792257-INS-102 |
| chr1-83829093-INS-1261 | chr1-83829094-INS-1261 |
| chr1-83859978-INS-58 | chr1-83859978-INS-73 |
| chr1-83870560-INS-66 | chr1-83870560-INS-56 |
| chr1-83883070-INS-57 | chr1-83883070-INS-68 |
| chr1-83885166-INS-78 | chr1-83885166-INS-78 |
| chr1-83885767-INS-6603 | chr1-83885768-INS-6605 |
| chr1-83889454-INS-3097 | chr1-83889455-INS-3098 |
| chr1-83907296-INS-55 | chr1-83907296-INS-55 |
| chr1-84052060-INS-6591 | chr1-84052061-INS-6591 |
| chr1-84061147-INS-74 | chr1-84061147-INS-76 |
| chr1-84080468-INS-806 | chr1-84080468-INS-818 |
| chr1-84092661-INS-112 | chr1-84092661-INS-140 |
| chr1-84095268-INS-182 | chr1-84095268-INS-182 |
| chr1-84098660-INS-528 | chr1-84098660-INS-528 |
| chr1-84139233-INS-53 | chr1-84139233-INS-53 |
| chr1-84163291-INS-1282 | chr1-84163292-INS-1282 |
| chr1-84168230-INS-52 | chr1-84168230-INS-52 |
| chr1-84168657-INS-2963 | chr1-84168658-INS-2963 |
| chr1-84235816-INS-63 | chr1-84235816-INS-75 |
| chr1-84255717-INS-109 | chr1-84255717-INS-72 |
| chr1-84307320-INS-98 | chr1-84307320-INS-98 |
| chr1-84319899-INS-516 | chr1-84319899-INS-516 |
| chr1-84334183-INS-119 | chr1-84334183-INS-119 |
| chr1-84345490-INS-54 | chr1-84345490-INS-54 |
| chr1-84360773-INS-498 | chr1-84360773-INS-498 |
| chr1-84384874-INS-243 | chr1-84384874-INS-243 |
| chr1-84402141-INS-166 | chr1-84402142-INS-170 |
| chr1-84413075-INS-52 | chr1-84413057-INS-67 |
| chr1-84416898-INS-580 | chr1-84416898-INS-595 |
| chr1-84428169-INS-83 | chr1-84428169-INS-88 |
| chr1-84439266-INS-832 | chr1-84439265-INS-882 |
| chr1-84441642-INS-140 | chr1-84441642-INS-140 |
| chr1-84450686-INS-51 | chr1-84450671-INS-66 |
| chr1-84460422-INS-145 | chr1-84460423-INS-143 |
| chr1-84465307-INS-142 | chr1-84465307-INS-126 |
| chr1-84475837-INS-64 | chr1-84475837-INS-97 |
| chr1-84482979-INS-1711 | chr1-84482980-INS-1711 |
| chr1-84554567-INS-51 | chr1-84554572-INS-56 |
| chr1-84563271-INS-54 | chr1-84563271-INS-54 |
| chr1-84571007-INS-66 | chr1-84571007-INS-66 |
| chr1-84574458-INS-51 | chr1-84574457-INS-61 |
| chr1-84598645-INS-112 | chr1-84598661-INS-142 |
| chr1-84634020-INS-218 | chr1-84634020-INS-222 |
| chr1-84638435-INS-213 | chr1-84638435-INS-214 |
| chr1-84658928-INS-1858 | chr1-84658929-INS-1858 |
| chr1-84672284-INS-52 | chr1-84672284-INS-58 |
| chr1-84689861-INS-76 | chr1-84689861-INS-79 |
| chr1-84804004-INS-218 | chr1-84804004-INS-219 |
| chr1-84808595-INS-252 | chr1-84808591-INS-266 |
| chr1-84822959-INS-58 | chr1-84822959-INS-62 |

| | |
|---|---|
| chr1-84844630-INS-206 | chr1-84844630-INS-206 |
| chr1-84865942-INS-80 | chr1-84865942-INS-80 |
| chr1-84966730-INS-85 | chr1-84966730-INS-85 |
| chr1-85013532-INS-71 | chr1-85013531-INS-64 |
| chr1-85039139-INS-2031 | chr1-85039140-INS-2032 |
| chr1-85169127-INS-434 | chr1-85169127-INS-434 |
| chr1-85421736-INS-277 | chr1-85421736-INS-277 |
| chr1-85521165-INS-129 | chr1-85521165-INS-129 |
| chr1-85595733-INS-234 | chr1-85595733-INS-234 |
| chr1-85621180-INS-439 | chr1-85621180-INS-440 |
| chr1-85656231-INS-76 | chr1-85656231-INS-96 |
| chr1-85691472-INS-60 | chr1-85691472-INS-70 |
| chr1-85702693-INS-498 | chr1-85702693-INS-498 |
| chr1-85742031-INS-198 | chr1-85742032-INS-204 |
| chr1-85807624-INS-219 | chr1-85807624-INS-219 |
| chr1-85843350-INS-105 | chr1-85843350-INS-129 |
| chr1-85900612-INS-66 | chr1-85900612-INS-66 |
| chr1-85903377-INS-130 | chr1-85903372-INS-135 |
| chr1-85905240-INS-137 | chr1-85905240-INS-149 |
| chr1-85915444-INS-557 | chr1-85915444-INS-564 |
| chr1-85923417-INS-205 | chr1-85923417-INS-204 |
| chr1-85942386-INS-252 | chr1-85942386-INS-252 |
| chr1-85943899-INS-72 | chr1-85943899-INS-72 |
| chr1-85963799-INS-2846 | chr1-85963800-INS-2846 |
| chr1-86013178-INS-175 | chr1-86013178-INS-210 |
| chr1-86018574-INS-210 | chr1-86018574-INS-210 |
| chr1-86075876-INS-64 | chr1-86075876-INS-54 |
| chr1-86076477-INS-181 | chr1-86076477-INS-182 |
| chr1-86078124-INS-423 | chr1-86078124-INS-423 |
| chr1-86079468-INS-59 | chr1-86079468-INS-59 |
| chr1-86082857-INS-105 | chr1-86082887-INS-120 |
| chr1-86094100-INS-196 | chr1-86094100-INS-196 |
| chr1-86106456-INS-1900 | chr1-86106457-INS-1900 |
| chr1-86141913-INS-236 | chr1-86141913-INS-236 |
| chr1-86143313-INS-62 | chr1-86143313-INS-76 |
| chr1-86168005-INS-116 | chr1-86168005-INS-116 |
| chr1-86193184-INS-60 | chr1-86193184-INS-61 |
| chr1-86196203-INS-206 | chr1-86196203-INS-207 |
| chr1-86206623-INS-78 | chr1-86206621-INS-80 |
| chr1-86243481-INS-178 | chr1-86243481-INS-178 |
| chr1-86254730-INS-221 | chr1-86254730-INS-221 |
| chr1-86293802-INS-196 | chr1-86293802-INS-196 |
| chr1-86297449-INS-60 | chr1-86297449-INS-60 |
| chr1-86301918-INS-196 | chr1-86301918-INS-196 |
| chr1-86339210-INS-184 | chr1-86339210-INS-185 |
| chr1-86350104-INS-205 | chr1-86350104-INS-205 |
| chr1-86372540-INS-191 | chr1-86372535-INS-198 |
| chr1-86377599-INS-89 | chr1-86377599-INS-111 |
| chr1-86388527-INS-140 | chr1-86388528-INS-175 |
| chr1-86390043-INS-199 | chr1-86390043-INS-199 |
| chr1-86393574-INS-65 | chr1-86393585-INS-60 |
| chr1-86402503-INS-205 | chr1-86402503-INS-205 |
| chr1-86413614-INS-60 | chr1-86413603-INS-56 |
| chr1-86435255-INS-73 | chr1-86435255-INS-73 |
| chr1-86464819-INS-220 | chr1-86464819-INS-220 |
| chr1-86476309-INS-211 | chr1-86476309-INS-212 |
| chr1-86482867-INS-217 | chr1-86482867-INS-214 |
| chr1-86491721-INS-218 | chr1-86491721-INS-218 |
| chr1-86500162-INS-83 | chr1-86500166-INS-127 |
| chr1-86505265-INS-109 | chr1-86505265-INS-109 |
| chr1-86507388-INS-52 | chr1-86507398-INS-56 |
| chr1-86512642-INS-6710 | chr1-86512643-INS-6713 |
| chr1-86515893-INS-191 | chr1-86515893-INS-194 |
| chr1-86517885-INS-827 | chr1-86517885-INS-784 |
| chr1-86535261-INS-58 | chr1-86535273-INS-64 |
| chr1-86572538-INS-207 | chr1-86572538-INS-209 |
| chr1-86613077-INS-248 | chr1-86613061-INS-253 |
| chr1-86695318-INS-4808 | chr1-86695319-INS-4808 |
| chr1-86724959-INS-1615 | chr1-86724960-INS-1631 |
| chr1-86806747-INS-869 | chr1-86806747-INS-870 |
| chr1-86817559-INS-197 | chr1-86817559-INS-198 |
| chr1-86902727-INS-73 | chr1-86902727-INS-57 |
| chr1-86916638-INS-388 | chr1-86916639-INS-386 |
| chr1-86941422-INS-99 | chr1-86941422-INS-99 |
| chr1-87034011-INS-128 | chr1-87034011-INS-128 |
| chr1-87124414-INS-79 | chr1-87124404-INS-82 |
| chr1-87147189-INS-134 | chr1-87147189-INS-134 |

| | |
|---|---|
| chr1-87169603-INS-226 | chr1-87169603-INS-228 |
| chr1-87174555-INS-102 | chr1-87174589-INS-112 |
| chr1-87181263-INS-92 | chr1-87181263-INS-92 |
| chr1-87194879-INS-55 | chr1-87194879-INS-55 |
| chr1-87202560-INS-197 | chr1-87202560-INS-197 |
| chr1-87219590-INS-96 | chr1-87219622-INS-104 |
| chr1-87231810-INS-116 | chr1-87231827-INS-129 |
| chr1-87235644-INS-219 | chr1-87235645-INS-230 |
| chr1-87281155-INS-72 | chr1-87281155-INS-75 |
| chr1-87288290-INS-66 | chr1-87288288-INS-55 |
| chr1-87290210-INS-120 | chr1-87290210-INS-90 |
| chr1-87298896-INS-200 | chr1-87298896-INS-199 |
| chr1-87311102-INS-56 | chr1-87311102-INS-56 |
| chr1-87319641-INS-59 | chr1-87319651-INS-59 |
| chr1-87336703-INS-126 | chr1-87336703-INS-126 |
| chr1-87358795-INS-192 | chr1-87358795-INS-192 |
| chr1-87373547-INS-320 | chr1-87373545-INS-330 |
| chr1-87395661-INS-63 | chr1-87395661-INS-63 |
| chr1-87421198-INS-84 | chr1-87421198-INS-52 |
| chr1-87440498-INS-125 | chr1-87440498-INS-125 |
| chr1-87446053-INS-53 | chr1-87446053-INS-56 |
| chr1-87462623-INS-5238 | chr1-87462624-INS-5238 |
| chr1-87577698-INS-52 | chr1-87577698-INS-52 |
| chr1-87581104-INS-99 | chr1-87581104-INS-93 |
| chr1-87588061-INS-58 | chr1-87588061-INS-53 |
| chr1-87592921-INS-306 | chr1-87592921-INS-306 |
| chr1-87618841-INS-203 | chr1-87618841-INS-203 |
| chr1-87656520-INS-65 | chr1-87656540-INS-50 |
| chr1-87666728-INS-176 | chr1-87666728-INS-176 |
| chr1-87676940-INS-683 | chr1-87676937-INS-681 |
| chr1-87689916-INS-164 | chr1-87689916-INS-164 |
| chr1-87711209-INS-1525 | chr1-87711210-INS-1525 |
| chr1-87738219-INS-130 | chr1-87738220-INS-130 |
| chr1-87758204-INS-59 | chr1-87758204-INS-59 |
| chr1-87763187-INS-80 | chr1-87763161-INS-76 |
| chr1-87764499-INS-95 | chr1-87764458-INS-106 |
| chr1-87800440-INS-58 | chr1-87800440-INS-58 |
| chr1-87818489-INS-96 | chr1-87818489-INS-96 |
| chr1-87918241-INS-255 | chr1-87918241-INS-255 |
| chr1-87977655-INS-149 | chr1-87977655-INS-149 |
| chr1-87992653-INS-147 | chr1-87992653-INS-147 |
| chr1-88012710-INS-651 | chr1-88012710-INS-651 |
| chr1-88036485-INS-100 | chr1-88036489-INS-112 |
| chr1-88107624-INS-822 | chr1-88107624-INS-822 |
| chr1-88178872-INS-219 | chr1-88178872-INS-219 |
| chr1-88201851-INS-103 | chr1-88201837-INS-77 |
| chr1-88225617-INS-92 | chr1-88225614-INS-76 |
| chr1-88268230-INS-113 | chr1-88268230-INS-68 |
| chr1-88272579-INS-52 | chr1-88272579-INS-54 |
| chr1-88342227-INS-169 | chr1-88342224-INS-126 |
| chr1-88349183-INS-67 | chr1-88349183-INS-67 |
| chr1-88370684-INS-186 | chr1-88370684-INS-175 |
| chr1-88383031-INS-169 | chr1-88383031-INS-169 |
| chr1-88395505-INS-399 | chr1-88395505-INS-399 |
| chr1-88397926-INS-95 | chr1-88397926-INS-95 |
| chr1-88430242-INS-60 | chr1-88430243-INS-60 |
| chr1-88450228-INS-194 | chr1-88450228-INS-194 |
| chr1-88460364-INS-80 | chr1-88460364-INS-80 |
| chr1-88461067-INS-283 | chr1-88461067-INS-285 |
| chr1-88472881-INS-2724 | chr1-88472882-INS-2724 |
| chr1-88552481-INS-61 | chr1-88552481-INS-63 |
| chr1-88599759-INS-65 | chr1-88599734-INS-79 |
| chr1-88605651-INS-90 | chr1-88605651-INS-90 |
| chr1-88619618-INS-58 | chr1-88619618-INS-58 |
| chr1-88641787-INS-321 | chr1-88641787-INS-321 |
| chr1-88649956-INS-132 | chr1-88649953-INS-138 |
| chr1-88659686-INS-417 | chr1-88659686-INS-417 |
| chr1-88689472-INS-199 | chr1-88689472-INS-199 |
| chr1-88724190-INS-301 | chr1-88724190-INS-301 |
| chr1-88801030-INS-196 | chr1-88801030-INS-197 |
| chr1-88806582-INS-110 | chr1-88806582-INS-110 |
| chr1-88811229-INS-76 | chr1-88811234-INS-72 |
| chr1-88818227-INS-52 | chr1-88818228-INS-60 |
| chr1-88848486-INS-53 | chr1-88848486-INS-53 |
| chr1-88867362-INS-73 | chr1-88867362-INS-78 |
| chr1-88882164-INS-61 | chr1-88882164-INS-61 |
| chr1-88901172-INS-60 | chr1-88901172-INS-60 |

| | |
|---|---|
| chr1-88914752-INS-498 | chr1-88914752-INS-498 |
| chr1-88917105-INS-64 | chr1-88917105-INS-64 |
| chr1-88936241-INS-77 | chr1-88936241-INS-67 |
| chr1-88947208-INS-170 | chr1-88947208-INS-202 |
| chr1-88950831-INS-52 | chr1-88950831-INS-53 |
| chr1-88956387-INS-202 | chr1-88956387-INS-203 |
| chr1-88972045-INS-60 | chr1-88972043-INS-84 |
| chr1-89023746-INS-103 | chr1-89023746-INS-87 |
| chr1-89047835-INS-58 | chr1-89047796-INS-58 |
| chr1-89060525-INS-64 | chr1-89060525-INS-56 |
| chr1-89079249-INS-128 | chr1-89079249-INS-128 |
| chr1-89091073-INS-166 | chr1-89091073-INS-167 |
| chr1-89106120-INS-66 | chr1-89106120-INS-66 |
| chr1-89113731-INS-75 | chr1-89113731-INS-75 |
| chr1-89123161-INS-6463 | chr1-89123162-INS-6463 |
| chr1-89235014-INS-63 | chr1-89235014-INS-63 |
| chr1-89239993-INS-457 | chr1-89239993-INS-457 |
| chr1-89242850-INS-60 | chr1-89242850-INS-54 |
| chr1-89254501-INS-203 | chr1-89254501-INS-203 |
| chr1-89259857-INS-489 | chr1-89259857-INS-489 |
| chr1-89298230-INS-74 | chr1-89298204-INS-66 |
| chr1-89336116-INS-205 | chr1-89336116-INS-206 |
| chr1-89339253-INS-56 | chr1-89339260-INS-84 |
| chr1-89353332-INS-115 | chr1-89353332-INS-115 |
| chr1-89373824-INS-70 | chr1-89373824-INS-70 |
| chr1-89415589-INS-404 | chr1-89415589-INS-404 |
| chr1-89434637-INS-621 | chr1-89434632-INS-623 |
| chr1-89436712-INS-106 | chr1-89436712-INS-107 |
| chr1-89464933-INS-427 | chr1-89464931-INS-467 |
| chr1-89467461-INS-53 | chr1-89467464-INS-55 |
| chr1-89498878-INS-71 | chr1-89498878-INS-71 |
| chr1-89511493-INS-88 | chr1-89511493-INS-116 |
| chr1-89517452-INS-428 | chr1-89517487-INS-400 |
| chr1-89579199-INS-51 | chr1-89579193-INS-51 |
| chr1-89585005-INS-233 | chr1-89585005-INS-233 |
| chr1-89608132-INS-140 | chr1-89608132-INS-140 |
| chr1-89612684-INS-5207 | chr1-89612685-INS-5207 |
| chr1-89637812-INS-477 | chr1-89637812-INS-477 |
| chr1-89756149-INS-60 | chr1-89756159-INS-57 |
| chr1-89783105-INS-200 | chr1-89783105-INS-206 |
| chr1-89868498-INS-141 | chr1-89868498-INS-132 |
| chr1-89874810-INS-7209 | chr1-89874811-INS-7209 |
| chr1-89884174-INS-65 | chr1-89884174-INS-65 |
| chr1-89885845-INS-200 | chr1-89885845-INS-212 |
| chr1-89887852-INS-83 | chr1-89887870-INS-68 |
| chr1-89894816-INS-63 | chr1-89894816-INS-58 |
| chr1-89904052-INS-63 | chr1-89904052-INS-63 |
| chr1-89925232-INS-534 | chr1-89925232-INS-534 |
| chr1-89936469-INS-98 | chr1-89936469-INS-98 |
| chr1-90009516-INS-60 | chr1-90009516-INS-60 |
| chr1-90052306-INS-166 | chr1-90052306-INS-165 |
| chr1-90075684-INS-71 | chr1-90075684-INS-71 |
| chr1-90090012-INS-172 | chr1-90090016-INS-174 |
| chr1-90108569-INS-63 | chr1-90108569-INS-85 |
| chr1-90120232-INS-112 | chr1-90120232-INS-112 |
| chr1-90125642-INS-68 | chr1-90125642-INS-68 |
| chr1-90201887-INS-70 | chr1-90201887-INS-60 |
| chr1-90216479-INS-57 | chr1-90216479-INS-63 |
| chr1-90238405-INS-155 | chr1-90238405-INS-155 |
| chr1-90252510-INS-82 | chr1-90252510-INS-82 |
| chr1-90277124-INS-52 | chr1-90277124-INS-52 |
| chr1-90295758-INS-133 | chr1-90295752-INS-144 |
| chr1-90298051-INS-51 | chr1-90298051-INS-51 |
| chr1-90299711-INS-60 | chr1-90299711-INS-60 |
| chr1-90348226-INS-79 | chr1-90348226-INS-79 |
| chr1-90348931-INS-2749 | chr1-90348932-INS-2749 |
| chr1-90374285-INS-52 | chr1-90374285-INS-52 |
| chr1-90375541-INS-457 | chr1-90375541-INS-457 |
| chr1-90387199-INS-297 | chr1-90387199-INS-297 |
| chr1-90413041-INS-56 | chr1-90413041-INS-56 |
| chr1-90415009-INS-1344 | chr1-90415010-INS-1345 |
| chr1-90425842-INS-5261 | chr1-90425843-INS-5261 |
| chr1-90439132-INS-7142 | chr1-90439133-INS-7142 |
| chr1-90441472-INS-56 | chr1-90441476-INS-58 |
| chr1-90442742-INS-184 | chr1-90442742-INS-184 |
| chr1-90451156-INS-85 | chr1-90451156-INS-85 |
| chr1-90462021-INS-75 | chr1-90462021-INS-60 |

| | |
|---|---|
| chr1-90466085-INS-152 | chr1-90466085-INS-152 |
| chr1-90471083-INS-184 | chr1-90471083-INS-184 |
| chr1-90476058-INS-105 | chr1-90476058-INS-105 |
| chr1-90487748-INS-71 | chr1-90487748-INS-71 |
| chr1-90489747-INS-156 | chr1-90489747-INS-156 |
| chr1-90490859-INS-72 | chr1-90490847-INS-78 |
| chr1-90512600-INS-133 | chr1-90512600-INS-133 |
| chr1-90526774-INS-58 | chr1-90526774-INS-60 |
| chr1-90549401-INS-59 | chr1-90549400-INS-62 |
| chr1-90558090-INS-3117 | chr1-90558091-INS-3118 |
| chr1-90564677-INS-80 | chr1-90564677-INS-96 |
| chr1-90573575-INS-80 | chr1-90573575-INS-80 |
| chr1-90596505-INS-1896 | chr1-90596509-INS-1894 |
| chr1-90602051-INS-52 | chr1-90602051-INS-50 |
| chr1-90615469-INS-6407 | chr1-90615470-INS-6407 |
| chr1-90617065-INS-64 | chr1-90617065-INS-64 |
| chr1-90663722-INS-52 | chr1-90663722-INS-52 |
| chr1-90664541-INS-7091 | chr1-90664542-INS-7091 |
| chr1-90675642-INS-61 | chr1-90675642-INS-61 |
| chr1-90677124-INS-2672 | chr1-90677125-INS-2672 |
| chr1-90690047-INS-59 | chr1-90690047-INS-59 |
| chr1-90696815-INS-143 | chr1-90696815-INS-143 |
| chr1-90702395-INS-64 | chr1-90702391-INS-64 |
| chr1-90710122-INS-409 | chr1-90710122-INS-408 |
| chr1-90734181-INS-70 | chr1-90734181-INS-70 |
| chr1-90760075-INS-420 | chr1-90760075-INS-420 |
| chr1-90768280-INS-204 | chr1-90768283-INS-202 |
| chr1-90777966-INS-819 | chr1-90777967-INS-818 |
| chr1-90841398-INS-204 | chr1-90841398-INS-204 |
| chr1-90844107-INS-60 | chr1-90844107-INS-60 |
| chr1-90850701-INS-52 | chr1-90850701-INS-52 |
| chr1-90856265-INS-99 | chr1-90856265-INS-106 |
| chr1-90870798-INS-712 | chr1-90870798-INS-709 |
| chr1-90877734-INS-89 | chr1-90877734-INS-89 |
| chr1-90881298-INS-138 | chr1-90881336-INS-138 |
| chr1-90883223-INS-66 | chr1-90883223-INS-66 |
| chr1-90901477-INS-98 | chr1-90901477-INS-107 |
| chr1-90941019-INS-88 | chr1-90941019-INS-88 |
| chr1-90946363-INS-80 | chr1-90946363-INS-76 |
| chr1-90946722-INS-1891 | chr1-90946723-INS-1891 |
| chr1-90953096-INS-82 | chr1-90953096-INS-90 |
| chr1-90967226-INS-146 | chr1-90967226-INS-146 |
| chr1-90973569-INS-60 | chr1-90973569-INS-65 |
| chr1-91003074-INS-202 | chr1-91003074-INS-202 |
| chr1-91013525-INS-123 | chr1-91013525-INS-124 |
| chr1-91022874-INS-261 | chr1-91022874-INS-261 |
| chr1-91034239-INS-72 | chr1-91034239-INS-62 |
| chr1-91067239-INS-993 | chr1-91067239-INS-971 |
| chr1-91070688-INS-68 | chr1-91070688-INS-69 |
| chr1-91086881-INS-387 | chr1-91086881-INS-387 |
| chr1-91104171-INS-98 | chr1-91104171-INS-102 |
| chr1-91147795-INS-96 | chr1-91147795-INS-96 |
| chr1-91153995-INS-504 | chr1-91153995-INS-505 |
| chr1-91173869-INS-155 | chr1-91173869-INS-155 |
| chr1-91208275-INS-6439 | chr1-91208276-INS-6439 |
| chr1-91240714-INS-200 | chr1-91240714-INS-200 |
| chr1-91241404-INS-611 | chr1-91241404-INS-627 |
| chr1-91326851-INS-221 | chr1-91326851-INS-222 |
| chr1-91346010-INS-62 | chr1-91346010-INS-62 |
| chr1-91359812-INS-165 | chr1-91359812-INS-207 |
| chr1-91365270-INS-228 | chr1-91365270-INS-228 |
| chr1-91410100-INS-65 | chr1-91410100-INS-65 |
| chr1-91433853-INS-401 | chr1-91433853-INS-401 |
| chr1-91436447-INS-216 | chr1-91436447-INS-208 |
| chr1-91439510-INS-60 | chr1-91439510-INS-64 |
| chr1-91441336-INS-145 | chr1-91441336-INS-145 |
| chr1-91443062-INS-53 | chr1-91443062-INS-53 |
| chr1-91483493-INS-216 | chr1-91483493-INS-216 |
| chr1-91486902-INS-70 | chr1-91486902-INS-70 |
| chr1-91487931-INS-51 | chr1-91487931-INS-51 |
| chr1-91514676-INS-328 | chr1-91514676-INS-328 |
| chr1-91527605-INS-82 | chr1-91527605-INS-123 |
| chr1-91541424-INS-183 | chr1-91541424-INS-184 |
| chr1-91563400-INS-77 | chr1-91563400-INS-81 |
| chr1-91590155-INS-66 | chr1-91590111-INS-108 |
| chr1-91591500-INS-67 | chr1-91591500-INS-63 |
| chr1-91595354-INS-61 | chr1-91595354-INS-61 |

| | |
|---|---|
| chr1-91633060-INS-54 | chr1-91633060-INS-51 |
| chr1-91644241-INS-311 | chr1-91644241-INS-300 |
| chr1-91649557-INS-76 | chr1-91649557-INS-72 |
| chr1-91672153-INS-60 | chr1-91672164-INS-61 |
| chr1-91689252-INS-55 | chr1-91689253-INS-65 |
| chr1-91714757-INS-107 | chr1-91714757-INS-105 |
| chr1-91718074-INS-226 | chr1-91718074-INS-227 |
| chr1-91734987-INS-82 | chr1-91734987-INS-123 |
| chr1-91739656-INS-66 | chr1-91739656-INS-66 |
| chr1-91749208-INS-415 | chr1-91749208-INS-419 |
| chr1-91774380-INS-163 | chr1-91774380-INS-163 |
| chr1-91790193-INS-200 | chr1-91790193-INS-201 |
| chr1-91794648-INS-67 | chr1-91794648-INS-67 |
| chr1-91804842-INS-96 | chr1-91804842-INS-96 |
| chr1-91816730-INS-1461 | chr1-91816731-INS-1461 |
| chr1-91823145-INS-52 | chr1-91823145-INS-52 |
| chr1-91828321-INS-62 | chr1-91828321-INS-62 |
| chr1-91830563-INS-584 | chr1-91830563-INS-587 |
| chr1-91838271-INS-1888 | chr1-91838272-INS-1888 |
| chr1-91842122-INS-104 | chr1-91842122-INS-108 |
| chr1-91843319-INS-59 | chr1-91843319-INS-59 |
| chr1-91909440-INS-81 | chr1-91909440-INS-81 |
| chr1-91956339-INS-57 | chr1-91956339-INS-51 |
| chr1-92063506-INS-56 | chr1-92063506-INS-56 |
| chr1-92068813-INS-132 | chr1-92068813-INS-132 |
| chr1-92080375-INS-113 | chr1-92080363-INS-118 |
| chr1-92084512-INS-387 | chr1-92084512-INS-387 |
| chr1-92104743-INS-208 | chr1-92104743-INS-208 |
| chr1-92126645-INS-116 | chr1-92126645-INS-116 |
| chr1-92153667-INS-53 | chr1-92153667-INS-81 |
| chr1-92178812-INS-64 | chr1-92178811-INS-64 |
| chr1-92201272-INS-52 | chr1-92201254-INS-57 |
| chr1-92231990-INS-51 | chr1-92231990-INS-55 |
| chr1-92235025-INS-389 | chr1-92235025-INS-397 |
| chr1-92245451-INS-481 | chr1-92245451-INS-481 |
| chr1-92247685-INS-136 | chr1-92247682-INS-136 |
| chr1-92264739-INS-55 | chr1-92264739-INS-57 |
| chr1-92290677-INS-90 | chr1-92290677-INS-90 |
| chr1-92318139-INS-118 | chr1-92318139-INS-118 |
| chr1-92355416-INS-203 | chr1-92355416-INS-231 |
| chr1-92387121-INS-214 | chr1-92387121-INS-214 |
| chr1-92404232-INS-70 | chr1-92404244-INS-77 |
| chr1-92405287-INS-170 | chr1-92405288-INS-170 |
| chr1-92445061-INS-214 | chr1-92445061-INS-214 |
| chr1-92447648-INS-90 | chr1-92447648-INS-72 |
| chr1-92448253-INS-376 | chr1-92448253-INS-376 |
| chr1-92448952-INS-59 | chr1-92448952-INS-59 |
| chr1-92452201-INS-52 | chr1-92452201-INS-52 |
| chr1-92463835-INS-611 | chr1-92463835-INS-621 |
| chr1-92476508-INS-580 | chr1-92476508-INS-579 |
| chr1-92485028-INS-2741 | chr1-92485029-INS-2741 |
| chr1-92491578-INS-199 | chr1-92491578-INS-199 |
| chr1-92495559-INS-247 | chr1-92495559-INS-247 |
| chr1-92555720-INS-98 | chr1-92555708-INS-74 |
| chr1-92567270-INS-156 | chr1-92567270-INS-156 |
| chr1-92585477-INS-1004 | chr1-92585478-INS-1005 |
| chr1-92588277-INS-138 | chr1-92588277-INS-154 |
| chr1-92736927-INS-334 | chr1-92736926-INS-348 |
| chr1-92742001-INS-54 | chr1-92742001-INS-54 |
| chr1-92819731-INS-78 | chr1-92819731-INS-78 |
| chr1-92877724-INS-7112 | chr1-92877725-INS-7112 |
| chr1-92889744-INS-58 | chr1-92889744-INS-90 |
| chr1-92903003-INS-53 | chr1-92903003-INS-54 |
| chr1-92909451-INS-139 | chr1-92909451-INS-139 |
| chr1-92914759-INS-498 | chr1-92914759-INS-498 |
| chr1-92917896-INS-2931 | chr1-92917897-INS-2931 |
| chr1-92930711-INS-179 | chr1-92930729-INS-140 |
| chr1-92937333-INS-1892 | chr1-92937334-INS-1892 |
| chr1-92940197-INS-65 | chr1-92940197-INS-65 |
| chr1-93006976-INS-1556 | chr1-93006977-INS-1556 |
| chr1-93130750-INS-188 | chr1-93130774-INS-141 |
| chr1-93132878-INS-94 | chr1-93132878-INS-94 |
| chr1-93181236-INS-69 | chr1-93181236-INS-55 |
| chr1-93252702-INS-209 | chr1-93252702-INS-209 |
| chr1-93307191-INS-169 | chr1-93307191-INS-168 |
| chr1-93316997-INS-96 | chr1-93317014-INS-96 |
| chr1-93363454-INS-1018 | chr1-93363455-INS-1019 |

| | |
|---|---|
| chr1-93395609-INS-218 | chr1-93395609-INS-219 |
| chr1-93430330-INS-86 | chr1-93430319-INS-90 |
| chr1-93464104-INS-207 | chr1-93464104-INS-208 |
| chr1-93475239-INS-179 | chr1-93475239-INS-181 |
| chr1-93478680-INS-190 | chr1-93478680-INS-190 |
| chr1-93492698-INS-205 | chr1-93492698-INS-204 |
| chr1-93530499-INS-220 | chr1-93530499-INS-192 |
| chr1-93554576-INS-225 | chr1-93554576-INS-225 |
| chr1-93571726-INS-64 | chr1-93571748-INS-64 |
| chr1-93573634-INS-74 | chr1-93573634-INS-60 |
| chr1-93606886-INS-64 | chr1-93606886-INS-64 |
| chr1-93684648-INS-158 | chr1-93684648-INS-160 |
| chr1-93692398-INS-95 | chr1-93692398-INS-105 |
| chr1-93703844-INS-127 | chr1-93703842-INS-126 |
| chr1-93725583-INS-144 | chr1-93725583-INS-144 |
| chr1-93921779-INS-173 | chr1-93921779-INS-172 |
| chr1-93965489-INS-85 | chr1-93965489-INS-80 |
| chr1-93997839-INS-196 | chr1-93997839-INS-196 |
| chr1-94009201-INS-60 | chr1-94009201-INS-60 |
| chr1-94057102-INS-603 | chr1-94057102-INS-602 |
| chr1-94057756-INS-207 | chr1-94057756-INS-205 |
| chr1-94066708-INS-62 | chr1-94066708-INS-63 |
| chr1-94101660-INS-800 | chr1-94101660-INS-800 |
| chr1-94104727-INS-60 | chr1-94104727-INS-60 |
| chr1-94139152-INS-212 | chr1-94139152-INS-200 |
| chr1-94161350-INS-677 | chr1-94161350-INS-677 |
| chr1-94177695-INS-69 | chr1-94177695-INS-69 |
| chr1-94181463-INS-313 | chr1-94181463-INS-317 |
| chr1-94184058-INS-864 | chr1-94184058-INS-865 |
| chr1-94230136-INS-4969 | chr1-94230137-INS-4998 |
| chr1-94233855-INS-83 | chr1-94233849-INS-89 |
| chr1-94237849-INS-175 | chr1-94237849-INS-175 |
| chr1-94243301-INS-202 | chr1-94243301-INS-202 |
| chr1-94246655-INS-817 | chr1-94246655-INS-817 |
| chr1-94249916-INS-523 | chr1-94249916-INS-522 |
| chr1-94308633-INS-2689 | chr1-94308634-INS-2689 |
| chr1-94365364-INS-65 | chr1-94365391-INS-79 |
| chr1-94375322-INS-253 | chr1-94375322-INS-253 |
| chr1-94381364-INS-91 | chr1-94381369-INS-92 |
| chr1-94384496-INS-124 | chr1-94384496-INS-124 |
| chr1-94410650-INS-81 | chr1-94410640-INS-78 |
| chr1-94443497-INS-7179 | chr1-94443498-INS-7179 |
| chr1-94450037-INS-657 | chr1-94450037-INS-657 |
| chr1-94456057-INS-132 | chr1-94456057-INS-137 |
| chr1-94478346-INS-861 | chr1-94478346-INS-861 |
| chr1-94486713-INS-54 | chr1-94486713-INS-66 |
| chr1-94496020-INS-940 | chr1-94496020-INS-939 |
| chr1-94497760-INS-6254 | chr1-94497761-INS-6258 |
| chr1-94515274-INS-620 | chr1-94515274-INS-623 |
| chr1-94515945-INS-6401 | chr1-94515946-INS-6402 |
| chr1-94561207-INS-954 | chr1-94561207-INS-964 |
| chr1-94585545-INS-471 | chr1-94585545-INS-471 |
| chr1-94598812-INS-89 | chr1-94598812-INS-85 |
| chr1-94600088-INS-54 | chr1-94600088-INS-62 |
| chr1-94613423-INS-109 | chr1-94613423-INS-109 |
| chr1-94668345-INS-64 | chr1-94668345-INS-72 |
| chr1-94686127-INS-944 | chr1-94686127-INS-950 |
| chr1-94689881-INS-414 | chr1-94689882-INS-415 |
| chr1-94693462-INS-143 | chr1-94693462-INS-144 |
| chr1-94703737-INS-59 | chr1-94703737-INS-61 |
| chr1-94720955-INS-424 | chr1-94720955-INS-423 |
| chr1-94753894-INS-60 | chr1-94753894-INS-60 |
| chr1-94762408-INS-82 | chr1-94762408-INS-82 |
| chr1-94763605-INS-69 | chr1-94763605-INS-68 |
| chr1-94777276-INS-82 | chr1-94777276-INS-86 |
| chr1-94790797-INS-60 | chr1-94790797-INS-50 |
| chr1-94799887-INS-828 | chr1-94799887-INS-828 |
| chr1-94832634-INS-150 | chr1-94832634-INS-131 |
| chr1-94854852-INS-1890 | chr1-94854853-INS-1890 |
| chr1-94876125-INS-6624 | chr1-94876126-INS-6625 |
| chr1-94882788-INS-414 | chr1-94882786-INS-422 |
| chr1-94885286-INS-478 | chr1-94885286-INS-478 |
| chr1-94890797-INS-212 | chr1-94890797-INS-210 |
| chr1-94922239-INS-601 | chr1-94922239-INS-601 |
| chr1-94934835-INS-3841 | chr1-94934836-INS-3841 |
| chr1-94980651-INS-4542 | chr1-94980652-INS-4542 |
| chr1-94985019-INS-6856 | chr1-94985020-INS-6856 |

| | |
|---|---|
| chr1-94985046-INS-353 | chr1-94985051-INS-357 |
| chr1-95049476-INS-53 | chr1-95049476-INS-53 |
| chr1-95069240-INS-6838 | chr1-95069241-INS-6839 |
| chr1-95089139-INS-125 | chr1-95089139-INS-125 |
| chr1-95093078-INS-197 | chr1-95093078-INS-197 |
| chr1-95133496-INS-78 | chr1-95133496-INS-78 |
| chr1-95139669-INS-96 | chr1-95139669-INS-100 |
| chr1-95143624-INS-459 | chr1-95143624-INS-457 |
| chr1-95176196-INS-55 | chr1-95176196-INS-54 |
| chr1-95236727-INS-88 | chr1-95236727-INS-82 |
| chr1-95273215-INS-1757 | chr1-95273216-INS-1757 |
| chr1-95304548-INS-67 | chr1-95304568-INS-73 |
| chr1-95331241-INS-5663 | chr1-95331242-INS-5663 |
| chr1-95344101-INS-95 | chr1-95344101-INS-95 |
| chr1-95353825-INS-81 | chr1-95353831-INS-84 |
| chr1-95371444-INS-4800 | chr1-95371445-INS-4800 |
| chr1-95372140-INS-255 | chr1-95372135-INS-254 |
| chr1-95377867-INS-57 | chr1-95377867-INS-57 |
| chr1-95397819-INS-595 | chr1-95397819-INS-595 |
| chr1-95420293-INS-192 | chr1-95420293-INS-192 |
| chr1-95438620-INS-1888 | chr1-95438621-INS-1888 |
| chr1-95469990-INS-1605 | chr1-95469991-INS-1609 |
| chr1-95504610-INS-390 | chr1-95504610-INS-390 |
| chr1-95530945-INS-527 | chr1-95530945-INS-527 |
| chr1-95540116-INS-176 | chr1-95540124-INS-183 |
| chr1-95580724-INS-59 | chr1-95580724-INS-59 |
| chr1-95643324-INS-7704 | chr1-95643325-INS-7705 |
| chr1-95655391-INS-6225 | chr1-95655392-INS-6225 |
| chr1-95655832-INS-222 | chr1-95655832-INS-222 |
| chr1-95658736-INS-2346 | chr1-95658737-INS-2346 |
| chr1-95682920-INS-419 | chr1-95682920-INS-419 |
| chr1-95695453-INS-4347 | chr1-95695454-INS-4347 |
| chr1-95722226-INS-297 | chr1-95722226-INS-282 |
| chr1-95743686-INS-266 | chr1-95743686-INS-266 |
| chr1-95753532-INS-353 | chr1-95753532-INS-353 |
| chr1-95785590-INS-50 | chr1-95785590-INS-78 |
| chr1-95801654-INS-6061 | chr1-95801655-INS-6067 |
| chr1-95802346-INS-6341 | chr1-95802347-INS-6341 |
| chr1-95818892-INS-111 | chr1-95818890-INS-114 |
| chr1-95844755-INS-185 | chr1-95844755-INS-185 |
| chr1-95854926-INS-72 | chr1-95854918-INS-60 |
| chr1-95868734-INS-74 | chr1-95868734-INS-75 |
| chr1-95874561-INS-1336 | chr1-95874562-INS-1336 |
| chr1-95889652-INS-1028 | chr1-95889653-INS-1028 |
| chr1-95908922-INS-86 | chr1-95908922-INS-86 |
| chr1-95913747-INS-112 | chr1-95913747-INS-112 |
| chr1-96024451-INS-4752 | chr1-96024452-INS-4752 |
| chr1-96035183-INS-89 | chr1-96035183-INS-98 |
| chr1-96055644-INS-120 | chr1-96055635-INS-89 |
| chr1-96060562-INS-58 | chr1-96060562-INS-58 |
| chr1-96089687-INS-500 | chr1-96089687-INS-500 |
| chr1-96146530-INS-1300 | chr1-96146531-INS-1300 |
| chr1-96157564-INS-437 | chr1-96157564-INS-458 |
| chr1-96159168-INS-91 | chr1-96159168-INS-84 |
| chr1-96182050-INS-90 | chr1-96182050-INS-90 |
| chr1-96188924-INS-65 | chr1-96188922-INS-62 |
| chr1-96194579-INS-161 | chr1-96194575-INS-154 |
| chr1-96206322-INS-1943 | chr1-96206321-INS-1924 |
| chr1-96268434-INS-112 | chr1-96268434-INS-112 |
| chr1-96305738-INS-2707 | chr1-96305739-INS-2707 |
| chr1-96312251-INS-55 | chr1-96312251-INS-55 |
| chr1-96376325-INS-93 | chr1-96376325-INS-93 |
| chr1-96381809-INS-325 | chr1-96381809-INS-325 |
| chr1-96424907-INS-753 | chr1-96424908-INS-753 |
| chr1-96432246-INS-506 | chr1-96432246-INS-506 |
| chr1-96436462-INS-64 | chr1-96436463-INS-64 |
| chr1-96460039-INS-975 | chr1-96460039-INS-975 |
| chr1-96473011-INS-103 | chr1-96473011-INS-99 |
| chr1-96480431-INS-503 | chr1-96480431-INS-504 |
| chr1-96483450-INS-471 | chr1-96483450-INS-471 |
| chr1-96498571-INS-60 | chr1-96498571-INS-60 |
| chr1-96517724-INS-88 | chr1-96517739-INS-86 |
| chr1-96538939-INS-187 | chr1-96538939-INS-192 |
| chr1-96565140-INS-5158 | chr1-96565141-INS-5158 |
| chr1-96571327-INS-3545 | chr1-96571328-INS-3547 |
| chr1-96580815-INS-139 | chr1-96580812-INS-163 |
| chr1-96582180-INS-107 | chr1-96582180-INS-107 |

| | |
|---|---|
| chr1-96611271-INS-205 | chr1-96611271-INS-206 |
| chr1-96617865-INS-60 | chr1-96617835-INS-54 |
| chr1-96648328-INS-4614 | chr1-96648329-INS-4614 |
| chr1-96660567-INS-276 | chr1-96660567-INS-312 |
| chr1-96666623-INS-844 | chr1-96666623-INS-847 |
| chr1-96668730-INS-2033 | chr1-96668731-INS-2034 |
| chr1-96693075-INS-6516 | chr1-96693076-INS-6516 |
| chr1-96695172-INS-195 | chr1-96695165-INS-238 |
| chr1-96697333-INS-109 | chr1-96697333-INS-99 |
| chr1-96702084-INS-374 | chr1-96702084-INS-376 |
| chr1-96747990-INS-56 | chr1-96747990-INS-56 |
| chr1-96751606-INS-181 | chr1-96751606-INS-182 |
| chr1-96754719-INS-53 | chr1-96754721-INS-53 |
| chr1-96783076-INS-52 | chr1-96783074-INS-58 |
| chr1-96784350-INS-52 | chr1-96784350-INS-56 |
| chr1-96787827-INS-592 | chr1-96787827-INS-593 |
| chr1-96791682-INS-52 | chr1-96791682-INS-54 |
| chr1-96826766-INS-6187 | chr1-96826767-INS-6187 |
| chr1-96858726-INS-52 | chr1-96858723-INS-78 |
| chr1-96894744-INS-60 | chr1-96894744-INS-60 |
| chr1-96929422-INS-7097 | chr1-96929423-INS-7097 |
| chr1-96933876-INS-54 | chr1-96933876-INS-54 |
| chr1-96934736-INS-1286 | chr1-96934737-INS-1286 |
| chr1-96949682-INS-75 | chr1-96949694-INS-57 |
| chr1-96966819-INS-232 | chr1-96966819-INS-232 |
| chr1-96983579-INS-196 | chr1-96983575-INS-208 |
| chr1-97030059-INS-685 | chr1-97030059-INS-686 |
| chr1-97114534-INS-179 | chr1-97114532-INS-179 |
| chr1-97202495-INS-458 | chr1-97202495-INS-458 |
| chr1-97209320-INS-574 | chr1-97209320-INS-574 |
| chr1-97257834-INS-7709 | chr1-97257835-INS-7709 |
| chr1-97270766-INS-563 | chr1-97270766-INS-563 |
| chr1-97274740-INS-98 | chr1-97274740-INS-98 |
| chr1-97280235-INS-61 | chr1-97280235-INS-56 |
| chr1-97325996-INS-199 | chr1-97325996-INS-199 |
| chr1-97361838-INS-4567 | chr1-97361839-INS-4567 |
| chr1-97370898-INS-418 | chr1-97370898-INS-418 |
| chr1-97405243-INS-284 | chr1-97405243-INS-278 |
| chr1-97407470-INS-165 | chr1-97407470-INS-157 |
| chr1-97412999-INS-326 | chr1-97412999-INS-326 |
| chr1-97435829-INS-1347 | chr1-97435830-INS-1348 |
| chr1-97485856-INS-58 | chr1-97485857-INS-88 |
| chr1-97493304-INS-6387 | chr1-97493305-INS-6391 |
| chr1-97512734-INS-125 | chr1-97512734-INS-125 |
| chr1-97545428-INS-560 | chr1-97545428-INS-560 |
| chr1-97604035-INS-192 | chr1-97604005-INS-189 |
| chr1-97622259-INS-196 | chr1-97622259-INS-198 |
| chr1-97656325-INS-182 | chr1-97656325-INS-182 |
| chr1-97702513-INS-875 | chr1-97702513-INS-875 |
| chr1-97717943-INS-152 | chr1-97717943-INS-114 |
| chr1-97742504-INS-433 | chr1-97742504-INS-431 |
| chr1-97750332-INS-69 | chr1-97750313-INS-72 |
| chr1-97764683-INS-207 | chr1-97764683-INS-207 |
| chr1-97765540-INS-213 | chr1-97765540-INS-213 |
| chr1-97783327-INS-800 | chr1-97783327-INS-810 |
| chr1-97792785-INS-68 | chr1-97792796-INS-58 |
| chr1-97800305-INS-113 | chr1-97800305-INS-113 |
| chr1-97802657-INS-324 | chr1-97802657-INS-292 |
| chr1-97831274-INS-68 | chr1-97831274-INS-90 |
| chr1-97833623-INS-6438 | chr1-97833624-INS-6440 |
| chr1-97839558-INS-59 | chr1-97839558-INS-59 |
| chr1-97842920-INS-339 | chr1-97842927-INS-308 |
| chr1-97855568-INS-212 | chr1-97855568-INS-213 |
| chr1-97900011-INS-80 | chr1-97900011-INS-105 |
| chr1-97902127-INS-52 | chr1-97902127-INS-52 |
| chr1-97928142-INS-313 | chr1-97928144-INS-330 |
| chr1-97938376-INS-85 | chr1-97938386-INS-95 |
| chr1-97958704-INS-2262 | chr1-97958705-INS-2262 |
| chr1-98054282-INS-2490 | chr1-98054283-INS-2490 |
| chr1-98054483-INS-401 | chr1-98054483-INS-401 |
| chr1-98126715-INS-105 | chr1-98126715-INS-105 |
| chr1-98201280-INS-415 | chr1-98201280-INS-413 |
| chr1-98253402-INS-74 | chr1-98253402-INS-74 |
| chr1-98260159-INS-810 | chr1-98260160-INS-800 |
| chr1-98262702-INS-554 | chr1-98262702-INS-554 |
| chr1-98264484-INS-52 | chr1-98264484-INS-52 |
| chr1-98282086-INS-51 | chr1-98282086-INS-51 |

| | |
|---|---|
| chr1-98346945-INS-201 | chr1-98346945-INS-201 |
| chr1-98410383-INS-1893 | chr1-98410384-INS-1893 |
| chr1-98450621-INS-7117 | chr1-98450622-INS-7117 |
| chr1-98461826-INS-74 | chr1-98461826-INS-71 |
| chr1-98463322-INS-72 | chr1-98463322-INS-52 |
| chr1-98464180-INS-6675 | chr1-98464181-INS-6675 |
| chr1-98469813-INS-1467 | chr1-98469814-INS-1467 |
| chr1-98594536-INS-63 | chr1-98594537-INS-89 |
| chr1-98607188-INS-78 | chr1-98607188-INS-82 |
| chr1-98613884-INS-825 | chr1-98613884-INS-825 |
| chr1-98618204-INS-827 | chr1-98618204-INS-849 |
| chr1-98625187-INS-124 | chr1-98625206-INS-136 |
| chr1-98630867-INS-735 | chr1-98630866-INS-735 |
| chr1-98637147-INS-106 | chr1-98637147-INS-106 |
| chr1-98638378-INS-324 | chr1-98638378-INS-324 |
| chr1-98655381-INS-7152 | chr1-98655382-INS-7153 |
| chr1-98685320-INS-52 | chr1-98685320-INS-56 |
| chr1-98687197-INS-156 | chr1-98687194-INS-156 |
| chr1-98690711-INS-72 | chr1-98690711-INS-104 |
| chr1-98729679-INS-75 | chr1-98729679-INS-85 |
| chr1-98738793-INS-53 | chr1-98738807-INS-53 |
| chr1-98756904-INS-60 | chr1-98756904-INS-60 |
| chr1-98759478-INS-593 | chr1-98759478-INS-595 |
| chr1-98773942-INS-201 | chr1-98773942-INS-201 |
| chr1-98780480-INS-2447 | chr1-98780481-INS-2444 |
| chr1-98803508-INS-396 | chr1-98803508-INS-396 |
| chr1-98844656-INS-229 | chr1-98844656-INS-229 |
| chr1-98848537-INS-77 | chr1-98848537-INS-77 |
| chr1-98856866-INS-3394 | chr1-98856867-INS-3394 |
| chr1-98892637-INS-64 | chr1-98892637-INS-64 |
| chr1-98893940-INS-104 | chr1-98893940-INS-107 |
| chr1-98902648-INS-75 | chr1-98902648-INS-75 |
| chr1-98908475-INS-1166 | chr1-98908476-INS-1168 |
| chr1-98947165-INS-569 | chr1-98947165-INS-569 |
| chr1-98949752-INS-401 | chr1-98949752-INS-401 |
| chr1-98962301-INS-55 | chr1-98962304-INS-60 |
| chr1-98964173-INS-707 | chr1-98964173-INS-707 |
| chr1-98973694-INS-120 | chr1-98973694-INS-120 |
| chr1-98979545-INS-51 | chr1-98979545-INS-51 |
| chr1-99053597-INS-300 | chr1-99053597-INS-300 |
| chr1-99080852-INS-82 | chr1-99080881-INS-53 |
| chr1-99082486-INS-56 | chr1-99082486-INS-58 |
| chr1-99097194-INS-57 | chr1-99097198-INS-59 |
| chr1-99099740-INS-189 | chr1-99099740-INS-186 |
| chr1-99100019-INS-995 | chr1-99100019-INS-995 |
| chr1-99157457-INS-4023 | chr1-99157458-INS-4023 |
| chr1-99187626-INS-2709 | chr1-99187627-INS-2709 |
| chr1-99224259-INS-201 | chr1-99224259-INS-201 |
| chr1-99269590-INS-522 | chr1-99269567-INS-512 |
| chr1-99279526-INS-208 | chr1-99279526-INS-208 |
| chr1-99282573-INS-1599 | chr1-99282574-INS-1599 |
| chr1-99282930-INS-285 | chr1-99282930-INS-299 |
| chr1-99295164-INS-74 | chr1-99295173-INS-65 |
| chr1-99317887-INS-2664 | chr1-99317888-INS-2664 |
| chr1-99327106-INS-52 | chr1-99327106-INS-54 |
| chr1-99327138-INS-520 | chr1-99327138-INS-526 |
| chr1-99354431-INS-63 | chr1-99354458-INS-50 |
| chr1-99356889-INS-105 | chr1-99356889-INS-105 |
| chr1-99368809-INS-89 | chr1-99368809-INS-89 |
| chr1-99371412-INS-57 | chr1-99371412-INS-57 |
| chr1-99379148-INS-133 | chr1-99379148-INS-140 |
| chr1-99399583-INS-401 | chr1-99399583-INS-401 |
| chr1-99406696-INS-484 | chr1-99406696-INS-484 |
| chr1-99412760-INS-6438 | chr1-99412761-INS-6438 |
| chr1-99431902-INS-60 | chr1-99431902-INS-60 |
| chr1-99444140-INS-69 | chr1-99444140-INS-69 |
| chr1-99451296-INS-58 | chr1-99451319-INS-68 |
| chr1-99459523-INS-215 | chr1-99459537-INS-234 |
| chr1-99482485-INS-89 | chr1-99482485-INS-89 |
| chr1-99502178-INS-1888 | chr1-99502179-INS-1888 |
| chr1-99537606-INS-64 | chr1-99537587-INS-67 |
| chr1-99552034-INS-130 | chr1-99552034-INS-130 |
| chr1-99553711-INS-1685 | chr1-99553712-INS-1685 |
| chr1-99601988-INS-261 | chr1-99601988-INS-261 |
| chr1-99607011-INS-68 | chr1-99607011-INS-66 |
| chr1-99608882-INS-323 | chr1-99608882-INS-323 |
| chr1-99615093-INS-390 | chr1-99615084-INS-404 |

| | |
|---|---|
| chr1-99642320-INS-6271 | chr1-99642321-INS-6271 |
| chr1-99647142-INS-55 | chr1-99647142-INS-55 |
| chr1-99667500-INS-1136 | chr1-99667501-INS-1136 |
| chr1-99675218-INS-54 | chr1-99675218-INS-54 |
| chr1-99705162-INS-566 | chr1-99705162-INS-566 |
| chr1-99715814-INS-64 | chr1-99715815-INS-68 |
| chr1-99717227-INS-876 | chr1-99717227-INS-875 |
| chr1-99721729-INS-639 | chr1-99721729-INS-639 |
| chr1-99725308-INS-64 | chr1-99725309-INS-56 |
| chr1-99726707-INS-369 | chr1-99726707-INS-374 |
| chr1-99728719-INS-1085 | chr1-99728719-INS-1084 |
| chr1-99751791-INS-200 | chr1-99751791-INS-200 |
| chr1-99758899-INS-56 | chr1-99758899-INS-54 |
| chr1-99815320-INS-52 | chr1-99815320-INS-52 |
| chr1-99835964-INS-230 | chr1-99835964-INS-229 |
| chr1-99857533-INS-59 | chr1-99857533-INS-56 |
| chr1-99886357-INS-54 | chr1-99886357-INS-54 |
| chr1-99908721-INS-54 | chr1-99908721-INS-54 |
| chr1-99915980-INS-2182 | chr1-99915981-INS-2182 |
| chr1-99915981-INS-324 | chr1-99915981-INS-323 |
| chr1-99922682-INS-7031 | chr1-99922683-INS-7038 |
| chr1-99935442-INS-393 | chr1-99935442-INS-393 |
| chr1-99937201-INS-60 | chr1-99937176-INS-105 |
| chr1-99975942-INS-62 | chr1-99975950-INS-61 |
| chr1-99977893-INS-1819 | chr1-99977894-INS-1818 |
| chr1-100010906-INS-56 | chr1-100010906-INS-56 |
| chr1-100012664-INS-76 | chr1-100012677-INS-60 |
| chr1-100018196-INS-875 | chr1-100018196-INS-875 |
| chr1-100036325-INS-604 | chr1-100036325-INS-604 |
| chr1-100065446-INS-432 | chr1-100065446-INS-439 |
| chr1-100072716-INS-2243 | chr1-100072718-INS-2240 |
| chr1-100090274-INS-4766 | chr1-100090275-INS-4768 |
| chr1-100094631-INS-76 | chr1-100094631-INS-80 |
| chr1-100174903-INS-87 | chr1-100174903-INS-84 |
| chr1-100184260-INS-491 | chr1-100184260-INS-491 |
| chr1-100194904-INS-54 | chr1-100194904-INS-52 |
| chr1-100228448-INS-166 | chr1-100228434-INS-174 |
| chr1-100255591-INS-290 | chr1-100255591-INS-290 |
| chr1-100257098-INS-62 | chr1-100257094-INS-66 |
| chr1-100268477-INS-72 | chr1-100268477-INS-52 |
| chr1-100281226-INS-445 | chr1-100281226-INS-445 |
| chr1-100288796-INS-212 | chr1-100288799-INS-213 |
| chr1-100289694-INS-102 | chr1-100289694-INS-106 |
| chr1-100306675-INS-112 | chr1-100306675-INS-118 |
| chr1-100327774-INS-96 | chr1-100327782-INS-60 |
| chr1-100350289-INS-54 | chr1-100350319-INS-51 |
| chr1-100354822-INS-875 | chr1-100354822-INS-873 |
| chr1-100365444-INS-613 | chr1-100365444-INS-613 |
| chr1-100377960-INS-111 | chr1-100377960-INS-123 |
| chr1-100420184-INS-417 | chr1-100420185-INS-415 |
| chr1-100424200-INS-762 | chr1-100424200-INS-763 |
| chr1-100427909-INS-54 | chr1-100427909-INS-54 |
| chr1-100459129-INS-285 | chr1-100459129-INS-288 |
| chr1-100484149-INS-535 | chr1-100484149-INS-536 |
| chr1-100515083-INS-1118 | chr1-100515084-INS-1115 |
| chr1-100535173-INS-6373 | chr1-100535174-INS-6372 |
| chr1-100560297-INS-51 | chr1-100560298-INS-81 |
| chr1-100590968-INS-54 | chr1-100590968-INS-54 |
| chr1-100600963-INS-152 | chr1-100600963-INS-152 |
| chr1-100611898-INS-6650 | chr1-100611899-INS-6650 |
| chr1-100642645-INS-6370 | chr1-100642648-INS-6368 |
| chr1-100648783-INS-131 | chr1-100648784-INS-91 |
| chr1-100656844-INS-74 | chr1-100656846-INS-60 |
| chr1-100658017-INS-651 | chr1-100658017-INS-671 |
| chr1-100677176-INS-1886 | chr1-100677177-INS-1886 |
| chr1-100718227-INS-623 | chr1-100718227-INS-620 |
| chr1-100737714-INS-74 | chr1-100737714-INS-74 |
| chr1-100753287-INS-52 | chr1-100753248-INS-62 |
| chr1-100768310-INS-1170 | chr1-100768311-INS-1176 |
| chr1-100832989-INS-367 | chr1-100832989-INS-367 |
| chr1-100839877-INS-85 | chr1-100839871-INS-66 |
| chr1-100870020-INS-630 | chr1-100870020-INS-630 |
| chr1-100879534-INS-986 | chr1-100879534-INS-987 |
| chr1-100887137-INS-405 | chr1-100887159-INS-374 |
| chr1-100920537-INS-1236 | chr1-100920538-INS-1236 |
| chr1-100922337-INS-208 | chr1-100922349-INS-208 |
| chr1-100933765-INS-287 | chr1-100933765-INS-283 |

| | |
|---|---|
| chr1-100955403-INS-55 | chr1-100955403-INS-55 |
| chr1-100957314-INS-69 | chr1-100957300-INS-93 |
| chr1-100966171-INS-52 | chr1-100966171-INS-52 |
| chr1-100990071-INS-1251 | chr1-100990072-INS-1253 |
| chr1-101006588-INS-328 | chr1-101006588-INS-330 |
| chr1-101045214-INS-784 | chr1-101045214-INS-784 |
| chr1-101068206-INS-100 | chr1-101068206-INS-104 |
| chr1-101104466-INS-6186 | chr1-101104467-INS-6185 |
| chr1-101140386-INS-1378 | chr1-101140387-INS-1376 |
| chr1-101148996-INS-7086 | chr1-101148997-INS-7086 |
| chr1-101201659-INS-6362 | chr1-101201660-INS-6362 |
| chr1-101239989-INS-146 | chr1-101239988-INS-174 |
| chr1-101293191-INS-64 | chr1-101293191-INS-64 |
| chr1-101379416-INS-7118 | chr1-101379417-INS-7118 |
| chr1-101381209-INS-665 | chr1-101381209-INS-665 |
| chr1-101389742-INS-100 | chr1-101389754-INS-92 |
| chr1-101397569-INS-7185 | chr1-101397570-INS-7185 |
| chr1-101409547-INS-3965 | chr1-101409517-INS-3973 |
| chr1-101449947-INS-3422 | chr1-101449948-INS-3428 |
| chr1-101491562-INS-1818 | chr1-101491563-INS-1814 |
| chr1-101494405-INS-1211 | chr1-101494406-INS-1212 |
| chr1-101506127-INS-56 | chr1-101506119-INS-68 |
| chr1-101515205-INS-60 | chr1-101515205-INS-60 |
| chr1-101521544-INS-199 | chr1-101521544-INS-199 |
| chr1-101523222-INS-200 | chr1-101523222-INS-200 |
| chr1-101568620-INS-1048 | chr1-101568621-INS-1054 |
| chr1-101575701-INS-86 | chr1-101575701-INS-62 |
| chr1-101615021-INS-328 | chr1-101615021-INS-328 |
| chr1-101621810-INS-195 | chr1-101621810-INS-200 |
| chr1-101625266-INS-102 | chr1-101625266-INS-68 |
| chr1-101638961-INS-212 | chr1-101638943-INS-212 |
| chr1-101649501-INS-211 | chr1-101649501-INS-211 |
| chr1-101651141-INS-6333 | chr1-101651142-INS-6322 |
| chr1-101666560-INS-2706 | chr1-101666561-INS-2705 |
| chr1-101673690-INS-1330 | chr1-101673691-INS-1330 |
| chr1-101675057-INS-421 | chr1-101675057-INS-421 |
| chr1-101711440-INS-56 | chr1-101711412-INS-96 |
| chr1-101727272-INS-484 | chr1-101727272-INS-484 |
| chr1-101738726-INS-88 | chr1-101738767-INS-76 |
| chr1-101751054-INS-100 | chr1-101751055-INS-98 |
| chr1-101791201-INS-141 | chr1-101791201-INS-141 |
| chr1-101796320-INS-6547 | chr1-101796321-INS-6547 |
| chr1-101816630-INS-417 | chr1-101816630-INS-417 |
| chr1-101869333-INS-109 | chr1-101869333-INS-67 |
| chr1-101873743-INS-432 | chr1-101873743-INS-432 |
| chr1-101898194-INS-70 | chr1-101898194-INS-70 |
| chr1-101927321-INS-148 | chr1-101927321-INS-148 |
| chr1-101933140-INS-97 | chr1-101933140-INS-97 |
| chr1-101965720-INS-55 | chr1-101965720-INS-56 |
| chr1-101972453-INS-6658 | chr1-101972454-INS-6658 |
| chr1-101985052-INS-111 | chr1-101985053-INS-118 |
| chr1-101996964-INS-99 | chr1-101996964-INS-99 |
| chr1-102035492-INS-74 | chr1-102035507-INS-80 |
| chr1-102038575-INS-253 | chr1-102038578-INS-253 |
| chr1-102050723-INS-409 | chr1-102050723-INS-409 |
| chr1-102086772-INS-50 | chr1-102086772-INS-56 |
| chr1-102088266-INS-6649 | chr1-102088267-INS-6649 |
| chr1-102098385-INS-71 | chr1-102098385-INS-71 |
| chr1-102127382-INS-507 | chr1-102127382-INS-507 |
| chr1-102140664-INS-62 | chr1-102140664-INS-65 |
| chr1-102151821-INS-144 | chr1-102151821-INS-144 |
| chr1-102207666-INS-174 | chr1-102207666-INS-174 |
| chr1-102215481-INS-134 | chr1-102215481-INS-146 |
| chr1-102240402-INS-6776 | chr1-102240403-INS-6776 |
| chr1-102244426-INS-74 | chr1-102244426-INS-74 |
| chr1-102249073-INS-157 | chr1-102249078-INS-199 |
| chr1-102300238-INS-686 | chr1-102300238-INS-686 |
| chr1-102324241-INS-447 | chr1-102324241-INS-443 |
| chr1-102337275-INS-64 | chr1-102337275-INS-68 |
| chr1-102344704-INS-66 | chr1-102344704-INS-69 |
| chr1-102374425-INS-84 | chr1-102374411-INS-72 |
| chr1-102379431-INS-224 | chr1-102379431-INS-233 |
| chr1-102387073-INS-106 | chr1-102387073-INS-106 |
| chr1-102398687-INS-72 | chr1-102398687-INS-72 |
| chr1-102409294-INS-1379 | chr1-102409295-INS-1379 |
| chr1-102410413-INS-6463 | chr1-102410414-INS-6463 |
| chr1-102431535-INS-88 | chr1-102431524-INS-64 |

| | |
|---|---|
| chr1-102436343-INS-72 | chr1-102436343-INS-72 |
| chr1-102499161-INS-6226 | chr1-102499162-INS-6225 |
| chr1-102504741-INS-156 | chr1-102504759-INS-146 |
| chr1-102513982-INS-92 | chr1-102513982-INS-92 |
| chr1-102522255-INS-2323 | chr1-102522256-INS-2325 |
| chr1-102537554-INS-52 | chr1-102537555-INS-52 |
| chr1-102546722-INS-306 | chr1-102546722-INS-306 |
| chr1-102548091-INS-2311 | chr1-102548092-INS-2311 |
| chr1-102559329-INS-56 | chr1-102559328-INS-59 |
| chr1-102561635-INS-98 | chr1-102561635-INS-96 |
| chr1-102568282-INS-615 | chr1-102568282-INS-615 |
| chr1-102575150-INS-52 | chr1-102575134-INS-52 |
| chr1-102582576-INS-54 | chr1-102582564-INS-72 |
| chr1-102600558-INS-102 | chr1-102600558-INS-90 |
| chr1-102611707-INS-52 | chr1-102611707-INS-52 |
| chr1-102663851-INS-2425 | chr1-102663854-INS-2422 |
| chr1-102675653-INS-68 | chr1-102675659-INS-96 |
| chr1-102714365-INS-51 | chr1-102714365-INS-66 |
| chr1-102722599-INS-105 | chr1-102722604-INS-106 |
| chr1-102744458-INS-452 | chr1-102744458-INS-451 |
| chr1-102749574-INS-227 | chr1-102749574-INS-227 |
| chr1-102784116-INS-587 | chr1-102784116-INS-588 |
| chr1-102789348-INS-52 | chr1-102789348-INS-52 |
| chr1-102801246-INS-397 | chr1-102801246-INS-397 |
| chr1-102823938-INS-98 | chr1-102823939-INS-116 |
| chr1-102829803-INS-225 | chr1-102829800-INS-226 |
| chr1-102862959-INS-627 | chr1-102862959-INS-627 |
| chr1-102914596-INS-657 | chr1-102914596-INS-657 |
| chr1-102920118-INS-754 | chr1-102920118-INS-752 |
| chr1-102976828-INS-132 | chr1-102976828-INS-132 |
| chr1-102977510-INS-900 | chr1-102977510-INS-864 |
| chr1-102990071-INS-275 | chr1-102990071-INS-275 |
| chr1-103036882-INS-81 | chr1-103036882-INS-89 |
| chr1-103062322-INS-802 | chr1-103062322-INS-784 |
| chr1-103074302-INS-162 | chr1-103074302-INS-162 |
| chr1-103097555-INS-226 | chr1-103097555-INS-204 |
| chr1-103116002-INS-364 | chr1-103116002-INS-365 |
| chr1-103166233-INS-80 | chr1-103166233-INS-82 |
| chr1-103172451-INS-75 | chr1-103172451-INS-50 |
| chr1-103173447-INS-52 | chr1-103173447-INS-66 |
| chr1-103179151-INS-376 | chr1-103179151-INS-376 |
| chr1-103181055-INS-67 | chr1-103181055-INS-67 |
| chr1-103192402-INS-243 | chr1-103192402-INS-242 |
| chr1-103194938-INS-1470 | chr1-103194939-INS-1470 |
| chr1-103201155-INS-199 | chr1-103201155-INS-199 |
| chr1-103263420-INS-1117 | chr1-103263421-INS-1117 |
| chr1-103268912-INS-53 | chr1-103268909-INS-97 |
| chr1-103269494-INS-1136 | chr1-103269495-INS-1137 |
| chr1-103279789-INS-6457 | chr1-103279785-INS-6458 |
| chr1-103296728-INS-174 | chr1-103296723-INS-189 |
| chr1-103299079-INS-104 | chr1-103299092-INS-95 |
| chr1-103316751-INS-166 | chr1-103316751-INS-116 |
| chr1-103320521-INS-286 | chr1-103320543-INS-244 |
| chr1-103328739-INS-55 | chr1-103328739-INS-74 |
| chr1-103416200-INS-1864 | chr1-103416201-INS-1864 |
| chr1-103497967-INS-676 | chr1-103497967-INS-676 |
| chr1-103501981-INS-110 | chr1-103501981-INS-110 |
| chr1-103527337-INS-64 | chr1-103527337-INS-64 |
| chr1-103540085-INS-76 | chr1-103540085-INS-76 |
| chr1-103557652-INS-58 | chr1-103557652-INS-66 |
| chr1-103559822-INS-149 | chr1-103559822-INS-123 |
| chr1-103575577-INS-52 | chr1-103575577-INS-52 |
| chr1-103587822-INS-56 | chr1-103587822-INS-56 |
| chr1-103589309-INS-55 | chr1-103589309-INS-55 |
| chr1-103627377-INS-93 | chr1-103627377-INS-99 |
| chr1-103659149-INS-554 | chr1-103659149-INS-553 |
| chr1-103661681-INS-72 | chr1-103661679-INS-74 |
| chr1-103662403-INS-328 | chr1-103662403-INS-328 |
| chr1-103690123-INS-245 | chr1-103690123-INS-245 |
| chr1-103702050-INS-60 | chr1-103702050-INS-60 |
| chr1-103707125-INS-193 | chr1-103707123-INS-204 |
| chr1-103710797-INS-84 | chr1-103710797-INS-75 |
| chr1-103725674-INS-204 | chr1-103725674-INS-216 |
| chr1-103730465-INS-400 | chr1-103730465-INS-400 |
| chr1-103731870-INS-167 | chr1-103731865-INS-181 |
| chr1-103734745-INS-78 | chr1-103734745-INS-68 |
| chr1-103748796-INS-6144 | chr1-103748797-INS-6144 |

| | |
|---|---|
| chr1-103776483-INS-62 | chr1-103776483-INS-62 |
| chr1-103786069-INS-6429 | chr1-103786070-INS-6429 |
| chr1-103872477-INS-123 | chr1-103872485-INS-132 |
| chr1-103879559-INS-76 | chr1-103879559-INS-76 |
| chr1-103905200-INS-7175 | chr1-103905201-INS-7174 |
| chr1-103955242-INS-6193 | chr1-103955243-INS-6193 |
| chr1-103959071-INS-5924 | chr1-103959072-INS-5925 |
| chr1-103969225-INS-5204 | chr1-103969226-INS-5204 |
| chr1-103977878-INS-64 | chr1-103977834-INS-96 |
| chr1-103989632-INS-170 | chr1-103989632-INS-170 |
| chr1-103992522-INS-71 | chr1-103992522-INS-71 |
| chr1-104051318-INS-75 | chr1-104051330-INS-103 |
| chr1-104132980-INS-3869 | chr1-104132987-INS-3856 |
| chr1-104190494-INS-94 | chr1-104190494-INS-94 |
| chr1-104259091-INS-55 | chr1-104259054-INS-82 |
| chr1-104261955-INS-116 | chr1-104261955-INS-117 |
| chr1-104338437-INS-343 | chr1-104338437-INS-343 |
| chr1-104345857-INS-64 | chr1-104345856-INS-100 |
| chr1-104374370-INS-110 | chr1-104374370-INS-102 |
| chr1-104377906-INS-52 | chr1-104377906-INS-52 |
| chr1-104388220-INS-912 | chr1-104388220-INS-912 |
| chr1-104406358-INS-94 | chr1-104406358-INS-64 |
| chr1-104426892-INS-96 | chr1-104426892-INS-96 |
| chr1-104448946-INS-382 | chr1-104448946-INS-382 |
| chr1-104488055-INS-53 | chr1-104488055-INS-53 |
| chr1-104495950-INS-664 | chr1-104495950-INS-672 |
| chr1-104518227-INS-184 | chr1-104518227-INS-184 |
| chr1-104522046-INS-139 | chr1-104522046-INS-139 |
| chr1-104531776-INS-6367 | chr1-104531777-INS-6367 |
| chr1-104561089-INS-8900 | chr1-104561090-INS-8900 |
| chr1-104595184-INS-57 | chr1-104595184-INS-60 |
| chr1-104596239-INS-716 | chr1-104596239-INS-716 |
| chr1-104597215-INS-280 | chr1-104597215-INS-275 |
| chr1-104603109-INS-87 | chr1-104603109-INS-89 |
| chr1-104626634-INS-104 | chr1-104626590-INS-98 |
| chr1-104688805-INS-54 | chr1-104688805-INS-50 |
| chr1-104713375-INS-50 | chr1-104713375-INS-58 |
| chr1-104739531-INS-124 | chr1-104739530-INS-135 |
| chr1-104741301-INS-111 | chr1-104741301-INS-111 |
| chr1-104761314-INS-1285 | chr1-104761315-INS-1284 |
| chr1-104791572-INS-1314 | chr1-104791573-INS-1314 |
| chr1-104800471-INS-175 | chr1-104800471-INS-176 |
| chr1-104807404-INS-367 | chr1-104807404-INS-367 |
| chr1-104828827-INS-219 | chr1-104828827-INS-219 |
| chr1-104836483-INS-141 | chr1-104836483-INS-139 |
| chr1-104847812-INS-57 | chr1-104847812-INS-58 |
| chr1-104887073-INS-123 | chr1-104887073-INS-110 |
| chr1-104899809-INS-202 | chr1-104899810-INS-203 |
| chr1-104909275-INS-53 | chr1-104909275-INS-53 |
| chr1-104914231-INS-915 | chr1-104914231-INS-915 |
| chr1-104960452-INS-60 | chr1-104960404-INS-62 |
| chr1-104967147-INS-4382 | chr1-104967148-INS-4382 |
| chr1-104967261-INS-83 | chr1-104967261-INS-83 |
| chr1-104992946-INS-86 | chr1-104992904-INS-110 |
| chr1-104997393-INS-53 | chr1-104997393-INS-53 |
| chr1-105001119-INS-342 | chr1-105001118-INS-304 |
| chr1-105007503-INS-7090 | chr1-105007504-INS-7090 |
| chr1-105008656-INS-53 | chr1-105008657-INS-82 |
| chr1-105017909-INS-146 | chr1-105017909-INS-146 |
| chr1-105029761-INS-74 | chr1-105029761-INS-75 |
| chr1-105079821-INS-53 | chr1-105079821-INS-55 |
| chr1-105083061-INS-3013 | chr1-105083062-INS-3013 |
| chr1-105089362-INS-6489 | chr1-105089363-INS-6489 |
| chr1-105091058-INS-60 | chr1-105091055-INS-58 |
| chr1-105107562-INS-224 | chr1-105107548-INS-210 |
| chr1-105125289-INS-6733 | chr1-105125290-INS-6743 |
| chr1-105175325-INS-6883 | chr1-105175326-INS-6884 |
| chr1-105191904-INS-64 | chr1-105191904-INS-64 |
| chr1-105203218-INS-72 | chr1-105203218-INS-74 |
| chr1-105220038-INS-203 | chr1-105220038-INS-203 |
| chr1-105235306-INS-5441 | chr1-105235307-INS-5440 |
| chr1-105250778-INS-6740 | chr1-105250779-INS-6741 |
| chr1-105257548-INS-55 | chr1-105257538-INS-50 |
| chr1-105292284-INS-531 | chr1-105292284-INS-531 |
| chr1-105296482-INS-401 | chr1-105296482-INS-393 |
| chr1-105327722-INS-55 | chr1-105327730-INS-54 |
| chr1-105340168-INS-80 | chr1-105340168-INS-100 |

| | |
|---|---|
| chr1-105364187-INS-91 | chr1-105364187-INS-91 |
| chr1-105372925-INS-9124 | chr1-105372926-INS-9125 |
| chr1-105383516-INS-166 | chr1-105383517-INS-166 |
| chr1-105388407-INS-58 | chr1-105388407-INS-82 |
| chr1-105409654-INS-151 | chr1-105409654-INS-139 |
| chr1-105413763-INS-177 | chr1-105413760-INS-193 |
| chr1-105421245-INS-190 | chr1-105421245-INS-192 |
| chr1-105422782-INS-79 | chr1-105422782-INS-79 |
| chr1-105435775-INS-56 | chr1-105435775-INS-56 |
| chr1-105440295-INS-133 | chr1-105440296-INS-143 |
| chr1-105482753-INS-92 | chr1-105482753-INS-108 |
| chr1-105483577-INS-1352 | chr1-105483578-INS-1357 |
| chr1-105524483-INS-837 | chr1-105524483-INS-837 |
| chr1-105528680-INS-57 | chr1-105528680-INS-59 |
| chr1-105550394-INS-52 | chr1-105550394-INS-52 |
| chr1-105570785-INS-168 | chr1-105570785-INS-170 |
| chr1-105587407-INS-58 | chr1-105587407-INS-58 |
| chr1-105620925-INS-242 | chr1-105620925-INS-244 |
| chr1-105644718-INS-328 | chr1-105644718-INS-328 |
| chr1-105672794-INS-82 | chr1-105672794-INS-54 |
| chr1-105695747-INS-230 | chr1-105695754-INS-220 |
| chr1-105698584-INS-400 | chr1-105698584-INS-399 |
| chr1-105740329-INS-565 | chr1-105740329-INS-564 |
| chr1-105757982-INS-60 | chr1-105757982-INS-68 |
| chr1-105772969-INS-165 | chr1-105772969-INS-167 |
| chr1-105781485-INS-164 | chr1-105781485-INS-170 |
| chr1-105837333-INS-89 | chr1-105837347-INS-73 |
| chr1-105849386-INS-81 | chr1-105849386-INS-81 |
| chr1-105856404-INS-126 | chr1-105856404-INS-138 |
| chr1-105862100-INS-374 | chr1-105862100-INS-374 |
| chr1-105890871-INS-72 | chr1-105890871-INS-72 |
| chr1-105904280-INS-750 | chr1-105904280-INS-750 |
| chr1-105965100-INS-74 | chr1-105965100-INS-74 |
| chr1-105978646-INS-98 | chr1-105978659-INS-62 |
| chr1-105983774-INS-6401 | chr1-105983775-INS-6401 |
| chr1-105999284-INS-673 | chr1-105999284-INS-673 |
| chr1-106009834-INS-90 | chr1-106009834-INS-90 |
| chr1-106089266-INS-67 | chr1-106089272-INS-64 |
| chr1-106096581-INS-222 | chr1-106096541-INS-230 |
| chr1-106114415-INS-173 | chr1-106114415-INS-192 |
| chr1-106137646-INS-55 | chr1-106137646-INS-64 |
| chr1-106139380-INS-66 | chr1-106139382-INS-54 |
| chr1-106192045-INS-86 | chr1-106192045-INS-86 |
| chr1-106258980-INS-267 | chr1-106258980-INS-269 |
| chr1-106271835-INS-52 | chr1-106271835-INS-52 |
| chr1-106306624-INS-107 | chr1-106306601-INS-88 |
| chr1-106352918-INS-1111 | chr1-106352919-INS-1112 |
| chr1-106365456-INS-401 | chr1-106365456-INS-401 |
| chr1-106378539-INS-656 | chr1-106378539-INS-656 |
| chr1-106386918-INS-64 | chr1-106386918-INS-64 |
| chr1-106400846-INS-80 | chr1-106400847-INS-80 |
| chr1-106418929-INS-95 | chr1-106418940-INS-80 |
| chr1-106422181-INS-4546 | chr1-106422182-INS-4546 |
| chr1-106435612-INS-117 | chr1-106435612-INS-117 |
| chr1-106438478-INS-94 | chr1-106438478-INS-94 |
| chr1-106446391-INS-54 | chr1-106446391-INS-54 |
| chr1-106553058-INS-60 | chr1-106553058-INS-54 |
| chr1-106562179-INS-70 | chr1-106562179-INS-68 |
| chr1-106570859-INS-1505 | chr1-106570860-INS-1505 |
| chr1-106612139-INS-1030 | chr1-106612153-INS-1060 |
| chr1-106624086-INS-91 | chr1-106624086-INS-91 |
| chr1-106657568-INS-60 | chr1-106657568-INS-58 |
| chr1-106666241-INS-2213 | chr1-106666242-INS-2213 |
| chr1-106683866-INS-140 | chr1-106683866-INS-188 |
| chr1-106689539-INS-94 | chr1-106689556-INS-99 |
| chr1-106715108-INS-4841 | chr1-106715109-INS-4841 |
| chr1-106721808-INS-275 | chr1-106721808-INS-242 |
| chr1-106733523-INS-148 | chr1-106733523-INS-148 |
| chr1-106746360-INS-57 | chr1-106746343-INS-67 |
| chr1-106749358-INS-1074 | chr1-106749359-INS-1075 |
| chr1-106765958-INS-6310 | chr1-106765959-INS-6312 |
| chr1-106837438-INS-163 | chr1-106837438-INS-156 |
| chr1-106848227-INS-882 | chr1-106848228-INS-882 |
| chr1-106872311-INS-571 | chr1-106872311-INS-568 |
| chr1-106873813-INS-131 | chr1-106873813-INS-106 |
| chr1-106889575-INS-74 | chr1-106889575-INS-75 |
| chr1-106915428-INS-101 | chr1-106915428-INS-101 |

| | |
|---|---|
| chr1-106931202-INS-1995 | chr1-106931209-INS-1989 |
| chr1-106953861-INS-6374 | chr1-106953862-INS-6374 |
| chr1-106963111-INS-197 | chr1-106963111-INS-197 |
| chr1-107001640-INS-805 | chr1-107001640-INS-805 |
| chr1-107005136-INS-271 | chr1-107005136-INS-271 |
| chr1-107018859-INS-109 | chr1-107018859-INS-109 |
| chr1-107054772-INS-204 | chr1-107054793-INS-204 |
| chr1-107087194-INS-498 | chr1-107087194-INS-498 |
| chr1-107117571-INS-300 | chr1-107117571-INS-274 |
| chr1-107120187-INS-995 | chr1-107120187-INS-995 |
| chr1-107169493-INS-243 | chr1-107169493-INS-243 |
| chr1-107192201-INS-3279 | chr1-107192202-INS-3296 |
| chr1-107197393-INS-2891 | chr1-107197394-INS-2883 |
| chr1-107208606-INS-72 | chr1-107208591-INS-50 |
| chr1-107242830-INS-56 | chr1-107242830-INS-58 |
| chr1-107246281-INS-60 | chr1-107246281-INS-60 |
| chr1-107265097-INS-2939 | chr1-107265098-INS-2939 |
| chr1-107319136-INS-199 | chr1-107319169-INS-199 |
| chr1-107333140-INS-59 | chr1-107333140-INS-59 |
| chr1-107371905-INS-55 | chr1-107371905-INS-55 |
| chr1-107382629-INS-116 | chr1-107382629-INS-117 |
| chr1-107393568-INS-115 | chr1-107393568-INS-115 |
| chr1-107418038-INS-674 | chr1-107418038-INS-674 |
| chr1-107419931-INS-70 | chr1-107419931-INS-68 |
| chr1-107438230-INS-281 | chr1-107438230-INS-282 |
| chr1-107451781-INS-361 | chr1-107451781-INS-362 |
| chr1-107459381-INS-2009 | chr1-107459382-INS-2009 |
| chr1-107461338-INS-92 | chr1-107461338-INS-90 |
| chr1-107463647-INS-77 | chr1-107463647-INS-77 |
| chr1-107508102-INS-652 | chr1-107508102-INS-652 |
| chr1-107536119-INS-199 | chr1-107536127-INS-182 |
| chr1-107543525-INS-3137 | chr1-107543526-INS-3137 |
| chr1-107547251-INS-6294 | chr1-107547252-INS-6293 |
| chr1-107575738-INS-81 | chr1-107575738-INS-81 |
| chr1-107576548-INS-1535 | chr1-107576549-INS-1533 |
| chr1-107599329-INS-7086 | chr1-107599330-INS-7086 |
| chr1-107631326-INS-6731 | chr1-107631327-INS-6729 |
| chr1-107650231-INS-4116 | chr1-107650232-INS-4116 |
| chr1-107654072-INS-460 | chr1-107654072-INS-462 |
| chr1-107658029-INS-5841 | chr1-107658030-INS-5841 |
| chr1-107672532-INS-84 | chr1-107672532-INS-70 |
| chr1-107691124-INS-106 | chr1-107691124-INS-106 |
| chr1-107702277-INS-63 | chr1-107702275-INS-53 |
| chr1-107732065-INS-105 | chr1-107732065-INS-105 |
| chr1-107737633-INS-64 | chr1-107737633-INS-64 |
| chr1-107818227-INS-268 | chr1-107818227-INS-268 |
| chr1-107822481-INS-154 | chr1-107822487-INS-106 |
| chr1-107830915-INS-675 | chr1-107830915-INS-675 |
| chr1-107843865-INS-110 | chr1-107843865-INS-112 |
| chr1-107878900-INS-528 | chr1-107878900-INS-516 |
| chr1-107902226-INS-64 | chr1-107902226-INS-64 |
| chr1-107920684-INS-61 | chr1-107920684-INS-61 |
| chr1-107963914-INS-54 | chr1-107963914-INS-75 |
| chr1-107980616-INS-6970 | chr1-107980617-INS-6970 |
| chr1-107981194-INS-965 | chr1-107981194-INS-965 |
| chr1-108000744-INS-503 | chr1-108000744-INS-503 |
| chr1-108012367-INS-1532 | chr1-108012368-INS-1534 |
| chr1-108021233-INS-2943 | chr1-108021234-INS-2943 |
| chr1-108024243-INS-172 | chr1-108024264-INS-188 |
| chr1-108025087-INS-6663 | chr1-108025088-INS-6664 |
| chr1-108037041-INS-100 | chr1-108037041-INS-104 |
| chr1-108044072-INS-327 | chr1-108044072-INS-327 |
| chr1-108051049-INS-238 | chr1-108051049-INS-238 |
| chr1-108055583-INS-158 | chr1-108055583-INS-168 |
| chr1-108063541-INS-395 | chr1-108063541-INS-395 |
| chr1-108084537-INS-347 | chr1-108084537-INS-347 |
| chr1-108089646-INS-73 | chr1-108089646-INS-53 |
| chr1-108091361-INS-326 | chr1-108091361-INS-326 |
| chr1-108119418-INS-95 | chr1-108119418-INS-50 |
| chr1-108145224-INS-64 | chr1-108145224-INS-66 |
| chr1-108146523-INS-5303 | chr1-108146524-INS-5303 |
| chr1-108163966-INS-6176 | chr1-108163967-INS-6175 |
| chr1-108173587-INS-54 | chr1-108173587-INS-54 |
| chr1-108179358-INS-66 | chr1-108179358-INS-61 |
| chr1-108188163-INS-6198 | chr1-108188164-INS-6198 |
| chr1-108234345-INS-60 | chr1-108234345-INS-60 |
| chr1-108236068-INS-389 | chr1-108236068-INS-391 |

| | |
|---|---|
| chr1-108268367-INS-3219 | chr1-108268368-INS-3220 |
| chr1-108274162-INS-90 | chr1-108274162-INS-90 |
| chr1-108312516-INS-522 | chr1-108312516-INS-496 |
| chr1-108319400-INS-549 | chr1-108319400-INS-549 |
| chr1-108330005-INS-82 | chr1-108330005-INS-82 |
| chr1-108346598-INS-320 | chr1-108346598-INS-322 |
| chr1-108357592-INS-10932 | chr1-108357593-INS-10932 |
| chr1-108376627-INS-78 | chr1-108376625-INS-74 |
| chr1-108382984-INS-71 | chr1-108382984-INS-71 |
| chr1-108417619-INS-6154 | chr1-108417620-INS-6154 |
| chr1-108439197-INS-130 | chr1-108439192-INS-180 |
| chr1-108442963-INS-634 | chr1-108442963-INS-634 |
| chr1-108464327-INS-5211 | chr1-108464324-INS-5207 |
| chr1-108466534-INS-182 | chr1-108466534-INS-172 |
| chr1-108485425-INS-220 | chr1-108485425-INS-211 |
| chr1-108494707-INS-7117 | chr1-108494708-INS-7117 |
| chr1-108524568-INS-5917 | chr1-108524569-INS-5919 |
| chr1-108533448-INS-198 | chr1-108533448-INS-197 |
| chr1-108535230-INS-1696 | chr1-108535231-INS-1698 |
| chr1-108537317-INS-7105 | chr1-108537318-INS-7105 |
| chr1-108567369-INS-207 | chr1-108567369-INS-207 |
| chr1-108568805-INS-459 | chr1-108568805-INS-459 |
| chr1-108569645-INS-8315 | chr1-108569646-INS-8316 |
| chr1-108572796-INS-96 | chr1-108572796-INS-96 |
| chr1-108576578-INS-61 | chr1-108576578-INS-68 |
| chr1-108591020-INS-479 | chr1-108591020-INS-479 |
| chr1-108602758-INS-6437 | chr1-108602759-INS-6436 |
| chr1-108604374-INS-119 | chr1-108604404-INS-95 |
| chr1-108605889-INS-178 | chr1-108605889-INS-178 |
| chr1-108677297-INS-2757 | chr1-108677298-INS-2757 |
| chr1-108680472-INS-55 | chr1-108680472-INS-55 |
| chr1-108684376-INS-128 | chr1-108684376-INS-128 |
| chr1-108697570-INS-92 | chr1-108697575-INS-83 |
| chr1-108702623-INS-52 | chr1-108702623-INS-52 |
| chr1-108705737-INS-76 | chr1-108705737-INS-79 |
| chr1-108765495-INS-52 | chr1-108765495-INS-68 |
| chr1-108770701-INS-401 | chr1-108770701-INS-401 |
| chr1-108777957-INS-72 | chr1-108777957-INS-72 |
| chr1-108783260-INS-173 | chr1-108783260-INS-173 |
| chr1-108802995-INS-56 | chr1-108802995-INS-56 |
| chr1-108811625-INS-502 | chr1-108811625-INS-502 |
| chr1-108824508-INS-385 | chr1-108824508-INS-385 |
| chr1-108842428-INS-124 | chr1-108842428-INS-124 |
| chr1-108856951-INS-820 | chr1-108856951-INS-820 |
| chr1-108880304-INS-60 | chr1-108880304-INS-60 |
| chr1-108906131-INS-611 | chr1-108906131-INS-606 |
| chr1-108923297-INS-52 | chr1-108923297-INS-52 |
| chr1-108928638-INS-57 | chr1-108928592-INS-58 |
| chr1-108935537-INS-328 | chr1-108935537-INS-328 |
| chr1-108936372-INS-572 | chr1-108936372-INS-572 |
| chr1-108955876-INS-108 | chr1-108955876-INS-108 |
| chr1-108978390-INS-7128 | chr1-108978391-INS-7128 |
| chr1-108980553-INS-381 | chr1-108980554-INS-381 |
| chr1-109010009-INS-59 | chr1-109010009-INS-59 |
| chr1-109012718-INS-475 | chr1-109012718-INS-475 |
| chr1-109022535-INS-180 | chr1-109022535-INS-140 |
| chr1-109031496-INS-56 | chr1-109031498-INS-56 |
| chr1-109065947-INS-101 | chr1-109065919-INS-92 |
| chr1-109092181-INS-159 | chr1-109092181-INS-159 |
| chr1-109101992-INS-1455 | chr1-109101993-INS-1455 |
| chr1-109104830-INS-238 | chr1-109104830-INS-237 |
| chr1-109115889-INS-351 | chr1-109115889-INS-351 |
| chr1-109127829-INS-63 | chr1-109127829-INS-63 |
| chr1-109149731-INS-1818 | chr1-109149732-INS-1818 |
| chr1-109151302-INS-73 | chr1-109151302-INS-68 |
| chr1-109165710-INS-71 | chr1-109165710-INS-73 |
| chr1-109178152-INS-6671 | chr1-109178153-INS-6671 |
| chr1-109211792-INS-431 | chr1-109211792-INS-431 |
| chr1-109243424-INS-121 | chr1-109243424-INS-121 |
| chr1-109267455-INS-80 | chr1-109267459-INS-84 |
| chr1-109279623-INS-55 | chr1-109279623-INS-55 |
| chr1-109299196-INS-7500 | chr1-109299197-INS-7500 |
| chr1-109323966-INS-68 | chr1-109323966-INS-62 |
| chr1-109325267-INS-111 | chr1-109325267-INS-110 |
| chr1-109329923-INS-81 | chr1-109329924-INS-61 |
| chr1-109363135-INS-6381 | chr1-109363136-INS-6364 |
| chr1-109376091-INS-459 | chr1-109376091-INS-459 |

| | |
|---|---|
| chr1-109378781-INS-64 | chr1-109378781-INS-92 |
| chr1-109379013-INS-3162 | chr1-109379014-INS-3162 |
| chr1-109383987-INS-154 | chr1-109383987-INS-154 |
| chr1-109390213-INS-60 | chr1-109390213-INS-56 |
| chr1-109402875-INS-64 | chr1-109402875-INS-64 |
| chr1-109427205-INS-84 | chr1-109427225-INS-94 |
| chr1-109461986-INS-164 | chr1-109461986-INS-164 |
| chr1-109484381-INS-114 | chr1-109484381-INS-114 |
| chr1-109516968-INS-109 | chr1-109516968-INS-117 |
| chr1-109527406-INS-77 | chr1-109527402-INS-85 |
| chr1-109540411-INS-137 | chr1-109540411-INS-137 |
| chr1-109550868-INS-145 | chr1-109550868-INS-145 |
| chr1-109612410-INS-128 | chr1-109612410-INS-128 |
| chr1-109614954-INS-4198 | chr1-109614955-INS-4232 |
| chr1-109630387-INS-58 | chr1-109630387-INS-56 |
| chr1-109635717-INS-6017 | chr1-109635718-INS-6017 |
| chr1-109647317-INS-1601 | chr1-109647318-INS-1604 |
| chr1-109668128-INS-82 | chr1-109668128-INS-82 |
| chr1-109681840-INS-5409 | chr1-109681841-INS-5409 |
| chr1-109713159-INS-58 | chr1-109713159-INS-74 |
| chr1-109723994-INS-7315 | chr1-109723995-INS-7317 |
| chr1-109736289-INS-312 | chr1-109736289-INS-312 |
| chr1-109741977-INS-166 | chr1-109741977-INS-165 |
| chr1-109768058-INS-114 | chr1-109768058-INS-114 |
| chr1-109812747-INS-635 | chr1-109812747-INS-643 |
| chr1-109814025-INS-3691 | chr1-109814026-INS-3691 |
| chr1-109825095-INS-7028 | chr1-109825096-INS-7029 |
| chr1-109861483-INS-97 | chr1-109861485-INS-73 |
| chr1-109862307-INS-6710 | chr1-109862308-INS-6711 |
| chr1-109883008-INS-743 | chr1-109883031-INS-743 |
| chr1-109894000-INS-400 | chr1-109894000-INS-404 |
| chr1-109917857-INS-78 | chr1-109917857-INS-94 |
| chr1-109937598-INS-231 | chr1-109937598-INS-233 |
| chr1-109949475-INS-56 | chr1-109949476-INS-66 |
| chr1-110001218-INS-55 | chr1-110001218-INS-55 |
| chr1-110002723-INS-450 | chr1-110002723-INS-450 |
| chr1-110006752-INS-527 | chr1-110006752-INS-526 |
| chr1-110120536-INS-845 | chr1-110120536-INS-845 |
| chr1-110128845-INS-57 | chr1-110128845-INS-57 |
| chr1-110148652-INS-495 | chr1-110148652-INS-495 |
| chr1-110150127-INS-627 | chr1-110150127-INS-627 |
| chr1-110155156-INS-5313 | chr1-110155157-INS-5313 |
| chr1-110169120-INS-4225 | chr1-110169121-INS-4225 |
| chr1-110174738-INS-54 | chr1-110174738-INS-54 |
| chr1-110176174-INS-835 | chr1-110176174-INS-833 |
| chr1-110205021-INS-429 | chr1-110205021-INS-429 |
| chr1-110227421-INS-67 | chr1-110227421-INS-67 |
| chr1-110229542-INS-87 | chr1-110229542-INS-87 |
| chr1-110265286-INS-180 | chr1-110265286-INS-180 |
| chr1-110283695-INS-68 | chr1-110283695-INS-70 |
| chr1-110287724-INS-72 | chr1-110287724-INS-72 |
| chr1-110302357-INS-84 | chr1-110302357-INS-96 |
| chr1-110308235-INS-585 | chr1-110308235-INS-585 |
| chr1-110319495-INS-198 | chr1-110319495-INS-198 |
| chr1-110321729-INS-6568 | chr1-110321730-INS-6568 |
| chr1-110325559-INS-6198 | chr1-110325560-INS-6199 |
| chr1-110330409-INS-4831 | chr1-110330410-INS-4832 |
| chr1-110333344-INS-53 | chr1-110333344-INS-53 |
| chr1-110337361-INS-7569 | chr1-110337362-INS-7569 |
| chr1-110349318-INS-2832 | chr1-110349319-INS-2832 |
| chr1-110350429-INS-3286 | chr1-110350430-INS-3286 |
| chr1-110352552-INS-463 | chr1-110352552-INS-463 |
| chr1-110361962-INS-1123 | chr1-110361963-INS-1123 |
| chr1-110369859-INS-7083 | chr1-110369860-INS-7084 |
| chr1-110407954-INS-80 | chr1-110407955-INS-120 |
| chr1-110437148-INS-95 | chr1-110437148-INS-95 |
| chr1-110450063-INS-88 | chr1-110450063-INS-88 |
| chr1-110476249-INS-2518 | chr1-110476250-INS-2515 |
| chr1-110480279-INS-991 | chr1-110480275-INS-992 |
| chr1-110517716-INS-62 | chr1-110517716-INS-62 |
| chr1-110528070-INS-4330 | chr1-110528071-INS-4330 |
| chr1-110545966-INS-116 | chr1-110545966-INS-70 |
| chr1-110549465-INS-3110 | chr1-110549466-INS-3112 |
| chr1-110565274-INS-204 | chr1-110565274-INS-252 |
| chr1-110588862-INS-202 | chr1-110588863-INS-214 |
| chr1-110604405-INS-2362 | chr1-110604406-INS-2362 |
| chr1-110605742-INS-67 | chr1-110605742-INS-67 |

| | |
|---|---|
| chr1-110607170-INS-149 | chr1-110607170-INS-149 |
| chr1-110625123-INS-6198 | chr1-110625124-INS-6198 |
| chr1-110629737-INS-936 | chr1-110629737-INS-936 |
| chr1-110634050-INS-61 | chr1-110634050-INS-63 |
| chr1-110636157-INS-54 | chr1-110636157-INS-54 |
| chr1-110641525-INS-255 | chr1-110641525-INS-257 |
| chr1-110657906-INS-612 | chr1-110657906-INS-612 |
| chr1-110662362-INS-66 | chr1-110662362-INS-68 |
| chr1-110686640-INS-7128 | chr1-110686641-INS-7128 |
| chr1-110710748-INS-104 | chr1-110710748-INS-102 |
| chr1-110717005-INS-52 | chr1-110717047-INS-52 |
| chr1-110728076-INS-1603 | chr1-110728077-INS-1603 |
| chr1-110728903-INS-171 | chr1-110728903-INS-171 |
| chr1-110739387-INS-319 | chr1-110739387-INS-319 |
| chr1-110740987-INS-211 | chr1-110740987-INS-211 |
| chr1-110746204-INS-361 | chr1-110746204-INS-361 |
| chr1-110749597-INS-208 | chr1-110749597-INS-208 |
| chr1-110760721-INS-114 | chr1-110760721-INS-102 |
| chr1-110765241-INS-52 | chr1-110765241-INS-52 |
| chr1-110838949-INS-739 | chr1-110838949-INS-739 |
| chr1-110888847-INS-64 | chr1-110888847-INS-64 |
| chr1-110956819-INS-72 | chr1-110956815-INS-72 |
| chr1-110985123-INS-207 | chr1-110985123-INS-207 |
| chr1-111004421-INS-358 | chr1-111004421-INS-358 |
| chr1-111021950-INS-73 | chr1-111021950-INS-73 |
| chr1-111023782-INS-693 | chr1-111023782-INS-694 |
| chr1-111028499-INS-5743 | chr1-111028503-INS-5782 |
| chr1-111128097-INS-6362 | chr1-111128098-INS-6362 |
| chr1-111129597-INS-562 | chr1-111129597-INS-562 |
| chr1-111165971-INS-105 | chr1-111165971-INS-104 |
| chr1-111241448-INS-98 | chr1-111241448-INS-97 |
| chr1-111241743-INS-929 | chr1-111241743-INS-925 |
| chr1-111256251-INS-52 | chr1-111256251-INS-52 |
| chr1-111269174-INS-4151 | chr1-111269175-INS-4151 |
| chr1-111273252-INS-212 | chr1-111273252-INS-212 |
| chr1-111277645-INS-1638 | chr1-111277646-INS-1638 |
| chr1-111290337-INS-4806 | chr1-111290338-INS-4806 |
| chr1-111306024-INS-438 | chr1-111306024-INS-438 |
| chr1-111307592-INS-82 | chr1-111307563-INS-64 |
| chr1-111320466-INS-84 | chr1-111320466-INS-82 |
| chr1-111321594-INS-55 | chr1-111321594-INS-54 |
| chr1-111338556-INS-460 | chr1-111338556-INS-462 |
| chr1-111342806-INS-53 | chr1-111342799-INS-64 |
| chr1-111345152-INS-490 | chr1-111345152-INS-490 |
| chr1-111354940-INS-54 | chr1-111354940-INS-62 |
| chr1-111390113-INS-2998 | chr1-111390114-INS-2998 |
| chr1-111435502-INS-267 | chr1-111435502-INS-267 |
| chr1-111453155-INS-81 | chr1-111453155-INS-81 |
| chr1-111483828-INS-324 | chr1-111483828-INS-324 |
| chr1-111498820-INS-64 | chr1-111498820-INS-70 |
| chr1-111528700-INS-769 | chr1-111528700-INS-770 |
| chr1-111550048-INS-353 | chr1-111550046-INS-357 |
| chr1-111554472-INS-268 | chr1-111554472-INS-268 |
| chr1-111562208-INS-2715 | chr1-111562209-INS-2716 |
| chr1-111578644-INS-507 | chr1-111578644-INS-507 |
| chr1-111611616-INS-164 | chr1-111611616-INS-164 |
| chr1-111674969-INS-58 | chr1-111674969-INS-54 |
| chr1-111681525-INS-74 | chr1-111681538-INS-60 |
| chr1-111690957-INS-943 | chr1-111690957-INS-940 |
| chr1-111693890-INS-116 | chr1-111693890-INS-116 |
| chr1-111696500-INS-4923 | chr1-111696501-INS-4923 |
| chr1-111718105-INS-112 | chr1-111718105-INS-112 |
| chr1-111730492-INS-2585 | chr1-111730493-INS-2585 |
| chr1-111730493-INS-358 | chr1-111730493-INS-358 |
| chr1-111755391-INS-192 | chr1-111755391-INS-195 |
| chr1-111761876-INS-8175 | chr1-111761877-INS-8177 |
| chr1-111770870-INS-154 | chr1-111770870-INS-148 |
| chr1-111804906-INS-245 | chr1-111804906-INS-245 |
| chr1-111807213-INS-278 | chr1-111807213-INS-277 |
| chr1-111829887-INS-80 | chr1-111829887-INS-88 |
| chr1-111843578-INS-488 | chr1-111843578-INS-478 |
| chr1-111883181-INS-693 | chr1-111883181-INS-689 |
| chr1-111908155-INS-272 | chr1-111908155-INS-273 |
| chr1-111927107-INS-60 | chr1-111927107-INS-66 |
| chr1-111932562-INS-204 | chr1-111932562-INS-204 |
| chr1-111935722-INS-58 | chr1-111935722-INS-58 |
| chr1-111956106-INS-56 | chr1-111956106-INS-55 |

| | |
|---|---|
| chr1-111960048-INS-60 | chr1-111960048-INS-64 |
| chr1-111963141-INS-6366 | chr1-111963142-INS-6366 |
| chr1-111973203-INS-54 | chr1-111973203-INS-54 |
| chr1-112008783-INS-6661 | chr1-112008784-INS-6661 |
| chr1-112019952-INS-3348 | chr1-112019953-INS-3356 |
| chr1-112020655-INS-69 | chr1-112020656-INS-69 |
| chr1-112036621-INS-126 | chr1-112036621-INS-152 |
| chr1-112040034-INS-53 | chr1-112040034-INS-52 |
| chr1-112048299-INS-57 | chr1-112048299-INS-57 |
| chr1-112055634-INS-154 | chr1-112055636-INS-136 |
| chr1-112066906-INS-96 | chr1-112066906-INS-96 |
| chr1-112074370-INS-385 | chr1-112074415-INS-385 |
| chr1-112105877-INS-427 | chr1-112105877-INS-464 |
| chr1-112116173-INS-6843 | chr1-112116174-INS-6843 |
| chr1-112124922-INS-6874 | chr1-112124923-INS-6875 |
| chr1-112129638-INS-60 | chr1-112129638-INS-60 |
| chr1-112136345-INS-1000 | chr1-112136345-INS-1004 |
| chr1-112182563-INS-294 | chr1-112182563-INS-282 |
| chr1-112188474-INS-142 | chr1-112188473-INS-149 |
| chr1-112239214-INS-80 | chr1-112239215-INS-60 |
| chr1-112246797-INS-74 | chr1-112246797-INS-72 |
| chr1-112259773-INS-773 | chr1-112259773-INS-774 |
| chr1-112268394-INS-351 | chr1-112268390-INS-357 |
| chr1-112277612-INS-524 | chr1-112277612-INS-524 |
| chr1-112287348-INS-199 | chr1-112287348-INS-199 |
| chr1-112288522-INS-73 | chr1-112288522-INS-73 |
| chr1-112305544-INS-191 | chr1-112305544-INS-191 |
| chr1-112313518-INS-228 | chr1-112313518-INS-228 |
| chr1-112327118-INS-59 | chr1-112327118-INS-59 |
| chr1-112333329-INS-608 | chr1-112333329-INS-606 |
| chr1-112335646-INS-202 | chr1-112335646-INS-202 |
| chr1-112341260-INS-253 | chr1-112341260-INS-254 |
| chr1-112342919-INS-63 | chr1-112342901-INS-66 |
| chr1-112358366-INS-73 | chr1-112358361-INS-80 |
| chr1-112374116-INS-358 | chr1-112374116-INS-358 |
| chr1-112379119-INS-88 | chr1-112379119-INS-66 |
| chr1-112384721-INS-51 | chr1-112384730-INS-57 |
| chr1-112395309-INS-336 | chr1-112395309-INS-338 |
| chr1-112411871-INS-444 | chr1-112411871-INS-444 |
| chr1-112419685-INS-52 | chr1-112419686-INS-56 |
| chr1-112421626-INS-70 | chr1-112421591-INS-60 |
| chr1-112428615-INS-115 | chr1-112428636-INS-116 |
| chr1-112444227-INS-137 | chr1-112444227-INS-137 |
| chr1-112445216-INS-1068 | chr1-112445217-INS-1066 |
| chr1-112446580-INS-3025 | chr1-112446581-INS-3025 |
| chr1-112450079-INS-1739 | chr1-112450078-INS-1738 |
| chr1-112451786-INS-66 | chr1-112451786-INS-60 |
| chr1-112461651-INS-635 | chr1-112461651-INS-635 |
| chr1-112467988-INS-710 | chr1-112467988-INS-710 |
| chr1-112559772-INS-149 | chr1-112559772-INS-152 |
| chr1-112563546-INS-164 | chr1-112563546-INS-164 |
| chr1-112574886-INS-6330 | chr1-112574887-INS-6330 |
| chr1-112581944-INS-4383 | chr1-112581945-INS-4383 |
| chr1-112588835-INS-3138 | chr1-112588836-INS-3138 |
| chr1-112595037-INS-646 | chr1-112595037-INS-648 |
| chr1-112612129-INS-5003 | chr1-112612130-INS-5003 |
| chr1-112623005-INS-938 | chr1-112623005-INS-936 |
| chr1-112636771-INS-804 | chr1-112636771-INS-804 |
| chr1-112662439-INS-120 | chr1-112662443-INS-96 |
| chr1-112667924-INS-482 | chr1-112667924-INS-482 |
| chr1-112671024-INS-64 | chr1-112671024-INS-64 |
| chr1-112677858-INS-100 | chr1-112677858-INS-100 |
| chr1-112680026-INS-715 | chr1-112680026-INS-716 |
| chr1-112685222-INS-123 | chr1-112685232-INS-96 |
| chr1-112694588-INS-70 | chr1-112694588-INS-74 |
| chr1-112711537-INS-77 | chr1-112711524-INS-103 |
| chr1-112713552-INS-68 | chr1-112713552-INS-68 |
| chr1-112716451-INS-7109 | chr1-112716452-INS-7109 |
| chr1-112721985-INS-789 | chr1-112721985-INS-788 |
| chr1-112741647-INS-7164 | chr1-112741648-INS-7164 |
| chr1-112762896-INS-96 | chr1-112762895-INS-126 |
| chr1-112765884-INS-424 | chr1-112765884-INS-424 |
| chr1-112787995-INS-305 | chr1-112787995-INS-306 |
| chr1-112814841-INS-6697 | chr1-112814842-INS-6697 |
| chr1-112840327-INS-1558 | chr1-112840328-INS-1558 |
| chr1-112844285-INS-52 | chr1-112844285-INS-68 |
| chr1-112864512-INS-656 | chr1-112864512-INS-656 |

| | |
|---|---|
| chr1-112885494-INS-424 | chr1-112885490-INS-424 |
| chr1-112900812-INS-5246 | chr1-112900813-INS-5246 |
| chr1-112932882-INS-343 | chr1-112932882-INS-343 |
| chr1-112968951-INS-60 | chr1-112968951-INS-64 |
| chr1-112999607-INS-3653 | chr1-112999608-INS-3655 |
| chr1-113039932-INS-110 | chr1-113039932-INS-66 |
| chr1-113060112-INS-747 | chr1-113060112-INS-746 |
| chr1-113075496-INS-53 | chr1-113075496-INS-53 |
| chr1-113082740-INS-52 | chr1-113082740-INS-52 |
| chr1-113110069-INS-65 | chr1-113110069-INS-73 |
| chr1-113118133-INS-73 | chr1-113118133-INS-73 |
| chr1-113121586-INS-74 | chr1-113121617-INS-74 |
| chr1-113122240-INS-52 | chr1-113122240-INS-52 |
| chr1-113147824-INS-191 | chr1-113147824-INS-191 |
| chr1-113166146-INS-56 | chr1-113166142-INS-63 |
| chr1-113179687-INS-85 | chr1-113179687-INS-68 |
| chr1-113184312-INS-396 | chr1-113184312-INS-398 |
| chr1-113192262-INS-71 | chr1-113192262-INS-60 |
| chr1-113195275-INS-80 | chr1-113195275-INS-80 |
| chr1-113201045-INS-109 | chr1-113201045-INS-109 |
| chr1-113210062-INS-94 | chr1-113210062-INS-52 |
| chr1-113244564-INS-89 | chr1-113244564-INS-89 |
| chr1-113247718-INS-289 | chr1-113247730-INS-267 |
| chr1-113277032-INS-1890 | chr1-113277033-INS-1890 |
| chr1-113299558-INS-144 | chr1-113299558-INS-144 |
| chr1-113308874-INS-7278 | chr1-113308875-INS-7278 |
| chr1-113336632-INS-402 | chr1-113336632-INS-403 |
| chr1-113341709-INS-3417 | chr1-113341710-INS-3418 |
| chr1-113358759-INS-216 | chr1-113358759-INS-216 |
| chr1-113369518-INS-584 | chr1-113369518-INS-584 |
| chr1-113371904-INS-98 | chr1-113371904-INS-98 |
| chr1-113406528-INS-68 | chr1-113406528-INS-68 |
| chr1-113420235-INS-78 | chr1-113420187-INS-116 |
| chr1-113437657-INS-4958 | chr1-113437658-INS-4958 |
| chr1-113450193-INS-58 | chr1-113450171-INS-59 |
| chr1-113452161-INS-63 | chr1-113452161-INS-63 |
| chr1-113453967-INS-3655 | chr1-113453968-INS-3667 |
| chr1-113454272-INS-161 | chr1-113454272-INS-161 |
| chr1-113454742-INS-1010 | chr1-113454743-INS-1010 |
| chr1-113463878-INS-187 | chr1-113463878-INS-187 |
| chr1-113466049-INS-202 | chr1-113466049-INS-211 |
| chr1-113472962-INS-682 | chr1-113472962-INS-685 |
| chr1-113475273-INS-515 | chr1-113475273-INS-516 |
| chr1-113476966-INS-92 | chr1-113476966-INS-92 |
| chr1-113502576-INS-153 | chr1-113502576-INS-153 |
| chr1-113513946-INS-153 | chr1-113513946-INS-153 |
| chr1-113518470-INS-7087 | chr1-113518471-INS-7087 |
| chr1-113536700-INS-197 | chr1-113536700-INS-197 |
| chr1-113555572-INS-75 | chr1-113555572-INS-75 |
| chr1-113561366-INS-510 | chr1-113561366-INS-511 |
| chr1-113562697-INS-68 | chr1-113562684-INS-100 |
| chr1-113564596-INS-66 | chr1-113564587-INS-74 |
| chr1-113584481-INS-473 | chr1-113584481-INS-473 |
| chr1-113588566-INS-332 | chr1-113588566-INS-332 |
| chr1-113593879-INS-4122 | chr1-113593880-INS-4122 |
| chr1-113596580-INS-59 | chr1-113596549-INS-65 |
| chr1-113600318-INS-479 | chr1-113600297-INS-486 |
| chr1-113628080-INS-69 | chr1-113628092-INS-104 |
| chr1-113646701-INS-283 | chr1-113646701-INS-283 |
| chr1-113671176-INS-97 | chr1-113671176-INS-97 |
| chr1-113696473-INS-212 | chr1-113696473-INS-206 |
| chr1-113712939-INS-62 | chr1-113712939-INS-62 |
| chr1-113713387-INS-5514 | chr1-113713388-INS-5514 |
| chr1-113719376-INS-50 | chr1-113719376-INS-50 |
| chr1-113799432-INS-56 | chr1-113799432-INS-56 |
| chr1-113819842-INS-1230 | chr1-113819843-INS-1230 |
| chr1-113897284-INS-173 | chr1-113897284-INS-173 |
| chr1-113921177-INS-94 | chr1-113921181-INS-100 |
| chr1-113924156-INS-139 | chr1-113924156-INS-140 |
| chr1-113926009-INS-309 | chr1-113926009-INS-308 |
| chr1-113944928-INS-6156 | chr1-113944929-INS-6156 |
| chr1-113992944-INS-254 | chr1-113992944-INS-254 |
| chr1-113998530-INS-91 | chr1-113998530-INS-91 |
| chr1-114031550-INS-850 | chr1-114031552-INS-849 |
| chr1-114077453-INS-54 | chr1-114077453-INS-56 |
| chr1-114143247-INS-7128 | chr1-114143248-INS-7128 |
| chr1-114149352-INS-2081 | chr1-114149353-INS-2081 |

| | |
|---|---|
| chr1-114159107-INS-67 | chr1-114159107-INS-67 |
| chr1-114190166-INS-70 | chr1-114190167-INS-72 |
| chr1-114192693-INS-59 | chr1-114192696-INS-54 |
| chr1-114198859-INS-6869 | chr1-114198860-INS-6869 |
| chr1-114220168-INS-64 | chr1-114220171-INS-79 |
| chr1-114223458-INS-851 | chr1-114223458-INS-851 |
| chr1-114227231-INS-377 | chr1-114227231-INS-371 |
| chr1-114258857-INS-76 | chr1-114258857-INS-76 |
| chr1-114277994-INS-734 | chr1-114277994-INS-734 |
| chr1-114315935-INS-6403 | chr1-114315936-INS-6403 |
| chr1-114317015-INS-2756 | chr1-114317016-INS-2756 |
| chr1-114331640-INS-823 | chr1-114331640-INS-823 |
| chr1-114346840-INS-212 | chr1-114346840-INS-212 |
| chr1-114354392-INS-225 | chr1-114354392-INS-225 |
| chr1-114387128-INS-485 | chr1-114387128-INS-485 |
| chr1-114392651-INS-12550 | chr1-114392652-INS-12550 |
| chr1-114394874-INS-70 | chr1-114394874-INS-52 |
| chr1-114414172-INS-66 | chr1-114414172-INS-70 |
| chr1-114423836-INS-77 | chr1-114423847-INS-89 |
| chr1-114427757-INS-210 | chr1-114427757-INS-210 |
| chr1-114445107-INS-54 | chr1-114445108-INS-69 |
| chr1-114461210-INS-146 | chr1-114461210-INS-138 |
| chr1-114465460-INS-69 | chr1-114465460-INS-69 |
| chr1-114475264-INS-78 | chr1-114475254-INS-78 |
| chr1-114503728-INS-164 | chr1-114503728-INS-164 |
| chr1-114511854-INS-54 | chr1-114511852-INS-59 |
| chr1-114521485-INS-53 | chr1-114521485-INS-53 |
| chr1-114530578-INS-81 | chr1-114530578-INS-81 |
| chr1-114543264-INS-58 | chr1-114543264-INS-58 |
| chr1-114552893-INS-577 | chr1-114552893-INS-577 |
| chr1-114556486-INS-515 | chr1-114556486-INS-515 |
| chr1-114573138-INS-152 | chr1-114573138-INS-152 |
| chr1-114579742-INS-85 | chr1-114579742-INS-85 |
| chr1-114608374-INS-65 | chr1-114608374-INS-67 |
| chr1-114609630-INS-1869 | chr1-114609631-INS-1870 |
| chr1-114615623-INS-62 | chr1-114615623-INS-62 |
| chr1-114623243-INS-671 | chr1-114623243-INS-682 |
| chr1-114636783-INS-6559 | chr1-114636784-INS-6559 |
| chr1-114639652-INS-88 | chr1-114639629-INS-87 |
| chr1-114646671-INS-54 | chr1-114646671-INS-56 |
| chr1-114679368-INS-143 | chr1-114679368-INS-143 |
| chr1-114727180-INS-105 | chr1-114727180-INS-105 |
| chr1-114797405-INS-1737 | chr1-114797406-INS-1737 |
| chr1-114798073-INS-90 | chr1-114798071-INS-92 |
| chr1-114798494-INS-472 | chr1-114798494-INS-472 |
| chr1-114807478-INS-54 | chr1-114807478-INS-56 |
| chr1-114842162-INS-6827 | chr1-114842163-INS-6827 |
| chr1-114847197-INS-5347 | chr1-114847198-INS-5347 |
| chr1-114862593-INS-4557 | chr1-114862594-INS-4557 |
| chr1-114875969-INS-498 | chr1-114875969-INS-498 |
| chr1-114877963-INS-52 | chr1-114877963-INS-54 |
| chr1-114883405-INS-6166 | chr1-114883406-INS-6166 |
| chr1-114902586-INS-1519 | chr1-114902587-INS-1519 |
| chr1-114911226-INS-56 | chr1-114911226-INS-56 |
| chr1-114914518-INS-376 | chr1-114914518-INS-376 |
| chr1-114915999-INS-1890 | chr1-114916000-INS-1890 |
| chr1-114962391-INS-185 | chr1-114962391-INS-184 |
| chr1-114966846-INS-4100 | chr1-114966847-INS-4100 |
| chr1-114973507-INS-6226 | chr1-114973508-INS-6226 |
| chr1-114980251-INS-291 | chr1-114980259-INS-297 |
| chr1-114981660-INS-4954 | chr1-114981661-INS-4954 |
| chr1-114989407-INS-236 | chr1-114989407-INS-236 |
| chr1-114991582-INS-2181 | chr1-114991583-INS-2181 |
| chr1-114995735-INS-6409 | chr1-114995736-INS-6410 |
| chr1-115008702-INS-1964 | chr1-115008703-INS-1964 |
| chr1-115028244-INS-402 | chr1-115028245-INS-402 |
| chr1-115042068-INS-54 | chr1-115042093-INS-56 |
| chr1-115050048-INS-7071 | chr1-115050049-INS-7071 |
| chr1-115066913-INS-639 | chr1-115066913-INS-639 |
| chr1-115097112-INS-512 | chr1-115097112-INS-512 |
| chr1-115106563-INS-444 | chr1-115106563-INS-439 |
| chr1-115115791-INS-56 | chr1-115115791-INS-56 |
| chr1-115117841-INS-504 | chr1-115117841-INS-504 |
| chr1-115124081-INS-1837 | chr1-115124084-INS-1843 |
| chr1-115177480-INS-873 | chr1-115177480-INS-872 |
| chr1-115179671-INS-9397 | chr1-115179672-INS-9402 |
| chr1-115202850-INS-462 | chr1-115202850-INS-462 |

| | |
|---|---|
| chr1-115223841-INS-774 | chr1-115223823-INS-775 |
| chr1-115226624-INS-198 | chr1-115226624-INS-198 |
| chr1-115236072-INS-2316 | chr1-115236073-INS-2318 |
| chr1-115237695-INS-470 | chr1-115237695-INS-471 |
| chr1-115239248-INS-54 | chr1-115239230-INS-50 |
| chr1-115245079-INS-6445 | chr1-115245080-INS-6447 |
| chr1-115257531-INS-60 | chr1-115257531-INS-60 |
| chr1-115260384-INS-52 | chr1-115260384-INS-52 |
| chr1-115279300-INS-203 | chr1-115279300-INS-200 |
| chr1-115303838-INS-5302 | chr1-115303839-INS-5302 |
| chr1-115305633-INS-6277 | chr1-115305634-INS-6277 |
| chr1-115312133-INS-85 | chr1-115312133-INS-87 |
| chr1-115340493-INS-68 | chr1-115340493-INS-68 |
| chr1-115343123-INS-181 | chr1-115343123-INS-181 |
| chr1-115344212-INS-774 | chr1-115344212-INS-774 |
| chr1-115384431-INS-709 | chr1-115384431-INS-713 |
| chr1-115452137-INS-827 | chr1-115452137-INS-827 |
| chr1-115471376-INS-116 | chr1-115471376-INS-84 |
| chr1-115472222-INS-52 | chr1-115472222-INS-50 |
| chr1-115482591-INS-780 | chr1-115482591-INS-780 |
| chr1-115497175-INS-2674 | chr1-115497176-INS-2674 |
| chr1-115502104-INS-99 | chr1-115502104-INS-90 |
| chr1-115529229-INS-641 | chr1-115529229-INS-651 |
| chr1-115532970-INS-798 | chr1-115532970-INS-798 |
| chr1-115543062-INS-520 | chr1-115543062-INS-520 |
| chr1-115567499-INS-56 | chr1-115567487-INS-56 |
| chr1-115568671-INS-2580 | chr1-115568672-INS-2580 |
| chr1-115571632-INS-98 | chr1-115571632-INS-78 |
| chr1-115579129-INS-70 | chr1-115579129-INS-74 |
| chr1-115580974-INS-54 | chr1-115580974-INS-60 |
| chr1-115592734-INS-55 | chr1-115592724-INS-50 |
| chr1-115694285-INS-56 | chr1-115694284-INS-60 |
| chr1-115695651-INS-58 | chr1-115695626-INS-53 |
| chr1-115702538-INS-51 | chr1-115702538-INS-52 |
| chr1-115713370-INS-1229 | chr1-115713371-INS-1229 |
| chr1-115713676-INS-97 | chr1-115713676-INS-77 |
| chr1-115719480-INS-235 | chr1-115719480-INS-234 |
| chr1-115725298-INS-64 | chr1-115725298-INS-74 |
| chr1-115738135-INS-122 | chr1-115738135-INS-121 |
| chr1-115751860-INS-6452 | chr1-115751861-INS-6453 |
| chr1-115763306-INS-524 | chr1-115763306-INS-524 |
| chr1-115767118-INS-119 | chr1-115767118-INS-121 |
| chr1-115775086-INS-1134 | chr1-115775087-INS-1134 |
| chr1-115811779-INS-267 | chr1-115811779-INS-267 |
| chr1-115820301-INS-64 | chr1-115820301-INS-74 |
| chr1-115827544-INS-114 | chr1-115827544-INS-114 |
| chr1-115844782-INS-178 | chr1-115844782-INS-179 |
| chr1-115900037-INS-392 | chr1-115900037-INS-392 |
| chr1-115920924-INS-70 | chr1-115920924-INS-116 |
| chr1-116029673-INS-62 | chr1-116029673-INS-62 |
| chr1-116030090-INS-2402 | chr1-116030106-INS-2426 |
| chr1-116033555-INS-504 | chr1-116033555-INS-504 |
| chr1-116048893-INS-56 | chr1-116048893-INS-58 |
| chr1-116063727-INS-54 | chr1-116063731-INS-60 |
| chr1-116078792-INS-467 | chr1-116078792-INS-467 |
| chr1-116080683-INS-967 | chr1-116080684-INS-961 |
| chr1-116122732-INS-2082 | chr1-116122733-INS-2082 |
| chr1-116154740-INS-72 | chr1-116154735-INS-62 |
| chr1-116158687-INS-59 | chr1-116158714-INS-52 |
| chr1-116166537-INS-52 | chr1-116166537-INS-52 |
| chr1-116171151-INS-6427 | chr1-116171152-INS-6427 |
| chr1-116172791-INS-246 | chr1-116172791-INS-246 |
| chr1-116173275-INS-3547 | chr1-116173276-INS-3548 |
| chr1-116175602-INS-78 | chr1-116175602-INS-80 |
| chr1-116179796-INS-96 | chr1-116179808-INS-128 |
| chr1-116206693-INS-193 | chr1-116206693-INS-193 |
| chr1-116220730-INS-7093 | chr1-116220731-INS-7093 |
| chr1-116230064-INS-80 | chr1-116230064-INS-81 |
| chr1-116236492-INS-64 | chr1-116236495-INS-70 |
| chr1-116250501-INS-68 | chr1-116250501-INS-68 |
| chr1-116252988-INS-325 | chr1-116252988-INS-324 |
| chr1-116261612-INS-54 | chr1-116261611-INS-52 |
| chr1-116263026-INS-120 | chr1-116263026-INS-132 |
| chr1-116270171-INS-77 | chr1-116270171-INS-77 |
| chr1-116280895-INS-54 | chr1-116280895-INS-56 |
| chr1-116297449-INS-56 | chr1-116297449-INS-66 |
| chr1-116315465-INS-1018 | chr1-116315466-INS-1018 |

| | |
|---|---|
| chr1-116328816-INS-126 | chr1-116328817-INS-126 |
| chr1-116329663-INS-52 | chr1-116329663-INS-68 |
| chr1-116344527-INS-6482 | chr1-116344528-INS-6483 |
| chr1-116362639-INS-73 | chr1-116362639-INS-73 |
| chr1-116381490-INS-777 | chr1-116381490-INS-776 |
| chr1-116393407-INS-82 | chr1-116393411-INS-75 |
| chr1-116398179-INS-67 | chr1-116398180-INS-64 |
| chr1-116412640-INS-67 | chr1-116412638-INS-115 |
| chr1-116425121-INS-835 | chr1-116425121-INS-851 |
| chr1-116431091-INS-6657 | chr1-116431092-INS-6657 |
| chr1-116442407-INS-411 | chr1-116442407-INS-411 |
| chr1-116453763-INS-60 | chr1-116453771-INS-73 |
| chr1-116455320-INS-456 | chr1-116455320-INS-456 |
| chr1-116460349-INS-455 | chr1-116460349-INS-455 |
| chr1-116464679-INS-51 | chr1-116464679-INS-55 |
| chr1-116480626-INS-2761 | chr1-116480627-INS-2762 |
| chr1-116483550-INS-1044 | chr1-116483552-INS-1044 |
| chr1-116524614-INS-400 | chr1-116524614-INS-400 |
| chr1-116556052-INS-98 | chr1-116556042-INS-96 |
| chr1-116561900-INS-2399 | chr1-116561901-INS-2399 |
| chr1-116602637-INS-791 | chr1-116602637-INS-791 |
| chr1-116611125-INS-4399 | chr1-116611126-INS-4408 |
| chr1-116614291-INS-78 | chr1-116614295-INS-81 |
| chr1-116616410-INS-74 | chr1-116616410-INS-74 |
| chr1-116627065-INS-100 | chr1-116627071-INS-103 |
| chr1-116644007-INS-115 | chr1-116644007-INS-115 |
| chr1-116652396-INS-434 | chr1-116652397-INS-434 |
| chr1-116654351-INS-80 | chr1-116654351-INS-80 |
| chr1-116674829-INS-59 | chr1-116674829-INS-97 |
| chr1-116680536-INS-450 | chr1-116680536-INS-450 |
| chr1-116693270-INS-66 | chr1-116693270-INS-66 |
| chr1-116714982-INS-84 | chr1-116714992-INS-114 |
| chr1-116718517-INS-595 | chr1-116718517-INS-595 |
| chr1-116721581-INS-103 | chr1-116721581-INS-103 |
| chr1-116725025-INS-229 | chr1-116725025-INS-229 |
| chr1-116756790-INS-6444 | chr1-116756791-INS-6443 |
| chr1-116758831-INS-156 | chr1-116758831-INS-156 |
| chr1-116773889-INS-528 | chr1-116773889-INS-528 |
| chr1-116802065-INS-54 | chr1-116802077-INS-70 |
| chr1-116807876-INS-7118 | chr1-116807877-INS-7118 |
| chr1-116825265-INS-575 | chr1-116825265-INS-575 |
| chr1-116867291-INS-51 | chr1-116867292-INS-58 |
| chr1-116882705-INS-67 | chr1-116882705-INS-96 |
| chr1-116908606-INS-55 | chr1-116908605-INS-53 |
| chr1-116912731-INS-1757 | chr1-116912732-INS-1757 |
| chr1-116920582-INS-152 | chr1-116920582-INS-152 |
| chr1-116944802-INS-6180 | chr1-116944803-INS-6183 |
| chr1-116956651-INS-107 | chr1-116956652-INS-82 |
| chr1-116957538-INS-441 | chr1-116957538-INS-441 |
| chr1-116966718-INS-504 | chr1-116966718-INS-504 |
| chr1-116988081-INS-54 | chr1-116988081-INS-52 |
| chr1-117003519-INS-2699 | chr1-117003520-INS-2699 |
| chr1-117014169-INS-75 | chr1-117014142-INS-58 |
| chr1-117023149-INS-57 | chr1-117023125-INS-53 |
| chr1-117044868-INS-148 | chr1-117044868-INS-148 |
| chr1-117049288-INS-963 | chr1-117049288-INS-963 |
| chr1-117075667-INS-82 | chr1-117075667-INS-82 |
| chr1-117087119-INS-258 | chr1-117087119-INS-258 |
| chr1-117095203-INS-212 | chr1-117095203-INS-212 |
| chr1-117147488-INS-133 | chr1-117147488-INS-133 |
| chr1-117154004-INS-67 | chr1-117154004-INS-67 |
| chr1-117199679-INS-129 | chr1-117199679-INS-129 |
| chr1-117233317-INS-4534 | chr1-117233318-INS-4534 |
| chr1-117234768-INS-97 | chr1-117234770-INS-117 |
| chr1-117236530-INS-312 | chr1-117236530-INS-312 |
| chr1-117244957-INS-163 | chr1-117244957-INS-163 |
| chr1-117257666-INS-502 | chr1-117257666-INS-503 |
| chr1-117268001-INS-3291 | chr1-117268002-INS-3303 |
| chr1-117280786-INS-62 | chr1-117280786-INS-62 |
| chr1-117293982-INS-3110 | chr1-117293983-INS-3110 |
| chr1-117308986-INS-303 | chr1-117308986-INS-305 |
| chr1-117310586-INS-60 | chr1-117310586-INS-60 |
| chr1-117322109-INS-93 | chr1-117322109-INS-129 |
| chr1-117328020-INS-72 | chr1-117328020-INS-73 |
| chr1-117338870-INS-58 | chr1-117338870-INS-58 |
| chr1-117471329-INS-275 | chr1-117471329-INS-275 |
| chr1-117474969-INS-64 | chr1-117474969-INS-66 |

| | |
|---|---|
| chr1-117488603-INS-123 | chr1-117488603-INS-123 |
| chr1-117495149-INS-211 | chr1-117495149-INS-215 |
| chr1-117522740-INS-8233 | chr1-117522741-INS-8253 |
| chr1-117543288-INS-52 | chr1-117543288-INS-56 |
| chr1-117620819-INS-2756 | chr1-117620820-INS-2753 |
| chr1-117664532-INS-2988 | chr1-117664533-INS-2991 |
| chr1-117740805-INS-74 | chr1-117740805-INS-76 |
| chr1-117813467-INS-4892 | chr1-117813468-INS-4892 |
| chr1-117855540-INS-1396 | chr1-117855541-INS-1396 |
| chr1-117903488-INS-6903 | chr1-117903489-INS-6903 |
| chr1-117911094-INS-2469 | chr1-117911095-INS-2469 |
| chr1-117921001-INS-7086 | chr1-117921002-INS-7086 |
| chr1-117923226-INS-203 | chr1-117923226-INS-203 |
| chr1-117952884-INS-1862 | chr1-117952885-INS-1862 |
| chr1-117952885-INS-312 | chr1-117952885-INS-309 |
| chr1-118031136-INS-534 | chr1-118031136-INS-536 |
| chr1-118042770-INS-202 | chr1-118042770-INS-202 |
| chr1-118106936-INS-6647 | chr1-118106937-INS-6647 |
| chr1-118115089-INS-52 | chr1-118115089-INS-54 |
| chr1-118121050-INS-227 | chr1-118121050-INS-227 |
| chr1-118203139-INS-6289 | chr1-118203140-INS-6289 |
| chr1-118211940-INS-897 | chr1-118211940-INS-897 |
| chr1-118222543-INS-196 | chr1-118222543-INS-199 |
| chr1-118253350-INS-206 | chr1-118253350-INS-207 |
| chr1-118268617-INS-416 | chr1-118268655-INS-416 |
| chr1-118270720-INS-109 | chr1-118270720-INS-109 |
| chr1-118275203-INS-445 | chr1-118275203-INS-444 |
| chr1-118282844-INS-513 | chr1-118282844-INS-511 |
| chr1-118286754-INS-88 | chr1-118286754-INS-103 |
| chr1-118288440-INS-56 | chr1-118288430-INS-95 |
| chr1-118291107-INS-108 | chr1-118291107-INS-108 |
| chr1-118298638-INS-52 | chr1-118298637-INS-77 |
| chr1-118303269-INS-52 | chr1-118303269-INS-52 |
| chr1-118306618-INS-101 | chr1-118306616-INS-110 |
| chr1-118340108-INS-314 | chr1-118340108-INS-314 |
| chr1-118349633-INS-116 | chr1-118349633-INS-114 |
| chr1-118368137-INS-191 | chr1-118368137-INS-191 |
| chr1-118387705-INS-99 | chr1-118387705-INS-99 |
| chr1-118394404-INS-96 | chr1-118394404-INS-96 |
| chr1-118404009-INS-68 | chr1-118404001-INS-86 |
| chr1-118484335-INS-122 | chr1-118484335-INS-122 |
| chr1-118500306-INS-319 | chr1-118500307-INS-316 |
| chr1-118503292-INS-209 | chr1-118503292-INS-209 |
| chr1-118508765-INS-204 | chr1-118508765-INS-204 |
| chr1-118534253-INS-78 | chr1-118534253-INS-78 |
| chr1-118541202-INS-204 | chr1-118541202-INS-213 |
| chr1-118551535-INS-53 | chr1-118551536-INS-53 |
| chr1-118565078-INS-202 | chr1-118565078-INS-199 |
| chr1-118572319-INS-158 | chr1-118572319-INS-111 |
| chr1-118575021-INS-81 | chr1-118575021-INS-81 |
| chr1-118580922-INS-82 | chr1-118580922-INS-106 |
| chr1-118588267-INS-90 | chr1-118588275-INS-60 |
| chr1-118595870-INS-71 | chr1-118595870-INS-71 |
| chr1-118598504-INS-757 | chr1-118598504-INS-758 |
| chr1-118608036-INS-70 | chr1-118608036-INS-70 |
| chr1-118618753-INS-150 | chr1-118618753-INS-150 |
| chr1-118627732-INS-202 | chr1-118627732-INS-201 |
| chr1-118628374-INS-5543 | chr1-118628375-INS-5543 |
| chr1-118630602-INS-220 | chr1-118630602-INS-220 |
| chr1-118637540-INS-91 | chr1-118637557-INS-93 |
| chr1-118642693-INS-132 | chr1-118642693-INS-112 |
| chr1-118651191-INS-1293 | chr1-118651192-INS-1293 |
| chr1-118658956-INS-64 | chr1-118658956-INS-64 |
| chr1-118668676-INS-94 | chr1-118668676-INS-84 |
| chr1-118677336-INS-59 | chr1-118677337-INS-63 |
| chr1-118681292-INS-66 | chr1-118681292-INS-50 |
| chr1-118698158-INS-719 | chr1-118698158-INS-718 |
| chr1-118710522-INS-6401 | chr1-118710523-INS-6401 |
| chr1-118722539-INS-319 | chr1-118722539-INS-341 |
| chr1-118727834-INS-62 | chr1-118727834-INS-65 |
| chr1-118739256-INS-2619 | chr1-118739257-INS-2619 |
| chr1-118739257-INS-420 | chr1-118739257-INS-420 |
| chr1-118768516-INS-489 | chr1-118768516-INS-493 |
| chr1-118786224-INS-60 | chr1-118786234-INS-62 |
| chr1-118810421-INS-212 | chr1-118810429-INS-212 |
| chr1-118833747-INS-52 | chr1-118833747-INS-54 |
| chr1-118846776-INS-270 | chr1-118846776-INS-262 |

| | |
|---|---|
| chr1-118875451-INS-68 | chr1-118875451-INS-68 |
| chr1-118880376-INS-292 | chr1-118880376-INS-292 |
| chr1-118899547-INS-51 | chr1-118899547-INS-51 |
| chr1-118907191-INS-54 | chr1-118907192-INS-68 |
| chr1-118941335-INS-51 | chr1-118941335-INS-51 |
| chr1-118942683-INS-68 | chr1-118942676-INS-60 |
| chr1-118947267-INS-508 | chr1-118947267-INS-508 |
| chr1-118975125-INS-52 | chr1-118975125-INS-51 |
| chr1-118984806-INS-86 | chr1-118984806-INS-97 |
| chr1-119016651-INS-1286 | chr1-119016652-INS-1289 |
| chr1-119019380-INS-239 | chr1-119019380-INS-242 |
| chr1-119033543-INS-408 | chr1-119033543-INS-408 |
| chr1-119035746-INS-59 | chr1-119035746-INS-59 |
| chr1-119041736-INS-6612 | chr1-119041737-INS-6612 |
| chr1-119051429-INS-72 | chr1-119051429-INS-76 |
| chr1-119056029-INS-58 | chr1-119056029-INS-58 |
| chr1-119101560-INS-4922 | chr1-119101561-INS-4922 |
| chr1-119148125-INS-179 | chr1-119148125-INS-180 |
| chr1-119183818-INS-68 | chr1-119183818-INS-68 |
| chr1-119192288-INS-156 | chr1-119192290-INS-160 |
| chr1-119225387-INS-82 | chr1-119225377-INS-129 |
| chr1-119227326-INS-638 | chr1-119227326-INS-637 |
| chr1-119231579-INS-73 | chr1-119231579-INS-63 |
| chr1-119238141-INS-77 | chr1-119238141-INS-55 |
| chr1-119239660-INS-272 | chr1-119239688-INS-284 |
| chr1-119268012-INS-60 | chr1-119268012-INS-74 |
| chr1-119286834-INS-207 | chr1-119286834-INS-208 |
| chr1-119290654-INS-60 | chr1-119290654-INS-56 |
| chr1-119302969-INS-6476 | chr1-119302970-INS-6476 |
| chr1-119310106-INS-53 | chr1-119310106-INS-53 |
| chr1-119329849-INS-55 | chr1-119329850-INS-50 |
| chr1-119409279-INS-58 | chr1-119409261-INS-76 |
| chr1-119503445-INS-228 | chr1-119503445-INS-228 |
| chr1-119580078-INS-207 | chr1-119580078-INS-207 |
| chr1-119658116-INS-100 | chr1-119658116-INS-78 |
| chr1-119743778-INS-52 | chr1-119743778-INS-52 |
| chr1-119769100-INS-87 | chr1-119769100-INS-87 |
| chr1-119787449-INS-3231 | chr1-119787450-INS-3231 |
| chr1-119794107-INS-59 | chr1-119794107-INS-59 |
| chr1-119803283-INS-72 | chr1-119803283-INS-74 |
| chr1-119805909-INS-62 | chr1-119805900-INS-52 |
| chr1-119811671-INS-620 | chr1-119811671-INS-609 |
| chr1-119826155-INS-863 | chr1-119826155-INS-875 |
| chr1-119851483-INS-52 | chr1-119851483-INS-52 |
| chr1-119891914-INS-54 | chr1-119891914-INS-54 |
| chr1-119911334-INS-180 | chr1-119911334-INS-180 |
| chr1-119920013-INS-96 | chr1-119920016-INS-81 |
| chr1-119930160-INS-1737 | chr1-119930161-INS-1737 |
| chr1-119933097-INS-111 | chr1-119933097-INS-111 |
| chr1-119954810-INS-60 | chr1-119954810-INS-60 |
| chr1-119969563-INS-58 | chr1-119969563-INS-59 |
| chr1-119980539-INS-53 | chr1-119980551-INS-64 |
| chr1-120017799-INS-128 | chr1-120017799-INS-136 |
| chr1-120034584-INS-202 | chr1-120034584-INS-202 |
| chr1-120084190-INS-73 | chr1-120084190-INS-73 |
| chr1-120124496-INS-52 | chr1-120124468-INS-58 |
| chr1-120144384-INS-361 | chr1-120144384-INS-361 |
| chr1-120161238-INS-112 | chr1-120161230-INS-158 |
| chr1-120189749-INS-212 | chr1-120189749-INS-212 |
| chr1-120205091-INS-190 | chr1-120205091-INS-192 |
| chr1-120206066-INS-72 | chr1-120206066-INS-68 |
| chr1-120210141-INS-84 | chr1-120210133-INS-120 |
| chr1-120211384-INS-55 | chr1-120211362-INS-56 |
| chr1-120211402-INS-672 | chr1-120211402-INS-672 |
| chr1-120227727-INS-72 | chr1-120227727-INS-78 |
| chr1-120235805-INS-6867 | chr1-120235806-INS-6868 |
| chr1-120236235-INS-90 | chr1-120236235-INS-60 |
| chr1-120244887-INS-635 | chr1-120244887-INS-635 |
| chr1-120247057-INS-170 | chr1-120247057-INS-130 |
| chr1-120254969-INS-105 | chr1-120255006-INS-92 |
| chr1-120283502-INS-135 | chr1-120283502-INS-135 |
| chr1-120300032-INS-1911 | chr1-120300033-INS-1914 |
| chr1-120334132-INS-115 | chr1-120334132-INS-115 |
| chr1-120370940-INS-108 | chr1-120370940-INS-134 |
| chr1-120399333-INS-88 | chr1-120399333-INS-88 |
| chr1-120433271-INS-58 | chr1-120433271-INS-58 |
| chr1-120451700-INS-56 | chr1-120451700-INS-58 |

| | |
|---|---|
| chr1-120457314-INS-193 | chr1-120457314-INS-208 |
| chr1-120465066-INS-79 | chr1-120465083-INS-52 |
| chr1-120468258-INS-58 | chr1-120468258-INS-62 |
| chr1-120471799-INS-6470 | chr1-120471800-INS-6471 |
| chr1-120513603-INS-60 | chr1-120513603-INS-51 |
| chr1-120518022-INS-78 | chr1-120518022-INS-64 |
| chr1-120555917-INS-212 | chr1-120555917-INS-212 |
| chr1-120565982-INS-66 | chr1-120565982-INS-66 |
| chr1-120580470-INS-113 | chr1-120580470-INS-114 |
| chr1-120582165-INS-220 | chr1-120582165-INS-220 |
| chr1-120605698-INS-220 | chr1-120605698-INS-220 |
| chr1-120615375-INS-144 | chr1-120615375-INS-132 |
| chr1-120636868-INS-138 | chr1-120636868-INS-139 |
| chr1-120658556-INS-117 | chr1-120658556-INS-117 |
| chr1-120754669-INS-106 | chr1-120754669-INS-106 |
| chr1-120760628-INS-72 | chr1-120760628-INS-64 |
| chr1-120793748-INS-1915 | chr1-120793749-INS-1915 |
| chr1-120808120-INS-399 | chr1-120808121-INS-399 |
| chr1-120820416-INS-66 | chr1-120820416-INS-84 |
| chr1-120841595-INS-120 | chr1-120841595-INS-126 |
| chr1-120850234-INS-493 | chr1-120850234-INS-493 |
| chr1-120866708-INS-6858 | chr1-120866709-INS-6858 |
| chr1-120879359-INS-51 | chr1-120879360-INS-54 |
| chr1-120894645-INS-79 | chr1-120894646-INS-84 |
| chr1-120906028-INS-64 | chr1-120906028-INS-64 |
| chr1-120912556-INS-90 | chr1-120912556-INS-90 |
| chr1-120922128-INS-212 | chr1-120922128-INS-212 |
| chr1-120926521-INS-1865 | chr1-120926521-INS-1865 |
| chr1-120928165-INS-424 | chr1-120928165-INS-423 |
| chr1-120940686-INS-790 | chr1-120940686-INS-790 |
| chr1-120977580-INS-55 | chr1-120977580-INS-55 |
| chr1-120989195-INS-58 | chr1-120989195-INS-58 |
| chr1-121004948-INS-132 | chr1-121004948-INS-132 |
| chr1-121005831-INS-58 | chr1-121005831-INS-58 |
| chr1-121011558-INS-94 | chr1-121011558-INS-86 |
| chr1-121018239-INS-282 | chr1-121018247-INS-270 |
| chr1-121033856-INS-100 | chr1-121033898-INS-52 |
| chr1-121035019-INS-103 | chr1-121035019-INS-103 |
| chr1-121036692-INS-204 | chr1-121036691-INS-226 |
| chr1-121049619-INS-6732 | chr1-121049620-INS-6734 |
| chr1-121081209-INS-325 | chr1-121081209-INS-325 |
| chr1-121085835-INS-68 | chr1-121085834-INS-62 |
| chr1-121087171-INS-56 | chr1-121087171-INS-56 |
| chr1-121112991-INS-274 | chr1-121112967-INS-253 |
| chr1-121166273-INS-76 | chr1-121166264-INS-66 |
| chr1-121198984-INS-5027 | chr1-121198985-INS-5027 |
| chr1-121199790-INS-81 | chr1-121199790-INS-81 |
| chr1-121204373-INS-400 | chr1-121204373-INS-400 |
| chr1-121235546-INS-56 | chr1-121235546-INS-64 |
| chr1-121285822-INS-58 | chr1-121285825-INS-100 |
| chr1-121286426-INS-282 | chr1-121286427-INS-282 |
| chr1-121287032-INS-880 | chr1-121287033-INS-880 |
| chr1-121289560-INS-66 | chr1-121289560-INS-63 |
| chr1-121303389-INS-57 | chr1-121303397-INS-59 |
| chr1-121329256-INS-106 | chr1-121329256-INS-108 |
| chr1-121334504-INS-68 | chr1-121334508-INS-72 |
| chr1-121346943-INS-416 | chr1-121346944-INS-416 |
| chr1-121352281-INS-99 | chr1-121352281-INS-99 |
| chr1-121354030-INS-60 | chr1-121354030-INS-74 |
| chr1-121366581-INS-90 | chr1-121366581-INS-92 |
| chr1-121369788-INS-74 | chr1-121369788-INS-74 |
| chr1-121386700-INS-241 | chr1-121386701-INS-242 |
| chr1-121395415-INS-137 | chr1-121395418-INS-178 |
| chr1-121399712-INS-433 | chr1-121399712-INS-432 |
| chr1-121400562-INS-91 | chr1-121400555-INS-50 |
| chr1-121405696-INS-360 | chr1-121405696-INS-360 |
| chr1-121412537-INS-78 | chr1-121412537-INS-78 |
| chr1-121436032-INS-58 | chr1-121436033-INS-60 |
| chr1-121439010-INS-249 | chr1-121439010-INS-249 |
| chr1-121442555-INS-60 | chr1-121442555-INS-60 |
| chr1-121448869-INS-114 | chr1-121448869-INS-138 |
| chr1-121450360-INS-72 | chr1-121450360-INS-72 |
| chr1-121459488-INS-68 | chr1-121459488-INS-68 |
| chr1-121474649-INS-76 | chr1-121474649-INS-76 |
| chr1-121479868-INS-57 | chr1-121479893-INS-55 |
| chr1-121485895-INS-77 | chr1-121485894-INS-77 |
| chr1-121502638-INS-498 | chr1-121502638-INS-498 |

| | |
|---|---|
| chr1-121505137-INS-291 | chr1-121505138-INS-288 |
| chr1-121507145-INS-113 | chr1-121507145-INS-113 |
| chr1-121542036-INS-347 | chr1-121542036-INS-347 |
| chr1-121562729-INS-504 | chr1-121562729-INS-506 |
| chr1-121596800-INS-3302 | chr1-121596801-INS-3302 |
| chr1-121598405-INS-427 | chr1-121598405-INS-427 |
| chr1-121599560-INS-155 | chr1-121599560-INS-174 |
| chr1-121605950-INS-738 | chr1-121605950-INS-738 |
| chr1-121616906-INS-7214 | chr1-121616907-INS-7215 |
| chr1-121636578-INS-72 | chr1-121636586-INS-54 |
| chr1-121701724-INS-127 | chr1-121701724-INS-127 |
| chr1-121703447-INS-103 | chr1-121703447-INS-103 |
| chr1-121715881-INS-1021 | chr1-121715882-INS-1021 |
| chr1-121717117-INS-7100 | chr1-121717118-INS-7100 |
| chr1-121726405-INS-1200 | chr1-121726406-INS-1200 |
| chr1-121727604-INS-87 | chr1-121727604-INS-87 |
| chr1-121728751-INS-52 | chr1-121728751-INS-56 |
| chr1-121739211-INS-2117 | chr1-121739212-INS-2117 |
| chr1-121756560-INS-5540 | chr1-121756561-INS-5540 |
| chr1-121770296-INS-62 | chr1-121770296-INS-80 |
| chr1-121776741-INS-66 | chr1-121776741-INS-66 |
| chr1-121784042-INS-72 | chr1-121784075-INS-79 |
| chr1-121814738-INS-85 | chr1-121814738-INS-85 |
| chr1-121839021-INS-75 | chr1-121839021-INS-50 |
| chr1-121906734-INS-102 | chr1-121906735-INS-105 |
| chr1-121920645-INS-1346 | chr1-121920646-INS-1347 |
| chr1-121923454-INS-95 | chr1-121923454-INS-89 |
| chr1-122008750-INS-62 | chr1-122008750-INS-58 |
| chr1-122014610-INS-212 | chr1-122014610-INS-212 |
| chr1-122024360-INS-55 | chr1-122024379-INS-56 |
| chr1-122025813-INS-60 | chr1-122025854-INS-54 |
| chr1-122050180-INS-58 | chr1-122050180-INS-58 |
| chr1-122087312-INS-52 | chr1-122087334-INS-52 |
| chr1-122104184-INS-66 | chr1-122104184-INS-68 |
| chr1-122109238-INS-945 | chr1-122109238-INS-945 |
| chr1-122124200-INS-6239 | chr1-122124201-INS-6239 |
| chr1-122160565-INS-62 | chr1-122160565-INS-62 |
| chr1-122162359-INS-2862 | chr1-122162360-INS-2862 |
| chr1-122168064-INS-69 | chr1-122168064-INS-74 |
| chr1-122178516-INS-54 | chr1-122178516-INS-59 |
| chr1-122184274-INS-395 | chr1-122184274-INS-395 |
| chr1-122186839-INS-65 | chr1-122186839-INS-65 |
| chr1-122193523-INS-6169 | chr1-122193524-INS-6169 |
| chr1-122194854-INS-241 | chr1-122194854-INS-241 |
| chr1-122197677-INS-52 | chr1-122197677-INS-66 |
| chr1-122206588-INS-52 | chr1-122206588-INS-64 |
| chr1-122228439-INS-3724 | chr1-122228440-INS-3724 |
| chr1-122296880-INS-78 | chr1-122296880-INS-77 |
| chr1-122309901-INS-67 | chr1-122309900-INS-81 |
| chr1-122323388-INS-120 | chr1-122323388-INS-108 |
| chr1-122371605-INS-200 | chr1-122371605-INS-201 |
| chr1-122383325-INS-537 | chr1-122383325-INS-533 |
| chr1-122398038-INS-6445 | chr1-122398039-INS-6445 |
| chr1-122415795-INS-7882 | chr1-122415796-INS-7883 |
| chr1-122454468-INS-120 | chr1-122454468-INS-97 |
| chr1-122457151-INS-52 | chr1-122457151-INS-54 |
| chr1-122464765-INS-68 | chr1-122464765-INS-68 |
| chr1-122475106-INS-51 | chr1-122475111-INS-51 |
| chr1-122498793-INS-501 | chr1-122498793-INS-503 |
| chr1-122503689-INS-64 | chr1-122503689-INS-64 |
| chr1-122521428-INS-707 | chr1-122521428-INS-707 |
| chr1-122544019-INS-224 | chr1-122544019-INS-224 |
| chr1-122554065-INS-77 | chr1-122554066-INS-63 |
| chr1-122554255-INS-541 | chr1-122554256-INS-541 |
| chr1-122567476-INS-7337 | chr1-122567477-INS-7337 |
| chr1-122582245-INS-6760 | chr1-122582246-INS-6759 |
| chr1-122590851-INS-6389 | chr1-122590852-INS-6390 |
| chr1-122621497-INS-2962 | chr1-122621498-INS-2962 |
| chr1-122659812-INS-474 | chr1-122659812-INS-473 |
| chr1-122677809-INS-109 | chr1-122677803-INS-95 |
| chr1-122716536-INS-79 | chr1-122716538-INS-82 |
| chr1-122724329-INS-52 | chr1-122724329-INS-52 |
| chr1-122725178-INS-313 | chr1-122725178-INS-313 |
| chr1-122739656-INS-1500 | chr1-122739657-INS-1500 |
| chr1-122740086-INS-662 | chr1-122740086-INS-663 |
| chr1-122809333-INS-1195 | chr1-122809350-INS-1171 |
| chr1-122823108-INS-109 | chr1-122823108-INS-66 |

| | |
|---|---|
| chr1-122827262-INS-6179 | chr1-122827263-INS-6180 |
| chr1-122847879-INS-396 | chr1-122847880-INS-396 |
| chr1-122850761-INS-515 | chr1-122850761-INS-515 |
| chr1-122876137-INS-1404 | chr1-122876138-INS-1404 |
| chr1-122880109-INS-51 | chr1-122880109-INS-51 |
| chr1-122907757-INS-6367 | chr1-122907758-INS-6367 |
| chr1-122910029-INS-3116 | chr1-122910030-INS-3116 |
| chr1-122914930-INS-1630 | chr1-122914931-INS-1630 |
| chr1-122915776-INS-111 | chr1-122915809-INS-86 |
| chr1-122926183-INS-6381 | chr1-122926184-INS-6381 |
| chr1-122929023-INS-87 | chr1-122929023-INS-79 |
| chr1-122933274-INS-68 | chr1-122933274-INS-67 |
| chr1-122959382-INS-56 | chr1-122959382-INS-54 |
| chr1-122974058-INS-284 | chr1-122974020-INS-312 |
| chr1-122994649-INS-1345 | chr1-122994650-INS-1342 |
| chr1-122998171-INS-54 | chr1-122998171-INS-87 |
| chr1-123002053-INS-1325 | chr1-123002054-INS-1324 |
| chr1-123004794-INS-61 | chr1-123004794-INS-54 |
| chr1-123005841-INS-6205 | chr1-123005842-INS-6203 |
| chr1-123016090-INS-1988 | chr1-123016091-INS-1988 |
| chr1-123035818-INS-3358 | chr1-123035819-INS-3358 |
| chr1-123039379-INS-61 | chr1-123039379-INS-62 |
| chr1-123045974-INS-64 | chr1-123045972-INS-71 |
| chr1-123115929-INS-744 | chr1-123115929-INS-745 |
| chr1-123132994-INS-72 | chr1-123132994-INS-73 |
| chr1-123144183-INS-7124 | chr1-123144184-INS-7124 |
| chr1-123157884-INS-84 | chr1-123157890-INS-78 |
| chr1-123167616-INS-72 | chr1-123167616-INS-72 |
| chr1-123209467-INS-615 | chr1-123209467-INS-615 |
| chr1-123221630-INS-4220 | chr1-123221631-INS-4220 |
| chr1-123245293-INS-72 | chr1-123245292-INS-65 |
| chr1-123267797-INS-70 | chr1-123267779-INS-65 |
| chr1-123274753-INS-57 | chr1-123274755-INS-57 |
| chr1-123285101-INS-96 | chr1-123285111-INS-110 |
| chr1-123287097-INS-474 | chr1-123287097-INS-474 |
| chr1-123291362-INS-88 | chr1-123291362-INS-92 |
| chr1-123327638-INS-84 | chr1-123327638-INS-84 |
| chr1-123337515-INS-260 | chr1-123337515-INS-262 |
| chr1-123342903-INS-6593 | chr1-123342904-INS-6595 |
| chr1-123344752-INS-84 | chr1-123344753-INS-83 |
| chr1-123351588-INS-58 | chr1-123351588-INS-60 |
| chr1-123369554-INS-819 | chr1-123369552-INS-821 |
| chr1-123379259-INS-222 | chr1-123379248-INS-236 |
| chr1-123380548-INS-50 | chr1-123380548-INS-52 |
| chr1-123417755-INS-6447 | chr1-123417756-INS-6439 |
| chr1-123426460-INS-610 | chr1-123426460-INS-608 |
| chr1-123430753-INS-51 | chr1-123430753-INS-63 |
| chr1-123436873-INS-52 | chr1-123436871-INS-54 |
| chr1-123457910-INS-56 | chr1-123457914-INS-102 |
| chr1-123458748-INS-655 | chr1-123458748-INS-662 |
| chr1-123464201-INS-52 | chr1-123464201-INS-52 |
| chr1-123476816-INS-113 | chr1-123476816-INS-114 |
| chr1-123489421-INS-116 | chr1-123489406-INS-108 |
| chr1-123513508-INS-66 | chr1-123513506-INS-64 |
| chr1-123525055-INS-63 | chr1-123525055-INS-63 |
| chr1-123551771-INS-4258 | chr1-123551772-INS-4271 |
| chr1-123562522-INS-64 | chr1-123562534-INS-50 |
| chr1-123562902-INS-7005 | chr1-123562903-INS-7005 |
| chr1-123584307-INS-53 | chr1-123584309-INS-55 |
| chr1-123587454-INS-114 | chr1-123587454-INS-94 |
| chr1-123597142-INS-60 | chr1-123597142-INS-60 |
| chr1-123599034-INS-201 | chr1-123599034-INS-201 |
| chr1-123633304-INS-693 | chr1-123633304-INS-694 |
| chr1-123646350-INS-97 | chr1-123646350-INS-97 |
| chr1-123664086-INS-112 | chr1-123664086-INS-112 |
| chr1-123669325-INS-112 | chr1-123669325-INS-118 |
| chr1-123672669-INS-1633 | chr1-123672670-INS-1633 |
| chr1-123674325-INS-60 | chr1-123674325-INS-60 |
| chr1-123676276-INS-316 | chr1-123676276-INS-316 |
| chr1-123682552-INS-479 | chr1-123682552-INS-479 |
| chr1-123689726-INS-261 | chr1-123689726-INS-258 |
| chr1-123707539-INS-689 | chr1-123707539-INS-689 |
| chr1-123710452-INS-843 | chr1-123710452-INS-843 |
| chr1-123724801-INS-1756 | chr1-123724802-INS-1756 |
| chr1-123727091-INS-130 | chr1-123727091-INS-130 |
| chr1-123733378-INS-263 | chr1-123733378-INS-266 |
| chr1-123738202-INS-54 | chr1-123738202-INS-54 |

| | |
|---|---|
| chr1-123777855-INS-93 | chr1-123777855-INS-93 |
| chr1-123819884-INS-6746 | chr1-123819885-INS-6734 |
| chr1-123841880-INS-536 | chr1-123841880-INS-536 |
| chr1-123860149-INS-126 | chr1-123860149-INS-108 |
| chr1-123905902-INS-110 | chr1-123905906-INS-122 |
| chr1-123911388-INS-401 | chr1-123911388-INS-401 |
| chr1-123922476-INS-408 | chr1-123922476-INS-408 |
| chr1-123964935-INS-3268 | chr1-123964936-INS-3289 |
| chr1-123968073-INS-59 | chr1-123968084-INS-60 |
| chr1-123969397-INS-53 | chr1-123969406-INS-54 |
| chr1-123984398-INS-59 | chr1-123984398-INS-63 |
| chr1-124010491-INS-55 | chr1-124010491-INS-56 |
| chr1-124039313-INS-51 | chr1-124039313-INS-57 |
| chr1-124047991-INS-187 | chr1-124047991-INS-157 |
| chr1-124052019-INS-7131 | chr1-124052020-INS-7131 |
| chr1-124086958-INS-97 | chr1-124086958-INS-106 |
| chr1-124112702-INS-102 | chr1-124112702-INS-52 |
| chr1-124204833-INS-84 | chr1-124204830-INS-87 |
| chr1-124254225-INS-365 | chr1-124254225-INS-365 |
| chr1-124265959-INS-66 | chr1-124265959-INS-60 |
| chr1-124322927-INS-83 | chr1-124322927-INS-83 |
| chr1-124351102-INS-110 | chr1-124351102-INS-95 |
| chr1-124377413-INS-67 | chr1-124377413-INS-96 |
| chr1-124394556-INS-68 | chr1-124394556-INS-56 |
| chr1-124412191-INS-69 | chr1-124412191-INS-72 |
| chr1-124435248-INS-221 | chr1-124435245-INS-219 |
| chr1-124450302-INS-73 | chr1-124450302-INS-73 |
| chr1-124476135-INS-56 | chr1-124476135-INS-52 |
| chr1-124483326-INS-1251 | chr1-124483339-INS-1246 |
| chr1-124500565-INS-302 | chr1-124500565-INS-302 |
| chr1-124501576-INS-1157 | chr1-124501577-INS-1179 |
| chr1-124514487-INS-76 | chr1-124514488-INS-66 |
| chr1-124517666-INS-60 | chr1-124517666-INS-60 |
| chr1-124534289-INS-52 | chr1-124534289-INS-52 |
| chr1-124572734-INS-53 | chr1-124572734-INS-51 |
| chr1-124591343-INS-84 | chr1-124591373-INS-54 |
| chr1-124592778-INS-7217 | chr1-124592779-INS-7217 |
| chr1-124653211-INS-76 | chr1-124653211-INS-76 |
| chr1-124663079-INS-464 | chr1-124663079-INS-464 |
| chr1-124678082-INS-178 | chr1-124678083-INS-179 |
| chr1-124682514-INS-81 | chr1-124682514-INS-81 |
| chr1-124686143-INS-55 | chr1-124686143-INS-55 |
| chr1-124694376-INS-79 | chr1-124694378-INS-76 |
| chr1-124697243-INS-70 | chr1-124697243-INS-52 |
| chr1-124700853-INS-189 | chr1-124700853-INS-189 |
| chr1-124707053-INS-78 | chr1-124707053-INS-54 |
| chr1-124711344-INS-66 | chr1-124711344-INS-66 |
| chr1-124714958-INS-79 | chr1-124714958-INS-79 |
| chr1-124718951-INS-363 | chr1-124718961-INS-374 |
| chr1-124740654-INS-245 | chr1-124740654-INS-245 |
| chr1-124754353-INS-115 | chr1-124754388-INS-82 |
| chr1-124765370-INS-245 | chr1-124765370-INS-246 |
| chr1-124778089-INS-842 | chr1-124778089-INS-841 |
| chr1-124782297-INS-64 | chr1-124782297-INS-54 |
| chr1-124792739-INS-401 | chr1-124792739-INS-401 |
| chr1-124796664-INS-216 | chr1-124796660-INS-220 |
| chr1-124811509-INS-59 | chr1-124811509-INS-58 |
| chr1-124823087-INS-80 | chr1-124823087-INS-76 |
| chr1-124830550-INS-7230 | chr1-124830551-INS-7225 |
| chr1-124883059-INS-276 | chr1-124883059-INS-264 |
| chr1-124901364-INS-140 | chr1-124901364-INS-140 |
| chr1-124905086-INS-129 | chr1-124905086-INS-131 |
| chr1-124920280-INS-56 | chr1-124920280-INS-56 |
| chr1-124928091-INS-68 | chr1-124928091-INS-70 |
| chr1-124931576-INS-82 | chr1-124931576-INS-82 |
| chr1-124955422-INS-6892 | chr1-124955423-INS-6892 |
| chr1-124969194-INS-374 | chr1-124969200-INS-376 |
| chr1-124981251-INS-81 | chr1-124981251-INS-83 |
| chr1-124988161-INS-73 | chr1-124988161-INS-73 |
| chr1-124991583-INS-157 | chr1-124991583-INS-156 |
| chr1-124998144-INS-315 | chr1-124998144-INS-315 |
| chr1-125015761-INS-63 | chr1-125015761-INS-65 |
| chr1-125045748-INS-454 | chr1-125045748-INS-475 |
| chr1-125047545-INS-132 | chr1-125047545-INS-132 |
| chr1-125076379-INS-63 | chr1-125076379-INS-63 |
| chr1-125090793-INS-169 | chr1-125090793-INS-197 |
| chr1-125095609-INS-213 | chr1-125095609-INS-213 |

| | |
|---|---|
| chr1-125126136-INS-99 | chr1-125126148-INS-63 |
| chr1-125145118-INS-95 | chr1-125145118-INS-95 |
| chr1-125159414-INS-267 | chr1-125159414-INS-266 |
| chr1-125188395-INS-113 | chr1-125188395-INS-113 |
| chr1-125199626-INS-99 | chr1-125199671-INS-60 |
| chr1-125210290-INS-405 | chr1-125210290-INS-405 |
| chr1-125217111-INS-74 | chr1-125217111-INS-74 |
| chr1-125225080-INS-381 | chr1-125225080-INS-381 |
| chr1-125229835-INS-366 | chr1-125229835-INS-366 |
| chr1-125240608-INS-56 | chr1-125240608-INS-60 |
| chr1-125270676-INS-268 | chr1-125270676-INS-268 |
| chr1-125302668-INS-648 | chr1-125302668-INS-648 |
| chr1-125315331-INS-520 | chr1-125315331-INS-520 |
| chr1-125351702-INS-82 | chr1-125351702-INS-83 |
| chr1-125354004-INS-497 | chr1-125354004-INS-497 |
| chr1-125407784-INS-87 | chr1-125407783-INS-94 |
| chr1-125417632-INS-62 | chr1-125417632-INS-60 |
| chr1-125432276-INS-59 | chr1-125432276-INS-59 |
| chr1-125435346-INS-246 | chr1-125435346-INS-261 |
| chr1-125465787-INS-494 | chr1-125465793-INS-487 |
| chr1-125471904-INS-62 | chr1-125471906-INS-63 |
| chr1-125480879-INS-68 | chr1-125480879-INS-68 |
| chr1-125486357-INS-68 | chr1-125486357-INS-78 |
| chr1-125489391-INS-65 | chr1-125489391-INS-67 |
| chr1-125496489-INS-316 | chr1-125496489-INS-316 |
| chr1-125507237-INS-386 | chr1-125507237-INS-388 |
| chr1-125518936-INS-56 | chr1-125518965-INS-60 |
| chr1-125546006-INS-452 | chr1-125546006-INS-452 |
| chr1-125574625-INS-110 | chr1-125574625-INS-110 |
| chr1-125610066-INS-493 | chr1-125610066-INS-493 |
| chr1-125623659-INS-750 | chr1-125623659-INS-750 |
| chr1-125634549-INS-3341 | chr1-125634550-INS-3341 |
| chr1-125656404-INS-86 | chr1-125656384-INS-104 |
| chr1-125659232-INS-86 | chr1-125659232-INS-86 |
| chr1-125736045-INS-52 | chr1-125736045-INS-52 |
| chr1-125739718-INS-500 | chr1-125739718-INS-500 |
| chr1-125757459-INS-204 | chr1-125757460-INS-203 |
| chr1-125770132-INS-58 | chr1-125770136-INS-56 |
| chr1-125781249-INS-105 | chr1-125781249-INS-105 |
| chr1-125783908-INS-60 | chr1-125783908-INS-64 |
| chr1-125794620-INS-600 | chr1-125794620-INS-550 |
| chr1-125812651-INS-51 | chr1-125812651-INS-51 |
| chr1-125819641-INS-225 | chr1-125819641-INS-226 |
| chr1-125820584-INS-54 | chr1-125820584-INS-52 |
| chr1-125822206-INS-52 | chr1-125822206-INS-56 |
| chr1-125825898-INS-78 | chr1-125825898-INS-52 |
| chr1-125851832-INS-6352 | chr1-125851833-INS-6352 |
| chr1-125855967-INS-126 | chr1-125855967-INS-126 |
| chr1-125881679-INS-955 | chr1-125881679-INS-955 |
| chr1-125883082-INS-75 | chr1-125883095-INS-94 |
| chr1-125889678-INS-64 | chr1-125889678-INS-60 |
| chr1-125892111-INS-619 | chr1-125892111-INS-619 |
| chr1-125897211-INS-6452 | chr1-125897212-INS-6452 |
| chr1-125937120-INS-455 | chr1-125937120-INS-455 |
| chr1-126000966-INS-2704 | chr1-126000967-INS-2704 |
| chr1-126008328-INS-156 | chr1-126008328-INS-139 |
| chr1-126076882-INS-6166 | chr1-126076883-INS-6166 |
| chr1-126077624-INS-65 | chr1-126077624-INS-62 |
| chr1-126084524-INS-52 | chr1-126084524-INS-50 |
| chr1-126108063-INS-58 | chr1-126108063-INS-58 |
| chr1-126187254-INS-524 | chr1-126187254-INS-524 |
| chr1-126189240-INS-105 | chr1-126189261-INS-131 |
| chr1-126191132-INS-464 | chr1-126191132-INS-465 |
| chr1-126194634-INS-71 | chr1-126194629-INS-75 |
| chr1-126198959-INS-80 | chr1-126198959-INS-82 |
| chr1-126203304-INS-158 | chr1-126203278-INS-150 |
| chr1-126214574-INS-265 | chr1-126214569-INS-232 |
| chr1-126215724-INS-70 | chr1-126215724-INS-58 |
| chr1-126221406-INS-70 | chr1-126221406-INS-70 |
| chr1-126282641-INS-52 | chr1-126282641-INS-52 |
| chr1-126301377-INS-64 | chr1-126301377-INS-62 |
| chr1-126311671-INS-819 | chr1-126311671-INS-819 |
| chr1-126313582-INS-60 | chr1-126313582-INS-60 |
| chr1-126336324-INS-107 | chr1-126336341-INS-106 |
| chr1-126367064-INS-499 | chr1-126367064-INS-505 |
| chr1-126428079-INS-665 | chr1-126428079-INS-666 |
| chr1-126470772-INS-94 | chr1-126470778-INS-126 |

| | |
|---|---|
| chr1-126479676-INS-52 | chr1-126479676-INS-52 |
| chr1-126494766-INS-124 | chr1-126494730-INS-132 |
| chr1-126499399-INS-481 | chr1-126499399-INS-481 |
| chr1-126511764-INS-58 | chr1-126511764-INS-58 |
| chr1-126528160-INS-94 | chr1-126528168-INS-94 |
| chr1-126546296-INS-105 | chr1-126546297-INS-105 |
| chr1-126547738-INS-1490 | chr1-126547739-INS-1490 |
| chr1-126566938-INS-52 | chr1-126566939-INS-50 |
| chr1-126581967-INS-90 | chr1-126581967-INS-66 |
| chr1-126613187-INS-188 | chr1-126613187-INS-167 |
| chr1-126620112-INS-87 | chr1-126620112-INS-87 |
| chr1-126636113-INS-102 | chr1-126636113-INS-102 |
| chr1-126665124-INS-60 | chr1-126665101-INS-60 |
| chr1-126669742-INS-52 | chr1-126669710-INS-64 |
| chr1-126677934-INS-51 | chr1-126677934-INS-60 |
| chr1-126709516-INS-81 | chr1-126709516-INS-73 |
| chr1-126775322-INS-144 | chr1-126775322-INS-118 |
| chr1-126780211-INS-7315 | chr1-126780212-INS-7291 |
| chr1-126786567-INS-52 | chr1-126786565-INS-60 |
| chr1-126823999-INS-56 | chr1-126823998-INS-66 |
| chr1-126824807-INS-109 | chr1-126824830-INS-110 |
| chr1-126830646-INS-486 | chr1-126830646-INS-486 |
| chr1-126865795-INS-328 | chr1-126865795-INS-328 |
| chr1-126892847-INS-388 | chr1-126892847-INS-388 |
| chr1-126894220-INS-291 | chr1-126894220-INS-291 |
| chr1-126901300-INS-56 | chr1-126901300-INS-56 |
| chr1-126911347-INS-6336 | chr1-126911348-INS-6337 |
| chr1-126936043-INS-61 | chr1-126936043-INS-64 |
| chr1-126948858-INS-58 | chr1-126948858-INS-58 |
| chr1-126957737-INS-748 | chr1-126957737-INS-748 |
| chr1-126978920-INS-56 | chr1-126978920-INS-58 |
| chr1-127006204-INS-84 | chr1-127006203-INS-86 |
| chr1-127021024-INS-55 | chr1-127021025-INS-55 |
| chr1-127043160-INS-138 | chr1-127043179-INS-160 |
| chr1-127048810-INS-172 | chr1-127048810-INS-172 |
| chr1-127077566-INS-578 | chr1-127077566-INS-608 |
| chr1-127083668-INS-54 | chr1-127083640-INS-58 |
| chr1-127095185-INS-6904 | chr1-127095186-INS-6904 |
| chr1-127099689-INS-78 | chr1-127099690-INS-54 |
| chr1-127107000-INS-1888 | chr1-127107001-INS-1888 |
| chr1-127118926-INS-77 | chr1-127118913-INS-69 |
| chr1-127143286-INS-81 | chr1-127143286-INS-87 |
| chr1-127149311-INS-195 | chr1-127149311-INS-195 |
| chr1-127171377-INS-152 | chr1-127171377-INS-173 |
| chr1-127211685-INS-92 | chr1-127211685-INS-64 |
| chr1-127217025-INS-133 | chr1-127217022-INS-128 |
| chr1-127222244-INS-183 | chr1-127222244-INS-184 |
| chr1-127229449-INS-7114 | chr1-127229450-INS-7114 |
| chr1-127243891-INS-58 | chr1-127243846-INS-58 |
| chr1-127254249-INS-77 | chr1-127254249-INS-77 |
| chr1-127261000-INS-110 | chr1-127261000-INS-110 |
| chr1-127269636-INS-65 | chr1-127269636-INS-65 |
| chr1-127286239-INS-69 | chr1-127286239-INS-54 |
| chr1-127337809-INS-210 | chr1-127337809-INS-210 |
| chr1-127340828-INS-227 | chr1-127340828-INS-227 |
| chr1-127346326-INS-74 | chr1-127346326-INS-76 |
| chr1-127356243-INS-94 | chr1-127356206-INS-85 |
| chr1-127363722-INS-62 | chr1-127363722-INS-56 |
| chr1-127371157-INS-1866 | chr1-127371158-INS-1866 |
| chr1-127395433-INS-87 | chr1-127395433-INS-87 |
| chr1-127424509-INS-70 | chr1-127424509-INS-65 |
| chr1-127435754-INS-52 | chr1-127435758-INS-61 |
| chr1-127443188-INS-2873 | chr1-127443189-INS-2873 |
| chr1-127453197-INS-55 | chr1-127453197-INS-63 |
| chr1-127455025-INS-1066 | chr1-127455026-INS-1064 |
| chr1-127479620-INS-54 | chr1-127479620-INS-54 |
| chr1-127494183-INS-108 | chr1-127494181-INS-112 |
| chr1-127499197-INS-57 | chr1-127499222-INS-72 |
| chr1-127507031-INS-112 | chr1-127507035-INS-63 |
| chr1-127532559-INS-56 | chr1-127532559-INS-56 |
| chr1-127535575-INS-202 | chr1-127535575-INS-202 |
| chr1-127541404-INS-293 | chr1-127541404-INS-286 |
| chr1-127544736-INS-322 | chr1-127544736-INS-324 |
| chr1-127546151-INS-58 | chr1-127546152-INS-67 |
| chr1-127547534-INS-84 | chr1-127547486-INS-98 |
| chr1-127556123-INS-500 | chr1-127556123-INS-496 |
| chr1-127560641-INS-56 | chr1-127560641-INS-60 |

| | |
|---|---|
| chr1-127565070-INS-68 | chr1-127565068-INS-72 |
| chr1-127576599-INS-1353 | chr1-127576600-INS-1353 |
| chr1-127611137-INS-1205 | chr1-127611138-INS-1206 |
| chr1-127615891-INS-281 | chr1-127615891-INS-281 |
| chr1-127667498-INS-80 | chr1-127667505-INS-50 |
| chr1-127766304-INS-242 | chr1-127766304-INS-242 |
| chr1-127775256-INS-54 | chr1-127775256-INS-83 |
| chr1-127783641-INS-92 | chr1-127783641-INS-92 |
| chr1-127790470-INS-207 | chr1-127790470-INS-206 |
| chr1-127860548-INS-600 | chr1-127860548-INS-603 |
| chr1-127878198-INS-114 | chr1-127878199-INS-110 |
| chr1-127900087-INS-194 | chr1-127900078-INS-171 |
| chr1-127930519-INS-208 | chr1-127930519-INS-208 |
| chr1-127967429-INS-218 | chr1-127967430-INS-221 |
| chr1-128022498-INS-62 | chr1-128022477-INS-68 |
| chr1-128049649-INS-211 | chr1-128049649-INS-211 |
| chr1-128087471-INS-216 | chr1-128087471-INS-217 |
| chr1-128138235-INS-208 | chr1-128138235-INS-208 |
| chr1-128175931-INS-200 | chr1-128175931-INS-198 |
| chr1-128221735-INS-765 | chr1-128221735-INS-776 |
| chr1-128241612-INS-401 | chr1-128241612-INS-403 |
| chr1-128245846-INS-129 | chr1-128245846-INS-129 |
| chr1-128264856-INS-51 | chr1-128264856-INS-51 |
| chr1-128267458-INS-239 | chr1-128267458-INS-239 |
| chr1-128272825-INS-249 | chr1-128272825-INS-250 |
| chr1-128289659-INS-204 | chr1-128289659-INS-204 |
| chr1-128340937-INS-187 | chr1-128340937-INS-187 |
| chr1-128390944-INS-141 | chr1-128390944-INS-141 |
| chr1-128394512-INS-52 | chr1-128394529-INS-52 |
| chr1-128406213-INS-62 | chr1-128406213-INS-60 |
| chr1-128407420-INS-405 | chr1-128407420-INS-405 |
| chr1-128413549-INS-203 | chr1-128413549-INS-204 |
| chr1-128433802-INS-77 | chr1-128433802-INS-67 |
| chr1-128448056-INS-7001 | chr1-128448057-INS-7001 |
| chr1-128491048-INS-65 | chr1-128491048-INS-87 |
| chr1-128492949-INS-89 | chr1-128492949-INS-89 |
| chr1-128503606-INS-202 | chr1-128503606-INS-193 |
| chr1-128555157-INS-78 | chr1-128555157-INS-58 |
| chr1-128556004-INS-60 | chr1-128556004-INS-60 |
| chr1-128566268-INS-243 | chr1-128566268-INS-244 |
| chr1-128585095-INS-88 | chr1-128585095-INS-68 |
| chr1-128587101-INS-56 | chr1-128587101-INS-50 |
| chr1-128598195-INS-77 | chr1-128598222-INS-77 |
| chr1-128616499-INS-60 | chr1-128616499-INS-62 |
| chr1-128618335-INS-53 | chr1-128618294-INS-59 |
| chr1-128664649-INS-800 | chr1-128664649-INS-800 |
| chr1-128682097-INS-354 | chr1-128682097-INS-354 |
| chr1-128683822-INS-64 | chr1-128683822-INS-83 |
| chr1-128696798-INS-156 | chr1-128696798-INS-156 |
| chr1-128707980-INS-63 | chr1-128707980-INS-61 |
| chr1-128738622-INS-100 | chr1-128738622-INS-100 |
| chr1-128790274-INS-216 | chr1-128790274-INS-216 |
| chr1-128793161-INS-51 | chr1-128793161-INS-51 |
| chr1-128800425-INS-110 | chr1-128800439-INS-68 |
| chr1-128821024-INS-64 | chr1-128821024-INS-62 |
| chr1-128834980-INS-135 | chr1-128834980-INS-135 |
| chr1-128843645-INS-5222 | chr1-128843646-INS-5222 |
| chr1-128850409-INS-80 | chr1-128850408-INS-96 |
| chr1-128874218-INS-267 | chr1-128874218-INS-267 |
| chr1-128931687-INS-52 | chr1-128931705-INS-66 |
| chr1-128947119-INS-59 | chr1-128947096-INS-70 |
| chr1-128952929-INS-2951 | chr1-128952930-INS-2951 |
| chr1-128956589-INS-78 | chr1-128956589-INS-78 |
| chr1-128964161-INS-3199 | chr1-128964162-INS-3200 |
| chr1-128966774-INS-285 | chr1-128966774-INS-294 |
| chr1-128968586-INS-6359 | chr1-128968587-INS-6359 |
| chr1-129050086-INS-6364 | chr1-129050087-INS-6364 |
| chr1-129058496-INS-54 | chr1-129058496-INS-54 |
| chr1-129095230-INS-3423 | chr1-129095231-INS-3424 |
| chr1-129104613-INS-58 | chr1-129104625-INS-54 |
| chr1-129123369-INS-78 | chr1-129123369-INS-84 |
| chr1-129125790-INS-6870 | chr1-129125791-INS-6870 |
| chr1-129127982-INS-60 | chr1-129127982-INS-50 |
| chr1-129130944-INS-77 | chr1-129130944-INS-77 |
| chr1-129140442-INS-124 | chr1-129140424-INS-96 |
| chr1-129175005-INS-80 | chr1-129175005-INS-75 |
| chr1-129264833-INS-214 | chr1-129264834-INS-214 |

| | |
|---|---|
| chr1-129314170-INS-83 | chr1-129314170-INS-83 |
| chr1-129333023-INS-72 | chr1-129333023-INS-76 |
| chr1-129340218-INS-504 | chr1-129340218-INS-502 |
| chr1-129385133-INS-52 | chr1-129385133-INS-54 |
| chr1-129406937-INS-58 | chr1-129406937-INS-52 |
| chr1-129416951-INS-395 | chr1-129416951-INS-395 |
| chr1-129427817-INS-82 | chr1-129427817-INS-82 |
| chr1-129441528-INS-62 | chr1-129441528-INS-64 |
| chr1-129453018-INS-80 | chr1-129453026-INS-58 |
| chr1-129490137-INS-493 | chr1-129490137-INS-493 |
| chr1-129531866-INS-56 | chr1-129531866-INS-65 |
| chr1-129551198-INS-1350 | chr1-129551199-INS-1350 |
| chr1-129574152-INS-245 | chr1-129574152-INS-245 |
| chr1-129582763-INS-142 | chr1-129582763-INS-142 |
| chr1-129643192-INS-119 | chr1-129643192-INS-119 |
| chr1-129671679-INS-154 | chr1-129671679-INS-154 |
| chr1-129705087-INS-426 | chr1-129705088-INS-429 |
| chr1-129714259-INS-54 | chr1-129714282-INS-55 |
| chr1-129815109-INS-232 | chr1-129815112-INS-248 |
| chr1-129833021-INS-129 | chr1-129833013-INS-138 |
| chr1-129840719-INS-145 | chr1-129840719-INS-122 |
| chr1-129855335-INS-6586 | chr1-129855336-INS-6585 |
| chr1-129859073-INS-180 | chr1-129859073-INS-180 |
| chr1-129914331-INS-503 | chr1-129914331-INS-504 |
| chr1-129915679-INS-54 | chr1-129915679-INS-54 |
| chr1-129925483-INS-54 | chr1-129925483-INS-54 |
| chr1-129946542-INS-185 | chr1-129946542-INS-185 |
| chr1-129984418-INS-6682 | chr1-129984419-INS-6689 |
| chr1-130030268-INS-81 | chr1-130030268-INS-84 |
| chr1-130033036-INS-360 | chr1-130033036-INS-360 |
| chr1-130047649-INS-52 | chr1-130047649-INS-54 |
| chr1-130050150-INS-170 | chr1-130050150-INS-176 |
| chr1-130103706-INS-7246 | chr1-130103707-INS-7247 |
| chr1-130104921-INS-99 | chr1-130104921-INS-99 |
| chr1-130106671-INS-448 | chr1-130106671-INS-447 |
| chr1-130125268-INS-84 | chr1-130125280-INS-72 |
| chr1-130126380-INS-150 | chr1-130126384-INS-129 |
| chr1-130166620-INS-94 | chr1-130166620-INS-89 |
| chr1-130166823-INS-806 | chr1-130166823-INS-806 |
| chr1-130184652-INS-906 | chr1-130184652-INS-903 |
| chr1-130194616-INS-782 | chr1-130194616-INS-782 |
| chr1-130201119-INS-88 | chr1-130201119-INS-96 |
| chr1-130206925-INS-216 | chr1-130206925-INS-216 |
| chr1-130211423-INS-231 | chr1-130211416-INS-231 |
| chr1-130219097-INS-350 | chr1-130219099-INS-353 |
| chr1-130224250-INS-1443 | chr1-130224251-INS-1443 |
| chr1-130280659-INS-69 | chr1-130280659-INS-69 |
| chr1-130324966-INS-363 | chr1-130324966-INS-363 |
| chr1-130338861-INS-390 | chr1-130338861-INS-390 |
| chr1-130376569-INS-195 | chr1-130376575-INS-195 |
| chr1-130397576-INS-199 | chr1-130397576-INS-198 |
| chr1-130447046-INS-56 | chr1-130447046-INS-56 |
| chr1-130450348-INS-70 | chr1-130450348-INS-70 |
| chr1-130456944-INS-341 | chr1-130456944-INS-342 |
| chr1-130471455-INS-325 | chr1-130471455-INS-325 |
| chr1-130476770-INS-218 | chr1-130476770-INS-218 |
| chr1-130479524-INS-1055 | chr1-130479525-INS-1055 |
| chr1-130545602-INS-60 | chr1-130545602-INS-60 |
| chr1-130575060-INS-225 | chr1-130575060-INS-225 |
| chr1-130612735-INS-171 | chr1-130612735-INS-170 |
| chr1-130613927-INS-3412 | chr1-130613928-INS-3412 |
| chr1-130634066-INS-52 | chr1-130634066-INS-72 |
| chr1-130639412-INS-61 | chr1-130639462-INS-61 |
| chr1-130673118-INS-203 | chr1-130673118-INS-210 |
| chr1-130679111-INS-105 | chr1-130679111-INS-105 |
| chr1-130727518-INS-72 | chr1-130727518-INS-60 |
| chr1-130743062-INS-104 | chr1-130743062-INS-104 |
| chr1-130779619-INS-57 | chr1-130779594-INS-66 |
| chr1-130807266-INS-83 | chr1-130807266-INS-83 |
| chr1-130846593-INS-6600 | chr1-130846594-INS-6600 |
| chr1-130849820-INS-355 | chr1-130849820-INS-355 |
| chr1-130859400-INS-62 | chr1-130859391-INS-52 |
| chr1-130861645-INS-402 | chr1-130861645-INS-402 |
| chr1-130864682-INS-52 | chr1-130864682-INS-52 |
| chr1-130874278-INS-366 | chr1-130874278-INS-366 |
| chr1-130881212-INS-53 | chr1-130881212-INS-53 |
| chr1-130884160-INS-323 | chr1-130884160-INS-306 |

| | |
|---|---|
| chr1-130895705-INS-90 | chr1-130895705-INS-93 |
| chr1-130964805-INS-64 | chr1-130964805-INS-64 |
| chr1-130970563-INS-162 | chr1-130970563-INS-162 |
| chr1-130972769-INS-180 | chr1-130972769-INS-136 |
| chr1-130994526-INS-203 | chr1-130994526-INS-203 |
| chr1-130996533-INS-2390 | chr1-130996534-INS-2390 |
| chr1-131003729-INS-56 | chr1-131003729-INS-56 |
| chr1-131004510-INS-1050 | chr1-131004511-INS-1049 |
| chr1-131006022-INS-1634 | chr1-131006023-INS-1634 |
| chr1-131016358-INS-78 | chr1-131016358-INS-78 |
| chr1-131026042-INS-203 | chr1-131026042-INS-203 |
| chr1-131041190-INS-1075 | chr1-131041191-INS-1075 |
| chr1-131047643-INS-202 | chr1-131047639-INS-199 |
| chr1-131053647-INS-64 | chr1-131053647-INS-64 |
| chr1-131078804-INS-85 | chr1-131078804-INS-63 |
| chr1-131087740-INS-146 | chr1-131087740-INS-186 |
| chr1-131136734-INS-201 | chr1-131136734-INS-203 |
| chr1-131151520-INS-78 | chr1-131151520-INS-79 |
| chr1-131153697-INS-106 | chr1-131153698-INS-106 |
| chr1-131213773-INS-232 | chr1-131213772-INS-248 |
| chr1-131225224-INS-124 | chr1-131225224-INS-125 |
| chr1-131276213-INS-331 | chr1-131276213-INS-331 |
| chr1-131284159-INS-84 | chr1-131284192-INS-68 |
| chr1-131381605-INS-1914 | chr1-131381606-INS-1869 |
| chr1-131431971-INS-133 | chr1-131431971-INS-123 |
| chr1-131480691-INS-510 | chr1-131480691-INS-514 |
| chr1-131500443-INS-54 | chr1-131500443-INS-54 |
| chr1-131520792-INS-503 | chr1-131520792-INS-503 |
| chr1-131526612-INS-91 | chr1-131526612-INS-109 |
| chr1-131532494-INS-208 | chr1-131532494-INS-209 |
| chr1-131540649-INS-1815 | chr1-131540650-INS-1815 |
| chr1-131543490-INS-2652 | chr1-131543491-INS-2652 |
| chr1-131622727-INS-271 | chr1-131622727-INS-271 |
| chr1-131640794-INS-410 | chr1-131640794-INS-410 |
| chr1-131661399-INS-7179 | chr1-131661400-INS-7179 |
| chr1-131669056-INS-100 | chr1-131669056-INS-100 |
| chr1-131680402-INS-76 | chr1-131680402-INS-76 |
| chr1-131705723-INS-220 | chr1-131705713-INS-219 |
| chr1-131722022-INS-56 | chr1-131722022-INS-80 |
| chr1-131727209-INS-106 | chr1-131727209-INS-156 |
| chr1-131754759-INS-84 | chr1-131754759-INS-84 |
| chr1-131763894-INS-203 | chr1-131763894-INS-203 |
| chr1-131806100-INS-78 | chr1-131806075-INS-71 |
| chr1-131809064-INS-152 | chr1-131809064-INS-152 |
| chr1-131816954-INS-71 | chr1-131816954-INS-89 |
| chr1-131817811-INS-519 | chr1-131817811-INS-519 |
| chr1-131818976-INS-175 | chr1-131818981-INS-168 |
| chr1-131834394-INS-51 | chr1-131834394-INS-69 |
| chr1-131841913-INS-173 | chr1-131841913-INS-173 |
| chr1-131867333-INS-307 | chr1-131867333-INS-315 |
| chr1-131871883-INS-200 | chr1-131871883-INS-200 |
| chr1-131894597-INS-218 | chr1-131894597-INS-218 |
| chr1-131911839-INS-207 | chr1-131911839-INS-208 |
| chr1-131918203-INS-186 | chr1-131918203-INS-186 |
| chr1-131960799-INS-54 | chr1-131960799-INS-58 |
| chr1-131971338-INS-100 | chr1-131971350-INS-121 |
| chr1-131972444-INS-224 | chr1-131972444-INS-224 |
| chr1-132016934-INS-125 | chr1-132016934-INS-135 |
| chr1-132085091-INS-208 | chr1-132085091-INS-208 |
| chr1-132091435-INS-406 | chr1-132091435-INS-406 |
| chr1-132111774-INS-174 | chr1-132111774-INS-175 |
| chr1-132144369-INS-206 | chr1-132144369-INS-207 |
| chr1-132153445-INS-120 | chr1-132153413-INS-102 |
| chr1-132199970-INS-212 | chr1-132199970-INS-212 |
| chr1-132239613-INS-80 | chr1-132239613-INS-80 |
| chr1-132280218-INS-126 | chr1-132280218-INS-126 |
| chr1-132283927-INS-90 | chr1-132283927-INS-84 |
| chr1-132336963-INS-63 | chr1-132336963-INS-64 |
| chr1-132396123-INS-4703 | chr1-132396124-INS-4703 |
| chr1-132398491-INS-2120 | chr1-132398492-INS-2120 |
| chr1-132430221-INS-1888 | chr1-132430222-INS-1888 |
| chr1-132451767-INS-66 | chr1-132451763-INS-90 |
| chr1-132475383-INS-151 | chr1-132475383-INS-156 |
| chr1-132480078-INS-180 | chr1-132480078-INS-178 |
| chr1-132483261-INS-66 | chr1-132483238-INS-95 |
| chr1-132501836-INS-194 | chr1-132501836-INS-194 |
| chr1-132530045-INS-231 | chr1-132530045-INS-231 |

| | |
|---|---|
| chr1-132563217-INS-131 | chr1-132563217-INS-132 |
| chr1-132574714-INS-66 | chr1-132574714-INS-66 |
| chr1-132602372-INS-67 | chr1-132602372-INS-66 |
| chr1-132623488-INS-297 | chr1-132623488-INS-297 |
| chr1-132641652-INS-66 | chr1-132641652-INS-66 |
| chr1-132683256-INS-207 | chr1-132683256-INS-206 |
| chr1-132685078-INS-155 | chr1-132685078-INS-151 |
| chr1-132695957-INS-1348 | chr1-132696001-INS-1349 |
| chr1-132749957-INS-74 | chr1-132749966-INS-74 |
| chr1-132802170-INS-106 | chr1-132802208-INS-65 |
| chr1-132824412-INS-1397 | chr1-132824413-INS-1432 |
| chr1-132864121-INS-85 | chr1-132864121-INS-85 |
| chr1-132889916-INS-138 | chr1-132889897-INS-161 |
| chr1-132896218-INS-64 | chr1-132896218-INS-80 |
| chr1-132906934-INS-86 | chr1-132906934-INS-86 |
| chr1-132927768-INS-189 | chr1-132927768-INS-189 |
| chr1-132935576-INS-208 | chr1-132935576-INS-208 |
| chr1-132945564-INS-239 | chr1-132945564-INS-239 |
| chr1-132953562-INS-250 | chr1-132953562-INS-250 |
| chr1-132965352-INS-206 | chr1-132965352-INS-206 |
| chr1-132984699-INS-59 | chr1-132984697-INS-50 |
| chr1-132990462-INS-87 | chr1-132990456-INS-96 |
| chr1-133006638-INS-207 | chr1-133006638-INS-208 |
| chr1-133017025-INS-201 | chr1-133017025-INS-205 |
| chr1-133019814-INS-205 | chr1-133019814-INS-205 |
| chr1-133030264-INS-68 | chr1-133030264-INS-68 |
| chr1-133040232-INS-141 | chr1-133040232-INS-145 |
| chr1-133043707-INS-86 | chr1-133043707-INS-50 |
| chr1-133053211-INS-204 | chr1-133053211-INS-204 |
| chr1-133065684-INS-190 | chr1-133065684-INS-190 |
| chr1-133070049-INS-122 | chr1-133070055-INS-148 |
| chr1-133074879-INS-54 | chr1-133074879-INS-59 |
| chr1-133079231-INS-3485 | chr1-133079232-INS-3485 |
| chr1-133082739-INS-159 | chr1-133082739-INS-159 |
| chr1-133087628-INS-78 | chr1-133087628-INS-73 |
| chr1-133095370-INS-86 | chr1-133095370-INS-86 |
| chr1-133107947-INS-6442 | chr1-133107948-INS-6442 |
| chr1-133112568-INS-394 | chr1-133112568-INS-394 |
| chr1-133114449-INS-84 | chr1-133114449-INS-113 |
| chr1-133115509-INS-56 | chr1-133115483-INS-64 |
| chr1-133151349-INS-113 | chr1-133151349-INS-113 |
| chr1-133156526-INS-231 | chr1-133156526-INS-230 |
| chr1-133173132-INS-73 | chr1-133173132-INS-73 |
| chr1-133218364-INS-72 | chr1-133218364-INS-70 |
| chr1-133269161-INS-278 | chr1-133269134-INS-274 |
| chr1-133272290-INS-99 | chr1-133272290-INS-99 |
| chr1-133329769-INS-60 | chr1-133329769-INS-58 |
| chr1-133333919-INS-277 | chr1-133333919-INS-279 |
| chr1-133352269-INS-56 | chr1-133352269-INS-57 |
| chr1-133355962-INS-222 | chr1-133355962-INS-223 |
| chr1-133385191-INS-150 | chr1-133385191-INS-150 |
| chr1-133414789-INS-387 | chr1-133414791-INS-391 |
| chr1-133417669-INS-72 | chr1-133417669-INS-72 |
| chr1-133423011-INS-116 | chr1-133423011-INS-146 |
| chr1-133429363-INS-1467 | chr1-133429364-INS-1467 |
| chr1-133434747-INS-7185 | chr1-133434748-INS-7185 |
| chr1-133459481-INS-80 | chr1-133459481-INS-70 |
| chr1-133465511-INS-467 | chr1-133465503-INS-474 |
| chr1-133467713-INS-427 | chr1-133467713-INS-427 |
| chr1-133469146-INS-166 | chr1-133469146-INS-145 |
| chr1-133472886-INS-128 | chr1-133472886-INS-128 |
| chr1-133494783-INS-205 | chr1-133494783-INS-205 |
| chr1-133508348-INS-205 | chr1-133508348-INS-205 |
| chr1-133509763-INS-58 | chr1-133509733-INS-63 |
| chr1-133512165-INS-70 | chr1-133512165-INS-76 |
| chr1-133516041-INS-218 | chr1-133516041-INS-218 |
| chr1-133554248-INS-208 | chr1-133554248-INS-208 |
| chr1-133557485-INS-205 | chr1-133557485-INS-205 |
| chr1-133569600-INS-228 | chr1-133569600-INS-228 |
| chr1-133573082-INS-206 | chr1-133573082-INS-206 |
| chr1-133574822-INS-442 | chr1-133574822-INS-441 |
| chr1-133590900-INS-217 | chr1-133590900-INS-218 |
| chr1-133615744-INS-203 | chr1-133615744-INS-203 |
| chr1-133626806-INS-239 | chr1-133626806-INS-240 |
| chr1-133629537-INS-70 | chr1-133629538-INS-74 |
| chr1-133640665-INS-534 | chr1-133640665-INS-535 |
| chr1-133653062-INS-610 | chr1-133653062-INS-610 |

| | |
|---|---|
| chr1-133661671-INS-70 | chr1-133661671-INS-68 |
| chr1-133719101-INS-72 | chr1-133719101-INS-74 |
| chr1-133760104-INS-90 | chr1-133760104-INS-90 |
| chr1-133766651-INS-134 | chr1-133766647-INS-140 |
| chr1-133786370-INS-688 | chr1-133786371-INS-686 |
| chr1-133826294-INS-74 | chr1-133826294-INS-74 |
| chr1-133832556-INS-416 | chr1-133832556-INS-416 |
| chr1-133845633-INS-286 | chr1-133845633-INS-289 |
| chr1-133851130-INS-82 | chr1-133851130-INS-81 |
| chr1-133904704-INS-492 | chr1-133904704-INS-457 |
| chr1-133943079-INS-174 | chr1-133943079-INS-174 |
| chr1-133950328-INS-106 | chr1-133950278-INS-133 |
| chr1-133984679-INS-363 | chr1-133984679-INS-363 |
| chr1-133985772-INS-170 | chr1-133985772-INS-163 |
| chr1-134031849-INS-60 | chr1-134031849-INS-84 |
| chr1-134032859-INS-65 | chr1-134032859-INS-65 |
| chr1-134038221-INS-8507 | chr1-134038222-INS-8507 |
| chr1-134059005-INS-168 | chr1-134059005-INS-169 |
| chr1-134065271-INS-822 | chr1-134065272-INS-822 |
| chr1-134066308-INS-115 | chr1-134066308-INS-120 |
| chr1-134097676-INS-55 | chr1-134097676-INS-60 |
| chr1-134100612-INS-113 | chr1-134100612-INS-70 |
| chr1-134135257-INS-156 | chr1-134135257-INS-156 |
| chr1-134165020-INS-61 | chr1-134165020-INS-55 |
| chr1-134170615-INS-63 | chr1-134170615-INS-63 |
| chr1-134175994-INS-60 | chr1-134175994-INS-63 |
| chr1-134195019-INS-573 | chr1-134195019-INS-573 |
| chr1-134197368-INS-199 | chr1-134197368-INS-199 |
| chr1-134200135-INS-88 | chr1-134200135-INS-88 |
| chr1-134229678-INS-203 | chr1-134229678-INS-202 |
| chr1-134231291-INS-1232 | chr1-134231292-INS-1234 |
| chr1-134248043-INS-125 | chr1-134248036-INS-100 |
| chr1-134258886-INS-225 | chr1-134258886-INS-226 |
| chr1-134285431-INS-119 | chr1-134285431-INS-85 |
| chr1-134291178-INS-205 | chr1-134291178-INS-202 |
| chr1-134294295-INS-242 | chr1-134294295-INS-238 |
| chr1-134296428-INS-117 | chr1-134296428-INS-114 |
| chr1-134297821-INS-864 | chr1-134297821-INS-862 |
| chr1-134300785-INS-74 | chr1-134300785-INS-76 |
| chr1-134303063-INS-66 | chr1-134303066-INS-63 |
| chr1-134303507-INS-237 | chr1-134303507-INS-237 |
| chr1-134315186-INS-4920 | chr1-134315187-INS-4920 |
| chr1-134316666-INS-191 | chr1-134316694-INS-183 |
| chr1-134320723-INS-54 | chr1-134320680-INS-69 |
| chr1-134325902-INS-218 | chr1-134325902-INS-218 |
| chr1-134335093-INS-128 | chr1-134335093-INS-128 |
| chr1-134406965-INS-208 | chr1-134406965-INS-208 |
| chr1-134427543-INS-63 | chr1-134427541-INS-62 |
| chr1-134429494-INS-684 | chr1-134429494-INS-682 |
| chr1-134439258-INS-206 | chr1-134439258-INS-206 |
| chr1-134459726-INS-225 | chr1-134459726-INS-210 |
| chr1-134464705-INS-239 | chr1-134464705-INS-239 |
| chr1-134471603-INS-55 | chr1-134471578-INS-74 |
| chr1-134476979-INS-69 | chr1-134476979-INS-69 |
| chr1-134492900-INS-206 | chr1-134492900-INS-206 |
| chr1-134494957-INS-223 | chr1-134494957-INS-223 |
| chr1-134502080-INS-73 | chr1-134502080-INS-71 |
| chr1-134504429-INS-1691 | chr1-134504430-INS-1691 |
| chr1-134504825-INS-202 | chr1-134504825-INS-202 |
| chr1-134507715-INS-211 | chr1-134507715-INS-211 |
| chr1-134508239-INS-222 | chr1-134508239-INS-222 |
| chr1-134527035-INS-341 | chr1-134527035-INS-341 |
| chr1-134577085-INS-182 | chr1-134577085-INS-183 |
| chr1-134583427-INS-556 | chr1-134583387-INS-560 |
| chr1-134593407-INS-388 | chr1-134593407-INS-388 |
| chr1-134597640-INS-6487 | chr1-134597641-INS-6487 |
| chr1-134603412-INS-55 | chr1-134603412-INS-55 |
| chr1-134643815-INS-55 | chr1-134643815-INS-59 |
| chr1-134658200-INS-1639 | chr1-134658201-INS-1659 |
| chr1-134706818-INS-53 | chr1-134706818-INS-53 |
| chr1-134725619-INS-52 | chr1-134725619-INS-86 |
| chr1-134741948-INS-229 | chr1-134741948-INS-229 |
| chr1-134743117-INS-66 | chr1-134743117-INS-66 |
| chr1-134765334-INS-254 | chr1-134765334-INS-254 |
| chr1-134775372-INS-65 | chr1-134775372-INS-65 |
| chr1-134784471-INS-1327 | chr1-134784472-INS-1327 |
| chr1-134811124-INS-93 | chr1-134811124-INS-93 |

| | |
|---|---|
| chr1-134842184-INS-403 | chr1-134842184-INS-403 |
| chr1-134846429-INS-96 | chr1-134846429-INS-75 |
| chr1-134848090-INS-545 | chr1-134848090-INS-523 |
| chr1-134849640-INS-532 | chr1-134849640-INS-530 |
| chr1-134863939-INS-56 | chr1-134863939-INS-59 |
| chr1-134877299-INS-218 | chr1-134877299-INS-225 |
| chr1-134878363-INS-6401 | chr1-134878364-INS-6401 |
| chr1-134881028-INS-68 | chr1-134881042-INS-75 |
| chr1-134882313-INS-60 | chr1-134882313-INS-52 |
| chr1-134890567-INS-148 | chr1-134890567-INS-148 |
| chr1-134902965-INS-1193 | chr1-134902966-INS-1193 |
| chr1-134951389-INS-56 | chr1-134951389-INS-63 |
| chr1-134972224-INS-212 | chr1-134972224-INS-211 |
| chr1-135003407-INS-148 | chr1-135003407-INS-144 |
| chr1-135027661-INS-83 | chr1-135027658-INS-100 |
| chr1-135043220-INS-211 | chr1-135043220-INS-212 |
| chr1-135046871-INS-52 | chr1-135046901-INS-60 |
| chr1-135068192-INS-58 | chr1-135068192-INS-61 |
| chr1-135077992-INS-63 | chr1-135077992-INS-58 |
| chr1-135080796-INS-57 | chr1-135080796-INS-66 |
| chr1-135093672-INS-78 | chr1-135093672-INS-78 |
| chr1-135100303-INS-180 | chr1-135100303-INS-180 |
| chr1-135104894-INS-65 | chr1-135104892-INS-60 |
| chr1-135113612-INS-201 | chr1-135113612-INS-201 |
| chr1-135115184-INS-173 | chr1-135115184-INS-177 |
| chr1-135133081-INS-242 | chr1-135133081-INS-244 |
| chr1-135136342-INS-95 | chr1-135136342-INS-95 |
| chr1-135138014-INS-82 | chr1-135138014-INS-84 |
| chr1-135143631-INS-197 | chr1-135143631-INS-197 |
| chr1-135201221-INS-78 | chr1-135201221-INS-78 |
| chr1-135203761-INS-200 | chr1-135203761-INS-199 |
| chr1-135206452-INS-55 | chr1-135206452-INS-55 |
| chr1-135210043-INS-220 | chr1-135210043-INS-220 |
| chr1-135214759-INS-804 | chr1-135214759-INS-805 |
| chr1-135216693-INS-273 | chr1-135216693-INS-274 |
| chr1-135222234-INS-94 | chr1-135222234-INS-94 |
| chr1-135226013-INS-98 | chr1-135226013-INS-98 |
| chr1-135231313-INS-82 | chr1-135231295-INS-56 |
| chr1-135234337-INS-188 | chr1-135234338-INS-190 |
| chr1-135244782-INS-84 | chr1-135244782-INS-84 |
| chr1-135247168-INS-235 | chr1-135247168-INS-235 |
| chr1-135260878-INS-201 | chr1-135260878-INS-201 |
| chr1-135281019-INS-223 | chr1-135281019-INS-223 |
| chr1-135290272-INS-198 | chr1-135290272-INS-199 |
| chr1-135336988-INS-237 | chr1-135336988-INS-240 |
| chr1-135343032-INS-177 | chr1-135343032-INS-177 |
| chr1-135355876-INS-960 | chr1-135355876-INS-970 |
| chr1-135370899-INS-200 | chr1-135370899-INS-200 |
| chr1-135375061-INS-212 | chr1-135375061-INS-211 |
| chr1-135387139-INS-80 | chr1-135387139-INS-56 |
| chr1-135389721-INS-330 | chr1-135389721-INS-330 |
| chr1-135411284-INS-211 | chr1-135411284-INS-210 |
| chr1-135412771-INS-193 | chr1-135412771-INS-193 |
| chr1-135422224-INS-66 | chr1-135422224-INS-66 |
| chr1-135423696-INS-1895 | chr1-135423697-INS-1895 |
| chr1-135450779-INS-238 | chr1-135450779-INS-237 |
| chr1-135454199-INS-177 | chr1-135454199-INS-181 |
| chr1-135455411-INS-57 | chr1-135455411-INS-57 |
| chr1-135462077-INS-140 | chr1-135462091-INS-142 |
| chr1-135467624-INS-214 | chr1-135467624-INS-215 |
| chr1-135471466-INS-444 | chr1-135471466-INS-444 |
| chr1-135472747-INS-141 | chr1-135472747-INS-141 |
| chr1-135481359-INS-134 | chr1-135481359-INS-126 |
| chr1-135493287-INS-66 | chr1-135493287-INS-66 |
| chr1-135495973-INS-90 | chr1-135495973-INS-90 |
| chr1-135497136-INS-197 | chr1-135497136-INS-197 |
| chr1-135528977-INS-196 | chr1-135528977-INS-196 |
| chr1-135583513-INS-170 | chr1-135583513-INS-171 |
| chr1-135585498-INS-90 | chr1-135585510-INS-74 |
| chr1-135633668-INS-209 | chr1-135633668-INS-209 |
| chr1-135634531-INS-70 | chr1-135634532-INS-66 |
| chr1-135634832-INS-523 | chr1-135634832-INS-523 |
| chr1-135665292-INS-341 | chr1-135665292-INS-354 |
| chr1-135668159-INS-107 | chr1-135668159-INS-107 |
| chr1-135673734-INS-58 | chr1-135673734-INS-60 |
| chr1-135676470-INS-219 | chr1-135676470-INS-220 |
| chr1-135681770-INS-260 | chr1-135681770-INS-260 |

| | |
|---|---|
| chr1-135717114-INS-196 | chr1-135717114-INS-196 |
| chr1-135726099-INS-278 | chr1-135726099-INS-270 |
| chr1-135740243-INS-56 | chr1-135740243-INS-58 |
| chr1-135753088-INS-77 | chr1-135753071-INS-87 |
| chr1-135756265-INS-55 | chr1-135756265-INS-58 |
| chr1-135787853-INS-162 | chr1-135787853-INS-162 |
| chr1-135850646-INS-218 | chr1-135850646-INS-218 |
| chr1-135862059-INS-6896 | chr1-135862060-INS-6896 |
| chr1-135876490-INS-54 | chr1-135876472-INS-56 |
| chr1-135908696-INS-62 | chr1-135908696-INS-76 |
| chr1-135921207-INS-209 | chr1-135921207-INS-210 |
| chr1-135927246-INS-297 | chr1-135927254-INS-272 |
| chr1-135987462-INS-86 | chr1-135987462-INS-86 |
| chr1-136006598-INS-291 | chr1-136006598-INS-287 |
| chr1-136008463-INS-169 | chr1-136008463-INS-169 |
| chr1-136014122-INS-87 | chr1-136014122-INS-87 |
| chr1-136019350-INS-6440 | chr1-136019351-INS-6440 |
| chr1-136036991-INS-57 | chr1-136036993-INS-69 |
| chr1-136050913-INS-203 | chr1-136050913-INS-203 |
| chr1-136074089-INS-85 | chr1-136074082-INS-97 |
| chr1-136097998-INS-80 | chr1-136097998-INS-80 |
| chr1-136117485-INS-53 | chr1-136117483-INS-63 |
| chr1-136123297-INS-210 | chr1-136123297-INS-209 |
| chr1-136123598-INS-1925 | chr1-136123595-INS-1921 |
| chr1-136129464-INS-199 | chr1-136129464-INS-198 |
| chr1-136130482-INS-151 | chr1-136130475-INS-152 |
| chr1-136132980-INS-200 | chr1-136132980-INS-200 |
| chr1-136200590-INS-125 | chr1-136200590-INS-102 |
| chr1-136218714-INS-226 | chr1-136218714-INS-224 |
| chr1-136222678-INS-89 | chr1-136222678-INS-89 |
| chr1-136259886-INS-205 | chr1-136259886-INS-210 |
| chr1-136269621-INS-78 | chr1-136269618-INS-51 |
| chr1-136298924-INS-155 | chr1-136298924-INS-155 |
| chr1-136314670-INS-57 | chr1-136314671-INS-52 |
| chr1-136331506-INS-117 | chr1-136331506-INS-129 |
| chr1-136351733-INS-75 | chr1-136351735-INS-75 |
| chr1-136364210-INS-173 | chr1-136364210-INS-173 |
| chr1-136379593-INS-330 | chr1-136379595-INS-328 |
| chr1-136393727-INS-209 | chr1-136393727-INS-209 |
| chr1-136411963-INS-58 | chr1-136411963-INS-60 |
| chr1-136413213-INS-72 | chr1-136413212-INS-72 |
| chr1-136422865-INS-220 | chr1-136422865-INS-220 |
| chr1-136434200-INS-120 | chr1-136434200-INS-120 |
| chr1-136434936-INS-1890 | chr1-136434937-INS-1890 |
| chr1-136436656-INS-219 | chr1-136436656-INS-219 |
| chr1-136447960-INS-217 | chr1-136447960-INS-218 |
| chr1-136448522-INS-194 | chr1-136448522-INS-194 |
| chr1-136453815-INS-6568 | chr1-136453816-INS-6568 |
| chr1-136460365-INS-372 | chr1-136460365-INS-354 |
| chr1-136461203-INS-547 | chr1-136461203-INS-552 |
| chr1-136462512-INS-198 | chr1-136462512-INS-198 |
| chr1-136464622-INS-119 | chr1-136464622-INS-119 |
| chr1-136466503-INS-202 | chr1-136466503-INS-204 |
| chr1-136472670-INS-56 | chr1-136472670-INS-56 |
| chr1-136482194-INS-829 | chr1-136482194-INS-822 |
| chr1-136502298-INS-92 | chr1-136502298-INS-93 |
| chr1-136543823-INS-419 | chr1-136543823-INS-419 |
| chr1-136545975-INS-240 | chr1-136545975-INS-240 |
| chr1-136561220-INS-5549 | chr1-136561221-INS-5549 |
| chr1-136573943-INS-160 | chr1-136573943-INS-160 |
| chr1-136575856-INS-108 | chr1-136575856-INS-108 |
| chr1-136579063-INS-172 | chr1-136579063-INS-173 |
| chr1-136583469-INS-104 | chr1-136583469-INS-104 |
| chr1-136584884-INS-68 | chr1-136584884-INS-68 |
| chr1-136595266-INS-82 | chr1-136595271-INS-52 |
| chr1-136610094-INS-216 | chr1-136610090-INS-230 |
| chr1-136614882-INS-182 | chr1-136614882-INS-183 |
| chr1-136616415-INS-463 | chr1-136616415-INS-463 |
| chr1-136625828-INS-169 | chr1-136625828-INS-166 |
| chr1-136627700-INS-141 | chr1-136627700-INS-141 |
| chr1-136670526-INS-2451 | chr1-136670527-INS-2451 |
| chr1-136675621-INS-115 | chr1-136675621-INS-125 |
| chr1-136680281-INS-241 | chr1-136680281-INS-241 |
| chr1-136681039-INS-223 | chr1-136681039-INS-224 |
| chr1-136692043-INS-51 | chr1-136692043-INS-53 |
| chr1-136695647-INS-54 | chr1-136695647-INS-54 |
| chr1-136697224-INS-184 | chr1-136697224-INS-186 |

| | |
|---|---|
| chr1-136700089-INS-59 | chr1-136700089-INS-59 |
| chr1-136709210-INS-810 | chr1-136709210-INS-810 |
| chr1-136715556-INS-96 | chr1-136715556-INS-100 |
| chr1-136724882-INS-77 | chr1-136724904-INS-104 |
| chr1-136735840-INS-84 | chr1-136735840-INS-84 |
| chr1-136737662-INS-75 | chr1-136737662-INS-75 |
| chr1-136742857-INS-62 | chr1-136742857-INS-54 |
| chr1-136801967-INS-182 | chr1-136801967-INS-187 |
| chr1-136806632-INS-343 | chr1-136806632-INS-343 |
| chr1-136819065-INS-54 | chr1-136819065-INS-54 |
| chr1-136819614-INS-1825 | chr1-136819615-INS-1825 |
| chr1-136822425-INS-79 | chr1-136822431-INS-52 |
| chr1-136826558-INS-207 | chr1-136826558-INS-207 |
| chr1-136828235-INS-191 | chr1-136828235-INS-191 |
| chr1-136830618-INS-88 | chr1-136830618-INS-88 |
| chr1-136837774-INS-123 | chr1-136837774-INS-119 |
| chr1-136865227-INS-54 | chr1-136865265-INS-53 |
| chr1-136865989-INS-203 | chr1-136865989-INS-203 |
| chr1-136870984-INS-88 | chr1-136870984-INS-88 |
| chr1-136889963-INS-181 | chr1-136889963-INS-181 |
| chr1-136916049-INS-185 | chr1-136916049-INS-185 |
| chr1-136923924-INS-193 | chr1-136923924-INS-193 |
| chr1-136925413-INS-85 | chr1-136925414-INS-85 |
| chr1-136935831-INS-221 | chr1-136935831-INS-221 |
| chr1-136941624-INS-207 | chr1-136941624-INS-208 |
| chr1-136948309-INS-55 | chr1-136948309-INS-60 |
| chr1-136949269-INS-340 | chr1-136949269-INS-340 |
| chr1-136953886-INS-68 | chr1-136953886-INS-72 |
| chr1-136956207-INS-59 | chr1-136956207-INS-64 |
| chr1-136978375-INS-190 | chr1-136978375-INS-190 |
| chr1-136982652-INS-131 | chr1-136982652-INS-131 |
| chr1-137014610-INS-220 | chr1-137014610-INS-220 |
| chr1-137023180-INS-621 | chr1-137023180-INS-624 |
| chr1-137031775-INS-65 | chr1-137031775-INS-65 |
| chr1-137049147-INS-84 | chr1-137049147-INS-79 |
| chr1-137095864-INS-55 | chr1-137095864-INS-55 |
| chr1-137139727-INS-90 | chr1-137139727-INS-90 |
| chr1-137143230-INS-56 | chr1-137143230-INS-56 |
| chr1-137144235-INS-6312 | chr1-137144236-INS-6313 |
| chr1-137195750-INS-6458 | chr1-137195751-INS-6458 |
| chr1-137196923-INS-285 | chr1-137196884-INS-309 |
| chr1-137231994-INS-808 | chr1-137231994-INS-808 |
| chr1-137244814-INS-75 | chr1-137244814-INS-75 |
| chr1-137298904-INS-54 | chr1-137298904-INS-52 |
| chr1-137302128-INS-71 | chr1-137302129-INS-69 |
| chr1-137324918-INS-67 | chr1-137324918-INS-67 |
| chr1-137356851-INS-167 | chr1-137356857-INS-147 |
| chr1-137363761-INS-64 | chr1-137363761-INS-64 |
| chr1-137386320-INS-60 | chr1-137386320-INS-70 |
| chr1-137390270-INS-2523 | chr1-137390271-INS-2523 |
| chr1-137394512-INS-7451 | chr1-137394513-INS-7451 |
| chr1-137395051-INS-64 | chr1-137395051-INS-64 |
| chr1-137397375-INS-267 | chr1-137397375-INS-267 |
| chr1-137404297-INS-87 | chr1-137404283-INS-60 |
| chr1-137411003-INS-103 | chr1-137411005-INS-103 |
| chr1-137411723-INS-531 | chr1-137411723-INS-531 |
| chr1-137425031-INS-206 | chr1-137425031-INS-223 |
| chr1-137460907-INS-65 | chr1-137460907-INS-65 |
| chr1-137472678-INS-126 | chr1-137472678-INS-126 |
| chr1-137481745-INS-6379 | chr1-137481746-INS-6382 |
| chr1-137508919-INS-53 | chr1-137508919-INS-53 |
| chr1-137512920-INS-220 | chr1-137512920-INS-220 |
| chr1-137514395-INS-89 | chr1-137514395-INS-89 |
| chr1-137556631-INS-777 | chr1-137556631-INS-777 |
| chr1-137562774-INS-70 | chr1-137562774-INS-70 |
| chr1-137567552-INS-1285 | chr1-137567553-INS-1285 |
| chr1-137571184-INS-80 | chr1-137571222-INS-82 |
| chr1-137580416-INS-328 | chr1-137580416-INS-330 |
| chr1-137599381-INS-87 | chr1-137599381-INS-94 |
| chr1-137615266-INS-96 | chr1-137615266-INS-99 |
| chr1-137635421-INS-56 | chr1-137635421-INS-56 |
| chr1-137694159-INS-298 | chr1-137694159-INS-300 |
| chr1-137724917-INS-741 | chr1-137724917-INS-734 |
| chr1-137738656-INS-1873 | chr1-137738657-INS-1872 |
| chr1-137759414-INS-119 | chr1-137759414-INS-88 |
| chr1-137786098-INS-4979 | chr1-137786099-INS-4979 |
| chr1-137806499-INS-644 | chr1-137806499-INS-644 |

| | |
|---|---|
| chr1-137810458-INS-228 | chr1-137810458-INS-228 |
| chr1-137847346-INS-241 | chr1-137847346-INS-248 |
| chr1-137853412-INS-6892 | chr1-137853413-INS-6892 |
| chr1-137871634-INS-74 | chr1-137871634-INS-74 |
| chr1-137882563-INS-448 | chr1-137882563-INS-448 |
| chr1-137899658-INS-60 | chr1-137899658-INS-60 |
| chr1-137922128-INS-6202 | chr1-137922129-INS-6202 |
| chr1-137930984-INS-117 | chr1-137930984-INS-117 |
| chr1-137938669-INS-172 | chr1-137938669-INS-173 |
| chr1-137946936-INS-1131 | chr1-137946937-INS-1131 |
| chr1-137949703-INS-121 | chr1-137949703-INS-128 |
| chr1-137963563-INS-98 | chr1-137963563-INS-98 |
| chr1-137967932-INS-241 | chr1-137967932-INS-241 |
| chr1-137973524-INS-54 | chr1-137973524-INS-56 |
| chr1-138005368-INS-476 | chr1-138005368-INS-476 |
| chr1-138017832-INS-96 | chr1-138017832-INS-96 |
| chr1-138050998-INS-72 | chr1-138050998-INS-69 |
| chr1-138057646-INS-52 | chr1-138057646-INS-56 |
| chr1-138061852-INS-105 | chr1-138061866-INS-111 |
| chr1-138084628-INS-189 | chr1-138084628-INS-189 |
| chr1-138136532-INS-233 | chr1-138136532-INS-233 |
| chr1-138138224-INS-58 | chr1-138138234-INS-62 |
| chr1-138193133-INS-172 | chr1-138193133-INS-172 |
| chr1-138218146-INS-59 | chr1-138218144-INS-56 |
| chr1-138218877-INS-77 | chr1-138218877-INS-89 |
| chr1-138239570-INS-424 | chr1-138239571-INS-424 |
| chr1-138260102-INS-4755 | chr1-138260103-INS-4755 |
| chr1-138293777-INS-106 | chr1-138293777-INS-104 |
| chr1-138350959-INS-78 | chr1-138350959-INS-77 |
| chr1-138354551-INS-395 | chr1-138354551-INS-395 |
| chr1-138375278-INS-7358 | chr1-138375279-INS-7359 |
| chr1-138443356-INS-88 | chr1-138443356-INS-88 |
| chr1-138460148-INS-61 | chr1-138460148-INS-61 |
| chr1-138487109-INS-56 | chr1-138487109-INS-56 |
| chr1-138506482-INS-113 | chr1-138506482-INS-113 |
| chr1-138511821-INS-60 | chr1-138511821-INS-60 |
| chr1-138516529-INS-170 | chr1-138516529-INS-170 |
| chr1-138521634-INS-899 | chr1-138521634-INS-899 |
| chr1-138525745-INS-50 | chr1-138525724-INS-50 |
| chr1-138533106-INS-72 | chr1-138533106-INS-68 |
| chr1-138551911-INS-83 | chr1-138551911-INS-83 |
| chr1-138591536-INS-79 | chr1-138591536-INS-82 |
| chr1-138598908-INS-53 | chr1-138598908-INS-53 |
| chr1-138613678-INS-416 | chr1-138613678-INS-422 |
| chr1-138614778-INS-73 | chr1-138614778-INS-73 |
| chr1-138627175-INS-645 | chr1-138627175-INS-646 |
| chr1-138638636-INS-400 | chr1-138638636-INS-400 |
| chr1-138674656-INS-2555 | chr1-138674657-INS-2555 |
| chr1-138678926-INS-1656 | chr1-138678927-INS-1656 |
| chr1-138682145-INS-80 | chr1-138682185-INS-71 |
| chr1-138684219-INS-825 | chr1-138684219-INS-826 |
| chr1-138736996-INS-67 | chr1-138736996-INS-67 |
| chr1-138743203-INS-425 | chr1-138743203-INS-425 |
| chr1-138743730-INS-202 | chr1-138743730-INS-206 |
| chr1-138747081-INS-4761 | chr1-138747082-INS-4761 |
| chr1-138804452-INS-66 | chr1-138804452-INS-60 |
| chr1-138818513-INS-676 | chr1-138818513-INS-675 |
| chr1-138828436-INS-84 | chr1-138828436-INS-88 |
| chr1-138851880-INS-185 | chr1-138851880-INS-185 |
| chr1-138858031-INS-191 | chr1-138858031-INS-191 |
| chr1-138870997-INS-480 | chr1-138870997-INS-480 |
| chr1-138924906-INS-65 | chr1-138924907-INS-65 |
| chr1-138981466-INS-123 | chr1-138981466-INS-123 |
| chr1-139017240-INS-297 | chr1-139017240-INS-297 |
| chr1-139073573-INS-60 | chr1-139073537-INS-52 |
| chr1-139106655-INS-65 | chr1-139106655-INS-65 |
| chr1-139124244-INS-173 | chr1-139124241-INS-140 |
| chr1-139129789-INS-61 | chr1-139129789-INS-65 |
| chr1-139199733-INS-122 | chr1-139199733-INS-122 |
| chr1-139200075-INS-7125 | chr1-139200076-INS-7125 |
| chr1-139205436-INS-7497 | chr1-139205437-INS-7496 |
| chr1-139207901-INS-908 | chr1-139207901-INS-908 |
| chr1-139211501-INS-56 | chr1-139211501-INS-56 |
| chr1-139216892-INS-6741 | chr1-139216893-INS-6741 |
| chr1-139220544-INS-298 | chr1-139220544-INS-300 |
| chr1-139242693-INS-1047 | chr1-139242694-INS-1047 |
| chr1-139249764-INS-201 | chr1-139249764-INS-200 |

| | |
|---|---|
| chr1-139250900-INS-56 | chr1-139250900-INS-56 |
| chr1-139254506-INS-56 | chr1-139254506-INS-56 |
| chr1-139259291-INS-200 | chr1-139259252-INS-200 |
| chr1-139261685-INS-240 | chr1-139261685-INS-270 |
| chr1-139268461-INS-72 | chr1-139268461-INS-100 |
| chr1-139279334-INS-116 | chr1-139279334-INS-117 |
| chr1-139304041-INS-199 | chr1-139304041-INS-200 |
| chr1-139306104-INS-54 | chr1-139306104-INS-56 |
| chr1-139307960-INS-96 | chr1-139307960-INS-96 |
| chr1-139309161-INS-51 | chr1-139309165-INS-54 |
| chr1-139323170-INS-995 | chr1-139323170-INS-995 |
| chr1-139323855-INS-51 | chr1-139323855-INS-54 |
| chr1-139325280-INS-1182 | chr1-139325281-INS-1184 |
| chr1-139329123-INS-195 | chr1-139329123-INS-195 |
| chr1-139337238-INS-168 | chr1-139337238-INS-168 |
| chr1-139346911-INS-1120 | chr1-139346912-INS-1120 |
| chr1-139399982-INS-63 | chr1-139399983-INS-58 |
| chr1-139410611-INS-279 | chr1-139410611-INS-280 |
| chr1-139433619-INS-64 | chr1-139433619-INS-64 |
| chr1-139435778-INS-224 | chr1-139435778-INS-224 |
| chr1-139446099-INS-56 | chr1-139446099-INS-60 |
| chr1-139461959-INS-2195 | chr1-139461960-INS-2195 |
| chr1-139514482-INS-313 | chr1-139514483-INS-313 |
| chr1-139532335-INS-845 | chr1-139532335-INS-843 |
| chr1-139536519-INS-227 | chr1-139536519-INS-235 |
| chr1-139556831-INS-141 | chr1-139556836-INS-142 |
| chr1-139621037-INS-54 | chr1-139621037-INS-54 |
| chr1-139633793-INS-188 | chr1-139633793-INS-194 |
| chr1-139640069-INS-437 | chr1-139640069-INS-437 |
| chr1-139664775-INS-52 | chr1-139664775-INS-52 |
| chr1-139709902-INS-136 | chr1-139709902-INS-136 |
| chr1-139745602-INS-298 | chr1-139745602-INS-298 |
| chr1-139747116-INS-190 | chr1-139747116-INS-190 |
| chr1-139756456-INS-486 | chr1-139756456-INS-486 |
| chr1-139776413-INS-198 | chr1-139776413-INS-198 |
| chr1-139777861-INS-2712 | chr1-139777862-INS-2709 |
| chr1-139860351-INS-90 | chr1-139860351-INS-99 |
| chr1-139950980-INS-3080 | chr1-139950981-INS-3080 |
| chr1-139962846-INS-2898 | chr1-139962847-INS-2898 |
| chr1-140052247-INS-510 | chr1-140052247-INS-510 |
| chr1-140062969-INS-6487 | chr1-140062970-INS-6490 |
| chr1-140146227-INS-51 | chr1-140146227-INS-51 |
| chr1-140153588-INS-73 | chr1-140153588-INS-85 |
| chr1-140160231-INS-1579 | chr1-140160232-INS-1580 |
| chr1-140170240-INS-264 | chr1-140170240-INS-264 |
| chr1-140182142-INS-131 | chr1-140182146-INS-132 |
| chr1-140240853-INS-730 | chr1-140240853-INS-730 |
| chr1-140241996-INS-1938 | chr1-140241997-INS-1938 |
| chr1-140254112-INS-6919 | chr1-140254113-INS-6920 |
| chr1-140267811-INS-154 | chr1-140267799-INS-163 |
| chr1-140268787-INS-55 | chr1-140268787-INS-55 |
| chr1-140330651-INS-1047 | chr1-140330652-INS-1049 |
| chr1-140348672-INS-6030 | chr1-140348673-INS-6030 |
| chr1-140353590-INS-189 | chr1-140353590-INS-189 |
| chr1-140370631-INS-6191 | chr1-140370632-INS-6191 |
| chr1-140388684-INS-1358 | chr1-140388685-INS-1358 |
| chr1-140398495-INS-317 | chr1-140398495-INS-315 |
| chr1-140408311-INS-464 | chr1-140408311-INS-464 |
| chr1-140426823-INS-251 | chr1-140426823-INS-250 |
| chr1-140504108-INS-206 | chr1-140504108-INS-206 |
| chr1-140524845-INS-6979 | chr1-140524846-INS-6979 |
| chr1-140527366-INS-519 | chr1-140527366-INS-519 |
| chr1-140559813-INS-107 | chr1-140559813-INS-107 |
| chr1-140591042-INS-111 | chr1-140590997-INS-126 |
| chr1-140603817-INS-216 | chr1-140603817-INS-216 |
| chr1-140614051-INS-5448 | chr1-140614052-INS-5448 |
| chr1-140621349-INS-201 | chr1-140621349-INS-201 |
| chr1-140683427-INS-201 | chr1-140683427-INS-201 |
| chr1-140727257-INS-51 | chr1-140727257-INS-51 |
| chr1-140781369-INS-2855 | chr1-140781370-INS-2856 |
| chr1-140789407-INS-51 | chr1-140789411-INS-62 |
| chr1-140795498-INS-189 | chr1-140795498-INS-192 |
| chr1-140810012-INS-792 | chr1-140810012-INS-792 |
| chr1-140860381-INS-66 | chr1-140860381-INS-66 |
| chr1-140867272-INS-1177 | chr1-140867273-INS-1175 |
| chr1-140870524-INS-969 | chr1-140870524-INS-969 |
| chr1-140903276-INS-154 | chr1-140903276-INS-154 |

| | |
|---|---|
| chr1-140932217-INS-171 | chr1-140932217-INS-171 |
| chr1-140972132-INS-936 | chr1-140972132-INS-938 |
| chr1-140986900-INS-110 | chr1-140986900-INS-112 |
| chr1-141080294-INS-84 | chr1-141080290-INS-84 |
| chr1-141084211-INS-68 | chr1-141084211-INS-68 |
| chr1-141099129-INS-63 | chr1-141099129-INS-67 |
| chr1-141103259-INS-843 | chr1-141103259-INS-845 |
| chr1-141112679-INS-163 | chr1-141112654-INS-139 |
| chr1-141179967-INS-414 | chr1-141179967-INS-416 |
| chr1-141200836-INS-374 | chr1-141200836-INS-373 |
| chr1-141222894-INS-512 | chr1-141222894-INS-512 |
| chr1-141258674-INS-59 | chr1-141258674-INS-59 |
| chr1-141286219-INS-53 | chr1-141286219-INS-52 |
| chr1-141310751-INS-86 | chr1-141310751-INS-82 |
| chr1-141358598-INS-328 | chr1-141358598-INS-328 |
| chr1-141361639-INS-8767 | chr1-141361640-INS-8771 |
| chr1-141400793-INS-62 | chr1-141400793-INS-62 |
| chr1-141403800-INS-69 | chr1-141403800-INS-68 |
| chr1-141404859-INS-192 | chr1-141404859-INS-192 |
| chr1-141412458-INS-69 | chr1-141412458-INS-69 |
| chr1-141421227-INS-121 | chr1-141421241-INS-155 |
| chr1-141452993-INS-54 | chr1-141452993-INS-69 |
| chr1-141474995-INS-390 | chr1-141474994-INS-391 |
| chr1-141518375-INS-873 | chr1-141518375-INS-873 |
| chr1-141522774-INS-127 | chr1-141522774-INS-133 |
| chr1-141535188-INS-209 | chr1-141535188-INS-210 |
| chr1-141536496-INS-592 | chr1-141536496-INS-592 |
| chr1-141553675-INS-64 | chr1-141553675-INS-72 |
| chr1-141586222-INS-6256 | chr1-141586223-INS-6256 |
| chr1-141604035-INS-62 | chr1-141604036-INS-63 |
| chr1-141604999-INS-244 | chr1-141605015-INS-251 |
| chr1-141606478-INS-520 | chr1-141606478-INS-520 |
| chr1-141610410-INS-826 | chr1-141610410-INS-826 |
| chr1-141619987-INS-183 | chr1-141619987-INS-183 |
| chr1-141672748-INS-74 | chr1-141672748-INS-74 |
| chr1-141681594-INS-62 | chr1-141681594-INS-62 |
| chr1-141686364-INS-108 | chr1-141686364-INS-99 |
| chr1-141700444-INS-451 | chr1-141700444-INS-450 |
| chr1-141707045-INS-470 | chr1-141707045-INS-470 |
| chr1-141736056-INS-305 | chr1-141736056-INS-306 |
| chr1-141763077-INS-63 | chr1-141763089-INS-82 |
| chr1-141796882-INS-690 | chr1-141796882-INS-692 |
| chr1-141802811-INS-6539 | chr1-141802812-INS-6539 |
| chr1-141809189-INS-6877 | chr1-141809190-INS-6877 |
| chr1-141809240-INS-634 | chr1-141809240-INS-634 |
| chr1-141809872-INS-915 | chr1-141809872-INS-936 |
| chr1-141876564-INS-1800 | chr1-141876565-INS-1800 |
| chr1-141881077-INS-86 | chr1-141881077-INS-88 |
| chr1-141913456-INS-52 | chr1-141913456-INS-53 |
| chr1-142006948-INS-241 | chr1-142006948-INS-241 |
| chr1-142007651-INS-1817 | chr1-142007652-INS-1816 |
| chr1-142015874-INS-4267 | chr1-142015875-INS-4267 |
| chr1-142028148-INS-1911 | chr1-142028135-INS-1911 |
| chr1-142047150-INS-397 | chr1-142047150-INS-399 |
| chr1-142050891-INS-8391 | chr1-142050892-INS-8391 |
| chr1-142069728-INS-1173 | chr1-142069730-INS-1167 |
| chr1-142070654-INS-74 | chr1-142070654-INS-74 |
| chr1-142074444-INS-6210 | chr1-142074445-INS-6210 |
| chr1-142079645-INS-6436 | chr1-142079646-INS-6436 |
| chr1-142089789-INS-6348 | chr1-142089790-INS-6351 |
| chr1-142097529-INS-512 | chr1-142097529-INS-512 |
| chr1-142104574-INS-155 | chr1-142104620-INS-147 |
| chr1-142111081-INS-119 | chr1-142111080-INS-123 |
| chr1-142112805-INS-74 | chr1-142112805-INS-74 |
| chr1-142122269-INS-385 | chr1-142122265-INS-374 |
| chr1-142191624-INS-175 | chr1-142191624-INS-175 |
| chr1-142320411-INS-82 | chr1-142320411-INS-82 |
| chr1-142322227-INS-2770 | chr1-142322228-INS-2755 |
| chr1-142346519-INS-155 | chr1-142346471-INS-176 |
| chr1-142375328-INS-68 | chr1-142375328-INS-83 |
| chr1-142392311-INS-52 | chr1-142392311-INS-66 |
| chr1-142401074-INS-59 | chr1-142401069-INS-63 |
| chr1-142410501-INS-114 | chr1-142410501-INS-111 |
| chr1-142428357-INS-208 | chr1-142428357-INS-208 |
| chr1-142429255-INS-6350 | chr1-142429256-INS-6350 |
| chr1-142432170-INS-485 | chr1-142432170-INS-485 |
| chr1-142460169-INS-1176 | chr1-142460170-INS-1176 |

| | |
|---|---|
| chr1-142470567-INS-3025 | chr1-142470568-INS-3025 |
| chr1-142483398-INS-496 | chr1-142483398-INS-496 |
| chr1-142495997-INS-395 | chr1-142495997-INS-397 |
| chr1-142525044-INS-99 | chr1-142525043-INS-87 |
| chr1-142530977-INS-237 | chr1-142530977-INS-237 |
| chr1-142533037-INS-141 | chr1-142533037-INS-141 |
| chr1-142545069-INS-123 | chr1-142545069-INS-123 |
| chr1-142584146-INS-494 | chr1-142584146-INS-493 |
| chr1-142590545-INS-57 | chr1-142590545-INS-56 |
| chr1-142595675-INS-6492 | chr1-142595676-INS-6493 |
| chr1-142612367-INS-110 | chr1-142612367-INS-128 |
| chr1-142621280-INS-54 | chr1-142621280-INS-66 |
| chr1-142627326-INS-2816 | chr1-142627327-INS-2816 |
| chr1-142649363-INS-54 | chr1-142649363-INS-54 |
| chr1-142651112-INS-51 | chr1-142651112-INS-51 |
| chr1-142672232-INS-313 | chr1-142672250-INS-311 |
| chr1-142678724-INS-202 | chr1-142678724-INS-178 |
| chr1-142694459-INS-969 | chr1-142694459-INS-969 |
| chr1-142720744-INS-278 | chr1-142720744-INS-280 |
| chr1-142722330-INS-179 | chr1-142722330-INS-179 |
| chr1-142752151-INS-931 | chr1-142752151-INS-931 |
| chr1-142759349-INS-147 | chr1-142759349-INS-147 |
| chr1-142764671-INS-192 | chr1-142764671-INS-192 |
| chr1-142773279-INS-799 | chr1-142773279-INS-799 |
| chr1-142774759-INS-6260 | chr1-142774760-INS-6260 |
| chr1-142785412-INS-1097 | chr1-142785413-INS-1099 |
| chr1-142805223-INS-6491 | chr1-142805224-INS-6493 |
| chr1-142810410-INS-3126 | chr1-142810411-INS-3126 |
| chr1-142813913-INS-5773 | chr1-142813914-INS-5773 |
| chr1-142816181-INS-7538 | chr1-142816182-INS-7537 |
| chr1-142822279-INS-141 | chr1-142822279-INS-137 |
| chr1-142867389-INS-1671 | chr1-142867390-INS-1671 |
| chr1-142870200-INS-51 | chr1-142870200-INS-51 |
| chr1-142889116-INS-436 | chr1-142889116-INS-436 |
| chr1-142897755-INS-54 | chr1-142897755-INS-56 |
| chr1-142926278-INS-62 | chr1-142926280-INS-64 |
| chr1-142955379-INS-83 | chr1-142955379-INS-83 |
| chr1-142956575-INS-136 | chr1-142956575-INS-136 |
| chr1-142959395-INS-789 | chr1-142959395-INS-789 |
| chr1-142959794-INS-1412 | chr1-142959795-INS-1412 |
| chr1-142985250-INS-55 | chr1-142985250-INS-65 |
| chr1-143001448-INS-116 | chr1-143001448-INS-116 |
| chr1-143014390-INS-6387 | chr1-143014391-INS-6387 |
| chr1-143040842-INS-2936 | chr1-143040843-INS-2936 |
| chr1-143044449-INS-959 | chr1-143044449-INS-959 |
| chr1-143059478-INS-1481 | chr1-143059479-INS-1481 |
| chr1-143108333-INS-66 | chr1-143108334-INS-66 |
| chr1-143108678-INS-903 | chr1-143108678-INS-919 |
| chr1-143122819-INS-6751 | chr1-143122820-INS-6751 |
| chr1-143130668-INS-305 | chr1-143130668-INS-305 |
| chr1-143131333-INS-424 | chr1-143131333-INS-424 |
| chr1-143171117-INS-123 | chr1-143171117-INS-123 |
| chr1-143185918-INS-52 | chr1-143185918-INS-52 |
| chr1-143211779-INS-6098 | chr1-143211780-INS-6098 |
| chr1-143219148-INS-274 | chr1-143219148-INS-274 |
| chr1-143220955-INS-51 | chr1-143220962-INS-51 |
| chr1-143233359-INS-51 | chr1-143233359-INS-66 |
| chr1-143234615-INS-6223 | chr1-143234616-INS-6223 |
| chr1-143240871-INS-362 | chr1-143240871-INS-362 |
| chr1-143251158-INS-52 | chr1-143251158-INS-54 |
| chr1-143264753-INS-1994 | chr1-143264754-INS-1985 |
| chr1-143277900-INS-994 | chr1-143277894-INS-972 |
| chr1-143294792-INS-6438 | chr1-143294793-INS-6438 |
| chr1-143300642-INS-99 | chr1-143300621-INS-140 |
| chr1-143376590-INS-56 | chr1-143376582-INS-56 |
| chr1-143379809-INS-1035 | chr1-143379810-INS-1036 |
| chr1-143387875-INS-214 | chr1-143387875-INS-214 |
| chr1-143392386-INS-56 | chr1-143392390-INS-72 |
| chr1-143401258-INS-559 | chr1-143401258-INS-559 |
| chr1-143402689-INS-405 | chr1-143402693-INS-405 |
| chr1-143416747-INS-106 | chr1-143416750-INS-72 |
| chr1-143418871-INS-228 | chr1-143418871-INS-198 |
| chr1-143426694-INS-6777 | chr1-143426695-INS-6777 |
| chr1-143430904-INS-4728 | chr1-143430905-INS-4728 |
| chr1-143461515-INS-210 | chr1-143461515-INS-226 |
| chr1-143469887-INS-300 | chr1-143469887-INS-300 |
| chr1-143491018-INS-6215 | chr1-143491019-INS-6215 |

| | |
|---|---|
| chr1-143544192-INS-59 | chr1-143544193-INS-52 |
| chr1-143548335-INS-3427 | chr1-143548336-INS-3427 |
| chr1-143589330-INS-2811 | chr1-143589331-INS-2811 |
| chr1-143592846-INS-214 | chr1-143592846-INS-214 |
| chr1-143596254-INS-212 | chr1-143596254-INS-213 |
| chr1-143602135-INS-883 | chr1-143602135-INS-873 |
| chr1-143604108-INS-197 | chr1-143604108-INS-196 |
| chr1-143611670-INS-497 | chr1-143611670-INS-497 |
| chr1-143625218-INS-218 | chr1-143625218-INS-218 |
| chr1-143630205-INS-199 | chr1-143630205-INS-199 |
| chr1-143643819-INS-200 | chr1-143643819-INS-199 |
| chr1-143671582-INS-257 | chr1-143671582-INS-257 |
| chr1-143682922-INS-218 | chr1-143682922-INS-218 |
| chr1-143752776-INS-347 | chr1-143752776-INS-357 |
| chr1-143765037-INS-53 | chr1-143765037-INS-53 |
| chr1-143785535-INS-213 | chr1-143785535-INS-213 |
| chr1-143817668-INS-135 | chr1-143817668-INS-135 |
| chr1-143823792-INS-6906 | chr1-143823793-INS-6906 |
| chr1-143863357-INS-157 | chr1-143863358-INS-136 |
| chr1-143912146-INS-82 | chr1-143912146-INS-82 |
| chr1-143930101-INS-6882 | chr1-143930102-INS-6884 |
| chr1-143936305-INS-232 | chr1-143936305-INS-232 |
| chr1-143961270-INS-79 | chr1-143961270-INS-79 |
| chr1-143973049-INS-7080 | chr1-143973050-INS-7080 |
| chr1-143983204-INS-58 | chr1-143983204-INS-58 |
| chr1-144017660-INS-646 | chr1-144017660-INS-646 |
| chr1-144021169-INS-248 | chr1-144021169-INS-248 |
| chr1-144026779-INS-792 | chr1-144026779-INS-793 |
| chr1-144066557-INS-146 | chr1-144066557-INS-146 |
| chr1-144068758-INS-210 | chr1-144068758-INS-210 |
| chr1-144069632-INS-891 | chr1-144069632-INS-891 |
| chr1-144075280-INS-168 | chr1-144075280-INS-156 |
| chr1-144083328-INS-95 | chr1-144083321-INS-60 |
| chr1-144095944-INS-2470 | chr1-144095945-INS-2470 |
| chr1-144115652-INS-132 | chr1-144115652-INS-131 |
| chr1-144134757-INS-54 | chr1-144134757-INS-60 |
| chr1-144136741-INS-61 | chr1-144136741-INS-70 |
| chr1-144164505-INS-70 | chr1-144164511-INS-70 |
| chr1-144217495-INS-68 | chr1-144217496-INS-89 |
| chr1-144241735-INS-179 | chr1-144241735-INS-179 |
| chr1-144270811-INS-107 | chr1-144270811-INS-108 |
| chr1-144293273-INS-4864 | chr1-144293274-INS-4864 |
| chr1-144387793-INS-140 | chr1-144387793-INS-144 |
| chr1-144395319-INS-54 | chr1-144395319-INS-50 |
| chr1-144455761-INS-52 | chr1-144455761-INS-60 |
| chr1-144462062-INS-115 | chr1-144462046-INS-107 |
| chr1-144477359-INS-54 | chr1-144477364-INS-54 |
| chr1-144481587-INS-61 | chr1-144481587-INS-61 |
| chr1-144492196-INS-5966 | chr1-144492197-INS-5966 |
| chr1-144492197-INS-344 | chr1-144492197-INS-344 |
| chr1-144526868-INS-392 | chr1-144526868-INS-391 |
| chr1-144533198-INS-2584 | chr1-144533199-INS-2584 |
| chr1-144574227-INS-91 | chr1-144574227-INS-91 |
| chr1-144574841-INS-606 | chr1-144574841-INS-601 |
| chr1-144579144-INS-456 | chr1-144579144-INS-456 |
| chr1-144591009-INS-185 | chr1-144591009-INS-185 |
| chr1-144595941-INS-5316 | chr1-144595942-INS-5316 |
| chr1-144616124-INS-5059 | chr1-144616125-INS-5059 |
| chr1-144654391-INS-7129 | chr1-144654392-INS-7129 |
| chr1-144673690-INS-399 | chr1-144673690-INS-399 |
| chr1-144706880-INS-1935 | chr1-144706881-INS-1935 |
| chr1-144717383-INS-6865 | chr1-144717384-INS-6865 |
| chr1-144719335-INS-482 | chr1-144719337-INS-473 |
| chr1-144760542-INS-52 | chr1-144760542-INS-56 |
| chr1-144764216-INS-64 | chr1-144764216-INS-59 |
| chr1-144766742-INS-109 | chr1-144766739-INS-108 |
| chr1-144783136-INS-125 | chr1-144783132-INS-128 |
| chr1-144826810-INS-1363 | chr1-144826811-INS-1364 |
| chr1-144842058-INS-54 | chr1-144842058-INS-66 |
| chr1-144846086-INS-316 | chr1-144846086-INS-316 |
| chr1-144864178-INS-121 | chr1-144864178-INS-121 |
| chr1-144873822-INS-88 | chr1-144873800-INS-64 |
| chr1-144878046-INS-458 | chr1-144878046-INS-458 |
| chr1-144889378-INS-51 | chr1-144889378-INS-51 |
| chr1-144904645-INS-277 | chr1-144904650-INS-278 |
| chr1-144908098-INS-2088 | chr1-144908099-INS-2088 |
| chr1-144946825-INS-146 | chr1-144946825-INS-146 |

| | |
|---|---|
| chr1-144949050-INS-133 | chr1-144949050-INS-133 |
| chr1-144969224-INS-206 | chr1-144969224-INS-205 |
| chr1-144985551-INS-141 | chr1-144985559-INS-141 |
| chr1-144991408-INS-7059 | chr1-144991409-INS-7059 |
| chr1-145000155-INS-56 | chr1-145000155-INS-50 |
| chr1-145055833-INS-204 | chr1-145055833-INS-204 |
| chr1-145075567-INS-52 | chr1-145075567-INS-68 |
| chr1-145079311-INS-83 | chr1-145079311-INS-59 |
| chr1-145094882-INS-51 | chr1-145094882-INS-51 |
| chr1-145114299-INS-58 | chr1-145114299-INS-57 |
| chr1-145138613-INS-174 | chr1-145138583-INS-192 |
| chr1-145139434-INS-5332 | chr1-145139435-INS-5333 |
| chr1-145154549-INS-56 | chr1-145154549-INS-60 |
| chr1-145177759-INS-3410 | chr1-145177760-INS-3410 |
| chr1-145182413-INS-4734 | chr1-145182414-INS-4735 |
| chr1-145184595-INS-452 | chr1-145184595-INS-454 |
| chr1-145185025-INS-7098 | chr1-145185026-INS-7098 |
| chr1-145189514-INS-88 | chr1-145189514-INS-88 |
| chr1-145261558-INS-4377 | chr1-145261559-INS-4377 |
| chr1-145265072-INS-152 | chr1-145265072-INS-170 |
| chr1-145329549-INS-64 | chr1-145329549-INS-57 |
| chr1-145333186-INS-80 | chr1-145333186-INS-68 |
| chr1-145336788-INS-964 | chr1-145336788-INS-964 |
| chr1-145352333-INS-75 | chr1-145352333-INS-55 |
| chr1-145352989-INS-274 | chr1-145352998-INS-271 |
| chr1-145380439-INS-185 | chr1-145380439-INS-185 |
| chr1-145399036-INS-400 | chr1-145399036-INS-400 |
| chr1-145402326-INS-66 | chr1-145402326-INS-70 |
| chr1-145418407-INS-1773 | chr1-145418408-INS-1780 |
| chr1-145484877-INS-51 | chr1-145484877-INS-51 |
| chr1-145492504-INS-56 | chr1-145492504-INS-62 |
| chr1-145493713-INS-56 | chr1-145493713-INS-58 |
| chr1-145500895-INS-1443 | chr1-145500896-INS-1438 |
| chr1-145508039-INS-114 | chr1-145508039-INS-114 |
| chr1-145513902-INS-56 | chr1-145513892-INS-57 |
| chr1-145525102-INS-389 | chr1-145525102-INS-389 |
| chr1-145537838-INS-52 | chr1-145537838-INS-52 |
| chr1-145544170-INS-4740 | chr1-145544171-INS-4740 |
| chr1-145584083-INS-393 | chr1-145584083-INS-393 |
| chr1-145617180-INS-590 | chr1-145617180-INS-589 |
| chr1-145724778-INS-196 | chr1-145724778-INS-196 |
| chr1-145733466-INS-4939 | chr1-145733467-INS-4939 |
| chr1-145743734-INS-374 | chr1-145743734-INS-370 |
| chr1-145758487-INS-529 | chr1-145758487-INS-533 |
| chr1-145761657-INS-408 | chr1-145761657-INS-413 |
| chr1-145769416-INS-63 | chr1-145769416-INS-66 |
| chr1-145770636-INS-724 | chr1-145770636-INS-727 |
| chr1-145811610-INS-84 | chr1-145811606-INS-94 |
| chr1-145812038-INS-569 | chr1-145812038-INS-569 |
| chr1-145815496-INS-7123 | chr1-145815497-INS-7123 |
| chr1-145828011-INS-98 | chr1-145828011-INS-98 |
| chr1-145854394-INS-640 | chr1-145854394-INS-640 |
| chr1-145862506-INS-56 | chr1-145862529-INS-68 |
| chr1-145904417-INS-1545 | chr1-145904418-INS-1544 |
| chr1-145935064-INS-4509 | chr1-145935065-INS-4509 |
| chr1-145949627-INS-889 | chr1-145949626-INS-907 |
| chr1-145957103-INS-124 | chr1-145957103-INS-148 |
| chr1-145990744-INS-77 | chr1-145990744-INS-77 |
| chr1-145996025-INS-55 | chr1-145996025-INS-60 |
| chr1-146021172-INS-74 | chr1-146021172-INS-74 |
| chr1-146048766-INS-60 | chr1-146048766-INS-62 |
| chr1-146060150-INS-57 | chr1-146060150-INS-72 |
| chr1-146122420-INS-68 | chr1-146122420-INS-52 |
| chr1-146125182-INS-5988 | chr1-146125183-INS-5988 |
| chr1-146128313-INS-420 | chr1-146128313-INS-420 |
| chr1-146131312-INS-211 | chr1-146131312-INS-211 |
| chr1-146135811-INS-410 | chr1-146135811-INS-410 |
| chr1-146153178-INS-140 | chr1-146153178-INS-145 |
| chr1-146154789-INS-54 | chr1-146154789-INS-54 |
| chr1-146181315-INS-72 | chr1-146181315-INS-72 |
| chr1-146199378-INS-1149 | chr1-146199379-INS-1149 |
| chr1-146220326-INS-363 | chr1-146220326-INS-363 |
| chr1-146226437-INS-7373 | chr1-146226438-INS-7373 |
| chr1-146241535-INS-78 | chr1-146241535-INS-78 |
| chr1-146243883-INS-390 | chr1-146243883-INS-390 |
| chr1-146307147-INS-440 | chr1-146307147-INS-440 |
| chr1-146308705-INS-260 | chr1-146308705-INS-260 |

| | |
|---|---|
| chr1-146315760-INS-53 | chr1-146315760-INS-53 |
| chr1-146338937-INS-1893 | chr1-146338938-INS-1893 |
| chr1-146387695-INS-549 | chr1-146387695-INS-549 |
| chr1-146419774-INS-5345 | chr1-146419775-INS-5345 |
| chr1-146431143-INS-52 | chr1-146431143-INS-56 |
| chr1-146503261-INS-68 | chr1-146503261-INS-64 |
| chr1-146514244-INS-207 | chr1-146514244-INS-208 |
| chr1-146638505-INS-107 | chr1-146638505-INS-117 |
| chr1-146668187-INS-60 | chr1-146668215-INS-61 |
| chr1-146674299-INS-52 | chr1-146674300-INS-54 |
| chr1-146676590-INS-505 | chr1-146676590-INS-505 |
| chr1-146701844-INS-117 | chr1-146701857-INS-115 |
| chr1-146726951-INS-64 | chr1-146726951-INS-56 |
| chr1-146733302-INS-203 | chr1-146733302-INS-203 |
| chr1-146748751-INS-100 | chr1-146748726-INS-86 |
| chr1-146756313-INS-212 | chr1-146756313-INS-212 |
| chr1-146764062-INS-481 | chr1-146764062-INS-481 |
| chr1-146775472-INS-986 | chr1-146775472-INS-986 |
| chr1-146783636-INS-52 | chr1-146783637-INS-80 |
| chr1-146793062-INS-68 | chr1-146793062-INS-68 |
| chr1-146803216-INS-163 | chr1-146803216-INS-163 |
| chr1-146834825-INS-311 | chr1-146834825-INS-311 |
| chr1-146852683-INS-382 | chr1-146852683-INS-382 |
| chr1-146854593-INS-150 | chr1-146854593-INS-150 |
| chr1-146873971-INS-69 | chr1-146874005-INS-57 |
| chr1-146882077-INS-390 | chr1-146882077-INS-390 |
| chr1-146885785-INS-900 | chr1-146885753-INS-887 |
| chr1-146900970-INS-78 | chr1-146900970-INS-56 |
| chr1-146911502-INS-115 | chr1-146911502-INS-113 |
| chr1-146922071-INS-53 | chr1-146922071-INS-53 |
| chr1-146963282-INS-400 | chr1-146963282-INS-400 |
| chr1-146981655-INS-6441 | chr1-146981656-INS-6442 |
| chr1-147014751-INS-52 | chr1-147014751-INS-52 |
| chr1-147067854-INS-61 | chr1-147067854-INS-57 |
| chr1-147068507-INS-2698 | chr1-147068508-INS-2698 |
| chr1-147077658-INS-176 | chr1-147077658-INS-176 |
| chr1-147128183-INS-268 | chr1-147128183-INS-268 |
| chr1-147132299-INS-508 | chr1-147132299-INS-508 |
| chr1-147133520-INS-6854 | chr1-147133521-INS-6854 |
| chr1-147148324-INS-54 | chr1-147148324-INS-50 |
| chr1-147157667-INS-52 | chr1-147157667-INS-50 |
| chr1-147172337-INS-55 | chr1-147172337-INS-51 |
| chr1-147179948-INS-328 | chr1-147179948-INS-328 |
| chr1-147181873-INS-1086 | chr1-147181874-INS-1086 |
| chr1-147184837-INS-76 | chr1-147184838-INS-69 |
| chr1-147191638-INS-82 | chr1-147191640-INS-60 |
| chr1-147218552-INS-190 | chr1-147218552-INS-191 |
| chr1-147231116-INS-4316 | chr1-147231117-INS-4316 |
| chr1-147233829-INS-396 | chr1-147233830-INS-426 |
| chr1-147236641-INS-6364 | chr1-147236642-INS-6364 |
| chr1-147273104-INS-78 | chr1-147273104-INS-84 |
| chr1-147277110-INS-136 | chr1-147277110-INS-136 |
| chr1-147279207-INS-56 | chr1-147279207-INS-56 |
| chr1-147281697-INS-573 | chr1-147281697-INS-573 |
| chr1-147297496-INS-56 | chr1-147297496-INS-56 |
| chr1-147331903-INS-94 | chr1-147331903-INS-74 |
| chr1-147346401-INS-4554 | chr1-147346402-INS-4554 |
| chr1-147350691-INS-95 | chr1-147350691-INS-104 |
| chr1-147353362-INS-58 | chr1-147353362-INS-58 |
| chr1-147386804-INS-1626 | chr1-147386805-INS-1627 |
| chr1-147406560-INS-2610 | chr1-147406561-INS-2611 |
| chr1-147443416-INS-114 | chr1-147443416-INS-114 |
| chr1-147466714-INS-56 | chr1-147466720-INS-51 |
| chr1-147467730-INS-208 | chr1-147467730-INS-208 |
| chr1-147491086-INS-842 | chr1-147491086-INS-842 |
| chr1-147494725-INS-159 | chr1-147494725-INS-114 |
| chr1-147500266-INS-109 | chr1-147500304-INS-115 |
| chr1-147513722-INS-429 | chr1-147513722-INS-429 |
| chr1-147516048-INS-939 | chr1-147516048-INS-939 |
| chr1-147528795-INS-4022 | chr1-147528799-INS-4019 |
| chr1-147584351-INS-106 | chr1-147584351-INS-106 |
| chr1-147594286-INS-3786 | chr1-147594287-INS-3786 |
| chr1-147597700-INS-2958 | chr1-147597702-INS-2959 |
| chr1-147624801-INS-7011 | chr1-147624802-INS-7011 |
| chr1-147627059-INS-1117 | chr1-147627061-INS-1109 |
| chr1-147635126-INS-59 | chr1-147635116-INS-51 |
| chr1-147696791-INS-53 | chr1-147696791-INS-53 |

| | |
|---|---|
| chr1-147753646-INS-72 | chr1-147753666-INS-70 |
| chr1-147770941-INS-6883 | chr1-147770942-INS-6883 |
| chr1-147772190-INS-731 | chr1-147772190-INS-731 |
| chr1-147785067-INS-6200 | chr1-147785068-INS-6200 |
| chr1-147805320-INS-68 | chr1-147805320-INS-68 |
| chr1-147826231-INS-282 | chr1-147826231-INS-282 |
| chr1-147837566-INS-2252 | chr1-147837567-INS-2255 |
| chr1-147871245-INS-51 | chr1-147871235-INS-60 |
| chr1-147872795-INS-121 | chr1-147872795-INS-121 |
| chr1-147875556-INS-89 | chr1-147875555-INS-86 |
| chr1-147880979-INS-314 | chr1-147880979-INS-318 |
| chr1-147893080-INS-90 | chr1-147893080-INS-90 |
| chr1-147898493-INS-347 | chr1-147898513-INS-357 |
| chr1-147906913-INS-124 | chr1-147906913-INS-124 |
| chr1-147913551-INS-52 | chr1-147913551-INS-54 |
| chr1-147921784-INS-6555 | chr1-147921785-INS-6555 |
| chr1-147964324-INS-515 | chr1-147964345-INS-534 |
| chr1-147978938-INS-2011 | chr1-147978939-INS-2011 |
| chr1-147987466-INS-332 | chr1-147987467-INS-332 |
| chr1-147993452-INS-62 | chr1-147993433-INS-106 |
| chr1-148005138-INS-2592 | chr1-148005139-INS-2592 |
| chr1-148016745-INS-52 | chr1-148016745-INS-52 |
| chr1-148029380-INS-3315 | chr1-148029381-INS-3291 |
| chr1-148034446-INS-80 | chr1-148034446-INS-82 |
| chr1-148044001-INS-52 | chr1-148043997-INS-54 |
| chr1-148054768-INS-93 | chr1-148054771-INS-84 |
| chr1-148077915-INS-73 | chr1-148077916-INS-67 |
| chr1-148078007-INS-6984 | chr1-148078008-INS-6984 |
| chr1-148094243-INS-214 | chr1-148094243-INS-214 |
| chr1-148117644-INS-1408 | chr1-148117645-INS-1408 |
| chr1-148127868-INS-400 | chr1-148127868-INS-400 |
| chr1-148130560-INS-6438 | chr1-148130561-INS-6438 |
| chr1-148134789-INS-3327 | chr1-148134790-INS-3327 |
| chr1-148163116-INS-147 | chr1-148163116-INS-143 |
| chr1-148279167-INS-68 | chr1-148279137-INS-102 |
| chr1-148295742-INS-52 | chr1-148295761-INS-55 |
| chr1-148313147-INS-52 | chr1-148313147-INS-52 |
| chr1-148319222-INS-86 | chr1-148319222-INS-60 |
| chr1-148355665-INS-56 | chr1-148355665-INS-54 |
| chr1-148358223-INS-56 | chr1-148358223-INS-56 |
| chr1-148448115-INS-57 | chr1-148448115-INS-57 |
| chr1-148478420-INS-6871 | chr1-148478421-INS-6870 |
| chr1-148482617-INS-340 | chr1-148482617-INS-340 |
| chr1-148507823-INS-1023 | chr1-148507824-INS-1023 |
| chr1-148508576-INS-923 | chr1-148508577-INS-923 |
| chr1-148546288-INS-1865 | chr1-148546289-INS-1865 |
| chr1-148556958-INS-109 | chr1-148556958-INS-109 |
| chr1-148557323-INS-873 | chr1-148557323-INS-873 |
| chr1-148560326-INS-595 | chr1-148560326-INS-596 |
| chr1-148563785-INS-66 | chr1-148563785-INS-64 |
| chr1-148575094-INS-52 | chr1-148575094-INS-85 |
| chr1-148580498-INS-7338 | chr1-148580499-INS-7338 |
| chr1-148630979-INS-78 | chr1-148631006-INS-64 |
| chr1-148644536-INS-261 | chr1-148644536-INS-261 |
| chr1-148652317-INS-102 | chr1-148652317-INS-116 |
| chr1-148659371-INS-414 | chr1-148659371-INS-414 |
| chr1-148661645-INS-1002 | chr1-148661646-INS-1002 |
| chr1-148664125-INS-54 | chr1-148664125-INS-62 |
| chr1-148666214-INS-558 | chr1-148666214-INS-558 |
| chr1-148669258-INS-72 | chr1-148669258-INS-72 |
| chr1-148675407-INS-110 | chr1-148675407-INS-100 |
| chr1-148688871-INS-6258 | chr1-148688872-INS-6260 |
| chr1-148732520-INS-3201 | chr1-148732521-INS-3201 |
| chr1-148749799-INS-1891 | chr1-148749800-INS-1891 |
| chr1-148759985-INS-416 | chr1-148759985-INS-416 |
| chr1-148773646-INS-305 | chr1-148773646-INS-305 |
| chr1-148787549-INS-81 | chr1-148787549-INS-81 |
| chr1-148789499-INS-561 | chr1-148789499-INS-561 |
| chr1-148805647-INS-150 | chr1-148805648-INS-104 |
| chr1-148827439-INS-80 | chr1-148827439-INS-80 |
| chr1-148848299-INS-120 | chr1-148848299-INS-120 |
| chr1-148850283-INS-6360 | chr1-148850284-INS-6360 |
| chr1-148853557-INS-54 | chr1-148853557-INS-57 |
| chr1-148867779-INS-367 | chr1-148867779-INS-367 |
| chr1-148915644-INS-52 | chr1-148915644-INS-52 |
| chr1-148916255-INS-4804 | chr1-148916256-INS-4804 |
| chr1-148952164-INS-62 | chr1-148952139-INS-56 |

| | |
|---|---|
| chr1-148954643-INS-593 | chr1-148954643-INS-594 |
| chr1-148986266-INS-82 | chr1-148986266-INS-82 |
| chr1-149005486-INS-54 | chr1-149005487-INS-53 |
| chr1-149010049-INS-146 | chr1-149010023-INS-171 |
| chr1-149042065-INS-56 | chr1-149042023-INS-56 |
| chr1-149093925-INS-6462 | chr1-149093926-INS-6462 |
| chr1-149102830-INS-200 | chr1-149102830-INS-200 |
| chr1-149137291-INS-4074 | chr1-149137292-INS-4070 |
| chr1-149138775-INS-1472 | chr1-149138776-INS-1472 |
| chr1-149159105-INS-107 | chr1-149159105-INS-113 |
| chr1-149162292-INS-219 | chr1-149162292-INS-219 |
| chr1-149186103-INS-141 | chr1-149186103-INS-142 |
| chr1-149220482-INS-79 | chr1-149220480-INS-59 |
| chr1-149263676-INS-90 | chr1-149263676-INS-96 |
| chr1-149277180-INS-573 | chr1-149277181-INS-573 |
| chr1-149280581-INS-110 | chr1-149280579-INS-114 |
| chr1-149283823-INS-274 | chr1-149283801-INS-295 |
| chr1-149293840-INS-72 | chr1-149293840-INS-72 |
| chr1-149299698-INS-190 | chr1-149299698-INS-189 |
| chr1-149322745-INS-60 | chr1-149322745-INS-56 |
| chr1-149334120-INS-7376 | chr1-149334121-INS-7377 |
| chr1-149359783-INS-794 | chr1-149359783-INS-795 |
| chr1-149374145-INS-131 | chr1-149374145-INS-131 |
| chr1-149414147-INS-120 | chr1-149414147-INS-90 |
| chr1-149434734-INS-2967 | chr1-149434735-INS-2967 |
| chr1-149440772-INS-52 | chr1-149440772-INS-54 |
| chr1-149442110-INS-579 | chr1-149442110-INS-578 |
| chr1-149443718-INS-85 | chr1-149443718-INS-90 |
| chr1-149454344-INS-1127 | chr1-149454345-INS-1135 |
| chr1-149454531-INS-202 | chr1-149454531-INS-206 |
| chr1-149486646-INS-4342 | chr1-149486647-INS-4342 |
| chr1-149499564-INS-60 | chr1-149499564-INS-60 |
| chr1-149514974-INS-66 | chr1-149514974-INS-66 |
| chr1-149523957-INS-630 | chr1-149523957-INS-631 |
| chr1-149527056-INS-75 | chr1-149527056-INS-50 |
| chr1-149544887-INS-204 | chr1-149544887-INS-205 |
| chr1-149579318-INS-3182 | chr1-149579319-INS-3183 |
| chr1-149596812-INS-152 | chr1-149596812-INS-152 |
| chr1-149622293-INS-52 | chr1-149622293-INS-52 |
| chr1-149644330-INS-439 | chr1-149644330-INS-439 |
| chr1-149661120-INS-70 | chr1-149661080-INS-87 |
| chr1-149662557-INS-62 | chr1-149662557-INS-58 |
| chr1-149665254-INS-131 | chr1-149665227-INS-141 |
| chr1-149672917-INS-1775 | chr1-149672918-INS-1776 |
| chr1-149681549-INS-6240 | chr1-149681550-INS-6241 |
| chr1-149686070-INS-82 | chr1-149686070-INS-81 |
| chr1-149700749-INS-450 | chr1-149700749-INS-450 |
| chr1-149708596-INS-101 | chr1-149708596-INS-112 |
| chr1-149734294-INS-213 | chr1-149734294-INS-214 |
| chr1-149740815-INS-56 | chr1-149740815-INS-56 |
| chr1-149763620-INS-359 | chr1-149763620-INS-359 |
| chr1-149788262-INS-70 | chr1-149788262-INS-72 |
| chr1-149796632-INS-54 | chr1-149796603-INS-54 |
| chr1-149814374-INS-6444 | chr1-149814375-INS-6444 |
| chr1-149828360-INS-53 | chr1-149828345-INS-63 |
| chr1-149845587-INS-56 | chr1-149845587-INS-80 |
| chr1-149847677-INS-84 | chr1-149847677-INS-90 |
| chr1-149849727-INS-74 | chr1-149849727-INS-102 |
| chr1-149868956-INS-96 | chr1-149868956-INS-80 |
| chr1-149910708-INS-94 | chr1-149910708-INS-94 |
| chr1-149925237-INS-98 | chr1-149925224-INS-141 |
| chr1-149948267-INS-230 | chr1-149948267-INS-238 |
| chr1-149965431-INS-153 | chr1-149965431-INS-183 |
| chr1-149978168-INS-83 | chr1-149978168-INS-83 |
| chr1-150017025-INS-407 | chr1-150017025-INS-407 |
| chr1-150024950-INS-2437 | chr1-150024951-INS-2436 |
| chr1-150030732-INS-904 | chr1-150030732-INS-904 |
| chr1-150055259-INS-130 | chr1-150055245-INS-142 |
| chr1-150056647-INS-546 | chr1-150056647-INS-546 |
| chr1-150076241-INS-1575 | chr1-150076242-INS-1575 |
| chr1-150084436-INS-6365 | chr1-150084437-INS-6366 |
| chr1-150098791-INS-56 | chr1-150098791-INS-64 |
| chr1-150120035-INS-204 | chr1-150120035-INS-204 |
| chr1-150122255-INS-696 | chr1-150122255-INS-695 |
| chr1-150188138-INS-757 | chr1-150188138-INS-759 |
| chr1-150193757-INS-106 | chr1-150193761-INS-58 |
| chr1-150205135-INS-312 | chr1-150205135-INS-313 |

| | |
|---|---|
| chr1-150210995-INS-7080 | chr1-150210996-INS-7080 |
| chr1-150211449-INS-252 | chr1-150211449-INS-267 |
| chr1-150212958-INS-64 | chr1-150212958-INS-78 |
| chr1-150213341-INS-203 | chr1-150213341-INS-203 |
| chr1-150250250-INS-167 | chr1-150250250-INS-168 |
| chr1-150260411-INS-200 | chr1-150260411-INS-215 |
| chr1-150288581-INS-221 | chr1-150288581-INS-220 |
| chr1-150331351-INS-90 | chr1-150331328-INS-72 |
| chr1-150351661-INS-370 | chr1-150351661-INS-370 |
| chr1-150366267-INS-117 | chr1-150366267-INS-95 |
| chr1-150367452-INS-233 | chr1-150367452-INS-233 |
| chr1-150368839-INS-62 | chr1-150368839-INS-56 |
| chr1-150371266-INS-178 | chr1-150371266-INS-191 |
| chr1-150392639-INS-401 | chr1-150392639-INS-401 |
| chr1-150403927-INS-828 | chr1-150403928-INS-827 |
| chr1-150445372-INS-122 | chr1-150445372-INS-122 |
| chr1-150542966-INS-3741 | chr1-150542967-INS-3740 |
| chr1-150571484-INS-67 | chr1-150571483-INS-71 |
| chr1-150604322-INS-6611 | chr1-150604323-INS-6611 |
| chr1-150606697-INS-56 | chr1-150606697-INS-54 |
| chr1-150616304-INS-668 | chr1-150616304-INS-668 |
| chr1-150625566-INS-382 | chr1-150625566-INS-391 |
| chr1-150670806-INS-68 | chr1-150670806-INS-70 |
| chr1-150691489-INS-96 | chr1-150691489-INS-84 |
| chr1-150703872-INS-86 | chr1-150703872-INS-90 |
| chr1-150723476-INS-57 | chr1-150723476-INS-57 |
| chr1-150760163-INS-98 | chr1-150760163-INS-98 |
| chr1-150768234-INS-51 | chr1-150768234-INS-51 |
| chr1-150795229-INS-92 | chr1-150795262-INS-91 |
| chr1-150795634-INS-738 | chr1-150795634-INS-738 |
| chr1-150826726-INS-592 | chr1-150826726-INS-591 |
| chr1-150843791-INS-1141 | chr1-150843792-INS-1141 |
| chr1-150862278-INS-58 | chr1-150862278-INS-58 |
| chr1-150888259-INS-1223 | chr1-150888260-INS-1223 |
| chr1-150894269-INS-57 | chr1-150894230-INS-60 |
| chr1-150905984-INS-138 | chr1-150905984-INS-138 |
| chr1-150907344-INS-210 | chr1-150907344-INS-210 |
| chr1-150908451-INS-6706 | chr1-150908452-INS-6708 |
| chr1-150913817-INS-124 | chr1-150913817-INS-134 |
| chr1-150920346-INS-146 | chr1-150920352-INS-129 |
| chr1-150921598-INS-1309 | chr1-150921603-INS-1304 |
| chr1-150933619-INS-78 | chr1-150933619-INS-78 |
| chr1-150940232-INS-418 | chr1-150940232-INS-418 |
| chr1-150942580-INS-95 | chr1-150942580-INS-95 |
| chr1-150950185-INS-73 | chr1-150950213-INS-53 |
| chr1-150952269-INS-193 | chr1-150952269-INS-193 |
| chr1-150989459-INS-505 | chr1-150989459-INS-504 |
| chr1-150995077-INS-400 | chr1-150995077-INS-400 |
| chr1-151002542-INS-73 | chr1-151002542-INS-72 |
| chr1-151023443-INS-75 | chr1-151023443-INS-51 |
| chr1-151028599-INS-60 | chr1-151028599-INS-60 |
| chr1-151079000-INS-275 | chr1-151079000-INS-276 |
| chr1-151094139-INS-2675 | chr1-151094140-INS-2675 |
| chr1-151121950-INS-208 | chr1-151121950-INS-201 |
| chr1-151153108-INS-240 | chr1-151153108-INS-240 |
| chr1-151153778-INS-493 | chr1-151153778-INS-495 |
| chr1-151172773-INS-199 | chr1-151172773-INS-199 |
| chr1-151174960-INS-347 | chr1-151174960-INS-347 |
| chr1-151180701-INS-4947 | chr1-151180702-INS-4948 |
| chr1-151192898-INS-316 | chr1-151192898-INS-316 |
| chr1-151212180-INS-231 | chr1-151212180-INS-231 |
| chr1-151225448-INS-190 | chr1-151225448-INS-228 |
| chr1-151225958-INS-226 | chr1-151225958-INS-226 |
| chr1-151266067-INS-198 | chr1-151266067-INS-198 |
| chr1-151280155-INS-277 | chr1-151280155-INS-277 |
| chr1-151282386-INS-192 | chr1-151282386-INS-190 |
| chr1-151283549-INS-92 | chr1-151283549-INS-92 |
| chr1-151290730-INS-202 | chr1-151290730-INS-203 |
| chr1-151298442-INS-208 | chr1-151298442-INS-208 |
| chr1-151310663-INS-191 | chr1-151310663-INS-191 |
| chr1-151314978-INS-98 | chr1-151314978-INS-98 |
| chr1-151323262-INS-70 | chr1-151323263-INS-65 |
| chr1-151351437-INS-367 | chr1-151351437-INS-377 |
| chr1-151381553-INS-124 | chr1-151381553-INS-124 |
| chr1-151403562-INS-125 | chr1-151403562-INS-125 |
| chr1-151413858-INS-171 | chr1-151413858-INS-171 |
| chr1-151420542-INS-563 | chr1-151420542-INS-563 |

| | |
|---|---|
| chr1-151428642-INS-103 | chr1-151428642-INS-111 |
| chr1-151434475-INS-88 | chr1-151434475-INS-79 |
| chr1-151444582-INS-380 | chr1-151444582-INS-382 |
| chr1-151469881-INS-68 | chr1-151469881-INS-70 |
| chr1-151475034-INS-127 | chr1-151475034-INS-127 |
| chr1-151480984-INS-83 | chr1-151480985-INS-84 |
| chr1-151482719-INS-162 | chr1-151482719-INS-162 |
| chr1-151497008-INS-7067 | chr1-151497009-INS-7067 |
| chr1-151504174-INS-78 | chr1-151504174-INS-78 |
| chr1-151513183-INS-70 | chr1-151513183-INS-72 |
| chr1-151527265-INS-68 | chr1-151527265-INS-68 |
| chr1-151542163-INS-511 | chr1-151542163-INS-463 |
| chr1-151547557-INS-201 | chr1-151547556-INS-201 |
| chr1-151550372-INS-130 | chr1-151550372-INS-130 |
| chr1-151556634-INS-268 | chr1-151556634-INS-268 |
| chr1-151588619-INS-3197 | chr1-151588620-INS-3197 |
| chr1-151604385-INS-510 | chr1-151604385-INS-510 |
| chr1-151613915-INS-4736 | chr1-151613916-INS-4736 |
| chr1-151616629-INS-433 | chr1-151616629-INS-430 |
| chr1-151617662-INS-492 | chr1-151617661-INS-487 |
| chr1-151624981-INS-136 | chr1-151624981-INS-136 |
| chr1-151651013-INS-418 | chr1-151651013-INS-418 |
| chr1-151672692-INS-780 | chr1-151672692-INS-780 |
| chr1-151672712-INS-108 | chr1-151672692-INS-106 |
| chr1-151686139-INS-207 | chr1-151686139-INS-207 |
| chr1-151699541-INS-248 | chr1-151699541-INS-243 |
| chr1-151706544-INS-76 | chr1-151706544-INS-84 |
| chr1-151707986-INS-1866 | chr1-151707987-INS-1866 |
| chr1-151723773-INS-56 | chr1-151723773-INS-58 |
| chr1-151726782-INS-119 | chr1-151726782-INS-119 |
| chr1-151729480-INS-56 | chr1-151729480-INS-60 |
| chr1-151734246-INS-156 | chr1-151734252-INS-129 |
| chr1-151749158-INS-722 | chr1-151749158-INS-684 |
| chr1-151749791-INS-825 | chr1-151749792-INS-825 |
| chr1-151803526-INS-487 | chr1-151803526-INS-493 |
| chr1-151817263-INS-168 | chr1-151817263-INS-168 |
| chr1-151820548-INS-86 | chr1-151820540-INS-86 |
| chr1-151847649-INS-5095 | chr1-151847650-INS-5095 |
| chr1-151855017-INS-996 | chr1-151855017-INS-996 |
| chr1-151872622-INS-109 | chr1-151872622-INS-109 |
| chr1-151901816-INS-52 | chr1-151901816-INS-56 |
| chr1-151905251-INS-228 | chr1-151905251-INS-228 |
| chr1-151966454-INS-54 | chr1-151966454-INS-54 |
| chr1-151978487-INS-214 | chr1-151978487-INS-214 |
| chr1-151983294-INS-54 | chr1-151983294-INS-54 |
| chr1-151985708-INS-90 | chr1-151985708-INS-94 |
| chr1-151988952-INS-444 | chr1-151988952-INS-444 |
| chr1-152028837-INS-373 | chr1-152028837-INS-323 |
| chr1-152036356-INS-60 | chr1-152036356-INS-106 |
| chr1-152042341-INS-183 | chr1-152042341-INS-183 |
| chr1-152066603-INS-325 | chr1-152066560-INS-374 |
| chr1-152080331-INS-60 | chr1-152080331-INS-60 |
| chr1-152089629-INS-1776 | chr1-152089633-INS-1770 |
| chr1-152107933-INS-208 | chr1-152107933-INS-208 |
| chr1-152115220-INS-576 | chr1-152115220-INS-575 |
| chr1-152116103-INS-98 | chr1-152116103-INS-99 |
| chr1-152122942-INS-498 | chr1-152122942-INS-498 |
| chr1-152144161-INS-1904 | chr1-152144162-INS-1905 |
| chr1-152157922-INS-138 | chr1-152157922-INS-138 |
| chr1-152161196-INS-76 | chr1-152161190-INS-50 |
| chr1-152176039-INS-86 | chr1-152176011-INS-110 |
| chr1-152201715-INS-58 | chr1-152201715-INS-58 |
| chr1-152219645-INS-84 | chr1-152219645-INS-84 |
| chr1-152223645-INS-156 | chr1-152223645-INS-156 |
| chr1-152260462-INS-129 | chr1-152260462-INS-129 |
| chr1-152260652-INS-532 | chr1-152260652-INS-533 |
| chr1-152273677-INS-227 | chr1-152273677-INS-227 |
| chr1-152277231-INS-66 | chr1-152277231-INS-66 |
| chr1-152284604-INS-138 | chr1-152284623-INS-135 |
| chr1-152300339-INS-331 | chr1-152300339-INS-331 |
| chr1-152334288-INS-60 | chr1-152334288-INS-62 |
| chr1-152348352-INS-179 | chr1-152348352-INS-179 |
| chr1-152377893-INS-64 | chr1-152377878-INS-112 |
| chr1-152400766-INS-55 | chr1-152400766-INS-50 |
| chr1-152415904-INS-65 | chr1-152415904-INS-65 |
| chr1-152423467-INS-74 | chr1-152423467-INS-54 |
| chr1-152429084-INS-273 | chr1-152429119-INS-238 |

| | |
|---|---|
| chr1-152438539-INS-4378 | chr1-152438540-INS-4378 |
| chr1-152448645-INS-76 | chr1-152448645-INS-54 |
| chr1-152459492-INS-2340 | chr1-152459515-INS-2335 |
| chr1-152479369-INS-52 | chr1-152479369-INS-50 |
| chr1-152480902-INS-58 | chr1-152480902-INS-60 |
| chr1-152500255-INS-60 | chr1-152500255-INS-50 |
| chr1-152509924-INS-103 | chr1-152509928-INS-107 |
| chr1-152519221-INS-4890 | chr1-152519222-INS-4896 |
| chr1-152529302-INS-368 | chr1-152529302-INS-368 |
| chr1-152529969-INS-67 | chr1-152529969-INS-67 |
| chr1-152531119-INS-199 | chr1-152531119-INS-199 |
| chr1-152535910-INS-62 | chr1-152535921-INS-62 |
| chr1-152581850-INS-929 | chr1-152581850-INS-929 |
| chr1-152584135-INS-233 | chr1-152584135-INS-233 |
| chr1-152590116-INS-52 | chr1-152590116-INS-52 |
| chr1-152593827-INS-170 | chr1-152593828-INS-128 |
| chr1-152595519-INS-55 | chr1-152595519-INS-55 |
| chr1-152602715-INS-87 | chr1-152602716-INS-87 |
| chr1-152607086-INS-100 | chr1-152607088-INS-72 |
| chr1-152610868-INS-210 | chr1-152610868-INS-160 |
| chr1-152616101-INS-220 | chr1-152616101-INS-220 |
| chr1-152617086-INS-629 | chr1-152617086-INS-629 |
| chr1-152630125-INS-317 | chr1-152630125-INS-319 |
| chr1-152635584-INS-60 | chr1-152635584-INS-60 |
| chr1-152637955-INS-58 | chr1-152637955-INS-58 |
| chr1-152667318-INS-400 | chr1-152667318-INS-400 |
| chr1-152678044-INS-80 | chr1-152678056-INS-86 |
| chr1-152680320-INS-105 | chr1-152680320-INS-105 |
| chr1-152681531-INS-202 | chr1-152681531-INS-202 |
| chr1-152695063-INS-60 | chr1-152695063-INS-56 |
| chr1-152697077-INS-213 | chr1-152697077-INS-213 |
| chr1-152698928-INS-1990 | chr1-152698929-INS-1990 |
| chr1-152701728-INS-348 | chr1-152701728-INS-354 |
| chr1-152742739-INS-826 | chr1-152742739-INS-826 |
| chr1-152749611-INS-189 | chr1-152749611-INS-189 |
| chr1-152801898-INS-233 | chr1-152801898-INS-233 |
| chr1-152803754-INS-206 | chr1-152803754-INS-206 |
| chr1-152805200-INS-735 | chr1-152805200-INS-738 |
| chr1-152815435-INS-390 | chr1-152815435-INS-390 |
| chr1-152816596-INS-489 | chr1-152816596-INS-464 |
| chr1-152821606-INS-218 | chr1-152821606-INS-218 |
| chr1-152835952-INS-221 | chr1-152835952-INS-222 |
| chr1-152853471-INS-63 | chr1-152853471-INS-72 |
| chr1-152877435-INS-213 | chr1-152877435-INS-213 |
| chr1-152897826-INS-110 | chr1-152897826-INS-105 |
| chr1-152899357-INS-191 | chr1-152899357-INS-192 |
| chr1-152902217-INS-479 | chr1-152902217-INS-479 |
| chr1-152902897-INS-61 | chr1-152902897-INS-61 |
| chr1-152943158-INS-52 | chr1-152943158-INS-56 |
| chr1-152947491-INS-341 | chr1-152947491-INS-341 |
| chr1-152949430-INS-205 | chr1-152949430-INS-205 |
| chr1-152973537-INS-51 | chr1-152973537-INS-51 |
| chr1-152993614-INS-95 | chr1-152993614-INS-95 |
| chr1-153000762-INS-58 | chr1-153000762-INS-60 |
| chr1-153022155-INS-228 | chr1-153022155-INS-228 |
| chr1-153031834-INS-210 | chr1-153031834-INS-210 |
| chr1-153040902-INS-158 | chr1-153040902-INS-143 |
| chr1-153050380-INS-462 | chr1-153050380-INS-462 |
| chr1-153056987-INS-56 | chr1-153056987-INS-56 |
| chr1-153083936-INS-50 | chr1-153083936-INS-54 |
| chr1-153086083-INS-72 | chr1-153086083-INS-72 |
| chr1-153087956-INS-59 | chr1-153087956-INS-59 |
| chr1-153091394-INS-68 | chr1-153091394-INS-68 |
| chr1-153100455-INS-147 | chr1-153100455-INS-147 |
| chr1-153110669-INS-217 | chr1-153110669-INS-219 |
| chr1-153111845-INS-493 | chr1-153111856-INS-515 |
| chr1-153152469-INS-226 | chr1-153152469-INS-227 |
| chr1-153154201-INS-187 | chr1-153154201-INS-188 |
| chr1-153171793-INS-104 | chr1-153171793-INS-104 |
| chr1-153178914-INS-66 | chr1-153178914-INS-66 |
| chr1-153241423-INS-137 | chr1-153241423-INS-142 |
| chr1-153262275-INS-90 | chr1-153262275-INS-91 |
| chr1-153298818-INS-4099 | chr1-153298819-INS-4099 |
| chr1-153350417-INS-74 | chr1-153350417-INS-74 |
| chr1-153366631-INS-176 | chr1-153366631-INS-161 |
| chr1-153376100-INS-202 | chr1-153376100-INS-202 |
| chr1-153389758-INS-282 | chr1-153389759-INS-282 |

| | |
|---|---|
| chr1-153395184-INS-301 | chr1-153395184-INS-303 |
| chr1-153404226-INS-203 | chr1-153404226-INS-203 |
| chr1-153417749-INS-209 | chr1-153417749-INS-209 |
| chr1-153436523-INS-104 | chr1-153436523-INS-104 |
| chr1-153439494-INS-217 | chr1-153439502-INS-222 |
| chr1-153441486-INS-93 | chr1-153441486-INS-93 |
| chr1-153473321-INS-213 | chr1-153473321-INS-213 |
| chr1-153488804-INS-341 | chr1-153488804-INS-341 |
| chr1-153535175-INS-111 | chr1-153535175-INS-119 |
| chr1-153565068-INS-107 | chr1-153565065-INS-72 |
| chr1-153633039-INS-72 | chr1-153633039-INS-72 |
| chr1-153638463-INS-74 | chr1-153638468-INS-60 |
| chr1-153670839-INS-1032 | chr1-153670840-INS-1034 |
| chr1-153704017-INS-1058 | chr1-153704018-INS-1058 |
| chr1-153724955-INS-6440 | chr1-153724956-INS-6440 |
| chr1-153740395-INS-65 | chr1-153740395-INS-72 |
| chr1-153753789-INS-54 | chr1-153753789-INS-54 |
| chr1-153776450-INS-51 | chr1-153776450-INS-51 |
| chr1-153795098-INS-69 | chr1-153795067-INS-70 |
| chr1-153799932-INS-94 | chr1-153799932-INS-94 |
| chr1-153808393-INS-212 | chr1-153808393-INS-212 |
| chr1-153814322-INS-220 | chr1-153814322-INS-231 |
| chr1-153815335-INS-591 | chr1-153815335-INS-591 |
| chr1-153818738-INS-150 | chr1-153818738-INS-150 |
| chr1-153828272-INS-229 | chr1-153828272-INS-231 |
| chr1-153832049-INS-64 | chr1-153832053-INS-63 |
| chr1-153856468-INS-59 | chr1-153856468-INS-59 |
| chr1-153865593-INS-94 | chr1-153865593-INS-94 |
| chr1-153881439-INS-75 | chr1-153881439-INS-75 |
| chr1-153886721-INS-245 | chr1-153886721-INS-245 |
| chr1-153900263-INS-701 | chr1-153900251-INS-715 |
| chr1-153916066-INS-52 | chr1-153916104-INS-51 |
| chr1-153984101-INS-93 | chr1-153984116-INS-99 |
| chr1-153990593-INS-64 | chr1-153990593-INS-64 |
| chr1-154000780-INS-205 | chr1-154000780-INS-205 |
| chr1-154002868-INS-189 | chr1-154002868-INS-189 |
| chr1-154014297-INS-81 | chr1-154014297-INS-81 |
| chr1-154031685-INS-361 | chr1-154031685-INS-361 |
| chr1-154041089-INS-169 | chr1-154041089-INS-170 |
| chr1-154071310-INS-113 | chr1-154071310-INS-113 |
| chr1-154083834-INS-447 | chr1-154083835-INS-447 |
| chr1-154104439-INS-54 | chr1-154104439-INS-72 |
| chr1-154189938-INS-106 | chr1-154189938-INS-106 |
| chr1-154210332-INS-68 | chr1-154210338-INS-54 |
| chr1-154236540-INS-576 | chr1-154236540-INS-576 |
| chr1-154340223-INS-1853 | chr1-154340224-INS-1854 |
| chr1-154385110-INS-2751 | chr1-154385109-INS-2743 |
| chr1-154419503-INS-58 | chr1-154419502-INS-59 |
| chr1-154487323-INS-116 | chr1-154487323-INS-116 |
| chr1-154488301-INS-7124 | chr1-154488302-INS-7124 |
| chr1-154503837-INS-148 | chr1-154503837-INS-144 |
| chr1-154524797-INS-228 | chr1-154524797-INS-228 |
| chr1-154529084-INS-260 | chr1-154529084-INS-260 |
| chr1-154553616-INS-397 | chr1-154553616-INS-397 |
| chr1-154565394-INS-111 | chr1-154565394-INS-115 |
| chr1-154579609-INS-59 | chr1-154579605-INS-59 |
| chr1-154610065-INS-72 | chr1-154610065-INS-67 |
| chr1-154637315-INS-74 | chr1-154637311-INS-76 |
| chr1-154643719-INS-72 | chr1-154643719-INS-72 |
| chr1-154665548-INS-100 | chr1-154665548-INS-101 |
| chr1-154674144-INS-131 | chr1-154674156-INS-119 |
| chr1-154680253-INS-1020 | chr1-154680254-INS-1019 |
| chr1-154702532-INS-6299 | chr1-154702533-INS-6299 |
| chr1-154720829-INS-990 | chr1-154720829-INS-990 |
| chr1-154720981-INS-52 | chr1-154720981-INS-52 |
| chr1-154735772-INS-843 | chr1-154735772-INS-847 |
| chr1-154751953-INS-589 | chr1-154751953-INS-589 |
| chr1-154754240-INS-100 | chr1-154754240-INS-67 |
| chr1-154761154-INS-59 | chr1-154761154-INS-50 |
| chr1-154795211-INS-100 | chr1-154795211-INS-100 |
| chr1-154804051-INS-1888 | chr1-154804052-INS-1888 |
| chr1-154847366-INS-58 | chr1-154847369-INS-52 |
| chr1-154851573-INS-466 | chr1-154851573-INS-466 |
| chr1-154862350-INS-63 | chr1-154862350-INS-63 |
| chr1-154868112-INS-204 | chr1-154868113-INS-204 |
| chr1-154894010-INS-128 | chr1-154894010-INS-134 |
| chr1-154922172-INS-7087 | chr1-154922173-INS-7087 |

| | |
|---|---|
| chr1-154929413-INS-110 | chr1-154929399-INS-126 |
| chr1-154951261-INS-76 | chr1-154951261-INS-76 |
| chr1-154983848-INS-227 | chr1-154983848-INS-228 |
| chr1-154986991-INS-221 | chr1-154986991-INS-221 |
| chr1-154995988-INS-6298 | chr1-154995989-INS-6303 |
| chr1-155025344-INS-2298 | chr1-155025345-INS-2251 |
| chr1-155050739-INS-87 | chr1-155050742-INS-54 |
| chr1-155052359-INS-52 | chr1-155052359-INS-56 |
| chr1-155060285-INS-170 | chr1-155060288-INS-170 |
| chr1-155114826-INS-65 | chr1-155114826-INS-65 |
| chr1-155124585-INS-60 | chr1-155124585-INS-63 |
| chr1-155131971-INS-213 | chr1-155131971-INS-213 |
| chr1-155171201-INS-70 | chr1-155171201-INS-75 |
| chr1-155191563-INS-59 | chr1-155191563-INS-55 |
| chr1-155233649-INS-926 | chr1-155233649-INS-927 |
| chr1-155249696-INS-5313 | chr1-155249697-INS-5313 |
| chr1-155264813-INS-52 | chr1-155264814-INS-60 |
| chr1-155270501-INS-174 | chr1-155270501-INS-174 |
| chr1-155298779-INS-142 | chr1-155298779-INS-135 |
| chr1-155302316-INS-66 | chr1-155302316-INS-52 |
| chr1-155311912-INS-60 | chr1-155311912-INS-68 |
| chr1-155326248-INS-358 | chr1-155326248-INS-372 |
| chr1-155336681-INS-118 | chr1-155336681-INS-118 |
| chr1-155360613-INS-519 | chr1-155360613-INS-519 |
| chr1-155383538-INS-217 | chr1-155383538-INS-218 |
| chr1-155384490-INS-211 | chr1-155384490-INS-211 |
| chr1-155430033-INS-60 | chr1-155430033-INS-64 |
| chr1-155431324-INS-74 | chr1-155431324-INS-74 |
| chr1-155545782-INS-3247 | chr1-155545783-INS-3247 |
| chr1-155574699-INS-96 | chr1-155574699-INS-96 |
| chr1-155577550-INS-123 | chr1-155577550-INS-124 |
| chr1-155585373-INS-100 | chr1-155585373-INS-100 |
| chr1-155621285-INS-52 | chr1-155621285-INS-56 |
| chr1-155624926-INS-190 | chr1-155624926-INS-191 |
| chr1-155633158-INS-84 | chr1-155633158-INS-80 |
| chr1-155637807-INS-737 | chr1-155637807-INS-737 |
| chr1-155657963-INS-114 | chr1-155657918-INS-135 |
| chr1-155677263-INS-52 | chr1-155677263-INS-52 |
| chr1-155685165-INS-82 | chr1-155685165-INS-87 |
| chr1-155695046-INS-575 | chr1-155695046-INS-575 |
| chr1-155712135-INS-218 | chr1-155712135-INS-218 |
| chr1-155730340-INS-57 | chr1-155730340-INS-57 |
| chr1-155774398-INS-215 | chr1-155774398-INS-215 |
| chr1-155787590-INS-60 | chr1-155787590-INS-68 |
| chr1-155819138-INS-61 | chr1-155819129-INS-61 |
| chr1-155849024-INS-52 | chr1-155849024-INS-50 |
| chr1-155857517-INS-128 | chr1-155857517-INS-120 |
| chr1-155896634-INS-218 | chr1-155896634-INS-218 |
| chr1-155900161-INS-148 | chr1-155900161-INS-148 |
| chr1-155903531-INS-52 | chr1-155903531-INS-52 |
| chr1-155923526-INS-96 | chr1-155923526-INS-96 |
| chr1-155936151-INS-54 | chr1-155936151-INS-55 |
| chr1-155937251-INS-60 | chr1-155937251-INS-62 |
| chr1-155975397-INS-203 | chr1-155975397-INS-203 |
| chr1-155982579-INS-52 | chr1-155982579-INS-52 |
| chr1-155994462-INS-98 | chr1-155994462-INS-96 |
| chr1-156004821-INS-210 | chr1-156004821-INS-210 |
| chr1-156022696-INS-54 | chr1-156022696-INS-54 |
| chr1-156024310-INS-201 | chr1-156024310-INS-201 |
| chr1-156036297-INS-59 | chr1-156036297-INS-66 |
| chr1-156051497-INS-1311 | chr1-156051498-INS-1307 |
| chr1-156096739-INS-105 | chr1-156096739-INS-105 |
| chr1-156102813-INS-62 | chr1-156102813-INS-62 |
| chr1-156123860-INS-65 | chr1-156123860-INS-65 |
| chr1-156126119-INS-423 | chr1-156126119-INS-423 |
| chr1-156126763-INS-203 | chr1-156126763-INS-202 |
| chr1-156133196-INS-1089 | chr1-156133197-INS-1089 |
| chr1-156142928-INS-60 | chr1-156142928-INS-62 |
| chr1-156149421-INS-130 | chr1-156149421-INS-130 |
| chr1-156152119-INS-151 | chr1-156152128-INS-161 |
| chr1-156178850-INS-59 | chr1-156178850-INS-63 |
| chr1-156185395-INS-796 | chr1-156185396-INS-796 |
| chr1-156216251-INS-198 | chr1-156216251-INS-198 |
| chr1-156225584-INS-207 | chr1-156225584-INS-207 |
| chr1-156227997-INS-211 | chr1-156227997-INS-211 |
| chr1-156239623-INS-162 | chr1-156239623-INS-160 |
| chr1-156263885-INS-52 | chr1-156263885-INS-61 |

| | |
|---|---|
| chr1-156267481-INS-244 | chr1-156267481-INS-244 |
| chr1-156282826-INS-62 | chr1-156282826-INS-62 |
| chr1-156288143-INS-145 | chr1-156288143-INS-145 |
| chr1-156289228-INS-86 | chr1-156289228-INS-86 |
| chr1-156318375-INS-207 | chr1-156318375-INS-207 |
| chr1-156338165-INS-192 | chr1-156338165-INS-192 |
| chr1-156339230-INS-211 | chr1-156339230-INS-211 |
| chr1-156360142-INS-331 | chr1-156360142-INS-331 |
| chr1-156360972-INS-96 | chr1-156360972-INS-96 |
| chr1-156376010-INS-112 | chr1-156376010-INS-104 |
| chr1-156378545-INS-56 | chr1-156378545-INS-56 |
| chr1-156383707-INS-104 | chr1-156383707-INS-104 |
| chr1-156385366-INS-577 | chr1-156385366-INS-576 |
| chr1-156387899-INS-174 | chr1-156387899-INS-174 |
| chr1-156414651-INS-294 | chr1-156414651-INS-294 |
| chr1-156425659-INS-206 | chr1-156425659-INS-206 |
| chr1-156427054-INS-62 | chr1-156427054-INS-62 |
| chr1-156450675-INS-103 | chr1-156450675-INS-103 |
| chr1-156456004-INS-206 | chr1-156456004-INS-206 |
| chr1-156484217-INS-185 | chr1-156484218-INS-178 |
| chr1-156509229-INS-208 | chr1-156509229-INS-208 |
| chr1-156518629-INS-888 | chr1-156518629-INS-888 |
| chr1-156597722-INS-185 | chr1-156597722-INS-185 |
| chr1-156666175-INS-61 | chr1-156666175-INS-63 |
| chr1-156689392-INS-135 | chr1-156689392-INS-90 |
| chr1-156693578-INS-207 | chr1-156693578-INS-207 |
| chr1-156700966-INS-243 | chr1-156700966-INS-243 |
| chr1-156719422-INS-239 | chr1-156719422-INS-240 |
| chr1-156721320-INS-200 | chr1-156721317-INS-184 |
| chr1-156730454-INS-210 | chr1-156730454-INS-211 |
| chr1-156740237-INS-88 | chr1-156740237-INS-88 |
| chr1-156774212-INS-6454 | chr1-156774213-INS-6454 |
| chr1-156777786-INS-99 | chr1-156777786-INS-76 |
| chr1-156785140-INS-293 | chr1-156785140-INS-247 |
| chr1-156786708-INS-56 | chr1-156786708-INS-50 |
| chr1-156790541-INS-52 | chr1-156790541-INS-56 |
| chr1-156795808-INS-301 | chr1-156795808-INS-301 |
| chr1-156798683-INS-229 | chr1-156798683-INS-229 |
| chr1-156803742-INS-58 | chr1-156803742-INS-58 |
| chr1-156811551-INS-183 | chr1-156811551-INS-184 |
| chr1-156829446-INS-54 | chr1-156829446-INS-60 |
| chr1-156845884-INS-132 | chr1-156845884-INS-126 |
| chr1-156854165-INS-170 | chr1-156854165-INS-160 |
| chr1-156857713-INS-79 | chr1-156857689-INS-54 |
| chr1-156866354-INS-745 | chr1-156866354-INS-745 |
| chr1-156896514-INS-95 | chr1-156896514-INS-93 |
| chr1-156897602-INS-507 | chr1-156897602-INS-507 |
| chr1-156903730-INS-335 | chr1-156903730-INS-335 |
| chr1-156914357-INS-549 | chr1-156914357-INS-549 |
| chr1-156915949-INS-62 | chr1-156915949-INS-64 |
| chr1-156918346-INS-318 | chr1-156918346-INS-318 |
| chr1-156923353-INS-216 | chr1-156923353-INS-216 |
| chr1-156936957-INS-140 | chr1-156936957-INS-162 |
| chr1-156978692-INS-648 | chr1-156978692-INS-649 |
| chr1-156989402-INS-4546 | chr1-156989403-INS-4546 |
| chr1-157011981-INS-137 | chr1-157011981-INS-137 |
| chr1-157013970-INS-222 | chr1-157013970-INS-222 |
| chr1-157021375-INS-109 | chr1-157021375-INS-109 |
| chr1-157042494-INS-80 | chr1-157042494-INS-80 |
| chr1-157055661-INS-6462 | chr1-157055662-INS-6461 |
| chr1-157057195-INS-1569 | chr1-157057196-INS-1569 |
| chr1-157065505-INS-1103 | chr1-157065506-INS-1103 |
| chr1-157077372-INS-546 | chr1-157077373-INS-547 |
| chr1-157096523-INS-98 | chr1-157096524-INS-93 |
| chr1-157098621-INS-7185 | chr1-157098622-INS-7185 |
| chr1-157114796-INS-204 | chr1-157114796-INS-204 |
| chr1-157116913-INS-210 | chr1-157116913-INS-209 |
| chr1-157121156-INS-226 | chr1-157121156-INS-227 |
| chr1-157130648-INS-220 | chr1-157130648-INS-220 |
| chr1-157137971-INS-200 | chr1-157137971-INS-201 |
| chr1-157177060-INS-60 | chr1-157177060-INS-60 |
| chr1-157185331-INS-75 | chr1-157185331-INS-57 |
| chr1-157223837-INS-209 | chr1-157223837-INS-202 |
| chr1-157250905-INS-66 | chr1-157250905-INS-82 |
| chr1-157263527-INS-284 | chr1-157263527-INS-284 |
| chr1-157269663-INS-99 | chr1-157269672-INS-105 |
| chr1-157306362-INS-52 | chr1-157306362-INS-54 |

| | |
|---|---|
| chr1-157306557-INS-773 | chr1-157306557-INS-773 |
| chr1-157325162-INS-64 | chr1-157325162-INS-64 |
| chr1-157326531-INS-145 | chr1-157326535-INS-157 |
| chr1-157328050-INS-142 | chr1-157328090-INS-142 |
| chr1-157347844-INS-7111 | chr1-157347845-INS-7111 |
| chr1-157357175-INS-471 | chr1-157357176-INS-471 |
| chr1-157358572-INS-440 | chr1-157358572-INS-440 |
| chr1-157366572-INS-62 | chr1-157366598-INS-63 |
| chr1-157376226-INS-56 | chr1-157376226-INS-52 |
| chr1-157377431-INS-222 | chr1-157377431-INS-222 |
| chr1-157392778-INS-75 | chr1-157392755-INS-58 |
| chr1-157424233-INS-244 | chr1-157424234-INS-244 |
| chr1-157428013-INS-6616 | chr1-157428014-INS-6618 |
| chr1-157428287-INS-58 | chr1-157428287-INS-58 |
| chr1-157445590-INS-65 | chr1-157445590-INS-55 |
| chr1-157448027-INS-407 | chr1-157448005-INS-420 |
| chr1-157452947-INS-51 | chr1-157452947-INS-51 |
| chr1-157469390-INS-439 | chr1-157469390-INS-441 |
| chr1-157481279-INS-72 | chr1-157481279-INS-68 |
| chr1-157486073-INS-231 | chr1-157486073-INS-212 |
| chr1-157494298-INS-491 | chr1-157494298-INS-491 |
| chr1-157513276-INS-116 | chr1-157513276-INS-146 |
| chr1-157525741-INS-66 | chr1-157525741-INS-66 |
| chr1-157542380-INS-132 | chr1-157542380-INS-136 |
| chr1-157546919-INS-122 | chr1-157546874-INS-136 |
| chr1-157594177-INS-673 | chr1-157594177-INS-673 |
| chr1-157602252-INS-3086 | chr1-157602253-INS-3086 |
| chr1-157613433-INS-6652 | chr1-157613434-INS-6652 |
| chr1-157635122-INS-99 | chr1-157635130-INS-72 |
| chr1-157637342-INS-95 | chr1-157637342-INS-95 |
| chr1-157644371-INS-57 | chr1-157644371-INS-67 |
| chr1-157656240-INS-82 | chr1-157656240-INS-82 |
| chr1-157686662-INS-115 | chr1-157686662-INS-115 |
| chr1-157690536-INS-60 | chr1-157690536-INS-76 |
| chr1-157694525-INS-90 | chr1-157694523-INS-106 |
| chr1-157726586-INS-97 | chr1-157726578-INS-77 |
| chr1-157738366-INS-92 | chr1-157738366-INS-80 |
| chr1-157783778-INS-56 | chr1-157783778-INS-56 |
| chr1-157796691-INS-6434 | chr1-157796692-INS-6441 |
| chr1-157804890-INS-1244 | chr1-157804885-INS-1245 |
| chr1-157804896-INS-98 | chr1-157804879-INS-97 |
| chr1-157833790-INS-188 | chr1-157833790-INS-218 |
| chr1-157839176-INS-61 | chr1-157839160-INS-60 |
| chr1-157857042-INS-62 | chr1-157857042-INS-62 |
| chr1-157912365-INS-474 | chr1-157912365-INS-474 |
| chr1-157952151-INS-58 | chr1-157952151-INS-60 |
| chr1-157974596-INS-126 | chr1-157974596-INS-126 |
| chr1-157976472-INS-214 | chr1-157976481-INS-224 |
| chr1-157986210-INS-451 | chr1-157986210-INS-451 |
| chr1-158002471-INS-1609 | chr1-158002472-INS-1609 |
| chr1-158022415-INS-1162 | chr1-158022416-INS-1161 |
| chr1-158050449-INS-293 | chr1-158050442-INS-304 |
| chr1-158051741-INS-307 | chr1-158051741-INS-308 |
| chr1-158055180-INS-52 | chr1-158055206-INS-64 |
| chr1-158055797-INS-1358 | chr1-158055798-INS-1369 |
| chr1-158059275-INS-108 | chr1-158059273-INS-84 |
| chr1-158075025-INS-200 | chr1-158075002-INS-200 |
| chr1-158126027-INS-78 | chr1-158126027-INS-78 |
| chr1-158213368-INS-64 | chr1-158213368-INS-82 |
| chr1-158220491-INS-81 | chr1-158220491-INS-84 |
| chr1-158238107-INS-1328 | chr1-158238098-INS-1314 |
| chr1-158239858-INS-51 | chr1-158239858-INS-63 |
| chr1-158286615-INS-874 | chr1-158286624-INS-877 |
| chr1-158290226-INS-751 | chr1-158290226-INS-754 |
| chr1-158316854-INS-58 | chr1-158316870-INS-58 |
| chr1-158337855-INS-102 | chr1-158337855-INS-102 |
| chr1-158368014-INS-134 | chr1-158368014-INS-126 |
| chr1-158403403-INS-74 | chr1-158403395-INS-51 |
| chr1-158415432-INS-6841 | chr1-158415433-INS-6842 |
| chr1-158431207-INS-60 | chr1-158431173-INS-95 |
| chr1-158484215-INS-137 | chr1-158484215-INS-138 |
| chr1-158509458-INS-235 | chr1-158509458-INS-235 |
| chr1-158532794-INS-864 | chr1-158532794-INS-865 |
| chr1-158545279-INS-169 | chr1-158545279-INS-169 |
| chr1-158577002-INS-60 | chr1-158577002-INS-52 |
| chr1-158582111-INS-57 | chr1-158582081-INS-59 |
| chr1-158586728-INS-59 | chr1-158586728-INS-54 |

| | |
|---|---|
| chr1-158591234-INS-64 | chr1-158591234-INS-54 |
| chr1-158593746-INS-57 | chr1-158593746-INS-105 |
| chr1-158606732-INS-484 | chr1-158606732-INS-484 |
| chr1-158637310-INS-75 | chr1-158637310-INS-75 |
| chr1-158650263-INS-56 | chr1-158650263-INS-75 |
| chr1-158656922-INS-6854 | chr1-158656923-INS-6853 |
| chr1-158656923-INS-167 | chr1-158656923-INS-167 |
| chr1-158665264-INS-58 | chr1-158665236-INS-58 |
| chr1-158670713-INS-77 | chr1-158670713-INS-77 |
| chr1-158690012-INS-242 | chr1-158690012-INS-242 |
| chr1-158706557-INS-1440 | chr1-158706551-INS-1430 |
| chr1-158707812-INS-424 | chr1-158707812-INS-424 |
| chr1-158727018-INS-194 | chr1-158727054-INS-208 |
| chr1-158729490-INS-52 | chr1-158729490-INS-50 |
| chr1-158789728-INS-60 | chr1-158789728-INS-60 |
| chr1-158809832-INS-58 | chr1-158809832-INS-58 |
| chr1-158810861-INS-56 | chr1-158810861-INS-56 |
| chr1-158841608-INS-401 | chr1-158841608-INS-401 |
| chr1-158880337-INS-5052 | chr1-158880338-INS-5052 |
| chr1-158886055-INS-158 | chr1-158886045-INS-158 |
| chr1-158895288-INS-52 | chr1-158895288-INS-54 |
| chr1-158904467-INS-6592 | chr1-158904468-INS-6592 |
| chr1-158930586-INS-237 | chr1-158930587-INS-238 |
| chr1-158954593-INS-108 | chr1-158954593-INS-60 |
| chr1-158975744-INS-1521 | chr1-158975745-INS-1521 |
| chr1-158980381-INS-100 | chr1-158980381-INS-84 |
| chr1-158984875-INS-127 | chr1-158984906-INS-90 |
| chr1-158999521-INS-206 | chr1-158999521-INS-206 |
| chr1-159017223-INS-4818 | chr1-159017224-INS-4818 |
| chr1-159040605-INS-56 | chr1-159040606-INS-54 |
| chr1-159041283-INS-597 | chr1-159041285-INS-590 |
| chr1-159042756-INS-543 | chr1-159042756-INS-544 |
| chr1-159055884-INS-5144 | chr1-159055885-INS-5144 |
| chr1-159063227-INS-151 | chr1-159063227-INS-151 |
| chr1-159066760-INS-182 | chr1-159066760-INS-180 |
| chr1-159080375-INS-168 | chr1-159080375-INS-137 |
| chr1-159098147-INS-233 | chr1-159098147-INS-233 |
| chr1-159110683-INS-148 | chr1-159110682-INS-150 |
| chr1-159133063-INS-58 | chr1-159133063-INS-99 |
| chr1-159159458-INS-64 | chr1-159159458-INS-60 |
| chr1-159169099-INS-70 | chr1-159169099-INS-70 |
| chr1-159169673-INS-1436 | chr1-159169674-INS-1448 |
| chr1-159171367-INS-1000 | chr1-159171367-INS-1000 |
| chr1-159208903-INS-461 | chr1-159208903-INS-461 |
| chr1-159210358-INS-117 | chr1-159210358-INS-117 |
| chr1-159218076-INS-77 | chr1-159218047-INS-70 |
| chr1-159223308-INS-190 | chr1-159223308-INS-190 |
| chr1-159225669-INS-232 | chr1-159225669-INS-239 |
| chr1-159249046-INS-66 | chr1-159249046-INS-60 |
| chr1-159271294-INS-80 | chr1-159271294-INS-60 |
| chr1-159285005-INS-126 | chr1-159285009-INS-84 |
| chr1-159291001-INS-62 | chr1-159291002-INS-54 |
| chr1-159304021-INS-62 | chr1-159304045-INS-67 |
| chr1-159306243-INS-266 | chr1-159306232-INS-254 |
| chr1-159337760-INS-52 | chr1-159337773-INS-86 |
| chr1-159410748-INS-319 | chr1-159410748-INS-319 |
| chr1-159416591-INS-65 | chr1-159416579-INS-55 |
| chr1-159420241-INS-163 | chr1-159420238-INS-154 |
| chr1-159442163-INS-106 | chr1-159442195-INS-62 |
| chr1-159454534-INS-1147 | chr1-159454535-INS-1157 |
| chr1-159465315-INS-66 | chr1-159465315-INS-64 |
| chr1-159493269-INS-56 | chr1-159493274-INS-56 |
| chr1-159495465-INS-72 | chr1-159495465-INS-74 |
| chr1-159527814-INS-2712 | chr1-159527815-INS-2712 |
| chr1-159546338-INS-104 | chr1-159546338-INS-76 |
| chr1-159586432-INS-68 | chr1-159586438-INS-54 |
| chr1-159587504-INS-138 | chr1-159587504-INS-138 |
| chr1-159632498-INS-243 | chr1-159632498-INS-243 |
| chr1-159641098-INS-51 | chr1-159641099-INS-50 |
| chr1-159651742-INS-6716 | chr1-159651743-INS-6717 |
| chr1-159707133-INS-71 | chr1-159707130-INS-95 |
| chr1-159712253-INS-51 | chr1-159712253-INS-57 |
| chr1-159725585-INS-400 | chr1-159725585-INS-400 |
| chr1-159753917-INS-108 | chr1-159753917-INS-63 |
| chr1-159793893-INS-54 | chr1-159793867-INS-61 |
| chr1-159806325-INS-61 | chr1-159806325-INS-61 |
| chr1-159822355-INS-385 | chr1-159822355-INS-385 |

| | |
|---|---|
| chr1-159840635-INS-368 | chr1-159840635-INS-368 |
| chr1-159845553-INS-795 | chr1-159845553-INS-795 |
| chr1-159879243-INS-54 | chr1-159879243-INS-60 |
| chr1-159918492-INS-6694 | chr1-159918493-INS-6693 |
| chr1-159930938-INS-98 | chr1-159930938-INS-98 |
| chr1-159957466-INS-98 | chr1-159957466-INS-75 |
| chr1-159962282-INS-71 | chr1-159962267-INS-50 |
| chr1-159973379-INS-326 | chr1-159973379-INS-326 |
| chr1-159991163-INS-6307 | chr1-159991164-INS-6307 |
| chr1-159992507-INS-186 | chr1-159992507-INS-187 |
| chr1-160009346-INS-60 | chr1-160009346-INS-51 |
| chr1-160010571-INS-7083 | chr1-160010572-INS-7084 |
| chr1-160026705-INS-2922 | chr1-160026706-INS-2922 |
| chr1-160026706-INS-463 | chr1-160026706-INS-463 |
| chr1-160055088-INS-210 | chr1-160055088-INS-210 |
| chr1-160062863-INS-226 | chr1-160062863-INS-226 |
| chr1-160066022-INS-54 | chr1-160066022-INS-58 |
| chr1-160077252-INS-78 | chr1-160077252-INS-78 |
| chr1-160093029-INS-1010 | chr1-160093030-INS-1006 |
| chr1-160100233-INS-143 | chr1-160100233-INS-143 |
| chr1-160120930-INS-53 | chr1-160120930-INS-54 |
| chr1-160147369-INS-55 | chr1-160147369-INS-55 |
| chr1-160157270-INS-58 | chr1-160157270-INS-58 |
| chr1-160160267-INS-54 | chr1-160160267-INS-50 |
| chr1-160219295-INS-168 | chr1-160219295-INS-167 |
| chr1-160239367-INS-400 | chr1-160239367-INS-400 |
| chr1-160252181-INS-7091 | chr1-160252181-INS-7091 |
| chr1-160256370-INS-219 | chr1-160256371-INS-219 |
| chr1-160301065-INS-6675 | chr1-160301066-INS-6675 |
| chr1-160352309-INS-249 | chr1-160352309-INS-250 |
| chr1-160381326-INS-78 | chr1-160381326-INS-66 |
| chr1-160396854-INS-54 | chr1-160396858-INS-65 |
| chr1-160401390-INS-115 | chr1-160401438-INS-108 |
| chr1-160434970-INS-3252 | chr1-160434971-INS-3252 |
| chr1-160472016-INS-97 | chr1-160472016-INS-97 |
| chr1-160504431-INS-202 | chr1-160504431-INS-201 |
| chr1-160511624-INS-200 | chr1-160511624-INS-200 |
| chr1-160518326-INS-6888 | chr1-160518327-INS-6911 |
| chr1-160556376-INS-113 | chr1-160556367-INS-146 |
| chr1-160579437-INS-74 | chr1-160579437-INS-50 |
| chr1-160588548-INS-237 | chr1-160588548-INS-237 |
| chr1-160605435-INS-72 | chr1-160605435-INS-99 |
| chr1-160607657-INS-186 | chr1-160607657-INS-182 |
| chr1-160630420-INS-91 | chr1-160630420-INS-67 |
| chr1-160648834-INS-236 | chr1-160648835-INS-236 |
| chr1-160682071-INS-80 | chr1-160682071-INS-74 |
| chr1-160686176-INS-1310 | chr1-160686177-INS-1310 |
| chr1-160689478-INS-169 | chr1-160689478-INS-174 |
| chr1-160690935-INS-64 | chr1-160690935-INS-60 |
| chr1-160719192-INS-70 | chr1-160719192-INS-70 |
| chr1-160750583-INS-196 | chr1-160750583-INS-196 |
| chr1-160783807-INS-57 | chr1-160783807-INS-57 |
| chr1-160787791-INS-197 | chr1-160787791-INS-197 |
| chr1-160816645-INS-202 | chr1-160816645-INS-205 |
| chr1-160817728-INS-138 | chr1-160817728-INS-140 |
| chr1-160827085-INS-324 | chr1-160827085-INS-324 |
| chr1-160835404-INS-211 | chr1-160835404-INS-206 |
| chr1-160850351-INS-216 | chr1-160850351-INS-216 |
| chr1-160865263-INS-218 | chr1-160865263-INS-218 |
| chr1-160867251-INS-704 | chr1-160867267-INS-667 |
| chr1-160875515-INS-230 | chr1-160875515-INS-230 |
| chr1-160878368-INS-189 | chr1-160878368-INS-191 |
| chr1-160886096-INS-218 | chr1-160886096-INS-219 |
| chr1-160891105-INS-301 | chr1-160891105-INS-302 |
| chr1-160917163-INS-80 | chr1-160917163-INS-120 |
| chr1-160925218-INS-1643 | chr1-160925219-INS-1650 |
| chr1-160940805-INS-375 | chr1-160940805-INS-376 |
| chr1-160941510-INS-7165 | chr1-160941511-INS-7165 |
| chr1-160948682-INS-212 | chr1-160948682-INS-212 |
| chr1-160990589-INS-58 | chr1-160990589-INS-58 |
| chr1-160999407-INS-190 | chr1-160999407-INS-190 |
| chr1-161004330-INS-57 | chr1-161004330-INS-89 |
| chr1-161005732-INS-305 | chr1-161005732-INS-305 |
| chr1-161014614-INS-197 | chr1-161014615-INS-197 |
| chr1-161018164-INS-76 | chr1-161018164-INS-76 |
| chr1-161024750-INS-52 | chr1-161024749-INS-54 |
| chr1-161026589-INS-494 | chr1-161026589-INS-515 |

| | |
|---|---|
| chr1-161032385-INS-66 | chr1-161032385-INS-66 |
| chr1-161036343-INS-252 | chr1-161036360-INS-253 |
| chr1-161041744-INS-196 | chr1-161041744-INS-196 |
| chr1-161053435-INS-66 | chr1-161053435-INS-66 |
| chr1-161057786-INS-191 | chr1-161057786-INS-191 |
| chr1-161060663-INS-200 | chr1-161060663-INS-200 |
| chr1-161061939-INS-146 | chr1-161061965-INS-149 |
| chr1-161063985-INS-6469 | chr1-161063986-INS-6469 |
| chr1-161090056-INS-121 | chr1-161090053-INS-124 |
| chr1-161093981-INS-174 | chr1-161093981-INS-174 |
| chr1-161100798-INS-68 | chr1-161100799-INS-66 |
| chr1-161113475-INS-54 | chr1-161113475-INS-54 |
| chr1-161120061-INS-60 | chr1-161120061-INS-60 |
| chr1-161129008-INS-161 | chr1-161129017-INS-126 |
| chr1-161133163-INS-72 | chr1-161133163-INS-56 |
| chr1-161134607-INS-227 | chr1-161134607-INS-228 |
| chr1-161137936-INS-235 | chr1-161137936-INS-235 |
| chr1-161144712-INS-56 | chr1-161144712-INS-58 |
| chr1-161152043-INS-65 | chr1-161152043-INS-59 |
| chr1-161162189-INS-366 | chr1-161162189-INS-367 |
| chr1-161164204-INS-747 | chr1-161164204-INS-745 |
| chr1-161183719-INS-158 | chr1-161183719-INS-159 |
| chr1-161267613-INS-8529 | chr1-161267614-INS-8529 |
| chr1-161288003-INS-74 | chr1-161287964-INS-104 |
| chr1-161290524-INS-58 | chr1-161290522-INS-60 |
| chr1-161298863-INS-5087 | chr1-161298864-INS-5087 |
| chr1-161315810-INS-330 | chr1-161315810-INS-330 |
| chr1-161349571-INS-306 | chr1-161349571-INS-306 |
| chr1-161371161-INS-344 | chr1-161371161-INS-344 |
| chr1-161396256-INS-92 | chr1-161396250-INS-82 |
| chr1-161419130-INS-2100 | chr1-161419131-INS-2100 |
| chr1-161420278-INS-58 | chr1-161420278-INS-62 |
| chr1-161432722-INS-7292 | chr1-161432724-INS-7287 |
| chr1-161468358-INS-504 | chr1-161468358-INS-509 |
| chr1-161476773-INS-56 | chr1-161476773-INS-56 |
| chr1-161499852-INS-94 | chr1-161499852-INS-94 |
| chr1-161512120-INS-71 | chr1-161512120-INS-71 |
| chr1-161557107-INS-165 | chr1-161557107-INS-136 |
| chr1-161594108-INS-54 | chr1-161594108-INS-56 |
| chr1-161700159-INS-64 | chr1-161700159-INS-64 |
| chr1-161705709-INS-426 | chr1-161705709-INS-426 |
| chr1-161706131-INS-4875 | chr1-161706133-INS-4876 |
| chr1-161712458-INS-1046 | chr1-161712459-INS-1051 |
| chr1-161724291-INS-60 | chr1-161724291-INS-56 |
| chr1-161751503-INS-342 | chr1-161751503-INS-342 |
| chr1-161756638-INS-70 | chr1-161756638-INS-63 |
| chr1-161758722-INS-102 | chr1-161758722-INS-106 |
| chr1-161760534-INS-212 | chr1-161760534-INS-212 |
| chr1-161783037-INS-87 | chr1-161783037-INS-67 |
| chr1-161803337-INS-143 | chr1-161803352-INS-145 |
| chr1-161828200-INS-285 | chr1-161828200-INS-285 |
| chr1-161838991-INS-62 | chr1-161839023-INS-54 |
| chr1-161849666-INS-294 | chr1-161849666-INS-294 |
| chr1-161861240-INS-661 | chr1-161861241-INS-661 |
| chr1-161866196-INS-82 | chr1-161866197-INS-90 |
| chr1-161873557-INS-4575 | chr1-161873558-INS-4575 |
| chr1-161920844-INS-68 | chr1-161920812-INS-52 |
| chr1-161926227-INS-56 | chr1-161926227-INS-56 |
| chr1-161931598-INS-961 | chr1-161931609-INS-958 |
| chr1-161933086-INS-64 | chr1-161933086-INS-64 |
| chr1-161973927-INS-6345 | chr1-161973928-INS-6345 |
| chr1-161994069-INS-126 | chr1-161994069-INS-126 |
| chr1-161999293-INS-81 | chr1-161999294-INS-81 |
| chr1-162031394-INS-140 | chr1-162031394-INS-142 |
| chr1-162054489-INS-128 | chr1-162054489-INS-116 |
| chr1-162084476-INS-641 | chr1-162084476-INS-641 |
| chr1-162089911-INS-60 | chr1-162089911-INS-60 |
| chr1-162104706-INS-146 | chr1-162104706-INS-146 |
| chr1-162115301-INS-91 | chr1-162115318-INS-88 |
| chr1-162136289-INS-57 | chr1-162136281-INS-102 |
| chr1-162160911-INS-498 | chr1-162160911-INS-498 |
| chr1-162166254-INS-206 | chr1-162166255-INS-206 |
| chr1-162178763-INS-593 | chr1-162178763-INS-593 |
| chr1-162183654-INS-79 | chr1-162183654-INS-79 |
| chr1-162190707-INS-175 | chr1-162190707-INS-175 |
| chr1-162223998-INS-803 | chr1-162223998-INS-803 |
| chr1-162243624-INS-278 | chr1-162243624-INS-278 |

| | |
|---|---|
| chr1-162249358-INS-58 | chr1-162249358-INS-58 |
| chr1-162265596-INS-56 | chr1-162265590-INS-60 |
| chr1-162283349-INS-100 | chr1-162283339-INS-86 |
| chr1-162283886-INS-60 | chr1-162283846-INS-110 |
| chr1-162312693-INS-247 | chr1-162312683-INS-248 |
| chr1-162316860-INS-176 | chr1-162316860-INS-177 |
| chr1-162325082-INS-376 | chr1-162325082-INS-370 |
| chr1-162338444-INS-480 | chr1-162338444-INS-480 |
| chr1-162343388-INS-65 | chr1-162343388-INS-66 |
| chr1-162345634-INS-96 | chr1-162345634-INS-78 |
| chr1-162365729-INS-99 | chr1-162365729-INS-106 |
| chr1-162380234-INS-234 | chr1-162380234-INS-234 |
| chr1-162391992-INS-7123 | chr1-162391993-INS-7123 |
| chr1-162412021-INS-167 | chr1-162412021-INS-167 |
| chr1-162435268-INS-561 | chr1-162435268-INS-561 |
| chr1-162454321-INS-401 | chr1-162454321-INS-401 |
| chr1-162463074-INS-63 | chr1-162463074-INS-63 |
| chr1-162470810-INS-241 | chr1-162470810-INS-240 |
| chr1-162475244-INS-74 | chr1-162475244-INS-74 |
| chr1-162556208-INS-173 | chr1-162556208-INS-173 |
| chr1-162590851-INS-356 | chr1-162590863-INS-359 |
| chr1-162616459-INS-562 | chr1-162616459-INS-562 |
| chr1-162683021-INS-696 | chr1-162683016-INS-695 |
| chr1-162695928-INS-1280 | chr1-162695929-INS-1280 |
| chr1-162701736-INS-441 | chr1-162701736-INS-441 |
| chr1-162717233-INS-181 | chr1-162717233-INS-181 |
| chr1-162742943-INS-52 | chr1-162742943-INS-65 |
| chr1-162757042-INS-168 | chr1-162757045-INS-203 |
| chr1-162758403-INS-3143 | chr1-162758404-INS-3177 |
| chr1-162772301-INS-617 | chr1-162772301-INS-619 |
| chr1-162784499-INS-59 | chr1-162784499-INS-59 |
| chr1-162798687-INS-104 | chr1-162798687-INS-72 |
| chr1-162819166-INS-939 | chr1-162819166-INS-939 |
| chr1-162825723-INS-3026 | chr1-162825724-INS-3026 |
| chr1-162828694-INS-831 | chr1-162828694-INS-831 |
| chr1-162869470-INS-5816 | chr1-162869471-INS-5816 |
| chr1-162874952-INS-128 | chr1-162874947-INS-135 |
| chr1-162897091-INS-214 | chr1-162897091-INS-214 |
| chr1-162898835-INS-233 | chr1-162898835-INS-233 |
| chr1-162907838-INS-124 | chr1-162907838-INS-124 |
| chr1-162923677-INS-74 | chr1-162923677-INS-74 |
| chr1-162981785-INS-76 | chr1-162981785-INS-62 |
| chr1-163016115-INS-103 | chr1-163016120-INS-101 |
| chr1-163062824-INS-245 | chr1-163062824-INS-245 |
| chr1-163068888-INS-58 | chr1-163068874-INS-98 |
| chr1-163100343-INS-107 | chr1-163100343-INS-108 |
| chr1-163133081-INS-91 | chr1-163133081-INS-91 |
| chr1-163138586-INS-51 | chr1-163138586-INS-51 |
| chr1-163144204-INS-204 | chr1-163144205-INS-213 |
| chr1-163152459-INS-68 | chr1-163152459-INS-56 |
| chr1-163156280-INS-390 | chr1-163156280-INS-390 |
| chr1-163170348-INS-104 | chr1-163170348-INS-106 |
| chr1-163186243-INS-54 | chr1-163186229-INS-50 |
| chr1-163190777-INS-58 | chr1-163190777-INS-74 |
| chr1-163205038-INS-104 | chr1-163205038-INS-112 |
| chr1-163212213-INS-74 | chr1-163212213-INS-74 |
| chr1-163264093-INS-6360 | chr1-163264094-INS-6361 |
| chr1-163282969-INS-144 | chr1-163282965-INS-124 |
| chr1-163349567-INS-56 | chr1-163349567-INS-56 |
| chr1-163354646-INS-175 | chr1-163354646-INS-177 |
| chr1-163379375-INS-245 | chr1-163379375-INS-210 |
| chr1-163382784-INS-51 | chr1-163382784-INS-50 |
| chr1-163405043-INS-663 | chr1-163405043-INS-664 |
| chr1-163413028-INS-870 | chr1-163413028-INS-872 |
| chr1-163432844-INS-358 | chr1-163432844-INS-358 |
| chr1-163434461-INS-754 | chr1-163434461-INS-755 |
| chr1-163436700-INS-52 | chr1-163436667-INS-80 |
| chr1-163467922-INS-1831 | chr1-163467923-INS-1831 |
| chr1-163476085-INS-294 | chr1-163476079-INS-290 |
| chr1-163519180-INS-77 | chr1-163519180-INS-77 |
| chr1-163532937-INS-54 | chr1-163532937-INS-54 |
| chr1-163544948-INS-55 | chr1-163544948-INS-55 |
| chr1-163566940-INS-773 | chr1-163566938-INS-766 |
| chr1-163568868-INS-55 | chr1-163568868-INS-53 |
| chr1-163582339-INS-107 | chr1-163582339-INS-107 |
| chr1-163599307-INS-201 | chr1-163599307-INS-201 |
| chr1-163622105-INS-818 | chr1-163622105-INS-818 |

| | |
|---|---|
| chr1-163641044-INS-60 | chr1-163641044-INS-60 |
| chr1-163669430-INS-111 | chr1-163669430-INS-111 |
| chr1-163682732-INS-308 | chr1-163682732-INS-308 |
| chr1-163695279-INS-144 | chr1-163695278-INS-148 |
| chr1-163716816-INS-67 | chr1-163716816-INS-67 |
| chr1-163745868-INS-223 | chr1-163745868-INS-223 |
| chr1-163749991-INS-51 | chr1-163749991-INS-51 |
| chr1-163757146-INS-52 | chr1-163757146-INS-52 |
| chr1-163771098-INS-84 | chr1-163771084-INS-84 |
| chr1-163780647-INS-234 | chr1-163780647-INS-235 |
| chr1-163790567-INS-106 | chr1-163790558-INS-155 |
| chr1-163793049-INS-177 | chr1-163793049-INS-179 |
| chr1-163802724-INS-124 | chr1-163802724-INS-124 |
| chr1-163805127-INS-53 | chr1-163805127-INS-53 |
| chr1-163820155-INS-87 | chr1-163820155-INS-87 |
| chr1-163830789-INS-8455 | chr1-163830790-INS-8455 |
| chr1-163832882-INS-97 | chr1-163832882-INS-100 |
| chr1-163839291-INS-64 | chr1-163839294-INS-96 |
| chr1-163882765-INS-58 | chr1-163882765-INS-60 |
| chr1-163911251-INS-6360 | chr1-163911252-INS-6360 |
| chr1-163947354-INS-89 | chr1-163947354-INS-95 |
| chr1-163952134-INS-51 | chr1-163952134-INS-53 |
| chr1-163954388-INS-576 | chr1-163954388-INS-576 |
| chr1-163963363-INS-140 | chr1-163963360-INS-97 |
| chr1-163979804-INS-62 | chr1-163979804-INS-57 |
| chr1-163988311-INS-145 | chr1-163988311-INS-149 |
| chr1-163997317-INS-54 | chr1-163997315-INS-56 |
| chr1-164006864-INS-65 | chr1-164006864-INS-65 |
| chr1-164014648-INS-283 | chr1-164014648-INS-288 |
| chr1-164038581-INS-193 | chr1-164038581-INS-193 |
| chr1-164045863-INS-55 | chr1-164045863-INS-55 |
| chr1-164071520-INS-212 | chr1-164071520-INS-223 |
| chr1-164074179-INS-213 | chr1-164074176-INS-197 |
| chr1-164094158-INS-67 | chr1-164094158-INS-67 |
| chr1-164106788-INS-54 | chr1-164106752-INS-56 |
| chr1-164136005-INS-5219 | chr1-164136006-INS-5219 |
| chr1-164145348-INS-65 | chr1-164145348-INS-65 |
| chr1-164148781-INS-147 | chr1-164148781-INS-147 |
| chr1-164156920-INS-80 | chr1-164156920-INS-99 |
| chr1-164166431-INS-75 | chr1-164166431-INS-75 |
| chr1-164171769-INS-68 | chr1-164171769-INS-68 |
| chr1-164177254-INS-74 | chr1-164177254-INS-62 |
| chr1-164180321-INS-207 | chr1-164180321-INS-207 |
| chr1-164199936-INS-51 | chr1-164199936-INS-56 |
| chr1-164221258-INS-84 | chr1-164221261-INS-88 |
| chr1-164224851-INS-63 | chr1-164224851-INS-71 |
| chr1-164238806-INS-74 | chr1-164238806-INS-74 |
| chr1-164239876-INS-133 | chr1-164239876-INS-133 |
| chr1-164257195-INS-425 | chr1-164257194-INS-426 |
| chr1-164259559-INS-64 | chr1-164259549-INS-78 |
| chr1-164267537-INS-55 | chr1-164267537-INS-77 |
| chr1-164275014-INS-68 | chr1-164275000-INS-54 |
| chr1-164276631-INS-329 | chr1-164276658-INS-323 |
| chr1-164307999-INS-60 | chr1-164307999-INS-60 |
| chr1-164320016-INS-57 | chr1-164320016-INS-62 |
| chr1-164338793-INS-95 | chr1-164338837-INS-65 |
| chr1-164361373-INS-598 | chr1-164361373-INS-598 |
| chr1-164372711-INS-80 | chr1-164372711-INS-60 |
| chr1-164403014-INS-68 | chr1-164403014-INS-68 |
| chr1-164415173-INS-51 | chr1-164415173-INS-51 |
| chr1-164420286-INS-54 | chr1-164420286-INS-58 |
| chr1-164426889-INS-395 | chr1-164426889-INS-388 |
| chr1-164445820-INS-56 | chr1-164445812-INS-60 |
| chr1-164459177-INS-57 | chr1-164459182-INS-81 |
| chr1-164460467-INS-186 | chr1-164460434-INS-186 |
| chr1-164461454-INS-56 | chr1-164461454-INS-58 |
| chr1-164464853-INS-113 | chr1-164464853-INS-113 |
| chr1-164468156-INS-165 | chr1-164468156-INS-165 |
| chr1-164489637-INS-499 | chr1-164489637-INS-500 |
| chr1-164495858-INS-127 | chr1-164495871-INS-116 |
| chr1-164533123-INS-116 | chr1-164533123-INS-123 |
| chr1-164578447-INS-193 | chr1-164578447-INS-187 |
| chr1-164585857-INS-629 | chr1-164585857-INS-629 |
| chr1-164592469-INS-102 | chr1-164592469-INS-102 |
| chr1-164599924-INS-83 | chr1-164599924-INS-93 |
| chr1-164627611-INS-539 | chr1-164627611-INS-539 |
| chr1-164628309-INS-248 | chr1-164628309-INS-247 |

| | |
|---|---|
| chr1-164644392-INS-134 | chr1-164644392-INS-134 |
| chr1-164645437-INS-6394 | chr1-164645438-INS-6393 |
| chr1-164645852-INS-1954 | chr1-164645853-INS-1962 |
| chr1-164664273-INS-142 | chr1-164664273-INS-150 |
| chr1-164672771-INS-55 | chr1-164672771-INS-97 |
| chr1-164694139-INS-100 | chr1-164694139-INS-125 |
| chr1-164719007-INS-147 | chr1-164719009-INS-132 |
| chr1-164725188-INS-56 | chr1-164725188-INS-56 |
| chr1-164734477-INS-58 | chr1-164734477-INS-56 |
| chr1-164765668-INS-3230 | chr1-164765669-INS-3230 |
| chr1-164766812-INS-58 | chr1-164766812-INS-60 |
| chr1-164774388-INS-69 | chr1-164774388-INS-77 |
| chr1-164797564-INS-6686 | chr1-164797565-INS-6664 |
| chr1-164807691-INS-95 | chr1-164807691-INS-95 |
| chr1-164822266-INS-735 | chr1-164822266-INS-732 |
| chr1-164824477-INS-4834 | chr1-164824478-INS-4834 |
| chr1-164837741-INS-3515 | chr1-164837742-INS-3515 |
| chr1-164839793-INS-51 | chr1-164839794-INS-70 |
| chr1-164872135-INS-114 | chr1-164872135-INS-114 |
| chr1-164927950-INS-104 | chr1-164927950-INS-104 |
| chr1-164933761-INS-357 | chr1-164933761-INS-363 |
| chr1-164955058-INS-74 | chr1-164955057-INS-88 |
| chr1-164959610-INS-52 | chr1-164959610-INS-52 |
| chr1-164965805-INS-68 | chr1-164965805-INS-68 |
| chr1-164990358-INS-169 | chr1-164990358-INS-170 |
| chr1-165014164-INS-720 | chr1-165014172-INS-733 |
| chr1-165016669-INS-248 | chr1-165016669-INS-248 |
| chr1-165025979-INS-221 | chr1-165025980-INS-216 |
| chr1-165031526-INS-538 | chr1-165031526-INS-538 |
| chr1-165032476-INS-74 | chr1-165032476-INS-74 |
| chr1-165033695-INS-397 | chr1-165033695-INS-397 |
| chr1-165035877-INS-149 | chr1-165035877-INS-157 |
| chr1-165037022-INS-155 | chr1-165037022-INS-157 |
| chr1-165052242-INS-64 | chr1-165052228-INS-64 |
| chr1-165060641-INS-354 | chr1-165060625-INS-382 |
| chr1-165065500-INS-400 | chr1-165065500-INS-400 |
| chr1-165067957-INS-79 | chr1-165067952-INS-80 |
| chr1-165102705-INS-168 | chr1-165102705-INS-168 |
| chr1-165120259-INS-102 | chr1-165120259-INS-102 |
| chr1-165137730-INS-135 | chr1-165137730-INS-137 |
| chr1-165139414-INS-56 | chr1-165139414-INS-56 |
| chr1-165144024-INS-568 | chr1-165144024-INS-571 |
| chr1-165145000-INS-120 | chr1-165145000-INS-120 |
| chr1-165152131-INS-56 | chr1-165152131-INS-54 |
| chr1-165169178-INS-222 | chr1-165169178-INS-222 |
| chr1-165177797-INS-196 | chr1-165177797-INS-197 |
| chr1-165178573-INS-75 | chr1-165178595-INS-75 |
| chr1-165187456-INS-1888 | chr1-165187457-INS-1888 |
| chr1-165193873-INS-204 | chr1-165193873-INS-204 |
| chr1-165197118-INS-226 | chr1-165197118-INS-226 |
| chr1-165207128-INS-369 | chr1-165207128-INS-369 |
| chr1-165231387-INS-283 | chr1-165231387-INS-283 |
| chr1-165232718-INS-212 | chr1-165232712-INS-212 |
| chr1-165246938-INS-173 | chr1-165246938-INS-173 |
| chr1-165253128-INS-180 | chr1-165253128-INS-180 |
| chr1-165258930-INS-52 | chr1-165258924-INS-64 |
| chr1-165273786-INS-224 | chr1-165273786-INS-224 |
| chr1-165291661-INS-159 | chr1-165291661-INS-159 |
| chr1-165323018-INS-206 | chr1-165323018-INS-206 |
| chr1-165324002-INS-605 | chr1-165324002-INS-605 |
| chr1-165342095-INS-63 | chr1-165342095-INS-63 |
| chr1-165361928-INS-82 | chr1-165361928-INS-82 |
| chr1-165368526-INS-227 | chr1-165368526-INS-227 |
| chr1-165391945-INS-105 | chr1-165391945-INS-105 |
| chr1-165403769-INS-100 | chr1-165403768-INS-116 |
| chr1-165408790-INS-5443 | chr1-165408791-INS-5444 |
| chr1-165409075-INS-195 | chr1-165409075-INS-195 |
| chr1-165411021-INS-108 | chr1-165411068-INS-68 |
| chr1-165412982-INS-4735 | chr1-165412983-INS-4735 |
| chr1-165424378-INS-99 | chr1-165424378-INS-99 |
| chr1-165465600-INS-54 | chr1-165465600-INS-63 |
| chr1-165466398-INS-441 | chr1-165466398-INS-441 |
| chr1-165488412-INS-56 | chr1-165488412-INS-56 |
| chr1-165492423-INS-63 | chr1-165492423-INS-63 |
| chr1-165493796-INS-80 | chr1-165493796-INS-86 |
| chr1-165556418-INS-200 | chr1-165556418-INS-200 |
| chr1-165557188-INS-440 | chr1-165557188-INS-422 |

| | |
|---|---|
| chr1-165572780-INS-492 | chr1-165572780-INS-492 |
| chr1-165583746-INS-52 | chr1-165583744-INS-64 |
| chr1-165587308-INS-252 | chr1-165587308-INS-252 |
| chr1-165588953-INS-71 | chr1-165588953-INS-71 |
| chr1-165597822-INS-816 | chr1-165597822-INS-821 |
| chr1-165635530-INS-78 | chr1-165635530-INS-71 |
| chr1-165688081-INS-72 | chr1-165688081-INS-72 |
| chr1-165731294-INS-191 | chr1-165731294-INS-191 |
| chr1-165743615-INS-614 | chr1-165743593-INS-650 |
| chr1-165745142-INS-89 | chr1-165745142-INS-89 |
| chr1-165748305-INS-82 | chr1-165748305-INS-82 |
| chr1-165754190-INS-64 | chr1-165754190-INS-72 |
| chr1-165798920-INS-459 | chr1-165798921-INS-456 |
| chr1-165822095-INS-172 | chr1-165822087-INS-177 |
| chr1-165842173-INS-83 | chr1-165842173-INS-83 |
| chr1-165853217-INS-499 | chr1-165853217-INS-499 |
| chr1-165943921-INS-200 | chr1-165943921-INS-200 |
| chr1-165946704-INS-202 | chr1-165946704-INS-203 |
| chr1-165947568-INS-52 | chr1-165947568-INS-52 |
| chr1-165950691-INS-201 | chr1-165950691-INS-201 |
| chr1-165954254-INS-138 | chr1-165954254-INS-138 |
| chr1-165963641-INS-199 | chr1-165963641-INS-199 |
| chr1-165978243-INS-99 | chr1-165978243-INS-100 |
| chr1-165982762-INS-4398 | chr1-165982763-INS-4398 |
| chr1-165997158-INS-145 | chr1-165997158-INS-121 |
| chr1-165999079-INS-446 | chr1-165999079-INS-471 |
| chr1-166019443-INS-369 | chr1-166019441-INS-372 |
| chr1-166025278-INS-4594 | chr1-166025279-INS-4593 |
| chr1-166032885-INS-91 | chr1-166032885-INS-91 |
| chr1-166056057-INS-84 | chr1-166056057-INS-84 |
| chr1-166079014-INS-8469 | chr1-166079015-INS-8469 |
| chr1-166079017-INS-60 | chr1-166079015-INS-66 |
| chr1-166080050-INS-1893 | chr1-166080051-INS-1893 |
| chr1-166081174-INS-66 | chr1-166081174-INS-66 |
| chr1-166090613-INS-109 | chr1-166090613-INS-109 |
| chr1-166095693-INS-112 | chr1-166095693-INS-112 |
| chr1-166096584-INS-436 | chr1-166096584-INS-436 |
| chr1-166107566-INS-54 | chr1-166107566-INS-50 |
| chr1-166159820-INS-69 | chr1-166159820-INS-69 |
| chr1-166164270-INS-226 | chr1-166164270-INS-226 |
| chr1-166169136-INS-56 | chr1-166169136-INS-56 |
| chr1-166198370-INS-204 | chr1-166198373-INS-208 |
| chr1-166201882-INS-72 | chr1-166201882-INS-72 |
| chr1-166230734-INS-306 | chr1-166230734-INS-306 |
| chr1-166266146-INS-61 | chr1-166266146-INS-61 |
| chr1-166268113-INS-367 | chr1-166268113-INS-372 |
| chr1-166302866-INS-212 | chr1-166302866-INS-212 |
| chr1-166321674-INS-124 | chr1-166321712-INS-120 |
| chr1-166340247-INS-624 | chr1-166340247-INS-623 |
| chr1-166358728-INS-85 | chr1-166358724-INS-50 |
| chr1-166383052-INS-175 | chr1-166383049-INS-146 |
| chr1-166404530-INS-219 | chr1-166404529-INS-219 |
| chr1-166413985-INS-75 | chr1-166413985-INS-75 |
| chr1-166505016-INS-60 | chr1-166505016-INS-64 |
| chr1-166527173-INS-7154 | chr1-166527174-INS-7154 |
| chr1-166531667-INS-66 | chr1-166531667-INS-62 |
| chr1-166551323-INS-187 | chr1-166551323-INS-187 |
| chr1-166558625-INS-276 | chr1-166558625-INS-233 |
| chr1-166598848-INS-172 | chr1-166598848-INS-198 |
| chr1-166606105-INS-263 | chr1-166606105-INS-247 |
| chr1-166649815-INS-62 | chr1-166649822-INS-66 |
| chr1-166651131-INS-210 | chr1-166651131-INS-210 |
| chr1-166654298-INS-54 | chr1-166654322-INS-54 |
| chr1-166656466-INS-533 | chr1-166656466-INS-532 |
| chr1-166663811-INS-60 | chr1-166663811-INS-62 |
| chr1-166675455-INS-59 | chr1-166675455-INS-59 |
| chr1-166706035-INS-255 | chr1-166706035-INS-255 |
| chr1-166749600-INS-530 | chr1-166749600-INS-530 |
| chr1-166755131-INS-166 | chr1-166755131-INS-166 |
| chr1-166762924-INS-637 | chr1-166762924-INS-638 |
| chr1-166772640-INS-210 | chr1-166772641-INS-210 |
| chr1-166781545-INS-76 | chr1-166781542-INS-74 |
| chr1-166801570-INS-299 | chr1-166801570-INS-299 |
| chr1-166809698-INS-60 | chr1-166809698-INS-60 |
| chr1-166827102-INS-59 | chr1-166827102-INS-59 |
| chr1-166847729-INS-6442 | chr1-166847730-INS-6442 |
| chr1-166857886-INS-162 | chr1-166857886-INS-160 |

| | |
|---|---|
| chr1-166872330-INS-400 | chr1-166872330-INS-400 |
| chr1-166885596-INS-76 | chr1-166885594-INS-84 |
| chr1-166885899-INS-1892 | chr1-166885900-INS-1892 |
| chr1-166914986-INS-736 | chr1-166914986-INS-736 |
| chr1-166935443-INS-69 | chr1-166935443-INS-69 |
| chr1-166945282-INS-65 | chr1-166945282-INS-55 |
| chr1-166952267-INS-497 | chr1-166952267-INS-497 |
| chr1-166957000-INS-65 | chr1-166956995-INS-51 |
| chr1-166958008-INS-486 | chr1-166958008-INS-486 |
| chr1-166963603-INS-189 | chr1-166963603-INS-189 |
| chr1-166973192-INS-96 | chr1-166973192-INS-96 |
| chr1-166974014-INS-96 | chr1-166974014-INS-88 |
| chr1-167073228-INS-53 | chr1-167073228-INS-51 |
| chr1-167079694-INS-67 | chr1-167079694-INS-67 |
| chr1-167083660-INS-121 | chr1-167083660-INS-121 |
| chr1-167094679-INS-62 | chr1-167094679-INS-62 |
| chr1-167144848-INS-410 | chr1-167144848-INS-411 |
| chr1-167168342-INS-506 | chr1-167168342-INS-506 |
| chr1-167173475-INS-206 | chr1-167173475-INS-206 |
| chr1-167174780-INS-72 | chr1-167174776-INS-70 |
| chr1-167212199-INS-144 | chr1-167212199-INS-144 |
| chr1-167231311-INS-2994 | chr1-167231312-INS-2994 |
| chr1-167240733-INS-222 | chr1-167240733-INS-222 |
| chr1-167246627-INS-213 | chr1-167246627-INS-213 |
| chr1-167251635-INS-177 | chr1-167251635-INS-177 |
| chr1-167274300-INS-51 | chr1-167274300-INS-68 |
| chr1-167277060-INS-198 | chr1-167277060-INS-201 |
| chr1-167277843-INS-6187 | chr1-167277844-INS-6187 |
| chr1-167279178-INS-892 | chr1-167279178-INS-887 |
| chr1-167290097-INS-62 | chr1-167290097-INS-62 |
| chr1-167291955-INS-86 | chr1-167291955-INS-86 |
| chr1-167308806-INS-6440 | chr1-167308807-INS-6440 |
| chr1-167315359-INS-282 | chr1-167315359-INS-282 |
| chr1-167329404-INS-256 | chr1-167329405-INS-242 |
| chr1-167330166-INS-56 | chr1-167330166-INS-52 |
| chr1-167347506-INS-459 | chr1-167347506-INS-459 |
| chr1-167358277-INS-66 | chr1-167358277-INS-66 |
| chr1-167364294-INS-280 | chr1-167364297-INS-276 |
| chr1-167428146-INS-60 | chr1-167428146-INS-60 |
| chr1-167430653-INS-64 | chr1-167430653-INS-64 |
| chr1-167473791-INS-330 | chr1-167473791-INS-335 |
| chr1-167487014-INS-57 | chr1-167487014-INS-57 |
| chr1-167522155-INS-52 | chr1-167522147-INS-88 |
| chr1-167529492-INS-159 | chr1-167529492-INS-163 |
| chr1-167552587-INS-88 | chr1-167552587-INS-88 |
| chr1-167553613-INS-490 | chr1-167553613-INS-491 |
| chr1-167575559-INS-76 | chr1-167575559-INS-76 |
| chr1-167579314-INS-53 | chr1-167579319-INS-84 |
| chr1-167586658-INS-59 | chr1-167586658-INS-59 |
| chr1-167596738-INS-71 | chr1-167596734-INS-76 |
| chr1-167623115-INS-740 | chr1-167623115-INS-739 |
| chr1-167694805-INS-54 | chr1-167694805-INS-54 |
| chr1-167700935-INS-608 | chr1-167700935-INS-608 |
| chr1-167711197-INS-51 | chr1-167711172-INS-56 |
| chr1-167714258-INS-427 | chr1-167714258-INS-427 |
| chr1-167733353-INS-200 | chr1-167733353-INS-200 |
| chr1-167740628-INS-114 | chr1-167740664-INS-123 |
| chr1-167742226-INS-157 | chr1-167742226-INS-157 |
| chr1-167743383-INS-84 | chr1-167743383-INS-70 |
| chr1-167752546-INS-400 | chr1-167752546-INS-400 |
| chr1-167754794-INS-63 | chr1-167754794-INS-68 |
| chr1-167764411-INS-4484 | chr1-167764412-INS-4484 |
| chr1-167779670-INS-64 | chr1-167779664-INS-58 |
| chr1-167862531-INS-58 | chr1-167862536-INS-60 |
| chr1-167868255-INS-399 | chr1-167868255-INS-399 |
| chr1-167916064-INS-60 | chr1-167916078-INS-82 |
| chr1-167917705-INS-52 | chr1-167917734-INS-69 |
| chr1-167940477-INS-61 | chr1-167940477-INS-71 |
| chr1-167944048-INS-90 | chr1-167944048-INS-56 |
| chr1-167952126-INS-61 | chr1-167952126-INS-81 |
| chr1-167953413-INS-626 | chr1-167953413-INS-627 |
| chr1-167957791-INS-140 | chr1-167957791-INS-140 |
| chr1-167962557-INS-364 | chr1-167962557-INS-364 |
| chr1-167991955-INS-88 | chr1-167991955-INS-90 |
| chr1-167999633-INS-160 | chr1-167999641-INS-155 |
| chr1-168035053-INS-93 | chr1-168035053-INS-93 |
| chr1-168049571-INS-92 | chr1-168049560-INS-53 |

| | |
|---|---|
| chr1-168050732-INS-68 | chr1-168050732-INS-50 |
| chr1-168058470-INS-56 | chr1-168058470-INS-56 |
| chr1-168087551-INS-95 | chr1-168087551-INS-95 |
| chr1-168091564-INS-210 | chr1-168091564-INS-210 |
| chr1-168107131-INS-68 | chr1-168107131-INS-68 |
| chr1-168134376-INS-78 | chr1-168134376-INS-78 |
| chr1-168144837-INS-56 | chr1-168144837-INS-56 |
| chr1-168189877-INS-64 | chr1-168189877-INS-64 |
| chr1-168206255-INS-55 | chr1-168206255-INS-57 |
| chr1-168208412-INS-90 | chr1-168208412-INS-86 |
| chr1-168225453-INS-54 | chr1-168225454-INS-52 |
| chr1-168236266-INS-102 | chr1-168236266-INS-102 |
| chr1-168266026-INS-79 | chr1-168266026-INS-82 |
| chr1-168275860-INS-126 | chr1-168275860-INS-126 |
| chr1-168286827-INS-52 | chr1-168286827-INS-50 |
| chr1-168287908-INS-66 | chr1-168287908-INS-66 |
| chr1-168306028-INS-246 | chr1-168306028-INS-246 |
| chr1-168311918-INS-222 | chr1-168311918-INS-221 |
| chr1-168336519-INS-177 | chr1-168336519-INS-177 |
| chr1-168343113-INS-79 | chr1-168343090-INS-60 |
| chr1-168347496-INS-262 | chr1-168347496-INS-262 |
| chr1-168385889-INS-506 | chr1-168385889-INS-506 |
| chr1-168401393-INS-58 | chr1-168401393-INS-50 |
| chr1-168406096-INS-504 | chr1-168406096-INS-504 |
| chr1-168477394-INS-205 | chr1-168477394-INS-205 |
| chr1-168496669-INS-88 | chr1-168496669-INS-88 |
| chr1-168505812-INS-113 | chr1-168505812-INS-113 |
| chr1-168524597-INS-196 | chr1-168524597-INS-196 |
| chr1-168537830-INS-80 | chr1-168537830-INS-80 |
| chr1-168542838-INS-139 | chr1-168542838-INS-139 |
| chr1-168566270-INS-64 | chr1-168566270-INS-64 |
| chr1-168589760-INS-70 | chr1-168589760-INS-70 |
| chr1-168609484-INS-58 | chr1-168609484-INS-62 |
| chr1-168648045-INS-70 | chr1-168648045-INS-70 |
| chr1-168657423-INS-3262 | chr1-168657424-INS-3262 |
| chr1-168664795-INS-52 | chr1-168664770-INS-56 |
| chr1-168665372-INS-133 | chr1-168665372-INS-140 |
| chr1-168700209-INS-68 | chr1-168700209-INS-70 |
| chr1-168722360-INS-67 | chr1-168722360-INS-69 |
| chr1-168727683-INS-208 | chr1-168727683-INS-162 |
| chr1-168743795-INS-139 | chr1-168743795-INS-140 |
| chr1-168772268-INS-1474 | chr1-168772269-INS-1474 |
| chr1-168783425-INS-384 | chr1-168783427-INS-394 |
| chr1-168820814-INS-86 | chr1-168820816-INS-72 |
| chr1-168825971-INS-130 | chr1-168825971-INS-127 |
| chr1-168830212-INS-60 | chr1-168830212-INS-60 |
| chr1-168860865-INS-80 | chr1-168860865-INS-62 |
| chr1-168895938-INS-53 | chr1-168895938-INS-53 |
| chr1-168905656-INS-498 | chr1-168905656-INS-498 |
| chr1-168907351-INS-51 | chr1-168907361-INS-64 |
| chr1-168918307-INS-178 | chr1-168918307-INS-177 |
| chr1-168929029-INS-118 | chr1-168929029-INS-116 |
| chr1-168930260-INS-54 | chr1-168930261-INS-54 |
| chr1-168955433-INS-54 | chr1-168955433-INS-56 |
| chr1-169098211-INS-333 | chr1-169098211-INS-318 |
| chr1-169128441-INS-535 | chr1-169128441-INS-535 |
| chr1-169168387-INS-82 | chr1-169168383-INS-86 |
| chr1-169169258-INS-68 | chr1-169169258-INS-68 |
| chr1-169184628-INS-68 | chr1-169184628-INS-78 |
| chr1-169201793-INS-54 | chr1-169201793-INS-50 |
| chr1-169203795-INS-83 | chr1-169203813-INS-76 |
| chr1-169260892-INS-572 | chr1-169260892-INS-572 |
| chr1-169263413-INS-85 | chr1-169263413-INS-95 |
| chr1-169277962-INS-469 | chr1-169277962-INS-469 |
| chr1-169285578-INS-64 | chr1-169285578-INS-64 |
| chr1-169325127-INS-422 | chr1-169325127-INS-422 |
| chr1-169327905-INS-209 | chr1-169327905-INS-209 |
| chr1-169328615-INS-95 | chr1-169328615-INS-99 |
| chr1-169329364-INS-88 | chr1-169329364-INS-88 |
| chr1-169333432-INS-52 | chr1-169333432-INS-64 |
| chr1-169334763-INS-80 | chr1-169334763-INS-80 |
| chr1-169353133-INS-85 | chr1-169353133-INS-85 |
| chr1-169366301-INS-269 | chr1-169366301-INS-269 |
| chr1-169380130-INS-74 | chr1-169380130-INS-74 |
| chr1-169390962-INS-62 | chr1-169390944-INS-88 |
| chr1-169433410-INS-172 | chr1-169433410-INS-172 |
| chr1-169441951-INS-207 | chr1-169441951-INS-204 |

| | |
|---|---|
| chr1-169467912-INS-2080 | chr1-169467913-INS-2059 |
| chr1-169478374-INS-1879 | chr1-169478375-INS-1879 |
| chr1-169489238-INS-70 | chr1-169489238-INS-51 |
| chr1-169492353-INS-74 | chr1-169492349-INS-50 |
| chr1-169497231-INS-112 | chr1-169497231-INS-108 |
| chr1-169531050-INS-59 | chr1-169531050-INS-77 |
| chr1-169534706-INS-54 | chr1-169534706-INS-54 |
| chr1-169535363-INS-765 | chr1-169535363-INS-766 |
| chr1-169555353-INS-60 | chr1-169555369-INS-58 |
| chr1-169576020-INS-53 | chr1-169576020-INS-53 |
| chr1-169594335-INS-60 | chr1-169594335-INS-62 |
| chr1-169601698-INS-6398 | chr1-169601699-INS-6397 |
| chr1-169607437-INS-1890 | chr1-169607438-INS-1890 |
| chr1-169646145-INS-1641 | chr1-169646146-INS-1649 |
| chr1-169648358-INS-63 | chr1-169648358-INS-63 |
| chr1-169650308-INS-3316 | chr1-169650309-INS-3299 |
| chr1-169674951-INS-205 | chr1-169674955-INS-214 |
| chr1-169679579-INS-6721 | chr1-169679580-INS-6721 |
| chr1-169683515-INS-402 | chr1-169683515-INS-402 |
| chr1-169687546-INS-388 | chr1-169687546-INS-382 |
| chr1-169692115-INS-6429 | chr1-169692116-INS-6429 |
| chr1-169693149-INS-794 | chr1-169693149-INS-795 |
| chr1-169718928-INS-58 | chr1-169718929-INS-58 |
| chr1-169771893-INS-91 | chr1-169771893-INS-91 |
| chr1-169793507-INS-140 | chr1-169793516-INS-144 |
| chr1-169800493-INS-167 | chr1-169800493-INS-167 |
| chr1-169843021-INS-419 | chr1-169843021-INS-420 |
| chr1-169869770-INS-185 | chr1-169869770-INS-189 |
| chr1-169871639-INS-60 | chr1-169871639-INS-54 |
| chr1-169877328-INS-498 | chr1-169877328-INS-498 |
| chr1-169906802-INS-404 | chr1-169906802-INS-404 |
| chr1-169935724-INS-59 | chr1-169935724-INS-59 |
| chr1-169961242-INS-58 | chr1-169961242-INS-70 |
| chr1-169998856-INS-304 | chr1-169998856-INS-304 |
| chr1-170004162-INS-68 | chr1-170004162-INS-68 |
| chr1-170009590-INS-69 | chr1-170009590-INS-69 |
| chr1-170035406-INS-226 | chr1-170035406-INS-226 |
| chr1-170045385-INS-527 | chr1-170045385-INS-530 |
| chr1-170063934-INS-72 | chr1-170063934-INS-72 |
| chr1-170067453-INS-401 | chr1-170067453-INS-401 |
| chr1-170071661-INS-59 | chr1-170071661-INS-55 |
| chr1-170119606-INS-211 | chr1-170119606-INS-211 |
| chr1-170124553-INS-72 | chr1-170124553-INS-72 |
| chr1-170163060-INS-63 | chr1-170163027-INS-57 |
| chr1-170164165-INS-64 | chr1-170164123-INS-56 |
| chr1-170169719-INS-198 | chr1-170169719-INS-200 |
| chr1-170185862-INS-246 | chr1-170185862-INS-235 |
| chr1-170199961-INS-84 | chr1-170199961-INS-94 |
| chr1-170205351-INS-527 | chr1-170205351-INS-526 |
| chr1-170225028-INS-221 | chr1-170225028-INS-221 |
| chr1-170228917-INS-1601 | chr1-170228918-INS-1601 |
| chr1-170238506-INS-185 | chr1-170238506-INS-185 |
| chr1-170246598-INS-2893 | chr1-170246599-INS-2877 |
| chr1-170253853-INS-7158 | chr1-170253854-INS-7158 |
| chr1-170290728-INS-58 | chr1-170290728-INS-58 |
| chr1-170295882-INS-60 | chr1-170295882-INS-60 |
| chr1-170336137-INS-60 | chr1-170336137-INS-60 |
| chr1-170341545-INS-191 | chr1-170341545-INS-191 |
| chr1-170342636-INS-51 | chr1-170342637-INS-58 |
| chr1-170388956-INS-60 | chr1-170388956-INS-50 |
| chr1-170395024-INS-6265 | chr1-170395025-INS-6264 |
| chr1-170437357-INS-74 | chr1-170437340-INS-110 |
| chr1-170450377-INS-68 | chr1-170450377-INS-68 |
| chr1-170452275-INS-185 | chr1-170452275-INS-185 |
| chr1-170454578-INS-504 | chr1-170454578-INS-504 |
| chr1-170501309-INS-55 | chr1-170501321-INS-58 |
| chr1-170510478-INS-818 | chr1-170510478-INS-818 |
| chr1-170542662-INS-52 | chr1-170542662-INS-52 |
| chr1-170548147-INS-201 | chr1-170548147-INS-203 |
| chr1-170558469-INS-84 | chr1-170558469-INS-84 |
| chr1-170574173-INS-6547 | chr1-170574174-INS-6549 |
| chr1-170601524-INS-235 | chr1-170601556-INS-244 |
| chr1-170621316-INS-204 | chr1-170621316-INS-202 |
| chr1-170625660-INS-756 | chr1-170625660-INS-756 |
| chr1-170626727-INS-780 | chr1-170626727-INS-780 |
| chr1-170642290-INS-183 | chr1-170642290-INS-183 |
| chr1-170662219-INS-223 | chr1-170662220-INS-223 |

| | |
|---|---|
| chr1-170681558-INS-155 | chr1-170681561-INS-156 |
| chr1-170683174-INS-73 | chr1-170683178-INS-74 |
| chr1-170687347-INS-70 | chr1-170687347-INS-70 |
| chr1-170708383-INS-190 | chr1-170708383-INS-190 |
| chr1-170709112-INS-202 | chr1-170709112-INS-202 |
| chr1-170716881-INS-68 | chr1-170716881-INS-52 |
| chr1-170720289-INS-63 | chr1-170720289-INS-61 |
| chr1-170721873-INS-199 | chr1-170721873-INS-200 |
| chr1-170744280-INS-173 | chr1-170744280-INS-173 |
| chr1-170768766-INS-487 | chr1-170768766-INS-487 |
| chr1-170805205-INS-210 | chr1-170805205-INS-210 |
| chr1-170808760-INS-112 | chr1-170808760-INS-112 |
| chr1-170818564-INS-66 | chr1-170818564-INS-53 |
| chr1-170820075-INS-72 | chr1-170820075-INS-68 |
| chr1-170871088-INS-235 | chr1-170871088-INS-235 |
| chr1-170872217-INS-373 | chr1-170872217-INS-373 |
| chr1-170875143-INS-506 | chr1-170875143-INS-506 |
| chr1-170877114-INS-434 | chr1-170877114-INS-434 |
| chr1-170885701-INS-264 | chr1-170885701-INS-266 |
| chr1-170889859-INS-203 | chr1-170889859-INS-203 |
| chr1-170891252-INS-980 | chr1-170891253-INS-981 |
| chr1-170910054-INS-358 | chr1-170910054-INS-358 |
| chr1-170925712-INS-56 | chr1-170925712-INS-56 |
| chr1-170927012-INS-453 | chr1-170927012-INS-453 |
| chr1-170927781-INS-208 | chr1-170927781-INS-208 |
| chr1-170930291-INS-130 | chr1-170930291-INS-130 |
| chr1-170933600-INS-57 | chr1-170933600-INS-60 |
| chr1-170935064-INS-51 | chr1-170935064-INS-51 |
| chr1-170936340-INS-409 | chr1-170936340-INS-409 |
| chr1-170947726-INS-84 | chr1-170947727-INS-84 |
| chr1-170951093-INS-930 | chr1-170951093-INS-931 |
| chr1-170951800-INS-59 | chr1-170951778-INS-83 |
| chr1-170968937-INS-187 | chr1-170968937-INS-188 |
| chr1-170989206-INS-212 | chr1-170989206-INS-212 |
| chr1-170999932-INS-214 | chr1-170999932-INS-214 |
| chr1-171003835-INS-52 | chr1-171003835-INS-51 |
| chr1-171014202-INS-59 | chr1-171014202-INS-70 |
| chr1-171040352-INS-218 | chr1-171040352-INS-218 |
| chr1-171085045-INS-161 | chr1-171085045-INS-162 |
| chr1-171092622-INS-7182 | chr1-171092623-INS-7182 |
| chr1-171128819-INS-205 | chr1-171128819-INS-205 |
| chr1-171129434-INS-224 | chr1-171129434-INS-224 |
| chr1-171131009-INS-62 | chr1-171131009-INS-62 |
| chr1-171145089-INS-239 | chr1-171145090-INS-209 |
| chr1-171150464-INS-225 | chr1-171150465-INS-211 |
| chr1-171175137-INS-207 | chr1-171175138-INS-203 |
| chr1-171175875-INS-212 | chr1-171175876-INS-212 |
| chr1-171176879-INS-137 | chr1-171176879-INS-142 |
| chr1-171191273-INS-201 | chr1-171191273-INS-201 |
| chr1-171242339-INS-181 | chr1-171242339-INS-180 |
| chr1-171257540-INS-178 | chr1-171257540-INS-178 |
| chr1-171260615-INS-125 | chr1-171260615-INS-124 |
| chr1-171261711-INS-59 | chr1-171261711-INS-63 |
| chr1-171262926-INS-205 | chr1-171262926-INS-206 |
| chr1-171271767-INS-215 | chr1-171271767-INS-214 |
| chr1-171276239-INS-204 | chr1-171276239-INS-204 |
| chr1-171281703-INS-103 | chr1-171281703-INS-103 |
| chr1-171285452-INS-221 | chr1-171285452-INS-222 |
| chr1-171289673-INS-537 | chr1-171289673-INS-538 |
| chr1-171305569-INS-194 | chr1-171305569-INS-194 |
| chr1-171306303-INS-195 | chr1-171306303-INS-196 |
| chr1-171320612-INS-70 | chr1-171320612-INS-68 |
| chr1-171323539-INS-186 | chr1-171323539-INS-190 |
| chr1-171329068-INS-222 | chr1-171329068-INS-222 |
| chr1-171329862-INS-197 | chr1-171329863-INS-209 |
| chr1-171358386-INS-66 | chr1-171358388-INS-74 |
| chr1-171460903-INS-65 | chr1-171460903-INS-65 |
| chr1-171501030-INS-1333 | chr1-171501031-INS-1333 |
| chr1-171518904-INS-69 | chr1-171518904-INS-69 |
| chr1-171529198-INS-6423 | chr1-171529199-INS-6423 |
| chr1-171539280-INS-69 | chr1-171539280-INS-74 |
| chr1-171568700-INS-79 | chr1-171568705-INS-84 |
| chr1-171588547-INS-62 | chr1-171588551-INS-62 |
| chr1-171601339-INS-59 | chr1-171601329-INS-64 |
| chr1-171635817-INS-67 | chr1-171635825-INS-59 |
| chr1-171646112-INS-124 | chr1-171646112-INS-122 |
| chr1-171672995-INS-98 | chr1-171672995-INS-98 |

| | |
|---|---|
| chr1-171708849-INS-54 | chr1-171708849-INS-54 |
| chr1-171727738-INS-3104 | chr1-171727739-INS-3104 |
| chr1-171729974-INS-88 | chr1-171729974-INS-88 |
| chr1-171734970-INS-50 | chr1-171734970-INS-50 |
| chr1-171738036-INS-899 | chr1-171738036-INS-904 |
| chr1-171742619-INS-51 | chr1-171742621-INS-50 |
| chr1-171751299-INS-63 | chr1-171751299-INS-63 |
| chr1-171768940-INS-157 | chr1-171768940-INS-157 |
| chr1-171780686-INS-208 | chr1-171780686-INS-209 |
| chr1-171809060-INS-60 | chr1-171809060-INS-60 |
| chr1-171811581-INS-97 | chr1-171811581-INS-97 |
| chr1-171815726-INS-74 | chr1-171815726-INS-74 |
| chr1-171845056-INS-56 | chr1-171845056-INS-54 |
| chr1-171867434-INS-51 | chr1-171867434-INS-51 |
| chr1-171888092-INS-60 | chr1-171888138-INS-56 |
| chr1-171914247-INS-240 | chr1-171914247-INS-240 |
| chr1-171941115-INS-175 | chr1-171941115-INS-175 |
| chr1-171943686-INS-260 | chr1-171943686-INS-260 |
| chr1-171949155-INS-57 | chr1-171949155-INS-57 |
| chr1-171953301-INS-208 | chr1-171953301-INS-208 |
| chr1-171959557-INS-251 | chr1-171959557-INS-252 |
| chr1-171967767-INS-303 | chr1-171967767-INS-303 |
| chr1-171977495-INS-80 | chr1-171977495-INS-80 |
| chr1-171983900-INS-228 | chr1-171983900-INS-210 |
| chr1-171998313-INS-51 | chr1-171998313-INS-51 |
| chr1-172004705-INS-100 | chr1-172004705-INS-90 |
| chr1-172048665-INS-448 | chr1-172048665-INS-448 |
| chr1-172054187-INS-882 | chr1-172054187-INS-882 |
| chr1-172066883-INS-55 | chr1-172066883-INS-55 |
| chr1-172166119-INS-223 | chr1-172166119-INS-224 |
| chr1-172181242-INS-75 | chr1-172181246-INS-81 |
| chr1-172182497-INS-83 | chr1-172182497-INS-83 |
| chr1-172184839-INS-212 | chr1-172184839-INS-212 |
| chr1-172195489-INS-59 | chr1-172195489-INS-59 |
| chr1-172227130-INS-54 | chr1-172227139-INS-66 |
| chr1-172228050-INS-202 | chr1-172228050-INS-203 |
| chr1-172250626-INS-60 | chr1-172250627-INS-67 |
| chr1-172270463-INS-478 | chr1-172270463-INS-478 |
| chr1-172271792-INS-170 | chr1-172271792-INS-170 |
| chr1-172278211-INS-132 | chr1-172278206-INS-127 |
| chr1-172279941-INS-53 | chr1-172279941-INS-95 |
| chr1-172296229-INS-60 | chr1-172296229-INS-60 |
| chr1-172298350-INS-229 | chr1-172298350-INS-229 |
| chr1-172349664-INS-1891 | chr1-172349665-INS-1891 |
| chr1-172364005-INS-204 | chr1-172364005-INS-204 |
| chr1-172387030-INS-110 | chr1-172387030-INS-110 |
| chr1-172405369-INS-63 | chr1-172405328-INS-64 |
| chr1-172441097-INS-1042 | chr1-172441098-INS-1059 |
| chr1-172455111-INS-1893 | chr1-172455112-INS-1893 |
| chr1-172464196-INS-60 | chr1-172464196-INS-52 |
| chr1-172485557-INS-77 | chr1-172485557-INS-50 |
| chr1-172499842-INS-834 | chr1-172499842-INS-834 |
| chr1-172506499-INS-60 | chr1-172506499-INS-66 |
| chr1-172526846-INS-54 | chr1-172526846-INS-50 |
| chr1-172529432-INS-358 | chr1-172529432-INS-358 |
| chr1-172541064-INS-71 | chr1-172541064-INS-87 |
| chr1-172544602-INS-624 | chr1-172544603-INS-624 |
| chr1-172586993-INS-126 | chr1-172586993-INS-126 |
| chr1-172588519-INS-58 | chr1-172588520-INS-54 |
| chr1-172592513-INS-139 | chr1-172592513-INS-139 |
| chr1-172595516-INS-57 | chr1-172595516-INS-59 |
| chr1-172605520-INS-106 | chr1-172605520-INS-104 |
| chr1-172624036-INS-999 | chr1-172624036-INS-1001 |
| chr1-172633925-INS-351 | chr1-172633925-INS-351 |
| chr1-172638267-INS-70 | chr1-172638267-INS-70 |
| chr1-172664317-INS-72 | chr1-172664318-INS-60 |
| chr1-172674314-INS-80 | chr1-172674314-INS-80 |
| chr1-172684859-INS-440 | chr1-172684859-INS-440 |
| chr1-172686264-INS-3094 | chr1-172686265-INS-3094 |
| chr1-172693915-INS-607 | chr1-172693915-INS-607 |
| chr1-172698235-INS-3165 | chr1-172698236-INS-3165 |
| chr1-172701981-INS-418 | chr1-172701981-INS-418 |
| chr1-172715633-INS-235 | chr1-172715633-INS-235 |
| chr1-172766536-INS-831 | chr1-172766536-INS-832 |
| chr1-172772831-INS-443 | chr1-172772831-INS-443 |
| chr1-172787785-INS-95 | chr1-172787790-INS-90 |
| chr1-172821532-INS-104 | chr1-172821515-INS-140 |

| | |
|---|---|
| chr1-172824457-INS-83 | chr1-172824457-INS-51 |
| chr1-172825758-INS-1014 | chr1-172825759-INS-1051 |
| chr1-172826538-INS-77 | chr1-172826554-INS-99 |
| chr1-172848274-INS-1408 | chr1-172848275-INS-1408 |
| chr1-172853072-INS-6362 | chr1-172853073-INS-6362 |
| chr1-172896459-INS-839 | chr1-172896459-INS-836 |
| chr1-172905444-INS-52 | chr1-172905444-INS-52 |
| chr1-172922997-INS-54 | chr1-172922997-INS-56 |
| chr1-172977518-INS-582 | chr1-172977518-INS-582 |
| chr1-172990505-INS-4709 | chr1-172990506-INS-4709 |
| chr1-172992210-INS-239 | chr1-172992210-INS-248 |
| chr1-172993498-INS-445 | chr1-172993498-INS-445 |
| chr1-173000640-INS-910 | chr1-173000640-INS-911 |
| chr1-173023450-INS-6704 | chr1-173023451-INS-6704 |
| chr1-173024556-INS-62 | chr1-173024556-INS-60 |
| chr1-173026617-INS-258 | chr1-173026617-INS-258 |
| chr1-173036454-INS-209 | chr1-173036454-INS-209 |
| chr1-173046945-INS-79 | chr1-173046945-INS-81 |
| chr1-173056321-INS-517 | chr1-173056321-INS-517 |
| chr1-173058622-INS-1904 | chr1-173058623-INS-1904 |
| chr1-173066518-INS-62 | chr1-173066510-INS-64 |
| chr1-173072144-INS-939 | chr1-173072144-INS-939 |
| chr1-173075227-INS-2291 | chr1-173075228-INS-2292 |
| chr1-173095442-INS-68 | chr1-173095442-INS-70 |
| chr1-173118711-INS-67 | chr1-173118717-INS-53 |
| chr1-173133553-INS-208 | chr1-173133553-INS-208 |
| chr1-173166098-INS-60 | chr1-173166084-INS-52 |
| chr1-173180448-INS-104 | chr1-173180448-INS-104 |
| chr1-173186051-INS-52 | chr1-173186051-INS-50 |
| chr1-173222116-INS-600 | chr1-173222116-INS-600 |
| chr1-173224568-INS-651 | chr1-173224568-INS-651 |
| chr1-173234276-INS-52 | chr1-173234276-INS-52 |
| chr1-173291952-INS-57 | chr1-173291940-INS-78 |
| chr1-173318695-INS-80 | chr1-173318695-INS-82 |
| chr1-173334803-INS-88 | chr1-173334803-INS-88 |
| chr1-173341473-INS-54 | chr1-173341473-INS-52 |
| chr1-173408768-INS-5241 | chr1-173408769-INS-5241 |
| chr1-173450802-INS-323 | chr1-173450802-INS-323 |
| chr1-173475426-INS-1330 | chr1-173475427-INS-1329 |
| chr1-173481771-INS-53 | chr1-173481771-INS-53 |
| chr1-173496864-INS-245 | chr1-173496864-INS-245 |
| chr1-173507892-INS-8981 | chr1-173507893-INS-8976 |
| chr1-173533322-INS-54 | chr1-173533323-INS-54 |
| chr1-173553907-INS-236 | chr1-173553907-INS-236 |
| chr1-173556779-INS-331 | chr1-173556779-INS-336 |
| chr1-173587714-INS-63 | chr1-173587714-INS-63 |
| chr1-173631040-INS-110 | chr1-173631040-INS-110 |
| chr1-173638455-INS-528 | chr1-173638455-INS-528 |
| chr1-173651620-INS-251 | chr1-173651620-INS-251 |
| chr1-173654267-INS-882 | chr1-173654267-INS-882 |
| chr1-173656838-INS-84 | chr1-173656840-INS-90 |
| chr1-173677584-INS-458 | chr1-173677584-INS-458 |
| chr1-173705835-INS-142 | chr1-173705835-INS-142 |
| chr1-173733734-INS-62 | chr1-173733716-INS-63 |
| chr1-173793217-INS-853 | chr1-173793218-INS-853 |
| chr1-173833593-INS-204 | chr1-173833593-INS-203 |
| chr1-173852527-INS-76 | chr1-173852527-INS-76 |
| chr1-173854754-INS-63 | chr1-173854754-INS-63 |
| chr1-173858300-INS-108 | chr1-173858300-INS-108 |
| chr1-173896677-INS-66 | chr1-173896677-INS-66 |
| chr1-173916201-INS-62 | chr1-173916201-INS-62 |
| chr1-173920176-INS-7232 | chr1-173920177-INS-7232 |
| chr1-173931937-INS-120 | chr1-173931937-INS-124 |
| chr1-173969825-INS-70 | chr1-173969825-INS-70 |
| chr1-173991998-INS-54 | chr1-173991998-INS-56 |
| chr1-173995151-INS-394 | chr1-173995151-INS-399 |
| chr1-174001536-INS-70 | chr1-174001536-INS-70 |
| chr1-174020795-INS-660 | chr1-174020795-INS-660 |
| chr1-174027945-INS-199 | chr1-174027945-INS-199 |
| chr1-174064171-INS-70 | chr1-174064171-INS-70 |
| chr1-174067367-INS-163 | chr1-174067353-INS-166 |
| chr1-174078603-INS-54 | chr1-174078603-INS-54 |
| chr1-174144689-INS-563 | chr1-174144689-INS-563 |
| chr1-174151955-INS-654 | chr1-174151953-INS-642 |
| chr1-174158436-INS-204 | chr1-174158436-INS-204 |
| chr1-174164652-INS-449 | chr1-174164652-INS-449 |
| chr1-174182903-INS-87 | chr1-174182903-INS-87 |

| | |
|---|---|
| chr1-174184028-INS-837 | chr1-174184028-INS-837 |
| chr1-174235255-INS-328 | chr1-174235255-INS-327 |
| chr1-174236834-INS-93 | chr1-174236834-INS-93 |
| chr1-174241873-INS-553 | chr1-174241873-INS-552 |
| chr1-174257671-INS-54 | chr1-174257671-INS-54 |
| chr1-174269162-INS-52 | chr1-174269162-INS-60 |
| chr1-174273602-INS-108 | chr1-174273602-INS-109 |
| chr1-174280835-INS-58 | chr1-174280821-INS-82 |
| chr1-174283566-INS-68 | chr1-174283566-INS-70 |
| chr1-174298377-INS-641 | chr1-174298377-INS-640 |
| chr1-174304050-INS-199 | chr1-174304050-INS-199 |
| chr1-174323814-INS-67 | chr1-174323814-INS-67 |
| chr1-174326677-INS-71 | chr1-174326677-INS-79 |
| chr1-174337545-INS-128 | chr1-174337545-INS-128 |
| chr1-174347606-INS-86 | chr1-174347606-INS-56 |
| chr1-174367886-INS-374 | chr1-174367887-INS-375 |
| chr1-174372789-INS-203 | chr1-174372790-INS-222 |
| chr1-174379353-INS-230 | chr1-174379353-INS-232 |
| chr1-174386463-INS-162 | chr1-174386463-INS-162 |
| chr1-174395492-INS-204 | chr1-174395492-INS-204 |
| chr1-174397998-INS-251 | chr1-174397998-INS-251 |
| chr1-174399165-INS-57 | chr1-174399165-INS-57 |
| chr1-174402071-INS-475 | chr1-174402071-INS-475 |
| chr1-174416089-INS-74 | chr1-174416089-INS-74 |
| chr1-174424133-INS-125 | chr1-174424133-INS-125 |
| chr1-174426334-INS-245 | chr1-174426334-INS-245 |
| chr1-174429899-INS-164 | chr1-174429900-INS-152 |
| chr1-174440797-INS-7127 | chr1-174440798-INS-7127 |
| chr1-174451166-INS-7026 | chr1-174451167-INS-7024 |
| chr1-174484563-INS-555 | chr1-174484563-INS-555 |
| chr1-174496185-INS-239 | chr1-174496188-INS-237 |
| chr1-174507722-INS-68 | chr1-174507722-INS-76 |
| chr1-174519116-INS-70 | chr1-174519108-INS-58 |
| chr1-174543419-INS-91 | chr1-174543419-INS-99 |
| chr1-174553329-INS-828 | chr1-174553329-INS-828 |
| chr1-174569275-INS-60 | chr1-174569276-INS-81 |
| chr1-174572985-INS-6585 | chr1-174572986-INS-6587 |
| chr1-174577313-INS-341 | chr1-174577311-INS-349 |
| chr1-174600063-INS-86 | chr1-174600063-INS-82 |
| chr1-174604415-INS-2758 | chr1-174604416-INS-2758 |
| chr1-174628408-INS-305 | chr1-174628408-INS-305 |
| chr1-174633698-INS-58 | chr1-174633728-INS-68 |
| chr1-174635622-INS-63 | chr1-174635609-INS-62 |
| chr1-174646090-INS-395 | chr1-174646090-INS-396 |
| chr1-174650334-INS-84 | chr1-174650334-INS-90 |
| chr1-174695530-INS-823 | chr1-174695530-INS-823 |
| chr1-174722154-INS-167 | chr1-174722154-INS-167 |
| chr1-174722794-INS-1257 | chr1-174722784-INS-1227 |
| chr1-174730947-INS-56 | chr1-174730948-INS-56 |
| chr1-174734750-INS-769 | chr1-174734750-INS-769 |
| chr1-174763608-INS-268 | chr1-174763608-INS-268 |
| chr1-174767899-INS-83 | chr1-174767897-INS-80 |
| chr1-174778946-INS-196 | chr1-174778946-INS-196 |
| chr1-174802015-INS-68 | chr1-174802015-INS-68 |
| chr1-174802687-INS-64 | chr1-174802687-INS-64 |
| chr1-174807302-INS-84 | chr1-174807302-INS-84 |
| chr1-174845634-INS-299 | chr1-174845634-INS-299 |
| chr1-174848882-INS-82 | chr1-174848882-INS-82 |
| chr1-174855882-INS-55 | chr1-174855875-INS-94 |
| chr1-174857154-INS-64 | chr1-174857154-INS-64 |
| chr1-174860753-INS-54 | chr1-174860753-INS-58 |
| chr1-174947470-INS-4520 | chr1-174947471-INS-4520 |
| chr1-174948675-INS-134 | chr1-174948675-INS-134 |
| chr1-174958182-INS-222 | chr1-174958182-INS-222 |
| chr1-174958887-INS-60 | chr1-174958887-INS-62 |
| chr1-174984745-INS-74 | chr1-174984745-INS-74 |
| chr1-175056132-INS-2831 | chr1-175056140-INS-2823 |
| chr1-175144267-INS-86 | chr1-175144267-INS-86 |
| chr1-175152993-INS-64 | chr1-175152974-INS-62 |
| chr1-175194281-INS-203 | chr1-175194281-INS-203 |
| chr1-175206307-INS-56 | chr1-175206307-INS-72 |
| chr1-175212783-INS-464 | chr1-175212783-INS-464 |
| chr1-175215696-INS-72 | chr1-175215698-INS-73 |
| chr1-175218479-INS-57 | chr1-175218467-INS-79 |
| chr1-175219386-INS-209 | chr1-175219386-INS-211 |
| chr1-175234132-INS-7115 | chr1-175234133-INS-7116 |
| chr1-175270495-INS-117 | chr1-175270495-INS-117 |

| | |
|---|---|
| chr1-175302785-INS-53 | chr1-175302776-INS-59 |
| chr1-175304571-INS-52 | chr1-175304571-INS-52 |
| chr1-175310220-INS-158 | chr1-175310220-INS-158 |
| chr1-175343288-INS-6179 | chr1-175343289-INS-6179 |
| chr1-175346134-INS-493 | chr1-175346134-INS-493 |
| chr1-175347381-INS-82 | chr1-175347381-INS-56 |
| chr1-175361442-INS-1204 | chr1-175361443-INS-1204 |
| chr1-175389484-INS-72 | chr1-175389484-INS-72 |
| chr1-175390647-INS-54 | chr1-175390647-INS-57 |
| chr1-175419484-INS-750 | chr1-175419484-INS-750 |
| chr1-175429406-INS-149 | chr1-175429406-INS-149 |
| chr1-175468877-INS-70 | chr1-175468877-INS-72 |
| chr1-175494041-INS-426 | chr1-175494041-INS-425 |
| chr1-175527686-INS-114 | chr1-175527686-INS-118 |
| chr1-175554316-INS-112 | chr1-175554316-INS-112 |
| chr1-175571881-INS-6440 | chr1-175571882-INS-6440 |
| chr1-175586380-INS-6867 | chr1-175586381-INS-6867 |
| chr1-175589415-INS-88 | chr1-175589415-INS-88 |
| chr1-175605522-INS-68 | chr1-175605522-INS-50 |
| chr1-175614231-INS-3851 | chr1-175614232-INS-3851 |
| chr1-175620141-INS-608 | chr1-175620141-INS-608 |
| chr1-175629455-INS-201 | chr1-175629455-INS-201 |
| chr1-175659668-INS-197 | chr1-175659668-INS-198 |
| chr1-175682400-INS-215 | chr1-175682403-INS-224 |
| chr1-175729329-INS-55 | chr1-175729329-INS-55 |
| chr1-175731496-INS-164 | chr1-175731490-INS-128 |
| chr1-175752127-INS-76 | chr1-175752127-INS-76 |
| chr1-175753935-INS-462 | chr1-175753935-INS-462 |
| chr1-175764555-INS-61 | chr1-175764513-INS-75 |
| chr1-175767911-INS-121 | chr1-175767912-INS-124 |
| chr1-175783009-INS-158 | chr1-175782959-INS-186 |
| chr1-175794979-INS-72 | chr1-175795015-INS-61 |
| chr1-175803781-INS-5305 | chr1-175803782-INS-5305 |
| chr1-175805702-INS-212 | chr1-175805702-INS-212 |
| chr1-175806965-INS-57 | chr1-175806965-INS-57 |
| chr1-175807612-INS-116 | chr1-175807612-INS-116 |
| chr1-175814949-INS-255 | chr1-175814961-INS-263 |
| chr1-175829816-INS-61 | chr1-175829810-INS-52 |
| chr1-175842460-INS-409 | chr1-175842460-INS-409 |
| chr1-175851318-INS-180 | chr1-175851318-INS-135 |
| chr1-175853262-INS-6660 | chr1-175853263-INS-6661 |
| chr1-175853402-INS-115 | chr1-175853403-INS-83 |
| chr1-175854896-INS-56 | chr1-175854896-INS-56 |
| chr1-175865742-INS-52 | chr1-175865742-INS-52 |
| chr1-175879599-INS-1500 | chr1-175879612-INS-1491 |
| chr1-175883290-INS-81 | chr1-175883290-INS-81 |
| chr1-175897032-INS-88 | chr1-175897030-INS-132 |
| chr1-175909678-INS-54 | chr1-175909678-INS-54 |
| chr1-175912985-INS-220 | chr1-175912985-INS-220 |
| chr1-175915469-INS-146 | chr1-175915473-INS-112 |
| chr1-175924724-INS-155 | chr1-175924723-INS-126 |
| chr1-175954072-INS-60 | chr1-175954072-INS-64 |
| chr1-175957732-INS-81 | chr1-175957732-INS-81 |
| chr1-176024167-INS-6454 | chr1-176024168-INS-6454 |
| chr1-176045302-INS-161 | chr1-176045326-INS-169 |
| chr1-176046899-INS-93 | chr1-176046899-INS-93 |
| chr1-176057973-INS-75 | chr1-176057974-INS-60 |
| chr1-176063748-INS-945 | chr1-176063748-INS-945 |
| chr1-176064440-INS-90 | chr1-176064440-INS-98 |
| chr1-176065147-INS-789 | chr1-176065147-INS-788 |
| chr1-176067990-INS-193 | chr1-176067990-INS-193 |
| chr1-176082030-INS-255 | chr1-176082030-INS-255 |
| chr1-176086967-INS-6483 | chr1-176086968-INS-6483 |
| chr1-176107606-INS-224 | chr1-176107606-INS-224 |
| chr1-176149623-INS-7127 | chr1-176149624-INS-7127 |
| chr1-176156023-INS-72 | chr1-176156055-INS-50 |
| chr1-176162032-INS-140 | chr1-176162032-INS-140 |
| chr1-176184031-INS-497 | chr1-176184031-INS-497 |
| chr1-176208684-INS-1889 | chr1-176208694-INS-1897 |
| chr1-176219782-INS-328 | chr1-176219782-INS-328 |
| chr1-176232415-INS-6730 | chr1-176232416-INS-6730 |
| chr1-176263238-INS-55 | chr1-176263239-INS-59 |
| chr1-176266807-INS-77 | chr1-176266807-INS-93 |
| chr1-176288932-INS-52 | chr1-176288932-INS-54 |
| chr1-176304164-INS-92 | chr1-176304164-INS-50 |
| chr1-176309140-INS-221 | chr1-176309145-INS-220 |
| chr1-176318892-INS-90 | chr1-176318897-INS-80 |

| | |
|---|---|
| chr1-176327707-INS-68 | chr1-176327707-INS-68 |
| chr1-176348807-INS-523 | chr1-176348807-INS-523 |
| chr1-176352818-INS-163 | chr1-176352818-INS-163 |
| chr1-176361911-INS-56 | chr1-176361911-INS-56 |
| chr1-176397025-INS-81 | chr1-176397025-INS-81 |
| chr1-176400526-INS-281 | chr1-176400525-INS-281 |
| chr1-176491706-INS-2384 | chr1-176491707-INS-2384 |
| chr1-176508327-INS-6615 | chr1-176508328-INS-6615 |
| chr1-176530822-INS-175 | chr1-176530822-INS-175 |
| chr1-176552606-INS-213 | chr1-176552606-INS-214 |
| chr1-176557952-INS-889 | chr1-176557952-INS-889 |
| chr1-176565838-INS-2920 | chr1-176565835-INS-2928 |
| chr1-176599003-INS-1676 | chr1-176599004-INS-1676 |
| chr1-176633889-INS-706 | chr1-176633889-INS-659 |
| chr1-176676070-INS-7113 | chr1-176676071-INS-7113 |
| chr1-176683962-INS-1484 | chr1-176683937-INS-1484 |
| chr1-176688936-INS-752 | chr1-176688936-INS-752 |
| chr1-176717795-INS-191 | chr1-176717796-INS-160 |
| chr1-176739617-INS-4369 | chr1-176739618-INS-4369 |
| chr1-176785219-INS-573 | chr1-176785219-INS-576 |
| chr1-176801956-INS-340 | chr1-176801956-INS-340 |
| chr1-176853902-INS-100 | chr1-176853902-INS-100 |
| chr1-176868325-INS-1395 | chr1-176868326-INS-1410 |
| chr1-176898807-INS-3675 | chr1-176898808-INS-3673 |
| chr1-176944893-INS-52 | chr1-176944893-INS-54 |
| chr1-176945239-INS-220 | chr1-176945239-INS-219 |
| chr1-177005636-INS-955 | chr1-177005636-INS-955 |
| chr1-177007962-INS-6854 | chr1-177007963-INS-6854 |
| chr1-177019393-INS-186 | chr1-177019393-INS-192 |
| chr1-177035568-INS-940 | chr1-177035553-INS-957 |
| chr1-177054410-INS-1942 | chr1-177054411-INS-1941 |
| chr1-177095630-INS-345 | chr1-177095630-INS-345 |
| chr1-177097860-INS-78 | chr1-177097860-INS-78 |
| chr1-177100697-INS-1075 | chr1-177100698-INS-1074 |
| chr1-177116515-INS-96 | chr1-177116515-INS-126 |
| chr1-177131389-INS-226 | chr1-177131389-INS-226 |
| chr1-177156997-INS-2298 | chr1-177156973-INS-2298 |
| chr1-177171164-INS-389 | chr1-177171164-INS-407 |
| chr1-177206867-INS-77 | chr1-177206867-INS-54 |
| chr1-177221513-INS-85 | chr1-177221513-INS-55 |
| chr1-177223467-INS-74 | chr1-177223467-INS-77 |
| chr1-177261547-INS-56 | chr1-177261547-INS-66 |
| chr1-177325285-INS-52 | chr1-177325285-INS-58 |
| chr1-177384778-INS-62 | chr1-177384778-INS-52 |
| chr1-177425160-INS-54 | chr1-177425150-INS-78 |
| chr1-177426447-INS-1195 | chr1-177426448-INS-1192 |
| chr1-177433436-INS-78 | chr1-177433436-INS-72 |
| chr1-177476272-INS-88 | chr1-177476272-INS-88 |
| chr1-177480009-INS-51 | chr1-177480009-INS-51 |
| chr1-177508262-INS-187 | chr1-177508262-INS-190 |
| chr1-177526735-INS-78 | chr1-177526735-INS-102 |
| chr1-177610492-INS-371 | chr1-177610492-INS-371 |
| chr1-177634430-INS-131 | chr1-177634430-INS-129 |
| chr1-177708269-INS-6439 | chr1-177708270-INS-6439 |
| chr1-177793247-INS-2478 | chr1-177793248-INS-2479 |
| chr1-177843026-INS-7098 | chr1-177843027-INS-7098 |
| chr1-177883846-INS-241 | chr1-177883846-INS-242 |
| chr1-177924655-INS-376 | chr1-177924655-INS-376 |
| chr1-177950200-INS-92 | chr1-177950200-INS-92 |
| chr1-177963294-INS-522 | chr1-177963294-INS-522 |
| chr1-177977765-INS-66 | chr1-177977765-INS-66 |
| chr1-178010800-INS-58 | chr1-178010800-INS-60 |
| chr1-178023051-INS-6440 | chr1-178023052-INS-6440 |
| chr1-178033387-INS-5087 | chr1-178033388-INS-5087 |
| chr1-178038882-INS-52 | chr1-178038882-INS-52 |
| chr1-178045653-INS-68 | chr1-178045653-INS-72 |
| chr1-178089249-INS-246 | chr1-178089249-INS-247 |
| chr1-178112180-INS-8326 | chr1-178112181-INS-8327 |
| chr1-178123718-INS-3286 | chr1-178123719-INS-3286 |
| chr1-178134889-INS-174 | chr1-178134889-INS-174 |
| chr1-178174156-INS-200 | chr1-178174156-INS-200 |
| chr1-178175520-INS-54 | chr1-178175547-INS-53 |
| chr1-178179288-INS-60 | chr1-178179288-INS-56 |
| chr1-178194296-INS-132 | chr1-178194284-INS-174 |
| chr1-178213198-INS-420 | chr1-178213198-INS-420 |
| chr1-178227008-INS-82 | chr1-178226970-INS-91 |
| chr1-178258873-INS-52 | chr1-178258873-INS-52 |

| | |
|---|---|
| chr1-178281151-INS-221 | chr1-178281152-INS-201 |
| chr1-178287870-INS-5272 | chr1-178287871-INS-5272 |
| chr1-178288825-INS-492 | chr1-178288825-INS-493 |
| chr1-178331675-INS-3661 | chr1-178331676-INS-3661 |
| chr1-178375481-INS-90 | chr1-178375481-INS-90 |
| chr1-178377639-INS-116 | chr1-178377641-INS-116 |
| chr1-178394241-INS-100 | chr1-178394241-INS-100 |
| chr1-178396069-INS-116 | chr1-178396073-INS-114 |
| chr1-178415865-INS-64 | chr1-178415865-INS-68 |
| chr1-178421510-INS-589 | chr1-178421510-INS-589 |
| chr1-178442927-INS-58 | chr1-178442928-INS-62 |
| chr1-178460643-INS-7299 | chr1-178460644-INS-7323 |
| chr1-178460866-INS-176 | chr1-178460866-INS-180 |
| chr1-178554397-INS-129 | chr1-178554397-INS-129 |
| chr1-178556360-INS-515 | chr1-178556360-INS-515 |
| chr1-178570631-INS-68 | chr1-178570631-INS-70 |
| chr1-178593581-INS-4453 | chr1-178593582-INS-4448 |
| chr1-178638681-INS-163 | chr1-178638681-INS-185 |
| chr1-178669703-INS-4605 | chr1-178669704-INS-4605 |
| chr1-178691104-INS-502 | chr1-178691120-INS-505 |
| chr1-178747215-INS-1643 | chr1-178747216-INS-1644 |
| chr1-178772803-INS-477 | chr1-178772803-INS-477 |
| chr1-178776409-INS-874 | chr1-178776409-INS-874 |
| chr1-178810029-INS-66 | chr1-178810029-INS-66 |
| chr1-178828717-INS-81 | chr1-178828717-INS-93 |
| chr1-178851036-INS-205 | chr1-178851036-INS-205 |
| chr1-178862220-INS-58 | chr1-178862220-INS-60 |
| chr1-178867292-INS-54 | chr1-178867263-INS-54 |
| chr1-178871044-INS-93 | chr1-178871044-INS-93 |
| chr1-178916937-INS-74 | chr1-178916937-INS-74 |
| chr1-178947615-INS-2022 | chr1-178947616-INS-2022 |
| chr1-178970434-INS-185 | chr1-178970434-INS-153 |
| chr1-178979630-INS-212 | chr1-178979605-INS-198 |
| chr1-178995803-INS-92 | chr1-178995803-INS-96 |
| chr1-179037197-INS-311 | chr1-179037197-INS-287 |
| chr1-179049488-INS-107 | chr1-179049488-INS-102 |
| chr1-179053799-INS-65 | chr1-179053799-INS-65 |
| chr1-179055689-INS-65 | chr1-179055692-INS-71 |
| chr1-179057124-INS-151 | chr1-179057124-INS-151 |
| chr1-179064054-INS-200 | chr1-179064054-INS-200 |
| chr1-179072297-INS-5445 | chr1-179072298-INS-5445 |
| chr1-179078351-INS-220 | chr1-179078351-INS-222 |
| chr1-179082106-INS-1041 | chr1-179082107-INS-1043 |
| chr1-179084899-INS-100 | chr1-179084909-INS-102 |
| chr1-179087348-INS-60 | chr1-179087348-INS-56 |
| chr1-179101648-INS-142 | chr1-179101647-INS-96 |
| chr1-179104516-INS-75 | chr1-179104516-INS-83 |
| chr1-179113184-INS-194 | chr1-179113184-INS-182 |
| chr1-179133053-INS-79 | chr1-179133053-INS-79 |
| chr1-179141622-INS-417 | chr1-179141622-INS-417 |
| chr1-179142667-INS-52 | chr1-179142651-INS-52 |
| chr1-179155641-INS-119 | chr1-179155645-INS-119 |
| chr1-179173990-INS-60 | chr1-179173990-INS-60 |
| chr1-179181626-INS-365 | chr1-179181626-INS-365 |
| chr1-179201375-INS-112 | chr1-179201417-INS-84 |
| chr1-179210541-INS-537 | chr1-179210541-INS-537 |
| chr1-179231035-INS-877 | chr1-179231035-INS-877 |
| chr1-179236802-INS-369 | chr1-179236799-INS-374 |
| chr1-179300798-INS-218 | chr1-179300798-INS-218 |
| chr1-179324631-INS-112 | chr1-179324631-INS-112 |
| chr1-179330066-INS-9049 | chr1-179330067-INS-9049 |
| chr1-179342639-INS-206 | chr1-179342639-INS-207 |
| chr1-179343895-INS-66 | chr1-179343895-INS-66 |
| chr1-179404661-INS-900 | chr1-179404677-INS-911 |
| chr1-179444119-INS-337 | chr1-179444119-INS-339 |
| chr1-179459235-INS-55 | chr1-179459235-INS-55 |
| chr1-179467792-INS-208 | chr1-179467792-INS-208 |
| chr1-179470799-INS-80 | chr1-179470800-INS-63 |
| chr1-179472570-INS-55 | chr1-179472570-INS-55 |
| chr1-179527285-INS-64 | chr1-179527285-INS-64 |
| chr1-179569814-INS-51 | chr1-179569814-INS-51 |
| chr1-179627507-INS-231 | chr1-179627507-INS-231 |
| chr1-179666175-INS-70 | chr1-179666175-INS-70 |
| chr1-179674398-INS-52 | chr1-179674398-INS-52 |
| chr1-179683573-INS-87 | chr1-179683573-INS-87 |
| chr1-179687364-INS-109 | chr1-179687364-INS-117 |
| chr1-179690063-INS-1895 | chr1-179690064-INS-1895 |

| | |
|---|---|
| chr1-179691659-INS-217 | chr1-179691659-INS-189 |
| chr1-179731126-INS-105 | chr1-179731126-INS-105 |
| chr1-179777586-INS-222 | chr1-179777586-INS-222 |
| chr1-179783528-INS-65 | chr1-179783528-INS-71 |
| chr1-179801300-INS-168 | chr1-179801300-INS-169 |
| chr1-179837669-INS-79 | chr1-179837669-INS-79 |
| chr1-179865697-INS-7101 | chr1-179865698-INS-7102 |
| chr1-179884274-INS-635 | chr1-179884274-INS-631 |
| chr1-179885279-INS-94 | chr1-179885279-INS-94 |
| chr1-179893565-INS-6465 | chr1-179893566-INS-6465 |
| chr1-179909321-INS-900 | chr1-179909321-INS-900 |
| chr1-179923733-INS-199 | chr1-179923733-INS-186 |
| chr1-179948773-INS-564 | chr1-179948774-INS-564 |
| chr1-180003104-INS-794 | chr1-180003104-INS-795 |
| chr1-180023997-INS-55 | chr1-180023997-INS-60 |
| chr1-180030498-INS-440 | chr1-180030498-INS-440 |
| chr1-180037209-INS-60 | chr1-180037209-INS-62 |
| chr1-180054455-INS-371 | chr1-180054456-INS-371 |
| chr1-180056381-INS-120 | chr1-180056381-INS-120 |
| chr1-180063702-INS-70 | chr1-180063702-INS-76 |
| chr1-180086683-INS-56 | chr1-180086683-INS-56 |
| chr1-180100559-INS-168 | chr1-180100559-INS-168 |
| chr1-180113543-INS-72 | chr1-180113543-INS-52 |
| chr1-180131604-INS-196 | chr1-180131604-INS-194 |
| chr1-180138830-INS-88 | chr1-180138830-INS-99 |
| chr1-180143481-INS-314 | chr1-180143481-INS-314 |
| chr1-180168675-INS-397 | chr1-180168675-INS-396 |
| chr1-180243589-INS-66 | chr1-180243589-INS-66 |
| chr1-180254627-INS-55 | chr1-180254627-INS-54 |
| chr1-180270576-INS-201 | chr1-180270588-INS-202 |
| chr1-180283275-INS-86 | chr1-180283275-INS-86 |
| chr1-180294412-INS-229 | chr1-180294412-INS-237 |
| chr1-180319592-INS-3107 | chr1-180319593-INS-3107 |
| chr1-180331935-INS-140 | chr1-180331935-INS-140 |
| chr1-180333236-INS-191 | chr1-180333236-INS-193 |
| chr1-180352289-INS-108 | chr1-180352289-INS-108 |
| chr1-180354092-INS-5544 | chr1-180354093-INS-5544 |
| chr1-180365018-INS-201 | chr1-180365018-INS-201 |
| chr1-180388450-INS-247 | chr1-180388450-INS-247 |
| chr1-180393462-INS-220 | chr1-180393462-INS-220 |
| chr1-180397763-INS-208 | chr1-180397763-INS-207 |
| chr1-180435669-INS-248 | chr1-180435669-INS-255 |
| chr1-180438999-INS-74 | chr1-180438999-INS-74 |
| chr1-180445021-INS-195 | chr1-180445021-INS-205 |
| chr1-180461456-INS-70 | chr1-180461456-INS-55 |
| chr1-180467209-INS-206 | chr1-180467209-INS-205 |
| chr1-180492077-INS-200 | chr1-180492077-INS-202 |
| chr1-180499825-INS-74 | chr1-180499787-INS-72 |
| chr1-180521856-INS-538 | chr1-180521856-INS-538 |
| chr1-180538400-INS-892 | chr1-180538400-INS-891 |
| chr1-180543418-INS-102 | chr1-180543414-INS-102 |
| chr1-180543961-INS-213 | chr1-180543961-INS-212 |
| chr1-180547133-INS-392 | chr1-180547133-INS-392 |
| chr1-180548211-INS-232 | chr1-180548212-INS-232 |
| chr1-180560428-INS-100 | chr1-180560459-INS-104 |
| chr1-180571236-INS-209 | chr1-180571236-INS-209 |
| chr1-180586480-INS-233 | chr1-180586480-INS-229 |
| chr1-180608881-INS-7178 | chr1-180608882-INS-7178 |
| chr1-180612038-INS-84 | chr1-180612038-INS-84 |
| chr1-180625178-INS-1501 | chr1-180625179-INS-1503 |
| chr1-180625458-INS-199 | chr1-180625458-INS-196 |
| chr1-180632077-INS-526 | chr1-180632077-INS-526 |
| chr1-180634427-INS-61 | chr1-180634427-INS-61 |
| chr1-180644668-INS-201 | chr1-180644668-INS-208 |
| chr1-180647797-INS-498 | chr1-180647797-INS-498 |
| chr1-180699218-INS-86 | chr1-180699218-INS-86 |
| chr1-180723616-INS-105 | chr1-180723658-INS-83 |
| chr1-180731300-INS-9050 | chr1-180731301-INS-9050 |
| chr1-180752989-INS-76 | chr1-180752989-INS-76 |
| chr1-180760406-INS-233 | chr1-180760437-INS-208 |
| chr1-180764395-INS-54 | chr1-180764395-INS-54 |
| chr1-180775121-INS-969 | chr1-180775121-INS-969 |
| chr1-180783115-INS-178 | chr1-180783115-INS-181 |
| chr1-180798317-INS-50 | chr1-180798317-INS-78 |
| chr1-180800567-INS-145 | chr1-180800567-INS-150 |
| chr1-180827929-INS-108 | chr1-180827929-INS-108 |
| chr1-180836841-INS-78 | chr1-180836841-INS-80 |

| | |
|---|---|
| chr1-180841257-INS-243 | chr1-180841257-INS-244 |
| chr1-180849434-INS-1515 | chr1-180849435-INS-1519 |
| chr1-180873744-INS-54 | chr1-180873744-INS-54 |
| chr1-180899048-INS-189 | chr1-180899048-INS-189 |
| chr1-180908823-INS-202 | chr1-180908827-INS-204 |
| chr1-180937882-INS-213 | chr1-180937882-INS-203 |
| chr1-180942265-INS-883 | chr1-180942265-INS-883 |
| chr1-180943685-INS-242 | chr1-180943685-INS-242 |
| chr1-180948578-INS-206 | chr1-180948578-INS-206 |
| chr1-180958266-INS-52 | chr1-180958266-INS-52 |
| chr1-180965480-INS-1080 | chr1-180965481-INS-1080 |
| chr1-180974069-INS-54 | chr1-180974069-INS-54 |
| chr1-180994765-INS-220 | chr1-180994765-INS-221 |
| chr1-181009276-INS-288 | chr1-181009276-INS-288 |
| chr1-181010126-INS-477 | chr1-181010127-INS-477 |
| chr1-181011666-INS-228 | chr1-181011666-INS-228 |
| chr1-181013461-INS-203 | chr1-181013461-INS-204 |
| chr1-181022276-INS-226 | chr1-181022276-INS-226 |
| chr1-181023639-INS-213 | chr1-181023639-INS-214 |
| chr1-181027464-INS-59 | chr1-181027464-INS-59 |
| chr1-181037673-INS-52 | chr1-181037673-INS-66 |
| chr1-181041262-INS-249 | chr1-181041262-INS-251 |
| chr1-181049138-INS-64 | chr1-181049138-INS-54 |
| chr1-181063412-INS-125 | chr1-181063412-INS-126 |
| chr1-181088991-INS-61 | chr1-181088991-INS-50 |
| chr1-181106038-INS-65 | chr1-181106004-INS-80 |
| chr1-181107318-INS-4656 | chr1-181107319-INS-4657 |
| chr1-181144196-INS-431 | chr1-181144196-INS-431 |
| chr1-181197717-INS-64 | chr1-181197718-INS-64 |
| chr1-181204707-INS-80 | chr1-181204674-INS-72 |
| chr1-181266601-INS-67 | chr1-181266601-INS-54 |
| chr1-181299712-INS-69 | chr1-181299686-INS-52 |
| chr1-181309387-INS-411 | chr1-181309387-INS-411 |
| chr1-181327199-INS-60 | chr1-181327200-INS-62 |
| chr1-181329515-INS-353 | chr1-181329515-INS-353 |
| chr1-181350927-INS-135 | chr1-181350927-INS-130 |
| chr1-181353106-INS-64 | chr1-181353106-INS-83 |
| chr1-181355494-INS-146 | chr1-181355494-INS-150 |
| chr1-181377996-INS-68 | chr1-181377997-INS-76 |
| chr1-181384543-INS-6796 | chr1-181384544-INS-6796 |
| chr1-181398036-INS-151 | chr1-181398036-INS-151 |
| chr1-181440128-INS-68 | chr1-181440152-INS-61 |
| chr1-181446652-INS-60 | chr1-181446652-INS-62 |
| chr1-181479461-INS-214 | chr1-181479461-INS-217 |
| chr1-181540612-INS-105 | chr1-181540612-INS-90 |
| chr1-181554787-INS-201 | chr1-181554787-INS-201 |
| chr1-181583326-INS-808 | chr1-181583326-INS-809 |
| chr1-181613729-INS-6342 | chr1-181613730-INS-6342 |
| chr1-181627042-INS-1893 | chr1-181627043-INS-1893 |
| chr1-181632663-INS-126 | chr1-181632663-INS-84 |
| chr1-181641885-INS-211 | chr1-181641885-INS-211 |
| chr1-181661537-INS-552 | chr1-181661537-INS-553 |
| chr1-181662415-INS-210 | chr1-181662415-INS-210 |
| chr1-181677192-INS-719 | chr1-181677192-INS-719 |
| chr1-181693725-INS-187 | chr1-181693725-INS-189 |
| chr1-181703750-INS-200 | chr1-181703750-INS-200 |
| chr1-181710299-INS-203 | chr1-181710299-INS-203 |
| chr1-181720869-INS-210 | chr1-181720869-INS-210 |
| chr1-181737894-INS-117 | chr1-181737894-INS-149 |
| chr1-181785199-INS-362 | chr1-181785199-INS-362 |
| chr1-181800349-INS-171 | chr1-181800349-INS-172 |
| chr1-181809945-INS-162 | chr1-181809945-INS-162 |
| chr1-181824172-INS-189 | chr1-181824174-INS-181 |
| chr1-181861294-INS-55 | chr1-181861294-INS-55 |
| chr1-181898701-INS-399 | chr1-181898701-INS-399 |
| chr1-181900593-INS-53 | chr1-181900593-INS-53 |
| chr1-181903304-INS-56 | chr1-181903304-INS-56 |
| chr1-181904388-INS-90 | chr1-181904388-INS-51 |
| chr1-181911795-INS-54 | chr1-181911795-INS-70 |
| chr1-181934542-INS-71 | chr1-181934542-INS-79 |
| chr1-181935927-INS-54 | chr1-181935927-INS-66 |
| chr1-181937731-INS-252 | chr1-181937731-INS-252 |
| chr1-181939442-INS-5495 | chr1-181939443-INS-5496 |
| chr1-181953960-INS-614 | chr1-181953965-INS-572 |
| chr1-181961903-INS-159 | chr1-181961904-INS-161 |
| chr1-181963432-INS-53 | chr1-181963422-INS-55 |
| chr1-181973075-INS-84 | chr1-181973060-INS-60 |

| | |
|---|---|
| chr1-181980324-INS-229 | chr1-181980324-INS-227 |
| chr1-181990170-INS-56 | chr1-181990170-INS-56 |
| chr1-181991654-INS-218 | chr1-181991654-INS-218 |
| chr1-181993726-INS-70 | chr1-181993740-INS-71 |
| chr1-181994673-INS-121 | chr1-181994673-INS-141 |
| chr1-181998467-INS-7621 | chr1-181998468-INS-7621 |
| chr1-181999801-INS-5507 | chr1-181999802-INS-5544 |
| chr1-182018248-INS-204 | chr1-182018248-INS-204 |
| chr1-182040327-INS-203 | chr1-182040327-INS-203 |
| chr1-182080723-INS-207 | chr1-182080723-INS-207 |
| chr1-182114218-INS-486 | chr1-182114218-INS-486 |
| chr1-182117668-INS-81 | chr1-182117660-INS-62 |
| chr1-182169824-INS-52 | chr1-182169824-INS-52 |
| chr1-182173073-INS-364 | chr1-182173073-INS-364 |
| chr1-182206113-INS-185 | chr1-182206113-INS-185 |
| chr1-182216483-INS-5441 | chr1-182216484-INS-5441 |
| chr1-182228570-INS-1901 | chr1-182228569-INS-1930 |
| chr1-182245883-INS-102 | chr1-182245883-INS-102 |
| chr1-182250889-INS-72 | chr1-182250886-INS-79 |
| chr1-182313529-INS-82 | chr1-182313537-INS-64 |
| chr1-182356949-INS-111 | chr1-182356949-INS-120 |
| chr1-182396946-INS-227 | chr1-182396946-INS-227 |
| chr1-182428980-INS-6401 | chr1-182428981-INS-6401 |
| chr1-182429229-INS-150 | chr1-182429229-INS-150 |
| chr1-182471959-INS-53 | chr1-182471959-INS-53 |
| chr1-182474138-INS-108 | chr1-182474138-INS-108 |
| chr1-182475568-INS-537 | chr1-182475568-INS-537 |
| chr1-182507092-INS-6937 | chr1-182507093-INS-6937 |
| chr1-182516525-INS-495 | chr1-182516525-INS-495 |
| chr1-182543828-INS-83 | chr1-182543827-INS-51 |
| chr1-182549650-INS-108 | chr1-182549654-INS-112 |
| chr1-182550679-INS-137 | chr1-182550679-INS-137 |
| chr1-182554756-INS-74 | chr1-182554759-INS-84 |
| chr1-182562773-INS-671 | chr1-182562773-INS-672 |
| chr1-182583088-INS-143 | chr1-182583122-INS-143 |
| chr1-182586957-INS-94 | chr1-182586954-INS-112 |
| chr1-182603471-INS-184 | chr1-182603493-INS-166 |
| chr1-182612681-INS-202 | chr1-182612681-INS-199 |
| chr1-182627820-INS-69 | chr1-182627820-INS-69 |
| chr1-182642431-INS-60 | chr1-182642431-INS-60 |
| chr1-182656034-INS-1888 | chr1-182656035-INS-1888 |
| chr1-182673155-INS-7057 | chr1-182673156-INS-7057 |
| chr1-182693641-INS-59 | chr1-182693641-INS-59 |
| chr1-182733202-INS-110 | chr1-182733202-INS-110 |
| chr1-182746595-INS-1341 | chr1-182746596-INS-1342 |
| chr1-182749329-INS-400 | chr1-182749329-INS-400 |
| chr1-182752462-INS-74 | chr1-182752462-INS-71 |
| chr1-182757274-INS-81 | chr1-182757274-INS-105 |
| chr1-182763418-INS-120 | chr1-182763428-INS-156 |
| chr1-182771788-INS-283 | chr1-182771788-INS-284 |
| chr1-182775012-INS-60 | chr1-182775012-INS-60 |
| chr1-182791650-INS-281 | chr1-182791656-INS-301 |
| chr1-182804859-INS-54 | chr1-182804859-INS-51 |
| chr1-182810102-INS-89 | chr1-182810102-INS-89 |
| chr1-182838588-INS-166 | chr1-182838588-INS-167 |
| chr1-182857376-INS-52 | chr1-182857389-INS-52 |
| chr1-182876795-INS-485 | chr1-182876795-INS-485 |
| chr1-182878053-INS-62 | chr1-182878053-INS-64 |
| chr1-182885452-INS-61 | chr1-182885452-INS-61 |
| chr1-182887835-INS-75 | chr1-182887835-INS-75 |
| chr1-182889192-INS-54 | chr1-182889192-INS-52 |
| chr1-182893578-INS-62 | chr1-182893578-INS-62 |
| chr1-182899183-INS-63 | chr1-182899184-INS-75 |
| chr1-182927237-INS-66 | chr1-182927237-INS-66 |
| chr1-182931148-INS-653 | chr1-182931148-INS-676 |
| chr1-182942867-INS-1888 | chr1-182942868-INS-1888 |
| chr1-182943790-INS-2240 | chr1-182943791-INS-2240 |
| chr1-182977504-INS-115 | chr1-182977507-INS-115 |
| chr1-183009215-INS-6893 | chr1-183009216-INS-6896 |
| chr1-183030148-INS-136 | chr1-183030150-INS-145 |
| chr1-183048271-INS-120 | chr1-183048271-INS-120 |
| chr1-183051525-INS-463 | chr1-183051525-INS-463 |
| chr1-183054726-INS-180 | chr1-183054726-INS-180 |
| chr1-183071995-INS-109 | chr1-183072006-INS-64 |
| chr1-183074472-INS-61 | chr1-183074472-INS-61 |
| chr1-183113658-INS-176 | chr1-183113658-INS-176 |
| chr1-183141413-INS-53 | chr1-183141413-INS-53 |

| | |
|---|---|
| chr1-183143230-INS-116 | chr1-183143230-INS-116 |
| chr1-183154925-INS-215 | chr1-183154925-INS-215 |
| chr1-183158417-INS-68 | chr1-183158417-INS-68 |
| chr1-183165409-INS-391 | chr1-183165409-INS-390 |
| chr1-183168206-INS-213 | chr1-183168206-INS-213 |
| chr1-183201677-INS-243 | chr1-183201664-INS-245 |
| chr1-183205075-INS-217 | chr1-183205075-INS-217 |
| chr1-183210118-INS-52 | chr1-183210118-INS-80 |
| chr1-183222272-INS-62 | chr1-183222272-INS-100 |
| chr1-183245945-INS-57 | chr1-183245945-INS-72 |
| chr1-183249230-INS-62 | chr1-183249220-INS-80 |
| chr1-183279666-INS-60 | chr1-183279666-INS-60 |
| chr1-183319786-INS-237 | chr1-183319787-INS-237 |
| chr1-183333433-INS-397 | chr1-183333433-INS-397 |
| chr1-183336059-INS-160 | chr1-183336059-INS-156 |
| chr1-183346400-INS-110 | chr1-183346403-INS-101 |
| chr1-183347455-INS-50 | chr1-183347455-INS-50 |
| chr1-183348974-INS-232 | chr1-183348974-INS-232 |
| chr1-183363341-INS-109 | chr1-183363341-INS-111 |
| chr1-183394243-INS-7159 | chr1-183394244-INS-7159 |
| chr1-183400542-INS-1888 | chr1-183400543-INS-1888 |
| chr1-183407863-INS-235 | chr1-183407863-INS-248 |
| chr1-183410411-INS-54 | chr1-183410411-INS-54 |
| chr1-183432231-INS-58 | chr1-183432231-INS-58 |
| chr1-183434281-INS-112 | chr1-183434281-INS-112 |
| chr1-183451173-INS-60 | chr1-183451173-INS-56 |
| chr1-183454118-INS-935 | chr1-183454118-INS-937 |
| chr1-183473152-INS-1141 | chr1-183473153-INS-1141 |
| chr1-183478302-INS-99 | chr1-183478302-INS-83 |
| chr1-183480347-INS-440 | chr1-183480347-INS-440 |
| chr1-183508173-INS-164 | chr1-183508173-INS-188 |
| chr1-183518473-INS-68 | chr1-183518473-INS-70 |
| chr1-183521510-INS-66 | chr1-183521510-INS-70 |
| chr1-183542983-INS-4780 | chr1-183542984-INS-4780 |
| chr1-183550154-INS-7059 | chr1-183550155-INS-7059 |
| chr1-183555921-INS-80 | chr1-183555921-INS-76 |
| chr1-183564385-INS-1904 | chr1-183564364-INS-1905 |
| chr1-183565169-INS-60 | chr1-183565169-INS-75 |
| chr1-183570781-INS-152 | chr1-183570781-INS-152 |
| chr1-183572426-INS-55 | chr1-183572426-INS-59 |
| chr1-183588620-INS-105 | chr1-183588620-INS-111 |
| chr1-183589221-INS-1794 | chr1-183589222-INS-1795 |
| chr1-183590421-INS-67 | chr1-183590421-INS-73 |
| chr1-183596487-INS-1435 | chr1-183596488-INS-1435 |
| chr1-183599156-INS-2866 | chr1-183599157-INS-2867 |
| chr1-183605158-INS-440 | chr1-183605158-INS-440 |
| chr1-183636813-INS-248 | chr1-183636813-INS-248 |
| chr1-183657002-INS-91 | chr1-183657040-INS-90 |
| chr1-183685853-INS-507 | chr1-183685853-INS-507 |
| chr1-183739040-INS-956 | chr1-183739040-INS-965 |
| chr1-183745027-INS-82 | chr1-183745027-INS-82 |
| chr1-183745739-INS-533 | chr1-183745736-INS-527 |
| chr1-183751291-INS-176 | chr1-183751291-INS-177 |
| chr1-183752569-INS-52 | chr1-183752569-INS-50 |
| chr1-183877631-INS-87 | chr1-183877631-INS-87 |
| chr1-183880695-INS-283 | chr1-183880695-INS-283 |
| chr1-183962166-INS-55 | chr1-183962166-INS-100 |
| chr1-183992191-INS-136 | chr1-183992190-INS-141 |
| chr1-184017526-INS-55 | chr1-184017487-INS-71 |
| chr1-184046629-INS-6626 | chr1-184046630-INS-6627 |
| chr1-184147670-INS-1895 | chr1-184147671-INS-1895 |
| chr1-184148963-INS-4811 | chr1-184148964-INS-4811 |
| chr1-184150807-INS-1534 | chr1-184150808-INS-1535 |
| chr1-184158754-INS-67 | chr1-184158754-INS-67 |
| chr1-184164523-INS-61 | chr1-184164523-INS-64 |
| chr1-184190134-INS-52 | chr1-184190134-INS-80 |
| chr1-184194002-INS-58 | chr1-184194028-INS-56 |
| chr1-184202680-INS-80 | chr1-184202680-INS-83 |
| chr1-184261821-INS-52 | chr1-184261821-INS-50 |
| chr1-184264306-INS-111 | chr1-184264303-INS-103 |
| chr1-184279406-INS-153 | chr1-184279406-INS-154 |
| chr1-184280932-INS-72 | chr1-184280932-INS-72 |
| chr1-184281736-INS-1094 | chr1-184281737-INS-1094 |
| chr1-184356967-INS-69 | chr1-184356967-INS-67 |
| chr1-184361505-INS-62 | chr1-184361505-INS-62 |
| chr1-184381078-INS-81 | chr1-184381078-INS-81 |
| chr1-184413385-INS-201 | chr1-184413385-INS-201 |

| | |
|---|---|
| chr1-184418238-INS-68 | chr1-184418238-INS-70 |
| chr1-184429222-INS-491 | chr1-184429222-INS-486 |
| chr1-184454879-INS-76 | chr1-184454900-INS-56 |
| chr1-184467089-INS-149 | chr1-184467089-INS-135 |
| chr1-184472327-INS-246 | chr1-184472327-INS-254 |
| chr1-184483445-INS-52 | chr1-184483441-INS-82 |
| chr1-184521450-INS-59 | chr1-184521450-INS-67 |
| chr1-184529077-INS-53 | chr1-184529079-INS-82 |
| chr1-184533130-INS-405 | chr1-184533130-INS-405 |
| chr1-184543548-INS-52 | chr1-184543548-INS-52 |
| chr1-184589403-INS-95 | chr1-184589403-INS-95 |
| chr1-184599783-INS-178 | chr1-184599783-INS-178 |
| chr1-184617015-INS-81 | chr1-184617010-INS-96 |
| chr1-184673142-INS-195 | chr1-184673143-INS-196 |
| chr1-184676604-INS-547 | chr1-184676604-INS-547 |
| chr1-184761024-INS-382 | chr1-184761025-INS-367 |
| chr1-184793540-INS-238 | chr1-184793540-INS-238 |
| chr1-184813621-INS-52 | chr1-184813621-INS-58 |
| chr1-184815098-INS-747 | chr1-184815098-INS-747 |
| chr1-184818103-INS-71 | chr1-184818101-INS-70 |
| chr1-184840874-INS-60 | chr1-184840874-INS-60 |
| chr1-184846524-INS-146 | chr1-184846524-INS-186 |
| chr1-184849068-INS-531 | chr1-184849068-INS-531 |
| chr1-184898991-INS-91 | chr1-184899002-INS-100 |
| chr1-184909951-INS-98 | chr1-184909951-INS-98 |
| chr1-184928491-INS-100 | chr1-184928491-INS-100 |
| chr1-184949255-INS-927 | chr1-184949255-INS-927 |
| chr1-184982486-INS-51 | chr1-184982486-INS-51 |
| chr1-184996150-INS-614 | chr1-184996150-INS-614 |
| chr1-185010224-INS-64 | chr1-185010224-INS-64 |
| chr1-185020695-INS-206 | chr1-185020695-INS-206 |
| chr1-185058476-INS-121 | chr1-185058476-INS-121 |
| chr1-185059460-INS-90 | chr1-185059460-INS-90 |
| chr1-185069137-INS-53 | chr1-185069137-INS-53 |
| chr1-185073116-INS-198 | chr1-185073116-INS-199 |
| chr1-185078466-INS-57 | chr1-185078485-INS-61 |
| chr1-185091326-INS-56 | chr1-185091326-INS-56 |
| chr1-185100870-INS-56 | chr1-185100863-INS-72 |
| chr1-185108041-INS-124 | chr1-185108041-INS-124 |
| chr1-185116002-INS-68 | chr1-185116002-INS-72 |
| chr1-185124854-INS-253 | chr1-185124838-INS-256 |
| chr1-185134690-INS-134 | chr1-185134690-INS-128 |
| chr1-185137590-INS-105 | chr1-185137590-INS-108 |
| chr1-185151702-INS-66 | chr1-185151702-INS-58 |
| chr1-185158386-INS-260 | chr1-185158386-INS-260 |
| chr1-185161486-INS-751 | chr1-185161486-INS-751 |
| chr1-185167594-INS-149 | chr1-185167594-INS-136 |
| chr1-185174582-INS-307 | chr1-185174582-INS-307 |
| chr1-185182477-INS-186 | chr1-185182477-INS-186 |
| chr1-185221341-INS-234 | chr1-185221341-INS-233 |
| chr1-185223932-INS-101 | chr1-185223953-INS-105 |
| chr1-185247749-INS-378 | chr1-185247749-INS-378 |
| chr1-185250091-INS-285 | chr1-185250091-INS-285 |
| chr1-185265241-INS-298 | chr1-185265241-INS-248 |
| chr1-185271065-INS-375 | chr1-185271065-INS-374 |
| chr1-185282715-INS-60 | chr1-185282715-INS-60 |
| chr1-185284082-INS-51 | chr1-185284092-INS-66 |
| chr1-185287922-INS-76 | chr1-185287922-INS-76 |
| chr1-185335775-INS-352 | chr1-185335775-INS-353 |
| chr1-185361925-INS-993 | chr1-185361925-INS-992 |
| chr1-185374327-INS-140 | chr1-185374327-INS-124 |
| chr1-185396417-INS-247 | chr1-185396417-INS-255 |
| chr1-185436726-INS-151 | chr1-185436726-INS-171 |
| chr1-185444871-INS-80 | chr1-185444871-INS-85 |
| chr1-185472305-INS-1058 | chr1-185472306-INS-1058 |
| chr1-185482708-INS-982 | chr1-185482708-INS-987 |
| chr1-185490847-INS-7771 | chr1-185490848-INS-7773 |
| chr1-185492148-INS-6278 | chr1-185492149-INS-6277 |
| chr1-185496130-INS-73 | chr1-185496130-INS-73 |
| chr1-185507651-INS-56 | chr1-185507651-INS-56 |
| chr1-185529208-INS-77 | chr1-185529208-INS-77 |
| chr1-185534935-INS-54 | chr1-185534935-INS-60 |
| chr1-185536351-INS-53 | chr1-185536351-INS-53 |
| chr1-185546009-INS-58 | chr1-185546009-INS-58 |
| chr1-185561944-INS-74 | chr1-185561944-INS-74 |
| chr1-185587656-INS-1253 | chr1-185587657-INS-1255 |
| chr1-185592933-INS-55 | chr1-185592933-INS-62 |

| | |
|---|---|
| chr1-185612978-INS-60 | chr1-185612978-INS-60 |
| chr1-185654239-INS-60 | chr1-185654239-INS-60 |
| chr1-185685741-INS-66 | chr1-185685741-INS-68 |
| chr1-185706097-INS-144 | chr1-185706097-INS-144 |
| chr1-185707777-INS-78 | chr1-185707777-INS-94 |
| chr1-185723801-INS-101 | chr1-185723821-INS-51 |
| chr1-185788987-INS-78 | chr1-185788987-INS-78 |
| chr1-185808564-INS-101 | chr1-185808564-INS-84 |
| chr1-185816303-INS-500 | chr1-185816303-INS-522 |
| chr1-185819340-INS-344 | chr1-185819340-INS-343 |
| chr1-185821092-INS-298 | chr1-185821092-INS-298 |
| chr1-185842492-INS-212 | chr1-185842492-INS-212 |
| chr1-185843238-INS-162 | chr1-185843238-INS-161 |
| chr1-185845778-INS-52 | chr1-185845778-INS-60 |
| chr1-185856252-INS-60 | chr1-185856252-INS-60 |
| chr1-185860106-INS-6572 | chr1-185860107-INS-6572 |
| chr1-185863622-INS-73 | chr1-185863643-INS-81 |
| chr1-185873970-INS-507 | chr1-185873970-INS-507 |
| chr1-185876571-INS-167 | chr1-185876571-INS-168 |
| chr1-185903445-INS-512 | chr1-185903460-INS-513 |
| chr1-185937035-INS-59 | chr1-185937035-INS-59 |
| chr1-185945417-INS-497 | chr1-185945417-INS-497 |
| chr1-185975731-INS-6576 | chr1-185975732-INS-6576 |
| chr1-185983806-INS-2780 | chr1-185983807-INS-2780 |
| chr1-186000678-INS-59 | chr1-186000678-INS-59 |
| chr1-186013967-INS-108 | chr1-186013967-INS-108 |
| chr1-186018407-INS-8274 | chr1-186018408-INS-8275 |
| chr1-186023904-INS-838 | chr1-186023904-INS-838 |
| chr1-186053188-INS-954 | chr1-186053188-INS-955 |
| chr1-186070586-INS-111 | chr1-186070586-INS-111 |
| chr1-186072270-INS-54 | chr1-186072268-INS-56 |
| chr1-186079078-INS-496 | chr1-186079078-INS-496 |
| chr1-186102099-INS-71 | chr1-186102099-INS-60 |
| chr1-186108813-INS-542 | chr1-186108814-INS-541 |
| chr1-186118554-INS-87 | chr1-186118554-INS-108 |
| chr1-186128439-INS-64 | chr1-186128433-INS-78 |
| chr1-186138691-INS-65 | chr1-186138691-INS-65 |
| chr1-186145157-INS-143 | chr1-186145155-INS-150 |
| chr1-186150310-INS-66 | chr1-186150310-INS-66 |
| chr1-186150973-INS-402 | chr1-186150973-INS-402 |
| chr1-186152876-INS-520 | chr1-186152877-INS-520 |
| chr1-186158300-INS-269 | chr1-186158300-INS-275 |
| chr1-186180860-INS-86 | chr1-186180860-INS-86 |
| chr1-186183233-INS-52 | chr1-186183233-INS-52 |
| chr1-186186454-INS-95 | chr1-186186454-INS-94 |
| chr1-186190021-INS-159 | chr1-186190021-INS-157 |
| chr1-186204652-INS-199 | chr1-186204652-INS-198 |
| chr1-186258309-INS-76 | chr1-186258309-INS-76 |
| chr1-186274158-INS-7370 | chr1-186274159-INS-7370 |
| chr1-186277708-INS-76 | chr1-186277708-INS-82 |
| chr1-186289555-INS-6574 | chr1-186289556-INS-6574 |
| chr1-186308589-INS-160 | chr1-186308589-INS-143 |
| chr1-186387050-INS-52 | chr1-186387050-INS-56 |
| chr1-186419998-INS-52 | chr1-186419998-INS-52 |
| chr1-186429338-INS-323 | chr1-186429338-INS-323 |
| chr1-186461811-INS-192 | chr1-186461812-INS-194 |
| chr1-186472548-INS-59 | chr1-186472548-INS-59 |
| chr1-186480105-INS-80 | chr1-186480106-INS-80 |
| chr1-186489663-INS-54 | chr1-186489663-INS-57 |
| chr1-186495976-INS-65 | chr1-186495976-INS-65 |
| chr1-186508690-INS-858 | chr1-186508690-INS-860 |
| chr1-186528714-INS-873 | chr1-186528714-INS-873 |
| chr1-186573271-INS-70 | chr1-186573259-INS-108 |
| chr1-186636375-INS-70 | chr1-186636375-INS-70 |
| chr1-186656996-INS-80 | chr1-186657012-INS-54 |
| chr1-186680431-INS-268 | chr1-186680431-INS-268 |
| chr1-186772127-INS-995 | chr1-186772127-INS-996 |
| chr1-186816307-INS-550 | chr1-186816307-INS-550 |
| chr1-186823438-INS-578 | chr1-186823438-INS-578 |
| chr1-186826982-INS-843 | chr1-186826982-INS-844 |
| chr1-186831375-INS-58 | chr1-186831372-INS-61 |
| chr1-186891271-INS-93 | chr1-186891271-INS-93 |
| chr1-186904027-INS-66 | chr1-186904028-INS-106 |
| chr1-186923723-INS-68 | chr1-186923723-INS-68 |
| chr1-186936320-INS-70 | chr1-186936330-INS-81 |
| chr1-186942444-INS-233 | chr1-186942444-INS-233 |
| chr1-186955392-INS-77 | chr1-186955384-INS-77 |

| | |
|---|---|
| chr1-186982019-INS-62 | chr1-186982019-INS-66 |
| chr1-186989763-INS-54 | chr1-186989763-INS-54 |
| chr1-186996005-INS-51 | chr1-186996005-INS-51 |
| chr1-187012333-INS-96 | chr1-187012333-INS-96 |
| chr1-187017132-INS-84 | chr1-187017132-INS-84 |
| chr1-187018148-INS-86 | chr1-187018148-INS-86 |
| chr1-187063152-INS-114 | chr1-187063152-INS-114 |
| chr1-187119432-INS-95 | chr1-187119454-INS-86 |
| chr1-187157953-INS-305 | chr1-187157953-INS-305 |
| chr1-187189900-INS-56 | chr1-187189900-INS-56 |
| chr1-187195283-INS-168 | chr1-187195281-INS-180 |
| chr1-187212639-INS-205 | chr1-187212639-INS-205 |
| chr1-187279244-INS-62 | chr1-187279244-INS-62 |
| chr1-187286368-INS-117 | chr1-187286368-INS-117 |
| chr1-187310867-INS-110 | chr1-187310867-INS-110 |
| chr1-187313723-INS-162 | chr1-187313723-INS-171 |
| chr1-187335990-INS-76 | chr1-187335988-INS-80 |
| chr1-187357703-INS-85 | chr1-187357703-INS-85 |
| chr1-187383050-INS-83 | chr1-187383050-INS-83 |
| chr1-187411758-INS-1982 | chr1-187411759-INS-1975 |
| chr1-187467039-INS-722 | chr1-187467037-INS-725 |
| chr1-187469590-INS-54 | chr1-187469590-INS-54 |
| chr1-187506076-INS-82 | chr1-187506103-INS-82 |
| chr1-187538627-INS-56 | chr1-187538627-INS-50 |
| chr1-187547189-INS-72 | chr1-187547189-INS-54 |
| chr1-187550598-INS-389 | chr1-187550598-INS-389 |
| chr1-187576717-INS-95 | chr1-187576717-INS-56 |
| chr1-187606241-INS-117 | chr1-187606219-INS-117 |
| chr1-187622982-INS-72 | chr1-187622983-INS-55 |
| chr1-187635118-INS-86 | chr1-187635111-INS-121 |
| chr1-187662820-INS-224 | chr1-187662820-INS-224 |
| chr1-187672877-INS-113 | chr1-187672877-INS-120 |
| chr1-187679225-INS-61 | chr1-187679225-INS-61 |
| chr1-187687873-INS-61 | chr1-187687873-INS-61 |
| chr1-187723438-INS-78 | chr1-187723438-INS-81 |
| chr1-187758466-INS-272 | chr1-187758466-INS-273 |
| chr1-187805254-INS-68 | chr1-187805254-INS-66 |
| chr1-187822228-INS-88 | chr1-187822241-INS-56 |
| chr1-187826655-INS-111 | chr1-187826666-INS-77 |
| chr1-187864050-INS-58 | chr1-187864050-INS-79 |
| chr1-187876308-INS-52 | chr1-187876308-INS-52 |
| chr1-187933485-INS-168 | chr1-187933485-INS-168 |
| chr1-187959221-INS-241 | chr1-187959221-INS-238 |
| chr1-187961711-INS-66 | chr1-187961711-INS-66 |
| chr1-187990112-INS-192 | chr1-187990112-INS-192 |
| chr1-188007644-INS-101 | chr1-188007644-INS-101 |
| chr1-188029983-INS-96 | chr1-188029983-INS-96 |
| chr1-188038862-INS-69 | chr1-188038862-INS-71 |
| chr1-188044989-INS-70 | chr1-188044989-INS-73 |
| chr1-188053681-INS-209 | chr1-188053681-INS-209 |
| chr1-188082084-INS-253 | chr1-188082118-INS-254 |
| chr1-188117088-INS-69 | chr1-188117089-INS-68 |
| chr1-188122813-INS-55 | chr1-188122813-INS-55 |
| chr1-188165424-INS-64 | chr1-188165425-INS-65 |
| chr1-188195609-INS-51 | chr1-188195609-INS-51 |
| chr1-188249672-INS-6334 | chr1-188249673-INS-6334 |
| chr1-188292796-INS-691 | chr1-188292796-INS-692 |
| chr1-188309301-INS-101 | chr1-188309323-INS-72 |
| chr1-188320977-INS-256 | chr1-188320977-INS-228 |
| chr1-188355287-INS-53 | chr1-188355287-INS-53 |
| chr1-188368208-INS-401 | chr1-188368208-INS-401 |
| chr1-188378257-INS-740 | chr1-188378257-INS-740 |
| chr1-188418690-INS-208 | chr1-188418690-INS-208 |
| chr1-188419176-INS-1360 | chr1-188419177-INS-1361 |
| chr1-188429007-INS-3249 | chr1-188429008-INS-3249 |
| chr1-188436685-INS-71 | chr1-188436685-INS-60 |
| chr1-188447776-INS-209 | chr1-188447776-INS-209 |
| chr1-188453593-INS-58 | chr1-188453593-INS-58 |
| chr1-188498022-INS-504 | chr1-188498022-INS-504 |
| chr1-188499807-INS-7309 | chr1-188499808-INS-7309 |
| chr1-188506399-INS-81 | chr1-188506399-INS-78 |
| chr1-188527768-INS-69 | chr1-188527768-INS-69 |
| chr1-188535140-INS-124 | chr1-188535140-INS-124 |
| chr1-188568927-INS-141 | chr1-188568927-INS-141 |
| chr1-188591967-INS-258 | chr1-188591967-INS-301 |
| chr1-188619870-INS-772 | chr1-188619870-INS-739 |
| chr1-188621037-INS-5219 | chr1-188621038-INS-5220 |

| | |
|---|---|
| chr1-188649910-INS-215 | chr1-188649910-INS-215 |
| chr1-188664058-INS-401 | chr1-188664058-INS-401 |
| chr1-188672338-INS-66 | chr1-188672338-INS-66 |
| chr1-188686233-INS-55 | chr1-188686242-INS-61 |
| chr1-188714537-INS-76 | chr1-188714537-INS-76 |
| chr1-188723885-INS-240 | chr1-188723885-INS-240 |
| chr1-188732073-INS-93 | chr1-188732073-INS-113 |
| chr1-188742087-INS-210 | chr1-188742087-INS-210 |
| chr1-188761923-INS-438 | chr1-188761923-INS-438 |
| chr1-188765063-INS-76 | chr1-188765063-INS-76 |
| chr1-188792219-INS-399 | chr1-188792219-INS-399 |
| chr1-188802416-INS-73 | chr1-188802416-INS-74 |
| chr1-188817530-INS-372 | chr1-188817530-INS-326 |
| chr1-188835571-INS-80 | chr1-188835571-INS-81 |
| chr1-188837039-INS-682 | chr1-188837039-INS-682 |
| chr1-188840520-INS-63 | chr1-188840520-INS-50 |
| chr1-188884237-INS-932 | chr1-188884238-INS-931 |
| chr1-188901440-INS-72 | chr1-188901440-INS-72 |
| chr1-188911454-INS-293 | chr1-188911458-INS-288 |
| chr1-188922350-INS-1889 | chr1-188922351-INS-1889 |
| chr1-188942882-INS-80 | chr1-188942882-INS-88 |
| chr1-188975007-INS-6814 | chr1-188975008-INS-6814 |
| chr1-188981433-INS-121 | chr1-188981433-INS-121 |
| chr1-188985755-INS-210 | chr1-188985754-INS-209 |
| chr1-189032883-INS-66 | chr1-189032852-INS-63 |
| chr1-189033435-INS-76 | chr1-189033435-INS-51 |
| chr1-189049541-INS-148 | chr1-189049500-INS-156 |
| chr1-189059016-INS-134 | chr1-189059016-INS-132 |
| chr1-189074354-INS-50 | chr1-189074354-INS-54 |
| chr1-189079510-INS-114 | chr1-189079510-INS-114 |
| chr1-189107783-INS-1238 | chr1-189107784-INS-1235 |
| chr1-189122693-INS-64 | chr1-189122693-INS-64 |
| chr1-189130682-INS-185 | chr1-189130682-INS-193 |
| chr1-189199468-INS-335 | chr1-189199468-INS-334 |
| chr1-189228417-INS-55 | chr1-189228417-INS-55 |
| chr1-189243819-INS-219 | chr1-189243820-INS-219 |
| chr1-189280630-INS-217 | chr1-189280630-INS-219 |
| chr1-189289878-INS-78 | chr1-189289878-INS-78 |
| chr1-189296262-INS-112 | chr1-189296262-INS-113 |
| chr1-189299114-INS-55 | chr1-189299114-INS-55 |
| chr1-189311534-INS-50 | chr1-189311534-INS-51 |
| chr1-189342258-INS-72 | chr1-189342254-INS-80 |
| chr1-189357385-INS-203 | chr1-189357385-INS-204 |
| chr1-189365933-INS-144 | chr1-189365933-INS-136 |
| chr1-189382273-INS-176 | chr1-189382265-INS-193 |
| chr1-189388069-INS-198 | chr1-189388069-INS-198 |
| chr1-189429983-INS-515 | chr1-189429983-INS-515 |
| chr1-189431220-INS-76 | chr1-189431184-INS-92 |
| chr1-189433955-INS-202 | chr1-189433955-INS-202 |
| chr1-189444428-INS-223 | chr1-189444428-INS-224 |
| chr1-189450034-INS-121 | chr1-189450034-INS-121 |
| chr1-189450708-INS-207 | chr1-189450708-INS-207 |
| chr1-189452612-INS-194 | chr1-189452628-INS-194 |
| chr1-189478745-INS-100 | chr1-189478766-INS-121 |
| chr1-189479886-INS-207 | chr1-189479886-INS-207 |
| chr1-189484141-INS-100 | chr1-189484142-INS-90 |
| chr1-189485758-INS-60 | chr1-189485759-INS-64 |
| chr1-189511056-INS-56 | chr1-189511056-INS-56 |
| chr1-189527149-INS-3000 | chr1-189527150-INS-3000 |
| chr1-189529026-INS-52 | chr1-189529026-INS-58 |
| chr1-189570212-INS-114 | chr1-189570208-INS-161 |
| chr1-189605316-INS-153 | chr1-189605316-INS-153 |
| chr1-189633918-INS-52 | chr1-189633918-INS-54 |
| chr1-189644220-INS-75 | chr1-189644226-INS-69 |
| chr1-189651358-INS-67 | chr1-189651358-INS-67 |
| chr1-189659739-INS-763 | chr1-189659739-INS-763 |
| chr1-189662805-INS-66 | chr1-189662805-INS-66 |
| chr1-189686101-INS-1506 | chr1-189686102-INS-1506 |
| chr1-189693562-INS-173 | chr1-189693589-INS-172 |
| chr1-189698491-INS-116 | chr1-189698491-INS-115 |
| chr1-189701640-INS-120 | chr1-189701640-INS-156 |
| chr1-189744599-INS-68 | chr1-189744599-INS-68 |
| chr1-189748514-INS-59 | chr1-189748514-INS-53 |
| chr1-189761084-INS-562 | chr1-189761084-INS-562 |
| chr1-189792564-INS-82 | chr1-189792564-INS-82 |
| chr1-189804890-INS-76 | chr1-189804890-INS-74 |
| chr1-189830485-INS-61 | chr1-189830485-INS-61 |

| | |
|---|---|
| chr1-189834035-INS-60 | chr1-189834035-INS-60 |
| chr1-189838050-INS-59 | chr1-189838044-INS-62 |
| chr1-189839763-INS-52 | chr1-189839763-INS-52 |
| chr1-189863491-INS-157 | chr1-189863495-INS-149 |
| chr1-189906805-INS-52 | chr1-189906805-INS-76 |
| chr1-189943996-INS-56 | chr1-189943996-INS-63 |
| chr1-189945148-INS-60 | chr1-189945148-INS-60 |
| chr1-189954089-INS-60 | chr1-189954089-INS-62 |
| chr1-190007982-INS-60 | chr1-190007982-INS-60 |
| chr1-190009831-INS-74 | chr1-190009831-INS-74 |
| chr1-190051301-INS-52 | chr1-190051301-INS-56 |
| chr1-190062945-INS-55 | chr1-190062945-INS-55 |
| chr1-190073722-INS-147 | chr1-190073722-INS-147 |
| chr1-190075242-INS-730 | chr1-190075242-INS-730 |
| chr1-190075947-INS-317 | chr1-190075947-INS-317 |
| chr1-190114852-INS-56 | chr1-190114851-INS-64 |
| chr1-190136878-INS-82 | chr1-190136878-INS-82 |
| chr1-190181480-INS-55 | chr1-190181485-INS-57 |
| chr1-190195495-INS-471 | chr1-190195490-INS-465 |
| chr1-190206927-INS-320 | chr1-190206927-INS-320 |
| chr1-190208837-INS-54 | chr1-190208837-INS-52 |
| chr1-190209951-INS-437 | chr1-190209947-INS-437 |
| chr1-190216310-INS-96 | chr1-190216310-INS-96 |
| chr1-190235732-INS-368 | chr1-190235732-INS-368 |
| chr1-190243108-INS-206 | chr1-190243108-INS-215 |
| chr1-190244330-INS-477 | chr1-190244328-INS-509 |
| chr1-190250950-INS-63 | chr1-190250950-INS-63 |
| chr1-190268867-INS-72 | chr1-190268867-INS-72 |
| chr1-190274140-INS-60 | chr1-190274140-INS-60 |
| chr1-190280268-INS-894 | chr1-190280268-INS-890 |
| chr1-190287165-INS-104 | chr1-190287165-INS-104 |
| chr1-190298334-INS-84 | chr1-190298334-INS-78 |
| chr1-190343960-INS-62 | chr1-190343960-INS-50 |
| chr1-190371404-INS-54 | chr1-190371404-INS-50 |
| chr1-190373937-INS-498 | chr1-190373937-INS-498 |
| chr1-190405255-INS-82 | chr1-190405255-INS-82 |
| chr1-190410985-INS-112 | chr1-190410985-INS-89 |
| chr1-190427732-INS-52 | chr1-190427732-INS-52 |
| chr1-190434161-INS-389 | chr1-190434161-INS-389 |
| chr1-190468982-INS-201 | chr1-190468982-INS-205 |
| chr1-190500907-INS-147 | chr1-190500892-INS-167 |
| chr1-190518788-INS-977 | chr1-190518788-INS-977 |
| chr1-190560151-INS-132 | chr1-190560151-INS-132 |
| chr1-190564686-INS-68 | chr1-190564686-INS-68 |
| chr1-190571076-INS-94 | chr1-190571076-INS-84 |
| chr1-190592741-INS-54 | chr1-190592704-INS-81 |
| chr1-190595853-INS-57 | chr1-190595853-INS-57 |
| chr1-190597241-INS-57 | chr1-190597241-INS-59 |
| chr1-190602426-INS-199 | chr1-190602426-INS-199 |
| chr1-190616586-INS-288 | chr1-190616586-INS-312 |
| chr1-190624499-INS-68 | chr1-190624499-INS-68 |
| chr1-190634962-INS-173 | chr1-190634962-INS-173 |
| chr1-190635222-INS-1853 | chr1-190635223-INS-1853 |
| chr1-190639441-INS-161 | chr1-190639438-INS-166 |
| chr1-190644844-INS-60 | chr1-190644844-INS-60 |
| chr1-190650172-INS-209 | chr1-190650172-INS-209 |
| chr1-190696409-INS-249 | chr1-190696409-INS-249 |
| chr1-190724766-INS-88 | chr1-190724766-INS-88 |
| chr1-190730361-INS-599 | chr1-190730361-INS-600 |
| chr1-190742232-INS-71 | chr1-190742232-INS-71 |
| chr1-190754330-INS-75 | chr1-190754330-INS-75 |
| chr1-190777678-INS-85 | chr1-190777678-INS-85 |
| chr1-190786635-INS-839 | chr1-190786635-INS-839 |
| chr1-190796987-INS-215 | chr1-190796987-INS-215 |
| chr1-190798590-INS-197 | chr1-190798590-INS-197 |
| chr1-190837932-INS-58 | chr1-190837932-INS-58 |
| chr1-190893954-INS-80 | chr1-190893921-INS-95 |
| chr1-190909120-INS-76 | chr1-190909122-INS-70 |
| chr1-190923263-INS-72 | chr1-190923258-INS-72 |
| chr1-190935764-INS-115 | chr1-190935764-INS-115 |
| chr1-190959739-INS-58 | chr1-190959748-INS-60 |
| chr1-190994842-INS-90 | chr1-190994842-INS-90 |
| chr1-190996149-INS-67 | chr1-190996149-INS-67 |
| chr1-191001170-INS-52 | chr1-191001170-INS-52 |
| chr1-191007943-INS-52 | chr1-191007943-INS-52 |
| chr1-191009535-INS-258 | chr1-191009535-INS-258 |
| chr1-191015895-INS-134 | chr1-191015895-INS-133 |

| | |
|---|---|
| chr1-191035638-INS-73 | chr1-191035638-INS-73 |
| chr1-191047836-INS-215 | chr1-191047836-INS-216 |
| chr1-191067657-INS-69 | chr1-191067657-INS-69 |
| chr1-191138793-INS-7102 | chr1-191138794-INS-7102 |
| chr1-191160224-INS-3105 | chr1-191160225-INS-3105 |
| chr1-191194930-INS-61 | chr1-191194930-INS-62 |
| chr1-191196459-INS-693 | chr1-191196459-INS-693 |
| chr1-191200705-INS-51 | chr1-191200705-INS-51 |
| chr1-191223396-INS-60 | chr1-191223396-INS-60 |
| chr1-191224386-INS-60 | chr1-191224356-INS-59 |
| chr1-191229988-INS-102 | chr1-191229988-INS-102 |
| chr1-191237627-INS-387 | chr1-191237627-INS-387 |
| chr1-191239484-INS-114 | chr1-191239484-INS-150 |
| chr1-191245200-INS-202 | chr1-191245200-INS-202 |
| chr1-191256193-INS-52 | chr1-191256193-INS-52 |
| chr1-191266869-INS-198 | chr1-191266869-INS-198 |
| chr1-191277456-INS-1253 | chr1-191277457-INS-1249 |
| chr1-191282038-INS-209 | chr1-191282038-INS-209 |
| chr1-191292781-INS-216 | chr1-191292781-INS-216 |
| chr1-191317804-INS-64 | chr1-191317804-INS-66 |
| chr1-191328662-INS-96 | chr1-191328654-INS-58 |
| chr1-191339521-INS-178 | chr1-191339521-INS-178 |
| chr1-191356557-INS-1590 | chr1-191356558-INS-1589 |
| chr1-191378953-INS-88 | chr1-191378953-INS-88 |
| chr1-191414480-INS-202 | chr1-191414480-INS-202 |
| chr1-191491126-INS-53 | chr1-191491126-INS-53 |
| chr1-191493040-INS-113 | chr1-191493047-INS-120 |
| chr1-191497572-INS-89 | chr1-191497572-INS-87 |
| chr1-191533727-INS-9047 | chr1-191533728-INS-9047 |
| chr1-191601540-INS-129 | chr1-191601540-INS-129 |
| chr1-191609215-INS-146 | chr1-191609217-INS-146 |
| chr1-191623298-INS-52 | chr1-191623298-INS-50 |
| chr1-191652538-INS-218 | chr1-191652538-INS-218 |
| chr1-191663493-INS-52 | chr1-191663493-INS-56 |
| chr1-191690971-INS-215 | chr1-191690971-INS-217 |
| chr1-191699548-INS-55 | chr1-191699548-INS-55 |
| chr1-191716843-INS-6452 | chr1-191716844-INS-6452 |
| chr1-191799800-INS-319 | chr1-191799800-INS-319 |
| chr1-191803301-INS-268 | chr1-191803301-INS-267 |
| chr1-191826929-INS-4938 | chr1-191826930-INS-4938 |
| chr1-191853927-INS-83 | chr1-191853927-INS-83 |
| chr1-191855708-INS-89 | chr1-191855715-INS-59 |
| chr1-191869167-INS-226 | chr1-191869167-INS-226 |
| chr1-191878143-INS-52 | chr1-191878143-INS-52 |
| chr1-191884950-INS-118 | chr1-191884951-INS-118 |
| chr1-191899436-INS-96 | chr1-191899436-INS-70 |
| chr1-191914093-INS-1488 | chr1-191914094-INS-1488 |
| chr1-191917337-INS-69 | chr1-191917337-INS-69 |
| chr1-191941409-INS-492 | chr1-191941409-INS-492 |
| chr1-191944257-INS-530 | chr1-191944257-INS-530 |
| chr1-191971981-INS-53 | chr1-191971981-INS-60 |
| chr1-191972568-INS-1286 | chr1-191972569-INS-1281 |
| chr1-191975042-INS-52 | chr1-191975042-INS-54 |
| chr1-192000225-INS-98 | chr1-192000225-INS-98 |
| chr1-192005226-INS-505 | chr1-192005226-INS-504 |
| chr1-192022635-INS-2956 | chr1-192022636-INS-2956 |
| chr1-192027153-INS-52 | chr1-192027154-INS-84 |
| chr1-192031086-INS-61 | chr1-192031086-INS-65 |
| chr1-192047555-INS-96 | chr1-192047555-INS-96 |
| chr1-192073943-INS-109 | chr1-192073944-INS-109 |
| chr1-192097063-INS-537 | chr1-192097063-INS-535 |
| chr1-192153774-INS-208 | chr1-192153774-INS-208 |
| chr1-192199572-INS-2820 | chr1-192199573-INS-2820 |
| chr1-192260071-INS-411 | chr1-192260071-INS-411 |
| chr1-192275621-INS-56 | chr1-192275621-INS-56 |
| chr1-192299934-INS-58 | chr1-192299934-INS-50 |
| chr1-192320772-INS-100 | chr1-192320772-INS-100 |
| chr1-192337552-INS-69 | chr1-192337552-INS-69 |
| chr1-192340655-INS-87 | chr1-192340671-INS-83 |
| chr1-192347921-INS-980 | chr1-192347922-INS-984 |
| chr1-192350031-INS-60 | chr1-192350031-INS-96 |
| chr1-192352866-INS-246 | chr1-192352866-INS-246 |
| chr1-192363592-INS-78 | chr1-192363592-INS-56 |
| chr1-192416921-INS-148 | chr1-192416921-INS-132 |
| chr1-192425868-INS-51 | chr1-192425868-INS-51 |
| chr1-192432836-INS-112 | chr1-192432880-INS-82 |
| chr1-192439399-INS-352 | chr1-192439406-INS-330 |

| | |
|---|---|
| chr1-192484138-INS-64 | chr1-192484108-INS-72 |
| chr1-192507934-INS-65 | chr1-192507934-INS-69 |
| chr1-192545180-INS-68 | chr1-192545180-INS-68 |
| chr1-192546425-INS-52 | chr1-192546425-INS-54 |
| chr1-192606372-INS-1688 | chr1-192606373-INS-1691 |
| chr1-192624586-INS-236 | chr1-192624586-INS-236 |
| chr1-192658800-INS-373 | chr1-192658800-INS-374 |
| chr1-192681550-INS-195 | chr1-192681550-INS-195 |
| chr1-192686312-INS-684 | chr1-192686312-INS-690 |
| chr1-192695338-INS-207 | chr1-192695338-INS-207 |
| chr1-192724056-INS-58 | chr1-192724057-INS-56 |
| chr1-192730725-INS-222 | chr1-192730725-INS-222 |
| chr1-192734771-INS-7255 | chr1-192734772-INS-7255 |
| chr1-192786967-INS-1023 | chr1-192786968-INS-1023 |
| chr1-192790700-INS-71 | chr1-192790721-INS-51 |
| chr1-192808512-INS-75 | chr1-192808530-INS-102 |
| chr1-192814426-INS-72 | chr1-192814456-INS-76 |
| chr1-192819730-INS-58 | chr1-192819728-INS-64 |
| chr1-192913317-INS-100 | chr1-192913344-INS-52 |
| chr1-192919152-INS-122 | chr1-192919152-INS-141 |
| chr1-192934123-INS-2112 | chr1-192934123-INS-2124 |
| chr1-192938243-INS-59 | chr1-192938243-INS-51 |
| chr1-192989372-INS-130 | chr1-192989372-INS-134 |
| chr1-192992155-INS-128 | chr1-192992155-INS-128 |
| chr1-192996306-INS-109 | chr1-192996306-INS-108 |
| chr1-193007651-INS-70 | chr1-193007651-INS-73 |
| chr1-193057187-INS-65 | chr1-193057187-INS-65 |
| chr1-193058358-INS-52 | chr1-193058358-INS-52 |
| chr1-193099710-INS-141 | chr1-193099710-INS-141 |
| chr1-193105237-INS-143 | chr1-193105238-INS-143 |
| chr1-193108913-INS-317 | chr1-193108913-INS-317 |
| chr1-193109869-INS-68 | chr1-193109869-INS-68 |
| chr1-193124010-INS-75 | chr1-193124009-INS-79 |
| chr1-193126312-INS-400 | chr1-193126312-INS-400 |
| chr1-193157768-INS-418 | chr1-193157768-INS-421 |
| chr1-193159703-INS-68 | chr1-193159703-INS-68 |
| chr1-193162664-INS-52 | chr1-193162681-INS-57 |
| chr1-193171294-INS-262 | chr1-193171294-INS-282 |
| chr1-193195699-INS-225 | chr1-193195699-INS-225 |
| chr1-193206353-INS-55 | chr1-193206354-INS-50 |
| chr1-193218945-INS-103 | chr1-193218945-INS-103 |
| chr1-193221022-INS-520 | chr1-193221022-INS-520 |
| chr1-193230422-INS-75 | chr1-193230422-INS-75 |
| chr1-193239748-INS-88 | chr1-193239748-INS-89 |
| chr1-193244105-INS-72 | chr1-193244105-INS-72 |
| chr1-193265888-INS-85 | chr1-193265888-INS-85 |
| chr1-193273145-INS-3111 | chr1-193273146-INS-3111 |
| chr1-193286922-INS-1895 | chr1-193286923-INS-1895 |
| chr1-193307322-INS-74 | chr1-193307322-INS-88 |
| chr1-193317012-INS-90 | chr1-193317012-INS-90 |
| chr1-193341434-INS-82 | chr1-193341434-INS-76 |
| chr1-193349303-INS-655 | chr1-193349298-INS-651 |
| chr1-193444679-INS-750 | chr1-193444679-INS-751 |
| chr1-193481585-INS-463 | chr1-193481585-INS-463 |
| chr1-193563094-INS-3037 | chr1-193563095-INS-3037 |
| chr1-193589959-INS-81 | chr1-193589959-INS-53 |
| chr1-193613081-INS-171 | chr1-193613081-INS-171 |
| chr1-193719585-INS-1854 | chr1-193719586-INS-1838 |
| chr1-193781128-INS-125 | chr1-193781128-INS-127 |
| chr1-193792511-INS-68 | chr1-193792511-INS-62 |
| chr1-193812328-INS-259 | chr1-193812328-INS-260 |
| chr1-193813693-INS-263 | chr1-193813693-INS-263 |
| chr1-193815854-INS-6418 | chr1-193815855-INS-6418 |
| chr1-193829273-INS-6737 | chr1-193829274-INS-6742 |
| chr1-193917683-INS-6207 | chr1-193917684-INS-6205 |
| chr1-193947698-INS-70 | chr1-193947659-INS-90 |
| chr1-194009949-INS-7095 | chr1-194009950-INS-7095 |
| chr1-194033012-INS-433 | chr1-194033012-INS-433 |
| chr1-194094875-INS-5143 | chr1-194094876-INS-5143 |
| chr1-194101381-INS-2449 | chr1-194101382-INS-2449 |
| chr1-194111518-INS-123 | chr1-194111518-INS-123 |
| chr1-194128201-INS-272 | chr1-194128201-INS-272 |
| chr1-194151696-INS-220 | chr1-194151696-INS-220 |
| chr1-194175793-INS-6399 | chr1-194175794-INS-6399 |
| chr1-194188476-INS-82 | chr1-194188476-INS-82 |
| chr1-194195851-INS-58 | chr1-194195851-INS-60 |
| chr1-194227383-INS-905 | chr1-194227397-INS-901 |

| | |
|---|---|
| chr1-194227414-INS-7170 | chr1-194227415-INS-7187 |
| chr1-194261954-INS-81 | chr1-194261954-INS-81 |
| chr1-194276213-INS-57 | chr1-194276213-INS-57 |
| chr1-194301070-INS-178 | chr1-194301070-INS-208 |
| chr1-194353248-INS-60 | chr1-194353248-INS-60 |
| chr1-194354825-INS-400 | chr1-194354825-INS-400 |
| chr1-194357202-INS-7102 | chr1-194357203-INS-7117 |
| chr1-194370927-INS-217 | chr1-194370927-INS-218 |
| chr1-194651032-INS-5164 | chr1-194651033-INS-5164 |
| chr1-194723637-INS-231 | chr1-194723637-INS-231 |
| chr1-194753561-INS-54 | chr1-194753561-INS-55 |
| chr1-194777153-INS-70 | chr1-194777153-INS-70 |
| chr1-194821520-INS-7986 | chr1-194821521-INS-7986 |
| chr1-194825308-INS-73 | chr1-194825308-INS-73 |
| chr1-194874694-INS-587 | chr1-194874694-INS-586 |
| chr1-194977110-INS-1317 | chr1-194977097-INS-1306 |
| chr1-194982108-INS-206 | chr1-194982108-INS-206 |
| chr1-194986956-INS-397 | chr1-194986956-INS-397 |
| chr1-195025703-INS-6268 | chr1-195025704-INS-6268 |
| chr10-3055502-INS-60 | chr10-3055502-INS-60 |
| chr10-3076815-INS-85 | chr10-3076815-INS-85 |
| chr10-3092925-INS-56 | chr10-3092922-INS-54 |
| chr10-3098243-INS-344 | chr10-3098243-INS-344 |
| chr10-3111849-INS-3542 | chr10-3111850-INS-3542 |
| chr10-3121075-INS-1045 | chr10-3121076-INS-1045 |
| chr10-3121258-INS-76 | chr10-3121258-INS-76 |
| chr10-3157244-INS-1200 | chr10-3157245-INS-1201 |
| chr10-3184065-INS-72 | chr10-3184065-INS-72 |
| chr10-3218230-INS-56 | chr10-3218230-INS-90 |
| chr10-3220842-INS-4270 | chr10-3220843-INS-4270 |
| chr10-3248540-INS-212 | chr10-3248540-INS-212 |
| chr10-3377206-INS-118 | chr10-3377206-INS-102 |
| chr10-3435644-INS-84 | chr10-3435644-INS-50 |
| chr10-3443239-INS-75 | chr10-3443239-INS-75 |
| chr10-3460965-INS-144 | chr10-3460965-INS-142 |
| chr10-3467192-INS-7088 | chr10-3467193-INS-7088 |
| chr10-3475967-INS-68 | chr10-3475978-INS-82 |
| chr10-3482475-INS-52 | chr10-3482475-INS-62 |
| chr10-3489893-INS-252 | chr10-3489893-INS-275 |
| chr10-3536941-INS-69 | chr10-3536941-INS-69 |
| chr10-3588101-INS-283 | chr10-3588109-INS-269 |
| chr10-3615769-INS-56 | chr10-3615769-INS-60 |
| chr10-3622242-INS-208 | chr10-3622242-INS-208 |
| chr10-3639271-INS-197 | chr10-3639271-INS-197 |
| chr10-3656818-INS-52 | chr10-3656818-INS-52 |
| chr10-3670957-INS-68 | chr10-3670957-INS-68 |
| chr10-3676668-INS-7129 | chr10-3676669-INS-7129 |
| chr10-3693365-INS-245 | chr10-3693365-INS-246 |
| chr10-3696658-INS-944 | chr10-3696658-INS-944 |
| chr10-3699557-INS-6477 | chr10-3699558-INS-6477 |
| chr10-3710980-INS-144 | chr10-3710964-INS-141 |
| chr10-3741081-INS-138 | chr10-3741081-INS-122 |
| chr10-3816588-INS-64 | chr10-3816588-INS-85 |
| chr10-3830381-INS-215 | chr10-3830381-INS-219 |
| chr10-3833032-INS-190 | chr10-3833032-INS-191 |
| chr10-3845418-INS-66 | chr10-3845418-INS-66 |
| chr10-3848447-INS-228 | chr10-3848447-INS-212 |
| chr10-3876716-INS-84 | chr10-3876716-INS-84 |
| chr10-3879173-INS-56 | chr10-3879140-INS-88 |
| chr10-3924417-INS-237 | chr10-3924417-INS-237 |
| chr10-3945811-INS-162 | chr10-3945811-INS-160 |
| chr10-3952409-INS-137 | chr10-3952409-INS-101 |
| chr10-3975012-INS-178 | chr10-3975012-INS-178 |
| chr10-3995989-INS-68 | chr10-3995990-INS-63 |
| chr10-4003210-INS-51 | chr10-4003210-INS-56 |
| chr10-4005240-INS-69 | chr10-4005240-INS-72 |
| chr10-4016842-INS-205 | chr10-4016842-INS-207 |
| chr10-4024303-INS-196 | chr10-4024269-INS-217 |
| chr10-4031779-INS-70 | chr10-4031779-INS-70 |
| chr10-4048142-INS-309 | chr10-4048142-INS-309 |
| chr10-4048738-INS-64 | chr10-4048734-INS-52 |
| chr10-4067449-INS-64 | chr10-4067473-INS-68 |
| chr10-4070384-INS-93 | chr10-4070384-INS-93 |
| chr10-4087231-INS-180 | chr10-4087228-INS-187 |
| chr10-4094003-INS-69 | chr10-4094003-INS-56 |
| chr10-4099568-INS-231 | chr10-4099568-INS-231 |
| chr10-4101372-INS-95 | chr10-4101372-INS-55 |

| | |
|---|---|
| chr10-4111271-INS-213 | chr10-4111271-INS-212 |
| chr10-4112022-INS-95 | chr10-4112020-INS-94 |
| chr10-4121100-INS-350 | chr10-4121100-INS-350 |
| chr10-4148326-INS-217 | chr10-4148326-INS-217 |
| chr10-4159055-INS-609 | chr10-4159055-INS-609 |
| chr10-4161416-INS-54 | chr10-4161416-INS-74 |
| chr10-4164475-INS-53 | chr10-4164475-INS-55 |
| chr10-4168348-INS-70 | chr10-4168348-INS-90 |
| chr10-4170988-INS-178 | chr10-4170991-INS-179 |
| chr10-4179618-INS-55 | chr10-4179618-INS-55 |
| chr10-4181732-INS-283 | chr10-4181733-INS-283 |
| chr10-4184727-INS-207 | chr10-4184727-INS-206 |
| chr10-4200855-INS-90 | chr10-4200855-INS-90 |
| chr10-4204270-INS-174 | chr10-4204270-INS-174 |
| chr10-4208277-INS-63 | chr10-4208277-INS-81 |
| chr10-4210122-INS-177 | chr10-4210121-INS-185 |
| chr10-4224934-INS-214 | chr10-4224934-INS-213 |
| chr10-4236726-INS-212 | chr10-4236729-INS-212 |
| chr10-4242764-INS-178 | chr10-4242764-INS-164 |
| chr10-4255921-INS-65 | chr10-4255921-INS-65 |
| chr10-4267260-INS-98 | chr10-4267260-INS-102 |
| chr10-4286906-INS-57 | chr10-4286880-INS-59 |
| chr10-4290206-INS-604 | chr10-4290214-INS-606 |
| chr10-4312522-INS-432 | chr10-4312521-INS-447 |
| chr10-4314861-INS-1873 | chr10-4314862-INS-1873 |
| chr10-4327183-INS-63 | chr10-4327183-INS-63 |
| chr10-4352193-INS-221 | chr10-4352193-INS-220 |
| chr10-4353384-INS-235 | chr10-4353384-INS-236 |
| chr10-4359818-INS-569 | chr10-4359818-INS-569 |
| chr10-4361747-INS-228 | chr10-4361747-INS-228 |
| chr10-4362686-INS-210 | chr10-4362686-INS-210 |
| chr10-4365408-INS-58 | chr10-4365408-INS-58 |
| chr10-4369033-INS-162 | chr10-4369033-INS-160 |
| chr10-4378016-INS-81 | chr10-4378016-INS-81 |
| chr10-4390181-INS-59 | chr10-4390181-INS-60 |
| chr10-4401750-INS-455 | chr10-4401750-INS-456 |
| chr10-4408680-INS-770 | chr10-4408680-INS-770 |
| chr10-4418634-INS-52 | chr10-4418634-INS-54 |
| chr10-4420412-INS-72 | chr10-4420412-INS-54 |
| chr10-4421059-INS-84 | chr10-4421059-INS-56 |
| chr10-4422521-INS-210 | chr10-4422521-INS-210 |
| chr10-4423117-INS-106 | chr10-4423117-INS-133 |
| chr10-4430182-INS-69 | chr10-4430182-INS-69 |
| chr10-4433524-INS-70 | chr10-4433524-INS-55 |
| chr10-4436667-INS-62 | chr10-4436649-INS-84 |
| chr10-4438049-INS-231 | chr10-4438049-INS-231 |
| chr10-4439810-INS-558 | chr10-4439810-INS-557 |
| chr10-4440789-INS-216 | chr10-4440789-INS-215 |
| chr10-4445419-INS-54 | chr10-4445419-INS-54 |
| chr10-4445970-INS-107 | chr10-4445970-INS-111 |
| chr10-4448611-INS-212 | chr10-4448611-INS-212 |
| chr10-4462047-INS-57 | chr10-4462047-INS-52 |
| chr10-4469735-INS-2255 | chr10-4469736-INS-2246 |
| chr10-4470454-INS-1864 | chr10-4470455-INS-1864 |
| chr10-4487985-INS-75 | chr10-4487985-INS-75 |
| chr10-4497495-INS-498 | chr10-4497495-INS-498 |
| chr10-4502564-INS-66 | chr10-4502550-INS-72 |
| chr10-4512211-INS-84 | chr10-4512211-INS-84 |
| chr10-4550042-INS-73 | chr10-4550042-INS-77 |
| chr10-4580792-INS-64 | chr10-4580792-INS-64 |
| chr10-4591302-INS-54 | chr10-4591302-INS-70 |
| chr10-4610248-INS-60 | chr10-4610248-INS-60 |
| chr10-4630746-INS-561 | chr10-4630746-INS-561 |
| chr10-4649850-INS-747 | chr10-4649850-INS-747 |
| chr10-4651041-INS-205 | chr10-4651041-INS-205 |
| chr10-4655002-INS-54 | chr10-4655002-INS-54 |
| chr10-4677587-INS-83 | chr10-4677588-INS-88 |
| chr10-4694669-INS-69 | chr10-4694669-INS-110 |
| chr10-4701177-INS-67 | chr10-4701177-INS-58 |
| chr10-4710834-INS-253 | chr10-4710834-INS-253 |
| chr10-4719733-INS-144 | chr10-4719737-INS-116 |
| chr10-4738545-INS-138 | chr10-4738545-INS-138 |
| chr10-4748918-INS-108 | chr10-4748918-INS-108 |
| chr10-4769086-INS-70 | chr10-4769086-INS-70 |
| chr10-4773961-INS-52 | chr10-4773936-INS-96 |
| chr10-4809344-INS-51 | chr10-4809344-INS-51 |
| chr10-4826633-INS-68 | chr10-4826633-INS-77 |

| | |
|---|---|
| chr10-4843601-INS-105 | chr10-4843601-INS-105 |
| chr10-4848176-INS-4824 | chr10-4848177-INS-4824 |
| chr10-4856977-INS-211 | chr10-4857023-INS-211 |
| chr10-4912221-INS-4383 | chr10-4912222-INS-4383 |
| chr10-4912410-INS-64 | chr10-4912410-INS-64 |
| chr10-4919898-INS-68 | chr10-4919898-INS-68 |
| chr10-4926286-INS-2340 | chr10-4926287-INS-2311 |
| chr10-5012259-INS-656 | chr10-5012259-INS-656 |
| chr10-5013622-INS-82 | chr10-5013622-INS-85 |
| chr10-5019955-INS-70 | chr10-5019955-INS-70 |
| chr10-5027516-INS-52 | chr10-5027516-INS-52 |
| chr10-5077509-INS-6962 | chr10-5077510-INS-6963 |
| chr10-5090486-INS-54 | chr10-5090486-INS-56 |
| chr10-5116533-INS-60 | chr10-5116529-INS-60 |
| chr10-5165155-INS-60 | chr10-5165110-INS-80 |
| chr10-5198008-INS-54 | chr10-5198018-INS-74 |
| chr10-5204549-INS-546 | chr10-5204549-INS-546 |
| chr10-5205747-INS-169 | chr10-5205747-INS-169 |
| chr10-5218471-INS-137 | chr10-5218454-INS-140 |
| chr10-5240501-INS-69 | chr10-5240501-INS-69 |
| chr10-5263988-INS-309 | chr10-5263988-INS-309 |
| chr10-5271250-INS-1888 | chr10-5271251-INS-1888 |
| chr10-5273620-INS-59 | chr10-5273619-INS-65 |
| chr10-5278642-INS-492 | chr10-5278642-INS-492 |
| chr10-5303203-INS-945 | chr10-5303203-INS-947 |
| chr10-5333457-INS-72 | chr10-5333457-INS-72 |
| chr10-5337608-INS-221 | chr10-5337608-INS-221 |
| chr10-5383226-INS-230 | chr10-5383226-INS-229 |
| chr10-5387379-INS-879 | chr10-5387375-INS-879 |
| chr10-5397860-INS-50 | chr10-5397860-INS-52 |
| chr10-5409090-INS-62 | chr10-5409086-INS-66 |
| chr10-5413470-INS-289 | chr10-5413470-INS-289 |
| chr10-5420537-INS-7108 | chr10-5420538-INS-7108 |
| chr10-5425159-INS-827 | chr10-5425159-INS-827 |
| chr10-5439120-INS-215 | chr10-5439120-INS-215 |
| chr10-5441092-INS-183 | chr10-5441092-INS-185 |
| chr10-5448373-INS-225 | chr10-5448374-INS-225 |
| chr10-5457572-INS-202 | chr10-5457572-INS-202 |
| chr10-5475561-INS-64 | chr10-5475561-INS-64 |
| chr10-5481012-INS-201 | chr10-5481012-INS-201 |
| chr10-5560694-INS-61 | chr10-5560705-INS-50 |
| chr10-5565102-INS-363 | chr10-5565102-INS-363 |
| chr10-5574894-INS-211 | chr10-5574894-INS-211 |
| chr10-5577184-INS-107 | chr10-5577188-INS-143 |
| chr10-5582284-INS-890 | chr10-5582284-INS-869 |
| chr10-5609859-INS-412 | chr10-5609859-INS-412 |
| chr10-5622409-INS-162 | chr10-5622409-INS-162 |
| chr10-5630022-INS-60 | chr10-5630022-INS-60 |
| chr10-5630840-INS-646 | chr10-5630840-INS-646 |
| chr10-5633569-INS-6403 | chr10-5633570-INS-6403 |
| chr10-5641842-INS-115 | chr10-5641837-INS-122 |
| chr10-5647602-INS-120 | chr10-5647602-INS-120 |
| chr10-5662425-INS-54 | chr10-5662425-INS-54 |
| chr10-5694800-INS-162 | chr10-5694800-INS-173 |
| chr10-5695719-INS-63 | chr10-5695719-INS-63 |
| chr10-5701607-INS-204 | chr10-5701607-INS-205 |
| chr10-5712352-INS-57 | chr10-5712320-INS-96 |
| chr10-5725514-INS-561 | chr10-5725514-INS-561 |
| chr10-5737124-INS-76 | chr10-5737124-INS-76 |
| chr10-5757099-INS-193 | chr10-5757099-INS-193 |
| chr10-5758616-INS-201 | chr10-5758616-INS-200 |
| chr10-5765998-INS-188 | chr10-5765998-INS-188 |
| chr10-5767922-INS-210 | chr10-5767922-INS-210 |
| chr10-5782225-INS-72 | chr10-5782225-INS-77 |
| chr10-5783624-INS-160 | chr10-5783624-INS-160 |
| chr10-5786005-INS-400 | chr10-5786005-INS-400 |
| chr10-5787611-INS-602 | chr10-5787611-INS-606 |
| chr10-5795655-INS-337 | chr10-5795655-INS-337 |
| chr10-5797696-INS-68 | chr10-5797696-INS-92 |
| chr10-5807303-INS-328 | chr10-5807303-INS-328 |
| chr10-5819375-INS-215 | chr10-5819375-INS-215 |
| chr10-5820745-INS-156 | chr10-5820697-INS-172 |
| chr10-5832691-INS-53 | chr10-5832691-INS-55 |
| chr10-5833304-INS-1238 | chr10-5833305-INS-1237 |
| chr10-5849096-INS-207 | chr10-5849101-INS-231 |
| chr10-5864397-INS-87 | chr10-5864397-INS-87 |
| chr10-5879874-INS-70 | chr10-5879874-INS-52 |

| | |
|---|---|
| chr10-5908639-INS-7091 | chr10-5908640-INS-7089 |
| chr10-5919446-INS-548 | chr10-5919446-INS-548 |
| chr10-5929900-INS-60 | chr10-5929901-INS-104 |
| chr10-5934702-INS-335 | chr10-5934702-INS-336 |
| chr10-5937970-INS-210 | chr10-5937970-INS-210 |
| chr10-5953425-INS-176 | chr10-5953425-INS-176 |
| chr10-5959664-INS-230 | chr10-5959664-INS-206 |
| chr10-5972754-INS-105 | chr10-5972775-INS-66 |
| chr10-6000710-INS-102 | chr10-6000710-INS-84 |
| chr10-6006643-INS-173 | chr10-6006643-INS-173 |
| chr10-6067551-INS-322 | chr10-6067551-INS-322 |
| chr10-6085757-INS-112 | chr10-6085757-INS-108 |
| chr10-6113325-INS-936 | chr10-6113325-INS-937 |
| chr10-6118013-INS-516 | chr10-6118013-INS-516 |
| chr10-6134268-INS-1888 | chr10-6134269-INS-1888 |
| chr10-6140868-INS-52 | chr10-6140868-INS-52 |
| chr10-6144543-INS-861 | chr10-6144543-INS-861 |
| chr10-6146421-INS-357 | chr10-6146421-INS-357 |
| chr10-6158428-INS-58 | chr10-6158428-INS-58 |
| chr10-6191789-INS-244 | chr10-6191789-INS-240 |
| chr10-6197186-INS-58 | chr10-6197186-INS-58 |
| chr10-6200109-INS-191 | chr10-6200109-INS-191 |
| chr10-6240815-INS-51 | chr10-6240815-INS-51 |
| chr10-6243510-INS-476 | chr10-6243510-INS-476 |
| chr10-6256745-INS-438 | chr10-6256745-INS-438 |
| chr10-6257955-INS-54 | chr10-6257955-INS-52 |
| chr10-6279065-INS-6169 | chr10-6279066-INS-6169 |
| chr10-6280721-INS-164 | chr10-6280721-INS-164 |
| chr10-6286613-INS-4335 | chr10-6286614-INS-4335 |
| chr10-6292712-INS-974 | chr10-6292712-INS-974 |
| chr10-6314711-INS-181 | chr10-6314711-INS-226 |
| chr10-6317558-INS-4588 | chr10-6317559-INS-4573 |
| chr10-6318177-INS-427 | chr10-6318177-INS-427 |
| chr10-6334249-INS-79 | chr10-6334249-INS-102 |
| chr10-6354994-INS-1885 | chr10-6354995-INS-1885 |
| chr10-6394647-INS-249 | chr10-6394647-INS-249 |
| chr10-6398046-INS-5108 | chr10-6398047-INS-5108 |
| chr10-6416904-INS-4350 | chr10-6416905-INS-4350 |
| chr10-6439114-INS-6442 | chr10-6439115-INS-6442 |
| chr10-6464803-INS-115 | chr10-6464769-INS-165 |
| chr10-6470054-INS-455 | chr10-6470054-INS-455 |
| chr10-6470884-INS-329 | chr10-6470884-INS-329 |
| chr10-6493913-INS-52 | chr10-6493903-INS-62 |
| chr10-6499823-INS-2913 | chr10-6499824-INS-2913 |
| chr10-6596994-INS-109 | chr10-6596949-INS-148 |
| chr10-6670006-INS-92 | chr10-6670006-INS-92 |
| chr10-6676602-INS-652 | chr10-6676602-INS-651 |
| chr10-6683838-INS-74 | chr10-6683838-INS-93 |
| chr10-6715456-INS-54 | chr10-6715456-INS-56 |
| chr10-6716687-INS-64 | chr10-6716674-INS-50 |
| chr10-6745575-INS-72 | chr10-6745575-INS-75 |
| chr10-6751548-INS-62 | chr10-6751548-INS-61 |
| chr10-6753175-INS-65 | chr10-6753176-INS-61 |
| chr10-6759147-INS-7107 | chr10-6759148-INS-7106 |
| chr10-6821167-INS-73 | chr10-6821167-INS-73 |
| chr10-6830442-INS-61 | chr10-6830429-INS-73 |
| chr10-6851339-INS-244 | chr10-6851339-INS-244 |
| chr10-6876314-INS-52 | chr10-6876324-INS-50 |
| chr10-6893999-INS-55 | chr10-6893999-INS-55 |
| chr10-6907575-INS-1135 | chr10-6907576-INS-1135 |
| chr10-6921984-INS-60 | chr10-6921984-INS-60 |
| chr10-6941270-INS-161 | chr10-6941270-INS-161 |
| chr10-6961943-INS-156 | chr10-6961932-INS-182 |
| chr10-6974216-INS-7034 | chr10-6974217-INS-7034 |
| chr10-6985616-INS-126 | chr10-6985616-INS-126 |
| chr10-7007069-INS-78 | chr10-7007069-INS-54 |
| chr10-7027198-INS-61 | chr10-7027199-INS-52 |
| chr10-7058714-INS-65 | chr10-7058709-INS-72 |
| chr10-7095518-INS-107 | chr10-7095518-INS-119 |
| chr10-7119243-INS-5668 | chr10-7119244-INS-5668 |
| chr10-7128066-INS-122 | chr10-7128066-INS-106 |
| chr10-7154685-INS-447 | chr10-7154685-INS-447 |
| chr10-7174705-INS-604 | chr10-7174705-INS-604 |
| chr10-7177038-INS-88 | chr10-7177038-INS-88 |
| chr10-7207551-INS-63 | chr10-7207551-INS-64 |
| chr10-7234640-INS-193 | chr10-7234665-INS-202 |
| chr10-7252081-INS-68 | chr10-7252081-INS-64 |

| | |
|---|---|
| chr10-7252876-INS-332 | chr10-7252876-INS-332 |
| chr10-7265758-INS-235 | chr10-7265758-INS-234 |
| chr10-7283124-INS-325 | chr10-7283124-INS-325 |
| chr10-7312999-INS-88 | chr10-7312999-INS-138 |
| chr10-7319006-INS-591 | chr10-7319016-INS-591 |
| chr10-7320156-INS-81 | chr10-7320156-INS-81 |
| chr10-7333605-INS-6597 | chr10-7333606-INS-6597 |
| chr10-7393517-INS-57 | chr10-7393517-INS-57 |
| chr10-7396895-INS-577 | chr10-7396895-INS-577 |
| chr10-7401938-INS-2383 | chr10-7401939-INS-2383 |
| chr10-7453810-INS-167 | chr10-7453810-INS-168 |
| chr10-7455731-INS-488 | chr10-7455733-INS-497 |
| chr10-7468327-INS-239 | chr10-7468327-INS-240 |
| chr10-7470276-INS-228 | chr10-7470276-INS-228 |
| chr10-7487573-INS-55 | chr10-7487573-INS-55 |
| chr10-7491073-INS-203 | chr10-7491073-INS-203 |
| chr10-7493969-INS-203 | chr10-7493969-INS-203 |
| chr10-7518794-INS-205 | chr10-7518794-INS-205 |
| chr10-7535104-INS-64 | chr10-7535104-INS-64 |
| chr10-7573159-INS-68 | chr10-7573159-INS-94 |
| chr10-7603510-INS-808 | chr10-7603510-INS-812 |
| chr10-7618332-INS-205 | chr10-7618332-INS-205 |
| chr10-7619018-INS-235 | chr10-7619018-INS-233 |
| chr10-7621729-INS-66 | chr10-7621729-INS-70 |
| chr10-7626798-INS-205 | chr10-7626798-INS-205 |
| chr10-7642278-INS-209 | chr10-7642278-INS-209 |
| chr10-7647439-INS-76 | chr10-7647439-INS-52 |
| chr10-7650240-INS-120 | chr10-7650240-INS-120 |
| chr10-7662646-INS-54 | chr10-7662646-INS-54 |
| chr10-7664179-INS-201 | chr10-7664179-INS-201 |
| chr10-7675927-INS-159 | chr10-7675917-INS-166 |
| chr10-7690620-INS-68 | chr10-7690620-INS-68 |
| chr10-7712726-INS-133 | chr10-7712726-INS-136 |
| chr10-7738027-INS-777 | chr10-7738027-INS-778 |
| chr10-7742211-INS-56 | chr10-7742214-INS-99 |
| chr10-7745520-INS-74 | chr10-7745520-INS-74 |
| chr10-7747811-INS-63 | chr10-7747841-INS-52 |
| chr10-7763607-INS-100 | chr10-7763602-INS-70 |
| chr10-7769850-INS-773 | chr10-7769850-INS-773 |
| chr10-7778764-INS-914 | chr10-7778765-INS-914 |
| chr10-7780411-INS-200 | chr10-7780411-INS-201 |
| chr10-7799567-INS-274 | chr10-7799567-INS-274 |
| chr10-7811575-INS-234 | chr10-7811575-INS-234 |
| chr10-7851885-INS-209 | chr10-7851885-INS-209 |
| chr10-7883709-INS-80 | chr10-7883755-INS-50 |
| chr10-7897428-INS-136 | chr10-7897441-INS-95 |
| chr10-7901851-INS-385 | chr10-7901824-INS-412 |
| chr10-7924963-INS-881 | chr10-7924965-INS-880 |
| chr10-7947542-INS-189 | chr10-7947546-INS-183 |
| chr10-7949365-INS-56 | chr10-7949365-INS-72 |
| chr10-7953081-INS-250 | chr10-7953082-INS-250 |
| chr10-7955079-INS-464 | chr10-7955079-INS-465 |
| chr10-7989507-INS-65 | chr10-7989507-INS-65 |
| chr10-7997414-INS-122 | chr10-7997413-INS-116 |
| chr10-8011116-INS-141 | chr10-8011116-INS-142 |
| chr10-8013258-INS-62 | chr10-8013258-INS-62 |
| chr10-8014216-INS-112 | chr10-8014216-INS-112 |
| chr10-8069681-INS-339 | chr10-8069681-INS-339 |
| chr10-8071626-INS-56 | chr10-8071612-INS-56 |
| chr10-8084349-INS-209 | chr10-8084349-INS-211 |
| chr10-8150998-INS-627 | chr10-8150998-INS-627 |
| chr10-8162476-INS-58 | chr10-8162476-INS-50 |
| chr10-8201039-INS-255 | chr10-8201041-INS-246 |
| chr10-8201436-INS-1008 | chr10-8201437-INS-1004 |
| chr10-8206294-INS-164 | chr10-8206294-INS-166 |
| chr10-8218202-INS-51 | chr10-8218202-INS-51 |
| chr10-8233767-INS-2242 | chr10-8233768-INS-2262 |
| chr10-8233929-INS-362 | chr10-8233929-INS-362 |
| chr10-8234705-INS-59 | chr10-8234736-INS-76 |
| chr10-8254253-INS-199 | chr10-8254253-INS-199 |
| chr10-8255752-INS-338 | chr10-8255752-INS-338 |
| chr10-8288402-INS-448 | chr10-8288402-INS-448 |
| chr10-8296415-INS-69 | chr10-8296415-INS-69 |
| chr10-8300687-INS-73 | chr10-8300687-INS-75 |
| chr10-8307193-INS-92 | chr10-8307193-INS-92 |
| chr10-8325664-INS-260 | chr10-8325664-INS-260 |
| chr10-8378535-INS-57 | chr10-8378535-INS-57 |

| | |
|---|---|
| chr10-8388567-INS-84 | chr10-8388604-INS-74 |
| chr10-8416113-INS-248 | chr10-8416132-INS-216 |
| chr10-8430519-INS-523 | chr10-8430519-INS-524 |
| chr10-8439758-INS-243 | chr10-8439758-INS-239 |
| chr10-8467562-INS-347 | chr10-8467562-INS-347 |
| chr10-8490397-INS-115 | chr10-8490397-INS-124 |
| chr10-8495916-INS-176 | chr10-8495916-INS-176 |
| chr10-8498275-INS-59 | chr10-8498275-INS-60 |
| chr10-8501436-INS-124 | chr10-8501436-INS-124 |
| chr10-8504290-INS-536 | chr10-8504284-INS-533 |
| chr10-8505400-INS-392 | chr10-8505400-INS-392 |
| chr10-8507430-INS-66 | chr10-8507430-INS-66 |
| chr10-8514948-INS-210 | chr10-8514921-INS-209 |
| chr10-8530836-INS-55 | chr10-8530836-INS-55 |
| chr10-8532823-INS-2988 | chr10-8532824-INS-2988 |
| chr10-8558956-INS-126 | chr10-8558956-INS-118 |
| chr10-8559895-INS-328 | chr10-8559895-INS-328 |
| chr10-8574477-INS-108 | chr10-8574477-INS-108 |
| chr10-8576274-INS-6400 | chr10-8576275-INS-6400 |
| chr10-8584362-INS-736 | chr10-8584362-INS-736 |
| chr10-8600307-INS-90 | chr10-8600307-INS-90 |
| chr10-8615022-INS-138 | chr10-8615042-INS-152 |
| chr10-8660834-INS-64 | chr10-8660834-INS-68 |
| chr10-8669647-INS-69 | chr10-8669647-INS-69 |
| chr10-8681883-INS-73 | chr10-8681894-INS-54 |
| chr10-8686938-INS-207 | chr10-8686937-INS-210 |
| chr10-8692000-INS-79 | chr10-8692000-INS-79 |
| chr10-8702919-INS-316 | chr10-8702919-INS-316 |
| chr10-8705369-INS-75 | chr10-8705342-INS-87 |
| chr10-8728635-INS-90 | chr10-8728635-INS-106 |
| chr10-8731653-INS-67 | chr10-8731653-INS-67 |
| chr10-8747561-INS-62 | chr10-8747561-INS-66 |
| chr10-8755674-INS-206 | chr10-8755674-INS-206 |
| chr10-8793106-INS-60 | chr10-8793106-INS-52 |
| chr10-8797865-INS-198 | chr10-8797865-INS-198 |
| chr10-8801683-INS-71 | chr10-8801636-INS-90 |
| chr10-8812764-INS-70 | chr10-8812748-INS-82 |
| chr10-8845876-INS-51 | chr10-8845876-INS-51 |
| chr10-8853877-INS-240 | chr10-8853877-INS-240 |
| chr10-8861792-INS-656 | chr10-8861792-INS-656 |
| chr10-8869885-INS-68 | chr10-8869885-INS-55 |
| chr10-8873578-INS-909 | chr10-8873578-INS-909 |
| chr10-8894000-INS-111 | chr10-8894000-INS-111 |
| chr10-8896340-INS-72 | chr10-8896340-INS-72 |
| chr10-8897628-INS-66 | chr10-8897629-INS-79 |
| chr10-8907226-INS-53 | chr10-8907226-INS-53 |
| chr10-8938285-INS-91 | chr10-8938273-INS-109 |
| chr10-8950118-INS-56 | chr10-8950113-INS-56 |
| chr10-8963094-INS-2667 | chr10-8963095-INS-2667 |
| chr10-8966618-INS-690 | chr10-8966615-INS-698 |
| chr10-8967555-INS-6586 | chr10-8967556-INS-6586 |
| chr10-9009356-INS-68 | chr10-9009356-INS-68 |
| chr10-9053878-INS-56 | chr10-9053878-INS-56 |
| chr10-9056357-INS-131 | chr10-9056357-INS-133 |
| chr10-9059686-INS-225 | chr10-9059686-INS-225 |
| chr10-9060126-INS-955 | chr10-9060126-INS-955 |
| chr10-9071842-INS-510 | chr10-9071842-INS-510 |
| chr10-9083112-INS-140 | chr10-9083112-INS-140 |
| chr10-9129485-INS-177 | chr10-9129485-INS-177 |
| chr10-9154738-INS-120 | chr10-9154738-INS-120 |
| chr10-9157284-INS-343 | chr10-9157284-INS-343 |
| chr10-9180848-INS-2929 | chr10-9180849-INS-2929 |
| chr10-9195782-INS-51 | chr10-9195782-INS-53 |
| chr10-9208225-INS-64 | chr10-9208223-INS-52 |
| chr10-9210749-INS-7101 | chr10-9210750-INS-7101 |
| chr10-9247009-INS-97 | chr10-9247027-INS-60 |
| chr10-9252802-INS-2878 | chr10-9252803-INS-2880 |
| chr10-9274365-INS-58 | chr10-9274365-INS-68 |
| chr10-9275606-INS-54 | chr10-9275606-INS-56 |
| chr10-9278785-INS-7159 | chr10-9278786-INS-7159 |
| chr10-9323620-INS-255 | chr10-9323620-INS-258 |
| chr10-9336256-INS-1864 | chr10-9336257-INS-1864 |
| chr10-9352355-INS-96 | chr10-9352361-INS-92 |
| chr10-9355914-INS-98 | chr10-9355914-INS-81 |
| chr10-9416284-INS-58 | chr10-9416284-INS-58 |
| chr10-9419004-INS-93 | chr10-9419004-INS-51 |
| chr10-9422357-INS-83 | chr10-9422362-INS-63 |

| | |
|---|---|
| chr10-9430445-INS-50 | chr10-9430445-INS-64 |
| chr10-9455229-INS-7124 | chr10-9455230-INS-7124 |
| chr10-9457263-INS-56 | chr10-9457263-INS-58 |
| chr10-9458399-INS-57 | chr10-9458393-INS-57 |
| chr10-9464453-INS-74 | chr10-9464444-INS-67 |
| chr10-9469301-INS-856 | chr10-9469301-INS-855 |
| chr10-9470345-INS-139 | chr10-9470347-INS-133 |
| chr10-9483509-INS-198 | chr10-9483509-INS-199 |
| chr10-9485755-INS-63 | chr10-9485716-INS-70 |
| chr10-9487756-INS-461 | chr10-9487757-INS-457 |
| chr10-9503431-INS-153 | chr10-9503431-INS-153 |
| chr10-9520014-INS-313 | chr10-9520014-INS-316 |
| chr10-9528644-INS-2910 | chr10-9528645-INS-2910 |
| chr10-9528645-INS-450 | chr10-9528645-INS-450 |
| chr10-9541414-INS-2845 | chr10-9541415-INS-2845 |
| chr10-9564951-INS-399 | chr10-9564951-INS-399 |
| chr10-9578827-INS-5411 | chr10-9578828-INS-5411 |
| chr10-9593326-INS-155 | chr10-9593326-INS-164 |
| chr10-9613252-INS-54 | chr10-9613245-INS-68 |
| chr10-9621369-INS-2470 | chr10-9621370-INS-2484 |
| chr10-9652249-INS-82 | chr10-9652249-INS-82 |
| chr10-9663213-INS-216 | chr10-9663213-INS-216 |
| chr10-9680163-INS-199 | chr10-9680163-INS-199 |
| chr10-9715568-INS-6440 | chr10-9715569-INS-6441 |
| chr10-9823161-INS-6524 | chr10-9823162-INS-6525 |
| chr10-9835245-INS-296 | chr10-9835243-INS-295 |
| chr10-9836609-INS-55 | chr10-9836609-INS-62 |
| chr10-9838304-INS-574 | chr10-9838302-INS-576 |
| chr10-9893361-INS-742 | chr10-9893361-INS-747 |
| chr10-9900975-INS-168 | chr10-9900975-INS-168 |
| chr10-9927254-INS-54 | chr10-9927254-INS-54 |
| chr10-9951187-INS-60 | chr10-9951187-INS-62 |
| chr10-9960724-INS-214 | chr10-9960724-INS-214 |
| chr10-9982970-INS-60 | chr10-9982970-INS-65 |
| chr10-9999591-INS-503 | chr10-9999591-INS-504 |
| chr10-10005764-INS-431 | chr10-10005764-INS-431 |
| chr10-10031996-INS-6660 | chr10-10031997-INS-6662 |
| chr10-10038473-INS-131 | chr10-10038511-INS-108 |
| chr10-10074953-INS-90 | chr10-10074953-INS-84 |
| chr10-10179512-INS-6308 | chr10-10179513-INS-6308 |
| chr10-10183797-INS-66 | chr10-10183797-INS-66 |
| chr10-10193453-INS-86 | chr10-10193453-INS-90 |
| chr10-10211291-INS-102 | chr10-10211312-INS-88 |
| chr10-10242610-INS-631 | chr10-10242608-INS-628 |
| chr10-10250257-INS-165 | chr10-10250257-INS-165 |
| chr10-10274531-INS-88 | chr10-10274531-INS-88 |
| chr10-10280549-INS-54 | chr10-10280549-INS-58 |
| chr10-10285780-INS-6486 | chr10-10285781-INS-6486 |
| chr10-10314604-INS-51 | chr10-10314598-INS-60 |
| chr10-10332245-INS-123 | chr10-10332245-INS-123 |
| chr10-10333891-INS-126 | chr10-10333891-INS-126 |
| chr10-10342304-INS-6445 | chr10-10342305-INS-6445 |
| chr10-10400744-INS-356 | chr10-10400744-INS-363 |
| chr10-10427278-INS-63 | chr10-10427278-INS-63 |
| chr10-10449298-INS-5495 | chr10-10449299-INS-5495 |
| chr10-10450319-INS-194 | chr10-10450319-INS-196 |
| chr10-10456848-INS-203 | chr10-10456848-INS-203 |
| chr10-10460620-INS-208 | chr10-10460620-INS-208 |
| chr10-10483037-INS-94 | chr10-10483043-INS-74 |
| chr10-10518148-INS-72 | chr10-10518139-INS-88 |
| chr10-10531131-INS-60 | chr10-10531131-INS-62 |
| chr10-10537028-INS-363 | chr10-10537028-INS-363 |
| chr10-10564500-INS-99 | chr10-10564500-INS-95 |
| chr10-10585241-INS-202 | chr10-10585250-INS-212 |
| chr10-10599084-INS-993 | chr10-10599084-INS-1004 |
| chr10-10609780-INS-58 | chr10-10609780-INS-62 |
| chr10-10657496-INS-66 | chr10-10657496-INS-62 |
| chr10-10664318-INS-55 | chr10-10664298-INS-66 |
| chr10-10684140-INS-54 | chr10-10684140-INS-54 |
| chr10-10697337-INS-52 | chr10-10697364-INS-55 |
| chr10-10715092-INS-6370 | chr10-10715093-INS-6369 |
| chr10-10720045-INS-268 | chr10-10720045-INS-268 |
| chr10-10742930-INS-69 | chr10-10742930-INS-69 |
| chr10-10756514-INS-96 | chr10-10756514-INS-80 |
| chr10-10776298-INS-7168 | chr10-10776299-INS-7169 |
| chr10-10781943-INS-4106 | chr10-10781944-INS-4107 |
| chr10-10782557-INS-945 | chr10-10782557-INS-945 |

| | |
|---|---|
| chr10-10811097-INS-51 | chr10-10811097-INS-51 |
| chr10-10839771-INS-1863 | chr10-10839772-INS-1863 |
| chr10-10862453-INS-214 | chr10-10862453-INS-212 |
| chr10-10897667-INS-232 | chr10-10897667-INS-232 |
| chr10-10900085-INS-540 | chr10-10900086-INS-540 |
| chr10-10913217-INS-6603 | chr10-10913218-INS-6604 |
| chr10-10914228-INS-99 | chr10-10914228-INS-99 |
| chr10-10927333-INS-68 | chr10-10927333-INS-68 |
| chr10-10931394-INS-102 | chr10-10931394-INS-54 |
| chr10-10943733-INS-73 | chr10-10943733-INS-72 |
| chr10-10998010-INS-7369 | chr10-10998011-INS-7369 |
| chr10-10999418-INS-110 | chr10-10999418-INS-110 |
| chr10-11008712-INS-62 | chr10-11008712-INS-70 |
| chr10-11047412-INS-129 | chr10-11047412-INS-135 |
| chr10-11069296-INS-211 | chr10-11069296-INS-211 |
| chr10-11071371-INS-1717 | chr10-11071372-INS-1717 |
| chr10-11072902-INS-85 | chr10-11072902-INS-85 |
| chr10-11115533-INS-214 | chr10-11115533-INS-210 |
| chr10-11127701-INS-4779 | chr10-11127702-INS-4779 |
| chr10-11142876-INS-244 | chr10-11142876-INS-244 |
| chr10-11146793-INS-7064 | chr10-11146794-INS-7064 |
| chr10-11147520-INS-245 | chr10-11147520-INS-243 |
| chr10-11161799-INS-215 | chr10-11161799-INS-215 |
| chr10-11188578-INS-2392 | chr10-11188579-INS-2392 |
| chr10-11189074-INS-89 | chr10-11189107-INS-96 |
| chr10-11191010-INS-198 | chr10-11191010-INS-198 |
| chr10-11192544-INS-199 | chr10-11192544-INS-202 |
| chr10-11206293-INS-58 | chr10-11206293-INS-58 |
| chr10-11209697-INS-196 | chr10-11209697-INS-196 |
| chr10-11226466-INS-302 | chr10-11226466-INS-310 |
| chr10-11230835-INS-5978 | chr10-11230836-INS-5978 |
| chr10-11244491-INS-60 | chr10-11244491-INS-81 |
| chr10-11264024-INS-73 | chr10-11264024-INS-73 |
| chr10-11278551-INS-1729 | chr10-11278552-INS-1726 |
| chr10-11282440-INS-569 | chr10-11282440-INS-568 |
| chr10-11285502-INS-1847 | chr10-11285503-INS-1847 |
| chr10-11285503-INS-386 | chr10-11285503-INS-386 |
| chr10-11286989-INS-695 | chr10-11286989-INS-679 |
| chr10-11289963-INS-250 | chr10-11289963-INS-250 |
| chr10-11331578-INS-63 | chr10-11331578-INS-63 |
| chr10-11334909-INS-58 | chr10-11334909-INS-56 |
| chr10-11406964-INS-87 | chr10-11406964-INS-87 |
| chr10-11411728-INS-64 | chr10-11411728-INS-64 |
| chr10-11414737-INS-51 | chr10-11414732-INS-59 |
| chr10-11431406-INS-769 | chr10-11431406-INS-767 |
| chr10-11437104-INS-58 | chr10-11437076-INS-66 |
| chr10-11447203-INS-54 | chr10-11447203-INS-56 |
| chr10-11512076-INS-425 | chr10-11512077-INS-425 |
| chr10-11565484-INS-61 | chr10-11565484-INS-61 |
| chr10-11632902-INS-73 | chr10-11632904-INS-70 |
| chr10-11637181-INS-84 | chr10-11637181-INS-120 |
| chr10-11653139-INS-532 | chr10-11653139-INS-532 |
| chr10-11654343-INS-114 | chr10-11654343-INS-108 |
| chr10-11669570-INS-7232 | chr10-11669571-INS-7232 |
| chr10-11685764-INS-536 | chr10-11685764-INS-536 |
| chr10-11693485-INS-75 | chr10-11693485-INS-76 |
| chr10-11713255-INS-168 | chr10-11713255-INS-171 |
| chr10-11724516-INS-78 | chr10-11724547-INS-77 |
| chr10-11760262-INS-68 | chr10-11760262-INS-68 |
| chr10-11780642-INS-891 | chr10-11780629-INS-896 |
| chr10-11788840-INS-60 | chr10-11788840-INS-61 |
| chr10-11808881-INS-514 | chr10-11808881-INS-514 |
| chr10-11857472-INS-199 | chr10-11857472-INS-199 |
| chr10-11859755-INS-6384 | chr10-11859756-INS-6383 |
| chr10-11875525-INS-626 | chr10-11875525-INS-626 |
| chr10-11890534-INS-707 | chr10-11890534-INS-698 |
| chr10-11892331-INS-1365 | chr10-11892332-INS-1373 |
| chr10-11914559-INS-5194 | chr10-11914560-INS-5205 |
| chr10-11919561-INS-954 | chr10-11919561-INS-956 |
| chr10-11926902-INS-572 | chr10-11926903-INS-571 |
| chr10-11935524-INS-1793 | chr10-11935525-INS-1804 |
| chr10-11938926-INS-7205 | chr10-11938927-INS-7205 |
| chr10-11944369-INS-55 | chr10-11944369-INS-55 |
| chr10-11946636-INS-73 | chr10-11946634-INS-76 |
| chr10-11968658-INS-72 | chr10-11968668-INS-89 |
| chr10-11976705-INS-56 | chr10-11976701-INS-52 |
| chr10-11979943-INS-7472 | chr10-11979944-INS-7473 |

| | |
|---|---|
| chr10-11981273-INS-6777 | chr10-11981274-INS-6777 |
| chr10-11981574-INS-341 | chr10-11981574-INS-340 |
| chr10-11985631-INS-95 | chr10-11985625-INS-82 |
| chr10-11989962-INS-105 | chr10-11989967-INS-74 |
| chr10-12004417-INS-211 | chr10-12004417-INS-211 |
| chr10-12021438-INS-352 | chr10-12021438-INS-352 |
| chr10-12030895-INS-5284 | chr10-12030896-INS-5284 |
| chr10-12046691-INS-70 | chr10-12046679-INS-64 |
| chr10-12096206-INS-80 | chr10-12096206-INS-88 |
| chr10-12096877-INS-442 | chr10-12096877-INS-442 |
| chr10-12123961-INS-3685 | chr10-12123962-INS-3686 |
| chr10-12125742-INS-54 | chr10-12125742-INS-54 |
| chr10-12150984-INS-56 | chr10-12150984-INS-56 |
| chr10-12176161-INS-148 | chr10-12176161-INS-148 |
| chr10-12206054-INS-204 | chr10-12206054-INS-204 |
| chr10-12212087-INS-235 | chr10-12212087-INS-189 |
| chr10-12218840-INS-98 | chr10-12218860-INS-125 |
| chr10-12221776-INS-64 | chr10-12221812-INS-69 |
| chr10-12272888-INS-179 | chr10-12272888-INS-194 |
| chr10-12322937-INS-80 | chr10-12322929-INS-62 |
| chr10-12333798-INS-219 | chr10-12333798-INS-222 |
| chr10-12376959-INS-276 | chr10-12376959-INS-276 |
| chr10-12379342-INS-68 | chr10-12379344-INS-74 |
| chr10-12382114-INS-215 | chr10-12382114-INS-216 |
| chr10-12404331-INS-88 | chr10-12404282-INS-98 |
| chr10-12414285-INS-100 | chr10-12414286-INS-100 |
| chr10-12433740-INS-654 | chr10-12433741-INS-654 |
| chr10-12438170-INS-215 | chr10-12438170-INS-217 |
| chr10-12443422-INS-138 | chr10-12443422-INS-150 |
| chr10-12447299-INS-119 | chr10-12447299-INS-121 |
| chr10-12456640-INS-175 | chr10-12456638-INS-177 |
| chr10-12472447-INS-258 | chr10-12472426-INS-260 |
| chr10-12515996-INS-1908 | chr10-12515997-INS-1906 |
| chr10-12527223-INS-88 | chr10-12527223-INS-88 |
| chr10-12530769-INS-226 | chr10-12530768-INS-226 |
| chr10-12545616-INS-68 | chr10-12545616-INS-68 |
| chr10-12563939-INS-56 | chr10-12563939-INS-56 |
| chr10-12582075-INS-76 | chr10-12582075-INS-82 |
| chr10-12584028-INS-74 | chr10-12584067-INS-57 |
| chr10-12614558-INS-197 | chr10-12614558-INS-198 |
| chr10-12624106-INS-420 | chr10-12624122-INS-404 |
| chr10-12660028-INS-204 | chr10-12660028-INS-204 |
| chr10-12682665-INS-98 | chr10-12682656-INS-95 |
| chr10-12697469-INS-70 | chr10-12697469-INS-70 |
| chr10-12720435-INS-207 | chr10-12720435-INS-207 |
| chr10-12722755-INS-497 | chr10-12722755-INS-497 |
| chr10-12727389-INS-90 | chr10-12727389-INS-90 |
| chr10-12739205-INS-56 | chr10-12739205-INS-56 |
| chr10-12742187-INS-75 | chr10-12742187-INS-85 |
| chr10-12749775-INS-3778 | chr10-12749776-INS-3778 |
| chr10-12754029-INS-192 | chr10-12754029-INS-195 |
| chr10-12757621-INS-138 | chr10-12757621-INS-138 |
| chr10-12783174-INS-252 | chr10-12783174-INS-252 |
| chr10-12789020-INS-507 | chr10-12789020-INS-507 |
| chr10-12795327-INS-56 | chr10-12795327-INS-56 |
| chr10-12796013-INS-64 | chr10-12796010-INS-74 |
| chr10-12808070-INS-65 | chr10-12808070-INS-70 |
| chr10-12813038-INS-128 | chr10-12813038-INS-124 |
| chr10-12827936-INS-124 | chr10-12827936-INS-124 |
| chr10-12831963-INS-583 | chr10-12831963-INS-585 |
| chr10-12845657-INS-147 | chr10-12845658-INS-146 |
| chr10-12859638-INS-54 | chr10-12859591-INS-55 |
| chr10-12866086-INS-53 | chr10-12866092-INS-56 |
| chr10-12885780-INS-224 | chr10-12885780-INS-224 |
| chr10-12904365-INS-71 | chr10-12904365-INS-68 |
| chr10-12908594-INS-123 | chr10-12908594-INS-123 |
| chr10-12912871-INS-166 | chr10-12912871-INS-177 |
| chr10-12914663-INS-105 | chr10-12914627-INS-102 |
| chr10-12929878-INS-443 | chr10-12929878-INS-442 |
| chr10-12932907-INS-58 | chr10-12932907-INS-58 |
| chr10-12940433-INS-63 | chr10-12940408-INS-55 |
| chr10-12943227-INS-309 | chr10-12943227-INS-310 |
| chr10-12945157-INS-66 | chr10-12945176-INS-69 |
| chr10-12947273-INS-426 | chr10-12947273-INS-426 |
| chr10-12955326-INS-119 | chr10-12955327-INS-115 |
| chr10-13005992-INS-58 | chr10-13005992-INS-58 |
| chr10-13009874-INS-6224 | chr10-13009875-INS-6217 |

| | |
|---|---|
| chr10-13014514-INS-54 | chr10-13014477-INS-51 |
| chr10-13020593-INS-259 | chr10-13020593-INS-247 |
| chr10-13083976-INS-96 | chr10-13083976-INS-105 |
| chr10-13145056-INS-611 | chr10-13145056-INS-612 |
| chr10-13157155-INS-69 | chr10-13157165-INS-70 |
| chr10-13177408-INS-3230 | chr10-13177409-INS-3233 |
| chr10-13181448-INS-63 | chr10-13181448-INS-63 |
| chr10-13242761-INS-66 | chr10-13242761-INS-54 |
| chr10-13318539-INS-114 | chr10-13318573-INS-129 |
| chr10-13321213-INS-86 | chr10-13321213-INS-86 |
| chr10-13326156-INS-107 | chr10-13326152-INS-105 |
| chr10-13344254-INS-160 | chr10-13344268-INS-174 |
| chr10-13358476-INS-403 | chr10-13358466-INS-430 |
| chr10-13361032-INS-699 | chr10-13361032-INS-702 |
| chr10-13364495-INS-75 | chr10-13364495-INS-87 |
| chr10-13372008-INS-1924 | chr10-13372009-INS-1924 |
| chr10-13377180-INS-82 | chr10-13377180-INS-84 |
| chr10-13380251-INS-885 | chr10-13380252-INS-881 |
| chr10-13426273-INS-210 | chr10-13426273-INS-210 |
| chr10-13439231-INS-7169 | chr10-13439232-INS-7169 |
| chr10-13449184-INS-341 | chr10-13449184-INS-341 |
| chr10-13454711-INS-221 | chr10-13454711-INS-221 |
| chr10-13456417-INS-76 | chr10-13456417-INS-76 |
| chr10-13468000-INS-53 | chr10-13468000-INS-53 |
| chr10-13470784-INS-76 | chr10-13470784-INS-76 |
| chr10-13490756-INS-6061 | chr10-13490757-INS-6058 |
| chr10-13522305-INS-1558 | chr10-13522306-INS-1560 |
| chr10-13560809-INS-435 | chr10-13560809-INS-435 |
| chr10-13582935-INS-408 | chr10-13582935-INS-408 |
| chr10-13620395-INS-70 | chr10-13620395-INS-70 |
| chr10-13636592-INS-117 | chr10-13636592-INS-117 |
| chr10-13639711-INS-192 | chr10-13639711-INS-192 |
| chr10-13648899-INS-88 | chr10-13648899-INS-87 |
| chr10-13654026-INS-1049 | chr10-13654027-INS-1045 |
| chr10-13660217-INS-55 | chr10-13660213-INS-62 |
| chr10-13694323-INS-931 | chr10-13694340-INS-908 |
| chr10-13706000-INS-82 | chr10-13705995-INS-80 |
| chr10-13725320-INS-293 | chr10-13725331-INS-298 |
| chr10-13727684-INS-94 | chr10-13727684-INS-96 |
| chr10-13747089-INS-101 | chr10-13747089-INS-101 |
| chr10-13763023-INS-3416 | chr10-13763024-INS-3427 |
| chr10-13790312-INS-491 | chr10-13790309-INS-475 |
| chr10-13811705-INS-64 | chr10-13811705-INS-64 |
| chr10-13814117-INS-107 | chr10-13814117-INS-112 |
| chr10-13875626-INS-69 | chr10-13875623-INS-57 |
| chr10-13876629-INS-71 | chr10-13876629-INS-71 |
| chr10-13879519-INS-56 | chr10-13879519-INS-56 |
| chr10-13901579-INS-96 | chr10-13901579-INS-96 |
| chr10-13930599-INS-221 | chr10-13930599-INS-221 |
| chr10-13935500-INS-56 | chr10-13935500-INS-60 |
| chr10-13945325-INS-78 | chr10-13945325-INS-52 |
| chr10-13962702-INS-90 | chr10-13962694-INS-90 |
| chr10-13964649-INS-167 | chr10-13964649-INS-167 |
| chr10-14016389-INS-54 | chr10-14016389-INS-54 |
| chr10-14035225-INS-105 | chr10-14035258-INS-105 |
| chr10-14047720-INS-60 | chr10-14047720-INS-52 |
| chr10-14049787-INS-84 | chr10-14049811-INS-83 |
| chr10-14054530-INS-67 | chr10-14054530-INS-67 |
| chr10-14056829-INS-52 | chr10-14056796-INS-82 |
| chr10-14067825-INS-383 | chr10-14067825-INS-383 |
| chr10-14077489-INS-61 | chr10-14077489-INS-90 |
| chr10-14078580-INS-6655 | chr10-14078581-INS-6655 |
| chr10-14099897-INS-55 | chr10-14099934-INS-66 |
| chr10-14102340-INS-51 | chr10-14102340-INS-51 |
| chr10-14120588-INS-6457 | chr10-14120591-INS-6445 |
| chr10-14159213-INS-621 | chr10-14159212-INS-625 |
| chr10-14172618-INS-81 | chr10-14172618-INS-82 |
| chr10-14183144-INS-62 | chr10-14183144-INS-66 |
| chr10-14184665-INS-108 | chr10-14184665-INS-120 |
| chr10-14213731-INS-285 | chr10-14213731-INS-285 |
| chr10-14215614-INS-199 | chr10-14215614-INS-199 |
| chr10-14229673-INS-2094 | chr10-14229674-INS-2098 |
| chr10-14250762-INS-51 | chr10-14250762-INS-94 |
| chr10-14298863-INS-51 | chr10-14298864-INS-54 |
| chr10-14302271-INS-69 | chr10-14302252-INS-72 |
| chr10-14311621-INS-64 | chr10-14311621-INS-64 |
| chr10-14329357-INS-86 | chr10-14329353-INS-66 |

| | |
|---|---|
| chr10-14355604-INS-445 | chr10-14355604-INS-474 |
| chr10-14367093-INS-100 | chr10-14367093-INS-102 |
| chr10-14371537-INS-75 | chr10-14371537-INS-77 |
| chr10-14372509-INS-62 | chr10-14372509-INS-62 |
| chr10-14402506-INS-546 | chr10-14402505-INS-560 |
| chr10-14404609-INS-171 | chr10-14404609-INS-171 |
| chr10-14432891-INS-385 | chr10-14432891-INS-396 |
| chr10-14456624-INS-199 | chr10-14456624-INS-199 |
| chr10-14456686-INS-1263 | chr10-14456687-INS-1263 |
| chr10-14462758-INS-199 | chr10-14462758-INS-199 |
| chr10-14491857-INS-1683 | chr10-14491856-INS-1697 |
| chr10-14502808-INS-208 | chr10-14502808-INS-208 |
| chr10-14522544-INS-378 | chr10-14522547-INS-376 |
| chr10-14621693-INS-3149 | chr10-14621694-INS-3149 |
| chr10-14628531-INS-56 | chr10-14628531-INS-56 |
| chr10-14634689-INS-6368 | chr10-14634690-INS-6368 |
| chr10-14663197-INS-92 | chr10-14663205-INS-97 |
| chr10-14676429-INS-331 | chr10-14676429-INS-331 |
| chr10-14720783-INS-105 | chr10-14720783-INS-106 |
| chr10-14744362-INS-94 | chr10-14744362-INS-94 |
| chr10-14788596-INS-55 | chr10-14788596-INS-53 |
| chr10-14805467-INS-148 | chr10-14805467-INS-147 |
| chr10-14816690-INS-156 | chr10-14816720-INS-126 |
| chr10-14837435-INS-56 | chr10-14837435-INS-56 |
| chr10-14867622-INS-506 | chr10-14867622-INS-506 |
| chr10-14870500-INS-83 | chr10-14870500-INS-85 |
| chr10-14874340-INS-7243 | chr10-14874341-INS-7243 |
| chr10-14876746-INS-328 | chr10-14876746-INS-328 |
| chr10-14897006-INS-257 | chr10-14897006-INS-257 |
| chr10-14908797-INS-202 | chr10-14908797-INS-202 |
| chr10-14974951-INS-158 | chr10-14974951-INS-158 |
| chr10-14994800-INS-3095 | chr10-14994801-INS-3096 |
| chr10-15004125-INS-417 | chr10-15004125-INS-416 |
| chr10-15020527-INS-114 | chr10-15020527-INS-114 |
| chr10-15032891-INS-134 | chr10-15032891-INS-134 |
| chr10-15037033-INS-6448 | chr10-15037034-INS-6450 |
| chr10-15099858-INS-54 | chr10-15099858-INS-58 |
| chr10-15119702-INS-55 | chr10-15119710-INS-60 |
| chr10-15127133-INS-6360 | chr10-15127134-INS-6360 |
| chr10-15139229-INS-6490 | chr10-15139230-INS-6491 |
| chr10-15165825-INS-97 | chr10-15165825-INS-97 |
| chr10-15190952-INS-76 | chr10-15190956-INS-72 |
| chr10-15230393-INS-111 | chr10-15230393-INS-91 |
| chr10-15239299-INS-86 | chr10-15239299-INS-54 |
| chr10-15280699-INS-68 | chr10-15280683-INS-69 |
| chr10-15301120-INS-6448 | chr10-15301121-INS-6450 |
| chr10-15302404-INS-2079 | chr10-15302405-INS-2079 |
| chr10-15317184-INS-6844 | chr10-15317185-INS-6845 |
| chr10-15358083-INS-56 | chr10-15358083-INS-56 |
| chr10-15403295-INS-80 | chr10-15403332-INS-75 |
| chr10-15420807-INS-1726 | chr10-15420808-INS-1693 |
| chr10-15432130-INS-2957 | chr10-15432131-INS-2957 |
| chr10-15485840-INS-428 | chr10-15485840-INS-428 |
| chr10-15491584-INS-55 | chr10-15491584-INS-50 |
| chr10-15534407-INS-271 | chr10-15534408-INS-273 |
| chr10-15574622-INS-452 | chr10-15574622-INS-452 |
| chr10-15586693-INS-184 | chr10-15586693-INS-134 |
| chr10-15602355-INS-218 | chr10-15602356-INS-217 |
| chr10-15613338-INS-66 | chr10-15613338-INS-56 |
| chr10-15645553-INS-6705 | chr10-15645554-INS-6707 |
| chr10-15657634-INS-341 | chr10-15657634-INS-340 |
| chr10-15664679-INS-390 | chr10-15664679-INS-386 |
| chr10-15686170-INS-6009 | chr10-15686171-INS-6009 |
| chr10-15688583-INS-230 | chr10-15688583-INS-230 |
| chr10-15692518-INS-87 | chr10-15692518-INS-66 |
| chr10-15704001-INS-132 | chr10-15704001-INS-147 |
| chr10-15712058-INS-541 | chr10-15712058-INS-541 |
| chr10-15722835-INS-74 | chr10-15722835-INS-74 |
| chr10-15744504-INS-541 | chr10-15744504-INS-541 |
| chr10-15745530-INS-6579 | chr10-15745531-INS-6579 |
| chr10-15772307-INS-214 | chr10-15772313-INS-214 |
| chr10-15794409-INS-57 | chr10-15794409-INS-57 |
| chr10-15804465-INS-5287 | chr10-15804466-INS-5287 |
| chr10-15818991-INS-60 | chr10-15818991-INS-70 |
| chr10-15836133-INS-56 | chr10-15836133-INS-60 |
| chr10-15855476-INS-6170 | chr10-15855477-INS-6170 |
| chr10-15856364-INS-6179 | chr10-15856365-INS-6178 |

| | |
|---|---|
| chr10-15857016-INS-800 | chr10-15857016-INS-800 |
| chr10-15859674-INS-243 | chr10-15859674-INS-243 |
| chr10-15889688-INS-202 | chr10-15889688-INS-202 |
| chr10-15903365-INS-371 | chr10-15903365-INS-371 |
| chr10-15942213-INS-53 | chr10-15942218-INS-52 |
| chr10-15969579-INS-610 | chr10-15969579-INS-610 |
| chr10-16001546-INS-85 | chr10-16001544-INS-55 |
| chr10-16002481-INS-235 | chr10-16002481-INS-235 |
| chr10-16032035-INS-311 | chr10-16032031-INS-293 |
| chr10-16037052-INS-510 | chr10-16037052-INS-510 |
| chr10-16063879-INS-52 | chr10-16063879-INS-52 |
| chr10-16091631-INS-493 | chr10-16091632-INS-495 |
| chr10-16108737-INS-100 | chr10-16108737-INS-97 |
| chr10-16125974-INS-925 | chr10-16125974-INS-927 |
| chr10-16142188-INS-1161 | chr10-16142189-INS-1150 |
| chr10-16145011-INS-56 | chr10-16145011-INS-60 |
| chr10-16152265-INS-75 | chr10-16152265-INS-75 |
| chr10-16154289-INS-113 | chr10-16154289-INS-120 |
| chr10-16200460-INS-713 | chr10-16200460-INS-713 |
| chr10-16212923-INS-608 | chr10-16212928-INS-610 |
| chr10-16214924-INS-68 | chr10-16214928-INS-60 |
| chr10-16221633-INS-115 | chr10-16221633-INS-115 |
| chr10-16247055-INS-445 | chr10-16247055-INS-445 |
| chr10-16254055-INS-95 | chr10-16254055-INS-60 |
| chr10-16268900-INS-6588 | chr10-16268901-INS-6591 |
| chr10-16280255-INS-181 | chr10-16280255-INS-197 |
| chr10-16298646-INS-762 | chr10-16298646-INS-769 |
| chr10-16315813-INS-6423 | chr10-16315814-INS-6423 |
| chr10-16320782-INS-58 | chr10-16320782-INS-58 |
| chr10-16350202-INS-4151 | chr10-16350203-INS-4150 |
| chr10-16354723-INS-5563 | chr10-16354724-INS-5563 |
| chr10-16387846-INS-88 | chr10-16387848-INS-70 |
| chr10-16409825-INS-1686 | chr10-16409826-INS-1677 |
| chr10-16432911-INS-450 | chr10-16432911-INS-450 |
| chr10-16441654-INS-904 | chr10-16441654-INS-904 |
| chr10-16447886-INS-58 | chr10-16447886-INS-58 |
| chr10-16528743-INS-6698 | chr10-16528744-INS-6701 |
| chr10-16555145-INS-1507 | chr10-16555141-INS-1504 |
| chr10-16603544-INS-3278 | chr10-16603545-INS-3278 |
| chr10-16612130-INS-281 | chr10-16612130-INS-281 |
| chr10-16642392-INS-58 | chr10-16642392-INS-66 |
| chr10-16709345-INS-141 | chr10-16709345-INS-141 |
| chr10-16714608-INS-142 | chr10-16714606-INS-173 |
| chr10-16718918-INS-6182 | chr10-16718919-INS-6182 |
| chr10-16750905-INS-4895 | chr10-16750906-INS-4895 |
| chr10-16778971-INS-905 | chr10-16778972-INS-906 |
| chr10-16793880-INS-157 | chr10-16793883-INS-149 |
| chr10-16805348-INS-52 | chr10-16805383-INS-58 |
| chr10-16806420-INS-3466 | chr10-16806421-INS-3463 |
| chr10-16808742-INS-6213 | chr10-16808743-INS-6214 |
| chr10-16820969-INS-105 | chr10-16820969-INS-105 |
| chr10-16851604-INS-115 | chr10-16851604-INS-115 |
| chr10-16880401-INS-4856 | chr10-16880402-INS-4856 |
| chr10-16896475-INS-54 | chr10-16896475-INS-88 |
| chr10-16905227-INS-253 | chr10-16905227-INS-253 |
| chr10-16909236-INS-79 | chr10-16909236-INS-72 |
| chr10-16957054-INS-56 | chr10-16957054-INS-74 |
| chr10-16959728-INS-56 | chr10-16959728-INS-77 |
| chr10-16967684-INS-296 | chr10-16967684-INS-296 |
| chr10-16967979-INS-1854 | chr10-16967980-INS-1854 |
| chr10-16981110-INS-1673 | chr10-16981111-INS-1673 |
| chr10-16986751-INS-127 | chr10-16986751-INS-94 |
| chr10-17035239-INS-59 | chr10-17035239-INS-60 |
| chr10-17042811-INS-209 | chr10-17042811-INS-210 |
| chr10-17091913-INS-889 | chr10-17091913-INS-877 |
| chr10-17091914-INS-166 | chr10-17091892-INS-182 |
| chr10-17104984-INS-52 | chr10-17104986-INS-52 |
| chr10-17126310-INS-56 | chr10-17126310-INS-58 |
| chr10-17169800-INS-211 | chr10-17169800-INS-216 |
| chr10-17186168-INS-188 | chr10-17186168-INS-184 |
| chr10-17200755-INS-177 | chr10-17200755-INS-187 |
| chr10-17221615-INS-57 | chr10-17221628-INS-61 |
| chr10-17227474-INS-64 | chr10-17227474-INS-96 |
| chr10-17260699-INS-262 | chr10-17260711-INS-216 |
| chr10-17285548-INS-52 | chr10-17285548-INS-52 |
| chr10-17303539-INS-310 | chr10-17303539-INS-310 |
| chr10-17305281-INS-67 | chr10-17305281-INS-67 |

| | |
|---|---|
| chr10-17315599-INS-161 | chr10-17315599-INS-161 |
| chr10-17322655-INS-72 | chr10-17322659-INS-84 |
| chr10-17324520-INS-171 | chr10-17324520-INS-171 |
| chr10-17330948-INS-83 | chr10-17330948-INS-83 |
| chr10-17336106-INS-197 | chr10-17336106-INS-194 |
| chr10-17348872-INS-98 | chr10-17348883-INS-88 |
| chr10-17352091-INS-3147 | chr10-17352092-INS-3147 |
| chr10-17541985-INS-652 | chr10-17541985-INS-652 |
| chr10-17642521-INS-175 | chr10-17642521-INS-175 |
| chr10-17643878-INS-1090 | chr10-17643879-INS-1090 |
| chr10-17645967-INS-83 | chr10-17645967-INS-83 |
| chr10-17675404-INS-62 | chr10-17675404-INS-72 |
| chr10-17707540-INS-86 | chr10-17707540-INS-68 |
| chr10-17726272-INS-97 | chr10-17726272-INS-99 |
| chr10-17759547-INS-91 | chr10-17759547-INS-75 |
| chr10-17826993-INS-334 | chr10-17826993-INS-343 |
| chr10-17849833-INS-52 | chr10-17849833-INS-52 |
| chr10-17860324-INS-1240 | chr10-17860325-INS-1240 |
| chr10-17867903-INS-106 | chr10-17867903-INS-106 |
| chr10-17915910-INS-82 | chr10-17915910-INS-82 |
| chr10-17951494-INS-1888 | chr10-17951495-INS-1888 |
| chr10-17952820-INS-105 | chr10-17952820-INS-105 |
| chr10-18011230-INS-191 | chr10-18011234-INS-195 |
| chr10-18014868-INS-4067 | chr10-18014869-INS-4067 |
| chr10-18025005-INS-270 | chr10-18025006-INS-269 |
| chr10-18062017-INS-54 | chr10-18062017-INS-54 |
| chr10-18070266-INS-211 | chr10-18070266-INS-211 |
| chr10-18076222-INS-193 | chr10-18076223-INS-193 |
| chr10-18082336-INS-1864 | chr10-18082337-INS-1864 |
| chr10-18084347-INS-81 | chr10-18084347-INS-81 |
| chr10-18085654-INS-197 | chr10-18085658-INS-197 |
| chr10-18109177-INS-68 | chr10-18109177-INS-68 |
| chr10-18114170-INS-52 | chr10-18114170-INS-74 |
| chr10-18118600-INS-143 | chr10-18118600-INS-143 |
| chr10-18127836-INS-91 | chr10-18127817-INS-90 |
| chr10-18140454-INS-7084 | chr10-18140455-INS-7084 |
| chr10-18157152-INS-142 | chr10-18157152-INS-148 |
| chr10-18164453-INS-133 | chr10-18164453-INS-131 |
| chr10-18167141-INS-64 | chr10-18167141-INS-64 |
| chr10-18185260-INS-65 | chr10-18185260-INS-59 |
| chr10-18188836-INS-595 | chr10-18188836-INS-595 |
| chr10-18203567-INS-63 | chr10-18203567-INS-63 |
| chr10-18213474-INS-56 | chr10-18213474-INS-57 |
| chr10-18223810-INS-207 | chr10-18223810-INS-206 |
| chr10-18227491-INS-200 | chr10-18227491-INS-200 |
| chr10-18242179-INS-1886 | chr10-18242187-INS-1879 |
| chr10-18249294-INS-53 | chr10-18249294-INS-57 |
| chr10-18250109-INS-2811 | chr10-18250110-INS-2811 |
| chr10-18257697-INS-198 | chr10-18257697-INS-198 |
| chr10-18263795-INS-83 | chr10-18263795-INS-88 |
| chr10-18270756-INS-216 | chr10-18270756-INS-217 |
| chr10-18271615-INS-68 | chr10-18271615-INS-64 |
| chr10-18297299-INS-178 | chr10-18297299-INS-179 |
| chr10-18299834-INS-97 | chr10-18299834-INS-96 |
| chr10-18310293-INS-207 | chr10-18310293-INS-207 |
| chr10-18321841-INS-209 | chr10-18321841-INS-209 |
| chr10-18352602-INS-177 | chr10-18352602-INS-177 |
| chr10-18360920-INS-223 | chr10-18360920-INS-224 |
| chr10-18369514-INS-55 | chr10-18369514-INS-54 |
| chr10-18375982-INS-175 | chr10-18375982-INS-175 |
| chr10-18383332-INS-198 | chr10-18383332-INS-198 |
| chr10-18395707-INS-56 | chr10-18395702-INS-61 |
| chr10-18396246-INS-52 | chr10-18396246-INS-50 |
| chr10-18406887-INS-218 | chr10-18406919-INS-208 |
| chr10-18435163-INS-57 | chr10-18435163-INS-57 |
| chr10-18448548-INS-196 | chr10-18448549-INS-158 |
| chr10-18449739-INS-604 | chr10-18449739-INS-604 |
| chr10-18450343-INS-6164 | chr10-18450344-INS-6164 |
| chr10-18455469-INS-170 | chr10-18455469-INS-170 |
| chr10-18478557-INS-401 | chr10-18478557-INS-401 |
| chr10-18489823-INS-68 | chr10-18489823-INS-64 |
| chr10-18493696-INS-63 | chr10-18493696-INS-63 |
| chr10-18558580-INS-84 | chr10-18558580-INS-84 |
| chr10-18580848-INS-1271 | chr10-18580849-INS-1273 |
| chr10-18583517-INS-54 | chr10-18583543-INS-75 |
| chr10-18611737-INS-56 | chr10-18611737-INS-60 |
| chr10-18637370-INS-393 | chr10-18637370-INS-393 |

| | |
|---|---|
| chr10-18686237-INS-54 | chr10-18686237-INS-56 |
| chr10-18698959-INS-548 | chr10-18698959-INS-548 |
| chr10-18702871-INS-73 | chr10-18702871-INS-52 |
| chr10-18711497-INS-52 | chr10-18711497-INS-54 |
| chr10-18723923-INS-71 | chr10-18723923-INS-55 |
| chr10-18737260-INS-99 | chr10-18737254-INS-146 |
| chr10-18737998-INS-56 | chr10-18737977-INS-63 |
| chr10-18759175-INS-74 | chr10-18759175-INS-74 |
| chr10-18775974-INS-68 | chr10-18775974-INS-69 |
| chr10-18783691-INS-1365 | chr10-18783692-INS-1368 |
| chr10-18796291-INS-52 | chr10-18796319-INS-50 |
| chr10-18827812-INS-72 | chr10-18827812-INS-75 |
| chr10-18846148-INS-126 | chr10-18846148-INS-126 |
| chr10-18847237-INS-60 | chr10-18847237-INS-60 |
| chr10-18848221-INS-194 | chr10-18848221-INS-195 |
| chr10-18934983-INS-347 | chr10-18934983-INS-347 |
| chr10-18940263-INS-160 | chr10-18940263-INS-132 |
| chr10-18949523-INS-555 | chr10-18949523-INS-556 |
| chr10-18970679-INS-508 | chr10-18970676-INS-504 |
| chr10-18984245-INS-400 | chr10-18984245-INS-400 |
| chr10-18989866-INS-900 | chr10-18989866-INS-900 |
| chr10-18993652-INS-95 | chr10-18993652-INS-128 |
| chr10-18999394-INS-142 | chr10-18999394-INS-140 |
| chr10-19022748-INS-54 | chr10-19022748-INS-56 |
| chr10-19023842-INS-122 | chr10-19023842-INS-122 |
| chr10-19042032-INS-70 | chr10-19042032-INS-66 |
| chr10-19082065-INS-624 | chr10-19082065-INS-624 |
| chr10-19083153-INS-237 | chr10-19083153-INS-237 |
| chr10-19127188-INS-206 | chr10-19127188-INS-212 |
| chr10-19139538-INS-283 | chr10-19139538-INS-283 |
| chr10-19157787-INS-76 | chr10-19157787-INS-76 |
| chr10-19164079-INS-92 | chr10-19164079-INS-92 |
| chr10-19164968-INS-3315 | chr10-19164969-INS-3291 |
| chr10-19169725-INS-52 | chr10-19169727-INS-54 |
| chr10-19183925-INS-257 | chr10-19183925-INS-247 |
| chr10-19194694-INS-230 | chr10-19194694-INS-226 |
| chr10-19239138-INS-66 | chr10-19239138-INS-66 |
| chr10-19279050-INS-59 | chr10-19279050-INS-54 |
| chr10-19305501-INS-101 | chr10-19305501-INS-101 |
| chr10-19307463-INS-100 | chr10-19307463-INS-123 |
| chr10-19313104-INS-51 | chr10-19313104-INS-51 |
| chr10-19346278-INS-1393 | chr10-19346279-INS-1397 |
| chr10-19360815-INS-88 | chr10-19360815-INS-76 |
| chr10-19390529-INS-854 | chr10-19390530-INS-854 |
| chr10-19413269-INS-1756 | chr10-19413276-INS-1726 |
| chr10-19433485-INS-348 | chr10-19433490-INS-353 |
| chr10-19435135-INS-390 | chr10-19435135-INS-390 |
| chr10-19452018-INS-6437 | chr10-19452019-INS-6437 |
| chr10-19454800-INS-55 | chr10-19454800-INS-60 |
| chr10-19479988-INS-208 | chr10-19479988-INS-208 |
| chr10-19484514-INS-73 | chr10-19484514-INS-73 |
| chr10-19487403-INS-264 | chr10-19487403-INS-263 |
| chr10-19540794-INS-78 | chr10-19540794-INS-84 |
| chr10-19545884-INS-66 | chr10-19545884-INS-72 |
| chr10-19565416-INS-108 | chr10-19565416-INS-91 |
| chr10-19601340-INS-63 | chr10-19601294-INS-75 |
| chr10-19612950-INS-52 | chr10-19612950-INS-63 |
| chr10-19630696-INS-1891 | chr10-19630697-INS-1891 |
| chr10-19642295-INS-121 | chr10-19642296-INS-121 |
| chr10-19645045-INS-258 | chr10-19645045-INS-257 |
| chr10-19678105-INS-304 | chr10-19678110-INS-259 |
| chr10-19686882-INS-1893 | chr10-19686883-INS-1892 |
| chr10-19688499-INS-73 | chr10-19688475-INS-84 |
| chr10-19696211-INS-52 | chr10-19696211-INS-52 |
| chr10-19717214-INS-538 | chr10-19717214-INS-538 |
| chr10-19722320-INS-205 | chr10-19722320-INS-206 |
| chr10-19742769-INS-58 | chr10-19742769-INS-58 |
| chr10-19759873-INS-73 | chr10-19759873-INS-73 |
| chr10-19769586-INS-84 | chr10-19769586-INS-86 |
| chr10-19799439-INS-639 | chr10-19799439-INS-637 |
| chr10-19808409-INS-200 | chr10-19808409-INS-200 |
| chr10-19819964-INS-179 | chr10-19819955-INS-154 |
| chr10-19843756-INS-91 | chr10-19843756-INS-91 |
| chr10-19878423-INS-55 | chr10-19878427-INS-96 |
| chr10-19929069-INS-1501 | chr10-19929070-INS-1501 |
| chr10-19933499-INS-120 | chr10-19933499-INS-109 |
| chr10-19955236-INS-146 | chr10-19955236-INS-152 |

| | |
|---|---|
| chr10-19988625-INS-203 | chr10-19988625-INS-204 |
| chr10-20036085-INS-181 | chr10-20036085-INS-182 |
| chr10-20044039-INS-74 | chr10-20044076-INS-74 |
| chr10-20063450-INS-712 | chr10-20063450-INS-712 |
| chr10-20068033-INS-55 | chr10-20068026-INS-55 |
| chr10-20081851-INS-113 | chr10-20081851-INS-113 |
| chr10-20087132-INS-5219 | chr10-20087133-INS-5219 |
| chr10-20091548-INS-274 | chr10-20091548-INS-274 |
| chr10-20094350-INS-52 | chr10-20094350-INS-58 |
| chr10-20119493-INS-641 | chr10-20119481-INS-632 |
| chr10-20140849-INS-51 | chr10-20140849-INS-50 |
| chr10-20142019-INS-216 | chr10-20142019-INS-216 |
| chr10-20174431-INS-209 | chr10-20174431-INS-210 |
| chr10-20178932-INS-225 | chr10-20178932-INS-226 |
| chr10-20280306-INS-426 | chr10-20280306-INS-426 |
| chr10-20284178-INS-54 | chr10-20284178-INS-51 |
| chr10-20298602-INS-3801 | chr10-20298603-INS-3793 |
| chr10-20304410-INS-81 | chr10-20304410-INS-81 |
| chr10-20306382-INS-72 | chr10-20306382-INS-76 |
| chr10-20328867-INS-227 | chr10-20328867-INS-227 |
| chr10-20334364-INS-170 | chr10-20334362-INS-172 |
| chr10-20353180-INS-69 | chr10-20353180-INS-69 |
| chr10-20354622-INS-6467 | chr10-20354623-INS-6467 |
| chr10-20356173-INS-66 | chr10-20356173-INS-54 |
| chr10-20357748-INS-59 | chr10-20357748-INS-59 |
| chr10-20418697-INS-89 | chr10-20418697-INS-88 |
| chr10-20420482-INS-256 | chr10-20420482-INS-256 |
| chr10-20429004-INS-115 | chr10-20429004-INS-115 |
| chr10-20501148-INS-498 | chr10-20501148-INS-498 |
| chr10-20511774-INS-76 | chr10-20511757-INS-104 |
| chr10-20517540-INS-328 | chr10-20517540-INS-328 |
| chr10-20564754-INS-186 | chr10-20564754-INS-186 |
| chr10-20570519-INS-211 | chr10-20570516-INS-220 |
| chr10-20575164-INS-80 | chr10-20575164-INS-80 |
| chr10-20579051-INS-88 | chr10-20579051-INS-88 |
| chr10-20581602-INS-54 | chr10-20581602-INS-50 |
| chr10-20582744-INS-128 | chr10-20582744-INS-128 |
| chr10-20585224-INS-400 | chr10-20585224-INS-400 |
| chr10-20596779-INS-72 | chr10-20596779-INS-72 |
| chr10-20616505-INS-60 | chr10-20616505-INS-60 |
| chr10-20630537-INS-83 | chr10-20630537-INS-85 |
| chr10-20633008-INS-328 | chr10-20633008-INS-328 |
| chr10-20635918-INS-53 | chr10-20635918-INS-59 |
| chr10-20641263-INS-129 | chr10-20641276-INS-144 |
| chr10-20642905-INS-96 | chr10-20642906-INS-55 |
| chr10-20664750-INS-95 | chr10-20664750-INS-95 |
| chr10-20698713-INS-126 | chr10-20698719-INS-132 |
| chr10-20700374-INS-64 | chr10-20700374-INS-64 |
| chr10-20720840-INS-528 | chr10-20720850-INS-496 |
| chr10-20736590-INS-6894 | chr10-20736591-INS-6896 |
| chr10-20759366-INS-198 | chr10-20759366-INS-200 |
| chr10-20857676-INS-515 | chr10-20857676-INS-515 |
| chr10-20923592-INS-2850 | chr10-20923593-INS-2838 |
| chr10-20928915-INS-72 | chr10-20928915-INS-72 |
| chr10-20970861-INS-154 | chr10-20970860-INS-116 |
| chr10-20984487-INS-62 | chr10-20984493-INS-102 |
| chr10-21006647-INS-83 | chr10-21006637-INS-55 |
| chr10-21010034-INS-60 | chr10-21010034-INS-60 |
| chr10-21035405-INS-75 | chr10-21035403-INS-70 |
| chr10-21128404-INS-162 | chr10-21128404-INS-162 |
| chr10-21142444-INS-234 | chr10-21142444-INS-224 |
| chr10-21257699-INS-75 | chr10-21257699-INS-60 |
| chr10-21269187-INS-111 | chr10-21269187-INS-111 |
| chr10-21304121-INS-160 | chr10-21304121-INS-160 |
| chr10-21335000-INS-357 | chr10-21335000-INS-357 |
| chr10-21368352-INS-4667 | chr10-21368353-INS-4667 |
| chr10-21386762-INS-220 | chr10-21386762-INS-218 |
| chr10-21434391-INS-133 | chr10-21434382-INS-166 |
| chr10-21440574-INS-299 | chr10-21440574-INS-299 |
| chr10-21444158-INS-84 | chr10-21444158-INS-84 |
| chr10-21446035-INS-111 | chr10-21446035-INS-111 |
| chr10-21447108-INS-2572 | chr10-21447109-INS-2572 |
| chr10-21469578-INS-250 | chr10-21469578-INS-250 |
| chr10-21481177-INS-78 | chr10-21481177-INS-78 |
| chr10-21495119-INS-368 | chr10-21495119-INS-368 |
| chr10-21527409-INS-199 | chr10-21527409-INS-199 |
| chr10-21532088-INS-204 | chr10-21532088-INS-204 |

| | |
|---|---|
| chr10-21540333-INS-322 | chr10-21540333-INS-322 |
| chr10-21574345-INS-400 | chr10-21574345-INS-399 |
| chr10-21595414-INS-328 | chr10-21595414-INS-328 |
| chr10-21619276-INS-3494 | chr10-21619277-INS-3494 |
| chr10-21632561-INS-107 | chr10-21632561-INS-107 |
| chr10-21646462-INS-343 | chr10-21646462-INS-343 |
| chr10-21657692-INS-2156 | chr10-21657693-INS-2156 |
| chr10-21684612-INS-113 | chr10-21684612-INS-138 |
| chr10-21708794-INS-55 | chr10-21708794-INS-55 |
| chr10-21717901-INS-201 | chr10-21717901-INS-201 |
| chr10-21741163-INS-217 | chr10-21741163-INS-217 |
| chr10-21764828-INS-56 | chr10-21764828-INS-58 |
| chr10-21780045-INS-51 | chr10-21780028-INS-61 |
| chr10-21798632-INS-115 | chr10-21798632-INS-117 |
| chr10-21812189-INS-51 | chr10-21812176-INS-87 |
| chr10-21828224-INS-275 | chr10-21828224-INS-299 |
| chr10-21835656-INS-227 | chr10-21835656-INS-227 |
| chr10-21877937-INS-103 | chr10-21877937-INS-108 |
| chr10-21893398-INS-127 | chr10-21893398-INS-120 |
| chr10-21897287-INS-838 | chr10-21897287-INS-838 |
| chr10-21910442-INS-61 | chr10-21910442-INS-60 |
| chr10-21915948-INS-499 | chr10-21915948-INS-501 |
| chr10-22045771-INS-300 | chr10-22045771-INS-308 |
| chr10-22075273-INS-177 | chr10-22075273-INS-177 |
| chr10-22146064-INS-102 | chr10-22146058-INS-112 |
| chr10-22169223-INS-102 | chr10-22169223-INS-102 |
| chr10-22214246-INS-4143 | chr10-22214247-INS-4143 |
| chr10-22264023-INS-70 | chr10-22264024-INS-93 |
| chr10-22330168-INS-97 | chr10-22330162-INS-100 |
| chr10-22332417-INS-415 | chr10-22332417-INS-415 |
| chr10-22333493-INS-160 | chr10-22333493-INS-161 |
| chr10-22337396-INS-66 | chr10-22337396-INS-60 |
| chr10-22346789-INS-2343 | chr10-22346790-INS-2343 |
| chr10-22355457-INS-180 | chr10-22355457-INS-180 |
| chr10-22358962-INS-62 | chr10-22358962-INS-70 |
| chr10-22377732-INS-4083 | chr10-22377733-INS-4087 |
| chr10-22416978-INS-221 | chr10-22416990-INS-225 |
| chr10-22454940-INS-630 | chr10-22454940-INS-631 |
| chr10-22467531-INS-412 | chr10-22467531-INS-444 |
| chr10-22476667-INS-175 | chr10-22476667-INS-178 |
| chr10-22529126-INS-207 | chr10-22529126-INS-205 |
| chr10-22593581-INS-219 | chr10-22593581-INS-219 |
| chr10-22621258-INS-96 | chr10-22621258-INS-96 |
| chr10-22672888-INS-246 | chr10-22672888-INS-252 |
| chr10-22691157-INS-54 | chr10-22691151-INS-72 |
| chr10-22693878-INS-63 | chr10-22693878-INS-63 |
| chr10-22701492-INS-410 | chr10-22701504-INS-391 |
| chr10-22724542-INS-161 | chr10-22724542-INS-164 |
| chr10-22735220-INS-212 | chr10-22735220-INS-212 |
| chr10-22767527-INS-497 | chr10-22767527-INS-497 |
| chr10-22778729-INS-240 | chr10-22778760-INS-257 |
| chr10-22791953-INS-82 | chr10-22791953-INS-82 |
| chr10-22798175-INS-52 | chr10-22798175-INS-52 |
| chr10-22806216-INS-82 | chr10-22806217-INS-84 |
| chr10-22807121-INS-81 | chr10-22807121-INS-81 |
| chr10-22808829-INS-57 | chr10-22808829-INS-57 |
| chr10-22819989-INS-366 | chr10-22819989-INS-366 |
| chr10-22835881-INS-400 | chr10-22835881-INS-400 |
| chr10-22836559-INS-1327 | chr10-22836560-INS-1327 |
| chr10-22841782-INS-2895 | chr10-22841783-INS-2895 |
| chr10-22849570-INS-96 | chr10-22849585-INS-95 |
| chr10-22852961-INS-108 | chr10-22852957-INS-116 |
| chr10-22884450-INS-266 | chr10-22884450-INS-266 |
| chr10-22887395-INS-51 | chr10-22887387-INS-51 |
| chr10-22892182-INS-66 | chr10-22892164-INS-86 |
| chr10-22912844-INS-92 | chr10-22912844-INS-86 |
| chr10-22924335-INS-211 | chr10-22924335-INS-211 |
| chr10-22946416-INS-663 | chr10-22946416-INS-667 |
| chr10-22973224-INS-64 | chr10-22973224-INS-64 |
| chr10-22976201-INS-7122 | chr10-22976202-INS-7122 |
| chr10-23015737-INS-55 | chr10-23015737-INS-67 |
| chr10-23083949-INS-61 | chr10-23083949-INS-63 |
| chr10-23105951-INS-59 | chr10-23105951-INS-71 |
| chr10-23113959-INS-62 | chr10-23113959-INS-62 |
| chr10-23125263-INS-96 | chr10-23125259-INS-100 |
| chr10-23132672-INS-123 | chr10-23132672-INS-99 |
| chr10-23156439-INS-568 | chr10-23156439-INS-568 |

| | |
|---|---|
| chr10-23170221-INS-498 | chr10-23170221-INS-498 |
| chr10-23178351-INS-408 | chr10-23178351-INS-420 |
| chr10-23195763-INS-105 | chr10-23195763-INS-105 |
| chr10-23262312-INS-74 | chr10-23262324-INS-64 |
| chr10-23267496-INS-540 | chr10-23267492-INS-533 |
| chr10-23286114-INS-58 | chr10-23286114-INS-58 |
| chr10-23292102-INS-74 | chr10-23292102-INS-78 |
| chr10-23306033-INS-304 | chr10-23306033-INS-304 |
| chr10-23307732-INS-3648 | chr10-23307733-INS-3648 |
| chr10-23327888-INS-819 | chr10-23327888-INS-819 |
| chr10-23332652-INS-73 | chr10-23332653-INS-70 |
| chr10-23364935-INS-70 | chr10-23364935-INS-56 |
| chr10-23376414-INS-65 | chr10-23376414-INS-65 |
| chr10-23385578-INS-578 | chr10-23385578-INS-579 |
| chr10-23397611-INS-69 | chr10-23397611-INS-69 |
| chr10-23449097-INS-172 | chr10-23449097-INS-173 |
| chr10-23455246-INS-81 | chr10-23455246-INS-123 |
| chr10-23460521-INS-68 | chr10-23460521-INS-68 |
| chr10-23532585-INS-72 | chr10-23532585-INS-76 |
| chr10-23549079-INS-105 | chr10-23549079-INS-105 |
| chr10-23562136-INS-176 | chr10-23562136-INS-176 |
| chr10-23568594-INS-101 | chr10-23568594-INS-101 |
| chr10-23608345-INS-78 | chr10-23608345-INS-68 |
| chr10-23610610-INS-55 | chr10-23610611-INS-53 |
| chr10-23616635-INS-229 | chr10-23616635-INS-229 |
| chr10-23617580-INS-438 | chr10-23617580-INS-438 |
| chr10-23622490-INS-175 | chr10-23622490-INS-168 |
| chr10-23639797-INS-76 | chr10-23639797-INS-76 |
| chr10-23645202-INS-219 | chr10-23645202-INS-219 |
| chr10-23645967-INS-986 | chr10-23645967-INS-986 |
| chr10-23648916-INS-607 | chr10-23648916-INS-610 |
| chr10-23652069-INS-204 | chr10-23652069-INS-204 |
| chr10-23659611-INS-87 | chr10-23659611-INS-84 |
| chr10-23668470-INS-60 | chr10-23668470-INS-60 |
| chr10-23672230-INS-233 | chr10-23672230-INS-233 |
| chr10-23679295-INS-206 | chr10-23679296-INS-203 |
| chr10-23717019-INS-53 | chr10-23717019-INS-53 |
| chr10-23726552-INS-306 | chr10-23726552-INS-305 |
| chr10-23728358-INS-78 | chr10-23728358-INS-78 |
| chr10-23760659-INS-66 | chr10-23760688-INS-50 |
| chr10-23820174-INS-211 | chr10-23820161-INS-186 |
| chr10-23824386-INS-55 | chr10-23824386-INS-55 |
| chr10-23825156-INS-875 | chr10-23825156-INS-875 |
| chr10-23833910-INS-335 | chr10-23833910-INS-338 |
| chr10-23838197-INS-169 | chr10-23838193-INS-168 |
| chr10-23839274-INS-989 | chr10-23839274-INS-995 |
| chr10-23840943-INS-81 | chr10-23840946-INS-81 |
| chr10-23845573-INS-62 | chr10-23845573-INS-68 |
| chr10-23923114-INS-344 | chr10-23923114-INS-344 |
| chr10-23964089-INS-60 | chr10-23964089-INS-72 |
| chr10-23981383-INS-72 | chr10-23981383-INS-70 |
| chr10-24016279-INS-150 | chr10-24016279-INS-150 |
| chr10-24039914-INS-70 | chr10-24039914-INS-70 |
| chr10-24084047-INS-340 | chr10-24084046-INS-336 |
| chr10-24089043-INS-53 | chr10-24089043-INS-87 |
| chr10-24092819-INS-60 | chr10-24092819-INS-60 |
| chr10-24097043-INS-52 | chr10-24097043-INS-52 |
| chr10-24116438-INS-76 | chr10-24116438-INS-76 |
| chr10-24128592-INS-184 | chr10-24128592-INS-188 |
| chr10-24134690-INS-399 | chr10-24134690-INS-400 |
| chr10-24149118-INS-52 | chr10-24149118-INS-50 |
| chr10-24152151-INS-8115 | chr10-24152152-INS-8116 |
| chr10-24161774-INS-57 | chr10-24161774-INS-57 |
| chr10-24192572-INS-66 | chr10-24192572-INS-66 |
| chr10-24200915-INS-64 | chr10-24200931-INS-76 |
| chr10-24231957-INS-52 | chr10-24231957-INS-52 |
| chr10-24246124-INS-138 | chr10-24246118-INS-132 |
| chr10-24252649-INS-491 | chr10-24252649-INS-491 |
| chr10-24287900-INS-93 | chr10-24287900-INS-112 |
| chr10-24290613-INS-6219 | chr10-24290614-INS-6220 |
| chr10-24293088-INS-1978 | chr10-24293089-INS-1996 |
| chr10-24294970-INS-121 | chr10-24294970-INS-121 |
| chr10-24320197-INS-3472 | chr10-24320198-INS-3472 |
| chr10-24336406-INS-593 | chr10-24336402-INS-593 |
| chr10-24349704-INS-54 | chr10-24349704-INS-54 |
| chr10-24356949-INS-6228 | chr10-24356950-INS-6228 |
| chr10-24391349-INS-387 | chr10-24391348-INS-386 |

| | |
|---|---|
| chr10-24413270-INS-794 | chr10-24413270-INS-794 |
| chr10-24419508-INS-91 | chr10-24419527-INS-52 |
| chr10-24425189-INS-76 | chr10-24425189-INS-76 |
| chr10-24479119-INS-98 | chr10-24479119-INS-98 |
| chr10-24481721-INS-207 | chr10-24481721-INS-207 |
| chr10-24488057-INS-88 | chr10-24488057-INS-88 |
| chr10-24518020-INS-319 | chr10-24518018-INS-323 |
| chr10-24522832-INS-86 | chr10-24522829-INS-86 |
| chr10-24525604-INS-107 | chr10-24525604-INS-107 |
| chr10-24531073-INS-78 | chr10-24531073-INS-78 |
| chr10-24541310-INS-57 | chr10-24541310-INS-57 |
| chr10-24549926-INS-61 | chr10-24549881-INS-70 |
| chr10-24591701-INS-52 | chr10-24591701-INS-52 |
| chr10-24611678-INS-70 | chr10-24611678-INS-68 |
| chr10-24614261-INS-140 | chr10-24614261-INS-140 |
| chr10-24681619-INS-139 | chr10-24681619-INS-135 |
| chr10-24690526-INS-177 | chr10-24690526-INS-181 |
| chr10-24700594-INS-88 | chr10-24700574-INS-94 |
| chr10-24706016-INS-60 | chr10-24706016-INS-94 |
| chr10-24733444-INS-7000 | chr10-24733445-INS-7000 |
| chr10-24733506-INS-497 | chr10-24733506-INS-497 |
| chr10-24738778-INS-64 | chr10-24738800-INS-63 |
| chr10-24748683-INS-399 | chr10-24748683-INS-399 |
| chr10-24753533-INS-164 | chr10-24753533-INS-205 |
| chr10-24768219-INS-170 | chr10-24768219-INS-172 |
| chr10-24845421-INS-62 | chr10-24845421-INS-68 |
| chr10-24855128-INS-92 | chr10-24855128-INS-92 |
| chr10-24868993-INS-90 | chr10-24868993-INS-90 |
| chr10-24879832-INS-133 | chr10-24879832-INS-133 |
| chr10-24885909-INS-555 | chr10-24885909-INS-555 |
| chr10-24937205-INS-250 | chr10-24937205-INS-248 |
| chr10-24944238-INS-102 | chr10-24944238-INS-102 |
| chr10-24965343-INS-229 | chr10-24965387-INS-235 |
| chr10-24976847-INS-81 | chr10-24976847-INS-81 |
| chr10-24989779-INS-211 | chr10-24989779-INS-211 |
| chr10-25000634-INS-767 | chr10-25000634-INS-767 |
| chr10-25010316-INS-1704 | chr10-25010317-INS-1705 |
| chr10-25014172-INS-129 | chr10-25014172-INS-129 |
| chr10-25025583-INS-499 | chr10-25025583-INS-499 |
| chr10-25029878-INS-123 | chr10-25029924-INS-83 |
| chr10-25057013-INS-98 | chr10-25057013-INS-100 |
| chr10-25060131-INS-209 | chr10-25060131-INS-209 |
| chr10-25084225-INS-699 | chr10-25084225-INS-667 |
| chr10-25090043-INS-108 | chr10-25090052-INS-147 |
| chr10-25103647-INS-154 | chr10-25103647-INS-154 |
| chr10-25108560-INS-447 | chr10-25108560-INS-447 |
| chr10-25112919-INS-58 | chr10-25112912-INS-72 |
| chr10-25112950-INS-708 | chr10-25112950-INS-708 |
| chr10-25120085-INS-203 | chr10-25120079-INS-186 |
| chr10-25127549-INS-624 | chr10-25127546-INS-635 |
| chr10-25141111-INS-562 | chr10-25141111-INS-562 |
| chr10-25143489-INS-130 | chr10-25143489-INS-129 |
| chr10-25150375-INS-195 | chr10-25150375-INS-197 |
| chr10-25166400-INS-155 | chr10-25166361-INS-160 |
| chr10-25173567-INS-330 | chr10-25173567-INS-330 |
| chr10-25183202-INS-514 | chr10-25183202-INS-514 |
| chr10-25188909-INS-76 | chr10-25188946-INS-54 |
| chr10-25198237-INS-774 | chr10-25198233-INS-763 |
| chr10-25201886-INS-217 | chr10-25201886-INS-217 |
| chr10-25206669-INS-206 | chr10-25206669-INS-205 |
| chr10-25210505-INS-326 | chr10-25210505-INS-326 |
| chr10-25213796-INS-365 | chr10-25213796-INS-365 |
| chr10-25215011-INS-150 | chr10-25215011-INS-150 |
| chr10-25217576-INS-56 | chr10-25217576-INS-56 |
| chr10-25226747-INS-332 | chr10-25226747-INS-332 |
| chr10-25232657-INS-147 | chr10-25232654-INS-139 |
| chr10-25232665-INS-1195 | chr10-25232653-INS-1216 |
| chr10-25233492-INS-4453 | chr10-25233493-INS-4453 |
| chr10-25251855-INS-206 | chr10-25251855-INS-206 |
| chr10-25258886-INS-320 | chr10-25258886-INS-321 |
| chr10-25265475-INS-66 | chr10-25265475-INS-66 |
| chr10-25267695-INS-102 | chr10-25267695-INS-101 |
| chr10-25270183-INS-909 | chr10-25270183-INS-909 |
| chr10-25274878-INS-77 | chr10-25274878-INS-79 |
| chr10-25276331-INS-154 | chr10-25276331-INS-154 |
| chr10-25293336-INS-272 | chr10-25293336-INS-272 |
| chr10-25309583-INS-93 | chr10-25309612-INS-62 |

| | |
|---|---|
| chr10-25318423-INS-119 | chr10-25318423-INS-161 |
| chr10-25322386-INS-70 | chr10-25322386-INS-70 |
| chr10-25346248-INS-5305 | chr10-25346249-INS-5305 |
| chr10-25347067-INS-71 | chr10-25347067-INS-108 |
| chr10-25360990-INS-56 | chr10-25360981-INS-92 |
| chr10-25408531-INS-187 | chr10-25408531-INS-187 |
| chr10-25409463-INS-221 | chr10-25409463-INS-221 |
| chr10-25410316-INS-62 | chr10-25410316-INS-62 |
| chr10-25425623-INS-95 | chr10-25425623-INS-95 |
| chr10-25450796-INS-204 | chr10-25450796-INS-205 |
| chr10-25455725-INS-393 | chr10-25455725-INS-441 |
| chr10-25466254-INS-514 | chr10-25466254-INS-514 |
| chr10-25475969-INS-507 | chr10-25475969-INS-507 |
| chr10-25481017-INS-168 | chr10-25481017-INS-192 |
| chr10-25484398-INS-743 | chr10-25484398-INS-743 |
| chr10-25538695-INS-55 | chr10-25538695-INS-55 |
| chr10-25560337-INS-5544 | chr10-25560338-INS-5544 |
| chr10-25561306-INS-120 | chr10-25561288-INS-156 |
| chr10-25600840-INS-70 | chr10-25600840-INS-70 |
| chr10-25602994-INS-393 | chr10-25602994-INS-393 |
| chr10-25611006-INS-85 | chr10-25611006-INS-85 |
| chr10-25621614-INS-59 | chr10-25621614-INS-59 |
| chr10-25629209-INS-70 | chr10-25629199-INS-62 |
| chr10-25639238-INS-470 | chr10-25639238-INS-471 |
| chr10-25643406-INS-52 | chr10-25643382-INS-76 |
| chr10-25653433-INS-264 | chr10-25653433-INS-264 |
| chr10-25712433-INS-8759 | chr10-25712434-INS-8760 |
| chr10-25783438-INS-210 | chr10-25783438-INS-210 |
| chr10-25787109-INS-500 | chr10-25787109-INS-501 |
| chr10-25800044-INS-94 | chr10-25800044-INS-94 |
| chr10-25803167-INS-129 | chr10-25803167-INS-132 |
| chr10-25807033-INS-52 | chr10-25807024-INS-62 |
| chr10-25829979-INS-64 | chr10-25829979-INS-76 |
| chr10-25872488-INS-79 | chr10-25872484-INS-50 |
| chr10-25878958-INS-545 | chr10-25878958-INS-545 |
| chr10-25884184-INS-118 | chr10-25884184-INS-118 |
| chr10-25906441-INS-54 | chr10-25906441-INS-54 |
| chr10-26002887-INS-201 | chr10-26002887-INS-201 |
| chr10-26003729-INS-386 | chr10-26003729-INS-386 |
| chr10-26011724-INS-117 | chr10-26011724-INS-117 |
| chr10-26049371-INS-102 | chr10-26049371-INS-98 |
| chr10-26050911-INS-90 | chr10-26050915-INS-126 |
| chr10-26072026-INS-7064 | chr10-26072027-INS-7064 |
| chr10-26077751-INS-603 | chr10-26077751-INS-601 |
| chr10-26081138-INS-168 | chr10-26081138-INS-168 |
| chr10-26090853-INS-76 | chr10-26090847-INS-50 |
| chr10-26092405-INS-69 | chr10-26092405-INS-66 |
| chr10-26098448-INS-69 | chr10-26098448-INS-65 |
| chr10-26110373-INS-796 | chr10-26110373-INS-803 |
| chr10-26125159-INS-64 | chr10-26125159-INS-68 |
| chr10-26145116-INS-60 | chr10-26145116-INS-60 |
| chr10-26149707-INS-82 | chr10-26149707-INS-84 |
| chr10-26197666-INS-135 | chr10-26197668-INS-123 |
| chr10-26209171-INS-54 | chr10-26209171-INS-58 |
| chr10-26210026-INS-195 | chr10-26210026-INS-202 |
| chr10-26236233-INS-113 | chr10-26236233-INS-119 |
| chr10-26262509-INS-54 | chr10-26262509-INS-56 |
| chr10-26265949-INS-222 | chr10-26265949-INS-222 |
| chr10-26278180-INS-208 | chr10-26278180-INS-208 |
| chr10-26298221-INS-5248 | chr10-26298222-INS-5248 |
| chr10-26323746-INS-100 | chr10-26323746-INS-100 |
| chr10-26350094-INS-97 | chr10-26350061-INS-104 |
| chr10-26388825-INS-115 | chr10-26388825-INS-115 |
| chr10-26400506-INS-408 | chr10-26400506-INS-408 |
| chr10-26432858-INS-114 | chr10-26432858-INS-114 |
| chr10-26445488-INS-498 | chr10-26445488-INS-498 |
| chr10-26451137-INS-84 | chr10-26451137-INS-84 |
| chr10-26458656-INS-57 | chr10-26458656-INS-57 |
| chr10-26466448-INS-82 | chr10-26466448-INS-82 |
| chr10-26477146-INS-1890 | chr10-26477147-INS-1890 |
| chr10-26492690-INS-342 | chr10-26492690-INS-342 |
| chr10-26559689-INS-311 | chr10-26559689-INS-311 |
| chr10-26560705-INS-603 | chr10-26560706-INS-611 |
| chr10-26561783-INS-109 | chr10-26561783-INS-109 |
| chr10-26574204-INS-124 | chr10-26574204-INS-124 |
| chr10-26590660-INS-6995 | chr10-26590661-INS-6996 |
| chr10-26612598-INS-79 | chr10-26612598-INS-79 |

| | |
|---|---|
| chr10-26617746-INS-102 | chr10-26617746-INS-102 |
| chr10-26634324-INS-91 | chr10-26634324-INS-91 |
| chr10-26642909-INS-74 | chr10-26642911-INS-78 |
| chr10-26646197-INS-168 | chr10-26646197-INS-169 |
| chr10-26656510-INS-201 | chr10-26656510-INS-201 |
| chr10-26694093-INS-415 | chr10-26694093-INS-415 |
| chr10-26718391-INS-119 | chr10-26718391-INS-122 |
| chr10-26725030-INS-103 | chr10-26725003-INS-102 |
| chr10-26729800-INS-400 | chr10-26729800-INS-400 |
| chr10-26755495-INS-60 | chr10-26755495-INS-60 |
| chr10-26763020-INS-97 | chr10-26763020-INS-97 |
| chr10-26788943-INS-138 | chr10-26788941-INS-174 |
| chr10-26791072-INS-300 | chr10-26791063-INS-295 |
| chr10-26822853-INS-4243 | chr10-26822854-INS-4245 |
| chr10-26831047-INS-6614 | chr10-26831048-INS-6614 |
| chr10-26831770-INS-219 | chr10-26831770-INS-219 |
| chr10-26833034-INS-383 | chr10-26833034-INS-382 |
| chr10-26834208-INS-91 | chr10-26834208-INS-91 |
| chr10-26838032-INS-5295 | chr10-26838033-INS-5296 |
| chr10-26842584-INS-221 | chr10-26842584-INS-222 |
| chr10-26888680-INS-795 | chr10-26888680-INS-795 |
| chr10-26901613-INS-6469 | chr10-26901618-INS-6466 |
| chr10-26942208-INS-80 | chr10-26942208-INS-80 |
| chr10-26948013-INS-126 | chr10-26948013-INS-126 |
| chr10-26975660-INS-287 | chr10-26975660-INS-287 |
| chr10-26984997-INS-99 | chr10-26985000-INS-120 |
| chr10-26992235-INS-6532 | chr10-26992236-INS-6500 |
| chr10-26994282-INS-106 | chr10-26994282-INS-108 |
| chr10-27070506-INS-356 | chr10-27070505-INS-368 |
| chr10-27097674-INS-6586 | chr10-27097675-INS-6586 |
| chr10-27116840-INS-66 | chr10-27116840-INS-69 |
| chr10-27141471-INS-2295 | chr10-27141472-INS-2295 |
| chr10-27205268-INS-6795 | chr10-27205269-INS-6795 |
| chr10-27218238-INS-76 | chr10-27218238-INS-72 |
| chr10-27252197-INS-222 | chr10-27252197-INS-222 |
| chr10-27327752-INS-66 | chr10-27327759-INS-63 |
| chr10-27341104-INS-140 | chr10-27341104-INS-140 |
| chr10-27354669-INS-226 | chr10-27354669-INS-226 |
| chr10-27391711-INS-270 | chr10-27391711-INS-268 |
| chr10-27411502-INS-84 | chr10-27411516-INS-62 |
| chr10-27417816-INS-927 | chr10-27417816-INS-928 |
| chr10-27421368-INS-358 | chr10-27421368-INS-358 |
| chr10-27424107-INS-135 | chr10-27424107-INS-112 |
| chr10-27468577-INS-60 | chr10-27468562-INS-60 |
| chr10-27509614-INS-4525 | chr10-27509615-INS-4525 |
| chr10-27528618-INS-60 | chr10-27528618-INS-60 |
| chr10-27538368-INS-65 | chr10-27538368-INS-65 |
| chr10-27575803-INS-6389 | chr10-27575804-INS-6389 |
| chr10-27598902-INS-477 | chr10-27598902-INS-477 |
| chr10-27635417-INS-1451 | chr10-27635418-INS-1451 |
| chr10-27643748-INS-1792 | chr10-27643749-INS-1793 |
| chr10-27649332-INS-401 | chr10-27649332-INS-401 |
| chr10-27681917-INS-330 | chr10-27681917-INS-327 |
| chr10-27730165-INS-1736 | chr10-27730168-INS-1774 |
| chr10-27739289-INS-60 | chr10-27739289-INS-60 |
| chr10-27741757-INS-106 | chr10-27741757-INS-108 |
| chr10-27763587-INS-126 | chr10-27763587-INS-126 |
| chr10-27798143-INS-564 | chr10-27798138-INS-557 |
| chr10-27800987-INS-6442 | chr10-27800988-INS-6442 |
| chr10-27805355-INS-63 | chr10-27805355-INS-63 |
| chr10-27816354-INS-3767 | chr10-27816355-INS-3767 |
| chr10-27829763-INS-1140 | chr10-27829764-INS-1134 |
| chr10-27860724-INS-6441 | chr10-27860725-INS-6441 |
| chr10-27860725-INS-498 | chr10-27860725-INS-498 |
| chr10-27865299-INS-167 | chr10-27865308-INS-156 |
| chr10-27873272-INS-78 | chr10-27873272-INS-84 |
| chr10-27891611-INS-72 | chr10-27891609-INS-74 |
| chr10-27900468-INS-111 | chr10-27900466-INS-119 |
| chr10-27909742-INS-63 | chr10-27909706-INS-55 |
| chr10-27914504-INS-240 | chr10-27914552-INS-225 |
| chr10-27916330-INS-60 | chr10-27916330-INS-66 |
| chr10-27933366-INS-64 | chr10-27933366-INS-64 |
| chr10-27956668-INS-66 | chr10-27956668-INS-57 |
| chr10-27959589-INS-103 | chr10-27959589-INS-103 |
| chr10-27999751-INS-62 | chr10-27999714-INS-63 |
| chr10-28001048-INS-952 | chr10-28001048-INS-952 |
| chr10-28030402-INS-54 | chr10-28030402-INS-60 |

| | |
|---|---|
| chr10-28042887-INS-4668 | chr10-28042902-INS-4659 |
| chr10-28051965-INS-118 | chr10-28051949-INS-136 |
| chr10-28058352-INS-203 | chr10-28058352-INS-202 |
| chr10-28078148-INS-401 | chr10-28078148-INS-401 |
| chr10-28081296-INS-96 | chr10-28081296-INS-97 |
| chr10-28092202-INS-71 | chr10-28092202-INS-70 |
| chr10-28096689-INS-68 | chr10-28096689-INS-108 |
| chr10-28114188-INS-72 | chr10-28114210-INS-65 |
| chr10-28209044-INS-784 | chr10-28209044-INS-812 |
| chr10-28235184-INS-112 | chr10-28235184-INS-112 |
| chr10-28274042-INS-150 | chr10-28274042-INS-147 |
| chr10-28275562-INS-66 | chr10-28275562-INS-60 |
| chr10-28278179-INS-54 | chr10-28278180-INS-50 |
| chr10-28280678-INS-632 | chr10-28280692-INS-607 |
| chr10-28285556-INS-60 | chr10-28285558-INS-76 |
| chr10-28286313-INS-72 | chr10-28286313-INS-74 |
| chr10-28290852-INS-215 | chr10-28290852-INS-216 |
| chr10-28293787-INS-84 | chr10-28293786-INS-82 |
| chr10-28296449-INS-162 | chr10-28296449-INS-170 |
| chr10-28307980-INS-214 | chr10-28307980-INS-214 |
| chr10-28317651-INS-1109 | chr10-28317665-INS-1087 |
| chr10-28342371-INS-3952 | chr10-28342372-INS-3952 |
| chr10-28352425-INS-214 | chr10-28352425-INS-212 |
| chr10-28429616-INS-55 | chr10-28429620-INS-60 |
| chr10-28453556-INS-107 | chr10-28453557-INS-107 |
| chr10-28486599-INS-81 | chr10-28486599-INS-77 |
| chr10-28506979-INS-189 | chr10-28506979-INS-189 |
| chr10-28522103-INS-58 | chr10-28522103-INS-58 |
| chr10-28576935-INS-3732 | chr10-28576936-INS-3732 |
| chr10-28608092-INS-203 | chr10-28608092-INS-204 |
| chr10-28632977-INS-422 | chr10-28632977-INS-427 |
| chr10-28644702-INS-201 | chr10-28644702-INS-201 |
| chr10-28663936-INS-7188 | chr10-28663937-INS-7188 |
| chr10-28680381-INS-1670 | chr10-28680382-INS-1684 |
| chr10-28690683-INS-162 | chr10-28690683-INS-162 |
| chr10-28704740-INS-252 | chr10-28704740-INS-260 |
| chr10-28721629-INS-74 | chr10-28721629-INS-74 |
| chr10-28838334-INS-378 | chr10-28838334-INS-378 |
| chr10-28855415-INS-6690 | chr10-28855416-INS-6690 |
| chr10-28889504-INS-81 | chr10-28889504-INS-81 |
| chr10-28933003-INS-198 | chr10-28933003-INS-198 |
| chr10-28938527-INS-58 | chr10-28938527-INS-62 |
| chr10-28953562-INS-6979 | chr10-28953563-INS-6986 |
| chr10-28960467-INS-749 | chr10-28960467-INS-753 |
| chr10-28974223-INS-71 | chr10-28974201-INS-66 |
| chr10-28998988-INS-470 | chr10-28998988-INS-470 |
| chr10-29014885-INS-179 | chr10-29014885-INS-181 |
| chr10-29038317-INS-484 | chr10-29038317-INS-484 |
| chr10-29050817-INS-400 | chr10-29050817-INS-390 |
| chr10-29070421-INS-7280 | chr10-29070422-INS-7280 |
| chr10-29079536-INS-156 | chr10-29079536-INS-156 |
| chr10-29102642-INS-181 | chr10-29102641-INS-183 |
| chr10-29138278-INS-54 | chr10-29138278-INS-54 |
| chr10-29140019-INS-423 | chr10-29140019-INS-423 |
| chr10-29147153-INS-239 | chr10-29147153-INS-239 |
| chr10-29152687-INS-905 | chr10-29152687-INS-909 |
| chr10-29155203-INS-826 | chr10-29155203-INS-826 |
| chr10-29160167-INS-56 | chr10-29160167-INS-58 |
| chr10-29179127-INS-89 | chr10-29179127-INS-88 |
| chr10-29184382-INS-5546 | chr10-29184383-INS-5546 |
| chr10-29194051-INS-62 | chr10-29194051-INS-52 |
| chr10-29202708-INS-122 | chr10-29202714-INS-104 |
| chr10-29221207-INS-68 | chr10-29221207-INS-68 |
| chr10-29244266-INS-225 | chr10-29244266-INS-225 |
| chr10-29258277-INS-6441 | chr10-29258278-INS-6441 |
| chr10-29267183-INS-299 | chr10-29267183-INS-299 |
| chr10-29269428-INS-60 | chr10-29269423-INS-72 |
| chr10-29272364-INS-321 | chr10-29272364-INS-321 |
| chr10-29295437-INS-749 | chr10-29295437-INS-749 |
| chr10-29303049-INS-69 | chr10-29303049-INS-54 |
| chr10-29335643-INS-5546 | chr10-29335644-INS-5546 |
| chr10-29337451-INS-83 | chr10-29337451-INS-83 |
| chr10-29369384-INS-86 | chr10-29369384-INS-86 |
| chr10-29374249-INS-1088 | chr10-29374250-INS-1088 |
| chr10-29424654-INS-138 | chr10-29424652-INS-132 |
| chr10-29476418-INS-1119 | chr10-29476389-INS-1108 |
| chr10-29492495-INS-1179 | chr10-29492496-INS-1180 |

| | |
|---|---|
| chr10-29552202-INS-378 | chr10-29552202-INS-378 |
| chr10-29560672-INS-62 | chr10-29560672-INS-74 |
| chr10-29579022-INS-184 | chr10-29579022-INS-184 |
| chr10-29589923-INS-399 | chr10-29589923-INS-399 |
| chr10-29604766-INS-102 | chr10-29604766-INS-102 |
| chr10-29621101-INS-72 | chr10-29621101-INS-54 |
| chr10-29675960-INS-83 | chr10-29675960-INS-109 |
| chr10-29752663-INS-1812 | chr10-29752664-INS-1817 |
| chr10-29783361-INS-102 | chr10-29783361-INS-102 |
| chr10-29785377-INS-192 | chr10-29785373-INS-153 |
| chr10-29810370-INS-1478 | chr10-29810371-INS-1478 |
| chr10-29822718-INS-428 | chr10-29822718-INS-428 |
| chr10-29848694-INS-69 | chr10-29848703-INS-63 |
| chr10-29916675-INS-60 | chr10-29916675-INS-60 |
| chr10-29919563-INS-224 | chr10-29919564-INS-225 |
| chr10-29943234-INS-62 | chr10-29943234-INS-66 |
| chr10-29958633-INS-70 | chr10-29958633-INS-74 |
| chr10-29976494-INS-112 | chr10-29976494-INS-112 |
| chr10-29990799-INS-489 | chr10-29990803-INS-501 |
| chr10-29994578-INS-1683 | chr10-29994579-INS-1683 |
| chr10-30017183-INS-264 | chr10-30017183-INS-264 |
| chr10-30080892-INS-63 | chr10-30080892-INS-63 |
| chr10-30118000-INS-298 | chr10-30118003-INS-292 |
| chr10-30151188-INS-209 | chr10-30151188-INS-209 |
| chr10-30154051-INS-1431 | chr10-30154052-INS-1431 |
| chr10-30162837-INS-90 | chr10-30162825-INS-122 |
| chr10-30180515-INS-163 | chr10-30180515-INS-165 |
| chr10-30182109-INS-343 | chr10-30182109-INS-343 |
| chr10-30182899-INS-76 | chr10-30182899-INS-76 |
| chr10-30186785-INS-3061 | chr10-30186786-INS-3106 |
| chr10-30209590-INS-64 | chr10-30209590-INS-64 |
| chr10-30230558-INS-3061 | chr10-30230559-INS-3061 |
| chr10-30245000-INS-243 | chr10-30245000-INS-243 |
| chr10-30249413-INS-118 | chr10-30249413-INS-118 |
| chr10-30319021-INS-89 | chr10-30319039-INS-85 |
| chr10-30325542-INS-978 | chr10-30325542-INS-977 |
| chr10-30381159-INS-4139 | chr10-30381160-INS-4139 |
| chr10-30383886-INS-299 | chr10-30383886-INS-307 |
| chr10-30403257-INS-164 | chr10-30403257-INS-164 |
| chr10-30418918-INS-188 | chr10-30418918-INS-188 |
| chr10-30437185-INS-508 | chr10-30437185-INS-524 |
| chr10-30445934-INS-89 | chr10-30445918-INS-70 |
| chr10-30464716-INS-883 | chr10-30464716-INS-885 |
| chr10-30472295-INS-144 | chr10-30472295-INS-144 |
| chr10-30483533-INS-87 | chr10-30483532-INS-87 |
| chr10-30484444-INS-7175 | chr10-30484445-INS-7175 |
| chr10-30484938-INS-64 | chr10-30484938-INS-60 |
| chr10-30491458-INS-235 | chr10-30491458-INS-231 |
| chr10-30529513-INS-186 | chr10-30529513-INS-193 |
| chr10-30553174-INS-7094 | chr10-30553175-INS-7094 |
| chr10-30562890-INS-62 | chr10-30562890-INS-62 |
| chr10-30587085-INS-354 | chr10-30587085-INS-371 |
| chr10-30596152-INS-82 | chr10-30596112-INS-82 |
| chr10-30604534-INS-88 | chr10-30604538-INS-90 |
| chr10-30629105-INS-179 | chr10-30629105-INS-175 |
| chr10-30661872-INS-52 | chr10-30661873-INS-52 |
| chr10-30699764-INS-69 | chr10-30699764-INS-94 |
| chr10-30731876-INS-52 | chr10-30731876-INS-50 |
| chr10-30801069-INS-242 | chr10-30801061-INS-222 |
| chr10-30806605-INS-80 | chr10-30806624-INS-72 |
| chr10-30808499-INS-52 | chr10-30808499-INS-53 |
| chr10-30833973-INS-54 | chr10-30833989-INS-88 |
| chr10-30866121-INS-232 | chr10-30866121-INS-227 |
| chr10-30918734-INS-119 | chr10-30918734-INS-116 |
| chr10-31021949-INS-584 | chr10-31021949-INS-585 |
| chr10-31029230-INS-2415 | chr10-31029231-INS-2415 |
| chr10-31038938-INS-6221 | chr10-31038939-INS-6215 |
| chr10-31044516-INS-1855 | chr10-31044517-INS-1855 |
| chr10-31053551-INS-396 | chr10-31053551-INS-396 |
| chr10-31095218-INS-456 | chr10-31095218-INS-456 |
| chr10-31104824-INS-216 | chr10-31104824-INS-215 |
| chr10-31120459-INS-486 | chr10-31120459-INS-486 |
| chr10-31127569-INS-350 | chr10-31127569-INS-354 |
| chr10-31132373-INS-448 | chr10-31132381-INS-458 |
| chr10-31139450-INS-56 | chr10-31139450-INS-58 |
| chr10-31153662-INS-167 | chr10-31153662-INS-167 |
| chr10-31160159-INS-166 | chr10-31160159-INS-166 |

| | |
|---|---|
| chr10-31192578-INS-78 | chr10-31192578-INS-78 |
| chr10-31222500-INS-625 | chr10-31222500-INS-624 |
| chr10-31234186-INS-91 | chr10-31234186-INS-92 |
| chr10-31261816-INS-219 | chr10-31261816-INS-220 |
| chr10-31293175-INS-968 | chr10-31293175-INS-969 |
| chr10-31313140-INS-56 | chr10-31313140-INS-56 |
| chr10-31340223-INS-200 | chr10-31340223-INS-200 |
| chr10-31346643-INS-168 | chr10-31346643-INS-171 |
| chr10-31350417-INS-207 | chr10-31350417-INS-211 |
| chr10-31354819-INS-207 | chr10-31354819-INS-211 |
| chr10-31389335-INS-603 | chr10-31389335-INS-610 |
| chr10-31414114-INS-86 | chr10-31414102-INS-108 |
| chr10-31419034-INS-187 | chr10-31419029-INS-182 |
| chr10-31457323-INS-52 | chr10-31457307-INS-56 |
| chr10-31474792-INS-56 | chr10-31474792-INS-56 |
| chr10-31481544-INS-102 | chr10-31481544-INS-102 |
| chr10-31487807-INS-6244 | chr10-31487808-INS-6244 |
| chr10-31503735-INS-87 | chr10-31503735-INS-87 |
| chr10-31510385-INS-6273 | chr10-31510386-INS-6273 |
| chr10-31521045-INS-56 | chr10-31521045-INS-52 |
| chr10-31527573-INS-557 | chr10-31527573-INS-558 |
| chr10-31618340-INS-112 | chr10-31618340-INS-122 |
| chr10-31639660-INS-6366 | chr10-31639661-INS-6366 |
| chr10-31687745-INS-243 | chr10-31687745-INS-250 |
| chr10-31702421-INS-58 | chr10-31702411-INS-70 |
| chr10-31714049-INS-59 | chr10-31714010-INS-59 |
| chr10-31745299-INS-2243 | chr10-31745300-INS-2238 |
| chr10-31772600-INS-129 | chr10-31772600-INS-135 |
| chr10-31773766-INS-423 | chr10-31773766-INS-423 |
| chr10-31792431-INS-856 | chr10-31792431-INS-855 |
| chr10-31795255-INS-901 | chr10-31795255-INS-901 |
| chr10-31812603-INS-98 | chr10-31812603-INS-96 |
| chr10-31813698-INS-1160 | chr10-31813699-INS-1160 |
| chr10-31835947-INS-2910 | chr10-31835948-INS-2908 |
| chr10-31851513-INS-170 | chr10-31851520-INS-181 |
| chr10-31925895-INS-1116 | chr10-31925877-INS-1109 |
| chr10-31933627-INS-262 | chr10-31933627-INS-262 |
| chr10-31952834-INS-52 | chr10-31952834-INS-54 |
| chr10-31961799-INS-1416 | chr10-31961800-INS-1416 |
| chr10-31964511-INS-1119 | chr10-31964512-INS-1121 |
| chr10-31967067-INS-199 | chr10-31967068-INS-199 |
| chr10-31986631-INS-211 | chr10-31986631-INS-211 |
| chr10-32000475-INS-65 | chr10-32000475-INS-65 |
| chr10-32006537-INS-1268 | chr10-32006541-INS-1264 |
| chr10-32007094-INS-81 | chr10-32007091-INS-76 |
| chr10-32038371-INS-1637 | chr10-32038372-INS-1637 |
| chr10-32078730-INS-55 | chr10-32078730-INS-55 |
| chr10-32079770-INS-160 | chr10-32079770-INS-140 |
| chr10-32174337-INS-392 | chr10-32174337-INS-392 |
| chr10-32181950-INS-656 | chr10-32181950-INS-656 |
| chr10-32219697-INS-195 | chr10-32219698-INS-196 |
| chr10-32235489-INS-84 | chr10-32235489-INS-84 |
| chr10-32285434-INS-70 | chr10-32285434-INS-50 |
| chr10-32289163-INS-125 | chr10-32289163-INS-125 |
| chr10-32333305-INS-477 | chr10-32333305-INS-477 |
| chr10-32338201-INS-80 | chr10-32338217-INS-72 |
| chr10-32353922-INS-828 | chr10-32353922-INS-828 |
| chr10-32366120-INS-148 | chr10-32366123-INS-151 |
| chr10-32367401-INS-317 | chr10-32367402-INS-318 |
| chr10-32405466-INS-811 | chr10-32405466-INS-811 |
| chr10-32406283-INS-1760 | chr10-32406284-INS-1761 |
| chr10-32442792-INS-83 | chr10-32442792-INS-83 |
| chr10-32470067-INS-135 | chr10-32470067-INS-134 |
| chr10-32490770-INS-83 | chr10-32490770-INS-70 |
| chr10-32506565-INS-56 | chr10-32506565-INS-56 |
| chr10-32532564-INS-53 | chr10-32532564-INS-53 |
| chr10-32540433-INS-2040 | chr10-32540434-INS-2040 |
| chr10-32579859-INS-58 | chr10-32579859-INS-60 |
| chr10-32619906-INS-90 | chr10-32619868-INS-100 |
| chr10-32627080-INS-1226 | chr10-32627085-INS-1215 |
| chr10-32636072-INS-381 | chr10-32636072-INS-382 |
| chr10-32638500-INS-175 | chr10-32638500-INS-177 |
| chr10-32663728-INS-335 | chr10-32663728-INS-335 |
| chr10-32669891-INS-57 | chr10-32669868-INS-56 |
| chr10-32728044-INS-6419 | chr10-32728045-INS-6419 |
| chr10-32821798-INS-57 | chr10-32821798-INS-57 |
| chr10-32848612-INS-212 | chr10-32848612-INS-212 |

| | |
|---|---|
| chr10-32866791-INS-68 | chr10-32866791-INS-68 |
| chr10-32902840-INS-204 | chr10-32902840-INS-203 |
| chr10-32928859-INS-60 | chr10-32928859-INS-60 |
| chr10-32948681-INS-390 | chr10-32948681-INS-390 |
| chr10-32951757-INS-115 | chr10-32951757-INS-115 |
| chr10-32953154-INS-58 | chr10-32953168-INS-54 |
| chr10-32965180-INS-130 | chr10-32965180-INS-130 |
| chr10-32977207-INS-115 | chr10-32977181-INS-96 |
| chr10-33040076-INS-180 | chr10-33040076-INS-198 |
| chr10-33059637-INS-59 | chr10-33059637-INS-60 |
| chr10-33070853-INS-700 | chr10-33070853-INS-700 |
| chr10-33102193-INS-446 | chr10-33102193-INS-448 |
| chr10-33107760-INS-1031 | chr10-33107761-INS-1032 |
| chr10-33135521-INS-119 | chr10-33135521-INS-119 |
| chr10-33138250-INS-219 | chr10-33138250-INS-219 |
| chr10-33140337-INS-1596 | chr10-33140338-INS-1596 |
| chr10-33161246-INS-1186 | chr10-33161249-INS-1178 |
| chr10-33165346-INS-199 | chr10-33165346-INS-200 |
| chr10-33171904-INS-53 | chr10-33171904-INS-88 |
| chr10-33178911-INS-114 | chr10-33178911-INS-114 |
| chr10-33189894-INS-128 | chr10-33189894-INS-128 |
| chr10-33212255-INS-171 | chr10-33212255-INS-173 |
| chr10-33226645-INS-4492 | chr10-33226646-INS-4492 |
| chr10-33232026-INS-90 | chr10-33232028-INS-89 |
| chr10-33261055-INS-400 | chr10-33261055-INS-400 |
| chr10-33296273-INS-53 | chr10-33296273-INS-62 |
| chr10-33316210-INS-50 | chr10-33316210-INS-52 |
| chr10-33333809-INS-82 | chr10-33333809-INS-87 |
| chr10-33347620-INS-890 | chr10-33347620-INS-891 |
| chr10-33350151-INS-62 | chr10-33350151-INS-96 |
| chr10-33430757-INS-390 | chr10-33430757-INS-389 |
| chr10-33571418-INS-5513 | chr10-33571419-INS-5514 |
| chr10-33612095-INS-71 | chr10-33612095-INS-71 |
| chr10-33615248-INS-123 | chr10-33615248-INS-123 |
| chr10-33629585-INS-558 | chr10-33629585-INS-558 |
| chr10-33694591-INS-190 | chr10-33694591-INS-187 |
| chr10-33697676-INS-424 | chr10-33697676-INS-424 |
| chr10-33723196-INS-107 | chr10-33723180-INS-96 |
| chr10-33745926-INS-163 | chr10-33745926-INS-163 |
| chr10-33830723-INS-199 | chr10-33830723-INS-200 |
| chr10-33835454-INS-508 | chr10-33835454-INS-509 |
| chr10-33841163-INS-205 | chr10-33841163-INS-205 |
| chr10-33847671-INS-194 | chr10-33847671-INS-194 |
| chr10-33951806-INS-97 | chr10-33951781-INS-77 |
| chr10-33965445-INS-132 | chr10-33965442-INS-160 |
| chr10-33983346-INS-105 | chr10-33983346-INS-105 |
| chr10-34038271-INS-116 | chr10-34038271-INS-116 |
| chr10-34045717-INS-76 | chr10-34045717-INS-76 |
| chr10-34054949-INS-59 | chr10-34054949-INS-70 |
| chr10-34077188-INS-209 | chr10-34077188-INS-209 |
| chr10-34077648-INS-468 | chr10-34077648-INS-466 |
| chr10-34101167-INS-86 | chr10-34101167-INS-58 |
| chr10-34109770-INS-6724 | chr10-34109771-INS-6725 |
| chr10-34126822-INS-531 | chr10-34126822-INS-538 |
| chr10-34132290-INS-1546 | chr10-34132291-INS-1555 |
| chr10-34133398-INS-78 | chr10-34133398-INS-78 |
| chr10-34170502-INS-103 | chr10-34170502-INS-107 |
| chr10-34178924-INS-284 | chr10-34178924-INS-285 |
| chr10-34268392-INS-131 | chr10-34268392-INS-131 |
| chr10-34284277-INS-99 | chr10-34284277-INS-92 |
| chr10-34285242-INS-392 | chr10-34285242-INS-392 |
| chr10-34297418-INS-95 | chr10-34297418-INS-95 |
| chr10-34341825-INS-64 | chr10-34341825-INS-64 |
| chr10-34379012-INS-436 | chr10-34378984-INS-439 |
| chr10-34383675-INS-70 | chr10-34383675-INS-70 |
| chr10-34445110-INS-196 | chr10-34445110-INS-196 |
| chr10-34512400-INS-1628 | chr10-34512401-INS-1625 |
| chr10-34590620-INS-166 | chr10-34590620-INS-166 |
| chr10-34633145-INS-106 | chr10-34633145-INS-102 |
| chr10-34637070-INS-400 | chr10-34637070-INS-400 |
| chr10-34654874-INS-794 | chr10-34654872-INS-795 |
| chr10-34666216-INS-88 | chr10-34666216-INS-132 |
| chr10-34676257-INS-577 | chr10-34676257-INS-577 |
| chr10-34739272-INS-62 | chr10-34739272-INS-82 |
| chr10-34741699-INS-52 | chr10-34741699-INS-54 |
| chr10-34744078-INS-55 | chr10-34744078-INS-63 |
| chr10-34794275-INS-899 | chr10-34794275-INS-899 |

| | |
|---|---|
| chr10-34808670-INS-636 | chr10-34808690-INS-655 |
| chr10-34832805-INS-97 | chr10-34832805-INS-93 |
| chr10-34840990-INS-55 | chr10-34840990-INS-55 |
| chr10-34862102-INS-205 | chr10-34862102-INS-205 |
| chr10-34902537-INS-568 | chr10-34902537-INS-568 |
| chr10-34918898-INS-77 | chr10-34918898-INS-77 |
| chr10-34977144-INS-210 | chr10-34977144-INS-210 |
| chr10-34988301-INS-52 | chr10-34988301-INS-52 |
| chr10-34993291-INS-5067 | chr10-34993292-INS-5067 |
| chr10-35013998-INS-773 | chr10-35013998-INS-773 |
| chr10-35021811-INS-309 | chr10-35021812-INS-309 |
| chr10-35026476-INS-127 | chr10-35026476-INS-77 |
| chr10-35046397-INS-53 | chr10-35046397-INS-53 |
| chr10-35053327-INS-212 | chr10-35053327-INS-212 |
| chr10-35053717-INS-661 | chr10-35053717-INS-662 |
| chr10-35085822-INS-6281 | chr10-35085823-INS-6281 |
| chr10-35097893-INS-119 | chr10-35097893-INS-119 |
| chr10-35117634-INS-959 | chr10-35117634-INS-959 |
| chr10-35124898-INS-64 | chr10-35124897-INS-71 |
| chr10-35134765-INS-60 | chr10-35134740-INS-92 |
| chr10-35146048-INS-80 | chr10-35146048-INS-88 |
| chr10-35186236-INS-416 | chr10-35186236-INS-416 |
| chr10-35216943-INS-455 | chr10-35216943-INS-455 |
| chr10-35242454-INS-5219 | chr10-35242455-INS-5219 |
| chr10-35281955-INS-605 | chr10-35281955-INS-605 |
| chr10-35302484-INS-51 | chr10-35302478-INS-60 |
| chr10-35306211-INS-63 | chr10-35306211-INS-63 |
| chr10-35372875-INS-6463 | chr10-35372876-INS-6463 |
| chr10-35375149-INS-401 | chr10-35375151-INS-401 |
| chr10-35382730-INS-1175 | chr10-35382731-INS-1177 |
| chr10-35403883-INS-2504 | chr10-35403884-INS-2504 |
| chr10-35412717-INS-68 | chr10-35412717-INS-52 |
| chr10-35454150-INS-102 | chr10-35454150-INS-102 |
| chr10-35461215-INS-52 | chr10-35461215-INS-72 |
| chr10-35465457-INS-8038 | chr10-35465458-INS-8039 |
| chr10-35484882-INS-778 | chr10-35484882-INS-778 |
| chr10-35536817-INS-85 | chr10-35536817-INS-85 |
| chr10-35563747-INS-117 | chr10-35563747-INS-117 |
| chr10-35591253-INS-93 | chr10-35591253-INS-94 |
| chr10-35619091-INS-63 | chr10-35619091-INS-63 |
| chr10-35629107-INS-124 | chr10-35629107-INS-124 |
| chr10-35689156-INS-63 | chr10-35689156-INS-63 |
| chr10-35689676-INS-613 | chr10-35689676-INS-613 |
| chr10-35710169-INS-54 | chr10-35710169-INS-58 |
| chr10-35723193-INS-219 | chr10-35723193-INS-214 |
| chr10-35770419-INS-6387 | chr10-35770420-INS-6387 |
| chr10-35822395-INS-82 | chr10-35822390-INS-84 |
| chr10-35831303-INS-2847 | chr10-35831304-INS-2847 |
| chr10-35852326-INS-166 | chr10-35852326-INS-166 |
| chr10-35857458-INS-96 | chr10-35857448-INS-90 |
| chr10-35866899-INS-60 | chr10-35866899-INS-58 |
| chr10-35910807-INS-939 | chr10-35910807-INS-940 |
| chr10-35919409-INS-701 | chr10-35919409-INS-701 |
| chr10-35937996-INS-52 | chr10-35937997-INS-68 |
| chr10-35942268-INS-304 | chr10-35942269-INS-304 |
| chr10-35954617-INS-1362 | chr10-35954618-INS-1361 |
| chr10-35967867-INS-310 | chr10-35967867-INS-324 |
| chr10-36003066-INS-6897 | chr10-36003067-INS-6897 |
| chr10-36030942-INS-160 | chr10-36030930-INS-187 |
| chr10-36050332-INS-3764 | chr10-36050333-INS-3774 |
| chr10-36078487-INS-331 | chr10-36078487-INS-331 |
| chr10-36122278-INS-1655 | chr10-36122279-INS-1655 |
| chr10-36123942-INS-487 | chr10-36123942-INS-487 |
| chr10-36169135-INS-64 | chr10-36169135-INS-89 |
| chr10-36203817-INS-64 | chr10-36203768-INS-88 |
| chr10-36206359-INS-368 | chr10-36206359-INS-368 |
| chr10-36224825-INS-3718 | chr10-36224826-INS-3718 |
| chr10-36263692-INS-4605 | chr10-36263693-INS-4596 |
| chr10-36282828-INS-6527 | chr10-36282829-INS-6527 |
| chr10-36286780-INS-480 | chr10-36286775-INS-478 |
| chr10-36309044-INS-105 | chr10-36309044-INS-95 |
| chr10-36318048-INS-103 | chr10-36318048-INS-92 |
| chr10-36320212-INS-2755 | chr10-36320196-INS-2735 |
| chr10-36357472-INS-54 | chr10-36357446-INS-54 |
| chr10-36391325-INS-101 | chr10-36391325-INS-51 |
| chr10-36397218-INS-78 | chr10-36397224-INS-57 |
| chr10-36432709-INS-83 | chr10-36432709-INS-81 |

| | |
|---|---|
| chr10-36507066-INS-60 | chr10-36507066-INS-60 |
| chr10-36531361-INS-228 | chr10-36531361-INS-229 |
| chr10-36607539-INS-77 | chr10-36607542-INS-71 |
| chr10-36610836-INS-75 | chr10-36610836-INS-76 |
| chr10-36613238-INS-280 | chr10-36613238-INS-281 |
| chr10-36664773-INS-71 | chr10-36664773-INS-74 |
| chr10-36666120-INS-829 | chr10-36666133-INS-832 |
| chr10-36713328-INS-123 | chr10-36713328-INS-129 |
| chr10-36770133-INS-129 | chr10-36770133-INS-131 |
| chr10-36791180-INS-85 | chr10-36791180-INS-52 |
| chr10-36799013-INS-100 | chr10-36799054-INS-93 |
| chr10-36829258-INS-236 | chr10-36829258-INS-236 |
| chr10-36844119-INS-73 | chr10-36844119-INS-73 |
| chr10-36847548-INS-6476 | chr10-36847549-INS-6477 |
| chr10-36865730-INS-243 | chr10-36865730-INS-244 |
| chr10-36876442-INS-53 | chr10-36876433-INS-53 |
| chr10-36883377-INS-234 | chr10-36883377-INS-235 |
| chr10-36885679-INS-53 | chr10-36885679-INS-53 |
| chr10-36899120-INS-204 | chr10-36899101-INS-177 |
| chr10-36930239-INS-540 | chr10-36930239-INS-540 |
| chr10-36956736-INS-1609 | chr10-36956737-INS-1616 |
| chr10-36961976-INS-274 | chr10-36961976-INS-274 |
| chr10-36971324-INS-80 | chr10-36971324-INS-72 |
| chr10-36998226-INS-1056 | chr10-36998227-INS-1056 |
| chr10-37000759-INS-88 | chr10-37000757-INS-90 |
| chr10-37021476-INS-335 | chr10-37021476-INS-311 |
| chr10-37026077-INS-6357 | chr10-37026078-INS-6359 |
| chr10-37055704-INS-126 | chr10-37055704-INS-126 |
| chr10-37097806-INS-165 | chr10-37097806-INS-165 |
| chr10-37165222-INS-493 | chr10-37165222-INS-493 |
| chr10-37195739-INS-6251 | chr10-37195740-INS-6252 |
| chr10-37198614-INS-394 | chr10-37198614-INS-379 |
| chr10-37257643-INS-51 | chr10-37257643-INS-51 |
| chr10-37267286-INS-56 | chr10-37267286-INS-60 |
| chr10-37269106-INS-78 | chr10-37269106-INS-76 |
| chr10-37272746-INS-6452 | chr10-37272747-INS-6452 |
| chr10-37292987-INS-6647 | chr10-37292988-INS-6648 |
| chr10-37312228-INS-1886 | chr10-37312229-INS-1886 |
| chr10-37333077-INS-6447 | chr10-37333078-INS-6445 |
| chr10-37334153-INS-91 | chr10-37334153-INS-78 |
| chr10-37363989-INS-200 | chr10-37363989-INS-200 |
| chr10-37403062-INS-697 | chr10-37403062-INS-697 |
| chr10-37407189-INS-248 | chr10-37407189-INS-210 |
| chr10-37414746-INS-69 | chr10-37414746-INS-69 |
| chr10-37422210-INS-6366 | chr10-37422211-INS-6367 |
| chr10-37445053-INS-7209 | chr10-37445054-INS-7209 |
| chr10-37475943-INS-91 | chr10-37475943-INS-91 |
| chr10-37478172-INS-56 | chr10-37478172-INS-56 |
| chr10-37479480-INS-55 | chr10-37479480-INS-61 |
| chr10-37482574-INS-68 | chr10-37482574-INS-62 |
| chr10-37486620-INS-115 | chr10-37486620-INS-70 |
| chr10-37516686-INS-64 | chr10-37516686-INS-72 |
| chr10-37522513-INS-62 | chr10-37522513-INS-58 |
| chr10-37524668-INS-521 | chr10-37524668-INS-520 |
| chr10-37530561-INS-70 | chr10-37530561-INS-90 |
| chr10-37539178-INS-6860 | chr10-37539179-INS-6861 |
| chr10-37572474-INS-348 | chr10-37572474-INS-348 |
| chr10-37594261-INS-100 | chr10-37594261-INS-138 |
| chr10-37601338-INS-2718 | chr10-37601339-INS-2718 |
| chr10-37631050-INS-937 | chr10-37631050-INS-937 |
| chr10-37634173-INS-205 | chr10-37634173-INS-206 |
| chr10-37640377-INS-70 | chr10-37640412-INS-52 |
| chr10-37652733-INS-64 | chr10-37652733-INS-68 |
| chr10-37666475-INS-2420 | chr10-37666476-INS-2420 |
| chr10-37671948-INS-6736 | chr10-37671949-INS-6737 |
| chr10-37673150-INS-90 | chr10-37673150-INS-60 |
| chr10-37710974-INS-6638 | chr10-37710975-INS-6638 |
| chr10-37713025-INS-208 | chr10-37713025-INS-206 |
| chr10-37737768-INS-54 | chr10-37737768-INS-54 |
| chr10-37743169-INS-111 | chr10-37743169-INS-111 |
| chr10-37753380-INS-717 | chr10-37753380-INS-717 |
| chr10-37771524-INS-713 | chr10-37771524-INS-712 |
| chr10-37774310-INS-6484 | chr10-37774311-INS-6485 |
| chr10-37794427-INS-705 | chr10-37794427-INS-705 |
| chr10-37807922-INS-2563 | chr10-37807897-INS-2563 |
| chr10-37855500-INS-56 | chr10-37855500-INS-56 |
| chr10-37873227-INS-4743 | chr10-37873228-INS-4743 |

| | |
|---|---|
| chr10-37889452-INS-51 | chr10-37889453-INS-51 |
| chr10-37892647-INS-82 | chr10-37892647-INS-90 |
| chr10-37899882-INS-89 | chr10-37899882-INS-89 |
| chr10-37909166-INS-4285 | chr10-37909167-INS-4286 |
| chr10-37913988-INS-518 | chr10-37913988-INS-518 |
| chr10-37915191-INS-60 | chr10-37915191-INS-60 |
| chr10-37917823-INS-335 | chr10-37917823-INS-335 |
| chr10-37958531-INS-72 | chr10-37958529-INS-58 |
| chr10-37973141-INS-721 | chr10-37973141-INS-723 |
| chr10-37980673-INS-6512 | chr10-37980674-INS-6508 |
| chr10-38008786-INS-143 | chr10-38008798-INS-144 |
| chr10-38013342-INS-107 | chr10-38013342-INS-107 |
| chr10-38017441-INS-56 | chr10-38017441-INS-60 |
| chr10-38018979-INS-209 | chr10-38018980-INS-213 |
| chr10-38025913-INS-66 | chr10-38025913-INS-72 |
| chr10-38043976-INS-482 | chr10-38043977-INS-483 |
| chr10-38057810-INS-2720 | chr10-38057811-INS-2721 |
| chr10-38062435-INS-1730 | chr10-38062436-INS-1716 |
| chr10-38064623-INS-1065 | chr10-38064624-INS-1067 |
| chr10-38075446-INS-2128 | chr10-38075447-INS-2128 |
| chr10-38153160-INS-1758 | chr10-38153161-INS-1758 |
| chr10-38170026-INS-422 | chr10-38170034-INS-423 |
| chr10-38174462-INS-482 | chr10-38174462-INS-483 |
| chr10-38176252-INS-169 | chr10-38176244-INS-208 |
| chr10-38189460-INS-1836 | chr10-38189461-INS-1837 |
| chr10-38199136-INS-246 | chr10-38199136-INS-246 |
| chr10-38209824-INS-438 | chr10-38209821-INS-440 |
| chr10-38211630-INS-111 | chr10-38211630-INS-111 |
| chr10-38226379-INS-1147 | chr10-38226380-INS-1143 |
| chr10-38239499-INS-1775 | chr10-38239500-INS-1781 |
| chr10-38284243-INS-84 | chr10-38284243-INS-86 |
| chr10-38293818-INS-6282 | chr10-38293819-INS-6288 |
| chr10-38295848-INS-136 | chr10-38295848-INS-137 |
| chr10-38317532-INS-2746 | chr10-38317533-INS-2746 |
| chr10-38327808-INS-6478 | chr10-38327809-INS-6478 |
| chr10-38338657-INS-3922 | chr10-38338658-INS-3922 |
| chr10-38339233-INS-1660 | chr10-38339234-INS-1661 |
| chr10-38341153-INS-55 | chr10-38341153-INS-55 |
| chr10-38348051-INS-54 | chr10-38348051-INS-54 |
| chr10-38351384-INS-6456 | chr10-38351385-INS-6456 |
| chr10-38387666-INS-78 | chr10-38387664-INS-78 |
| chr10-38395758-INS-261 | chr10-38395758-INS-261 |
| chr10-38400265-INS-75 | chr10-38400265-INS-74 |
| chr10-38411385-INS-91 | chr10-38411385-INS-91 |
| chr10-38418216-INS-401 | chr10-38418216-INS-401 |
| chr10-38419068-INS-647 | chr10-38419068-INS-647 |
| chr10-38420116-INS-708 | chr10-38420116-INS-708 |
| chr10-38436586-INS-6643 | chr10-38436587-INS-6643 |
| chr10-38487277-INS-498 | chr10-38487277-INS-498 |
| chr10-38495125-INS-148 | chr10-38495125-INS-148 |
| chr10-38566859-INS-599 | chr10-38566859-INS-600 |
| chr10-38570291-INS-86 | chr10-38570333-INS-94 |
| chr10-38590348-INS-117 | chr10-38590305-INS-166 |
| chr10-38595963-INS-57 | chr10-38595960-INS-60 |
| chr10-38606803-INS-385 | chr10-38606803-INS-385 |
| chr10-38607978-INS-93 | chr10-38607978-INS-99 |
| chr10-38611674-INS-374 | chr10-38611705-INS-381 |
| chr10-38611679-INS-93 | chr10-38611679-INS-93 |
| chr10-38649215-INS-212 | chr10-38649215-INS-212 |
| chr10-38656969-INS-780 | chr10-38656969-INS-780 |
| chr10-38664740-INS-186 | chr10-38664740-INS-186 |
| chr10-38664943-INS-6741 | chr10-38664944-INS-6740 |
| chr10-38669524-INS-758 | chr10-38669524-INS-759 |
| chr10-38685947-INS-398 | chr10-38685947-INS-398 |
| chr10-38698286-INS-82 | chr10-38698286-INS-52 |
| chr10-38702504-INS-6433 | chr10-38702505-INS-6431 |
| chr10-38710265-INS-213 | chr10-38710265-INS-212 |
| chr10-38716272-INS-168 | chr10-38716272-INS-168 |
| chr10-38726579-INS-61 | chr10-38726579-INS-61 |
| chr10-38728655-INS-875 | chr10-38728655-INS-878 |
| chr10-38733887-INS-1737 | chr10-38733888-INS-1739 |
| chr10-38739809-INS-1217 | chr10-38739810-INS-1247 |
| chr10-38750192-INS-6709 | chr10-38750193-INS-6709 |
| chr10-38755696-INS-625 | chr10-38755696-INS-669 |
| chr10-38788685-INS-227 | chr10-38788685-INS-225 |
| chr10-38797798-INS-735 | chr10-38797798-INS-733 |
| chr10-38798602-INS-458 | chr10-38798602-INS-458 |

| | |
|---|---|
| chr10-38848269-INS-58 | chr10-38848269-INS-58 |
| chr10-38902015-INS-218 | chr10-38902016-INS-252 |
| chr10-38906903-INS-341 | chr10-38906903-INS-341 |
| chr10-38926647-INS-6444 | chr10-38926648-INS-6445 |
| chr10-38937847-INS-832 | chr10-38937847-INS-832 |
| chr10-38947207-INS-113 | chr10-38947207-INS-74 |
| chr10-38968561-INS-217 | chr10-38968561-INS-217 |
| chr10-38969880-INS-355 | chr10-38969880-INS-356 |
| chr10-39028629-INS-79 | chr10-39028631-INS-66 |
| chr10-39089186-INS-92 | chr10-39089204-INS-84 |
| chr10-39115317-INS-148 | chr10-39115317-INS-148 |
| chr10-39119694-INS-90 | chr10-39119694-INS-90 |
| chr10-39145105-INS-51 | chr10-39145105-INS-68 |
| chr10-39160475-INS-5176 | chr10-39160476-INS-5176 |
| chr10-39189460-INS-6662 | chr10-39189461-INS-6662 |
| chr10-39196958-INS-90 | chr10-39196958-INS-80 |
| chr10-39208652-INS-1704 | chr10-39208653-INS-1699 |
| chr10-39218503-INS-104 | chr10-39218503-INS-104 |
| chr10-39225731-INS-68 | chr10-39225731-INS-68 |
| chr10-39237563-INS-215 | chr10-39237564-INS-189 |
| chr10-39243485-INS-116 | chr10-39243468-INS-89 |
| chr10-39292309-INS-164 | chr10-39292309-INS-164 |
| chr10-39326042-INS-58 | chr10-39326042-INS-58 |
| chr10-39336436-INS-1864 | chr10-39336437-INS-1864 |
| chr10-39356374-INS-210 | chr10-39356374-INS-210 |
| chr10-39363018-INS-152 | chr10-39363017-INS-154 |
| chr10-39408226-INS-102 | chr10-39408210-INS-68 |
| chr10-39435332-INS-1890 | chr10-39435333-INS-1890 |
| chr10-39450237-INS-62 | chr10-39450237-INS-62 |
| chr10-39455658-INS-208 | chr10-39455658-INS-208 |
| chr10-39490589-INS-120 | chr10-39490589-INS-152 |
| chr10-39501346-INS-157 | chr10-39501346-INS-157 |
| chr10-39512233-INS-200 | chr10-39512233-INS-202 |
| chr10-39519756-INS-92 | chr10-39519787-INS-92 |
| chr10-39520956-INS-99 | chr10-39520954-INS-103 |
| chr10-39533673-INS-245 | chr10-39533673-INS-245 |
| chr10-39558528-INS-406 | chr10-39558528-INS-406 |
| chr10-39632098-INS-434 | chr10-39632098-INS-438 |
| chr10-39642808-INS-66 | chr10-39642808-INS-56 |
| chr10-39654317-INS-63 | chr10-39654317-INS-84 |
| chr10-39762831-INS-80 | chr10-39762831-INS-80 |
| chr10-39764137-INS-196 | chr10-39764137-INS-196 |
| chr10-39794910-INS-64 | chr10-39794922-INS-58 |
| chr10-39799310-INS-643 | chr10-39799311-INS-643 |
| chr10-39818006-INS-56 | chr10-39818006-INS-56 |
| chr10-39829694-INS-2689 | chr10-39829695-INS-2691 |
| chr10-39864009-INS-100 | chr10-39864009-INS-100 |
| chr10-39871977-INS-117 | chr10-39871977-INS-145 |
| chr10-39882540-INS-7073 | chr10-39882547-INS-7066 |
| chr10-39897291-INS-106 | chr10-39897291-INS-108 |
| chr10-39908276-INS-4754 | chr10-39908277-INS-4752 |
| chr10-40025433-INS-177 | chr10-40025433-INS-176 |
| chr10-40045067-INS-374 | chr10-40045067-INS-416 |
| chr10-40047546-INS-1505 | chr10-40047551-INS-1500 |
| chr10-40093161-INS-314 | chr10-40093161-INS-314 |
| chr10-40095662-INS-1017 | chr10-40095663-INS-1017 |
| chr10-40121572-INS-68 | chr10-40121572-INS-68 |
| chr10-40124320-INS-619 | chr10-40124320-INS-615 |
| chr10-40125002-INS-108 | chr10-40125002-INS-111 |
| chr10-40141453-INS-223 | chr10-40141457-INS-195 |
| chr10-40149330-INS-139 | chr10-40149331-INS-137 |
| chr10-40150337-INS-176 | chr10-40150337-INS-181 |
| chr10-40185101-INS-5544 | chr10-40185102-INS-5544 |
| chr10-40194066-INS-77 | chr10-40194076-INS-69 |
| chr10-40209596-INS-110 | chr10-40209596-INS-110 |
| chr10-40211539-INS-212 | chr10-40211539-INS-211 |
| chr10-40230921-INS-204 | chr10-40230921-INS-204 |
| chr10-40271972-INS-70 | chr10-40271972-INS-80 |
| chr10-40274117-INS-270 | chr10-40274117-INS-270 |
| chr10-40280458-INS-2942 | chr10-40280459-INS-2942 |
| chr10-40287777-INS-242 | chr10-40287777-INS-242 |
| chr10-40290403-INS-70 | chr10-40290401-INS-75 |
| chr10-40303426-INS-192 | chr10-40303426-INS-193 |
| chr10-40307318-INS-207 | chr10-40307318-INS-207 |
| chr10-40316025-INS-181 | chr10-40316025-INS-181 |
| chr10-40318284-INS-53 | chr10-40318284-INS-53 |
| chr10-40321465-INS-102 | chr10-40321465-INS-102 |

| | |
|---|---|
| chr10-40325707-INS-64 | chr10-40325707-INS-64 |
| chr10-40365659-INS-375 | chr10-40365659-INS-375 |
| chr10-40372621-INS-169 | chr10-40372597-INS-173 |
| chr10-40374338-INS-4085 | chr10-40374339-INS-4085 |
| chr10-40394774-INS-64 | chr10-40394768-INS-65 |
| chr10-40420377-INS-57 | chr10-40420377-INS-57 |
| chr10-40496690-INS-211 | chr10-40496690-INS-211 |
| chr10-40501403-INS-288 | chr10-40501403-INS-288 |
| chr10-40531276-INS-253 | chr10-40531276-INS-253 |
| chr10-40542178-INS-7233 | chr10-40542179-INS-7232 |
| chr10-40564114-INS-52 | chr10-40564114-INS-54 |
| chr10-40576069-INS-53 | chr10-40576074-INS-50 |
| chr10-40577493-INS-132 | chr10-40577493-INS-132 |
| chr10-40584201-INS-80 | chr10-40584201-INS-80 |
| chr10-40591930-INS-63 | chr10-40591930-INS-54 |
| chr10-40602040-INS-82 | chr10-40602038-INS-52 |
| chr10-40612399-INS-58 | chr10-40612399-INS-60 |
| chr10-40645884-INS-1388 | chr10-40645910-INS-1385 |
| chr10-40657144-INS-245 | chr10-40657144-INS-254 |
| chr10-40666415-INS-346 | chr10-40666415-INS-346 |
| chr10-40672055-INS-217 | chr10-40672055-INS-217 |
| chr10-40692475-INS-211 | chr10-40692475-INS-212 |
| chr10-40698502-INS-542 | chr10-40698502-INS-542 |
| chr10-40700139-INS-77 | chr10-40700139-INS-77 |
| chr10-40715593-INS-60 | chr10-40715617-INS-80 |
| chr10-40727440-INS-6415 | chr10-40727441-INS-6415 |
| chr10-40739290-INS-97 | chr10-40739290-INS-97 |
| chr10-40753361-INS-498 | chr10-40753361-INS-498 |
| chr10-40771299-INS-199 | chr10-40771299-INS-199 |
| chr10-40772914-INS-67 | chr10-40772920-INS-54 |
| chr10-40785541-INS-52 | chr10-40785541-INS-54 |
| chr10-40787465-INS-153 | chr10-40787474-INS-156 |
| chr10-40819347-INS-355 | chr10-40819347-INS-356 |
| chr10-40846268-INS-119 | chr10-40846268-INS-119 |
| chr10-40862136-INS-173 | chr10-40862136-INS-175 |
| chr10-40891438-INS-111 | chr10-40891438-INS-111 |
| chr10-40894397-INS-364 | chr10-40894350-INS-372 |
| chr10-40898605-INS-54 | chr10-40898605-INS-56 |
| chr10-40971080-INS-439 | chr10-40971081-INS-444 |
| chr10-40977562-INS-215 | chr10-40977562-INS-215 |
| chr10-40990460-INS-52 | chr10-40990460-INS-52 |
| chr10-41015035-INS-92 | chr10-41015035-INS-100 |
| chr10-41051939-INS-68 | chr10-41051901-INS-64 |
| chr10-41120372-INS-92 | chr10-41120366-INS-92 |
| chr10-41130959-INS-64 | chr10-41130959-INS-64 |
| chr10-41157197-INS-214 | chr10-41157201-INS-217 |
| chr10-41226284-INS-88 | chr10-41226284-INS-84 |
| chr10-41227496-INS-208 | chr10-41227496-INS-208 |
| chr10-41249208-INS-65 | chr10-41249208-INS-65 |
| chr10-41280008-INS-198 | chr10-41280008-INS-198 |
| chr10-41297657-INS-55 | chr10-41297657-INS-53 |
| chr10-41301585-INS-53 | chr10-41301585-INS-53 |
| chr10-41310473-INS-216 | chr10-41310473-INS-218 |
| chr10-41325499-INS-248 | chr10-41325499-INS-298 |
| chr10-41341303-INS-70 | chr10-41341303-INS-70 |
| chr10-41342954-INS-209 | chr10-41342954-INS-209 |
| chr10-41351890-INS-62 | chr10-41351892-INS-69 |
| chr10-41370193-INS-243 | chr10-41370193-INS-243 |
| chr10-41411679-INS-186 | chr10-41411679-INS-186 |
| chr10-41458280-INS-68 | chr10-41458280-INS-50 |
| chr10-41462731-INS-57 | chr10-41462731-INS-57 |
| chr10-41469657-INS-398 | chr10-41469657-INS-398 |
| chr10-41484030-INS-78 | chr10-41484030-INS-78 |
| chr10-41507294-INS-205 | chr10-41507294-INS-205 |
| chr10-41518997-INS-64 | chr10-41518997-INS-76 |
| chr10-41527518-INS-81 | chr10-41527510-INS-82 |
| chr10-41573453-INS-65 | chr10-41573453-INS-65 |
| chr10-41575010-INS-63 | chr10-41575012-INS-52 |
| chr10-41614383-INS-6452 | chr10-41614384-INS-6452 |
| chr10-41615915-INS-81 | chr10-41615915-INS-52 |
| chr10-41617714-INS-107 | chr10-41617714-INS-111 |
| chr10-41637816-INS-344 | chr10-41637816-INS-344 |
| chr10-41644242-INS-52 | chr10-41644242-INS-50 |
| chr10-41695054-INS-54 | chr10-41695051-INS-102 |
| chr10-41703232-INS-8958 | chr10-41703233-INS-8958 |
| chr10-41737011-INS-61 | chr10-41737011-INS-61 |
| chr10-41742559-INS-7141 | chr10-41742560-INS-7141 |

| | |
|---|---|
| chr10-41789142-INS-718 | chr10-41789156-INS-732 |
| chr10-41798727-INS-99 | chr10-41798727-INS-99 |
| chr10-41810356-INS-88 | chr10-41810356-INS-90 |
| chr10-41832518-INS-123 | chr10-41832518-INS-123 |
| chr10-41836400-INS-65 | chr10-41836400-INS-65 |
| chr10-41844861-INS-59 | chr10-41844861-INS-61 |
| chr10-41864487-INS-69 | chr10-41864487-INS-60 |
| chr10-41871548-INS-88 | chr10-41871548-INS-88 |
| chr10-41888638-INS-96 | chr10-41888639-INS-94 |
| chr10-41898920-INS-52 | chr10-41898932-INS-85 |
| chr10-41916367-INS-97 | chr10-41916371-INS-101 |
| chr10-41940642-INS-893 | chr10-41940642-INS-893 |
| chr10-41941718-INS-52 | chr10-41941718-INS-71 |
| chr10-41957510-INS-60 | chr10-41957548-INS-68 |
| chr10-41970983-INS-526 | chr10-41970984-INS-526 |
| chr10-41976769-INS-52 | chr10-41976746-INS-72 |
| chr10-41981193-INS-108 | chr10-41981193-INS-108 |
| chr10-41982739-INS-108 | chr10-41982739-INS-108 |
| chr10-41989819-INS-108 | chr10-41989819-INS-108 |
| chr10-42024506-INS-999 | chr10-42024506-INS-1000 |
| chr10-42039814-INS-77 | chr10-42039814-INS-84 |
| chr10-42062796-INS-58 | chr10-42062796-INS-54 |
| chr10-42074942-INS-56 | chr10-42074942-INS-58 |
| chr10-42104526-INS-64 | chr10-42104526-INS-64 |
| chr10-42112278-INS-74 | chr10-42112278-INS-74 |
| chr10-42122168-INS-208 | chr10-42122168-INS-209 |
| chr10-42171091-INS-86 | chr10-42171085-INS-87 |
| chr10-42177712-INS-55 | chr10-42177667-INS-56 |
| chr10-42178813-INS-369 | chr10-42178813-INS-369 |
| chr10-42184195-INS-134 | chr10-42184195-INS-134 |
| chr10-42190099-INS-122 | chr10-42190095-INS-78 |
| chr10-42237393-INS-371 | chr10-42237393-INS-379 |
| chr10-42288436-INS-138 | chr10-42288439-INS-130 |
| chr10-42302727-INS-104 | chr10-42302727-INS-104 |
| chr10-42308506-INS-62 | chr10-42308506-INS-62 |
| chr10-42331639-INS-51 | chr10-42331639-INS-51 |
| chr10-42333174-INS-59 | chr10-42333174-INS-59 |
| chr10-42336988-INS-61 | chr10-42336988-INS-61 |
| chr10-42338014-INS-203 | chr10-42338014-INS-205 |
| chr10-42345523-INS-140 | chr10-42345523-INS-139 |
| chr10-42366257-INS-183 | chr10-42366257-INS-183 |
| chr10-42371392-INS-182 | chr10-42371392-INS-199 |
| chr10-42381099-INS-56 | chr10-42381099-INS-58 |
| chr10-42413020-INS-839 | chr10-42413020-INS-839 |
| chr10-42421485-INS-221 | chr10-42421485-INS-221 |
| chr10-42457649-INS-68 | chr10-42457629-INS-72 |
| chr10-42483429-INS-1217 | chr10-42483430-INS-1217 |
| chr10-42488451-INS-200 | chr10-42488452-INS-200 |
| chr10-42513452-INS-204 | chr10-42513452-INS-204 |
| chr10-42514416-INS-217 | chr10-42514416-INS-217 |
| chr10-42522021-INS-172 | chr10-42522021-INS-172 |
| chr10-42527606-INS-52 | chr10-42527607-INS-60 |
| chr10-42531929-INS-58 | chr10-42531929-INS-58 |
| chr10-42569357-INS-55 | chr10-42569357-INS-55 |
| chr10-42584872-INS-98 | chr10-42584892-INS-110 |
| chr10-42588436-INS-86 | chr10-42588436-INS-86 |
| chr10-42590206-INS-82 | chr10-42590218-INS-102 |
| chr10-42605833-INS-200 | chr10-42605833-INS-199 |
| chr10-42607189-INS-54 | chr10-42607175-INS-54 |
| chr10-42627055-INS-52 | chr10-42627055-INS-52 |
| chr10-42640125-INS-103 | chr10-42640125-INS-109 |
| chr10-42643388-INS-5908 | chr10-42643390-INS-5907 |
| chr10-42658288-INS-2033 | chr10-42658289-INS-2034 |
| chr10-42726639-INS-59 | chr10-42726639-INS-59 |
| chr10-42744436-INS-201 | chr10-42744436-INS-201 |
| chr10-42753654-INS-62 | chr10-42753654-INS-55 |
| chr10-42782112-INS-72 | chr10-42782112-INS-72 |
| chr10-42787441-INS-167 | chr10-42787441-INS-167 |
| chr10-42791363-INS-135 | chr10-42791363-INS-135 |
| chr10-42805271-INS-422 | chr10-42805270-INS-423 |
| chr10-42811659-INS-166 | chr10-42811659-INS-166 |
| chr10-42822602-INS-54 | chr10-42822590-INS-71 |
| chr10-42824015-INS-441 | chr10-42824015-INS-418 |
| chr10-42826589-INS-102 | chr10-42826589-INS-104 |
| chr10-42832561-INS-96 | chr10-42832561-INS-132 |
| chr10-42859644-INS-201 | chr10-42859644-INS-201 |
| chr10-42921366-INS-56 | chr10-42921366-INS-76 |

| | |
|---|---|
| chr10-42934884-INS-140 | chr10-42934884-INS-154 |
| chr10-42940258-INS-56 | chr10-42940258-INS-96 |
| chr10-42944824-INS-183 | chr10-42944824-INS-183 |
| chr10-42955175-INS-56 | chr10-42955175-INS-56 |
| chr10-42980559-INS-242 | chr10-42980559-INS-242 |
| chr10-43018875-INS-107 | chr10-43018873-INS-115 |
| chr10-43025204-INS-70 | chr10-43025204-INS-71 |
| chr10-43031775-INS-3939 | chr10-43031776-INS-3891 |
| chr10-43075864-INS-315 | chr10-43075864-INS-315 |
| chr10-43084087-INS-64 | chr10-43084087-INS-62 |
| chr10-43099865-INS-112 | chr10-43099865-INS-112 |
| chr10-43130315-INS-84 | chr10-43130315-INS-88 |
| chr10-43180344-INS-147 | chr10-43180346-INS-146 |
| chr10-43211425-INS-62 | chr10-43211425-INS-62 |
| chr10-43249715-INS-59 | chr10-43249734-INS-59 |
| chr10-43267545-INS-221 | chr10-43267545-INS-221 |
| chr10-43270657-INS-203 | chr10-43270657-INS-203 |
| chr10-43294599-INS-72 | chr10-43294599-INS-76 |
| chr10-43296617-INS-394 | chr10-43296617-INS-395 |
| chr10-43309113-INS-221 | chr10-43309112-INS-220 |
| chr10-43311723-INS-391 | chr10-43311723-INS-391 |
| chr10-43351709-INS-194 | chr10-43351709-INS-196 |
| chr10-43372725-INS-203 | chr10-43372725-INS-207 |
| chr10-43395625-INS-190 | chr10-43395625-INS-190 |
| chr10-43421266-INS-206 | chr10-43421266-INS-207 |
| chr10-43432841-INS-210 | chr10-43432842-INS-210 |
| chr10-43435799-INS-569 | chr10-43435799-INS-569 |
| chr10-43438573-INS-1083 | chr10-43438577-INS-1051 |
| chr10-43445446-INS-60 | chr10-43445447-INS-60 |
| chr10-43451395-INS-367 | chr10-43451395-INS-367 |
| chr10-43477162-INS-95 | chr10-43477153-INS-55 |
| chr10-43483153-INS-110 | chr10-43483153-INS-112 |
| chr10-43485753-INS-74 | chr10-43485753-INS-74 |
| chr10-43498777-INS-943 | chr10-43498777-INS-943 |
| chr10-43507077-INS-153 | chr10-43507071-INS-112 |
| chr10-43518227-INS-109 | chr10-43518227-INS-109 |
| chr10-43523399-INS-407 | chr10-43523423-INS-399 |
| chr10-43589241-INS-1074 | chr10-43589243-INS-1075 |
| chr10-43604609-INS-108 | chr10-43604620-INS-140 |
| chr10-43623060-INS-174 | chr10-43623061-INS-174 |
| chr10-43627554-INS-69 | chr10-43627561-INS-98 |
| chr10-43628707-INS-64 | chr10-43628716-INS-77 |
| chr10-43635791-INS-145 | chr10-43635791-INS-151 |
| chr10-43639070-INS-62 | chr10-43639070-INS-66 |
| chr10-43668819-INS-217 | chr10-43668819-INS-221 |
| chr10-43671987-INS-397 | chr10-43671987-INS-397 |
| chr10-43681708-INS-60 | chr10-43681708-INS-92 |
| chr10-43692375-INS-94 | chr10-43692375-INS-94 |
| chr10-43692450-INS-548 | chr10-43692450-INS-548 |
| chr10-43705234-INS-5330 | chr10-43705235-INS-5330 |
| chr10-43715012-INS-513 | chr10-43715012-INS-513 |
| chr10-43724813-INS-180 | chr10-43724813-INS-182 |
| chr10-43734342-INS-7526 | chr10-43734343-INS-7526 |
| chr10-43737099-INS-390 | chr10-43737095-INS-377 |
| chr10-43739413-INS-1472 | chr10-43739414-INS-1472 |
| chr10-43751058-INS-74 | chr10-43751058-INS-74 |
| chr10-43810732-INS-325 | chr10-43810732-INS-325 |
| chr10-43817106-INS-673 | chr10-43817106-INS-720 |
| chr10-43848770-INS-56 | chr10-43848770-INS-56 |
| chr10-43852956-INS-148 | chr10-43852956-INS-148 |
| chr10-43868243-INS-140 | chr10-43868243-INS-140 |
| chr10-43871249-INS-172 | chr10-43871245-INS-196 |
| chr10-43882753-INS-134 | chr10-43882752-INS-123 |
| chr10-43908315-INS-82 | chr10-43908315-INS-58 |
| chr10-43976321-INS-111 | chr10-43976321-INS-111 |
| chr10-43985926-INS-351 | chr10-43985926-INS-354 |
| chr10-44018724-INS-167 | chr10-44018726-INS-177 |
| chr10-44028044-INS-52 | chr10-44028044-INS-60 |
| chr10-44072206-INS-5314 | chr10-44072207-INS-5314 |
| chr10-44082328-INS-547 | chr10-44082328-INS-547 |
| chr10-44149352-INS-2345 | chr10-44149366-INS-2348 |
| chr10-44153726-INS-1796 | chr10-44153727-INS-1796 |
| chr10-44191107-INS-70 | chr10-44191107-INS-70 |
| chr10-44249126-INS-299 | chr10-44249126-INS-299 |
| chr10-44271394-INS-331 | chr10-44271394-INS-334 |
| chr10-44296722-INS-50 | chr10-44296722-INS-64 |
| chr10-44311640-INS-58 | chr10-44311640-INS-60 |

| | |
|---|---|
| chr10-44434091-INS-353 | chr10-44434091-INS-353 |
| chr10-44469983-INS-453 | chr10-44469983-INS-453 |
| chr10-44480711-INS-233 | chr10-44480711-INS-233 |
| chr10-44487238-INS-204 | chr10-44487238-INS-204 |
| chr10-44559573-INS-55 | chr10-44559573-INS-55 |
| chr10-44586326-INS-56 | chr10-44586330-INS-102 |
| chr10-44606231-INS-1197 | chr10-44606232-INS-1206 |
| chr10-44633620-INS-54 | chr10-44633620-INS-54 |
| chr10-44638851-INS-51 | chr10-44638852-INS-74 |
| chr10-44652842-INS-2986 | chr10-44652843-INS-2986 |
| chr10-44655435-INS-486 | chr10-44655435-INS-486 |
| chr10-44677916-INS-84 | chr10-44677916-INS-84 |
| chr10-44685491-INS-80 | chr10-44685491-INS-80 |
| chr10-44687445-INS-59 | chr10-44687445-INS-73 |
| chr10-44765062-INS-124 | chr10-44765062-INS-88 |
| chr10-44777104-INS-92 | chr10-44777104-INS-92 |
| chr10-44796408-INS-604 | chr10-44796368-INS-604 |
| chr10-44837983-INS-60 | chr10-44837983-INS-64 |
| chr10-44856622-INS-112 | chr10-44856621-INS-110 |
| chr10-44861853-INS-5241 | chr10-44861854-INS-5241 |
| chr10-44883599-INS-265 | chr10-44883599-INS-231 |
| chr10-44893314-INS-129 | chr10-44893314-INS-129 |
| chr10-44909468-INS-58 | chr10-44909468-INS-58 |
| chr10-44928843-INS-90 | chr10-44928827-INS-95 |
| chr10-44930862-INS-159 | chr10-44930862-INS-159 |
| chr10-44934364-INS-181 | chr10-44934364-INS-173 |
| chr10-44967320-INS-70 | chr10-44967320-INS-62 |
| chr10-44973736-INS-217 | chr10-44973736-INS-217 |
| chr10-44980108-INS-603 | chr10-44980108-INS-581 |
| chr10-44986921-INS-152 | chr10-44986933-INS-152 |
| chr10-45012025-INS-476 | chr10-45012025-INS-476 |
| chr10-45033282-INS-173 | chr10-45033282-INS-173 |
| chr10-45041543-INS-856 | chr10-45041543-INS-848 |
| chr10-45064283-INS-59 | chr10-45064289-INS-72 |
| chr10-45076908-INS-52 | chr10-45076908-INS-52 |
| chr10-45089359-INS-835 | chr10-45089359-INS-834 |
| chr10-45097885-INS-294 | chr10-45097885-INS-294 |
| chr10-45102202-INS-786 | chr10-45102195-INS-787 |
| chr10-45118647-INS-101 | chr10-45118647-INS-78 |
| chr10-45120556-INS-57 | chr10-45120556-INS-57 |
| chr10-45135325-INS-52 | chr10-45135325-INS-52 |
| chr10-45152046-INS-439 | chr10-45152046-INS-439 |
| chr10-45162308-INS-55 | chr10-45162308-INS-50 |
| chr10-45164326-INS-60 | chr10-45164326-INS-59 |
| chr10-45168934-INS-105 | chr10-45168934-INS-70 |
| chr10-45196938-INS-87 | chr10-45196938-INS-80 |
| chr10-45205913-INS-191 | chr10-45205913-INS-191 |
| chr10-45207183-INS-169 | chr10-45207183-INS-169 |
| chr10-45222741-INS-208 | chr10-45222741-INS-200 |
| chr10-45226521-INS-825 | chr10-45226521-INS-825 |
| chr10-45275921-INS-855 | chr10-45275921-INS-850 |
| chr10-45299639-INS-105 | chr10-45299639-INS-92 |
| chr10-45322575-INS-112 | chr10-45322575-INS-112 |
| chr10-45364822-INS-203 | chr10-45364822-INS-203 |
| chr10-45372496-INS-68 | chr10-45372464-INS-52 |
| chr10-45375540-INS-89 | chr10-45375540-INS-101 |
| chr10-45454736-INS-72 | chr10-45454736-INS-72 |
| chr10-45482537-INS-76 | chr10-45482543-INS-111 |
| chr10-45513422-INS-70 | chr10-45513425-INS-112 |
| chr10-45516513-INS-74 | chr10-45516502-INS-54 |
| chr10-45541277-INS-113 | chr10-45541277-INS-113 |
| chr10-45598216-INS-166 | chr10-45598216-INS-166 |
| chr10-45621216-INS-210 | chr10-45621216-INS-210 |
| chr10-45695388-INS-6549 | chr10-45695389-INS-6549 |
| chr10-45698843-INS-57 | chr10-45698843-INS-57 |
| chr10-45712606-INS-2201 | chr10-45712607-INS-2202 |
| chr10-45732290-INS-63 | chr10-45732290-INS-62 |
| chr10-45737780-INS-51 | chr10-45737780-INS-51 |
| chr10-45760784-INS-6432 | chr10-45760785-INS-6432 |
| chr10-45773223-INS-423 | chr10-45773222-INS-410 |
| chr10-45784344-INS-80 | chr10-45784346-INS-122 |
| chr10-45789380-INS-60 | chr10-45789380-INS-60 |
| chr10-45802977-INS-57 | chr10-45802977-INS-51 |
| chr10-45804697-INS-67 | chr10-45804697-INS-69 |
| chr10-45827303-INS-199 | chr10-45827303-INS-199 |
| chr10-45850087-INS-164 | chr10-45850087-INS-164 |
| chr10-45867551-INS-7133 | chr10-45867552-INS-7133 |

| | |
|---|---|
| chr10-45877415-INS-102 | chr10-45877415-INS-102 |
| chr10-45889373-INS-287 | chr10-45889373-INS-287 |
| chr10-45911016-INS-622 | chr10-45911016-INS-622 |
| chr10-45946083-INS-88 | chr10-45946080-INS-70 |
| chr10-45951323-INS-76 | chr10-45951323-INS-77 |
| chr10-45966513-INS-59 | chr10-45966513-INS-72 |
| chr10-45971726-INS-74 | chr10-45971726-INS-74 |
| chr10-45973327-INS-310 | chr10-45973327-INS-310 |
| chr10-45998560-INS-87 | chr10-45998600-INS-92 |
| chr10-46013153-INS-2839 | chr10-46013154-INS-2839 |
| chr10-46018843-INS-300 | chr10-46018839-INS-306 |
| chr10-46051948-INS-6037 | chr10-46051949-INS-6037 |
| chr10-46075680-INS-388 | chr10-46075680-INS-388 |
| chr10-46077337-INS-4373 | chr10-46077338-INS-4366 |
| chr10-46127154-INS-689 | chr10-46127154-INS-697 |
| chr10-46128982-INS-936 | chr10-46128982-INS-921 |
| chr10-46132539-INS-62 | chr10-46132540-INS-62 |
| chr10-46135833-INS-85 | chr10-46135833-INS-70 |
| chr10-46140074-INS-70 | chr10-46140035-INS-86 |
| chr10-46148512-INS-2201 | chr10-46148513-INS-2201 |
| chr10-46198506-INS-54 | chr10-46198506-INS-67 |
| chr10-46201842-INS-743 | chr10-46201842-INS-743 |
| chr10-46268291-INS-430 | chr10-46268291-INS-430 |
| chr10-46271268-INS-118 | chr10-46271266-INS-118 |
| chr10-46283028-INS-64 | chr10-46283028-INS-78 |
| chr10-46335276-INS-148 | chr10-46335318-INS-152 |
| chr10-46350980-INS-303 | chr10-46350980-INS-303 |
| chr10-46371116-INS-390 | chr10-46371116-INS-390 |
| chr10-46403281-INS-387 | chr10-46403281-INS-388 |
| chr10-46420459-INS-71 | chr10-46420459-INS-66 |
| chr10-46437079-INS-62 | chr10-46437079-INS-66 |
| chr10-46437703-INS-600 | chr10-46437703-INS-600 |
| chr10-46502472-INS-184 | chr10-46502470-INS-192 |
| chr10-46544175-INS-446 | chr10-46544175-INS-446 |
| chr10-46574913-INS-530 | chr10-46574913-INS-529 |
| chr10-46580464-INS-3139 | chr10-46580465-INS-3139 |
| chr10-46595179-INS-1132 | chr10-46595180-INS-1132 |
| chr10-46607814-INS-492 | chr10-46607814-INS-492 |
| chr10-46608905-INS-668 | chr10-46608919-INS-678 |
| chr10-46640222-INS-162 | chr10-46640222-INS-144 |
| chr10-46651409-INS-74 | chr10-46651409-INS-68 |
| chr10-46667683-INS-2824 | chr10-46667684-INS-2824 |
| chr10-46667752-INS-6401 | chr10-46667753-INS-6401 |
| chr10-46678200-INS-294 | chr10-46678200-INS-294 |
| chr10-46710139-INS-230 | chr10-46710139-INS-230 |
| chr10-46755542-INS-62 | chr10-46755542-INS-62 |
| chr10-46758459-INS-945 | chr10-46758459-INS-942 |
| chr10-46765924-INS-174 | chr10-46765924-INS-158 |
| chr10-46802302-INS-75 | chr10-46802290-INS-60 |
| chr10-46816608-INS-150 | chr10-46816609-INS-101 |
| chr10-46822435-INS-221 | chr10-46822435-INS-221 |
| chr10-46866538-INS-1699 | chr10-46866539-INS-1699 |
| chr10-46875508-INS-72 | chr10-46875507-INS-68 |
| chr10-46901451-INS-109 | chr10-46901451-INS-109 |
| chr10-46925697-INS-285 | chr10-46925697-INS-285 |
| chr10-46970590-INS-969 | chr10-46970590-INS-967 |
| chr10-46976483-INS-969 | chr10-46976483-INS-970 |
| chr10-47021291-INS-60 | chr10-47021292-INS-61 |
| chr10-47060309-INS-497 | chr10-47060309-INS-459 |
| chr10-47063149-INS-210 | chr10-47063149-INS-210 |
| chr10-47082637-INS-161 | chr10-47082641-INS-149 |
| chr10-47092656-INS-419 | chr10-47092656-INS-419 |
| chr10-47104886-INS-2183 | chr10-47104924-INS-2183 |
| chr10-47108938-INS-52 | chr10-47108938-INS-52 |
| chr10-47110830-INS-576 | chr10-47110830-INS-576 |
| chr10-47155839-INS-3996 | chr10-47155840-INS-3996 |
| chr10-47172784-INS-151 | chr10-47172784-INS-152 |
| chr10-47229074-INS-201 | chr10-47229074-INS-201 |
| chr10-47232289-INS-56 | chr10-47232285-INS-68 |
| chr10-47236201-INS-5081 | chr10-47236202-INS-5081 |
| chr10-47270150-INS-675 | chr10-47270150-INS-675 |
| chr10-47276575-INS-1284 | chr10-47276576-INS-1283 |
| chr10-47301749-INS-52 | chr10-47301750-INS-52 |
| chr10-47381641-INS-57 | chr10-47381641-INS-57 |
| chr10-47411520-INS-529 | chr10-47411520-INS-529 |
| chr10-47431899-INS-347 | chr10-47431899-INS-370 |
| chr10-47499039-INS-1864 | chr10-47499040-INS-1864 |

| | |
|---|---|
| chr10-47523028-INS-94 | chr10-47523031-INS-95 |
| chr10-47568371-INS-52 | chr10-47568371-INS-52 |
| chr10-47570653-INS-638 | chr10-47570653-INS-654 |
| chr10-47586844-INS-4778 | chr10-47586845-INS-4778 |
| chr10-47644169-INS-72 | chr10-47644169-INS-54 |
| chr10-47651248-INS-1097 | chr10-47651249-INS-1096 |
| chr10-47668137-INS-62 | chr10-47668137-INS-64 |
| chr10-47695948-INS-90 | chr10-47695952-INS-94 |
| chr10-47701845-INS-660 | chr10-47701845-INS-663 |
| chr10-47720985-INS-481 | chr10-47720987-INS-485 |
| chr10-47726476-INS-76 | chr10-47726476-INS-76 |
| chr10-47740880-INS-1214 | chr10-47740881-INS-1214 |
| chr10-47781795-INS-83 | chr10-47781795-INS-84 |
| chr10-47792012-INS-64 | chr10-47792000-INS-56 |
| chr10-47802952-INS-114 | chr10-47802964-INS-99 |
| chr10-47835597-INS-248 | chr10-47835597-INS-248 |
| chr10-47857934-INS-121 | chr10-47857934-INS-121 |
| chr10-47861239-INS-299 | chr10-47861241-INS-299 |
| chr10-47877175-INS-834 | chr10-47877175-INS-833 |
| chr10-47884248-INS-203 | chr10-47884248-INS-206 |
| chr10-47888594-INS-270 | chr10-47888594-INS-274 |
| chr10-47916011-INS-55 | chr10-47916011-INS-55 |
| chr10-47945457-INS-54 | chr10-47945457-INS-68 |
| chr10-47963971-INS-762 | chr10-47963971-INS-762 |
| chr10-47976452-INS-325 | chr10-47976452-INS-325 |
| chr10-47988454-INS-56 | chr10-47988454-INS-72 |
| chr10-47995888-INS-6397 | chr10-47995889-INS-6397 |
| chr10-48005732-INS-184 | chr10-48005732-INS-184 |
| chr10-48017848-INS-7091 | chr10-48017849-INS-7091 |
| chr10-48021123-INS-72 | chr10-48021123-INS-84 |
| chr10-48022679-INS-311 | chr10-48022679-INS-295 |
| chr10-48030520-INS-175 | chr10-48030520-INS-175 |
| chr10-48034395-INS-130 | chr10-48034395-INS-130 |
| chr10-48041122-INS-832 | chr10-48041122-INS-835 |
| chr10-48042720-INS-74 | chr10-48042721-INS-78 |
| chr10-48068183-INS-779 | chr10-48068184-INS-776 |
| chr10-48085938-INS-183 | chr10-48085926-INS-203 |
| chr10-48092487-INS-72 | chr10-48092488-INS-53 |
| chr10-48101078-INS-52 | chr10-48101078-INS-62 |
| chr10-48179026-INS-2265 | chr10-48179027-INS-2265 |
| chr10-48191730-INS-410 | chr10-48191730-INS-410 |
| chr10-48202819-INS-351 | chr10-48202819-INS-349 |
| chr10-48247332-INS-1433 | chr10-48247333-INS-1433 |
| chr10-48273453-INS-1609 | chr10-48273456-INS-1613 |
| chr10-48313567-INS-333 | chr10-48313567-INS-333 |
| chr10-48374157-INS-8808 | chr10-48374158-INS-8808 |
| chr10-48378249-INS-63 | chr10-48378249-INS-63 |
| chr10-48398423-INS-58 | chr10-48398423-INS-58 |
| chr10-48404293-INS-85 | chr10-48404293-INS-85 |
| chr10-48408568-INS-72 | chr10-48408568-INS-70 |
| chr10-48422952-INS-7171 | chr10-48422953-INS-7171 |
| chr10-48446779-INS-7171 | chr10-48446779-INS-7171 |
| chr10-48484927-INS-214 | chr10-48484927-INS-220 |
| chr10-48487542-INS-54 | chr10-48487530-INS-56 |
| chr10-48504714-INS-421 | chr10-48504714-INS-434 |
| chr10-48538450-INS-94 | chr10-48538470-INS-110 |
| chr10-48542705-INS-60 | chr10-48542706-INS-64 |
| chr10-48545536-INS-2547 | chr10-48545537-INS-2548 |
| chr10-48570881-INS-1888 | chr10-48570882-INS-1888 |
| chr10-48597559-INS-1114 | chr10-48597560-INS-1114 |
| chr10-48607810-INS-1905 | chr10-48607811-INS-1950 |
| chr10-48632772-INS-6960 | chr10-48632773-INS-6961 |
| chr10-48651099-INS-116 | chr10-48651095-INS-124 |
| chr10-48657528-INS-56 | chr10-48657528-INS-54 |
| chr10-48699335-INS-77 | chr10-48699353-INS-50 |
| chr10-48743671-INS-146 | chr10-48743671-INS-146 |
| chr10-48765171-INS-7320 | chr10-48765172-INS-7319 |
| chr10-48779170-INS-73 | chr10-48779170-INS-73 |
| chr10-48793015-INS-153 | chr10-48793015-INS-153 |
| chr10-48814738-INS-75 | chr10-48814738-INS-75 |
| chr10-48839620-INS-6261 | chr10-48839621-INS-6251 |
| chr10-48857237-INS-55 | chr10-48857237-INS-55 |
| chr10-48862278-INS-55 | chr10-48862278-INS-65 |
| chr10-48871597-INS-78 | chr10-48871597-INS-78 |
| chr10-48924009-INS-51 | chr10-48924010-INS-50 |
| chr10-48943243-INS-4799 | chr10-48943244-INS-4799 |
| chr10-48950766-INS-212 | chr10-48950766-INS-212 |

| | |
|---|---|
| chr10-49014481-INS-8020 | chr10-49014482-INS-8024 |
| chr10-49017069-INS-1459 | chr10-49017070-INS-1458 |
| chr10-49018882-INS-78 | chr10-49018882-INS-60 |
| chr10-49039653-INS-76 | chr10-49039624-INS-56 |
| chr10-49050118-INS-693 | chr10-49050117-INS-692 |
| chr10-49058576-INS-56 | chr10-49058576-INS-68 |
| chr10-49068373-INS-6179 | chr10-49068374-INS-6180 |
| chr10-49073161-INS-313 | chr10-49073161-INS-291 |
| chr10-49073702-INS-6401 | chr10-49073703-INS-6386 |
| chr10-49074664-INS-1938 | chr10-49074665-INS-1935 |
| chr10-49090747-INS-498 | chr10-49090747-INS-498 |
| chr10-49177266-INS-170 | chr10-49177266-INS-170 |
| chr10-49184193-INS-1007 | chr10-49184194-INS-1007 |
| chr10-49211490-INS-909 | chr10-49211490-INS-910 |
| chr10-49223533-INS-52 | chr10-49223533-INS-52 |
| chr10-49231214-INS-384 | chr10-49231214-INS-387 |
| chr10-49232376-INS-1472 | chr10-49232377-INS-1474 |
| chr10-49240431-INS-54 | chr10-49240431-INS-56 |
| chr10-49268638-INS-414 | chr10-49268638-INS-435 |
| chr10-49270786-INS-53 | chr10-49270776-INS-63 |
| chr10-49288144-INS-179 | chr10-49288144-INS-179 |
| chr10-49322818-INS-51 | chr10-49322804-INS-63 |
| chr10-49327436-INS-54 | chr10-49327436-INS-56 |
| chr10-49378408-INS-52 | chr10-49378367-INS-54 |
| chr10-49394536-INS-72 | chr10-49394542-INS-96 |
| chr10-49432355-INS-219 | chr10-49432355-INS-219 |
| chr10-49441465-INS-62 | chr10-49441465-INS-56 |
| chr10-49472971-INS-387 | chr10-49472971-INS-387 |
| chr10-49483717-INS-667 | chr10-49483717-INS-662 |
| chr10-49488135-INS-1933 | chr10-49488136-INS-1932 |
| chr10-49495760-INS-91 | chr10-49495760-INS-91 |
| chr10-49513107-INS-89 | chr10-49513107-INS-90 |
| chr10-49520869-INS-872 | chr10-49520869-INS-873 |
| chr10-49592723-INS-188 | chr10-49592723-INS-188 |
| chr10-49606530-INS-68 | chr10-49606530-INS-78 |
| chr10-49631376-INS-3083 | chr10-49631377-INS-3083 |
| chr10-49633415-INS-954 | chr10-49633415-INS-932 |
| chr10-49643209-INS-3345 | chr10-49643210-INS-3346 |
| chr10-49688836-INS-71 | chr10-49688836-INS-63 |
| chr10-49708032-INS-159 | chr10-49708032-INS-158 |
| chr10-49724563-INS-1807 | chr10-49724564-INS-1807 |
| chr10-49731000-INS-859 | chr10-49731000-INS-858 |
| chr10-49741569-INS-612 | chr10-49741569-INS-612 |
| chr10-49748469-INS-435 | chr10-49748469-INS-434 |
| chr10-49764975-INS-3178 | chr10-49764976-INS-3178 |
| chr10-49774658-INS-78 | chr10-49774658-INS-78 |
| chr10-49797257-INS-463 | chr10-49797257-INS-449 |
| chr10-49800635-INS-52 | chr10-49800635-INS-58 |
| chr10-49801602-INS-67 | chr10-49801602-INS-67 |
| chr10-49812007-INS-898 | chr10-49812007-INS-898 |
| chr10-49841316-INS-648 | chr10-49841316-INS-648 |
| chr10-49856307-INS-109 | chr10-49856307-INS-109 |
| chr10-49873713-INS-336 | chr10-49873713-INS-332 |
| chr10-49898442-INS-56 | chr10-49898442-INS-60 |
| chr10-49916201-INS-2601 | chr10-49916202-INS-2590 |
| chr10-49921385-INS-342 | chr10-49921385-INS-342 |
| chr10-49949513-INS-4782 | chr10-49949514-INS-4782 |
| chr10-50062592-INS-98 | chr10-50062592-INS-95 |
| chr10-50084658-INS-290 | chr10-50084658-INS-296 |
| chr10-50095778-INS-762 | chr10-50095778-INS-762 |
| chr10-50102358-INS-51 | chr10-50102369-INS-53 |
| chr10-50110416-INS-140 | chr10-50110416-INS-140 |
| chr10-50114623-INS-371 | chr10-50114623-INS-371 |
| chr10-50116268-INS-108 | chr10-50116280-INS-82 |
| chr10-50117195-INS-216 | chr10-50117195-INS-216 |
| chr10-50155076-INS-6781 | chr10-50155077-INS-6781 |
| chr10-50173441-INS-382 | chr10-50173441-INS-382 |
| chr10-50179992-INS-198 | chr10-50180019-INS-199 |
| chr10-50184731-INS-78 | chr10-50184731-INS-78 |
| chr10-50191696-INS-270 | chr10-50191696-INS-270 |
| chr10-50198933-INS-108 | chr10-50198929-INS-114 |
| chr10-50222570-INS-6781 | chr10-50222571-INS-6798 |
| chr10-50228848-INS-257 | chr10-50228848-INS-269 |
| chr10-50232372-INS-82 | chr10-50232372-INS-83 |
| chr10-50275700-INS-4779 | chr10-50275701-INS-4779 |
| chr10-50279739-INS-120 | chr10-50279739-INS-80 |
| chr10-50304512-INS-390 | chr10-50304512-INS-390 |

| | |
|---|---|
| chr10-50321300-INS-88 | chr10-50321327-INS-107 |
| chr10-50325030-INS-414 | chr10-50325030-INS-414 |
| chr10-50344593-INS-234 | chr10-50344593-INS-236 |
| chr10-50360599-INS-1193 | chr10-50360600-INS-1193 |
| chr10-50377136-INS-486 | chr10-50377136-INS-486 |
| chr10-50384166-INS-3074 | chr10-50384167-INS-3080 |
| chr10-50399638-INS-63 | chr10-50399643-INS-57 |
| chr10-50402811-INS-1811 | chr10-50402812-INS-1809 |
| chr10-50412685-INS-74 | chr10-50412685-INS-74 |
| chr10-50413614-INS-204 | chr10-50413614-INS-204 |
| chr10-50438215-INS-497 | chr10-50438215-INS-497 |
| chr10-50446248-INS-70 | chr10-50446242-INS-50 |
| chr10-50458629-INS-3474 | chr10-50458630-INS-3474 |
| chr10-50525572-INS-2297 | chr10-50525573-INS-2293 |
| chr10-50536036-INS-120 | chr10-50536036-INS-120 |
| chr10-50538063-INS-1695 | chr10-50538064-INS-1698 |
| chr10-50542299-INS-54 | chr10-50542299-INS-54 |
| chr10-50577719-INS-617 | chr10-50577719-INS-619 |
| chr10-50634043-INS-2366 | chr10-50634044-INS-2366 |
| chr10-50641953-INS-6233 | chr10-50641954-INS-6233 |
| chr10-50645456-INS-201 | chr10-50645462-INS-196 |
| chr10-50647921-INS-149 | chr10-50647921-INS-150 |
| chr10-50652924-INS-478 | chr10-50652924-INS-480 |
| chr10-50656342-INS-495 | chr10-50656342-INS-497 |
| chr10-50665094-INS-908 | chr10-50665094-INS-908 |
| chr10-50690107-INS-7026 | chr10-50690108-INS-7027 |
| chr10-50697972-INS-84 | chr10-50697972-INS-84 |
| chr10-50707493-INS-6359 | chr10-50707494-INS-6361 |
| chr10-50729972-INS-112 | chr10-50729972-INS-112 |
| chr10-50748471-INS-80 | chr10-50748471-INS-90 |
| chr10-50792690-INS-131 | chr10-50792680-INS-124 |
| chr10-50798212-INS-258 | chr10-50798212-INS-260 |
| chr10-50816392-INS-62 | chr10-50816392-INS-102 |
| chr10-50838720-INS-1292 | chr10-50838721-INS-1278 |
| chr10-50880742-INS-6629 | chr10-50880743-INS-6630 |
| chr10-50896917-INS-221 | chr10-50896917-INS-223 |
| chr10-50903575-INS-54 | chr10-50903580-INS-56 |
| chr10-50922051-INS-202 | chr10-50922051-INS-202 |
| chr10-50954825-INS-355 | chr10-50954825-INS-355 |
| chr10-50969960-INS-3275 | chr10-50969961-INS-3275 |
| chr10-50975377-INS-103 | chr10-50975378-INS-148 |
| chr10-50977326-INS-6479 | chr10-50977327-INS-6479 |
| chr10-51046409-INS-7088 | chr10-51046410-INS-7088 |
| chr10-51049287-INS-58 | chr10-51049287-INS-60 |
| chr10-51097475-INS-53 | chr10-51097475-INS-53 |
| chr10-51114332-INS-1369 | chr10-51114333-INS-1374 |
| chr10-51123246-INS-123 | chr10-51123246-INS-123 |
| chr10-51123724-INS-1159 | chr10-51123725-INS-1159 |
| chr10-51129789-INS-58 | chr10-51129789-INS-52 |
| chr10-51152174-INS-2240 | chr10-51152175-INS-2241 |
| chr10-51155261-INS-64 | chr10-51155261-INS-56 |
| chr10-51172476-INS-52 | chr10-51172479-INS-91 |
| chr10-51197417-INS-6320 | chr10-51197418-INS-6320 |
| chr10-51215231-INS-2061 | chr10-51215232-INS-2062 |
| chr10-51286928-INS-6725 | chr10-51286929-INS-6722 |
| chr10-51338485-INS-942 | chr10-51338485-INS-944 |
| chr10-51384607-INS-3069 | chr10-51384608-INS-3069 |
| chr10-51395614-INS-6209 | chr10-51395615-INS-6209 |
| chr10-51401365-INS-245 | chr10-51401364-INS-250 |
| chr10-51417382-INS-118 | chr10-51417382-INS-118 |
| chr10-51429218-INS-214 | chr10-51429218-INS-225 |
| chr10-51438798-INS-113 | chr10-51438798-INS-113 |
| chr10-51451244-INS-397 | chr10-51451241-INS-400 |
| chr10-51472672-INS-814 | chr10-51472672-INS-814 |
| chr10-51475112-INS-64 | chr10-51475100-INS-68 |
| chr10-51487235-INS-936 | chr10-51487235-INS-936 |
| chr10-51515803-INS-452 | chr10-51515803-INS-452 |
| chr10-51519599-INS-66 | chr10-51519599-INS-56 |
| chr10-51533686-INS-383 | chr10-51533686-INS-383 |
| chr10-51537046-INS-3424 | chr10-51537047-INS-3424 |
| chr10-51574395-INS-228 | chr10-51574395-INS-229 |
| chr10-51586773-INS-174 | chr10-51586773-INS-173 |
| chr10-51605684-INS-82 | chr10-51605684-INS-82 |
| chr10-51618546-INS-199 | chr10-51618556-INS-199 |
| chr10-51626878-INS-81 | chr10-51626878-INS-78 |
| chr10-51634447-INS-130 | chr10-51634447-INS-130 |
| chr10-51652085-INS-315 | chr10-51652085-INS-314 |

| | |
|---|---|
| chr10-51654101-INS-60 | chr10-51654101-INS-60 |
| chr10-51658677-INS-602 | chr10-51658677-INS-602 |
| chr10-51673015-INS-231 | chr10-51673015-INS-229 |
| chr10-51675539-INS-297 | chr10-51675539-INS-297 |
| chr10-51678792-INS-214 | chr10-51678792-INS-214 |
| chr10-51714817-INS-86 | chr10-51714817-INS-86 |
| chr10-51719590-INS-126 | chr10-51719590-INS-126 |
| chr10-51778521-INS-226 | chr10-51778521-INS-229 |
| chr10-51797982-INS-2581 | chr10-51797983-INS-2581 |
| chr10-51803943-INS-140 | chr10-51803943-INS-140 |
| chr10-51808726-INS-52 | chr10-51808726-INS-50 |
| chr10-51823754-INS-212 | chr10-51823765-INS-216 |
| chr10-51826846-INS-161 | chr10-51826846-INS-175 |
| chr10-51830812-INS-61 | chr10-51830812-INS-78 |
| chr10-51868004-INS-68 | chr10-51868005-INS-60 |
| chr10-51908887-INS-459 | chr10-51908887-INS-459 |
| chr10-51913201-INS-197 | chr10-51913201-INS-197 |
| chr10-51991111-INS-214 | chr10-51991111-INS-214 |
| chr10-52053396-INS-178 | chr10-52053396-INS-178 |
| chr10-52074422-INS-56 | chr10-52074426-INS-90 |
| chr10-52115417-INS-186 | chr10-52115417-INS-186 |
| chr10-52172299-INS-190 | chr10-52172299-INS-188 |
| chr10-52177800-INS-300 | chr10-52177800-INS-300 |
| chr10-52245612-INS-252 | chr10-52245612-INS-252 |
| chr10-52308773-INS-810 | chr10-52308773-INS-809 |
| chr10-52330838-INS-94 | chr10-52330838-INS-94 |
| chr10-52344533-INS-58 | chr10-52344533-INS-60 |
| chr10-52379185-INS-214 | chr10-52379185-INS-214 |
| chr10-52390117-INS-302 | chr10-52390117-INS-302 |
| chr10-52398258-INS-7456 | chr10-52398259-INS-7456 |
| chr10-52483366-INS-1091 | chr10-52483367-INS-1092 |
| chr10-52489715-INS-762 | chr10-52489715-INS-762 |
| chr10-52552045-INS-531 | chr10-52552045-INS-531 |
| chr10-52576597-INS-101 | chr10-52576597-INS-101 |
| chr10-52578218-INS-160 | chr10-52578218-INS-160 |
| chr10-52579494-INS-54 | chr10-52579494-INS-54 |
| chr10-52614820-INS-52 | chr10-52614820-INS-54 |
| chr10-52631474-INS-879 | chr10-52631474-INS-879 |
| chr10-52652581-INS-3378 | chr10-52652582-INS-3378 |
| chr10-52670734-INS-586 | chr10-52670734-INS-586 |
| chr10-52675497-INS-372 | chr10-52675497-INS-363 |
| chr10-52699986-INS-60 | chr10-52699986-INS-60 |
| chr10-52725014-INS-245 | chr10-52725014-INS-245 |
| chr10-52727563-INS-80 | chr10-52727563-INS-80 |
| chr10-52768845-INS-97 | chr10-52768871-INS-66 |
| chr10-52808394-INS-54 | chr10-52808394-INS-54 |
| chr10-52831424-INS-56 | chr10-52831424-INS-56 |
| chr10-52855775-INS-56 | chr10-52855775-INS-56 |
| chr10-52866803-INS-56 | chr10-52866801-INS-58 |
| chr10-52891025-INS-6432 | chr10-52891026-INS-6432 |
| chr10-52902394-INS-79 | chr10-52902394-INS-81 |
| chr10-52910237-INS-72 | chr10-52910237-INS-68 |
| chr10-52921725-INS-78 | chr10-52921725-INS-78 |
| chr10-52927848-INS-332 | chr10-52927848-INS-336 |
| chr10-52929910-INS-195 | chr10-52929910-INS-197 |
| chr10-52940456-INS-96 | chr10-52940451-INS-101 |
| chr10-52943217-INS-81 | chr10-52943217-INS-87 |
| chr10-52956784-INS-578 | chr10-52956784-INS-577 |
| chr10-52963358-INS-100 | chr10-52963358-INS-52 |
| chr10-52983729-INS-6370 | chr10-52983730-INS-6370 |
| chr10-53003391-INS-70 | chr10-53003391-INS-70 |
| chr10-53004991-INS-60 | chr10-53004991-INS-56 |
| chr10-53014466-INS-839 | chr10-53014466-INS-843 |
| chr10-53063508-INS-551 | chr10-53063508-INS-554 |
| chr10-53085857-INS-1887 | chr10-53085858-INS-1887 |
| chr10-53123408-INS-1985 | chr10-53123409-INS-1988 |
| chr10-53133010-INS-206 | chr10-53133010-INS-208 |
| chr10-53166119-INS-140 | chr10-53166119-INS-140 |
| chr10-53179941-INS-63 | chr10-53179933-INS-60 |
| chr10-53189202-INS-71 | chr10-53189202-INS-71 |
| chr10-53230725-INS-2633 | chr10-53230726-INS-2645 |
| chr10-53230870-INS-6478 | chr10-53230871-INS-6479 |
| chr10-53246861-INS-523 | chr10-53246861-INS-523 |
| chr10-53249683-INS-116 | chr10-53249683-INS-112 |
| chr10-53257250-INS-83 | chr10-53257250-INS-70 |
| chr10-53312023-INS-1197 | chr10-53312024-INS-1168 |
| chr10-53354715-INS-146 | chr10-53354715-INS-149 |

| | |
|---|---|
| chr10-53433048-INS-78 | chr10-53433048-INS-78 |
| chr10-53475366-INS-202 | chr10-53475366-INS-195 |
| chr10-53496665-INS-54 | chr10-53496657-INS-50 |
| chr10-53504521-INS-58 | chr10-53504476-INS-88 |
| chr10-53608993-INS-179 | chr10-53608993-INS-169 |
| chr10-53646877-INS-68 | chr10-53646874-INS-59 |
| chr10-53683321-INS-64 | chr10-53683321-INS-68 |
| chr10-53736230-INS-6569 | chr10-53736231-INS-6569 |
| chr10-53752379-INS-400 | chr10-53752379-INS-450 |
| chr10-53755674-INS-68 | chr10-53755674-INS-64 |
| chr10-53814642-INS-1898 | chr10-53814643-INS-1898 |
| chr10-53816462-INS-466 | chr10-53816462-INS-485 |
| chr10-53827142-INS-229 | chr10-53827142-INS-229 |
| chr10-53858914-INS-72 | chr10-53858914-INS-72 |
| chr10-53884207-INS-6508 | chr10-53884215-INS-6510 |
| chr10-53899882-INS-53 | chr10-53899882-INS-53 |
| chr10-53931147-INS-194 | chr10-53931169-INS-198 |
| chr10-53933246-INS-423 | chr10-53933246-INS-423 |
| chr10-53943388-INS-347 | chr10-53943388-INS-312 |
| chr10-53973448-INS-1322 | chr10-53973449-INS-1319 |
| chr10-54031138-INS-52 | chr10-54031139-INS-76 |
| chr10-54042513-INS-190 | chr10-54042513-INS-190 |
| chr10-54043685-INS-2005 | chr10-54043686-INS-2005 |
| chr10-54052184-INS-358 | chr10-54052184-INS-358 |
| chr10-54056811-INS-127 | chr10-54056811-INS-127 |
| chr10-54070184-INS-52 | chr10-54070180-INS-51 |
| chr10-54081085-INS-1345 | chr10-54081086-INS-1345 |
| chr10-54109904-INS-7108 | chr10-54109905-INS-7108 |
| chr10-54125328-INS-105 | chr10-54125317-INS-122 |
| chr10-54128672-INS-456 | chr10-54128672-INS-456 |
| chr10-54154647-INS-66 | chr10-54154661-INS-99 |
| chr10-54155299-INS-6875 | chr10-54155300-INS-6877 |
| chr10-54165589-INS-56 | chr10-54165589-INS-56 |
| chr10-54182022-INS-931 | chr10-54182022-INS-931 |
| chr10-54182499-INS-1272 | chr10-54182500-INS-1271 |
| chr10-54195963-INS-639 | chr10-54195962-INS-641 |
| chr10-54211816-INS-6570 | chr10-54211817-INS-6570 |
| chr10-54227596-INS-107 | chr10-54227596-INS-107 |
| chr10-54231009-INS-1017 | chr10-54231010-INS-1000 |
| chr10-54248087-INS-50 | chr10-54248087-INS-85 |
| chr10-54253537-INS-105 | chr10-54253537-INS-105 |
| chr10-54256098-INS-55 | chr10-54256098-INS-70 |
| chr10-54260061-INS-2980 | chr10-54260062-INS-2980 |
| chr10-54261635-INS-53 | chr10-54261635-INS-55 |
| chr10-54298722-INS-56 | chr10-54298722-INS-82 |
| chr10-54306937-INS-68 | chr10-54306937-INS-62 |
| chr10-54320103-INS-600 | chr10-54320103-INS-600 |
| chr10-54359994-INS-223 | chr10-54359994-INS-223 |
| chr10-54415221-INS-62 | chr10-54415221-INS-52 |
| chr10-54446718-INS-345 | chr10-54446721-INS-349 |
| chr10-54516234-INS-112 | chr10-54516214-INS-92 |
| chr10-54524906-INS-119 | chr10-54524906-INS-135 |
| chr10-54528036-INS-72 | chr10-54528036-INS-74 |
| chr10-54531066-INS-4756 | chr10-54531067-INS-4756 |
| chr10-54534824-INS-179 | chr10-54534824-INS-184 |
| chr10-54584877-INS-608 | chr10-54584877-INS-608 |
| chr10-54622057-INS-334 | chr10-54622057-INS-334 |
| chr10-54646857-INS-1871 | chr10-54646858-INS-1876 |
| chr10-54647607-INS-257 | chr10-54647607-INS-257 |
| chr10-54652394-INS-52 | chr10-54652395-INS-52 |
| chr10-54683025-INS-368 | chr10-54683025-INS-369 |
| chr10-54727997-INS-409 | chr10-54727997-INS-409 |
| chr10-54736500-INS-2733 | chr10-54736501-INS-2733 |
| chr10-54742421-INS-315 | chr10-54742421-INS-315 |
| chr10-54751601-INS-1301 | chr10-54751602-INS-1301 |
| chr10-54783821-INS-64 | chr10-54783821-INS-64 |
| chr10-54800706-INS-108 | chr10-54800706-INS-108 |
| chr10-54816207-INS-110 | chr10-54816207-INS-99 |
| chr10-54869162-INS-68 | chr10-54869158-INS-106 |
| chr10-54871180-INS-74 | chr10-54871180-INS-74 |
| chr10-54874355-INS-561 | chr10-54874355-INS-560 |
| chr10-54875198-INS-6646 | chr10-54875199-INS-6653 |
| chr10-54889724-INS-199 | chr10-54889724-INS-199 |
| chr10-54891303-INS-87 | chr10-54891332-INS-75 |
| chr10-54906949-INS-3485 | chr10-54906950-INS-3475 |
| chr10-54920968-INS-80 | chr10-54920938-INS-60 |
| chr10-54986146-INS-621 | chr10-54986146-INS-621 |

| | |
|---|---|
| chr10-55026891-INS-673 | chr10-55026891-INS-672 |
| chr10-55078331-INS-2027 | chr10-55078332-INS-2027 |
| chr10-55126033-INS-59 | chr10-55126033-INS-57 |
| chr10-55150535-INS-260 | chr10-55150535-INS-260 |
| chr10-55180786-INS-60 | chr10-55180786-INS-54 |
| chr10-55183651-INS-2817 | chr10-55183652-INS-2817 |
| chr10-55210880-INS-6531 | chr10-55210881-INS-6531 |
| chr10-55218187-INS-90 | chr10-55218187-INS-90 |
| chr10-55223107-INS-214 | chr10-55223107-INS-216 |
| chr10-55238496-INS-529 | chr10-55238496-INS-529 |
| chr10-55243252-INS-57 | chr10-55243252-INS-51 |
| chr10-55244386-INS-85 | chr10-55244386-INS-83 |
| chr10-55254494-INS-68 | chr10-55254494-INS-102 |
| chr10-55297591-INS-198 | chr10-55297591-INS-199 |
| chr10-55299841-INS-377 | chr10-55299841-INS-377 |
| chr10-55368036-INS-87 | chr10-55368036-INS-88 |
| chr10-55433910-INS-111 | chr10-55433910-INS-113 |
| chr10-55443732-INS-371 | chr10-55443732-INS-371 |
| chr10-55465914-INS-1733 | chr10-55465915-INS-1733 |
| chr10-55492828-INS-54 | chr10-55492826-INS-64 |
| chr10-55519705-INS-7463 | chr10-55519706-INS-7463 |
| chr10-55547130-INS-102 | chr10-55547124-INS-96 |
| chr10-55554398-INS-91 | chr10-55554396-INS-91 |
| chr10-55563228-INS-63 | chr10-55563228-INS-51 |
| chr10-55578356-INS-53 | chr10-55578356-INS-53 |
| chr10-55606553-INS-73 | chr10-55606553-INS-71 |
| chr10-55650103-INS-6245 | chr10-55650104-INS-6245 |
| chr10-55663471-INS-987 | chr10-55663471-INS-987 |
| chr10-55687213-INS-1981 | chr10-55687214-INS-1980 |
| chr10-55689235-INS-180 | chr10-55689235-INS-177 |
| chr10-55708067-INS-64 | chr10-55708067-INS-52 |
| chr10-55708580-INS-2394 | chr10-55708581-INS-2394 |
| chr10-55735587-INS-2845 | chr10-55735588-INS-2845 |
| chr10-55758122-INS-208 | chr10-55758122-INS-209 |
| chr10-55790573-INS-54 | chr10-55790573-INS-60 |
| chr10-55793448-INS-931 | chr10-55793448-INS-931 |
| chr10-55802014-INS-56 | chr10-55802014-INS-60 |
| chr10-55814150-INS-70 | chr10-55814150-INS-50 |
| chr10-55814413-INS-2816 | chr10-55814414-INS-2816 |
| chr10-55819850-INS-70 | chr10-55819850-INS-70 |
| chr10-55822499-INS-72 | chr10-55822499-INS-72 |
| chr10-55846762-INS-91 | chr10-55846762-INS-60 |
| chr10-55848454-INS-92 | chr10-55848454-INS-92 |
| chr10-55850192-INS-7231 | chr10-55850193-INS-7234 |
| chr10-55872032-INS-798 | chr10-55872032-INS-798 |
| chr10-55873475-INS-274 | chr10-55873475-INS-292 |
| chr10-55880527-INS-4093 | chr10-55880528-INS-4093 |
| chr10-55892160-INS-72 | chr10-55892156-INS-60 |
| chr10-55903268-INS-65 | chr10-55903268-INS-73 |
| chr10-55919234-INS-98 | chr10-55919234-INS-98 |
| chr10-55934440-INS-736 | chr10-55934440-INS-736 |
| chr10-55946626-INS-401 | chr10-55946626-INS-401 |
| chr10-55970658-INS-513 | chr10-55970658-INS-513 |
| chr10-55978974-INS-65 | chr10-55978974-INS-65 |
| chr10-56020288-INS-208 | chr10-56020288-INS-208 |
| chr10-56023950-INS-810 | chr10-56023950-INS-810 |
| chr10-56035254-INS-1356 | chr10-56035255-INS-1352 |
| chr10-56037863-INS-60 | chr10-56037863-INS-66 |
| chr10-56063026-INS-268 | chr10-56063026-INS-270 |
| chr10-56077874-INS-88 | chr10-56077874-INS-64 |
| chr10-56096850-INS-64 | chr10-56096850-INS-64 |
| chr10-56101809-INS-114 | chr10-56101809-INS-65 |
| chr10-56132979-INS-636 | chr10-56132979-INS-636 |
| chr10-56133812-INS-190 | chr10-56133812-INS-230 |
| chr10-56175117-INS-132 | chr10-56175117-INS-163 |
| chr10-56180830-INS-75 | chr10-56180830-INS-76 |
| chr10-56188521-INS-685 | chr10-56188521-INS-685 |
| chr10-56195904-INS-821 | chr10-56195920-INS-844 |
| chr10-56211061-INS-101 | chr10-56211061-INS-101 |
| chr10-56212226-INS-3071 | chr10-56212227-INS-3071 |
| chr10-56216833-INS-64 | chr10-56216833-INS-66 |
| chr10-56223462-INS-414 | chr10-56223462-INS-394 |
| chr10-56239639-INS-744 | chr10-56239642-INS-754 |
| chr10-56244810-INS-66 | chr10-56244810-INS-52 |
| chr10-56269783-INS-128 | chr10-56269783-INS-122 |
| chr10-56270027-INS-812 | chr10-56270027-INS-812 |
| chr10-56271087-INS-172 | chr10-56271090-INS-171 |

| | |
|---|---|
| chr10-56276014-INS-195 | chr10-56276015-INS-195 |
| chr10-56279792-INS-130 | chr10-56279822-INS-91 |
| chr10-56284636-INS-171 | chr10-56284636-INS-187 |
| chr10-56295746-INS-191 | chr10-56295746-INS-191 |
| chr10-56301933-INS-66 | chr10-56301933-INS-66 |
| chr10-56323753-INS-979 | chr10-56323753-INS-979 |
| chr10-56351544-INS-58 | chr10-56351545-INS-56 |
| chr10-56353084-INS-68 | chr10-56353056-INS-77 |
| chr10-56353811-INS-63 | chr10-56353811-INS-66 |
| chr10-56370708-INS-64 | chr10-56370735-INS-59 |
| chr10-56387823-INS-289 | chr10-56387823-INS-289 |
| chr10-56394759-INS-82 | chr10-56394759-INS-82 |
| chr10-56397602-INS-205 | chr10-56397602-INS-205 |
| chr10-56400836-INS-185 | chr10-56400836-INS-184 |
| chr10-56423960-INS-57 | chr10-56423959-INS-80 |
| chr10-56443000-INS-53 | chr10-56443000-INS-53 |
| chr10-56448878-INS-123 | chr10-56448899-INS-95 |
| chr10-56457130-INS-88 | chr10-56457128-INS-92 |
| chr10-56459830-INS-1457 | chr10-56459831-INS-1457 |
| chr10-56466042-INS-398 | chr10-56466042-INS-398 |
| chr10-56467538-INS-54 | chr10-56467566-INS-55 |
| chr10-56480121-INS-6817 | chr10-56480122-INS-6817 |
| chr10-56491923-INS-7071 | chr10-56491924-INS-7071 |
| chr10-56546929-INS-56 | chr10-56546929-INS-64 |
| chr10-56566334-INS-225 | chr10-56566334-INS-225 |
| chr10-56568086-INS-65 | chr10-56568095-INS-70 |
| chr10-56571686-INS-300 | chr10-56571686-INS-300 |
| chr10-56574063-INS-54 | chr10-56574063-INS-54 |
| chr10-56591342-INS-2532 | chr10-56591343-INS-2527 |
| chr10-56617943-INS-129 | chr10-56617944-INS-134 |
| chr10-56623922-INS-186 | chr10-56623922-INS-184 |
| chr10-56627168-INS-55 | chr10-56627167-INS-55 |
| chr10-56632725-INS-501 | chr10-56632725-INS-501 |
| chr10-56644087-INS-58 | chr10-56644087-INS-62 |
| chr10-56645991-INS-79 | chr10-56645991-INS-85 |
| chr10-56676551-INS-54 | chr10-56676551-INS-92 |
| chr10-56739914-INS-2175 | chr10-56739915-INS-2176 |
| chr10-56743572-INS-714 | chr10-56743572-INS-691 |
| chr10-56769238-INS-227 | chr10-56769238-INS-229 |
| chr10-56785770-INS-68 | chr10-56785770-INS-68 |
| chr10-56791410-INS-605 | chr10-56791410-INS-605 |
| chr10-56822419-INS-1897 | chr10-56822420-INS-1897 |
| chr10-56851341-INS-7877 | chr10-56851342-INS-7877 |
| chr10-56907103-INS-102 | chr10-56907103-INS-102 |
| chr10-56920316-INS-6822 | chr10-56920317-INS-6822 |
| chr10-56951322-INS-6733 | chr10-56951323-INS-6733 |
| chr10-57103998-INS-103 | chr10-57103998-INS-104 |
| chr10-57106629-INS-123 | chr10-57106629-INS-133 |
| chr10-57126345-INS-5153 | chr10-57126346-INS-5153 |
| chr10-57157206-INS-74 | chr10-57157206-INS-54 |
| chr10-57165700-INS-681 | chr10-57165700-INS-681 |
| chr10-57175347-INS-123 | chr10-57175347-INS-123 |
| chr10-57179693-INS-72 | chr10-57179698-INS-77 |
| chr10-57196940-INS-791 | chr10-57196940-INS-791 |
| chr10-57220849-INS-576 | chr10-57220849-INS-576 |
| chr10-57249188-INS-81 | chr10-57249188-INS-81 |
| chr10-57262266-INS-206 | chr10-57262271-INS-210 |
| chr10-57286073-INS-1590 | chr10-57286074-INS-1592 |
| chr10-57289287-INS-7171 | chr10-57289288-INS-7171 |
| chr10-57302973-INS-628 | chr10-57302973-INS-627 |
| chr10-57311708-INS-357 | chr10-57311714-INS-351 |
| chr10-57313734-INS-58 | chr10-57313758-INS-64 |
| chr10-57323738-INS-307 | chr10-57323738-INS-307 |
| chr10-57325016-INS-213 | chr10-57325016-INS-213 |
| chr10-57326845-INS-213 | chr10-57326824-INS-214 |
| chr10-57397820-INS-54 | chr10-57397820-INS-54 |
| chr10-57418860-INS-119 | chr10-57418860-INS-119 |
| chr10-57420591-INS-204 | chr10-57420591-INS-209 |
| chr10-57425280-INS-76 | chr10-57425280-INS-82 |
| chr10-57428119-INS-372 | chr10-57428119-INS-372 |
| chr10-57431608-INS-53 | chr10-57431608-INS-72 |
| chr10-57465410-INS-97 | chr10-57465394-INS-108 |
| chr10-57477413-INS-3041 | chr10-57477414-INS-3041 |
| chr10-57478949-INS-54 | chr10-57478949-INS-54 |
| chr10-57490679-INS-54 | chr10-57490722-INS-60 |
| chr10-57514225-INS-610 | chr10-57514225-INS-644 |
| chr10-57534806-INS-400 | chr10-57534806-INS-400 |

| | |
|---|---|
| chr10-57562116-INS-54 | chr10-57562116-INS-56 |
| chr10-57563234-INS-281 | chr10-57563234-INS-295 |
| chr10-57609491-INS-250 | chr10-57609491-INS-250 |
| chr10-57618511-INS-92 | chr10-57618508-INS-54 |
| chr10-57621918-INS-834 | chr10-57621918-INS-834 |
| chr10-57629881-INS-56 | chr10-57629878-INS-92 |
| chr10-57650522-INS-1884 | chr10-57650523-INS-1884 |
| chr10-57713037-INS-58 | chr10-57713037-INS-60 |
| chr10-58025147-INS-468 | chr10-58025147-INS-468 |
| chr10-58032385-INS-95 | chr10-58032385-INS-69 |
| chr10-58078218-INS-178 | chr10-58078218-INS-177 |
| chr10-58118784-INS-82 | chr10-58118790-INS-66 |
| chr10-58130484-INS-394 | chr10-58130484-INS-394 |
| chr10-58143382-INS-349 | chr10-58143382-INS-350 |
| chr10-58152595-INS-76 | chr10-58152604-INS-117 |
| chr10-58156595-INS-91 | chr10-58156595-INS-91 |
| chr10-58164293-INS-66 | chr10-58164293-INS-66 |
| chr10-58169721-INS-62 | chr10-58169721-INS-62 |
| chr10-58237048-INS-65 | chr10-58237048-INS-65 |
| chr10-58272985-INS-99 | chr10-58272985-INS-56 |
| chr10-58324206-INS-54 | chr10-58324206-INS-50 |
| chr10-58325980-INS-60 | chr10-58326013-INS-73 |
| chr10-58392198-INS-7279 | chr10-58392199-INS-7280 |
| chr10-58516995-INS-347 | chr10-58516995-INS-340 |
| chr10-58519833-INS-154 | chr10-58519833-INS-157 |
| chr10-58525547-INS-91 | chr10-58525595-INS-81 |
| chr10-58583268-INS-56 | chr10-58583282-INS-56 |
| chr10-58592421-INS-717 | chr10-58592421-INS-721 |
| chr10-58658066-INS-68 | chr10-58658066-INS-68 |
| chr10-58679370-INS-75 | chr10-58679370-INS-75 |
| chr10-58685300-INS-100 | chr10-58685300-INS-108 |
| chr10-58696649-INS-115 | chr10-58696650-INS-115 |
| chr10-58703994-INS-178 | chr10-58703994-INS-178 |
| chr10-58726893-INS-75 | chr10-58726893-INS-75 |
| chr10-58732248-INS-78 | chr10-58732248-INS-78 |
| chr10-58746259-INS-2961 | chr10-58746260-INS-2962 |
| chr10-58755965-INS-88 | chr10-58755965-INS-110 |
| chr10-58759528-INS-72 | chr10-58759528-INS-68 |
| chr10-58801381-INS-69 | chr10-58801363-INS-114 |
| chr10-58812940-INS-6205 | chr10-58812941-INS-6206 |
| chr10-58839713-INS-428 | chr10-58839713-INS-428 |
| chr10-58898350-INS-75 | chr10-58898350-INS-73 |
| chr10-58902156-INS-175 | chr10-58902160-INS-169 |
| chr10-58917458-INS-374 | chr10-58917458-INS-374 |
| chr10-58919869-INS-957 | chr10-58919869-INS-957 |
| chr10-58928194-INS-704 | chr10-58928195-INS-706 |
| chr10-58949844-INS-63 | chr10-58949844-INS-63 |
| chr10-58957987-INS-797 | chr10-58957987-INS-803 |
| chr10-58982715-INS-64 | chr10-58982715-INS-64 |
| chr10-58984682-INS-750 | chr10-58984682-INS-758 |
| chr10-59010025-INS-356 | chr10-59010025-INS-356 |
| chr10-59043308-INS-504 | chr10-59043308-INS-504 |
| chr10-59072994-INS-519 | chr10-59072994-INS-519 |
| chr10-59086546-INS-86 | chr10-59086546-INS-58 |
| chr10-59098867-INS-174 | chr10-59098867-INS-182 |
| chr10-59130683-INS-494 | chr10-59130683-INS-494 |
| chr10-59170480-INS-102 | chr10-59170480-INS-140 |
| chr10-59205304-INS-220 | chr10-59205304-INS-220 |
| chr10-59216514-INS-164 | chr10-59216514-INS-164 |
| chr10-59238059-INS-206 | chr10-59238059-INS-207 |
| chr10-59281642-INS-525 | chr10-59281640-INS-537 |
| chr10-59288892-INS-94 | chr10-59288892-INS-94 |
| chr10-59290027-INS-63 | chr10-59290027-INS-63 |
| chr10-59292187-INS-201 | chr10-59292187-INS-195 |
| chr10-59347647-INS-71 | chr10-59347647-INS-71 |
| chr10-59363179-INS-57 | chr10-59363179-INS-55 |
| chr10-59408589-INS-80 | chr10-59408589-INS-80 |
| chr10-59418902-INS-360 | chr10-59418902-INS-360 |
| chr10-59440047-INS-149 | chr10-59440047-INS-132 |
| chr10-59440666-INS-72 | chr10-59440666-INS-72 |
| chr10-59448025-INS-356 | chr10-59448025-INS-357 |
| chr10-59456951-INS-133 | chr10-59456934-INS-131 |
| chr10-59494519-INS-236 | chr10-59494519-INS-236 |
| chr10-59508263-INS-253 | chr10-59508263-INS-253 |
| chr10-59509127-INS-1989 | chr10-59509128-INS-1989 |
| chr10-59518068-INS-133 | chr10-59518068-INS-133 |
| chr10-59527649-INS-199 | chr10-59527649-INS-199 |

| | |
|---|---|
| chr10-59540387-INS-60 | chr10-59540387-INS-74 |
| chr10-59542788-INS-96 | chr10-59542788-INS-96 |
| chr10-59552041-INS-94 | chr10-59552046-INS-50 |
| chr10-59569818-INS-75 | chr10-59569818-INS-50 |
| chr10-59571206-INS-55 | chr10-59571206-INS-56 |
| chr10-59590378-INS-57 | chr10-59590378-INS-68 |
| chr10-59661756-INS-51 | chr10-59661756-INS-54 |
| chr10-59669519-INS-208 | chr10-59669519-INS-208 |
| chr10-59679412-INS-231 | chr10-59679412-INS-231 |
| chr10-59708503-INS-161 | chr10-59708503-INS-149 |
| chr10-59724990-INS-208 | chr10-59724990-INS-209 |
| chr10-59762205-INS-203 | chr10-59762205-INS-203 |
| chr10-59767461-INS-64 | chr10-59767461-INS-81 |
| chr10-59797187-INS-71 | chr10-59797187-INS-71 |
| chr10-59804174-INS-207 | chr10-59804174-INS-207 |
| chr10-59820324-INS-894 | chr10-59820324-INS-895 |
| chr10-59825540-INS-60 | chr10-59825540-INS-60 |
| chr10-59836168-INS-200 | chr10-59836168-INS-200 |
| chr10-59854328-INS-350 | chr10-59854328-INS-353 |
| chr10-59855250-INS-200 | chr10-59855250-INS-200 |
| chr10-59856682-INS-62 | chr10-59856682-INS-53 |
| chr10-59857884-INS-201 | chr10-59857884-INS-201 |
| chr10-59890509-INS-493 | chr10-59890509-INS-493 |
| chr10-59899008-INS-1891 | chr10-59899009-INS-1891 |
| chr10-59922552-INS-52 | chr10-59922552-INS-52 |
| chr10-59926186-INS-58 | chr10-59926186-INS-66 |
| chr10-59930973-INS-186 | chr10-59930973-INS-161 |
| chr10-59952828-INS-183 | chr10-59952828-INS-173 |
| chr10-59980563-INS-74 | chr10-59980563-INS-74 |
| chr10-59994445-INS-182 | chr10-59994445-INS-156 |
| chr10-60011557-INS-63 | chr10-60011561-INS-92 |
| chr10-60012919-INS-82 | chr10-60012919-INS-82 |
| chr10-60028586-INS-210 | chr10-60028586-INS-210 |
| chr10-60085504-INS-138 | chr10-60085504-INS-142 |
| chr10-60090221-INS-158 | chr10-60090221-INS-162 |
| chr10-60094601-INS-135 | chr10-60094597-INS-161 |
| chr10-60098825-INS-222 | chr10-60098825-INS-204 |
| chr10-60100166-INS-91 | chr10-60100166-INS-84 |
| chr10-60101419-INS-555 | chr10-60101419-INS-555 |
| chr10-60154266-INS-179 | chr10-60154266-INS-179 |
| chr10-60176891-INS-75 | chr10-60176881-INS-52 |
| chr10-60202607-INS-91 | chr10-60202607-INS-91 |
| chr10-60220120-INS-5566 | chr10-60220121-INS-5566 |
| chr10-60312944-INS-741 | chr10-60312944-INS-741 |
| chr10-60339356-INS-52 | chr10-60339354-INS-53 |
| chr10-60343796-INS-743 | chr10-60343796-INS-749 |
| chr10-60348462-INS-2031 | chr10-60348463-INS-2037 |
| chr10-60353405-INS-56 | chr10-60353405-INS-56 |
| chr10-60392893-INS-108 | chr10-60392893-INS-84 |
| chr10-60403776-INS-126 | chr10-60403776-INS-126 |
| chr10-60453619-INS-68 | chr10-60453619-INS-76 |
| chr10-60462767-INS-52 | chr10-60462767-INS-52 |
| chr10-60486388-INS-400 | chr10-60486388-INS-400 |
| chr10-60498141-INS-119 | chr10-60498156-INS-131 |
| chr10-60531169-INS-874 | chr10-60531169-INS-871 |
| chr10-60556356-INS-796 | chr10-60556356-INS-792 |
| chr10-60591407-INS-9046 | chr10-60591408-INS-9046 |
| chr10-60605720-INS-65 | chr10-60605720-INS-65 |
| chr10-60683218-INS-53 | chr10-60683218-INS-53 |
| chr10-60694345-INS-54 | chr10-60694330-INS-58 |
| chr10-60736703-INS-131 | chr10-60736703-INS-120 |
| chr10-60739831-INS-56 | chr10-60739829-INS-83 |
| chr10-60807817-INS-200 | chr10-60807817-INS-201 |
| chr10-60824567-INS-59 | chr10-60824575-INS-53 |
| chr10-60844860-INS-77 | chr10-60844860-INS-56 |
| chr10-60911840-INS-203 | chr10-60911840-INS-197 |
| chr10-60925885-INS-96 | chr10-60925885-INS-102 |
| chr10-60934281-INS-56 | chr10-60934283-INS-92 |
| chr10-60957402-INS-175 | chr10-60957402-INS-176 |
| chr10-60972817-INS-216 | chr10-60972817-INS-217 |
| chr10-60981828-INS-211 | chr10-60981828-INS-211 |
| chr10-60982942-INS-68 | chr10-60982942-INS-68 |
| chr10-61018509-INS-109 | chr10-61018485-INS-109 |
| chr10-61058534-INS-868 | chr10-61058532-INS-867 |
| chr10-61066553-INS-54 | chr10-61066553-INS-54 |
| chr10-61101336-INS-67 | chr10-61101337-INS-75 |
| chr10-61115142-INS-210 | chr10-61115142-INS-210 |

| | |
|---|---|
| chr10-61130486-INS-86 | chr10-61130486-INS-86 |
| chr10-61135016-INS-110 | chr10-61135013-INS-94 |
| chr10-61165198-INS-205 | chr10-61165198-INS-204 |
| chr10-61171095-INS-190 | chr10-61171095-INS-186 |
| chr10-61187845-INS-83 | chr10-61187846-INS-83 |
| chr10-61224509-INS-198 | chr10-61224509-INS-198 |
| chr10-61229978-INS-68 | chr10-61229978-INS-68 |
| chr10-61234899-INS-248 | chr10-61234899-INS-248 |
| chr10-61237036-INS-716 | chr10-61237045-INS-699 |
| chr10-61238560-INS-86 | chr10-61238560-INS-112 |
| chr10-61242714-INS-123 | chr10-61242714-INS-123 |
| chr10-61249768-INS-213 | chr10-61249768-INS-213 |
| chr10-61256716-INS-208 | chr10-61256716-INS-209 |
| chr10-61273618-INS-217 | chr10-61273618-INS-218 |
| chr10-61295498-INS-195 | chr10-61295497-INS-196 |
| chr10-61387449-INS-74 | chr10-61387449-INS-74 |
| chr10-61393500-INS-209 | chr10-61393500-INS-209 |
| chr10-61402938-INS-57 | chr10-61402938-INS-57 |
| chr10-61431676-INS-58 | chr10-61431714-INS-52 |
| chr10-61438057-INS-96 | chr10-61438057-INS-72 |
| chr10-61472704-INS-56 | chr10-61472700-INS-79 |
| chr10-61475929-INS-7097 | chr10-61475930-INS-7097 |
| chr10-61477331-INS-237 | chr10-61477331-INS-237 |
| chr10-61492361-INS-688 | chr10-61492360-INS-698 |
| chr10-61517813-INS-193 | chr10-61517813-INS-192 |
| chr10-61546255-INS-208 | chr10-61546255-INS-209 |
| chr10-61547681-INS-102 | chr10-61547681-INS-100 |
| chr10-61555817-INS-105 | chr10-61555817-INS-60 |
| chr10-61568619-INS-66 | chr10-61568619-INS-62 |
| chr10-61579762-INS-74 | chr10-61579762-INS-74 |
| chr10-61600178-INS-176 | chr10-61600178-INS-176 |
| chr10-61629634-INS-106 | chr10-61629634-INS-106 |
| chr10-61636486-INS-441 | chr10-61636486-INS-441 |
| chr10-61651572-INS-84 | chr10-61651596-INS-105 |
| chr10-61653806-INS-73 | chr10-61653806-INS-73 |
| chr10-61658419-INS-54 | chr10-61658419-INS-52 |
| chr10-61664058-INS-62 | chr10-61664058-INS-62 |
| chr10-61701079-INS-75 | chr10-61701079-INS-75 |
| chr10-61733154-INS-860 | chr10-61733156-INS-855 |
| chr10-61793627-INS-226 | chr10-61793627-INS-226 |
| chr10-61808268-INS-69 | chr10-61808268-INS-69 |
| chr10-61834757-INS-170 | chr10-61834757-INS-157 |
| chr10-61864491-INS-70 | chr10-61864494-INS-108 |
| chr10-61921903-INS-323 | chr10-61921903-INS-323 |
| chr10-61933240-INS-52 | chr10-61933211-INS-67 |
| chr10-61958162-INS-54 | chr10-61958142-INS-65 |
| chr10-61978478-INS-102 | chr10-61978479-INS-103 |
| chr10-62008617-INS-56 | chr10-62008617-INS-58 |
| chr10-62011914-INS-5253 | chr10-62011915-INS-5253 |
| chr10-62025814-INS-372 | chr10-62025814-INS-372 |
| chr10-62034565-INS-61 | chr10-62034569-INS-66 |
| chr10-62047205-INS-141 | chr10-62047178-INS-137 |
| chr10-62059627-INS-414 | chr10-62059627-INS-414 |
| chr10-62063828-INS-54 | chr10-62063828-INS-54 |
| chr10-62069190-INS-228 | chr10-62069190-INS-228 |
| chr10-62070712-INS-55 | chr10-62070712-INS-55 |
| chr10-62097212-INS-642 | chr10-62097212-INS-642 |
| chr10-62104030-INS-99 | chr10-62104030-INS-99 |
| chr10-62134070-INS-178 | chr10-62134070-INS-178 |
| chr10-62144172-INS-68 | chr10-62144172-INS-52 |
| chr10-62146447-INS-72 | chr10-62146447-INS-72 |
| chr10-62149594-INS-51 | chr10-62149576-INS-51 |
| chr10-62150975-INS-56 | chr10-62150975-INS-56 |
| chr10-62167559-INS-179 | chr10-62167546-INS-181 |
| chr10-62171189-INS-186 | chr10-62171186-INS-195 |
| chr10-62186811-INS-208 | chr10-62186812-INS-218 |
| chr10-62225563-INS-195 | chr10-62225582-INS-217 |
| chr10-62237042-INS-80 | chr10-62237042-INS-80 |
| chr10-62248035-INS-66 | chr10-62248035-INS-50 |
| chr10-62275278-INS-76 | chr10-62275278-INS-78 |
| chr10-62296218-INS-161 | chr10-62296182-INS-180 |
| chr10-62299041-INS-221 | chr10-62299041-INS-221 |
| chr10-62304844-INS-72 | chr10-62304844-INS-75 |
| chr10-62356304-INS-72 | chr10-62356304-INS-81 |
| chr10-62388546-INS-55 | chr10-62388546-INS-60 |
| chr10-62408789-INS-2007 | chr10-62408790-INS-2007 |
| chr10-62409539-INS-112 | chr10-62409539-INS-112 |

| | |
|---|---|
| chr10-62419142-INS-219 | chr10-62419142-INS-192 |
| chr10-62433970-INS-53 | chr10-62433973-INS-50 |
| chr10-62442125-INS-190 | chr10-62442125-INS-183 |
| chr10-62497761-INS-52 | chr10-62497761-INS-52 |
| chr10-62503966-INS-211 | chr10-62503966-INS-211 |
| chr10-62510211-INS-54 | chr10-62510211-INS-63 |
| chr10-62533678-INS-480 | chr10-62533678-INS-496 |
| chr10-62557233-INS-181 | chr10-62557233-INS-132 |
| chr10-62574072-INS-999 | chr10-62574072-INS-998 |
| chr10-62584112-INS-112 | chr10-62584112-INS-108 |
| chr10-62617514-INS-218 | chr10-62617514-INS-220 |
| chr10-62619138-INS-201 | chr10-62619138-INS-203 |
| chr10-62645032-INS-52 | chr10-62645032-INS-52 |
| chr10-62649853-INS-55 | chr10-62649853-INS-55 |
| chr10-62657583-INS-62 | chr10-62657583-INS-79 |
| chr10-62663889-INS-221 | chr10-62663889-INS-221 |
| chr10-62673028-INS-86 | chr10-62673028-INS-94 |
| chr10-62688190-INS-200 | chr10-62688190-INS-202 |
| chr10-62694249-INS-80 | chr10-62694254-INS-89 |
| chr10-62713065-INS-248 | chr10-62713075-INS-251 |
| chr10-62717283-INS-408 | chr10-62717284-INS-408 |
| chr10-62725493-INS-229 | chr10-62725493-INS-229 |
| chr10-62727200-INS-53 | chr10-62727200-INS-54 |
| chr10-62749922-INS-76 | chr10-62749923-INS-88 |
| chr10-62752079-INS-335 | chr10-62752086-INS-329 |
| chr10-62760864-INS-78 | chr10-62760864-INS-78 |
| chr10-62768028-INS-96 | chr10-62768028-INS-110 |
| chr10-62775905-INS-183 | chr10-62775905-INS-189 |
| chr10-62782456-INS-199 | chr10-62782456-INS-198 |
| chr10-62801009-INS-144 | chr10-62801009-INS-144 |
| chr10-62803166-INS-382 | chr10-62803164-INS-396 |
| chr10-62810515-INS-1848 | chr10-62810516-INS-1848 |
| chr10-62813127-INS-114 | chr10-62813163-INS-95 |
| chr10-62831294-INS-54 | chr10-62831294-INS-56 |
| chr10-62871655-INS-175 | chr10-62871669-INS-173 |
| chr10-62876997-INS-165 | chr10-62876997-INS-167 |
| chr10-62906123-INS-722 | chr10-62906123-INS-723 |
| chr10-62923994-INS-505 | chr10-62923993-INS-507 |
| chr10-62936780-INS-188 | chr10-62936780-INS-187 |
| chr10-62977326-INS-195 | chr10-62977326-INS-195 |
| chr10-63004492-INS-105 | chr10-63004496-INS-99 |
| chr10-63009525-INS-183 | chr10-63009562-INS-164 |
| chr10-63013148-INS-58 | chr10-63013148-INS-58 |
| chr10-63033698-INS-82 | chr10-63033711-INS-60 |
| chr10-63036908-INS-183 | chr10-63036935-INS-175 |
| chr10-63047750-INS-76 | chr10-63047750-INS-76 |
| chr10-63056240-INS-199 | chr10-63056244-INS-199 |
| chr10-63058502-INS-180 | chr10-63058502-INS-181 |
| chr10-63093929-INS-54 | chr10-63093929-INS-52 |
| chr10-63107839-INS-163 | chr10-63107839-INS-163 |
| chr10-63135315-INS-513 | chr10-63135315-INS-515 |
| chr10-63158415-INS-199 | chr10-63158415-INS-201 |
| chr10-63176997-INS-4515 | chr10-63176998-INS-4515 |
| chr10-63182157-INS-614 | chr10-63182157-INS-614 |
| chr10-63189194-INS-225 | chr10-63189194-INS-226 |
| chr10-63205105-INS-201 | chr10-63205105-INS-201 |
| chr10-63211784-INS-184 | chr10-63211787-INS-231 |
| chr10-63215989-INS-102 | chr10-63215994-INS-62 |
| chr10-63242023-INS-70 | chr10-63241997-INS-67 |
| chr10-63251848-INS-224 | chr10-63251848-INS-227 |
| chr10-63273076-INS-211 | chr10-63273076-INS-211 |
| chr10-63275578-INS-62 | chr10-63275578-INS-62 |
| chr10-63285697-INS-70 | chr10-63285697-INS-70 |
| chr10-63292623-INS-57 | chr10-63292622-INS-67 |
| chr10-63297895-INS-933 | chr10-63297895-INS-938 |
| chr10-63325098-INS-105 | chr10-63325098-INS-105 |
| chr10-63330569-INS-117 | chr10-63330569-INS-117 |
| chr10-63334130-INS-85 | chr10-63334130-INS-58 |
| chr10-63357791-INS-62 | chr10-63357791-INS-63 |
| chr10-63361532-INS-60 | chr10-63361532-INS-84 |
| chr10-63377474-INS-52 | chr10-63377474-INS-52 |
| chr10-63380373-INS-634 | chr10-63380373-INS-638 |
| chr10-63382075-INS-229 | chr10-63382076-INS-229 |
| chr10-63387200-INS-73 | chr10-63387210-INS-93 |
| chr10-63408525-INS-476 | chr10-63408525-INS-465 |
| chr10-63426676-INS-60 | chr10-63426676-INS-60 |
| chr10-63429333-INS-406 | chr10-63429333-INS-406 |

| | |
|---|---|
| chr10-63460687-INS-7261 | chr10-63460688-INS-7261 |
| chr10-63461262-INS-58 | chr10-63461280-INS-52 |
| chr10-63509040-INS-58 | chr10-63509040-INS-60 |
| chr10-63528158-INS-213 | chr10-63528158-INS-215 |
| chr10-63543837-INS-54 | chr10-63543836-INS-56 |
| chr10-63600197-INS-508 | chr10-63600197-INS-508 |
| chr10-63628441-INS-498 | chr10-63628441-INS-505 |
| chr10-63629873-INS-4781 | chr10-63629874-INS-4781 |
| chr10-63659818-INS-60 | chr10-63659818-INS-74 |
| chr10-63673091-INS-631 | chr10-63673091-INS-632 |
| chr10-63687483-INS-544 | chr10-63687483-INS-562 |
| chr10-63694904-INS-445 | chr10-63694904-INS-446 |
| chr10-63707627-INS-108 | chr10-63707659-INS-70 |
| chr10-63708127-INS-931 | chr10-63708127-INS-931 |
| chr10-63709517-INS-116 | chr10-63709517-INS-117 |
| chr10-63721339-INS-129 | chr10-63721320-INS-120 |
| chr10-63729905-INS-62 | chr10-63729905-INS-62 |
| chr10-63734746-INS-52 | chr10-63734746-INS-52 |
| chr10-63750497-INS-534 | chr10-63750497-INS-535 |
| chr10-63787436-INS-60 | chr10-63787412-INS-62 |
| chr10-63791876-INS-2861 | chr10-63791877-INS-2865 |
| chr10-63808493-INS-296 | chr10-63808493-INS-280 |
| chr10-63821877-INS-60 | chr10-63821853-INS-83 |
| chr10-63882361-INS-81 | chr10-63882362-INS-107 |
| chr10-63895662-INS-136 | chr10-63895664-INS-119 |
| chr10-63899103-INS-74 | chr10-63899109-INS-55 |
| chr10-63931085-INS-1166 | chr10-63931086-INS-1166 |
| chr10-63938726-INS-284 | chr10-63938726-INS-291 |
| chr10-63958713-INS-5051 | chr10-63958714-INS-5051 |
| chr10-63959413-INS-60 | chr10-63959413-INS-60 |
| chr10-63975421-INS-97 | chr10-63975421-INS-97 |
| chr10-63977358-INS-92 | chr10-63977359-INS-120 |
| chr10-63978474-INS-1072 | chr10-63978475-INS-1056 |
| chr10-63991490-INS-143 | chr10-63991492-INS-148 |
| chr10-64043814-INS-214 | chr10-64043814-INS-214 |
| chr10-64056627-INS-80 | chr10-64056627-INS-52 |
| chr10-64057605-INS-172 | chr10-64057587-INS-172 |
| chr10-64182522-INS-68 | chr10-64182522-INS-68 |
| chr10-64186279-INS-6578 | chr10-64186280-INS-6578 |
| chr10-64210446-INS-400 | chr10-64210446-INS-400 |
| chr10-64227711-INS-54 | chr10-64227711-INS-52 |
| chr10-64238958-INS-203 | chr10-64238951-INS-212 |
| chr10-64243337-INS-110 | chr10-64243331-INS-77 |
| chr10-64313253-INS-126 | chr10-64313253-INS-141 |
| chr10-64316014-INS-60 | chr10-64316014-INS-59 |
| chr10-64328738-INS-76 | chr10-64328728-INS-60 |
| chr10-64338806-INS-403 | chr10-64338806-INS-403 |
| chr10-64346354-INS-392 | chr10-64346354-INS-394 |
| chr10-64354073-INS-95 | chr10-64354066-INS-58 |
| chr10-64378733-INS-212 | chr10-64378733-INS-212 |
| chr10-64425841-INS-51 | chr10-64425841-INS-51 |
| chr10-64461502-INS-125 | chr10-64461502-INS-125 |
| chr10-64490519-INS-93 | chr10-64490519-INS-92 |
| chr10-64510570-INS-91 | chr10-64510567-INS-93 |
| chr10-64524803-INS-737 | chr10-64524803-INS-734 |
| chr10-64568259-INS-6219 | chr10-64568260-INS-6222 |
| chr10-64583098-INS-61 | chr10-64583098-INS-50 |
| chr10-64597142-INS-2694 | chr10-64597149-INS-2689 |
| chr10-64644542-INS-86 | chr10-64644542-INS-86 |
| chr10-64665139-INS-54 | chr10-64665139-INS-54 |
| chr10-64721812-INS-191 | chr10-64721812-INS-191 |
| chr10-64734365-INS-108 | chr10-64734365-INS-120 |
| chr10-64795161-INS-64 | chr10-64795162-INS-80 |
| chr10-64806411-INS-1214 | chr10-64806412-INS-1214 |
| chr10-64856040-INS-90 | chr10-64856040-INS-135 |
| chr10-64862242-INS-1177 | chr10-64862243-INS-1178 |
| chr10-64862243-INS-391 | chr10-64862243-INS-391 |
| chr10-64868957-INS-51 | chr10-64868957-INS-55 |
| chr10-64892622-INS-8541 | chr10-64892623-INS-8541 |
| chr10-64893516-INS-434 | chr10-64893516-INS-434 |
| chr10-64943260-INS-80 | chr10-64943260-INS-78 |
| chr10-64953489-INS-6356 | chr10-64953490-INS-6357 |
| chr10-64957247-INS-503 | chr10-64957247-INS-503 |
| chr10-64964798-INS-1738 | chr10-64964798-INS-1732 |
| chr10-64978278-INS-521 | chr10-64978278-INS-521 |
| chr10-65022974-INS-247 | chr10-65022974-INS-247 |
| chr10-65045273-INS-162 | chr10-65045273-INS-162 |

| | |
|---|---|
| chr10-65085792-INS-1046 | chr10-65085804-INS-1026 |
| chr10-65111889-INS-87 | chr10-65111889-INS-87 |
| chr10-65129404-INS-96 | chr10-65129367-INS-85 |
| chr10-65205422-INS-5285 | chr10-65205422-INS-5286 |
| chr10-65274137-INS-51 | chr10-65274137-INS-51 |
| chr10-65276707-INS-93 | chr10-65276705-INS-117 |
| chr10-65279803-INS-62 | chr10-65279804-INS-54 |
| chr10-65281843-INS-54 | chr10-65281843-INS-52 |
| chr10-65295710-INS-56 | chr10-65295710-INS-56 |
| chr10-65306039-INS-59 | chr10-65306066-INS-59 |
| chr10-65333414-INS-1064 | chr10-65333415-INS-1065 |
| chr10-65345668-INS-885 | chr10-65345668-INS-884 |
| chr10-65348069-INS-6808 | chr10-65348070-INS-6807 |
| chr10-65351500-INS-408 | chr10-65351500-INS-408 |
| chr10-65376237-INS-52 | chr10-65376237-INS-64 |
| chr10-65398134-INS-323 | chr10-65398134-INS-323 |
| chr10-65414787-INS-74 | chr10-65414787-INS-77 |
| chr10-65439300-INS-51 | chr10-65439300-INS-51 |
| chr10-65472179-INS-124 | chr10-65472179-INS-154 |
| chr10-65472750-INS-953 | chr10-65472750-INS-953 |
| chr10-65518002-INS-52 | chr10-65517988-INS-56 |
| chr10-65533587-INS-68 | chr10-65533606-INS-61 |
| chr10-65550743-INS-561 | chr10-65550743-INS-561 |
| chr10-65567277-INS-68 | chr10-65567277-INS-68 |
| chr10-65595848-INS-845 | chr10-65595847-INS-849 |
| chr10-65604366-INS-102 | chr10-65604366-INS-134 |
| chr10-65644150-INS-65 | chr10-65644150-INS-92 |
| chr10-65646905-INS-204 | chr10-65646905-INS-204 |
| chr10-65706200-INS-76 | chr10-65706200-INS-76 |
| chr10-65807725-INS-138 | chr10-65807725-INS-140 |
| chr10-65810189-INS-904 | chr10-65810189-INS-906 |
| chr10-65845850-INS-54 | chr10-65845850-INS-56 |
| chr10-65865767-INS-202 | chr10-65865767-INS-203 |
| chr10-65869028-INS-52 | chr10-65869029-INS-72 |
| chr10-65894316-INS-58 | chr10-65894316-INS-78 |
| chr10-65904890-INS-54 | chr10-65904900-INS-52 |
| chr10-65908233-INS-88 | chr10-65908229-INS-54 |
| chr10-65921236-INS-101 | chr10-65921240-INS-121 |
| chr10-65960891-INS-1079 | chr10-65960892-INS-1079 |
| chr10-65972303-INS-317 | chr10-65972303-INS-317 |
| chr10-65987450-INS-56 | chr10-65987450-INS-56 |
| chr10-66011543-INS-6378 | chr10-66011543-INS-6388 |
| chr10-66017936-INS-525 | chr10-66017936-INS-525 |
| chr10-66099237-INS-894 | chr10-66099237-INS-893 |
| chr10-66110426-INS-64 | chr10-66110464-INS-50 |
| chr10-66115088-INS-97 | chr10-66115088-INS-97 |
| chr10-66121982-INS-134 | chr10-66121982-INS-146 |
| chr10-66135502-INS-64 | chr10-66135502-INS-64 |
| chr10-66156825-INS-54 | chr10-66156825-INS-54 |
| chr10-66201777-INS-70 | chr10-66201777-INS-105 |
| chr10-66225290-INS-92 | chr10-66225290-INS-92 |
| chr10-66227480-INS-4495 | chr10-66227481-INS-4495 |
| chr10-66231148-INS-74 | chr10-66231148-INS-88 |
| chr10-66278549-INS-2982 | chr10-66278550-INS-2982 |
| chr10-66295468-INS-63 | chr10-66295483-INS-68 |
| chr10-66316194-INS-56 | chr10-66316194-INS-60 |
| chr10-66322804-INS-120 | chr10-66322804-INS-120 |
| chr10-66326379-INS-160 | chr10-66326379-INS-160 |
| chr10-66332913-INS-68 | chr10-66332913-INS-68 |
| chr10-66346286-INS-74 | chr10-66346286-INS-67 |
| chr10-66347548-INS-234 | chr10-66347548-INS-234 |
| chr10-66367724-INS-418 | chr10-66367724-INS-418 |
| chr10-66389063-INS-124 | chr10-66389063-INS-124 |
| chr10-66405024-INS-52 | chr10-66405052-INS-63 |
| chr10-66412630-INS-90 | chr10-66412620-INS-90 |
| chr10-66434558-INS-66 | chr10-66434558-INS-52 |
| chr10-66446401-INS-73 | chr10-66446401-INS-73 |
| chr10-66469930-INS-64 | chr10-66469930-INS-72 |
| chr10-66544211-INS-175 | chr10-66544211-INS-174 |
| chr10-66557099-INS-55 | chr10-66557099-INS-65 |
| chr10-66568877-INS-72 | chr10-66568861-INS-71 |
| chr10-66635144-INS-159 | chr10-66635144-INS-169 |
| chr10-66642514-INS-526 | chr10-66642514-INS-527 |
| chr10-66649060-INS-172 | chr10-66649060-INS-177 |
| chr10-66657102-INS-197 | chr10-66657102-INS-196 |
| chr10-66666083-INS-80 | chr10-66666083-INS-80 |
| chr10-66671703-INS-77 | chr10-66671705-INS-58 |

| | |
|---|---|
| chr10-66674315-INS-60 | chr10-66674330-INS-75 |
| chr10-66700868-INS-393 | chr10-66700868-INS-392 |
| chr10-66703249-INS-198 | chr10-66703250-INS-148 |
| chr10-66710973-INS-288 | chr10-66710973-INS-288 |
| chr10-66727312-INS-71 | chr10-66727314-INS-66 |
| chr10-66737299-INS-206 | chr10-66737299-INS-205 |
| chr10-66739785-INS-244 | chr10-66739785-INS-258 |
| chr10-66752073-INS-204 | chr10-66752062-INS-228 |
| chr10-66777141-INS-55 | chr10-66777141-INS-55 |
| chr10-66809607-INS-126 | chr10-66809607-INS-126 |
| chr10-66850045-INS-328 | chr10-66850045-INS-320 |
| chr10-66851888-INS-187 | chr10-66851887-INS-188 |
| chr10-66872182-INS-206 | chr10-66872182-INS-206 |
| chr10-66890318-INS-85 | chr10-66890318-INS-85 |
| chr10-66909377-INS-169 | chr10-66909377-INS-172 |
| chr10-66917023-INS-203 | chr10-66917023-INS-205 |
| chr10-66926355-INS-210 | chr10-66926355-INS-211 |
| chr10-66927654-INS-80 | chr10-66927654-INS-84 |
| chr10-66981473-INS-217 | chr10-66981473-INS-217 |
| chr10-67020476-INS-56 | chr10-67020481-INS-74 |
| chr10-67031385-INS-205 | chr10-67031385-INS-205 |
| chr10-67036626-INS-246 | chr10-67036626-INS-198 |
| chr10-67111759-INS-127 | chr10-67111759-INS-127 |
| chr10-67138177-INS-97 | chr10-67138177-INS-97 |
| chr10-67139207-INS-55 | chr10-67139207-INS-58 |
| chr10-67143431-INS-382 | chr10-67143432-INS-382 |
| chr10-67170323-INS-92 | chr10-67170323-INS-92 |
| chr10-67181657-INS-82 | chr10-67181657-INS-82 |
| chr10-67189525-INS-123 | chr10-67189525-INS-119 |
| chr10-67191182-INS-60 | chr10-67191198-INS-60 |
| chr10-67202930-INS-80 | chr10-67202939-INS-80 |
| chr10-67205776-INS-52 | chr10-67205776-INS-56 |
| chr10-67264332-INS-219 | chr10-67264352-INS-222 |
| chr10-67267412-INS-400 | chr10-67267412-INS-400 |
| chr10-67339117-INS-167 | chr10-67339117-INS-153 |
| chr10-67356548-INS-102 | chr10-67356548-INS-102 |
| chr10-67395400-INS-67 | chr10-67395400-INS-65 |
| chr10-67396557-INS-6701 | chr10-67396558-INS-6701 |
| chr10-67405817-INS-51 | chr10-67405817-INS-51 |
| chr10-67410133-INS-119 | chr10-67410131-INS-131 |
| chr10-67419809-INS-58 | chr10-67419824-INS-57 |
| chr10-67422997-INS-52 | chr10-67422997-INS-76 |
| chr10-67430167-INS-52 | chr10-67430167-INS-52 |
| chr10-67448064-INS-68 | chr10-67448064-INS-70 |
| chr10-67469352-INS-212 | chr10-67469352-INS-212 |
| chr10-67486446-INS-6442 | chr10-67486447-INS-6442 |
| chr10-67511583-INS-118 | chr10-67511583-INS-118 |
| chr10-67532942-INS-99 | chr10-67532942-INS-66 |
| chr10-67553941-INS-76 | chr10-67553907-INS-76 |
| chr10-67590470-INS-69 | chr10-67590470-INS-69 |
| chr10-67631406-INS-97 | chr10-67631406-INS-81 |
| chr10-67673511-INS-58 | chr10-67673495-INS-80 |
| chr10-67700819-INS-54 | chr10-67700819-INS-54 |
| chr10-67702712-INS-397 | chr10-67702712-INS-397 |
| chr10-67710723-INS-73 | chr10-67710723-INS-73 |
| chr10-67717328-INS-1149 | chr10-67717329-INS-1149 |
| chr10-67731066-INS-232 | chr10-67731066-INS-232 |
| chr10-67739102-INS-204 | chr10-67739102-INS-204 |
| chr10-67744345-INS-53 | chr10-67744345-INS-53 |
| chr10-67754846-INS-6594 | chr10-67754847-INS-6594 |
| chr10-67814205-INS-56 | chr10-67814205-INS-56 |
| chr10-67825594-INS-54 | chr10-67825594-INS-54 |
| chr10-67837007-INS-110 | chr10-67837007-INS-110 |
| chr10-67842980-INS-212 | chr10-67842986-INS-213 |
| chr10-67846940-INS-80 | chr10-67846940-INS-80 |
| chr10-67859001-INS-90 | chr10-67859005-INS-85 |
| chr10-67860697-INS-138 | chr10-67860697-INS-138 |
| chr10-67885559-INS-611 | chr10-67885559-INS-611 |
| chr10-67901152-INS-110 | chr10-67901152-INS-110 |
| chr10-67911988-INS-696 | chr10-67911988-INS-696 |
| chr10-67915579-INS-78 | chr10-67915579-INS-78 |
| chr10-67917032-INS-91 | chr10-67917032-INS-91 |
| chr10-67945557-INS-100 | chr10-67945557-INS-100 |
| chr10-67970267-INS-227 | chr10-67970267-INS-227 |
| chr10-67983421-INS-203 | chr10-67983421-INS-203 |
| chr10-67985106-INS-175 | chr10-67985106-INS-175 |
| chr10-67995135-INS-199 | chr10-67995135-INS-199 |

| | |
|---|---|
| chr10-68008074-INS-205 | chr10-68008074-INS-205 |
| chr10-68015243-INS-109 | chr10-68015243-INS-109 |
| chr10-68032191-INS-207 | chr10-68032191-INS-207 |
| chr10-68065197-INS-373 | chr10-68065197-INS-373 |
| chr10-68101149-INS-298 | chr10-68101149-INS-301 |
| chr10-68136676-INS-191 | chr10-68136676-INS-191 |
| chr10-68204198-INS-93 | chr10-68204198-INS-93 |
| chr10-68208846-INS-198 | chr10-68208846-INS-198 |
| chr10-68227336-INS-90 | chr10-68227336-INS-91 |
| chr10-68234510-INS-52 | chr10-68234510-INS-54 |
| chr10-68268119-INS-454 | chr10-68268119-INS-456 |
| chr10-68273420-INS-92 | chr10-68273420-INS-72 |
| chr10-68309375-INS-70 | chr10-68309375-INS-68 |
| chr10-68362180-INS-90 | chr10-68362180-INS-104 |
| chr10-68388311-INS-6225 | chr10-68388312-INS-6225 |
| chr10-68402821-INS-89 | chr10-68402821-INS-89 |
| chr10-68439748-INS-116 | chr10-68439748-INS-124 |
| chr10-68446318-INS-58 | chr10-68446306-INS-56 |
| chr10-68451206-INS-57 | chr10-68451206-INS-57 |
| chr10-68456928-INS-64 | chr10-68456928-INS-64 |
| chr10-68510937-INS-203 | chr10-68510937-INS-203 |
| chr10-68515083-INS-52 | chr10-68515079-INS-55 |
| chr10-68530054-INS-52 | chr10-68530054-INS-52 |
| chr10-68539774-INS-1481 | chr10-68539775-INS-1482 |
| chr10-68549173-INS-62 | chr10-68549139-INS-66 |
| chr10-68571239-INS-313 | chr10-68571239-INS-313 |
| chr10-68576617-INS-63 | chr10-68576617-INS-70 |
| chr10-68637433-INS-60 | chr10-68637433-INS-60 |
| chr10-68665690-INS-5234 | chr10-68665691-INS-5234 |
| chr10-68666892-INS-139 | chr10-68666892-INS-139 |
| chr10-68735465-INS-101 | chr10-68735472-INS-63 |
| chr10-68740817-INS-62 | chr10-68740817-INS-62 |
| chr10-68808165-INS-56 | chr10-68808168-INS-57 |
| chr10-68839651-INS-219 | chr10-68839651-INS-239 |
| chr10-68854582-INS-4208 | chr10-68854583-INS-4208 |
| chr10-68855560-INS-63 | chr10-68855560-INS-63 |
| chr10-68877479-INS-229 | chr10-68877479-INS-229 |
| chr10-68905002-INS-63 | chr10-68905007-INS-67 |
| chr10-68929698-INS-60 | chr10-68929698-INS-62 |
| chr10-68935842-INS-125 | chr10-68935842-INS-127 |
| chr10-68944560-INS-60 | chr10-68944560-INS-60 |
| chr10-68947737-INS-64 | chr10-68947737-INS-64 |
| chr10-68965127-INS-123 | chr10-68965127-INS-123 |
| chr10-69092996-INS-365 | chr10-69092996-INS-366 |
| chr10-69116378-INS-166 | chr10-69116378-INS-166 |
| chr10-69128629-INS-67 | chr10-69128629-INS-75 |
| chr10-69147461-INS-164 | chr10-69147461-INS-161 |
| chr10-69153777-INS-94 | chr10-69153777-INS-94 |
| chr10-69211346-INS-73 | chr10-69211346-INS-73 |
| chr10-69224404-INS-51 | chr10-69224358-INS-81 |
| chr10-69239648-INS-68 | chr10-69239648-INS-71 |
| chr10-69242660-INS-74 | chr10-69242660-INS-116 |
| chr10-69244921-INS-568 | chr10-69244921-INS-568 |
| chr10-69257850-INS-141 | chr10-69257850-INS-125 |
| chr10-69273892-INS-201 | chr10-69273892-INS-213 |
| chr10-69279661-INS-194 | chr10-69279660-INS-188 |
| chr10-69283565-INS-6442 | chr10-69283566-INS-6442 |
| chr10-69284577-INS-6797 | chr10-69284578-INS-6797 |
| chr10-69305823-INS-53 | chr10-69305823-INS-51 |
| chr10-69311926-INS-150 | chr10-69311927-INS-124 |
| chr10-69319949-INS-112 | chr10-69319949-INS-112 |
| chr10-69362640-INS-603 | chr10-69362640-INS-603 |
| chr10-69454526-INS-231 | chr10-69454526-INS-189 |
| chr10-69469934-INS-326 | chr10-69469910-INS-301 |
| chr10-69556804-INS-60 | chr10-69556800-INS-58 |
| chr10-69570959-INS-88 | chr10-69570959-INS-88 |
| chr10-69589093-INS-86 | chr10-69589093-INS-54 |
| chr10-69602890-INS-100 | chr10-69602890-INS-100 |
| chr10-69612207-INS-56 | chr10-69612207-INS-62 |
| chr10-69682817-INS-811 | chr10-69682818-INS-812 |
| chr10-69737899-INS-176 | chr10-69737899-INS-182 |
| chr10-69746716-INS-133 | chr10-69746716-INS-133 |
| chr10-69748923-INS-98 | chr10-69748926-INS-117 |
| chr10-69753558-INS-53 | chr10-69753558-INS-75 |
| chr10-69770582-INS-84 | chr10-69770582-INS-68 |
| chr10-69788472-INS-58 | chr10-69788472-INS-66 |
| chr10-69795022-INS-95 | chr10-69795022-INS-95 |

| | |
|---|---|
| chr10-69849924-INS-78 | chr10-69849970-INS-78 |
| chr10-69904827-INS-92 | chr10-69904827-INS-80 |
| chr10-69917943-INS-204 | chr10-69917943-INS-204 |
| chr10-69929795-INS-140 | chr10-69929795-INS-140 |
| chr10-69945068-INS-114 | chr10-69945068-INS-114 |
| chr10-69950303-INS-270 | chr10-69950303-INS-269 |
| chr10-69992384-INS-53 | chr10-69992384-INS-53 |
| chr10-69993555-INS-67 | chr10-69993555-INS-67 |
| chr10-69996392-INS-67 | chr10-69996377-INS-56 |
| chr10-70030892-INS-136 | chr10-70030892-INS-183 |
| chr10-70035322-INS-169 | chr10-70035322-INS-171 |
| chr10-70065629-INS-99 | chr10-70065648-INS-69 |
| chr10-70072440-INS-81 | chr10-70072463-INS-81 |
| chr10-70075923-INS-60 | chr10-70075923-INS-58 |
| chr10-70078046-INS-59 | chr10-70078022-INS-52 |
| chr10-70087133-INS-74 | chr10-70087113-INS-66 |
| chr10-70095892-INS-54 | chr10-70095885-INS-57 |
| chr10-70130432-INS-53 | chr10-70130432-INS-53 |
| chr10-70144933-INS-58 | chr10-70144933-INS-62 |
| chr10-70147853-INS-324 | chr10-70147854-INS-324 |
| chr10-70156723-INS-830 | chr10-70156723-INS-831 |
| chr10-70157804-INS-58 | chr10-70157804-INS-58 |
| chr10-70173902-INS-309 | chr10-70173902-INS-307 |
| chr10-70180551-INS-5539 | chr10-70180552-INS-5539 |
| chr10-70186128-INS-343 | chr10-70186128-INS-343 |
| chr10-70194336-INS-442 | chr10-70194336-INS-442 |
| chr10-70240153-INS-216 | chr10-70240153-INS-215 |
| chr10-70283578-INS-674 | chr10-70283578-INS-678 |
| chr10-70305772-INS-160 | chr10-70305774-INS-110 |
| chr10-70323963-INS-213 | chr10-70323963-INS-213 |
| chr10-70335668-INS-54 | chr10-70335668-INS-54 |
| chr10-70415892-INS-57 | chr10-70415850-INS-57 |
| chr10-70416470-INS-717 | chr10-70416470-INS-717 |
| chr10-70421563-INS-1872 | chr10-70421564-INS-1872 |
| chr10-70441354-INS-6401 | chr10-70441355-INS-6401 |
| chr10-70471668-INS-102 | chr10-70471668-INS-102 |
| chr10-70504639-INS-98 | chr10-70504639-INS-98 |
| chr10-70507474-INS-138 | chr10-70507474-INS-150 |
| chr10-70523233-INS-3091 | chr10-70523234-INS-3091 |
| chr10-70526457-INS-455 | chr10-70526457-INS-455 |
| chr10-70542107-INS-74 | chr10-70542110-INS-90 |
| chr10-70550421-INS-7080 | chr10-70550422-INS-7080 |
| chr10-70608410-INS-169 | chr10-70608411-INS-125 |
| chr10-70660960-INS-62 | chr10-70660960-INS-78 |
| chr10-70666795-INS-56 | chr10-70666795-INS-62 |
| chr10-70671308-INS-636 | chr10-70671308-INS-636 |
| chr10-70683793-INS-60 | chr10-70683793-INS-60 |
| chr10-70688987-INS-87 | chr10-70688987-INS-87 |
| chr10-70729015-INS-203 | chr10-70729015-INS-203 |
| chr10-70736650-INS-116 | chr10-70736650-INS-92 |
| chr10-70745450-INS-104 | chr10-70745450-INS-104 |
| chr10-70749377-INS-448 | chr10-70749377-INS-452 |
| chr10-70782205-INS-52 | chr10-70782205-INS-52 |
| chr10-70803525-INS-208 | chr10-70803525-INS-209 |
| chr10-70866255-INS-112 | chr10-70866255-INS-112 |
| chr10-70876069-INS-78 | chr10-70876069-INS-78 |
| chr10-70882111-INS-520 | chr10-70882111-INS-518 |
| chr10-70885864-INS-366 | chr10-70885864-INS-366 |
| chr10-70889987-INS-108 | chr10-70889987-INS-112 |
| chr10-70896926-INS-200 | chr10-70896926-INS-196 |
| chr10-70905959-INS-52 | chr10-70905959-INS-54 |
| chr10-70909012-INS-68 | chr10-70909012-INS-76 |
| chr10-70911414-INS-826 | chr10-70911414-INS-826 |
| chr10-70931555-INS-143 | chr10-70931546-INS-138 |
| chr10-70986373-INS-52 | chr10-70986365-INS-52 |
| chr10-70998573-INS-220 | chr10-70998573-INS-221 |
| chr10-71002503-INS-360 | chr10-71002503-INS-360 |
| chr10-71012053-INS-166 | chr10-71012053-INS-166 |
| chr10-71023089-INS-64 | chr10-71023104-INS-63 |
| chr10-71035719-INS-402 | chr10-71035719-INS-402 |
| chr10-71050349-INS-400 | chr10-71050349-INS-400 |
| chr10-71051103-INS-256 | chr10-71051103-INS-256 |
| chr10-71076672-INS-60 | chr10-71076672-INS-60 |
| chr10-71080376-INS-216 | chr10-71080376-INS-215 |
| chr10-71085798-INS-217 | chr10-71085798-INS-217 |
| chr10-71092390-INS-86 | chr10-71092390-INS-118 |
| chr10-71132562-INS-164 | chr10-71132568-INS-180 |

| | |
|---|---|
| chr10-71142680-INS-343 | chr10-71142680-INS-343 |
| chr10-71161140-INS-191 | chr10-71161140-INS-191 |
| chr10-71189484-INS-2893 | chr10-71189490-INS-2899 |
| chr10-71193179-INS-162 | chr10-71193179-INS-164 |
| chr10-71202601-INS-7100 | chr10-71202602-INS-7100 |
| chr10-71265457-INS-325 | chr10-71265457-INS-325 |
| chr10-71277034-INS-85 | chr10-71277013-INS-115 |
| chr10-71279766-INS-58 | chr10-71279766-INS-58 |
| chr10-71293927-INS-353 | chr10-71293927-INS-354 |
| chr10-71321873-INS-70 | chr10-71321873-INS-70 |
| chr10-71351643-INS-205 | chr10-71351643-INS-205 |
| chr10-71430923-INS-52 | chr10-71430909-INS-51 |
| chr10-71505120-INS-70 | chr10-71505134-INS-64 |
| chr10-71516621-INS-556 | chr10-71516621-INS-557 |
| chr10-71518284-INS-103 | chr10-71518284-INS-103 |
| chr10-71521323-INS-64 | chr10-71521323-INS-64 |
| chr10-71554502-INS-127 | chr10-71554502-INS-128 |
| chr10-71587487-INS-600 | chr10-71587487-INS-600 |
| chr10-71595042-INS-77 | chr10-71595042-INS-77 |
| chr10-71622785-INS-52 | chr10-71622785-INS-52 |
| chr10-71670960-INS-6835 | chr10-71670961-INS-6835 |
| chr10-71677102-INS-201 | chr10-71677130-INS-201 |
| chr10-71697920-INS-74 | chr10-71697920-INS-70 |
| chr10-71705401-INS-60 | chr10-71705401-INS-52 |
| chr10-71728395-INS-702 | chr10-71728395-INS-702 |
| chr10-71760939-INS-662 | chr10-71760939-INS-668 |
| chr10-71765540-INS-125 | chr10-71765540-INS-125 |
| chr10-71775188-INS-73 | chr10-71775188-INS-55 |
| chr10-71795396-INS-63 | chr10-71795396-INS-60 |
| chr10-71826807-INS-216 | chr10-71826807-INS-216 |
| chr10-71844099-INS-416 | chr10-71844099-INS-416 |
| chr10-71878358-INS-58 | chr10-71878358-INS-60 |
| chr10-71898877-INS-580 | chr10-71898877-INS-580 |
| chr10-71922496-INS-1552 | chr10-71922497-INS-1552 |
| chr10-71926563-INS-131 | chr10-71926563-INS-130 |
| chr10-71930109-INS-1027 | chr10-71930110-INS-1027 |
| chr10-71939905-INS-89 | chr10-71939903-INS-85 |
| chr10-71941845-INS-6594 | chr10-71941846-INS-6594 |
| chr10-71968891-INS-84 | chr10-71968935-INS-58 |
| chr10-71982378-INS-54 | chr10-71982378-INS-56 |
| chr10-71992004-INS-145 | chr10-71992033-INS-129 |
| chr10-72017028-INS-245 | chr10-72017028-INS-247 |
| chr10-72019889-INS-965 | chr10-72019889-INS-954 |
| chr10-72023166-INS-211 | chr10-72023166-INS-222 |
| chr10-72029206-INS-922 | chr10-72029206-INS-916 |
| chr10-72031361-INS-817 | chr10-72031361-INS-817 |
| chr10-72068304-INS-2826 | chr10-72068305-INS-2826 |
| chr10-72202216-INS-6569 | chr10-72202217-INS-6569 |
| chr10-72208895-INS-238 | chr10-72208881-INS-232 |
| chr10-72238938-INS-60 | chr10-72238937-INS-52 |
| chr10-72258674-INS-6628 | chr10-72258675-INS-6628 |
| chr10-72277228-INS-6445 | chr10-72277229-INS-6445 |
| chr10-72324539-INS-811 | chr10-72324539-INS-843 |
| chr10-72359278-INS-54 | chr10-72359278-INS-56 |
| chr10-72372369-INS-4702 | chr10-72372370-INS-4702 |
| chr10-72386902-INS-2537 | chr10-72386903-INS-2537 |
| chr10-72392741-INS-168 | chr10-72392746-INS-178 |
| chr10-72400351-INS-64 | chr10-72400357-INS-68 |
| chr10-72407684-INS-64 | chr10-72407687-INS-54 |
| chr10-72419423-INS-527 | chr10-72419423-INS-527 |
| chr10-72430238-INS-111 | chr10-72430238-INS-112 |
| chr10-72458730-INS-94 | chr10-72458730-INS-95 |
| chr10-72480717-INS-6356 | chr10-72480718-INS-6356 |
| chr10-72611781-INS-467 | chr10-72611781-INS-467 |
| chr10-72612379-INS-6504 | chr10-72612380-INS-6504 |
| chr10-72638925-INS-3704 | chr10-72638926-INS-3702 |
| chr10-72648386-INS-1529 | chr10-72648387-INS-1541 |
| chr10-72675561-INS-232 | chr10-72675561-INS-232 |
| chr10-72731025-INS-120 | chr10-72731057-INS-128 |
| chr10-72737075-INS-52 | chr10-72737075-INS-58 |
| chr10-72781925-INS-268 | chr10-72781925-INS-238 |
| chr10-72835200-INS-297 | chr10-72835200-INS-297 |
| chr10-72855281-INS-6207 | chr10-72855282-INS-6208 |
| chr10-72887351-INS-145 | chr10-72887351-INS-145 |
| chr10-72915737-INS-74 | chr10-72915737-INS-70 |
| chr10-72941657-INS-154 | chr10-72941657-INS-154 |
| chr10-72983666-INS-614 | chr10-72983666-INS-614 |

| | |
|---|---|
| chr10-72984375-INS-116 | chr10-72984375-INS-116 |
| chr10-73000543-INS-380 | chr10-73000543-INS-380 |
| chr10-73031173-INS-821 | chr10-73031173-INS-821 |
| chr10-73095543-INS-78 | chr10-73095543-INS-78 |
| chr10-73135902-INS-3812 | chr10-73135903-INS-3812 |
| chr10-73145194-INS-192 | chr10-73145194-INS-192 |
| chr10-73162871-INS-56 | chr10-73162871-INS-56 |
| chr10-73198280-INS-4887 | chr10-73198281-INS-4887 |
| chr10-73216868-INS-6126 | chr10-73216869-INS-6126 |
| chr10-73233259-INS-556 | chr10-73233259-INS-557 |
| chr10-73238638-INS-234 | chr10-73238638-INS-237 |
| chr10-73269971-INS-93 | chr10-73269971-INS-122 |
| chr10-73283125-INS-156 | chr10-73283125-INS-186 |
| chr10-73292720-INS-59 | chr10-73292716-INS-59 |
| chr10-73331460-INS-71 | chr10-73331460-INS-78 |
| chr10-73334267-INS-60 | chr10-73334273-INS-63 |
| chr10-73344226-INS-66 | chr10-73344226-INS-66 |
| chr10-73360135-INS-100 | chr10-73360135-INS-100 |
| chr10-73363827-INS-80 | chr10-73363827-INS-80 |
| chr10-73391144-INS-190 | chr10-73391144-INS-191 |
| chr10-73396953-INS-374 | chr10-73396953-INS-374 |
| chr10-73440770-INS-116 | chr10-73440770-INS-116 |
| chr10-73459504-INS-88 | chr10-73459504-INS-88 |
| chr10-73464471-INS-735 | chr10-73464471-INS-736 |
| chr10-73536429-INS-861 | chr10-73536429-INS-862 |
| chr10-73540123-INS-60 | chr10-73540123-INS-62 |
| chr10-73565800-INS-62 | chr10-73565800-INS-60 |
| chr10-73628325-INS-52 | chr10-73628325-INS-52 |
| chr10-73650412-INS-1936 | chr10-73650413-INS-1937 |
| chr10-73655075-INS-75 | chr10-73655075-INS-75 |
| chr10-73668795-INS-59 | chr10-73668795-INS-61 |
| chr10-73690171-INS-1248 | chr10-73690172-INS-1248 |
| chr10-73777840-INS-66 | chr10-73777836-INS-58 |
| chr10-73794013-INS-70 | chr10-73794008-INS-113 |
| chr10-73797558-INS-936 | chr10-73797558-INS-936 |
| chr10-73854982-INS-138 | chr10-73854982-INS-138 |
| chr10-73877833-INS-208 | chr10-73877833-INS-208 |
| chr10-73884412-INS-64 | chr10-73884402-INS-61 |
| chr10-73927850-INS-51 | chr10-73927850-INS-89 |
| chr10-73963283-INS-401 | chr10-73963283-INS-401 |
| chr10-74040090-INS-5332 | chr10-74040091-INS-5332 |
| chr10-74049151-INS-236 | chr10-74049151-INS-236 |
| chr10-74068093-INS-64 | chr10-74068093-INS-64 |
| chr10-74072206-INS-1377 | chr10-74072207-INS-1377 |
| chr10-74101475-INS-288 | chr10-74101475-INS-289 |
| chr10-74137597-INS-332 | chr10-74137597-INS-332 |
| chr10-74147753-INS-55 | chr10-74147753-INS-55 |
| chr10-74158993-INS-73 | chr10-74158993-INS-74 |
| chr10-74179410-INS-5583 | chr10-74179411-INS-5583 |
| chr10-74296023-INS-61 | chr10-74296023-INS-62 |
| chr10-74304561-INS-90 | chr10-74304561-INS-88 |
| chr10-74313051-INS-975 | chr10-74313051-INS-974 |
| chr10-74319395-INS-70 | chr10-74319395-INS-70 |
| chr10-74333043-INS-186 | chr10-74333043-INS-186 |
| chr10-74345930-INS-6215 | chr10-74345931-INS-6216 |
| chr10-74355351-INS-62 | chr10-74355351-INS-62 |
| chr10-74384019-INS-87 | chr10-74384019-INS-87 |
| chr10-74389117-INS-116 | chr10-74389117-INS-116 |
| chr10-74446639-INS-6449 | chr10-74446640-INS-6451 |
| chr10-74455376-INS-518 | chr10-74455378-INS-518 |
| chr10-74459863-INS-1061 | chr10-74459864-INS-1061 |
| chr10-74486634-INS-52 | chr10-74486634-INS-52 |
| chr10-74522725-INS-54 | chr10-74522725-INS-54 |
| chr10-74531468-INS-74 | chr10-74531468-INS-56 |
| chr10-74532380-INS-200 | chr10-74532380-INS-200 |
| chr10-74538669-INS-2000 | chr10-74538670-INS-2000 |
| chr10-74538972-INS-176 | chr10-74538972-INS-176 |
| chr10-74579193-INS-8310 | chr10-74579194-INS-8310 |
| chr10-74597049-INS-1188 | chr10-74597050-INS-1187 |
| chr10-74597607-INS-2925 | chr10-74597608-INS-2925 |
| chr10-74615017-INS-8916 | chr10-74615018-INS-8922 |
| chr10-74616698-INS-486 | chr10-74616693-INS-482 |
| chr10-74623660-INS-1611 | chr10-74623649-INS-1608 |
| chr10-74653786-INS-6264 | chr10-74653787-INS-6264 |
| chr10-74706955-INS-498 | chr10-74706955-INS-498 |
| chr10-74723811-INS-56 | chr10-74723811-INS-56 |
| chr10-74730478-INS-62 | chr10-74730478-INS-62 |

| | |
|---|---|
| chr10-74744426-INS-5254 | chr10-74744427-INS-5253 |
| chr10-74762447-INS-171 | chr10-74762451-INS-171 |
| chr10-74763707-INS-64 | chr10-74763707-INS-64 |
| chr10-74766137-INS-68 | chr10-74766141-INS-69 |
| chr10-74772133-INS-400 | chr10-74772133-INS-400 |
| chr10-74844064-INS-77 | chr10-74844064-INS-81 |
| chr10-74875089-INS-209 | chr10-74875089-INS-211 |
| chr10-74895173-INS-62 | chr10-74895175-INS-66 |
| chr10-74903281-INS-181 | chr10-74903281-INS-138 |
| chr10-74914159-INS-62 | chr10-74914159-INS-62 |
| chr10-74918577-INS-195 | chr10-74918577-INS-195 |
| chr10-74948076-INS-57 | chr10-74948077-INS-84 |
| chr10-74968369-INS-412 | chr10-74968369-INS-418 |
| chr10-74995059-INS-159 | chr10-74995059-INS-163 |
| chr10-74997237-INS-471 | chr10-74997237-INS-470 |
| chr10-75000268-INS-150 | chr10-75000268-INS-146 |
| chr10-75024062-INS-109 | chr10-75024062-INS-109 |
| chr10-75031016-INS-765 | chr10-75031016-INS-767 |
| chr10-75037940-INS-155 | chr10-75037940-INS-155 |
| chr10-75039788-INS-276 | chr10-75039788-INS-276 |
| chr10-75051122-INS-288 | chr10-75051122-INS-276 |
| chr10-75114756-INS-140 | chr10-75114754-INS-145 |
| chr10-75119635-INS-2052 | chr10-75119636-INS-2052 |
| chr10-75184669-INS-71 | chr10-75184669-INS-71 |
| chr10-75197464-INS-337 | chr10-75197464-INS-351 |
| chr10-75199124-INS-62 | chr10-75199149-INS-65 |
| chr10-75214525-INS-186 | chr10-75214525-INS-193 |
| chr10-75221180-INS-70 | chr10-75221180-INS-94 |
| chr10-75230927-INS-6403 | chr10-75230928-INS-6403 |
| chr10-75268543-INS-201 | chr10-75268543-INS-201 |
| chr10-75273695-INS-838 | chr10-75273695-INS-839 |
| chr10-75296568-INS-85 | chr10-75296568-INS-80 |
| chr10-75308834-INS-543 | chr10-75308834-INS-563 |
| chr10-75309823-INS-308 | chr10-75309823-INS-308 |
| chr10-75318734-INS-206 | chr10-75318734-INS-207 |
| chr10-75329304-INS-200 | chr10-75329304-INS-200 |
| chr10-75331142-INS-389 | chr10-75331142-INS-389 |
| chr10-75363498-INS-245 | chr10-75363498-INS-245 |
| chr10-75373721-INS-793 | chr10-75373721-INS-816 |
| chr10-75394770-INS-65 | chr10-75394770-INS-84 |
| chr10-75404831-INS-247 | chr10-75404831-INS-249 |
| chr10-75411069-INS-200 | chr10-75411070-INS-202 |
| chr10-75421617-INS-212 | chr10-75421618-INS-212 |
| chr10-75431217-INS-207 | chr10-75431217-INS-207 |
| chr10-75444240-INS-197 | chr10-75444240-INS-197 |
| chr10-75451077-INS-402 | chr10-75451077-INS-402 |
| chr10-75453307-INS-148 | chr10-75453307-INS-148 |
| chr10-75458748-INS-97 | chr10-75458747-INS-112 |
| chr10-75518338-INS-100 | chr10-75518338-INS-100 |
| chr10-75572861-INS-56 | chr10-75572861-INS-56 |
| chr10-75576701-INS-260 | chr10-75576701-INS-260 |
| chr10-75600776-INS-208 | chr10-75600776-INS-208 |
| chr10-75603479-INS-60 | chr10-75603480-INS-61 |
| chr10-75621345-INS-210 | chr10-75621345-INS-210 |
| chr10-75623107-INS-232 | chr10-75623107-INS-233 |
| chr10-75625235-INS-425 | chr10-75625235-INS-425 |
| chr10-75628867-INS-68 | chr10-75628867-INS-68 |
| chr10-75644527-INS-97 | chr10-75644527-INS-102 |
| chr10-75660592-INS-399 | chr10-75660592-INS-399 |
| chr10-75680469-INS-212 | chr10-75680469-INS-212 |
| chr10-75706828-INS-58 | chr10-75706828-INS-52 |
| chr10-75724045-INS-89 | chr10-75724045-INS-92 |
| chr10-75754001-INS-578 | chr10-75754001-INS-578 |
| chr10-75774846-INS-82 | chr10-75774846-INS-82 |
| chr10-75778527-INS-151 | chr10-75778527-INS-155 |
| chr10-75779621-INS-62 | chr10-75779619-INS-64 |
| chr10-75796368-INS-866 | chr10-75796343-INS-868 |
| chr10-75803417-INS-1176 | chr10-75803418-INS-1166 |
| chr10-75822601-INS-58 | chr10-75822601-INS-58 |
| chr10-75827339-INS-179 | chr10-75827339-INS-179 |
| chr10-75830775-INS-79 | chr10-75830775-INS-79 |
| chr10-75837819-INS-192 | chr10-75837820-INS-144 |
| chr10-75847417-INS-54 | chr10-75847417-INS-54 |
| chr10-75853514-INS-96 | chr10-75853514-INS-88 |
| chr10-75879149-INS-189 | chr10-75879149-INS-189 |
| chr10-75886226-INS-66 | chr10-75886226-INS-66 |
| chr10-75897537-INS-202 | chr10-75897537-INS-202 |

| | |
|---|---|
| chr10-75919419-INS-943 | chr10-75919419-INS-940 |
| chr10-75956302-INS-205 | chr10-75956302-INS-207 |
| chr10-76020116-INS-57 | chr10-76020116-INS-57 |
| chr10-76033686-INS-143 | chr10-76033686-INS-152 |
| chr10-76061559-INS-104 | chr10-76061592-INS-104 |
| chr10-76069638-INS-105 | chr10-76069622-INS-147 |
| chr10-76085941-INS-57 | chr10-76085941-INS-57 |
| chr10-76109388-INS-81 | chr10-76109389-INS-77 |
| chr10-76125634-INS-131 | chr10-76125634-INS-135 |
| chr10-76128418-INS-7178 | chr10-76128419-INS-7178 |
| chr10-76129289-INS-387 | chr10-76129289-INS-387 |
| chr10-76131051-INS-98 | chr10-76131051-INS-98 |
| chr10-76148409-INS-69 | chr10-76148387-INS-88 |
| chr10-76163683-INS-56 | chr10-76163683-INS-54 |
| chr10-76174662-INS-1857 | chr10-76174663-INS-1857 |
| chr10-76180002-INS-51 | chr10-76180010-INS-53 |
| chr10-76231886-INS-51 | chr10-76231886-INS-51 |
| chr10-76232437-INS-237 | chr10-76232437-INS-237 |
| chr10-76261225-INS-384 | chr10-76261225-INS-383 |
| chr10-76272038-INS-72 | chr10-76272038-INS-72 |
| chr10-76273109-INS-5399 | chr10-76273110-INS-5400 |
| chr10-76286039-INS-95 | chr10-76286039-INS-98 |
| chr10-76309214-INS-69 | chr10-76309214-INS-69 |
| chr10-76329210-INS-206 | chr10-76329210-INS-205 |
| chr10-76373428-INS-7118 | chr10-76373429-INS-7118 |
| chr10-76417785-INS-335 | chr10-76417785-INS-350 |
| chr10-76454966-INS-164 | chr10-76454966-INS-179 |
| chr10-76476306-INS-55 | chr10-76476306-INS-55 |
| chr10-76526536-INS-77 | chr10-76526536-INS-78 |
| chr10-76541221-INS-4448 | chr10-76541222-INS-4448 |
| chr10-76547895-INS-62 | chr10-76547895-INS-62 |
| chr10-76557016-INS-57 | chr10-76557018-INS-58 |
| chr10-76582800-INS-195 | chr10-76582800-INS-195 |
| chr10-76686496-INS-59 | chr10-76686496-INS-63 |
| chr10-76712667-INS-114 | chr10-76712667-INS-114 |
| chr10-76739150-INS-254 | chr10-76739150-INS-256 |
| chr10-76755039-INS-233 | chr10-76755039-INS-233 |
| chr10-76815284-INS-309 | chr10-76815285-INS-309 |
| chr10-76837432-INS-58 | chr10-76837412-INS-76 |
| chr10-76840491-INS-52 | chr10-76840491-INS-52 |
| chr10-76848636-INS-62 | chr10-76848636-INS-77 |
| chr10-76853317-INS-76 | chr10-76853317-INS-78 |
| chr10-76900765-INS-84 | chr10-76900765-INS-84 |
| chr10-76904176-INS-167 | chr10-76904176-INS-165 |
| chr10-76944253-INS-152 | chr10-76944253-INS-152 |
| chr10-76951889-INS-119 | chr10-76951911-INS-86 |
| chr10-76959069-INS-196 | chr10-76959069-INS-196 |
| chr10-76971587-INS-79 | chr10-76971587-INS-79 |
| chr10-77007190-INS-240 | chr10-77007190-INS-220 |
| chr10-77034055-INS-52 | chr10-77034055-INS-52 |
| chr10-77043832-INS-60 | chr10-77043832-INS-60 |
| chr10-77046018-INS-80 | chr10-77046034-INS-50 |
| chr10-77050417-INS-213 | chr10-77050417-INS-213 |
| chr10-77051331-INS-1504 | chr10-77051332-INS-1504 |
| chr10-77077965-INS-70 | chr10-77077965-INS-70 |
| chr10-77088543-INS-172 | chr10-77088543-INS-171 |
| chr10-77092515-INS-86 | chr10-77092515-INS-57 |
| chr10-77093835-INS-199 | chr10-77093835-INS-214 |
| chr10-77103256-INS-81 | chr10-77103256-INS-81 |
| chr10-77113349-INS-61 | chr10-77113349-INS-61 |
| chr10-77114707-INS-2149 | chr10-77114708-INS-2162 |
| chr10-77144277-INS-175 | chr10-77144277-INS-152 |
| chr10-77150244-INS-59 | chr10-77150259-INS-58 |
| chr10-77150332-INS-1018 | chr10-77150333-INS-1019 |
| chr10-77154473-INS-60 | chr10-77154473-INS-60 |
| chr10-77186372-INS-315 | chr10-77186372-INS-319 |
| chr10-77214811-INS-202 | chr10-77214811-INS-207 |
| chr10-77240925-INS-202 | chr10-77240925-INS-199 |
| chr10-77296919-INS-236 | chr10-77296918-INS-224 |
| chr10-77319464-INS-1006 | chr10-77319465-INS-1006 |
| chr10-77325322-INS-4491 | chr10-77325323-INS-4491 |
| chr10-77344383-INS-328 | chr10-77344383-INS-326 |
| chr10-77347336-INS-146 | chr10-77347336-INS-146 |
| chr10-77351201-INS-674 | chr10-77351201-INS-678 |
| chr10-77352260-INS-53 | chr10-77352260-INS-53 |
| chr10-77363979-INS-183 | chr10-77363979-INS-183 |
| chr10-77377512-INS-72 | chr10-77377512-INS-60 |

| | |
|---|---|
| chr10-77387978-INS-199 | chr10-77387978-INS-199 |
| chr10-77388842-INS-207 | chr10-77388842-INS-207 |
| chr10-77404335-INS-107 | chr10-77404335-INS-114 |
| chr10-77420492-INS-80 | chr10-77420492-INS-76 |
| chr10-77427145-INS-179 | chr10-77427145-INS-179 |
| chr10-77431417-INS-201 | chr10-77431417-INS-200 |
| chr10-77434852-INS-363 | chr10-77434852-INS-363 |
| chr10-77444916-INS-200 | chr10-77444916-INS-201 |
| chr10-77471250-INS-212 | chr10-77471250-INS-211 |
| chr10-77478575-INS-252 | chr10-77478575-INS-252 |
| chr10-77496811-INS-202 | chr10-77496811-INS-200 |
| chr10-77546769-INS-52 | chr10-77546764-INS-54 |
| chr10-77565429-INS-560 | chr10-77565429-INS-559 |
| chr10-77568360-INS-80 | chr10-77568360-INS-80 |
| chr10-77578270-INS-75 | chr10-77578270-INS-75 |
| chr10-77583379-INS-474 | chr10-77583379-INS-475 |
| chr10-77639738-INS-405 | chr10-77639738-INS-446 |
| chr10-77649317-INS-76 | chr10-77649302-INS-60 |
| chr10-77670819-INS-90 | chr10-77670819-INS-90 |
| chr10-77688053-INS-1043 | chr10-77688054-INS-1043 |
| chr10-77689990-INS-113 | chr10-77689990-INS-113 |
| chr10-77690532-INS-56 | chr10-77690532-INS-56 |
| chr10-77711951-INS-756 | chr10-77711953-INS-752 |
| chr10-77764514-INS-144 | chr10-77764514-INS-144 |
| chr10-77780600-INS-120 | chr10-77780600-INS-90 |
| chr10-77784706-INS-408 | chr10-77784706-INS-408 |
| chr10-77803422-INS-159 | chr10-77803422-INS-117 |
| chr10-77808153-INS-872 | chr10-77808153-INS-872 |
| chr10-77838321-INS-1885 | chr10-77838322-INS-1885 |
| chr10-77846214-INS-145 | chr10-77846170-INS-146 |
| chr10-77863400-INS-208 | chr10-77863407-INS-212 |
| chr10-77900011-INS-88 | chr10-77900011-INS-88 |
| chr10-77943822-INS-356 | chr10-77943822-INS-351 |
| chr10-77953706-INS-120 | chr10-77953698-INS-95 |
| chr10-77994815-INS-102 | chr10-77994815-INS-102 |
| chr10-78001068-INS-64 | chr10-78001068-INS-64 |
| chr10-78038409-INS-211 | chr10-78038409-INS-213 |
| chr10-78085839-INS-52 | chr10-78085840-INS-60 |
| chr10-78087315-INS-126 | chr10-78087315-INS-126 |
| chr10-78090326-INS-124 | chr10-78090326-INS-128 |
| chr10-78112160-INS-92 | chr10-78112160-INS-64 |
| chr10-78140713-INS-210 | chr10-78140713-INS-210 |
| chr10-78197305-INS-215 | chr10-78197305-INS-215 |
| chr10-78216233-INS-149 | chr10-78216233-INS-149 |
| chr10-78217137-INS-445 | chr10-78217137-INS-449 |
| chr10-78233290-INS-201 | chr10-78233290-INS-201 |
| chr10-78247020-INS-212 | chr10-78247020-INS-213 |
| chr10-78286548-INS-145 | chr10-78286548-INS-145 |
| chr10-78288984-INS-53 | chr10-78288984-INS-53 |
| chr10-78289832-INS-628 | chr10-78289832-INS-625 |
| chr10-78342038-INS-963 | chr10-78342038-INS-963 |
| chr10-78358100-INS-90 | chr10-78358100-INS-92 |
| chr10-78388023-INS-2974 | chr10-78388024-INS-2952 |
| chr10-78402041-INS-120 | chr10-78402057-INS-72 |
| chr10-78405499-INS-166 | chr10-78405499-INS-162 |
| chr10-78424018-INS-83 | chr10-78424039-INS-72 |
| chr10-78429438-INS-83 | chr10-78429444-INS-124 |
| chr10-78433089-INS-70 | chr10-78433089-INS-70 |
| chr10-78475463-INS-984 | chr10-78475463-INS-984 |
| chr10-78483409-INS-65 | chr10-78483409-INS-68 |
| chr10-78492459-INS-218 | chr10-78492459-INS-218 |
| chr10-78525876-INS-7443 | chr10-78525877-INS-7455 |
| chr10-78535838-INS-118 | chr10-78535852-INS-118 |
| chr10-78557386-INS-108 | chr10-78557386-INS-112 |
| chr10-78586995-INS-257 | chr10-78586995-INS-258 |
| chr10-78597914-INS-5349 | chr10-78597915-INS-5350 |
| chr10-78607061-INS-60 | chr10-78607061-INS-60 |
| chr10-78611274-INS-1892 | chr10-78611275-INS-1892 |
| chr10-78621555-INS-574 | chr10-78621555-INS-574 |
| chr10-78624154-INS-1173 | chr10-78624155-INS-1173 |
| chr10-78630507-INS-82 | chr10-78630503-INS-58 |
| chr10-78672053-INS-52 | chr10-78672029-INS-102 |
| chr10-78681181-INS-293 | chr10-78681197-INS-291 |
| chr10-78803142-INS-56 | chr10-78803138-INS-52 |
| chr10-78810248-INS-5095 | chr10-78810249-INS-5095 |
| chr10-78898445-INS-6317 | chr10-78898446-INS-6318 |
| chr10-78910256-INS-97 | chr10-78910256-INS-97 |

| | |
|---|---|
| chr10-78915136-INS-398 | chr10-78915136-INS-396 |
| chr10-78916167-INS-218 | chr10-78916167-INS-219 |
| chr10-78927098-INS-65 | chr10-78927098-INS-50 |
| chr10-78933756-INS-90 | chr10-78933756-INS-90 |
| chr10-78936353-INS-58 | chr10-78936385-INS-50 |
| chr10-78949833-INS-50 | chr10-78949833-INS-60 |
| chr10-78953497-INS-431 | chr10-78953497-INS-440 |
| chr10-78964197-INS-1198 | chr10-78964198-INS-1210 |
| chr10-78966808-INS-444 | chr10-78966808-INS-445 |
| chr10-78968681-INS-6399 | chr10-78968682-INS-6399 |
| chr10-79022699-INS-330 | chr10-79022699-INS-340 |
| chr10-79042709-INS-6264 | chr10-79042710-INS-6264 |
| chr10-79181557-INS-327 | chr10-79181557-INS-324 |
| chr10-79222737-INS-6891 | chr10-79222738-INS-6891 |
| chr10-79242491-INS-63 | chr10-79242491-INS-67 |
| chr10-79281477-INS-52 | chr10-79281477-INS-54 |
| chr10-79359108-INS-53 | chr10-79359108-INS-53 |
| chr10-79427616-INS-200 | chr10-79427616-INS-200 |
| chr10-79460334-INS-59 | chr10-79460336-INS-57 |
| chr10-79477807-INS-408 | chr10-79477807-INS-408 |
| chr10-79487042-INS-84 | chr10-79487042-INS-84 |
| chr10-79512671-INS-90 | chr10-79512671-INS-90 |
| chr10-79519613-INS-129 | chr10-79519613-INS-130 |
| chr10-79550853-INS-53 | chr10-79550853-INS-53 |
| chr10-79566326-INS-93 | chr10-79566326-INS-93 |
| chr10-79604896-INS-205 | chr10-79604896-INS-205 |
| chr10-79624069-INS-56 | chr10-79624069-INS-56 |
| chr10-79627767-INS-201 | chr10-79627767-INS-201 |
| chr10-79631307-INS-202 | chr10-79631307-INS-202 |
| chr10-79668611-INS-182 | chr10-79668611-INS-182 |
| chr10-79670148-INS-207 | chr10-79670148-INS-207 |
| chr10-79706243-INS-188 | chr10-79706243-INS-188 |
| chr10-79714876-INS-73 | chr10-79714876-INS-80 |
| chr10-79740207-INS-73 | chr10-79740207-INS-73 |
| chr10-79795178-INS-198 | chr10-79795196-INS-159 |
| chr10-79826564-INS-96 | chr10-79826599-INS-102 |
| chr10-79851492-INS-63 | chr10-79851492-INS-63 |
| chr10-79876332-INS-77 | chr10-79876336-INS-77 |
| chr10-79926121-INS-399 | chr10-79926121-INS-399 |
| chr10-79934068-INS-219 | chr10-79934068-INS-239 |
| chr10-79944306-INS-143 | chr10-79944304-INS-145 |
| chr10-79977586-INS-56 | chr10-79977586-INS-56 |
| chr10-79980578-INS-212 | chr10-79980578-INS-211 |
| chr10-80037164-INS-202 | chr10-80037164-INS-202 |
| chr10-80045594-INS-51 | chr10-80045594-INS-66 |
| chr10-80083577-INS-67 | chr10-80083575-INS-86 |
| chr10-80084352-INS-688 | chr10-80084352-INS-688 |
| chr10-80139232-INS-174 | chr10-80139232-INS-168 |
| chr10-80158140-INS-64 | chr10-80158140-INS-64 |
| chr10-80174792-INS-213 | chr10-80174788-INS-204 |
| chr10-80204156-INS-2374 | chr10-80204157-INS-2374 |
| chr10-80207629-INS-51 | chr10-80207630-INS-51 |
| chr10-80212354-INS-261 | chr10-80212354-INS-261 |
| chr10-80230743-INS-342 | chr10-80230743-INS-342 |
| chr10-80270426-INS-65 | chr10-80270426-INS-64 |
| chr10-80276042-INS-54 | chr10-80276042-INS-54 |
| chr10-80323486-INS-113 | chr10-80323486-INS-93 |
| chr10-80337856-INS-64 | chr10-80337856-INS-62 |
| chr10-80339261-INS-193 | chr10-80339261-INS-193 |
| chr10-80340516-INS-85 | chr10-80340516-INS-85 |
| chr10-80438452-INS-61 | chr10-80438452-INS-61 |
| chr10-80492976-INS-267 | chr10-80492976-INS-274 |
| chr10-80538223-INS-216 | chr10-80538223-INS-217 |
| chr10-80567165-INS-232 | chr10-80567165-INS-233 |
| chr10-80596874-INS-85 | chr10-80596874-INS-85 |
| chr10-80613047-INS-215 | chr10-80613047-INS-210 |
| chr10-80650054-INS-295 | chr10-80650035-INS-295 |
| chr10-80650954-INS-166 | chr10-80650954-INS-166 |
| chr10-80651588-INS-1640 | chr10-80651588-INS-1621 |
| chr10-80652047-INS-363 | chr10-80652055-INS-392 |
| chr10-80654150-INS-96 | chr10-80654150-INS-96 |
| chr10-80657581-INS-171 | chr10-80657581-INS-173 |
| chr10-80692196-INS-228 | chr10-80692196-INS-230 |
| chr10-80743553-INS-210 | chr10-80743553-INS-214 |
| chr10-80747654-INS-210 | chr10-80747655-INS-214 |
| chr10-80758372-INS-196 | chr10-80758372-INS-196 |
| chr10-80760949-INS-295 | chr10-80760949-INS-295 |

| | |
|---|---|
| chr10-80782225-INS-91 | chr10-80782225-INS-96 |
| chr10-80819460-INS-473 | chr10-80819460-INS-473 |
| chr10-80882631-INS-60 | chr10-80882631-INS-80 |
| chr10-80889440-INS-237 | chr10-80889440-INS-237 |
| chr10-80898762-INS-220 | chr10-80898762-INS-217 |
| chr10-80912577-INS-209 | chr10-80912577-INS-211 |
| chr10-80933698-INS-53 | chr10-80933700-INS-53 |
| chr10-80936364-INS-96 | chr10-80936362-INS-119 |
| chr10-80962826-INS-243 | chr10-80962865-INS-247 |
| chr10-80971067-INS-69 | chr10-80971067-INS-77 |
| chr10-81007416-INS-175 | chr10-81007420-INS-182 |
| chr10-81034466-INS-193 | chr10-81034466-INS-193 |
| chr10-81045019-INS-74 | chr10-81045019-INS-74 |
| chr10-81047144-INS-208 | chr10-81047144-INS-209 |
| chr10-81048245-INS-148 | chr10-81048245-INS-148 |
| chr10-81054611-INS-85 | chr10-81054611-INS-85 |
| chr10-81055770-INS-198 | chr10-81055770-INS-200 |
| chr10-81103155-INS-200 | chr10-81103155-INS-200 |
| chr10-81150497-INS-91 | chr10-81150497-INS-91 |
| chr10-81188303-INS-185 | chr10-81188282-INS-163 |
| chr10-81189278-INS-214 | chr10-81189278-INS-214 |
| chr10-81193749-INS-60 | chr10-81193749-INS-60 |
| chr10-81240305-INS-109 | chr10-81240305-INS-109 |
| chr10-81280325-INS-60 | chr10-81280325-INS-63 |
| chr10-81346839-INS-52 | chr10-81346867-INS-55 |
| chr10-81355201-INS-128 | chr10-81355201-INS-130 |
| chr10-81372306-INS-337 | chr10-81372306-INS-337 |
| chr10-81383900-INS-186 | chr10-81383888-INS-194 |
| chr10-81419626-INS-53 | chr10-81419626-INS-53 |
| chr10-81449669-INS-2832 | chr10-81449670-INS-2832 |
| chr10-81449670-INS-500 | chr10-81449670-INS-500 |
| chr10-81464871-INS-80 | chr10-81464872-INS-100 |
| chr10-81487119-INS-214 | chr10-81487119-INS-214 |
| chr10-81492417-INS-144 | chr10-81492429-INS-120 |
| chr10-81518961-INS-217 | chr10-81518961-INS-216 |
| chr10-81854541-INS-5548 | chr10-81854542-INS-5548 |
| chr10-81872139-INS-181 | chr10-81872139-INS-181 |
| chr10-81878091-INS-152 | chr10-81878091-INS-152 |
| chr10-81893227-INS-230 | chr10-81893227-INS-230 |
| chr10-81935213-INS-6060 | chr10-81935214-INS-6060 |
| chr10-82027593-INS-6244 | chr10-82027594-INS-6245 |
| chr10-82054274-INS-196 | chr10-82054274-INS-195 |
| chr10-82085009-INS-84 | chr10-82085009-INS-84 |
| chr10-82091364-INS-1806 | chr10-82091365-INS-1815 |
| chr10-82093432-INS-1892 | chr10-82093433-INS-1888 |
| chr10-82095859-INS-7356 | chr10-82095860-INS-7358 |
| chr10-82104824-INS-107 | chr10-82104822-INS-123 |
| chr10-82116727-INS-6407 | chr10-82116728-INS-6392 |
| chr10-82117285-INS-1438 | chr10-82117286-INS-1430 |
| chr10-82136745-INS-82 | chr10-82136745-INS-82 |
| chr10-82150383-INS-76 | chr10-82150395-INS-106 |
| chr10-82163815-INS-143 | chr10-82163815-INS-140 |
| chr10-82165349-INS-194 | chr10-82165349-INS-194 |
| chr10-82168882-INS-7111 | chr10-82168883-INS-7111 |
| chr10-82170569-INS-220 | chr10-82170569-INS-220 |
| chr10-82199514-INS-140 | chr10-82199514-INS-140 |
| chr10-82247609-INS-204 | chr10-82247609-INS-204 |
| chr10-82266540-INS-682 | chr10-82266535-INS-694 |
| chr10-82274269-INS-129 | chr10-82274269-INS-129 |
| chr10-82286316-INS-153 | chr10-82286316-INS-149 |
| chr10-82298253-INS-6772 | chr10-82298254-INS-6771 |
| chr10-82307318-INS-6458 | chr10-82307319-INS-6458 |
| chr10-82307515-INS-343 | chr10-82307515-INS-343 |
| chr10-82313136-INS-7160 | chr10-82313137-INS-7160 |
| chr10-82314558-INS-51 | chr10-82314558-INS-51 |
| chr10-82328957-INS-3800 | chr10-82328958-INS-3800 |
| chr10-82352727-INS-132 | chr10-82352727-INS-132 |
| chr10-82362249-INS-237 | chr10-82362247-INS-261 |
| chr10-82370133-INS-199 | chr10-82370133-INS-199 |
| chr10-82388788-INS-212 | chr10-82388788-INS-212 |
| chr10-82400601-INS-74 | chr10-82400601-INS-74 |
| chr10-82430400-INS-993 | chr10-82430400-INS-1005 |
| chr10-82453874-INS-171 | chr10-82453874-INS-171 |
| chr10-82478657-INS-177 | chr10-82478657-INS-177 |
| chr10-82480888-INS-70 | chr10-82480888-INS-70 |
| chr10-82487743-INS-251 | chr10-82487743-INS-251 |
| chr10-82490967-INS-136 | chr10-82490967-INS-136 |

| | |
|---|---|
| chr10-82511845-INS-1337 | chr10-82511846-INS-1338 |
| chr10-82513612-INS-130 | chr10-82513652-INS-88 |
| chr10-82542457-INS-80 | chr10-82542457-INS-80 |
| chr10-82577481-INS-62 | chr10-82577481-INS-62 |
| chr10-82582787-INS-836 | chr10-82582787-INS-836 |
| chr10-82605525-INS-147 | chr10-82605525-INS-169 |
| chr10-82625196-INS-57 | chr10-82625196-INS-57 |
| chr10-82627847-INS-201 | chr10-82627847-INS-201 |
| chr10-82644021-INS-82 | chr10-82644021-INS-72 |
| chr10-82663557-INS-7070 | chr10-82663558-INS-7070 |
| chr10-82720481-INS-136 | chr10-82720481-INS-136 |
| chr10-82725034-INS-203 | chr10-82725034-INS-203 |
| chr10-82733216-INS-84 | chr10-82733216-INS-84 |
| chr10-82750384-INS-172 | chr10-82750384-INS-173 |
| chr10-82771897-INS-78 | chr10-82771897-INS-82 |
| chr10-82784192-INS-310 | chr10-82784192-INS-310 |
| chr10-82795048-INS-54 | chr10-82795048-INS-52 |
| chr10-82799969-INS-52 | chr10-82799969-INS-54 |
| chr10-82811303-INS-590 | chr10-82811303-INS-590 |
| chr10-82812749-INS-231 | chr10-82812749-INS-231 |
| chr10-82864857-INS-83 | chr10-82864857-INS-83 |
| chr10-82875315-INS-265 | chr10-82875315-INS-267 |
| chr10-82891657-INS-50 | chr10-82891616-INS-85 |
| chr10-82901327-INS-197 | chr10-82901327-INS-191 |
| chr10-82910931-INS-69 | chr10-82910931-INS-61 |
| chr10-82928392-INS-540 | chr10-82928392-INS-510 |
| chr10-82941382-INS-261 | chr10-82941411-INS-260 |
| chr10-82944507-INS-5217 | chr10-82944508-INS-5217 |
| chr10-82986780-INS-98 | chr10-82986780-INS-98 |
| chr10-82999652-INS-51 | chr10-82999642-INS-75 |
| chr10-83016772-INS-67 | chr10-83016772-INS-67 |
| chr10-83033371-INS-54 | chr10-83033371-INS-54 |
| chr10-83084999-INS-91 | chr10-83084999-INS-91 |
| chr10-83095850-INS-236 | chr10-83095853-INS-253 |
| chr10-83099843-INS-75 | chr10-83099822-INS-51 |
| chr10-83118114-INS-401 | chr10-83118114-INS-401 |
| chr10-83178080-INS-7152 | chr10-83178081-INS-7152 |
| chr10-83187340-INS-88 | chr10-83187350-INS-67 |
| chr10-83403619-INS-202 | chr10-83403619-INS-203 |
| chr10-83496799-INS-103 | chr10-83496799-INS-105 |
| chr10-83501464-INS-92 | chr10-83501464-INS-92 |
| chr10-83514459-INS-72 | chr10-83514443-INS-68 |
| chr10-83518567-INS-345 | chr10-83518567-INS-345 |
| chr10-83551284-INS-88 | chr10-83551284-INS-102 |
| chr10-83572260-INS-57 | chr10-83572260-INS-77 |
| chr10-83590592-INS-70 | chr10-83590582-INS-77 |
| chr10-83599419-INS-241 | chr10-83599419-INS-240 |
| chr10-83614692-INS-121 | chr10-83614692-INS-121 |
| chr10-83659123-INS-60 | chr10-83659123-INS-60 |
| chr10-83707550-INS-120 | chr10-83707550-INS-124 |
| chr10-83721091-INS-63 | chr10-83721113-INS-66 |
| chr10-83732330-INS-208 | chr10-83732330-INS-208 |
| chr10-83737643-INS-310 | chr10-83737644-INS-310 |
| chr10-83748519-INS-55 | chr10-83748534-INS-50 |
| chr10-83750312-INS-384 | chr10-83750312-INS-378 |
| chr10-83791007-INS-90 | chr10-83791007-INS-50 |
| chr10-83796434-INS-301 | chr10-83796434-INS-301 |
| chr10-83810132-INS-53 | chr10-83810168-INS-53 |
| chr10-83811315-INS-143 | chr10-83811319-INS-134 |
| chr10-83816943-INS-57 | chr10-83816937-INS-76 |
| chr10-83817785-INS-2359 | chr10-83817786-INS-2359 |
| chr10-83823115-INS-56 | chr10-83823115-INS-60 |
| chr10-83839070-INS-77 | chr10-83839070-INS-82 |
| chr10-83839850-INS-7044 | chr10-83839851-INS-7044 |
| chr10-83841443-INS-337 | chr10-83841441-INS-340 |
| chr10-83862789-INS-631 | chr10-83862789-INS-630 |
| chr10-83874966-INS-53 | chr10-83874966-INS-66 |
| chr10-83877438-INS-81 | chr10-83877426-INS-85 |
| chr10-83883165-INS-51 | chr10-83883165-INS-51 |
| chr10-83887543-INS-331 | chr10-83887544-INS-332 |
| chr10-83907618-INS-148 | chr10-83907618-INS-116 |
| chr10-83912260-INS-106 | chr10-83912260-INS-108 |
| chr10-83951888-INS-151 | chr10-83951888-INS-151 |
| chr10-83957041-INS-196 | chr10-83957041-INS-224 |
| chr10-83961243-INS-58 | chr10-83961241-INS-62 |
| chr10-83967596-INS-86 | chr10-83967588-INS-68 |
| chr10-83972281-INS-2877 | chr10-83972282-INS-2877 |

| | |
|---|---|
| chr10-83976684-INS-294 | chr10-83976684-INS-291 |
| chr10-84031524-INS-54 | chr10-84031524-INS-54 |
| chr10-84032367-INS-54 | chr10-84032367-INS-54 |
| chr10-84098780-INS-56 | chr10-84098773-INS-76 |
| chr10-84149544-INS-971 | chr10-84149544-INS-971 |
| chr10-84153150-INS-56 | chr10-84153150-INS-56 |
| chr10-84172838-INS-263 | chr10-84172838-INS-262 |
| chr10-84175829-INS-60 | chr10-84175833-INS-57 |
| chr10-84189117-INS-142 | chr10-84189117-INS-141 |
| chr10-84190268-INS-6513 | chr10-84190269-INS-6513 |
| chr10-84204607-INS-94 | chr10-84204607-INS-94 |
| chr10-84206438-INS-119 | chr10-84206438-INS-119 |
| chr10-84232317-INS-151 | chr10-84232322-INS-156 |
| chr10-84248642-INS-118 | chr10-84248642-INS-118 |
| chr10-84266110-INS-244 | chr10-84266110-INS-256 |
| chr10-84291425-INS-66 | chr10-84291433-INS-73 |
| chr10-84311180-INS-56 | chr10-84311180-INS-56 |
| chr10-84338425-INS-93 | chr10-84338425-INS-93 |
| chr10-84342064-INS-53 | chr10-84342064-INS-53 |
| chr10-84351803-INS-151 | chr10-84351803-INS-151 |
| chr10-84377430-INS-507 | chr10-84377430-INS-507 |
| chr10-84428164-INS-197 | chr10-84428164-INS-197 |
| chr10-84447738-INS-181 | chr10-84447738-INS-181 |
| chr10-84450983-INS-694 | chr10-84450983-INS-694 |
| chr10-84489748-INS-202 | chr10-84489748-INS-205 |
| chr10-84491107-INS-174 | chr10-84491107-INS-174 |
| chr10-84517220-INS-66 | chr10-84517220-INS-68 |
| chr10-84519403-INS-51 | chr10-84519403-INS-51 |
| chr10-84523225-INS-161 | chr10-84523225-INS-172 |
| chr10-84527845-INS-58 | chr10-84527845-INS-58 |
| chr10-84553384-INS-136 | chr10-84553384-INS-136 |
| chr10-84560522-INS-131 | chr10-84560522-INS-131 |
| chr10-84589656-INS-56 | chr10-84589649-INS-76 |
| chr10-84595878-INS-68 | chr10-84595878-INS-66 |
| chr10-84605489-INS-151 | chr10-84605491-INS-143 |
| chr10-84623376-INS-6498 | chr10-84623377-INS-6498 |
| chr10-84646921-INS-88 | chr10-84646921-INS-72 |
| chr10-84654409-INS-82 | chr10-84654413-INS-68 |
| chr10-84655149-INS-213 | chr10-84655149-INS-213 |
| chr10-84687744-INS-1890 | chr10-84687745-INS-1890 |
| chr10-84702894-INS-133 | chr10-84702893-INS-149 |
| chr10-84765377-INS-231 | chr10-84765377-INS-231 |
| chr10-84778235-INS-276 | chr10-84778235-INS-277 |
| chr10-84861768-INS-166 | chr10-84861768-INS-168 |
| chr10-84881729-INS-170 | chr10-84881729-INS-170 |
| chr10-84885389-INS-1496 | chr10-84885390-INS-1495 |
| chr10-84893099-INS-75 | chr10-84893099-INS-75 |
| chr10-84893966-INS-1118 | chr10-84893967-INS-1125 |
| chr10-84894505-INS-2198 | chr10-84894506-INS-2195 |
| chr10-84948395-INS-209 | chr10-84948395-INS-209 |
| chr10-84949690-INS-757 | chr10-84949690-INS-744 |
| chr10-84958193-INS-56 | chr10-84958193-INS-58 |
| chr10-84959906-INS-57 | chr10-84959906-INS-57 |
| chr10-84982836-INS-72 | chr10-84982836-INS-50 |
| chr10-84994050-INS-93 | chr10-84994050-INS-93 |
| chr10-85001079-INS-86 | chr10-85001079-INS-86 |
| chr10-85014895-INS-207 | chr10-85014895-INS-208 |
| chr10-85019282-INS-56 | chr10-85019282-INS-56 |
| chr10-85024741-INS-205 | chr10-85024741-INS-208 |
| chr10-85031208-INS-54 | chr10-85031210-INS-57 |
| chr10-85032721-INS-197 | chr10-85032721-INS-197 |
| chr10-85060152-INS-7551 | chr10-85060153-INS-7550 |
| chr10-85060153-INS-210 | chr10-85060153-INS-210 |
| chr10-85065590-INS-56 | chr10-85065590-INS-56 |
| chr10-85074189-INS-1225 | chr10-85074190-INS-1229 |
| chr10-85075955-INS-66 | chr10-85075944-INS-68 |
| chr10-85078616-INS-6401 | chr10-85078617-INS-6401 |
| chr10-85080097-INS-332 | chr10-85080097-INS-332 |
| chr10-85082387-INS-163 | chr10-85082387-INS-163 |
| chr10-85086200-INS-203 | chr10-85086200-INS-203 |
| chr10-85101309-INS-68 | chr10-85101309-INS-68 |
| chr10-85103688-INS-528 | chr10-85103688-INS-528 |
| chr10-85111836-INS-106 | chr10-85111836-INS-106 |
| chr10-85180111-INS-62 | chr10-85180111-INS-62 |
| chr10-85183778-INS-4129 | chr10-85183779-INS-4133 |
| chr10-85189961-INS-72 | chr10-85189961-INS-72 |
| chr10-85191674-INS-475 | chr10-85191674-INS-475 |

| | |
|---|---|
| chr10-85192722-INS-62 | chr10-85192722-INS-62 |
| chr10-85195759-INS-445 | chr10-85195759-INS-445 |
| chr10-85203699-INS-84 | chr10-85203699-INS-84 |
| chr10-85213533-INS-64 | chr10-85213533-INS-64 |
| chr10-85219237-INS-6324 | chr10-85219238-INS-6324 |
| chr10-85229071-INS-110 | chr10-85229071-INS-82 |
| chr10-85230096-INS-191 | chr10-85230096-INS-207 |
| chr10-85235548-INS-118 | chr10-85235548-INS-118 |
| chr10-85264998-INS-315 | chr10-85264998-INS-280 |
| chr10-85265768-INS-81 | chr10-85265768-INS-80 |
| chr10-85363493-INS-54 | chr10-85363493-INS-60 |
| chr10-85365245-INS-81 | chr10-85365245-INS-81 |
| chr10-85371380-INS-81 | chr10-85371377-INS-56 |
| chr10-85373554-INS-418 | chr10-85373554-INS-418 |
| chr10-85379993-INS-7007 | chr10-85379994-INS-7009 |
| chr10-85381280-INS-88 | chr10-85381233-INS-99 |
| chr10-85399294-INS-54 | chr10-85399294-INS-64 |
| chr10-85406092-INS-60 | chr10-85406091-INS-64 |
| chr10-85414519-INS-1040 | chr10-85414520-INS-1040 |
| chr10-85431223-INS-52 | chr10-85431223-INS-50 |
| chr10-85439780-INS-228 | chr10-85439777-INS-227 |
| chr10-85455554-INS-270 | chr10-85455554-INS-270 |
| chr10-85472525-INS-81 | chr10-85472525-INS-75 |
| chr10-85474703-INS-121 | chr10-85474703-INS-111 |
| chr10-85489186-INS-204 | chr10-85489149-INS-227 |
| chr10-85508655-INS-2046 | chr10-85508656-INS-2046 |
| chr10-85541905-INS-285 | chr10-85541905-INS-285 |
| chr10-85567340-INS-126 | chr10-85567340-INS-92 |
| chr10-85575161-INS-4455 | chr10-85575162-INS-4455 |
| chr10-85580432-INS-70 | chr10-85580432-INS-70 |
| chr10-85648392-INS-342 | chr10-85648387-INS-342 |
| chr10-85701153-INS-498 | chr10-85701153-INS-498 |
| chr10-85720896-INS-250 | chr10-85720896-INS-250 |
| chr10-85735951-INS-55 | chr10-85735951-INS-55 |
| chr10-85740397-INS-132 | chr10-85740397-INS-132 |
| chr10-85745835-INS-3147 | chr10-85745836-INS-3147 |
| chr10-85772354-INS-6489 | chr10-85772355-INS-6489 |
| chr10-85802085-INS-56 | chr10-85802085-INS-94 |
| chr10-85805322-INS-130 | chr10-85805322-INS-131 |
| chr10-85806066-INS-204 | chr10-85806076-INS-205 |
| chr10-85818682-INS-623 | chr10-85818682-INS-624 |
| chr10-85831056-INS-1213 | chr10-85831057-INS-1205 |
| chr10-85852980-INS-792 | chr10-85852983-INS-793 |
| chr10-85873092-INS-6431 | chr10-85873093-INS-6431 |
| chr10-85937169-INS-63 | chr10-85937169-INS-63 |
| chr10-85942474-INS-104 | chr10-85942474-INS-96 |
| chr10-85951451-INS-78 | chr10-85951463-INS-92 |
| chr10-85979543-INS-158 | chr10-85979544-INS-163 |
| chr10-85991979-INS-96 | chr10-85991979-INS-61 |
| chr10-86010425-INS-52 | chr10-86010425-INS-54 |
| chr10-86018093-INS-874 | chr10-86018093-INS-874 |
| chr10-86027494-INS-2064 | chr10-86027495-INS-2064 |
| chr10-86064248-INS-81 | chr10-86064248-INS-81 |
| chr10-86110423-INS-51 | chr10-86110423-INS-51 |
| chr10-86111315-INS-68 | chr10-86111315-INS-68 |
| chr10-86119815-INS-73 | chr10-86119797-INS-74 |
| chr10-86128971-INS-108 | chr10-86128971-INS-108 |
| chr10-86162531-INS-78 | chr10-86162531-INS-78 |
| chr10-86203280-INS-94 | chr10-86203280-INS-94 |
| chr10-86218605-INS-88 | chr10-86218605-INS-88 |
| chr10-86225534-INS-182 | chr10-86225534-INS-187 |
| chr10-86240691-INS-69 | chr10-86240691-INS-69 |
| chr10-86256876-INS-5263 | chr10-86256877-INS-5263 |
| chr10-86295854-INS-64 | chr10-86295854-INS-56 |
| chr10-86311399-INS-74 | chr10-86311399-INS-74 |
| chr10-86315446-INS-64 | chr10-86315430-INS-83 |
| chr10-86323743-INS-1076 | chr10-86323744-INS-1072 |
| chr10-86342044-INS-217 | chr10-86342057-INS-211 |
| chr10-86358117-INS-70 | chr10-86358117-INS-70 |
| chr10-86387707-INS-333 | chr10-86387707-INS-333 |
| chr10-86496492-INS-171 | chr10-86496492-INS-171 |
| chr10-86522177-INS-190 | chr10-86522177-INS-178 |
| chr10-86538583-INS-201 | chr10-86538583-INS-214 |
| chr10-86553722-INS-78 | chr10-86553722-INS-81 |
| chr10-86554506-INS-662 | chr10-86554506-INS-662 |
| chr10-86556020-INS-203 | chr10-86556020-INS-203 |
| chr10-86558134-INS-447 | chr10-86558134-INS-455 |

| | |
|---|---|
| chr10-86568201-INS-1379 | chr10-86568202-INS-1379 |
| chr10-86576187-INS-623 | chr10-86576187-INS-624 |
| chr10-86582240-INS-199 | chr10-86582240-INS-195 |
| chr10-86583364-INS-221 | chr10-86583364-INS-221 |
| chr10-86588169-INS-183 | chr10-86588169-INS-183 |
| chr10-86590742-INS-149 | chr10-86590764-INS-117 |
| chr10-86601181-INS-81 | chr10-86601167-INS-83 |
| chr10-86610522-INS-207 | chr10-86610522-INS-207 |
| chr10-86616458-INS-202 | chr10-86616458-INS-212 |
| chr10-86618650-INS-183 | chr10-86618650-INS-183 |
| chr10-86624006-INS-70 | chr10-86624006-INS-70 |
| chr10-86625757-INS-399 | chr10-86625757-INS-399 |
| chr10-86640422-INS-684 | chr10-86640430-INS-691 |
| chr10-86647925-INS-3509 | chr10-86647926-INS-3508 |
| chr10-86663735-INS-382 | chr10-86663735-INS-382 |
| chr10-86691796-INS-80 | chr10-86691796-INS-80 |
| chr10-86712414-INS-722 | chr10-86712414-INS-722 |
| chr10-86714970-INS-60 | chr10-86714970-INS-62 |
| chr10-86738998-INS-67 | chr10-86738998-INS-94 |
| chr10-86784127-INS-82 | chr10-86784123-INS-65 |
| chr10-86785499-INS-383 | chr10-86785499-INS-383 |
| chr10-86786091-INS-727 | chr10-86786091-INS-727 |
| chr10-86797052-INS-64 | chr10-86797052-INS-64 |
| chr10-86806461-INS-67 | chr10-86806461-INS-67 |
| chr10-86834913-INS-54 | chr10-86834913-INS-58 |
| chr10-86842628-INS-132 | chr10-86842628-INS-136 |
| chr10-86845993-INS-51 | chr10-86845993-INS-51 |
| chr10-86852393-INS-58 | chr10-86852393-INS-70 |
| chr10-86860738-INS-65 | chr10-86860738-INS-83 |
| chr10-86863520-INS-778 | chr10-86863520-INS-803 |
| chr10-86899390-INS-489 | chr10-86899390-INS-489 |
| chr10-86907582-INS-104 | chr10-86907582-INS-104 |
| chr10-86919025-INS-60 | chr10-86919025-INS-60 |
| chr10-87010001-INS-52 | chr10-87010001-INS-54 |
| chr10-87032873-INS-79 | chr10-87032873-INS-85 |
| chr10-87058587-INS-88 | chr10-87058587-INS-88 |
| chr10-87099155-INS-565 | chr10-87099155-INS-565 |
| chr10-87107258-INS-85 | chr10-87107278-INS-112 |
| chr10-87142911-INS-201 | chr10-87142911-INS-201 |
| chr10-87190676-INS-639 | chr10-87190676-INS-639 |
| chr10-87191569-INS-86 | chr10-87191555-INS-70 |
| chr10-87195506-INS-96 | chr10-87195506-INS-96 |
| chr10-87214694-INS-2583 | chr10-87214696-INS-2585 |
| chr10-87257006-INS-74 | chr10-87257000-INS-120 |
| chr10-87264017-INS-57 | chr10-87264017-INS-62 |
| chr10-87266767-INS-52 | chr10-87266767-INS-58 |
| chr10-87291972-INS-256 | chr10-87291972-INS-256 |
| chr10-87303198-INS-385 | chr10-87303199-INS-386 |
| chr10-87307438-INS-72 | chr10-87307410-INS-50 |
| chr10-87327557-INS-100 | chr10-87327557-INS-102 |
| chr10-87335466-INS-155 | chr10-87335466-INS-161 |
| chr10-87339470-INS-102 | chr10-87339470-INS-102 |
| chr10-87346322-INS-201 | chr10-87346322-INS-201 |
| chr10-87355231-INS-212 | chr10-87355231-INS-236 |
| chr10-87374932-INS-384 | chr10-87374932-INS-384 |
| chr10-87383448-INS-51 | chr10-87383448-INS-54 |
| chr10-87466504-INS-85 | chr10-87466496-INS-84 |
| chr10-87494767-INS-72 | chr10-87494767-INS-104 |
| chr10-87500039-INS-64 | chr10-87500039-INS-65 |
| chr10-87511194-INS-51 | chr10-87511184-INS-87 |
| chr10-87530147-INS-856 | chr10-87530147-INS-856 |
| chr10-87531899-INS-463 | chr10-87531899-INS-463 |
| chr10-87532741-INS-7100 | chr10-87532742-INS-7100 |
| chr10-87539364-INS-975 | chr10-87539364-INS-975 |
| chr10-87595499-INS-117 | chr10-87595499-INS-117 |
| chr10-87628192-INS-51 | chr10-87628193-INS-55 |
| chr10-87661747-INS-56 | chr10-87661747-INS-56 |
| chr10-87663408-INS-2810 | chr10-87663409-INS-2810 |
| chr10-87674171-INS-272 | chr10-87674171-INS-272 |
| chr10-87714977-INS-262 | chr10-87714977-INS-262 |
| chr10-87778668-INS-1223 | chr10-87778669-INS-1226 |
| chr10-87790534-INS-1170 | chr10-87790535-INS-1170 |
| chr10-87790943-INS-189 | chr10-87790943-INS-189 |
| chr10-87798538-INS-3135 | chr10-87798539-INS-3135 |
| chr10-87799168-INS-964 | chr10-87799168-INS-964 |
| chr10-87804432-INS-141 | chr10-87804432-INS-141 |
| chr10-87840772-INS-313 | chr10-87840772-INS-313 |

| | |
|---|---|
| chr10-87846595-INS-271 | chr10-87846596-INS-271 |
| chr10-87920152-INS-65 | chr10-87920152-INS-65 |
| chr10-87977395-INS-117 | chr10-87977415-INS-126 |
| chr10-87985875-INS-84 | chr10-87985875-INS-122 |
| chr10-87992150-INS-264 | chr10-87992150-INS-264 |
| chr10-88014461-INS-825 | chr10-88014461-INS-825 |
| chr10-88017123-INS-76 | chr10-88017137-INS-54 |
| chr10-88037500-INS-57 | chr10-88037500-INS-85 |
| chr10-88050903-INS-247 | chr10-88050903-INS-251 |
| chr10-88095658-INS-54 | chr10-88095658-INS-81 |
| chr10-88099980-INS-6403 | chr10-88099981-INS-6403 |
| chr10-88118432-INS-203 | chr10-88118432-INS-195 |
| chr10-88152382-INS-72 | chr10-88152401-INS-80 |
| chr10-88154377-INS-74 | chr10-88154350-INS-62 |
| chr10-88163897-INS-194 | chr10-88163897-INS-194 |
| chr10-88173754-INS-170 | chr10-88173754-INS-170 |
| chr10-88195508-INS-83 | chr10-88195508-INS-83 |
| chr10-88205981-INS-174 | chr10-88205981-INS-166 |
| chr10-88218249-INS-4548 | chr10-88218250-INS-4548 |
| chr10-88237339-INS-293 | chr10-88237339-INS-293 |
| chr10-88254697-INS-190 | chr10-88254697-INS-190 |
| chr10-88260422-INS-68 | chr10-88260422-INS-68 |
| chr10-88262569-INS-217 | chr10-88262569-INS-217 |
| chr10-88268829-INS-171 | chr10-88268829-INS-171 |
| chr10-88270968-INS-464 | chr10-88270968-INS-464 |
| chr10-88272911-INS-226 | chr10-88272911-INS-178 |
| chr10-88278874-INS-970 | chr10-88278874-INS-970 |
| chr10-88279595-INS-141 | chr10-88279595-INS-141 |
| chr10-88283903-INS-60 | chr10-88283903-INS-60 |
| chr10-88284612-INS-1661 | chr10-88284613-INS-1665 |
| chr10-88288862-INS-219 | chr10-88288862-INS-220 |
| chr10-88323722-INS-214 | chr10-88323722-INS-214 |
| chr10-88328787-INS-199 | chr10-88328787-INS-198 |
| chr10-88329387-INS-80 | chr10-88329387-INS-92 |
| chr10-88330630-INS-546 | chr10-88330630-INS-546 |
| chr10-88336661-INS-166 | chr10-88336661-INS-166 |
| chr10-88346145-INS-179 | chr10-88346145-INS-179 |
| chr10-88352924-INS-266 | chr10-88352924-INS-261 |
| chr10-88371039-INS-171 | chr10-88371039-INS-171 |
| chr10-88374236-INS-160 | chr10-88374236-INS-160 |
| chr10-88382217-INS-205 | chr10-88382217-INS-205 |
| chr10-88390198-INS-218 | chr10-88390198-INS-219 |
| chr10-88400370-INS-52 | chr10-88400370-INS-66 |
| chr10-88411344-INS-56 | chr10-88411344-INS-52 |
| chr10-88419069-INS-212 | chr10-88419069-INS-210 |
| chr10-88424812-INS-102 | chr10-88424812-INS-102 |
| chr10-88430142-INS-68 | chr10-88430142-INS-68 |
| chr10-88447386-INS-68 | chr10-88447386-INS-70 |
| chr10-88459357-INS-72 | chr10-88459357-INS-72 |
| chr10-88467905-INS-68 | chr10-88467905-INS-68 |
| chr10-88470629-INS-52 | chr10-88470630-INS-52 |
| chr10-88485164-INS-1371 | chr10-88485165-INS-1384 |
| chr10-88498466-INS-2887 | chr10-88498515-INS-2886 |
| chr10-88512868-INS-244 | chr10-88512868-INS-244 |
| chr10-88527542-INS-172 | chr10-88527542-INS-157 |
| chr10-88529253-INS-60 | chr10-88529253-INS-54 |
| chr10-88530805-INS-55 | chr10-88530805-INS-55 |
| chr10-88556401-INS-92 | chr10-88556386-INS-60 |
| chr10-88564938-INS-8326 | chr10-88564939-INS-8322 |
| chr10-88669272-INS-52 | chr10-88669272-INS-52 |
| chr10-88711546-INS-210 | chr10-88711546-INS-210 |
| chr10-88727690-INS-978 | chr10-88727690-INS-981 |
| chr10-88747399-INS-7567 | chr10-88747400-INS-7570 |
| chr10-88749915-INS-52 | chr10-88749919-INS-76 |
| chr10-88758372-INS-621 | chr10-88758372-INS-628 |
| chr10-88760985-INS-82 | chr10-88760940-INS-118 |
| chr10-88809891-INS-167 | chr10-88809891-INS-167 |
| chr10-88811574-INS-123 | chr10-88811574-INS-123 |
| chr10-88815930-INS-70 | chr10-88815934-INS-66 |
| chr10-88825811-INS-146 | chr10-88825847-INS-128 |
| chr10-88834297-INS-331 | chr10-88834297-INS-331 |
| chr10-88836067-INS-3650 | chr10-88836068-INS-3676 |
| chr10-88836858-INS-69 | chr10-88836852-INS-58 |
| chr10-88857466-INS-385 | chr10-88857466-INS-384 |
| chr10-88860734-INS-117 | chr10-88860734-INS-142 |
| chr10-88877586-INS-216 | chr10-88877586-INS-188 |
| chr10-88879947-INS-197 | chr10-88879947-INS-197 |

| | |
|---|---|
| chr10-88880586-INS-282 | chr10-88880586-INS-282 |
| chr10-88881941-INS-73 | chr10-88881941-INS-72 |
| chr10-88889837-INS-708 | chr10-88889837-INS-709 |
| chr10-88912080-INS-6408 | chr10-88912081-INS-6408 |
| chr10-88924874-INS-801 | chr10-88924874-INS-803 |
| chr10-88980034-INS-799 | chr10-88980035-INS-803 |
| chr10-88996820-INS-6658 | chr10-88996821-INS-6659 |
| chr10-89001330-INS-400 | chr10-89001330-INS-400 |
| chr10-89018339-INS-65 | chr10-89018339-INS-65 |
| chr10-89048187-INS-588 | chr10-89048187-INS-588 |
| chr10-89061589-INS-206 | chr10-89061589-INS-223 |
| chr10-89067429-INS-65 | chr10-89067429-INS-90 |
| chr10-89071459-INS-68 | chr10-89071441-INS-58 |
| chr10-89092232-INS-395 | chr10-89092233-INS-395 |
| chr10-89097539-INS-160 | chr10-89097539-INS-160 |
| chr10-89106216-INS-680 | chr10-89106217-INS-680 |
| chr10-89139872-INS-1243 | chr10-89139873-INS-1243 |
| chr10-89142375-INS-1541 | chr10-89142376-INS-1536 |
| chr10-89183400-INS-6413 | chr10-89183401-INS-6413 |
| chr10-89190343-INS-104 | chr10-89190343-INS-104 |
| chr10-89192508-INS-54 | chr10-89192508-INS-50 |
| chr10-89222977-INS-84 | chr10-89222977-INS-70 |
| chr10-89283205-INS-3318 | chr10-89283210-INS-3317 |
| chr10-89331913-INS-124 | chr10-89331913-INS-130 |
| chr10-89346189-INS-114 | chr10-89346189-INS-114 |
| chr10-89366504-INS-135 | chr10-89366500-INS-129 |
| chr10-89383651-INS-7155 | chr10-89383652-INS-7155 |
| chr10-89404613-INS-306 | chr10-89404613-INS-306 |
| chr10-89415600-INS-1974 | chr10-89415601-INS-1974 |
| chr10-89429313-INS-63 | chr10-89429313-INS-63 |
| chr10-89436626-INS-455 | chr10-89436626-INS-455 |
| chr10-89438076-INS-62 | chr10-89438076-INS-62 |
| chr10-89453933-INS-56 | chr10-89453934-INS-56 |
| chr10-89456113-INS-91 | chr10-89456135-INS-72 |
| chr10-89459042-INS-299 | chr10-89459042-INS-299 |
| chr10-89500040-INS-360 | chr10-89500040-INS-360 |
| chr10-89532441-INS-65 | chr10-89532440-INS-72 |
| chr10-89561926-INS-204 | chr10-89561926-INS-203 |
| chr10-89566000-INS-442 | chr10-89566000-INS-459 |
| chr10-89591269-INS-869 | chr10-89591269-INS-869 |
| chr10-89594708-INS-98 | chr10-89594710-INS-106 |
| chr10-89610396-INS-87 | chr10-89610373-INS-93 |
| chr10-89621497-INS-144 | chr10-89621497-INS-156 |
| chr10-89638564-INS-104 | chr10-89638564-INS-84 |
| chr10-89642708-INS-2159 | chr10-89642709-INS-2159 |
| chr10-89670575-INS-315 | chr10-89670575-INS-270 |
| chr10-89677733-INS-497 | chr10-89677733-INS-498 |
| chr10-89686938-INS-66 | chr10-89686938-INS-66 |
| chr10-89695954-INS-62 | chr10-89695924-INS-83 |
| chr10-89703234-INS-196 | chr10-89703235-INS-196 |
| chr10-89706429-INS-97 | chr10-89706430-INS-97 |
| chr10-89715855-INS-82 | chr10-89715855-INS-82 |
| chr10-89736738-INS-57 | chr10-89736732-INS-50 |
| chr10-89765430-INS-167 | chr10-89765428-INS-172 |
| chr10-89767398-INS-6453 | chr10-89767399-INS-6453 |
| chr10-89768847-INS-178 | chr10-89768847-INS-178 |
| chr10-89800497-INS-199 | chr10-89800497-INS-199 |
| chr10-89820133-INS-4572 | chr10-89820134-INS-4571 |
| chr10-89839845-INS-571 | chr10-89839845-INS-573 |
| chr10-89841391-INS-54 | chr10-89841391-INS-50 |
| chr10-89845473-INS-52 | chr10-89845491-INS-51 |
| chr10-89870126-INS-56 | chr10-89870126-INS-56 |
| chr10-89886753-INS-1275 | chr10-89886754-INS-1308 |
| chr10-89888229-INS-202 | chr10-89888229-INS-187 |
| chr10-89905174-INS-54 | chr10-89905174-INS-54 |
| chr10-89910574-INS-188 | chr10-89910574-INS-189 |
| chr10-89911421-INS-89 | chr10-89911397-INS-88 |
| chr10-89916854-INS-58 | chr10-89916855-INS-66 |
| chr10-89924236-INS-168 | chr10-89924236-INS-128 |
| chr10-89940753-INS-598 | chr10-89940753-INS-600 |
| chr10-89949093-INS-199 | chr10-89949093-INS-199 |
| chr10-89950011-INS-207 | chr10-89950011-INS-204 |
| chr10-89951299-INS-55 | chr10-89951299-INS-55 |
| chr10-89953663-INS-55 | chr10-89953663-INS-55 |
| chr10-89965837-INS-71 | chr10-89965837-INS-70 |
| chr10-89976847-INS-3252 | chr10-89976848-INS-3252 |
| chr10-89977833-INS-6412 | chr10-89977834-INS-6412 |

| | |
|---|---|
| chr10-89985739-INS-7119 | chr10-89985740-INS-7119 |
| chr10-90033297-INS-514 | chr10-90033298-INS-513 |
| chr10-90037089-INS-57 | chr10-90037051-INS-78 |
| chr10-90051435-INS-226 | chr10-90051436-INS-225 |
| chr10-90092073-INS-100 | chr10-90092073-INS-64 |
| chr10-90112803-INS-890 | chr10-90112799-INS-892 |
| chr10-90118898-INS-346 | chr10-90118898-INS-346 |
| chr10-90120521-INS-333 | chr10-90120521-INS-333 |
| chr10-90140915-INS-130 | chr10-90140915-INS-130 |
| chr10-90189760-INS-63 | chr10-90189760-INS-61 |
| chr10-90190603-INS-735 | chr10-90190603-INS-736 |
| chr10-90201386-INS-119 | chr10-90201386-INS-119 |
| chr10-90208420-INS-77 | chr10-90208420-INS-77 |
| chr10-90215376-INS-241 | chr10-90215376-INS-242 |
| chr10-90219059-INS-923 | chr10-90219059-INS-923 |
| chr10-90231502-INS-466 | chr10-90231502-INS-467 |
| chr10-90260135-INS-167 | chr10-90260135-INS-167 |
| chr10-90263902-INS-1278 | chr10-90263903-INS-1277 |
| chr10-90265108-INS-117 | chr10-90265113-INS-121 |
| chr10-90272703-INS-367 | chr10-90272703-INS-367 |
| chr10-90288441-INS-6234 | chr10-90288442-INS-6234 |
| chr10-90291504-INS-295 | chr10-90291504-INS-295 |
| chr10-90302392-INS-64 | chr10-90302392-INS-72 |
| chr10-90315911-INS-204 | chr10-90315911-INS-204 |
| chr10-90333682-INS-461 | chr10-90333682-INS-461 |
| chr10-90343372-INS-57 | chr10-90343372-INS-57 |
| chr10-90352273-INS-80 | chr10-90352268-INS-85 |
| chr10-90359060-INS-593 | chr10-90359060-INS-593 |
| chr10-90392699-INS-94 | chr10-90392699-INS-96 |
| chr10-90421857-INS-56 | chr10-90421863-INS-88 |
| chr10-90464824-INS-71 | chr10-90464824-INS-83 |
| chr10-90505350-INS-471 | chr10-90505350-INS-471 |
| chr10-90529113-INS-70 | chr10-90529113-INS-70 |
| chr10-90535745-INS-52 | chr10-90535745-INS-52 |
| chr10-90536559-INS-258 | chr10-90536559-INS-259 |
| chr10-90569573-INS-128 | chr10-90569573-INS-88 |
| chr10-90585263-INS-657 | chr10-90585263-INS-659 |
| chr10-90602862-INS-54 | chr10-90602862-INS-51 |
| chr10-90664386-INS-207 | chr10-90664386-INS-207 |
| chr10-90680828-INS-101 | chr10-90680828-INS-101 |
| chr10-90681849-INS-126 | chr10-90681849-INS-131 |
| chr10-90703434-INS-83 | chr10-90703433-INS-79 |
| chr10-90710996-INS-144 | chr10-90710996-INS-144 |
| chr10-90712524-INS-550 | chr10-90712524-INS-549 |
| chr10-90768552-INS-128 | chr10-90768552-INS-128 |
| chr10-90787021-INS-6455 | chr10-90787022-INS-6455 |
| chr10-90820063-INS-5418 | chr10-90820064-INS-5418 |
| chr10-90821377-INS-79 | chr10-90821378-INS-79 |
| chr10-90822183-INS-855 | chr10-90822184-INS-853 |
| chr10-90822354-INS-108 | chr10-90822355-INS-74 |
| chr10-90823711-INS-80 | chr10-90823711-INS-80 |
| chr10-90832043-INS-179 | chr10-90832043-INS-173 |
| chr10-90842885-INS-121 | chr10-90842880-INS-152 |
| chr10-90855355-INS-188 | chr10-90855355-INS-188 |
| chr10-90865691-INS-73 | chr10-90865691-INS-74 |
| chr10-90882456-INS-2444 | chr10-90882457-INS-2448 |
| chr10-90892921-INS-201 | chr10-90892921-INS-201 |
| chr10-90903303-INS-186 | chr10-90903303-INS-186 |
| chr10-90908714-INS-108 | chr10-90908714-INS-108 |
| chr10-90922209-INS-2652 | chr10-90922210-INS-2652 |
| chr10-90925466-INS-116 | chr10-90925466-INS-116 |
| chr10-90936712-INS-236 | chr10-90936712-INS-236 |
| chr10-90938737-INS-210 | chr10-90938737-INS-211 |
| chr10-90941037-INS-187 | chr10-90941038-INS-182 |
| chr10-90941560-INS-193 | chr10-90941561-INS-192 |
| chr10-90942151-INS-224 | chr10-90942152-INS-200 |
| chr10-90942705-INS-159 | chr10-90942706-INS-159 |
| chr10-90943234-INS-246 | chr10-90943235-INS-250 |
| chr10-90944253-INS-209 | chr10-90944253-INS-209 |
| chr10-90961578-INS-7264 | chr10-90961579-INS-7264 |
| chr10-90962393-INS-55 | chr10-90962393-INS-63 |
| chr10-90963985-INS-61 | chr10-90963985-INS-65 |
| chr10-90970184-INS-205 | chr10-90970184-INS-205 |
| chr10-90975803-INS-204 | chr10-90975803-INS-199 |
| chr10-90989835-INS-498 | chr10-90989835-INS-498 |
| chr10-91009182-INS-7366 | chr10-91009183-INS-7366 |
| chr10-91022589-INS-61 | chr10-91022589-INS-61 |

| | |
|---|---|
| chr10-91024588-INS-233 | chr10-91024588-INS-232 |
| chr10-91085190-INS-81 | chr10-91085190-INS-81 |
| chr10-91104948-INS-64 | chr10-91104948-INS-70 |
| chr10-91111846-INS-56 | chr10-91111864-INS-50 |
| chr10-91117633-INS-57 | chr10-91117633-INS-57 |
| chr10-91132026-INS-66 | chr10-91132025-INS-70 |
| chr10-91197865-INS-66 | chr10-91197865-INS-66 |
| chr10-91212902-INS-155 | chr10-91212881-INS-152 |
| chr10-91227944-INS-110 | chr10-91227953-INS-76 |
| chr10-91230551-INS-97 | chr10-91230551-INS-97 |
| chr10-91243713-INS-6421 | chr10-91243714-INS-6421 |
| chr10-91294763-INS-70 | chr10-91294764-INS-58 |
| chr10-91334965-INS-6999 | chr10-91334966-INS-6997 |
| chr10-91339504-INS-162 | chr10-91339502-INS-154 |
| chr10-91347290-INS-1101 | chr10-91347291-INS-1101 |
| chr10-91390340-INS-265 | chr10-91390340-INS-265 |
| chr10-91409672-INS-63 | chr10-91409672-INS-63 |
| chr10-91443806-INS-104 | chr10-91443806-INS-87 |
| chr10-91466748-INS-83 | chr10-91466740-INS-85 |
| chr10-91492795-INS-58 | chr10-91492757-INS-52 |
| chr10-91500185-INS-88 | chr10-91500185-INS-88 |
| chr10-91507756-INS-104 | chr10-91507763-INS-92 |
| chr10-91515456-INS-64 | chr10-91515456-INS-76 |
| chr10-91519875-INS-106 | chr10-91519911-INS-110 |
| chr10-91563383-INS-51 | chr10-91563383-INS-57 |
| chr10-91566909-INS-68 | chr10-91566918-INS-100 |
| chr10-91574208-INS-58 | chr10-91574208-INS-58 |
| chr10-91592716-INS-169 | chr10-91592716-INS-161 |
| chr10-91647081-INS-222 | chr10-91647081-INS-220 |
| chr10-91675186-INS-65 | chr10-91675186-INS-65 |
| chr10-91692063-INS-68 | chr10-91692063-INS-74 |
| chr10-91712620-INS-386 | chr10-91712620-INS-386 |
| chr10-91785184-INS-60 | chr10-91785184-INS-58 |
| chr10-91794698-INS-72 | chr10-91794665-INS-81 |
| chr10-91814343-INS-87 | chr10-91814325-INS-126 |
| chr10-91821148-INS-108 | chr10-91821148-INS-96 |
| chr10-91829352-INS-56 | chr10-91829352-INS-52 |
| chr10-91851943-INS-68 | chr10-91851943-INS-66 |
| chr10-91855800-INS-4579 | chr10-91855801-INS-4579 |
| chr10-91857053-INS-54 | chr10-91857053-INS-54 |
| chr10-91867580-INS-79 | chr10-91867580-INS-79 |
| chr10-91878678-INS-77 | chr10-91878677-INS-70 |
| chr10-91883455-INS-206 | chr10-91883455-INS-206 |
| chr10-91955503-INS-463 | chr10-91955503-INS-463 |
| chr10-91956404-INS-53 | chr10-91956404-INS-53 |
| chr10-91960605-INS-54 | chr10-91960605-INS-64 |
| chr10-91962227-INS-110 | chr10-91962215-INS-128 |
| chr10-91972270-INS-68 | chr10-91972280-INS-73 |
| chr10-91990482-INS-99 | chr10-91990490-INS-100 |
| chr10-92014935-INS-173 | chr10-92014936-INS-185 |
| chr10-92057629-INS-147 | chr10-92057629-INS-163 |
| chr10-92075451-INS-3623 | chr10-92075452-INS-3623 |
| chr10-92086842-INS-498 | chr10-92086842-INS-498 |
| chr10-92087920-INS-51 | chr10-92087920-INS-51 |
| chr10-92104318-INS-82 | chr10-92104318-INS-82 |
| chr10-92123331-INS-66 | chr10-92123331-INS-68 |
| chr10-92124656-INS-65 | chr10-92124656-INS-65 |
| chr10-92135259-INS-60 | chr10-92135259-INS-57 |
| chr10-92139408-INS-154 | chr10-92139408-INS-154 |
| chr10-92167851-INS-64 | chr10-92167851-INS-60 |
| chr10-92171454-INS-66 | chr10-92171454-INS-58 |
| chr10-92189925-INS-54 | chr10-92189925-INS-54 |
| chr10-92198597-INS-394 | chr10-92198597-INS-402 |
| chr10-92254454-INS-434 | chr10-92254454-INS-434 |
| chr10-92270893-INS-60 | chr10-92270893-INS-60 |
| chr10-92295289-INS-101 | chr10-92295292-INS-104 |
| chr10-92303467-INS-2648 | chr10-92303468-INS-2651 |
| chr10-92305087-INS-56 | chr10-92305087-INS-50 |
| chr10-92308887-INS-273 | chr10-92308887-INS-273 |
| chr10-92314159-INS-466 | chr10-92314159-INS-466 |
| chr10-92319958-INS-58 | chr10-92319958-INS-70 |
| chr10-92341215-INS-84 | chr10-92341215-INS-78 |
| chr10-92354202-INS-190 | chr10-92354202-INS-198 |
| chr10-92394780-INS-7152 | chr10-92394781-INS-7153 |
| chr10-92417005-INS-102 | chr10-92417005-INS-104 |
| chr10-92429762-INS-76 | chr10-92429762-INS-84 |
| chr10-92481516-INS-142 | chr10-92481516-INS-142 |

| | |
|---|---|
| chr10-92497720-INS-66 | chr10-92497720-INS-60 |
| chr10-92503579-INS-269 | chr10-92503579-INS-269 |
| chr10-92566761-INS-119 | chr10-92566757-INS-131 |
| chr10-92575142-INS-73 | chr10-92575142-INS-89 |
| chr10-92576353-INS-227 | chr10-92576345-INS-274 |
| chr10-92597312-INS-250 | chr10-92597312-INS-268 |
| chr10-92634243-INS-141 | chr10-92634243-INS-141 |
| chr10-92655911-INS-787 | chr10-92655911-INS-787 |
| chr10-92699739-INS-94 | chr10-92699739-INS-141 |
| chr10-92705903-INS-132 | chr10-92705927-INS-132 |
| chr10-92731111-INS-598 | chr10-92731121-INS-598 |
| chr10-92732402-INS-60 | chr10-92732402-INS-59 |
| chr10-92737014-INS-208 | chr10-92737014-INS-209 |
| chr10-92769365-INS-1434 | chr10-92769366-INS-1435 |
| chr10-92786903-INS-2872 | chr10-92786904-INS-2872 |
| chr10-92786904-INS-508 | chr10-92786904-INS-508 |
| chr10-92803522-INS-52 | chr10-92803522-INS-50 |
| chr10-92805040-INS-136 | chr10-92805040-INS-136 |
| chr10-92824426-INS-152 | chr10-92824427-INS-152 |
| chr10-92829126-INS-328 | chr10-92829130-INS-338 |
| chr10-92851390-INS-1116 | chr10-92851391-INS-1117 |
| chr10-92855264-INS-414 | chr10-92855264-INS-414 |
| chr10-92861169-INS-58 | chr10-92861170-INS-58 |
| chr10-92918537-INS-214 | chr10-92918537-INS-215 |
| chr10-92934926-INS-62 | chr10-92934926-INS-57 |
| chr10-92940008-INS-483 | chr10-92940008-INS-494 |
| chr10-92957381-INS-455 | chr10-92957381-INS-455 |
| chr10-92960512-INS-639 | chr10-92960512-INS-640 |
| chr10-92981579-INS-50 | chr10-92981579-INS-52 |
| chr10-92986500-INS-317 | chr10-92986500-INS-344 |
| chr10-92989557-INS-165 | chr10-92989553-INS-167 |
| chr10-92994090-INS-58 | chr10-92994085-INS-51 |
| chr10-93010422-INS-64 | chr10-93010422-INS-64 |
| chr10-93023285-INS-78 | chr10-93023285-INS-108 |
| chr10-93025496-INS-54 | chr10-93025496-INS-54 |
| chr10-93035858-INS-198 | chr10-93035858-INS-198 |
| chr10-93067506-INS-143 | chr10-93067506-INS-143 |
| chr10-93078074-INS-487 | chr10-93078075-INS-501 |
| chr10-93106840-INS-100 | chr10-93106840-INS-100 |
| chr10-93159396-INS-223 | chr10-93159396-INS-223 |
| chr10-93167282-INS-434 | chr10-93167282-INS-434 |
| chr10-93190850-INS-2632 | chr10-93190851-INS-2632 |
| chr10-93207933-INS-6460 | chr10-93207934-INS-6457 |
| chr10-93228018-INS-105 | chr10-93228018-INS-81 |
| chr10-93243018-INS-60 | chr10-93243018-INS-60 |
| chr10-93277105-INS-97 | chr10-93277105-INS-97 |
| chr10-93281692-INS-52 | chr10-93281692-INS-56 |
| chr10-93315829-INS-131 | chr10-93315829-INS-129 |
| chr10-93319874-INS-80 | chr10-93319874-INS-80 |
| chr10-93349025-INS-807 | chr10-93349025-INS-807 |
| chr10-93356099-INS-60 | chr10-93356099-INS-60 |
| chr10-93366230-INS-604 | chr10-93366230-INS-604 |
| chr10-93427573-INS-55 | chr10-93427573-INS-56 |
| chr10-93430917-INS-457 | chr10-93430917-INS-457 |
| chr10-93457126-INS-57 | chr10-93457126-INS-55 |
| chr10-93468195-INS-1183 | chr10-93468196-INS-1166 |
| chr10-93469240-INS-54 | chr10-93469240-INS-84 |
| chr10-93489160-INS-72 | chr10-93489160-INS-72 |
| chr10-93494915-INS-80 | chr10-93494903-INS-67 |
| chr10-93501098-INS-59 | chr10-93501098-INS-55 |
| chr10-93533913-INS-84 | chr10-93533913-INS-84 |
| chr10-93535239-INS-110 | chr10-93535256-INS-115 |
| chr10-93550172-INS-93 | chr10-93550172-INS-93 |
| chr10-93551637-INS-55 | chr10-93551637-INS-61 |
| chr10-93554480-INS-98 | chr10-93554479-INS-100 |
| chr10-93565729-INS-258 | chr10-93565728-INS-256 |
| chr10-93572638-INS-60 | chr10-93572638-INS-81 |
| chr10-93584802-INS-212 | chr10-93584802-INS-212 |
| chr10-93598636-INS-179 | chr10-93598650-INS-184 |
| chr10-93605508-INS-154 | chr10-93605502-INS-152 |
| chr10-93612099-INS-176 | chr10-93612099-INS-176 |
| chr10-93613821-INS-211 | chr10-93613785-INS-215 |
| chr10-93615495-INS-214 | chr10-93615495-INS-214 |
| chr10-93620310-INS-223 | chr10-93620310-INS-223 |
| chr10-93632152-INS-52 | chr10-93632153-INS-54 |
| chr10-93652083-INS-84 | chr10-93652083-INS-72 |
| chr10-93657727-INS-112 | chr10-93657727-INS-125 |

| | |
|---|---|
| chr10-93674977-INS-194 | chr10-93674977-INS-197 |
| chr10-93675669-INS-509 | chr10-93675669-INS-509 |
| chr10-93700774-INS-209 | chr10-93700774-INS-210 |
| chr10-93730083-INS-2435 | chr10-93730084-INS-2435 |
| chr10-93730946-INS-61 | chr10-93730946-INS-61 |
| chr10-93731598-INS-221 | chr10-93731598-INS-223 |
| chr10-93746283-INS-226 | chr10-93746283-INS-214 |
| chr10-93747257-INS-164 | chr10-93747257-INS-176 |
| chr10-93789131-INS-52 | chr10-93789131-INS-53 |
| chr10-93791980-INS-65 | chr10-93791980-INS-65 |
| chr10-93803102-INS-60 | chr10-93803102-INS-60 |
| chr10-93818854-INS-93 | chr10-93818854-INS-93 |
| chr10-93852145-INS-285 | chr10-93852145-INS-285 |
| chr10-93852466-INS-496 | chr10-93852466-INS-485 |
| chr10-93853578-INS-238 | chr10-93853578-INS-238 |
| chr10-93858999-INS-209 | chr10-93858999-INS-209 |
| chr10-93896668-INS-89 | chr10-93896667-INS-80 |
| chr10-93899982-INS-203 | chr10-93899982-INS-202 |
| chr10-93938488-INS-63 | chr10-93938488-INS-63 |
| chr10-93949098-INS-5219 | chr10-93949099-INS-5219 |
| chr10-93954942-INS-80 | chr10-93954936-INS-68 |
| chr10-93957679-INS-1210 | chr10-93957680-INS-1211 |
| chr10-93974760-INS-203 | chr10-93974760-INS-203 |
| chr10-94006592-INS-56 | chr10-94006592-INS-56 |
| chr10-94020051-INS-182 | chr10-94020051-INS-182 |
| chr10-94055907-INS-160 | chr10-94055907-INS-158 |
| chr10-94056874-INS-53 | chr10-94056874-INS-53 |
| chr10-94058255-INS-93 | chr10-94058255-INS-93 |
| chr10-94069698-INS-69 | chr10-94069698-INS-57 |
| chr10-94113105-INS-52 | chr10-94113105-INS-52 |
| chr10-94119418-INS-74 | chr10-94119418-INS-74 |
| chr10-94129086-INS-55 | chr10-94129086-INS-57 |
| chr10-94132498-INS-260 | chr10-94132498-INS-260 |
| chr10-94139060-INS-827 | chr10-94139060-INS-829 |
| chr10-94174955-INS-165 | chr10-94174955-INS-165 |
| chr10-94190621-INS-78 | chr10-94190621-INS-78 |
| chr10-94237819-INS-4763 | chr10-94237820-INS-4763 |
| chr10-94271526-INS-126 | chr10-94271526-INS-126 |
| chr10-94316962-INS-192 | chr10-94316962-INS-198 |
| chr10-94334412-INS-64 | chr10-94334412-INS-64 |
| chr10-94364338-INS-56 | chr10-94364338-INS-52 |
| chr10-94399308-INS-174 | chr10-94399308-INS-174 |
| chr10-94400396-INS-416 | chr10-94400396-INS-416 |
| chr10-94409904-INS-242 | chr10-94409904-INS-242 |
| chr10-94429272-INS-84 | chr10-94429266-INS-66 |
| chr10-94452495-INS-2859 | chr10-94452496-INS-2859 |
| chr10-94452540-INS-60 | chr10-94452540-INS-56 |
| chr10-94460728-INS-85 | chr10-94460726-INS-90 |
| chr10-94470176-INS-355 | chr10-94470176-INS-355 |
| chr10-94505208-INS-613 | chr10-94505208-INS-613 |
| chr10-94521168-INS-4707 | chr10-94521169-INS-4707 |
| chr10-94568800-INS-54 | chr10-94568800-INS-54 |
| chr10-94578008-INS-1073 | chr10-94578009-INS-1073 |
| chr10-94578823-INS-73 | chr10-94578819-INS-55 |
| chr10-94600168-INS-521 | chr10-94600168-INS-522 |
| chr10-94623428-INS-107 | chr10-94623428-INS-111 |
| chr10-94624765-INS-414 | chr10-94624765-INS-417 |
| chr10-94707134-INS-80 | chr10-94707134-INS-80 |
| chr10-94711721-INS-85 | chr10-94711721-INS-52 |
| chr10-94729400-INS-854 | chr10-94729414-INS-856 |
| chr10-94737916-INS-150 | chr10-94737916-INS-150 |
| chr10-94777207-INS-64 | chr10-94777207-INS-64 |
| chr10-94811662-INS-963 | chr10-94811662-INS-963 |
| chr10-94835105-INS-79 | chr10-94835105-INS-79 |
| chr10-94840217-INS-56 | chr10-94840217-INS-64 |
| chr10-94875295-INS-78 | chr10-94875281-INS-59 |
| chr10-94921560-INS-321 | chr10-94921560-INS-321 |
| chr10-94927949-INS-270 | chr10-94927949-INS-270 |
| chr10-94930094-INS-401 | chr10-94930094-INS-401 |
| chr10-94970801-INS-60 | chr10-94970801-INS-64 |
| chr10-94989477-INS-3002 | chr10-94989478-INS-3002 |
| chr10-95010667-INS-1625 | chr10-95010668-INS-1611 |
| chr10-95032784-INS-66 | chr10-95032784-INS-66 |
| chr10-95037115-INS-302 | chr10-95037115-INS-304 |
| chr10-95043088-INS-75 | chr10-95043088-INS-79 |
| chr10-95046389-INS-60 | chr10-95046389-INS-56 |
| chr10-95049807-INS-426 | chr10-95049807-INS-426 |

| | |
|---|---|
| chr10-95051768-INS-1473 | chr10-95051769-INS-1474 |
| chr10-95073185-INS-7116 | chr10-95073186-INS-7116 |
| chr10-95104665-INS-114 | chr10-95104665-INS-114 |
| chr10-95125121-INS-56 | chr10-95125121-INS-56 |
| chr10-95134136-INS-245 | chr10-95134136-INS-245 |
| chr10-95136956-INS-69 | chr10-95136956-INS-70 |
| chr10-95178560-INS-52 | chr10-95178585-INS-64 |
| chr10-95188217-INS-284 | chr10-95188217-INS-284 |
| chr10-95205878-INS-59 | chr10-95205884-INS-51 |
| chr10-95216724-INS-59 | chr10-95216724-INS-59 |
| chr10-95242183-INS-67 | chr10-95242183-INS-78 |
| chr10-95269873-INS-200 | chr10-95269873-INS-202 |
| chr10-95280617-INS-196 | chr10-95280617-INS-203 |
| chr10-95282378-INS-210 | chr10-95282378-INS-210 |
| chr10-95283438-INS-86 | chr10-95283433-INS-92 |
| chr10-95335052-INS-92 | chr10-95335052-INS-62 |
| chr10-95338994-INS-99 | chr10-95338994-INS-95 |
| chr10-95340065-INS-185 | chr10-95340065-INS-185 |
| chr10-95359480-INS-212 | chr10-95359480-INS-212 |
| chr10-95373843-INS-983 | chr10-95373843-INS-983 |
| chr10-95386548-INS-212 | chr10-95386548-INS-212 |
| chr10-95387064-INS-62 | chr10-95387064-INS-62 |
| chr10-95391526-INS-78 | chr10-95391526-INS-78 |
| chr10-95410373-INS-6401 | chr10-95410374-INS-6401 |
| chr10-95425961-INS-5549 | chr10-95425962-INS-5549 |
| chr10-95434894-INS-71 | chr10-95434871-INS-69 |
| chr10-95449941-INS-811 | chr10-95449941-INS-820 |
| chr10-95474665-INS-70 | chr10-95474698-INS-82 |
| chr10-95487639-INS-63 | chr10-95487635-INS-66 |
| chr10-95509910-INS-73 | chr10-95509910-INS-70 |
| chr10-95535209-INS-52 | chr10-95535209-INS-52 |
| chr10-95541408-INS-68 | chr10-95541408-INS-74 |
| chr10-95547231-INS-54 | chr10-95547231-INS-72 |
| chr10-95555317-INS-54 | chr10-95555317-INS-54 |
| chr10-95619223-INS-70 | chr10-95619223-INS-62 |
| chr10-95621242-INS-110 | chr10-95621228-INS-102 |
| chr10-95636115-INS-183 | chr10-95636101-INS-216 |
| chr10-95650655-INS-400 | chr10-95650655-INS-400 |
| chr10-95671613-INS-52 | chr10-95671613-INS-56 |
| chr10-95673994-INS-202 | chr10-95673994-INS-202 |
| chr10-95692569-INS-56 | chr10-95692569-INS-56 |
| chr10-95695686-INS-187 | chr10-95695686-INS-188 |
| chr10-95702856-INS-62 | chr10-95702856-INS-62 |
| chr10-95711369-INS-230 | chr10-95711369-INS-231 |
| chr10-95740211-INS-894 | chr10-95740210-INS-849 |
| chr10-95769211-INS-399 | chr10-95769211-INS-399 |
| chr10-95785902-INS-56 | chr10-95785902-INS-58 |
| chr10-95789669-INS-116 | chr10-95789669-INS-120 |
| chr10-95796105-INS-6484 | chr10-95796106-INS-6479 |
| chr10-95805766-INS-64 | chr10-95805766-INS-64 |
| chr10-95812589-INS-214 | chr10-95812589-INS-214 |
| chr10-95911413-INS-1444 | chr10-95911414-INS-1433 |
| chr10-95926838-INS-417 | chr10-95926838-INS-423 |
| chr10-95952382-INS-6019 | chr10-95952383-INS-6021 |
| chr10-96055758-INS-58 | chr10-96055758-INS-60 |
| chr10-96060838-INS-62 | chr10-96060838-INS-62 |
| chr10-96069991-INS-54 | chr10-96069991-INS-56 |
| chr10-96075576-INS-918 | chr10-96075576-INS-918 |
| chr10-96090138-INS-96 | chr10-96090138-INS-96 |
| chr10-96133217-INS-138 | chr10-96133217-INS-92 |
| chr10-96143716-INS-100 | chr10-96143716-INS-63 |
| chr10-96150489-INS-76 | chr10-96150489-INS-76 |
| chr10-96153326-INS-97 | chr10-96153344-INS-112 |
| chr10-96171434-INS-498 | chr10-96171435-INS-498 |
| chr10-96185706-INS-506 | chr10-96185707-INS-506 |
| chr10-96213440-INS-184 | chr10-96213440-INS-184 |
| chr10-96243144-INS-61 | chr10-96243144-INS-61 |
| chr10-96256244-INS-122 | chr10-96256244-INS-124 |
| chr10-96259364-INS-57 | chr10-96259364-INS-57 |
| chr10-96268784-INS-634 | chr10-96268784-INS-633 |
| chr10-96270300-INS-76 | chr10-96270300-INS-76 |
| chr10-96326281-INS-258 | chr10-96326281-INS-214 |
| chr10-96376709-INS-132 | chr10-96376709-INS-110 |
| chr10-96388647-INS-173 | chr10-96388647-INS-173 |
| chr10-96408882-INS-351 | chr10-96408882-INS-352 |
| chr10-96411829-INS-346 | chr10-96411807-INS-368 |
| chr10-96429600-INS-51 | chr10-96429600-INS-51 |

| | |
|---|---|
| chr10-96437278-INS-210 | chr10-96437278-INS-210 |
| chr10-96513227-INS-58 | chr10-96513227-INS-58 |
| chr10-96546508-INS-130 | chr10-96546508-INS-138 |
| chr10-96554998-INS-250 | chr10-96554979-INS-247 |
| chr10-96563019-INS-71 | chr10-96563019-INS-71 |
| chr10-96586582-INS-223 | chr10-96586582-INS-223 |
| chr10-96602275-INS-156 | chr10-96602275-INS-108 |
| chr10-96610917-INS-167 | chr10-96610917-INS-167 |
| chr10-96629813-INS-61 | chr10-96629813-INS-61 |
| chr10-96631829-INS-408 | chr10-96631829-INS-418 |
| chr10-96645903-INS-52 | chr10-96645903-INS-52 |
| chr10-96657114-INS-391 | chr10-96657114-INS-391 |
| chr10-96724326-INS-66 | chr10-96724326-INS-66 |
| chr10-96755612-INS-79 | chr10-96755612-INS-66 |
| chr10-96800371-INS-7058 | chr10-96800372-INS-7058 |
| chr10-96888566-INS-52 | chr10-96888568-INS-52 |
| chr10-96895605-INS-986 | chr10-96895608-INS-991 |
| chr10-96982605-INS-635 | chr10-96982603-INS-644 |
| chr10-96985856-INS-6190 | chr10-96985857-INS-6190 |
| chr10-96989141-INS-6505 | chr10-96989142-INS-6505 |
| chr10-96993741-INS-79 | chr10-96993741-INS-62 |
| chr10-97026282-INS-903 | chr10-97026282-INS-889 |
| chr10-97034685-INS-122 | chr10-97034685-INS-106 |
| chr10-97039033-INS-865 | chr10-97039033-INS-864 |
| chr10-97040775-INS-6164 | chr10-97040776-INS-6164 |
| chr10-97093010-INS-1491 | chr10-97093011-INS-1491 |
| chr10-97152778-INS-105 | chr10-97152762-INS-134 |
| chr10-97170659-INS-85 | chr10-97170659-INS-86 |
| chr10-97176113-INS-80 | chr10-97176113-INS-80 |
| chr10-97190250-INS-4958 | chr10-97190251-INS-4958 |
| chr10-97198952-INS-498 | chr10-97198952-INS-492 |
| chr10-97216135-INS-83 | chr10-97216135-INS-83 |
| chr10-97218651-INS-52 | chr10-97218669-INS-52 |
| chr10-97220481-INS-836 | chr10-97220480-INS-849 |
| chr10-97227030-INS-173 | chr10-97227030-INS-173 |
| chr10-97249921-INS-967 | chr10-97249921-INS-966 |
| chr10-97251117-INS-168 | chr10-97251108-INS-169 |
| chr10-97259889-INS-69 | chr10-97259889-INS-53 |
| chr10-97267346-INS-565 | chr10-97267346-INS-565 |
| chr10-97269196-INS-80 | chr10-97269196-INS-80 |
| chr10-97273082-INS-327 | chr10-97273082-INS-327 |
| chr10-97276515-INS-393 | chr10-97276515-INS-359 |
| chr10-97333426-INS-57 | chr10-97333426-INS-80 |
| chr10-97336439-INS-60 | chr10-97336439-INS-58 |
| chr10-97383272-INS-276 | chr10-97383272-INS-276 |
| chr10-97389589-INS-63 | chr10-97389589-INS-59 |
| chr10-97438709-INS-3785 | chr10-97438710-INS-3785 |
| chr10-97450986-INS-190 | chr10-97450986-INS-226 |
| chr10-97472438-INS-335 | chr10-97472438-INS-335 |
| chr10-97526754-INS-72 | chr10-97526754-INS-60 |
| chr10-97564884-INS-154 | chr10-97564900-INS-182 |
| chr10-97572581-INS-505 | chr10-97572575-INS-516 |
| chr10-97579629-INS-520 | chr10-97579637-INS-520 |
| chr10-97582101-INS-54 | chr10-97582101-INS-52 |
| chr10-97605686-INS-1371 | chr10-97605687-INS-1371 |
| chr10-97606326-INS-2293 | chr10-97606327-INS-2293 |
| chr10-97616857-INS-828 | chr10-97616857-INS-828 |
| chr10-97620241-INS-6460 | chr10-97620242-INS-6462 |
| chr10-97621173-INS-51 | chr10-97621173-INS-55 |
| chr10-97624090-INS-114 | chr10-97624105-INS-111 |
| chr10-97638805-INS-423 | chr10-97638805-INS-429 |
| chr10-97652153-INS-52 | chr10-97652153-INS-52 |
| chr10-97686134-INS-92 | chr10-97686134-INS-92 |
| chr10-97697288-INS-334 | chr10-97697288-INS-334 |
| chr10-97708332-INS-69 | chr10-97708332-INS-71 |
| chr10-97740691-INS-63 | chr10-97740691-INS-63 |
| chr10-97794605-INS-77 | chr10-97794605-INS-59 |
| chr10-97803218-INS-76 | chr10-97803170-INS-106 |
| chr10-97808656-INS-111 | chr10-97808656-INS-111 |
| chr10-97844770-INS-554 | chr10-97844770-INS-554 |
| chr10-97863465-INS-56 | chr10-97863465-INS-60 |
| chr10-97866896-INS-59 | chr10-97866896-INS-59 |
| chr10-97873372-INS-54 | chr10-97873372-INS-60 |
| chr10-97879008-INS-6592 | chr10-97879009-INS-6593 |
| chr10-97956314-INS-251 | chr10-97956314-INS-251 |
| chr10-97960110-INS-108 | chr10-97960108-INS-103 |
| chr10-98050119-INS-234 | chr10-98050119-INS-235 |

| | |
|---|---|
| chr10-98088204-INS-61 | chr10-98088204-INS-60 |
| chr10-98133892-INS-203 | chr10-98133860-INS-195 |
| chr10-98135535-INS-62 | chr10-98135535-INS-62 |
| chr10-98138560-INS-60 | chr10-98138560-INS-64 |
| chr10-98149295-INS-78 | chr10-98149295-INS-81 |
| chr10-98158557-INS-97 | chr10-98158557-INS-97 |
| chr10-98166859-INS-122 | chr10-98166858-INS-132 |
| chr10-98250277-INS-54 | chr10-98250277-INS-64 |
| chr10-98290648-INS-58 | chr10-98290648-INS-58 |
| chr10-98304551-INS-318 | chr10-98304551-INS-318 |
| chr10-98321259-INS-6207 | chr10-98321260-INS-6207 |
| chr10-98327811-INS-63 | chr10-98327832-INS-70 |
| chr10-98347186-INS-58 | chr10-98347186-INS-58 |
| chr10-98436276-INS-214 | chr10-98436276-INS-214 |
| chr10-98477641-INS-194 | chr10-98477641-INS-194 |
| chr10-98480099-INS-109 | chr10-98480099-INS-110 |
| chr10-98557499-INS-96 | chr10-98557499-INS-100 |
| chr10-98628463-INS-859 | chr10-98628463-INS-859 |
| chr10-98641797-INS-343 | chr10-98641797-INS-343 |
| chr10-98648955-INS-74 | chr10-98648955-INS-74 |
| chr10-98686840-INS-177 | chr10-98686840-INS-176 |
| chr10-98691081-INS-91 | chr10-98691078-INS-88 |
| chr10-98699569-INS-135 | chr10-98699569-INS-141 |
| chr10-98721885-INS-54 | chr10-98721885-INS-52 |
| chr10-98726259-INS-62 | chr10-98726259-INS-66 |
| chr10-98728062-INS-56 | chr10-98728062-INS-58 |
| chr10-98856083-INS-54 | chr10-98856081-INS-56 |
| chr10-98890819-INS-112 | chr10-98890819-INS-120 |
| chr10-98962834-INS-69 | chr10-98962834-INS-69 |
| chr10-98973666-INS-493 | chr10-98973666-INS-493 |
| chr10-98991656-INS-85 | chr10-98991656-INS-85 |
| chr10-99002772-INS-75 | chr10-99002772-INS-75 |
| chr10-99006902-INS-154 | chr10-99006902-INS-154 |
| chr10-99044577-INS-95 | chr10-99044577-INS-95 |
| chr10-99148684-INS-155 | chr10-99148664-INS-176 |
| chr10-99171558-INS-90 | chr10-99171558-INS-90 |
| chr10-99198309-INS-192 | chr10-99198309-INS-192 |
| chr10-99217366-INS-73 | chr10-99217366-INS-73 |
| chr10-99310709-INS-65 | chr10-99310713-INS-51 |
| chr10-99344119-INS-126 | chr10-99344119-INS-126 |
| chr10-99371545-INS-462 | chr10-99371545-INS-462 |
| chr10-99393084-INS-54 | chr10-99393065-INS-60 |
| chr10-99399886-INS-102 | chr10-99399850-INS-89 |
| chr10-99407806-INS-84 | chr10-99407806-INS-84 |
| chr10-99415824-INS-132 | chr10-99415824-INS-140 |
| chr10-99418007-INS-744 | chr10-99418007-INS-744 |
| chr10-99418987-INS-228 | chr10-99418987-INS-229 |
| chr10-99442047-INS-69 | chr10-99442048-INS-71 |
| chr10-99447484-INS-138 | chr10-99447484-INS-122 |
| chr10-99457043-INS-6670 | chr10-99457044-INS-6670 |
| chr10-99470859-INS-52 | chr10-99470859-INS-52 |
| chr10-99481591-INS-2092 | chr10-99481592-INS-2093 |
| chr10-99510517-INS-237 | chr10-99510517-INS-246 |
| chr10-99554402-INS-84 | chr10-99554402-INS-84 |
| chr10-99587459-INS-68 | chr10-99587459-INS-56 |
| chr10-99591460-INS-73 | chr10-99591460-INS-73 |
| chr10-99603898-INS-61 | chr10-99603898-INS-52 |
| chr10-99612768-INS-56 | chr10-99612768-INS-54 |
| chr10-99615456-INS-123 | chr10-99615456-INS-123 |
| chr10-99632910-INS-939 | chr10-99632910-INS-939 |
| chr10-99650834-INS-139 | chr10-99650834-INS-139 |
| chr10-99654873-INS-6474 | chr10-99654874-INS-6471 |
| chr10-99667857-INS-72 | chr10-99667857-INS-86 |
| chr10-99669405-INS-68 | chr10-99669405-INS-68 |
| chr10-99714361-INS-88 | chr10-99714361-INS-88 |
| chr10-99722797-INS-1862 | chr10-99722798-INS-1862 |
| chr10-99764545-INS-71 | chr10-99764560-INS-52 |
| chr10-99798526-INS-56 | chr10-99798526-INS-56 |
| chr10-99805291-INS-3603 | chr10-99805292-INS-3603 |
| chr10-99806852-INS-94 | chr10-99806852-INS-94 |
| chr10-99814318-INS-268 | chr10-99814341-INS-284 |
| chr10-99817702-INS-92 | chr10-99817702-INS-92 |
| chr10-99832103-INS-82 | chr10-99832103-INS-82 |
| chr10-99845486-INS-444 | chr10-99845486-INS-443 |
| chr10-99852709-INS-70 | chr10-99852708-INS-111 |
| chr10-99859543-INS-68 | chr10-99859542-INS-102 |
| chr10-99861117-INS-789 | chr10-99861117-INS-781 |

| | |
|---|---|
| chr10-99886972-INS-57 | chr10-99886983-INS-58 |
| chr10-99904571-INS-535 | chr10-99904572-INS-546 |
| chr10-99908988-INS-119 | chr10-99908981-INS-80 |
| chr10-99940634-INS-292 | chr10-99940634-INS-292 |
| chr10-99943481-INS-76 | chr10-99943461-INS-62 |
| chr10-99985497-INS-307 | chr10-99985497-INS-307 |
| chr10-100005972-INS-312 | chr10-100005972-INS-312 |
| chr10-100075763-INS-201 | chr10-100075763-INS-201 |
| chr10-100100862-INS-995 | chr10-100100862-INS-995 |
| chr10-100128168-INS-349 | chr10-100128168-INS-349 |
| chr10-100293508-INS-57 | chr10-100293508-INS-57 |
| chr10-100298854-INS-246 | chr10-100298854-INS-270 |
| chr10-100307456-INS-88 | chr10-100307456-INS-87 |
| chr10-100309872-INS-68 | chr10-100309872-INS-68 |
| chr10-100330409-INS-208 | chr10-100330396-INS-211 |
| chr10-100426046-INS-56 | chr10-100426046-INS-56 |
| chr10-100434073-INS-198 | chr10-100434073-INS-198 |
| chr10-100438989-INS-5450 | chr10-100438990-INS-5450 |
| chr10-100444365-INS-80 | chr10-100444365-INS-80 |
| chr10-100478326-INS-412 | chr10-100478326-INS-412 |
| chr10-100505113-INS-3658 | chr10-100505114-INS-3657 |
| chr10-100542786-INS-51 | chr10-100542786-INS-52 |
| chr10-100616031-INS-564 | chr10-100616031-INS-564 |
| chr10-100622867-INS-292 | chr10-100622867-INS-292 |
| chr10-100657770-INS-453 | chr10-100657770-INS-453 |
| chr10-100707422-INS-4710 | chr10-100707428-INS-4703 |
| chr10-100724550-INS-8831 | chr10-100724551-INS-8832 |
| chr10-100726633-INS-2818 | chr10-100726634-INS-2818 |
| chr10-100739337-INS-58 | chr10-100739337-INS-55 |
| chr10-100755363-INS-56 | chr10-100755363-INS-58 |
| chr10-100756169-INS-60 | chr10-100756181-INS-63 |
| chr10-100766318-INS-6456 | chr10-100766319-INS-6456 |
| chr10-100796411-INS-418 | chr10-100796411-INS-418 |
| chr10-100804800-INS-6210 | chr10-100804801-INS-6210 |
| chr10-100819125-INS-63 | chr10-100819125-INS-65 |
| chr10-100826851-INS-54 | chr10-100826851-INS-54 |
| chr10-100846438-INS-170 | chr10-100846438-INS-170 |
| chr10-100864817-INS-3568 | chr10-100864818-INS-3594 |
| chr10-100874077-INS-56 | chr10-100874077-INS-52 |
| chr10-100875746-INS-481 | chr10-100875746-INS-481 |
| chr10-100878475-INS-56 | chr10-100878475-INS-56 |
| chr10-100905295-INS-64 | chr10-100905326-INS-50 |
| chr10-100910326-INS-51 | chr10-100910326-INS-51 |
| chr10-100928116-INS-3536 | chr10-100928117-INS-3537 |
| chr10-100956667-INS-56 | chr10-100956667-INS-58 |
| chr10-100974336-INS-50 | chr10-100974344-INS-56 |
| chr10-100984066-INS-57 | chr10-100984066-INS-57 |
| chr10-101006826-INS-522 | chr10-101006826-INS-522 |
| chr10-101055204-INS-249 | chr10-101055204-INS-249 |
| chr10-101058041-INS-1482 | chr10-101058042-INS-1481 |
| chr10-101128641-INS-3931 | chr10-101128642-INS-3931 |
| chr10-101134112-INS-408 | chr10-101134112-INS-372 |
| chr10-101142379-INS-1062 | chr10-101142380-INS-1062 |
| chr10-101272051-INS-65 | chr10-101272051-INS-77 |
| chr10-101293640-INS-56 | chr10-101293640-INS-56 |
| chr10-101297660-INS-106 | chr10-101297658-INS-110 |
| chr10-101321279-INS-327 | chr10-101321279-INS-307 |
| chr10-101340011-INS-94 | chr10-101340011-INS-94 |
| chr10-101397649-INS-391 | chr10-101397649-INS-391 |
| chr10-101484341-INS-383 | chr10-101484341-INS-383 |
| chr10-101534830-INS-138 | chr10-101534830-INS-136 |
| chr10-101537819-INS-711 | chr10-101537826-INS-719 |
| chr10-101538533-INS-63 | chr10-101538543-INS-85 |
| chr10-101616883-INS-92 | chr10-101616883-INS-92 |
| chr10-101618349-INS-76 | chr10-101618349-INS-78 |
| chr10-101662591-INS-505 | chr10-101662591-INS-505 |
| chr10-101689717-INS-60 | chr10-101689717-INS-60 |
| chr10-101717063-INS-7138 | chr10-101717064-INS-7138 |
| chr10-101718157-INS-222 | chr10-101718157-INS-222 |
| chr10-101774697-INS-360 | chr10-101774697-INS-360 |
| chr10-101782640-INS-95 | chr10-101782640-INS-95 |
| chr10-101795921-INS-227 | chr10-101795921-INS-227 |
| chr10-101803128-INS-2710 | chr10-101803129-INS-2710 |
| chr10-101805978-INS-721 | chr10-101805973-INS-729 |
| chr10-101816545-INS-52 | chr10-101816545-INS-58 |
| chr10-101819847-INS-160 | chr10-101819847-INS-160 |
| chr10-101836317-INS-725 | chr10-101836317-INS-726 |

| | |
|---|---|
| chr10-101843894-INS-1332 | chr10-101843895-INS-1332 |
| chr10-101897794-INS-454 | chr10-101897794-INS-462 |
| chr10-101905206-INS-116 | chr10-101905206-INS-118 |
| chr10-101933082-INS-87 | chr10-101933105-INS-51 |
| chr10-101944909-INS-6389 | chr10-101944910-INS-6390 |
| chr10-101953767-INS-79 | chr10-101953767-INS-79 |
| chr10-101955382-INS-83 | chr10-101955382-INS-71 |
| chr10-101961717-INS-248 | chr10-101961717-INS-248 |
| chr10-102000496-INS-3100 | chr10-102000497-INS-3100 |
| chr10-102012277-INS-107 | chr10-102012260-INS-80 |
| chr10-102019851-INS-119 | chr10-102019851-INS-121 |
| chr10-102050866-INS-134 | chr10-102050866-INS-140 |
| chr10-102079583-INS-57 | chr10-102079583-INS-59 |
| chr10-102085665-INS-408 | chr10-102085665-INS-408 |
| chr10-102093285-INS-983 | chr10-102093285-INS-974 |
| chr10-102094965-INS-135 | chr10-102094965-INS-137 |
| chr10-102122088-INS-54 | chr10-102122099-INS-51 |
| chr10-102130422-INS-252 | chr10-102130422-INS-258 |
| chr10-102146887-INS-396 | chr10-102146887-INS-396 |
| chr10-102178355-INS-54 | chr10-102178355-INS-70 |
| chr10-102199200-INS-2745 | chr10-102199201-INS-2745 |
| chr10-102214374-INS-62 | chr10-102214332-INS-82 |
| chr10-102262275-INS-98 | chr10-102262275-INS-100 |
| chr10-102284883-INS-52 | chr10-102284886-INS-52 |
| chr10-102308399-INS-668 | chr10-102308399-INS-668 |
| chr10-102310651-INS-6794 | chr10-102310652-INS-6796 |
| chr10-102315405-INS-54 | chr10-102315429-INS-60 |
| chr10-102321376-INS-80 | chr10-102321376-INS-80 |
| chr10-102353774-INS-58 | chr10-102353774-INS-58 |
| chr10-102376913-INS-423 | chr10-102376913-INS-423 |
| chr10-102383797-INS-78 | chr10-102383797-INS-78 |
| chr10-102421728-INS-74 | chr10-102421716-INS-59 |
| chr10-102432907-INS-99 | chr10-102432904-INS-102 |
| chr10-102454006-INS-80 | chr10-102454007-INS-76 |
| chr10-102456626-INS-3758 | chr10-102456627-INS-3758 |
| chr10-102462657-INS-55 | chr10-102462657-INS-57 |
| chr10-102492566-INS-78 | chr10-102492566-INS-76 |
| chr10-102498643-INS-134 | chr10-102498643-INS-134 |
| chr10-102523720-INS-135 | chr10-102523720-INS-131 |
| chr10-102525372-INS-78 | chr10-102525372-INS-78 |
| chr10-102536463-INS-98 | chr10-102536463-INS-106 |
| chr10-102552601-INS-90 | chr10-102552601-INS-90 |
| chr10-102556472-INS-66 | chr10-102556472-INS-70 |
| chr10-102579033-INS-54 | chr10-102579033-INS-54 |
| chr10-102590266-INS-3413 | chr10-102590267-INS-3414 |
| chr10-102594631-INS-483 | chr10-102594631-INS-483 |
| chr10-102617142-INS-204 | chr10-102617142-INS-204 |
| chr10-102636393-INS-115 | chr10-102636393-INS-115 |
| chr10-102650953-INS-1252 | chr10-102650954-INS-1252 |
| chr10-102653191-INS-85 | chr10-102653191-INS-85 |
| chr10-102655847-INS-89 | chr10-102655847-INS-91 |
| chr10-102670547-INS-5318 | chr10-102670548-INS-5318 |
| chr10-102672499-INS-112 | chr10-102672456-INS-115 |
| chr10-102679880-INS-450 | chr10-102679883-INS-454 |
| chr10-102680561-INS-52 | chr10-102680561-INS-52 |
| chr10-102741210-INS-107 | chr10-102741210-INS-107 |
| chr10-102741660-INS-3105 | chr10-102741661-INS-3105 |
| chr10-102743704-INS-258 | chr10-102743704-INS-260 |
| chr10-102767884-INS-54 | chr10-102767884-INS-58 |
| chr10-102823381-INS-718 | chr10-102823381-INS-718 |
| chr10-102850812-INS-123 | chr10-102850812-INS-123 |
| chr10-102884489-INS-127 | chr10-102884489-INS-135 |
| chr10-102890358-INS-56 | chr10-102890334-INS-94 |
| chr10-102894267-INS-280 | chr10-102894258-INS-288 |
| chr10-102906524-INS-203 | chr10-102906524-INS-203 |
| chr10-102929885-INS-556 | chr10-102929885-INS-556 |
| chr10-102931194-INS-322 | chr10-102931194-INS-323 |
| chr10-102936919-INS-139 | chr10-102936919-INS-139 |
| chr10-102941092-INS-283 | chr10-102941092-INS-283 |
| chr10-102977673-INS-397 | chr10-102977673-INS-397 |
| chr10-102989637-INS-68 | chr10-102989637-INS-70 |
| chr10-103011440-INS-607 | chr10-103011440-INS-607 |
| chr10-103013311-INS-254 | chr10-103013311-INS-221 |
| chr10-103015822-INS-478 | chr10-103015822-INS-478 |
| chr10-103052469-INS-707 | chr10-103052469-INS-706 |
| chr10-103057616-INS-145 | chr10-103057616-INS-145 |
| chr10-103080452-INS-224 | chr10-103080452-INS-226 |

| | |
|---|---|
| chr10-103081051-INS-636 | chr10-103081051-INS-636 |
| chr10-103085144-INS-583 | chr10-103085144-INS-583 |
| chr10-103086568-INS-53 | chr10-103086568-INS-53 |
| chr10-103103753-INS-205 | chr10-103103753-INS-205 |
| chr10-103116626-INS-529 | chr10-103116626-INS-529 |
| chr10-103120950-INS-52 | chr10-103120950-INS-54 |
| chr10-103132045-INS-485 | chr10-103132045-INS-486 |
| chr10-103136354-INS-80 | chr10-103136354-INS-103 |
| chr10-103138438-INS-52 | chr10-103138445-INS-66 |
| chr10-103140754-INS-1544 | chr10-103140755-INS-1545 |
| chr10-103155112-INS-1218 | chr10-103155113-INS-1218 |
| chr10-103163055-INS-945 | chr10-103163055-INS-946 |
| chr10-103179226-INS-451 | chr10-103179226-INS-451 |
| chr10-103198181-INS-94 | chr10-103198181-INS-94 |
| chr10-103200492-INS-133 | chr10-103200492-INS-133 |
| chr10-103237597-INS-100 | chr10-103237597-INS-100 |
| chr10-103261262-INS-185 | chr10-103261262-INS-186 |
| chr10-103267573-INS-390 | chr10-103267573-INS-390 |
| chr10-103279717-INS-136 | chr10-103279717-INS-136 |
| chr10-103294717-INS-856 | chr10-103294717-INS-857 |
| chr10-103317310-INS-130 | chr10-103317294-INS-130 |
| chr10-103320567-INS-126 | chr10-103320567-INS-126 |
| chr10-103322001-INS-229 | chr10-103322001-INS-229 |
| chr10-103338159-INS-665 | chr10-103338159-INS-665 |
| chr10-103360978-INS-229 | chr10-103360979-INS-229 |
| chr10-103384210-INS-410 | chr10-103384210-INS-410 |
| chr10-103392729-INS-552 | chr10-103392729-INS-552 |
| chr10-103393565-INS-72 | chr10-103393565-INS-66 |
| chr10-103400787-INS-66 | chr10-103400787-INS-66 |
| chr10-103405303-INS-359 | chr10-103405303-INS-359 |
| chr10-103414904-INS-151 | chr10-103414904-INS-146 |
| chr10-103417550-INS-520 | chr10-103417550-INS-520 |
| chr10-103427266-INS-787 | chr10-103427266-INS-787 |
| chr10-103519890-INS-66 | chr10-103519890-INS-66 |
| chr10-103522313-INS-2354 | chr10-103522314-INS-2351 |
| chr10-103534809-INS-2252 | chr10-103534810-INS-2252 |
| chr10-103557651-INS-7048 | chr10-103557652-INS-7050 |
| chr10-103568817-INS-545 | chr10-103568817-INS-545 |
| chr10-103572969-INS-368 | chr10-103572969-INS-368 |
| chr10-103590841-INS-86 | chr10-103590841-INS-86 |
| chr10-103605117-INS-7099 | chr10-103605118-INS-7099 |
| chr10-103619609-INS-58 | chr10-103619609-INS-58 |
| chr10-103629096-INS-53 | chr10-103629093-INS-60 |
| chr10-103638567-INS-3356 | chr10-103638568-INS-3356 |
| chr10-103638993-INS-56 | chr10-103638993-INS-55 |
| chr10-103644858-INS-55 | chr10-103644858-INS-100 |
| chr10-103662588-INS-189 | chr10-103662588-INS-189 |
| chr10-103664928-INS-5564 | chr10-103664929-INS-5565 |
| chr10-103665451-INS-52 | chr10-103665451-INS-52 |
| chr10-103666429-INS-375 | chr10-103666429-INS-377 |
| chr10-103667378-INS-907 | chr10-103667378-INS-909 |
| chr10-103678313-INS-198 | chr10-103678313-INS-204 |
| chr10-103699507-INS-175 | chr10-103699507-INS-175 |
| chr10-103726018-INS-128 | chr10-103726018-INS-128 |
| chr10-103729256-INS-175 | chr10-103729256-INS-189 |
| chr10-103754210-INS-59 | chr10-103754208-INS-52 |
| chr10-103763545-INS-53 | chr10-103763545-INS-57 |
| chr10-103767735-INS-76 | chr10-103767735-INS-64 |
| chr10-103787219-INS-52 | chr10-103787219-INS-52 |
| chr10-103817436-INS-54 | chr10-103817436-INS-57 |
| chr10-103851848-INS-51 | chr10-103851847-INS-51 |
| chr10-103869722-INS-58 | chr10-103869722-INS-54 |
| chr10-103888707-INS-61 | chr10-103888707-INS-63 |
| chr10-103895343-INS-199 | chr10-103895343-INS-199 |
| chr10-103917518-INS-123 | chr10-103917503-INS-92 |
| chr10-103922095-INS-585 | chr10-103922095-INS-581 |
| chr10-103953926-INS-235 | chr10-103953963-INS-238 |
| chr10-103975985-INS-795 | chr10-103975985-INS-796 |
| chr10-104041034-INS-105 | chr10-104041034-INS-105 |
| chr10-104053649-INS-6532 | chr10-104053650-INS-6532 |
| chr10-104088549-INS-314 | chr10-104088548-INS-310 |
| chr10-104103137-INS-544 | chr10-104103137-INS-548 |
| chr10-104138850-INS-1014 | chr10-104138851-INS-1008 |
| chr10-104142152-INS-133 | chr10-104142154-INS-134 |
| chr10-104192670-INS-6731 | chr10-104192671-INS-6732 |
| chr10-104221485-INS-304 | chr10-104221485-INS-304 |
| chr10-104231621-INS-52 | chr10-104231621-INS-52 |

| | |
|---|---|
| chr10-104243822-INS-84 | chr10-104243822-INS-84 |
| chr10-104274484-INS-96 | chr10-104274484-INS-98 |
| chr10-104282047-INS-66 | chr10-104282047-INS-66 |
| chr10-104300691-INS-125 | chr10-104300691-INS-125 |
| chr10-104301358-INS-58 | chr10-104301358-INS-62 |
| chr10-104314963-INS-1077 | chr10-104314964-INS-1077 |
| chr10-104402461-INS-443 | chr10-104402461-INS-443 |
| chr10-104419245-INS-60 | chr10-104419245-INS-62 |
| chr10-104424154-INS-593 | chr10-104424154-INS-593 |
| chr10-104448762-INS-75 | chr10-104448762-INS-75 |
| chr10-104502007-INS-282 | chr10-104502008-INS-282 |
| chr10-104535826-INS-7099 | chr10-104535827-INS-7099 |
| chr10-104543794-INS-68 | chr10-104543794-INS-68 |
| chr10-104552523-INS-72 | chr10-104552523-INS-70 |
| chr10-104614247-INS-80 | chr10-104614247-INS-80 |
| chr10-104630787-INS-132 | chr10-104630787-INS-136 |
| chr10-104692497-INS-1919 | chr10-104692498-INS-1922 |
| chr10-104713773-INS-75 | chr10-104713773-INS-75 |
| chr10-104716107-INS-69 | chr10-104716107-INS-69 |
| chr10-104716618-INS-676 | chr10-104716618-INS-676 |
| chr10-104740234-INS-53 | chr10-104740234-INS-51 |
| chr10-104748913-INS-112 | chr10-104748913-INS-112 |
| chr10-104777316-INS-2064 | chr10-104777317-INS-2064 |
| chr10-104816516-INS-461 | chr10-104816516-INS-461 |
| chr10-104818025-INS-74 | chr10-104818025-INS-74 |
| chr10-104819928-INS-526 | chr10-104819928-INS-526 |
| chr10-104840160-INS-103 | chr10-104840160-INS-90 |
| chr10-104850532-INS-357 | chr10-104850535-INS-358 |
| chr10-104853480-INS-64 | chr10-104853480-INS-56 |
| chr10-104863318-INS-72 | chr10-104863318-INS-72 |
| chr10-104867023-INS-55 | chr10-104867023-INS-50 |
| chr10-104903567-INS-103 | chr10-104903567-INS-103 |
| chr10-104972499-INS-223 | chr10-104972499-INS-223 |
| chr10-104991565-INS-109 | chr10-104991563-INS-115 |
| chr10-105009493-INS-88 | chr10-105009493-INS-92 |
| chr10-105028908-INS-80 | chr10-105028908-INS-80 |
| chr10-105036538-INS-58 | chr10-105036541-INS-50 |
| chr10-105042263-INS-86 | chr10-105042263-INS-86 |
| chr10-105053793-INS-79 | chr10-105053836-INS-54 |
| chr10-105067859-INS-1094 | chr10-105067859-INS-1109 |
| chr10-105097617-INS-85 | chr10-105097617-INS-100 |
| chr10-105106189-INS-161 | chr10-105106189-INS-161 |
| chr10-105121027-INS-91 | chr10-105121020-INS-101 |
| chr10-105131330-INS-56 | chr10-105131330-INS-60 |
| chr10-105169940-INS-509 | chr10-105169940-INS-484 |
| chr10-105171124-INS-167 | chr10-105171128-INS-165 |
| chr10-105230826-INS-385 | chr10-105230826-INS-378 |
| chr10-105282420-INS-83 | chr10-105282420-INS-83 |
| chr10-105288339-INS-64 | chr10-105288339-INS-68 |
| chr10-105290140-INS-60 | chr10-105290140-INS-60 |
| chr10-105295154-INS-222 | chr10-105295154-INS-214 |
| chr10-105313046-INS-300 | chr10-105313046-INS-300 |
| chr10-105325552-INS-264 | chr10-105325550-INS-282 |
| chr10-105332922-INS-409 | chr10-105332922-INS-409 |
| chr10-105336851-INS-214 | chr10-105336851-INS-214 |
| chr10-105348590-INS-164 | chr10-105348588-INS-166 |
| chr10-105354758-INS-67 | chr10-105354770-INS-71 |
| chr10-105361192-INS-146 | chr10-105361192-INS-146 |
| chr10-105369596-INS-64 | chr10-105369596-INS-60 |
| chr10-105371758-INS-92 | chr10-105371758-INS-92 |
| chr10-105372583-INS-64 | chr10-105372583-INS-68 |
| chr10-105422926-INS-112 | chr10-105422926-INS-112 |
| chr10-105424009-INS-6699 | chr10-105424010-INS-6699 |
| chr10-105432391-INS-120 | chr10-105432391-INS-120 |
| chr10-105443604-INS-372 | chr10-105443604-INS-372 |
| chr10-105451656-INS-202 | chr10-105451656-INS-202 |
| chr10-105459077-INS-212 | chr10-105459075-INS-214 |
| chr10-105460491-INS-213 | chr10-105460491-INS-213 |
| chr10-105462264-INS-55 | chr10-105462264-INS-55 |
| chr10-105484844-INS-54 | chr10-105484844-INS-56 |
| chr10-105496298-INS-125 | chr10-105496298-INS-122 |
| chr10-105551808-INS-60 | chr10-105551808-INS-60 |
| chr10-105555209-INS-62 | chr10-105555219-INS-72 |
| chr10-105555425-INS-6162 | chr10-105555426-INS-6162 |
| chr10-105564101-INS-436 | chr10-105564101-INS-436 |
| chr10-105570598-INS-1890 | chr10-105570599-INS-1889 |
| chr10-105586177-INS-68 | chr10-105586177-INS-68 |

| | |
|---|---|
| chr10-105603275-INS-6397 | chr10-105603276-INS-6397 |
| chr10-105604072-INS-77 | chr10-105604057-INS-79 |
| chr10-105609273-INS-78 | chr10-105609273-INS-60 |
| chr10-105610652-INS-59 | chr10-105610652-INS-59 |
| chr10-105619409-INS-62 | chr10-105619407-INS-58 |
| chr10-105621189-INS-1864 | chr10-105621190-INS-1864 |
| chr10-105670360-INS-199 | chr10-105670360-INS-196 |
| chr10-105676407-INS-115 | chr10-105676407-INS-118 |
| chr10-105697594-INS-297 | chr10-105697594-INS-297 |
| chr10-105704735-INS-274 | chr10-105704735-INS-274 |
| chr10-105711233-INS-184 | chr10-105711233-INS-184 |
| chr10-105717140-INS-351 | chr10-105717140-INS-351 |
| chr10-105736409-INS-703 | chr10-105736409-INS-698 |
| chr10-105766647-INS-90 | chr10-105766645-INS-116 |
| chr10-105773199-INS-70 | chr10-105773199-INS-68 |
| chr10-105814332-INS-81 | chr10-105814332-INS-81 |
| chr10-105832236-INS-7087 | chr10-105832237-INS-7087 |
| chr10-105842949-INS-485 | chr10-105842949-INS-485 |
| chr10-105850557-INS-2809 | chr10-105850558-INS-2809 |
| chr10-105851193-INS-835 | chr10-105851193-INS-835 |
| chr10-105860932-INS-379 | chr10-105860918-INS-383 |
| chr10-105865386-INS-58 | chr10-105865388-INS-58 |
| chr10-105866579-INS-572 | chr10-105866579-INS-572 |
| chr10-105868871-INS-352 | chr10-105868871-INS-362 |
| chr10-105875225-INS-401 | chr10-105875225-INS-402 |
| chr10-105915619-INS-56 | chr10-105915619-INS-70 |
| chr10-105935046-INS-151 | chr10-105935046-INS-151 |
| chr10-105968206-INS-2591 | chr10-105968207-INS-2591 |
| chr10-105996815-INS-3784 | chr10-105996816-INS-3784 |
| chr10-106005719-INS-115 | chr10-106005719-INS-99 |
| chr10-106037526-INS-140 | chr10-106037487-INS-189 |
| chr10-106066832-INS-90 | chr10-106066827-INS-99 |
| chr10-106093556-INS-1127 | chr10-106093545-INS-1141 |
| chr10-106098075-INS-66 | chr10-106098075-INS-66 |
| chr10-106110084-INS-100 | chr10-106110084-INS-105 |
| chr10-106119875-INS-56 | chr10-106119876-INS-56 |
| chr10-106158380-INS-54 | chr10-106158380-INS-54 |
| chr10-106160650-INS-664 | chr10-106160650-INS-670 |
| chr10-106186797-INS-129 | chr10-106186797-INS-129 |
| chr10-106187889-INS-62 | chr10-106187889-INS-68 |
| chr10-106210790-INS-190 | chr10-106210781-INS-209 |
| chr10-106243306-INS-418 | chr10-106243304-INS-423 |
| chr10-106263022-INS-61 | chr10-106262974-INS-60 |
| chr10-106270062-INS-125 | chr10-106270062-INS-128 |
| chr10-106280246-INS-60 | chr10-106280246-INS-60 |
| chr10-106389467-INS-52 | chr10-106389467-INS-60 |
| chr10-106390585-INS-55 | chr10-106390585-INS-89 |
| chr10-106391638-INS-4420 | chr10-106391639-INS-4459 |
| chr10-106412472-INS-66 | chr10-106412508-INS-56 |
| chr10-106458070-INS-191 | chr10-106458070-INS-195 |
| chr10-106477674-INS-134 | chr10-106477674-INS-134 |
| chr10-106504056-INS-122 | chr10-106504057-INS-125 |
| chr10-106508287-INS-60 | chr10-106508283-INS-59 |
| chr10-106527429-INS-54 | chr10-106527464-INS-50 |
| chr10-106544637-INS-658 | chr10-106544637-INS-658 |
| chr10-106554386-INS-76 | chr10-106554387-INS-62 |
| chr10-106562768-INS-467 | chr10-106562768-INS-467 |
| chr10-106565281-INS-56 | chr10-106565281-INS-56 |
| chr10-106580814-INS-2758 | chr10-106580815-INS-2758 |
| chr10-106587077-INS-52 | chr10-106587077-INS-52 |
| chr10-106605955-INS-805 | chr10-106605955-INS-805 |
| chr10-106620769-INS-726 | chr10-106620769-INS-725 |
| chr10-106697423-INS-482 | chr10-106697423-INS-483 |
| chr10-106699009-INS-778 | chr10-106699009-INS-778 |
| chr10-106705084-INS-71 | chr10-106705084-INS-71 |
| chr10-106739611-INS-66 | chr10-106739613-INS-55 |
| chr10-106773894-INS-80 | chr10-106773894-INS-84 |
| chr10-106779741-INS-58 | chr10-106779747-INS-54 |
| chr10-106797816-INS-98 | chr10-106797816-INS-98 |
| chr10-106814326-INS-6196 | chr10-106814327-INS-6196 |
| chr10-106815535-INS-97 | chr10-106815535-INS-106 |
| chr10-106820027-INS-107 | chr10-106820027-INS-103 |
| chr10-106839269-INS-171 | chr10-106839269-INS-170 |
| chr10-106858156-INS-68 | chr10-106858155-INS-50 |
| chr10-106879446-INS-56 | chr10-106879446-INS-56 |
| chr10-106895908-INS-53 | chr10-106895908-INS-57 |
| chr10-106996137-INS-51 | chr10-106996135-INS-55 |

| | |
|---|---|
| chr10-107018917-INS-117 | chr10-107018962-INS-67 |
| chr10-107042632-INS-138 | chr10-107042632-INS-142 |
| chr10-107135766-INS-56 | chr10-107135727-INS-92 |
| chr10-107156335-INS-67 | chr10-107156335-INS-67 |
| chr10-107191318-INS-279 | chr10-107191318-INS-279 |
| chr10-107235297-INS-64 | chr10-107235300-INS-53 |
| chr10-107239936-INS-62 | chr10-107239936-INS-62 |
| chr10-107246719-INS-607 | chr10-107246719-INS-609 |
| chr10-107249833-INS-266 | chr10-107249833-INS-268 |
| chr10-107262337-INS-82 | chr10-107262337-INS-77 |
| chr10-107266073-INS-226 | chr10-107266073-INS-227 |
| chr10-107269025-INS-151 | chr10-107269025-INS-131 |
| chr10-107354064-INS-6444 | chr10-107354065-INS-6445 |
| chr10-107354199-INS-142 | chr10-107354199-INS-142 |
| chr10-107452564-INS-63 | chr10-107452564-INS-63 |
| chr10-107499511-INS-512 | chr10-107499511-INS-512 |
| chr10-107534062-INS-227 | chr10-107534062-INS-227 |
| chr10-107556446-INS-79 | chr10-107556447-INS-80 |
| chr10-107557759-INS-56 | chr10-107557759-INS-56 |
| chr10-107601704-INS-510 | chr10-107601704-INS-511 |
| chr10-107638970-INS-60 | chr10-107638979-INS-65 |
| chr10-107661351-INS-706 | chr10-107661351-INS-706 |
| chr10-107666458-INS-422 | chr10-107666458-INS-422 |
| chr10-107669198-INS-58 | chr10-107669198-INS-58 |
| chr10-107671769-INS-52 | chr10-107671769-INS-52 |
| chr10-107677633-INS-458 | chr10-107677633-INS-464 |
| chr10-107681511-INS-208 | chr10-107681512-INS-208 |
| chr10-107692211-INS-58 | chr10-107692211-INS-50 |
| chr10-107724493-INS-81 | chr10-107724493-INS-79 |
| chr10-107731725-INS-62 | chr10-107731725-INS-62 |
| chr10-107745237-INS-62 | chr10-107745233-INS-50 |
| chr10-107761858-INS-90 | chr10-107761858-INS-90 |
| chr10-107763214-INS-155 | chr10-107763214-INS-165 |
| chr10-107767749-INS-83 | chr10-107767749-INS-72 |
| chr10-107773280-INS-577 | chr10-107773280-INS-577 |
| chr10-107791090-INS-1265 | chr10-107791091-INS-1237 |
| chr10-107801148-INS-1088 | chr10-107801149-INS-1088 |
| chr10-107805519-INS-190 | chr10-107805519-INS-190 |
| chr10-107884752-INS-422 | chr10-107884752-INS-465 |
| chr10-107921654-INS-5975 | chr10-107921655-INS-5975 |
| chr10-107937115-INS-68 | chr10-107937111-INS-76 |
| chr10-107990375-INS-228 | chr10-107990375-INS-228 |
| chr10-108010005-INS-54 | chr10-108010001-INS-76 |
| chr10-108019271-INS-289 | chr10-108019271-INS-289 |
| chr10-108037486-INS-324 | chr10-108037486-INS-324 |
| chr10-108039354-INS-6236 | chr10-108039355-INS-6237 |
| chr10-108046136-INS-524 | chr10-108046136-INS-524 |
| chr10-108058520-INS-70 | chr10-108058520-INS-70 |
| chr10-108064452-INS-51 | chr10-108064437-INS-63 |
| chr10-108068927-INS-234 | chr10-108068927-INS-236 |
| chr10-108075346-INS-68 | chr10-108075346-INS-68 |
| chr10-108082906-INS-101 | chr10-108082938-INS-52 |
| chr10-108101337-INS-215 | chr10-108101337-INS-216 |
| chr10-108120668-INS-59 | chr10-108120668-INS-59 |
| chr10-108134830-INS-248 | chr10-108134830-INS-250 |
| chr10-108138762-INS-64 | chr10-108138786-INS-77 |
| chr10-108145967-INS-6440 | chr10-108145968-INS-6440 |
| chr10-108151036-INS-234 | chr10-108151036-INS-234 |
| chr10-108193793-INS-92 | chr10-108193793-INS-92 |
| chr10-108205149-INS-52 | chr10-108205149-INS-52 |
| chr10-108234384-INS-81 | chr10-108234384-INS-81 |
| chr10-108236173-INS-337 | chr10-108236173-INS-337 |
| chr10-108277519-INS-184 | chr10-108277519-INS-184 |
| chr10-108294873-INS-56 | chr10-108294873-INS-59 |
| chr10-108298309-INS-498 | chr10-108298309-INS-498 |
| chr10-108306923-INS-58 | chr10-108306923-INS-58 |
| chr10-108308494-INS-6401 | chr10-108308495-INS-6403 |
| chr10-108320924-INS-103 | chr10-108320924-INS-118 |
| chr10-108323583-INS-55 | chr10-108323583-INS-55 |
| chr10-108325307-INS-150 | chr10-108325301-INS-175 |
| chr10-108328417-INS-323 | chr10-108328417-INS-341 |
| chr10-108341424-INS-124 | chr10-108341424-INS-124 |
| chr10-108348086-INS-86 | chr10-108348086-INS-132 |
| chr10-108360002-INS-94 | chr10-108360002-INS-94 |
| chr10-108381150-INS-711 | chr10-108381150-INS-711 |
| chr10-108387155-INS-5307 | chr10-108387156-INS-5307 |
| chr10-108396230-INS-687 | chr10-108396230-INS-687 |

| | |
|---|---|
| chr10-108443539-INS-516 | chr10-108443539-INS-516 |
| chr10-108447418-INS-133 | chr10-108447418-INS-121 |
| chr10-108469916-INS-180 | chr10-108469916-INS-180 |
| chr10-108534657-INS-6442 | chr10-108534658-INS-6442 |
| chr10-108590512-INS-64 | chr10-108590512-INS-64 |
| chr10-108609834-INS-66 | chr10-108609834-INS-66 |
| chr10-108628505-INS-1062 | chr10-108628506-INS-1060 |
| chr10-108763844-INS-67 | chr10-108763855-INS-94 |
| chr10-108790001-INS-614 | chr10-108789991-INS-613 |
| chr10-108794188-INS-3658 | chr10-108794189-INS-3658 |
| chr10-108795937-INS-75 | chr10-108795937-INS-75 |
| chr10-108809943-INS-254 | chr10-108809943-INS-250 |
| chr10-108851560-INS-60 | chr10-108851560-INS-60 |
| chr10-108858348-INS-56 | chr10-108858348-INS-56 |
| chr10-108895302-INS-96 | chr10-108895302-INS-96 |
| chr10-108907551-INS-80 | chr10-108907551-INS-50 |
| chr10-108919500-INS-204 | chr10-108919500-INS-204 |
| chr10-108937315-INS-61 | chr10-108937319-INS-66 |
| chr10-108960974-INS-581 | chr10-108960974-INS-574 |
| chr10-108977122-INS-360 | chr10-108977122-INS-360 |
| chr10-109004862-INS-304 | chr10-109004862-INS-304 |
| chr10-109014852-INS-87 | chr10-109014852-INS-87 |
| chr10-109023430-INS-4760 | chr10-109023431-INS-4760 |
| chr10-109110264-INS-55 | chr10-109110254-INS-57 |
| chr10-109113422-INS-141 | chr10-109113422-INS-141 |
| chr10-109120025-INS-2084 | chr10-109120026-INS-2081 |
| chr10-109130372-INS-62 | chr10-109130372-INS-62 |
| chr10-109135230-INS-64 | chr10-109135230-INS-58 |
| chr10-109136665-INS-81 | chr10-109136665-INS-87 |
| chr10-109198864-INS-53 | chr10-109198864-INS-54 |
| chr10-109204886-INS-74 | chr10-109204886-INS-70 |
| chr10-109209018-INS-56 | chr10-109209018-INS-56 |
| chr10-109213588-INS-96 | chr10-109213588-INS-70 |
| chr10-109223979-INS-56 | chr10-109223979-INS-60 |
| chr10-109240004-INS-148 | chr10-109239990-INS-129 |
| chr10-109257410-INS-71 | chr10-109257410-INS-71 |
| chr10-109308306-INS-67 | chr10-109308306-INS-51 |
| chr10-109335211-INS-292 | chr10-109335211-INS-292 |
| chr10-109340439-INS-141 | chr10-109340439-INS-142 |
| chr10-109373535-INS-6755 | chr10-109373536-INS-6763 |
| chr10-109403445-INS-52 | chr10-109403445-INS-52 |
| chr10-109413184-INS-1396 | chr10-109413185-INS-1398 |
| chr10-109427310-INS-1717 | chr10-109427311-INS-1765 |
| chr10-109440434-INS-60 | chr10-109440482-INS-55 |
| chr10-109445782-INS-52 | chr10-109445782-INS-58 |
| chr10-109446617-INS-394 | chr10-109446617-INS-393 |
| chr10-109461807-INS-425 | chr10-109461807-INS-424 |
| chr10-109477097-INS-208 | chr10-109477097-INS-208 |
| chr10-109493921-INS-76 | chr10-109493921-INS-76 |
| chr10-109495078-INS-234 | chr10-109495078-INS-234 |
| chr10-109498090-INS-64 | chr10-109498090-INS-64 |
| chr10-109499324-INS-82 | chr10-109499324-INS-83 |
| chr10-109506707-INS-5554 | chr10-109506708-INS-5554 |
| chr10-109529803-INS-71 | chr10-109529794-INS-96 |
| chr10-109543005-INS-232 | chr10-109543005-INS-232 |
| chr10-109550988-INS-56 | chr10-109550988-INS-56 |
| chr10-109569514-INS-60 | chr10-109569514-INS-60 |
| chr10-109585279-INS-581 | chr10-109585284-INS-537 |
| chr10-109624114-INS-65 | chr10-109624113-INS-62 |
| chr10-109666055-INS-198 | chr10-109666055-INS-198 |
| chr10-109693662-INS-154 | chr10-109693662-INS-154 |
| chr10-109716064-INS-156 | chr10-109716064-INS-156 |
| chr10-109726461-INS-176 | chr10-109726461-INS-176 |
| chr10-109730091-INS-53 | chr10-109730076-INS-50 |
| chr10-109772695-INS-427 | chr10-109772677-INS-396 |
| chr10-109807456-INS-667 | chr10-109807456-INS-673 |
| chr10-109818760-INS-89 | chr10-109818760-INS-89 |
| chr10-109913945-INS-125 | chr10-109913948-INS-128 |
| chr10-109917219-INS-400 | chr10-109917219-INS-400 |
| chr10-109929674-INS-90 | chr10-109929627-INS-132 |
| chr10-109956372-INS-171 | chr10-109956378-INS-157 |
| chr10-110007190-INS-248 | chr10-110007190-INS-250 |
| chr10-110050623-INS-75 | chr10-110050622-INS-80 |
| chr10-110104271-INS-56 | chr10-110104271-INS-56 |
| chr10-110106143-INS-58 | chr10-110106143-INS-58 |
| chr10-110139645-INS-506 | chr10-110139645-INS-506 |
| chr10-110140106-INS-1314 | chr10-110140107-INS-1309 |

| | |
|---|---|
| chr10-110150981-INS-84 | chr10-110151009-INS-64 |
| chr10-110174141-INS-68 | chr10-110174141-INS-68 |
| chr10-110176802-INS-438 | chr10-110176802-INS-438 |
| chr10-110180423-INS-62 | chr10-110180408-INS-62 |
| chr10-110239952-INS-468 | chr10-110239968-INS-462 |
| chr10-110276927-INS-55 | chr10-110276927-INS-55 |
| chr10-110287228-INS-85 | chr10-110287228-INS-86 |
| chr10-110309723-INS-450 | chr10-110309723-INS-450 |
| chr10-110364422-INS-6440 | chr10-110364423-INS-6441 |
| chr10-110368109-INS-405 | chr10-110368109-INS-405 |
| chr10-110392281-INS-93 | chr10-110392281-INS-93 |
| chr10-110414370-INS-62 | chr10-110414370-INS-64 |
| chr10-110433621-INS-86 | chr10-110433623-INS-106 |
| chr10-110438868-INS-984 | chr10-110438868-INS-981 |
| chr10-110496900-INS-289 | chr10-110496900-INS-289 |
| chr10-110498123-INS-555 | chr10-110498123-INS-555 |
| chr10-110499668-INS-347 | chr10-110499668-INS-339 |
| chr10-110512127-INS-54 | chr10-110512101-INS-54 |
| chr10-110547033-INS-138 | chr10-110547033-INS-151 |
| chr10-110570985-INS-104 | chr10-110570986-INS-81 |
| chr10-110593530-INS-156 | chr10-110593530-INS-146 |
| chr10-110636629-INS-52 | chr10-110636629-INS-52 |
| chr10-110639657-INS-76 | chr10-110639655-INS-65 |
| chr10-110644055-INS-73 | chr10-110644055-INS-73 |
| chr10-110661751-INS-51 | chr10-110661751-INS-57 |
| chr10-110698874-INS-3675 | chr10-110698875-INS-3675 |
| chr10-110706113-INS-262 | chr10-110706113-INS-262 |
| chr10-110723272-INS-194 | chr10-110723272-INS-194 |
| chr10-110739995-INS-194 | chr10-110739995-INS-194 |
| chr10-110747486-INS-66 | chr10-110747486-INS-61 |
| chr10-110795092-INS-284 | chr10-110795092-INS-284 |
| chr10-110806406-INS-204 | chr10-110806406-INS-205 |
| chr10-110836563-INS-81 | chr10-110836563-INS-81 |
| chr10-110848039-INS-109 | chr10-110848023-INS-88 |
| chr10-110875541-INS-188 | chr10-110875542-INS-188 |
| chr10-110882270-INS-52 | chr10-110882270-INS-52 |
| chr10-110887932-INS-1896 | chr10-110887933-INS-1896 |
| chr10-110895277-INS-7329 | chr10-110895278-INS-7329 |
| chr10-110906114-INS-120 | chr10-110906114-INS-120 |
| chr10-110908181-INS-219 | chr10-110908181-INS-231 |
| chr10-110911459-INS-183 | chr10-110911459-INS-195 |
| chr10-110923825-INS-7143 | chr10-110923826-INS-7143 |
| chr10-110935873-INS-215 | chr10-110935873-INS-214 |
| chr10-110943348-INS-68 | chr10-110943348-INS-66 |
| chr10-110957790-INS-66 | chr10-110957790-INS-74 |
| chr10-110966672-INS-7144 | chr10-110966673-INS-7144 |
| chr10-110998823-INS-199 | chr10-110998823-INS-198 |
| chr10-111019071-INS-88 | chr10-111019071-INS-88 |
| chr10-111031880-INS-4961 | chr10-111031881-INS-4961 |
| chr10-111047832-INS-52 | chr10-111047832-INS-52 |
| chr10-111072390-INS-2064 | chr10-111072391-INS-2065 |
| chr10-111089377-INS-58 | chr10-111089377-INS-58 |
| chr10-111127587-INS-497 | chr10-111127587-INS-491 |
| chr10-111144561-INS-92 | chr10-111144565-INS-68 |
| chr10-111156574-INS-86 | chr10-111156574-INS-86 |
| chr10-111162201-INS-325 | chr10-111162189-INS-326 |
| chr10-111214453-INS-200 | chr10-111214453-INS-200 |
| chr10-111246830-INS-100 | chr10-111246830-INS-90 |
| chr10-111249297-INS-377 | chr10-111249297-INS-368 |
| chr10-111254749-INS-230 | chr10-111254749-INS-230 |
| chr10-111265172-INS-208 | chr10-111265172-INS-212 |
| chr10-111275894-INS-74 | chr10-111275894-INS-74 |
| chr10-111289564-INS-64 | chr10-111289610-INS-62 |
| chr10-111290985-INS-218 | chr10-111290985-INS-218 |
| chr10-111294567-INS-80 | chr10-111294567-INS-80 |
| chr10-111361857-INS-141 | chr10-111361857-INS-141 |
| chr10-111369871-INS-78 | chr10-111369871-INS-99 |
| chr10-111369883-INS-585 | chr10-111369872-INS-589 |
| chr10-111443133-INS-173 | chr10-111443133-INS-178 |
| chr10-111444534-INS-1895 | chr10-111444535-INS-1895 |
| chr10-111450892-INS-343 | chr10-111450892-INS-343 |
| chr10-111453790-INS-109 | chr10-111453790-INS-113 |
| chr10-111458387-INS-126 | chr10-111458387-INS-110 |
| chr10-111477216-INS-991 | chr10-111477212-INS-984 |
| chr10-111479086-INS-143 | chr10-111479090-INS-143 |
| chr10-111491693-INS-54 | chr10-111491684-INS-62 |
| chr10-111495196-INS-99 | chr10-111495196-INS-52 |

| | |
|---|---|
| chr10-111513596-INS-52 | chr10-111513596-INS-52 |
| chr10-111518178-INS-101 | chr10-111518179-INS-75 |
| chr10-111520823-INS-56 | chr10-111520831-INS-54 |
| chr10-111573596-INS-57 | chr10-111573596-INS-57 |
| chr10-111583380-INS-247 | chr10-111583380-INS-265 |
| chr10-111632468-INS-66 | chr10-111632469-INS-62 |
| chr10-111643238-INS-51 | chr10-111643238-INS-51 |
| chr10-111649021-INS-229 | chr10-111649021-INS-229 |
| chr10-111658066-INS-75 | chr10-111658033-INS-86 |
| chr10-111682729-INS-190 | chr10-111682729-INS-190 |
| chr10-111688759-INS-549 | chr10-111688759-INS-549 |
| chr10-111690896-INS-164 | chr10-111690896-INS-164 |
| chr10-111702910-INS-66 | chr10-111702910-INS-51 |
| chr10-111713922-INS-56 | chr10-111713897-INS-50 |
| chr10-111736372-INS-76 | chr10-111736372-INS-56 |
| chr10-111776186-INS-541 | chr10-111776186-INS-538 |
| chr10-111795764-INS-85 | chr10-111795764-INS-134 |
| chr10-111800119-INS-604 | chr10-111800119-INS-604 |
| chr10-111802294-INS-94 | chr10-111802294-INS-141 |
| chr10-111807546-INS-80 | chr10-111807546-INS-80 |
| chr10-111841054-INS-173 | chr10-111841054-INS-172 |
| chr10-111858902-INS-86 | chr10-111858902-INS-54 |
| chr10-111864697-INS-790 | chr10-111864697-INS-790 |
| chr10-111887602-INS-277 | chr10-111887598-INS-283 |
| chr10-111909246-INS-61 | chr10-111909261-INS-59 |
| chr10-111917218-INS-148 | chr10-111917218-INS-148 |
| chr10-111926526-INS-398 | chr10-111926526-INS-397 |
| chr10-111947708-INS-735 | chr10-111947674-INS-756 |
| chr10-111949922-INS-345 | chr10-111949922-INS-345 |
| chr10-111974895-INS-803 | chr10-111974895-INS-803 |
| chr10-112002655-INS-52 | chr10-112002649-INS-52 |
| chr10-112012181-INS-214 | chr10-112012181-INS-215 |
| chr10-112042887-INS-524 | chr10-112042887-INS-524 |
| chr10-112061054-INS-2941 | chr10-112061055-INS-2941 |
| chr10-112061442-INS-525 | chr10-112061441-INS-526 |
| chr10-112074863-INS-1127 | chr10-112074864-INS-1127 |
| chr10-112085008-INS-52 | chr10-112085008-INS-74 |
| chr10-112116860-INS-990 | chr10-112116860-INS-988 |
| chr10-112134282-INS-107 | chr10-112134282-INS-109 |
| chr10-112181245-INS-6634 | chr10-112181246-INS-6635 |
| chr10-112193905-INS-123 | chr10-112193905-INS-123 |
| chr10-112204809-INS-158 | chr10-112204809-INS-159 |
| chr10-112246517-INS-71 | chr10-112246512-INS-120 |
| chr10-112281730-INS-1063 | chr10-112281731-INS-1063 |
| chr10-112288563-INS-247 | chr10-112288563-INS-247 |
| chr10-112319780-INS-228 | chr10-112319780-INS-228 |
| chr10-112332547-INS-427 | chr10-112332551-INS-408 |
| chr10-112335197-INS-514 | chr10-112335197-INS-509 |
| chr10-112372605-INS-54 | chr10-112372603-INS-60 |
| chr10-112404222-INS-212 | chr10-112404222-INS-212 |
| chr10-112405620-INS-241 | chr10-112405620-INS-241 |
| chr10-112416499-INS-380 | chr10-112416499-INS-380 |
| chr10-112440638-INS-201 | chr10-112440638-INS-201 |
| chr10-112454013-INS-369 | chr10-112454013-INS-370 |
| chr10-112486231-INS-131 | chr10-112486231-INS-128 |
| chr10-112513606-INS-288 | chr10-112513606-INS-301 |
| chr10-112524908-INS-282 | chr10-112524908-INS-286 |
| chr10-112529659-INS-133 | chr10-112529659-INS-133 |
| chr10-112530599-INS-147 | chr10-112530606-INS-138 |
| chr10-112546667-INS-66 | chr10-112546667-INS-66 |
| chr10-112566755-INS-76 | chr10-112566755-INS-76 |
| chr10-112574507-INS-96 | chr10-112574477-INS-84 |
| chr10-112579520-INS-82 | chr10-112579538-INS-74 |
| chr10-112629553-INS-206 | chr10-112629553-INS-207 |
| chr10-112640154-INS-220 | chr10-112640166-INS-230 |
| chr10-112649335-INS-54 | chr10-112649335-INS-56 |
| chr10-112655042-INS-89 | chr10-112655048-INS-95 |
| chr10-112656664-INS-181 | chr10-112656664-INS-182 |
| chr10-112660904-INS-55 | chr10-112660897-INS-55 |
| chr10-112678057-INS-68 | chr10-112678059-INS-78 |
| chr10-112679160-INS-6158 | chr10-112679161-INS-6158 |
| chr10-112686277-INS-90 | chr10-112686277-INS-96 |
| chr10-112693738-INS-478 | chr10-112693738-INS-478 |
| chr10-112716896-INS-256 | chr10-112716896-INS-233 |
| chr10-112719626-INS-130 | chr10-112719630-INS-146 |
| chr10-112768835-INS-398 | chr10-112768835-INS-398 |
| chr10-112851465-INS-146 | chr10-112851465-INS-146 |

| | |
|---|---|
| chr10-112953737-INS-137 | chr10-112953737-INS-137 |
| chr10-112978649-INS-77 | chr10-112978649-INS-78 |
| chr10-113034157-INS-89 | chr10-113034157-INS-89 |
| chr10-113035614-INS-54 | chr10-113035625-INS-55 |
| chr10-113056865-INS-67 | chr10-113056899-INS-67 |
| chr10-113074972-INS-139 | chr10-113074978-INS-176 |
| chr10-113085311-INS-57 | chr10-113085361-INS-56 |
| chr10-113085798-INS-277 | chr10-113085800-INS-281 |
| chr10-113097428-INS-230 | chr10-113097428-INS-232 |
| chr10-113115486-INS-66 | chr10-113115486-INS-66 |
| chr10-113144885-INS-549 | chr10-113144885-INS-549 |
| chr10-113196896-INS-98 | chr10-113196896-INS-98 |
| chr10-113229907-INS-70 | chr10-113229907-INS-50 |
| chr10-113233819-INS-1721 | chr10-113233820-INS-1721 |
| chr10-113264303-INS-682 | chr10-113264303-INS-681 |
| chr10-113282093-INS-62 | chr10-113282093-INS-50 |
| chr10-113322116-INS-66 | chr10-113322116-INS-66 |
| chr10-113346976-INS-943 | chr10-113346976-INS-943 |
| chr10-113355346-INS-6427 | chr10-113355347-INS-6427 |
| chr10-113390190-INS-6179 | chr10-113390191-INS-6179 |
| chr10-113452600-INS-1890 | chr10-113452601-INS-1890 |
| chr10-113465148-INS-556 | chr10-113465148-INS-560 |
| chr10-113488140-INS-5357 | chr10-113488141-INS-5357 |
| chr10-113501644-INS-6589 | chr10-113501645-INS-6589 |
| chr10-113513624-INS-299 | chr10-113513624-INS-294 |
| chr10-113518654-INS-76 | chr10-113518679-INS-80 |
| chr10-113555474-INS-81 | chr10-113555474-INS-81 |
| chr10-113573879-INS-84 | chr10-113573862-INS-51 |
| chr10-113575762-INS-74 | chr10-113575762-INS-74 |
| chr10-113588802-INS-942 | chr10-113588802-INS-943 |
| chr10-113643511-INS-59 | chr10-113643511-INS-63 |
| chr10-113663431-INS-231 | chr10-113663431-INS-244 |
| chr10-113670037-INS-56 | chr10-113670025-INS-58 |
| chr10-113685248-INS-3028 | chr10-113685249-INS-3028 |
| chr10-113692572-INS-445 | chr10-113692572-INS-445 |
| chr10-113723848-INS-74 | chr10-113723844-INS-80 |
| chr10-113725799-INS-6514 | chr10-113725800-INS-6514 |
| chr10-113734490-INS-1388 | chr10-113734491-INS-1388 |
| chr10-113770635-INS-1179 | chr10-113770636-INS-1179 |
| chr10-113872929-INS-55 | chr10-113872929-INS-55 |
| chr10-113881689-INS-333 | chr10-113881689-INS-333 |
| chr10-113902480-INS-293 | chr10-113902480-INS-294 |
| chr10-113918735-INS-958 | chr10-113918735-INS-959 |
| chr10-113922316-INS-56 | chr10-113922316-INS-58 |
| chr10-113924734-INS-88 | chr10-113924734-INS-80 |
| chr10-113938809-INS-218 | chr10-113938809-INS-227 |
| chr10-113944814-INS-6390 | chr10-113944815-INS-6390 |
| chr10-113952264-INS-577 | chr10-113952264-INS-577 |
| chr10-113984987-INS-554 | chr10-113984987-INS-554 |
| chr10-114067038-INS-6464 | chr10-114067039-INS-6461 |
| chr10-114085063-INS-54 | chr10-114085063-INS-54 |
| chr10-114089040-INS-882 | chr10-114089040-INS-882 |
| chr10-114103928-INS-410 | chr10-114103928-INS-410 |
| chr10-114123480-INS-680 | chr10-114123480-INS-683 |
| chr10-114132515-INS-193 | chr10-114132515-INS-192 |
| chr10-114143637-INS-74 | chr10-114143643-INS-58 |
| chr10-114145922-INS-134 | chr10-114145922-INS-134 |
| chr10-114173164-INS-83 | chr10-114173164-INS-83 |
| chr10-114185203-INS-292 | chr10-114185203-INS-292 |
| chr10-114192496-INS-99 | chr10-114192512-INS-85 |
| chr10-114208514-INS-2810 | chr10-114208515-INS-2810 |
| chr10-114243433-INS-397 | chr10-114243433-INS-397 |
| chr10-114270932-INS-518 | chr10-114270932-INS-518 |
| chr10-114273115-INS-54 | chr10-114273115-INS-53 |
| chr10-114287124-INS-105 | chr10-114287121-INS-120 |
| chr10-114291063-INS-6121 | chr10-114291064-INS-6122 |
| chr10-114304389-INS-139 | chr10-114304385-INS-147 |
| chr10-114319440-INS-53 | chr10-114319440-INS-53 |
| chr10-114336743-INS-184 | chr10-114336743-INS-185 |
| chr10-114379287-INS-110 | chr10-114379287-INS-110 |
| chr10-114388067-INS-56 | chr10-114388068-INS-72 |
| chr10-114435415-INS-102 | chr10-114435411-INS-72 |
| chr10-114437067-INS-476 | chr10-114437067-INS-477 |
| chr10-114450853-INS-147 | chr10-114450861-INS-163 |
| chr10-114476803-INS-528 | chr10-114476803-INS-531 |
| chr10-114483015-INS-51 | chr10-114483015-INS-51 |
| chr10-114519151-INS-188 | chr10-114519151-INS-188 |

| | |
|---|---|
| chr10-114543164-INS-187 | chr10-114543164-INS-186 |
| chr10-114548506-INS-102 | chr10-114548506-INS-53 |
| chr10-114554280-INS-3022 | chr10-114554283-INS-3046 |
| chr10-114556779-INS-439 | chr10-114556780-INS-439 |
| chr10-114559057-INS-64 | chr10-114559079-INS-68 |
| chr10-114573197-INS-63 | chr10-114573185-INS-75 |
| chr10-114584339-INS-92 | chr10-114584339-INS-92 |
| chr10-114640320-INS-55 | chr10-114640294-INS-60 |
| chr10-114662067-INS-324 | chr10-114662046-INS-342 |
| chr10-114668408-INS-167 | chr10-114668408-INS-165 |
| chr10-114678593-INS-62 | chr10-114678583-INS-78 |
| chr10-114703689-INS-53 | chr10-114703689-INS-53 |
| chr10-114725995-INS-65 | chr10-114725989-INS-59 |
| chr10-114745490-INS-6319 | chr10-114745491-INS-6320 |
| chr10-114758290-INS-78 | chr10-114758290-INS-122 |
| chr10-114766797-INS-242 | chr10-114766797-INS-242 |
| chr10-114799003-INS-130 | chr10-114799003-INS-130 |
| chr10-114810315-INS-462 | chr10-114810315-INS-434 |
| chr10-114859824-INS-126 | chr10-114859824-INS-108 |
| chr10-114865740-INS-80 | chr10-114865740-INS-80 |
| chr10-114868323-INS-94 | chr10-114868323-INS-94 |
| chr10-114870009-INS-209 | chr10-114870009-INS-210 |
| chr10-114890949-INS-51 | chr10-114890949-INS-51 |
| chr10-114894564-INS-10307 | chr10-114894565-INS-10308 |
| chr10-114923057-INS-56 | chr10-114923057-INS-56 |
| chr10-114925983-INS-63 | chr10-114925983-INS-63 |
| chr10-114935174-INS-460 | chr10-114935174-INS-459 |
| chr10-114936125-INS-219 | chr10-114936120-INS-213 |
| chr10-114937203-INS-96 | chr10-114937203-INS-112 |
| chr10-114938051-INS-579 | chr10-114938051-INS-577 |
| chr10-114945946-INS-80 | chr10-114945902-INS-108 |
| chr10-114978180-INS-106 | chr10-114978180-INS-120 |
| chr10-114993610-INS-132 | chr10-114993609-INS-135 |
| chr10-115011388-INS-59 | chr10-115011388-INS-59 |
| chr10-115023719-INS-199 | chr10-115023719-INS-199 |
| chr10-115039621-INS-90 | chr10-115039621-INS-88 |
| chr10-115074292-INS-222 | chr10-115074292-INS-234 |
| chr10-115081583-INS-68 | chr10-115081583-INS-68 |
| chr10-115090754-INS-189 | chr10-115090754-INS-189 |
| chr10-115095032-INS-1891 | chr10-115095033-INS-1891 |
| chr10-115096336-INS-64 | chr10-115096336-INS-52 |
| chr10-115096667-INS-298 | chr10-115096667-INS-298 |
| chr10-115109506-INS-347 | chr10-115109507-INS-316 |
| chr10-115111028-INS-83 | chr10-115111028-INS-83 |
| chr10-115122384-INS-76 | chr10-115122387-INS-77 |
| chr10-115134890-INS-76 | chr10-115134890-INS-66 |
| chr10-115138162-INS-357 | chr10-115138162-INS-358 |
| chr10-115155404-INS-515 | chr10-115155404-INS-515 |
| chr10-115165001-INS-99 | chr10-115165001-INS-119 |
| chr10-115173872-INS-206 | chr10-115173872-INS-206 |
| chr10-115175348-INS-74 | chr10-115175348-INS-74 |
| chr10-115178772-INS-66 | chr10-115178772-INS-66 |
| chr10-115182773-INS-110 | chr10-115182781-INS-104 |
| chr10-115190913-INS-215 | chr10-115190913-INS-215 |
| chr10-115191961-INS-52 | chr10-115191961-INS-50 |
| chr10-115213212-INS-7086 | chr10-115213213-INS-7086 |
| chr10-115273007-INS-54 | chr10-115273007-INS-54 |
| chr10-115282005-INS-242 | chr10-115282005-INS-242 |
| chr10-115303211-INS-196 | chr10-115303211-INS-198 |
| chr10-115317810-INS-72 | chr10-115317810-INS-72 |
| chr10-115347036-INS-133 | chr10-115347036-INS-114 |
| chr10-115359202-INS-72 | chr10-115359212-INS-72 |
| chr10-115363968-INS-1588 | chr10-115363969-INS-1588 |
| chr10-115366386-INS-92 | chr10-115366386-INS-101 |
| chr10-115386573-INS-148 | chr10-115386573-INS-166 |
| chr10-115393487-INS-1125 | chr10-115393488-INS-1125 |
| chr10-115432379-INS-51 | chr10-115432395-INS-51 |
| chr10-115434874-INS-64 | chr10-115434874-INS-65 |
| chr10-115449461-INS-495 | chr10-115449461-INS-495 |
| chr10-115454499-INS-60 | chr10-115454498-INS-80 |
| chr10-115491158-INS-117 | chr10-115491153-INS-122 |
| chr10-115541847-INS-114 | chr10-115541847-INS-114 |
| chr10-115551876-INS-176 | chr10-115551876-INS-177 |
| chr10-115561014-INS-70 | chr10-115561014-INS-94 |
| chr10-115563836-INS-199 | chr10-115563836-INS-185 |
| chr10-115575865-INS-944 | chr10-115575865-INS-943 |
| chr10-115584883-INS-266 | chr10-115584883-INS-310 |

| | |
|---|---|
| chr10-115596801-INS-173 | chr10-115596801-INS-173 |
| chr10-115612880-INS-108 | chr10-115612891-INS-82 |
| chr10-115644998-INS-52 | chr10-115644998-INS-52 |
| chr10-115659144-INS-197 | chr10-115659144-INS-197 |
| chr10-115666260-INS-52 | chr10-115666260-INS-52 |
| chr10-115667464-INS-1027 | chr10-115667465-INS-1027 |
| chr10-115672967-INS-59 | chr10-115672967-INS-75 |
| chr10-115684966-INS-173 | chr10-115684966-INS-173 |
| chr10-115690072-INS-774 | chr10-115690072-INS-771 |
| chr10-115705646-INS-133 | chr10-115705646-INS-122 |
| chr10-115747318-INS-304 | chr10-115747318-INS-304 |
| chr10-115752699-INS-75 | chr10-115752699-INS-75 |
| chr10-115763208-INS-76 | chr10-115763208-INS-78 |
| chr10-115809377-INS-1124 | chr10-115809378-INS-1124 |
| chr10-115824964-INS-120 | chr10-115824964-INS-120 |
| chr10-115839469-INS-76 | chr10-115839469-INS-76 |
| chr10-115843239-INS-90 | chr10-115843239-INS-90 |
| chr10-115877278-INS-168 | chr10-115877278-INS-168 |
| chr10-115899949-INS-56 | chr10-115899949-INS-56 |
| chr10-115917238-INS-202 | chr10-115917238-INS-206 |
| chr10-115960561-INS-84 | chr10-115960574-INS-60 |
| chr10-115966944-INS-86 | chr10-115966944-INS-90 |
| chr10-116025128-INS-1022 | chr10-116025129-INS-1022 |
| chr10-116029575-INS-529 | chr10-116029575-INS-526 |
| chr10-116046596-INS-236 | chr10-116046596-INS-249 |
| chr10-116061227-INS-110 | chr10-116061227-INS-110 |
| chr10-116062272-INS-192 | chr10-116062272-INS-192 |
| chr10-116067194-INS-149 | chr10-116067194-INS-149 |
| chr10-116084630-INS-414 | chr10-116084630-INS-414 |
| chr10-116095772-INS-57 | chr10-116095772-INS-72 |
| chr10-116096542-INS-752 | chr10-116096542-INS-753 |
| chr10-116098140-INS-409 | chr10-116098140-INS-409 |
| chr10-116131868-INS-60 | chr10-116131868-INS-60 |
| chr10-116152281-INS-68 | chr10-116152281-INS-68 |
| chr10-116170993-INS-103 | chr10-116170993-INS-71 |
| chr10-116202572-INS-339 | chr10-116202572-INS-339 |
| chr10-116204986-INS-113 | chr10-116204986-INS-113 |
| chr10-116210436-INS-141 | chr10-116210431-INS-180 |
| chr10-116211440-INS-87 | chr10-116211417-INS-107 |
| chr10-116217596-INS-156 | chr10-116217596-INS-156 |
| chr10-116229314-INS-62 | chr10-116229316-INS-65 |
| chr10-116231825-INS-55 | chr10-116231825-INS-55 |
| chr10-116241660-INS-202 | chr10-116241660-INS-203 |
| chr10-116266901-INS-276 | chr10-116266901-INS-292 |
| chr10-116269385-INS-60 | chr10-116269382-INS-63 |
| chr10-116281947-INS-162 | chr10-116281948-INS-169 |
| chr10-116292861-INS-72 | chr10-116292862-INS-74 |
| chr10-116295660-INS-116 | chr10-116295664-INS-118 |
| chr10-116311001-INS-680 | chr10-116311001-INS-680 |
| chr10-116314076-INS-211 | chr10-116314077-INS-213 |
| chr10-116374271-INS-84 | chr10-116374271-INS-76 |
| chr10-116388972-INS-178 | chr10-116388973-INS-180 |
| chr10-116390687-INS-252 | chr10-116390726-INS-256 |
| chr10-116403170-INS-482 | chr10-116403170-INS-482 |
| chr10-116426073-INS-161 | chr10-116426073-INS-161 |
| chr10-116431071-INS-231 | chr10-116431071-INS-229 |
| chr10-116436525-INS-508 | chr10-116436525-INS-508 |
| chr10-116463142-INS-137 | chr10-116463142-INS-107 |
| chr10-116492946-INS-198 | chr10-116492940-INS-202 |
| chr10-116495830-INS-56 | chr10-116495830-INS-60 |
| chr10-116518211-INS-184 | chr10-116518211-INS-184 |
| chr10-116533062-INS-319 | chr10-116533062-INS-321 |
| chr10-116552287-INS-284 | chr10-116552287-INS-282 |
| chr10-116585655-INS-60 | chr10-116585662-INS-52 |
| chr10-116640465-INS-129 | chr10-116640465-INS-129 |
| chr10-116665254-INS-7111 | chr10-116665255-INS-7113 |
| chr10-116673103-INS-1399 | chr10-116673104-INS-1399 |
| chr10-116687111-INS-76 | chr10-116687111-INS-86 |
| chr10-116696771-INS-63 | chr10-116696768-INS-51 |
| chr10-116718971-INS-68 | chr10-116718971-INS-60 |
| chr10-116728338-INS-54 | chr10-116728338-INS-54 |
| chr10-116731348-INS-86 | chr10-116731324-INS-86 |
| chr10-116748327-INS-80 | chr10-116748327-INS-81 |
| chr10-116761611-INS-51 | chr10-116761611-INS-51 |
| chr10-116787199-INS-222 | chr10-116787199-INS-222 |
| chr10-116821351-INS-364 | chr10-116821351-INS-364 |
| chr10-116824293-INS-2190 | chr10-116824294-INS-2190 |

| | |
|---|---|
| chr10-116839581-INS-72 | chr10-116839581-INS-72 |
| chr10-116861552-INS-297 | chr10-116861552-INS-297 |
| chr10-116871814-INS-63 | chr10-116871854-INS-57 |
| chr10-116895946-INS-52 | chr10-116895946-INS-50 |
| chr10-116937336-INS-202 | chr10-116937336-INS-202 |
| chr10-116964602-INS-219 | chr10-116964602-INS-219 |
| chr10-116965661-INS-185 | chr10-116965661-INS-186 |
| chr10-117003669-INS-151 | chr10-117003668-INS-191 |
| chr10-117011106-INS-202 | chr10-117011106-INS-198 |
| chr10-117012988-INS-56 | chr10-117012988-INS-58 |
| chr10-117019675-INS-114 | chr10-117019675-INS-114 |
| chr10-117034799-INS-201 | chr10-117034799-INS-201 |
| chr10-117035968-INS-80 | chr10-117035968-INS-60 |
| chr10-117047452-INS-396 | chr10-117047452-INS-396 |
| chr10-117063566-INS-100 | chr10-117063566-INS-100 |
| chr10-117138383-INS-212 | chr10-117138383-INS-212 |
| chr10-117149470-INS-58 | chr10-117149465-INS-70 |
| chr10-117155733-INS-288 | chr10-117155733-INS-308 |
| chr10-117194809-INS-87 | chr10-117194809-INS-85 |
| chr10-117217831-INS-57 | chr10-117217831-INS-57 |
| chr10-117218512-INS-486 | chr10-117218512-INS-486 |
| chr10-117220421-INS-436 | chr10-117220421-INS-436 |
| chr10-117223509-INS-51 | chr10-117223502-INS-78 |
| chr10-117236542-INS-302 | chr10-117236542-INS-302 |
| chr10-117248530-INS-54 | chr10-117248530-INS-54 |
| chr10-117257964-INS-213 | chr10-117257964-INS-215 |
| chr10-117266272-INS-141 | chr10-117266270-INS-147 |
| chr10-117282062-INS-86 | chr10-117282062-INS-129 |
| chr10-117283451-INS-549 | chr10-117283441-INS-552 |
| chr10-117288067-INS-126 | chr10-117288065-INS-130 |
| chr10-117295438-INS-54 | chr10-117295439-INS-54 |
| chr10-117296812-INS-108 | chr10-117296813-INS-110 |
| chr10-117298654-INS-70 | chr10-117298622-INS-65 |
| chr10-117300592-INS-506 | chr10-117300593-INS-506 |
| chr10-117302103-INS-58 | chr10-117302106-INS-68 |
| chr10-117311106-INS-568 | chr10-117311106-INS-568 |
| chr10-117331590-INS-101 | chr10-117331590-INS-107 |
| chr10-117335465-INS-171 | chr10-117335465-INS-170 |
| chr10-117338442-INS-55 | chr10-117338442-INS-57 |
| chr10-117358569-INS-70 | chr10-117358570-INS-55 |
| chr10-117386517-INS-51 | chr10-117386517-INS-51 |
| chr10-117388588-INS-740 | chr10-117388588-INS-740 |
| chr10-117390755-INS-481 | chr10-117390755-INS-481 |
| chr10-117400915-INS-70 | chr10-117400936-INS-75 |
| chr10-117403994-INS-420 | chr10-117403994-INS-419 |
| chr10-117412672-INS-216 | chr10-117412672-INS-217 |
| chr10-117420369-INS-197 | chr10-117420369-INS-196 |
| chr10-117451651-INS-75 | chr10-117451648-INS-56 |
| chr10-117468823-INS-66 | chr10-117468823-INS-66 |
| chr10-117475071-INS-334 | chr10-117475071-INS-353 |
| chr10-117550084-INS-195 | chr10-117550084-INS-200 |
| chr10-117568156-INS-400 | chr10-117568156-INS-400 |
| chr10-117576107-INS-196 | chr10-117576107-INS-191 |
| chr10-117590155-INS-1303 | chr10-117590156-INS-1303 |
| chr10-117616864-INS-347 | chr10-117616864-INS-350 |
| chr10-117636970-INS-65 | chr10-117636970-INS-70 |
| chr10-117689537-INS-446 | chr10-117689537-INS-439 |
| chr10-117701503-INS-55 | chr10-117701503-INS-55 |
| chr10-117702290-INS-401 | chr10-117702290-INS-405 |
| chr10-117706270-INS-62 | chr10-117706270-INS-56 |
| chr10-117713017-INS-71 | chr10-117713017-INS-68 |
| chr10-117751186-INS-7111 | chr10-117751187-INS-7111 |
| chr10-117778053-INS-116 | chr10-117778053-INS-116 |
| chr10-117781267-INS-86 | chr10-117781267-INS-94 |
| chr10-117804323-INS-175 | chr10-117804323-INS-176 |
| chr10-117813974-INS-432 | chr10-117813974-INS-432 |
| chr10-117820400-INS-124 | chr10-117820404-INS-124 |
| chr10-117837296-INS-7125 | chr10-117837297-INS-7125 |
| chr10-117841695-INS-78 | chr10-117841683-INS-68 |
| chr10-117850806-INS-95 | chr10-117850807-INS-95 |
| chr10-117861558-INS-468 | chr10-117861558-INS-468 |
| chr10-117865409-INS-178 | chr10-117865450-INS-167 |
| chr10-117884740-INS-76 | chr10-117884740-INS-66 |
| chr10-117894262-INS-250 | chr10-117894269-INS-250 |
| chr10-117900092-INS-478 | chr10-117900092-INS-431 |
| chr10-117918107-INS-60 | chr10-117918107-INS-60 |
| chr10-117924719-INS-58 | chr10-117924719-INS-57 |

| | |
|---|---|
| chr10-117927152-INS-111 | chr10-117927173-INS-122 |
| chr10-117931386-INS-133 | chr10-117931386-INS-133 |
| chr10-117938268-INS-55 | chr10-117938268-INS-57 |
| chr10-117952256-INS-112 | chr10-117952263-INS-71 |
| chr10-117971524-INS-68 | chr10-117971536-INS-69 |
| chr10-117973292-INS-206 | chr10-117973292-INS-206 |
| chr10-117975949-INS-51 | chr10-117975950-INS-50 |
| chr10-117986583-INS-74 | chr10-117986583-INS-74 |
| chr10-117996406-INS-560 | chr10-117996406-INS-560 |
| chr10-118009466-INS-57 | chr10-118009466-INS-57 |
| chr10-118025954-INS-3207 | chr10-118025955-INS-3207 |
| chr10-118026828-INS-135 | chr10-118026828-INS-94 |
| chr10-118036077-INS-156 | chr10-118036077-INS-156 |
| chr10-118051255-INS-365 | chr10-118051255-INS-365 |
| chr10-118088945-INS-53 | chr10-118088925-INS-53 |
| chr10-118126841-INS-61 | chr10-118126841-INS-56 |
| chr10-118204354-INS-62 | chr10-118204354-INS-62 |
| chr10-118214894-INS-58 | chr10-118214894-INS-58 |
| chr10-118216788-INS-72 | chr10-118216788-INS-72 |
| chr10-118222293-INS-135 | chr10-118222293-INS-135 |
| chr10-118242230-INS-120 | chr10-118242230-INS-120 |
| chr10-118258271-INS-54 | chr10-118258271-INS-54 |
| chr10-118267387-INS-70 | chr10-118267387-INS-65 |
| chr10-118292264-INS-658 | chr10-118292264-INS-656 |
| chr10-118334550-INS-122 | chr10-118334550-INS-126 |
| chr10-118360369-INS-97 | chr10-118360369-INS-97 |
| chr10-118368773-INS-70 | chr10-118368773-INS-70 |
| chr10-118376400-INS-63 | chr10-118376401-INS-63 |
| chr10-118384460-INS-65 | chr10-118384460-INS-65 |
| chr10-118402490-INS-77 | chr10-118402481-INS-66 |
| chr10-118416868-INS-72 | chr10-118416886-INS-66 |
| chr10-118444752-INS-54 | chr10-118444752-INS-54 |
| chr10-118448603-INS-65 | chr10-118448603-INS-65 |
| chr10-118450004-INS-82 | chr10-118450004-INS-82 |
| chr10-118478754-INS-56 | chr10-118478754-INS-56 |
| chr10-118483125-INS-147 | chr10-118483125-INS-147 |
| chr10-118496976-INS-146 | chr10-118496976-INS-146 |
| chr10-118538599-INS-95 | chr10-118538599-INS-95 |
| chr10-118542398-INS-329 | chr10-118542398-INS-329 |
| chr10-118544609-INS-58 | chr10-118544581-INS-64 |
| chr10-118567561-INS-111 | chr10-118567561-INS-111 |
| chr10-118593291-INS-76 | chr10-118593291-INS-76 |
| chr10-118595249-INS-1097 | chr10-118595250-INS-1097 |
| chr10-118597042-INS-110 | chr10-118597042-INS-104 |
| chr10-118603413-INS-76 | chr10-118603425-INS-59 |
| chr10-118646381-INS-72 | chr10-118646381-INS-70 |
| chr10-118650455-INS-93 | chr10-118650459-INS-54 |
| chr10-118653601-INS-538 | chr10-118653601-INS-539 |
| chr10-118655497-INS-58 | chr10-118655500-INS-88 |
| chr10-118658288-INS-341 | chr10-118658288-INS-342 |
| chr10-118665332-INS-91 | chr10-118665338-INS-85 |
| chr10-118668121-INS-183 | chr10-118668121-INS-177 |
| chr10-118687851-INS-53 | chr10-118687851-INS-53 |
| chr10-118790881-INS-400 | chr10-118790881-INS-400 |
| chr10-118804795-INS-100 | chr10-118804795-INS-118 |
| chr10-118808003-INS-70 | chr10-118808003-INS-60 |
| chr10-118813603-INS-155 | chr10-118813603-INS-134 |
| chr10-118831909-INS-75 | chr10-118831909-INS-81 |
| chr10-118836174-INS-5373 | chr10-118836175-INS-5373 |
| chr10-118845872-INS-75 | chr10-118845872-INS-75 |
| chr10-118850184-INS-59 | chr10-118850184-INS-59 |
| chr10-118883552-INS-979 | chr10-118883552-INS-980 |
| chr10-118883796-INS-201 | chr10-118883796-INS-203 |
| chr10-118904977-INS-54 | chr10-118904999-INS-57 |
| chr10-118925613-INS-199 | chr10-118925613-INS-199 |
| chr10-118932740-INS-7504 | chr10-118932741-INS-7504 |
| chr10-118934118-INS-163 | chr10-118934118-INS-163 |
| chr10-118935450-INS-186 | chr10-118935450-INS-192 |
| chr10-118974124-INS-64 | chr10-118974124-INS-64 |
| chr10-118978541-INS-213 | chr10-118978541-INS-212 |
| chr10-119011006-INS-322 | chr10-119011005-INS-278 |
| chr10-119013625-INS-497 | chr10-119013625-INS-497 |
| chr10-119016518-INS-5441 | chr10-119016519-INS-5441 |
| chr10-119021712-INS-91 | chr10-119021712-INS-71 |
| chr10-119024212-INS-54 | chr10-119024209-INS-51 |
| chr10-119132459-INS-77 | chr10-119132459-INS-93 |
| chr10-119144632-INS-282 | chr10-119144632-INS-280 |

| | |
|---|---|
| chr10-119148825-INS-72 | chr10-119148825-INS-72 |
| chr10-119171422-INS-50 | chr10-119171422-INS-55 |
| chr10-119184217-INS-319 | chr10-119184217-INS-319 |
| chr10-119192540-INS-321 | chr10-119192540-INS-321 |
| chr10-119202398-INS-55 | chr10-119202398-INS-55 |
| chr10-119207858-INS-120 | chr10-119207847-INS-132 |
| chr10-119216549-INS-609 | chr10-119216549-INS-611 |
| chr10-119273023-INS-161 | chr10-119273024-INS-161 |
| chr10-119315048-INS-60 | chr10-119315048-INS-62 |
| chr10-119320068-INS-91 | chr10-119320067-INS-85 |
| chr10-119330682-INS-203 | chr10-119330682-INS-203 |
| chr10-119369407-INS-2601 | chr10-119369408-INS-2602 |
| chr10-119376919-INS-4775 | chr10-119376920-INS-4775 |
| chr10-119378104-INS-55 | chr10-119378104-INS-55 |
| chr10-119389782-INS-312 | chr10-119389782-INS-312 |
| chr10-119402382-INS-162 | chr10-119402382-INS-162 |
| chr10-119422612-INS-62 | chr10-119422612-INS-80 |
| chr10-119477549-INS-325 | chr10-119477549-INS-330 |
| chr10-119491283-INS-54 | chr10-119491283-INS-54 |
| chr10-119501673-INS-208 | chr10-119501673-INS-202 |
| chr10-119509042-INS-148 | chr10-119509060-INS-166 |
| chr10-119549276-INS-70 | chr10-119549276-INS-70 |
| chr10-119583751-INS-84 | chr10-119583751-INS-84 |
| chr10-119592473-INS-203 | chr10-119592473-INS-203 |
| chr10-119637470-INS-878 | chr10-119637470-INS-880 |
| chr10-119653602-INS-85 | chr10-119653602-INS-85 |
| chr10-119686207-INS-58 | chr10-119686205-INS-60 |
| chr10-119715515-INS-101 | chr10-119715515-INS-101 |
| chr10-119716603-INS-534 | chr10-119716603-INS-544 |
| chr10-119745275-INS-6575 | chr10-119745276-INS-6576 |
| chr10-119749016-INS-62 | chr10-119749016-INS-62 |
| chr10-119753051-INS-96 | chr10-119753082-INS-98 |
| chr10-119757713-INS-64 | chr10-119757713-INS-64 |
| chr10-119761572-INS-134 | chr10-119761572-INS-93 |
| chr10-119767470-INS-61 | chr10-119767469-INS-58 |
| chr10-119770617-INS-82 | chr10-119770655-INS-90 |
| chr10-119789889-INS-63 | chr10-119789889-INS-63 |
| chr10-119794960-INS-114 | chr10-119794960-INS-114 |
| chr10-119800842-INS-175 | chr10-119800843-INS-175 |
| chr10-119808617-INS-202 | chr10-119808617-INS-202 |
| chr10-119839067-INS-267 | chr10-119839067-INS-267 |
| chr10-119860361-INS-162 | chr10-119860361-INS-162 |
| chr10-119865394-INS-96 | chr10-119865390-INS-60 |
| chr10-119868905-INS-52 | chr10-119868905-INS-51 |
| chr10-119907263-INS-79 | chr10-119907263-INS-70 |
| chr10-119909941-INS-172 | chr10-119909941-INS-172 |
| chr10-119922101-INS-69 | chr10-119922101-INS-69 |
| chr10-119957601-INS-197 | chr10-119957601-INS-197 |
| chr10-119960475-INS-196 | chr10-119960475-INS-195 |
| chr10-119967029-INS-109 | chr10-119967029-INS-109 |
| chr10-119968383-INS-96 | chr10-119968383-INS-96 |
| chr10-119985353-INS-74 | chr10-119985353-INS-74 |
| chr10-120041207-INS-333 | chr10-120041207-INS-333 |
| chr10-120046729-INS-1892 | chr10-120046730-INS-1892 |
| chr10-120189788-INS-268 | chr10-120189788-INS-268 |
| chr10-120202174-INS-60 | chr10-120202174-INS-60 |
| chr10-120229740-INS-54 | chr10-120229740-INS-54 |
| chr10-120231466-INS-215 | chr10-120231463-INS-230 |
| chr10-120255720-INS-89 | chr10-120255720-INS-89 |
| chr10-120287246-INS-51 | chr10-120287247-INS-63 |
| chr10-120291281-INS-95 | chr10-120291281-INS-95 |
| chr10-120327107-INS-217 | chr10-120327107-INS-215 |
| chr10-120331355-INS-209 | chr10-120331355-INS-209 |
| chr10-120352935-INS-220 | chr10-120352935-INS-217 |
| chr10-120354149-INS-155 | chr10-120354161-INS-111 |
| chr10-120365450-INS-122 | chr10-120365448-INS-112 |
| chr10-120367326-INS-120 | chr10-120367326-INS-120 |
| chr10-120427717-INS-56 | chr10-120427717-INS-56 |
| chr10-120508963-INS-102 | chr10-120508963-INS-104 |
| chr10-120512404-INS-52 | chr10-120512389-INS-67 |
| chr10-120534426-INS-66 | chr10-120534426-INS-50 |
| chr10-120541556-INS-223 | chr10-120541556-INS-224 |
| chr10-120542070-INS-6468 | chr10-120542071-INS-6468 |
| chr10-120566940-INS-157 | chr10-120566940-INS-130 |
| chr10-120568575-INS-89 | chr10-120568565-INS-95 |
| chr10-120577108-INS-70 | chr10-120577108-INS-50 |
| chr10-120582856-INS-91 | chr10-120582856-INS-55 |

| | |
|---|---|
| chr10-120604196-INS-82 | chr10-120604196-INS-82 |
| chr10-120618611-INS-76 | chr10-120618611-INS-76 |
| chr10-120619799-INS-117 | chr10-120619796-INS-82 |
| chr10-120626105-INS-63 | chr10-120626105-INS-63 |
| chr10-120631440-INS-442 | chr10-120631440-INS-442 |
| chr10-120636526-INS-100 | chr10-120636526-INS-100 |
| chr10-120638935-INS-82 | chr10-120638935-INS-76 |
| chr10-120654261-INS-62 | chr10-120654261-INS-62 |
| chr10-120668780-INS-92 | chr10-120668780-INS-92 |
| chr10-120670306-INS-141 | chr10-120670306-INS-141 |
| chr10-120682403-INS-208 | chr10-120682403-INS-207 |
| chr10-120736670-INS-69 | chr10-120736670-INS-69 |
| chr10-120743003-INS-114 | chr10-120743005-INS-94 |
| chr10-120757632-INS-68 | chr10-120757632-INS-60 |
| chr10-120772734-INS-845 | chr10-120772734-INS-846 |
| chr10-120793613-INS-63 | chr10-120793613-INS-63 |
| chr10-120823441-INS-103 | chr10-120823441-INS-96 |
| chr10-120824840-INS-184 | chr10-120824846-INS-195 |
| chr10-120832254-INS-828 | chr10-120832255-INS-825 |
| chr10-120860155-INS-148 | chr10-120860155-INS-146 |
| chr10-120885239-INS-164 | chr10-120885239-INS-164 |
| chr10-120905932-INS-131 | chr10-120905932-INS-131 |
| chr10-120908763-INS-120 | chr10-120908763-INS-120 |
| chr10-120946136-INS-55 | chr10-120946136-INS-54 |
| chr10-120970625-INS-81 | chr10-120970625-INS-81 |
| chr10-121008671-INS-80 | chr10-121008671-INS-78 |
| chr10-121020108-INS-72 | chr10-121020108-INS-72 |
| chr10-121036623-INS-57 | chr10-121036611-INS-78 |
| chr10-121051615-INS-1570 | chr10-121051616-INS-1570 |
| chr10-121067039-INS-120 | chr10-121067039-INS-120 |
| chr10-121069831-INS-2408 | chr10-121069832-INS-2408 |
| chr10-121081261-INS-65 | chr10-121081261-INS-65 |
| chr10-121083804-INS-54 | chr10-121083804-INS-54 |
| chr10-121088558-INS-204 | chr10-121088566-INS-207 |
| chr10-121091410-INS-78 | chr10-121091410-INS-78 |
| chr10-121099018-INS-58 | chr10-121099018-INS-58 |
| chr10-121104158-INS-93 | chr10-121104158-INS-93 |
| chr10-121110106-INS-60 | chr10-121110106-INS-52 |
| chr10-121112936-INS-657 | chr10-121112936-INS-659 |
| chr10-121117820-INS-936 | chr10-121117819-INS-943 |
| chr10-121125161-INS-380 | chr10-121125161-INS-380 |
| chr10-121131965-INS-342 | chr10-121131965-INS-342 |
| chr10-121143023-INS-72 | chr10-121143023-INS-72 |
| chr10-121165379-INS-66 | chr10-121165366-INS-64 |
| chr10-121168940-INS-765 | chr10-121168940-INS-765 |
| chr10-121180756-INS-110 | chr10-121180754-INS-126 |
| chr10-121193383-INS-431 | chr10-121193383-INS-431 |
| chr10-121198818-INS-52 | chr10-121198818-INS-52 |
| chr10-121212151-INS-144 | chr10-121212151-INS-144 |
| chr10-121220372-INS-232 | chr10-121220372-INS-261 |
| chr10-121221963-INS-248 | chr10-121221963-INS-248 |
| chr10-121227793-INS-139 | chr10-121227790-INS-147 |
| chr10-121237565-INS-108 | chr10-121237565-INS-60 |
| chr10-121249602-INS-63 | chr10-121249602-INS-63 |
| chr10-121251607-INS-368 | chr10-121251607-INS-368 |
| chr10-121260751-INS-52 | chr10-121260751-INS-52 |
| chr10-121263080-INS-90 | chr10-121263080-INS-102 |
| chr10-121281072-INS-70 | chr10-121281072-INS-64 |
| chr10-121285129-INS-170 | chr10-121285129-INS-170 |
| chr10-121294402-INS-392 | chr10-121294402-INS-392 |
| chr10-121301943-INS-65 | chr10-121301939-INS-64 |
| chr10-121323878-INS-52 | chr10-121323878-INS-52 |
| chr10-121334951-INS-133 | chr10-121334958-INS-162 |
| chr10-121360737-INS-229 | chr10-121360737-INS-229 |
| chr10-121386270-INS-52 | chr10-121386270-INS-52 |
| chr10-121394812-INS-241 | chr10-121394812-INS-240 |
| chr10-121427758-INS-2491 | chr10-121427759-INS-2491 |
| chr10-121432572-INS-63 | chr10-121432572-INS-63 |
| chr10-121435213-INS-54 | chr10-121435213-INS-54 |
| chr10-121463639-INS-220 | chr10-121463639-INS-220 |
| chr10-121485731-INS-347 | chr10-121485731-INS-347 |
| chr10-121498326-INS-52 | chr10-121498326-INS-52 |
| chr10-121499806-INS-2828 | chr10-121499807-INS-2828 |
| chr10-121505874-INS-751 | chr10-121505874-INS-751 |
| chr10-121512481-INS-194 | chr10-121512481-INS-196 |
| chr10-121524441-INS-198 | chr10-121524441-INS-198 |
| chr10-121525084-INS-87 | chr10-121525084-INS-87 |

| | |
|---|---|
| chr10-121536471-INS-52 | chr10-121536471-INS-53 |
| chr10-121559667-INS-62 | chr10-121559620-INS-63 |
| chr10-121567337-INS-82 | chr10-121567337-INS-82 |
| chr10-121569516-INS-60 | chr10-121569476-INS-55 |
| chr10-121589905-INS-378 | chr10-121589905-INS-378 |
| chr10-121606369-INS-52 | chr10-121606369-INS-52 |
| chr10-121623516-INS-203 | chr10-121623516-INS-162 |
| chr10-121626879-INS-173 | chr10-121626879-INS-183 |
| chr10-121648250-INS-156 | chr10-121648250-INS-156 |
| chr10-121683290-INS-70 | chr10-121683290-INS-62 |
| chr10-121693758-INS-72 | chr10-121693758-INS-72 |
| chr10-121721836-INS-105 | chr10-121721836-INS-84 |
| chr10-121728306-INS-58 | chr10-121728337-INS-57 |
| chr10-121733264-INS-566 | chr10-121733264-INS-566 |
| chr10-121737774-INS-308 | chr10-121737783-INS-309 |
| chr10-121771951-INS-1429 | chr10-121771952-INS-1429 |
| chr10-121780559-INS-212 | chr10-121780559-INS-212 |
| chr10-121828026-INS-66 | chr10-121828022-INS-110 |
| chr10-121830298-INS-1127 | chr10-121830299-INS-1131 |
| chr10-121891076-INS-53 | chr10-121891104-INS-50 |
| chr10-121916389-INS-74 | chr10-121916393-INS-78 |
| chr10-121935318-INS-105 | chr10-121935338-INS-70 |
| chr10-121937978-INS-88 | chr10-121937978-INS-102 |
| chr10-121972395-INS-73 | chr10-121972395-INS-73 |
| chr10-122062029-INS-296 | chr10-122062029-INS-296 |
| chr10-122064792-INS-52 | chr10-122064792-INS-52 |
| chr10-122066792-INS-421 | chr10-122066792-INS-420 |
| chr10-122088208-INS-100 | chr10-122088208-INS-128 |
| chr10-122096146-INS-75 | chr10-122096146-INS-89 |
| chr10-122097745-INS-505 | chr10-122097745-INS-498 |
| chr10-122105100-INS-6619 | chr10-122105101-INS-6618 |
| chr10-122114023-INS-52 | chr10-122114019-INS-60 |
| chr10-122132344-INS-167 | chr10-122132339-INS-195 |
| chr10-122171329-INS-6763 | chr10-122171330-INS-6762 |
| chr10-122180537-INS-51 | chr10-122180537-INS-51 |
| chr10-122195522-INS-310 | chr10-122195522-INS-310 |
| chr10-122200406-INS-198 | chr10-122200406-INS-198 |
| chr10-122209076-INS-3232 | chr10-122209077-INS-3232 |
| chr10-122211172-INS-377 | chr10-122211170-INS-393 |
| chr10-122214157-INS-79 | chr10-122214157-INS-85 |
| chr10-122272791-INS-169 | chr10-122272789-INS-180 |
| chr10-122273476-INS-919 | chr10-122273466-INS-928 |
| chr10-122275127-INS-98 | chr10-122275127-INS-98 |
| chr10-122295279-INS-66 | chr10-122295279-INS-66 |
| chr10-122311387-INS-180 | chr10-122311387-INS-172 |
| chr10-122313262-INS-554 | chr10-122313262-INS-569 |
| chr10-122317347-INS-172 | chr10-122317347-INS-172 |
| chr10-122329885-INS-334 | chr10-122329885-INS-334 |
| chr10-122334347-INS-126 | chr10-122334347-INS-126 |
| chr10-122356944-INS-255 | chr10-122356944-INS-255 |
| chr10-122408325-INS-59 | chr10-122408346-INS-59 |
| chr10-122416562-INS-64 | chr10-122416562-INS-64 |
| chr10-122425575-INS-116 | chr10-122425575-INS-116 |
| chr10-122427194-INS-50 | chr10-122427194-INS-56 |
| chr10-122439011-INS-253 | chr10-122439010-INS-259 |
| chr10-122445922-INS-1089 | chr10-122445923-INS-1089 |
| chr10-122463678-INS-120 | chr10-122463678-INS-120 |
| chr10-122465585-INS-82 | chr10-122465585-INS-82 |
| chr10-122486878-INS-54 | chr10-122486878-INS-54 |
| chr10-122493074-INS-649 | chr10-122493074-INS-653 |
| chr10-122496340-INS-6402 | chr10-122496341-INS-6403 |
| chr10-122584348-INS-1888 | chr10-122584349-INS-1888 |
| chr10-122589029-INS-91 | chr10-122589025-INS-120 |
| chr10-122607493-INS-62 | chr10-122607496-INS-64 |
| chr10-122633172-INS-76 | chr10-122633172-INS-114 |
| chr10-122665971-INS-127 | chr10-122665971-INS-127 |
| chr10-122678655-INS-787 | chr10-122678655-INS-787 |
| chr10-122680106-INS-144 | chr10-122680156-INS-142 |
| chr10-122696415-INS-260 | chr10-122696463-INS-247 |
| chr10-122707962-INS-2887 | chr10-122707963-INS-2887 |
| chr10-122722158-INS-51 | chr10-122722158-INS-67 |
| chr10-122735591-INS-68 | chr10-122735588-INS-56 |
| chr10-122745127-INS-200 | chr10-122745127-INS-200 |
| chr10-122757825-INS-6383 | chr10-122757826-INS-6383 |
| chr10-122761395-INS-225 | chr10-122761395-INS-225 |
| chr10-122765044-INS-144 | chr10-122765044-INS-144 |
| chr10-122767010-INS-70 | chr10-122766980-INS-58 |

| | |
|---|---|
| chr10-122784932-INS-91 | chr10-122784934-INS-94 |
| chr10-122786506-INS-1862 | chr10-122786507-INS-1862 |
| chr10-122815304-INS-89 | chr10-122815304-INS-60 |
| chr10-122827876-INS-6441 | chr10-122827877-INS-6441 |
| chr10-122844968-INS-139 | chr10-122844968-INS-138 |
| chr10-122849910-INS-57 | chr10-122849910-INS-57 |
| chr10-122858146-INS-52 | chr10-122858145-INS-52 |
| chr10-122899693-INS-58 | chr10-122899693-INS-58 |
| chr10-122918022-INS-207 | chr10-122918022-INS-207 |
| chr10-122928137-INS-70 | chr10-122928141-INS-65 |
| chr10-122958693-INS-180 | chr10-122958693-INS-180 |
| chr10-122973158-INS-5403 | chr10-122973159-INS-5415 |
| chr10-122974208-INS-94 | chr10-122974208-INS-94 |
| chr10-122981300-INS-146 | chr10-122981300-INS-146 |
| chr10-122988355-INS-90 | chr10-122988355-INS-54 |
| chr10-123026085-INS-4245 | chr10-123026086-INS-4245 |
| chr10-123029164-INS-525 | chr10-123029164-INS-522 |
| chr10-123031715-INS-100 | chr10-123031730-INS-74 |
| chr10-123048129-INS-57 | chr10-123048091-INS-63 |
| chr10-123049971-INS-6369 | chr10-123049972-INS-6369 |
| chr10-123057741-INS-53 | chr10-123057741-INS-53 |
| chr10-123088886-INS-393 | chr10-123088884-INS-391 |
| chr10-123141539-INS-871 | chr10-123141539-INS-871 |
| chr10-123144834-INS-1262 | chr10-123144835-INS-1262 |
| chr10-123166299-INS-639 | chr10-123166299-INS-639 |
| chr10-123175579-INS-3797 | chr10-123175580-INS-3798 |
| chr10-123178152-INS-209 | chr10-123178152-INS-210 |
| chr10-123181579-INS-105 | chr10-123181579-INS-101 |
| chr10-123192367-INS-57 | chr10-123192328-INS-56 |
| chr10-123199316-INS-60 | chr10-123199315-INS-69 |
| chr10-123210008-INS-56 | chr10-123210008-INS-66 |
| chr10-123250149-INS-149 | chr10-123250149-INS-149 |
| chr10-123262093-INS-849 | chr10-123262114-INS-849 |
| chr10-123268746-INS-216 | chr10-123268746-INS-216 |
| chr10-123305610-INS-325 | chr10-123305603-INS-324 |
| chr10-123320236-INS-390 | chr10-123320245-INS-399 |
| chr10-123326347-INS-76 | chr10-123326347-INS-76 |
| chr10-123353516-INS-94 | chr10-123353516-INS-94 |
| chr10-123358440-INS-156 | chr10-123358434-INS-124 |
| chr10-123363406-INS-765 | chr10-123363406-INS-763 |
| chr10-123366181-INS-2037 | chr10-123366182-INS-2040 |
| chr10-123372253-INS-201 | chr10-123372274-INS-192 |
| chr10-123383746-INS-221 | chr10-123383724-INS-230 |
| chr10-123405691-INS-714 | chr10-123405689-INS-714 |
| chr10-123410544-INS-254 | chr10-123410544-INS-268 |
| chr10-123420684-INS-943 | chr10-123420684-INS-973 |
| chr10-123453600-INS-52 | chr10-123453600-INS-52 |
| chr10-123457800-INS-2224 | chr10-123457801-INS-2224 |
| chr10-123469010-INS-151 | chr10-123469010-INS-105 |
| chr10-123477366-INS-60 | chr10-123477366-INS-60 |
| chr10-123480868-INS-64 | chr10-123480870-INS-68 |
| chr10-123513569-INS-90 | chr10-123513569-INS-90 |
| chr10-123516257-INS-152 | chr10-123516257-INS-152 |
| chr10-123592781-INS-100 | chr10-123592781-INS-96 |
| chr10-123612344-INS-446 | chr10-123612344-INS-442 |
| chr10-123613255-INS-437 | chr10-123613255-INS-429 |
| chr10-123633621-INS-312 | chr10-123633622-INS-313 |
| chr10-123642617-INS-111 | chr10-123642617-INS-111 |
| chr10-123649609-INS-85 | chr10-123649609-INS-56 |
| chr10-123654638-INS-56 | chr10-123654638-INS-56 |
| chr10-123673037-INS-1051 | chr10-123673038-INS-1052 |
| chr10-123695828-INS-75 | chr10-123695828-INS-75 |
| chr10-123705517-INS-105 | chr10-123705518-INS-105 |
| chr10-123736030-INS-721 | chr10-123736030-INS-721 |
| chr10-123754769-INS-400 | chr10-123754769-INS-400 |
| chr10-123755550-INS-2670 | chr10-123755551-INS-2679 |
| chr10-123781717-INS-62 | chr10-123781717-INS-62 |
| chr10-123801927-INS-289 | chr10-123801927-INS-289 |
| chr10-123817066-INS-435 | chr10-123817066-INS-435 |
| chr10-123852029-INS-80 | chr10-123852035-INS-99 |
| chr10-123876404-INS-617 | chr10-123876404-INS-618 |
| chr10-123878021-INS-54 | chr10-123878021-INS-56 |
| chr10-123887052-INS-56 | chr10-123887052-INS-60 |
| chr10-123888079-INS-80 | chr10-123888079-INS-92 |
| chr10-123914374-INS-112 | chr10-123914374-INS-112 |
| chr10-123919078-INS-173 | chr10-123919078-INS-173 |
| chr10-123929485-INS-78 | chr10-123929483-INS-82 |

| | |
|---|---|
| chr10-123944897-INS-73 | chr10-123944897-INS-73 |
| chr10-123984770-INS-549 | chr10-123984770-INS-549 |
| chr10-124006826-INS-4699 | chr10-124006827-INS-4699 |
| chr10-124013496-INS-64 | chr10-124013496-INS-52 |
| chr10-124025023-INS-103 | chr10-124025023-INS-139 |
| chr10-124034633-INS-252 | chr10-124034633-INS-252 |
| chr10-124087037-INS-101 | chr10-124087037-INS-101 |
| chr10-124096912-INS-78 | chr10-124096912-INS-78 |
| chr10-124099803-INS-203 | chr10-124099803-INS-201 |
| chr10-124112902-INS-84 | chr10-124112902-INS-78 |
| chr10-124131370-INS-870 | chr10-124131370-INS-870 |
| chr10-124140260-INS-7195 | chr10-124140261-INS-7195 |
| chr10-124155927-INS-827 | chr10-124155927-INS-826 |
| chr10-124184273-INS-1799 | chr10-124184274-INS-1810 |
| chr10-124219374-INS-136 | chr10-124219374-INS-136 |
| chr10-124296808-INS-574 | chr10-124296808-INS-567 |
| chr10-124333686-INS-212 | chr10-124333686-INS-212 |
| chr10-124381249-INS-59 | chr10-124381249-INS-59 |
| chr10-124395252-INS-60 | chr10-124395256-INS-56 |
| chr10-124431387-INS-401 | chr10-124431387-INS-401 |
| chr10-124464823-INS-78 | chr10-124464821-INS-86 |
| chr10-124486433-INS-7285 | chr10-124486434-INS-7285 |
| chr10-124509955-INS-134 | chr10-124509955-INS-135 |
| chr10-124543389-INS-55 | chr10-124543389-INS-55 |
| chr10-124547440-INS-564 | chr10-124547440-INS-563 |
| chr10-124592659-INS-63 | chr10-124592660-INS-69 |
| chr10-124619763-INS-456 | chr10-124619763-INS-448 |
| chr10-124649452-INS-122 | chr10-124649452-INS-77 |
| chr10-124675012-INS-193 | chr10-124675034-INS-204 |
| chr10-124684696-INS-52 | chr10-124684696-INS-52 |
| chr10-124707659-INS-413 | chr10-124707659-INS-413 |
| chr10-124721679-INS-134 | chr10-124721679-INS-129 |
| chr10-124733429-INS-60 | chr10-124733429-INS-52 |
| chr10-124739594-INS-536 | chr10-124739594-INS-535 |
| chr10-124768561-INS-829 | chr10-124768561-INS-829 |
| chr10-124786064-INS-90 | chr10-124786064-INS-96 |
| chr10-124801116-INS-166 | chr10-124801116-INS-166 |
| chr10-124812731-INS-82 | chr10-124812731-INS-104 |
| chr10-124826920-INS-326 | chr10-124826920-INS-326 |
| chr10-124844623-INS-6568 | chr10-124844624-INS-6568 |
| chr10-124857938-INS-3474 | chr10-124857939-INS-3475 |
| chr10-124860046-INS-472 | chr10-124860046-INS-472 |
| chr10-124871023-INS-134 | chr10-124871023-INS-134 |
| chr10-124894689-INS-1291 | chr10-124894690-INS-1293 |
| chr10-124919416-INS-1506 | chr10-124919417-INS-1503 |
| chr10-124948807-INS-892 | chr10-124948807-INS-893 |
| chr10-124953729-INS-119 | chr10-124953729-INS-119 |
| chr10-124966153-INS-56 | chr10-124966153-INS-56 |
| chr10-124978630-INS-61 | chr10-124978630-INS-61 |
| chr10-124998690-INS-422 | chr10-124998691-INS-420 |
| chr10-125033910-INS-53 | chr10-125033910-INS-51 |
| chr10-125049369-INS-118 | chr10-125049369-INS-118 |
| chr10-125096768-INS-53 | chr10-125096768-INS-53 |
| chr10-125110223-INS-76 | chr10-125110223-INS-78 |
| chr10-125116380-INS-93 | chr10-125116381-INS-93 |
| chr10-125118944-INS-52 | chr10-125118948-INS-50 |
| chr10-125132495-INS-99 | chr10-125132527-INS-55 |
| chr10-125145560-INS-290 | chr10-125145560-INS-290 |
| chr10-125187400-INS-358 | chr10-125187400-INS-358 |
| chr10-125209594-INS-53 | chr10-125209617-INS-57 |
| chr10-125232580-INS-77 | chr10-125232580-INS-82 |
| chr10-125239147-INS-55 | chr10-125239146-INS-102 |
| chr10-125240811-INS-94 | chr10-125240811-INS-94 |
| chr10-125243253-INS-52 | chr10-125243253-INS-54 |
| chr10-125260419-INS-51 | chr10-125260422-INS-61 |
| chr10-125264663-INS-154 | chr10-125264663-INS-154 |
| chr10-125275812-INS-121 | chr10-125275858-INS-75 |
| chr10-125289846-INS-116 | chr10-125289846-INS-121 |
| chr10-125297785-INS-585 | chr10-125297785-INS-585 |
| chr10-125301320-INS-164 | chr10-125301320-INS-164 |
| chr10-125312552-INS-156 | chr10-125312552-INS-156 |
| chr10-125322407-INS-6828 | chr10-125322408-INS-6828 |
| chr10-125328258-INS-58 | chr10-125328258-INS-64 |
| chr10-125345168-INS-85 | chr10-125345167-INS-123 |
| chr10-125375712-INS-97 | chr10-125375712-INS-97 |
| chr10-125378610-INS-276 | chr10-125378610-INS-276 |
| chr10-125382288-INS-627 | chr10-125382288-INS-628 |

| | |
|---|---|
| chr10-125385161-INS-54 | chr10-125385161-INS-60 |
| chr10-125392304-INS-942 | chr10-125392304-INS-941 |
| chr10-125394880-INS-76 | chr10-125394865-INS-78 |
| chr10-125398219-INS-2211 | chr10-125398220-INS-2213 |
| chr10-125437632-INS-66 | chr10-125437638-INS-108 |
| chr10-125439078-INS-144 | chr10-125439078-INS-144 |
| chr10-125455671-INS-110 | chr10-125455671-INS-110 |
| chr10-125461636-INS-245 | chr10-125461636-INS-257 |
| chr10-125471720-INS-1869 | chr10-125471721-INS-1869 |
| chr10-125484116-INS-63 | chr10-125484141-INS-77 |
| chr10-125485159-INS-517 | chr10-125485161-INS-543 |
| chr10-125488034-INS-139 | chr10-125488034-INS-139 |
| chr10-125494690-INS-61 | chr10-125494701-INS-70 |
| chr10-125533532-INS-95 | chr10-125533531-INS-72 |
| chr10-125537258-INS-74 | chr10-125537259-INS-74 |
| chr10-125564064-INS-60 | chr10-125564064-INS-60 |
| chr10-125575012-INS-3601 | chr10-125575013-INS-3601 |
| chr10-125578367-INS-7127 | chr10-125578368-INS-7127 |
| chr10-125582610-INS-54 | chr10-125582610-INS-54 |
| chr10-125593422-INS-143 | chr10-125593422-INS-143 |
| chr10-125608067-INS-6234 | chr10-125608068-INS-6236 |
| chr10-125614424-INS-293 | chr10-125614424-INS-292 |
| chr10-125634360-INS-54 | chr10-125634363-INS-62 |
| chr10-125648575-INS-111 | chr10-125648567-INS-114 |
| chr10-125652617-INS-52 | chr10-125652617-INS-62 |
| chr10-125704490-INS-488 | chr10-125704490-INS-488 |
| chr10-125715006-INS-198 | chr10-125715013-INS-160 |
| chr10-125729098-INS-202 | chr10-125729098-INS-202 |
| chr10-125737206-INS-52 | chr10-125737206-INS-56 |
| chr10-125774280-INS-61 | chr10-125774278-INS-79 |
| chr10-125811445-INS-933 | chr10-125811445-INS-944 |
| chr10-125816503-INS-60 | chr10-125816503-INS-60 |
| chr10-125821799-INS-228 | chr10-125821799-INS-230 |
| chr10-125856196-INS-52 | chr10-125856200-INS-80 |
| chr10-125860847-INS-214 | chr10-125860847-INS-214 |
| chr10-125869730-INS-3915 | chr10-125869731-INS-3915 |
| chr10-125873357-INS-1015 | chr10-125873358-INS-1015 |
| chr10-125882346-INS-7080 | chr10-125882347-INS-7081 |
| chr10-125888844-INS-60 | chr10-125888844-INS-60 |
| chr10-125918299-INS-6757 | chr10-125918300-INS-6777 |
| chr10-125926980-INS-60 | chr10-125926980-INS-60 |
| chr10-125972305-INS-737 | chr10-125972305-INS-737 |
| chr10-125975921-INS-186 | chr10-125975922-INS-186 |
| chr10-125984837-INS-50 | chr10-125984796-INS-75 |
| chr10-125985574-INS-134 | chr10-125985579-INS-127 |
| chr10-126008173-INS-1132 | chr10-126008174-INS-1132 |
| chr10-126046525-INS-99 | chr10-126046528-INS-112 |
| chr10-126047836-INS-238 | chr10-126047836-INS-238 |
| chr10-126055917-INS-53 | chr10-126055911-INS-53 |
| chr10-126082222-INS-6086 | chr10-126082223-INS-6086 |
| chr10-126113959-INS-88 | chr10-126113917-INS-88 |
| chr10-126156305-INS-53 | chr10-126156305-INS-53 |
| chr10-126180914-INS-207 | chr10-126180914-INS-207 |
| chr10-126183777-INS-884 | chr10-126183786-INS-886 |
| chr10-126201223-INS-7140 | chr10-126201224-INS-7140 |
| chr10-126222567-INS-425 | chr10-126222567-INS-424 |
| chr10-126226435-INS-127 | chr10-126226435-INS-127 |
| chr10-126228499-INS-102 | chr10-126228459-INS-120 |
| chr10-126231576-INS-70 | chr10-126231576-INS-70 |
| chr10-126259957-INS-125 | chr10-126259957-INS-125 |
| chr10-126272481-INS-144 | chr10-126272481-INS-112 |
| chr10-126273349-INS-7242 | chr10-126273350-INS-7242 |
| chr10-126277248-INS-167 | chr10-126277248-INS-169 |
| chr10-126283195-INS-6579 | chr10-126283196-INS-6579 |
| chr10-126287268-INS-131 | chr10-126287266-INS-139 |
| chr10-126292526-INS-74 | chr10-126292544-INS-70 |
| chr10-126304941-INS-58 | chr10-126304941-INS-58 |
| chr10-126305533-INS-56 | chr10-126305533-INS-56 |
| chr10-126306599-INS-213 | chr10-126306599-INS-213 |
| chr10-126309068-INS-376 | chr10-126309068-INS-400 |
| chr10-126313722-INS-437 | chr10-126313722-INS-437 |
| chr10-126338784-INS-66 | chr10-126338755-INS-50 |
| chr10-126364834-INS-126 | chr10-126364834-INS-155 |
| chr10-126367034-INS-69 | chr10-126367028-INS-65 |
| chr10-126380269-INS-56 | chr10-126380227-INS-64 |
| chr10-126383266-INS-72 | chr10-126383287-INS-60 |
| chr10-126397764-INS-59 | chr10-126397764-INS-88 |

| | |
|---|---|
| chr10-126410640-INS-57 | chr10-126410631-INS-70 |
| chr10-126423940-INS-136 | chr10-126423940-INS-137 |
| chr10-126440128-INS-8976 | chr10-126440129-INS-8976 |
| chr10-126444732-INS-1373 | chr10-126444748-INS-1358 |
| chr10-126459400-INS-1729 | chr10-126459401-INS-1729 |
| chr10-126482140-INS-6214 | chr10-126482141-INS-6233 |
| chr10-126483731-INS-863 | chr10-126483731-INS-863 |
| chr10-126499909-INS-6898 | chr10-126499910-INS-6899 |
| chr10-126518497-INS-51 | chr10-126518498-INS-53 |
| chr10-126534855-INS-203 | chr10-126534855-INS-224 |
| chr10-126552439-INS-117 | chr10-126552442-INS-113 |
| chr10-126566396-INS-52 | chr10-126566396-INS-52 |
| chr10-126586387-INS-127 | chr10-126586387-INS-127 |
| chr10-126592555-INS-745 | chr10-126592555-INS-744 |
| chr10-126612716-INS-80 | chr10-126612716-INS-80 |
| chr10-126620042-INS-294 | chr10-126620006-INS-298 |
| chr10-126654979-INS-181 | chr10-126654979-INS-181 |
| chr10-126678161-INS-400 | chr10-126678161-INS-400 |
| chr10-126688688-INS-173 | chr10-126688688-INS-180 |
| chr10-126694778-INS-3667 | chr10-126694779-INS-3666 |
| chr10-126702659-INS-87 | chr10-126702659-INS-112 |
| chr10-126705196-INS-111 | chr10-126705204-INS-139 |
| chr10-126712877-INS-217 | chr10-126712877-INS-217 |
| chr10-126715000-INS-215 | chr10-126715000-INS-214 |
| chr10-126719708-INS-181 | chr10-126719708-INS-178 |
| chr10-126722246-INS-85 | chr10-126722248-INS-51 |
| chr10-126727076-INS-90 | chr10-126727082-INS-83 |
| chr10-126737548-INS-222 | chr10-126737548-INS-222 |
| chr10-126760528-INS-97 | chr10-126760492-INS-84 |
| chr10-126778558-INS-86 | chr10-126778554-INS-78 |
| chr10-126794263-INS-117 | chr10-126794263-INS-112 |
| chr10-126807294-INS-84 | chr10-126807298-INS-96 |
| chr10-126811562-INS-244 | chr10-126811562-INS-244 |
| chr10-126846496-INS-83 | chr10-126846496-INS-83 |
| chr10-126847902-INS-54 | chr10-126847902-INS-62 |
| chr10-126850990-INS-91 | chr10-126851005-INS-75 |
| chr10-126853471-INS-55 | chr10-126853472-INS-55 |
| chr10-126878534-INS-206 | chr10-126878534-INS-205 |
| chr10-126934088-INS-59 | chr10-126934068-INS-63 |
| chr10-126937586-INS-270 | chr10-126937586-INS-273 |
| chr10-126945015-INS-433 | chr10-126945015-INS-443 |
| chr10-126965646-INS-66 | chr10-126965646-INS-66 |
| chr10-126969657-INS-200 | chr10-126969657-INS-200 |
| chr10-126976855-INS-58 | chr10-126976854-INS-58 |
| chr10-126977710-INS-208 | chr10-126977710-INS-208 |
| chr10-126980192-INS-468 | chr10-126980192-INS-468 |
| chr10-126989756-INS-5446 | chr10-126989757-INS-5406 |
| chr10-126995246-INS-64 | chr10-126995246-INS-64 |
| chr10-127041427-INS-220 | chr10-127041427-INS-220 |
| chr10-127045572-INS-63 | chr10-127045572-INS-63 |
| chr10-127054371-INS-50 | chr10-127054371-INS-50 |
| chr10-127057386-INS-62 | chr10-127057399-INS-57 |
| chr10-127070223-INS-121 | chr10-127070224-INS-121 |
| chr10-127072906-INS-144 | chr10-127072906-INS-144 |
| chr10-127079152-INS-182 | chr10-127079152-INS-182 |
| chr10-127080703-INS-87 | chr10-127080703-INS-87 |
| chr10-127085592-INS-209 | chr10-127085592-INS-221 |
| chr10-127114494-INS-460 | chr10-127114494-INS-460 |
| chr10-127151535-INS-156 | chr10-127151531-INS-186 |
| chr10-127174589-INS-209 | chr10-127174590-INS-209 |
| chr10-127181782-INS-4747 | chr10-127181783-INS-4747 |
| chr10-127196125-INS-68 | chr10-127196125-INS-68 |
| chr10-127203010-INS-83 | chr10-127203012-INS-84 |
| chr10-127207143-INS-72 | chr10-127207157-INS-61 |
| chr10-127210107-INS-224 | chr10-127210109-INS-203 |
| chr10-127211455-INS-203 | chr10-127211455-INS-204 |
| chr10-127212250-INS-305 | chr10-127212250-INS-320 |
| chr10-127215549-INS-57 | chr10-127215549-INS-52 |
| chr10-127255933-INS-67 | chr10-127255932-INS-74 |
| chr10-127283933-INS-72 | chr10-127283933-INS-72 |
| chr10-127301340-INS-74 | chr10-127301341-INS-74 |
| chr10-127331629-INS-238 | chr10-127331629-INS-239 |
| chr10-127341286-INS-262 | chr10-127341286-INS-260 |
| chr10-127343187-INS-60 | chr10-127343187-INS-55 |
| chr10-127395732-INS-94 | chr10-127395776-INS-90 |
| chr10-127438474-INS-175 | chr10-127438474-INS-175 |
| chr10-127463805-INS-66 | chr10-127463805-INS-62 |

| | |
|---|---|
| chr10-127468455-INS-104 | chr10-127468455-INS-104 |
| chr10-127489828-INS-217 | chr10-127489828-INS-221 |
| chr10-127503841-INS-63 | chr10-127503846-INS-63 |
| chr10-127506496-INS-6395 | chr10-127506497-INS-6395 |
| chr10-127510790-INS-61 | chr10-127510790-INS-61 |
| chr10-127533459-INS-54 | chr10-127533459-INS-54 |
| chr10-127542126-INS-90 | chr10-127542126-INS-90 |
| chr10-127561049-INS-70 | chr10-127561056-INS-58 |
| chr10-127570563-INS-52 | chr10-127570563-INS-64 |
| chr10-127578251-INS-51 | chr10-127578251-INS-51 |
| chr10-127581270-INS-52 | chr10-127581292-INS-54 |
| chr10-127602899-INS-223 | chr10-127602899-INS-224 |
| chr10-127613423-INS-6443 | chr10-127613424-INS-6443 |
| chr10-127615042-INS-227 | chr10-127615042-INS-227 |
| chr10-127656755-INS-6401 | chr10-127656756-INS-6401 |
| chr10-127683355-INS-105 | chr10-127683355-INS-105 |
| chr10-127706624-INS-213 | chr10-127706624-INS-213 |
| chr10-127764762-INS-101 | chr10-127764762-INS-53 |
| chr10-127777822-INS-4987 | chr10-127777823-INS-4997 |
| chr10-127826416-INS-497 | chr10-127826416-INS-485 |
| chr10-127827651-INS-198 | chr10-127827651-INS-198 |
| chr10-127835094-INS-211 | chr10-127835094-INS-211 |
| chr10-127870327-INS-56 | chr10-127870327-INS-105 |
| chr10-127886998-INS-54 | chr10-127886998-INS-54 |
| chr10-127889694-INS-234 | chr10-127889694-INS-231 |
| chr10-127903218-INS-51 | chr10-127903218-INS-51 |
| chr10-127906163-INS-216 | chr10-127906163-INS-216 |
| chr10-127916018-INS-229 | chr10-127916018-INS-190 |
| chr10-127932015-INS-99 | chr10-127932015-INS-99 |
| chr10-127957649-INS-237 | chr10-127957649-INS-237 |
| chr10-127977626-INS-220 | chr10-127977626-INS-218 |
| chr10-127985413-INS-65 | chr10-127985413-INS-60 |
| chr10-127999864-INS-240 | chr10-127999864-INS-240 |
| chr10-128000919-INS-199 | chr10-128000919-INS-199 |
| chr10-128004029-INS-59 | chr10-128004025-INS-52 |
| chr10-128005722-INS-239 | chr10-128005722-INS-239 |
| chr10-128017981-INS-170 | chr10-128017981-INS-170 |
| chr10-128041603-INS-225 | chr10-128041602-INS-221 |
| chr10-128048458-INS-160 | chr10-128048458-INS-160 |
| chr10-128056388-INS-109 | chr10-128056388-INS-109 |
| chr10-128063756-INS-62 | chr10-128063756-INS-62 |
| chr10-128065587-INS-204 | chr10-128065587-INS-205 |
| chr10-128069501-INS-72 | chr10-128069501-INS-108 |
| chr10-128083726-INS-209 | chr10-128083726-INS-224 |
| chr10-128095401-INS-52 | chr10-128095401-INS-52 |
| chr10-128097028-INS-195 | chr10-128097028-INS-199 |
| chr10-128102633-INS-67 | chr10-128102633-INS-67 |
| chr10-128148489-INS-167 | chr10-128148489-INS-167 |
| chr10-128151995-INS-185 | chr10-128151995-INS-191 |
| chr10-128184730-INS-201 | chr10-128184730-INS-201 |
| chr10-128193078-INS-208 | chr10-128193078-INS-208 |
| chr10-128215768-INS-202 | chr10-128215768-INS-202 |
| chr10-128246401-INS-229 | chr10-128246401-INS-229 |
| chr10-128270427-INS-64 | chr10-128270427-INS-76 |
| chr10-128281828-INS-145 | chr10-128281828-INS-145 |
| chr10-128284514-INS-74 | chr10-128284508-INS-54 |
| chr10-128285538-INS-194 | chr10-128285538-INS-194 |
| chr10-128303416-INS-106 | chr10-128303416-INS-106 |
| chr10-128343608-INS-70 | chr10-128343612-INS-74 |
| chr10-128357819-INS-224 | chr10-128357819-INS-226 |
| chr10-128362758-INS-427 | chr10-128362758-INS-427 |
| chr10-128363540-INS-68 | chr10-128363540-INS-68 |
| chr10-128367741-INS-215 | chr10-128367741-INS-215 |
| chr10-128373447-INS-465 | chr10-128373447-INS-465 |
| chr10-128378812-INS-174 | chr10-128378812-INS-175 |
| chr10-128388517-INS-131 | chr10-128388504-INS-115 |
| chr10-128390095-INS-173 | chr10-128390095-INS-175 |
| chr10-128391102-INS-5546 | chr10-128391103-INS-5546 |
| chr10-128392534-INS-183 | chr10-128392535-INS-187 |
| chr10-128402569-INS-167 | chr10-128402578-INS-168 |
| chr10-128406510-INS-73 | chr10-128406510-INS-53 |
| chr10-128427733-INS-209 | chr10-128427733-INS-208 |
| chr10-128432171-INS-197 | chr10-128432169-INS-198 |
| chr10-128438891-INS-203 | chr10-128438891-INS-203 |
| chr10-128442218-INS-201 | chr10-128442218-INS-208 |
| chr10-128443172-INS-484 | chr10-128443161-INS-507 |
| chr10-128444512-INS-268 | chr10-128444512-INS-268 |

| | |
|---|---|
| chr10-128453649-INS-171 | chr10-128453650-INS-169 |
| chr10-128473748-INS-117 | chr10-128473771-INS-76 |
| chr10-128475062-INS-229 | chr10-128475062-INS-220 |
| chr10-128548934-INS-225 | chr10-128548935-INS-210 |
| chr10-128561074-INS-81 | chr10-128561074-INS-81 |
| chr10-128581727-INS-182 | chr10-128581727-INS-182 |
| chr10-128589470-INS-213 | chr10-128589470-INS-214 |
| chr10-128593196-INS-178 | chr10-128593196-INS-178 |
| chr10-128607716-INS-70 | chr10-128607716-INS-70 |
| chr10-128609369-INS-125 | chr10-128609369-INS-134 |
| chr10-128623591-INS-112 | chr10-128623591-INS-88 |
| chr10-128635147-INS-280 | chr10-128635179-INS-272 |
| chr10-128645077-INS-300 | chr10-128645078-INS-296 |
| chr10-128667498-INS-54 | chr10-128667499-INS-54 |
| chr10-128692522-INS-936 | chr10-128692522-INS-935 |
| chr10-128696179-INS-66 | chr10-128696179-INS-64 |
| chr10-128712962-INS-62 | chr10-128712962-INS-62 |
| chr10-128716357-INS-105 | chr10-128716351-INS-77 |
| chr10-128725926-INS-54 | chr10-128725926-INS-54 |
| chr10-128727878-INS-60 | chr10-128727878-INS-74 |
| chr10-128733444-INS-181 | chr10-128733406-INS-213 |
| chr10-128742190-INS-217 | chr10-128742190-INS-217 |
| chr10-128767915-INS-677 | chr10-128767915-INS-677 |
| chr10-128784157-INS-63 | chr10-128784157-INS-61 |
| chr10-128787156-INS-56 | chr10-128787150-INS-68 |
| chr10-128788449-INS-126 | chr10-128788450-INS-151 |
| chr10-128790170-INS-68 | chr10-128790170-INS-70 |
| chr10-128795228-INS-82 | chr10-128795217-INS-131 |
| chr10-128805129-INS-121 | chr10-128805135-INS-94 |
| chr10-128823989-INS-900 | chr10-128823989-INS-900 |
| chr10-128825545-INS-93 | chr10-128825545-INS-93 |
| chr10-128827796-INS-162 | chr10-128827797-INS-121 |
| chr10-128832722-INS-131 | chr10-128832719-INS-136 |
| chr10-128849456-INS-4341 | chr10-128849457-INS-4341 |
| chr10-128863905-INS-3350 | chr10-128863899-INS-3362 |
| chr10-128865464-INS-4075 | chr10-128865465-INS-4075 |
| chr10-128883292-INS-318 | chr10-128883292-INS-318 |
| chr10-128886482-INS-63 | chr10-128886482-INS-63 |
| chr10-128888636-INS-1869 | chr10-128888637-INS-1869 |
| chr10-128911048-INS-2021 | chr10-128911049-INS-2021 |
| chr10-128912802-INS-174 | chr10-128912802-INS-174 |
| chr10-128938800-INS-138 | chr10-128938800-INS-138 |
| chr10-128949731-INS-55 | chr10-128949756-INS-52 |
| chr10-129005078-INS-186 | chr10-129005078-INS-186 |
| chr10-129009102-INS-6671 | chr10-129009103-INS-6671 |
| chr10-129068457-INS-75 | chr10-129068457-INS-75 |
| chr10-129068816-INS-6368 | chr10-129068817-INS-6368 |
| chr10-129080912-INS-115 | chr10-129080912-INS-115 |
| chr10-129114909-INS-204 | chr10-129114874-INS-204 |
| chr10-129148478-INS-81 | chr10-129148478-INS-129 |
| chr10-129173741-INS-1888 | chr10-129173742-INS-1888 |
| chr10-129189518-INS-1575 | chr10-129189519-INS-1570 |
| chr10-129206921-INS-101 | chr10-129206925-INS-108 |
| chr10-129274265-INS-74 | chr10-129274265-INS-74 |
| chr10-129295857-INS-72 | chr10-129295857-INS-68 |
| chr10-129297529-INS-63 | chr10-129297529-INS-63 |
| chr10-129303015-INS-63 | chr10-129302998-INS-65 |
| chr10-129316593-INS-69 | chr10-129316593-INS-69 |
| chr10-129342392-INS-81 | chr10-129342392-INS-81 |
| chr10-129369162-INS-1228 | chr10-129369163-INS-1228 |
| chr10-129379673-INS-100 | chr10-129379674-INS-54 |
| chr10-129386071-INS-124 | chr10-129386071-INS-88 |
| chr10-129396187-INS-3095 | chr10-129396188-INS-3095 |
| chr10-129406147-INS-1064 | chr10-129406148-INS-1064 |
| chr10-129453925-INS-57 | chr10-129453925-INS-81 |
| chr10-129488811-INS-192 | chr10-129488811-INS-149 |
| chr10-129535684-INS-80 | chr10-129535696-INS-76 |
| chr10-129579649-INS-182 | chr10-129579649-INS-183 |
| chr10-129582155-INS-154 | chr10-129582155-INS-157 |
| chr10-129584422-INS-6229 | chr10-129584408-INS-6216 |
| chr10-129600987-INS-6392 | chr10-129600988-INS-6393 |
| chr10-129624944-INS-6199 | chr10-129624929-INS-6200 |
| chr10-129702606-INS-54 | chr10-129702606-INS-85 |
| chr10-129793541-INS-1445 | chr10-129793542-INS-1449 |
| chr10-129796781-INS-379 | chr10-129796781-INS-379 |
| chr10-129835269-INS-358 | chr10-129835269-INS-358 |
| chr10-129849167-INS-58 | chr10-129849167-INS-60 |

| | |
|---|---|
| chr10-129868294-INS-64 | chr10-129868294-INS-52 |
| chr10-129897922-INS-1006 | chr10-129897923-INS-999 |
| chr10-129944825-INS-446 | chr10-129944825-INS-446 |
| chr10-130010895-INS-127 | chr10-130010895-INS-127 |
| chr10-130024098-INS-1303 | chr10-130024099-INS-1301 |
| chr10-130169030-INS-443 | chr10-130169030-INS-437 |
| chr10-130232841-INS-52 | chr10-130232841-INS-52 |
| chr10-130246089-INS-506 | chr10-130246089-INS-513 |
| chr10-130302141-INS-1489 | chr10-130302142-INS-1489 |
| chr10-130362050-INS-52 | chr10-130362074-INS-50 |
| chr10-130372260-INS-5500 | chr10-130372261-INS-5500 |
| chr10-130416587-INS-51 | chr10-130416587-INS-52 |
| chr10-130427894-INS-181 | chr10-130427894-INS-185 |
| chr11-3068846-INS-362 | chr11-3068847-INS-366 |
| chr11-3101893-INS-248 | chr11-3101893-INS-248 |
| chr11-3131900-INS-168 | chr11-3131900-INS-168 |
| chr11-3160787-INS-61 | chr11-3160787-INS-55 |
| chr11-3175897-INS-54 | chr11-3175897-INS-54 |
| chr11-3187699-INS-54 | chr11-3187704-INS-62 |
| chr11-3192972-INS-428 | chr11-3192967-INS-433 |
| chr11-3201015-INS-62 | chr11-3201015-INS-62 |
| chr11-3220736-INS-56 | chr11-3220736-INS-58 |
| chr11-3236388-INS-53 | chr11-3236388-INS-53 |
| chr11-3268020-INS-77 | chr11-3268063-INS-56 |
| chr11-3284232-INS-182 | chr11-3284232-INS-187 |
| chr11-3290606-INS-68 | chr11-3290578-INS-50 |
| chr11-3293932-INS-90 | chr11-3293932-INS-90 |
| chr11-3309319-INS-61 | chr11-3309319-INS-61 |
| chr11-3336030-INS-231 | chr11-3336030-INS-230 |
| chr11-3349601-INS-240 | chr11-3349601-INS-240 |
| chr11-3351817-INS-96 | chr11-3351817-INS-96 |
| chr11-3380781-INS-3986 | chr11-3380782-INS-4012 |
| chr11-3380782-INS-66 | chr11-3380782-INS-60 |
| chr11-3416139-INS-62 | chr11-3416139-INS-62 |
| chr11-3433099-INS-92 | chr11-3433107-INS-78 |
| chr11-3455983-INS-60 | chr11-3455983-INS-78 |
| chr11-3483043-INS-90 | chr11-3483074-INS-106 |
| chr11-3507319-INS-63 | chr11-3507319-INS-63 |
| chr11-3552555-INS-132 | chr11-3552555-INS-117 |
| chr11-3561210-INS-281 | chr11-3561210-INS-281 |
| chr11-3570593-INS-83 | chr11-3570593-INS-80 |
| chr11-3573262-INS-59 | chr11-3573262-INS-51 |
| chr11-3674286-INS-150 | chr11-3674286-INS-150 |
| chr11-3776702-INS-202 | chr11-3776702-INS-202 |
| chr11-3786780-INS-218 | chr11-3786780-INS-210 |
| chr11-3789041-INS-167 | chr11-3789041-INS-146 |
| chr11-3824638-INS-74 | chr11-3824616-INS-58 |
| chr11-3830456-INS-69 | chr11-3830456-INS-69 |
| chr11-3836537-INS-132 | chr11-3836537-INS-132 |
| chr11-3845848-INS-400 | chr11-3845848-INS-402 |
| chr11-3858376-INS-81 | chr11-3858374-INS-84 |
| chr11-3866370-INS-304 | chr11-3866370-INS-304 |
| chr11-3868308-INS-255 | chr11-3868307-INS-256 |
| chr11-3882397-INS-61 | chr11-3882397-INS-70 |
| chr11-3892087-INS-205 | chr11-3892087-INS-205 |
| chr11-3904960-INS-61 | chr11-3904956-INS-61 |
| chr11-3912662-INS-1284 | chr11-3912663-INS-1284 |
| chr11-3963551-INS-159 | chr11-3963551-INS-159 |
| chr11-3975027-INS-74 | chr11-3975027-INS-79 |
| chr11-4005086-INS-414 | chr11-4005086-INS-414 |
| chr11-4007841-INS-54 | chr11-4007841-INS-54 |
| chr11-4099482-INS-218 | chr11-4099482-INS-217 |
| chr11-4105450-INS-70 | chr11-4105450-INS-60 |
| chr11-4108127-INS-191 | chr11-4108127-INS-185 |
| chr11-4116251-INS-70 | chr11-4116251-INS-70 |
| chr11-4149584-INS-219 | chr11-4149584-INS-222 |
| chr11-4213397-INS-321 | chr11-4213397-INS-323 |
| chr11-4225768-INS-207 | chr11-4225768-INS-206 |
| chr11-4227630-INS-100 | chr11-4227628-INS-103 |
| chr11-4275193-INS-427 | chr11-4275193-INS-425 |
| chr11-4275994-INS-62 | chr11-4275994-INS-90 |
| chr11-4285228-INS-58 | chr11-4285228-INS-52 |
| chr11-4292882-INS-62 | chr11-4292882-INS-98 |
| chr11-4305454-INS-331 | chr11-4305454-INS-331 |
| chr11-4306418-INS-108 | chr11-4306418-INS-108 |
| chr11-4317590-INS-74 | chr11-4317590-INS-76 |
| chr11-4328964-INS-98 | chr11-4328933-INS-96 |

| | |
|---|---|
| chr11-4352453-INS-73 | chr11-4352463-INS-54 |
| chr11-4409439-INS-91 | chr11-4409439-INS-91 |
| chr11-4427354-INS-144 | chr11-4427354-INS-144 |
| chr11-4464035-INS-209 | chr11-4464035-INS-210 |
| chr11-4548043-INS-356 | chr11-4548043-INS-386 |
| chr11-4600863-INS-400 | chr11-4600863-INS-400 |
| chr11-4606595-INS-175 | chr11-4606595-INS-175 |
| chr11-4610928-INS-167 | chr11-4610928-INS-167 |
| chr11-4629610-INS-66 | chr11-4629610-INS-62 |
| chr11-4654932-INS-205 | chr11-4654932-INS-206 |
| chr11-4657673-INS-204 | chr11-4657673-INS-204 |
| chr11-4666407-INS-54 | chr11-4666407-INS-54 |
| chr11-4689457-INS-72 | chr11-4689457-INS-72 |
| chr11-4740046-INS-231 | chr11-4740046-INS-232 |
| chr11-4746339-INS-196 | chr11-4746339-INS-196 |
| chr11-4746831-INS-2921 | chr11-4746832-INS-2921 |
| chr11-4771917-INS-211 | chr11-4771917-INS-211 |
| chr11-4782115-INS-444 | chr11-4782109-INS-445 |
| chr11-4790779-INS-116 | chr11-4790779-INS-116 |
| chr11-4792781-INS-57 | chr11-4792781-INS-61 |
| chr11-4801118-INS-230 | chr11-4801118-INS-211 |
| chr11-4813602-INS-248 | chr11-4813602-INS-248 |
| chr11-4816115-INS-184 | chr11-4816115-INS-184 |
| chr11-4823512-INS-57 | chr11-4823512-INS-59 |
| chr11-4828982-INS-4492 | chr11-4828983-INS-4492 |
| chr11-4857401-INS-58 | chr11-4857401-INS-58 |
| chr11-4860394-INS-208 | chr11-4860394-INS-209 |
| chr11-4879502-INS-203 | chr11-4879502-INS-215 |
| chr11-4909216-INS-78 | chr11-4909216-INS-78 |
| chr11-4909334-INS-679 | chr11-4909334-INS-679 |
| chr11-4917339-INS-406 | chr11-4917346-INS-406 |
| chr11-4921641-INS-226 | chr11-4921641-INS-226 |
| chr11-4941716-INS-209 | chr11-4941716-INS-208 |
| chr11-4961805-INS-70 | chr11-4961805-INS-80 |
| chr11-4971732-INS-1033 | chr11-4971733-INS-1033 |
| chr11-5034782-INS-231 | chr11-5034782-INS-205 |
| chr11-5045249-INS-199 | chr11-5045249-INS-199 |
| chr11-5063179-INS-60 | chr11-5063179-INS-71 |
| chr11-5073724-INS-55 | chr11-5073724-INS-55 |
| chr11-5074859-INS-194 | chr11-5074859-INS-194 |
| chr11-5085922-INS-228 | chr11-5085922-INS-228 |
| chr11-5093131-INS-88 | chr11-5093131-INS-88 |
| chr11-5119702-INS-2556 | chr11-5119703-INS-2556 |
| chr11-5129325-INS-59 | chr11-5129325-INS-50 |
| chr11-5145203-INS-303 | chr11-5145203-INS-303 |
| chr11-5156523-INS-2365 | chr11-5156524-INS-2365 |
| chr11-5165122-INS-54 | chr11-5165122-INS-58 |
| chr11-5172892-INS-282 | chr11-5172892-INS-282 |
| chr11-5186670-INS-128 | chr11-5186686-INS-144 |
| chr11-5196079-INS-105 | chr11-5196079-INS-105 |
| chr11-5202966-INS-200 | chr11-5202966-INS-200 |
| chr11-5277152-INS-200 | chr11-5277152-INS-200 |
| chr11-5336067-INS-126 | chr11-5336067-INS-126 |
| chr11-5337565-INS-3599 | chr11-5337566-INS-3593 |
| chr11-5351219-INS-927 | chr11-5351219-INS-936 |
| chr11-5390520-INS-51 | chr11-5390554-INS-53 |
| chr11-5399883-INS-53 | chr11-5399883-INS-53 |
| chr11-5401522-INS-4313 | chr11-5401523-INS-4303 |
| chr11-5419377-INS-200 | chr11-5419377-INS-200 |
| chr11-5425471-INS-172 | chr11-5425471-INS-172 |
| chr11-5426523-INS-210 | chr11-5426523-INS-210 |
| chr11-5457311-INS-136 | chr11-5457311-INS-87 |
| chr11-5459413-INS-58 | chr11-5459413-INS-58 |
| chr11-5492952-INS-196 | chr11-5492952-INS-196 |
| chr11-5532136-INS-71 | chr11-5532136-INS-71 |
| chr11-5538801-INS-171 | chr11-5538801-INS-171 |
| chr11-5547133-INS-82 | chr11-5547133-INS-82 |
| chr11-5567978-INS-7172 | chr11-5567979-INS-7172 |
| chr11-5574876-INS-1919 | chr11-5574877-INS-1919 |
| chr11-5575778-INS-85 | chr11-5575767-INS-114 |
| chr11-5577200-INS-208 | chr11-5577200-INS-208 |
| chr11-5580885-INS-7087 | chr11-5580886-INS-7087 |
| chr11-5620901-INS-102 | chr11-5620901-INS-121 |
| chr11-5638308-INS-104 | chr11-5638308-INS-104 |
| chr11-5659683-INS-60 | chr11-5659661-INS-50 |
| chr11-5673029-INS-98 | chr11-5673029-INS-98 |
| chr11-5675267-INS-55 | chr11-5675267-INS-55 |

| | |
|---|---|
| chr11-5683371-INS-131 | chr11-5683371-INS-131 |
| chr11-5692335-INS-445 | chr11-5692335-INS-445 |
| chr11-5692946-INS-84 | chr11-5692946-INS-84 |
| chr11-5697699-INS-51 | chr11-5697699-INS-55 |
| chr11-5707657-INS-504 | chr11-5707657-INS-504 |
| chr11-5712615-INS-209 | chr11-5712615-INS-209 |
| chr11-5714969-INS-175 | chr11-5714969-INS-175 |
| chr11-5723931-INS-212 | chr11-5723931-INS-212 |
| chr11-5740915-INS-204 | chr11-5740915-INS-204 |
| chr11-5769198-INS-55 | chr11-5769198-INS-55 |
| chr11-5770659-INS-556 | chr11-5770659-INS-556 |
| chr11-5771357-INS-171 | chr11-5771357-INS-171 |
| chr11-5773911-INS-168 | chr11-5773911-INS-168 |
| chr11-5808980-INS-5362 | chr11-5808981-INS-5362 |
| chr11-5813511-INS-213 | chr11-5813511-INS-210 |
| chr11-5845449-INS-222 | chr11-5845449-INS-222 |
| chr11-5854541-INS-57 | chr11-5854541-INS-57 |
| chr11-5855912-INS-1014 | chr11-5855913-INS-1014 |
| chr11-5855948-INS-70 | chr11-5855906-INS-70 |
| chr11-5866219-INS-66 | chr11-5866219-INS-78 |
| chr11-5875022-INS-122 | chr11-5875022-INS-122 |
| chr11-5889944-INS-2252 | chr11-5889948-INS-2250 |
| chr11-5907436-INS-246 | chr11-5907436-INS-246 |
| chr11-5917874-INS-1829 | chr11-5917875-INS-1829 |
| chr11-5951626-INS-126 | chr11-5951626-INS-126 |
| chr11-5955403-INS-177 | chr11-5955403-INS-158 |
| chr11-5963201-INS-249 | chr11-5963201-INS-248 |
| chr11-5966441-INS-96 | chr11-5966419-INS-68 |
| chr11-5989465-INS-132 | chr11-5989465-INS-132 |
| chr11-5997062-INS-78 | chr11-5997062-INS-78 |
| chr11-5999366-INS-54 | chr11-5999360-INS-84 |
| chr11-6036504-INS-214 | chr11-6036504-INS-213 |
| chr11-6131347-INS-255 | chr11-6131347-INS-237 |
| chr11-6152963-INS-422 | chr11-6152963-INS-422 |
| chr11-6157551-INS-5254 | chr11-6157552-INS-5254 |
| chr11-6163150-INS-116 | chr11-6163150-INS-124 |
| chr11-6199892-INS-236 | chr11-6199892-INS-236 |
| chr11-6233866-INS-84 | chr11-6233888-INS-54 |
| chr11-6236350-INS-7128 | chr11-6236351-INS-7128 |
| chr11-6246671-INS-227 | chr11-6246671-INS-226 |
| chr11-6291396-INS-1144 | chr11-6291397-INS-1144 |
| chr11-6303226-INS-216 | chr11-6303226-INS-216 |
| chr11-6305448-INS-181 | chr11-6305448-INS-182 |
| chr11-6314027-INS-68 | chr11-6314027-INS-60 |
| chr11-6315461-INS-204 | chr11-6315461-INS-204 |
| chr11-6319550-INS-179 | chr11-6319550-INS-179 |
| chr11-6336706-INS-158 | chr11-6336707-INS-158 |
| chr11-6338447-INS-211 | chr11-6338438-INS-213 |
| chr11-6341998-INS-216 | chr11-6341998-INS-221 |
| chr11-6372414-INS-194 | chr11-6372414-INS-195 |
| chr11-6379956-INS-71 | chr11-6379956-INS-71 |
| chr11-6385676-INS-60 | chr11-6385686-INS-100 |
| chr11-6408874-INS-71 | chr11-6408874-INS-75 |
| chr11-6411135-INS-221 | chr11-6411135-INS-220 |
| chr11-6412481-INS-198 | chr11-6412481-INS-199 |
| chr11-6415269-INS-66 | chr11-6415276-INS-71 |
| chr11-6464359-INS-202 | chr11-6464359-INS-202 |
| chr11-6477370-INS-136 | chr11-6477370-INS-139 |
| chr11-6510433-INS-205 | chr11-6510433-INS-205 |
| chr11-6514552-INS-76 | chr11-6514552-INS-76 |
| chr11-6526769-INS-202 | chr11-6526769-INS-202 |
| chr11-6533169-INS-216 | chr11-6533169-INS-216 |
| chr11-6549338-INS-66 | chr11-6549338-INS-70 |
| chr11-6577115-INS-239 | chr11-6577115-INS-239 |
| chr11-6587563-INS-99 | chr11-6587564-INS-99 |
| chr11-6599310-INS-687 | chr11-6599310-INS-687 |
| chr11-6640095-INS-66 | chr11-6640095-INS-66 |
| chr11-6660623-INS-77 | chr11-6660623-INS-77 |
| chr11-6689393-INS-146 | chr11-6689393-INS-146 |
| chr11-6698192-INS-208 | chr11-6698192-INS-209 |
| chr11-6713873-INS-228 | chr11-6713873-INS-229 |
| chr11-6720159-INS-58 | chr11-6720159-INS-58 |
| chr11-6730973-INS-529 | chr11-6730973-INS-529 |
| chr11-6734159-INS-1187 | chr11-6734160-INS-1212 |
| chr11-6755445-INS-51 | chr11-6755445-INS-51 |
| chr11-6760215-INS-64 | chr11-6760215-INS-68 |
| chr11-6768716-INS-930 | chr11-6768716-INS-933 |

| | |
|---|---|
| chr11-6770949-INS-327 | chr11-6770949-INS-329 |
| chr11-6773259-INS-90 | chr11-6773259-INS-90 |
| chr11-6777510-INS-256 | chr11-6777510-INS-256 |
| chr11-6778630-INS-64 | chr11-6778630-INS-62 |
| chr11-6809282-INS-6403 | chr11-6809283-INS-6403 |
| chr11-6820041-INS-110 | chr11-6820041-INS-110 |
| chr11-6829617-INS-74 | chr11-6829617-INS-74 |
| chr11-6836179-INS-411 | chr11-6836179-INS-411 |
| chr11-6838218-INS-894 | chr11-6838218-INS-877 |
| chr11-6852203-INS-486 | chr11-6852203-INS-486 |
| chr11-6867740-INS-79 | chr11-6867740-INS-79 |
| chr11-6878201-INS-58 | chr11-6878201-INS-53 |
| chr11-6880392-INS-52 | chr11-6880392-INS-54 |
| chr11-6890299-INS-87 | chr11-6890324-INS-86 |
| chr11-6920128-INS-4856 | chr11-6920129-INS-4856 |
| chr11-6939803-INS-75 | chr11-6939797-INS-81 |
| chr11-6969152-INS-6195 | chr11-6969153-INS-6197 |
| chr11-6971779-INS-515 | chr11-6971779-INS-515 |
| chr11-6977800-INS-711 | chr11-6977800-INS-712 |
| chr11-6993802-INS-67 | chr11-6993802-INS-67 |
| chr11-7035812-INS-115 | chr11-7035812-INS-115 |
| chr11-7050994-INS-7111 | chr11-7050995-INS-7111 |
| chr11-7082208-INS-59 | chr11-7082209-INS-76 |
| chr11-7135520-INS-325 | chr11-7135520-INS-325 |
| chr11-7136837-INS-4838 | chr11-7136838-INS-4838 |
| chr11-7145922-INS-227 | chr11-7145922-INS-228 |
| chr11-7193916-INS-3654 | chr11-7193915-INS-3656 |
| chr11-7282578-INS-767 | chr11-7282578-INS-765 |
| chr11-7285039-INS-1124 | chr11-7285040-INS-1124 |
| chr11-7285606-INS-248 | chr11-7285606-INS-248 |
| chr11-7287369-INS-70 | chr11-7287369-INS-70 |
| chr11-7298421-INS-52 | chr11-7298421-INS-53 |
| chr11-7342944-INS-179 | chr11-7342944-INS-188 |
| chr11-7359473-INS-1079 | chr11-7359474-INS-1079 |
| chr11-7362238-INS-6440 | chr11-7362239-INS-6440 |
| chr11-7366532-INS-58 | chr11-7366532-INS-58 |
| chr11-7411025-INS-440 | chr11-7411025-INS-440 |
| chr11-7420810-INS-68 | chr11-7420810-INS-72 |
| chr11-7427300-INS-80 | chr11-7427300-INS-80 |
| chr11-7439137-INS-1263 | chr11-7439138-INS-1263 |
| chr11-7442996-INS-52 | chr11-7442996-INS-52 |
| chr11-7476574-INS-97 | chr11-7476574-INS-97 |
| chr11-7478405-INS-559 | chr11-7478405-INS-562 |
| chr11-7489163-INS-6291 | chr11-7489164-INS-6293 |
| chr11-7491248-INS-76 | chr11-7491240-INS-124 |
| chr11-7493854-INS-70 | chr11-7493854-INS-70 |
| chr11-7509437-INS-70 | chr11-7509437-INS-73 |
| chr11-7564492-INS-58 | chr11-7564492-INS-52 |
| chr11-7588068-INS-6933 | chr11-7588069-INS-6930 |
| chr11-7588430-INS-110 | chr11-7588394-INS-108 |
| chr11-7603883-INS-204 | chr11-7603883-INS-204 |
| chr11-7621668-INS-92 | chr11-7621668-INS-92 |
| chr11-7626251-INS-126 | chr11-7626251-INS-126 |
| chr11-7633233-INS-182 | chr11-7633278-INS-151 |
| chr11-7661216-INS-7338 | chr11-7661217-INS-7338 |
| chr11-7685459-INS-78 | chr11-7685459-INS-80 |
| chr11-7696407-INS-215 | chr11-7696407-INS-215 |
| chr11-7698240-INS-57 | chr11-7698240-INS-56 |
| chr11-7700105-INS-64 | chr11-7700105-INS-68 |
| chr11-7727635-INS-6172 | chr11-7727636-INS-6172 |
| chr11-7770565-INS-57 | chr11-7770565-INS-57 |
| chr11-7784134-INS-53 | chr11-7784134-INS-53 |
| chr11-7787987-INS-94 | chr11-7787987-INS-94 |
| chr11-7795481-INS-97 | chr11-7795481-INS-93 |
| chr11-7848612-INS-85 | chr11-7848612-INS-61 |
| chr11-7921831-INS-579 | chr11-7921831-INS-579 |
| chr11-7944998-INS-6691 | chr11-7944999-INS-6691 |
| chr11-7963050-INS-351 | chr11-7963050-INS-351 |
| chr11-7963659-INS-87 | chr11-7963658-INS-54 |
| chr11-7968140-INS-328 | chr11-7968140-INS-328 |
| chr11-8002689-INS-148 | chr11-8002689-INS-148 |
| chr11-8005044-INS-53 | chr11-8005044-INS-53 |
| chr11-8009789-INS-1045 | chr11-8009790-INS-1046 |
| chr11-8022125-INS-58 | chr11-8022125-INS-78 |
| chr11-8041298-INS-60 | chr11-8041298-INS-60 |
| chr11-8053684-INS-327 | chr11-8053684-INS-327 |
| chr11-8067990-INS-401 | chr11-8067990-INS-401 |

| | |
|---|---|
| chr11-8068202-INS-6586 | chr11-8068203-INS-6587 |
| chr11-8072833-INS-128 | chr11-8072845-INS-157 |
| chr11-8089172-INS-1607 | chr11-8089173-INS-1626 |
| chr11-8092480-INS-102 | chr11-8092511-INS-87 |
| chr11-8098940-INS-80 | chr11-8098940-INS-51 |
| chr11-8123286-INS-69 | chr11-8123286-INS-69 |
| chr11-8134156-INS-6257 | chr11-8134157-INS-6262 |
| chr11-8140021-INS-3260 | chr11-8140023-INS-3256 |
| chr11-8164999-INS-60 | chr11-8164999-INS-55 |
| chr11-8171160-INS-65 | chr11-8171160-INS-57 |
| chr11-8201063-INS-52 | chr11-8201063-INS-65 |
| chr11-8205033-INS-389 | chr11-8205033-INS-389 |
| chr11-8268457-INS-198 | chr11-8268418-INS-198 |
| chr11-8281403-INS-217 | chr11-8281403-INS-217 |
| chr11-8333846-INS-133 | chr11-8333846-INS-133 |
| chr11-8335097-INS-389 | chr11-8335097-INS-389 |
| chr11-8338289-INS-68 | chr11-8338289-INS-68 |
| chr11-8356116-INS-681 | chr11-8356116-INS-681 |
| chr11-8404690-INS-116 | chr11-8404687-INS-106 |
| chr11-8411067-INS-52 | chr11-8411067-INS-62 |
| chr11-8474585-INS-232 | chr11-8474585-INS-232 |
| chr11-8505208-INS-211 | chr11-8505208-INS-211 |
| chr11-8518265-INS-80 | chr11-8518235-INS-83 |
| chr11-8541791-INS-191 | chr11-8541791-INS-191 |
| chr11-8548747-INS-78 | chr11-8548745-INS-72 |
| chr11-8561082-INS-678 | chr11-8561082-INS-679 |
| chr11-8563825-INS-80 | chr11-8563825-INS-66 |
| chr11-8595696-INS-65 | chr11-8595696-INS-65 |
| chr11-8598160-INS-5547 | chr11-8598161-INS-5547 |
| chr11-8619080-INS-236 | chr11-8619080-INS-236 |
| chr11-8621748-INS-145 | chr11-8621748-INS-145 |
| chr11-8623358-INS-249 | chr11-8623370-INS-249 |
| chr11-8632330-INS-6079 | chr11-8632331-INS-6079 |
| chr11-8634158-INS-56 | chr11-8634158-INS-56 |
| chr11-8637301-INS-76 | chr11-8637301-INS-76 |
| chr11-8656133-INS-52 | chr11-8656133-INS-52 |
| chr11-8659527-INS-1583 | chr11-8659528-INS-1576 |
| chr11-8660241-INS-7162 | chr11-8660242-INS-7162 |
| chr11-8663850-INS-81 | chr11-8663850-INS-81 |
| chr11-8665473-INS-206 | chr11-8665473-INS-206 |
| chr11-8666970-INS-130 | chr11-8666970-INS-130 |
| chr11-8696042-INS-66 | chr11-8696071-INS-76 |
| chr11-8698983-INS-59 | chr11-8698984-INS-57 |
| chr11-8709343-INS-70 | chr11-8709343-INS-54 |
| chr11-8710829-INS-261 | chr11-8710829-INS-262 |
| chr11-8717321-INS-51 | chr11-8717331-INS-51 |
| chr11-8723416-INS-470 | chr11-8723416-INS-472 |
| chr11-8735561-INS-84 | chr11-8735561-INS-84 |
| chr11-8756702-INS-904 | chr11-8756702-INS-904 |
| chr11-8756726-INS-295 | chr11-8756702-INS-304 |
| chr11-8762743-INS-216 | chr11-8762743-INS-216 |
| chr11-8824819-INS-54 | chr11-8824819-INS-54 |
| chr11-8843227-INS-287 | chr11-8843233-INS-291 |
| chr11-8846833-INS-190 | chr11-8846833-INS-189 |
| chr11-8849120-INS-159 | chr11-8849120-INS-159 |
| chr11-8851561-INS-211 | chr11-8851561-INS-211 |
| chr11-8861499-INS-212 | chr11-8861499-INS-212 |
| chr11-8863049-INS-1890 | chr11-8863050-INS-1890 |
| chr11-8902831-INS-58 | chr11-8902831-INS-58 |
| chr11-8935027-INS-71 | chr11-8935027-INS-72 |
| chr11-8936442-INS-68 | chr11-8936442-INS-70 |
| chr11-8957367-INS-338 | chr11-8957367-INS-338 |
| chr11-8972231-INS-96 | chr11-8972231-INS-96 |
| chr11-9005853-INS-82 | chr11-9005853-INS-84 |
| chr11-9056746-INS-1894 | chr11-9056747-INS-1894 |
| chr11-9059661-INS-208 | chr11-9059661-INS-207 |
| chr11-9089185-INS-1831 | chr11-9089186-INS-1831 |
| chr11-9121417-INS-1040 | chr11-9121418-INS-1039 |
| chr11-9123003-INS-274 | chr11-9123005-INS-275 |
| chr11-9126854-INS-120 | chr11-9126854-INS-120 |
| chr11-9153301-INS-214 | chr11-9153301-INS-214 |
| chr11-9159300-INS-154 | chr11-9159261-INS-146 |
| chr11-9164941-INS-71 | chr11-9164941-INS-71 |
| chr11-9168097-INS-382 | chr11-9168097-INS-382 |
| chr11-9190372-INS-93 | chr11-9190372-INS-113 |
| chr11-9192327-INS-53 | chr11-9192305-INS-50 |
| chr11-9255940-INS-102 | chr11-9255940-INS-84 |

| | |
|---|---|
| chr11-9256077-INS-6581 | chr11-9256078-INS-6581 |
| chr11-9259523-INS-219 | chr11-9259523-INS-219 |
| chr11-9277463-INS-557 | chr11-9277463-INS-557 |
| chr11-9279459-INS-707 | chr11-9279459-INS-707 |
| chr11-9282484-INS-50 | chr11-9282484-INS-74 |
| chr11-9339676-INS-75 | chr11-9339676-INS-75 |
| chr11-9343024-INS-7385 | chr11-9343025-INS-7385 |
| chr11-9343205-INS-92 | chr11-9343205-INS-92 |
| chr11-9352011-INS-458 | chr11-9352011-INS-459 |
| chr11-9356154-INS-401 | chr11-9356154-INS-401 |
| chr11-9401949-INS-6442 | chr11-9401950-INS-6442 |
| chr11-9417756-INS-55 | chr11-9417778-INS-61 |
| chr11-9438309-INS-55 | chr11-9438309-INS-54 |
| chr11-9451658-INS-147 | chr11-9451687-INS-147 |
| chr11-9452323-INS-6803 | chr11-9452324-INS-6804 |
| chr11-9520039-INS-55 | chr11-9520039-INS-55 |
| chr11-9532511-INS-79 | chr11-9532511-INS-79 |
| chr11-9535738-INS-564 | chr11-9535738-INS-564 |
| chr11-9542332-INS-65 | chr11-9542332-INS-75 |
| chr11-9549109-INS-109 | chr11-9549109-INS-109 |
| chr11-9553699-INS-180 | chr11-9553700-INS-183 |
| chr11-9568524-INS-60 | chr11-9568524-INS-60 |
| chr11-9616258-INS-74 | chr11-9616258-INS-74 |
| chr11-9647208-INS-201 | chr11-9647208-INS-201 |
| chr11-9668692-INS-116 | chr11-9668692-INS-110 |
| chr11-9686806-INS-84 | chr11-9686806-INS-84 |
| chr11-9705201-INS-2709 | chr11-9705202-INS-2708 |
| chr11-9709526-INS-832 | chr11-9709526-INS-833 |
| chr11-9723104-INS-1491 | chr11-9723105-INS-1491 |
| chr11-9731044-INS-488 | chr11-9731044-INS-488 |
| chr11-9733691-INS-104 | chr11-9733691-INS-104 |
| chr11-9735539-INS-56 | chr11-9735539-INS-58 |
| chr11-9789736-INS-163 | chr11-9789736-INS-163 |
| chr11-9796679-INS-56 | chr11-9796679-INS-51 |
| chr11-9806602-INS-325 | chr11-9806602-INS-325 |
| chr11-9844144-INS-390 | chr11-9844144-INS-390 |
| chr11-9852093-INS-92 | chr11-9852090-INS-125 |
| chr11-9859543-INS-75 | chr11-9859543-INS-75 |
| chr11-9875392-INS-7968 | chr11-9875393-INS-7968 |
| chr11-9894741-INS-51 | chr11-9894741-INS-51 |
| chr11-9911460-INS-562 | chr11-9911460-INS-562 |
| chr11-9929964-INS-59 | chr11-9929964-INS-60 |
| chr11-9968507-INS-200 | chr11-9968507-INS-200 |
| chr11-9975367-INS-2278 | chr11-9975368-INS-2280 |
| chr11-9982907-INS-67 | chr11-9982907-INS-67 |
| chr11-10003460-INS-52 | chr11-10003422-INS-58 |
| chr11-10003609-INS-1240 | chr11-10003610-INS-1241 |
| chr11-10011965-INS-52 | chr11-10011965-INS-50 |
| chr11-10028181-INS-369 | chr11-10028181-INS-369 |
| chr11-10030180-INS-56 | chr11-10030169-INS-68 |
| chr11-10053512-INS-4763 | chr11-10053513-INS-4763 |
| chr11-10062794-INS-8809 | chr11-10062795-INS-8809 |
| chr11-10082751-INS-51 | chr11-10082751-INS-57 |
| chr11-10102381-INS-4962 | chr11-10102382-INS-4962 |
| chr11-10112104-INS-4305 | chr11-10112105-INS-4305 |
| chr11-10125750-INS-6442 | chr11-10125742-INS-6438 |
| chr11-10128798-INS-64 | chr11-10128798-INS-64 |
| chr11-10132833-INS-1724 | chr11-10132834-INS-1723 |
| chr11-10138824-INS-1889 | chr11-10138825-INS-1889 |
| chr11-10152478-INS-72 | chr11-10152478-INS-72 |
| chr11-10154970-INS-51 | chr11-10154980-INS-86 |
| chr11-10161317-INS-61 | chr11-10161318-INS-61 |
| chr11-10219076-INS-66 | chr11-10219076-INS-66 |
| chr11-10263434-INS-276 | chr11-10263434-INS-276 |
| chr11-10264363-INS-386 | chr11-10264397-INS-339 |
| chr11-10268783-INS-231 | chr11-10268783-INS-231 |
| chr11-10290186-INS-585 | chr11-10290192-INS-626 |
| chr11-10330887-INS-166 | chr11-10330887-INS-184 |
| chr11-10332054-INS-541 | chr11-10332054-INS-541 |
| chr11-10336359-INS-474 | chr11-10336359-INS-474 |
| chr11-10342479-INS-57 | chr11-10342479-INS-60 |
| chr11-10346797-INS-67 | chr11-10346797-INS-67 |
| chr11-10361613-INS-74 | chr11-10361613-INS-82 |
| chr11-10417480-INS-674 | chr11-10417480-INS-674 |
| chr11-10447103-INS-122 | chr11-10447116-INS-136 |
| chr11-10449132-INS-3848 | chr11-10449133-INS-3808 |
| chr11-10454806-INS-576 | chr11-10454806-INS-593 |

| | |
|---|---|
| chr11-10479734-INS-208 | chr11-10479734-INS-208 |
| chr11-10492697-INS-125 | chr11-10492697-INS-125 |
| chr11-10503024-INS-84 | chr11-10503024-INS-98 |
| chr11-10559253-INS-420 | chr11-10559253-INS-420 |
| chr11-10609991-INS-5981 | chr11-10609992-INS-5981 |
| chr11-10620302-INS-104 | chr11-10620302-INS-104 |
| chr11-10671135-INS-375 | chr11-10671135-INS-375 |
| chr11-10678248-INS-547 | chr11-10678248-INS-547 |
| chr11-10696828-INS-195 | chr11-10696828-INS-195 |
| chr11-10697761-INS-68 | chr11-10697761-INS-70 |
| chr11-10708526-INS-7162 | chr11-10708527-INS-7162 |
| chr11-10746790-INS-57 | chr11-10746790-INS-59 |
| chr11-10850444-INS-211 | chr11-10850444-INS-211 |
| chr11-10851379-INS-63 | chr11-10851379-INS-76 |
| chr11-10856158-INS-894 | chr11-10856158-INS-894 |
| chr11-10873462-INS-103 | chr11-10873462-INS-96 |
| chr11-10937203-INS-4053 | chr11-10937204-INS-4053 |
| chr11-10940152-INS-399 | chr11-10940153-INS-399 |
| chr11-10945878-INS-197 | chr11-10945878-INS-203 |
| chr11-10952168-INS-7042 | chr11-10952169-INS-7042 |
| chr11-10958513-INS-1901 | chr11-10958514-INS-1901 |
| chr11-10965065-INS-82 | chr11-10965065-INS-82 |
| chr11-10967194-INS-71 | chr11-10967194-INS-73 |
| chr11-10970672-INS-131 | chr11-10970699-INS-130 |
| chr11-11029741-INS-7106 | chr11-11029742-INS-7106 |
| chr11-11050286-INS-2585 | chr11-11050289-INS-2581 |
| chr11-11053253-INS-105 | chr11-11053248-INS-104 |
| chr11-11059684-INS-1890 | chr11-11059685-INS-1890 |
| chr11-11079893-INS-66 | chr11-11079893-INS-68 |
| chr11-11088412-INS-116 | chr11-11088412-INS-116 |
| chr11-11110374-INS-201 | chr11-11110374-INS-201 |
| chr11-11128279-INS-140 | chr11-11128279-INS-140 |
| chr11-11133949-INS-52 | chr11-11133949-INS-50 |
| chr11-11194022-INS-53 | chr11-11194041-INS-57 |
| chr11-11199365-INS-88 | chr11-11199365-INS-74 |
| chr11-11237379-INS-212 | chr11-11237379-INS-212 |
| chr11-11257005-INS-289 | chr11-11257005-INS-289 |
| chr11-11288690-INS-7175 | chr11-11288691-INS-7175 |
| chr11-11343888-INS-76 | chr11-11343888-INS-76 |
| chr11-11350568-INS-127 | chr11-11350568-INS-127 |
| chr11-11364034-INS-61 | chr11-11364032-INS-67 |
| chr11-11403812-INS-94 | chr11-11403812-INS-101 |
| chr11-11404801-INS-267 | chr11-11404801-INS-267 |
| chr11-11451171-INS-87 | chr11-11451171-INS-87 |
| chr11-11461070-INS-64 | chr11-11461070-INS-64 |
| chr11-11465267-INS-219 | chr11-11465267-INS-218 |
| chr11-11495292-INS-84 | chr11-11495292-INS-84 |
| chr11-11502308-INS-86 | chr11-11502308-INS-97 |
| chr11-11503918-INS-5202 | chr11-11503919-INS-5202 |
| chr11-11515427-INS-130 | chr11-11515427-INS-130 |
| chr11-11544093-INS-300 | chr11-11544093-INS-300 |
| chr11-11549722-INS-325 | chr11-11549722-INS-325 |
| chr11-11628424-INS-618 | chr11-11628424-INS-613 |
| chr11-11645532-INS-67 | chr11-11645532-INS-67 |
| chr11-11678663-INS-702 | chr11-11678663-INS-703 |
| chr11-11691510-INS-56 | chr11-11691520-INS-68 |
| chr11-11694761-INS-52 | chr11-11694761-INS-52 |
| chr11-11716095-INS-52 | chr11-11716095-INS-52 |
| chr11-11738207-INS-260 | chr11-11738207-INS-260 |
| chr11-11740384-INS-194 | chr11-11740384-INS-193 |
| chr11-11742318-INS-86 | chr11-11742318-INS-104 |
| chr11-11766971-INS-69 | chr11-11766975-INS-60 |
| chr11-11767819-INS-199 | chr11-11767819-INS-199 |
| chr11-11785153-INS-216 | chr11-11785153-INS-216 |
| chr11-11792035-INS-199 | chr11-11792035-INS-202 |
| chr11-11797423-INS-67 | chr11-11797412-INS-95 |
| chr11-11812040-INS-219 | chr11-11812040-INS-219 |
| chr11-11817131-INS-7102 | chr11-11817132-INS-7102 |
| chr11-11819680-INS-62 | chr11-11819680-INS-60 |
| chr11-11849215-INS-61 | chr11-11849186-INS-107 |
| chr11-11850986-INS-118 | chr11-11850986-INS-118 |
| chr11-11860798-INS-118 | chr11-11860799-INS-118 |
| chr11-11945340-INS-241 | chr11-11945340-INS-241 |
| chr11-12030412-INS-3790 | chr11-12030413-INS-3790 |
| chr11-12042588-INS-146 | chr11-12042588-INS-146 |
| chr11-12065042-INS-96 | chr11-12065042-INS-96 |
| chr11-12066954-INS-384 | chr11-12066954-INS-384 |

| | |
|---|---|
| chr11-12068890-INS-1473 | chr11-12068891-INS-1473 |
| chr11-12115945-INS-58 | chr11-12115945-INS-58 |
| chr11-12143531-INS-5396 | chr11-12143532-INS-5396 |
| chr11-12146291-INS-835 | chr11-12146291-INS-839 |
| chr11-12153883-INS-54 | chr11-12153883-INS-56 |
| chr11-12155183-INS-1290 | chr11-12155184-INS-1292 |
| chr11-12192250-INS-7299 | chr11-12192251-INS-7300 |
| chr11-12242767-INS-6648 | chr11-12242768-INS-6649 |
| chr11-12254644-INS-267 | chr11-12254644-INS-267 |
| chr11-12263785-INS-348 | chr11-12263785-INS-348 |
| chr11-12374648-INS-69 | chr11-12374648-INS-69 |
| chr11-12381777-INS-6489 | chr11-12381771-INS-6477 |
| chr11-12418399-INS-85 | chr11-12418390-INS-70 |
| chr11-12419261-INS-269 | chr11-12419261-INS-267 |
| chr11-12420066-INS-72 | chr11-12420066-INS-76 |
| chr11-12433329-INS-73 | chr11-12433329-INS-76 |
| chr11-12436329-INS-248 | chr11-12436329-INS-248 |
| chr11-12445651-INS-74 | chr11-12445651-INS-74 |
| chr11-12447453-INS-67 | chr11-12447453-INS-67 |
| chr11-12448439-INS-60 | chr11-12448439-INS-68 |
| chr11-12465542-INS-233 | chr11-12465542-INS-233 |
| chr11-12481920-INS-54 | chr11-12481920-INS-54 |
| chr11-12494191-INS-76 | chr11-12494191-INS-75 |
| chr11-12507599-INS-342 | chr11-12507599-INS-354 |
| chr11-12536363-INS-53 | chr11-12536363-INS-54 |
| chr11-12548714-INS-130 | chr11-12548714-INS-130 |
| chr11-12572840-INS-6406 | chr11-12572841-INS-6407 |
| chr11-12599757-INS-3060 | chr11-12599758-INS-3060 |
| chr11-12605882-INS-4327 | chr11-12605883-INS-4329 |
| chr11-12605901-INS-227 | chr11-12605901-INS-227 |
| chr11-12607059-INS-1956 | chr11-12607060-INS-1956 |
| chr11-12644971-INS-82 | chr11-12644969-INS-84 |
| chr11-12665589-INS-56 | chr11-12665589-INS-56 |
| chr11-12681214-INS-1628 | chr11-12681215-INS-1628 |
| chr11-12683987-INS-75 | chr11-12683987-INS-80 |
| chr11-12694347-INS-282 | chr11-12694347-INS-284 |
| chr11-12728371-INS-203 | chr11-12728371-INS-203 |
| chr11-12764607-INS-1539 | chr11-12764608-INS-1539 |
| chr11-12779792-INS-7120 | chr11-12779793-INS-7120 |
| chr11-12803880-INS-6869 | chr11-12803881-INS-6870 |
| chr11-12819753-INS-3338 | chr11-12819754-INS-3338 |
| chr11-12823340-INS-249 | chr11-12823340-INS-241 |
| chr11-12826279-INS-52 | chr11-12826279-INS-54 |
| chr11-12827137-INS-751 | chr11-12827137-INS-734 |
| chr11-12831731-INS-57 | chr11-12831731-INS-54 |
| chr11-12835896-INS-71 | chr11-12835897-INS-57 |
| chr11-12856269-INS-198 | chr11-12856302-INS-198 |
| chr11-12863570-INS-2190 | chr11-12863573-INS-2186 |
| chr11-12918261-INS-629 | chr11-12918261-INS-629 |
| chr11-12984348-INS-592 | chr11-12984348-INS-592 |
| chr11-12985751-INS-78 | chr11-12985784-INS-62 |
| chr11-13002705-INS-763 | chr11-13002705-INS-763 |
| chr11-13030593-INS-3350 | chr11-13030594-INS-3350 |
| chr11-13032943-INS-81 | chr11-13032944-INS-80 |
| chr11-13048554-INS-6222 | chr11-13048555-INS-6222 |
| chr11-13052984-INS-749 | chr11-13052984-INS-749 |
| chr11-13060599-INS-69 | chr11-13060599-INS-69 |
| chr11-13075339-INS-56 | chr11-13075339-INS-56 |
| chr11-13076546-INS-74 | chr11-13076546-INS-78 |
| chr11-13079843-INS-393 | chr11-13079843-INS-393 |
| chr11-13093110-INS-7057 | chr11-13093111-INS-7058 |
| chr11-13158159-INS-185 | chr11-13158159-INS-185 |
| chr11-13179958-INS-60 | chr11-13179958-INS-60 |
| chr11-13183297-INS-94 | chr11-13183297-INS-94 |
| chr11-13201661-INS-52 | chr11-13201661-INS-54 |
| chr11-13204089-INS-92 | chr11-13204089-INS-84 |
| chr11-13215689-INS-103 | chr11-13215689-INS-103 |
| chr11-13245992-INS-245 | chr11-13245991-INS-242 |
| chr11-13248129-INS-518 | chr11-13248129-INS-519 |
| chr11-13313635-INS-600 | chr11-13313635-INS-600 |
| chr11-13338259-INS-97 | chr11-13338242-INS-126 |
| chr11-13356416-INS-68 | chr11-13356416-INS-56 |
| chr11-13372017-INS-413 | chr11-13372017-INS-449 |
| chr11-13407976-INS-288 | chr11-13407964-INS-268 |
| chr11-13413878-INS-52 | chr11-13413878-INS-52 |
| chr11-13462023-INS-193 | chr11-13462023-INS-193 |
| chr11-13463222-INS-445 | chr11-13463222-INS-445 |

| | |
|---|---|
| chr11-13478266-INS-589 | chr11-13478266-INS-589 |
| chr11-13507236-INS-343 | chr11-13507236-INS-343 |
| chr11-13512089-INS-941 | chr11-13512089-INS-941 |
| chr11-13529348-INS-708 | chr11-13529348-INS-708 |
| chr11-13545119-INS-501 | chr11-13545104-INS-520 |
| chr11-13590165-INS-886 | chr11-13590165-INS-886 |
| chr11-13626515-INS-319 | chr11-13626515-INS-319 |
| chr11-13664436-INS-1888 | chr11-13664437-INS-1888 |
| chr11-13675054-INS-668 | chr11-13675054-INS-669 |
| chr11-13688961-INS-1893 | chr11-13688962-INS-1893 |
| chr11-13702556-INS-4878 | chr11-13702557-INS-4878 |
| chr11-13724791-INS-64 | chr11-13724791-INS-64 |
| chr11-13842567-INS-54 | chr11-13842567-INS-54 |
| chr11-13849451-INS-59 | chr11-13849451-INS-59 |
| chr11-13856556-INS-70 | chr11-13856556-INS-74 |
| chr11-13887397-INS-54 | chr11-13887353-INS-56 |
| chr11-13947234-INS-120 | chr11-13947234-INS-120 |
| chr11-13953571-INS-902 | chr11-13953571-INS-902 |
| chr11-13957089-INS-84 | chr11-13957089-INS-120 |
| chr11-13972952-INS-51 | chr11-13972953-INS-51 |
| chr11-13991431-INS-1515 | chr11-13991432-INS-1514 |
| chr11-14003938-INS-199 | chr11-14003938-INS-199 |
| chr11-14004808-INS-101 | chr11-14004808-INS-90 |
| chr11-14008486-INS-2499 | chr11-14008487-INS-2498 |
| chr11-14036147-INS-366 | chr11-14036147-INS-366 |
| chr11-14042799-INS-1435 | chr11-14042800-INS-1435 |
| chr11-14092430-INS-4470 | chr11-14092431-INS-4470 |
| chr11-14104571-INS-6566 | chr11-14104572-INS-6597 |
| chr11-14117995-INS-6156 | chr11-14117996-INS-6159 |
| chr11-14122371-INS-79 | chr11-14122371-INS-79 |
| chr11-14139475-INS-401 | chr11-14139475-INS-401 |
| chr11-14146669-INS-6502 | chr11-14146670-INS-6502 |
| chr11-14151146-INS-67 | chr11-14151146-INS-67 |
| chr11-14217064-INS-211 | chr11-14217064-INS-210 |
| chr11-14246873-INS-60 | chr11-14246873-INS-60 |
| chr11-14251678-INS-509 | chr11-14251678-INS-531 |
| chr11-14278941-INS-321 | chr11-14278941-INS-305 |
| chr11-14285004-INS-805 | chr11-14285004-INS-805 |
| chr11-14302620-INS-53 | chr11-14302620-INS-56 |
| chr11-14348298-INS-66 | chr11-14348298-INS-92 |
| chr11-14402567-INS-275 | chr11-14402567-INS-275 |
| chr11-14423269-INS-3257 | chr11-14423270-INS-3257 |
| chr11-14427378-INS-721 | chr11-14427378-INS-721 |
| chr11-14469718-INS-215 | chr11-14469718-INS-215 |
| chr11-14474493-INS-218 | chr11-14474493-INS-217 |
| chr11-14508643-INS-529 | chr11-14508643-INS-531 |
| chr11-14509311-INS-4268 | chr11-14509312-INS-4268 |
| chr11-14515169-INS-69 | chr11-14515169-INS-73 |
| chr11-14524860-INS-52 | chr11-14524860-INS-61 |
| chr11-14541345-INS-749 | chr11-14541345-INS-752 |
| chr11-14586897-INS-3062 | chr11-14586898-INS-3056 |
| chr11-14642098-INS-62 | chr11-14642098-INS-62 |
| chr11-14753202-INS-70 | chr11-14753209-INS-74 |
| chr11-14762407-INS-88 | chr11-14762407-INS-88 |
| chr11-14783144-INS-1913 | chr11-14783145-INS-1914 |
| chr11-14806143-INS-580 | chr11-14806151-INS-603 |
| chr11-14816373-INS-436 | chr11-14816373-INS-436 |
| chr11-14830632-INS-6406 | chr11-14830633-INS-6407 |
| chr11-14833843-INS-74 | chr11-14833843-INS-74 |
| chr11-14874868-INS-312 | chr11-14874868-INS-312 |
| chr11-14897857-INS-6274 | chr11-14897858-INS-6279 |
| chr11-14900989-INS-419 | chr11-14900989-INS-419 |
| chr11-14901215-INS-7075 | chr11-14901216-INS-7075 |
| chr11-14920980-INS-99 | chr11-14920980-INS-99 |
| chr11-14936364-INS-77 | chr11-14936364-INS-77 |
| chr11-14979217-INS-3012 | chr11-14979218-INS-3012 |
| chr11-15068197-INS-233 | chr11-15068197-INS-233 |
| chr11-15071205-INS-466 | chr11-15071205-INS-466 |
| chr11-15106815-INS-76 | chr11-15106815-INS-76 |
| chr11-15117295-INS-197 | chr11-15117282-INS-197 |
| chr11-15123715-INS-282 | chr11-15123715-INS-282 |
| chr11-15158574-INS-84 | chr11-15158574-INS-84 |
| chr11-15168756-INS-1288 | chr11-15168757-INS-1296 |
| chr11-15206673-INS-51 | chr11-15206673-INS-95 |
| chr11-15263528-INS-2871 | chr11-15263529-INS-2871 |
| chr11-15266138-INS-187 | chr11-15266138-INS-187 |
| chr11-15268490-INS-56 | chr11-15268457-INS-50 |

| | |
|---|---|
| chr11-15273128-INS-523 | chr11-15273128-INS-523 |
| chr11-15278488-INS-60 | chr11-15278479-INS-86 |
| chr11-15284296-INS-3002 | chr11-15284297-INS-3002 |
| chr11-15290133-INS-53 | chr11-15290133-INS-59 |
| chr11-15312699-INS-264 | chr11-15312698-INS-264 |
| chr11-15332814-INS-390 | chr11-15332814-INS-390 |
| chr11-15335706-INS-73 | chr11-15335706-INS-104 |
| chr11-15337609-INS-1864 | chr11-15337610-INS-1864 |
| chr11-15339749-INS-54 | chr11-15339749-INS-54 |
| chr11-15358532-INS-54 | chr11-15358488-INS-54 |
| chr11-15373559-INS-711 | chr11-15373559-INS-748 |
| chr11-15386084-INS-7130 | chr11-15386085-INS-7130 |
| chr11-15395822-INS-6421 | chr11-15395823-INS-6423 |
| chr11-15405499-INS-346 | chr11-15405499-INS-346 |
| chr11-15422665-INS-58 | chr11-15422665-INS-58 |
| chr11-15431311-INS-288 | chr11-15431311-INS-288 |
| chr11-15451104-INS-303 | chr11-15451104-INS-303 |
| chr11-15471308-INS-592 | chr11-15471308-INS-592 |
| chr11-15473929-INS-116 | chr11-15473901-INS-123 |
| chr11-15479710-INS-56 | chr11-15479710-INS-56 |
| chr11-15526586-INS-883 | chr11-15526585-INS-882 |
| chr11-15549853-INS-387 | chr11-15549853-INS-387 |
| chr11-15578983-INS-153 | chr11-15578983-INS-127 |
| chr11-15611409-INS-971 | chr11-15611409-INS-971 |
| chr11-15620132-INS-865 | chr11-15620132-INS-865 |
| chr11-15674803-INS-88 | chr11-15674803-INS-88 |
| chr11-15764038-INS-83 | chr11-15764038-INS-83 |
| chr11-15792805-INS-72 | chr11-15792803-INS-72 |
| chr11-15825151-INS-6391 | chr11-15825152-INS-6392 |
| chr11-15832394-INS-93 | chr11-15832394-INS-57 |
| chr11-15845123-INS-93 | chr11-15845123-INS-62 |
| chr11-15863514-INS-56 | chr11-15863514-INS-56 |
| chr11-15892564-INS-973 | chr11-15892564-INS-974 |
| chr11-15895399-INS-201 | chr11-15895399-INS-201 |
| chr11-15904914-INS-3856 | chr11-15904915-INS-3856 |
| chr11-15916779-INS-171 | chr11-15916779-INS-171 |
| chr11-15952447-INS-2076 | chr11-15952448-INS-2076 |
| chr11-15975731-INS-214 | chr11-15975731-INS-214 |
| chr11-16016305-INS-3273 | chr11-16016306-INS-3273 |
| chr11-16026499-INS-364 | chr11-16026499-INS-364 |
| chr11-16033868-INS-56 | chr11-16033868-INS-52 |
| chr11-16063319-INS-675 | chr11-16063332-INS-677 |
| chr11-16065615-INS-64 | chr11-16065615-INS-64 |
| chr11-16160108-INS-511 | chr11-16160108-INS-511 |
| chr11-16244836-INS-58 | chr11-16244836-INS-58 |
| chr11-16288430-INS-328 | chr11-16288429-INS-330 |
| chr11-16297619-INS-51 | chr11-16297619-INS-51 |
| chr11-16315800-INS-124 | chr11-16315800-INS-125 |
| chr11-16321644-INS-100 | chr11-16321643-INS-128 |
| chr11-16322923-INS-81 | chr11-16322923-INS-87 |
| chr11-16329172-INS-104 | chr11-16329172-INS-104 |
| chr11-16331581-INS-67 | chr11-16331573-INS-72 |
| chr11-16365367-INS-58 | chr11-16365367-INS-62 |
| chr11-16392792-INS-3234 | chr11-16392793-INS-3232 |
| chr11-16400712-INS-176 | chr11-16400712-INS-175 |
| chr11-16416826-INS-444 | chr11-16416826-INS-424 |
| chr11-16443470-INS-91 | chr11-16443470-INS-91 |
| chr11-16471976-INS-83 | chr11-16471976-INS-55 |
| chr11-16475683-INS-88 | chr11-16475683-INS-58 |
| chr11-16487644-INS-89 | chr11-16487644-INS-59 |
| chr11-16502913-INS-414 | chr11-16502904-INS-422 |
| chr11-16507534-INS-56 | chr11-16507532-INS-60 |
| chr11-16508367-INS-55 | chr11-16508367-INS-57 |
| chr11-16511537-INS-201 | chr11-16511537-INS-201 |
| chr11-16524369-INS-53 | chr11-16524369-INS-56 |
| chr11-16583652-INS-84 | chr11-16583650-INS-103 |
| chr11-16586662-INS-66 | chr11-16586662-INS-66 |
| chr11-16623554-INS-53 | chr11-16623554-INS-53 |
| chr11-16630106-INS-237 | chr11-16630106-INS-237 |
| chr11-16634980-INS-68 | chr11-16634980-INS-68 |
| chr11-16645727-INS-327 | chr11-16645727-INS-327 |
| chr11-16669779-INS-90 | chr11-16669779-INS-90 |
| chr11-16671432-INS-64 | chr11-16671432-INS-64 |
| chr11-16676373-INS-6640 | chr11-16676374-INS-6641 |
| chr11-16678206-INS-61 | chr11-16678205-INS-66 |
| chr11-16726062-INS-70 | chr11-16726062-INS-70 |
| chr11-16761011-INS-81 | chr11-16761010-INS-77 |

| | |
|---|---|
| chr11-16766185-INS-184 | chr11-16766185-INS-183 |
| chr11-16781175-INS-440 | chr11-16781175-INS-440 |
| chr11-16814475-INS-171 | chr11-16814475-INS-171 |
| chr11-16867412-INS-94 | chr11-16867412-INS-96 |
| chr11-16875323-INS-125 | chr11-16875323-INS-126 |
| chr11-16890779-INS-129 | chr11-16890779-INS-129 |
| chr11-16916651-INS-87 | chr11-16916651-INS-87 |
| chr11-16976759-INS-368 | chr11-16976759-INS-369 |
| chr11-16980947-INS-76 | chr11-16980947-INS-88 |
| chr11-16981514-INS-80 | chr11-16981514-INS-80 |
| chr11-16991560-INS-56 | chr11-16991561-INS-56 |
| chr11-17025294-INS-501 | chr11-17025294-INS-501 |
| chr11-17037301-INS-93 | chr11-17037313-INS-128 |
| chr11-17053904-INS-597 | chr11-17053904-INS-598 |
| chr11-17098230-INS-114 | chr11-17098276-INS-110 |
| chr11-17128137-INS-194 | chr11-17128137-INS-194 |
| chr11-17133636-INS-114 | chr11-17133636-INS-118 |
| chr11-17163148-INS-84 | chr11-17163148-INS-84 |
| chr11-17164569-INS-140 | chr11-17164569-INS-140 |
| chr11-17169888-INS-81 | chr11-17169884-INS-79 |
| chr11-17173525-INS-64 | chr11-17173515-INS-96 |
| chr11-17201361-INS-74 | chr11-17201361-INS-74 |
| chr11-17219821-INS-56 | chr11-17219821-INS-60 |
| chr11-17247045-INS-426 | chr11-17247046-INS-426 |
| chr11-17247570-INS-98 | chr11-17247571-INS-98 |
| chr11-17258469-INS-60 | chr11-17258459-INS-56 |
| chr11-17300809-INS-1888 | chr11-17300810-INS-1888 |
| chr11-17310904-INS-442 | chr11-17310904-INS-434 |
| chr11-17319346-INS-196 | chr11-17319346-INS-196 |
| chr11-17322396-INS-328 | chr11-17322396-INS-333 |
| chr11-17333196-INS-658 | chr11-17333196-INS-669 |
| chr11-17350844-INS-252 | chr11-17350844-INS-249 |
| chr11-17358772-INS-92 | chr11-17358772-INS-52 |
| chr11-17366334-INS-72 | chr11-17366334-INS-66 |
| chr11-17393192-INS-88 | chr11-17393192-INS-88 |
| chr11-17422056-INS-179 | chr11-17422056-INS-166 |
| chr11-17431353-INS-96 | chr11-17431353-INS-98 |
| chr11-17439698-INS-64 | chr11-17439698-INS-64 |
| chr11-17443192-INS-7130 | chr11-17443193-INS-7130 |
| chr11-17456755-INS-196 | chr11-17456739-INS-197 |
| chr11-17466902-INS-5297 | chr11-17466903-INS-5297 |
| chr11-17517438-INS-68 | chr11-17517414-INS-96 |
| chr11-17528575-INS-59 | chr11-17528571-INS-58 |
| chr11-17540809-INS-62 | chr11-17540809-INS-70 |
| chr11-17551765-INS-56 | chr11-17551729-INS-71 |
| chr11-17579310-INS-161 | chr11-17579310-INS-161 |
| chr11-17588098-INS-97 | chr11-17588098-INS-97 |
| chr11-17612581-INS-72 | chr11-17612581-INS-72 |
| chr11-17628112-INS-73 | chr11-17628112-INS-74 |
| chr11-17633891-INS-53 | chr11-17633857-INS-82 |
| chr11-17649576-INS-100 | chr11-17649576-INS-100 |
| chr11-17650067-INS-684 | chr11-17650067-INS-684 |
| chr11-17660194-INS-52 | chr11-17660194-INS-54 |
| chr11-17664544-INS-708 | chr11-17664544-INS-708 |
| chr11-17678546-INS-333 | chr11-17678546-INS-336 |
| chr11-17757171-INS-1976 | chr11-17757157-INS-1964 |
| chr11-17767422-INS-199 | chr11-17767422-INS-199 |
| chr11-17772420-INS-127 | chr11-17772421-INS-135 |
| chr11-17784773-INS-204 | chr11-17784773-INS-204 |
| chr11-17789994-INS-6343 | chr11-17789995-INS-6343 |
| chr11-17813735-INS-160 | chr11-17813742-INS-169 |
| chr11-17824291-INS-341 | chr11-17824291-INS-341 |
| chr11-17840322-INS-84 | chr11-17840322-INS-84 |
| chr11-17861806-INS-2801 | chr11-17861806-INS-2801 |
| chr11-17894977-INS-52 | chr11-17894977-INS-52 |
| chr11-17904907-INS-54 | chr11-17904907-INS-54 |
| chr11-17905758-INS-6235 | chr11-17905759-INS-6235 |
| chr11-17912627-INS-448 | chr11-17912627-INS-448 |
| chr11-17914720-INS-66 | chr11-17914720-INS-66 |
| chr11-17925872-INS-514 | chr11-17925872-INS-519 |
| chr11-17928918-INS-71 | chr11-17928918-INS-71 |
| chr11-17936122-INS-485 | chr11-17936122-INS-485 |
| chr11-17958938-INS-75 | chr11-17958938-INS-75 |
| chr11-17990212-INS-54 | chr11-17990244-INS-53 |
| chr11-17996006-INS-226 | chr11-17996006-INS-227 |
| chr11-18050600-INS-400 | chr11-18050600-INS-400 |
| chr11-18053410-INS-1516 | chr11-18053411-INS-1515 |

| | |
|---|---|
| chr11-18055194-INS-125 | chr11-18055194-INS-126 |
| chr11-18067790-INS-198 | chr11-18067790-INS-197 |
| chr11-18088188-INS-105 | chr11-18088188-INS-137 |
| chr11-18149585-INS-55 | chr11-18149585-INS-56 |
| chr11-18172193-INS-486 | chr11-18172193-INS-486 |
| chr11-18191562-INS-2768 | chr11-18191563-INS-2768 |
| chr11-18243042-INS-468 | chr11-18243042-INS-468 |
| chr11-18244072-INS-456 | chr11-18244072-INS-456 |
| chr11-18247896-INS-848 | chr11-18247896-INS-848 |
| chr11-18262488-INS-64 | chr11-18262488-INS-64 |
| chr11-18270854-INS-88 | chr11-18270854-INS-92 |
| chr11-18275472-INS-240 | chr11-18275472-INS-249 |
| chr11-18281406-INS-102 | chr11-18281406-INS-102 |
| chr11-18282006-INS-2904 | chr11-18282007-INS-2928 |
| chr11-18296754-INS-70 | chr11-18296759-INS-70 |
| chr11-18317787-INS-131 | chr11-18317787-INS-131 |
| chr11-18333535-INS-351 | chr11-18333535-INS-339 |
| chr11-18337613-INS-54 | chr11-18337613-INS-54 |
| chr11-18350279-INS-56 | chr11-18350279-INS-56 |
| chr11-18354198-INS-76 | chr11-18354195-INS-52 |
| chr11-18374658-INS-101 | chr11-18374658-INS-53 |
| chr11-18434061-INS-3040 | chr11-18434062-INS-3040 |
| chr11-18484015-INS-62 | chr11-18484015-INS-60 |
| chr11-18514161-INS-75 | chr11-18514161-INS-75 |
| chr11-18529380-INS-3238 | chr11-18529381-INS-3238 |
| chr11-18529537-INS-54 | chr11-18529537-INS-50 |
| chr11-18661292-INS-88 | chr11-18661292-INS-90 |
| chr11-18768779-INS-83 | chr11-18768773-INS-108 |
| chr11-18850373-INS-54 | chr11-18850373-INS-51 |
| chr11-18886880-INS-110 | chr11-18886880-INS-105 |
| chr11-18899920-INS-56 | chr11-18899910-INS-72 |
| chr11-18919477-INS-207 | chr11-18919477-INS-207 |
| chr11-18947435-INS-193 | chr11-18947435-INS-214 |
| chr11-18986750-INS-315 | chr11-18986750-INS-316 |
| chr11-18992869-INS-2174 | chr11-18992870-INS-2177 |
| chr11-19045336-INS-53 | chr11-19045336-INS-61 |
| chr11-19051110-INS-469 | chr11-19051111-INS-463 |
| chr11-19073365-INS-6492 | chr11-19073366-INS-6493 |
| chr11-19078447-INS-193 | chr11-19078447-INS-194 |
| chr11-19111206-INS-76 | chr11-19111206-INS-76 |
| chr11-19132928-INS-96 | chr11-19132928-INS-90 |
| chr11-19137739-INS-110 | chr11-19137735-INS-70 |
| chr11-19146075-INS-268 | chr11-19146075-INS-255 |
| chr11-19164438-INS-308 | chr11-19164438-INS-308 |
| chr11-19171472-INS-155 | chr11-19171469-INS-131 |
| chr11-19244425-INS-318 | chr11-19244425-INS-321 |
| chr11-19264948-INS-523 | chr11-19264948-INS-523 |
| chr11-19268669-INS-2568 | chr11-19268670-INS-2568 |
| chr11-19279759-INS-6236 | chr11-19279760-INS-6236 |
| chr11-19280622-INS-142 | chr11-19280622-INS-173 |
| chr11-19289212-INS-90 | chr11-19289207-INS-66 |
| chr11-19326066-INS-216 | chr11-19326066-INS-216 |
| chr11-19335472-INS-6165 | chr11-19335473-INS-6165 |
| chr11-19376526-INS-58 | chr11-19376520-INS-56 |
| chr11-19388014-INS-52 | chr11-19388002-INS-59 |
| chr11-19397058-INS-473 | chr11-19397058-INS-474 |
| chr11-19398312-INS-435 | chr11-19398312-INS-435 |
| chr11-19407857-INS-59 | chr11-19407857-INS-59 |
| chr11-19443946-INS-7192 | chr11-19443947-INS-7192 |
| chr11-19499473-INS-222 | chr11-19499473-INS-222 |
| chr11-19513577-INS-205 | chr11-19513577-INS-205 |
| chr11-19556322-INS-5208 | chr11-19556323-INS-5208 |
| chr11-19565691-INS-135 | chr11-19565691-INS-100 |
| chr11-19588069-INS-123 | chr11-19588069-INS-123 |
| chr11-19589867-INS-331 | chr11-19589867-INS-331 |
| chr11-19597395-INS-974 | chr11-19597395-INS-973 |
| chr11-19619517-INS-175 | chr11-19619516-INS-198 |
| chr11-19653017-INS-84 | chr11-19653027-INS-56 |
| chr11-19683084-INS-1890 | chr11-19683085-INS-1890 |
| chr11-19684735-INS-64 | chr11-19684735-INS-64 |
| chr11-19690294-INS-68 | chr11-19690278-INS-78 |
| chr11-19711072-INS-205 | chr11-19711072-INS-205 |
| chr11-19729589-INS-100 | chr11-19729589-INS-100 |
| chr11-19734676-INS-140 | chr11-19734676-INS-133 |
| chr11-19746015-INS-161 | chr11-19746020-INS-176 |
| chr11-19759615-INS-88 | chr11-19759573-INS-84 |
| chr11-19770241-INS-52 | chr11-19770241-INS-52 |

| | |
|---|---|
| chr11-19797572-INS-203 | chr11-19797572-INS-205 |
| chr11-19801490-INS-162 | chr11-19801490-INS-162 |
| chr11-19805288-INS-318 | chr11-19805288-INS-318 |
| chr11-19812333-INS-58 | chr11-19812333-INS-58 |
| chr11-19830361-INS-225 | chr11-19830361-INS-226 |
| chr11-19833146-INS-2937 | chr11-19833147-INS-2937 |
| chr11-19834332-INS-1991 | chr11-19834333-INS-1991 |
| chr11-19838778-INS-151 | chr11-19838772-INS-118 |
| chr11-19842377-INS-204 | chr11-19842377-INS-204 |
| chr11-19855260-INS-55 | chr11-19855260-INS-54 |
| chr11-19863084-INS-224 | chr11-19863084-INS-224 |
| chr11-19868038-INS-69 | chr11-19868038-INS-51 |
| chr11-19887015-INS-207 | chr11-19887015-INS-207 |
| chr11-19902294-INS-183 | chr11-19902294-INS-183 |
| chr11-19919352-INS-130 | chr11-19919347-INS-112 |
| chr11-19933127-INS-58 | chr11-19933127-INS-58 |
| chr11-19967794-INS-6403 | chr11-19967795-INS-6403 |
| chr11-19980854-INS-215 | chr11-19980854-INS-215 |
| chr11-19994049-INS-200 | chr11-19994049-INS-200 |
| chr11-19994681-INS-216 | chr11-19994681-INS-216 |
| chr11-20006636-INS-273 | chr11-20006636-INS-264 |
| chr11-20010200-INS-201 | chr11-20010200-INS-202 |
| chr11-20024127-INS-175 | chr11-20024127-INS-175 |
| chr11-20029921-INS-50 | chr11-20029921-INS-52 |
| chr11-20032642-INS-222 | chr11-20032635-INS-219 |
| chr11-20033921-INS-361 | chr11-20033922-INS-368 |
| chr11-20039435-INS-210 | chr11-20039435-INS-210 |
| chr11-20040815-INS-332 | chr11-20040815-INS-332 |
| chr11-20054153-INS-208 | chr11-20054153-INS-208 |
| chr11-20065848-INS-516 | chr11-20065848-INS-516 |
| chr11-20073508-INS-53 | chr11-20073508-INS-53 |
| chr11-20075244-INS-201 | chr11-20075244-INS-201 |
| chr11-20077372-INS-102 | chr11-20077372-INS-101 |
| chr11-20078139-INS-55 | chr11-20078139-INS-56 |
| chr11-20079175-INS-205 | chr11-20079175-INS-206 |
| chr11-20085587-INS-196 | chr11-20085602-INS-205 |
| chr11-20094437-INS-182 | chr11-20094437-INS-183 |
| chr11-20097225-INS-198 | chr11-20097225-INS-198 |
| chr11-20104927-INS-204 | chr11-20104927-INS-204 |
| chr11-20108079-INS-55 | chr11-20108079-INS-55 |
| chr11-20118574-INS-169 | chr11-20118574-INS-169 |
| chr11-20130917-INS-205 | chr11-20130917-INS-202 |
| chr11-20132787-INS-73 | chr11-20132787-INS-68 |
| chr11-20135567-INS-8339 | chr11-20135568-INS-8339 |
| chr11-20150596-INS-154 | chr11-20150596-INS-154 |
| chr11-20169456-INS-197 | chr11-20169456-INS-198 |
| chr11-20173827-INS-199 | chr11-20173827-INS-202 |
| chr11-20211755-INS-60 | chr11-20211755-INS-99 |
| chr11-20220523-INS-325 | chr11-20220523-INS-325 |
| chr11-20223733-INS-154 | chr11-20223720-INS-168 |
| chr11-20225152-INS-212 | chr11-20225152-INS-212 |
| chr11-20226425-INS-201 | chr11-20226425-INS-215 |
| chr11-20227686-INS-441 | chr11-20227686-INS-441 |
| chr11-20236129-INS-56 | chr11-20236129-INS-56 |
| chr11-20256150-INS-509 | chr11-20256150-INS-509 |
| chr11-20270280-INS-64 | chr11-20270280-INS-64 |
| chr11-20288726-INS-249 | chr11-20288726-INS-249 |
| chr11-20292173-INS-215 | chr11-20292173-INS-215 |
| chr11-20302488-INS-71 | chr11-20302488-INS-67 |
| chr11-20321171-INS-88 | chr11-20321171-INS-88 |
| chr11-20326852-INS-52 | chr11-20326852-INS-52 |
| chr11-20329693-INS-1854 | chr11-20329694-INS-1854 |
| chr11-20338616-INS-70 | chr11-20338616-INS-52 |
| chr11-20342754-INS-231 | chr11-20342754-INS-231 |
| chr11-20351253-INS-211 | chr11-20351253-INS-212 |
| chr11-20355376-INS-52 | chr11-20355376-INS-52 |
| chr11-20356706-INS-443 | chr11-20356706-INS-437 |
| chr11-20372944-INS-197 | chr11-20372944-INS-197 |
| chr11-20380462-INS-59 | chr11-20380462-INS-59 |
| chr11-20416913-INS-52 | chr11-20416913-INS-52 |
| chr11-20418488-INS-198 | chr11-20418488-INS-200 |
| chr11-20424074-INS-54 | chr11-20424074-INS-66 |
| chr11-20442508-INS-506 | chr11-20442508-INS-506 |
| chr11-20447280-INS-176 | chr11-20447280-INS-182 |
| chr11-20452723-INS-62 | chr11-20452723-INS-97 |
| chr11-20453898-INS-114 | chr11-20453898-INS-136 |
| chr11-20484423-INS-69 | chr11-20484430-INS-69 |

| | |
|---|---|
| chr11-20487837-INS-189 | chr11-20487837-INS-189 |
| chr11-20504120-INS-216 | chr11-20504120-INS-216 |
| chr11-20506547-INS-61 | chr11-20506524-INS-70 |
| chr11-20507824-INS-269 | chr11-20507827-INS-266 |
| chr11-20532033-INS-97 | chr11-20532037-INS-123 |
| chr11-20539968-INS-81 | chr11-20539968-INS-81 |
| chr11-20562985-INS-86 | chr11-20563016-INS-72 |
| chr11-20617280-INS-441 | chr11-20617280-INS-470 |
| chr11-20618814-INS-56 | chr11-20618830-INS-50 |
| chr11-20620384-INS-202 | chr11-20620384-INS-202 |
| chr11-20624539-INS-168 | chr11-20624539-INS-168 |
| chr11-20721649-INS-6659 | chr11-20721618-INS-6660 |
| chr11-20737574-INS-7153 | chr11-20737575-INS-7153 |
| chr11-20746644-INS-51 | chr11-20746602-INS-78 |
| chr11-20749962-INS-60 | chr11-20749962-INS-72 |
| chr11-20755856-INS-454 | chr11-20755856-INS-454 |
| chr11-20760681-INS-433 | chr11-20760682-INS-433 |
| chr11-20762549-INS-58 | chr11-20762549-INS-74 |
| chr11-20773220-INS-165 | chr11-20773220-INS-165 |
| chr11-20788338-INS-98 | chr11-20788348-INS-75 |
| chr11-20859732-INS-540 | chr11-20859732-INS-542 |
| chr11-20865016-INS-67 | chr11-20865016-INS-67 |
| chr11-20877695-INS-146 | chr11-20877695-INS-154 |
| chr11-20883410-INS-160 | chr11-20883410-INS-160 |
| chr11-20895938-INS-51 | chr11-20895950-INS-77 |
| chr11-20922913-INS-206 | chr11-20922913-INS-206 |
| chr11-20926029-INS-94 | chr11-20926029-INS-75 |
| chr11-20928555-INS-4303 | chr11-20928556-INS-4303 |
| chr11-20967676-INS-105 | chr11-20967676-INS-105 |
| chr11-20980867-INS-60 | chr11-20980867-INS-60 |
| chr11-20997848-INS-84 | chr11-20997848-INS-84 |
| chr11-21006494-INS-639 | chr11-21006494-INS-659 |
| chr11-21050338-INS-1115 | chr11-21050339-INS-1115 |
| chr11-21116384-INS-62 | chr11-21116384-INS-63 |
| chr11-21118152-INS-92 | chr11-21118152-INS-92 |
| chr11-21119517-INS-400 | chr11-21119517-INS-400 |
| chr11-21136822-INS-173 | chr11-21136822-INS-173 |
| chr11-21142339-INS-200 | chr11-21142339-INS-199 |
| chr11-21159995-INS-54 | chr11-21159995-INS-62 |
| chr11-21254426-INS-72 | chr11-21254426-INS-72 |
| chr11-21257124-INS-110 | chr11-21257124-INS-112 |
| chr11-21280675-INS-202 | chr11-21280675-INS-202 |
| chr11-21313182-INS-7211 | chr11-21313183-INS-7211 |
| chr11-21325798-INS-129 | chr11-21325799-INS-130 |
| chr11-21371189-INS-92 | chr11-21371189-INS-92 |
| chr11-21404171-INS-1273 | chr11-21404172-INS-1273 |
| chr11-21412534-INS-6983 | chr11-21412535-INS-6983 |
| chr11-21428618-INS-640 | chr11-21428618-INS-640 |
| chr11-21439800-INS-1738 | chr11-21439801-INS-1738 |
| chr11-21450558-INS-1862 | chr11-21450559-INS-1862 |
| chr11-21456247-INS-2644 | chr11-21456248-INS-2644 |
| chr11-21473171-INS-341 | chr11-21473171-INS-341 |
| chr11-21475715-INS-226 | chr11-21475715-INS-226 |
| chr11-21526943-INS-120 | chr11-21526943-INS-120 |
| chr11-21571848-INS-3884 | chr11-21571849-INS-3885 |
| chr11-21577303-INS-5579 | chr11-21577304-INS-5579 |
| chr11-21579557-INS-481 | chr11-21579557-INS-482 |
| chr11-21584147-INS-550 | chr11-21584147-INS-550 |
| chr11-21585723-INS-711 | chr11-21585726-INS-717 |
| chr11-21587918-INS-1862 | chr11-21587919-INS-1862 |
| chr11-21592157-INS-209 | chr11-21592157-INS-209 |
| chr11-21634770-INS-5762 | chr11-21634771-INS-5762 |
| chr11-21694180-INS-4553 | chr11-21694181-INS-4553 |
| chr11-21749090-INS-1162 | chr11-21749091-INS-1163 |
| chr11-21758147-INS-128 | chr11-21758147-INS-128 |
| chr11-21764320-INS-522 | chr11-21764320-INS-522 |
| chr11-21768275-INS-54 | chr11-21768265-INS-54 |
| chr11-21801103-INS-207 | chr11-21801103-INS-207 |
| chr11-21818987-INS-146 | chr11-21818987-INS-146 |
| chr11-21847878-INS-59 | chr11-21847878-INS-59 |
| chr11-21895966-INS-112 | chr11-21895966-INS-112 |
| chr11-21897490-INS-201 | chr11-21897490-INS-201 |
| chr11-21922406-INS-54 | chr11-21922406-INS-54 |
| chr11-21929436-INS-391 | chr11-21929436-INS-390 |
| chr11-21969788-INS-1012 | chr11-21969789-INS-1013 |
| chr11-21973629-INS-75 | chr11-21973629-INS-75 |
| chr11-22051888-INS-76 | chr11-22051888-INS-76 |

| | |
|---|---|
| chr11-22060054-INS-199 | chr11-22060054-INS-199 |
| chr11-22066741-INS-186 | chr11-22066741-INS-186 |
| chr11-22080375-INS-70 | chr11-22080375-INS-74 |
| chr11-22139313-INS-114 | chr11-22139313-INS-114 |
| chr11-22150343-INS-7306 | chr11-22150344-INS-7306 |
| chr11-22177809-INS-62 | chr11-22177809-INS-62 |
| chr11-22181402-INS-110 | chr11-22181378-INS-118 |
| chr11-22191973-INS-1891 | chr11-22191974-INS-1891 |
| chr11-22217935-INS-74 | chr11-22217933-INS-60 |
| chr11-22269706-INS-57 | chr11-22269706-INS-57 |
| chr11-22271911-INS-221 | chr11-22271911-INS-222 |
| chr11-22296597-INS-419 | chr11-22296597-INS-419 |
| chr11-22297289-INS-73 | chr11-22297289-INS-73 |
| chr11-22321733-INS-6901 | chr11-22321734-INS-6902 |
| chr11-22322346-INS-51 | chr11-22322344-INS-51 |
| chr11-22334400-INS-91 | chr11-22334400-INS-91 |
| chr11-22364375-INS-1869 | chr11-22364376-INS-1869 |
| chr11-22416477-INS-53 | chr11-22416477-INS-53 |
| chr11-22419423-INS-186 | chr11-22419423-INS-186 |
| chr11-22452604-INS-217 | chr11-22452604-INS-217 |
| chr11-22476903-INS-50 | chr11-22476903-INS-54 |
| chr11-22495847-INS-238 | chr11-22495835-INS-246 |
| chr11-22503298-INS-62 | chr11-22503298-INS-64 |
| chr11-22547003-INS-7215 | chr11-22547004-INS-7215 |
| chr11-22551313-INS-58 | chr11-22551313-INS-56 |
| chr11-22566194-INS-72 | chr11-22566194-INS-80 |
| chr11-22569557-INS-119 | chr11-22569557-INS-118 |
| chr11-22571406-INS-197 | chr11-22571406-INS-198 |
| chr11-22578466-INS-69 | chr11-22578466-INS-69 |
| chr11-22585483-INS-93 | chr11-22585483-INS-120 |
| chr11-22589486-INS-81 | chr11-22589486-INS-81 |
| chr11-22598069-INS-1531 | chr11-22598070-INS-1531 |
| chr11-22600834-INS-64 | chr11-22600843-INS-91 |
| chr11-22610449-INS-195 | chr11-22610449-INS-195 |
| chr11-22625509-INS-252 | chr11-22625509-INS-252 |
| chr11-22635284-INS-130 | chr11-22635284-INS-136 |
| chr11-22656515-INS-197 | chr11-22656515-INS-197 |
| chr11-22674768-INS-56 | chr11-22674768-INS-50 |
| chr11-22676715-INS-53 | chr11-22676715-INS-53 |
| chr11-22688284-INS-933 | chr11-22688284-INS-933 |
| chr11-22696767-INS-71 | chr11-22696767-INS-81 |
| chr11-22701379-INS-61 | chr11-22701379-INS-63 |
| chr11-22704278-INS-64 | chr11-22704289-INS-62 |
| chr11-22717608-INS-217 | chr11-22717608-INS-217 |
| chr11-22741566-INS-57 | chr11-22741572-INS-52 |
| chr11-22774197-INS-169 | chr11-22774197-INS-169 |
| chr11-22788813-INS-6713 | chr11-22788814-INS-6716 |
| chr11-22791981-INS-110 | chr11-22791981-INS-110 |
| chr11-22796708-INS-69 | chr11-22796708-INS-69 |
| chr11-22799262-INS-208 | chr11-22799262-INS-208 |
| chr11-22805903-INS-88 | chr11-22805951-INS-57 |
| chr11-22839096-INS-7140 | chr11-22839097-INS-7140 |
| chr11-22840114-INS-142 | chr11-22840114-INS-140 |
| chr11-22857069-INS-365 | chr11-22857069-INS-365 |
| chr11-22866307-INS-140 | chr11-22866307-INS-145 |
| chr11-22871192-INS-6401 | chr11-22871193-INS-6401 |
| chr11-22884399-INS-212 | chr11-22884399-INS-212 |
| chr11-22907488-INS-72 | chr11-22907488-INS-72 |
| chr11-22913869-INS-72 | chr11-22913845-INS-76 |
| chr11-22941417-INS-108 | chr11-22941417-INS-108 |
| chr11-22951721-INS-78 | chr11-22951721-INS-78 |
| chr11-22991532-INS-468 | chr11-22991532-INS-468 |
| chr11-22994053-INS-206 | chr11-22994053-INS-206 |
| chr11-23002898-INS-56 | chr11-23002895-INS-89 |
| chr11-23012249-INS-2311 | chr11-23012250-INS-2311 |
| chr11-23040381-INS-218 | chr11-23040381-INS-218 |
| chr11-23054307-INS-315 | chr11-23054307-INS-315 |
| chr11-23057421-INS-204 | chr11-23057421-INS-203 |
| chr11-23059895-INS-188 | chr11-23059895-INS-188 |
| chr11-23111714-INS-299 | chr11-23111714-INS-302 |
| chr11-23122710-INS-98 | chr11-23122710-INS-98 |
| chr11-23123768-INS-252 | chr11-23123768-INS-252 |
| chr11-23125070-INS-5298 | chr11-23125071-INS-5298 |
| chr11-23129833-INS-84 | chr11-23129833-INS-84 |
| chr11-23137009-INS-201 | chr11-23137009-INS-201 |
| chr11-23157607-INS-135 | chr11-23157607-INS-134 |
| chr11-23158308-INS-227 | chr11-23158308-INS-227 |

| | |
|---|---|
| chr11-23160939-INS-190 | chr11-23160939-INS-190 |
| chr11-23162502-INS-201 | chr11-23162502-INS-203 |
| chr11-23164256-INS-324 | chr11-23164256-INS-324 |
| chr11-23177226-INS-213 | chr11-23177226-INS-225 |
| chr11-23179197-INS-92 | chr11-23179197-INS-92 |
| chr11-23243374-INS-66 | chr11-23243374-INS-66 |
| chr11-23260723-INS-71 | chr11-23260723-INS-72 |
| chr11-23366154-INS-86 | chr11-23366154-INS-78 |
| chr11-23391388-INS-60 | chr11-23391388-INS-60 |
| chr11-23399979-INS-79 | chr11-23399979-INS-67 |
| chr11-23445545-INS-377 | chr11-23445545-INS-375 |
| chr11-23478865-INS-400 | chr11-23478865-INS-431 |
| chr11-23486097-INS-100 | chr11-23486097-INS-102 |
| chr11-23505995-INS-61 | chr11-23505975-INS-77 |
| chr11-23510034-INS-116 | chr11-23510011-INS-116 |
| chr11-23547794-INS-91 | chr11-23547794-INS-91 |
| chr11-23552723-INS-94 | chr11-23552723-INS-94 |
| chr11-23573596-INS-197 | chr11-23573596-INS-197 |
| chr11-23596671-INS-225 | chr11-23596671-INS-226 |
| chr11-23630521-INS-136 | chr11-23630521-INS-150 |
| chr11-23661392-INS-688 | chr11-23661392-INS-688 |
| chr11-23665094-INS-183 | chr11-23665094-INS-185 |
| chr11-23670059-INS-369 | chr11-23670059-INS-369 |
| chr11-23705561-INS-66 | chr11-23705561-INS-64 |
| chr11-23724508-INS-351 | chr11-23724508-INS-351 |
| chr11-23748555-INS-136 | chr11-23748555-INS-128 |
| chr11-23779099-INS-174 | chr11-23779099-INS-174 |
| chr11-23791057-INS-848 | chr11-23791058-INS-848 |
| chr11-23847249-INS-129 | chr11-23847251-INS-132 |
| chr11-23848377-INS-128 | chr11-23848376-INS-125 |
| chr11-23852555-INS-149 | chr11-23852555-INS-121 |
| chr11-23858010-INS-1050 | chr11-23858011-INS-1050 |
| chr11-23860932-INS-78 | chr11-23860932-INS-88 |
| chr11-23868619-INS-58 | chr11-23868619-INS-74 |
| chr11-23957670-INS-204 | chr11-23957670-INS-204 |
| chr11-23972787-INS-69 | chr11-23972787-INS-92 |
| chr11-24085108-INS-59 | chr11-24085104-INS-55 |
| chr11-24107418-INS-66 | chr11-24107418-INS-66 |
| chr11-24109100-INS-51 | chr11-24109096-INS-52 |
| chr11-24188329-INS-66 | chr11-24188329-INS-66 |
| chr11-24210794-INS-78 | chr11-24210794-INS-66 |
| chr11-24266720-INS-54 | chr11-24266720-INS-54 |
| chr11-24302544-INS-165 | chr11-24302544-INS-158 |
| chr11-24313757-INS-5536 | chr11-24313758-INS-5536 |
| chr11-24315202-INS-64 | chr11-24315202-INS-56 |
| chr11-24340089-INS-163 | chr11-24340089-INS-163 |
| chr11-24346465-INS-6029 | chr11-24346466-INS-6029 |
| chr11-24386430-INS-5473 | chr11-24386431-INS-5473 |
| chr11-24402897-INS-1893 | chr11-24402898-INS-1893 |
| chr11-24426285-INS-53 | chr11-24426285-INS-69 |
| chr11-24456965-INS-311 | chr11-24456965-INS-311 |
| chr11-24491338-INS-81 | chr11-24491338-INS-81 |
| chr11-24515195-INS-51 | chr11-24515195-INS-51 |
| chr11-24522925-INS-646 | chr11-24522925-INS-647 |
| chr11-24538923-INS-97 | chr11-24538923-INS-97 |
| chr11-24539309-INS-488 | chr11-24539309-INS-488 |
| chr11-24558765-INS-72 | chr11-24558765-INS-72 |
| chr11-24570461-INS-58 | chr11-24570485-INS-80 |
| chr11-24571408-INS-87 | chr11-24571415-INS-77 |
| chr11-24616180-INS-68 | chr11-24616180-INS-71 |
| chr11-24636870-INS-60 | chr11-24636871-INS-76 |
| chr11-24675549-INS-58 | chr11-24675549-INS-64 |
| chr11-24724881-INS-74 | chr11-24724881-INS-78 |
| chr11-24775298-INS-78 | chr11-24775298-INS-78 |
| chr11-24806238-INS-6448 | chr11-24806239-INS-6449 |
| chr11-24849136-INS-504 | chr11-24849136-INS-504 |
| chr11-24851613-INS-53 | chr11-24851613-INS-53 |
| chr11-24860592-INS-112 | chr11-24860592-INS-118 |
| chr11-24865519-INS-1888 | chr11-24865520-INS-1888 |
| chr11-24872992-INS-434 | chr11-24872992-INS-434 |
| chr11-24885160-INS-640 | chr11-24885160-INS-640 |
| chr11-24888716-INS-155 | chr11-24888716-INS-176 |
| chr11-24909050-INS-3192 | chr11-24909051-INS-3192 |
| chr11-24909471-INS-503 | chr11-24909471-INS-503 |
| chr11-24923267-INS-98 | chr11-24923267-INS-98 |
| chr11-24926426-INS-106 | chr11-24926426-INS-106 |
| chr11-24964215-INS-201 | chr11-24964215-INS-201 |

| | |
|---|---|
| chr11-24968677-INS-63 | chr11-24968677-INS-63 |
| chr11-24988377-INS-4966 | chr11-24988378-INS-4966 |
| chr11-25015880-INS-94 | chr11-25015884-INS-98 |
| chr11-25042010-INS-964 | chr11-25042010-INS-964 |
| chr11-25050155-INS-75 | chr11-25050155-INS-70 |
| chr11-25062658-INS-52 | chr11-25062658-INS-52 |
| chr11-25065966-INS-54 | chr11-25065962-INS-50 |
| chr11-25076582-INS-102 | chr11-25076574-INS-102 |
| chr11-25077747-INS-252 | chr11-25077747-INS-252 |
| chr11-25114420-INS-223 | chr11-25114420-INS-223 |
| chr11-25125711-INS-911 | chr11-25125711-INS-911 |
| chr11-25130774-INS-52 | chr11-25130776-INS-50 |
| chr11-25158054-INS-52 | chr11-25158054-INS-52 |
| chr11-25165507-INS-402 | chr11-25165514-INS-401 |
| chr11-25192680-INS-292 | chr11-25192680-INS-292 |
| chr11-25198178-INS-138 | chr11-25198174-INS-140 |
| chr11-25203145-INS-484 | chr11-25203145-INS-480 |
| chr11-25234195-INS-136 | chr11-25234195-INS-136 |
| chr11-25235917-INS-67 | chr11-25235917-INS-67 |
| chr11-25242419-INS-89 | chr11-25242419-INS-87 |
| chr11-25244761-INS-162 | chr11-25244761-INS-162 |
| chr11-25246837-INS-225 | chr11-25246837-INS-225 |
| chr11-25248869-INS-111 | chr11-25248869-INS-117 |
| chr11-25272118-INS-178 | chr11-25272118-INS-170 |
| chr11-25277138-INS-82 | chr11-25277138-INS-88 |
| chr11-25299416-INS-82 | chr11-25299416-INS-98 |
| chr11-25328438-INS-6385 | chr11-25328439-INS-6385 |
| chr11-25330667-INS-96 | chr11-25330667-INS-94 |
| chr11-25335059-INS-207 | chr11-25335059-INS-207 |
| chr11-25345729-INS-1053 | chr11-25345730-INS-1053 |
| chr11-25397353-INS-180 | chr11-25397353-INS-180 |
| chr11-25400305-INS-5380 | chr11-25400306-INS-5380 |
| chr11-25402254-INS-189 | chr11-25402254-INS-189 |
| chr11-25408330-INS-445 | chr11-25408330-INS-445 |
| chr11-25433633-INS-200 | chr11-25433633-INS-200 |
| chr11-25434221-INS-75 | chr11-25434221-INS-85 |
| chr11-25436624-INS-54 | chr11-25436624-INS-54 |
| chr11-25457686-INS-1896 | chr11-25457687-INS-1896 |
| chr11-25470588-INS-56 | chr11-25470588-INS-64 |
| chr11-25483113-INS-64 | chr11-25483113-INS-68 |
| chr11-25486649-INS-58 | chr11-25486650-INS-58 |
| chr11-25501955-INS-86 | chr11-25501955-INS-90 |
| chr11-25527590-INS-81 | chr11-25527587-INS-80 |
| chr11-25534372-INS-621 | chr11-25534372-INS-622 |
| chr11-25535358-INS-181 | chr11-25535358-INS-181 |
| chr11-25542805-INS-128 | chr11-25542805-INS-128 |
| chr11-25548071-INS-213 | chr11-25548071-INS-213 |
| chr11-25570356-INS-60 | chr11-25570356-INS-60 |
| chr11-25590958-INS-64 | chr11-25590918-INS-56 |
| chr11-25592026-INS-52 | chr11-25592027-INS-56 |
| chr11-25619577-INS-80 | chr11-25619577-INS-80 |
| chr11-25658019-INS-6265 | chr11-25658020-INS-6262 |
| chr11-25658020-INS-59 | chr11-25658020-INS-59 |
| chr11-25666889-INS-194 | chr11-25666862-INS-183 |
| chr11-25691650-INS-79 | chr11-25691650-INS-60 |
| chr11-25695330-INS-173 | chr11-25695330-INS-173 |
| chr11-25703895-INS-54 | chr11-25703895-INS-55 |
| chr11-25704633-INS-670 | chr11-25704633-INS-670 |
| chr11-25728041-INS-115 | chr11-25728041-INS-115 |
| chr11-25742313-INS-51 | chr11-25742314-INS-57 |
| chr11-25815737-INS-78 | chr11-25815737-INS-78 |
| chr11-25823944-INS-62 | chr11-25823946-INS-66 |
| chr11-25833822-INS-59 | chr11-25833822-INS-53 |
| chr11-25835152-INS-58 | chr11-25835152-INS-60 |
| chr11-25847171-INS-87 | chr11-25847199-INS-69 |
| chr11-25849551-INS-109 | chr11-25849552-INS-107 |
| chr11-25850852-INS-2498 | chr11-25850853-INS-2498 |
| chr11-25881706-INS-1092 | chr11-25881707-INS-1093 |
| chr11-25902259-INS-95 | chr11-25902259-INS-95 |
| chr11-25911268-INS-91 | chr11-25911270-INS-70 |
| chr11-25922861-INS-6607 | chr11-25922862-INS-6607 |
| chr11-25969765-INS-888 | chr11-25969765-INS-888 |
| chr11-26014023-INS-1178 | chr11-26014024-INS-1178 |
| chr11-26015967-INS-646 | chr11-26015967-INS-649 |
| chr11-26019626-INS-56 | chr11-26019626-INS-60 |
| chr11-26049260-INS-52 | chr11-26049260-INS-52 |
| chr11-26111310-INS-91 | chr11-26111310-INS-91 |

| | |
|---|---|
| chr11-26126388-INS-61 | chr11-26126388-INS-73 |
| chr11-26151162-INS-55 | chr11-26151162-INS-60 |
| chr11-26157456-INS-76 | chr11-26157456-INS-76 |
| chr11-26195934-INS-102 | chr11-26195937-INS-75 |
| chr11-26198930-INS-641 | chr11-26198930-INS-641 |
| chr11-26207083-INS-536 | chr11-26207083-INS-536 |
| chr11-26243669-INS-153 | chr11-26243665-INS-193 |
| chr11-26249656-INS-186 | chr11-26249651-INS-149 |
| chr11-26251985-INS-71 | chr11-26251985-INS-71 |
| chr11-26257064-INS-84 | chr11-26257064-INS-84 |
| chr11-26258535-INS-342 | chr11-26258535-INS-342 |
| chr11-26259266-INS-134 | chr11-26259266-INS-134 |
| chr11-26271073-INS-75 | chr11-26271073-INS-75 |
| chr11-26283377-INS-224 | chr11-26283377-INS-224 |
| chr11-26289869-INS-104 | chr11-26289869-INS-104 |
| chr11-26304032-INS-5266 | chr11-26304033-INS-5266 |
| chr11-26350870-INS-932 | chr11-26350864-INS-906 |
| chr11-26353662-INS-6398 | chr11-26353663-INS-6398 |
| chr11-26367781-INS-380 | chr11-26367781-INS-380 |
| chr11-26376053-INS-80 | chr11-26376053-INS-52 |
| chr11-26387452-INS-266 | chr11-26387452-INS-301 |
| chr11-26395358-INS-557 | chr11-26395358-INS-557 |
| chr11-26407572-INS-207 | chr11-26407572-INS-207 |
| chr11-26440786-INS-79 | chr11-26440786-INS-79 |
| chr11-26459584-INS-999 | chr11-26459584-INS-1002 |
| chr11-26528205-INS-68 | chr11-26528205-INS-68 |
| chr11-26531834-INS-52 | chr11-26531834-INS-60 |
| chr11-26544539-INS-62 | chr11-26544530-INS-55 |
| chr11-26555677-INS-533 | chr11-26555686-INS-535 |
| chr11-26570097-INS-154 | chr11-26570097-INS-154 |
| chr11-26571597-INS-52 | chr11-26571597-INS-52 |
| chr11-26572812-INS-6446 | chr11-26572813-INS-6447 |
| chr11-26602452-INS-981 | chr11-26602452-INS-981 |
| chr11-26612965-INS-126 | chr11-26612965-INS-126 |
| chr11-26613833-INS-1144 | chr11-26613834-INS-1144 |
| chr11-26616872-INS-869 | chr11-26616872-INS-869 |
| chr11-26668692-INS-185 | chr11-26668692-INS-185 |
| chr11-26677392-INS-54 | chr11-26677392-INS-54 |
| chr11-26708718-INS-131 | chr11-26708721-INS-122 |
| chr11-26722572-INS-58 | chr11-26722590-INS-61 |
| chr11-26727472-INS-54 | chr11-26727470-INS-82 |
| chr11-26730030-INS-4080 | chr11-26730031-INS-4080 |
| chr11-26755710-INS-54 | chr11-26755711-INS-54 |
| chr11-26763954-INS-55 | chr11-26763954-INS-55 |
| chr11-26781166-INS-7140 | chr11-26781167-INS-7139 |
| chr11-26830076-INS-6838 | chr11-26830077-INS-6838 |
| chr11-26836316-INS-55 | chr11-26836316-INS-50 |
| chr11-26843563-INS-241 | chr11-26843551-INS-244 |
| chr11-26892643-INS-2800 | chr11-26892644-INS-2801 |
| chr11-26904788-INS-75 | chr11-26904788-INS-80 |
| chr11-26924415-INS-96 | chr11-26924415-INS-96 |
| chr11-26972443-INS-199 | chr11-26972443-INS-199 |
| chr11-26982538-INS-2483 | chr11-26982539-INS-2483 |
| chr11-26984226-INS-70 | chr11-26984226-INS-72 |
| chr11-26989152-INS-173 | chr11-26989143-INS-180 |
| chr11-27002159-INS-165 | chr11-27002159-INS-159 |
| chr11-27043992-INS-458 | chr11-27043992-INS-458 |
| chr11-27047046-INS-71 | chr11-27047046-INS-52 |
| chr11-27052952-INS-165 | chr11-27052952-INS-165 |
| chr11-27119557-INS-72 | chr11-27119557-INS-80 |
| chr11-27122511-INS-186 | chr11-27122511-INS-186 |
| chr11-27190490-INS-521 | chr11-27190490-INS-525 |
| chr11-27197234-INS-6442 | chr11-27197235-INS-6442 |
| chr11-27204516-INS-478 | chr11-27204516-INS-477 |
| chr11-27217101-INS-63 | chr11-27217101-INS-87 |
| chr11-27218912-INS-634 | chr11-27218912-INS-634 |
| chr11-27221402-INS-817 | chr11-27221402-INS-818 |
| chr11-27255162-INS-128 | chr11-27255162-INS-128 |
| chr11-27272631-INS-759 | chr11-27272631-INS-759 |
| chr11-27287840-INS-6442 | chr11-27287841-INS-6442 |
| chr11-27318727-INS-346 | chr11-27318727-INS-346 |
| chr11-27336596-INS-5406 | chr11-27336596-INS-5406 |
| chr11-27356409-INS-52 | chr11-27356419-INS-56 |
| chr11-27378245-INS-730 | chr11-27378245-INS-729 |
| chr11-27390256-INS-195 | chr11-27390256-INS-195 |
| chr11-27394609-INS-956 | chr11-27394609-INS-956 |
| chr11-27406459-INS-202 | chr11-27406459-INS-202 |

| | |
|---|---|
| chr11-27415709-INS-85 | chr11-27415709-INS-85 |
| chr11-27420015-INS-5564 | chr11-27420016-INS-5564 |
| chr11-27424097-INS-144 | chr11-27424097-INS-144 |
| chr11-27429425-INS-95 | chr11-27429425-INS-95 |
| chr11-27474747-INS-76 | chr11-27474747-INS-76 |
| chr11-27476754-INS-52 | chr11-27476754-INS-52 |
| chr11-27539989-INS-883 | chr11-27539989-INS-883 |
| chr11-27557776-INS-52 | chr11-27557778-INS-55 |
| chr11-27574312-INS-122 | chr11-27574312-INS-122 |
| chr11-27587112-INS-212 | chr11-27587095-INS-212 |
| chr11-27600788-INS-2370 | chr11-27600789-INS-2369 |
| chr11-27602975-INS-13939 | chr11-27602976-INS-13939 |
| chr11-27617554-INS-466 | chr11-27617554-INS-466 |
| chr11-27623553-INS-449 | chr11-27623553-INS-449 |
| chr11-27646490-INS-82 | chr11-27646490-INS-82 |
| chr11-27666942-INS-52 | chr11-27666942-INS-52 |
| chr11-27671811-INS-199 | chr11-27671811-INS-195 |
| chr11-27680810-INS-2214 | chr11-27680811-INS-2217 |
| chr11-27687126-INS-52 | chr11-27687126-INS-60 |
| chr11-27689934-INS-306 | chr11-27689934-INS-306 |
| chr11-27694033-INS-58 | chr11-27694033-INS-58 |
| chr11-27699235-INS-5300 | chr11-27699236-INS-5300 |
| chr11-27702598-INS-2989 | chr11-27702599-INS-2989 |
| chr11-27730693-INS-83 | chr11-27730691-INS-66 |
| chr11-27742273-INS-54 | chr11-27742273-INS-54 |
| chr11-27747796-INS-6164 | chr11-27747797-INS-6164 |
| chr11-27760247-INS-52 | chr11-27760287-INS-52 |
| chr11-27785660-INS-70 | chr11-27785660-INS-70 |
| chr11-27796897-INS-59 | chr11-27796897-INS-59 |
| chr11-27810094-INS-497 | chr11-27810094-INS-496 |
| chr11-27813438-INS-60 | chr11-27813441-INS-65 |
| chr11-27893471-INS-135 | chr11-27893471-INS-135 |
| chr11-27914959-INS-95 | chr11-27914942-INS-67 |
| chr11-27933982-INS-507 | chr11-27933982-INS-506 |
| chr11-27951360-INS-52 | chr11-27951360-INS-52 |
| chr11-27972627-INS-187 | chr11-27972627-INS-187 |
| chr11-27995310-INS-7486 | chr11-27995311-INS-7486 |
| chr11-28012131-INS-96 | chr11-28012155-INS-61 |
| chr11-28017085-INS-209 | chr11-28017085-INS-209 |
| chr11-28043722-INS-51 | chr11-28043722-INS-51 |
| chr11-28102838-INS-135 | chr11-28102838-INS-135 |
| chr11-28103814-INS-413 | chr11-28103814-INS-413 |
| chr11-28105828-INS-54 | chr11-28105828-INS-54 |
| chr11-28154020-INS-268 | chr11-28154026-INS-265 |
| chr11-28160907-INS-70 | chr11-28160909-INS-64 |
| chr11-28162531-INS-72 | chr11-28162535-INS-68 |
| chr11-28166628-INS-249 | chr11-28166628-INS-249 |
| chr11-28190240-INS-213 | chr11-28190240-INS-213 |
| chr11-28201893-INS-287 | chr11-28201893-INS-287 |
| chr11-28210937-INS-50 | chr11-28210963-INS-65 |
| chr11-28214605-INS-150 | chr11-28214605-INS-150 |
| chr11-28238871-INS-3724 | chr11-28238872-INS-3724 |
| chr11-28269212-INS-157 | chr11-28269196-INS-144 |
| chr11-28285544-INS-980 | chr11-28285544-INS-980 |
| chr11-28285562-INS-398 | chr11-28285562-INS-398 |
| chr11-28294531-INS-134 | chr11-28294531-INS-134 |
| chr11-28317293-INS-56 | chr11-28317293-INS-56 |
| chr11-28319478-INS-56 | chr11-28319478-INS-51 |
| chr11-28330764-INS-204 | chr11-28330764-INS-205 |
| chr11-28335199-INS-7090 | chr11-28335200-INS-7090 |
| chr11-28335491-INS-144 | chr11-28335491-INS-149 |
| chr11-28348395-INS-279 | chr11-28348395-INS-288 |
| chr11-28353808-INS-6498 | chr11-28353809-INS-6498 |
| chr11-28356160-INS-530 | chr11-28356160-INS-526 |
| chr11-28356919-INS-146 | chr11-28356897-INS-162 |
| chr11-28371366-INS-66 | chr11-28371366-INS-66 |
| chr11-28379826-INS-186 | chr11-28379826-INS-187 |
| chr11-28400598-INS-73 | chr11-28400598-INS-73 |
| chr11-28401304-INS-63 | chr11-28401304-INS-52 |
| chr11-28404670-INS-226 | chr11-28404670-INS-226 |
| chr11-28420840-INS-2022 | chr11-28420841-INS-2022 |
| chr11-28437931-INS-52 | chr11-28437932-INS-52 |
| chr11-28440096-INS-54 | chr11-28440096-INS-54 |
| chr11-28440497-INS-1953 | chr11-28440498-INS-1953 |
| chr11-28446141-INS-5398 | chr11-28446142-INS-5398 |
| chr11-28460212-INS-70 | chr11-28460212-INS-70 |
| chr11-28487351-INS-228 | chr11-28487351-INS-267 |

| | |
|---|---|
| chr11-28488853-INS-64 | chr11-28488853-INS-64 |
| chr11-28490771-INS-52 | chr11-28490771-INS-55 |
| chr11-28505528-INS-55 | chr11-28505528-INS-55 |
| chr11-28512438-INS-64 | chr11-28512438-INS-72 |
| chr11-28517355-INS-665 | chr11-28517355-INS-665 |
| chr11-28518287-INS-426 | chr11-28518287-INS-429 |
| chr11-28528146-INS-66 | chr11-28528146-INS-66 |
| chr11-28538967-INS-2547 | chr11-28538968-INS-2547 |
| chr11-28547879-INS-270 | chr11-28547879-INS-270 |
| chr11-28561026-INS-228 | chr11-28561026-INS-240 |
| chr11-28564260-INS-60 | chr11-28564260-INS-60 |
| chr11-28579165-INS-269 | chr11-28579166-INS-269 |
| chr11-28610937-INS-3428 | chr11-28610938-INS-3428 |
| chr11-28618576-INS-185 | chr11-28618576-INS-185 |
| chr11-28623738-INS-71 | chr11-28623738-INS-71 |
| chr11-28626307-INS-106 | chr11-28626307-INS-106 |
| chr11-28685864-INS-1891 | chr11-28685865-INS-1891 |
| chr11-28713458-INS-68 | chr11-28713451-INS-74 |
| chr11-28771520-INS-6151 | chr11-28771521-INS-6151 |
| chr11-28787457-INS-3210 | chr11-28787458-INS-3210 |
| chr11-28806625-INS-1944 | chr11-28806626-INS-1944 |
| chr11-28945321-INS-64 | chr11-28945329-INS-52 |
| chr11-28956076-INS-6421 | chr11-28956077-INS-6421 |
| chr11-28971013-INS-72 | chr11-28971013-INS-77 |
| chr11-29013015-INS-55 | chr11-29013015-INS-61 |
| chr11-29051238-INS-6362 | chr11-29051239-INS-6361 |
| chr11-29083935-INS-134 | chr11-29083935-INS-134 |
| chr11-29086238-INS-140 | chr11-29086238-INS-144 |
| chr11-29097773-INS-748 | chr11-29097773-INS-748 |
| chr11-29098541-INS-1890 | chr11-29098542-INS-1890 |
| chr11-29104848-INS-92 | chr11-29104848-INS-92 |
| chr11-29108439-INS-229 | chr11-29108439-INS-229 |
| chr11-29110419-INS-105 | chr11-29110419-INS-107 |
| chr11-29130228-INS-1333 | chr11-29130229-INS-1333 |
| chr11-29165878-INS-1435 | chr11-29165879-INS-1435 |
| chr11-29171710-INS-202 | chr11-29171710-INS-203 |
| chr11-29184775-INS-200 | chr11-29184775-INS-200 |
| chr11-29190594-INS-122 | chr11-29190594-INS-134 |
| chr11-29197912-INS-151 | chr11-29197912-INS-151 |
| chr11-29223499-INS-396 | chr11-29223499-INS-396 |
| chr11-29243737-INS-125 | chr11-29243737-INS-117 |
| chr11-29245806-INS-4538 | chr11-29245807-INS-4538 |
| chr11-29311326-INS-1023 | chr11-29311327-INS-1023 |
| chr11-29318764-INS-326 | chr11-29318763-INS-366 |
| chr11-29343107-INS-1793 | chr11-29343108-INS-1794 |
| chr11-29345590-INS-75 | chr11-29345590-INS-79 |
| chr11-29359239-INS-111 | chr11-29359240-INS-111 |
| chr11-29368098-INS-63 | chr11-29368100-INS-95 |
| chr11-29381508-INS-1091 | chr11-29381509-INS-1091 |
| chr11-29388714-INS-2267 | chr11-29388715-INS-2267 |
| chr11-29407205-INS-56 | chr11-29407205-INS-52 |
| chr11-29432930-INS-58 | chr11-29432924-INS-66 |
| chr11-29434716-INS-200 | chr11-29434716-INS-200 |
| chr11-29439393-INS-201 | chr11-29439393-INS-201 |
| chr11-29442595-INS-137 | chr11-29442595-INS-137 |
| chr11-29478408-INS-64 | chr11-29478408-INS-52 |
| chr11-29503069-INS-74 | chr11-29503091-INS-56 |
| chr11-29506341-INS-176 | chr11-29506341-INS-176 |
| chr11-29525870-INS-61 | chr11-29525870-INS-61 |
| chr11-29549630-INS-204 | chr11-29549630-INS-204 |
| chr11-29591035-INS-692 | chr11-29591035-INS-693 |
| chr11-29592874-INS-120 | chr11-29592875-INS-78 |
| chr11-29599527-INS-229 | chr11-29599554-INS-213 |
| chr11-29599559-INS-1081 | chr11-29599554-INS-1088 |
| chr11-29606991-INS-275 | chr11-29606992-INS-275 |
| chr11-29608565-INS-380 | chr11-29608566-INS-379 |
| chr11-29610372-INS-52 | chr11-29610374-INS-52 |
| chr11-29614810-INS-316 | chr11-29614810-INS-316 |
| chr11-29618893-INS-74 | chr11-29618893-INS-56 |
| chr11-29624898-INS-198 | chr11-29624898-INS-198 |
| chr11-29628537-INS-203 | chr11-29628537-INS-203 |
| chr11-29641837-INS-86 | chr11-29641837-INS-88 |
| chr11-29651579-INS-115 | chr11-29651579-INS-110 |
| chr11-29665952-INS-78 | chr11-29665972-INS-86 |
| chr11-29672832-INS-210 | chr11-29672832-INS-210 |
| chr11-29674738-INS-198 | chr11-29674738-INS-198 |
| chr11-29698385-INS-88 | chr11-29698385-INS-88 |

| | |
|---|---|
| chr11-29703788-INS-1215 | chr11-29703789-INS-1215 |
| chr11-29708778-INS-137 | chr11-29708778-INS-137 |
| chr11-29725788-INS-1579 | chr11-29725789-INS-1580 |
| chr11-29732255-INS-120 | chr11-29732256-INS-120 |
| chr11-29743289-INS-567 | chr11-29743289-INS-567 |
| chr11-29757209-INS-123 | chr11-29757209-INS-119 |
| chr11-29782951-INS-53 | chr11-29782927-INS-87 |
| chr11-29810595-INS-108 | chr11-29810595-INS-120 |
| chr11-29812266-INS-485 | chr11-29812266-INS-485 |
| chr11-29813159-INS-80 | chr11-29813159-INS-64 |
| chr11-29825923-INS-465 | chr11-29825923-INS-465 |
| chr11-29862271-INS-92 | chr11-29862269-INS-122 |
| chr11-29866902-INS-58 | chr11-29866902-INS-58 |
| chr11-29869279-INS-6427 | chr11-29869280-INS-6427 |
| chr11-29878820-INS-70 | chr11-29878820-INS-105 |
| chr11-29896385-INS-6438 | chr11-29896386-INS-6438 |
| chr11-29900674-INS-694 | chr11-29900674-INS-694 |
| chr11-29901198-INS-62 | chr11-29901198-INS-96 |
| chr11-29932395-INS-797 | chr11-29932395-INS-797 |
| chr11-29958244-INS-75 | chr11-29958266-INS-78 |
| chr11-29961524-INS-160 | chr11-29961524-INS-160 |
| chr11-29965774-INS-82 | chr11-29965774-INS-83 |
| chr11-29967954-INS-199 | chr11-29967954-INS-199 |
| chr11-29989073-INS-2479 | chr11-29989074-INS-2484 |
| chr11-30003576-INS-201 | chr11-30003576-INS-202 |
| chr11-30010571-INS-2958 | chr11-30010572-INS-2954 |
| chr11-30028282-INS-3105 | chr11-30028283-INS-3105 |
| chr11-30028794-INS-70 | chr11-30028794-INS-50 |
| chr11-30078868-INS-88 | chr11-30078869-INS-81 |
| chr11-30084240-INS-166 | chr11-30084240-INS-168 |
| chr11-30104789-INS-1865 | chr11-30104790-INS-1884 |
| chr11-30113033-INS-682 | chr11-30113033-INS-682 |
| chr11-30126816-INS-60 | chr11-30126816-INS-52 |
| chr11-30129275-INS-6878 | chr11-30129276-INS-6878 |
| chr11-30138916-INS-172 | chr11-30138916-INS-172 |
| chr11-30142887-INS-93 | chr11-30142887-INS-93 |
| chr11-30163454-INS-88 | chr11-30163454-INS-82 |
| chr11-30170510-INS-226 | chr11-30170510-INS-224 |
| chr11-30176166-INS-198 | chr11-30176166-INS-196 |
| chr11-30229909-INS-2269 | chr11-30229910-INS-2270 |
| chr11-30255791-INS-826 | chr11-30255791-INS-826 |
| chr11-30279696-INS-871 | chr11-30279696-INS-871 |
| chr11-30311071-INS-468 | chr11-30311071-INS-484 |
| chr11-30329178-INS-662 | chr11-30329178-INS-662 |
| chr11-30332958-INS-55 | chr11-30332958-INS-55 |
| chr11-30359097-INS-5203 | chr11-30359098-INS-5203 |
| chr11-30391210-INS-56 | chr11-30391182-INS-54 |
| chr11-30401617-INS-57 | chr11-30401617-INS-57 |
| chr11-30403965-INS-92 | chr11-30403966-INS-92 |
| chr11-30409196-INS-6204 | chr11-30409197-INS-6204 |
| chr11-30414381-INS-218 | chr11-30414381-INS-217 |
| chr11-30417163-INS-210 | chr11-30417163-INS-210 |
| chr11-30436063-INS-227 | chr11-30436063-INS-229 |
| chr11-30437207-INS-51 | chr11-30437207-INS-53 |
| chr11-30455606-INS-418 | chr11-30455606-INS-417 |
| chr11-30459919-INS-65 | chr11-30459921-INS-77 |
| chr11-30474370-INS-56 | chr11-30474370-INS-54 |
| chr11-30475323-INS-237 | chr11-30475323-INS-238 |
| chr11-30479136-INS-103 | chr11-30479136-INS-108 |
| chr11-30484593-INS-54 | chr11-30484593-INS-54 |
| chr11-30492885-INS-89 | chr11-30492885-INS-89 |
| chr11-30506028-INS-53 | chr11-30506039-INS-67 |
| chr11-30510716-INS-54 | chr11-30510716-INS-54 |
| chr11-30512621-INS-615 | chr11-30512621-INS-615 |
| chr11-30552067-INS-72 | chr11-30552061-INS-77 |
| chr11-30555634-INS-51 | chr11-30555633-INS-53 |
| chr11-30566670-INS-91 | chr11-30566670-INS-90 |
| chr11-30583526-INS-451 | chr11-30583527-INS-451 |
| chr11-30604719-INS-148 | chr11-30604719-INS-148 |
| chr11-30653897-INS-66 | chr11-30653897-INS-72 |
| chr11-30656222-INS-56 | chr11-30656223-INS-51 |
| chr11-30659726-INS-208 | chr11-30659726-INS-210 |
| chr11-30680320-INS-54 | chr11-30680320-INS-54 |
| chr11-30689808-INS-828 | chr11-30689808-INS-830 |
| chr11-30700538-INS-70 | chr11-30700538-INS-54 |
| chr11-30708113-INS-1890 | chr11-30708114-INS-1890 |
| chr11-30720119-INS-209 | chr11-30720119-INS-209 |

| | |
|---|---|
| chr11-30726931-INS-2323 | chr11-30726932-INS-2322 |
| chr11-30734063-INS-6442 | chr11-30734064-INS-6442 |
| chr11-30737379-INS-196 | chr11-30737379-INS-188 |
| chr11-30748951-INS-99 | chr11-30748951-INS-132 |
| chr11-30762697-INS-60 | chr11-30762743-INS-55 |
| chr11-30788667-INS-78 | chr11-30788667-INS-72 |
| chr11-30794348-INS-6921 | chr11-30794349-INS-6921 |
| chr11-30816687-INS-87 | chr11-30816687-INS-87 |
| chr11-30827722-INS-57 | chr11-30827722-INS-57 |
| chr11-30831810-INS-124 | chr11-30831810-INS-130 |
| chr11-30879671-INS-135 | chr11-30879671-INS-136 |
| chr11-30918001-INS-125 | chr11-30918001-INS-125 |
| chr11-30929533-INS-208 | chr11-30929533-INS-208 |
| chr11-30977503-INS-92 | chr11-30977503-INS-95 |
| chr11-31113428-INS-1241 | chr11-31113429-INS-1241 |
| chr11-31120465-INS-104 | chr11-31120465-INS-104 |
| chr11-31139375-INS-400 | chr11-31139375-INS-400 |
| chr11-31140435-INS-6725 | chr11-31140436-INS-6725 |
| chr11-31141715-INS-213 | chr11-31141715-INS-213 |
| chr11-31154185-INS-215 | chr11-31154187-INS-202 |
| chr11-31155110-INS-59 | chr11-31155110-INS-61 |
| chr11-31190300-INS-430 | chr11-31190300-INS-430 |
| chr11-31230112-INS-7301 | chr11-31230113-INS-7301 |
| chr11-31230649-INS-401 | chr11-31230649-INS-401 |
| chr11-31252044-INS-56 | chr11-31252044-INS-56 |
| chr11-31256156-INS-1679 | chr11-31256157-INS-1683 |
| chr11-31259903-INS-6331 | chr11-31259904-INS-6331 |
| chr11-31271402-INS-229 | chr11-31271415-INS-182 |
| chr11-31278210-INS-58 | chr11-31278210-INS-58 |
| chr11-31295805-INS-1790 | chr11-31295806-INS-1790 |
| chr11-31343719-INS-92 | chr11-31343719-INS-92 |
| chr11-31386406-INS-689 | chr11-31386406-INS-689 |
| chr11-31395440-INS-132 | chr11-31395440-INS-133 |
| chr11-31427496-INS-58 | chr11-31427496-INS-60 |
| chr11-31429175-INS-328 | chr11-31429175-INS-328 |
| chr11-31445609-INS-82 | chr11-31445609-INS-84 |
| chr11-31453016-INS-119 | chr11-31453016-INS-121 |
| chr11-31459218-INS-871 | chr11-31459216-INS-874 |
| chr11-31473764-INS-74 | chr11-31473764-INS-74 |
| chr11-31481725-INS-168 | chr11-31481725-INS-168 |
| chr11-31488949-INS-54 | chr11-31488947-INS-72 |
| chr11-31568828-INS-186 | chr11-31568828-INS-192 |
| chr11-31600884-INS-498 | chr11-31600884-INS-498 |
| chr11-31602182-INS-4086 | chr11-31602183-INS-4086 |
| chr11-31621229-INS-99 | chr11-31621233-INS-91 |
| chr11-31705310-INS-85 | chr11-31705310-INS-85 |
| chr11-31737848-INS-161 | chr11-31737848-INS-162 |
| chr11-31746195-INS-352 | chr11-31746191-INS-351 |
| chr11-31755912-INS-61 | chr11-31755912-INS-61 |
| chr11-31759054-INS-175 | chr11-31759054-INS-175 |
| chr11-31783239-INS-60 | chr11-31783239-INS-56 |
| chr11-31807386-INS-73 | chr11-31807386-INS-73 |
| chr11-31836854-INS-86 | chr11-31836854-INS-86 |
| chr11-31899635-INS-237 | chr11-31899635-INS-237 |
| chr11-31903467-INS-239 | chr11-31903468-INS-264 |
| chr11-31904695-INS-4506 | chr11-31904696-INS-4506 |
| chr11-31913476-INS-586 | chr11-31913476-INS-586 |
| chr11-31940849-INS-65 | chr11-31940849-INS-66 |
| chr11-31944640-INS-70 | chr11-31944640-INS-70 |
| chr11-31945860-INS-79 | chr11-31945860-INS-56 |
| chr11-31958319-INS-55 | chr11-31958319-INS-55 |
| chr11-31964491-INS-4367 | chr11-31964492-INS-4367 |
| chr11-31975325-INS-316 | chr11-31975329-INS-316 |
| chr11-31986207-INS-114 | chr11-31986207-INS-114 |
| chr11-32003698-INS-199 | chr11-32003698-INS-200 |
| chr11-32017653-INS-213 | chr11-32017653-INS-213 |
| chr11-32019789-INS-1257 | chr11-32019790-INS-1257 |
| chr11-32022728-INS-52 | chr11-32022728-INS-52 |
| chr11-32024042-INS-199 | chr11-32024042-INS-195 |
| chr11-32027629-INS-144 | chr11-32027629-INS-146 |
| chr11-32043854-INS-112 | chr11-32043843-INS-112 |
| chr11-32064145-INS-200 | chr11-32064145-INS-200 |
| chr11-32072925-INS-81 | chr11-32072925-INS-81 |
| chr11-32074494-INS-209 | chr11-32074494-INS-209 |
| chr11-32075376-INS-108 | chr11-32075376-INS-108 |
| chr11-32081569-INS-56 | chr11-32081569-INS-70 |
| chr11-32086398-INS-2443 | chr11-32086399-INS-2443 |

| | |
|---|---|
| chr11-32094202-INS-2241 | chr11-32094203-INS-2244 |
| chr11-32105552-INS-252 | chr11-32105552-INS-252 |
| chr11-32126112-INS-218 | chr11-32126112-INS-218 |
| chr11-32136032-INS-61 | chr11-32136013-INS-71 |
| chr11-32145266-INS-154 | chr11-32145287-INS-155 |
| chr11-32153718-INS-87 | chr11-32153718-INS-87 |
| chr11-32153719-INS-1896 | chr11-32153720-INS-1896 |
| chr11-32194077-INS-97 | chr11-32194077-INS-97 |
| chr11-32208551-INS-164 | chr11-32208551-INS-164 |
| chr11-32211470-INS-92 | chr11-32211470-INS-60 |
| chr11-32222716-INS-345 | chr11-32222716-INS-345 |
| chr11-32223593-INS-138 | chr11-32223593-INS-138 |
| chr11-32224671-INS-198 | chr11-32224671-INS-198 |
| chr11-32238134-INS-64 | chr11-32238134-INS-68 |
| chr11-32247762-INS-72 | chr11-32247752-INS-70 |
| chr11-32248569-INS-76 | chr11-32248569-INS-60 |
| chr11-32293364-INS-93 | chr11-32293381-INS-52 |
| chr11-32309474-INS-59 | chr11-32309470-INS-60 |
| chr11-32318149-INS-69 | chr11-32318149-INS-69 |
| chr11-32323212-INS-192 | chr11-32323212-INS-192 |
| chr11-32360251-INS-52 | chr11-32360251-INS-52 |
| chr11-32371307-INS-386 | chr11-32371307-INS-386 |
| chr11-32408898-INS-214 | chr11-32408898-INS-213 |
| chr11-32421087-INS-84 | chr11-32421087-INS-120 |
| chr11-32435638-INS-592 | chr11-32435638-INS-596 |
| chr11-32445948-INS-380 | chr11-32445948-INS-381 |
| chr11-32457958-INS-125 | chr11-32457958-INS-125 |
| chr11-32477638-INS-204 | chr11-32477638-INS-208 |
| chr11-32495168-INS-72 | chr11-32495168-INS-78 |
| chr11-32519898-INS-170 | chr11-32519898-INS-170 |
| chr11-32530255-INS-85 | chr11-32530245-INS-94 |
| chr11-32534340-INS-80 | chr11-32534340-INS-80 |
| chr11-32569504-INS-325 | chr11-32569504-INS-325 |
| chr11-32577052-INS-81 | chr11-32577053-INS-93 |
| chr11-32577949-INS-79 | chr11-32577950-INS-79 |
| chr11-32580232-INS-65 | chr11-32580232-INS-98 |
| chr11-32601533-INS-78 | chr11-32601533-INS-78 |
| chr11-32606705-INS-81 | chr11-32606705-INS-63 |
| chr11-32609948-INS-216 | chr11-32609948-INS-216 |
| chr11-32611113-INS-516 | chr11-32611113-INS-519 |
| chr11-32664969-INS-718 | chr11-32664969-INS-718 |
| chr11-32685888-INS-350 | chr11-32685888-INS-355 |
| chr11-32687011-INS-6440 | chr11-32687012-INS-6471 |
| chr11-32703153-INS-808 | chr11-32703187-INS-817 |
| chr11-32712370-INS-132 | chr11-32712370-INS-132 |
| chr11-32752361-INS-190 | chr11-32752361-INS-190 |
| chr11-32767806-INS-102 | chr11-32767806-INS-104 |
| chr11-32777226-INS-64 | chr11-32777223-INS-92 |
| chr11-32782792-INS-230 | chr11-32782792-INS-230 |
| chr11-32838262-INS-64 | chr11-32838258-INS-60 |
| chr11-32845109-INS-220 | chr11-32845109-INS-219 |
| chr11-32863531-INS-68 | chr11-32863531-INS-68 |
| chr11-32888907-INS-114 | chr11-32888907-INS-114 |
| chr11-32904361-INS-75 | chr11-32904358-INS-91 |
| chr11-32906614-INS-190 | chr11-32906614-INS-191 |
| chr11-32935808-INS-67 | chr11-32935808-INS-67 |
| chr11-32965538-INS-496 | chr11-32965538-INS-495 |
| chr11-32984133-INS-77 | chr11-32984133-INS-77 |
| chr11-32986124-INS-175 | chr11-32986124-INS-175 |
| chr11-32986886-INS-217 | chr11-32986885-INS-220 |
| chr11-32995339-INS-383 | chr11-32995339-INS-383 |
| chr11-32996952-INS-393 | chr11-32996952-INS-393 |
| chr11-32996955-INS-1860 | chr11-32996952-INS-1860 |
| chr11-33023598-INS-199 | chr11-33023598-INS-203 |
| chr11-33037409-INS-62 | chr11-33037411-INS-50 |
| chr11-33043430-INS-62 | chr11-33043430-INS-68 |
| chr11-33046274-INS-167 | chr11-33046248-INS-177 |
| chr11-33052262-INS-995 | chr11-33052262-INS-994 |
| chr11-33056241-INS-226 | chr11-33056241-INS-227 |
| chr11-33079016-INS-60 | chr11-33079016-INS-60 |
| chr11-33105545-INS-115 | chr11-33105545-INS-114 |
| chr11-33115613-INS-205 | chr11-33115613-INS-206 |
| chr11-33126166-INS-55 | chr11-33126166-INS-60 |
| chr11-33139891-INS-1929 | chr11-33139892-INS-1929 |
| chr11-33164355-INS-58 | chr11-33164355-INS-54 |
| chr11-33199947-INS-199 | chr11-33199947-INS-199 |
| chr11-33213134-INS-192 | chr11-33213134-INS-192 |

| | |
|---|---|
| chr11-33225895-INS-2733 | chr11-33225896-INS-2733 |
| chr11-33229263-INS-1893 | chr11-33229264-INS-1893 |
| chr11-33232266-INS-1864 | chr11-33232267-INS-1864 |
| chr11-33233530-INS-130 | chr11-33233530-INS-130 |
| chr11-33254873-INS-513 | chr11-33254873-INS-513 |
| chr11-33354701-INS-218 | chr11-33354702-INS-218 |
| chr11-33362745-INS-1745 | chr11-33362746-INS-1745 |
| chr11-33385459-INS-239 | chr11-33385460-INS-217 |
| chr11-33401928-INS-410 | chr11-33401928-INS-410 |
| chr11-33411037-INS-949 | chr11-33411037-INS-949 |
| chr11-33412465-INS-208 | chr11-33412465-INS-208 |
| chr11-33418147-INS-212 | chr11-33418147-INS-212 |
| chr11-33425596-INS-6248 | chr11-33425597-INS-6248 |
| chr11-33447585-INS-61 | chr11-33447585-INS-57 |
| chr11-33468475-INS-202 | chr11-33468475-INS-202 |
| chr11-33476100-INS-178 | chr11-33476100-INS-178 |
| chr11-33490589-INS-233 | chr11-33490589-INS-235 |
| chr11-33492711-INS-187 | chr11-33492711-INS-187 |
| chr11-33494102-INS-72 | chr11-33494102-INS-72 |
| chr11-33505135-INS-112 | chr11-33505135-INS-112 |
| chr11-33509844-INS-60 | chr11-33509844-INS-56 |
| chr11-33511813-INS-120 | chr11-33511813-INS-120 |
| chr11-33542232-INS-65 | chr11-33542223-INS-74 |
| chr11-33548182-INS-1893 | chr11-33548183-INS-1893 |
| chr11-33554443-INS-70 | chr11-33554441-INS-71 |
| chr11-33573446-INS-81 | chr11-33573490-INS-54 |
| chr11-33594594-INS-72 | chr11-33594578-INS-50 |
| chr11-33610482-INS-100 | chr11-33610482-INS-100 |
| chr11-33618640-INS-102 | chr11-33618640-INS-102 |
| chr11-33631240-INS-84 | chr11-33631240-INS-86 |
| chr11-33632904-INS-126 | chr11-33632904-INS-126 |
| chr11-33639939-INS-284 | chr11-33639939-INS-284 |
| chr11-33639975-INS-766 | chr11-33639975-INS-760 |
| chr11-33648437-INS-58 | chr11-33648437-INS-60 |
| chr11-33652109-INS-80 | chr11-33652109-INS-80 |
| chr11-33659014-INS-222 | chr11-33659014-INS-222 |
| chr11-33660548-INS-437 | chr11-33660548-INS-432 |
| chr11-33673020-INS-110 | chr11-33673020-INS-110 |
| chr11-33675270-INS-148 | chr11-33675287-INS-145 |
| chr11-33682646-INS-52 | chr11-33682646-INS-52 |
| chr11-33688310-INS-280 | chr11-33688310-INS-280 |
| chr11-33703103-INS-55 | chr11-33703103-INS-57 |
| chr11-33715066-INS-7742 | chr11-33715067-INS-7743 |
| chr11-33726673-INS-189 | chr11-33726678-INS-200 |
| chr11-33764615-INS-88 | chr11-33764615-INS-121 |
| chr11-33805547-INS-60 | chr11-33805547-INS-68 |
| chr11-33865145-INS-82 | chr11-33865145-INS-87 |
| chr11-33868230-INS-57 | chr11-33868230-INS-57 |
| chr11-33874288-INS-76 | chr11-33874288-INS-76 |
| chr11-33881615-INS-442 | chr11-33881615-INS-442 |
| chr11-33894364-INS-160 | chr11-33894364-INS-166 |
| chr11-33951407-INS-2578 | chr11-33951408-INS-2578 |
| chr11-33964031-INS-214 | chr11-33964031-INS-214 |
| chr11-33995427-INS-121 | chr11-33995428-INS-121 |
| chr11-33997038-INS-133 | chr11-33997038-INS-133 |
| chr11-34038656-INS-51 | chr11-34038655-INS-66 |
| chr11-34062532-INS-196 | chr11-34062532-INS-196 |
| chr11-34073903-INS-57 | chr11-34073903-INS-58 |
| chr11-34077145-INS-623 | chr11-34077145-INS-623 |
| chr11-34095940-INS-60 | chr11-34095928-INS-88 |
| chr11-34104734-INS-54 | chr11-34104734-INS-56 |
| chr11-34113099-INS-305 | chr11-34113099-INS-305 |
| chr11-34116871-INS-106 | chr11-34116871-INS-110 |
| chr11-34120566-INS-125 | chr11-34120564-INS-126 |
| chr11-34151146-INS-3271 | chr11-34151147-INS-3271 |
| chr11-34158721-INS-52 | chr11-34158722-INS-54 |
| chr11-34169538-INS-52 | chr11-34169538-INS-52 |
| chr11-34219718-INS-55 | chr11-34219718-INS-55 |
| chr11-34225679-INS-53 | chr11-34225679-INS-53 |
| chr11-34229743-INS-62 | chr11-34229743-INS-50 |
| chr11-34242015-INS-52 | chr11-34242015-INS-54 |
| chr11-34246833-INS-187 | chr11-34246833-INS-187 |
| chr11-34285994-INS-54 | chr11-34285994-INS-54 |
| chr11-34300967-INS-101 | chr11-34300967-INS-101 |
| chr11-34321877-INS-206 | chr11-34321877-INS-206 |
| chr11-34385375-INS-152 | chr11-34385375-INS-176 |
| chr11-34395992-INS-186 | chr11-34395992-INS-186 |

| | |
|---|---|
| chr11-34403161-INS-139 | chr11-34403163-INS-135 |
| chr11-34417178-INS-635 | chr11-34417178-INS-635 |
| chr11-34447091-INS-56 | chr11-34447101-INS-66 |
| chr11-34465726-INS-102 | chr11-34465726-INS-108 |
| chr11-34468025-INS-52 | chr11-34468025-INS-62 |
| chr11-34480433-INS-500 | chr11-34480433-INS-500 |
| chr11-34493536-INS-237 | chr11-34493536-INS-237 |
| chr11-34552881-INS-837 | chr11-34552882-INS-837 |
| chr11-34586799-INS-56 | chr11-34586800-INS-56 |
| chr11-34635535-INS-55 | chr11-34635535-INS-55 |
| chr11-34674978-INS-95 | chr11-34674978-INS-96 |
| chr11-34684307-INS-3774 | chr11-34684308-INS-3776 |
| chr11-34701665-INS-185 | chr11-34701665-INS-185 |
| chr11-34713739-INS-64 | chr11-34713760-INS-76 |
| chr11-34714800-INS-244 | chr11-34714800-INS-232 |
| chr11-34719451-INS-63 | chr11-34719451-INS-63 |
| chr11-34732218-INS-178 | chr11-34732218-INS-145 |
| chr11-34740779-INS-212 | chr11-34740779-INS-212 |
| chr11-34804298-INS-93 | chr11-34804290-INS-96 |
| chr11-34806657-INS-132 | chr11-34806688-INS-88 |
| chr11-34849197-INS-102 | chr11-34849198-INS-60 |
| chr11-34849934-INS-58 | chr11-34849932-INS-58 |
| chr11-34856012-INS-52 | chr11-34856012-INS-56 |
| chr11-34916120-INS-168 | chr11-34916117-INS-169 |
| chr11-34925750-INS-141 | chr11-34925750-INS-142 |
| chr11-34931178-INS-62 | chr11-34931178-INS-66 |
| chr11-34950765-INS-378 | chr11-34950765-INS-402 |
| chr11-34955651-INS-195 | chr11-34955651-INS-196 |
| chr11-34985356-INS-54 | chr11-34985356-INS-58 |
| chr11-34987221-INS-66 | chr11-34987221-INS-66 |
| chr11-35003630-INS-101 | chr11-35003630-INS-101 |
| chr11-35043727-INS-65 | chr11-35043727-INS-65 |
| chr11-35047274-INS-163 | chr11-35047274-INS-167 |
| chr11-35078512-INS-52 | chr11-35078512-INS-52 |
| chr11-35092118-INS-75 | chr11-35092118-INS-75 |
| chr11-35095166-INS-117 | chr11-35095166-INS-117 |
| chr11-35098059-INS-58 | chr11-35098060-INS-64 |
| chr11-35127137-INS-163 | chr11-35127137-INS-163 |
| chr11-35130763-INS-174 | chr11-35130775-INS-176 |
| chr11-35178766-INS-76 | chr11-35178766-INS-76 |
| chr11-35192562-INS-78 | chr11-35192562-INS-80 |
| chr11-35205917-INS-98 | chr11-35205917-INS-98 |
| chr11-35236250-INS-215 | chr11-35236250-INS-216 |
| chr11-35238837-INS-101 | chr11-35238829-INS-107 |
| chr11-35242572-INS-86 | chr11-35242577-INS-85 |
| chr11-35248689-INS-144 | chr11-35248719-INS-148 |
| chr11-35250885-INS-70 | chr11-35250885-INS-72 |
| chr11-35256979-INS-1088 | chr11-35257001-INS-1045 |
| chr11-35268220-INS-145 | chr11-35268220-INS-145 |
| chr11-35271230-INS-78 | chr11-35271218-INS-108 |
| chr11-35272616-INS-54 | chr11-35272615-INS-54 |
| chr11-35280519-INS-4927 | chr11-35280522-INS-4928 |
| chr11-35283753-INS-53 | chr11-35283753-INS-53 |
| chr11-35311947-INS-123 | chr11-35311947-INS-123 |
| chr11-35336364-INS-207 | chr11-35336364-INS-206 |
| chr11-35355769-INS-52 | chr11-35355769-INS-54 |
| chr11-35410377-INS-732 | chr11-35410377-INS-732 |
| chr11-35471437-INS-400 | chr11-35471437-INS-400 |
| chr11-35491142-INS-354 | chr11-35491142-INS-354 |
| chr11-35532972-INS-376 | chr11-35532972-INS-376 |
| chr11-35564311-INS-157 | chr11-35564311-INS-157 |
| chr11-35567960-INS-52 | chr11-35567960-INS-52 |
| chr11-35586339-INS-387 | chr11-35586339-INS-387 |
| chr11-35594894-INS-100 | chr11-35594894-INS-100 |
| chr11-35620902-INS-249 | chr11-35620902-INS-249 |
| chr11-35624089-INS-91 | chr11-35624082-INS-92 |
| chr11-35630112-INS-204 | chr11-35630112-INS-204 |
| chr11-35643795-INS-434 | chr11-35643795-INS-440 |
| chr11-35647723-INS-198 | chr11-35647723-INS-198 |
| chr11-35652919-INS-101 | chr11-35652919-INS-101 |
| chr11-35700774-INS-244 | chr11-35700774-INS-248 |
| chr11-35739845-INS-52 | chr11-35739845-INS-52 |
| chr11-35740736-INS-198 | chr11-35740736-INS-198 |
| chr11-35741302-INS-88 | chr11-35741302-INS-88 |
| chr11-35744976-INS-58 | chr11-35744976-INS-58 |
| chr11-35745151-INS-503 | chr11-35745151-INS-503 |
| chr11-35801925-INS-59 | chr11-35801925-INS-75 |

| | |
|---|---|
| chr11-35827498-INS-577 | chr11-35827498-INS-577 |
| chr11-35832920-INS-104 | chr11-35832920-INS-105 |
| chr11-35856493-INS-75 | chr11-35856501-INS-56 |
| chr11-35886774-INS-359 | chr11-35886774-INS-359 |
| chr11-35906599-INS-80 | chr11-35906599-INS-80 |
| chr11-35962869-INS-58 | chr11-35962869-INS-56 |
| chr11-35973868-INS-120 | chr11-35973868-INS-114 |
| chr11-36038552-INS-56 | chr11-36038552-INS-64 |
| chr11-36040345-INS-908 | chr11-36040347-INS-905 |
| chr11-36047097-INS-108 | chr11-36047097-INS-108 |
| chr11-36082963-INS-220 | chr11-36082943-INS-204 |
| chr11-36088833-INS-182 | chr11-36088833-INS-182 |
| chr11-36131075-INS-52 | chr11-36131075-INS-52 |
| chr11-36164560-INS-63 | chr11-36164560-INS-71 |
| chr11-36173045-INS-51 | chr11-36173042-INS-56 |
| chr11-36205535-INS-70 | chr11-36205535-INS-50 |
| chr11-36218870-INS-922 | chr11-36218870-INS-923 |
| chr11-36278493-INS-200 | chr11-36278493-INS-199 |
| chr11-36293228-INS-56 | chr11-36293230-INS-64 |
| chr11-36301069-INS-55 | chr11-36301070-INS-55 |
| chr11-36320813-INS-92 | chr11-36320858-INS-50 |
| chr11-36330742-INS-87 | chr11-36330742-INS-76 |
| chr11-36343467-INS-1890 | chr11-36343468-INS-1890 |
| chr11-36366103-INS-143 | chr11-36366079-INS-154 |
| chr11-36389432-INS-56 | chr11-36389414-INS-60 |
| chr11-36426653-INS-92 | chr11-36426624-INS-92 |
| chr11-36450346-INS-104 | chr11-36450311-INS-104 |
| chr11-36472973-INS-54 | chr11-36472973-INS-56 |
| chr11-36499910-INS-1051 | chr11-36499911-INS-1052 |
| chr11-36519387-INS-120 | chr11-36519389-INS-146 |
| chr11-36537646-INS-81 | chr11-36537634-INS-70 |
| chr11-36606078-INS-104 | chr11-36606078-INS-104 |
| chr11-36662530-INS-52 | chr11-36662508-INS-57 |
| chr11-36676127-INS-66 | chr11-36676089-INS-64 |
| chr11-36689714-INS-1620 | chr11-36689715-INS-1611 |
| chr11-36698045-INS-54 | chr11-36698050-INS-57 |
| chr11-36710548-INS-58 | chr11-36710549-INS-62 |
| chr11-36820172-INS-188 | chr11-36820172-INS-188 |
| chr11-36827926-INS-91 | chr11-36827926-INS-118 |
| chr11-36885723-INS-120 | chr11-36885723-INS-119 |
| chr11-36890885-INS-64 | chr11-36890868-INS-54 |
| chr11-36899881-INS-58 | chr11-36899881-INS-51 |
| chr11-36998496-INS-54 | chr11-36998496-INS-56 |
| chr11-37036155-INS-58 | chr11-37036155-INS-62 |
| chr11-37069471-INS-6684 | chr11-37069472-INS-6684 |
| chr11-37084726-INS-157 | chr11-37084726-INS-157 |
| chr11-37141009-INS-424 | chr11-37141009-INS-424 |
| chr11-37153798-INS-347 | chr11-37153798-INS-336 |
| chr11-37156453-INS-2902 | chr11-37156454-INS-2887 |
| chr11-37159164-INS-360 | chr11-37159165-INS-341 |
| chr11-37167411-INS-60 | chr11-37167411-INS-60 |
| chr11-37176644-INS-480 | chr11-37176644-INS-480 |
| chr11-37184200-INS-717 | chr11-37184200-INS-717 |
| chr11-37193438-INS-58 | chr11-37193438-INS-62 |
| chr11-37207919-INS-81 | chr11-37207919-INS-51 |
| chr11-37226316-INS-95 | chr11-37226316-INS-143 |
| chr11-37232855-INS-88 | chr11-37232856-INS-88 |
| chr11-37257958-INS-6628 | chr11-37257959-INS-6629 |
| chr11-37268714-INS-2809 | chr11-37268715-INS-2809 |
| chr11-37297009-INS-324 | chr11-37297009-INS-325 |
| chr11-37307964-INS-199 | chr11-37307964-INS-199 |
| chr11-37312759-INS-229 | chr11-37312759-INS-229 |
| chr11-37327867-INS-1684 | chr11-37327868-INS-1684 |
| chr11-37336312-INS-64 | chr11-37336312-INS-62 |
| chr11-37349102-INS-96 | chr11-37349102-INS-96 |
| chr11-37357916-INS-122 | chr11-37357916-INS-122 |
| chr11-37362403-INS-52 | chr11-37362403-INS-54 |
| chr11-37393246-INS-84 | chr11-37393246-INS-70 |
| chr11-37411382-INS-54 | chr11-37411380-INS-58 |
| chr11-37479176-INS-92 | chr11-37479176-INS-92 |
| chr11-37481256-INS-75 | chr11-37481257-INS-95 |
| chr11-37484324-INS-54 | chr11-37484324-INS-56 |
| chr11-37489275-INS-469 | chr11-37489275-INS-487 |
| chr11-37499413-INS-56 | chr11-37499413-INS-56 |
| chr11-37503317-INS-54 | chr11-37503317-INS-54 |
| chr11-37505810-INS-1037 | chr11-37505811-INS-1037 |
| chr11-37515001-INS-497 | chr11-37515001-INS-497 |

| | |
|---|---|
| chr11-37520243-INS-217 | chr11-37520243-INS-217 |
| chr11-37523877-INS-111 | chr11-37523877-INS-111 |
| chr11-37547957-INS-76 | chr11-37547979-INS-50 |
| chr11-37550377-INS-54 | chr11-37550377-INS-54 |
| chr11-37555007-INS-75 | chr11-37555008-INS-52 |
| chr11-37577934-INS-54 | chr11-37577934-INS-54 |
| chr11-37585619-INS-2209 | chr11-37585602-INS-2189 |
| chr11-37597297-INS-60 | chr11-37597297-INS-60 |
| chr11-37607606-INS-6317 | chr11-37607607-INS-6317 |
| chr11-37610574-INS-301 | chr11-37610574-INS-300 |
| chr11-37613112-INS-105 | chr11-37613114-INS-103 |
| chr11-37620379-INS-6472 | chr11-37620380-INS-6472 |
| chr11-37633415-INS-95 | chr11-37633415-INS-100 |
| chr11-37656537-INS-356 | chr11-37656537-INS-356 |
| chr11-37674865-INS-6385 | chr11-37674866-INS-6385 |
| chr11-37693967-INS-55 | chr11-37693969-INS-59 |
| chr11-37704912-INS-1902 | chr11-37704913-INS-1901 |
| chr11-37731376-INS-52 | chr11-37731376-INS-56 |
| chr11-37781044-INS-53 | chr11-37781044-INS-65 |
| chr11-37797899-INS-159 | chr11-37797899-INS-159 |
| chr11-37863145-INS-88 | chr11-37863145-INS-88 |
| chr11-37881895-INS-56 | chr11-37881895-INS-56 |
| chr11-37899468-INS-685 | chr11-37899468-INS-685 |
| chr11-37930436-INS-71 | chr11-37930436-INS-71 |
| chr11-37956269-INS-318 | chr11-37956269-INS-315 |
| chr11-37961893-INS-51 | chr11-37961893-INS-54 |
| chr11-37973955-INS-66 | chr11-37973955-INS-66 |
| chr11-37980272-INS-60 | chr11-37980272-INS-60 |
| chr11-37994301-INS-58 | chr11-37994272-INS-50 |
| chr11-37999436-INS-103 | chr11-37999431-INS-97 |
| chr11-38005996-INS-60 | chr11-38005996-INS-60 |
| chr11-38012534-INS-52 | chr11-38012534-INS-54 |
| chr11-38056480-INS-62 | chr11-38056480-INS-56 |
| chr11-38065025-INS-180 | chr11-38065025-INS-180 |
| chr11-38082769-INS-642 | chr11-38082775-INS-657 |
| chr11-38088828-INS-1829 | chr11-38088829-INS-1829 |
| chr11-38099665-INS-57 | chr11-38099669-INS-53 |
| chr11-38148102-INS-290 | chr11-38148102-INS-290 |
| chr11-38182973-INS-147 | chr11-38182976-INS-157 |
| chr11-38206054-INS-52 | chr11-38206054-INS-60 |
| chr11-38217475-INS-84 | chr11-38217475-INS-84 |
| chr11-38253363-INS-189 | chr11-38253360-INS-190 |
| chr11-38258107-INS-59 | chr11-38258110-INS-60 |
| chr11-38273768-INS-179 | chr11-38273768-INS-160 |
| chr11-38308753-INS-144 | chr11-38308753-INS-162 |
| chr11-38317930-INS-55 | chr11-38317930-INS-54 |
| chr11-38356951-INS-105 | chr11-38356951-INS-105 |
| chr11-38393583-INS-196 | chr11-38393583-INS-197 |
| chr11-38427487-INS-152 | chr11-38427487-INS-152 |
| chr11-38431922-INS-81 | chr11-38431922-INS-81 |
| chr11-38433611-INS-240 | chr11-38433611-INS-246 |
| chr11-38434621-INS-108 | chr11-38434621-INS-84 |
| chr11-38439075-INS-82 | chr11-38439075-INS-86 |
| chr11-38440644-INS-52 | chr11-38440644-INS-52 |
| chr11-38486321-INS-64 | chr11-38486329-INS-60 |
| chr11-38486872-INS-6365 | chr11-38486873-INS-6365 |
| chr11-38490125-INS-618 | chr11-38490125-INS-618 |
| chr11-38497980-INS-3703 | chr11-38497974-INS-3700 |
| chr11-38510835-INS-60 | chr11-38510832-INS-64 |
| chr11-38511996-INS-115 | chr11-38511996-INS-116 |
| chr11-38515262-INS-52 | chr11-38515214-INS-60 |
| chr11-38529304-INS-56 | chr11-38529304-INS-56 |
| chr11-38557885-INS-51 | chr11-38557885-INS-51 |
| chr11-38564291-INS-91 | chr11-38564291-INS-113 |
| chr11-38568002-INS-342 | chr11-38568008-INS-351 |
| chr11-38628076-INS-60 | chr11-38628046-INS-62 |
| chr11-38641816-INS-623 | chr11-38641816-INS-623 |
| chr11-38674845-INS-72 | chr11-38674845-INS-72 |
| chr11-38676780-INS-134 | chr11-38676780-INS-134 |
| chr11-38699430-INS-71 | chr11-38699420-INS-76 |
| chr11-38699610-INS-3937 | chr11-38699611-INS-3937 |
| chr11-38727881-INS-7253 | chr11-38727882-INS-7253 |
| chr11-38758394-INS-489 | chr11-38758397-INS-486 |
| chr11-38890439-INS-155 | chr11-38890439-INS-150 |
| chr11-38891326-INS-107 | chr11-38891326-INS-108 |
| chr11-38895568-INS-4164 | chr11-38895569-INS-4166 |
| chr11-38901129-INS-6243 | chr11-38901130-INS-6243 |

| | |
|---|---|
| chr11-38926448-INS-1329 | chr11-38926449-INS-1329 |
| chr11-38963027-INS-59 | chr11-38963027-INS-59 |
| chr11-38963649-INS-934 | chr11-38963649-INS-934 |
| chr11-38968521-INS-60 | chr11-38968521-INS-66 |
| chr11-38988616-INS-2192 | chr11-38988617-INS-2192 |
| chr11-39003148-INS-213 | chr11-39003148-INS-217 |
| chr11-39008832-INS-52 | chr11-39008834-INS-56 |
| chr11-39016557-INS-1882 | chr11-39016558-INS-1882 |
| chr11-39029570-INS-141 | chr11-39029546-INS-138 |
| chr11-39032982-INS-730 | chr11-39032982-INS-745 |
| chr11-39059399-INS-75 | chr11-39059399-INS-75 |
| chr11-39061060-INS-75 | chr11-39061060-INS-75 |
| chr11-39079279-INS-61 | chr11-39079280-INS-56 |
| chr11-39080067-INS-114 | chr11-39080068-INS-71 |
| chr11-39110535-INS-6790 | chr11-39110536-INS-6790 |
| chr11-39149399-INS-51 | chr11-39149376-INS-86 |
| chr11-39186408-INS-7307 | chr11-39186409-INS-7308 |
| chr11-39213892-INS-2415 | chr11-39213893-INS-2419 |
| chr11-39263029-INS-2816 | chr11-39263030-INS-2816 |
| chr11-39267067-INS-2512 | chr11-39267068-INS-2514 |
| chr11-39280549-INS-3154 | chr11-39280550-INS-3155 |
| chr11-39286418-INS-66 | chr11-39286418-INS-66 |
| chr11-39326080-INS-60 | chr11-39326080-INS-70 |
| chr11-39348322-INS-8296 | chr11-39348323-INS-8297 |
| chr11-39352557-INS-56 | chr11-39352557-INS-56 |
| chr11-39354419-INS-60 | chr11-39354419-INS-63 |
| chr11-39410362-INS-5328 | chr11-39410363-INS-5327 |
| chr11-39425373-INS-72 | chr11-39425373-INS-78 |
| chr11-39443292-INS-272 | chr11-39443292-INS-272 |
| chr11-39471327-INS-54 | chr11-39471327-INS-54 |
| chr11-39508235-INS-145 | chr11-39508235-INS-145 |
| chr11-39534567-INS-52 | chr11-39534567-INS-56 |
| chr11-39540551-INS-414 | chr11-39540551-INS-414 |
| chr11-39543707-INS-52 | chr11-39543707-INS-50 |
| chr11-39556921-INS-500 | chr11-39556921-INS-500 |
| chr11-39581221-INS-65 | chr11-39581221-INS-65 |
| chr11-39650821-INS-6455 | chr11-39650822-INS-6456 |
| chr11-39654767-INS-103 | chr11-39654767-INS-103 |
| chr11-39704519-INS-88 | chr11-39704517-INS-90 |
| chr11-39730806-INS-52 | chr11-39730806-INS-56 |
| chr11-39750916-INS-753 | chr11-39750916-INS-753 |
| chr11-39756394-INS-7162 | chr11-39756395-INS-7162 |
| chr11-39759915-INS-76 | chr11-39759915-INS-78 |
| chr11-39768600-INS-234 | chr11-39768600-INS-234 |
| chr11-39781193-INS-263 | chr11-39781193-INS-263 |
| chr11-39796910-INS-68 | chr11-39796899-INS-64 |
| chr11-39807151-INS-70 | chr11-39807151-INS-70 |
| chr11-39818738-INS-6203 | chr11-39818739-INS-6203 |
| chr11-39818846-INS-573 | chr11-39818847-INS-573 |
| chr11-39866147-INS-351 | chr11-39866147-INS-349 |
| chr11-39868486-INS-550 | chr11-39868486-INS-551 |
| chr11-39875843-INS-59 | chr11-39875843-INS-59 |
| chr11-39896338-INS-98 | chr11-39896338-INS-98 |
| chr11-39899688-INS-200 | chr11-39899688-INS-200 |
| chr11-39901584-INS-153 | chr11-39901633-INS-112 |
| chr11-39928707-INS-164 | chr11-39928707-INS-164 |
| chr11-39935810-INS-66 | chr11-39935810-INS-66 |
| chr11-39938163-INS-1866 | chr11-39938164-INS-1866 |
| chr11-39938857-INS-503 | chr11-39938857-INS-503 |
| chr11-39940255-INS-3138 | chr11-39940256-INS-3138 |
| chr11-39949450-INS-70 | chr11-39949448-INS-88 |
| chr11-39980728-INS-418 | chr11-39980728-INS-418 |
| chr11-40006819-INS-256 | chr11-40006819-INS-256 |
| chr11-40014858-INS-244 | chr11-40014859-INS-244 |
| chr11-40035655-INS-1704 | chr11-40035656-INS-1704 |
| chr11-40042257-INS-52 | chr11-40042257-INS-60 |
| chr11-40059231-INS-214 | chr11-40059231-INS-214 |
| chr11-40105727-INS-7085 | chr11-40105728-INS-7087 |
| chr11-40111398-INS-1959 | chr11-40111399-INS-1959 |
| chr11-40116967-INS-69 | chr11-40116967-INS-105 |
| chr11-40127534-INS-208 | chr11-40127534-INS-208 |
| chr11-40144558-INS-63 | chr11-40144558-INS-63 |
| chr11-40155310-INS-56 | chr11-40155310-INS-54 |
| chr11-40166619-INS-82 | chr11-40166615-INS-74 |
| chr11-40171724-INS-84 | chr11-40171730-INS-78 |
| chr11-40178130-INS-116 | chr11-40178130-INS-117 |
| chr11-40191508-INS-78 | chr11-40191508-INS-88 |

| | |
|---|---|
| chr11-40210433-INS-1003 | chr11-40210434-INS-1023 |
| chr11-40222696-INS-512 | chr11-40222696-INS-513 |
| chr11-40243022-INS-88 | chr11-40243022-INS-92 |
| chr11-40259798-INS-69 | chr11-40259839-INS-72 |
| chr11-40269954-INS-64 | chr11-40269954-INS-68 |
| chr11-40302776-INS-76 | chr11-40302755-INS-113 |
| chr11-40311745-INS-116 | chr11-40311751-INS-128 |
| chr11-40412893-INS-176 | chr11-40412933-INS-140 |
| chr11-40432709-INS-96 | chr11-40432709-INS-94 |
| chr11-40461331-INS-424 | chr11-40461331-INS-424 |
| chr11-40474656-INS-60 | chr11-40474656-INS-60 |
| chr11-40501310-INS-200 | chr11-40501310-INS-200 |
| chr11-40506036-INS-62 | chr11-40506031-INS-50 |
| chr11-40514057-INS-66 | chr11-40514077-INS-65 |
| chr11-40562584-INS-538 | chr11-40562585-INS-538 |
| chr11-40601035-INS-126 | chr11-40601035-INS-126 |
| chr11-40620209-INS-175 | chr11-40620210-INS-175 |
| chr11-40622506-INS-206 | chr11-40622506-INS-206 |
| chr11-40638187-INS-171 | chr11-40638187-INS-171 |
| chr11-40659070-INS-124 | chr11-40659076-INS-136 |
| chr11-40662601-INS-68 | chr11-40662573-INS-112 |
| chr11-40666293-INS-112 | chr11-40666293-INS-112 |
| chr11-40673426-INS-607 | chr11-40673426-INS-607 |
| chr11-40678746-INS-52 | chr11-40678746-INS-50 |
| chr11-40723462-INS-208 | chr11-40723462-INS-210 |
| chr11-40747959-INS-94 | chr11-40747959-INS-94 |
| chr11-40759113-INS-79 | chr11-40759113-INS-71 |
| chr11-40763091-INS-6599 | chr11-40763092-INS-6600 |
| chr11-40769290-INS-65 | chr11-40769289-INS-54 |
| chr11-40770314-INS-54 | chr11-40770314-INS-51 |
| chr11-40774365-INS-72 | chr11-40774365-INS-72 |
| chr11-40825464-INS-55 | chr11-40825464-INS-55 |
| chr11-40889902-INS-52 | chr11-40889917-INS-66 |
| chr11-40899613-INS-3963 | chr11-40899614-INS-3963 |
| chr11-40921310-INS-64 | chr11-40921360-INS-50 |
| chr11-40943394-INS-72 | chr11-40943394-INS-76 |
| chr11-40965250-INS-52 | chr11-40965205-INS-51 |
| chr11-41028903-INS-100 | chr11-41028903-INS-75 |
| chr11-41072869-INS-73 | chr11-41072869-INS-74 |
| chr11-41091876-INS-432 | chr11-41091876-INS-432 |
| chr11-41093491-INS-894 | chr11-41093491-INS-894 |
| chr11-41094073-INS-101 | chr11-41094069-INS-109 |
| chr11-41148794-INS-1070 | chr11-41148802-INS-1090 |
| chr11-41175188-INS-209 | chr11-41175188-INS-208 |
| chr11-41177967-INS-925 | chr11-41177967-INS-926 |
| chr11-41188523-INS-253 | chr11-41188523-INS-253 |
| chr11-41220563-INS-536 | chr11-41220563-INS-535 |
| chr11-41235085-INS-3108 | chr11-41235086-INS-3108 |
| chr11-41263429-INS-75 | chr11-41263429-INS-77 |
| chr11-41284867-INS-5352 | chr11-41284868-INS-5352 |
| chr11-41288514-INS-174 | chr11-41288514-INS-174 |
| chr11-41291782-INS-74 | chr11-41291784-INS-74 |
| chr11-41299825-INS-243 | chr11-41299817-INS-224 |
| chr11-41326386-INS-52 | chr11-41326386-INS-50 |
| chr11-41336104-INS-133 | chr11-41336104-INS-138 |
| chr11-41386287-INS-506 | chr11-41386284-INS-490 |
| chr11-41421563-INS-118 | chr11-41421563-INS-122 |
| chr11-41423459-INS-264 | chr11-41423459-INS-264 |
| chr11-41434112-INS-52 | chr11-41434112-INS-52 |
| chr11-41459932-INS-135 | chr11-41459954-INS-137 |
| chr11-41470793-INS-69 | chr11-41470793-INS-74 |
| chr11-41472364-INS-74 | chr11-41472364-INS-80 |
| chr11-41492501-INS-69 | chr11-41492501-INS-113 |
| chr11-41518413-INS-583 | chr11-41518413-INS-582 |
| chr11-41523440-INS-1752 | chr11-41523441-INS-1752 |
| chr11-41523535-INS-176 | chr11-41523535-INS-176 |
| chr11-41525177-INS-6571 | chr11-41525178-INS-6572 |
| chr11-41562101-INS-1262 | chr11-41562102-INS-1263 |
| chr11-41579983-INS-6902 | chr11-41579984-INS-6902 |
| chr11-41649506-INS-62 | chr11-41649507-INS-70 |
| chr11-41668788-INS-54 | chr11-41668788-INS-54 |
| chr11-41671162-INS-575 | chr11-41671162-INS-578 |
| chr11-41687384-INS-454 | chr11-41687384-INS-454 |
| chr11-41695764-INS-3789 | chr11-41695765-INS-3789 |
| chr11-41737012-INS-4121 | chr11-41737013-INS-4121 |
| chr11-41741917-INS-92 | chr11-41741917-INS-92 |
| chr11-41744985-INS-110 | chr11-41744985-INS-106 |

| | |
|---|---|
| chr11-41751932-INS-2923 | chr11-41751933-INS-2923 |
| chr11-41759903-INS-243 | chr11-41759903-INS-241 |
| chr11-41764392-INS-56 | chr11-41764392-INS-58 |
| chr11-41797333-INS-413 | chr11-41797333-INS-413 |
| chr11-41798087-INS-6488 | chr11-41798088-INS-6489 |
| chr11-41800316-INS-62 | chr11-41800317-INS-66 |
| chr11-41833726-INS-66 | chr11-41833726-INS-66 |
| chr11-41835088-INS-175 | chr11-41835084-INS-163 |
| chr11-41838628-INS-4458 | chr11-41838629-INS-4458 |
| chr11-41846785-INS-63 | chr11-41846785-INS-63 |
| chr11-41874881-INS-768 | chr11-41874881-INS-768 |
| chr11-41876412-INS-615 | chr11-41876412-INS-615 |
| chr11-41894194-INS-51 | chr11-41894194-INS-51 |
| chr11-41920659-INS-300 | chr11-41920659-INS-301 |
| chr11-41933496-INS-54 | chr11-41933496-INS-54 |
| chr11-41974932-INS-3644 | chr11-41974933-INS-3643 |
| chr11-41978677-INS-60 | chr11-41978665-INS-57 |
| chr11-41985621-INS-343 | chr11-41985621-INS-383 |
| chr11-42001466-INS-5276 | chr11-42001467-INS-5276 |
| chr11-42030670-INS-171 | chr11-42030670-INS-171 |
| chr11-42057178-INS-72 | chr11-42057178-INS-72 |
| chr11-42062947-INS-65 | chr11-42062947-INS-65 |
| chr11-42074628-INS-7078 | chr11-42074629-INS-7078 |
| chr11-42093103-INS-60 | chr11-42093103-INS-61 |
| chr11-42094347-INS-6401 | chr11-42094348-INS-6401 |
| chr11-42097667-INS-1918 | chr11-42097668-INS-1918 |
| chr11-42106687-INS-63 | chr11-42106687-INS-63 |
| chr11-42120443-INS-490 | chr11-42120439-INS-491 |
| chr11-42128051-INS-180 | chr11-42128051-INS-166 |
| chr11-42140536-INS-403 | chr11-42140536-INS-418 |
| chr11-42161497-INS-804 | chr11-42161497-INS-804 |
| chr11-42163423-INS-95 | chr11-42163423-INS-91 |
| chr11-42217638-INS-76 | chr11-42217638-INS-78 |
| chr11-42229091-INS-55 | chr11-42229091-INS-55 |
| chr11-42240612-INS-55 | chr11-42240612-INS-56 |
| chr11-42253662-INS-51 | chr11-42253662-INS-51 |
| chr11-42289476-INS-132 | chr11-42289466-INS-152 |
| chr11-42291058-INS-5746 | chr11-42291059-INS-5745 |
| chr11-42296391-INS-2839 | chr11-42296393-INS-2840 |
| chr11-42297865-INS-71 | chr11-42297865-INS-75 |
| chr11-42299584-INS-128 | chr11-42299584-INS-128 |
| chr11-42311780-INS-53 | chr11-42311780-INS-56 |
| chr11-42318523-INS-107 | chr11-42318523-INS-107 |
| chr11-42343622-INS-60 | chr11-42343622-INS-52 |
| chr11-42359894-INS-86 | chr11-42359894-INS-88 |
| chr11-42413639-INS-107 | chr11-42413639-INS-91 |
| chr11-42420929-INS-87 | chr11-42420929-INS-63 |
| chr11-42430444-INS-75 | chr11-42430444-INS-79 |
| chr11-42432635-INS-72 | chr11-42432610-INS-94 |
| chr11-42449963-INS-69 | chr11-42449963-INS-69 |
| chr11-42467949-INS-254 | chr11-42467949-INS-254 |
| chr11-42486855-INS-778 | chr11-42486855-INS-778 |
| chr11-42492284-INS-92 | chr11-42492284-INS-92 |
| chr11-42498982-INS-1675 | chr11-42498983-INS-1676 |
| chr11-42525228-INS-114 | chr11-42525249-INS-72 |
| chr11-42550967-INS-113 | chr11-42550967-INS-99 |
| chr11-42554110-INS-3311 | chr11-42554111-INS-3311 |
| chr11-42562376-INS-601 | chr11-42562377-INS-601 |
| chr11-42566292-INS-485 | chr11-42566292-INS-484 |
| chr11-42590292-INS-62 | chr11-42590292-INS-62 |
| chr11-42621060-INS-146 | chr11-42621060-INS-146 |
| chr11-42625994-INS-52 | chr11-42625994-INS-54 |
| chr11-42630997-INS-747 | chr11-42630997-INS-769 |
| chr11-42654931-INS-68 | chr11-42654924-INS-64 |
| chr11-42666875-INS-112 | chr11-42666875-INS-112 |
| chr11-42691296-INS-58 | chr11-42691274-INS-96 |
| chr11-42698710-INS-84 | chr11-42698691-INS-67 |
| chr11-42700461-INS-4316 | chr11-42700462-INS-4312 |
| chr11-42701040-INS-525 | chr11-42701040-INS-525 |
| chr11-42703052-INS-54 | chr11-42703060-INS-57 |
| chr11-42705225-INS-54 | chr11-42705225-INS-58 |
| chr11-42715664-INS-1894 | chr11-42715665-INS-1894 |
| chr11-42720115-INS-226 | chr11-42720116-INS-225 |
| chr11-42725630-INS-68 | chr11-42725630-INS-68 |
| chr11-42727477-INS-57 | chr11-42727477-INS-57 |
| chr11-42732659-INS-170 | chr11-42732659-INS-170 |
| chr11-42733506-INS-398 | chr11-42733511-INS-401 |

| | |
|---|---|
| chr11-42742267-INS-66 | chr11-42742267-INS-56 |
| chr11-42754625-INS-56 | chr11-42754625-INS-56 |
| chr11-42777002-INS-7075 | chr11-42777003-INS-7075 |
| chr11-42806747-INS-252 | chr11-42806747-INS-252 |
| chr11-42819088-INS-87 | chr11-42819085-INS-69 |
| chr11-42825470-INS-186 | chr11-42825469-INS-189 |
| chr11-42833534-INS-6442 | chr11-42833535-INS-6442 |
| chr11-42846069-INS-541 | chr11-42846072-INS-545 |
| chr11-42850966-INS-1413 | chr11-42850971-INS-1422 |
| chr11-42854989-INS-2446 | chr11-42854990-INS-2446 |
| chr11-42857453-INS-55 | chr11-42857451-INS-70 |
| chr11-42893927-INS-91 | chr11-42893927-INS-91 |
| chr11-42899233-INS-217 | chr11-42899233-INS-217 |
| chr11-42948491-INS-1888 | chr11-42948492-INS-1888 |
| chr11-42964902-INS-355 | chr11-42964902-INS-355 |
| chr11-42975885-INS-1505 | chr11-42975886-INS-1505 |
| chr11-42997097-INS-6502 | chr11-42997098-INS-6502 |
| chr11-43003104-INS-1476 | chr11-43003105-INS-1474 |
| chr11-43018708-INS-90 | chr11-43018708-INS-91 |
| chr11-43027245-INS-74 | chr11-43027245-INS-70 |
| chr11-43055705-INS-53 | chr11-43055705-INS-53 |
| chr11-43061587-INS-129 | chr11-43061587-INS-129 |
| chr11-43083134-INS-52 | chr11-43083134-INS-54 |
| chr11-43109993-INS-68 | chr11-43109993-INS-78 |
| chr11-43120151-INS-1224 | chr11-43120152-INS-1224 |
| chr11-43153870-INS-78 | chr11-43153870-INS-78 |
| chr11-43157328-INS-68 | chr11-43157328-INS-70 |
| chr11-43164041-INS-497 | chr11-43164041-INS-497 |
| chr11-43164919-INS-107 | chr11-43164917-INS-118 |
| chr11-43173262-INS-58 | chr11-43173262-INS-58 |
| chr11-43208093-INS-459 | chr11-43208093-INS-459 |
| chr11-43225349-INS-7541 | chr11-43225350-INS-7542 |
| chr11-43225799-INS-127 | chr11-43225816-INS-86 |
| chr11-43230284-INS-189 | chr11-43230283-INS-186 |
| chr11-43237979-INS-56 | chr11-43237979-INS-58 |
| chr11-43274648-INS-54 | chr11-43274649-INS-54 |
| chr11-43278510-INS-60 | chr11-43278510-INS-64 |
| chr11-43280840-INS-78 | chr11-43280843-INS-87 |
| chr11-43281365-INS-55 | chr11-43281365-INS-55 |
| chr11-43285379-INS-75 | chr11-43285383-INS-71 |
| chr11-43316858-INS-576 | chr11-43316858-INS-576 |
| chr11-43318938-INS-69 | chr11-43318938-INS-66 |
| chr11-43322537-INS-172 | chr11-43322537-INS-172 |
| chr11-43329327-INS-122 | chr11-43329327-INS-101 |
| chr11-43341788-INS-56 | chr11-43341788-INS-52 |
| chr11-43354865-INS-66 | chr11-43354866-INS-63 |
| chr11-43361851-INS-63 | chr11-43361867-INS-56 |
| chr11-43373018-INS-86 | chr11-43373012-INS-50 |
| chr11-43375509-INS-293 | chr11-43375509-INS-293 |
| chr11-43392694-INS-289 | chr11-43392694-INS-287 |
| chr11-43409837-INS-56 | chr11-43409837-INS-56 |
| chr11-43411154-INS-352 | chr11-43411172-INS-375 |
| chr11-43431287-INS-50 | chr11-43431271-INS-56 |
| chr11-43445884-INS-60 | chr11-43445880-INS-70 |
| chr11-43463084-INS-108 | chr11-43463073-INS-112 |
| chr11-43465308-INS-237 | chr11-43465308-INS-237 |
| chr11-43471596-INS-123 | chr11-43471596-INS-123 |
| chr11-43487943-INS-86 | chr11-43487943-INS-66 |
| chr11-43492129-INS-108 | chr11-43492129-INS-108 |
| chr11-43502476-INS-182 | chr11-43502476-INS-182 |
| chr11-43505518-INS-68 | chr11-43505518-INS-68 |
| chr11-43524867-INS-5400 | chr11-43524868-INS-5400 |
| chr11-43524868-INS-701 | chr11-43524868-INS-701 |
| chr11-43531997-INS-534 | chr11-43531997-INS-544 |
| chr11-43533433-INS-57 | chr11-43533433-INS-72 |
| chr11-43534948-INS-99 | chr11-43534948-INS-86 |
| chr11-43536048-INS-76 | chr11-43536048-INS-76 |
| chr11-43557208-INS-189 | chr11-43557208-INS-189 |
| chr11-43566864-INS-84 | chr11-43566867-INS-90 |
| chr11-43571954-INS-56 | chr11-43571923-INS-96 |
| chr11-43586000-INS-75 | chr11-43586000-INS-75 |
| chr11-43588622-INS-68 | chr11-43588622-INS-74 |
| chr11-43605610-INS-283 | chr11-43605610-INS-283 |
| chr11-43638079-INS-54 | chr11-43638104-INS-52 |
| chr11-43647266-INS-650 | chr11-43647266-INS-647 |
| chr11-43654778-INS-213 | chr11-43654778-INS-186 |
| chr11-43656876-INS-390 | chr11-43656876-INS-390 |

| | |
|---|---|
| chr11-43674550-INS-55 | chr11-43674550-INS-55 |
| chr11-43675945-INS-123 | chr11-43675945-INS-123 |
| chr11-43694372-INS-545 | chr11-43694372-INS-545 |
| chr11-43703436-INS-72 | chr11-43703436-INS-84 |
| chr11-43713559-INS-72 | chr11-43713559-INS-79 |
| chr11-43728092-INS-3017 | chr11-43728093-INS-3017 |
| chr11-43729178-INS-58 | chr11-43729178-INS-58 |
| chr11-43731758-INS-66 | chr11-43731770-INS-102 |
| chr11-43737645-INS-164 | chr11-43737645-INS-140 |
| chr11-43763219-INS-498 | chr11-43763219-INS-498 |
| chr11-43789121-INS-220 | chr11-43789140-INS-208 |
| chr11-43792648-INS-105 | chr11-43792648-INS-105 |
| chr11-43799625-INS-249 | chr11-43799625-INS-249 |
| chr11-43839090-INS-80 | chr11-43839088-INS-118 |
| chr11-43872929-INS-80 | chr11-43872913-INS-68 |
| chr11-43877526-INS-56 | chr11-43877489-INS-54 |
| chr11-43880500-INS-58 | chr11-43880500-INS-58 |
| chr11-43883643-INS-2057 | chr11-43883644-INS-2075 |
| chr11-43907708-INS-1888 | chr11-43907709-INS-1888 |
| chr11-43951522-INS-92 | chr11-43951522-INS-90 |
| chr11-43972540-INS-6211 | chr11-43972541-INS-6211 |
| chr11-43994918-INS-74 | chr11-43994918-INS-64 |
| chr11-44009973-INS-1834 | chr11-44009974-INS-1834 |
| chr11-44019148-INS-93 | chr11-44019148-INS-93 |
| chr11-44027716-INS-65 | chr11-44027704-INS-66 |
| chr11-44029599-INS-54 | chr11-44029599-INS-54 |
| chr11-44042969-INS-510 | chr11-44042969-INS-510 |
| chr11-44065162-INS-92 | chr11-44065162-INS-136 |
| chr11-44096869-INS-75 | chr11-44096870-INS-90 |
| chr11-44100053-INS-7285 | chr11-44100054-INS-7285 |
| chr11-44105998-INS-237 | chr11-44105998-INS-232 |
| chr11-44114207-INS-380 | chr11-44114230-INS-402 |
| chr11-44123335-INS-350 | chr11-44123335-INS-350 |
| chr11-44137327-INS-5617 | chr11-44137328-INS-5618 |
| chr11-44144927-INS-55 | chr11-44144927-INS-55 |
| chr11-44158980-INS-64 | chr11-44158980-INS-59 |
| chr11-44170432-INS-170 | chr11-44170409-INS-146 |
| chr11-44173600-INS-361 | chr11-44173600-INS-361 |
| chr11-44195741-INS-180 | chr11-44195741-INS-180 |
| chr11-44206245-INS-124 | chr11-44206251-INS-142 |
| chr11-44207516-INS-92 | chr11-44207516-INS-92 |
| chr11-44227073-INS-103 | chr11-44227073-INS-85 |
| chr11-44237815-INS-204 | chr11-44237815-INS-204 |
| chr11-44243197-INS-70 | chr11-44243203-INS-110 |
| chr11-44245238-INS-68 | chr11-44245233-INS-60 |
| chr11-44246895-INS-81 | chr11-44246895-INS-81 |
| chr11-44321938-INS-216 | chr11-44321938-INS-216 |
| chr11-44380862-INS-163 | chr11-44380862-INS-146 |
| chr11-44388678-INS-102 | chr11-44388668-INS-80 |
| chr11-44393614-INS-160 | chr11-44393615-INS-150 |
| chr11-44397273-INS-2665 | chr11-44397274-INS-2665 |
| chr11-44432732-INS-645 | chr11-44432732-INS-626 |
| chr11-44456647-INS-96 | chr11-44456647-INS-56 |
| chr11-44466912-INS-84 | chr11-44466922-INS-86 |
| chr11-44493024-INS-60 | chr11-44493046-INS-74 |
| chr11-44528841-INS-120 | chr11-44528841-INS-126 |
| chr11-44562585-INS-69 | chr11-44562585-INS-69 |
| chr11-44610473-INS-247 | chr11-44610473-INS-206 |
| chr11-44623049-INS-6477 | chr11-44623050-INS-6477 |
| chr11-44645211-INS-77 | chr11-44645192-INS-56 |
| chr11-44646891-INS-60 | chr11-44646885-INS-80 |
| chr11-44647892-INS-599 | chr11-44647892-INS-599 |
| chr11-44664292-INS-56 | chr11-44664292-INS-56 |
| chr11-44701275-INS-66 | chr11-44701275-INS-66 |
| chr11-44747079-INS-151 | chr11-44747077-INS-124 |
| chr11-44805044-INS-59 | chr11-44805043-INS-90 |
| chr11-44826759-INS-132 | chr11-44826759-INS-135 |
| chr11-44841092-INS-72 | chr11-44841092-INS-72 |
| chr11-44874367-INS-68 | chr11-44874373-INS-74 |
| chr11-44907077-INS-60 | chr11-44907077-INS-68 |
| chr11-44911857-INS-53 | chr11-44911857-INS-55 |
| chr11-44914544-INS-240 | chr11-44914535-INS-240 |
| chr11-44919929-INS-56 | chr11-44919927-INS-58 |
| chr11-44936097-INS-81 | chr11-44936094-INS-84 |
| chr11-44938696-INS-499 | chr11-44938696-INS-513 |
| chr11-44947557-INS-986 | chr11-44947559-INS-988 |
| chr11-44953828-INS-51 | chr11-44953843-INS-50 |

| | |
|---|---|
| chr11-44997861-INS-111 | chr11-44997860-INS-118 |
| chr11-45016750-INS-60 | chr11-45016750-INS-60 |
| chr11-45102422-INS-116 | chr11-45102429-INS-99 |
| chr11-45146514-INS-2480 | chr11-45146515-INS-2480 |
| chr11-45195803-INS-197 | chr11-45195803-INS-197 |
| chr11-45203415-INS-60 | chr11-45203401-INS-60 |
| chr11-45210121-INS-95 | chr11-45210108-INS-99 |
| chr11-45216042-INS-891 | chr11-45216042-INS-870 |
| chr11-45218049-INS-5345 | chr11-45218050-INS-5345 |
| chr11-45238849-INS-1863 | chr11-45238850-INS-1863 |
| chr11-45239631-INS-52 | chr11-45239645-INS-80 |
| chr11-45247455-INS-80 | chr11-45247455-INS-72 |
| chr11-45284559-INS-56 | chr11-45284559-INS-64 |
| chr11-45298090-INS-1667 | chr11-45298091-INS-1668 |
| chr11-45301264-INS-3052 | chr11-45301265-INS-3052 |
| chr11-45328060-INS-254 | chr11-45328060-INS-254 |
| chr11-45344476-INS-55 | chr11-45344476-INS-55 |
| chr11-45348936-INS-68 | chr11-45348946-INS-96 |
| chr11-45363814-INS-55 | chr11-45363808-INS-83 |
| chr11-45369102-INS-403 | chr11-45369102-INS-403 |
| chr11-45405153-INS-363 | chr11-45405153-INS-365 |
| chr11-45407892-INS-8136 | chr11-45407893-INS-8138 |
| chr11-45467482-INS-97 | chr11-45467476-INS-108 |
| chr11-45480343-INS-7165 | chr11-45480344-INS-7166 |
| chr11-45504637-INS-197 | chr11-45504637-INS-197 |
| chr11-45513350-INS-613 | chr11-45513350-INS-613 |
| chr11-45515398-INS-271 | chr11-45515398-INS-271 |
| chr11-45519452-INS-198 | chr11-45519452-INS-201 |
| chr11-45536214-INS-100 | chr11-45536183-INS-100 |
| chr11-45540509-INS-111 | chr11-45540469-INS-154 |
| chr11-45555871-INS-236 | chr11-45555871-INS-204 |
| chr11-45559670-INS-196 | chr11-45559670-INS-188 |
| chr11-45582494-INS-402 | chr11-45582494-INS-402 |
| chr11-45584960-INS-6275 | chr11-45584961-INS-6275 |
| chr11-45634538-INS-80 | chr11-45634531-INS-56 |
| chr11-45639521-INS-59 | chr11-45639521-INS-60 |
| chr11-45670028-INS-193 | chr11-45670028-INS-239 |
| chr11-45684443-INS-156 | chr11-45684443-INS-179 |
| chr11-45695437-INS-58 | chr11-45695438-INS-55 |
| chr11-45699438-INS-85 | chr11-45699443-INS-91 |
| chr11-45718447-INS-199 | chr11-45718447-INS-199 |
| chr11-45727148-INS-70 | chr11-45727149-INS-58 |
| chr11-45730892-INS-204 | chr11-45730890-INS-216 |
| chr11-45747418-INS-75 | chr11-45747417-INS-60 |
| chr11-45755820-INS-205 | chr11-45755820-INS-205 |
| chr11-45760150-INS-56 | chr11-45760150-INS-56 |
| chr11-45766468-INS-200 | chr11-45766468-INS-200 |
| chr11-45804393-INS-104 | chr11-45804393-INS-101 |
| chr11-45817282-INS-203 | chr11-45817282-INS-203 |
| chr11-45821507-INS-174 | chr11-45821507-INS-177 |
| chr11-45838219-INS-52 | chr11-45838219-INS-54 |
| chr11-45853413-INS-170 | chr11-45853413-INS-179 |
| chr11-45856491-INS-108 | chr11-45856490-INS-88 |
| chr11-45859056-INS-55 | chr11-45859056-INS-57 |
| chr11-45869797-INS-196 | chr11-45869797-INS-197 |
| chr11-45880927-INS-206 | chr11-45880927-INS-206 |
| chr11-45883777-INS-198 | chr11-45883777-INS-198 |
| chr11-45919395-INS-82 | chr11-45919395-INS-78 |
| chr11-45939895-INS-60 | chr11-45939895-INS-68 |
| chr11-45947291-INS-173 | chr11-45947291-INS-138 |
| chr11-45949870-INS-168 | chr11-45949870-INS-168 |
| chr11-45959235-INS-60 | chr11-45959235-INS-60 |
| chr11-45963261-INS-91 | chr11-45963260-INS-79 |
| chr11-45978976-INS-192 | chr11-45978950-INS-198 |
| chr11-45989071-INS-1893 | chr11-45989072-INS-1893 |
| chr11-45994630-INS-227 | chr11-45994630-INS-228 |
| chr11-46015761-INS-76 | chr11-46015777-INS-95 |
| chr11-46024554-INS-51 | chr11-46024554-INS-51 |
| chr11-46032456-INS-200 | chr11-46032456-INS-202 |
| chr11-46043573-INS-219 | chr11-46043573-INS-220 |
| chr11-46077489-INS-105 | chr11-46077489-INS-60 |
| chr11-46079181-INS-93 | chr11-46079181-INS-118 |
| chr11-46087832-INS-211 | chr11-46087833-INS-203 |
| chr11-46139916-INS-85 | chr11-46139916-INS-85 |
| chr11-46141887-INS-130 | chr11-46141887-INS-130 |
| chr11-46144727-INS-939 | chr11-46144727-INS-939 |
| chr11-46192237-INS-54 | chr11-46192237-INS-52 |

| | |
|---|---|
| chr11-46207257-INS-65 | chr11-46207257-INS-54 |
| chr11-46232160-INS-99 | chr11-46232160-INS-99 |
| chr11-46284996-INS-60 | chr11-46284996-INS-60 |
| chr11-46288917-INS-200 | chr11-46288917-INS-201 |
| chr11-46308135-INS-170 | chr11-46308135-INS-175 |
| chr11-46321719-INS-219 | chr11-46321719-INS-217 |
| chr11-46340796-INS-114 | chr11-46340796-INS-136 |
| chr11-46351061-INS-1030 | chr11-46351062-INS-1030 |
| chr11-46375566-INS-57 | chr11-46375566-INS-61 |
| chr11-46417477-INS-227 | chr11-46417477-INS-227 |
| chr11-46423669-INS-188 | chr11-46423669-INS-183 |
| chr11-46431919-INS-85 | chr11-46431919-INS-75 |
| chr11-46452278-INS-455 | chr11-46452278-INS-455 |
| chr11-46457171-INS-175 | chr11-46457182-INS-142 |
| chr11-46464180-INS-408 | chr11-46464180-INS-414 |
| chr11-46464884-INS-257 | chr11-46464884-INS-258 |
| chr11-46473128-INS-58 | chr11-46473128-INS-74 |
| chr11-46475804-INS-219 | chr11-46475805-INS-202 |
| chr11-46481552-INS-172 | chr11-46481552-INS-173 |
| chr11-46483798-INS-85 | chr11-46483808-INS-50 |
| chr11-46492552-INS-5023 | chr11-46492552-INS-5023 |
| chr11-46515210-INS-218 | chr11-46515230-INS-230 |
| chr11-46541117-INS-959 | chr11-46541117-INS-979 |
| chr11-46541812-INS-107 | chr11-46541812-INS-108 |
| chr11-46543855-INS-6699 | chr11-46543856-INS-6699 |
| chr11-46544319-INS-57 | chr11-46544319-INS-57 |
| chr11-46546372-INS-199 | chr11-46546372-INS-214 |
| chr11-46546716-INS-1812 | chr11-46546717-INS-1812 |
| chr11-46560312-INS-199 | chr11-46560312-INS-199 |
| chr11-46571685-INS-202 | chr11-46571685-INS-201 |
| chr11-46581590-INS-6360 | chr11-46581591-INS-6363 |
| chr11-46590240-INS-1566 | chr11-46590241-INS-1566 |
| chr11-46591796-INS-227 | chr11-46591796-INS-226 |
| chr11-46595723-INS-58 | chr11-46595723-INS-58 |
| chr11-46602428-INS-2496 | chr11-46602429-INS-2497 |
| chr11-46604361-INS-172 | chr11-46604361-INS-172 |
| chr11-46607823-INS-614 | chr11-46607823-INS-614 |
| chr11-46609059-INS-108 | chr11-46609059-INS-138 |
| chr11-46630139-INS-258 | chr11-46630139-INS-258 |
| chr11-46634073-INS-54 | chr11-46634073-INS-54 |
| chr11-46672628-INS-104 | chr11-46672628-INS-104 |
| chr11-46723396-INS-1116 | chr11-46723397-INS-1116 |
| chr11-46725962-INS-56 | chr11-46725962-INS-60 |
| chr11-46735500-INS-401 | chr11-46735500-INS-401 |
| chr11-46737203-INS-51 | chr11-46737192-INS-52 |
| chr11-46739263-INS-3136 | chr11-46739264-INS-3136 |
| chr11-46741822-INS-117 | chr11-46741871-INS-74 |
| chr11-46742501-INS-1919 | chr11-46742502-INS-1904 |
| chr11-46745112-INS-58 | chr11-46745113-INS-58 |
| chr11-46756954-INS-244 | chr11-46756943-INS-264 |
| chr11-46757850-INS-79 | chr11-46757850-INS-96 |
| chr11-46760838-INS-182 | chr11-46760838-INS-182 |
| chr11-46761392-INS-124 | chr11-46761393-INS-132 |
| chr11-46768043-INS-67 | chr11-46768043-INS-63 |
| chr11-46787039-INS-83 | chr11-46787039-INS-80 |
| chr11-46804709-INS-473 | chr11-46804710-INS-471 |
| chr11-46805308-INS-60 | chr11-46805309-INS-56 |
| chr11-46814209-INS-1102 | chr11-46814210-INS-1102 |
| chr11-46825129-INS-2110 | chr11-46825130-INS-2114 |
| chr11-46836347-INS-3552 | chr11-46836348-INS-3552 |
| chr11-46841540-INS-470 | chr11-46841540-INS-471 |
| chr11-46844458-INS-6452 | chr11-46844459-INS-6452 |
| chr11-46878912-INS-71 | chr11-46878936-INS-78 |
| chr11-46879837-INS-1213 | chr11-46879838-INS-1213 |
| chr11-46881182-INS-2603 | chr11-46881183-INS-2602 |
| chr11-46898540-INS-1661 | chr11-46898541-INS-1661 |
| chr11-46938127-INS-52 | chr11-46938127-INS-82 |
| chr11-46942246-INS-7624 | chr11-46942247-INS-7625 |
| chr11-46951908-INS-206 | chr11-46951908-INS-206 |
| chr11-46968540-INS-52 | chr11-46968540-INS-52 |
| chr11-47006022-INS-215 | chr11-47006026-INS-258 |
| chr11-47069827-INS-1009 | chr11-47069828-INS-1011 |
| chr11-47088312-INS-202 | chr11-47088312-INS-200 |
| chr11-47092106-INS-63 | chr11-47092106-INS-60 |
| chr11-47134729-INS-7087 | chr11-47134730-INS-7087 |
| chr11-47174557-INS-68 | chr11-47174557-INS-68 |
| chr11-47269597-INS-53 | chr11-47269597-INS-54 |

| | |
|---|---|
| chr11-47297788-INS-618 | chr11-47297788-INS-617 |
| chr11-47347963-INS-241 | chr11-47347963-INS-237 |
| chr11-47351615-INS-51 | chr11-47351623-INS-58 |
| chr11-47359008-INS-214 | chr11-47359008-INS-214 |
| chr11-47391338-INS-413 | chr11-47391338-INS-409 |
| chr11-47397079-INS-6449 | chr11-47397080-INS-6449 |
| chr11-47420350-INS-132 | chr11-47420351-INS-132 |
| chr11-47445660-INS-561 | chr11-47445660-INS-561 |
| chr11-47460577-INS-762 | chr11-47460577-INS-762 |
| chr11-47489903-INS-162 | chr11-47489903-INS-162 |
| chr11-47546759-INS-63 | chr11-47546759-INS-63 |
| chr11-47585298-INS-78 | chr11-47585298-INS-80 |
| chr11-47613978-INS-52 | chr11-47613978-INS-52 |
| chr11-47614824-INS-208 | chr11-47614824-INS-208 |
| chr11-47709826-INS-56 | chr11-47709802-INS-62 |
| chr11-47715902-INS-61 | chr11-47715906-INS-61 |
| chr11-47729813-INS-1446 | chr11-47729814-INS-1429 |
| chr11-47756651-INS-54 | chr11-47756651-INS-54 |
| chr11-47842325-INS-60 | chr11-47842325-INS-60 |
| chr11-47848263-INS-242 | chr11-47848263-INS-260 |
| chr11-47856766-INS-62 | chr11-47856766-INS-50 |
| chr11-47860697-INS-241 | chr11-47860697-INS-271 |
| chr11-47884106-INS-128 | chr11-47884106-INS-83 |
| chr11-47893562-INS-385 | chr11-47893562-INS-385 |
| chr11-47898619-INS-51 | chr11-47898619-INS-58 |
| chr11-47910594-INS-52 | chr11-47910594-INS-52 |
| chr11-47919391-INS-67 | chr11-47919384-INS-60 |
| chr11-47969285-INS-509 | chr11-47969285-INS-508 |
| chr11-47988935-INS-6479 | chr11-47988936-INS-6479 |
| chr11-47996900-INS-75 | chr11-47996899-INS-70 |
| chr11-48050717-INS-82 | chr11-48050717-INS-82 |
| chr11-48105143-INS-52 | chr11-48105152-INS-72 |
| chr11-48116524-INS-821 | chr11-48116524-INS-821 |
| chr11-48118187-INS-7218 | chr11-48118188-INS-7220 |
| chr11-48123072-INS-104 | chr11-48123072-INS-104 |
| chr11-48126627-INS-4967 | chr11-48126628-INS-4967 |
| chr11-48128493-INS-259 | chr11-48128493-INS-259 |
| chr11-48157303-INS-117 | chr11-48157304-INS-129 |
| chr11-48159215-INS-8455 | chr11-48159216-INS-8455 |
| chr11-48182588-INS-109 | chr11-48182588-INS-109 |
| chr11-48193520-INS-399 | chr11-48193520-INS-399 |
| chr11-48246827-INS-64 | chr11-48246827-INS-72 |
| chr11-48276228-INS-129 | chr11-48276228-INS-128 |
| chr11-48283326-INS-58 | chr11-48283293-INS-57 |
| chr11-48300927-INS-66 | chr11-48300933-INS-91 |
| chr11-48309251-INS-53 | chr11-48309251-INS-53 |
| chr11-48322258-INS-72 | chr11-48322257-INS-80 |
| chr11-48327192-INS-60 | chr11-48327186-INS-69 |
| chr11-48336697-INS-1491 | chr11-48336698-INS-1491 |
| chr11-48352141-INS-555 | chr11-48352141-INS-556 |
| chr11-48358060-INS-103 | chr11-48358060-INS-103 |
| chr11-48383112-INS-94 | chr11-48383112-INS-51 |
| chr11-48386808-INS-7085 | chr11-48386809-INS-7085 |
| chr11-48388753-INS-394 | chr11-48388753-INS-400 |
| chr11-48390111-INS-90 | chr11-48390111-INS-102 |
| chr11-48391346-INS-217 | chr11-48391346-INS-195 |
| chr11-48410247-INS-479 | chr11-48410247-INS-477 |
| chr11-48410891-INS-117 | chr11-48410891-INS-121 |
| chr11-48433011-INS-342 | chr11-48433011-INS-342 |
| chr11-48437852-INS-52 | chr11-48437878-INS-72 |
| chr11-48450773-INS-79 | chr11-48450773-INS-78 |
| chr11-48471288-INS-1996 | chr11-48471289-INS-1996 |
| chr11-48483539-INS-1711 | chr11-48483540-INS-1709 |
| chr11-48487109-INS-371 | chr11-48487109-INS-372 |
| chr11-48488848-INS-327 | chr11-48488848-INS-327 |
| chr11-48542683-INS-108 | chr11-48542683-INS-64 |
| chr11-48548504-INS-130 | chr11-48548504-INS-129 |
| chr11-48568173-INS-168 | chr11-48568150-INS-159 |
| chr11-48580291-INS-726 | chr11-48580291-INS-727 |
| chr11-48597046-INS-64 | chr11-48597046-INS-76 |
| chr11-48629313-INS-1079 | chr11-48629318-INS-1080 |
| chr11-48632097-INS-195 | chr11-48632097-INS-194 |
| chr11-48633474-INS-252 | chr11-48633474-INS-252 |
| chr11-48639605-INS-5420 | chr11-48639606-INS-5420 |
| chr11-48651994-INS-54 | chr11-48651994-INS-54 |
| chr11-48662139-INS-130 | chr11-48662110-INS-120 |
| chr11-48680153-INS-199 | chr11-48680153-INS-200 |

| | |
|---|---|
| chr11-48694893-INS-210 | chr11-48694893-INS-222 |
| chr11-48709459-INS-110 | chr11-48709459-INS-110 |
| chr11-48712120-INS-126 | chr11-48712120-INS-126 |
| chr11-48720221-INS-63 | chr11-48720221-INS-60 |
| chr11-48758983-INS-96 | chr11-48758983-INS-60 |
| chr11-48769197-INS-100 | chr11-48769206-INS-101 |
| chr11-48775811-INS-305 | chr11-48775810-INS-268 |
| chr11-48782278-INS-189 | chr11-48782277-INS-179 |
| chr11-48801140-INS-68 | chr11-48801140-INS-68 |
| chr11-48881907-INS-81 | chr11-48881907-INS-81 |
| chr11-48882695-INS-540 | chr11-48882695-INS-542 |
| chr11-48883791-INS-332 | chr11-48883790-INS-332 |
| chr11-48972078-INS-184 | chr11-48972078-INS-179 |
| chr11-48978296-INS-54 | chr11-48978302-INS-59 |
| chr11-48992109-INS-220 | chr11-48992109-INS-220 |
| chr11-48997864-INS-193 | chr11-48997864-INS-193 |
| chr11-48998463-INS-59 | chr11-48998463-INS-53 |
| chr11-49008011-INS-147 | chr11-49008011-INS-143 |
| chr11-49018042-INS-532 | chr11-49018042-INS-533 |
| chr11-49049116-INS-6864 | chr11-49049117-INS-6864 |
| chr11-49054698-INS-723 | chr11-49054698-INS-723 |
| chr11-49063172-INS-550 | chr11-49063172-INS-552 |
| chr11-49148389-INS-5460 | chr11-49148390-INS-5460 |
| chr11-49157116-INS-167 | chr11-49157113-INS-168 |
| chr11-49164587-INS-109 | chr11-49164540-INS-109 |
| chr11-49241289-INS-184 | chr11-49241289-INS-184 |
| chr11-49246777-INS-60 | chr11-49246777-INS-60 |
| chr11-49264271-INS-6545 | chr11-49264272-INS-6545 |
| chr11-49289328-INS-3024 | chr11-49289329-INS-3024 |
| chr11-49299195-INS-57 | chr11-49299195-INS-57 |
| chr11-49306715-INS-769 | chr11-49306715-INS-769 |
| chr11-49319761-INS-51 | chr11-49319761-INS-51 |
| chr11-49359195-INS-672 | chr11-49359195-INS-672 |
| chr11-49375486-INS-730 | chr11-49375486-INS-730 |
| chr11-49389271-INS-75 | chr11-49389271-INS-78 |
| chr11-49413732-INS-483 | chr11-49413732-INS-483 |
| chr11-49433516-INS-195 | chr11-49433516-INS-194 |
| chr11-49441807-INS-143 | chr11-49441813-INS-120 |
| chr11-49451893-INS-296 | chr11-49451893-INS-293 |
| chr11-49460246-INS-201 | chr11-49460246-INS-199 |
| chr11-49494276-INS-464 | chr11-49494276-INS-464 |
| chr11-49510972-INS-66 | chr11-49510958-INS-74 |
| chr11-49529386-INS-208 | chr11-49529386-INS-208 |
| chr11-49591027-INS-264 | chr11-49591027-INS-264 |
| chr11-49628896-INS-50 | chr11-49628900-INS-68 |
| chr11-49651363-INS-3217 | chr11-49651364-INS-3217 |
| chr11-49664249-INS-55 | chr11-49664249-INS-61 |
| chr11-49682830-INS-201 | chr11-49682830-INS-199 |
| chr11-49718224-INS-202 | chr11-49718224-INS-202 |
| chr11-49733264-INS-54 | chr11-49733264-INS-54 |
| chr11-49740064-INS-444 | chr11-49740064-INS-443 |
| chr11-49742948-INS-197 | chr11-49742948-INS-196 |
| chr11-49788143-INS-78 | chr11-49788143-INS-52 |
| chr11-49834349-INS-642 | chr11-49834349-INS-642 |
| chr11-49844782-INS-115 | chr11-49844782-INS-85 |
| chr11-49853130-INS-209 | chr11-49853130-INS-212 |
| chr11-49863372-INS-195 | chr11-49863372-INS-195 |
| chr11-49889477-INS-63 | chr11-49889474-INS-56 |
| chr11-49897051-INS-155 | chr11-49897054-INS-157 |
| chr11-49900388-INS-3322 | chr11-49900389-INS-3322 |
| chr11-49915102-INS-189 | chr11-49915102-INS-189 |
| chr11-49946333-INS-52 | chr11-49946333-INS-52 |
| chr11-50020792-INS-56 | chr11-50020792-INS-68 |
| chr11-50025851-INS-59 | chr11-50025851-INS-59 |
| chr11-50055883-INS-52 | chr11-50055883-INS-52 |
| chr11-50068641-INS-1755 | chr11-50068642-INS-1757 |
| chr11-50083017-INS-198 | chr11-50083016-INS-195 |
| chr11-50087478-INS-201 | chr11-50087478-INS-201 |
| chr11-50103316-INS-57 | chr11-50103316-INS-57 |
| chr11-50104389-INS-218 | chr11-50104389-INS-217 |
| chr11-50110604-INS-194 | chr11-50110604-INS-194 |
| chr11-50142026-INS-60 | chr11-50142026-INS-60 |
| chr11-50151414-INS-215 | chr11-50151414-INS-215 |
| chr11-50173374-INS-902 | chr11-50173373-INS-940 |
| chr11-50173549-INS-51 | chr11-50173549-INS-51 |
| chr11-50197103-INS-149 | chr11-50197115-INS-139 |
| chr11-50229268-INS-210 | chr11-50229268-INS-212 |

| | |
|---|---|
| chr11-50234017-INS-202 | chr11-50234017-INS-204 |
| chr11-50260433-INS-71 | chr11-50260433-INS-71 |
| chr11-50264489-INS-188 | chr11-50264489-INS-188 |
| chr11-50279011-INS-1886 | chr11-50279012-INS-1886 |
| chr11-50286765-INS-151 | chr11-50286765-INS-151 |
| chr11-50296927-INS-233 | chr11-50296938-INS-202 |
| chr11-50298298-INS-68 | chr11-50298298-INS-68 |
| chr11-50314301-INS-61 | chr11-50314301-INS-61 |
| chr11-50326193-INS-74 | chr11-50326225-INS-71 |
| chr11-50344131-INS-227 | chr11-50344131-INS-227 |
| chr11-50353494-INS-115 | chr11-50353494-INS-115 |
| chr11-50370664-INS-471 | chr11-50370674-INS-472 |
| chr11-50374486-INS-228 | chr11-50374486-INS-228 |
| chr11-50386239-INS-309 | chr11-50386237-INS-312 |
| chr11-50387628-INS-80 | chr11-50387628-INS-81 |
| chr11-50390904-INS-79 | chr11-50390904-INS-79 |
| chr11-50428725-INS-70 | chr11-50428725-INS-70 |
| chr11-50430987-INS-6275 | chr11-50430988-INS-6275 |
| chr11-50463181-INS-60 | chr11-50463181-INS-75 |
| chr11-50473138-INS-92 | chr11-50473138-INS-95 |
| chr11-50474515-INS-3263 | chr11-50474516-INS-3261 |
| chr11-50479943-INS-105 | chr11-50479977-INS-60 |
| chr11-50505443-INS-187 | chr11-50505443-INS-193 |
| chr11-50510836-INS-6310 | chr11-50510820-INS-6304 |
| chr11-50517258-INS-60 | chr11-50517258-INS-66 |
| chr11-50541737-INS-62 | chr11-50541737-INS-64 |
| chr11-50585287-INS-184 | chr11-50585288-INS-171 |
| chr11-50588684-INS-937 | chr11-50588684-INS-937 |
| chr11-50590696-INS-64 | chr11-50590696-INS-64 |
| chr11-50602989-INS-93 | chr11-50602989-INS-93 |
| chr11-50632155-INS-71 | chr11-50632155-INS-75 |
| chr11-50651055-INS-218 | chr11-50651067-INS-184 |
| chr11-50669723-INS-63 | chr11-50669723-INS-63 |
| chr11-50680623-INS-64 | chr11-50680623-INS-70 |
| chr11-50698626-INS-265 | chr11-50698626-INS-265 |
| chr11-50707214-INS-60 | chr11-50707214-INS-60 |
| chr11-50711621-INS-5541 | chr11-50711622-INS-5541 |
| chr11-50734078-INS-121 | chr11-50734078-INS-124 |
| chr11-50747021-INS-107 | chr11-50747021-INS-107 |
| chr11-50760166-INS-84 | chr11-50760166-INS-81 |
| chr11-50767848-INS-889 | chr11-50767848-INS-890 |
| chr11-50774502-INS-60 | chr11-50774502-INS-60 |
| chr11-50786327-INS-108 | chr11-50786327-INS-108 |
| chr11-50787879-INS-59 | chr11-50787879-INS-59 |
| chr11-50796782-INS-85 | chr11-50796783-INS-63 |
| chr11-50811054-INS-54 | chr11-50811054-INS-54 |
| chr11-50836487-INS-75 | chr11-50836484-INS-78 |
| chr11-50848623-INS-63 | chr11-50848623-INS-63 |
| chr11-50853730-INS-63 | chr11-50853734-INS-60 |
| chr11-50856660-INS-72 | chr11-50856660-INS-72 |
| chr11-50865657-INS-72 | chr11-50865657-INS-72 |
| chr11-50904582-INS-399 | chr11-50904582-INS-399 |
| chr11-50952193-INS-54 | chr11-50952193-INS-55 |
| chr11-50957922-INS-205 | chr11-50957922-INS-204 |
| chr11-50977434-INS-164 | chr11-50977434-INS-162 |
| chr11-50985375-INS-358 | chr11-50985375-INS-358 |
| chr11-50996225-INS-60 | chr11-50996225-INS-60 |
| chr11-51067131-INS-193 | chr11-51067118-INS-183 |
| chr11-51082637-INS-84 | chr11-51082637-INS-84 |
| chr11-51094026-INS-71 | chr11-51094026-INS-71 |
| chr11-51117867-INS-60 | chr11-51117867-INS-65 |
| chr11-51139375-INS-429 | chr11-51139375-INS-429 |
| chr11-51140071-INS-390 | chr11-51140072-INS-390 |
| chr11-51200531-INS-52 | chr11-51200531-INS-52 |
| chr11-51212209-INS-184 | chr11-51212209-INS-184 |
| chr11-51217688-INS-79 | chr11-51217688-INS-79 |
| chr11-51218691-INS-72 | chr11-51218691-INS-54 |
| chr11-51223132-INS-419 | chr11-51223132-INS-419 |
| chr11-51227118-INS-134 | chr11-51227118-INS-138 |
| chr11-51231002-INS-229 | chr11-51231002-INS-227 |
| chr11-51231354-INS-2707 | chr11-51231355-INS-2707 |
| chr11-51238996-INS-52 | chr11-51238996-INS-51 |
| chr11-51251633-INS-1020 | chr11-51251634-INS-1027 |
| chr11-51256149-INS-62 | chr11-51256149-INS-58 |
| chr11-51261054-INS-490 | chr11-51261054-INS-490 |
| chr11-51263818-INS-80 | chr11-51263818-INS-80 |
| chr11-51291025-INS-94 | chr11-51291024-INS-104 |

| | |
|---|---|
| chr11-51293383-INS-55 | chr11-51293383-INS-56 |
| chr11-51304458-INS-205 | chr11-51304458-INS-208 |
| chr11-51365012-INS-56 | chr11-51365012-INS-56 |
| chr11-51370117-INS-1514 | chr11-51370118-INS-1527 |
| chr11-51445598-INS-191 | chr11-51445598-INS-166 |
| chr11-51452549-INS-66 | chr11-51452585-INS-68 |
| chr11-51465341-INS-109 | chr11-51465341-INS-109 |
| chr11-51466014-INS-1406 | chr11-51466015-INS-1406 |
| chr11-51501329-INS-52 | chr11-51501329-INS-52 |
| chr11-51502213-INS-168 | chr11-51502213-INS-167 |
| chr11-51503507-INS-98 | chr11-51503525-INS-50 |
| chr11-51530961-INS-197 | chr11-51530961-INS-197 |
| chr11-51547434-INS-195 | chr11-51547434-INS-195 |
| chr11-51563361-INS-211 | chr11-51563378-INS-188 |
| chr11-51565609-INS-784 | chr11-51565609-INS-784 |
| chr11-51574681-INS-209 | chr11-51574681-INS-209 |
| chr11-51588884-INS-82 | chr11-51588884-INS-84 |
| chr11-51590115-INS-214 | chr11-51590115-INS-215 |
| chr11-51592498-INS-222 | chr11-51592498-INS-223 |
| chr11-51594439-INS-221 | chr11-51594439-INS-221 |
| chr11-51598257-INS-56 | chr11-51598253-INS-78 |
| chr11-51602972-INS-169 | chr11-51602972-INS-169 |
| chr11-51608285-INS-512 | chr11-51608285-INS-512 |
| chr11-51618547-INS-115 | chr11-51618547-INS-115 |
| chr11-51629121-INS-71 | chr11-51629121-INS-71 |
| chr11-51638082-INS-181 | chr11-51638082-INS-181 |
| chr11-51641629-INS-139 | chr11-51641629-INS-139 |
| chr11-51651451-INS-652 | chr11-51651451-INS-654 |
| chr11-51656530-INS-202 | chr11-51656530-INS-202 |
| chr11-51673338-INS-53 | chr11-51673338-INS-83 |
| chr11-51689221-INS-242 | chr11-51689221-INS-242 |
| chr11-51694318-INS-176 | chr11-51694304-INS-154 |
| chr11-51702847-INS-211 | chr11-51702847-INS-211 |
| chr11-51727391-INS-158 | chr11-51727392-INS-136 |
| chr11-51755729-INS-213 | chr11-51755729-INS-213 |
| chr11-51769943-INS-52 | chr11-51769943-INS-62 |
| chr11-51780397-INS-11194 | chr11-51780398-INS-11194 |
| chr11-51792930-INS-59 | chr11-51792932-INS-64 |
| chr11-51814014-INS-369 | chr11-51814014-INS-369 |
| chr11-51832811-INS-126 | chr11-51832811-INS-126 |
| chr11-51839309-INS-65 | chr11-51839309-INS-53 |
| chr11-51851652-INS-86 | chr11-51851652-INS-50 |
| chr11-51854086-INS-208 | chr11-51854081-INS-185 |
| chr11-51860114-INS-209 | chr11-51860114-INS-209 |
| chr11-51864634-INS-219 | chr11-51864634-INS-220 |
| chr11-51872307-INS-60 | chr11-51872319-INS-86 |
| chr11-51904775-INS-212 | chr11-51904775-INS-215 |
| chr11-51910166-INS-175 | chr11-51910166-INS-175 |
| chr11-51912961-INS-206 | chr11-51912961-INS-206 |
| chr11-51945913-INS-4527 | chr11-51945914-INS-4527 |
| chr11-51959062-INS-197 | chr11-51959063-INS-198 |
| chr11-51964774-INS-212 | chr11-51964774-INS-211 |
| chr11-51984354-INS-220 | chr11-51984354-INS-220 |
| chr11-52006977-INS-223 | chr11-52006977-INS-223 |
| chr11-52012044-INS-580 | chr11-52012044-INS-580 |
| chr11-52065117-INS-209 | chr11-52065117-INS-209 |
| chr11-52076757-INS-52 | chr11-52076757-INS-54 |
| chr11-52079423-INS-144 | chr11-52079423-INS-144 |
| chr11-52088516-INS-64 | chr11-52088524-INS-64 |
| chr11-52105175-INS-62 | chr11-52105175-INS-62 |
| chr11-52122332-INS-62 | chr11-52122332-INS-58 |
| chr11-52179691-INS-131 | chr11-52179691-INS-131 |
| chr11-52197741-INS-90 | chr11-52197741-INS-90 |
| chr11-52202615-INS-211 | chr11-52202618-INS-230 |
| chr11-52216783-INS-236 | chr11-52216783-INS-235 |
| chr11-52235863-INS-6442 | chr11-52235864-INS-6442 |
| chr11-52236684-INS-222 | chr11-52236684-INS-249 |
| chr11-52238026-INS-75 | chr11-52238026-INS-78 |
| chr11-52241263-INS-52 | chr11-52241263-INS-56 |
| chr11-52244331-INS-1215 | chr11-52244332-INS-1215 |
| chr11-52244609-INS-236 | chr11-52244609-INS-230 |
| chr11-52256224-INS-133 | chr11-52256224-INS-101 |
| chr11-52258485-INS-210 | chr11-52258485-INS-211 |
| chr11-52260522-INS-54 | chr11-52260522-INS-54 |
| chr11-52264349-INS-53 | chr11-52264349-INS-65 |
| chr11-52267862-INS-150 | chr11-52267862-INS-150 |
| chr11-52271005-INS-73 | chr11-52271005-INS-73 |

| | |
|---|---|
| chr11-52273098-INS-76 | chr11-52273098-INS-60 |
| chr11-52274603-INS-205 | chr11-52274603-INS-205 |
| chr11-52278387-INS-217 | chr11-52278387-INS-217 |
| chr11-52285264-INS-58 | chr11-52285264-INS-90 |
| chr11-52315234-INS-212 | chr11-52315234-INS-211 |
| chr11-52320150-INS-59 | chr11-52320150-INS-59 |
| chr11-52320612-INS-193 | chr11-52320611-INS-204 |
| chr11-52339860-INS-51 | chr11-52339861-INS-50 |
| chr11-52343789-INS-199 | chr11-52343789-INS-182 |
| chr11-52352348-INS-138 | chr11-52352348-INS-138 |
| chr11-52358357-INS-465 | chr11-52358357-INS-466 |
| chr11-52361165-INS-58 | chr11-52361165-INS-62 |
| chr11-52374957-INS-4015 | chr11-52374958-INS-4015 |
| chr11-52403906-INS-325 | chr11-52403906-INS-325 |
| chr11-52428327-INS-63 | chr11-52428327-INS-79 |
| chr11-52435896-INS-55 | chr11-52435896-INS-95 |
| chr11-52477033-INS-3043 | chr11-52477034-INS-3046 |
| chr11-52489735-INS-84 | chr11-52489735-INS-108 |
| chr11-52534027-INS-71 | chr11-52534018-INS-54 |
| chr11-52548132-INS-62 | chr11-52548132-INS-63 |
| chr11-52551349-INS-54 | chr11-52551349-INS-54 |
| chr11-52566698-INS-71 | chr11-52566698-INS-73 |
| chr11-52591537-INS-84 | chr11-52591537-INS-63 |
| chr11-52622875-INS-60 | chr11-52622875-INS-61 |
| chr11-52635037-INS-1493 | chr11-52635038-INS-1493 |
| chr11-52644710-INS-676 | chr11-52644710-INS-677 |
| chr11-52696177-INS-185 | chr11-52696177-INS-185 |
| chr11-52721843-INS-142 | chr11-52721843-INS-141 |
| chr11-52733964-INS-3981 | chr11-52733965-INS-3983 |
| chr11-52740608-INS-109 | chr11-52740608-INS-109 |
| chr11-52743299-INS-182 | chr11-52743299-INS-182 |
| chr11-52776067-INS-5207 | chr11-52776068-INS-5207 |
| chr11-52803259-INS-56 | chr11-52803259-INS-56 |
| chr11-52811950-INS-68 | chr11-52811950-INS-56 |
| chr11-52873901-INS-82 | chr11-52873901-INS-82 |
| chr11-52902250-INS-54 | chr11-52902250-INS-54 |
| chr11-52903073-INS-6245 | chr11-52903074-INS-6246 |
| chr11-52920380-INS-94 | chr11-52920380-INS-94 |
| chr11-52928638-INS-84 | chr11-52928638-INS-86 |
| chr11-53016615-INS-113 | chr11-53016613-INS-124 |
| chr11-53029322-INS-201 | chr11-53029322-INS-201 |
| chr11-53034683-INS-78 | chr11-53034683-INS-82 |
| chr11-53055016-INS-183 | chr11-53055016-INS-183 |
| chr11-53059180-INS-171 | chr11-53059180-INS-171 |
| chr11-53071202-INS-651 | chr11-53071202-INS-651 |
| chr11-53095715-INS-74 | chr11-53095715-INS-80 |
| chr11-53112394-INS-71 | chr11-53112394-INS-68 |
| chr11-53153043-INS-62 | chr11-53153001-INS-56 |
| chr11-53154667-INS-209 | chr11-53154667-INS-210 |
| chr11-53177082-INS-210 | chr11-53177082-INS-234 |
| chr11-53193986-INS-377 | chr11-53193986-INS-377 |
| chr11-53195311-INS-382 | chr11-53195311-INS-375 |
| chr11-53196499-INS-397 | chr11-53196499-INS-391 |
| chr11-53212254-INS-57 | chr11-53212254-INS-57 |
| chr11-53218489-INS-202 | chr11-53218489-INS-205 |
| chr11-53227176-INS-174 | chr11-53227192-INS-159 |
| chr11-53228950-INS-198 | chr11-53228950-INS-198 |
| chr11-53232310-INS-188 | chr11-53232310-INS-187 |
| chr11-53250892-INS-153 | chr11-53250892-INS-153 |
| chr11-53254250-INS-216 | chr11-53254250-INS-212 |
| chr11-53274495-INS-214 | chr11-53274495-INS-214 |
| chr11-53286305-INS-221 | chr11-53286305-INS-220 |
| chr11-53339292-INS-203 | chr11-53339292-INS-202 |
| chr11-53393204-INS-311 | chr11-53393204-INS-311 |
| chr11-53461310-INS-359 | chr11-53461310-INS-359 |
| chr11-53470723-INS-97 | chr11-53470723-INS-97 |
| chr11-53474589-INS-111 | chr11-53474589-INS-110 |
| chr11-53497264-INS-81 | chr11-53497298-INS-56 |
| chr11-53519032-INS-76 | chr11-53519032-INS-79 |
| chr11-53571445-INS-207 | chr11-53571445-INS-207 |
| chr11-53588321-INS-215 | chr11-53588321-INS-215 |
| chr11-53594465-INS-73 | chr11-53594465-INS-73 |
| chr11-53607268-INS-204 | chr11-53607268-INS-204 |
| chr11-53630002-INS-227 | chr11-53630002-INS-227 |
| chr11-53631511-INS-208 | chr11-53631511-INS-208 |
| chr11-53632856-INS-363 | chr11-53632856-INS-363 |
| chr11-53644710-INS-205 | chr11-53644710-INS-205 |

| | |
|---|---|
| chr11-53663032-INS-56 | chr11-53663031-INS-69 |
| chr11-53691608-INS-211 | chr11-53691610-INS-211 |
| chr11-53708186-INS-206 | chr11-53708186-INS-208 |
| chr11-53720523-INS-69 | chr11-53720523-INS-69 |
| chr11-53720687-INS-3668 | chr11-53720688-INS-3668 |
| chr11-53737516-INS-237 | chr11-53737516-INS-237 |
| chr11-53739555-INS-175 | chr11-53739555-INS-175 |
| chr11-53768450-INS-248 | chr11-53768450-INS-248 |
| chr11-53784585-INS-205 | chr11-53784585-INS-205 |
| chr11-53828222-INS-520 | chr11-53828222-INS-530 |
| chr11-53832128-INS-316 | chr11-53832124-INS-315 |
| chr11-53855210-INS-54 | chr11-53855210-INS-90 |
| chr11-53858401-INS-225 | chr11-53858401-INS-225 |
| chr11-53887533-INS-5024 | chr11-53887534-INS-5024 |
| chr11-53908356-INS-119 | chr11-53908356-INS-119 |
| chr11-53910835-INS-57 | chr11-53910835-INS-63 |
| chr11-53930860-INS-727 | chr11-53930860-INS-727 |
| chr11-53940886-INS-239 | chr11-53940886-INS-240 |
| chr11-53971817-INS-136 | chr11-53971813-INS-146 |
| chr11-53972975-INS-239 | chr11-53972975-INS-239 |
| chr11-54031544-INS-70 | chr11-54031544-INS-78 |
| chr11-54052166-INS-109 | chr11-54052166-INS-109 |
| chr11-54057499-INS-378 | chr11-54057499-INS-389 |
| chr11-54070262-INS-89 | chr11-54070257-INS-53 |
| chr11-54087384-INS-210 | chr11-54087384-INS-217 |
| chr11-54116647-INS-63 | chr11-54116647-INS-64 |
| chr11-54128340-INS-92 | chr11-54128340-INS-94 |
| chr11-54148645-INS-137 | chr11-54148633-INS-128 |
| chr11-54150458-INS-409 | chr11-54150458-INS-409 |
| chr11-54173813-INS-204 | chr11-54173813-INS-204 |
| chr11-54175637-INS-74 | chr11-54175632-INS-79 |
| chr11-54189019-INS-358 | chr11-54189019-INS-358 |
| chr11-54199433-INS-119 | chr11-54199433-INS-127 |
| chr11-54208958-INS-56 | chr11-54208958-INS-58 |
| chr11-54211450-INS-64 | chr11-54211450-INS-64 |
| chr11-54256590-INS-55 | chr11-54256585-INS-85 |
| chr11-54257530-INS-215 | chr11-54257530-INS-216 |
| chr11-54290817-INS-211 | chr11-54290817-INS-211 |
| chr11-54297645-INS-162 | chr11-54297645-INS-162 |
| chr11-54333444-INS-95 | chr11-54333444-INS-95 |
| chr11-54355742-INS-205 | chr11-54355742-INS-205 |
| chr11-54360149-INS-112 | chr11-54360149-INS-112 |
| chr11-54380809-INS-939 | chr11-54380809-INS-938 |
| chr11-54418413-INS-75 | chr11-54418413-INS-75 |
| chr11-54432032-INS-2010 | chr11-54432033-INS-2010 |
| chr11-54449324-INS-420 | chr11-54449324-INS-420 |
| chr11-54450032-INS-6396 | chr11-54450033-INS-6396 |
| chr11-54456640-INS-1239 | chr11-54456641-INS-1239 |
| chr11-54503590-INS-53 | chr11-54503590-INS-53 |
| chr11-54523526-INS-199 | chr11-54523526-INS-199 |
| chr11-54525158-INS-429 | chr11-54525158-INS-429 |
| chr11-54599779-INS-204 | chr11-54599779-INS-204 |
| chr11-54628270-INS-192 | chr11-54628270-INS-180 |
| chr11-54633272-INS-210 | chr11-54633272-INS-208 |
| chr11-54635869-INS-73 | chr11-54635869-INS-73 |
| chr11-54656910-INS-220 | chr11-54656910-INS-220 |
| chr11-54696683-INS-185 | chr11-54696683-INS-144 |
| chr11-54705155-INS-152 | chr11-54705155-INS-148 |
| chr11-54719981-INS-66 | chr11-54719981-INS-66 |
| chr11-54731188-INS-54 | chr11-54731188-INS-56 |
| chr11-54740466-INS-144 | chr11-54740466-INS-164 |
| chr11-54751762-INS-5305 | chr11-54751763-INS-5306 |
| chr11-54763770-INS-401 | chr11-54763769-INS-402 |
| chr11-54783383-INS-226 | chr11-54783383-INS-226 |
| chr11-54784415-INS-243 | chr11-54784415-INS-243 |
| chr11-54786678-INS-201 | chr11-54786678-INS-201 |
| chr11-54794703-INS-55 | chr11-54794703-INS-58 |
| chr11-54830809-INS-503 | chr11-54830809-INS-503 |
| chr11-54833347-INS-218 | chr11-54833347-INS-219 |
| chr11-54848873-INS-52 | chr11-54848873-INS-54 |
| chr11-54856487-INS-203 | chr11-54856487-INS-203 |
| chr11-54862410-INS-53 | chr11-54862410-INS-55 |
| chr11-54873850-INS-193 | chr11-54873850-INS-193 |
| chr11-54891652-INS-60 | chr11-54891653-INS-64 |
| chr11-54892986-INS-70 | chr11-54892972-INS-56 |
| chr11-54896345-INS-56 | chr11-54896345-INS-58 |
| chr11-54899325-INS-251 | chr11-54899325-INS-250 |

| | |
|---|---|
| chr11-54927385-INS-90 | chr11-54927385-INS-81 |
| chr11-54930040-INS-346 | chr11-54930040-INS-362 |
| chr11-54964951-INS-196 | chr11-54964951-INS-196 |
| chr11-54966942-INS-58 | chr11-54966942-INS-58 |
| chr11-54978469-INS-262 | chr11-54978469-INS-262 |
| chr11-54980336-INS-110 | chr11-54980336-INS-116 |
| chr11-54988489-INS-209 | chr11-54988489-INS-212 |
| chr11-54996989-INS-165 | chr11-54996989-INS-166 |
| chr11-55005562-INS-55 | chr11-55005562-INS-57 |
| chr11-55008351-INS-715 | chr11-55008351-INS-727 |
| chr11-55011124-INS-60 | chr11-55011124-INS-58 |
| chr11-55014161-INS-498 | chr11-55014161-INS-498 |
| chr11-55034657-INS-134 | chr11-55034674-INS-121 |
| chr11-55076147-INS-56 | chr11-55076148-INS-100 |
| chr11-55081841-INS-60 | chr11-55081841-INS-68 |
| chr11-55100478-INS-78 | chr11-55100478-INS-78 |
| chr11-55118776-INS-57 | chr11-55118776-INS-57 |
| chr11-55129517-INS-95 | chr11-55129517-INS-95 |
| chr11-55131355-INS-128 | chr11-55131364-INS-141 |
| chr11-55145627-INS-80 | chr11-55145613-INS-130 |
| chr11-55155338-INS-62 | chr11-55155338-INS-62 |
| chr11-55170260-INS-76 | chr11-55170260-INS-76 |
| chr11-55179133-INS-52 | chr11-55179133-INS-66 |
| chr11-55188131-INS-51 | chr11-55188131-INS-54 |
| chr11-55196305-INS-84 | chr11-55196305-INS-84 |
| chr11-55234074-INS-60 | chr11-55234074-INS-62 |
| chr11-55242288-INS-66 | chr11-55242288-INS-66 |
| chr11-55245914-INS-63 | chr11-55245914-INS-63 |
| chr11-55250090-INS-165 | chr11-55250090-INS-166 |
| chr11-55270942-INS-441 | chr11-55270942-INS-441 |
| chr11-55276226-INS-85 | chr11-55276226-INS-78 |
| chr11-55284761-INS-432 | chr11-55284761-INS-439 |
| chr11-55298258-INS-193 | chr11-55298258-INS-193 |
| chr11-55316879-INS-107 | chr11-55316879-INS-107 |
| chr11-55318340-INS-138 | chr11-55318349-INS-139 |
| chr11-55324413-INS-468 | chr11-55324413-INS-468 |
| chr11-55325327-INS-197 | chr11-55325327-INS-197 |
| chr11-55341551-INS-69 | chr11-55341551-INS-69 |
| chr11-55344562-INS-187 | chr11-55344562-INS-188 |
| chr11-55355164-INS-137 | chr11-55355164-INS-137 |
| chr11-55358386-INS-206 | chr11-55358386-INS-205 |
| chr11-55382636-INS-63 | chr11-55382629-INS-55 |
| chr11-55407706-INS-197 | chr11-55407706-INS-197 |
| chr11-55408562-INS-52 | chr11-55408562-INS-52 |
| chr11-55410979-INS-248 | chr11-55410979-INS-240 |
| chr11-55412193-INS-675 | chr11-55412193-INS-675 |
| chr11-55412756-INS-195 | chr11-55412743-INS-196 |
| chr11-55413969-INS-415 | chr11-55413969-INS-425 |
| chr11-55416760-INS-88 | chr11-55416760-INS-88 |
| chr11-55430020-INS-226 | chr11-55430020-INS-227 |
| chr11-55456020-INS-54 | chr11-55456042-INS-52 |
| chr11-55460127-INS-211 | chr11-55460127-INS-212 |
| chr11-55462434-INS-52 | chr11-55462434-INS-52 |
| chr11-55477173-INS-211 | chr11-55477173-INS-211 |
| chr11-55489892-INS-86 | chr11-55489896-INS-90 |
| chr11-55518799-INS-66 | chr11-55518799-INS-50 |
| chr11-55525200-INS-126 | chr11-55525200-INS-119 |
| chr11-55567631-INS-55 | chr11-55567626-INS-105 |
| chr11-55568698-INS-80 | chr11-55568745-INS-62 |
| chr11-55581508-INS-3233 | chr11-55581509-INS-3233 |
| chr11-55586690-INS-295 | chr11-55586690-INS-267 |
| chr11-55628358-INS-56 | chr11-55628358-INS-54 |
| chr11-55632794-INS-165 | chr11-55632810-INS-180 |
| chr11-55643591-INS-54 | chr11-55643591-INS-54 |
| chr11-55651432-INS-239 | chr11-55651432-INS-239 |
| chr11-55654216-INS-914 | chr11-55654216-INS-914 |
| chr11-55673705-INS-78 | chr11-55673705-INS-78 |
| chr11-55697377-INS-80 | chr11-55697375-INS-82 |
| chr11-55698695-INS-270 | chr11-55698695-INS-270 |
| chr11-55710652-INS-150 | chr11-55710652-INS-150 |
| chr11-55712024-INS-56 | chr11-55712024-INS-56 |
| chr11-55736663-INS-66 | chr11-55736664-INS-53 |
| chr11-55739546-INS-145 | chr11-55739546-INS-145 |
| chr11-55753643-INS-316 | chr11-55753643-INS-315 |
| chr11-55779674-INS-400 | chr11-55779674-INS-400 |
| chr11-55786619-INS-155 | chr11-55786619-INS-153 |
| chr11-55789375-INS-53 | chr11-55789375-INS-53 |

| | |
|---|---|
| chr11-55793831-INS-81 | chr11-55793831-INS-79 |
| chr11-55801597-INS-85 | chr11-55801597-INS-86 |
| chr11-55807431-INS-139 | chr11-55807431-INS-139 |
| chr11-55814464-INS-76 | chr11-55814466-INS-88 |
| chr11-55816486-INS-228 | chr11-55816486-INS-248 |
| chr11-55834722-INS-84 | chr11-55834722-INS-78 |
| chr11-55836531-INS-68 | chr11-55836531-INS-82 |
| chr11-55875200-INS-767 | chr11-55875200-INS-768 |
| chr11-55886884-INS-52 | chr11-55886884-INS-58 |
| chr11-55899737-INS-291 | chr11-55899737-INS-302 |
| chr11-55900900-INS-57 | chr11-55900917-INS-81 |
| chr11-55908344-INS-53 | chr11-55908331-INS-56 |
| chr11-55915061-INS-2980 | chr11-55915062-INS-2980 |
| chr11-55915062-INS-74 | chr11-55915062-INS-74 |
| chr11-55930311-INS-80 | chr11-55930312-INS-78 |
| chr11-55939612-INS-539 | chr11-55939612-INS-539 |
| chr11-55943641-INS-54 | chr11-55943641-INS-54 |
| chr11-55949556-INS-104 | chr11-55949533-INS-90 |
| chr11-55952056-INS-78 | chr11-55952056-INS-78 |
| chr11-55967857-INS-1662 | chr11-55967858-INS-1662 |
| chr11-55974594-INS-55 | chr11-55974600-INS-65 |
| chr11-55979853-INS-280 | chr11-55979854-INS-281 |
| chr11-55995007-INS-51 | chr11-55995007-INS-51 |
| chr11-55999481-INS-6701 | chr11-55999482-INS-6699 |
| chr11-56011197-INS-52 | chr11-56011194-INS-58 |
| chr11-56021102-INS-107 | chr11-56021102-INS-106 |
| chr11-56026498-INS-75 | chr11-56026498-INS-75 |
| chr11-56034547-INS-1807 | chr11-56034548-INS-1808 |
| chr11-56038214-INS-734 | chr11-56038214-INS-734 |
| chr11-56038265-INS-60 | chr11-56038265-INS-64 |
| chr11-56039745-INS-68 | chr11-56039762-INS-62 |
| chr11-56064110-INS-56 | chr11-56064111-INS-57 |
| chr11-56067131-INS-196 | chr11-56067131-INS-196 |
| chr11-56158969-INS-148 | chr11-56158969-INS-148 |
| chr11-56165020-INS-6095 | chr11-56165021-INS-6095 |
| chr11-56170389-INS-52 | chr11-56170389-INS-52 |
| chr11-56189087-INS-68 | chr11-56189087-INS-68 |
| chr11-56198744-INS-1864 | chr11-56198745-INS-1864 |
| chr11-56198797-INS-194 | chr11-56198797-INS-198 |
| chr11-56202678-INS-78 | chr11-56202678-INS-78 |
| chr11-56212581-INS-411 | chr11-56212581-INS-411 |
| chr11-56241912-INS-276 | chr11-56241912-INS-276 |
| chr11-56246599-INS-2280 | chr11-56246600-INS-2280 |
| chr11-56278732-INS-124 | chr11-56278740-INS-128 |
| chr11-56291130-INS-428 | chr11-56291130-INS-428 |
| chr11-56293312-INS-64 | chr11-56293316-INS-80 |
| chr11-56309093-INS-55 | chr11-56309093-INS-55 |
| chr11-56314480-INS-66 | chr11-56314480-INS-70 |
| chr11-56326102-INS-59 | chr11-56326102-INS-59 |
| chr11-56359083-INS-80 | chr11-56359083-INS-81 |
| chr11-56375421-INS-369 | chr11-56375432-INS-370 |
| chr11-56387650-INS-62 | chr11-56387650-INS-60 |
| chr11-56419807-INS-71 | chr11-56419807-INS-71 |
| chr11-56463009-INS-52 | chr11-56463009-INS-52 |
| chr11-56467530-INS-6739 | chr11-56467531-INS-6741 |
| chr11-56476532-INS-70 | chr11-56476532-INS-70 |
| chr11-56495611-INS-379 | chr11-56495611-INS-379 |
| chr11-56507513-INS-631 | chr11-56507513-INS-645 |
| chr11-56536853-INS-826 | chr11-56536853-INS-830 |
| chr11-56542219-INS-857 | chr11-56542219-INS-857 |
| chr11-56556175-INS-77 | chr11-56556174-INS-96 |
| chr11-56560673-INS-65 | chr11-56560673-INS-65 |
| chr11-56561210-INS-2163 | chr11-56561211-INS-2163 |
| chr11-56562537-INS-6831 | chr11-56562540-INS-6835 |
| chr11-56565141-INS-200 | chr11-56565173-INS-212 |
| chr11-56572515-INS-159 | chr11-56572515-INS-161 |
| chr11-56573694-INS-6899 | chr11-56573695-INS-6898 |
| chr11-56585026-INS-6442 | chr11-56585027-INS-6442 |
| chr11-56601709-INS-155 | chr11-56601709-INS-155 |
| chr11-56607147-INS-68 | chr11-56607147-INS-68 |
| chr11-56607764-INS-52 | chr11-56607754-INS-58 |
| chr11-56631673-INS-57 | chr11-56631674-INS-54 |
| chr11-56645840-INS-1886 | chr11-56645841-INS-1887 |
| chr11-56662583-INS-105 | chr11-56662595-INS-58 |
| chr11-56667074-INS-60 | chr11-56667074-INS-58 |
| chr11-56687749-INS-499 | chr11-56687749-INS-499 |
| chr11-56710576-INS-381 | chr11-56710572-INS-427 |

| | |
|---|---|
| chr11-56736333-INS-235 | chr11-56736333-INS-239 |
| chr11-56761898-INS-310 | chr11-56761903-INS-322 |
| chr11-56776494-INS-64 | chr11-56776494-INS-65 |
| chr11-56789951-INS-60 | chr11-56789951-INS-62 |
| chr11-56849220-INS-146 | chr11-56849220-INS-146 |
| chr11-56858668-INS-54 | chr11-56858668-INS-54 |
| chr11-56858993-INS-276 | chr11-56858993-INS-276 |
| chr11-56865744-INS-6941 | chr11-56865745-INS-6940 |
| chr11-56874653-INS-66 | chr11-56874653-INS-68 |
| chr11-56897486-INS-104 | chr11-56897486-INS-104 |
| chr11-56898640-INS-54 | chr11-56898640-INS-56 |
| chr11-56944945-INS-52 | chr11-56944946-INS-54 |
| chr11-56950214-INS-66 | chr11-56950214-INS-66 |
| chr11-56959826-INS-59 | chr11-56959826-INS-53 |
| chr11-56978285-INS-4309 | chr11-56978286-INS-4310 |
| chr11-56979295-INS-56 | chr11-56979295-INS-62 |
| chr11-56982954-INS-437 | chr11-56982954-INS-437 |
| chr11-56993132-INS-666 | chr11-56993132-INS-666 |
| chr11-56999311-INS-6403 | chr11-56999312-INS-6403 |
| chr11-57009972-INS-56 | chr11-57010007-INS-58 |
| chr11-57014031-INS-66 | chr11-57014031-INS-66 |
| chr11-57015796-INS-182 | chr11-57015799-INS-181 |
| chr11-57027670-INS-58 | chr11-57027670-INS-60 |
| chr11-57028446-INS-58 | chr11-57028424-INS-59 |
| chr11-57051468-INS-707 | chr11-57051473-INS-719 |
| chr11-57071548-INS-1888 | chr11-57071549-INS-1888 |
| chr11-57077723-INS-147 | chr11-57077723-INS-147 |
| chr11-57087880-INS-59 | chr11-57087880-INS-60 |
| chr11-57111235-INS-72 | chr11-57111235-INS-72 |
| chr11-57114461-INS-248 | chr11-57114461-INS-248 |
| chr11-57144384-INS-239 | chr11-57144384-INS-243 |
| chr11-57173416-INS-411 | chr11-57173416-INS-411 |
| chr11-57178742-INS-52 | chr11-57178742-INS-54 |
| chr11-57184928-INS-64 | chr11-57184911-INS-60 |
| chr11-57224090-INS-88 | chr11-57224120-INS-75 |
| chr11-57243814-INS-66 | chr11-57243814-INS-69 |
| chr11-57260705-INS-520 | chr11-57260705-INS-520 |
| chr11-57268838-INS-55 | chr11-57268838-INS-55 |
| chr11-57272048-INS-56 | chr11-57272046-INS-58 |
| chr11-57328630-INS-230 | chr11-57328630-INS-226 |
| chr11-57342611-INS-224 | chr11-57342611-INS-223 |
| chr11-57344799-INS-171 | chr11-57344799-INS-171 |
| chr11-57351199-INS-1333 | chr11-57351200-INS-1345 |
| chr11-57355317-INS-71 | chr11-57355317-INS-71 |
| chr11-57367126-INS-82 | chr11-57367126-INS-82 |
| chr11-57368198-INS-248 | chr11-57368198-INS-255 |
| chr11-57384312-INS-72 | chr11-57384312-INS-72 |
| chr11-57385027-INS-214 | chr11-57385027-INS-214 |
| chr11-57428118-INS-75 | chr11-57428118-INS-67 |
| chr11-57436984-INS-68 | chr11-57436960-INS-76 |
| chr11-57442632-INS-648 | chr11-57442632-INS-648 |
| chr11-57488757-INS-82 | chr11-57488757-INS-82 |
| chr11-57489864-INS-70 | chr11-57489864-INS-70 |
| chr11-57501330-INS-155 | chr11-57501330-INS-155 |
| chr11-57513634-INS-80 | chr11-57513634-INS-70 |
| chr11-57528741-INS-59 | chr11-57528741-INS-59 |
| chr11-57537123-INS-75 | chr11-57537137-INS-67 |
| chr11-57546379-INS-153 | chr11-57546405-INS-153 |
| chr11-57559116-INS-51 | chr11-57559116-INS-51 |
| chr11-57573826-INS-52 | chr11-57573826-INS-52 |
| chr11-57582220-INS-202 | chr11-57582221-INS-187 |
| chr11-57585740-INS-54 | chr11-57585740-INS-54 |
| chr11-57602674-INS-86 | chr11-57602674-INS-86 |
| chr11-57605684-INS-54 | chr11-57605684-INS-54 |
| chr11-57625211-INS-62 | chr11-57625211-INS-62 |
| chr11-57634647-INS-131 | chr11-57634647-INS-134 |
| chr11-57635992-INS-211 | chr11-57635992-INS-211 |
| chr11-57639616-INS-51 | chr11-57639618-INS-61 |
| chr11-57641500-INS-193 | chr11-57641500-INS-194 |
| chr11-57657953-INS-54 | chr11-57657940-INS-90 |
| chr11-57664959-INS-52 | chr11-57664959-INS-52 |
| chr11-57674823-INS-237 | chr11-57674823-INS-237 |
| chr11-57685025-INS-178 | chr11-57685025-INS-178 |
| chr11-57688567-INS-235 | chr11-57688567-INS-234 |
| chr11-57696639-INS-76 | chr11-57696639-INS-76 |
| chr11-57699596-INS-208 | chr11-57699584-INS-226 |
| chr11-57707010-INS-52 | chr11-57707051-INS-52 |

| | |
|---|---|
| chr11-57707954-INS-725 | chr11-57707954-INS-725 |
| chr11-57749412-INS-249 | chr11-57749412-INS-245 |
| chr11-57755437-INS-99 | chr11-57755437-INS-74 |
| chr11-57763204-INS-491 | chr11-57763204-INS-491 |
| chr11-57764463-INS-120 | chr11-57764463-INS-120 |
| chr11-57784669-INS-128 | chr11-57784669-INS-122 |
| chr11-57786231-INS-257 | chr11-57786231-INS-258 |
| chr11-57827965-INS-100 | chr11-57827965-INS-100 |
| chr11-57833087-INS-136 | chr11-57833086-INS-147 |
| chr11-57836011-INS-54 | chr11-57836011-INS-54 |
| chr11-57838934-INS-204 | chr11-57838929-INS-198 |
| chr11-57841129-INS-218 | chr11-57841129-INS-218 |
| chr11-57851352-INS-218 | chr11-57851352-INS-224 |
| chr11-57855446-INS-55 | chr11-57855446-INS-80 |
| chr11-57869013-INS-64 | chr11-57869013-INS-64 |
| chr11-57883902-INS-209 | chr11-57883902-INS-203 |
| chr11-57896598-INS-57 | chr11-57896598-INS-51 |
| chr11-57903940-INS-200 | chr11-57903940-INS-205 |
| chr11-57918569-INS-183 | chr11-57918569-INS-180 |
| chr11-57953040-INS-206 | chr11-57953040-INS-206 |
| chr11-57972534-INS-56 | chr11-57972534-INS-56 |
| chr11-57976355-INS-179 | chr11-57976355-INS-179 |
| chr11-57997791-INS-190 | chr11-57997787-INS-194 |
| chr11-57999801-INS-198 | chr11-57999801-INS-198 |
| chr11-58029546-INS-123 | chr11-58029546-INS-123 |
| chr11-58039954-INS-58 | chr11-58039929-INS-68 |
| chr11-58075698-INS-238 | chr11-58075698-INS-239 |
| chr11-58085357-INS-214 | chr11-58085357-INS-214 |
| chr11-58102896-INS-400 | chr11-58102896-INS-400 |
| chr11-58107587-INS-106 | chr11-58107587-INS-106 |
| chr11-58119299-INS-117 | chr11-58119299-INS-87 |
| chr11-58165051-INS-275 | chr11-58165052-INS-275 |
| chr11-58187299-INS-211 | chr11-58187299-INS-211 |
| chr11-58190583-INS-198 | chr11-58190583-INS-200 |
| chr11-58225990-INS-76 | chr11-58225990-INS-76 |
| chr11-58241782-INS-55 | chr11-58241782-INS-57 |
| chr11-58294463-INS-152 | chr11-58294463-INS-191 |
| chr11-58303415-INS-375 | chr11-58303414-INS-391 |
| chr11-58319654-INS-105 | chr11-58319654-INS-105 |
| chr11-58333509-INS-72 | chr11-58333509-INS-72 |
| chr11-58376271-INS-76 | chr11-58376271-INS-76 |
| chr11-58426238-INS-6217 | chr11-58426239-INS-6216 |
| chr11-58461084-INS-76 | chr11-58461084-INS-82 |
| chr11-58490275-INS-62 | chr11-58490275-INS-62 |
| chr11-58500300-INS-69 | chr11-58500317-INS-52 |
| chr11-58523572-INS-90 | chr11-58523574-INS-82 |
| chr11-58539640-INS-563 | chr11-58539640-INS-562 |
| chr11-58540422-INS-3118 | chr11-58540423-INS-3118 |
| chr11-58550093-INS-91 | chr11-58550093-INS-99 |
| chr11-58551078-INS-56 | chr11-58551078-INS-52 |
| chr11-58552287-INS-60 | chr11-58552287-INS-54 |
| chr11-58599759-INS-68 | chr11-58599759-INS-72 |
| chr11-58613198-INS-52 | chr11-58613198-INS-52 |
| chr11-58620451-INS-200 | chr11-58620463-INS-205 |
| chr11-58633490-INS-7127 | chr11-58633491-INS-7126 |
| chr11-58634455-INS-929 | chr11-58634455-INS-929 |
| chr11-58658772-INS-109 | chr11-58658772-INS-109 |
| chr11-58680657-INS-82 | chr11-58680653-INS-81 |
| chr11-58685492-INS-181 | chr11-58685492-INS-181 |
| chr11-58696241-INS-161 | chr11-58696230-INS-173 |
| chr11-58699976-INS-57 | chr11-58699976-INS-61 |
| chr11-58702772-INS-80 | chr11-58702772-INS-80 |
| chr11-58706890-INS-91 | chr11-58706890-INS-91 |
| chr11-58730851-INS-122 | chr11-58730852-INS-147 |
| chr11-58790564-INS-207 | chr11-58790564-INS-210 |
| chr11-58801836-INS-213 | chr11-58801836-INS-213 |
| chr11-58818025-INS-211 | chr11-58818025-INS-213 |
| chr11-58825514-INS-216 | chr11-58825514-INS-216 |
| chr11-58843440-INS-216 | chr11-58843440-INS-216 |
| chr11-58858548-INS-201 | chr11-58858548-INS-201 |
| chr11-58871224-INS-180 | chr11-58871224-INS-180 |
| chr11-58951870-INS-619 | chr11-58951824-INS-629 |
| chr11-59004201-INS-71 | chr11-59004201-INS-79 |
| chr11-59015938-INS-206 | chr11-59015938-INS-207 |
| chr11-59017052-INS-75 | chr11-59017052-INS-75 |
| chr11-59017950-INS-542 | chr11-59017950-INS-542 |
| chr11-59042847-INS-202 | chr11-59042847-INS-202 |

| | |
|---|---|
| chr11-59084372-INS-60 | chr11-59084372-INS-60 |
| chr11-59091268-INS-208 | chr11-59091268-INS-208 |
| chr11-59093896-INS-214 | chr11-59093896-INS-214 |
| chr11-59144247-INS-51 | chr11-59144211-INS-78 |
| chr11-59145808-INS-55 | chr11-59145813-INS-90 |
| chr11-59160834-INS-209 | chr11-59160834-INS-209 |
| chr11-59166328-INS-150 | chr11-59166328-INS-150 |
| chr11-59182208-INS-215 | chr11-59182208-INS-214 |
| chr11-59215122-INS-353 | chr11-59215122-INS-354 |
| chr11-59236020-INS-88 | chr11-59236020-INS-52 |
| chr11-59259318-INS-64 | chr11-59259318-INS-64 |
| chr11-59264639-INS-77 | chr11-59264596-INS-77 |
| chr11-59268013-INS-73 | chr11-59268014-INS-73 |
| chr11-59269767-INS-63 | chr11-59269767-INS-63 |
| chr11-59283097-INS-51 | chr11-59283099-INS-92 |
| chr11-59286765-INS-56 | chr11-59286765-INS-52 |
| chr11-59315024-INS-12380 | chr11-59315025-INS-12380 |
| chr11-59359039-INS-96 | chr11-59359085-INS-97 |
| chr11-59359850-INS-1871 | chr11-59359851-INS-1871 |
| chr11-59381364-INS-204 | chr11-59381364-INS-204 |
| chr11-59411127-INS-71 | chr11-59411117-INS-94 |
| chr11-59413414-INS-125 | chr11-59413414-INS-126 |
| chr11-59458377-INS-218 | chr11-59458378-INS-190 |
| chr11-59471494-INS-145 | chr11-59471494-INS-144 |
| chr11-59476628-INS-204 | chr11-59476628-INS-207 |
| chr11-59498730-INS-84 | chr11-59498730-INS-84 |
| chr11-59501033-INS-59 | chr11-59501033-INS-75 |
| chr11-59521367-INS-80 | chr11-59521368-INS-78 |
| chr11-59542375-INS-208 | chr11-59542375-INS-208 |
| chr11-59581523-INS-110 | chr11-59581523-INS-110 |
| chr11-59592874-INS-6447 | chr11-59592875-INS-6447 |
| chr11-59609277-INS-983 | chr11-59609277-INS-982 |
| chr11-59624422-INS-197 | chr11-59624422-INS-198 |
| chr11-59659396-INS-214 | chr11-59659396-INS-214 |
| chr11-59699553-INS-78 | chr11-59699553-INS-65 |
| chr11-59705570-INS-110 | chr11-59705570-INS-110 |
| chr11-59706696-INS-130 | chr11-59706651-INS-128 |
| chr11-59732634-INS-764 | chr11-59732634-INS-762 |
| chr11-59745512-INS-66 | chr11-59745512-INS-66 |
| chr11-59762411-INS-205 | chr11-59762411-INS-204 |
| chr11-59774264-INS-430 | chr11-59774262-INS-448 |
| chr11-59777889-INS-252 | chr11-59777889-INS-252 |
| chr11-59793748-INS-62 | chr11-59793748-INS-62 |
| chr11-59794282-INS-633 | chr11-59794282-INS-629 |
| chr11-59804983-INS-1433 | chr11-59804984-INS-1437 |
| chr11-59805936-INS-186 | chr11-59805936-INS-186 |
| chr11-59809474-INS-200 | chr11-59809474-INS-200 |
| chr11-59818119-INS-4371 | chr11-59818120-INS-4371 |
| chr11-59864629-INS-292 | chr11-59864629-INS-292 |
| chr11-59871187-INS-95 | chr11-59871187-INS-103 |
| chr11-59881647-INS-76 | chr11-59881647-INS-85 |
| chr11-59903558-INS-61 | chr11-59903558-INS-51 |
| chr11-59908493-INS-165 | chr11-59908493-INS-174 |
| chr11-59916027-INS-256 | chr11-59916027-INS-256 |
| chr11-59932954-INS-52 | chr11-59932952-INS-52 |
| chr11-59948723-INS-99 | chr11-59948765-INS-105 |
| chr11-59953274-INS-55 | chr11-59953274-INS-55 |
| chr11-59957719-INS-144 | chr11-59957720-INS-152 |
| chr11-59958972-INS-207 | chr11-59958972-INS-207 |
| chr11-59970860-INS-547 | chr11-59970860-INS-543 |
| chr11-59973626-INS-167 | chr11-59973620-INS-165 |
| chr11-59994170-INS-57 | chr11-59994171-INS-60 |
| chr11-60003499-INS-74 | chr11-60003499-INS-74 |
| chr11-60010141-INS-59 | chr11-60010142-INS-53 |
| chr11-60061034-INS-1085 | chr11-60061039-INS-1081 |
| chr11-60110521-INS-226 | chr11-60110521-INS-226 |
| chr11-60125857-INS-174 | chr11-60125857-INS-174 |
| chr11-60131025-INS-96 | chr11-60131025-INS-50 |
| chr11-60137964-INS-72 | chr11-60137925-INS-96 |
| chr11-60163391-INS-204 | chr11-60163391-INS-204 |
| chr11-60167992-INS-140 | chr11-60167992-INS-140 |
| chr11-60172863-INS-209 | chr11-60172863-INS-209 |
| chr11-60182291-INS-192 | chr11-60182291-INS-194 |
| chr11-60186972-INS-83 | chr11-60186972-INS-85 |
| chr11-60194791-INS-235 | chr11-60194791-INS-235 |
| chr11-60222610-INS-150 | chr11-60222619-INS-152 |
| chr11-60229207-INS-212 | chr11-60229207-INS-216 |

| | |
|---|---|
| chr11-60247391-INS-123 | chr11-60247391-INS-123 |
| chr11-60251213-INS-209 | chr11-60251213-INS-209 |
| chr11-60267491-INS-285 | chr11-60267491-INS-285 |
| chr11-60286620-INS-201 | chr11-60286620-INS-201 |
| chr11-60301903-INS-209 | chr11-60301903-INS-211 |
| chr11-60320584-INS-213 | chr11-60320584-INS-213 |
| chr11-60322659-INS-497 | chr11-60322659-INS-497 |
| chr11-60325980-INS-69 | chr11-60325980-INS-69 |
| chr11-60329860-INS-64 | chr11-60329860-INS-52 |
| chr11-60338898-INS-80 | chr11-60338898-INS-80 |
| chr11-60342006-INS-808 | chr11-60341980-INS-846 |
| chr11-60350821-INS-57 | chr11-60350821-INS-57 |
| chr11-60360582-INS-110 | chr11-60360582-INS-110 |
| chr11-60403116-INS-183 | chr11-60403116-INS-183 |
| chr11-60410114-INS-201 | chr11-60410114-INS-201 |
| chr11-60449932-INS-634 | chr11-60449932-INS-634 |
| chr11-60523905-INS-66 | chr11-60523905-INS-66 |
| chr11-60578796-INS-56 | chr11-60578796-INS-50 |
| chr11-60628177-INS-333 | chr11-60628177-INS-333 |
| chr11-60630174-INS-67 | chr11-60630174-INS-67 |
| chr11-60647300-INS-52 | chr11-60647300-INS-56 |
| chr11-60661804-INS-215 | chr11-60661792-INS-225 |
| chr11-60663949-INS-204 | chr11-60663949-INS-204 |
| chr11-60671945-INS-110 | chr11-60671945-INS-110 |
| chr11-60679177-INS-229 | chr11-60679177-INS-228 |
| chr11-60696252-INS-65 | chr11-60696253-INS-106 |
| chr11-60708775-INS-159 | chr11-60708775-INS-163 |
| chr11-60725535-INS-72 | chr11-60725535-INS-72 |
| chr11-60731565-INS-222 | chr11-60731565-INS-222 |
| chr11-60733065-INS-84 | chr11-60733069-INS-85 |
| chr11-60738226-INS-205 | chr11-60738226-INS-205 |
| chr11-60739075-INS-423 | chr11-60739075-INS-423 |
| chr11-60771091-INS-205 | chr11-60771091-INS-209 |
| chr11-60782738-INS-193 | chr11-60782738-INS-193 |
| chr11-60785111-INS-203 | chr11-60785111-INS-203 |
| chr11-60788264-INS-175 | chr11-60788264-INS-175 |
| chr11-60857734-INS-195 | chr11-60857734-INS-195 |
| chr11-60870439-INS-6442 | chr11-60870440-INS-6442 |
| chr11-60896909-INS-69 | chr11-60896902-INS-72 |
| chr11-60909718-INS-70 | chr11-60909718-INS-74 |
| chr11-60936736-INS-124 | chr11-60936736-INS-134 |
| chr11-60942145-INS-264 | chr11-60942145-INS-264 |
| chr11-60952278-INS-56 | chr11-60952278-INS-56 |
| chr11-60978334-INS-69 | chr11-60978334-INS-79 |
| chr11-60986068-INS-180 | chr11-60986068-INS-185 |
| chr11-60992452-INS-5380 | chr11-60992453-INS-5385 |
| chr11-60998427-INS-58 | chr11-60998427-INS-53 |
| chr11-61012501-INS-56 | chr11-61012501-INS-80 |
| chr11-61026470-INS-100 | chr11-61026470-INS-63 |
| chr11-61031844-INS-111 | chr11-61031844-INS-96 |
| chr11-61077176-INS-68 | chr11-61077176-INS-56 |
| chr11-61097983-INS-293 | chr11-61097983-INS-293 |
| chr11-61123624-INS-193 | chr11-61123624-INS-189 |
| chr11-61145742-INS-99 | chr11-61145742-INS-99 |
| chr11-61229447-INS-55 | chr11-61229447-INS-55 |
| chr11-61249916-INS-52 | chr11-61249916-INS-52 |
| chr11-61271052-INS-198 | chr11-61271052-INS-198 |
| chr11-61319141-INS-60 | chr11-61319141-INS-60 |
| chr11-61324345-INS-135 | chr11-61324345-INS-88 |
| chr11-61353644-INS-690 | chr11-61353644-INS-690 |
| chr11-61363722-INS-227 | chr11-61363683-INS-256 |
| chr11-61374783-INS-56 | chr11-61374783-INS-65 |
| chr11-61429997-INS-270 | chr11-61429997-INS-270 |
| chr11-61473533-INS-184 | chr11-61473540-INS-181 |
| chr11-61498247-INS-201 | chr11-61498247-INS-201 |
| chr11-61499599-INS-68 | chr11-61499599-INS-72 |
| chr11-61513514-INS-506 | chr11-61513514-INS-506 |
| chr11-61518650-INS-206 | chr11-61518650-INS-206 |
| chr11-61521904-INS-157 | chr11-61521904-INS-155 |
| chr11-61548721-INS-232 | chr11-61548721-INS-252 |
| chr11-61656502-INS-223 | chr11-61656502-INS-223 |
| chr11-61696122-INS-55 | chr11-61696122-INS-55 |
| chr11-61702945-INS-58 | chr11-61702945-INS-52 |
| chr11-61712601-INS-64 | chr11-61712605-INS-60 |
| chr11-61717685-INS-108 | chr11-61717685-INS-108 |
| chr11-61721693-INS-52 | chr11-61721693-INS-58 |
| chr11-61724492-INS-198 | chr11-61724492-INS-198 |

| | |
|---|---|
| chr11-61729453-INS-102 | chr11-61729453-INS-102 |
| chr11-61733930-INS-52 | chr11-61733930-INS-58 |
| chr11-61736910-INS-135 | chr11-61736910-INS-135 |
| chr11-61741336-INS-81 | chr11-61741336-INS-75 |
| chr11-61752411-INS-355 | chr11-61752411-INS-359 |
| chr11-61765490-INS-80 | chr11-61765489-INS-68 |
| chr11-61766378-INS-210 | chr11-61766378-INS-210 |
| chr11-61773319-INS-62 | chr11-61773319-INS-62 |
| chr11-61782497-INS-199 | chr11-61782497-INS-199 |
| chr11-61801944-INS-134 | chr11-61801944-INS-134 |
| chr11-61802383-INS-209 | chr11-61802383-INS-209 |
| chr11-61809563-INS-167 | chr11-61809563-INS-167 |
| chr11-61822121-INS-2243 | chr11-61822122-INS-2245 |
| chr11-61832718-INS-1041 | chr11-61832730-INS-1000 |
| chr11-61836446-INS-399 | chr11-61836446-INS-399 |
| chr11-61843910-INS-95 | chr11-61843910-INS-95 |
| chr11-61844730-INS-541 | chr11-61844730-INS-543 |
| chr11-61847051-INS-440 | chr11-61847051-INS-440 |
| chr11-61863390-INS-184 | chr11-61863390-INS-184 |
| chr11-61866018-INS-121 | chr11-61865999-INS-147 |
| chr11-61871132-INS-7217 | chr11-61871133-INS-7217 |
| chr11-61877166-INS-502 | chr11-61877157-INS-512 |
| chr11-61908753-INS-188 | chr11-61908755-INS-206 |
| chr11-61949415-INS-65 | chr11-61949415-INS-65 |
| chr11-61956781-INS-88 | chr11-61956781-INS-89 |
| chr11-61983156-INS-224 | chr11-61983156-INS-224 |
| chr11-61992425-INS-580 | chr11-61992425-INS-581 |
| chr11-61996927-INS-63 | chr11-61996927-INS-64 |
| chr11-62026076-INS-92 | chr11-62026076-INS-93 |
| chr11-62051123-INS-193 | chr11-62051123-INS-193 |
| chr11-62066793-INS-225 | chr11-62066793-INS-224 |
| chr11-62067961-INS-66 | chr11-62067961-INS-66 |
| chr11-62121819-INS-133 | chr11-62121819-INS-145 |
| chr11-62125292-INS-78 | chr11-62125292-INS-72 |
| chr11-62128792-INS-68 | chr11-62128792-INS-68 |
| chr11-62132219-INS-770 | chr11-62132219-INS-769 |
| chr11-62136975-INS-80 | chr11-62136975-INS-80 |
| chr11-62152047-INS-216 | chr11-62152047-INS-211 |
| chr11-62192976-INS-220 | chr11-62192976-INS-221 |
| chr11-62213272-INS-207 | chr11-62213272-INS-208 |
| chr11-62258635-INS-1893 | chr11-62258636-INS-1893 |
| chr11-62263214-INS-68 | chr11-62263174-INS-73 |
| chr11-62275253-INS-55 | chr11-62275253-INS-55 |
| chr11-62280907-INS-54 | chr11-62280907-INS-52 |
| chr11-62282447-INS-183 | chr11-62282447-INS-183 |
| chr11-62290103-INS-406 | chr11-62290103-INS-406 |
| chr11-62301149-INS-1344 | chr11-62301150-INS-1350 |
| chr11-62320925-INS-214 | chr11-62320925-INS-214 |
| chr11-62325242-INS-50 | chr11-62325242-INS-80 |
| chr11-62344399-INS-333 | chr11-62344399-INS-348 |
| chr11-62353987-INS-54 | chr11-62353987-INS-54 |
| chr11-62359185-INS-74 | chr11-62359175-INS-62 |
| chr11-62362900-INS-166 | chr11-62362900-INS-210 |
| chr11-62369015-INS-190 | chr11-62369015-INS-194 |
| chr11-62370922-INS-51 | chr11-62370922-INS-51 |
| chr11-62376055-INS-209 | chr11-62376055-INS-210 |
| chr11-62390450-INS-62 | chr11-62390450-INS-62 |
| chr11-62395155-INS-6454 | chr11-62395156-INS-6454 |
| chr11-62401349-INS-232 | chr11-62401350-INS-232 |
| chr11-62404327-INS-60 | chr11-62404327-INS-60 |
| chr11-62408980-INS-260 | chr11-62408980-INS-260 |
| chr11-62414823-INS-7138 | chr11-62414824-INS-7138 |
| chr11-62438265-INS-1343 | chr11-62438266-INS-1343 |
| chr11-62440176-INS-58 | chr11-62440176-INS-58 |
| chr11-62457634-INS-110 | chr11-62457634-INS-110 |
| chr11-62460624-INS-70 | chr11-62460621-INS-70 |
| chr11-62462924-INS-473 | chr11-62462924-INS-473 |
| chr11-62468675-INS-96 | chr11-62468688-INS-62 |
| chr11-62471697-INS-97 | chr11-62471697-INS-97 |
| chr11-62497295-INS-201 | chr11-62497295-INS-203 |
| chr11-62540929-INS-427 | chr11-62540929-INS-429 |
| chr11-62542952-INS-204 | chr11-62542953-INS-204 |
| chr11-62544687-INS-193 | chr11-62544687-INS-193 |
| chr11-62572075-INS-213 | chr11-62572075-INS-213 |
| chr11-62576277-INS-73 | chr11-62576277-INS-62 |
| chr11-62585290-INS-167 | chr11-62585290-INS-167 |
| chr11-62586435-INS-222 | chr11-62586435-INS-217 |

| | |
|---|---|
| chr11-62586435-INS-919 | chr11-62586435-INS-920 |
| chr11-62590975-INS-209 | chr11-62590975-INS-209 |
| chr11-62615214-INS-110 | chr11-62615206-INS-82 |
| chr11-62617965-INS-225 | chr11-62617965-INS-225 |
| chr11-62635732-INS-64 | chr11-62635732-INS-66 |
| chr11-62636087-INS-829 | chr11-62636087-INS-830 |
| chr11-62640023-INS-545 | chr11-62640023-INS-545 |
| chr11-62657184-INS-217 | chr11-62657184-INS-217 |
| chr11-62665288-INS-175 | chr11-62665288-INS-175 |
| chr11-62667225-INS-52 | chr11-62667225-INS-52 |
| chr11-62685814-INS-64 | chr11-62685814-INS-64 |
| chr11-62691118-INS-66 | chr11-62691118-INS-66 |
| chr11-62706439-INS-64 | chr11-62706439-INS-64 |
| chr11-62712362-INS-54 | chr11-62712362-INS-70 |
| chr11-62715610-INS-62 | chr11-62715610-INS-72 |
| chr11-62718341-INS-58 | chr11-62718339-INS-80 |
| chr11-62721147-INS-72 | chr11-62721137-INS-52 |
| chr11-62735612-INS-53 | chr11-62735612-INS-53 |
| chr11-62740861-INS-93 | chr11-62740861-INS-111 |
| chr11-62763015-INS-104 | chr11-62763047-INS-78 |
| chr11-62769818-INS-216 | chr11-62769818-INS-215 |
| chr11-62774982-INS-179 | chr11-62774982-INS-179 |
| chr11-62788450-INS-52 | chr11-62788450-INS-52 |
| chr11-62820049-INS-1075 | chr11-62820050-INS-1075 |
| chr11-62824719-INS-791 | chr11-62824719-INS-791 |
| chr11-62831093-INS-6242 | chr11-62831094-INS-6241 |
| chr11-62865259-INS-208 | chr11-62865259-INS-208 |
| chr11-62901561-INS-75 | chr11-62901558-INS-79 |
| chr11-62902897-INS-405 | chr11-62902897-INS-405 |
| chr11-62925902-INS-259 | chr11-62925905-INS-259 |
| chr11-62932583-INS-5884 | chr11-62932584-INS-5884 |
| chr11-62944532-INS-703 | chr11-62944532-INS-703 |
| chr11-62947483-INS-57 | chr11-62947480-INS-60 |
| chr11-62950265-INS-57 | chr11-62950261-INS-51 |
| chr11-62988020-INS-756 | chr11-62988020-INS-755 |
| chr11-62991174-INS-62 | chr11-62991174-INS-70 |
| chr11-63009185-INS-72 | chr11-63009147-INS-114 |
| chr11-63051786-INS-79 | chr11-63051786-INS-80 |
| chr11-63063354-INS-547 | chr11-63063354-INS-548 |
| chr11-63070148-INS-192 | chr11-63070148-INS-180 |
| chr11-63074015-INS-51 | chr11-63074008-INS-59 |
| chr11-63086843-INS-5192 | chr11-63086844-INS-5192 |
| chr11-63116070-INS-57 | chr11-63116070-INS-58 |
| chr11-63116678-INS-475 | chr11-63116678-INS-475 |
| chr11-63121429-INS-420 | chr11-63121439-INS-428 |
| chr11-63130205-INS-281 | chr11-63130206-INS-289 |
| chr11-63142059-INS-99 | chr11-63142059-INS-124 |
| chr11-63154644-INS-749 | chr11-63154644-INS-749 |
| chr11-63172527-INS-387 | chr11-63172527-INS-387 |
| chr11-63197881-INS-51 | chr11-63197881-INS-51 |
| chr11-63209580-INS-910 | chr11-63209580-INS-876 |
| chr11-63213915-INS-89 | chr11-63213915-INS-84 |
| chr11-63222664-INS-152 | chr11-63222658-INS-158 |
| chr11-63254921-INS-7121 | chr11-63254922-INS-7121 |
| chr11-63263821-INS-54 | chr11-63263830-INS-62 |
| chr11-63306635-INS-549 | chr11-63306635-INS-549 |
| chr11-63313611-INS-54 | chr11-63313611-INS-51 |
| chr11-63317781-INS-437 | chr11-63317781-INS-437 |
| chr11-63338409-INS-274 | chr11-63338409-INS-274 |
| chr11-63340188-INS-150 | chr11-63340188-INS-150 |
| chr11-63367930-INS-209 | chr11-63367930-INS-207 |
| chr11-63369563-INS-384 | chr11-63369563-INS-384 |
| chr11-63370895-INS-92 | chr11-63370929-INS-106 |
| chr11-63375665-INS-385 | chr11-63375665-INS-381 |
| chr11-63401862-INS-913 | chr11-63401862-INS-913 |
| chr11-63410990-INS-69 | chr11-63410993-INS-66 |
| chr11-63421878-INS-62 | chr11-63421885-INS-50 |
| chr11-63442516-INS-104 | chr11-63442516-INS-78 |
| chr11-63476333-INS-650 | chr11-63476333-INS-647 |
| chr11-63479936-INS-192 | chr11-63479936-INS-193 |
| chr11-63483334-INS-440 | chr11-63483334-INS-440 |
| chr11-63493692-INS-78 | chr11-63493692-INS-78 |
| chr11-63498345-INS-172 | chr11-63498345-INS-174 |
| chr11-63504917-INS-54 | chr11-63504917-INS-54 |
| chr11-63522603-INS-53 | chr11-63522603-INS-53 |
| chr11-63528725-INS-302 | chr11-63528725-INS-305 |
| chr11-63531757-INS-274 | chr11-63531757-INS-272 |

| | |
|---|---|
| chr11-63551060-INS-226 | chr11-63551060-INS-228 |
| chr11-63557186-INS-6262 | chr11-63557187-INS-6262 |
| chr11-63583475-INS-82 | chr11-63583475-INS-60 |
| chr11-63590774-INS-210 | chr11-63590780-INS-214 |
| chr11-63593117-INS-204 | chr11-63593108-INS-218 |
| chr11-63602057-INS-115 | chr11-63602057-INS-125 |
| chr11-63606477-INS-146 | chr11-63606465-INS-152 |
| chr11-63620296-INS-109 | chr11-63620296-INS-93 |
| chr11-63650831-INS-51 | chr11-63650845-INS-72 |
| chr11-63651673-INS-1979 | chr11-63651674-INS-1979 |
| chr11-63655308-INS-58 | chr11-63655308-INS-60 |
| chr11-63659357-INS-13625 | chr11-63659358-INS-13625 |
| chr11-63672016-INS-5262 | chr11-63672017-INS-5262 |
| chr11-63702288-INS-58 | chr11-63702288-INS-58 |
| chr11-63710497-INS-374 | chr11-63710498-INS-380 |
| chr11-63715566-INS-51 | chr11-63715531-INS-52 |
| chr11-63717831-INS-105 | chr11-63717851-INS-105 |
| chr11-63740546-INS-128 | chr11-63740546-INS-128 |
| chr11-63753545-INS-136 | chr11-63753529-INS-172 |
| chr11-63760741-INS-157 | chr11-63760741-INS-153 |
| chr11-63764070-INS-88 | chr11-63764070-INS-88 |
| chr11-63790760-INS-78 | chr11-63790760-INS-81 |
| chr11-63820620-INS-476 | chr11-63820617-INS-470 |
| chr11-63861006-INS-52 | chr11-63861006-INS-52 |
| chr11-63866865-INS-152 | chr11-63866866-INS-152 |
| chr11-63883823-INS-54 | chr11-63883823-INS-58 |
| chr11-63943275-INS-496 | chr11-63943275-INS-496 |
| chr11-63982676-INS-68 | chr11-63982676-INS-64 |
| chr11-63991685-INS-53 | chr11-63991668-INS-53 |
| chr11-64040775-INS-543 | chr11-64040775-INS-543 |
| chr11-64057907-INS-52 | chr11-64057896-INS-52 |
| chr11-64094254-INS-78 | chr11-64094254-INS-78 |
| chr11-64094522-INS-5340 | chr11-64094523-INS-5340 |
| chr11-64114137-INS-54 | chr11-64114121-INS-66 |
| chr11-64118939-INS-116 | chr11-64118939-INS-108 |
| chr11-64126098-INS-222 | chr11-64126098-INS-222 |
| chr11-64137786-INS-60 | chr11-64137783-INS-60 |
| chr11-64147971-INS-65 | chr11-64147971-INS-59 |
| chr11-64160510-INS-440 | chr11-64160510-INS-440 |
| chr11-64199129-INS-786 | chr11-64199129-INS-786 |
| chr11-64210569-INS-6455 | chr11-64210570-INS-6456 |
| chr11-64213461-INS-84 | chr11-64213449-INS-54 |
| chr11-64220605-INS-234 | chr11-64220587-INS-231 |
| chr11-64235291-INS-52 | chr11-64235291-INS-54 |
| chr11-64235965-INS-301 | chr11-64235965-INS-305 |
| chr11-64241998-INS-210 | chr11-64241998-INS-177 |
| chr11-64268630-INS-3100 | chr11-64268631-INS-3100 |
| chr11-64280031-INS-99 | chr11-64280031-INS-99 |
| chr11-64293847-INS-828 | chr11-64293847-INS-838 |
| chr11-64294523-INS-80 | chr11-64294523-INS-80 |
| chr11-64298027-INS-110 | chr11-64298042-INS-84 |
| chr11-64308929-INS-6155 | chr11-64308930-INS-6155 |
| chr11-64323943-INS-58 | chr11-64323953-INS-50 |
| chr11-64333321-INS-176 | chr11-64333321-INS-201 |
| chr11-64352387-INS-59 | chr11-64352392-INS-51 |
| chr11-64356837-INS-327 | chr11-64356837-INS-327 |
| chr11-64362484-INS-218 | chr11-64362454-INS-198 |
| chr11-64365620-INS-201 | chr11-64365620-INS-183 |
| chr11-64367398-INS-149 | chr11-64367398-INS-148 |
| chr11-64399148-INS-170 | chr11-64399148-INS-170 |
| chr11-64414897-INS-79 | chr11-64414896-INS-128 |
| chr11-64422549-INS-80 | chr11-64422534-INS-98 |
| chr11-64439476-INS-133 | chr11-64439476-INS-134 |
| chr11-64454858-INS-7498 | chr11-64454859-INS-7498 |
| chr11-64484334-INS-1444 | chr11-64484335-INS-1445 |
| chr11-64510375-INS-188 | chr11-64510375-INS-194 |
| chr11-64528396-INS-383 | chr11-64528396-INS-381 |
| chr11-64544560-INS-52 | chr11-64544561-INS-52 |
| chr11-64574606-INS-195 | chr11-64574606-INS-195 |
| chr11-64597462-INS-102 | chr11-64597462-INS-106 |
| chr11-64616781-INS-152 | chr11-64616781-INS-158 |
| chr11-64664946-INS-65 | chr11-64664946-INS-65 |
| chr11-64667068-INS-86 | chr11-64667068-INS-82 |
| chr11-64670549-INS-52 | chr11-64670549-INS-52 |
| chr11-64672483-INS-53 | chr11-64672483-INS-53 |
| chr11-64705321-INS-116 | chr11-64705321-INS-114 |
| chr11-64723680-INS-86 | chr11-64723680-INS-60 |

| | |
|---|---|
| chr11-64728065-INS-192 | chr11-64728065-INS-192 |
| chr11-64745010-INS-114 | chr11-64745010-INS-118 |
| chr11-64755854-INS-53 | chr11-64755855-INS-71 |
| chr11-64818059-INS-8455 | chr11-64818060-INS-8456 |
| chr11-64827380-INS-106 | chr11-64827382-INS-120 |
| chr11-64882302-INS-61 | chr11-64882302-INS-61 |
| chr11-64886549-INS-113 | chr11-64886549-INS-113 |
| chr11-64901565-INS-483 | chr11-64901565-INS-483 |
| chr11-64955813-INS-206 | chr11-64955826-INS-198 |
| chr11-64958995-INS-4054 | chr11-64958996-INS-4058 |
| chr11-64968886-INS-70 | chr11-64968886-INS-70 |
| chr11-64981962-INS-53 | chr11-64981962-INS-53 |
| chr11-65009247-INS-85 | chr11-65009247-INS-85 |
| chr11-65043439-INS-57 | chr11-65043439-INS-59 |
| chr11-65048791-INS-93 | chr11-65048791-INS-93 |
| chr11-65049970-INS-144 | chr11-65049970-INS-144 |
| chr11-65071969-INS-6419 | chr11-65071970-INS-6419 |
| chr11-65152394-INS-109 | chr11-65152394-INS-111 |
| chr11-65172998-INS-69 | chr11-65172998-INS-76 |
| chr11-65177791-INS-64 | chr11-65177791-INS-64 |
| chr11-65200341-INS-93 | chr11-65200341-INS-93 |
| chr11-65228142-INS-87 | chr11-65228147-INS-50 |
| chr11-65249342-INS-112 | chr11-65249364-INS-77 |
| chr11-65289128-INS-5249 | chr11-65289129-INS-5249 |
| chr11-65298883-INS-105 | chr11-65298883-INS-84 |
| chr11-65307565-INS-286 | chr11-65307565-INS-283 |
| chr11-65329533-INS-6647 | chr11-65329534-INS-6647 |
| chr11-65359879-INS-56 | chr11-65359893-INS-56 |
| chr11-65369652-INS-171 | chr11-65369652-INS-210 |
| chr11-65376776-INS-75 | chr11-65376776-INS-80 |
| chr11-65386926-INS-50 | chr11-65386926-INS-63 |
| chr11-65395159-INS-67 | chr11-65395159-INS-66 |
| chr11-65397949-INS-95 | chr11-65397946-INS-92 |
| chr11-65420213-INS-58 | chr11-65420215-INS-59 |
| chr11-65434005-INS-55 | chr11-65434005-INS-55 |
| chr11-65440913-INS-203 | chr11-65440911-INS-231 |
| chr11-65443877-INS-130 | chr11-65443875-INS-136 |
| chr11-65457999-INS-399 | chr11-65457999-INS-405 |
| chr11-65469688-INS-6379 | chr11-65469698-INS-6362 |
| chr11-65474809-INS-6582 | chr11-65474810-INS-6583 |
| chr11-65483035-INS-118 | chr11-65483041-INS-86 |
| chr11-65508630-INS-69 | chr11-65508630-INS-69 |
| chr11-65564665-INS-90 | chr11-65564665-INS-96 |
| chr11-65566748-INS-123 | chr11-65566748-INS-123 |
| chr11-65639496-INS-55 | chr11-65639496-INS-54 |
| chr11-65657411-INS-6222 | chr11-65657412-INS-6222 |
| chr11-65683448-INS-60 | chr11-65683456-INS-97 |
| chr11-65685993-INS-58 | chr11-65685996-INS-57 |
| chr11-65722828-INS-920 | chr11-65722828-INS-920 |
| chr11-65739476-INS-75 | chr11-65739476-INS-60 |
| chr11-65768891-INS-57 | chr11-65768933-INS-62 |
| chr11-65772374-INS-376 | chr11-65772374-INS-377 |
| chr11-65782181-INS-128 | chr11-65782181-INS-126 |
| chr11-65806623-INS-401 | chr11-65806623-INS-400 |
| chr11-65806625-INS-6198 | chr11-65806626-INS-6198 |
| chr11-65816325-INS-205 | chr11-65816324-INS-203 |
| chr11-65825219-INS-55 | chr11-65825219-INS-55 |
| chr11-65836536-INS-1393 | chr11-65836537-INS-1393 |
| chr11-65844936-INS-104 | chr11-65844915-INS-110 |
| chr11-65904718-INS-51 | chr11-65904718-INS-51 |
| chr11-65906853-INS-6230 | chr11-65906854-INS-6230 |
| chr11-65942955-INS-303 | chr11-65942955-INS-303 |
| chr11-65945022-INS-54 | chr11-65945022-INS-65 |
| chr11-65946832-INS-60 | chr11-65946832-INS-56 |
| chr11-65954715-INS-62 | chr11-65954762-INS-56 |
| chr11-65961254-INS-69 | chr11-65961254-INS-85 |
| chr11-65980110-INS-104 | chr11-65980110-INS-104 |
| chr11-66025583-INS-56 | chr11-66025583-INS-58 |
| chr11-66035112-INS-100 | chr11-66035112-INS-98 |
| chr11-66041138-INS-96 | chr11-66041138-INS-72 |
| chr11-66061263-INS-103 | chr11-66061263-INS-103 |
| chr11-66073766-INS-178 | chr11-66073766-INS-176 |
| chr11-66079820-INS-528 | chr11-66079820-INS-545 |
| chr11-66127626-INS-787 | chr11-66127626-INS-801 |
| chr11-66133026-INS-76 | chr11-66133024-INS-74 |
| chr11-66146256-INS-236 | chr11-66146256-INS-242 |
| chr11-66149923-INS-1373 | chr11-66149924-INS-1374 |

| | |
|---|---|
| chr11-66150778-INS-88 | chr11-66150778-INS-88 |
| chr11-66166661-INS-68 | chr11-66166634-INS-89 |
| chr11-66188136-INS-222 | chr11-66188102-INS-211 |
| chr11-66211346-INS-147 | chr11-66211346-INS-147 |
| chr11-66227586-INS-57 | chr11-66227586-INS-55 |
| chr11-66235775-INS-527 | chr11-66235775-INS-527 |
| chr11-66258016-INS-307 | chr11-66258002-INS-331 |
| chr11-66263671-INS-55 | chr11-66263671-INS-55 |
| chr11-66286599-INS-6444 | chr11-66286586-INS-6437 |
| chr11-66300914-INS-182 | chr11-66300914-INS-183 |
| chr11-66313410-INS-59 | chr11-66313410-INS-60 |
| chr11-66323201-INS-76 | chr11-66323201-INS-76 |
| chr11-66395998-INS-235 | chr11-66395998-INS-235 |
| chr11-66403895-INS-71 | chr11-66403895-INS-71 |
| chr11-66441798-INS-301 | chr11-66441798-INS-302 |
| chr11-66447119-INS-72 | chr11-66447116-INS-106 |
| chr11-66455631-INS-699 | chr11-66455631-INS-699 |
| chr11-66457900-INS-205 | chr11-66457900-INS-204 |
| chr11-66461707-INS-7403 | chr11-66461708-INS-7403 |
| chr11-66476274-INS-206 | chr11-66476274-INS-206 |
| chr11-66479675-INS-964 | chr11-66479675-INS-965 |
| chr11-66489985-INS-425 | chr11-66489985-INS-425 |
| chr11-66515581-INS-382 | chr11-66515581-INS-382 |
| chr11-66517024-INS-164 | chr11-66517024-INS-165 |
| chr11-66529038-INS-505 | chr11-66529038-INS-505 |
| chr11-66551411-INS-70 | chr11-66551411-INS-70 |
| chr11-66573708-INS-54 | chr11-66573708-INS-60 |
| chr11-66645986-INS-6460 | chr11-66645987-INS-6459 |
| chr11-66649678-INS-51 | chr11-66649676-INS-67 |
| chr11-66663348-INS-106 | chr11-66663348-INS-106 |
| chr11-66706473-INS-7536 | chr11-66706474-INS-7536 |
| chr11-66720064-INS-89 | chr11-66720071-INS-93 |
| chr11-66722453-INS-106 | chr11-66722453-INS-107 |
| chr11-66732273-INS-137 | chr11-66732273-INS-137 |
| chr11-66753828-INS-1893 | chr11-66753829-INS-1893 |
| chr11-66779666-INS-497 | chr11-66779666-INS-497 |
| chr11-66788962-INS-200 | chr11-66788962-INS-200 |
| chr11-66811207-INS-388 | chr11-66811207-INS-369 |
| chr11-66829997-INS-68 | chr11-66829997-INS-68 |
| chr11-66868216-INS-300 | chr11-66868231-INS-319 |
| chr11-66871871-INS-141 | chr11-66871871-INS-141 |
| chr11-66886049-INS-494 | chr11-66886049-INS-495 |
| chr11-66905356-INS-121 | chr11-66905356-INS-128 |
| chr11-66906438-INS-105 | chr11-66906453-INS-105 |
| chr11-66913927-INS-52 | chr11-66913927-INS-56 |
| chr11-66918393-INS-112 | chr11-66918382-INS-88 |
| chr11-66926835-INS-192 | chr11-66926835-INS-192 |
| chr11-66928571-INS-93 | chr11-66928550-INS-81 |
| chr11-67045542-INS-82 | chr11-67045542-INS-82 |
| chr11-67048795-INS-77 | chr11-67048793-INS-53 |
| chr11-67117910-INS-55 | chr11-67117910-INS-65 |
| chr11-67124668-INS-145 | chr11-67124668-INS-146 |
| chr11-67126405-INS-1295 | chr11-67126406-INS-1296 |
| chr11-67138346-INS-60 | chr11-67138346-INS-82 |
| chr11-67164628-INS-66 | chr11-67164628-INS-70 |
| chr11-67210400-INS-54 | chr11-67210400-INS-66 |
| chr11-67269474-INS-103 | chr11-67269474-INS-102 |
| chr11-67284295-INS-56 | chr11-67284311-INS-73 |
| chr11-67302267-INS-1091 | chr11-67302268-INS-1084 |
| chr11-67302268-INS-315 | chr11-67302268-INS-301 |
| chr11-67321356-INS-56 | chr11-67321356-INS-60 |
| chr11-67353679-INS-256 | chr11-67353679-INS-256 |
| chr11-67356126-INS-339 | chr11-67356126-INS-339 |
| chr11-67364566-INS-611 | chr11-67364566-INS-615 |
| chr11-67378931-INS-550 | chr11-67378931-INS-552 |
| chr11-67404286-INS-56 | chr11-67404286-INS-56 |
| chr11-67431912-INS-56 | chr11-67431912-INS-56 |
| chr11-67453032-INS-71 | chr11-67453032-INS-67 |
| chr11-67461161-INS-269 | chr11-67461161-INS-269 |
| chr11-67465030-INS-187 | chr11-67465037-INS-192 |
| chr11-67497349-INS-165 | chr11-67497351-INS-165 |
| chr11-67501071-INS-58 | chr11-67501071-INS-50 |
| chr11-67506911-INS-64 | chr11-67506868-INS-86 |
| chr11-67526461-INS-63 | chr11-67526461-INS-53 |
| chr11-67530023-INS-94 | chr11-67530023-INS-92 |
| chr11-67536560-INS-59 | chr11-67536560-INS-58 |
| chr11-67603429-INS-55 | chr11-67603429-INS-55 |

| | |
|---|---|
| chr11-67606684-INS-51 | chr11-67606685-INS-51 |
| chr11-67619425-INS-60 | chr11-67619425-INS-52 |
| chr11-67626045-INS-281 | chr11-67626045-INS-281 |
| chr11-67628083-INS-814 | chr11-67628083-INS-814 |
| chr11-67657858-INS-95 | chr11-67657858-INS-95 |
| chr11-67700010-INS-455 | chr11-67700011-INS-458 |
| chr11-67705162-INS-7125 | chr11-67705163-INS-7125 |
| chr11-67712986-INS-54 | chr11-67712986-INS-54 |
| chr11-67726665-INS-458 | chr11-67726665-INS-458 |
| chr11-67737514-INS-77 | chr11-67737514-INS-56 |
| chr11-67740795-INS-66 | chr11-67740795-INS-74 |
| chr11-67756169-INS-58 | chr11-67756169-INS-58 |
| chr11-67768264-INS-80 | chr11-67768264-INS-86 |
| chr11-67771467-INS-206 | chr11-67771467-INS-206 |
| chr11-67776411-INS-51 | chr11-67776411-INS-51 |
| chr11-67797805-INS-6812 | chr11-67797806-INS-6813 |
| chr11-67819317-INS-80 | chr11-67819305-INS-72 |
| chr11-67825700-INS-166 | chr11-67825701-INS-167 |
| chr11-67828797-INS-95 | chr11-67828797-INS-103 |
| chr11-67854776-INS-62 | chr11-67854776-INS-61 |
| chr11-67873543-INS-89 | chr11-67873544-INS-73 |
| chr11-67890915-INS-111 | chr11-67890915-INS-99 |
| chr11-67910723-INS-51 | chr11-67910746-INS-51 |
| chr11-67917885-INS-68 | chr11-67917885-INS-76 |
| chr11-67923597-INS-55 | chr11-67923597-INS-55 |
| chr11-67947414-INS-344 | chr11-67947415-INS-344 |
| chr11-67987496-INS-271 | chr11-67987496-INS-271 |
| chr11-68007754-INS-62 | chr11-68007754-INS-62 |
| chr11-68023737-INS-51 | chr11-68023737-INS-51 |
| chr11-68075307-INS-102 | chr11-68075307-INS-105 |
| chr11-68095918-INS-56 | chr11-68095918-INS-56 |
| chr11-68113962-INS-56 | chr11-68113963-INS-56 |
| chr11-68129029-INS-208 | chr11-68129029-INS-205 |
| chr11-68148728-INS-530 | chr11-68148726-INS-533 |
| chr11-68158610-INS-216 | chr11-68158610-INS-224 |
| chr11-68163780-INS-60 | chr11-68163780-INS-60 |
| chr11-68254141-INS-977 | chr11-68254141-INS-977 |
| chr11-68298440-INS-212 | chr11-68298440-INS-212 |
| chr11-68300343-INS-197 | chr11-68300343-INS-197 |
| chr11-68301911-INS-66 | chr11-68301926-INS-76 |
| chr11-68319329-INS-275 | chr11-68319329-INS-275 |
| chr11-68339894-INS-65 | chr11-68339878-INS-100 |
| chr11-68375073-INS-7115 | chr11-68375074-INS-7115 |
| chr11-68387617-INS-182 | chr11-68387617-INS-182 |
| chr11-68393081-INS-54 | chr11-68393081-INS-54 |
| chr11-68440547-INS-125 | chr11-68440580-INS-77 |
| chr11-68444262-INS-213 | chr11-68444262-INS-216 |
| chr11-68446533-INS-96 | chr11-68446533-INS-96 |
| chr11-68457378-INS-74 | chr11-68457378-INS-76 |
| chr11-68513502-INS-52 | chr11-68513502-INS-54 |
| chr11-68516815-INS-173 | chr11-68516815-INS-173 |
| chr11-68530080-INS-60 | chr11-68530081-INS-60 |
| chr11-68551259-INS-94 | chr11-68551259-INS-94 |
| chr11-68559773-INS-56 | chr11-68559773-INS-64 |
| chr11-68560675-INS-400 | chr11-68560675-INS-400 |
| chr11-68562733-INS-177 | chr11-68562733-INS-177 |
| chr11-68597290-INS-204 | chr11-68597290-INS-204 |
| chr11-68603305-INS-83 | chr11-68603305-INS-85 |
| chr11-68609191-INS-90 | chr11-68609191-INS-90 |
| chr11-68611064-INS-343 | chr11-68611064-INS-327 |
| chr11-68656874-INS-93 | chr11-68656874-INS-93 |
| chr11-68667990-INS-52 | chr11-68667990-INS-52 |
| chr11-68669455-INS-209 | chr11-68669455-INS-203 |
| chr11-68699376-INS-2028 | chr11-68699377-INS-2028 |
| chr11-68719955-INS-83 | chr11-68719955-INS-75 |
| chr11-68732602-INS-171 | chr11-68732564-INS-171 |
| chr11-68738684-INS-57 | chr11-68738684-INS-62 |
| chr11-68740116-INS-194 | chr11-68740116-INS-195 |
| chr11-68751851-INS-61 | chr11-68751844-INS-55 |
| chr11-68752665-INS-67 | chr11-68752666-INS-55 |
| chr11-68758355-INS-69 | chr11-68758355-INS-69 |
| chr11-68764503-INS-54 | chr11-68764503-INS-54 |
| chr11-68787197-INS-438 | chr11-68787198-INS-438 |
| chr11-68787759-INS-154 | chr11-68787759-INS-154 |
| chr11-68794222-INS-172 | chr11-68794222-INS-172 |
| chr11-68801200-INS-100 | chr11-68801179-INS-80 |
| chr11-68805830-INS-61 | chr11-68805830-INS-60 |

| | |
|---|---|
| chr11-68806756-INS-216 | chr11-68806757-INS-215 |
| chr11-68813529-INS-55 | chr11-68813529-INS-55 |
| chr11-68814958-INS-64 | chr11-68814958-INS-90 |
| chr11-68821826-INS-218 | chr11-68821826-INS-211 |
| chr11-68824225-INS-201 | chr11-68824225-INS-202 |
| chr11-68826248-INS-61 | chr11-68826248-INS-61 |
| chr11-68834800-INS-53 | chr11-68834800-INS-57 |
| chr11-68848924-INS-61 | chr11-68848922-INS-73 |
| chr11-68853102-INS-96 | chr11-68853102-INS-93 |
| chr11-68856107-INS-210 | chr11-68856108-INS-211 |
| chr11-68869984-INS-211 | chr11-68869984-INS-217 |
| chr11-68872588-INS-65 | chr11-68872588-INS-98 |
| chr11-68900222-INS-222 | chr11-68900222-INS-222 |
| chr11-68906042-INS-61 | chr11-68906050-INS-75 |
| chr11-68917627-INS-87 | chr11-68917627-INS-87 |
| chr11-68930208-INS-136 | chr11-68930208-INS-138 |
| chr11-68947218-INS-7225 | chr11-68947219-INS-7225 |
| chr11-68956500-INS-174 | chr11-68956500-INS-174 |
| chr11-68963909-INS-115 | chr11-68963909-INS-115 |
| chr11-69021079-INS-72 | chr11-69021079-INS-56 |
| chr11-69046863-INS-114 | chr11-69046863-INS-126 |
| chr11-69050682-INS-92 | chr11-69050682-INS-92 |
| chr11-69062833-INS-162 | chr11-69062833-INS-158 |
| chr11-69074332-INS-383 | chr11-69074332-INS-395 |
| chr11-69079225-INS-85 | chr11-69079215-INS-85 |
| chr11-69082152-INS-205 | chr11-69082152-INS-205 |
| chr11-69086101-INS-64 | chr11-69086101-INS-64 |
| chr11-69095336-INS-7081 | chr11-69095337-INS-7081 |
| chr11-69106941-INS-200 | chr11-69106941-INS-200 |
| chr11-69108039-INS-88 | chr11-69108054-INS-68 |
| chr11-69111732-INS-63 | chr11-69111732-INS-69 |
| chr11-69124579-INS-66 | chr11-69124579-INS-67 |
| chr11-69132071-INS-120 | chr11-69132071-INS-120 |
| chr11-69133506-INS-336 | chr11-69133502-INS-323 |
| chr11-69155959-INS-549 | chr11-69155959-INS-549 |
| chr11-69169057-INS-81 | chr11-69169065-INS-74 |
| chr11-69172936-INS-259 | chr11-69172936-INS-259 |
| chr11-69174955-INS-139 | chr11-69174955-INS-164 |
| chr11-69207306-INS-242 | chr11-69207306-INS-242 |
| chr11-69340417-INS-63 | chr11-69340418-INS-66 |
| chr11-69357841-INS-284 | chr11-69357841-INS-271 |
| chr11-69360033-INS-178 | chr11-69360033-INS-190 |
| chr11-69369353-INS-73 | chr11-69369363-INS-70 |
| chr11-69398470-INS-194 | chr11-69398470-INS-173 |
| chr11-69400986-INS-187 | chr11-69400986-INS-188 |
| chr11-69401520-INS-209 | chr11-69401520-INS-209 |
| chr11-69402460-INS-78 | chr11-69402460-INS-80 |
| chr11-69410214-INS-93 | chr11-69410214-INS-93 |
| chr11-69419138-INS-198 | chr11-69419138-INS-199 |
| chr11-69423133-INS-92 | chr11-69423133-INS-92 |
| chr11-69431993-INS-52 | chr11-69431993-INS-52 |
| chr11-69435144-INS-72 | chr11-69435144-INS-72 |
| chr11-69440937-INS-53 | chr11-69440937-INS-57 |
| chr11-69460174-INS-59 | chr11-69460174-INS-59 |
| chr11-69489528-INS-211 | chr11-69489528-INS-211 |
| chr11-69504060-INS-263 | chr11-69504060-INS-264 |
| chr11-69506183-INS-189 | chr11-69506183-INS-189 |
| chr11-69514502-INS-167 | chr11-69514502-INS-167 |
| chr11-69516829-INS-363 | chr11-69516834-INS-358 |
| chr11-69517876-INS-197 | chr11-69517876-INS-199 |
| chr11-69529009-INS-208 | chr11-69529009-INS-216 |
| chr11-69531585-INS-225 | chr11-69531585-INS-224 |
| chr11-69532752-INS-221 | chr11-69532752-INS-222 |
| chr11-69537632-INS-431 | chr11-69537632-INS-418 |
| chr11-69544847-INS-210 | chr11-69544862-INS-243 |
| chr11-69569927-INS-175 | chr11-69569927-INS-175 |
| chr11-69570659-INS-213 | chr11-69570659-INS-213 |
| chr11-69592500-INS-424 | chr11-69592500-INS-413 |
| chr11-69603159-INS-191 | chr11-69603159-INS-192 |
| chr11-69603648-INS-212 | chr11-69603648-INS-212 |
| chr11-69603822-INS-1015 | chr11-69603823-INS-1007 |
| chr11-69607136-INS-204 | chr11-69607136-INS-205 |
| chr11-69607770-INS-53 | chr11-69607770-INS-53 |
| chr11-69611575-INS-226 | chr11-69611575-INS-226 |
| chr11-69615404-INS-62 | chr11-69615404-INS-62 |
| chr11-69639195-INS-213 | chr11-69639182-INS-206 |
| chr11-69640241-INS-204 | chr11-69640241-INS-211 |

| | |
|---|---|
| chr11-69645658-INS-202 | chr11-69645659-INS-201 |
| chr11-69653399-INS-64 | chr11-69653399-INS-88 |
| chr11-69659869-INS-202 | chr11-69659869-INS-202 |
| chr11-69683219-INS-71 | chr11-69683219-INS-57 |
| chr11-69687918-INS-68 | chr11-69687918-INS-64 |
| chr11-69700028-INS-426 | chr11-69700028-INS-428 |
| chr11-69748646-INS-64 | chr11-69748646-INS-60 |
| chr11-69757087-INS-278 | chr11-69757087-INS-278 |
| chr11-69763892-INS-1355 | chr11-69763888-INS-1345 |
| chr11-69779896-INS-221 | chr11-69779896-INS-221 |
| chr11-69787087-INS-219 | chr11-69787087-INS-219 |
| chr11-69823375-INS-196 | chr11-69823376-INS-195 |
| chr11-69840155-INS-215 | chr11-69840155-INS-215 |
| chr11-69869595-INS-205 | chr11-69869595-INS-202 |
| chr11-69951831-INS-73 | chr11-69951850-INS-78 |
| chr11-69960516-INS-52 | chr11-69960504-INS-52 |
| chr11-69971222-INS-212 | chr11-69971222-INS-212 |
| chr11-69974036-INS-75 | chr11-69974036-INS-70 |
| chr11-69980135-INS-80 | chr11-69980135-INS-92 |
| chr11-69982101-INS-129 | chr11-69982136-INS-124 |
| chr11-70002172-INS-50 | chr11-70002172-INS-50 |
| chr11-70028268-INS-221 | chr11-70028288-INS-221 |
| chr11-70033897-INS-52 | chr11-70033897-INS-80 |
| chr11-70038610-INS-99 | chr11-70038610-INS-99 |
| chr11-70053771-INS-519 | chr11-70053772-INS-548 |
| chr11-70060404-INS-53 | chr11-70060404-INS-53 |
| chr11-70066894-INS-51 | chr11-70066894-INS-52 |
| chr11-70088589-INS-218 | chr11-70088589-INS-218 |
| chr11-70114137-INS-253 | chr11-70114137-INS-253 |
| chr11-70151303-INS-382 | chr11-70151303-INS-382 |
| chr11-70166690-INS-71 | chr11-70166690-INS-64 |
| chr11-70189851-INS-175 | chr11-70189851-INS-179 |
| chr11-70216833-INS-68 | chr11-70216833-INS-70 |
| chr11-70230373-INS-168 | chr11-70230373-INS-169 |
| chr11-70235307-INS-68 | chr11-70235307-INS-68 |
| chr11-70240296-INS-71 | chr11-70240296-INS-71 |
| chr11-70262711-INS-91 | chr11-70262711-INS-91 |
| chr11-70284338-INS-194 | chr11-70284338-INS-196 |
| chr11-70291996-INS-67 | chr11-70291996-INS-67 |
| chr11-70301728-INS-213 | chr11-70301728-INS-212 |
| chr11-70304340-INS-87 | chr11-70304340-INS-87 |
| chr11-70307347-INS-82 | chr11-70307347-INS-120 |
| chr11-70315823-INS-1091 | chr11-70315824-INS-1095 |
| chr11-70316559-INS-125 | chr11-70316559-INS-130 |
| chr11-70361554-INS-115 | chr11-70361531-INS-106 |
| chr11-70384731-INS-66 | chr11-70384730-INS-86 |
| chr11-70394523-INS-60 | chr11-70394523-INS-64 |
| chr11-70402086-INS-52 | chr11-70402086-INS-50 |
| chr11-70424791-INS-183 | chr11-70424791-INS-183 |
| chr11-70450262-INS-833 | chr11-70450262-INS-832 |
| chr11-70450782-INS-62 | chr11-70450782-INS-60 |
| chr11-70461642-INS-169 | chr11-70461642-INS-169 |
| chr11-70491357-INS-191 | chr11-70491358-INS-194 |
| chr11-70500649-INS-1017 | chr11-70500650-INS-978 |
| chr11-70517201-INS-65 | chr11-70517201-INS-65 |
| chr11-70617505-INS-69 | chr11-70617505-INS-69 |
| chr11-70629207-INS-71 | chr11-70629207-INS-71 |
| chr11-70632860-INS-213 | chr11-70632860-INS-213 |
| chr11-70648605-INS-70 | chr11-70648605-INS-75 |
| chr11-70651952-INS-222 | chr11-70651952-INS-224 |
| chr11-70655654-INS-60 | chr11-70655654-INS-54 |
| chr11-70658420-INS-165 | chr11-70658420-INS-162 |
| chr11-70663908-INS-54 | chr11-70663908-INS-54 |
| chr11-70681335-INS-203 | chr11-70681335-INS-203 |
| chr11-70682750-INS-78 | chr11-70682750-INS-78 |
| chr11-70684184-INS-67 | chr11-70684184-INS-67 |
| chr11-70689611-INS-53 | chr11-70689611-INS-50 |
| chr11-70700832-INS-172 | chr11-70700836-INS-137 |
| chr11-70701212-INS-656 | chr11-70701212-INS-656 |
| chr11-70710688-INS-210 | chr11-70710688-INS-210 |
| chr11-70713305-INS-3309 | chr11-70713306-INS-3309 |
| chr11-70719651-INS-54 | chr11-70719652-INS-50 |
| chr11-70725296-INS-189 | chr11-70725296-INS-189 |
| chr11-70772676-INS-397 | chr11-70772676-INS-397 |
| chr11-70786870-INS-189 | chr11-70786870-INS-189 |
| chr11-70798043-INS-201 | chr11-70798043-INS-239 |
| chr11-70800571-INS-211 | chr11-70800571-INS-211 |

| | |
|---|---|
| chr11-70824425-INS-56 | chr11-70824425-INS-66 |
| chr11-70859110-INS-64 | chr11-70859110-INS-58 |
| chr11-70864015-INS-511 | chr11-70864015-INS-514 |
| chr11-70891806-INS-53 | chr11-70891806-INS-53 |
| chr11-70892583-INS-218 | chr11-70892583-INS-220 |
| chr11-70924870-INS-1154 | chr11-70924866-INS-1158 |
| chr11-70927855-INS-482 | chr11-70927855-INS-482 |
| chr11-70936068-INS-205 | chr11-70936068-INS-205 |
| chr11-70947019-INS-120 | chr11-70947019-INS-120 |
| chr11-70961819-INS-218 | chr11-70961819-INS-230 |
| chr11-70968421-INS-204 | chr11-70968421-INS-204 |
| chr11-71004811-INS-7102 | chr11-71004812-INS-7102 |
| chr11-71015171-INS-64 | chr11-71015171-INS-52 |
| chr11-71024854-INS-7076 | chr11-71024855-INS-7076 |
| chr11-71038302-INS-341 | chr11-71038302-INS-350 |
| chr11-71050678-INS-658 | chr11-71050678-INS-659 |
| chr11-71067330-INS-7501 | chr11-71067331-INS-7501 |
| chr11-71153200-INS-206 | chr11-71153220-INS-232 |
| chr11-71165956-INS-124 | chr11-71165956-INS-124 |
| chr11-71214005-INS-89 | chr11-71214005-INS-89 |
| chr11-71217103-INS-66 | chr11-71217077-INS-60 |
| chr11-71227348-INS-256 | chr11-71227348-INS-240 |
| chr11-71228483-INS-71 | chr11-71228483-INS-73 |
| chr11-71235079-INS-62 | chr11-71235079-INS-62 |
| chr11-71247021-INS-500 | chr11-71247021-INS-500 |
| chr11-71255860-INS-612 | chr11-71255860-INS-612 |
| chr11-71258545-INS-88 | chr11-71258545-INS-88 |
| chr11-71265322-INS-61 | chr11-71265322-INS-61 |
| chr11-71274221-INS-1186 | chr11-71274222-INS-1186 |
| chr11-71296118-INS-358 | chr11-71296118-INS-357 |
| chr11-71297798-INS-490 | chr11-71297798-INS-490 |
| chr11-71338039-INS-73 | chr11-71338015-INS-77 |
| chr11-71379137-INS-162 | chr11-71379137-INS-160 |
| chr11-71379665-INS-6425 | chr11-71379666-INS-6425 |
| chr11-71388972-INS-92 | chr11-71388972-INS-87 |
| chr11-71419226-INS-76 | chr11-71419226-INS-56 |
| chr11-71455289-INS-125 | chr11-71455305-INS-125 |
| chr11-71479656-INS-93 | chr11-71479656-INS-93 |
| chr11-71486811-INS-78 | chr11-71486811-INS-72 |
| chr11-71492471-INS-51 | chr11-71492471-INS-51 |
| chr11-71497884-INS-324 | chr11-71497884-INS-324 |
| chr11-71510145-INS-52 | chr11-71510145-INS-50 |
| chr11-71514320-INS-100 | chr11-71514320-INS-80 |
| chr11-71527461-INS-71 | chr11-71527428-INS-68 |
| chr11-71564078-INS-283 | chr11-71564078-INS-283 |
| chr11-71573326-INS-52 | chr11-71573326-INS-54 |
| chr11-71582908-INS-8631 | chr11-71582909-INS-8649 |
| chr11-71590474-INS-79 | chr11-71590436-INS-96 |
| chr11-71600894-INS-69 | chr11-71600894-INS-69 |
| chr11-71602455-INS-88 | chr11-71602455-INS-60 |
| chr11-71612775-INS-58 | chr11-71612775-INS-52 |
| chr11-71613353-INS-75 | chr11-71613353-INS-75 |
| chr11-71618933-INS-53 | chr11-71618933-INS-53 |
| chr11-71621523-INS-77 | chr11-71621523-INS-59 |
| chr11-71628099-INS-54 | chr11-71628099-INS-54 |
| chr11-71652481-INS-63 | chr11-71652481-INS-63 |
| chr11-71701777-INS-1892 | chr11-71701778-INS-1892 |
| chr11-71704209-INS-56 | chr11-71704209-INS-58 |
| chr11-71723227-INS-732 | chr11-71723227-INS-732 |
| chr11-71725271-INS-2823 | chr11-71725272-INS-2823 |
| chr11-71736675-INS-419 | chr11-71736671-INS-420 |
| chr11-71744530-INS-7118 | chr11-71744531-INS-7118 |
| chr11-71746985-INS-85 | chr11-71746969-INS-77 |
| chr11-71758214-INS-463 | chr11-71758214-INS-463 |
| chr11-71759925-INS-60 | chr11-71759887-INS-60 |
| chr11-71763325-INS-360 | chr11-71763325-INS-375 |
| chr11-71764296-INS-115 | chr11-71764296-INS-115 |
| chr11-71774305-INS-73 | chr11-71774322-INS-76 |
| chr11-71809237-INS-119 | chr11-71809243-INS-120 |
| chr11-71820325-INS-65 | chr11-71820325-INS-60 |
| chr11-71823356-INS-54 | chr11-71823358-INS-60 |
| chr11-71838497-INS-6169 | chr11-71838498-INS-6171 |
| chr11-71854271-INS-329 | chr11-71854271-INS-329 |
| chr11-71868517-INS-219 | chr11-71868517-INS-219 |
| chr11-71878476-INS-76 | chr11-71878474-INS-74 |
| chr11-71912421-INS-53 | chr11-71912421-INS-54 |
| chr11-71993522-INS-89 | chr11-71993522-INS-89 |

| | |
|---|---|
| chr11-71998417-INS-95 | chr11-71998417-INS-100 |
| chr11-72060784-INS-52 | chr11-72060784-INS-52 |
| chr11-72075424-INS-219 | chr11-72075424-INS-219 |
| chr11-72087790-INS-212 | chr11-72087790-INS-208 |
| chr11-72091219-INS-199 | chr11-72091219-INS-198 |
| chr11-72160416-INS-592 | chr11-72160416-INS-592 |
| chr11-72198844-INS-401 | chr11-72198844-INS-401 |
| chr11-72253838-INS-68 | chr11-72253838-INS-72 |
| chr11-72267300-INS-208 | chr11-72267300-INS-208 |
| chr11-72274579-INS-99 | chr11-72274579-INS-149 |
| chr11-72301543-INS-52 | chr11-72301541-INS-50 |
| chr11-72324173-INS-268 | chr11-72324173-INS-268 |
| chr11-72441891-INS-207 | chr11-72441891-INS-204 |
| chr11-72443498-INS-350 | chr11-72443498-INS-350 |
| chr11-72462189-INS-207 | chr11-72462189-INS-207 |
| chr11-72478303-INS-56 | chr11-72478303-INS-64 |
| chr11-72484065-INS-212 | chr11-72484065-INS-212 |
| chr11-72512607-INS-1106 | chr11-72512608-INS-1105 |
| chr11-72527146-INS-1070 | chr11-72527147-INS-1071 |
| chr11-72532615-INS-206 | chr11-72532615-INS-206 |
| chr11-72542744-INS-52 | chr11-72542744-INS-56 |
| chr11-72549374-INS-219 | chr11-72549374-INS-202 |
| chr11-72550424-INS-153 | chr11-72550424-INS-153 |
| chr11-72554292-INS-54 | chr11-72554292-INS-68 |
| chr11-72573444-INS-4816 | chr11-72573445-INS-4816 |
| chr11-72574014-INS-92 | chr11-72574014-INS-92 |
| chr11-72583838-INS-80 | chr11-72583845-INS-58 |
| chr11-72587961-INS-6442 | chr11-72587962-INS-6442 |
| chr11-72597900-INS-215 | chr11-72597908-INS-218 |
| chr11-72623819-INS-217 | chr11-72623819-INS-217 |
| chr11-72670021-INS-51 | chr11-72670021-INS-51 |
| chr11-72675541-INS-214 | chr11-72675541-INS-214 |
| chr11-72708176-INS-148 | chr11-72708176-INS-163 |
| chr11-72724874-INS-53 | chr11-72724874-INS-53 |
| chr11-72740504-INS-61 | chr11-72740494-INS-69 |
| chr11-72759865-INS-93 | chr11-72759866-INS-94 |
| chr11-72762440-INS-189 | chr11-72762440-INS-189 |
| chr11-72825056-INS-60 | chr11-72825056-INS-60 |
| chr11-72837877-INS-56 | chr11-72837877-INS-50 |
| chr11-72872278-INS-57 | chr11-72872288-INS-54 |
| chr11-72886021-INS-56 | chr11-72886021-INS-56 |
| chr11-72945595-INS-179 | chr11-72945595-INS-178 |
| chr11-72947920-INS-293 | chr11-72947920-INS-294 |
| chr11-72954765-INS-60 | chr11-72954765-INS-60 |
| chr11-73000228-INS-108 | chr11-73000228-INS-108 |
| chr11-73012840-INS-337 | chr11-73012844-INS-335 |
| chr11-73043726-INS-3048 | chr11-73043727-INS-3048 |
| chr11-73044967-INS-80 | chr11-73044967-INS-88 |
| chr11-73100807-INS-547 | chr11-73100807-INS-547 |
| chr11-73102303-INS-63 | chr11-73102303-INS-65 |
| chr11-73107530-INS-107 | chr11-73107518-INS-130 |
| chr11-73116723-INS-198 | chr11-73116723-INS-199 |
| chr11-73150060-INS-3343 | chr11-73150061-INS-3343 |
| chr11-73213351-INS-108 | chr11-73213381-INS-84 |
| chr11-73225789-INS-62 | chr11-73225789-INS-60 |
| chr11-73240207-INS-60 | chr11-73240213-INS-88 |
| chr11-73260251-INS-198 | chr11-73260212-INS-198 |
| chr11-73269055-INS-1117 | chr11-73269056-INS-1117 |
| chr11-73269697-INS-4689 | chr11-73269698-INS-4689 |
| chr11-73312122-INS-6902 | chr11-73312123-INS-6902 |
| chr11-73319865-INS-100 | chr11-73319869-INS-139 |
| chr11-73334136-INS-78 | chr11-73334136-INS-78 |
| chr11-73350804-INS-351 | chr11-73350805-INS-340 |
| chr11-73392241-INS-135 | chr11-73392241-INS-132 |
| chr11-73474087-INS-70 | chr11-73474064-INS-58 |
| chr11-73496664-INS-65 | chr11-73496664-INS-66 |
| chr11-73523406-INS-94 | chr11-73523399-INS-53 |
| chr11-73551711-INS-172 | chr11-73551711-INS-172 |
| chr11-73563952-INS-2975 | chr11-73563953-INS-2975 |
| chr11-73592668-INS-7087 | chr11-73592669-INS-7087 |
| chr11-73656267-INS-51 | chr11-73656267-INS-51 |
| chr11-73659626-INS-57 | chr11-73659626-INS-52 |
| chr11-73662396-INS-150 | chr11-73662396-INS-132 |
| chr11-73666641-INS-6692 | chr11-73666642-INS-6694 |
| chr11-73673458-INS-6486 | chr11-73673459-INS-6486 |
| chr11-73681961-INS-84 | chr11-73681961-INS-84 |
| chr11-73698542-INS-72 | chr11-73698542-INS-74 |

| | |
|---|---|
| chr11-73703017-INS-256 | chr11-73703017-INS-256 |
| chr11-73772953-INS-522 | chr11-73772953-INS-522 |
| chr11-73788880-INS-191 | chr11-73788880-INS-191 |
| chr11-73815852-INS-634 | chr11-73815852-INS-635 |
| chr11-73829158-INS-4310 | chr11-73829159-INS-4310 |
| chr11-73849582-INS-73 | chr11-73849536-INS-98 |
| chr11-73966513-INS-55 | chr11-73966513-INS-50 |
| chr11-73975983-INS-252 | chr11-73975983-INS-252 |
| chr11-74001718-INS-339 | chr11-74001718-INS-342 |
| chr11-74035214-INS-66 | chr11-74035214-INS-66 |
| chr11-74074999-INS-6895 | chr11-74075000-INS-6895 |
| chr11-74099565-INS-58 | chr11-74099565-INS-58 |
| chr11-74150362-INS-66 | chr11-74150362-INS-66 |
| chr11-74162504-INS-111 | chr11-74162504-INS-114 |
| chr11-74174509-INS-134 | chr11-74174497-INS-142 |
| chr11-74226763-INS-54 | chr11-74226763-INS-54 |
| chr11-74231693-INS-7088 | chr11-74231694-INS-7088 |
| chr11-74251224-INS-590 | chr11-74251224-INS-593 |
| chr11-74260282-INS-75 | chr11-74260282-INS-105 |
| chr11-74283529-INS-79 | chr11-74283529-INS-79 |
| chr11-74313355-INS-54 | chr11-74313355-INS-57 |
| chr11-74429561-INS-1282 | chr11-74429562-INS-1282 |
| chr11-74459170-INS-189 | chr11-74459170-INS-191 |
| chr11-74460589-INS-190 | chr11-74460589-INS-190 |
| chr11-74469060-INS-86 | chr11-74469060-INS-86 |
| chr11-74470262-INS-189 | chr11-74470262-INS-190 |
| chr11-74489889-INS-191 | chr11-74489889-INS-191 |
| chr11-74493573-INS-635 | chr11-74493573-INS-635 |
| chr11-74496274-INS-58 | chr11-74496274-INS-56 |
| chr11-74499370-INS-400 | chr11-74499370-INS-400 |
| chr11-74500541-INS-80 | chr11-74500541-INS-84 |
| chr11-74524475-INS-194 | chr11-74524475-INS-194 |
| chr11-74527693-INS-65 | chr11-74527693-INS-65 |
| chr11-74549570-INS-208 | chr11-74549570-INS-208 |
| chr11-74586100-INS-209 | chr11-74586100-INS-209 |
| chr11-74607649-INS-185 | chr11-74607649-INS-184 |
| chr11-74609778-INS-184 | chr11-74609778-INS-184 |
| chr11-74618992-INS-52 | chr11-74618992-INS-52 |
| chr11-74624704-INS-170 | chr11-74624704-INS-135 |
| chr11-74627452-INS-197 | chr11-74627452-INS-197 |
| chr11-74629126-INS-67 | chr11-74629126-INS-67 |
| chr11-74633671-INS-1203 | chr11-74633672-INS-1203 |
| chr11-74635913-INS-51 | chr11-74635913-INS-51 |
| chr11-74646471-INS-298 | chr11-74646471-INS-298 |
| chr11-74649094-INS-58 | chr11-74649094-INS-58 |
| chr11-74651671-INS-199 | chr11-74651671-INS-202 |
| chr11-74660810-INS-190 | chr11-74660810-INS-193 |
| chr11-74667611-INS-56 | chr11-74667611-INS-56 |
| chr11-74690217-INS-214 | chr11-74690217-INS-214 |
| chr11-74694189-INS-116 | chr11-74694189-INS-116 |
| chr11-74769861-INS-617 | chr11-74769858-INS-635 |
| chr11-74805634-INS-7269 | chr11-74805635-INS-7269 |
| chr11-74808156-INS-72 | chr11-74808197-INS-76 |
| chr11-74832636-INS-145 | chr11-74832653-INS-135 |
| chr11-74891245-INS-60 | chr11-74891245-INS-58 |
| chr11-74893621-INS-134 | chr11-74893621-INS-134 |
| chr11-74895151-INS-201 | chr11-74895151-INS-196 |
| chr11-74915766-INS-414 | chr11-74915766-INS-414 |
| chr11-74954723-INS-301 | chr11-74954723-INS-302 |
| chr11-74989066-INS-209 | chr11-74989066-INS-209 |
| chr11-74989999-INS-6487 | chr11-74990000-INS-6487 |
| chr11-74990598-INS-187 | chr11-74990598-INS-187 |
| chr11-75013396-INS-54 | chr11-75013396-INS-54 |
| chr11-75026686-INS-201 | chr11-75026686-INS-201 |
| chr11-75049841-INS-218 | chr11-75049841-INS-218 |
| chr11-75080155-INS-212 | chr11-75080155-INS-212 |
| chr11-75095223-INS-2848 | chr11-75095224-INS-2848 |
| chr11-75149248-INS-97 | chr11-75149241-INS-97 |
| chr11-75206393-INS-169 | chr11-75206393-INS-169 |
| chr11-75208246-INS-99 | chr11-75208246-INS-99 |
| chr11-75285258-INS-499 | chr11-75285258-INS-499 |
| chr11-75325079-INS-196 | chr11-75325079-INS-196 |
| chr11-75349032-INS-940 | chr11-75349032-INS-950 |
| chr11-75359969-INS-83 | chr11-75359969-INS-69 |
| chr11-75362303-INS-1612 | chr11-75362304-INS-1612 |
| chr11-75406429-INS-5537 | chr11-75406430-INS-5537 |
| chr11-75407358-INS-129 | chr11-75407358-INS-129 |

| | |
|---|---|
| chr11-75472424-INS-205 | chr11-75472424-INS-205 |
| chr11-75484562-INS-64 | chr11-75484562-INS-68 |
| chr11-75500535-INS-92 | chr11-75500535-INS-92 |
| chr11-75507419-INS-235 | chr11-75507419-INS-235 |
| chr11-75567622-INS-1000 | chr11-75567622-INS-1001 |
| chr11-75575420-INS-157 | chr11-75575420-INS-157 |
| chr11-75586450-INS-486 | chr11-75586450-INS-492 |
| chr11-75593144-INS-200 | chr11-75593144-INS-200 |
| chr11-75645982-INS-204 | chr11-75645982-INS-204 |
| chr11-75675285-INS-98 | chr11-75675286-INS-98 |
| chr11-75681931-INS-125 | chr11-75681950-INS-77 |
| chr11-75695619-INS-197 | chr11-75695619-INS-197 |
| chr11-75700846-INS-183 | chr11-75700846-INS-183 |
| chr11-75709094-INS-218 | chr11-75709094-INS-218 |
| chr11-75716302-INS-192 | chr11-75716301-INS-197 |
| chr11-75726212-INS-57 | chr11-75726213-INS-58 |
| chr11-75742096-INS-52 | chr11-75742108-INS-66 |
| chr11-75765842-INS-207 | chr11-75765842-INS-207 |
| chr11-75810417-INS-201 | chr11-75810417-INS-201 |
| chr11-75819127-INS-75 | chr11-75819121-INS-79 |
| chr11-75836233-INS-88 | chr11-75836233-INS-88 |
| chr11-75840811-INS-1890 | chr11-75840812-INS-1890 |
| chr11-75860790-INS-360 | chr11-75860790-INS-360 |
| chr11-75870166-INS-67 | chr11-75870170-INS-50 |
| chr11-75873090-INS-55 | chr11-75873090-INS-82 |
| chr11-75881259-INS-229 | chr11-75881259-INS-229 |
| chr11-75886709-INS-56 | chr11-75886709-INS-60 |
| chr11-75906411-INS-127 | chr11-75906411-INS-100 |
| chr11-75920578-INS-219 | chr11-75920578-INS-219 |
| chr11-75934521-INS-142 | chr11-75934521-INS-142 |
| chr11-75955154-INS-112 | chr11-75955155-INS-112 |
| chr11-75961338-INS-5661 | chr11-75961339-INS-5661 |
| chr11-75994906-INS-167 | chr11-75994906-INS-167 |
| chr11-76033419-INS-53 | chr11-76033419-INS-53 |
| chr11-76033975-INS-206 | chr11-76033975-INS-206 |
| chr11-76040674-INS-5755 | chr11-76040675-INS-5755 |
| chr11-76041813-INS-8116 | chr11-76041814-INS-8116 |
| chr11-76055755-INS-210 | chr11-76055756-INS-203 |
| chr11-76075858-INS-213 | chr11-76075858-INS-213 |
| chr11-76077519-INS-62 | chr11-76077519-INS-62 |
| chr11-76082264-INS-216 | chr11-76082264-INS-214 |
| chr11-76091907-INS-208 | chr11-76091907-INS-208 |
| chr11-76094465-INS-62 | chr11-76094465-INS-62 |
| chr11-76125429-INS-199 | chr11-76125429-INS-199 |
| chr11-76129976-INS-218 | chr11-76129976-INS-219 |
| chr11-76168926-INS-64 | chr11-76168926-INS-68 |
| chr11-76171313-INS-54 | chr11-76171313-INS-55 |
| chr11-76197625-INS-60 | chr11-76197638-INS-80 |
| chr11-76263762-INS-96 | chr11-76263762-INS-96 |
| chr11-76284866-INS-169 | chr11-76284866-INS-169 |
| chr11-76297227-INS-181 | chr11-76297239-INS-184 |
| chr11-76344500-INS-200 | chr11-76344500-INS-200 |
| chr11-76353408-INS-112 | chr11-76353409-INS-112 |
| chr11-76393251-INS-176 | chr11-76393251-INS-176 |
| chr11-76394169-INS-213 | chr11-76394169-INS-213 |
| chr11-76431560-INS-52 | chr11-76431560-INS-52 |
| chr11-76452335-INS-52 | chr11-76452335-INS-52 |
| chr11-76460698-INS-58 | chr11-76460698-INS-87 |
| chr11-76471308-INS-559 | chr11-76471308-INS-583 |
| chr11-76473564-INS-76 | chr11-76473567-INS-77 |
| chr11-76511680-INS-207 | chr11-76511680-INS-208 |
| chr11-76521027-INS-120 | chr11-76521049-INS-117 |
| chr11-76532311-INS-4747 | chr11-76532323-INS-4739 |
| chr11-76541641-INS-74 | chr11-76541615-INS-60 |
| chr11-76559168-INS-51 | chr11-76559139-INS-68 |
| chr11-76589555-INS-7077 | chr11-76589556-INS-7077 |
| chr11-76594629-INS-93 | chr11-76594629-INS-93 |
| chr11-76603514-INS-200 | chr11-76603514-INS-200 |
| chr11-76605124-INS-766 | chr11-76605124-INS-754 |
| chr11-76606462-INS-54 | chr11-76606469-INS-78 |
| chr11-76621411-INS-54 | chr11-76621412-INS-54 |
| chr11-76674405-INS-214 | chr11-76674405-INS-214 |
| chr11-76691074-INS-173 | chr11-76691074-INS-173 |
| chr11-76694239-INS-58 | chr11-76694239-INS-58 |
| chr11-76706447-INS-3127 | chr11-76706448-INS-3151 |
| chr11-76748600-INS-84 | chr11-76748600-INS-88 |
| chr11-76762920-INS-164 | chr11-76762920-INS-156 |

| | |
|---|---|
| chr11-76769699-INS-200 | chr11-76769699-INS-200 |
| chr11-76848059-INS-216 | chr11-76848059-INS-222 |
| chr11-76850130-INS-350 | chr11-76850130-INS-350 |
| chr11-76879151-INS-209 | chr11-76879151-INS-214 |
| chr11-76893115-INS-99 | chr11-76893115-INS-96 |
| chr11-76928386-INS-64 | chr11-76928351-INS-72 |
| chr11-76930852-INS-457 | chr11-76930852-INS-457 |
| chr11-76979210-INS-177 | chr11-76979210-INS-179 |
| chr11-77011834-INS-156 | chr11-77011834-INS-156 |
| chr11-77020251-INS-92 | chr11-77020251-INS-100 |
| chr11-77027729-INS-133 | chr11-77027729-INS-135 |
| chr11-77028661-INS-119 | chr11-77028644-INS-120 |
| chr11-77059056-INS-177 | chr11-77059056-INS-182 |
| chr11-77119599-INS-118 | chr11-77119599-INS-113 |
| chr11-77137523-INS-60 | chr11-77137523-INS-56 |
| chr11-77141974-INS-194 | chr11-77141974-INS-194 |
| chr11-77155407-INS-680 | chr11-77155407-INS-680 |
| chr11-77160676-INS-171 | chr11-77160676-INS-193 |
| chr11-77172760-INS-134 | chr11-77172760-INS-106 |
| chr11-77185382-INS-625 | chr11-77185382-INS-626 |
| chr11-77280428-INS-202 | chr11-77280428-INS-202 |
| chr11-77322680-INS-133 | chr11-77322680-INS-127 |
| chr11-77417461-INS-236 | chr11-77417461-INS-236 |
| chr11-77443741-INS-134 | chr11-77443741-INS-134 |
| chr11-77463612-INS-230 | chr11-77463612-INS-234 |
| chr11-77473450-INS-208 | chr11-77473450-INS-208 |
| chr11-77513995-INS-322 | chr11-77513995-INS-308 |
| chr11-77515007-INS-83 | chr11-77515007-INS-83 |
| chr11-77523178-INS-172 | chr11-77523178-INS-170 |
| chr11-77542999-INS-213 | chr11-77542999-INS-212 |
| chr11-77576583-INS-58 | chr11-77576561-INS-94 |
| chr11-77586585-INS-174 | chr11-77586585-INS-175 |
| chr11-77589507-INS-193 | chr11-77589507-INS-197 |
| chr11-77590818-INS-209 | chr11-77590770-INS-210 |
| chr11-77608133-INS-89 | chr11-77608133-INS-86 |
| chr11-77617686-INS-178 | chr11-77617686-INS-178 |
| chr11-77623633-INS-64 | chr11-77623633-INS-78 |
| chr11-77627533-INS-64 | chr11-77627533-INS-64 |
| chr11-77666040-INS-219 | chr11-77666040-INS-219 |
| chr11-77670596-INS-393 | chr11-77670596-INS-393 |
| chr11-77689420-INS-1204 | chr11-77689421-INS-1214 |
| chr11-77689969-INS-59 | chr11-77689969-INS-59 |
| chr11-77722783-INS-52 | chr11-77722783-INS-54 |
| chr11-77743652-INS-115 | chr11-77743630-INS-96 |
| chr11-77768656-INS-210 | chr11-77768656-INS-210 |
| chr11-77769474-INS-54 | chr11-77769474-INS-54 |
| chr11-77780745-INS-98 | chr11-77780745-INS-68 |
| chr11-77790153-INS-52 | chr11-77790153-INS-52 |
| chr11-77797549-INS-58 | chr11-77797549-INS-62 |
| chr11-77838542-INS-57 | chr11-77838542-INS-57 |
| chr11-77839739-INS-78 | chr11-77839739-INS-76 |
| chr11-77843574-INS-81 | chr11-77843574-INS-93 |
| chr11-77866807-INS-103 | chr11-77866807-INS-103 |
| chr11-77885824-INS-196 | chr11-77885824-INS-196 |
| chr11-77903117-INS-60 | chr11-77903117-INS-58 |
| chr11-77909763-INS-56 | chr11-77909763-INS-56 |
| chr11-77930185-INS-63 | chr11-77930185-INS-63 |
| chr11-77938971-INS-223 | chr11-77938971-INS-223 |
| chr11-77942568-INS-192 | chr11-77942568-INS-185 |
| chr11-77953996-INS-995 | chr11-77953996-INS-995 |
| chr11-77956140-INS-57 | chr11-77956140-INS-60 |
| chr11-77957805-INS-246 | chr11-77957805-INS-246 |
| chr11-77964371-INS-169 | chr11-77964371-INS-169 |
| chr11-77972867-INS-903 | chr11-77972868-INS-876 |
| chr11-77973420-INS-377 | chr11-77973420-INS-378 |
| chr11-78000352-INS-176 | chr11-78000352-INS-176 |
| chr11-78020820-INS-208 | chr11-78020820-INS-208 |
| chr11-78077201-INS-204 | chr11-78077201-INS-206 |
| chr11-78090076-INS-775 | chr11-78090076-INS-775 |
| chr11-78093641-INS-199 | chr11-78093641-INS-199 |
| chr11-78115967-INS-190 | chr11-78115967-INS-182 |
| chr11-78129397-INS-398 | chr11-78129397-INS-398 |
| chr11-78142103-INS-52 | chr11-78142103-INS-52 |
| chr11-78161450-INS-516 | chr11-78161450-INS-516 |
| chr11-78188357-INS-206 | chr11-78188357-INS-205 |
| chr11-78230373-INS-205 | chr11-78230373-INS-210 |
| chr11-78296559-INS-204 | chr11-78296559-INS-205 |

| | |
|---|---|
| chr11-78302684-INS-169 | chr11-78302684-INS-182 |
| chr11-78333514-INS-459 | chr11-78333514-INS-459 |
| chr11-78335185-INS-197 | chr11-78335186-INS-161 |
| chr11-78337863-INS-199 | chr11-78337863-INS-198 |
| chr11-78339644-INS-495 | chr11-78339644-INS-499 |
| chr11-78344583-INS-272 | chr11-78344583-INS-294 |
| chr11-78347898-INS-173 | chr11-78347898-INS-174 |
| chr11-78434273-INS-193 | chr11-78434273-INS-195 |
| chr11-78479835-INS-56 | chr11-78479835-INS-60 |
| chr11-78510990-INS-52 | chr11-78510990-INS-56 |
| chr11-78564296-INS-62 | chr11-78564297-INS-62 |
| chr11-78576456-INS-69 | chr11-78576456-INS-78 |
| chr11-78591041-INS-105 | chr11-78591041-INS-105 |
| chr11-78593022-INS-76 | chr11-78593022-INS-76 |
| chr11-78618741-INS-335 | chr11-78618741-INS-321 |
| chr11-78635616-INS-64 | chr11-78635616-INS-62 |
| chr11-78636759-INS-226 | chr11-78636759-INS-225 |
| chr11-78643122-INS-59 | chr11-78643122-INS-72 |
| chr11-78682845-INS-84 | chr11-78682845-INS-84 |
| chr11-78703098-INS-67 | chr11-78703098-INS-81 |
| chr11-78744240-INS-120 | chr11-78744240-INS-120 |
| chr11-78750305-INS-53 | chr11-78750305-INS-53 |
| chr11-78762523-INS-938 | chr11-78762523-INS-929 |
| chr11-78827590-INS-82 | chr11-78827590-INS-82 |
| chr11-78861692-INS-67 | chr11-78861693-INS-81 |
| chr11-78871257-INS-53 | chr11-78871244-INS-53 |
| chr11-78874302-INS-196 | chr11-78874302-INS-201 |
| chr11-78901858-INS-197 | chr11-78901858-INS-197 |
| chr11-78947414-INS-401 | chr11-78947414-INS-401 |
| chr11-78971671-INS-59 | chr11-78971671-INS-51 |
| chr11-78988642-INS-105 | chr11-78988642-INS-105 |
| chr11-78991712-INS-163 | chr11-78991712-INS-163 |
| chr11-79001906-INS-94 | chr11-79001895-INS-58 |
| chr11-79027914-INS-266 | chr11-79027914-INS-243 |
| chr11-79030635-INS-312 | chr11-79030635-INS-312 |
| chr11-79035851-INS-79 | chr11-79035851-INS-79 |
| chr11-79038140-INS-200 | chr11-79038140-INS-200 |
| chr11-79038758-INS-73 | chr11-79038726-INS-74 |
| chr11-79047463-INS-81 | chr11-79047463-INS-102 |
| chr11-79054499-INS-84 | chr11-79054499-INS-84 |
| chr11-79056070-INS-166 | chr11-79056070-INS-168 |
| chr11-79064276-INS-174 | chr11-79064276-INS-174 |
| chr11-79067866-INS-192 | chr11-79067866-INS-194 |
| chr11-79081041-INS-202 | chr11-79081041-INS-205 |
| chr11-79088561-INS-57 | chr11-79088561-INS-57 |
| chr11-79103535-INS-70 | chr11-79103535-INS-72 |
| chr11-79119559-INS-75 | chr11-79119559-INS-93 |
| chr11-79121618-INS-110 | chr11-79121618-INS-111 |
| chr11-79130574-INS-203 | chr11-79130574-INS-203 |
| chr11-79156955-INS-185 | chr11-79156955-INS-170 |
| chr11-79157930-INS-51 | chr11-79157939-INS-51 |
| chr11-79165047-INS-123 | chr11-79165043-INS-116 |
| chr11-79176699-INS-52 | chr11-79176699-INS-64 |
| chr11-79187290-INS-169 | chr11-79187317-INS-183 |
| chr11-79203358-INS-2394 | chr11-79203359-INS-2394 |
| chr11-79213347-INS-645 | chr11-79213352-INS-644 |
| chr11-79222378-INS-177 | chr11-79222378-INS-178 |
| chr11-79225854-INS-148 | chr11-79225860-INS-125 |
| chr11-79247746-INS-83 | chr11-79247746-INS-83 |
| chr11-79267795-INS-207 | chr11-79267795-INS-207 |
| chr11-79287184-INS-155 | chr11-79287188-INS-152 |
| chr11-79289969-INS-56 | chr11-79289969-INS-59 |
| chr11-79310456-INS-79 | chr11-79310456-INS-79 |
| chr11-79310750-INS-964 | chr11-79310750-INS-964 |
| chr11-79311257-INS-210 | chr11-79311257-INS-210 |
| chr11-79342535-INS-715 | chr11-79342536-INS-728 |
| chr11-79392356-INS-214 | chr11-79392356-INS-214 |
| chr11-79398071-INS-211 | chr11-79398071-INS-210 |
| chr11-79408699-INS-174 | chr11-79408699-INS-174 |
| chr11-79416597-INS-84 | chr11-79416597-INS-84 |
| chr11-79425919-INS-394 | chr11-79425919-INS-397 |
| chr11-79444047-INS-175 | chr11-79444047-INS-168 |
| chr11-79488860-INS-187 | chr11-79488860-INS-185 |
| chr11-79494087-INS-91 | chr11-79494087-INS-91 |
| chr11-79507791-INS-199 | chr11-79507791-INS-199 |
| chr11-79511726-INS-205 | chr11-79511726-INS-199 |
| chr11-79513390-INS-51 | chr11-79513390-INS-51 |

| | |
|---|---|
| chr11-79523198-INS-212 | chr11-79523198-INS-212 |
| chr11-79526437-INS-196 | chr11-79526429-INS-186 |
| chr11-79528124-INS-154 | chr11-79528124-INS-159 |
| chr11-79563585-INS-434 | chr11-79563609-INS-414 |
| chr11-79625236-INS-162 | chr11-79625236-INS-162 |
| chr11-79629185-INS-211 | chr11-79629185-INS-204 |
| chr11-79641593-INS-79 | chr11-79641583-INS-53 |
| chr11-79647815-INS-422 | chr11-79647799-INS-434 |
| chr11-79671478-INS-60 | chr11-79671483-INS-54 |
| chr11-79682140-INS-570 | chr11-79682159-INS-590 |
| chr11-79689078-INS-73 | chr11-79689126-INS-50 |
| chr11-79719995-INS-363 | chr11-79719995-INS-363 |
| chr11-79779587-INS-222 | chr11-79779587-INS-222 |
| chr11-79791926-INS-703 | chr11-79791921-INS-706 |
| chr11-79793074-INS-64 | chr11-79793072-INS-70 |
| chr11-79794703-INS-199 | chr11-79794703-INS-197 |
| chr11-79800442-INS-234 | chr11-79800442-INS-215 |
| chr11-79812864-INS-108 | chr11-79812866-INS-132 |
| chr11-79822352-INS-202 | chr11-79822352-INS-167 |
| chr11-79827935-INS-194 | chr11-79827900-INS-194 |
| chr11-79832655-INS-190 | chr11-79832655-INS-188 |
| chr11-79843673-INS-182 | chr11-79843673-INS-183 |
| chr11-79851841-INS-181 | chr11-79851841-INS-181 |
| chr11-79867408-INS-216 | chr11-79867408-INS-216 |
| chr11-79875700-INS-207 | chr11-79875700-INS-208 |
| chr11-79886916-INS-2427 | chr11-79886917-INS-2427 |
| chr11-79891294-INS-62 | chr11-79891297-INS-69 |
| chr11-79920510-INS-216 | chr11-79920510-INS-226 |
| chr11-79921509-INS-213 | chr11-79921509-INS-213 |
| chr11-79936345-INS-197 | chr11-79936345-INS-197 |
| chr11-79945971-INS-206 | chr11-79945971-INS-206 |
| chr11-79952834-INS-84 | chr11-79952834-INS-60 |
| chr11-79957484-INS-185 | chr11-79957484-INS-185 |
| chr11-79964069-INS-210 | chr11-79964069-INS-210 |
| chr11-79966411-INS-188 | chr11-79966411-INS-188 |
| chr11-79973964-INS-212 | chr11-79973965-INS-204 |
| chr11-79976338-INS-203 | chr11-79976338-INS-203 |
| chr11-79979987-INS-213 | chr11-79979987-INS-213 |
| chr11-80005967-INS-343 | chr11-80005967-INS-343 |
| chr11-80009153-INS-218 | chr11-80009153-INS-218 |
| chr11-80014288-INS-187 | chr11-80014288-INS-188 |
| chr11-80027223-INS-421 | chr11-80027222-INS-433 |
| chr11-80028591-INS-58 | chr11-80028591-INS-70 |
| chr11-80050377-INS-101 | chr11-80050377-INS-101 |
| chr11-80077179-INS-217 | chr11-80077179-INS-217 |
| chr11-80079649-INS-198 | chr11-80079649-INS-198 |
| chr11-80088311-INS-1377 | chr11-80088292-INS-1379 |
| chr11-80093537-INS-65 | chr11-80093537-INS-65 |
| chr11-80105982-INS-214 | chr11-80105982-INS-214 |
| chr11-80114693-INS-220 | chr11-80114693-INS-221 |
| chr11-80120170-INS-149 | chr11-80120170-INS-149 |
| chr11-80121972-INS-391 | chr11-80121973-INS-411 |
| chr11-80122472-INS-178 | chr11-80122440-INS-215 |
| chr11-80144148-INS-248 | chr11-80144148-INS-248 |
| chr11-80161260-INS-222 | chr11-80161260-INS-223 |
| chr11-80171685-INS-397 | chr11-80171686-INS-396 |
| chr11-80176174-INS-202 | chr11-80176174-INS-212 |
| chr11-80177214-INS-181 | chr11-80177214-INS-176 |
| chr11-80179044-INS-296 | chr11-80179044-INS-296 |
| chr11-80187596-INS-213 | chr11-80187593-INS-217 |
| chr11-80188912-INS-178 | chr11-80188912-INS-178 |
| chr11-80225947-INS-195 | chr11-80225951-INS-199 |
| chr11-80240374-INS-212 | chr11-80240373-INS-210 |
| chr11-80255141-INS-194 | chr11-80255141-INS-194 |
| chr11-80281251-INS-200 | chr11-80281251-INS-203 |
| chr11-80308023-INS-532 | chr11-80308023-INS-526 |
| chr11-80312181-INS-655 | chr11-80312182-INS-653 |
| chr11-80314756-INS-197 | chr11-80314757-INS-197 |
| chr11-80316007-INS-70 | chr11-80316007-INS-70 |
| chr11-80326853-INS-775 | chr11-80326853-INS-746 |
| chr11-80354594-INS-672 | chr11-80354594-INS-675 |
| chr11-80371768-INS-439 | chr11-80371768-INS-441 |
| chr11-80383768-INS-200 | chr11-80383768-INS-202 |
| chr11-80428650-INS-224 | chr11-80428650-INS-224 |
| chr11-80449154-INS-457 | chr11-80449154-INS-457 |
| chr11-80459136-INS-587 | chr11-80459136-INS-590 |
| chr11-80476626-INS-104 | chr11-80476592-INS-105 |

| | |
|---|---|
| chr11-80489733-INS-214 | chr11-80489733-INS-214 |
| chr11-80530219-INS-60 | chr11-80530205-INS-60 |
| chr11-80566283-INS-414 | chr11-80566283-INS-414 |
| chr11-80567954-INS-166 | chr11-80567954-INS-166 |
| chr11-80570372-INS-122 | chr11-80570372-INS-117 |
| chr11-80576876-INS-83 | chr11-80576875-INS-75 |
| chr11-80580365-INS-238 | chr11-80580365-INS-238 |
| chr11-80595836-INS-164 | chr11-80595798-INS-164 |
| chr11-80607267-INS-5257 | chr11-80607268-INS-5257 |
| chr11-80634403-INS-256 | chr11-80634403-INS-235 |
| chr11-80639372-INS-54 | chr11-80639372-INS-54 |
| chr11-80655506-INS-120 | chr11-80655504-INS-81 |
| chr11-80670045-INS-120 | chr11-80670045-INS-123 |
| chr11-80697861-INS-75 | chr11-80697861-INS-75 |
| chr11-80715052-INS-52 | chr11-80715052-INS-53 |
| chr11-80750289-INS-67 | chr11-80750289-INS-67 |
| chr11-80796749-INS-94 | chr11-80796750-INS-79 |
| chr11-80802441-INS-50 | chr11-80802441-INS-50 |
| chr11-80853111-INS-98 | chr11-80853111-INS-98 |
| chr11-80872023-INS-535 | chr11-80872023-INS-535 |
| chr11-80880810-INS-1077 | chr11-80880811-INS-1073 |
| chr11-80887045-INS-59 | chr11-80887045-INS-60 |
| chr11-80947284-INS-5311 | chr11-80947285-INS-5311 |
| chr11-81004722-INS-498 | chr11-81004722-INS-498 |
| chr11-81015192-INS-78 | chr11-81015205-INS-82 |
| chr11-81063399-INS-227 | chr11-81063399-INS-227 |
| chr11-81083519-INS-399 | chr11-81083519-INS-399 |
| chr11-81086494-INS-75 | chr11-81086498-INS-76 |
| chr11-81088355-INS-52 | chr11-81088355-INS-52 |
| chr11-81093139-INS-62 | chr11-81093139-INS-62 |
| chr11-81107942-INS-86 | chr11-81107942-INS-82 |
| chr11-81125349-INS-189 | chr11-81125394-INS-144 |
| chr11-81133314-INS-54 | chr11-81133314-INS-52 |
| chr11-81141309-INS-325 | chr11-81141309-INS-327 |
| chr11-81157361-INS-66 | chr11-81157361-INS-66 |
| chr11-81173031-INS-90 | chr11-81173031-INS-66 |
| chr11-81234757-INS-1203 | chr11-81234755-INS-1216 |
| chr11-81244394-INS-7302 | chr11-81244395-INS-7302 |
| chr11-81247935-INS-83 | chr11-81247936-INS-83 |
| chr11-81282049-INS-365 | chr11-81282049-INS-365 |
| chr11-81288469-INS-74 | chr11-81288469-INS-74 |
| chr11-81301058-INS-80 | chr11-81301072-INS-56 |
| chr11-81361220-INS-137 | chr11-81361228-INS-152 |
| chr11-81362295-INS-91 | chr11-81362317-INS-104 |
| chr11-81393652-INS-75 | chr11-81393651-INS-103 |
| chr11-81401606-INS-90 | chr11-81401606-INS-56 |
| chr11-81452337-INS-294 | chr11-81452337-INS-294 |
| chr11-81467922-INS-486 | chr11-81467922-INS-486 |
| chr11-81477473-INS-58 | chr11-81477473-INS-58 |
| chr11-81520371-INS-216 | chr11-81520371-INS-216 |
| chr11-81531960-INS-76 | chr11-81531960-INS-118 |
| chr11-81536761-INS-57 | chr11-81536761-INS-56 |
| chr11-81538769-INS-76 | chr11-81538730-INS-72 |
| chr11-81563038-INS-71 | chr11-81563038-INS-71 |
| chr11-81566351-INS-52 | chr11-81566349-INS-64 |
| chr11-81574873-INS-382 | chr11-81574873-INS-382 |
| chr11-81608909-INS-58 | chr11-81608909-INS-61 |
| chr11-81615817-INS-55 | chr11-81615817-INS-55 |
| chr11-81617903-INS-66 | chr11-81617903-INS-60 |
| chr11-81636422-INS-207 | chr11-81636422-INS-211 |
| chr11-81654838-INS-52 | chr11-81654791-INS-78 |
| chr11-81666021-INS-701 | chr11-81666021-INS-702 |
| chr11-81702056-INS-449 | chr11-81702055-INS-450 |
| chr11-81727980-INS-518 | chr11-81727980-INS-519 |
| chr11-81743618-INS-53 | chr11-81743616-INS-56 |
| chr11-81772152-INS-86 | chr11-81772156-INS-78 |
| chr11-81811040-INS-170 | chr11-81811041-INS-180 |
| chr11-81830361-INS-695 | chr11-81830361-INS-695 |
| chr11-81858048-INS-66 | chr11-81858048-INS-52 |
| chr11-81889556-INS-81 | chr11-81889563-INS-79 |
| chr11-81904544-INS-53 | chr11-81904544-INS-53 |
| chr11-81905537-INS-245 | chr11-81905539-INS-255 |
| chr11-81907928-INS-256 | chr11-81907928-INS-256 |
| chr11-81949879-INS-54 | chr11-81949879-INS-54 |
| chr11-81984569-INS-82 | chr11-81984569-INS-108 |
| chr11-82029195-INS-218 | chr11-82029195-INS-218 |
| chr11-82042250-INS-62 | chr11-82042213-INS-89 |

| | |
|---|---|
| chr11-82052417-INS-90 | chr11-82052417-INS-76 |
| chr11-82088385-INS-126 | chr11-82088385-INS-126 |
| chr11-82090912-INS-126 | chr11-82090912-INS-128 |
| chr11-82119551-INS-109 | chr11-82119559-INS-101 |
| chr11-82123819-INS-58 | chr11-82123819-INS-60 |
| chr11-82151577-INS-75 | chr11-82151577-INS-75 |
| chr11-82212624-INS-218 | chr11-82212624-INS-218 |
| chr11-82218252-INS-62 | chr11-82218252-INS-62 |
| chr11-82220587-INS-3057 | chr11-82220588-INS-3057 |
| chr11-82296103-INS-184 | chr11-82296101-INS-168 |
| chr11-82350132-INS-7141 | chr11-82350133-INS-7142 |
| chr11-82352934-INS-134 | chr11-82352934-INS-130 |
| chr11-82357698-INS-263 | chr11-82357698-INS-263 |
| chr11-82391086-INS-204 | chr11-82391086-INS-204 |
| chr11-82398150-INS-64 | chr11-82398150-INS-64 |
| chr11-82408317-INS-75 | chr11-82408280-INS-93 |
| chr11-82411393-INS-59 | chr11-82411393-INS-59 |
| chr11-82421274-INS-1270 | chr11-82421275-INS-1270 |
| chr11-82452801-INS-418 | chr11-82452801-INS-418 |
| chr11-82475703-INS-74 | chr11-82475703-INS-74 |
| chr11-82483021-INS-4530 | chr11-82483022-INS-4530 |
| chr11-82488192-INS-432 | chr11-82488192-INS-432 |
| chr11-82534234-INS-135 | chr11-82534234-INS-148 |
| chr11-82597698-INS-6441 | chr11-82597699-INS-6442 |
| chr11-82615276-INS-344 | chr11-82615276-INS-344 |
| chr11-82616280-INS-301 | chr11-82616280-INS-301 |
| chr11-82621488-INS-52 | chr11-82621488-INS-56 |
| chr11-82631524-INS-185 | chr11-82631524-INS-185 |
| chr11-82633802-INS-91 | chr11-82633806-INS-60 |
| chr11-82635221-INS-317 | chr11-82635221-INS-275 |
| chr11-82643490-INS-202 | chr11-82643490-INS-199 |
| chr11-82656783-INS-405 | chr11-82656783-INS-405 |
| chr11-82659696-INS-204 | chr11-82659696-INS-204 |
| chr11-82678915-INS-207 | chr11-82678915-INS-207 |
| chr11-82682623-INS-231 | chr11-82682623-INS-230 |
| chr11-82687157-INS-205 | chr11-82687158-INS-204 |
| chr11-82711460-INS-197 | chr11-82711460-INS-202 |
| chr11-82727180-INS-362 | chr11-82727180-INS-362 |
| chr11-82732078-INS-175 | chr11-82732062-INS-174 |
| chr11-82746010-INS-195 | chr11-82746010-INS-195 |
| chr11-82760154-INS-147 | chr11-82760154-INS-124 |
| chr11-82777898-INS-200 | chr11-82777898-INS-201 |
| chr11-82793324-INS-73 | chr11-82793324-INS-61 |
| chr11-82808726-INS-196 | chr11-82808726-INS-196 |
| chr11-82815080-INS-68 | chr11-82815082-INS-52 |
| chr11-82833264-INS-106 | chr11-82833264-INS-107 |
| chr11-82835176-INS-66 | chr11-82835176-INS-96 |
| chr11-82847008-INS-325 | chr11-82847008-INS-328 |
| chr11-82856953-INS-85 | chr11-82856953-INS-60 |
| chr11-82859502-INS-197 | chr11-82859502-INS-197 |
| chr11-82860574-INS-211 | chr11-82860574-INS-211 |
| chr11-82874786-INS-508 | chr11-82874786-INS-508 |
| chr11-82921515-INS-52 | chr11-82921515-INS-50 |
| chr11-82929967-INS-53 | chr11-82929967-INS-53 |
| chr11-82932666-INS-123 | chr11-82932667-INS-75 |
| chr11-82938769-INS-418 | chr11-82938769-INS-418 |
| chr11-82971150-INS-61 | chr11-82971150-INS-61 |
| chr11-83003675-INS-4850 | chr11-83003676-INS-4850 |
| chr11-83068070-INS-107 | chr11-83068070-INS-107 |
| chr11-83090109-INS-88 | chr11-83090109-INS-88 |
| chr11-83104616-INS-135 | chr11-83104616-INS-135 |
| chr11-83115365-INS-60 | chr11-83115365-INS-64 |
| chr11-83153711-INS-51 | chr11-83153711-INS-52 |
| chr11-83159341-INS-114 | chr11-83159341-INS-114 |
| chr11-83166239-INS-375 | chr11-83166235-INS-377 |
| chr11-83171088-INS-68 | chr11-83171071-INS-84 |
| chr11-83177136-INS-177 | chr11-83177136-INS-167 |
| chr11-83178177-INS-204 | chr11-83178177-INS-204 |
| chr11-83185853-INS-207 | chr11-83185853-INS-207 |
| chr11-83214621-INS-214 | chr11-83214621-INS-214 |
| chr11-83230433-INS-214 | chr11-83230433-INS-214 |
| chr11-83241040-INS-173 | chr11-83241040-INS-173 |
| chr11-83244616-INS-377 | chr11-83244616-INS-377 |
| chr11-83272800-INS-51 | chr11-83272800-INS-51 |
| chr11-83298108-INS-100 | chr11-83298108-INS-100 |
| chr11-83317493-INS-53 | chr11-83317493-INS-53 |
| chr11-83322882-INS-51 | chr11-83322882-INS-51 |

| | |
|---|---|
| chr11-83327125-INS-56 | chr11-83327125-INS-56 |
| chr11-83352131-INS-57 | chr11-83352131-INS-57 |
| chr11-83360449-INS-7108 | chr11-83360450-INS-7108 |
| chr11-83362040-INS-254 | chr11-83362040-INS-211 |
| chr11-83366284-INS-174 | chr11-83366284-INS-174 |
| chr11-83381343-INS-91 | chr11-83381343-INS-91 |
| chr11-83386754-INS-221 | chr11-83386754-INS-221 |
| chr11-83408347-INS-81 | chr11-83408339-INS-125 |
| chr11-83426510-INS-772 | chr11-83426510-INS-772 |
| chr11-83442049-INS-110 | chr11-83442024-INS-133 |
| chr11-83471171-INS-586 | chr11-83471171-INS-586 |
| chr11-83473160-INS-243 | chr11-83473160-INS-243 |
| chr11-83478121-INS-220 | chr11-83478121-INS-220 |
| chr11-83561531-INS-731 | chr11-83561531-INS-731 |
| chr11-83567560-INS-54 | chr11-83567561-INS-58 |
| chr11-83568102-INS-6318 | chr11-83568103-INS-6319 |
| chr11-83611295-INS-79 | chr11-83611295-INS-79 |
| chr11-83630874-INS-365 | chr11-83630874-INS-365 |
| chr11-83633480-INS-52 | chr11-83633480-INS-64 |
| chr11-83663832-INS-186 | chr11-83663832-INS-186 |
| chr11-83692146-INS-99 | chr11-83692146-INS-99 |
| chr11-83693180-INS-215 | chr11-83693180-INS-215 |
| chr11-83726207-INS-66 | chr11-83726207-INS-66 |
| chr11-83727822-INS-104 | chr11-83727822-INS-104 |
| chr11-83733485-INS-55 | chr11-83733485-INS-69 |
| chr11-83751643-INS-224 | chr11-83751643-INS-226 |
| chr11-83757460-INS-111 | chr11-83757460-INS-111 |
| chr11-83827538-INS-257 | chr11-83827539-INS-255 |
| chr11-83837480-INS-170 | chr11-83837480-INS-168 |
| chr11-83839050-INS-102 | chr11-83839051-INS-70 |
| chr11-83847744-INS-165 | chr11-83847744-INS-165 |
| chr11-83853926-INS-180 | chr11-83853926-INS-182 |
| chr11-83868868-INS-218 | chr11-83868868-INS-219 |
| chr11-83893204-INS-220 | chr11-83893204-INS-220 |
| chr11-83897007-INS-212 | chr11-83897007-INS-210 |
| chr11-83920048-INS-1891 | chr11-83920049-INS-1891 |
| chr11-83925771-INS-199 | chr11-83925771-INS-200 |
| chr11-83947430-INS-325 | chr11-83947430-INS-289 |
| chr11-83952725-INS-197 | chr11-83952725-INS-197 |
| chr11-83976645-INS-54 | chr11-83976645-INS-54 |
| chr11-83979893-INS-222 | chr11-83979893-INS-230 |
| chr11-84009343-INS-214 | chr11-84009344-INS-165 |
| chr11-84022265-INS-134 | chr11-84022265-INS-140 |
| chr11-84033384-INS-214 | chr11-84033384-INS-212 |
| chr11-84042827-INS-122 | chr11-84042828-INS-78 |
| chr11-84045649-INS-224 | chr11-84045649-INS-224 |
| chr11-84052878-INS-184 | chr11-84052878-INS-179 |
| chr11-84064567-INS-77 | chr11-84064567-INS-95 |
| chr11-84066377-INS-183 | chr11-84066377-INS-183 |
| chr11-84068217-INS-91 | chr11-84068217-INS-91 |
| chr11-84092961-INS-171 | chr11-84092961-INS-171 |
| chr11-84111870-INS-196 | chr11-84111870-INS-197 |
| chr11-84143953-INS-165 | chr11-84143953-INS-172 |
| chr11-84145215-INS-847 | chr11-84145215-INS-848 |
| chr11-84175431-INS-201 | chr11-84175431-INS-202 |
| chr11-84217843-INS-210 | chr11-84217843-INS-210 |
| chr11-84237298-INS-203 | chr11-84237298-INS-205 |
| chr11-84238451-INS-326 | chr11-84238451-INS-325 |
| chr11-84255919-INS-126 | chr11-84255919-INS-126 |
| chr11-84265514-INS-217 | chr11-84265514-INS-217 |
| chr11-84285528-INS-59 | chr11-84285528-INS-53 |
| chr11-84299109-INS-87 | chr11-84299105-INS-60 |
| chr11-84300622-INS-183 | chr11-84300622-INS-183 |
| chr11-84311089-INS-328 | chr11-84311089-INS-329 |
| chr11-84333771-INS-52 | chr11-84333771-INS-52 |
| chr11-84352447-INS-672 | chr11-84352447-INS-672 |
| chr11-84384316-INS-206 | chr11-84384316-INS-206 |
| chr11-84392422-INS-195 | chr11-84392422-INS-194 |
| chr11-84396619-INS-212 | chr11-84396619-INS-212 |
| chr11-84397839-INS-189 | chr11-84397839-INS-189 |
| chr11-84436206-INS-91 | chr11-84436207-INS-99 |
| chr11-84441497-INS-1641 | chr11-84441498-INS-1639 |
| chr11-84494597-INS-75 | chr11-84494597-INS-75 |
| chr11-84505972-INS-58 | chr11-84505972-INS-58 |
| chr11-84525044-INS-82 | chr11-84525048-INS-55 |
| chr11-84542402-INS-94 | chr11-84542395-INS-121 |
| chr11-84544085-INS-60 | chr11-84544085-INS-66 |

| | |
|---|---|
| chr11-84587507-INS-88 | chr11-84587507-INS-88 |
| chr11-84593855-INS-87 | chr11-84593855-INS-87 |
| chr11-84607798-INS-226 | chr11-84607792-INS-220 |
| chr11-84631127-INS-54 | chr11-84631127-INS-56 |
| chr11-84636653-INS-199 | chr11-84636653-INS-198 |
| chr11-84659913-INS-131 | chr11-84659909-INS-136 |
| chr11-84665329-INS-211 | chr11-84665329-INS-211 |
| chr11-84668112-INS-73 | chr11-84668112-INS-51 |
| chr11-84669659-INS-57 | chr11-84669659-INS-57 |
| chr11-84687479-INS-82 | chr11-84687511-INS-72 |
| chr11-84702330-INS-5101 | chr11-84702331-INS-5101 |
| chr11-84705446-INS-60 | chr11-84705446-INS-50 |
| chr11-84736396-INS-426 | chr11-84736396-INS-433 |
| chr11-84744592-INS-76 | chr11-84744592-INS-81 |
| chr11-84752216-INS-84 | chr11-84752216-INS-88 |
| chr11-84766427-INS-207 | chr11-84766427-INS-207 |
| chr11-84770418-INS-221 | chr11-84770418-INS-222 |
| chr11-84781477-INS-174 | chr11-84781477-INS-174 |
| chr11-84804487-INS-220 | chr11-84804487-INS-220 |
| chr11-84806104-INS-226 | chr11-84806104-INS-227 |
| chr11-84812363-INS-58 | chr11-84812363-INS-58 |
| chr11-84816015-INS-6434 | chr11-84816016-INS-6434 |
| chr11-84822399-INS-303 | chr11-84822400-INS-324 |
| chr11-84826145-INS-174 | chr11-84826145-INS-197 |
| chr11-84829171-INS-203 | chr11-84829171-INS-203 |
| chr11-84857378-INS-59 | chr11-84857379-INS-59 |
| chr11-84864021-INS-184 | chr11-84864021-INS-184 |
| chr11-84886879-INS-229 | chr11-84886879-INS-229 |
| chr11-84893777-INS-84 | chr11-84893777-INS-88 |
| chr11-84908805-INS-212 | chr11-84908805-INS-209 |
| chr11-84910359-INS-7111 | chr11-84910360-INS-7111 |
| chr11-84927044-INS-54 | chr11-84927044-INS-59 |
| chr11-84931338-INS-130 | chr11-84931338-INS-116 |
| chr11-84935394-INS-200 | chr11-84935394-INS-201 |
| chr11-84950950-INS-5379 | chr11-84950951-INS-5379 |
| chr11-84974364-INS-251 | chr11-84974364-INS-246 |
| chr11-85003752-INS-2882 | chr11-85003753-INS-2881 |
| chr11-85021917-INS-212 | chr11-85021917-INS-213 |
| chr11-85055272-INS-54 | chr11-85055283-INS-50 |
| chr11-85056441-INS-1316 | chr11-85056442-INS-1322 |
| chr11-85068524-INS-72 | chr11-85068524-INS-72 |
| chr11-85087973-INS-217 | chr11-85087973-INS-217 |
| chr11-85089784-INS-199 | chr11-85089784-INS-199 |
| chr11-85102638-INS-202 | chr11-85102638-INS-203 |
| chr11-85122836-INS-772 | chr11-85122836-INS-754 |
| chr11-85134802-INS-228 | chr11-85134802-INS-229 |
| chr11-85136105-INS-7638 | chr11-85136106-INS-7638 |
| chr11-85137618-INS-365 | chr11-85137618-INS-366 |
| chr11-85155436-INS-209 | chr11-85155436-INS-209 |
| chr11-85158325-INS-210 | chr11-85158325-INS-210 |
| chr11-85165448-INS-234 | chr11-85165448-INS-234 |
| chr11-85170451-INS-60 | chr11-85170451-INS-60 |
| chr11-85171776-INS-78 | chr11-85171776-INS-68 |
| chr11-85173863-INS-214 | chr11-85173863-INS-214 |
| chr11-85189234-INS-62 | chr11-85189206-INS-70 |
| chr11-85209626-INS-1012 | chr11-85209627-INS-1007 |
| chr11-85210275-INS-93 | chr11-85210275-INS-93 |
| chr11-85211169-INS-227 | chr11-85211169-INS-227 |
| chr11-85219423-INS-64 | chr11-85219423-INS-64 |
| chr11-85219937-INS-208 | chr11-85219937-INS-209 |
| chr11-85225326-INS-58 | chr11-85225326-INS-54 |
| chr11-85227855-INS-207 | chr11-85227855-INS-206 |
| chr11-85230667-INS-140 | chr11-85230660-INS-150 |
| chr11-85245347-INS-223 | chr11-85245347-INS-224 |
| chr11-85255268-INS-204 | chr11-85255268-INS-204 |
| chr11-85289016-INS-327 | chr11-85289016-INS-332 |
| chr11-85311380-INS-207 | chr11-85311380-INS-207 |
| chr11-85317291-INS-735 | chr11-85317297-INS-731 |
| chr11-85345847-INS-95 | chr11-85345847-INS-95 |
| chr11-85349987-INS-607 | chr11-85349969-INS-615 |
| chr11-85394421-INS-204 | chr11-85394422-INS-203 |
| chr11-85409871-INS-90 | chr11-85409871-INS-90 |
| chr11-85470011-INS-177 | chr11-85470011-INS-177 |
| chr11-85484035-INS-72 | chr11-85484035-INS-72 |
| chr11-85490915-INS-124 | chr11-85490915-INS-124 |
| chr11-85497349-INS-188 | chr11-85497349-INS-188 |
| chr11-85509766-INS-244 | chr11-85509766-INS-196 |

| | |
|---|---|
| chr11-85539473-INS-277 | chr11-85539503-INS-241 |
| chr11-85542254-INS-71 | chr11-85542254-INS-71 |
| chr11-85595809-INS-108 | chr11-85595809-INS-72 |
| chr11-85693464-INS-282 | chr11-85693472-INS-288 |
| chr11-85719255-INS-115 | chr11-85719255-INS-123 |
| chr11-85740325-INS-100 | chr11-85740325-INS-100 |
| chr11-85773182-INS-216 | chr11-85773182-INS-213 |
| chr11-85790407-INS-66 | chr11-85790407-INS-71 |
| chr11-85816769-INS-162 | chr11-85816769-INS-158 |
| chr11-85853373-INS-240 | chr11-85853373-INS-230 |
| chr11-85910573-INS-340 | chr11-85910573-INS-340 |
| chr11-85934208-INS-142 | chr11-85934208-INS-113 |
| chr11-85958115-INS-88 | chr11-85958115-INS-88 |
| chr11-85967534-INS-90 | chr11-85967534-INS-114 |
| chr11-85977398-INS-409 | chr11-85977398-INS-422 |
| chr11-85988225-INS-79 | chr11-85988225-INS-79 |
| chr11-86011095-INS-64 | chr11-86011049-INS-80 |
| chr11-86027594-INS-114 | chr11-86027594-INS-114 |
| chr11-86033146-INS-80 | chr11-86033147-INS-59 |
| chr11-86040458-INS-89 | chr11-86040458-INS-89 |
| chr11-86085073-INS-201 | chr11-86085073-INS-202 |
| chr11-86149489-INS-177 | chr11-86149489-INS-179 |
| chr11-86187692-INS-216 | chr11-86187692-INS-216 |
| chr11-86191420-INS-56 | chr11-86191420-INS-56 |
| chr11-86249548-INS-224 | chr11-86249548-INS-224 |
| chr11-86254221-INS-52 | chr11-86254221-INS-52 |
| chr11-86260385-INS-68 | chr11-86260385-INS-68 |
| chr11-86284255-INS-200 | chr11-86284255-INS-200 |
| chr11-86289067-INS-208 | chr11-86289067-INS-208 |
| chr11-86299934-INS-62 | chr11-86299934-INS-62 |
| chr11-86307231-INS-68 | chr11-86307229-INS-72 |
| chr11-86340513-INS-87 | chr11-86340505-INS-74 |
| chr11-86358506-INS-6442 | chr11-86358507-INS-6442 |
| chr11-86370174-INS-213 | chr11-86370175-INS-213 |
| chr11-86372050-INS-92 | chr11-86372050-INS-92 |
| chr11-86372779-INS-4816 | chr11-86372780-INS-4816 |
| chr11-86389501-INS-202 | chr11-86389501-INS-202 |
| chr11-86397935-INS-213 | chr11-86397935-INS-213 |
| chr11-86444535-INS-58 | chr11-86444535-INS-58 |
| chr11-86458448-INS-216 | chr11-86458448-INS-216 |
| chr11-86467011-INS-210 | chr11-86467011-INS-210 |
| chr11-86471948-INS-202 | chr11-86471948-INS-195 |
| chr11-86488837-INS-201 | chr11-86488837-INS-201 |
| chr11-86498356-INS-200 | chr11-86498356-INS-200 |
| chr11-86508290-INS-52 | chr11-86508290-INS-52 |
| chr11-86512133-INS-78 | chr11-86512133-INS-78 |
| chr11-86539535-INS-208 | chr11-86539535-INS-208 |
| chr11-86557625-INS-196 | chr11-86557625-INS-196 |
| chr11-86562558-INS-196 | chr11-86562558-INS-196 |
| chr11-86575718-INS-411 | chr11-86575718-INS-402 |
| chr11-86612637-INS-536 | chr11-86612637-INS-536 |
| chr11-86657848-INS-90 | chr11-86657848-INS-90 |
| chr11-86682213-INS-239 | chr11-86682213-INS-241 |
| chr11-86689560-INS-208 | chr11-86689560-INS-208 |
| chr11-86711396-INS-194 | chr11-86711396-INS-194 |
| chr11-86713092-INS-75 | chr11-86713092-INS-81 |
| chr11-86731240-INS-52 | chr11-86731240-INS-54 |
| chr11-86735043-INS-54 | chr11-86735043-INS-54 |
| chr11-86751814-INS-179 | chr11-86751814-INS-167 |
| chr11-86754428-INS-60 | chr11-86754428-INS-60 |
| chr11-86791932-INS-178 | chr11-86791932-INS-178 |
| chr11-86803123-INS-54 | chr11-86803118-INS-55 |
| chr11-86832179-INS-66 | chr11-86832177-INS-54 |
| chr11-86866050-INS-140 | chr11-86866050-INS-140 |
| chr11-86875441-INS-73 | chr11-86875441-INS-76 |
| chr11-86907587-INS-439 | chr11-86907587-INS-439 |
| chr11-86919494-INS-183 | chr11-86919494-INS-183 |
| chr11-86921294-INS-52 | chr11-86921294-INS-52 |
| chr11-86931910-INS-94 | chr11-86931910-INS-94 |
| chr11-86937035-INS-258 | chr11-86937035-INS-258 |
| chr11-86945819-INS-63 | chr11-86945819-INS-63 |
| chr11-86953743-INS-286 | chr11-86953743-INS-286 |
| chr11-86974684-INS-173 | chr11-86974684-INS-188 |
| chr11-86980703-INS-242 | chr11-86980721-INS-241 |
| chr11-86988211-INS-176 | chr11-86988211-INS-178 |
| chr11-87008150-INS-77 | chr11-87008150-INS-77 |
| chr11-87009894-INS-190 | chr11-87009894-INS-190 |

| | |
|---|---|
| chr11-87021176-INS-64 | chr11-87021176-INS-62 |
| chr11-87050869-INS-100 | chr11-87050873-INS-92 |
| chr11-87063459-INS-320 | chr11-87063459-INS-320 |
| chr11-87065644-INS-890 | chr11-87065644-INS-890 |
| chr11-87152447-INS-56 | chr11-87152447-INS-56 |
| chr11-87184220-INS-76 | chr11-87184220-INS-76 |
| chr11-87242064-INS-210 | chr11-87242064-INS-211 |
| chr11-87253473-INS-109 | chr11-87253466-INS-111 |
| chr11-87262220-INS-207 | chr11-87262220-INS-204 |
| chr11-87264003-INS-68 | chr11-87264003-INS-51 |
| chr11-87281238-INS-208 | chr11-87281238-INS-208 |
| chr11-87287120-INS-209 | chr11-87287120-INS-209 |
| chr11-87326372-INS-69 | chr11-87326372-INS-67 |
| chr11-87335579-INS-565 | chr11-87335579-INS-565 |
| chr11-87348159-INS-207 | chr11-87348159-INS-207 |
| chr11-87354572-INS-77 | chr11-87354572-INS-83 |
| chr11-87366970-INS-213 | chr11-87366970-INS-212 |
| chr11-87407473-INS-469 | chr11-87407473-INS-469 |
| chr11-87428284-INS-201 | chr11-87428284-INS-201 |
| chr11-87429494-INS-1264 | chr11-87429495-INS-1265 |
| chr11-87578154-INS-5547 | chr11-87578155-INS-5548 |
| chr11-87597915-INS-72 | chr11-87597914-INS-88 |
| chr11-87604079-INS-56 | chr11-87604076-INS-60 |
| chr11-87640598-INS-61 | chr11-87640598-INS-55 |
| chr11-87649799-INS-178 | chr11-87649799-INS-178 |
| chr11-87683355-INS-50 | chr11-87683355-INS-64 |
| chr11-87689124-INS-65 | chr11-87689132-INS-91 |
| chr11-87691642-INS-70 | chr11-87691642-INS-52 |
| chr11-87693069-INS-183 | chr11-87693069-INS-183 |
| chr11-87696646-INS-76 | chr11-87696646-INS-76 |
| chr11-87718089-INS-150 | chr11-87718089-INS-100 |
| chr11-87766719-INS-204 | chr11-87766719-INS-204 |
| chr11-87790214-INS-54 | chr11-87790178-INS-72 |
| chr11-87799095-INS-246 | chr11-87799090-INS-226 |
| chr11-87812633-INS-4825 | chr11-87812634-INS-4826 |
| chr11-87815804-INS-183 | chr11-87815804-INS-183 |
| chr11-87816518-INS-538 | chr11-87816518-INS-539 |
| chr11-87819162-INS-512 | chr11-87819162-INS-520 |
| chr11-87824711-INS-82 | chr11-87824712-INS-82 |
| chr11-87826605-INS-4445 | chr11-87826606-INS-4445 |
| chr11-87833578-INS-346 | chr11-87833578-INS-346 |
| chr11-87842593-INS-80 | chr11-87842593-INS-80 |
| chr11-87843989-INS-111 | chr11-87843989-INS-111 |
| chr11-87846031-INS-209 | chr11-87846031-INS-210 |
| chr11-87848383-INS-215 | chr11-87848383-INS-216 |
| chr11-87855908-INS-53 | chr11-87855908-INS-53 |
| chr11-87856964-INS-218 | chr11-87856964-INS-218 |
| chr11-87894227-INS-62 | chr11-87894227-INS-62 |
| chr11-87915486-INS-88 | chr11-87915498-INS-68 |
| chr11-87926374-INS-210 | chr11-87926374-INS-210 |
| chr11-87932838-INS-1151 | chr11-87932839-INS-1151 |
| chr11-87935051-INS-186 | chr11-87935051-INS-186 |
| chr11-87936152-INS-207 | chr11-87936152-INS-207 |
| chr11-87947035-INS-197 | chr11-87947035-INS-196 |
| chr11-87952258-INS-102 | chr11-87952258-INS-102 |
| chr11-87953804-INS-187 | chr11-87953804-INS-187 |
| chr11-87956778-INS-74 | chr11-87956778-INS-75 |
| chr11-87984326-INS-219 | chr11-87984326-INS-216 |
| chr11-87987611-INS-70 | chr11-87987611-INS-70 |
| chr11-88034237-INS-131 | chr11-88034237-INS-131 |
| chr11-88052391-INS-62 | chr11-88052391-INS-54 |
| chr11-88076448-INS-188 | chr11-88076457-INS-198 |
| chr11-88101140-INS-56 | chr11-88101140-INS-58 |
| chr11-88129476-INS-421 | chr11-88129476-INS-421 |
| chr11-88135020-INS-792 | chr11-88135020-INS-792 |
| chr11-88137467-INS-51 | chr11-88137468-INS-51 |
| chr11-88138676-INS-61 | chr11-88138676-INS-64 |
| chr11-88154555-INS-52 | chr11-88154552-INS-98 |
| chr11-88194131-INS-271 | chr11-88194131-INS-270 |
| chr11-88219216-INS-68 | chr11-88219216-INS-65 |
| chr11-88277763-INS-83 | chr11-88277763-INS-83 |
| chr11-88288167-INS-196 | chr11-88288167-INS-195 |
| chr11-88312568-INS-68 | chr11-88312568-INS-70 |
| chr11-88323758-INS-200 | chr11-88323752-INS-195 |
| chr11-88339776-INS-85 | chr11-88339776-INS-85 |
| chr11-88350364-INS-204 | chr11-88350364-INS-204 |
| chr11-88382403-INS-90 | chr11-88382393-INS-86 |

| | |
|---|---|
| chr11-88386167-INS-72 | chr11-88386162-INS-64 |
| chr11-88390477-INS-84 | chr11-88390477-INS-82 |
| chr11-88398670-INS-176 | chr11-88398670-INS-176 |
| chr11-88471649-INS-247 | chr11-88471649-INS-248 |
| chr11-88554108-INS-222 | chr11-88554108-INS-222 |
| chr11-88579397-INS-64 | chr11-88579397-INS-55 |
| chr11-88615582-INS-593 | chr11-88615582-INS-593 |
| chr11-88621612-INS-198 | chr11-88621612-INS-198 |
| chr11-88633658-INS-504 | chr11-88633658-INS-505 |
| chr11-88638693-INS-205 | chr11-88638693-INS-205 |
| chr11-88651278-INS-499 | chr11-88651278-INS-499 |
| chr11-88667372-INS-186 | chr11-88667372-INS-185 |
| chr11-88669936-INS-56 | chr11-88669936-INS-56 |
| chr11-88675801-INS-826 | chr11-88675801-INS-820 |
| chr11-88686285-INS-54 | chr11-88686285-INS-54 |
| chr11-88691728-INS-88 | chr11-88691728-INS-88 |
| chr11-88703323-INS-262 | chr11-88703323-INS-262 |
| chr11-88709137-INS-224 | chr11-88709137-INS-223 |
| chr11-88715426-INS-179 | chr11-88715426-INS-180 |
| chr11-88741203-INS-50 | chr11-88741201-INS-52 |
| chr11-88767472-INS-161 | chr11-88767472-INS-161 |
| chr11-88773069-INS-52 | chr11-88773069-INS-54 |
| chr11-88777477-INS-177 | chr11-88777477-INS-177 |
| chr11-88789903-INS-189 | chr11-88789903-INS-189 |
| chr11-88792840-INS-229 | chr11-88792840-INS-218 |
| chr11-88794729-INS-959 | chr11-88794729-INS-947 |
| chr11-88796056-INS-58 | chr11-88796056-INS-60 |
| chr11-88800039-INS-67 | chr11-88800039-INS-67 |
| chr11-88812900-INS-201 | chr11-88812900-INS-201 |
| chr11-88826798-INS-201 | chr11-88826798-INS-203 |
| chr11-88842351-INS-166 | chr11-88842351-INS-166 |
| chr11-88843548-INS-165 | chr11-88843548-INS-165 |
| chr11-88846756-INS-199 | chr11-88846756-INS-199 |
| chr11-88848368-INS-106 | chr11-88848368-INS-106 |
| chr11-88851124-INS-211 | chr11-88851124-INS-211 |
| chr11-88858694-INS-379 | chr11-88858694-INS-379 |
| chr11-88871120-INS-189 | chr11-88871120-INS-189 |
| chr11-88880443-INS-420 | chr11-88880443-INS-420 |
| chr11-88886072-INS-213 | chr11-88886072-INS-213 |
| chr11-88889467-INS-88 | chr11-88889467-INS-88 |
| chr11-88898854-INS-225 | chr11-88898854-INS-225 |
| chr11-88912620-INS-223 | chr11-88912620-INS-223 |
| chr11-88917267-INS-214 | chr11-88917267-INS-224 |
| chr11-88917818-INS-58 | chr11-88917818-INS-54 |
| chr11-88918868-INS-209 | chr11-88918868-INS-209 |
| chr11-88925900-INS-201 | chr11-88925900-INS-202 |
| chr11-88930820-INS-176 | chr11-88930820-INS-159 |
| chr11-88934251-INS-65 | chr11-88934251-INS-65 |
| chr11-88935566-INS-620 | chr11-88935566-INS-625 |
| chr11-88939238-INS-171 | chr11-88939237-INS-172 |
| chr11-88944176-INS-386 | chr11-88944176-INS-395 |
| chr11-88949897-INS-222 | chr11-88949897-INS-211 |
| chr11-88965498-INS-56 | chr11-88965498-INS-56 |
| chr11-88987376-INS-7139 | chr11-88987377-INS-7139 |
| chr11-88997084-INS-202 | chr11-88997084-INS-205 |
| chr11-89030240-INS-88 | chr11-89030240-INS-92 |
| chr11-89037607-INS-207 | chr11-89037604-INS-211 |
| chr11-89043291-INS-62 | chr11-89043291-INS-62 |
| chr11-89055734-INS-457 | chr11-89055734-INS-457 |
| chr11-89111992-INS-57 | chr11-89111992-INS-57 |
| chr11-89122553-INS-180 | chr11-89122553-INS-188 |
| chr11-89149214-INS-75 | chr11-89149214-INS-75 |
| chr11-89159210-INS-81 | chr11-89159210-INS-81 |
| chr11-89188333-INS-78 | chr11-89188333-INS-72 |
| chr11-89204737-INS-199 | chr11-89204737-INS-199 |
| chr11-89267168-INS-205 | chr11-89267168-INS-205 |
| chr11-89268386-INS-426 | chr11-89268386-INS-423 |
| chr11-89268959-INS-58 | chr11-89268957-INS-58 |
| chr11-89288982-INS-7415 | chr11-89288983-INS-7415 |
| chr11-89294923-INS-425 | chr11-89294923-INS-434 |
| chr11-89294923-INS-949 | chr11-89294923-INS-902 |
| chr11-89352676-INS-233 | chr11-89352674-INS-239 |
| chr11-89355146-INS-147 | chr11-89355158-INS-153 |
| chr11-89434408-INS-51 | chr11-89434408-INS-51 |
| chr11-89442806-INS-149 | chr11-89442806-INS-149 |
| chr11-89452033-INS-57 | chr11-89452033-INS-87 |
| chr11-89454853-INS-64 | chr11-89454850-INS-67 |

| | |
|---|---|
| chr11-89461779-INS-51 | chr11-89461779-INS-56 |
| chr11-89466806-INS-61 | chr11-89466802-INS-64 |
| chr11-89469666-INS-52 | chr11-89469666-INS-50 |
| chr11-89481182-INS-54 | chr11-89481182-INS-58 |
| chr11-89547213-INS-57 | chr11-89547213-INS-56 |
| chr11-89551680-INS-54 | chr11-89551680-INS-54 |
| chr11-89569083-INS-82 | chr11-89569083-INS-80 |
| chr11-89577323-INS-56 | chr11-89577323-INS-56 |
| chr11-89580553-INS-100 | chr11-89580555-INS-98 |
| chr11-89592459-INS-172 | chr11-89592460-INS-172 |
| chr11-89601461-INS-301 | chr11-89601471-INS-307 |
| chr11-89624754-INS-74 | chr11-89624755-INS-50 |
| chr11-89674318-INS-1933 | chr11-89674319-INS-1933 |
| chr11-89675937-INS-62 | chr11-89675937-INS-64 |
| chr11-89688874-INS-146 | chr11-89688877-INS-147 |
| chr11-89694362-INS-89 | chr11-89694362-INS-72 |
| chr11-89697186-INS-56 | chr11-89697186-INS-56 |
| chr11-89698965-INS-400 | chr11-89698965-INS-400 |
| chr11-89737512-INS-816 | chr11-89737512-INS-816 |
| chr11-89748334-INS-907 | chr11-89748334-INS-907 |
| chr11-89752169-INS-55 | chr11-89752169-INS-55 |
| chr11-89758406-INS-52 | chr11-89758404-INS-54 |
| chr11-89777936-INS-249 | chr11-89777936-INS-246 |
| chr11-89784209-INS-52 | chr11-89784209-INS-56 |
| chr11-89793386-INS-767 | chr11-89793386-INS-767 |
| chr11-89803242-INS-76 | chr11-89803242-INS-51 |
| chr11-89818356-INS-300 | chr11-89818400-INS-296 |
| chr11-89821669-INS-163 | chr11-89821669-INS-163 |
| chr11-89842247-INS-205 | chr11-89842253-INS-206 |
| chr11-89849208-INS-975 | chr11-89849208-INS-975 |
| chr11-89850113-INS-73 | chr11-89850113-INS-73 |
| chr11-89879861-INS-365 | chr11-89879861-INS-365 |
| chr11-89883392-INS-522 | chr11-89883392-INS-523 |
| chr11-89890676-INS-107 | chr11-89890676-INS-117 |
| chr11-89894138-INS-100 | chr11-89894138-INS-100 |
| chr11-89903788-INS-70 | chr11-89903788-INS-65 |
| chr11-89929619-INS-84 | chr11-89929619-INS-84 |
| chr11-89954532-INS-725 | chr11-89954532-INS-725 |
| chr11-89970452-INS-126 | chr11-89970453-INS-126 |
| chr11-90018430-INS-75 | chr11-90018430-INS-70 |
| chr11-90039635-INS-127 | chr11-90039635-INS-131 |
| chr11-90061304-INS-53 | chr11-90061324-INS-69 |
| chr11-90071529-INS-76 | chr11-90071529-INS-74 |
| chr11-90074577-INS-150 | chr11-90074577-INS-171 |
| chr11-90078363-INS-102 | chr11-90078349-INS-70 |
| chr11-90087238-INS-204 | chr11-90087238-INS-204 |
| chr11-90097823-INS-55 | chr11-90097823-INS-52 |
| chr11-90114881-INS-202 | chr11-90114881-INS-202 |
| chr11-90127097-INS-57 | chr11-90127097-INS-57 |
| chr11-90132323-INS-224 | chr11-90132323-INS-225 |
| chr11-90177206-INS-66 | chr11-90177206-INS-66 |
| chr11-90186265-INS-170 | chr11-90186266-INS-177 |
| chr11-90205365-INS-634 | chr11-90205365-INS-635 |
| chr11-90225874-INS-81 | chr11-90225874-INS-71 |
| chr11-90254123-INS-60 | chr11-90254123-INS-60 |
| chr11-90265854-INS-102 | chr11-90265854-INS-106 |
| chr11-90292454-INS-338 | chr11-90292454-INS-338 |
| chr11-90294860-INS-73 | chr11-90294863-INS-63 |
| chr11-90311124-INS-670 | chr11-90311124-INS-670 |
| chr11-90311650-INS-79 | chr11-90311650-INS-79 |
| chr11-90316583-INS-105 | chr11-90316583-INS-105 |
| chr11-90337222-INS-251 | chr11-90337222-INS-251 |
| chr11-90361551-INS-144 | chr11-90361570-INS-142 |
| chr11-90374302-INS-60 | chr11-90374302-INS-60 |
| chr11-90382856-INS-64 | chr11-90382856-INS-69 |
| chr11-90433802-INS-6996 | chr11-90433803-INS-6996 |
| chr11-90507180-INS-52 | chr11-90507180-INS-56 |
| chr11-90517958-INS-359 | chr11-90517959-INS-359 |
| chr11-90519394-INS-1348 | chr11-90519395-INS-1348 |
| chr11-90572036-INS-56 | chr11-90572036-INS-54 |
| chr11-90588924-INS-364 | chr11-90588924-INS-363 |
| chr11-90594460-INS-102 | chr11-90594460-INS-73 |
| chr11-90598552-INS-6790 | chr11-90598553-INS-6790 |
| chr11-90602078-INS-53 | chr11-90602078-INS-53 |
| chr11-90609435-INS-257 | chr11-90609435-INS-257 |
| chr11-90611535-INS-4843 | chr11-90611536-INS-4844 |
| chr11-90656588-INS-6588 | chr11-90656589-INS-6589 |

| | |
|---|---|
| chr11-90658472-INS-424 | chr11-90658473-INS-424 |
| chr11-90709934-INS-352 | chr11-90709934-INS-344 |
| chr11-90727627-INS-257 | chr11-90727627-INS-252 |
| chr11-90733274-INS-362 | chr11-90733274-INS-362 |
| chr11-90741923-INS-74 | chr11-90741920-INS-98 |
| chr11-90745016-INS-183 | chr11-90745016-INS-167 |
| chr11-90770789-INS-64 | chr11-90770789-INS-64 |
| chr11-90809304-INS-95 | chr11-90809304-INS-95 |
| chr11-90811779-INS-284 | chr11-90811779-INS-283 |
| chr11-90824686-INS-66 | chr11-90824686-INS-98 |
| chr11-90830288-INS-70 | chr11-90830288-INS-72 |
| chr11-90841507-INS-52 | chr11-90841507-INS-54 |
| chr11-90867326-INS-146 | chr11-90867326-INS-151 |
| chr11-90883497-INS-63 | chr11-90883484-INS-72 |
| chr11-90884821-INS-81 | chr11-90884810-INS-81 |
| chr11-90921447-INS-80 | chr11-90921453-INS-80 |
| chr11-90921616-INS-1135 | chr11-90921617-INS-1147 |
| chr11-90928966-INS-106 | chr11-90928966-INS-106 |
| chr11-90929960-INS-204 | chr11-90929960-INS-208 |
| chr11-90943066-INS-60 | chr11-90943067-INS-60 |
| chr11-90949156-INS-92 | chr11-90949158-INS-116 |
| chr11-90980711-INS-110 | chr11-90980711-INS-92 |
| chr11-91053621-INS-90 | chr11-91053621-INS-90 |
| chr11-91054483-INS-240 | chr11-91054483-INS-240 |
| chr11-91059207-INS-50 | chr11-91059207-INS-50 |
| chr11-91090397-INS-56 | chr11-91090397-INS-54 |
| chr11-91091336-INS-420 | chr11-91091336-INS-420 |
| chr11-91118437-INS-407 | chr11-91118437-INS-407 |
| chr11-91136141-INS-1351 | chr11-91136142-INS-1351 |
| chr11-91157572-INS-184 | chr11-91157572-INS-184 |
| chr11-91190851-INS-88 | chr11-91190820-INS-128 |
| chr11-91194791-INS-724 | chr11-91194791-INS-717 |
| chr11-91211857-INS-203 | chr11-91211857-INS-203 |
| chr11-91214468-INS-112 | chr11-91214468-INS-112 |
| chr11-91230922-INS-232 | chr11-91230922-INS-233 |
| chr11-91264349-INS-56 | chr11-91264350-INS-57 |
| chr11-91265243-INS-271 | chr11-91265243-INS-271 |
| chr11-91272333-INS-58 | chr11-91272333-INS-62 |
| chr11-91280370-INS-62 | chr11-91280370-INS-62 |
| chr11-91299993-INS-220 | chr11-91299993-INS-235 |
| chr11-91330214-INS-69 | chr11-91330214-INS-72 |
| chr11-91348894-INS-61 | chr11-91348894-INS-61 |
| chr11-91351803-INS-82 | chr11-91351803-INS-86 |
| chr11-91416401-INS-598 | chr11-91416401-INS-598 |
| chr11-91422972-INS-84 | chr11-91422972-INS-66 |
| chr11-91473991-INS-606 | chr11-91473991-INS-604 |
| chr11-91474623-INS-95 | chr11-91474623-INS-95 |
| chr11-91491601-INS-116 | chr11-91491601-INS-97 |
| chr11-91506960-INS-199 | chr11-91506960-INS-199 |
| chr11-91536093-INS-157 | chr11-91536093-INS-156 |
| chr11-91570279-INS-737 | chr11-91570279-INS-737 |
| chr11-91572900-INS-131 | chr11-91572900-INS-135 |
| chr11-91587420-INS-52 | chr11-91587420-INS-50 |
| chr11-91597034-INS-58 | chr11-91597034-INS-58 |
| chr11-91600585-INS-6523 | chr11-91600586-INS-6523 |
| chr11-91619953-INS-150 | chr11-91619974-INS-172 |
| chr11-91643157-INS-53 | chr11-91643157-INS-76 |
| chr11-91645377-INS-113 | chr11-91645377-INS-109 |
| chr11-91672759-INS-52 | chr11-91672782-INS-66 |
| chr11-91680813-INS-72 | chr11-91680813-INS-60 |
| chr11-91685636-INS-70 | chr11-91685636-INS-86 |
| chr11-91693085-INS-56 | chr11-91693085-INS-56 |
| chr11-91693813-INS-64 | chr11-91693813-INS-50 |
| chr11-91703542-INS-341 | chr11-91703544-INS-338 |
| chr11-91711711-INS-215 | chr11-91711711-INS-215 |
| chr11-91734190-INS-56 | chr11-91734190-INS-54 |
| chr11-91742044-INS-439 | chr11-91742044-INS-439 |
| chr11-91745532-INS-965 | chr11-91745532-INS-965 |
| chr11-91747146-INS-1892 | chr11-91747147-INS-1892 |
| chr11-91754509-INS-52 | chr11-91754463-INS-66 |
| chr11-91755879-INS-54 | chr11-91755879-INS-80 |
| chr11-91762432-INS-62 | chr11-91762432-INS-66 |
| chr11-91766694-INS-497 | chr11-91766694-INS-497 |
| chr11-91768424-INS-88 | chr11-91768424-INS-90 |
| chr11-91785787-INS-64 | chr11-91785782-INS-80 |
| chr11-91797244-INS-58 | chr11-91797245-INS-59 |
| chr11-91860196-INS-175 | chr11-91860196-INS-175 |

| | |
|---|---|
| chr11-91887294-INS-66 | chr11-91887317-INS-64 |
| chr11-91906383-INS-1687 | chr11-91906384-INS-1687 |
| chr11-91907382-INS-58 | chr11-91907382-INS-76 |
| chr11-92005949-INS-215 | chr11-92005950-INS-207 |
| chr11-92023715-INS-5363 | chr11-92023716-INS-5363 |
| chr11-92080704-INS-2027 | chr11-92080705-INS-2027 |
| chr11-92101042-INS-54 | chr11-92101018-INS-76 |
| chr11-92130118-INS-98 | chr11-92130118-INS-115 |
| chr11-92169062-INS-86 | chr11-92169062-INS-86 |
| chr11-92170356-INS-60 | chr11-92170356-INS-60 |
| chr11-92189320-INS-6171 | chr11-92189321-INS-6171 |
| chr11-92193141-INS-64 | chr11-92193141-INS-52 |
| chr11-92205610-INS-562 | chr11-92205610-INS-546 |
| chr11-92220899-INS-71 | chr11-92220897-INS-80 |
| chr11-92270795-INS-352 | chr11-92270795-INS-352 |
| chr11-92297004-INS-119 | chr11-92297008-INS-87 |
| chr11-92327032-INS-72 | chr11-92327032-INS-72 |
| chr11-92350154-INS-400 | chr11-92350154-INS-400 |
| chr11-92365979-INS-116 | chr11-92365979-INS-118 |
| chr11-92391814-INS-149 | chr11-92391814-INS-150 |
| chr11-92394047-INS-3889 | chr11-92394048-INS-3889 |
| chr11-92403554-INS-108 | chr11-92403554-INS-105 |
| chr11-92411695-INS-937 | chr11-92411695-INS-937 |
| chr11-92422984-INS-92 | chr11-92422996-INS-84 |
| chr11-92441100-INS-147 | chr11-92441100-INS-147 |
| chr11-92448449-INS-107 | chr11-92448449-INS-107 |
| chr11-92474071-INS-96 | chr11-92474071-INS-96 |
| chr11-92515723-INS-52 | chr11-92515723-INS-50 |
| chr11-92520723-INS-130 | chr11-92520723-INS-130 |
| chr11-92526522-INS-244 | chr11-92526522-INS-244 |
| chr11-92533384-INS-471 | chr11-92533384-INS-471 |
| chr11-92535156-INS-1488 | chr11-92535157-INS-1490 |
| chr11-92539888-INS-7143 | chr11-92539889-INS-7143 |
| chr11-92547255-INS-79 | chr11-92547255-INS-77 |
| chr11-92557404-INS-574 | chr11-92557404-INS-576 |
| chr11-92563731-INS-164 | chr11-92563731-INS-164 |
| chr11-92564802-INS-7115 | chr11-92564803-INS-7115 |
| chr11-92576692-INS-205 | chr11-92576692-INS-205 |
| chr11-92577681-INS-81 | chr11-92577681-INS-81 |
| chr11-92580509-INS-118 | chr11-92580505-INS-112 |
| chr11-92598908-INS-73 | chr11-92598908-INS-78 |
| chr11-92620635-INS-52 | chr11-92620635-INS-54 |
| chr11-92641061-INS-645 | chr11-92641061-INS-645 |
| chr11-92673248-INS-3059 | chr11-92673249-INS-3059 |
| chr11-92697803-INS-131 | chr11-92697803-INS-115 |
| chr11-92710767-INS-325 | chr11-92710767-INS-325 |
| chr11-92724874-INS-6453 | chr11-92724875-INS-6453 |
| chr11-92725466-INS-56 | chr11-92725466-INS-56 |
| chr11-92736146-INS-52 | chr11-92736146-INS-52 |
| chr11-92757209-INS-325 | chr11-92757209-INS-325 |
| chr11-92766789-INS-64 | chr11-92766789-INS-64 |
| chr11-92767372-INS-504 | chr11-92767372-INS-489 |
| chr11-92775088-INS-52 | chr11-92775053-INS-97 |
| chr11-92792559-INS-208 | chr11-92792559-INS-208 |
| chr11-92797671-INS-4363 | chr11-92797672-INS-4365 |
| chr11-92800992-INS-216 | chr11-92800984-INS-213 |
| chr11-92803280-INS-8457 | chr11-92803281-INS-8457 |
| chr11-92805085-INS-58 | chr11-92805085-INS-64 |
| chr11-92820308-INS-156 | chr11-92820308-INS-118 |
| chr11-92823111-INS-135 | chr11-92823111-INS-135 |
| chr11-92825462-INS-64 | chr11-92825462-INS-79 |
| chr11-92827950-INS-87 | chr11-92827961-INS-96 |
| chr11-92831474-INS-6909 | chr11-92831475-INS-6909 |
| chr11-92838689-INS-1672 | chr11-92838690-INS-1670 |
| chr11-92849359-INS-286 | chr11-92849359-INS-286 |
| chr11-92861320-INS-58 | chr11-92861320-INS-58 |
| chr11-92878430-INS-88 | chr11-92878441-INS-73 |
| chr11-92919897-INS-71 | chr11-92919897-INS-71 |
| chr11-92937463-INS-1770 | chr11-92937464-INS-1770 |
| chr11-92940016-INS-216 | chr11-92940016-INS-216 |
| chr11-92959945-INS-505 | chr11-92959945-INS-503 |
| chr11-92961357-INS-185 | chr11-92961357-INS-185 |
| chr11-93005391-INS-7141 | chr11-93005392-INS-7141 |
| chr11-93013216-INS-410 | chr11-93013216-INS-409 |
| chr11-93035472-INS-1011 | chr11-93035473-INS-1017 |
| chr11-93062890-INS-759 | chr11-93062890-INS-758 |
| chr11-93073783-INS-6357 | chr11-93073784-INS-6357 |

| | |
|---|---|
| chr11-93100279-INS-104 | chr11-93100276-INS-104 |
| chr11-93102467-INS-100 | chr11-93102467-INS-100 |
| chr11-93106355-INS-68 | chr11-93106355-INS-69 |
| chr11-93113654-INS-644 | chr11-93113654-INS-644 |
| chr11-93124465-INS-86 | chr11-93124465-INS-86 |
| chr11-93156252-INS-437 | chr11-93156252-INS-437 |
| chr11-93162046-INS-54 | chr11-93162069-INS-76 |
| chr11-93178683-INS-110 | chr11-93178681-INS-112 |
| chr11-93202247-INS-70 | chr11-93202247-INS-70 |
| chr11-93213972-INS-68 | chr11-93213968-INS-68 |
| chr11-93219134-INS-962 | chr11-93219134-INS-970 |
| chr11-93228162-INS-3120 | chr11-93228163-INS-3120 |
| chr11-93238753-INS-56 | chr11-93238759-INS-50 |
| chr11-93244969-INS-63 | chr11-93244969-INS-67 |
| chr11-93249497-INS-56 | chr11-93249498-INS-68 |
| chr11-93252540-INS-73 | chr11-93252540-INS-73 |
| chr11-93282479-INS-77 | chr11-93282479-INS-63 |
| chr11-93283282-INS-54 | chr11-93283284-INS-68 |
| chr11-93294176-INS-55 | chr11-93294176-INS-60 |
| chr11-93298050-INS-52 | chr11-93298050-INS-52 |
| chr11-93309810-INS-1075 | chr11-93309811-INS-1072 |
| chr11-93318130-INS-471 | chr11-93318130-INS-471 |
| chr11-93332498-INS-89 | chr11-93332498-INS-108 |
| chr11-93348748-INS-220 | chr11-93348748-INS-219 |
| chr11-93356193-INS-128 | chr11-93356186-INS-138 |
| chr11-93376078-INS-338 | chr11-93376078-INS-338 |
| chr11-93376900-INS-104 | chr11-93376900-INS-108 |
| chr11-93384072-INS-52 | chr11-93384073-INS-87 |
| chr11-93386937-INS-257 | chr11-93386937-INS-256 |
| chr11-93395167-INS-7081 | chr11-93395168-INS-7081 |
| chr11-93399671-INS-390 | chr11-93399681-INS-420 |
| chr11-93409550-INS-139 | chr11-93409550-INS-139 |
| chr11-93448088-INS-85 | chr11-93448091-INS-101 |
| chr11-93452379-INS-69 | chr11-93452344-INS-55 |
| chr11-93471482-INS-521 | chr11-93471482-INS-521 |
| chr11-93507252-INS-72 | chr11-93507252-INS-72 |
| chr11-93508420-INS-75 | chr11-93508420-INS-81 |
| chr11-93518622-INS-66 | chr11-93518622-INS-66 |
| chr11-93518622-INS-843 | chr11-93518622-INS-843 |
| chr11-93533129-INS-60 | chr11-93533129-INS-60 |
| chr11-93540903-INS-212 | chr11-93540903-INS-212 |
| chr11-93545478-INS-56 | chr11-93545478-INS-56 |
| chr11-93554320-INS-321 | chr11-93554320-INS-321 |
| chr11-93559633-INS-74 | chr11-93559633-INS-63 |
| chr11-93590928-INS-65 | chr11-93590929-INS-61 |
| chr11-93596643-INS-211 | chr11-93596629-INS-220 |
| chr11-93600166-INS-136 | chr11-93600166-INS-134 |
| chr11-93646907-INS-64 | chr11-93646907-INS-74 |
| chr11-93663858-INS-55 | chr11-93663852-INS-50 |
| chr11-93668741-INS-569 | chr11-93668741-INS-570 |
| chr11-93697650-INS-51 | chr11-93697650-INS-52 |
| chr11-93711551-INS-216 | chr11-93711549-INS-218 |
| chr11-93712129-INS-704 | chr11-93712129-INS-704 |
| chr11-93734172-INS-52 | chr11-93734172-INS-56 |
| chr11-93743378-INS-100 | chr11-93743378-INS-107 |
| chr11-93768619-INS-534 | chr11-93768619-INS-536 |
| chr11-93769584-INS-94 | chr11-93769578-INS-95 |
| chr11-93782875-INS-215 | chr11-93782875-INS-215 |
| chr11-93791884-INS-74 | chr11-93791884-INS-76 |
| chr11-93799370-INS-56 | chr11-93799370-INS-56 |
| chr11-93814439-INS-218 | chr11-93814439-INS-219 |
| chr11-93819482-INS-196 | chr11-93819482-INS-196 |
| chr11-93838914-INS-72 | chr11-93838914-INS-72 |
| chr11-93850600-INS-325 | chr11-93850600-INS-326 |
| chr11-93862528-INS-203 | chr11-93862528-INS-203 |
| chr11-93866687-INS-80 | chr11-93866687-INS-72 |
| chr11-93872361-INS-693 | chr11-93872361-INS-693 |
| chr11-93877626-INS-114 | chr11-93877626-INS-126 |
| chr11-93877739-INS-7164 | chr11-93877740-INS-7164 |
| chr11-93883721-INS-96 | chr11-93883721-INS-96 |
| chr11-93885196-INS-1343 | chr11-93885197-INS-1343 |
| chr11-93896601-INS-211 | chr11-93896601-INS-210 |
| chr11-93906823-INS-212 | chr11-93906823-INS-212 |
| chr11-93910470-INS-206 | chr11-93910470-INS-206 |
| chr11-93917935-INS-605 | chr11-93917935-INS-605 |
| chr11-93919637-INS-82 | chr11-93919637-INS-87 |
| chr11-93926078-INS-183 | chr11-93926078-INS-183 |

| | |
|---|---|
| chr11-93994574-INS-88 | chr11-93994574-INS-120 |
| chr11-94024038-INS-206 | chr11-94024038-INS-206 |
| chr11-94034379-INS-170 | chr11-94034379-INS-171 |
| chr11-94060216-INS-57 | chr11-94060217-INS-66 |
| chr11-94078273-INS-174 | chr11-94078273-INS-174 |
| chr11-94079467-INS-84 | chr11-94079478-INS-69 |
| chr11-94095607-INS-90 | chr11-94095607-INS-90 |
| chr11-94157023-INS-112 | chr11-94157019-INS-84 |
| chr11-94160461-INS-4663 | chr11-94160462-INS-4663 |
| chr11-94168757-INS-113 | chr11-94168757-INS-105 |
| chr11-94173833-INS-169 | chr11-94173833-INS-169 |
| chr11-94177123-INS-78 | chr11-94177123-INS-78 |
| chr11-94180656-INS-64 | chr11-94180656-INS-64 |
| chr11-94186651-INS-76 | chr11-94186651-INS-64 |
| chr11-94196485-INS-210 | chr11-94196485-INS-208 |
| chr11-94221872-INS-201 | chr11-94221872-INS-201 |
| chr11-94240465-INS-5202 | chr11-94240466-INS-5204 |
| chr11-94257965-INS-72 | chr11-94257965-INS-68 |
| chr11-94269841-INS-51 | chr11-94269841-INS-51 |
| chr11-94272570-INS-66 | chr11-94272570-INS-66 |
| chr11-94278549-INS-308 | chr11-94278549-INS-308 |
| chr11-94283017-INS-131 | chr11-94283017-INS-131 |
| chr11-94404208-INS-105 | chr11-94404202-INS-111 |
| chr11-94426958-INS-64 | chr11-94426958-INS-52 |
| chr11-94430679-INS-56 | chr11-94430679-INS-56 |
| chr11-94451089-INS-75 | chr11-94451089-INS-80 |
| chr11-94452803-INS-71 | chr11-94452803-INS-71 |
| chr11-94454245-INS-208 | chr11-94454245-INS-209 |
| chr11-94471661-INS-76 | chr11-94471661-INS-76 |
| chr11-94472756-INS-212 | chr11-94472756-INS-212 |
| chr11-94477236-INS-168 | chr11-94477236-INS-168 |
| chr11-94483351-INS-205 | chr11-94483310-INS-211 |
| chr11-94495692-INS-56 | chr11-94495692-INS-56 |
| chr11-94496932-INS-1258 | chr11-94496933-INS-1258 |
| chr11-94504019-INS-212 | chr11-94504019-INS-213 |
| chr11-94507651-INS-100 | chr11-94507655-INS-72 |
| chr11-94542985-INS-126 | chr11-94542985-INS-127 |
| chr11-94544225-INS-171 | chr11-94544225-INS-172 |
| chr11-94564537-INS-103 | chr11-94564537-INS-103 |
| chr11-94573735-INS-205 | chr11-94573735-INS-208 |
| chr11-94599597-INS-180 | chr11-94599600-INS-183 |
| chr11-94612033-INS-52 | chr11-94612033-INS-52 |
| chr11-94621580-INS-166 | chr11-94621580-INS-165 |
| chr11-94625985-INS-214 | chr11-94625985-INS-214 |
| chr11-94664893-INS-56 | chr11-94664893-INS-51 |
| chr11-94683396-INS-180 | chr11-94683399-INS-176 |
| chr11-94745647-INS-90 | chr11-94745647-INS-90 |
| chr11-94774716-INS-53 | chr11-94774716-INS-52 |
| chr11-94782971-INS-171 | chr11-94782971-INS-171 |
| chr11-94796629-INS-180 | chr11-94796629-INS-180 |
| chr11-94798353-INS-52 | chr11-94798353-INS-52 |
| chr11-94814467-INS-66 | chr11-94814467-INS-66 |
| chr11-94831217-INS-62 | chr11-94831217-INS-62 |
| chr11-94849890-INS-179 | chr11-94849890-INS-179 |
| chr11-94851627-INS-230 | chr11-94851627-INS-230 |
| chr11-94868261-INS-186 | chr11-94868261-INS-186 |
| chr11-94872351-INS-217 | chr11-94872351-INS-216 |
| chr11-94873424-INS-225 | chr11-94873425-INS-225 |
| chr11-94902173-INS-78 | chr11-94902173-INS-80 |
| chr11-94907817-INS-69 | chr11-94907820-INS-70 |
| chr11-94932751-INS-75 | chr11-94932751-INS-72 |
| chr11-94941734-INS-76 | chr11-94941734-INS-70 |
| chr11-94973189-INS-400 | chr11-94973189-INS-400 |
| chr11-94980841-INS-2984 | chr11-94980842-INS-2939 |
| chr11-94998210-INS-204 | chr11-94998210-INS-204 |
| chr11-95003403-INS-216 | chr11-95003403-INS-216 |
| chr11-95006965-INS-121 | chr11-95006965-INS-128 |
| chr11-95022949-INS-230 | chr11-95022949-INS-231 |
| chr11-95038053-INS-84 | chr11-95038053-INS-105 |
| chr11-95040842-INS-73 | chr11-95040842-INS-73 |
| chr11-95064825-INS-201 | chr11-95064825-INS-201 |
| chr11-95067675-INS-238 | chr11-95067675-INS-239 |
| chr11-95087000-INS-192 | chr11-95087000-INS-192 |
| chr11-95098026-INS-54 | chr11-95098048-INS-62 |
| chr11-95103319-INS-95 | chr11-95103319-INS-80 |
| chr11-95112591-INS-153 | chr11-95112591-INS-153 |
| chr11-95116846-INS-467 | chr11-95116846-INS-467 |

| | |
|---|---|
| chr11-95143945-INS-173 | chr11-95143945-INS-174 |
| chr11-95147365-INS-68 | chr11-95147365-INS-68 |
| chr11-95148778-INS-404 | chr11-95148778-INS-404 |
| chr11-95154200-INS-52 | chr11-95154200-INS-52 |
| chr11-95163837-INS-61 | chr11-95163837-INS-61 |
| chr11-95170313-INS-220 | chr11-95170313-INS-220 |
| chr11-95178496-INS-211 | chr11-95178496-INS-211 |
| chr11-95183817-INS-209 | chr11-95183817-INS-210 |
| chr11-95190238-INS-221 | chr11-95190238-INS-225 |
| chr11-95217705-INS-170 | chr11-95217705-INS-170 |
| chr11-95221854-INS-203 | chr11-95221854-INS-203 |
| chr11-95223804-INS-62 | chr11-95223816-INS-82 |
| chr11-95252829-INS-205 | chr11-95252829-INS-205 |
| chr11-95259070-INS-70 | chr11-95259070-INS-70 |
| chr11-95262022-INS-94 | chr11-95262022-INS-90 |
| chr11-95274590-INS-207 | chr11-95274590-INS-207 |
| chr11-95281046-INS-77 | chr11-95281046-INS-77 |
| chr11-95287219-INS-205 | chr11-95287219-INS-206 |
| chr11-95288274-INS-68 | chr11-95288274-INS-68 |
| chr11-95315671-INS-277 | chr11-95315671-INS-277 |
| chr11-95342454-INS-213 | chr11-95342454-INS-213 |
| chr11-95361292-INS-58 | chr11-95361292-INS-58 |
| chr11-95408512-INS-52 | chr11-95408512-INS-52 |
| chr11-95414707-INS-116 | chr11-95414707-INS-116 |
| chr11-95435710-INS-213 | chr11-95435706-INS-213 |
| chr11-95442097-INS-206 | chr11-95442097-INS-203 |
| chr11-95444516-INS-65 | chr11-95444516-INS-65 |
| chr11-95456979-INS-175 | chr11-95456979-INS-178 |
| chr11-95482421-INS-204 | chr11-95482421-INS-204 |
| chr11-95499304-INS-182 | chr11-95499304-INS-182 |
| chr11-95499874-INS-1472 | chr11-95499875-INS-1472 |
| chr11-95506861-INS-66 | chr11-95506861-INS-68 |
| chr11-95558665-INS-169 | chr11-95558665-INS-169 |
| chr11-95566251-INS-219 | chr11-95566251-INS-219 |
| chr11-95569713-INS-152 | chr11-95569713-INS-152 |
| chr11-95580907-INS-63 | chr11-95580928-INS-84 |
| chr11-95584773-INS-208 | chr11-95584773-INS-208 |
| chr11-95593501-INS-201 | chr11-95593501-INS-201 |
| chr11-95602601-INS-88 | chr11-95602601-INS-92 |
| chr11-95612955-INS-70 | chr11-95612955-INS-70 |
| chr11-95616475-INS-209 | chr11-95616475-INS-209 |
| chr11-95618437-INS-199 | chr11-95618437-INS-199 |
| chr11-95648377-INS-211 | chr11-95648377-INS-211 |
| chr11-95660269-INS-214 | chr11-95660269-INS-214 |
| chr11-95664662-INS-58 | chr11-95664662-INS-58 |
| chr11-95677900-INS-1385 | chr11-95677901-INS-1385 |
| chr11-95686205-INS-222 | chr11-95686205-INS-222 |
| chr11-95688470-INS-233 | chr11-95688470-INS-233 |
| chr11-95690195-INS-357 | chr11-95690195-INS-357 |
| chr11-95697501-INS-7365 | chr11-95697502-INS-7365 |
| chr11-95710838-INS-51 | chr11-95710837-INS-51 |
| chr11-95712335-INS-73 | chr11-95712335-INS-108 |
| chr11-95717149-INS-58 | chr11-95717149-INS-58 |
| chr11-95727668-INS-56 | chr11-95727666-INS-60 |
| chr11-95736095-INS-72 | chr11-95736095-INS-96 |
| chr11-95736908-INS-7262 | chr11-95736909-INS-7262 |
| chr11-95738490-INS-206 | chr11-95738490-INS-206 |
| chr11-95741152-INS-82 | chr11-95741152-INS-88 |
| chr11-95753483-INS-7258 | chr11-95753484-INS-7258 |
| chr11-95755509-INS-66 | chr11-95755509-INS-73 |
| chr11-95767701-INS-52 | chr11-95767701-INS-66 |
| chr11-95779080-INS-84 | chr11-95779080-INS-84 |
| chr11-95819741-INS-514 | chr11-95819741-INS-514 |
| chr11-95822185-INS-63 | chr11-95822186-INS-66 |
| chr11-95823044-INS-466 | chr11-95823044-INS-466 |
| chr11-95846857-INS-474 | chr11-95846857-INS-473 |
| chr11-95862518-INS-180 | chr11-95862511-INS-230 |
| chr11-95873656-INS-240 | chr11-95873656-INS-240 |
| chr11-95891917-INS-97 | chr11-95891917-INS-97 |
| chr11-95911105-INS-100 | chr11-95911101-INS-72 |
| chr11-95916464-INS-220 | chr11-95916464-INS-221 |
| chr11-95923474-INS-302 | chr11-95923474-INS-298 |
| chr11-95960534-INS-52 | chr11-95960534-INS-58 |
| chr11-95979420-INS-212 | chr11-95979420-INS-212 |
| chr11-95983281-INS-203 | chr11-95983281-INS-203 |
| chr11-95986925-INS-218 | chr11-95986925-INS-219 |
| chr11-95992948-INS-196 | chr11-95992948-INS-196 |

| | |
|---|---|
| chr11-95996741-INS-148 | chr11-95996739-INS-150 |
| chr11-96014347-INS-148 | chr11-96014347-INS-148 |
| chr11-96050513-INS-244 | chr11-96050513-INS-243 |
| chr11-96093729-INS-141 | chr11-96093729-INS-115 |
| chr11-96104361-INS-154 | chr11-96104361-INS-123 |
| chr11-96106831-INS-77 | chr11-96106831-INS-55 |
| chr11-96107708-INS-114 | chr11-96107708-INS-114 |
| chr11-96110614-INS-146 | chr11-96110614-INS-146 |
| chr11-96137441-INS-1731 | chr11-96137442-INS-1731 |
| chr11-96247081-INS-165 | chr11-96247081-INS-165 |
| chr11-96275079-INS-91 | chr11-96275079-INS-91 |
| chr11-96292371-INS-72 | chr11-96292377-INS-66 |
| chr11-96294355-INS-74 | chr11-96294387-INS-86 |
| chr11-96315376-INS-51 | chr11-96315376-INS-51 |
| chr11-96335232-INS-209 | chr11-96335232-INS-209 |
| chr11-96354489-INS-52 | chr11-96354489-INS-68 |
| chr11-96360496-INS-109 | chr11-96360454-INS-110 |
| chr11-96394747-INS-103 | chr11-96394736-INS-109 |
| chr11-96400311-INS-61 | chr11-96400311-INS-61 |
| chr11-96404667-INS-82 | chr11-96404667-INS-82 |
| chr11-96412941-INS-54 | chr11-96412912-INS-93 |
| chr11-96462131-INS-56 | chr11-96462131-INS-56 |
| chr11-96503242-INS-62 | chr11-96503242-INS-53 |
| chr11-96503931-INS-75 | chr11-96503932-INS-75 |
| chr11-96511252-INS-251 | chr11-96511252-INS-252 |
| chr11-96534535-INS-58 | chr11-96534536-INS-94 |
| chr11-96541427-INS-180 | chr11-96541434-INS-181 |
| chr11-96543306-INS-70 | chr11-96543306-INS-70 |
| chr11-96562699-INS-6399 | chr11-96562700-INS-6399 |
| chr11-96563895-INS-95 | chr11-96563895-INS-95 |
| chr11-96592862-INS-195 | chr11-96592862-INS-195 |
| chr11-96613636-INS-56 | chr11-96613636-INS-92 |
| chr11-96619412-INS-198 | chr11-96619413-INS-184 |
| chr11-96625606-INS-59 | chr11-96625606-INS-53 |
| chr11-96646760-INS-198 | chr11-96646760-INS-198 |
| chr11-96651117-INS-178 | chr11-96651117-INS-178 |
| chr11-96670600-INS-221 | chr11-96670600-INS-221 |
| chr11-96674306-INS-50 | chr11-96674277-INS-81 |
| chr11-96677633-INS-208 | chr11-96677633-INS-208 |
| chr11-96678311-INS-212 | chr11-96678311-INS-212 |
| chr11-96681888-INS-212 | chr11-96681888-INS-212 |
| chr11-96685962-INS-184 | chr11-96685962-INS-184 |
| chr11-96703192-INS-201 | chr11-96703192-INS-201 |
| chr11-96704717-INS-213 | chr11-96704717-INS-213 |
| chr11-96735151-INS-208 | chr11-96735151-INS-208 |
| chr11-96737527-INS-199 | chr11-96737527-INS-242 |
| chr11-96750811-INS-183 | chr11-96750811-INS-199 |
| chr11-96761859-INS-202 | chr11-96761859-INS-202 |
| chr11-96782536-INS-139 | chr11-96782536-INS-139 |
| chr11-96788931-INS-164 | chr11-96788931-INS-164 |
| chr11-96812731-INS-68 | chr11-96812731-INS-68 |
| chr11-96818867-INS-1888 | chr11-96818868-INS-1888 |
| chr11-96831538-INS-71 | chr11-96831538-INS-71 |
| chr11-96841221-INS-194 | chr11-96841221-INS-194 |
| chr11-96855411-INS-233 | chr11-96855411-INS-197 |
| chr11-96900363-INS-188 | chr11-96900363-INS-181 |
| chr11-96936973-INS-60 | chr11-96936973-INS-52 |
| chr11-96949426-INS-132 | chr11-96949426-INS-132 |
| chr11-96965353-INS-88 | chr11-96965353-INS-88 |
| chr11-96971216-INS-57 | chr11-96971216-INS-61 |
| chr11-96982322-INS-196 | chr11-96982322-INS-162 |
| chr11-96984883-INS-71 | chr11-96984879-INS-72 |
| chr11-96996330-INS-134 | chr11-96996330-INS-122 |
| chr11-97068290-INS-54 | chr11-97068290-INS-54 |
| chr11-97092760-INS-190 | chr11-97092760-INS-192 |
| chr11-97094663-INS-181 | chr11-97094663-INS-181 |
| chr11-97096626-INS-212 | chr11-97096626-INS-212 |
| chr11-97104963-INS-214 | chr11-97104963-INS-213 |
| chr11-97109143-INS-273 | chr11-97109143-INS-273 |
| chr11-97110103-INS-234 | chr11-97110085-INS-267 |
| chr11-97113050-INS-281 | chr11-97113050-INS-281 |
| chr11-97117000-INS-201 | chr11-97117000-INS-201 |
| chr11-97118194-INS-110 | chr11-97118194-INS-110 |
| chr11-97133250-INS-208 | chr11-97133250-INS-209 |
| chr11-97146481-INS-210 | chr11-97146481-INS-208 |
| chr11-97151561-INS-204 | chr11-97151561-INS-248 |
| chr11-97159791-INS-229 | chr11-97159791-INS-228 |

| | |
|---|---|
| chr11-97174056-INS-237 | chr11-97174056-INS-237 |
| chr11-97179672-INS-7114 | chr11-97179693-INS-7122 |
| chr11-97188394-INS-205 | chr11-97188394-INS-205 |
| chr11-97196260-INS-215 | chr11-97196260-INS-216 |
| chr11-97204675-INS-62 | chr11-97204661-INS-55 |
| chr11-97222305-INS-52 | chr11-97222305-INS-54 |
| chr11-97235809-INS-88 | chr11-97235809-INS-88 |
| chr11-97267022-INS-64 | chr11-97267022-INS-64 |
| chr11-97281515-INS-379 | chr11-97281516-INS-379 |
| chr11-97291138-INS-532 | chr11-97291156-INS-527 |
| chr11-97300170-INS-225 | chr11-97300170-INS-225 |
| chr11-97304642-INS-105 | chr11-97304642-INS-105 |
| chr11-97306181-INS-53 | chr11-97306181-INS-53 |
| chr11-97312993-INS-73 | chr11-97312993-INS-72 |
| chr11-97316089-INS-99 | chr11-97316108-INS-59 |
| chr11-97348419-INS-53 | chr11-97348420-INS-59 |
| chr11-97361013-INS-581 | chr11-97361013-INS-582 |
| chr11-97378814-INS-575 | chr11-97378814-INS-575 |
| chr11-97397744-INS-427 | chr11-97397745-INS-427 |
| chr11-97415715-INS-59 | chr11-97415715-INS-61 |
| chr11-97481720-INS-60 | chr11-97481720-INS-68 |
| chr11-97508317-INS-250 | chr11-97508326-INS-225 |
| chr11-97527112-INS-52 | chr11-97527112-INS-52 |
| chr11-97531762-INS-165 | chr11-97531762-INS-171 |
| chr11-97533692-INS-7098 | chr11-97533693-INS-7098 |
| chr11-97598737-INS-216 | chr11-97598737-INS-212 |
| chr11-97669343-INS-209 | chr11-97669343-INS-210 |
| chr11-97677196-INS-330 | chr11-97677196-INS-333 |
| chr11-97678024-INS-62 | chr11-97678024-INS-66 |
| chr11-97679130-INS-247 | chr11-97679130-INS-249 |
| chr11-97680479-INS-84 | chr11-97680479-INS-56 |
| chr11-97708818-INS-214 | chr11-97708818-INS-218 |
| chr11-97716823-INS-174 | chr11-97716823-INS-174 |
| chr11-97731274-INS-200 | chr11-97731274-INS-200 |
| chr11-97747268-INS-207 | chr11-97747268-INS-207 |
| chr11-97754390-INS-168 | chr11-97754390-INS-168 |
| chr11-97773544-INS-210 | chr11-97773544-INS-210 |
| chr11-97778035-INS-174 | chr11-97778035-INS-174 |
| chr11-97786169-INS-127 | chr11-97786169-INS-127 |
| chr11-97800519-INS-375 | chr11-97800519-INS-374 |
| chr11-97817132-INS-118 | chr11-97817132-INS-118 |
| chr11-97822597-INS-115 | chr11-97822597-INS-115 |
| chr11-97834016-INS-167 | chr11-97834016-INS-167 |
| chr11-97835818-INS-176 | chr11-97835820-INS-178 |
| chr11-97840258-INS-120 | chr11-97840280-INS-110 |
| chr11-97868941-INS-206 | chr11-97868941-INS-206 |
| chr11-97884768-INS-77 | chr11-97884768-INS-77 |
| chr11-97905556-INS-248 | chr11-97905556-INS-236 |
| chr11-97922536-INS-182 | chr11-97922536-INS-183 |
| chr11-97933202-INS-76 | chr11-97933202-INS-76 |
| chr11-97955305-INS-181 | chr11-97955305-INS-181 |
| chr11-97956812-INS-279 | chr11-97956813-INS-278 |
| chr11-97969577-INS-74 | chr11-97969577-INS-78 |
| chr11-97975264-INS-54 | chr11-97975264-INS-54 |
| chr11-97979155-INS-52 | chr11-97979155-INS-52 |
| chr11-97985554-INS-2177 | chr11-97985555-INS-2151 |
| chr11-97995221-INS-204 | chr11-97995221-INS-204 |
| chr11-98003577-INS-94 | chr11-98003577-INS-94 |
| chr11-98007207-INS-87 | chr11-98007207-INS-87 |
| chr11-98007238-INS-927 | chr11-98007238-INS-927 |
| chr11-98015477-INS-270 | chr11-98015478-INS-270 |
| chr11-98020160-INS-141 | chr11-98020160-INS-141 |
| chr11-98025372-INS-90 | chr11-98025372-INS-90 |
| chr11-98038113-INS-228 | chr11-98038113-INS-229 |
| chr11-98049728-INS-70 | chr11-98049728-INS-70 |
| chr11-98053421-INS-179 | chr11-98053421-INS-180 |
| chr11-98056818-INS-211 | chr11-98056818-INS-205 |
| chr11-98076525-INS-212 | chr11-98076525-INS-212 |
| chr11-98078694-INS-230 | chr11-98078694-INS-230 |
| chr11-98086087-INS-172 | chr11-98086087-INS-172 |
| chr11-98098306-INS-212 | chr11-98098306-INS-214 |
| chr11-98116021-INS-229 | chr11-98116021-INS-233 |
| chr11-98120156-INS-197 | chr11-98120156-INS-202 |
| chr11-98120664-INS-218 | chr11-98120664-INS-219 |
| chr11-98121684-INS-248 | chr11-98121684-INS-248 |
| chr11-98125658-INS-201 | chr11-98125659-INS-200 |
| chr11-98129546-INS-175 | chr11-98129546-INS-175 |

| | |
|---|---|
| chr11-98133131-INS-212 | chr11-98133131-INS-211 |
| chr11-98165708-INS-223 | chr11-98165728-INS-244 |
| chr11-98184953-INS-202 | chr11-98184953-INS-202 |
| chr11-98192672-INS-93 | chr11-98192672-INS-93 |
| chr11-98215763-INS-74 | chr11-98215743-INS-54 |
| chr11-98226822-INS-106 | chr11-98226822-INS-106 |
| chr11-98234668-INS-200 | chr11-98234668-INS-200 |
| chr11-98256115-INS-67 | chr11-98256086-INS-76 |
| chr11-98295034-INS-174 | chr11-98295034-INS-177 |
| chr11-98306833-INS-942 | chr11-98306833-INS-942 |
| chr11-98362641-INS-174 | chr11-98362641-INS-174 |
| chr11-98380625-INS-71 | chr11-98380625-INS-71 |
| chr11-98396496-INS-51 | chr11-98396492-INS-56 |
| chr11-98410214-INS-75 | chr11-98410214-INS-75 |
| chr11-98412610-INS-303 | chr11-98412610-INS-303 |
| chr11-98453901-INS-115 | chr11-98453901-INS-115 |
| chr11-98456106-INS-76 | chr11-98456093-INS-56 |
| chr11-98485784-INS-78 | chr11-98485784-INS-81 |
| chr11-98486813-INS-71 | chr11-98486813-INS-71 |
| chr11-98492015-INS-212 | chr11-98492015-INS-214 |
| chr11-98534042-INS-4571 | chr11-98534043-INS-4571 |
| chr11-98538899-INS-66 | chr11-98538899-INS-63 |
| chr11-98559532-INS-168 | chr11-98559530-INS-176 |
| chr11-98567803-INS-498 | chr11-98567803-INS-498 |
| chr11-98614256-INS-971 | chr11-98614256-INS-987 |
| chr11-98619905-INS-61 | chr11-98619905-INS-55 |
| chr11-98631659-INS-80 | chr11-98631659-INS-82 |
| chr11-98649993-INS-360 | chr11-98649993-INS-361 |
| chr11-98671531-INS-52 | chr11-98671531-INS-54 |
| chr11-98672596-INS-214 | chr11-98672596-INS-216 |
| chr11-98678095-INS-218 | chr11-98678095-INS-219 |
| chr11-98679104-INS-86 | chr11-98679104-INS-86 |
| chr11-98698686-INS-6452 | chr11-98698687-INS-6452 |
| chr11-98738611-INS-190 | chr11-98738611-INS-187 |
| chr11-98748398-INS-207 | chr11-98748396-INS-208 |
| chr11-98750600-INS-675 | chr11-98750600-INS-675 |
| chr11-98750935-INS-5305 | chr11-98750936-INS-5305 |
| chr11-98759323-INS-207 | chr11-98759323-INS-207 |
| chr11-98772809-INS-218 | chr11-98772809-INS-218 |
| chr11-98811002-INS-171 | chr11-98811004-INS-179 |
| chr11-98817980-INS-60 | chr11-98817980-INS-60 |
| chr11-98842869-INS-200 | chr11-98842873-INS-203 |
| chr11-98874862-INS-198 | chr11-98874862-INS-198 |
| chr11-98883517-INS-172 | chr11-98883517-INS-212 |
| chr11-98895517-INS-179 | chr11-98895517-INS-179 |
| chr11-98907142-INS-6483 | chr11-98907143-INS-6483 |
| chr11-98934338-INS-99 | chr11-98934338-INS-79 |
| chr11-98975098-INS-66 | chr11-98975098-INS-66 |
| chr11-99006134-INS-66 | chr11-99006134-INS-60 |
| chr11-99009622-INS-56 | chr11-99009622-INS-56 |
| chr11-99024673-INS-54 | chr11-99024673-INS-54 |
| chr11-99046530-INS-231 | chr11-99046530-INS-235 |
| chr11-99087762-INS-203 | chr11-99087762-INS-203 |
| chr11-99126310-INS-63 | chr11-99126327-INS-64 |
| chr11-99138083-INS-497 | chr11-99138083-INS-497 |
| chr11-99140717-INS-78 | chr11-99140717-INS-78 |
| chr11-99157159-INS-124 | chr11-99157159-INS-124 |
| chr11-99159161-INS-64 | chr11-99159157-INS-52 |
| chr11-99165894-INS-52 | chr11-99165894-INS-52 |
| chr11-99166981-INS-176 | chr11-99166981-INS-176 |
| chr11-99181400-INS-183 | chr11-99181400-INS-183 |
| chr11-99182934-INS-531 | chr11-99182934-INS-531 |
| chr11-99183349-INS-7218 | chr11-99183350-INS-7218 |
| chr11-99185488-INS-62 | chr11-99185488-INS-60 |
| chr11-99191805-INS-504 | chr11-99191805-INS-508 |
| chr11-99216657-INS-52 | chr11-99216657-INS-52 |
| chr11-99234701-INS-58 | chr11-99234701-INS-54 |
| chr11-99245429-INS-122 | chr11-99245441-INS-74 |
| chr11-99251113-INS-399 | chr11-99251113-INS-399 |
| chr11-99269090-INS-6882 | chr11-99269091-INS-6882 |
| chr11-99290435-INS-176 | chr11-99290435-INS-176 |
| chr11-99295566-INS-66 | chr11-99295566-INS-66 |
| chr11-99297878-INS-112 | chr11-99297878-INS-122 |
| chr11-99300753-INS-480 | chr11-99300753-INS-480 |
| chr11-99314147-INS-784 | chr11-99314147-INS-768 |
| chr11-99325874-INS-82 | chr11-99325874-INS-64 |
| chr11-99336794-INS-79 | chr11-99336794-INS-79 |

| | |
|---|---|
| chr11-99347211-INS-92 | chr11-99347211-INS-92 |
| chr11-99408844-INS-1893 | chr11-99408845-INS-1893 |
| chr11-99425196-INS-65 | chr11-99425196-INS-65 |
| chr11-99446066-INS-91 | chr11-99446066-INS-116 |
| chr11-99452064-INS-356 | chr11-99452064-INS-356 |
| chr11-99476268-INS-56 | chr11-99476286-INS-52 |
| chr11-99479826-INS-64 | chr11-99479826-INS-64 |
| chr11-99496584-INS-122 | chr11-99496584-INS-121 |
| chr11-99527270-INS-230 | chr11-99527270-INS-230 |
| chr11-99531019-INS-2171 | chr11-99531020-INS-2171 |
| chr11-99532213-INS-55 | chr11-99532213-INS-64 |
| chr11-99544553-INS-114 | chr11-99544553-INS-96 |
| chr11-99617860-INS-886 | chr11-99617860-INS-886 |
| chr11-99627372-INS-532 | chr11-99627372-INS-541 |
| chr11-99630096-INS-445 | chr11-99630096-INS-445 |
| chr11-99633440-INS-91 | chr11-99633440-INS-95 |
| chr11-99642889-INS-91 | chr11-99642880-INS-53 |
| chr11-99645424-INS-56 | chr11-99645424-INS-72 |
| chr11-99649442-INS-5436 | chr11-99649443-INS-5436 |
| chr11-99683648-INS-75 | chr11-99683648-INS-75 |
| chr11-99693177-INS-107 | chr11-99693131-INS-112 |
| chr11-99700282-INS-90 | chr11-99700282-INS-90 |
| chr11-99730130-INS-200 | chr11-99730130-INS-200 |
| chr11-99808753-INS-1164 | chr11-99808754-INS-1164 |
| chr11-99830287-INS-66 | chr11-99830287-INS-66 |
| chr11-99833509-INS-55 | chr11-99833491-INS-50 |
| chr11-99835825-INS-216 | chr11-99835825-INS-216 |
| chr11-99861695-INS-197 | chr11-99861707-INS-197 |
| chr11-99885804-INS-63 | chr11-99885804-INS-63 |
| chr11-99888250-INS-74 | chr11-99888257-INS-60 |
| chr11-99890198-INS-98 | chr11-99890245-INS-98 |
| chr11-99913528-INS-7570 | chr11-99913529-INS-7572 |
| chr11-99969394-INS-123 | chr11-99969394-INS-123 |
| chr11-99971240-INS-6494 | chr11-99971241-INS-6494 |
| chr11-100001304-INS-60 | chr11-100001304-INS-60 |
| chr11-100031201-INS-124 | chr11-100031201-INS-126 |
| chr11-100034009-INS-81 | chr11-100034009-INS-73 |
| chr11-100051900-INS-322 | chr11-100051900-INS-355 |
| chr11-100056261-INS-60 | chr11-100056261-INS-60 |
| chr11-100061652-INS-146 | chr11-100061652-INS-156 |
| chr11-100064983-INS-83 | chr11-100064983-INS-83 |
| chr11-100074865-INS-64 | chr11-100074865-INS-62 |
| chr11-100085213-INS-145 | chr11-100085213-INS-145 |
| chr11-100093251-INS-96 | chr11-100093251-INS-69 |
| chr11-100094604-INS-73 | chr11-100094604-INS-59 |
| chr11-100122904-INS-897 | chr11-100122904-INS-882 |
| chr11-100128324-INS-411 | chr11-100128325-INS-410 |
| chr11-100163331-INS-95 | chr11-100163332-INS-95 |
| chr11-100178822-INS-51 | chr11-100178828-INS-95 |
| chr11-100189774-INS-210 | chr11-100189774-INS-210 |
| chr11-100192194-INS-56 | chr11-100192194-INS-59 |
| chr11-100194072-INS-208 | chr11-100194072-INS-208 |
| chr11-100207922-INS-78 | chr11-100207922-INS-80 |
| chr11-100214097-INS-60 | chr11-100214097-INS-60 |
| chr11-100221691-INS-195 | chr11-100221691-INS-195 |
| chr11-100257358-INS-81 | chr11-100257358-INS-54 |
| chr11-100262806-INS-87 | chr11-100262806-INS-87 |
| chr11-100270725-INS-201 | chr11-100270725-INS-201 |
| chr11-100280488-INS-210 | chr11-100280488-INS-215 |
| chr11-100339732-INS-203 | chr11-100339737-INS-215 |
| chr11-100357871-INS-82 | chr11-100357871-INS-82 |
| chr11-100399342-INS-204 | chr11-100399342-INS-204 |
| chr11-100465235-INS-184 | chr11-100465235-INS-186 |
| chr11-100487209-INS-180 | chr11-100487211-INS-181 |
| chr11-100501940-INS-212 | chr11-100501940-INS-212 |
| chr11-100505341-INS-79 | chr11-100505361-INS-72 |
| chr11-100546183-INS-271 | chr11-100546183-INS-265 |
| chr11-100574834-INS-51 | chr11-100574834-INS-63 |
| chr11-100616915-INS-79 | chr11-100616906-INS-52 |
| chr11-100624011-INS-203 | chr11-100624011-INS-202 |
| chr11-100654816-INS-54 | chr11-100654816-INS-50 |
| chr11-100657125-INS-62 | chr11-100657125-INS-62 |
| chr11-100698185-INS-319 | chr11-100698186-INS-319 |
| chr11-100703423-INS-144 | chr11-100703423-INS-144 |
| chr11-100706350-INS-213 | chr11-100706350-INS-237 |
| chr11-100709209-INS-54 | chr11-100709207-INS-56 |
| chr11-100770673-INS-55 | chr11-100770673-INS-55 |

| | |
|---|---|
| chr11-100786152-INS-80 | chr11-100786152-INS-64 |
| chr11-100790971-INS-177 | chr11-100790971-INS-177 |
| chr11-100824198-INS-52 | chr11-100824206-INS-77 |
| chr11-100856503-INS-54 | chr11-100856503-INS-54 |
| chr11-100857901-INS-146 | chr11-100857897-INS-151 |
| chr11-100864842-INS-227 | chr11-100864840-INS-230 |
| chr11-100866130-INS-2992 | chr11-100866131-INS-2992 |
| chr11-100873119-INS-86 | chr11-100873119-INS-86 |
| chr11-100874889-INS-107 | chr11-100874889-INS-108 |
| chr11-100875621-INS-56 | chr11-100875582-INS-66 |
| chr11-100884890-INS-231 | chr11-100884888-INS-230 |
| chr11-100887336-INS-283 | chr11-100887336-INS-282 |
| chr11-100922412-INS-128 | chr11-100922400-INS-120 |
| chr11-100932858-INS-172 | chr11-100932858-INS-172 |
| chr11-101007256-INS-221 | chr11-101007256-INS-208 |
| chr11-101036907-INS-193 | chr11-101036907-INS-194 |
| chr11-101042527-INS-60 | chr11-101042527-INS-60 |
| chr11-101045268-INS-85 | chr11-101045268-INS-81 |
| chr11-101093563-INS-59 | chr11-101093563-INS-80 |
| chr11-101094940-INS-204 | chr11-101094940-INS-210 |
| chr11-101113120-INS-215 | chr11-101113120-INS-216 |
| chr11-101127256-INS-392 | chr11-101127257-INS-392 |
| chr11-101131740-INS-52 | chr11-101131740-INS-52 |
| chr11-101134386-INS-174 | chr11-101134386-INS-174 |
| chr11-101155485-INS-54 | chr11-101155485-INS-54 |
| chr11-101163724-INS-158 | chr11-101163725-INS-158 |
| chr11-101175930-INS-1892 | chr11-101175931-INS-1892 |
| chr11-101180311-INS-215 | chr11-101180311-INS-215 |
| chr11-101183263-INS-211 | chr11-101183263-INS-208 |
| chr11-101184205-INS-167 | chr11-101184205-INS-166 |
| chr11-101204573-INS-245 | chr11-101204573-INS-245 |
| chr11-101205726-INS-119 | chr11-101205726-INS-119 |
| chr11-101209488-INS-184 | chr11-101209488-INS-181 |
| chr11-101210084-INS-66 | chr11-101210084-INS-76 |
| chr11-101232610-INS-97 | chr11-101232610-INS-97 |
| chr11-101239283-INS-383 | chr11-101239283-INS-383 |
| chr11-101257756-INS-217 | chr11-101257756-INS-203 |
| chr11-101262871-INS-141 | chr11-101262871-INS-141 |
| chr11-101264849-INS-64 | chr11-101264838-INS-80 |
| chr11-101277168-INS-54 | chr11-101277168-INS-54 |
| chr11-101295918-INS-203 | chr11-101295918-INS-203 |
| chr11-101296833-INS-171 | chr11-101296833-INS-171 |
| chr11-101323493-INS-207 | chr11-101323493-INS-192 |
| chr11-101346352-INS-195 | chr11-101346352-INS-195 |
| chr11-101347889-INS-208 | chr11-101347889-INS-207 |
| chr11-101374233-INS-207 | chr11-101374233-INS-207 |
| chr11-101403077-INS-201 | chr11-101403077-INS-203 |
| chr11-101404835-INS-205 | chr11-101404836-INS-213 |
| chr11-101433363-INS-609 | chr11-101433363-INS-610 |
| chr11-101436544-INS-275 | chr11-101436544-INS-275 |
| chr11-101482776-INS-190 | chr11-101482776-INS-189 |
| chr11-101489318-INS-179 | chr11-101489318-INS-179 |
| chr11-101490539-INS-199 | chr11-101490539-INS-199 |
| chr11-101492332-INS-174 | chr11-101492332-INS-222 |
| chr11-101494015-INS-952 | chr11-101494015-INS-972 |
| chr11-101497209-INS-191 | chr11-101497209-INS-193 |
| chr11-101498879-INS-349 | chr11-101498882-INS-348 |
| chr11-101503481-INS-54 | chr11-101503458-INS-54 |
| chr11-101505224-INS-206 | chr11-101505224-INS-206 |
| chr11-101510173-INS-204 | chr11-101510173-INS-205 |
| chr11-101516238-INS-480 | chr11-101516238-INS-486 |
| chr11-101534074-INS-154 | chr11-101534074-INS-154 |
| chr11-101544771-INS-650 | chr11-101544771-INS-680 |
| chr11-101547348-INS-207 | chr11-101547348-INS-207 |
| chr11-101577024-INS-56 | chr11-101577024-INS-51 |
| chr11-101597236-INS-263 | chr11-101597236-INS-251 |
| chr11-101600813-INS-74 | chr11-101600813-INS-74 |
| chr11-101602141-INS-123 | chr11-101602141-INS-123 |
| chr11-101611687-INS-62 | chr11-101611681-INS-105 |
| chr11-101621578-INS-395 | chr11-101621578-INS-396 |
| chr11-101639727-INS-51 | chr11-101639717-INS-55 |
| chr11-101641498-INS-84 | chr11-101641498-INS-82 |
| chr11-101686537-INS-51 | chr11-101686543-INS-52 |
| chr11-101693831-INS-57 | chr11-101693831-INS-57 |
| chr11-101699790-INS-188 | chr11-101699790-INS-204 |
| chr11-101702351-INS-165 | chr11-101702351-INS-165 |
| chr11-101704541-INS-82 | chr11-101704541-INS-84 |

| | |
|---|---|
| chr11-101711612-INS-7331 | chr11-101711613-INS-7331 |
| chr11-101718889-INS-178 | chr11-101718889-INS-183 |
| chr11-101722440-INS-207 | chr11-101722440-INS-207 |
| chr11-101727131-INS-59 | chr11-101727131-INS-59 |
| chr11-101731302-INS-208 | chr11-101731302-INS-208 |
| chr11-101750645-INS-442 | chr11-101750645-INS-442 |
| chr11-101769784-INS-1189 | chr11-101769785-INS-1189 |
| chr11-101787281-INS-254 | chr11-101787281-INS-254 |
| chr11-101790424-INS-213 | chr11-101790425-INS-213 |
| chr11-101792747-INS-176 | chr11-101792747-INS-176 |
| chr11-101802080-INS-68 | chr11-101802074-INS-53 |
| chr11-101809294-INS-150 | chr11-101809294-INS-171 |
| chr11-101810113-INS-89 | chr11-101810113-INS-89 |
| chr11-101821440-INS-81 | chr11-101821407-INS-68 |
| chr11-101825957-INS-216 | chr11-101825958-INS-240 |
| chr11-101845400-INS-56 | chr11-101845400-INS-54 |
| chr11-101861820-INS-66 | chr11-101861820-INS-66 |
| chr11-101862617-INS-221 | chr11-101862617-INS-222 |
| chr11-101870076-INS-268 | chr11-101870076-INS-268 |
| chr11-101902703-INS-78 | chr11-101902703-INS-80 |
| chr11-101947011-INS-106 | chr11-101947004-INS-82 |
| chr11-101959241-INS-187 | chr11-101959241-INS-188 |
| chr11-102003713-INS-238 | chr11-102003713-INS-240 |
| chr11-102008109-INS-56 | chr11-102008109-INS-56 |
| chr11-102013727-INS-82 | chr11-102013727-INS-82 |
| chr11-102017278-INS-180 | chr11-102017278-INS-172 |
| chr11-102029810-INS-786 | chr11-102029810-INS-786 |
| chr11-102042736-INS-198 | chr11-102042736-INS-200 |
| chr11-102048534-INS-51 | chr11-102048534-INS-51 |
| chr11-102049651-INS-215 | chr11-102049651-INS-214 |
| chr11-102063276-INS-211 | chr11-102063276-INS-211 |
| chr11-102116090-INS-398 | chr11-102116090-INS-397 |
| chr11-102124557-INS-181 | chr11-102124557-INS-181 |
| chr11-102131093-INS-178 | chr11-102131093-INS-198 |
| chr11-102165185-INS-186 | chr11-102165185-INS-185 |
| chr11-102175332-INS-208 | chr11-102175332-INS-208 |
| chr11-102216577-INS-238 | chr11-102216577-INS-221 |
| chr11-102219952-INS-199 | chr11-102219952-INS-199 |
| chr11-102228822-INS-175 | chr11-102228822-INS-175 |
| chr11-102230542-INS-172 | chr11-102230542-INS-172 |
| chr11-102232747-INS-8985 | chr11-102232748-INS-8985 |
| chr11-102233024-INS-382 | chr11-102233024-INS-382 |
| chr11-102234476-INS-52 | chr11-102234476-INS-54 |
| chr11-102237557-INS-146 | chr11-102237557-INS-147 |
| chr11-102267944-INS-65 | chr11-102267944-INS-53 |
| chr11-102269368-INS-200 | chr11-102269368-INS-200 |
| chr11-102272155-INS-213 | chr11-102272155-INS-209 |
| chr11-102273726-INS-64 | chr11-102273755-INS-54 |
| chr11-102277042-INS-2683 | chr11-102277043-INS-2683 |
| chr11-102277347-INS-189 | chr11-102277347-INS-189 |
| chr11-102279509-INS-112 | chr11-102279509-INS-113 |
| chr11-102342096-INS-6466 | chr11-102342097-INS-6466 |
| chr11-102359828-INS-211 | chr11-102359828-INS-212 |
| chr11-102435500-INS-196 | chr11-102435500-INS-196 |
| chr11-102453049-INS-455 | chr11-102453046-INS-449 |
| chr11-102455574-INS-197 | chr11-102455574-INS-196 |
| chr11-102466119-INS-184 | chr11-102466119-INS-185 |
| chr11-102480932-INS-88 | chr11-102480932-INS-88 |
| chr11-102491398-INS-200 | chr11-102491399-INS-200 |
| chr11-102493527-INS-215 | chr11-102493527-INS-216 |
| chr11-102503092-INS-87 | chr11-102503097-INS-79 |
| chr11-102516850-INS-56 | chr11-102516853-INS-54 |
| chr11-102520314-INS-1890 | chr11-102520315-INS-1890 |
| chr11-102524926-INS-183 | chr11-102524926-INS-185 |
| chr11-102525445-INS-1867 | chr11-102525446-INS-1867 |
| chr11-102545555-INS-201 | chr11-102545555-INS-202 |
| chr11-102553607-INS-93 | chr11-102553599-INS-80 |
| chr11-102596073-INS-198 | chr11-102596073-INS-196 |
| chr11-102606227-INS-235 | chr11-102606227-INS-235 |
| chr11-102627727-INS-115 | chr11-102627728-INS-94 |
| chr11-102631238-INS-120 | chr11-102631238-INS-120 |
| chr11-102672370-INS-164 | chr11-102672370-INS-172 |
| chr11-102692881-INS-185 | chr11-102692881-INS-185 |
| chr11-102711949-INS-138 | chr11-102711948-INS-138 |
| chr11-102717835-INS-76 | chr11-102717823-INS-75 |
| chr11-102720666-INS-195 | chr11-102720666-INS-195 |
| chr11-102723944-INS-79 | chr11-102723944-INS-79 |

| | |
|---|---|
| chr11-102738565-INS-56 | chr11-102738565-INS-56 |
| chr11-102740224-INS-296 | chr11-102740224-INS-299 |
| chr11-102748484-INS-204 | chr11-102748484-INS-204 |
| chr11-102754129-INS-104 | chr11-102754129-INS-64 |
| chr11-102757597-INS-275 | chr11-102757597-INS-275 |
| chr11-102763658-INS-85 | chr11-102763658-INS-86 |
| chr11-102782430-INS-96 | chr11-102782428-INS-97 |
| chr11-102783007-INS-56 | chr11-102783006-INS-78 |
| chr11-102808392-INS-109 | chr11-102808392-INS-108 |
| chr11-102809896-INS-224 | chr11-102809896-INS-223 |
| chr11-102823687-INS-214 | chr11-102823687-INS-214 |
| chr11-102844765-INS-64 | chr11-102844765-INS-68 |
| chr11-102850474-INS-74 | chr11-102850474-INS-77 |
| chr11-102869936-INS-52 | chr11-102869936-INS-52 |
| chr11-102888650-INS-213 | chr11-102888650-INS-213 |
| chr11-102909520-INS-153 | chr11-102909520-INS-153 |
| chr11-102916607-INS-231 | chr11-102916607-INS-231 |
| chr11-102922543-INS-217 | chr11-102922543-INS-217 |
| chr11-102924445-INS-69 | chr11-102924442-INS-109 |
| chr11-102925266-INS-187 | chr11-102925266-INS-187 |
| chr11-102942086-INS-219 | chr11-102942086-INS-220 |
| chr11-102948787-INS-3689 | chr11-102948788-INS-3689 |
| chr11-102952286-INS-218 | chr11-102952286-INS-218 |
| chr11-102963183-INS-102 | chr11-102963183-INS-102 |
| chr11-102999335-INS-62 | chr11-102999335-INS-62 |
| chr11-103054505-INS-201 | chr11-103054505-INS-201 |
| chr11-103058674-INS-59 | chr11-103058674-INS-52 |
| chr11-103074693-INS-137 | chr11-103074703-INS-120 |
| chr11-103113953-INS-67 | chr11-103113953-INS-71 |
| chr11-103115417-INS-90 | chr11-103115417-INS-78 |
| chr11-103133366-INS-60 | chr11-103133390-INS-66 |
| chr11-103138232-INS-136 | chr11-103138232-INS-136 |
| chr11-103152055-INS-72 | chr11-103152053-INS-50 |
| chr11-103175102-INS-156 | chr11-103175102-INS-192 |
| chr11-103177063-INS-161 | chr11-103177068-INS-134 |
| chr11-103182689-INS-66 | chr11-103182683-INS-72 |
| chr11-103184338-INS-189 | chr11-103184338-INS-179 |
| chr11-103189891-INS-156 | chr11-103189891-INS-192 |
| chr11-103202313-INS-112 | chr11-103202313-INS-116 |
| chr11-103205541-INS-304 | chr11-103205573-INS-298 |
| chr11-103214179-INS-60 | chr11-103214168-INS-52 |
| chr11-103255689-INS-151 | chr11-103255671-INS-166 |
| chr11-103269088-INS-220 | chr11-103269088-INS-220 |
| chr11-103275710-INS-206 | chr11-103275710-INS-206 |
| chr11-103277427-INS-67 | chr11-103277427-INS-67 |
| chr11-103282091-INS-57 | chr11-103282091-INS-57 |
| chr11-103296265-INS-119 | chr11-103296265-INS-118 |
| chr11-103300554-INS-70 | chr11-103300554-INS-70 |
| chr11-103307260-INS-205 | chr11-103307260-INS-205 |
| chr11-103311928-INS-61 | chr11-103311928-INS-59 |
| chr11-103312426-INS-77 | chr11-103312426-INS-75 |
| chr11-103322442-INS-56 | chr11-103322442-INS-52 |
| chr11-103339117-INS-56 | chr11-103339117-INS-50 |
| chr11-103371239-INS-105 | chr11-103371258-INS-66 |
| chr11-103444611-INS-209 | chr11-103444611-INS-209 |
| chr11-103453987-INS-1893 | chr11-103453988-INS-1893 |
| chr11-103501943-INS-578 | chr11-103501943-INS-578 |
| chr11-103512961-INS-316 | chr11-103512961-INS-316 |
| chr11-103530674-INS-69 | chr11-103530674-INS-69 |
| chr11-103533162-INS-200 | chr11-103533164-INS-198 |
| chr11-103599269-INS-191 | chr11-103599269-INS-203 |
| chr11-103610023-INS-55 | chr11-103610023-INS-55 |
| chr11-103611450-INS-375 | chr11-103611450-INS-375 |
| chr11-103611577-INS-5219 | chr11-103611578-INS-5219 |
| chr11-103619021-INS-88 | chr11-103619021-INS-88 |
| chr11-103711021-INS-102 | chr11-103711021-INS-102 |
| chr11-103734933-INS-72 | chr11-103734933-INS-72 |
| chr11-103769160-INS-218 | chr11-103769160-INS-218 |
| chr11-103778065-INS-64 | chr11-103778065-INS-64 |
| chr11-103798530-INS-339 | chr11-103798527-INS-357 |
| chr11-103806546-INS-89 | chr11-103806546-INS-89 |
| chr11-103820522-INS-74 | chr11-103820522-INS-74 |
| chr11-103827748-INS-73 | chr11-103827748-INS-73 |
| chr11-103853259-INS-319 | chr11-103853259-INS-320 |
| chr11-103856079-INS-59 | chr11-103856079-INS-59 |
| chr11-103868406-INS-111 | chr11-103868406-INS-112 |
| chr11-103887156-INS-553 | chr11-103887157-INS-555 |

| | |
|---|---|
| chr11-103914847-INS-64 | chr11-103914847-INS-64 |
| chr11-103921366-INS-5466 | chr11-103921367-INS-5466 |
| chr11-103934893-INS-118 | chr11-103934893-INS-118 |
| chr11-103973310-INS-91 | chr11-103973306-INS-68 |
| chr11-103977754-INS-244 | chr11-103977754-INS-244 |
| chr11-103980580-INS-91 | chr11-103980580-INS-91 |
| chr11-104086979-INS-204 | chr11-104086979-INS-211 |
| chr11-104090968-INS-70 | chr11-104090968-INS-66 |
| chr11-104092693-INS-59 | chr11-104092693-INS-67 |
| chr11-104094953-INS-990 | chr11-104094953-INS-986 |
| chr11-104121705-INS-177 | chr11-104121705-INS-181 |
| chr11-104143713-INS-653 | chr11-104143713-INS-653 |
| chr11-104144232-INS-67 | chr11-104144232-INS-67 |
| chr11-104172201-INS-1374 | chr11-104172202-INS-1374 |
| chr11-104200803-INS-90 | chr11-104200808-INS-80 |
| chr11-104201482-INS-942 | chr11-104201482-INS-942 |
| chr11-104204051-INS-52 | chr11-104204051-INS-54 |
| chr11-104206351-INS-213 | chr11-104206352-INS-213 |
| chr11-104247769-INS-202 | chr11-104247769-INS-202 |
| chr11-104259264-INS-182 | chr11-104259264-INS-182 |
| chr11-104261716-INS-156 | chr11-104261716-INS-156 |
| chr11-104278426-INS-57 | chr11-104278426-INS-75 |
| chr11-104312689-INS-58 | chr11-104312689-INS-58 |
| chr11-104349836-INS-220 | chr11-104349836-INS-220 |
| chr11-104357827-INS-228 | chr11-104357827-INS-228 |
| chr11-104363287-INS-402 | chr11-104363287-INS-423 |
| chr11-104367045-INS-196 | chr11-104367045-INS-196 |
| chr11-104373260-INS-510 | chr11-104373260-INS-508 |
| chr11-104381397-INS-95 | chr11-104381397-INS-95 |
| chr11-104397101-INS-216 | chr11-104397101-INS-216 |
| chr11-104400649-INS-182 | chr11-104400649-INS-183 |
| chr11-104421948-INS-62 | chr11-104421948-INS-62 |
| chr11-104423946-INS-54 | chr11-104423946-INS-56 |
| chr11-104429307-INS-222 | chr11-104429307-INS-199 |
| chr11-104432615-INS-65 | chr11-104432615-INS-69 |
| chr11-104445908-INS-53 | chr11-104445908-INS-53 |
| chr11-104450183-INS-59 | chr11-104450183-INS-60 |
| chr11-104451261-INS-103 | chr11-104451252-INS-103 |
| chr11-104467421-INS-443 | chr11-104467421-INS-443 |
| chr11-104475584-INS-52 | chr11-104475584-INS-65 |
| chr11-104495053-INS-328 | chr11-104495053-INS-329 |
| chr11-104496553-INS-56 | chr11-104496553-INS-56 |
| chr11-104512046-INS-6440 | chr11-104512047-INS-6440 |
| chr11-104520150-INS-194 | chr11-104520150-INS-194 |
| chr11-104548981-INS-59 | chr11-104548981-INS-59 |
| chr11-104566315-INS-55 | chr11-104566315-INS-58 |
| chr11-104568226-INS-78 | chr11-104568193-INS-81 |
| chr11-104572103-INS-50 | chr11-104572139-INS-57 |
| chr11-104588545-INS-92 | chr11-104588546-INS-100 |
| chr11-104597573-INS-74 | chr11-104597579-INS-50 |
| chr11-104605781-INS-262 | chr11-104605781-INS-264 |
| chr11-104621307-INS-132 | chr11-104621307-INS-132 |
| chr11-104638438-INS-7248 | chr11-104638439-INS-7248 |
| chr11-104659517-INS-179 | chr11-104659517-INS-179 |
| chr11-104660743-INS-4983 | chr11-104660744-INS-4983 |
| chr11-104696831-INS-535 | chr11-104696831-INS-537 |
| chr11-104699844-INS-1370 | chr11-104699845-INS-1370 |
| chr11-104712628-INS-590 | chr11-104712628-INS-590 |
| chr11-104716783-INS-61 | chr11-104716784-INS-52 |
| chr11-104772148-INS-52 | chr11-104772148-INS-68 |
| chr11-104785753-INS-78 | chr11-104785753-INS-78 |
| chr11-104809410-INS-1185 | chr11-104809411-INS-1185 |
| chr11-104815836-INS-6604 | chr11-104815837-INS-6604 |
| chr11-104824342-INS-141 | chr11-104824342-INS-141 |
| chr11-104842749-INS-98 | chr11-104842749-INS-98 |
| chr11-104845805-INS-1066 | chr11-104845806-INS-1066 |
| chr11-104874292-INS-202 | chr11-104874292-INS-202 |
| chr11-104921077-INS-56 | chr11-104921077-INS-56 |
| chr11-104935251-INS-6578 | chr11-104935252-INS-6578 |
| chr11-104938098-INS-840 | chr11-104938073-INS-847 |
| chr11-104950431-INS-238 | chr11-104950431-INS-238 |
| chr11-104978715-INS-124 | chr11-104978683-INS-138 |
| chr11-104993880-INS-76 | chr11-104993880-INS-80 |
| chr11-104996068-INS-192 | chr11-104996068-INS-194 |
| chr11-105029342-INS-82 | chr11-105029342-INS-82 |
| chr11-105033577-INS-54 | chr11-105033577-INS-54 |
| chr11-105058614-INS-5723 | chr11-105058615-INS-5724 |

| | |
|---|---|
| chr11-105083695-INS-197 | chr11-105083695-INS-196 |
| chr11-105094447-INS-72 | chr11-105094447-INS-78 |
| chr11-105112867-INS-205 | chr11-105112867-INS-205 |
| chr11-105129178-INS-58 | chr11-105129178-INS-58 |
| chr11-105164264-INS-206 | chr11-105164264-INS-205 |
| chr11-105185168-INS-55 | chr11-105185168-INS-61 |
| chr11-105190504-INS-136 | chr11-105190504-INS-136 |
| chr11-105192354-INS-123 | chr11-105192393-INS-88 |
| chr11-105195940-INS-71 | chr11-105195940-INS-71 |
| chr11-105217134-INS-60 | chr11-105217134-INS-66 |
| chr11-105225493-INS-104 | chr11-105225493-INS-104 |
| chr11-105233292-INS-276 | chr11-105233243-INS-275 |
| chr11-105248664-INS-198 | chr11-105248664-INS-195 |
| chr11-105304720-INS-83 | chr11-105304720-INS-84 |
| chr11-105316031-INS-62 | chr11-105316031-INS-68 |
| chr11-105332052-INS-88 | chr11-105332052-INS-88 |
| chr11-105332455-INS-764 | chr11-105332455-INS-764 |
| chr11-105333676-INS-68 | chr11-105333676-INS-68 |
| chr11-105336277-INS-63 | chr11-105336234-INS-83 |
| chr11-105374814-INS-74 | chr11-105374814-INS-74 |
| chr11-105383617-INS-232 | chr11-105383642-INS-246 |
| chr11-105442644-INS-75 | chr11-105442644-INS-75 |
| chr11-105445223-INS-156 | chr11-105445223-INS-156 |
| chr11-105453049-INS-593 | chr11-105453049-INS-593 |
| chr11-105472494-INS-1813 | chr11-105472495-INS-1814 |
| chr11-105488488-INS-52 | chr11-105488488-INS-50 |
| chr11-105508093-INS-139 | chr11-105508093-INS-139 |
| chr11-105560177-INS-74 | chr11-105560156-INS-50 |
| chr11-105595922-INS-6847 | chr11-105595923-INS-6851 |
| chr11-105609654-INS-1206 | chr11-105609655-INS-1207 |
| chr11-105651623-INS-2310 | chr11-105651624-INS-2310 |
| chr11-105668398-INS-216 | chr11-105668398-INS-216 |
| chr11-105673350-INS-6838 | chr11-105673351-INS-6839 |
| chr11-105697445-INS-3339 | chr11-105697446-INS-3339 |
| chr11-105719647-INS-63 | chr11-105719647-INS-63 |
| chr11-105752794-INS-58 | chr11-105752794-INS-58 |
| chr11-105783823-INS-80 | chr11-105783823-INS-80 |
| chr11-105852758-INS-887 | chr11-105852758-INS-887 |
| chr11-105929648-INS-4210 | chr11-105929649-INS-4210 |
| chr11-105931215-INS-277 | chr11-105931216-INS-284 |
| chr11-105940368-INS-62 | chr11-105940368-INS-64 |
| chr11-105950469-INS-213 | chr11-105950469-INS-213 |
| chr11-105952547-INS-176 | chr11-105952547-INS-158 |
| chr11-105956286-INS-138 | chr11-105956286-INS-138 |
| chr11-105964542-INS-183 | chr11-105964542-INS-184 |
| chr11-105965631-INS-187 | chr11-105965631-INS-188 |
| chr11-105966934-INS-104 | chr11-105966925-INS-67 |
| chr11-105980882-INS-212 | chr11-105980882-INS-212 |
| chr11-105998813-INS-211 | chr11-105998813-INS-211 |
| chr11-106074870-INS-60 | chr11-106074870-INS-64 |
| chr11-106078345-INS-65 | chr11-106078345-INS-65 |
| chr11-106087810-INS-181 | chr11-106087810-INS-181 |
| chr11-106172424-INS-203 | chr11-106172437-INS-199 |
| chr11-106183253-INS-207 | chr11-106183253-INS-206 |
| chr11-106198098-INS-88 | chr11-106198098-INS-104 |
| chr11-106231682-INS-175 | chr11-106231678-INS-186 |
| chr11-106237038-INS-105 | chr11-106237037-INS-94 |
| chr11-106255607-INS-94 | chr11-106255607-INS-94 |
| chr11-106305841-INS-205 | chr11-106305841-INS-206 |
| chr11-106345175-INS-233 | chr11-106345175-INS-234 |
| chr11-106356263-INS-206 | chr11-106356263-INS-206 |
| chr11-106364799-INS-187 | chr11-106364797-INS-199 |
| chr11-106396186-INS-265 | chr11-106396195-INS-275 |
| chr11-106398202-INS-967 | chr11-106398202-INS-968 |
| chr11-106398491-INS-55 | chr11-106398491-INS-55 |
| chr11-106416157-INS-149 | chr11-106416157-INS-113 |
| chr11-106422465-INS-226 | chr11-106422465-INS-226 |
| chr11-106432625-INS-200 | chr11-106432625-INS-200 |
| chr11-106443225-INS-205 | chr11-106443225-INS-219 |
| chr11-106462385-INS-60 | chr11-106462385-INS-60 |
| chr11-106470820-INS-57 | chr11-106470820-INS-59 |
| chr11-106494985-INS-181 | chr11-106494985-INS-175 |
| chr11-106510317-INS-108 | chr11-106510319-INS-91 |
| chr11-106517462-INS-59 | chr11-106517458-INS-55 |
| chr11-106523608-INS-217 | chr11-106523608-INS-217 |
| chr11-106526195-INS-216 | chr11-106526195-INS-228 |
| chr11-106532598-INS-370 | chr11-106532598-INS-370 |

| | |
|---|---|
| chr11-106535972-INS-464 | chr11-106535972-INS-464 |
| chr11-106537161-INS-223 | chr11-106537161-INS-222 |
| chr11-106541990-INS-59 | chr11-106542001-INS-60 |
| chr11-106553138-INS-156 | chr11-106553129-INS-154 |
| chr11-106558573-INS-243 | chr11-106558579-INS-213 |
| chr11-106577019-INS-484 | chr11-106577019-INS-484 |
| chr11-106602326-INS-51 | chr11-106602326-INS-51 |
| chr11-106612590-INS-58 | chr11-106612590-INS-58 |
| chr11-106631525-INS-60 | chr11-106631525-INS-53 |
| chr11-106632647-INS-67 | chr11-106632642-INS-96 |
| chr11-106633380-INS-79 | chr11-106633380-INS-95 |
| chr11-106654989-INS-166 | chr11-106654989-INS-166 |
| chr11-106668882-INS-171 | chr11-106668882-INS-171 |
| chr11-106700912-INS-159 | chr11-106700912-INS-160 |
| chr11-106740340-INS-240 | chr11-106740340-INS-232 |
| chr11-106741101-INS-163 | chr11-106741101-INS-185 |
| chr11-106776352-INS-54 | chr11-106776352-INS-54 |
| chr11-106785144-INS-209 | chr11-106785144-INS-202 |
| chr11-106832100-INS-116 | chr11-106832083-INS-114 |
| chr11-106841365-INS-106 | chr11-106841373-INS-113 |
| chr11-106859245-INS-65 | chr11-106859255-INS-52 |
| chr11-106865528-INS-217 | chr11-106865528-INS-217 |
| chr11-106870590-INS-198 | chr11-106870590-INS-200 |
| chr11-106876657-INS-202 | chr11-106876657-INS-202 |
| chr11-106877424-INS-215 | chr11-106877424-INS-215 |
| chr11-106922260-INS-155 | chr11-106922260-INS-163 |
| chr11-106948228-INS-62 | chr11-106948228-INS-62 |
| chr11-106980502-INS-226 | chr11-106980502-INS-226 |
| chr11-106993075-INS-183 | chr11-106993075-INS-185 |
| chr11-107005391-INS-168 | chr11-107005400-INS-166 |
| chr11-107009806-INS-293 | chr11-107009806-INS-294 |
| chr11-107016197-INS-72 | chr11-107016197-INS-72 |
| chr11-107017302-INS-82 | chr11-107017302-INS-82 |
| chr11-107028168-INS-55 | chr11-107028168-INS-54 |
| chr11-107035964-INS-59 | chr11-107035964-INS-59 |
| chr11-107040492-INS-77 | chr11-107040492-INS-77 |
| chr11-107050381-INS-205 | chr11-107050381-INS-203 |
| chr11-107051320-INS-150 | chr11-107051333-INS-114 |
| chr11-107061347-INS-185 | chr11-107061347-INS-186 |
| chr11-107069048-INS-200 | chr11-107069065-INS-202 |
| chr11-107120486-INS-211 | chr11-107120486-INS-210 |
| chr11-107143601-INS-258 | chr11-107143601-INS-258 |
| chr11-107148024-INS-385 | chr11-107148024-INS-383 |
| chr11-107173002-INS-95 | chr11-107173002-INS-109 |
| chr11-107177372-INS-212 | chr11-107177372-INS-212 |
| chr11-107187886-INS-3698 | chr11-107187887-INS-3698 |
| chr11-107202538-INS-195 | chr11-107202538-INS-192 |
| chr11-107227284-INS-61 | chr11-107227284-INS-61 |
| chr11-107234022-INS-230 | chr11-107234022-INS-230 |
| chr11-107238053-INS-137 | chr11-107238053-INS-153 |
| chr11-107243263-INS-108 | chr11-107243263-INS-156 |
| chr11-107244894-INS-209 | chr11-107244894-INS-208 |
| chr11-107247904-INS-80 | chr11-107247904-INS-80 |
| chr11-107268904-INS-115 | chr11-107268904-INS-120 |
| chr11-107270814-INS-79 | chr11-107270814-INS-78 |
| chr11-107272551-INS-53 | chr11-107272551-INS-53 |
| chr11-107274873-INS-65 | chr11-107274873-INS-68 |
| chr11-107283239-INS-98 | chr11-107283239-INS-100 |
| chr11-107291270-INS-125 | chr11-107291270-INS-126 |
| chr11-107297440-INS-242 | chr11-107297440-INS-197 |
| chr11-107305459-INS-72 | chr11-107305459-INS-61 |
| chr11-107314126-INS-170 | chr11-107314126-INS-170 |
| chr11-107326115-INS-9045 | chr11-107326116-INS-9046 |
| chr11-107337427-INS-71 | chr11-107337427-INS-77 |
| chr11-107345382-INS-51 | chr11-107345403-INS-60 |
| chr11-107373740-INS-188 | chr11-107373740-INS-189 |
| chr11-107374982-INS-82 | chr11-107375004-INS-88 |
| chr11-107380221-INS-206 | chr11-107380221-INS-207 |
| chr11-107397254-INS-233 | chr11-107397254-INS-233 |
| chr11-107406854-INS-60 | chr11-107406854-INS-56 |
| chr11-107408299-INS-208 | chr11-107408299-INS-207 |
| chr11-107414386-INS-63 | chr11-107414386-INS-63 |
| chr11-107417835-INS-197 | chr11-107417865-INS-208 |
| chr11-107419598-INS-104 | chr11-107419599-INS-77 |
| chr11-107422609-INS-206 | chr11-107422609-INS-207 |
| chr11-107449839-INS-206 | chr11-107449839-INS-206 |
| chr11-107462385-INS-211 | chr11-107462385-INS-212 |

| | |
|---|---|
| chr11-107471832-INS-248 | chr11-107471832-INS-248 |
| chr11-107472898-INS-164 | chr11-107472898-INS-156 |
| chr11-107474409-INS-196 | chr11-107474409-INS-196 |
| chr11-107501038-INS-205 | chr11-107501038-INS-205 |
| chr11-107548391-INS-175 | chr11-107548391-INS-175 |
| chr11-107588254-INS-79 | chr11-107588260-INS-79 |
| chr11-107618776-INS-236 | chr11-107618776-INS-236 |
| chr11-107638683-INS-77 | chr11-107638656-INS-110 |
| chr11-107643081-INS-91 | chr11-107643081-INS-91 |
| chr11-107649159-INS-63 | chr11-107649160-INS-63 |
| chr11-107656840-INS-63 | chr11-107656840-INS-63 |
| chr11-107677727-INS-60 | chr11-107677727-INS-56 |
| chr11-107721770-INS-94 | chr11-107721770-INS-94 |
| chr11-107733137-INS-61 | chr11-107733116-INS-65 |
| chr11-107737956-INS-100 | chr11-107737956-INS-104 |
| chr11-107745146-INS-84 | chr11-107745146-INS-84 |
| chr11-107749999-INS-136 | chr11-107749999-INS-130 |
| chr11-107759236-INS-185 | chr11-107759235-INS-182 |
| chr11-107794447-INS-576 | chr11-107794447-INS-576 |
| chr11-107797769-INS-160 | chr11-107797769-INS-160 |
| chr11-107805540-INS-52 | chr11-107805540-INS-58 |
| chr11-107810237-INS-219 | chr11-107810237-INS-209 |
| chr11-107823717-INS-146 | chr11-107823717-INS-120 |
| chr11-107837253-INS-111 | chr11-107837253-INS-111 |
| chr11-107839193-INS-293 | chr11-107839193-INS-293 |
| chr11-107848394-INS-7111 | chr11-107848395-INS-7111 |
| chr11-107878152-INS-68 | chr11-107878152-INS-68 |
| chr11-107895295-INS-1893 | chr11-107895296-INS-1893 |
| chr11-107898095-INS-51 | chr11-107898101-INS-61 |
| chr11-107908891-INS-216 | chr11-107908891-INS-216 |
| chr11-107914989-INS-56 | chr11-107914989-INS-54 |
| chr11-107943076-INS-69 | chr11-107943076-INS-53 |
| chr11-107949445-INS-50 | chr11-107949445-INS-50 |
| chr11-107971759-INS-72 | chr11-107971799-INS-80 |
| chr11-107980988-INS-498 | chr11-107980988-INS-498 |
| chr11-107983242-INS-116 | chr11-107983242-INS-116 |
| chr11-107985295-INS-71 | chr11-107985280-INS-52 |
| chr11-107986530-INS-202 | chr11-107986530-INS-202 |
| chr11-107987141-INS-55 | chr11-107987148-INS-63 |
| chr11-108004930-INS-94 | chr11-108004930-INS-98 |
| chr11-108038192-INS-64 | chr11-108038192-INS-66 |
| chr11-108040624-INS-106 | chr11-108040664-INS-64 |
| chr11-108057963-INS-104 | chr11-108057957-INS-66 |
| chr11-108081802-INS-56 | chr11-108081802-INS-57 |
| chr11-108089032-INS-72 | chr11-108089032-INS-80 |
| chr11-108097286-INS-58 | chr11-108097286-INS-58 |
| chr11-108119715-INS-92 | chr11-108119715-INS-92 |
| chr11-108154017-INS-60 | chr11-108154017-INS-64 |
| chr11-108191164-INS-58 | chr11-108191164-INS-60 |
| chr11-108245231-INS-206 | chr11-108245226-INS-198 |
| chr11-108249318-INS-202 | chr11-108249318-INS-199 |
| chr11-108277891-INS-90 | chr11-108277891-INS-67 |
| chr11-108278534-INS-179 | chr11-108278534-INS-179 |
| chr11-108282518-INS-69 | chr11-108282514-INS-60 |
| chr11-108295857-INS-55 | chr11-108295857-INS-55 |
| chr11-108303610-INS-131 | chr11-108303610-INS-131 |
| chr11-108326861-INS-253 | chr11-108326861-INS-253 |
| chr11-108347726-INS-204 | chr11-108347726-INS-205 |
| chr11-108369359-INS-76 | chr11-108369359-INS-56 |
| chr11-108372361-INS-612 | chr11-108372361-INS-612 |
| chr11-108374181-INS-65 | chr11-108374181-INS-65 |
| chr11-108405701-INS-60 | chr11-108405701-INS-60 |
| chr11-108416719-INS-100 | chr11-108416719-INS-100 |
| chr11-108422637-INS-84 | chr11-108422637-INS-84 |
| chr11-108442496-INS-56 | chr11-108442497-INS-56 |
| chr11-108446860-INS-266 | chr11-108446860-INS-266 |
| chr11-108523474-INS-1491 | chr11-108523475-INS-1491 |
| chr11-108525581-INS-6638 | chr11-108525582-INS-6637 |
| chr11-108532105-INS-1790 | chr11-108532106-INS-1789 |
| chr11-108536211-INS-127 | chr11-108536211-INS-124 |
| chr11-108547128-INS-58 | chr11-108547128-INS-64 |
| chr11-108551619-INS-103 | chr11-108551619-INS-103 |
| chr11-108571050-INS-1743 | chr11-108571051-INS-1744 |
| chr11-108577267-INS-5292 | chr11-108577268-INS-5292 |
| chr11-108634504-INS-237 | chr11-108634504-INS-237 |
| chr11-108636185-INS-183 | chr11-108636170-INS-175 |
| chr11-108639701-INS-102 | chr11-108639701-INS-102 |

| | |
|---|---|
| chr11-108680934-INS-55 | chr11-108680934-INS-55 |
| chr11-108702293-INS-67 | chr11-108702293-INS-105 |
| chr11-108728848-INS-440 | chr11-108728848-INS-440 |
| chr11-108839904-INS-54 | chr11-108839904-INS-54 |
| chr11-108869773-INS-7111 | chr11-108869774-INS-7111 |
| chr11-108880034-INS-114 | chr11-108880034-INS-114 |
| chr11-108904949-INS-731 | chr11-108904949-INS-712 |
| chr11-108909025-INS-86 | chr11-108909025-INS-86 |
| chr11-108920650-INS-453 | chr11-108920650-INS-453 |
| chr11-108924365-INS-130 | chr11-108924365-INS-130 |
| chr11-108936312-INS-69 | chr11-108936312-INS-87 |
| chr11-108953112-INS-70 | chr11-108953112-INS-70 |
| chr11-108958168-INS-188 | chr11-108958168-INS-188 |
| chr11-108960660-INS-103 | chr11-108960660-INS-103 |
| chr11-108986273-INS-117 | chr11-108986273-INS-117 |
| chr11-108996846-INS-82 | chr11-108996848-INS-106 |
| chr11-109025602-INS-63 | chr11-109025602-INS-57 |
| chr11-109045987-INS-68 | chr11-109045987-INS-68 |
| chr11-109048574-INS-257 | chr11-109048576-INS-298 |
| chr11-109054538-INS-219 | chr11-109054537-INS-195 |
| chr11-109081145-INS-126 | chr11-109081146-INS-134 |
| chr11-109106315-INS-150 | chr11-109106315-INS-150 |
| chr11-109127824-INS-363 | chr11-109127824-INS-372 |
| chr11-109147074-INS-70 | chr11-109147086-INS-70 |
| chr11-109153092-INS-72 | chr11-109153093-INS-96 |
| chr11-109159410-INS-223 | chr11-109159410-INS-223 |
| chr11-109165600-INS-92 | chr11-109165600-INS-92 |
| chr11-109186336-INS-102 | chr11-109186288-INS-142 |
| chr11-109200598-INS-56 | chr11-109200598-INS-56 |
| chr11-109223925-INS-100 | chr11-109223925-INS-98 |
| chr11-109245567-INS-116 | chr11-109245567-INS-120 |
| chr11-109259372-INS-53 | chr11-109259372-INS-64 |
| chr11-109283688-INS-92 | chr11-109283688-INS-92 |
| chr11-109293261-INS-205 | chr11-109293261-INS-206 |
| chr11-109304503-INS-87 | chr11-109304503-INS-87 |
| chr11-109315060-INS-100 | chr11-109315063-INS-92 |
| chr11-109322386-INS-114 | chr11-109322384-INS-159 |
| chr11-109330209-INS-76 | chr11-109330209-INS-84 |
| chr11-109338921-INS-96 | chr11-109338892-INS-104 |
| chr11-109402099-INS-64 | chr11-109402099-INS-70 |
| chr11-109409164-INS-183 | chr11-109409164-INS-183 |
| chr11-109420099-INS-73 | chr11-109420099-INS-73 |
| chr11-109446787-INS-164 | chr11-109446787-INS-164 |
| chr11-109460546-INS-101 | chr11-109460548-INS-97 |
| chr11-109464601-INS-54 | chr11-109464563-INS-58 |
| chr11-109488810-INS-312 | chr11-109488810-INS-312 |
| chr11-109496650-INS-50 | chr11-109496650-INS-52 |
| chr11-109498918-INS-136 | chr11-109498918-INS-136 |
| chr11-109515553-INS-149 | chr11-109515553-INS-149 |
| chr11-109517761-INS-385 | chr11-109517761-INS-385 |
| chr11-109539090-INS-66 | chr11-109539090-INS-68 |
| chr11-109574334-INS-114 | chr11-109574334-INS-114 |
| chr11-109598304-INS-150 | chr11-109598302-INS-152 |
| chr11-109605051-INS-62 | chr11-109605051-INS-62 |
| chr11-109614191-INS-70 | chr11-109614209-INS-96 |
| chr11-109618714-INS-84 | chr11-109618714-INS-84 |
| chr11-109628818-INS-227 | chr11-109628818-INS-227 |
| chr11-109634590-INS-64 | chr11-109634633-INS-58 |
| chr11-109641063-INS-70 | chr11-109641063-INS-70 |
| chr11-109645853-INS-203 | chr11-109645853-INS-203 |
| chr11-109649679-INS-6401 | chr11-109649680-INS-6401 |
| chr11-109650864-INS-163 | chr11-109650864-INS-163 |
| chr11-109665874-INS-64 | chr11-109665874-INS-62 |
| chr11-109678425-INS-5423 | chr11-109678426-INS-5423 |
| chr11-109695721-INS-109 | chr11-109695721-INS-109 |
| chr11-109703863-INS-271 | chr11-109703863-INS-271 |
| chr11-109706510-INS-80 | chr11-109706510-INS-87 |
| chr11-109714346-INS-90 | chr11-109714348-INS-96 |
| chr11-109719361-INS-64 | chr11-109719361-INS-64 |
| chr11-109724580-INS-62 | chr11-109724598-INS-62 |
| chr11-109742885-INS-66 | chr11-109742885-INS-68 |
| chr11-109749918-INS-480 | chr11-109749918-INS-480 |
| chr11-109757383-INS-232 | chr11-109757381-INS-236 |
| chr11-109761023-INS-64 | chr11-109761023-INS-67 |
| chr11-109766230-INS-124 | chr11-109766230-INS-124 |
| chr11-109770790-INS-90 | chr11-109770790-INS-90 |
| chr11-109807736-INS-176 | chr11-109807736-INS-177 |

| | |
|---|---|
| chr11-109826207-INS-69 | chr11-109826207-INS-87 |
| chr11-109838781-INS-71 | chr11-109838781-INS-71 |
| chr11-109841140-INS-916 | chr11-109841140-INS-913 |
| chr11-109971082-INS-311 | chr11-109971082-INS-311 |
| chr11-109976425-INS-76 | chr11-109976425-INS-76 |
| chr11-109981418-INS-687 | chr11-109981419-INS-693 |
| chr11-109985863-INS-51 | chr11-109985863-INS-51 |
| chr11-109987572-INS-122 | chr11-109987572-INS-126 |
| chr11-109992711-INS-59 | chr11-109992711-INS-61 |
| chr11-110020847-INS-533 | chr11-110020847-INS-533 |
| chr11-110044456-INS-56 | chr11-110044459-INS-62 |
| chr11-110055603-INS-125 | chr11-110055603-INS-125 |
| chr11-110092842-INS-69 | chr11-110092842-INS-69 |
| chr11-110097845-INS-68 | chr11-110097845-INS-68 |
| chr11-110099363-INS-52 | chr11-110099364-INS-52 |
| chr11-110112901-INS-97 | chr11-110112911-INS-109 |
| chr11-110114759-INS-110 | chr11-110114759-INS-110 |
| chr11-110139962-INS-76 | chr11-110139962-INS-64 |
| chr11-110142597-INS-60 | chr11-110142597-INS-64 |
| chr11-110195414-INS-111 | chr11-110195405-INS-102 |
| chr11-110236391-INS-54 | chr11-110236425-INS-50 |
| chr11-110237464-INS-65 | chr11-110237464-INS-65 |
| chr11-110248079-INS-241 | chr11-110248079-INS-241 |
| chr11-110248871-INS-6674 | chr11-110248872-INS-6675 |
| chr11-110250583-INS-177 | chr11-110250583-INS-177 |
| chr11-110264829-INS-209 | chr11-110264829-INS-209 |
| chr11-110266800-INS-149 | chr11-110266780-INS-174 |
| chr11-110278706-INS-112 | chr11-110278700-INS-88 |
| chr11-110307043-INS-85 | chr11-110307043-INS-76 |
| chr11-110308263-INS-94 | chr11-110308263-INS-94 |
| chr11-110316394-INS-162 | chr11-110316406-INS-165 |
| chr11-110327418-INS-830 | chr11-110327415-INS-832 |
| chr11-110327498-INS-210 | chr11-110327459-INS-256 |
| chr11-110341243-INS-3446 | chr11-110341244-INS-3446 |
| chr11-110360161-INS-94 | chr11-110360157-INS-96 |
| chr11-110368076-INS-51 | chr11-110368076-INS-78 |
| chr11-110374765-INS-965 | chr11-110374765-INS-967 |
| chr11-110386830-INS-117 | chr11-110386830-INS-109 |
| chr11-110423186-INS-105 | chr11-110423186-INS-106 |
| chr11-110429631-INS-66 | chr11-110429631-INS-51 |
| chr11-110430923-INS-81 | chr11-110430923-INS-81 |
| chr11-110434514-INS-507 | chr11-110434514-INS-507 |
| chr11-110439787-INS-58 | chr11-110439787-INS-54 |
| chr11-110450879-INS-456 | chr11-110450879-INS-456 |
| chr11-110458712-INS-238 | chr11-110458722-INS-236 |
| chr11-110476621-INS-113 | chr11-110476621-INS-113 |
| chr11-110497367-INS-207 | chr11-110497339-INS-220 |
| chr11-110512814-INS-435 | chr11-110512814-INS-435 |
| chr11-110516805-INS-52 | chr11-110516805-INS-52 |
| chr11-110557157-INS-232 | chr11-110557157-INS-236 |
| chr11-110561399-INS-183 | chr11-110561399-INS-137 |
| chr11-110567847-INS-149 | chr11-110567862-INS-138 |
| chr11-110584604-INS-52 | chr11-110584604-INS-52 |
| chr11-110589652-INS-117 | chr11-110589636-INS-139 |
| chr11-110596453-INS-360 | chr11-110596453-INS-360 |
| chr11-110620231-INS-130 | chr11-110620231-INS-130 |
| chr11-110624993-INS-950 | chr11-110624993-INS-951 |
| chr11-110653467-INS-513 | chr11-110653467-INS-513 |
| chr11-110676418-INS-393 | chr11-110676418-INS-393 |
| chr11-110741829-INS-90 | chr11-110741819-INS-59 |
| chr11-110751782-INS-149 | chr11-110751782-INS-145 |
| chr11-110755933-INS-63 | chr11-110755923-INS-66 |
| chr11-110764383-INS-82 | chr11-110764383-INS-86 |
| chr11-110774364-INS-63 | chr11-110774348-INS-71 |
| chr11-110792776-INS-168 | chr11-110792776-INS-172 |
| chr11-110832688-INS-98 | chr11-110832706-INS-104 |
| chr11-110836362-INS-134 | chr11-110836362-INS-133 |
| chr11-110861720-INS-167 | chr11-110861720-INS-144 |
| chr11-110862320-INS-120 | chr11-110862337-INS-95 |
| chr11-110869816-INS-304 | chr11-110869816-INS-305 |
| chr11-110874843-INS-365 | chr11-110874851-INS-372 |
| chr11-110878996-INS-198 | chr11-110878996-INS-198 |
| chr11-110895481-INS-77 | chr11-110895481-INS-81 |
| chr11-110913829-INS-54 | chr11-110913855-INS-54 |
| chr11-110927389-INS-57 | chr11-110927389-INS-57 |
| chr11-110941188-INS-634 | chr11-110941188-INS-634 |
| chr11-110945607-INS-68 | chr11-110945607-INS-68 |

| | |
|---|---|
| chr11-110993223-INS-52 | chr11-110993223-INS-56 |
| chr11-111009156-INS-2118 | chr11-111009157-INS-2119 |
| chr11-111025497-INS-90 | chr11-111025497-INS-90 |
| chr11-111046548-INS-72 | chr11-111046539-INS-120 |
| chr11-111051117-INS-64 | chr11-111051117-INS-64 |
| chr11-111099576-INS-77 | chr11-111099576-INS-77 |
| chr11-111101890-INS-56 | chr11-111101893-INS-54 |
| chr11-111121488-INS-59 | chr11-111121488-INS-59 |
| chr11-111185070-INS-71 | chr11-111185095-INS-60 |
| chr11-111186241-INS-2241 | chr11-111186246-INS-2225 |
| chr11-111216207-INS-6863 | chr11-111216208-INS-6863 |
| chr11-111231000-INS-77 | chr11-111231000-INS-77 |
| chr11-111240029-INS-115 | chr11-111240029-INS-115 |
| chr11-111284827-INS-62 | chr11-111284827-INS-62 |
| chr11-111299085-INS-488 | chr11-111299085-INS-488 |
| chr11-111308428-INS-52 | chr11-111308430-INS-50 |
| chr11-111319167-INS-55 | chr11-111319167-INS-60 |
| chr11-111333125-INS-54 | chr11-111333125-INS-55 |
| chr11-111343960-INS-60 | chr11-111343960-INS-60 |
| chr11-111375731-INS-56 | chr11-111375731-INS-57 |
| chr11-111377103-INS-73 | chr11-111377103-INS-73 |
| chr11-111399988-INS-186 | chr11-111399988-INS-186 |
| chr11-111408834-INS-220 | chr11-111408834-INS-219 |
| chr11-111419283-INS-108 | chr11-111419283-INS-108 |
| chr11-111445927-INS-59 | chr11-111445927-INS-59 |
| chr11-111453676-INS-96 | chr11-111453676-INS-95 |
| chr11-111460678-INS-167 | chr11-111460678-INS-167 |
| chr11-111470132-INS-83 | chr11-111470132-INS-83 |
| chr11-111493339-INS-51 | chr11-111493339-INS-52 |
| chr11-111507493-INS-84 | chr11-111507495-INS-88 |
| chr11-111509845-INS-69 | chr11-111509845-INS-69 |
| chr11-111553355-INS-605 | chr11-111553355-INS-611 |
| chr11-111567002-INS-127 | chr11-111567006-INS-128 |
| chr11-111578477-INS-74 | chr11-111578477-INS-74 |
| chr11-111597270-INS-74 | chr11-111597270-INS-74 |
| chr11-111614389-INS-493 | chr11-111614372-INS-488 |
| chr11-111644112-INS-390 | chr11-111644112-INS-390 |
| chr11-111663037-INS-53 | chr11-111663037-INS-55 |
| chr11-111671808-INS-70 | chr11-111671808-INS-70 |
| chr11-111681814-INS-85 | chr11-111681808-INS-58 |
| chr11-111682830-INS-114 | chr11-111682830-INS-114 |
| chr11-111690964-INS-108 | chr11-111690964-INS-104 |
| chr11-111699888-INS-101 | chr11-111699932-INS-66 |
| chr11-111714189-INS-56 | chr11-111714189-INS-56 |
| chr11-111721691-INS-59 | chr11-111721691-INS-59 |
| chr11-111741651-INS-56 | chr11-111741651-INS-56 |
| chr11-111745262-INS-84 | chr11-111745265-INS-52 |
| chr11-111772441-INS-538 | chr11-111772461-INS-532 |
| chr11-111800302-INS-101 | chr11-111800342-INS-100 |
| chr11-111828141-INS-679 | chr11-111828141-INS-679 |
| chr11-111923041-INS-513 | chr11-111923041-INS-512 |
| chr11-111926631-INS-946 | chr11-111926632-INS-947 |
| chr11-111934063-INS-57 | chr11-111934063-INS-61 |
| chr11-111937375-INS-56 | chr11-111937375-INS-56 |
| chr11-111941023-INS-400 | chr11-111941023-INS-396 |
| chr11-111957545-INS-72 | chr11-111957545-INS-72 |
| chr11-111964148-INS-90 | chr11-111964148-INS-90 |
| chr11-111979857-INS-199 | chr11-111979857-INS-199 |
| chr11-111991513-INS-100 | chr11-111991520-INS-90 |
| chr11-112018492-INS-73 | chr11-112018492-INS-73 |
| chr11-112039627-INS-66 | chr11-112039627-INS-68 |
| chr11-112078749-INS-67 | chr11-112078712-INS-114 |
| chr11-112080718-INS-54 | chr11-112080718-INS-50 |
| chr11-112085829-INS-352 | chr11-112085829-INS-352 |
| chr11-112087788-INS-434 | chr11-112087789-INS-434 |
| chr11-112102328-INS-95 | chr11-112102328-INS-98 |
| chr11-112106975-INS-373 | chr11-112106975-INS-323 |
| chr11-112112723-INS-64 | chr11-112112723-INS-67 |
| chr11-112151372-INS-65 | chr11-112151405-INS-64 |
| chr11-112154384-INS-432 | chr11-112154384-INS-413 |
| chr11-112159650-INS-58 | chr11-112159650-INS-52 |
| chr11-112168473-INS-54 | chr11-112168455-INS-74 |
| chr11-112170141-INS-54 | chr11-112170145-INS-50 |
| chr11-112174992-INS-77 | chr11-112174945-INS-112 |
| chr11-112175206-INS-5357 | chr11-112175207-INS-5357 |
| chr11-112194560-INS-56 | chr11-112194561-INS-54 |
| chr11-112199311-INS-56 | chr11-112199311-INS-60 |

| | |
|---|---|
| chr11-112202942-INS-141 | chr11-112202969-INS-118 |
| chr11-112209740-INS-124 | chr11-112209740-INS-128 |
| chr11-112231299-INS-304 | chr11-112231299-INS-310 |
| chr11-112280846-INS-3145 | chr11-112280847-INS-3145 |
| chr11-112307555-INS-134 | chr11-112307559-INS-141 |
| chr11-112308876-INS-66 | chr11-112308876-INS-66 |
| chr11-112315430-INS-80 | chr11-112315430-INS-88 |
| chr11-112327855-INS-62 | chr11-112327830-INS-54 |
| chr11-112342284-INS-1909 | chr11-112342285-INS-1913 |
| chr11-112355142-INS-6394 | chr11-112355143-INS-6394 |
| chr11-112367573-INS-54 | chr11-112367573-INS-54 |
| chr11-112368727-INS-56 | chr11-112368727-INS-52 |
| chr11-112378766-INS-54 | chr11-112378766-INS-81 |
| chr11-112394700-INS-99 | chr11-112394700-INS-54 |
| chr11-112427624-INS-52 | chr11-112427624-INS-54 |
| chr11-112456848-INS-133 | chr11-112456848-INS-140 |
| chr11-112459906-INS-108 | chr11-112459932-INS-73 |
| chr11-112467002-INS-1933 | chr11-112467003-INS-1933 |
| chr11-112478576-INS-63 | chr11-112478576-INS-63 |
| chr11-112485183-INS-108 | chr11-112485183-INS-86 |
| chr11-112493510-INS-122 | chr11-112493510-INS-122 |
| chr11-112512514-INS-55 | chr11-112512512-INS-56 |
| chr11-112523907-INS-166 | chr11-112523907-INS-182 |
| chr11-112529622-INS-50 | chr11-112529622-INS-57 |
| chr11-112556417-INS-83 | chr11-112556390-INS-105 |
| chr11-112564934-INS-87 | chr11-112564934-INS-91 |
| chr11-112570691-INS-615 | chr11-112570691-INS-615 |
| chr11-112576338-INS-386 | chr11-112576329-INS-416 |
| chr11-112578399-INS-55 | chr11-112578399-INS-55 |
| chr11-112580527-INS-62 | chr11-112580527-INS-66 |
| chr11-112587915-INS-129 | chr11-112587915-INS-129 |
| chr11-112589585-INS-133 | chr11-112589585-INS-133 |
| chr11-112621015-INS-76 | chr11-112621015-INS-76 |
| chr11-112627672-INS-83 | chr11-112627671-INS-83 |
| chr11-112629641-INS-71 | chr11-112629641-INS-71 |
| chr11-112631392-INS-1242 | chr11-112631393-INS-1249 |
| chr11-112642263-INS-59 | chr11-112642263-INS-59 |
| chr11-112653790-INS-7392 | chr11-112653791-INS-7387 |
| chr11-112653791-INS-594 | chr11-112653791-INS-594 |
| chr11-112683984-INS-96 | chr11-112683984-INS-96 |
| chr11-112707714-INS-55 | chr11-112707700-INS-66 |
| chr11-112711022-INS-51 | chr11-112711022-INS-51 |
| chr11-112728598-INS-168 | chr11-112728598-INS-168 |
| chr11-112730065-INS-52 | chr11-112730065-INS-50 |
| chr11-112765577-INS-52 | chr11-112765534-INS-52 |
| chr11-112777551-INS-82 | chr11-112777532-INS-77 |
| chr11-112800585-INS-87 | chr11-112800585-INS-92 |
| chr11-112825102-INS-128 | chr11-112825102-INS-128 |
| chr11-112830310-INS-111 | chr11-112830310-INS-111 |
| chr11-112853112-INS-108 | chr11-112853112-INS-108 |
| chr11-112868530-INS-52 | chr11-112868531-INS-52 |
| chr11-112876723-INS-399 | chr11-112876723-INS-399 |
| chr11-112885512-INS-59 | chr11-112885513-INS-60 |
| chr11-112891419-INS-120 | chr11-112891419-INS-120 |
| chr11-112938595-INS-160 | chr11-112938595-INS-160 |
| chr11-112940076-INS-67 | chr11-112940076-INS-50 |
| chr11-112946416-INS-56 | chr11-112946439-INS-60 |
| chr11-112963833-INS-92 | chr11-112963833-INS-96 |
| chr11-112990428-INS-193 | chr11-112990428-INS-202 |
| chr11-113026405-INS-58 | chr11-113026411-INS-60 |
| chr11-113053900-INS-215 | chr11-113053900-INS-203 |
| chr11-113057263-INS-57 | chr11-113057281-INS-56 |
| chr11-113064623-INS-314 | chr11-113064623-INS-312 |
| chr11-113093241-INS-139 | chr11-113093241-INS-141 |
| chr11-113107532-INS-64 | chr11-113107532-INS-66 |
| chr11-113120613-INS-195 | chr11-113120613-INS-195 |
| chr11-113123281-INS-2040 | chr11-113123282-INS-2040 |
| chr11-113123383-INS-173 | chr11-113123383-INS-173 |
| chr11-113125920-INS-423 | chr11-113125920-INS-423 |
| chr11-113135164-INS-61 | chr11-113135164-INS-61 |
| chr11-113149264-INS-56 | chr11-113149266-INS-56 |
| chr11-113159738-INS-651 | chr11-113159738-INS-654 |
| chr11-113175705-INS-69 | chr11-113175688-INS-80 |
| chr11-113179985-INS-72 | chr11-113179985-INS-72 |
| chr11-113185941-INS-2002 | chr11-113185942-INS-1999 |
| chr11-113193580-INS-118 | chr11-113193580-INS-118 |
| chr11-113194671-INS-67 | chr11-113194670-INS-75 |

| | |
|---|---|
| chr11-113221514-INS-68 | chr11-113221514-INS-68 |
| chr11-113230314-INS-7438 | chr11-113230315-INS-7438 |
| chr11-113249206-INS-183 | chr11-113249206-INS-183 |
| chr11-113261476-INS-526 | chr11-113261476-INS-526 |
| chr11-113291224-INS-918 | chr11-113291227-INS-922 |
| chr11-113312771-INS-437 | chr11-113312771-INS-439 |
| chr11-113318838-INS-104 | chr11-113318836-INS-112 |
| chr11-113322129-INS-123 | chr11-113322129-INS-150 |
| chr11-113345663-INS-69 | chr11-113345663-INS-54 |
| chr11-113359529-INS-299 | chr11-113359529-INS-299 |
| chr11-113367348-INS-62 | chr11-113367346-INS-74 |
| chr11-113368660-INS-79 | chr11-113368660-INS-53 |
| chr11-113370322-INS-351 | chr11-113370323-INS-335 |
| chr11-113391964-INS-228 | chr11-113391964-INS-228 |
| chr11-113392565-INS-54 | chr11-113392565-INS-56 |
| chr11-113397970-INS-899 | chr11-113397970-INS-899 |
| chr11-113400918-INS-68 | chr11-113400918-INS-68 |
| chr11-113403199-INS-77 | chr11-113403199-INS-77 |
| chr11-113406370-INS-60 | chr11-113406370-INS-68 |
| chr11-113409561-INS-54 | chr11-113409561-INS-54 |
| chr11-113425173-INS-215 | chr11-113425173-INS-214 |
| chr11-113428458-INS-195 | chr11-113428458-INS-195 |
| chr11-113434935-INS-84 | chr11-113434942-INS-112 |
| chr11-113442435-INS-58 | chr11-113442435-INS-58 |
| chr11-113451394-INS-113 | chr11-113451390-INS-115 |
| chr11-113466312-INS-78 | chr11-113466305-INS-78 |
| chr11-113469229-INS-54 | chr11-113469230-INS-75 |
| chr11-113475596-INS-54 | chr11-113475596-INS-54 |
| chr11-113501670-INS-72 | chr11-113501666-INS-112 |
| chr11-113505687-INS-73 | chr11-113505659-INS-73 |
| chr11-113532497-INS-182 | chr11-113532497-INS-183 |
| chr11-113543791-INS-719 | chr11-113543791-INS-719 |
| chr11-113548957-INS-232 | chr11-113548957-INS-232 |
| chr11-113551927-INS-58 | chr11-113551927-INS-58 |
| chr11-113557799-INS-236 | chr11-113557799-INS-236 |
| chr11-113582221-INS-79 | chr11-113582221-INS-79 |
| chr11-113584553-INS-216 | chr11-113584553-INS-216 |
| chr11-113587356-INS-402 | chr11-113587356-INS-402 |
| chr11-113603884-INS-223 | chr11-113603884-INS-223 |
| chr11-113611028-INS-81 | chr11-113611028-INS-96 |
| chr11-113632914-INS-324 | chr11-113632914-INS-324 |
| chr11-113639034-INS-372 | chr11-113639034-INS-358 |
| chr11-113654472-INS-440 | chr11-113654472-INS-440 |
| chr11-113655662-INS-201 | chr11-113655662-INS-201 |
| chr11-113663573-INS-172 | chr11-113663573-INS-178 |
| chr11-113680412-INS-307 | chr11-113680412-INS-283 |
| chr11-113696421-INS-61 | chr11-113696421-INS-61 |
| chr11-113699889-INS-76 | chr11-113699889-INS-76 |
| chr11-113702873-INS-266 | chr11-113702873-INS-266 |
| chr11-113704690-INS-116 | chr11-113704690-INS-116 |
| chr11-113705701-INS-64 | chr11-113705734-INS-69 |
| chr11-113730028-INS-74 | chr11-113730028-INS-74 |
| chr11-113741071-INS-209 | chr11-113741071-INS-210 |
| chr11-113756081-INS-104 | chr11-113756081-INS-104 |
| chr11-113825918-INS-83 | chr11-113825918-INS-83 |
| chr11-113832571-INS-80 | chr11-113832571-INS-116 |
| chr11-113866832-INS-54 | chr11-113866832-INS-54 |
| chr11-113869563-INS-56 | chr11-113869563-INS-56 |
| chr11-113893154-INS-94 | chr11-113893154-INS-94 |
| chr11-113920469-INS-352 | chr11-113920469-INS-352 |
| chr11-113932052-INS-55 | chr11-113932052-INS-55 |
| chr11-113937059-INS-189 | chr11-113937059-INS-189 |
| chr11-113970345-INS-400 | chr11-113970345-INS-400 |
| chr11-113981274-INS-133 | chr11-113981274-INS-143 |
| chr11-113985241-INS-6442 | chr11-113985242-INS-6442 |
| chr11-113986854-INS-431 | chr11-113986854-INS-431 |
| chr11-113994960-INS-144 | chr11-113994950-INS-120 |
| chr11-113998363-INS-63 | chr11-113998363-INS-63 |
| chr11-114018141-INS-65 | chr11-114018141-INS-65 |
| chr11-114050019-INS-56 | chr11-114050019-INS-56 |
| chr11-114051662-INS-132 | chr11-114051662-INS-132 |
| chr11-114079092-INS-116 | chr11-114079092-INS-116 |
| chr11-114091551-INS-52 | chr11-114091551-INS-56 |
| chr11-114113299-INS-401 | chr11-114113299-INS-401 |
| chr11-114117286-INS-74 | chr11-114117286-INS-84 |
| chr11-114125732-INS-88 | chr11-114125732-INS-92 |
| chr11-114170955-INS-112 | chr11-114170951-INS-68 |

| | |
|---|---|
| chr11-114171996-INS-52 | chr11-114171996-INS-50 |
| chr11-114174381-INS-67 | chr11-114174381-INS-74 |
| chr11-114184088-INS-68 | chr11-114184092-INS-85 |
| chr11-114195726-INS-73 | chr11-114195726-INS-118 |
| chr11-114205586-INS-121 | chr11-114205570-INS-124 |
| chr11-114226127-INS-53 | chr11-114226128-INS-53 |
| chr11-114236148-INS-416 | chr11-114236148-INS-406 |
| chr11-114239411-INS-72 | chr11-114239412-INS-62 |
| chr11-114245450-INS-74 | chr11-114245450-INS-74 |
| chr11-114252478-INS-57 | chr11-114252478-INS-55 |
| chr11-114260511-INS-155 | chr11-114260514-INS-191 |
| chr11-114276859-INS-96 | chr11-114276842-INS-64 |
| chr11-114291310-INS-131 | chr11-114291310-INS-131 |
| chr11-114317009-INS-99 | chr11-114317009-INS-96 |
| chr11-114318866-INS-91 | chr11-114318866-INS-91 |
| chr11-114329201-INS-817 | chr11-114329201-INS-820 |
| chr11-114427715-INS-77 | chr11-114427715-INS-77 |
| chr11-114428733-INS-524 | chr11-114428733-INS-492 |
| chr11-114438252-INS-51 | chr11-114438252-INS-51 |
| chr11-114441354-INS-394 | chr11-114441354-INS-394 |
| chr11-114459883-INS-135 | chr11-114459892-INS-144 |
| chr11-114466198-INS-243 | chr11-114466198-INS-243 |
| chr11-114483611-INS-132 | chr11-114483604-INS-113 |
| chr11-114489789-INS-220 | chr11-114489780-INS-216 |
| chr11-114493396-INS-62 | chr11-114493396-INS-62 |
| chr11-114495498-INS-5514 | chr11-114495499-INS-5514 |
| chr11-114497681-INS-217 | chr11-114497681-INS-217 |
| chr11-114508698-INS-120 | chr11-114508696-INS-120 |
| chr11-114534169-INS-396 | chr11-114534169-INS-396 |
| chr11-114543898-INS-212 | chr11-114543898-INS-213 |
| chr11-114554135-INS-91 | chr11-114554146-INS-91 |
| chr11-114563658-INS-217 | chr11-114563658-INS-217 |
| chr11-114578359-INS-114 | chr11-114578359-INS-97 |
| chr11-114593730-INS-201 | chr11-114593731-INS-201 |
| chr11-114622659-INS-137 | chr11-114622657-INS-152 |
| chr11-114624367-INS-102 | chr11-114624327-INS-103 |
| chr11-114632377-INS-1353 | chr11-114632378-INS-1355 |
| chr11-114635752-INS-138 | chr11-114635752-INS-100 |
| chr11-114638293-INS-149 | chr11-114638293-INS-149 |
| chr11-114644712-INS-6442 | chr11-114644713-INS-6442 |
| chr11-114665434-INS-162 | chr11-114665434-INS-160 |
| chr11-114694431-INS-69 | chr11-114694431-INS-69 |
| chr11-114702991-INS-100 | chr11-114702991-INS-100 |
| chr11-114707304-INS-67 | chr11-114707323-INS-65 |
| chr11-114708573-INS-769 | chr11-114708573-INS-770 |
| chr11-114713030-INS-51 | chr11-114713031-INS-54 |
| chr11-114718487-INS-345 | chr11-114718469-INS-362 |
| chr11-114721952-INS-56 | chr11-114721952-INS-54 |
| chr11-114729868-INS-257 | chr11-114729868-INS-257 |
| chr11-114730395-INS-118 | chr11-114730395-INS-118 |
| chr11-114745250-INS-209 | chr11-114745250-INS-209 |
| chr11-114746257-INS-232 | chr11-114746257-INS-231 |
| chr11-114747249-INS-103 | chr11-114747249-INS-103 |
| chr11-114748857-INS-88 | chr11-114748857-INS-88 |
| chr11-114753266-INS-72 | chr11-114753266-INS-72 |
| chr11-114758714-INS-98 | chr11-114758714-INS-100 |
| chr11-114769765-INS-89 | chr11-114769765-INS-89 |
| chr11-114771892-INS-90 | chr11-114771889-INS-90 |
| chr11-114781803-INS-50 | chr11-114781803-INS-54 |
| chr11-114788517-INS-52 | chr11-114788517-INS-52 |
| chr11-114797767-INS-1888 | chr11-114797768-INS-1888 |
| chr11-114837885-INS-135 | chr11-114837885-INS-135 |
| chr11-114848211-INS-111 | chr11-114848212-INS-111 |
| chr11-114852591-INS-476 | chr11-114852591-INS-476 |
| chr11-114857304-INS-414 | chr11-114857305-INS-414 |
| chr11-114861486-INS-504 | chr11-114861486-INS-505 |
| chr11-114866247-INS-417 | chr11-114866247-INS-417 |
| chr11-114889960-INS-276 | chr11-114889960-INS-282 |
| chr11-114899358-INS-69 | chr11-114899359-INS-81 |
| chr11-114901079-INS-356 | chr11-114901036-INS-383 |
| chr11-114911473-INS-172 | chr11-114911473-INS-178 |
| chr11-114920477-INS-66 | chr11-114920470-INS-66 |
| chr11-114932924-INS-271 | chr11-114932931-INS-244 |
| chr11-114935657-INS-68 | chr11-114935657-INS-68 |
| chr11-114951270-INS-71 | chr11-114951271-INS-71 |
| chr11-114974617-INS-80 | chr11-114974617-INS-80 |
| chr11-115006529-INS-169 | chr11-115006529-INS-151 |

| | |
|---|---|
| chr11-115008932-INS-68 | chr11-115008932-INS-68 |
| chr11-115022389-INS-54 | chr11-115022390-INS-54 |
| chr11-115065467-INS-98 | chr11-115065469-INS-101 |
| chr11-115068384-INS-233 | chr11-115068384-INS-233 |
| chr11-115082612-INS-211 | chr11-115082612-INS-211 |
| chr11-115110898-INS-448 | chr11-115110898-INS-448 |
| chr11-115125900-INS-81 | chr11-115125900-INS-81 |
| chr11-115132142-INS-268 | chr11-115132142-INS-268 |
| chr11-115154209-INS-62 | chr11-115154209-INS-68 |
| chr11-115210559-INS-64 | chr11-115210559-INS-96 |
| chr11-115217018-INS-81 | chr11-115217014-INS-55 |
| chr11-115242623-INS-220 | chr11-115242623-INS-233 |
| chr11-115267416-INS-102 | chr11-115267407-INS-78 |
| chr11-115267425-INS-738 | chr11-115267425-INS-738 |
| chr11-115281872-INS-99 | chr11-115281872-INS-93 |
| chr11-115297691-INS-187 | chr11-115297691-INS-187 |
| chr11-115314271-INS-198 | chr11-115314271-INS-198 |
| chr11-115318743-INS-90 | chr11-115318743-INS-90 |
| chr11-115334953-INS-60 | chr11-115334954-INS-88 |
| chr11-115339667-INS-4994 | chr11-115339668-INS-4994 |
| chr11-115343528-INS-80 | chr11-115343528-INS-104 |
| chr11-115345253-INS-103 | chr11-115345253-INS-99 |
| chr11-115346442-INS-177 | chr11-115346442-INS-178 |
| chr11-115359509-INS-200 | chr11-115359509-INS-201 |
| chr11-115394524-INS-96 | chr11-115394524-INS-96 |
| chr11-115408825-INS-219 | chr11-115408828-INS-216 |
| chr11-115431343-INS-65 | chr11-115431343-INS-65 |
| chr11-115437989-INS-58 | chr11-115437989-INS-51 |
| chr11-115447089-INS-215 | chr11-115447089-INS-215 |
| chr11-115456314-INS-220 | chr11-115456314-INS-221 |
| chr11-115460466-INS-932 | chr11-115460466-INS-932 |
| chr11-115484012-INS-596 | chr11-115484012-INS-596 |
| chr11-115489117-INS-224 | chr11-115489117-INS-225 |
| chr11-115509565-INS-206 | chr11-115509565-INS-213 |
| chr11-115519482-INS-60 | chr11-115519482-INS-60 |
| chr11-115520051-INS-52 | chr11-115520051-INS-50 |
| chr11-115527514-INS-52 | chr11-115527514-INS-54 |
| chr11-115531800-INS-203 | chr11-115531800-INS-203 |
| chr11-115533641-INS-226 | chr11-115533641-INS-227 |
| chr11-115555585-INS-354 | chr11-115555585-INS-356 |
| chr11-115612218-INS-83 | chr11-115612218-INS-100 |
| chr11-115613308-INS-53 | chr11-115613308-INS-70 |
| chr11-115615661-INS-239 | chr11-115615661-INS-213 |
| chr11-115625813-INS-249 | chr11-115625813-INS-250 |
| chr11-115626680-INS-70 | chr11-115626680-INS-51 |
| chr11-115659268-INS-51 | chr11-115659268-INS-51 |
| chr11-115659596-INS-218 | chr11-115659597-INS-209 |
| chr11-115662537-INS-73 | chr11-115662538-INS-76 |
| chr11-115665282-INS-153 | chr11-115665282-INS-153 |
| chr11-115669252-INS-69 | chr11-115669250-INS-85 |
| chr11-115672121-INS-76 | chr11-115672121-INS-78 |
| chr11-115675073-INS-1400 | chr11-115675074-INS-1400 |
| chr11-115681244-INS-354 | chr11-115681240-INS-333 |
| chr11-115696299-INS-72 | chr11-115696299-INS-72 |
| chr11-115736917-INS-210 | chr11-115736917-INS-210 |
| chr11-115757796-INS-51 | chr11-115757796-INS-51 |
| chr11-115759549-INS-202 | chr11-115759549-INS-202 |
| chr11-115763829-INS-80 | chr11-115763829-INS-82 |
| chr11-115772913-INS-160 | chr11-115772909-INS-165 |
| chr11-115779365-INS-116 | chr11-115779365-INS-116 |
| chr11-115779665-INS-963 | chr11-115779665-INS-961 |
| chr11-115801070-INS-174 | chr11-115801070-INS-176 |
| chr11-115802833-INS-78 | chr11-115802833-INS-78 |
| chr11-115830408-INS-106 | chr11-115830408-INS-108 |
| chr11-115833928-INS-463 | chr11-115833928-INS-464 |
| chr11-115861374-INS-66 | chr11-115861375-INS-67 |
| chr11-115863908-INS-131 | chr11-115863868-INS-134 |
| chr11-115885865-INS-53 | chr11-115885865-INS-53 |
| chr11-115889580-INS-184 | chr11-115889580-INS-184 |
| chr11-115910633-INS-222 | chr11-115910633-INS-218 |
| chr11-115919904-INS-56 | chr11-115919903-INS-59 |
| chr11-115926121-INS-204 | chr11-115926121-INS-204 |
| chr11-115927962-INS-199 | chr11-115927962-INS-199 |
| chr11-115935883-INS-207 | chr11-115935883-INS-207 |
| chr11-115940698-INS-55 | chr11-115940679-INS-52 |
| chr11-115979044-INS-364 | chr11-115979044-INS-368 |
| chr11-115985647-INS-58 | chr11-115985647-INS-66 |

| | |
|---|---|
| chr11-115987148-INS-62 | chr11-115987154-INS-62 |
| chr11-116005467-INS-198 | chr11-116005470-INS-207 |
| chr11-116008965-INS-88 | chr11-116008965-INS-84 |
| chr11-116011137-INS-198 | chr11-116011137-INS-200 |
| chr11-116013202-INS-199 | chr11-116013202-INS-200 |
| chr11-116021363-INS-51 | chr11-116021363-INS-51 |
| chr11-116031059-INS-197 | chr11-116031059-INS-198 |
| chr11-116044927-INS-61 | chr11-116044927-INS-61 |
| chr11-116057178-INS-207 | chr11-116057178-INS-209 |
| chr11-116059649-INS-206 | chr11-116059650-INS-200 |
| chr11-116078493-INS-51 | chr11-116078493-INS-51 |
| chr11-116084810-INS-51 | chr11-116084791-INS-69 |
| chr11-116087818-INS-185 | chr11-116087818-INS-185 |
| chr11-116098037-INS-61 | chr11-116098037-INS-61 |
| chr11-116100005-INS-210 | chr11-116100005-INS-210 |
| chr11-116105463-INS-62 | chr11-116105463-INS-64 |
| chr11-116110382-INS-124 | chr11-116110382-INS-104 |
| chr11-116116793-INS-152 | chr11-116116794-INS-134 |
| chr11-116117718-INS-215 | chr11-116117718-INS-215 |
| chr11-116129995-INS-76 | chr11-116129994-INS-73 |
| chr11-116140946-INS-74 | chr11-116140946-INS-74 |
| chr11-116143069-INS-62 | chr11-116143069-INS-62 |
| chr11-116209095-INS-56 | chr11-116209095-INS-56 |
| chr11-116231868-INS-222 | chr11-116231868-INS-224 |
| chr11-116250222-INS-60 | chr11-116250221-INS-66 |
| chr11-116257614-INS-105 | chr11-116257614-INS-105 |
| chr11-116270347-INS-222 | chr11-116270347-INS-225 |
| chr11-116275729-INS-64 | chr11-116275729-INS-68 |
| chr11-116293282-INS-228 | chr11-116293282-INS-228 |
| chr11-116296731-INS-197 | chr11-116296731-INS-196 |
| chr11-116343956-INS-88 | chr11-116343956-INS-88 |
| chr11-116377086-INS-838 | chr11-116377086-INS-838 |
| chr11-116383444-INS-957 | chr11-116383443-INS-963 |
| chr11-116386806-INS-52 | chr11-116386810-INS-53 |
| chr11-116395025-INS-217 | chr11-116395025-INS-218 |
| chr11-116396265-INS-220 | chr11-116396265-INS-220 |
| chr11-116405636-INS-54 | chr11-116405627-INS-60 |
| chr11-116408729-INS-576 | chr11-116408729-INS-576 |
| chr11-116460667-INS-58 | chr11-116460667-INS-58 |
| chr11-116474880-INS-87 | chr11-116474880-INS-87 |
| chr11-116482571-INS-63 | chr11-116482569-INS-68 |
| chr11-116495586-INS-68 | chr11-116495585-INS-90 |
| chr11-116532064-INS-324 | chr11-116532064-INS-324 |
| chr11-116533229-INS-134 | chr11-116533229-INS-134 |
| chr11-116564159-INS-187 | chr11-116564150-INS-201 |
| chr11-116566129-INS-282 | chr11-116566129-INS-278 |
| chr11-116591475-INS-197 | chr11-116591475-INS-197 |
| chr11-116599382-INS-191 | chr11-116599382-INS-192 |
| chr11-116617182-INS-400 | chr11-116617182-INS-400 |
| chr11-116621054-INS-211 | chr11-116621054-INS-211 |
| chr11-116622703-INS-1137 | chr11-116622704-INS-1141 |
| chr11-116625047-INS-96 | chr11-116625047-INS-86 |
| chr11-116640378-INS-895 | chr11-116640378-INS-895 |
| chr11-116666936-INS-84 | chr11-116666936-INS-86 |
| chr11-116674010-INS-206 | chr11-116674010-INS-206 |
| chr11-116688928-INS-107 | chr11-116688928-INS-107 |
| chr11-116693163-INS-54 | chr11-116693137-INS-56 |
| chr11-116700043-INS-215 | chr11-116700043-INS-206 |
| chr11-116712743-INS-53 | chr11-116712743-INS-53 |
| chr11-116714758-INS-165 | chr11-116714758-INS-144 |
| chr11-116724745-INS-205 | chr11-116724745-INS-205 |
| chr11-116725262-INS-339 | chr11-116725262-INS-341 |
| chr11-116752246-INS-205 | chr11-116752246-INS-205 |
| chr11-116770317-INS-96 | chr11-116770300-INS-72 |
| chr11-116799201-INS-6413 | chr11-116799202-INS-6413 |
| chr11-116803253-INS-145 | chr11-116803252-INS-130 |
| chr11-116806124-INS-189 | chr11-116806124-INS-189 |
| chr11-116834046-INS-54 | chr11-116834046-INS-62 |
| chr11-116860009-INS-192 | chr11-116860041-INS-192 |
| chr11-116880949-INS-198 | chr11-116880949-INS-198 |
| chr11-116886849-INS-97 | chr11-116886849-INS-96 |
| chr11-116896171-INS-302 | chr11-116896171-INS-302 |
| chr11-116930758-INS-169 | chr11-116930758-INS-169 |
| chr11-116944801-INS-222 | chr11-116944801-INS-230 |
| chr11-116949219-INS-84 | chr11-116949219-INS-66 |
| chr11-116978257-INS-368 | chr11-116978257-INS-355 |
| chr11-116980101-INS-107 | chr11-116980101-INS-107 |

| | |
|---|---|
| chr11-116984303-INS-544 | chr11-116984310-INS-526 |
| chr11-117001934-INS-67 | chr11-117001934-INS-55 |
| chr11-117004765-INS-84 | chr11-117004765-INS-84 |
| chr11-117148063-INS-210 | chr11-117148063-INS-210 |
| chr11-117165684-INS-204 | chr11-117165684-INS-204 |
| chr11-117260071-INS-209 | chr11-117260072-INS-200 |
| chr11-117264830-INS-52 | chr11-117264830-INS-52 |
| chr11-117267637-INS-218 | chr11-117267637-INS-218 |
| chr11-117269303-INS-68 | chr11-117269303-INS-68 |
| chr11-117278449-INS-65 | chr11-117278449-INS-65 |
| chr11-117279671-INS-58 | chr11-117279669-INS-62 |
| chr11-117290624-INS-64 | chr11-117290624-INS-64 |
| chr11-117308331-INS-101 | chr11-117308331-INS-101 |
| chr11-117324078-INS-201 | chr11-117324078-INS-200 |
| chr11-117330443-INS-80 | chr11-117330443-INS-80 |
| chr11-117334234-INS-62 | chr11-117334234-INS-62 |
| chr11-117340067-INS-232 | chr11-117340067-INS-236 |
| chr11-117349129-INS-52 | chr11-117349129-INS-52 |
| chr11-117351056-INS-354 | chr11-117351056-INS-356 |
| chr11-117353595-INS-79 | chr11-117353595-INS-90 |
| chr11-117356697-INS-98 | chr11-117356697-INS-102 |
| chr11-117359052-INS-61 | chr11-117359052-INS-61 |
| chr11-117366463-INS-74 | chr11-117366463-INS-74 |
| chr11-117373965-INS-51 | chr11-117373965-INS-51 |
| chr11-117380979-INS-59 | chr11-117380979-INS-59 |
| chr11-117404689-INS-763 | chr11-117404689-INS-746 |
| chr11-117415647-INS-6442 | chr11-117415648-INS-6442 |
| chr11-117435764-INS-73 | chr11-117435764-INS-73 |
| chr11-117440481-INS-257 | chr11-117440481-INS-257 |
| chr11-117446388-INS-83 | chr11-117446388-INS-83 |
| chr11-117447037-INS-52 | chr11-117447037-INS-52 |
| chr11-117467096-INS-86 | chr11-117467096-INS-90 |
| chr11-117474767-INS-4155 | chr11-117474768-INS-4155 |
| chr11-117476297-INS-197 | chr11-117476297-INS-197 |
| chr11-117506769-INS-370 | chr11-117506769-INS-370 |
| chr11-117515802-INS-210 | chr11-117515802-INS-210 |
| chr11-117525348-INS-179 | chr11-117525348-INS-176 |
| chr11-117528515-INS-222 | chr11-117528515-INS-222 |
| chr11-117584416-INS-63 | chr11-117584416-INS-63 |
| chr11-117585620-INS-100 | chr11-117585620-INS-100 |
| chr11-117643163-INS-219 | chr11-117643163-INS-219 |
| chr11-117648078-INS-250 | chr11-117648078-INS-254 |
| chr11-117662435-INS-208 | chr11-117662435-INS-208 |
| chr11-117676322-INS-175 | chr11-117676322-INS-175 |
| chr11-117694567-INS-64 | chr11-117694567-INS-53 |
| chr11-117696191-INS-1068 | chr11-117696192-INS-1068 |
| chr11-117728185-INS-54 | chr11-117728185-INS-52 |
| chr11-117764405-INS-962 | chr11-117764405-INS-962 |
| chr11-117771350-INS-64 | chr11-117771350-INS-64 |
| chr11-117775984-INS-439 | chr11-117775984-INS-439 |
| chr11-117792230-INS-237 | chr11-117792228-INS-197 |
| chr11-117794652-INS-63 | chr11-117794688-INS-54 |
| chr11-117796960-INS-133 | chr11-117796960-INS-133 |
| chr11-117812902-INS-70 | chr11-117812902-INS-70 |
| chr11-117824688-INS-70 | chr11-117824688-INS-70 |
| chr11-117826966-INS-213 | chr11-117826966-INS-213 |
| chr11-117830417-INS-75 | chr11-117830412-INS-70 |
| chr11-117848278-INS-71 | chr11-117848278-INS-73 |
| chr11-117875574-INS-76 | chr11-117875574-INS-56 |
| chr11-117884614-INS-225 | chr11-117884614-INS-225 |
| chr11-117888033-INS-213 | chr11-117888033-INS-213 |
| chr11-117897967-INS-210 | chr11-117897967-INS-212 |
| chr11-117935240-INS-267 | chr11-117935240-INS-268 |
| chr11-117950540-INS-156 | chr11-117950540-INS-156 |
| chr11-117952656-INS-88 | chr11-117952656-INS-88 |
| chr11-117990603-INS-213 | chr11-117990603-INS-213 |
| chr11-118012188-INS-95 | chr11-118012188-INS-61 |
| chr11-118018589-INS-238 | chr11-118018589-INS-238 |
| chr11-118036012-INS-175 | chr11-118036012-INS-225 |
| chr11-118042698-INS-209 | chr11-118042698-INS-209 |
| chr11-118076447-INS-222 | chr11-118076447-INS-223 |
| chr11-118111117-INS-211 | chr11-118111117-INS-211 |
| chr11-118116710-INS-62 | chr11-118116708-INS-64 |
| chr11-118143909-INS-222 | chr11-118143909-INS-222 |
| chr11-118165019-INS-56 | chr11-118165019-INS-56 |
| chr11-118172162-INS-93 | chr11-118172162-INS-93 |
| chr11-118204085-INS-83 | chr11-118204069-INS-83 |

| | |
|---|---|
| chr11-118236215-INS-61 | chr11-118236216-INS-61 |
| chr11-118308122-INS-113 | chr11-118308122-INS-113 |
| chr11-118330554-INS-52 | chr11-118330554-INS-56 |
| chr11-118341278-INS-194 | chr11-118341278-INS-194 |
| chr11-118343060-INS-62 | chr11-118343060-INS-66 |
| chr11-118369213-INS-60 | chr11-118369201-INS-61 |
| chr11-118391508-INS-63 | chr11-118391508-INS-63 |
| chr11-118408356-INS-204 | chr11-118408356-INS-205 |
| chr11-118423698-INS-214 | chr11-118423698-INS-213 |
| chr11-118446661-INS-259 | chr11-118446661-INS-222 |
| chr11-118495462-INS-75 | chr11-118495462-INS-125 |
| chr11-118592388-INS-390 | chr11-118592388-INS-390 |
| chr11-118628957-INS-123 | chr11-118628957-INS-86 |
| chr11-118633052-INS-76 | chr11-118633040-INS-70 |
| chr11-118648838-INS-96 | chr11-118648838-INS-96 |
| chr11-118659213-INS-54 | chr11-118659213-INS-54 |
| chr11-118681179-INS-223 | chr11-118681179-INS-224 |
| chr11-118687212-INS-66 | chr11-118687207-INS-56 |
| chr11-118690706-INS-87 | chr11-118690688-INS-78 |
| chr11-118699514-INS-74 | chr11-118699505-INS-88 |
| chr11-118778112-INS-528 | chr11-118778112-INS-528 |
| chr11-118806717-INS-159 | chr11-118806717-INS-159 |
| chr11-118824862-INS-58 | chr11-118824862-INS-60 |
| chr11-118826950-INS-74 | chr11-118826950-INS-78 |
| chr11-118839689-INS-54 | chr11-118839689-INS-54 |
| chr11-118878832-INS-334 | chr11-118878832-INS-334 |
| chr11-118889261-INS-7185 | chr11-118889262-INS-7185 |
| chr11-118892605-INS-95 | chr11-118892582-INS-116 |
| chr11-118918039-INS-72 | chr11-118918042-INS-62 |
| chr11-118923841-INS-70 | chr11-118923841-INS-70 |
| chr11-118986595-INS-130 | chr11-118986601-INS-83 |
| chr11-118997337-INS-159 | chr11-118997333-INS-155 |
| chr11-119003978-INS-68 | chr11-119003990-INS-72 |
| chr11-119023896-INS-59 | chr11-119023896-INS-60 |
| chr11-119029925-INS-59 | chr11-119029925-INS-59 |
| chr11-119095994-INS-199 | chr11-119095979-INS-232 |
| chr11-119102197-INS-194 | chr11-119102197-INS-194 |
| chr11-119111626-INS-98 | chr11-119111626-INS-98 |
| chr11-119118709-INS-52 | chr11-119118709-INS-60 |
| chr11-119140426-INS-243 | chr11-119140426-INS-193 |
| chr11-119163447-INS-943 | chr11-119163447-INS-943 |
| chr11-119167020-INS-80 | chr11-119167020-INS-80 |
| chr11-119173537-INS-487 | chr11-119173538-INS-487 |
| chr11-119199468-INS-70 | chr11-119199468-INS-71 |
| chr11-119207110-INS-106 | chr11-119207110-INS-106 |
| chr11-119210275-INS-61 | chr11-119210275-INS-79 |
| chr11-119222665-INS-56 | chr11-119222665-INS-56 |
| chr11-119223561-INS-106 | chr11-119223561-INS-112 |
| chr11-119224065-INS-225 | chr11-119224065-INS-225 |
| chr11-119238375-INS-158 | chr11-119238375-INS-158 |
| chr11-119239745-INS-499 | chr11-119239745-INS-499 |
| chr11-119251669-INS-113 | chr11-119251669-INS-113 |
| chr11-119283467-INS-72 | chr11-119283467-INS-67 |
| chr11-119292364-INS-176 | chr11-119292364-INS-176 |
| chr11-119294182-INS-221 | chr11-119294183-INS-205 |
| chr11-119299403-INS-773 | chr11-119299403-INS-775 |
| chr11-119301052-INS-72 | chr11-119301052-INS-108 |
| chr11-119303583-INS-192 | chr11-119303583-INS-192 |
| chr11-119305725-INS-57 | chr11-119305725-INS-57 |
| chr11-119330637-INS-221 | chr11-119330637-INS-218 |
| chr11-119349004-INS-327 | chr11-119349004-INS-327 |
| chr11-119355245-INS-99 | chr11-119355245-INS-99 |
| chr11-119402322-INS-206 | chr11-119402322-INS-206 |
| chr11-119415271-INS-56 | chr11-119415271-INS-58 |
| chr11-119424235-INS-120 | chr11-119424235-INS-120 |
| chr11-119426268-INS-115 | chr11-119426268-INS-115 |
| chr11-119440834-INS-56 | chr11-119440834-INS-105 |
| chr11-119452837-INS-459 | chr11-119452837-INS-459 |
| chr11-119465369-INS-65 | chr11-119465369-INS-65 |
| chr11-119466615-INS-226 | chr11-119466615-INS-216 |
| chr11-119468986-INS-198 | chr11-119468986-INS-199 |
| chr11-119470345-INS-196 | chr11-119470345-INS-166 |
| chr11-119475670-INS-216 | chr11-119475670-INS-225 |
| chr11-119487175-INS-215 | chr11-119487175-INS-215 |
| chr11-119502060-INS-56 | chr11-119502060-INS-56 |
| chr11-119504240-INS-65 | chr11-119504240-INS-76 |
| chr11-119505885-INS-64 | chr11-119505885-INS-65 |

| | |
|---|---|
| chr11-119528917-INS-7111 | chr11-119528918-INS-7111 |
| chr11-119530689-INS-102 | chr11-119530677-INS-81 |
| chr11-119571366-INS-401 | chr11-119571366-INS-401 |
| chr11-119602424-INS-82 | chr11-119602382-INS-92 |
| chr11-119617399-INS-110 | chr11-119617400-INS-115 |
| chr11-119647527-INS-235 | chr11-119647527-INS-235 |
| chr11-119654233-INS-180 | chr11-119654233-INS-180 |
| chr11-119663397-INS-91 | chr11-119663401-INS-52 |
| chr11-119679637-INS-384 | chr11-119679637-INS-352 |
| chr11-119700848-INS-62 | chr11-119700868-INS-50 |
| chr11-119703970-INS-895 | chr11-119703970-INS-895 |
| chr11-119711174-INS-67 | chr11-119711141-INS-71 |
| chr11-119720443-INS-74 | chr11-119720436-INS-111 |
| chr11-119751833-INS-96 | chr11-119751833-INS-73 |
| chr11-119773279-INS-203 | chr11-119773279-INS-203 |
| chr11-119793830-INS-204 | chr11-119793830-INS-205 |
| chr11-119840077-INS-207 | chr11-119840077-INS-208 |
| chr11-119918407-INS-120 | chr11-119918407-INS-126 |
| chr11-119933182-INS-204 | chr11-119933182-INS-204 |
| chr11-119941632-INS-105 | chr11-119941681-INS-105 |
| chr11-119947834-INS-437 | chr11-119947834-INS-437 |
| chr11-119961546-INS-76 | chr11-119961561-INS-76 |
| chr11-120014944-INS-232 | chr11-120014944-INS-188 |
| chr11-120028647-INS-383 | chr11-120028659-INS-384 |
| chr11-120030501-INS-145 | chr11-120030501-INS-146 |
| chr11-120031783-INS-105 | chr11-120031783-INS-105 |
| chr11-120043536-INS-298 | chr11-120043536-INS-298 |
| chr11-120047193-INS-134 | chr11-120047193-INS-134 |
| chr11-120049355-INS-198 | chr11-120049355-INS-198 |
| chr11-120085996-INS-63 | chr11-120085972-INS-81 |
| chr11-120112855-INS-58 | chr11-120112855-INS-60 |
| chr11-120115024-INS-58 | chr11-120115022-INS-56 |
| chr11-120140689-INS-107 | chr11-120140689-INS-107 |
| chr11-120144954-INS-72 | chr11-120144982-INS-94 |
| chr11-120154164-INS-59 | chr11-120154164-INS-59 |
| chr11-120169775-INS-740 | chr11-120169775-INS-751 |
| chr11-120194960-INS-57 | chr11-120194960-INS-57 |
| chr11-120199972-INS-214 | chr11-120199972-INS-214 |
| chr11-120206664-INS-85 | chr11-120206664-INS-75 |
| chr11-120210136-INS-400 | chr11-120210136-INS-400 |
| chr11-120225791-INS-56 | chr11-120225791-INS-56 |
| chr11-120227908-INS-170 | chr11-120227908-INS-177 |
| chr11-120234229-INS-214 | chr11-120234229-INS-214 |
| chr11-120290039-INS-445 | chr11-120290039-INS-445 |
| chr11-120306039-INS-551 | chr11-120306039-INS-561 |
| chr11-120309499-INS-56 | chr11-120309499-INS-62 |
| chr11-120330327-INS-555 | chr11-120330328-INS-555 |
| chr11-120342626-INS-416 | chr11-120342626-INS-417 |
| chr11-120346504-INS-216 | chr11-120346504-INS-217 |
| chr11-120352447-INS-99 | chr11-120352447-INS-99 |
| chr11-120356316-INS-211 | chr11-120356316-INS-214 |
| chr11-120367561-INS-138 | chr11-120367561-INS-138 |
| chr11-120401936-INS-53 | chr11-120401936-INS-57 |
| chr11-120402994-INS-205 | chr11-120402994-INS-194 |
| chr11-120405055-INS-132 | chr11-120405055-INS-131 |
| chr11-120414161-INS-206 | chr11-120414161-INS-205 |
| chr11-120449647-INS-185 | chr11-120449647-INS-185 |
| chr11-120458230-INS-218 | chr11-120458230-INS-219 |
| chr11-120474206-INS-220 | chr11-120474206-INS-221 |
| chr11-120501231-INS-54 | chr11-120501231-INS-54 |
| chr11-120526498-INS-220 | chr11-120526498-INS-221 |
| chr11-120554920-INS-96 | chr11-120554920-INS-96 |
| chr11-120560766-INS-56 | chr11-120560766-INS-56 |
| chr11-120566655-INS-190 | chr11-120566655-INS-189 |
| chr11-120578909-INS-78 | chr11-120578909-INS-78 |
| chr11-120627194-INS-165 | chr11-120627194-INS-179 |
| chr11-120644211-INS-213 | chr11-120644211-INS-213 |
| chr11-120681126-INS-216 | chr11-120681126-INS-215 |
| chr11-120689843-INS-90 | chr11-120689837-INS-99 |
| chr11-120728399-INS-62 | chr11-120728399-INS-74 |
| chr11-120731477-INS-70 | chr11-120731477-INS-50 |
| chr11-120735600-INS-6415 | chr11-120735601-INS-6415 |
| chr11-120739359-INS-118 | chr11-120739359-INS-118 |
| chr11-120748874-INS-340 | chr11-120748861-INS-345 |
| chr11-120807083-INS-209 | chr11-120807083-INS-229 |
| chr11-120820091-INS-240 | chr11-120820087-INS-243 |
| chr11-120825791-INS-7083 | chr11-120825792-INS-7083 |

| | |
|---|---|
| chr11-120857807-INS-249 | chr11-120857807-INS-238 |
| chr11-120894556-INS-125 | chr11-120894556-INS-125 |
| chr11-120897628-INS-184 | chr11-120897628-INS-184 |
| chr11-120910075-INS-586 | chr11-120910075-INS-577 |
| chr11-120944083-INS-1877 | chr11-120944084-INS-1877 |
| chr11-120951857-INS-152 | chr11-120951857-INS-156 |
| chr11-121000181-INS-189 | chr11-121000181-INS-189 |
| chr11-121000603-INS-52 | chr11-121000603-INS-52 |
| chr11-121044155-INS-192 | chr11-121044155-INS-198 |
| chr11-121071952-INS-273 | chr11-121071953-INS-242 |
| chr11-121086112-INS-214 | chr11-121086112-INS-214 |
| chr11-121092661-INS-233 | chr11-121092661-INS-233 |
| chr11-121123166-INS-7160 | chr11-121123167-INS-7160 |
| chr11-121130277-INS-208 | chr11-121130277-INS-208 |
| chr11-121158146-INS-203 | chr11-121158146-INS-203 |
| chr11-121163405-INS-214 | chr11-121163405-INS-218 |
| chr11-121167088-INS-456 | chr11-121167089-INS-460 |
| chr11-121178000-INS-220 | chr11-121178000-INS-220 |
| chr11-121216229-INS-211 | chr11-121216229-INS-211 |
| chr11-121222586-INS-480 | chr11-121222586-INS-480 |
| chr11-121226903-INS-201 | chr11-121226903-INS-201 |
| chr11-121230141-INS-51 | chr11-121230141-INS-51 |
| chr11-121246530-INS-184 | chr11-121246530-INS-184 |
| chr11-121252580-INS-177 | chr11-121252580-INS-177 |
| chr11-121260276-INS-102 | chr11-121260276-INS-102 |
| chr11-121270025-INS-713 | chr11-121270025-INS-713 |
| chr11-121302013-INS-438 | chr11-121302013-INS-444 |
| chr11-121303168-INS-206 | chr11-121303171-INS-206 |
| chr11-121337777-INS-75 | chr11-121337777-INS-52 |
| chr11-121347711-INS-177 | chr11-121347711-INS-177 |
| chr11-121351884-INS-75 | chr11-121351900-INS-99 |
| chr11-121364932-INS-62 | chr11-121364932-INS-62 |
| chr11-121473226-INS-2047 | chr11-121473227-INS-2064 |
| chr11-121478063-INS-76 | chr11-121478063-INS-92 |
| chr11-121533814-INS-108 | chr11-121533815-INS-90 |
| chr11-121551561-INS-82 | chr11-121551561-INS-116 |
| chr11-121573761-INS-277 | chr11-121573761-INS-277 |
| chr11-121584671-INS-3695 | chr11-121584672-INS-3694 |
| chr11-121592692-INS-75 | chr11-121592692-INS-58 |
| chr11-121596308-INS-68 | chr11-121596308-INS-70 |
| chr11-121618415-INS-6404 | chr11-121618416-INS-6404 |
| chr11-121621216-INS-184 | chr11-121621216-INS-193 |
| chr11-121645089-INS-421 | chr11-121645089-INS-421 |
| chr11-121781710-INS-400 | chr11-121781710-INS-401 |
| chr11-121794134-INS-6883 | chr11-121794135-INS-6883 |
| chr11-121809022-INS-201 | chr11-121809022-INS-201 |
| chr11-121809873-INS-203 | chr11-121809873-INS-203 |
| chr11-121840718-INS-114 | chr11-121840731-INS-122 |
| chr11-121846652-INS-188 | chr11-121846652-INS-189 |
| chr11-121849472-INS-203 | chr11-121849472-INS-199 |
| chr12-3135754-INS-339 | chr12-3135757-INS-341 |
| chr12-3136789-INS-186 | chr12-3136816-INS-201 |
| chr12-3182328-INS-68 | chr12-3182296-INS-52 |
| chr12-3208816-INS-212 | chr12-3208816-INS-215 |
| chr12-3217348-INS-90 | chr12-3217390-INS-64 |
| chr12-3240113-INS-637 | chr12-3240113-INS-637 |
| chr12-3270512-INS-498 | chr12-3270512-INS-498 |
| chr12-3273493-INS-75 | chr12-3273493-INS-75 |
| chr12-3275059-INS-68 | chr12-3275059-INS-68 |
| chr12-3284401-INS-185 | chr12-3284401-INS-185 |
| chr12-3290591-INS-114 | chr12-3290591-INS-114 |
| chr12-3291558-INS-432 | chr12-3291563-INS-432 |
| chr12-3347773-INS-135 | chr12-3347775-INS-155 |
| chr12-3354403-INS-179 | chr12-3354406-INS-184 |
| chr12-3361776-INS-212 | chr12-3361776-INS-215 |
| chr12-3370246-INS-74 | chr12-3370243-INS-65 |
| chr12-3372842-INS-66 | chr12-3372842-INS-66 |
| chr12-3374008-INS-74 | chr12-3374008-INS-54 |
| chr12-3394026-INS-205 | chr12-3394026-INS-205 |
| chr12-3396210-INS-360 | chr12-3396210-INS-360 |
| chr12-3435779-INS-57 | chr12-3435779-INS-51 |
| chr12-3447629-INS-60 | chr12-3447629-INS-60 |
| chr12-3457027-INS-108 | chr12-3457027-INS-108 |
| chr12-3460661-INS-5282 | chr12-3460662-INS-5282 |
| chr12-3463948-INS-75 | chr12-3463948-INS-74 |
| chr12-3485884-INS-113 | chr12-3485884-INS-95 |
| chr12-3496538-INS-215 | chr12-3496538-INS-214 |

| | |
|---|---|
| chr12-3510151-INS-55 | chr12-3510148-INS-59 |
| chr12-3531437-INS-60 | chr12-3531437-INS-60 |
| chr12-3566408-INS-147 | chr12-3566408-INS-167 |
| chr12-3577783-INS-209 | chr12-3577783-INS-209 |
| chr12-3580761-INS-62 | chr12-3580761-INS-57 |
| chr12-3582389-INS-54 | chr12-3582389-INS-84 |
| chr12-3601778-INS-61 | chr12-3601778-INS-61 |
| chr12-3604923-INS-214 | chr12-3604923-INS-214 |
| chr12-3616683-INS-80 | chr12-3616684-INS-60 |
| chr12-3631063-INS-603 | chr12-3631063-INS-584 |
| chr12-3652471-INS-202 | chr12-3652471-INS-203 |
| chr12-3679012-INS-260 | chr12-3679012-INS-260 |
| chr12-3685020-INS-192 | chr12-3685053-INS-159 |
| chr12-3690532-INS-295 | chr12-3690532-INS-295 |
| chr12-3691823-INS-176 | chr12-3691823-INS-180 |
| chr12-3700921-INS-244 | chr12-3700921-INS-231 |
| chr12-3709569-INS-106 | chr12-3709581-INS-94 |
| chr12-3738470-INS-116 | chr12-3738470-INS-120 |
| chr12-3763022-INS-7079 | chr12-3763023-INS-7077 |
| chr12-3838391-INS-188 | chr12-3838391-INS-188 |
| chr12-3897328-INS-54 | chr12-3897328-INS-54 |
| chr12-3913776-INS-198 | chr12-3913776-INS-198 |
| chr12-3918248-INS-129 | chr12-3918248-INS-129 |
| chr12-3922465-INS-197 | chr12-3922465-INS-243 |
| chr12-3967513-INS-2625 | chr12-3967514-INS-2625 |
| chr12-3969022-INS-434 | chr12-3969022-INS-434 |
| chr12-3969597-INS-76 | chr12-3969597-INS-76 |
| chr12-3983966-INS-70 | chr12-3983966-INS-70 |
| chr12-3993201-INS-75 | chr12-3993201-INS-51 |
| chr12-3996141-INS-56 | chr12-3996147-INS-66 |
| chr12-4017441-INS-120 | chr12-4017441-INS-120 |
| chr12-4036762-INS-230 | chr12-4036762-INS-230 |
| chr12-4055054-INS-1882 | chr12-4055052-INS-1870 |
| chr12-4057302-INS-82 | chr12-4057302-INS-82 |
| chr12-4067743-INS-51 | chr12-4067762-INS-60 |
| chr12-4069163-INS-52 | chr12-4069155-INS-64 |
| chr12-4069983-INS-196 | chr12-4069982-INS-198 |
| chr12-4096596-INS-148 | chr12-4096596-INS-148 |
| chr12-4100416-INS-52 | chr12-4100416-INS-56 |
| chr12-4116309-INS-649 | chr12-4116309-INS-645 |
| chr12-4117446-INS-563 | chr12-4117446-INS-565 |
| chr12-4125143-INS-203 | chr12-4125143-INS-203 |
| chr12-4135649-INS-499 | chr12-4135649-INS-499 |
| chr12-4160837-INS-108 | chr12-4160838-INS-108 |
| chr12-4161619-INS-4000 | chr12-4161620-INS-3999 |
| chr12-4167318-INS-104 | chr12-4167318-INS-56 |
| chr12-4172025-INS-801 | chr12-4172025-INS-801 |
| chr12-4173752-INS-189 | chr12-4173753-INS-189 |
| chr12-4178678-INS-311 | chr12-4178678-INS-311 |
| chr12-4192913-INS-198 | chr12-4192913-INS-198 |
| chr12-4193820-INS-100 | chr12-4193820-INS-100 |
| chr12-4199086-INS-53 | chr12-4199086-INS-53 |
| chr12-4215762-INS-50 | chr12-4215762-INS-58 |
| chr12-4219410-INS-173 | chr12-4219410-INS-180 |
| chr12-4235891-INS-300 | chr12-4235881-INS-292 |
| chr12-4237852-INS-63 | chr12-4237852-INS-60 |
| chr12-4282033-INS-211 | chr12-4282033-INS-211 |
| chr12-4290695-INS-76 | chr12-4290701-INS-56 |
| chr12-4314802-INS-482 | chr12-4314803-INS-482 |
| chr12-4315367-INS-67 | chr12-4315352-INS-70 |
| chr12-4326720-INS-141 | chr12-4326720-INS-141 |
| chr12-4343355-INS-88 | chr12-4343355-INS-88 |
| chr12-4356807-INS-2468 | chr12-4356808-INS-2469 |
| chr12-4375741-INS-412 | chr12-4375741-INS-411 |
| chr12-4384354-INS-69 | chr12-4384354-INS-69 |
| chr12-4403149-INS-1254 | chr12-4403150-INS-1257 |
| chr12-4430289-INS-203 | chr12-4430289-INS-205 |
| chr12-4433086-INS-4555 | chr12-4433087-INS-4555 |
| chr12-4438321-INS-520 | chr12-4438321-INS-520 |
| chr12-4442027-INS-114 | chr12-4442027-INS-114 |
| chr12-4450690-INS-322 | chr12-4450692-INS-351 |
| chr12-4452848-INS-279 | chr12-4452850-INS-280 |
| chr12-4474638-INS-194 | chr12-4474638-INS-194 |
| chr12-4476989-INS-199 | chr12-4476989-INS-199 |
| chr12-4477773-INS-59 | chr12-4477776-INS-64 |
| chr12-4483053-INS-172 | chr12-4483053-INS-172 |
| chr12-4487754-INS-144 | chr12-4487754-INS-144 |

| | |
|---|---|
| chr12-4493829-INS-3336 | chr12-4493825-INS-3337 |
| chr12-4495713-INS-146 | chr12-4495713-INS-146 |
| chr12-4500348-INS-302 | chr12-4500348-INS-292 |
| chr12-4516721-INS-476 | chr12-4516753-INS-442 |
| chr12-4527964-INS-171 | chr12-4527964-INS-173 |
| chr12-4541078-INS-196 | chr12-4541062-INS-210 |
| chr12-4567200-INS-86 | chr12-4567178-INS-102 |
| chr12-4579211-INS-180 | chr12-4579211-INS-181 |
| chr12-4604637-INS-506 | chr12-4604637-INS-506 |
| chr12-4619633-INS-175 | chr12-4619633-INS-126 |
| chr12-4626383-INS-166 | chr12-4626383-INS-166 |
| chr12-4632038-INS-162 | chr12-4632038-INS-162 |
| chr12-4660376-INS-151 | chr12-4660376-INS-151 |
| chr12-4664816-INS-226 | chr12-4664816-INS-226 |
| chr12-4684452-INS-55 | chr12-4684457-INS-50 |
| chr12-4701108-INS-350 | chr12-4701108-INS-350 |
| chr12-4714985-INS-72 | chr12-4714997-INS-71 |
| chr12-4732569-INS-66 | chr12-4732569-INS-88 |
| chr12-4740993-INS-185 | chr12-4740993-INS-185 |
| chr12-4767585-INS-54 | chr12-4767585-INS-54 |
| chr12-4772390-INS-210 | chr12-4772390-INS-208 |
| chr12-4800957-INS-1905 | chr12-4800958-INS-1905 |
| chr12-4845223-INS-732 | chr12-4845221-INS-737 |
| chr12-4863632-INS-213 | chr12-4863632-INS-213 |
| chr12-4872127-INS-201 | chr12-4872127-INS-201 |
| chr12-4877441-INS-171 | chr12-4877441-INS-170 |
| chr12-4878174-INS-1075 | chr12-4878175-INS-1075 |
| chr12-4880536-INS-78 | chr12-4880535-INS-95 |
| chr12-4889851-INS-204 | chr12-4889856-INS-199 |
| chr12-4902860-INS-122 | chr12-4902860-INS-120 |
| chr12-4913913-INS-181 | chr12-4913914-INS-173 |
| chr12-4935998-INS-65 | chr12-4935998-INS-68 |
| chr12-4949632-INS-5381 | chr12-4949633-INS-5382 |
| chr12-4949774-INS-204 | chr12-4949775-INS-211 |
| chr12-4952868-INS-2629 | chr12-4952899-INS-2627 |
| chr12-4958738-INS-94 | chr12-4958738-INS-124 |
| chr12-4961295-INS-217 | chr12-4961296-INS-179 |
| chr12-4971528-INS-168 | chr12-4971528-INS-168 |
| chr12-4975764-INS-232 | chr12-4975764-INS-233 |
| chr12-4978548-INS-140 | chr12-4978548-INS-148 |
| chr12-4983137-INS-60 | chr12-4983137-INS-74 |
| chr12-5037097-INS-212 | chr12-5037097-INS-205 |
| chr12-5073290-INS-559 | chr12-5073290-INS-559 |
| chr12-5076674-INS-56 | chr12-5076674-INS-62 |
| chr12-5125207-INS-59 | chr12-5125207-INS-59 |
| chr12-5168008-INS-90 | chr12-5168008-INS-90 |
| chr12-5203861-INS-120 | chr12-5203861-INS-120 |
| chr12-5272149-INS-69 | chr12-5272111-INS-69 |
| chr12-5324027-INS-215 | chr12-5324027-INS-215 |
| chr12-5363967-INS-126 | chr12-5363967-INS-126 |
| chr12-5375387-INS-58 | chr12-5375387-INS-58 |
| chr12-5398863-INS-63 | chr12-5398908-INS-63 |
| chr12-5412809-INS-210 | chr12-5412844-INS-175 |
| chr12-5430100-INS-91 | chr12-5430099-INS-92 |
| chr12-5441439-INS-117 | chr12-5441439-INS-117 |
| chr12-5453095-INS-70 | chr12-5453095-INS-52 |
| chr12-5462639-INS-54 | chr12-5462639-INS-54 |
| chr12-5513330-INS-438 | chr12-5513330-INS-435 |
| chr12-5519695-INS-73 | chr12-5519695-INS-73 |
| chr12-5562985-INS-137 | chr12-5562985-INS-137 |
| chr12-5571852-INS-6457 | chr12-5571853-INS-6457 |
| chr12-5585973-INS-6884 | chr12-5585974-INS-6885 |
| chr12-5585982-INS-175 | chr12-5585982-INS-175 |
| chr12-5605033-INS-1064 | chr12-5605034-INS-1064 |
| chr12-5617485-INS-78 | chr12-5617485-INS-84 |
| chr12-5624758-INS-825 | chr12-5624778-INS-840 |
| chr12-5630606-INS-160 | chr12-5630606-INS-152 |
| chr12-5676958-INS-6888 | chr12-5676959-INS-6861 |
| chr12-5682750-INS-209 | chr12-5682750-INS-209 |
| chr12-5698292-INS-84 | chr12-5698293-INS-84 |
| chr12-5738169-INS-122 | chr12-5738169-INS-123 |
| chr12-5741910-INS-85 | chr12-5741896-INS-76 |
| chr12-5754734-INS-54 | chr12-5754735-INS-76 |
| chr12-5777489-INS-3668 | chr12-5777490-INS-3662 |
| chr12-5786531-INS-3373 | chr12-5786532-INS-3375 |
| chr12-5807120-INS-86 | chr12-5807120-INS-86 |
| chr12-5822530-INS-56 | chr12-5822530-INS-60 |

| | |
|---|---|
| chr12-5842751-INS-210 | chr12-5842751-INS-210 |
| chr12-5843555-INS-55 | chr12-5843554-INS-61 |
| chr12-5886574-INS-425 | chr12-5886574-INS-421 |
| chr12-5902924-INS-1534 | chr12-5902925-INS-1535 |
| chr12-5915217-INS-95 | chr12-5915209-INS-126 |
| chr12-5939968-INS-400 | chr12-5939968-INS-400 |
| chr12-5946475-INS-152 | chr12-5946475-INS-153 |
| chr12-6001601-INS-1889 | chr12-6001602-INS-1889 |
| chr12-6012809-INS-318 | chr12-6012809-INS-318 |
| chr12-6024580-INS-60 | chr12-6024538-INS-60 |
| chr12-6196286-INS-349 | chr12-6196286-INS-349 |
| chr12-6210765-INS-527 | chr12-6210765-INS-527 |
| chr12-6221083-INS-1683 | chr12-6221084-INS-1684 |
| chr12-6223790-INS-331 | chr12-6223790-INS-350 |
| chr12-6280374-INS-938 | chr12-6280374-INS-938 |
| chr12-6289186-INS-101 | chr12-6289186-INS-102 |
| chr12-6324742-INS-141 | chr12-6324736-INS-168 |
| chr12-6328137-INS-4420 | chr12-6328138-INS-4421 |
| chr12-6361654-INS-62 | chr12-6361654-INS-93 |
| chr12-6395116-INS-2039 | chr12-6395117-INS-2043 |
| chr12-6456146-INS-3103 | chr12-6456147-INS-3104 |
| chr12-6462464-INS-866 | chr12-6462463-INS-867 |
| chr12-6466655-INS-238 | chr12-6466655-INS-238 |
| chr12-6501899-INS-6444 | chr12-6501900-INS-6445 |
| chr12-6515200-INS-215 | chr12-6515200-INS-213 |
| chr12-6519712-INS-82 | chr12-6519711-INS-90 |
| chr12-6528656-INS-530 | chr12-6528656-INS-530 |
| chr12-6559669-INS-1890 | chr12-6559670-INS-1890 |
| chr12-6559670-INS-399 | chr12-6559670-INS-399 |
| chr12-6602319-INS-6240 | chr12-6602320-INS-6238 |
| chr12-6603356-INS-104 | chr12-6603356-INS-104 |
| chr12-6652585-INS-52 | chr12-6652541-INS-56 |
| chr12-6733241-INS-512 | chr12-6733241-INS-512 |
| chr12-6764119-INS-3359 | chr12-6764120-INS-3359 |
| chr12-6767361-INS-102 | chr12-6767361-INS-102 |
| chr12-6774725-INS-54 | chr12-6774725-INS-56 |
| chr12-6775961-INS-64 | chr12-6775961-INS-64 |
| chr12-6781340-INS-6692 | chr12-6781341-INS-6692 |
| chr12-6791211-INS-6680 | chr12-6791212-INS-6680 |
| chr12-6793350-INS-72 | chr12-6793336-INS-82 |
| chr12-6804326-INS-6610 | chr12-6804327-INS-6610 |
| chr12-6821353-INS-208 | chr12-6821353-INS-208 |
| chr12-6850339-INS-4220 | chr12-6850340-INS-4220 |
| chr12-6852689-INS-88 | chr12-6852693-INS-56 |
| chr12-6873547-INS-1200 | chr12-6873548-INS-1201 |
| chr12-6874348-INS-963 | chr12-6874348-INS-963 |
| chr12-6937496-INS-60 | chr12-6937496-INS-61 |
| chr12-6964621-INS-377 | chr12-6964621-INS-377 |
| chr12-7009661-INS-221 | chr12-7009660-INS-229 |
| chr12-7013690-INS-455 | chr12-7013690-INS-455 |
| chr12-7033146-INS-220 | chr12-7033146-INS-222 |
| chr12-7036400-INS-118 | chr12-7036378-INS-101 |
| chr12-7040549-INS-155 | chr12-7040526-INS-158 |
| chr12-7052284-INS-143 | chr12-7052284-INS-143 |
| chr12-7059493-INS-68 | chr12-7059493-INS-68 |
| chr12-7082878-INS-834 | chr12-7082878-INS-834 |
| chr12-7110732-INS-440 | chr12-7110732-INS-441 |
| chr12-7129744-INS-75 | chr12-7129744-INS-60 |
| chr12-7139999-INS-52 | chr12-7140001-INS-56 |
| chr12-7162002-INS-416 | chr12-7162002-INS-416 |
| chr12-7199772-INS-214 | chr12-7199772-INS-214 |
| chr12-7215066-INS-197 | chr12-7215066-INS-197 |
| chr12-7226544-INS-321 | chr12-7226544-INS-321 |
| chr12-7235821-INS-6511 | chr12-7235822-INS-6511 |
| chr12-7253364-INS-62 | chr12-7253354-INS-66 |
| chr12-7389242-INS-68 | chr12-7389242-INS-68 |
| chr12-7470411-INS-189 | chr12-7470411-INS-189 |
| chr12-7476034-INS-75 | chr12-7476026-INS-71 |
| chr12-7490709-INS-81 | chr12-7490709-INS-95 |
| chr12-7497423-INS-808 | chr12-7497423-INS-808 |
| chr12-7535650-INS-198 | chr12-7535650-INS-198 |
| chr12-7552725-INS-1500 | chr12-7552726-INS-1500 |
| chr12-7560030-INS-6203 | chr12-7560031-INS-6203 |
| chr12-7616297-INS-6803 | chr12-7616298-INS-6804 |
| chr12-7618770-INS-57 | chr12-7618770-INS-57 |
| chr12-7680189-INS-58 | chr12-7680189-INS-58 |
| chr12-7687295-INS-67 | chr12-7687300-INS-70 |

| | |
|---|---|
| chr12-7696244-INS-233 | chr12-7696244-INS-212 |
| chr12-7706204-INS-1749 | chr12-7706205-INS-1749 |
| chr12-7747117-INS-106 | chr12-7747117-INS-106 |
| chr12-7769014-INS-126 | chr12-7769010-INS-100 |
| chr12-7796115-INS-55 | chr12-7796110-INS-54 |
| chr12-7808240-INS-66 | chr12-7808240-INS-66 |
| chr12-7828335-INS-1412 | chr12-7828336-INS-1413 |
| chr12-7836230-INS-56 | chr12-7836230-INS-72 |
| chr12-7863063-INS-553 | chr12-7863096-INS-509 |
| chr12-7871852-INS-109 | chr12-7871870-INS-110 |
| chr12-7890120-INS-62 | chr12-7890120-INS-67 |
| chr12-7908425-INS-1240 | chr12-7908410-INS-1219 |
| chr12-7910112-INS-503 | chr12-7910112-INS-504 |
| chr12-7917461-INS-1887 | chr12-7917462-INS-1887 |
| chr12-7924589-INS-78 | chr12-7924597-INS-93 |
| chr12-7947129-INS-110 | chr12-7947129-INS-116 |
| chr12-7950630-INS-68 | chr12-7950630-INS-68 |
| chr12-7971598-INS-77 | chr12-7971632-INS-68 |
| chr12-7988282-INS-4746 | chr12-7988283-INS-4746 |
| chr12-7999345-INS-6596 | chr12-7999346-INS-6596 |
| chr12-8013675-INS-68 | chr12-8013675-INS-54 |
| chr12-8032701-INS-503 | chr12-8032701-INS-503 |
| chr12-8068760-INS-233 | chr12-8068760-INS-234 |
| chr12-8084222-INS-85 | chr12-8084222-INS-83 |
| chr12-8095818-INS-51 | chr12-8095818-INS-85 |
| chr12-8112930-INS-411 | chr12-8112938-INS-411 |
| chr12-8114248-INS-5224 | chr12-8114249-INS-5224 |
| chr12-8123557-INS-168 | chr12-8123557-INS-175 |
| chr12-8131956-INS-59 | chr12-8131956-INS-55 |
| chr12-8139504-INS-8985 | chr12-8139505-INS-8985 |
| chr12-8164768-INS-243 | chr12-8164768-INS-256 |
| chr12-8171749-INS-102 | chr12-8171749-INS-102 |
| chr12-8182583-INS-358 | chr12-8182583-INS-358 |
| chr12-8187592-INS-525 | chr12-8187592-INS-525 |
| chr12-8200874-INS-737 | chr12-8200874-INS-737 |
| chr12-8202451-INS-229 | chr12-8202451-INS-230 |
| chr12-8238246-INS-51 | chr12-8238246-INS-50 |
| chr12-8241232-INS-298 | chr12-8241232-INS-289 |
| chr12-8243115-INS-294 | chr12-8243115-INS-294 |
| chr12-8260034-INS-52 | chr12-8260027-INS-88 |
| chr12-8265111-INS-172 | chr12-8265111-INS-172 |
| chr12-8275246-INS-377 | chr12-8275246-INS-377 |
| chr12-8276920-INS-1095 | chr12-8276918-INS-1085 |
| chr12-8283343-INS-1537 | chr12-8283344-INS-1538 |
| chr12-8288638-INS-56 | chr12-8288638-INS-56 |
| chr12-8359641-INS-114 | chr12-8359641-INS-114 |
| chr12-8373349-INS-52 | chr12-8373349-INS-52 |
| chr12-8393916-INS-90 | chr12-8393915-INS-110 |
| chr12-8402690-INS-246 | chr12-8402690-INS-246 |
| chr12-8403847-INS-66 | chr12-8403847-INS-72 |
| chr12-8415184-INS-52 | chr12-8415184-INS-52 |
| chr12-8415246-INS-7140 | chr12-8415247-INS-7140 |
| chr12-8416643-INS-384 | chr12-8416643-INS-385 |
| chr12-8426572-INS-56 | chr12-8426581-INS-70 |
| chr12-8441236-INS-57 | chr12-8441236-INS-60 |
| chr12-8447770-INS-506 | chr12-8447770-INS-519 |
| chr12-8464446-INS-142 | chr12-8464451-INS-139 |
| chr12-8494717-INS-64 | chr12-8494718-INS-68 |
| chr12-8530081-INS-60 | chr12-8530081-INS-60 |
| chr12-8544748-INS-232 | chr12-8544748-INS-232 |
| chr12-8555195-INS-222 | chr12-8555195-INS-222 |
| chr12-8558912-INS-7213 | chr12-8558913-INS-7213 |
| chr12-8566661-INS-207 | chr12-8566661-INS-207 |
| chr12-8576458-INS-101 | chr12-8576458-INS-101 |
| chr12-8580496-INS-497 | chr12-8580493-INS-498 |
| chr12-8584176-INS-66 | chr12-8584176-INS-66 |
| chr12-8587808-INS-56 | chr12-8587808-INS-56 |
| chr12-8594134-INS-62 | chr12-8594134-INS-62 |
| chr12-8604553-INS-102 | chr12-8604536-INS-120 |
| chr12-8609317-INS-104 | chr12-8609288-INS-104 |
| chr12-8620702-INS-64 | chr12-8620702-INS-59 |
| chr12-8657307-INS-58 | chr12-8657307-INS-58 |
| chr12-8658217-INS-89 | chr12-8658217-INS-89 |
| chr12-8668071-INS-161 | chr12-8668096-INS-164 |
| chr12-8699935-INS-177 | chr12-8699935-INS-183 |
| chr12-8721871-INS-55 | chr12-8721871-INS-55 |
| chr12-8741102-INS-53 | chr12-8741102-INS-53 |

| | |
|---|---|
| chr12-8851169-INS-162 | chr12-8851169-INS-162 |
| chr12-8867425-INS-58 | chr12-8867425-INS-60 |
| chr12-8934819-INS-195 | chr12-8934819-INS-195 |
| chr12-8951564-INS-207 | chr12-8951564-INS-213 |
| chr12-8978365-INS-58 | chr12-8978365-INS-70 |
| chr12-8980036-INS-240 | chr12-8980036-INS-243 |
| chr12-9018434-INS-76 | chr12-9018434-INS-76 |
| chr12-9035166-INS-85 | chr12-9035166-INS-120 |
| chr12-9041715-INS-533 | chr12-9041716-INS-541 |
| chr12-9089395-INS-227 | chr12-9089395-INS-227 |
| chr12-9115343-INS-238 | chr12-9115349-INS-239 |
| chr12-9135136-INS-102 | chr12-9135136-INS-88 |
| chr12-9149822-INS-327 | chr12-9149822-INS-323 |
| chr12-9174940-INS-201 | chr12-9174940-INS-201 |
| chr12-9197111-INS-52 | chr12-9197111-INS-60 |
| chr12-9234184-INS-109 | chr12-9234184-INS-97 |
| chr12-9268798-INS-512 | chr12-9268798-INS-512 |
| chr12-9304523-INS-133 | chr12-9304523-INS-133 |
| chr12-9313023-INS-56 | chr12-9313012-INS-55 |
| chr12-9347576-INS-1363 | chr12-9347577-INS-1363 |
| chr12-9356813-INS-50 | chr12-9356813-INS-50 |
| chr12-9367317-INS-58 | chr12-9367317-INS-74 |
| chr12-9406610-INS-59 | chr12-9406610-INS-54 |
| chr12-9454332-INS-58 | chr12-9454332-INS-60 |
| chr12-9491872-INS-56 | chr12-9491872-INS-54 |
| chr12-9502094-INS-253 | chr12-9502100-INS-262 |
| chr12-9506756-INS-57 | chr12-9506756-INS-60 |
| chr12-9533091-INS-265 | chr12-9533090-INS-266 |
| chr12-9535489-INS-70 | chr12-9535489-INS-70 |
| chr12-9570604-INS-934 | chr12-9570605-INS-935 |
| chr12-9576218-INS-328 | chr12-9576218-INS-328 |
| chr12-9594767-INS-6387 | chr12-9594768-INS-6387 |
| chr12-9595441-INS-1749 | chr12-9595442-INS-1749 |
| chr12-9608683-INS-80 | chr12-9608683-INS-80 |
| chr12-9617426-INS-105 | chr12-9617426-INS-114 |
| chr12-9633930-INS-52 | chr12-9633895-INS-78 |
| chr12-9649952-INS-6029 | chr12-9649953-INS-6029 |
| chr12-9661990-INS-62 | chr12-9661990-INS-62 |
| chr12-9735276-INS-108 | chr12-9735264-INS-92 |
| chr12-9738064-INS-4738 | chr12-9738065-INS-4738 |
| chr12-9762039-INS-6907 | chr12-9762040-INS-6907 |
| chr12-9763280-INS-544 | chr12-9763280-INS-544 |
| chr12-9793101-INS-5924 | chr12-9793102-INS-5924 |
| chr12-9799048-INS-1727 | chr12-9799049-INS-1728 |
| chr12-9806454-INS-86 | chr12-9806454-INS-86 |
| chr12-9832642-INS-6406 | chr12-9832643-INS-6406 |
| chr12-9837750-INS-51 | chr12-9837750-INS-51 |
| chr12-9851380-INS-107 | chr12-9851380-INS-75 |
| chr12-9853843-INS-889 | chr12-9853843-INS-890 |
| chr12-9862861-INS-52 | chr12-9862861-INS-52 |
| chr12-9871937-INS-59 | chr12-9871937-INS-59 |
| chr12-9880913-INS-2320 | chr12-9880944-INS-2322 |
| chr12-9917994-INS-1103 | chr12-9917995-INS-1103 |
| chr12-9933753-INS-92 | chr12-9933755-INS-92 |
| chr12-9943838-INS-1287 | chr12-9943839-INS-1290 |
| chr12-9959231-INS-319 | chr12-9959231-INS-319 |
| chr12-10061112-INS-7001 | chr12-10061113-INS-7002 |
| chr12-10081105-INS-6440 | chr12-10081106-INS-6440 |
| chr12-10120456-INS-737 | chr12-10120456-INS-741 |
| chr12-10135417-INS-1012 | chr12-10135418-INS-1012 |
| chr12-10135645-INS-117 | chr12-10135645-INS-117 |
| chr12-10145619-INS-6325 | chr12-10145620-INS-6325 |
| chr12-10155816-INS-654 | chr12-10155816-INS-653 |
| chr12-10157598-INS-52 | chr12-10157599-INS-54 |
| chr12-10158536-INS-65 | chr12-10158536-INS-75 |
| chr12-10221610-INS-60 | chr12-10221610-INS-60 |
| chr12-10224917-INS-142 | chr12-10224917-INS-188 |
| chr12-10228689-INS-4354 | chr12-10228690-INS-4354 |
| chr12-10231842-INS-488 | chr12-10231842-INS-489 |
| chr12-10247463-INS-82 | chr12-10247463-INS-82 |
| chr12-10247712-INS-844 | chr12-10247712-INS-846 |
| chr12-10256992-INS-78 | chr12-10256992-INS-92 |
| chr12-10294368-INS-204 | chr12-10294368-INS-204 |
| chr12-10302803-INS-63 | chr12-10302794-INS-52 |
| chr12-10309071-INS-201 | chr12-10309071-INS-222 |
| chr12-10310321-INS-54 | chr12-10310321-INS-56 |
| chr12-10327849-INS-86 | chr12-10327872-INS-52 |

| | |
|---|---|
| chr12-10340600-INS-83 | chr12-10340600-INS-85 |
| chr12-10357536-INS-220 | chr12-10357536-INS-220 |
| chr12-10369591-INS-5666 | chr12-10369576-INS-5665 |
| chr12-10408713-INS-204 | chr12-10408713-INS-204 |
| chr12-10412472-INS-7169 | chr12-10412473-INS-7170 |
| chr12-10421141-INS-208 | chr12-10421141-INS-208 |
| chr12-10457862-INS-80 | chr12-10457868-INS-78 |
| chr12-10460014-INS-59 | chr12-10460014-INS-66 |
| chr12-10466625-INS-159 | chr12-10466623-INS-135 |
| chr12-10482434-INS-90 | chr12-10482436-INS-76 |
| chr12-10487322-INS-568 | chr12-10487322-INS-572 |
| chr12-10489157-INS-51 | chr12-10489157-INS-51 |
| chr12-10523987-INS-74 | chr12-10523987-INS-74 |
| chr12-10527427-INS-130 | chr12-10527424-INS-178 |
| chr12-10538221-INS-107 | chr12-10538221-INS-107 |
| chr12-10559206-INS-975 | chr12-10559206-INS-974 |
| chr12-10588887-INS-6786 | chr12-10588888-INS-6786 |
| chr12-10593651-INS-483 | chr12-10593651-INS-483 |
| chr12-10605696-INS-69 | chr12-10605696-INS-69 |
| chr12-10618452-INS-6398 | chr12-10618453-INS-6398 |
| chr12-10644097-INS-6442 | chr12-10644098-INS-6442 |
| chr12-10664931-INS-7474 | chr12-10664932-INS-7486 |
| chr12-10682311-INS-385 | chr12-10682311-INS-385 |
| chr12-10725152-INS-8538 | chr12-10725153-INS-8538 |
| chr12-10776784-INS-798 | chr12-10776784-INS-798 |
| chr12-10780328-INS-963 | chr12-10780328-INS-963 |
| chr12-10784256-INS-80 | chr12-10784256-INS-82 |
| chr12-10809871-INS-7526 | chr12-10809872-INS-7528 |
| chr12-10820487-INS-588 | chr12-10820487-INS-588 |
| chr12-10867234-INS-83 | chr12-10867234-INS-78 |
| chr12-10889998-INS-207 | chr12-10889998-INS-209 |
| chr12-10910473-INS-59 | chr12-10910503-INS-68 |
| chr12-10915632-INS-949 | chr12-10915632-INS-949 |
| chr12-10930439-INS-283 | chr12-10930439-INS-284 |
| chr12-10932650-INS-499 | chr12-10932650-INS-500 |
| chr12-10934463-INS-56 | chr12-10934463-INS-56 |
| chr12-10987881-INS-5152 | chr12-10987882-INS-5152 |
| chr12-10992407-INS-104 | chr12-10992407-INS-109 |
| chr12-11003130-INS-3933 | chr12-11003131-INS-3910 |
| chr12-11021496-INS-208 | chr12-11021496-INS-208 |
| chr12-11081599-INS-204 | chr12-11081599-INS-204 |
| chr12-11106307-INS-401 | chr12-11106307-INS-401 |
| chr12-11127644-INS-6709 | chr12-11127645-INS-6709 |
| chr12-11146764-INS-214 | chr12-11146764-INS-214 |
| chr12-11170601-INS-67 | chr12-11170601-INS-67 |
| chr12-11194195-INS-583 | chr12-11194195-INS-583 |
| chr12-11207301-INS-202 | chr12-11207301-INS-202 |
| chr12-11226055-INS-194 | chr12-11226055-INS-194 |
| chr12-11227239-INS-198 | chr12-11227238-INS-193 |
| chr12-11229961-INS-53 | chr12-11229961-INS-54 |
| chr12-11260327-INS-54 | chr12-11260320-INS-50 |
| chr12-11270403-INS-206 | chr12-11270409-INS-200 |
| chr12-11271317-INS-177 | chr12-11271317-INS-177 |
| chr12-11273990-INS-212 | chr12-11273990-INS-212 |
| chr12-11277124-INS-5284 | chr12-11277125-INS-5285 |
| chr12-11282088-INS-52 | chr12-11282088-INS-62 |
| chr12-11304055-INS-440 | chr12-11304055-INS-440 |
| chr12-11306538-INS-211 | chr12-11306538-INS-211 |
| chr12-11320357-INS-56 | chr12-11320357-INS-56 |
| chr12-11345446-INS-171 | chr12-11345446-INS-172 |
| chr12-11355054-INS-210 | chr12-11355054-INS-210 |
| chr12-11357697-INS-1005 | chr12-11357698-INS-998 |
| chr12-11379091-INS-298 | chr12-11379091-INS-298 |
| chr12-11398146-INS-3182 | chr12-11398147-INS-3182 |
| chr12-11402200-INS-122 | chr12-11402200-INS-122 |
| chr12-11415869-INS-91 | chr12-11415867-INS-121 |
| chr12-11429851-INS-62 | chr12-11429851-INS-64 |
| chr12-11441370-INS-85 | chr12-11441370-INS-85 |
| chr12-11464222-INS-200 | chr12-11464222-INS-200 |
| chr12-11497446-INS-295 | chr12-11497446-INS-296 |
| chr12-11518031-INS-728 | chr12-11518031-INS-728 |
| chr12-11531790-INS-55 | chr12-11531790-INS-55 |
| chr12-11569113-INS-3800 | chr12-11569114-INS-3800 |
| chr12-11570818-INS-52 | chr12-11570818-INS-60 |
| chr12-11590843-INS-100 | chr12-11590843-INS-103 |
| chr12-11622555-INS-52 | chr12-11622555-INS-52 |
| chr12-11650175-INS-434 | chr12-11650175-INS-434 |

| | |
|---|---|
| chr12-11651282-INS-65 | chr12-11651282-INS-65 |
| chr12-11663310-INS-70 | chr12-11663301-INS-120 |
| chr12-11666783-INS-69 | chr12-11666783-INS-69 |
| chr12-11670779-INS-6489 | chr12-11670780-INS-6489 |
| chr12-11675406-INS-2624 | chr12-11675407-INS-2632 |
| chr12-11682150-INS-52 | chr12-11682150-INS-58 |
| chr12-11727995-INS-544 | chr12-11728004-INS-545 |
| chr12-11737262-INS-6332 | chr12-11737263-INS-6332 |
| chr12-11744082-INS-447 | chr12-11744082-INS-447 |
| chr12-11755701-INS-2326 | chr12-11755702-INS-2328 |
| chr12-11779598-INS-383 | chr12-11779598-INS-383 |
| chr12-11787575-INS-56 | chr12-11787575-INS-56 |
| chr12-11815379-INS-634 | chr12-11815379-INS-634 |
| chr12-11833900-INS-628 | chr12-11833900-INS-630 |
| chr12-11838922-INS-463 | chr12-11838876-INS-475 |
| chr12-11856325-INS-80 | chr12-11856325-INS-80 |
| chr12-11859138-INS-525 | chr12-11859138-INS-547 |
| chr12-11868858-INS-5214 | chr12-11868859-INS-5214 |
| chr12-11870451-INS-558 | chr12-11870451-INS-556 |
| chr12-11895075-INS-81 | chr12-11895075-INS-81 |
| chr12-11897956-INS-1136 | chr12-11897957-INS-1146 |
| chr12-11902784-INS-72 | chr12-11902784-INS-80 |
| chr12-11917789-INS-176 | chr12-11917790-INS-145 |
| chr12-11938405-INS-6270 | chr12-11938406-INS-6264 |
| chr12-11947091-INS-371 | chr12-11947091-INS-375 |
| chr12-11948624-INS-6574 | chr12-11948625-INS-6574 |
| chr12-11965677-INS-1130 | chr12-11965678-INS-1130 |
| chr12-11975467-INS-208 | chr12-11975511-INS-202 |
| chr12-12049370-INS-1630 | chr12-12049371-INS-1630 |
| chr12-12062119-INS-90 | chr12-12062104-INS-96 |
| chr12-12070273-INS-503 | chr12-12070273-INS-502 |
| chr12-12088538-INS-218 | chr12-12088538-INS-218 |
| chr12-12097194-INS-2449 | chr12-12097195-INS-2449 |
| chr12-12102080-INS-6435 | chr12-12102081-INS-6436 |
| chr12-12102234-INS-122 | chr12-12102234-INS-122 |
| chr12-12120466-INS-85 | chr12-12120466-INS-85 |
| chr12-12140410-INS-92 | chr12-12140410-INS-92 |
| chr12-12151216-INS-54 | chr12-12151216-INS-60 |
| chr12-12155073-INS-575 | chr12-12155073-INS-575 |
| chr12-12217631-INS-341 | chr12-12217631-INS-341 |
| chr12-12221045-INS-372 | chr12-12221045-INS-372 |
| chr12-12227429-INS-113 | chr12-12227435-INS-108 |
| chr12-12232468-INS-531 | chr12-12232468-INS-531 |
| chr12-12256782-INS-165 | chr12-12256782-INS-165 |
| chr12-12284021-INS-56 | chr12-12284021-INS-68 |
| chr12-12297621-INS-96 | chr12-12297604-INS-100 |
| chr12-12299353-INS-52 | chr12-12299353-INS-52 |
| chr12-12301283-INS-3486 | chr12-12301284-INS-3487 |
| chr12-12359799-INS-108 | chr12-12359799-INS-108 |
| chr12-12372586-INS-74 | chr12-12372586-INS-74 |
| chr12-12390056-INS-228 | chr12-12390056-INS-228 |
| chr12-12411231-INS-3337 | chr12-12411232-INS-3337 |
| chr12-12421035-INS-92 | chr12-12421035-INS-76 |
| chr12-12432263-INS-55 | chr12-12432263-INS-53 |
| chr12-12432854-INS-60 | chr12-12432854-INS-60 |
| chr12-12486304-INS-154 | chr12-12486302-INS-160 |
| chr12-12510135-INS-108 | chr12-12510135-INS-109 |
| chr12-12512726-INS-7137 | chr12-12512727-INS-7137 |
| chr12-12525691-INS-302 | chr12-12525691-INS-302 |
| chr12-12561289-INS-58 | chr12-12561297-INS-68 |
| chr12-12571444-INS-55 | chr12-12571444-INS-55 |
| chr12-12580709-INS-60 | chr12-12580709-INS-60 |
| chr12-12587342-INS-1683 | chr12-12587343-INS-1683 |
| chr12-12596684-INS-865 | chr12-12596641-INS-867 |
| chr12-12597969-INS-164 | chr12-12597969-INS-163 |
| chr12-12637863-INS-227 | chr12-12637826-INS-255 |
| chr12-12666326-INS-7091 | chr12-12666327-INS-7092 |
| chr12-12682555-INS-54 | chr12-12682555-INS-56 |
| chr12-12684769-INS-497 | chr12-12684769-INS-497 |
| chr12-12689118-INS-72 | chr12-12689118-INS-64 |
| chr12-12707577-INS-65 | chr12-12707577-INS-65 |
| chr12-12748788-INS-62 | chr12-12748788-INS-66 |
| chr12-12749847-INS-452 | chr12-12749847-INS-452 |
| chr12-12814041-INS-194 | chr12-12814037-INS-193 |
| chr12-12819805-INS-61 | chr12-12819800-INS-72 |
| chr12-12825091-INS-199 | chr12-12825091-INS-199 |
| chr12-12844658-INS-78 | chr12-12844659-INS-90 |

| | |
|---|---|
| chr12-12845487-INS-194 | chr12-12845488-INS-197 |
| chr12-12848394-INS-108 | chr12-12848394-INS-108 |
| chr12-12854640-INS-53 | chr12-12854640-INS-65 |
| chr12-12865886-INS-437 | chr12-12865886-INS-437 |
| chr12-12875303-INS-60 | chr12-12875305-INS-73 |
| chr12-12876937-INS-63 | chr12-12876937-INS-50 |
| chr12-12886731-INS-215 | chr12-12886731-INS-215 |
| chr12-12905988-INS-193 | chr12-12905984-INS-208 |
| chr12-12915997-INS-52 | chr12-12915997-INS-52 |
| chr12-12924846-INS-358 | chr12-12924846-INS-358 |
| chr12-12937862-INS-93 | chr12-12937861-INS-74 |
| chr12-12943697-INS-6456 | chr12-12943698-INS-6457 |
| chr12-12948821-INS-68 | chr12-12948821-INS-70 |
| chr12-12957771-INS-99 | chr12-12957785-INS-111 |
| chr12-12961608-INS-367 | chr12-12961608-INS-367 |
| chr12-13019063-INS-57 | chr12-13019089-INS-54 |
| chr12-13038336-INS-100 | chr12-13038336-INS-110 |
| chr12-13044306-INS-62 | chr12-13044306-INS-64 |
| chr12-13052756-INS-96 | chr12-13052760-INS-68 |
| chr12-13058558-INS-59 | chr12-13058558-INS-59 |
| chr12-13059848-INS-159 | chr12-13059853-INS-114 |
| chr12-13060801-INS-52 | chr12-13060789-INS-68 |
| chr12-13063695-INS-95 | chr12-13063695-INS-103 |
| chr12-13089406-INS-161 | chr12-13089406-INS-161 |
| chr12-13098140-INS-173 | chr12-13098140-INS-173 |
| chr12-13101363-INS-52 | chr12-13101363-INS-52 |
| chr12-13102787-INS-54 | chr12-13102787-INS-58 |
| chr12-13111640-INS-53 | chr12-13111640-INS-52 |
| chr12-13111979-INS-418 | chr12-13111979-INS-418 |
| chr12-13116150-INS-152 | chr12-13116199-INS-106 |
| chr12-13118674-INS-51 | chr12-13118674-INS-51 |
| chr12-13121224-INS-214 | chr12-13121224-INS-214 |
| chr12-13129448-INS-238 | chr12-13129448-INS-238 |
| chr12-13138202-INS-63 | chr12-13138202-INS-61 |
| chr12-13175988-INS-70 | chr12-13175988-INS-70 |
| chr12-13184155-INS-120 | chr12-13184155-INS-120 |
| chr12-13219931-INS-91 | chr12-13219931-INS-103 |
| chr12-13247255-INS-55 | chr12-13247255-INS-55 |
| chr12-13254013-INS-82 | chr12-13254013-INS-92 |
| chr12-13255157-INS-189 | chr12-13255157-INS-189 |
| chr12-13301367-INS-1298 | chr12-13301368-INS-1298 |
| chr12-13313066-INS-72 | chr12-13313066-INS-88 |
| chr12-13323484-INS-74 | chr12-13323484-INS-74 |
| chr12-13340736-INS-1466 | chr12-13340737-INS-1467 |
| chr12-13342664-INS-88 | chr12-13342658-INS-100 |
| chr12-13356685-INS-526 | chr12-13356685-INS-526 |
| chr12-13363605-INS-242 | chr12-13363605-INS-242 |
| chr12-13386785-INS-100 | chr12-13386785-INS-88 |
| chr12-13388158-INS-222 | chr12-13388152-INS-198 |
| chr12-13441765-INS-4031 | chr12-13441766-INS-4034 |
| chr12-13448967-INS-1889 | chr12-13448968-INS-1889 |
| chr12-13479698-INS-226 | chr12-13479698-INS-226 |
| chr12-13489117-INS-395 | chr12-13489117-INS-373 |
| chr12-13516899-INS-56 | chr12-13516899-INS-56 |
| chr12-13519254-INS-71 | chr12-13519254-INS-71 |
| chr12-13530853-INS-492 | chr12-13530853-INS-492 |
| chr12-13534836-INS-6401 | chr12-13534837-INS-6401 |
| chr12-13536932-INS-76 | chr12-13536932-INS-76 |
| chr12-13537902-INS-51 | chr12-13537902-INS-60 |
| chr12-13563850-INS-394 | chr12-13563850-INS-413 |
| chr12-13613409-INS-76 | chr12-13613409-INS-76 |
| chr12-13692117-INS-3025 | chr12-13692118-INS-3022 |
| chr12-13731863-INS-7525 | chr12-13731864-INS-7525 |
| chr12-13764656-INS-6427 | chr12-13764657-INS-6428 |
| chr12-13783181-INS-78 | chr12-13783228-INS-50 |
| chr12-13786349-INS-66 | chr12-13786356-INS-50 |
| chr12-13790048-INS-3318 | chr12-13790049-INS-3318 |
| chr12-13800831-INS-533 | chr12-13800831-INS-526 |
| chr12-13813824-INS-212 | chr12-13813824-INS-212 |
| chr12-13835819-INS-6611 | chr12-13835820-INS-6614 |
| chr12-13857038-INS-695 | chr12-13857049-INS-683 |
| chr12-13872116-INS-958 | chr12-13872140-INS-967 |
| chr12-13879283-INS-186 | chr12-13879283-INS-185 |
| chr12-13880767-INS-54 | chr12-13880767-INS-50 |
| chr12-13883533-INS-78 | chr12-13883533-INS-52 |
| chr12-13885366-INS-111 | chr12-13885357-INS-119 |
| chr12-13906658-INS-1427 | chr12-13906659-INS-1427 |

| | |
|---|---|
| chr12-13917688-INS-464 | chr12-13917688-INS-464 |
| chr12-13941927-INS-187 | chr12-13941927-INS-187 |
| chr12-13961153-INS-50 | chr12-13961137-INS-64 |
| chr12-13976800-INS-215 | chr12-13976800-INS-220 |
| chr12-13980864-INS-1893 | chr12-13980865-INS-1893 |
| chr12-14009982-INS-52 | chr12-14009982-INS-52 |
| chr12-14041035-INS-117 | chr12-14041035-INS-122 |
| chr12-14049322-INS-80 | chr12-14049323-INS-79 |
| chr12-14089274-INS-72 | chr12-14089274-INS-72 |
| chr12-14116118-INS-188 | chr12-14116118-INS-189 |
| chr12-14121118-INS-6230 | chr12-14121119-INS-6228 |
| chr12-14122325-INS-341 | chr12-14122325-INS-340 |
| chr12-14143710-INS-6351 | chr12-14143711-INS-6349 |
| chr12-14151350-INS-362 | chr12-14151350-INS-329 |
| chr12-14155502-INS-246 | chr12-14155502-INS-246 |
| chr12-14185507-INS-4150 | chr12-14185508-INS-4150 |
| chr12-14193574-INS-100 | chr12-14193617-INS-100 |
| chr12-14202456-INS-60 | chr12-14202456-INS-60 |
| chr12-14211126-INS-136 | chr12-14211126-INS-133 |
| chr12-14227226-INS-1063 | chr12-14227227-INS-1063 |
| chr12-14227785-INS-1024 | chr12-14227786-INS-1024 |
| chr12-14234041-INS-1091 | chr12-14234049-INS-1086 |
| chr12-14235336-INS-7084 | chr12-14235337-INS-7094 |
| chr12-14289149-INS-51 | chr12-14289150-INS-51 |
| chr12-14306515-INS-261 | chr12-14306515-INS-264 |
| chr12-14348393-INS-180 | chr12-14348393-INS-183 |
| chr12-14368813-INS-401 | chr12-14368813-INS-400 |
| chr12-14370467-INS-104 | chr12-14370482-INS-120 |
| chr12-14371580-INS-3342 | chr12-14371581-INS-3342 |
| chr12-14382375-INS-6305 | chr12-14382376-INS-6305 |
| chr12-14387888-INS-228 | chr12-14387888-INS-228 |
| chr12-14406924-INS-3446 | chr12-14406925-INS-3447 |
| chr12-14448407-INS-4552 | chr12-14448408-INS-4552 |
| chr12-14482261-INS-59 | chr12-14482261-INS-58 |
| chr12-14488589-INS-204 | chr12-14488589-INS-204 |
| chr12-14510929-INS-166 | chr12-14510929-INS-166 |
| chr12-14511963-INS-360 | chr12-14511963-INS-360 |
| chr12-14516379-INS-56 | chr12-14516360-INS-63 |
| chr12-14522693-INS-88 | chr12-14522693-INS-76 |
| chr12-14539041-INS-472 | chr12-14539043-INS-459 |
| chr12-14549666-INS-400 | chr12-14549666-INS-400 |
| chr12-14565437-INS-448 | chr12-14565428-INS-485 |
| chr12-14611837-INS-734 | chr12-14611837-INS-734 |
| chr12-14624908-INS-145 | chr12-14624904-INS-148 |
| chr12-14642689-INS-51 | chr12-14642690-INS-74 |
| chr12-14682515-INS-451 | chr12-14682525-INS-445 |
| chr12-14745449-INS-719 | chr12-14745449-INS-719 |
| chr12-14759778-INS-79 | chr12-14759778-INS-53 |
| chr12-14770046-INS-517 | chr12-14770046-INS-516 |
| chr12-14812663-INS-77 | chr12-14812663-INS-103 |
| chr12-14840447-INS-469 | chr12-14840447-INS-430 |
| chr12-14863286-INS-1271 | chr12-14863287-INS-1278 |
| chr12-14881205-INS-143 | chr12-14881205-INS-143 |
| chr12-14911711-INS-52 | chr12-14911711-INS-52 |
| chr12-14941364-INS-70 | chr12-14941364-INS-88 |
| chr12-14949591-INS-71 | chr12-14949591-INS-50 |
| chr12-14950834-INS-261 | chr12-14950833-INS-271 |
| chr12-14963205-INS-583 | chr12-14963205-INS-583 |
| chr12-14971952-INS-120 | chr12-14971952-INS-120 |
| chr12-14975927-INS-52 | chr12-14975927-INS-52 |
| chr12-15056508-INS-5587 | chr12-15056509-INS-5587 |
| chr12-15103842-INS-68 | chr12-15103842-INS-68 |
| chr12-15121720-INS-62 | chr12-15121720-INS-77 |
| chr12-15133856-INS-52 | chr12-15133856-INS-52 |
| chr12-15136786-INS-209 | chr12-15136786-INS-209 |
| chr12-15137385-INS-3359 | chr12-15137386-INS-3359 |
| chr12-15150689-INS-3647 | chr12-15150690-INS-3647 |
| chr12-15154932-INS-82 | chr12-15154932-INS-82 |
| chr12-15166562-INS-66 | chr12-15166562-INS-69 |
| chr12-15177707-INS-56 | chr12-15177676-INS-50 |
| chr12-15211556-INS-203 | chr12-15211556-INS-201 |
| chr12-15234848-INS-56 | chr12-15234820-INS-68 |
| chr12-15256032-INS-66 | chr12-15256042-INS-55 |
| chr12-15272917-INS-73 | chr12-15272917-INS-73 |
| chr12-15295312-INS-156 | chr12-15295312-INS-156 |
| chr12-15306008-INS-139 | chr12-15306008-INS-139 |
| chr12-15317879-INS-410 | chr12-15317879-INS-410 |

| | |
|---|---|
| chr12-15326164-INS-7850 | chr12-15326165-INS-7850 |
| chr12-15332168-INS-1977 | chr12-15332169-INS-1977 |
| chr12-15360297-INS-265 | chr12-15360297-INS-265 |
| chr12-15373706-INS-358 | chr12-15373706-INS-358 |
| chr12-15382440-INS-267 | chr12-15382440-INS-267 |
| chr12-15444119-INS-64 | chr12-15444119-INS-64 |
| chr12-15448336-INS-4974 | chr12-15448337-INS-4976 |
| chr12-15449853-INS-6241 | chr12-15449854-INS-6241 |
| chr12-15452723-INS-755 | chr12-15452723-INS-754 |
| chr12-15466576-INS-5482 | chr12-15466577-INS-5482 |
| chr12-15470934-INS-297 | chr12-15470934-INS-299 |
| chr12-15473424-INS-209 | chr12-15473424-INS-209 |
| chr12-15510316-INS-81 | chr12-15510312-INS-82 |
| chr12-15578019-INS-95 | chr12-15578020-INS-64 |
| chr12-15583809-INS-73 | chr12-15583825-INS-58 |
| chr12-15589364-INS-433 | chr12-15589364-INS-433 |
| chr12-15595154-INS-56 | chr12-15595154-INS-56 |
| chr12-15607320-INS-69 | chr12-15607320-INS-77 |
| chr12-15616804-INS-470 | chr12-15616804-INS-470 |
| chr12-15619217-INS-189 | chr12-15619217-INS-189 |
| chr12-15662980-INS-85 | chr12-15662980-INS-111 |
| chr12-15688230-INS-67 | chr12-15688230-INS-67 |
| chr12-15710433-INS-4045 | chr12-15710434-INS-4046 |
| chr12-15748226-INS-120 | chr12-15748226-INS-88 |
| chr12-15756207-INS-890 | chr12-15756207-INS-891 |
| chr12-15779208-INS-194 | chr12-15779208-INS-194 |
| chr12-15787705-INS-77 | chr12-15787705-INS-103 |
| chr12-15824983-INS-400 | chr12-15824983-INS-400 |
| chr12-15868849-INS-71 | chr12-15868811-INS-59 |
| chr12-15869965-INS-76 | chr12-15869965-INS-51 |
| chr12-15888928-INS-52 | chr12-15888879-INS-56 |
| chr12-15896196-INS-200 | chr12-15896196-INS-200 |
| chr12-15899512-INS-64 | chr12-15899546-INS-68 |
| chr12-15941761-INS-72 | chr12-15941761-INS-72 |
| chr12-15954129-INS-138 | chr12-15954167-INS-116 |
| chr12-15978374-INS-117 | chr12-15978374-INS-84 |
| chr12-15995145-INS-7109 | chr12-15995146-INS-7109 |
| chr12-16057422-INS-1888 | chr12-16057423-INS-1888 |
| chr12-16069073-INS-56 | chr12-16069073-INS-56 |
| chr12-16117976-INS-101 | chr12-16117976-INS-102 |
| chr12-16128756-INS-6295 | chr12-16128757-INS-6294 |
| chr12-16128775-INS-419 | chr12-16128775-INS-418 |
| chr12-16133238-INS-1140 | chr12-16133239-INS-1139 |
| chr12-16152061-INS-56 | chr12-16152061-INS-56 |
| chr12-16172750-INS-62 | chr12-16172750-INS-64 |
| chr12-16178638-INS-51 | chr12-16178639-INS-51 |
| chr12-16206669-INS-82 | chr12-16206647-INS-54 |
| chr12-16254747-INS-1393 | chr12-16254748-INS-1393 |
| chr12-16261456-INS-156 | chr12-16261489-INS-156 |
| chr12-16286599-INS-891 | chr12-16286599-INS-893 |
| chr12-16292066-INS-78 | chr12-16292066-INS-86 |
| chr12-16308898-INS-59 | chr12-16308898-INS-51 |
| chr12-16313535-INS-63 | chr12-16313535-INS-63 |
| chr12-16358686-INS-104 | chr12-16358686-INS-104 |
| chr12-16372982-INS-110 | chr12-16372983-INS-96 |
| chr12-16400512-INS-76 | chr12-16400512-INS-84 |
| chr12-16450877-INS-51 | chr12-16450895-INS-62 |
| chr12-16472956-INS-218 | chr12-16472956-INS-218 |
| chr12-16496835-INS-112 | chr12-16496835-INS-112 |
| chr12-16502640-INS-78 | chr12-16502644-INS-90 |
| chr12-16546906-INS-56 | chr12-16546900-INS-60 |
| chr12-16549970-INS-513 | chr12-16549971-INS-513 |
| chr12-16570308-INS-60 | chr12-16570306-INS-50 |
| chr12-16574105-INS-467 | chr12-16574105-INS-467 |
| chr12-16614101-INS-84 | chr12-16614148-INS-50 |
| chr12-16633821-INS-917 | chr12-16633821-INS-917 |
| chr12-16656968-INS-544 | chr12-16656968-INS-544 |
| chr12-16665198-INS-319 | chr12-16665198-INS-319 |
| chr12-16668868-INS-52 | chr12-16668894-INS-60 |
| chr12-16698062-INS-76 | chr12-16698062-INS-78 |
| chr12-16700717-INS-127 | chr12-16700717-INS-115 |
| chr12-16707406-INS-84 | chr12-16707406-INS-84 |
| chr12-16728984-INS-62 | chr12-16728984-INS-62 |
| chr12-16730980-INS-175 | chr12-16730980-INS-174 |
| chr12-16754343-INS-1328 | chr12-16754359-INS-1310 |
| chr12-16778870-INS-2229 | chr12-16778871-INS-2229 |
| chr12-16792142-INS-62 | chr12-16792130-INS-60 |

| | |
|---|---|
| chr12-16793925-INS-52 | chr12-16793925-INS-51 |
| chr12-16812205-INS-182 | chr12-16812206-INS-186 |
| chr12-16835806-INS-88 | chr12-16835806-INS-79 |
| chr12-16837899-INS-57 | chr12-16837899-INS-57 |
| chr12-16839464-INS-52 | chr12-16839465-INS-56 |
| chr12-16849708-INS-93 | chr12-16849709-INS-97 |
| chr12-16851490-INS-69 | chr12-16851469-INS-72 |
| chr12-16858754-INS-553 | chr12-16858754-INS-553 |
| chr12-16871509-INS-215 | chr12-16871509-INS-215 |
| chr12-16936726-INS-209 | chr12-16936726-INS-209 |
| chr12-16959172-INS-6452 | chr12-16959173-INS-6452 |
| chr12-16995257-INS-52 | chr12-16995257-INS-68 |
| chr12-17067756-INS-79 | chr12-17067756-INS-79 |
| chr12-17068964-INS-156 | chr12-17068964-INS-156 |
| chr12-17078673-INS-498 | chr12-17078673-INS-498 |
| chr12-17079405-INS-53 | chr12-17079405-INS-53 |
| chr12-17080726-INS-228 | chr12-17080726-INS-228 |
| chr12-17090842-INS-206 | chr12-17090842-INS-206 |
| chr12-17116426-INS-82 | chr12-17116426-INS-82 |
| chr12-17138338-INS-51 | chr12-17138339-INS-54 |
| chr12-17169598-INS-58 | chr12-17169597-INS-57 |
| chr12-17185550-INS-54 | chr12-17185550-INS-54 |
| chr12-17201485-INS-70 | chr12-17201485-INS-70 |
| chr12-17211956-INS-52 | chr12-17211956-INS-52 |
| chr12-17218017-INS-441 | chr12-17218017-INS-438 |
| chr12-17230712-INS-57 | chr12-17230712-INS-57 |
| chr12-17254291-INS-410 | chr12-17254291-INS-402 |
| chr12-17275566-INS-175 | chr12-17275566-INS-175 |
| chr12-17283351-INS-195 | chr12-17283351-INS-190 |
| chr12-17292639-INS-787 | chr12-17292639-INS-787 |
| chr12-17307723-INS-171 | chr12-17307690-INS-179 |
| chr12-17336319-INS-59 | chr12-17336319-INS-59 |
| chr12-17357772-INS-66 | chr12-17357772-INS-66 |
| chr12-17360703-INS-51 | chr12-17360703-INS-63 |
| chr12-17363845-INS-94 | chr12-17363845-INS-94 |
| chr12-17420954-INS-90 | chr12-17420954-INS-91 |
| chr12-17464206-INS-72 | chr12-17464210-INS-104 |
| chr12-17475210-INS-76 | chr12-17475195-INS-61 |
| chr12-17484712-INS-270 | chr12-17484743-INS-266 |
| chr12-17487520-INS-51 | chr12-17487505-INS-58 |
| chr12-17496143-INS-56 | chr12-17496103-INS-63 |
| chr12-17511982-INS-6471 | chr12-17511983-INS-6472 |
| chr12-17520948-INS-65 | chr12-17520948-INS-68 |
| chr12-17537106-INS-58 | chr12-17537106-INS-70 |
| chr12-17547548-INS-343 | chr12-17547548-INS-343 |
| chr12-17560984-INS-54 | chr12-17560984-INS-54 |
| chr12-17564877-INS-96 | chr12-17564878-INS-80 |
| chr12-17580965-INS-435 | chr12-17580957-INS-398 |
| chr12-17652535-INS-215 | chr12-17652535-INS-215 |
| chr12-17665836-INS-207 | chr12-17665793-INS-246 |
| chr12-17704735-INS-208 | chr12-17704735-INS-207 |
| chr12-17718576-INS-343 | chr12-17718576-INS-343 |
| chr12-17746105-INS-52 | chr12-17746105-INS-54 |
| chr12-17766323-INS-81 | chr12-17766323-INS-81 |
| chr12-17770173-INS-199 | chr12-17770173-INS-200 |
| chr12-17773731-INS-200 | chr12-17773731-INS-201 |
| chr12-17787180-INS-253 | chr12-17787175-INS-258 |
| chr12-17798605-INS-1319 | chr12-17798606-INS-1319 |
| chr12-17801376-INS-971 | chr12-17801376-INS-971 |
| chr12-17803235-INS-54 | chr12-17803235-INS-60 |
| chr12-17810438-INS-341 | chr12-17810438-INS-341 |
| chr12-17826017-INS-216 | chr12-17826017-INS-217 |
| chr12-17836807-INS-215 | chr12-17836807-INS-231 |
| chr12-17936623-INS-455 | chr12-17936623-INS-455 |
| chr12-18047317-INS-115 | chr12-18047317-INS-115 |
| chr12-18076600-INS-168 | chr12-18076600-INS-168 |
| chr12-18246198-INS-4994 | chr12-18246199-INS-4994 |
| chr12-18277926-INS-546 | chr12-18277931-INS-549 |
| chr12-18286959-INS-547 | chr12-18286959-INS-547 |
| chr12-18370307-INS-57 | chr12-18370307-INS-57 |
| chr12-18416949-INS-114 | chr12-18416949-INS-114 |
| chr12-18441819-INS-52 | chr12-18441819-INS-54 |
| chr12-18443023-INS-230 | chr12-18443023-INS-230 |
| chr12-18584005-INS-167 | chr12-18584005-INS-167 |
| chr12-18627474-INS-65 | chr12-18627474-INS-75 |
| chr12-18951706-INS-603 | chr12-18951706-INS-603 |
| chr12-18999341-INS-206 | chr12-18999341-INS-204 |

| | |
|---|---|
| chr12-19248943-INS-126 | chr12-19248943-INS-84 |
| chr12-19255848-INS-68 | chr12-19255849-INS-64 |
| chr12-19487902-INS-726 | chr12-19487903-INS-724 |
| chr12-19614143-INS-208 | chr12-19614143-INS-208 |
| chr12-19620336-INS-711 | chr12-19620336-INS-711 |
| chr12-19621492-INS-86 | chr12-19621492-INS-88 |
| chr12-19980823-INS-51 | chr12-19980823-INS-71 |
| chr12-20147199-INS-65 | chr12-20147199-INS-65 |
| chr12-20336495-INS-4827 | chr12-20336496-INS-4827 |
| chr12-20354143-INS-144 | chr12-20354152-INS-123 |
| chr12-20485766-INS-84 | chr12-20485767-INS-84 |
| chr12-20529724-INS-338 | chr12-20529768-INS-341 |
| chr12-20546540-INS-132 | chr12-20546540-INS-132 |
| chr12-20573988-INS-82 | chr12-20573988-INS-86 |
| chr12-20712055-INS-4909 | chr12-20712056-INS-4909 |
| chr12-20712056-INS-565 | chr12-20712056-INS-565 |
| chr12-20719593-INS-77 | chr12-20719594-INS-77 |
| chr12-20820697-INS-84 | chr12-20820697-INS-84 |
| chr12-20824936-INS-385 | chr12-20824936-INS-385 |
| chr12-20857212-INS-55 | chr12-20857212-INS-50 |
| chr12-20862346-INS-1892 | chr12-20862347-INS-1895 |
| chr12-20875625-INS-72 | chr12-20875625-INS-50 |
| chr12-20881036-INS-118 | chr12-20881036-INS-99 |
| chr12-20884844-INS-1476 | chr12-20884845-INS-1490 |
| chr12-20916796-INS-66 | chr12-20916796-INS-66 |
| chr12-20942574-INS-5736 | chr12-20942574-INS-5725 |
| chr12-20950118-INS-336 | chr12-20950119-INS-336 |
| chr12-20950118-INS-923 | chr12-20950116-INS-924 |
| chr12-20958676-INS-569 | chr12-20958676-INS-569 |
| chr12-21113289-INS-68 | chr12-21113289-INS-69 |
| chr12-21116410-INS-56 | chr12-21116410-INS-50 |
| chr12-21136868-INS-1302 | chr12-21136869-INS-1304 |
| chr12-21156514-INS-52 | chr12-21156512-INS-52 |
| chr12-21157650-INS-199 | chr12-21157650-INS-199 |
| chr12-21160057-INS-52 | chr12-21160057-INS-52 |
| chr12-21197255-INS-138 | chr12-21197255-INS-138 |
| chr12-21208487-INS-74 | chr12-21208487-INS-80 |
| chr12-21226860-INS-99 | chr12-21226860-INS-102 |
| chr12-21228289-INS-89 | chr12-21228285-INS-96 |
| chr12-21232193-INS-52 | chr12-21232193-INS-52 |
| chr12-21240457-INS-462 | chr12-21240457-INS-462 |
| chr12-21246334-INS-100 | chr12-21246334-INS-100 |
| chr12-21263029-INS-120 | chr12-21263029-INS-112 |
| chr12-21338733-INS-294 | chr12-21338733-INS-294 |
| chr12-21358242-INS-80 | chr12-21358242-INS-80 |
| chr12-21394757-INS-136 | chr12-21394757-INS-134 |
| chr12-21428059-INS-51 | chr12-21428060-INS-51 |
| chr12-21434745-INS-210 | chr12-21434745-INS-210 |
| chr12-21435816-INS-57 | chr12-21435840-INS-57 |
| chr12-21449208-INS-208 | chr12-21449222-INS-214 |
| chr12-21455420-INS-124 | chr12-21455444-INS-123 |
| chr12-21473777-INS-183 | chr12-21473777-INS-184 |
| chr12-21477564-INS-156 | chr12-21477564-INS-163 |
| chr12-21479688-INS-105 | chr12-21479688-INS-105 |
| chr12-21482986-INS-209 | chr12-21482986-INS-209 |
| chr12-21492520-INS-175 | chr12-21492505-INS-173 |
| chr12-21502916-INS-304 | chr12-21502912-INS-308 |
| chr12-21503607-INS-211 | chr12-21503607-INS-211 |
| chr12-21514013-INS-159 | chr12-21514013-INS-159 |
| chr12-21516947-INS-199 | chr12-21516947-INS-199 |
| chr12-21517607-INS-6401 | chr12-21517608-INS-6401 |
| chr12-21535669-INS-208 | chr12-21535669-INS-208 |
| chr12-21591143-INS-565 | chr12-21591143-INS-565 |
| chr12-21591880-INS-52 | chr12-21591881-INS-52 |
| chr12-21595536-INS-211 | chr12-21595536-INS-213 |
| chr12-21602547-INS-66 | chr12-21602547-INS-66 |
| chr12-21607673-INS-8846 | chr12-21607674-INS-8846 |
| chr12-21626594-INS-66 | chr12-21626594-INS-66 |
| chr12-21631862-INS-75 | chr12-21631862-INS-84 |
| chr12-21639547-INS-4380 | chr12-21639548-INS-4412 |
| chr12-21672536-INS-3250 | chr12-21672537-INS-3254 |
| chr12-21687651-INS-66 | chr12-21687651-INS-66 |
| chr12-21823189-INS-58 | chr12-21823187-INS-57 |
| chr12-21867072-INS-448 | chr12-21867072-INS-448 |
| chr12-21900309-INS-4314 | chr12-21900310-INS-4314 |
| chr12-21919015-INS-78 | chr12-21919028-INS-78 |
| chr12-22497484-INS-84 | chr12-22497485-INS-84 |

| | |
|---|---|
| chr12-23028580-INS-244 | chr12-23028580-INS-244 |
| chr12-23132873-INS-57 | chr12-23132873-INS-57 |
| chr12-23692883-INS-76 | chr12-23692883-INS-114 |
| chr12-23786408-INS-523 | chr12-23786414-INS-521 |
| chr12-23965501-INS-56 | chr12-23965502-INS-52 |
| chr12-24122543-INS-84 | chr12-24122544-INS-84 |
| chr12-24188956-INS-73 | chr12-24188955-INS-73 |
| chr12-24491278-INS-201 | chr12-24491278-INS-201 |
| chr12-24501855-INS-7068 | chr12-24501856-INS-7068 |
| chr12-24537576-INS-54 | chr12-24537577-INS-57 |
| chr12-24567897-INS-215 | chr12-24567897-INS-215 |
| chr12-24571840-INS-4920 | chr12-24571841-INS-4920 |
| chr12-24580485-INS-2530 | chr12-24580486-INS-2529 |
| chr12-24590796-INS-495 | chr12-24590796-INS-495 |
| chr12-24592645-INS-52 | chr12-24592645-INS-50 |
| chr12-24611865-INS-56 | chr12-24611865-INS-56 |
| chr12-24649972-INS-166 | chr12-24649972-INS-166 |
| chr12-24659410-INS-186 | chr12-24659410-INS-186 |
| chr12-24696901-INS-482 | chr12-24696902-INS-483 |
| chr12-24698369-INS-85 | chr12-24698369-INS-85 |
| chr12-24713731-INS-52 | chr12-24713733-INS-50 |
| chr12-24716919-INS-58 | chr12-24716919-INS-50 |
| chr12-24722146-INS-246 | chr12-24722146-INS-233 |
| chr12-24732460-INS-154 | chr12-24732460-INS-154 |
| chr12-24749314-INS-202 | chr12-24749314-INS-202 |
| chr12-24755857-INS-57 | chr12-24755857-INS-57 |
| chr12-24782492-INS-203 | chr12-24782492-INS-203 |
| chr12-24782953-INS-4793 | chr12-24782954-INS-4793 |
| chr12-24789240-INS-1006 | chr12-24789241-INS-1006 |
| chr12-24791570-INS-577 | chr12-24791570-INS-577 |
| chr12-24808063-INS-54 | chr12-24808063-INS-54 |
| chr12-24810129-INS-400 | chr12-24810129-INS-400 |
| chr12-24812024-INS-56 | chr12-24812024-INS-56 |
| chr12-24824647-INS-209 | chr12-24824647-INS-209 |
| chr12-24825790-INS-59 | chr12-24825792-INS-57 |
| chr12-24827742-INS-211 | chr12-24827742-INS-212 |
| chr12-24832087-INS-211 | chr12-24832084-INS-204 |
| chr12-24851728-INS-132 | chr12-24851728-INS-132 |
| chr12-24903486-INS-211 | chr12-24903486-INS-211 |
| chr12-24912566-INS-219 | chr12-24912566-INS-218 |
| chr12-24917119-INS-634 | chr12-24917117-INS-633 |
| chr12-24925905-INS-59 | chr12-24925902-INS-74 |
| chr12-24943641-INS-56 | chr12-24943641-INS-56 |
| chr12-24946154-INS-176 | chr12-24946154-INS-172 |
| chr12-24959018-INS-90 | chr12-24959007-INS-110 |
| chr12-24965014-INS-122 | chr12-24964995-INS-155 |
| chr12-24979541-INS-59 | chr12-24979493-INS-72 |
| chr12-24985475-INS-172 | chr12-24985475-INS-172 |
| chr12-24990968-INS-519 | chr12-24990968-INS-527 |
| chr12-24996801-INS-72 | chr12-24996802-INS-76 |
| chr12-24998467-INS-66 | chr12-24998467-INS-50 |
| chr12-25007296-INS-134 | chr12-25007296-INS-134 |
| chr12-25020793-INS-323 | chr12-25020793-INS-323 |
| chr12-25045322-INS-207 | chr12-25045322-INS-207 |
| chr12-25086733-INS-92 | chr12-25086765-INS-76 |
| chr12-25128842-INS-175 | chr12-25128842-INS-175 |
| chr12-25136068-INS-2208 | chr12-25136069-INS-2227 |
| chr12-25136788-INS-70 | chr12-25136788-INS-75 |
| chr12-25140023-INS-58 | chr12-25140037-INS-68 |
| chr12-25172523-INS-55 | chr12-25172523-INS-55 |
| chr12-25203674-INS-205 | chr12-25203674-INS-205 |
| chr12-25213166-INS-53 | chr12-25213166-INS-53 |
| chr12-25219110-INS-58 | chr12-25219116-INS-52 |
| chr12-25220252-INS-6405 | chr12-25220253-INS-6405 |
| chr12-25247649-INS-63 | chr12-25247649-INS-75 |
| chr12-25266673-INS-86 | chr12-25266673-INS-86 |
| chr12-25292032-INS-6224 | chr12-25292033-INS-6225 |
| chr12-25301278-INS-54 | chr12-25301278-INS-55 |
| chr12-25313469-INS-81 | chr12-25313469-INS-80 |
| chr12-25318178-INS-253 | chr12-25318178-INS-256 |
| chr12-25339664-INS-192 | chr12-25339664-INS-193 |
| chr12-25386806-INS-76 | chr12-25386807-INS-50 |
| chr12-25393407-INS-57 | chr12-25393407-INS-57 |
| chr12-25395456-INS-159 | chr12-25395456-INS-159 |
| chr12-25410419-INS-1266 | chr12-25410420-INS-1267 |
| chr12-25419992-INS-125 | chr12-25419990-INS-145 |
| chr12-25446210-INS-54 | chr12-25446210-INS-66 |

| | |
|---|---|
| chr12-25456043-INS-54 | chr12-25456059-INS-70 |
| chr12-25471337-INS-59 | chr12-25471337-INS-59 |
| chr12-25531954-INS-83 | chr12-25531962-INS-92 |
| chr12-25560021-INS-3855 | chr12-25560022-INS-3855 |
| chr12-25572498-INS-4399 | chr12-25572499-INS-4399 |
| chr12-25601520-INS-6430 | chr12-25601521-INS-6430 |
| chr12-25638362-INS-225 | chr12-25638362-INS-221 |
| chr12-25648367-INS-58 | chr12-25648367-INS-56 |
| chr12-25657499-INS-98 | chr12-25657499-INS-110 |
| chr12-25657889-INS-775 | chr12-25657889-INS-775 |
| chr12-25684601-INS-60 | chr12-25684601-INS-60 |
| chr12-25696146-INS-161 | chr12-25696146-INS-161 |
| chr12-25711109-INS-63 | chr12-25711109-INS-69 |
| chr12-25728325-INS-58 | chr12-25728325-INS-58 |
| chr12-25736552-INS-58 | chr12-25736550-INS-56 |
| chr12-25742079-INS-395 | chr12-25742079-INS-387 |
| chr12-25759345-INS-182 | chr12-25759345-INS-183 |
| chr12-25779517-INS-159 | chr12-25779517-INS-142 |
| chr12-25846803-INS-6510 | chr12-25846804-INS-6510 |
| chr12-25860239-INS-77 | chr12-25860239-INS-77 |
| chr12-25893045-INS-60 | chr12-25893045-INS-62 |
| chr12-25908762-INS-439 | chr12-25908762-INS-439 |
| chr12-25930657-INS-67 | chr12-25930657-INS-67 |
| chr12-26000456-INS-6503 | chr12-26000457-INS-6503 |
| chr12-26010784-INS-104 | chr12-26010784-INS-104 |
| chr12-26054652-INS-384 | chr12-26054652-INS-384 |
| chr12-26081710-INS-90 | chr12-26081710-INS-108 |
| chr12-26113717-INS-929 | chr12-26113717-INS-928 |
| chr12-26126945-INS-80 | chr12-26126945-INS-80 |
| chr12-26132786-INS-206 | chr12-26132786-INS-205 |
| chr12-26146340-INS-508 | chr12-26146340-INS-508 |
| chr12-26150852-INS-54 | chr12-26150856-INS-50 |
| chr12-26159915-INS-59 | chr12-26159915-INS-90 |
| chr12-26203975-INS-70 | chr12-26203975-INS-70 |
| chr12-26230649-INS-54 | chr12-26230649-INS-60 |
| chr12-26243989-INS-290 | chr12-26243989-INS-301 |
| chr12-26245376-INS-375 | chr12-26245358-INS-351 |
| chr12-26265108-INS-202 | chr12-26265108-INS-201 |
| chr12-26286538-INS-190 | chr12-26286538-INS-186 |
| chr12-26310804-INS-75 | chr12-26310814-INS-80 |
| chr12-26353509-INS-224 | chr12-26353509-INS-224 |
| chr12-26363979-INS-58 | chr12-26363979-INS-60 |
| chr12-26371724-INS-65 | chr12-26371734-INS-78 |
| chr12-26376853-INS-2032 | chr12-26376854-INS-2032 |
| chr12-26388662-INS-54 | chr12-26388668-INS-52 |
| chr12-26428756-INS-93 | chr12-26428764-INS-132 |
| chr12-26446443-INS-493 | chr12-26446444-INS-507 |
| chr12-26462377-INS-750 | chr12-26462377-INS-751 |
| chr12-26489208-INS-158 | chr12-26489208-INS-158 |
| chr12-26543096-INS-54 | chr12-26543070-INS-52 |
| chr12-26568028-INS-91 | chr12-26568028-INS-96 |
| chr12-26585667-INS-66 | chr12-26585667-INS-66 |
| chr12-26587346-INS-118 | chr12-26587346-INS-118 |
| chr12-26591746-INS-80 | chr12-26591746-INS-80 |
| chr12-26608471-INS-62 | chr12-26608471-INS-62 |
| chr12-26619056-INS-198 | chr12-26619056-INS-198 |
| chr12-26630461-INS-572 | chr12-26630461-INS-572 |
| chr12-26632040-INS-65 | chr12-26632040-INS-65 |
| chr12-26645163-INS-943 | chr12-26645163-INS-943 |
| chr12-26649869-INS-76 | chr12-26649863-INS-92 |
| chr12-26678379-INS-243 | chr12-26678379-INS-243 |
| chr12-26690430-INS-6395 | chr12-26690431-INS-6395 |
| chr12-26701947-INS-54 | chr12-26701947-INS-56 |
| chr12-26705828-INS-78 | chr12-26705829-INS-71 |
| chr12-26707935-INS-5541 | chr12-26707936-INS-5541 |
| chr12-26708264-INS-173 | chr12-26708265-INS-151 |
| chr12-26721459-INS-96 | chr12-26721459-INS-96 |
| chr12-26735192-INS-259 | chr12-26735192-INS-259 |
| chr12-26755157-INS-64 | chr12-26755139-INS-66 |
| chr12-26791845-INS-120 | chr12-26791845-INS-120 |
| chr12-26794496-INS-52 | chr12-26794496-INS-50 |
| chr12-26805633-INS-2163 | chr12-26805634-INS-2166 |
| chr12-26810124-INS-100 | chr12-26810124-INS-100 |
| chr12-26817545-INS-188 | chr12-26817546-INS-189 |
| chr12-26830307-INS-181 | chr12-26830307-INS-181 |
| chr12-26844613-INS-59 | chr12-26844613-INS-59 |
| chr12-26858530-INS-109 | chr12-26858530-INS-73 |

| | |
|---|---|
| chr12-26872673-INS-6327 | chr12-26872674-INS-6327 |
| chr12-26929885-INS-470 | chr12-26929885-INS-470 |
| chr12-26959509-INS-620 | chr12-26959509-INS-620 |
| chr12-26977555-INS-240 | chr12-26977555-INS-240 |
| chr12-26985812-INS-2637 | chr12-26985813-INS-2636 |
| chr12-26990862-INS-84 | chr12-26990862-INS-84 |
| chr12-26998780-INS-1505 | chr12-26998781-INS-1505 |
| chr12-27028065-INS-487 | chr12-27028065-INS-488 |
| chr12-27061829-INS-6859 | chr12-27061830-INS-6859 |
| chr12-27066592-INS-137 | chr12-27066592-INS-137 |
| chr12-27071185-INS-52 | chr12-27071185-INS-52 |
| chr12-27086625-INS-58 | chr12-27086615-INS-52 |
| chr12-27107928-INS-52 | chr12-27107929-INS-52 |
| chr12-27120680-INS-136 | chr12-27120680-INS-139 |
| chr12-27136453-INS-59 | chr12-27136454-INS-69 |
| chr12-27160414-INS-51 | chr12-27160414-INS-51 |
| chr12-27183015-INS-120 | chr12-27183015-INS-96 |
| chr12-27183136-INS-742 | chr12-27183136-INS-743 |
| chr12-27231358-INS-52 | chr12-27231358-INS-52 |
| chr12-27255508-INS-342 | chr12-27255502-INS-345 |
| chr12-27265956-INS-6323 | chr12-27265957-INS-6325 |
| chr12-27301955-INS-58 | chr12-27301951-INS-68 |
| chr12-27305286-INS-256 | chr12-27305286-INS-254 |
| chr12-27309676-INS-52 | chr12-27309676-INS-52 |
| chr12-27312845-INS-70 | chr12-27312833-INS-64 |
| chr12-27368156-INS-68 | chr12-27368156-INS-60 |
| chr12-27396353-INS-945 | chr12-27396353-INS-947 |
| chr12-27405063-INS-92 | chr12-27405063-INS-92 |
| chr12-27408473-INS-226 | chr12-27408473-INS-225 |
| chr12-27416794-INS-779 | chr12-27416794-INS-779 |
| chr12-27428087-INS-698 | chr12-27428087-INS-699 |
| chr12-27437024-INS-280 | chr12-27437024-INS-280 |
| chr12-27447657-INS-84 | chr12-27447657-INS-84 |
| chr12-27506654-INS-61 | chr12-27506654-INS-61 |
| chr12-27519990-INS-63 | chr12-27519990-INS-63 |
| chr12-27521070-INS-387 | chr12-27521070-INS-387 |
| chr12-27530450-INS-1892 | chr12-27530451-INS-1893 |
| chr12-27543099-INS-52 | chr12-27543099-INS-53 |
| chr12-27546467-INS-52 | chr12-27546467-INS-52 |
| chr12-27558866-INS-73 | chr12-27558864-INS-59 |
| chr12-27598601-INS-104 | chr12-27598607-INS-117 |
| chr12-27607880-INS-69 | chr12-27607880-INS-63 |
| chr12-27613122-INS-68 | chr12-27613122-INS-68 |
| chr12-27646024-INS-393 | chr12-27646024-INS-393 |
| chr12-27666276-INS-100 | chr12-27666276-INS-102 |
| chr12-27687040-INS-221 | chr12-27687040-INS-222 |
| chr12-27689779-INS-2644 | chr12-27689780-INS-2644 |
| chr12-27704099-INS-52 | chr12-27704099-INS-52 |
| chr12-27723115-INS-805 | chr12-27723165-INS-804 |
| chr12-27725719-INS-6621 | chr12-27725720-INS-6622 |
| chr12-27752614-INS-544 | chr12-27752614-INS-544 |
| chr12-27758249-INS-100 | chr12-27758249-INS-100 |
| chr12-27815177-INS-65 | chr12-27815177-INS-65 |
| chr12-27831801-INS-6649 | chr12-27831802-INS-6649 |
| chr12-27833730-INS-531 | chr12-27833730-INS-533 |
| chr12-27839640-INS-874 | chr12-27839640-INS-874 |
| chr12-27855729-INS-60 | chr12-27855729-INS-60 |
| chr12-27857766-INS-83 | chr12-27857766-INS-87 |
| chr12-27864734-INS-106 | chr12-27864734-INS-106 |
| chr12-27938527-INS-4466 | chr12-27938528-INS-4466 |
| chr12-27955668-INS-52 | chr12-27955668-INS-52 |
| chr12-27971894-INS-6387 | chr12-27971895-INS-6388 |
| chr12-27991746-INS-759 | chr12-27991746-INS-759 |
| chr12-28049588-INS-220 | chr12-28049588-INS-220 |
| chr12-28070336-INS-78 | chr12-28070336-INS-98 |
| chr12-28072466-INS-1890 | chr12-28072467-INS-1890 |
| chr12-28103028-INS-74 | chr12-28103021-INS-56 |
| chr12-28127721-INS-100 | chr12-28127709-INS-102 |
| chr12-28139291-INS-701 | chr12-28139291-INS-703 |
| chr12-28282361-INS-1660 | chr12-28282362-INS-1660 |
| chr12-28310735-INS-6276 | chr12-28310736-INS-6276 |
| chr12-28323239-INS-78 | chr12-28323239-INS-78 |
| chr12-28396416-INS-96 | chr12-28396416-INS-96 |
| chr12-28409434-INS-144 | chr12-28409434-INS-143 |
| chr12-28424378-INS-442 | chr12-28424378-INS-444 |
| chr12-28466013-INS-71 | chr12-28466013-INS-71 |
| chr12-28497148-INS-764 | chr12-28497148-INS-768 |

| | |
|---|---|
| chr12-28516175-INS-66 | chr12-28516194-INS-96 |
| chr12-28538588-INS-884 | chr12-28538588-INS-885 |
| chr12-28563053-INS-352 | chr12-28563053-INS-352 |
| chr12-28610158-INS-250 | chr12-28610158-INS-225 |
| chr12-28611848-INS-350 | chr12-28611848-INS-350 |
| chr12-28615951-INS-291 | chr12-28615952-INS-288 |
| chr12-28626806-INS-487 | chr12-28626806-INS-487 |
| chr12-28643508-INS-143 | chr12-28643508-INS-169 |
| chr12-28660241-INS-413 | chr12-28660249-INS-429 |
| chr12-28665172-INS-287 | chr12-28665172-INS-288 |
| chr12-28670798-INS-53 | chr12-28670771-INS-52 |
| chr12-28677050-INS-207 | chr12-28677050-INS-207 |
| chr12-28683882-INS-249 | chr12-28683882-INS-249 |
| chr12-28687842-INS-79 | chr12-28687842-INS-120 |
| chr12-28694397-INS-147 | chr12-28694397-INS-155 |
| chr12-28698631-INS-223 | chr12-28698631-INS-223 |
| chr12-28712134-INS-133 | chr12-28712134-INS-134 |
| chr12-28723940-INS-162 | chr12-28723940-INS-164 |
| chr12-28744384-INS-243 | chr12-28744384-INS-243 |
| chr12-28747318-INS-52 | chr12-28747319-INS-54 |
| chr12-28779827-INS-6442 | chr12-28779828-INS-6442 |
| chr12-28780348-INS-204 | chr12-28780348-INS-203 |
| chr12-28814875-INS-183 | chr12-28814875-INS-183 |
| chr12-28827998-INS-237 | chr12-28827998-INS-237 |
| chr12-28850635-INS-555 | chr12-28850635-INS-555 |
| chr12-28891945-INS-57 | chr12-28891945-INS-57 |
| chr12-28919861-INS-55 | chr12-28919861-INS-75 |
| chr12-28924376-INS-108 | chr12-28924376-INS-108 |
| chr12-28931236-INS-400 | chr12-28931236-INS-400 |
| chr12-28936206-INS-315 | chr12-28936206-INS-319 |
| chr12-28960937-INS-97 | chr12-28960937-INS-101 |
| chr12-28987668-INS-221 | chr12-28987668-INS-221 |
| chr12-28995260-INS-83 | chr12-28995262-INS-60 |
| chr12-29000413-INS-251 | chr12-29000413-INS-251 |
| chr12-29002194-INS-53 | chr12-29002194-INS-53 |
| chr12-29003291-INS-55 | chr12-29003291-INS-54 |
| chr12-29003960-INS-6752 | chr12-29003961-INS-6752 |
| chr12-29055826-INS-2113 | chr12-29055827-INS-2107 |
| chr12-29061425-INS-95 | chr12-29061425-INS-95 |
| chr12-29082380-INS-143 | chr12-29082381-INS-143 |
| chr12-29106636-INS-6546 | chr12-29106637-INS-6547 |
| chr12-29123920-INS-663 | chr12-29123920-INS-663 |
| chr12-29129766-INS-92 | chr12-29129766-INS-92 |
| chr12-29138107-INS-6378 | chr12-29138108-INS-6378 |
| chr12-29148957-INS-1258 | chr12-29148958-INS-1258 |
| chr12-29192358-INS-103 | chr12-29192358-INS-103 |
| chr12-29208001-INS-969 | chr12-29208001-INS-968 |
| chr12-29209097-INS-82 | chr12-29209097-INS-84 |
| chr12-29272595-INS-112 | chr12-29272595-INS-114 |
| chr12-29277659-INS-795 | chr12-29277659-INS-795 |
| chr12-29287949-INS-59 | chr12-29287949-INS-59 |
| chr12-29306513-INS-60 | chr12-29306513-INS-58 |
| chr12-29316283-INS-162 | chr12-29316284-INS-136 |
| chr12-29357910-INS-99 | chr12-29357910-INS-51 |
| chr12-29386611-INS-80 | chr12-29386611-INS-60 |
| chr12-29398045-INS-60 | chr12-29398045-INS-100 |
| chr12-29457568-INS-56 | chr12-29457568-INS-56 |
| chr12-29465405-INS-61 | chr12-29465405-INS-69 |
| chr12-29487037-INS-55 | chr12-29487037-INS-59 |
| chr12-29491973-INS-692 | chr12-29491973-INS-693 |
| chr12-29520395-INS-108 | chr12-29520389-INS-110 |
| chr12-29552914-INS-2793 | chr12-29552915-INS-2808 |
| chr12-29552915-INS-401 | chr12-29552915-INS-401 |
| chr12-29571387-INS-408 | chr12-29571388-INS-372 |
| chr12-29594837-INS-341 | chr12-29594837-INS-341 |
| chr12-29608749-INS-78 | chr12-29608749-INS-78 |
| chr12-29615239-INS-65 | chr12-29615255-INS-78 |
| chr12-29625469-INS-68 | chr12-29625469-INS-72 |
| chr12-29644148-INS-135 | chr12-29644148-INS-121 |
| chr12-29660067-INS-69 | chr12-29660067-INS-69 |
| chr12-29667101-INS-218 | chr12-29667101-INS-218 |
| chr12-29671804-INS-96 | chr12-29671804-INS-96 |
| chr12-29676275-INS-330 | chr12-29676275-INS-330 |
| chr12-29681751-INS-66 | chr12-29681751-INS-66 |
| chr12-29683751-INS-53 | chr12-29683751-INS-53 |
| chr12-29745097-INS-52 | chr12-29745097-INS-52 |
| chr12-29766651-INS-611 | chr12-29766651-INS-611 |

| | |
|---|---|
| chr12-29795288-INS-1859 | chr12-29795289-INS-1859 |
| chr12-29808948-INS-62 | chr12-29808908-INS-70 |
| chr12-29829761-INS-53 | chr12-29829761-INS-53 |
| chr12-29855487-INS-157 | chr12-29855487-INS-167 |
| chr12-29926691-INS-54 | chr12-29926692-INS-93 |
| chr12-29944201-INS-52 | chr12-29944201-INS-64 |
| chr12-29998273-INS-191 | chr12-29998273-INS-192 |
| chr12-30015449-INS-168 | chr12-30015449-INS-168 |
| chr12-30022440-INS-58 | chr12-30022440-INS-58 |
| chr12-30024110-INS-400 | chr12-30024110-INS-400 |
| chr12-30054349-INS-52 | chr12-30054349-INS-52 |
| chr12-30085050-INS-230 | chr12-30085050-INS-232 |
| chr12-30114618-INS-66 | chr12-30114618-INS-88 |
| chr12-30118384-INS-71 | chr12-30118384-INS-62 |
| chr12-30165475-INS-3438 | chr12-30165476-INS-3438 |
| chr12-30193162-INS-92 | chr12-30193162-INS-92 |
| chr12-30212829-INS-54 | chr12-30212829-INS-54 |
| chr12-30216532-INS-52 | chr12-30216536-INS-56 |
| chr12-30223779-INS-67 | chr12-30223778-INS-71 |
| chr12-30233375-INS-76 | chr12-30233375-INS-75 |
| chr12-30235902-INS-115 | chr12-30235902-INS-114 |
| chr12-30267927-INS-59 | chr12-30267927-INS-59 |
| chr12-30321853-INS-188 | chr12-30321853-INS-203 |
| chr12-30325056-INS-53 | chr12-30325056-INS-53 |
| chr12-30330665-INS-326 | chr12-30330665-INS-325 |
| chr12-30361807-INS-90 | chr12-30361807-INS-90 |
| chr12-30369926-INS-6323 | chr12-30369927-INS-6323 |
| chr12-30372188-INS-212 | chr12-30372188-INS-212 |
| chr12-30399091-INS-6577 | chr12-30399092-INS-6577 |
| chr12-30406150-INS-665 | chr12-30406148-INS-661 |
| chr12-30406851-INS-225 | chr12-30406851-INS-225 |
| chr12-30439602-INS-76 | chr12-30439602-INS-76 |
| chr12-30448572-INS-54 | chr12-30448572-INS-50 |
| chr12-30453333-INS-84 | chr12-30453333-INS-84 |
| chr12-30505443-INS-52 | chr12-30505443-INS-54 |
| chr12-30515738-INS-118 | chr12-30515738-INS-119 |
| chr12-30524061-INS-1073 | chr12-30524062-INS-1073 |
| chr12-30533779-INS-6452 | chr12-30533780-INS-6452 |
| chr12-30549774-INS-242 | chr12-30549774-INS-242 |
| chr12-30552414-INS-2050 | chr12-30552415-INS-2050 |
| chr12-30584995-INS-6297 | chr12-30584996-INS-6297 |
| chr12-30600442-INS-394 | chr12-30600442-INS-394 |
| chr12-30608013-INS-7129 | chr12-30608014-INS-7130 |
| chr12-30610080-INS-1462 | chr12-30610081-INS-1462 |
| chr12-30613775-INS-54 | chr12-30613775-INS-56 |
| chr12-30616521-INS-534 | chr12-30616521-INS-532 |
| chr12-30623272-INS-1077 | chr12-30623273-INS-1077 |
| chr12-30631431-INS-68 | chr12-30631431-INS-68 |
| chr12-30646430-INS-72 | chr12-30646430-INS-72 |
| chr12-30647262-INS-523 | chr12-30647262-INS-523 |
| chr12-30699652-INS-97 | chr12-30699650-INS-127 |
| chr12-30704274-INS-894 | chr12-30704272-INS-896 |
| chr12-30712032-INS-623 | chr12-30712032-INS-623 |
| chr12-30717708-INS-53 | chr12-30717708-INS-53 |
| chr12-30718306-INS-723 | chr12-30718305-INS-722 |
| chr12-30727313-INS-107 | chr12-30727313-INS-101 |
| chr12-30752985-INS-6216 | chr12-30752986-INS-6216 |
| chr12-30762933-INS-66 | chr12-30762933-INS-66 |
| chr12-30771748-INS-104 | chr12-30771752-INS-54 |
| chr12-30773317-INS-88 | chr12-30773317-INS-88 |
| chr12-30793888-INS-908 | chr12-30793888-INS-908 |
| chr12-30815390-INS-207 | chr12-30815390-INS-207 |
| chr12-30822581-INS-52 | chr12-30822581-INS-52 |
| chr12-30835553-INS-7117 | chr12-30835554-INS-7117 |
| chr12-30917440-INS-103 | chr12-30917444-INS-95 |
| chr12-30923100-INS-358 | chr12-30923100-INS-358 |
| chr12-30928318-INS-70 | chr12-30928318-INS-70 |
| chr12-30942688-INS-140 | chr12-30942688-INS-160 |
| chr12-30979170-INS-445 | chr12-30979170-INS-445 |
| chr12-31007374-INS-6391 | chr12-31007375-INS-6391 |
| chr12-31041775-INS-424 | chr12-31041776-INS-424 |
| chr12-31070716-INS-101 | chr12-31070716-INS-101 |
| chr12-31083925-INS-58 | chr12-31083925-INS-58 |
| chr12-31091919-INS-1720 | chr12-31091920-INS-1720 |
| chr12-31143174-INS-89 | chr12-31143174-INS-86 |
| chr12-31154893-INS-105 | chr12-31154893-INS-133 |
| chr12-31170581-INS-150 | chr12-31170581-INS-149 |

| | |
|---|---|
| chr12-31172962-INS-55 | chr12-31172962-INS-55 |
| chr12-31188603-INS-959 | chr12-31188603-INS-970 |
| chr12-31191895-INS-402 | chr12-31191895-INS-404 |
| chr12-31192964-INS-99 | chr12-31192964-INS-100 |
| chr12-31210226-INS-76 | chr12-31210224-INS-50 |
| chr12-31247732-INS-63 | chr12-31247732-INS-63 |
| chr12-31254841-INS-196 | chr12-31254841-INS-175 |
| chr12-31260038-INS-183 | chr12-31260038-INS-181 |
| chr12-31285042-INS-198 | chr12-31285043-INS-201 |
| chr12-31285874-INS-58 | chr12-31285874-INS-56 |
| chr12-31290598-INS-82 | chr12-31290598-INS-82 |
| chr12-31336107-INS-120 | chr12-31336107-INS-120 |
| chr12-31350676-INS-105 | chr12-31350676-INS-105 |
| chr12-31369353-INS-167 | chr12-31369353-INS-168 |
| chr12-31382595-INS-96 | chr12-31382580-INS-79 |
| chr12-31426293-INS-72 | chr12-31426293-INS-72 |
| chr12-31439075-INS-120 | chr12-31439075-INS-120 |
| chr12-31460582-INS-195 | chr12-31460590-INS-148 |
| chr12-31473095-INS-1096 | chr12-31473096-INS-1097 |
| chr12-31483333-INS-160 | chr12-31483310-INS-188 |
| chr12-31521866-INS-275 | chr12-31521866-INS-275 |
| chr12-31524011-INS-217 | chr12-31524011-INS-217 |
| chr12-31547602-INS-172 | chr12-31547602-INS-181 |
| chr12-31571247-INS-65 | chr12-31571248-INS-62 |
| chr12-31576061-INS-89 | chr12-31576061-INS-83 |
| chr12-31580155-INS-83 | chr12-31580155-INS-98 |
| chr12-31589640-INS-209 | chr12-31589640-INS-210 |
| chr12-31602957-INS-57 | chr12-31602931-INS-82 |
| chr12-31610193-INS-80 | chr12-31610193-INS-78 |
| chr12-31619008-INS-113 | chr12-31619014-INS-76 |
| chr12-31635807-INS-74 | chr12-31635807-INS-74 |
| chr12-31650354-INS-198 | chr12-31650354-INS-198 |
| chr12-31655971-INS-143 | chr12-31655971-INS-136 |
| chr12-31656632-INS-55 | chr12-31656632-INS-55 |
| chr12-31661703-INS-353 | chr12-31661703-INS-353 |
| chr12-31667590-INS-122 | chr12-31667590-INS-122 |
| chr12-31722510-INS-7556 | chr12-31722511-INS-7581 |
| chr12-31729862-INS-90 | chr12-31729862-INS-90 |
| chr12-31773924-INS-7425 | chr12-31773925-INS-7425 |
| chr12-31786898-INS-105 | chr12-31786898-INS-105 |
| chr12-31799635-INS-489 | chr12-31799635-INS-489 |
| chr12-31818664-INS-6442 | chr12-31818665-INS-6442 |
| chr12-31819607-INS-55 | chr12-31819607-INS-55 |
| chr12-31821652-INS-210 | chr12-31821652-INS-211 |
| chr12-31846065-INS-203 | chr12-31846065-INS-203 |
| chr12-31853888-INS-54 | chr12-31853888-INS-50 |
| chr12-31854314-INS-207 | chr12-31854314-INS-208 |
| chr12-31876029-INS-184 | chr12-31875994-INS-187 |
| chr12-31919925-INS-159 | chr12-31919925-INS-159 |
| chr12-31923777-INS-207 | chr12-31923777-INS-208 |
| chr12-31950587-INS-74 | chr12-31950587-INS-74 |
| chr12-31994548-INS-155 | chr12-31994548-INS-152 |
| chr12-31996972-INS-63 | chr12-31996972-INS-63 |
| chr12-32014283-INS-59 | chr12-32014283-INS-59 |
| chr12-32017185-INS-117 | chr12-32017185-INS-117 |
| chr12-32024584-INS-108 | chr12-32024584-INS-108 |
| chr12-32043083-INS-56 | chr12-32043081-INS-52 |
| chr12-32098870-INS-964 | chr12-32098868-INS-954 |
| chr12-32118607-INS-1123 | chr12-32118613-INS-1114 |
| chr12-32148925-INS-8457 | chr12-32148926-INS-8458 |
| chr12-32186267-INS-71 | chr12-32186267-INS-71 |
| chr12-32198104-INS-225 | chr12-32198104-INS-225 |
| chr12-32201650-INS-92 | chr12-32201650-INS-92 |
| chr12-32206581-INS-119 | chr12-32206581-INS-119 |
| chr12-32207965-INS-245 | chr12-32207965-INS-245 |
| chr12-32214286-INS-3144 | chr12-32214287-INS-3144 |
| chr12-32214710-INS-1128 | chr12-32214711-INS-1126 |
| chr12-32216159-INS-601 | chr12-32216160-INS-565 |
| chr12-32239452-INS-1587 | chr12-32239453-INS-1582 |
| chr12-32242774-INS-346 | chr12-32242774-INS-346 |
| chr12-32293250-INS-340 | chr12-32293250-INS-340 |
| chr12-32295966-INS-1116 | chr12-32295967-INS-1116 |
| chr12-32310105-INS-484 | chr12-32310105-INS-485 |
| chr12-32313987-INS-578 | chr12-32313987-INS-569 |
| chr12-32323787-INS-2800 | chr12-32323788-INS-2800 |
| chr12-32343529-INS-72 | chr12-32343530-INS-72 |
| chr12-32355333-INS-67 | chr12-32355334-INS-66 |

| | |
|---|---|
| chr12-32377104-INS-833 | chr12-32377104-INS-833 |
| chr12-32405907-INS-3037 | chr12-32405908-INS-3037 |
| chr12-32418104-INS-275 | chr12-32418104-INS-272 |
| chr12-32426150-INS-7155 | chr12-32426151-INS-7155 |
| chr12-32455323-INS-111 | chr12-32455324-INS-110 |
| chr12-32465439-INS-6178 | chr12-32465439-INS-6178 |
| chr12-32477678-INS-70 | chr12-32477678-INS-70 |
| chr12-32489989-INS-84 | chr12-32489989-INS-96 |
| chr12-32490550-INS-6608 | chr12-32490551-INS-6609 |
| chr12-32614706-INS-326 | chr12-32614706-INS-326 |
| chr12-32619491-INS-3143 | chr12-32619492-INS-3162 |
| chr12-32620339-INS-110 | chr12-32620346-INS-146 |
| chr12-32630390-INS-176 | chr12-32630390-INS-176 |
| chr12-32661669-INS-450 | chr12-32661671-INS-443 |
| chr12-32696167-INS-464 | chr12-32696167-INS-464 |
| chr12-32715555-INS-292 | chr12-32715555-INS-292 |
| chr12-32728647-INS-451 | chr12-32728647-INS-451 |
| chr12-32733283-INS-52 | chr12-32733283-INS-58 |
| chr12-32774116-INS-86 | chr12-32774116-INS-80 |
| chr12-32810070-INS-6325 | chr12-32810071-INS-6326 |
| chr12-32820306-INS-1265 | chr12-32820307-INS-1267 |
| chr12-32843042-INS-88 | chr12-32843043-INS-88 |
| chr12-32855646-INS-622 | chr12-32855646-INS-620 |
| chr12-32867957-INS-196 | chr12-32867957-INS-196 |
| chr12-32882732-INS-55 | chr12-32882732-INS-55 |
| chr12-32910267-INS-57 | chr12-32910267-INS-57 |
| chr12-32911357-INS-1885 | chr12-32911358-INS-1886 |
| chr12-32924882-INS-106 | chr12-32924882-INS-114 |
| chr12-32942339-INS-56 | chr12-32942339-INS-56 |
| chr12-32946092-INS-163 | chr12-32946089-INS-177 |
| chr12-32970403-INS-381 | chr12-32970403-INS-380 |
| chr12-32980342-INS-434 | chr12-32980342-INS-434 |
| chr12-33000916-INS-125 | chr12-33000916-INS-125 |
| chr12-33012881-INS-182 | chr12-33012881-INS-182 |
| chr12-33031292-INS-259 | chr12-33031292-INS-259 |
| chr12-33032908-INS-60 | chr12-33032906-INS-62 |
| chr12-33039256-INS-71 | chr12-33039256-INS-77 |
| chr12-33041941-INS-207 | chr12-33041941-INS-205 |
| chr12-33043596-INS-357 | chr12-33043597-INS-332 |
| chr12-33046667-INS-333 | chr12-33046667-INS-331 |
| chr12-33057974-INS-223 | chr12-33057974-INS-223 |
| chr12-33116188-INS-80 | chr12-33116194-INS-83 |
| chr12-33123264-INS-161 | chr12-33123264-INS-161 |
| chr12-33131685-INS-144 | chr12-33131685-INS-140 |
| chr12-33133197-INS-86 | chr12-33133192-INS-121 |
| chr12-33140919-INS-451 | chr12-33140919-INS-451 |
| chr12-33147974-INS-60 | chr12-33147970-INS-64 |
| chr12-33149089-INS-6352 | chr12-33149090-INS-6352 |
| chr12-33149914-INS-207 | chr12-33149915-INS-207 |
| chr12-33157655-INS-66 | chr12-33157655-INS-68 |
| chr12-33164344-INS-175 | chr12-33164344-INS-175 |
| chr12-33180024-INS-60 | chr12-33180024-INS-60 |
| chr12-33236651-INS-188 | chr12-33236651-INS-189 |
| chr12-33264863-INS-399 | chr12-33264863-INS-399 |
| chr12-33329998-INS-53 | chr12-33329998-INS-53 |
| chr12-33335140-INS-69 | chr12-33335140-INS-69 |
| chr12-33341919-INS-89 | chr12-33341919-INS-89 |
| chr12-33360883-INS-441 | chr12-33360883-INS-443 |
| chr12-33383631-INS-105 | chr12-33383631-INS-107 |
| chr12-33437999-INS-66 | chr12-33437999-INS-72 |
| chr12-33453759-INS-580 | chr12-33453759-INS-580 |
| chr12-33461680-INS-186 | chr12-33461680-INS-187 |
| chr12-33466230-INS-392 | chr12-33466230-INS-392 |
| chr12-33496697-INS-213 | chr12-33496697-INS-225 |
| chr12-33499021-INS-429 | chr12-33499021-INS-428 |
| chr12-33511779-INS-198 | chr12-33511779-INS-198 |
| chr12-33533922-INS-6459 | chr12-33533923-INS-6459 |
| chr12-33535263-INS-210 | chr12-33535263-INS-211 |
| chr12-33536069-INS-66 | chr12-33536069-INS-66 |
| chr12-33560834-INS-449 | chr12-33560834-INS-450 |
| chr12-33562895-INS-132 | chr12-33562895-INS-145 |
| chr12-33563394-INS-1091 | chr12-33563395-INS-1092 |
| chr12-33570061-INS-164 | chr12-33570061-INS-168 |
| chr12-33571778-INS-132 | chr12-33571778-INS-116 |
| chr12-33580880-INS-6481 | chr12-33580881-INS-6482 |
| chr12-33607940-INS-596 | chr12-33607940-INS-596 |
| chr12-33622428-INS-1815 | chr12-33622429-INS-1815 |

| | |
|---|---|
| chr12-33639892-INS-401 | chr12-33639892-INS-401 |
| chr12-33693904-INS-57 | chr12-33693904-INS-61 |
| chr12-33709098-INS-52 | chr12-33709098-INS-50 |
| chr12-33711659-INS-55 | chr12-33711661-INS-59 |
| chr12-33713643-INS-72 | chr12-33713643-INS-72 |
| chr12-33746930-INS-1786 | chr12-33746931-INS-1786 |
| chr12-33769854-INS-6482 | chr12-33769855-INS-6482 |
| chr12-33776498-INS-5444 | chr12-33776499-INS-5461 |
| chr12-33782552-INS-424 | chr12-33782552-INS-424 |
| chr12-33791576-INS-134 | chr12-33791576-INS-134 |
| chr12-33793299-INS-429 | chr12-33793299-INS-429 |
| chr12-33802927-INS-52 | chr12-33802927-INS-52 |
| chr12-33816725-INS-214 | chr12-33816725-INS-214 |
| chr12-33824781-INS-61 | chr12-33824781-INS-62 |
| chr12-33827182-INS-119 | chr12-33827182-INS-119 |
| chr12-33827873-INS-126 | chr12-33827873-INS-134 |
| chr12-33849851-INS-403 | chr12-33849851-INS-403 |
| chr12-33857851-INS-99 | chr12-33857845-INS-87 |
| chr12-33864470-INS-2727 | chr12-33864471-INS-2727 |
| chr12-33866894-INS-52 | chr12-33866894-INS-52 |
| chr12-33882973-INS-123 | chr12-33882974-INS-132 |
| chr12-33887420-INS-68 | chr12-33887420-INS-70 |
| chr12-33896054-INS-240 | chr12-33896054-INS-248 |
| chr12-33948985-INS-120 | chr12-33948985-INS-100 |
| chr12-33959663-INS-451 | chr12-33959663-INS-452 |
| chr12-33960673-INS-1244 | chr12-33960674-INS-1244 |
| chr12-33966290-INS-277 | chr12-33966290-INS-277 |
| chr12-33972355-INS-7046 | chr12-33972356-INS-7048 |
| chr12-33974123-INS-791 | chr12-33974123-INS-791 |
| chr12-33989820-INS-134 | chr12-33989825-INS-94 |
| chr12-34015635-INS-96 | chr12-34015635-INS-98 |
| chr12-34039104-INS-56 | chr12-34039104-INS-58 |
| chr12-34056214-INS-641 | chr12-34056214-INS-641 |
| chr12-34070173-INS-336 | chr12-34070173-INS-339 |
| chr12-34071865-INS-184 | chr12-34071865-INS-184 |
| chr12-34130479-INS-4753 | chr12-34130480-INS-4753 |
| chr12-34143462-INS-78 | chr12-34143464-INS-50 |
| chr12-34147463-INS-81 | chr12-34147440-INS-57 |
| chr12-34176854-INS-6983 | chr12-34176855-INS-6983 |
| chr12-34221516-INS-68 | chr12-34221516-INS-69 |
| chr12-34241319-INS-254 | chr12-34241319-INS-254 |
| chr12-34267369-INS-96 | chr12-34267369-INS-96 |
| chr12-34283967-INS-136 | chr12-34283967-INS-136 |
| chr12-34304924-INS-94 | chr12-34304924-INS-94 |
| chr12-34305370-INS-1866 | chr12-34305371-INS-1866 |
| chr12-34308622-INS-902 | chr12-34308622-INS-908 |
| chr12-34317152-INS-105 | chr12-34317152-INS-105 |
| chr12-34399266-INS-54 | chr12-34399266-INS-54 |
| chr12-34455149-INS-194 | chr12-34455149-INS-194 |
| chr12-34463529-INS-68 | chr12-34463558-INS-50 |
| chr12-34465588-INS-89 | chr12-34465588-INS-89 |
| chr12-34473301-INS-363 | chr12-34473301-INS-364 |
| chr12-34494863-INS-3315 | chr12-34494864-INS-3315 |
| chr12-34630933-INS-69 | chr12-34630933-INS-68 |
| chr12-34681487-INS-64 | chr12-34681448-INS-64 |
| chr12-34759668-INS-101 | chr12-34759663-INS-64 |
| chr12-34865041-INS-56 | chr12-34865041-INS-58 |
| chr12-34897155-INS-81 | chr12-34897149-INS-96 |
| chr12-34929631-INS-3438 | chr12-34929632-INS-3434 |
| chr12-34944290-INS-52 | chr12-34944290-INS-60 |
| chr12-34957964-INS-245 | chr12-34957964-INS-245 |
| chr12-34975463-INS-58 | chr12-34975463-INS-63 |
| chr12-34979010-INS-1433 | chr12-34979013-INS-1431 |
| chr12-34982703-INS-54 | chr12-34982703-INS-60 |
| chr12-34987352-INS-178 | chr12-34987352-INS-178 |
| chr12-34988325-INS-658 | chr12-34988325-INS-666 |
| chr12-34992209-INS-897 | chr12-34992209-INS-897 |
| chr12-34997791-INS-179 | chr12-34997791-INS-179 |
| chr12-35014071-INS-6436 | chr12-35014072-INS-6436 |
| chr12-35025330-INS-506 | chr12-35025330-INS-506 |
| chr12-35036912-INS-1499 | chr12-35036913-INS-1499 |
| chr12-35052419-INS-204 | chr12-35052419-INS-204 |
| chr12-35053712-INS-65 | chr12-35053712-INS-65 |
| chr12-35061500-INS-60 | chr12-35061500-INS-57 |
| chr12-35070560-INS-147 | chr12-35070560-INS-149 |
| chr12-35083450-INS-692 | chr12-35083450-INS-692 |
| chr12-35109742-INS-222 | chr12-35109742-INS-221 |

| | |
|---|---|
| chr12-35124879-INS-96 | chr12-35124876-INS-84 |
| chr12-35190467-INS-52 | chr12-35190468-INS-84 |
| chr12-35215369-INS-498 | chr12-35215369-INS-498 |
| chr12-35223689-INS-588 | chr12-35223689-INS-588 |
| chr12-35254307-INS-101 | chr12-35254301-INS-113 |
| chr12-35258495-INS-118 | chr12-35258495-INS-117 |
| chr12-35274688-INS-1768 | chr12-35274689-INS-1769 |
| chr12-35276795-INS-145 | chr12-35276791-INS-162 |
| chr12-35290611-INS-52 | chr12-35290617-INS-53 |
| chr12-35310416-INS-91 | chr12-35310417-INS-76 |
| chr12-35310921-INS-343 | chr12-35310922-INS-343 |
| chr12-35323443-INS-131 | chr12-35323443-INS-131 |
| chr12-35356069-INS-50 | chr12-35356069-INS-58 |
| chr12-35357465-INS-74 | chr12-35357465-INS-72 |
| chr12-35367968-INS-100 | chr12-35367968-INS-100 |
| chr12-35375777-INS-6672 | chr12-35375761-INS-6664 |
| chr12-35416896-INS-6570 | chr12-35416897-INS-6570 |
| chr12-35417659-INS-934 | chr12-35417662-INS-929 |
| chr12-35424624-INS-68 | chr12-35424630-INS-56 |
| chr12-35445895-INS-208 | chr12-35445895-INS-206 |
| chr12-35457078-INS-50 | chr12-35457078-INS-60 |
| chr12-35471459-INS-88 | chr12-35471467-INS-84 |
| chr12-35481974-INS-176 | chr12-35481974-INS-176 |
| chr12-35516893-INS-329 | chr12-35516893-INS-329 |
| chr12-35542272-INS-125 | chr12-35542267-INS-160 |
| chr12-35545705-INS-51 | chr12-35545705-INS-51 |
| chr12-35545894-INS-813 | chr12-35545894-INS-818 |
| chr12-35557189-INS-349 | chr12-35557189-INS-348 |
| chr12-35579379-INS-70 | chr12-35579379-INS-70 |
| chr12-35590397-INS-52 | chr12-35590397-INS-52 |
| chr12-35591624-INS-54 | chr12-35591624-INS-54 |
| chr12-35632927-INS-467 | chr12-35632927-INS-468 |
| chr12-35685883-INS-82 | chr12-35685883-INS-89 |
| chr12-35705285-INS-341 | chr12-35705285-INS-341 |
| chr12-35710621-INS-256 | chr12-35710621-INS-256 |
| chr12-35723910-INS-224 | chr12-35723910-INS-224 |
| chr12-35758633-INS-57 | chr12-35758633-INS-55 |
| chr12-35764902-INS-61 | chr12-35764904-INS-66 |
| chr12-35768556-INS-6694 | chr12-35768557-INS-6695 |
| chr12-35779347-INS-2437 | chr12-35779348-INS-2437 |
| chr12-35791680-INS-6588 | chr12-35791681-INS-6588 |
| chr12-35805764-INS-127 | chr12-35805765-INS-130 |
| chr12-35840699-INS-77 | chr12-35840699-INS-75 |
| chr12-35868926-INS-51 | chr12-35868926-INS-52 |
| chr12-35873493-INS-2077 | chr12-35873494-INS-2076 |
| chr12-35880542-INS-58 | chr12-35880572-INS-60 |
| chr12-35884000-INS-52 | chr12-35884000-INS-57 |
| chr12-35887473-INS-67 | chr12-35887473-INS-81 |
| chr12-35894129-INS-1443 | chr12-35894130-INS-1443 |
| chr12-35896304-INS-201 | chr12-35896304-INS-201 |
| chr12-35901970-INS-51 | chr12-35901993-INS-55 |
| chr12-35943077-INS-96 | chr12-35943103-INS-100 |
| chr12-35947289-INS-261 | chr12-35947290-INS-262 |
| chr12-35966207-INS-64 | chr12-35966207-INS-64 |
| chr12-35974353-INS-834 | chr12-35974353-INS-834 |
| chr12-35983250-INS-189 | chr12-35983250-INS-189 |
| chr12-35983959-INS-893 | chr12-35983959-INS-894 |
| chr12-35989961-INS-691 | chr12-35989961-INS-686 |
| chr12-35992810-INS-52 | chr12-35992810-INS-52 |
| chr12-36000837-INS-6384 | chr12-36000838-INS-6384 |
| chr12-36007458-INS-573 | chr12-36007458-INS-550 |
| chr12-36017996-INS-548 | chr12-36017996-INS-548 |
| chr12-36034631-INS-624 | chr12-36034631-INS-574 |
| chr12-36036311-INS-80 | chr12-36036307-INS-68 |
| chr12-36037342-INS-108 | chr12-36037342-INS-84 |
| chr12-36063198-INS-58 | chr12-36063183-INS-64 |
| chr12-36076212-INS-83 | chr12-36076212-INS-83 |
| chr12-36095290-INS-681 | chr12-36095290-INS-691 |
| chr12-36108492-INS-86 | chr12-36108485-INS-92 |
| chr12-36113167-INS-297 | chr12-36113167-INS-297 |
| chr12-36119524-INS-401 | chr12-36119524-INS-401 |
| chr12-36124001-INS-2915 | chr12-36124002-INS-2928 |
| chr12-36124806-INS-497 | chr12-36124806-INS-497 |
| chr12-36127876-INS-136 | chr12-36127876-INS-138 |
| chr12-36155936-INS-74 | chr12-36155937-INS-74 |
| chr12-36197091-INS-162 | chr12-36197091-INS-162 |
| chr12-36226922-INS-174 | chr12-36226922-INS-174 |

| | |
|---|---|
| chr12-36228248-INS-7102 | chr12-36228249-INS-7102 |
| chr12-36244662-INS-214 | chr12-36244662-INS-214 |
| chr12-36248192-INS-325 | chr12-36248192-INS-324 |
| chr12-36258774-INS-90 | chr12-36258774-INS-90 |
| chr12-36260759-INS-140 | chr12-36260759-INS-140 |
| chr12-36289037-INS-2867 | chr12-36289038-INS-2868 |
| chr12-36291543-INS-248 | chr12-36291543-INS-212 |
| chr12-36293814-INS-60 | chr12-36293814-INS-100 |
| chr12-36308513-INS-197 | chr12-36308513-INS-197 |
| chr12-36319025-INS-52 | chr12-36319026-INS-52 |
| chr12-36319797-INS-7613 | chr12-36319798-INS-7616 |
| chr12-36338234-INS-93 | chr12-36338235-INS-80 |
| chr12-36340296-INS-78 | chr12-36340296-INS-82 |
| chr12-36347021-INS-99 | chr12-36347021-INS-99 |
| chr12-36379743-INS-183 | chr12-36379743-INS-177 |
| chr12-36410506-INS-78 | chr12-36410506-INS-78 |
| chr12-36439118-INS-55 | chr12-36439118-INS-55 |
| chr12-36448175-INS-2467 | chr12-36448176-INS-2452 |
| chr12-36449697-INS-6637 | chr12-36449698-INS-6637 |
| chr12-36451836-INS-538 | chr12-36451836-INS-538 |
| chr12-36455908-INS-56 | chr12-36455908-INS-56 |
| chr12-36493689-INS-533 | chr12-36493695-INS-533 |
| chr12-36509053-INS-257 | chr12-36509053-INS-258 |
| chr12-36514545-INS-52 | chr12-36514545-INS-58 |
| chr12-36520663-INS-57 | chr12-36520663-INS-57 |
| chr12-36536162-INS-206 | chr12-36536163-INS-206 |
| chr12-36561636-INS-134 | chr12-36561636-INS-132 |
| chr12-36564910-INS-6483 | chr12-36564911-INS-6484 |
| chr12-36613842-INS-76 | chr12-36613841-INS-75 |
| chr12-36618590-INS-231 | chr12-36618590-INS-232 |
| chr12-36642095-INS-437 | chr12-36642095-INS-440 |
| chr12-36649807-INS-90 | chr12-36649807-INS-90 |
| chr12-36663752-INS-400 | chr12-36663752-INS-400 |
| chr12-36670902-INS-1190 | chr12-36670896-INS-1184 |
| chr12-36694638-INS-3085 | chr12-36694639-INS-3085 |
| chr12-36706515-INS-218 | chr12-36706515-INS-217 |
| chr12-36713664-INS-1556 | chr12-36713665-INS-1557 |
| chr12-36720660-INS-69 | chr12-36720660-INS-69 |
| chr12-36732833-INS-66 | chr12-36732833-INS-66 |
| chr12-36756915-INS-57 | chr12-36756915-INS-59 |
| chr12-36758433-INS-789 | chr12-36758433-INS-789 |
| chr12-36782808-INS-3165 | chr12-36782809-INS-3165 |
| chr12-36785386-INS-57 | chr12-36785383-INS-55 |
| chr12-36791280-INS-826 | chr12-36791280-INS-826 |
| chr12-36797929-INS-55 | chr12-36797922-INS-59 |
| chr12-36827416-INS-185 | chr12-36827416-INS-185 |
| chr12-36827904-INS-6990 | chr12-36827905-INS-6990 |
| chr12-36830012-INS-130 | chr12-36830012-INS-128 |
| chr12-36850966-INS-390 | chr12-36850966-INS-401 |
| chr12-36874183-INS-86 | chr12-36874183-INS-72 |
| chr12-36916932-INS-399 | chr12-36916932-INS-404 |
| chr12-36920296-INS-330 | chr12-36920296-INS-330 |
| chr12-36942299-INS-1887 | chr12-36942300-INS-1887 |
| chr12-36943142-INS-187 | chr12-36943142-INS-184 |
| chr12-36952618-INS-51 | chr12-36952618-INS-61 |
| chr12-37018692-INS-59 | chr12-37018692-INS-63 |
| chr12-37036739-INS-392 | chr12-37036739-INS-392 |
| chr12-37053544-INS-109 | chr12-37053544-INS-147 |
| chr12-37085659-INS-1471 | chr12-37085628-INS-1471 |
| chr12-37124556-INS-1089 | chr12-37124557-INS-1089 |
| chr12-37135286-INS-86 | chr12-37135286-INS-86 |
| chr12-37135435-INS-699 | chr12-37135435-INS-699 |
| chr12-37148521-INS-1176 | chr12-37148522-INS-1176 |
| chr12-37151348-INS-1576 | chr12-37151349-INS-1575 |
| chr12-37155674-INS-51 | chr12-37155674-INS-51 |
| chr12-37159850-INS-71 | chr12-37159850-INS-71 |
| chr12-37164320-INS-106 | chr12-37164320-INS-130 |
| chr12-37185460-INS-6689 | chr12-37185461-INS-6690 |
| chr12-37226087-INS-64 | chr12-37226047-INS-98 |
| chr12-37227341-INS-60 | chr12-37227341-INS-66 |
| chr12-37233650-INS-89 | chr12-37233650-INS-89 |
| chr12-37240882-INS-416 | chr12-37240882-INS-416 |
| chr12-37270100-INS-136 | chr12-37270100-INS-136 |
| chr12-37272373-INS-849 | chr12-37272373-INS-861 |
| chr12-37298333-INS-1033 | chr12-37298334-INS-1034 |
| chr12-37313046-INS-288 | chr12-37313046-INS-287 |
| chr12-37329679-INS-7083 | chr12-37329680-INS-7083 |

| | |
|---|---|
| chr12-37355054-INS-90 | chr12-37355051-INS-93 |
| chr12-37366724-INS-1953 | chr12-37366725-INS-1953 |
| chr12-37424373-INS-82 | chr12-37424373-INS-82 |
| chr12-37437704-INS-99 | chr12-37437704-INS-99 |
| chr12-37490942-INS-400 | chr12-37490942-INS-399 |
| chr12-37497554-INS-7086 | chr12-37497555-INS-7086 |
| chr12-37503088-INS-67 | chr12-37503088-INS-67 |
| chr12-37527078-INS-844 | chr12-37527063-INS-843 |
| chr12-37528490-INS-66 | chr12-37528481-INS-80 |
| chr12-37537600-INS-6344 | chr12-37537601-INS-6344 |
| chr12-37564998-INS-52 | chr12-37564998-INS-52 |
| chr12-37617016-INS-1665 | chr12-37617039-INS-1649 |
| chr12-37617676-INS-6406 | chr12-37617677-INS-6406 |
| chr12-37628917-INS-452 | chr12-37628917-INS-452 |
| chr12-37740840-INS-91 | chr12-37740840-INS-104 |
| chr12-37744582-INS-1034 | chr12-37744583-INS-1034 |
| chr12-37750494-INS-1363 | chr12-37750495-INS-1363 |
| chr12-37757434-INS-2181 | chr12-37757435-INS-2180 |
| chr12-37763824-INS-2832 | chr12-37763825-INS-2839 |
| chr12-37790225-INS-1699 | chr12-37790226-INS-1699 |
| chr12-37816403-INS-104 | chr12-37816403-INS-105 |
| chr12-37849524-INS-6632 | chr12-37849525-INS-6633 |
| chr12-37876507-INS-72 | chr12-37876490-INS-60 |
| chr12-37879751-INS-347 | chr12-37879751-INS-348 |
| chr12-37926969-INS-133 | chr12-37926969-INS-133 |
| chr12-37929335-INS-72 | chr12-37929371-INS-72 |
| chr12-37941280-INS-58 | chr12-37941280-INS-60 |
| chr12-37957001-INS-64 | chr12-37957001-INS-68 |
| chr12-37960330-INS-52 | chr12-37960331-INS-56 |
| chr12-37977741-INS-856 | chr12-37977741-INS-856 |
| chr12-38011526-INS-79 | chr12-38011494-INS-66 |
| chr12-38032221-INS-56 | chr12-38032221-INS-56 |
| chr12-38043961-INS-115 | chr12-38043961-INS-122 |
| chr12-38055556-INS-75 | chr12-38055556-INS-64 |
| chr12-38061927-INS-166 | chr12-38061927-INS-166 |
| chr12-38069819-INS-1043 | chr12-38069820-INS-1042 |
| chr12-38082450-INS-792 | chr12-38082450-INS-794 |
| chr12-38106246-INS-454 | chr12-38106246-INS-454 |
| chr12-38110090-INS-166 | chr12-38110090-INS-166 |
| chr12-38114407-INS-90 | chr12-38114407-INS-92 |
| chr12-38118871-INS-56 | chr12-38118871-INS-56 |
| chr12-38133852-INS-78 | chr12-38133817-INS-78 |
| chr12-38154547-INS-705 | chr12-38154547-INS-710 |
| chr12-38163602-INS-96 | chr12-38163602-INS-88 |
| chr12-38167706-INS-73 | chr12-38167706-INS-73 |
| chr12-38179439-INS-55 | chr12-38179430-INS-52 |
| chr12-38241382-INS-64 | chr12-38241384-INS-64 |
| chr12-38246769-INS-307 | chr12-38246769-INS-303 |
| chr12-38266672-INS-196 | chr12-38266672-INS-196 |
| chr12-38271097-INS-324 | chr12-38271097-INS-324 |
| chr12-38279163-INS-6442 | chr12-38279164-INS-6442 |
| chr12-38279690-INS-401 | chr12-38279690-INS-401 |
| chr12-38281945-INS-58 | chr12-38281945-INS-78 |
| chr12-38294244-INS-119 | chr12-38294244-INS-119 |
| chr12-38298849-INS-673 | chr12-38298849-INS-673 |
| chr12-38310714-INS-509 | chr12-38310714-INS-509 |
| chr12-38311312-INS-322 | chr12-38311312-INS-322 |
| chr12-38341163-INS-285 | chr12-38341163-INS-267 |
| chr12-38345761-INS-661 | chr12-38345761-INS-661 |
| chr12-38363453-INS-1002 | chr12-38363454-INS-993 |
| chr12-38371730-INS-405 | chr12-38371730-INS-405 |
| chr12-38382424-INS-6611 | chr12-38382425-INS-6611 |
| chr12-38387393-INS-6333 | chr12-38387394-INS-6331 |
| chr12-38388744-INS-4244 | chr12-38388744-INS-4237 |
| chr12-38401919-INS-197 | chr12-38401917-INS-195 |
| chr12-38410557-INS-117 | chr12-38410557-INS-118 |
| chr12-38415837-INS-84 | chr12-38415837-INS-84 |
| chr12-38421976-INS-401 | chr12-38421976-INS-401 |
| chr12-38432591-INS-7136 | chr12-38432592-INS-7137 |
| chr12-38440803-INS-66 | chr12-38440791-INS-74 |
| chr12-38491439-INS-72 | chr12-38491452-INS-72 |
| chr12-38498681-INS-56 | chr12-38498681-INS-56 |
| chr12-38541278-INS-142 | chr12-38541278-INS-142 |
| chr12-38587253-INS-257 | chr12-38587253-INS-257 |
| chr12-38593195-INS-80 | chr12-38593203-INS-70 |
| chr12-38617709-INS-160 | chr12-38617709-INS-155 |
| chr12-38635653-INS-111 | chr12-38635653-INS-111 |

| | |
|---|---|
| chr12-38656294-INS-54 | chr12-38656247-INS-54 |
| chr12-38749271-INS-7598 | chr12-38749272-INS-7600 |
| chr12-38788950-INS-61 | chr12-38788985-INS-59 |
| chr12-38793085-INS-724 | chr12-38793085-INS-724 |
| chr12-38807488-INS-6239 | chr12-38807489-INS-6240 |
| chr12-38809285-INS-59 | chr12-38809269-INS-82 |
| chr12-38810066-INS-156 | chr12-38810034-INS-182 |
| chr12-38811076-INS-4682 | chr12-38811084-INS-4669 |
| chr12-38819569-INS-468 | chr12-38819570-INS-471 |
| chr12-38856371-INS-102 | chr12-38856371-INS-102 |
| chr12-38867185-INS-86 | chr12-38867185-INS-86 |
| chr12-38927410-INS-106 | chr12-38927410-INS-106 |
| chr12-38929834-INS-877 | chr12-38929834-INS-878 |
| chr12-38950290-INS-68 | chr12-38950290-INS-70 |
| chr12-38980411-INS-400 | chr12-38980411-INS-400 |
| chr12-38984632-INS-70 | chr12-38984653-INS-66 |
| chr12-38991089-INS-441 | chr12-38991089-INS-441 |
| chr12-39023100-INS-200 | chr12-39023100-INS-200 |
| chr12-39062053-INS-264 | chr12-39062053-INS-263 |
| chr12-39097186-INS-541 | chr12-39097186-INS-542 |
| chr12-39108175-INS-178 | chr12-39108175-INS-180 |
| chr12-39153514-INS-400 | chr12-39153514-INS-400 |
| chr12-39170655-INS-255 | chr12-39170655-INS-256 |
| chr12-39170965-INS-6231 | chr12-39170966-INS-6231 |
| chr12-39184062-INS-55 | chr12-39184073-INS-81 |
| chr12-39228481-INS-592 | chr12-39228481-INS-592 |
| chr12-39231109-INS-53 | chr12-39231109-INS-53 |
| chr12-39272938-INS-55 | chr12-39272938-INS-55 |
| chr12-39288969-INS-3074 | chr12-39288970-INS-3077 |
| chr12-39293982-INS-6578 | chr12-39293983-INS-6578 |
| chr12-39293983-INS-228 | chr12-39293984-INS-228 |
| chr12-39300587-INS-75 | chr12-39300587-INS-51 |
| chr12-39345415-INS-78 | chr12-39345424-INS-64 |
| chr12-39352644-INS-56 | chr12-39352644-INS-52 |
| chr12-39397298-INS-402 | chr12-39397298-INS-402 |
| chr12-39429341-INS-3087 | chr12-39429342-INS-3087 |
| chr12-39489773-INS-80 | chr12-39489773-INS-108 |
| chr12-39508002-INS-1665 | chr12-39508003-INS-1665 |
| chr12-39524988-INS-517 | chr12-39524989-INS-497 |
| chr12-39567422-INS-6685 | chr12-39567423-INS-6685 |
| chr12-39638741-INS-8400 | chr12-39638742-INS-8402 |
| chr12-39648737-INS-228 | chr12-39648737-INS-228 |
| chr12-39660228-INS-138 | chr12-39660228-INS-138 |
| chr12-39665533-INS-927 | chr12-39665533-INS-929 |
| chr12-39687712-INS-84 | chr12-39687712-INS-85 |
| chr12-39720164-INS-56 | chr12-39720168-INS-70 |
| chr12-39721380-INS-2532 | chr12-39721381-INS-2532 |
| chr12-39740982-INS-150 | chr12-39740982-INS-150 |
| chr12-39757195-INS-277 | chr12-39757195-INS-277 |
| chr12-39777643-INS-6696 | chr12-39777644-INS-6702 |
| chr12-39798299-INS-283 | chr12-39798281-INS-326 |
| chr12-39830315-INS-6075 | chr12-39830316-INS-6075 |
| chr12-39841790-INS-293 | chr12-39841790-INS-293 |
| chr12-39843794-INS-709 | chr12-39843794-INS-709 |
| chr12-39847180-INS-396 | chr12-39847180-INS-396 |
| chr12-39872338-INS-1346 | chr12-39872339-INS-1366 |
| chr12-39873731-INS-411 | chr12-39873731-INS-411 |
| chr12-39884732-INS-90 | chr12-39884732-INS-90 |
| chr12-39886375-INS-5516 | chr12-39886376-INS-5508 |
| chr12-39896851-INS-168 | chr12-39896857-INS-166 |
| chr12-39909877-INS-146 | chr12-39909878-INS-146 |
| chr12-39913728-INS-422 | chr12-39913728-INS-422 |
| chr12-39938829-INS-952 | chr12-39938829-INS-952 |
| chr12-39962700-INS-110 | chr12-39962700-INS-110 |
| chr12-40004433-INS-356 | chr12-40004434-INS-357 |
| chr12-40010628-INS-406 | chr12-40010628-INS-406 |
| chr12-40031169-INS-5345 | chr12-40031170-INS-5345 |
| chr12-40034784-INS-63 | chr12-40034784-INS-63 |
| chr12-40036308-INS-6615 | chr12-40036309-INS-6615 |
| chr12-40050946-INS-732 | chr12-40050948-INS-722 |
| chr12-40060244-INS-141 | chr12-40060246-INS-144 |
| chr12-40070806-INS-72 | chr12-40070806-INS-66 |
| chr12-40075118-INS-139 | chr12-40075118-INS-139 |
| chr12-40125084-INS-74 | chr12-40125091-INS-60 |
| chr12-40139910-INS-54 | chr12-40139910-INS-56 |
| chr12-40187330-INS-130 | chr12-40187355-INS-107 |
| chr12-40211185-INS-108 | chr12-40211185-INS-104 |

| | |
|---|---|
| chr12-40243955-INS-197 | chr12-40243955-INS-195 |
| chr12-40293473-INS-202 | chr12-40293473-INS-202 |
| chr12-40314664-INS-69 | chr12-40314633-INS-52 |
| chr12-40341956-INS-366 | chr12-40341956-INS-366 |
| chr12-40362932-INS-480 | chr12-40362932-INS-480 |
| chr12-40372904-INS-52 | chr12-40372904-INS-56 |
| chr12-40409200-INS-53 | chr12-40409200-INS-53 |
| chr12-40443792-INS-163 | chr12-40443789-INS-160 |
| chr12-40451171-INS-107 | chr12-40451171-INS-74 |
| chr12-40493034-INS-221 | chr12-40493034-INS-221 |
| chr12-40501377-INS-68 | chr12-40501377-INS-68 |
| chr12-40531525-INS-229 | chr12-40531525-INS-216 |
| chr12-40559466-INS-704 | chr12-40559466-INS-704 |
| chr12-40587119-INS-604 | chr12-40587119-INS-606 |
| chr12-40600436-INS-289 | chr12-40600437-INS-290 |
| chr12-40623340-INS-54 | chr12-40623340-INS-54 |
| chr12-40623724-INS-4325 | chr12-40623725-INS-4326 |
| chr12-40627680-INS-55 | chr12-40627680-INS-55 |
| chr12-40629069-INS-208 | chr12-40629069-INS-208 |
| chr12-40690911-INS-163 | chr12-40690911-INS-163 |
| chr12-40721520-INS-65 | chr12-40721520-INS-65 |
| chr12-40733294-INS-213 | chr12-40733294-INS-212 |
| chr12-40734629-INS-140 | chr12-40734623-INS-172 |
| chr12-40771089-INS-58 | chr12-40771089-INS-58 |
| chr12-40795091-INS-74 | chr12-40795080-INS-62 |
| chr12-40814186-INS-208 | chr12-40814186-INS-208 |
| chr12-40852877-INS-72 | chr12-40852877-INS-85 |
| chr12-40863771-INS-91 | chr12-40863771-INS-64 |
| chr12-40877884-INS-6544 | chr12-40877885-INS-6544 |
| chr12-40905916-INS-108 | chr12-40905916-INS-112 |
| chr12-40936646-INS-222 | chr12-40936646-INS-222 |
| chr12-40964410-INS-697 | chr12-40964411-INS-673 |
| chr12-40974712-INS-220 | chr12-40974712-INS-221 |
| chr12-40979856-INS-330 | chr12-40979856-INS-330 |
| chr12-40994232-INS-7022 | chr12-40994233-INS-7022 |
| chr12-41000827-INS-55 | chr12-41000827-INS-55 |
| chr12-41002282-INS-128 | chr12-41002282-INS-128 |
| chr12-41004924-INS-441 | chr12-41004924-INS-441 |
| chr12-41009489-INS-2062 | chr12-41009490-INS-2062 |
| chr12-41023749-INS-352 | chr12-41023749-INS-352 |
| chr12-41034781-INS-470 | chr12-41034781-INS-470 |
| chr12-41037996-INS-509 | chr12-41037993-INS-508 |
| chr12-41051347-INS-188 | chr12-41051347-INS-189 |
| chr12-41057358-INS-172 | chr12-41057358-INS-207 |
| chr12-41059302-INS-6440 | chr12-41059303-INS-6440 |
| chr12-41072603-INS-52 | chr12-41072603-INS-54 |
| chr12-41079511-INS-70 | chr12-41079506-INS-51 |
| chr12-41117243-INS-205 | chr12-41117236-INS-208 |
| chr12-41119206-INS-1197 | chr12-41119207-INS-1197 |
| chr12-41155803-INS-61 | chr12-41155799-INS-67 |
| chr12-41156953-INS-70 | chr12-41156955-INS-64 |
| chr12-41163584-INS-82 | chr12-41163584-INS-108 |
| chr12-41166227-INS-531 | chr12-41166227-INS-531 |
| chr12-41182935-INS-551 | chr12-41182935-INS-551 |
| chr12-41196337-INS-5143 | chr12-41196338-INS-5143 |
| chr12-41224528-INS-2037 | chr12-41224529-INS-2037 |
| chr12-41233667-INS-56 | chr12-41233667-INS-56 |
| chr12-41250027-INS-200 | chr12-41250027-INS-200 |
| chr12-41253089-INS-861 | chr12-41253089-INS-861 |
| chr12-41254476-INS-76 | chr12-41254476-INS-74 |
| chr12-41259009-INS-199 | chr12-41259009-INS-200 |
| chr12-41282772-INS-78 | chr12-41282772-INS-78 |
| chr12-41300419-INS-458 | chr12-41300420-INS-457 |
| chr12-41312257-INS-152 | chr12-41312257-INS-152 |
| chr12-41335900-INS-236 | chr12-41335900-INS-236 |
| chr12-41349536-INS-796 | chr12-41349536-INS-796 |
| chr12-41371872-INS-94 | chr12-41371872-INS-94 |
| chr12-41374018-INS-54 | chr12-41374018-INS-55 |
| chr12-41378150-INS-52 | chr12-41378151-INS-52 |
| chr12-41394686-INS-96 | chr12-41394686-INS-96 |
| chr12-41399357-INS-118 | chr12-41399357-INS-104 |
| chr12-41467643-INS-523 | chr12-41467643-INS-523 |
| chr12-41472066-INS-326 | chr12-41472066-INS-326 |
| chr12-41476116-INS-439 | chr12-41476116-INS-439 |
| chr12-41482077-INS-58 | chr12-41482077-INS-52 |
| chr12-41521708-INS-7091 | chr12-41521709-INS-7091 |
| chr12-41607679-INS-197 | chr12-41607679-INS-197 |

| | |
|---|---|
| chr12-41636889-INS-6181 | chr12-41636890-INS-6182 |
| chr12-41641778-INS-1544 | chr12-41641779-INS-1544 |
| chr12-41671447-INS-760 | chr12-41671447-INS-760 |
| chr12-41683475-INS-56 | chr12-41683475-INS-82 |
| chr12-41685267-INS-5264 | chr12-41685268-INS-5264 |
| chr12-41717649-INS-2111 | chr12-41717650-INS-2109 |
| chr12-41743219-INS-211 | chr12-41743219-INS-210 |
| chr12-41749537-INS-156 | chr12-41749537-INS-178 |
| chr12-41784717-INS-293 | chr12-41784719-INS-265 |
| chr12-41801743-INS-114 | chr12-41801743-INS-114 |
| chr12-41810516-INS-6730 | chr12-41810517-INS-6730 |
| chr12-41830062-INS-1887 | chr12-41830063-INS-1903 |
| chr12-41834360-INS-148 | chr12-41834361-INS-119 |
| chr12-41838869-INS-375 | chr12-41838870-INS-370 |
| chr12-41873791-INS-325 | chr12-41873791-INS-307 |
| chr12-41875882-INS-503 | chr12-41875882-INS-500 |
| chr12-41884400-INS-131 | chr12-41884397-INS-114 |
| chr12-41931248-INS-56 | chr12-41931248-INS-56 |
| chr12-41955277-INS-6349 | chr12-41955278-INS-6349 |
| chr12-41966667-INS-213 | chr12-41966667-INS-213 |
| chr12-41972829-INS-157 | chr12-41972829-INS-157 |
| chr12-41976434-INS-514 | chr12-41976434-INS-515 |
| chr12-42018617-INS-152 | chr12-42018617-INS-152 |
| chr12-42062770-INS-58 | chr12-42062770-INS-58 |
| chr12-42070850-INS-7822 | chr12-42070851-INS-7822 |
| chr12-42072192-INS-193 | chr12-42072192-INS-193 |
| chr12-42085538-INS-171 | chr12-42085538-INS-171 |
| chr12-42088677-INS-166 | chr12-42088677-INS-166 |
| chr12-42094484-INS-696 | chr12-42094484-INS-712 |
| chr12-42098863-INS-106 | chr12-42098863-INS-107 |
| chr12-42127775-INS-1203 | chr12-42127734-INS-1236 |
| chr12-42132343-INS-3100 | chr12-42132344-INS-3100 |
| chr12-42141716-INS-232 | chr12-42141716-INS-230 |
| chr12-42153940-INS-372 | chr12-42153940-INS-372 |
| chr12-42161789-INS-53 | chr12-42161789-INS-56 |
| chr12-42175935-INS-59 | chr12-42175935-INS-59 |
| chr12-42181081-INS-59 | chr12-42181047-INS-54 |
| chr12-42230186-INS-6576 | chr12-42230187-INS-6576 |
| chr12-42235238-INS-1667 | chr12-42235239-INS-1667 |
| chr12-42253149-INS-1932 | chr12-42253150-INS-1932 |
| chr12-42253915-INS-251 | chr12-42253915-INS-251 |
| chr12-42296852-INS-56 | chr12-42296852-INS-56 |
| chr12-42305486-INS-355 | chr12-42305486-INS-360 |
| chr12-42308941-INS-1461 | chr12-42308942-INS-1460 |
| chr12-42317619-INS-399 | chr12-42317619-INS-399 |
| chr12-42335720-INS-691 | chr12-42335720-INS-690 |
| chr12-42359475-INS-6066 | chr12-42359476-INS-6066 |
| chr12-42366095-INS-390 | chr12-42366099-INS-386 |
| chr12-42404219-INS-1890 | chr12-42404220-INS-1890 |
| chr12-42411586-INS-164 | chr12-42411586-INS-164 |
| chr12-42433604-INS-78 | chr12-42433631-INS-56 |
| chr12-42439592-INS-350 | chr12-42439592-INS-350 |
| chr12-42464313-INS-51 | chr12-42464270-INS-58 |
| chr12-42470241-INS-151 | chr12-42470241-INS-154 |
| chr12-42507011-INS-143 | chr12-42507011-INS-143 |
| chr12-42527649-INS-361 | chr12-42527649-INS-361 |
| chr12-42539986-INS-555 | chr12-42539986-INS-555 |
| chr12-42561598-INS-66 | chr12-42561598-INS-66 |
| chr12-42564273-INS-54 | chr12-42564273-INS-54 |
| chr12-42568698-INS-4441 | chr12-42568699-INS-4476 |
| chr12-42602168-INS-161 | chr12-42602168-INS-161 |
| chr12-42641829-INS-4075 | chr12-42641830-INS-4075 |
| chr12-42653399-INS-4926 | chr12-42653400-INS-4926 |
| chr12-42703992-INS-108 | chr12-42704003-INS-109 |
| chr12-42731050-INS-52 | chr12-42731050-INS-52 |
| chr12-42753962-INS-65 | chr12-42753962-INS-65 |
| chr12-42778616-INS-472 | chr12-42778616-INS-472 |
| chr12-42779553-INS-554 | chr12-42779553-INS-554 |
| chr12-42792161-INS-424 | chr12-42792161-INS-424 |
| chr12-42820921-INS-74 | chr12-42820921-INS-80 |
| chr12-42844741-INS-73 | chr12-42844741-INS-74 |
| chr12-42852438-INS-220 | chr12-42852438-INS-220 |
| chr12-42853945-INS-338 | chr12-42853945-INS-338 |
| chr12-42903814-INS-596 | chr12-42903814-INS-596 |
| chr12-42914156-INS-150 | chr12-42914150-INS-118 |
| chr12-42926330-INS-2671 | chr12-42926320-INS-2672 |
| chr12-42936622-INS-423 | chr12-42936622-INS-426 |

| | |
|---|---|
| chr12-42947452-INS-191 | chr12-42947452-INS-192 |
| chr12-42961803-INS-116 | chr12-42961821-INS-70 |
| chr12-42975850-INS-6483 | chr12-42975851-INS-6483 |
| chr12-42986188-INS-7084 | chr12-42986189-INS-7084 |
| chr12-43016647-INS-90 | chr12-43016647-INS-104 |
| chr12-43021416-INS-210 | chr12-43021416-INS-210 |
| chr12-43036054-INS-53 | chr12-43036050-INS-83 |
| chr12-43080382-INS-68 | chr12-43080382-INS-76 |
| chr12-43099014-INS-58 | chr12-43099013-INS-55 |
| chr12-43100371-INS-92 | chr12-43100376-INS-86 |
| chr12-43107119-INS-115 | chr12-43107119-INS-115 |
| chr12-43175185-INS-6469 | chr12-43175186-INS-6470 |
| chr12-43186932-INS-593 | chr12-43186932-INS-591 |
| chr12-43207498-INS-51 | chr12-43207499-INS-80 |
| chr12-43237885-INS-452 | chr12-43237885-INS-452 |
| chr12-43247307-INS-54 | chr12-43247307-INS-54 |
| chr12-43252227-INS-56 | chr12-43252227-INS-60 |
| chr12-43271915-INS-418 | chr12-43271915-INS-418 |
| chr12-43276172-INS-119 | chr12-43276172-INS-119 |
| chr12-43379686-INS-56 | chr12-43379686-INS-50 |
| chr12-43385168-INS-259 | chr12-43385168-INS-260 |
| chr12-43398015-INS-216 | chr12-43398017-INS-218 |
| chr12-43401812-INS-161 | chr12-43401808-INS-162 |
| chr12-43405021-INS-106 | chr12-43405021-INS-68 |
| chr12-43454138-INS-51 | chr12-43454138-INS-51 |
| chr12-43483518-INS-56 | chr12-43483518-INS-62 |
| chr12-43497213-INS-50 | chr12-43497182-INS-64 |
| chr12-43512086-INS-940 | chr12-43512086-INS-940 |
| chr12-43512879-INS-5262 | chr12-43512880-INS-5262 |
| chr12-43517460-INS-287 | chr12-43517460-INS-287 |
| chr12-43596112-INS-96 | chr12-43596112-INS-108 |
| chr12-43603031-INS-1035 | chr12-43603032-INS-1035 |
| chr12-43603710-INS-847 | chr12-43603710-INS-847 |
| chr12-43612846-INS-101 | chr12-43612846-INS-100 |
| chr12-43634638-INS-62 | chr12-43634638-INS-62 |
| chr12-43687593-INS-85 | chr12-43687591-INS-95 |
| chr12-43700752-INS-73 | chr12-43700752-INS-73 |
| chr12-43754193-INS-62 | chr12-43754193-INS-76 |
| chr12-43811471-INS-2721 | chr12-43811472-INS-2721 |
| chr12-43819148-INS-961 | chr12-43819148-INS-963 |
| chr12-43850762-INS-2675 | chr12-43850763-INS-2675 |
| chr12-43865361-INS-918 | chr12-43865361-INS-918 |
| chr12-43880748-INS-756 | chr12-43880748-INS-776 |
| chr12-43895106-INS-422 | chr12-43895106-INS-425 |
| chr12-43922336-INS-110 | chr12-43922336-INS-110 |
| chr12-43944396-INS-61 | chr12-43944396-INS-63 |
| chr12-43972686-INS-228 | chr12-43972686-INS-228 |
| chr12-43975713-INS-214 | chr12-43975713-INS-214 |
| chr12-43983026-INS-4254 | chr12-43983027-INS-4253 |
| chr12-44046002-INS-220 | chr12-44046002-INS-220 |
| chr12-44051642-INS-66 | chr12-44051642-INS-66 |
| chr12-44065353-INS-5981 | chr12-44065354-INS-5981 |
| chr12-44097064-INS-236 | chr12-44097067-INS-257 |
| chr12-44102004-INS-156 | chr12-44102004-INS-156 |
| chr12-44108641-INS-767 | chr12-44108641-INS-770 |
| chr12-44111091-INS-214 | chr12-44111091-INS-214 |
| chr12-44123416-INS-322 | chr12-44123416-INS-323 |
| chr12-44143179-INS-849 | chr12-44143179-INS-850 |
| chr12-44168062-INS-205 | chr12-44168062-INS-205 |
| chr12-44211368-INS-509 | chr12-44211368-INS-515 |
| chr12-44216141-INS-84 | chr12-44216136-INS-62 |
| chr12-44249184-INS-746 | chr12-44249184-INS-748 |
| chr12-44272831-INS-103 | chr12-44272830-INS-85 |
| chr12-44289101-INS-678 | chr12-44289100-INS-678 |
| chr12-44289720-INS-463 | chr12-44289720-INS-464 |
| chr12-44325401-INS-283 | chr12-44325401-INS-283 |
| chr12-44333934-INS-60 | chr12-44333934-INS-66 |
| chr12-44335792-INS-52 | chr12-44335792-INS-52 |
| chr12-44361112-INS-94 | chr12-44361112-INS-94 |
| chr12-44388189-INS-175 | chr12-44388189-INS-178 |
| chr12-44389793-INS-2786 | chr12-44389794-INS-2788 |
| chr12-44391482-INS-212 | chr12-44391482-INS-212 |
| chr12-44397019-INS-254 | chr12-44397019-INS-255 |
| chr12-44426005-INS-441 | chr12-44426005-INS-441 |
| chr12-44449431-INS-3012 | chr12-44449432-INS-3012 |
| chr12-44470441-INS-158 | chr12-44470392-INS-169 |
| chr12-44474926-INS-6767 | chr12-44474927-INS-6767 |

| | |
|---|---|
| chr12-44478700-INS-205 | chr12-44478700-INS-205 |
| chr12-44491813-INS-113 | chr12-44491813-INS-113 |
| chr12-44493426-INS-540 | chr12-44493426-INS-541 |
| chr12-44501064-INS-74 | chr12-44501064-INS-60 |
| chr12-44507968-INS-54 | chr12-44507952-INS-52 |
| chr12-44524527-INS-74 | chr12-44524527-INS-74 |
| chr12-44526044-INS-389 | chr12-44526044-INS-389 |
| chr12-44595283-INS-842 | chr12-44595283-INS-842 |
| chr12-44597825-INS-414 | chr12-44597825-INS-414 |
| chr12-44611614-INS-958 | chr12-44611615-INS-957 |
| chr12-44634577-INS-62 | chr12-44634570-INS-72 |
| chr12-44635655-INS-62 | chr12-44635656-INS-62 |
| chr12-44638390-INS-112 | chr12-44638390-INS-112 |
| chr12-44648958-INS-76 | chr12-44648958-INS-94 |
| chr12-44652778-INS-105 | chr12-44652779-INS-135 |
| chr12-44657621-INS-503 | chr12-44657621-INS-503 |
| chr12-44659644-INS-475 | chr12-44659644-INS-475 |
| chr12-44661381-INS-93 | chr12-44661423-INS-54 |
| chr12-44663324-INS-3073 | chr12-44663325-INS-3071 |
| chr12-44673726-INS-657 | chr12-44673727-INS-621 |
| chr12-44679298-INS-58 | chr12-44679298-INS-60 |
| chr12-44686984-INS-537 | chr12-44686984-INS-564 |
| chr12-44691523-INS-1310 | chr12-44691524-INS-1309 |
| chr12-44703666-INS-386 | chr12-44703666-INS-386 |
| chr12-44730836-INS-72 | chr12-44730836-INS-82 |
| chr12-44744659-INS-1162 | chr12-44744668-INS-1148 |
| chr12-44756634-INS-546 | chr12-44756634-INS-546 |
| chr12-44757835-INS-170 | chr12-44757835-INS-170 |
| chr12-44787900-INS-134 | chr12-44787900-INS-98 |
| chr12-44800570-INS-694 | chr12-44800570-INS-694 |
| chr12-44808007-INS-54 | chr12-44808007-INS-56 |
| chr12-44850380-INS-204 | chr12-44850380-INS-204 |
| chr12-44865961-INS-347 | chr12-44865961-INS-347 |
| chr12-44897730-INS-368 | chr12-44897730-INS-368 |
| chr12-44925375-INS-532 | chr12-44925375-INS-532 |
| chr12-44931230-INS-147 | chr12-44931230-INS-147 |
| chr12-44931652-INS-443 | chr12-44931652-INS-443 |
| chr12-44966001-INS-320 | chr12-44966001-INS-326 |
| chr12-44980426-INS-240 | chr12-44980426-INS-252 |
| chr12-44997574-INS-578 | chr12-44997574-INS-578 |
| chr12-45056697-INS-207 | chr12-45056681-INS-212 |
| chr12-45086179-INS-58 | chr12-45086179-INS-62 |
| chr12-45101642-INS-336 | chr12-45101642-INS-336 |
| chr12-45116766-INS-72 | chr12-45116809-INS-57 |
| chr12-45127152-INS-200 | chr12-45127152-INS-200 |
| chr12-45147967-INS-721 | chr12-45147967-INS-721 |
| chr12-45153867-INS-90 | chr12-45153867-INS-120 |
| chr12-45159629-INS-875 | chr12-45159629-INS-875 |
| chr12-45166508-INS-117 | chr12-45166475-INS-130 |
| chr12-45178477-INS-50 | chr12-45178477-INS-66 |
| chr12-45220493-INS-994 | chr12-45220493-INS-994 |
| chr12-45269101-INS-119 | chr12-45269096-INS-144 |
| chr12-45280302-INS-69 | chr12-45280295-INS-117 |
| chr12-45297349-INS-82 | chr12-45297334-INS-130 |
| chr12-45302571-INS-60 | chr12-45302571-INS-60 |
| chr12-45315003-INS-84 | chr12-45315003-INS-72 |
| chr12-45320807-INS-73 | chr12-45320788-INS-62 |
| chr12-45322622-INS-56 | chr12-45322614-INS-88 |
| chr12-45324790-INS-68 | chr12-45324790-INS-60 |
| chr12-45330931-INS-52 | chr12-45330931-INS-52 |
| chr12-45355131-INS-359 | chr12-45355131-INS-351 |
| chr12-45357845-INS-52 | chr12-45357845-INS-52 |
| chr12-45381257-INS-68 | chr12-45381257-INS-68 |
| chr12-45385443-INS-420 | chr12-45385443-INS-420 |
| chr12-45404906-INS-1565 | chr12-45404907-INS-1565 |
| chr12-45416847-INS-123 | chr12-45416847-INS-125 |
| chr12-45430979-INS-343 | chr12-45430979-INS-350 |
| chr12-45459395-INS-89 | chr12-45459395-INS-89 |
| chr12-45469224-INS-6377 | chr12-45469225-INS-6377 |
| chr12-45472407-INS-336 | chr12-45472407-INS-336 |
| chr12-45544240-INS-6401 | chr12-45544241-INS-6401 |
| chr12-45552596-INS-112 | chr12-45552597-INS-98 |
| chr12-45558363-INS-69 | chr12-45558363-INS-68 |
| chr12-45558637-INS-1408 | chr12-45558638-INS-1407 |
| chr12-45577979-INS-475 | chr12-45577985-INS-473 |
| chr12-45596262-INS-2533 | chr12-45596263-INS-2533 |
| chr12-45603318-INS-52 | chr12-45603319-INS-52 |

| | |
|---|---|
| chr12-45619075-INS-66 | chr12-45619075-INS-66 |
| chr12-45627573-INS-76 | chr12-45627593-INS-60 |
| chr12-45644046-INS-419 | chr12-45644046-INS-419 |
| chr12-45646847-INS-53 | chr12-45646847-INS-53 |
| chr12-45654212-INS-158 | chr12-45654212-INS-158 |
| chr12-45669555-INS-197 | chr12-45669555-INS-197 |
| chr12-45702456-INS-154 | chr12-45702452-INS-158 |
| chr12-45712072-INS-3328 | chr12-45712073-INS-3328 |
| chr12-45713979-INS-316 | chr12-45713979-INS-316 |
| chr12-45719238-INS-2763 | chr12-45719239-INS-2763 |
| chr12-45733216-INS-66 | chr12-45733216-INS-66 |
| chr12-45772142-INS-505 | chr12-45772142-INS-509 |
| chr12-45786764-INS-56 | chr12-45786783-INS-84 |
| chr12-45791874-INS-126 | chr12-45791874-INS-128 |
| chr12-45796225-INS-108 | chr12-45796225-INS-108 |
| chr12-45803858-INS-400 | chr12-45803859-INS-400 |
| chr12-45815041-INS-585 | chr12-45815041-INS-585 |
| chr12-45829033-INS-641 | chr12-45829033-INS-641 |
| chr12-45836849-INS-6609 | chr12-45836850-INS-6610 |
| chr12-45852837-INS-184 | chr12-45852837-INS-184 |
| chr12-45874079-INS-72 | chr12-45874079-INS-56 |
| chr12-45886635-INS-109 | chr12-45886634-INS-74 |
| chr12-45898145-INS-620 | chr12-45898144-INS-622 |
| chr12-45907184-INS-591 | chr12-45907184-INS-591 |
| chr12-45940100-INS-931 | chr12-45940101-INS-932 |
| chr12-45946714-INS-669 | chr12-45946714-INS-669 |
| chr12-45993094-INS-62 | chr12-45993078-INS-70 |
| chr12-46022452-INS-56 | chr12-46022452-INS-60 |
| chr12-46028519-INS-2684 | chr12-46028520-INS-2684 |
| chr12-46045066-INS-56 | chr12-46045066-INS-58 |
| chr12-46069324-INS-479 | chr12-46069325-INS-477 |
| chr12-46078617-INS-103 | chr12-46078633-INS-60 |
| chr12-46114775-INS-68 | chr12-46114775-INS-90 |
| chr12-46117590-INS-947 | chr12-46117590-INS-962 |
| chr12-46121428-INS-62 | chr12-46121427-INS-81 |
| chr12-46163349-INS-433 | chr12-46163349-INS-432 |
| chr12-46169465-INS-76 | chr12-46169465-INS-76 |
| chr12-46210796-INS-240 | chr12-46210797-INS-240 |
| chr12-46212782-INS-127 | chr12-46212782-INS-127 |
| chr12-46251621-INS-59 | chr12-46251621-INS-63 |
| chr12-46303954-INS-6197 | chr12-46303955-INS-6197 |
| chr12-46352669-INS-64 | chr12-46352669-INS-62 |
| chr12-46354337-INS-7327 | chr12-46354338-INS-7328 |
| chr12-46364281-INS-4433 | chr12-46364282-INS-4433 |
| chr12-46377865-INS-156 | chr12-46377865-INS-156 |
| chr12-46381672-INS-551 | chr12-46381672-INS-551 |
| chr12-46383779-INS-147 | chr12-46383779-INS-147 |
| chr12-46397602-INS-128 | chr12-46397602-INS-128 |
| chr12-46409424-INS-192 | chr12-46409424-INS-146 |
| chr12-46422563-INS-59 | chr12-46422563-INS-59 |
| chr12-46428640-INS-66 | chr12-46428624-INS-90 |
| chr12-46446850-INS-78 | chr12-46446850-INS-54 |
| chr12-46448706-INS-7186 | chr12-46448707-INS-7186 |
| chr12-46463084-INS-110 | chr12-46463084-INS-70 |
| chr12-46469789-INS-62 | chr12-46469789-INS-62 |
| chr12-46482203-INS-58 | chr12-46482203-INS-58 |
| chr12-46483235-INS-56 | chr12-46483235-INS-56 |
| chr12-46490780-INS-207 | chr12-46490780-INS-199 |
| chr12-46532728-INS-277 | chr12-46532728-INS-277 |
| chr12-46541983-INS-52 | chr12-46541983-INS-56 |
| chr12-46560855-INS-7588 | chr12-46560856-INS-7589 |
| chr12-46564497-INS-72 | chr12-46564497-INS-79 |
| chr12-46572151-INS-72 | chr12-46572151-INS-72 |
| chr12-46575082-INS-6672 | chr12-46575083-INS-6671 |
| chr12-46584647-INS-104 | chr12-46584647-INS-106 |
| chr12-46590992-INS-400 | chr12-46590992-INS-400 |
| chr12-46592732-INS-1538 | chr12-46592733-INS-1538 |
| chr12-46632073-INS-8482 | chr12-46632073-INS-8481 |
| chr12-46637379-INS-55 | chr12-46637346-INS-88 |
| chr12-46640446-INS-176 | chr12-46640446-INS-176 |
| chr12-46647308-INS-52 | chr12-46647308-INS-50 |
| chr12-46657463-INS-110 | chr12-46657475-INS-96 |
| chr12-46665619-INS-6662 | chr12-46665620-INS-6662 |
| chr12-46703229-INS-72 | chr12-46703229-INS-74 |
| chr12-46711057-INS-80 | chr12-46711057-INS-80 |
| chr12-46726461-INS-455 | chr12-46726461-INS-455 |
| chr12-46754796-INS-1742 | chr12-46754797-INS-1742 |

| | |
|---|---|
| chr12-46778766-INS-77 | chr12-46778770-INS-61 |
| chr12-46780089-INS-52 | chr12-46780089-INS-52 |
| chr12-46823156-INS-737 | chr12-46823156-INS-737 |
| chr12-46835147-INS-651 | chr12-46835147-INS-651 |
| chr12-46841187-INS-81 | chr12-46841187-INS-87 |
| chr12-46872903-INS-57 | chr12-46872903-INS-63 |
| chr12-46880180-INS-389 | chr12-46880180-INS-389 |
| chr12-46900077-INS-1567 | chr12-46900078-INS-1567 |
| chr12-46921443-INS-1034 | chr12-46921444-INS-1034 |
| chr12-46924226-INS-52 | chr12-46924234-INS-51 |
| chr12-46943472-INS-498 | chr12-46943472-INS-498 |
| chr12-47024051-INS-6777 | chr12-47024052-INS-6777 |
| chr12-47030613-INS-80 | chr12-47030622-INS-85 |
| chr12-47033707-INS-78 | chr12-47033708-INS-76 |
| chr12-47046483-INS-388 | chr12-47046483-INS-402 |
| chr12-47073973-INS-199 | chr12-47073973-INS-199 |
| chr12-47081666-INS-1541 | chr12-47081667-INS-1541 |
| chr12-47086158-INS-124 | chr12-47086158-INS-124 |
| chr12-47090631-INS-7231 | chr12-47090632-INS-7231 |
| chr12-47129803-INS-75 | chr12-47129803-INS-75 |
| chr12-47152706-INS-94 | chr12-47152706-INS-94 |
| chr12-47175274-INS-54 | chr12-47175270-INS-90 |
| chr12-47213823-INS-108 | chr12-47213823-INS-72 |
| chr12-47225661-INS-1790 | chr12-47225662-INS-1817 |
| chr12-47229834-INS-86 | chr12-47229835-INS-96 |
| chr12-47257146-INS-102 | chr12-47257146-INS-106 |
| chr12-47264777-INS-6403 | chr12-47264778-INS-6403 |
| chr12-47269032-INS-108 | chr12-47269032-INS-108 |
| chr12-47304535-INS-801 | chr12-47304535-INS-792 |
| chr12-47315603-INS-78 | chr12-47315603-INS-78 |
| chr12-47320198-INS-238 | chr12-47320198-INS-238 |
| chr12-47324368-INS-51 | chr12-47324369-INS-84 |
| chr12-47357849-INS-52 | chr12-47357849-INS-54 |
| chr12-47370307-INS-54 | chr12-47370307-INS-58 |
| chr12-47383424-INS-711 | chr12-47383424-INS-711 |
| chr12-47413090-INS-13271 | chr12-47413091-INS-13272 |
| chr12-47449899-INS-636 | chr12-47449899-INS-640 |
| chr12-47456849-INS-82 | chr12-47456847-INS-93 |
| chr12-47458231-INS-318 | chr12-47458231-INS-318 |
| chr12-47545209-INS-58 | chr12-47545209-INS-58 |
| chr12-47552844-INS-264 | chr12-47552844-INS-264 |
| chr12-47553958-INS-69 | chr12-47553958-INS-72 |
| chr12-47644273-INS-199 | chr12-47644273-INS-200 |
| chr12-47662709-INS-92 | chr12-47662709-INS-94 |
| chr12-47676799-INS-70 | chr12-47676812-INS-58 |
| chr12-47688841-INS-80 | chr12-47688841-INS-114 |
| chr12-47794334-INS-633 | chr12-47794334-INS-633 |
| chr12-47844992-INS-809 | chr12-47844989-INS-843 |
| chr12-47857573-INS-823 | chr12-47857573-INS-818 |
| chr12-47896701-INS-60 | chr12-47896701-INS-60 |
| chr12-47918591-INS-945 | chr12-47918591-INS-946 |
| chr12-47925281-INS-86 | chr12-47925282-INS-73 |
| chr12-47997758-INS-65 | chr12-47997758-INS-69 |
| chr12-48066676-INS-1214 | chr12-48066677-INS-1215 |
| chr12-48078559-INS-70 | chr12-48078559-INS-100 |
| chr12-48101243-INS-216 | chr12-48101208-INS-216 |
| chr12-48106889-INS-290 | chr12-48106889-INS-290 |
| chr12-48166539-INS-2362 | chr12-48166540-INS-2362 |
| chr12-48168436-INS-52 | chr12-48168436-INS-52 |
| chr12-48207180-INS-691 | chr12-48207180-INS-693 |
| chr12-48218408-INS-2581 | chr12-48218409-INS-2581 |
| chr12-48233959-INS-52 | chr12-48233959-INS-52 |
| chr12-48239763-INS-53 | chr12-48239763-INS-61 |
| chr12-48242660-INS-6454 | chr12-48242661-INS-6454 |
| chr12-48266785-INS-78 | chr12-48266773-INS-68 |
| chr12-48301114-INS-8056 | chr12-48301115-INS-8056 |
| chr12-48305040-INS-778 | chr12-48305040-INS-779 |
| chr12-48325859-INS-506 | chr12-48325859-INS-506 |
| chr12-48345664-INS-203 | chr12-48345664-INS-203 |
| chr12-48352726-INS-51 | chr12-48352726-INS-57 |
| chr12-48367797-INS-328 | chr12-48367797-INS-328 |
| chr12-48380575-INS-2421 | chr12-48380576-INS-2421 |
| chr12-48392740-INS-322 | chr12-48392740-INS-322 |
| chr12-48448026-INS-74 | chr12-48448026-INS-76 |
| chr12-48450644-INS-1778 | chr12-48450652-INS-1780 |
| chr12-48463315-INS-78 | chr12-48463315-INS-78 |
| chr12-48494501-INS-121 | chr12-48494504-INS-121 |

| | |
|---|---|
| chr12-48495893-INS-174 | chr12-48495893-INS-167 |
| chr12-48508522-INS-98 | chr12-48508522-INS-98 |
| chr12-48514098-INS-1845 | chr12-48514099-INS-1846 |
| chr12-48518970-INS-506 | chr12-48518970-INS-493 |
| chr12-48524262-INS-7131 | chr12-48524263-INS-7131 |
| chr12-48545215-INS-172 | chr12-48545215-INS-188 |
| chr12-48611967-INS-61 | chr12-48611967-INS-61 |
| chr12-48624293-INS-61 | chr12-48624291-INS-86 |
| chr12-48628624-INS-3958 | chr12-48628625-INS-3920 |
| chr12-48642748-INS-6765 | chr12-48642749-INS-6765 |
| chr12-48646597-INS-441 | chr12-48646597-INS-441 |
| chr12-48649254-INS-325 | chr12-48649254-INS-325 |
| chr12-48663550-INS-56 | chr12-48663550-INS-54 |
| chr12-48684942-INS-165 | chr12-48684955-INS-173 |
| chr12-48697381-INS-780 | chr12-48697381-INS-779 |
| chr12-48726500-INS-68 | chr12-48726500-INS-54 |
| chr12-48766219-INS-611 | chr12-48766219-INS-612 |
| chr12-48786254-INS-466 | chr12-48786254-INS-466 |
| chr12-48788345-INS-6545 | chr12-48788346-INS-6545 |
| chr12-48836788-INS-1812 | chr12-48836789-INS-1810 |
| chr12-48851611-INS-3347 | chr12-48851612-INS-3333 |
| chr12-48859113-INS-6380 | chr12-48859114-INS-6383 |
| chr12-48914286-INS-7960 | chr12-48914287-INS-7961 |
| chr12-48921523-INS-53 | chr12-48921516-INS-102 |
| chr12-48937075-INS-231 | chr12-48937075-INS-231 |
| chr12-48954499-INS-174 | chr12-48954499-INS-174 |
| chr12-48957649-INS-190 | chr12-48957649-INS-199 |
| chr12-48958609-INS-6718 | chr12-48958610-INS-6719 |
| chr12-48982102-INS-1066 | chr12-48982103-INS-1067 |
| chr12-48986158-INS-419 | chr12-48986158-INS-419 |
| chr12-49005902-INS-51 | chr12-49005902-INS-51 |
| chr12-49088941-INS-158 | chr12-49088941-INS-158 |
| chr12-49154872-INS-315 | chr12-49154873-INS-315 |
| chr12-49169415-INS-215 | chr12-49169415-INS-214 |
| chr12-49189478-INS-1376 | chr12-49189479-INS-1376 |
| chr12-49190219-INS-68 | chr12-49190219-INS-68 |
| chr12-49224983-INS-67 | chr12-49224983-INS-66 |
| chr12-49278934-INS-71 | chr12-49278955-INS-71 |
| chr12-49337213-INS-634 | chr12-49337213-INS-634 |
| chr12-49388338-INS-1734 | chr12-49388339-INS-1736 |
| chr12-49404689-INS-62 | chr12-49404689-INS-62 |
| chr12-49472399-INS-196 | chr12-49472399-INS-197 |
| chr12-49497917-INS-648 | chr12-49497917-INS-648 |
| chr12-49503113-INS-53 | chr12-49503113-INS-65 |
| chr12-49506361-INS-329 | chr12-49506361-INS-331 |
| chr12-49618949-INS-108 | chr12-49618933-INS-94 |
| chr12-49667750-INS-1925 | chr12-49667751-INS-1925 |
| chr12-49703888-INS-830 | chr12-49703888-INS-840 |
| chr12-49727387-INS-6269 | chr12-49727388-INS-6270 |
| chr12-49734153-INS-52 | chr12-49734153-INS-52 |
| chr12-49763341-INS-72 | chr12-49763341-INS-54 |
| chr12-49792784-INS-63 | chr12-49792785-INS-71 |
| chr12-49795748-INS-63 | chr12-49795748-INS-63 |
| chr12-49832922-INS-88 | chr12-49832922-INS-64 |
| chr12-49879564-INS-104 | chr12-49879564-INS-104 |
| chr12-49904391-INS-2142 | chr12-49904392-INS-2149 |
| chr12-49915456-INS-641 | chr12-49915454-INS-634 |
| chr12-49952856-INS-2466 | chr12-49952857-INS-2467 |
| chr12-49988844-INS-130 | chr12-49988888-INS-80 |
| chr12-50002549-INS-80 | chr12-50002553-INS-64 |
| chr12-50015998-INS-62 | chr12-50015998-INS-62 |
| chr12-50059965-INS-164 | chr12-50059966-INS-166 |
| chr12-50067812-INS-572 | chr12-50067812-INS-574 |
| chr12-50149024-INS-4556 | chr12-50149025-INS-4556 |
| chr12-50188505-INS-343 | chr12-50188505-INS-343 |
| chr12-50196420-INS-125 | chr12-50196444-INS-82 |
| chr12-50198879-INS-77 | chr12-50198879-INS-78 |
| chr12-50248933-INS-94 | chr12-50248933-INS-94 |
| chr12-50251721-INS-899 | chr12-50251721-INS-899 |
| chr12-50352591-INS-875 | chr12-50352591-INS-875 |
| chr12-50368005-INS-64 | chr12-50368009-INS-88 |
| chr12-50373824-INS-719 | chr12-50373824-INS-720 |
| chr12-50381793-INS-124 | chr12-50381793-INS-146 |
| chr12-50393957-INS-52 | chr12-50393957-INS-52 |
| chr12-50421840-INS-52 | chr12-50421840-INS-52 |
| chr12-50435539-INS-54 | chr12-50435553-INS-50 |
| chr12-50436653-INS-826 | chr12-50436653-INS-826 |

| | |
|---|---|
| chr12-50441196-INS-55 | chr12-50441196-INS-55 |
| chr12-50462125-INS-100 | chr12-50462128-INS-90 |
| chr12-50482598-INS-60 | chr12-50482598-INS-60 |
| chr12-50488257-INS-172 | chr12-50488233-INS-192 |
| chr12-50502021-INS-132 | chr12-50502021-INS-140 |
| chr12-50514415-INS-61 | chr12-50514415-INS-61 |
| chr12-50525796-INS-1210 | chr12-50525797-INS-1210 |
| chr12-50546151-INS-88 | chr12-50546149-INS-90 |
| chr12-50582352-INS-231 | chr12-50582352-INS-231 |
| chr12-50599519-INS-487 | chr12-50599519-INS-492 |
| chr12-50601832-INS-465 | chr12-50601832-INS-465 |
| chr12-50664444-INS-688 | chr12-50664444-INS-688 |
| chr12-50711257-INS-6744 | chr12-50711258-INS-6744 |
| chr12-50716184-INS-6598 | chr12-50716185-INS-6598 |
| chr12-50726932-INS-60 | chr12-50726932-INS-60 |
| chr12-50731638-INS-1635 | chr12-50731639-INS-1635 |
| chr12-50744802-INS-99 | chr12-50744830-INS-58 |
| chr12-50770906-INS-70 | chr12-50770906-INS-70 |
| chr12-50776224-INS-298 | chr12-50776224-INS-298 |
| chr12-50800050-INS-2757 | chr12-50800055-INS-2774 |
| chr12-50817878-INS-120 | chr12-50817878-INS-128 |
| chr12-50853902-INS-400 | chr12-50853902-INS-400 |
| chr12-50863620-INS-56 | chr12-50863618-INS-55 |
| chr12-50883995-INS-3004 | chr12-50883996-INS-3003 |
| chr12-50890981-INS-6587 | chr12-50890982-INS-6587 |
| chr12-50911702-INS-178 | chr12-50911702-INS-178 |
| chr12-50916165-INS-275 | chr12-50916165-INS-275 |
| chr12-50946434-INS-80 | chr12-50946436-INS-54 |
| chr12-50980374-INS-500 | chr12-50980374-INS-500 |
| chr12-50984209-INS-120 | chr12-50984229-INS-88 |
| chr12-51061077-INS-52 | chr12-51061077-INS-52 |
| chr12-51065712-INS-447 | chr12-51065711-INS-435 |
| chr12-51087485-INS-92 | chr12-51087485-INS-80 |
| chr12-51093903-INS-273 | chr12-51093903-INS-273 |
| chr12-51119134-INS-71 | chr12-51119135-INS-68 |
| chr12-51119263-INS-1491 | chr12-51119264-INS-1490 |
| chr12-51144174-INS-6502 | chr12-51144175-INS-6500 |
| chr12-51180660-INS-310 | chr12-51180660-INS-316 |
| chr12-51197556-INS-59 | chr12-51197556-INS-61 |
| chr12-51202469-INS-87 | chr12-51202469-INS-87 |
| chr12-51206300-INS-701 | chr12-51206300-INS-701 |
| chr12-51212960-INS-78 | chr12-51212960-INS-66 |
| chr12-51225725-INS-79 | chr12-51225689-INS-105 |
| chr12-51229583-INS-60 | chr12-51229583-INS-58 |
| chr12-51247775-INS-58 | chr12-51247755-INS-101 |
| chr12-51248787-INS-64 | chr12-51248787-INS-60 |
| chr12-51260439-INS-215 | chr12-51260439-INS-215 |
| chr12-51275076-INS-1244 | chr12-51275077-INS-1244 |
| chr12-51278251-INS-6146 | chr12-51278252-INS-6146 |
| chr12-51290927-INS-386 | chr12-51290927-INS-384 |
| chr12-51313166-INS-7067 | chr12-51313167-INS-7067 |
| chr12-51315992-INS-220 | chr12-51315992-INS-220 |
| chr12-51317139-INS-200 | chr12-51317139-INS-202 |
| chr12-51334947-INS-56 | chr12-51334933-INS-58 |
| chr12-51373062-INS-219 | chr12-51373062-INS-219 |
| chr12-51374509-INS-71 | chr12-51374509-INS-92 |
| chr12-51380485-INS-823 | chr12-51380485-INS-823 |
| chr12-51390909-INS-206 | chr12-51390909-INS-206 |
| chr12-51399525-INS-198 | chr12-51399525-INS-199 |
| chr12-51401945-INS-203 | chr12-51401945-INS-201 |
| chr12-51441976-INS-220 | chr12-51441976-INS-220 |
| chr12-51459915-INS-203 | chr12-51459915-INS-204 |
| chr12-51474035-INS-794 | chr12-51474035-INS-794 |
| chr12-51489746-INS-248 | chr12-51489746-INS-247 |
| chr12-51512132-INS-648 | chr12-51512132-INS-648 |
| chr12-51527036-INS-748 | chr12-51527036-INS-748 |
| chr12-51558570-INS-80 | chr12-51558579-INS-59 |
| chr12-51565052-INS-121 | chr12-51565052-INS-121 |
| chr12-51575739-INS-6442 | chr12-51575740-INS-6442 |
| chr12-51579534-INS-162 | chr12-51579534-INS-166 |
| chr12-51583072-INS-99 | chr12-51583073-INS-108 |
| chr12-51584818-INS-73 | chr12-51584818-INS-75 |
| chr12-51603169-INS-305 | chr12-51603175-INS-310 |
| chr12-51615714-INS-119 | chr12-51615714-INS-119 |
| chr12-51627556-INS-808 | chr12-51627556-INS-807 |
| chr12-51634214-INS-125 | chr12-51634214-INS-155 |
| chr12-51660375-INS-87 | chr12-51660398-INS-87 |

| | |
|---|---|
| chr12-51678698-INS-75 | chr12-51678698-INS-79 |
| chr12-51694277-INS-209 | chr12-51694277-INS-209 |
| chr12-51700066-INS-178 | chr12-51700066-INS-216 |
| chr12-51709523-INS-69 | chr12-51709530-INS-71 |
| chr12-51718585-INS-1600 | chr12-51718586-INS-1600 |
| chr12-51732391-INS-209 | chr12-51732391-INS-208 |
| chr12-51743072-INS-203 | chr12-51743072-INS-203 |
| chr12-51745020-INS-196 | chr12-51745020-INS-196 |
| chr12-51755101-INS-85 | chr12-51755101-INS-73 |
| chr12-51775617-INS-188 | chr12-51775617-INS-187 |
| chr12-51780561-INS-63 | chr12-51780561-INS-63 |
| chr12-51781730-INS-198 | chr12-51781730-INS-193 |
| chr12-51795973-INS-98 | chr12-51795973-INS-98 |
| chr12-51797451-INS-132 | chr12-51797451-INS-132 |
| chr12-51804831-INS-208 | chr12-51804831-INS-208 |
| chr12-51851137-INS-185 | chr12-51851137-INS-185 |
| chr12-51864672-INS-64 | chr12-51864672-INS-64 |
| chr12-51867455-INS-59 | chr12-51867455-INS-68 |
| chr12-51896997-INS-108 | chr12-51896997-INS-109 |
| chr12-51906343-INS-175 | chr12-51906343-INS-175 |
| chr12-51913886-INS-82 | chr12-51913886-INS-82 |
| chr12-51915785-INS-434 | chr12-51915785-INS-434 |
| chr12-51918104-INS-60 | chr12-51918150-INS-50 |
| chr12-51928207-INS-52 | chr12-51928207-INS-50 |
| chr12-51932686-INS-76 | chr12-51932686-INS-76 |
| chr12-51947096-INS-492 | chr12-51947097-INS-493 |
| chr12-51947856-INS-416 | chr12-51947857-INS-417 |
| chr12-51956042-INS-176 | chr12-51956042-INS-174 |
| chr12-51962201-INS-218 | chr12-51962201-INS-217 |
| chr12-51969209-INS-164 | chr12-51969209-INS-164 |
| chr12-51995135-INS-204 | chr12-51995135-INS-204 |
| chr12-51997495-INS-77 | chr12-51997495-INS-77 |
| chr12-52000116-INS-400 | chr12-52000116-INS-400 |
| chr12-52004151-INS-96 | chr12-52004151-INS-80 |
| chr12-52023154-INS-140 | chr12-52023154-INS-141 |
| chr12-52035390-INS-206 | chr12-52035390-INS-206 |
| chr12-52039896-INS-390 | chr12-52039896-INS-390 |
| chr12-52044760-INS-368 | chr12-52044760-INS-371 |
| chr12-52047397-INS-183 | chr12-52047397-INS-183 |
| chr12-52050590-INS-167 | chr12-52050590-INS-167 |
| chr12-52068997-INS-65 | chr12-52068997-INS-67 |
| chr12-52071177-INS-214 | chr12-52071177-INS-214 |
| chr12-52072033-INS-52 | chr12-52072033-INS-53 |
| chr12-52074081-INS-196 | chr12-52074081-INS-195 |
| chr12-52100038-INS-99 | chr12-52100038-INS-99 |
| chr12-52105598-INS-54 | chr12-52105584-INS-50 |
| chr12-52108462-INS-123 | chr12-52108462-INS-123 |
| chr12-52113345-INS-97 | chr12-52113338-INS-116 |
| chr12-52205790-INS-1158 | chr12-52205791-INS-1169 |
| chr12-52223704-INS-3012 | chr12-52223705-INS-3012 |
| chr12-52276106-INS-52 | chr12-52276106-INS-50 |
| chr12-52294009-INS-52 | chr12-52294009-INS-54 |
| chr12-52297440-INS-4438 | chr12-52297441-INS-4438 |
| chr12-52329111-INS-5262 | chr12-52329112-INS-5265 |
| chr12-52385574-INS-398 | chr12-52385574-INS-398 |
| chr12-52395866-INS-877 | chr12-52395867-INS-886 |
| chr12-52397720-INS-588 | chr12-52397720-INS-588 |
| chr12-52406461-INS-2059 | chr12-52406462-INS-2059 |
| chr12-52420957-INS-180 | chr12-52420957-INS-180 |
| chr12-52455327-INS-136 | chr12-52455327-INS-136 |
| chr12-52456764-INS-170 | chr12-52456764-INS-171 |
| chr12-52460436-INS-206 | chr12-52460434-INS-210 |
| chr12-52484425-INS-185 | chr12-52484425-INS-185 |
| chr12-52496534-INS-60 | chr12-52496535-INS-83 |
| chr12-52516282-INS-55 | chr12-52516282-INS-56 |
| chr12-52522989-INS-135 | chr12-52522989-INS-135 |
| chr12-52536318-INS-108 | chr12-52536318-INS-78 |
| chr12-52576103-INS-210 | chr12-52576103-INS-210 |
| chr12-52594606-INS-668 | chr12-52594606-INS-668 |
| chr12-52603121-INS-58 | chr12-52603121-INS-56 |
| chr12-52658950-INS-612 | chr12-52658950-INS-612 |
| chr12-52664436-INS-87 | chr12-52664436-INS-87 |
| chr12-52678819-INS-52 | chr12-52678819-INS-52 |
| chr12-52684197-INS-58 | chr12-52684197-INS-58 |
| chr12-52700764-INS-6668 | chr12-52700765-INS-6668 |
| chr12-52725184-INS-61 | chr12-52725184-INS-73 |
| chr12-52734005-INS-150 | chr12-52734005-INS-157 |

| | |
|---|---|
| chr12-52735584-INS-67 | chr12-52735584-INS-67 |
| chr12-52770067-INS-108 | chr12-52770067-INS-136 |
| chr12-52791775-INS-93 | chr12-52791775-INS-92 |
| chr12-52796238-INS-63 | chr12-52796238-INS-57 |
| chr12-52806301-INS-201 | chr12-52806301-INS-201 |
| chr12-52814189-INS-67 | chr12-52814190-INS-68 |
| chr12-52815324-INS-214 | chr12-52815321-INS-223 |
| chr12-52824864-INS-217 | chr12-52824864-INS-218 |
| chr12-52840570-INS-74 | chr12-52840570-INS-74 |
| chr12-52860587-INS-143 | chr12-52860567-INS-123 |
| chr12-52866981-INS-521 | chr12-52866981-INS-522 |
| chr12-52885144-INS-55 | chr12-52885144-INS-74 |
| chr12-52886463-INS-56 | chr12-52886463-INS-56 |
| chr12-52898631-INS-68 | chr12-52898631-INS-70 |
| chr12-52905417-INS-52 | chr12-52905417-INS-52 |
| chr12-52910299-INS-204 | chr12-52910299-INS-204 |
| chr12-52943151-INS-149 | chr12-52943151-INS-154 |
| chr12-52949919-INS-60 | chr12-52949918-INS-102 |
| chr12-52954740-INS-135 | chr12-52954740-INS-135 |
| chr12-52993128-INS-3701 | chr12-52993129-INS-3701 |
| chr12-52993129-INS-497 | chr12-52993129-INS-497 |
| chr12-53001876-INS-458 | chr12-53001876-INS-458 |
| chr12-53017156-INS-431 | chr12-53017156-INS-435 |
| chr12-53048997-INS-347 | chr12-53048997-INS-347 |
| chr12-53055852-INS-5544 | chr12-53055853-INS-5544 |
| chr12-53124657-INS-52 | chr12-53124657-INS-54 |
| chr12-53186117-INS-83 | chr12-53186117-INS-91 |
| chr12-53204585-INS-51 | chr12-53204585-INS-51 |
| chr12-53238163-INS-64 | chr12-53238163-INS-64 |
| chr12-53252304-INS-52 | chr12-53252304-INS-52 |
| chr12-53266803-INS-2771 | chr12-53266804-INS-2771 |
| chr12-53271296-INS-227 | chr12-53271296-INS-227 |
| chr12-53273927-INS-149 | chr12-53273902-INS-164 |
| chr12-53293980-INS-114 | chr12-53293980-INS-102 |
| chr12-53306492-INS-56 | chr12-53306492-INS-56 |
| chr12-53323703-INS-86 | chr12-53323703-INS-86 |
| chr12-53332683-INS-595 | chr12-53332683-INS-595 |
| chr12-53377169-INS-144 | chr12-53377149-INS-188 |
| chr12-53385687-INS-78 | chr12-53385687-INS-68 |
| chr12-53405125-INS-233 | chr12-53405125-INS-233 |
| chr12-53411783-INS-181 | chr12-53411783-INS-135 |
| chr12-53428898-INS-55 | chr12-53428898-INS-55 |
| chr12-53440304-INS-698 | chr12-53440304-INS-707 |
| chr12-53446025-INS-4907 | chr12-53446026-INS-4907 |
| chr12-53488446-INS-143 | chr12-53488446-INS-143 |
| chr12-53538364-INS-154 | chr12-53538364-INS-183 |
| chr12-53560372-INS-54 | chr12-53560372-INS-54 |
| chr12-53575485-INS-64 | chr12-53575485-INS-66 |
| chr12-53579110-INS-321 | chr12-53579110-INS-321 |
| chr12-53582486-INS-4415 | chr12-53582487-INS-4415 |
| chr12-53586829-INS-76 | chr12-53586829-INS-76 |
| chr12-53601368-INS-73 | chr12-53601368-INS-73 |
| chr12-53612201-INS-3602 | chr12-53612202-INS-3602 |
| chr12-53615437-INS-1087 | chr12-53615438-INS-1087 |
| chr12-53616081-INS-1413 | chr12-53616082-INS-1413 |
| chr12-53644187-INS-65 | chr12-53644173-INS-72 |
| chr12-53659599-INS-296 | chr12-53659599-INS-295 |
| chr12-53709987-INS-126 | chr12-53710003-INS-88 |
| chr12-53728610-INS-202 | chr12-53728610-INS-202 |
| chr12-53755867-INS-52 | chr12-53755899-INS-63 |
| chr12-53790292-INS-74 | chr12-53790292-INS-74 |
| chr12-53807148-INS-52 | chr12-53807148-INS-56 |
| chr12-53838968-INS-54 | chr12-53838968-INS-56 |
| chr12-53847763-INS-117 | chr12-53847775-INS-80 |
| chr12-53858166-INS-531 | chr12-53858166-INS-513 |
| chr12-53875067-INS-52 | chr12-53875067-INS-52 |
| chr12-53916068-INS-52 | chr12-53916068-INS-52 |
| chr12-53980027-INS-51 | chr12-53980027-INS-51 |
| chr12-54003510-INS-51 | chr12-54003510-INS-53 |
| chr12-54013802-INS-130 | chr12-54013802-INS-100 |
| chr12-54038364-INS-62 | chr12-54038364-INS-62 |
| chr12-54041249-INS-66 | chr12-54041249-INS-66 |
| chr12-54048546-INS-138 | chr12-54048546-INS-135 |
| chr12-54062764-INS-67 | chr12-54062764-INS-70 |
| chr12-54068827-INS-657 | chr12-54068827-INS-658 |
| chr12-54129420-INS-158 | chr12-54129420-INS-158 |
| chr12-54135655-INS-88 | chr12-54135663-INS-89 |

| | |
|---|---|
| chr12-54147432-INS-327 | chr12-54147432-INS-327 |
| chr12-54180277-INS-397 | chr12-54180277-INS-397 |
| chr12-54192089-INS-352 | chr12-54192089-INS-352 |
| chr12-54192888-INS-69 | chr12-54192888-INS-75 |
| chr12-54201827-INS-332 | chr12-54201828-INS-332 |
| chr12-54204712-INS-7062 | chr12-54204713-INS-7062 |
| chr12-54229796-INS-204 | chr12-54229796-INS-204 |
| chr12-54278629-INS-343 | chr12-54278629-INS-343 |
| chr12-54281532-INS-52 | chr12-54281539-INS-50 |
| chr12-54301300-INS-10012 | chr12-54301301-INS-10012 |
| chr12-54303027-INS-1753 | chr12-54303028-INS-1753 |
| chr12-54305162-INS-52 | chr12-54305162-INS-52 |
| chr12-54317740-INS-198 | chr12-54317740-INS-197 |
| chr12-54325806-INS-78 | chr12-54325792-INS-76 |
| chr12-54329610-INS-69 | chr12-54329610-INS-57 |
| chr12-54341967-INS-58 | chr12-54341967-INS-56 |
| chr12-54353703-INS-203 | chr12-54353703-INS-203 |
| chr12-54357282-INS-96 | chr12-54357256-INS-88 |
| chr12-54364686-INS-154 | chr12-54364683-INS-157 |
| chr12-54367732-INS-100 | chr12-54367732-INS-106 |
| chr12-54416824-INS-188 | chr12-54416824-INS-188 |
| chr12-54423040-INS-195 | chr12-54423040-INS-197 |
| chr12-54455626-INS-437 | chr12-54455626-INS-437 |
| chr12-54465923-INS-60 | chr12-54465924-INS-60 |
| chr12-54495637-INS-89 | chr12-54495621-INS-125 |
| chr12-54509294-INS-51 | chr12-54509281-INS-60 |
| chr12-54514517-INS-100 | chr12-54514517-INS-80 |
| chr12-54523235-INS-52 | chr12-54523224-INS-51 |
| chr12-54536309-INS-6428 | chr12-54536310-INS-6429 |
| chr12-54554115-INS-134 | chr12-54554115-INS-134 |
| chr12-54558114-INS-170 | chr12-54558114-INS-170 |
| chr12-54571535-INS-52 | chr12-54571535-INS-52 |
| chr12-54583078-INS-705 | chr12-54583078-INS-705 |
| chr12-54594926-INS-71 | chr12-54594926-INS-101 |
| chr12-54607271-INS-86 | chr12-54607271-INS-86 |
| chr12-54634409-INS-98 | chr12-54634409-INS-121 |
| chr12-54635707-INS-168 | chr12-54635707-INS-177 |
| chr12-54651793-INS-103 | chr12-54651789-INS-72 |
| chr12-54653241-INS-102 | chr12-54653241-INS-102 |
| chr12-54657000-INS-323 | chr12-54657000-INS-323 |
| chr12-54670046-INS-60 | chr12-54670046-INS-52 |
| chr12-54690007-INS-441 | chr12-54690007-INS-441 |
| chr12-54692546-INS-112 | chr12-54692546-INS-113 |
| chr12-54697558-INS-167 | chr12-54697558-INS-167 |
| chr12-54699175-INS-56 | chr12-54699175-INS-52 |
| chr12-54714481-INS-194 | chr12-54714481-INS-191 |
| chr12-54719914-INS-244 | chr12-54719914-INS-245 |
| chr12-54734632-INS-205 | chr12-54734633-INS-205 |
| chr12-54758543-INS-55 | chr12-54758510-INS-74 |
| chr12-54794127-INS-201 | chr12-54794127-INS-201 |
| chr12-54800527-INS-425 | chr12-54800527-INS-425 |
| chr12-54802538-INS-200 | chr12-54802538-INS-200 |
| chr12-54803663-INS-168 | chr12-54803663-INS-168 |
| chr12-54807103-INS-60 | chr12-54807103-INS-60 |
| chr12-54846854-INS-215 | chr12-54846854-INS-216 |
| chr12-54849014-INS-236 | chr12-54849014-INS-236 |
| chr12-54870287-INS-182 | chr12-54870287-INS-182 |
| chr12-54872383-INS-373 | chr12-54872383-INS-373 |
| chr12-54875911-INS-163 | chr12-54875911-INS-163 |
| chr12-54887178-INS-260 | chr12-54887179-INS-218 |
| chr12-54898375-INS-203 | chr12-54898375-INS-203 |
| chr12-54902433-INS-214 | chr12-54902433-INS-214 |
| chr12-54921340-INS-51 | chr12-54921340-INS-56 |
| chr12-54933983-INS-209 | chr12-54933983-INS-209 |
| chr12-54946360-INS-670 | chr12-54946361-INS-623 |
| chr12-54987903-INS-256 | chr12-54987903-INS-256 |
| chr12-54990186-INS-52 | chr12-54990174-INS-64 |
| chr12-54991052-INS-209 | chr12-54991052-INS-209 |
| chr12-54992715-INS-223 | chr12-54992715-INS-223 |
| chr12-55002503-INS-214 | chr12-55002503-INS-210 |
| chr12-55010718-INS-59 | chr12-55010718-INS-72 |
| chr12-55014665-INS-157 | chr12-55014666-INS-157 |
| chr12-55018441-INS-139 | chr12-55018441-INS-139 |
| chr12-55031382-INS-124 | chr12-55031382-INS-124 |
| chr12-55053479-INS-54 | chr12-55053479-INS-54 |
| chr12-55067433-INS-219 | chr12-55067433-INS-217 |
| chr12-55073613-INS-57 | chr12-55073613-INS-57 |

| | |
|---|---|
| chr12-55095320-INS-325 | chr12-55095320-INS-326 |
| chr12-55102076-INS-139 | chr12-55102066-INS-118 |
| chr12-55112171-INS-82 | chr12-55112171-INS-82 |
| chr12-55369658-INS-1271 | chr12-55369659-INS-1271 |
| chr12-55393822-INS-6399 | chr12-55393823-INS-6399 |
| chr12-55426988-INS-79 | chr12-55426988-INS-79 |
| chr12-55429957-INS-134 | chr12-55429957-INS-134 |
| chr12-55442397-INS-67 | chr12-55442397-INS-55 |
| chr12-55475968-INS-1049 | chr12-55475969-INS-1063 |
| chr12-55478468-INS-238 | chr12-55478451-INS-235 |
| chr12-55496203-INS-212 | chr12-55496203-INS-210 |
| chr12-55548906-INS-183 | chr12-55548906-INS-183 |
| chr12-55552942-INS-184 | chr12-55552942-INS-178 |
| chr12-55554752-INS-52 | chr12-55554752-INS-52 |
| chr12-55557527-INS-110 | chr12-55557518-INS-75 |
| chr12-55572681-INS-1887 | chr12-55572682-INS-1887 |
| chr12-55585231-INS-842 | chr12-55585231-INS-842 |
| chr12-55598567-INS-80 | chr12-55598544-INS-64 |
| chr12-55603039-INS-95 | chr12-55603033-INS-70 |
| chr12-55642396-INS-156 | chr12-55642396-INS-156 |
| chr12-55693822-INS-606 | chr12-55693822-INS-607 |
| chr12-55714464-INS-84 | chr12-55714464-INS-84 |
| chr12-55725510-INS-1746 | chr12-55725511-INS-1747 |
| chr12-55779698-INS-6451 | chr12-55779699-INS-6451 |
| chr12-55783172-INS-84 | chr12-55783172-INS-84 |
| chr12-55802811-INS-397 | chr12-55802811-INS-397 |
| chr12-55805294-INS-2708 | chr12-55805295-INS-2708 |
| chr12-55809840-INS-6239 | chr12-55809841-INS-6239 |
| chr12-55813460-INS-510 | chr12-55813460-INS-510 |
| chr12-55849269-INS-180 | chr12-55849269-INS-171 |
| chr12-55851752-INS-176 | chr12-55851752-INS-180 |
| chr12-55871984-INS-149 | chr12-55871989-INS-144 |
| chr12-55912369-INS-94 | chr12-55912369-INS-94 |
| chr12-55931668-INS-68 | chr12-55931668-INS-67 |
| chr12-55958466-INS-52 | chr12-55958466-INS-56 |
| chr12-55969078-INS-76 | chr12-55969078-INS-76 |
| chr12-55970120-INS-2350 | chr12-55970121-INS-2350 |
| chr12-55975116-INS-116 | chr12-55975116-INS-134 |
| chr12-55995862-INS-55 | chr12-55995862-INS-56 |
| chr12-56007926-INS-6440 | chr12-56007927-INS-6440 |
| chr12-56014473-INS-1435 | chr12-56014474-INS-1435 |
| chr12-56017404-INS-903 | chr12-56017404-INS-910 |
| chr12-56031477-INS-52 | chr12-56031477-INS-56 |
| chr12-56048560-INS-207 | chr12-56048560-INS-207 |
| chr12-56062489-INS-54 | chr12-56062489-INS-54 |
| chr12-56077162-INS-785 | chr12-56077162-INS-786 |
| chr12-56077833-INS-6700 | chr12-56077834-INS-6701 |
| chr12-56104689-INS-151 | chr12-56104680-INS-157 |
| chr12-56111895-INS-60 | chr12-56111895-INS-60 |
| chr12-56122917-INS-60 | chr12-56122921-INS-63 |
| chr12-56146418-INS-57 | chr12-56146418-INS-57 |
| chr12-56164422-INS-76 | chr12-56164417-INS-56 |
| chr12-56170377-INS-54 | chr12-56170377-INS-54 |
| chr12-56190148-INS-77 | chr12-56190148-INS-75 |
| chr12-56199218-INS-6509 | chr12-56199219-INS-6511 |
| chr12-56200407-INS-311 | chr12-56200408-INS-310 |
| chr12-56209845-INS-56 | chr12-56209843-INS-50 |
| chr12-56226499-INS-561 | chr12-56226499-INS-567 |
| chr12-56234379-INS-70 | chr12-56234363-INS-86 |
| chr12-56247316-INS-215 | chr12-56247316-INS-215 |
| chr12-56270006-INS-193 | chr12-56270006-INS-229 |
| chr12-56295940-INS-1115 | chr12-56295941-INS-1115 |
| chr12-56297009-INS-193 | chr12-56297009-INS-192 |
| chr12-56298361-INS-1185 | chr12-56298362-INS-1186 |
| chr12-56333459-INS-1078 | chr12-56333460-INS-1074 |
| chr12-56333941-INS-63 | chr12-56333941-INS-63 |
| chr12-56368116-INS-100 | chr12-56368116-INS-106 |
| chr12-56386705-INS-95 | chr12-56386705-INS-95 |
| chr12-56403502-INS-72 | chr12-56403460-INS-89 |
| chr12-56417170-INS-6227 | chr12-56417171-INS-6227 |
| chr12-56420221-INS-61 | chr12-56420217-INS-58 |
| chr12-56431692-INS-817 | chr12-56431692-INS-817 |
| chr12-56439095-INS-96 | chr12-56439095-INS-98 |
| chr12-56443715-INS-57 | chr12-56443715-INS-74 |
| chr12-56458517-INS-4559 | chr12-56458518-INS-4559 |
| chr12-56468406-INS-70 | chr12-56468406-INS-50 |
| chr12-56516319-INS-87 | chr12-56516319-INS-87 |

| | |
|---|---|
| chr12-56519087-INS-108 | chr12-56519087-INS-106 |
| chr12-56570825-INS-51 | chr12-56570825-INS-51 |
| chr12-56579572-INS-100 | chr12-56579572-INS-144 |
| chr12-56589942-INS-75 | chr12-56589942-INS-78 |
| chr12-56596529-INS-94 | chr12-56596529-INS-96 |
| chr12-56611343-INS-136 | chr12-56611342-INS-136 |
| chr12-56614440-INS-166 | chr12-56614440-INS-166 |
| chr12-56617941-INS-358 | chr12-56617941-INS-358 |
| chr12-56622989-INS-6288 | chr12-56622990-INS-6287 |
| chr12-56642872-INS-66 | chr12-56642860-INS-57 |
| chr12-56663923-INS-78 | chr12-56663940-INS-86 |
| chr12-56692060-INS-203 | chr12-56692060-INS-203 |
| chr12-56694853-INS-714 | chr12-56694853-INS-714 |
| chr12-56696842-INS-4577 | chr12-56696840-INS-4576 |
| chr12-56697036-INS-90 | chr12-56697036-INS-89 |
| chr12-56701249-INS-211 | chr12-56701249-INS-211 |
| chr12-56716064-INS-64 | chr12-56716064-INS-56 |
| chr12-56827536-INS-58 | chr12-56827536-INS-58 |
| chr12-56832372-INS-219 | chr12-56832372-INS-217 |
| chr12-56841320-INS-108 | chr12-56841320-INS-152 |
| chr12-56863338-INS-198 | chr12-56863338-INS-187 |
| chr12-56868098-INS-953 | chr12-56868098-INS-952 |
| chr12-56879796-INS-66 | chr12-56879796-INS-69 |
| chr12-56884524-INS-141 | chr12-56884553-INS-142 |
| chr12-56902916-INS-76 | chr12-56902916-INS-78 |
| chr12-56922204-INS-875 | chr12-56922204-INS-875 |
| chr12-56927473-INS-96 | chr12-56927473-INS-69 |
| chr12-56956273-INS-51 | chr12-56956273-INS-51 |
| chr12-56974110-INS-81 | chr12-56974110-INS-81 |
| chr12-56977909-INS-56 | chr12-56977913-INS-54 |
| chr12-56979698-INS-88 | chr12-56979674-INS-84 |
| chr12-57017474-INS-51 | chr12-57017474-INS-51 |
| chr12-57019434-INS-233 | chr12-57019434-INS-233 |
| chr12-57021592-INS-193 | chr12-57021592-INS-193 |
| chr12-57030308-INS-60 | chr12-57030271-INS-55 |
| chr12-57075694-INS-52 | chr12-57075695-INS-52 |
| chr12-57140804-INS-53 | chr12-57140804-INS-53 |
| chr12-57143616-INS-79 | chr12-57143616-INS-79 |
| chr12-57198386-INS-100 | chr12-57198386-INS-100 |
| chr12-57203119-INS-98 | chr12-57203119-INS-102 |
| chr12-57216646-INS-2577 | chr12-57216647-INS-2578 |
| chr12-57222135-INS-66 | chr12-57222135-INS-66 |
| chr12-57245891-INS-710 | chr12-57245891-INS-711 |
| chr12-57253687-INS-58 | chr12-57253687-INS-58 |
| chr12-57255873-INS-78 | chr12-57255873-INS-78 |
| chr12-57258727-INS-54 | chr12-57258727-INS-54 |
| chr12-57264779-INS-493 | chr12-57264779-INS-493 |
| chr12-57290132-INS-96 | chr12-57290133-INS-98 |
| chr12-57342010-INS-553 | chr12-57342010-INS-554 |
| chr12-57351049-INS-299 | chr12-57351049-INS-299 |
| chr12-57370118-INS-85 | chr12-57370118-INS-85 |
| chr12-57371479-INS-185 | chr12-57371479-INS-180 |
| chr12-57419274-INS-64 | chr12-57419229-INS-95 |
| chr12-57422531-INS-2862 | chr12-57422532-INS-2862 |
| chr12-57433327-INS-209 | chr12-57433327-INS-209 |
| chr12-57441377-INS-1642 | chr12-57441378-INS-1642 |
| chr12-57450355-INS-56 | chr12-57450355-INS-72 |
| chr12-57461810-INS-667 | chr12-57461810-INS-668 |
| chr12-57467076-INS-394 | chr12-57467076-INS-394 |
| chr12-57478289-INS-6603 | chr12-57478290-INS-6604 |
| chr12-57481709-INS-63 | chr12-57481702-INS-79 |
| chr12-57517524-INS-134 | chr12-57517524-INS-134 |
| chr12-57532521-INS-149 | chr12-57532518-INS-117 |
| chr12-57539248-INS-1058 | chr12-57539249-INS-1058 |
| chr12-57595047-INS-66 | chr12-57595047-INS-68 |
| chr12-57634243-INS-395 | chr12-57634243-INS-395 |
| chr12-57652942-INS-214 | chr12-57652942-INS-214 |
| chr12-57672792-INS-54 | chr12-57672792-INS-54 |
| chr12-57694173-INS-6777 | chr12-57694174-INS-6778 |
| chr12-57726655-INS-2773 | chr12-57726656-INS-2773 |
| chr12-57762405-INS-68 | chr12-57762405-INS-66 |
| chr12-57811931-INS-63 | chr12-57811931-INS-50 |
| chr12-57834010-INS-1211 | chr12-57834011-INS-1230 |
| chr12-57840649-INS-6639 | chr12-57840650-INS-6639 |
| chr12-57853423-INS-602 | chr12-57853423-INS-602 |
| chr12-57880439-INS-2894 | chr12-57880440-INS-2894 |
| chr12-57880440-INS-441 | chr12-57880440-INS-441 |

| | |
|---|---|
| chr12-57920512-INS-698 | chr12-57920512-INS-698 |
| chr12-57921083-INS-3225 | chr12-57921084-INS-3226 |
| chr12-57933603-INS-199 | chr12-57933603-INS-199 |
| chr12-57962010-INS-74 | chr12-57962010-INS-74 |
| chr12-57963601-INS-416 | chr12-57963601-INS-416 |
| chr12-57965611-INS-701 | chr12-57965611-INS-700 |
| chr12-57983616-INS-1259 | chr12-57983626-INS-1261 |
| chr12-57986921-INS-108 | chr12-57986921-INS-108 |
| chr12-57998239-INS-55 | chr12-57998239-INS-56 |
| chr12-57999519-INS-357 | chr12-57999519-INS-359 |
| chr12-58012569-INS-751 | chr12-58012569-INS-746 |
| chr12-58019275-INS-78 | chr12-58019275-INS-74 |
| chr12-58046901-INS-264 | chr12-58046901-INS-264 |
| chr12-58048704-INS-871 | chr12-58048704-INS-873 |
| chr12-58050533-INS-106 | chr12-58050533-INS-110 |
| chr12-58064300-INS-65 | chr12-58064300-INS-65 |
| chr12-58076148-INS-170 | chr12-58076148-INS-170 |
| chr12-58078918-INS-1368 | chr12-58078920-INS-1365 |
| chr12-58081842-INS-6860 | chr12-58081843-INS-6861 |
| chr12-58109155-INS-407 | chr12-58109155-INS-407 |
| chr12-58139042-INS-69 | chr12-58139022-INS-69 |
| chr12-58163952-INS-68 | chr12-58163903-INS-78 |
| chr12-58273599-INS-67 | chr12-58273599-INS-67 |
| chr12-58289223-INS-1888 | chr12-58289224-INS-1888 |
| chr12-58291660-INS-254 | chr12-58291660-INS-254 |
| chr12-58330152-INS-322 | chr12-58330152-INS-323 |
| chr12-58346598-INS-1500 | chr12-58346599-INS-1500 |
| chr12-58391707-INS-481 | chr12-58391707-INS-481 |
| chr12-58392516-INS-2453 | chr12-58392534-INS-2439 |
| chr12-58393385-INS-233 | chr12-58393385-INS-229 |
| chr12-58399756-INS-52 | chr12-58399756-INS-52 |
| chr12-58405226-INS-412 | chr12-58405226-INS-413 |
| chr12-58419116-INS-57 | chr12-58419105-INS-50 |
| chr12-58425759-INS-562 | chr12-58425759-INS-562 |
| chr12-58451177-INS-562 | chr12-58451177-INS-562 |
| chr12-58464856-INS-73 | chr12-58464856-INS-83 |
| chr12-58475987-INS-149 | chr12-58475982-INS-149 |
| chr12-58488716-INS-282 | chr12-58488695-INS-280 |
| chr12-58490644-INS-56 | chr12-58490638-INS-60 |
| chr12-58499592-INS-117 | chr12-58499592-INS-114 |
| chr12-58540281-INS-214 | chr12-58540281-INS-214 |
| chr12-58567378-INS-80 | chr12-58567378-INS-82 |
| chr12-58568142-INS-2610 | chr12-58568143-INS-2611 |
| chr12-58611776-INS-167 | chr12-58611776-INS-167 |
| chr12-58622336-INS-251 | chr12-58622336-INS-252 |
| chr12-58639228-INS-56 | chr12-58639228-INS-58 |
| chr12-58671014-INS-6403 | chr12-58671015-INS-6404 |
| chr12-58716178-INS-2819 | chr12-58716179-INS-2819 |
| chr12-58722590-INS-98 | chr12-58722590-INS-105 |
| chr12-58738855-INS-945 | chr12-58738855-INS-945 |
| chr12-58765867-INS-6678 | chr12-58765868-INS-6678 |
| chr12-58767012-INS-50 | chr12-58767012-INS-55 |
| chr12-58784080-INS-732 | chr12-58784080-INS-732 |
| chr12-58800768-INS-63 | chr12-58800768-INS-63 |
| chr12-58811697-INS-682 | chr12-58811697-INS-681 |
| chr12-58816102-INS-201 | chr12-58816102-INS-201 |
| chr12-58841695-INS-1074 | chr12-58841696-INS-1074 |
| chr12-58841700-INS-222 | chr12-58841700-INS-222 |
| chr12-58867583-INS-2181 | chr12-58867584-INS-2179 |
| chr12-58867929-INS-186 | chr12-58867929-INS-186 |
| chr12-58871877-INS-58 | chr12-58871877-INS-62 |
| chr12-58878764-INS-1781 | chr12-58878765-INS-1782 |
| chr12-58879606-INS-3407 | chr12-58879607-INS-3404 |
| chr12-58913791-INS-249 | chr12-58913791-INS-249 |
| chr12-58918650-INS-3480 | chr12-58918651-INS-3480 |
| chr12-58922205-INS-63 | chr12-58922205-INS-51 |
| chr12-58926381-INS-1314 | chr12-58926382-INS-1314 |
| chr12-58927452-INS-52 | chr12-58927452-INS-53 |
| chr12-58960074-INS-115 | chr12-58960074-INS-115 |
| chr12-58983866-INS-1022 | chr12-58983867-INS-1022 |
| chr12-58984591-INS-120 | chr12-58984588-INS-76 |
| chr12-58985516-INS-206 | chr12-58985516-INS-206 |
| chr12-58996521-INS-87 | chr12-58996522-INS-51 |
| chr12-58998398-INS-66 | chr12-58998396-INS-66 |
| chr12-59001841-INS-141 | chr12-59001841-INS-141 |
| chr12-59003602-INS-1892 | chr12-59003603-INS-1892 |
| chr12-59009292-INS-336 | chr12-59009292-INS-332 |

| | |
|---|---|
| chr12-59041229-INS-1892 | chr12-59041230-INS-1892 |
| chr12-59057081-INS-199 | chr12-59057081-INS-199 |
| chr12-59073548-INS-476 | chr12-59073549-INS-520 |
| chr12-59086148-INS-53 | chr12-59086148-INS-53 |
| chr12-59091967-INS-135 | chr12-59091967-INS-138 |
| chr12-59096121-INS-87 | chr12-59096120-INS-89 |
| chr12-59109292-INS-55 | chr12-59109295-INS-56 |
| chr12-59111943-INS-336 | chr12-59111942-INS-355 |
| chr12-59134318-INS-148 | chr12-59134318-INS-160 |
| chr12-59134777-INS-207 | chr12-59134777-INS-208 |
| chr12-59142989-INS-179 | chr12-59142987-INS-171 |
| chr12-59147077-INS-7088 | chr12-59147078-INS-7088 |
| chr12-59157021-INS-498 | chr12-59157021-INS-498 |
| chr12-59172325-INS-314 | chr12-59172325-INS-313 |
| chr12-59175016-INS-5345 | chr12-59175017-INS-5345 |
| chr12-59179715-INS-210 | chr12-59179715-INS-214 |
| chr12-59187662-INS-164 | chr12-59187698-INS-177 |
| chr12-59199760-INS-51 | chr12-59199760-INS-51 |
| chr12-59227651-INS-199 | chr12-59227651-INS-199 |
| chr12-59234540-INS-89 | chr12-59234540-INS-89 |
| chr12-59238846-INS-53 | chr12-59238846-INS-58 |
| chr12-59243554-INS-218 | chr12-59243554-INS-219 |
| chr12-59250653-INS-196 | chr12-59250653-INS-196 |
| chr12-59255475-INS-818 | chr12-59255475-INS-818 |
| chr12-59335776-INS-111 | chr12-59335776-INS-120 |
| chr12-59340613-INS-96 | chr12-59340613-INS-108 |
| chr12-59352183-INS-210 | chr12-59352183-INS-200 |
| chr12-59360657-INS-82 | chr12-59360657-INS-85 |
| chr12-59371848-INS-126 | chr12-59371848-INS-128 |
| chr12-59389624-INS-300 | chr12-59389624-INS-300 |
| chr12-59401975-INS-94 | chr12-59401975-INS-94 |
| chr12-59426316-INS-116 | chr12-59426316-INS-93 |
| chr12-59431339-INS-6582 | chr12-59431340-INS-6584 |
| chr12-59434812-INS-59 | chr12-59434812-INS-55 |
| chr12-59447991-INS-336 | chr12-59447991-INS-336 |
| chr12-59464348-INS-6412 | chr12-59464349-INS-6412 |
| chr12-59469179-INS-114 | chr12-59469179-INS-114 |
| chr12-59472493-INS-7170 | chr12-59472494-INS-7170 |
| chr12-59501605-INS-179 | chr12-59501605-INS-181 |
| chr12-59509629-INS-70 | chr12-59509629-INS-76 |
| chr12-59537493-INS-85 | chr12-59537490-INS-50 |
| chr12-59542335-INS-2772 | chr12-59542336-INS-2772 |
| chr12-59554319-INS-719 | chr12-59554319-INS-719 |
| chr12-59565352-INS-1888 | chr12-59565353-INS-1888 |
| chr12-59572529-INS-58 | chr12-59572529-INS-58 |
| chr12-59592850-INS-3372 | chr12-59592851-INS-3370 |
| chr12-59597355-INS-6688 | chr12-59597356-INS-6688 |
| chr12-59597380-INS-437 | chr12-59597380-INS-439 |
| chr12-59614554-INS-54 | chr12-59614554-INS-60 |
| chr12-59630752-INS-460 | chr12-59630752-INS-462 |
| chr12-59631046-INS-6564 | chr12-59631047-INS-6564 |
| chr12-59643129-INS-2124 | chr12-59643130-INS-2125 |
| chr12-59654562-INS-1026 | chr12-59654563-INS-1026 |
| chr12-59705507-INS-98 | chr12-59705507-INS-96 |
| chr12-59707278-INS-70 | chr12-59707278-INS-70 |
| chr12-59712993-INS-1389 | chr12-59712994-INS-1390 |
| chr12-59716221-INS-424 | chr12-59716221-INS-424 |
| chr12-59746128-INS-174 | chr12-59746128-INS-178 |
| chr12-59775487-INS-178 | chr12-59775485-INS-181 |
| chr12-59799408-INS-51 | chr12-59799408-INS-51 |
| chr12-59810395-INS-5286 | chr12-59810396-INS-5286 |
| chr12-59817530-INS-7140 | chr12-59817531-INS-7140 |
| chr12-59840221-INS-217 | chr12-59840221-INS-218 |
| chr12-59845809-INS-102 | chr12-59845810-INS-64 |
| chr12-59851912-INS-148 | chr12-59851912-INS-149 |
| chr12-59861870-INS-4803 | chr12-59861871-INS-4803 |
| chr12-59884287-INS-7299 | chr12-59884288-INS-7299 |
| chr12-59894178-INS-84 | chr12-59894178-INS-84 |
| chr12-59907613-INS-4407 | chr12-59907614-INS-4407 |
| chr12-59931856-INS-7104 | chr12-59931857-INS-7104 |
| chr12-59940021-INS-743 | chr12-59940021-INS-744 |
| chr12-59955714-INS-75 | chr12-59955714-INS-75 |
| chr12-59959281-INS-7149 | chr12-59959282-INS-7149 |
| chr12-59966996-INS-2339 | chr12-59966997-INS-2342 |
| chr12-59974758-INS-94 | chr12-59974758-INS-94 |
| chr12-59977089-INS-526 | chr12-59977089-INS-527 |
| chr12-59978579-INS-58 | chr12-59978579-INS-60 |

| | |
|---|---|
| chr12-60000088-INS-3788 | chr12-60000089-INS-3788 |
| chr12-60020051-INS-1589 | chr12-60020052-INS-1589 |
| chr12-60021469-INS-1092 | chr12-60021470-INS-1092 |
| chr12-60023917-INS-499 | chr12-60023917-INS-499 |
| chr12-60028119-INS-180 | chr12-60028105-INS-178 |
| chr12-60043352-INS-139 | chr12-60043352-INS-129 |
| chr12-60046093-INS-162 | chr12-60046095-INS-174 |
| chr12-60048896-INS-182 | chr12-60048896-INS-182 |
| chr12-60050233-INS-1206 | chr12-60050234-INS-1206 |
| chr12-60069475-INS-1395 | chr12-60069465-INS-1395 |
| chr12-60085679-INS-456 | chr12-60085679-INS-459 |
| chr12-60092469-INS-623 | chr12-60092469-INS-630 |
| chr12-60096408-INS-80 | chr12-60096400-INS-54 |
| chr12-60098504-INS-51 | chr12-60098504-INS-54 |
| chr12-60100532-INS-478 | chr12-60100532-INS-486 |
| chr12-60101543-INS-79 | chr12-60101543-INS-79 |
| chr12-60118652-INS-83 | chr12-60118652-INS-83 |
| chr12-60135347-INS-72 | chr12-60135347-INS-74 |
| chr12-60147155-INS-809 | chr12-60147152-INS-808 |
| chr12-60159879-INS-806 | chr12-60159879-INS-808 |
| chr12-60160962-INS-151 | chr12-60160962-INS-151 |
| chr12-60165910-INS-3030 | chr12-60165911-INS-3030 |
| chr12-60167028-INS-6453 | chr12-60167029-INS-6454 |
| chr12-60181893-INS-440 | chr12-60181893-INS-440 |
| chr12-60184586-INS-59 | chr12-60184586-INS-67 |
| chr12-60204008-INS-59 | chr12-60204008-INS-60 |
| chr12-60214506-INS-57 | chr12-60214506-INS-57 |
| chr12-60220336-INS-7009 | chr12-60220337-INS-7010 |
| chr12-60238921-INS-871 | chr12-60238921-INS-871 |
| chr12-60246775-INS-158 | chr12-60246775-INS-165 |
| chr12-60265496-INS-237 | chr12-60265497-INS-239 |
| chr12-60279341-INS-786 | chr12-60279344-INS-785 |
| chr12-60281990-INS-399 | chr12-60281990-INS-400 |
| chr12-60295012-INS-7023 | chr12-60295013-INS-7019 |
| chr12-60316388-INS-499 | chr12-60316388-INS-498 |
| chr12-60341416-INS-117 | chr12-60341416-INS-117 |
| chr12-60350424-INS-420 | chr12-60350424-INS-420 |
| chr12-60374104-INS-507 | chr12-60374104-INS-503 |
| chr12-60375136-INS-640 | chr12-60375136-INS-616 |
| chr12-60390355-INS-52 | chr12-60390352-INS-84 |
| chr12-60405347-INS-233 | chr12-60405347-INS-232 |
| chr12-60422909-INS-58 | chr12-60422913-INS-58 |
| chr12-60438766-INS-56 | chr12-60438766-INS-72 |
| chr12-60461982-INS-93 | chr12-60461982-INS-124 |
| chr12-60478815-INS-81 | chr12-60478815-INS-81 |
| chr12-60520747-INS-171 | chr12-60520738-INS-185 |
| chr12-60524029-INS-74 | chr12-60524049-INS-58 |
| chr12-60524295-INS-397 | chr12-60524295-INS-397 |
| chr12-60571053-INS-116 | chr12-60571053-INS-116 |
| chr12-60624652-INS-1726 | chr12-60624653-INS-1716 |
| chr12-60628115-INS-2733 | chr12-60628116-INS-2742 |
| chr12-60635211-INS-2084 | chr12-60635212-INS-2083 |
| chr12-60661576-INS-51 | chr12-60661576-INS-53 |
| chr12-60664074-INS-52 | chr12-60664074-INS-56 |
| chr12-60680586-INS-190 | chr12-60680586-INS-190 |
| chr12-60684692-INS-400 | chr12-60684692-INS-400 |
| chr12-60718380-INS-481 | chr12-60718380-INS-481 |
| chr12-60728543-INS-80 | chr12-60728543-INS-80 |
| chr12-60808432-INS-1821 | chr12-60808433-INS-1821 |
| chr12-60808595-INS-831 | chr12-60808595-INS-831 |
| chr12-60810737-INS-85 | chr12-60810737-INS-85 |
| chr12-60815037-INS-159 | chr12-60815037-INS-158 |
| chr12-60816758-INS-1656 | chr12-60816759-INS-1658 |
| chr12-60842247-INS-394 | chr12-60842247-INS-394 |
| chr12-60851726-INS-887 | chr12-60851726-INS-887 |
| chr12-60858576-INS-746 | chr12-60858576-INS-746 |
| chr12-60866502-INS-52 | chr12-60866502-INS-52 |
| chr12-60891606-INS-55 | chr12-60891606-INS-63 |
| chr12-60943367-INS-6446 | chr12-60943368-INS-6446 |
| chr12-60943566-INS-351 | chr12-60943566-INS-351 |
| chr12-60948127-INS-219 | chr12-60948127-INS-218 |
| chr12-60952786-INS-207 | chr12-60952786-INS-206 |
| chr12-60975705-INS-6568 | chr12-60975706-INS-6569 |
| chr12-60988168-INS-216 | chr12-60988168-INS-216 |
| chr12-60993685-INS-203 | chr12-60993685-INS-203 |
| chr12-61008777-INS-6936 | chr12-61008778-INS-6927 |
| chr12-61013176-INS-5622 | chr12-61013177-INS-5620 |

| | |
|---|---|
| chr12-61018409-INS-1885 | chr12-61018410-INS-1877 |
| chr12-61029435-INS-290 | chr12-61029435-INS-290 |
| chr12-61052809-INS-434 | chr12-61052809-INS-434 |
| chr12-61090361-INS-55 | chr12-61090357-INS-54 |
| chr12-61094416-INS-343 | chr12-61094416-INS-343 |
| chr12-61094715-INS-6957 | chr12-61094716-INS-6958 |
| chr12-61114241-INS-547 | chr12-61114241-INS-547 |
| chr12-61119004-INS-1034 | chr12-61119005-INS-1034 |
| chr12-61120831-INS-3061 | chr12-61120832-INS-3061 |
| chr12-61122886-INS-531 | chr12-61122886-INS-531 |
| chr12-61151616-INS-6194 | chr12-61151617-INS-6195 |
| chr12-61166382-INS-382 | chr12-61166382-INS-382 |
| chr12-61168961-INS-57 | chr12-61168961-INS-51 |
| chr12-61179616-INS-80 | chr12-61179589-INS-80 |
| chr12-61185414-INS-54 | chr12-61185414-INS-54 |
| chr12-61186524-INS-208 | chr12-61186524-INS-208 |
| chr12-61213252-INS-1330 | chr12-61213253-INS-1330 |
| chr12-61226953-INS-2067 | chr12-61226954-INS-2067 |
| chr12-61294319-INS-755 | chr12-61294319-INS-763 |
| chr12-61342912-INS-1186 | chr12-61342913-INS-1187 |
| chr12-61355955-INS-70 | chr12-61355955-INS-70 |
| chr12-61361149-INS-53 | chr12-61361149-INS-81 |
| chr12-61381288-INS-867 | chr12-61381288-INS-867 |
| chr12-61419689-INS-143 | chr12-61419689-INS-143 |
| chr12-61432223-INS-667 | chr12-61432223-INS-667 |
| chr12-61448424-INS-75 | chr12-61448424-INS-80 |
| chr12-61453356-INS-92 | chr12-61453356-INS-90 |
| chr12-61517291-INS-58 | chr12-61517291-INS-58 |
| chr12-61579662-INS-68 | chr12-61579662-INS-50 |
| chr12-61596971-INS-344 | chr12-61596971-INS-344 |
| chr12-61615269-INS-1764 | chr12-61615270-INS-1764 |
| chr12-61632840-INS-6659 | chr12-61632841-INS-6659 |
| chr12-61636486-INS-848 | chr12-61636486-INS-848 |
| chr12-61644732-INS-555 | chr12-61644732-INS-555 |
| chr12-61650647-INS-57 | chr12-61650647-INS-53 |
| chr12-61653907-INS-6859 | chr12-61653908-INS-6858 |
| chr12-61687413-INS-764 | chr12-61687413-INS-734 |
| chr12-61694401-INS-234 | chr12-61694401-INS-234 |
| chr12-61709575-INS-340 | chr12-61709575-INS-340 |
| chr12-61710638-INS-104 | chr12-61710638-INS-104 |
| chr12-61712724-INS-216 | chr12-61712724-INS-216 |
| chr12-61715018-INS-54 | chr12-61715019-INS-61 |
| chr12-61721320-INS-2309 | chr12-61721321-INS-2307 |
| chr12-61766463-INS-93 | chr12-61766459-INS-93 |
| chr12-61782413-INS-235 | chr12-61782413-INS-235 |
| chr12-61793450-INS-540 | chr12-61793450-INS-538 |
| chr12-61823673-INS-6669 | chr12-61823674-INS-6670 |
| chr12-61846614-INS-1005 | chr12-61846615-INS-1005 |
| chr12-61865334-INS-9313 | chr12-61865336-INS-9295 |
| chr12-61866188-INS-885 | chr12-61866188-INS-886 |
| chr12-61899985-INS-54 | chr12-61899971-INS-66 |
| chr12-61903312-INS-146 | chr12-61903316-INS-139 |
| chr12-61911715-INS-385 | chr12-61911716-INS-385 |
| chr12-61913261-INS-721 | chr12-61913261-INS-721 |
| chr12-61918511-INS-55 | chr12-61918511-INS-65 |
| chr12-61921401-INS-52 | chr12-61921421-INS-52 |
| chr12-61927827-INS-6796 | chr12-61927828-INS-6796 |
| chr12-61939275-INS-4394 | chr12-61939276-INS-4394 |
| chr12-61948797-INS-56 | chr12-61948797-INS-77 |
| chr12-61976037-INS-58 | chr12-61976037-INS-58 |
| chr12-61993912-INS-82 | chr12-61993912-INS-82 |
| chr12-62033603-INS-6281 | chr12-62033604-INS-6281 |
| chr12-62037526-INS-849 | chr12-62037526-INS-849 |
| chr12-62137184-INS-68 | chr12-62137184-INS-68 |
| chr12-62164906-INS-1293 | chr12-62164907-INS-1284 |
| chr12-62173734-INS-2395 | chr12-62173745-INS-2386 |
| chr12-62188887-INS-72 | chr12-62188887-INS-78 |
| chr12-62204608-INS-328 | chr12-62204608-INS-328 |
| chr12-62242360-INS-405 | chr12-62242360-INS-405 |
| chr12-62256599-INS-594 | chr12-62256599-INS-594 |
| chr12-62268439-INS-108 | chr12-62268439-INS-108 |
| chr12-62316813-INS-72 | chr12-62316813-INS-72 |
| chr12-62345668-INS-313 | chr12-62345668-INS-313 |
| chr12-62362142-INS-328 | chr12-62362142-INS-328 |
| chr12-62372585-INS-294 | chr12-62372585-INS-294 |
| chr12-62377207-INS-72 | chr12-62377196-INS-93 |
| chr12-62504758-INS-75 | chr12-62504758-INS-75 |

| | |
|---|---|
| chr12-62515373-INS-336 | chr12-62515373-INS-336 |
| chr12-62574412-INS-152 | chr12-62574412-INS-152 |
| chr12-62587731-INS-86 | chr12-62587731-INS-86 |
| chr12-62621360-INS-68 | chr12-62621340-INS-72 |
| chr12-62626658-INS-8767 | chr12-62626651-INS-8765 |
| chr12-62627899-INS-75 | chr12-62627894-INS-60 |
| chr12-62648482-INS-398 | chr12-62648482-INS-368 |
| chr12-62652284-INS-257 | chr12-62652284-INS-257 |
| chr12-62659538-INS-148 | chr12-62659532-INS-110 |
| chr12-62684596-INS-68 | chr12-62684596-INS-70 |
| chr12-62688173-INS-67 | chr12-62688173-INS-66 |
| chr12-62702030-INS-121 | chr12-62702030-INS-114 |
| chr12-62707805-INS-175 | chr12-62707805-INS-176 |
| chr12-62711966-INS-226 | chr12-62711966-INS-221 |
| chr12-62739104-INS-146 | chr12-62739104-INS-146 |
| chr12-62742805-INS-181 | chr12-62742805-INS-181 |
| chr12-62771276-INS-59 | chr12-62771276-INS-56 |
| chr12-62792970-INS-138 | chr12-62792970-INS-138 |
| chr12-62797412-INS-151 | chr12-62797412-INS-151 |
| chr12-62832272-INS-81 | chr12-62832287-INS-56 |
| chr12-62840879-INS-322 | chr12-62840879-INS-322 |
| chr12-62846838-INS-161 | chr12-62846838-INS-159 |
| chr12-62865493-INS-491 | chr12-62865493-INS-491 |
| chr12-62896745-INS-58 | chr12-62896745-INS-58 |
| chr12-62901243-INS-66 | chr12-62901241-INS-64 |
| chr12-62913309-INS-344 | chr12-62913309-INS-344 |
| chr12-62932028-INS-104 | chr12-62932058-INS-72 |
| chr12-62980084-INS-64 | chr12-62980084-INS-54 |
| chr12-62990037-INS-165 | chr12-62990037-INS-170 |
| chr12-62996877-INS-99 | chr12-62996877-INS-105 |
| chr12-63042651-INS-58 | chr12-63042651-INS-58 |
| chr12-63106375-INS-122 | chr12-63106375-INS-118 |
| chr12-63126411-INS-52 | chr12-63126411-INS-52 |
| chr12-63141608-INS-56 | chr12-63141608-INS-56 |
| chr12-63181721-INS-54 | chr12-63181721-INS-58 |
| chr12-63192783-INS-70 | chr12-63192783-INS-70 |
| chr12-63201957-INS-333 | chr12-63201957-INS-333 |
| chr12-63241738-INS-290 | chr12-63241738-INS-290 |
| chr12-63320341-INS-236 | chr12-63320341-INS-232 |
| chr12-63329023-INS-74 | chr12-63329029-INS-96 |
| chr12-63375457-INS-52 | chr12-63375457-INS-52 |
| chr12-63377063-INS-56 | chr12-63377085-INS-84 |
| chr12-63384351-INS-213 | chr12-63384351-INS-213 |
| chr12-63386103-INS-53 | chr12-63386103-INS-52 |
| chr12-63389048-INS-922 | chr12-63389048-INS-922 |
| chr12-63396885-INS-2594 | chr12-63396886-INS-2594 |
| chr12-63427727-INS-56 | chr12-63427727-INS-56 |
| chr12-63450665-INS-357 | chr12-63450665-INS-360 |
| chr12-63480545-INS-204 | chr12-63480554-INS-204 |
| chr12-63493299-INS-91 | chr12-63493299-INS-93 |
| chr12-63518261-INS-1310 | chr12-63518262-INS-1308 |
| chr12-63518332-INS-180 | chr12-63518332-INS-180 |
| chr12-63520666-INS-482 | chr12-63520666-INS-482 |
| chr12-63540377-INS-65 | chr12-63540377-INS-55 |
| chr12-63550316-INS-109 | chr12-63550327-INS-104 |
| chr12-63577373-INS-219 | chr12-63577377-INS-251 |
| chr12-63583525-INS-58 | chr12-63583525-INS-58 |
| chr12-63598702-INS-6460 | chr12-63598703-INS-6461 |
| chr12-63648615-INS-6694 | chr12-63648616-INS-6694 |
| chr12-63651751-INS-108 | chr12-63651751-INS-75 |
| chr12-63714456-INS-435 | chr12-63714456-INS-428 |
| chr12-63717097-INS-4348 | chr12-63717098-INS-4347 |
| chr12-63717978-INS-122 | chr12-63717978-INS-140 |
| chr12-63723260-INS-340 | chr12-63723274-INS-323 |
| chr12-63757345-INS-110 | chr12-63757345-INS-92 |
| chr12-63807958-INS-1251 | chr12-63807956-INS-1240 |
| chr12-63822179-INS-717 | chr12-63822179-INS-725 |
| chr12-63827216-INS-938 | chr12-63827216-INS-940 |
| chr12-63832705-INS-96 | chr12-63832705-INS-100 |
| chr12-63834463-INS-84 | chr12-63834442-INS-68 |
| chr12-63859301-INS-1017 | chr12-63859302-INS-1017 |
| chr12-63864274-INS-74 | chr12-63864274-INS-78 |
| chr12-63887405-INS-58 | chr12-63887405-INS-60 |
| chr12-63931165-INS-477 | chr12-63931165-INS-444 |
| chr12-63955330-INS-2814 | chr12-63955331-INS-2814 |
| chr12-63964572-INS-136 | chr12-63964572-INS-136 |
| chr12-63972373-INS-55 | chr12-63972374-INS-62 |

| | |
|---|---|
| chr12-63992661-INS-89 | chr12-63992661-INS-84 |
| chr12-63994565-INS-317 | chr12-63994565-INS-315 |
| chr12-63996120-INS-6255 | chr12-63996121-INS-6256 |
| chr12-64028004-INS-995 | chr12-64028004-INS-995 |
| chr12-64032440-INS-1109 | chr12-64032441-INS-1110 |
| chr12-64041134-INS-74 | chr12-64041134-INS-74 |
| chr12-64058293-INS-1421 | chr12-64058294-INS-1422 |
| chr12-64069205-INS-7275 | chr12-64069206-INS-7275 |
| chr12-64072968-INS-55 | chr12-64072968-INS-55 |
| chr12-64087008-INS-100 | chr12-64087008-INS-96 |
| chr12-64091715-INS-5637 | chr12-64091716-INS-5637 |
| chr12-64141413-INS-4699 | chr12-64141414-INS-4699 |
| chr12-64143797-INS-54 | chr12-64143797-INS-52 |
| chr12-64147570-INS-940 | chr12-64147570-INS-939 |
| chr12-64159449-INS-219 | chr12-64159449-INS-219 |
| chr12-64180324-INS-1596 | chr12-64180325-INS-1596 |
| chr12-64219778-INS-627 | chr12-64219778-INS-627 |
| chr12-64230260-INS-99 | chr12-64230300-INS-52 |
| chr12-64272701-INS-76 | chr12-64272701-INS-107 |
| chr12-64280535-INS-60 | chr12-64280535-INS-55 |
| chr12-64293190-INS-64 | chr12-64293190-INS-67 |
| chr12-64309350-INS-2459 | chr12-64309351-INS-2457 |
| chr12-64309616-INS-104 | chr12-64309616-INS-109 |
| chr12-64321473-INS-86 | chr12-64321473-INS-86 |
| chr12-64336398-INS-64 | chr12-64336398-INS-80 |
| chr12-64375617-INS-52 | chr12-64375617-INS-52 |
| chr12-64390313-INS-1113 | chr12-64390317-INS-1103 |
| chr12-64395131-INS-1159 | chr12-64395132-INS-1159 |
| chr12-64430336-INS-627 | chr12-64430336-INS-627 |
| chr12-64472525-INS-52 | chr12-64472527-INS-56 |
| chr12-64491525-INS-58 | chr12-64491525-INS-60 |
| chr12-64574287-INS-3811 | chr12-64574288-INS-3795 |
| chr12-64583891-INS-68 | chr12-64583891-INS-66 |
| chr12-64610805-INS-70 | chr12-64610805-INS-70 |
| chr12-64622030-INS-944 | chr12-64622030-INS-944 |
| chr12-64643121-INS-152 | chr12-64643121-INS-149 |
| chr12-64655768-INS-208 | chr12-64655768-INS-208 |
| chr12-64699245-INS-106 | chr12-64699245-INS-101 |
| chr12-64700239-INS-190 | chr12-64700239-INS-212 |
| chr12-64718596-INS-178 | chr12-64718596-INS-178 |
| chr12-64737978-INS-1000 | chr12-64737979-INS-1000 |
| chr12-64769100-INS-419 | chr12-64769100-INS-419 |
| chr12-64771538-INS-53 | chr12-64771538-INS-59 |
| chr12-64819916-INS-71 | chr12-64819913-INS-75 |
| chr12-64825382-INS-6235 | chr12-64825363-INS-6230 |
| chr12-64852283-INS-5946 | chr12-64852284-INS-5945 |
| chr12-64865464-INS-178 | chr12-64865471-INS-156 |
| chr12-64889371-INS-53 | chr12-64889371-INS-53 |
| chr12-64914387-INS-5002 | chr12-64914388-INS-5002 |
| chr12-64945991-INS-164 | chr12-64945991-INS-164 |
| chr12-64948639-INS-66 | chr12-64948645-INS-73 |
| chr12-64981421-INS-239 | chr12-64981421-INS-239 |
| chr12-65001136-INS-108 | chr12-65001146-INS-108 |
| chr12-65024213-INS-7367 | chr12-65024214-INS-7369 |
| chr12-65045138-INS-143 | chr12-65045138-INS-143 |
| chr12-65073174-INS-175 | chr12-65073174-INS-174 |
| chr12-65092999-INS-59 | chr12-65092999-INS-59 |
| chr12-65114520-INS-252 | chr12-65114520-INS-247 |
| chr12-65118799-INS-208 | chr12-65118799-INS-205 |
| chr12-65128637-INS-92 | chr12-65128637-INS-88 |
| chr12-65134196-INS-2090 | chr12-65134197-INS-2090 |
| chr12-65138352-INS-51 | chr12-65138352-INS-51 |
| chr12-65166889-INS-211 | chr12-65166889-INS-210 |
| chr12-65192704-INS-68 | chr12-65192704-INS-64 |
| chr12-65209849-INS-1764 | chr12-65209850-INS-1765 |
| chr12-65221893-INS-64 | chr12-65221893-INS-66 |
| chr12-65233476-INS-54 | chr12-65233476-INS-56 |
| chr12-65253597-INS-69 | chr12-65253597-INS-67 |
| chr12-65270358-INS-7309 | chr12-65270359-INS-7311 |
| chr12-65281652-INS-141 | chr12-65281652-INS-140 |
| chr12-65330448-INS-290 | chr12-65330448-INS-290 |
| chr12-65387503-INS-616 | chr12-65387502-INS-615 |
| chr12-65389142-INS-72 | chr12-65389151-INS-90 |
| chr12-65415401-INS-8831 | chr12-65415402-INS-8831 |
| chr12-65416956-INS-72 | chr12-65416956-INS-68 |
| chr12-65433891-INS-52 | chr12-65433891-INS-52 |
| chr12-65439930-INS-891 | chr12-65439930-INS-891 |

| | |
|---|---|
| chr12-65452578-INS-169 | chr12-65452576-INS-144 |
| chr12-65501822-INS-451 | chr12-65501822-INS-451 |
| chr12-65555543-INS-73 | chr12-65555543-INS-73 |
| chr12-65557701-INS-190 | chr12-65557701-INS-190 |
| chr12-65572905-INS-427 | chr12-65572905-INS-397 |
| chr12-65581736-INS-720 | chr12-65581736-INS-720 |
| chr12-65590314-INS-74 | chr12-65590314-INS-72 |
| chr12-65620416-INS-6627 | chr12-65620417-INS-6626 |
| chr12-65632831-INS-4518 | chr12-65632832-INS-4519 |
| chr12-65648246-INS-242 | chr12-65648246-INS-279 |
| chr12-65655136-INS-85 | chr12-65655136-INS-77 |
| chr12-65661642-INS-441 | chr12-65661642-INS-441 |
| chr12-65673392-INS-320 | chr12-65673392-INS-320 |
| chr12-65674994-INS-1982 | chr12-65674995-INS-1979 |
| chr12-65675396-INS-893 | chr12-65675396-INS-894 |
| chr12-65699806-INS-91 | chr12-65699806-INS-60 |
| chr12-65719871-INS-92 | chr12-65719886-INS-76 |
| chr12-65758211-INS-68 | chr12-65758211-INS-68 |
| chr12-65769986-INS-6966 | chr12-65769987-INS-6966 |
| chr12-65874543-INS-64 | chr12-65874543-INS-69 |
| chr12-65897062-INS-1514 | chr12-65897063-INS-1514 |
| chr12-65940521-INS-637 | chr12-65940521-INS-636 |
| chr12-65943929-INS-363 | chr12-65943929-INS-363 |
| chr12-65951601-INS-64 | chr12-65951637-INS-56 |
| chr12-65973298-INS-1085 | chr12-65973299-INS-1085 |
| chr12-65975067-INS-136 | chr12-65975067-INS-136 |
| chr12-65978622-INS-551 | chr12-65978622-INS-551 |
| chr12-65983581-INS-2266 | chr12-65983582-INS-2270 |
| chr12-66022438-INS-1156 | chr12-66022429-INS-1155 |
| chr12-66044501-INS-114 | chr12-66044501-INS-110 |
| chr12-66069424-INS-586 | chr12-66069424-INS-586 |
| chr12-66081718-INS-102 | chr12-66081718-INS-102 |
| chr12-66105024-INS-56 | chr12-66105024-INS-68 |
| chr12-66125298-INS-216 | chr12-66125298-INS-216 |
| chr12-66139218-INS-1563 | chr12-66139219-INS-1563 |
| chr12-66150311-INS-200 | chr12-66150311-INS-200 |
| chr12-66192338-INS-5713 | chr12-66192339-INS-5713 |
| chr12-66222955-INS-1674 | chr12-66222956-INS-1670 |
| chr12-66244749-INS-343 | chr12-66244749-INS-343 |
| chr12-66251373-INS-148 | chr12-66251373-INS-148 |
| chr12-66265056-INS-104 | chr12-66265054-INS-90 |
| chr12-66297688-INS-274 | chr12-66297679-INS-275 |
| chr12-66377821-INS-122 | chr12-66377821-INS-122 |
| chr12-66406917-INS-304 | chr12-66406917-INS-303 |
| chr12-66426955-INS-58 | chr12-66426919-INS-50 |
| chr12-66457580-INS-497 | chr12-66457580-INS-497 |
| chr12-66545967-INS-57 | chr12-66545967-INS-57 |
| chr12-66561352-INS-56 | chr12-66561352-INS-60 |
| chr12-66573736-INS-105 | chr12-66573736-INS-104 |
| chr12-66586493-INS-522 | chr12-66586493-INS-519 |
| chr12-66590784-INS-2031 | chr12-66590785-INS-2031 |
| chr12-66642739-INS-1032 | chr12-66642740-INS-1032 |
| chr12-66679221-INS-6970 | chr12-66679222-INS-6971 |
| chr12-66682545-INS-389 | chr12-66682545-INS-389 |
| chr12-66684926-INS-66 | chr12-66684927-INS-66 |
| chr12-66712013-INS-186 | chr12-66712007-INS-187 |
| chr12-66726979-INS-58 | chr12-66726979-INS-58 |
| chr12-66733842-INS-198 | chr12-66733842-INS-198 |
| chr12-66749348-INS-6400 | chr12-66749349-INS-6400 |
| chr12-66800866-INS-457 | chr12-66800866-INS-460 |
| chr12-66895405-INS-389 | chr12-66895405-INS-390 |
| chr12-66909563-INS-1767 | chr12-66909564-INS-1765 |
| chr12-66921621-INS-549 | chr12-66921621-INS-549 |
| chr12-66956303-INS-56 | chr12-66956303-INS-84 |
| chr12-67031057-INS-471 | chr12-67031057-INS-466 |
| chr12-67067734-INS-224 | chr12-67067734-INS-224 |
| chr12-67093586-INS-54 | chr12-67093587-INS-54 |
| chr12-67106376-INS-118 | chr12-67106376-INS-119 |
| chr12-67128701-INS-447 | chr12-67128701-INS-449 |
| chr12-67169116-INS-53 | chr12-67169126-INS-87 |
| chr12-67204837-INS-54 | chr12-67204837-INS-65 |
| chr12-67219877-INS-594 | chr12-67219878-INS-620 |
| chr12-67234594-INS-62 | chr12-67234592-INS-74 |
| chr12-67236258-INS-64 | chr12-67236258-INS-52 |
| chr12-67240626-INS-112 | chr12-67240628-INS-102 |
| chr12-67251497-INS-1093 | chr12-67251498-INS-1093 |
| chr12-67287582-INS-3357 | chr12-67287583-INS-3369 |

| | |
|---|---|
| chr12-67306283-INS-57 | chr12-67306283-INS-61 |
| chr12-67329420-INS-60 | chr12-67329375-INS-90 |
| chr12-67333555-INS-79 | chr12-67333555-INS-79 |
| chr12-67351007-INS-1329 | chr12-67351008-INS-1330 |
| chr12-67351243-INS-573 | chr12-67351243-INS-573 |
| chr12-67413100-INS-7085 | chr12-67413101-INS-7084 |
| chr12-67430682-INS-140 | chr12-67430673-INS-170 |
| chr12-67433389-INS-58 | chr12-67433389-INS-58 |
| chr12-67441285-INS-134 | chr12-67441285-INS-135 |
| chr12-67447567-INS-3177 | chr12-67447568-INS-3177 |
| chr12-67448061-INS-1880 | chr12-67448062-INS-1877 |
| chr12-67458524-INS-52 | chr12-67458524-INS-62 |
| chr12-67475230-INS-1464 | chr12-67475231-INS-1464 |
| chr12-67482321-INS-4333 | chr12-67482322-INS-4333 |
| chr12-67484395-INS-87 | chr12-67484395-INS-87 |
| chr12-67505493-INS-78 | chr12-67505493-INS-78 |
| chr12-67513900-INS-84 | chr12-67513900-INS-84 |
| chr12-67520590-INS-413 | chr12-67520590-INS-413 |
| chr12-67543197-INS-4740 | chr12-67543198-INS-4742 |
| chr12-67544448-INS-1376 | chr12-67544449-INS-1375 |
| chr12-67548102-INS-78 | chr12-67548101-INS-84 |
| chr12-67550208-INS-6577 | chr12-67550209-INS-6577 |
| chr12-67576057-INS-76 | chr12-67576057-INS-78 |
| chr12-67579953-INS-201 | chr12-67579953-INS-201 |
| chr12-67656009-INS-94 | chr12-67656012-INS-94 |
| chr12-67657526-INS-87 | chr12-67657526-INS-96 |
| chr12-67664222-INS-68 | chr12-67664222-INS-68 |
| chr12-67745246-INS-56 | chr12-67745246-INS-60 |
| chr12-67789447-INS-101 | chr12-67789417-INS-150 |
| chr12-67792306-INS-7891 | chr12-67792307-INS-7889 |
| chr12-67803103-INS-51 | chr12-67803103-INS-51 |
| chr12-67826961-INS-7374 | chr12-67826962-INS-7374 |
| chr12-67842325-INS-1445 | chr12-67842326-INS-1445 |
| chr12-67851472-INS-1662 | chr12-67851473-INS-1687 |
| chr12-67857435-INS-799 | chr12-67857435-INS-799 |
| chr12-67867912-INS-344 | chr12-67867912-INS-344 |
| chr12-67871306-INS-8967 | chr12-67871307-INS-8968 |
| chr12-67907845-INS-6823 | chr12-67907846-INS-6824 |
| chr12-67933979-INS-171 | chr12-67933969-INS-156 |
| chr12-67945622-INS-59 | chr12-67945622-INS-59 |
| chr12-67948315-INS-249 | chr12-67948315-INS-249 |
| chr12-67952135-INS-6969 | chr12-67952136-INS-6972 |
| chr12-68020535-INS-1304 | chr12-68020536-INS-1304 |
| chr12-68039551-INS-842 | chr12-68039551-INS-844 |
| chr12-68045161-INS-66 | chr12-68045161-INS-70 |
| chr12-68093793-INS-6470 | chr12-68093794-INS-6469 |
| chr12-68111646-INS-1815 | chr12-68111647-INS-1817 |
| chr12-68124487-INS-76 | chr12-68124498-INS-50 |
| chr12-68156608-INS-400 | chr12-68156608-INS-400 |
| chr12-68165534-INS-2642 | chr12-68165532-INS-2642 |
| chr12-68178844-INS-1645 | chr12-68178845-INS-1638 |
| chr12-68235464-INS-1127 | chr12-68235465-INS-1128 |
| chr12-68245003-INS-606 | chr12-68245003-INS-606 |
| chr12-68249824-INS-124 | chr12-68249827-INS-128 |
| chr12-68258348-INS-1232 | chr12-68258349-INS-1232 |
| chr12-68260753-INS-335 | chr12-68260753-INS-335 |
| chr12-68261501-INS-106 | chr12-68261521-INS-68 |
| chr12-68272158-INS-62 | chr12-68272156-INS-60 |
| chr12-68284369-INS-96 | chr12-68284369-INS-64 |
| chr12-68287183-INS-1841 | chr12-68287184-INS-1841 |
| chr12-68293921-INS-5316 | chr12-68293922-INS-5316 |
| chr12-68296986-INS-1411 | chr12-68296987-INS-1411 |
| chr12-68305238-INS-239 | chr12-68305238-INS-239 |
| chr12-68311578-INS-74 | chr12-68311579-INS-74 |
| chr12-68313698-INS-128 | chr12-68313698-INS-128 |
| chr12-68384640-INS-582 | chr12-68384640-INS-582 |
| chr12-68426837-INS-1736 | chr12-68426838-INS-1736 |
| chr12-68439556-INS-59 | chr12-68439556-INS-59 |
| chr12-68477335-INS-418 | chr12-68477336-INS-420 |
| chr12-68501748-INS-670 | chr12-68501748-INS-670 |
| chr12-68512318-INS-120 | chr12-68512318-INS-120 |
| chr12-68539411-INS-103 | chr12-68539409-INS-94 |
| chr12-68544313-INS-2217 | chr12-68544314-INS-2217 |
| chr12-68562922-INS-186 | chr12-68562922-INS-184 |
| chr12-68570115-INS-98 | chr12-68570115-INS-102 |
| chr12-68573893-INS-6213 | chr12-68573894-INS-6211 |
| chr12-68586407-INS-131 | chr12-68586407-INS-131 |

| | |
|---|---|
| chr12-68661497-INS-56 | chr12-68661498-INS-84 |
| chr12-68662894-INS-265 | chr12-68662894-INS-269 |
| chr12-68686376-INS-507 | chr12-68686376-INS-507 |
| chr12-68687645-INS-169 | chr12-68687645-INS-169 |
| chr12-68715729-INS-58 | chr12-68715717-INS-70 |
| chr12-68730613-INS-255 | chr12-68730613-INS-263 |
| chr12-68777890-INS-98 | chr12-68777890-INS-98 |
| chr12-68787628-INS-76 | chr12-68787628-INS-50 |
| chr12-68828432-INS-9346 | chr12-68828433-INS-9346 |
| chr12-68847005-INS-106 | chr12-68847005-INS-104 |
| chr12-68863052-INS-2767 | chr12-68863053-INS-2767 |
| chr12-68868030-INS-55 | chr12-68868030-INS-56 |
| chr12-68880713-INS-142 | chr12-68880713-INS-144 |
| chr12-68883867-INS-74 | chr12-68883879-INS-50 |
| chr12-68884549-INS-250 | chr12-68884549-INS-251 |
| chr12-68911965-INS-179 | chr12-68911965-INS-179 |
| chr12-68991420-INS-149 | chr12-68991420-INS-139 |
| chr12-68997874-INS-72 | chr12-68997874-INS-74 |
| chr12-69021648-INS-492 | chr12-69021649-INS-492 |
| chr12-69031649-INS-511 | chr12-69031649-INS-511 |
| chr12-69039591-INS-54 | chr12-69039591-INS-54 |
| chr12-69061176-INS-107 | chr12-69061172-INS-108 |
| chr12-69064726-INS-200 | chr12-69064726-INS-201 |
| chr12-69074797-INS-63 | chr12-69074797-INS-63 |
| chr12-69084462-INS-169 | chr12-69084462-INS-169 |
| chr12-69087933-INS-76 | chr12-69087933-INS-76 |
| chr12-69092737-INS-7444 | chr12-69092738-INS-7444 |
| chr12-69118065-INS-654 | chr12-69118075-INS-651 |
| chr12-69135639-INS-207 | chr12-69135639-INS-207 |
| chr12-69138512-INS-85 | chr12-69138513-INS-62 |
| chr12-69150293-INS-215 | chr12-69150293-INS-215 |
| chr12-69161636-INS-242 | chr12-69161605-INS-232 |
| chr12-69163251-INS-68 | chr12-69163251-INS-98 |
| chr12-69181270-INS-78 | chr12-69181271-INS-80 |
| chr12-69187441-INS-209 | chr12-69187441-INS-209 |
| chr12-69191514-INS-198 | chr12-69191515-INS-198 |
| chr12-69193762-INS-6778 | chr12-69193763-INS-6778 |
| chr12-69237390-INS-202 | chr12-69237399-INS-203 |
| chr12-69247346-INS-2431 | chr12-69247347-INS-2445 |
| chr12-69268492-INS-70 | chr12-69268496-INS-94 |
| chr12-69284074-INS-256 | chr12-69284074-INS-256 |
| chr12-69296285-INS-168 | chr12-69296285-INS-168 |
| chr12-69304911-INS-192 | chr12-69304911-INS-216 |
| chr12-69306467-INS-80 | chr12-69306467-INS-80 |
| chr12-69347671-INS-448 | chr12-69347671-INS-462 |
| chr12-69349976-INS-188 | chr12-69349976-INS-188 |
| chr12-69352309-INS-232 | chr12-69352309-INS-232 |
| chr12-69356153-INS-153 | chr12-69356153-INS-153 |
| chr12-69360259-INS-60 | chr12-69360259-INS-60 |
| chr12-69385729-INS-667 | chr12-69385728-INS-670 |
| chr12-69387429-INS-220 | chr12-69387429-INS-220 |
| chr12-69390058-INS-214 | chr12-69390058-INS-215 |
| chr12-69402993-INS-54 | chr12-69402993-INS-50 |
| chr12-69409312-INS-52 | chr12-69409312-INS-52 |
| chr12-69433073-INS-156 | chr12-69433107-INS-152 |
| chr12-69444429-INS-56 | chr12-69444429-INS-57 |
| chr12-69448498-INS-92 | chr12-69448493-INS-72 |
| chr12-69461924-INS-159 | chr12-69461924-INS-159 |
| chr12-69463055-INS-148 | chr12-69463055-INS-146 |
| chr12-69487811-INS-82 | chr12-69487811-INS-83 |
| chr12-69506585-INS-111 | chr12-69506585-INS-111 |
| chr12-69517184-INS-171 | chr12-69517184-INS-172 |
| chr12-69522900-INS-209 | chr12-69522900-INS-209 |
| chr12-69532219-INS-53 | chr12-69532219-INS-56 |
| chr12-69541783-INS-81 | chr12-69541783-INS-101 |
| chr12-69557051-INS-58 | chr12-69557051-INS-58 |
| chr12-69559633-INS-58 | chr12-69559633-INS-68 |
| chr12-69577337-INS-225 | chr12-69577337-INS-225 |
| chr12-69583461-INS-52 | chr12-69583461-INS-52 |
| chr12-69604107-INS-1194 | chr12-69604108-INS-1194 |
| chr12-69605821-INS-319 | chr12-69605821-INS-319 |
| chr12-69613497-INS-109 | chr12-69613497-INS-109 |
| chr12-69633925-INS-64 | chr12-69633926-INS-64 |
| chr12-69648552-INS-206 | chr12-69648552-INS-207 |
| chr12-69688653-INS-366 | chr12-69688653-INS-368 |
| chr12-69713022-INS-212 | chr12-69713023-INS-210 |
| chr12-69735067-INS-112 | chr12-69735067-INS-100 |

| | |
|---|---|
| chr12-69736363-INS-78 | chr12-69736363-INS-78 |
| chr12-69745004-INS-206 | chr12-69745004-INS-207 |
| chr12-69750973-INS-241 | chr12-69750973-INS-240 |
| chr12-69758228-INS-358 | chr12-69758228-INS-318 |
| chr12-69777850-INS-481 | chr12-69777850-INS-481 |
| chr12-69809256-INS-58 | chr12-69809256-INS-52 |
| chr12-69830421-INS-55 | chr12-69830385-INS-93 |
| chr12-69894502-INS-207 | chr12-69894502-INS-209 |
| chr12-69905629-INS-224 | chr12-69905629-INS-218 |
| chr12-69911092-INS-529 | chr12-69911092-INS-513 |
| chr12-69912832-INS-215 | chr12-69912832-INS-221 |
| chr12-69920999-INS-7122 | chr12-69921000-INS-7122 |
| chr12-69935022-INS-326 | chr12-69935022-INS-326 |
| chr12-69948065-INS-201 | chr12-69948065-INS-201 |
| chr12-69969737-INS-486 | chr12-69969737-INS-486 |
| chr12-69979309-INS-80 | chr12-69979309-INS-80 |
| chr12-69988521-INS-175 | chr12-69988521-INS-175 |
| chr12-70001874-INS-207 | chr12-70001874-INS-207 |
| chr12-70002818-INS-186 | chr12-70002819-INS-186 |
| chr12-70004649-INS-6454 | chr12-70004650-INS-6454 |
| chr12-70009972-INS-56 | chr12-70009972-INS-56 |
| chr12-70023509-INS-227 | chr12-70023509-INS-227 |
| chr12-70084335-INS-128 | chr12-70084326-INS-128 |
| chr12-70109984-INS-56 | chr12-70109984-INS-56 |
| chr12-70116699-INS-206 | chr12-70116699-INS-203 |
| chr12-70138983-INS-72 | chr12-70138983-INS-90 |
| chr12-70143904-INS-199 | chr12-70143904-INS-209 |
| chr12-70150750-INS-120 | chr12-70150750-INS-84 |
| chr12-70186765-INS-186 | chr12-70186795-INS-196 |
| chr12-70191769-INS-52 | chr12-70191769-INS-52 |
| chr12-70209449-INS-66 | chr12-70209449-INS-66 |
| chr12-70211760-INS-57 | chr12-70211760-INS-57 |
| chr12-70219423-INS-198 | chr12-70219423-INS-198 |
| chr12-70237228-INS-202 | chr12-70237228-INS-202 |
| chr12-70239802-INS-136 | chr12-70239802-INS-136 |
| chr12-70245526-INS-73 | chr12-70245526-INS-73 |
| chr12-70249002-INS-54 | chr12-70248973-INS-66 |
| chr12-70259903-INS-150 | chr12-70259903-INS-150 |
| chr12-70265087-INS-129 | chr12-70265087-INS-129 |
| chr12-70268944-INS-136 | chr12-70268944-INS-130 |
| chr12-70325214-INS-925 | chr12-70325214-INS-925 |
| chr12-70329806-INS-284 | chr12-70329806-INS-288 |
| chr12-70335151-INS-102 | chr12-70335151-INS-104 |
| chr12-70343828-INS-114 | chr12-70343828-INS-116 |
| chr12-70386850-INS-194 | chr12-70386850-INS-193 |
| chr12-70389518-INS-515 | chr12-70389518-INS-517 |
| chr12-70403752-INS-273 | chr12-70403752-INS-273 |
| chr12-70446924-INS-412 | chr12-70446925-INS-413 |
| chr12-70453270-INS-76 | chr12-70453270-INS-74 |
| chr12-70466288-INS-58 | chr12-70466285-INS-54 |
| chr12-70494742-INS-372 | chr12-70494742-INS-372 |
| chr12-70550489-INS-58 | chr12-70550489-INS-58 |
| chr12-70555822-INS-84 | chr12-70555822-INS-84 |
| chr12-70573665-INS-497 | chr12-70573665-INS-497 |
| chr12-70577794-INS-1443 | chr12-70577795-INS-1442 |
| chr12-70595674-INS-106 | chr12-70595710-INS-84 |
| chr12-70600351-INS-299 | chr12-70600351-INS-299 |
| chr12-70610029-INS-218 | chr12-70610029-INS-218 |
| chr12-70620841-INS-93 | chr12-70620846-INS-93 |
| chr12-70630417-INS-103 | chr12-70630417-INS-103 |
| chr12-70664585-INS-60 | chr12-70664585-INS-60 |
| chr12-70666095-INS-79 | chr12-70666107-INS-55 |
| chr12-70668422-INS-90 | chr12-70668422-INS-104 |
| chr12-70718262-INS-60 | chr12-70718262-INS-53 |
| chr12-70736179-INS-64 | chr12-70736179-INS-64 |
| chr12-70794622-INS-118 | chr12-70794622-INS-102 |
| chr12-70813410-INS-54 | chr12-70813410-INS-54 |
| chr12-70833085-INS-51 | chr12-70833085-INS-51 |
| chr12-70844645-INS-83 | chr12-70844638-INS-60 |
| chr12-70903151-INS-142 | chr12-70903151-INS-142 |
| chr12-70916016-INS-76 | chr12-70916017-INS-76 |
| chr12-70924539-INS-93 | chr12-70924541-INS-70 |
| chr12-70941939-INS-386 | chr12-70941939-INS-386 |
| chr12-70952607-INS-7134 | chr12-70952608-INS-7134 |
| chr12-70978655-INS-72 | chr12-70978655-INS-76 |
| chr12-70993800-INS-92 | chr12-70993802-INS-60 |
| chr12-70997594-INS-408 | chr12-70997594-INS-409 |

| | |
|---|---|
| chr12-71001359-INS-67 | chr12-71001359-INS-66 |
| chr12-71026960-INS-139 | chr12-71026960-INS-139 |
| chr12-71030817-INS-151 | chr12-71030817-INS-152 |
| chr12-71035011-INS-169 | chr12-71035011-INS-169 |
| chr12-71047420-INS-64 | chr12-71047420-INS-68 |
| chr12-71133609-INS-61 | chr12-71133609-INS-62 |
| chr12-71147400-INS-604 | chr12-71147400-INS-603 |
| chr12-71152339-INS-210 | chr12-71152339-INS-206 |
| chr12-71159997-INS-199 | chr12-71159997-INS-200 |
| chr12-71162048-INS-199 | chr12-71162048-INS-199 |
| chr12-71164014-INS-65 | chr12-71164015-INS-62 |
| chr12-71165206-INS-201 | chr12-71165206-INS-200 |
| chr12-71177133-INS-51 | chr12-71177133-INS-51 |
| chr12-71177518-INS-723 | chr12-71177518-INS-724 |
| chr12-71186523-INS-65 | chr12-71186523-INS-65 |
| chr12-71203186-INS-218 | chr12-71203186-INS-218 |
| chr12-71222491-INS-96 | chr12-71222491-INS-52 |
| chr12-71252931-INS-860 | chr12-71252931-INS-864 |
| chr12-71273164-INS-1303 | chr12-71273165-INS-1303 |
| chr12-71283869-INS-68 | chr12-71283869-INS-68 |
| chr12-71300517-INS-225 | chr12-71300517-INS-225 |
| chr12-71349343-INS-96 | chr12-71349343-INS-96 |
| chr12-71361225-INS-52 | chr12-71361225-INS-52 |
| chr12-71367565-INS-1346 | chr12-71367566-INS-1346 |
| chr12-71367974-INS-193 | chr12-71367974-INS-193 |
| chr12-71377216-INS-116 | chr12-71377216-INS-133 |
| chr12-71389129-INS-57 | chr12-71389129-INS-51 |
| chr12-71396899-INS-328 | chr12-71396899-INS-328 |
| chr12-71398042-INS-508 | chr12-71398042-INS-508 |
| chr12-71406101-INS-1327 | chr12-71406102-INS-1330 |
| chr12-71409750-INS-106 | chr12-71409765-INS-110 |
| chr12-71425376-INS-3272 | chr12-71425377-INS-3272 |
| chr12-71431666-INS-65 | chr12-71431657-INS-101 |
| chr12-71433640-INS-250 | chr12-71433640-INS-248 |
| chr12-71437439-INS-66 | chr12-71437439-INS-64 |
| chr12-71439196-INS-58 | chr12-71439243-INS-56 |
| chr12-71481680-INS-117 | chr12-71481680-INS-117 |
| chr12-71509837-INS-61 | chr12-71509837-INS-61 |
| chr12-71512736-INS-54 | chr12-71512736-INS-54 |
| chr12-71522047-INS-5548 | chr12-71522048-INS-5548 |
| chr12-71534231-INS-283 | chr12-71534231-INS-283 |
| chr12-71563679-INS-408 | chr12-71563679-INS-408 |
| chr12-71569036-INS-490 | chr12-71569036-INS-490 |
| chr12-71594299-INS-112 | chr12-71594300-INS-79 |
| chr12-71651578-INS-55 | chr12-71651578-INS-57 |
| chr12-71654083-INS-576 | chr12-71654083-INS-576 |
| chr12-71668228-INS-60 | chr12-71668228-INS-50 |
| chr12-71678715-INS-131 | chr12-71678718-INS-135 |
| chr12-71687704-INS-401 | chr12-71687704-INS-401 |
| chr12-71690795-INS-115 | chr12-71690790-INS-92 |
| chr12-71698681-INS-64 | chr12-71698681-INS-64 |
| chr12-71707594-INS-56 | chr12-71707594-INS-60 |
| chr12-71708508-INS-221 | chr12-71708508-INS-211 |
| chr12-71726046-INS-425 | chr12-71726046-INS-428 |
| chr12-71733525-INS-65 | chr12-71733525-INS-60 |
| chr12-71758210-INS-84 | chr12-71758210-INS-86 |
| chr12-71793671-INS-116 | chr12-71793666-INS-118 |
| chr12-71795394-INS-65 | chr12-71795399-INS-70 |
| chr12-71797109-INS-53 | chr12-71797080-INS-63 |
| chr12-71801340-INS-192 | chr12-71801340-INS-192 |
| chr12-71811836-INS-107 | chr12-71811836-INS-107 |
| chr12-71818811-INS-111 | chr12-71818811-INS-111 |
| chr12-71829801-INS-63 | chr12-71829801-INS-63 |
| chr12-71846494-INS-90 | chr12-71846494-INS-90 |
| chr12-71858580-INS-78 | chr12-71858581-INS-80 |
| chr12-71859091-INS-54 | chr12-71859091-INS-54 |
| chr12-71892819-INS-60 | chr12-71892819-INS-60 |
| chr12-71909189-INS-161 | chr12-71909189-INS-161 |
| chr12-71918011-INS-209 | chr12-71918011-INS-209 |
| chr12-71940871-INS-290 | chr12-71940871-INS-290 |
| chr12-71953733-INS-116 | chr12-71953733-INS-114 |
| chr12-71968839-INS-68 | chr12-71968840-INS-56 |
| chr12-71995315-INS-179 | chr12-71995315-INS-184 |
| chr12-72006335-INS-216 | chr12-72006335-INS-216 |
| chr12-72044563-INS-88 | chr12-72044564-INS-100 |
| chr12-72052817-INS-128 | chr12-72052820-INS-121 |
| chr12-72060146-INS-4758 | chr12-72060147-INS-4758 |

| | |
|---|---|
| chr12-72082033-INS-219 | chr12-72082033-INS-219 |
| chr12-72101745-INS-177 | chr12-72101745-INS-180 |
| chr12-72111546-INS-401 | chr12-72111546-INS-401 |
| chr12-72118584-INS-511 | chr12-72118584-INS-511 |
| chr12-72120169-INS-80 | chr12-72120186-INS-57 |
| chr12-72126131-INS-174 | chr12-72126131-INS-173 |
| chr12-72131038-INS-600 | chr12-72131038-INS-600 |
| chr12-72186312-INS-68 | chr12-72186312-INS-68 |
| chr12-72193238-INS-100 | chr12-72193238-INS-100 |
| chr12-72224909-INS-2054 | chr12-72224910-INS-2054 |
| chr12-72242510-INS-93 | chr12-72242503-INS-67 |
| chr12-72245435-INS-224 | chr12-72245435-INS-226 |
| chr12-72247488-INS-225 | chr12-72247488-INS-225 |
| chr12-72248803-INS-332 | chr12-72248803-INS-332 |
| chr12-72260939-INS-72 | chr12-72260939-INS-72 |
| chr12-72286922-INS-51 | chr12-72286922-INS-52 |
| chr12-72329691-INS-6176 | chr12-72329692-INS-6173 |
| chr12-72358312-INS-65 | chr12-72358312-INS-65 |
| chr12-72359137-INS-60 | chr12-72359137-INS-60 |
| chr12-72370325-INS-406 | chr12-72370325-INS-406 |
| chr12-72375518-INS-6475 | chr12-72375519-INS-6475 |
| chr12-72422446-INS-203 | chr12-72422446-INS-203 |
| chr12-72427714-INS-359 | chr12-72427714-INS-359 |
| chr12-72445235-INS-96 | chr12-72445236-INS-97 |
| chr12-72450072-INS-64 | chr12-72450075-INS-60 |
| chr12-72465531-INS-113 | chr12-72465537-INS-118 |
| chr12-72472903-INS-57 | chr12-72472903-INS-53 |
| chr12-72525686-INS-377 | chr12-72525686-INS-377 |
| chr12-72538740-INS-232 | chr12-72538723-INS-236 |
| chr12-72586554-INS-1645 | chr12-72586555-INS-1645 |
| chr12-72628015-INS-114 | chr12-72628029-INS-68 |
| chr12-72643759-INS-572 | chr12-72643759-INS-572 |
| chr12-72671090-INS-50 | chr12-72671090-INS-54 |
| chr12-72689510-INS-254 | chr12-72689510-INS-257 |
| chr12-72698620-INS-65 | chr12-72698595-INS-63 |
| chr12-72703606-INS-226 | chr12-72703609-INS-192 |
| chr12-72705393-INS-76 | chr12-72705380-INS-51 |
| chr12-72706563-INS-204 | chr12-72706563-INS-216 |
| chr12-72747330-INS-53 | chr12-72747330-INS-53 |
| chr12-72778382-INS-52 | chr12-72778382-INS-52 |
| chr12-72782262-INS-349 | chr12-72782262-INS-352 |
| chr12-72789120-INS-59 | chr12-72789120-INS-74 |
| chr12-72812270-INS-213 | chr12-72812270-INS-213 |
| chr12-72813063-INS-231 | chr12-72813063-INS-221 |
| chr12-72818930-INS-434 | chr12-72818930-INS-435 |
| chr12-72828295-INS-65 | chr12-72828295-INS-65 |
| chr12-72852003-INS-59 | chr12-72852003-INS-50 |
| chr12-72853584-INS-54 | chr12-72853584-INS-54 |
| chr12-72870243-INS-445 | chr12-72870243-INS-445 |
| chr12-72889223-INS-4756 | chr12-72889224-INS-4756 |
| chr12-72908340-INS-3235 | chr12-72908341-INS-3229 |
| chr12-72919836-INS-2004 | chr12-72919837-INS-2004 |
| chr12-72937599-INS-124 | chr12-72937599-INS-124 |
| chr12-72953515-INS-65 | chr12-72953515-INS-65 |
| chr12-72954269-INS-209 | chr12-72954269-INS-210 |
| chr12-73002086-INS-202 | chr12-73002086-INS-202 |
| chr12-73023273-INS-55 | chr12-73023256-INS-52 |
| chr12-73024319-INS-112 | chr12-73024319-INS-111 |
| chr12-73040827-INS-162 | chr12-73040823-INS-171 |
| chr12-73049151-INS-97 | chr12-73049151-INS-96 |
| chr12-73054548-INS-7110 | chr12-73054549-INS-7111 |
| chr12-73074017-INS-95 | chr12-73074017-INS-100 |
| chr12-73080084-INS-230 | chr12-73080084-INS-230 |
| chr12-73107095-INS-67 | chr12-73107095-INS-77 |
| chr12-73128275-INS-68 | chr12-73128276-INS-92 |
| chr12-73144820-INS-229 | chr12-73144820-INS-229 |
| chr12-73161502-INS-132 | chr12-73161505-INS-110 |
| chr12-73179386-INS-56 | chr12-73179386-INS-56 |
| chr12-73180377-INS-202 | chr12-73180377-INS-202 |
| chr12-73192737-INS-74 | chr12-73192737-INS-74 |
| chr12-73219746-INS-113 | chr12-73219746-INS-113 |
| chr12-73232823-INS-65 | chr12-73232823-INS-65 |
| chr12-73243602-INS-68 | chr12-73243602-INS-70 |
| chr12-73261414-INS-101 | chr12-73261414-INS-101 |
| chr12-73310335-INS-202 | chr12-73310335-INS-203 |
| chr12-73325152-INS-214 | chr12-73325152-INS-214 |
| chr12-73338531-INS-295 | chr12-73338531-INS-295 |

| | |
|---|---|
| chr12-73344800-INS-2665 | chr12-73344801-INS-2665 |
| chr12-73347658-INS-1283 | chr12-73347659-INS-1283 |
| chr12-73370134-INS-72 | chr12-73370134-INS-72 |
| chr12-73386941-INS-175 | chr12-73386938-INS-184 |
| chr12-73397983-INS-151 | chr12-73397983-INS-156 |
| chr12-73403150-INS-255 | chr12-73403183-INS-224 |
| chr12-73403923-INS-365 | chr12-73403923-INS-365 |
| chr12-73424497-INS-113 | chr12-73424498-INS-68 |
| chr12-73430948-INS-465 | chr12-73430949-INS-464 |
| chr12-73454035-INS-60 | chr12-73454035-INS-60 |
| chr12-73464917-INS-86 | chr12-73464935-INS-58 |
| chr12-73468311-INS-7092 | chr12-73468312-INS-7091 |
| chr12-73496313-INS-67 | chr12-73496313-INS-68 |
| chr12-73501004-INS-82 | chr12-73501004-INS-80 |
| chr12-73508660-INS-61 | chr12-73508661-INS-85 |
| chr12-73522225-INS-161 | chr12-73522225-INS-161 |
| chr12-73537900-INS-8985 | chr12-73537901-INS-8985 |
| chr12-73544622-INS-52 | chr12-73544622-INS-50 |
| chr12-73575074-INS-122 | chr12-73575074-INS-122 |
| chr12-73586683-INS-336 | chr12-73586683-INS-336 |
| chr12-73587781-INS-52 | chr12-73587781-INS-52 |
| chr12-73597111-INS-221 | chr12-73597111-INS-222 |
| chr12-73601503-INS-234 | chr12-73601503-INS-234 |
| chr12-73607173-INS-60 | chr12-73607173-INS-63 |
| chr12-73618249-INS-2883 | chr12-73618250-INS-2883 |
| chr12-73633630-INS-172 | chr12-73633630-INS-169 |
| chr12-73642583-INS-99 | chr12-73642565-INS-102 |
| chr12-73644127-INS-201 | chr12-73644127-INS-201 |
| chr12-73686590-INS-6403 | chr12-73686591-INS-6403 |
| chr12-73688278-INS-1321 | chr12-73688279-INS-1307 |
| chr12-73690508-INS-178 | chr12-73690508-INS-178 |
| chr12-73741260-INS-52 | chr12-73741260-INS-52 |
| chr12-73751805-INS-64 | chr12-73751805-INS-64 |
| chr12-73753905-INS-56 | chr12-73753905-INS-72 |
| chr12-73756353-INS-149 | chr12-73756353-INS-138 |
| chr12-73768394-INS-76 | chr12-73768394-INS-76 |
| chr12-73800708-INS-52 | chr12-73800708-INS-54 |
| chr12-73813501-INS-60 | chr12-73813499-INS-101 |
| chr12-73830729-INS-3576 | chr12-73830730-INS-3576 |
| chr12-73848626-INS-62 | chr12-73848626-INS-62 |
| chr12-73861616-INS-52 | chr12-73861617-INS-67 |
| chr12-73880768-INS-6758 | chr12-73880769-INS-6759 |
| chr12-73911160-INS-53 | chr12-73911160-INS-68 |
| chr12-73997072-INS-54 | chr12-73997072-INS-56 |
| chr12-73997818-INS-75 | chr12-73997818-INS-75 |
| chr12-74042511-INS-95 | chr12-74042511-INS-109 |
| chr12-74045597-INS-118 | chr12-74045597-INS-116 |
| chr12-74075285-INS-3063 | chr12-74075286-INS-3063 |
| chr12-74081380-INS-78 | chr12-74081380-INS-78 |
| chr12-74082865-INS-76 | chr12-74082865-INS-64 |
| chr12-74087088-INS-119 | chr12-74087088-INS-120 |
| chr12-74119091-INS-260 | chr12-74119095-INS-277 |
| chr12-74148922-INS-170 | chr12-74148892-INS-205 |
| chr12-74169936-INS-52 | chr12-74169936-INS-58 |
| chr12-74196023-INS-6642 | chr12-74196024-INS-6642 |
| chr12-74254088-INS-51 | chr12-74254115-INS-54 |
| chr12-74254679-INS-477 | chr12-74254679-INS-480 |
| chr12-74292604-INS-162 | chr12-74292604-INS-162 |
| chr12-74333978-INS-461 | chr12-74333980-INS-462 |
| chr12-74358311-INS-176 | chr12-74358289-INS-203 |
| chr12-74376175-INS-525 | chr12-74376175-INS-525 |
| chr12-74402999-INS-221 | chr12-74402999-INS-222 |
| chr12-74405422-INS-378 | chr12-74405422-INS-379 |
| chr12-74409065-INS-54 | chr12-74409065-INS-54 |
| chr12-74414042-INS-2321 | chr12-74414043-INS-2335 |
| chr12-74449546-INS-232 | chr12-74449533-INS-209 |
| chr12-74477140-INS-117 | chr12-74477141-INS-132 |
| chr12-74478855-INS-428 | chr12-74478835-INS-423 |
| chr12-74496290-INS-225 | chr12-74496290-INS-225 |
| chr12-74508363-INS-125 | chr12-74508363-INS-125 |
| chr12-74519035-INS-5060 | chr12-74519036-INS-5061 |
| chr12-74545689-INS-486 | chr12-74545689-INS-486 |
| chr12-74549166-INS-3657 | chr12-74549167-INS-3657 |
| chr12-74562551-INS-732 | chr12-74562551-INS-715 |
| chr12-74570276-INS-492 | chr12-74570276-INS-492 |
| chr12-74600567-INS-104 | chr12-74600567-INS-124 |
| chr12-74611575-INS-4594 | chr12-74611576-INS-4609 |

| | |
|---|---|
| chr12-74675427-INS-6669 | chr12-74675428-INS-6669 |
| chr12-74719983-INS-6685 | chr12-74719984-INS-6683 |
| chr12-74741955-INS-68 | chr12-74741955-INS-68 |
| chr12-74745658-INS-162 | chr12-74745658-INS-162 |
| chr12-74757801-INS-53 | chr12-74757801-INS-53 |
| chr12-74808629-INS-311 | chr12-74808629-INS-311 |
| chr12-74815536-INS-183 | chr12-74815536-INS-191 |
| chr12-74838796-INS-51 | chr12-74838786-INS-50 |
| chr12-74864233-INS-346 | chr12-74864233-INS-351 |
| chr12-74888157-INS-5046 | chr12-74888158-INS-5046 |
| chr12-74949517-INS-92 | chr12-74949515-INS-114 |
| chr12-75025201-INS-120 | chr12-75025201-INS-123 |
| chr12-75050222-INS-66 | chr12-75050222-INS-66 |
| chr12-75186910-INS-60 | chr12-75186910-INS-60 |
| chr12-75198487-INS-71 | chr12-75198487-INS-56 |
| chr12-75206314-INS-91 | chr12-75206308-INS-54 |
| chr12-75216337-INS-182 | chr12-75216337-INS-182 |
| chr12-75322655-INS-442 | chr12-75322655-INS-440 |
| chr12-75337232-INS-7119 | chr12-75337233-INS-7119 |
| chr12-75372285-INS-55 | chr12-75372285-INS-50 |
| chr12-75382537-INS-948 | chr12-75382537-INS-947 |
| chr12-75393780-INS-111 | chr12-75393780-INS-107 |
| chr12-75396658-INS-485 | chr12-75396658-INS-485 |
| chr12-75403233-INS-72 | chr12-75403233-INS-72 |
| chr12-75422360-INS-6444 | chr12-75422361-INS-6444 |
| chr12-75521706-INS-54 | chr12-75521706-INS-66 |
| chr12-75538796-INS-498 | chr12-75538796-INS-498 |
| chr12-75562661-INS-181 | chr12-75562661-INS-182 |
| chr12-75644997-INS-207 | chr12-75644997-INS-207 |
| chr12-75669134-INS-6401 | chr12-75669135-INS-6401 |
| chr12-75742511-INS-52 | chr12-75742494-INS-52 |
| chr12-75792354-INS-202 | chr12-75792354-INS-203 |
| chr12-75804383-INS-147 | chr12-75804383-INS-147 |
| chr12-75825924-INS-165 | chr12-75825924-INS-170 |
| chr12-75828542-INS-56 | chr12-75828542-INS-68 |
| chr12-75838186-INS-192 | chr12-75838186-INS-192 |
| chr12-75847456-INS-222 | chr12-75847456-INS-216 |
| chr12-75850313-INS-194 | chr12-75850313-INS-195 |
| chr12-75904840-INS-50 | chr12-75904840-INS-55 |
| chr12-75937484-INS-874 | chr12-75937484-INS-871 |
| chr12-75942616-INS-311 | chr12-75942600-INS-315 |
| chr12-75947581-INS-600 | chr12-75947584-INS-587 |
| chr12-75986319-INS-235 | chr12-75986319-INS-232 |
| chr12-75990997-INS-97 | chr12-75990997-INS-96 |
| chr12-76020665-INS-198 | chr12-76020665-INS-198 |
| chr12-76024349-INS-160 | chr12-76024349-INS-162 |
| chr12-76025540-INS-219 | chr12-76025540-INS-219 |
| chr12-76042265-INS-60 | chr12-76042265-INS-60 |
| chr12-76090729-INS-60 | chr12-76090729-INS-60 |
| chr12-76105262-INS-66 | chr12-76105262-INS-66 |
| chr12-76114244-INS-174 | chr12-76114244-INS-176 |
| chr12-76119287-INS-65 | chr12-76119287-INS-105 |
| chr12-76138474-INS-108 | chr12-76138474-INS-113 |
| chr12-76140654-INS-54 | chr12-76140654-INS-54 |
| chr12-76148318-INS-65 | chr12-76148319-INS-97 |
| chr12-76150220-INS-273 | chr12-76150220-INS-273 |
| chr12-76154504-INS-78 | chr12-76154504-INS-78 |
| chr12-76159656-INS-116 | chr12-76159656-INS-116 |
| chr12-76190278-INS-101 | chr12-76190279-INS-101 |
| chr12-76195715-INS-403 | chr12-76195726-INS-387 |
| chr12-76199975-INS-6453 | chr12-76199976-INS-6453 |
| chr12-76224624-INS-54 | chr12-76224617-INS-70 |
| chr12-76234038-INS-52 | chr12-76234038-INS-52 |
| chr12-76258686-INS-117 | chr12-76258686-INS-117 |
| chr12-76261924-INS-86 | chr12-76261924-INS-90 |
| chr12-76264076-INS-222 | chr12-76264076-INS-222 |
| chr12-76282833-INS-205 | chr12-76282833-INS-205 |
| chr12-76284548-INS-286 | chr12-76284548-INS-286 |
| chr12-76294007-INS-54 | chr12-76294007-INS-56 |
| chr12-76326604-INS-203 | chr12-76326604-INS-203 |
| chr12-76333559-INS-53 | chr12-76333559-INS-54 |
| chr12-76343007-INS-921 | chr12-76343007-INS-871 |
| chr12-76364676-INS-57 | chr12-76364676-INS-57 |
| chr12-76369484-INS-57 | chr12-76369484-INS-57 |
| chr12-76381250-INS-80 | chr12-76381254-INS-84 |
| chr12-76401311-INS-56 | chr12-76401311-INS-68 |
| chr12-76413508-INS-197 | chr12-76413508-INS-197 |

| | |
|---|---|
| chr12-76456152-INS-204 | chr12-76456152-INS-204 |
| chr12-76463571-INS-56 | chr12-76463571-INS-64 |
| chr12-76471620-INS-164 | chr12-76471620-INS-168 |
| chr12-76484148-INS-1210 | chr12-76484149-INS-1234 |
| chr12-76493893-INS-7138 | chr12-76493894-INS-7139 |
| chr12-76501779-INS-76 | chr12-76501771-INS-112 |
| chr12-76509439-INS-204 | chr12-76509439-INS-206 |
| chr12-76522681-INS-202 | chr12-76522681-INS-202 |
| chr12-76524088-INS-202 | chr12-76524086-INS-201 |
| chr12-76531826-INS-293 | chr12-76531826-INS-293 |
| chr12-76544164-INS-68 | chr12-76544164-INS-86 |
| chr12-76545393-INS-87 | chr12-76545404-INS-78 |
| chr12-76551708-INS-208 | chr12-76551708-INS-206 |
| chr12-76558041-INS-114 | chr12-76558042-INS-118 |
| chr12-76565154-INS-99 | chr12-76565157-INS-67 |
| chr12-76566913-INS-77 | chr12-76566902-INS-118 |
| chr12-76568092-INS-117 | chr12-76568089-INS-133 |
| chr12-76579425-INS-198 | chr12-76579425-INS-198 |
| chr12-76632172-INS-392 | chr12-76632172-INS-392 |
| chr12-76654447-INS-50 | chr12-76654447-INS-50 |
| chr12-76672699-INS-75 | chr12-76672699-INS-75 |
| chr12-76679743-INS-140 | chr12-76679743-INS-140 |
| chr12-76715189-INS-84 | chr12-76715153-INS-90 |
| chr12-76720575-INS-132 | chr12-76720575-INS-132 |
| chr12-76729037-INS-100 | chr12-76729037-INS-100 |
| chr12-76741299-INS-109 | chr12-76741299-INS-130 |
| chr12-76743657-INS-90 | chr12-76743657-INS-90 |
| chr12-76768161-INS-187 | chr12-76768156-INS-192 |
| chr12-76778500-INS-53 | chr12-76778500-INS-59 |
| chr12-76786155-INS-4311 | chr12-76786156-INS-4311 |
| chr12-76810788-INS-63 | chr12-76810806-INS-63 |
| chr12-76816728-INS-896 | chr12-76816728-INS-896 |
| chr12-76821470-INS-67 | chr12-76821470-INS-50 |
| chr12-76827029-INS-207 | chr12-76827029-INS-207 |
| chr12-76832385-INS-207 | chr12-76832385-INS-206 |
| chr12-76871597-INS-220 | chr12-76871597-INS-220 |
| chr12-76876223-INS-6474 | chr12-76876224-INS-6474 |
| chr12-76907963-INS-230 | chr12-76907963-INS-229 |
| chr12-76914927-INS-216 | chr12-76914927-INS-216 |
| chr12-76924898-INS-92 | chr12-76924898-INS-100 |
| chr12-76929961-INS-97 | chr12-76929961-INS-98 |
| chr12-76960064-INS-203 | chr12-76960064-INS-203 |
| chr12-76969722-INS-190 | chr12-76969722-INS-190 |
| chr12-76976219-INS-166 | chr12-76976219-INS-198 |
| chr12-76995046-INS-72 | chr12-76995046-INS-72 |
| chr12-77015160-INS-200 | chr12-77015160-INS-200 |
| chr12-77032716-INS-56 | chr12-77032717-INS-90 |
| chr12-77034205-INS-63 | chr12-77034206-INS-62 |
| chr12-77046047-INS-79 | chr12-77046019-INS-98 |
| chr12-77063514-INS-278 | chr12-77063514-INS-278 |
| chr12-77073714-INS-51 | chr12-77073715-INS-57 |
| chr12-77083189-INS-764 | chr12-77083208-INS-773 |
| chr12-77104953-INS-87 | chr12-77104934-INS-75 |
| chr12-77118833-INS-60 | chr12-77118833-INS-60 |
| chr12-77120785-INS-140 | chr12-77120785-INS-142 |
| chr12-77123361-INS-338 | chr12-77123361-INS-338 |
| chr12-77138231-INS-74 | chr12-77138231-INS-122 |
| chr12-77159815-INS-152 | chr12-77159815-INS-153 |
| chr12-77172705-INS-112 | chr12-77172702-INS-125 |
| chr12-77207416-INS-347 | chr12-77207416-INS-346 |
| chr12-77228426-INS-190 | chr12-77228426-INS-166 |
| chr12-77243837-INS-78 | chr12-77243837-INS-78 |
| chr12-77262670-INS-134 | chr12-77262670-INS-135 |
| chr12-77266781-INS-52 | chr12-77266781-INS-52 |
| chr12-77267180-INS-396 | chr12-77267180-INS-397 |
| chr12-77293697-INS-98 | chr12-77293697-INS-98 |
| chr12-77310936-INS-101 | chr12-77310936-INS-101 |
| chr12-77327099-INS-1158 | chr12-77327100-INS-1158 |
| chr12-77332336-INS-161 | chr12-77332336-INS-171 |
| chr12-77336704-INS-134 | chr12-77336704-INS-94 |
| chr12-77344599-INS-837 | chr12-77344599-INS-838 |
| chr12-77352776-INS-114 | chr12-77352776-INS-124 |
| chr12-77381432-INS-1436 | chr12-77381433-INS-1436 |
| chr12-77407974-INS-6570 | chr12-77407975-INS-6570 |
| chr12-77451437-INS-68 | chr12-77451437-INS-58 |
| chr12-77471427-INS-107 | chr12-77471419-INS-96 |
| chr12-77486378-INS-114 | chr12-77486378-INS-114 |

| | |
|---|---|
| chr12-77519639-INS-303 | chr12-77519639-INS-299 |
| chr12-77521638-INS-50 | chr12-77521616-INS-84 |
| chr12-77539553-INS-81 | chr12-77539555-INS-80 |
| chr12-77567411-INS-67 | chr12-77567411-INS-67 |
| chr12-77576758-INS-60 | chr12-77576760-INS-64 |
| chr12-77579984-INS-55 | chr12-77579984-INS-55 |
| chr12-77616634-INS-58 | chr12-77616634-INS-58 |
| chr12-77617701-INS-116 | chr12-77617701-INS-144 |
| chr12-77642475-INS-63 | chr12-77642475-INS-63 |
| chr12-77656841-INS-1263 | chr12-77656842-INS-1263 |
| chr12-77667329-INS-71 | chr12-77667329-INS-58 |
| chr12-77674332-INS-105 | chr12-77674332-INS-89 |
| chr12-77696954-INS-968 | chr12-77696954-INS-962 |
| chr12-77722958-INS-90 | chr12-77722958-INS-114 |
| chr12-77732854-INS-2645 | chr12-77732855-INS-2644 |
| chr12-77757220-INS-57 | chr12-77757220-INS-64 |
| chr12-77778968-INS-60 | chr12-77778968-INS-60 |
| chr12-77791186-INS-66 | chr12-77791186-INS-114 |
| chr12-77824713-INS-630 | chr12-77824713-INS-639 |
| chr12-77842548-INS-401 | chr12-77842548-INS-401 |
| chr12-77845148-INS-84 | chr12-77845148-INS-84 |
| chr12-77846964-INS-76 | chr12-77846966-INS-76 |
| chr12-77880366-INS-106 | chr12-77880366-INS-154 |
| chr12-77908290-INS-4553 | chr12-77908291-INS-4553 |
| chr12-77924915-INS-1669 | chr12-77924916-INS-1669 |
| chr12-77925867-INS-214 | chr12-77925867-INS-214 |
| chr12-77941911-INS-56 | chr12-77941911-INS-50 |
| chr12-77944423-INS-72 | chr12-77944423-INS-75 |
| chr12-77947254-INS-809 | chr12-77947254-INS-824 |
| chr12-77951243-INS-56 | chr12-77951243-INS-53 |
| chr12-77959431-INS-78 | chr12-77959431-INS-91 |
| chr12-77963901-INS-212 | chr12-77963901-INS-212 |
| chr12-77969311-INS-203 | chr12-77969311-INS-203 |
| chr12-77995054-INS-130 | chr12-77995047-INS-138 |
| chr12-78035815-INS-6800 | chr12-78035816-INS-6800 |
| chr12-78042269-INS-51 | chr12-78042269-INS-51 |
| chr12-78060932-INS-90 | chr12-78060932-INS-90 |
| chr12-78069051-INS-78 | chr12-78069051-INS-56 |
| chr12-78070569-INS-51 | chr12-78070569-INS-67 |
| chr12-78080153-INS-110 | chr12-78080153-INS-135 |
| chr12-78092493-INS-52 | chr12-78092493-INS-60 |
| chr12-78107496-INS-6163 | chr12-78107496-INS-6120 |
| chr12-78118715-INS-1762 | chr12-78118716-INS-1762 |
| chr12-78122235-INS-60 | chr12-78122235-INS-60 |
| chr12-78137521-INS-6468 | chr12-78137522-INS-6468 |
| chr12-78140962-INS-6629 | chr12-78140963-INS-6629 |
| chr12-78146623-INS-221 | chr12-78146623-INS-220 |
| chr12-78183315-INS-56 | chr12-78183315-INS-50 |
| chr12-78205536-INS-98 | chr12-78205536-INS-108 |
| chr12-78216557-INS-6569 | chr12-78216558-INS-6569 |
| chr12-78267615-INS-52 | chr12-78267590-INS-68 |
| chr12-78272489-INS-105 | chr12-78272494-INS-141 |
| chr12-78292400-INS-61 | chr12-78292400-INS-51 |
| chr12-78304956-INS-356 | chr12-78304953-INS-333 |
| chr12-78313710-INS-6363 | chr12-78313711-INS-6363 |
| chr12-78356609-INS-71 | chr12-78356609-INS-71 |
| chr12-78363421-INS-1738 | chr12-78363422-INS-1736 |
| chr12-78417306-INS-1288 | chr12-78417307-INS-1288 |
| chr12-78447172-INS-3653 | chr12-78447173-INS-3653 |
| chr12-78493753-INS-1763 | chr12-78493754-INS-1764 |
| chr12-78499899-INS-519 | chr12-78499899-INS-520 |
| chr12-78507428-INS-2879 | chr12-78507429-INS-2881 |
| chr12-78527311-INS-53 | chr12-78527311-INS-53 |
| chr12-78546951-INS-745 | chr12-78546951-INS-745 |
| chr12-78549535-INS-1263 | chr12-78549536-INS-1264 |
| chr12-78647603-INS-1502 | chr12-78647604-INS-1515 |
| chr12-78689095-INS-504 | chr12-78689095-INS-504 |
| chr12-78692543-INS-82 | chr12-78692562-INS-50 |
| chr12-78708766-INS-552 | chr12-78708800-INS-526 |
| chr12-78709430-INS-114 | chr12-78709430-INS-114 |
| chr12-78736802-INS-110 | chr12-78736802-INS-110 |
| chr12-78763395-INS-66 | chr12-78763395-INS-66 |
| chr12-78770097-INS-60 | chr12-78770097-INS-64 |
| chr12-78782183-INS-461 | chr12-78782183-INS-461 |
| chr12-78827443-INS-396 | chr12-78827443-INS-396 |
| chr12-78841712-INS-70 | chr12-78841712-INS-70 |
| chr12-78850917-INS-1822 | chr12-78850918-INS-1822 |

| | |
|---|---|
| chr12-78895917-INS-62 | chr12-78895917-INS-64 |
| chr12-78900225-INS-53 | chr12-78900219-INS-69 |
| chr12-78903751-INS-199 | chr12-78903751-INS-199 |
| chr12-78935217-INS-195 | chr12-78935217-INS-191 |
| chr12-78939519-INS-54 | chr12-78939519-INS-66 |
| chr12-78949217-INS-502 | chr12-78949217-INS-502 |
| chr12-78955618-INS-209 | chr12-78955618-INS-208 |
| chr12-78960751-INS-201 | chr12-78960751-INS-200 |
| chr12-78966305-INS-57 | chr12-78966305-INS-58 |
| chr12-78973178-INS-305 | chr12-78973178-INS-305 |
| chr12-78976372-INS-211 | chr12-78976372-INS-211 |
| chr12-78978039-INS-102 | chr12-78978039-INS-92 |
| chr12-78986091-INS-459 | chr12-78986091-INS-458 |
| chr12-78995000-INS-64 | chr12-78995013-INS-67 |
| chr12-78998644-INS-62 | chr12-78998644-INS-62 |
| chr12-79020622-INS-74 | chr12-79020618-INS-58 |
| chr12-79022138-INS-57 | chr12-79022138-INS-57 |
| chr12-79026769-INS-197 | chr12-79026769-INS-197 |
| chr12-79038897-INS-521 | chr12-79038897-INS-515 |
| chr12-79040251-INS-74 | chr12-79040251-INS-74 |
| chr12-79060595-INS-254 | chr12-79060595-INS-254 |
| chr12-79066485-INS-226 | chr12-79066485-INS-227 |
| chr12-79075246-INS-75 | chr12-79075246-INS-100 |
| chr12-79087516-INS-86 | chr12-79087514-INS-74 |
| chr12-79094466-INS-244 | chr12-79094466-INS-244 |
| chr12-79110037-INS-58 | chr12-79110037-INS-58 |
| chr12-79122539-INS-211 | chr12-79122539-INS-218 |
| chr12-79138982-INS-203 | chr12-79138982-INS-203 |
| chr12-79148555-INS-136 | chr12-79148559-INS-108 |
| chr12-79156701-INS-239 | chr12-79156701-INS-239 |
| chr12-79163039-INS-2271 | chr12-79163040-INS-2271 |
| chr12-79163350-INS-339 | chr12-79163350-INS-339 |
| chr12-79181153-INS-144 | chr12-79181153-INS-144 |
| chr12-79184596-INS-1110 | chr12-79184597-INS-1110 |
| chr12-79194148-INS-215 | chr12-79194148-INS-215 |
| chr12-79199365-INS-213 | chr12-79199365-INS-212 |
| chr12-79217724-INS-183 | chr12-79217724-INS-183 |
| chr12-79219835-INS-289 | chr12-79219835-INS-288 |
| chr12-79222324-INS-874 | chr12-79222324-INS-872 |
| chr12-79232780-INS-263 | chr12-79232780-INS-263 |
| chr12-79240712-INS-2365 | chr12-79240713-INS-2384 |
| chr12-79242136-INS-4797 | chr12-79242137-INS-4797 |
| chr12-79249072-INS-183 | chr12-79249072-INS-182 |
| chr12-79251431-INS-198 | chr12-79251431-INS-199 |
| chr12-79258999-INS-91 | chr12-79259019-INS-70 |
| chr12-79278813-INS-98 | chr12-79278820-INS-94 |
| chr12-79289937-INS-58 | chr12-79289937-INS-58 |
| chr12-79292380-INS-199 | chr12-79292380-INS-200 |
| chr12-79294780-INS-83 | chr12-79294780-INS-93 |
| chr12-79300066-INS-60 | chr12-79300066-INS-60 |
| chr12-79302160-INS-185 | chr12-79302160-INS-186 |
| chr12-79324414-INS-53 | chr12-79324414-INS-52 |
| chr12-79328692-INS-113 | chr12-79328670-INS-136 |
| chr12-79346715-INS-76 | chr12-79346715-INS-88 |
| chr12-79364995-INS-77 | chr12-79364991-INS-105 |
| chr12-79368682-INS-60 | chr12-79368682-INS-60 |
| chr12-79369749-INS-766 | chr12-79369749-INS-767 |
| chr12-79372957-INS-192 | chr12-79372957-INS-193 |
| chr12-79378347-INS-911 | chr12-79378347-INS-910 |
| chr12-79385726-INS-57 | chr12-79385776-INS-57 |
| chr12-79414225-INS-51 | chr12-79414225-INS-51 |
| chr12-79420431-INS-7181 | chr12-79420432-INS-7181 |
| chr12-79429583-INS-91 | chr12-79429585-INS-89 |
| chr12-79434413-INS-175 | chr12-79434413-INS-175 |
| chr12-79439883-INS-83 | chr12-79439883-INS-81 |
| chr12-79455245-INS-216 | chr12-79455245-INS-216 |
| chr12-79458425-INS-133 | chr12-79458425-INS-133 |
| chr12-79481206-INS-118 | chr12-79481206-INS-76 |
| chr12-79507608-INS-60 | chr12-79507608-INS-88 |
| chr12-79509939-INS-1145 | chr12-79509940-INS-1145 |
| chr12-79517199-INS-189 | chr12-79517199-INS-189 |
| chr12-79518169-INS-6808 | chr12-79518170-INS-6808 |
| chr12-79544446-INS-51 | chr12-79544446-INS-51 |
| chr12-79565940-INS-59 | chr12-79565940-INS-54 |
| chr12-79592717-INS-219 | chr12-79592717-INS-219 |
| chr12-79618671-INS-135 | chr12-79618671-INS-135 |
| chr12-79623744-INS-4724 | chr12-79623745-INS-4724 |

| | |
|---|---|
| chr12-79629906-INS-111 | chr12-79629904-INS-114 |
| chr12-79650080-INS-4998 | chr12-79650081-INS-4998 |
| chr12-79662131-INS-62 | chr12-79662131-INS-67 |
| chr12-79680097-INS-72 | chr12-79680097-INS-72 |
| chr12-79686583-INS-158 | chr12-79686613-INS-168 |
| chr12-79691820-INS-769 | chr12-79691820-INS-769 |
| chr12-79695221-INS-52 | chr12-79695221-INS-54 |
| chr12-79698389-INS-82 | chr12-79698389-INS-82 |
| chr12-79761318-INS-112 | chr12-79761318-INS-112 |
| chr12-79802909-INS-54 | chr12-79802909-INS-64 |
| chr12-79804229-INS-58 | chr12-79804229-INS-56 |
| chr12-79808631-INS-399 | chr12-79808631-INS-399 |
| chr12-79840050-INS-114 | chr12-79840050-INS-114 |
| chr12-79866050-INS-112 | chr12-79866050-INS-114 |
| chr12-79871623-INS-108 | chr12-79871623-INS-108 |
| chr12-79878777-INS-210 | chr12-79878776-INS-196 |
| chr12-79899798-INS-58 | chr12-79899798-INS-54 |
| chr12-79919360-INS-80 | chr12-79919360-INS-80 |
| chr12-79947594-INS-65 | chr12-79947594-INS-69 |
| chr12-79948941-INS-67 | chr12-79948941-INS-69 |
| chr12-79966551-INS-211 | chr12-79966551-INS-216 |
| chr12-79977901-INS-1695 | chr12-79977902-INS-1699 |
| chr12-80029445-INS-83 | chr12-80029445-INS-102 |
| chr12-80062283-INS-67 | chr12-80062283-INS-69 |
| chr12-80076725-INS-104 | chr12-80076725-INS-148 |
| chr12-80084028-INS-786 | chr12-80084028-INS-785 |
| chr12-80086259-INS-59 | chr12-80086260-INS-63 |
| chr12-80123041-INS-72 | chr12-80123041-INS-62 |
| chr12-80137291-INS-70 | chr12-80137291-INS-52 |
| chr12-80141154-INS-55 | chr12-80141154-INS-55 |
| chr12-80144820-INS-141 | chr12-80144822-INS-140 |
| chr12-80148019-INS-828 | chr12-80148019-INS-828 |
| chr12-80152827-INS-60 | chr12-80152827-INS-64 |
| chr12-80171991-INS-56 | chr12-80171991-INS-62 |
| chr12-80180350-INS-129 | chr12-80180350-INS-129 |
| chr12-80209390-INS-456 | chr12-80209391-INS-456 |
| chr12-80246868-INS-208 | chr12-80246868-INS-208 |
| chr12-80261439-INS-317 | chr12-80261462-INS-336 |
| chr12-80272298-INS-212 | chr12-80272298-INS-211 |
| chr12-80284515-INS-791 | chr12-80284515-INS-787 |
| chr12-80300642-INS-230 | chr12-80300642-INS-229 |
| chr12-80338728-INS-90 | chr12-80338728-INS-96 |
| chr12-80354823-INS-336 | chr12-80354823-INS-336 |
| chr12-80356701-INS-52 | chr12-80356701-INS-56 |
| chr12-80389269-INS-209 | chr12-80389269-INS-207 |
| chr12-80399740-INS-74 | chr12-80399740-INS-74 |
| chr12-80410416-INS-221 | chr12-80410416-INS-221 |
| chr12-80418536-INS-421 | chr12-80418536-INS-425 |
| chr12-80439469-INS-212 | chr12-80439469-INS-212 |
| chr12-80440707-INS-200 | chr12-80440707-INS-200 |
| chr12-80443623-INS-181 | chr12-80443623-INS-170 |
| chr12-80457017-INS-216 | chr12-80457017-INS-216 |
| chr12-80512682-INS-243 | chr12-80512682-INS-243 |
| chr12-80518361-INS-95 | chr12-80518361-INS-95 |
| chr12-80529670-INS-1524 | chr12-80529671-INS-1525 |
| chr12-80553320-INS-214 | chr12-80553320-INS-215 |
| chr12-80586150-INS-63 | chr12-80586150-INS-63 |
| chr12-80616014-INS-224 | chr12-80616014-INS-224 |
| chr12-80623931-INS-53 | chr12-80623931-INS-59 |
| chr12-80628127-INS-60 | chr12-80628127-INS-50 |
| chr12-80640028-INS-62 | chr12-80640034-INS-50 |
| chr12-80658578-INS-212 | chr12-80658578-INS-212 |
| chr12-80668853-INS-217 | chr12-80668853-INS-217 |
| chr12-80676414-INS-206 | chr12-80676414-INS-206 |
| chr12-80712702-INS-202 | chr12-80712702-INS-202 |
| chr12-80734919-INS-53 | chr12-80734919-INS-53 |
| chr12-80737598-INS-201 | chr12-80737598-INS-202 |
| chr12-80753455-INS-57 | chr12-80753455-INS-57 |
| chr12-80756350-INS-312 | chr12-80756350-INS-312 |
| chr12-80782576-INS-294 | chr12-80782577-INS-294 |
| chr12-80785407-INS-58 | chr12-80785407-INS-61 |
| chr12-80792349-INS-52 | chr12-80792349-INS-56 |
| chr12-80812211-INS-81 | chr12-80812211-INS-81 |
| chr12-80844475-INS-3388 | chr12-80844476-INS-3388 |
| chr12-80916558-INS-184 | chr12-80916558-INS-185 |
| chr12-80918668-INS-137 | chr12-80918668-INS-137 |
| chr12-80931393-INS-52 | chr12-80931393-INS-50 |

| | |
|---|---|
| chr12-80933327-INS-188 | chr12-80933327-INS-188 |
| chr12-80935453-INS-148 | chr12-80935453-INS-148 |
| chr12-80999206-INS-222 | chr12-80999206-INS-222 |
| chr12-81009266-INS-92 | chr12-81009266-INS-92 |
| chr12-81066655-INS-109 | chr12-81066655-INS-104 |
| chr12-81092703-INS-51 | chr12-81092706-INS-55 |
| chr12-81107248-INS-60 | chr12-81107248-INS-60 |
| chr12-81126395-INS-176 | chr12-81126388-INS-154 |
| chr12-81138180-INS-456 | chr12-81138180-INS-442 |
| chr12-81143677-INS-52 | chr12-81143677-INS-52 |
| chr12-81145834-INS-100 | chr12-81145874-INS-76 |
| chr12-81159910-INS-80 | chr12-81159910-INS-52 |
| chr12-81160500-INS-54 | chr12-81160501-INS-99 |
| chr12-81172207-INS-56 | chr12-81172207-INS-54 |
| chr12-81193541-INS-64 | chr12-81193541-INS-64 |
| chr12-81208981-INS-59 | chr12-81208981-INS-59 |
| chr12-81237897-INS-51 | chr12-81237897-INS-51 |
| chr12-81238769-INS-4704 | chr12-81238770-INS-4704 |
| chr12-81266766-INS-116 | chr12-81266764-INS-118 |
| chr12-81270054-INS-88 | chr12-81270054-INS-88 |
| chr12-81271091-INS-62 | chr12-81271091-INS-62 |
| chr12-81331347-INS-58 | chr12-81331347-INS-58 |
| chr12-81421134-INS-63 | chr12-81421134-INS-57 |
| chr12-81477077-INS-2907 | chr12-81477078-INS-2907 |
| chr12-81499638-INS-7559 | chr12-81499639-INS-7559 |
| chr12-81530590-INS-67 | chr12-81530590-INS-67 |
| chr12-81580277-INS-62 | chr12-81580277-INS-84 |
| chr12-81601721-INS-104 | chr12-81601721-INS-104 |
| chr12-81601923-INS-8897 | chr12-81601924-INS-8901 |
| chr12-81656504-INS-80 | chr12-81656504-INS-71 |
| chr12-81662154-INS-71 | chr12-81662157-INS-88 |
| chr12-81697327-INS-105 | chr12-81697327-INS-102 |
| chr12-81730970-INS-173 | chr12-81730970-INS-173 |
| chr12-81747651-INS-401 | chr12-81747651-INS-401 |
| chr12-81758621-INS-54 | chr12-81758621-INS-60 |
| chr12-81760863-INS-54 | chr12-81760878-INS-54 |
| chr12-81797526-INS-839 | chr12-81797527-INS-883 |
| chr12-81798034-INS-54 | chr12-81798034-INS-60 |
| chr12-81820875-INS-160 | chr12-81820875-INS-160 |
| chr12-81830055-INS-69 | chr12-81830054-INS-79 |
| chr12-81895161-INS-434 | chr12-81895161-INS-459 |
| chr12-81897769-INS-248 | chr12-81897769-INS-249 |
| chr12-81903138-INS-79 | chr12-81903138-INS-79 |
| chr12-81913332-INS-70 | chr12-81913332-INS-69 |
| chr12-81917181-INS-600 | chr12-81917170-INS-601 |
| chr12-81929061-INS-197 | chr12-81929061-INS-197 |
| chr12-81932089-INS-5722 | chr12-81932090-INS-5722 |
| chr12-81939481-INS-52 | chr12-81939481-INS-52 |
| chr12-82035836-INS-75 | chr12-82035836-INS-75 |
| chr12-82059824-INS-146 | chr12-82059824-INS-146 |
| chr12-82123327-INS-115 | chr12-82123327-INS-115 |
| chr12-82143119-INS-64 | chr12-82143115-INS-53 |
| chr12-82145104-INS-381 | chr12-82145104-INS-381 |
| chr12-82159396-INS-4119 | chr12-82159397-INS-4137 |
| chr12-82162137-INS-253 | chr12-82162137-INS-253 |
| chr12-82167596-INS-94 | chr12-82167596-INS-94 |
| chr12-82168707-INS-370 | chr12-82168707-INS-370 |
| chr12-82173817-INS-75 | chr12-82173817-INS-75 |
| chr12-82176775-INS-51 | chr12-82176775-INS-51 |
| chr12-82190364-INS-6494 | chr12-82190365-INS-6494 |
| chr12-82236689-INS-174 | chr12-82236690-INS-176 |
| chr12-82257930-INS-86 | chr12-82257930-INS-131 |
| chr12-82261306-INS-96 | chr12-82261306-INS-96 |
| chr12-82274874-INS-215 | chr12-82274874-INS-216 |
| chr12-82286312-INS-76 | chr12-82286334-INS-68 |
| chr12-82307733-INS-54 | chr12-82307733-INS-54 |
| chr12-82331901-INS-90 | chr12-82331901-INS-135 |
| chr12-82337382-INS-65 | chr12-82337382-INS-61 |
| chr12-82338644-INS-63 | chr12-82338644-INS-63 |
| chr12-82358713-INS-101 | chr12-82358713-INS-101 |
| chr12-82411548-INS-1639 | chr12-82411549-INS-1639 |
| chr12-82419993-INS-396 | chr12-82419993-INS-386 |
| chr12-82427238-INS-58 | chr12-82427238-INS-58 |
| chr12-82437467-INS-212 | chr12-82437467-INS-212 |
| chr12-82438406-INS-65 | chr12-82438406-INS-60 |
| chr12-82446971-INS-70 | chr12-82446971-INS-70 |
| chr12-82451832-INS-59 | chr12-82451832-INS-73 |

| | |
|---|---|
| chr12-82464883-INS-70 | chr12-82464883-INS-70 |
| chr12-82472317-INS-177 | chr12-82472317-INS-177 |
| chr12-82500772-INS-72 | chr12-82500772-INS-72 |
| chr12-82521062-INS-7116 | chr12-82521063-INS-7118 |
| chr12-82524153-INS-90 | chr12-82524198-INS-52 |
| chr12-82555346-INS-1854 | chr12-82555347-INS-1854 |
| chr12-82561367-INS-633 | chr12-82561367-INS-633 |
| chr12-82564422-INS-511 | chr12-82564422-INS-511 |
| chr12-82610741-INS-51 | chr12-82610741-INS-51 |
| chr12-82628652-INS-980 | chr12-82628652-INS-975 |
| chr12-82636503-INS-149 | chr12-82636503-INS-154 |
| chr12-82670448-INS-179 | chr12-82670482-INS-159 |
| chr12-82689711-INS-51 | chr12-82689711-INS-51 |
| chr12-82691202-INS-64 | chr12-82691202-INS-64 |
| chr12-82694367-INS-119 | chr12-82694367-INS-107 |
| chr12-82723634-INS-108 | chr12-82723644-INS-125 |
| chr12-82736204-INS-135 | chr12-82736204-INS-135 |
| chr12-82738509-INS-74 | chr12-82738511-INS-75 |
| chr12-82760483-INS-140 | chr12-82760483-INS-140 |
| chr12-82786001-INS-54 | chr12-82786001-INS-56 |
| chr12-82808306-INS-976 | chr12-82808306-INS-975 |
| chr12-82852918-INS-3303 | chr12-82852919-INS-3303 |
| chr12-82855979-INS-2376 | chr12-82855980-INS-2377 |
| chr12-82915211-INS-646 | chr12-82915211-INS-646 |
| chr12-82929861-INS-64 | chr12-82929856-INS-58 |
| chr12-82945194-INS-96 | chr12-82945194-INS-100 |
| chr12-82952952-INS-72 | chr12-82952952-INS-56 |
| chr12-82963235-INS-480 | chr12-82963235-INS-480 |
| chr12-82974807-INS-76 | chr12-82974807-INS-76 |
| chr12-82988056-INS-199 | chr12-82988057-INS-199 |
| chr12-82998622-INS-225 | chr12-82998622-INS-230 |
| chr12-83013354-INS-496 | chr12-83013354-INS-496 |
| chr12-83058971-INS-55 | chr12-83058972-INS-79 |
| chr12-83061649-INS-92 | chr12-83061684-INS-51 |
| chr12-83076600-INS-320 | chr12-83076600-INS-320 |
| chr12-83078574-INS-65 | chr12-83078574-INS-65 |
| chr12-83101190-INS-64 | chr12-83101186-INS-68 |
| chr12-83102780-INS-277 | chr12-83102781-INS-278 |
| chr12-83109088-INS-124 | chr12-83109088-INS-122 |
| chr12-83109638-INS-71 | chr12-83109629-INS-68 |
| chr12-83130290-INS-209 | chr12-83130290-INS-209 |
| chr12-83165712-INS-66 | chr12-83165712-INS-60 |
| chr12-83236712-INS-66 | chr12-83236712-INS-68 |
| chr12-83258694-INS-162 | chr12-83258692-INS-162 |
| chr12-83286094-INS-231 | chr12-83286094-INS-231 |
| chr12-83296259-INS-906 | chr12-83296259-INS-906 |
| chr12-83299016-INS-190 | chr12-83299052-INS-154 |
| chr12-83326420-INS-162 | chr12-83326420-INS-162 |
| chr12-83369856-INS-67 | chr12-83369856-INS-67 |
| chr12-83375111-INS-52 | chr12-83375111-INS-54 |
| chr12-83376695-INS-83 | chr12-83376695-INS-89 |
| chr12-83410909-INS-96 | chr12-83410909-INS-78 |
| chr12-83414795-INS-147 | chr12-83414795-INS-147 |
| chr12-83425172-INS-251 | chr12-83425174-INS-295 |
| chr12-83477932-INS-68 | chr12-83477906-INS-114 |
| chr12-83497477-INS-60 | chr12-83497477-INS-60 |
| chr12-83513905-INS-71 | chr12-83513904-INS-74 |
| chr12-83545591-INS-206 | chr12-83545591-INS-206 |
| chr12-83554032-INS-86 | chr12-83554032-INS-81 |
| chr12-83560843-INS-116 | chr12-83560812-INS-136 |
| chr12-83564237-INS-202 | chr12-83564237-INS-221 |
| chr12-83565119-INS-322 | chr12-83565113-INS-323 |
| chr12-83570713-INS-6463 | chr12-83570714-INS-6463 |
| chr12-83573569-INS-201 | chr12-83573569-INS-201 |
| chr12-83583732-INS-78 | chr12-83583734-INS-78 |
| chr12-83613270-INS-207 | chr12-83613270-INS-207 |
| chr12-83617306-INS-51 | chr12-83617306-INS-51 |
| chr12-83620071-INS-234 | chr12-83620071-INS-234 |
| chr12-83623135-INS-182 | chr12-83623135-INS-182 |
| chr12-83628349-INS-198 | chr12-83628349-INS-198 |
| chr12-83638332-INS-266 | chr12-83638332-INS-266 |
| chr12-83655211-INS-54 | chr12-83655211-INS-54 |
| chr12-83661443-INS-57 | chr12-83661443-INS-58 |
| chr12-83682925-INS-407 | chr12-83682925-INS-407 |
| chr12-83688394-INS-192 | chr12-83688394-INS-193 |
| chr12-83696455-INS-54 | chr12-83696455-INS-52 |
| chr12-83700911-INS-221 | chr12-83700911-INS-220 |

| | |
|---|---|
| chr12-83715404-INS-60 | chr12-83715404-INS-64 |
| chr12-83733366-INS-200 | chr12-83733366-INS-203 |
| chr12-83736412-INS-196 | chr12-83736412-INS-200 |
| chr12-83743525-INS-4786 | chr12-83743526-INS-4786 |
| chr12-83774403-INS-108 | chr12-83774403-INS-108 |
| chr12-83789491-INS-224 | chr12-83789491-INS-224 |
| chr12-83802829-INS-185 | chr12-83802829-INS-219 |
| chr12-83806477-INS-197 | chr12-83806477-INS-197 |
| chr12-83807910-INS-137 | chr12-83807910-INS-137 |
| chr12-83826858-INS-77 | chr12-83826858-INS-51 |
| chr12-83838080-INS-520 | chr12-83838080-INS-524 |
| chr12-83855140-INS-60 | chr12-83855140-INS-60 |
| chr12-83865377-INS-1045 | chr12-83865378-INS-1046 |
| chr12-83882258-INS-207 | chr12-83882258-INS-208 |
| chr12-83899681-INS-143 | chr12-83899681-INS-143 |
| chr12-83905504-INS-114 | chr12-83905504-INS-114 |
| chr12-83908127-INS-539 | chr12-83908127-INS-538 |
| chr12-83943505-INS-611 | chr12-83943505-INS-619 |
| chr12-83943523-INS-86 | chr12-83943530-INS-60 |
| chr12-83950417-INS-109 | chr12-83950417-INS-109 |
| chr12-83964913-INS-220 | chr12-83964913-INS-231 |
| chr12-83980118-INS-179 | chr12-83980118-INS-200 |
| chr12-83985502-INS-118 | chr12-83985502-INS-102 |
| chr12-84018226-INS-366 | chr12-84018226-INS-366 |
| chr12-84024059-INS-95 | chr12-84024059-INS-90 |
| chr12-84032894-INS-492 | chr12-84032894-INS-493 |
| chr12-84033541-INS-106 | chr12-84033541-INS-98 |
| chr12-84039155-INS-409 | chr12-84039127-INS-451 |
| chr12-84040013-INS-54 | chr12-84040019-INS-59 |
| chr12-84066103-INS-205 | chr12-84066103-INS-206 |
| chr12-84096383-INS-3863 | chr12-84096384-INS-3863 |
| chr12-84099253-INS-88 | chr12-84099253-INS-52 |
| chr12-84102685-INS-185 | chr12-84102685-INS-185 |
| chr12-84107362-INS-502 | chr12-84107359-INS-512 |
| chr12-84126337-INS-263 | chr12-84126337-INS-263 |
| chr12-84135508-INS-6866 | chr12-84135509-INS-6866 |
| chr12-84145041-INS-200 | chr12-84145041-INS-201 |
| chr12-84174227-INS-220 | chr12-84174230-INS-220 |
| chr12-84180430-INS-51 | chr12-84180430-INS-51 |
| chr12-84183495-INS-60 | chr12-84183495-INS-60 |
| chr12-84259088-INS-51 | chr12-84259067-INS-53 |
| chr12-84302448-INS-208 | chr12-84302448-INS-208 |
| chr12-84309891-INS-212 | chr12-84309886-INS-221 |
| chr12-84347279-INS-218 | chr12-84347279-INS-219 |
| chr12-84353935-INS-199 | chr12-84353935-INS-199 |
| chr12-84358083-INS-79 | chr12-84358083-INS-73 |
| chr12-84402498-INS-116 | chr12-84402498-INS-116 |
| chr12-84440008-INS-248 | chr12-84440031-INS-272 |
| chr12-84441618-INS-368 | chr12-84441628-INS-371 |
| chr12-84465859-INS-626 | chr12-84465860-INS-626 |
| chr12-84491641-INS-199 | chr12-84491641-INS-195 |
| chr12-84492676-INS-127 | chr12-84492676-INS-127 |
| chr12-84508960-INS-124 | chr12-84508968-INS-135 |
| chr12-84525437-INS-493 | chr12-84525436-INS-492 |
| chr12-84543687-INS-174 | chr12-84543687-INS-174 |
| chr12-84545476-INS-221 | chr12-84545476-INS-221 |
| chr12-84549834-INS-213 | chr12-84549834-INS-213 |
| chr12-84565445-INS-68 | chr12-84565443-INS-70 |
| chr12-84574488-INS-65 | chr12-84574488-INS-111 |
| chr12-84583656-INS-194 | chr12-84583656-INS-194 |
| chr12-84601416-INS-56 | chr12-84601409-INS-53 |
| chr12-84622281-INS-66 | chr12-84622251-INS-70 |
| chr12-84635733-INS-171 | chr12-84635733-INS-182 |
| chr12-84643438-INS-60 | chr12-84643438-INS-60 |
| chr12-84648011-INS-64 | chr12-84648011-INS-63 |
| chr12-84657354-INS-264 | chr12-84657354-INS-264 |
| chr12-84693566-INS-216 | chr12-84693566-INS-217 |
| chr12-84694220-INS-628 | chr12-84694220-INS-624 |
| chr12-84709127-INS-215 | chr12-84709127-INS-215 |
| chr12-84731889-INS-237 | chr12-84731889-INS-233 |
| chr12-84736401-INS-6413 | chr12-84736402-INS-6413 |
| chr12-84737999-INS-93 | chr12-84738022-INS-92 |
| chr12-84739317-INS-205 | chr12-84739317-INS-205 |
| chr12-84746455-INS-61 | chr12-84746455-INS-61 |
| chr12-84757710-INS-214 | chr12-84757710-INS-214 |
| chr12-84764077-INS-87 | chr12-84764077-INS-73 |
| chr12-84789290-INS-402 | chr12-84789290-INS-402 |

| | |
|---|---|
| chr12-84795827-INS-60 | chr12-84795827-INS-60 |
| chr12-84796697-INS-217 | chr12-84796698-INS-206 |
| chr12-84804925-INS-400 | chr12-84804925-INS-401 |
| chr12-84808717-INS-202 | chr12-84808717-INS-207 |
| chr12-84815410-INS-207 | chr12-84815413-INS-184 |
| chr12-84818272-INS-116 | chr12-84818272-INS-116 |
| chr12-84874888-INS-393 | chr12-84874888-INS-345 |
| chr12-84897053-INS-439 | chr12-84897053-INS-439 |
| chr12-84900150-INS-116 | chr12-84900150-INS-116 |
| chr12-84906947-INS-68 | chr12-84906947-INS-71 |
| chr12-84911197-INS-55 | chr12-84911197-INS-62 |
| chr12-84913582-INS-210 | chr12-84913590-INS-179 |
| chr12-84923948-INS-5254 | chr12-84923949-INS-5254 |
| chr12-84926330-INS-78 | chr12-84926330-INS-62 |
| chr12-84933566-INS-78 | chr12-84933566-INS-66 |
| chr12-85017138-INS-56 | chr12-85017114-INS-66 |
| chr12-85094094-INS-253 | chr12-85094094-INS-237 |
| chr12-85139015-INS-63 | chr12-85139015-INS-63 |
| chr12-85149108-INS-134 | chr12-85149108-INS-132 |
| chr12-85154424-INS-68 | chr12-85154424-INS-68 |
| chr12-85177583-INS-124 | chr12-85177583-INS-124 |
| chr12-85184472-INS-122 | chr12-85184472-INS-116 |
| chr12-85198990-INS-76 | chr12-85198990-INS-50 |
| chr12-85200180-INS-72 | chr12-85200180-INS-52 |
| chr12-85230436-INS-117 | chr12-85230436-INS-116 |
| chr12-85252223-INS-6413 | chr12-85252224-INS-6413 |
| chr12-85256207-INS-131 | chr12-85256210-INS-142 |
| chr12-85263559-INS-156 | chr12-85263569-INS-182 |
| chr12-85268844-INS-60 | chr12-85268841-INS-60 |
| chr12-85273781-INS-74 | chr12-85273781-INS-74 |
| chr12-85288589-INS-114 | chr12-85288589-INS-64 |
| chr12-85325048-INS-107 | chr12-85325048-INS-96 |
| chr12-85335267-INS-108 | chr12-85335267-INS-72 |
| chr12-85341032-INS-123 | chr12-85341032-INS-123 |
| chr12-85387248-INS-4076 | chr12-85387249-INS-4066 |
| chr12-85389422-INS-75 | chr12-85389422-INS-93 |
| chr12-85406269-INS-99 | chr12-85406269-INS-99 |
| chr12-85416864-INS-178 | chr12-85416864-INS-178 |
| chr12-85428155-INS-60 | chr12-85428155-INS-60 |
| chr12-85429335-INS-498 | chr12-85429335-INS-498 |
| chr12-85433118-INS-96 | chr12-85433120-INS-67 |
| chr12-85438158-INS-177 | chr12-85438158-INS-176 |
| chr12-85442474-INS-64 | chr12-85442474-INS-68 |
| chr12-85473469-INS-224 | chr12-85473469-INS-226 |
| chr12-85481971-INS-154 | chr12-85481971-INS-154 |
| chr12-85504296-INS-56 | chr12-85504296-INS-52 |
| chr12-85512814-INS-68 | chr12-85512814-INS-84 |
| chr12-85546209-INS-67 | chr12-85546209-INS-98 |
| chr12-85551606-INS-101 | chr12-85551618-INS-75 |
| chr12-85581630-INS-52 | chr12-85581630-INS-52 |
| chr12-85638669-INS-64 | chr12-85638684-INS-51 |
| chr12-85673328-INS-78 | chr12-85673328-INS-56 |
| chr12-85694069-INS-80 | chr12-85694069-INS-84 |
| chr12-85696464-INS-1333 | chr12-85696469-INS-1363 |
| chr12-85757976-INS-60 | chr12-85757976-INS-58 |
| chr12-85777101-INS-190 | chr12-85777101-INS-187 |
| chr12-85787112-INS-150 | chr12-85787112-INS-155 |
| chr12-85789498-INS-71 | chr12-85789498-INS-71 |
| chr12-85817892-INS-123 | chr12-85817893-INS-76 |
| chr12-85853974-INS-498 | chr12-85853974-INS-498 |
| chr12-85857031-INS-164 | chr12-85857031-INS-132 |
| chr12-85864173-INS-60 | chr12-85864173-INS-60 |
| chr12-85947212-INS-140 | chr12-85947212-INS-140 |
| chr12-85995498-INS-2878 | chr12-85995499-INS-2878 |
| chr12-86054969-INS-848 | chr12-86054969-INS-848 |
| chr12-86077376-INS-51 | chr12-86077375-INS-71 |
| chr12-86081797-INS-204 | chr12-86081797-INS-204 |
| chr12-86118978-INS-193 | chr12-86118978-INS-192 |
| chr12-86119817-INS-199 | chr12-86119817-INS-199 |
| chr12-86170182-INS-68 | chr12-86170182-INS-68 |
| chr12-86171752-INS-82 | chr12-86171752-INS-82 |
| chr12-86175165-INS-57 | chr12-86175165-INS-57 |
| chr12-86185370-INS-54 | chr12-86185370-INS-54 |
| chr12-86212442-INS-7088 | chr12-86212443-INS-7088 |
| chr12-86216603-INS-184 | chr12-86216603-INS-184 |
| chr12-86232553-INS-82 | chr12-86232573-INS-62 |
| chr12-86251148-INS-70 | chr12-86251148-INS-52 |

| | |
|---|---|
| chr12-86268147-INS-68 | chr12-86268147-INS-68 |
| chr12-86284362-INS-103 | chr12-86284362-INS-113 |
| chr12-86322549-INS-452 | chr12-86322549-INS-451 |
| chr12-86329743-INS-190 | chr12-86329743-INS-189 |
| chr12-86346514-INS-403 | chr12-86346514-INS-403 |
| chr12-86351480-INS-136 | chr12-86351480-INS-135 |
| chr12-86354688-INS-181 | chr12-86354687-INS-195 |
| chr12-86366766-INS-72 | chr12-86366751-INS-72 |
| chr12-86372891-INS-626 | chr12-86372889-INS-626 |
| chr12-86374703-INS-293 | chr12-86374703-INS-300 |
| chr12-86376069-INS-67 | chr12-86376069-INS-81 |
| chr12-86378698-INS-93 | chr12-86378698-INS-93 |
| chr12-86382102-INS-55 | chr12-86382102-INS-55 |
| chr12-86383655-INS-57 | chr12-86383659-INS-57 |
| chr12-86388049-INS-76 | chr12-86388010-INS-76 |
| chr12-86400159-INS-60 | chr12-86400159-INS-60 |
| chr12-86428695-INS-69 | chr12-86428695-INS-69 |
| chr12-86447140-INS-277 | chr12-86447140-INS-276 |
| chr12-86458474-INS-200 | chr12-86458474-INS-200 |
| chr12-86491294-INS-208 | chr12-86491307-INS-201 |
| chr12-86510672-INS-198 | chr12-86510672-INS-200 |
| chr12-86526543-INS-204 | chr12-86526543-INS-204 |
| chr12-86529841-INS-8679 | chr12-86529842-INS-8680 |
| chr12-86556325-INS-401 | chr12-86556325-INS-397 |
| chr12-86575588-INS-498 | chr12-86575588-INS-494 |
| chr12-86580399-INS-67 | chr12-86580399-INS-67 |
| chr12-86589852-INS-178 | chr12-86589852-INS-181 |
| chr12-86593524-INS-65 | chr12-86593524-INS-65 |
| chr12-86599810-INS-138 | chr12-86599853-INS-131 |
| chr12-86603820-INS-63 | chr12-86603820-INS-61 |
| chr12-86610270-INS-122 | chr12-86610270-INS-106 |
| chr12-86667681-INS-432 | chr12-86667681-INS-432 |
| chr12-86673044-INS-52 | chr12-86673044-INS-52 |
| chr12-86673826-INS-314 | chr12-86673826-INS-314 |
| chr12-86680451-INS-745 | chr12-86680451-INS-753 |
| chr12-86697822-INS-73 | chr12-86697822-INS-73 |
| chr12-86721718-INS-2911 | chr12-86721719-INS-2911 |
| chr12-86724968-INS-54 | chr12-86724968-INS-58 |
| chr12-86791632-INS-52 | chr12-86791632-INS-54 |
| chr12-86812327-INS-208 | chr12-86812327-INS-208 |
| chr12-86824913-INS-55 | chr12-86824913-INS-55 |
| chr12-86827990-INS-78 | chr12-86827990-INS-78 |
| chr12-86828918-INS-182 | chr12-86828918-INS-183 |
| chr12-86834831-INS-54 | chr12-86834831-INS-51 |
| chr12-86836597-INS-207 | chr12-86836597-INS-207 |
| chr12-86857110-INS-52 | chr12-86857110-INS-56 |
| chr12-86888816-INS-211 | chr12-86888816-INS-211 |
| chr12-86901975-INS-213 | chr12-86901975-INS-213 |
| chr12-86912728-INS-100 | chr12-86912728-INS-52 |
| chr12-86938090-INS-104 | chr12-86938101-INS-88 |
| chr12-86945901-INS-147 | chr12-86945901-INS-133 |
| chr12-86954411-INS-141 | chr12-86954417-INS-117 |
| chr12-86966673-INS-374 | chr12-86966673-INS-374 |
| chr12-86970101-INS-3083 | chr12-86970102-INS-3083 |
| chr12-86975777-INS-84 | chr12-86975777-INS-84 |
| chr12-87019537-INS-740 | chr12-87019537-INS-740 |
| chr12-87023823-INS-203 | chr12-87023823-INS-203 |
| chr12-87048933-INS-425 | chr12-87048944-INS-426 |
| chr12-87087274-INS-52 | chr12-87087274-INS-57 |
| chr12-87092063-INS-98 | chr12-87092061-INS-68 |
| chr12-87092449-INS-497 | chr12-87092449-INS-497 |
| chr12-87095259-INS-331 | chr12-87095259-INS-329 |
| chr12-87120937-INS-698 | chr12-87120936-INS-703 |
| chr12-87120971-INS-167 | chr12-87120971-INS-168 |
| chr12-87147333-INS-99 | chr12-87147354-INS-67 |
| chr12-87148345-INS-52 | chr12-87148346-INS-52 |
| chr12-87160162-INS-74 | chr12-87160162-INS-78 |
| chr12-87173614-INS-363 | chr12-87173614-INS-363 |
| chr12-87252872-INS-55 | chr12-87252871-INS-50 |
| chr12-87256269-INS-364 | chr12-87256270-INS-364 |
| chr12-87257756-INS-207 | chr12-87257756-INS-206 |
| chr12-87275893-INS-53 | chr12-87275893-INS-53 |
| chr12-87278906-INS-1070 | chr12-87278907-INS-1082 |
| chr12-87283876-INS-205 | chr12-87283876-INS-205 |
| chr12-87294562-INS-226 | chr12-87294562-INS-218 |
| chr12-87298111-INS-209 | chr12-87298111-INS-209 |
| chr12-87309565-INS-222 | chr12-87309565-INS-220 |

| | |
|---|---|
| chr12-87340509-INS-58 | chr12-87340470-INS-52 |
| chr12-87341403-INS-864 | chr12-87341447-INS-815 |
| chr12-87366546-INS-216 | chr12-87366546-INS-216 |
| chr12-87368405-INS-209 | chr12-87368405-INS-209 |
| chr12-87369671-INS-81 | chr12-87369671-INS-81 |
| chr12-87379018-INS-115 | chr12-87379018-INS-131 |
| chr12-87411106-INS-254 | chr12-87411106-INS-254 |
| chr12-87412850-INS-203 | chr12-87412850-INS-202 |
| chr12-87443241-INS-182 | chr12-87443241-INS-182 |
| chr12-87447531-INS-5185 | chr12-87447532-INS-5185 |
| chr12-87452001-INS-64 | chr12-87452001-INS-64 |
| chr12-87458121-INS-208 | chr12-87458121-INS-210 |
| chr12-87459677-INS-1673 | chr12-87459683-INS-1683 |
| chr12-87469531-INS-163 | chr12-87469532-INS-118 |
| chr12-87471807-INS-57 | chr12-87471782-INS-60 |
| chr12-87482021-INS-52 | chr12-87482021-INS-62 |
| chr12-87489620-INS-51 | chr12-87489620-INS-51 |
| chr12-87514969-INS-207 | chr12-87514969-INS-207 |
| chr12-87531333-INS-104 | chr12-87531333-INS-104 |
| chr12-87540098-INS-377 | chr12-87540098-INS-377 |
| chr12-87610960-INS-200 | chr12-87610960-INS-200 |
| chr12-87620832-INS-570 | chr12-87620832-INS-570 |
| chr12-87666030-INS-64 | chr12-87666026-INS-60 |
| chr12-87922500-INS-58 | chr12-87922500-INS-58 |
| chr12-87994165-INS-64 | chr12-87994165-INS-64 |
| chr12-88010966-INS-55 | chr12-88010959-INS-58 |
| chr12-88012970-INS-171 | chr12-88012970-INS-171 |
| chr12-88042911-INS-3482 | chr12-88042912-INS-3481 |
| chr12-88093629-INS-109 | chr12-88093632-INS-109 |
| chr12-88101745-INS-104 | chr12-88101744-INS-107 |
| chr12-88208391-INS-3094 | chr12-88208392-INS-3093 |
| chr12-88213860-INS-80 | chr12-88213860-INS-80 |
| chr12-88281838-INS-700 | chr12-88281844-INS-699 |
| chr12-88298491-INS-60 | chr12-88298491-INS-60 |
| chr12-88318555-INS-4682 | chr12-88318556-INS-4682 |
| chr12-88332766-INS-139 | chr12-88332768-INS-140 |
| chr12-88349287-INS-80 | chr12-88349287-INS-80 |
| chr12-88353571-INS-135 | chr12-88353571-INS-131 |
| chr12-88366274-INS-426 | chr12-88366274-INS-422 |
| chr12-88369629-INS-52 | chr12-88369629-INS-54 |
| chr12-88398600-INS-1889 | chr12-88398601-INS-1889 |
| chr12-88415183-INS-351 | chr12-88415183-INS-351 |
| chr12-88437193-INS-6170 | chr12-88437194-INS-6170 |
| chr12-88440511-INS-96 | chr12-88440508-INS-110 |
| chr12-88458827-INS-92 | chr12-88458843-INS-58 |
| chr12-88463765-INS-51 | chr12-88463764-INS-53 |
| chr12-88475323-INS-3085 | chr12-88475324-INS-3085 |
| chr12-88490046-INS-68 | chr12-88490046-INS-68 |
| chr12-88503380-INS-63 | chr12-88503380-INS-50 |
| chr12-88504953-INS-528 | chr12-88504953-INS-528 |
| chr12-88507580-INS-196 | chr12-88507580-INS-196 |
| chr12-88526581-INS-116 | chr12-88526580-INS-120 |
| chr12-88533734-INS-7586 | chr12-88533735-INS-7585 |
| chr12-88588972-INS-87 | chr12-88588972-INS-101 |
| chr12-88592170-INS-52 | chr12-88592152-INS-56 |
| chr12-88597017-INS-6030 | chr12-88597018-INS-6030 |
| chr12-88599394-INS-70 | chr12-88599394-INS-72 |
| chr12-88616244-INS-3780 | chr12-88616245-INS-3779 |
| chr12-88616958-INS-172 | chr12-88616958-INS-173 |
| chr12-88632005-INS-1017 | chr12-88632006-INS-1017 |
| chr12-88676277-INS-52 | chr12-88676277-INS-52 |
| chr12-88712053-INS-68 | chr12-88712057-INS-104 |
| chr12-88723548-INS-6437 | chr12-88723548-INS-6438 |
| chr12-88742498-INS-52 | chr12-88742498-INS-52 |
| chr12-88752801-INS-103 | chr12-88752801-INS-103 |
| chr12-88777302-INS-51 | chr12-88777302-INS-51 |
| chr12-88791901-INS-1960 | chr12-88791902-INS-1960 |
| chr12-88794851-INS-111 | chr12-88794851-INS-111 |
| chr12-88815760-INS-84 | chr12-88815765-INS-85 |
| chr12-88857065-INS-84 | chr12-88857065-INS-84 |
| chr12-88912268-INS-3916 | chr12-88912269-INS-3919 |
| chr12-88913318-INS-2563 | chr12-88913319-INS-2546 |
| chr12-88945042-INS-528 | chr12-88945042-INS-528 |
| chr12-88985414-INS-95 | chr12-88985408-INS-113 |
| chr12-88994255-INS-91 | chr12-88994255-INS-91 |
| chr12-89004340-INS-55 | chr12-89004359-INS-70 |
| chr12-89007480-INS-85 | chr12-89007480-INS-85 |

| | |
|---|---|
| chr12-89037280-INS-1893 | chr12-89037281-INS-1893 |
| chr12-89055277-INS-221 | chr12-89055278-INS-212 |
| chr12-89065515-INS-222 | chr12-89065515-INS-222 |
| chr12-89072008-INS-7136 | chr12-89072009-INS-7136 |
| chr12-89080965-INS-147 | chr12-89080965-INS-144 |
| chr12-89083559-INS-95 | chr12-89083605-INS-74 |
| chr12-89122207-INS-6685 | chr12-89122208-INS-6685 |
| chr12-89137888-INS-92 | chr12-89137888-INS-84 |
| chr12-89167177-INS-1054 | chr12-89167178-INS-1054 |
| chr12-89183641-INS-503 | chr12-89183641-INS-503 |
| chr12-89200960-INS-82 | chr12-89200960-INS-82 |
| chr12-89211945-INS-133 | chr12-89211946-INS-137 |
| chr12-89220979-INS-6030 | chr12-89220980-INS-6030 |
| chr12-89272245-INS-2186 | chr12-89272246-INS-2188 |
| chr12-89365122-INS-4483 | chr12-89365126-INS-4473 |
| chr12-89375286-INS-539 | chr12-89375286-INS-543 |
| chr12-89389101-INS-249 | chr12-89389101-INS-249 |
| chr12-89396408-INS-239 | chr12-89396408-INS-227 |
| chr12-89397938-INS-135 | chr12-89397938-INS-135 |
| chr12-89414520-INS-161 | chr12-89414520-INS-162 |
| chr12-89419408-INS-7072 | chr12-89419409-INS-7072 |
| chr12-89431500-INS-791 | chr12-89431500-INS-775 |
| chr12-89457046-INS-64 | chr12-89457046-INS-64 |
| chr12-89471326-INS-155 | chr12-89471326-INS-155 |
| chr12-89494564-INS-7186 | chr12-89494565-INS-7186 |
| chr12-89506148-INS-635 | chr12-89506148-INS-635 |
| chr12-89525333-INS-200 | chr12-89525333-INS-200 |
| chr12-89556607-INS-156 | chr12-89556619-INS-121 |
| chr12-89585586-INS-120 | chr12-89585578-INS-141 |
| chr12-89593734-INS-54 | chr12-89593734-INS-80 |
| chr12-89596451-INS-60 | chr12-89596450-INS-108 |
| chr12-89605763-INS-93 | chr12-89605754-INS-114 |
| chr12-89607499-INS-80 | chr12-89607504-INS-68 |
| chr12-89617044-INS-100 | chr12-89617044-INS-100 |
| chr12-89633522-INS-54 | chr12-89633524-INS-60 |
| chr12-89642695-INS-234 | chr12-89642695-INS-234 |
| chr12-89664794-INS-76 | chr12-89664790-INS-82 |
| chr12-89668949-INS-200 | chr12-89668949-INS-200 |
| chr12-89707583-INS-52 | chr12-89707583-INS-52 |
| chr12-89718584-INS-80 | chr12-89718630-INS-81 |
| chr12-89724965-INS-103 | chr12-89724965-INS-105 |
| chr12-89794094-INS-188 | chr12-89794094-INS-188 |
| chr12-89803336-INS-342 | chr12-89803336-INS-342 |
| chr12-89873048-INS-51 | chr12-89873048-INS-81 |
| chr12-89910564-INS-98 | chr12-89910564-INS-98 |
| chr12-89964527-INS-687 | chr12-89964527-INS-687 |
| chr12-90052208-INS-280 | chr12-90052208-INS-280 |
| chr12-90199225-INS-432 | chr12-90199233-INS-430 |
| chr12-90204015-INS-182 | chr12-90204015-INS-222 |
| chr12-90258526-INS-530 | chr12-90258547-INS-549 |
| chr12-90276686-INS-57 | chr12-90276686-INS-57 |
| chr12-90290418-INS-56 | chr12-90290418-INS-60 |
| chr12-90358328-INS-1589 | chr12-90358329-INS-1587 |
| chr12-90405137-INS-984 | chr12-90405137-INS-984 |
| chr12-90437653-INS-1184 | chr12-90437654-INS-1191 |
| chr12-90438871-INS-73 | chr12-90438871-INS-73 |
| chr12-90452757-INS-60 | chr12-90452757-INS-56 |
| chr12-90459072-INS-53 | chr12-90459072-INS-57 |
| chr12-90467572-INS-442 | chr12-90467572-INS-451 |
| chr12-90487475-INS-375 | chr12-90487475-INS-375 |
| chr12-90495810-INS-6887 | chr12-90495811-INS-6887 |
| chr12-90526037-INS-80 | chr12-90526037-INS-82 |
| chr12-90529433-INS-290 | chr12-90529426-INS-325 |
| chr12-90558302-INS-62 | chr12-90558302-INS-62 |
| chr12-90573399-INS-54 | chr12-90573429-INS-60 |
| chr12-90574641-INS-56 | chr12-90574641-INS-54 |
| chr12-90593723-INS-6482 | chr12-90593724-INS-6482 |
| chr12-90594475-INS-528 | chr12-90594475-INS-528 |
| chr12-90622426-INS-178 | chr12-90622428-INS-173 |
| chr12-90650069-INS-272 | chr12-90650069-INS-272 |
| chr12-90653892-INS-169 | chr12-90653892-INS-169 |
| chr12-90683377-INS-6246 | chr12-90683378-INS-6244 |
| chr12-90747398-INS-84 | chr12-90747398-INS-76 |
| chr12-90763722-INS-595 | chr12-90763722-INS-555 |
| chr12-90797420-INS-68 | chr12-90797420-INS-68 |
| chr12-90812154-INS-505 | chr12-90812154-INS-505 |
| chr12-90816322-INS-54 | chr12-90816322-INS-54 |

| | |
|---|---|
| chr12-90831344-INS-238 | chr12-90831344-INS-232 |
| chr12-90837321-INS-69 | chr12-90837321-INS-87 |
| chr12-90840591-INS-193 | chr12-90840591-INS-193 |
| chr12-90865861-INS-138 | chr12-90865861-INS-138 |
| chr12-90883603-INS-56 | chr12-90883603-INS-56 |
| chr12-90947730-INS-376 | chr12-90947730-INS-376 |
| chr12-90951222-INS-249 | chr12-90951222-INS-249 |
| chr12-90956861-INS-373 | chr12-90956860-INS-383 |
| chr12-90996416-INS-64 | chr12-90996416-INS-66 |
| chr12-91049445-INS-116 | chr12-91049440-INS-86 |
| chr12-91054673-INS-73 | chr12-91054661-INS-78 |
| chr12-91104962-INS-60 | chr12-91104965-INS-62 |
| chr12-91121809-INS-60 | chr12-91121807-INS-58 |
| chr12-91123021-INS-68 | chr12-91122994-INS-72 |
| chr12-91125000-INS-133 | chr12-91125000-INS-133 |
| chr12-91169162-INS-64 | chr12-91169162-INS-56 |
| chr12-91181059-INS-124 | chr12-91181060-INS-124 |
| chr12-91194988-INS-75 | chr12-91194988-INS-75 |
| chr12-91206275-INS-412 | chr12-91206275-INS-411 |
| chr12-91238280-INS-82 | chr12-91238280-INS-82 |
| chr12-91244035-INS-142 | chr12-91244048-INS-94 |
| chr12-91244915-INS-199 | chr12-91244915-INS-199 |
| chr12-91254906-INS-104 | chr12-91254906-INS-117 |
| chr12-91265216-INS-60 | chr12-91265216-INS-60 |
| chr12-91268065-INS-2265 | chr12-91268065-INS-2250 |
| chr12-91285484-INS-1403 | chr12-91285485-INS-1402 |
| chr12-91303301-INS-167 | chr12-91303301-INS-141 |
| chr12-91347608-INS-647 | chr12-91347608-INS-648 |
| chr12-91364474-INS-134 | chr12-91364474-INS-137 |
| chr12-91384325-INS-169 | chr12-91384325-INS-169 |
| chr12-91391065-INS-61 | chr12-91391073-INS-98 |
| chr12-91398601-INS-189 | chr12-91398601-INS-189 |
| chr12-91400695-INS-151 | chr12-91400695-INS-151 |
| chr12-91450658-INS-178 | chr12-91450658-INS-172 |
| chr12-91455014-INS-52 | chr12-91455014-INS-62 |
| chr12-91469046-INS-66 | chr12-91469046-INS-66 |
| chr12-91478270-INS-132 | chr12-91478270-INS-130 |
| chr12-91508173-INS-496 | chr12-91508173-INS-524 |
| chr12-91512399-INS-236 | chr12-91512399-INS-236 |
| chr12-91520280-INS-2363 | chr12-91520281-INS-2359 |
| chr12-91538349-INS-959 | chr12-91538349-INS-960 |
| chr12-91550324-INS-58 | chr12-91550324-INS-58 |
| chr12-91567349-INS-213 | chr12-91567349-INS-213 |
| chr12-91576713-INS-100 | chr12-91576678-INS-105 |
| chr12-91581742-INS-77 | chr12-91581742-INS-108 |
| chr12-91583891-INS-6702 | chr12-91583892-INS-6707 |
| chr12-91595298-INS-120 | chr12-91595298-INS-120 |
| chr12-91604862-INS-58 | chr12-91604853-INS-52 |
| chr12-91626514-INS-203 | chr12-91626514-INS-203 |
| chr12-91628925-INS-83 | chr12-91628925-INS-82 |
| chr12-91633366-INS-114 | chr12-91633366-INS-114 |
| chr12-91641651-INS-433 | chr12-91641651-INS-432 |
| chr12-91648523-INS-499 | chr12-91648523-INS-499 |
| chr12-91649971-INS-153 | chr12-91649940-INS-170 |
| chr12-91654211-INS-73 | chr12-91654211-INS-73 |
| chr12-91666884-INS-65 | chr12-91666852-INS-77 |
| chr12-91674564-INS-201 | chr12-91674564-INS-201 |
| chr12-91675732-INS-205 | chr12-91675736-INS-205 |
| chr12-91686922-INS-76 | chr12-91686922-INS-78 |
| chr12-91687924-INS-66 | chr12-91687924-INS-66 |
| chr12-91704564-INS-1890 | chr12-91704565-INS-1890 |
| chr12-91715808-INS-108 | chr12-91715808-INS-64 |
| chr12-91749127-INS-176 | chr12-91749127-INS-196 |
| chr12-91757991-INS-161 | chr12-91757991-INS-160 |
| chr12-91766207-INS-765 | chr12-91766207-INS-765 |
| chr12-91779127-INS-202 | chr12-91779127-INS-202 |
| chr12-91796386-INS-148 | chr12-91796386-INS-152 |
| chr12-91806072-INS-121 | chr12-91806110-INS-120 |
| chr12-91809046-INS-503 | chr12-91809046-INS-503 |
| chr12-91819985-INS-375 | chr12-91819985-INS-373 |
| chr12-91822466-INS-72 | chr12-91822466-INS-88 |
| chr12-91827455-INS-51 | chr12-91827455-INS-51 |
| chr12-91841762-INS-82 | chr12-91841746-INS-96 |
| chr12-91850716-INS-131 | chr12-91850736-INS-137 |
| chr12-91853373-INS-68 | chr12-91853393-INS-52 |
| chr12-91858138-INS-52 | chr12-91858138-INS-52 |
| chr12-91859442-INS-418 | chr12-91859437-INS-423 |

| | |
|---|---|
| chr12-91867649-INS-6268 | chr12-91867650-INS-6269 |
| chr12-91868413-INS-484 | chr12-91868413-INS-480 |
| chr12-91877678-INS-587 | chr12-91877678-INS-585 |
| chr12-91931856-INS-136 | chr12-91931820-INS-182 |
| chr12-91942632-INS-469 | chr12-91942640-INS-478 |
| chr12-91948610-INS-498 | chr12-91948610-INS-498 |
| chr12-91962914-INS-62 | chr12-91962905-INS-72 |
| chr12-91963537-INS-765 | chr12-91963537-INS-764 |
| chr12-91976039-INS-571 | chr12-91976039-INS-571 |
| chr12-91979713-INS-66 | chr12-91979713-INS-66 |
| chr12-91981166-INS-6259 | chr12-91981167-INS-6259 |
| chr12-91982608-INS-780 | chr12-91982608-INS-780 |
| chr12-91983353-INS-93 | chr12-91983347-INS-59 |
| chr12-91986077-INS-588 | chr12-91986077-INS-588 |
| chr12-91999960-INS-682 | chr12-91999960-INS-682 |
| chr12-92017782-INS-52 | chr12-92017782-INS-72 |
| chr12-92022051-INS-603 | chr12-92022051-INS-604 |
| chr12-92022878-INS-526 | chr12-92022878-INS-526 |
| chr12-92035020-INS-251 | chr12-92035020-INS-261 |
| chr12-92037500-INS-6198 | chr12-92037501-INS-6198 |
| chr12-92042781-INS-84 | chr12-92042781-INS-50 |
| chr12-92047150-INS-390 | chr12-92047150-INS-390 |
| chr12-92048516-INS-349 | chr12-92048516-INS-352 |
| chr12-92052533-INS-1884 | chr12-92052534-INS-1884 |
| chr12-92059545-INS-6509 | chr12-92059546-INS-6509 |
| chr12-92065737-INS-52 | chr12-92065737-INS-74 |
| chr12-92069684-INS-2260 | chr12-92069685-INS-2260 |
| chr12-92083049-INS-54 | chr12-92083049-INS-54 |
| chr12-92091070-INS-131 | chr12-92091070-INS-131 |
| chr12-92104002-INS-534 | chr12-92104004-INS-513 |
| chr12-92114387-INS-202 | chr12-92114387-INS-202 |
| chr12-92125975-INS-220 | chr12-92125967-INS-225 |
| chr12-92131952-INS-255 | chr12-92131952-INS-260 |
| chr12-92179289-INS-64 | chr12-92179319-INS-52 |
| chr12-92186070-INS-108 | chr12-92186070-INS-108 |
| chr12-92187999-INS-80 | chr12-92187999-INS-80 |
| chr12-92205085-INS-53 | chr12-92205083-INS-52 |
| chr12-92217910-INS-2811 | chr12-92217911-INS-2811 |
| chr12-92295139-INS-5292 | chr12-92295140-INS-5292 |
| chr12-92301033-INS-56 | chr12-92301033-INS-58 |
| chr12-92302712-INS-77 | chr12-92302690-INS-52 |
| chr12-92339672-INS-174 | chr12-92339672-INS-177 |
| chr12-92349337-INS-2218 | chr12-92349338-INS-2216 |
| chr12-92351866-INS-54 | chr12-92351866-INS-54 |
| chr12-92368641-INS-66 | chr12-92368650-INS-56 |
| chr12-92386290-INS-52 | chr12-92386290-INS-52 |
| chr12-92388245-INS-121 | chr12-92388241-INS-117 |
| chr12-92399653-INS-7072 | chr12-92399654-INS-7073 |
| chr12-92409818-INS-636 | chr12-92409811-INS-641 |
| chr12-92416768-INS-78 | chr12-92416768-INS-80 |
| chr12-92420748-INS-343 | chr12-92420748-INS-343 |
| chr12-92427540-INS-55 | chr12-92427540-INS-55 |
| chr12-92439169-INS-77 | chr12-92439176-INS-66 |
| chr12-92446361-INS-6674 | chr12-92446362-INS-6674 |
| chr12-92457028-INS-7767 | chr12-92457029-INS-7780 |
| chr12-92459594-INS-287 | chr12-92459594-INS-287 |
| chr12-92479305-INS-871 | chr12-92479305-INS-871 |
| chr12-92482986-INS-243 | chr12-92482986-INS-227 |
| chr12-92497217-INS-52 | chr12-92497217-INS-52 |
| chr12-92529159-INS-684 | chr12-92529159-INS-685 |
| chr12-92536863-INS-5409 | chr12-92536864-INS-5409 |
| chr12-92557097-INS-6413 | chr12-92557098-INS-6413 |
| chr12-92575614-INS-72 | chr12-92575614-INS-72 |
| chr12-92580886-INS-517 | chr12-92580886-INS-516 |
| chr12-92586875-INS-99 | chr12-92586875-INS-99 |
| chr12-92596226-INS-53 | chr12-92596226-INS-55 |
| chr12-92601894-INS-209 | chr12-92601895-INS-200 |
| chr12-92663818-INS-122 | chr12-92663818-INS-128 |
| chr12-92666192-INS-1518 | chr12-92666193-INS-1519 |
| chr12-92670567-INS-67 | chr12-92670567-INS-67 |
| chr12-92698203-INS-53 | chr12-92698203-INS-53 |
| chr12-92703586-INS-102 | chr12-92703586-INS-103 |
| chr12-92746588-INS-1895 | chr12-92746589-INS-1895 |
| chr12-92747226-INS-91 | chr12-92747226-INS-109 |
| chr12-92753122-INS-2069 | chr12-92753123-INS-2069 |
| chr12-92760954-INS-131 | chr12-92760954-INS-131 |
| chr12-92789827-INS-559 | chr12-92789827-INS-562 |

| | |
|---|---|
| chr12-92803637-INS-262 | chr12-92803637-INS-246 |
| chr12-92839954-INS-354 | chr12-92839954-INS-354 |
| chr12-92846393-INS-57 | chr12-92846392-INS-52 |
| chr12-92862293-INS-2833 | chr12-92862294-INS-2833 |
| chr12-92866107-INS-96 | chr12-92866107-INS-96 |
| chr12-92878717-INS-80 | chr12-92878717-INS-80 |
| chr12-92911953-INS-5737 | chr12-92911954-INS-5743 |
| chr12-92947010-INS-326 | chr12-92947010-INS-326 |
| chr12-92966778-INS-170 | chr12-92966778-INS-167 |
| chr12-92970533-INS-180 | chr12-92970533-INS-180 |
| chr12-92979776-INS-51 | chr12-92979777-INS-53 |
| chr12-92986162-INS-182 | chr12-92986162-INS-184 |
| chr12-92991357-INS-53 | chr12-92991357-INS-61 |
| chr12-93010587-INS-62 | chr12-93010587-INS-63 |
| chr12-93014978-INS-92 | chr12-93014978-INS-64 |
| chr12-93018348-INS-531 | chr12-93018348-INS-532 |
| chr12-93098125-INS-538 | chr12-93098125-INS-538 |
| chr12-93100994-INS-506 | chr12-93100994-INS-506 |
| chr12-93108552-INS-237 | chr12-93108552-INS-237 |
| chr12-93110949-INS-1293 | chr12-93110950-INS-1291 |
| chr12-93134549-INS-105 | chr12-93134549-INS-105 |
| chr12-93145174-INS-395 | chr12-93145174-INS-395 |
| chr12-93163699-INS-154 | chr12-93163699-INS-159 |
| chr12-93175360-INS-69 | chr12-93175360-INS-57 |
| chr12-93182597-INS-492 | chr12-93182597-INS-491 |
| chr12-93207281-INS-106 | chr12-93207281-INS-108 |
| chr12-93212011-INS-6574 | chr12-93212012-INS-6575 |
| chr12-93219404-INS-102 | chr12-93219404-INS-102 |
| chr12-93237043-INS-346 | chr12-93237043-INS-346 |
| chr12-93276418-INS-701 | chr12-93276418-INS-696 |
| chr12-93326224-INS-52 | chr12-93326224-INS-52 |
| chr12-93351041-INS-110 | chr12-93351043-INS-75 |
| chr12-93353504-INS-56 | chr12-93353504-INS-64 |
| chr12-93358441-INS-71 | chr12-93358423-INS-52 |
| chr12-93364086-INS-150 | chr12-93364086-INS-145 |
| chr12-93373125-INS-439 | chr12-93373125-INS-439 |
| chr12-93376804-INS-278 | chr12-93376806-INS-282 |
| chr12-93382377-INS-64 | chr12-93382409-INS-74 |
| chr12-93387586-INS-219 | chr12-93387615-INS-201 |
| chr12-93411896-INS-3779 | chr12-93411889-INS-3783 |
| chr12-93533481-INS-60 | chr12-93533481-INS-60 |
| chr12-93592493-INS-605 | chr12-93592493-INS-607 |
| chr12-93601106-INS-344 | chr12-93601109-INS-339 |
| chr12-93608064-INS-78 | chr12-93608089-INS-52 |
| chr12-93693048-INS-108 | chr12-93693048-INS-108 |
| chr12-93710121-INS-79 | chr12-93710121-INS-79 |
| chr12-93726086-INS-263 | chr12-93726086-INS-263 |
| chr12-93732184-INS-60 | chr12-93732184-INS-68 |
| chr12-93740019-INS-688 | chr12-93740019-INS-704 |
| chr12-93782104-INS-6248 | chr12-93782105-INS-6248 |
| chr12-93792871-INS-499 | chr12-93792871-INS-499 |
| chr12-93829564-INS-114 | chr12-93829565-INS-114 |
| chr12-93836074-INS-71 | chr12-93836074-INS-71 |
| chr12-93840457-INS-72 | chr12-93840457-INS-72 |
| chr12-93877295-INS-72 | chr12-93877295-INS-60 |
| chr12-93878070-INS-582 | chr12-93878070-INS-580 |
| chr12-93903522-INS-304 | chr12-93903522-INS-323 |
| chr12-93969055-INS-305 | chr12-93969055-INS-305 |
| chr12-93993302-INS-576 | chr12-93993301-INS-576 |
| chr12-93999149-INS-724 | chr12-93999149-INS-724 |
| chr12-94011335-INS-459 | chr12-94011335-INS-456 |
| chr12-94026129-INS-193 | chr12-94026129-INS-193 |
| chr12-94035724-INS-629 | chr12-94035727-INS-633 |
| chr12-94035724-INS-88 | chr12-94035727-INS-79 |
| chr12-94049543-INS-6666 | chr12-94049544-INS-6666 |
| chr12-94055379-INS-306 | chr12-94055379-INS-306 |
| chr12-94061791-INS-214 | chr12-94061791-INS-214 |
| chr12-94074140-INS-51 | chr12-94074135-INS-51 |
| chr12-94079788-INS-599 | chr12-94079788-INS-577 |
| chr12-94097407-INS-793 | chr12-94097407-INS-791 |
| chr12-94106296-INS-58 | chr12-94106297-INS-56 |
| chr12-94107492-INS-64 | chr12-94107494-INS-53 |
| chr12-94117736-INS-503 | chr12-94117736-INS-514 |
| chr12-94136213-INS-4345 | chr12-94136214-INS-4345 |
| chr12-94142152-INS-89 | chr12-94142174-INS-66 |
| chr12-94149723-INS-95 | chr12-94149723-INS-95 |
| chr12-94186124-INS-184 | chr12-94186124-INS-184 |

| | |
|---|---|
| chr12-94197228-INS-91 | chr12-94197189-INS-120 |
| chr12-94218136-INS-56 | chr12-94218138-INS-52 |
| chr12-94245438-INS-65 | chr12-94245429-INS-102 |
| chr12-94263025-INS-443 | chr12-94263025-INS-443 |
| chr12-94287415-INS-74 | chr12-94287415-INS-54 |
| chr12-94334155-INS-509 | chr12-94334155-INS-509 |
| chr12-94337218-INS-72 | chr12-94337227-INS-96 |
| chr12-94351753-INS-54 | chr12-94351753-INS-56 |
| chr12-94375559-INS-80 | chr12-94375559-INS-80 |
| chr12-94393703-INS-53 | chr12-94393703-INS-53 |
| chr12-94397456-INS-189 | chr12-94397456-INS-190 |
| chr12-94399146-INS-91 | chr12-94399146-INS-55 |
| chr12-94421774-INS-243 | chr12-94421774-INS-240 |
| chr12-94427996-INS-203 | chr12-94427996-INS-203 |
| chr12-94429512-INS-92 | chr12-94429512-INS-103 |
| chr12-94431149-INS-104 | chr12-94431149-INS-135 |
| chr12-94461574-INS-378 | chr12-94461575-INS-378 |
| chr12-94468089-INS-64 | chr12-94468089-INS-70 |
| chr12-94478311-INS-55 | chr12-94478311-INS-57 |
| chr12-94529813-INS-1124 | chr12-94529814-INS-1108 |
| chr12-94535905-INS-89 | chr12-94535905-INS-89 |
| chr12-94602832-INS-562 | chr12-94602832-INS-562 |
| chr12-94633821-INS-332 | chr12-94633821-INS-332 |
| chr12-94675006-INS-75 | chr12-94675006-INS-69 |
| chr12-94676541-INS-386 | chr12-94676534-INS-390 |
| chr12-94700693-INS-150 | chr12-94700693-INS-150 |
| chr12-94709056-INS-51 | chr12-94709056-INS-55 |
| chr12-94721011-INS-209 | chr12-94721011-INS-210 |
| chr12-94725534-INS-86 | chr12-94725534-INS-86 |
| chr12-94790908-INS-637 | chr12-94790908-INS-637 |
| chr12-94837726-INS-88 | chr12-94837726-INS-88 |
| chr12-94847675-INS-98 | chr12-94847675-INS-100 |
| chr12-94854114-INS-72 | chr12-94854112-INS-72 |
| chr12-94864989-INS-223 | chr12-94864989-INS-223 |
| chr12-94874120-INS-54 | chr12-94874120-INS-57 |
| chr12-94894978-INS-80 | chr12-94894978-INS-80 |
| chr12-94997761-INS-452 | chr12-94997761-INS-452 |
| chr12-95006439-INS-80 | chr12-95006440-INS-52 |
| chr12-95019364-INS-468 | chr12-95019364-INS-473 |
| chr12-95028821-INS-66 | chr12-95028821-INS-66 |
| chr12-95038787-INS-87 | chr12-95038787-INS-51 |
| chr12-95045705-INS-62 | chr12-95045705-INS-62 |
| chr12-95056538-INS-2400 | chr12-95056539-INS-2400 |
| chr12-95101078-INS-54 | chr12-95101078-INS-54 |
| chr12-95150114-INS-6871 | chr12-95150115-INS-6871 |
| chr12-95151659-INS-318 | chr12-95151659-INS-302 |
| chr12-95170309-INS-212 | chr12-95170309-INS-212 |
| chr12-95178801-INS-539 | chr12-95178801-INS-538 |
| chr12-95181752-INS-88 | chr12-95181752-INS-88 |
| chr12-95183684-INS-213 | chr12-95183684-INS-217 |
| chr12-95211541-INS-79 | chr12-95211541-INS-88 |
| chr12-95215482-INS-214 | chr12-95215482-INS-214 |
| chr12-95242367-INS-82 | chr12-95242367-INS-82 |
| chr12-95246390-INS-5421 | chr12-95246391-INS-5421 |
| chr12-95251658-INS-381 | chr12-95251658-INS-382 |
| chr12-95273267-INS-1046 | chr12-95273268-INS-1046 |
| chr12-95286818-INS-273 | chr12-95286818-INS-273 |
| chr12-95324785-INS-6287 | chr12-95324786-INS-6287 |
| chr12-95325127-INS-52 | chr12-95325127-INS-50 |
| chr12-95326135-INS-63 | chr12-95326139-INS-77 |
| chr12-95349341-INS-77 | chr12-95349341-INS-73 |
| chr12-95369116-INS-98 | chr12-95369116-INS-98 |
| chr12-95376845-INS-171 | chr12-95376845-INS-171 |
| chr12-95379132-INS-69 | chr12-95379132-INS-69 |
| chr12-95387628-INS-73 | chr12-95387628-INS-73 |
| chr12-95391707-INS-122 | chr12-95391707-INS-123 |
| chr12-95409260-INS-59 | chr12-95409260-INS-59 |
| chr12-95412793-INS-51 | chr12-95412793-INS-54 |
| chr12-95462906-INS-348 | chr12-95462906-INS-347 |
| chr12-95493782-INS-1000 | chr12-95493782-INS-1001 |
| chr12-95499069-INS-67 | chr12-95499076-INS-72 |
| chr12-95523644-INS-502 | chr12-95523644-INS-502 |
| chr12-95555559-INS-607 | chr12-95555559-INS-607 |
| chr12-95604432-INS-53 | chr12-95604418-INS-91 |
| chr12-95605762-INS-242 | chr12-95605762-INS-242 |
| chr12-95621504-INS-805 | chr12-95621503-INS-805 |
| chr12-95642169-INS-58 | chr12-95642169-INS-58 |

| | |
|---|---|
| chr12-95695459-INS-52 | chr12-95695453-INS-58 |
| chr12-95725095-INS-84 | chr12-95725095-INS-84 |
| chr12-95743477-INS-1385 | chr12-95743478-INS-1381 |
| chr12-95772293-INS-166 | chr12-95772293-INS-201 |
| chr12-95866693-INS-79 | chr12-95866690-INS-54 |
| chr12-95876286-INS-87 | chr12-95876286-INS-68 |
| chr12-95888871-INS-197 | chr12-95888871-INS-197 |
| chr12-95895745-INS-190 | chr12-95895745-INS-190 |
| chr12-95918660-INS-93 | chr12-95918665-INS-50 |
| chr12-95939986-INS-56 | chr12-95939986-INS-56 |
| chr12-95943789-INS-6656 | chr12-95943790-INS-6657 |
| chr12-95968319-INS-1290 | chr12-95968320-INS-1290 |
| chr12-95980864-INS-104 | chr12-95980864-INS-116 |
| chr12-96014169-INS-497 | chr12-96014169-INS-497 |
| chr12-96027318-INS-219 | chr12-96027318-INS-219 |
| chr12-96037990-INS-120 | chr12-96037990-INS-121 |
| chr12-96052225-INS-641 | chr12-96052225-INS-641 |
| chr12-96059088-INS-52 | chr12-96059088-INS-52 |
| chr12-96084018-INS-77 | chr12-96083996-INS-101 |
| chr12-96090102-INS-472 | chr12-96090102-INS-473 |
| chr12-96092944-INS-71 | chr12-96092901-INS-88 |
| chr12-96094416-INS-183 | chr12-96094416-INS-183 |
| chr12-96095923-INS-75 | chr12-96095920-INS-55 |
| chr12-96109752-INS-67 | chr12-96109752-INS-75 |
| chr12-96129340-INS-54 | chr12-96129340-INS-59 |
| chr12-96136922-INS-52 | chr12-96136922-INS-52 |
| chr12-96152492-INS-52 | chr12-96152492-INS-52 |
| chr12-96168036-INS-54 | chr12-96168036-INS-52 |
| chr12-96188195-INS-247 | chr12-96188195-INS-267 |
| chr12-96220563-INS-111 | chr12-96220563-INS-114 |
| chr12-96220736-INS-6050 | chr12-96220736-INS-6050 |
| chr12-96225519-INS-1842 | chr12-96225520-INS-1843 |
| chr12-96239872-INS-214 | chr12-96239869-INS-218 |
| chr12-96274351-INS-78 | chr12-96274350-INS-79 |
| chr12-96304294-INS-52 | chr12-96304294-INS-52 |
| chr12-96327789-INS-4739 | chr12-96327790-INS-4739 |
| chr12-96328916-INS-116 | chr12-96328916-INS-69 |
| chr12-96367206-INS-74 | chr12-96367206-INS-76 |
| chr12-96374601-INS-4983 | chr12-96374602-INS-4983 |
| chr12-96402086-INS-77 | chr12-96402086-INS-77 |
| chr12-96405456-INS-128 | chr12-96405451-INS-119 |
| chr12-96410535-INS-129 | chr12-96410534-INS-139 |
| chr12-96411776-INS-299 | chr12-96411776-INS-301 |
| chr12-96429910-INS-193 | chr12-96429910-INS-193 |
| chr12-96431013-INS-75 | chr12-96431013-INS-71 |
| chr12-96444011-INS-61 | chr12-96444011-INS-61 |
| chr12-96449823-INS-605 | chr12-96449823-INS-605 |
| chr12-96454048-INS-112 | chr12-96454050-INS-62 |
| chr12-96457648-INS-66 | chr12-96457648-INS-74 |
| chr12-96474004-INS-919 | chr12-96474005-INS-920 |
| chr12-96494589-INS-58 | chr12-96494589-INS-64 |
| chr12-96515760-INS-62 | chr12-96515760-INS-54 |
| chr12-96564873-INS-277 | chr12-96564873-INS-275 |
| chr12-96587253-INS-149 | chr12-96587253-INS-150 |
| chr12-96603566-INS-2152 | chr12-96603567-INS-2150 |
| chr12-96616838-INS-77 | chr12-96616838-INS-77 |
| chr12-96636014-INS-66 | chr12-96636014-INS-66 |
| chr12-96650553-INS-686 | chr12-96650553-INS-685 |
| chr12-96660495-INS-292 | chr12-96660504-INS-274 |
| chr12-96671986-INS-422 | chr12-96671986-INS-422 |
| chr12-96673370-INS-197 | chr12-96673370-INS-197 |
| chr12-96690436-INS-63 | chr12-96690436-INS-63 |
| chr12-96691818-INS-79 | chr12-96691818-INS-81 |
| chr12-96704702-INS-126 | chr12-96704702-INS-126 |
| chr12-96710393-INS-1227 | chr12-96710394-INS-1225 |
| chr12-96716018-INS-56 | chr12-96716018-INS-50 |
| chr12-96739617-INS-446 | chr12-96739617-INS-449 |
| chr12-96754847-INS-86 | chr12-96754816-INS-74 |
| chr12-96764730-INS-1506 | chr12-96764722-INS-1518 |
| chr12-96765431-INS-362 | chr12-96765431-INS-362 |
| chr12-96786902-INS-84 | chr12-96786902-INS-54 |
| chr12-96896534-INS-56 | chr12-96896534-INS-89 |
| chr12-96904085-INS-75 | chr12-96904130-INS-75 |
| chr12-96905178-INS-77 | chr12-96905178-INS-77 |
| chr12-96925968-INS-60 | chr12-96925968-INS-60 |
| chr12-96926862-INS-7125 | chr12-96926863-INS-7125 |
| chr12-96928082-INS-5543 | chr12-96928083-INS-5543 |

| | |
|---|---|
| chr12-96988359-INS-63 | chr12-96988359-INS-63 |
| chr12-96990235-INS-915 | chr12-96990235-INS-912 |
| chr12-96995726-INS-294 | chr12-96995727-INS-278 |
| chr12-97002683-INS-193 | chr12-97002683-INS-192 |
| chr12-97013950-INS-1204 | chr12-97013951-INS-1204 |
| chr12-97065886-INS-1482 | chr12-97065887-INS-1484 |
| chr12-97099452-INS-54 | chr12-97099455-INS-66 |
| chr12-97105583-INS-130 | chr12-97105583-INS-130 |
| chr12-97137926-INS-104 | chr12-97137926-INS-104 |
| chr12-97144824-INS-52 | chr12-97144824-INS-54 |
| chr12-97178730-INS-102 | chr12-97178730-INS-104 |
| chr12-97204324-INS-827 | chr12-97204333-INS-830 |
| chr12-97207976-INS-1047 | chr12-97207974-INS-1041 |
| chr12-97277035-INS-81 | chr12-97277035-INS-81 |
| chr12-97290511-INS-176 | chr12-97290491-INS-181 |
| chr12-97291505-INS-106 | chr12-97291505-INS-113 |
| chr12-97317387-INS-197 | chr12-97317387-INS-197 |
| chr12-97330188-INS-912 | chr12-97330188-INS-912 |
| chr12-97335731-INS-60 | chr12-97335732-INS-60 |
| chr12-97370826-INS-522 | chr12-97370825-INS-530 |
| chr12-97375126-INS-52 | chr12-97375118-INS-78 |
| chr12-97408525-INS-406 | chr12-97408525-INS-409 |
| chr12-97417236-INS-1088 | chr12-97417237-INS-1087 |
| chr12-97427641-INS-145 | chr12-97427641-INS-145 |
| chr12-97504322-INS-6196 | chr12-97504323-INS-6196 |
| chr12-97654552-INS-54 | chr12-97654552-INS-52 |
| chr12-97668220-INS-6678 | chr12-97668221-INS-6678 |
| chr12-97701938-INS-95 | chr12-97701938-INS-95 |
| chr12-97722056-INS-964 | chr12-97722056-INS-964 |
| chr12-97737868-INS-283 | chr12-97737868-INS-288 |
| chr12-97809883-INS-50 | chr12-97809843-INS-50 |
| chr12-97839257-INS-183 | chr12-97839257-INS-184 |
| chr12-97851507-INS-68 | chr12-97851507-INS-51 |
| chr12-97881305-INS-89 | chr12-97881305-INS-97 |
| chr12-97934825-INS-56 | chr12-97934825-INS-52 |
| chr12-97948845-INS-6502 | chr12-97948846-INS-6502 |
| chr12-97955301-INS-55 | chr12-97955300-INS-55 |
| chr12-97994142-INS-160 | chr12-97994142-INS-170 |
| chr12-98004690-INS-108 | chr12-98004664-INS-100 |
| chr12-98007126-INS-70 | chr12-98007126-INS-70 |
| chr12-98007889-INS-498 | chr12-98007889-INS-498 |
| chr12-98019793-INS-192 | chr12-98019793-INS-192 |
| chr12-98027909-INS-61 | chr12-98027909-INS-55 |
| chr12-98028904-INS-563 | chr12-98028907-INS-550 |
| chr12-98089489-INS-966 | chr12-98089489-INS-966 |
| chr12-98102734-INS-90 | chr12-98102736-INS-85 |
| chr12-98129113-INS-2018 | chr12-98129090-INS-1997 |
| chr12-98152407-INS-991 | chr12-98152410-INS-989 |
| chr12-98155659-INS-2908 | chr12-98155660-INS-2908 |
| chr12-98168154-INS-85 | chr12-98168147-INS-50 |
| chr12-98179209-INS-132 | chr12-98179209-INS-132 |
| chr12-98191130-INS-60 | chr12-98191130-INS-60 |
| chr12-98208727-INS-1431 | chr12-98208728-INS-1430 |
| chr12-98223404-INS-622 | chr12-98223404-INS-622 |
| chr12-98264144-INS-736 | chr12-98264144-INS-736 |
| chr12-98266602-INS-7119 | chr12-98266603-INS-7119 |
| chr12-98270482-INS-92 | chr12-98270482-INS-78 |
| chr12-98271619-INS-86 | chr12-98271619-INS-86 |
| chr12-98276556-INS-145 | chr12-98276556-INS-145 |
| chr12-98299202-INS-273 | chr12-98299202-INS-273 |
| chr12-98301865-INS-146 | chr12-98301865-INS-149 |
| chr12-98320879-INS-1537 | chr12-98320880-INS-1537 |
| chr12-98320896-INS-416 | chr12-98320896-INS-416 |
| chr12-98338017-INS-60 | chr12-98338047-INS-50 |
| chr12-98379610-INS-77 | chr12-98379610-INS-77 |
| chr12-98393096-INS-85 | chr12-98393096-INS-85 |
| chr12-98411695-INS-58 | chr12-98411695-INS-87 |
| chr12-98420569-INS-271 | chr12-98420569-INS-268 |
| chr12-98437073-INS-182 | chr12-98437073-INS-186 |
| chr12-98457384-INS-208 | chr12-98457384-INS-208 |
| chr12-98461225-INS-102 | chr12-98461207-INS-107 |
| chr12-98464262-INS-363 | chr12-98464262-INS-365 |
| chr12-98471205-INS-51 | chr12-98471205-INS-51 |
| chr12-98492596-INS-94 | chr12-98492592-INS-54 |
| chr12-98496364-INS-13649 | chr12-98496365-INS-13649 |
| chr12-98497672-INS-76 | chr12-98497672-INS-84 |
| chr12-98524725-INS-75 | chr12-98524725-INS-101 |

| | |
|---|---|
| chr12-98530625-INS-52 | chr12-98530626-INS-64 |
| chr12-98533312-INS-113 | chr12-98533312-INS-113 |
| chr12-98547933-INS-81 | chr12-98547933-INS-81 |
| chr12-98576780-INS-6541 | chr12-98576781-INS-6543 |
| chr12-98584627-INS-1005 | chr12-98584628-INS-1005 |
| chr12-98609660-INS-205 | chr12-98609660-INS-205 |
| chr12-98613139-INS-71 | chr12-98613139-INS-73 |
| chr12-98630376-INS-180 | chr12-98630376-INS-181 |
| chr12-98651278-INS-105 | chr12-98651278-INS-105 |
| chr12-98718709-INS-54 | chr12-98718709-INS-56 |
| chr12-98729521-INS-243 | chr12-98729521-INS-243 |
| chr12-98763968-INS-109 | chr12-98763991-INS-91 |
| chr12-98817410-INS-179 | chr12-98817410-INS-181 |
| chr12-98842081-INS-59 | chr12-98842081-INS-59 |
| chr12-98851015-INS-790 | chr12-98851015-INS-790 |
| chr12-98863597-INS-209 | chr12-98863598-INS-209 |
| chr12-98869583-INS-87 | chr12-98869587-INS-80 |
| chr12-98876864-INS-75 | chr12-98876862-INS-77 |
| chr12-98883693-INS-119 | chr12-98883707-INS-69 |
| chr12-98897544-INS-136 | chr12-98897544-INS-136 |
| chr12-98941809-INS-168 | chr12-98941809-INS-168 |
| chr12-98947780-INS-60 | chr12-98947780-INS-60 |
| chr12-98949223-INS-188 | chr12-98949223-INS-188 |
| chr12-98964099-INS-173 | chr12-98964099-INS-172 |
| chr12-98980207-INS-88 | chr12-98980207-INS-82 |
| chr12-98987204-INS-60 | chr12-98987203-INS-60 |
| chr12-99040973-INS-105 | chr12-99040973-INS-105 |
| chr12-99046737-INS-2034 | chr12-99046738-INS-2047 |
| chr12-99072658-INS-56 | chr12-99072701-INS-55 |
| chr12-99084850-INS-196 | chr12-99084850-INS-196 |
| chr12-99088192-INS-869 | chr12-99088192-INS-869 |
| chr12-99142458-INS-159 | chr12-99142458-INS-189 |
| chr12-99153908-INS-70 | chr12-99153908-INS-70 |
| chr12-99168069-INS-218 | chr12-99168069-INS-218 |
| chr12-99217011-INS-89 | chr12-99217039-INS-53 |
| chr12-99240815-INS-86 | chr12-99240815-INS-100 |
| chr12-99262104-INS-76 | chr12-99262102-INS-76 |
| chr12-99278359-INS-247 | chr12-99278359-INS-247 |
| chr12-99287476-INS-52 | chr12-99287476-INS-58 |
| chr12-99293084-INS-99 | chr12-99293084-INS-99 |
| chr12-99343575-INS-100 | chr12-99343574-INS-100 |
| chr12-99360373-INS-62 | chr12-99360373-INS-58 |
| chr12-99372278-INS-52 | chr12-99372283-INS-54 |
| chr12-99377366-INS-89 | chr12-99377366-INS-89 |
| chr12-99386381-INS-504 | chr12-99386381-INS-504 |
| chr12-99390722-INS-214 | chr12-99390722-INS-214 |
| chr12-99402844-INS-205 | chr12-99402847-INS-206 |
| chr12-99412827-INS-179 | chr12-99412827-INS-160 |
| chr12-99418353-INS-129 | chr12-99418354-INS-86 |
| chr12-99422453-INS-82 | chr12-99422452-INS-118 |
| chr12-99429796-INS-66 | chr12-99429757-INS-96 |
| chr12-99433335-INS-64 | chr12-99433335-INS-96 |
| chr12-99444143-INS-59 | chr12-99444146-INS-67 |
| chr12-99447532-INS-102 | chr12-99447532-INS-102 |
| chr12-99455624-INS-3712 | chr12-99455625-INS-3712 |
| chr12-99457041-INS-201 | chr12-99457041-INS-201 |
| chr12-99504130-INS-89 | chr12-99504111-INS-72 |
| chr12-99550611-INS-199 | chr12-99550612-INS-199 |
| chr12-99557979-INS-60 | chr12-99557980-INS-70 |
| chr12-99574412-INS-66 | chr12-99574412-INS-66 |
| chr12-99613626-INS-889 | chr12-99613626-INS-925 |
| chr12-99614261-INS-319 | chr12-99614262-INS-319 |
| chr12-99649578-INS-1001 | chr12-99649579-INS-1009 |
| chr12-99698949-INS-1230 | chr12-99698950-INS-1230 |
| chr12-99716038-INS-6094 | chr12-99716039-INS-6094 |
| chr12-99716546-INS-246 | chr12-99716550-INS-246 |
| chr12-99770359-INS-68 | chr12-99770359-INS-68 |
| chr12-99801742-INS-403 | chr12-99801742-INS-403 |
| chr12-99804556-INS-79 | chr12-99804556-INS-79 |
| chr12-99811087-INS-59 | chr12-99811087-INS-64 |
| chr12-99815890-INS-77 | chr12-99815873-INS-80 |
| chr12-99853534-INS-91 | chr12-99853534-INS-91 |
| chr12-99855752-INS-199 | chr12-99855752-INS-199 |
| chr12-99871538-INS-1727 | chr12-99871539-INS-1729 |
| chr12-99888373-INS-341 | chr12-99888373-INS-341 |
| chr12-99920526-INS-126 | chr12-99920526-INS-126 |
| chr12-99955523-INS-136 | chr12-99955530-INS-148 |

| | |
|---|---|
| chr12-99975812-INS-127 | chr12-99975812-INS-127 |
| chr12-100007298-INS-234 | chr12-100007298-INS-234 |
| chr12-100049047-INS-342 | chr12-100049047-INS-342 |
| chr12-100066526-INS-194 | chr12-100066565-INS-194 |
| chr12-100079754-INS-56 | chr12-100079757-INS-63 |
| chr12-100102346-INS-53 | chr12-100102346-INS-53 |
| chr12-100103626-INS-90 | chr12-100103618-INS-72 |
| chr12-100130110-INS-81 | chr12-100130110-INS-54 |
| chr12-100137081-INS-4708 | chr12-100137082-INS-4709 |
| chr12-100150034-INS-996 | chr12-100150034-INS-1000 |
| chr12-100200919-INS-74 | chr12-100200919-INS-74 |
| chr12-100202975-INS-191 | chr12-100202975-INS-191 |
| chr12-100205282-INS-65 | chr12-100205282-INS-65 |
| chr12-100207437-INS-109 | chr12-100207437-INS-109 |
| chr12-100208463-INS-69 | chr12-100208463-INS-55 |
| chr12-100217518-INS-177 | chr12-100217518-INS-198 |
| chr12-100241314-INS-58 | chr12-100241304-INS-72 |
| chr12-100254924-INS-54 | chr12-100254924-INS-54 |
| chr12-100289089-INS-147 | chr12-100289089-INS-157 |
| chr12-100309706-INS-80 | chr12-100309706-INS-121 |
| chr12-100357353-INS-87 | chr12-100357353-INS-89 |
| chr12-100405429-INS-209 | chr12-100405429-INS-209 |
| chr12-100410863-INS-54 | chr12-100410863-INS-64 |
| chr12-100420753-INS-58 | chr12-100420775-INS-58 |
| chr12-100424374-INS-5883 | chr12-100424375-INS-5883 |
| chr12-100434164-INS-58 | chr12-100434164-INS-60 |
| chr12-100445304-INS-69 | chr12-100445305-INS-69 |
| chr12-100449509-INS-62 | chr12-100449509-INS-62 |
| chr12-100457395-INS-594 | chr12-100457395-INS-594 |
| chr12-100472376-INS-6438 | chr12-100472377-INS-6438 |
| chr12-100491840-INS-180 | chr12-100491840-INS-179 |
| chr12-100503748-INS-72 | chr12-100503734-INS-57 |
| chr12-100505372-INS-199 | chr12-100505372-INS-200 |
| chr12-100539359-INS-62 | chr12-100539340-INS-93 |
| chr12-100545875-INS-52 | chr12-100545875-INS-52 |
| chr12-100550449-INS-65 | chr12-100550449-INS-50 |
| chr12-100557729-INS-215 | chr12-100557729-INS-213 |
| chr12-100567381-INS-97 | chr12-100567378-INS-66 |
| chr12-100570619-INS-120 | chr12-100570619-INS-110 |
| chr12-100605120-INS-69 | chr12-100605120-INS-69 |
| chr12-100610997-INS-500 | chr12-100610997-INS-500 |
| chr12-100626479-INS-127 | chr12-100626479-INS-127 |
| chr12-100653573-INS-63 | chr12-100653573-INS-55 |
| chr12-100665306-INS-535 | chr12-100665306-INS-535 |
| chr12-100696271-INS-81 | chr12-100696276-INS-94 |
| chr12-100700589-INS-140 | chr12-100700570-INS-159 |
| chr12-100713492-INS-100 | chr12-100713492-INS-100 |
| chr12-100716140-INS-58 | chr12-100716140-INS-58 |
| chr12-100738020-INS-7085 | chr12-100738021-INS-7085 |
| chr12-100739322-INS-78 | chr12-100739322-INS-84 |
| chr12-100741595-INS-92 | chr12-100741595-INS-92 |
| chr12-100759493-INS-205 | chr12-100759493-INS-205 |
| chr12-100812664-INS-1261 | chr12-100812665-INS-1261 |
| chr12-100833916-INS-1204 | chr12-100833917-INS-1216 |
| chr12-100859013-INS-212 | chr12-100859013-INS-211 |
| chr12-100882286-INS-64 | chr12-100882286-INS-64 |
| chr12-100889244-INS-228 | chr12-100889244-INS-228 |
| chr12-100897208-INS-1706 | chr12-100897215-INS-1705 |
| chr12-100898600-INS-53 | chr12-100898584-INS-77 |
| chr12-100905141-INS-59 | chr12-100905141-INS-83 |
| chr12-100912200-INS-156 | chr12-100912200-INS-156 |
| chr12-100929690-INS-273 | chr12-100929690-INS-273 |
| chr12-100940777-INS-215 | chr12-100940777-INS-215 |
| chr12-100945206-INS-77 | chr12-100945206-INS-77 |
| chr12-100945775-INS-285 | chr12-100945775-INS-285 |
| chr12-100953326-INS-165 | chr12-100953326-INS-117 |
| chr12-100976151-INS-204 | chr12-100976151-INS-215 |
| chr12-100979852-INS-261 | chr12-100979870-INS-263 |
| chr12-101069741-INS-220 | chr12-101069741-INS-206 |
| chr12-101091661-INS-490 | chr12-101091652-INS-493 |
| chr12-101098588-INS-54 | chr12-101098590-INS-56 |
| chr12-101099118-INS-1891 | chr12-101099119-INS-1891 |
| chr12-101158884-INS-198 | chr12-101158884-INS-198 |
| chr12-101162641-INS-719 | chr12-101162641-INS-719 |
| chr12-101163721-INS-217 | chr12-101163721-INS-217 |
| chr12-101199643-INS-203 | chr12-101199643-INS-203 |
| chr12-101241996-INS-455 | chr12-101241998-INS-459 |

| | |
|---|---|
| chr12-101243738-INS-2217 | chr12-101243739-INS-2217 |
| chr12-101270235-INS-6213 | chr12-101270236-INS-6213 |
| chr12-101324305-INS-6431 | chr12-101324306-INS-6431 |
| chr12-101324656-INS-197 | chr12-101324656-INS-197 |
| chr12-101400212-INS-409 | chr12-101400243-INS-409 |
| chr12-101424223-INS-174 | chr12-101424223-INS-174 |
| chr12-101425485-INS-896 | chr12-101425485-INS-902 |
| chr12-101449360-INS-75 | chr12-101449362-INS-66 |
| chr12-101457504-INS-1961 | chr12-101457505-INS-1961 |
| chr12-101486055-INS-3013 | chr12-101486056-INS-3013 |
| chr12-101506107-INS-6452 | chr12-101506108-INS-6451 |
| chr12-101556222-INS-164 | chr12-101556222-INS-164 |
| chr12-101600347-INS-390 | chr12-101600347-INS-390 |
| chr12-101652776-INS-52 | chr12-101652776-INS-72 |
| chr12-101666722-INS-203 | chr12-101666722-INS-202 |
| chr12-101691913-INS-5120 | chr12-101691914-INS-5120 |
| chr12-101704288-INS-413 | chr12-101704288-INS-413 |
| chr12-101718556-INS-847 | chr12-101718556-INS-847 |
| chr12-101739615-INS-115 | chr12-101739615-INS-92 |
| chr12-101763094-INS-861 | chr12-101763094-INS-862 |
| chr12-101772137-INS-92 | chr12-101772137-INS-92 |
| chr12-101794795-INS-6190 | chr12-101794796-INS-6190 |
| chr12-101800769-INS-138 | chr12-101800769-INS-93 |
| chr12-101833818-INS-114 | chr12-101833857-INS-70 |
| chr12-101843629-INS-273 | chr12-101843629-INS-273 |
| chr12-101858824-INS-67 | chr12-101858824-INS-87 |
| chr12-101872917-INS-81 | chr12-101872919-INS-84 |
| chr12-101887123-INS-207 | chr12-101887123-INS-207 |
| chr12-101927590-INS-6585 | chr12-101927591-INS-6585 |
| chr12-101931036-INS-549 | chr12-101931037-INS-548 |
| chr12-101932126-INS-78 | chr12-101932126-INS-76 |
| chr12-101937644-INS-144 | chr12-101937644-INS-144 |
| chr12-101944416-INS-68 | chr12-101944417-INS-105 |
| chr12-101945327-INS-195 | chr12-101945328-INS-198 |
| chr12-101952451-INS-186 | chr12-101952451-INS-186 |
| chr12-101954300-INS-62 | chr12-101954300-INS-62 |
| chr12-101965962-INS-2306 | chr12-101965963-INS-2315 |
| chr12-101979706-INS-65 | chr12-101979706-INS-69 |
| chr12-101998510-INS-52 | chr12-101998510-INS-52 |
| chr12-102010254-INS-62 | chr12-102010265-INS-56 |
| chr12-102032378-INS-6403 | chr12-102032379-INS-6403 |
| chr12-102046474-INS-503 | chr12-102046474-INS-506 |
| chr12-102060125-INS-1708 | chr12-102060126-INS-1708 |
| chr12-102060755-INS-63 | chr12-102060755-INS-63 |
| chr12-102061902-INS-65 | chr12-102061890-INS-79 |
| chr12-102070294-INS-1562 | chr12-102070295-INS-1562 |
| chr12-102073542-INS-1166 | chr12-102073546-INS-1162 |
| chr12-102137772-INS-62 | chr12-102137772-INS-62 |
| chr12-102184132-INS-93 | chr12-102184159-INS-58 |
| chr12-102188050-INS-68 | chr12-102188050-INS-68 |
| chr12-102213422-INS-102 | chr12-102213422-INS-68 |
| chr12-102265451-INS-145 | chr12-102265451-INS-145 |
| chr12-102290820-INS-194 | chr12-102290826-INS-187 |
| chr12-102304180-INS-78 | chr12-102304180-INS-114 |
| chr12-102313033-INS-109 | chr12-102313033-INS-109 |
| chr12-102314629-INS-98 | chr12-102314629-INS-98 |
| chr12-102319149-INS-444 | chr12-102319149-INS-446 |
| chr12-102321421-INS-52 | chr12-102321421-INS-54 |
| chr12-102399029-INS-189 | chr12-102399030-INS-192 |
| chr12-102410311-INS-127 | chr12-102410311-INS-150 |
| chr12-102414128-INS-229 | chr12-102414138-INS-189 |
| chr12-102421454-INS-181 | chr12-102421454-INS-182 |
| chr12-102423994-INS-222 | chr12-102423994-INS-222 |
| chr12-102433215-INS-792 | chr12-102433220-INS-801 |
| chr12-102452084-INS-213 | chr12-102452084-INS-213 |
| chr12-102459017-INS-199 | chr12-102459017-INS-199 |
| chr12-102474135-INS-215 | chr12-102474135-INS-215 |
| chr12-102483495-INS-51 | chr12-102483495-INS-51 |
| chr12-102485478-INS-6401 | chr12-102485479-INS-6401 |
| chr12-102496658-INS-204 | chr12-102496653-INS-209 |
| chr12-102511371-INS-202 | chr12-102511371-INS-202 |
| chr12-102528465-INS-54 | chr12-102528462-INS-52 |
| chr12-102536101-INS-62 | chr12-102536102-INS-75 |
| chr12-102538529-INS-63 | chr12-102538529-INS-63 |
| chr12-102593675-INS-205 | chr12-102593675-INS-205 |
| chr12-102596637-INS-54 | chr12-102596639-INS-69 |
| chr12-102660452-INS-180 | chr12-102660452-INS-180 |

| | |
|---|---|
| chr12-102674954-INS-95 | chr12-102674955-INS-96 |
| chr12-102676664-INS-77 | chr12-102676664-INS-77 |
| chr12-102682541-INS-185 | chr12-102682541-INS-184 |
| chr12-102718299-INS-52 | chr12-102718299-INS-52 |
| chr12-102727504-INS-208 | chr12-102727504-INS-210 |
| chr12-102729634-INS-203 | chr12-102729634-INS-204 |
| chr12-102736772-INS-181 | chr12-102736773-INS-171 |
| chr12-102765516-INS-118 | chr12-102765562-INS-101 |
| chr12-102791905-INS-326 | chr12-102791905-INS-325 |
| chr12-102817118-INS-1024 | chr12-102817119-INS-1024 |
| chr12-102830896-INS-230 | chr12-102830896-INS-231 |
| chr12-102873664-INS-58 | chr12-102873664-INS-62 |
| chr12-102876341-INS-403 | chr12-102876341-INS-389 |
| chr12-102927073-INS-78 | chr12-102927055-INS-52 |
| chr12-102930533-INS-84 | chr12-102930533-INS-84 |
| chr12-102932688-INS-58 | chr12-102932688-INS-58 |
| chr12-102952098-INS-79 | chr12-102952099-INS-63 |
| chr12-103018894-INS-448 | chr12-103018894-INS-448 |
| chr12-103057162-INS-243 | chr12-103057162-INS-243 |
| chr12-103066823-INS-62 | chr12-103066823-INS-66 |
| chr12-103105282-INS-123 | chr12-103105282-INS-76 |
| chr12-103196260-INS-168 | chr12-103196260-INS-182 |
| chr12-103230499-INS-60 | chr12-103230499-INS-79 |
| chr12-103246108-INS-94 | chr12-103246108-INS-94 |
| chr12-103289860-INS-102 | chr12-103289860-INS-103 |
| chr12-103312834-INS-80 | chr12-103312834-INS-80 |
| chr12-103346710-INS-405 | chr12-103346710-INS-405 |
| chr12-103348523-INS-64 | chr12-103348523-INS-69 |
| chr12-103349529-INS-64 | chr12-103349529-INS-66 |
| chr12-103396122-INS-6442 | chr12-103396123-INS-6442 |
| chr12-103417476-INS-81 | chr12-103417476-INS-81 |
| chr12-103454456-INS-88 | chr12-103454466-INS-53 |
| chr12-103468982-INS-52 | chr12-103468990-INS-62 |
| chr12-103489391-INS-6227 | chr12-103489392-INS-6227 |
| chr12-103505917-INS-68 | chr12-103505917-INS-68 |
| chr12-103522628-INS-219 | chr12-103522628-INS-218 |
| chr12-103544394-INS-152 | chr12-103544394-INS-120 |
| chr12-103559633-INS-57 | chr12-103559633-INS-57 |
| chr12-103563493-INS-261 | chr12-103563493-INS-263 |
| chr12-103575768-INS-85 | chr12-103575768-INS-85 |
| chr12-103617052-INS-74 | chr12-103617052-INS-84 |
| chr12-103633634-INS-874 | chr12-103633634-INS-874 |
| chr12-103694341-INS-84 | chr12-103694345-INS-54 |
| chr12-103803975-INS-187 | chr12-103803975-INS-183 |
| chr12-103807508-INS-559 | chr12-103807508-INS-559 |
| chr12-103910197-INS-60 | chr12-103910166-INS-60 |
| chr12-103924960-INS-98 | chr12-103924960-INS-98 |
| chr12-103952809-INS-179 | chr12-103952809-INS-187 |
| chr12-103953843-INS-54 | chr12-103953843-INS-54 |
| chr12-103962610-INS-137 | chr12-103962610-INS-114 |
| chr12-103967838-INS-214 | chr12-103967838-INS-241 |
| chr12-103990399-INS-81 | chr12-103990399-INS-89 |
| chr12-104000824-INS-360 | chr12-104000824-INS-360 |
| chr12-104028918-INS-98 | chr12-104028921-INS-118 |
| chr12-104037874-INS-78 | chr12-104037874-INS-78 |
| chr12-104049230-INS-51 | chr12-104049231-INS-51 |
| chr12-104073321-INS-12297 | chr12-104073321-INS-12292 |
| chr12-104079293-INS-58 | chr12-104079293-INS-58 |
| chr12-104086502-INS-110 | chr12-104086502-INS-110 |
| chr12-104108683-INS-56 | chr12-104108683-INS-62 |
| chr12-104125597-INS-50 | chr12-104125597-INS-54 |
| chr12-104130867-INS-632 | chr12-104130867-INS-632 |
| chr12-104134813-INS-59 | chr12-104134813-INS-59 |
| chr12-104156970-INS-109 | chr12-104156970-INS-109 |
| chr12-104197848-INS-75 | chr12-104197848-INS-74 |
| chr12-104204054-INS-420 | chr12-104204055-INS-431 |
| chr12-104206689-INS-871 | chr12-104206689-INS-871 |
| chr12-104282549-INS-84 | chr12-104282530-INS-84 |
| chr12-104298592-INS-59 | chr12-104298553-INS-78 |
| chr12-104321345-INS-1727 | chr12-104321346-INS-1730 |
| chr12-104338483-INS-92 | chr12-104338524-INS-90 |
| chr12-104339737-INS-571 | chr12-104339737-INS-573 |
| chr12-104342966-INS-536 | chr12-104342966-INS-536 |
| chr12-104363433-INS-725 | chr12-104363433-INS-725 |
| chr12-104368040-INS-63 | chr12-104368040-INS-63 |
| chr12-104381176-INS-118 | chr12-104381179-INS-120 |
| chr12-104421986-INS-134 | chr12-104421986-INS-134 |

| | |
|---|---|
| chr12-104424986-INS-206 | chr12-104424986-INS-205 |
| chr12-104430325-INS-90 | chr12-104430325-INS-92 |
| chr12-104471246-INS-56 | chr12-104471277-INS-63 |
| chr12-104505591-INS-508 | chr12-104505591-INS-508 |
| chr12-104510182-INS-65 | chr12-104510231-INS-60 |
| chr12-104524611-INS-486 | chr12-104524611-INS-486 |
| chr12-104577664-INS-60 | chr12-104577665-INS-52 |
| chr12-104578924-INS-735 | chr12-104578924-INS-735 |
| chr12-104584190-INS-78 | chr12-104584190-INS-98 |
| chr12-104586584-INS-54 | chr12-104586578-INS-58 |
| chr12-104588554-INS-89 | chr12-104588554-INS-99 |
| chr12-104605335-INS-183 | chr12-104605348-INS-180 |
| chr12-104634262-INS-56 | chr12-104634262-INS-56 |
| chr12-104649728-INS-241 | chr12-104649728-INS-237 |
| chr12-104650867-INS-3076 | chr12-104650868-INS-3076 |
| chr12-104685443-INS-264 | chr12-104685443-INS-264 |
| chr12-104690671-INS-89 | chr12-104690671-INS-87 |
| chr12-104698303-INS-86 | chr12-104698303-INS-129 |
| chr12-104706860-INS-147 | chr12-104706890-INS-108 |
| chr12-104720417-INS-63 | chr12-104720417-INS-63 |
| chr12-104726002-INS-104 | chr12-104726002-INS-68 |
| chr12-104735101-INS-274 | chr12-104735101-INS-256 |
| chr12-104760382-INS-70 | chr12-104760382-INS-60 |
| chr12-104781419-INS-53 | chr12-104781419-INS-65 |
| chr12-104784915-INS-194 | chr12-104784915-INS-194 |
| chr12-104787862-INS-235 | chr12-104787862-INS-235 |
| chr12-104792212-INS-66 | chr12-104792212-INS-70 |
| chr12-104802706-INS-1944 | chr12-104802707-INS-1944 |
| chr12-104809020-INS-70 | chr12-104809020-INS-70 |
| chr12-104820546-INS-96 | chr12-104820546-INS-96 |
| chr12-104830411-INS-371 | chr12-104830411-INS-375 |
| chr12-104834736-INS-75 | chr12-104834736-INS-75 |
| chr12-104849981-INS-145 | chr12-104849981-INS-145 |
| chr12-104870957-INS-77 | chr12-104870957-INS-76 |
| chr12-104872345-INS-62 | chr12-104872345-INS-52 |
| chr12-104872803-INS-695 | chr12-104872803-INS-695 |
| chr12-104884627-INS-156 | chr12-104884623-INS-168 |
| chr12-104928034-INS-167 | chr12-104928034-INS-167 |
| chr12-104928822-INS-950 | chr12-104928822-INS-949 |
| chr12-104941972-INS-128 | chr12-104941972-INS-128 |
| chr12-104956256-INS-65 | chr12-104956256-INS-73 |
| chr12-104963529-INS-58 | chr12-104963529-INS-58 |
| chr12-104980274-INS-62 | chr12-104980276-INS-50 |
| chr12-104991985-INS-124 | chr12-104991985-INS-124 |
| chr12-104996135-INS-275 | chr12-104996135-INS-275 |
| chr12-105012517-INS-77 | chr12-105012517-INS-77 |
| chr12-105020379-INS-213 | chr12-105020379-INS-213 |
| chr12-105048793-INS-62 | chr12-105048793-INS-62 |
| chr12-105050471-INS-198 | chr12-105050471-INS-198 |
| chr12-105051911-INS-168 | chr12-105051911-INS-168 |
| chr12-105063448-INS-79 | chr12-105063448-INS-78 |
| chr12-105071559-INS-90 | chr12-105071559-INS-90 |
| chr12-105092164-INS-459 | chr12-105092164-INS-459 |
| chr12-105105236-INS-70 | chr12-105105236-INS-102 |
| chr12-105126503-INS-109 | chr12-105126503-INS-61 |
| chr12-105127801-INS-149 | chr12-105127801-INS-149 |
| chr12-105162498-INS-96 | chr12-105162498-INS-84 |
| chr12-105188550-INS-52 | chr12-105188550-INS-52 |
| chr12-105195000-INS-162 | chr12-105195000-INS-135 |
| chr12-105220075-INS-190 | chr12-105220076-INS-190 |
| chr12-105245224-INS-64 | chr12-105245224-INS-60 |
| chr12-105255574-INS-133 | chr12-105255574-INS-165 |
| chr12-105260548-INS-63 | chr12-105260548-INS-63 |
| chr12-105265831-INS-112 | chr12-105265831-INS-112 |
| chr12-105286417-INS-68 | chr12-105286417-INS-68 |
| chr12-105290978-INS-417 | chr12-105290978-INS-443 |
| chr12-105323866-INS-61 | chr12-105323866-INS-67 |
| chr12-105330721-INS-64 | chr12-105330724-INS-68 |
| chr12-105338887-INS-87 | chr12-105338887-INS-105 |
| chr12-105351172-INS-52 | chr12-105351173-INS-52 |
| chr12-105389038-INS-5227 | chr12-105389039-INS-5227 |
| chr12-105391522-INS-54 | chr12-105391522-INS-58 |
| chr12-105392507-INS-254 | chr12-105392507-INS-281 |
| chr12-105402436-INS-2698 | chr12-105402437-INS-2698 |
| chr12-105417814-INS-564 | chr12-105417814-INS-564 |
| chr12-105425842-INS-92 | chr12-105425842-INS-60 |
| chr12-105439289-INS-344 | chr12-105439289-INS-344 |

| | |
|---|---|
| chr12-105451281-INS-62 | chr12-105451281-INS-58 |
| chr12-105453718-INS-5313 | chr12-105453719-INS-5313 |
| chr12-105462050-INS-54 | chr12-105462050-INS-54 |
| chr12-105511637-INS-143 | chr12-105511636-INS-170 |
| chr12-105512894-INS-54 | chr12-105512874-INS-50 |
| chr12-105514583-INS-209 | chr12-105514583-INS-207 |
| chr12-105537360-INS-56 | chr12-105537360-INS-72 |
| chr12-105544684-INS-72 | chr12-105544684-INS-72 |
| chr12-105557071-INS-172 | chr12-105557071-INS-172 |
| chr12-105569804-INS-60 | chr12-105569804-INS-62 |
| chr12-105579257-INS-56 | chr12-105579269-INS-50 |
| chr12-105621707-INS-51 | chr12-105621707-INS-51 |
| chr12-105640680-INS-208 | chr12-105640680-INS-208 |
| chr12-105649825-INS-211 | chr12-105649825-INS-210 |
| chr12-105675536-INS-139 | chr12-105675536-INS-151 |
| chr12-105704233-INS-86 | chr12-105704233-INS-59 |
| chr12-105709982-INS-194 | chr12-105709983-INS-191 |
| chr12-105712490-INS-86 | chr12-105712491-INS-106 |
| chr12-105728246-INS-79 | chr12-105728246-INS-79 |
| chr12-105736284-INS-403 | chr12-105736284-INS-417 |
| chr12-105741125-INS-186 | chr12-105741125-INS-186 |
| chr12-105745316-INS-59 | chr12-105745332-INS-60 |
| chr12-105749770-INS-114 | chr12-105749770-INS-107 |
| chr12-105759177-INS-200 | chr12-105759177-INS-200 |
| chr12-105835777-INS-54 | chr12-105835754-INS-54 |
| chr12-105842818-INS-51 | chr12-105842819-INS-52 |
| chr12-105853128-INS-94 | chr12-105853128-INS-84 |
| chr12-105857038-INS-7359 | chr12-105857039-INS-7359 |
| chr12-105863114-INS-4962 | chr12-105863115-INS-4962 |
| chr12-105867971-INS-52 | chr12-105867971-INS-52 |
| chr12-105872570-INS-91 | chr12-105872570-INS-91 |
| chr12-105879418-INS-59 | chr12-105879420-INS-58 |
| chr12-105886009-INS-84 | chr12-105885978-INS-84 |
| chr12-105909457-INS-208 | chr12-105909456-INS-213 |
| chr12-105920276-INS-418 | chr12-105920261-INS-418 |
| chr12-105925924-INS-157 | chr12-105925932-INS-184 |
| chr12-105930774-INS-76 | chr12-105930787-INS-51 |
| chr12-105948541-INS-3416 | chr12-105948542-INS-3416 |
| chr12-105949453-INS-134 | chr12-105949453-INS-134 |
| chr12-105965373-INS-66 | chr12-105965373-INS-70 |
| chr12-105990951-INS-2683 | chr12-105990981-INS-2669 |
| chr12-106018888-INS-200 | chr12-106018888-INS-200 |
| chr12-106025410-INS-56 | chr12-106025399-INS-56 |
| chr12-106036245-INS-84 | chr12-106036245-INS-84 |
| chr12-106056923-INS-216 | chr12-106056923-INS-217 |
| chr12-106070723-INS-62 | chr12-106070723-INS-62 |
| chr12-106075936-INS-51 | chr12-106075936-INS-53 |
| chr12-106086248-INS-157 | chr12-106086249-INS-124 |
| chr12-106087698-INS-82 | chr12-106087698-INS-82 |
| chr12-106100230-INS-1517 | chr12-106100231-INS-1517 |
| chr12-106122137-INS-52 | chr12-106122137-INS-52 |
| chr12-106144589-INS-72 | chr12-106144589-INS-72 |
| chr12-106153250-INS-59 | chr12-106153250-INS-56 |
| chr12-106167687-INS-156 | chr12-106167683-INS-139 |
| chr12-106170639-INS-105 | chr12-106170639-INS-138 |
| chr12-106176759-INS-64 | chr12-106176759-INS-72 |
| chr12-106186214-INS-93 | chr12-106186214-INS-82 |
| chr12-106206559-INS-62 | chr12-106206559-INS-62 |
| chr12-106213258-INS-90 | chr12-106213258-INS-51 |
| chr12-106219196-INS-64 | chr12-106219190-INS-68 |
| chr12-106230891-INS-120 | chr12-106230891-INS-126 |
| chr12-106237405-INS-103 | chr12-106237405-INS-103 |
| chr12-106241332-INS-63 | chr12-106241333-INS-65 |
| chr12-106269787-INS-133 | chr12-106269786-INS-174 |
| chr12-106297680-INS-898 | chr12-106297680-INS-898 |
| chr12-106334434-INS-357 | chr12-106334434-INS-347 |
| chr12-106378946-INS-110 | chr12-106378934-INS-116 |
| chr12-106392742-INS-64 | chr12-106392742-INS-64 |
| chr12-106400146-INS-71 | chr12-106400146-INS-71 |
| chr12-106405599-INS-68 | chr12-106405579-INS-74 |
| chr12-106407343-INS-185 | chr12-106407339-INS-186 |
| chr12-106410680-INS-218 | chr12-106410656-INS-235 |
| chr12-106416507-INS-100 | chr12-106416507-INS-108 |
| chr12-106423530-INS-64 | chr12-106423530-INS-72 |
| chr12-106433152-INS-130 | chr12-106433122-INS-150 |
| chr12-106464920-INS-104 | chr12-106464918-INS-59 |
| chr12-106486397-INS-56 | chr12-106486429-INS-68 |

| | |
|---|---|
| chr12-106499362-INS-54 | chr12-106499363-INS-94 |
| chr12-106503050-INS-60 | chr12-106503050-INS-72 |
| chr12-106518693-INS-52 | chr12-106518693-INS-53 |
| chr12-106539028-INS-52 | chr12-106539028-INS-53 |
| chr12-106551619-INS-194 | chr12-106551619-INS-194 |
| chr12-106565723-INS-57 | chr12-106565723-INS-54 |
| chr12-106578266-INS-165 | chr12-106578266-INS-165 |
| chr12-106592484-INS-68 | chr12-106592484-INS-68 |
| chr12-106597938-INS-106 | chr12-106597976-INS-106 |
| chr12-106607019-INS-53 | chr12-106607030-INS-89 |
| chr12-106612756-INS-1038 | chr12-106612757-INS-1038 |
| chr12-106619906-INS-137 | chr12-106619906-INS-102 |
| chr12-106620880-INS-74 | chr12-106620880-INS-90 |
| chr12-106634546-INS-86 | chr12-106634546-INS-86 |
| chr12-106638431-INS-203 | chr12-106638431-INS-189 |
| chr12-106652816-INS-80 | chr12-106652816-INS-76 |
| chr12-106656407-INS-84 | chr12-106656407-INS-84 |
| chr12-106741996-INS-56 | chr12-106741996-INS-50 |
| chr12-106747076-INS-130 | chr12-106747076-INS-130 |
| chr12-106821755-INS-68 | chr12-106821755-INS-76 |
| chr12-106885485-INS-76 | chr12-106885485-INS-76 |
| chr12-106897161-INS-72 | chr12-106897161-INS-78 |
| chr12-106898332-INS-66 | chr12-106898332-INS-66 |
| chr12-106902811-INS-125 | chr12-106902811-INS-125 |
| chr12-106912400-INS-71 | chr12-106912400-INS-77 |
| chr12-106921531-INS-51 | chr12-106921532-INS-72 |
| chr12-106969710-INS-78 | chr12-106969717-INS-62 |
| chr12-106979967-INS-110 | chr12-106979972-INS-105 |
| chr12-106992619-INS-103 | chr12-106992630-INS-90 |
| chr12-107010022-INS-83 | chr12-107010022-INS-64 |
| chr12-107025325-INS-1890 | chr12-107025326-INS-1890 |
| chr12-107028090-INS-412 | chr12-107028090-INS-412 |
| chr12-107040454-INS-5547 | chr12-107040455-INS-5547 |
| chr12-107055041-INS-93 | chr12-107055025-INS-113 |
| chr12-107136764-INS-70 | chr12-107136764-INS-70 |
| chr12-107167982-INS-439 | chr12-107167984-INS-436 |
| chr12-107180286-INS-60 | chr12-107180286-INS-60 |
| chr12-107203326-INS-2774 | chr12-107203327-INS-2774 |
| chr12-107227291-INS-401 | chr12-107227291-INS-401 |
| chr12-107235808-INS-102 | chr12-107235795-INS-92 |
| chr12-107241114-INS-82 | chr12-107241114-INS-82 |
| chr12-107263757-INS-66 | chr12-107263757-INS-63 |
| chr12-107278009-INS-55 | chr12-107278010-INS-55 |
| chr12-107291769-INS-78 | chr12-107291768-INS-78 |
| chr12-107295690-INS-66 | chr12-107295690-INS-65 |
| chr12-107297356-INS-52 | chr12-107297339-INS-76 |
| chr12-107312064-INS-76 | chr12-107312064-INS-78 |
| chr12-107314103-INS-82 | chr12-107314103-INS-72 |
| chr12-107325165-INS-60 | chr12-107325167-INS-56 |
| chr12-107337061-INS-77 | chr12-107337061-INS-72 |
| chr12-107340136-INS-51 | chr12-107340136-INS-51 |
| chr12-107350810-INS-66 | chr12-107350834-INS-56 |
| chr12-107361814-INS-58 | chr12-107361814-INS-58 |
| chr12-107365069-INS-88 | chr12-107365067-INS-96 |
| chr12-107372439-INS-114 | chr12-107372439-INS-114 |
| chr12-107399054-INS-100 | chr12-107399054-INS-100 |
| chr12-107413678-INS-55 | chr12-107413719-INS-64 |
| chr12-107418741-INS-353 | chr12-107418742-INS-348 |
| chr12-107525355-INS-110 | chr12-107525355-INS-127 |
| chr12-107565999-INS-52 | chr12-107565999-INS-60 |
| chr12-107581486-INS-69 | chr12-107581503-INS-72 |
| chr12-107600365-INS-59 | chr12-107600365-INS-57 |
| chr12-107615731-INS-220 | chr12-107615731-INS-220 |
| chr12-107634998-INS-387 | chr12-107634998-INS-384 |
| chr12-107638035-INS-54 | chr12-107638029-INS-60 |
| chr12-107650622-INS-150 | chr12-107650622-INS-160 |
| chr12-107659678-INS-1640 | chr12-107659679-INS-1668 |
| chr12-107699503-INS-717 | chr12-107699503-INS-717 |
| chr12-107704958-INS-240 | chr12-107704958-INS-240 |
| chr12-107705827-INS-399 | chr12-107705827-INS-399 |
| chr12-107713881-INS-60 | chr12-107713881-INS-58 |
| chr12-107738859-INS-52 | chr12-107738857-INS-54 |
| chr12-107764204-INS-86 | chr12-107764204-INS-94 |
| chr12-107784292-INS-60 | chr12-107784292-INS-60 |
| chr12-107812507-INS-77 | chr12-107812495-INS-72 |
| chr12-107813678-INS-67 | chr12-107813678-INS-66 |
| chr12-107848864-INS-62 | chr12-107848864-INS-62 |

| | |
|---|---|
| chr12-107877986-INS-56 | chr12-107877941-INS-84 |
| chr12-107912148-INS-60 | chr12-107912148-INS-60 |
| chr12-107924162-INS-75 | chr12-107924162-INS-80 |
| chr12-107934405-INS-213 | chr12-107934405-INS-213 |
| chr12-107935023-INS-56 | chr12-107935023-INS-58 |
| chr12-107953860-INS-101 | chr12-107953860-INS-89 |
| chr12-107963126-INS-60 | chr12-107963126-INS-56 |
| chr12-107976171-INS-50 | chr12-107976171-INS-53 |
| chr12-107988406-INS-150 | chr12-107988406-INS-150 |
| chr12-107991246-INS-197 | chr12-107991246-INS-187 |
| chr12-108025442-INS-131 | chr12-108025442-INS-131 |
| chr12-108026748-INS-60 | chr12-108026748-INS-60 |
| chr12-108031752-INS-55 | chr12-108031753-INS-56 |
| chr12-108107006-INS-174 | chr12-108107006-INS-174 |
| chr12-108149835-INS-7094 | chr12-108149836-INS-7094 |
| chr12-108195089-INS-96 | chr12-108195089-INS-68 |
| chr12-108197944-INS-140 | chr12-108197944-INS-141 |
| chr12-108212317-INS-85 | chr12-108212312-INS-62 |
| chr12-108236710-INS-117 | chr12-108236704-INS-81 |
| chr12-108254510-INS-56 | chr12-108254510-INS-55 |
| chr12-108258866-INS-120 | chr12-108258866-INS-120 |
| chr12-108261634-INS-111 | chr12-108261634-INS-111 |
| chr12-108266725-INS-302 | chr12-108266725-INS-296 |
| chr12-108279409-INS-6439 | chr12-108279410-INS-6439 |
| chr12-108281683-INS-60 | chr12-108281683-INS-60 |
| chr12-108283723-INS-494 | chr12-108283723-INS-494 |
| chr12-108292252-INS-213 | chr12-108292252-INS-213 |
| chr12-108311519-INS-55 | chr12-108311519-INS-55 |
| chr12-108345937-INS-53 | chr12-108345937-INS-54 |
| chr12-108354977-INS-129 | chr12-108354977-INS-117 |
| chr12-108368283-INS-58 | chr12-108368283-INS-60 |
| chr12-108420764-INS-69 | chr12-108420764-INS-57 |
| chr12-108422917-INS-498 | chr12-108422917-INS-498 |
| chr12-108446628-INS-52 | chr12-108446628-INS-56 |
| chr12-108455701-INS-86 | chr12-108455701-INS-87 |
| chr12-108471309-INS-218 | chr12-108471309-INS-218 |
| chr12-108490313-INS-56 | chr12-108490317-INS-50 |
| chr12-108500783-INS-182 | chr12-108500783-INS-192 |
| chr12-108511028-INS-75 | chr12-108511028-INS-74 |
| chr12-108516384-INS-576 | chr12-108516384-INS-576 |
| chr12-108524720-INS-217 | chr12-108524720-INS-217 |
| chr12-108531880-INS-196 | chr12-108531880-INS-196 |
| chr12-108544536-INS-7107 | chr12-108544537-INS-7107 |
| chr12-108562524-INS-505 | chr12-108562524-INS-506 |
| chr12-108575620-INS-470 | chr12-108575620-INS-470 |
| chr12-108577594-INS-339 | chr12-108577594-INS-345 |
| chr12-108594768-INS-54 | chr12-108594768-INS-56 |
| chr12-108601784-INS-89 | chr12-108601784-INS-89 |
| chr12-108614181-INS-449 | chr12-108614181-INS-449 |
| chr12-108620297-INS-71 | chr12-108620294-INS-71 |
| chr12-108640246-INS-80 | chr12-108640246-INS-75 |
| chr12-108648615-INS-55 | chr12-108648615-INS-55 |
| chr12-108661204-INS-57 | chr12-108661199-INS-90 |
| chr12-108673745-INS-51 | chr12-108673745-INS-51 |
| chr12-108684663-INS-60 | chr12-108684663-INS-90 |
| chr12-108700065-INS-205 | chr12-108700065-INS-162 |
| chr12-108716024-INS-332 | chr12-108716024-INS-332 |
| chr12-108723647-INS-191 | chr12-108723647-INS-171 |
| chr12-108734025-INS-192 | chr12-108734025-INS-192 |
| chr12-108735579-INS-283 | chr12-108735579-INS-285 |
| chr12-108736638-INS-54 | chr12-108736638-INS-56 |
| chr12-108738413-INS-267 | chr12-108738413-INS-267 |
| chr12-108745017-INS-516 | chr12-108745017-INS-516 |
| chr12-108752100-INS-66 | chr12-108752100-INS-74 |
| chr12-108764100-INS-780 | chr12-108764100-INS-780 |
| chr12-108814793-INS-489 | chr12-108814793-INS-489 |
| chr12-108828594-INS-250 | chr12-108828594-INS-250 |
| chr12-108875444-INS-96 | chr12-108875444-INS-96 |
| chr12-108887843-INS-81 | chr12-108887843-INS-81 |
| chr12-108904724-INS-410 | chr12-108904724-INS-410 |
| chr12-108925758-INS-54 | chr12-108925758-INS-54 |
| chr12-108966252-INS-82 | chr12-108966257-INS-102 |
| chr12-108984265-INS-522 | chr12-108984265-INS-522 |
| chr12-109008636-INS-77 | chr12-109008636-INS-77 |
| chr12-109022376-INS-51 | chr12-109022376-INS-51 |
| chr12-109032684-INS-73 | chr12-109032699-INS-51 |
| chr12-109061390-INS-60 | chr12-109061390-INS-60 |

| | |
|---|---|
| chr12-109067581-INS-78 | chr12-109067581-INS-78 |
| chr12-109069597-INS-152 | chr12-109069579-INS-154 |
| chr12-109091487-INS-399 | chr12-109091487-INS-399 |
| chr12-109108041-INS-70 | chr12-109108041-INS-70 |
| chr12-109121543-INS-365 | chr12-109121543-INS-365 |
| chr12-109138762-INS-807 | chr12-109138762-INS-807 |
| chr12-109139549-INS-336 | chr12-109139549-INS-336 |
| chr12-109147126-INS-498 | chr12-109147126-INS-498 |
| chr12-109149998-INS-333 | chr12-109149998-INS-333 |
| chr12-109158939-INS-1891 | chr12-109158940-INS-1892 |
| chr12-109165201-INS-73 | chr12-109165201-INS-73 |
| chr12-109174555-INS-105 | chr12-109174555-INS-105 |
| chr12-109183488-INS-56 | chr12-109183476-INS-76 |
| chr12-109186657-INS-87 | chr12-109186657-INS-79 |
| chr12-109221670-INS-82 | chr12-109221698-INS-64 |
| chr12-109247867-INS-6342 | chr12-109247868-INS-6342 |
| chr12-109254656-INS-277 | chr12-109254657-INS-278 |
| chr12-109278180-INS-53 | chr12-109278203-INS-54 |
| chr12-109299909-INS-368 | chr12-109299909-INS-368 |
| chr12-109312360-INS-569 | chr12-109312360-INS-569 |
| chr12-109314644-INS-1730 | chr12-109314645-INS-1730 |
| chr12-109321386-INS-76 | chr12-109321387-INS-84 |
| chr12-109327950-INS-6489 | chr12-109327951-INS-6490 |
| chr12-109330084-INS-166 | chr12-109330099-INS-193 |
| chr12-109338653-INS-176 | chr12-109338653-INS-169 |
| chr12-109342520-INS-202 | chr12-109342520-INS-202 |
| chr12-109346983-INS-151 | chr12-109346983-INS-151 |
| chr12-109356107-INS-56 | chr12-109356107-INS-64 |
| chr12-109393437-INS-54 | chr12-109393440-INS-51 |
| chr12-109400901-INS-64 | chr12-109400902-INS-64 |
| chr12-109415752-INS-142 | chr12-109415752-INS-142 |
| chr12-109423284-INS-62 | chr12-109423284-INS-62 |
| chr12-109459624-INS-96 | chr12-109459624-INS-96 |
| chr12-109465521-INS-66 | chr12-109465521-INS-64 |
| chr12-109531554-INS-156 | chr12-109531554-INS-156 |
| chr12-109540605-INS-142 | chr12-109540606-INS-142 |
| chr12-109548605-INS-66 | chr12-109548605-INS-66 |
| chr12-109561092-INS-66 | chr12-109561092-INS-66 |
| chr12-109603001-INS-89 | chr12-109603001-INS-89 |
| chr12-109620489-INS-104 | chr12-109620489-INS-104 |
| chr12-109623330-INS-220 | chr12-109623330-INS-221 |
| chr12-109652726-INS-99 | chr12-109652726-INS-145 |
| chr12-109660663-INS-111 | chr12-109660663-INS-111 |
| chr12-109665862-INS-66 | chr12-109665862-INS-68 |
| chr12-109674191-INS-82 | chr12-109674191-INS-97 |
| chr12-109770227-INS-58 | chr12-109770227-INS-58 |
| chr12-109784714-INS-429 | chr12-109784714-INS-429 |
| chr12-109796747-INS-52 | chr12-109796747-INS-70 |
| chr12-109804214-INS-241 | chr12-109804215-INS-241 |
| chr12-109844673-INS-221 | chr12-109844673-INS-219 |
| chr12-109845562-INS-197 | chr12-109845562-INS-197 |
| chr12-109854495-INS-330 | chr12-109854496-INS-324 |
| chr12-109866408-INS-198 | chr12-109866408-INS-198 |
| chr12-109887378-INS-60 | chr12-109887378-INS-82 |
| chr12-109905066-INS-52 | chr12-109905066-INS-52 |
| chr12-109910102-INS-9045 | chr12-109910103-INS-9046 |
| chr12-109911529-INS-200 | chr12-109911529-INS-199 |
| chr12-109924156-INS-56 | chr12-109924156-INS-78 |
| chr12-109931423-INS-98 | chr12-109931423-INS-98 |
| chr12-109966641-INS-7306 | chr12-109966642-INS-7306 |
| chr12-109970707-INS-51 | chr12-109970686-INS-55 |
| chr12-110004250-INS-599 | chr12-110004250-INS-617 |
| chr12-110020956-INS-60 | chr12-110020956-INS-60 |
| chr12-110041009-INS-54 | chr12-110041009-INS-54 |
| chr12-110084961-INS-50 | chr12-110084961-INS-56 |
| chr12-110104529-INS-386 | chr12-110104529-INS-385 |
| chr12-110109567-INS-62 | chr12-110109545-INS-82 |
| chr12-110116954-INS-63 | chr12-110116954-INS-63 |
| chr12-110129027-INS-61 | chr12-110129027-INS-61 |
| chr12-110157768-INS-78 | chr12-110157768-INS-82 |
| chr12-110159131-INS-77 | chr12-110159151-INS-63 |
| chr12-110229334-INS-109 | chr12-110229334-INS-115 |
| chr12-110253041-INS-54 | chr12-110253054-INS-55 |
| chr12-110269494-INS-60 | chr12-110269495-INS-56 |
| chr12-110279359-INS-3105 | chr12-110279360-INS-3103 |
| chr12-110281057-INS-134 | chr12-110281057-INS-134 |
| chr12-110310134-INS-57 | chr12-110310134-INS-57 |

| | |
|---|---|
| chr12-110322020-INS-145 | chr12-110322020-INS-144 |
| chr12-110332987-INS-74 | chr12-110332987-INS-64 |
| chr12-110348225-INS-677 | chr12-110348247-INS-668 |
| chr12-110358895-INS-374 | chr12-110358895-INS-374 |
| chr12-110374485-INS-213 | chr12-110374485-INS-213 |
| chr12-110375806-INS-179 | chr12-110375806-INS-177 |
| chr12-110378294-INS-177 | chr12-110378290-INS-158 |
| chr12-110382379-INS-65 | chr12-110382379-INS-57 |
| chr12-110394066-INS-3545 | chr12-110394067-INS-3545 |
| chr12-110397474-INS-189 | chr12-110397474-INS-197 |
| chr12-110405961-INS-212 | chr12-110405961-INS-213 |
| chr12-110449863-INS-462 | chr12-110449863-INS-480 |
| chr12-110456384-INS-80 | chr12-110456384-INS-80 |
| chr12-110485845-INS-84 | chr12-110485845-INS-83 |
| chr12-110505547-INS-63 | chr12-110505547-INS-82 |
| chr12-110521799-INS-207 | chr12-110521799-INS-203 |
| chr12-110533168-INS-51 | chr12-110533169-INS-63 |
| chr12-110564366-INS-120 | chr12-110564366-INS-120 |
| chr12-110608166-INS-213 | chr12-110608150-INS-219 |
| chr12-110609437-INS-96 | chr12-110609437-INS-96 |
| chr12-110611126-INS-221 | chr12-110611126-INS-221 |
| chr12-110620008-INS-202 | chr12-110620008-INS-200 |
| chr12-110624052-INS-143 | chr12-110624052-INS-143 |
| chr12-110626189-INS-94 | chr12-110626189-INS-94 |
| chr12-110626682-INS-258 | chr12-110626682-INS-258 |
| chr12-110639695-INS-220 | chr12-110639681-INS-201 |
| chr12-110644989-INS-197 | chr12-110644989-INS-197 |
| chr12-110656991-INS-199 | chr12-110656991-INS-199 |
| chr12-110664799-INS-221 | chr12-110664799-INS-221 |
| chr12-110670657-INS-196 | chr12-110670657-INS-196 |
| chr12-110675055-INS-168 | chr12-110675055-INS-168 |
| chr12-110677800-INS-197 | chr12-110677800-INS-197 |
| chr12-110682125-INS-51 | chr12-110682125-INS-63 |
| chr12-110682827-INS-212 | chr12-110682827-INS-211 |
| chr12-110684487-INS-78 | chr12-110684457-INS-88 |
| chr12-110686150-INS-1431 | chr12-110686150-INS-1440 |
| chr12-110689969-INS-170 | chr12-110689968-INS-171 |
| chr12-110695513-INS-203 | chr12-110695513-INS-204 |
| chr12-110712855-INS-195 | chr12-110712855-INS-195 |
| chr12-110716020-INS-217 | chr12-110716013-INS-222 |
| chr12-110718521-INS-203 | chr12-110718521-INS-204 |
| chr12-110729252-INS-375 | chr12-110729252-INS-376 |
| chr12-110734483-INS-209 | chr12-110734483-INS-209 |
| chr12-110781963-INS-199 | chr12-110781963-INS-198 |
| chr12-110784758-INS-156 | chr12-110784758-INS-195 |
| chr12-110814928-INS-327 | chr12-110814928-INS-327 |
| chr12-110820357-INS-366 | chr12-110820382-INS-355 |
| chr12-110823262-INS-4732 | chr12-110823263-INS-4732 |
| chr12-110825906-INS-701 | chr12-110825906-INS-701 |
| chr12-110829605-INS-80 | chr12-110829605-INS-80 |
| chr12-110842181-INS-205 | chr12-110842182-INS-174 |
| chr12-110857289-INS-213 | chr12-110857289-INS-213 |
| chr12-110861265-INS-2215 | chr12-110861266-INS-2215 |
| chr12-110870567-INS-177 | chr12-110870567-INS-178 |
| chr12-110898281-INS-175 | chr12-110898281-INS-175 |
| chr12-110917092-INS-6604 | chr12-110917093-INS-6604 |
| chr12-110920744-INS-230 | chr12-110920744-INS-184 |
| chr12-110924061-INS-68 | chr12-110924061-INS-72 |
| chr12-110931035-INS-60 | chr12-110931035-INS-60 |
| chr12-110933298-INS-211 | chr12-110933298-INS-212 |
| chr12-110934918-INS-54 | chr12-110934918-INS-57 |
| chr12-110963452-INS-54 | chr12-110963452-INS-60 |
| chr12-110964039-INS-103 | chr12-110964039-INS-93 |
| chr12-110984486-INS-194 | chr12-110984486-INS-195 |
| chr12-111014051-INS-134 | chr12-111014051-INS-134 |
| chr12-111020504-INS-110 | chr12-111020504-INS-72 |
| chr12-111025749-INS-200 | chr12-111025745-INS-203 |
| chr12-111031324-INS-1823 | chr12-111031325-INS-1823 |
| chr12-111043676-INS-60 | chr12-111043676-INS-83 |
| chr12-111074341-INS-62 | chr12-111074336-INS-60 |
| chr12-111092328-INS-237 | chr12-111092328-INS-252 |
| chr12-111123774-INS-217 | chr12-111123774-INS-217 |
| chr12-111144312-INS-52 | chr12-111144312-INS-52 |
| chr12-111145299-INS-282 | chr12-111145337-INS-274 |
| chr12-111151514-INS-200 | chr12-111151514-INS-200 |
| chr12-111165515-INS-84 | chr12-111165551-INS-84 |
| chr12-111174267-INS-133 | chr12-111174267-INS-93 |

| | |
|---|---|
| chr12-111175412-INS-206 | chr12-111175412-INS-205 |
| chr12-111190586-INS-211 | chr12-111190586-INS-211 |
| chr12-111202406-INS-782 | chr12-111202406-INS-783 |
| chr12-111246967-INS-284 | chr12-111246967-INS-277 |
| chr12-111258171-INS-109 | chr12-111258159-INS-72 |
| chr12-111279746-INS-59 | chr12-111279746-INS-59 |
| chr12-111283461-INS-73 | chr12-111283461-INS-73 |
| chr12-111291683-INS-159 | chr12-111291700-INS-160 |
| chr12-111295892-INS-173 | chr12-111295892-INS-173 |
| chr12-111312429-INS-219 | chr12-111312429-INS-217 |
| chr12-111314695-INS-144 | chr12-111314694-INS-146 |
| chr12-111326584-INS-224 | chr12-111326584-INS-238 |
| chr12-111332247-INS-1039 | chr12-111332248-INS-1039 |
| chr12-111367621-INS-203 | chr12-111367621-INS-203 |
| chr12-111419285-INS-136 | chr12-111419285-INS-132 |
| chr12-111442570-INS-132 | chr12-111442578-INS-140 |
| chr12-111445036-INS-176 | chr12-111445036-INS-176 |
| chr12-111459625-INS-122 | chr12-111459625-INS-115 |
| chr12-111464276-INS-288 | chr12-111464307-INS-306 |
| chr12-111472824-INS-319 | chr12-111472825-INS-319 |
| chr12-111480103-INS-60 | chr12-111480115-INS-63 |
| chr12-111481654-INS-204 | chr12-111481654-INS-204 |
| chr12-111495325-INS-95 | chr12-111495329-INS-69 |
| chr12-111520569-INS-178 | chr12-111520569-INS-177 |
| chr12-111521694-INS-77 | chr12-111521732-INS-67 |
| chr12-111527227-INS-100 | chr12-111527220-INS-101 |
| chr12-111528188-INS-56 | chr12-111528188-INS-56 |
| chr12-111532154-INS-200 | chr12-111532154-INS-200 |
| chr12-111533160-INS-252 | chr12-111533160-INS-251 |
| chr12-111550452-INS-58 | chr12-111550452-INS-58 |
| chr12-111576151-INS-219 | chr12-111576151-INS-219 |
| chr12-111577234-INS-397 | chr12-111577234-INS-397 |
| chr12-111583742-INS-213 | chr12-111583742-INS-213 |
| chr12-111589769-INS-213 | chr12-111589769-INS-214 |
| chr12-111602562-INS-170 | chr12-111602562-INS-170 |
| chr12-111607508-INS-7483 | chr12-111607509-INS-7483 |
| chr12-111608207-INS-207 | chr12-111608207-INS-207 |
| chr12-111611834-INS-228 | chr12-111611834-INS-266 |
| chr12-111616091-INS-860 | chr12-111616091-INS-860 |
| chr12-111641338-INS-192 | chr12-111641338-INS-178 |
| chr12-111675549-INS-78 | chr12-111675549-INS-78 |
| chr12-111685881-INS-63 | chr12-111685881-INS-63 |
| chr12-111702591-INS-52 | chr12-111702590-INS-62 |
| chr12-111705027-INS-68 | chr12-111705027-INS-74 |
| chr12-111707124-INS-59 | chr12-111707118-INS-63 |
| chr12-111714634-INS-63 | chr12-111714634-INS-64 |
| chr12-111720267-INS-55 | chr12-111720267-INS-51 |
| chr12-111747360-INS-91 | chr12-111747360-INS-91 |
| chr12-111750328-INS-203 | chr12-111750344-INS-214 |
| chr12-111751248-INS-400 | chr12-111751249-INS-400 |
| chr12-111757161-INS-267 | chr12-111757161-INS-267 |
| chr12-111758513-INS-217 | chr12-111758513-INS-217 |
| chr12-111776586-INS-80 | chr12-111776586-INS-82 |
| chr12-111778592-INS-73 | chr12-111778592-INS-73 |
| chr12-111788630-INS-1722 | chr12-111788631-INS-1721 |
| chr12-111804632-INS-624 | chr12-111804632-INS-624 |
| chr12-111807786-INS-216 | chr12-111807786-INS-217 |
| chr12-111812118-INS-82 | chr12-111812118-INS-50 |
| chr12-111816005-INS-62 | chr12-111816025-INS-60 |
| chr12-111846936-INS-285 | chr12-111846930-INS-242 |
| chr12-111848428-INS-95 | chr12-111848428-INS-95 |
| chr12-111852762-INS-68 | chr12-111852762-INS-70 |
| chr12-111887304-INS-79 | chr12-111887304-INS-79 |
| chr12-111906839-INS-54 | chr12-111906841-INS-62 |
| chr12-111935894-INS-161 | chr12-111935894-INS-161 |
| chr12-111969815-INS-65 | chr12-111969815-INS-65 |
| chr12-111990191-INS-66 | chr12-111990191-INS-66 |
| chr12-112004008-INS-213 | chr12-112004008-INS-213 |
| chr12-112033119-INS-212 | chr12-112033119-INS-214 |
| chr12-112034534-INS-203 | chr12-112034534-INS-203 |
| chr12-112045685-INS-101 | chr12-112045685-INS-101 |
| chr12-112076636-INS-196 | chr12-112076636-INS-197 |
| chr12-112121589-INS-166 | chr12-112121589-INS-166 |
| chr12-112128600-INS-61 | chr12-112128600-INS-61 |
| chr12-112132213-INS-172 | chr12-112132213-INS-172 |
| chr12-112135720-INS-592 | chr12-112135720-INS-592 |
| chr12-112158648-INS-112 | chr12-112158648-INS-108 |

| | |
|---|---|
| chr12-112211839-INS-170 | chr12-112211839-INS-170 |
| chr12-112213483-INS-268 | chr12-112213482-INS-266 |
| chr12-112219035-INS-70 | chr12-112219035-INS-70 |
| chr12-112236936-INS-406 | chr12-112236936-INS-406 |
| chr12-112247547-INS-99 | chr12-112247547-INS-99 |
| chr12-112260862-INS-56 | chr12-112260893-INS-56 |
| chr12-112280939-INS-1858 | chr12-112280940-INS-1858 |
| chr12-112282431-INS-115 | chr12-112282431-INS-115 |
| chr12-112298349-INS-66 | chr12-112298350-INS-65 |
| chr12-112306098-INS-79 | chr12-112306098-INS-79 |
| chr12-112319538-INS-61 | chr12-112319539-INS-55 |
| chr12-112333707-INS-58 | chr12-112333707-INS-58 |
| chr12-112348637-INS-88 | chr12-112348637-INS-90 |
| chr12-112363047-INS-54 | chr12-112363047-INS-54 |
| chr12-112374157-INS-158 | chr12-112374157-INS-195 |
| chr12-112405719-INS-85 | chr12-112405719-INS-67 |
| chr12-112411763-INS-57 | chr12-112411764-INS-57 |
| chr12-112416829-INS-378 | chr12-112416829-INS-378 |
| chr12-112423487-INS-105 | chr12-112423487-INS-153 |
| chr12-112439477-INS-237 | chr12-112439477-INS-228 |
| chr12-112492493-INS-201 | chr12-112492493-INS-202 |
| chr12-112498149-INS-54 | chr12-112498149-INS-52 |
| chr12-112527297-INS-55 | chr12-112527299-INS-86 |
| chr12-112531541-INS-205 | chr12-112531541-INS-205 |
| chr12-112552763-INS-230 | chr12-112552763-INS-230 |
| chr12-112567083-INS-225 | chr12-112567083-INS-228 |
| chr12-112573031-INS-93 | chr12-112573031-INS-93 |
| chr12-112597583-INS-215 | chr12-112597583-INS-235 |
| chr12-112608678-INS-210 | chr12-112608679-INS-210 |
| chr12-112633478-INS-56 | chr12-112633478-INS-56 |
| chr12-112643431-INS-82 | chr12-112643431-INS-82 |
| chr12-112651371-INS-96 | chr12-112651371-INS-96 |
| chr12-112651583-INS-5311 | chr12-112651584-INS-5311 |
| chr12-112679875-INS-61 | chr12-112679853-INS-80 |
| chr12-112710691-INS-80 | chr12-112710722-INS-60 |
| chr12-112718312-INS-440 | chr12-112718312-INS-440 |
| chr12-112766487-INS-59 | chr12-112766489-INS-56 |
| chr12-112778904-INS-55 | chr12-112778904-INS-55 |
| chr12-112796831-INS-225 | chr12-112796831-INS-214 |
| chr12-112803044-INS-121 | chr12-112803044-INS-121 |
| chr12-112804834-INS-270 | chr12-112804834-INS-284 |
| chr12-112822747-INS-63 | chr12-112822747-INS-63 |
| chr12-112833534-INS-84 | chr12-112833534-INS-84 |
| chr12-112846571-INS-203 | chr12-112846571-INS-203 |
| chr12-112857222-INS-257 | chr12-112857222-INS-257 |
| chr12-112860421-INS-54 | chr12-112860421-INS-56 |
| chr12-112863688-INS-63 | chr12-112863688-INS-63 |
| chr12-112869380-INS-67 | chr12-112869380-INS-67 |
| chr12-112883810-INS-57 | chr12-112883810-INS-57 |
| chr12-112935868-INS-102 | chr12-112935868-INS-101 |
| chr12-112949885-INS-66 | chr12-112949931-INS-70 |
| chr12-112954829-INS-188 | chr12-112954828-INS-187 |
| chr12-112986334-INS-184 | chr12-112986332-INS-190 |
| chr12-113017299-INS-137 | chr12-113017299-INS-143 |
| chr12-113040798-INS-100 | chr12-113040798-INS-97 |
| chr12-113058829-INS-144 | chr12-113058829-INS-192 |
| chr12-113085353-INS-61 | chr12-113085353-INS-61 |
| chr12-113130431-INS-325 | chr12-113130431-INS-325 |
| chr12-113137960-INS-3148 | chr12-113137961-INS-3165 |
| chr12-113155819-INS-333 | chr12-113155819-INS-333 |
| chr12-113165798-INS-744 | chr12-113165798-INS-744 |
| chr12-113168751-INS-56 | chr12-113168751-INS-56 |
| chr12-113171730-INS-210 | chr12-113171730-INS-210 |
| chr12-113177661-INS-199 | chr12-113177661-INS-199 |
| chr12-113178730-INS-199 | chr12-113178730-INS-203 |
| chr12-113188962-INS-238 | chr12-113188962-INS-221 |
| chr12-113196607-INS-117 | chr12-113196607-INS-113 |
| chr12-113199628-INS-75 | chr12-113199628-INS-50 |
| chr12-113203546-INS-76 | chr12-113203546-INS-76 |
| chr12-113207740-INS-70 | chr12-113207744-INS-60 |
| chr12-113212504-INS-257 | chr12-113212503-INS-223 |
| chr12-113226172-INS-70 | chr12-113226173-INS-70 |
| chr12-113228133-INS-70 | chr12-113228133-INS-70 |
| chr12-113255755-INS-122 | chr12-113255755-INS-122 |
| chr12-113266711-INS-121 | chr12-113266711-INS-122 |
| chr12-113282154-INS-135 | chr12-113282154-INS-183 |
| chr12-113313447-INS-311 | chr12-113313447-INS-311 |

| | |
|---|---|
| chr12-113327219-INS-722 | chr12-113327219-INS-722 |
| chr12-113337425-INS-128 | chr12-113337425-INS-128 |
| chr12-113346305-INS-958 | chr12-113346290-INS-962 |
| chr12-113356307-INS-74 | chr12-113356307-INS-74 |
| chr12-113359381-INS-1800 | chr12-113359382-INS-1800 |
| chr12-113361044-INS-400 | chr12-113361044-INS-400 |
| chr12-113388011-INS-3149 | chr12-113388012-INS-3149 |
| chr12-113400353-INS-6192 | chr12-113400354-INS-6192 |
| chr12-113401127-INS-140 | chr12-113401127-INS-140 |
| chr12-113403018-INS-582 | chr12-113403018-INS-584 |
| chr12-113411615-INS-54 | chr12-113411615-INS-54 |
| chr12-113430655-INS-327 | chr12-113430655-INS-327 |
| chr12-113437648-INS-61 | chr12-113437648-INS-81 |
| chr12-113442008-INS-315 | chr12-113442008-INS-315 |
| chr12-113468363-INS-6434 | chr12-113468364-INS-6434 |
| chr12-113488002-INS-60 | chr12-113488006-INS-56 |
| chr12-113500735-INS-207 | chr12-113500735-INS-213 |
| chr12-113512191-INS-184 | chr12-113512191-INS-182 |
| chr12-113514320-INS-6482 | chr12-113514321-INS-6483 |
| chr12-113529448-INS-585 | chr12-113529453-INS-587 |
| chr12-113542034-INS-404 | chr12-113542032-INS-407 |
| chr12-113546624-INS-806 | chr12-113546624-INS-806 |
| chr12-113547966-INS-60 | chr12-113547964-INS-50 |
| chr12-113610594-INS-496 | chr12-113610594-INS-489 |
| chr12-113660711-INS-106 | chr12-113660694-INS-92 |
| chr12-113711577-INS-72 | chr12-113711577-INS-70 |
| chr12-113747881-INS-6718 | chr12-113747882-INS-6717 |
| chr12-113771899-INS-115 | chr12-113771899-INS-80 |
| chr12-113789786-INS-65 | chr12-113789835-INS-65 |
| chr12-113797214-INS-95 | chr12-113797214-INS-95 |
| chr12-113820890-INS-54 | chr12-113820910-INS-64 |
| chr12-113833713-INS-6430 | chr12-113833714-INS-6430 |
| chr12-113866678-INS-166 | chr12-113866678-INS-166 |
| chr12-113875382-INS-194 | chr12-113875382-INS-194 |
| chr12-113887723-INS-51 | chr12-113887723-INS-51 |
| chr12-113895048-INS-406 | chr12-113895049-INS-406 |
| chr12-113984766-INS-243 | chr12-113984766-INS-243 |
| chr12-114002541-INS-404 | chr12-114002541-INS-404 |
| chr12-114021697-INS-764 | chr12-114021697-INS-764 |
| chr12-114046239-INS-341 | chr12-114046239-INS-341 |
| chr12-114047962-INS-411 | chr12-114047985-INS-429 |
| chr12-114110228-INS-86 | chr12-114110228-INS-69 |
| chr12-114127859-INS-181 | chr12-114127859-INS-179 |
| chr12-114153242-INS-168 | chr12-114153242-INS-167 |
| chr12-114167464-INS-870 | chr12-114167464-INS-870 |
| chr12-114178276-INS-498 | chr12-114178276-INS-498 |
| chr12-114181346-INS-52 | chr12-114181346-INS-58 |
| chr12-114182418-INS-359 | chr12-114182418-INS-359 |
| chr12-114234255-INS-83 | chr12-114234255-INS-83 |
| chr12-114235076-INS-331 | chr12-114235076-INS-331 |
| chr12-114236618-INS-737 | chr12-114236618-INS-737 |
| chr12-114288239-INS-283 | chr12-114288239-INS-283 |
| chr12-114362324-INS-1100 | chr12-114362325-INS-1100 |
| chr12-114374192-INS-64 | chr12-114374192-INS-64 |
| chr12-114410229-INS-476 | chr12-114410229-INS-476 |
| chr12-114416683-INS-214 | chr12-114416683-INS-216 |
| chr12-114428812-INS-503 | chr12-114428812-INS-503 |
| chr12-114484938-INS-273 | chr12-114484938-INS-273 |
| chr12-114486240-INS-64 | chr12-114486240-INS-64 |
| chr12-114510318-INS-74 | chr12-114510310-INS-63 |
| chr12-114538673-INS-102 | chr12-114538663-INS-80 |
| chr12-114540126-INS-115 | chr12-114540126-INS-115 |
| chr12-114551313-INS-83 | chr12-114551314-INS-83 |
| chr12-114607934-INS-433 | chr12-114607934-INS-433 |
| chr12-114613819-INS-74 | chr12-114613820-INS-63 |
| chr12-114614280-INS-818 | chr12-114614277-INS-824 |
| chr12-114616288-INS-89 | chr12-114616288-INS-89 |
| chr12-114622297-INS-806 | chr12-114622297-INS-806 |
| chr12-114631333-INS-171 | chr12-114631336-INS-199 |
| chr12-114635446-INS-54 | chr12-114635446-INS-53 |
| chr12-114689972-INS-218 | chr12-114689972-INS-218 |
| chr12-114869448-INS-420 | chr12-114869448-INS-420 |
| chr12-114875692-INS-190 | chr12-114875692-INS-190 |
| chr12-114880763-INS-697 | chr12-114880766-INS-698 |
| chr12-114892834-INS-974 | chr12-114892834-INS-974 |
| chr12-114899351-INS-81 | chr12-114899351-INS-81 |
| chr12-114903937-INS-56 | chr12-114903937-INS-60 |

| | |
|---|---|
| chr12-114930456-INS-56 | chr12-114930456-INS-50 |
| chr12-114931634-INS-68 | chr12-114931634-INS-66 |
| chr12-114940654-INS-7302 | chr12-114940655-INS-7302 |
| chr12-114961254-INS-416 | chr12-114961254-INS-416 |
| chr12-114963874-INS-87 | chr12-114963874-INS-87 |
| chr12-114964789-INS-315 | chr12-114964789-INS-324 |
| chr12-114973451-INS-799 | chr12-114973451-INS-799 |
| chr12-114975748-INS-56 | chr12-114975748-INS-56 |
| chr12-114982428-INS-563 | chr12-114982428-INS-563 |
| chr12-114998853-INS-483 | chr12-114998853-INS-483 |
| chr12-115105418-INS-90 | chr12-115105423-INS-104 |
| chr12-115107887-INS-83 | chr12-115107887-INS-83 |
| chr12-115148416-INS-225 | chr12-115148416-INS-225 |
| chr12-115329999-INS-79 | chr12-115329999-INS-80 |
| chr12-115343325-INS-62 | chr12-115343334-INS-69 |
| chr12-115409857-INS-188 | chr12-115409857-INS-188 |
| chr12-115422164-INS-318 | chr12-115422164-INS-318 |
| chr12-115433824-INS-307 | chr12-115433824-INS-306 |
| chr12-115509385-INS-942 | chr12-115509385-INS-944 |
| chr12-115668285-INS-141 | chr12-115668285-INS-141 |
| chr12-115677276-INS-351 | chr12-115677241-INS-345 |
| chr12-115729272-INS-131 | chr12-115729272-INS-131 |
| chr12-115803973-INS-61 | chr12-115803973-INS-56 |
| chr12-115817966-INS-60 | chr12-115817966-INS-61 |
| chr12-115854735-INS-66 | chr12-115854735-INS-66 |
| chr12-115872974-INS-100 | chr12-115872974-INS-100 |
| chr12-115899216-INS-2538 | chr12-115899217-INS-2539 |
| chr12-115964688-INS-68 | chr12-115964672-INS-86 |
| chr12-115988451-INS-53 | chr12-115988448-INS-50 |
| chr12-115999628-INS-400 | chr12-115999628-INS-400 |
| chr12-116014517-INS-106 | chr12-116014517-INS-106 |
| chr12-116034038-INS-3154 | chr12-116034039-INS-3154 |
| chr12-116036639-INS-72 | chr12-116036639-INS-72 |
| chr12-116039950-INS-258 | chr12-116039950-INS-261 |
| chr12-116040761-INS-478 | chr12-116040761-INS-478 |
| chr12-116044509-INS-170 | chr12-116044509-INS-171 |
| chr12-116048783-INS-508 | chr12-116048783-INS-508 |
| chr12-116052590-INS-254 | chr12-116052590-INS-254 |
| chr12-116054298-INS-51 | chr12-116054298-INS-51 |
| chr12-116055986-INS-1416 | chr12-116055987-INS-1416 |
| chr12-116060283-INS-7087 | chr12-116060284-INS-7087 |
| chr12-116061081-INS-71 | chr12-116061081-INS-71 |
| chr12-116079710-INS-1061 | chr12-116079711-INS-1061 |
| chr12-116080596-INS-184 | chr12-116080596-INS-184 |
| chr12-116093608-INS-719 | chr12-116093608-INS-719 |
| chr12-116114744-INS-116 | chr12-116114744-INS-116 |
| chr12-116116225-INS-338 | chr12-116116225-INS-338 |
| chr12-116122471-INS-230 | chr12-116122471-INS-223 |
| chr12-116128959-INS-186 | chr12-116128959-INS-186 |
| chr12-116140836-INS-185 | chr12-116140836-INS-185 |
| chr12-116166492-INS-187 | chr12-116166492-INS-187 |
| chr12-116167143-INS-79 | chr12-116167143-INS-79 |
| chr12-116175241-INS-204 | chr12-116175241-INS-204 |
| chr12-116183710-INS-86 | chr12-116183710-INS-86 |
| chr12-116187912-INS-54 | chr12-116187912-INS-54 |
| chr12-116189237-INS-60 | chr12-116189237-INS-60 |
| chr12-116190880-INS-110 | chr12-116190878-INS-116 |
| chr12-116217608-INS-64 | chr12-116217608-INS-50 |
| chr12-116237961-INS-67 | chr12-116237961-INS-67 |
| chr12-116253503-INS-198 | chr12-116253503-INS-198 |
| chr12-116258006-INS-122 | chr12-116258006-INS-122 |
| chr12-116270243-INS-80 | chr12-116270278-INS-50 |
| chr12-116276338-INS-224 | chr12-116276336-INS-227 |
| chr12-116308461-INS-86 | chr12-116308461-INS-86 |
| chr12-116324389-INS-120 | chr12-116324389-INS-120 |
| chr12-116329895-INS-241 | chr12-116329895-INS-241 |
| chr12-116338091-INS-97 | chr12-116338091-INS-97 |
| chr12-116339458-INS-348 | chr12-116339458-INS-348 |
| chr12-116346140-INS-221 | chr12-116346140-INS-221 |
| chr12-116355283-INS-196 | chr12-116355283-INS-196 |
| chr12-116395324-INS-61 | chr12-116395324-INS-61 |
| chr12-116415078-INS-1463 | chr12-116415079-INS-1463 |
| chr12-116442105-INS-53 | chr12-116442110-INS-57 |
| chr12-116467701-INS-88 | chr12-116467701-INS-88 |
| chr12-116472669-INS-83 | chr12-116472669-INS-83 |
| chr12-116477101-INS-128 | chr12-116477057-INS-132 |
| chr12-116478884-INS-1223 | chr12-116478885-INS-1224 |

| | |
|---|---|
| chr12-116489454-INS-79 | chr12-116489454-INS-79 |
| chr12-116503671-INS-120 | chr12-116503671-INS-120 |
| chr12-116533197-INS-67 | chr12-116533197-INS-67 |
| chr12-116553827-INS-114 | chr12-116553827-INS-116 |
| chr12-116568544-INS-84 | chr12-116568544-INS-84 |
| chr12-116570349-INS-60 | chr12-116570349-INS-60 |
| chr12-116581022-INS-231 | chr12-116581022-INS-231 |
| chr12-116589546-INS-985 | chr12-116589546-INS-985 |
| chr12-116593706-INS-283 | chr12-116593706-INS-283 |
| chr12-116598911-INS-116 | chr12-116598911-INS-116 |
| chr12-116602675-INS-86 | chr12-116602675-INS-86 |
| chr12-116613237-INS-150 | chr12-116613255-INS-163 |
| chr12-116618177-INS-66 | chr12-116618177-INS-67 |
| chr12-116624422-INS-71 | chr12-116624422-INS-73 |
| chr12-116626973-INS-1915 | chr12-116626974-INS-1915 |
| chr12-116629083-INS-61 | chr12-116629083-INS-73 |
| chr12-116632690-INS-503 | chr12-116632690-INS-503 |
| chr12-116637282-INS-823 | chr12-116637282-INS-819 |
| chr12-116642118-INS-1463 | chr12-116642119-INS-1438 |
| chr12-116649996-INS-469 | chr12-116649996-INS-469 |
| chr12-116667462-INS-6665 | chr12-116667468-INS-6665 |
| chr12-116671859-INS-356 | chr12-116671859-INS-356 |
| chr12-116690583-INS-86 | chr12-116690583-INS-86 |
| chr12-116699870-INS-203 | chr12-116699870-INS-203 |
| chr12-116700573-INS-56 | chr12-116700573-INS-94 |
| chr12-116732551-INS-138 | chr12-116732551-INS-140 |
| chr12-116750877-INS-1187 | chr12-116750878-INS-1187 |
| chr12-116781191-INS-153 | chr12-116781191-INS-150 |
| chr12-116805829-INS-70 | chr12-116805829-INS-81 |
| chr12-116816463-INS-66 | chr12-116816463-INS-66 |
| chr12-116825466-INS-419 | chr12-116825466-INS-420 |
| chr12-116844292-INS-80 | chr12-116844292-INS-80 |
| chr12-116899304-INS-200 | chr12-116899304-INS-202 |
| chr12-116900406-INS-2348 | chr12-116900407-INS-2349 |
| chr12-116905468-INS-6175 | chr12-116905469-INS-6176 |
| chr12-116912395-INS-65 | chr12-116912417-INS-80 |
| chr12-116917187-INS-5652 | chr12-116917188-INS-5652 |
| chr12-116960228-INS-64 | chr12-116960228-INS-66 |
| chr12-116965755-INS-51 | chr12-116965755-INS-51 |
| chr12-116981293-INS-534 | chr12-116981293-INS-534 |
| chr12-117049202-INS-147 | chr12-117049202-INS-147 |
| chr12-117058330-INS-1443 | chr12-117058331-INS-1443 |
| chr12-117059215-INS-134 | chr12-117059215-INS-134 |
| chr12-117068312-INS-1461 | chr12-117068269-INS-1461 |
| chr12-117080952-INS-144 | chr12-117080952-INS-144 |
| chr12-117103696-INS-55 | chr12-117103697-INS-55 |
| chr12-117109315-INS-192 | chr12-117109315-INS-192 |
| chr12-117142322-INS-341 | chr12-117142322-INS-340 |
| chr12-117178412-INS-70 | chr12-117178412-INS-72 |
| chr12-117206164-INS-81 | chr12-117206164-INS-81 |
| chr12-117223249-INS-64 | chr12-117223249-INS-76 |
| chr12-117243952-INS-327 | chr12-117243952-INS-327 |
| chr12-117252514-INS-1866 | chr12-117252515-INS-1866 |
| chr12-117275212-INS-58 | chr12-117275212-INS-60 |
| chr12-117279531-INS-57 | chr12-117279531-INS-57 |
| chr12-117284920-INS-2756 | chr12-117284921-INS-2756 |
| chr12-117286925-INS-225 | chr12-117286925-INS-232 |
| chr12-117299728-INS-482 | chr12-117299728-INS-482 |
| chr12-117306159-INS-63 | chr12-117306159-INS-63 |
| chr12-117330266-INS-52 | chr12-117330229-INS-58 |
| chr12-117331077-INS-68 | chr12-117331077-INS-70 |
| chr12-117349066-INS-318 | chr12-117349067-INS-320 |
| chr12-117355300-INS-163 | chr12-117355300-INS-163 |
| chr12-117362455-INS-92 | chr12-117362455-INS-92 |
| chr12-117363794-INS-52 | chr12-117363794-INS-54 |
| chr12-117382368-INS-153 | chr12-117382338-INS-161 |
| chr12-117399098-INS-69 | chr12-117399095-INS-99 |
| chr12-117421051-INS-141 | chr12-117421051-INS-137 |
| chr12-117449696-INS-498 | chr12-117449696-INS-498 |
| chr12-117455842-INS-4362 | chr12-117455843-INS-4362 |
| chr12-117457564-INS-149 | chr12-117457564-INS-149 |
| chr12-117484502-INS-263 | chr12-117484502-INS-263 |
| chr12-117525670-INS-58 | chr12-117525670-INS-58 |
| chr12-117528048-INS-90 | chr12-117528046-INS-92 |
| chr12-117551973-INS-2380 | chr12-117551974-INS-2380 |
| chr12-117564319-INS-177 | chr12-117564319-INS-179 |
| chr12-117575042-INS-90 | chr12-117575042-INS-90 |

| | |
|---|---|
| chr12-117579756-INS-138 | chr12-117579756-INS-138 |
| chr12-117588917-INS-104 | chr12-117588917-INS-104 |
| chr12-117643123-INS-6814 | chr12-117643124-INS-6813 |
| chr12-117651741-INS-3481 | chr12-117651742-INS-3481 |
| chr12-117669100-INS-80 | chr12-117669100-INS-80 |
| chr12-117677202-INS-205 | chr12-117677202-INS-205 |
| chr12-117680579-INS-653 | chr12-117680579-INS-653 |
| chr12-117689653-INS-56 | chr12-117689653-INS-56 |
| chr12-117699705-INS-181 | chr12-117699705-INS-181 |
| chr12-117709436-INS-168 | chr12-117709424-INS-178 |
| chr12-117748840-INS-7280 | chr12-117748841-INS-7280 |
| chr12-117754881-INS-492 | chr12-117754881-INS-492 |
| chr12-117771716-INS-70 | chr12-117771715-INS-71 |
| chr12-117779035-INS-60 | chr12-117779035-INS-60 |
| chr12-117779545-INS-1182 | chr12-117779547-INS-1186 |
| chr12-117791798-INS-560 | chr12-117791798-INS-560 |
| chr12-117800817-INS-121 | chr12-117800791-INS-97 |
| chr12-117801406-INS-1504 | chr12-117801406-INS-1490 |
| chr12-117805065-INS-128 | chr12-117805068-INS-175 |
| chr12-117823118-INS-254 | chr12-117823118-INS-255 |
| chr12-117825567-INS-1742 | chr12-117825568-INS-1772 |
| chr12-117831541-INS-245 | chr12-117831541-INS-253 |
| chr12-117833450-INS-64 | chr12-117833450-INS-69 |
| chr12-117840134-INS-54 | chr12-117840135-INS-54 |
| chr12-117844486-INS-55 | chr12-117844486-INS-55 |
| chr12-117848089-INS-63 | chr12-117848089-INS-63 |
| chr12-117851595-INS-199 | chr12-117851595-INS-197 |
| chr12-117879350-INS-630 | chr12-117879349-INS-631 |
| chr12-117880908-INS-431 | chr12-117880908-INS-431 |
| chr12-117881765-INS-124 | chr12-117881778-INS-132 |
| chr12-117889545-INS-220 | chr12-117889545-INS-220 |
| chr12-117909465-INS-168 | chr12-117909503-INS-122 |
| chr12-117922931-INS-218 | chr12-117922931-INS-224 |
| chr12-117940890-INS-79 | chr12-117940890-INS-79 |
| chr12-117954971-INS-52 | chr12-117954971-INS-52 |
| chr12-117989067-INS-80 | chr12-117989067-INS-80 |
| chr12-117991722-INS-5338 | chr12-117991723-INS-5338 |
| chr12-118008340-INS-229 | chr12-118008340-INS-229 |
| chr12-118034523-INS-6843 | chr12-118034524-INS-6844 |
| chr12-118035779-INS-106 | chr12-118035764-INS-120 |
| chr12-118043481-INS-115 | chr12-118043465-INS-84 |
| chr12-118072252-INS-120 | chr12-118072252-INS-120 |
| chr12-118074985-INS-6969 | chr12-118074986-INS-6969 |
| chr12-118078256-INS-107 | chr12-118078263-INS-96 |
| chr12-118078532-INS-1554 | chr12-118078533-INS-1558 |
| chr12-118138657-INS-6853 | chr12-118138658-INS-6853 |
| chr12-118140578-INS-114 | chr12-118140578-INS-112 |
| chr12-118146853-INS-6438 | chr12-118146854-INS-6438 |
| chr12-118155269-INS-415 | chr12-118155269-INS-415 |
| chr12-118173026-INS-92 | chr12-118173026-INS-96 |
| chr12-118190154-INS-62 | chr12-118190154-INS-52 |
| chr12-118245851-INS-6403 | chr12-118245852-INS-6403 |
| chr12-118252698-INS-262 | chr12-118252698-INS-254 |
| chr12-118304720-INS-255 | chr12-118304720-INS-280 |
| chr12-118326229-INS-440 | chr12-118326229-INS-461 |
| chr12-118339476-INS-1011 | chr12-118339477-INS-1011 |
| chr12-118340816-INS-140 | chr12-118340815-INS-177 |
| chr12-118366163-INS-3801 | chr12-118366164-INS-3847 |
| chr12-118393919-INS-57 | chr12-118393919-INS-57 |
| chr12-118396958-INS-164 | chr12-118396958-INS-166 |
| chr12-118430226-INS-2144 | chr12-118430227-INS-2144 |
| chr12-118461074-INS-5259 | chr12-118461075-INS-5259 |
| chr12-118462092-INS-98 | chr12-118462055-INS-114 |
| chr12-118464730-INS-206 | chr12-118464730-INS-208 |
| chr12-118471372-INS-3355 | chr12-118471373-INS-3350 |
| chr12-118481585-INS-531 | chr12-118481585-INS-531 |
| chr12-118483149-INS-100 | chr12-118483149-INS-132 |
| chr12-118517198-INS-97 | chr12-118517191-INS-103 |
| chr12-118550863-INS-222 | chr12-118550863-INS-222 |
| chr12-118556777-INS-5149 | chr12-118556778-INS-5149 |
| chr12-118561970-INS-132 | chr12-118561970-INS-120 |
| chr12-118620804-INS-483 | chr12-118620804-INS-483 |
| chr12-118673557-INS-307 | chr12-118673557-INS-308 |
| chr12-118692103-INS-90 | chr12-118692103-INS-90 |
| chr12-118734102-INS-5156 | chr12-118734103-INS-5156 |
| chr12-118772381-INS-76 | chr12-118772381-INS-77 |
| chr12-118778692-INS-77 | chr12-118778692-INS-59 |

| | |
|---|---|
| chr12-118820706-INS-212 | chr12-118820706-INS-212 |
| chr12-118860088-INS-53 | chr12-118860088-INS-52 |
| chr12-118869275-INS-65 | chr12-118869279-INS-51 |
| chr12-118888491-INS-59 | chr12-118888491-INS-59 |
| chr12-118892140-INS-73 | chr12-118892140-INS-73 |
| chr12-118898313-INS-111 | chr12-118898313-INS-117 |
| chr12-118911457-INS-92 | chr12-118911457-INS-50 |
| chr12-118923634-INS-223 | chr12-118923634-INS-225 |
| chr12-118926125-INS-503 | chr12-118926125-INS-501 |
| chr12-118942432-INS-208 | chr12-118942432-INS-208 |
| chr12-118996033-INS-453 | chr12-118996033-INS-453 |
| chr12-119014420-INS-4527 | chr12-119014421-INS-4527 |
| chr12-119018821-INS-6634 | chr12-119018822-INS-6634 |
| chr12-119051218-INS-6056 | chr12-119051219-INS-6057 |
| chr12-119071824-INS-225 | chr12-119071824-INS-226 |
| chr12-119085716-INS-72 | chr12-119085717-INS-106 |
| chr12-119089636-INS-3340 | chr12-119089637-INS-3340 |
| chr12-119104934-INS-302 | chr12-119104933-INS-297 |
| chr12-119114994-INS-86 | chr12-119114994-INS-86 |
| chr12-119148500-INS-200 | chr12-119148500-INS-200 |
| chr12-119153282-INS-90 | chr12-119153282-INS-87 |
| chr12-119163815-INS-91 | chr12-119163815-INS-91 |
| chr12-119185158-INS-65 | chr12-119185158-INS-65 |
| chr12-119214706-INS-7128 | chr12-119214707-INS-7128 |
| chr12-119220555-INS-173 | chr12-119220555-INS-173 |
| chr12-119221441-INS-448 | chr12-119221441-INS-448 |
| chr12-119245700-INS-2028 | chr12-119245701-INS-2028 |
| chr12-119266702-INS-54 | chr12-119266702-INS-84 |
| chr12-119278976-INS-6368 | chr12-119278977-INS-6368 |
| chr12-119285344-INS-371 | chr12-119285344-INS-371 |
| chr12-119288750-INS-78 | chr12-119288750-INS-78 |
| chr12-119300218-INS-53 | chr12-119300218-INS-53 |
| chr12-119301497-INS-560 | chr12-119301496-INS-561 |
| chr12-119322327-INS-6278 | chr12-119322328-INS-6278 |
| chr12-119348175-INS-64 | chr12-119348175-INS-59 |
| chr12-119352688-INS-56 | chr12-119352666-INS-53 |
| chr12-119373972-INS-6751 | chr12-119373973-INS-6752 |
| chr12-119420015-INS-7341 | chr12-119420016-INS-7343 |
| chr12-119459724-INS-1866 | chr12-119459725-INS-1866 |
| chr12-119472728-INS-2576 | chr12-119472729-INS-2576 |
| chr12-119511893-INS-66 | chr12-119511893-INS-66 |
| chr12-119551927-INS-2268 | chr12-119551928-INS-2268 |
| chr12-119579145-INS-226 | chr12-119579145-INS-226 |
| chr12-119581841-INS-66 | chr12-119581841-INS-66 |
| chr12-119608002-INS-227 | chr12-119608002-INS-234 |
| chr12-119608803-INS-1042 | chr12-119608803-INS-1023 |
| chr12-119622370-INS-205 | chr12-119622370-INS-205 |
| chr12-119641385-INS-1123 | chr12-119641386-INS-1123 |
| chr12-119650757-INS-698 | chr12-119650757-INS-698 |
| chr12-119654819-INS-64 | chr12-119654819-INS-64 |
| chr12-119674824-INS-589 | chr12-119674824-INS-589 |
| chr12-119689842-INS-62 | chr12-119689842-INS-62 |
| chr12-119697051-INS-159 | chr12-119697051-INS-159 |
| chr12-119708402-INS-9181 | chr12-119708403-INS-9178 |
| chr12-119728477-INS-7227 | chr12-119728478-INS-7227 |
| chr12-119738993-INS-84 | chr12-119738993-INS-84 |
| chr12-119744090-INS-81 | chr12-119744090-INS-85 |
| chr12-119756442-INS-882 | chr12-119756442-INS-883 |
| chr12-119776471-INS-569 | chr12-119776471-INS-569 |
| chr12-119802161-INS-58 | chr12-119802161-INS-70 |
| chr12-119809931-INS-52 | chr12-119809931-INS-52 |
| chr12-119818979-INS-139 | chr12-119818975-INS-150 |
| chr12-119837206-INS-534 | chr12-119837206-INS-537 |
| chr12-119853410-INS-583 | chr12-119853410-INS-583 |
| chr12-119869825-INS-6398 | chr12-119869826-INS-6398 |
| chr12-119885108-INS-6490 | chr12-119885109-INS-6492 |
| chr12-119901553-INS-6792 | chr12-119901553-INS-6792 |
| chr12-119919569-INS-128 | chr12-119919565-INS-136 |
| chr12-119920576-INS-122 | chr12-119920576-INS-122 |
| chr12-119967354-INS-332 | chr12-119967354-INS-332 |
| chr12-119971093-INS-4964 | chr12-119971094-INS-5001 |
| chr12-119973274-INS-661 | chr12-119973274-INS-661 |
| chr12-119976798-INS-169 | chr12-119976798-INS-169 |
| chr13-3084980-INS-71 | chr13-3084981-INS-50 |
| chr13-3089980-INS-117 | chr13-3089980-INS-117 |
| chr13-3108465-INS-80 | chr13-3108470-INS-65 |
| chr13-3117664-INS-81 | chr13-3117664-INS-81 |

| | |
|---|---|
| chr13-3137940-INS-100 | chr13-3137940-INS-50 |
| chr13-3138925-INS-71 | chr13-3138925-INS-71 |
| chr13-3141560-INS-54 | chr13-3141560-INS-60 |
| chr13-3204754-INS-449 | chr13-3204754-INS-449 |
| chr13-3264536-INS-54 | chr13-3264536-INS-54 |
| chr13-3266846-INS-88 | chr13-3266846-INS-91 |
| chr13-3270544-INS-463 | chr13-3270544-INS-463 |
| chr13-3288965-INS-440 | chr13-3288966-INS-440 |
| chr13-3325160-INS-56 | chr13-3325160-INS-56 |
| chr13-3343357-INS-2680 | chr13-3343358-INS-2680 |
| chr13-3348426-INS-72 | chr13-3348426-INS-78 |
| chr13-3354483-INS-739 | chr13-3354483-INS-739 |
| chr13-3357988-INS-1193 | chr13-3357989-INS-1193 |
| chr13-3406035-INS-81 | chr13-3406035-INS-81 |
| chr13-3422590-INS-609 | chr13-3422590-INS-609 |
| chr13-3429629-INS-7116 | chr13-3429630-INS-7126 |
| chr13-3498954-INS-1891 | chr13-3498955-INS-1890 |
| chr13-3499537-INS-65 | chr13-3499537-INS-53 |
| chr13-3501573-INS-100 | chr13-3501573-INS-100 |
| chr13-3524567-INS-6622 | chr13-3524568-INS-6622 |
| chr13-3526716-INS-496 | chr13-3526716-INS-496 |
| chr13-3532957-INS-481 | chr13-3532957-INS-481 |
| chr13-3571248-INS-56 | chr13-3571248-INS-56 |
| chr13-3574212-INS-184 | chr13-3574212-INS-184 |
| chr13-3628348-INS-70 | chr13-3628350-INS-88 |
| chr13-3629750-INS-224 | chr13-3629750-INS-228 |
| chr13-3641359-INS-218 | chr13-3641359-INS-218 |
| chr13-3664240-INS-103 | chr13-3664241-INS-103 |
| chr13-3696872-INS-859 | chr13-3696872-INS-859 |
| chr13-3703272-INS-75 | chr13-3703272-INS-75 |
| chr13-3710806-INS-78 | chr13-3710806-INS-78 |
| chr13-3769952-INS-112 | chr13-3769952-INS-112 |
| chr13-3792594-INS-58 | chr13-3792594-INS-60 |
| chr13-3794758-INS-75 | chr13-3794758-INS-81 |
| chr13-3830367-INS-81 | chr13-3830366-INS-53 |
| chr13-3833113-INS-203 | chr13-3833122-INS-206 |
| chr13-3875155-INS-282 | chr13-3875155-INS-284 |
| chr13-3880074-INS-54 | chr13-3880074-INS-54 |
| chr13-3890032-INS-212 | chr13-3890032-INS-212 |
| chr13-3923340-INS-1893 | chr13-3923341-INS-1893 |
| chr13-3928500-INS-536 | chr13-3928500-INS-536 |
| chr13-3997396-INS-114 | chr13-3997397-INS-86 |
| chr13-4022796-INS-473 | chr13-4022796-INS-472 |
| chr13-4030075-INS-63 | chr13-4030063-INS-75 |
| chr13-4067721-INS-66 | chr13-4067721-INS-66 |
| chr13-4069728-INS-2019 | chr13-4069729-INS-2019 |
| chr13-4075428-INS-6550 | chr13-4075429-INS-6550 |
| chr13-4085134-INS-96 | chr13-4085134-INS-93 |
| chr13-4103053-INS-6403 | chr13-4103054-INS-6403 |
| chr13-4114512-INS-188 | chr13-4114512-INS-185 |
| chr13-4136346-INS-57 | chr13-4136346-INS-57 |
| chr13-4162084-INS-198 | chr13-4162084-INS-198 |
| chr13-4162620-INS-61 | chr13-4162596-INS-76 |
| chr13-4173799-INS-6160 | chr13-4173800-INS-6161 |
| chr13-4177814-INS-197 | chr13-4177814-INS-197 |
| chr13-4204784-INS-237 | chr13-4204781-INS-245 |
| chr13-4229453-INS-445 | chr13-4229453-INS-445 |
| chr13-4230752-INS-524 | chr13-4230752-INS-524 |
| chr13-4257108-INS-56 | chr13-4257108-INS-87 |
| chr13-4263107-INS-508 | chr13-4263107-INS-508 |
| chr13-4268137-INS-56 | chr13-4268137-INS-52 |
| chr13-4269933-INS-285 | chr13-4269933-INS-283 |
| chr13-4283388-INS-324 | chr13-4283388-INS-324 |
| chr13-4290855-INS-431 | chr13-4290855-INS-431 |
| chr13-4299322-INS-1406 | chr13-4299323-INS-1406 |
| chr13-4299323-INS-682 | chr13-4299323-INS-681 |
| chr13-4318845-INS-73 | chr13-4318845-INS-73 |
| chr13-4319898-INS-2033 | chr13-4319899-INS-2029 |
| chr13-4332847-INS-6746 | chr13-4332848-INS-6746 |
| chr13-4365802-INS-1258 | chr13-4365803-INS-1256 |
| chr13-4401952-INS-64 | chr13-4401947-INS-64 |
| chr13-4453996-INS-4817 | chr13-4453997-INS-4817 |
| chr13-4467744-INS-52 | chr13-4467745-INS-54 |
| chr13-4492704-INS-111 | chr13-4492704-INS-111 |
| chr13-4502591-INS-58 | chr13-4502591-INS-60 |
| chr13-4506183-INS-152 | chr13-4506183-INS-146 |
| chr13-4523176-INS-50 | chr13-4523181-INS-59 |

| | |
|---|---|
| chr13-4536457-INS-399 | chr13-4536457-INS-399 |
| chr13-4541358-INS-92 | chr13-4541358-INS-93 |
| chr13-4543856-INS-57 | chr13-4543856-INS-62 |
| chr13-4544512-INS-1629 | chr13-4544513-INS-1629 |
| chr13-4572909-INS-85 | chr13-4572913-INS-85 |
| chr13-4637718-INS-829 | chr13-4637716-INS-834 |
| chr13-4642247-INS-8265 | chr13-4642248-INS-8265 |
| chr13-4653522-INS-64 | chr13-4653522-INS-75 |
| chr13-4654672-INS-129 | chr13-4654672-INS-129 |
| chr13-4676748-INS-567 | chr13-4676748-INS-567 |
| chr13-4679562-INS-283 | chr13-4679573-INS-314 |
| chr13-4728994-INS-2941 | chr13-4728995-INS-2941 |
| chr13-4730843-INS-1889 | chr13-4730844-INS-1889 |
| chr13-4735939-INS-226 | chr13-4735939-INS-226 |
| chr13-4763747-INS-66 | chr13-4763747-INS-66 |
| chr13-4772787-INS-152 | chr13-4772784-INS-167 |
| chr13-4803496-INS-483 | chr13-4803496-INS-483 |
| chr13-4804859-INS-6930 | chr13-4804860-INS-6930 |
| chr13-4806931-INS-235 | chr13-4806931-INS-223 |
| chr13-4809546-INS-60 | chr13-4809546-INS-72 |
| chr13-4814480-INS-7201 | chr13-4814481-INS-7201 |
| chr13-4828418-INS-756 | chr13-4828419-INS-756 |
| chr13-4839372-INS-6418 | chr13-4839373-INS-6417 |
| chr13-4839536-INS-328 | chr13-4839536-INS-328 |
| chr13-4854630-INS-6417 | chr13-4854632-INS-6415 |
| chr13-4860219-INS-1064 | chr13-4860220-INS-1063 |
| chr13-4875350-INS-54 | chr13-4875350-INS-56 |
| chr13-4885961-INS-1625 | chr13-4885962-INS-1625 |
| chr13-4904016-INS-54 | chr13-4904016-INS-64 |
| chr13-4909826-INS-1754 | chr13-4909827-INS-1754 |
| chr13-4914380-INS-290 | chr13-4914380-INS-293 |
| chr13-4991276-INS-74 | chr13-4991276-INS-74 |
| chr13-5011428-INS-6475 | chr13-5011429-INS-6475 |
| chr13-5013858-INS-1449 | chr13-5013859-INS-1449 |
| chr13-5082965-INS-93 | chr13-5083000-INS-93 |
| chr13-5103555-INS-105 | chr13-5103555-INS-59 |
| chr13-5133310-INS-132 | chr13-5133310-INS-132 |
| chr13-5162799-INS-1894 | chr13-5162800-INS-1894 |
| chr13-5254642-INS-86 | chr13-5254644-INS-76 |
| chr13-5263136-INS-55 | chr13-5263137-INS-55 |
| chr13-5285052-INS-184 | chr13-5285080-INS-197 |
| chr13-5297017-INS-64 | chr13-5297017-INS-64 |
| chr13-5311011-INS-5298 | chr13-5311012-INS-5298 |
| chr13-5325442-INS-211 | chr13-5325442-INS-213 |
| chr13-5344522-INS-52 | chr13-5344522-INS-52 |
| chr13-5381903-INS-83 | chr13-5381903-INS-83 |
| chr13-5421630-INS-181 | chr13-5421630-INS-181 |
| chr13-5499576-INS-65 | chr13-5499576-INS-65 |
| chr13-5560835-INS-55 | chr13-5560835-INS-52 |
| chr13-5576195-INS-62 | chr13-5576195-INS-60 |
| chr13-5613293-INS-3437 | chr13-5613294-INS-3437 |
| chr13-5627727-INS-56 | chr13-5627727-INS-50 |
| chr13-5641640-INS-289 | chr13-5641650-INS-280 |
| chr13-5652905-INS-114 | chr13-5652905-INS-120 |
| chr13-5663577-INS-54 | chr13-5663577-INS-54 |
| chr13-5668477-INS-58 | chr13-5668477-INS-58 |
| chr13-5695514-INS-2183 | chr13-5695515-INS-2187 |
| chr13-5719665-INS-65 | chr13-5719665-INS-65 |
| chr13-5727816-INS-1270 | chr13-5727817-INS-1270 |
| chr13-5751532-INS-72 | chr13-5751532-INS-80 |
| chr13-5753639-INS-60 | chr13-5753660-INS-80 |
| chr13-5778480-INS-52 | chr13-5778480-INS-52 |
| chr13-5803471-INS-53 | chr13-5803471-INS-53 |
| chr13-5810895-INS-211 | chr13-5810895-INS-211 |
| chr13-5848014-INS-471 | chr13-5848000-INS-487 |
| chr13-5902716-INS-86 | chr13-5902735-INS-93 |
| chr13-5910269-INS-195 | chr13-5910269-INS-195 |
| chr13-5938057-INS-1289 | chr13-5938058-INS-1298 |
| chr13-5940637-INS-246 | chr13-5940637-INS-246 |
| chr13-5953978-INS-3000 | chr13-5953979-INS-3000 |
| chr13-5979218-INS-80 | chr13-5979218-INS-84 |
| chr13-5979688-INS-1742 | chr13-5979689-INS-1742 |
| chr13-5983274-INS-99 | chr13-5983271-INS-81 |
| chr13-5996095-INS-6509 | chr13-5996096-INS-6509 |
| chr13-6004502-INS-67 | chr13-6004502-INS-67 |
| chr13-6037340-INS-52 | chr13-6037340-INS-52 |
| chr13-6042859-INS-595 | chr13-6042859-INS-593 |

| | |
|---|---|
| chr13-6069317-INS-149 | chr13-6069317-INS-149 |
| chr13-6071826-INS-65 | chr13-6071826-INS-66 |
| chr13-6081289-INS-66 | chr13-6081289-INS-66 |
| chr13-6099217-INS-73 | chr13-6099217-INS-73 |
| chr13-6108793-INS-82 | chr13-6108793-INS-68 |
| chr13-6126544-INS-129 | chr13-6126587-INS-90 |
| chr13-6130884-INS-93 | chr13-6130884-INS-93 |
| chr13-6134965-INS-186 | chr13-6134965-INS-186 |
| chr13-6168275-INS-80 | chr13-6168275-INS-81 |
| chr13-6178383-INS-794 | chr13-6178383-INS-794 |
| chr13-6192479-INS-56 | chr13-6192475-INS-52 |
| chr13-6199942-INS-654 | chr13-6199944-INS-654 |
| chr13-6230513-INS-251 | chr13-6230513-INS-251 |
| chr13-6263502-INS-6149 | chr13-6263503-INS-6149 |
| chr13-6282194-INS-60 | chr13-6282194-INS-60 |
| chr13-6297233-INS-65 | chr13-6297233-INS-65 |
| chr13-6310490-INS-50 | chr13-6310488-INS-54 |
| chr13-6334271-INS-6260 | chr13-6334272-INS-6261 |
| chr13-6344983-INS-58 | chr13-6344983-INS-62 |
| chr13-6346094-INS-139 | chr13-6346094-INS-139 |
| chr13-6436731-INS-51 | chr13-6436731-INS-50 |
| chr13-6458380-INS-124 | chr13-6458380-INS-124 |
| chr13-6514709-INS-235 | chr13-6514710-INS-228 |
| chr13-6535303-INS-761 | chr13-6535303-INS-758 |
| chr13-6557438-INS-84 | chr13-6557465-INS-92 |
| chr13-6593432-INS-70 | chr13-6593432-INS-72 |
| chr13-6604065-INS-217 | chr13-6604065-INS-221 |
| chr13-6630709-INS-85 | chr13-6630709-INS-85 |
| chr13-6634852-INS-84 | chr13-6634852-INS-84 |
| chr13-6638114-INS-60 | chr13-6638114-INS-62 |
| chr13-6654401-INS-220 | chr13-6654401-INS-221 |
| chr13-6667740-INS-1887 | chr13-6667741-INS-1887 |
| chr13-6674428-INS-162 | chr13-6674428-INS-162 |
| chr13-6820655-INS-231 | chr13-6820655-INS-231 |
| chr13-6822279-INS-136 | chr13-6822279-INS-120 |
| chr13-6825897-INS-545 | chr13-6825897-INS-549 |
| chr13-6826947-INS-55 | chr13-6826947-INS-55 |
| chr13-6837339-INS-70 | chr13-6837339-INS-70 |
| chr13-6843127-INS-1269 | chr13-6843128-INS-1269 |
| chr13-6843351-INS-52 | chr13-6843352-INS-63 |
| chr13-6849487-INS-52 | chr13-6849487-INS-84 |
| chr13-6864080-INS-194 | chr13-6864080-INS-193 |
| chr13-6880523-INS-128 | chr13-6880523-INS-128 |
| chr13-6888170-INS-54 | chr13-6888170-INS-54 |
| chr13-6909189-INS-356 | chr13-6909189-INS-356 |
| chr13-6940020-INS-553 | chr13-6940020-INS-553 |
| chr13-6943107-INS-4413 | chr13-6943108-INS-4413 |
| chr13-6953136-INS-58 | chr13-6953136-INS-60 |
| chr13-6963277-INS-66 | chr13-6963277-INS-66 |
| chr13-6972796-INS-72 | chr13-6972796-INS-72 |
| chr13-6978255-INS-55 | chr13-6978256-INS-54 |
| chr13-6986436-INS-79 | chr13-6986436-INS-82 |
| chr13-7031695-INS-64 | chr13-7031695-INS-64 |
| chr13-7044803-INS-64 | chr13-7044803-INS-64 |
| chr13-7046343-INS-550 | chr13-7046343-INS-550 |
| chr13-7078635-INS-208 | chr13-7078635-INS-208 |
| chr13-7087191-INS-3143 | chr13-7087192-INS-3143 |
| chr13-7130381-INS-1135 | chr13-7130382-INS-1135 |
| chr13-7151328-INS-358 | chr13-7151328-INS-358 |
| chr13-7184408-INS-688 | chr13-7184408-INS-688 |
| chr13-7193236-INS-78 | chr13-7193236-INS-78 |
| chr13-7231871-INS-117 | chr13-7231871-INS-120 |
| chr13-7248402-INS-731 | chr13-7248402-INS-731 |
| chr13-7257979-INS-51 | chr13-7257979-INS-51 |
| chr13-7294164-INS-720 | chr13-7294164-INS-720 |
| chr13-7340279-INS-54 | chr13-7340279-INS-54 |
| chr13-7379445-INS-7182 | chr13-7379446-INS-7182 |
| chr13-7394243-INS-6337 | chr13-7394244-INS-6337 |
| chr13-7402707-INS-522 | chr13-7402707-INS-523 |
| chr13-7404302-INS-1459 | chr13-7404303-INS-1459 |
| chr13-7437842-INS-492 | chr13-7437842-INS-492 |
| chr13-7449061-INS-78 | chr13-7449061-INS-71 |
| chr13-7473576-INS-4502 | chr13-7473577-INS-4503 |
| chr13-7484904-INS-2546 | chr13-7484905-INS-2546 |
| chr13-7503102-INS-60 | chr13-7503102-INS-72 |
| chr13-7537439-INS-58 | chr13-7537407-INS-101 |
| chr13-7542182-INS-72 | chr13-7542174-INS-78 |

| | |
|---|---|
| chr13-7557367-INS-84 | chr13-7557367-INS-84 |
| chr13-7559954-INS-94 | chr13-7559954-INS-94 |
| chr13-7570459-INS-52 | chr13-7570459-INS-52 |
| chr13-7621369-INS-417 | chr13-7621370-INS-423 |
| chr13-7636635-INS-160 | chr13-7636651-INS-140 |
| chr13-7640432-INS-2307 | chr13-7640433-INS-2307 |
| chr13-7666682-INS-266 | chr13-7666689-INS-289 |
| chr13-7677796-INS-453 | chr13-7677796-INS-451 |
| chr13-7680997-INS-74 | chr13-7680997-INS-111 |
| chr13-7684117-INS-53 | chr13-7684117-INS-53 |
| chr13-7690297-INS-395 | chr13-7690297-INS-395 |
| chr13-7695753-INS-100 | chr13-7695753-INS-100 |
| chr13-7722058-INS-1891 | chr13-7722059-INS-1887 |
| chr13-7727038-INS-201 | chr13-7727038-INS-199 |
| chr13-7730633-INS-1751 | chr13-7730634-INS-1752 |
| chr13-7739785-INS-7128 | chr13-7739786-INS-7128 |
| chr13-7746900-INS-56 | chr13-7746900-INS-56 |
| chr13-7763331-INS-308 | chr13-7763331-INS-308 |
| chr13-7785212-INS-128 | chr13-7785207-INS-127 |
| chr13-7792727-INS-128 | chr13-7792727-INS-128 |
| chr13-7818956-INS-3638 | chr13-7818957-INS-3638 |
| chr13-7845843-INS-312 | chr13-7845843-INS-311 |
| chr13-7862162-INS-214 | chr13-7862162-INS-214 |
| chr13-7885837-INS-89 | chr13-7885836-INS-134 |
| chr13-7918142-INS-110 | chr13-7918142-INS-150 |
| chr13-7945069-INS-1940 | chr13-7945070-INS-1940 |
| chr13-7949664-INS-214 | chr13-7949664-INS-214 |
| chr13-7995339-INS-651 | chr13-7995339-INS-651 |
| chr13-8035668-INS-578 | chr13-8035668-INS-579 |
| chr13-8056108-INS-2400 | chr13-8056109-INS-2400 |
| chr13-8063006-INS-534 | chr13-8063006-INS-565 |
| chr13-8067947-INS-6357 | chr13-8067948-INS-6357 |
| chr13-8097001-INS-60 | chr13-8097001-INS-60 |
| chr13-8106097-INS-57 | chr13-8106099-INS-62 |
| chr13-8121429-INS-6162 | chr13-8121430-INS-6162 |
| chr13-8148626-INS-51 | chr13-8148626-INS-57 |
| chr13-8188945-INS-61 | chr13-8188940-INS-58 |
| chr13-8210307-INS-390 | chr13-8210307-INS-390 |
| chr13-8211664-INS-88 | chr13-8211665-INS-92 |
| chr13-8215608-INS-100 | chr13-8215645-INS-70 |
| chr13-8236091-INS-54 | chr13-8236091-INS-56 |
| chr13-8238210-INS-434 | chr13-8238196-INS-410 |
| chr13-8238987-INS-931 | chr13-8238987-INS-931 |
| chr13-8245734-INS-220 | chr13-8245732-INS-222 |
| chr13-8279661-INS-51 | chr13-8279661-INS-55 |
| chr13-8288495-INS-58 | chr13-8288495-INS-60 |
| chr13-8293341-INS-66 | chr13-8293341-INS-66 |
| chr13-8325484-INS-129 | chr13-8325484-INS-129 |
| chr13-8330018-INS-85 | chr13-8330021-INS-86 |
| chr13-8368485-INS-102 | chr13-8368492-INS-114 |
| chr13-8376853-INS-216 | chr13-8376853-INS-216 |
| chr13-8389146-INS-66 | chr13-8389146-INS-60 |
| chr13-8407551-INS-203 | chr13-8407551-INS-173 |
| chr13-8428191-INS-79 | chr13-8428191-INS-79 |
| chr13-8446810-INS-64 | chr13-8446810-INS-64 |
| chr13-8495961-INS-119 | chr13-8495973-INS-121 |
| chr13-8510678-INS-61 | chr13-8510678-INS-61 |
| chr13-8540115-INS-63 | chr13-8540115-INS-63 |
| chr13-8544668-INS-100 | chr13-8544669-INS-59 |
| chr13-8554538-INS-507 | chr13-8554538-INS-507 |
| chr13-8562609-INS-99 | chr13-8562609-INS-99 |
| chr13-8567218-INS-59 | chr13-8567218-INS-59 |
| chr13-8581347-INS-68 | chr13-8581347-INS-80 |
| chr13-8608605-INS-52 | chr13-8608605-INS-54 |
| chr13-8610423-INS-195 | chr13-8610423-INS-190 |
| chr13-8677145-INS-152 | chr13-8677146-INS-152 |
| chr13-8703837-INS-62 | chr13-8703837-INS-56 |
| chr13-8706669-INS-543 | chr13-8706669-INS-544 |
| chr13-8710934-INS-136 | chr13-8710934-INS-134 |
| chr13-8733495-INS-199 | chr13-8733495-INS-187 |
| chr13-8735218-INS-498 | chr13-8735218-INS-499 |
| chr13-8739386-INS-52 | chr13-8739383-INS-57 |
| chr13-8741457-INS-2893 | chr13-8741458-INS-2900 |
| chr13-8775092-INS-115 | chr13-8775070-INS-107 |
| chr13-8788185-INS-122 | chr13-8788187-INS-153 |
| chr13-8792284-INS-149 | chr13-8792284-INS-149 |
| chr13-8806823-INS-55 | chr13-8806823-INS-55 |

| | |
|---|---|
| chr13-8836130-INS-175 | chr13-8836130-INS-175 |
| chr13-8849435-INS-50 | chr13-8849435-INS-52 |
| chr13-8849919-INS-80 | chr13-8849919-INS-84 |
| chr13-8879548-INS-70 | chr13-8879548-INS-70 |
| chr13-8883306-INS-98 | chr13-8883306-INS-98 |
| chr13-8887365-INS-314 | chr13-8887365-INS-314 |
| chr13-8910804-INS-64 | chr13-8910804-INS-64 |
| chr13-8919034-INS-222 | chr13-8919034-INS-222 |
| chr13-8926167-INS-58 | chr13-8926167-INS-58 |
| chr13-8971036-INS-126 | chr13-8971036-INS-126 |
| chr13-8976003-INS-207 | chr13-8976003-INS-207 |
| chr13-8988191-INS-471 | chr13-8988191-INS-471 |
| chr13-8991405-INS-216 | chr13-8991405-INS-216 |
| chr13-9001585-INS-596 | chr13-9001585-INS-600 |
| chr13-9019470-INS-111 | chr13-9019471-INS-111 |
| chr13-9022214-INS-284 | chr13-9022205-INS-285 |
| chr13-9030662-INS-677 | chr13-9030662-INS-678 |
| chr13-9039447-INS-80 | chr13-9039485-INS-50 |
| chr13-9052262-INS-186 | chr13-9052262-INS-186 |
| chr13-9055551-INS-401 | chr13-9055551-INS-401 |
| chr13-9059188-INS-90 | chr13-9059188-INS-90 |
| chr13-9069351-INS-74 | chr13-9069351-INS-82 |
| chr13-9074676-INS-201 | chr13-9074676-INS-201 |
| chr13-9085088-INS-176 | chr13-9085068-INS-165 |
| chr13-9088843-INS-400 | chr13-9088843-INS-400 |
| chr13-9092644-INS-54 | chr13-9092644-INS-54 |
| chr13-9099646-INS-558 | chr13-9099646-INS-558 |
| chr13-9100274-INS-205 | chr13-9100302-INS-177 |
| chr13-9103938-INS-5231 | chr13-9103939-INS-5231 |
| chr13-9112420-INS-58 | chr13-9112420-INS-58 |
| chr13-9134773-INS-160 | chr13-9134796-INS-160 |
| chr13-9143123-INS-60 | chr13-9143123-INS-50 |
| chr13-9162501-INS-195 | chr13-9162501-INS-195 |
| chr13-9168023-INS-135 | chr13-9168023-INS-135 |
| chr13-9175357-INS-578 | chr13-9175357-INS-576 |
| chr13-9185837-INS-243 | chr13-9185837-INS-246 |
| chr13-9199429-INS-120 | chr13-9199429-INS-160 |
| chr13-9201832-INS-152 | chr13-9201832-INS-152 |
| chr13-9214214-INS-239 | chr13-9214214-INS-239 |
| chr13-9226093-INS-197 | chr13-9226093-INS-197 |
| chr13-9232935-INS-56 | chr13-9232935-INS-56 |
| chr13-9255406-INS-221 | chr13-9255406-INS-221 |
| chr13-9278057-INS-438 | chr13-9278057-INS-438 |
| chr13-9281911-INS-69 | chr13-9281911-INS-71 |
| chr13-9283348-INS-83 | chr13-9283348-INS-81 |
| chr13-9286461-INS-566 | chr13-9286461-INS-567 |
| chr13-9288893-INS-62 | chr13-9288893-INS-56 |
| chr13-9290221-INS-70 | chr13-9290221-INS-70 |
| chr13-9293420-INS-198 | chr13-9293420-INS-211 |
| chr13-9298765-INS-6178 | chr13-9298766-INS-6178 |
| chr13-9308109-INS-211 | chr13-9308110-INS-213 |
| chr13-9311753-INS-96 | chr13-9311759-INS-94 |
| chr13-9313500-INS-69 | chr13-9313500-INS-69 |
| chr13-9317310-INS-84 | chr13-9317310-INS-84 |
| chr13-9327595-INS-51 | chr13-9327595-INS-51 |
| chr13-9329214-INS-177 | chr13-9329228-INS-205 |
| chr13-9344645-INS-78 | chr13-9344645-INS-78 |
| chr13-9360079-INS-191 | chr13-9360079-INS-193 |
| chr13-9361540-INS-195 | chr13-9361540-INS-195 |
| chr13-9363763-INS-62 | chr13-9363763-INS-62 |
| chr13-9377353-INS-420 | chr13-9377353-INS-420 |
| chr13-9388415-INS-57 | chr13-9388415-INS-68 |
| chr13-9448433-INS-80 | chr13-9448433-INS-105 |
| chr13-9491612-INS-96 | chr13-9491611-INS-96 |
| chr13-9522763-INS-209 | chr13-9522763-INS-209 |
| chr13-9557645-INS-160 | chr13-9557645-INS-160 |
| chr13-9559796-INS-420 | chr13-9559796-INS-420 |
| chr13-9574579-INS-2965 | chr13-9574585-INS-2963 |
| chr13-9579194-INS-1917 | chr13-9579195-INS-1917 |
| chr13-9585714-INS-4771 | chr13-9585715-INS-4773 |
| chr13-9649927-INS-335 | chr13-9649927-INS-335 |
| chr13-9678296-INS-79 | chr13-9678296-INS-70 |
| chr13-9696932-INS-70 | chr13-9696932-INS-70 |
| chr13-9702233-INS-456 | chr13-9702233-INS-459 |
| chr13-9703044-INS-78 | chr13-9703080-INS-78 |
| chr13-9703132-INS-832 | chr13-9703132-INS-832 |
| chr13-9757945-INS-214 | chr13-9757945-INS-213 |

| | |
|---|---|
| chr13-9777330-INS-62 | chr13-9777330-INS-62 |
| chr13-9792807-INS-1878 | chr13-9792808-INS-1878 |
| chr13-9807139-INS-93 | chr13-9807139-INS-93 |
| chr13-9823213-INS-70 | chr13-9823213-INS-100 |
| chr13-9830543-INS-92 | chr13-9830543-INS-92 |
| chr13-9849650-INS-96 | chr13-9849650-INS-96 |
| chr13-9851884-INS-72 | chr13-9851884-INS-84 |
| chr13-9878881-INS-192 | chr13-9878930-INS-192 |
| chr13-9924106-INS-600 | chr13-9924136-INS-614 |
| chr13-9929881-INS-140 | chr13-9929881-INS-140 |
| chr13-9934916-INS-245 | chr13-9934916-INS-245 |
| chr13-9947836-INS-56 | chr13-9947826-INS-72 |
| chr13-9949067-INS-101 | chr13-9949067-INS-101 |
| chr13-9961543-INS-52 | chr13-9961543-INS-50 |
| chr13-9967408-INS-70 | chr13-9967408-INS-72 |
| chr13-9969750-INS-941 | chr13-9969750-INS-899 |
| chr13-9994775-INS-56 | chr13-9994773-INS-64 |
| chr13-10010515-INS-123 | chr13-10010515-INS-155 |
| chr13-10018352-INS-69 | chr13-10018352-INS-69 |
| chr13-10038792-INS-130 | chr13-10038792-INS-136 |
| chr13-10058011-INS-781 | chr13-10058011-INS-782 |
| chr13-10064222-INS-7126 | chr13-10064223-INS-7126 |
| chr13-10075706-INS-68 | chr13-10075706-INS-60 |
| chr13-10077777-INS-6705 | chr13-10077778-INS-6706 |
| chr13-10090659-INS-6862 | chr13-10090660-INS-6864 |
| chr13-10095014-INS-53 | chr13-10095015-INS-58 |
| chr13-10097123-INS-1274 | chr13-10097124-INS-1283 |
| chr13-10152903-INS-8121 | chr13-10152904-INS-8121 |
| chr13-10153211-INS-133 | chr13-10153211-INS-133 |
| chr13-10179308-INS-5207 | chr13-10179309-INS-5207 |
| chr13-10216712-INS-243 | chr13-10216712-INS-243 |
| chr13-10222587-INS-499 | chr13-10222587-INS-498 |
| chr13-10250583-INS-66 | chr13-10250583-INS-78 |
| chr13-10252675-INS-952 | chr13-10252661-INS-951 |
| chr13-10253021-INS-102 | chr13-10253021-INS-102 |
| chr13-10256414-INS-194 | chr13-10256414-INS-194 |
| chr13-10265855-INS-134 | chr13-10265855-INS-134 |
| chr13-10268940-INS-221 | chr13-10268940-INS-221 |
| chr13-10343681-INS-96 | chr13-10343681-INS-96 |
| chr13-10369446-INS-7116 | chr13-10369447-INS-7116 |
| chr13-10377933-INS-52 | chr13-10377953-INS-66 |
| chr13-10395470-INS-65 | chr13-10395470-INS-56 |
| chr13-10419332-INS-51 | chr13-10419332-INS-51 |
| chr13-10449960-INS-75 | chr13-10449960-INS-81 |
| chr13-10452197-INS-100 | chr13-10452197-INS-100 |
| chr13-10457799-INS-65 | chr13-10457799-INS-65 |
| chr13-10465511-INS-2941 | chr13-10465512-INS-2925 |
| chr13-10470073-INS-84 | chr13-10470080-INS-64 |
| chr13-10499936-INS-76 | chr13-10499960-INS-72 |
| chr13-10507230-INS-52 | chr13-10507230-INS-64 |
| chr13-10509524-INS-210 | chr13-10509524-INS-210 |
| chr13-10513968-INS-92 | chr13-10513960-INS-96 |
| chr13-10518347-INS-504 | chr13-10518347-INS-529 |
| chr13-10530023-INS-54 | chr13-10530040-INS-53 |
| chr13-10549978-INS-686 | chr13-10549978-INS-686 |
| chr13-10579791-INS-129 | chr13-10579791-INS-129 |
| chr13-10596333-INS-52 | chr13-10596333-INS-54 |
| chr13-10604511-INS-90 | chr13-10604510-INS-64 |
| chr13-10605418-INS-5698 | chr13-10605419-INS-5698 |
| chr13-10610467-INS-8407 | chr13-10610468-INS-8407 |
| chr13-10623075-INS-577 | chr13-10623075-INS-581 |
| chr13-10626776-INS-70 | chr13-10626776-INS-62 |
| chr13-10649925-INS-655 | chr13-10649925-INS-655 |
| chr13-10651759-INS-89 | chr13-10651759-INS-90 |
| chr13-10670973-INS-167 | chr13-10670973-INS-158 |
| chr13-10681363-INS-76 | chr13-10681389-INS-56 |
| chr13-10702172-INS-60 | chr13-10702172-INS-62 |
| chr13-10702815-INS-56 | chr13-10702815-INS-56 |
| chr13-10739319-INS-885 | chr13-10739319-INS-885 |
| chr13-10740757-INS-56 | chr13-10740757-INS-60 |
| chr13-10753606-INS-6491 | chr13-10753607-INS-6491 |
| chr13-10761972-INS-322 | chr13-10761972-INS-322 |
| chr13-10782758-INS-138 | chr13-10782792-INS-100 |
| chr13-10785814-INS-6574 | chr13-10785815-INS-6574 |
| chr13-10786687-INS-174 | chr13-10786687-INS-174 |
| chr13-10838488-INS-51 | chr13-10838490-INS-57 |
| chr13-10838667-INS-5496 | chr13-10838668-INS-5497 |

| | |
|---|---|
| chr13-10865412-INS-54 | chr13-10865412-INS-54 |
| chr13-10875376-INS-55 | chr13-10875327-INS-76 |
| chr13-10892751-INS-119 | chr13-10892751-INS-119 |
| chr13-10905484-INS-95 | chr13-10905486-INS-89 |
| chr13-10912620-INS-170 | chr13-10912620-INS-170 |
| chr13-10914732-INS-157 | chr13-10914732-INS-157 |
| chr13-10927277-INS-538 | chr13-10927277-INS-536 |
| chr13-10932238-INS-128 | chr13-10932238-INS-128 |
| chr13-10965072-INS-61 | chr13-10965072-INS-61 |
| chr13-10966225-INS-484 | chr13-10966225-INS-486 |
| chr13-10969123-INS-71 | chr13-10969123-INS-71 |
| chr13-10984408-INS-7384 | chr13-10984409-INS-7384 |
| chr13-11043006-INS-1596 | chr13-11043007-INS-1596 |
| chr13-11046173-INS-61 | chr13-11046173-INS-59 |
| chr13-11052347-INS-118 | chr13-11052347-INS-118 |
| chr13-11053955-INS-6243 | chr13-11053956-INS-6243 |
| chr13-11055052-INS-175 | chr13-11055052-INS-170 |
| chr13-11067266-INS-316 | chr13-11067266-INS-322 |
| chr13-11079912-INS-150 | chr13-11079912-INS-108 |
| chr13-11091146-INS-200 | chr13-11091146-INS-200 |
| chr13-11101289-INS-224 | chr13-11101289-INS-224 |
| chr13-11102924-INS-60 | chr13-11102924-INS-60 |
| chr13-11117062-INS-317 | chr13-11117062-INS-317 |
| chr13-11137538-INS-57 | chr13-11137538-INS-62 |
| chr13-11147481-INS-85 | chr13-11147464-INS-54 |
| chr13-11147843-INS-1192 | chr13-11147844-INS-1192 |
| chr13-11150463-INS-397 | chr13-11150463-INS-397 |
| chr13-11181650-INS-60 | chr13-11181650-INS-60 |
| chr13-11186475-INS-2207 | chr13-11186476-INS-2206 |
| chr13-11202785-INS-416 | chr13-11202785-INS-416 |
| chr13-11206054-INS-56 | chr13-11206046-INS-64 |
| chr13-11222469-INS-327 | chr13-11222469-INS-298 |
| chr13-11243335-INS-80 | chr13-11243335-INS-80 |
| chr13-11245142-INS-52 | chr13-11245147-INS-55 |
| chr13-11284903-INS-612 | chr13-11284903-INS-613 |
| chr13-11342542-INS-161 | chr13-11342542-INS-161 |
| chr13-11351272-INS-53 | chr13-11351272-INS-60 |
| chr13-11357017-INS-160 | chr13-11357018-INS-159 |
| chr13-11373049-INS-66 | chr13-11373049-INS-70 |
| chr13-11398703-INS-203 | chr13-11398703-INS-203 |
| chr13-11411030-INS-928 | chr13-11411030-INS-928 |
| chr13-11415342-INS-68 | chr13-11415342-INS-72 |
| chr13-11434642-INS-403 | chr13-11434642-INS-402 |
| chr13-11444664-INS-6342 | chr13-11444665-INS-6342 |
| chr13-11459746-INS-137 | chr13-11459746-INS-139 |
| chr13-11464643-INS-68 | chr13-11464643-INS-68 |
| chr13-11465583-INS-2899 | chr13-11465584-INS-2894 |
| chr13-11473842-INS-6525 | chr13-11473843-INS-6525 |
| chr13-11524598-INS-606 | chr13-11524598-INS-606 |
| chr13-11531764-INS-754 | chr13-11531764-INS-754 |
| chr13-11544089-INS-1097 | chr13-11544090-INS-1097 |
| chr13-11548612-INS-172 | chr13-11548612-INS-172 |
| chr13-11559511-INS-440 | chr13-11559511-INS-440 |
| chr13-11572009-INS-54 | chr13-11572009-INS-54 |
| chr13-11579767-INS-86 | chr13-11579767-INS-87 |
| chr13-11600467-INS-115 | chr13-11600467-INS-115 |
| chr13-11604204-INS-175 | chr13-11604204-INS-175 |
| chr13-11607930-INS-763 | chr13-11607930-INS-763 |
| chr13-11609487-INS-113 | chr13-11609481-INS-119 |
| chr13-11619711-INS-200 | chr13-11619726-INS-208 |
| chr13-11646082-INS-2106 | chr13-11646083-INS-2107 |
| chr13-11656964-INS-336 | chr13-11656964-INS-341 |
| chr13-11669840-INS-743 | chr13-11669840-INS-743 |
| chr13-11674657-INS-62 | chr13-11674630-INS-72 |
| chr13-11675845-INS-1094 | chr13-11675846-INS-1094 |
| chr13-11678743-INS-51 | chr13-11678741-INS-98 |
| chr13-11693705-INS-54 | chr13-11693703-INS-65 |
| chr13-11696209-INS-52 | chr13-11696209-INS-60 |
| chr13-11696503-INS-296 | chr13-11696503-INS-296 |
| chr13-11701993-INS-56 | chr13-11701993-INS-63 |
| chr13-11716357-INS-231 | chr13-11716357-INS-234 |
| chr13-11751460-INS-51 | chr13-11751460-INS-53 |
| chr13-11763080-INS-55 | chr13-11763080-INS-51 |
| chr13-11775105-INS-148 | chr13-11775105-INS-168 |
| chr13-11823690-INS-83 | chr13-11823688-INS-87 |
| chr13-11824679-INS-57 | chr13-11824679-INS-64 |
| chr13-11841292-INS-62 | chr13-11841300-INS-70 |

| | |
|---|---|
| chr13-11849383-INS-53 | chr13-11849383-INS-53 |
| chr13-11854064-INS-120 | chr13-11854064-INS-120 |
| chr13-11857997-INS-940 | chr13-11857997-INS-936 |
| chr13-11859613-INS-66 | chr13-11859613-INS-66 |
| chr13-11880008-INS-308 | chr13-11880008-INS-311 |
| chr13-11901545-INS-200 | chr13-11901545-INS-200 |
| chr13-11904716-INS-80 | chr13-11904717-INS-80 |
| chr13-11928803-INS-55 | chr13-11928803-INS-55 |
| chr13-11946254-INS-74 | chr13-11946254-INS-76 |
| chr13-11958634-INS-166 | chr13-11958634-INS-166 |
| chr13-11968006-INS-88 | chr13-11968006-INS-88 |
| chr13-11970011-INS-484 | chr13-11970011-INS-484 |
| chr13-11974558-INS-54 | chr13-11974558-INS-72 |
| chr13-11984457-INS-619 | chr13-11984457-INS-619 |
| chr13-11996398-INS-228 | chr13-11996398-INS-229 |
| chr13-12006645-INS-54 | chr13-12006669-INS-54 |
| chr13-12018121-INS-364 | chr13-12018121-INS-393 |
| chr13-12043073-INS-3273 | chr13-12043074-INS-3273 |
| chr13-12062620-INS-429 | chr13-12062620-INS-429 |
| chr13-12090316-INS-711 | chr13-12090316-INS-711 |
| chr13-12097268-INS-108 | chr13-12097268-INS-114 |
| chr13-12112656-INS-219 | chr13-12112641-INS-202 |
| chr13-12131616-INS-7127 | chr13-12131617-INS-7127 |
| chr13-12142122-INS-2739 | chr13-12142123-INS-2739 |
| chr13-12151402-INS-201 | chr13-12151402-INS-202 |
| chr13-12153643-INS-52 | chr13-12153643-INS-50 |
| chr13-12166353-INS-5466 | chr13-12166354-INS-5465 |
| chr13-12173736-INS-574 | chr13-12173736-INS-574 |
| chr13-12180089-INS-148 | chr13-12180089-INS-118 |
| chr13-12204968-INS-210 | chr13-12204968-INS-210 |
| chr13-12217997-INS-99 | chr13-12217993-INS-88 |
| chr13-12236758-INS-55 | chr13-12236758-INS-55 |
| chr13-12256626-INS-480 | chr13-12256626-INS-481 |
| chr13-12269806-INS-213 | chr13-12269806-INS-213 |
| chr13-12276001-INS-212 | chr13-12276001-INS-214 |
| chr13-12278466-INS-246 | chr13-12278466-INS-251 |
| chr13-12292993-INS-172 | chr13-12292993-INS-172 |
| chr13-12294871-INS-206 | chr13-12294871-INS-211 |
| chr13-12331970-INS-54 | chr13-12331970-INS-54 |
| chr13-12333569-INS-52 | chr13-12333569-INS-58 |
| chr13-12347761-INS-213 | chr13-12347761-INS-213 |
| chr13-12361707-INS-5469 | chr13-12361708-INS-5469 |
| chr13-12377011-INS-401 | chr13-12377011-INS-401 |
| chr13-12398082-INS-184 | chr13-12398082-INS-183 |
| chr13-12401984-INS-54 | chr13-12401966-INS-54 |
| chr13-12404796-INS-147 | chr13-12404796-INS-147 |
| chr13-12412690-INS-207 | chr13-12412690-INS-213 |
| chr13-12417844-INS-99 | chr13-12417844-INS-99 |
| chr13-12435265-INS-170 | chr13-12435265-INS-171 |
| chr13-12463221-INS-135 | chr13-12463221-INS-135 |
| chr13-12477204-INS-879 | chr13-12477204-INS-849 |
| chr13-12485276-INS-78 | chr13-12485276-INS-52 |
| chr13-12502962-INS-251 | chr13-12502962-INS-221 |
| chr13-12513855-INS-113 | chr13-12513833-INS-99 |
| chr13-12528634-INS-64 | chr13-12528634-INS-64 |
| chr13-12531142-INS-204 | chr13-12531143-INS-205 |
| chr13-12537129-INS-71 | chr13-12537129-INS-66 |
| chr13-12547792-INS-314 | chr13-12547792-INS-314 |
| chr13-12553527-INS-211 | chr13-12553527-INS-209 |
| chr13-12575641-INS-145 | chr13-12575641-INS-145 |
| chr13-12592781-INS-187 | chr13-12592781-INS-188 |
| chr13-12595785-INS-62 | chr13-12595785-INS-62 |
| chr13-12597549-INS-85 | chr13-12597549-INS-85 |
| chr13-12600179-INS-396 | chr13-12600179-INS-410 |
| chr13-12609573-INS-207 | chr13-12609573-INS-206 |
| chr13-12616431-INS-219 | chr13-12616432-INS-213 |
| chr13-12755738-INS-372 | chr13-12755738-INS-372 |
| chr13-13145269-INS-177 | chr13-13145293-INS-156 |
| chr13-13521013-INS-97 | chr13-13521013-INS-97 |
| chr13-13543947-INS-55 | chr13-13543947-INS-55 |
| chr13-13635414-INS-242 | chr13-13635414-INS-241 |
| chr13-13644278-INS-52 | chr13-13644278-INS-54 |
| chr13-13720536-INS-102 | chr13-13720561-INS-116 |
| chr13-13742074-INS-73 | chr13-13742074-INS-73 |
| chr13-13775021-INS-216 | chr13-13775021-INS-224 |
| chr13-13816423-INS-57 | chr13-13816423-INS-57 |
| chr13-13832961-INS-220 | chr13-13832945-INS-250 |

| | |
|---|---|
| chr13-13843917-INS-63 | chr13-13843937-INS-81 |
| chr13-13867599-INS-1001 | chr13-13867600-INS-1001 |
| chr13-13870655-INS-412 | chr13-13870655-INS-401 |
| chr13-13927758-INS-814 | chr13-13927770-INS-816 |
| chr13-13929072-INS-275 | chr13-13929095-INS-275 |
| chr13-13984805-INS-69 | chr13-13984805-INS-69 |
| chr13-14007272-INS-150 | chr13-14007272-INS-165 |
| chr13-14022884-INS-164 | chr13-14022884-INS-164 |
| chr13-14039922-INS-187 | chr13-14039922-INS-188 |
| chr13-14068495-INS-3321 | chr13-14068496-INS-3321 |
| chr13-14090220-INS-3375 | chr13-14090221-INS-3375 |
| chr13-14099118-INS-398 | chr13-14099118-INS-398 |
| chr13-14122936-INS-93 | chr13-14122937-INS-93 |
| chr13-14177634-INS-200 | chr13-14177634-INS-200 |
| chr13-14245878-INS-206 | chr13-14245878-INS-206 |
| chr13-14276771-INS-90 | chr13-14276771-INS-89 |
| chr13-14283991-INS-51 | chr13-14283991-INS-51 |
| chr13-14290319-INS-234 | chr13-14290319-INS-234 |
| chr13-14304325-INS-117 | chr13-14304327-INS-125 |
| chr13-14336416-INS-475 | chr13-14336416-INS-473 |
| chr13-14429851-INS-211 | chr13-14429851-INS-211 |
| chr13-14475987-INS-1000 | chr13-14475987-INS-1000 |
| chr13-14511402-INS-427 | chr13-14511398-INS-428 |
| chr13-14520094-INS-71 | chr13-14520094-INS-71 |
| chr13-14540168-INS-6809 | chr13-14540169-INS-6809 |
| chr13-14541221-INS-1753 | chr13-14541222-INS-1753 |
| chr13-14544021-INS-82 | chr13-14544019-INS-82 |
| chr13-14549427-INS-106 | chr13-14549427-INS-106 |
| chr13-14595235-INS-58 | chr13-14595235-INS-62 |
| chr13-14598574-INS-66 | chr13-14598574-INS-68 |
| chr13-14626608-INS-496 | chr13-14626608-INS-496 |
| chr13-14635687-INS-91 | chr13-14635687-INS-92 |
| chr13-14648654-INS-174 | chr13-14648654-INS-174 |
| chr13-14652751-INS-1060 | chr13-14652746-INS-1063 |
| chr13-14687734-INS-192 | chr13-14687734-INS-192 |
| chr13-14700964-INS-160 | chr13-14700964-INS-160 |
| chr13-14706239-INS-6588 | chr13-14706240-INS-6592 |
| chr13-14707089-INS-234 | chr13-14707089-INS-234 |
| chr13-14713947-INS-7506 | chr13-14713948-INS-7506 |
| chr13-14771826-INS-127 | chr13-14771826-INS-127 |
| chr13-14780675-INS-441 | chr13-14780675-INS-441 |
| chr13-14796896-INS-54 | chr13-14796896-INS-54 |
| chr13-14806366-INS-54 | chr13-14806366-INS-50 |
| chr13-14890364-INS-4256 | chr13-14890365-INS-4257 |
| chr13-14897126-INS-432 | chr13-14897126-INS-432 |
| chr13-14908778-INS-55 | chr13-14908782-INS-65 |
| chr13-14916357-INS-212 | chr13-14916357-INS-211 |
| chr13-14925634-INS-647 | chr13-14925634-INS-647 |
| chr13-14985410-INS-224 | chr13-14985410-INS-224 |
| chr13-15018583-INS-832 | chr13-15018583-INS-831 |
| chr13-15034192-INS-3628 | chr13-15034193-INS-3628 |
| chr13-15043004-INS-362 | chr13-15043004-INS-354 |
| chr13-15051661-INS-6541 | chr13-15051662-INS-6542 |
| chr13-15064078-INS-73 | chr13-15064075-INS-77 |
| chr13-15073681-INS-51 | chr13-15073684-INS-50 |
| chr13-15075461-INS-64 | chr13-15075461-INS-60 |
| chr13-15090539-INS-3095 | chr13-15090540-INS-3095 |
| chr13-15126037-INS-123 | chr13-15126035-INS-126 |
| chr13-15128700-INS-125 | chr13-15128700-INS-114 |
| chr13-15144410-INS-83 | chr13-15144406-INS-71 |
| chr13-15188711-INS-601 | chr13-15188711-INS-601 |
| chr13-15201844-INS-428 | chr13-15201844-INS-428 |
| chr13-15219574-INS-68 | chr13-15219574-INS-64 |
| chr13-15222909-INS-59 | chr13-15222909-INS-59 |
| chr13-15241363-INS-372 | chr13-15241363-INS-374 |
| chr13-15253060-INS-649 | chr13-15253060-INS-653 |
| chr13-15263426-INS-2232 | chr13-15263428-INS-2233 |
| chr13-15290300-INS-88 | chr13-15290300-INS-92 |
| chr13-15292966-INS-6244 | chr13-15292967-INS-6244 |
| chr13-15323779-INS-70 | chr13-15323780-INS-71 |
| chr13-15324640-INS-693 | chr13-15324640-INS-693 |
| chr13-15325592-INS-2459 | chr13-15325595-INS-2455 |
| chr13-15398117-INS-687 | chr13-15398117-INS-687 |
| chr13-15406191-INS-53 | chr13-15406191-INS-53 |
| chr13-15426552-INS-204 | chr13-15426552-INS-204 |
| chr13-15493991-INS-6277 | chr13-15493992-INS-6277 |
| chr13-15502529-INS-7081 | chr13-15502530-INS-7081 |

| | |
|---|---|
| chr13-15526620-INS-498 | chr13-15526620-INS-498 |
| chr13-15571091-INS-120 | chr13-15571091-INS-112 |
| chr13-15603319-INS-141 | chr13-15603319-INS-108 |
| chr13-15613541-INS-111 | chr13-15613535-INS-79 |
| chr13-15649593-INS-187 | chr13-15649593-INS-187 |
| chr13-15653591-INS-394 | chr13-15653591-INS-394 |
| chr13-15658717-INS-249 | chr13-15658717-INS-250 |
| chr13-15673998-INS-85 | chr13-15674044-INS-85 |
| chr13-15708108-INS-128 | chr13-15708108-INS-88 |
| chr13-15741662-INS-51 | chr13-15741663-INS-75 |
| chr13-15798572-INS-271 | chr13-15798572-INS-271 |
| chr13-15802914-INS-68 | chr13-15802916-INS-98 |
| chr13-15851413-INS-6247 | chr13-15851414-INS-6249 |
| chr13-15876369-INS-52 | chr13-15876361-INS-51 |
| chr13-15905767-INS-3175 | chr13-15905768-INS-3175 |
| chr13-15920139-INS-6459 | chr13-15920140-INS-6459 |
| chr13-15924080-INS-1173 | chr13-15924081-INS-1173 |
| chr13-15929052-INS-6234 | chr13-15929053-INS-6234 |
| chr13-15932879-INS-78 | chr13-15932879-INS-54 |
| chr13-15935091-INS-107 | chr13-15935091-INS-111 |
| chr13-15945044-INS-243 | chr13-15945058-INS-243 |
| chr13-15963224-INS-508 | chr13-15963224-INS-508 |
| chr13-15965985-INS-1376 | chr13-15965986-INS-1377 |
| chr13-15991358-INS-542 | chr13-15991358-INS-539 |
| chr13-16017173-INS-325 | chr13-16017173-INS-325 |
| chr13-16033522-INS-174 | chr13-16033522-INS-174 |
| chr13-16055860-INS-7163 | chr13-16055864-INS-7164 |
| chr13-16104950-INS-52 | chr13-16104950-INS-56 |
| chr13-16109400-INS-54 | chr13-16109400-INS-54 |
| chr13-16118931-INS-1449 | chr13-16118932-INS-1432 |
| chr13-16119513-INS-55 | chr13-16119513-INS-55 |
| chr13-16145559-INS-2756 | chr13-16145560-INS-2756 |
| chr13-16146049-INS-580 | chr13-16146049-INS-580 |
| chr13-16150810-INS-64 | chr13-16150810-INS-70 |
| chr13-16151126-INS-543 | chr13-16151126-INS-543 |
| chr13-16152449-INS-1063 | chr13-16152450-INS-1061 |
| chr13-16167224-INS-4825 | chr13-16167225-INS-4825 |
| chr13-16167972-INS-183 | chr13-16167972-INS-183 |
| chr13-16181406-INS-63 | chr13-16181406-INS-63 |
| chr13-16220901-INS-78 | chr13-16220892-INS-77 |
| chr13-16272792-INS-56 | chr13-16272791-INS-58 |
| chr13-16297154-INS-238 | chr13-16297154-INS-238 |
| chr13-16298866-INS-200 | chr13-16298866-INS-200 |
| chr13-16307610-INS-175 | chr13-16307610-INS-175 |
| chr13-16327728-INS-52 | chr13-16327728-INS-54 |
| chr13-16333470-INS-78 | chr13-16333470-INS-86 |
| chr13-16343477-INS-54 | chr13-16343477-INS-56 |
| chr13-16356801-INS-214 | chr13-16356801-INS-214 |
| chr13-16391103-INS-1466 | chr13-16391104-INS-1466 |
| chr13-16406440-INS-388 | chr13-16406421-INS-378 |
| chr13-16474091-INS-404 | chr13-16474086-INS-413 |
| chr13-16509747-INS-6309 | chr13-16509748-INS-6276 |
| chr13-16512154-INS-130 | chr13-16512154-INS-130 |
| chr13-16555805-INS-1162 | chr13-16555806-INS-1161 |
| chr13-16599872-INS-434 | chr13-16599872-INS-434 |
| chr13-16609346-INS-1072 | chr13-16609347-INS-1073 |
| chr13-16613760-INS-2618 | chr13-16613761-INS-2618 |
| chr13-16623967-INS-84 | chr13-16623961-INS-88 |
| chr13-16644681-INS-440 | chr13-16644681-INS-440 |
| chr13-16647724-INS-66 | chr13-16647724-INS-78 |
| chr13-16661095-INS-276 | chr13-16661095-INS-276 |
| chr13-16705117-INS-308 | chr13-16705117-INS-308 |
| chr13-16737241-INS-52 | chr13-16737213-INS-80 |
| chr13-16738378-INS-53 | chr13-16738378-INS-51 |
| chr13-16752417-INS-412 | chr13-16752417-INS-409 |
| chr13-16762695-INS-4415 | chr13-16762704-INS-4406 |
| chr13-16788171-INS-5540 | chr13-16788172-INS-5540 |
| chr13-16794398-INS-63 | chr13-16794410-INS-66 |
| chr13-16804578-INS-1521 | chr13-16804579-INS-1521 |
| chr13-16896389-INS-936 | chr13-16896389-INS-936 |
| chr13-16903462-INS-513 | chr13-16903462-INS-513 |
| chr13-16906844-INS-1321 | chr13-16906845-INS-1322 |
| chr13-16916855-INS-907 | chr13-16916855-INS-908 |
| chr13-16919776-INS-72 | chr13-16919776-INS-72 |
| chr13-16957927-INS-110 | chr13-16957928-INS-96 |
| chr13-16975112-INS-491 | chr13-16975112-INS-502 |
| chr13-16984303-INS-359 | chr13-16984303-INS-359 |

| | |
|---|---|
| chr13-16995108-INS-52 | chr13-16995108-INS-52 |
| chr13-17010123-INS-766 | chr13-17010123-INS-767 |
| chr13-17034086-INS-168 | chr13-17034084-INS-164 |
| chr13-17063336-INS-140 | chr13-17063312-INS-188 |
| chr13-17067856-INS-3659 | chr13-17067857-INS-3676 |
| chr13-17086626-INS-2487 | chr13-17086627-INS-2487 |
| chr13-17111319-INS-470 | chr13-17111319-INS-470 |
| chr13-17114789-INS-424 | chr13-17114789-INS-424 |
| chr13-17134418-INS-541 | chr13-17134418-INS-510 |
| chr13-17171088-INS-323 | chr13-17171088-INS-322 |
| chr13-17186950-INS-51 | chr13-17186950-INS-56 |
| chr13-17188618-INS-83 | chr13-17188618-INS-83 |
| chr13-17217972-INS-64 | chr13-17217970-INS-56 |
| chr13-17235750-INS-504 | chr13-17235750-INS-504 |
| chr13-17271534-INS-3238 | chr13-17271535-INS-3233 |
| chr13-17280840-INS-88 | chr13-17280840-INS-88 |
| chr13-17374521-INS-151 | chr13-17374501-INS-163 |
| chr13-17386897-INS-7047 | chr13-17386898-INS-7053 |
| chr13-17461860-INS-193 | chr13-17461860-INS-193 |
| chr13-17475898-INS-53 | chr13-17475892-INS-50 |
| chr13-17480730-INS-593 | chr13-17480730-INS-593 |
| chr13-17498994-INS-217 | chr13-17498994-INS-218 |
| chr13-17512379-INS-6903 | chr13-17512380-INS-6903 |
| chr13-17512425-INS-92 | chr13-17512425-INS-92 |
| chr13-17554674-INS-79 | chr13-17554674-INS-55 |
| chr13-17573864-INS-6366 | chr13-17573865-INS-6366 |
| chr13-17584714-INS-62 | chr13-17584714-INS-62 |
| chr13-17613034-INS-5402 | chr13-17613035-INS-5402 |
| chr13-17654262-INS-449 | chr13-17654262-INS-449 |
| chr13-17684695-INS-64 | chr13-17684695-INS-58 |
| chr13-17693451-INS-633 | chr13-17693451-INS-632 |
| chr13-17728665-INS-209 | chr13-17728665-INS-208 |
| chr13-17750226-INS-60 | chr13-17750210-INS-68 |
| chr13-17754040-INS-7275 | chr13-17754040-INS-7275 |
| chr13-17782284-INS-64 | chr13-17782284-INS-50 |
| chr13-17789023-INS-1711 | chr13-17789024-INS-1711 |
| chr13-17794261-INS-3131 | chr13-17794262-INS-3131 |
| chr13-17795955-INS-64 | chr13-17795955-INS-60 |
| chr13-17813530-INS-241 | chr13-17813530-INS-241 |
| chr13-17823269-INS-118 | chr13-17823269-INS-118 |
| chr13-17824259-INS-4835 | chr13-17824260-INS-4835 |
| chr13-17825789-INS-192 | chr13-17825789-INS-192 |
| chr13-17835846-INS-6483 | chr13-17835847-INS-6483 |
| chr13-17841652-INS-543 | chr13-17841652-INS-543 |
| chr13-17847887-INS-59 | chr13-17847887-INS-59 |
| chr13-17881864-INS-68 | chr13-17881864-INS-72 |
| chr13-17902145-INS-6455 | chr13-17902146-INS-6455 |
| chr13-17928517-INS-63 | chr13-17928517-INS-63 |
| chr13-17939293-INS-86 | chr13-17939293-INS-86 |
| chr13-17949094-INS-85 | chr13-17949094-INS-83 |
| chr13-17957822-INS-66 | chr13-17957826-INS-51 |
| chr13-17962101-INS-7090 | chr13-17962102-INS-7090 |
| chr13-17964890-INS-61 | chr13-17964890-INS-65 |
| chr13-18003941-INS-1607 | chr13-18003942-INS-1607 |
| chr13-18025448-INS-159 | chr13-18025448-INS-159 |
| chr13-18032542-INS-53 | chr13-18032578-INS-59 |
| chr13-18036244-INS-250 | chr13-18036246-INS-251 |
| chr13-18084941-INS-56 | chr13-18084939-INS-58 |
| chr13-18086302-INS-80 | chr13-18086302-INS-82 |
| chr13-18135054-INS-1240 | chr13-18135055-INS-1240 |
| chr13-18141108-INS-7119 | chr13-18141109-INS-7119 |
| chr13-18146336-INS-687 | chr13-18146336-INS-692 |
| chr13-18173600-INS-74 | chr13-18173600-INS-74 |
| chr13-18203461-INS-185 | chr13-18203461-INS-185 |
| chr13-18218573-INS-334 | chr13-18218573-INS-333 |
| chr13-18244138-INS-399 | chr13-18244138-INS-399 |
| chr13-18245909-INS-89 | chr13-18245911-INS-69 |
| chr13-18246350-INS-435 | chr13-18246350-INS-435 |
| chr13-18261512-INS-6452 | chr13-18261513-INS-6453 |
| chr13-18272184-INS-380 | chr13-18272184-INS-344 |
| chr13-18286018-INS-587 | chr13-18286018-INS-587 |
| chr13-18294565-INS-57 | chr13-18294539-INS-60 |
| chr13-18307412-INS-78 | chr13-18307392-INS-102 |
| chr13-18310129-INS-52 | chr13-18310144-INS-54 |
| chr13-18317803-INS-507 | chr13-18317803-INS-507 |
| chr13-18325513-INS-78 | chr13-18325513-INS-78 |
| chr13-18354507-INS-243 | chr13-18354507-INS-243 |

| | |
|---|---|
| chr13-18357089-INS-208 | chr13-18357089-INS-208 |
| chr13-18392889-INS-80 | chr13-18392889-INS-80 |
| chr13-18401966-INS-154 | chr13-18401966-INS-154 |
| chr13-18441484-INS-783 | chr13-18441484-INS-783 |
| chr13-18469037-INS-437 | chr13-18469037-INS-437 |
| chr13-18535531-INS-4373 | chr13-18535532-INS-4373 |
| chr13-18536510-INS-126 | chr13-18536510-INS-125 |
| chr13-18556905-INS-54 | chr13-18556903-INS-54 |
| chr13-18562602-INS-108 | chr13-18562565-INS-98 |
| chr13-18586317-INS-6869 | chr13-18586318-INS-6869 |
| chr13-18596896-INS-78 | chr13-18596896-INS-78 |
| chr13-18609868-INS-1161 | chr13-18609869-INS-1165 |
| chr13-18621025-INS-84 | chr13-18621025-INS-84 |
| chr13-18623371-INS-6662 | chr13-18623372-INS-6651 |
| chr13-18631363-INS-4318 | chr13-18631364-INS-4318 |
| chr13-18633497-INS-159 | chr13-18633497-INS-169 |
| chr13-18657060-INS-214 | chr13-18657060-INS-218 |
| chr13-18658732-INS-6406 | chr13-18658733-INS-6406 |
| chr13-18690194-INS-1972 | chr13-18690195-INS-1974 |
| chr13-18696416-INS-3330 | chr13-18696417-INS-3330 |
| chr13-18707684-INS-215 | chr13-18707701-INS-217 |
| chr13-18719252-INS-6761 | chr13-18719253-INS-6761 |
| chr13-18728527-INS-100 | chr13-18728533-INS-72 |
| chr13-18740437-INS-111 | chr13-18740437-INS-111 |
| chr13-18780029-INS-488 | chr13-18780029-INS-488 |
| chr13-18802252-INS-440 | chr13-18802252-INS-440 |
| chr13-18803880-INS-6712 | chr13-18803881-INS-6712 |
| chr13-18805252-INS-6631 | chr13-18805253-INS-6631 |
| chr13-18867732-INS-390 | chr13-18867732-INS-390 |
| chr13-18890614-INS-211 | chr13-18890614-INS-211 |
| chr13-18893420-INS-220 | chr13-18893420-INS-220 |
| chr13-18894288-INS-874 | chr13-18894284-INS-879 |
| chr13-18898267-INS-704 | chr13-18898267-INS-704 |
| chr13-18903272-INS-88 | chr13-18903272-INS-88 |
| chr13-18918861-INS-3556 | chr13-18918862-INS-3558 |
| chr13-18925524-INS-2190 | chr13-18925525-INS-2190 |
| chr13-18926447-INS-60 | chr13-18926447-INS-68 |
| chr13-18934329-INS-5201 | chr13-18934330-INS-5202 |
| chr13-18970072-INS-615 | chr13-18970081-INS-622 |
| chr13-18987624-INS-345 | chr13-18987624-INS-345 |
| chr13-18989798-INS-489 | chr13-18989798-INS-489 |
| chr13-19011542-INS-70 | chr13-19011542-INS-70 |
| chr13-19016086-INS-944 | chr13-19016085-INS-948 |
| chr13-19031244-INS-450 | chr13-19031247-INS-447 |
| chr13-19064362-INS-6588 | chr13-19064363-INS-6588 |
| chr13-19073327-INS-56 | chr13-19073323-INS-68 |
| chr13-19073511-INS-8755 | chr13-19073512-INS-8759 |
| chr13-19077094-INS-730 | chr13-19077094-INS-731 |
| chr13-19086948-INS-1896 | chr13-19086934-INS-1874 |
| chr13-19094320-INS-760 | chr13-19094320-INS-761 |
| chr13-19096504-INS-281 | chr13-19096504-INS-281 |
| chr13-19103423-INS-59 | chr13-19103423-INS-59 |
| chr13-19112076-INS-566 | chr13-19112076-INS-569 |
| chr13-19155787-INS-54 | chr13-19155787-INS-54 |
| chr13-19160283-INS-200 | chr13-19160283-INS-195 |
| chr13-19168891-INS-58 | chr13-19168864-INS-52 |
| chr13-19202660-INS-440 | chr13-19202694-INS-451 |
| chr13-19216009-INS-157 | chr13-19216009-INS-157 |
| chr13-19238347-INS-640 | chr13-19238347-INS-641 |
| chr13-19264189-INS-293 | chr13-19264189-INS-291 |
| chr13-19276238-INS-4948 | chr13-19276239-INS-4948 |
| chr13-19277821-INS-74 | chr13-19277821-INS-64 |
| chr13-19325905-INS-60 | chr13-19325905-INS-60 |
| chr13-19330287-INS-407 | chr13-19330287-INS-407 |
| chr13-19360223-INS-2392 | chr13-19360224-INS-2393 |
| chr13-19364098-INS-95 | chr13-19364098-INS-95 |
| chr13-19373133-INS-278 | chr13-19373133-INS-272 |
| chr13-19507815-INS-79 | chr13-19507815-INS-79 |
| chr13-19527374-INS-1310 | chr13-19527375-INS-1310 |
| chr13-19532961-INS-543 | chr13-19532961-INS-543 |
| chr13-19534094-INS-97 | chr13-19534094-INS-96 |
| chr13-19550401-INS-121 | chr13-19550401-INS-121 |
| chr13-19557474-INS-58 | chr13-19557474-INS-54 |
| chr13-19577169-INS-7271 | chr13-19577170-INS-7271 |
| chr13-19595694-INS-87 | chr13-19595694-INS-87 |
| chr13-19633376-INS-110 | chr13-19633376-INS-110 |
| chr13-19645903-INS-100 | chr13-19645903-INS-100 |

| | |
|---|---|
| chr13-19649674-INS-327 | chr13-19649681-INS-319 |
| chr13-19670535-INS-64 | chr13-19670535-INS-64 |
| chr13-19676257-INS-5614 | chr13-19676258-INS-5614 |
| chr13-19681650-INS-198 | chr13-19681657-INS-208 |
| chr13-19695233-INS-524 | chr13-19695233-INS-525 |
| chr13-19704564-INS-76 | chr13-19704579-INS-70 |
| chr13-19711868-INS-223 | chr13-19711863-INS-222 |
| chr13-19712040-INS-2864 | chr13-19712041-INS-2864 |
| chr13-19726657-INS-59 | chr13-19726657-INS-59 |
| chr13-19732329-INS-141 | chr13-19732329-INS-141 |
| chr13-19737905-INS-5483 | chr13-19737906-INS-5483 |
| chr13-19742167-INS-560 | chr13-19742167-INS-560 |
| chr13-19755362-INS-434 | chr13-19755362-INS-434 |
| chr13-19758701-INS-58 | chr13-19758716-INS-62 |
| chr13-19761723-INS-578 | chr13-19761723-INS-578 |
| chr13-19766041-INS-222 | chr13-19766041-INS-222 |
| chr13-19815987-INS-219 | chr13-19815987-INS-219 |
| chr13-19862801-INS-85 | chr13-19862801-INS-55 |
| chr13-19868079-INS-484 | chr13-19868079-INS-484 |
| chr13-19876609-INS-63 | chr13-19876607-INS-57 |
| chr13-19881458-INS-56 | chr13-19881458-INS-52 |
| chr13-19909502-INS-51 | chr13-19909502-INS-51 |
| chr13-19922174-INS-511 | chr13-19922174-INS-511 |
| chr13-19938914-INS-64 | chr13-19938915-INS-63 |
| chr13-19943527-INS-52 | chr13-19943527-INS-56 |
| chr13-19952429-INS-55 | chr13-19952428-INS-55 |
| chr13-19955685-INS-52 | chr13-19955685-INS-52 |
| chr13-19976769-INS-99 | chr13-19976784-INS-53 |
| chr13-19988856-INS-498 | chr13-19988856-INS-498 |
| chr13-19990233-INS-142 | chr13-19990232-INS-145 |
| chr13-19998861-INS-102 | chr13-19998861-INS-96 |
| chr13-20003642-INS-152 | chr13-20003642-INS-152 |
| chr13-20017428-INS-141 | chr13-20017428-INS-163 |
| chr13-20023679-INS-7897 | chr13-20023680-INS-7898 |
| chr13-20044287-INS-72 | chr13-20044288-INS-76 |
| chr13-20045764-INS-62 | chr13-20045775-INS-64 |
| chr13-20050626-INS-422 | chr13-20050626-INS-422 |
| chr13-20059450-INS-322 | chr13-20059450-INS-322 |
| chr13-20062384-INS-296 | chr13-20062384-INS-296 |
| chr13-20064966-INS-148 | chr13-20064966-INS-148 |
| chr13-20073720-INS-73 | chr13-20073720-INS-73 |
| chr13-20076116-INS-630 | chr13-20076116-INS-630 |
| chr13-20093972-INS-200 | chr13-20093975-INS-200 |
| chr13-20099499-INS-73 | chr13-20099499-INS-73 |
| chr13-20112319-INS-66 | chr13-20112319-INS-80 |
| chr13-20126353-INS-2941 | chr13-20126354-INS-2941 |
| chr13-20129732-INS-56 | chr13-20129732-INS-56 |
| chr13-20153794-INS-66 | chr13-20153794-INS-66 |
| chr13-20158194-INS-407 | chr13-20158194-INS-407 |
| chr13-20188866-INS-209 | chr13-20188866-INS-209 |
| chr13-20192852-INS-688 | chr13-20192852-INS-687 |
| chr13-20195811-INS-151 | chr13-20195811-INS-151 |
| chr13-20212477-INS-116 | chr13-20212477-INS-116 |
| chr13-20237371-INS-50 | chr13-20237373-INS-68 |
| chr13-20239854-INS-65 | chr13-20239854-INS-65 |
| chr13-20242971-INS-50 | chr13-20242961-INS-77 |
| chr13-20319583-INS-7991 | chr13-20319583-INS-7983 |
| chr13-20334610-INS-71 | chr13-20334610-INS-71 |
| chr13-20366035-INS-143 | chr13-20366035-INS-107 |
| chr13-20377465-INS-80 | chr13-20377454-INS-80 |
| chr13-20400045-INS-1992 | chr13-20400046-INS-1994 |
| chr13-20402693-INS-52 | chr13-20402695-INS-62 |
| chr13-20403285-INS-343 | chr13-20403285-INS-343 |
| chr13-20406901-INS-158 | chr13-20406901-INS-197 |
| chr13-20411369-INS-56 | chr13-20411366-INS-64 |
| chr13-20413223-INS-5838 | chr13-20413224-INS-5842 |
| chr13-20423175-INS-554 | chr13-20423175-INS-553 |
| chr13-20428191-INS-288 | chr13-20428191-INS-286 |
| chr13-20437735-INS-110 | chr13-20437747-INS-84 |
| chr13-20442350-INS-507 | chr13-20442350-INS-506 |
| chr13-20459969-INS-62 | chr13-20459969-INS-52 |
| chr13-20463240-INS-84 | chr13-20463240-INS-68 |
| chr13-20488500-INS-75 | chr13-20488475-INS-98 |
| chr13-20508140-INS-64 | chr13-20508140-INS-64 |
| chr13-20509327-INS-7167 | chr13-20509328-INS-7167 |
| chr13-20530780-INS-375 | chr13-20530760-INS-376 |
| chr13-20548169-INS-119 | chr13-20548169-INS-108 |

| | |
|---|---|
| chr13-20553888-INS-186 | chr13-20553888-INS-183 |
| chr13-20566146-INS-1892 | chr13-20566147-INS-1892 |
| chr13-20614729-INS-122 | chr13-20614729-INS-122 |
| chr13-20625966-INS-222 | chr13-20625966-INS-216 |
| chr13-20626308-INS-441 | chr13-20626308-INS-440 |
| chr13-20638309-INS-507 | chr13-20638309-INS-507 |
| chr13-20646672-INS-54 | chr13-20646672-INS-56 |
| chr13-20662145-INS-252 | chr13-20662096-INS-282 |
| chr13-20672032-INS-134 | chr13-20672031-INS-162 |
| chr13-20687004-INS-112 | chr13-20687004-INS-112 |
| chr13-20747481-INS-545 | chr13-20747481-INS-545 |
| chr13-20750458-INS-83 | chr13-20750458-INS-83 |
| chr13-20753908-INS-52 | chr13-20753908-INS-52 |
| chr13-20764190-INS-105 | chr13-20764190-INS-105 |
| chr13-20793037-INS-51 | chr13-20793037-INS-54 |
| chr13-20796317-INS-490 | chr13-20796317-INS-505 |
| chr13-20798477-INS-59 | chr13-20798477-INS-59 |
| chr13-20845038-INS-63 | chr13-20845038-INS-63 |
| chr13-20865178-INS-151 | chr13-20865178-INS-151 |
| chr13-20878322-INS-2589 | chr13-20878323-INS-2589 |
| chr13-20880103-INS-1077 | chr13-20880104-INS-1077 |
| chr13-20912166-INS-826 | chr13-20912166-INS-826 |
| chr13-20913195-INS-657 | chr13-20913195-INS-657 |
| chr13-20921861-INS-906 | chr13-20921861-INS-907 |
| chr13-20933332-INS-487 | chr13-20933332-INS-487 |
| chr13-20950370-INS-6844 | chr13-20950371-INS-6844 |
| chr13-20953591-INS-607 | chr13-20953591-INS-608 |
| chr13-20965947-INS-990 | chr13-20965947-INS-991 |
| chr13-20966651-INS-7098 | chr13-20966652-INS-7098 |
| chr13-20980735-INS-1699 | chr13-20980736-INS-1699 |
| chr13-20988138-INS-75 | chr13-20988138-INS-83 |
| chr13-21000131-INS-609 | chr13-21000131-INS-609 |
| chr13-21016885-INS-77 | chr13-21016881-INS-81 |
| chr13-21030687-INS-54 | chr13-21030713-INS-66 |
| chr13-21034287-INS-60 | chr13-21034287-INS-60 |
| chr13-21038254-INS-118 | chr13-21038268-INS-141 |
| chr13-21069635-INS-150 | chr13-21069635-INS-180 |
| chr13-21084297-INS-57 | chr13-21084297-INS-87 |
| chr13-21092321-INS-6307 | chr13-21092322-INS-6306 |
| chr13-21135371-INS-274 | chr13-21135371-INS-274 |
| chr13-21182850-INS-187 | chr13-21182850-INS-187 |
| chr13-21183794-INS-55 | chr13-21183798-INS-79 |
| chr13-21185178-INS-90 | chr13-21185178-INS-53 |
| chr13-21191848-INS-206 | chr13-21191848-INS-207 |
| chr13-21211804-INS-761 | chr13-21211804-INS-761 |
| chr13-21214902-INS-175 | chr13-21214902-INS-176 |
| chr13-21242073-INS-499 | chr13-21242073-INS-499 |
| chr13-21254785-INS-205 | chr13-21254785-INS-205 |
| chr13-21265621-INS-136 | chr13-21265657-INS-99 |
| chr13-21279073-INS-7086 | chr13-21279074-INS-7086 |
| chr13-21282075-INS-62 | chr13-21282075-INS-78 |
| chr13-21293871-INS-72 | chr13-21293871-INS-72 |
| chr13-21304386-INS-95 | chr13-21304386-INS-95 |
| chr13-21306384-INS-439 | chr13-21306385-INS-439 |
| chr13-21324048-INS-199 | chr13-21324048-INS-198 |
| chr13-21327604-INS-89 | chr13-21327604-INS-89 |
| chr13-21332630-INS-135 | chr13-21332631-INS-132 |
| chr13-21334974-INS-513 | chr13-21334974-INS-513 |
| chr13-21353774-INS-62 | chr13-21353774-INS-51 |
| chr13-21375244-INS-288 | chr13-21375244-INS-240 |
| chr13-21400420-INS-221 | chr13-21400420-INS-228 |
| chr13-21411621-INS-160 | chr13-21411621-INS-170 |
| chr13-21415199-INS-80 | chr13-21415206-INS-119 |
| chr13-21419690-INS-271 | chr13-21419690-INS-313 |
| chr13-21427790-INS-66 | chr13-21427790-INS-66 |
| chr13-21431507-INS-56 | chr13-21431507-INS-56 |
| chr13-21441755-INS-177 | chr13-21441755-INS-177 |
| chr13-21455706-INS-209 | chr13-21455706-INS-211 |
| chr13-21480206-INS-175 | chr13-21480206-INS-183 |
| chr13-21494069-INS-265 | chr13-21494074-INS-268 |
| chr13-21523420-INS-82 | chr13-21523424-INS-82 |
| chr13-21534584-INS-151 | chr13-21534584-INS-151 |
| chr13-21602849-INS-113 | chr13-21602849-INS-102 |
| chr13-21610900-INS-7113 | chr13-21610901-INS-7113 |
| chr13-21613489-INS-66 | chr13-21613489-INS-66 |
| chr13-21745721-INS-4743 | chr13-21745722-INS-4743 |
| chr13-21749978-INS-395 | chr13-21749978-INS-395 |

| | |
|---|---|
| chr13-21769526-INS-142 | chr13-21769526-INS-146 |
| chr13-21773021-INS-318 | chr13-21773021-INS-318 |
| chr13-21775215-INS-830 | chr13-21775215-INS-830 |
| chr13-21800519-INS-72 | chr13-21800519-INS-96 |
| chr13-21801990-INS-75 | chr13-21801990-INS-75 |
| chr13-21841863-INS-128 | chr13-21841863-INS-108 |
| chr13-21843846-INS-58 | chr13-21843846-INS-57 |
| chr13-21866099-INS-766 | chr13-21866099-INS-766 |
| chr13-21884034-INS-188 | chr13-21884034-INS-190 |
| chr13-21891935-INS-83 | chr13-21891935-INS-83 |
| chr13-21905193-INS-64 | chr13-21905211-INS-63 |
| chr13-21911009-INS-3308 | chr13-21911010-INS-3308 |
| chr13-22067225-INS-66 | chr13-22067225-INS-66 |
| chr13-22093436-INS-52 | chr13-22093436-INS-56 |
| chr13-22101917-INS-139 | chr13-22101917-INS-128 |
| chr13-22137927-INS-201 | chr13-22137927-INS-201 |
| chr13-22151446-INS-93 | chr13-22151446-INS-68 |
| chr13-22162853-INS-185 | chr13-22162853-INS-186 |
| chr13-22169560-INS-119 | chr13-22169560-INS-119 |
| chr13-22173658-INS-100 | chr13-22173658-INS-100 |
| chr13-22177618-INS-656 | chr13-22177616-INS-660 |
| chr13-22197242-INS-369 | chr13-22197242-INS-363 |
| chr13-22211914-INS-6454 | chr13-22211915-INS-6454 |
| chr13-22215757-INS-64 | chr13-22215757-INS-52 |
| chr13-22227946-INS-53 | chr13-22227946-INS-53 |
| chr13-22235910-INS-64 | chr13-22235924-INS-52 |
| chr13-22240775-INS-192 | chr13-22240778-INS-191 |
| chr13-22246310-INS-123 | chr13-22246310-INS-123 |
| chr13-22267441-INS-69 | chr13-22267441-INS-72 |
| chr13-22278750-INS-115 | chr13-22278750-INS-115 |
| chr13-22330153-INS-852 | chr13-22330153-INS-852 |
| chr13-22368063-INS-66 | chr13-22368063-INS-52 |
| chr13-22395337-INS-203 | chr13-22395337-INS-203 |
| chr13-22411853-INS-87 | chr13-22411853-INS-87 |
| chr13-22448729-INS-817 | chr13-22448729-INS-816 |
| chr13-22457222-INS-78 | chr13-22457222-INS-56 |
| chr13-22471725-INS-6617 | chr13-22471726-INS-6617 |
| chr13-22481573-INS-6448 | chr13-22481574-INS-6448 |
| chr13-22629150-INS-5251 | chr13-22629151-INS-5251 |
| chr13-22660663-INS-6597 | chr13-22660664-INS-6597 |
| chr13-22665709-INS-98 | chr13-22665710-INS-100 |
| chr13-22673418-INS-6436 | chr13-22673419-INS-6436 |
| chr13-22696032-INS-354 | chr13-22696032-INS-354 |
| chr13-22698052-INS-327 | chr13-22698052-INS-327 |
| chr13-22700275-INS-54 | chr13-22700279-INS-54 |
| chr13-22738212-INS-104 | chr13-22738212-INS-105 |
| chr13-22782606-INS-208 | chr13-22782606-INS-208 |
| chr13-22818038-INS-2975 | chr13-22818039-INS-2975 |
| chr13-22834432-INS-66 | chr13-22834432-INS-66 |
| chr13-22855398-INS-747 | chr13-22855437-INS-749 |
| chr13-22899244-INS-57 | chr13-22899223-INS-78 |
| chr13-22901017-INS-219 | chr13-22901017-INS-219 |
| chr13-22911092-INS-52 | chr13-22911092-INS-50 |
| chr13-23001671-INS-1888 | chr13-23001672-INS-1888 |
| chr13-23016725-INS-6622 | chr13-23016726-INS-6618 |
| chr13-23049459-INS-6248 | chr13-23049460-INS-6248 |
| chr13-23160910-INS-109 | chr13-23160910-INS-110 |
| chr13-23178911-INS-80 | chr13-23178911-INS-80 |
| chr13-23232783-INS-68 | chr13-23232783-INS-72 |
| chr13-23251639-INS-50 | chr13-23251639-INS-54 |
| chr13-23265609-INS-6206 | chr13-23265610-INS-6206 |
| chr13-23276339-INS-299 | chr13-23276339-INS-299 |
| chr13-23289831-INS-6586 | chr13-23289832-INS-6586 |
| chr13-23323506-INS-91 | chr13-23323506-INS-91 |
| chr13-23361996-INS-4786 | chr13-23361997-INS-4786 |
| chr13-23425139-INS-57 | chr13-23425139-INS-60 |
| chr13-23443407-INS-583 | chr13-23443407-INS-583 |
| chr13-23445329-INS-80 | chr13-23445329-INS-80 |
| chr13-23464261-INS-623 | chr13-23464261-INS-623 |
| chr13-23477168-INS-300 | chr13-23477168-INS-300 |
| chr13-23505130-INS-362 | chr13-23505130-INS-362 |
| chr13-23507332-INS-214 | chr13-23507332-INS-214 |
| chr13-23537034-INS-417 | chr13-23537034-INS-417 |
| chr13-23558906-INS-178 | chr13-23558906-INS-178 |
| chr13-23589906-INS-80 | chr13-23589906-INS-80 |
| chr13-23620991-INS-167 | chr13-23620992-INS-165 |
| chr13-23627695-INS-6601 | chr13-23627696-INS-6601 |

| | |
|---|---|
| chr13-23638558-INS-576 | chr13-23638558-INS-589 |
| chr13-23677740-INS-538 | chr13-23677740-INS-539 |
| chr13-23687013-INS-59 | chr13-23687014-INS-58 |
| chr13-23762612-INS-141 | chr13-23762592-INS-178 |
| chr13-23788058-INS-130 | chr13-23788058-INS-169 |
| chr13-23810338-INS-90 | chr13-23810338-INS-90 |
| chr13-23840296-INS-4465 | chr13-23840297-INS-4465 |
| chr13-23841775-INS-261 | chr13-23841775-INS-261 |
| chr13-23850837-INS-52 | chr13-23850838-INS-54 |
| chr13-23874610-INS-184 | chr13-23874610-INS-185 |
| chr13-23894117-INS-54 | chr13-23894117-INS-57 |
| chr13-23941788-INS-109 | chr13-23941788-INS-107 |
| chr13-23993042-INS-443 | chr13-23993040-INS-450 |
| chr13-23997520-INS-61 | chr13-23997520-INS-61 |
| chr13-24000444-INS-205 | chr13-24000444-INS-206 |
| chr13-24014051-INS-50 | chr13-24014051-INS-66 |
| chr13-24019576-INS-106 | chr13-24019576-INS-112 |
| chr13-24025860-INS-55 | chr13-24025860-INS-57 |
| chr13-24028201-INS-78 | chr13-24028201-INS-78 |
| chr13-24071599-INS-99 | chr13-24071599-INS-99 |
| chr13-24077761-INS-60 | chr13-24077761-INS-60 |
| chr13-24142230-INS-56 | chr13-24142230-INS-56 |
| chr13-24144670-INS-647 | chr13-24144670-INS-647 |
| chr13-24161422-INS-64 | chr13-24161422-INS-72 |
| chr13-24177853-INS-275 | chr13-24177853-INS-275 |
| chr13-24191608-INS-7139 | chr13-24191609-INS-7139 |
| chr13-24193972-INS-942 | chr13-24193972-INS-944 |
| chr13-24203117-INS-58 | chr13-24203117-INS-58 |
| chr13-24241210-INS-76 | chr13-24241210-INS-90 |
| chr13-24242278-INS-64 | chr13-24242278-INS-70 |
| chr13-24247264-INS-126 | chr13-24247264-INS-130 |
| chr13-24252411-INS-100 | chr13-24252412-INS-60 |
| chr13-24258582-INS-1124 | chr13-24258583-INS-1124 |
| chr13-24313831-INS-93 | chr13-24313831-INS-97 |
| chr13-24324005-INS-368 | chr13-24324031-INS-372 |
| chr13-24333399-INS-64 | chr13-24333389-INS-87 |
| chr13-24367271-INS-85 | chr13-24367271-INS-51 |
| chr13-24374743-INS-170 | chr13-24374743-INS-170 |
| chr13-24392038-INS-63 | chr13-24392037-INS-65 |
| chr13-24405720-INS-52 | chr13-24405720-INS-52 |
| chr13-24448910-INS-2181 | chr13-24448915-INS-2147 |
| chr13-24457787-INS-75 | chr13-24457787-INS-78 |
| chr13-24473752-INS-61 | chr13-24473715-INS-62 |
| chr13-24478866-INS-251 | chr13-24478866-INS-259 |
| chr13-24500924-INS-84 | chr13-24500924-INS-120 |
| chr13-24549780-INS-1132 | chr13-24549781-INS-1132 |
| chr13-24588551-INS-62 | chr13-24588551-INS-62 |
| chr13-24611931-INS-76 | chr13-24611931-INS-76 |
| chr13-24612243-INS-399 | chr13-24612243-INS-399 |
| chr13-24640027-INS-80 | chr13-24640027-INS-75 |
| chr13-24641713-INS-166 | chr13-24641714-INS-166 |
| chr13-24665952-INS-1923 | chr13-24665953-INS-1923 |
| chr13-24670478-INS-4844 | chr13-24670479-INS-4844 |
| chr13-24699596-INS-112 | chr13-24699596-INS-112 |
| chr13-24707552-INS-154 | chr13-24707550-INS-116 |
| chr13-24721904-INS-56 | chr13-24721904-INS-60 |
| chr13-24725765-INS-444 | chr13-24725765-INS-444 |
| chr13-24767730-INS-139 | chr13-24767731-INS-139 |
| chr13-24784674-INS-217 | chr13-24784674-INS-217 |
| chr13-24788582-INS-53 | chr13-24788582-INS-68 |
| chr13-24825061-INS-64 | chr13-24825061-INS-64 |
| chr13-24869131-INS-163 | chr13-24869132-INS-191 |
| chr13-24870237-INS-408 | chr13-24870203-INS-408 |
| chr13-24873686-INS-212 | chr13-24873686-INS-212 |
| chr13-24878056-INS-59 | chr13-24878056-INS-59 |
| chr13-24926559-INS-77 | chr13-24926559-INS-77 |
| chr13-24934755-INS-314 | chr13-24934758-INS-315 |
| chr13-24938168-INS-238 | chr13-24938168-INS-238 |
| chr13-24961524-INS-104 | chr13-24961524-INS-84 |
| chr13-25005287-INS-206 | chr13-25005287-INS-206 |
| chr13-25006585-INS-261 | chr13-25006585-INS-261 |
| chr13-25009117-INS-240 | chr13-25009117-INS-240 |
| chr13-25011629-INS-203 | chr13-25011629-INS-203 |
| chr13-25019671-INS-246 | chr13-25019671-INS-246 |
| chr13-25053878-INS-477 | chr13-25053878-INS-481 |
| chr13-25068868-INS-3918 | chr13-25068869-INS-3923 |
| chr13-25075117-INS-147 | chr13-25075154-INS-147 |

| | |
|---|---|
| chr13-25081646-INS-965 | chr13-25081646-INS-965 |
| chr13-25086312-INS-114 | chr13-25086312-INS-114 |
| chr13-25101144-INS-358 | chr13-25101144-INS-362 |
| chr13-25106517-INS-132 | chr13-25106517-INS-132 |
| chr13-25117623-INS-84 | chr13-25117623-INS-84 |
| chr13-25118573-INS-979 | chr13-25118573-INS-982 |
| chr13-25120986-INS-180 | chr13-25120986-INS-162 |
| chr13-25148819-INS-96 | chr13-25148819-INS-96 |
| chr13-25162369-INS-173 | chr13-25162369-INS-173 |
| chr13-25187531-INS-68 | chr13-25187531-INS-60 |
| chr13-25189092-INS-87 | chr13-25189092-INS-87 |
| chr13-25201218-INS-290 | chr13-25201218-INS-290 |
| chr13-25234393-INS-3142 | chr13-25234380-INS-3150 |
| chr13-25238673-INS-131 | chr13-25238673-INS-134 |
| chr13-25249918-INS-101 | chr13-25249918-INS-102 |
| chr13-25271125-INS-53 | chr13-25271125-INS-59 |
| chr13-25278659-INS-53 | chr13-25278662-INS-54 |
| chr13-25287393-INS-200 | chr13-25287393-INS-200 |
| chr13-25300520-INS-553 | chr13-25300520-INS-504 |
| chr13-25352540-INS-86 | chr13-25352562-INS-86 |
| chr13-25358418-INS-81 | chr13-25358426-INS-97 |
| chr13-25362440-INS-84 | chr13-25362440-INS-84 |
| chr13-25374221-INS-106 | chr13-25374221-INS-106 |
| chr13-25376234-INS-54 | chr13-25376234-INS-54 |
| chr13-25380954-INS-176 | chr13-25380954-INS-176 |
| chr13-25387425-INS-114 | chr13-25387423-INS-145 |
| chr13-25393522-INS-6719 | chr13-25393523-INS-6719 |
| chr13-25394649-INS-6443 | chr13-25394650-INS-6443 |
| chr13-25426920-INS-199 | chr13-25426920-INS-199 |
| chr13-25441051-INS-6403 | chr13-25441051-INS-6403 |
| chr13-25488865-INS-229 | chr13-25488865-INS-229 |
| chr13-25501292-INS-75 | chr13-25501292-INS-105 |
| chr13-25523786-INS-80 | chr13-25523786-INS-86 |
| chr13-25533378-INS-213 | chr13-25533378-INS-218 |
| chr13-25563338-INS-143 | chr13-25563338-INS-146 |
| chr13-25577636-INS-388 | chr13-25577649-INS-400 |
| chr13-25648036-INS-431 | chr13-25648036-INS-431 |
| chr13-25653418-INS-68 | chr13-25653426-INS-88 |
| chr13-25663864-INS-111 | chr13-25663864-INS-74 |
| chr13-25670828-INS-83 | chr13-25670828-INS-52 |
| chr13-25676383-INS-380 | chr13-25676383-INS-380 |
| chr13-25705223-INS-5548 | chr13-25705223-INS-5548 |
| chr13-25732420-INS-274 | chr13-25732420-INS-274 |
| chr13-25736297-INS-317 | chr13-25736297-INS-317 |
| chr13-25744850-INS-61 | chr13-25744850-INS-61 |
| chr13-25753095-INS-842 | chr13-25753095-INS-845 |
| chr13-25803104-INS-425 | chr13-25803104-INS-425 |
| chr13-25809293-INS-61 | chr13-25809293-INS-61 |
| chr13-25817679-INS-131 | chr13-25817679-INS-147 |
| chr13-25821860-INS-480 | chr13-25821860-INS-480 |
| chr13-25822666-INS-188 | chr13-25822666-INS-188 |
| chr13-25835612-INS-8475 | chr13-25835613-INS-8475 |
| chr13-25837401-INS-105 | chr13-25837396-INS-108 |
| chr13-25843593-INS-55 | chr13-25843593-INS-56 |
| chr13-25851701-INS-2984 | chr13-25851702-INS-2984 |
| chr13-25853681-INS-488 | chr13-25853681-INS-488 |
| chr13-25855043-INS-828 | chr13-25855043-INS-828 |
| chr13-25872702-INS-657 | chr13-25872702-INS-657 |
| chr13-25883586-INS-64 | chr13-25883537-INS-66 |
| chr13-25914717-INS-83 | chr13-25914717-INS-85 |
| chr13-25918626-INS-578 | chr13-25918619-INS-581 |
| chr13-25926035-INS-222 | chr13-25926035-INS-223 |
| chr13-25931338-INS-138 | chr13-25931338-INS-140 |
| chr13-25952534-INS-57 | chr13-25952534-INS-52 |
| chr13-25961987-INS-214 | chr13-25961987-INS-213 |
| chr13-25967413-INS-530 | chr13-25967413-INS-530 |
| chr13-26003137-INS-52 | chr13-26003137-INS-54 |
| chr13-26020983-INS-54 | chr13-26020983-INS-54 |
| chr13-26037972-INS-54 | chr13-26037972-INS-54 |
| chr13-26062964-INS-67 | chr13-26062964-INS-73 |
| chr13-26064463-INS-6461 | chr13-26064464-INS-6462 |
| chr13-26108240-INS-607 | chr13-26108240-INS-607 |
| chr13-26109408-INS-88 | chr13-26109408-INS-88 |
| chr13-26114294-INS-64 | chr13-26114294-INS-56 |
| chr13-26147280-INS-85 | chr13-26147281-INS-54 |
| chr13-26156643-INS-74 | chr13-26156643-INS-74 |
| chr13-26201867-INS-700 | chr13-26201867-INS-700 |

| | |
|---|---|
| chr13-26224239-INS-64 | chr13-26224239-INS-64 |
| chr13-26229888-INS-527 | chr13-26229888-INS-528 |
| chr13-26244656-INS-476 | chr13-26244652-INS-474 |
| chr13-26247625-INS-62 | chr13-26247631-INS-50 |
| chr13-26281316-INS-56 | chr13-26281309-INS-76 |
| chr13-26285243-INS-63 | chr13-26285243-INS-63 |
| chr13-26292097-INS-6322 | chr13-26292098-INS-6322 |
| chr13-26299765-INS-415 | chr13-26299765-INS-415 |
| chr13-26301086-INS-127 | chr13-26301086-INS-127 |
| chr13-26307858-INS-53 | chr13-26307859-INS-53 |
| chr13-26317550-INS-92 | chr13-26317532-INS-102 |
| chr13-26331729-INS-6621 | chr13-26331730-INS-6621 |
| chr13-26410392-INS-60 | chr13-26410394-INS-51 |
| chr13-26419891-INS-56 | chr13-26419933-INS-52 |
| chr13-26431047-INS-60 | chr13-26431047-INS-62 |
| chr13-26458688-INS-116 | chr13-26458688-INS-116 |
| chr13-26492528-INS-68 | chr13-26492528-INS-64 |
| chr13-26504602-INS-7582 | chr13-26504603-INS-7583 |
| chr13-26507044-INS-549 | chr13-26507045-INS-548 |
| chr13-26564112-INS-708 | chr13-26564112-INS-709 |
| chr13-26613602-INS-167 | chr13-26613609-INS-151 |
| chr13-26619792-INS-96 | chr13-26619792-INS-70 |
| chr13-26622461-INS-56 | chr13-26622449-INS-55 |
| chr13-26641487-INS-922 | chr13-26641487-INS-923 |
| chr13-26656831-INS-259 | chr13-26656831-INS-259 |
| chr13-26660343-INS-70 | chr13-26660343-INS-72 |
| chr13-26675081-INS-53 | chr13-26675081-INS-53 |
| chr13-26685403-INS-52 | chr13-26685403-INS-50 |
| chr13-26751738-INS-2033 | chr13-26751739-INS-2034 |
| chr13-26796364-INS-58 | chr13-26796364-INS-68 |
| chr13-26804119-INS-473 | chr13-26804113-INS-474 |
| chr13-26832080-INS-1315 | chr13-26832081-INS-1315 |
| chr13-26835135-INS-1933 | chr13-26835136-INS-1949 |
| chr13-26836189-INS-165 | chr13-26836189-INS-185 |
| chr13-26836466-INS-7006 | chr13-26836467-INS-7006 |
| chr13-26838544-INS-703 | chr13-26838544-INS-705 |
| chr13-26872592-INS-86 | chr13-26872592-INS-86 |
| chr13-26913367-INS-69 | chr13-26913363-INS-73 |
| chr13-26941064-INS-76 | chr13-26941064-INS-80 |
| chr13-26947951-INS-6281 | chr13-26947952-INS-6278 |
| chr13-26953497-INS-102 | chr13-26953497-INS-102 |
| chr13-26964738-INS-54 | chr13-26964738-INS-52 |
| chr13-26971028-INS-77 | chr13-26971035-INS-69 |
| chr13-27021284-INS-213 | chr13-27021284-INS-213 |
| chr13-27047713-INS-66 | chr13-27047713-INS-70 |
| chr13-27075335-INS-72 | chr13-27075335-INS-68 |
| chr13-27082697-INS-65 | chr13-27082697-INS-65 |
| chr13-27083670-INS-243 | chr13-27083670-INS-243 |
| chr13-27091599-INS-314 | chr13-27091600-INS-316 |
| chr13-27095720-INS-52 | chr13-27095721-INS-59 |
| chr13-27103284-INS-54 | chr13-27103238-INS-60 |
| chr13-27108130-INS-58 | chr13-27108118-INS-55 |
| chr13-27179206-INS-1317 | chr13-27179207-INS-1320 |
| chr13-27194566-INS-6487 | chr13-27194567-INS-6487 |
| chr13-27207537-INS-7507 | chr13-27207538-INS-7507 |
| chr13-27213424-INS-124 | chr13-27213450-INS-114 |
| chr13-27237539-INS-6440 | chr13-27237540-INS-6440 |
| chr13-27262085-INS-97 | chr13-27262095-INS-78 |
| chr13-27288355-INS-91 | chr13-27288355-INS-91 |
| chr13-27343095-INS-170 | chr13-27343095-INS-170 |
| chr13-27351132-INS-910 | chr13-27351132-INS-910 |
| chr13-27374986-INS-6308 | chr13-27374987-INS-6321 |
| chr13-27399922-INS-52 | chr13-27399923-INS-64 |
| chr13-27410373-INS-55 | chr13-27410332-INS-59 |
| chr13-27419903-INS-80 | chr13-27419899-INS-84 |
| chr13-27451593-INS-187 | chr13-27451593-INS-187 |
| chr13-27453329-INS-593 | chr13-27453329-INS-592 |
| chr13-27506338-INS-61 | chr13-27506338-INS-88 |
| chr13-27565047-INS-66 | chr13-27565047-INS-68 |
| chr13-27565797-INS-7079 | chr13-27565798-INS-7084 |
| chr13-27568792-INS-68 | chr13-27568792-INS-70 |
| chr13-27602395-INS-60 | chr13-27602395-INS-56 |
| chr13-27631630-INS-1232 | chr13-27631631-INS-1233 |
| chr13-27702194-INS-1476 | chr13-27702195-INS-1476 |
| chr13-27715227-INS-119 | chr13-27715260-INS-113 |
| chr13-27773898-INS-5260 | chr13-27773899-INS-5260 |
| chr13-27785108-INS-6586 | chr13-27785109-INS-6586 |

| | |
|---|---|
| chr13-27815779-INS-89 | chr13-27815779-INS-89 |
| chr13-27888010-INS-206 | chr13-27888010-INS-206 |
| chr13-27918031-INS-485 | chr13-27918030-INS-496 |
| chr13-27920408-INS-64 | chr13-27920408-INS-64 |
| chr13-27995398-INS-4906 | chr13-27995399-INS-4906 |
| chr13-28015784-INS-164 | chr13-28015784-INS-169 |
| chr13-28055503-INS-88 | chr13-28055503-INS-88 |
| chr13-28065826-INS-199 | chr13-28065826-INS-199 |
| chr13-28108760-INS-659 | chr13-28108760-INS-659 |
| chr13-28111810-INS-984 | chr13-28111810-INS-985 |
| chr13-28180058-INS-3985 | chr13-28180059-INS-3991 |
| chr13-28209838-INS-146 | chr13-28209838-INS-130 |
| chr13-28215710-INS-7040 | chr13-28215711-INS-7041 |
| chr13-28232313-INS-139 | chr13-28232313-INS-145 |
| chr13-28236064-INS-2787 | chr13-28236065-INS-2787 |
| chr13-28257051-INS-138 | chr13-28257006-INS-149 |
| chr13-28336031-INS-6224 | chr13-28336032-INS-6224 |
| chr13-28346867-INS-522 | chr13-28346867-INS-523 |
| chr13-28350924-INS-711 | chr13-28350924-INS-711 |
| chr13-28428663-INS-495 | chr13-28428663-INS-496 |
| chr13-28455199-INS-161 | chr13-28455199-INS-163 |
| chr13-28526910-INS-6414 | chr13-28526911-INS-6414 |
| chr13-28542626-INS-648 | chr13-28542626-INS-686 |
| chr13-28546695-INS-486 | chr13-28546695-INS-488 |
| chr13-28621101-INS-7124 | chr13-28621102-INS-7124 |
| chr13-28647401-INS-3289 | chr13-28647402-INS-3289 |
| chr13-28671504-INS-73 | chr13-28671504-INS-69 |
| chr13-28678973-INS-166 | chr13-28678973-INS-174 |
| chr13-28731487-INS-201 | chr13-28731487-INS-201 |
| chr13-28736643-INS-281 | chr13-28736643-INS-286 |
| chr13-28743779-INS-275 | chr13-28743779-INS-275 |
| chr13-28781663-INS-58 | chr13-28781663-INS-58 |
| chr13-28795695-INS-223 | chr13-28795695-INS-242 |
| chr13-28811194-INS-136 | chr13-28811190-INS-116 |
| chr13-28815936-INS-138 | chr13-28815933-INS-127 |
| chr13-28844443-INS-460 | chr13-28844439-INS-460 |
| chr13-28850858-INS-134 | chr13-28850858-INS-140 |
| chr13-28881583-INS-321 | chr13-28881583-INS-316 |
| chr13-28926019-INS-763 | chr13-28926019-INS-763 |
| chr13-28928383-INS-60 | chr13-28928383-INS-62 |
| chr13-28930189-INS-88 | chr13-28930189-INS-87 |
| chr13-28933141-INS-66 | chr13-28933141-INS-66 |
| chr13-28937515-INS-67 | chr13-28937515-INS-95 |
| chr13-28940365-INS-819 | chr13-28940365-INS-820 |
| chr13-28947406-INS-367 | chr13-28947406-INS-367 |
| chr13-28977127-INS-68 | chr13-28977128-INS-68 |
| chr13-29012391-INS-80 | chr13-29012391-INS-96 |
| chr13-29016407-INS-2125 | chr13-29016396-INS-2116 |
| chr13-29027162-INS-63 | chr13-29027162-INS-51 |
| chr13-29053911-INS-108 | chr13-29053911-INS-108 |
| chr13-29058987-INS-222 | chr13-29058987-INS-228 |
| chr13-29064585-INS-210 | chr13-29064585-INS-210 |
| chr13-29075625-INS-60 | chr13-29075638-INS-52 |
| chr13-29109041-INS-180 | chr13-29109041-INS-171 |
| chr13-29115260-INS-968 | chr13-29115260-INS-970 |
| chr13-29122478-INS-55 | chr13-29122478-INS-55 |
| chr13-29151455-INS-436 | chr13-29151455-INS-436 |
| chr13-29155080-INS-201 | chr13-29155080-INS-201 |
| chr13-29156279-INS-938 | chr13-29156279-INS-943 |
| chr13-29159951-INS-67 | chr13-29159951-INS-69 |
| chr13-29163583-INS-187 | chr13-29163583-INS-187 |
| chr13-29184854-INS-1776 | chr13-29184855-INS-1776 |
| chr13-29192362-INS-68 | chr13-29192362-INS-57 |
| chr13-29194540-INS-52 | chr13-29194540-INS-97 |
| chr13-29196215-INS-68 | chr13-29196239-INS-70 |
| chr13-29204522-INS-832 | chr13-29204521-INS-838 |
| chr13-29211597-INS-52 | chr13-29211598-INS-84 |
| chr13-29235388-INS-406 | chr13-29235388-INS-405 |
| chr13-29246264-INS-184 | chr13-29246264-INS-180 |
| chr13-29302778-INS-6907 | chr13-29302779-INS-6907 |
| chr13-29326479-INS-217 | chr13-29326479-INS-216 |
| chr13-29366104-INS-53 | chr13-29366065-INS-60 |
| chr13-29395680-INS-126 | chr13-29395701-INS-117 |
| chr13-29397913-INS-81 | chr13-29397923-INS-58 |
| chr13-29401712-INS-61 | chr13-29401712-INS-60 |
| chr13-29466741-INS-332 | chr13-29466741-INS-332 |
| chr13-29473717-INS-6590 | chr13-29473718-INS-6591 |

| | |
|---|---|
| chr13-29479463-INS-162 | chr13-29479461-INS-160 |
| chr13-29485232-INS-54 | chr13-29485232-INS-58 |
| chr13-29502166-INS-401 | chr13-29502166-INS-401 |
| chr13-29513549-INS-105 | chr13-29513549-INS-105 |
| chr13-29535662-INS-3456 | chr13-29535663-INS-3456 |
| chr13-29546790-INS-331 | chr13-29546790-INS-331 |
| chr13-29550228-INS-52 | chr13-29550228-INS-52 |
| chr13-29566242-INS-482 | chr13-29566242-INS-482 |
| chr13-29583329-INS-76 | chr13-29583329-INS-64 |
| chr13-29588450-INS-110 | chr13-29588448-INS-112 |
| chr13-29606454-INS-51 | chr13-29606454-INS-51 |
| chr13-29606613-INS-672 | chr13-29606613-INS-672 |
| chr13-29615657-INS-64 | chr13-29615657-INS-76 |
| chr13-29619425-INS-78 | chr13-29619425-INS-78 |
| chr13-29627371-INS-55 | chr13-29627371-INS-55 |
| chr13-29680040-INS-812 | chr13-29680040-INS-812 |
| chr13-29689981-INS-65 | chr13-29689981-INS-57 |
| chr13-29702877-INS-169 | chr13-29702877-INS-169 |
| chr13-29728205-INS-84 | chr13-29728205-INS-84 |
| chr13-29744171-INS-1284 | chr13-29744172-INS-1284 |
| chr13-29748932-INS-244 | chr13-29748932-INS-247 |
| chr13-29761450-INS-6479 | chr13-29761451-INS-6453 |
| chr13-29797387-INS-125 | chr13-29797387-INS-100 |
| chr13-29869306-INS-62 | chr13-29869352-INS-64 |
| chr13-29876858-INS-499 | chr13-29876858-INS-499 |
| chr13-29894238-INS-65 | chr13-29894236-INS-66 |
| chr13-29896232-INS-249 | chr13-29896232-INS-274 |
| chr13-29971683-INS-676 | chr13-29971681-INS-652 |
| chr13-29972733-INS-329 | chr13-29972733-INS-311 |
| chr13-30004974-INS-415 | chr13-30004974-INS-439 |
| chr13-30010973-INS-214 | chr13-30010973-INS-215 |
| chr13-30042449-INS-148 | chr13-30042449-INS-138 |
| chr13-30133258-INS-73 | chr13-30133258-INS-73 |
| chr13-30135552-INS-87 | chr13-30135552-INS-87 |
| chr13-30154822-INS-68 | chr13-30154822-INS-70 |
| chr13-30177456-INS-128 | chr13-30177493-INS-102 |
| chr13-30208228-INS-72 | chr13-30208239-INS-76 |
| chr13-30214650-INS-66 | chr13-30214636-INS-65 |
| chr13-30218342-INS-121 | chr13-30218348-INS-104 |
| chr13-30224125-INS-78 | chr13-30224125-INS-80 |
| chr13-30259913-INS-401 | chr13-30259913-INS-401 |
| chr13-30278237-INS-57 | chr13-30278237-INS-57 |
| chr13-30292952-INS-382 | chr13-30292952-INS-382 |
| chr13-30293845-INS-84 | chr13-30293845-INS-96 |
| chr13-30312345-INS-257 | chr13-30312345-INS-243 |
| chr13-30356777-INS-1234 | chr13-30356783-INS-1215 |
| chr13-30358839-INS-70 | chr13-30358839-INS-70 |
| chr13-30365817-INS-152 | chr13-30365817-INS-152 |
| chr13-30376828-INS-57 | chr13-30376828-INS-57 |
| chr13-30398629-INS-62 | chr13-30398666-INS-50 |
| chr13-30418803-INS-115 | chr13-30418803-INS-107 |
| chr13-30434180-INS-235 | chr13-30434180-INS-235 |
| chr13-30465311-INS-51 | chr13-30465311-INS-51 |
| chr13-30497221-INS-177 | chr13-30497221-INS-177 |
| chr13-30506892-INS-96 | chr13-30506892-INS-96 |
| chr13-30507485-INS-906 | chr13-30507485-INS-909 |
| chr13-30515777-INS-103 | chr13-30515777-INS-103 |
| chr13-30516214-INS-961 | chr13-30516214-INS-992 |
| chr13-30519770-INS-71 | chr13-30519734-INS-66 |
| chr13-30534000-INS-120 | chr13-30534000-INS-136 |
| chr13-30534454-INS-1054 | chr13-30534455-INS-1046 |
| chr13-30585095-INS-96 | chr13-30585095-INS-96 |
| chr13-30609855-INS-446 | chr13-30609855-INS-446 |
| chr13-30637216-INS-7088 | chr13-30637217-INS-7088 |
| chr13-30653287-INS-605 | chr13-30653287-INS-609 |
| chr13-30685643-INS-187 | chr13-30685643-INS-187 |
| chr13-30691000-INS-451 | chr13-30691000-INS-451 |
| chr13-30696539-INS-489 | chr13-30696539-INS-489 |
| chr13-30710634-INS-240 | chr13-30710634-INS-239 |
| chr13-30715826-INS-1390 | chr13-30715827-INS-1388 |
| chr13-30744452-INS-3193 | chr13-30744453-INS-3193 |
| chr13-30759291-INS-211 | chr13-30759291-INS-211 |
| chr13-30764821-INS-2802 | chr13-30764822-INS-2802 |
| chr13-30773275-INS-58 | chr13-30773275-INS-60 |
| chr13-30791831-INS-56 | chr13-30791831-INS-56 |
| chr13-30827084-INS-76 | chr13-30827084-INS-86 |
| chr13-30856104-INS-594 | chr13-30856104-INS-592 |

| | |
|---|---|
| chr13-30883250-INS-76 | chr13-30883250-INS-76 |
| chr13-30885412-INS-100 | chr13-30885375-INS-100 |
| chr13-30888644-INS-75 | chr13-30888644-INS-65 |
| chr13-30968310-INS-75 | chr13-30968310-INS-76 |
| chr13-30971720-INS-93 | chr13-30971720-INS-93 |
| chr13-30991062-INS-70 | chr13-30991078-INS-85 |
| chr13-31005861-INS-80 | chr13-31005861-INS-80 |
| chr13-31016937-INS-338 | chr13-31016936-INS-339 |
| chr13-31030837-INS-6548 | chr13-31030838-INS-6548 |
| chr13-31060669-INS-2498 | chr13-31060670-INS-2502 |
| chr13-31062637-INS-57 | chr13-31062637-INS-57 |
| chr13-31065599-INS-7095 | chr13-31065600-INS-7095 |
| chr13-31069327-INS-460 | chr13-31069327-INS-459 |
| chr13-31076848-INS-154 | chr13-31076849-INS-142 |
| chr13-31093814-INS-570 | chr13-31093814-INS-570 |
| chr13-31110058-INS-7144 | chr13-31110059-INS-7145 |
| chr13-31136419-INS-227 | chr13-31136419-INS-227 |
| chr13-31153916-INS-68 | chr13-31153916-INS-68 |
| chr13-31175806-INS-60 | chr13-31175806-INS-60 |
| chr13-31193494-INS-413 | chr13-31193491-INS-418 |
| chr13-31200910-INS-50 | chr13-31200910-INS-52 |
| chr13-31253012-INS-72 | chr13-31253012-INS-57 |
| chr13-31254369-INS-159 | chr13-31254369-INS-206 |
| chr13-31269249-INS-103 | chr13-31269246-INS-90 |
| chr13-31288768-INS-3676 | chr13-31288769-INS-3683 |
| chr13-31306365-INS-541 | chr13-31306365-INS-541 |
| chr13-31307142-INS-104 | chr13-31307142-INS-104 |
| chr13-31311288-INS-132 | chr13-31311288-INS-132 |
| chr13-31340492-INS-77 | chr13-31340492-INS-77 |
| chr13-31357643-INS-1037 | chr13-31357644-INS-1037 |
| chr13-31381992-INS-92 | chr13-31381992-INS-100 |
| chr13-31408852-INS-706 | chr13-31408852-INS-706 |
| chr13-31435295-INS-132 | chr13-31435295-INS-138 |
| chr13-31449036-INS-6715 | chr13-31449037-INS-6713 |
| chr13-31481096-INS-197 | chr13-31481120-INS-212 |
| chr13-31506458-INS-54 | chr13-31506442-INS-80 |
| chr13-31596868-INS-320 | chr13-31596868-INS-320 |
| chr13-31599201-INS-196 | chr13-31599201-INS-195 |
| chr13-31608705-INS-158 | chr13-31608705-INS-158 |
| chr13-31663650-INS-80 | chr13-31663671-INS-56 |
| chr13-31666412-INS-72 | chr13-31666412-INS-72 |
| chr13-31668459-INS-67 | chr13-31668459-INS-67 |
| chr13-31687275-INS-64 | chr13-31687275-INS-64 |
| chr13-31721733-INS-6912 | chr13-31721734-INS-6913 |
| chr13-31747990-INS-62 | chr13-31747989-INS-66 |
| chr13-31757561-INS-216 | chr13-31757561-INS-216 |
| chr13-31770735-INS-72 | chr13-31770735-INS-72 |
| chr13-31783285-INS-156 | chr13-31783285-INS-156 |
| chr13-31813395-INS-233 | chr13-31813395-INS-233 |
| chr13-31841008-INS-64 | chr13-31841014-INS-82 |
| chr13-31846153-INS-192 | chr13-31846153-INS-192 |
| chr13-31853184-INS-1727 | chr13-31853182-INS-1719 |
| chr13-31865001-INS-102 | chr13-31865001-INS-102 |
| chr13-31865479-INS-5549 | chr13-31865480-INS-5549 |
| chr13-31900204-INS-115 | chr13-31900204-INS-145 |
| chr13-31920172-INS-115 | chr13-31920172-INS-115 |
| chr13-31960151-INS-86 | chr13-31960151-INS-88 |
| chr13-32016867-INS-134 | chr13-32016886-INS-84 |
| chr13-32032899-INS-66 | chr13-32032899-INS-66 |
| chr13-32061565-INS-136 | chr13-32061565-INS-136 |
| chr13-32076713-INS-262 | chr13-32076713-INS-263 |
| chr13-32077779-INS-1150 | chr13-32077780-INS-1152 |
| chr13-32092151-INS-211 | chr13-32092151-INS-211 |
| chr13-32125501-INS-62 | chr13-32125501-INS-62 |
| chr13-32130433-INS-2151 | chr13-32130434-INS-2151 |
| chr13-32133804-INS-5882 | chr13-32133805-INS-5882 |
| chr13-32190354-INS-64 | chr13-32190354-INS-64 |
| chr13-32222800-INS-79 | chr13-32222800-INS-79 |
| chr13-32236040-INS-461 | chr13-32236040-INS-461 |
| chr13-32250833-INS-64 | chr13-32250833-INS-64 |
| chr13-32268188-INS-52 | chr13-32268188-INS-50 |
| chr13-32278609-INS-155 | chr13-32278609-INS-159 |
| chr13-32295116-INS-1302 | chr13-32295117-INS-1303 |
| chr13-32319955-INS-4252 | chr13-32319956-INS-4253 |
| chr13-32398588-INS-68 | chr13-32398588-INS-68 |
| chr13-32454624-INS-6795 | chr13-32454625-INS-6795 |
| chr13-32520873-INS-211 | chr13-32520874-INS-224 |

| | |
|---|---|
| chr13-32557844-INS-2192 | chr13-32557831-INS-2181 |
| chr13-32568365-INS-104 | chr13-32568358-INS-78 |
| chr13-32589555-INS-60 | chr13-32589555-INS-56 |
| chr13-32590546-INS-179 | chr13-32590546-INS-186 |
| chr13-32607460-INS-80 | chr13-32607460-INS-80 |
| chr13-32622815-INS-64 | chr13-32622815-INS-68 |
| chr13-32659309-INS-6393 | chr13-32659310-INS-6394 |
| chr13-32660953-INS-65 | chr13-32660953-INS-70 |
| chr13-32741243-INS-6399 | chr13-32741244-INS-6400 |
| chr13-32806766-INS-109 | chr13-32806766-INS-109 |
| chr13-32949907-INS-621 | chr13-32949907-INS-617 |
| chr13-32999408-INS-206 | chr13-32999408-INS-205 |
| chr13-33003930-INS-52 | chr13-33003930-INS-52 |
| chr13-33036199-INS-337 | chr13-33036199-INS-337 |
| chr13-33054574-INS-420 | chr13-33054574-INS-420 |
| chr13-33070741-INS-109 | chr13-33070741-INS-130 |
| chr13-33085406-INS-527 | chr13-33085406-INS-528 |
| chr13-33097852-INS-223 | chr13-33097852-INS-176 |
| chr13-33130007-INS-107 | chr13-33130007-INS-107 |
| chr13-33142280-INS-2964 | chr13-33142281-INS-2960 |
| chr13-33195125-INS-68 | chr13-33195125-INS-68 |
| chr13-33235608-INS-55 | chr13-33235608-INS-63 |
| chr13-33265623-INS-530 | chr13-33265623-INS-529 |
| chr13-33269906-INS-90 | chr13-33269906-INS-96 |
| chr13-33314720-INS-481 | chr13-33314720-INS-481 |
| chr13-33323096-INS-719 | chr13-33323096-INS-718 |
| chr13-33329570-INS-390 | chr13-33329570-INS-390 |
| chr13-33355202-INS-4522 | chr13-33355203-INS-4522 |
| chr13-33374248-INS-1197 | chr13-33374249-INS-1197 |
| chr13-33387115-INS-1251 | chr13-33387116-INS-1251 |
| chr13-33400052-INS-3889 | chr13-33400053-INS-3889 |
| chr13-33420286-INS-200 | chr13-33420286-INS-200 |
| chr13-33455988-INS-62 | chr13-33455988-INS-62 |
| chr13-33467116-INS-76 | chr13-33467114-INS-76 |
| chr13-33469946-INS-2018 | chr13-33469947-INS-2018 |
| chr13-33497094-INS-273 | chr13-33497094-INS-273 |
| chr13-33534510-INS-54 | chr13-33534510-INS-57 |
| chr13-33544585-INS-1999 | chr13-33544586-INS-1999 |
| chr13-33691218-INS-942 | chr13-33691218-INS-942 |
| chr13-33693400-INS-475 | chr13-33693400-INS-475 |
| chr13-33819474-INS-52 | chr13-33819477-INS-51 |
| chr13-33824840-INS-547 | chr13-33824840-INS-547 |
| chr13-33847265-INS-243 | chr13-33847265-INS-260 |
| chr13-33856580-INS-344 | chr13-33856580-INS-344 |
| chr13-33928864-INS-2133 | chr13-33928865-INS-2146 |
| chr13-33932225-INS-2927 | chr13-33932226-INS-2927 |
| chr13-33998784-INS-2674 | chr13-33998785-INS-2674 |
| chr13-34007800-INS-400 | chr13-34007800-INS-400 |
| chr13-34023410-INS-403 | chr13-34023420-INS-403 |
| chr13-34027877-INS-253 | chr13-34027863-INS-262 |
| chr13-34042500-INS-83 | chr13-34042500-INS-61 |
| chr13-34067120-INS-196 | chr13-34067120-INS-196 |
| chr13-34067639-INS-7051 | chr13-34067640-INS-7053 |
| chr13-34122064-INS-965 | chr13-34122065-INS-966 |
| chr13-34139117-INS-4915 | chr13-34139118-INS-4915 |
| chr13-34142238-INS-243 | chr13-34142238-INS-243 |
| chr13-34208343-INS-595 | chr13-34208341-INS-604 |
| chr13-34273949-INS-343 | chr13-34273949-INS-346 |
| chr13-34296860-INS-60 | chr13-34296860-INS-61 |
| chr13-34298029-INS-64 | chr13-34298021-INS-56 |
| chr13-34303031-INS-204 | chr13-34303031-INS-230 |
| chr13-34328312-INS-112 | chr13-34328312-INS-112 |
| chr13-34338917-INS-60 | chr13-34338917-INS-52 |
| chr13-34384347-INS-72 | chr13-34384335-INS-94 |
| chr13-34411333-INS-66 | chr13-34411333-INS-68 |
| chr13-34534118-INS-221 | chr13-34534118-INS-221 |
| chr13-34538020-INS-443 | chr13-34538020-INS-456 |
| chr13-34545437-INS-59 | chr13-34545434-INS-63 |
| chr13-34555076-INS-121 | chr13-34555076-INS-121 |
| chr13-34580158-INS-70 | chr13-34580158-INS-70 |
| chr13-34595531-INS-520 | chr13-34595531-INS-520 |
| chr13-34610750-INS-76 | chr13-34610750-INS-66 |
| chr13-34663887-INS-85 | chr13-34663887-INS-75 |
| chr13-34692845-INS-680 | chr13-34692845-INS-680 |
| chr13-34710637-INS-50 | chr13-34710637-INS-54 |
| chr13-34711936-INS-84 | chr13-34711936-INS-83 |
| chr13-34718652-INS-347 | chr13-34718649-INS-350 |

| | |
|---|---|
| chr13-34724430-INS-228 | chr13-34724430-INS-242 |
| chr13-34731768-INS-399 | chr13-34731768-INS-399 |
| chr13-34778704-INS-60 | chr13-34778704-INS-60 |
| chr13-34789765-INS-124 | chr13-34789787-INS-92 |
| chr13-34805103-INS-385 | chr13-34805103-INS-385 |
| chr13-34816068-INS-186 | chr13-34816068-INS-186 |
| chr13-34847541-INS-370 | chr13-34847541-INS-370 |
| chr13-34849488-INS-62 | chr13-34849488-INS-62 |
| chr13-34885285-INS-108 | chr13-34885285-INS-108 |
| chr13-34890994-INS-58 | chr13-34890990-INS-88 |
| chr13-34892733-INS-1103 | chr13-34892734-INS-1103 |
| chr13-34901139-INS-199 | chr13-34901151-INS-204 |
| chr13-34903534-INS-60 | chr13-34903534-INS-56 |
| chr13-34986140-INS-145 | chr13-34986147-INS-159 |
| chr13-35027216-INS-991 | chr13-35027216-INS-989 |
| chr13-35030188-INS-83 | chr13-35030174-INS-65 |
| chr13-35043862-INS-74 | chr13-35043862-INS-91 |
| chr13-35044998-INS-56 | chr13-35044998-INS-58 |
| chr13-35051403-INS-54 | chr13-35051403-INS-54 |
| chr13-35053576-INS-400 | chr13-35053576-INS-400 |
| chr13-35064598-INS-72 | chr13-35064598-INS-73 |
| chr13-35126527-INS-422 | chr13-35126527-INS-422 |
| chr13-35134886-INS-429 | chr13-35134886-INS-429 |
| chr13-35145999-INS-197 | chr13-35145999-INS-198 |
| chr13-35158339-INS-565 | chr13-35158339-INS-565 |
| chr13-35169071-INS-146 | chr13-35169092-INS-172 |
| chr13-35225104-INS-56 | chr13-35225124-INS-60 |
| chr13-35251840-INS-115 | chr13-35251840-INS-115 |
| chr13-35284724-INS-60 | chr13-35284724-INS-60 |
| chr13-35289744-INS-3222 | chr13-35289745-INS-3221 |
| chr13-35290462-INS-54 | chr13-35290462-INS-54 |
| chr13-35336808-INS-605 | chr13-35336808-INS-605 |
| chr13-35383988-INS-69 | chr13-35383988-INS-69 |
| chr13-35393890-INS-134 | chr13-35393886-INS-109 |
| chr13-35422693-INS-52 | chr13-35422691-INS-52 |
| chr13-35442208-INS-56 | chr13-35442208-INS-56 |
| chr13-35443447-INS-68 | chr13-35443447-INS-76 |
| chr13-35444910-INS-1888 | chr13-35444911-INS-1888 |
| chr13-35468670-INS-53 | chr13-35468670-INS-53 |
| chr13-35474796-INS-7067 | chr13-35474797-INS-7067 |
| chr13-35521249-INS-68 | chr13-35521249-INS-70 |
| chr13-35541424-INS-1982 | chr13-35541425-INS-1984 |
| chr13-35544454-INS-132 | chr13-35544444-INS-124 |
| chr13-35545926-INS-75 | chr13-35545926-INS-81 |
| chr13-35546787-INS-7344 | chr13-35546788-INS-7344 |
| chr13-35610872-INS-52 | chr13-35610854-INS-50 |
| chr13-35649744-INS-4774 | chr13-35649745-INS-4774 |
| chr13-35650905-INS-1624 | chr13-35650906-INS-1626 |
| chr13-35665461-INS-107 | chr13-35665461-INS-109 |
| chr13-35711217-INS-6410 | chr13-35711218-INS-6410 |
| chr13-35711860-INS-6312 | chr13-35711861-INS-6307 |
| chr13-35735649-INS-203 | chr13-35735649-INS-203 |
| chr13-35862490-INS-60 | chr13-35862490-INS-70 |
| chr13-35872034-INS-80 | chr13-35872034-INS-80 |
| chr13-35879000-INS-80 | chr13-35879000-INS-80 |
| chr13-35906335-INS-51 | chr13-35906335-INS-51 |
| chr13-35918238-INS-80 | chr13-35918238-INS-84 |
| chr13-35951878-INS-239 | chr13-35951878-INS-239 |
| chr13-35964428-INS-1030 | chr13-35964429-INS-1028 |
| chr13-35984554-INS-1090 | chr13-35984555-INS-1103 |
| chr13-36008575-INS-99 | chr13-36008574-INS-102 |
| chr13-36055940-INS-194 | chr13-36055940-INS-194 |
| chr13-36095360-INS-262 | chr13-36095360-INS-262 |
| chr13-36100968-INS-296 | chr13-36100968-INS-295 |
| chr13-36112839-INS-72 | chr13-36112841-INS-72 |
| chr13-36115922-INS-342 | chr13-36115922-INS-342 |
| chr13-36120167-INS-645 | chr13-36120167-INS-645 |
| chr13-36166422-INS-125 | chr13-36166422-INS-125 |
| chr13-36178670-INS-182 | chr13-36178664-INS-182 |
| chr13-36198906-INS-1122 | chr13-36198907-INS-1122 |
| chr13-36209347-INS-138 | chr13-36209347-INS-138 |
| chr13-36233550-INS-427 | chr13-36233550-INS-427 |
| chr13-36286526-INS-214 | chr13-36286526-INS-214 |
| chr13-36318833-INS-66 | chr13-36318833-INS-66 |
| chr13-36340390-INS-1695 | chr13-36340391-INS-1680 |
| chr13-36397077-INS-828 | chr13-36397077-INS-828 |
| chr13-36401681-INS-162 | chr13-36401681-INS-162 |

| | |
|---|---|
| chr13-36410431-INS-1991 | chr13-36410432-INS-1998 |
| chr13-36445119-INS-701 | chr13-36445119-INS-701 |
| chr13-36513418-INS-1037 | chr13-36513419-INS-1037 |
| chr13-36530872-INS-72 | chr13-36530870-INS-62 |
| chr13-36549362-INS-468 | chr13-36549362-INS-468 |
| chr13-36563090-INS-126 | chr13-36563090-INS-126 |
| chr13-36577831-INS-2303 | chr13-36577832-INS-2304 |
| chr13-36581260-INS-564 | chr13-36581260-INS-561 |
| chr13-36588463-INS-70 | chr13-36588467-INS-88 |
| chr13-36602120-INS-201 | chr13-36602120-INS-201 |
| chr13-36609684-INS-1774 | chr13-36609685-INS-1775 |
| chr13-36609928-INS-332 | chr13-36609928-INS-332 |
| chr13-36628430-INS-84 | chr13-36628477-INS-52 |
| chr13-36656441-INS-989 | chr13-36656441-INS-989 |
| chr13-36698338-INS-1299 | chr13-36698339-INS-1299 |
| chr13-36710820-INS-6500 | chr13-36710821-INS-6500 |
| chr13-36734632-INS-3238 | chr13-36734633-INS-3238 |
| chr13-36737658-INS-96 | chr13-36737658-INS-94 |
| chr13-36742383-INS-107 | chr13-36742383-INS-107 |
| chr13-36744776-INS-56 | chr13-36744776-INS-54 |
| chr13-36757378-INS-50 | chr13-36757378-INS-52 |
| chr13-36768898-INS-55 | chr13-36768898-INS-55 |
| chr13-36772397-INS-6243 | chr13-36772398-INS-6241 |
| chr13-36778114-INS-64 | chr13-36778097-INS-52 |
| chr13-36806483-INS-138 | chr13-36806483-INS-135 |
| chr13-36810127-INS-1084 | chr13-36810128-INS-1083 |
| chr13-36828158-INS-7259 | chr13-36828159-INS-7259 |
| chr13-36833286-INS-84 | chr13-36833287-INS-100 |
| chr13-36841582-INS-56 | chr13-36841582-INS-56 |
| chr13-36880460-INS-1888 | chr13-36880461-INS-1888 |
| chr13-36886419-INS-148 | chr13-36886419-INS-148 |
| chr13-36890596-INS-80 | chr13-36890596-INS-80 |
| chr13-36894897-INS-7291 | chr13-36894898-INS-7291 |
| chr13-36903511-INS-176 | chr13-36903513-INS-182 |
| chr13-36921752-INS-414 | chr13-36921752-INS-412 |
| chr13-36927511-INS-79 | chr13-36927511-INS-79 |
| chr13-37013445-INS-64 | chr13-37013445-INS-68 |
| chr13-37015515-INS-62 | chr13-37015515-INS-62 |
| chr13-37019795-INS-576 | chr13-37019795-INS-577 |
| chr13-37040761-INS-213 | chr13-37040761-INS-215 |
| chr13-37061510-INS-68 | chr13-37061483-INS-60 |
| chr13-37068388-INS-76 | chr13-37068375-INS-65 |
| chr13-37084411-INS-77 | chr13-37084421-INS-88 |
| chr13-37085772-INS-7589 | chr13-37085773-INS-7590 |
| chr13-37086135-INS-55 | chr13-37086139-INS-50 |
| chr13-37088419-INS-288 | chr13-37088419-INS-288 |
| chr13-37117410-INS-172 | chr13-37117410-INS-173 |
| chr13-37142446-INS-54 | chr13-37142446-INS-56 |
| chr13-37149095-INS-60 | chr13-37149095-INS-60 |
| chr13-37201460-INS-343 | chr13-37201460-INS-343 |
| chr13-37238319-INS-116 | chr13-37238360-INS-112 |
| chr13-37323889-INS-788 | chr13-37323889-INS-787 |
| chr13-37332940-INS-67 | chr13-37332924-INS-76 |
| chr13-37338331-INS-1120 | chr13-37338332-INS-1120 |
| chr13-37391823-INS-51 | chr13-37391788-INS-52 |
| chr13-37403342-INS-607 | chr13-37403342-INS-607 |
| chr13-37418833-INS-66 | chr13-37418833-INS-66 |
| chr13-37422436-INS-4274 | chr13-37422437-INS-4274 |
| chr13-37447006-INS-642 | chr13-37447006-INS-642 |
| chr13-37456355-INS-1479 | chr13-37456356-INS-1479 |
| chr13-37464395-INS-58 | chr13-37464395-INS-60 |
| chr13-37490134-INS-673 | chr13-37490134-INS-673 |
| chr13-37501185-INS-376 | chr13-37501185-INS-376 |
| chr13-37502655-INS-3458 | chr13-37502656-INS-3458 |
| chr13-37503072-INS-6452 | chr13-37503073-INS-6452 |
| chr13-37559596-INS-1888 | chr13-37559597-INS-1888 |
| chr13-37563501-INS-84 | chr13-37563501-INS-84 |
| chr13-37584197-INS-88 | chr13-37584197-INS-112 |
| chr13-37621916-INS-1236 | chr13-37621917-INS-1236 |
| chr13-37634494-INS-99 | chr13-37634494-INS-76 |
| chr13-37653849-INS-209 | chr13-37653849-INS-209 |
| chr13-37655135-INS-53 | chr13-37655135-INS-53 |
| chr13-37657963-INS-89 | chr13-37657974-INS-74 |
| chr13-37665150-INS-75 | chr13-37665150-INS-75 |
| chr13-37670898-INS-207 | chr13-37670898-INS-207 |
| chr13-37700027-INS-52 | chr13-37700027-INS-52 |
| chr13-37711023-INS-207 | chr13-37711023-INS-207 |

| | |
|---|---|
| chr13-37725681-INS-1500 | chr13-37725682-INS-1500 |
| chr13-37734585-INS-116 | chr13-37734585-INS-116 |
| chr13-37842309-INS-90 | chr13-37842309-INS-76 |
| chr13-37874113-INS-89 | chr13-37874114-INS-53 |
| chr13-37879554-INS-553 | chr13-37879554-INS-553 |
| chr13-37883896-INS-190 | chr13-37883896-INS-190 |
| chr13-37884814-INS-845 | chr13-37884814-INS-845 |
| chr13-37904703-INS-719 | chr13-37904703-INS-719 |
| chr13-37914612-INS-132 | chr13-37914620-INS-142 |
| chr13-37938046-INS-406 | chr13-37938046-INS-406 |
| chr13-37973617-INS-213 | chr13-37973617-INS-213 |
| chr13-37984909-INS-64 | chr13-37984909-INS-69 |
| chr13-37986710-INS-119 | chr13-37986710-INS-119 |
| chr13-37990273-INS-60 | chr13-37990265-INS-76 |
| chr13-37999569-INS-205 | chr13-37999569-INS-206 |
| chr13-38003261-INS-52 | chr13-38003261-INS-52 |
| chr13-38067991-INS-56 | chr13-38067991-INS-56 |
| chr13-38137858-INS-218 | chr13-38137858-INS-218 |
| chr13-38139206-INS-92 | chr13-38139206-INS-92 |
| chr13-38141209-INS-152 | chr13-38141239-INS-146 |
| chr13-38157485-INS-1009 | chr13-38157486-INS-1015 |
| chr13-38157733-INS-67 | chr13-38157733-INS-68 |
| chr13-38164947-INS-207 | chr13-38164947-INS-207 |
| chr13-38172942-INS-72 | chr13-38172958-INS-55 |
| chr13-38173719-INS-62 | chr13-38173719-INS-62 |
| chr13-38181295-INS-209 | chr13-38181295-INS-209 |
| chr13-38184803-INS-201 | chr13-38184803-INS-199 |
| chr13-38190350-INS-72 | chr13-38190322-INS-104 |
| chr13-38194327-INS-7113 | chr13-38194328-INS-7113 |
| chr13-38200035-INS-74 | chr13-38200035-INS-80 |
| chr13-38216170-INS-181 | chr13-38216181-INS-191 |
| chr13-38225292-INS-54 | chr13-38225292-INS-64 |
| chr13-38234387-INS-249 | chr13-38234387-INS-243 |
| chr13-38235099-INS-2720 | chr13-38235100-INS-2720 |
| chr13-38250332-INS-191 | chr13-38250332-INS-172 |
| chr13-38255982-INS-195 | chr13-38255982-INS-195 |
| chr13-38262592-INS-71 | chr13-38262592-INS-50 |
| chr13-38276668-INS-66 | chr13-38276668-INS-66 |
| chr13-38287409-INS-51 | chr13-38287409-INS-55 |
| chr13-38344019-INS-102 | chr13-38344019-INS-98 |
| chr13-38360667-INS-834 | chr13-38360673-INS-836 |
| chr13-38364343-INS-54 | chr13-38364342-INS-54 |
| chr13-38365666-INS-82 | chr13-38365666-INS-74 |
| chr13-38368373-INS-104 | chr13-38368373-INS-108 |
| chr13-38437510-INS-104 | chr13-38437516-INS-72 |
| chr13-38451380-INS-7098 | chr13-38451381-INS-7098 |
| chr13-38471821-INS-401 | chr13-38471811-INS-410 |
| chr13-38481646-INS-58 | chr13-38481646-INS-62 |
| chr13-38493977-INS-52 | chr13-38493977-INS-52 |
| chr13-38522503-INS-3833 | chr13-38522504-INS-3881 |
| chr13-38565285-INS-1542 | chr13-38565286-INS-1520 |
| chr13-38579388-INS-80 | chr13-38579388-INS-100 |
| chr13-38690535-INS-439 | chr13-38690535-INS-425 |
| chr13-38721461-INS-58 | chr13-38721461-INS-58 |
| chr13-38774091-INS-154 | chr13-38774103-INS-159 |
| chr13-38776423-INS-1316 | chr13-38776424-INS-1316 |
| chr13-38785013-INS-1992 | chr13-38785014-INS-1992 |
| chr13-38792245-INS-167 | chr13-38792245-INS-167 |
| chr13-38813886-INS-51 | chr13-38813878-INS-55 |
| chr13-38832259-INS-228 | chr13-38832259-INS-226 |
| chr13-38849534-INS-72 | chr13-38849501-INS-62 |
| chr13-38855760-INS-52 | chr13-38855767-INS-58 |
| chr13-38859186-INS-60 | chr13-38859186-INS-52 |
| chr13-38919696-INS-80 | chr13-38919696-INS-80 |
| chr13-38947300-INS-82 | chr13-38947300-INS-92 |
| chr13-38947873-INS-3066 | chr13-38947874-INS-3066 |
| chr13-38954164-INS-7223 | chr13-38954165-INS-7223 |
| chr13-38989097-INS-7315 | chr13-38989098-INS-7316 |
| chr13-39006245-INS-97 | chr13-39006245-INS-97 |
| chr13-39007851-INS-70 | chr13-39007851-INS-70 |
| chr13-39037721-INS-154 | chr13-39037721-INS-153 |
| chr13-39047939-INS-266 | chr13-39047939-INS-267 |
| chr13-39048340-INS-6606 | chr13-39048341-INS-6606 |
| chr13-39058828-INS-457 | chr13-39058828-INS-457 |
| chr13-39075570-INS-222 | chr13-39075570-INS-222 |
| chr13-39079815-INS-218 | chr13-39079815-INS-218 |
| chr13-39097864-INS-81 | chr13-39097864-INS-80 |

| | |
|---|---|
| chr13-39136467-INS-643 | chr13-39136467-INS-643 |
| chr13-39154246-INS-402 | chr13-39154246-INS-402 |
| chr13-39158515-INS-7111 | chr13-39158516-INS-7109 |
| chr13-39167247-INS-254 | chr13-39167247-INS-254 |
| chr13-39192332-INS-1888 | chr13-39192333-INS-1888 |
| chr13-39207401-INS-136 | chr13-39207401-INS-144 |
| chr13-39220308-INS-83 | chr13-39220308-INS-94 |
| chr13-39244401-INS-191 | chr13-39244401-INS-191 |
| chr13-39265324-INS-74 | chr13-39265324-INS-70 |
| chr13-39271947-INS-4040 | chr13-39271948-INS-4040 |
| chr13-39274926-INS-76 | chr13-39274926-INS-84 |
| chr13-39276421-INS-97 | chr13-39276421-INS-90 |
| chr13-39315594-INS-2053 | chr13-39315595-INS-2053 |
| chr13-39334666-INS-96 | chr13-39334666-INS-96 |
| chr13-39339856-INS-120 | chr13-39339856-INS-120 |
| chr13-39345846-INS-369 | chr13-39345846-INS-369 |
| chr13-39356897-INS-79 | chr13-39356897-INS-79 |
| chr13-39358545-INS-60 | chr13-39358545-INS-60 |
| chr13-39378770-INS-51 | chr13-39378770-INS-59 |
| chr13-39382726-INS-1576 | chr13-39382727-INS-1580 |
| chr13-39397709-INS-6548 | chr13-39397710-INS-6548 |
| chr13-39414757-INS-122 | chr13-39414757-INS-160 |
| chr13-39436519-INS-64 | chr13-39436519-INS-68 |
| chr13-39462661-INS-175 | chr13-39462661-INS-201 |
| chr13-39478366-INS-61 | chr13-39478402-INS-65 |
| chr13-39486237-INS-207 | chr13-39486237-INS-207 |
| chr13-39489042-INS-88 | chr13-39489042-INS-78 |
| chr13-39507862-INS-52 | chr13-39507862-INS-52 |
| chr13-39516598-INS-76 | chr13-39516606-INS-82 |
| chr13-39519791-INS-59 | chr13-39519791-INS-65 |
| chr13-39526029-INS-116 | chr13-39526025-INS-109 |
| chr13-39539662-INS-74 | chr13-39539662-INS-76 |
| chr13-39559614-INS-76 | chr13-39559614-INS-76 |
| chr13-39566480-INS-156 | chr13-39566480-INS-156 |
| chr13-39576661-INS-77 | chr13-39576686-INS-53 |
| chr13-39580648-INS-57 | chr13-39580648-INS-96 |
| chr13-39604837-INS-684 | chr13-39604837-INS-684 |
| chr13-39606399-INS-1852 | chr13-39606400-INS-1852 |
| chr13-39611611-INS-73 | chr13-39611562-INS-74 |
| chr13-39622620-INS-166 | chr13-39622621-INS-130 |
| chr13-39652679-INS-62 | chr13-39652679-INS-62 |
| chr13-39653985-INS-680 | chr13-39653985-INS-679 |
| chr13-39654725-INS-6239 | chr13-39654726-INS-6240 |
| chr13-39654827-INS-130 | chr13-39654827-INS-130 |
| chr13-39664885-INS-60 | chr13-39664903-INS-65 |
| chr13-39671311-INS-68 | chr13-39671312-INS-104 |
| chr13-39672021-INS-3668 | chr13-39672022-INS-3677 |
| chr13-39677795-INS-2052 | chr13-39677796-INS-2051 |
| chr13-39703322-INS-53 | chr13-39703322-INS-53 |
| chr13-39712486-INS-54 | chr13-39712486-INS-58 |
| chr13-39718863-INS-74 | chr13-39718843-INS-50 |
| chr13-39816312-INS-61 | chr13-39816312-INS-61 |
| chr13-39834582-INS-176 | chr13-39834582-INS-176 |
| chr13-39842993-INS-60 | chr13-39842993-INS-60 |
| chr13-39844509-INS-996 | chr13-39844509-INS-996 |
| chr13-39856095-INS-400 | chr13-39856095-INS-400 |
| chr13-39873522-INS-70 | chr13-39873507-INS-58 |
| chr13-39879512-INS-55 | chr13-39879512-INS-55 |
| chr13-39887898-INS-1564 | chr13-39887899-INS-1565 |
| chr13-39891469-INS-72 | chr13-39891469-INS-74 |
| chr13-39919991-INS-62 | chr13-39919990-INS-54 |
| chr13-39921292-INS-140 | chr13-39921304-INS-104 |
| chr13-39938280-INS-291 | chr13-39938280-INS-291 |
| chr13-39939908-INS-54 | chr13-39939908-INS-54 |
| chr13-39986766-INS-72 | chr13-39986766-INS-72 |
| chr13-39997947-INS-96 | chr13-39997947-INS-72 |
| chr13-40007206-INS-64 | chr13-40007206-INS-64 |
| chr13-40023323-INS-90 | chr13-40023323-INS-94 |
| chr13-40026949-INS-120 | chr13-40026949-INS-98 |
| chr13-40038981-INS-80 | chr13-40038999-INS-70 |
| chr13-40066803-INS-850 | chr13-40066803-INS-846 |
| chr13-40119926-INS-88 | chr13-40119926-INS-74 |
| chr13-40172695-INS-84 | chr13-40172695-INS-82 |
| chr13-40174193-INS-1127 | chr13-40174194-INS-1129 |
| chr13-40247202-INS-4683 | chr13-40247203-INS-4683 |
| chr13-40249578-INS-99 | chr13-40249578-INS-99 |
| chr13-40262633-INS-87 | chr13-40262639-INS-92 |

| | |
|---|---|
| chr13-40263611-INS-136 | chr13-40263611-INS-118 |
| chr13-40264258-INS-960 | chr13-40264258-INS-959 |
| chr13-40271847-INS-496 | chr13-40271847-INS-496 |
| chr13-40285120-INS-61 | chr13-40285157-INS-55 |
| chr13-40286768-INS-66 | chr13-40286768-INS-72 |
| chr13-40291984-INS-119 | chr13-40291984-INS-120 |
| chr13-40303336-INS-109 | chr13-40303336-INS-115 |
| chr13-40318724-INS-551 | chr13-40318724-INS-551 |
| chr13-40324643-INS-72 | chr13-40324643-INS-81 |
| chr13-40362389-INS-53 | chr13-40362389-INS-56 |
| chr13-40375870-INS-2695 | chr13-40375871-INS-2695 |
| chr13-40377226-INS-53 | chr13-40377226-INS-64 |
| chr13-40379600-INS-60 | chr13-40379600-INS-60 |
| chr13-40387670-INS-68 | chr13-40387670-INS-60 |
| chr13-40388643-INS-62 | chr13-40388643-INS-62 |
| chr13-40390003-INS-1158 | chr13-40390004-INS-1158 |
| chr13-40391880-INS-56 | chr13-40391880-INS-62 |
| chr13-40394938-INS-65 | chr13-40394973-INS-56 |
| chr13-40401606-INS-60 | chr13-40401606-INS-60 |
| chr13-40410032-INS-7087 | chr13-40410033-INS-7087 |
| chr13-40417556-INS-90 | chr13-40417557-INS-90 |
| chr13-40459439-INS-400 | chr13-40459439-INS-400 |
| chr13-40466142-INS-75 | chr13-40466142-INS-79 |
| chr13-40488662-INS-1827 | chr13-40488663-INS-1827 |
| chr13-40513498-INS-66 | chr13-40513498-INS-70 |
| chr13-40539630-INS-127 | chr13-40539633-INS-165 |
| chr13-40540273-INS-1206 | chr13-40540274-INS-1206 |
| chr13-40593722-INS-63 | chr13-40593722-INS-57 |
| chr13-40612942-INS-95 | chr13-40612942-INS-95 |
| chr13-40633758-INS-229 | chr13-40633758-INS-228 |
| chr13-40636625-INS-64 | chr13-40636625-INS-64 |
| chr13-40668089-INS-54 | chr13-40668089-INS-58 |
| chr13-40694207-INS-50 | chr13-40694207-INS-58 |
| chr13-40718314-INS-108 | chr13-40718314-INS-132 |
| chr13-40727289-INS-1834 | chr13-40727290-INS-1817 |
| chr13-40763578-INS-54 | chr13-40763578-INS-56 |
| chr13-40814445-INS-85 | chr13-40814440-INS-85 |
| chr13-40831972-INS-6820 | chr13-40831978-INS-6819 |
| chr13-40846771-INS-57 | chr13-40846771-INS-56 |
| chr13-40853802-INS-126 | chr13-40853802-INS-81 |
| chr13-40869382-INS-70 | chr13-40869382-INS-56 |
| chr13-40888750-INS-51 | chr13-40888750-INS-51 |
| chr13-40891042-INS-52 | chr13-40891042-INS-50 |
| chr13-40901208-INS-64 | chr13-40901208-INS-66 |
| chr13-40917505-INS-63 | chr13-40917505-INS-65 |
| chr13-40919135-INS-169 | chr13-40919135-INS-181 |
| chr13-40920969-INS-69 | chr13-40920949-INS-78 |
| chr13-40929544-INS-510 | chr13-40929544-INS-510 |
| chr13-40951490-INS-111 | chr13-40951490-INS-110 |
| chr13-40998750-INS-101 | chr13-40998750-INS-101 |
| chr13-41005310-INS-76 | chr13-41005310-INS-80 |
| chr13-41005390-INS-6442 | chr13-41005391-INS-6442 |
| chr13-41024461-INS-53 | chr13-41024461-INS-95 |
| chr13-41025579-INS-55 | chr13-41025587-INS-64 |
| chr13-41057100-INS-53 | chr13-41057100-INS-53 |
| chr13-41064623-INS-192 | chr13-41064623-INS-192 |
| chr13-41085390-INS-60 | chr13-41085390-INS-60 |
| chr13-41099943-INS-6401 | chr13-41099944-INS-6401 |
| chr13-41135248-INS-4174 | chr13-41135249-INS-4174 |
| chr13-41135580-INS-87 | chr13-41135580-INS-87 |
| chr13-41175012-INS-179 | chr13-41175012-INS-184 |
| chr13-41181633-INS-2999 | chr13-41181634-INS-2999 |
| chr13-41189394-INS-274 | chr13-41189394-INS-274 |
| chr13-41225953-INS-56 | chr13-41225953-INS-56 |
| chr13-41238452-INS-145 | chr13-41238452-INS-151 |
| chr13-41262833-INS-70 | chr13-41262833-INS-70 |
| chr13-41264541-INS-100 | chr13-41264541-INS-100 |
| chr13-41287886-INS-276 | chr13-41287886-INS-276 |
| chr13-41298482-INS-112 | chr13-41298500-INS-119 |
| chr13-41314884-INS-7558 | chr13-41314885-INS-7559 |
| chr13-41387526-INS-70 | chr13-41387526-INS-70 |
| chr13-41418236-INS-86 | chr13-41418236-INS-54 |
| chr13-41424774-INS-708 | chr13-41424771-INS-712 |
| chr13-41437803-INS-327 | chr13-41437803-INS-327 |
| chr13-41458662-INS-55 | chr13-41458662-INS-55 |
| chr13-41485944-INS-93 | chr13-41485944-INS-98 |
| chr13-41528088-INS-162 | chr13-41528088-INS-168 |

| | |
|---|---|
| chr13-41540748-INS-54 | chr13-41540748-INS-54 |
| chr13-41551743-INS-59 | chr13-41551743-INS-66 |
| chr13-41576606-INS-98 | chr13-41576617-INS-122 |
| chr13-41589507-INS-51 | chr13-41589504-INS-57 |
| chr13-41611808-INS-741 | chr13-41611808-INS-742 |
| chr13-41632400-INS-70 | chr13-41632400-INS-70 |
| chr13-41653955-INS-57 | chr13-41653955-INS-51 |
| chr13-41663055-INS-56 | chr13-41663055-INS-58 |
| chr13-41675757-INS-104 | chr13-41675757-INS-104 |
| chr13-41680601-INS-105 | chr13-41680601-INS-105 |
| chr13-41682700-INS-518 | chr13-41682700-INS-518 |
| chr13-41693009-INS-60 | chr13-41693015-INS-80 |
| chr13-41720807-INS-163 | chr13-41720807-INS-163 |
| chr13-41758036-INS-60 | chr13-41758036-INS-63 |
| chr13-41802945-INS-133 | chr13-41802946-INS-137 |
| chr13-41807025-INS-403 | chr13-41807025-INS-403 |
| chr13-41831269-INS-251 | chr13-41831269-INS-251 |
| chr13-41876119-INS-133 | chr13-41876139-INS-108 |
| chr13-41888415-INS-61 | chr13-41888415-INS-61 |
| chr13-41906766-INS-3479 | chr13-41906767-INS-3479 |
| chr13-41909698-INS-104 | chr13-41909687-INS-126 |
| chr13-41958863-INS-240 | chr13-41958863-INS-240 |
| chr13-41972161-INS-215 | chr13-41972161-INS-215 |
| chr13-41978736-INS-214 | chr13-41978736-INS-215 |
| chr13-41989349-INS-53 | chr13-41989345-INS-53 |
| chr13-41997317-INS-64 | chr13-41997317-INS-66 |
| chr13-42003399-INS-189 | chr13-42003399-INS-202 |
| chr13-42026831-INS-6845 | chr13-42026829-INS-6842 |
| chr13-42055322-INS-580 | chr13-42055322-INS-580 |
| chr13-42056623-INS-250 | chr13-42056623-INS-244 |
| chr13-42072811-INS-676 | chr13-42072811-INS-677 |
| chr13-42096703-INS-54 | chr13-42096703-INS-54 |
| chr13-42120912-INS-80 | chr13-42120912-INS-74 |
| chr13-42164701-INS-691 | chr13-42164701-INS-691 |
| chr13-42181474-INS-8452 | chr13-42181475-INS-8452 |
| chr13-42185842-INS-58 | chr13-42185842-INS-60 |
| chr13-42191709-INS-712 | chr13-42191709-INS-712 |
| chr13-42241570-INS-248 | chr13-42241570-INS-248 |
| chr13-42246915-INS-112 | chr13-42246929-INS-116 |
| chr13-42286847-INS-505 | chr13-42286847-INS-505 |
| chr13-42294001-INS-359 | chr13-42294001-INS-359 |
| chr13-42298414-INS-95 | chr13-42298412-INS-80 |
| chr13-42352112-INS-86 | chr13-42352141-INS-68 |
| chr13-42372616-INS-893 | chr13-42372616-INS-893 |
| chr13-42386731-INS-66 | chr13-42386729-INS-96 |
| chr13-42415504-INS-52 | chr13-42415532-INS-64 |
| chr13-42421442-INS-56 | chr13-42421453-INS-54 |
| chr13-42459908-INS-60 | chr13-42459908-INS-60 |
| chr13-42507233-INS-63 | chr13-42507233-INS-63 |
| chr13-42535962-INS-73 | chr13-42535962-INS-68 |
| chr13-42560611-INS-56 | chr13-42560611-INS-60 |
| chr13-42592239-INS-382 | chr13-42592239-INS-382 |
| chr13-42628190-INS-63 | chr13-42628195-INS-63 |
| chr13-42659291-INS-202 | chr13-42659291-INS-202 |
| chr13-42664603-INS-227 | chr13-42664603-INS-228 |
| chr13-42718421-INS-75 | chr13-42718421-INS-82 |
| chr13-42737366-INS-303 | chr13-42737366-INS-296 |
| chr13-42768157-INS-193 | chr13-42768167-INS-176 |
| chr13-42780735-INS-4538 | chr13-42780736-INS-4538 |
| chr13-42787421-INS-147 | chr13-42787421-INS-147 |
| chr13-42870204-INS-401 | chr13-42870204-INS-401 |
| chr13-42871062-INS-66 | chr13-42871062-INS-66 |
| chr13-42872123-INS-52 | chr13-42872123-INS-52 |
| chr13-42886625-INS-206 | chr13-42886625-INS-206 |
| chr13-42907219-INS-65 | chr13-42907219-INS-60 |
| chr13-42937647-INS-52 | chr13-42937647-INS-52 |
| chr13-42950103-INS-172 | chr13-42950095-INS-182 |
| chr13-42969833-INS-189 | chr13-42969836-INS-201 |
| chr13-42973388-INS-51 | chr13-42973385-INS-76 |
| chr13-43072828-INS-964 | chr13-43072828-INS-964 |
| chr13-43121603-INS-502 | chr13-43121603-INS-512 |
| chr13-43136910-INS-54 | chr13-43136910-INS-54 |
| chr13-43146865-INS-88 | chr13-43146865-INS-94 |
| chr13-43151039-INS-276 | chr13-43151058-INS-252 |
| chr13-43163512-INS-86 | chr13-43163512-INS-86 |
| chr13-43184330-INS-52 | chr13-43184330-INS-74 |
| chr13-43193196-INS-52 | chr13-43193196-INS-52 |

| | |
|---|---|
| chr13-43201100-INS-143 | chr13-43201104-INS-147 |
| chr13-43218859-INS-59 | chr13-43218863-INS-53 |
| chr13-43228034-INS-266 | chr13-43228035-INS-269 |
| chr13-43233383-INS-212 | chr13-43233383-INS-212 |
| chr13-43245656-INS-59 | chr13-43245676-INS-75 |
| chr13-43253901-INS-240 | chr13-43253901-INS-232 |
| chr13-43263654-INS-406 | chr13-43263654-INS-412 |
| chr13-43282208-INS-56 | chr13-43282206-INS-55 |
| chr13-43298082-INS-99 | chr13-43298082-INS-99 |
| chr13-43309916-INS-51 | chr13-43309916-INS-51 |
| chr13-43320819-INS-214 | chr13-43320819-INS-214 |
| chr13-43327876-INS-122 | chr13-43327876-INS-104 |
| chr13-43334568-INS-78 | chr13-43334587-INS-88 |
| chr13-43340709-INS-197 | chr13-43340709-INS-205 |
| chr13-43376628-INS-85 | chr13-43376628-INS-83 |
| chr13-43411073-INS-102 | chr13-43411073-INS-102 |
| chr13-43418514-INS-267 | chr13-43418514-INS-270 |
| chr13-43504906-INS-203 | chr13-43504903-INS-189 |
| chr13-43513198-INS-78 | chr13-43513198-INS-80 |
| chr13-43551118-INS-211 | chr13-43551118-INS-211 |
| chr13-43562722-INS-1873 | chr13-43562723-INS-1873 |
| chr13-43565750-INS-68 | chr13-43565750-INS-68 |
| chr13-43621813-INS-62 | chr13-43621813-INS-62 |
| chr13-43648047-INS-1891 | chr13-43648048-INS-1891 |
| chr13-43663966-INS-7098 | chr13-43663967-INS-7098 |
| chr13-43689855-INS-311 | chr13-43689855-INS-314 |
| chr13-43703802-INS-65 | chr13-43703802-INS-67 |
| chr13-43705388-INS-54 | chr13-43705388-INS-54 |
| chr13-43728785-INS-69 | chr13-43728785-INS-69 |
| chr13-43734616-INS-384 | chr13-43734618-INS-376 |
| chr13-43777017-INS-120 | chr13-43777017-INS-120 |
| chr13-43788057-INS-662 | chr13-43788057-INS-662 |
| chr13-43795435-INS-56 | chr13-43795462-INS-60 |
| chr13-43855736-INS-55 | chr13-43855736-INS-55 |
| chr13-43919946-INS-93 | chr13-43919946-INS-93 |
| chr13-43929169-INS-107 | chr13-43929169-INS-141 |
| chr13-43993367-INS-56 | chr13-43993367-INS-56 |
| chr13-44021245-INS-158 | chr13-44021245-INS-163 |
| chr13-44087144-INS-77 | chr13-44087144-INS-75 |
| chr13-44089693-INS-60 | chr13-44089693-INS-60 |
| chr13-44100761-INS-60 | chr13-44100761-INS-66 |
| chr13-44186999-INS-126 | chr13-44187009-INS-148 |
| chr13-44194682-INS-56 | chr13-44194683-INS-56 |
| chr13-44204595-INS-4675 | chr13-44204596-INS-4675 |
| chr13-44234556-INS-63 | chr13-44234556-INS-63 |
| chr13-44235692-INS-75 | chr13-44235692-INS-115 |
| chr13-44249011-INS-466 | chr13-44249011-INS-466 |
| chr13-44265506-INS-391 | chr13-44265506-INS-393 |
| chr13-44346675-INS-409 | chr13-44346675-INS-409 |
| chr13-44348773-INS-257 | chr13-44348773-INS-257 |
| chr13-44370832-INS-108 | chr13-44370832-INS-108 |
| chr13-44396413-INS-78 | chr13-44396413-INS-78 |
| chr13-44446749-INS-70 | chr13-44446726-INS-100 |
| chr13-44448373-INS-159 | chr13-44448373-INS-169 |
| chr13-44450165-INS-95 | chr13-44450171-INS-87 |
| chr13-44452533-INS-52 | chr13-44452534-INS-50 |
| chr13-44487940-INS-59 | chr13-44487936-INS-70 |
| chr13-44497713-INS-404 | chr13-44497713-INS-404 |
| chr13-44511253-INS-60 | chr13-44511253-INS-56 |
| chr13-44526944-INS-148 | chr13-44526944-INS-148 |
| chr13-44534316-INS-204 | chr13-44534316-INS-204 |
| chr13-44538133-INS-52 | chr13-44538133-INS-56 |
| chr13-44561058-INS-61 | chr13-44561058-INS-63 |
| chr13-44568615-INS-56 | chr13-44568615-INS-55 |
| chr13-44586154-INS-271 | chr13-44586167-INS-286 |
| chr13-44608164-INS-72 | chr13-44608164-INS-72 |
| chr13-44617103-INS-593 | chr13-44617103-INS-583 |
| chr13-44619590-INS-66 | chr13-44619573-INS-84 |
| chr13-44671749-INS-83 | chr13-44671749-INS-65 |
| chr13-44681975-INS-64 | chr13-44681975-INS-69 |
| chr13-44692623-INS-231 | chr13-44692623-INS-231 |
| chr13-44701479-INS-1888 | chr13-44701480-INS-1888 |
| chr13-44748667-INS-66 | chr13-44748667-INS-63 |
| chr13-44749041-INS-249 | chr13-44749069-INS-251 |
| chr13-44752614-INS-72 | chr13-44752614-INS-72 |
| chr13-44763619-INS-203 | chr13-44763620-INS-200 |
| chr13-44768693-INS-64 | chr13-44768664-INS-55 |

| | |
|---|---|
| chr13-44772499-INS-113 | chr13-44772489-INS-86 |
| chr13-44779099-INS-116 | chr13-44779099-INS-116 |
| chr13-44788990-INS-62 | chr13-44788990-INS-62 |
| chr13-44811637-INS-220 | chr13-44811637-INS-219 |
| chr13-44822499-INS-214 | chr13-44822499-INS-215 |
| chr13-44832011-INS-231 | chr13-44832011-INS-231 |
| chr13-44911106-INS-215 | chr13-44911106-INS-216 |
| chr13-44916602-INS-203 | chr13-44916602-INS-203 |
| chr13-44944425-INS-115 | chr13-44944425-INS-115 |
| chr13-44993294-INS-58 | chr13-44993294-INS-60 |
| chr13-45001515-INS-88 | chr13-45001515-INS-88 |
| chr13-45124013-INS-199 | chr13-45124013-INS-199 |
| chr13-45178297-INS-3528 | chr13-45178298-INS-3530 |
| chr13-45178884-INS-60 | chr13-45178884-INS-58 |
| chr13-45185233-INS-83 | chr13-45185233-INS-83 |
| chr13-45226044-INS-1955 | chr13-45226045-INS-1955 |
| chr13-45228192-INS-874 | chr13-45228192-INS-874 |
| chr13-45230025-INS-100 | chr13-45230025-INS-100 |
| chr13-45239891-INS-55 | chr13-45239891-INS-54 |
| chr13-45274919-INS-515 | chr13-45274919-INS-516 |
| chr13-45279489-INS-875 | chr13-45279489-INS-876 |
| chr13-45336837-INS-61 | chr13-45336837-INS-61 |
| chr13-45339003-INS-441 | chr13-45339003-INS-441 |
| chr13-45342042-INS-51 | chr13-45342042-INS-51 |
| chr13-45402346-INS-63 | chr13-45402346-INS-63 |
| chr13-45417390-INS-3794 | chr13-45417391-INS-3806 |
| chr13-45423827-INS-607 | chr13-45423827-INS-607 |
| chr13-45446940-INS-192 | chr13-45446940-INS-188 |
| chr13-45463219-INS-737 | chr13-45463219-INS-738 |
| chr13-45485275-INS-934 | chr13-45485275-INS-935 |
| chr13-45489566-INS-198 | chr13-45489566-INS-198 |
| chr13-45499598-INS-7133 | chr13-45499599-INS-7133 |
| chr13-45506777-INS-94 | chr13-45506777-INS-94 |
| chr13-45530861-INS-208 | chr13-45530861-INS-208 |
| chr13-45563660-INS-92 | chr13-45563662-INS-80 |
| chr13-45598803-INS-219 | chr13-45598803-INS-220 |
| chr13-45671994-INS-76 | chr13-45671994-INS-60 |
| chr13-45673596-INS-177 | chr13-45673578-INS-205 |
| chr13-45722059-INS-66 | chr13-45722059-INS-81 |
| chr13-45744012-INS-202 | chr13-45744012-INS-202 |
| chr13-45819665-INS-6635 | chr13-45819666-INS-6638 |
| chr13-45836978-INS-56 | chr13-45836978-INS-104 |
| chr13-45866201-INS-98 | chr13-45866156-INS-96 |
| chr13-45873796-INS-147 | chr13-45873796-INS-147 |
| chr13-45910727-INS-163 | chr13-45910727-INS-163 |
| chr13-45933586-INS-75 | chr13-45933586-INS-75 |
| chr13-45939906-INS-51 | chr13-45939907-INS-62 |
| chr13-45942295-INS-68 | chr13-45942295-INS-50 |
| chr13-45944831-INS-227 | chr13-45944831-INS-227 |
| chr13-45990584-INS-57 | chr13-45990584-INS-60 |
| chr13-46056721-INS-110 | chr13-46056722-INS-110 |
| chr13-46065717-INS-883 | chr13-46065717-INS-890 |
| chr13-46076553-INS-71 | chr13-46076553-INS-71 |
| chr13-46091066-INS-76 | chr13-46091066-INS-85 |
| chr13-46164196-INS-52 | chr13-46164196-INS-52 |
| chr13-46179605-INS-137 | chr13-46179605-INS-131 |
| chr13-46235889-INS-928 | chr13-46235889-INS-928 |
| chr13-46236034-INS-2194 | chr13-46236035-INS-2146 |
| chr13-46245760-INS-56 | chr13-46245760-INS-58 |
| chr13-46269210-INS-303 | chr13-46269210-INS-302 |
| chr13-46276238-INS-515 | chr13-46276238-INS-515 |
| chr13-46280499-INS-82 | chr13-46280491-INS-80 |
| chr13-46304889-INS-54 | chr13-46304894-INS-59 |
| chr13-46316746-INS-196 | chr13-46316746-INS-196 |
| chr13-46325285-INS-101 | chr13-46325327-INS-60 |
| chr13-46340422-INS-131 | chr13-46340418-INS-120 |
| chr13-46349368-INS-72 | chr13-46349364-INS-76 |
| chr13-46354141-INS-60 | chr13-46354141-INS-60 |
| chr13-46381867-INS-401 | chr13-46381867-INS-401 |
| chr13-46387465-INS-54 | chr13-46387465-INS-78 |
| chr13-46403125-INS-68 | chr13-46403125-INS-70 |
| chr13-46404606-INS-52 | chr13-46404606-INS-52 |
| chr13-46421053-INS-593 | chr13-46421053-INS-581 |
| chr13-46432677-INS-52 | chr13-46432702-INS-70 |
| chr13-46460927-INS-61 | chr13-46460927-INS-61 |
| chr13-46470030-INS-211 | chr13-46470031-INS-211 |
| chr13-46473172-INS-357 | chr13-46473172-INS-361 |

| | |
|---|---|
| chr13-46489541-INS-78 | chr13-46489541-INS-96 |
| chr13-46498631-INS-7086 | chr13-46498632-INS-7086 |
| chr13-46524141-INS-174 | chr13-46524141-INS-175 |
| chr13-46553532-INS-1051 | chr13-46553533-INS-1051 |
| chr13-46583232-INS-66 | chr13-46583232-INS-66 |
| chr13-46598885-INS-169 | chr13-46598885-INS-185 |
| chr13-46652925-INS-64 | chr13-46652925-INS-64 |
| chr13-46680694-INS-59 | chr13-46680692-INS-54 |
| chr13-46717021-INS-53 | chr13-46717021-INS-53 |
| chr13-46769118-INS-54 | chr13-46769118-INS-54 |
| chr13-46772756-INS-58 | chr13-46772756-INS-58 |
| chr13-46775877-INS-85 | chr13-46775877-INS-95 |
| chr13-46778758-INS-5249 | chr13-46778759-INS-5249 |
| chr13-46781160-INS-216 | chr13-46781160-INS-216 |
| chr13-46803780-INS-468 | chr13-46803780-INS-469 |
| chr13-46850891-INS-76 | chr13-46850891-INS-76 |
| chr13-46874011-INS-106 | chr13-46874011-INS-92 |
| chr13-46897398-INS-353 | chr13-46897398-INS-353 |
| chr13-46923011-INS-204 | chr13-46923011-INS-204 |
| chr13-46943262-INS-72 | chr13-46943262-INS-73 |
| chr13-46945502-INS-79 | chr13-46945493-INS-81 |
| chr13-46963138-INS-82 | chr13-46963138-INS-84 |
| chr13-46967878-INS-55 | chr13-46967878-INS-55 |
| chr13-46974319-INS-69 | chr13-46974319-INS-69 |
| chr13-46978996-INS-72 | chr13-46978996-INS-56 |
| chr13-46984007-INS-188 | chr13-46984007-INS-188 |
| chr13-46993128-INS-59 | chr13-46993128-INS-63 |
| chr13-47001417-INS-51 | chr13-47001417-INS-51 |
| chr13-47014109-INS-72 | chr13-47014109-INS-72 |
| chr13-47042239-INS-86 | chr13-47042239-INS-86 |
| chr13-47080982-INS-217 | chr13-47080982-INS-217 |
| chr13-47089500-INS-7107 | chr13-47089501-INS-7107 |
| chr13-47098044-INS-92 | chr13-47098044-INS-104 |
| chr13-47124981-INS-3848 | chr13-47124982-INS-3848 |
| chr13-47206527-INS-273 | chr13-47206527-INS-272 |
| chr13-47269694-INS-293 | chr13-47269694-INS-293 |
| chr13-47280701-INS-259 | chr13-47280701-INS-259 |
| chr13-47295698-INS-169 | chr13-47295698-INS-148 |
| chr13-47303669-INS-93 | chr13-47303669-INS-93 |
| chr13-47309020-INS-51 | chr13-47309020-INS-51 |
| chr13-47310663-INS-112 | chr13-47310674-INS-62 |
| chr13-47320557-INS-216 | chr13-47320557-INS-210 |
| chr13-47323943-INS-94 | chr13-47323943-INS-94 |
| chr13-47386677-INS-64 | chr13-47386677-INS-64 |
| chr13-47403451-INS-67 | chr13-47403451-INS-70 |
| chr13-47409106-INS-344 | chr13-47409106-INS-339 |
| chr13-47426017-INS-136 | chr13-47426013-INS-120 |
| chr13-47442715-INS-159 | chr13-47442715-INS-159 |
| chr13-47448035-INS-168 | chr13-47448019-INS-156 |
| chr13-47452132-INS-482 | chr13-47452132-INS-482 |
| chr13-47474554-INS-283 | chr13-47474554-INS-283 |
| chr13-47493481-INS-56 | chr13-47493481-INS-56 |
| chr13-47507182-INS-53 | chr13-47507182-INS-54 |
| chr13-47532610-INS-549 | chr13-47532610-INS-549 |
| chr13-47554844-INS-60 | chr13-47554858-INS-58 |
| chr13-47563918-INS-123 | chr13-47563918-INS-123 |
| chr13-47577318-INS-51 | chr13-47577318-INS-51 |
| chr13-47595528-INS-202 | chr13-47595528-INS-201 |
| chr13-47601346-INS-122 | chr13-47601346-INS-122 |
| chr13-47604564-INS-58 | chr13-47604564-INS-50 |
| chr13-47606395-INS-701 | chr13-47606396-INS-702 |
| chr13-47607125-INS-2159 | chr13-47607126-INS-2157 |
| chr13-47620005-INS-1891 | chr13-47620006-INS-1891 |
| chr13-47663822-INS-90 | chr13-47663776-INS-86 |
| chr13-47699701-INS-68 | chr13-47699701-INS-56 |
| chr13-47701336-INS-105 | chr13-47701333-INS-99 |
| chr13-47715710-INS-425 | chr13-47715710-INS-429 |
| chr13-47746857-INS-6576 | chr13-47746858-INS-6576 |
| chr13-47750846-INS-73 | chr13-47750847-INS-61 |
| chr13-47757948-INS-55 | chr13-47757948-INS-57 |
| chr13-47797444-INS-86 | chr13-47797444-INS-86 |
| chr13-47831571-INS-787 | chr13-47831571-INS-787 |
| chr13-47851619-INS-604 | chr13-47851619-INS-607 |
| chr13-47863789-INS-79 | chr13-47863839-INS-70 |
| chr13-47866653-INS-255 | chr13-47866653-INS-255 |
| chr13-47900691-INS-109 | chr13-47900691-INS-109 |
| chr13-47910429-INS-295 | chr13-47910427-INS-274 |

| | |
|---|---|
| chr13-47926533-INS-129 | chr13-47926537-INS-130 |
| chr13-47939215-INS-84 | chr13-47939211-INS-120 |
| chr13-48003928-INS-84 | chr13-48003928-INS-86 |
| chr13-48018948-INS-358 | chr13-48018948-INS-358 |
| chr13-48066834-INS-182 | chr13-48066834-INS-186 |
| chr13-48105751-INS-100 | chr13-48105751-INS-80 |
| chr13-48106608-INS-490 | chr13-48106608-INS-481 |
| chr13-48134606-INS-77 | chr13-48134606-INS-79 |
| chr13-48141429-INS-70 | chr13-48141429-INS-70 |
| chr13-48164529-INS-115 | chr13-48164529-INS-102 |
| chr13-48204929-INS-6562 | chr13-48204930-INS-6562 |
| chr13-48206258-INS-60 | chr13-48206262-INS-62 |
| chr13-48252506-INS-6440 | chr13-48252507-INS-6440 |
| chr13-48290631-INS-5294 | chr13-48290632-INS-5294 |
| chr13-48311550-INS-86 | chr13-48311550-INS-86 |
| chr13-48327805-INS-65 | chr13-48327806-INS-70 |
| chr13-48337662-INS-679 | chr13-48337662-INS-679 |
| chr13-48365838-INS-72 | chr13-48365863-INS-60 |
| chr13-48375726-INS-125 | chr13-48375703-INS-138 |
| chr13-48392173-INS-156 | chr13-48392173-INS-156 |
| chr13-48405022-INS-129 | chr13-48405022-INS-132 |
| chr13-48429073-INS-198 | chr13-48429073-INS-198 |
| chr13-48479293-INS-78 | chr13-48479293-INS-78 |
| chr13-48508450-INS-6362 | chr13-48508451-INS-6362 |
| chr13-48574815-INS-136 | chr13-48574815-INS-135 |
| chr13-48596588-INS-177 | chr13-48596588-INS-178 |
| chr13-48608976-INS-92 | chr13-48608976-INS-90 |
| chr13-48612745-INS-344 | chr13-48612745-INS-344 |
| chr13-48615789-INS-52 | chr13-48615785-INS-100 |
| chr13-48655006-INS-2345 | chr13-48655007-INS-2345 |
| chr13-48701061-INS-175 | chr13-48701061-INS-175 |
| chr13-48706815-INS-656 | chr13-48706815-INS-656 |
| chr13-48783435-INS-201 | chr13-48783435-INS-201 |
| chr13-48784473-INS-111 | chr13-48784473-INS-111 |
| chr13-48801815-INS-588 | chr13-48801815-INS-588 |
| chr13-48825653-INS-58 | chr13-48825653-INS-58 |
| chr13-48836967-INS-59 | chr13-48836968-INS-60 |
| chr13-48851508-INS-426 | chr13-48851508-INS-427 |
| chr13-48857010-INS-70 | chr13-48857010-INS-80 |
| chr13-48866691-INS-52 | chr13-48866691-INS-52 |
| chr13-48898181-INS-80 | chr13-48898181-INS-81 |
| chr13-48938954-INS-361 | chr13-48938954-INS-361 |
| chr13-48944636-INS-222 | chr13-48944636-INS-222 |
| chr13-48978919-INS-51 | chr13-48978919-INS-51 |
| chr13-48980608-INS-61 | chr13-48980608-INS-61 |
| chr13-48993052-INS-52 | chr13-48993052-INS-54 |
| chr13-48999704-INS-116 | chr13-48999700-INS-89 |
| chr13-49037862-INS-52 | chr13-49037862-INS-54 |
| chr13-49069508-INS-213 | chr13-49069508-INS-214 |
| chr13-49111459-INS-175 | chr13-49111459-INS-175 |
| chr13-49112873-INS-138 | chr13-49112873-INS-138 |
| chr13-49144016-INS-149 | chr13-49144016-INS-149 |
| chr13-49149486-INS-59 | chr13-49149486-INS-59 |
| chr13-49200906-INS-58 | chr13-49200906-INS-58 |
| chr13-49219041-INS-51 | chr13-49219041-INS-57 |
| chr13-49295337-INS-173 | chr13-49295337-INS-175 |
| chr13-49306876-INS-50 | chr13-49306871-INS-66 |
| chr13-49314204-INS-62 | chr13-49314204-INS-62 |
| chr13-49325559-INS-195 | chr13-49325559-INS-197 |
| chr13-49329818-INS-71 | chr13-49329836-INS-71 |
| chr13-49347768-INS-203 | chr13-49347768-INS-203 |
| chr13-49390515-INS-71 | chr13-49390515-INS-71 |
| chr13-49449527-INS-92 | chr13-49449530-INS-56 |
| chr13-49464592-INS-64 | chr13-49464592-INS-87 |
| chr13-49468873-INS-439 | chr13-49468860-INS-465 |
| chr13-49472678-INS-171 | chr13-49472678-INS-200 |
| chr13-49475355-INS-6459 | chr13-49475356-INS-6459 |
| chr13-49479359-INS-366 | chr13-49479357-INS-332 |
| chr13-49500938-INS-408 | chr13-49500938-INS-409 |
| chr13-49505332-INS-74 | chr13-49505332-INS-74 |
| chr13-49527318-INS-207 | chr13-49527318-INS-206 |
| chr13-49550538-INS-213 | chr13-49550538-INS-214 |
| chr13-49553146-INS-105 | chr13-49553146-INS-105 |
| chr13-49563290-INS-69 | chr13-49563290-INS-68 |
| chr13-49581050-INS-206 | chr13-49581050-INS-215 |
| chr13-49591554-INS-116 | chr13-49591553-INS-117 |
| chr13-49624117-INS-206 | chr13-49624117-INS-205 |

| | |
|---|---|
| chr13-49721511-INS-1075 | chr13-49721512-INS-1072 |
| chr13-49857494-INS-851 | chr13-49857494-INS-852 |
| chr13-49861343-INS-187 | chr13-49861343-INS-192 |
| chr13-49869966-INS-400 | chr13-49869966-INS-400 |
| chr13-49895657-INS-68 | chr13-49895650-INS-60 |
| chr13-49908808-INS-518 | chr13-49908808-INS-518 |
| chr13-49909860-INS-479 | chr13-49909860-INS-482 |
| chr13-49924916-INS-699 | chr13-49924916-INS-699 |
| chr13-49949010-INS-52 | chr13-49949010-INS-60 |
| chr13-49981462-INS-411 | chr13-49981462-INS-411 |
| chr13-49988096-INS-202 | chr13-49988096-INS-202 |
| chr13-50034956-INS-514 | chr13-50034956-INS-503 |
| chr13-50091964-INS-6368 | chr13-50091965-INS-6368 |
| chr13-50106812-INS-84 | chr13-50106821-INS-96 |
| chr13-50115679-INS-112 | chr13-50115679-INS-113 |
| chr13-50211647-INS-60 | chr13-50211647-INS-72 |
| chr13-50244637-INS-2670 | chr13-50244638-INS-2677 |
| chr13-50245099-INS-91 | chr13-50245099-INS-91 |
| chr13-50516817-INS-205 | chr13-50516817-INS-205 |
| chr13-50618223-INS-74 | chr13-50618223-INS-76 |
| chr13-50646416-INS-76 | chr13-50646416-INS-80 |
| chr13-50681626-INS-60 | chr13-50681626-INS-60 |
| chr13-50713108-INS-78 | chr13-50713108-INS-78 |
| chr13-51023987-INS-201 | chr13-51023987-INS-201 |
| chr13-51096715-INS-149 | chr13-51096715-INS-149 |
| chr13-51124630-INS-60 | chr13-51124630-INS-60 |
| chr13-51137404-INS-170 | chr13-51137404-INS-170 |
| chr13-51143506-INS-60 | chr13-51143506-INS-60 |
| chr13-51160171-INS-330 | chr13-51160171-INS-330 |
| chr13-51163341-INS-511 | chr13-51163341-INS-511 |
| chr13-51163924-INS-212 | chr13-51163924-INS-211 |
| chr13-51169951-INS-996 | chr13-51169951-INS-996 |
| chr13-51182071-INS-99 | chr13-51182071-INS-98 |
| chr13-51183540-INS-604 | chr13-51183540-INS-604 |
| chr13-51224176-INS-358 | chr13-51224177-INS-368 |
| chr13-51240644-INS-71 | chr13-51240644-INS-61 |
| chr13-51247316-INS-171 | chr13-51247316-INS-167 |
| chr13-51301055-INS-241 | chr13-51301055-INS-241 |
| chr13-51312305-INS-212 | chr13-51312305-INS-212 |
| chr13-51334610-INS-202 | chr13-51334610-INS-196 |
| chr13-51364275-INS-52 | chr13-51364275-INS-54 |
| chr13-51366701-INS-63 | chr13-51366701-INS-63 |
| chr13-51368527-INS-308 | chr13-51368523-INS-304 |
| chr13-51406770-INS-52 | chr13-51406770-INS-52 |
| chr13-51433256-INS-74 | chr13-51433256-INS-93 |
| chr13-51448930-INS-122 | chr13-51448930-INS-122 |
| chr13-51476084-INS-75 | chr13-51476084-INS-57 |
| chr13-51482190-INS-59 | chr13-51482196-INS-96 |
| chr13-51483580-INS-888 | chr13-51483579-INS-898 |
| chr13-51487231-INS-52 | chr13-51487232-INS-52 |
| chr13-51492214-INS-60 | chr13-51492214-INS-62 |
| chr13-51497878-INS-395 | chr13-51497878-INS-395 |
| chr13-51502730-INS-62 | chr13-51502730-INS-64 |
| chr13-51535970-INS-374 | chr13-51535970-INS-374 |
| chr13-51556764-INS-79 | chr13-51556767-INS-68 |
| chr13-51564363-INS-54 | chr13-51564328-INS-94 |
| chr13-51566897-INS-58 | chr13-51566897-INS-56 |
| chr13-51574313-INS-60 | chr13-51574313-INS-62 |
| chr13-51596136-INS-91 | chr13-51596125-INS-92 |
| chr13-51633436-INS-121 | chr13-51633436-INS-125 |
| chr13-51639739-INS-56 | chr13-51639738-INS-52 |
| chr13-51649064-INS-207 | chr13-51649064-INS-208 |
| chr13-51661093-INS-195 | chr13-51661093-INS-206 |
| chr13-51666708-INS-59 | chr13-51666708-INS-59 |
| chr13-51676053-INS-59 | chr13-51676053-INS-59 |
| chr13-51707414-INS-69 | chr13-51707414-INS-69 |
| chr13-51711137-INS-54 | chr13-51711137-INS-54 |
| chr13-51742297-INS-97 | chr13-51742322-INS-85 |
| chr13-51766161-INS-98 | chr13-51766170-INS-50 |
| chr13-51773040-INS-74 | chr13-51773040-INS-100 |
| chr13-51797522-INS-497 | chr13-51797523-INS-500 |
| chr13-51819651-INS-173 | chr13-51819651-INS-174 |
| chr13-51847989-INS-76 | chr13-51848007-INS-53 |
| chr13-51860504-INS-997 | chr13-51860504-INS-997 |
| chr13-51868378-INS-175 | chr13-51868378-INS-174 |
| chr13-51884658-INS-113 | chr13-51884658-INS-79 |
| chr13-51892362-INS-69 | chr13-51892362-INS-72 |

| | |
|---|---|
| chr13-51893132-INS-9049 | chr13-51893133-INS-9049 |
| chr13-51899505-INS-1874 | chr13-51899506-INS-1874 |
| chr13-51910052-INS-133 | chr13-51910052-INS-133 |
| chr13-51923357-INS-81 | chr13-51923372-INS-71 |
| chr13-51941882-INS-60 | chr13-51941879-INS-70 |
| chr13-51982583-INS-56 | chr13-51982583-INS-56 |
| chr13-51991675-INS-98 | chr13-51991675-INS-98 |
| chr13-52022140-INS-88 | chr13-52022140-INS-58 |
| chr13-52043941-INS-328 | chr13-52043941-INS-329 |
| chr13-52068836-INS-62 | chr13-52068836-INS-62 |
| chr13-52080731-INS-3100 | chr13-52080732-INS-3100 |
| chr13-52110547-INS-53 | chr13-52110547-INS-58 |
| chr13-52134117-INS-81 | chr13-52134117-INS-79 |
| chr13-52157093-INS-92 | chr13-52157043-INS-133 |
| chr13-52194614-INS-440 | chr13-52194615-INS-440 |
| chr13-52204399-INS-234 | chr13-52204399-INS-234 |
| chr13-52210577-INS-92 | chr13-52210577-INS-92 |
| chr13-52245764-INS-198 | chr13-52245760-INS-202 |
| chr13-52256695-INS-349 | chr13-52256695-INS-347 |
| chr13-52315594-INS-70 | chr13-52315594-INS-84 |
| chr13-52328712-INS-1817 | chr13-52328713-INS-1818 |
| chr13-52333752-INS-59 | chr13-52333752-INS-61 |
| chr13-52334621-INS-270 | chr13-52334641-INS-271 |
| chr13-52358590-INS-1034 | chr13-52358591-INS-1034 |
| chr13-52402163-INS-69 | chr13-52402139-INS-94 |
| chr13-52416740-INS-6182 | chr13-52416739-INS-6184 |
| chr13-52420107-INS-493 | chr13-52420107-INS-493 |
| chr13-52425643-INS-78 | chr13-52425644-INS-72 |
| chr13-52434702-INS-371 | chr13-52434717-INS-402 |
| chr13-52455185-INS-251 | chr13-52455185-INS-251 |
| chr13-52461579-INS-110 | chr13-52461579-INS-110 |
| chr13-52464253-INS-1498 | chr13-52464254-INS-1499 |
| chr13-52475976-INS-286 | chr13-52475976-INS-285 |
| chr13-52480404-INS-54 | chr13-52480404-INS-54 |
| chr13-52492104-INS-235 | chr13-52492104-INS-236 |
| chr13-52494019-INS-195 | chr13-52494019-INS-195 |
| chr13-52501026-INS-85 | chr13-52501026-INS-86 |
| chr13-52522585-INS-169 | chr13-52522587-INS-169 |
| chr13-52524520-INS-145 | chr13-52524520-INS-145 |
| chr13-52546657-INS-106 | chr13-52546657-INS-118 |
| chr13-52567294-INS-92 | chr13-52567302-INS-102 |
| chr13-52572644-INS-96 | chr13-52572644-INS-96 |
| chr13-52578491-INS-6592 | chr13-52578492-INS-6592 |
| chr13-52603841-INS-202 | chr13-52603841-INS-200 |
| chr13-52610028-INS-401 | chr13-52610028-INS-401 |
| chr13-52626368-INS-144 | chr13-52626368-INS-144 |
| chr13-52649684-INS-186 | chr13-52649684-INS-186 |
| chr13-52651670-INS-57 | chr13-52651670-INS-64 |
| chr13-52712982-INS-1355 | chr13-52712983-INS-1355 |
| chr13-52717505-INS-63 | chr13-52717505-INS-63 |
| chr13-52725520-INS-52 | chr13-52725520-INS-56 |
| chr13-52760238-INS-565 | chr13-52760238-INS-565 |
| chr13-52775389-INS-206 | chr13-52775389-INS-206 |
| chr13-52777050-INS-175 | chr13-52777050-INS-176 |
| chr13-52789617-INS-56 | chr13-52789617-INS-58 |
| chr13-52797282-INS-66 | chr13-52797282-INS-66 |
| chr13-52798931-INS-96 | chr13-52798931-INS-96 |
| chr13-52830304-INS-693 | chr13-52830304-INS-693 |
| chr13-52851057-INS-55 | chr13-52851057-INS-55 |
| chr13-52853463-INS-104 | chr13-52853463-INS-114 |
| chr13-52892964-INS-56 | chr13-52892964-INS-56 |
| chr13-52935383-INS-54 | chr13-52935383-INS-54 |
| chr13-52942356-INS-4558 | chr13-52942357-INS-4558 |
| chr13-52960649-INS-64 | chr13-52960642-INS-94 |
| chr13-52980405-INS-76 | chr13-52980403-INS-96 |
| chr13-53021647-INS-52 | chr13-53021647-INS-52 |
| chr13-53024914-INS-51 | chr13-53024914-INS-51 |
| chr13-53049188-INS-99 | chr13-53049188-INS-99 |
| chr13-53085195-INS-80 | chr13-53085195-INS-80 |
| chr13-53089212-INS-497 | chr13-53089212-INS-497 |
| chr13-53145132-INS-60 | chr13-53145118-INS-56 |
| chr13-53171561-INS-72 | chr13-53171547-INS-70 |
| chr13-53183126-INS-203 | chr13-53183126-INS-203 |
| chr13-53227303-INS-5557 | chr13-53227304-INS-5557 |
| chr13-53246205-INS-278 | chr13-53246207-INS-292 |
| chr13-53255156-INS-208 | chr13-53255156-INS-210 |
| chr13-53267983-INS-60 | chr13-53267983-INS-69 |

| | |
|---|---|
| chr13-53305036-INS-90 | chr13-53305036-INS-90 |
| chr13-53327187-INS-68 | chr13-53327187-INS-68 |
| chr13-53333218-INS-206 | chr13-53333218-INS-206 |
| chr13-53357394-INS-214 | chr13-53357394-INS-182 |
| chr13-53362283-INS-154 | chr13-53362283-INS-170 |
| chr13-53369381-INS-217 | chr13-53369384-INS-173 |
| chr13-53377978-INS-62 | chr13-53377990-INS-81 |
| chr13-53381045-INS-73 | chr13-53381045-INS-83 |
| chr13-53383635-INS-214 | chr13-53383635-INS-214 |
| chr13-53423234-INS-67 | chr13-53423234-INS-67 |
| chr13-53442689-INS-416 | chr13-53442689-INS-416 |
| chr13-53455044-INS-60 | chr13-53455044-INS-56 |
| chr13-53469314-INS-219 | chr13-53469314-INS-219 |
| chr13-53474157-INS-197 | chr13-53474126-INS-179 |
| chr13-53500583-INS-120 | chr13-53500583-INS-120 |
| chr13-53543055-INS-86 | chr13-53543055-INS-82 |
| chr13-53552963-INS-191 | chr13-53552963-INS-166 |
| chr13-53599917-INS-97 | chr13-53599917-INS-97 |
| chr13-53614582-INS-401 | chr13-53614582-INS-401 |
| chr13-53633810-INS-278 | chr13-53633806-INS-241 |
| chr13-53636417-INS-436 | chr13-53636417-INS-436 |
| chr13-53650343-INS-64 | chr13-53650322-INS-50 |
| chr13-53657334-INS-74 | chr13-53657334-INS-50 |
| chr13-53662230-INS-60 | chr13-53662214-INS-86 |
| chr13-53747093-INS-134 | chr13-53747093-INS-134 |
| chr13-53756799-INS-5313 | chr13-53756800-INS-5313 |
| chr13-53785921-INS-96 | chr13-53785916-INS-62 |
| chr13-53793554-INS-236 | chr13-53793554-INS-236 |
| chr13-53810610-INS-537 | chr13-53810610-INS-535 |
| chr13-53813141-INS-932 | chr13-53813141-INS-934 |
| chr13-53843453-INS-91 | chr13-53843453-INS-96 |
| chr13-53847203-INS-68 | chr13-53847203-INS-68 |
| chr13-53914185-INS-530 | chr13-53914185-INS-530 |
| chr13-53940084-INS-166 | chr13-53940084-INS-178 |
| chr13-53951976-INS-6562 | chr13-53951977-INS-6563 |
| chr13-53954655-INS-56 | chr13-53954670-INS-55 |
| chr13-54009661-INS-58 | chr13-54009661-INS-64 |
| chr13-54023941-INS-117 | chr13-54023985-INS-70 |
| chr13-54030996-INS-146 | chr13-54030996-INS-146 |
| chr13-54038237-INS-53 | chr13-54038237-INS-53 |
| chr13-54042375-INS-92 | chr13-54042375-INS-96 |
| chr13-54055208-INS-463 | chr13-54055208-INS-455 |
| chr13-54063638-INS-56 | chr13-54063638-INS-56 |
| chr13-54085026-INS-7210 | chr13-54085027-INS-7210 |
| chr13-54097676-INS-72 | chr13-54097648-INS-82 |
| chr13-54122666-INS-3291 | chr13-54122667-INS-3291 |
| chr13-54140948-INS-885 | chr13-54140948-INS-885 |
| chr13-54150307-INS-4897 | chr13-54150308-INS-4896 |
| chr13-54155432-INS-537 | chr13-54155432-INS-537 |
| chr13-54243099-INS-54 | chr13-54243099-INS-54 |
| chr13-54305994-INS-479 | chr13-54305994-INS-479 |
| chr13-54307370-INS-81 | chr13-54307370-INS-81 |
| chr13-54391254-INS-211 | chr13-54391254-INS-211 |
| chr13-54398928-INS-4076 | chr13-54398929-INS-4092 |
| chr13-54402875-INS-68 | chr13-54402875-INS-66 |
| chr13-54426612-INS-345 | chr13-54426612-INS-345 |
| chr13-54428279-INS-97 | chr13-54428279-INS-75 |
| chr13-54442226-INS-54 | chr13-54442214-INS-78 |
| chr13-54454814-INS-610 | chr13-54454814-INS-610 |
| chr13-54462661-INS-67 | chr13-54462673-INS-56 |
| chr13-54488397-INS-74 | chr13-54488393-INS-123 |
| chr13-54494028-INS-666 | chr13-54494041-INS-666 |
| chr13-54527805-INS-73 | chr13-54527805-INS-63 |
| chr13-54543970-INS-52 | chr13-54543970-INS-52 |
| chr13-54574562-INS-200 | chr13-54574562-INS-200 |
| chr13-54580186-INS-126 | chr13-54580186-INS-158 |
| chr13-54593187-INS-69 | chr13-54593191-INS-52 |
| chr13-54598047-INS-258 | chr13-54598047-INS-260 |
| chr13-54630325-INS-232 | chr13-54630325-INS-232 |
| chr13-54638028-INS-233 | chr13-54638030-INS-216 |
| chr13-54641951-INS-76 | chr13-54641951-INS-72 |
| chr13-54644540-INS-211 | chr13-54644540-INS-213 |
| chr13-54675243-INS-139 | chr13-54675243-INS-139 |
| chr13-54684025-INS-303 | chr13-54684025-INS-303 |
| chr13-54685933-INS-198 | chr13-54685934-INS-161 |
| chr13-54691837-INS-217 | chr13-54691837-INS-217 |
| chr13-54695601-INS-216 | chr13-54695601-INS-218 |

| | |
|---|---|
| chr13-54702854-INS-122 | chr13-54702858-INS-100 |
| chr13-54711809-INS-211 | chr13-54711809-INS-212 |
| chr13-54717666-INS-55 | chr13-54717666-INS-55 |
| chr13-54718534-INS-212 | chr13-54718534-INS-212 |
| chr13-54721228-INS-257 | chr13-54721228-INS-258 |
| chr13-54731133-INS-111 | chr13-54731133-INS-111 |
| chr13-54737344-INS-68 | chr13-54737344-INS-52 |
| chr13-54742090-INS-173 | chr13-54742090-INS-173 |
| chr13-54752779-INS-213 | chr13-54752779-INS-213 |
| chr13-54764261-INS-208 | chr13-54764261-INS-208 |
| chr13-54776463-INS-184 | chr13-54776463-INS-184 |
| chr13-54807217-INS-206 | chr13-54807217-INS-207 |
| chr13-54823908-INS-1299 | chr13-54823909-INS-1291 |
| chr13-54843241-INS-106 | chr13-54843241-INS-108 |
| chr13-54854277-INS-205 | chr13-54854277-INS-205 |
| chr13-54864681-INS-215 | chr13-54864681-INS-211 |
| chr13-54870639-INS-223 | chr13-54870639-INS-223 |
| chr13-54882899-INS-93 | chr13-54882904-INS-51 |
| chr13-54903622-INS-76 | chr13-54903622-INS-69 |
| chr13-54949491-INS-263 | chr13-54949491-INS-262 |
| chr13-54957710-INS-206 | chr13-54957710-INS-207 |
| chr13-54989221-INS-196 | chr13-54989221-INS-196 |
| chr13-55007177-INS-78 | chr13-55007193-INS-76 |
| chr13-55008478-INS-3523 | chr13-55008479-INS-3523 |
| chr13-55039089-INS-213 | chr13-55039089-INS-213 |
| chr13-55050567-INS-50 | chr13-55050567-INS-50 |
| chr13-55062423-INS-72 | chr13-55062423-INS-64 |
| chr13-55164978-INS-62 | chr13-55164963-INS-58 |
| chr13-55167438-INS-265 | chr13-55167438-INS-265 |
| chr13-55178405-INS-70 | chr13-55178390-INS-66 |
| chr13-55192431-INS-221 | chr13-55192431-INS-221 |
| chr13-55198471-INS-57 | chr13-55198472-INS-53 |
| chr13-55215766-INS-161 | chr13-55215766-INS-132 |
| chr13-55237747-INS-65 | chr13-55237747-INS-65 |
| chr13-55240307-INS-2820 | chr13-55240308-INS-2820 |
| chr13-55274698-INS-70 | chr13-55274698-INS-70 |
| chr13-55279289-INS-271 | chr13-55279289-INS-239 |
| chr13-55317833-INS-212 | chr13-55317833-INS-212 |
| chr13-55319061-INS-203 | chr13-55319061-INS-206 |
| chr13-55343468-INS-277 | chr13-55343468-INS-241 |
| chr13-55345061-INS-68 | chr13-55345061-INS-60 |
| chr13-55347787-INS-139 | chr13-55347789-INS-138 |
| chr13-55366086-INS-62 | chr13-55366086-INS-62 |
| chr13-55403162-INS-212 | chr13-55403162-INS-212 |
| chr13-55408012-INS-200 | chr13-55408012-INS-200 |
| chr13-55471424-INS-1892 | chr13-55471425-INS-1892 |
| chr13-55501414-INS-285 | chr13-55501417-INS-276 |
| chr13-55503720-INS-82 | chr13-55503720-INS-82 |
| chr13-55507464-INS-51 | chr13-55507464-INS-51 |
| chr13-55513370-INS-214 | chr13-55513370-INS-215 |
| chr13-55513766-INS-330 | chr13-55513766-INS-331 |
| chr13-55536657-INS-111 | chr13-55536657-INS-111 |
| chr13-55542324-INS-206 | chr13-55542324-INS-206 |
| chr13-55565773-INS-223 | chr13-55565773-INS-224 |
| chr13-55572732-INS-157 | chr13-55572732-INS-163 |
| chr13-55589200-INS-175 | chr13-55589192-INS-177 |
| chr13-55603184-INS-187 | chr13-55603184-INS-187 |
| chr13-55652354-INS-52 | chr13-55652309-INS-71 |
| chr13-55715946-INS-194 | chr13-55715946-INS-194 |
| chr13-55725226-INS-175 | chr13-55725226-INS-185 |
| chr13-55732228-INS-935 | chr13-55732228-INS-939 |
| chr13-55733099-INS-172 | chr13-55733099-INS-172 |
| chr13-55767138-INS-391 | chr13-55767138-INS-392 |
| chr13-55768791-INS-187 | chr13-55768785-INS-193 |
| chr13-55769335-INS-67 | chr13-55769327-INS-60 |
| chr13-55774760-INS-168 | chr13-55774760-INS-168 |
| chr13-55815121-INS-242 | chr13-55815121-INS-243 |
| chr13-55820911-INS-184 | chr13-55820911-INS-184 |
| chr13-55831219-INS-206 | chr13-55831219-INS-206 |
| chr13-55839091-INS-450 | chr13-55839091-INS-456 |
| chr13-55848635-INS-96 | chr13-55848635-INS-92 |
| chr13-55852557-INS-195 | chr13-55852557-INS-195 |
| chr13-55860305-INS-203 | chr13-55860303-INS-207 |
| chr13-55885075-INS-206 | chr13-55885075-INS-207 |
| chr13-55907737-INS-74 | chr13-55907765-INS-84 |
| chr13-55920677-INS-77 | chr13-55920646-INS-63 |
| chr13-55926569-INS-359 | chr13-55926569-INS-360 |

| | |
|---|---|
| chr13-55933317-INS-60 | chr13-55933317-INS-60 |
| chr13-55947400-INS-104 | chr13-55947400-INS-96 |
| chr13-55955108-INS-70 | chr13-55955108-INS-60 |
| chr13-55961424-INS-60 | chr13-55961424-INS-56 |
| chr13-55997924-INS-112 | chr13-55997924-INS-126 |
| chr13-56009568-INS-233 | chr13-56009568-INS-262 |
| chr13-56014593-INS-74 | chr13-56014593-INS-74 |
| chr13-56016641-INS-589 | chr13-56016641-INS-593 |
| chr13-56039800-INS-63 | chr13-56039800-INS-63 |
| chr13-56047197-INS-64 | chr13-56047239-INS-70 |
| chr13-56143334-INS-5544 | chr13-56143335-INS-5544 |
| chr13-56169359-INS-54 | chr13-56169389-INS-54 |
| chr13-56182630-INS-202 | chr13-56182630-INS-202 |
| chr13-56188153-INS-78 | chr13-56188153-INS-84 |
| chr13-56190213-INS-52 | chr13-56190213-INS-52 |
| chr13-56195089-INS-1074 | chr13-56195090-INS-1074 |
| chr13-56227588-INS-121 | chr13-56227588-INS-121 |
| chr13-56237051-INS-132 | chr13-56237051-INS-132 |
| chr13-56245941-INS-90 | chr13-56245941-INS-92 |
| chr13-56247216-INS-52 | chr13-56247216-INS-52 |
| chr13-56254750-INS-62 | chr13-56254750-INS-62 |
| chr13-56288119-INS-71 | chr13-56288119-INS-71 |
| chr13-56289421-INS-66 | chr13-56289415-INS-90 |
| chr13-56303000-INS-292 | chr13-56303000-INS-293 |
| chr13-56316016-INS-61 | chr13-56316016-INS-62 |
| chr13-56333975-INS-91 | chr13-56333975-INS-62 |
| chr13-56335709-INS-358 | chr13-56335709-INS-358 |
| chr13-56336316-INS-5233 | chr13-56336317-INS-5233 |
| chr13-56337843-INS-1898 | chr13-56337844-INS-1898 |
| chr13-56348353-INS-626 | chr13-56348353-INS-635 |
| chr13-56353160-INS-89 | chr13-56353131-INS-75 |
| chr13-56357681-INS-404 | chr13-56357687-INS-399 |
| chr13-56400485-INS-133 | chr13-56400478-INS-132 |
| chr13-56409486-INS-66 | chr13-56409486-INS-66 |
| chr13-56410677-INS-5142 | chr13-56410678-INS-5142 |
| chr13-56412021-INS-223 | chr13-56412021-INS-223 |
| chr13-56422979-INS-155 | chr13-56422979-INS-185 |
| chr13-56433621-INS-556 | chr13-56433621-INS-559 |
| chr13-56435756-INS-109 | chr13-56435756-INS-109 |
| chr13-56446093-INS-230 | chr13-56446093-INS-230 |
| chr13-56461823-INS-60 | chr13-56461823-INS-60 |
| chr13-56486384-INS-59 | chr13-56486384-INS-59 |
| chr13-56491071-INS-5348 | chr13-56491072-INS-5348 |
| chr13-56509184-INS-1606 | chr13-56509185-INS-1634 |
| chr13-56556314-INS-731 | chr13-56556314-INS-731 |
| chr13-56557667-INS-56 | chr13-56557647-INS-77 |
| chr13-56579105-INS-74 | chr13-56579105-INS-74 |
| chr13-56604847-INS-414 | chr13-56604843-INS-423 |
| chr13-56631666-INS-62 | chr13-56631666-INS-62 |
| chr13-56639955-INS-3564 | chr13-56639956-INS-3564 |
| chr13-56665789-INS-65 | chr13-56665785-INS-57 |
| chr13-56683730-INS-106 | chr13-56683739-INS-72 |
| chr13-56694295-INS-76 | chr13-56694343-INS-53 |
| chr13-56698118-INS-1891 | chr13-56698119-INS-1891 |
| chr13-56739788-INS-59 | chr13-56739788-INS-59 |
| chr13-56750102-INS-76 | chr13-56750102-INS-76 |
| chr13-56797559-INS-194 | chr13-56797559-INS-196 |
| chr13-56817700-INS-62 | chr13-56817700-INS-64 |
| chr13-56838117-INS-74 | chr13-56838118-INS-62 |
| chr13-56839589-INS-749 | chr13-56839589-INS-743 |
| chr13-56842010-INS-69 | chr13-56842010-INS-69 |
| chr13-56879721-INS-76 | chr13-56879721-INS-76 |
| chr13-56894982-INS-366 | chr13-56894982-INS-366 |
| chr13-56930973-INS-202 | chr13-56930973-INS-202 |
| chr13-56947616-INS-190 | chr13-56947617-INS-190 |
| chr13-56965743-INS-61 | chr13-56965748-INS-69 |
| chr13-56983608-INS-185 | chr13-56983608-INS-185 |
| chr13-56998285-INS-64 | chr13-56998285-INS-64 |
| chr13-57021415-INS-213 | chr13-57021415-INS-204 |
| chr13-57053155-INS-922 | chr13-57053155-INS-922 |
| chr13-57079672-INS-5569 | chr13-57079673-INS-5569 |
| chr13-57080666-INS-51 | chr13-57080666-INS-51 |
| chr13-57111588-INS-104 | chr13-57111588-INS-104 |
| chr13-57137225-INS-57 | chr13-57137225-INS-57 |
| chr13-57224531-INS-7098 | chr13-57224532-INS-7098 |
| chr13-57224787-INS-476 | chr13-57224787-INS-476 |
| chr13-57240826-INS-542 | chr13-57240826-INS-542 |

| | |
|---|---|
| chr13-57244889-INS-722 | chr13-57244889-INS-723 |
| chr13-57248932-INS-326 | chr13-57248932-INS-326 |
| chr13-57266761-INS-6440 | chr13-57266762-INS-6440 |
| chr13-57277184-INS-72 | chr13-57277184-INS-84 |
| chr13-57292997-INS-979 | chr13-57292997-INS-979 |
| chr13-57311577-INS-218 | chr13-57311577-INS-218 |
| chr13-57312363-INS-6735 | chr13-57312364-INS-6739 |
| chr13-57323464-INS-1017 | chr13-57323465-INS-1018 |
| chr13-57326059-INS-1164 | chr13-57326060-INS-1164 |
| chr13-57329956-INS-51 | chr13-57329956-INS-54 |
| chr13-57340621-INS-57 | chr13-57340621-INS-57 |
| chr13-57345217-INS-62 | chr13-57345217-INS-62 |
| chr13-57369411-INS-433 | chr13-57369411-INS-433 |
| chr13-57373990-INS-58 | chr13-57373990-INS-60 |
| chr13-57404314-INS-53 | chr13-57404314-INS-53 |
| chr13-57412927-INS-1850 | chr13-57412929-INS-1845 |
| chr13-57469908-INS-56 | chr13-57469908-INS-56 |
| chr13-57474661-INS-68 | chr13-57474661-INS-70 |
| chr13-57475676-INS-140 | chr13-57475676-INS-140 |
| chr13-57500417-INS-523 | chr13-57500417-INS-523 |
| chr13-57504873-INS-4776 | chr13-57504874-INS-4775 |
| chr13-57522792-INS-51 | chr13-57522792-INS-51 |
| chr13-57530445-INS-177 | chr13-57530395-INS-181 |
| chr13-57539135-INS-2056 | chr13-57539136-INS-2056 |
| chr13-57539261-INS-98 | chr13-57539261-INS-98 |
| chr13-57556950-INS-62 | chr13-57556950-INS-62 |
| chr13-57593789-INS-6262 | chr13-57593790-INS-6262 |
| chr13-57601953-INS-98 | chr13-57601953-INS-122 |
| chr13-57614101-INS-506 | chr13-57614101-INS-506 |
| chr13-57616392-INS-89 | chr13-57616392-INS-89 |
| chr13-57643109-INS-79 | chr13-57643109-INS-79 |
| chr13-57652965-INS-126 | chr13-57652965-INS-126 |
| chr13-57680395-INS-64 | chr13-57680395-INS-64 |
| chr13-57687748-INS-52 | chr13-57687748-INS-52 |
| chr13-57689236-INS-654 | chr13-57689236-INS-679 |
| chr13-57693127-INS-54 | chr13-57693127-INS-54 |
| chr13-57699388-INS-80 | chr13-57699388-INS-80 |
| chr13-57726080-INS-174 | chr13-57726077-INS-173 |
| chr13-57779210-INS-998 | chr13-57779210-INS-996 |
| chr13-57801506-INS-53 | chr13-57801506-INS-53 |
| chr13-57858412-INS-85 | chr13-57858431-INS-114 |
| chr13-57902660-INS-1890 | chr13-57902661-INS-1890 |
| chr13-57909446-INS-52 | chr13-57909446-INS-52 |
| chr13-57967169-INS-115 | chr13-57967169-INS-115 |
| chr13-58020023-INS-287 | chr13-58020023-INS-287 |
| chr13-58043915-INS-68 | chr13-58043916-INS-68 |
| chr13-58058069-INS-60 | chr13-58058069-INS-54 |
| chr13-58077685-INS-156 | chr13-58077685-INS-174 |
| chr13-58084687-INS-145 | chr13-58084688-INS-138 |
| chr13-58124094-INS-429 | chr13-58124094-INS-429 |
| chr13-58127536-INS-130 | chr13-58127536-INS-120 |
| chr13-58170236-INS-8458 | chr13-58170237-INS-8458 |
| chr13-58210004-INS-90 | chr13-58210016-INS-90 |
| chr13-58220725-INS-55 | chr13-58220725-INS-56 |
| chr13-58222368-INS-104 | chr13-58222368-INS-104 |
| chr13-58250812-INS-58 | chr13-58250812-INS-58 |
| chr13-58258495-INS-152 | chr13-58258495-INS-114 |
| chr13-58272365-INS-344 | chr13-58272365-INS-344 |
| chr13-58286690-INS-51 | chr13-58286690-INS-51 |
| chr13-58288558-INS-467 | chr13-58288558-INS-473 |
| chr13-58290204-INS-208 | chr13-58290204-INS-208 |
| chr13-58294314-INS-441 | chr13-58294314-INS-441 |
| chr13-58297979-INS-51 | chr13-58297960-INS-62 |
| chr13-58306121-INS-76 | chr13-58306121-INS-58 |
| chr13-58328895-INS-67 | chr13-58328895-INS-51 |
| chr13-58331459-INS-205 | chr13-58331459-INS-205 |
| chr13-58335362-INS-208 | chr13-58335362-INS-206 |
| chr13-58337573-INS-214 | chr13-58337573-INS-214 |
| chr13-58358075-INS-119 | chr13-58358075-INS-89 |
| chr13-58360969-INS-89 | chr13-58360969-INS-90 |
| chr13-58364648-INS-72 | chr13-58364648-INS-74 |
| chr13-58368501-INS-182 | chr13-58368501-INS-151 |
| chr13-58419602-INS-62 | chr13-58419602-INS-62 |
| chr13-58444330-INS-212 | chr13-58444330-INS-212 |
| chr13-58446243-INS-124 | chr13-58446278-INS-93 |
| chr13-58478508-INS-289 | chr13-58478503-INS-296 |
| chr13-58503640-INS-6415 | chr13-58503641-INS-6415 |

| | |
|---|---|
| chr13-58511658-INS-2757 | chr13-58511659-INS-2757 |
| chr13-58516165-INS-165 | chr13-58516165-INS-175 |
| chr13-58588191-INS-440 | chr13-58588191-INS-400 |
| chr13-58622905-INS-7329 | chr13-58622906-INS-7329 |
| chr13-58654368-INS-51 | chr13-58654368-INS-51 |
| chr13-58657415-INS-58 | chr13-58657415-INS-58 |
| chr13-58658570-INS-439 | chr13-58658570-INS-439 |
| chr13-58671450-INS-729 | chr13-58671450-INS-729 |
| chr13-58672134-INS-161 | chr13-58672134-INS-185 |
| chr13-58675147-INS-68 | chr13-58675147-INS-70 |
| chr13-58676128-INS-202 | chr13-58676128-INS-203 |
| chr13-58682893-INS-65 | chr13-58682893-INS-65 |
| chr13-58688731-INS-419 | chr13-58688731-INS-419 |
| chr13-58715270-INS-722 | chr13-58715270-INS-722 |
| chr13-58728035-INS-248 | chr13-58728035-INS-241 |
| chr13-58750929-INS-124 | chr13-58750929-INS-124 |
| chr13-58759318-INS-98 | chr13-58759318-INS-95 |
| chr13-58802300-INS-60 | chr13-58802301-INS-60 |
| chr13-58813958-INS-52 | chr13-58813958-INS-52 |
| chr13-58835863-INS-56 | chr13-58835863-INS-64 |
| chr13-58837299-INS-63 | chr13-58837299-INS-63 |
| chr13-58850622-INS-58 | chr13-58850616-INS-56 |
| chr13-58860915-INS-312 | chr13-58860915-INS-310 |
| chr13-58872435-INS-119 | chr13-58872435-INS-119 |
| chr13-58874435-INS-55 | chr13-58874435-INS-55 |
| chr13-58882290-INS-60 | chr13-58882287-INS-63 |
| chr13-58905065-INS-74 | chr13-58905062-INS-76 |
| chr13-58917790-INS-75 | chr13-58917790-INS-113 |
| chr13-58918794-INS-444 | chr13-58918794-INS-442 |
| chr13-59005269-INS-62 | chr13-59005269-INS-62 |
| chr13-59015541-INS-125 | chr13-59015541-INS-100 |
| chr13-59033792-INS-74 | chr13-59033792-INS-79 |
| chr13-59053286-INS-57 | chr13-59053286-INS-57 |
| chr13-59072954-INS-179 | chr13-59072954-INS-179 |
| chr13-59147818-INS-269 | chr13-59147818-INS-269 |
| chr13-59149130-INS-5292 | chr13-59149131-INS-5292 |
| chr13-59167922-INS-64 | chr13-59167922-INS-64 |
| chr13-59184956-INS-50 | chr13-59184963-INS-62 |
| chr13-59203697-INS-87 | chr13-59203697-INS-87 |
| chr13-59216907-INS-68 | chr13-59216907-INS-68 |
| chr13-59255428-INS-212 | chr13-59255426-INS-212 |
| chr13-59287254-INS-360 | chr13-59287255-INS-360 |
| chr13-59298105-INS-1893 | chr13-59298106-INS-1893 |
| chr13-59303822-INS-180 | chr13-59303822-INS-178 |
| chr13-59322132-INS-54 | chr13-59322133-INS-73 |
| chr13-59336402-INS-309 | chr13-59336402-INS-310 |
| chr13-59343860-INS-5201 | chr13-59343861-INS-5201 |
| chr13-59355082-INS-54 | chr13-59355082-INS-54 |
| chr13-59372415-INS-64 | chr13-59372412-INS-61 |
| chr13-59373014-INS-513 | chr13-59373014-INS-515 |
| chr13-59389704-INS-1104 | chr13-59389705-INS-1104 |
| chr13-59391978-INS-188 | chr13-59391978-INS-200 |
| chr13-59397353-INS-64 | chr13-59397379-INS-70 |
| chr13-59400547-INS-222 | chr13-59400547-INS-222 |
| chr13-59404748-INS-509 | chr13-59404751-INS-522 |
| chr13-59408573-INS-54 | chr13-59408566-INS-51 |
| chr13-59436463-INS-80 | chr13-59436463-INS-50 |
| chr13-59438842-INS-118 | chr13-59438842-INS-162 |
| chr13-59457449-INS-60 | chr13-59457401-INS-92 |
| chr13-59490491-INS-763 | chr13-59490460-INS-763 |
| chr13-59492590-INS-2317 | chr13-59492591-INS-2317 |
| chr13-59530831-INS-159 | chr13-59530831-INS-159 |
| chr13-59535201-INS-612 | chr13-59535201-INS-612 |
| chr13-59538415-INS-378 | chr13-59538416-INS-376 |
| chr13-59544441-INS-72 | chr13-59544435-INS-62 |
| chr13-59565100-INS-71 | chr13-59565074-INS-62 |
| chr13-59606308-INS-92 | chr13-59606308-INS-92 |
| chr13-59607000-INS-171 | chr13-59607000-INS-171 |
| chr13-59612012-INS-227 | chr13-59612013-INS-220 |
| chr13-59628493-INS-510 | chr13-59628493-INS-509 |
| chr13-59644482-INS-70 | chr13-59644482-INS-70 |
| chr13-59652678-INS-56 | chr13-59652708-INS-56 |
| chr13-59657822-INS-493 | chr13-59657822-INS-489 |
| chr13-59672535-INS-135 | chr13-59672535-INS-135 |
| chr13-59686564-INS-174 | chr13-59686564-INS-174 |
| chr13-59715653-INS-922 | chr13-59715653-INS-923 |
| chr13-59720203-INS-62 | chr13-59720203-INS-62 |

| | |
|---|---|
| chr13-59838075-INS-199 | chr13-59838075-INS-199 |
| chr13-59844158-INS-3143 | chr13-59844159-INS-3143 |
| chr13-59860853-INS-73 | chr13-59860853-INS-73 |
| chr13-59885925-INS-71 | chr13-59885921-INS-73 |
| chr13-59895158-INS-203 | chr13-59895158-INS-233 |
| chr13-59897011-INS-1004 | chr13-59897012-INS-1004 |
| chr13-59904415-INS-75 | chr13-59904415-INS-75 |
| chr13-59907834-INS-204 | chr13-59907834-INS-204 |
| chr13-59909412-INS-204 | chr13-59909412-INS-215 |
| chr13-59912281-INS-2773 | chr13-59912282-INS-2772 |
| chr13-60008073-INS-76 | chr13-60008073-INS-86 |
| chr13-60032218-INS-1718 | chr13-60032219-INS-1718 |
| chr13-60043713-INS-60 | chr13-60043713-INS-60 |
| chr13-60052136-INS-76 | chr13-60052136-INS-76 |
| chr13-60060523-INS-498 | chr13-60060523-INS-498 |
| chr13-60080759-INS-56 | chr13-60080760-INS-52 |
| chr13-60092949-INS-205 | chr13-60092949-INS-202 |
| chr13-60096907-INS-6381 | chr13-60096908-INS-6381 |
| chr13-60102605-INS-121 | chr13-60102605-INS-121 |
| chr13-60116171-INS-197 | chr13-60116171-INS-197 |
| chr13-60133207-INS-64 | chr13-60133206-INS-78 |
| chr13-60154020-INS-56 | chr13-60154020-INS-56 |
| chr13-60176971-INS-118 | chr13-60176943-INS-120 |
| chr13-60187871-INS-623 | chr13-60187871-INS-623 |
| chr13-60194500-INS-68 | chr13-60194500-INS-64 |
| chr13-60196435-INS-91 | chr13-60196434-INS-91 |
| chr13-60209365-INS-118 | chr13-60209365-INS-119 |
| chr13-60218400-INS-515 | chr13-60218400-INS-515 |
| chr13-60223387-INS-52 | chr13-60223387-INS-54 |
| chr13-60237764-INS-136 | chr13-60237765-INS-136 |
| chr13-60240196-INS-412 | chr13-60240196-INS-412 |
| chr13-60241262-INS-100 | chr13-60241262-INS-60 |
| chr13-60244574-INS-69 | chr13-60244575-INS-50 |
| chr13-60254418-INS-139 | chr13-60254419-INS-132 |
| chr13-60256119-INS-539 | chr13-60256119-INS-539 |
| chr13-60260678-INS-1894 | chr13-60260679-INS-1894 |
| chr13-60289166-INS-104 | chr13-60289166-INS-104 |
| chr13-60300156-INS-154 | chr13-60300153-INS-107 |
| chr13-60356783-INS-804 | chr13-60356783-INS-804 |
| chr13-60364070-INS-212 | chr13-60364070-INS-212 |
| chr13-60441735-INS-410 | chr13-60441735-INS-410 |
| chr13-60445772-INS-68 | chr13-60445773-INS-84 |
| chr13-60447397-INS-59 | chr13-60447397-INS-71 |
| chr13-60471823-INS-175 | chr13-60471823-INS-175 |
| chr13-60481723-INS-65 | chr13-60481711-INS-59 |
| chr13-60502908-INS-52 | chr13-60502908-INS-58 |
| chr13-60507943-INS-92 | chr13-60507944-INS-92 |
| chr13-60519922-INS-58 | chr13-60519922-INS-54 |
| chr13-60522984-INS-161 | chr13-60522984-INS-156 |
| chr13-60534837-INS-577 | chr13-60534837-INS-577 |
| chr13-60546315-INS-1886 | chr13-60546316-INS-1886 |
| chr13-60552504-INS-75 | chr13-60552536-INS-77 |
| chr13-60574988-INS-108 | chr13-60574988-INS-75 |
| chr13-60576464-INS-517 | chr13-60576464-INS-511 |
| chr13-60586565-INS-97 | chr13-60586561-INS-96 |
| chr13-60591558-INS-173 | chr13-60591558-INS-124 |
| chr13-60618114-INS-215 | chr13-60618114-INS-215 |
| chr13-60622026-INS-158 | chr13-60622011-INS-199 |
| chr13-60631279-INS-82 | chr13-60631286-INS-76 |
| chr13-60652666-INS-60 | chr13-60652666-INS-60 |
| chr13-60657341-INS-7260 | chr13-60657342-INS-7260 |
| chr13-60673242-INS-487 | chr13-60673242-INS-487 |
| chr13-60675178-INS-71 | chr13-60675178-INS-72 |
| chr13-60682232-INS-164 | chr13-60682232-INS-164 |
| chr13-60684215-INS-432 | chr13-60684215-INS-432 |
| chr13-60690409-INS-211 | chr13-60690409-INS-213 |
| chr13-60731933-INS-98 | chr13-60731933-INS-78 |
| chr13-60740657-INS-729 | chr13-60740657-INS-724 |
| chr13-60778925-INS-227 | chr13-60778925-INS-227 |
| chr13-60782648-INS-839 | chr13-60782648-INS-839 |
| chr13-60788033-INS-208 | chr13-60788033-INS-202 |
| chr13-60789237-INS-199 | chr13-60789237-INS-174 |
| chr13-60805533-INS-207 | chr13-60805533-INS-199 |
| chr13-60846386-INS-104 | chr13-60846386-INS-104 |
| chr13-60875523-INS-153 | chr13-60875523-INS-153 |
| chr13-60900300-INS-108 | chr13-60900300-INS-108 |
| chr13-60902198-INS-92 | chr13-60902198-INS-62 |

| | |
|---|---|
| chr13-60903363-INS-89 | chr13-60903364-INS-89 |
| chr13-60917100-INS-52 | chr13-60917100-INS-52 |
| chr13-60941691-INS-72 | chr13-60941691-INS-84 |
| chr13-61064477-INS-6002 | chr13-61064478-INS-6003 |
| chr13-61090905-INS-3530 | chr13-61090906-INS-3530 |
| chr13-61145699-INS-94 | chr13-61145701-INS-75 |
| chr13-61181679-INS-390 | chr13-61181679-INS-391 |
| chr13-61186489-INS-71 | chr13-61186490-INS-90 |
| chr13-61192598-INS-51 | chr13-61192596-INS-84 |
| chr13-61292571-INS-70 | chr13-61292571-INS-70 |
| chr13-61305899-INS-195 | chr13-61305899-INS-195 |
| chr13-61311826-INS-56 | chr13-61311826-INS-56 |
| chr13-61373478-INS-4354 | chr13-61373479-INS-4355 |
| chr13-61389126-INS-181 | chr13-61389126-INS-181 |
| chr13-61390342-INS-358 | chr13-61390342-INS-358 |
| chr13-61392959-INS-2497 | chr13-61392960-INS-2496 |
| chr13-61420149-INS-144 | chr13-61420149-INS-145 |
| chr13-61421549-INS-199 | chr13-61421549-INS-199 |
| chr13-61445253-INS-2302 | chr13-61445254-INS-2315 |
| chr13-61453452-INS-1178 | chr13-61453453-INS-1178 |
| chr13-61456629-INS-180 | chr13-61456629-INS-194 |
| chr13-61475712-INS-6552 | chr13-61475713-INS-6552 |
| chr13-61476319-INS-1082 | chr13-61476320-INS-1081 |
| chr13-61520149-INS-221 | chr13-61520149-INS-224 |
| chr13-61527950-INS-626 | chr13-61527950-INS-626 |
| chr13-61533429-INS-60 | chr13-61533429-INS-60 |
| chr13-61585124-INS-183 | chr13-61585124-INS-183 |
| chr13-61593953-INS-876 | chr13-61593953-INS-876 |
| chr13-61608950-INS-1134 | chr13-61608951-INS-1134 |
| chr13-61620284-INS-507 | chr13-61620284-INS-477 |
| chr13-61622306-INS-72 | chr13-61622306-INS-72 |
| chr13-61629348-INS-74 | chr13-61629349-INS-74 |
| chr13-61640672-INS-311 | chr13-61640672-INS-311 |
| chr13-61643694-INS-194 | chr13-61643694-INS-194 |
| chr13-61648477-INS-765 | chr13-61648477-INS-763 |
| chr13-61652286-INS-571 | chr13-61652286-INS-571 |
| chr13-61654561-INS-193 | chr13-61654561-INS-193 |
| chr13-61720727-INS-6023 | chr13-61720724-INS-6023 |
| chr13-61729504-INS-6202 | chr13-61729505-INS-6202 |
| chr13-61751426-INS-629 | chr13-61751426-INS-629 |
| chr13-61754604-INS-51 | chr13-61754586-INS-62 |
| chr13-61760701-INS-282 | chr13-61760701-INS-282 |
| chr13-61763000-INS-470 | chr13-61763000-INS-446 |
| chr13-61777809-INS-178 | chr13-61777809-INS-178 |
| chr13-61783672-INS-91 | chr13-61783672-INS-91 |
| chr13-61793682-INS-493 | chr13-61793682-INS-493 |
| chr13-61907018-INS-187 | chr13-61907019-INS-187 |
| chr13-62166442-INS-72 | chr13-62166442-INS-72 |
| chr13-62202154-INS-179 | chr13-62202154-INS-179 |
| chr13-62366299-INS-87 | chr13-62366299-INS-87 |
| chr13-62385772-INS-52 | chr13-62385772-INS-52 |
| chr13-62389720-INS-202 | chr13-62389720-INS-202 |
| chr13-62404205-INS-1026 | chr13-62404206-INS-1026 |
| chr13-62408341-INS-7110 | chr13-62408342-INS-7110 |
| chr13-62413306-INS-51 | chr13-62413308-INS-50 |
| chr13-62422346-INS-84 | chr13-62422346-INS-84 |
| chr13-62438333-INS-4912 | chr13-62438334-INS-4912 |
| chr13-62453214-INS-863 | chr13-62453214-INS-863 |
| chr13-62456572-INS-4246 | chr13-62456573-INS-4246 |
| chr13-62467108-INS-1864 | chr13-62467109-INS-1864 |
| chr13-62484520-INS-222 | chr13-62484520-INS-223 |
| chr13-62499891-INS-84 | chr13-62499891-INS-84 |
| chr13-62517890-INS-168 | chr13-62517890-INS-168 |
| chr13-62520157-INS-174 | chr13-62520157-INS-174 |
| chr13-62534294-INS-241 | chr13-62534294-INS-237 |
| chr13-62582891-INS-55 | chr13-62582891-INS-105 |
| chr13-62617613-INS-206 | chr13-62617613-INS-218 |
| chr13-62690084-INS-590 | chr13-62690084-INS-606 |
| chr13-62690795-INS-5391 | chr13-62690796-INS-5391 |
| chr13-62710973-INS-80 | chr13-62710950-INS-66 |
| chr13-62715846-INS-81 | chr13-62715846-INS-81 |
| chr13-62718461-INS-203 | chr13-62718462-INS-198 |
| chr13-62747004-INS-558 | chr13-62747004-INS-559 |
| chr13-62752055-INS-84 | chr13-62752055-INS-84 |
| chr13-62770110-INS-62 | chr13-62770110-INS-62 |
| chr13-62771116-INS-68 | chr13-62771116-INS-68 |
| chr13-62806459-INS-172 | chr13-62806459-INS-172 |

| | |
|---|---|
| chr13-62824767-INS-764 | chr13-62824767-INS-764 |
| chr13-62838776-INS-122 | chr13-62838776-INS-122 |
| chr13-62907799-INS-202 | chr13-62907799-INS-203 |
| chr13-62918656-INS-54 | chr13-62918656-INS-54 |
| chr13-62927201-INS-838 | chr13-62927201-INS-838 |
| chr13-62935727-INS-190 | chr13-62935727-INS-197 |
| chr13-62949782-INS-6500 | chr13-62949783-INS-6502 |
| chr13-62973862-INS-220 | chr13-62973862-INS-220 |
| chr13-62979588-INS-63 | chr13-62979587-INS-99 |
| chr13-62986466-INS-80 | chr13-62986466-INS-80 |
| chr13-62992775-INS-219 | chr13-62992775-INS-219 |
| chr13-62998419-INS-136 | chr13-62998419-INS-136 |
| chr13-63008225-INS-82 | chr13-63008225-INS-82 |
| chr13-63014424-INS-57 | chr13-63014424-INS-59 |
| chr13-63054137-INS-72 | chr13-63054137-INS-76 |
| chr13-63061906-INS-169 | chr13-63061906-INS-169 |
| chr13-63065178-INS-218 | chr13-63065178-INS-218 |
| chr13-63068771-INS-59 | chr13-63068783-INS-50 |
| chr13-63079477-INS-533 | chr13-63079477-INS-533 |
| chr13-63093367-INS-2185 | chr13-63093368-INS-2185 |
| chr13-63096418-INS-7332 | chr13-63096419-INS-7332 |
| chr13-63096839-INS-79 | chr13-63096843-INS-81 |
| chr13-63131704-INS-7279 | chr13-63131705-INS-7279 |
| chr13-63149186-INS-4775 | chr13-63149187-INS-4775 |
| chr13-63166368-INS-64 | chr13-63166368-INS-64 |
| chr13-63176337-INS-645 | chr13-63176337-INS-645 |
| chr13-63177691-INS-504 | chr13-63177691-INS-503 |
| chr13-63197457-INS-69 | chr13-63197457-INS-69 |
| chr13-63206311-INS-460 | chr13-63206311-INS-464 |
| chr13-63214794-INS-400 | chr13-63214794-INS-400 |
| chr13-63293573-INS-193 | chr13-63293573-INS-193 |
| chr13-63301639-INS-84 | chr13-63301639-INS-84 |
| chr13-63313578-INS-208 | chr13-63313578-INS-208 |
| chr13-63331012-INS-135 | chr13-63331003-INS-118 |
| chr13-63331803-INS-7126 | chr13-63331804-INS-7126 |
| chr13-63340863-INS-59 | chr13-63340874-INS-59 |
| chr13-63343761-INS-254 | chr13-63343768-INS-272 |
| chr13-63345531-INS-207 | chr13-63345531-INS-207 |
| chr13-63400065-INS-116 | chr13-63400065-INS-116 |
| chr13-63424913-INS-51 | chr13-63424906-INS-85 |
| chr13-63440767-INS-90 | chr13-63440767-INS-90 |
| chr13-63476056-INS-195 | chr13-63476056-INS-195 |
| chr13-63487007-INS-212 | chr13-63487007-INS-170 |
| chr13-63489785-INS-195 | chr13-63489785-INS-196 |
| chr13-63492998-INS-84 | chr13-63493004-INS-80 |
| chr13-63497538-INS-337 | chr13-63497538-INS-337 |
| chr13-63498565-INS-52 | chr13-63498565-INS-52 |
| chr13-63500488-INS-8986 | chr13-63500489-INS-8986 |
| chr13-63508449-INS-358 | chr13-63508449-INS-358 |
| chr13-63528687-INS-89 | chr13-63528687-INS-89 |
| chr13-63530039-INS-73 | chr13-63530047-INS-81 |
| chr13-63548181-INS-68 | chr13-63548181-INS-68 |
| chr13-63559377-INS-95 | chr13-63559370-INS-134 |
| chr13-63577859-INS-491 | chr13-63577859-INS-491 |
| chr13-63600348-INS-211 | chr13-63600348-INS-201 |
| chr13-63606331-INS-1890 | chr13-63606332-INS-1890 |
| chr13-63611932-INS-183 | chr13-63611946-INS-197 |
| chr13-63622966-INS-1904 | chr13-63622966-INS-1898 |
| chr13-63648951-INS-662 | chr13-63648951-INS-666 |
| chr13-63657237-INS-72 | chr13-63657231-INS-50 |
| chr13-63668971-INS-60 | chr13-63668971-INS-60 |
| chr13-63686773-INS-52 | chr13-63686773-INS-52 |
| chr13-63725314-INS-71 | chr13-63725314-INS-93 |
| chr13-63738290-INS-52 | chr13-63738290-INS-52 |
| chr13-63765171-INS-7076 | chr13-63765172-INS-7076 |
| chr13-63770453-INS-56 | chr13-63770416-INS-52 |
| chr13-63811315-INS-149 | chr13-63811315-INS-149 |
| chr13-63861329-INS-159 | chr13-63861329-INS-159 |
| chr13-63869484-INS-450 | chr13-63869484-INS-425 |
| chr13-63883070-INS-186 | chr13-63883070-INS-136 |
| chr13-63899625-INS-411 | chr13-63899625-INS-411 |
| chr13-63902265-INS-6263 | chr13-63902266-INS-6263 |
| chr13-63912195-INS-209 | chr13-63912195-INS-210 |
| chr13-63924974-INS-425 | chr13-63924974-INS-407 |
| chr13-63956838-INS-98 | chr13-63956838-INS-98 |
| chr13-64000359-INS-53 | chr13-64000359-INS-53 |
| chr13-64048485-INS-64 | chr13-64048485-INS-58 |

| | |
|---|---|
| chr13-64065909-INS-58 | chr13-64065912-INS-55 |
| chr13-64073658-INS-58 | chr13-64073658-INS-58 |
| chr13-64083812-INS-54 | chr13-64083812-INS-54 |
| chr13-64120375-INS-182 | chr13-64120375-INS-174 |
| chr13-64132061-INS-50 | chr13-64132061-INS-52 |
| chr13-64136725-INS-213 | chr13-64136725-INS-215 |
| chr13-64141556-INS-100 | chr13-64141556-INS-100 |
| chr13-64143738-INS-62 | chr13-64143723-INS-81 |
| chr13-64145284-INS-80 | chr13-64145284-INS-80 |
| chr13-64176652-INS-62 | chr13-64176652-INS-60 |
| chr13-64191951-INS-69 | chr13-64191951-INS-69 |
| chr13-64204476-INS-2820 | chr13-64204477-INS-2820 |
| chr13-64232230-INS-81 | chr13-64232230-INS-95 |
| chr13-64253440-INS-161 | chr13-64253440-INS-161 |
| chr13-64261500-INS-182 | chr13-64261500-INS-183 |
| chr13-64284893-INS-200 | chr13-64284893-INS-200 |
| chr13-64323522-INS-62 | chr13-64323522-INS-67 |
| chr13-64329141-INS-189 | chr13-64329141-INS-189 |
| chr13-64341493-INS-219 | chr13-64341493-INS-219 |
| chr13-64352088-INS-244 | chr13-64352088-INS-244 |
| chr13-64354716-INS-63 | chr13-64354742-INS-69 |
| chr13-64360258-INS-79 | chr13-64360258-INS-79 |
| chr13-64363142-INS-51 | chr13-64363142-INS-51 |
| chr13-64379089-INS-187 | chr13-64379089-INS-187 |
| chr13-64399908-INS-402 | chr13-64399908-INS-402 |
| chr13-64426461-INS-213 | chr13-64426461-INS-213 |
| chr13-64453962-INS-70 | chr13-64453962-INS-70 |
| chr13-64457058-INS-56 | chr13-64457058-INS-56 |
| chr13-64461379-INS-257 | chr13-64461379-INS-257 |
| chr13-64471913-INS-203 | chr13-64471913-INS-203 |
| chr13-64484889-INS-207 | chr13-64484889-INS-209 |
| chr13-64496038-INS-55 | chr13-64496038-INS-65 |
| chr13-64520966-INS-212 | chr13-64520966-INS-212 |
| chr13-64526271-INS-69 | chr13-64526271-INS-78 |
| chr13-64529401-INS-279 | chr13-64529401-INS-279 |
| chr13-64554589-INS-82 | chr13-64554617-INS-77 |
| chr13-64556104-INS-57 | chr13-64556104-INS-57 |
| chr13-64570882-INS-134 | chr13-64570882-INS-134 |
| chr13-64574416-INS-501 | chr13-64574416-INS-501 |
| chr13-64592809-INS-919 | chr13-64592809-INS-919 |
| chr13-64610261-INS-228 | chr13-64610261-INS-244 |
| chr13-64641271-INS-218 | chr13-64641272-INS-217 |
| chr13-64662598-INS-85 | chr13-64662598-INS-69 |
| chr13-64667354-INS-209 | chr13-64667354-INS-211 |
| chr13-64670162-INS-2798 | chr13-64670163-INS-2798 |
| chr13-64718240-INS-64 | chr13-64718240-INS-64 |
| chr13-64720127-INS-66 | chr13-64720127-INS-66 |
| chr13-64765878-INS-186 | chr13-64765878-INS-186 |
| chr13-64770358-INS-626 | chr13-64770358-INS-626 |
| chr13-64788792-INS-72 | chr13-64788792-INS-78 |
| chr13-64812122-INS-272 | chr13-64812121-INS-271 |
| chr13-64838261-INS-77 | chr13-64838261-INS-105 |
| chr13-64861906-INS-199 | chr13-64861906-INS-199 |
| chr13-64898344-INS-203 | chr13-64898342-INS-202 |
| chr13-64908278-INS-54 | chr13-64908278-INS-54 |
| chr13-64922968-INS-1692 | chr13-64922969-INS-1692 |
| chr13-64937563-INS-97 | chr13-64937563-INS-73 |
| chr13-64957906-INS-307 | chr13-64957906-INS-307 |
| chr13-65003605-INS-9045 | chr13-65003606-INS-9045 |
| chr13-65006723-INS-56 | chr13-65006723-INS-54 |
| chr13-65064673-INS-57 | chr13-65064673-INS-59 |
| chr13-65067078-INS-70 | chr13-65067078-INS-72 |
| chr13-65092632-INS-366 | chr13-65092632-INS-366 |
| chr13-65111736-INS-129 | chr13-65111736-INS-129 |
| chr13-65152418-INS-67 | chr13-65152418-INS-57 |
| chr13-65188841-INS-371 | chr13-65188841-INS-371 |
| chr13-65200916-INS-99 | chr13-65200922-INS-102 |
| chr13-65218088-INS-94 | chr13-65218088-INS-94 |
| chr13-65224433-INS-191 | chr13-65224426-INS-204 |
| chr13-65232480-INS-222 | chr13-65232480-INS-222 |
| chr13-65241394-INS-2417 | chr13-65241395-INS-2417 |
| chr13-65249619-INS-202 | chr13-65249619-INS-203 |
| chr13-65256321-INS-809 | chr13-65256321-INS-810 |
| chr13-65274222-INS-56 | chr13-65274222-INS-56 |
| chr13-65277160-INS-3339 | chr13-65277161-INS-3339 |
| chr13-65289205-INS-68 | chr13-65289205-INS-68 |
| chr13-65294067-INS-69 | chr13-65294067-INS-69 |

| | |
|---|---|
| chr13-65303139-INS-200 | chr13-65303139-INS-202 |
| chr13-65364101-INS-652 | chr13-65364111-INS-653 |
| chr13-65390066-INS-545 | chr13-65390066-INS-545 |
| chr13-65415234-INS-287 | chr13-65415234-INS-288 |
| chr13-65419977-INS-504 | chr13-65419977-INS-504 |
| chr13-65423328-INS-2081 | chr13-65423329-INS-2082 |
| chr13-65494357-INS-72 | chr13-65494375-INS-71 |
| chr13-65528833-INS-260 | chr13-65528833-INS-260 |
| chr13-65733746-INS-65 | chr13-65733746-INS-65 |
| chr13-65737690-INS-74 | chr13-65737681-INS-83 |
| chr13-65907282-INS-92 | chr13-65907314-INS-80 |
| chr13-66048732-INS-261 | chr13-66048732-INS-228 |
| chr13-66103355-INS-874 | chr13-66103342-INS-874 |
| chr13-66104568-INS-50 | chr13-66104568-INS-50 |
| chr13-66106065-INS-84 | chr13-66106066-INS-84 |
| chr13-66287176-INS-199 | chr13-66287176-INS-199 |
| chr13-66294231-INS-256 | chr13-66294231-INS-256 |
| chr13-66464500-INS-72 | chr13-66464488-INS-101 |
| chr13-66465776-INS-180 | chr13-66465782-INS-181 |
| chr13-66818726-INS-420 | chr13-66818727-INS-420 |
| chr13-67006450-INS-260 | chr13-67006451-INS-259 |
| chr13-67008019-INS-72 | chr13-67008019-INS-72 |
| chr13-67055422-INS-56 | chr13-67055395-INS-54 |
| chr13-67121669-INS-359 | chr13-67121669-INS-359 |
| chr13-67142148-INS-100 | chr13-67142148-INS-100 |
| chr13-67152265-INS-142 | chr13-67152265-INS-157 |
| chr13-67180695-INS-208 | chr13-67180695-INS-208 |
| chr13-67183398-INS-194 | chr13-67183398-INS-194 |
| chr13-67194626-INS-52 | chr13-67194626-INS-58 |
| chr13-67223612-INS-82 | chr13-67223612-INS-82 |
| chr13-67236638-INS-351 | chr13-67236634-INS-334 |
| chr13-67246775-INS-84 | chr13-67246775-INS-84 |
| chr13-67313441-INS-223 | chr13-67313448-INS-227 |
| chr13-67325926-INS-52 | chr13-67325926-INS-52 |
| chr13-67345023-INS-57 | chr13-67345023-INS-57 |
| chr13-67346953-INS-226 | chr13-67346953-INS-225 |
| chr13-67357985-INS-455 | chr13-67357985-INS-454 |
| chr13-67362492-INS-6867 | chr13-67362493-INS-6868 |
| chr13-67402146-INS-56 | chr13-67402146-INS-58 |
| chr13-67415135-INS-202 | chr13-67415135-INS-202 |
| chr13-67441178-INS-84 | chr13-67441178-INS-84 |
| chr13-67450457-INS-76 | chr13-67450457-INS-77 |
| chr13-67474234-INS-616 | chr13-67474234-INS-616 |
| chr13-67478754-INS-60 | chr13-67478754-INS-60 |
| chr13-67484978-INS-1889 | chr13-67484979-INS-1889 |
| chr13-67512180-INS-1936 | chr13-67512181-INS-1937 |
| chr13-67538581-INS-3287 | chr13-67538582-INS-3287 |
| chr13-67541876-INS-244 | chr13-67541876-INS-243 |
| chr13-67545129-INS-156 | chr13-67545129-INS-156 |
| chr13-67548480-INS-3096 | chr13-67548481-INS-3096 |
| chr13-67556946-INS-198 | chr13-67556946-INS-199 |
| chr13-67570349-INS-2936 | chr13-67570350-INS-2936 |
| chr13-67570958-INS-159 | chr13-67570958-INS-159 |
| chr13-67572141-INS-157 | chr13-67572141-INS-198 |
| chr13-67576561-INS-327 | chr13-67576561-INS-327 |
| chr13-67590027-INS-272 | chr13-67590027-INS-271 |
| chr13-67593783-INS-194 | chr13-67593783-INS-194 |
| chr13-67641092-INS-225 | chr13-67641092-INS-225 |
| chr13-67651532-INS-6779 | chr13-67651533-INS-6779 |
| chr13-67677815-INS-1893 | chr13-67677816-INS-1893 |
| chr13-67681052-INS-198 | chr13-67681052-INS-198 |
| chr13-67691531-INS-187 | chr13-67691531-INS-189 |
| chr13-67713670-INS-7107 | chr13-67713671-INS-7107 |
| chr13-67729084-INS-215 | chr13-67729084-INS-215 |
| chr13-67736822-INS-62 | chr13-67736822-INS-63 |
| chr13-67746586-INS-254 | chr13-67746586-INS-254 |
| chr13-67751838-INS-200 | chr13-67751838-INS-200 |
| chr13-67753115-INS-6257 | chr13-67753088-INS-6253 |
| chr13-67777330-INS-208 | chr13-67777330-INS-209 |
| chr13-67853736-INS-189 | chr13-67853736-INS-189 |
| chr13-67863540-INS-60 | chr13-67863540-INS-60 |
| chr13-67870839-INS-213 | chr13-67870839-INS-214 |
| chr13-67873813-INS-163 | chr13-67873813-INS-163 |
| chr13-67900379-INS-52 | chr13-67900379-INS-52 |
| chr13-67907538-INS-565 | chr13-67907538-INS-565 |
| chr13-67914078-INS-212 | chr13-67914078-INS-212 |
| chr13-67917906-INS-8815 | chr13-67917907-INS-8815 |

| | |
|---|---|
| chr13-67942803-INS-3259 | chr13-67942812-INS-3255 |
| chr13-67946464-INS-4371 | chr13-67946465-INS-4385 |
| chr13-67954905-INS-54 | chr13-67954905-INS-52 |
| chr13-67980151-INS-66 | chr13-67980151-INS-70 |
| chr13-68002389-INS-782 | chr13-68002389-INS-782 |
| chr13-68007769-INS-57 | chr13-68007769-INS-50 |
| chr13-68009118-INS-159 | chr13-68009116-INS-161 |
| chr13-68053381-INS-830 | chr13-68053381-INS-830 |
| chr13-68084073-INS-214 | chr13-68084076-INS-239 |
| chr13-68103441-INS-143 | chr13-68103441-INS-143 |
| chr13-68232974-INS-195 | chr13-68232974-INS-195 |
| chr13-68236026-INS-505 | chr13-68236024-INS-506 |
| chr13-68278243-INS-384 | chr13-68278243-INS-385 |
| chr13-68283183-INS-659 | chr13-68283184-INS-660 |
| chr13-68335525-INS-59 | chr13-68335546-INS-59 |
| chr13-68337387-INS-51 | chr13-68337388-INS-51 |
| chr13-68532977-INS-132 | chr13-68532976-INS-132 |
| chr13-68547694-INS-235 | chr13-68547695-INS-235 |
| chr13-68744703-INS-69 | chr13-68744705-INS-90 |
| chr13-68762423-INS-208 | chr13-68762423-INS-208 |
| chr13-68776919-INS-6452 | chr13-68776920-INS-6452 |
| chr13-68842553-INS-6219 | chr13-68842554-INS-6218 |
| chr13-68849559-INS-263 | chr13-68849559-INS-263 |
| chr13-68879859-INS-436 | chr13-68879859-INS-436 |
| chr13-68894497-INS-97 | chr13-68894468-INS-130 |
| chr13-68902469-INS-1025 | chr13-68902467-INS-1019 |
| chr13-68919802-INS-402 | chr13-68919802-INS-402 |
| chr13-68975364-INS-1292 | chr13-68975365-INS-1292 |
| chr13-68987601-INS-73 | chr13-68987601-INS-83 |
| chr13-68991146-INS-53 | chr13-68991146-INS-52 |
| chr13-68996367-INS-108 | chr13-68996367-INS-108 |
| chr13-69001613-INS-52 | chr13-69001613-INS-52 |
| chr13-69018161-INS-110 | chr13-69018161-INS-114 |
| chr13-69021128-INS-1453 | chr13-69021129-INS-1453 |
| chr13-69050874-INS-74 | chr13-69050874-INS-74 |
| chr13-69062714-INS-66 | chr13-69062715-INS-94 |
| chr13-69067507-INS-437 | chr13-69067507-INS-437 |
| chr13-69077894-INS-528 | chr13-69077894-INS-528 |
| chr13-69084144-INS-277 | chr13-69084144-INS-277 |
| chr13-69091847-INS-50 | chr13-69091847-INS-56 |
| chr13-69106034-INS-6440 | chr13-69106035-INS-6440 |
| chr13-69107033-INS-58 | chr13-69107033-INS-58 |
| chr13-69157606-INS-140 | chr13-69157606-INS-137 |
| chr13-69180968-INS-86 | chr13-69180968-INS-86 |
| chr13-69186033-INS-53 | chr13-69186033-INS-61 |
| chr13-69187224-INS-491 | chr13-69187224-INS-497 |
| chr13-69212542-INS-56 | chr13-69212543-INS-58 |
| chr13-69215692-INS-50 | chr13-69215692-INS-54 |
| chr13-69238177-INS-400 | chr13-69238177-INS-400 |
| chr13-69241840-INS-107 | chr13-69241841-INS-107 |
| chr13-69274985-INS-400 | chr13-69274985-INS-400 |
| chr13-69345201-INS-83 | chr13-69345201-INS-83 |
| chr13-69358733-INS-740 | chr13-69358733-INS-738 |
| chr13-69363664-INS-100 | chr13-69363664-INS-100 |
| chr13-69392359-INS-76 | chr13-69392356-INS-72 |
| chr13-69417393-INS-1187 | chr13-69417394-INS-1187 |
| chr13-69449079-INS-90 | chr13-69449079-INS-90 |
| chr13-69456194-INS-172 | chr13-69456194-INS-172 |
| chr13-69468761-INS-53 | chr13-69468761-INS-53 |
| chr13-69497154-INS-58 | chr13-69497154-INS-58 |
| chr13-69500935-INS-207 | chr13-69500935-INS-200 |
| chr13-69518681-INS-500 | chr13-69518670-INS-510 |
| chr13-69562447-INS-405 | chr13-69562447-INS-405 |
| chr13-69585269-INS-360 | chr13-69585270-INS-364 |
| chr13-69587243-INS-4743 | chr13-69587244-INS-4743 |
| chr13-69595247-INS-118 | chr13-69595247-INS-118 |
| chr13-69599059-INS-401 | chr13-69599059-INS-401 |
| chr13-69602902-INS-57 | chr13-69602902-INS-57 |
| chr13-69605643-INS-52 | chr13-69605643-INS-56 |
| chr13-69613814-INS-157 | chr13-69613814-INS-169 |
| chr13-69614871-INS-52 | chr13-69614871-INS-52 |
| chr13-69645174-INS-64 | chr13-69645176-INS-86 |
| chr13-69677809-INS-80 | chr13-69677809-INS-80 |
| chr13-69694891-INS-394 | chr13-69694891-INS-394 |
| chr13-69699750-INS-55 | chr13-69699752-INS-77 |
| chr13-69699870-INS-578 | chr13-69699870-INS-578 |
| chr13-69702036-INS-52 | chr13-69702036-INS-60 |

| | |
|---|---|
| chr13-69705158-INS-93 | chr13-69705159-INS-73 |
| chr13-69729551-INS-58 | chr13-69729551-INS-66 |
| chr13-69746644-INS-450 | chr13-69746644-INS-450 |
| chr13-69750900-INS-215 | chr13-69750900-INS-215 |
| chr13-69771440-INS-108 | chr13-69771440-INS-108 |
| chr13-69795496-INS-58 | chr13-69795496-INS-58 |
| chr13-69811307-INS-70 | chr13-69811307-INS-70 |
| chr13-69828839-INS-195 | chr13-69828839-INS-195 |
| chr13-69831519-INS-255 | chr13-69831519-INS-255 |
| chr13-69836382-INS-76 | chr13-69836382-INS-76 |
| chr13-69855678-INS-86 | chr13-69855684-INS-113 |
| chr13-69929162-INS-53 | chr13-69929162-INS-56 |
| chr13-69936700-INS-1891 | chr13-69936701-INS-1891 |
| chr13-69944042-INS-216 | chr13-69944042-INS-217 |
| chr13-69972743-INS-57 | chr13-69972743-INS-55 |
| chr13-69984302-INS-282 | chr13-69984302-INS-282 |
| chr13-69986879-INS-6991 | chr13-69986880-INS-6992 |
| chr13-69991457-INS-1216 | chr13-69991458-INS-1216 |
| chr13-70009243-INS-52 | chr13-70009243-INS-52 |
| chr13-70041954-INS-63 | chr13-70041954-INS-63 |
| chr13-70046448-INS-483 | chr13-70046448-INS-484 |
| chr13-70049532-INS-348 | chr13-70049532-INS-348 |
| chr13-70052163-INS-224 | chr13-70052163-INS-224 |
| chr13-70060620-INS-504 | chr13-70060620-INS-504 |
| chr13-70066929-INS-7149 | chr13-70066930-INS-7149 |
| chr13-70075587-INS-64 | chr13-70075587-INS-64 |
| chr13-70096341-INS-64 | chr13-70096341-INS-60 |
| chr13-70115585-INS-1984 | chr13-70115586-INS-1984 |
| chr13-70120862-INS-82 | chr13-70120862-INS-66 |
| chr13-70133382-INS-62 | chr13-70133382-INS-62 |
| chr13-70153858-INS-874 | chr13-70153858-INS-874 |
| chr13-70170996-INS-51 | chr13-70171003-INS-91 |
| chr13-70178556-INS-189 | chr13-70178556-INS-189 |
| chr13-70199506-INS-52 | chr13-70199506-INS-54 |
| chr13-70205971-INS-75 | chr13-70205971-INS-75 |
| chr13-70211879-INS-113 | chr13-70211879-INS-113 |
| chr13-70247066-INS-80 | chr13-70247066-INS-68 |
| chr13-70252362-INS-209 | chr13-70252362-INS-209 |
| chr13-70276343-INS-7154 | chr13-70276344-INS-7154 |
| chr13-70335084-INS-65 | chr13-70335085-INS-60 |
| chr13-70351165-INS-5939 | chr13-70351165-INS-5939 |
| chr13-70405322-INS-89 | chr13-70405322-INS-89 |
| chr13-70418764-INS-458 | chr13-70418764-INS-454 |
| chr13-70419779-INS-63 | chr13-70419779-INS-63 |
| chr13-70425264-INS-75 | chr13-70425224-INS-81 |
| chr13-70435518-INS-59 | chr13-70435519-INS-63 |
| chr13-70441903-INS-2255 | chr13-70441904-INS-2255 |
| chr13-70449429-INS-59 | chr13-70449429-INS-63 |
| chr13-70476323-INS-56 | chr13-70476323-INS-54 |
| chr13-70491716-INS-246 | chr13-70491716-INS-240 |
| chr13-70519124-INS-176 | chr13-70519124-INS-160 |
| chr13-70521430-INS-58 | chr13-70521434-INS-52 |
| chr13-70527603-INS-82 | chr13-70527603-INS-82 |
| chr13-70546078-INS-249 | chr13-70546076-INS-230 |
| chr13-70569973-INS-60 | chr13-70569973-INS-60 |
| chr13-70619992-INS-6211 | chr13-70619993-INS-6211 |
| chr13-70625016-INS-272 | chr13-70625016-INS-277 |
| chr13-70638104-INS-265 | chr13-70638101-INS-250 |
| chr13-70653291-INS-109 | chr13-70653291-INS-109 |
| chr13-70690624-INS-476 | chr13-70690624-INS-476 |
| chr13-70691977-INS-106 | chr13-70691996-INS-60 |
| chr13-70693751-INS-100 | chr13-70693743-INS-116 |
| chr13-70700138-INS-386 | chr13-70700138-INS-386 |
| chr13-70702301-INS-58 | chr13-70702301-INS-60 |
| chr13-70719392-INS-166 | chr13-70719392-INS-169 |
| chr13-70771648-INS-54 | chr13-70771648-INS-54 |
| chr13-70800079-INS-549 | chr13-70800079-INS-549 |
| chr13-70807356-INS-56 | chr13-70807356-INS-56 |
| chr13-70820475-INS-506 | chr13-70820476-INS-506 |
| chr13-70829473-INS-83 | chr13-70829473-INS-82 |
| chr13-70837983-INS-1891 | chr13-70837984-INS-1891 |
| chr13-70843305-INS-411 | chr13-70843305-INS-412 |
| chr13-70845919-INS-430 | chr13-70845921-INS-447 |
| chr13-70847640-INS-358 | chr13-70847640-INS-358 |
| chr13-70859716-INS-80 | chr13-70859728-INS-66 |
| chr13-70900086-INS-106 | chr13-70900086-INS-106 |
| chr13-70933680-INS-1474 | chr13-70933681-INS-1474 |

| | |
|---|---|
| chr13-70943187-INS-60 | chr13-70943179-INS-60 |
| chr13-70951749-INS-85 | chr13-70951749-INS-85 |
| chr13-70986321-INS-78 | chr13-70986321-INS-62 |
| chr13-70995146-INS-302 | chr13-70995146-INS-301 |
| chr13-71001743-INS-53 | chr13-71001743-INS-54 |
| chr13-71024020-INS-51 | chr13-71024020-INS-51 |
| chr13-71028504-INS-67 | chr13-71028494-INS-66 |
| chr13-71037113-INS-165 | chr13-71037113-INS-165 |
| chr13-71048575-INS-156 | chr13-71048595-INS-156 |
| chr13-71049701-INS-68 | chr13-71049701-INS-68 |
| chr13-71068554-INS-70 | chr13-71068507-INS-118 |
| chr13-71155109-INS-66 | chr13-71155109-INS-66 |
| chr13-71214069-INS-114 | chr13-71214069-INS-114 |
| chr13-71216102-INS-3298 | chr13-71216103-INS-3298 |
| chr13-71224025-INS-233 | chr13-71224025-INS-233 |
| chr13-71227046-INS-95 | chr13-71227046-INS-95 |
| chr13-71277089-INS-76 | chr13-71277089-INS-76 |
| chr13-71290809-INS-1106 | chr13-71290810-INS-1106 |
| chr13-71302223-INS-66 | chr13-71302236-INS-76 |
| chr13-71303318-INS-76 | chr13-71303336-INS-80 |
| chr13-71348354-INS-268 | chr13-71348354-INS-268 |
| chr13-71363492-INS-2442 | chr13-71363495-INS-2438 |
| chr13-71365181-INS-488 | chr13-71365181-INS-488 |
| chr13-71382045-INS-62 | chr13-71382045-INS-62 |
| chr13-71397958-INS-52 | chr13-71397958-INS-52 |
| chr13-71423979-INS-59 | chr13-71423979-INS-69 |
| chr13-71425724-INS-1118 | chr13-71425725-INS-1106 |
| chr13-71436574-INS-95 | chr13-71436574-INS-89 |
| chr13-71471728-INS-391 | chr13-71471728-INS-435 |
| chr13-71478228-INS-59 | chr13-71478228-INS-59 |
| chr13-71505698-INS-389 | chr13-71505698-INS-389 |
| chr13-71509436-INS-58 | chr13-71509436-INS-60 |
| chr13-71511833-INS-305 | chr13-71511833-INS-305 |
| chr13-71621270-INS-71 | chr13-71621270-INS-71 |
| chr13-71676917-INS-62 | chr13-71676917-INS-62 |
| chr13-71680315-INS-122 | chr13-71680315-INS-114 |
| chr13-71705530-INS-80 | chr13-71705530-INS-80 |
| chr13-71717825-INS-84 | chr13-71717825-INS-78 |
| chr13-71763351-INS-140 | chr13-71763351-INS-140 |
| chr13-71766313-INS-8453 | chr13-71766314-INS-8453 |
| chr13-71766541-INS-97 | chr13-71766541-INS-97 |
| chr13-71769868-INS-224 | chr13-71769868-INS-224 |
| chr13-71772159-INS-113 | chr13-71772159-INS-84 |
| chr13-71780617-INS-78 | chr13-71780617-INS-78 |
| chr13-71796450-INS-52 | chr13-71796450-INS-57 |
| chr13-71842056-INS-85 | chr13-71842056-INS-85 |
| chr13-71862681-INS-773 | chr13-71862681-INS-769 |
| chr13-71885750-INS-58 | chr13-71885750-INS-54 |
| chr13-71887432-INS-68 | chr13-71887464-INS-71 |
| chr13-71898550-INS-52 | chr13-71898550-INS-56 |
| chr13-71930779-INS-6936 | chr13-71930780-INS-6936 |
| chr13-71938539-INS-586 | chr13-71938539-INS-586 |
| chr13-71943394-INS-408 | chr13-71943394-INS-408 |
| chr13-71959078-INS-72 | chr13-71959078-INS-72 |
| chr13-71960573-INS-80 | chr13-71960562-INS-88 |
| chr13-71961285-INS-467 | chr13-71961285-INS-467 |
| chr13-71962234-INS-6597 | chr13-71962235-INS-6597 |
| chr13-71968446-INS-225 | chr13-71968446-INS-218 |
| chr13-71988161-INS-594 | chr13-71988161-INS-594 |
| chr13-72029362-INS-227 | chr13-72029362-INS-227 |
| chr13-72125031-INS-52 | chr13-72125007-INS-58 |
| chr13-72131789-INS-68 | chr13-72131789-INS-68 |
| chr13-72149407-INS-57 | chr13-72149407-INS-57 |
| chr13-72151964-INS-66 | chr13-72151964-INS-66 |
| chr13-72155617-INS-1862 | chr13-72155618-INS-1862 |
| chr13-72195042-INS-240 | chr13-72195042-INS-240 |
| chr13-72198970-INS-529 | chr13-72198970-INS-529 |
| chr13-72203426-INS-78 | chr13-72203426-INS-78 |
| chr13-72216448-INS-496 | chr13-72216448-INS-496 |
| chr13-72232264-INS-73 | chr13-72232264-INS-73 |
| chr13-72255939-INS-84 | chr13-72255939-INS-84 |
| chr13-72257560-INS-60 | chr13-72257560-INS-60 |
| chr13-72260496-INS-63 | chr13-72260496-INS-63 |
| chr13-72267685-INS-92 | chr13-72267683-INS-92 |
| chr13-72285665-INS-95 | chr13-72285665-INS-98 |
| chr13-72301850-INS-6488 | chr13-72301851-INS-6488 |
| chr13-72308959-INS-80 | chr13-72308959-INS-80 |

| | |
|---|---|
| chr13-72317953-INS-2613 | chr13-72317954-INS-2613 |
| chr13-72322094-INS-98 | chr13-72322118-INS-63 |
| chr13-72329926-INS-551 | chr13-72329926-INS-551 |
| chr13-72334838-INS-70 | chr13-72334838-INS-70 |
| chr13-72345735-INS-51 | chr13-72345735-INS-63 |
| chr13-72348376-INS-412 | chr13-72348376-INS-373 |
| chr13-72359469-INS-802 | chr13-72359469-INS-804 |
| chr13-72363626-INS-117 | chr13-72363626-INS-117 |
| chr13-72374521-INS-112 | chr13-72374521-INS-86 |
| chr13-72398359-INS-79 | chr13-72398357-INS-91 |
| chr13-72404254-INS-59 | chr13-72404254-INS-59 |
| chr13-72482682-INS-579 | chr13-72482682-INS-593 |
| chr13-72487532-INS-72 | chr13-72487532-INS-72 |
| chr13-72499567-INS-596 | chr13-72499582-INS-606 |
| chr13-72502171-INS-70 | chr13-72502171-INS-92 |
| chr13-72512855-INS-52 | chr13-72512855-INS-52 |
| chr13-72522135-INS-79 | chr13-72522135-INS-85 |
| chr13-72527462-INS-114 | chr13-72527469-INS-146 |
| chr13-72562893-INS-64 | chr13-72562893-INS-58 |
| chr13-72573927-INS-148 | chr13-72573927-INS-111 |
| chr13-72584418-INS-6214 | chr13-72584419-INS-6214 |
| chr13-72613506-INS-54 | chr13-72613506-INS-54 |
| chr13-72618100-INS-54 | chr13-72618098-INS-50 |
| chr13-72619742-INS-112 | chr13-72619742-INS-112 |
| chr13-72648820-INS-72 | chr13-72648820-INS-72 |
| chr13-72658690-INS-96 | chr13-72658653-INS-104 |
| chr13-72659993-INS-81 | chr13-72660018-INS-68 |
| chr13-72677102-INS-1595 | chr13-72677103-INS-1596 |
| chr13-72695660-INS-116 | chr13-72695660-INS-116 |
| chr13-72753498-INS-58 | chr13-72753498-INS-64 |
| chr13-72755929-INS-2066 | chr13-72755930-INS-2067 |
| chr13-72760225-INS-608 | chr13-72760226-INS-608 |
| chr13-72775064-INS-130 | chr13-72775064-INS-145 |
| chr13-72804875-INS-877 | chr13-72804875-INS-878 |
| chr13-72921055-INS-68 | chr13-72921054-INS-102 |
| chr13-72925433-INS-69 | chr13-72925433-INS-73 |
| chr13-72942331-INS-440 | chr13-72942331-INS-440 |
| chr13-72947593-INS-55 | chr13-72947593-INS-56 |
| chr13-72953498-INS-201 | chr13-72953498-INS-201 |
| chr13-72962146-INS-552 | chr13-72962146-INS-553 |
| chr13-72963627-INS-75 | chr13-72963627-INS-55 |
| chr13-73011351-INS-114 | chr13-73011351-INS-93 |
| chr13-73029121-INS-648 | chr13-73029121-INS-647 |
| chr13-73051572-INS-213 | chr13-73051572-INS-212 |
| chr13-73055518-INS-60 | chr13-73055518-INS-60 |
| chr13-73067785-INS-93 | chr13-73067781-INS-105 |
| chr13-73070673-INS-344 | chr13-73070673-INS-344 |
| chr13-73086297-INS-65 | chr13-73086297-INS-65 |
| chr13-73101251-INS-1348 | chr13-73101252-INS-1358 |
| chr13-73119844-INS-158 | chr13-73119844-INS-162 |
| chr13-73120958-INS-326 | chr13-73120958-INS-326 |
| chr13-73143082-INS-76 | chr13-73143082-INS-114 |
| chr13-73154529-INS-67 | chr13-73154535-INS-67 |
| chr13-73180715-INS-107 | chr13-73180715-INS-118 |
| chr13-73184488-INS-114 | chr13-73184488-INS-125 |
| chr13-73201575-INS-197 | chr13-73201575-INS-196 |
| chr13-73224522-INS-7256 | chr13-73224523-INS-7256 |
| chr13-73235949-INS-54 | chr13-73235949-INS-55 |
| chr13-73309948-INS-192 | chr13-73309948-INS-192 |
| chr13-73322648-INS-51 | chr13-73322648-INS-57 |
| chr13-73337393-INS-51 | chr13-73337393-INS-63 |
| chr13-73343140-INS-54 | chr13-73343140-INS-54 |
| chr13-73371010-INS-396 | chr13-73371010-INS-410 |
| chr13-73405141-INS-56 | chr13-73405137-INS-58 |
| chr13-73425522-INS-68 | chr13-73425514-INS-56 |
| chr13-73426762-INS-53 | chr13-73426762-INS-52 |
| chr13-73429279-INS-881 | chr13-73429279-INS-881 |
| chr13-73437576-INS-1261 | chr13-73437577-INS-1277 |
| chr13-73443064-INS-79 | chr13-73443064-INS-79 |
| chr13-73473516-INS-202 | chr13-73473516-INS-202 |
| chr13-73487738-INS-6440 | chr13-73487739-INS-6440 |
| chr13-73491209-INS-57 | chr13-73491209-INS-57 |
| chr13-73528877-INS-230 | chr13-73528877-INS-230 |
| chr13-73541387-INS-363 | chr13-73541387-INS-363 |
| chr13-73575839-INS-202 | chr13-73575839-INS-202 |
| chr13-73585949-INS-54 | chr13-73585949-INS-54 |
| chr13-73591119-INS-140 | chr13-73591119-INS-144 |

| | |
|---|---|
| chr13-73601892-INS-200 | chr13-73601892-INS-200 |
| chr13-73641794-INS-52 | chr13-73641794-INS-52 |
| chr13-73668883-INS-556 | chr13-73668883-INS-558 |
| chr13-73675706-INS-405 | chr13-73675706-INS-405 |
| chr13-73678704-INS-178 | chr13-73678704-INS-183 |
| chr13-73702545-INS-83 | chr13-73702545-INS-83 |
| chr13-73721638-INS-62 | chr13-73721638-INS-62 |
| chr13-73729428-INS-106 | chr13-73729428-INS-58 |
| chr13-73738599-INS-80 | chr13-73738599-INS-80 |
| chr13-73740121-INS-208 | chr13-73740121-INS-208 |
| chr13-73780387-INS-95 | chr13-73780387-INS-97 |
| chr13-73793402-INS-241 | chr13-73793402-INS-247 |
| chr13-73842515-INS-95 | chr13-73842515-INS-95 |
| chr13-73853081-INS-58 | chr13-73853081-INS-58 |
| chr13-73873996-INS-202 | chr13-73873996-INS-201 |
| chr13-73890308-INS-157 | chr13-73890308-INS-157 |
| chr13-73916463-INS-209 | chr13-73916463-INS-210 |
| chr13-73926328-INS-80 | chr13-73926343-INS-52 |
| chr13-73934734-INS-104 | chr13-73934734-INS-104 |
| chr13-74057807-INS-305 | chr13-74057806-INS-331 |
| chr13-74093083-INS-111 | chr13-74093083-INS-111 |
| chr13-74115432-INS-68 | chr13-74115432-INS-60 |
| chr13-74144978-INS-102 | chr13-74144978-INS-103 |
| chr13-74219782-INS-56 | chr13-74219775-INS-53 |
| chr13-74224630-INS-70 | chr13-74224642-INS-76 |
| chr13-74229432-INS-1085 | chr13-74229433-INS-1085 |
| chr13-74246091-INS-60 | chr13-74246099-INS-54 |
| chr13-74295351-INS-388 | chr13-74295354-INS-388 |
| chr13-74352218-INS-83 | chr13-74352218-INS-83 |
| chr13-74374235-INS-55 | chr13-74374235-INS-60 |
| chr13-74394208-INS-685 | chr13-74394207-INS-685 |
| chr13-74399256-INS-176 | chr13-74399247-INS-206 |
| chr13-74404963-INS-137 | chr13-74404963-INS-137 |
| chr13-74420386-INS-64 | chr13-74420374-INS-79 |
| chr13-74433624-INS-201 | chr13-74433624-INS-201 |
| chr13-74435145-INS-214 | chr13-74435145-INS-214 |
| chr13-74461496-INS-54 | chr13-74461496-INS-54 |
| chr13-74478857-INS-54 | chr13-74478857-INS-54 |
| chr13-74519204-INS-196 | chr13-74519204-INS-194 |
| chr13-74523296-INS-409 | chr13-74523295-INS-412 |
| chr13-74527199-INS-990 | chr13-74527199-INS-990 |
| chr13-74527916-INS-350 | chr13-74527917-INS-350 |
| chr13-74550293-INS-192 | chr13-74550289-INS-191 |
| chr13-74552292-INS-216 | chr13-74552292-INS-220 |
| chr13-74555780-INS-212 | chr13-74555780-INS-212 |
| chr13-74616398-INS-232 | chr13-74616398-INS-203 |
| chr13-74638557-INS-328 | chr13-74638557-INS-328 |
| chr13-74706530-INS-5436 | chr13-74706531-INS-5436 |
| chr13-74713986-INS-150 | chr13-74713986-INS-146 |
| chr13-74775347-INS-550 | chr13-74775347-INS-550 |
| chr13-74802349-INS-208 | chr13-74802349-INS-208 |
| chr13-74807964-INS-442 | chr13-74807964-INS-442 |
| chr13-74811834-INS-65 | chr13-74811834-INS-65 |
| chr13-74846036-INS-976 | chr13-74846036-INS-979 |
| chr13-74869016-INS-742 | chr13-74869016-INS-742 |
| chr13-74883154-INS-200 | chr13-74883154-INS-200 |
| chr13-74884615-INS-53 | chr13-74884615-INS-59 |
| chr13-74887186-INS-242 | chr13-74887186-INS-242 |
| chr13-74913344-INS-175 | chr13-74913344-INS-197 |
| chr13-74914805-INS-239 | chr13-74914809-INS-247 |
| chr13-74933629-INS-58 | chr13-74933629-INS-58 |
| chr13-74937260-INS-75 | chr13-74937260-INS-75 |
| chr13-74937974-INS-2864 | chr13-74937975-INS-2864 |
| chr13-74946178-INS-181 | chr13-74946178-INS-181 |
| chr13-74950145-INS-53 | chr13-74950145-INS-53 |
| chr13-74951921-INS-105 | chr13-74951921-INS-107 |
| chr13-74967829-INS-91 | chr13-74967829-INS-91 |
| chr13-75010205-INS-65 | chr13-75010205-INS-60 |
| chr13-75099052-INS-54 | chr13-75099063-INS-53 |
| chr13-75113040-INS-3143 | chr13-75113041-INS-3143 |
| chr13-75134954-INS-6377 | chr13-75134955-INS-6377 |
| chr13-75157865-INS-56 | chr13-75157865-INS-56 |
| chr13-75167249-INS-6590 | chr13-75167250-INS-6590 |
| chr13-75173044-INS-1292 | chr13-75173045-INS-1292 |
| chr13-75174518-INS-67 | chr13-75174518-INS-68 |
| chr13-75203188-INS-52 | chr13-75203188-INS-52 |
| chr13-75221646-INS-899 | chr13-75221646-INS-899 |

| | |
|---|---|
| chr13-75225482-INS-280 | chr13-75225482-INS-250 |
| chr13-75271853-INS-110 | chr13-75271845-INS-130 |
| chr13-75300388-INS-428 | chr13-75300388-INS-428 |
| chr13-75331413-INS-247 | chr13-75331413-INS-245 |
| chr13-75336615-INS-81 | chr13-75336615-INS-63 |
| chr13-75406210-INS-633 | chr13-75406210-INS-633 |
| chr13-75407387-INS-292 | chr13-75407387-INS-244 |
| chr13-75430113-INS-51 | chr13-75430113-INS-51 |
| chr13-75445740-INS-2142 | chr13-75445741-INS-2142 |
| chr13-75528674-INS-1129 | chr13-75528675-INS-1129 |
| chr13-75595257-INS-81 | chr13-75595257-INS-85 |
| chr13-75601268-INS-7474 | chr13-75601269-INS-7474 |
| chr13-75649462-INS-488 | chr13-75649462-INS-488 |
| chr13-75695563-INS-218 | chr13-75695563-INS-221 |
| chr13-75700276-INS-135 | chr13-75700276-INS-110 |
| chr13-75721132-INS-80 | chr13-75721109-INS-69 |
| chr13-75728122-INS-60 | chr13-75728122-INS-56 |
| chr13-75760814-INS-7102 | chr13-75760815-INS-7102 |
| chr13-75768391-INS-189 | chr13-75768391-INS-189 |
| chr13-75775036-INS-6583 | chr13-75775037-INS-6583 |
| chr13-75782443-INS-1208 | chr13-75782444-INS-1210 |
| chr13-75795910-INS-232 | chr13-75795910-INS-232 |
| chr13-75805022-INS-202 | chr13-75805022-INS-202 |
| chr13-75919518-INS-57 | chr13-75919518-INS-53 |
| chr13-75947557-INS-191 | chr13-75947557-INS-191 |
| chr13-75968198-INS-211 | chr13-75968198-INS-211 |
| chr13-75983359-INS-1609 | chr13-75983360-INS-1609 |
| chr13-75992132-INS-211 | chr13-75992132-INS-211 |
| chr13-76020334-INS-183 | chr13-76020334-INS-183 |
| chr13-76056058-INS-930 | chr13-76056058-INS-929 |
| chr13-76065663-INS-1057 | chr13-76065664-INS-1057 |
| chr13-76073737-INS-218 | chr13-76073737-INS-219 |
| chr13-76109382-INS-60 | chr13-76109382-INS-60 |
| chr13-76144242-INS-412 | chr13-76144242-INS-412 |
| chr13-76157661-INS-212 | chr13-76157661-INS-212 |
| chr13-76179192-INS-71 | chr13-76179192-INS-75 |
| chr13-76234755-INS-858 | chr13-76234755-INS-858 |
| chr13-76240856-INS-440 | chr13-76240856-INS-440 |
| chr13-76310882-INS-209 | chr13-76310882-INS-210 |
| chr13-76334565-INS-5123 | chr13-76334566-INS-5123 |
| chr13-76371462-INS-60 | chr13-76371462-INS-60 |
| chr13-76404230-INS-77 | chr13-76404230-INS-79 |
| chr13-76469323-INS-162 | chr13-76469323-INS-162 |
| chr13-76471717-INS-68 | chr13-76471719-INS-72 |
| chr13-76479534-INS-72 | chr13-76479534-INS-72 |
| chr13-76485131-INS-4671 | chr13-76485132-INS-4671 |
| chr13-76487311-INS-222 | chr13-76487313-INS-197 |
| chr13-76492030-INS-605 | chr13-76492030-INS-605 |
| chr13-76523134-INS-134 | chr13-76523134-INS-134 |
| chr13-76524504-INS-710 | chr13-76524504-INS-708 |
| chr13-76535201-INS-400 | chr13-76535201-INS-400 |
| chr13-76539076-INS-176 | chr13-76539076-INS-176 |
| chr13-76551328-INS-62 | chr13-76551332-INS-66 |
| chr13-76585675-INS-110 | chr13-76585679-INS-110 |
| chr13-76591128-INS-605 | chr13-76591128-INS-605 |
| chr13-76593437-INS-52 | chr13-76593437-INS-52 |
| chr13-76605100-INS-289 | chr13-76605100-INS-289 |
| chr13-76663976-INS-434 | chr13-76663976-INS-434 |
| chr13-76671175-INS-386 | chr13-76671175-INS-386 |
| chr13-76677128-INS-52 | chr13-76677128-INS-54 |
| chr13-76679847-INS-225 | chr13-76679847-INS-225 |
| chr13-76715073-INS-51 | chr13-76715073-INS-55 |
| chr13-76721614-INS-654 | chr13-76721614-INS-654 |
| chr13-76725733-INS-872 | chr13-76725733-INS-872 |
| chr13-76726609-INS-196 | chr13-76726609-INS-196 |
| chr13-76739177-INS-74 | chr13-76739193-INS-58 |
| chr13-76789081-INS-53 | chr13-76789081-INS-55 |
| chr13-76846869-INS-85 | chr13-76846869-INS-75 |
| chr13-76980061-INS-153 | chr13-76980102-INS-114 |
| chr13-76989187-INS-1519 | chr13-76989191-INS-1522 |
| chr13-77011258-INS-759 | chr13-77011259-INS-752 |
| chr13-77021214-INS-886 | chr13-77021214-INS-887 |
| chr13-77048504-INS-1793 | chr13-77048505-INS-1801 |
| chr13-77049486-INS-6927 | chr13-77049487-INS-6928 |
| chr13-77064044-INS-6661 | chr13-77064045-INS-6661 |
| chr13-77096012-INS-68 | chr13-77096012-INS-60 |
| chr13-77103093-INS-6324 | chr13-77103094-INS-6330 |

| | |
|---|---|
| chr13-77118918-INS-8059 | chr13-77118919-INS-8048 |
| chr13-77146726-INS-54 | chr13-77146726-INS-54 |
| chr13-77169345-INS-6477 | chr13-77169346-INS-6476 |
| chr13-77213467-INS-54 | chr13-77213467-INS-54 |
| chr13-77261230-INS-98 | chr13-77261230-INS-52 |
| chr13-77267225-INS-3002 | chr13-77267226-INS-3000 |
| chr13-77298096-INS-6597 | chr13-77298097-INS-6601 |
| chr13-77299908-INS-58 | chr13-77299904-INS-53 |
| chr13-77304410-INS-53 | chr13-77304410-INS-53 |
| chr13-77311991-INS-1314 | chr13-77311992-INS-1314 |
| chr13-77316039-INS-229 | chr13-77316039-INS-230 |
| chr13-77337894-INS-373 | chr13-77337894-INS-375 |
| chr13-77340272-INS-56 | chr13-77340272-INS-50 |
| chr13-77350375-INS-505 | chr13-77350375-INS-505 |
| chr13-77384895-INS-285 | chr13-77384897-INS-286 |
| chr13-77390882-INS-445 | chr13-77390888-INS-445 |
| chr13-77409696-INS-108 | chr13-77409696-INS-108 |
| chr13-77425753-INS-65 | chr13-77425751-INS-75 |
| chr13-77428774-INS-191 | chr13-77428774-INS-191 |
| chr13-77438010-INS-4368 | chr13-77438011-INS-4368 |
| chr13-77507162-INS-1100 | chr13-77507160-INS-1100 |
| chr13-77520884-INS-214 | chr13-77520884-INS-214 |
| chr13-77529653-INS-214 | chr13-77529653-INS-214 |
| chr13-77555828-INS-1065 | chr13-77555829-INS-1066 |
| chr13-77562249-INS-6443 | chr13-77562250-INS-6443 |
| chr13-77577003-INS-214 | chr13-77577003-INS-214 |
| chr13-77608868-INS-498 | chr13-77608868-INS-498 |
| chr13-77648028-INS-360 | chr13-77648028-INS-360 |
| chr13-77682539-INS-59 | chr13-77682539-INS-59 |
| chr13-77698794-INS-620 | chr13-77698794-INS-620 |
| chr13-77816745-INS-66 | chr13-77816707-INS-74 |
| chr13-77818336-INS-906 | chr13-77818336-INS-914 |
| chr13-77837087-INS-80 | chr13-77837087-INS-52 |
| chr13-77872271-INS-72 | chr13-77872271-INS-63 |
| chr13-77875930-INS-312 | chr13-77875930-INS-312 |
| chr13-77878833-INS-293 | chr13-77878833-INS-293 |
| chr13-77898017-INS-140 | chr13-77898017-INS-175 |
| chr13-77942728-INS-151 | chr13-77942728-INS-151 |
| chr13-77975175-INS-537 | chr13-77975175-INS-538 |
| chr13-77996459-INS-246 | chr13-77996459-INS-246 |
| chr13-78005275-INS-485 | chr13-78005275-INS-484 |
| chr13-78007133-INS-123 | chr13-78007133-INS-123 |
| chr13-78111491-INS-61 | chr13-78111491-INS-61 |
| chr13-78113817-INS-60 | chr13-78113813-INS-67 |
| chr13-78120042-INS-7921 | chr13-78120043-INS-7922 |
| chr13-78167762-INS-143 | chr13-78167762-INS-140 |
| chr13-78183309-INS-6401 | chr13-78183310-INS-6403 |
| chr13-78213470-INS-6493 | chr13-78213471-INS-6471 |
| chr13-78220437-INS-133 | chr13-78220437-INS-133 |
| chr13-78240762-INS-2249 | chr13-78240763-INS-2249 |
| chr13-78241255-INS-6601 | chr13-78241256-INS-6605 |
| chr13-78257477-INS-131 | chr13-78257477-INS-131 |
| chr13-78265785-INS-105 | chr13-78265787-INS-107 |
| chr13-78268362-INS-268 | chr13-78268362-INS-268 |
| chr13-78292233-INS-102 | chr13-78292233-INS-119 |
| chr13-78349602-INS-51 | chr13-78349602-INS-51 |
| chr13-78359667-INS-71 | chr13-78359667-INS-71 |
| chr13-78380328-INS-418 | chr13-78380328-INS-418 |
| chr13-78434015-INS-120 | chr13-78433983-INS-105 |
| chr13-78450368-INS-388 | chr13-78450368-INS-388 |
| chr13-78455127-INS-7552 | chr13-78455128-INS-7554 |
| chr13-78464966-INS-6474 | chr13-78464967-INS-6474 |
| chr13-78468758-INS-401 | chr13-78468758-INS-401 |
| chr13-78484011-INS-6255 | chr13-78484012-INS-6255 |
| chr13-78498132-INS-69 | chr13-78498098-INS-53 |
| chr13-78507833-INS-52 | chr13-78507833-INS-52 |
| chr13-78532636-INS-139 | chr13-78532636-INS-139 |
| chr13-78541938-INS-175 | chr13-78541938-INS-178 |
| chr13-78545338-INS-74 | chr13-78545359-INS-54 |
| chr13-78546010-INS-208 | chr13-78546010-INS-208 |
| chr13-78559268-INS-526 | chr13-78559268-INS-526 |
| chr13-78560978-INS-60 | chr13-78560977-INS-60 |
| chr13-78581790-INS-524 | chr13-78581796-INS-526 |
| chr13-78584122-INS-148 | chr13-78584122-INS-148 |
| chr13-78589870-INS-203 | chr13-78589876-INS-202 |
| chr13-78604207-INS-88 | chr13-78604207-INS-88 |
| chr13-78616552-INS-78 | chr13-78616552-INS-78 |

| | |
|---|---|
| chr13-78619538-INS-56 | chr13-78619538-INS-56 |
| chr13-78623405-INS-6197 | chr13-78623406-INS-6197 |
| chr13-78628638-INS-74 | chr13-78628638-INS-71 |
| chr13-78643095-INS-224 | chr13-78643107-INS-205 |
| chr13-78655272-INS-139 | chr13-78655272-INS-139 |
| chr13-78667323-INS-1891 | chr13-78667324-INS-1891 |
| chr13-78677317-INS-52 | chr13-78677330-INS-57 |
| chr13-78699507-INS-6396 | chr13-78699508-INS-6396 |
| chr13-78705167-INS-174 | chr13-78705167-INS-179 |
| chr13-78752198-INS-65 | chr13-78752199-INS-52 |
| chr13-78798855-INS-1112 | chr13-78798856-INS-1100 |
| chr13-78799426-INS-60 | chr13-78799426-INS-60 |
| chr13-78806326-INS-230 | chr13-78806334-INS-234 |
| chr13-78828832-INS-2787 | chr13-78828833-INS-2786 |
| chr13-78830731-INS-5799 | chr13-78830732-INS-5800 |
| chr13-78837412-INS-318 | chr13-78837412-INS-318 |
| chr13-78886289-INS-321 | chr13-78886289-INS-321 |
| chr13-78927972-INS-458 | chr13-78927972-INS-458 |
| chr13-78940954-INS-6787 | chr13-78940955-INS-6802 |
| chr13-78967468-INS-508 | chr13-78967468-INS-508 |
| chr13-78989834-INS-1118 | chr13-78989835-INS-1118 |
| chr13-78991268-INS-1350 | chr13-78991269-INS-1350 |
| chr13-78991594-INS-51 | chr13-78991591-INS-55 |
| chr13-79011653-INS-223 | chr13-79011659-INS-227 |
| chr13-79015582-INS-1953 | chr13-79015583-INS-1954 |
| chr13-79043942-INS-66 | chr13-79043945-INS-54 |
| chr13-79079437-INS-1044 | chr13-79079438-INS-1044 |
| chr13-79091322-INS-192 | chr13-79091322-INS-192 |
| chr13-79114049-INS-158 | chr13-79114049-INS-158 |
| chr13-79115634-INS-652 | chr13-79115635-INS-652 |
| chr13-79141727-INS-1501 | chr13-79141728-INS-1500 |
| chr13-79198057-INS-51 | chr13-79198058-INS-63 |
| chr13-79203962-INS-1523 | chr13-79203963-INS-1523 |
| chr13-79219898-INS-418 | chr13-79219898-INS-418 |
| chr13-79242138-INS-201 | chr13-79242138-INS-201 |
| chr13-79262685-INS-596 | chr13-79262685-INS-597 |
| chr13-79267743-INS-4676 | chr13-79267744-INS-4676 |
| chr13-79269540-INS-339 | chr13-79269533-INS-338 |
| chr13-79270679-INS-699 | chr13-79270679-INS-697 |
| chr13-79334154-INS-2085 | chr13-79334155-INS-2088 |
| chr13-79355504-INS-94 | chr13-79355512-INS-50 |
| chr13-79371733-INS-7091 | chr13-79371734-INS-7091 |
| chr13-79390714-INS-66 | chr13-79390714-INS-70 |
| chr13-79396800-INS-56 | chr13-79396800-INS-50 |
| chr13-79482448-INS-1786 | chr13-79482449-INS-1786 |
| chr13-79543502-INS-52 | chr13-79543502-INS-52 |
| chr13-79582111-INS-73 | chr13-79582111-INS-77 |
| chr13-79603178-INS-5261 | chr13-79603179-INS-5261 |
| chr13-79610465-INS-74 | chr13-79610465-INS-74 |
| chr13-79639185-INS-58 | chr13-79639185-INS-58 |
| chr13-79653803-INS-63 | chr13-79653803-INS-63 |
| chr13-79657874-INS-92 | chr13-79657854-INS-87 |
| chr13-79678334-INS-106 | chr13-79678334-INS-91 |
| chr13-79680241-INS-405 | chr13-79680236-INS-405 |
| chr13-79701302-INS-165 | chr13-79701327-INS-116 |
| chr13-79802717-INS-393 | chr13-79802717-INS-393 |
| chr13-79857116-INS-2009 | chr13-79857118-INS-1990 |
| chr13-79887607-INS-544 | chr13-79887607-INS-544 |
| chr13-79908843-INS-204 | chr13-79908843-INS-204 |
| chr13-79964083-INS-389 | chr13-79964083-INS-389 |
| chr13-79967933-INS-210 | chr13-79967933-INS-211 |
| chr13-79987416-INS-93 | chr13-79987404-INS-99 |
| chr13-79988446-INS-492 | chr13-79988446-INS-492 |
| chr13-79990302-INS-6178 | chr13-79990303-INS-6177 |
| chr13-80009565-INS-53 | chr13-80009561-INS-60 |
| chr13-80021437-INS-201 | chr13-80021437-INS-198 |
| chr13-80026252-INS-62 | chr13-80026242-INS-84 |
| chr13-80056080-INS-82 | chr13-80056080-INS-82 |
| chr13-80058459-INS-63 | chr13-80058459-INS-53 |
| chr13-80099475-INS-766 | chr13-80099475-INS-766 |
| chr13-80153301-INS-52 | chr13-80153301-INS-62 |
| chr13-80172616-INS-211 | chr13-80172616-INS-212 |
| chr13-80197608-INS-268 | chr13-80197608-INS-268 |
| chr13-80237041-INS-51 | chr13-80237041-INS-54 |
| chr13-80248101-INS-1098 | chr13-80248102-INS-1099 |
| chr13-80263542-INS-68 | chr13-80263540-INS-76 |
| chr13-80297314-INS-51 | chr13-80297314-INS-55 |

| | |
|---|---|
| chr13-80308111-INS-52 | chr13-80308111-INS-52 |
| chr13-80310326-INS-2885 | chr13-80310327-INS-2885 |
| chr13-80322431-INS-142 | chr13-80322431-INS-142 |
| chr13-80337417-INS-96 | chr13-80337413-INS-54 |
| chr13-80351107-INS-86 | chr13-80351107-INS-86 |
| chr13-80367057-INS-80 | chr13-80367059-INS-74 |
| chr13-80391754-INS-96 | chr13-80391754-INS-96 |
| chr13-80404054-INS-56 | chr13-80404054-INS-56 |
| chr13-80411143-INS-1007 | chr13-80411144-INS-1007 |
| chr13-80448687-INS-53 | chr13-80448687-INS-53 |
| chr13-80473742-INS-76 | chr13-80473742-INS-78 |
| chr13-80477598-INS-55 | chr13-80477598-INS-55 |
| chr13-80478527-INS-133 | chr13-80478527-INS-108 |
| chr13-80487655-INS-74 | chr13-80487655-INS-74 |
| chr13-80489632-INS-1576 | chr13-80489633-INS-1556 |
| chr13-80500398-INS-79 | chr13-80500398-INS-79 |
| chr13-80570041-INS-454 | chr13-80570041-INS-455 |
| chr13-80578283-INS-4164 | chr13-80578284-INS-4164 |
| chr13-80578284-INS-390 | chr13-80578284-INS-390 |
| chr13-80608586-INS-55 | chr13-80608585-INS-64 |
| chr13-80675546-INS-67 | chr13-80675512-INS-52 |
| chr13-80691853-INS-483 | chr13-80691853-INS-517 |
| chr13-80698942-INS-95 | chr13-80698939-INS-103 |
| chr13-80715943-INS-52 | chr13-80715943-INS-50 |
| chr13-80720005-INS-90 | chr13-80719995-INS-65 |
| chr13-80732251-INS-50 | chr13-80732251-INS-98 |
| chr13-80734095-INS-1791 | chr13-80734096-INS-1778 |
| chr13-80753997-INS-340 | chr13-80753997-INS-340 |
| chr13-80776507-INS-6208 | chr13-80776508-INS-6210 |
| chr13-80777698-INS-390 | chr13-80777698-INS-390 |
| chr13-80784169-INS-4235 | chr13-80784170-INS-4226 |
| chr13-80870659-INS-60 | chr13-80870705-INS-64 |
| chr13-80895442-INS-78 | chr13-80895442-INS-84 |
| chr13-80907257-INS-102 | chr13-80907257-INS-102 |
| chr13-80946629-INS-330 | chr13-80946629-INS-334 |
| chr13-80949653-INS-583 | chr13-80949653-INS-599 |
| chr13-80968944-INS-66 | chr13-80968944-INS-68 |
| chr13-80978352-INS-230 | chr13-80978356-INS-232 |
| chr13-80994257-INS-100 | chr13-80994257-INS-100 |
| chr13-80997850-INS-2804 | chr13-80997851-INS-2804 |
| chr13-81014748-INS-106 | chr13-81014748-INS-107 |
| chr13-81020951-INS-695 | chr13-81020951-INS-705 |
| chr13-81023640-INS-76 | chr13-81023640-INS-68 |
| chr13-81081432-INS-161 | chr13-81081432-INS-161 |
| chr13-81092896-INS-81 | chr13-81092896-INS-81 |
| chr13-81101747-INS-284 | chr13-81101745-INS-288 |
| chr13-81131582-INS-229 | chr13-81131581-INS-224 |
| chr13-81186450-INS-456 | chr13-81186450-INS-464 |
| chr13-81207902-INS-311 | chr13-81207902-INS-311 |
| chr13-81295377-INS-2213 | chr13-81295378-INS-2213 |
| chr13-81295434-INS-72 | chr13-81295434-INS-72 |
| chr13-81306709-INS-712 | chr13-81306710-INS-709 |
| chr13-81310571-INS-80 | chr13-81310571-INS-120 |
| chr13-81311863-INS-68 | chr13-81311863-INS-70 |
| chr13-81323819-INS-230 | chr13-81323819-INS-230 |
| chr13-81324675-INS-56 | chr13-81324675-INS-54 |
| chr13-81343063-INS-214 | chr13-81343063-INS-214 |
| chr13-81358298-INS-6442 | chr13-81358299-INS-6442 |
| chr13-81365146-INS-818 | chr13-81365146-INS-818 |
| chr13-81373873-INS-83 | chr13-81373873-INS-71 |
| chr13-81374545-INS-53 | chr13-81374546-INS-52 |
| chr13-81386735-INS-965 | chr13-81386753-INS-961 |
| chr13-81420809-INS-84 | chr13-81420812-INS-63 |
| chr13-81440507-INS-68 | chr13-81440505-INS-51 |
| chr13-81462318-INS-406 | chr13-81462318-INS-406 |
| chr13-81472618-INS-107 | chr13-81472618-INS-107 |
| chr13-81524774-INS-68 | chr13-81524774-INS-68 |
| chr13-81535264-INS-260 | chr13-81535264-INS-260 |
| chr13-81564259-INS-59 | chr13-81564259-INS-61 |
| chr13-81590051-INS-90 | chr13-81590051-INS-120 |
| chr13-81635126-INS-66 | chr13-81635126-INS-66 |
| chr13-81639131-INS-73 | chr13-81639131-INS-90 |
| chr13-81643659-INS-149 | chr13-81643631-INS-164 |
| chr13-81653551-INS-71 | chr13-81653551-INS-71 |
| chr13-81655041-INS-52 | chr13-81655031-INS-73 |
| chr13-81694677-INS-520 | chr13-81694677-INS-520 |
| chr13-81716925-INS-834 | chr13-81716925-INS-834 |

| | |
|---|---|
| chr13-81742192-INS-294 | chr13-81742192-INS-294 |
| chr13-81759477-INS-67 | chr13-81759477-INS-82 |
| chr13-81764681-INS-126 | chr13-81764681-INS-98 |
| chr13-81765897-INS-318 | chr13-81765898-INS-318 |
| chr13-81794339-INS-51 | chr13-81794340-INS-52 |
| chr13-81799413-INS-154 | chr13-81799419-INS-105 |
| chr13-81831539-INS-72 | chr13-81831539-INS-70 |
| chr13-81845391-INS-80 | chr13-81845393-INS-52 |
| chr13-81867083-INS-743 | chr13-81867083-INS-744 |
| chr13-81907404-INS-57 | chr13-81907404-INS-57 |
| chr13-81920521-INS-493 | chr13-81920521-INS-493 |
| chr13-81929299-INS-213 | chr13-81929299-INS-217 |
| chr13-81941968-INS-58 | chr13-81941968-INS-56 |
| chr13-81953970-INS-1171 | chr13-81953971-INS-1171 |
| chr13-81971660-INS-366 | chr13-81971660-INS-372 |
| chr13-82015942-INS-100 | chr13-82015942-INS-104 |
| chr13-82022913-INS-614 | chr13-82022913-INS-614 |
| chr13-82033218-INS-63 | chr13-82033218-INS-63 |
| chr13-82049355-INS-127 | chr13-82049355-INS-135 |
| chr13-82054355-INS-215 | chr13-82054355-INS-215 |
| chr13-82059292-INS-6421 | chr13-82059293-INS-6421 |
| chr13-82072758-INS-839 | chr13-82072758-INS-840 |
| chr13-82080039-INS-67 | chr13-82080016-INS-69 |
| chr13-82102017-INS-66 | chr13-82102017-INS-66 |
| chr13-82121784-INS-990 | chr13-82121784-INS-990 |
| chr13-82131108-INS-87 | chr13-82131073-INS-110 |
| chr13-82197670-INS-74 | chr13-82197670-INS-82 |
| chr13-82228454-INS-124 | chr13-82228454-INS-124 |
| chr13-82282832-INS-64 | chr13-82282832-INS-64 |
| chr13-82287715-INS-201 | chr13-82287715-INS-201 |
| chr13-82318727-INS-52 | chr13-82318727-INS-52 |
| chr13-82347493-INS-1269 | chr13-82347494-INS-1269 |
| chr13-82394645-INS-856 | chr13-82394646-INS-861 |
| chr13-82432343-INS-112 | chr13-82432343-INS-112 |
| chr13-82438286-INS-74 | chr13-82438266-INS-81 |
| chr13-82450609-INS-330 | chr13-82450609-INS-332 |
| chr13-82478243-INS-100 | chr13-82478236-INS-64 |
| chr13-82510806-INS-6888 | chr13-82510807-INS-6891 |
| chr13-82569959-INS-88 | chr13-82569959-INS-88 |
| chr13-82578243-INS-55 | chr13-82578243-INS-55 |
| chr13-82596258-INS-621 | chr13-82596258-INS-624 |
| chr13-82604087-INS-201 | chr13-82604081-INS-202 |
| chr13-82608353-INS-57 | chr13-82608338-INS-60 |
| chr13-82613837-INS-64 | chr13-82613837-INS-64 |
| chr13-82626995-INS-641 | chr13-82626995-INS-639 |
| chr13-82646888-INS-503 | chr13-82646888-INS-503 |
| chr13-82651619-INS-1213 | chr13-82651620-INS-1214 |
| chr13-82656306-INS-1659 | chr13-82656307-INS-1659 |
| chr13-82671100-INS-138 | chr13-82671100-INS-138 |
| chr13-82679154-INS-6991 | chr13-82679155-INS-6991 |
| chr13-82681175-INS-938 | chr13-82681175-INS-941 |
| chr13-82688998-INS-78 | chr13-82688998-INS-54 |
| chr13-82713959-INS-55 | chr13-82713963-INS-53 |
| chr13-82721194-INS-55 | chr13-82721157-INS-55 |
| chr13-82732862-INS-66 | chr13-82732872-INS-64 |
| chr13-82754615-INS-2261 | chr13-82754616-INS-2261 |
| chr13-82757557-INS-478 | chr13-82757557-INS-478 |
| chr13-82830914-INS-358 | chr13-82830914-INS-356 |
| chr13-82857776-INS-60 | chr13-82857776-INS-60 |
| chr13-82867096-INS-62 | chr13-82867096-INS-72 |
| chr13-82882343-INS-6592 | chr13-82882344-INS-6592 |
| chr13-82931865-INS-2674 | chr13-82931866-INS-2674 |
| chr13-82959219-INS-997 | chr13-82959219-INS-997 |
| chr13-83005852-INS-6351 | chr13-83005853-INS-6351 |
| chr13-83033882-INS-64 | chr13-83033882-INS-66 |
| chr13-83065139-INS-737 | chr13-83065139-INS-737 |
| chr13-83079600-INS-72 | chr13-83079600-INS-71 |
| chr13-83113551-INS-56 | chr13-83113551-INS-56 |
| chr13-83128785-INS-1931 | chr13-83128786-INS-1931 |
| chr13-83135397-INS-66 | chr13-83135397-INS-76 |
| chr13-83143730-INS-60 | chr13-83143730-INS-60 |
| chr13-83147769-INS-180 | chr13-83147769-INS-144 |
| chr13-83171195-INS-152 | chr13-83171196-INS-128 |
| chr13-83182953-INS-237 | chr13-83182953-INS-237 |
| chr13-83201727-INS-223 | chr13-83201740-INS-210 |
| chr13-83203573-INS-245 | chr13-83203573-INS-245 |
| chr13-83210352-INS-287 | chr13-83210352-INS-287 |

| | |
|---|---|
| chr13-83223061-INS-529 | chr13-83223061-INS-529 |
| chr13-83229925-INS-448 | chr13-83229929-INS-449 |
| chr13-83245553-INS-100 | chr13-83245553-INS-82 |
| chr13-83249722-INS-5313 | chr13-83249723-INS-5313 |
| chr13-83256126-INS-158 | chr13-83256132-INS-180 |
| chr13-83313361-INS-1630 | chr13-83313362-INS-1655 |
| chr13-83317215-INS-72 | chr13-83317215-INS-66 |
| chr13-83357817-INS-358 | chr13-83357817-INS-358 |
| chr13-83485377-INS-112 | chr13-83485377-INS-112 |
| chr13-83486509-INS-197 | chr13-83486509-INS-197 |
| chr13-83509390-INS-54 | chr13-83509390-INS-54 |
| chr13-83547415-INS-156 | chr13-83547415-INS-160 |
| chr13-83555676-INS-126 | chr13-83555676-INS-118 |
| chr13-83606192-INS-88 | chr13-83606192-INS-88 |
| chr13-83640343-INS-374 | chr13-83640343-INS-374 |
| chr13-83650304-INS-56 | chr13-83650304-INS-60 |
| chr13-83728112-INS-6484 | chr13-83728113-INS-6485 |
| chr13-83733146-INS-3502 | chr13-83733147-INS-3502 |
| chr13-83739656-INS-63 | chr13-83739656-INS-63 |
| chr13-83772874-INS-109 | chr13-83772874-INS-109 |
| chr13-83798365-INS-58 | chr13-83798353-INS-50 |
| chr13-83846120-INS-342 | chr13-83846120-INS-342 |
| chr13-83850545-INS-89 | chr13-83850543-INS-80 |
| chr13-83892149-INS-112 | chr13-83892149-INS-112 |
| chr13-83910492-INS-298 | chr13-83910492-INS-252 |
| chr13-83933357-INS-1776 | chr13-83933358-INS-1769 |
| chr13-83943026-INS-267 | chr13-83943020-INS-268 |
| chr13-83949185-INS-80 | chr13-83949185-INS-80 |
| chr13-83968002-INS-82 | chr13-83968002-INS-86 |
| chr13-83986776-INS-2237 | chr13-83986777-INS-2237 |
| chr13-83990496-INS-360 | chr13-83990496-INS-358 |
| chr13-83991782-INS-100 | chr13-83991797-INS-90 |
| chr13-84007140-INS-6416 | chr13-84007141-INS-6416 |
| chr13-84041054-INS-362 | chr13-84041054-INS-362 |
| chr13-84143986-INS-62 | chr13-84143987-INS-50 |
| chr13-84226284-INS-89 | chr13-84226284-INS-80 |
| chr13-84272542-INS-68 | chr13-84272542-INS-68 |
| chr13-84297839-INS-50 | chr13-84297854-INS-53 |
| chr13-84298403-INS-550 | chr13-84298403-INS-550 |
| chr13-84323326-INS-83 | chr13-84323326-INS-84 |
| chr13-84378312-INS-234 | chr13-84378312-INS-234 |
| chr13-84503134-INS-202 | chr13-84503134-INS-202 |
| chr13-84513716-INS-73 | chr13-84513716-INS-73 |
| chr13-84524535-INS-137 | chr13-84524533-INS-133 |
| chr13-84560642-INS-180 | chr13-84560642-INS-175 |
| chr13-84579435-INS-1821 | chr13-84579436-INS-1813 |
| chr13-84589373-INS-424 | chr13-84589373-INS-424 |
| chr13-84612337-INS-412 | chr13-84612337-INS-429 |
| chr13-84655983-INS-403 | chr13-84655983-INS-403 |
| chr13-84658666-INS-1222 | chr13-84658667-INS-1229 |
| chr13-84665080-INS-1026 | chr13-84665081-INS-1026 |
| chr13-84670392-INS-8628 | chr13-84670393-INS-8629 |
| chr13-84682745-INS-533 | chr13-84682745-INS-534 |
| chr13-84698947-INS-3710 | chr13-84698948-INS-3711 |
| chr13-84735694-INS-52 | chr13-84735694-INS-53 |
| chr13-84743729-INS-7174 | chr13-84743730-INS-7174 |
| chr13-84746759-INS-1096 | chr13-84746760-INS-1085 |
| chr13-84780859-INS-731 | chr13-84780859-INS-731 |
| chr13-84799847-INS-71 | chr13-84799864-INS-103 |
| chr13-84823604-INS-72 | chr13-84823604-INS-72 |
| chr13-84830609-INS-166 | chr13-84830610-INS-183 |
| chr13-84839375-INS-55 | chr13-84839375-INS-55 |
| chr13-84850211-INS-54 | chr13-84850211-INS-54 |
| chr13-84879075-INS-223 | chr13-84879075-INS-223 |
| chr13-84906936-INS-81 | chr13-84906936-INS-63 |
| chr13-84935071-INS-577 | chr13-84935075-INS-577 |
| chr13-84971828-INS-1714 | chr13-84971829-INS-1716 |
| chr13-84972446-INS-61 | chr13-84972432-INS-71 |
| chr13-84974134-INS-3076 | chr13-84974135-INS-3076 |
| chr13-85008616-INS-322 | chr13-85008616-INS-322 |
| chr13-85025791-INS-320 | chr13-85025791-INS-325 |
| chr13-85033517-INS-66 | chr13-85033517-INS-66 |
| chr13-85036689-INS-54 | chr13-85036689-INS-50 |
| chr13-85044511-INS-7318 | chr13-85044512-INS-7318 |
| chr13-85055785-INS-145 | chr13-85055785-INS-145 |
| chr13-85074427-INS-81 | chr13-85074427-INS-81 |
| chr13-85113992-INS-54 | chr13-85113992-INS-56 |

| | |
|---|---|
| chr13-85147084-INS-112 | chr13-85147084-INS-136 |
| chr13-85187740-INS-214 | chr13-85187740-INS-214 |
| chr13-85219344-INS-6491 | chr13-85219345-INS-6491 |
| chr13-85238427-INS-6198 | chr13-85238428-INS-6198 |
| chr13-85270837-INS-824 | chr13-85270837-INS-824 |
| chr13-85285001-INS-138 | chr13-85285001-INS-138 |
| chr13-85290282-INS-55 | chr13-85290282-INS-55 |
| chr13-85299256-INS-204 | chr13-85299247-INS-206 |
| chr13-85333047-INS-140 | chr13-85333090-INS-140 |
| chr13-85334586-INS-207 | chr13-85334586-INS-206 |
| chr13-85337157-INS-204 | chr13-85337157-INS-196 |
| chr13-85342617-INS-175 | chr13-85342617-INS-175 |
| chr13-85395228-INS-112 | chr13-85395229-INS-99 |
| chr13-85399042-INS-387 | chr13-85399042-INS-388 |
| chr13-85402195-INS-6442 | chr13-85402196-INS-6442 |
| chr13-85431667-INS-55 | chr13-85431656-INS-70 |
| chr13-85458058-INS-52 | chr13-85458058-INS-54 |
| chr13-85479390-INS-112 | chr13-85479390-INS-112 |
| chr13-85511306-INS-53 | chr13-85511306-INS-53 |
| chr13-85543226-INS-164 | chr13-85543190-INS-192 |
| chr13-85568752-INS-1321 | chr13-85568753-INS-1337 |
| chr13-85575243-INS-58 | chr13-85575243-INS-58 |
| chr13-85589313-INS-94 | chr13-85589313-INS-94 |
| chr13-85594876-INS-200 | chr13-85594876-INS-200 |
| chr13-85643727-INS-609 | chr13-85643727-INS-608 |
| chr13-85658324-INS-60 | chr13-85658324-INS-56 |
| chr13-85669903-INS-180 | chr13-85669903-INS-180 |
| chr13-85672880-INS-80 | chr13-85672880-INS-120 |
| chr13-85674806-INS-60 | chr13-85674806-INS-60 |
| chr13-85709997-INS-61 | chr13-85709993-INS-62 |
| chr13-85710947-INS-188 | chr13-85710950-INS-202 |
| chr13-85715179-INS-112 | chr13-85715179-INS-117 |
| chr13-85718164-INS-461 | chr13-85718164-INS-461 |
| chr13-85726748-INS-495 | chr13-85726748-INS-495 |
| chr13-85737759-INS-1221 | chr13-85737760-INS-1221 |
| chr13-85795195-INS-58 | chr13-85795196-INS-58 |
| chr13-85823109-INS-93 | chr13-85823109-INS-93 |
| chr13-85842175-INS-6037 | chr13-85842176-INS-6037 |
| chr13-85859494-INS-186 | chr13-85859494-INS-185 |
| chr13-85861357-INS-65 | chr13-85861357-INS-65 |
| chr13-85888385-INS-742 | chr13-85888385-INS-745 |
| chr13-85889030-INS-401 | chr13-85889030-INS-419 |
| chr13-85892207-INS-825 | chr13-85892207-INS-826 |
| chr13-85901113-INS-6221 | chr13-85901114-INS-6222 |
| chr13-85942199-INS-7305 | chr13-85942200-INS-7305 |
| chr13-85952550-INS-508 | chr13-85952550-INS-508 |
| chr13-85968932-INS-6472 | chr13-85968933-INS-6473 |
| chr13-85997692-INS-716 | chr13-85997692-INS-714 |
| chr13-86011292-INS-164 | chr13-86011292-INS-164 |
| chr13-86022951-INS-102 | chr13-86022951-INS-102 |
| chr13-86037970-INS-101 | chr13-86037970-INS-101 |
| chr13-86046641-INS-2011 | chr13-86046642-INS-2005 |
| chr13-86060705-INS-12828 | chr13-86060706-INS-12828 |
| chr13-86073831-INS-524 | chr13-86073832-INS-526 |
| chr13-86127618-INS-354 | chr13-86127618-INS-356 |
| chr13-86129883-INS-308 | chr13-86129883-INS-308 |
| chr13-86134176-INS-858 | chr13-86134176-INS-858 |
| chr13-86137374-INS-55 | chr13-86137374-INS-55 |
| chr13-86162315-INS-127 | chr13-86162307-INS-107 |
| chr13-86164672-INS-61 | chr13-86164669-INS-58 |
| chr13-86180690-INS-832 | chr13-86180690-INS-832 |
| chr13-86180869-INS-4582 | chr13-86180870-INS-4582 |
| chr13-86182968-INS-173 | chr13-86182968-INS-180 |
| chr13-86187513-INS-659 | chr13-86187513-INS-663 |
| chr13-86202743-INS-52 | chr13-86202743-INS-66 |
| chr13-86204314-INS-679 | chr13-86204314-INS-681 |
| chr13-86206961-INS-83 | chr13-86206961-INS-83 |
| chr13-86210334-INS-415 | chr13-86210334-INS-415 |
| chr13-86224826-INS-6432 | chr13-86224827-INS-6432 |
| chr13-86229388-INS-6442 | chr13-86229389-INS-6442 |
| chr13-86248589-INS-724 | chr13-86248589-INS-724 |
| chr13-86249503-INS-215 | chr13-86249503-INS-221 |
| chr13-86257388-INS-103 | chr13-86257388-INS-103 |
| chr13-86258446-INS-588 | chr13-86258446-INS-588 |
| chr13-86283358-INS-2744 | chr13-86283359-INS-2744 |
| chr13-86292906-INS-1864 | chr13-86292907-INS-1864 |
| chr13-86297068-INS-104 | chr13-86297068-INS-104 |

| | |
|---|---|
| chr13-86322954-INS-168 | chr13-86322954-INS-168 |
| chr13-86384692-INS-609 | chr13-86384692-INS-609 |
| chr13-86397929-INS-458 | chr13-86397929-INS-458 |
| chr13-86461715-INS-2006 | chr13-86461716-INS-2006 |
| chr13-86497599-INS-208 | chr13-86497599-INS-208 |
| chr13-86505192-INS-422 | chr13-86505192-INS-424 |
| chr13-86511164-INS-414 | chr13-86511164-INS-420 |
| chr13-86525009-INS-351 | chr13-86525009-INS-349 |
| chr13-86536648-INS-637 | chr13-86536648-INS-642 |
| chr13-86541888-INS-1862 | chr13-86541889-INS-1862 |
| chr13-86568345-INS-52 | chr13-86568347-INS-52 |
| chr13-86593043-INS-91 | chr13-86593043-INS-91 |
| chr13-86596477-INS-59 | chr13-86596478-INS-63 |
| chr13-86674317-INS-308 | chr13-86674317-INS-308 |
| chr13-86677361-INS-146 | chr13-86677362-INS-146 |
| chr13-86702080-INS-74 | chr13-86702088-INS-81 |
| chr13-86709864-INS-80 | chr13-86709864-INS-72 |
| chr13-86711477-INS-246 | chr13-86711478-INS-247 |
| chr13-86715956-INS-579 | chr13-86715955-INS-582 |
| chr13-86739931-INS-800 | chr13-86739931-INS-800 |
| chr13-86748197-INS-3049 | chr13-86748198-INS-3084 |
| chr13-86749567-INS-89 | chr13-86749567-INS-89 |
| chr13-86751276-INS-83 | chr13-86751251-INS-70 |
| chr13-86766752-INS-444 | chr13-86766752-INS-453 |
| chr13-86820140-INS-70 | chr13-86820140-INS-74 |
| chr13-86827960-INS-497 | chr13-86827960-INS-497 |
| chr13-86833167-INS-6457 | chr13-86833168-INS-6457 |
| chr13-86854671-INS-107 | chr13-86854671-INS-107 |
| chr13-86855628-INS-795 | chr13-86855629-INS-797 |
| chr13-86860033-INS-5291 | chr13-86860034-INS-5291 |
| chr13-86875992-INS-1870 | chr13-86875993-INS-1871 |
| chr13-86909960-INS-54 | chr13-86909959-INS-58 |
| chr13-86939972-INS-257 | chr13-86939972-INS-257 |
| chr13-86957021-INS-5913 | chr13-86957022-INS-5896 |
| chr13-86974125-INS-1801 | chr13-86974126-INS-1801 |
| chr13-86974851-INS-267 | chr13-86974851-INS-267 |
| chr13-86981543-INS-147 | chr13-86981546-INS-124 |
| chr13-86983880-INS-754 | chr13-86983880-INS-754 |
| chr13-87012815-INS-91 | chr13-87012815-INS-91 |
| chr13-87026392-INS-1894 | chr13-87026393-INS-1892 |
| chr13-87066194-INS-399 | chr13-87066194-INS-399 |
| chr13-87083474-INS-6251 | chr13-87083475-INS-6251 |
| chr13-87101443-INS-6237 | chr13-87101444-INS-6237 |
| chr13-87105669-INS-6517 | chr13-87105670-INS-6514 |
| chr13-87132532-INS-5468 | chr13-87132533-INS-5468 |
| chr13-87140999-INS-79 | chr13-87140999-INS-79 |
| chr13-87153862-INS-58 | chr13-87153862-INS-66 |
| chr13-87159962-INS-595 | chr13-87159962-INS-597 |
| chr13-87165687-INS-1395 | chr13-87165688-INS-1391 |
| chr13-87174419-INS-6191 | chr13-87174420-INS-6192 |
| chr13-87188635-INS-137 | chr13-87188635-INS-137 |
| chr13-87193516-INS-436 | chr13-87193516-INS-436 |
| chr13-87193896-INS-5914 | chr13-87193897-INS-5917 |
| chr13-87219914-INS-564 | chr13-87219914-INS-563 |
| chr13-87240032-INS-161 | chr13-87240032-INS-166 |
| chr13-87247875-INS-57 | chr13-87247875-INS-56 |
| chr13-87257336-INS-1864 | chr13-87257337-INS-1864 |
| chr13-87273682-INS-89 | chr13-87273682-INS-61 |
| chr13-87294639-INS-214 | chr13-87294639-INS-214 |
| chr13-87349412-INS-79 | chr13-87349412-INS-77 |
| chr13-87351109-INS-486 | chr13-87351109-INS-486 |
| chr13-87354481-INS-74 | chr13-87354481-INS-72 |
| chr13-87355189-INS-64 | chr13-87355189-INS-64 |
| chr13-87357185-INS-184 | chr13-87357185-INS-175 |
| chr13-87403710-INS-8579 | chr13-87403711-INS-8579 |
| chr13-87404435-INS-1892 | chr13-87404436-INS-1892 |
| chr13-87407901-INS-124 | chr13-87407901-INS-124 |
| chr13-87437113-INS-433 | chr13-87437123-INS-434 |
| chr13-87443321-INS-52 | chr13-87443321-INS-50 |
| chr13-87448545-INS-2154 | chr13-87448546-INS-2154 |
| chr13-87451787-INS-7091 | chr13-87451788-INS-7091 |
| chr13-87453335-INS-52 | chr13-87453335-INS-67 |
| chr13-87476488-INS-66 | chr13-87476488-INS-68 |
| chr13-87482855-INS-142 | chr13-87482855-INS-142 |
| chr13-87537386-INS-168 | chr13-87537386-INS-168 |
| chr13-87555126-INS-50 | chr13-87555126-INS-78 |
| chr13-87571576-INS-7428 | chr13-87571577-INS-7428 |

| | |
|---|---|
| chr13-87574488-INS-297 | chr13-87574488-INS-298 |
| chr13-87576742-INS-122 | chr13-87576742-INS-122 |
| chr13-87636699-INS-1099 | chr13-87636700-INS-1089 |
| chr13-87639129-INS-252 | chr13-87639129-INS-246 |
| chr13-87646608-INS-451 | chr13-87646608-INS-451 |
| chr13-87675850-INS-64 | chr13-87675850-INS-64 |
| chr13-87675890-INS-1060 | chr13-87675891-INS-1059 |
| chr13-87678045-INS-2754 | chr13-87678046-INS-2754 |
| chr13-87701152-INS-556 | chr13-87701152-INS-556 |
| chr13-87732113-INS-53 | chr13-87732081-INS-101 |
| chr13-87734025-INS-74 | chr13-87734025-INS-74 |
| chr13-87756133-INS-6439 | chr13-87756134-INS-6439 |
| chr13-87760921-INS-138 | chr13-87760921-INS-138 |
| chr13-87776279-INS-102 | chr13-87776279-INS-102 |
| chr13-87809545-INS-499 | chr13-87809545-INS-499 |
| chr13-87819284-INS-1352 | chr13-87819285-INS-1352 |
| chr13-87889242-INS-63 | chr13-87889242-INS-109 |
| chr13-87918730-INS-62 | chr13-87918730-INS-64 |
| chr13-87985438-INS-798 | chr13-87985438-INS-795 |
| chr13-87990559-INS-981 | chr13-87990559-INS-970 |
| chr13-88035801-INS-52 | chr13-88035801-INS-52 |
| chr13-88059196-INS-61 | chr13-88059196-INS-64 |
| chr13-88096953-INS-138 | chr13-88096914-INS-150 |
| chr13-88120717-INS-198 | chr13-88120717-INS-198 |
| chr13-88227882-INS-2722 | chr13-88227883-INS-2727 |
| chr13-88237254-INS-112 | chr13-88237254-INS-112 |
| chr13-88271875-INS-1502 | chr13-88271876-INS-1502 |
| chr13-88284204-INS-6205 | chr13-88284205-INS-6205 |
| chr13-88329430-INS-62 | chr13-88329392-INS-90 |
| chr13-88333865-INS-66 | chr13-88333865-INS-66 |
| chr13-88339802-INS-133 | chr13-88339802-INS-133 |
| chr13-88410376-INS-203 | chr13-88410376-INS-202 |
| chr13-88422865-INS-386 | chr13-88422865-INS-386 |
| chr13-88461061-INS-145 | chr13-88461051-INS-151 |
| chr13-88465741-INS-419 | chr13-88465741-INS-419 |
| chr13-88485703-INS-51 | chr13-88485703-INS-51 |
| chr13-88491883-INS-75 | chr13-88491883-INS-73 |
| chr13-88497173-INS-234 | chr13-88497174-INS-234 |
| chr13-88513855-INS-90 | chr13-88513855-INS-88 |
| chr13-88551913-INS-343 | chr13-88551913-INS-342 |
| chr13-88573405-INS-4714 | chr13-88573406-INS-4714 |
| chr13-88576023-INS-7109 | chr13-88576024-INS-7109 |
| chr13-88579344-INS-2408 | chr13-88579345-INS-2411 |
| chr13-88587670-INS-52 | chr13-88587670-INS-54 |
| chr13-88634646-INS-335 | chr13-88634646-INS-347 |
| chr13-88636782-INS-765 | chr13-88636782-INS-765 |
| chr13-88649752-INS-926 | chr13-88649752-INS-926 |
| chr13-88652977-INS-236 | chr13-88652968-INS-220 |
| chr13-88655343-INS-381 | chr13-88655343-INS-381 |
| chr13-88661258-INS-928 | chr13-88661258-INS-922 |
| chr13-88684743-INS-81 | chr13-88684740-INS-72 |
| chr13-88696792-INS-278 | chr13-88696792-INS-320 |
| chr13-88720571-INS-78 | chr13-88720571-INS-70 |
| chr13-88754588-INS-52 | chr13-88754588-INS-56 |
| chr13-88759233-INS-90 | chr13-88759233-INS-78 |
| chr13-88825170-INS-1346 | chr13-88825171-INS-1345 |
| chr13-88834626-INS-237 | chr13-88834626-INS-200 |
| chr13-88881375-INS-142 | chr13-88881385-INS-140 |
| chr13-88900536-INS-142 | chr13-88900536-INS-142 |
| chr13-88902559-INS-137 | chr13-88902559-INS-137 |
| chr13-88916027-INS-5612 | chr13-88916028-INS-5614 |
| chr13-88964503-INS-104 | chr13-88964503-INS-84 |
| chr13-88993588-INS-188 | chr13-88993588-INS-188 |
| chr13-88997850-INS-79 | chr13-88997850-INS-79 |
| chr13-89027910-INS-58 | chr13-89027910-INS-58 |
| chr13-89048762-INS-6818 | chr13-89048763-INS-6819 |
| chr13-89057484-INS-85 | chr13-89057485-INS-55 |
| chr13-89072330-INS-54 | chr13-89072330-INS-54 |
| chr13-89116110-INS-57 | chr13-89116110-INS-57 |
| chr13-89117351-INS-149 | chr13-89117351-INS-149 |
| chr13-89120554-INS-158 | chr13-89120554-INS-158 |
| chr13-89126785-INS-137 | chr13-89126785-INS-137 |
| chr13-89143803-INS-3041 | chr13-89143804-INS-3041 |
| chr13-89163980-INS-61 | chr13-89163980-INS-61 |
| chr13-89168067-INS-1466 | chr13-89168068-INS-1466 |
| chr13-89204201-INS-1365 | chr13-89204202-INS-1373 |
| chr13-89207837-INS-60 | chr13-89207829-INS-50 |

| | |
|---|---|
| chr13-89223598-INS-6473 | chr13-89223599-INS-6473 |
| chr13-89233771-INS-69 | chr13-89233771-INS-69 |
| chr13-89235838-INS-108 | chr13-89235838-INS-108 |
| chr13-89242450-INS-60 | chr13-89242450-INS-54 |
| chr13-89283101-INS-66 | chr13-89283101-INS-66 |
| chr13-89287428-INS-895 | chr13-89287428-INS-920 |
| chr13-89288753-INS-362 | chr13-89288753-INS-362 |
| chr13-89305416-INS-208 | chr13-89305417-INS-208 |
| chr13-89327836-INS-131 | chr13-89327852-INS-125 |
| chr13-89389761-INS-435 | chr13-89389761-INS-436 |
| chr13-89423596-INS-199 | chr13-89423596-INS-199 |
| chr13-89425447-INS-349 | chr13-89425447-INS-349 |
| chr13-89437752-INS-602 | chr13-89437753-INS-603 |
| chr13-89441825-INS-77 | chr13-89441804-INS-81 |
| chr13-89475645-INS-1157 | chr13-89475646-INS-1157 |
| chr13-89507314-INS-542 | chr13-89507314-INS-542 |
| chr13-89535151-INS-95 | chr13-89535151-INS-93 |
| chr13-89551866-INS-1089 | chr13-89551867-INS-1089 |
| chr13-89555413-INS-6877 | chr13-89555414-INS-6877 |
| chr13-89584052-INS-58 | chr13-89584052-INS-68 |
| chr13-89590089-INS-114 | chr13-89590089-INS-114 |
| chr13-89626452-INS-7086 | chr13-89626453-INS-7086 |
| chr13-89631159-INS-185 | chr13-89631159-INS-185 |
| chr13-89634519-INS-6676 | chr13-89634520-INS-6676 |
| chr13-89639900-INS-400 | chr13-89639900-INS-400 |
| chr13-89653301-INS-3107 | chr13-89653302-INS-3107 |
| chr13-89660017-INS-180 | chr13-89660017-INS-189 |
| chr13-89665149-INS-52 | chr13-89665150-INS-51 |
| chr13-89671286-INS-118 | chr13-89671286-INS-152 |
| chr13-89672609-INS-75 | chr13-89672606-INS-55 |
| chr13-89678220-INS-699 | chr13-89678220-INS-700 |
| chr13-89697142-INS-60 | chr13-89697142-INS-60 |
| chr13-89712396-INS-367 | chr13-89712396-INS-367 |
| chr13-89723881-INS-55 | chr13-89723881-INS-102 |
| chr13-89780852-INS-184 | chr13-89780852-INS-183 |
| chr13-89781941-INS-6178 | chr13-89781940-INS-6171 |
| chr13-89798636-INS-2313 | chr13-89798637-INS-2319 |
| chr13-89801264-INS-3314 | chr13-89801265-INS-3314 |
| chr13-89864162-INS-79 | chr13-89864143-INS-57 |
| chr13-89899059-INS-5813 | chr13-89899060-INS-5813 |
| chr13-89904342-INS-298 | chr13-89904342-INS-298 |
| chr13-89906848-INS-256 | chr13-89906848-INS-256 |
| chr13-89910279-INS-145 | chr13-89910279-INS-172 |
| chr13-89962008-INS-186 | chr13-89962008-INS-187 |
| chr13-89986544-INS-58 | chr13-89986544-INS-58 |
| chr13-89991060-INS-942 | chr13-89991060-INS-946 |
| chr13-89992647-INS-2641 | chr13-89992648-INS-2641 |
| chr13-89996880-INS-4525 | chr13-89996874-INS-4526 |
| chr13-90020299-INS-80 | chr13-90020299-INS-80 |
| chr13-90036699-INS-1890 | chr13-90036700-INS-1890 |
| chr13-90090754-INS-155 | chr13-90090755-INS-156 |
| chr13-90114467-INS-164 | chr13-90114468-INS-171 |
| chr13-90121692-INS-4806 | chr13-90121693-INS-4806 |
| chr13-90133524-INS-459 | chr13-90133524-INS-459 |
| chr13-90141450-INS-415 | chr13-90141450-INS-414 |
| chr13-90191213-INS-212 | chr13-90191213-INS-212 |
| chr13-90203147-INS-118 | chr13-90203147-INS-117 |
| chr13-90207124-INS-243 | chr13-90207124-INS-243 |
| chr13-90213765-INS-298 | chr13-90213753-INS-303 |
| chr13-90231511-INS-51 | chr13-90231511-INS-51 |
| chr13-90278201-INS-377 | chr13-90278204-INS-372 |
| chr13-90315199-INS-714 | chr13-90315199-INS-696 |
| chr13-90325598-INS-4486 | chr13-90325599-INS-4486 |
| chr13-90340998-INS-2627 | chr13-90340999-INS-2627 |
| chr13-90348859-INS-93 | chr13-90348859-INS-130 |
| chr13-90357884-INS-207 | chr13-90357884-INS-198 |
| chr13-90364021-INS-88 | chr13-90364021-INS-103 |
| chr13-90392613-INS-6434 | chr13-90392614-INS-6434 |
| chr13-90408396-INS-6269 | chr13-90408397-INS-6268 |
| chr13-90415999-INS-57 | chr13-90416002-INS-52 |
| chr13-90417192-INS-6291 | chr13-90417193-INS-6291 |
| chr13-90447015-INS-64 | chr13-90447007-INS-72 |
| chr13-90454882-INS-64 | chr13-90454903-INS-59 |
| chr13-90475559-INS-312 | chr13-90475559-INS-312 |
| chr13-90502757-INS-6455 | chr13-90502758-INS-6454 |
| chr13-90525756-INS-66 | chr13-90525796-INS-76 |
| chr13-90535158-INS-1582 | chr13-90535159-INS-1582 |

| | |
|---|---|
| chr13-90538662-INS-7288 | chr13-90538663-INS-7288 |
| chr13-90567760-INS-58 | chr13-90567760-INS-52 |
| chr13-90591696-INS-416 | chr13-90591696-INS-416 |
| chr13-90608146-INS-454 | chr13-90608146-INS-458 |
| chr13-90637914-INS-7115 | chr13-90637915-INS-7115 |
| chr13-90653677-INS-327 | chr13-90653677-INS-312 |
| chr13-90679360-INS-1161 | chr13-90679361-INS-1161 |
| chr13-90691136-INS-140 | chr13-90691136-INS-139 |
| chr13-90735978-INS-343 | chr13-90735978-INS-343 |
| chr13-90830913-INS-114 | chr13-90830913-INS-114 |
| chr13-90839504-INS-693 | chr13-90839504-INS-693 |
| chr13-90842777-INS-6939 | chr13-90842778-INS-6939 |
| chr13-90920208-INS-395 | chr13-90920208-INS-394 |
| chr13-90928382-INS-176 | chr13-90928382-INS-201 |
| chr13-90929764-INS-86 | chr13-90929765-INS-126 |
| chr13-90944390-INS-53 | chr13-90944390-INS-53 |
| chr13-90952504-INS-64 | chr13-90952504-INS-64 |
| chr13-90955419-INS-6284 | chr13-90955420-INS-6280 |
| chr13-90989289-INS-110 | chr13-90989289-INS-110 |
| chr13-91013369-INS-224 | chr13-91013368-INS-258 |
| chr13-91016922-INS-485 | chr13-91016922-INS-485 |
| chr13-91021077-INS-158 | chr13-91021077-INS-173 |
| chr13-91024563-INS-200 | chr13-91024563-INS-199 |
| chr13-91030998-INS-659 | chr13-91030998-INS-658 |
| chr13-91039905-INS-72 | chr13-91039901-INS-62 |
| chr13-91041204-INS-1831 | chr13-91041205-INS-1831 |
| chr13-91042390-INS-217 | chr13-91042390-INS-209 |
| chr13-91062683-INS-6475 | chr13-91062684-INS-6475 |
| chr13-91073800-INS-382 | chr13-91073801-INS-382 |
| chr13-91083114-INS-893 | chr13-91083114-INS-893 |
| chr13-91098379-INS-56 | chr13-91098379-INS-56 |
| chr13-91100770-INS-770 | chr13-91100770-INS-769 |
| chr13-91121422-INS-682 | chr13-91121422-INS-682 |
| chr13-91131789-INS-112 | chr13-91131793-INS-98 |
| chr13-91155593-INS-2791 | chr13-91155594-INS-2792 |
| chr13-91165583-INS-110 | chr13-91165583-INS-92 |
| chr13-91210800-INS-6374 | chr13-91210801-INS-6374 |
| chr13-91226135-INS-486 | chr13-91226135-INS-486 |
| chr13-91237773-INS-99 | chr13-91237807-INS-53 |
| chr13-91238801-INS-90 | chr13-91238801-INS-94 |
| chr13-91242177-INS-60 | chr13-91242177-INS-62 |
| chr13-91271542-INS-202 | chr13-91271542-INS-202 |
| chr13-91366335-INS-61 | chr13-91366335-INS-61 |
| chr13-91381309-INS-492 | chr13-91381309-INS-493 |
| chr13-91384241-INS-69 | chr13-91384225-INS-112 |
| chr13-91392866-INS-68 | chr13-91392866-INS-112 |
| chr13-91419079-INS-199 | chr13-91419079-INS-199 |
| chr13-91422289-INS-89 | chr13-91422289-INS-97 |
| chr13-91423154-INS-694 | chr13-91423154-INS-694 |
| chr13-91448195-INS-453 | chr13-91448195-INS-448 |
| chr13-91452167-INS-55 | chr13-91452168-INS-55 |
| chr13-91464253-INS-324 | chr13-91464253-INS-324 |
| chr13-91466372-INS-92 | chr13-91466375-INS-58 |
| chr13-91506412-INS-52 | chr13-91506412-INS-52 |
| chr13-91506482-INS-536 | chr13-91506482-INS-534 |
| chr13-91507065-INS-56 | chr13-91507064-INS-70 |
| chr13-91530531-INS-138 | chr13-91530531-INS-100 |
| chr13-91555657-INS-205 | chr13-91555657-INS-205 |
| chr13-91563971-INS-73 | chr13-91564006-INS-75 |
| chr13-91578859-INS-205 | chr13-91578860-INS-175 |
| chr13-91581050-INS-2850 | chr13-91581051-INS-2850 |
| chr13-91584280-INS-660 | chr13-91584280-INS-660 |
| chr13-91613219-INS-73 | chr13-91613204-INS-68 |
| chr13-91621329-INS-510 | chr13-91621329-INS-510 |
| chr13-91628598-INS-79 | chr13-91628598-INS-79 |
| chr13-91629332-INS-6385 | chr13-91629333-INS-6386 |
| chr13-91640144-INS-161 | chr13-91640140-INS-157 |
| chr13-91642656-INS-113 | chr13-91642656-INS-113 |
| chr13-91650623-INS-54 | chr13-91650623-INS-56 |
| chr13-91650828-INS-644 | chr13-91650828-INS-644 |
| chr13-91667902-INS-206 | chr13-91667902-INS-202 |
| chr13-91681063-INS-53 | chr13-91681045-INS-76 |
| chr13-91681945-INS-588 | chr13-91681945-INS-588 |
| chr13-91688907-INS-169 | chr13-91688907-INS-173 |
| chr13-91763233-INS-58 | chr13-91763233-INS-58 |
| chr13-91763903-INS-1757 | chr13-91763904-INS-1757 |
| chr13-91779049-INS-600 | chr13-91779049-INS-601 |

| | |
|---|---|
| chr13-91806785-INS-53 | chr13-91806785-INS-53 |
| chr13-91817273-INS-51 | chr13-91817273-INS-54 |
| chr13-91843551-INS-104 | chr13-91843551-INS-104 |
| chr13-91848741-INS-63 | chr13-91848741-INS-63 |
| chr13-91864821-INS-100 | chr13-91864822-INS-63 |
| chr13-91871328-INS-62 | chr13-91871328-INS-50 |
| chr13-91893427-INS-50 | chr13-91893427-INS-52 |
| chr13-91915599-INS-56 | chr13-91915605-INS-61 |
| chr13-91924706-INS-103 | chr13-91924687-INS-115 |
| chr13-91943656-INS-80 | chr13-91943656-INS-80 |
| chr13-91988665-INS-154 | chr13-91988665-INS-157 |
| chr13-91994541-INS-126 | chr13-91994541-INS-125 |
| chr13-91996659-INS-3631 | chr13-91996660-INS-3631 |
| chr13-92015230-INS-164 | chr13-92015230-INS-164 |
| chr13-92037266-INS-52 | chr13-92037279-INS-51 |
| chr13-92053294-INS-198 | chr13-92053294-INS-200 |
| chr13-92053453-INS-6401 | chr13-92053454-INS-6401 |
| chr13-92075776-INS-467 | chr13-92075776-INS-467 |
| chr13-92081529-INS-2896 | chr13-92081530-INS-2896 |
| chr13-92083314-INS-1871 | chr13-92083315-INS-1871 |
| chr13-92104898-INS-643 | chr13-92104898-INS-643 |
| chr13-92138303-INS-602 | chr13-92138303-INS-594 |
| chr13-92160006-INS-456 | chr13-92160006-INS-456 |
| chr13-92199775-INS-87 | chr13-92199779-INS-86 |
| chr13-92216103-INS-6335 | chr13-92216104-INS-6334 |
| chr13-92226391-INS-194 | chr13-92226391-INS-195 |
| chr13-92226937-INS-54 | chr13-92226937-INS-54 |
| chr13-92232878-INS-257 | chr13-92232878-INS-244 |
| chr13-92253795-INS-57 | chr13-92253795-INS-57 |
| chr13-92291813-INS-99 | chr13-92291813-INS-99 |
| chr13-92293322-INS-262 | chr13-92293322-INS-262 |
| chr13-92332014-INS-554 | chr13-92332014-INS-554 |
| chr13-92339573-INS-138 | chr13-92339573-INS-154 |
| chr13-92339973-INS-2904 | chr13-92339974-INS-2904 |
| chr13-92344679-INS-64 | chr13-92344683-INS-60 |
| chr13-92353678-INS-78 | chr13-92353660-INS-103 |
| chr13-92375732-INS-258 | chr13-92375732-INS-258 |
| chr13-92402259-INS-52 | chr13-92402259-INS-57 |
| chr13-92429918-INS-124 | chr13-92429918-INS-124 |
| chr13-92452486-INS-400 | chr13-92452486-INS-400 |
| chr13-92456836-INS-477 | chr13-92456836-INS-477 |
| chr13-92466424-INS-299 | chr13-92466424-INS-299 |
| chr13-92468363-INS-2443 | chr13-92468364-INS-2445 |
| chr13-92479562-INS-4851 | chr13-92479563-INS-4851 |
| chr13-92483693-INS-56 | chr13-92483693-INS-60 |
| chr13-92497670-INS-71 | chr13-92497670-INS-73 |
| chr13-92530472-INS-56 | chr13-92530472-INS-56 |
| chr13-92533479-INS-270 | chr13-92533479-INS-234 |
| chr13-92551946-INS-107 | chr13-92551940-INS-106 |
| chr13-92557676-INS-346 | chr13-92557676-INS-346 |
| chr13-92587326-INS-126 | chr13-92587326-INS-126 |
| chr13-92597491-INS-4846 | chr13-92597492-INS-4846 |
| chr13-92597841-INS-60 | chr13-92597841-INS-62 |
| chr13-92609021-INS-341 | chr13-92609021-INS-341 |
| chr13-92632949-INS-244 | chr13-92632949-INS-244 |
| chr13-92637717-INS-585 | chr13-92637717-INS-597 |
| chr13-92648744-INS-86 | chr13-92648744-INS-129 |
| chr13-92674620-INS-1378 | chr13-92674621-INS-1379 |
| chr13-92678784-INS-66 | chr13-92678784-INS-66 |
| chr13-92687850-INS-6466 | chr13-92687851-INS-6466 |
| chr13-92691857-INS-86 | chr13-92691857-INS-86 |
| chr13-92698395-INS-878 | chr13-92698395-INS-878 |
| chr13-92708173-INS-67 | chr13-92708173-INS-79 |
| chr13-92723076-INS-376 | chr13-92723076-INS-379 |
| chr13-92732276-INS-350 | chr13-92732287-INS-357 |
| chr13-92734728-INS-62 | chr13-92734728-INS-80 |
| chr13-92750143-INS-351 | chr13-92750143-INS-356 |
| chr13-92777498-INS-1653 | chr13-92777499-INS-1654 |
| chr13-92806490-INS-135 | chr13-92806490-INS-136 |
| chr13-92814396-INS-222 | chr13-92814396-INS-223 |
| chr13-92823354-INS-3333 | chr13-92823355-INS-3333 |
| chr13-92829041-INS-96 | chr13-92829041-INS-96 |
| chr13-92834339-INS-92 | chr13-92834339-INS-92 |
| chr13-92843344-INS-77 | chr13-92843344-INS-77 |
| chr13-92853243-INS-200 | chr13-92853243-INS-200 |
| chr13-92854346-INS-110 | chr13-92854346-INS-110 |
| chr13-92874419-INS-71 | chr13-92874419-INS-73 |

| | |
|---|---|
| chr13-92887069-INS-186 | chr13-92887069-INS-186 |
| chr13-92894335-INS-77 | chr13-92894335-INS-77 |
| chr13-92904687-INS-54 | chr13-92904687-INS-54 |
| chr13-92908811-INS-166 | chr13-92908811-INS-166 |
| chr13-93029489-INS-5970 | chr13-93029490-INS-5970 |
| chr13-93039287-INS-185 | chr13-93039287-INS-156 |
| chr13-93070463-INS-108 | chr13-93070500-INS-102 |
| chr13-93084929-INS-1179 | chr13-93084930-INS-1182 |
| chr13-93104898-INS-76 | chr13-93104892-INS-71 |
| chr13-93105653-INS-1891 | chr13-93105654-INS-1892 |
| chr13-93132413-INS-80 | chr13-93132417-INS-76 |
| chr13-93134318-INS-58 | chr13-93134318-INS-53 |
| chr13-93150289-INS-169 | chr13-93150289-INS-168 |
| chr13-93191371-INS-175 | chr13-93191366-INS-186 |
| chr13-93200529-INS-384 | chr13-93200529-INS-384 |
| chr13-93236087-INS-399 | chr13-93236087-INS-399 |
| chr13-93256725-INS-2372 | chr13-93256726-INS-2372 |
| chr13-93267727-INS-68 | chr13-93267727-INS-68 |
| chr13-93274302-INS-72 | chr13-93274302-INS-68 |
| chr13-93288981-INS-176 | chr13-93288981-INS-176 |
| chr13-93331163-INS-228 | chr13-93331163-INS-228 |
| chr13-93380856-INS-120 | chr13-93380855-INS-115 |
| chr13-93383919-INS-300 | chr13-93383919-INS-300 |
| chr13-93397958-INS-195 | chr13-93397958-INS-196 |
| chr13-93419472-INS-7221 | chr13-93419473-INS-7221 |
| chr13-93429180-INS-401 | chr13-93429180-INS-401 |
| chr13-93429588-INS-493 | chr13-93429588-INS-493 |
| chr13-93512395-INS-80 | chr13-93512395-INS-80 |
| chr13-93515775-INS-95 | chr13-93515775-INS-96 |
| chr13-93520154-INS-964 | chr13-93520154-INS-964 |
| chr13-93529263-INS-118 | chr13-93529289-INS-112 |
| chr13-93593844-INS-70 | chr13-93593854-INS-92 |
| chr13-93607631-INS-445 | chr13-93607631-INS-445 |
| chr13-93614983-INS-208 | chr13-93614983-INS-208 |
| chr13-93619851-INS-7432 | chr13-93619852-INS-7432 |
| chr13-93624811-INS-304 | chr13-93624811-INS-278 |
| chr13-93631996-INS-399 | chr13-93631996-INS-401 |
| chr13-93643642-INS-212 | chr13-93643642-INS-210 |
| chr13-93647731-INS-203 | chr13-93647731-INS-169 |
| chr13-93652083-INS-206 | chr13-93652083-INS-203 |
| chr13-93655796-INS-75 | chr13-93655796-INS-77 |
| chr13-93658787-INS-127 | chr13-93658787-INS-127 |
| chr13-93666390-INS-904 | chr13-93666390-INS-857 |
| chr13-93677037-INS-123 | chr13-93677037-INS-123 |
| chr13-93689458-INS-52 | chr13-93689470-INS-52 |
| chr13-93692010-INS-6950 | chr13-93692011-INS-6950 |
| chr13-93706462-INS-113 | chr13-93706462-INS-119 |
| chr13-93707622-INS-51 | chr13-93707622-INS-54 |
| chr13-93729819-INS-208 | chr13-93729819-INS-208 |
| chr13-93731221-INS-121 | chr13-93731221-INS-124 |
| chr13-93741918-INS-152 | chr13-93741918-INS-152 |
| chr13-93750450-INS-1572 | chr13-93750451-INS-1572 |
| chr13-93751935-INS-235 | chr13-93751935-INS-237 |
| chr13-93762645-INS-201 | chr13-93762644-INS-196 |
| chr13-93778016-INS-228 | chr13-93778016-INS-228 |
| chr13-93781012-INS-204 | chr13-93781012-INS-204 |
| chr13-93783921-INS-59 | chr13-93783925-INS-55 |
| chr13-93806846-INS-119 | chr13-93806846-INS-134 |
| chr13-93838818-INS-102 | chr13-93838818-INS-102 |
| chr13-93853479-INS-808 | chr13-93853479-INS-829 |
| chr13-93897244-INS-418 | chr13-93897244-INS-418 |
| chr13-93907201-INS-244 | chr13-93907201-INS-244 |
| chr13-93933005-INS-400 | chr13-93933005-INS-400 |
| chr13-93942838-INS-201 | chr13-93942838-INS-201 |
| chr13-93978486-INS-198 | chr13-93978486-INS-198 |
| chr13-93991411-INS-96 | chr13-93991411-INS-96 |
| chr13-94003019-INS-526 | chr13-94003019-INS-527 |
| chr13-94007502-INS-68 | chr13-94007502-INS-84 |
| chr13-94013839-INS-7119 | chr13-94013840-INS-7119 |
| chr13-94022223-INS-286 | chr13-94022223-INS-250 |
| chr13-94031289-INS-383 | chr13-94031290-INS-377 |
| chr13-94035467-INS-80 | chr13-94035467-INS-92 |
| chr13-94071296-INS-97 | chr13-94071295-INS-60 |
| chr13-94107390-INS-73 | chr13-94107390-INS-73 |
| chr13-94108673-INS-57 | chr13-94108673-INS-59 |
| chr13-94160346-INS-152 | chr13-94160346-INS-200 |
| chr13-94163682-INS-2484 | chr13-94163683-INS-2493 |

| | |
|---|---|
| chr13-94168310-INS-55 | chr13-94168310-INS-55 |
| chr13-94214970-INS-53 | chr13-94214970-INS-53 |
| chr13-94228306-INS-201 | chr13-94228290-INS-194 |
| chr13-94232355-INS-85 | chr13-94232352-INS-50 |
| chr13-94249193-INS-195 | chr13-94249193-INS-195 |
| chr13-94252256-INS-75 | chr13-94252256-INS-75 |
| chr13-94256118-INS-438 | chr13-94256118-INS-438 |
| chr13-94275030-INS-167 | chr13-94275030-INS-159 |
| chr13-94278792-INS-53 | chr13-94278792-INS-53 |
| chr13-94288832-INS-203 | chr13-94288832-INS-202 |
| chr13-94294908-INS-199 | chr13-94294908-INS-200 |
| chr13-94343737-INS-211 | chr13-94343737-INS-219 |
| chr13-94345046-INS-693 | chr13-94345046-INS-707 |
| chr13-94363041-INS-52 | chr13-94363041-INS-52 |
| chr13-94374599-INS-301 | chr13-94374579-INS-301 |
| chr13-94385235-INS-215 | chr13-94385235-INS-216 |
| chr13-94418201-INS-244 | chr13-94418201-INS-244 |
| chr13-94429025-INS-131 | chr13-94429025-INS-131 |
| chr13-94439411-INS-106 | chr13-94439406-INS-102 |
| chr13-94445299-INS-118 | chr13-94445299-INS-118 |
| chr13-94463033-INS-79 | chr13-94463033-INS-84 |
| chr13-94468284-INS-7083 | chr13-94468285-INS-7083 |
| chr13-94473461-INS-53 | chr13-94473461-INS-65 |
| chr13-94480752-INS-502 | chr13-94480756-INS-503 |
| chr13-94482093-INS-60 | chr13-94482094-INS-80 |
| chr13-94498128-INS-136 | chr13-94498128-INS-120 |
| chr13-94506286-INS-3198 | chr13-94506287-INS-3198 |
| chr13-94526676-INS-401 | chr13-94526676-INS-400 |
| chr13-94538748-INS-231 | chr13-94538745-INS-230 |
| chr13-94546440-INS-1906 | chr13-94546441-INS-1906 |
| chr13-94573431-INS-428 | chr13-94573431-INS-427 |
| chr13-94574184-INS-198 | chr13-94574184-INS-197 |
| chr13-94580757-INS-117 | chr13-94580757-INS-117 |
| chr13-94600649-INS-66 | chr13-94600649-INS-62 |
| chr13-94603626-INS-1809 | chr13-94603627-INS-1811 |
| chr13-94604586-INS-71 | chr13-94604586-INS-71 |
| chr13-94621947-INS-72 | chr13-94621947-INS-70 |
| chr13-94631133-INS-177 | chr13-94631133-INS-179 |
| chr13-94633797-INS-1895 | chr13-94633798-INS-1895 |
| chr13-94636989-INS-86 | chr13-94636989-INS-84 |
| chr13-94645911-INS-56 | chr13-94645911-INS-56 |
| chr13-94649395-INS-112 | chr13-94649395-INS-112 |
| chr13-94671830-INS-199 | chr13-94671830-INS-199 |
| chr13-94673792-INS-468 | chr13-94673792-INS-468 |
| chr13-94695623-INS-65 | chr13-94695634-INS-98 |
| chr13-94706387-INS-1080 | chr13-94706388-INS-1081 |
| chr13-94708847-INS-63 | chr13-94708868-INS-56 |
| chr13-94734981-INS-62 | chr13-94734981-INS-60 |
| chr13-94748105-INS-77 | chr13-94748095-INS-77 |
| chr13-94757769-INS-237 | chr13-94757756-INS-239 |
| chr13-94783252-INS-79 | chr13-94783252-INS-108 |
| chr13-94800004-INS-213 | chr13-94800004-INS-215 |
| chr13-94815657-INS-53 | chr13-94815657-INS-79 |
| chr13-94888939-INS-589 | chr13-94888939-INS-589 |
| chr13-94902071-INS-51 | chr13-94902071-INS-95 |
| chr13-94933183-INS-207 | chr13-94933183-INS-207 |
| chr13-94993674-INS-220 | chr13-94993674-INS-220 |
| chr13-95001677-INS-2682 | chr13-95001678-INS-2682 |
| chr13-95036283-INS-120 | chr13-95036283-INS-120 |
| chr13-95057486-INS-207 | chr13-95057486-INS-207 |
| chr13-95059676-INS-178 | chr13-95059676-INS-178 |
| chr13-95065977-INS-62 | chr13-95065977-INS-52 |
| chr13-95087694-INS-94 | chr13-95087695-INS-90 |
| chr13-95095226-INS-74 | chr13-95095226-INS-74 |
| chr13-95139955-INS-254 | chr13-95139955-INS-256 |
| chr13-95175710-INS-175 | chr13-95175710-INS-175 |
| chr13-95178137-INS-82 | chr13-95178137-INS-92 |
| chr13-95182383-INS-54 | chr13-95182383-INS-54 |
| chr13-95184918-INS-74 | chr13-95184903-INS-84 |
| chr13-95195268-INS-66 | chr13-95195268-INS-66 |
| chr13-95198822-INS-57 | chr13-95198822-INS-54 |
| chr13-95201216-INS-56 | chr13-95201216-INS-54 |
| chr13-95240940-INS-68 | chr13-95240940-INS-68 |
| chr13-95246825-INS-2965 | chr13-95246826-INS-2965 |
| chr13-95276838-INS-1115 | chr13-95276839-INS-1115 |
| chr13-95293173-INS-172 | chr13-95293173-INS-171 |
| chr13-95378534-INS-301 | chr13-95378535-INS-301 |

| | |
|---|---|
| chr13-95382052-INS-96 | chr13-95382052-INS-93 |
| chr13-95385488-INS-54 | chr13-95385488-INS-72 |
| chr13-95389478-INS-76 | chr13-95389478-INS-52 |
| chr13-95416589-INS-52 | chr13-95416589-INS-68 |
| chr13-95436960-INS-1891 | chr13-95436961-INS-1891 |
| chr13-95448106-INS-124 | chr13-95448098-INS-116 |
| chr13-95456875-INS-197 | chr13-95456876-INS-202 |
| chr13-95483435-INS-144 | chr13-95483435-INS-156 |
| chr13-95495819-INS-169 | chr13-95495819-INS-184 |
| chr13-95502431-INS-94 | chr13-95502431-INS-94 |
| chr13-95514369-INS-710 | chr13-95514369-INS-696 |
| chr13-95528937-INS-148 | chr13-95528933-INS-102 |
| chr13-95532693-INS-52 | chr13-95532693-INS-52 |
| chr13-95539045-INS-78 | chr13-95539036-INS-84 |
| chr13-95541736-INS-958 | chr13-95541736-INS-968 |
| chr13-95546992-INS-98 | chr13-95546992-INS-98 |
| chr13-95548687-INS-143 | chr13-95548713-INS-111 |
| chr13-95560214-INS-117 | chr13-95560214-INS-117 |
| chr13-95566938-INS-582 | chr13-95566938-INS-567 |
| chr13-95571601-INS-69 | chr13-95571601-INS-69 |
| chr13-95583121-INS-173 | chr13-95583112-INS-203 |
| chr13-95597624-INS-180 | chr13-95597602-INS-175 |
| chr13-95605924-INS-865 | chr13-95605925-INS-865 |
| chr13-95606125-INS-63 | chr13-95606136-INS-79 |
| chr13-95625277-INS-113 | chr13-95625280-INS-114 |
| chr13-95637411-INS-974 | chr13-95637411-INS-974 |
| chr13-95679908-INS-205 | chr13-95679908-INS-205 |
| chr13-95698501-INS-118 | chr13-95698501-INS-114 |
| chr13-95704150-INS-52 | chr13-95704148-INS-60 |
| chr13-95706471-INS-104 | chr13-95706471-INS-116 |
| chr13-95711810-INS-214 | chr13-95711810-INS-214 |
| chr13-95717238-INS-238 | chr13-95717238-INS-238 |
| chr13-95718749-INS-83 | chr13-95718749-INS-83 |
| chr13-95719828-INS-241 | chr13-95719828-INS-241 |
| chr13-95731363-INS-124 | chr13-95731356-INS-118 |
| chr13-95812081-INS-68 | chr13-95812081-INS-102 |
| chr13-95823094-INS-58 | chr13-95823094-INS-58 |
| chr13-95828313-INS-58 | chr13-95828313-INS-62 |
| chr13-95831372-INS-136 | chr13-95831381-INS-152 |
| chr13-95849956-INS-52 | chr13-95849956-INS-50 |
| chr13-95853586-INS-80 | chr13-95853586-INS-80 |
| chr13-95892147-INS-7601 | chr13-95892148-INS-7601 |
| chr13-95908288-INS-71 | chr13-95908285-INS-56 |
| chr13-95933863-INS-193 | chr13-95933863-INS-193 |
| chr13-95942889-INS-74 | chr13-95942888-INS-56 |
| chr13-95981730-INS-6172 | chr13-95981731-INS-6172 |
| chr13-95981908-INS-74 | chr13-95981908-INS-74 |
| chr13-95988510-INS-304 | chr13-95988510-INS-318 |
| chr13-95998651-INS-70 | chr13-95998651-INS-70 |
| chr13-96004022-INS-78 | chr13-96004022-INS-78 |
| chr13-96004738-INS-1182 | chr13-96004739-INS-1182 |
| chr13-96014597-INS-6440 | chr13-96014598-INS-6440 |
| chr13-96019972-INS-234 | chr13-96019972-INS-234 |
| chr13-96030905-INS-125 | chr13-96030905-INS-125 |
| chr13-96043904-INS-239 | chr13-96043904-INS-239 |
| chr13-96045138-INS-55 | chr13-96045138-INS-56 |
| chr13-96052297-INS-209 | chr13-96052297-INS-209 |
| chr13-96054270-INS-68 | chr13-96054270-INS-68 |
| chr13-96105792-INS-68 | chr13-96105792-INS-53 |
| chr13-96110310-INS-799 | chr13-96110310-INS-803 |
| chr13-96114273-INS-52 | chr13-96114265-INS-56 |
| chr13-96126909-INS-175 | chr13-96126909-INS-175 |
| chr13-96137723-INS-56 | chr13-96137728-INS-58 |
| chr13-96148878-INS-56 | chr13-96148878-INS-56 |
| chr13-96168990-INS-1219 | chr13-96168990-INS-1193 |
| chr13-96229876-INS-5141 | chr13-96229877-INS-5141 |
| chr13-96230204-INS-247 | chr13-96230204-INS-247 |
| chr13-96255092-INS-96 | chr13-96255092-INS-96 |
| chr13-96257965-INS-66 | chr13-96257965-INS-66 |
| chr13-96269790-INS-58 | chr13-96269790-INS-62 |
| chr13-96298201-INS-80 | chr13-96298201-INS-80 |
| chr13-96304648-INS-78 | chr13-96304648-INS-78 |
| chr13-96312817-INS-6465 | chr13-96312818-INS-6467 |
| chr13-96330638-INS-80 | chr13-96330638-INS-84 |
| chr13-96343022-INS-1488 | chr13-96343023-INS-1488 |
| chr13-96423578-INS-86 | chr13-96423578-INS-86 |
| chr13-96431079-INS-6583 | chr13-96431080-INS-6583 |

| | |
|---|---|
| chr13-96433936-INS-1587 | chr13-96433937-INS-1588 |
| chr13-96439898-INS-7117 | chr13-96439899-INS-7117 |
| chr13-96444236-INS-63 | chr13-96444236-INS-63 |
| chr13-96445162-INS-7004 | chr13-96445163-INS-7005 |
| chr13-96453864-INS-104 | chr13-96453864-INS-104 |
| chr13-96462663-INS-1481 | chr13-96462664-INS-1481 |
| chr13-96468419-INS-74 | chr13-96468419-INS-74 |
| chr13-96479052-INS-692 | chr13-96479052-INS-692 |
| chr13-96480116-INS-689 | chr13-96480116-INS-704 |
| chr13-96482315-INS-505 | chr13-96482315-INS-505 |
| chr13-96483159-INS-1030 | chr13-96483160-INS-1030 |
| chr13-96492355-INS-440 | chr13-96492355-INS-440 |
| chr13-96517464-INS-427 | chr13-96517464-INS-423 |
| chr13-96523871-INS-102 | chr13-96523871-INS-102 |
| chr13-96530034-INS-1251 | chr13-96530035-INS-1251 |
| chr13-96540106-INS-264 | chr13-96540106-INS-271 |
| chr13-96563071-INS-216 | chr13-96563054-INS-196 |
| chr13-96564284-INS-471 | chr13-96564284-INS-471 |
| chr13-96566092-INS-120 | chr13-96566092-INS-124 |
| chr13-96575539-INS-1074 | chr13-96575540-INS-1075 |
| chr13-96584891-INS-180 | chr13-96584892-INS-181 |
| chr13-96590038-INS-235 | chr13-96590038-INS-252 |
| chr13-96601388-INS-57 | chr13-96601388-INS-58 |
| chr13-96609958-INS-152 | chr13-96609958-INS-186 |
| chr13-96655523-INS-1386 | chr13-96655524-INS-1386 |
| chr13-96680596-INS-52 | chr13-96680596-INS-52 |
| chr13-96698646-INS-211 | chr13-96698646-INS-211 |
| chr13-96716406-INS-84 | chr13-96716406-INS-84 |
| chr13-96722033-INS-1862 | chr13-96722034-INS-1862 |
| chr13-96744188-INS-232 | chr13-96744188-INS-233 |
| chr13-96752775-INS-117 | chr13-96752775-INS-117 |
| chr13-96756605-INS-6440 | chr13-96756606-INS-6440 |
| chr13-96764411-INS-218 | chr13-96764411-INS-217 |
| chr13-96781478-INS-70 | chr13-96781478-INS-70 |
| chr13-96789378-INS-126 | chr13-96789378-INS-126 |
| chr13-96851150-INS-52 | chr13-96851150-INS-52 |
| chr13-96853811-INS-212 | chr13-96853811-INS-212 |
| chr13-96914184-INS-183 | chr13-96914184-INS-184 |
| chr13-96917390-INS-903 | chr13-96917390-INS-903 |
| chr13-96925538-INS-59 | chr13-96925538-INS-59 |
| chr13-96928951-INS-5609 | chr13-96928952-INS-5610 |
| chr13-96928952-INS-548 | chr13-96928952-INS-548 |
| chr13-96949649-INS-269 | chr13-96949649-INS-270 |
| chr13-96952930-INS-60 | chr13-96952930-INS-60 |
| chr13-96957065-INS-220 | chr13-96957065-INS-210 |
| chr13-97008182-INS-941 | chr13-97008182-INS-941 |
| chr13-97011079-INS-735 | chr13-97011079-INS-735 |
| chr13-97021444-INS-126 | chr13-97021444-INS-128 |
| chr13-97056314-INS-55 | chr13-97056314-INS-55 |
| chr13-97076689-INS-208 | chr13-97076689-INS-208 |
| chr13-97095976-INS-499 | chr13-97095976-INS-499 |
| chr13-97102341-INS-58 | chr13-97102341-INS-58 |
| chr13-97114332-INS-592 | chr13-97114332-INS-592 |
| chr13-97123566-INS-208 | chr13-97123566-INS-208 |
| chr13-97128872-INS-84 | chr13-97128872-INS-85 |
| chr13-97136410-INS-3310 | chr13-97136411-INS-3309 |
| chr13-97137034-INS-53 | chr13-97137029-INS-54 |
| chr13-97139966-INS-69 | chr13-97139966-INS-69 |
| chr13-97142524-INS-57 | chr13-97142524-INS-57 |
| chr13-97149529-INS-5390 | chr13-97149530-INS-5390 |
| chr13-97161898-INS-94 | chr13-97161898-INS-94 |
| chr13-97169729-INS-200 | chr13-97169729-INS-200 |
| chr13-97183309-INS-396 | chr13-97183331-INS-398 |
| chr13-97198351-INS-7106 | chr13-97198352-INS-7103 |
| chr13-97198352-INS-440 | chr13-97198352-INS-440 |
| chr13-97212643-INS-200 | chr13-97212643-INS-200 |
| chr13-97217070-INS-145 | chr13-97217070-INS-145 |
| chr13-97221406-INS-214 | chr13-97221406-INS-214 |
| chr13-97226244-INS-1218 | chr13-97226245-INS-1218 |
| chr13-97228194-INS-251 | chr13-97228194-INS-251 |
| chr13-97236097-INS-51 | chr13-97236097-INS-51 |
| chr13-97255819-INS-209 | chr13-97255820-INS-200 |
| chr13-97265288-INS-255 | chr13-97265288-INS-255 |
| chr13-97266755-INS-206 | chr13-97266755-INS-206 |
| chr13-97277571-INS-108 | chr13-97277571-INS-108 |
| chr13-97280833-INS-206 | chr13-97280833-INS-206 |
| chr13-97284984-INS-136 | chr13-97284982-INS-153 |

| | |
|---|---|
| chr13-97301139-INS-58 | chr13-97301139-INS-58 |
| chr13-97305848-INS-218 | chr13-97305848-INS-219 |
| chr13-97321264-INS-105 | chr13-97321265-INS-115 |
| chr13-97322455-INS-246 | chr13-97322455-INS-246 |
| chr13-97335152-INS-306 | chr13-97335152-INS-308 |
| chr13-97348436-INS-409 | chr13-97348436-INS-409 |
| chr13-97351606-INS-70 | chr13-97351606-INS-70 |
| chr13-97353512-INS-208 | chr13-97353512-INS-208 |
| chr13-97355035-INS-814 | chr13-97355035-INS-804 |
| chr13-97361436-INS-60 | chr13-97361436-INS-50 |
| chr13-97375180-INS-284 | chr13-97375180-INS-279 |
| chr13-97391364-INS-172 | chr13-97391364-INS-172 |
| chr13-97394224-INS-70 | chr13-97394225-INS-50 |
| chr13-97400812-INS-67 | chr13-97400812-INS-67 |
| chr13-97402587-INS-107 | chr13-97402586-INS-94 |
| chr13-97406118-INS-7127 | chr13-97406119-INS-7127 |
| chr13-97415294-INS-201 | chr13-97415320-INS-187 |
| chr13-97415776-INS-1268 | chr13-97415777-INS-1268 |
| chr13-97419835-INS-56 | chr13-97419835-INS-58 |
| chr13-97423794-INS-78 | chr13-97423772-INS-52 |
| chr13-97432656-INS-78 | chr13-97432656-INS-102 |
| chr13-97434292-INS-220 | chr13-97434292-INS-222 |
| chr13-97441835-INS-198 | chr13-97441835-INS-198 |
| chr13-97448305-INS-53 | chr13-97448305-INS-54 |
| chr13-97457385-INS-8421 | chr13-97457386-INS-8422 |
| chr13-97487806-INS-72 | chr13-97487806-INS-72 |
| chr13-97523150-INS-387 | chr13-97523150-INS-387 |
| chr13-97541310-INS-64 | chr13-97541310-INS-66 |
| chr13-97564701-INS-188 | chr13-97564701-INS-188 |
| chr13-97588381-INS-221 | chr13-97588381-INS-221 |
| chr13-97595570-INS-463 | chr13-97595567-INS-458 |
| chr13-97597205-INS-60 | chr13-97597201-INS-52 |
| chr13-97601991-INS-210 | chr13-97601991-INS-210 |
| chr13-97604978-INS-73 | chr13-97604978-INS-73 |
| chr13-97605167-INS-561 | chr13-97605167-INS-561 |
| chr13-97614473-INS-7195 | chr13-97614474-INS-7195 |
| chr13-97637908-INS-59 | chr13-97637899-INS-71 |
| chr13-97657945-INS-114 | chr13-97657946-INS-105 |
| chr13-97679874-INS-108 | chr13-97679874-INS-108 |
| chr13-97720568-INS-7120 | chr13-97720569-INS-7120 |
| chr13-97736105-INS-87 | chr13-97736103-INS-90 |
| chr13-97749998-INS-5547 | chr13-97749999-INS-5547 |
| chr13-97765763-INS-5148 | chr13-97765764-INS-5148 |
| chr13-97795847-INS-51 | chr13-97795847-INS-51 |
| chr13-97798248-INS-1539 | chr13-97798249-INS-1539 |
| chr13-97828555-INS-57 | chr13-97828547-INS-72 |
| chr13-97875916-INS-204 | chr13-97875916-INS-204 |
| chr13-97903992-INS-80 | chr13-97903992-INS-80 |
| chr13-97920245-INS-56 | chr13-97920245-INS-56 |
| chr13-97937772-INS-75 | chr13-97937757-INS-68 |
| chr13-97958211-INS-174 | chr13-97958212-INS-168 |
| chr13-97960507-INS-72 | chr13-97960507-INS-73 |
| chr13-97965648-INS-237 | chr13-97965648-INS-238 |
| chr13-97973106-INS-109 | chr13-97973106-INS-133 |
| chr13-98030707-INS-52 | chr13-98030702-INS-55 |
| chr13-98063714-INS-63 | chr13-98063714-INS-63 |
| chr13-98087069-INS-6462 | chr13-98087070-INS-6462 |
| chr13-98104574-INS-60 | chr13-98104574-INS-60 |
| chr13-98113166-INS-83 | chr13-98113166-INS-87 |
| chr13-98139035-INS-84 | chr13-98139040-INS-98 |
| chr13-98149347-INS-57 | chr13-98149347-INS-57 |
| chr13-98196023-INS-159 | chr13-98196023-INS-139 |
| chr13-98243258-INS-364 | chr13-98243258-INS-364 |
| chr13-98262173-INS-65 | chr13-98262173-INS-65 |
| chr13-98265444-INS-4805 | chr13-98265445-INS-4805 |
| chr13-98292875-INS-66 | chr13-98292875-INS-64 |
| chr13-98328211-INS-100 | chr13-98328211-INS-100 |
| chr13-98355496-INS-225 | chr13-98355496-INS-225 |
| chr13-98367726-INS-203 | chr13-98367726-INS-203 |
| chr13-98372584-INS-207 | chr13-98372584-INS-207 |
| chr13-98395498-INS-104 | chr13-98395498-INS-104 |
| chr13-98424045-INS-96 | chr13-98424045-INS-54 |
| chr13-98460530-INS-60 | chr13-98460530-INS-60 |
| chr13-98475412-INS-96 | chr13-98475412-INS-96 |
| chr13-98485016-INS-125 | chr13-98485016-INS-148 |
| chr13-98512436-INS-62 | chr13-98512436-INS-62 |
| chr13-98546515-INS-76 | chr13-98546494-INS-57 |

| | |
|---|---|
| chr13-98557556-INS-192 | chr13-98557556-INS-192 |
| chr13-98614069-INS-129 | chr13-98614068-INS-126 |
| chr13-98617278-INS-56 | chr13-98617278-INS-56 |
| chr13-98654057-INS-111 | chr13-98654057-INS-111 |
| chr13-98658871-INS-482 | chr13-98658871-INS-482 |
| chr13-98662151-INS-5668 | chr13-98662152-INS-5668 |
| chr13-98676206-INS-280 | chr13-98676206-INS-280 |
| chr13-98677297-INS-58 | chr13-98677297-INS-58 |
| chr13-98697371-INS-96 | chr13-98697371-INS-96 |
| chr13-98699783-INS-82 | chr13-98699759-INS-60 |
| chr13-98705332-INS-76 | chr13-98705332-INS-76 |
| chr13-98707419-INS-141 | chr13-98707419-INS-141 |
| chr13-98733124-INS-98 | chr13-98733124-INS-70 |
| chr13-98747377-INS-66 | chr13-98747371-INS-66 |
| chr13-98757560-INS-79 | chr13-98757560-INS-60 |
| chr13-98757970-INS-9609 | chr13-98757971-INS-9593 |
| chr13-98766258-INS-262 | chr13-98766258-INS-262 |
| chr13-98771154-INS-130 | chr13-98771154-INS-125 |
| chr13-98782445-INS-203 | chr13-98782445-INS-203 |
| chr13-98785294-INS-175 | chr13-98785294-INS-175 |
| chr13-98807962-INS-205 | chr13-98807962-INS-205 |
| chr13-98818269-INS-66 | chr13-98818269-INS-66 |
| chr13-98838141-INS-497 | chr13-98838141-INS-497 |
| chr13-98840632-INS-89 | chr13-98840632-INS-94 |
| chr13-98848048-INS-105 | chr13-98848048-INS-105 |
| chr13-98873678-INS-161 | chr13-98873677-INS-162 |
| chr13-98881167-INS-206 | chr13-98881167-INS-206 |
| chr13-98901493-INS-200 | chr13-98901493-INS-200 |
| chr13-98919442-INS-469 | chr13-98919442-INS-469 |
| chr13-98942146-INS-233 | chr13-98942146-INS-233 |
| chr13-98954532-INS-5545 | chr13-98954533-INS-5545 |
| chr13-98999169-INS-103 | chr13-98999169-INS-105 |
| chr13-99064927-INS-103 | chr13-99064923-INS-93 |
| chr13-99094796-INS-1999 | chr13-99094797-INS-1999 |
| chr13-99106149-INS-83 | chr13-99106149-INS-83 |
| chr13-99109652-INS-659 | chr13-99109652-INS-659 |
| chr13-99152987-INS-236 | chr13-99152987-INS-236 |
| chr13-99172633-INS-385 | chr13-99172633-INS-385 |
| chr13-99198347-INS-380 | chr13-99198344-INS-378 |
| chr13-99200888-INS-103 | chr13-99200888-INS-106 |
| chr13-99204600-INS-1116 | chr13-99204601-INS-1116 |
| chr13-99212247-INS-96 | chr13-99212247-INS-96 |
| chr13-99220025-INS-61 | chr13-99220025-INS-61 |
| chr13-99232378-INS-60 | chr13-99232378-INS-60 |
| chr13-99249913-INS-95 | chr13-99249919-INS-127 |
| chr13-99251273-INS-5544 | chr13-99251274-INS-5544 |
| chr13-99254556-INS-62 | chr13-99254556-INS-62 |
| chr13-99260265-INS-104 | chr13-99260265-INS-104 |
| chr13-99266770-INS-56 | chr13-99266776-INS-61 |
| chr13-99276744-INS-92 | chr13-99276744-INS-92 |
| chr13-99301293-INS-209 | chr13-99301293-INS-209 |
| chr13-99303537-INS-168 | chr13-99303537-INS-169 |
| chr13-99316855-INS-98 | chr13-99316855-INS-112 |
| chr13-99362503-INS-5515 | chr13-99362504-INS-5517 |
| chr13-99363167-INS-170 | chr13-99363167-INS-170 |
| chr13-99394486-INS-498 | chr13-99394486-INS-498 |
| chr13-99395691-INS-50 | chr13-99395691-INS-80 |
| chr13-99411438-INS-263 | chr13-99411439-INS-276 |
| chr13-99415507-INS-106 | chr13-99415507-INS-153 |
| chr13-99425657-INS-82 | chr13-99425657-INS-82 |
| chr13-99442363-INS-65 | chr13-99442363-INS-68 |
| chr13-99453026-INS-231 | chr13-99453026-INS-231 |
| chr13-99482487-INS-166 | chr13-99482487-INS-166 |
| chr13-99490364-INS-91 | chr13-99490364-INS-119 |
| chr13-99496757-INS-128 | chr13-99496757-INS-130 |
| chr13-99512949-INS-54 | chr13-99512949-INS-54 |
| chr13-99516525-INS-92 | chr13-99516531-INS-124 |
| chr13-99518771-INS-169 | chr13-99518771-INS-168 |
| chr13-99522461-INS-66 | chr13-99522461-INS-62 |
| chr13-99543704-INS-56 | chr13-99543704-INS-56 |
| chr13-99548644-INS-623 | chr13-99548644-INS-599 |
| chr13-99556579-INS-6440 | chr13-99556580-INS-6440 |
| chr13-99599879-INS-52 | chr13-99599879-INS-52 |
| chr13-99605797-INS-56 | chr13-99605797-INS-56 |
| chr13-99622738-INS-93 | chr13-99622732-INS-117 |
| chr13-99636132-INS-56 | chr13-99636132-INS-56 |
| chr13-99639910-INS-65 | chr13-99639910-INS-69 |

| | |
|---|---|
| chr13-99688530-INS-181 | chr13-99688530-INS-182 |
| chr13-99699377-INS-52 | chr13-99699366-INS-80 |
| chr13-99717078-INS-60 | chr13-99717078-INS-60 |
| chr13-99722806-INS-103 | chr13-99722807-INS-66 |
| chr13-99724923-INS-563 | chr13-99724923-INS-563 |
| chr13-99742838-INS-52 | chr13-99742839-INS-52 |
| chr13-99764816-INS-362 | chr13-99764816-INS-362 |
| chr13-99766951-INS-73 | chr13-99766951-INS-73 |
| chr13-99771581-INS-203 | chr13-99771581-INS-205 |
| chr13-99780060-INS-52 | chr13-99780061-INS-52 |
| chr13-99781844-INS-57 | chr13-99781844-INS-57 |
| chr13-99786414-INS-108 | chr13-99786414-INS-79 |
| chr13-99797840-INS-88 | chr13-99797842-INS-80 |
| chr13-99830011-INS-213 | chr13-99830011-INS-213 |
| chr13-99831425-INS-67 | chr13-99831425-INS-67 |
| chr13-99834871-INS-279 | chr13-99834871-INS-279 |
| chr13-99840833-INS-410 | chr13-99840833-INS-410 |
| chr13-99871143-INS-544 | chr13-99871143-INS-541 |
| chr13-99881231-INS-61 | chr13-99881238-INS-55 |
| chr13-99914794-INS-822 | chr13-99914794-INS-822 |
| chr13-99921915-INS-115 | chr13-99921915-INS-115 |
| chr13-99929323-INS-205 | chr13-99929323-INS-205 |
| chr13-99950163-INS-200 | chr13-99950163-INS-200 |
| chr13-99959873-INS-69 | chr13-99959873-INS-87 |
| chr13-99963465-INS-117 | chr13-99963465-INS-117 |
| chr13-99973193-INS-54 | chr13-99973193-INS-57 |
| chr13-99986018-INS-338 | chr13-99986017-INS-320 |
| chr13-99987511-INS-195 | chr13-99987511-INS-195 |
| chr13-100007787-INS-76 | chr13-100007787-INS-84 |
| chr13-100019085-INS-197 | chr13-100019085-INS-197 |
| chr13-100020665-INS-65 | chr13-100020665-INS-72 |
| chr13-100023861-INS-60 | chr13-100023861-INS-60 |
| chr13-100030904-INS-3699 | chr13-100030902-INS-3700 |
| chr13-100031058-INS-203 | chr13-100031047-INS-200 |
| chr13-100039438-INS-130 | chr13-100039438-INS-130 |
| chr13-100046725-INS-73 | chr13-100046725-INS-73 |
| chr13-100051407-INS-60 | chr13-100051375-INS-108 |
| chr13-100055324-INS-52 | chr13-100055324-INS-54 |
| chr13-100065811-INS-737 | chr13-100065811-INS-737 |
| chr13-100071380-INS-52 | chr13-100071373-INS-54 |
| chr13-100106938-INS-411 | chr13-100106938-INS-382 |
| chr13-100141245-INS-200 | chr13-100141245-INS-200 |
| chr13-100147963-INS-58 | chr13-100147961-INS-62 |
| chr13-100155479-INS-302 | chr13-100155482-INS-303 |
| chr13-100165060-INS-197 | chr13-100165060-INS-197 |
| chr13-100209733-INS-69 | chr13-100209754-INS-81 |
| chr13-100301741-INS-72 | chr13-100301741-INS-72 |
| chr13-100343629-INS-1315 | chr13-100343630-INS-1315 |
| chr13-100349053-INS-205 | chr13-100349053-INS-205 |
| chr13-100349872-INS-176 | chr13-100349872-INS-176 |
| chr13-100364531-INS-57 | chr13-100364532-INS-57 |
| chr13-100420849-INS-223 | chr13-100420849-INS-223 |
| chr13-100474732-INS-4985 | chr13-100474733-INS-4985 |
| chr13-100531827-INS-92 | chr13-100531827-INS-92 |
| chr13-100544218-INS-190 | chr13-100544218-INS-190 |
| chr13-100552806-INS-219 | chr13-100552806-INS-220 |
| chr13-100556298-INS-66 | chr13-100556298-INS-66 |
| chr13-100569471-INS-267 | chr13-100569471-INS-267 |
| chr13-100576608-INS-57 | chr13-100576608-INS-57 |
| chr13-100581453-INS-210 | chr13-100581453-INS-210 |
| chr13-100586928-INS-233 | chr13-100586928-INS-236 |
| chr13-100604435-INS-62 | chr13-100604435-INS-62 |
| chr13-100632558-INS-72 | chr13-100632558-INS-72 |
| chr13-100635615-INS-209 | chr13-100635615-INS-209 |
| chr13-100636844-INS-197 | chr13-100636850-INS-198 |
| chr13-100640083-INS-387 | chr13-100640084-INS-387 |
| chr13-100641244-INS-162 | chr13-100641247-INS-151 |
| chr13-100698773-INS-193 | chr13-100698773-INS-195 |
| chr13-100705424-INS-91 | chr13-100705424-INS-91 |
| chr13-100718822-INS-68 | chr13-100718822-INS-56 |
| chr13-100745756-INS-56 | chr13-100745756-INS-56 |
| chr13-100749218-INS-180 | chr13-100749218-INS-183 |
| chr13-100750223-INS-236 | chr13-100750223-INS-236 |
| chr13-100774653-INS-60 | chr13-100774653-INS-60 |
| chr13-100776049-INS-462 | chr13-100776049-INS-462 |
| chr13-100817632-INS-64 | chr13-100817632-INS-68 |
| chr13-100818841-INS-203 | chr13-100818841-INS-203 |

| | |
|---|---|
| chr13-100852353-INS-473 | chr13-100852353-INS-473 |
| chr13-100862108-INS-118 | chr13-100862108-INS-118 |
| chr13-100862892-INS-232 | chr13-100862893-INS-266 |
| chr13-100869676-INS-214 | chr13-100869676-INS-214 |
| chr13-100871461-INS-202 | chr13-100871461-INS-202 |
| chr13-100875293-INS-248 | chr13-100875293-INS-214 |
| chr13-100884973-INS-132 | chr13-100884973-INS-126 |
| chr13-100888847-INS-502 | chr13-100888847-INS-502 |
| chr13-100894076-INS-93 | chr13-100894090-INS-93 |
| chr13-100897580-INS-172 | chr13-100897580-INS-172 |
| chr13-100898875-INS-872 | chr13-100898875-INS-872 |
| chr13-100899828-INS-210 | chr13-100899828-INS-210 |
| chr13-100941553-INS-212 | chr13-100941553-INS-212 |
| chr13-100942925-INS-209 | chr13-100942925-INS-209 |
| chr13-100946970-INS-75 | chr13-100946970-INS-81 |
| chr13-100958194-INS-89 | chr13-100958194-INS-91 |
| chr13-100979783-INS-5549 | chr13-100979784-INS-5549 |
| chr13-101016813-INS-93 | chr13-101016801-INS-58 |
| chr13-101023368-INS-140 | chr13-101023368-INS-140 |
| chr13-101036837-INS-276 | chr13-101036843-INS-252 |
| chr13-101038767-INS-637 | chr13-101038767-INS-637 |
| chr13-101044504-INS-132 | chr13-101044504-INS-132 |
| chr13-101108982-INS-604 | chr13-101108983-INS-604 |
| chr13-101109554-INS-84 | chr13-101109555-INS-92 |
| chr13-101114140-INS-346 | chr13-101114140-INS-350 |
| chr13-101128804-INS-146 | chr13-101128804-INS-146 |
| chr13-101138639-INS-64 | chr13-101138639-INS-72 |
| chr13-101187022-INS-157 | chr13-101187022-INS-149 |
| chr13-101188532-INS-52 | chr13-101188530-INS-92 |
| chr13-101190611-INS-54 | chr13-101190611-INS-50 |
| chr13-101211434-INS-335 | chr13-101211435-INS-328 |
| chr13-101212846-INS-84 | chr13-101212846-INS-84 |
| chr13-101213625-INS-507 | chr13-101213625-INS-507 |
| chr13-101237021-INS-521 | chr13-101237021-INS-515 |
| chr13-101295764-INS-379 | chr13-101295764-INS-379 |
| chr13-101316007-INS-507 | chr13-101316007-INS-507 |
| chr13-101328837-INS-2003 | chr13-101328838-INS-2004 |
| chr13-101356622-INS-99 | chr13-101356623-INS-75 |
| chr13-101362558-INS-79 | chr13-101362558-INS-80 |
| chr13-101401860-INS-76 | chr13-101401860-INS-66 |
| chr13-101406586-INS-378 | chr13-101406586-INS-372 |
| chr13-101411815-INS-56 | chr13-101411779-INS-64 |
| chr13-101424047-INS-56 | chr13-101424021-INS-104 |
| chr13-101436448-INS-241 | chr13-101436448-INS-241 |
| chr13-101477215-INS-213 | chr13-101477215-INS-213 |
| chr13-101490768-INS-387 | chr13-101490768-INS-387 |
| chr13-101496802-INS-185 | chr13-101496802-INS-185 |
| chr13-101499484-INS-200 | chr13-101499484-INS-200 |
| chr13-101518613-INS-795 | chr13-101518608-INS-798 |
| chr13-101528629-INS-220 | chr13-101528629-INS-220 |
| chr13-101541640-INS-265 | chr13-101541640-INS-265 |
| chr13-101573729-INS-6464 | chr13-101573730-INS-6465 |
| chr13-101583784-INS-187 | chr13-101583784-INS-187 |
| chr13-101588237-INS-50 | chr13-101588237-INS-50 |
| chr13-101597297-INS-514 | chr13-101597297-INS-514 |
| chr13-101643696-INS-200 | chr13-101643696-INS-204 |
| chr13-101645419-INS-52 | chr13-101645419-INS-60 |
| chr13-101652236-INS-914 | chr13-101652236-INS-914 |
| chr13-101653918-INS-6676 | chr13-101653919-INS-6676 |
| chr13-101687323-INS-88 | chr13-101687323-INS-86 |
| chr13-101697524-INS-127 | chr13-101697539-INS-108 |
| chr13-101714537-INS-218 | chr13-101714537-INS-218 |
| chr13-101730034-INS-1776 | chr13-101730035-INS-1776 |
| chr13-101731987-INS-218 | chr13-101731987-INS-224 |
| chr13-101776388-INS-239 | chr13-101776388-INS-261 |
| chr13-101787444-INS-67 | chr13-101787444-INS-67 |
| chr13-101796540-INS-68 | chr13-101796540-INS-68 |
| chr13-101805469-INS-7139 | chr13-101805470-INS-7140 |
| chr13-101867381-INS-215 | chr13-101867381-INS-216 |
| chr13-101892195-INS-51 | chr13-101892196-INS-51 |
| chr13-101930032-INS-97 | chr13-101930032-INS-97 |
| chr13-101937856-INS-117 | chr13-101937856-INS-111 |
| chr13-101970269-INS-70 | chr13-101970269-INS-70 |
| chr13-102004212-INS-77 | chr13-102004212-INS-77 |
| chr13-102007794-INS-84 | chr13-102007798-INS-74 |
| chr13-102043916-INS-85 | chr13-102043920-INS-55 |
| chr13-102053289-INS-695 | chr13-102053289-INS-714 |

| | |
|---|---|
| chr13-102053297-INS-51 | chr13-102053288-INS-63 |
| chr13-102077925-INS-511 | chr13-102077931-INS-511 |
| chr13-102085065-INS-2862 | chr13-102085066-INS-2862 |
| chr13-102114114-INS-76 | chr13-102114114-INS-96 |
| chr13-102187502-INS-165 | chr13-102187502-INS-115 |
| chr13-102200162-INS-71 | chr13-102200205-INS-52 |
| chr13-102322213-INS-6817 | chr13-102322214-INS-6818 |
| chr13-102342727-INS-456 | chr13-102342727-INS-456 |
| chr13-102364355-INS-76 | chr13-102364355-INS-76 |
| chr13-102365515-INS-213 | chr13-102365515-INS-212 |
| chr13-102368664-INS-68 | chr13-102368664-INS-71 |
| chr13-102376972-INS-1627 | chr13-102376963-INS-1625 |
| chr13-102437765-INS-80 | chr13-102437765-INS-80 |
| chr13-102516298-INS-185 | chr13-102516298-INS-185 |
| chr13-102557942-INS-212 | chr13-102557942-INS-212 |
| chr13-102596721-INS-151 | chr13-102596721-INS-153 |
| chr13-102601367-INS-58 | chr13-102601367-INS-50 |
| chr13-102643891-INS-64 | chr13-102643891-INS-64 |
| chr13-102664607-INS-960 | chr13-102664607-INS-959 |
| chr13-102676191-INS-6640 | chr13-102676192-INS-6637 |
| chr13-102695143-INS-54 | chr13-102695154-INS-52 |
| chr13-102695378-INS-343 | chr13-102695378-INS-343 |
| chr13-102698877-INS-807 | chr13-102698877-INS-808 |
| chr13-102790486-INS-58 | chr13-102790491-INS-59 |
| chr13-102817751-INS-81 | chr13-102817751-INS-51 |
| chr13-102910943-INS-84 | chr13-102910943-INS-84 |
| chr13-102912152-INS-3658 | chr13-102912153-INS-3658 |
| chr13-102978568-INS-51 | chr13-102978566-INS-57 |
| chr13-102985209-INS-102 | chr13-102985209-INS-102 |
| chr13-102998170-INS-72 | chr13-102998180-INS-80 |
| chr13-103001854-INS-52 | chr13-103001854-INS-57 |
| chr13-103058874-INS-861 | chr13-103058874-INS-862 |
| chr13-103115740-INS-84 | chr13-103115764-INS-76 |
| chr13-103118462-INS-96 | chr13-103118462-INS-96 |
| chr13-103132307-INS-110 | chr13-103132307-INS-100 |
| chr13-103134140-INS-59 | chr13-103134140-INS-61 |
| chr13-103194959-INS-85 | chr13-103194950-INS-60 |
| chr13-103208216-INS-551 | chr13-103208216-INS-551 |
| chr13-103213889-INS-92 | chr13-103213889-INS-92 |
| chr13-103239549-INS-7098 | chr13-103239550-INS-7098 |
| chr13-103272968-INS-100 | chr13-103272968-INS-50 |
| chr13-103276459-INS-207 | chr13-103276459-INS-207 |
| chr13-103283094-INS-202 | chr13-103283094-INS-201 |
| chr13-103288835-INS-74 | chr13-103288835-INS-76 |
| chr13-103327358-INS-72 | chr13-103327358-INS-72 |
| chr13-103333220-INS-134 | chr13-103333220-INS-134 |
| chr13-103351396-INS-86 | chr13-103351434-INS-62 |
| chr13-103361511-INS-109 | chr13-103361554-INS-112 |
| chr13-103364621-INS-1006 | chr13-103364622-INS-1006 |
| chr13-103372114-INS-151 | chr13-103372114-INS-151 |
| chr13-103395181-INS-57 | chr13-103395181-INS-59 |
| chr13-103398985-INS-132 | chr13-103398985-INS-131 |
| chr13-103421217-INS-3997 | chr13-103421218-INS-3997 |
| chr13-103437466-INS-208 | chr13-103437466-INS-208 |
| chr13-103452264-INS-92 | chr13-103452264-INS-56 |
| chr13-103459659-INS-197 | chr13-103459659-INS-198 |
| chr13-103504896-INS-58 | chr13-103504896-INS-58 |
| chr13-103510963-INS-54 | chr13-103510963-INS-56 |
| chr13-103521908-INS-240 | chr13-103521908-INS-240 |
| chr13-103562223-INS-258 | chr13-103562249-INS-273 |
| chr13-103578856-INS-2169 | chr13-103578857-INS-2169 |
| chr13-103607267-INS-5050 | chr13-103607268-INS-5056 |
| chr13-103607466-INS-56 | chr13-103607466-INS-56 |
| chr13-103618286-INS-6833 | chr13-103618287-INS-6833 |
| chr13-103632374-INS-92 | chr13-103632374-INS-94 |
| chr13-103633762-INS-108 | chr13-103633767-INS-109 |
| chr13-103642843-INS-74 | chr13-103642843-INS-50 |
| chr13-103667911-INS-52 | chr13-103667911-INS-60 |
| chr13-103680075-INS-195 | chr13-103680075-INS-195 |
| chr13-103688368-INS-1140 | chr13-103688369-INS-1140 |
| chr13-103693433-INS-388 | chr13-103693433-INS-388 |
| chr13-103733549-INS-6176 | chr13-103733550-INS-6176 |
| chr13-103733844-INS-79 | chr13-103733844-INS-79 |
| chr13-103745213-INS-115 | chr13-103745213-INS-120 |
| chr13-103757714-INS-304 | chr13-103757714-INS-304 |
| chr13-103759415-INS-69 | chr13-103759415-INS-63 |
| chr13-103764648-INS-377 | chr13-103764648-INS-377 |

| | |
|---|---|
| chr13-103771457-INS-6370 | chr13-103771458-INS-6370 |
| chr13-103778187-INS-7104 | chr13-103778188-INS-7105 |
| chr13-103779331-INS-84 | chr13-103779331-INS-84 |
| chr13-103783078-INS-64 | chr13-103783078-INS-64 |
| chr13-103792176-INS-138 | chr13-103792176-INS-146 |
| chr13-103809870-INS-400 | chr13-103809870-INS-400 |
| chr13-103826383-INS-75 | chr13-103826379-INS-74 |
| chr13-103843802-INS-73 | chr13-103843802-INS-57 |
| chr13-103859230-INS-200 | chr13-103859230-INS-215 |
| chr13-103872715-INS-207 | chr13-103872715-INS-207 |
| chr13-103886073-INS-210 | chr13-103886073-INS-210 |
| chr13-103918739-INS-156 | chr13-103918739-INS-156 |
| chr13-103927365-INS-81 | chr13-103927365-INS-86 |
| chr13-103929042-INS-1007 | chr13-103929043-INS-1007 |
| chr13-103935533-INS-198 | chr13-103935533-INS-198 |
| chr13-103939877-INS-149 | chr13-103939877-INS-149 |
| chr13-103944463-INS-207 | chr13-103944463-INS-207 |
| chr13-103945814-INS-3810 | chr13-103945815-INS-3810 |
| chr13-103947742-INS-6442 | chr13-103947743-INS-6442 |
| chr13-103958419-INS-78 | chr13-103958419-INS-78 |
| chr13-103966395-INS-171 | chr13-103966395-INS-171 |
| chr13-103997048-INS-52 | chr13-103997048-INS-50 |
| chr13-103998243-INS-68 | chr13-103998260-INS-57 |
| chr13-104003739-INS-216 | chr13-104003739-INS-216 |
| chr13-104014521-INS-110 | chr13-104014517-INS-91 |
| chr13-104022350-INS-90 | chr13-104022350-INS-90 |
| chr13-104032686-INS-376 | chr13-104032686-INS-376 |
| chr13-104037433-INS-2652 | chr13-104037434-INS-2652 |
| chr13-104088902-INS-60 | chr13-104088902-INS-60 |
| chr13-104095658-INS-211 | chr13-104095658-INS-211 |
| chr13-104111328-INS-229 | chr13-104111328-INS-229 |
| chr13-104131431-INS-116 | chr13-104131431-INS-117 |
| chr13-104145218-INS-319 | chr13-104145218-INS-319 |
| chr13-104150299-INS-168 | chr13-104150299-INS-168 |
| chr13-104155778-INS-213 | chr13-104155778-INS-213 |
| chr13-104184344-INS-68 | chr13-104184344-INS-68 |
| chr13-104186934-INS-175 | chr13-104186925-INS-176 |
| chr13-104214263-INS-379 | chr13-104214263-INS-379 |
| chr13-104216248-INS-92 | chr13-104216242-INS-90 |
| chr13-104231040-INS-98 | chr13-104231040-INS-94 |
| chr13-104255167-INS-218 | chr13-104255167-INS-218 |
| chr13-104269295-INS-173 | chr13-104269295-INS-132 |
| chr13-104285838-INS-68 | chr13-104285838-INS-68 |
| chr13-104298108-INS-56 | chr13-104298108-INS-56 |
| chr13-104352942-INS-240 | chr13-104352942-INS-241 |
| chr13-104355720-INS-112 | chr13-104355720-INS-112 |
| chr13-104373526-INS-6616 | chr13-104373527-INS-6616 |
| chr13-104387648-INS-225 | chr13-104387648-INS-225 |
| chr13-104394011-INS-6877 | chr13-104394012-INS-6877 |
| chr13-104397239-INS-599 | chr13-104397239-INS-599 |
| chr13-104403872-INS-508 | chr13-104403872-INS-508 |
| chr13-104408929-INS-2285 | chr13-104408930-INS-2287 |
| chr13-104418095-INS-206 | chr13-104418095-INS-206 |
| chr13-104424339-INS-55 | chr13-104424339-INS-52 |
| chr13-104426887-INS-197 | chr13-104426887-INS-197 |
| chr13-104455988-INS-68 | chr13-104455988-INS-64 |
| chr13-104461161-INS-6712 | chr13-104461162-INS-6712 |
| chr13-104487261-INS-147 | chr13-104487249-INS-120 |
| chr13-104568571-INS-94 | chr13-104568571-INS-94 |
| chr13-104578950-INS-90 | chr13-104578951-INS-57 |
| chr13-104591083-INS-249 | chr13-104591083-INS-246 |
| chr13-104604933-INS-6783 | chr13-104604934-INS-6783 |
| chr13-104615517-INS-60 | chr13-104615517-INS-60 |
| chr13-104623994-INS-286 | chr13-104623994-INS-286 |
| chr13-104637503-INS-60 | chr13-104637503-INS-60 |
| chr13-104683721-INS-66 | chr13-104683721-INS-68 |
| chr13-104702542-INS-7199 | chr13-104702543-INS-7199 |
| chr13-104709919-INS-84 | chr13-104709909-INS-95 |
| chr13-104784565-INS-202 | chr13-104784565-INS-202 |
| chr13-104788675-INS-60 | chr13-104788675-INS-56 |
| chr13-104790669-INS-90 | chr13-104790669-INS-88 |
| chr13-104794078-INS-57 | chr13-104794075-INS-61 |
| chr13-104810575-INS-70 | chr13-104810566-INS-60 |
| chr13-104877607-INS-60 | chr13-104877609-INS-80 |
| chr13-104879578-INS-2028 | chr13-104879579-INS-2028 |
| chr13-104928566-INS-6147 | chr13-104928567-INS-6149 |
| chr13-104932085-INS-70 | chr13-104932080-INS-105 |

| | |
|---|---|
| chr13-104977805-INS-1039 | chr13-104977806-INS-1033 |
| chr13-104981178-INS-166 | chr13-104981222-INS-166 |
| chr13-104994425-INS-2373 | chr13-104994426-INS-2373 |
| chr13-105006849-INS-60 | chr13-105006849-INS-72 |
| chr13-105027399-INS-52 | chr13-105027399-INS-54 |
| chr13-105028350-INS-346 | chr13-105028350-INS-346 |
| chr13-105053607-INS-264 | chr13-105053607-INS-264 |
| chr13-105059079-INS-52 | chr13-105059080-INS-52 |
| chr13-105066228-INS-6404 | chr13-105066229-INS-6404 |
| chr13-105066486-INS-105 | chr13-105066486-INS-99 |
| chr13-105080943-INS-3129 | chr13-105080943-INS-3111 |
| chr13-105091560-INS-132 | chr13-105091561-INS-108 |
| chr13-105099557-INS-72 | chr13-105099557-INS-60 |
| chr13-105102830-INS-182 | chr13-105102830-INS-182 |
| chr13-105126165-INS-115 | chr13-105126165-INS-115 |
| chr13-105180402-INS-139 | chr13-105180402-INS-138 |
| chr13-105194283-INS-4728 | chr13-105194284-INS-4728 |
| chr13-105202884-INS-69 | chr13-105202884-INS-76 |
| chr13-105214679-INS-333 | chr13-105214679-INS-333 |
| chr13-105221759-INS-179 | chr13-105221759-INS-218 |
| chr13-105235439-INS-320 | chr13-105235439-INS-320 |
| chr13-105237565-INS-1210 | chr13-105237566-INS-1210 |
| chr13-105303331-INS-63 | chr13-105303331-INS-53 |
| chr13-105330532-INS-51 | chr13-105330533-INS-85 |
| chr13-105360033-INS-328 | chr13-105360033-INS-329 |
| chr13-105371332-INS-799 | chr13-105371332-INS-799 |
| chr13-105453067-INS-419 | chr13-105453067-INS-419 |
| chr13-105466823-INS-69 | chr13-105466823-INS-69 |
| chr13-105471747-INS-85 | chr13-105471747-INS-106 |
| chr13-105486894-INS-241 | chr13-105486894-INS-241 |
| chr13-105511411-INS-51 | chr13-105511411-INS-51 |
| chr13-105523195-INS-183 | chr13-105523195-INS-183 |
| chr13-105526206-INS-60 | chr13-105526216-INS-57 |
| chr13-105541136-INS-54 | chr13-105541136-INS-52 |
| chr13-105546018-INS-670 | chr13-105546018-INS-669 |
| chr13-105560091-INS-57 | chr13-105560073-INS-60 |
| chr13-105564110-INS-313 | chr13-105564110-INS-313 |
| chr13-105579846-INS-490 | chr13-105579846-INS-491 |
| chr13-105591727-INS-591 | chr13-105591727-INS-592 |
| chr13-105602496-INS-99 | chr13-105602496-INS-99 |
| chr13-105606778-INS-52 | chr13-105606778-INS-52 |
| chr13-105658361-INS-58 | chr13-105658361-INS-58 |
| chr13-105662182-INS-4365 | chr13-105662183-INS-4365 |
| chr13-105676099-INS-70 | chr13-105676099-INS-70 |
| chr13-105692777-INS-7110 | chr13-105692778-INS-7110 |
| chr13-105694811-INS-412 | chr13-105694811-INS-367 |
| chr13-105705629-INS-419 | chr13-105705629-INS-419 |
| chr13-105710157-INS-1893 | chr13-105710158-INS-1893 |
| chr13-105721294-INS-604 | chr13-105721294-INS-604 |
| chr13-105728662-INS-490 | chr13-105728662-INS-490 |
| chr13-105730971-INS-8594 | chr13-105730972-INS-8595 |
| chr13-105739647-INS-109 | chr13-105739647-INS-109 |
| chr13-105741122-INS-693 | chr13-105741122-INS-692 |
| chr13-105767401-INS-87 | chr13-105767389-INS-71 |
| chr13-105785648-INS-5212 | chr13-105785649-INS-5212 |
| chr13-105814575-INS-100 | chr13-105814575-INS-64 |
| chr13-105820942-INS-56 | chr13-105820939-INS-50 |
| chr13-105823363-INS-6575 | chr13-105823364-INS-6575 |
| chr13-105823617-INS-399 | chr13-105823617-INS-399 |
| chr13-105833456-INS-56 | chr13-105833456-INS-64 |
| chr13-105874891-INS-937 | chr13-105874891-INS-937 |
| chr13-105884427-INS-5861 | chr13-105884428-INS-5861 |
| chr13-105900123-INS-52 | chr13-105900123-INS-67 |
| chr13-105901020-INS-269 | chr13-105901020-INS-273 |
| chr13-105911948-INS-766 | chr13-105911948-INS-766 |
| chr13-105944325-INS-74 | chr13-105944325-INS-72 |
| chr13-105971485-INS-57 | chr13-105971485-INS-63 |
| chr13-106003274-INS-6684 | chr13-106003275-INS-6684 |
| chr13-106010015-INS-236 | chr13-106010015-INS-235 |
| chr13-106029588-INS-88 | chr13-106029589-INS-54 |
| chr13-106032807-INS-788 | chr13-106032807-INS-788 |
| chr13-106047838-INS-2685 | chr13-106047839-INS-2685 |
| chr13-106050773-INS-2077 | chr13-106050774-INS-2075 |
| chr13-106051413-INS-154 | chr13-106051413-INS-154 |
| chr13-106062647-INS-124 | chr13-106062647-INS-88 |
| chr13-106072256-INS-109 | chr13-106072281-INS-128 |
| chr13-106101424-INS-6842 | chr13-106101425-INS-6843 |

| | |
|---|---|
| chr13-106124126-INS-52 | chr13-106124126-INS-52 |
| chr13-106130945-INS-1480 | chr13-106130946-INS-1480 |
| chr13-106143620-INS-4750 | chr13-106143621-INS-4750 |
| chr13-106150041-INS-110 | chr13-106150053-INS-64 |
| chr13-106153342-INS-62 | chr13-106153342-INS-62 |
| chr13-106156774-INS-77 | chr13-106156790-INS-59 |
| chr13-106167337-INS-86 | chr13-106167337-INS-70 |
| chr13-106179568-INS-82 | chr13-106179568-INS-82 |
| chr13-106185680-INS-165 | chr13-106185680-INS-156 |
| chr13-106192435-INS-64 | chr13-106192435-INS-64 |
| chr13-106199023-INS-6694 | chr13-106199024-INS-6697 |
| chr13-106205463-INS-55 | chr13-106205463-INS-57 |
| chr13-106209469-INS-588 | chr13-106209469-INS-588 |
| chr13-106221777-INS-1453 | chr13-106221778-INS-1453 |
| chr13-106221912-INS-123 | chr13-106221912-INS-127 |
| chr13-106254692-INS-650 | chr13-106254692-INS-651 |
| chr13-106272496-INS-7915 | chr13-106272497-INS-7915 |
| chr13-106324929-INS-4899 | chr13-106324930-INS-4894 |
| chr13-106326140-INS-60 | chr13-106326140-INS-64 |
| chr13-106341302-INS-160 | chr13-106341302-INS-116 |
| chr13-106351359-INS-76 | chr13-106351359-INS-76 |
| chr13-106369496-INS-5421 | chr13-106369497-INS-5421 |
| chr13-106385056-INS-278 | chr13-106385056-INS-278 |
| chr13-106399251-INS-199 | chr13-106399296-INS-196 |
| chr13-106418926-INS-258 | chr13-106418926-INS-278 |
| chr13-106452135-INS-53 | chr13-106452171-INS-66 |
| chr13-106472934-INS-533 | chr13-106472934-INS-533 |
| chr13-106474328-INS-7039 | chr13-106474329-INS-7040 |
| chr13-106483900-INS-120 | chr13-106483900-INS-80 |
| chr13-106484372-INS-300 | chr13-106484372-INS-301 |
| chr13-106488024-INS-86 | chr13-106488024-INS-86 |
| chr13-106492478-INS-95 | chr13-106492478-INS-95 |
| chr13-106496282-INS-633 | chr13-106496282-INS-633 |
| chr13-106500100-INS-559 | chr13-106500100-INS-559 |
| chr13-106531608-INS-62 | chr13-106531608-INS-62 |
| chr13-106532638-INS-55 | chr13-106532638-INS-65 |
| chr13-106549417-INS-6668 | chr13-106549418-INS-6669 |
| chr13-106563527-INS-7078 | chr13-106563528-INS-7078 |
| chr13-106572500-INS-54 | chr13-106572506-INS-60 |
| chr13-106586215-INS-175 | chr13-106586215-INS-175 |
| chr13-106594483-INS-78 | chr13-106594483-INS-65 |
| chr13-106596880-INS-356 | chr13-106596880-INS-356 |
| chr13-106607091-INS-71 | chr13-106607091-INS-76 |
| chr13-106613703-INS-5307 | chr13-106613704-INS-5308 |
| chr13-106639337-INS-68 | chr13-106639337-INS-68 |
| chr13-106644414-INS-6806 | chr13-106644415-INS-6806 |
| chr13-106669795-INS-207 | chr13-106669795-INS-207 |
| chr13-106670212-INS-6442 | chr13-106670213-INS-6442 |
| chr13-106678473-INS-3868 | chr13-106678474-INS-3869 |
| chr13-106682935-INS-59 | chr13-106682937-INS-58 |
| chr13-106695429-INS-560 | chr13-106695429-INS-560 |
| chr13-106762559-INS-92 | chr13-106762559-INS-104 |
| chr13-106769315-INS-214 | chr13-106769315-INS-214 |
| chr13-106830283-INS-188 | chr13-106830283-INS-188 |
| chr13-106838052-INS-2645 | chr13-106838053-INS-2636 |
| chr13-106838052-INS-403 | chr13-106838052-INS-411 |
| chr13-106851404-INS-197 | chr13-106851402-INS-191 |
| chr13-106858029-INS-88 | chr13-106858031-INS-78 |
| chr13-106893538-INS-207 | chr13-106893538-INS-207 |
| chr13-106905322-INS-66 | chr13-106905322-INS-66 |
| chr13-106919142-INS-1890 | chr13-106919143-INS-1890 |
| chr13-106934225-INS-3917 | chr13-106934226-INS-3917 |
| chr13-106954495-INS-6401 | chr13-106954496-INS-6401 |
| chr13-107036433-INS-200 | chr13-107036433-INS-200 |
| chr13-107058326-INS-76 | chr13-107058326-INS-76 |
| chr13-107075304-INS-71 | chr13-107075304-INS-71 |
| chr13-107076332-INS-257 | chr13-107076332-INS-257 |
| chr13-107080080-INS-428 | chr13-107080128-INS-424 |
| chr13-107088740-INS-206 | chr13-107088740-INS-206 |
| chr13-107109393-INS-211 | chr13-107109393-INS-211 |
| chr13-107129133-INS-234 | chr13-107129133-INS-207 |
| chr13-107154026-INS-635 | chr13-107154026-INS-671 |
| chr13-107161352-INS-50 | chr13-107161352-INS-50 |
| chr13-107173481-INS-193 | chr13-107173481-INS-193 |
| chr13-107174314-INS-54 | chr13-107174314-INS-102 |
| chr13-107179912-INS-60 | chr13-107179912-INS-60 |
| chr13-107183767-INS-82 | chr13-107183767-INS-81 |

| | |
|---|---|
| chr13-107190929-INS-60 | chr13-107190929-INS-66 |
| chr13-107216059-INS-56 | chr13-107216059-INS-68 |
| chr13-107218804-INS-557 | chr13-107218804-INS-557 |
| chr13-107224233-INS-382 | chr13-107224233-INS-383 |
| chr13-107240012-INS-75 | chr13-107240012-INS-75 |
| chr13-107243315-INS-213 | chr13-107243315-INS-213 |
| chr13-107249955-INS-198 | chr13-107249955-INS-198 |
| chr13-107252060-INS-66 | chr13-107252060-INS-66 |
| chr13-107258262-INS-87 | chr13-107258262-INS-87 |
| chr13-107264685-INS-60 | chr13-107264691-INS-50 |
| chr13-107277257-INS-64 | chr13-107277257-INS-64 |
| chr13-107279240-INS-52 | chr13-107279240-INS-64 |
| chr13-107296056-INS-53 | chr13-107296056-INS-53 |
| chr13-107302317-INS-100 | chr13-107302349-INS-98 |
| chr13-107312035-INS-116 | chr13-107312035-INS-116 |
| chr13-107324649-INS-118 | chr13-107324657-INS-102 |
| chr13-107362213-INS-176 | chr13-107362213-INS-176 |
| chr13-107368810-INS-195 | chr13-107368810-INS-195 |
| chr13-107382009-INS-6489 | chr13-107382010-INS-6489 |
| chr13-107402066-INS-636 | chr13-107402066-INS-636 |
| chr13-107405437-INS-54 | chr13-107405437-INS-54 |
| chr13-107416939-INS-82 | chr13-107416939-INS-78 |
| chr13-107425400-INS-305 | chr13-107425400-INS-328 |
| chr13-107444121-INS-60 | chr13-107444101-INS-98 |
| chr13-107453721-INS-60 | chr13-107453721-INS-60 |
| chr13-107458298-INS-91 | chr13-107458298-INS-91 |
| chr13-107482640-INS-158 | chr13-107482640-INS-162 |
| chr13-107498125-INS-58 | chr13-107498125-INS-58 |
| chr13-107528800-INS-74 | chr13-107528800-INS-74 |
| chr13-107536122-INS-3665 | chr13-107536123-INS-3655 |
| chr13-107605549-INS-60 | chr13-107605549-INS-64 |
| chr13-107618541-INS-176 | chr13-107618541-INS-177 |
| chr13-107623843-INS-819 | chr13-107623843-INS-819 |
| chr13-107638165-INS-107 | chr13-107638165-INS-105 |
| chr13-107645584-INS-387 | chr13-107645584-INS-387 |
| chr13-107646097-INS-104 | chr13-107646097-INS-104 |
| chr13-107668503-INS-713 | chr13-107668503-INS-721 |
| chr13-107671282-INS-52 | chr13-107671293-INS-50 |
| chr13-107712734-INS-214 | chr13-107712781-INS-214 |
| chr13-107724042-INS-144 | chr13-107724034-INS-178 |
| chr13-107726043-INS-94 | chr13-107726043-INS-82 |
| chr13-107733699-INS-52 | chr13-107733699-INS-52 |
| chr13-107758249-INS-361 | chr13-107758249-INS-366 |
| chr13-107776514-INS-60 | chr13-107776514-INS-60 |
| chr13-107779393-INS-202 | chr13-107779393-INS-202 |
| chr13-107785155-INS-123 | chr13-107785155-INS-123 |
| chr13-107796682-INS-4828 | chr13-107796683-INS-4828 |
| chr13-107798765-INS-56 | chr13-107798766-INS-58 |
| chr13-107869390-INS-54 | chr13-107869390-INS-52 |
| chr13-107877195-INS-62 | chr13-107877195-INS-62 |
| chr13-107885056-INS-96 | chr13-107885056-INS-96 |
| chr13-107918761-INS-114 | chr13-107918761-INS-114 |
| chr13-107920878-INS-72 | chr13-107920879-INS-111 |
| chr13-107989472-INS-206 | chr13-107989472-INS-206 |
| chr13-107993680-INS-88 | chr13-107993680-INS-88 |
| chr13-108014618-INS-105 | chr13-108014618-INS-150 |
| chr13-108035563-INS-217 | chr13-108035563-INS-218 |
| chr13-108037301-INS-104 | chr13-108037311-INS-58 |
| chr13-108037912-INS-202 | chr13-108037912-INS-202 |
| chr13-108044950-INS-56 | chr13-108044950-INS-60 |
| chr13-108049842-INS-69 | chr13-108049848-INS-54 |
| chr13-108057121-INS-59 | chr13-108057121-INS-60 |
| chr13-108064775-INS-143 | chr13-108064775-INS-143 |
| chr13-108077834-INS-68 | chr13-108077834-INS-60 |
| chr13-108080926-INS-319 | chr13-108080926-INS-319 |
| chr13-108083149-INS-86 | chr13-108083135-INS-69 |
| chr13-108087665-INS-62 | chr13-108087643-INS-51 |
| chr13-108095380-INS-216 | chr13-108095380-INS-216 |
| chr13-108098021-INS-77 | chr13-108098021-INS-78 |
| chr13-108105162-INS-56 | chr13-108105162-INS-56 |
| chr13-108107051-INS-68 | chr13-108107051-INS-60 |
| chr13-108118286-INS-55 | chr13-108118287-INS-55 |
| chr13-108123552-INS-103 | chr13-108123552-INS-103 |
| chr13-108127224-INS-72 | chr13-108127224-INS-72 |
| chr13-108143577-INS-54 | chr13-108143577-INS-68 |
| chr13-108146490-INS-406 | chr13-108146490-INS-406 |
| chr13-108154610-INS-6590 | chr13-108154611-INS-6618 |

| | |
|---|---|
| chr13-108177071-INS-57 | chr13-108177071-INS-57 |
| chr13-108192864-INS-1490 | chr13-108192865-INS-1490 |
| chr13-108232770-INS-89 | chr13-108232770-INS-92 |
| chr13-108250415-INS-232 | chr13-108250415-INS-232 |
| chr13-108251439-INS-141 | chr13-108251439-INS-141 |
| chr13-108254456-INS-135 | chr13-108254456-INS-135 |
| chr13-108286042-INS-910 | chr13-108286042-INS-910 |
| chr13-108287231-INS-201 | chr13-108287231-INS-201 |
| chr13-108303623-INS-68 | chr13-108303653-INS-70 |
| chr13-108306662-INS-82 | chr13-108306662-INS-82 |
| chr13-108310797-INS-65 | chr13-108310797-INS-65 |
| chr13-108312705-INS-129 | chr13-108312705-INS-129 |
| chr13-108338532-INS-52 | chr13-108338532-INS-56 |
| chr13-108355441-INS-60 | chr13-108355441-INS-60 |
| chr13-108357897-INS-52 | chr13-108357897-INS-52 |
| chr13-108366504-INS-60 | chr13-108366509-INS-53 |
| chr13-108382027-INS-52 | chr13-108382027-INS-66 |
| chr13-108384781-INS-118 | chr13-108384781-INS-123 |
| chr13-108424056-INS-194 | chr13-108424056-INS-194 |
| chr13-108429915-INS-486 | chr13-108429915-INS-486 |
| chr13-108431676-INS-2827 | chr13-108431677-INS-2827 |
| chr13-108436544-INS-68 | chr13-108436544-INS-68 |
| chr13-108444741-INS-80 | chr13-108444741-INS-70 |
| chr13-108454578-INS-152 | chr13-108454578-INS-152 |
| chr13-108493337-INS-437 | chr13-108493337-INS-437 |
| chr13-108500658-INS-747 | chr13-108500658-INS-744 |
| chr13-108503424-INS-51 | chr13-108503424-INS-51 |
| chr13-108522439-INS-564 | chr13-108522439-INS-564 |
| chr13-108522464-INS-5268 | chr13-108522465-INS-5268 |
| chr13-108526345-INS-177 | chr13-108526345-INS-177 |
| chr13-108527874-INS-462 | chr13-108527874-INS-466 |
| chr13-108537099-INS-416 | chr13-108537099-INS-416 |
| chr13-108553216-INS-3302 | chr13-108553217-INS-3302 |
| chr13-108572785-INS-70 | chr13-108572785-INS-70 |
| chr13-108574871-INS-95 | chr13-108574871-INS-95 |
| chr13-108589120-INS-536 | chr13-108589120-INS-536 |
| chr13-108591505-INS-348 | chr13-108591480-INS-372 |
| chr13-108596204-INS-67 | chr13-108596200-INS-51 |
| chr13-108596867-INS-1019 | chr13-108596868-INS-1019 |
| chr13-108597625-INS-2830 | chr13-108597626-INS-2830 |
| chr13-108619229-INS-65 | chr13-108619229-INS-69 |
| chr13-108659083-INS-65 | chr13-108659083-INS-65 |
| chr13-108696068-INS-4992 | chr13-108696069-INS-4992 |
| chr13-108706684-INS-1028 | chr13-108706685-INS-1032 |
| chr13-108717588-INS-80 | chr13-108717624-INS-55 |
| chr13-108721885-INS-384 | chr13-108721886-INS-380 |
| chr13-108751531-INS-74 | chr13-108751531-INS-82 |
| chr13-108757858-INS-273 | chr13-108757869-INS-251 |
| chr13-108759536-INS-394 | chr13-108759536-INS-394 |
| chr13-108779976-INS-77 | chr13-108779976-INS-77 |
| chr13-108782865-INS-51 | chr13-108782865-INS-50 |
| chr13-108788922-INS-212 | chr13-108788922-INS-197 |
| chr13-108811966-INS-800 | chr13-108811966-INS-792 |
| chr13-108841346-INS-52 | chr13-108841346-INS-52 |
| chr13-108845943-INS-1950 | chr13-108845944-INS-1950 |
| chr13-108848218-INS-54 | chr13-108848218-INS-54 |
| chr13-108850376-INS-304 | chr13-108850358-INS-314 |
| chr13-108858183-INS-75 | chr13-108858183-INS-75 |
| chr13-108869859-INS-6401 | chr13-108869860-INS-6401 |
| chr13-108881249-INS-881 | chr13-108881249-INS-880 |
| chr13-108883437-INS-853 | chr13-108883437-INS-853 |
| chr13-108889474-INS-51 | chr13-108889474-INS-55 |
| chr13-108891774-INS-56 | chr13-108891774-INS-56 |
| chr13-108914206-INS-77 | chr13-108914206-INS-69 |
| chr13-108934214-INS-138 | chr13-108934192-INS-122 |
| chr13-108936210-INS-74 | chr13-108936208-INS-74 |
| chr13-108940289-INS-1831 | chr13-108940290-INS-1831 |
| chr13-108955970-INS-228 | chr13-108955970-INS-228 |
| chr13-108965809-INS-200 | chr13-108965809-INS-200 |
| chr13-108971627-INS-185 | chr13-108971628-INS-186 |
| chr13-108984356-INS-6332 | chr13-108984357-INS-6332 |
| chr13-109002936-INS-52 | chr13-109002936-INS-52 |
| chr13-109020230-INS-648 | chr13-109020230-INS-648 |
| chr13-109028690-INS-995 | chr13-109028690-INS-995 |
| chr13-109030922-INS-68 | chr13-109030912-INS-50 |
| chr13-109032866-INS-66 | chr13-109032866-INS-66 |
| chr13-109047291-INS-227 | chr13-109047291-INS-227 |

| | |
|---|---|
| chr13-109056018-INS-51 | chr13-109056018-INS-56 |
| chr13-109059368-INS-550 | chr13-109059368-INS-550 |
| chr13-109090409-INS-54 | chr13-109090409-INS-54 |
| chr13-109105338-INS-64 | chr13-109105338-INS-68 |
| chr13-109108936-INS-329 | chr13-109108930-INS-332 |
| chr13-109114979-INS-160 | chr13-109114979-INS-160 |
| chr13-109185664-INS-178 | chr13-109185664-INS-182 |
| chr13-109192365-INS-70 | chr13-109192365-INS-70 |
| chr13-109213118-INS-1816 | chr13-109213119-INS-1816 |
| chr13-109236690-INS-160 | chr13-109236690-INS-160 |
| chr13-109239529-INS-160 | chr13-109239529-INS-164 |
| chr13-109254658-INS-67 | chr13-109254658-INS-71 |
| chr13-109266478-INS-126 | chr13-109266478-INS-126 |
| chr13-109307753-INS-72 | chr13-109307753-INS-70 |
| chr13-109343091-INS-669 | chr13-109343091-INS-620 |
| chr13-109416871-INS-60 | chr13-109416871-INS-85 |
| chr13-109435864-INS-56 | chr13-109435864-INS-56 |
| chr13-109438623-INS-103 | chr13-109438623-INS-96 |
| chr13-109444115-INS-87 | chr13-109444115-INS-87 |
| chr13-109457269-INS-241 | chr13-109457269-INS-241 |
| chr13-109466670-INS-68 | chr13-109466670-INS-68 |
| chr13-109467767-INS-1126 | chr13-109467768-INS-1128 |
| chr13-109488857-INS-67 | chr13-109488856-INS-68 |
| chr13-109494898-INS-55 | chr13-109494891-INS-54 |
| chr13-109517265-INS-66 | chr13-109517277-INS-80 |
| chr13-109520554-INS-56 | chr13-109520554-INS-60 |
| chr13-109522199-INS-358 | chr13-109522199-INS-358 |
| chr13-109537866-INS-56 | chr13-109537866-INS-56 |
| chr13-109615578-INS-100 | chr13-109615578-INS-100 |
| chr13-109616471-INS-1704 | chr13-109616472-INS-1701 |
| chr13-109648983-INS-177 | chr13-109648988-INS-168 |
| chr13-109651068-INS-170 | chr13-109651068-INS-172 |
| chr13-109667065-INS-94 | chr13-109667065-INS-94 |
| chr13-109709272-INS-201 | chr13-109709272-INS-219 |
| chr13-109712389-INS-71 | chr13-109712389-INS-58 |
| chr13-109717046-INS-259 | chr13-109717046-INS-259 |
| chr13-109736881-INS-68 | chr13-109736909-INS-55 |
| chr13-109743629-INS-90 | chr13-109743594-INS-101 |
| chr13-109744615-INS-218 | chr13-109744615-INS-219 |
| chr13-109754384-INS-546 | chr13-109754384-INS-546 |
| chr13-109758634-INS-88 | chr13-109758643-INS-114 |
| chr13-109780345-INS-221 | chr13-109780345-INS-221 |
| chr13-109784863-INS-61 | chr13-109784863-INS-61 |
| chr13-109790804-INS-293 | chr13-109790804-INS-293 |
| chr13-109796692-INS-210 | chr13-109796697-INS-215 |
| chr13-109815031-INS-99 | chr13-109815031-INS-98 |
| chr13-109821131-INS-55 | chr13-109821131-INS-55 |
| chr13-109837036-INS-65 | chr13-109837036-INS-65 |
| chr13-109849569-INS-211 | chr13-109849569-INS-212 |
| chr13-109865889-INS-333 | chr13-109865889-INS-333 |
| chr13-109891211-INS-53 | chr13-109891211-INS-53 |
| chr13-109891259-INS-890 | chr13-109891259-INS-891 |
| chr13-109907923-INS-68 | chr13-109907923-INS-82 |
| chr13-109923308-INS-74 | chr13-109923356-INS-70 |
| chr13-109930722-INS-109 | chr13-109930722-INS-109 |
| chr13-109938526-INS-56 | chr13-109938526-INS-56 |
| chr13-109953289-INS-254 | chr13-109953289-INS-250 |
| chr13-109984794-INS-96 | chr13-109984794-INS-98 |
| chr13-109998543-INS-56 | chr13-109998543-INS-56 |
| chr13-110028520-INS-188 | chr13-110028520-INS-196 |
| chr13-110032538-INS-55 | chr13-110032539-INS-50 |
| chr13-110039292-INS-56 | chr13-110039292-INS-58 |
| chr13-110041298-INS-68 | chr13-110041298-INS-68 |
| chr13-110065535-INS-58 | chr13-110065535-INS-60 |
| chr13-110069927-INS-822 | chr13-110069924-INS-816 |
| chr13-110076577-INS-445 | chr13-110076577-INS-446 |
| chr13-110127735-INS-56 | chr13-110127729-INS-82 |
| chr13-110143424-INS-57 | chr13-110143424-INS-57 |
| chr13-110149128-INS-455 | chr13-110149128-INS-455 |
| chr13-110152055-INS-503 | chr13-110152055-INS-503 |
| chr13-110165375-INS-86 | chr13-110165375-INS-86 |
| chr13-110167846-INS-190 | chr13-110167846-INS-190 |
| chr13-110176007-INS-63 | chr13-110176007-INS-64 |
| chr13-110206609-INS-92 | chr13-110206609-INS-92 |
| chr13-110240600-INS-81 | chr13-110240600-INS-81 |
| chr13-110246052-INS-76 | chr13-110246070-INS-72 |
| chr13-110250689-INS-183 | chr13-110250689-INS-183 |

| | |
|---|---|
| chr13-110254065-INS-241 | chr13-110254065-INS-241 |
| chr13-110270046-INS-82 | chr13-110270048-INS-66 |
| chr13-110293203-INS-6657 | chr13-110293204-INS-6658 |
| chr13-110309181-INS-94 | chr13-110309181-INS-96 |
| chr13-110341482-INS-129 | chr13-110341482-INS-132 |
| chr13-110365712-INS-65 | chr13-110365712-INS-65 |
| chr13-110370616-INS-214 | chr13-110370616-INS-214 |
| chr13-110387257-INS-184 | chr13-110387257-INS-184 |
| chr13-110396598-INS-52 | chr13-110396598-INS-64 |
| chr13-110418685-INS-63 | chr13-110418685-INS-84 |
| chr13-110432473-INS-316 | chr13-110432473-INS-318 |
| chr13-110437542-INS-73 | chr13-110437542-INS-73 |
| chr13-110438825-INS-173 | chr13-110438825-INS-173 |
| chr13-110461406-INS-58 | chr13-110461406-INS-63 |
| chr13-110462877-INS-61 | chr13-110462877-INS-61 |
| chr13-110467359-INS-62 | chr13-110467359-INS-62 |
| chr13-110489382-INS-64 | chr13-110489404-INS-70 |
| chr13-110519896-INS-242 | chr13-110519896-INS-242 |
| chr13-110575563-INS-79 | chr13-110575563-INS-60 |
| chr13-110590336-INS-54 | chr13-110590336-INS-54 |
| chr13-110592624-INS-60 | chr13-110592625-INS-60 |
| chr13-110604647-INS-70 | chr13-110604647-INS-70 |
| chr13-110605376-INS-54 | chr13-110605376-INS-54 |
| chr13-110648882-INS-88 | chr13-110648882-INS-90 |
| chr13-110652274-INS-53 | chr13-110652274-INS-53 |
| chr13-110652342-INS-6440 | chr13-110652343-INS-6440 |
| chr13-110657378-INS-691 | chr13-110657378-INS-692 |
| chr13-110666959-INS-682 | chr13-110666959-INS-682 |
| chr13-110671164-INS-64 | chr13-110671164-INS-64 |
| chr13-110676102-INS-881 | chr13-110676102-INS-881 |
| chr13-110688709-INS-72 | chr13-110688722-INS-84 |
| chr13-110728915-INS-851 | chr13-110728914-INS-846 |
| chr13-110737518-INS-64 | chr13-110737518-INS-66 |
| chr13-110745999-INS-241 | chr13-110745999-INS-242 |
| chr13-110754202-INS-440 | chr13-110754202-INS-440 |
| chr13-110794058-INS-1223 | chr13-110794059-INS-1223 |
| chr13-110818744-INS-56 | chr13-110818744-INS-56 |
| chr13-110825487-INS-3371 | chr13-110825488-INS-3372 |
| chr13-110837709-INS-64 | chr13-110837709-INS-64 |
| chr13-110853221-INS-204 | chr13-110853221-INS-204 |
| chr13-110884164-INS-208 | chr13-110884164-INS-208 |
| chr13-110889362-INS-6690 | chr13-110889363-INS-6690 |
| chr13-110899206-INS-328 | chr13-110899206-INS-328 |
| chr13-110904457-INS-353 | chr13-110904457-INS-359 |
| chr13-110908064-INS-489 | chr13-110908064-INS-489 |
| chr13-110917961-INS-68 | chr13-110917996-INS-53 |
| chr13-110927029-INS-51 | chr13-110927038-INS-50 |
| chr13-110931268-INS-450 | chr13-110931268-INS-450 |
| chr13-110970124-INS-60 | chr13-110970124-INS-78 |
| chr13-110999195-INS-122 | chr13-110999193-INS-118 |
| chr13-111018634-INS-62 | chr13-111018634-INS-62 |
| chr13-111042012-INS-203 | chr13-111042012-INS-203 |
| chr13-111049100-INS-644 | chr13-111049100-INS-643 |
| chr13-111070807-INS-69 | chr13-111070807-INS-72 |
| chr13-111086203-INS-920 | chr13-111086203-INS-921 |
| chr13-111097096-INS-422 | chr13-111097077-INS-416 |
| chr13-111117306-INS-53 | chr13-111117306-INS-53 |
| chr13-111119514-INS-109 | chr13-111119492-INS-110 |
| chr13-111126722-INS-83 | chr13-111126722-INS-84 |
| chr13-111134276-INS-64 | chr13-111134276-INS-59 |
| chr13-111135220-INS-52 | chr13-111135220-INS-51 |
| chr13-111141361-INS-395 | chr13-111141361-INS-395 |
| chr13-111153513-INS-453 | chr13-111153513-INS-453 |
| chr13-111164220-INS-4970 | chr13-111164221-INS-4970 |
| chr13-111166607-INS-178 | chr13-111166607-INS-178 |
| chr13-111182868-INS-139 | chr13-111182869-INS-139 |
| chr13-111222757-INS-88 | chr13-111222757-INS-72 |
| chr13-111237048-INS-197 | chr13-111237048-INS-147 |
| chr13-111239366-INS-100 | chr13-111239366-INS-100 |
| chr13-111243415-INS-504 | chr13-111243448-INS-500 |
| chr13-111303703-INS-487 | chr13-111303703-INS-485 |
| chr13-111317493-INS-124 | chr13-111317493-INS-124 |
| chr13-111323249-INS-164 | chr13-111323249-INS-164 |
| chr13-111339959-INS-777 | chr13-111339959-INS-778 |
| chr13-111346309-INS-69 | chr13-111346309-INS-75 |
| chr13-111349675-INS-55 | chr13-111349667-INS-66 |
| chr13-111398110-INS-71 | chr13-111398079-INS-118 |

| | |
|---|---|
| chr13-111400565-INS-54 | chr13-111400565-INS-54 |
| chr13-111404744-INS-55 | chr13-111404744-INS-50 |
| chr13-111405988-INS-64 | chr13-111405988-INS-64 |
| chr13-111410265-INS-96 | chr13-111410265-INS-95 |
| chr13-111420437-INS-152 | chr13-111420437-INS-154 |
| chr13-111437720-INS-326 | chr13-111437720-INS-328 |
| chr13-111450333-INS-490 | chr13-111450333-INS-490 |
| chr13-111456391-INS-63 | chr13-111456391-INS-80 |
| chr13-111471739-INS-676 | chr13-111471739-INS-676 |
| chr13-111516663-INS-65 | chr13-111516663-INS-68 |
| chr13-111522184-INS-90 | chr13-111522200-INS-76 |
| chr13-111523960-INS-60 | chr13-111523960-INS-60 |
| chr13-111529129-INS-74 | chr13-111529129-INS-74 |
| chr13-111530279-INS-102 | chr13-111530279-INS-102 |
| chr13-111539978-INS-213 | chr13-111539978-INS-213 |
| chr13-111544647-INS-360 | chr13-111544649-INS-358 |
| chr13-111547299-INS-62 | chr13-111547299-INS-62 |
| chr13-111556265-INS-1676 | chr13-111556266-INS-1676 |
| chr13-111561519-INS-78 | chr13-111561519-INS-78 |
| chr13-111571064-INS-692 | chr13-111571064-INS-692 |
| chr13-111586980-INS-136 | chr13-111586980-INS-141 |
| chr13-111602134-INS-211 | chr13-111602134-INS-211 |
| chr13-111602761-INS-177 | chr13-111602761-INS-177 |
| chr13-111616457-INS-87 | chr13-111616457-INS-50 |
| chr13-111623457-INS-176 | chr13-111623457-INS-177 |
| chr13-111628891-INS-100 | chr13-111628891-INS-100 |
| chr13-111645621-INS-75 | chr13-111645621-INS-79 |
| chr13-111685972-INS-125 | chr13-111685980-INS-142 |
| chr13-111688134-INS-155 | chr13-111688134-INS-155 |
| chr13-111699758-INS-687 | chr13-111699759-INS-679 |
| chr13-111733402-INS-253 | chr13-111733403-INS-253 |
| chr13-111742330-INS-196 | chr13-111742324-INS-196 |
| chr13-111766049-INS-52 | chr13-111766049-INS-52 |
| chr13-111767504-INS-204 | chr13-111767504-INS-204 |
| chr13-111773917-INS-206 | chr13-111773918-INS-208 |
| chr13-111777755-INS-52 | chr13-111777753-INS-50 |
| chr13-111781121-INS-321 | chr13-111781121-INS-321 |
| chr13-111797681-INS-52 | chr13-111797681-INS-60 |
| chr13-111800096-INS-54 | chr13-111800096-INS-54 |
| chr13-111803822-INS-78 | chr13-111803822-INS-78 |
| chr13-111805185-INS-187 | chr13-111805210-INS-196 |
| chr13-111881069-INS-135 | chr13-111881069-INS-136 |
| chr13-111895997-INS-212 | chr13-111895997-INS-212 |
| chr13-111899848-INS-68 | chr13-111899848-INS-68 |
| chr13-111905656-INS-136 | chr13-111905656-INS-136 |
| chr13-111936121-INS-52 | chr13-111936121-INS-52 |
| chr13-111941870-INS-52 | chr13-111941871-INS-56 |
| chr13-111947231-INS-54 | chr13-111947231-INS-54 |
| chr13-111949067-INS-180 | chr13-111949067-INS-174 |
| chr13-111955953-INS-5575 | chr13-111955954-INS-5575 |
| chr13-111955964-INS-206 | chr13-111955964-INS-206 |
| chr13-111961256-INS-54 | chr13-111961256-INS-50 |
| chr13-111962331-INS-64 | chr13-111962331-INS-64 |
| chr13-111964364-INS-149 | chr13-111964364-INS-145 |
| chr13-111977980-INS-52 | chr13-111977980-INS-51 |
| chr13-111992514-INS-5382 | chr13-111992515-INS-5382 |
| chr13-112015468-INS-66 | chr13-112015468-INS-66 |
| chr13-112037190-INS-132 | chr13-112037190-INS-132 |
| chr13-112050270-INS-683 | chr13-112050275-INS-689 |
| chr13-112085450-INS-60 | chr13-112085450-INS-60 |
| chr13-112095425-INS-85 | chr13-112095425-INS-85 |
| chr13-112098393-INS-98 | chr13-112098393-INS-104 |
| chr13-112117450-INS-201 | chr13-112117450-INS-201 |
| chr13-112124266-INS-52 | chr13-112124266-INS-52 |
| chr13-112131323-INS-98 | chr13-112131323-INS-94 |
| chr13-112137371-INS-315 | chr13-112137371-INS-315 |
| chr13-112152898-INS-969 | chr13-112152898-INS-969 |
| chr13-112157142-INS-833 | chr13-112157142-INS-833 |
| chr13-112163691-INS-188 | chr13-112163691-INS-189 |
| chr13-112167238-INS-527 | chr13-112167238-INS-537 |
| chr13-112170712-INS-53 | chr13-112170713-INS-53 |
| chr13-112186523-INS-70 | chr13-112186523-INS-72 |
| chr13-112198171-INS-61 | chr13-112198171-INS-61 |
| chr13-112204453-INS-150 | chr13-112204453-INS-150 |
| chr13-112211745-INS-65 | chr13-112211745-INS-67 |
| chr13-112223125-INS-54 | chr13-112223125-INS-56 |
| chr13-112239027-INS-60 | chr13-112239027-INS-51 |

| | |
|---|---|
| chr13-112241658-INS-114 | chr13-112241665-INS-105 |
| chr13-112274403-INS-87 | chr13-112274403-INS-87 |
| chr13-112274893-INS-598 | chr13-112274890-INS-609 |
| chr13-112278639-INS-145 | chr13-112278639-INS-146 |
| chr13-112286472-INS-60 | chr13-112286472-INS-60 |
| chr13-112287198-INS-288 | chr13-112287198-INS-288 |
| chr13-112295950-INS-55 | chr13-112295943-INS-60 |
| chr13-112321915-INS-56 | chr13-112321915-INS-56 |
| chr13-112340011-INS-1459 | chr13-112340015-INS-1443 |
| chr13-112348117-INS-62 | chr13-112348117-INS-54 |
| chr13-112352247-INS-56 | chr13-112352247-INS-61 |
| chr13-112402934-INS-116 | chr13-112402934-INS-116 |
| chr13-112403231-INS-549 | chr13-112403247-INS-541 |
| chr13-112436251-INS-83 | chr13-112436251-INS-85 |
| chr13-112463505-INS-54 | chr13-112463506-INS-56 |
| chr13-112469707-INS-104 | chr13-112469704-INS-104 |
| chr13-112486690-INS-66 | chr13-112486690-INS-66 |
| chr13-112487946-INS-55 | chr13-112487928-INS-63 |
| chr13-112537180-INS-320 | chr13-112537180-INS-320 |
| chr13-112553236-INS-56 | chr13-112553236-INS-61 |
| chr13-112589195-INS-5311 | chr13-112589196-INS-5311 |
| chr13-112639757-INS-56 | chr13-112639757-INS-56 |
| chr13-112703046-INS-194 | chr13-112703046-INS-194 |
| chr13-112728288-INS-95 | chr13-112728288-INS-96 |
| chr13-112737535-INS-97 | chr13-112737537-INS-93 |
| chr13-112744851-INS-191 | chr13-112744851-INS-191 |
| chr13-112754135-INS-214 | chr13-112754135-INS-214 |
| chr13-112802205-INS-2039 | chr13-112802206-INS-2039 |
| chr13-112815563-INS-150 | chr13-112815561-INS-160 |
| chr13-112825430-INS-221 | chr13-112825430-INS-223 |
| chr13-112836435-INS-329 | chr13-112836435-INS-376 |
| chr13-112854822-INS-496 | chr13-112854822-INS-496 |
| chr13-112860184-INS-75 | chr13-112860184-INS-90 |
| chr13-112866730-INS-78 | chr13-112866730-INS-78 |
| chr13-112871130-INS-66 | chr13-112871130-INS-66 |
| chr13-112877032-INS-80 | chr13-112877032-INS-80 |
| chr13-112884043-INS-1005 | chr13-112884044-INS-1005 |
| chr13-112912026-INS-170 | chr13-112912026-INS-177 |
| chr13-112926087-INS-52 | chr13-112926094-INS-64 |
| chr13-112930796-INS-144 | chr13-112930796-INS-144 |
| chr13-112936081-INS-236 | chr13-112936081-INS-236 |
| chr13-112943167-INS-191 | chr13-112943167-INS-191 |
| chr13-112944668-INS-56 | chr13-112944668-INS-56 |
| chr13-112953324-INS-56 | chr13-112953324-INS-58 |
| chr13-112962710-INS-234 | chr13-112962710-INS-234 |
| chr13-112963917-INS-94 | chr13-112963917-INS-94 |
| chr13-112988987-INS-565 | chr13-112988978-INS-539 |
| chr13-112989907-INS-51 | chr13-112989917-INS-58 |
| chr13-113002884-INS-86 | chr13-113002884-INS-86 |
| chr13-113008066-INS-493 | chr13-113008066-INS-493 |
| chr13-113012692-INS-210 | chr13-113012692-INS-210 |
| chr13-113075282-INS-153 | chr13-113075282-INS-155 |
| chr13-113078617-INS-60 | chr13-113078617-INS-50 |
| chr13-113087416-INS-246 | chr13-113087416-INS-246 |
| chr13-113089159-INS-161 | chr13-113089159-INS-161 |
| chr13-113091112-INS-203 | chr13-113091112-INS-203 |
| chr13-113100759-INS-342 | chr13-113100759-INS-342 |
| chr13-113103420-INS-422 | chr13-113103420-INS-422 |
| chr13-113115879-INS-245 | chr13-113115879-INS-245 |
| chr13-113117113-INS-70 | chr13-113117113-INS-70 |
| chr13-113125900-INS-97 | chr13-113125900-INS-104 |
| chr13-113147510-INS-82 | chr13-113147510-INS-82 |
| chr13-113163979-INS-418 | chr13-113163979-INS-418 |
| chr13-113179219-INS-500 | chr13-113179221-INS-497 |
| chr13-113184771-INS-192 | chr13-113184771-INS-201 |
| chr13-113186659-INS-77 | chr13-113186659-INS-87 |
| chr13-113194645-INS-76 | chr13-113194645-INS-84 |
| chr13-113199869-INS-94 | chr13-113199869-INS-96 |
| chr13-113216848-INS-62 | chr13-113216849-INS-70 |
| chr13-113220083-INS-96 | chr13-113220083-INS-96 |
| chr13-113256463-INS-439 | chr13-113256463-INS-439 |
| chr13-113258418-INS-161 | chr13-113258447-INS-112 |
| chr13-113265231-INS-60 | chr13-113265231-INS-60 |
| chr13-113274697-INS-64 | chr13-113274697-INS-64 |
| chr13-113296918-INS-105 | chr13-113296918-INS-105 |
| chr13-113308791-INS-74 | chr13-113308791-INS-74 |
| chr13-113323768-INS-101 | chr13-113323768-INS-101 |

| | |
|---|---|
| chr13-113352440-INS-85 | chr13-113352437-INS-88 |
| chr13-113360093-INS-91 | chr13-113360093-INS-88 |
| chr13-113361224-INS-76 | chr13-113361224-INS-76 |
| chr13-113366302-INS-53 | chr13-113366302-INS-55 |
| chr13-113390632-INS-63 | chr13-113390616-INS-50 |
| chr13-113440873-INS-53 | chr13-113440873-INS-63 |
| chr13-113473201-INS-80 | chr13-113473201-INS-82 |
| chr13-113476847-INS-58 | chr13-113476847-INS-52 |
| chr13-113534461-INS-121 | chr13-113534460-INS-119 |
| chr13-113550456-INS-107 | chr13-113550432-INS-91 |
| chr13-113563699-INS-54 | chr13-113563695-INS-55 |
| chr13-113608266-INS-163 | chr13-113608265-INS-124 |
| chr13-113608318-INS-602 | chr13-113608283-INS-627 |
| chr13-113639584-INS-62 | chr13-113639586-INS-78 |
| chr13-113641276-INS-70 | chr13-113641276-INS-70 |
| chr13-113644192-INS-66 | chr13-113644192-INS-66 |
| chr13-113659819-INS-3163 | chr13-113659820-INS-3163 |
| chr13-113660558-INS-721 | chr13-113660558-INS-721 |
| chr13-113661608-INS-2814 | chr13-113661609-INS-2814 |
| chr13-113677579-INS-182 | chr13-113677579-INS-182 |
| chr13-113682916-INS-62 | chr13-113682916-INS-64 |
| chr13-113690384-INS-6186 | chr13-113690385-INS-6186 |
| chr13-113695396-INS-84 | chr13-113695396-INS-84 |
| chr13-113706061-INS-156 | chr13-113706061-INS-156 |
| chr13-113707944-INS-85 | chr13-113707941-INS-100 |
| chr13-113725230-INS-759 | chr13-113725230-INS-802 |
| chr13-113762597-INS-424 | chr13-113762597-INS-424 |
| chr13-113775190-INS-893 | chr13-113775190-INS-893 |
| chr13-113781663-INS-66 | chr13-113781663-INS-66 |
| chr13-113826094-INS-89 | chr13-113826094-INS-91 |
| chr13-113828760-INS-114 | chr13-113828756-INS-132 |
| chr13-113839743-INS-188 | chr13-113839743-INS-189 |
| chr13-113847546-INS-128 | chr13-113847535-INS-144 |
| chr13-113863485-INS-3573 | chr13-113863486-INS-3567 |
| chr13-113864311-INS-476 | chr13-113864311-INS-476 |
| chr13-113868184-INS-168 | chr13-113868184-INS-168 |
| chr13-113878062-INS-58 | chr13-113878062-INS-52 |
| chr13-113882841-INS-62 | chr13-113882841-INS-62 |
| chr13-113886350-INS-1697 | chr13-113886351-INS-1699 |
| chr13-113905537-INS-88 | chr13-113905537-INS-106 |
| chr13-113907979-INS-193 | chr13-113907979-INS-194 |
| chr13-113915448-INS-131 | chr13-113915448-INS-131 |
| chr13-113935115-INS-111 | chr13-113935115-INS-98 |
| chr13-113944119-INS-205 | chr13-113944119-INS-201 |
| chr13-113968686-INS-306 | chr13-113968686-INS-310 |
| chr13-113978539-INS-729 | chr13-113978541-INS-730 |
| chr13-113979759-INS-5293 | chr13-113979760-INS-5293 |
| chr13-113995016-INS-102 | chr13-113995055-INS-90 |
| chr13-114013317-INS-230 | chr13-114013317-INS-191 |
| chr13-114019307-INS-54 | chr13-114019307-INS-52 |
| chr13-114020515-INS-60 | chr13-114020515-INS-60 |
| chr13-114021192-INS-204 | chr13-114021192-INS-204 |
| chr13-114031953-INS-66 | chr13-114031953-INS-68 |
| chr13-114035273-INS-58 | chr13-114035273-INS-58 |
| chr13-114039116-INS-56 | chr13-114039116-INS-100 |
| chr13-114042854-INS-201 | chr13-114042854-INS-201 |
| chr13-114050604-INS-72 | chr13-114050604-INS-72 |
| chr13-114068024-INS-223 | chr13-114068044-INS-205 |
| chr13-114073755-INS-52 | chr13-114073755-INS-52 |
| chr13-114080327-INS-1687 | chr13-114080328-INS-1685 |
| chr13-114088784-INS-68 | chr13-114088784-INS-68 |
| chr13-114089940-INS-214 | chr13-114089940-INS-211 |
| chr13-114090345-INS-2683 | chr13-114090347-INS-2680 |
| chr13-114100533-INS-4822 | chr13-114100534-INS-4838 |
| chr13-114129576-INS-201 | chr13-114129576-INS-201 |
| chr13-114145632-INS-1368 | chr13-114145633-INS-1368 |
| chr13-114150816-INS-54 | chr13-114150824-INS-54 |
| chr13-114213823-INS-72 | chr13-114213827-INS-75 |
| chr13-114220776-INS-101 | chr13-114220776-INS-101 |
| chr13-114225613-INS-123 | chr13-114225613-INS-123 |
| chr13-114228588-INS-253 | chr13-114228588-INS-265 |
| chr13-114238891-INS-54 | chr13-114238909-INS-50 |
| chr13-114246430-INS-100 | chr13-114246430-INS-94 |
| chr13-114247912-INS-69 | chr13-114247912-INS-69 |
| chr13-114265643-INS-60 | chr13-114265643-INS-64 |
| chr13-114289592-INS-160 | chr13-114289592-INS-160 |
| chr13-114305142-INS-198 | chr13-114305142-INS-198 |

| | |
|---|---|
| chr13-114371534-INS-69 | chr13-114371534-INS-57 |
| chr13-114399181-INS-414 | chr13-114399182-INS-414 |
| chr13-114417579-INS-59 | chr13-114417579-INS-59 |
| chr13-114422103-INS-53 | chr13-114422103-INS-53 |
| chr13-114446038-INS-1730 | chr13-114446039-INS-1731 |
| chr13-114452222-INS-69 | chr13-114452222-INS-75 |
| chr13-114482805-INS-368 | chr13-114482805-INS-368 |
| chr13-114489314-INS-52 | chr13-114489314-INS-52 |
| chr13-114492232-INS-83 | chr13-114492207-INS-112 |
| chr13-114501056-INS-245 | chr13-114501056-INS-245 |
| chr13-114517798-INS-156 | chr13-114517810-INS-135 |
| chr13-114525505-INS-145 | chr13-114525505-INS-175 |
| chr13-114545594-INS-72 | chr13-114545564-INS-60 |
| chr13-114556604-INS-50 | chr13-114556604-INS-74 |
| chr13-114564112-INS-56 | chr13-114564112-INS-56 |
| chr13-114581479-INS-199 | chr13-114581479-INS-199 |
| chr13-114593276-INS-56 | chr13-114593272-INS-58 |
| chr13-114602552-INS-74 | chr13-114602552-INS-76 |
| chr13-114604949-INS-113 | chr13-114604943-INS-89 |
| chr13-114620773-INS-82 | chr13-114620797-INS-55 |
| chr13-114642223-INS-107 | chr13-114642223-INS-81 |
| chr13-114656603-INS-83 | chr13-114656559-INS-88 |
| chr13-114667500-INS-486 | chr13-114667500-INS-486 |
| chr13-114701097-INS-473 | chr13-114701097-INS-473 |
| chr13-114716659-INS-393 | chr13-114716659-INS-393 |
| chr13-114718312-INS-69 | chr13-114718313-INS-97 |
| chr13-114732640-INS-108 | chr13-114732640-INS-120 |
| chr13-114743925-INS-655 | chr13-114743925-INS-655 |
| chr13-114779241-INS-3213 | chr13-114779242-INS-3213 |
| chr13-114838641-INS-63 | chr13-114838641-INS-63 |
| chr13-114841045-INS-60 | chr13-114841045-INS-64 |
| chr13-114861220-INS-364 | chr13-114861220-INS-364 |
| chr13-114866829-INS-56 | chr13-114866829-INS-58 |
| chr13-114889068-INS-380 | chr13-114889068-INS-380 |
| chr13-114981547-INS-78 | chr13-114981535-INS-72 |
| chr13-114992607-INS-113 | chr13-114992619-INS-142 |
| chr13-115006856-INS-149 | chr13-115006856-INS-149 |
| chr13-115054182-INS-86 | chr13-115054182-INS-92 |
| chr13-115059300-INS-144 | chr13-115059307-INS-102 |
| chr13-115061734-INS-113 | chr13-115061734-INS-115 |
| chr13-115077915-INS-439 | chr13-115077915-INS-439 |
| chr13-115087490-INS-197 | chr13-115087490-INS-197 |
| chr13-115108017-INS-77 | chr13-115108008-INS-72 |
| chr13-115136250-INS-87 | chr13-115136250-INS-87 |
| chr13-115154405-INS-200 | chr13-115154405-INS-201 |
| chr13-115160307-INS-410 | chr13-115160307-INS-410 |
| chr13-115178446-INS-131 | chr13-115178451-INS-108 |
| chr13-115187143-INS-266 | chr13-115187143-INS-266 |
| chr13-115198199-INS-403 | chr13-115198199-INS-403 |
| chr13-115200992-INS-522 | chr13-115200992-INS-526 |
| chr13-115208252-INS-344 | chr13-115208252-INS-342 |
| chr13-115213581-INS-61 | chr13-115213581-INS-63 |
| chr13-115224468-INS-208 | chr13-115224468-INS-204 |
| chr13-115242135-INS-135 | chr13-115242135-INS-139 |
| chr13-115260702-INS-6515 | chr13-115260703-INS-6515 |
| chr13-115271579-INS-266 | chr13-115271579-INS-266 |
| chr13-115294463-INS-60 | chr13-115294463-INS-60 |
| chr13-115348987-INS-6383 | chr13-115348988-INS-6383 |
| chr13-115361519-INS-206 | chr13-115361519-INS-247 |
| chr13-115362859-INS-129 | chr13-115362859-INS-129 |
| chr13-115377903-INS-53 | chr13-115377903-INS-55 |
| chr13-115379995-INS-498 | chr13-115379995-INS-498 |
| chr13-115396739-INS-5532 | chr13-115396740-INS-5532 |
| chr13-115407052-INS-135 | chr13-115407052-INS-113 |
| chr13-115432020-INS-70 | chr13-115432020-INS-70 |
| chr13-115462450-INS-70 | chr13-115462450-INS-82 |
| chr13-115466534-INS-2648 | chr13-115466525-INS-2648 |
| chr13-115472941-INS-56 | chr13-115472941-INS-56 |
| chr13-115481213-INS-7197 | chr13-115481214-INS-7197 |
| chr13-115486246-INS-358 | chr13-115486246-INS-358 |
| chr13-115529306-INS-134 | chr13-115529306-INS-134 |
| chr13-115531643-INS-400 | chr13-115531643-INS-400 |
| chr13-115543915-INS-84 | chr13-115543915-INS-82 |
| chr13-115572020-INS-238 | chr13-115572012-INS-246 |
| chr13-115578505-INS-613 | chr13-115578505-INS-614 |
| chr13-115618305-INS-3242 | chr13-115618306-INS-3242 |
| chr13-115620288-INS-56 | chr13-115620288-INS-54 |

| | |
|---|---|
| chr13-115629507-INS-5276 | chr13-115629508-INS-5275 |
| chr13-115653086-INS-108 | chr13-115653086-INS-100 |
| chr13-115656701-INS-451 | chr13-115656701-INS-450 |
| chr13-115682906-INS-62 | chr13-115682906-INS-62 |
| chr13-115703496-INS-991 | chr13-115703496-INS-991 |
| chr13-115730099-INS-92 | chr13-115730099-INS-82 |
| chr13-115755718-INS-992 | chr13-115755718-INS-992 |
| chr13-115756291-INS-107 | chr13-115756291-INS-106 |
| chr13-115764843-INS-126 | chr13-115764843-INS-105 |
| chr13-115771239-INS-61 | chr13-115771239-INS-61 |
| chr13-115781038-INS-7197 | chr13-115781039-INS-7197 |
| chr13-115797412-INS-130 | chr13-115797412-INS-130 |
| chr13-115803805-INS-72 | chr13-115803805-INS-76 |
| chr13-115836363-INS-876 | chr13-115836363-INS-876 |
| chr13-115837411-INS-415 | chr13-115837411-INS-415 |
| chr13-115854074-INS-507 | chr13-115854074-INS-507 |
| chr13-115872681-INS-586 | chr13-115872681-INS-586 |
| chr13-115882317-INS-60 | chr13-115882317-INS-58 |
| chr13-115890841-INS-900 | chr13-115890841-INS-901 |
| chr13-115922636-INS-51 | chr13-115922636-INS-51 |
| chr13-115950717-INS-6375 | chr13-115950718-INS-6375 |
| chr13-115976997-INS-74 | chr13-115976997-INS-55 |
| chr13-115980924-INS-205 | chr13-115980924-INS-205 |
| chr13-115986362-INS-557 | chr13-115986362-INS-557 |
| chr13-115992923-INS-1133 | chr13-115992924-INS-1133 |
| chr13-116004229-INS-305 | chr13-116004229-INS-305 |
| chr13-116016118-INS-86 | chr13-116016114-INS-102 |
| chr13-116018443-INS-415 | chr13-116018443-INS-418 |
| chr13-116127915-INS-6612 | chr13-116127916-INS-6613 |
| chr13-116136381-INS-160 | chr13-116136381-INS-170 |
| chr13-116146121-INS-1468 | chr13-116146122-INS-1443 |
| chr13-116178383-INS-159 | chr13-116178383-INS-158 |
| chr13-116179474-INS-70 | chr13-116179474-INS-89 |
| chr13-116182694-INS-327 | chr13-116182694-INS-327 |
| chr13-116195348-INS-586 | chr13-116195348-INS-586 |
| chr13-116200419-INS-116 | chr13-116200420-INS-162 |
| chr13-116214726-INS-209 | chr13-116214726-INS-216 |
| chr13-116219281-INS-70 | chr13-116219281-INS-70 |
| chr13-116258539-INS-1625 | chr13-116258540-INS-1629 |
| chr13-116267212-INS-1518 | chr13-116267213-INS-1518 |
| chr13-116277873-INS-130 | chr13-116277874-INS-105 |
| chr13-116284985-INS-66 | chr13-116284970-INS-74 |
| chr13-116293260-INS-361 | chr13-116293260-INS-362 |
| chr13-116329012-INS-7115 | chr13-116329013-INS-7115 |
| chr13-116333286-INS-83 | chr13-116333286-INS-83 |
| chr13-116344459-INS-261 | chr13-116344459-INS-261 |
| chr13-116361506-INS-352 | chr13-116361506-INS-352 |
| chr13-116374094-INS-6521 | chr13-116374095-INS-6521 |
| chr13-116390681-INS-705 | chr13-116390681-INS-720 |
| chr13-116393495-INS-100 | chr13-116393495-INS-104 |
| chr13-116408017-INS-6347 | chr13-116408018-INS-6346 |
| chr13-116419022-INS-5544 | chr13-116419023-INS-5544 |
| chr13-116419353-INS-162 | chr13-116419353-INS-162 |
| chr13-116422240-INS-78 | chr13-116422223-INS-65 |
| chr13-116429499-INS-82 | chr13-116429499-INS-94 |
| chr13-116435508-INS-60 | chr13-116435508-INS-60 |
| chr13-116468437-INS-72 | chr13-116468388-INS-76 |
| chr13-116470266-INS-78 | chr13-116470266-INS-74 |
| chr13-116476523-INS-3189 | chr13-116476524-INS-3189 |
| chr13-116503486-INS-7174 | chr13-116503487-INS-7173 |
| chr13-116508627-INS-6776 | chr13-116508628-INS-6777 |
| chr13-116518087-INS-80 | chr13-116518090-INS-74 |
| chr13-116520173-INS-6628 | chr13-116520174-INS-6630 |
| chr13-116525864-INS-692 | chr13-116525864-INS-692 |
| chr13-116530127-INS-6355 | chr13-116530128-INS-6355 |
| chr13-116547743-INS-318 | chr13-116547743-INS-317 |
| chr13-116551852-INS-68 | chr13-116551852-INS-58 |
| chr13-116552776-INS-106 | chr13-116552776-INS-100 |
| chr13-116571020-INS-2639 | chr13-116571027-INS-2633 |
| chr13-116574992-INS-400 | chr13-116574992-INS-400 |
| chr13-116578836-INS-468 | chr13-116578836-INS-469 |
| chr13-116583200-INS-133 | chr13-116583200-INS-131 |
| chr13-116585796-INS-96 | chr13-116585796-INS-96 |
| chr13-116601604-INS-58 | chr13-116601604-INS-58 |
| chr13-116616610-INS-336 | chr13-116616610-INS-335 |
| chr13-116626541-INS-82 | chr13-116626541-INS-80 |
| chr13-116664158-INS-52 | chr13-116664158-INS-52 |

| | |
|---|---|
| chr13-116676482-INS-52 | chr13-116676482-INS-54 |
| chr13-116680960-INS-526 | chr13-116680960-INS-526 |
| chr13-116689898-INS-60 | chr13-116689898-INS-60 |
| chr13-116708078-INS-232 | chr13-116708078-INS-232 |
| chr13-116726100-INS-83 | chr13-116726074-INS-93 |
| chr13-116728623-INS-52 | chr13-116728623-INS-52 |
| chr13-116757116-INS-219 | chr13-116757116-INS-219 |
| chr13-116759313-INS-61 | chr13-116759313-INS-60 |
| chr13-116760189-INS-68 | chr13-116760189-INS-68 |
| chr13-116779693-INS-80 | chr13-116779693-INS-80 |
| chr13-116794077-INS-995 | chr13-116794078-INS-1043 |
| chr13-116806739-INS-84 | chr13-116806739-INS-84 |
| chr13-116815334-INS-52 | chr13-116815334-INS-54 |
| chr13-116837315-INS-383 | chr13-116837315-INS-383 |
| chr13-116839984-INS-164 | chr13-116839984-INS-164 |
| chr13-116863350-INS-1132 | chr13-116863351-INS-1132 |
| chr13-116872606-INS-6443 | chr13-116872607-INS-6444 |
| chr13-116913067-INS-105 | chr13-116913067-INS-95 |
| chr13-116921946-INS-53 | chr13-116921949-INS-54 |
| chr13-116929964-INS-675 | chr13-116929964-INS-677 |
| chr13-116930527-INS-1996 | chr13-116930528-INS-1996 |
| chr13-116943911-INS-65 | chr13-116943911-INS-60 |
| chr13-116965190-INS-2337 | chr13-116965191-INS-2338 |
| chr13-116969861-INS-879 | chr13-116969861-INS-879 |
| chr13-116981462-INS-1631 | chr13-116981463-INS-1631 |
| chr13-116993295-INS-275 | chr13-116993295-INS-271 |
| chr13-117010161-INS-135 | chr13-117010161-INS-135 |
| chr13-117013646-INS-798 | chr13-117013646-INS-799 |
| chr13-117025115-INS-453 | chr13-117025115-INS-453 |
| chr13-117028702-INS-54 | chr13-117028688-INS-50 |
| chr13-117054043-INS-54 | chr13-117054043-INS-58 |
| chr13-117060387-INS-6656 | chr13-117060388-INS-6656 |
| chr13-117066464-INS-206 | chr13-117066464-INS-206 |
| chr13-117097372-INS-209 | chr13-117097372-INS-209 |
| chr13-117123276-INS-161 | chr13-117123276-INS-165 |
| chr13-117125855-INS-821 | chr13-117125855-INS-821 |
| chr13-117137355-INS-96 | chr13-117137337-INS-131 |
| chr13-117146375-INS-56 | chr13-117146358-INS-76 |
| chr13-117147012-INS-1567 | chr13-117147013-INS-1567 |
| chr13-117168588-INS-57 | chr13-117168588-INS-63 |
| chr13-117175255-INS-987 | chr13-117175255-INS-987 |
| chr13-117186712-INS-184 | chr13-117186712-INS-184 |
| chr13-117204038-INS-263 | chr13-117204038-INS-263 |
| chr13-117221582-INS-112 | chr13-117221582-INS-112 |
| chr13-117224698-INS-145 | chr13-117224698-INS-139 |
| chr13-117238531-INS-5306 | chr13-117238532-INS-5306 |
| chr13-117248554-INS-360 | chr13-117248554-INS-360 |
| chr13-117251541-INS-7088 | chr13-117251542-INS-7088 |
| chr13-117256701-INS-78 | chr13-117256701-INS-78 |
| chr13-117263280-INS-107 | chr13-117263261-INS-112 |
| chr13-117281944-INS-205 | chr13-117281944-INS-205 |
| chr13-117282701-INS-441 | chr13-117282701-INS-441 |
| chr13-117301157-INS-78 | chr13-117301157-INS-82 |
| chr13-117314147-INS-209 | chr13-117314147-INS-209 |
| chr13-117333421-INS-210 | chr13-117333421-INS-210 |
| chr13-117335662-INS-345 | chr13-117335662-INS-345 |
| chr13-117337723-INS-60 | chr13-117337723-INS-60 |
| chr13-117343651-INS-208 | chr13-117343651-INS-208 |
| chr13-117350008-INS-97 | chr13-117350008-INS-97 |
| chr13-117417949-INS-52 | chr13-117417949-INS-52 |
| chr13-117420496-INS-350 | chr13-117420496-INS-350 |
| chr13-117421451-INS-118 | chr13-117421455-INS-112 |
| chr13-117423174-INS-808 | chr13-117423174-INS-808 |
| chr13-117429305-INS-64 | chr13-117429301-INS-92 |
| chr13-117447751-INS-113 | chr13-117447751-INS-113 |
| chr13-117456168-INS-168 | chr13-117456168-INS-169 |
| chr13-117464363-INS-64 | chr13-117464363-INS-64 |
| chr13-117466984-INS-96 | chr13-117466984-INS-116 |
| chr13-117472859-INS-202 | chr13-117472859-INS-202 |
| chr13-117490460-INS-180 | chr13-117490460-INS-180 |
| chr13-117491159-INS-182 | chr13-117491159-INS-182 |
| chr13-117506466-INS-66 | chr13-117506466-INS-66 |
| chr13-117533920-INS-132 | chr13-117533920-INS-132 |
| chr13-117535304-INS-504 | chr13-117535304-INS-504 |
| chr13-117537447-INS-84 | chr13-117537447-INS-84 |
| chr13-117549276-INS-68 | chr13-117549276-INS-70 |
| chr13-117566400-INS-74 | chr13-117566400-INS-74 |

| | |
|---|---|
| chr13-117589901-INS-64 | chr13-117589901-INS-64 |
| chr13-117613869-INS-379 | chr13-117613869-INS-379 |
| chr13-117623964-INS-507 | chr13-117623964-INS-507 |
| chr13-117648979-INS-58 | chr13-117648979-INS-61 |
| chr13-117663062-INS-943 | chr13-117663062-INS-943 |
| chr13-117691087-INS-7183 | chr13-117691088-INS-7183 |
| chr13-117719063-INS-805 | chr13-117719063-INS-805 |
| chr13-117729833-INS-331 | chr13-117729833-INS-331 |
| chr13-117743127-INS-84 | chr13-117743127-INS-80 |
| chr13-117743653-INS-151 | chr13-117743652-INS-153 |
| chr13-117752385-INS-555 | chr13-117752385-INS-555 |
| chr13-117752993-INS-126 | chr13-117752993-INS-126 |
| chr13-117756670-INS-538 | chr13-117756670-INS-538 |
| chr13-117792771-INS-6724 | chr13-117792772-INS-6728 |
| chr13-117808102-INS-290 | chr13-117808080-INS-278 |
| chr13-117858073-INS-77 | chr13-117858073-INS-95 |
| chr13-117866513-INS-854 | chr13-117866513-INS-854 |
| chr13-117884031-INS-79 | chr13-117884031-INS-84 |
| chr13-117910893-INS-52 | chr13-117910893-INS-52 |
| chr13-117916486-INS-437 | chr13-117916486-INS-437 |
| chr13-117920050-INS-59 | chr13-117920008-INS-55 |
| chr13-117940923-INS-801 | chr13-117940923-INS-802 |
| chr13-117945226-INS-62 | chr13-117945228-INS-76 |
| chr13-117946310-INS-99 | chr13-117946310-INS-100 |
| chr13-118052904-INS-54 | chr13-118052904-INS-56 |
| chr13-118073923-INS-6240 | chr13-118073924-INS-6240 |
| chr13-118147357-INS-198 | chr13-118147357-INS-198 |
| chr13-118163173-INS-52 | chr13-118163170-INS-55 |
| chr13-118170404-INS-6678 | chr13-118170405-INS-6671 |
| chr13-118181462-INS-60 | chr13-118181462-INS-58 |
| chr13-118202227-INS-673 | chr13-118202227-INS-673 |
| chr13-118213359-INS-68 | chr13-118213359-INS-68 |
| chr13-118237966-INS-308 | chr13-118237966-INS-308 |
| chr13-118243301-INS-148 | chr13-118243301-INS-151 |
| chr13-118274237-INS-423 | chr13-118274237-INS-423 |
| chr13-118278878-INS-457 | chr13-118278878-INS-470 |
| chr13-118281307-INS-542 | chr13-118281308-INS-542 |
| chr13-118281798-INS-814 | chr13-118281798-INS-813 |
| chr13-118304047-INS-52 | chr13-118304057-INS-54 |
| chr13-118327569-INS-52 | chr13-118327569-INS-52 |
| chr13-118329653-INS-237 | chr13-118329653-INS-237 |
| chr13-118370596-INS-73 | chr13-118370596-INS-73 |
| chr13-118443274-INS-52 | chr13-118443274-INS-54 |
| chr13-118453295-INS-63 | chr13-118453295-INS-63 |
| chr13-118464829-INS-818 | chr13-118464829-INS-818 |
| chr13-118465114-INS-2993 | chr13-118465115-INS-2993 |
| chr13-118467000-INS-399 | chr13-118467000-INS-399 |
| chr13-118484604-INS-84 | chr13-118484604-INS-84 |
| chr13-118499581-INS-7050 | chr13-118499582-INS-7050 |
| chr13-118513478-INS-84 | chr13-118513478-INS-84 |
| chr13-118520528-INS-120 | chr13-118520528-INS-120 |
| chr13-118525205-INS-459 | chr13-118525205-INS-459 |
| chr13-118533673-INS-197 | chr13-118533674-INS-197 |
| chr13-118539191-INS-65 | chr13-118539191-INS-77 |
| chr13-118546332-INS-52 | chr13-118546322-INS-62 |
| chr13-118553161-INS-52 | chr13-118553162-INS-55 |
| chr13-118574819-INS-82 | chr13-118574819-INS-83 |
| chr13-118577490-INS-636 | chr13-118577490-INS-636 |
| chr13-118579764-INS-427 | chr13-118579764-INS-427 |
| chr13-118581304-INS-769 | chr13-118581304-INS-770 |
| chr13-118587152-INS-762 | chr13-118587152-INS-763 |
| chr13-118588087-INS-144 | chr13-118588087-INS-144 |
| chr13-118599288-INS-295 | chr13-118599288-INS-295 |
| chr13-118677514-INS-223 | chr13-118677514-INS-223 |
| chr13-118687998-INS-51 | chr13-118687998-INS-51 |
| chr13-118692948-INS-129 | chr13-118692948-INS-133 |
| chr13-118694347-INS-90 | chr13-118694347-INS-72 |
| chr13-118741516-INS-184 | chr13-118741516-INS-184 |
| chr13-118759010-INS-714 | chr13-118759012-INS-716 |
| chr13-118781101-INS-270 | chr13-118781101-INS-270 |
| chr13-118783324-INS-67 | chr13-118783324-INS-67 |
| chr13-118798192-INS-63 | chr13-118798192-INS-63 |
| chr13-118836151-INS-140 | chr13-118836151-INS-140 |
| chr13-118844348-INS-65 | chr13-118844350-INS-112 |
| chr13-118864446-INS-170 | chr13-118864446-INS-174 |
| chr13-118897173-INS-324 | chr13-118897173-INS-324 |
| chr13-118911823-INS-180 | chr13-118911823-INS-180 |

| | |
|---|---|
| chr13-118921245-INS-66 | chr13-118921245-INS-66 |
| chr13-118972917-INS-200 | chr13-118972917-INS-200 |
| chr13-118975821-INS-6403 | chr13-118975822-INS-6403 |
| chr13-118985625-INS-6389 | chr13-118985626-INS-6389 |
| chr13-118996077-INS-268 | chr13-118996077-INS-266 |
| chr13-119013075-INS-3297 | chr13-119013076-INS-3297 |
| chr13-119060085-INS-197 | chr13-119060085-INS-197 |
| chr13-119062502-INS-68 | chr13-119062502-INS-68 |
| chr13-119142217-INS-94 | chr13-119142217-INS-98 |
| chr13-119151549-INS-132 | chr13-119151549-INS-147 |
| chr13-119158253-INS-103 | chr13-119158253-INS-103 |
| chr13-119177332-INS-212 | chr13-119177332-INS-212 |
| chr13-119182629-INS-52 | chr13-119182629-INS-54 |
| chr13-119220955-INS-1167 | chr13-119220956-INS-1167 |
| chr13-119221989-INS-114 | chr13-119221986-INS-118 |
| chr13-119280965-INS-90 | chr13-119280965-INS-90 |
| chr13-119283356-INS-158 | chr13-119283356-INS-158 |
| chr13-119296588-INS-10518 | chr13-119296589-INS-10518 |
| chr13-119354853-INS-6490 | chr13-119354854-INS-6490 |
| chr13-119406266-INS-71 | chr13-119406266-INS-70 |
| chr13-119413689-INS-80 | chr13-119413689-INS-80 |
| chr13-119446909-INS-181 | chr13-119446909-INS-181 |
| chr13-119447960-INS-335 | chr13-119447960-INS-335 |
| chr13-119450405-INS-129 | chr13-119450387-INS-134 |
| chr13-119456368-INS-209 | chr13-119456368-INS-210 |
| chr13-119456568-INS-7101 | chr13-119456569-INS-7101 |
| chr13-119519281-INS-199 | chr13-119519266-INS-206 |
| chr13-119520435-INS-199 | chr13-119520435-INS-199 |
| chr13-119528878-INS-220 | chr13-119528878-INS-221 |
| chr13-119532964-INS-199 | chr13-119532964-INS-199 |
| chr13-119548959-INS-4337 | chr13-119548956-INS-4313 |
| chr13-119550093-INS-58 | chr13-119550093-INS-58 |
| chr13-119551247-INS-213 | chr13-119551247-INS-209 |
| chr13-119553651-INS-108 | chr13-119553651-INS-94 |
| chr13-119576436-INS-421 | chr13-119576436-INS-421 |
| chr13-119625844-INS-184 | chr13-119625843-INS-196 |
| chr13-120072732-INS-55 | chr13-120072723-INS-54 |
| chr13-120093005-INS-82 | chr13-120093000-INS-82 |
| chr13-120096310-INS-203 | chr13-120096310-INS-207 |
| chr13-120109659-INS-104 | chr13-120109659-INS-104 |
| chr13-120114279-INS-268 | chr13-120114279-INS-266 |
| chr13-120138640-INS-105 | chr13-120138640-INS-103 |
| chr13-120145925-INS-444 | chr13-120145925-INS-444 |
| chr13-120149789-INS-98 | chr13-120149789-INS-98 |
| chr13-120181040-INS-61 | chr13-120181040-INS-61 |
| chr13-120188043-INS-182 | chr13-120188043-INS-182 |
| chr13-120202727-INS-55 | chr13-120202744-INS-53 |
| chr13-120211725-INS-58 | chr13-120211725-INS-64 |
| chr13-120247882-INS-205 | chr13-120247882-INS-204 |
| chr13-120261783-INS-64 | chr13-120261783-INS-64 |
| chr13-120264539-INS-658 | chr13-120264539-INS-670 |
| chr13-120268927-INS-535 | chr13-120268927-INS-535 |
| chr13-120278347-INS-66 | chr13-120278348-INS-58 |
| chr13-120280521-INS-664 | chr13-120280522-INS-658 |
| chr13-120281909-INS-179 | chr13-120281909-INS-179 |
| chr13-120282918-INS-114 | chr13-120282923-INS-132 |
| chr13-120285749-INS-202 | chr13-120285749-INS-202 |
| chr13-120294143-INS-54 | chr13-120294142-INS-55 |
| chr13-120297125-INS-840 | chr13-120297125-INS-840 |
| chr13-120342239-INS-343 | chr13-120342240-INS-343 |
| chr13-120363317-INS-141 | chr13-120363317-INS-134 |
| chr13-120367109-INS-60 | chr13-120367107-INS-60 |
| chr13-120389250-INS-1038 | chr13-120389251-INS-1038 |
| chr13-120499050-INS-54 | chr13-120499050-INS-54 |
| chr13-120526174-INS-58 | chr13-120526180-INS-54 |
| chr13-120635732-INS-492 | chr13-120635732-INS-492 |
| chr13-120663439-INS-323 | chr13-120663439-INS-323 |
| chr13-120699852-INS-89 | chr13-120699835-INS-96 |
| chr14-3075393-INS-6475 | chr14-3075394-INS-6475 |
| chr14-3135752-INS-6303 | chr14-3135753-INS-6303 |
| chr14-3145561-INS-76 | chr14-3145561-INS-80 |
| chr14-3269194-INS-424 | chr14-3269194-INS-424 |
| chr14-3293461-INS-241 | chr14-3293461-INS-242 |
| chr14-3369603-INS-3043 | chr14-3369603-INS-3043 |
| chr14-3382038-INS-148 | chr14-3382038-INS-168 |
| chr14-3393641-INS-6011 | chr14-3393642-INS-6011 |
| chr14-3397891-INS-3687 | chr14-3397892-INS-3688 |

| | |
|---|---|
| chr14-3403899-INS-146 | chr14-3403899-INS-146 |
| chr14-3429362-INS-57 | chr14-3429362-INS-53 |
| chr14-3432616-INS-60 | chr14-3432602-INS-64 |
| chr14-3448873-INS-68 | chr14-3448873-INS-76 |
| chr14-3478145-INS-73 | chr14-3478145-INS-123 |
| chr14-3498497-INS-6194 | chr14-3498498-INS-6194 |
| chr14-3525482-INS-180 | chr14-3525482-INS-188 |
| chr14-3554279-INS-313 | chr14-3554279-INS-319 |
| chr14-3555518-INS-54 | chr14-3555554-INS-52 |
| chr14-3589490-INS-53 | chr14-3589490-INS-66 |
| chr14-3591388-INS-1639 | chr14-3591389-INS-1639 |
| chr14-3646900-INS-106 | chr14-3646900-INS-106 |
| chr14-3659090-INS-298 | chr14-3659090-INS-298 |
| chr14-3663855-INS-84 | chr14-3663856-INS-92 |
| chr14-3690222-INS-188 | chr14-3690222-INS-188 |
| chr14-3701081-INS-2677 | chr14-3701082-INS-2677 |
| chr14-3756315-INS-7000 | chr14-3756316-INS-7000 |
| chr14-3762440-INS-7218 | chr14-3762441-INS-7218 |
| chr14-3771269-INS-3593 | chr14-3771270-INS-3593 |
| chr14-3810512-INS-5021 | chr14-3810513-INS-5021 |
| chr14-3849227-INS-349 | chr14-3849227-INS-349 |
| chr14-3858992-INS-64 | chr14-3858992-INS-60 |
| chr14-3875319-INS-70 | chr14-3875319-INS-69 |
| chr14-3878872-INS-196 | chr14-3878866-INS-216 |
| chr14-3888210-INS-97 | chr14-3888210-INS-97 |
| chr14-3892539-INS-74 | chr14-3892538-INS-76 |
| chr14-3962052-INS-177 | chr14-3962046-INS-182 |
| chr14-4005886-INS-198 | chr14-4005886-INS-198 |
| chr14-4067502-INS-210 | chr14-4067502-INS-166 |
| chr14-4082640-INS-521 | chr14-4082640-INS-521 |
| chr14-4092138-INS-200 | chr14-4092153-INS-202 |
| chr14-4095292-INS-58 | chr14-4095292-INS-58 |
| chr14-4098058-INS-64 | chr14-4098058-INS-64 |
| chr14-4103268-INS-352 | chr14-4103268-INS-352 |
| chr14-4116902-INS-2508 | chr14-4116903-INS-2508 |
| chr14-4128764-INS-445 | chr14-4128764-INS-445 |
| chr14-4132910-INS-52 | chr14-4132910-INS-52 |
| chr14-4137449-INS-61 | chr14-4137449-INS-61 |
| chr14-4144476-INS-50 | chr14-4144476-INS-52 |
| chr14-4162225-INS-216 | chr14-4162225-INS-216 |
| chr14-4165725-INS-212 | chr14-4165725-INS-212 |
| chr14-4203598-INS-470 | chr14-4203598-INS-470 |
| chr14-4208129-INS-209 | chr14-4208129-INS-209 |
| chr14-4211400-INS-75 | chr14-4211397-INS-79 |
| chr14-4220877-INS-74 | chr14-4220877-INS-74 |
| chr14-4228588-INS-128 | chr14-4228620-INS-144 |
| chr14-4284129-INS-52 | chr14-4284129-INS-52 |
| chr14-4332211-INS-328 | chr14-4332211-INS-328 |
| chr14-4356313-INS-4378 | chr14-4356314-INS-4378 |
| chr14-4388784-INS-2919 | chr14-4388785-INS-2919 |
| chr14-4440153-INS-2441 | chr14-4440154-INS-2441 |
| chr14-4500471-INS-1520 | chr14-4500472-INS-1520 |
| chr14-4505472-INS-218 | chr14-4505472-INS-217 |
| chr14-4512749-INS-649 | chr14-4512737-INS-645 |
| chr14-4518822-INS-62 | chr14-4518822-INS-56 |
| chr14-4534193-INS-435 | chr14-4534193-INS-435 |
| chr14-4546230-INS-835 | chr14-4546230-INS-835 |
| chr14-4560243-INS-1446 | chr14-4560244-INS-1446 |
| chr14-4613564-INS-52 | chr14-4613562-INS-52 |
| chr14-4632063-INS-185 | chr14-4632063-INS-185 |
| chr14-4639669-INS-92 | chr14-4639669-INS-80 |
| chr14-4668061-INS-64 | chr14-4668029-INS-72 |
| chr14-4678741-INS-63 | chr14-4678741-INS-63 |
| chr14-4721732-INS-73 | chr14-4721703-INS-58 |
| chr14-4789606-INS-2193 | chr14-4789607-INS-2193 |
| chr14-4804963-INS-59 | chr14-4804963-INS-59 |
| chr14-4820142-INS-4953 | chr14-4820143-INS-4953 |
| chr14-4828950-INS-52 | chr14-4828951-INS-52 |
| chr14-4834024-INS-58 | chr14-4834024-INS-58 |
| chr14-4860441-INS-72 | chr14-4860441-INS-72 |
| chr14-4881180-INS-63 | chr14-4881180-INS-55 |
| chr14-4896620-INS-141 | chr14-4896620-INS-141 |
| chr14-4915195-INS-349 | chr14-4915195-INS-348 |
| chr14-4925083-INS-204 | chr14-4925083-INS-204 |
| chr14-4930758-INS-64 | chr14-4930758-INS-68 |
| chr14-4963954-INS-252 | chr14-4963952-INS-220 |
| chr14-4977080-INS-7478 | chr14-4977081-INS-7478 |

| | |
|---|---|
| chr14-4977474-INS-78 | chr14-4977474-INS-78 |
| chr14-5002365-INS-7684 | chr14-5002366-INS-7684 |
| chr14-5012470-INS-105 | chr14-5012470-INS-105 |
| chr14-5025536-INS-1492 | chr14-5025537-INS-1492 |
| chr14-5032473-INS-434 | chr14-5032473-INS-434 |
| chr14-5042454-INS-195 | chr14-5042454-INS-195 |
| chr14-5050806-INS-4716 | chr14-5050807-INS-4716 |
| chr14-5055494-INS-459 | chr14-5055494-INS-459 |
| chr14-5083061-INS-6266 | chr14-5083062-INS-6266 |
| chr14-5096994-INS-51 | chr14-5096962-INS-100 |
| chr14-5132027-INS-61 | chr14-5132027-INS-53 |
| chr14-5206897-INS-4872 | chr14-5206898-INS-4872 |
| chr14-5213808-INS-78 | chr14-5213808-INS-78 |
| chr14-5216438-INS-4447 | chr14-5216439-INS-4445 |
| chr14-5238777-INS-53 | chr14-5238777-INS-53 |
| chr14-5249088-INS-57 | chr14-5249088-INS-51 |
| chr14-5268335-INS-79 | chr14-5268335-INS-52 |
| chr14-5272128-INS-400 | chr14-5272128-INS-400 |
| chr14-5273627-INS-157 | chr14-5273627-INS-157 |
| chr14-5307079-INS-233 | chr14-5307087-INS-230 |
| chr14-5322588-INS-1418 | chr14-5322589-INS-1418 |
| chr14-5345283-INS-90 | chr14-5345283-INS-90 |
| chr14-5355339-INS-60 | chr14-5355339-INS-60 |
| chr14-5397358-INS-498 | chr14-5397358-INS-498 |
| chr14-5415554-INS-316 | chr14-5415554-INS-316 |
| chr14-5421457-INS-690 | chr14-5421457-INS-690 |
| chr14-5434741-INS-211 | chr14-5434742-INS-211 |
| chr14-5456312-INS-258 | chr14-5456312-INS-258 |
| chr14-5473754-INS-56 | chr14-5473754-INS-52 |
| chr14-5475186-INS-76 | chr14-5475186-INS-76 |
| chr14-5488793-INS-78 | chr14-5488793-INS-78 |
| chr14-5535101-INS-67 | chr14-5535101-INS-66 |
| chr14-5549741-INS-56 | chr14-5549742-INS-56 |
| chr14-5555127-INS-947 | chr14-5555127-INS-949 |
| chr14-5561771-INS-214 | chr14-5561771-INS-214 |
| chr14-5595923-INS-532 | chr14-5595963-INS-532 |
| chr14-5602503-INS-208 | chr14-5602503-INS-208 |
| chr14-5640403-INS-6389 | chr14-5640404-INS-6389 |
| chr14-5653011-INS-3260 | chr14-5653012-INS-3260 |
| chr14-5653681-INS-62 | chr14-5653681-INS-62 |
| chr14-5688753-INS-81 | chr14-5688771-INS-96 |
| chr14-5693805-INS-1142 | chr14-5693806-INS-1142 |
| chr14-5708956-INS-52 | chr14-5708956-INS-52 |
| chr14-5738014-INS-2120 | chr14-5738015-INS-2120 |
| chr14-5825344-INS-1888 | chr14-5825345-INS-1888 |
| chr14-5841683-INS-6663 | chr14-5841684-INS-6665 |
| chr14-5866830-INS-56 | chr14-5866830-INS-50 |
| chr14-5881181-INS-156 | chr14-5881181-INS-156 |
| chr14-5916239-INS-66 | chr14-5916239-INS-65 |
| chr14-5923249-INS-51 | chr14-5923249-INS-51 |
| chr14-5932526-INS-127 | chr14-5932526-INS-125 |
| chr14-5935039-INS-58 | chr14-5935039-INS-58 |
| chr14-5942236-INS-80 | chr14-5942226-INS-58 |
| chr14-5955310-INS-70 | chr14-5955310-INS-70 |
| chr14-5966652-INS-89 | chr14-5966653-INS-92 |
| chr14-5981459-INS-63 | chr14-5981459-INS-63 |
| chr14-6058365-INS-106 | chr14-6058359-INS-104 |
| chr14-6072386-INS-96 | chr14-6072386-INS-96 |
| chr14-6076340-INS-95 | chr14-6076340-INS-95 |
| chr14-6171257-INS-60 | chr14-6171257-INS-55 |
| chr14-6189577-INS-60 | chr14-6189577-INS-60 |
| chr14-6207293-INS-221 | chr14-6207293-INS-221 |
| chr14-6236737-INS-181 | chr14-6236737-INS-188 |
| chr14-6248007-INS-100 | chr14-6248007-INS-112 |
| chr14-6249490-INS-60 | chr14-6249490-INS-54 |
| chr14-6286845-INS-478 | chr14-6286845-INS-478 |
| chr14-6298978-INS-87 | chr14-6298977-INS-91 |
| chr14-6325617-INS-674 | chr14-6325617-INS-674 |
| chr14-6330710-INS-846 | chr14-6330710-INS-846 |
| chr14-6375890-INS-213 | chr14-6375891-INS-216 |
| chr14-6394365-INS-919 | chr14-6394365-INS-919 |
| chr14-6408780-INS-54 | chr14-6408780-INS-54 |
| chr14-6442317-INS-199 | chr14-6442317-INS-199 |
| chr14-6445886-INS-222 | chr14-6445886-INS-222 |
| chr14-6460065-INS-210 | chr14-6460065-INS-210 |
| chr14-6464957-INS-60 | chr14-6464957-INS-60 |
| chr14-6471769-INS-54 | chr14-6471769-INS-54 |

| | |
|---|---|
| chr14-6489526-INS-160 | chr14-6489526-INS-160 |
| chr14-6534522-INS-406 | chr14-6534522-INS-406 |
| chr14-6581053-INS-75 | chr14-6581053-INS-75 |
| chr14-6596467-INS-2249 | chr14-6596468-INS-2249 |
| chr14-6673071-INS-82 | chr14-6673071-INS-82 |
| chr14-6685052-INS-71 | chr14-6685057-INS-67 |
| chr14-6703580-INS-248 | chr14-6703584-INS-240 |
| chr14-6718282-INS-223 | chr14-6718282-INS-223 |
| chr14-6724285-INS-74 | chr14-6724285-INS-74 |
| chr14-6729810-INS-449 | chr14-6729810-INS-449 |
| chr14-6736970-INS-55 | chr14-6736970-INS-50 |
| chr14-6756574-INS-141 | chr14-6756574-INS-137 |
| chr14-6779978-INS-283 | chr14-6779978-INS-283 |
| chr14-6781626-INS-6701 | chr14-6781627-INS-6701 |
| chr14-6781671-INS-101 | chr14-6781671-INS-101 |
| chr14-6783128-INS-69 | chr14-6783128-INS-69 |
| chr14-6841221-INS-65 | chr14-6841221-INS-65 |
| chr14-6883260-INS-2252 | chr14-6883261-INS-2252 |
| chr14-6902855-INS-120 | chr14-6902855-INS-122 |
| chr14-6922699-INS-52 | chr14-6922699-INS-52 |
| chr14-6947703-INS-1455 | chr14-6947704-INS-1455 |
| chr14-6968338-INS-85 | chr14-6968315-INS-66 |
| chr14-7075971-INS-976 | chr14-7075971-INS-978 |
| chr14-7117886-INS-87 | chr14-7117886-INS-87 |
| chr14-7125609-INS-5953 | chr14-7125610-INS-5953 |
| chr14-7144853-INS-1011 | chr14-7144854-INS-1011 |
| chr14-7154311-INS-266 | chr14-7154311-INS-266 |
| chr14-7157414-INS-4641 | chr14-7157415-INS-4641 |
| chr14-7166580-INS-106 | chr14-7166580-INS-108 |
| chr14-7166913-INS-2020 | chr14-7166914-INS-2016 |
| chr14-7176866-INS-92 | chr14-7176867-INS-93 |
| chr14-7192719-INS-207 | chr14-7192719-INS-207 |
| chr14-7194936-INS-63 | chr14-7194936-INS-57 |
| chr14-7246476-INS-61 | chr14-7246476-INS-63 |
| chr14-7252743-INS-1887 | chr14-7252744-INS-1887 |
| chr14-7261320-INS-176 | chr14-7261320-INS-176 |
| chr14-7261920-INS-1556 | chr14-7261921-INS-1548 |
| chr14-7262561-INS-7107 | chr14-7262562-INS-7107 |
| chr14-7265002-INS-82 | chr14-7265002-INS-82 |
| chr14-7285848-INS-7212 | chr14-7285849-INS-7212 |
| chr14-7288183-INS-454 | chr14-7288183-INS-454 |
| chr14-7293830-INS-913 | chr14-7293830-INS-913 |
| chr14-7297963-INS-115 | chr14-7297963-INS-115 |
| chr14-7324457-INS-259 | chr14-7324457-INS-259 |
| chr14-7331865-INS-299 | chr14-7331865-INS-299 |
| chr14-7334624-INS-79 | chr14-7334609-INS-57 |
| chr14-7339686-INS-2668 | chr14-7339687-INS-2668 |
| chr14-7361181-INS-52 | chr14-7361181-INS-54 |
| chr14-7367223-INS-3253 | chr14-7367224-INS-3277 |
| chr14-7372124-INS-57 | chr14-7372136-INS-65 |
| chr14-7373468-INS-67 | chr14-7373468-INS-62 |
| chr14-7375119-INS-120 | chr14-7375119-INS-120 |
| chr14-7388133-INS-68 | chr14-7388133-INS-66 |
| chr14-7413450-INS-106 | chr14-7413450-INS-106 |
| chr14-7417694-INS-65 | chr14-7417694-INS-80 |
| chr14-7418755-INS-71 | chr14-7418755-INS-79 |
| chr14-7419941-INS-147 | chr14-7419982-INS-97 |
| chr14-7422964-INS-91 | chr14-7422964-INS-89 |
| chr14-7438154-INS-774 | chr14-7438154-INS-774 |
| chr14-7500494-INS-147 | chr14-7500494-INS-147 |
| chr14-7501605-INS-127 | chr14-7501605-INS-127 |
| chr14-7504006-INS-129 | chr14-7504006-INS-129 |
| chr14-7511674-INS-78 | chr14-7511674-INS-78 |
| chr14-7520020-INS-209 | chr14-7520020-INS-210 |
| chr14-7530007-INS-145 | chr14-7530007-INS-149 |
| chr14-7543263-INS-324 | chr14-7543263-INS-333 |
| chr14-7559866-INS-452 | chr14-7559867-INS-452 |
| chr14-7566326-INS-543 | chr14-7566326-INS-543 |
| chr14-7573474-INS-73 | chr14-7573474-INS-73 |
| chr14-7585982-INS-1670 | chr14-7585983-INS-1670 |
| chr14-7600937-INS-66 | chr14-7600937-INS-65 |
| chr14-7616177-INS-274 | chr14-7616177-INS-274 |
| chr14-7624092-INS-122 | chr14-7624092-INS-122 |
| chr14-7630725-INS-50 | chr14-7630725-INS-50 |
| chr14-7635353-INS-744 | chr14-7635353-INS-744 |
| chr14-7675074-INS-60 | chr14-7675074-INS-64 |
| chr14-7695395-INS-623 | chr14-7695395-INS-623 |

| | |
|---|---|
| chr14-7714160-INS-490 | chr14-7714160-INS-490 |
| chr14-7735429-INS-101 | chr14-7735475-INS-52 |
| chr14-7740993-INS-75 | chr14-7740993-INS-93 |
| chr14-7762815-INS-52 | chr14-7762816-INS-52 |
| chr14-7783857-INS-55 | chr14-7783857-INS-63 |
| chr14-7788633-INS-69 | chr14-7788633-INS-69 |
| chr14-7793908-INS-114 | chr14-7793908-INS-126 |
| chr14-7802872-INS-6242 | chr14-7802873-INS-6242 |
| chr14-7816069-INS-387 | chr14-7816069-INS-387 |
| chr14-7824679-INS-156 | chr14-7824679-INS-159 |
| chr14-7828841-INS-2046 | chr14-7828842-INS-2048 |
| chr14-7862252-INS-161 | chr14-7862252-INS-161 |
| chr14-7927013-INS-1072 | chr14-7927014-INS-1095 |
| chr14-7935980-INS-58 | chr14-7935980-INS-50 |
| chr14-7942749-INS-52 | chr14-7942749-INS-52 |
| chr14-7976513-INS-302 | chr14-7976517-INS-320 |
| chr14-8000641-INS-203 | chr14-8000641-INS-203 |
| chr14-8022567-INS-59 | chr14-8022567-INS-59 |
| chr14-8046030-INS-6401 | chr14-8046031-INS-6401 |
| chr14-8104846-INS-2730 | chr14-8104847-INS-2730 |
| chr14-8173399-INS-328 | chr14-8173399-INS-328 |
| chr14-8188649-INS-403 | chr14-8188649-INS-403 |
| chr14-8191668-INS-296 | chr14-8191668-INS-294 |
| chr14-8221084-INS-124 | chr14-8221084-INS-124 |
| chr14-8227229-INS-252 | chr14-8227230-INS-252 |
| chr14-8241678-INS-6441 | chr14-8241679-INS-6441 |
| chr14-8249141-INS-826 | chr14-8249141-INS-826 |
| chr14-8296045-INS-60 | chr14-8296045-INS-60 |
| chr14-8301767-INS-202 | chr14-8301767-INS-201 |
| chr14-8305985-INS-3099 | chr14-8305986-INS-3099 |
| chr14-8316169-INS-64 | chr14-8316169-INS-64 |
| chr14-8316663-INS-75 | chr14-8316663-INS-75 |
| chr14-8349776-INS-273 | chr14-8349776-INS-273 |
| chr14-8417534-INS-96 | chr14-8417534-INS-96 |
| chr14-8427171-INS-6686 | chr14-8427172-INS-6687 |
| chr14-8431818-INS-118 | chr14-8431818-INS-118 |
| chr14-8437431-INS-215 | chr14-8437381-INS-252 |
| chr14-8447894-INS-186 | chr14-8447894-INS-186 |
| chr14-8483675-INS-72 | chr14-8483675-INS-92 |
| chr14-8485889-INS-52 | chr14-8485879-INS-58 |
| chr14-8504994-INS-4371 | chr14-8504995-INS-4371 |
| chr14-8505972-INS-203 | chr14-8505972-INS-203 |
| chr14-8509487-INS-74 | chr14-8509487-INS-72 |
| chr14-8520344-INS-176 | chr14-8520344-INS-176 |
| chr14-8523340-INS-197 | chr14-8523340-INS-197 |
| chr14-8525264-INS-339 | chr14-8525264-INS-339 |
| chr14-8527167-INS-222 | chr14-8527167-INS-222 |
| chr14-8537964-INS-112 | chr14-8537965-INS-112 |
| chr14-8539585-INS-209 | chr14-8539585-INS-209 |
| chr14-8542549-INS-529 | chr14-8542549-INS-529 |
| chr14-8565547-INS-177 | chr14-8565547-INS-177 |
| chr14-8570016-INS-295 | chr14-8570016-INS-296 |
| chr14-8599033-INS-193 | chr14-8599033-INS-193 |
| chr14-8600814-INS-61 | chr14-8600814-INS-61 |
| chr14-8606230-INS-95 | chr14-8606230-INS-95 |
| chr14-8610359-INS-6464 | chr14-8610360-INS-6464 |
| chr14-8616107-INS-7370 | chr14-8616108-INS-7371 |
| chr14-8622886-INS-972 | chr14-8622887-INS-972 |
| chr14-8628129-INS-82 | chr14-8628129-INS-82 |
| chr14-8629773-INS-424 | chr14-8629773-INS-424 |
| chr14-8650888-INS-67 | chr14-8650888-INS-67 |
| chr14-8655932-INS-109 | chr14-8655932-INS-109 |
| chr14-8669716-INS-64 | chr14-8669716-INS-64 |
| chr14-8680607-INS-122 | chr14-8680607-INS-116 |
| chr14-8706176-INS-57 | chr14-8706147-INS-57 |
| chr14-8708696-INS-137 | chr14-8708696-INS-137 |
| chr14-8719256-INS-356 | chr14-8719256-INS-356 |
| chr14-8724045-INS-183 | chr14-8724045-INS-183 |
| chr14-8724457-INS-498 | chr14-8724457-INS-498 |
| chr14-8732574-INS-1803 | chr14-8732575-INS-1803 |
| chr14-8868500-INS-221 | chr14-8868500-INS-221 |
| chr14-8880816-INS-188 | chr14-8880816-INS-188 |
| chr14-8882560-INS-4172 | chr14-8882561-INS-4172 |
| chr14-8882561-INS-390 | chr14-8882561-INS-390 |
| chr14-8890173-INS-68 | chr14-8890173-INS-68 |
| chr14-8910901-INS-51 | chr14-8910901-INS-51 |
| chr14-8932070-INS-307 | chr14-8932070-INS-307 |

| | |
|---|---|
| chr14-8940376-INS-6595 | chr14-8940377-INS-6595 |
| chr14-8946814-INS-63 | chr14-8946814-INS-69 |
| chr14-8958244-INS-140 | chr14-8958244-INS-140 |
| chr14-8961468-INS-1169 | chr14-8961469-INS-1169 |
| chr14-8962222-INS-6442 | chr14-8962223-INS-6442 |
| chr14-8968412-INS-220 | chr14-8968408-INS-224 |
| chr14-8971042-INS-177 | chr14-8971042-INS-177 |
| chr14-8978261-INS-935 | chr14-8978261-INS-934 |
| chr14-8988409-INS-201 | chr14-8988409-INS-201 |
| chr14-8999430-INS-7159 | chr14-8999431-INS-7159 |
| chr14-9060370-INS-69 | chr14-9060370-INS-73 |
| chr14-9083160-INS-2223 | chr14-9083161-INS-2223 |
| chr14-9112594-INS-318 | chr14-9112594-INS-320 |
| chr14-9142717-INS-72 | chr14-9142717-INS-66 |
| chr14-9146325-INS-201 | chr14-9146325-INS-201 |
| chr14-9148105-INS-582 | chr14-9148105-INS-582 |
| chr14-9158783-INS-942 | chr14-9158783-INS-942 |
| chr14-9164499-INS-500 | chr14-9164499-INS-500 |
| chr14-9169319-INS-897 | chr14-9169317-INS-897 |
| chr14-9170219-INS-62 | chr14-9170219-INS-60 |
| chr14-9177095-INS-706 | chr14-9177095-INS-706 |
| chr14-9178352-INS-100 | chr14-9178352-INS-100 |
| chr14-9193359-INS-212 | chr14-9193359-INS-212 |
| chr14-9210146-INS-1775 | chr14-9210147-INS-1775 |
| chr14-9221041-INS-183 | chr14-9221033-INS-174 |
| chr14-9227890-INS-116 | chr14-9227890-INS-116 |
| chr14-9229956-INS-401 | chr14-9229956-INS-401 |
| chr14-9238638-INS-507 | chr14-9238591-INS-508 |
| chr14-9240213-INS-578 | chr14-9240213-INS-578 |
| chr14-9267024-INS-64 | chr14-9267024-INS-72 |
| chr14-9268880-INS-442 | chr14-9268881-INS-442 |
| chr14-9270648-INS-55 | chr14-9270648-INS-70 |
| chr14-9319362-INS-467 | chr14-9319362-INS-467 |
| chr14-9329854-INS-148 | chr14-9329854-INS-148 |
| chr14-9358561-INS-126 | chr14-9358561-INS-126 |
| chr14-9374227-INS-60 | chr14-9374228-INS-55 |
| chr14-9388510-INS-53 | chr14-9388510-INS-53 |
| chr14-9390837-INS-145 | chr14-9390837-INS-145 |
| chr14-9397651-INS-94 | chr14-9397651-INS-55 |
| chr14-9402916-INS-554 | chr14-9402916-INS-554 |
| chr14-9407592-INS-66 | chr14-9407592-INS-66 |
| chr14-9420513-INS-264 | chr14-9420513-INS-264 |
| chr14-9429895-INS-61 | chr14-9429895-INS-61 |
| chr14-9433530-INS-66 | chr14-9433530-INS-66 |
| chr14-9440609-INS-419 | chr14-9440609-INS-419 |
| chr14-9461059-INS-480 | chr14-9461059-INS-480 |
| chr14-9466219-INS-7373 | chr14-9466220-INS-7373 |
| chr14-9496632-INS-299 | chr14-9496632-INS-298 |
| chr14-9504994-INS-2556 | chr14-9504995-INS-2556 |
| chr14-9515567-INS-353 | chr14-9515567-INS-353 |
| chr14-9516731-INS-70 | chr14-9516731-INS-70 |
| chr14-9523557-INS-200 | chr14-9523557-INS-200 |
| chr14-9527877-INS-214 | chr14-9527877-INS-214 |
| chr14-9530825-INS-341 | chr14-9530825-INS-341 |
| chr14-9531408-INS-53 | chr14-9531408-INS-53 |
| chr14-9539411-INS-4053 | chr14-9539412-INS-4053 |
| chr14-9543427-INS-179 | chr14-9543427-INS-179 |
| chr14-9557302-INS-1471 | chr14-9557303-INS-1472 |
| chr14-9566786-INS-7315 | chr14-9566787-INS-7318 |
| chr14-9580688-INS-125 | chr14-9580688-INS-105 |
| chr14-9587994-INS-2638 | chr14-9587995-INS-2638 |
| chr14-9588185-INS-208 | chr14-9588185-INS-208 |
| chr14-9628342-INS-363 | chr14-9628342-INS-363 |
| chr14-9654093-INS-85 | chr14-9654093-INS-79 |
| chr14-9698110-INS-6314 | chr14-9698111-INS-6314 |
| chr14-9716821-INS-69 | chr14-9716821-INS-69 |
| chr14-9723949-INS-51 | chr14-9723949-INS-52 |
| chr14-9727743-INS-86 | chr14-9727743-INS-86 |
| chr14-9735581-INS-400 | chr14-9735581-INS-400 |
| chr14-9739749-INS-62 | chr14-9739749-INS-72 |
| chr14-9749299-INS-3058 | chr14-9749300-INS-3058 |
| chr14-9755779-INS-6713 | chr14-9755780-INS-6715 |
| chr14-9761757-INS-271 | chr14-9761757-INS-272 |
| chr14-9779598-INS-2255 | chr14-9779599-INS-2255 |
| chr14-9786299-INS-238 | chr14-9786299-INS-241 |
| chr14-9820720-INS-287 | chr14-9820720-INS-287 |
| chr14-9835729-INS-53 | chr14-9835729-INS-53 |

| | |
|---|---|
| chr14-9841352-INS-80 | chr14-9841352-INS-80 |
| chr14-9843048-INS-52 | chr14-9843048-INS-52 |
| chr14-9851393-INS-213 | chr14-9851393-INS-213 |
| chr14-9863507-INS-64 | chr14-9863507-INS-64 |
| chr14-9871088-INS-434 | chr14-9871088-INS-438 |
| chr14-9873415-INS-73 | chr14-9873415-INS-73 |
| chr14-9879805-INS-85 | chr14-9879805-INS-85 |
| chr14-9902393-INS-495 | chr14-9902393-INS-495 |
| chr14-9911310-INS-63 | chr14-9911310-INS-63 |
| chr14-9919724-INS-102 | chr14-9919725-INS-78 |
| chr14-9924232-INS-178 | chr14-9924232-INS-178 |
| chr14-9924977-INS-7135 | chr14-9924978-INS-7136 |
| chr14-9925659-INS-122 | chr14-9925659-INS-126 |
| chr14-9928703-INS-158 | chr14-9928703-INS-160 |
| chr14-9932017-INS-66 | chr14-9932017-INS-66 |
| chr14-10024924-INS-66 | chr14-10024924-INS-66 |
| chr14-10038122-INS-324 | chr14-10038122-INS-303 |
| chr14-10059446-INS-80 | chr14-10059446-INS-80 |
| chr14-10062582-INS-134 | chr14-10062536-INS-157 |
| chr14-10086249-INS-77 | chr14-10086265-INS-50 |
| chr14-10110065-INS-54 | chr14-10110065-INS-54 |
| chr14-10115106-INS-84 | chr14-10115118-INS-69 |
| chr14-10139111-INS-107 | chr14-10139113-INS-112 |
| chr14-10157105-INS-56 | chr14-10157105-INS-56 |
| chr14-10202809-INS-524 | chr14-10202804-INS-516 |
| chr14-10240458-INS-1934 | chr14-10240459-INS-1932 |
| chr14-10259832-INS-724 | chr14-10259832-INS-724 |
| chr14-10264023-INS-98 | chr14-10264023-INS-98 |
| chr14-10316303-INS-391 | chr14-10316303-INS-391 |
| chr14-10334318-INS-201 | chr14-10334318-INS-202 |
| chr14-10338882-INS-124 | chr14-10338882-INS-124 |
| chr14-10355125-INS-70 | chr14-10355125-INS-70 |
| chr14-10360882-INS-56 | chr14-10360882-INS-56 |
| chr14-10362307-INS-280 | chr14-10362307-INS-280 |
| chr14-10373717-INS-212 | chr14-10373717-INS-212 |
| chr14-10379999-INS-52 | chr14-10379999-INS-56 |
| chr14-10396577-INS-8448 | chr14-10396578-INS-8448 |
| chr14-10408822-INS-1876 | chr14-10408823-INS-1876 |
| chr14-10427480-INS-5308 | chr14-10427481-INS-5308 |
| chr14-10430975-INS-55 | chr14-10430975-INS-55 |
| chr14-10452668-INS-400 | chr14-10452668-INS-400 |
| chr14-10478683-INS-205 | chr14-10478683-INS-205 |
| chr14-10483550-INS-51 | chr14-10483550-INS-53 |
| chr14-10485146-INS-54 | chr14-10485146-INS-54 |
| chr14-10498426-INS-212 | chr14-10498426-INS-212 |
| chr14-10509655-INS-204 | chr14-10509655-INS-204 |
| chr14-10514507-INS-84 | chr14-10514507-INS-84 |
| chr14-10532784-INS-462 | chr14-10532784-INS-462 |
| chr14-10546140-INS-5545 | chr14-10546141-INS-5546 |
| chr14-10579916-INS-1040 | chr14-10579917-INS-1040 |
| chr14-10587259-INS-203 | chr14-10587259-INS-206 |
| chr14-10639049-INS-256 | chr14-10639048-INS-256 |
| chr14-10647018-INS-180 | chr14-10647018-INS-180 |
| chr14-10652201-INS-200 | chr14-10652201-INS-205 |
| chr14-10655153-INS-60 | chr14-10655153-INS-72 |
| chr14-10656824-INS-74 | chr14-10656825-INS-66 |
| chr14-10663294-INS-70 | chr14-10663294-INS-70 |
| chr14-10682209-INS-92 | chr14-10682209-INS-92 |
| chr14-10695894-INS-81 | chr14-10695894-INS-77 |
| chr14-10708023-INS-257 | chr14-10708023-INS-257 |
| chr14-10721595-INS-88 | chr14-10721595-INS-88 |
| chr14-10789198-INS-185 | chr14-10789198-INS-185 |
| chr14-10794154-INS-136 | chr14-10794154-INS-136 |
| chr14-10836132-INS-55 | chr14-10836132-INS-70 |
| chr14-10851682-INS-184 | chr14-10851699-INS-184 |
| chr14-10859261-INS-54 | chr14-10859261-INS-50 |
| chr14-10862525-INS-1254 | chr14-10862526-INS-1254 |
| chr14-10866181-INS-204 | chr14-10866181-INS-204 |
| chr14-10883495-INS-78 | chr14-10883490-INS-80 |
| chr14-10884073-INS-804 | chr14-10884073-INS-804 |
| chr14-10906446-INS-217 | chr14-10906446-INS-216 |
| chr14-10908906-INS-75 | chr14-10908906-INS-75 |
| chr14-10911451-INS-221 | chr14-10911451-INS-221 |
| chr14-10918748-INS-53 | chr14-10918748-INS-53 |
| chr14-10926380-INS-132 | chr14-10926380-INS-151 |
| chr14-10941433-INS-112 | chr14-10941433-INS-113 |
| chr14-10946881-INS-55 | chr14-10946881-INS-55 |

| | |
|---|---|
| chr14-10949814-INS-7099 | chr14-10949815-INS-7100 |
| chr14-10953674-INS-304 | chr14-10953674-INS-304 |
| chr14-10958359-INS-211 | chr14-10958359-INS-212 |
| chr14-10985026-INS-54 | chr14-10985026-INS-87 |
| chr14-10988356-INS-373 | chr14-10988356-INS-370 |
| chr14-11002996-INS-146 | chr14-11002996-INS-146 |
| chr14-11005060-INS-188 | chr14-11005060-INS-188 |
| chr14-11012875-INS-54 | chr14-11012875-INS-54 |
| chr14-11029713-INS-120 | chr14-11029713-INS-120 |
| chr14-11047862-INS-646 | chr14-11047862-INS-622 |
| chr14-11065287-INS-291 | chr14-11065287-INS-291 |
| chr14-11103506-INS-338 | chr14-11103506-INS-346 |
| chr14-11105798-INS-151 | chr14-11105798-INS-151 |
| chr14-11119078-INS-69 | chr14-11119039-INS-78 |
| chr14-11126379-INS-1920 | chr14-11126380-INS-1929 |
| chr14-11184568-INS-801 | chr14-11184568-INS-801 |
| chr14-11201281-INS-55 | chr14-11201281-INS-54 |
| chr14-11205237-INS-92 | chr14-11205240-INS-51 |
| chr14-11271996-INS-149 | chr14-11271996-INS-145 |
| chr14-11284797-INS-6186 | chr14-11284798-INS-6186 |
| chr14-11312530-INS-56 | chr14-11312530-INS-54 |
| chr14-11387923-INS-358 | chr14-11387923-INS-358 |
| chr14-11417066-INS-88 | chr14-11417066-INS-56 |
| chr14-11493930-INS-6938 | chr14-11493931-INS-6938 |
| chr14-11539972-INS-54 | chr14-11539972-INS-50 |
| chr14-11592412-INS-856 | chr14-11592412-INS-856 |
| chr14-11608067-INS-91 | chr14-11608067-INS-91 |
| chr14-11609694-INS-80 | chr14-11609694-INS-82 |
| chr14-11612906-INS-57 | chr14-11612906-INS-57 |
| chr14-11613424-INS-236 | chr14-11613424-INS-236 |
| chr14-11615484-INS-498 | chr14-11615492-INS-498 |
| chr14-11626171-INS-2403 | chr14-11626172-INS-2407 |
| chr14-11649831-INS-117 | chr14-11649831-INS-147 |
| chr14-11702990-INS-2716 | chr14-11702991-INS-2716 |
| chr14-11733518-INS-235 | chr14-11733518-INS-248 |
| chr14-11744308-INS-56 | chr14-11744308-INS-56 |
| chr14-11788476-INS-67 | chr14-11788476-INS-73 |
| chr14-11798871-INS-145 | chr14-11798871-INS-145 |
| chr14-11835597-INS-107 | chr14-11835595-INS-109 |
| chr14-11846182-INS-2345 | chr14-11846183-INS-2345 |
| chr14-11866894-INS-52 | chr14-11866894-INS-52 |
| chr14-11893518-INS-89 | chr14-11893518-INS-50 |
| chr14-11935648-INS-301 | chr14-11935648-INS-301 |
| chr14-11939855-INS-1727 | chr14-11939856-INS-1733 |
| chr14-11948695-INS-1289 | chr14-11948696-INS-1289 |
| chr14-11951651-INS-84 | chr14-11951652-INS-84 |
| chr14-11952188-INS-64 | chr14-11952189-INS-64 |
| chr14-11954498-INS-53 | chr14-11954510-INS-54 |
| chr14-11958164-INS-681 | chr14-11958164-INS-669 |
| chr14-11964493-INS-410 | chr14-11964493-INS-410 |
| chr14-11972250-INS-318 | chr14-11972250-INS-318 |
| chr14-11985723-INS-111 | chr14-11985723-INS-110 |
| chr14-12004922-INS-1870 | chr14-12004928-INS-1869 |
| chr14-12007325-INS-327 | chr14-12007325-INS-327 |
| chr14-12010219-INS-458 | chr14-12010219-INS-462 |
| chr14-12081401-INS-498 | chr14-12081401-INS-498 |
| chr14-12097094-INS-262 | chr14-12097094-INS-262 |
| chr14-12113362-INS-6717 | chr14-12113363-INS-6718 |
| chr14-12113865-INS-55 | chr14-12113865-INS-55 |
| chr14-12116578-INS-63 | chr14-12116578-INS-63 |
| chr14-12118039-INS-573 | chr14-12118037-INS-573 |
| chr14-12142528-INS-414 | chr14-12142528-INS-414 |
| chr14-12155445-INS-105 | chr14-12155445-INS-125 |
| chr14-12183687-INS-817 | chr14-12183687-INS-817 |
| chr14-12238212-INS-60 | chr14-12238212-INS-60 |
| chr14-12262345-INS-53 | chr14-12262345-INS-51 |
| chr14-12301570-INS-80 | chr14-12301570-INS-80 |
| chr14-12304942-INS-90 | chr14-12304942-INS-92 |
| chr14-12313501-INS-50 | chr14-12313501-INS-62 |
| chr14-12327443-INS-355 | chr14-12327444-INS-355 |
| chr14-12336830-INS-480 | chr14-12336830-INS-481 |
| chr14-12411050-INS-52 | chr14-12411050-INS-52 |
| chr14-12430678-INS-260 | chr14-12430678-INS-253 |
| chr14-12492219-INS-170 | chr14-12492219-INS-177 |
| chr14-12505348-INS-51 | chr14-12505348-INS-51 |
| chr14-12515024-INS-52 | chr14-12515023-INS-52 |
| chr14-12535106-INS-68 | chr14-12535106-INS-66 |

| | |
|---|---|
| chr14-12568079-INS-52 | chr14-12568079-INS-52 |
| chr14-12584085-INS-248 | chr14-12584085-INS-248 |
| chr14-12597581-INS-86 | chr14-12597581-INS-86 |
| chr14-12652001-INS-7196 | chr14-12652002-INS-7196 |
| chr14-12685330-INS-88 | chr14-12685340-INS-64 |
| chr14-12688903-INS-86 | chr14-12688905-INS-88 |
| chr14-12700733-INS-560 | chr14-12700733-INS-560 |
| chr14-12709027-INS-421 | chr14-12709027-INS-422 |
| chr14-12748581-INS-191 | chr14-12748581-INS-191 |
| chr14-12749848-INS-66 | chr14-12749848-INS-72 |
| chr14-12750977-INS-418 | chr14-12750977-INS-423 |
| chr14-12752765-INS-125 | chr14-12752765-INS-150 |
| chr14-12758348-INS-90 | chr14-12758348-INS-90 |
| chr14-12762033-INS-178 | chr14-12762033-INS-178 |
| chr14-12763893-INS-86 | chr14-12763893-INS-86 |
| chr14-12769007-INS-6887 | chr14-12769008-INS-6887 |
| chr14-12772413-INS-479 | chr14-12772413-INS-479 |
| chr14-12780169-INS-56 | chr14-12780169-INS-60 |
| chr14-12784654-INS-172 | chr14-12784655-INS-172 |
| chr14-12798326-INS-72 | chr14-12798326-INS-72 |
| chr14-12813089-INS-7127 | chr14-12813090-INS-7127 |
| chr14-12829540-INS-5467 | chr14-12829541-INS-5467 |
| chr14-12832231-INS-199 | chr14-12832231-INS-199 |
| chr14-12847441-INS-59 | chr14-12847409-INS-109 |
| chr14-12849494-INS-56 | chr14-12849494-INS-56 |
| chr14-12850378-INS-6700 | chr14-12850379-INS-6702 |
| chr14-12871241-INS-52 | chr14-12871241-INS-52 |
| chr14-12883768-INS-61 | chr14-12883768-INS-63 |
| chr14-12915071-INS-179 | chr14-12915061-INS-185 |
| chr14-12927335-INS-999 | chr14-12927335-INS-999 |
| chr14-12946714-INS-467 | chr14-12946714-INS-467 |
| chr14-12950148-INS-258 | chr14-12950148-INS-258 |
| chr14-12957171-INS-400 | chr14-12957171-INS-400 |
| chr14-12971429-INS-119 | chr14-12971429-INS-119 |
| chr14-12973594-INS-3854 | chr14-12973595-INS-3854 |
| chr14-12979517-INS-111 | chr14-12979517-INS-112 |
| chr14-12987501-INS-234 | chr14-12987501-INS-234 |
| chr14-12994972-INS-629 | chr14-12994972-INS-630 |
| chr14-13002707-INS-78 | chr14-13002707-INS-80 |
| chr14-13005094-INS-104 | chr14-13005094-INS-104 |
| chr14-13022917-INS-80 | chr14-13022917-INS-80 |
| chr14-13027723-INS-6276 | chr14-13027724-INS-6277 |
| chr14-13028130-INS-108 | chr14-13028130-INS-108 |
| chr14-13029260-INS-56 | chr14-13029260-INS-56 |
| chr14-13082791-INS-116 | chr14-13082791-INS-116 |
| chr14-13084795-INS-1787 | chr14-13084796-INS-1787 |
| chr14-13089455-INS-2517 | chr14-13089456-INS-2517 |
| chr14-13089456-INS-357 | chr14-13089456-INS-357 |
| chr14-13098494-INS-184 | chr14-13098494-INS-184 |
| chr14-13099529-INS-66 | chr14-13099529-INS-66 |
| chr14-13116779-INS-87 | chr14-13116779-INS-86 |
| chr14-13147649-INS-616 | chr14-13147650-INS-617 |
| chr14-13165020-INS-56 | chr14-13165013-INS-86 |
| chr14-13205666-INS-708 | chr14-13205666-INS-708 |
| chr14-13215400-INS-56 | chr14-13215400-INS-60 |
| chr14-13231832-INS-2832 | chr14-13231833-INS-2832 |
| chr14-13265760-INS-762 | chr14-13265761-INS-759 |
| chr14-13267697-INS-52 | chr14-13267697-INS-54 |
| chr14-13278638-INS-58 | chr14-13278638-INS-64 |
| chr14-13280319-INS-517 | chr14-13280319-INS-517 |
| chr14-13297578-INS-57 | chr14-13297604-INS-71 |
| chr14-13345572-INS-52 | chr14-13345578-INS-50 |
| chr14-13353920-INS-1069 | chr14-13353921-INS-1070 |
| chr14-13357217-INS-310 | chr14-13357217-INS-307 |
| chr14-13386495-INS-64 | chr14-13386495-INS-64 |
| chr14-13393789-INS-6467 | chr14-13393790-INS-6467 |
| chr14-13411444-INS-63 | chr14-13411443-INS-67 |
| chr14-13437231-INS-57 | chr14-13437231-INS-57 |
| chr14-13463582-INS-53 | chr14-13463582-INS-55 |
| chr14-13470169-INS-3022 | chr14-13470170-INS-3022 |
| chr14-13492569-INS-498 | chr14-13492569-INS-498 |
| chr14-13512420-INS-490 | chr14-13512405-INS-482 |
| chr14-13515895-INS-126 | chr14-13515895-INS-126 |
| chr14-13531473-INS-68 | chr14-13531473-INS-74 |
| chr14-13532779-INS-54 | chr14-13532779-INS-56 |
| chr14-13534730-INS-54 | chr14-13534730-INS-66 |
| chr14-13540854-INS-56 | chr14-13540854-INS-54 |

| | |
|---|---|
| chr14-13564704-INS-88 | chr14-13564704-INS-88 |
| chr14-13565816-INS-3995 | chr14-13565817-INS-3995 |
| chr14-13574409-INS-547 | chr14-13574408-INS-574 |
| chr14-13574762-INS-4953 | chr14-13574763-INS-4953 |
| chr14-13654529-INS-98 | chr14-13654529-INS-98 |
| chr14-13659003-INS-558 | chr14-13659003-INS-558 |
| chr14-13665623-INS-68 | chr14-13665623-INS-72 |
| chr14-13665802-INS-6599 | chr14-13665803-INS-6599 |
| chr14-13687992-INS-6140 | chr14-13687993-INS-6144 |
| chr14-13695324-INS-57 | chr14-13695291-INS-51 |
| chr14-13703057-INS-279 | chr14-13703057-INS-277 |
| chr14-13733223-INS-66 | chr14-13733223-INS-66 |
| chr14-13740281-INS-60 | chr14-13740258-INS-54 |
| chr14-13741441-INS-186 | chr14-13741441-INS-186 |
| chr14-13755825-INS-57 | chr14-13755825-INS-57 |
| chr14-13760923-INS-5539 | chr14-13760924-INS-5539 |
| chr14-13761980-INS-205 | chr14-13761980-INS-205 |
| chr14-13767665-INS-200 | chr14-13767665-INS-200 |
| chr14-13781802-INS-70 | chr14-13781802-INS-70 |
| chr14-13782229-INS-989 | chr14-13782229-INS-989 |
| chr14-13792362-INS-520 | chr14-13792362-INS-520 |
| chr14-13794649-INS-206 | chr14-13794649-INS-206 |
| chr14-13820289-INS-99 | chr14-13820289-INS-99 |
| chr14-13840545-INS-97 | chr14-13840545-INS-101 |
| chr14-13844282-INS-7115 | chr14-13844283-INS-7115 |
| chr14-13931699-INS-168 | chr14-13931728-INS-168 |
| chr14-13942330-INS-56 | chr14-13942330-INS-56 |
| chr14-13942358-INS-6440 | chr14-13942343-INS-6428 |
| chr14-13947347-INS-186 | chr14-13947347-INS-186 |
| chr14-13967892-INS-840 | chr14-13967892-INS-840 |
| chr14-13979924-INS-709 | chr14-13979924-INS-708 |
| chr14-14006002-INS-153 | chr14-14006002-INS-153 |
| chr14-14020596-INS-114 | chr14-14020615-INS-138 |
| chr14-14064342-INS-1277 | chr14-14064343-INS-1293 |
| chr14-14082822-INS-64 | chr14-14082822-INS-64 |
| chr14-14095312-INS-422 | chr14-14095312-INS-422 |
| chr14-14097626-INS-309 | chr14-14097626-INS-301 |
| chr14-14103672-INS-60 | chr14-14103672-INS-60 |
| chr14-14104178-INS-64 | chr14-14104178-INS-64 |
| chr14-14105595-INS-73 | chr14-14105595-INS-73 |
| chr14-14112441-INS-218 | chr14-14112441-INS-218 |
| chr14-14139507-INS-66 | chr14-14139501-INS-68 |
| chr14-14162479-INS-64 | chr14-14162479-INS-64 |
| chr14-14181000-INS-2922 | chr14-14181001-INS-2922 |
| chr14-14185166-INS-361 | chr14-14185166-INS-361 |
| chr14-14187322-INS-528 | chr14-14187322-INS-528 |
| chr14-14201900-INS-57 | chr14-14201900-INS-53 |
| chr14-14206536-INS-54 | chr14-14206536-INS-52 |
| chr14-14217828-INS-54 | chr14-14217828-INS-56 |
| chr14-14229423-INS-160 | chr14-14229423-INS-160 |
| chr14-14233062-INS-56 | chr14-14233049-INS-63 |
| chr14-14236059-INS-98 | chr14-14236059-INS-104 |
| chr14-14276525-INS-154 | chr14-14276525-INS-154 |
| chr14-14280976-INS-78 | chr14-14280976-INS-66 |
| chr14-14286178-INS-3053 | chr14-14286179-INS-3053 |
| chr14-14310258-INS-67 | chr14-14310249-INS-70 |
| chr14-14337675-INS-64 | chr14-14337675-INS-62 |
| chr14-14372476-INS-60 | chr14-14372476-INS-60 |
| chr14-14374298-INS-211 | chr14-14374298-INS-211 |
| chr14-14375529-INS-6465 | chr14-14375530-INS-6465 |
| chr14-14376788-INS-56 | chr14-14376788-INS-56 |
| chr14-14383974-INS-204 | chr14-14383974-INS-204 |
| chr14-14393651-INS-86 | chr14-14393651-INS-63 |
| chr14-14394279-INS-214 | chr14-14394279-INS-214 |
| chr14-14404391-INS-228 | chr14-14404391-INS-228 |
| chr14-14410474-INS-174 | chr14-14410474-INS-172 |
| chr14-14420586-INS-218 | chr14-14420586-INS-218 |
| chr14-14425989-INS-177 | chr14-14425989-INS-178 |
| chr14-14430803-INS-62 | chr14-14430803-INS-62 |
| chr14-14468111-INS-60 | chr14-14468103-INS-100 |
| chr14-14487599-INS-181 | chr14-14487599-INS-181 |
| chr14-14490175-INS-66 | chr14-14490175-INS-74 |
| chr14-14498506-INS-204 | chr14-14498506-INS-204 |
| chr14-14534326-INS-98 | chr14-14534326-INS-98 |
| chr14-14541073-INS-884 | chr14-14541073-INS-880 |
| chr14-14576688-INS-141 | chr14-14576688-INS-141 |
| chr14-14590321-INS-57 | chr14-14590321-INS-57 |

| | |
|---|---|
| chr14-14594913-INS-63 | chr14-14594913-INS-63 |
| chr14-14612590-INS-52 | chr14-14612590-INS-52 |
| chr14-14623856-INS-84 | chr14-14623873-INS-60 |
| chr14-14625008-INS-176 | chr14-14625008-INS-176 |
| chr14-14627178-INS-128 | chr14-14627178-INS-131 |
| chr14-14628978-INS-60 | chr14-14628978-INS-73 |
| chr14-14633630-INS-388 | chr14-14633630-INS-388 |
| chr14-14649700-INS-81 | chr14-14649700-INS-81 |
| chr14-14668644-INS-147 | chr14-14668644-INS-147 |
| chr14-14674486-INS-207 | chr14-14674486-INS-213 |
| chr14-14701595-INS-201 | chr14-14701595-INS-201 |
| chr14-14706321-INS-73 | chr14-14706321-INS-73 |
| chr14-14708150-INS-62 | chr14-14708150-INS-79 |
| chr14-14715173-INS-654 | chr14-14715173-INS-654 |
| chr14-14719778-INS-81 | chr14-14719778-INS-85 |
| chr14-14723931-INS-105 | chr14-14723934-INS-75 |
| chr14-14762481-INS-76 | chr14-14762481-INS-72 |
| chr14-14776795-INS-306 | chr14-14776795-INS-306 |
| chr14-14795374-INS-409 | chr14-14795374-INS-409 |
| chr14-14806896-INS-155 | chr14-14806909-INS-171 |
| chr14-14808928-INS-4977 | chr14-14808929-INS-4977 |
| chr14-15184262-INS-54 | chr14-15184259-INS-57 |
| chr14-15491721-INS-52 | chr14-15491721-INS-52 |
| chr14-15512004-INS-180 | chr14-15512004-INS-181 |
| chr14-16733087-INS-108 | chr14-16733088-INS-107 |
| chr14-18235123-INS-214 | chr14-18235123-INS-214 |
| chr14-19713488-INS-644 | chr14-19713488-INS-644 |
| chr14-19725877-INS-250 | chr14-19725851-INS-293 |
| chr14-19728275-INS-2902 | chr14-19728276-INS-2902 |
| chr14-19743962-INS-498 | chr14-19743962-INS-498 |
| chr14-19756240-INS-83 | chr14-19756240-INS-84 |
| chr14-19762179-INS-74 | chr14-19762173-INS-88 |
| chr14-19764578-INS-56 | chr14-19764578-INS-56 |
| chr14-19766060-INS-152 | chr14-19766060-INS-152 |
| chr14-19775280-INS-230 | chr14-19775280-INS-213 |
| chr14-19783101-INS-151 | chr14-19783103-INS-151 |
| chr14-19791208-INS-111 | chr14-19791222-INS-134 |
| chr14-19797477-INS-2189 | chr14-19797478-INS-2192 |
| chr14-19817467-INS-108 | chr14-19817466-INS-131 |
| chr14-19823844-INS-100 | chr14-19823844-INS-110 |
| chr14-19833483-INS-162 | chr14-19833483-INS-162 |
| chr14-19835153-INS-61 | chr14-19835153-INS-61 |
| chr14-19853322-INS-63 | chr14-19853322-INS-63 |
| chr14-19880326-INS-401 | chr14-19880326-INS-401 |
| chr14-19891835-INS-204 | chr14-19891835-INS-202 |
| chr14-19897462-INS-76 | chr14-19897462-INS-76 |
| chr14-19935707-INS-86 | chr14-19935707-INS-86 |
| chr14-19940428-INS-206 | chr14-19940428-INS-206 |
| chr14-19964093-INS-54 | chr14-19964093-INS-54 |
| chr14-19979431-INS-178 | chr14-19979431-INS-178 |
| chr14-19994764-INS-228 | chr14-19994764-INS-228 |
| chr14-20002239-INS-111 | chr14-20002239-INS-117 |
| chr14-20004333-INS-60 | chr14-20004333-INS-64 |
| chr14-20028802-INS-780 | chr14-20028802-INS-780 |
| chr14-20031662-INS-115 | chr14-20031662-INS-91 |
| chr14-20039683-INS-212 | chr14-20039683-INS-212 |
| chr14-20046261-INS-538 | chr14-20046261-INS-538 |
| chr14-20053886-INS-400 | chr14-20053886-INS-400 |
| chr14-20059833-INS-69 | chr14-20059852-INS-66 |
| chr14-20068033-INS-96 | chr14-20068033-INS-96 |
| chr14-20075223-INS-454 | chr14-20075222-INS-452 |
| chr14-20075817-INS-107 | chr14-20075813-INS-84 |
| chr14-20120352-INS-59 | chr14-20120352-INS-73 |
| chr14-20130950-INS-219 | chr14-20130951-INS-222 |
| chr14-20154607-INS-514 | chr14-20154607-INS-514 |
| chr14-20188883-INS-249 | chr14-20188883-INS-249 |
| chr14-20198662-INS-1480 | chr14-20198663-INS-1502 |
| chr14-20237937-INS-63 | chr14-20237937-INS-50 |
| chr14-20252846-INS-212 | chr14-20252846-INS-212 |
| chr14-20268629-INS-2251 | chr14-20268630-INS-2251 |
| chr14-20273842-INS-54 | chr14-20273844-INS-88 |
| chr14-20292476-INS-340 | chr14-20292476-INS-351 |
| chr14-20297700-INS-150 | chr14-20297706-INS-163 |
| chr14-20324291-INS-199 | chr14-20324291-INS-200 |
| chr14-20338917-INS-421 | chr14-20338917-INS-421 |
| chr14-20342270-INS-52 | chr14-20342270-INS-52 |
| chr14-20346968-INS-215 | chr14-20346968-INS-215 |

| | |
|---|---|
| chr14-20358823-INS-122 | chr14-20358823-INS-83 |
| chr14-20368017-INS-90 | chr14-20368017-INS-90 |
| chr14-20384860-INS-68 | chr14-20384845-INS-97 |
| chr14-20406070-INS-195 | chr14-20406071-INS-165 |
| chr14-20462021-INS-62 | chr14-20462021-INS-64 |
| chr14-20481569-INS-200 | chr14-20481570-INS-198 |
| chr14-20483589-INS-63 | chr14-20483589-INS-67 |
| chr14-20490461-INS-94 | chr14-20490461-INS-94 |
| chr14-20495932-INS-765 | chr14-20495932-INS-765 |
| chr14-20516933-INS-218 | chr14-20516933-INS-218 |
| chr14-20595345-INS-62 | chr14-20595345-INS-50 |
| chr14-20641318-INS-222 | chr14-20641301-INS-258 |
| chr14-20643551-INS-6407 | chr14-20643552-INS-6407 |
| chr14-20652468-INS-216 | chr14-20652468-INS-217 |
| chr14-20680999-INS-84 | chr14-20680999-INS-84 |
| chr14-20694095-INS-236 | chr14-20694095-INS-236 |
| chr14-20698554-INS-161 | chr14-20698554-INS-165 |
| chr14-20704482-INS-100 | chr14-20704482-INS-100 |
| chr14-20740725-INS-60 | chr14-20740693-INS-57 |
| chr14-20765432-INS-206 | chr14-20765432-INS-210 |
| chr14-20787077-INS-206 | chr14-20787077-INS-206 |
| chr14-20793386-INS-52 | chr14-20793386-INS-52 |
| chr14-20811117-INS-264 | chr14-20811117-INS-263 |
| chr14-20812749-INS-273 | chr14-20812749-INS-273 |
| chr14-20818064-INS-66 | chr14-20818064-INS-66 |
| chr14-20848081-INS-178 | chr14-20848081-INS-173 |
| chr14-20870869-INS-189 | chr14-20870872-INS-145 |
| chr14-20886600-INS-87 | chr14-20886580-INS-106 |
| chr14-20902842-INS-218 | chr14-20902842-INS-218 |
| chr14-20906604-INS-1289 | chr14-20906605-INS-1289 |
| chr14-20928288-INS-1895 | chr14-20928289-INS-1895 |
| chr14-20933504-INS-511 | chr14-20933504-INS-511 |
| chr14-20967875-INS-129 | chr14-20967876-INS-102 |
| chr14-20999331-INS-58 | chr14-20999331-INS-58 |
| chr14-21036973-INS-159 | chr14-21036973-INS-157 |
| chr14-21061950-INS-75 | chr14-21061950-INS-83 |
| chr14-21068735-INS-680 | chr14-21068735-INS-682 |
| chr14-21090956-INS-135 | chr14-21090956-INS-134 |
| chr14-21093837-INS-302 | chr14-21093837-INS-304 |
| chr14-21119973-INS-166 | chr14-21119973-INS-166 |
| chr14-21147691-INS-954 | chr14-21147691-INS-953 |
| chr14-21174882-INS-197 | chr14-21174882-INS-197 |
| chr14-21176392-INS-1864 | chr14-21176393-INS-1864 |
| chr14-21198399-INS-52 | chr14-21198370-INS-56 |
| chr14-21218700-INS-209 | chr14-21218700-INS-209 |
| chr14-21219257-INS-6455 | chr14-21219258-INS-6456 |
| chr14-21240500-INS-60 | chr14-21240500-INS-60 |
| chr14-21246903-INS-189 | chr14-21246903-INS-189 |
| chr14-21264952-INS-6605 | chr14-21264953-INS-6606 |
| chr14-21266422-INS-262 | chr14-21266422-INS-262 |
| chr14-21298192-INS-137 | chr14-21298186-INS-168 |
| chr14-21313137-INS-221 | chr14-21313137-INS-221 |
| chr14-21327321-INS-555 | chr14-21327321-INS-555 |
| chr14-21329883-INS-54 | chr14-21329924-INS-50 |
| chr14-21354132-INS-85 | chr14-21354132-INS-85 |
| chr14-21359150-INS-75 | chr14-21359150-INS-75 |
| chr14-21369174-INS-155 | chr14-21369174-INS-155 |
| chr14-21375604-INS-351 | chr14-21375604-INS-351 |
| chr14-21377381-INS-413 | chr14-21377381-INS-413 |
| chr14-21408192-INS-183 | chr14-21408193-INS-164 |
| chr14-21428559-INS-64 | chr14-21428559-INS-64 |
| chr14-21452032-INS-204 | chr14-21452032-INS-204 |
| chr14-21456866-INS-204 | chr14-21456833-INS-195 |
| chr14-21478765-INS-1864 | chr14-21478766-INS-1864 |
| chr14-21508339-INS-163 | chr14-21508339-INS-163 |
| chr14-21516283-INS-80 | chr14-21516283-INS-87 |
| chr14-21527286-INS-7761 | chr14-21527287-INS-7761 |
| chr14-21559671-INS-95 | chr14-21559671-INS-95 |
| chr14-21564026-INS-175 | chr14-21564026-INS-191 |
| chr14-21571920-INS-213 | chr14-21571920-INS-214 |
| chr14-21581311-INS-171 | chr14-21581311-INS-171 |
| chr14-21587565-INS-115 | chr14-21587605-INS-76 |
| chr14-21602270-INS-213 | chr14-21602270-INS-210 |
| chr14-21615307-INS-81 | chr14-21615307-INS-84 |
| chr14-21616999-INS-180 | chr14-21616999-INS-181 |
| chr14-21620269-INS-291 | chr14-21620269-INS-291 |
| chr14-21621210-INS-63 | chr14-21621210-INS-63 |

| | |
|---|---|
| chr14-21623246-INS-62 | chr14-21623246-INS-66 |
| chr14-21629004-INS-203 | chr14-21629004-INS-195 |
| chr14-21690251-INS-86 | chr14-21690251-INS-86 |
| chr14-21715569-INS-487 | chr14-21715569-INS-487 |
| chr14-21728371-INS-113 | chr14-21728371-INS-117 |
| chr14-21731889-INS-106 | chr14-21731863-INS-102 |
| chr14-21742849-INS-60 | chr14-21742849-INS-60 |
| chr14-21743950-INS-153 | chr14-21743950-INS-153 |
| chr14-21748712-INS-130 | chr14-21748720-INS-88 |
| chr14-21749915-INS-56 | chr14-21749915-INS-71 |
| chr14-21765928-INS-134 | chr14-21765928-INS-135 |
| chr14-21778796-INS-224 | chr14-21778796-INS-224 |
| chr14-21795703-INS-98 | chr14-21795678-INS-90 |
| chr14-21819342-INS-56 | chr14-21819342-INS-58 |
| chr14-21823318-INS-109 | chr14-21823318-INS-109 |
| chr14-21824143-INS-203 | chr14-21824143-INS-202 |
| chr14-21850302-INS-199 | chr14-21850302-INS-199 |
| chr14-21855579-INS-58 | chr14-21855579-INS-58 |
| chr14-21859244-INS-149 | chr14-21859244-INS-147 |
| chr14-21860111-INS-52 | chr14-21860117-INS-59 |
| chr14-21867439-INS-90 | chr14-21867432-INS-118 |
| chr14-21877229-INS-216 | chr14-21877229-INS-216 |
| chr14-21919301-INS-105 | chr14-21919301-INS-102 |
| chr14-21921233-INS-202 | chr14-21921233-INS-202 |
| chr14-21926615-INS-504 | chr14-21926615-INS-505 |
| chr14-21951457-INS-317 | chr14-21951457-INS-315 |
| chr14-22032031-INS-80 | chr14-22032031-INS-80 |
| chr14-22059801-INS-188 | chr14-22059802-INS-188 |
| chr14-22063703-INS-67 | chr14-22063703-INS-67 |
| chr14-22078294-INS-3247 | chr14-22078295-INS-3247 |
| chr14-22093873-INS-215 | chr14-22093873-INS-215 |
| chr14-22133734-INS-137 | chr14-22133734-INS-137 |
| chr14-22140739-INS-61 | chr14-22140739-INS-58 |
| chr14-22170688-INS-76 | chr14-22170688-INS-76 |
| chr14-22190563-INS-54 | chr14-22190532-INS-58 |
| chr14-22193508-INS-1891 | chr14-22193509-INS-1891 |
| chr14-22209309-INS-693 | chr14-22209309-INS-694 |
| chr14-22223985-INS-2448 | chr14-22223986-INS-2448 |
| chr14-22227051-INS-70 | chr14-22227051-INS-50 |
| chr14-22236877-INS-99 | chr14-22236874-INS-93 |
| chr14-22240621-INS-67 | chr14-22240621-INS-68 |
| chr14-22256450-INS-174 | chr14-22256450-INS-174 |
| chr14-22286993-INS-74 | chr14-22286993-INS-74 |
| chr14-22294010-INS-6685 | chr14-22294011-INS-6689 |
| chr14-22329431-INS-872 | chr14-22329431-INS-872 |
| chr14-22369090-INS-54 | chr14-22369090-INS-54 |
| chr14-22380498-INS-4008 | chr14-22380499-INS-4008 |
| chr14-22381326-INS-100 | chr14-22381315-INS-74 |
| chr14-22412015-INS-6442 | chr14-22412016-INS-6442 |
| chr14-22415650-INS-200 | chr14-22415650-INS-200 |
| chr14-22438400-INS-216 | chr14-22438400-INS-216 |
| chr14-22451419-INS-52 | chr14-22451419-INS-52 |
| chr14-22481244-INS-62 | chr14-22481244-INS-62 |
| chr14-22561416-INS-280 | chr14-22561416-INS-288 |
| chr14-22564205-INS-6456 | chr14-22564206-INS-6456 |
| chr14-22620856-INS-56 | chr14-22620856-INS-56 |
| chr14-22624697-INS-78 | chr14-22624666-INS-82 |
| chr14-22650039-INS-52 | chr14-22650041-INS-60 |
| chr14-22651962-INS-101 | chr14-22651962-INS-95 |
| chr14-22660938-INS-144 | chr14-22660951-INS-127 |
| chr14-22664838-INS-1864 | chr14-22664839-INS-1864 |
| chr14-22668731-INS-158 | chr14-22668731-INS-154 |
| chr14-22684336-INS-51 | chr14-22684337-INS-51 |
| chr14-22687321-INS-109 | chr14-22687312-INS-155 |
| chr14-22707924-INS-84 | chr14-22707924-INS-86 |
| chr14-22727976-INS-524 | chr14-22727976-INS-522 |
| chr14-22728892-INS-261 | chr14-22728892-INS-261 |
| chr14-22739198-INS-52 | chr14-22739198-INS-52 |
| chr14-22766361-INS-55 | chr14-22766362-INS-53 |
| chr14-22781615-INS-835 | chr14-22781615-INS-835 |
| chr14-22799178-INS-75 | chr14-22799178-INS-77 |
| chr14-22818981-INS-102 | chr14-22818981-INS-102 |
| chr14-22828360-INS-86 | chr14-22828357-INS-109 |
| chr14-22870651-INS-80 | chr14-22870651-INS-82 |
| chr14-22918659-INS-6382 | chr14-22918660-INS-6382 |
| chr14-22920784-INS-51 | chr14-22920786-INS-87 |
| chr14-22958001-INS-76 | chr14-22958001-INS-84 |

| | |
|---|---|
| chr14-22965540-INS-244 | chr14-22965540-INS-240 |
| chr14-22970134-INS-95 | chr14-22970134-INS-95 |
| chr14-23038840-INS-54 | chr14-23038840-INS-54 |
| chr14-23043983-INS-271 | chr14-23043983-INS-271 |
| chr14-23094949-INS-606 | chr14-23094949-INS-606 |
| chr14-23126936-INS-52 | chr14-23126952-INS-57 |
| chr14-23169968-INS-74 | chr14-23169968-INS-80 |
| chr14-23178910-INS-156 | chr14-23178910-INS-153 |
| chr14-23212004-INS-245 | chr14-23212004-INS-220 |
| chr14-23220627-INS-7796 | chr14-23220628-INS-7800 |
| chr14-23222738-INS-4512 | chr14-23222739-INS-4512 |
| chr14-23253128-INS-4364 | chr14-23253129-INS-4364 |
| chr14-23270839-INS-6507 | chr14-23270840-INS-6507 |
| chr14-23276079-INS-283 | chr14-23276079-INS-283 |
| chr14-23288248-INS-6690 | chr14-23288249-INS-6690 |
| chr14-23327971-INS-2061 | chr14-23327967-INS-2063 |
| chr14-23366162-INS-124 | chr14-23366162-INS-144 |
| chr14-23407237-INS-77 | chr14-23407237-INS-82 |
| chr14-23460405-INS-214 | chr14-23460405-INS-214 |
| chr14-23484630-INS-79 | chr14-23484627-INS-69 |
| chr14-23489992-INS-292 | chr14-23489992-INS-304 |
| chr14-23508517-INS-169 | chr14-23508502-INS-214 |
| chr14-23513895-INS-52 | chr14-23513895-INS-52 |
| chr14-23518580-INS-56 | chr14-23518580-INS-64 |
| chr14-23528778-INS-118 | chr14-23528778-INS-118 |
| chr14-23545657-INS-63 | chr14-23545649-INS-58 |
| chr14-23582678-INS-91 | chr14-23582678-INS-91 |
| chr14-23594641-INS-82 | chr14-23594629-INS-58 |
| chr14-23596304-INS-184 | chr14-23596304-INS-178 |
| chr14-23597556-INS-51 | chr14-23597556-INS-51 |
| chr14-23635373-INS-76 | chr14-23635373-INS-76 |
| chr14-23651747-INS-124 | chr14-23651747-INS-140 |
| chr14-23655650-INS-64 | chr14-23655650-INS-67 |
| chr14-23672384-INS-84 | chr14-23672384-INS-84 |
| chr14-23678655-INS-73 | chr14-23678655-INS-73 |
| chr14-23689337-INS-292 | chr14-23689337-INS-292 |
| chr14-23691666-INS-78 | chr14-23691649-INS-52 |
| chr14-23719183-INS-65 | chr14-23719183-INS-97 |
| chr14-23733825-INS-216 | chr14-23733825-INS-217 |
| chr14-23766604-INS-59 | chr14-23766604-INS-61 |
| chr14-23768545-INS-7161 | chr14-23768546-INS-7161 |
| chr14-23781900-INS-610 | chr14-23781900-INS-609 |
| chr14-23815391-INS-87 | chr14-23815391-INS-84 |
| chr14-23843649-INS-188 | chr14-23843649-INS-188 |
| chr14-23847446-INS-60 | chr14-23847451-INS-81 |
| chr14-23852794-INS-54 | chr14-23852794-INS-54 |
| chr14-23855725-INS-695 | chr14-23855725-INS-695 |
| chr14-23861650-INS-125 | chr14-23861650-INS-125 |
| chr14-23874385-INS-60 | chr14-23874385-INS-84 |
| chr14-23922946-INS-168 | chr14-23922946-INS-178 |
| chr14-23937858-INS-334 | chr14-23937858-INS-337 |
| chr14-23943568-INS-96 | chr14-23943568-INS-104 |
| chr14-23945471-INS-119 | chr14-23945471-INS-119 |
| chr14-23967615-INS-62 | chr14-23967615-INS-62 |
| chr14-23975256-INS-67 | chr14-23975256-INS-67 |
| chr14-23977040-INS-52 | chr14-23976992-INS-95 |
| chr14-23990573-INS-116 | chr14-23990573-INS-117 |
| chr14-24010742-INS-242 | chr14-24010742-INS-242 |
| chr14-24029445-INS-480 | chr14-24029445-INS-479 |
| chr14-24040574-INS-94 | chr14-24040574-INS-94 |
| chr14-24095078-INS-6351 | chr14-24095079-INS-6347 |
| chr14-24105374-INS-511 | chr14-24105374-INS-511 |
| chr14-24128880-INS-848 | chr14-24128881-INS-849 |
| chr14-24141441-INS-56 | chr14-24141441-INS-58 |
| chr14-24175589-INS-108 | chr14-24175589-INS-131 |
| chr14-24178005-INS-54 | chr14-24178005-INS-54 |
| chr14-24181892-INS-218 | chr14-24181892-INS-218 |
| chr14-24193140-INS-5129 | chr14-24193141-INS-5129 |
| chr14-24200504-INS-75 | chr14-24200507-INS-69 |
| chr14-24205073-INS-120 | chr14-24205073-INS-119 |
| chr14-24242306-INS-52 | chr14-24242306-INS-52 |
| chr14-24246242-INS-68 | chr14-24246242-INS-68 |
| chr14-24310355-INS-263 | chr14-24310355-INS-263 |
| chr14-24325854-INS-54 | chr14-24325848-INS-86 |
| chr14-24336309-INS-398 | chr14-24336309-INS-400 |
| chr14-24338260-INS-259 | chr14-24338260-INS-266 |
| chr14-24341843-INS-68 | chr14-24341843-INS-68 |

| | |
|---|---|
| chr14-24351695-INS-177 | chr14-24351695-INS-159 |
| chr14-24355019-INS-166 | chr14-24355019-INS-166 |
| chr14-24357879-INS-60 | chr14-24357879-INS-62 |
| chr14-24373854-INS-212 | chr14-24373854-INS-212 |
| chr14-24391589-INS-198 | chr14-24391589-INS-199 |
| chr14-24396783-INS-289 | chr14-24396785-INS-287 |
| chr14-24414895-INS-60 | chr14-24414908-INS-51 |
| chr14-24422981-INS-90 | chr14-24422991-INS-54 |
| chr14-24424910-INS-510 | chr14-24424910-INS-510 |
| chr14-24449690-INS-185 | chr14-24449690-INS-185 |
| chr14-24469023-INS-63 | chr14-24469023-INS-63 |
| chr14-24501505-INS-278 | chr14-24501505-INS-279 |
| chr14-24511167-INS-84 | chr14-24511167-INS-52 |
| chr14-24547753-INS-255 | chr14-24547753-INS-227 |
| chr14-24550098-INS-573 | chr14-24550098-INS-572 |
| chr14-24561772-INS-86 | chr14-24561773-INS-82 |
| chr14-24564501-INS-52 | chr14-24564495-INS-60 |
| chr14-24569033-INS-200 | chr14-24569033-INS-201 |
| chr14-24587268-INS-328 | chr14-24587268-INS-323 |
| chr14-24616626-INS-122 | chr14-24616626-INS-122 |
| chr14-24619919-INS-121 | chr14-24619919-INS-121 |
| chr14-24658219-INS-61 | chr14-24658219-INS-61 |
| chr14-24662136-INS-940 | chr14-24662136-INS-941 |
| chr14-24679027-INS-549 | chr14-24679027-INS-549 |
| chr14-24691571-INS-52 | chr14-24691571-INS-68 |
| chr14-24699499-INS-90 | chr14-24699497-INS-90 |
| chr14-24713469-INS-79 | chr14-24713469-INS-105 |
| chr14-24715503-INS-1862 | chr14-24715504-INS-1862 |
| chr14-24748941-INS-51 | chr14-24748938-INS-50 |
| chr14-24788672-INS-96 | chr14-24788642-INS-76 |
| chr14-24843790-INS-103 | chr14-24843790-INS-97 |
| chr14-24863241-INS-55 | chr14-24863241-INS-55 |
| chr14-24873220-INS-54 | chr14-24873220-INS-54 |
| chr14-24939643-INS-56 | chr14-24939643-INS-56 |
| chr14-24946107-INS-67 | chr14-24946107-INS-65 |
| chr14-24963239-INS-200 | chr14-24963239-INS-200 |
| chr14-24971274-INS-69 | chr14-24971300-INS-71 |
| chr14-25071577-INS-56 | chr14-25071577-INS-60 |
| chr14-25090088-INS-7163 | chr14-25090089-INS-7164 |
| chr14-25112163-INS-1415 | chr14-25112164-INS-1415 |
| chr14-25177750-INS-58 | chr14-25177750-INS-62 |
| chr14-25216799-INS-55 | chr14-25216799-INS-55 |
| chr14-25242894-INS-332 | chr14-25242890-INS-288 |
| chr14-25248754-INS-172 | chr14-25248790-INS-124 |
| chr14-25251921-INS-205 | chr14-25251921-INS-205 |
| chr14-25291011-INS-160 | chr14-25291011-INS-162 |
| chr14-25293837-INS-56 | chr14-25293837-INS-56 |
| chr14-25302854-INS-553 | chr14-25302854-INS-553 |
| chr14-25319500-INS-672 | chr14-25319529-INS-671 |
| chr14-25323165-INS-59 | chr14-25323153-INS-72 |
| chr14-25332098-INS-68 | chr14-25332127-INS-66 |
| chr14-25410521-INS-103 | chr14-25410521-INS-103 |
| chr14-25417014-INS-64 | chr14-25417014-INS-64 |
| chr14-25436809-INS-429 | chr14-25436809-INS-431 |
| chr14-25513265-INS-54 | chr14-25513265-INS-54 |
| chr14-25532085-INS-64 | chr14-25532085-INS-68 |
| chr14-25543400-INS-209 | chr14-25543400-INS-209 |
| chr14-25562449-INS-72 | chr14-25562449-INS-70 |
| chr14-25589095-INS-572 | chr14-25589095-INS-580 |
| chr14-25642463-INS-301 | chr14-25642463-INS-301 |
| chr14-25671386-INS-103 | chr14-25671386-INS-91 |
| chr14-25673646-INS-196 | chr14-25673646-INS-196 |
| chr14-25678339-INS-58 | chr14-25678362-INS-60 |
| chr14-25686487-INS-400 | chr14-25686487-INS-400 |
| chr14-25705849-INS-126 | chr14-25705846-INS-136 |
| chr14-25710329-INS-89 | chr14-25710329-INS-87 |
| chr14-25717486-INS-52 | chr14-25717486-INS-52 |
| chr14-25788670-INS-218 | chr14-25788670-INS-219 |
| chr14-25793136-INS-165 | chr14-25793131-INS-166 |
| chr14-25823056-INS-211 | chr14-25823056-INS-211 |
| chr14-25840759-INS-211 | chr14-25840737-INS-239 |
| chr14-25855588-INS-82 | chr14-25855626-INS-52 |
| chr14-25869451-INS-177 | chr14-25869451-INS-177 |
| chr14-25881495-INS-96 | chr14-25881495-INS-100 |
| chr14-25897232-INS-123 | chr14-25897240-INS-74 |
| chr14-25922091-INS-75 | chr14-25922091-INS-50 |
| chr14-25929897-INS-125 | chr14-25929897-INS-125 |

| | |
|---|---|
| chr14-25936315-INS-515 | chr14-25936315-INS-514 |
| chr14-25961904-INS-196 | chr14-25961904-INS-196 |
| chr14-25966524-INS-209 | chr14-25966524-INS-199 |
| chr14-25968990-INS-319 | chr14-25968990-INS-319 |
| chr14-25982264-INS-233 | chr14-25982264-INS-234 |
| chr14-25998603-INS-73 | chr14-25998600-INS-95 |
| chr14-26011799-INS-122 | chr14-26011799-INS-122 |
| chr14-26019609-INS-126 | chr14-26019609-INS-127 |
| chr14-26058125-INS-442 | chr14-26058125-INS-441 |
| chr14-26069623-INS-54 | chr14-26069623-INS-54 |
| chr14-26095726-INS-124 | chr14-26095726-INS-124 |
| chr14-26105585-INS-146 | chr14-26105585-INS-146 |
| chr14-26113939-INS-206 | chr14-26113939-INS-211 |
| chr14-26125355-INS-80 | chr14-26125336-INS-78 |
| chr14-26176040-INS-64 | chr14-26176040-INS-96 |
| chr14-26213735-INS-960 | chr14-26213735-INS-987 |
| chr14-26214636-INS-53 | chr14-26214617-INS-51 |
| chr14-26218582-INS-270 | chr14-26218582-INS-270 |
| chr14-26260893-INS-64 | chr14-26260864-INS-59 |
| chr14-26279936-INS-225 | chr14-26279936-INS-225 |
| chr14-26286881-INS-353 | chr14-26286881-INS-352 |
| chr14-26299230-INS-170 | chr14-26299230-INS-169 |
| chr14-26318550-INS-1104 | chr14-26318551-INS-1104 |
| chr14-26322166-INS-1888 | chr14-26322167-INS-1888 |
| chr14-26326912-INS-188 | chr14-26326912-INS-190 |
| chr14-26339677-INS-58 | chr14-26339677-INS-58 |
| chr14-26349169-INS-3097 | chr14-26349170-INS-3083 |
| chr14-26362727-INS-237 | chr14-26362727-INS-237 |
| chr14-26371343-INS-363 | chr14-26371343-INS-364 |
| chr14-26376443-INS-123 | chr14-26376443-INS-141 |
| chr14-26406169-INS-480 | chr14-26406169-INS-480 |
| chr14-26416563-INS-172 | chr14-26416563-INS-172 |
| chr14-26467149-INS-54 | chr14-26467149-INS-54 |
| chr14-26475813-INS-53 | chr14-26475814-INS-53 |
| chr14-26478058-INS-109 | chr14-26478058-INS-109 |
| chr14-26503174-INS-377 | chr14-26503174-INS-377 |
| chr14-26555419-INS-169 | chr14-26555428-INS-185 |
| chr14-26557813-INS-120 | chr14-26557820-INS-160 |
| chr14-26585781-INS-86 | chr14-26585745-INS-87 |
| chr14-26600483-INS-76 | chr14-26600483-INS-76 |
| chr14-26634405-INS-144 | chr14-26634405-INS-144 |
| chr14-26680815-INS-171 | chr14-26680815-INS-172 |
| chr14-26681815-INS-206 | chr14-26681815-INS-206 |
| chr14-26694933-INS-66 | chr14-26694933-INS-66 |
| chr14-26698737-INS-143 | chr14-26698737-INS-144 |
| chr14-26702536-INS-87 | chr14-26702537-INS-87 |
| chr14-26710536-INS-197 | chr14-26710536-INS-197 |
| chr14-26712772-INS-61 | chr14-26712772-INS-76 |
| chr14-26726073-INS-176 | chr14-26726073-INS-176 |
| chr14-26729008-INS-52 | chr14-26729008-INS-52 |
| chr14-26730429-INS-216 | chr14-26730441-INS-207 |
| chr14-26731713-INS-215 | chr14-26731713-INS-215 |
| chr14-26733784-INS-171 | chr14-26733784-INS-171 |
| chr14-26752849-INS-213 | chr14-26752850-INS-213 |
| chr14-26771911-INS-210 | chr14-26771911-INS-210 |
| chr14-26778180-INS-69 | chr14-26778180-INS-69 |
| chr14-26817524-INS-51 | chr14-26817524-INS-51 |
| chr14-26878595-INS-142 | chr14-26878595-INS-146 |
| chr14-26885450-INS-117 | chr14-26885450-INS-117 |
| chr14-26911581-INS-218 | chr14-26911581-INS-216 |
| chr14-26912842-INS-197 | chr14-26912840-INS-157 |
| chr14-26924154-INS-70 | chr14-26924154-INS-85 |
| chr14-26938735-INS-1216 | chr14-26938736-INS-1173 |
| chr14-26940419-INS-167 | chr14-26940419-INS-167 |
| chr14-26942712-INS-2575 | chr14-26942713-INS-2577 |
| chr14-26987303-INS-65 | chr14-26987303-INS-64 |
| chr14-27003216-INS-175 | chr14-27003216-INS-175 |
| chr14-27010795-INS-208 | chr14-27010795-INS-208 |
| chr14-27014464-INS-106 | chr14-27014502-INS-102 |
| chr14-27022175-INS-401 | chr14-27022175-INS-401 |
| chr14-27023198-INS-202 | chr14-27023198-INS-202 |
| chr14-27038272-INS-401 | chr14-27038273-INS-383 |
| chr14-27043872-INS-207 | chr14-27043866-INS-207 |
| chr14-27056551-INS-173 | chr14-27056552-INS-178 |
| chr14-27116630-INS-206 | chr14-27116630-INS-206 |
| chr14-27118616-INS-77 | chr14-27118610-INS-80 |
| chr14-27145653-INS-209 | chr14-27145653-INS-209 |

| | |
|---|---|
| chr14-27169295-INS-1235 | chr14-27169296-INS-1235 |
| chr14-27261445-INS-351 | chr14-27261445-INS-351 |
| chr14-27279585-INS-56 | chr14-27279585-INS-60 |
| chr14-27281413-INS-178 | chr14-27281413-INS-179 |
| chr14-27300583-INS-66 | chr14-27300583-INS-66 |
| chr14-27306705-INS-1085 | chr14-27306706-INS-1085 |
| chr14-27307476-INS-60 | chr14-27307476-INS-56 |
| chr14-27337548-INS-56 | chr14-27337548-INS-56 |
| chr14-27342002-INS-494 | chr14-27342002-INS-494 |
| chr14-27366859-INS-432 | chr14-27366859-INS-432 |
| chr14-27372616-INS-830 | chr14-27372617-INS-832 |
| chr14-27417167-INS-88 | chr14-27417165-INS-68 |
| chr14-27423340-INS-159 | chr14-27423316-INS-155 |
| chr14-27427570-INS-54 | chr14-27427570-INS-60 |
| chr14-27442513-INS-68 | chr14-27442477-INS-68 |
| chr14-27471786-INS-208 | chr14-27471786-INS-208 |
| chr14-27513571-INS-71 | chr14-27513571-INS-104 |
| chr14-27514883-INS-54 | chr14-27514883-INS-58 |
| chr14-27547688-INS-54 | chr14-27547688-INS-54 |
| chr14-27551136-INS-530 | chr14-27551136-INS-530 |
| chr14-27582255-INS-196 | chr14-27582255-INS-196 |
| chr14-27587203-INS-52 | chr14-27587203-INS-58 |
| chr14-27610022-INS-59 | chr14-27610022-INS-60 |
| chr14-27648140-INS-6590 | chr14-27648141-INS-6590 |
| chr14-27661567-INS-69 | chr14-27661567-INS-69 |
| chr14-27673070-INS-704 | chr14-27673074-INS-705 |
| chr14-27676139-INS-249 | chr14-27676141-INS-250 |
| chr14-27715383-INS-4509 | chr14-27715384-INS-4509 |
| chr14-27718643-INS-427 | chr14-27718643-INS-427 |
| chr14-27727488-INS-1757 | chr14-27727489-INS-1736 |
| chr14-27773112-INS-68 | chr14-27773112-INS-77 |
| chr14-27807824-INS-63 | chr14-27807828-INS-75 |
| chr14-27823388-INS-981 | chr14-27823388-INS-981 |
| chr14-27834968-INS-96 | chr14-27834969-INS-90 |
| chr14-27859906-INS-92 | chr14-27859906-INS-92 |
| chr14-27876956-INS-6372 | chr14-27876957-INS-6372 |
| chr14-27879109-INS-52 | chr14-27879140-INS-50 |
| chr14-27966636-INS-441 | chr14-27966636-INS-454 |
| chr14-28018929-INS-6837 | chr14-28018930-INS-6838 |
| chr14-28074867-INS-96 | chr14-28074867-INS-50 |
| chr14-28088086-INS-55 | chr14-28088086-INS-50 |
| chr14-28089221-INS-1323 | chr14-28089222-INS-1323 |
| chr14-28147893-INS-401 | chr14-28147893-INS-401 |
| chr14-28148037-INS-6487 | chr14-28148038-INS-6487 |
| chr14-28173436-INS-207 | chr14-28173436-INS-207 |
| chr14-28179693-INS-74 | chr14-28179693-INS-80 |
| chr14-28203978-INS-80 | chr14-28203972-INS-60 |
| chr14-28207190-INS-67 | chr14-28207185-INS-54 |
| chr14-28244132-INS-54 | chr14-28244132-INS-58 |
| chr14-28252796-INS-92 | chr14-28252817-INS-60 |
| chr14-28283144-INS-56 | chr14-28283144-INS-58 |
| chr14-28286099-INS-60 | chr14-28286102-INS-70 |
| chr14-28304829-INS-467 | chr14-28304829-INS-467 |
| chr14-28353521-INS-221 | chr14-28353521-INS-220 |
| chr14-28365394-INS-56 | chr14-28365347-INS-54 |
| chr14-28376473-INS-178 | chr14-28376451-INS-162 |
| chr14-28380756-INS-123 | chr14-28380756-INS-125 |
| chr14-28391333-INS-128 | chr14-28391320-INS-91 |
| chr14-28400963-INS-83 | chr14-28400963-INS-83 |
| chr14-28414788-INS-228 | chr14-28414788-INS-225 |
| chr14-28415916-INS-59 | chr14-28415924-INS-69 |
| chr14-28427153-INS-70 | chr14-28427153-INS-72 |
| chr14-28439357-INS-311 | chr14-28439380-INS-322 |
| chr14-28460189-INS-4681 | chr14-28460190-INS-4681 |
| chr14-28492484-INS-363 | chr14-28492484-INS-363 |
| chr14-28516225-INS-66 | chr14-28516225-INS-62 |
| chr14-28523101-INS-1029 | chr14-28523102-INS-1029 |
| chr14-28530686-INS-179 | chr14-28530685-INS-194 |
| chr14-28532128-INS-2796 | chr14-28532129-INS-2796 |
| chr14-28662850-INS-102 | chr14-28662857-INS-102 |
| chr14-28672177-INS-90 | chr14-28672177-INS-90 |
| chr14-28692570-INS-60 | chr14-28692550-INS-83 |
| chr14-28698152-INS-579 | chr14-28698152-INS-579 |
| chr14-28716457-INS-73 | chr14-28716471-INS-88 |
| chr14-28735574-INS-96 | chr14-28735574-INS-96 |
| chr14-28806344-INS-80 | chr14-28806344-INS-80 |
| chr14-28858555-INS-56 | chr14-28858555-INS-58 |

| | |
|---|---|
| chr14-28900491-INS-55 | chr14-28900491-INS-55 |
| chr14-28949711-INS-57 | chr14-28949716-INS-59 |
| chr14-28962330-INS-326 | chr14-28962330-INS-326 |
| chr14-29000932-INS-199 | chr14-29000932-INS-199 |
| chr14-29041650-INS-74 | chr14-29041662-INS-50 |
| chr14-29043894-INS-986 | chr14-29043896-INS-981 |
| chr14-29103309-INS-84 | chr14-29103309-INS-84 |
| chr14-29159216-INS-66 | chr14-29159215-INS-76 |
| chr14-29160903-INS-356 | chr14-29160903-INS-356 |
| chr14-29168280-INS-61 | chr14-29168262-INS-80 |
| chr14-29172560-INS-226 | chr14-29172560-INS-231 |
| chr14-29200108-INS-6340 | chr14-29200109-INS-6340 |
| chr14-29213554-INS-80 | chr14-29213554-INS-80 |
| chr14-29232137-INS-71 | chr14-29232137-INS-71 |
| chr14-29234863-INS-77 | chr14-29234863-INS-55 |
| chr14-29242110-INS-51 | chr14-29242110-INS-51 |
| chr14-29244197-INS-200 | chr14-29244188-INS-171 |
| chr14-29287566-INS-280 | chr14-29287566-INS-280 |
| chr14-29295401-INS-74 | chr14-29295401-INS-78 |
| chr14-29325934-INS-432 | chr14-29325934-INS-432 |
| chr14-29342803-INS-506 | chr14-29342803-INS-513 |
| chr14-29350725-INS-1891 | chr14-29350726-INS-1891 |
| chr14-29384543-INS-136 | chr14-29384543-INS-128 |
| chr14-29405667-INS-66 | chr14-29405667-INS-68 |
| chr14-29475711-INS-91 | chr14-29475711-INS-67 |
| chr14-29482461-INS-864 | chr14-29482461-INS-865 |
| chr14-29636154-INS-80 | chr14-29636154-INS-80 |
| chr14-29637698-INS-87 | chr14-29637668-INS-113 |
| chr14-29648576-INS-1685 | chr14-29648577-INS-1685 |
| chr14-29652005-INS-93 | chr14-29652005-INS-93 |
| chr14-29665119-INS-127 | chr14-29665119-INS-127 |
| chr14-29670979-INS-54 | chr14-29670971-INS-52 |
| chr14-29692084-INS-203 | chr14-29692084-INS-203 |
| chr14-29746205-INS-50 | chr14-29746201-INS-54 |
| chr14-29747194-INS-96 | chr14-29747227-INS-99 |
| chr14-29749583-INS-522 | chr14-29749583-INS-507 |
| chr14-29771243-INS-58 | chr14-29771243-INS-52 |
| chr14-29806569-INS-270 | chr14-29806569-INS-270 |
| chr14-29833485-INS-902 | chr14-29833485-INS-901 |
| chr14-29859551-INS-823 | chr14-29859551-INS-823 |
| chr14-29900810-INS-61 | chr14-29900810-INS-69 |
| chr14-29933688-INS-189 | chr14-29933688-INS-168 |
| chr14-29935636-INS-138 | chr14-29935637-INS-138 |
| chr14-29938362-INS-1131 | chr14-29938363-INS-1131 |
| chr14-29993843-INS-781 | chr14-29993843-INS-781 |
| chr14-30034496-INS-83 | chr14-30034496-INS-83 |
| chr14-30056855-INS-55 | chr14-30056855-INS-55 |
| chr14-30095429-INS-77 | chr14-30095429-INS-77 |
| chr14-30142151-INS-78 | chr14-30142151-INS-78 |
| chr14-30189366-INS-540 | chr14-30189366-INS-540 |
| chr14-30211831-INS-164 | chr14-30211831-INS-164 |
| chr14-30216255-INS-207 | chr14-30216255-INS-207 |
| chr14-30229341-INS-52 | chr14-30229341-INS-52 |
| chr14-30236950-INS-56 | chr14-30236950-INS-52 |
| chr14-30252579-INS-97 | chr14-30252579-INS-97 |
| chr14-30255817-INS-223 | chr14-30255817-INS-223 |
| chr14-30258785-INS-53 | chr14-30258785-INS-81 |
| chr14-30265333-INS-79 | chr14-30265333-INS-93 |
| chr14-30293988-INS-221 | chr14-30293988-INS-221 |
| chr14-30299968-INS-7091 | chr14-30299969-INS-7091 |
| chr14-30311032-INS-189 | chr14-30311032-INS-189 |
| chr14-30326812-INS-62 | chr14-30326816-INS-50 |
| chr14-30361355-INS-769 | chr14-30361355-INS-769 |
| chr14-30419144-INS-54 | chr14-30419144-INS-54 |
| chr14-30458521-INS-1732 | chr14-30458522-INS-1732 |
| chr14-30512069-INS-201 | chr14-30512069-INS-201 |
| chr14-30530412-INS-56 | chr14-30530412-INS-50 |
| chr14-30559634-INS-120 | chr14-30559634-INS-120 |
| chr14-30569617-INS-56 | chr14-30569629-INS-64 |
| chr14-30625069-INS-58 | chr14-30625069-INS-50 |
| chr14-30627095-INS-63 | chr14-30627095-INS-72 |
| chr14-30632504-INS-622 | chr14-30632504-INS-634 |
| chr14-30648339-INS-517 | chr14-30648325-INS-523 |
| chr14-30662323-INS-57 | chr14-30662365-INS-57 |
| chr14-30665630-INS-54 | chr14-30665630-INS-52 |
| chr14-30669335-INS-210 | chr14-30669335-INS-209 |
| chr14-30674227-INS-290 | chr14-30674227-INS-296 |

| | |
|---|---|
| chr14-30708053-INS-222 | chr14-30708053-INS-223 |
| chr14-30726128-INS-92 | chr14-30726122-INS-57 |
| chr14-30805294-INS-209 | chr14-30805294-INS-209 |
| chr14-30825735-INS-94 | chr14-30825735-INS-94 |
| chr14-30877105-INS-226 | chr14-30877105-INS-226 |
| chr14-30949302-INS-52 | chr14-30949302-INS-56 |
| chr14-30956226-INS-1889 | chr14-30956227-INS-1889 |
| chr14-30965970-INS-163 | chr14-30965970-INS-164 |
| chr14-30977311-INS-216 | chr14-30977311-INS-216 |
| chr14-30990596-INS-51 | chr14-30990596-INS-51 |
| chr14-31050047-INS-52 | chr14-31050072-INS-54 |
| chr14-31054360-INS-405 | chr14-31054360-INS-399 |
| chr14-31106427-INS-440 | chr14-31106427-INS-440 |
| chr14-31112726-INS-62 | chr14-31112692-INS-62 |
| chr14-31116676-INS-60 | chr14-31116676-INS-60 |
| chr14-31138505-INS-582 | chr14-31138505-INS-583 |
| chr14-31145841-INS-281 | chr14-31145841-INS-281 |
| chr14-31152979-INS-58 | chr14-31152979-INS-58 |
| chr14-31154339-INS-58 | chr14-31154339-INS-78 |
| chr14-31161569-INS-56 | chr14-31161564-INS-62 |
| chr14-31169369-INS-59 | chr14-31169369-INS-53 |
| chr14-31175648-INS-464 | chr14-31175648-INS-464 |
| chr14-31186859-INS-133 | chr14-31186859-INS-133 |
| chr14-31189759-INS-2503 | chr14-31189760-INS-2503 |
| chr14-31193396-INS-208 | chr14-31193396-INS-252 |
| chr14-31237886-INS-466 | chr14-31237886-INS-466 |
| chr14-31245136-INS-228 | chr14-31245136-INS-229 |
| chr14-31256546-INS-203 | chr14-31256546-INS-203 |
| chr14-31262996-INS-164 | chr14-31262996-INS-166 |
| chr14-31310535-INS-70 | chr14-31310521-INS-80 |
| chr14-31402002-INS-123 | chr14-31402002-INS-120 |
| chr14-31406666-INS-540 | chr14-31406666-INS-540 |
| chr14-31411130-INS-6434 | chr14-31411131-INS-6434 |
| chr14-31457193-INS-170 | chr14-31457193-INS-170 |
| chr14-31487047-INS-612 | chr14-31487047-INS-610 |
| chr14-31491145-INS-205 | chr14-31491145-INS-205 |
| chr14-31512876-INS-60 | chr14-31512876-INS-60 |
| chr14-31516589-INS-90 | chr14-31516589-INS-88 |
| chr14-31581571-INS-50 | chr14-31581571-INS-50 |
| chr14-31599590-INS-229 | chr14-31599590-INS-229 |
| chr14-31647155-INS-65 | chr14-31647155-INS-65 |
| chr14-31651477-INS-76 | chr14-31651477-INS-94 |
| chr14-31655140-INS-77 | chr14-31655140-INS-77 |
| chr14-31690981-INS-100 | chr14-31690981-INS-100 |
| chr14-31699301-INS-73 | chr14-31699332-INS-89 |
| chr14-31708065-INS-277 | chr14-31708065-INS-280 |
| chr14-31728439-INS-208 | chr14-31728439-INS-233 |
| chr14-31763747-INS-148 | chr14-31763780-INS-148 |
| chr14-31766186-INS-1054 | chr14-31766187-INS-1054 |
| chr14-31772011-INS-130 | chr14-31772011-INS-130 |
| chr14-31775831-INS-76 | chr14-31775832-INS-77 |
| chr14-31798015-INS-138 | chr14-31798034-INS-135 |
| chr14-31834750-INS-172 | chr14-31834750-INS-169 |
| chr14-31849372-INS-202 | chr14-31849372-INS-202 |
| chr14-31851089-INS-405 | chr14-31851089-INS-405 |
| chr14-31872642-INS-124 | chr14-31872642-INS-120 |
| chr14-31876019-INS-6493 | chr14-31876020-INS-6493 |
| chr14-31910992-INS-240 | chr14-31910992-INS-240 |
| chr14-31925062-INS-108 | chr14-31925062-INS-108 |
| chr14-31940465-INS-215 | chr14-31940465-INS-214 |
| chr14-31985520-INS-55 | chr14-31985520-INS-60 |
| chr14-32007003-INS-340 | chr14-32007003-INS-341 |
| chr14-32022489-INS-154 | chr14-32022489-INS-156 |
| chr14-32033199-INS-118 | chr14-32033199-INS-127 |
| chr14-32035763-INS-5516 | chr14-32035764-INS-5517 |
| chr14-32036162-INS-60 | chr14-32036162-INS-60 |
| chr14-32037521-INS-194 | chr14-32037521-INS-180 |
| chr14-32052187-INS-52 | chr14-32052187-INS-56 |
| chr14-32077415-INS-78 | chr14-32077415-INS-81 |
| chr14-32093039-INS-213 | chr14-32093039-INS-214 |
| chr14-32094136-INS-155 | chr14-32094136-INS-130 |
| chr14-32114259-INS-195 | chr14-32114259-INS-195 |
| chr14-32128260-INS-58 | chr14-32128266-INS-72 |
| chr14-32130165-INS-80 | chr14-32130165-INS-84 |
| chr14-32138866-INS-1501 | chr14-32138867-INS-1501 |
| chr14-32139508-INS-152 | chr14-32139508-INS-152 |
| chr14-32141979-INS-88 | chr14-32141979-INS-88 |

| | |
|---|---|
| chr14-32160916-INS-62 | chr14-32160916-INS-56 |
| chr14-32175945-INS-60 | chr14-32175941-INS-60 |
| chr14-32193387-INS-174 | chr14-32193387-INS-165 |
| chr14-32211086-INS-67 | chr14-32211086-INS-66 |
| chr14-32228669-INS-4094 | chr14-32228670-INS-4094 |
| chr14-32233509-INS-7119 | chr14-32233510-INS-7119 |
| chr14-32245897-INS-658 | chr14-32245897-INS-658 |
| chr14-32290467-INS-60 | chr14-32290467-INS-60 |
| chr14-32291209-INS-1727 | chr14-32291199-INS-1715 |
| chr14-32335731-INS-70 | chr14-32335731-INS-70 |
| chr14-32344095-INS-124 | chr14-32344091-INS-132 |
| chr14-32361262-INS-60 | chr14-32361262-INS-92 |
| chr14-32418260-INS-103 | chr14-32418260-INS-104 |
| chr14-32423959-INS-207 | chr14-32423959-INS-207 |
| chr14-32428448-INS-57 | chr14-32428450-INS-53 |
| chr14-32447184-INS-184 | chr14-32447156-INS-222 |
| chr14-32469277-INS-72 | chr14-32469277-INS-72 |
| chr14-32488160-INS-55 | chr14-32488160-INS-55 |
| chr14-32499344-INS-76 | chr14-32499344-INS-88 |
| chr14-32507428-INS-168 | chr14-32507428-INS-168 |
| chr14-32541871-INS-66 | chr14-32541872-INS-60 |
| chr14-32546009-INS-3879 | chr14-32546010-INS-3879 |
| chr14-32553197-INS-688 | chr14-32553197-INS-688 |
| chr14-32589101-INS-63 | chr14-32589101-INS-62 |
| chr14-32600710-INS-114 | chr14-32600705-INS-134 |
| chr14-32602723-INS-95 | chr14-32602723-INS-95 |
| chr14-32615174-INS-69 | chr14-32615165-INS-59 |
| chr14-32632465-INS-6631 | chr14-32632466-INS-6631 |
| chr14-32674959-INS-103 | chr14-32674959-INS-103 |
| chr14-32694540-INS-119 | chr14-32694540-INS-114 |
| chr14-32734137-INS-56 | chr14-32734137-INS-52 |
| chr14-32759349-INS-172 | chr14-32759349-INS-172 |
| chr14-32762385-INS-158 | chr14-32762385-INS-156 |
| chr14-32806439-INS-112 | chr14-32806421-INS-142 |
| chr14-32841083-INS-479 | chr14-32841083-INS-479 |
| chr14-32841622-INS-72 | chr14-32841634-INS-96 |
| chr14-32849943-INS-82 | chr14-32849943-INS-82 |
| chr14-32856620-INS-99 | chr14-32856620-INS-101 |
| chr14-32863749-INS-154 | chr14-32863749-INS-154 |
| chr14-32872193-INS-59 | chr14-32872163-INS-53 |
| chr14-32900331-INS-80 | chr14-32900331-INS-79 |
| chr14-32910852-INS-74 | chr14-32910852-INS-74 |
| chr14-32912575-INS-66 | chr14-32912575-INS-62 |
| chr14-32920291-INS-198 | chr14-32920291-INS-198 |
| chr14-32938117-INS-80 | chr14-32938117-INS-110 |
| chr14-32959162-INS-52 | chr14-32959166-INS-52 |
| chr14-32992963-INS-175 | chr14-32992963-INS-175 |
| chr14-33001985-INS-112 | chr14-33001985-INS-136 |
| chr14-33013045-INS-373 | chr14-33013045-INS-377 |
| chr14-33063867-INS-1292 | chr14-33063868-INS-1292 |
| chr14-33079668-INS-583 | chr14-33079668-INS-584 |
| chr14-33093628-INS-212 | chr14-33093628-INS-212 |
| chr14-33139534-INS-6633 | chr14-33139535-INS-6633 |
| chr14-33144147-INS-689 | chr14-33144147-INS-695 |
| chr14-33152508-INS-88 | chr14-33152508-INS-80 |
| chr14-33174661-INS-2917 | chr14-33174662-INS-2929 |
| chr14-33194202-INS-372 | chr14-33194202-INS-372 |
| chr14-33197790-INS-64 | chr14-33197790-INS-66 |
| chr14-33220746-INS-133 | chr14-33220746-INS-133 |
| chr14-33248587-INS-54 | chr14-33248587-INS-56 |
| chr14-33251655-INS-73 | chr14-33251655-INS-57 |
| chr14-33252179-INS-399 | chr14-33252179-INS-399 |
| chr14-33277020-INS-88 | chr14-33277020-INS-88 |
| chr14-33288610-INS-61 | chr14-33288610-INS-59 |
| chr14-33349092-INS-78 | chr14-33349092-INS-84 |
| chr14-33351917-INS-480 | chr14-33351917-INS-480 |
| chr14-33375024-INS-365 | chr14-33375024-INS-365 |
| chr14-33382235-INS-354 | chr14-33382235-INS-351 |
| chr14-33423896-INS-103 | chr14-33423896-INS-110 |
| chr14-33427294-INS-129 | chr14-33427294-INS-129 |
| chr14-33438694-INS-85 | chr14-33438694-INS-85 |
| chr14-33478072-INS-73 | chr14-33478072-INS-73 |
| chr14-33487685-INS-88 | chr14-33487685-INS-138 |
| chr14-33492277-INS-211 | chr14-33492283-INS-181 |
| chr14-33493575-INS-110 | chr14-33493575-INS-110 |
| chr14-33498394-INS-150 | chr14-33498394-INS-150 |
| chr14-33527502-INS-6509 | chr14-33527503-INS-6509 |

| | |
|---|---|
| chr14-33559655-INS-499 | chr14-33559655-INS-499 |
| chr14-33572720-INS-173 | chr14-33572720-INS-173 |
| chr14-33592133-INS-163 | chr14-33592133-INS-163 |
| chr14-33619486-INS-54 | chr14-33619487-INS-52 |
| chr14-33623182-INS-2874 | chr14-33623183-INS-2874 |
| chr14-33633411-INS-6728 | chr14-33633412-INS-6729 |
| chr14-33633581-INS-229 | chr14-33633581-INS-264 |
| chr14-33634850-INS-550 | chr14-33634850-INS-550 |
| chr14-33649469-INS-398 | chr14-33649469-INS-398 |
| chr14-33654929-INS-65 | chr14-33654929-INS-50 |
| chr14-33664542-INS-52 | chr14-33664542-INS-52 |
| chr14-33731346-INS-239 | chr14-33731346-INS-239 |
| chr14-33740423-INS-464 | chr14-33740423-INS-464 |
| chr14-33754143-INS-66 | chr14-33754143-INS-66 |
| chr14-33771487-INS-69 | chr14-33771487-INS-71 |
| chr14-33777712-INS-60 | chr14-33777712-INS-60 |
| chr14-33827762-INS-54 | chr14-33827762-INS-52 |
| chr14-33841673-INS-339 | chr14-33841673-INS-339 |
| chr14-33861821-INS-54 | chr14-33861821-INS-54 |
| chr14-33876471-INS-69 | chr14-33876471-INS-69 |
| chr14-33890626-INS-95 | chr14-33890626-INS-55 |
| chr14-33914887-INS-834 | chr14-33914887-INS-834 |
| chr14-33937418-INS-64 | chr14-33937418-INS-64 |
| chr14-33939396-INS-109 | chr14-33939396-INS-112 |
| chr14-33945173-INS-78 | chr14-33945173-INS-78 |
| chr14-33956296-INS-95 | chr14-33956277-INS-104 |
| chr14-33970033-INS-194 | chr14-33970033-INS-194 |
| chr14-33977574-INS-92 | chr14-33977574-INS-84 |
| chr14-33978013-INS-230 | chr14-33978014-INS-209 |
| chr14-33986025-INS-207 | chr14-33986025-INS-207 |
| chr14-33999152-INS-7165 | chr14-33999153-INS-7165 |
| chr14-34001767-INS-185 | chr14-34001768-INS-184 |
| chr14-34022712-INS-220 | chr14-34022712-INS-221 |
| chr14-34029162-INS-192 | chr14-34029162-INS-190 |
| chr14-34030711-INS-75 | chr14-34030711-INS-75 |
| chr14-34038825-INS-709 | chr14-34038852-INS-712 |
| chr14-34040660-INS-175 | chr14-34040660-INS-175 |
| chr14-34046089-INS-78 | chr14-34046089-INS-78 |
| chr14-34053272-INS-210 | chr14-34053272-INS-212 |
| chr14-34056595-INS-206 | chr14-34056595-INS-206 |
| chr14-34079280-INS-54 | chr14-34079280-INS-56 |
| chr14-34101400-INS-52 | chr14-34101400-INS-52 |
| chr14-34128182-INS-207 | chr14-34128182-INS-207 |
| chr14-34150839-INS-128 | chr14-34150839-INS-128 |
| chr14-34153235-INS-207 | chr14-34153235-INS-207 |
| chr14-34153869-INS-222 | chr14-34153869-INS-222 |
| chr14-34185942-INS-67 | chr14-34185941-INS-72 |
| chr14-34229719-INS-92 | chr14-34229720-INS-72 |
| chr14-34231209-INS-66 | chr14-34231209-INS-66 |
| chr14-34231990-INS-190 | chr14-34231990-INS-190 |
| chr14-34238103-INS-213 | chr14-34238103-INS-214 |
| chr14-34257608-INS-52 | chr14-34257608-INS-52 |
| chr14-34326769-INS-4936 | chr14-34326770-INS-4936 |
| chr14-34336090-INS-180 | chr14-34336090-INS-180 |
| chr14-34341402-INS-397 | chr14-34341402-INS-397 |
| chr14-34353793-INS-194 | chr14-34353793-INS-194 |
| chr14-34361157-INS-231 | chr14-34361157-INS-229 |
| chr14-34364762-INS-72 | chr14-34364762-INS-72 |
| chr14-34376596-INS-211 | chr14-34376596-INS-211 |
| chr14-34377063-INS-5262 | chr14-34377064-INS-5262 |
| chr14-34378564-INS-177 | chr14-34378564-INS-177 |
| chr14-34380934-INS-85 | chr14-34380934-INS-78 |
| chr14-34385285-INS-164 | chr14-34385261-INS-151 |
| chr14-34393482-INS-201 | chr14-34393482-INS-200 |
| chr14-34407326-INS-212 | chr14-34407326-INS-213 |
| chr14-34422784-INS-167 | chr14-34422784-INS-167 |
| chr14-34437490-INS-299 | chr14-34437490-INS-289 |
| chr14-34445129-INS-106 | chr14-34445129-INS-122 |
| chr14-34485603-INS-88 | chr14-34485603-INS-88 |
| chr14-34530630-INS-840 | chr14-34530630-INS-816 |
| chr14-34535744-INS-999 | chr14-34535744-INS-996 |
| chr14-34553508-INS-99 | chr14-34553508-INS-99 |
| chr14-34614774-INS-320 | chr14-34614774-INS-316 |
| chr14-34642496-INS-60 | chr14-34642490-INS-58 |
| chr14-34649568-INS-79 | chr14-34649568-INS-79 |
| chr14-34677847-INS-358 | chr14-34677847-INS-358 |
| chr14-34682754-INS-368 | chr14-34682754-INS-368 |

| | |
|---|---|
| chr14-34695250-INS-60 | chr14-34695250-INS-58 |
| chr14-34711563-INS-68 | chr14-34711563-INS-60 |
| chr14-34739892-INS-641 | chr14-34739892-INS-641 |
| chr14-34750717-INS-117 | chr14-34750717-INS-117 |
| chr14-34768347-INS-448 | chr14-34768347-INS-448 |
| chr14-34790220-INS-7972 | chr14-34790221-INS-7972 |
| chr14-34822873-INS-526 | chr14-34822874-INS-517 |
| chr14-34861010-INS-56 | chr14-34861010-INS-56 |
| chr14-34862988-INS-64 | chr14-34862978-INS-62 |
| chr14-34888225-INS-76 | chr14-34888204-INS-71 |
| chr14-34894565-INS-119 | chr14-34894565-INS-119 |
| chr14-34904603-INS-6666 | chr14-34904604-INS-6666 |
| chr14-34911990-INS-179 | chr14-34911984-INS-161 |
| chr14-34931321-INS-124 | chr14-34931306-INS-142 |
| chr14-34944567-INS-1891 | chr14-34944568-INS-1892 |
| chr14-34949956-INS-56 | chr14-34949956-INS-58 |
| chr14-34952136-INS-67 | chr14-34952136-INS-69 |
| chr14-34954806-INS-195 | chr14-34954806-INS-195 |
| chr14-34956669-INS-83 | chr14-34956657-INS-64 |
| chr14-34973341-INS-5497 | chr14-34973342-INS-5497 |
| chr14-34990598-INS-81 | chr14-34990598-INS-81 |
| chr14-34990898-INS-2844 | chr14-34990899-INS-2844 |
| chr14-35005985-INS-484 | chr14-35005980-INS-487 |
| chr14-35019704-INS-2077 | chr14-35019705-INS-2078 |
| chr14-35049999-INS-56 | chr14-35049999-INS-56 |
| chr14-35064873-INS-1894 | chr14-35064874-INS-1894 |
| chr14-35103253-INS-80 | chr14-35103216-INS-90 |
| chr14-35178387-INS-75 | chr14-35178386-INS-71 |
| chr14-35180016-INS-1187 | chr14-35180017-INS-1187 |
| chr14-35189056-INS-510 | chr14-35189056-INS-509 |
| chr14-35217213-INS-957 | chr14-35217213-INS-957 |
| chr14-35260624-INS-223 | chr14-35260624-INS-228 |
| chr14-35266488-INS-939 | chr14-35266488-INS-939 |
| chr14-35282719-INS-84 | chr14-35282724-INS-50 |
| chr14-35310099-INS-7117 | chr14-35310100-INS-7118 |
| chr14-35328984-INS-6671 | chr14-35328985-INS-6671 |
| chr14-35348712-INS-60 | chr14-35348712-INS-60 |
| chr14-35363544-INS-221 | chr14-35363544-INS-221 |
| chr14-35375424-INS-182 | chr14-35375424-INS-152 |
| chr14-35376748-INS-65 | chr14-35376748-INS-65 |
| chr14-35383146-INS-84 | chr14-35383146-INS-60 |
| chr14-35384623-INS-60 | chr14-35384651-INS-75 |
| chr14-35389834-INS-825 | chr14-35389834-INS-825 |
| chr14-35414632-INS-90 | chr14-35414604-INS-93 |
| chr14-35444791-INS-59 | chr14-35444791-INS-59 |
| chr14-35472441-INS-82 | chr14-35472447-INS-67 |
| chr14-35475365-INS-146 | chr14-35475365-INS-146 |
| chr14-35477104-INS-602 | chr14-35477104-INS-601 |
| chr14-35478198-INS-6474 | chr14-35478199-INS-6474 |
| chr14-35488051-INS-53 | chr14-35488051-INS-53 |
| chr14-35501020-INS-70 | chr14-35501020-INS-70 |
| chr14-35516471-INS-80 | chr14-35516471-INS-93 |
| chr14-35550393-INS-67 | chr14-35550393-INS-79 |
| chr14-35554360-INS-6431 | chr14-35554361-INS-6431 |
| chr14-35561816-INS-84 | chr14-35561816-INS-78 |
| chr14-35565678-INS-9696 | chr14-35565679-INS-9696 |
| chr14-35569720-INS-61 | chr14-35569720-INS-78 |
| chr14-35575308-INS-80 | chr14-35575301-INS-100 |
| chr14-35580687-INS-55 | chr14-35580687-INS-58 |
| chr14-35586243-INS-498 | chr14-35586243-INS-498 |
| chr14-35591615-INS-54 | chr14-35591615-INS-86 |
| chr14-35594581-INS-62 | chr14-35594581-INS-62 |
| chr14-35632906-INS-54 | chr14-35632906-INS-58 |
| chr14-35650683-INS-76 | chr14-35650668-INS-70 |
| chr14-35657929-INS-65 | chr14-35657929-INS-65 |
| chr14-35676320-INS-92 | chr14-35676325-INS-137 |
| chr14-35694827-INS-807 | chr14-35694827-INS-806 |
| chr14-35695519-INS-52 | chr14-35695519-INS-56 |
| chr14-35703739-INS-107 | chr14-35703739-INS-108 |
| chr14-35710074-INS-836 | chr14-35710074-INS-837 |
| chr14-35795396-INS-5373 | chr14-35795401-INS-5358 |
| chr14-35833670-INS-65 | chr14-35833670-INS-60 |
| chr14-35839495-INS-681 | chr14-35839495-INS-681 |
| chr14-35852609-INS-58 | chr14-35852591-INS-68 |
| chr14-35853295-INS-940 | chr14-35853295-INS-940 |
| chr14-35871136-INS-67 | chr14-35871136-INS-89 |
| chr14-35884985-INS-10872 | chr14-35884986-INS-10872 |

| | |
|---|---|
| chr14-35886534-INS-7434 | chr14-35886535-INS-7434 |
| chr14-35898174-INS-94 | chr14-35898162-INS-82 |
| chr14-35939233-INS-232 | chr14-35939232-INS-250 |
| chr14-35968645-INS-68 | chr14-35968646-INS-76 |
| chr14-35997269-INS-75 | chr14-35997269-INS-83 |
| chr14-36005026-INS-4763 | chr14-36005027-INS-4763 |
| chr14-36037484-INS-118 | chr14-36037484-INS-114 |
| chr14-36042153-INS-54 | chr14-36042153-INS-54 |
| chr14-36059591-INS-52 | chr14-36059591-INS-54 |
| chr14-36070085-INS-465 | chr14-36070082-INS-468 |
| chr14-36076534-INS-60 | chr14-36076534-INS-60 |
| chr14-36080846-INS-444 | chr14-36080846-INS-450 |
| chr14-36092715-INS-6377 | chr14-36092716-INS-6377 |
| chr14-36130916-INS-277 | chr14-36130916-INS-277 |
| chr14-36162391-INS-127 | chr14-36162391-INS-130 |
| chr14-36213424-INS-116 | chr14-36213424-INS-116 |
| chr14-36215113-INS-77 | chr14-36215113-INS-77 |
| chr14-36222911-INS-80 | chr14-36222911-INS-82 |
| chr14-36230677-INS-659 | chr14-36230678-INS-659 |
| chr14-36237402-INS-399 | chr14-36237402-INS-399 |
| chr14-36281144-INS-55 | chr14-36281144-INS-55 |
| chr14-36286559-INS-429 | chr14-36286559-INS-429 |
| chr14-36291012-INS-70 | chr14-36291012-INS-70 |
| chr14-36307481-INS-84 | chr14-36307481-INS-84 |
| chr14-36318496-INS-208 | chr14-36318496-INS-208 |
| chr14-36326741-INS-811 | chr14-36326741-INS-812 |
| chr14-36354303-INS-408 | chr14-36354303-INS-364 |
| chr14-36362844-INS-60 | chr14-36362844-INS-62 |
| chr14-36368072-INS-271 | chr14-36368072-INS-272 |
| chr14-36381711-INS-924 | chr14-36381711-INS-920 |
| chr14-36384277-INS-53 | chr14-36384277-INS-53 |
| chr14-36394062-INS-180 | chr14-36394062-INS-181 |
| chr14-36399866-INS-68 | chr14-36399866-INS-70 |
| chr14-36414621-INS-1133 | chr14-36414622-INS-1133 |
| chr14-36420222-INS-137 | chr14-36420222-INS-137 |
| chr14-36426232-INS-133 | chr14-36426238-INS-133 |
| chr14-36432846-INS-51 | chr14-36432846-INS-51 |
| chr14-36435161-INS-495 | chr14-36435161-INS-495 |
| chr14-36450091-INS-78 | chr14-36450091-INS-65 |
| chr14-36463038-INS-3020 | chr14-36463039-INS-3020 |
| chr14-36467606-INS-1894 | chr14-36467607-INS-1894 |
| chr14-36484446-INS-6199 | chr14-36484447-INS-6210 |
| chr14-36526532-INS-7226 | chr14-36526533-INS-7226 |
| chr14-36531646-INS-6575 | chr14-36531647-INS-6575 |
| chr14-36534207-INS-88 | chr14-36534207-INS-92 |
| chr14-36535875-INS-1312 | chr14-36535876-INS-1321 |
| chr14-36538686-INS-6449 | chr14-36538687-INS-6449 |
| chr14-36539745-INS-1013 | chr14-36539746-INS-1014 |
| chr14-36554508-INS-60 | chr14-36554508-INS-58 |
| chr14-36561196-INS-6423 | chr14-36561197-INS-6423 |
| chr14-36574875-INS-2650 | chr14-36574876-INS-2650 |
| chr14-36575945-INS-3352 | chr14-36575946-INS-3352 |
| chr14-36581277-INS-7049 | chr14-36581278-INS-7049 |
| chr14-36586853-INS-5548 | chr14-36586854-INS-5548 |
| chr14-36590005-INS-676 | chr14-36590005-INS-677 |
| chr14-36591431-INS-850 | chr14-36591431-INS-851 |
| chr14-36593372-INS-64 | chr14-36593372-INS-64 |
| chr14-36624433-INS-85 | chr14-36624433-INS-85 |
| chr14-36628333-INS-140 | chr14-36628333-INS-140 |
| chr14-36628572-INS-644 | chr14-36628572-INS-644 |
| chr14-36673661-INS-201 | chr14-36673661-INS-201 |
| chr14-36677709-INS-416 | chr14-36677709-INS-416 |
| chr14-36689083-INS-307 | chr14-36689083-INS-308 |
| chr14-36696886-INS-295 | chr14-36696886-INS-295 |
| chr14-36742214-INS-6591 | chr14-36742215-INS-6591 |
| chr14-36746921-INS-1368 | chr14-36746922-INS-1367 |
| chr14-36762067-INS-115 | chr14-36762067-INS-108 |
| chr14-36768640-INS-91 | chr14-36768640-INS-71 |
| chr14-36792206-INS-175 | chr14-36792206-INS-178 |
| chr14-36810732-INS-79 | chr14-36810732-INS-72 |
| chr14-36822005-INS-89 | chr14-36822005-INS-89 |
| chr14-36834242-INS-85 | chr14-36834242-INS-85 |
| chr14-36846280-INS-170 | chr14-36846280-INS-170 |
| chr14-36847545-INS-366 | chr14-36847545-INS-366 |
| chr14-36858513-INS-158 | chr14-36858513-INS-123 |
| chr14-36863848-INS-935 | chr14-36863848-INS-942 |
| chr14-36874880-INS-50 | chr14-36874878-INS-50 |

| | |
|---|---|
| chr14-36895310-INS-6413 | chr14-36895311-INS-6413 |
| chr14-36900496-INS-70 | chr14-36900496-INS-70 |
| chr14-36906741-INS-76 | chr14-36906741-INS-76 |
| chr14-36910274-INS-112 | chr14-36910269-INS-111 |
| chr14-36959582-INS-98 | chr14-36959582-INS-98 |
| chr14-36967096-INS-78 | chr14-36967097-INS-69 |
| chr14-36982897-INS-214 | chr14-36982897-INS-214 |
| chr14-36985637-INS-2877 | chr14-36985638-INS-2878 |
| chr14-36998812-INS-636 | chr14-36998812-INS-636 |
| chr14-37004409-INS-6357 | chr14-37004410-INS-6357 |
| chr14-37020502-INS-186 | chr14-37020502-INS-186 |
| chr14-37022661-INS-368 | chr14-37022661-INS-365 |
| chr14-37026548-INS-60 | chr14-37026548-INS-67 |
| chr14-37054225-INS-399 | chr14-37054225-INS-399 |
| chr14-37056373-INS-78 | chr14-37056381-INS-98 |
| chr14-37099254-INS-51 | chr14-37099254-INS-53 |
| chr14-37110795-INS-56 | chr14-37110795-INS-91 |
| chr14-37132205-INS-51 | chr14-37132205-INS-51 |
| chr14-37132519-INS-401 | chr14-37132519-INS-401 |
| chr14-37140561-INS-2343 | chr14-37140562-INS-2343 |
| chr14-37148604-INS-1084 | chr14-37148605-INS-1085 |
| chr14-37163650-INS-6197 | chr14-37163651-INS-6197 |
| chr14-37165506-INS-475 | chr14-37165505-INS-491 |
| chr14-37167376-INS-3154 | chr14-37167377-INS-3154 |
| chr14-37212559-INS-70 | chr14-37212533-INS-100 |
| chr14-37216528-INS-428 | chr14-37216528-INS-428 |
| chr14-37235891-INS-1255 | chr14-37235892-INS-1255 |
| chr14-37237343-INS-1320 | chr14-37237344-INS-1320 |
| chr14-37241084-INS-54 | chr14-37241084-INS-52 |
| chr14-37242716-INS-219 | chr14-37242716-INS-220 |
| chr14-37244456-INS-549 | chr14-37244456-INS-550 |
| chr14-37248086-INS-6209 | chr14-37248087-INS-6209 |
| chr14-37279585-INS-76 | chr14-37279585-INS-76 |
| chr14-37281348-INS-68 | chr14-37281348-INS-72 |
| chr14-37322592-INS-62 | chr14-37322592-INS-62 |
| chr14-37334608-INS-617 | chr14-37334608-INS-617 |
| chr14-37362820-INS-61 | chr14-37362820-INS-51 |
| chr14-37370040-INS-82 | chr14-37370040-INS-82 |
| chr14-37396439-INS-6715 | chr14-37396440-INS-6715 |
| chr14-37404150-INS-113 | chr14-37404150-INS-113 |
| chr14-37410629-INS-245 | chr14-37410650-INS-202 |
| chr14-37449062-INS-55 | chr14-37449062-INS-55 |
| chr14-37469691-INS-5584 | chr14-37469692-INS-5584 |
| chr14-37476991-INS-497 | chr14-37476991-INS-497 |
| chr14-37479146-INS-1827 | chr14-37479147-INS-1826 |
| chr14-37494640-INS-51 | chr14-37494640-INS-51 |
| chr14-37506469-INS-624 | chr14-37506469-INS-624 |
| chr14-37520097-INS-6546 | chr14-37520098-INS-6546 |
| chr14-37529910-INS-1421 | chr14-37529911-INS-1416 |
| chr14-37537883-INS-60 | chr14-37537883-INS-72 |
| chr14-37541934-INS-105 | chr14-37541934-INS-110 |
| chr14-37565805-INS-416 | chr14-37565805-INS-416 |
| chr14-37601152-INS-285 | chr14-37601152-INS-289 |
| chr14-37635355-INS-2929 | chr14-37635356-INS-2929 |
| chr14-37641499-INS-1057 | chr14-37641500-INS-1057 |
| chr14-37656630-INS-130 | chr14-37656630-INS-130 |
| chr14-37698140-INS-53 | chr14-37698140-INS-51 |
| chr14-37702404-INS-476 | chr14-37702402-INS-478 |
| chr14-37709997-INS-99 | chr14-37710006-INS-100 |
| chr14-37720508-INS-5034 | chr14-37720509-INS-5035 |
| chr14-37721041-INS-1728 | chr14-37721042-INS-1729 |
| chr14-37731053-INS-76 | chr14-37731068-INS-65 |
| chr14-37735455-INS-6817 | chr14-37735456-INS-6818 |
| chr14-37767752-INS-52 | chr14-37767752-INS-50 |
| chr14-37768805-INS-6473 | chr14-37768806-INS-6473 |
| chr14-37777341-INS-94 | chr14-37777341-INS-68 |
| chr14-37798604-INS-3018 | chr14-37798605-INS-3018 |
| chr14-37807781-INS-82 | chr14-37807781-INS-82 |
| chr14-37823967-INS-6586 | chr14-37823968-INS-6586 |
| chr14-37846335-INS-91 | chr14-37846335-INS-95 |
| chr14-37854665-INS-120 | chr14-37854623-INS-120 |
| chr14-37875507-INS-56 | chr14-37875507-INS-58 |
| chr14-37877447-INS-67 | chr14-37877438-INS-67 |
| chr14-37885427-INS-52 | chr14-37885427-INS-52 |
| chr14-37906157-INS-3453 | chr14-37906158-INS-3453 |
| chr14-37906905-INS-55 | chr14-37906905-INS-55 |
| chr14-37924132-INS-180 | chr14-37924132-INS-180 |

| | |
|---|---|
| chr14-37930884-INS-93 | chr14-37930884-INS-56 |
| chr14-37940309-INS-54 | chr14-37940309-INS-54 |
| chr14-37953238-INS-157 | chr14-37953238-INS-157 |
| chr14-37956994-INS-4594 | chr14-37956995-INS-4594 |
| chr14-37995336-INS-74 | chr14-37995337-INS-73 |
| chr14-37998557-INS-7028 | chr14-37998558-INS-7029 |
| chr14-38002175-INS-576 | chr14-38002175-INS-576 |
| chr14-38058095-INS-78 | chr14-38058095-INS-78 |
| chr14-38069477-INS-52 | chr14-38069476-INS-52 |
| chr14-38098746-INS-699 | chr14-38098746-INS-699 |
| chr14-38101354-INS-54 | chr14-38101354-INS-54 |
| chr14-38122777-INS-53 | chr14-38122777-INS-53 |
| chr14-38124316-INS-52 | chr14-38124316-INS-52 |
| chr14-38148369-INS-317 | chr14-38148369-INS-317 |
| chr14-38209097-INS-109 | chr14-38209091-INS-118 |
| chr14-38213786-INS-136 | chr14-38213786-INS-136 |
| chr14-38234377-INS-667 | chr14-38234377-INS-667 |
| chr14-38235286-INS-5070 | chr14-38235287-INS-5070 |
| chr14-38268282-INS-82 | chr14-38268271-INS-55 |
| chr14-38270578-INS-193 | chr14-38270579-INS-193 |
| chr14-38290810-INS-120 | chr14-38290810-INS-124 |
| chr14-38310867-INS-53 | chr14-38310867-INS-57 |
| chr14-38315102-INS-7105 | chr14-38315103-INS-7105 |
| chr14-38320225-INS-67 | chr14-38320225-INS-67 |
| chr14-38336602-INS-276 | chr14-38336604-INS-285 |
| chr14-38387959-INS-78 | chr14-38387953-INS-66 |
| chr14-38401321-INS-708 | chr14-38401318-INS-717 |
| chr14-38404246-INS-256 | chr14-38404210-INS-242 |
| chr14-38413016-INS-1915 | chr14-38413017-INS-1915 |
| chr14-38425114-INS-54 | chr14-38425114-INS-54 |
| chr14-38469790-INS-101 | chr14-38469790-INS-97 |
| chr14-38485777-INS-500 | chr14-38485777-INS-500 |
| chr14-38494005-INS-696 | chr14-38494005-INS-696 |
| chr14-38518196-INS-90 | chr14-38518189-INS-50 |
| chr14-38527254-INS-4691 | chr14-38527255-INS-4691 |
| chr14-38558173-INS-64 | chr14-38558173-INS-68 |
| chr14-38567631-INS-56 | chr14-38567631-INS-62 |
| chr14-38588170-INS-66 | chr14-38588170-INS-66 |
| chr14-38597479-INS-58 | chr14-38597479-INS-58 |
| chr14-38605144-INS-66 | chr14-38605144-INS-72 |
| chr14-38629764-INS-390 | chr14-38629764-INS-390 |
| chr14-38639549-INS-104 | chr14-38639549-INS-104 |
| chr14-38656238-INS-66 | chr14-38656238-INS-66 |
| chr14-38659191-INS-811 | chr14-38659191-INS-811 |
| chr14-38660411-INS-271 | chr14-38660412-INS-266 |
| chr14-38670481-INS-605 | chr14-38670481-INS-597 |
| chr14-38672289-INS-978 | chr14-38672289-INS-978 |
| chr14-38673880-INS-194 | chr14-38673880-INS-195 |
| chr14-38701761-INS-204 | chr14-38701761-INS-204 |
| chr14-38738818-INS-136 | chr14-38738818-INS-136 |
| chr14-38746382-INS-54 | chr14-38746382-INS-54 |
| chr14-38752894-INS-66 | chr14-38752894-INS-58 |
| chr14-38784425-INS-72 | chr14-38784425-INS-71 |
| chr14-38804497-INS-54 | chr14-38804497-INS-62 |
| chr14-38811129-INS-125 | chr14-38811129-INS-125 |
| chr14-38821460-INS-450 | chr14-38821460-INS-451 |
| chr14-38833239-INS-869 | chr14-38833239-INS-869 |
| chr14-38854347-INS-56 | chr14-38854347-INS-56 |
| chr14-38862650-INS-124 | chr14-38862651-INS-104 |
| chr14-38865584-INS-69 | chr14-38865584-INS-69 |
| chr14-38907333-INS-60 | chr14-38907333-INS-60 |
| chr14-38909084-INS-176 | chr14-38909084-INS-176 |
| chr14-38942616-INS-710 | chr14-38942616-INS-710 |
| chr14-38960084-INS-103 | chr14-38960084-INS-102 |
| chr14-38995499-INS-340 | chr14-38995491-INS-335 |
| chr14-39005252-INS-208 | chr14-39005252-INS-208 |
| chr14-39059589-INS-60 | chr14-39059589-INS-62 |
| chr14-39062896-INS-1889 | chr14-39062897-INS-1889 |
| chr14-39082807-INS-93 | chr14-39082807-INS-125 |
| chr14-39110106-INS-61 | chr14-39110106-INS-57 |
| chr14-39129105-INS-54 | chr14-39129105-INS-58 |
| chr14-39132751-INS-720 | chr14-39132752-INS-720 |
| chr14-39138179-INS-202 | chr14-39138179-INS-200 |
| chr14-39139284-INS-2772 | chr14-39139285-INS-2772 |
| chr14-39160575-INS-138 | chr14-39160586-INS-96 |
| chr14-39202622-INS-70 | chr14-39202620-INS-70 |
| chr14-39259253-INS-358 | chr14-39259253-INS-358 |

| | |
|---|---|
| chr14-39266421-INS-2481 | chr14-39266422-INS-2481 |
| chr14-39359394-INS-548 | chr14-39359395-INS-548 |
| chr14-39378741-INS-773 | chr14-39378741-INS-777 |
| chr14-39393599-INS-120 | chr14-39393635-INS-85 |
| chr14-39402244-INS-52 | chr14-39402244-INS-52 |
| chr14-39406990-INS-341 | chr14-39406990-INS-341 |
| chr14-39421097-INS-55 | chr14-39421097-INS-55 |
| chr14-39468020-INS-60 | chr14-39468020-INS-60 |
| chr14-39498161-INS-121 | chr14-39498157-INS-133 |
| chr14-39547244-INS-252 | chr14-39547244-INS-252 |
| chr14-39547572-INS-6382 | chr14-39547573-INS-6382 |
| chr14-39550494-INS-70 | chr14-39550494-INS-72 |
| chr14-39566456-INS-607 | chr14-39566456-INS-609 |
| chr14-39573062-INS-860 | chr14-39573062-INS-860 |
| chr14-39580202-INS-2284 | chr14-39580203-INS-2284 |
| chr14-39589329-INS-377 | chr14-39589329-INS-377 |
| chr14-39592342-INS-108 | chr14-39592320-INS-111 |
| chr14-39619640-INS-451 | chr14-39619641-INS-440 |
| chr14-39627134-INS-214 | chr14-39627134-INS-214 |
| chr14-39637909-INS-64 | chr14-39637894-INS-75 |
| chr14-39649112-INS-766 | chr14-39649112-INS-772 |
| chr14-39660422-INS-171 | chr14-39660422-INS-171 |
| chr14-39680333-INS-76 | chr14-39680333-INS-76 |
| chr14-39720529-INS-88 | chr14-39720529-INS-90 |
| chr14-39745955-INS-54 | chr14-39745955-INS-52 |
| chr14-39758635-INS-97 | chr14-39758635-INS-101 |
| chr14-39761374-INS-445 | chr14-39761374-INS-445 |
| chr14-39768476-INS-56 | chr14-39768476-INS-58 |
| chr14-39769791-INS-100 | chr14-39769795-INS-97 |
| chr14-39776192-INS-64 | chr14-39776192-INS-52 |
| chr14-39879533-INS-64 | chr14-39879533-INS-60 |
| chr14-39887804-INS-51 | chr14-39887804-INS-51 |
| chr14-39890091-INS-7634 | chr14-39890092-INS-7635 |
| chr14-39896615-INS-120 | chr14-39896615-INS-120 |
| chr14-39905835-INS-212 | chr14-39905835-INS-212 |
| chr14-39927106-INS-633 | chr14-39927105-INS-633 |
| chr14-39930686-INS-116 | chr14-39930686-INS-116 |
| chr14-39935333-INS-68 | chr14-39935333-INS-68 |
| chr14-39936053-INS-52 | chr14-39936011-INS-53 |
| chr14-39938424-INS-6590 | chr14-39938425-INS-6590 |
| chr14-39942333-INS-1717 | chr14-39942334-INS-1717 |
| chr14-39976957-INS-5792 | chr14-39976946-INS-5792 |
| chr14-39990109-INS-341 | chr14-39990109-INS-341 |
| chr14-39997988-INS-440 | chr14-39997988-INS-440 |
| chr14-40012712-INS-56 | chr14-40012718-INS-59 |
| chr14-40020159-INS-70 | chr14-40020177-INS-54 |
| chr14-40058840-INS-77 | chr14-40058840-INS-77 |
| chr14-40120682-INS-6403 | chr14-40120683-INS-6403 |
| chr14-40126708-INS-66 | chr14-40126708-INS-66 |
| chr14-40141246-INS-7138 | chr14-40141247-INS-7138 |
| chr14-40165078-INS-4620 | chr14-40165079-INS-4620 |
| chr14-40168521-INS-5205 | chr14-40168522-INS-5205 |
| chr14-40169814-INS-5650 | chr14-40169815-INS-5650 |
| chr14-40197715-INS-846 | chr14-40197715-INS-846 |
| chr14-40235706-INS-386 | chr14-40235706-INS-396 |
| chr14-40263499-INS-108 | chr14-40263499-INS-108 |
| chr14-40269460-INS-424 | chr14-40269460-INS-424 |
| chr14-40293970-INS-65 | chr14-40293971-INS-65 |
| chr14-40297470-INS-260 | chr14-40297470-INS-260 |
| chr14-40309196-INS-397 | chr14-40309196-INS-397 |
| chr14-40322221-INS-2961 | chr14-40322222-INS-2961 |
| chr14-40338570-INS-87 | chr14-40338571-INS-91 |
| chr14-40351404-INS-67 | chr14-40351404-INS-60 |
| chr14-40357136-INS-509 | chr14-40357136-INS-509 |
| chr14-40365715-INS-270 | chr14-40365712-INS-270 |
| chr14-40378117-INS-470 | chr14-40378117-INS-470 |
| chr14-40378392-INS-1077 | chr14-40378393-INS-1079 |
| chr14-40386079-INS-201 | chr14-40386079-INS-201 |
| chr14-40403064-INS-129 | chr14-40403064-INS-138 |
| chr14-40405431-INS-130 | chr14-40405431-INS-130 |
| chr14-40426515-INS-395 | chr14-40426515-INS-395 |
| chr14-40431770-INS-50 | chr14-40431770-INS-54 |
| chr14-40445206-INS-579 | chr14-40445206-INS-579 |
| chr14-40448865-INS-203 | chr14-40448865-INS-203 |
| chr14-40450015-INS-200 | chr14-40450045-INS-200 |
| chr14-40455839-INS-141 | chr14-40455832-INS-156 |
| chr14-40482244-INS-66 | chr14-40482244-INS-54 |

| | |
|---|---|
| chr14-40493111-INS-153 | chr14-40493128-INS-162 |
| chr14-40516805-INS-99 | chr14-40516805-INS-99 |
| chr14-40537957-INS-504 | chr14-40537957-INS-504 |
| chr14-40541319-INS-59 | chr14-40541319-INS-55 |
| chr14-40558328-INS-60 | chr14-40558328-INS-66 |
| chr14-40591876-INS-260 | chr14-40591876-INS-259 |
| chr14-40593956-INS-65 | chr14-40593956-INS-80 |
| chr14-40598198-INS-95 | chr14-40598198-INS-95 |
| chr14-40600320-INS-197 | chr14-40600320-INS-156 |
| chr14-40607346-INS-56 | chr14-40607346-INS-56 |
| chr14-40622234-INS-383 | chr14-40622229-INS-390 |
| chr14-40662154-INS-176 | chr14-40662154-INS-179 |
| chr14-40693561-INS-203 | chr14-40693586-INS-174 |
| chr14-40703566-INS-148 | chr14-40703566-INS-147 |
| chr14-40705951-INS-216 | chr14-40705951-INS-215 |
| chr14-40715321-INS-108 | chr14-40715323-INS-84 |
| chr14-40721770-INS-216 | chr14-40721770-INS-217 |
| chr14-40778166-INS-56 | chr14-40778166-INS-56 |
| chr14-40785035-INS-58 | chr14-40785035-INS-58 |
| chr14-40847723-INS-5707 | chr14-40847724-INS-5707 |
| chr14-40851414-INS-112 | chr14-40851390-INS-108 |
| chr14-40851831-INS-172 | chr14-40851824-INS-203 |
| chr14-40859237-INS-812 | chr14-40859237-INS-811 |
| chr14-40872809-INS-54 | chr14-40872809-INS-54 |
| chr14-40918774-INS-71 | chr14-40918774-INS-75 |
| chr14-40920813-INS-199 | chr14-40920813-INS-199 |
| chr14-40938433-INS-1427 | chr14-40938434-INS-1428 |
| chr14-41334832-INS-56 | chr14-41334852-INS-60 |
| chr14-41364941-INS-106 | chr14-41364907-INS-132 |
| chr14-41382648-INS-62 | chr14-41382648-INS-62 |
| chr14-41851208-INS-79 | chr14-41851216-INS-70 |
| chr14-42184964-INS-84 | chr14-42184964-INS-91 |
| chr14-42510997-INS-104 | chr14-42510997-INS-106 |
| chr14-42646022-INS-170 | chr14-42646022-INS-179 |
| chr14-43090906-INS-58 | chr14-43090890-INS-58 |
| chr14-43145387-INS-559 | chr14-43145386-INS-560 |
| chr14-44040226-INS-187 | chr14-44040233-INS-189 |
| chr14-44327751-INS-62 | chr14-44327720-INS-50 |
| chr14-44348155-INS-77 | chr14-44348155-INS-61 |
| chr14-44683832-INS-529 | chr14-44683832-INS-515 |
| chr14-44964727-INS-62 | chr14-44964727-INS-64 |
| chr14-45037605-INS-344 | chr14-45037605-INS-344 |
| chr14-45046694-INS-497 | chr14-45046694-INS-497 |
| chr14-45047260-INS-210 | chr14-45047261-INS-212 |
| chr14-45068900-INS-132 | chr14-45068900-INS-117 |
| chr14-45099283-INS-62 | chr14-45099283-INS-57 |
| chr14-45139017-INS-222 | chr14-45139017-INS-225 |
| chr14-45208879-INS-77 | chr14-45208879-INS-60 |
| chr14-45214131-INS-60 | chr14-45214131-INS-60 |
| chr14-45222933-INS-190 | chr14-45222929-INS-180 |
| chr14-45293642-INS-6677 | chr14-45293643-INS-6677 |
| chr14-45317176-INS-76 | chr14-45317176-INS-84 |
| chr14-45361484-INS-352 | chr14-45361484-INS-352 |
| chr14-45366442-INS-197 | chr14-45366442-INS-197 |
| chr14-45375094-INS-87 | chr14-45375094-INS-89 |
| chr14-45391106-INS-392 | chr14-45391106-INS-392 |
| chr14-45454872-INS-54 | chr14-45454872-INS-54 |
| chr14-45524542-INS-180 | chr14-45524542-INS-180 |
| chr14-45527506-INS-98 | chr14-45527506-INS-92 |
| chr14-45533187-INS-133 | chr14-45533227-INS-119 |
| chr14-45604188-INS-129 | chr14-45604188-INS-129 |
| chr14-45632515-INS-339 | chr14-45632515-INS-340 |
| chr14-45635618-INS-477 | chr14-45635618-INS-478 |
| chr14-45670292-INS-220 | chr14-45670292-INS-218 |
| chr14-45672499-INS-4923 | chr14-45672500-INS-4923 |
| chr14-45674991-INS-193 | chr14-45674991-INS-194 |
| chr14-45686801-INS-7102 | chr14-45686802-INS-7102 |
| chr14-45688381-INS-211 | chr14-45688381-INS-210 |
| chr14-45716404-INS-264 | chr14-45716404-INS-265 |
| chr14-45718288-INS-198 | chr14-45718288-INS-204 |
| chr14-45723095-INS-198 | chr14-45723095-INS-198 |
| chr14-45726333-INS-80 | chr14-45726333-INS-80 |
| chr14-45776026-INS-67 | chr14-45776026-INS-63 |
| chr14-45787209-INS-62 | chr14-45787193-INS-72 |
| chr14-45790291-INS-70 | chr14-45790323-INS-74 |
| chr14-45815313-INS-55 | chr14-45815313-INS-51 |
| chr14-45819580-INS-170 | chr14-45819580-INS-161 |

| | |
|---|---|
| chr14-45849204-INS-668 | chr14-45849204-INS-670 |
| chr14-45851320-INS-230 | chr14-45851320-INS-226 |
| chr14-45869727-INS-240 | chr14-45869727-INS-240 |
| chr14-45893665-INS-6413 | chr14-45893666-INS-6413 |
| chr14-45908863-INS-181 | chr14-45908863-INS-183 |
| chr14-45911795-INS-60 | chr14-45911795-INS-62 |
| chr14-45917549-INS-205 | chr14-45917549-INS-206 |
| chr14-45933927-INS-5342 | chr14-45933928-INS-5342 |
| chr14-45941802-INS-825 | chr14-45941802-INS-825 |
| chr14-45957926-INS-162 | chr14-45957927-INS-162 |
| chr14-45959083-INS-175 | chr14-45959083-INS-176 |
| chr14-45968919-INS-53 | chr14-45968919-INS-53 |
| chr14-45975960-INS-65 | chr14-45975960-INS-65 |
| chr14-45981150-INS-253 | chr14-45981150-INS-254 |
| chr14-45982096-INS-169 | chr14-45982096-INS-169 |
| chr14-45992898-INS-50 | chr14-45992898-INS-50 |
| chr14-45994005-INS-154 | chr14-45994005-INS-154 |
| chr14-46004663-INS-73 | chr14-46004650-INS-52 |
| chr14-46005353-INS-666 | chr14-46005353-INS-666 |
| chr14-46016293-INS-202 | chr14-46016293-INS-204 |
| chr14-46024141-INS-176 | chr14-46024141-INS-178 |
| chr14-46026343-INS-6185 | chr14-46026344-INS-6183 |
| chr14-46047257-INS-139 | chr14-46047257-INS-139 |
| chr14-46060675-INS-133 | chr14-46060675-INS-133 |
| chr14-46091567-INS-176 | chr14-46091567-INS-176 |
| chr14-46105854-INS-54 | chr14-46105854-INS-58 |
| chr14-46115789-INS-63 | chr14-46115783-INS-69 |
| chr14-46134180-INS-58 | chr14-46134180-INS-58 |
| chr14-46144238-INS-201 | chr14-46144238-INS-201 |
| chr14-46153748-INS-6225 | chr14-46153749-INS-6248 |
| chr14-46156963-INS-96 | chr14-46156963-INS-102 |
| chr14-46162266-INS-76 | chr14-46162266-INS-76 |
| chr14-46181190-INS-393 | chr14-46181190-INS-393 |
| chr14-46190672-INS-69 | chr14-46190672-INS-69 |
| chr14-46194741-INS-1001 | chr14-46194742-INS-992 |
| chr14-46208601-INS-80 | chr14-46208591-INS-50 |
| chr14-46218706-INS-175 | chr14-46218706-INS-175 |
| chr14-46249929-INS-3082 | chr14-46249930-INS-3082 |
| chr14-46277814-INS-200 | chr14-46277814-INS-200 |
| chr14-46284854-INS-159 | chr14-46284855-INS-159 |
| chr14-46313836-INS-71 | chr14-46313836-INS-71 |
| chr14-46314537-INS-210 | chr14-46314537-INS-210 |
| chr14-46391556-INS-6637 | chr14-46391557-INS-6637 |
| chr14-46435701-INS-5462 | chr14-46435702-INS-5462 |
| chr14-46482132-INS-56 | chr14-46482132-INS-56 |
| chr14-46498330-INS-80 | chr14-46498330-INS-80 |
| chr14-46504803-INS-65 | chr14-46504803-INS-65 |
| chr14-46513735-INS-203 | chr14-46513735-INS-203 |
| chr14-46541246-INS-74 | chr14-46541246-INS-74 |
| chr14-46548406-INS-720 | chr14-46548406-INS-720 |
| chr14-46620327-INS-54 | chr14-46620334-INS-55 |
| chr14-46648786-INS-54 | chr14-46648786-INS-69 |
| chr14-46657729-INS-63 | chr14-46657729-INS-63 |
| chr14-46664439-INS-214 | chr14-46664439-INS-214 |
| chr14-46674589-INS-487 | chr14-46674589-INS-487 |
| chr14-46684374-INS-70 | chr14-46684374-INS-70 |
| chr14-46732220-INS-199 | chr14-46732220-INS-201 |
| chr14-46746837-INS-2223 | chr14-46746838-INS-2223 |
| chr14-46748796-INS-60 | chr14-46748796-INS-60 |
| chr14-46787288-INS-507 | chr14-46787288-INS-507 |
| chr14-46803049-INS-51 | chr14-46803050-INS-63 |
| chr14-46810539-INS-56 | chr14-46810539-INS-62 |
| chr14-46834492-INS-336 | chr14-46834492-INS-336 |
| chr14-46905052-INS-7016 | chr14-46905053-INS-7016 |
| chr14-46915222-INS-56 | chr14-46915222-INS-56 |
| chr14-46918033-INS-52 | chr14-46918033-INS-52 |
| chr14-46951605-INS-76 | chr14-46951606-INS-52 |
| chr14-46955003-INS-153 | chr14-46955003-INS-135 |
| chr14-47033015-INS-185 | chr14-47033015-INS-186 |
| chr14-47080809-INS-62 | chr14-47080809-INS-93 |
| chr14-47105359-INS-78 | chr14-47105359-INS-80 |
| chr14-47134229-INS-432 | chr14-47134229-INS-432 |
| chr14-47143062-INS-200 | chr14-47143062-INS-200 |
| chr14-47145158-INS-69 | chr14-47145158-INS-69 |
| chr14-47146879-INS-75 | chr14-47146879-INS-75 |
| chr14-47168997-INS-55 | chr14-47169015-INS-55 |
| chr14-47180518-INS-242 | chr14-47180518-INS-242 |

| | |
|---|---|
| chr14-47201718-INS-86 | chr14-47201718-INS-86 |
| chr14-47220252-INS-487 | chr14-47220252-INS-487 |
| chr14-47233808-INS-63 | chr14-47233808-INS-63 |
| chr14-47254597-INS-138 | chr14-47254597-INS-150 |
| chr14-47264164-INS-71 | chr14-47264164-INS-76 |
| chr14-47313820-INS-1710 | chr14-47313821-INS-1710 |
| chr14-47344786-INS-98 | chr14-47344786-INS-126 |
| chr14-47346328-INS-55 | chr14-47346328-INS-55 |
| chr14-47358406-INS-210 | chr14-47358406-INS-210 |
| chr14-47382803-INS-212 | chr14-47382803-INS-212 |
| chr14-47394992-INS-158 | chr14-47394992-INS-158 |
| chr14-47398266-INS-202 | chr14-47398266-INS-202 |
| chr14-47402674-INS-144 | chr14-47402674-INS-144 |
| chr14-47431587-INS-51 | chr14-47431584-INS-58 |
| chr14-47460159-INS-136 | chr14-47460167-INS-177 |
| chr14-47461201-INS-201 | chr14-47461201-INS-201 |
| chr14-47475704-INS-202 | chr14-47475704-INS-200 |
| chr14-47510273-INS-204 | chr14-47510273-INS-203 |
| chr14-47519064-INS-196 | chr14-47519064-INS-197 |
| chr14-47524450-INS-72 | chr14-47524450-INS-72 |
| chr14-47541732-INS-177 | chr14-47541732-INS-177 |
| chr14-47542211-INS-211 | chr14-47542211-INS-211 |
| chr14-47546489-INS-92 | chr14-47546489-INS-94 |
| chr14-47554647-INS-172 | chr14-47554647-INS-172 |
| chr14-47561828-INS-205 | chr14-47561828-INS-205 |
| chr14-47577306-INS-5049 | chr14-47577307-INS-5049 |
| chr14-47577720-INS-215 | chr14-47577720-INS-217 |
| chr14-47584063-INS-189 | chr14-47584062-INS-190 |
| chr14-47596049-INS-486 | chr14-47596049-INS-486 |
| chr14-47608607-INS-370 | chr14-47608626-INS-380 |
| chr14-47633563-INS-228 | chr14-47633563-INS-227 |
| chr14-47636154-INS-53 | chr14-47636154-INS-83 |
| chr14-47669325-INS-212 | chr14-47669325-INS-211 |
| chr14-47676927-INS-199 | chr14-47676927-INS-199 |
| chr14-47684964-INS-160 | chr14-47684964-INS-160 |
| chr14-47691910-INS-54 | chr14-47691910-INS-57 |
| chr14-47708477-INS-56 | chr14-47708473-INS-52 |
| chr14-47716015-INS-209 | chr14-47716015-INS-209 |
| chr14-47725468-INS-132 | chr14-47725468-INS-132 |
| chr14-47753374-INS-132 | chr14-47753374-INS-132 |
| chr14-47769471-INS-496 | chr14-47769471-INS-496 |
| chr14-47790345-INS-1276 | chr14-47790346-INS-1276 |
| chr14-47809797-INS-198 | chr14-47809797-INS-198 |
| chr14-47819980-INS-70 | chr14-47819980-INS-75 |
| chr14-47838422-INS-175 | chr14-47838422-INS-178 |
| chr14-47868497-INS-303 | chr14-47868497-INS-304 |
| chr14-47876386-INS-56 | chr14-47876386-INS-56 |
| chr14-47884424-INS-95 | chr14-47884424-INS-95 |
| chr14-47892001-INS-72 | chr14-47892005-INS-94 |
| chr14-47895927-INS-177 | chr14-47895928-INS-177 |
| chr14-47899785-INS-80 | chr14-47899785-INS-80 |
| chr14-47906243-INS-51 | chr14-47906254-INS-56 |
| chr14-47922502-INS-206 | chr14-47922503-INS-205 |
| chr14-47931786-INS-132 | chr14-47931786-INS-108 |
| chr14-47938201-INS-230 | chr14-47938201-INS-231 |
| chr14-47946790-INS-207 | chr14-47946790-INS-206 |
| chr14-47959495-INS-52 | chr14-47959495-INS-51 |
| chr14-48004007-INS-655 | chr14-48004015-INS-665 |
| chr14-48008321-INS-6552 | chr14-48008322-INS-6552 |
| chr14-48036274-INS-83 | chr14-48036274-INS-83 |
| chr14-48039458-INS-247 | chr14-48039453-INS-236 |
| chr14-48053929-INS-192 | chr14-48053929-INS-192 |
| chr14-48061361-INS-164 | chr14-48061361-INS-164 |
| chr14-48079108-INS-77 | chr14-48079108-INS-77 |
| chr14-48082662-INS-1156 | chr14-48082663-INS-1156 |
| chr14-48132848-INS-58 | chr14-48132848-INS-58 |
| chr14-48160735-INS-1903 | chr14-48160732-INS-1900 |
| chr14-48198851-INS-191 | chr14-48198851-INS-190 |
| chr14-48202233-INS-407 | chr14-48202233-INS-407 |
| chr14-48207849-INS-436 | chr14-48207850-INS-436 |
| chr14-48241904-INS-59 | chr14-48241904-INS-63 |
| chr14-48251613-INS-64 | chr14-48251613-INS-50 |
| chr14-48258033-INS-56 | chr14-48258033-INS-56 |
| chr14-48269228-INS-339 | chr14-48269228-INS-339 |
| chr14-48274430-INS-55 | chr14-48274430-INS-55 |
| chr14-48299415-INS-129 | chr14-48299415-INS-129 |
| chr14-48332625-INS-212 | chr14-48332625-INS-212 |

| | |
|---|---|
| chr14-48337388-INS-96 | chr14-48337388-INS-96 |
| chr14-48347491-INS-111 | chr14-48347480-INS-78 |
| chr14-48380833-INS-88 | chr14-48380833-INS-88 |
| chr14-48390993-INS-471 | chr14-48390993-INS-471 |
| chr14-48394020-INS-275 | chr14-48394020-INS-275 |
| chr14-48421706-INS-280 | chr14-48421706-INS-280 |
| chr14-48474586-INS-333 | chr14-48474586-INS-333 |
| chr14-48483076-INS-250 | chr14-48483076-INS-247 |
| chr14-48501004-INS-52 | chr14-48501005-INS-54 |
| chr14-48561137-INS-220 | chr14-48561137-INS-220 |
| chr14-48562433-INS-98 | chr14-48562433-INS-112 |
| chr14-48581658-INS-72 | chr14-48581658-INS-60 |
| chr14-48586213-INS-155 | chr14-48586213-INS-155 |
| chr14-48608910-INS-54 | chr14-48608910-INS-54 |
| chr14-48637393-INS-78 | chr14-48637393-INS-87 |
| chr14-48642883-INS-62 | chr14-48642883-INS-60 |
| chr14-48646070-INS-829 | chr14-48646070-INS-830 |
| chr14-48700134-INS-288 | chr14-48700134-INS-288 |
| chr14-48713159-INS-125 | chr14-48713159-INS-125 |
| chr14-48728218-INS-55 | chr14-48728218-INS-59 |
| chr14-48740238-INS-115 | chr14-48740238-INS-115 |
| chr14-48775142-INS-70 | chr14-48775146-INS-58 |
| chr14-48798375-INS-177 | chr14-48798375-INS-156 |
| chr14-48801033-INS-63 | chr14-48801033-INS-63 |
| chr14-48802089-INS-204 | chr14-48802089-INS-204 |
| chr14-48806099-INS-838 | chr14-48806099-INS-838 |
| chr14-48811480-INS-206 | chr14-48811480-INS-207 |
| chr14-48841417-INS-425 | chr14-48841417-INS-425 |
| chr14-48843488-INS-80 | chr14-48843488-INS-80 |
| chr14-48881227-INS-493 | chr14-48881212-INS-494 |
| chr14-48918942-INS-69 | chr14-48918923-INS-69 |
| chr14-48926321-INS-170 | chr14-48926321-INS-181 |
| chr14-48934178-INS-54 | chr14-48934178-INS-57 |
| chr14-48954187-INS-230 | chr14-48954187-INS-230 |
| chr14-48964953-INS-96 | chr14-48964953-INS-128 |
| chr14-48971163-INS-84 | chr14-48971163-INS-84 |
| chr14-48983498-INS-70 | chr14-48983496-INS-94 |
| chr14-48998305-INS-59 | chr14-48998306-INS-72 |
| chr14-49005525-INS-59 | chr14-49005556-INS-66 |
| chr14-49007196-INS-59 | chr14-49007196-INS-64 |
| chr14-49015000-INS-270 | chr14-49015000-INS-268 |
| chr14-49016716-INS-206 | chr14-49016716-INS-206 |
| chr14-49041274-INS-116 | chr14-49041274-INS-126 |
| chr14-49044037-INS-497 | chr14-49044037-INS-497 |
| chr14-49068106-INS-88 | chr14-49068106-INS-88 |
| chr14-49078579-INS-54 | chr14-49078579-INS-52 |
| chr14-49092849-INS-59 | chr14-49092848-INS-53 |
| chr14-49100977-INS-713 | chr14-49100977-INS-716 |
| chr14-49110071-INS-400 | chr14-49110071-INS-400 |
| chr14-49134798-INS-1216 | chr14-49134799-INS-1216 |
| chr14-49134799-INS-418 | chr14-49134799-INS-418 |
| chr14-49190138-INS-54 | chr14-49190138-INS-60 |
| chr14-49192808-INS-68 | chr14-49192808-INS-68 |
| chr14-49199431-INS-205 | chr14-49199431-INS-205 |
| chr14-49218433-INS-215 | chr14-49218433-INS-216 |
| chr14-49222077-INS-66 | chr14-49222077-INS-62 |
| chr14-49349098-INS-116 | chr14-49349098-INS-116 |
| chr14-49351846-INS-664 | chr14-49351846-INS-664 |
| chr14-49364216-INS-214 | chr14-49364216-INS-214 |
| chr14-49409229-INS-153 | chr14-49409243-INS-164 |
| chr14-49414217-INS-204 | chr14-49414217-INS-204 |
| chr14-49451616-INS-83 | chr14-49451616-INS-83 |
| chr14-49454494-INS-63 | chr14-49454494-INS-65 |
| chr14-49456600-INS-68 | chr14-49456593-INS-69 |
| chr14-49483368-INS-1280 | chr14-49483369-INS-1281 |
| chr14-49503720-INS-189 | chr14-49503720-INS-190 |
| chr14-49506980-INS-104 | chr14-49506980-INS-106 |
| chr14-49509302-INS-272 | chr14-49509302-INS-277 |
| chr14-49525087-INS-143 | chr14-49525087-INS-143 |
| chr14-49528222-INS-1388 | chr14-49528223-INS-1388 |
| chr14-49589165-INS-573 | chr14-49589165-INS-578 |
| chr14-49608127-INS-6441 | chr14-49608128-INS-6441 |
| chr14-49620165-INS-6925 | chr14-49620166-INS-6925 |
| chr14-49646384-INS-53 | chr14-49646384-INS-53 |
| chr14-49669611-INS-64 | chr14-49669611-INS-64 |
| chr14-49686887-INS-109 | chr14-49686887-INS-120 |
| chr14-49695430-INS-148 | chr14-49695448-INS-154 |

| | |
|---|---|
| chr14-49704749-INS-55 | chr14-49704743-INS-56 |
| chr14-49711240-INS-208 | chr14-49711240-INS-208 |
| chr14-49719382-INS-62 | chr14-49719382-INS-62 |
| chr14-49729208-INS-363 | chr14-49729208-INS-378 |
| chr14-49747188-INS-127 | chr14-49747188-INS-127 |
| chr14-49771425-INS-171 | chr14-49771425-INS-171 |
| chr14-49808994-INS-4363 | chr14-49808995-INS-4363 |
| chr14-49809601-INS-64 | chr14-49809601-INS-63 |
| chr14-49810128-INS-6543 | chr14-49810129-INS-6543 |
| chr14-49813105-INS-242 | chr14-49813105-INS-240 |
| chr14-49860121-INS-194 | chr14-49860114-INS-194 |
| chr14-49875952-INS-2009 | chr14-49875953-INS-2009 |
| chr14-49890520-INS-7092 | chr14-49890521-INS-7092 |
| chr14-49901242-INS-55 | chr14-49901242-INS-65 |
| chr14-49950972-INS-553 | chr14-49950972-INS-553 |
| chr14-49966421-INS-66 | chr14-49966422-INS-56 |
| chr14-49974978-INS-82 | chr14-49974978-INS-88 |
| chr14-49991929-INS-56 | chr14-49991941-INS-71 |
| chr14-49995744-INS-156 | chr14-49995744-INS-156 |
| chr14-50048609-INS-267 | chr14-50048609-INS-267 |
| chr14-50077373-INS-173 | chr14-50077373-INS-173 |
| chr14-50093536-INS-199 | chr14-50093537-INS-199 |
| chr14-50094567-INS-211 | chr14-50094551-INS-230 |
| chr14-50144539-INS-985 | chr14-50144539-INS-985 |
| chr14-50182721-INS-1947 | chr14-50182722-INS-1947 |
| chr14-50188517-INS-6396 | chr14-50188518-INS-6396 |
| chr14-50189736-INS-553 | chr14-50189737-INS-551 |
| chr14-50200588-INS-52 | chr14-50200588-INS-52 |
| chr14-50201215-INS-7072 | chr14-50201216-INS-7072 |
| chr14-50210544-INS-72 | chr14-50210544-INS-75 |
| chr14-50212933-INS-221 | chr14-50212969-INS-235 |
| chr14-50256214-INS-204 | chr14-50256214-INS-204 |
| chr14-50266461-INS-231 | chr14-50266461-INS-225 |
| chr14-50289240-INS-214 | chr14-50289240-INS-214 |
| chr14-50382781-INS-921 | chr14-50382781-INS-920 |
| chr14-50427307-INS-4853 | chr14-50427308-INS-4853 |
| chr14-50437661-INS-116 | chr14-50437661-INS-116 |
| chr14-50471627-INS-51 | chr14-50471627-INS-51 |
| chr14-50555329-INS-51 | chr14-50555329-INS-52 |
| chr14-50578277-INS-52 | chr14-50578277-INS-52 |
| chr14-50579537-INS-3204 | chr14-50579538-INS-3184 |
| chr14-50584005-INS-1095 | chr14-50584006-INS-1096 |
| chr14-50586581-INS-1203 | chr14-50586582-INS-1203 |
| chr14-50590651-INS-102 | chr14-50590651-INS-102 |
| chr14-50593411-INS-534 | chr14-50593411-INS-534 |
| chr14-50593957-INS-6497 | chr14-50593958-INS-6497 |
| chr14-50620461-INS-69 | chr14-50620461-INS-75 |
| chr14-50627244-INS-846 | chr14-50627244-INS-846 |
| chr14-50647687-INS-102 | chr14-50647687-INS-102 |
| chr14-50663298-INS-66 | chr14-50663328-INS-51 |
| chr14-50666471-INS-6348 | chr14-50666472-INS-6345 |
| chr14-50681723-INS-10820 | chr14-50681724-INS-10821 |
| chr14-50692959-INS-196 | chr14-50692959-INS-196 |
| chr14-50701362-INS-498 | chr14-50701362-INS-498 |
| chr14-50712475-INS-230 | chr14-50712475-INS-231 |
| chr14-50768453-INS-492 | chr14-50768453-INS-492 |
| chr14-50771843-INS-115 | chr14-50771843-INS-115 |
| chr14-50782313-INS-435 | chr14-50782313-INS-435 |
| chr14-50793844-INS-1116 | chr14-50793845-INS-1116 |
| chr14-50802491-INS-60 | chr14-50802491-INS-60 |
| chr14-50833143-INS-202 | chr14-50833143-INS-202 |
| chr14-50874138-INS-11845 | chr14-50874139-INS-11815 |
| chr14-50880883-INS-588 | chr14-50880883-INS-588 |
| chr14-50885194-INS-57 | chr14-50885194-INS-60 |
| chr14-50913680-INS-353 | chr14-50913680-INS-353 |
| chr14-50984781-INS-66 | chr14-50984781-INS-66 |
| chr14-50995101-INS-346 | chr14-50995090-INS-355 |
| chr14-51001975-INS-85 | chr14-51001975-INS-85 |
| chr14-51006337-INS-55 | chr14-51006337-INS-80 |
| chr14-51016983-INS-211 | chr14-51016983-INS-211 |
| chr14-51035813-INS-3828 | chr14-51035814-INS-3828 |
| chr14-51037169-INS-895 | chr14-51037169-INS-895 |
| chr14-51047346-INS-205 | chr14-51047346-INS-203 |
| chr14-51083368-INS-214 | chr14-51083368-INS-214 |
| chr14-51094597-INS-73 | chr14-51094597-INS-73 |
| chr14-51100903-INS-6442 | chr14-51100904-INS-6442 |
| chr14-51110286-INS-655 | chr14-51110286-INS-666 |

| | |
|---|---|
| chr14-51128283-INS-60 | chr14-51128262-INS-92 |
| chr14-51136606-INS-193 | chr14-51136606-INS-195 |
| chr14-51137654-INS-4847 | chr14-51137655-INS-4847 |
| chr14-51140532-INS-271 | chr14-51140532-INS-268 |
| chr14-51159579-INS-6595 | chr14-51159580-INS-6595 |
| chr14-51173102-INS-169 | chr14-51173107-INS-173 |
| chr14-51195256-INS-278 | chr14-51195256-INS-278 |
| chr14-51197215-INS-216 | chr14-51197215-INS-216 |
| chr14-51220143-INS-204 | chr14-51220143-INS-204 |
| chr14-51233681-INS-104 | chr14-51233682-INS-108 |
| chr14-51236539-INS-445 | chr14-51236539-INS-445 |
| chr14-51252872-INS-73 | chr14-51252872-INS-74 |
| chr14-51262365-INS-234 | chr14-51262365-INS-234 |
| chr14-51266450-INS-195 | chr14-51266450-INS-196 |
| chr14-51274811-INS-93 | chr14-51274811-INS-93 |
| chr14-51282337-INS-174 | chr14-51282337-INS-174 |
| chr14-51296005-INS-411 | chr14-51296005-INS-411 |
| chr14-51316132-INS-8066 | chr14-51316133-INS-8068 |
| chr14-51322698-INS-133 | chr14-51322698-INS-133 |
| chr14-51338617-INS-200 | chr14-51338617-INS-200 |
| chr14-51359646-INS-141 | chr14-51359691-INS-108 |
| chr14-51370574-INS-223 | chr14-51370566-INS-230 |
| chr14-51406620-INS-6403 | chr14-51406621-INS-6402 |
| chr14-51449987-INS-6779 | chr14-51449988-INS-6779 |
| chr14-51455729-INS-78 | chr14-51455729-INS-78 |
| chr14-51467922-INS-621 | chr14-51467922-INS-621 |
| chr14-51470649-INS-6159 | chr14-51470650-INS-6160 |
| chr14-51502382-INS-922 | chr14-51502382-INS-922 |
| chr14-51547404-INS-79 | chr14-51547404-INS-80 |
| chr14-51561965-INS-932 | chr14-51561965-INS-932 |
| chr14-51611362-INS-191 | chr14-51611362-INS-191 |
| chr14-51635309-INS-149 | chr14-51635309-INS-149 |
| chr14-51636914-INS-5157 | chr14-51636913-INS-5158 |
| chr14-51674317-INS-56 | chr14-51674312-INS-56 |
| chr14-51675650-INS-261 | chr14-51675650-INS-270 |
| chr14-51692875-INS-66 | chr14-51692847-INS-62 |
| chr14-51712023-INS-50 | chr14-51712023-INS-52 |
| chr14-51713810-INS-6485 | chr14-51713811-INS-6485 |
| chr14-51756285-INS-64 | chr14-51756285-INS-66 |
| chr14-51775101-INS-805 | chr14-51775101-INS-808 |
| chr14-51795599-INS-485 | chr14-51795599-INS-485 |
| chr14-51813005-INS-1110 | chr14-51813006-INS-1106 |
| chr14-51867356-INS-206 | chr14-51867356-INS-206 |
| chr14-51951435-INS-498 | chr14-51951435-INS-498 |
| chr14-52030579-INS-195 | chr14-52030579-INS-195 |
| chr14-52041933-INS-1839 | chr14-52041934-INS-1839 |
| chr14-52081149-INS-196 | chr14-52081150-INS-190 |
| chr14-52096295-INS-341 | chr14-52096294-INS-340 |
| chr14-52110070-INS-135 | chr14-52110070-INS-135 |
| chr14-52121934-INS-69 | chr14-52121934-INS-69 |
| chr14-52137192-INS-235 | chr14-52137192-INS-235 |
| chr14-52142544-INS-415 | chr14-52142588-INS-415 |
| chr14-52163273-INS-400 | chr14-52163273-INS-400 |
| chr14-52170892-INS-200 | chr14-52170892-INS-200 |
| chr14-52172941-INS-218 | chr14-52172941-INS-218 |
| chr14-52185308-INS-497 | chr14-52185308-INS-497 |
| chr14-52186677-INS-202 | chr14-52186677-INS-202 |
| chr14-52196480-INS-117 | chr14-52196461-INS-92 |
| chr14-52200492-INS-204 | chr14-52200492-INS-204 |
| chr14-52210803-INS-66 | chr14-52210796-INS-90 |
| chr14-52213036-INS-53 | chr14-52213036-INS-53 |
| chr14-52214670-INS-120 | chr14-52214668-INS-118 |
| chr14-52226042-INS-52 | chr14-52226042-INS-56 |
| chr14-52235859-INS-54 | chr14-52235859-INS-54 |
| chr14-52259318-INS-287 | chr14-52259318-INS-287 |
| chr14-52271194-INS-98 | chr14-52271194-INS-98 |
| chr14-52292802-INS-195 | chr14-52292810-INS-195 |
| chr14-52297074-INS-207 | chr14-52297087-INS-208 |
| chr14-52304315-INS-538 | chr14-52304315-INS-538 |
| chr14-52328010-INS-208 | chr14-52328010-INS-208 |
| chr14-52331683-INS-172 | chr14-52331683-INS-172 |
| chr14-52365315-INS-193 | chr14-52365315-INS-193 |
| chr14-52368476-INS-212 | chr14-52368476-INS-212 |
| chr14-52370615-INS-176 | chr14-52370615-INS-176 |
| chr14-52375183-INS-4221 | chr14-52375184-INS-4221 |
| chr14-52386323-INS-181 | chr14-52386323-INS-182 |
| chr14-52393310-INS-217 | chr14-52393310-INS-216 |

| | |
|---|---|
| chr14-52395131-INS-58 | chr14-52395131-INS-58 |
| chr14-52401051-INS-220 | chr14-52401051-INS-220 |
| chr14-52420212-INS-209 | chr14-52420212-INS-209 |
| chr14-52439253-INS-255 | chr14-52439253-INS-255 |
| chr14-52477197-INS-988 | chr14-52477197-INS-997 |
| chr14-52478354-INS-255 | chr14-52478354-INS-255 |
| chr14-52498008-INS-60 | chr14-52498008-INS-60 |
| chr14-52506026-INS-218 | chr14-52506026-INS-218 |
| chr14-52512804-INS-187 | chr14-52512804-INS-187 |
| chr14-52513506-INS-218 | chr14-52513506-INS-218 |
| chr14-52522968-INS-61 | chr14-52522974-INS-62 |
| chr14-52572120-INS-176 | chr14-52572120-INS-176 |
| chr14-52584800-INS-223 | chr14-52584799-INS-196 |
| chr14-52589930-INS-402 | chr14-52589930-INS-401 |
| chr14-52594563-INS-532 | chr14-52594563-INS-532 |
| chr14-52598713-INS-219 | chr14-52598713-INS-218 |
| chr14-52600185-INS-186 | chr14-52600185-INS-186 |
| chr14-52625418-INS-187 | chr14-52625418-INS-187 |
| chr14-52642183-INS-106 | chr14-52642178-INS-109 |
| chr14-52656411-INS-214 | chr14-52656411-INS-214 |
| chr14-52690587-INS-79 | chr14-52690587-INS-79 |
| chr14-52717436-INS-70 | chr14-52717436-INS-68 |
| chr14-52719451-INS-177 | chr14-52719451-INS-177 |
| chr14-52721704-INS-288 | chr14-52721704-INS-288 |
| chr14-52726001-INS-508 | chr14-52726001-INS-508 |
| chr14-52734146-INS-483 | chr14-52734146-INS-483 |
| chr14-52801778-INS-86 | chr14-52801780-INS-114 |
| chr14-52817402-INS-54 | chr14-52817402-INS-55 |
| chr14-52821847-INS-117 | chr14-52821847-INS-117 |
| chr14-52841557-INS-2612 | chr14-52841558-INS-2612 |
| chr14-52850779-INS-9063 | chr14-52850780-INS-9063 |
| chr14-52850887-INS-75 | chr14-52850887-INS-75 |
| chr14-52918436-INS-639 | chr14-52918435-INS-637 |
| chr14-52935825-INS-351 | chr14-52935825-INS-351 |
| chr14-52946906-INS-70 | chr14-52946906-INS-70 |
| chr14-52948285-INS-68 | chr14-52948279-INS-78 |
| chr14-52959090-INS-1060 | chr14-52959091-INS-1060 |
| chr14-52989899-INS-613 | chr14-52989899-INS-613 |
| chr14-53016834-INS-68 | chr14-53016834-INS-66 |
| chr14-53031697-INS-219 | chr14-53031697-INS-221 |
| chr14-53044738-INS-374 | chr14-53044738-INS-374 |
| chr14-53062893-INS-348 | chr14-53062893-INS-349 |
| chr14-53280426-INS-267 | chr14-53280427-INS-267 |
| chr14-53549679-INS-212 | chr14-53549679-INS-211 |
| chr14-53553112-INS-206 | chr14-53553112-INS-205 |
| chr14-53560147-INS-154 | chr14-53560147-INS-160 |
| chr14-53561153-INS-70 | chr14-53561153-INS-72 |
| chr14-53561938-INS-667 | chr14-53561938-INS-668 |
| chr14-53568093-INS-54 | chr14-53568093-INS-54 |
| chr14-53590261-INS-705 | chr14-53590261-INS-705 |
| chr14-53592279-INS-127 | chr14-53592279-INS-120 |
| chr14-53610893-INS-97 | chr14-53610893-INS-97 |
| chr14-53723623-INS-71 | chr14-53723623-INS-75 |
| chr14-53724688-INS-53 | chr14-53724663-INS-99 |
| chr14-53733841-INS-263 | chr14-53733841-INS-225 |
| chr14-53751335-INS-187 | chr14-53751335-INS-188 |
| chr14-53834978-INS-56 | chr14-53834978-INS-56 |
| chr14-53838869-INS-60 | chr14-53838848-INS-90 |
| chr14-53852945-INS-432 | chr14-53852945-INS-432 |
| chr14-53863672-INS-83 | chr14-53863626-INS-116 |
| chr14-53996433-INS-114 | chr14-53996433-INS-83 |
| chr14-53997763-INS-345 | chr14-53997763-INS-347 |
| chr14-54002736-INS-97 | chr14-54002736-INS-97 |
| chr14-54027619-INS-361 | chr14-54027619-INS-361 |
| chr14-54031544-INS-196 | chr14-54031544-INS-196 |
| chr14-54042285-INS-1306 | chr14-54042286-INS-1306 |
| chr14-54074414-INS-350 | chr14-54074414-INS-353 |
| chr14-54084919-INS-52 | chr14-54084919-INS-52 |
| chr14-54088602-INS-458 | chr14-54088639-INS-458 |
| chr14-54089877-INS-542 | chr14-54089877-INS-542 |
| chr14-54115730-INS-445 | chr14-54115730-INS-445 |
| chr14-54216831-INS-7128 | chr14-54216832-INS-7128 |
| chr14-54239946-INS-84 | chr14-54239946-INS-76 |
| chr14-54246262-INS-360 | chr14-54246262-INS-360 |
| chr14-54253021-INS-691 | chr14-54253021-INS-691 |
| chr14-54261540-INS-531 | chr14-54261540-INS-531 |
| chr14-54276269-INS-6672 | chr14-54276270-INS-6672 |

| | |
|---|---|
| chr14-54277206-INS-1492 | chr14-54277207-INS-1492 |
| chr14-54314830-INS-98 | chr14-54314830-INS-64 |
| chr14-54326823-INS-204 | chr14-54326823-INS-204 |
| chr14-54347337-INS-74 | chr14-54347330-INS-77 |
| chr14-54382018-INS-80 | chr14-54382018-INS-80 |
| chr14-54395378-INS-60 | chr14-54395378-INS-60 |
| chr14-54399460-INS-53 | chr14-54399460-INS-53 |
| chr14-54426245-INS-78 | chr14-54426245-INS-78 |
| chr14-54487174-INS-56 | chr14-54487174-INS-56 |
| chr14-54504173-INS-66 | chr14-54504173-INS-66 |
| chr14-54514921-INS-184 | chr14-54514921-INS-184 |
| chr14-54517221-INS-183 | chr14-54517221-INS-183 |
| chr14-54523191-INS-59 | chr14-54523191-INS-59 |
| chr14-54527053-INS-291 | chr14-54527053-INS-291 |
| chr14-54540034-INS-199 | chr14-54540034-INS-199 |
| chr14-54560488-INS-92 | chr14-54560488-INS-86 |
| chr14-54602709-INS-204 | chr14-54602709-INS-204 |
| chr14-54609370-INS-54 | chr14-54609370-INS-58 |
| chr14-54618890-INS-83 | chr14-54618890-INS-83 |
| chr14-54622786-INS-107 | chr14-54622786-INS-100 |
| chr14-54647082-INS-166 | chr14-54647082-INS-166 |
| chr14-54655896-INS-217 | chr14-54655896-INS-217 |
| chr14-54656895-INS-203 | chr14-54656895-INS-203 |
| chr14-54684817-INS-54 | chr14-54684865-INS-52 |
| chr14-54694958-INS-219 | chr14-54694958-INS-219 |
| chr14-54698822-INS-298 | chr14-54698822-INS-302 |
| chr14-54711356-INS-124 | chr14-54711356-INS-126 |
| chr14-54727063-INS-52 | chr14-54727063-INS-64 |
| chr14-54727635-INS-97 | chr14-54727635-INS-99 |
| chr14-54729905-INS-66 | chr14-54729905-INS-66 |
| chr14-54739351-INS-182 | chr14-54739351-INS-184 |
| chr14-54742431-INS-214 | chr14-54742431-INS-214 |
| chr14-54751617-INS-200 | chr14-54751617-INS-200 |
| chr14-54753073-INS-51 | chr14-54753073-INS-51 |
| chr14-54756783-INS-381 | chr14-54756830-INS-353 |
| chr14-54760925-INS-104 | chr14-54760926-INS-104 |
| chr14-54789529-INS-179 | chr14-54789529-INS-177 |
| chr14-54792315-INS-209 | chr14-54792315-INS-209 |
| chr14-54795944-INS-62 | chr14-54795944-INS-62 |
| chr14-54818017-INS-188 | chr14-54818017-INS-188 |
| chr14-54827003-INS-330 | chr14-54827003-INS-338 |
| chr14-54894996-INS-68 | chr14-54894996-INS-68 |
| chr14-54909576-INS-52 | chr14-54909576-INS-54 |
| chr14-54913706-INS-54 | chr14-54913706-INS-54 |
| chr14-54935508-INS-51 | chr14-54935508-INS-59 |
| chr14-54938551-INS-210 | chr14-54938551-INS-210 |
| chr14-54942804-INS-204 | chr14-54942804-INS-204 |
| chr14-54960250-INS-222 | chr14-54960250-INS-222 |
| chr14-54965379-INS-55 | chr14-54965379-INS-55 |
| chr14-54982823-INS-101 | chr14-54982823-INS-101 |
| chr14-55027531-INS-81 | chr14-55027534-INS-72 |
| chr14-55045735-INS-184 | chr14-55045735-INS-184 |
| chr14-55078829-INS-217 | chr14-55078829-INS-217 |
| chr14-55080512-INS-219 | chr14-55080512-INS-219 |
| chr14-55082625-INS-1149 | chr14-55082626-INS-1149 |
| chr14-55083330-INS-98 | chr14-55083330-INS-98 |
| chr14-55085032-INS-202 | chr14-55085033-INS-177 |
| chr14-55088076-INS-176 | chr14-55088076-INS-176 |
| chr14-55110163-INS-2775 | chr14-55110164-INS-2775 |
| chr14-55112292-INS-67 | chr14-55112292-INS-67 |
| chr14-55118918-INS-7114 | chr14-55118919-INS-7114 |
| chr14-55127093-INS-151 | chr14-55127093-INS-149 |
| chr14-55187691-INS-237 | chr14-55187691-INS-236 |
| chr14-55245374-INS-205 | chr14-55245374-INS-191 |
| chr14-55249163-INS-206 | chr14-55249164-INS-206 |
| chr14-55249884-INS-217 | chr14-55249884-INS-218 |
| chr14-55263713-INS-52 | chr14-55263713-INS-54 |
| chr14-55281965-INS-60 | chr14-55281965-INS-50 |
| chr14-55316621-INS-66 | chr14-55316621-INS-66 |
| chr14-55324927-INS-202 | chr14-55324927-INS-202 |
| chr14-55370407-INS-184 | chr14-55370407-INS-184 |
| chr14-55393664-INS-75 | chr14-55393684-INS-75 |
| chr14-55399268-INS-53 | chr14-55399264-INS-57 |
| chr14-55429314-INS-2885 | chr14-55429315-INS-2885 |
| chr14-55433383-INS-150 | chr14-55433383-INS-150 |
| chr14-55475616-INS-60 | chr14-55475592-INS-80 |
| chr14-55479650-INS-465 | chr14-55479650-INS-465 |

| | |
|---|---|
| chr14-55491708-INS-62 | chr14-55491708-INS-62 |
| chr14-55530708-INS-245 | chr14-55530708-INS-245 |
| chr14-55541257-INS-6402 | chr14-55541258-INS-6402 |
| chr14-55545620-INS-4292 | chr14-55545621-INS-4299 |
| chr14-55577288-INS-591 | chr14-55577288-INS-592 |
| chr14-55591163-INS-95 | chr14-55591163-INS-95 |
| chr14-55596390-INS-7505 | chr14-55596391-INS-7505 |
| chr14-55598416-INS-498 | chr14-55598416-INS-498 |
| chr14-55649478-INS-62 | chr14-55649478-INS-56 |
| chr14-55660398-INS-64 | chr14-55660398-INS-74 |
| chr14-55740527-INS-66 | chr14-55740527-INS-86 |
| chr14-55770124-INS-53 | chr14-55770124-INS-53 |
| chr14-55808929-INS-209 | chr14-55808929-INS-200 |
| chr14-55820088-INS-230 | chr14-55820088-INS-218 |
| chr14-55820713-INS-223 | chr14-55820713-INS-224 |
| chr14-55883278-INS-469 | chr14-55883272-INS-426 |
| chr14-55891188-INS-183 | chr14-55891188-INS-184 |
| chr14-55891744-INS-116 | chr14-55891744-INS-129 |
| chr14-55964372-INS-219 | chr14-55964372-INS-207 |
| chr14-55969305-INS-69 | chr14-55969305-INS-75 |
| chr14-55974506-INS-220 | chr14-55974506-INS-267 |
| chr14-55989905-INS-232 | chr14-55989905-INS-233 |
| chr14-56030559-INS-240 | chr14-56030559-INS-242 |
| chr14-56072214-INS-288 | chr14-56072214-INS-288 |
| chr14-56073441-INS-214 | chr14-56073441-INS-214 |
| chr14-56078805-INS-52 | chr14-56078819-INS-50 |
| chr14-56086603-INS-196 | chr14-56086603-INS-196 |
| chr14-56114610-INS-7125 | chr14-56114611-INS-7110 |
| chr14-56184265-INS-60 | chr14-56184265-INS-60 |
| chr14-56212234-INS-460 | chr14-56212234-INS-460 |
| chr14-56213721-INS-325 | chr14-56213721-INS-325 |
| chr14-56227597-INS-60 | chr14-56227597-INS-64 |
| chr14-56305534-INS-6195 | chr14-56305535-INS-6195 |
| chr14-56306042-INS-604 | chr14-56306042-INS-607 |
| chr14-56313220-INS-495 | chr14-56313220-INS-495 |
| chr14-56327380-INS-6549 | chr14-56327381-INS-6549 |
| chr14-56328014-INS-1717 | chr14-56328015-INS-1717 |
| chr14-56335086-INS-54 | chr14-56335086-INS-54 |
| chr14-56353817-INS-62 | chr14-56353817-INS-58 |
| chr14-56370350-INS-51 | chr14-56370350-INS-55 |
| chr14-56411064-INS-58 | chr14-56411064-INS-58 |
| chr14-56418625-INS-51 | chr14-56418625-INS-51 |
| chr14-56420642-INS-61 | chr14-56420642-INS-61 |
| chr14-56450713-INS-495 | chr14-56450713-INS-495 |
| chr14-56511957-INS-109 | chr14-56511957-INS-105 |
| chr14-56514221-INS-82 | chr14-56514221-INS-85 |
| chr14-56517548-INS-139 | chr14-56517548-INS-144 |
| chr14-56520390-INS-52 | chr14-56520390-INS-56 |
| chr14-56530763-INS-517 | chr14-56530763-INS-520 |
| chr14-56549653-INS-60 | chr14-56549686-INS-60 |
| chr14-56552366-INS-74 | chr14-56552366-INS-102 |
| chr14-56553406-INS-77 | chr14-56553406-INS-77 |
| chr14-56560217-INS-53 | chr14-56560217-INS-53 |
| chr14-56565233-INS-52 | chr14-56565230-INS-53 |
| chr14-56567439-INS-65 | chr14-56567439-INS-65 |
| chr14-56571034-INS-50 | chr14-56571030-INS-68 |
| chr14-56579660-INS-277 | chr14-56579660-INS-291 |
| chr14-56584733-INS-1862 | chr14-56584734-INS-1862 |
| chr14-56601467-INS-73 | chr14-56601467-INS-73 |
| chr14-56611405-INS-92 | chr14-56611405-INS-93 |
| chr14-56612750-INS-94 | chr14-56612750-INS-94 |
| chr14-56634723-INS-224 | chr14-56634723-INS-230 |
| chr14-56685972-INS-436 | chr14-56685972-INS-436 |
| chr14-56711567-INS-120 | chr14-56711567-INS-120 |
| chr14-56756107-INS-284 | chr14-56756107-INS-280 |
| chr14-56781960-INS-59 | chr14-56781960-INS-59 |
| chr14-56784303-INS-120 | chr14-56784303-INS-120 |
| chr14-56794257-INS-423 | chr14-56794257-INS-424 |
| chr14-56797790-INS-150 | chr14-56797790-INS-120 |
| chr14-56832250-INS-217 | chr14-56832250-INS-218 |
| chr14-56876596-INS-188 | chr14-56876584-INS-188 |
| chr14-56887521-INS-59 | chr14-56887522-INS-59 |
| chr14-56889309-INS-6769 | chr14-56889310-INS-6770 |
| chr14-56896070-INS-219 | chr14-56896070-INS-220 |
| chr14-56905130-INS-122 | chr14-56905130-INS-135 |
| chr14-56917434-INS-287 | chr14-56917434-INS-287 |
| chr14-56946248-INS-96 | chr14-56946271-INS-72 |

| | |
|---|---|
| chr14-56974405-INS-95 | chr14-56974405-INS-91 |
| chr14-57003440-INS-192 | chr14-57003440-INS-192 |
| chr14-57006937-INS-219 | chr14-57006937-INS-219 |
| chr14-57029250-INS-207 | chr14-57029250-INS-207 |
| chr14-57037919-INS-56 | chr14-57037919-INS-52 |
| chr14-57074469-INS-7182 | chr14-57074470-INS-7182 |
| chr14-57085083-INS-124 | chr14-57085083-INS-122 |
| chr14-57092724-INS-213 | chr14-57092724-INS-214 |
| chr14-57106612-INS-2916 | chr14-57106613-INS-2927 |
| chr14-57211610-INS-265 | chr14-57211610-INS-265 |
| chr14-57347141-INS-824 | chr14-57347141-INS-821 |
| chr14-57354819-INS-83 | chr14-57354819-INS-131 |
| chr14-57400376-INS-207 | chr14-57400376-INS-207 |
| chr14-57409946-INS-354 | chr14-57409946-INS-354 |
| chr14-57417602-INS-132 | chr14-57417602-INS-108 |
| chr14-57422042-INS-72 | chr14-57422034-INS-90 |
| chr14-57434892-INS-663 | chr14-57434892-INS-664 |
| chr14-57467066-INS-64 | chr14-57467066-INS-64 |
| chr14-57475654-INS-1819 | chr14-57475655-INS-1820 |
| chr14-57479271-INS-68 | chr14-57479271-INS-68 |
| chr14-57490577-INS-52 | chr14-57490577-INS-52 |
| chr14-57496185-INS-88 | chr14-57496186-INS-91 |
| chr14-57496712-INS-114 | chr14-57496702-INS-76 |
| chr14-57497689-INS-56 | chr14-57497689-INS-56 |
| chr14-57519481-INS-52 | chr14-57519440-INS-80 |
| chr14-57546125-INS-365 | chr14-57546134-INS-369 |
| chr14-57626488-INS-121 | chr14-57626488-INS-121 |
| chr14-57695075-INS-198 | chr14-57695075-INS-203 |
| chr14-57738095-INS-116 | chr14-57738095-INS-118 |
| chr14-57740358-INS-205 | chr14-57740358-INS-205 |
| chr14-57769763-INS-128 | chr14-57769763-INS-124 |
| chr14-57779499-INS-81 | chr14-57779499-INS-81 |
| chr14-57782982-INS-1248 | chr14-57782983-INS-1252 |
| chr14-57795363-INS-107 | chr14-57795363-INS-107 |
| chr14-57808364-INS-88 | chr14-57808364-INS-96 |
| chr14-57809326-INS-204 | chr14-57809326-INS-204 |
| chr14-57811669-INS-233 | chr14-57811669-INS-233 |
| chr14-57831527-INS-88 | chr14-57831527-INS-86 |
| chr14-57833142-INS-256 | chr14-57833142-INS-256 |
| chr14-57845745-INS-74 | chr14-57845745-INS-74 |
| chr14-57886847-INS-84 | chr14-57886845-INS-58 |
| chr14-57900370-INS-180 | chr14-57900370-INS-180 |
| chr14-57908108-INS-210 | chr14-57908108-INS-210 |
| chr14-57952058-INS-62 | chr14-57952056-INS-66 |
| chr14-57958265-INS-204 | chr14-57958265-INS-204 |
| chr14-57960131-INS-470 | chr14-57960133-INS-469 |
| chr14-57970665-INS-531 | chr14-57970665-INS-531 |
| chr14-57980327-INS-93 | chr14-57980327-INS-93 |
| chr14-58010481-INS-210 | chr14-58010481-INS-211 |
| chr14-58012600-INS-221 | chr14-58012600-INS-221 |
| chr14-58012868-INS-529 | chr14-58012868-INS-529 |
| chr14-58019131-INS-3204 | chr14-58019132-INS-3202 |
| chr14-58023245-INS-111 | chr14-58023245-INS-111 |
| chr14-58030857-INS-207 | chr14-58030857-INS-207 |
| chr14-58034830-INS-69 | chr14-58034830-INS-83 |
| chr14-58053301-INS-70 | chr14-58053328-INS-52 |
| chr14-58077839-INS-213 | chr14-58077839-INS-213 |
| chr14-58091130-INS-210 | chr14-58091130-INS-210 |
| chr14-58099091-INS-214 | chr14-58099091-INS-215 |
| chr14-58106065-INS-202 | chr14-58106065-INS-202 |
| chr14-58116027-INS-935 | chr14-58116027-INS-935 |
| chr14-58118369-INS-64 | chr14-58118369-INS-64 |
| chr14-58119278-INS-2402 | chr14-58119279-INS-2402 |
| chr14-58120186-INS-73 | chr14-58120186-INS-73 |
| chr14-58140317-INS-54 | chr14-58140317-INS-56 |
| chr14-58152865-INS-75 | chr14-58152865-INS-75 |
| chr14-58165783-INS-6444 | chr14-58165784-INS-6444 |
| chr14-58171699-INS-2003 | chr14-58171700-INS-2004 |
| chr14-58178213-INS-208 | chr14-58178213-INS-207 |
| chr14-58192660-INS-64 | chr14-58192660-INS-64 |
| chr14-58205325-INS-156 | chr14-58205325-INS-156 |
| chr14-58230064-INS-72 | chr14-58230064-INS-72 |
| chr14-58260865-INS-6369 | chr14-58260866-INS-6370 |
| chr14-58267095-INS-2812 | chr14-58267096-INS-2812 |
| chr14-58285890-INS-1415 | chr14-58285891-INS-1415 |
| chr14-58342041-INS-54 | chr14-58342041-INS-54 |
| chr14-58367082-INS-98 | chr14-58367068-INS-64 |

| | |
|---|---|
| chr14-58388573-INS-7078 | chr14-58388574-INS-7079 |
| chr14-58399631-INS-542 | chr14-58399631-INS-542 |
| chr14-58421580-INS-72 | chr14-58421576-INS-60 |
| chr14-58428631-INS-134 | chr14-58428631-INS-134 |
| chr14-58449739-INS-54 | chr14-58449739-INS-56 |
| chr14-58498191-INS-492 | chr14-58498191-INS-482 |
| chr14-58505737-INS-918 | chr14-58505737-INS-947 |
| chr14-58550138-INS-80 | chr14-58550138-INS-80 |
| chr14-58562284-INS-112 | chr14-58562284-INS-112 |
| chr14-58618215-INS-84 | chr14-58618215-INS-84 |
| chr14-58652940-INS-80 | chr14-58652940-INS-80 |
| chr14-58790903-INS-338 | chr14-58790903-INS-338 |
| chr14-58795628-INS-72 | chr14-58795628-INS-72 |
| chr14-58798918-INS-1892 | chr14-58798919-INS-1892 |
| chr14-58855142-INS-76 | chr14-58855142-INS-76 |
| chr14-58863069-INS-65 | chr14-58863073-INS-55 |
| chr14-58863990-INS-2833 | chr14-58863991-INS-2833 |
| chr14-58967847-INS-153 | chr14-58967847-INS-153 |
| chr14-58968943-INS-64 | chr14-58968940-INS-68 |
| chr14-58989378-INS-544 | chr14-58989373-INS-547 |
| chr14-59055149-INS-51 | chr14-59055149-INS-51 |
| chr14-59057200-INS-6692 | chr14-59057201-INS-6692 |
| chr14-59065741-INS-1111 | chr14-59065742-INS-1119 |
| chr14-59075226-INS-172 | chr14-59075226-INS-166 |
| chr14-59089043-INS-1183 | chr14-59089044-INS-1183 |
| chr14-59134136-INS-97 | chr14-59134136-INS-97 |
| chr14-59136618-INS-59 | chr14-59136618-INS-59 |
| chr14-59160375-INS-7088 | chr14-59160376-INS-7088 |
| chr14-59194294-INS-3051 | chr14-59194295-INS-3051 |
| chr14-59204336-INS-713 | chr14-59204336-INS-709 |
| chr14-59222806-INS-52 | chr14-59222806-INS-68 |
| chr14-59234355-INS-6645 | chr14-59234356-INS-6645 |
| chr14-59276773-INS-135 | chr14-59276760-INS-133 |
| chr14-59298264-INS-53 | chr14-59298263-INS-53 |
| chr14-59309232-INS-1147 | chr14-59309221-INS-1148 |
| chr14-59329418-INS-640 | chr14-59329418-INS-640 |
| chr14-59344177-INS-7068 | chr14-59344178-INS-7068 |
| chr14-59350366-INS-399 | chr14-59350366-INS-399 |
| chr14-59361362-INS-260 | chr14-59361362-INS-265 |
| chr14-59388740-INS-60 | chr14-59388740-INS-61 |
| chr14-59463362-INS-52 | chr14-59463362-INS-52 |
| chr14-59476474-INS-4429 | chr14-59476475-INS-4415 |
| chr14-59490538-INS-5385 | chr14-59490539-INS-5385 |
| chr14-59499201-INS-162 | chr14-59499201-INS-162 |
| chr14-59560113-INS-172 | chr14-59560113-INS-176 |
| chr14-59645425-INS-511 | chr14-59645425-INS-466 |
| chr14-59685120-INS-348 | chr14-59685120-INS-348 |
| chr14-59740172-INS-203 | chr14-59740172-INS-203 |
| chr14-59785333-INS-430 | chr14-59785333-INS-430 |
| chr14-59902092-INS-128 | chr14-59902092-INS-94 |
| chr14-59910485-INS-7110 | chr14-59910486-INS-7110 |
| chr14-60035313-INS-379 | chr14-60035304-INS-364 |
| chr14-60048041-INS-77 | chr14-60048041-INS-77 |
| chr14-60089967-INS-1244 | chr14-60089968-INS-1245 |
| chr14-60127041-INS-6905 | chr14-60127042-INS-6906 |
| chr14-60131200-INS-63 | chr14-60131200-INS-85 |
| chr14-60184315-INS-54 | chr14-60184315-INS-54 |
| chr14-60217542-INS-602 | chr14-60217542-INS-602 |
| chr14-60258864-INS-157 | chr14-60258864-INS-157 |
| chr14-60262640-INS-1216 | chr14-60262641-INS-1216 |
| chr14-60274996-INS-179 | chr14-60274996-INS-135 |
| chr14-60305324-INS-299 | chr14-60305324-INS-297 |
| chr14-60325292-INS-58 | chr14-60325292-INS-62 |
| chr14-60341222-INS-318 | chr14-60341222-INS-318 |
| chr14-60343975-INS-78 | chr14-60343975-INS-104 |
| chr14-60359684-INS-55 | chr14-60359684-INS-55 |
| chr14-60361097-INS-671 | chr14-60361097-INS-689 |
| chr14-60421691-INS-7189 | chr14-60421692-INS-7188 |
| chr14-60424612-INS-158 | chr14-60424589-INS-132 |
| chr14-60431333-INS-63 | chr14-60431333-INS-63 |
| chr14-60447273-INS-63 | chr14-60447273-INS-63 |
| chr14-60571489-INS-212 | chr14-60571489-INS-212 |
| chr14-60671527-INS-113 | chr14-60671527-INS-113 |
| chr14-60714109-INS-53 | chr14-60714109-INS-53 |
| chr14-60717568-INS-53 | chr14-60717568-INS-53 |
| chr14-60738479-INS-222 | chr14-60738479-INS-220 |
| chr14-60744630-INS-71 | chr14-60744630-INS-71 |

| | |
|---|---|
| chr14-60758061-INS-205 | chr14-60758061-INS-205 |
| chr14-60832485-INS-83 | chr14-60832485-INS-80 |
| chr14-60880008-INS-174 | chr14-60880008-INS-172 |
| chr14-60905657-INS-132 | chr14-60905657-INS-128 |
| chr14-60922638-INS-72 | chr14-60922638-INS-72 |
| chr14-60953923-INS-709 | chr14-60953923-INS-709 |
| chr14-60958044-INS-203 | chr14-60958044-INS-202 |
| chr14-60967929-INS-66 | chr14-60967900-INS-74 |
| chr14-61004672-INS-51 | chr14-61004672-INS-51 |
| chr14-61020309-INS-103 | chr14-61020309-INS-104 |
| chr14-61038903-INS-60 | chr14-61038903-INS-60 |
| chr14-61075421-INS-51 | chr14-61075421-INS-54 |
| chr14-61096440-INS-62 | chr14-61096440-INS-64 |
| chr14-61138751-INS-56 | chr14-61138751-INS-58 |
| chr14-61171473-INS-4983 | chr14-61171474-INS-4983 |
| chr14-61181374-INS-60 | chr14-61181380-INS-61 |
| chr14-61195841-INS-143 | chr14-61195841-INS-143 |
| chr14-61200982-INS-105 | chr14-61200982-INS-105 |
| chr14-61225009-INS-207 | chr14-61225009-INS-207 |
| chr14-61234988-INS-397 | chr14-61234988-INS-403 |
| chr14-61261488-INS-186 | chr14-61261488-INS-186 |
| chr14-61266313-INS-683 | chr14-61266313-INS-683 |
| chr14-61297584-INS-203 | chr14-61297584-INS-210 |
| chr14-61301912-INS-56 | chr14-61301889-INS-60 |
| chr14-61326322-INS-52 | chr14-61326322-INS-52 |
| chr14-61331066-INS-67 | chr14-61331066-INS-67 |
| chr14-61335072-INS-94 | chr14-61335073-INS-80 |
| chr14-61337379-INS-4527 | chr14-61337380-INS-4527 |
| chr14-61339910-INS-567 | chr14-61339910-INS-567 |
| chr14-61341652-INS-2832 | chr14-61341653-INS-2844 |
| chr14-61377540-INS-819 | chr14-61377540-INS-819 |
| chr14-61400864-INS-62 | chr14-61400864-INS-66 |
| chr14-61416545-INS-55 | chr14-61416555-INS-56 |
| chr14-61425229-INS-540 | chr14-61425229-INS-541 |
| chr14-61454401-INS-78 | chr14-61454408-INS-60 |
| chr14-61475650-INS-498 | chr14-61475650-INS-498 |
| chr14-61486445-INS-396 | chr14-61486445-INS-396 |
| chr14-61494517-INS-57 | chr14-61494518-INS-54 |
| chr14-61501167-INS-6440 | chr14-61501168-INS-6440 |
| chr14-61506078-INS-479 | chr14-61506078-INS-479 |
| chr14-61506559-INS-661 | chr14-61506559-INS-661 |
| chr14-61511657-INS-93 | chr14-61511658-INS-94 |
| chr14-61515961-INS-1230 | chr14-61515962-INS-1230 |
| chr14-61521691-INS-436 | chr14-61521691-INS-436 |
| chr14-61550747-INS-158 | chr14-61550747-INS-162 |
| chr14-61576057-INS-52 | chr14-61576057-INS-52 |
| chr14-61583855-INS-217 | chr14-61583855-INS-217 |
| chr14-61638184-INS-200 | chr14-61638184-INS-200 |
| chr14-61694132-INS-66 | chr14-61694132-INS-66 |
| chr14-61700336-INS-846 | chr14-61700336-INS-848 |
| chr14-61703288-INS-204 | chr14-61703288-INS-204 |
| chr14-61707210-INS-71 | chr14-61707210-INS-73 |
| chr14-61731097-INS-163 | chr14-61731097-INS-163 |
| chr14-61734024-INS-59 | chr14-61734025-INS-59 |
| chr14-61735521-INS-502 | chr14-61735521-INS-501 |
| chr14-61755321-INS-81 | chr14-61755321-INS-82 |
| chr14-61755901-INS-57 | chr14-61755901-INS-53 |
| chr14-61779317-INS-128 | chr14-61779317-INS-128 |
| chr14-61782993-INS-304 | chr14-61782993-INS-304 |
| chr14-61800136-INS-58 | chr14-61800136-INS-58 |
| chr14-61893208-INS-108 | chr14-61893208-INS-108 |
| chr14-61907847-INS-190 | chr14-61907847-INS-190 |
| chr14-61952355-INS-98 | chr14-61952355-INS-98 |
| chr14-61956864-INS-108 | chr14-61956864-INS-104 |
| chr14-61968174-INS-105 | chr14-61968174-INS-105 |
| chr14-61982154-INS-51 | chr14-61982121-INS-65 |
| chr14-62015312-INS-146 | chr14-62015294-INS-146 |
| chr14-62049492-INS-113 | chr14-62049492-INS-113 |
| chr14-62117487-INS-7120 | chr14-62117488-INS-7123 |
| chr14-62141715-INS-108 | chr14-62141715-INS-108 |
| chr14-62149345-INS-200 | chr14-62149345-INS-200 |
| chr14-62192628-INS-616 | chr14-62192628-INS-636 |
| chr14-62215023-INS-57 | chr14-62215023-INS-57 |
| chr14-62244453-INS-76 | chr14-62244453-INS-80 |
| chr14-62280660-INS-56 | chr14-62280660-INS-56 |
| chr14-62301656-INS-73 | chr14-62301656-INS-73 |
| chr14-62305791-INS-276 | chr14-62305791-INS-276 |

| | |
|---|---|
| chr14-62307129-INS-524 | chr14-62307129-INS-524 |
| chr14-62370862-INS-70 | chr14-62370862-INS-66 |
| chr14-62395810-INS-163 | chr14-62395809-INS-209 |
| chr14-62403043-INS-159 | chr14-62403030-INS-126 |
| chr14-62408248-INS-213 | chr14-62408248-INS-211 |
| chr14-62411881-INS-72 | chr14-62411881-INS-82 |
| chr14-62422488-INS-98 | chr14-62422488-INS-98 |
| chr14-62448255-INS-74 | chr14-62448255-INS-79 |
| chr14-62458575-INS-134 | chr14-62458575-INS-134 |
| chr14-62470962-INS-188 | chr14-62470962-INS-188 |
| chr14-62478413-INS-82 | chr14-62478413-INS-98 |
| chr14-62508842-INS-5370 | chr14-62508843-INS-5370 |
| chr14-62537885-INS-440 | chr14-62537885-INS-440 |
| chr14-62541894-INS-185 | chr14-62541894-INS-185 |
| chr14-62558942-INS-193 | chr14-62558943-INS-193 |
| chr14-62581750-INS-415 | chr14-62581750-INS-404 |
| chr14-62585182-INS-59 | chr14-62585182-INS-59 |
| chr14-62593257-INS-73 | chr14-62593257-INS-71 |
| chr14-62601987-INS-442 | chr14-62601987-INS-442 |
| chr14-62614820-INS-180 | chr14-62614820-INS-180 |
| chr14-62616571-INS-96 | chr14-62616579-INS-120 |
| chr14-62618928-INS-504 | chr14-62618928-INS-504 |
| chr14-62622998-INS-222 | chr14-62622998-INS-222 |
| chr14-62631948-INS-805 | chr14-62631948-INS-810 |
| chr14-62641680-INS-127 | chr14-62641683-INS-160 |
| chr14-62642564-INS-109 | chr14-62642565-INS-93 |
| chr14-62654190-INS-9049 | chr14-62654191-INS-9049 |
| chr14-62662432-INS-70 | chr14-62662432-INS-72 |
| chr14-62665894-INS-56 | chr14-62665894-INS-56 |
| chr14-62674396-INS-95 | chr14-62674396-INS-95 |
| chr14-62679979-INS-54 | chr14-62679993-INS-60 |
| chr14-62697430-INS-120 | chr14-62697430-INS-160 |
| chr14-62699919-INS-79 | chr14-62699919-INS-80 |
| chr14-62732976-INS-117 | chr14-62732976-INS-117 |
| chr14-62747609-INS-772 | chr14-62747615-INS-784 |
| chr14-62750476-INS-1381 | chr14-62750477-INS-1382 |
| chr14-62769479-INS-105 | chr14-62769506-INS-65 |
| chr14-62787989-INS-54 | chr14-62787990-INS-76 |
| chr14-62790664-INS-106 | chr14-62790649-INS-75 |
| chr14-62807374-INS-52 | chr14-62807374-INS-62 |
| chr14-62816571-INS-100 | chr14-62816565-INS-102 |
| chr14-62824396-INS-101 | chr14-62824396-INS-101 |
| chr14-62826583-INS-104 | chr14-62826583-INS-107 |
| chr14-62841863-INS-245 | chr14-62841863-INS-244 |
| chr14-62846118-INS-63 | chr14-62846091-INS-66 |
| chr14-62849269-INS-256 | chr14-62849248-INS-286 |
| chr14-62851269-INS-313 | chr14-62851269-INS-358 |
| chr14-62855310-INS-71 | chr14-62855310-INS-72 |
| chr14-62856488-INS-266 | chr14-62856488-INS-266 |
| chr14-62866774-INS-212 | chr14-62866774-INS-212 |
| chr14-62877234-INS-238 | chr14-62877234-INS-238 |
| chr14-62890194-INS-196 | chr14-62890194-INS-196 |
| chr14-62892586-INS-409 | chr14-62892586-INS-409 |
| chr14-62895466-INS-503 | chr14-62895466-INS-503 |
| chr14-62939949-INS-127 | chr14-62939949-INS-127 |
| chr14-62957670-INS-333 | chr14-62957670-INS-334 |
| chr14-63066550-INS-51 | chr14-63066552-INS-52 |
| chr14-63091351-INS-202 | chr14-63091351-INS-206 |
| chr14-63096194-INS-52 | chr14-63096194-INS-52 |
| chr14-63102758-INS-224 | chr14-63102758-INS-224 |
| chr14-63104547-INS-2595 | chr14-63104548-INS-2600 |
| chr14-63121798-INS-218 | chr14-63121798-INS-218 |
| chr14-63128013-INS-209 | chr14-63128013-INS-209 |
| chr14-63128592-INS-219 | chr14-63128592-INS-219 |
| chr14-63154091-INS-157 | chr14-63154091-INS-117 |
| chr14-63220680-INS-239 | chr14-63220680-INS-241 |
| chr14-63222849-INS-5803 | chr14-63222850-INS-5803 |
| chr14-63222850-INS-653 | chr14-63222850-INS-653 |
| chr14-63232840-INS-74 | chr14-63232804-INS-80 |
| chr14-63238773-INS-1684 | chr14-63238774-INS-1684 |
| chr14-63250539-INS-50 | chr14-63250539-INS-76 |
| chr14-63253809-INS-1493 | chr14-63253810-INS-1493 |
| chr14-63256282-INS-84 | chr14-63256282-INS-81 |
| chr14-63291328-INS-448 | chr14-63291328-INS-448 |
| chr14-63292954-INS-102 | chr14-63292954-INS-102 |
| chr14-63299458-INS-131 | chr14-63299448-INS-150 |
| chr14-63334746-INS-94 | chr14-63334746-INS-106 |

| | |
|---|---|
| chr14-63342296-INS-380 | chr14-63342296-INS-378 |
| chr14-63347543-INS-60 | chr14-63347536-INS-57 |
| chr14-63356535-INS-54 | chr14-63356535-INS-78 |
| chr14-63462892-INS-103 | chr14-63462886-INS-141 |
| chr14-63476446-INS-406 | chr14-63476446-INS-407 |
| chr14-63504961-INS-236 | chr14-63504961-INS-235 |
| chr14-63545408-INS-54 | chr14-63545408-INS-54 |
| chr14-63548385-INS-137 | chr14-63548385-INS-116 |
| chr14-63589170-INS-66 | chr14-63589170-INS-66 |
| chr14-63602160-INS-7170 | chr14-63602161-INS-7170 |
| chr14-63615282-INS-108 | chr14-63615280-INS-152 |
| chr14-63629807-INS-60 | chr14-63629807-INS-60 |
| chr14-63664318-INS-125 | chr14-63664316-INS-127 |
| chr14-63677620-INS-1888 | chr14-63677621-INS-1888 |
| chr14-63712448-INS-203 | chr14-63712448-INS-203 |
| chr14-63723703-INS-120 | chr14-63723703-INS-120 |
| chr14-63727879-INS-169 | chr14-63727879-INS-169 |
| chr14-63749742-INS-90 | chr14-63749746-INS-84 |
| chr14-63774964-INS-172 | chr14-63774956-INS-185 |
| chr14-63780041-INS-1269 | chr14-63780042-INS-1269 |
| chr14-63801967-INS-792 | chr14-63801967-INS-794 |
| chr14-63805351-INS-96 | chr14-63805351-INS-96 |
| chr14-63825766-INS-58 | chr14-63825766-INS-62 |
| chr14-63835107-INS-2234 | chr14-63835108-INS-2189 |
| chr14-63860902-INS-112 | chr14-63860920-INS-88 |
| chr14-63899445-INS-74 | chr14-63899445-INS-62 |
| chr14-63962361-INS-78 | chr14-63962361-INS-78 |
| chr14-64034711-INS-59 | chr14-64034711-INS-63 |
| chr14-64133253-INS-53 | chr14-64133253-INS-53 |
| chr14-64178889-INS-72 | chr14-64178889-INS-60 |
| chr14-64195760-INS-160 | chr14-64195760-INS-160 |
| chr14-64197692-INS-75 | chr14-64197692-INS-90 |
| chr14-64200717-INS-110 | chr14-64200737-INS-60 |
| chr14-64261598-INS-7317 | chr14-64261599-INS-7317 |
| chr14-64312431-INS-3098 | chr14-64312432-INS-3098 |
| chr14-64330046-INS-80 | chr14-64330034-INS-89 |
| chr14-64353321-INS-162 | chr14-64353322-INS-162 |
| chr14-64364489-INS-389 | chr14-64364486-INS-391 |
| chr14-64422624-INS-60 | chr14-64422624-INS-60 |
| chr14-64446951-INS-139 | chr14-64446951-INS-141 |
| chr14-64483234-INS-524 | chr14-64483234-INS-524 |
| chr14-64516463-INS-59 | chr14-64516463-INS-56 |
| chr14-64516912-INS-431 | chr14-64516912-INS-431 |
| chr14-64535371-INS-115 | chr14-64535386-INS-85 |
| chr14-64545120-INS-474 | chr14-64545120-INS-473 |
| chr14-64580184-INS-62 | chr14-64580184-INS-50 |
| chr14-64594327-INS-211 | chr14-64594327-INS-207 |
| chr14-64609935-INS-1189 | chr14-64609936-INS-1189 |
| chr14-64621309-INS-646 | chr14-64621309-INS-647 |
| chr14-64636436-INS-60 | chr14-64636436-INS-60 |
| chr14-64647708-INS-52 | chr14-64647708-INS-52 |
| chr14-64684659-INS-81 | chr14-64684659-INS-81 |
| chr14-64759039-INS-192 | chr14-64759039-INS-201 |
| chr14-64765046-INS-411 | chr14-64765046-INS-411 |
| chr14-64790129-INS-246 | chr14-64790129-INS-248 |
| chr14-64792012-INS-242 | chr14-64792012-INS-245 |
| chr14-64822433-INS-68 | chr14-64822435-INS-61 |
| chr14-64853181-INS-217 | chr14-64853181-INS-217 |
| chr14-64871288-INS-205 | chr14-64871288-INS-205 |
| chr14-64876663-INS-204 | chr14-64876674-INS-207 |
| chr14-64891051-INS-240 | chr14-64891051-INS-241 |
| chr14-64907931-INS-199 | chr14-64907931-INS-199 |
| chr14-64921831-INS-182 | chr14-64921831-INS-184 |
| chr14-64935863-INS-190 | chr14-64935867-INS-192 |
| chr14-64936983-INS-187 | chr14-64936983-INS-187 |
| chr14-64954206-INS-74 | chr14-64954206-INS-74 |
| chr14-64963280-INS-80 | chr14-64963283-INS-79 |
| chr14-64979315-INS-56 | chr14-64979315-INS-60 |
| chr14-65023567-INS-86 | chr14-65023563-INS-74 |
| chr14-65027835-INS-58 | chr14-65027835-INS-58 |
| chr14-65039045-INS-57 | chr14-65039045-INS-97 |
| chr14-65065114-INS-184 | chr14-65065134-INS-175 |
| chr14-65076927-INS-66 | chr14-65076927-INS-62 |
| chr14-65100849-INS-410 | chr14-65100825-INS-424 |
| chr14-65108220-INS-102 | chr14-65108220-INS-101 |
| chr14-65111379-INS-56 | chr14-65111379-INS-56 |
| chr14-65121549-INS-193 | chr14-65121552-INS-193 |

| | |
|---|---|
| chr14-65133029-INS-204 | chr14-65133029-INS-206 |
| chr14-65162538-INS-64 | chr14-65162538-INS-62 |
| chr14-65174140-INS-200 | chr14-65174141-INS-201 |
| chr14-65182918-INS-185 | chr14-65182918-INS-186 |
| chr14-65204979-INS-2936 | chr14-65204980-INS-2936 |
| chr14-65223912-INS-74 | chr14-65223912-INS-74 |
| chr14-65230964-INS-520 | chr14-65230964-INS-520 |
| chr14-65259328-INS-220 | chr14-65259328-INS-220 |
| chr14-65260877-INS-76 | chr14-65260877-INS-84 |
| chr14-65262902-INS-504 | chr14-65262902-INS-504 |
| chr14-65266774-INS-174 | chr14-65266774-INS-174 |
| chr14-65273444-INS-64 | chr14-65273444-INS-64 |
| chr14-65325620-INS-214 | chr14-65325620-INS-222 |
| chr14-65329610-INS-438 | chr14-65329610-INS-438 |
| chr14-65330540-INS-58 | chr14-65330540-INS-58 |
| chr14-65343504-INS-77 | chr14-65343504-INS-77 |
| chr14-65392718-INS-340 | chr14-65392718-INS-340 |
| chr14-65393595-INS-75 | chr14-65393595-INS-75 |
| chr14-65408361-INS-214 | chr14-65408361-INS-207 |
| chr14-65444630-INS-63 | chr14-65444611-INS-72 |
| chr14-65510162-INS-61 | chr14-65510162-INS-51 |
| chr14-65535191-INS-184 | chr14-65535191-INS-185 |
| chr14-65551049-INS-213 | chr14-65551049-INS-214 |
| chr14-65563532-INS-55 | chr14-65563532-INS-55 |
| chr14-65574058-INS-179 | chr14-65574058-INS-178 |
| chr14-65576008-INS-177 | chr14-65576008-INS-174 |
| chr14-65579690-INS-56 | chr14-65579690-INS-56 |
| chr14-65608363-INS-58 | chr14-65608363-INS-58 |
| chr14-65617208-INS-340 | chr14-65617208-INS-343 |
| chr14-65634248-INS-54 | chr14-65634245-INS-56 |
| chr14-65655690-INS-52 | chr14-65655689-INS-56 |
| chr14-65696227-INS-85 | chr14-65696227-INS-85 |
| chr14-65704098-INS-62 | chr14-65704098-INS-62 |
| chr14-65725423-INS-236 | chr14-65725423-INS-237 |
| chr14-65734679-INS-80 | chr14-65734679-INS-75 |
| chr14-65771961-INS-331 | chr14-65771961-INS-329 |
| chr14-65844598-INS-68 | chr14-65844598-INS-68 |
| chr14-65863315-INS-96 | chr14-65863315-INS-96 |
| chr14-65864945-INS-96 | chr14-65864945-INS-96 |
| chr14-65888775-INS-152 | chr14-65888776-INS-154 |
| chr14-65892130-INS-62 | chr14-65892130-INS-62 |
| chr14-65907893-INS-537 | chr14-65907893-INS-538 |
| chr14-65910652-INS-84 | chr14-65910652-INS-64 |
| chr14-65912946-INS-57 | chr14-65912946-INS-59 |
| chr14-65920833-INS-68 | chr14-65920833-INS-68 |
| chr14-65938208-INS-97 | chr14-65938240-INS-99 |
| chr14-65961198-INS-111 | chr14-65961198-INS-111 |
| chr14-65963053-INS-58 | chr14-65963053-INS-58 |
| chr14-65966267-INS-206 | chr14-65966267-INS-210 |
| chr14-65982727-INS-60 | chr14-65982704-INS-64 |
| chr14-65985121-INS-264 | chr14-65985148-INS-230 |
| chr14-65986302-INS-90 | chr14-65986306-INS-94 |
| chr14-66013721-INS-371 | chr14-66013721-INS-371 |
| chr14-66040995-INS-903 | chr14-66041023-INS-910 |
| chr14-66068178-INS-110 | chr14-66068178-INS-110 |
| chr14-66069988-INS-163 | chr14-66069988-INS-158 |
| chr14-66074576-INS-200 | chr14-66074576-INS-201 |
| chr14-66109241-INS-67 | chr14-66109241-INS-67 |
| chr14-66117209-INS-763 | chr14-66117209-INS-763 |
| chr14-66132872-INS-163 | chr14-66132877-INS-204 |
| chr14-66136836-INS-319 | chr14-66136836-INS-320 |
| chr14-66159687-INS-89 | chr14-66159687-INS-89 |
| chr14-66172439-INS-81 | chr14-66172439-INS-81 |
| chr14-66276976-INS-51 | chr14-66276976-INS-53 |
| chr14-66280142-INS-111 | chr14-66280138-INS-113 |
| chr14-66312896-INS-677 | chr14-66312896-INS-677 |
| chr14-66316542-INS-97 | chr14-66316542-INS-57 |
| chr14-66323045-INS-73 | chr14-66323060-INS-69 |
| chr14-66327993-INS-955 | chr14-66327993-INS-954 |
| chr14-66370016-INS-64 | chr14-66370016-INS-76 |
| chr14-66397018-INS-1899 | chr14-66397019-INS-1899 |
| chr14-66425288-INS-161 | chr14-66425288-INS-164 |
| chr14-66426309-INS-95 | chr14-66426309-INS-62 |
| chr14-66434000-INS-978 | chr14-66434000-INS-978 |
| chr14-66465163-INS-113 | chr14-66465163-INS-111 |
| chr14-66487917-INS-805 | chr14-66487917-INS-806 |
| chr14-66507107-INS-137 | chr14-66507107-INS-133 |

| | |
|---|---|
| chr14-66511908-INS-7174 | chr14-66511909-INS-7174 |
| chr14-66528009-INS-1335 | chr14-66528010-INS-1338 |
| chr14-66569083-INS-199 | chr14-66569083-INS-199 |
| chr14-66597226-INS-63 | chr14-66597226-INS-105 |
| chr14-66607625-INS-508 | chr14-66607622-INS-508 |
| chr14-66640420-INS-205 | chr14-66640420-INS-204 |
| chr14-66663757-INS-88 | chr14-66663757-INS-88 |
| chr14-66673519-INS-85 | chr14-66673534-INS-50 |
| chr14-66711153-INS-876 | chr14-66711153-INS-897 |
| chr14-66730431-INS-100 | chr14-66730431-INS-80 |
| chr14-66747259-INS-327 | chr14-66747259-INS-327 |
| chr14-66758546-INS-170 | chr14-66758546-INS-170 |
| chr14-66767868-INS-64 | chr14-66767868-INS-64 |
| chr14-66798395-INS-381 | chr14-66798395-INS-381 |
| chr14-66801315-INS-81 | chr14-66801315-INS-86 |
| chr14-66819048-INS-223 | chr14-66819059-INS-173 |
| chr14-66826766-INS-629 | chr14-66826770-INS-629 |
| chr14-66855846-INS-62 | chr14-66855846-INS-64 |
| chr14-66863059-INS-401 | chr14-66863059-INS-401 |
| chr14-66900002-INS-58 | chr14-66900002-INS-58 |
| chr14-66924240-INS-65 | chr14-66924240-INS-65 |
| chr14-66944457-INS-311 | chr14-66944457-INS-316 |
| chr14-66964020-INS-154 | chr14-66964020-INS-140 |
| chr14-66978064-INS-579 | chr14-66978064-INS-579 |
| chr14-66989334-INS-62 | chr14-66989332-INS-80 |
| chr14-66993462-INS-60 | chr14-66993476-INS-89 |
| chr14-67025198-INS-58 | chr14-67025198-INS-54 |
| chr14-67030548-INS-58 | chr14-67030548-INS-58 |
| chr14-67050794-INS-99 | chr14-67050794-INS-68 |
| chr14-67053285-INS-138 | chr14-67053285-INS-138 |
| chr14-67060941-INS-150 | chr14-67060941-INS-150 |
| chr14-67063420-INS-203 | chr14-67063420-INS-202 |
| chr14-67065257-INS-119 | chr14-67065257-INS-119 |
| chr14-67098197-INS-66 | chr14-67098197-INS-50 |
| chr14-67125764-INS-201 | chr14-67125764-INS-201 |
| chr14-67143459-INS-62 | chr14-67143459-INS-62 |
| chr14-67161460-INS-6476 | chr14-67161461-INS-6476 |
| chr14-67191639-INS-86 | chr14-67191639-INS-80 |
| chr14-67251877-INS-225 | chr14-67251877-INS-227 |
| chr14-67255848-INS-80 | chr14-67255844-INS-72 |
| chr14-67262396-INS-128 | chr14-67262396-INS-132 |
| chr14-67312422-INS-60 | chr14-67312422-INS-60 |
| chr14-67316190-INS-63 | chr14-67316197-INS-67 |
| chr14-67319964-INS-57 | chr14-67319964-INS-57 |
| chr14-67324364-INS-62 | chr14-67324365-INS-60 |
| chr14-67335926-INS-269 | chr14-67335926-INS-271 |
| chr14-67353072-INS-141 | chr14-67353072-INS-141 |
| chr14-67407594-INS-356 | chr14-67407594-INS-356 |
| chr14-67430002-INS-82 | chr14-67430009-INS-98 |
| chr14-67436304-INS-85 | chr14-67436304-INS-85 |
| chr14-67441440-INS-64 | chr14-67441441-INS-64 |
| chr14-67444077-INS-53 | chr14-67444077-INS-58 |
| chr14-67446871-INS-57 | chr14-67446871-INS-52 |
| chr14-67450695-INS-98 | chr14-67450695-INS-98 |
| chr14-67455147-INS-366 | chr14-67455147-INS-366 |
| chr14-67480357-INS-82 | chr14-67480357-INS-72 |
| chr14-67488117-INS-60 | chr14-67488094-INS-58 |
| chr14-67506628-INS-55 | chr14-67506628-INS-60 |
| chr14-67514819-INS-78 | chr14-67514819-INS-62 |
| chr14-67568546-INS-6401 | chr14-67568547-INS-6401 |
| chr14-67574386-INS-80 | chr14-67574386-INS-80 |
| chr14-67577167-INS-194 | chr14-67577209-INS-190 |
| chr14-67591385-INS-372 | chr14-67591385-INS-373 |
| chr14-67595645-INS-852 | chr14-67595645-INS-854 |
| chr14-67597760-INS-196 | chr14-67597765-INS-204 |
| chr14-67651109-INS-88 | chr14-67651109-INS-90 |
| chr14-67671545-INS-216 | chr14-67671545-INS-217 |
| chr14-67674341-INS-62 | chr14-67674341-INS-68 |
| chr14-67679773-INS-87 | chr14-67679778-INS-96 |
| chr14-67685581-INS-113 | chr14-67685581-INS-72 |
| chr14-67689758-INS-170 | chr14-67689758-INS-170 |
| chr14-67709766-INS-75 | chr14-67709766-INS-55 |
| chr14-67726274-INS-53 | chr14-67726274-INS-53 |
| chr14-67730849-INS-51 | chr14-67730832-INS-51 |
| chr14-67735167-INS-90 | chr14-67735167-INS-85 |
| chr14-67741817-INS-56 | chr14-67741817-INS-56 |
| chr14-67744022-INS-3490 | chr14-67744023-INS-3490 |

| | |
|---|---|
| chr14-67747017-INS-79 | chr14-67747017-INS-79 |
| chr14-67748026-INS-106 | chr14-67748026-INS-106 |
| chr14-67755053-INS-54 | chr14-67755068-INS-51 |
| chr14-67758684-INS-74 | chr14-67758685-INS-60 |
| chr14-67759181-INS-92 | chr14-67759181-INS-92 |
| chr14-67803951-INS-256 | chr14-67803951-INS-256 |
| chr14-67834917-INS-128 | chr14-67834897-INS-166 |
| chr14-67855613-INS-112 | chr14-67855585-INS-115 |
| chr14-67876560-INS-486 | chr14-67876560-INS-497 |
| chr14-67885803-INS-386 | chr14-67885797-INS-396 |
| chr14-67896212-INS-66 | chr14-67896212-INS-66 |
| chr14-67926004-INS-66 | chr14-67926004-INS-66 |
| chr14-67934079-INS-76 | chr14-67934079-INS-78 |
| chr14-67935862-INS-4199 | chr14-67935863-INS-4199 |
| chr14-67943509-INS-55 | chr14-67943509-INS-55 |
| chr14-67963737-INS-180 | chr14-67963737-INS-180 |
| chr14-67977570-INS-200 | chr14-67977570-INS-200 |
| chr14-67992901-INS-88 | chr14-67992901-INS-62 |
| chr14-67999566-INS-647 | chr14-67999567-INS-647 |
| chr14-68034301-INS-113 | chr14-68034345-INS-71 |
| chr14-68038815-INS-214 | chr14-68038815-INS-214 |
| chr14-68042085-INS-127 | chr14-68042085-INS-125 |
| chr14-68045647-INS-64 | chr14-68045647-INS-64 |
| chr14-68089830-INS-1103 | chr14-68089831-INS-1103 |
| chr14-68141285-INS-90 | chr14-68141285-INS-54 |
| chr14-68160303-INS-71 | chr14-68160307-INS-100 |
| chr14-68184240-INS-201 | chr14-68184240-INS-202 |
| chr14-68206961-INS-421 | chr14-68206961-INS-421 |
| chr14-68213351-INS-70 | chr14-68213351-INS-68 |
| chr14-68215901-INS-205 | chr14-68215901-INS-206 |
| chr14-68222617-INS-278 | chr14-68222617-INS-278 |
| chr14-68224333-INS-6382 | chr14-68224334-INS-6379 |
| chr14-68229394-INS-219 | chr14-68229394-INS-220 |
| chr14-68243758-INS-168 | chr14-68243759-INS-166 |
| chr14-68287255-INS-187 | chr14-68287253-INS-194 |
| chr14-68291617-INS-768 | chr14-68291617-INS-769 |
| chr14-68305481-INS-2590 | chr14-68305482-INS-2594 |
| chr14-68335538-INS-51 | chr14-68335538-INS-51 |
| chr14-68337306-INS-211 | chr14-68337306-INS-211 |
| chr14-68339666-INS-52 | chr14-68339666-INS-52 |
| chr14-68345761-INS-563 | chr14-68345761-INS-563 |
| chr14-68391314-INS-77 | chr14-68391314-INS-77 |
| chr14-68411551-INS-89 | chr14-68411551-INS-89 |
| chr14-68440960-INS-78 | chr14-68440960-INS-78 |
| chr14-68465861-INS-211 | chr14-68465861-INS-211 |
| chr14-68533846-INS-968 | chr14-68533846-INS-969 |
| chr14-68536480-INS-132 | chr14-68536480-INS-132 |
| chr14-68581399-INS-73 | chr14-68581399-INS-73 |
| chr14-68589698-INS-56 | chr14-68589698-INS-56 |
| chr14-68592016-INS-5542 | chr14-68592017-INS-5542 |
| chr14-68598314-INS-3715 | chr14-68598315-INS-3717 |
| chr14-68603996-INS-60 | chr14-68603995-INS-68 |
| chr14-68629972-INS-7481 | chr14-68629973-INS-7481 |
| chr14-68631769-INS-224 | chr14-68631769-INS-226 |
| chr14-68633654-INS-547 | chr14-68633654-INS-547 |
| chr14-68634889-INS-476 | chr14-68634889-INS-477 |
| chr14-68640356-INS-155 | chr14-68640356-INS-158 |
| chr14-68643526-INS-84 | chr14-68643526-INS-84 |
| chr14-68666738-INS-4228 | chr14-68666739-INS-4228 |
| chr14-68669964-INS-50 | chr14-68669926-INS-60 |
| chr14-68683557-INS-122 | chr14-68683557-INS-122 |
| chr14-68718215-INS-2713 | chr14-68718216-INS-2713 |
| chr14-68734660-INS-6281 | chr14-68734661-INS-6282 |
| chr14-68784415-INS-87 | chr14-68784415-INS-93 |
| chr14-68794916-INS-89 | chr14-68794916-INS-91 |
| chr14-68821402-INS-213 | chr14-68821402-INS-213 |
| chr14-68830177-INS-84 | chr14-68830177-INS-84 |
| chr14-68847102-INS-200 | chr14-68847102-INS-200 |
| chr14-68847879-INS-233 | chr14-68847879-INS-233 |
| chr14-68850075-INS-1458 | chr14-68850084-INS-1453 |
| chr14-68861572-INS-387 | chr14-68861572-INS-387 |
| chr14-68864025-INS-135 | chr14-68864031-INS-138 |
| chr14-68864934-INS-4165 | chr14-68864935-INS-4165 |
| chr14-68884564-INS-159 | chr14-68884564-INS-156 |
| chr14-68895285-INS-152 | chr14-68895289-INS-159 |
| chr14-68907905-INS-70 | chr14-68907905-INS-54 |
| chr14-68912718-INS-79 | chr14-68912717-INS-127 |

| | |
|---|---|
| chr14-68942531-INS-56 | chr14-68942531-INS-52 |
| chr14-68944500-INS-91 | chr14-68944500-INS-91 |
| chr14-68957740-INS-55 | chr14-68957740-INS-71 |
| chr14-68960115-INS-116 | chr14-68960115-INS-116 |
| chr14-68981404-INS-94 | chr14-68981404-INS-90 |
| chr14-69020565-INS-60 | chr14-69020565-INS-60 |
| chr14-69041186-INS-367 | chr14-69041186-INS-367 |
| chr14-69052997-INS-86 | chr14-69052997-INS-78 |
| chr14-69081545-INS-383 | chr14-69081545-INS-383 |
| chr14-69125184-INS-56 | chr14-69125184-INS-66 |
| chr14-69130664-INS-63 | chr14-69130690-INS-68 |
| chr14-69164548-INS-688 | chr14-69164548-INS-688 |
| chr14-69190544-INS-796 | chr14-69190544-INS-796 |
| chr14-69195809-INS-54 | chr14-69195809-INS-54 |
| chr14-69201697-INS-84 | chr14-69201669-INS-64 |
| chr14-69231081-INS-52 | chr14-69231081-INS-52 |
| chr14-69233692-INS-88 | chr14-69233716-INS-60 |
| chr14-69253098-INS-228 | chr14-69253098-INS-230 |
| chr14-69272886-INS-212 | chr14-69272886-INS-212 |
| chr14-69293483-INS-5326 | chr14-69293483-INS-5333 |
| chr14-69313537-INS-73 | chr14-69313537-INS-73 |
| chr14-69339144-INS-84 | chr14-69339144-INS-68 |
| chr14-69407310-INS-97 | chr14-69407310-INS-96 |
| chr14-69409811-INS-51 | chr14-69409809-INS-55 |
| chr14-69416093-INS-62 | chr14-69416093-INS-62 |
| chr14-69941745-INS-68 | chr14-69941746-INS-54 |
| chr14-69947458-INS-54 | chr14-69947458-INS-54 |
| chr14-69959561-INS-194 | chr14-69959561-INS-195 |
| chr14-69975295-INS-498 | chr14-69975295-INS-498 |
| chr14-69977961-INS-189 | chr14-69977941-INS-199 |
| chr14-69982436-INS-215 | chr14-69982436-INS-215 |
| chr14-70004127-INS-62 | chr14-70004127-INS-64 |
| chr14-70006633-INS-60 | chr14-70006633-INS-64 |
| chr14-70013032-INS-68 | chr14-70013032-INS-64 |
| chr14-70019102-INS-169 | chr14-70019102-INS-169 |
| chr14-70032339-INS-95 | chr14-70032339-INS-95 |
| chr14-70035291-INS-54 | chr14-70035291-INS-56 |
| chr14-70036432-INS-194 | chr14-70036432-INS-194 |
| chr14-70044642-INS-66 | chr14-70044639-INS-94 |
| chr14-70062791-INS-54 | chr14-70062764-INS-86 |
| chr14-70066161-INS-77 | chr14-70066161-INS-75 |
| chr14-70087377-INS-56 | chr14-70087379-INS-58 |
| chr14-70124290-INS-56 | chr14-70124290-INS-56 |
| chr14-70136563-INS-151 | chr14-70136563-INS-151 |
| chr14-70140411-INS-76 | chr14-70140411-INS-76 |
| chr14-70146056-INS-362 | chr14-70146062-INS-385 |
| chr14-70152561-INS-179 | chr14-70152561-INS-179 |
| chr14-70157848-INS-99 | chr14-70157858-INS-73 |
| chr14-70161083-INS-128 | chr14-70161083-INS-130 |
| chr14-70164099-INS-52 | chr14-70164099-INS-58 |
| chr14-70174001-INS-72 | chr14-70174001-INS-72 |
| chr14-70201905-INS-76 | chr14-70201902-INS-52 |
| chr14-70219228-INS-113 | chr14-70219228-INS-113 |
| chr14-70255428-INS-77 | chr14-70255428-INS-77 |
| chr14-70257097-INS-62 | chr14-70257097-INS-66 |
| chr14-70325796-INS-216 | chr14-70325796-INS-218 |
| chr14-70348436-INS-840 | chr14-70348436-INS-819 |
| chr14-70353378-INS-80 | chr14-70353378-INS-80 |
| chr14-70370894-INS-112 | chr14-70370894-INS-111 |
| chr14-70420305-INS-52 | chr14-70420305-INS-56 |
| chr14-70489204-INS-223 | chr14-70489204-INS-224 |
| chr14-70491373-INS-121 | chr14-70491373-INS-121 |
| chr14-70496882-INS-85 | chr14-70496882-INS-93 |
| chr14-70512223-INS-56 | chr14-70512223-INS-56 |
| chr14-70520735-INS-56 | chr14-70520735-INS-68 |
| chr14-70533462-INS-65 | chr14-70533462-INS-54 |
| chr14-70536674-INS-221 | chr14-70536674-INS-218 |
| chr14-70562302-INS-64 | chr14-70562302-INS-70 |
| chr14-70563367-INS-201 | chr14-70563367-INS-201 |
| chr14-70568194-INS-60 | chr14-70568194-INS-50 |
| chr14-70593719-INS-54 | chr14-70593719-INS-54 |
| chr14-70595570-INS-263 | chr14-70595570-INS-262 |
| chr14-70605617-INS-215 | chr14-70605617-INS-215 |
| chr14-70616176-INS-207 | chr14-70616176-INS-207 |
| chr14-70641013-INS-212 | chr14-70641013-INS-212 |
| chr14-70647364-INS-201 | chr14-70647364-INS-201 |
| chr14-70649227-INS-213 | chr14-70649227-INS-213 |

| | |
|---|---|
| chr14-70655368-INS-614 | chr14-70655368-INS-614 |
| chr14-70661666-INS-205 | chr14-70661666-INS-205 |
| chr14-70663414-INS-56 | chr14-70663414-INS-56 |
| chr14-70708462-INS-80 | chr14-70708462-INS-80 |
| chr14-70712965-INS-151 | chr14-70712959-INS-113 |
| chr14-70734124-INS-209 | chr14-70734124-INS-209 |
| chr14-70736178-INS-121 | chr14-70736187-INS-128 |
| chr14-70762842-INS-87 | chr14-70762842-INS-87 |
| chr14-70820980-INS-75 | chr14-70820966-INS-51 |
| chr14-70824441-INS-55 | chr14-70824441-INS-60 |
| chr14-70892733-INS-259 | chr14-70892733-INS-260 |
| chr14-70924276-INS-138 | chr14-70924276-INS-126 |
| chr14-70925768-INS-66 | chr14-70925768-INS-66 |
| chr14-70964278-INS-135 | chr14-70964299-INS-156 |
| chr14-70974287-INS-52 | chr14-70974296-INS-60 |
| chr14-71017126-INS-52 | chr14-71017126-INS-52 |
| chr14-71019529-INS-214 | chr14-71019529-INS-199 |
| chr14-71027889-INS-358 | chr14-71027889-INS-358 |
| chr14-71051713-INS-140 | chr14-71051713-INS-140 |
| chr14-71060251-INS-64 | chr14-71060251-INS-64 |
| chr14-71064337-INS-214 | chr14-71064337-INS-213 |
| chr14-71071887-INS-246 | chr14-71071887-INS-246 |
| chr14-71098527-INS-132 | chr14-71098527-INS-132 |
| chr14-71116485-INS-350 | chr14-71116485-INS-350 |
| chr14-71117159-INS-180 | chr14-71117159-INS-180 |
| chr14-71146950-INS-73 | chr14-71146950-INS-77 |
| chr14-71160883-INS-215 | chr14-71160883-INS-216 |
| chr14-71184862-INS-230 | chr14-71184862-INS-230 |
| chr14-71189879-INS-2568 | chr14-71189880-INS-2568 |
| chr14-71253083-INS-207 | chr14-71253083-INS-205 |
| chr14-71270816-INS-1892 | chr14-71270817-INS-1892 |
| chr14-71278534-INS-81 | chr14-71278534-INS-81 |
| chr14-71314098-INS-96 | chr14-71314098-INS-96 |
| chr14-71326660-INS-240 | chr14-71326660-INS-240 |
| chr14-71351140-INS-119 | chr14-71351140-INS-119 |
| chr14-71371824-INS-52 | chr14-71371824-INS-52 |
| chr14-71386885-INS-6213 | chr14-71386886-INS-6213 |
| chr14-71402669-INS-6359 | chr14-71402670-INS-6359 |
| chr14-71427970-INS-1112 | chr14-71427971-INS-1112 |
| chr14-71448558-INS-52 | chr14-71448558-INS-52 |
| chr14-71508940-INS-118 | chr14-71508940-INS-78 |
| chr14-71509727-INS-193 | chr14-71509693-INS-178 |
| chr14-71544618-INS-68 | chr14-71544616-INS-56 |
| chr14-71584920-INS-52 | chr14-71584920-INS-50 |
| chr14-71606835-INS-350 | chr14-71606835-INS-350 |
| chr14-71622959-INS-66 | chr14-71622959-INS-66 |
| chr14-71627105-INS-97 | chr14-71627105-INS-97 |
| chr14-71684496-INS-58 | chr14-71684496-INS-58 |
| chr14-71715187-INS-829 | chr14-71715187-INS-829 |
| chr14-71724101-INS-155 | chr14-71724101-INS-155 |
| chr14-71726769-INS-150 | chr14-71726742-INS-144 |
| chr14-71732280-INS-82 | chr14-71732280-INS-74 |
| chr14-71755363-INS-52 | chr14-71755363-INS-74 |
| chr14-71771143-INS-81 | chr14-71771178-INS-59 |
| chr14-71774257-INS-1325 | chr14-71774258-INS-1327 |
| chr14-71786668-INS-6488 | chr14-71786669-INS-6488 |
| chr14-71799890-INS-59 | chr14-71799890-INS-61 |
| chr14-71814164-INS-6376 | chr14-71814165-INS-6376 |
| chr14-71837182-INS-142 | chr14-71837182-INS-142 |
| chr14-71848875-INS-410 | chr14-71848875-INS-402 |
| chr14-71874274-INS-386 | chr14-71874267-INS-386 |
| chr14-71885116-INS-6569 | chr14-71885117-INS-6569 |
| chr14-71889546-INS-6407 | chr14-71889547-INS-6407 |
| chr14-71891773-INS-973 | chr14-71891773-INS-973 |
| chr14-71893132-INS-76 | chr14-71893132-INS-76 |
| chr14-71900984-INS-408 | chr14-71900984-INS-408 |
| chr14-71921472-INS-684 | chr14-71921459-INS-717 |
| chr14-71921481-INS-124 | chr14-71921481-INS-125 |
| chr14-71967978-INS-79 | chr14-71967978-INS-82 |
| chr14-72008642-INS-66 | chr14-72008644-INS-62 |
| chr14-72061434-INS-51 | chr14-72061433-INS-53 |
| chr14-72062189-INS-129 | chr14-72062189-INS-132 |
| chr14-72077392-INS-6601 | chr14-72077393-INS-6601 |
| chr14-72092972-INS-56 | chr14-72092972-INS-56 |
| chr14-72094046-INS-63 | chr14-72094046-INS-63 |
| chr14-72096449-INS-54 | chr14-72096406-INS-56 |
| chr14-72099421-INS-329 | chr14-72099421-INS-329 |

| | |
|---|---|
| chr14-72115723-INS-63 | chr14-72115731-INS-62 |
| chr14-72123677-INS-81 | chr14-72123677-INS-121 |
| chr14-72125484-INS-1060 | chr14-72125485-INS-1060 |
| chr14-72163454-INS-272 | chr14-72163454-INS-271 |
| chr14-72179875-INS-61 | chr14-72179874-INS-69 |
| chr14-72186148-INS-7125 | chr14-72186149-INS-7125 |
| chr14-72187202-INS-71 | chr14-72187202-INS-74 |
| chr14-72206881-INS-76 | chr14-72206881-INS-76 |
| chr14-72216492-INS-60 | chr14-72216501-INS-56 |
| chr14-72217578-INS-499 | chr14-72217578-INS-498 |
| chr14-72222327-INS-4085 | chr14-72222328-INS-4096 |
| chr14-72231113-INS-52 | chr14-72231113-INS-50 |
| chr14-72236382-INS-77 | chr14-72236382-INS-83 |
| chr14-72240976-INS-60 | chr14-72240976-INS-58 |
| chr14-72250018-INS-452 | chr14-72250018-INS-452 |
| chr14-72265997-INS-6456 | chr14-72265998-INS-6456 |
| chr14-72274976-INS-85 | chr14-72274976-INS-85 |
| chr14-72276173-INS-222 | chr14-72276173-INS-226 |
| chr14-72284801-INS-426 | chr14-72284801-INS-426 |
| chr14-72303242-INS-60 | chr14-72303242-INS-60 |
| chr14-72310306-INS-249 | chr14-72310306-INS-249 |
| chr14-72317997-INS-483 | chr14-72317997-INS-483 |
| chr14-72323765-INS-9323 | chr14-72323766-INS-9323 |
| chr14-72337046-INS-72 | chr14-72337046-INS-86 |
| chr14-72382593-INS-359 | chr14-72382594-INS-365 |
| chr14-72389419-INS-494 | chr14-72389419-INS-496 |
| chr14-72393432-INS-50 | chr14-72393413-INS-66 |
| chr14-72395157-INS-225 | chr14-72395157-INS-225 |
| chr14-72477156-INS-70 | chr14-72477156-INS-72 |
| chr14-72480910-INS-1861 | chr14-72480911-INS-1861 |
| chr14-72488693-INS-7083 | chr14-72488694-INS-7083 |
| chr14-72495879-INS-85 | chr14-72495879-INS-86 |
| chr14-72515928-INS-89 | chr14-72515928-INS-89 |
| chr14-72517114-INS-397 | chr14-72517114-INS-397 |
| chr14-72542114-INS-78 | chr14-72542114-INS-60 |
| chr14-72561355-INS-1891 | chr14-72561356-INS-1891 |
| chr14-72564263-INS-114 | chr14-72564263-INS-110 |
| chr14-72585086-INS-60 | chr14-72585086-INS-62 |
| chr14-72586417-INS-254 | chr14-72586435-INS-220 |
| chr14-72609954-INS-56 | chr14-72609954-INS-54 |
| chr14-72614673-INS-87 | chr14-72614670-INS-74 |
| chr14-72621639-INS-699 | chr14-72621649-INS-724 |
| chr14-72624928-INS-65 | chr14-72624928-INS-106 |
| chr14-72627931-INS-82 | chr14-72627929-INS-86 |
| chr14-72636073-INS-58 | chr14-72636073-INS-60 |
| chr14-72638240-INS-63 | chr14-72638240-INS-97 |
| chr14-72656699-INS-74 | chr14-72656699-INS-80 |
| chr14-72663248-INS-203 | chr14-72663248-INS-203 |
| chr14-72681391-INS-78 | chr14-72681430-INS-79 |
| chr14-72683513-INS-69 | chr14-72683513-INS-79 |
| chr14-72701862-INS-67 | chr14-72701862-INS-67 |
| chr14-72717318-INS-144 | chr14-72717318-INS-158 |
| chr14-72873857-INS-65 | chr14-72873857-INS-50 |
| chr14-72895524-INS-57 | chr14-72895511-INS-71 |
| chr14-72915953-INS-729 | chr14-72915953-INS-729 |
| chr14-72917160-INS-77 | chr14-72917160-INS-77 |
| chr14-72939915-INS-4716 | chr14-72939916-INS-4716 |
| chr14-72952268-INS-1108 | chr14-72952269-INS-1108 |
| chr14-72985916-INS-1664 | chr14-72985917-INS-1665 |
| chr14-72993986-INS-58 | chr14-72993986-INS-52 |
| chr14-73002481-INS-6517 | chr14-73002482-INS-6517 |
| chr14-73020599-INS-103 | chr14-73020599-INS-98 |
| chr14-73024741-INS-92 | chr14-73024741-INS-70 |
| chr14-73118313-INS-61 | chr14-73118313-INS-57 |
| chr14-73119525-INS-6570 | chr14-73119526-INS-6570 |
| chr14-73127400-INS-3278 | chr14-73127401-INS-3303 |
| chr14-73175489-INS-516 | chr14-73175483-INS-516 |
| chr14-73214121-INS-306 | chr14-73214122-INS-306 |
| chr14-73226228-INS-335 | chr14-73226228-INS-335 |
| chr14-73232197-INS-7099 | chr14-73232198-INS-7099 |
| chr14-73237275-INS-547 | chr14-73237275-INS-547 |
| chr14-73245754-INS-413 | chr14-73245754-INS-413 |
| chr14-73255794-INS-992 | chr14-73255794-INS-991 |
| chr14-73268545-INS-6676 | chr14-73268544-INS-6680 |
| chr14-73282763-INS-132 | chr14-73282763-INS-132 |
| chr14-73290794-INS-3847 | chr14-73290795-INS-3847 |
| chr14-73295969-INS-156 | chr14-73295969-INS-146 |

| | |
|---|---|
| chr14-73296543-INS-70 | chr14-73296543-INS-70 |
| chr14-73298694-INS-814 | chr14-73298694-INS-814 |
| chr14-73310737-INS-6582 | chr14-73310738-INS-6582 |
| chr14-73327916-INS-67 | chr14-73327916-INS-71 |
| chr14-73374529-INS-199 | chr14-73374529-INS-201 |
| chr14-73386000-INS-55 | chr14-73386000-INS-67 |
| chr14-73389607-INS-81 | chr14-73389607-INS-85 |
| chr14-73395514-INS-80 | chr14-73395515-INS-80 |
| chr14-73412454-INS-348 | chr14-73412455-INS-372 |
| chr14-73451490-INS-7087 | chr14-73451491-INS-7088 |
| chr14-73472741-INS-4841 | chr14-73472742-INS-4841 |
| chr14-73477904-INS-221 | chr14-73477904-INS-222 |
| chr14-73521528-INS-200 | chr14-73521528-INS-200 |
| chr14-73535401-INS-184 | chr14-73535401-INS-184 |
| chr14-73537412-INS-371 | chr14-73537412-INS-371 |
| chr14-73538455-INS-84 | chr14-73538455-INS-84 |
| chr14-73556212-INS-3105 | chr14-73556213-INS-3105 |
| chr14-73558233-INS-112 | chr14-73558233-INS-112 |
| chr14-73575749-INS-78 | chr14-73575749-INS-78 |
| chr14-73628927-INS-56 | chr14-73628927-INS-56 |
| chr14-73631757-INS-2235 | chr14-73631758-INS-2241 |
| chr14-73640105-INS-63 | chr14-73640105-INS-63 |
| chr14-73645445-INS-52 | chr14-73645433-INS-60 |
| chr14-73665055-INS-65 | chr14-73665055-INS-50 |
| chr14-73679052-INS-51 | chr14-73679053-INS-57 |
| chr14-73679949-INS-293 | chr14-73679950-INS-331 |
| chr14-73685376-INS-650 | chr14-73685377-INS-650 |
| chr14-73717285-INS-417 | chr14-73717286-INS-417 |
| chr14-73731602-INS-84 | chr14-73731602-INS-76 |
| chr14-73750435-INS-6161 | chr14-73750436-INS-6161 |
| chr14-73769946-INS-58 | chr14-73769946-INS-58 |
| chr14-73780940-INS-130 | chr14-73780924-INS-142 |
| chr14-73781589-INS-1451 | chr14-73781590-INS-1455 |
| chr14-73808225-INS-150 | chr14-73808225-INS-150 |
| chr14-73823028-INS-196 | chr14-73823028-INS-196 |
| chr14-73829494-INS-52 | chr14-73829494-INS-58 |
| chr14-73830202-INS-1622 | chr14-73830237-INS-1617 |
| chr14-73838889-INS-6664 | chr14-73838890-INS-6665 |
| chr14-73855388-INS-83 | chr14-73855388-INS-83 |
| chr14-73870721-INS-52 | chr14-73870721-INS-50 |
| chr14-73875160-INS-938 | chr14-73875160-INS-941 |
| chr14-73889185-INS-6746 | chr14-73889186-INS-6746 |
| chr14-73891548-INS-316 | chr14-73891548-INS-316 |
| chr14-73899801-INS-200 | chr14-73899801-INS-200 |
| chr14-73906265-INS-147 | chr14-73906265-INS-127 |
| chr14-73927532-INS-1194 | chr14-73927533-INS-1194 |
| chr14-73972422-INS-1627 | chr14-73972424-INS-1621 |
| chr14-73982121-INS-153 | chr14-73982121-INS-153 |
| chr14-74018021-INS-56 | chr14-74018022-INS-64 |
| chr14-74037889-INS-713 | chr14-74037890-INS-714 |
| chr14-74043096-INS-64 | chr14-74043097-INS-65 |
| chr14-74056543-INS-53 | chr14-74056543-INS-53 |
| chr14-74058238-INS-52 | chr14-74058238-INS-60 |
| chr14-74072006-INS-4555 | chr14-74072007-INS-4556 |
| chr14-74084805-INS-64 | chr14-74084805-INS-64 |
| chr14-74086623-INS-230 | chr14-74086623-INS-230 |
| chr14-74089085-INS-56 | chr14-74089045-INS-72 |
| chr14-74113385-INS-3081 | chr14-74113386-INS-3089 |
| chr14-74142833-INS-95 | chr14-74142827-INS-113 |
| chr14-74144853-INS-73 | chr14-74144853-INS-72 |
| chr14-74195202-INS-4802 | chr14-74195203-INS-4802 |
| chr14-74202509-INS-151 | chr14-74202509-INS-151 |
| chr14-74214655-INS-7205 | chr14-74214656-INS-7205 |
| chr14-74248377-INS-245 | chr14-74248377-INS-246 |
| chr14-74256960-INS-76 | chr14-74256960-INS-76 |
| chr14-74257516-INS-7326 | chr14-74257517-INS-7326 |
| chr14-74280047-INS-1584 | chr14-74280048-INS-1585 |
| chr14-74301591-INS-756 | chr14-74301591-INS-756 |
| chr14-74304434-INS-385 | chr14-74304431-INS-361 |
| chr14-74313948-INS-6920 | chr14-74313949-INS-6920 |
| chr14-74331331-INS-4368 | chr14-74331332-INS-4368 |
| chr14-74342913-INS-72 | chr14-74342913-INS-72 |
| chr14-74360263-INS-100 | chr14-74360263-INS-90 |
| chr14-74361982-INS-208 | chr14-74361990-INS-209 |
| chr14-74381343-INS-1329 | chr14-74381344-INS-1328 |
| chr14-74403566-INS-82 | chr14-74403566-INS-68 |
| chr14-74420460-INS-169 | chr14-74420474-INS-177 |

| | |
|---|---|
| chr14-74431602-INS-62 | chr14-74431602-INS-62 |
| chr14-74435588-INS-6506 | chr14-74435589-INS-6504 |
| chr14-74440273-INS-65 | chr14-74440273-INS-56 |
| chr14-74470156-INS-291 | chr14-74470156-INS-270 |
| chr14-74477209-INS-53 | chr14-74477209-INS-51 |
| chr14-74492769-INS-52 | chr14-74492773-INS-60 |
| chr14-74535409-INS-750 | chr14-74535409-INS-750 |
| chr14-74536114-INS-167 | chr14-74536114-INS-167 |
| chr14-74569004-INS-66 | chr14-74568993-INS-80 |
| chr14-74608999-INS-362 | chr14-74608999-INS-362 |
| chr14-74616999-INS-177 | chr14-74617032-INS-177 |
| chr14-74634620-INS-310 | chr14-74634620-INS-310 |
| chr14-74662624-INS-51 | chr14-74662624-INS-52 |
| chr14-74692415-INS-6276 | chr14-74692416-INS-6278 |
| chr14-74704913-INS-60 | chr14-74704913-INS-62 |
| chr14-74712046-INS-564 | chr14-74712046-INS-564 |
| chr14-74726743-INS-70 | chr14-74726743-INS-70 |
| chr14-74761069-INS-301 | chr14-74761080-INS-281 |
| chr14-74802503-INS-282 | chr14-74802503-INS-282 |
| chr14-74808971-INS-202 | chr14-74808971-INS-207 |
| chr14-74813948-INS-77 | chr14-74813948-INS-125 |
| chr14-74855462-INS-349 | chr14-74855462-INS-349 |
| chr14-74871087-INS-540 | chr14-74871087-INS-540 |
| chr14-74888112-INS-52 | chr14-74888112-INS-52 |
| chr14-74927700-INS-6451 | chr14-74927701-INS-6451 |
| chr14-74949776-INS-84 | chr14-74949776-INS-84 |
| chr14-74974086-INS-59 | chr14-74974086-INS-59 |
| chr14-74985469-INS-91 | chr14-74985469-INS-91 |
| chr14-74988260-INS-59 | chr14-74988260-INS-61 |
| chr14-74996302-INS-163 | chr14-74996302-INS-155 |
| chr14-75021439-INS-62 | chr14-75021439-INS-86 |
| chr14-75028133-INS-83 | chr14-75028133-INS-83 |
| chr14-75030174-INS-641 | chr14-75030174-INS-641 |
| chr14-75091323-INS-324 | chr14-75091322-INS-327 |
| chr14-75168379-INS-334 | chr14-75168420-INS-340 |
| chr14-75169840-INS-138 | chr14-75169840-INS-139 |
| chr14-75179275-INS-59 | chr14-75179275-INS-59 |
| chr14-75197568-INS-267 | chr14-75197568-INS-267 |
| chr14-75198560-INS-52 | chr14-75198560-INS-58 |
| chr14-75215753-INS-68 | chr14-75215723-INS-79 |
| chr14-75220050-INS-78 | chr14-75220050-INS-56 |
| chr14-75222748-INS-85 | chr14-75222748-INS-92 |
| chr14-75226117-INS-291 | chr14-75226117-INS-292 |
| chr14-75228581-INS-72 | chr14-75228581-INS-76 |
| chr14-75233739-INS-56 | chr14-75233689-INS-83 |
| chr14-75236220-INS-2409 | chr14-75236221-INS-2408 |
| chr14-75246167-INS-70 | chr14-75246170-INS-69 |
| chr14-75272368-INS-142 | chr14-75272368-INS-152 |
| chr14-75280410-INS-131 | chr14-75280410-INS-131 |
| chr14-75281044-INS-141 | chr14-75281057-INS-99 |
| chr14-75299554-INS-52 | chr14-75299554-INS-56 |
| chr14-75316182-INS-748 | chr14-75316182-INS-754 |
| chr14-75349357-INS-216 | chr14-75349357-INS-218 |
| chr14-75396324-INS-68 | chr14-75396324-INS-54 |
| chr14-75431995-INS-59 | chr14-75431989-INS-57 |
| chr14-75437656-INS-197 | chr14-75437657-INS-197 |
| chr14-75475533-INS-226 | chr14-75475533-INS-234 |
| chr14-75477966-INS-193 | chr14-75477966-INS-193 |
| chr14-75483175-INS-2015 | chr14-75483176-INS-2015 |
| chr14-75495560-INS-54 | chr14-75495560-INS-54 |
| chr14-75503727-INS-341 | chr14-75503727-INS-341 |
| chr14-75519967-INS-57 | chr14-75519967-INS-54 |
| chr14-75535932-INS-204 | chr14-75535932-INS-204 |
| chr14-75603841-INS-521 | chr14-75603841-INS-521 |
| chr14-75609929-INS-1971 | chr14-75609930-INS-1951 |
| chr14-75611470-INS-203 | chr14-75611470-INS-203 |
| chr14-75621714-INS-174 | chr14-75621716-INS-175 |
| chr14-75624663-INS-181 | chr14-75624663-INS-181 |
| chr14-75627306-INS-328 | chr14-75627304-INS-368 |
| chr14-75642608-INS-3233 | chr14-75642609-INS-3233 |
| chr14-75650477-INS-458 | chr14-75650477-INS-457 |
| chr14-75655291-INS-130 | chr14-75655291-INS-108 |
| chr14-75703652-INS-101 | chr14-75703652-INS-101 |
| chr14-75733285-INS-421 | chr14-75733285-INS-420 |
| chr14-75770788-INS-163 | chr14-75770788-INS-163 |
| chr14-75791176-INS-62 | chr14-75791176-INS-101 |
| chr14-75797724-INS-134 | chr14-75797727-INS-138 |

| | |
|---|---|
| chr14-75804646-INS-162 | chr14-75804646-INS-162 |
| chr14-75808020-INS-253 | chr14-75808020-INS-248 |
| chr14-75854660-INS-5090 | chr14-75854661-INS-5090 |
| chr14-75898797-INS-1888 | chr14-75898798-INS-1888 |
| chr14-75938904-INS-89 | chr14-75938904-INS-89 |
| chr14-75948438-INS-272 | chr14-75948438-INS-272 |
| chr14-75984938-INS-52 | chr14-75984907-INS-83 |
| chr14-76009876-INS-1866 | chr14-76009877-INS-1866 |
| chr14-76039499-INS-582 | chr14-76039499-INS-581 |
| chr14-76040527-INS-210 | chr14-76040530-INS-210 |
| chr14-76054110-INS-196 | chr14-76054110-INS-197 |
| chr14-76065634-INS-203 | chr14-76065634-INS-204 |
| chr14-76096956-INS-63 | chr14-76096956-INS-63 |
| chr14-76099541-INS-72 | chr14-76099541-INS-72 |
| chr14-76103667-INS-192 | chr14-76103667-INS-198 |
| chr14-76106418-INS-82 | chr14-76106418-INS-82 |
| chr14-76127471-INS-68 | chr14-76127471-INS-68 |
| chr14-76151910-INS-110 | chr14-76151910-INS-110 |
| chr14-76178447-INS-65 | chr14-76178447-INS-65 |
| chr14-76216374-INS-179 | chr14-76216374-INS-182 |
| chr14-76220241-INS-4019 | chr14-76220242-INS-4019 |
| chr14-76230095-INS-724 | chr14-76230099-INS-725 |
| chr14-76251729-INS-54 | chr14-76251729-INS-70 |
| chr14-76257149-INS-486 | chr14-76257149-INS-486 |
| chr14-76275982-INS-129 | chr14-76275978-INS-147 |
| chr14-76289106-INS-198 | chr14-76289106-INS-198 |
| chr14-76311310-INS-72 | chr14-76311310-INS-72 |
| chr14-76315267-INS-203 | chr14-76315267-INS-203 |
| chr14-76316151-INS-732 | chr14-76316151-INS-732 |
| chr14-76329532-INS-203 | chr14-76329532-INS-203 |
| chr14-76334250-INS-212 | chr14-76334250-INS-212 |
| chr14-76343380-INS-199 | chr14-76343380-INS-199 |
| chr14-76362200-INS-52 | chr14-76362200-INS-52 |
| chr14-76380777-INS-54 | chr14-76380777-INS-56 |
| chr14-76381783-INS-83 | chr14-76381783-INS-83 |
| chr14-76382373-INS-56 | chr14-76382373-INS-56 |
| chr14-76383955-INS-56 | chr14-76383955-INS-64 |
| chr14-76398454-INS-124 | chr14-76398454-INS-104 |
| chr14-76411257-INS-197 | chr14-76411257-INS-197 |
| chr14-76433510-INS-115 | chr14-76433510-INS-115 |
| chr14-76436452-INS-94 | chr14-76436453-INS-70 |
| chr14-76436900-INS-215 | chr14-76436900-INS-215 |
| chr14-76438826-INS-249 | chr14-76438826-INS-203 |
| chr14-76441709-INS-264 | chr14-76441709-INS-264 |
| chr14-76447860-INS-99 | chr14-76447852-INS-91 |
| chr14-76477817-INS-106 | chr14-76477817-INS-106 |
| chr14-76482580-INS-206 | chr14-76482580-INS-213 |
| chr14-76488402-INS-123 | chr14-76488402-INS-83 |
| chr14-76488748-INS-1334 | chr14-76488749-INS-1334 |
| chr14-76508632-INS-5223 | chr14-76508633-INS-5223 |
| chr14-76511783-INS-104 | chr14-76511783-INS-114 |
| chr14-76549985-INS-181 | chr14-76549985-INS-184 |
| chr14-76564948-INS-326 | chr14-76564948-INS-331 |
| chr14-76583068-INS-2937 | chr14-76583069-INS-2937 |
| chr14-76583937-INS-196 | chr14-76583937-INS-196 |
| chr14-76599647-INS-226 | chr14-76599647-INS-230 |
| chr14-76620015-INS-116 | chr14-76620015-INS-116 |
| chr14-76623445-INS-342 | chr14-76623493-INS-337 |
| chr14-76632497-INS-3992 | chr14-76632498-INS-3992 |
| chr14-76637606-INS-1864 | chr14-76637607-INS-1864 |
| chr14-76671339-INS-176 | chr14-76671339-INS-176 |
| chr14-76696769-INS-216 | chr14-76696769-INS-216 |
| chr14-76704750-INS-69 | chr14-76704750-INS-69 |
| chr14-76721234-INS-215 | chr14-76721234-INS-219 |
| chr14-76724680-INS-60 | chr14-76724681-INS-56 |
| chr14-76733199-INS-109 | chr14-76733199-INS-109 |
| chr14-76735025-INS-120 | chr14-76735047-INS-120 |
| chr14-76842298-INS-134 | chr14-76842298-INS-136 |
| chr14-76848310-INS-62 | chr14-76848310-INS-64 |
| chr14-76863471-INS-802 | chr14-76863471-INS-803 |
| chr14-76870392-INS-54 | chr14-76870392-INS-54 |
| chr14-76906132-INS-95 | chr14-76906132-INS-70 |
| chr14-76907708-INS-3806 | chr14-76907709-INS-3806 |
| chr14-76912100-INS-106 | chr14-76912100-INS-80 |
| chr14-76913972-INS-71 | chr14-76913972-INS-71 |
| chr14-76914763-INS-327 | chr14-76914763-INS-295 |
| chr14-76917950-INS-533 | chr14-76917950-INS-533 |

| | |
|---|---|
| chr14-76922284-INS-55 | chr14-76922284-INS-59 |
| chr14-76925050-INS-329 | chr14-76925050-INS-341 |
| chr14-76988148-INS-86 | chr14-76988148-INS-86 |
| chr14-77028711-INS-53 | chr14-77028711-INS-59 |
| chr14-77081984-INS-95 | chr14-77082009-INS-83 |
| chr14-77097805-INS-6660 | chr14-77097806-INS-6661 |
| chr14-77107122-INS-88 | chr14-77107122-INS-103 |
| chr14-77119762-INS-56 | chr14-77119762-INS-56 |
| chr14-77126328-INS-661 | chr14-77126328-INS-661 |
| chr14-77136923-INS-51 | chr14-77136923-INS-52 |
| chr14-77166850-INS-787 | chr14-77166850-INS-787 |
| chr14-77171149-INS-64 | chr14-77171149-INS-64 |
| chr14-77192918-INS-1077 | chr14-77192919-INS-1075 |
| chr14-77209638-INS-86 | chr14-77209628-INS-84 |
| chr14-77210803-INS-163 | chr14-77210803-INS-163 |
| chr14-77250859-INS-70 | chr14-77250859-INS-70 |
| chr14-77254278-INS-885 | chr14-77254278-INS-885 |
| chr14-77269405-INS-1103 | chr14-77269406-INS-1103 |
| chr14-77269575-INS-246 | chr14-77269581-INS-257 |
| chr14-77277598-INS-90 | chr14-77277598-INS-90 |
| chr14-77287273-INS-275 | chr14-77287273-INS-275 |
| chr14-77348301-INS-140 | chr14-77348301-INS-140 |
| chr14-77361386-INS-548 | chr14-77361386-INS-561 |
| chr14-77363475-INS-122 | chr14-77363475-INS-122 |
| chr14-77377088-INS-235 | chr14-77377134-INS-228 |
| chr14-77383664-INS-60 | chr14-77383662-INS-79 |
| chr14-77407141-INS-6704 | chr14-77407142-INS-6704 |
| chr14-77430708-INS-6792 | chr14-77430709-INS-6792 |
| chr14-77445867-INS-167 | chr14-77445867-INS-167 |
| chr14-77448729-INS-94 | chr14-77448743-INS-85 |
| chr14-77453807-INS-6897 | chr14-77453808-INS-6895 |
| chr14-77462176-INS-61 | chr14-77462176-INS-61 |
| chr14-77468762-INS-141 | chr14-77468766-INS-150 |
| chr14-77502001-INS-425 | chr14-77502001-INS-425 |
| chr14-77569761-INS-200 | chr14-77569761-INS-200 |
| chr14-77605891-INS-1110 | chr14-77605892-INS-1109 |
| chr14-77607710-INS-94 | chr14-77607710-INS-94 |
| chr14-77725501-INS-150 | chr14-77725501-INS-150 |
| chr14-77743109-INS-514 | chr14-77743109-INS-514 |
| chr14-77757437-INS-576 | chr14-77757437-INS-576 |
| chr14-77758375-INS-120 | chr14-77758375-INS-120 |
| chr14-77759723-INS-56 | chr14-77759723-INS-56 |
| chr14-77770749-INS-1138 | chr14-77770750-INS-1138 |
| chr14-77773373-INS-87 | chr14-77773373-INS-87 |
| chr14-77782599-INS-110 | chr14-77782599-INS-110 |
| chr14-77826203-INS-87 | chr14-77826203-INS-100 |
| chr14-77833641-INS-954 | chr14-77833641-INS-955 |
| chr14-77851448-INS-75 | chr14-77851448-INS-75 |
| chr14-77857809-INS-61 | chr14-77857809-INS-61 |
| chr14-77860897-INS-553 | chr14-77860897-INS-552 |
| chr14-77866291-INS-558 | chr14-77866291-INS-531 |
| chr14-77900869-INS-54 | chr14-77900869-INS-54 |
| chr14-77903195-INS-4760 | chr14-77903196-INS-4760 |
| chr14-77911903-INS-50 | chr14-77911903-INS-52 |
| chr14-77912931-INS-204 | chr14-77912931-INS-204 |
| chr14-77921229-INS-564 | chr14-77921229-INS-564 |
| chr14-77962087-INS-360 | chr14-77962087-INS-360 |
| chr14-77965160-INS-196 | chr14-77965160-INS-197 |
| chr14-77966853-INS-739 | chr14-77966853-INS-739 |
| chr14-77970738-INS-58 | chr14-77970738-INS-62 |
| chr14-77981678-INS-87 | chr14-77981678-INS-87 |
| chr14-77984494-INS-72 | chr14-77984494-INS-76 |
| chr14-77995570-INS-96 | chr14-77995558-INS-104 |
| chr14-78000313-INS-7072 | chr14-78000314-INS-7071 |
| chr14-78001398-INS-64 | chr14-78001398-INS-64 |
| chr14-78005032-INS-129 | chr14-78005032-INS-131 |
| chr14-78012304-INS-342 | chr14-78012304-INS-342 |
| chr14-78016468-INS-72 | chr14-78016467-INS-74 |
| chr14-78036023-INS-83 | chr14-78036023-INS-101 |
| chr14-78039398-INS-182 | chr14-78039398-INS-182 |
| chr14-78043440-INS-80 | chr14-78043440-INS-86 |
| chr14-78051151-INS-110 | chr14-78051151-INS-110 |
| chr14-78085489-INS-200 | chr14-78085489-INS-200 |
| chr14-78087146-INS-117 | chr14-78087146-INS-117 |
| chr14-78119280-INS-58 | chr14-78119280-INS-69 |
| chr14-78128754-INS-90 | chr14-78128754-INS-90 |
| chr14-78160975-INS-640 | chr14-78160975-INS-640 |

| | |
|---|---|
| chr14-78170592-INS-507 | chr14-78170592-INS-522 |
| chr14-78203763-INS-443 | chr14-78203764-INS-443 |
| chr14-78230208-INS-211 | chr14-78230208-INS-219 |
| chr14-78238107-INS-441 | chr14-78238107-INS-441 |
| chr14-78251085-INS-208 | chr14-78251085-INS-208 |
| chr14-78260033-INS-52 | chr14-78259992-INS-97 |
| chr14-78293932-INS-217 | chr14-78293932-INS-217 |
| chr14-78296923-INS-120 | chr14-78296923-INS-120 |
| chr14-78304759-INS-389 | chr14-78304771-INS-406 |
| chr14-78318844-INS-225 | chr14-78318844-INS-225 |
| chr14-78323329-INS-59 | chr14-78323330-INS-71 |
| chr14-78334434-INS-57 | chr14-78334434-INS-57 |
| chr14-78347769-INS-51 | chr14-78347769-INS-53 |
| chr14-78351676-INS-4764 | chr14-78351677-INS-4764 |
| chr14-78392930-INS-52 | chr14-78392931-INS-52 |
| chr14-78399733-INS-64 | chr14-78399733-INS-64 |
| chr14-78416829-INS-63 | chr14-78416829-INS-59 |
| chr14-78466169-INS-79 | chr14-78466169-INS-81 |
| chr14-78473399-INS-54 | chr14-78473399-INS-63 |
| chr14-78484024-INS-172 | chr14-78484030-INS-160 |
| chr14-78485876-INS-96 | chr14-78485876-INS-97 |
| chr14-78521648-INS-95 | chr14-78521648-INS-95 |
| chr14-78565361-INS-62 | chr14-78565361-INS-62 |
| chr14-78611870-INS-64 | chr14-78611870-INS-64 |
| chr14-78631270-INS-79 | chr14-78631270-INS-79 |
| chr14-78640649-INS-6333 | chr14-78640650-INS-6333 |
| chr14-78643977-INS-3096 | chr14-78643978-INS-3096 |
| chr14-78703102-INS-58 | chr14-78703102-INS-60 |
| chr14-78712150-INS-76 | chr14-78712150-INS-68 |
| chr14-78725488-INS-65 | chr14-78725488-INS-66 |
| chr14-78731272-INS-98 | chr14-78731272-INS-98 |
| chr14-78743370-INS-60 | chr14-78743371-INS-52 |
| chr14-78775891-INS-60 | chr14-78775891-INS-50 |
| chr14-78808204-INS-53 | chr14-78808204-INS-53 |
| chr14-78812094-INS-51 | chr14-78812094-INS-51 |
| chr14-78820969-INS-72 | chr14-78820969-INS-72 |
| chr14-78827660-INS-781 | chr14-78827660-INS-765 |
| chr14-78853423-INS-1347 | chr14-78853424-INS-1370 |
| chr14-78853433-INS-174 | chr14-78853433-INS-174 |
| chr14-78882120-INS-70 | chr14-78882120-INS-70 |
| chr14-78914271-INS-52 | chr14-78914271-INS-52 |
| chr14-78914725-INS-99 | chr14-78914726-INS-69 |
| chr14-78919414-INS-1109 | chr14-78919415-INS-1096 |
| chr14-78921098-INS-146 | chr14-78921103-INS-174 |
| chr14-78923990-INS-285 | chr14-78923990-INS-285 |
| chr14-78937918-INS-294 | chr14-78937961-INS-295 |
| chr14-78945130-INS-5075 | chr14-78945131-INS-5075 |
| chr14-78945840-INS-524 | chr14-78945840-INS-540 |
| chr14-78955604-INS-549 | chr14-78955604-INS-549 |
| chr14-78957972-INS-64 | chr14-78957958-INS-76 |
| chr14-78976234-INS-381 | chr14-78976234-INS-383 |
| chr14-78981382-INS-81 | chr14-78981385-INS-50 |
| chr14-78998961-INS-316 | chr14-78998967-INS-285 |
| chr14-79038326-INS-451 | chr14-79038326-INS-451 |
| chr14-79060824-INS-358 | chr14-79060824-INS-358 |
| chr14-79062414-INS-375 | chr14-79062414-INS-375 |
| chr14-79068312-INS-211 | chr14-79068312-INS-211 |
| chr14-79070832-INS-7487 | chr14-79070833-INS-7491 |
| chr14-79073136-INS-424 | chr14-79073136-INS-424 |
| chr14-79078546-INS-445 | chr14-79078546-INS-445 |
| chr14-79094253-INS-6644 | chr14-79094254-INS-6644 |
| chr14-79136925-INS-556 | chr14-79136925-INS-556 |
| chr14-79143718-INS-59 | chr14-79143718-INS-63 |
| chr14-79151855-INS-9088 | chr14-79151856-INS-9089 |
| chr14-79161490-INS-107 | chr14-79161490-INS-119 |
| chr14-79167614-INS-1783 | chr14-79167615-INS-1783 |
| chr14-79196742-INS-60 | chr14-79196742-INS-75 |
| chr14-79211589-INS-216 | chr14-79211589-INS-216 |
| chr14-79217357-INS-6539 | chr14-79217358-INS-6544 |
| chr14-79242254-INS-64 | chr14-79242254-INS-64 |
| chr14-79250575-INS-297 | chr14-79250575-INS-329 |
| chr14-79255421-INS-382 | chr14-79255421-INS-382 |
| chr14-79262709-INS-211 | chr14-79262709-INS-213 |
| chr14-79293171-INS-211 | chr14-79293171-INS-211 |
| chr14-79299396-INS-7047 | chr14-79299397-INS-7048 |
| chr14-79304794-INS-66 | chr14-79304794-INS-66 |
| chr14-79305502-INS-600 | chr14-79305502-INS-600 |

| | |
|---|---|
| chr14-79307339-INS-121 | chr14-79307339-INS-123 |
| chr14-79336595-INS-191 | chr14-79336595-INS-191 |
| chr14-79349789-INS-52 | chr14-79349789-INS-54 |
| chr14-79381671-INS-781 | chr14-79381671-INS-781 |
| chr14-79395966-INS-58 | chr14-79395966-INS-58 |
| chr14-79437529-INS-66 | chr14-79437529-INS-66 |
| chr14-79448585-INS-742 | chr14-79448585-INS-742 |
| chr14-79455061-INS-113 | chr14-79455012-INS-131 |
| chr14-79456041-INS-54 | chr14-79456041-INS-50 |
| chr14-79462316-INS-104 | chr14-79462316-INS-112 |
| chr14-79471087-INS-87 | chr14-79471087-INS-87 |
| chr14-79498005-INS-51 | chr14-79498005-INS-51 |
| chr14-79524806-INS-655 | chr14-79524806-INS-656 |
| chr14-79528133-INS-206 | chr14-79528133-INS-206 |
| chr14-79533227-INS-64 | chr14-79533227-INS-62 |
| chr14-79534417-INS-65 | chr14-79534417-INS-65 |
| chr14-79537278-INS-100 | chr14-79537278-INS-106 |
| chr14-79539848-INS-51 | chr14-79539848-INS-51 |
| chr14-79542682-INS-88 | chr14-79542682-INS-88 |
| chr14-79558905-INS-1325 | chr14-79558906-INS-1326 |
| chr14-79562420-INS-58 | chr14-79562410-INS-56 |
| chr14-79566221-INS-2153 | chr14-79566222-INS-2154 |
| chr14-79582593-INS-209 | chr14-79582593-INS-209 |
| chr14-79599016-INS-404 | chr14-79599016-INS-404 |
| chr14-79600685-INS-70 | chr14-79600685-INS-64 |
| chr14-79615713-INS-961 | chr14-79615714-INS-961 |
| chr14-79618026-INS-56 | chr14-79618026-INS-58 |
| chr14-79625257-INS-196 | chr14-79625257-INS-196 |
| chr14-79675241-INS-434 | chr14-79675241-INS-434 |
| chr14-79680807-INS-90 | chr14-79680822-INS-78 |
| chr14-79700142-INS-390 | chr14-79700142-INS-390 |
| chr14-79735607-INS-77 | chr14-79735607-INS-77 |
| chr14-79742704-INS-211 | chr14-79742704-INS-211 |
| chr14-79808666-INS-135 | chr14-79808666-INS-135 |
| chr14-79812432-INS-208 | chr14-79812432-INS-208 |
| chr14-79859303-INS-62 | chr14-79859303-INS-64 |
| chr14-79871495-INS-59 | chr14-79871495-INS-65 |
| chr14-79900559-INS-1149 | chr14-79900560-INS-1149 |
| chr14-79905507-INS-67 | chr14-79905507-INS-67 |
| chr14-79906620-INS-171 | chr14-79906620-INS-171 |
| chr14-79928092-INS-91 | chr14-79928092-INS-91 |
| chr14-79937797-INS-348 | chr14-79937797-INS-350 |
| chr14-79975128-INS-82 | chr14-79975102-INS-58 |
| chr14-79997791-INS-232 | chr14-79997792-INS-233 |
| chr14-80076374-INS-72 | chr14-80076374-INS-72 |
| chr14-80089805-INS-88 | chr14-80089805-INS-89 |
| chr14-80142486-INS-7517 | chr14-80142487-INS-7520 |
| chr14-80148540-INS-1409 | chr14-80148541-INS-1416 |
| chr14-80257558-INS-173 | chr14-80257559-INS-144 |
| chr14-80311937-INS-75 | chr14-80311955-INS-74 |
| chr14-80379365-INS-127 | chr14-80379369-INS-154 |
| chr14-80423069-INS-299 | chr14-80423069-INS-299 |
| chr14-80465335-INS-1920 | chr14-80465336-INS-1953 |
| chr14-80493787-INS-68 | chr14-80493787-INS-67 |
| chr14-80508548-INS-84 | chr14-80508550-INS-51 |
| chr14-80516936-INS-478 | chr14-80516936-INS-478 |
| chr14-80521236-INS-3177 | chr14-80521237-INS-3178 |
| chr14-80634287-INS-51 | chr14-80634287-INS-51 |
| chr14-80651989-INS-2731 | chr14-80651990-INS-2731 |
| chr14-80654175-INS-4432 | chr14-80654176-INS-4433 |
| chr14-80668018-INS-60 | chr14-80668055-INS-65 |
| chr14-80702029-INS-7039 | chr14-80702030-INS-7040 |
| chr14-80734052-INS-183 | chr14-80734052-INS-183 |
| chr14-80738583-INS-935 | chr14-80738583-INS-893 |
| chr14-80745254-INS-1468 | chr14-80745255-INS-1468 |
| chr14-80770111-INS-72 | chr14-80770111-INS-72 |
| chr14-80770583-INS-2934 | chr14-80770584-INS-2934 |
| chr14-80787236-INS-393 | chr14-80787236-INS-393 |
| chr14-80791869-INS-1348 | chr14-80791870-INS-1356 |
| chr14-80804011-INS-151 | chr14-80804019-INS-151 |
| chr14-80813971-INS-220 | chr14-80813971-INS-221 |
| chr14-80845841-INS-67 | chr14-80845841-INS-63 |
| chr14-80858575-INS-560 | chr14-80858575-INS-560 |
| chr14-80871347-INS-51 | chr14-80871347-INS-54 |
| chr14-80893061-INS-113 | chr14-80893061-INS-113 |
| chr14-80909005-INS-699 | chr14-80909005-INS-704 |
| chr14-80953325-INS-68 | chr14-80953325-INS-54 |

| | |
|---|---|
| chr14-80967876-INS-202 | chr14-80967876-INS-202 |
| chr14-80985301-INS-638 | chr14-80985301-INS-638 |
| chr14-80987853-INS-56 | chr14-80987853-INS-56 |
| chr14-81012857-INS-134 | chr14-81012857-INS-134 |
| chr14-81102980-INS-529 | chr14-81102980-INS-528 |
| chr14-81129029-INS-775 | chr14-81129029-INS-789 |
| chr14-81190629-INS-7390 | chr14-81190630-INS-7390 |
| chr14-81205322-INS-2396 | chr14-81205323-INS-2407 |
| chr14-81236768-INS-230 | chr14-81236768-INS-230 |
| chr14-81265065-INS-196 | chr14-81265065-INS-196 |
| chr14-81269795-INS-69 | chr14-81269795-INS-68 |
| chr14-81297930-INS-54 | chr14-81297930-INS-54 |
| chr14-81303239-INS-59 | chr14-81303239-INS-67 |
| chr14-81344078-INS-114 | chr14-81344078-INS-116 |
| chr14-81345033-INS-3149 | chr14-81345034-INS-3149 |
| chr14-81353856-INS-57 | chr14-81353856-INS-57 |
| chr14-81390833-INS-141 | chr14-81390833-INS-143 |
| chr14-81423196-INS-220 | chr14-81423196-INS-221 |
| chr14-81425451-INS-210 | chr14-81425451-INS-210 |
| chr14-81438211-INS-52 | chr14-81438211-INS-58 |
| chr14-81442003-INS-76 | chr14-81442002-INS-80 |
| chr14-81453265-INS-453 | chr14-81453265-INS-454 |
| chr14-81542607-INS-6341 | chr14-81542608-INS-6341 |
| chr14-81543788-INS-1892 | chr14-81543789-INS-1892 |
| chr14-81551824-INS-4445 | chr14-81551825-INS-4445 |
| chr14-81557915-INS-64 | chr14-81557916-INS-54 |
| chr14-81563409-INS-199 | chr14-81563409-INS-199 |
| chr14-81593107-INS-78 | chr14-81593107-INS-93 |
| chr14-81705827-INS-527 | chr14-81705827-INS-528 |
| chr14-81765824-INS-566 | chr14-81765824-INS-566 |
| chr14-81774110-INS-1861 | chr14-81774111-INS-1861 |
| chr14-81776214-INS-1232 | chr14-81776215-INS-1232 |
| chr14-81794787-INS-67 | chr14-81794787-INS-67 |
| chr14-81811135-INS-60 | chr14-81811135-INS-65 |
| chr14-81820008-INS-52 | chr14-81820008-INS-66 |
| chr14-81827816-INS-56 | chr14-81827816-INS-56 |
| chr14-81837573-INS-4146 | chr14-81837574-INS-4146 |
| chr14-81840514-INS-80 | chr14-81840537-INS-72 |
| chr14-81860478-INS-61 | chr14-81860478-INS-67 |
| chr14-81868527-INS-2900 | chr14-81868528-INS-2900 |
| chr14-81870709-INS-6598 | chr14-81870710-INS-6598 |
| chr14-81875449-INS-2141 | chr14-81875450-INS-2141 |
| chr14-81887815-INS-3187 | chr14-81887816-INS-3187 |
| chr14-81897024-INS-6818 | chr14-81897025-INS-6819 |
| chr14-81906733-INS-50 | chr14-81906733-INS-50 |
| chr14-81938939-INS-66 | chr14-81938948-INS-64 |
| chr14-81952374-INS-6251 | chr14-81952375-INS-6251 |
| chr14-81973112-INS-65 | chr14-81973112-INS-71 |
| chr14-81976403-INS-482 | chr14-81976403-INS-483 |
| chr14-81988217-INS-312 | chr14-81988217-INS-312 |
| chr14-82007113-INS-80 | chr14-82007113-INS-80 |
| chr14-82025463-INS-760 | chr14-82025463-INS-760 |
| chr14-82050700-INS-56 | chr14-82050700-INS-56 |
| chr14-82063912-INS-239 | chr14-82063912-INS-239 |
| chr14-82094201-INS-64 | chr14-82094201-INS-64 |
| chr14-82098862-INS-6386 | chr14-82098863-INS-6386 |
| chr14-82101426-INS-636 | chr14-82101426-INS-637 |
| chr14-82104907-INS-479 | chr14-82104907-INS-479 |
| chr14-82106209-INS-657 | chr14-82106209-INS-657 |
| chr14-82114116-INS-56 | chr14-82114116-INS-56 |
| chr14-82134122-INS-5311 | chr14-82134123-INS-5311 |
| chr14-82134533-INS-925 | chr14-82134533-INS-925 |
| chr14-82145664-INS-390 | chr14-82145664-INS-381 |
| chr14-82150011-INS-103 | chr14-82150011-INS-103 |
| chr14-82151006-INS-50 | chr14-82151006-INS-50 |
| chr14-82153892-INS-267 | chr14-82153892-INS-267 |
| chr14-82163129-INS-57 | chr14-82163129-INS-57 |
| chr14-82175981-INS-303 | chr14-82175981-INS-303 |
| chr14-82177106-INS-51 | chr14-82177106-INS-51 |
| chr14-82196311-INS-1654 | chr14-82196312-INS-1655 |
| chr14-82199365-INS-331 | chr14-82199365-INS-331 |
| chr14-82201526-INS-428 | chr14-82201526-INS-428 |
| chr14-82206274-INS-332 | chr14-82206274-INS-332 |
| chr14-82213653-INS-3201 | chr14-82213654-INS-3201 |
| chr14-82213781-INS-62 | chr14-82213770-INS-78 |
| chr14-82225641-INS-175 | chr14-82225641-INS-173 |
| chr14-82237219-INS-1324 | chr14-82237220-INS-1324 |

| | |
|---|---|
| chr14-82253460-INS-211 | chr14-82253460-INS-211 |
| chr14-82268616-INS-78 | chr14-82268613-INS-76 |
| chr14-82281205-INS-204 | chr14-82281205-INS-204 |
| chr14-82286075-INS-216 | chr14-82286072-INS-212 |
| chr14-82289674-INS-82 | chr14-82289674-INS-54 |
| chr14-82307465-INS-512 | chr14-82307465-INS-512 |
| chr14-82321845-INS-424 | chr14-82321846-INS-424 |
| chr14-82326382-INS-144 | chr14-82326382-INS-144 |
| chr14-82328585-INS-395 | chr14-82328585-INS-395 |
| chr14-82333713-INS-62 | chr14-82333713-INS-64 |
| chr14-82334676-INS-449 | chr14-82334676-INS-448 |
| chr14-82337466-INS-54 | chr14-82337466-INS-56 |
| chr14-82378985-INS-251 | chr14-82378985-INS-251 |
| chr14-82380487-INS-483 | chr14-82380487-INS-483 |
| chr14-82383044-INS-89 | chr14-82383044-INS-89 |
| chr14-82388918-INS-922 | chr14-82388918-INS-922 |
| chr14-82394826-INS-60 | chr14-82394826-INS-60 |
| chr14-82398894-INS-464 | chr14-82398894-INS-464 |
| chr14-82406369-INS-1614 | chr14-82406360-INS-1647 |
| chr14-82423156-INS-1864 | chr14-82423157-INS-1864 |
| chr14-82437923-INS-6385 | chr14-82437924-INS-6386 |
| chr14-82455316-INS-379 | chr14-82455316-INS-380 |
| chr14-82456715-INS-617 | chr14-82456715-INS-617 |
| chr14-82469224-INS-51 | chr14-82469224-INS-59 |
| chr14-82517011-INS-974 | chr14-82517011-INS-974 |
| chr14-82521696-INS-3117 | chr14-82521697-INS-3117 |
| chr14-82528970-INS-94 | chr14-82528970-INS-94 |
| chr14-82541187-INS-507 | chr14-82541187-INS-507 |
| chr14-82624535-INS-102 | chr14-82624535-INS-108 |
| chr14-82644981-INS-111 | chr14-82644981-INS-112 |
| chr14-82670028-INS-6233 | chr14-82670029-INS-6235 |
| chr14-82678655-INS-527 | chr14-82678655-INS-527 |
| chr14-82710184-INS-6272 | chr14-82710185-INS-6274 |
| chr14-82711900-INS-1202 | chr14-82711898-INS-1196 |
| chr14-82712745-INS-86 | chr14-82712758-INS-51 |
| chr14-82751762-INS-822 | chr14-82751762-INS-822 |
| chr14-82753474-INS-55 | chr14-82753474-INS-55 |
| chr14-82766144-INS-1050 | chr14-82766145-INS-1050 |
| chr14-82767906-INS-53 | chr14-82767906-INS-54 |
| chr14-82776882-INS-57 | chr14-82776882-INS-57 |
| chr14-82790321-INS-3876 | chr14-82790324-INS-3859 |
| chr14-82795124-INS-93 | chr14-82795124-INS-83 |
| chr14-82806375-INS-52 | chr14-82806375-INS-58 |
| chr14-82846894-INS-468 | chr14-82846894-INS-475 |
| chr14-82857397-INS-52 | chr14-82857397-INS-56 |
| chr14-82873243-INS-220 | chr14-82873245-INS-225 |
| chr14-82884761-INS-652 | chr14-82884761-INS-652 |
| chr14-82921682-INS-87 | chr14-82921682-INS-75 |
| chr14-82934977-INS-51 | chr14-82934977-INS-54 |
| chr14-82947028-INS-95 | chr14-82947028-INS-97 |
| chr14-82947544-INS-4921 | chr14-82947545-INS-4921 |
| chr14-82969527-INS-60 | chr14-82969527-INS-60 |
| chr14-82978941-INS-79 | chr14-82978941-INS-65 |
| chr14-82979987-INS-201 | chr14-82979987-INS-201 |
| chr14-82980103-INS-7131 | chr14-82980104-INS-7131 |
| chr14-82984992-INS-58 | chr14-82984992-INS-58 |
| chr14-83003538-INS-2059 | chr14-83003539-INS-2063 |
| chr14-83013299-INS-56 | chr14-83013299-INS-68 |
| chr14-83040095-INS-7114 | chr14-83040096-INS-7114 |
| chr14-83070645-INS-87 | chr14-83070645-INS-89 |
| chr14-83110696-INS-178 | chr14-83110696-INS-178 |
| chr14-83112757-INS-77 | chr14-83112757-INS-83 |
| chr14-83146563-INS-4553 | chr14-83146564-INS-4553 |
| chr14-83157981-INS-71 | chr14-83157981-INS-71 |
| chr14-83248130-INS-189 | chr14-83248130-INS-185 |
| chr14-83250554-INS-2820 | chr14-83250555-INS-2820 |
| chr14-83255426-INS-70 | chr14-83255426-INS-70 |
| chr14-83256702-INS-51 | chr14-83256702-INS-51 |
| chr14-83268975-INS-66 | chr14-83268975-INS-99 |
| chr14-83296381-INS-183 | chr14-83296381-INS-186 |
| chr14-83317953-INS-6423 | chr14-83317954-INS-6424 |
| chr14-83328164-INS-62 | chr14-83328163-INS-50 |
| chr14-83332631-INS-79 | chr14-83332631-INS-79 |
| chr14-83334832-INS-78 | chr14-83334832-INS-78 |
| chr14-83336851-INS-108 | chr14-83336851-INS-128 |
| chr14-83340254-INS-168 | chr14-83340254-INS-188 |
| chr14-83354149-INS-6747 | chr14-83354150-INS-6749 |

| | |
|---|---|
| chr14-83370144-INS-63 | chr14-83370144-INS-67 |
| chr14-83384159-INS-52 | chr14-83384159-INS-50 |
| chr14-83385516-INS-265 | chr14-83385516-INS-314 |
| chr14-83404436-INS-1056 | chr14-83404437-INS-1056 |
| chr14-83426881-INS-4012 | chr14-83426882-INS-4012 |
| chr14-83445249-INS-88 | chr14-83445249-INS-88 |
| chr14-83448287-INS-1131 | chr14-83448288-INS-1133 |
| chr14-83456497-INS-1811 | chr14-83456498-INS-1813 |
| chr14-83456622-INS-102 | chr14-83456622-INS-64 |
| chr14-83469236-INS-134 | chr14-83469236-INS-158 |
| chr14-83494160-INS-222 | chr14-83494147-INS-226 |
| chr14-83509404-INS-224 | chr14-83509404-INS-224 |
| chr14-83514577-INS-1016 | chr14-83514578-INS-1016 |
| chr14-83529794-INS-58 | chr14-83529794-INS-54 |
| chr14-83536406-INS-636 | chr14-83536406-INS-636 |
| chr14-83573444-INS-6983 | chr14-83573445-INS-6987 |
| chr14-83577502-INS-6310 | chr14-83577503-INS-6311 |
| chr14-83599353-INS-5455 | chr14-83599342-INS-5472 |
| chr14-83609508-INS-7264 | chr14-83609509-INS-7264 |
| chr14-83615319-INS-109 | chr14-83615320-INS-120 |
| chr14-83632031-INS-2637 | chr14-83632032-INS-2637 |
| chr14-83634257-INS-79 | chr14-83634258-INS-85 |
| chr14-83678086-INS-82 | chr14-83678092-INS-66 |
| chr14-83694438-INS-142 | chr14-83694438-INS-142 |
| chr14-83696074-INS-78 | chr14-83696074-INS-78 |
| chr14-83697929-INS-329 | chr14-83697929-INS-329 |
| chr14-83700182-INS-4327 | chr14-83700183-INS-4327 |
| chr14-83708357-INS-1384 | chr14-83708358-INS-1384 |
| chr14-83725305-INS-6424 | chr14-83725306-INS-6424 |
| chr14-83735730-INS-52 | chr14-83735731-INS-52 |
| chr14-83771248-INS-72 | chr14-83771248-INS-64 |
| chr14-83821817-INS-6460 | chr14-83821818-INS-6461 |
| chr14-83870829-INS-83 | chr14-83870829-INS-83 |
| chr14-83883206-INS-1341 | chr14-83883207-INS-1341 |
| chr14-83884617-INS-77 | chr14-83884617-INS-62 |
| chr14-83885857-INS-492 | chr14-83885857-INS-492 |
| chr14-83890496-INS-51 | chr14-83890497-INS-69 |
| chr14-83915250-INS-6497 | chr14-83915251-INS-6497 |
| chr14-83960533-INS-104 | chr14-83960533-INS-104 |
| chr14-83972404-INS-1260 | chr14-83972413-INS-1251 |
| chr14-83980396-INS-332 | chr14-83980396-INS-332 |
| chr14-83981907-INS-788 | chr14-83981907-INS-787 |
| chr14-83986374-INS-54 | chr14-83986374-INS-54 |
| chr14-83993525-INS-800 | chr14-83993525-INS-800 |
| chr14-84000088-INS-1124 | chr14-84000089-INS-1124 |
| chr14-84001194-INS-54 | chr14-84001194-INS-60 |
| chr14-84006952-INS-9749 | chr14-84006953-INS-9749 |
| chr14-84012634-INS-466 | chr14-84012634-INS-466 |
| chr14-84039547-INS-434 | chr14-84039547-INS-434 |
| chr14-84067453-INS-104 | chr14-84067453-INS-104 |
| chr14-84072863-INS-188 | chr14-84072863-INS-188 |
| chr14-84091350-INS-500 | chr14-84091350-INS-501 |
| chr14-84098593-INS-739 | chr14-84098593-INS-739 |
| chr14-84100808-INS-184 | chr14-84100808-INS-184 |
| chr14-84131125-INS-6202 | chr14-84131126-INS-6202 |
| chr14-84140740-INS-3496 | chr14-84140741-INS-3504 |
| chr14-84159862-INS-260 | chr14-84159862-INS-302 |
| chr14-84182778-INS-71 | chr14-84182751-INS-78 |
| chr14-84192189-INS-66 | chr14-84192189-INS-61 |
| chr14-84203488-INS-60 | chr14-84203488-INS-67 |
| chr14-84210266-INS-4859 | chr14-84210267-INS-4859 |
| chr14-84216301-INS-6646 | chr14-84216302-INS-6646 |
| chr14-84233510-INS-133 | chr14-84233510-INS-133 |
| chr14-84248256-INS-62 | chr14-84248256-INS-66 |
| chr14-84255543-INS-390 | chr14-84255543-INS-390 |
| chr14-84268947-INS-489 | chr14-84268968-INS-486 |
| chr14-84273096-INS-68 | chr14-84273096-INS-68 |
| chr14-84288059-INS-2579 | chr14-84288060-INS-2579 |
| chr14-84290587-INS-6514 | chr14-84290588-INS-6514 |
| chr14-84299875-INS-52 | chr14-84299895-INS-70 |
| chr14-84309228-INS-6137 | chr14-84309229-INS-6137 |
| chr14-84312809-INS-1113 | chr14-84312810-INS-1113 |
| chr14-84327540-INS-6526 | chr14-84327541-INS-6527 |
| chr14-84333170-INS-92 | chr14-84333184-INS-50 |
| chr14-84354058-INS-2760 | chr14-84354059-INS-2754 |
| chr14-84358331-INS-401 | chr14-84358331-INS-401 |
| chr14-84407415-INS-1493 | chr14-84407416-INS-1492 |

| | |
|---|---|
| chr14-84479714-INS-1495 | chr14-84479715-INS-1495 |
| chr14-84492813-INS-305 | chr14-84492813-INS-305 |
| chr14-84497792-INS-92 | chr14-84497792-INS-92 |
| chr14-84503662-INS-71 | chr14-84503662-INS-71 |
| chr14-84514628-INS-400 | chr14-84514628-INS-400 |
| chr14-84516415-INS-347 | chr14-84516415-INS-347 |
| chr14-84527043-INS-111 | chr14-84527043-INS-111 |
| chr14-84538590-INS-88 | chr14-84538588-INS-92 |
| chr14-84558149-INS-230 | chr14-84558149-INS-230 |
| chr14-84561711-INS-344 | chr14-84561711-INS-344 |
| chr14-84568853-INS-1243 | chr14-84568854-INS-1244 |
| chr14-84570810-INS-684 | chr14-84570810-INS-682 |
| chr14-84573217-INS-63 | chr14-84573217-INS-61 |
| chr14-84578015-INS-1269 | chr14-84578016-INS-1271 |
| chr14-84592966-INS-536 | chr14-84592966-INS-536 |
| chr14-84595905-INS-53 | chr14-84595905-INS-53 |
| chr14-84637654-INS-1259 | chr14-84637655-INS-1259 |
| chr14-84648051-INS-53 | chr14-84648051-INS-53 |
| chr14-84650782-INS-202 | chr14-84650782-INS-202 |
| chr14-84665131-INS-51 | chr14-84665131-INS-52 |
| chr14-84738751-INS-4828 | chr14-84738752-INS-4828 |
| chr14-84751464-INS-102 | chr14-84751464-INS-96 |
| chr14-84767056-INS-64 | chr14-84767056-INS-64 |
| chr14-84791231-INS-417 | chr14-84791232-INS-417 |
| chr14-84794970-INS-112 | chr14-84794970-INS-111 |
| chr14-84808812-INS-66 | chr14-84808812-INS-66 |
| chr14-84810380-INS-4801 | chr14-84810381-INS-4801 |
| chr14-84831892-INS-4704 | chr14-84831893-INS-4704 |
| chr14-84854454-INS-60 | chr14-84854454-INS-60 |
| chr14-84857010-INS-58 | chr14-84857010-INS-50 |
| chr14-84910767-INS-79 | chr14-84910771-INS-74 |
| chr14-84912228-INS-74 | chr14-84912229-INS-92 |
| chr14-84928848-INS-1201 | chr14-84928849-INS-1201 |
| chr14-84944299-INS-1708 | chr14-84944300-INS-1708 |
| chr14-84946783-INS-188 | chr14-84946783-INS-186 |
| chr14-84955877-INS-199 | chr14-84955877-INS-199 |
| chr14-84964617-INS-235 | chr14-84964618-INS-225 |
| chr14-84977595-INS-90 | chr14-84977595-INS-90 |
| chr14-84983021-INS-60 | chr14-84983021-INS-54 |
| chr14-84985859-INS-66 | chr14-84985859-INS-52 |
| chr14-84994790-INS-184 | chr14-84994794-INS-186 |
| chr14-85017787-INS-231 | chr14-85017787-INS-225 |
| chr14-85022218-INS-7087 | chr14-85022219-INS-7087 |
| chr14-85027336-INS-80 | chr14-85027336-INS-80 |
| chr14-85039286-INS-75 | chr14-85039286-INS-99 |
| chr14-85061331-INS-146 | chr14-85061331-INS-146 |
| chr14-85063202-INS-57 | chr14-85063202-INS-69 |
| chr14-85071209-INS-505 | chr14-85071209-INS-505 |
| chr14-85079134-INS-80 | chr14-85079134-INS-80 |
| chr14-85081908-INS-237 | chr14-85081908-INS-237 |
| chr14-85087485-INS-455 | chr14-85087485-INS-455 |
| chr14-85100014-INS-52 | chr14-85100007-INS-68 |
| chr14-85107741-INS-79 | chr14-85107741-INS-79 |
| chr14-85128201-INS-2779 | chr14-85128202-INS-2780 |
| chr14-85128729-INS-54 | chr14-85128709-INS-58 |
| chr14-85133704-INS-500 | chr14-85133704-INS-500 |
| chr14-85137484-INS-800 | chr14-85137484-INS-804 |
| chr14-85143951-INS-231 | chr14-85143951-INS-231 |
| chr14-85146710-INS-1866 | chr14-85146711-INS-1866 |
| chr14-85156814-INS-277 | chr14-85156814-INS-277 |
| chr14-85213346-INS-527 | chr14-85213346-INS-527 |
| chr14-85217521-INS-4123 | chr14-85217522-INS-4123 |
| chr14-85229813-INS-501 | chr14-85229813-INS-501 |
| chr14-85230616-INS-162 | chr14-85230616-INS-166 |
| chr14-85235134-INS-80 | chr14-85235134-INS-80 |
| chr14-85262006-INS-553 | chr14-85262006-INS-553 |
| chr14-85280397-INS-202 | chr14-85280397-INS-202 |
| chr14-85294565-INS-906 | chr14-85294565-INS-906 |
| chr14-85307417-INS-62 | chr14-85307424-INS-64 |
| chr14-85331759-INS-179 | chr14-85331759-INS-181 |
| chr14-85341780-INS-328 | chr14-85341780-INS-328 |
| chr14-85359654-INS-1442 | chr14-85359655-INS-1442 |
| chr14-85365886-INS-66 | chr14-85365886-INS-64 |
| chr14-85395464-INS-1114 | chr14-85395465-INS-1114 |
| chr14-85407412-INS-55 | chr14-85407412-INS-57 |
| chr14-85412521-INS-78 | chr14-85412521-INS-88 |
| chr14-85422738-INS-108 | chr14-85422737-INS-80 |

| | |
|---|---|
| chr14-85451643-INS-52 | chr14-85451643-INS-56 |
| chr14-85463109-INS-5152 | chr14-85463110-INS-5152 |
| chr14-85494729-INS-58 | chr14-85494729-INS-54 |
| chr14-85518462-INS-123 | chr14-85518462-INS-123 |
| chr14-85526012-INS-96 | chr14-85525991-INS-84 |
| chr14-85559077-INS-521 | chr14-85559077-INS-521 |
| chr14-85565435-INS-478 | chr14-85565435-INS-479 |
| chr14-85566282-INS-105 | chr14-85566282-INS-101 |
| chr14-85577201-INS-401 | chr14-85577201-INS-401 |
| chr14-85598675-INS-63 | chr14-85598675-INS-63 |
| chr14-85603846-INS-498 | chr14-85603846-INS-498 |
| chr14-85621590-INS-2059 | chr14-85621591-INS-2060 |
| chr14-85644322-INS-84 | chr14-85644330-INS-50 |
| chr14-85648788-INS-7104 | chr14-85648789-INS-7104 |
| chr14-85654412-INS-60 | chr14-85654412-INS-60 |
| chr14-85668080-INS-60 | chr14-85668080-INS-56 |
| chr14-85691578-INS-77 | chr14-85691578-INS-84 |
| chr14-85715848-INS-65 | chr14-85715848-INS-55 |
| chr14-85742880-INS-109 | chr14-85742880-INS-109 |
| chr14-85748068-INS-75 | chr14-85748068-INS-75 |
| chr14-85749272-INS-3277 | chr14-85749273-INS-3277 |
| chr14-85749980-INS-369 | chr14-85749980-INS-369 |
| chr14-85774230-INS-119 | chr14-85774230-INS-93 |
| chr14-85788745-INS-2493 | chr14-85788740-INS-2490 |
| chr14-85797207-INS-59 | chr14-85797207-INS-51 |
| chr14-85802187-INS-134 | chr14-85802187-INS-134 |
| chr14-85830957-INS-4761 | chr14-85830958-INS-4761 |
| chr14-85833610-INS-6655 | chr14-85833611-INS-6655 |
| chr14-85850884-INS-5551 | chr14-85850885-INS-5551 |
| chr14-85859708-INS-8849 | chr14-85859709-INS-8855 |
| chr14-85863977-INS-82 | chr14-85863976-INS-81 |
| chr14-85876496-INS-2628 | chr14-85876497-INS-2628 |
| chr14-85890726-INS-262 | chr14-85890726-INS-262 |
| chr14-85892113-INS-6803 | chr14-85892114-INS-6803 |
| chr14-85906491-INS-516 | chr14-85906491-INS-516 |
| chr14-85920506-INS-404 | chr14-85920506-INS-404 |
| chr14-85942421-INS-60 | chr14-85942421-INS-60 |
| chr14-85959904-INS-312 | chr14-85959904-INS-313 |
| chr14-85970753-INS-54 | chr14-85970755-INS-56 |
| chr14-85980114-INS-1526 | chr14-85980115-INS-1538 |
| chr14-86038014-INS-646 | chr14-86038017-INS-672 |
| chr14-86051816-INS-126 | chr14-86051824-INS-122 |
| chr14-86062579-INS-7120 | chr14-86062580-INS-7120 |
| chr14-86073520-INS-1465 | chr14-86073521-INS-1465 |
| chr14-86141510-INS-188 | chr14-86141510-INS-189 |
| chr14-86146106-INS-60 | chr14-86146106-INS-62 |
| chr14-86147711-INS-261 | chr14-86147711-INS-261 |
| chr14-86150427-INS-6766 | chr14-86150428-INS-6765 |
| chr14-86182067-INS-216 | chr14-86182067-INS-216 |
| chr14-86225511-INS-64 | chr14-86225511-INS-68 |
| chr14-86228935-INS-56 | chr14-86228939-INS-50 |
| chr14-86256356-INS-66 | chr14-86256321-INS-106 |
| chr14-86267426-INS-60 | chr14-86267426-INS-62 |
| chr14-86275569-INS-199 | chr14-86275569-INS-199 |
| chr14-86275993-INS-2884 | chr14-86275994-INS-2884 |
| chr14-86280418-INS-331 | chr14-86280418-INS-331 |
| chr14-86287087-INS-206 | chr14-86287087-INS-205 |
| chr14-86290368-INS-52 | chr14-86290368-INS-53 |
| chr14-86308847-INS-430 | chr14-86308847-INS-430 |
| chr14-86326911-INS-53 | chr14-86326911-INS-53 |
| chr14-86353896-INS-76 | chr14-86353896-INS-80 |
| chr14-86363288-INS-204 | chr14-86363288-INS-198 |
| chr14-86365170-INS-479 | chr14-86365170-INS-479 |
| chr14-86381843-INS-52 | chr14-86381843-INS-56 |
| chr14-86402298-INS-60 | chr14-86402298-INS-85 |
| chr14-86412700-INS-2575 | chr14-86412701-INS-2575 |
| chr14-86420081-INS-89 | chr14-86420081-INS-116 |
| chr14-86427214-INS-4579 | chr14-86427215-INS-4579 |
| chr14-86430424-INS-500 | chr14-86430424-INS-500 |
| chr14-86433003-INS-208 | chr14-86433003-INS-208 |
| chr14-86438036-INS-90 | chr14-86438036-INS-90 |
| chr14-86441165-INS-58 | chr14-86441167-INS-82 |
| chr14-86475801-INS-71 | chr14-86475794-INS-78 |
| chr14-86488528-INS-6386 | chr14-86488551-INS-6382 |
| chr14-86495968-INS-64 | chr14-86495968-INS-62 |
| chr14-86517810-INS-409 | chr14-86517810-INS-413 |
| chr14-86553507-INS-5183 | chr14-86553508-INS-5183 |

| | |
|---|---|
| chr14-86591105-INS-219 | chr14-86591105-INS-219 |
| chr14-86604042-INS-60 | chr14-86604056-INS-84 |
| chr14-86627995-INS-427 | chr14-86627995-INS-430 |
| chr14-86643952-INS-61 | chr14-86643952-INS-61 |
| chr14-86674764-INS-96 | chr14-86674764-INS-100 |
| chr14-86708531-INS-141 | chr14-86708559-INS-138 |
| chr14-86712758-INS-4593 | chr14-86712759-INS-4591 |
| chr14-86720450-INS-97 | chr14-86720448-INS-105 |
| chr14-86753931-INS-798 | chr14-86753931-INS-798 |
| chr14-86758588-INS-140 | chr14-86758588-INS-140 |
| chr14-86787337-INS-52 | chr14-86787337-INS-50 |
| chr14-86789217-INS-485 | chr14-86789217-INS-485 |
| chr14-86791978-INS-6214 | chr14-86791979-INS-6213 |
| chr14-86793476-INS-6457 | chr14-86793477-INS-6457 |
| chr14-86802615-INS-1866 | chr14-86802616-INS-1869 |
| chr14-86852893-INS-92 | chr14-86852893-INS-92 |
| chr14-86868607-INS-76 | chr14-86868607-INS-76 |
| chr14-86879268-INS-493 | chr14-86879268-INS-494 |
| chr14-86888394-INS-100 | chr14-86888394-INS-120 |
| chr14-86905411-INS-300 | chr14-86905411-INS-300 |
| chr14-86914276-INS-7223 | chr14-86914277-INS-7223 |
| chr14-86916285-INS-67 | chr14-86916276-INS-62 |
| chr14-86958174-INS-58 | chr14-86958174-INS-62 |
| chr14-86962088-INS-275 | chr14-86962088-INS-275 |
| chr14-86971634-INS-407 | chr14-86971634-INS-407 |
| chr14-86977382-INS-212 | chr14-86977382-INS-212 |
| chr14-86999436-INS-209 | chr14-86999436-INS-209 |
| chr14-87007888-INS-2807 | chr14-87007889-INS-2804 |
| chr14-87038705-INS-80 | chr14-87038705-INS-50 |
| chr14-87052339-INS-241 | chr14-87052339-INS-240 |
| chr14-87055649-INS-495 | chr14-87055649-INS-495 |
| chr14-87062014-INS-328 | chr14-87062015-INS-328 |
| chr14-87073480-INS-104 | chr14-87073480-INS-108 |
| chr14-87081677-INS-326 | chr14-87081678-INS-310 |
| chr14-87116547-INS-3420 | chr14-87116548-INS-3420 |
| chr14-87119452-INS-376 | chr14-87119452-INS-376 |
| chr14-87128545-INS-476 | chr14-87128545-INS-476 |
| chr14-87137422-INS-64 | chr14-87137422-INS-64 |
| chr14-87168499-INS-81 | chr14-87168498-INS-66 |
| chr14-87169422-INS-64 | chr14-87169434-INS-72 |
| chr14-87194107-INS-200 | chr14-87194107-INS-200 |
| chr14-87210323-INS-420 | chr14-87210323-INS-414 |
| chr14-87211735-INS-140 | chr14-87211689-INS-148 |
| chr14-87215995-INS-126 | chr14-87215995-INS-104 |
| chr14-87224700-INS-87 | chr14-87224700-INS-87 |
| chr14-87248735-INS-82 | chr14-87248735-INS-66 |
| chr14-87250430-INS-86 | chr14-87250441-INS-81 |
| chr14-87267656-INS-157 | chr14-87267656-INS-158 |
| chr14-87298287-INS-52 | chr14-87298287-INS-52 |
| chr14-87306795-INS-436 | chr14-87306795-INS-436 |
| chr14-87321975-INS-3520 | chr14-87321976-INS-3520 |
| chr14-87353726-INS-217 | chr14-87353726-INS-217 |
| chr14-87377505-INS-60 | chr14-87377505-INS-60 |
| chr14-87385411-INS-6684 | chr14-87385412-INS-6684 |
| chr14-87429479-INS-67 | chr14-87429479-INS-67 |
| chr14-87455356-INS-98 | chr14-87455356-INS-98 |
| chr14-87504246-INS-71 | chr14-87504246-INS-70 |
| chr14-87515494-INS-110 | chr14-87515494-INS-116 |
| chr14-87520612-INS-71 | chr14-87520612-INS-75 |
| chr14-87524408-INS-73 | chr14-87524414-INS-74 |
| chr14-87531838-INS-62 | chr14-87531811-INS-57 |
| chr14-87572338-INS-57 | chr14-87572338-INS-57 |
| chr14-87587838-INS-210 | chr14-87587838-INS-210 |
| chr14-87590276-INS-75 | chr14-87590281-INS-79 |
| chr14-87604378-INS-58 | chr14-87604378-INS-50 |
| chr14-87623525-INS-930 | chr14-87623525-INS-937 |
| chr14-87638406-INS-430 | chr14-87638406-INS-430 |
| chr14-87664410-INS-210 | chr14-87664410-INS-210 |
| chr14-87707695-INS-344 | chr14-87707696-INS-344 |
| chr14-87713484-INS-209 | chr14-87713484-INS-210 |
| chr14-87719819-INS-58 | chr14-87719819-INS-62 |
| chr14-87742233-INS-53 | chr14-87742234-INS-53 |
| chr14-87770523-INS-62 | chr14-87770523-INS-62 |
| chr14-87814294-INS-160 | chr14-87814294-INS-160 |
| chr14-87816049-INS-6629 | chr14-87816050-INS-6632 |
| chr14-87828836-INS-6802 | chr14-87828837-INS-6805 |
| chr14-87830638-INS-114 | chr14-87830639-INS-111 |

| | |
|---|---|
| chr14-87833650-INS-56 | chr14-87833650-INS-56 |
| chr14-87834617-INS-261 | chr14-87834592-INS-266 |
| chr14-87838351-INS-615 | chr14-87838351-INS-631 |
| chr14-87841108-INS-218 | chr14-87841108-INS-218 |
| chr14-87848098-INS-105 | chr14-87848098-INS-106 |
| chr14-87875891-INS-63 | chr14-87875891-INS-63 |
| chr14-87880161-INS-440 | chr14-87880161-INS-439 |
| chr14-87890704-INS-374 | chr14-87890704-INS-374 |
| chr14-87900889-INS-55 | chr14-87900889-INS-55 |
| chr14-87929511-INS-108 | chr14-87929527-INS-102 |
| chr14-87957601-INS-60 | chr14-87957604-INS-64 |
| chr14-87974774-INS-78 | chr14-87974774-INS-114 |
| chr14-87988493-INS-3568 | chr14-87988494-INS-3576 |
| chr14-88017756-INS-56 | chr14-88017756-INS-52 |
| chr14-88024407-INS-162 | chr14-88024407-INS-162 |
| chr14-88034947-INS-54 | chr14-88034947-INS-54 |
| chr14-88064656-INS-112 | chr14-88064656-INS-112 |
| chr14-88065918-INS-55 | chr14-88065918-INS-55 |
| chr14-88069589-INS-199 | chr14-88069589-INS-209 |
| chr14-88073020-INS-496 | chr14-88073026-INS-497 |
| chr14-88079641-INS-262 | chr14-88079639-INS-251 |
| chr14-88101226-INS-55 | chr14-88101226-INS-59 |
| chr14-88135321-INS-51 | chr14-88135321-INS-51 |
| chr14-88142059-INS-1493 | chr14-88142060-INS-1494 |
| chr14-88143502-INS-494 | chr14-88143503-INS-494 |
| chr14-88214997-INS-54 | chr14-88214997-INS-62 |
| chr14-88221250-INS-63 | chr14-88221252-INS-71 |
| chr14-88241742-INS-78 | chr14-88241742-INS-78 |
| chr14-88243955-INS-224 | chr14-88243955-INS-227 |
| chr14-88245038-INS-541 | chr14-88245038-INS-538 |
| chr14-88282835-INS-2001 | chr14-88282841-INS-2012 |
| chr14-88293721-INS-54 | chr14-88293721-INS-54 |
| chr14-88299319-INS-239 | chr14-88299319-INS-239 |
| chr14-88311843-INS-464 | chr14-88311843-INS-465 |
| chr14-88313278-INS-1485 | chr14-88313281-INS-1466 |
| chr14-88320564-INS-7050 | chr14-88320565-INS-7051 |
| chr14-88344387-INS-56 | chr14-88344381-INS-58 |
| chr14-88347828-INS-58 | chr14-88347828-INS-58 |
| chr14-88370876-INS-74 | chr14-88370884-INS-68 |
| chr14-88388584-INS-3059 | chr14-88388585-INS-3059 |
| chr14-88388650-INS-1019 | chr14-88388651-INS-1019 |
| chr14-88421386-INS-662 | chr14-88421386-INS-662 |
| chr14-88437018-INS-492 | chr14-88437018-INS-492 |
| chr14-88442090-INS-127 | chr14-88442090-INS-127 |
| chr14-88465660-INS-55 | chr14-88465660-INS-55 |
| chr14-88470189-INS-60 | chr14-88470189-INS-60 |
| chr14-88502442-INS-299 | chr14-88502442-INS-299 |
| chr14-88511848-INS-5896 | chr14-88511849-INS-5894 |
| chr14-88535514-INS-129 | chr14-88535514-INS-130 |
| chr14-88538211-INS-87 | chr14-88538211-INS-87 |
| chr14-88545892-INS-64 | chr14-88545892-INS-66 |
| chr14-88553035-INS-56 | chr14-88553031-INS-64 |
| chr14-88557549-INS-575 | chr14-88557549-INS-575 |
| chr14-88585201-INS-6722 | chr14-88585202-INS-6722 |
| chr14-88590098-INS-917 | chr14-88590098-INS-929 |
| chr14-88592465-INS-53 | chr14-88592465-INS-53 |
| chr14-88600775-INS-76 | chr14-88600775-INS-82 |
| chr14-88623479-INS-52 | chr14-88623479-INS-60 |
| chr14-88625741-INS-71 | chr14-88625741-INS-75 |
| chr14-88630802-INS-894 | chr14-88630802-INS-894 |
| chr14-88633004-INS-51 | chr14-88633009-INS-51 |
| chr14-88645310-INS-122 | chr14-88645310-INS-121 |
| chr14-88658517-INS-6149 | chr14-88658518-INS-6149 |
| chr14-88660395-INS-333 | chr14-88660397-INS-334 |
| chr14-88678733-INS-523 | chr14-88678733-INS-523 |
| chr14-88683682-INS-58 | chr14-88683655-INS-64 |
| chr14-88690280-INS-257 | chr14-88690280-INS-257 |
| chr14-88692310-INS-436 | chr14-88692310-INS-436 |
| chr14-88716814-INS-66 | chr14-88716814-INS-66 |
| chr14-88742087-INS-6395 | chr14-88742088-INS-6402 |
| chr14-88757033-INS-250 | chr14-88757007-INS-288 |
| chr14-88769815-INS-66 | chr14-88769815-INS-50 |
| chr14-88771398-INS-488 | chr14-88771398-INS-488 |
| chr14-88775679-INS-52 | chr14-88775680-INS-50 |
| chr14-88807961-INS-816 | chr14-88807961-INS-816 |
| chr14-88810073-INS-92 | chr14-88810073-INS-92 |
| chr14-88826362-INS-52 | chr14-88826362-INS-52 |

| | |
|---|---|
| chr14-88843307-INS-84 | chr14-88843315-INS-64 |
| chr14-88845864-INS-325 | chr14-88845868-INS-321 |
| chr14-88846959-INS-159 | chr14-88846960-INS-159 |
| chr14-88865093-INS-60 | chr14-88865121-INS-64 |
| chr14-88869685-INS-56 | chr14-88869685-INS-56 |
| chr14-88872808-INS-390 | chr14-88872808-INS-390 |
| chr14-88881956-INS-392 | chr14-88881956-INS-393 |
| chr14-88904895-INS-60 | chr14-88904895-INS-60 |
| chr14-88923377-INS-1892 | chr14-88923378-INS-1892 |
| chr14-88945379-INS-195 | chr14-88945379-INS-195 |
| chr14-88964497-INS-57 | chr14-88964497-INS-57 |
| chr14-88981561-INS-833 | chr14-88981561-INS-833 |
| chr14-88982289-INS-56 | chr14-88982289-INS-58 |
| chr14-88991318-INS-166 | chr14-88991318-INS-166 |
| chr14-89001431-INS-6254 | chr14-89001432-INS-6256 |
| chr14-89007207-INS-1625 | chr14-89007208-INS-1623 |
| chr14-89007951-INS-52 | chr14-89007951-INS-58 |
| chr14-89008459-INS-78 | chr14-89008465-INS-105 |
| chr14-89012371-INS-75 | chr14-89012371-INS-75 |
| chr14-89014954-INS-56 | chr14-89014954-INS-56 |
| chr14-89017022-INS-367 | chr14-89017022-INS-367 |
| chr14-89036702-INS-58 | chr14-89036702-INS-60 |
| chr14-89040265-INS-330 | chr14-89040265-INS-331 |
| chr14-89044204-INS-58 | chr14-89044204-INS-58 |
| chr14-89067512-INS-484 | chr14-89067512-INS-484 |
| chr14-89087129-INS-2628 | chr14-89087130-INS-2630 |
| chr14-89098227-INS-56 | chr14-89098190-INS-80 |
| chr14-89132191-INS-484 | chr14-89132193-INS-483 |
| chr14-89139655-INS-219 | chr14-89139655-INS-219 |
| chr14-89145095-INS-233 | chr14-89145095-INS-231 |
| chr14-89158060-INS-231 | chr14-89158060-INS-238 |
| chr14-89162651-INS-169 | chr14-89162657-INS-159 |
| chr14-89162869-INS-4614 | chr14-89162870-INS-4614 |
| chr14-89166463-INS-6338 | chr14-89166464-INS-6322 |
| chr14-89169500-INS-346 | chr14-89169500-INS-346 |
| chr14-89171230-INS-953 | chr14-89171230-INS-952 |
| chr14-89200751-INS-6388 | chr14-89200752-INS-6389 |
| chr14-89211740-INS-54 | chr14-89211740-INS-54 |
| chr14-89219674-INS-178 | chr14-89219674-INS-179 |
| chr14-89240111-INS-586 | chr14-89240111-INS-586 |
| chr14-89251658-INS-60 | chr14-89251658-INS-62 |
| chr14-89315962-INS-60 | chr14-89315964-INS-58 |
| chr14-89319461-INS-7102 | chr14-89319462-INS-7102 |
| chr14-89388603-INS-76 | chr14-89388603-INS-74 |
| chr14-89424032-INS-82 | chr14-89424032-INS-88 |
| chr14-89427035-INS-294 | chr14-89427030-INS-292 |
| chr14-89433743-INS-856 | chr14-89433743-INS-856 |
| chr14-89444300-INS-7100 | chr14-89444301-INS-7100 |
| chr14-89447722-INS-183 | chr14-89447722-INS-185 |
| chr14-89447927-INS-510 | chr14-89447927-INS-510 |
| chr14-89461155-INS-799 | chr14-89461155-INS-799 |
| chr14-89478214-INS-87 | chr14-89478212-INS-103 |
| chr14-89488813-INS-121 | chr14-89488813-INS-121 |
| chr14-89490140-INS-135 | chr14-89490140-INS-135 |
| chr14-89512096-INS-66 | chr14-89512096-INS-66 |
| chr14-89618706-INS-100 | chr14-89618706-INS-50 |
| chr14-89642759-INS-4690 | chr14-89642760-INS-4690 |
| chr14-89665915-INS-100 | chr14-89665915-INS-72 |
| chr14-89671834-INS-73 | chr14-89671871-INS-56 |
| chr14-89673727-INS-106 | chr14-89673727-INS-106 |
| chr14-89692558-INS-549 | chr14-89692549-INS-550 |
| chr14-89693862-INS-52 | chr14-89693862-INS-52 |
| chr14-89764790-INS-179 | chr14-89764790-INS-179 |
| chr14-89771780-INS-148 | chr14-89771780-INS-148 |
| chr14-89773733-INS-880 | chr14-89773733-INS-879 |
| chr14-89780362-INS-326 | chr14-89780362-INS-326 |
| chr14-89781087-INS-273 | chr14-89781091-INS-276 |
| chr14-89832271-INS-182 | chr14-89832271-INS-182 |
| chr14-89848173-INS-329 | chr14-89848173-INS-329 |
| chr14-89880304-INS-55 | chr14-89880310-INS-57 |
| chr14-89903406-INS-66 | chr14-89903406-INS-66 |
| chr14-89958595-INS-662 | chr14-89958595-INS-662 |
| chr14-89965351-INS-354 | chr14-89965351-INS-354 |
| chr14-89981219-INS-171 | chr14-89981219-INS-174 |
| chr14-89992097-INS-328 | chr14-89992097-INS-328 |
| chr14-89993232-INS-96 | chr14-89993232-INS-98 |
| chr14-90009842-INS-872 | chr14-90009842-INS-872 |

| | |
|---|---|
| chr14-90010718-INS-136 | chr14-90010718-INS-136 |
| chr14-90014604-INS-1024 | chr14-90014615-INS-1016 |
| chr14-90029507-INS-408 | chr14-90029507-INS-408 |
| chr14-90043853-INS-60 | chr14-90043853-INS-60 |
| chr14-90055575-INS-1426 | chr14-90055576-INS-1426 |
| chr14-90061254-INS-1562 | chr14-90061255-INS-1563 |
| chr14-90063461-INS-99 | chr14-90063461-INS-99 |
| chr14-90100970-INS-212 | chr14-90100970-INS-212 |
| chr14-90121346-INS-51 | chr14-90121346-INS-51 |
| chr14-90143343-INS-91 | chr14-90143343-INS-91 |
| chr14-90151699-INS-3275 | chr14-90151716-INS-3272 |
| chr14-90153969-INS-2048 | chr14-90153970-INS-2049 |
| chr14-90160135-INS-65 | chr14-90160135-INS-63 |
| chr14-90186371-INS-454 | chr14-90186372-INS-454 |
| chr14-90199697-INS-748 | chr14-90199697-INS-748 |
| chr14-90224858-INS-152 | chr14-90224854-INS-151 |
| chr14-90233770-INS-488 | chr14-90233770-INS-488 |
| chr14-90255653-INS-199 | chr14-90255653-INS-199 |
| chr14-90273225-INS-60 | chr14-90273225-INS-84 |
| chr14-90279614-INS-440 | chr14-90279614-INS-440 |
| chr14-90288212-INS-7094 | chr14-90288213-INS-7094 |
| chr14-90296177-INS-584 | chr14-90296177-INS-584 |
| chr14-90346950-INS-53 | chr14-90346950-INS-53 |
| chr14-90351082-INS-138 | chr14-90351082-INS-137 |
| chr14-90366300-INS-136 | chr14-90366300-INS-137 |
| chr14-90372448-INS-53 | chr14-90372448-INS-83 |
| chr14-90381586-INS-123 | chr14-90381592-INS-78 |
| chr14-90388025-INS-54 | chr14-90388007-INS-57 |
| chr14-90471230-INS-150 | chr14-90471230-INS-150 |
| chr14-90475678-INS-111 | chr14-90475678-INS-111 |
| chr14-90505013-INS-50 | chr14-90505025-INS-60 |
| chr14-90525484-INS-87 | chr14-90525479-INS-109 |
| chr14-90525834-INS-406 | chr14-90525821-INS-407 |
| chr14-90528750-INS-227 | chr14-90528750-INS-227 |
| chr14-90569725-INS-60 | chr14-90569726-INS-60 |
| chr14-90628888-INS-1741 | chr14-90628888-INS-1745 |
| chr14-90632796-INS-52 | chr14-90632796-INS-62 |
| chr14-90641966-INS-125 | chr14-90641966-INS-121 |
| chr14-90642737-INS-89 | chr14-90642737-INS-89 |
| chr14-90657431-INS-54 | chr14-90657431-INS-54 |
| chr14-90658298-INS-205 | chr14-90658298-INS-205 |
| chr14-90666018-INS-107 | chr14-90666017-INS-110 |
| chr14-90671203-INS-135 | chr14-90671203-INS-135 |
| chr14-90673686-INS-757 | chr14-90673686-INS-757 |
| chr14-90677635-INS-6488 | chr14-90677636-INS-6488 |
| chr14-90688465-INS-203 | chr14-90688465-INS-203 |
| chr14-90693331-INS-86 | chr14-90693331-INS-126 |
| chr14-90715744-INS-58 | chr14-90715744-INS-56 |
| chr14-90718374-INS-7127 | chr14-90718375-INS-7127 |
| chr14-90719562-INS-7280 | chr14-90719563-INS-7280 |
| chr14-90741061-INS-426 | chr14-90741061-INS-426 |
| chr14-90797473-INS-479 | chr14-90797473-INS-479 |
| chr14-90822130-INS-66 | chr14-90822130-INS-66 |
| chr14-90829158-INS-62 | chr14-90829158-INS-64 |
| chr14-90837795-INS-6163 | chr14-90837796-INS-6163 |
| chr14-90839066-INS-384 | chr14-90839066-INS-384 |
| chr14-90853024-INS-59 | chr14-90853024-INS-59 |
| chr14-90895842-INS-54 | chr14-90895796-INS-74 |
| chr14-90897099-INS-390 | chr14-90897099-INS-390 |
| chr14-90898805-INS-220 | chr14-90898805-INS-215 |
| chr14-90899744-INS-1209 | chr14-90899745-INS-1209 |
| chr14-90902061-INS-6645 | chr14-90902062-INS-6645 |
| chr14-90910810-INS-440 | chr14-90910810-INS-440 |
| chr14-90913639-INS-55 | chr14-90913639-INS-55 |
| chr14-90919235-INS-199 | chr14-90919235-INS-199 |
| chr14-90932037-INS-495 | chr14-90932037-INS-495 |
| chr14-90936109-INS-206 | chr14-90936109-INS-206 |
| chr14-90951562-INS-68 | chr14-90951562-INS-68 |
| chr14-90960775-INS-60 | chr14-90960775-INS-50 |
| chr14-90977138-INS-325 | chr14-90977138-INS-325 |
| chr14-90993222-INS-91 | chr14-90993222-INS-79 |
| chr14-90997435-INS-89 | chr14-90997435-INS-90 |
| chr14-91009685-INS-59 | chr14-91009688-INS-82 |
| chr14-91062815-INS-524 | chr14-91062815-INS-521 |
| chr14-91071181-INS-184 | chr14-91071181-INS-184 |
| chr14-91080753-INS-64 | chr14-91080803-INS-64 |
| chr14-91081017-INS-4366 | chr14-91081018-INS-4366 |

| | |
|---|---|
| chr14-91095291-INS-4315 | chr14-91095292-INS-4315 |
| chr14-91101530-INS-6330 | chr14-91101531-INS-6330 |
| chr14-91122147-INS-95 | chr14-91122147-INS-95 |
| chr14-91125098-INS-7127 | chr14-91125099-INS-7127 |
| chr14-91159022-INS-203 | chr14-91159022-INS-201 |
| chr14-91164853-INS-152 | chr14-91164853-INS-153 |
| chr14-91198304-INS-79 | chr14-91198304-INS-78 |
| chr14-91199560-INS-109 | chr14-91199557-INS-76 |
| chr14-91260823-INS-6214 | chr14-91260824-INS-6213 |
| chr14-91287139-INS-8535 | chr14-91287140-INS-8535 |
| chr14-91291050-INS-55 | chr14-91291050-INS-55 |
| chr14-91296275-INS-58 | chr14-91296275-INS-87 |
| chr14-91314730-INS-83 | chr14-91314721-INS-76 |
| chr14-91352338-INS-54 | chr14-91352338-INS-56 |
| chr14-91371359-INS-52 | chr14-91371340-INS-63 |
| chr14-91372516-INS-145 | chr14-91372515-INS-142 |
| chr14-91423552-INS-109 | chr14-91423552-INS-100 |
| chr14-91427700-INS-2331 | chr14-91427701-INS-2329 |
| chr14-91443404-INS-490 | chr14-91443404-INS-490 |
| chr14-91469383-INS-430 | chr14-91469383-INS-430 |
| chr14-91484998-INS-93 | chr14-91484998-INS-93 |
| chr14-91486898-INS-87 | chr14-91486904-INS-91 |
| chr14-91489387-INS-501 | chr14-91489387-INS-501 |
| chr14-91491374-INS-538 | chr14-91491374-INS-538 |
| chr14-91493063-INS-706 | chr14-91493063-INS-707 |
| chr14-91505632-INS-52 | chr14-91505647-INS-68 |
| chr14-91507451-INS-286 | chr14-91507451-INS-286 |
| chr14-91534777-INS-328 | chr14-91534760-INS-335 |
| chr14-91540755-INS-213 | chr14-91540755-INS-213 |
| chr14-91549628-INS-495 | chr14-91549628-INS-502 |
| chr14-91560470-INS-52 | chr14-91560470-INS-50 |
| chr14-91580439-INS-496 | chr14-91580439-INS-496 |
| chr14-91581384-INS-200 | chr14-91581384-INS-200 |
| chr14-91615616-INS-6332 | chr14-91615617-INS-6332 |
| chr14-91617842-INS-896 | chr14-91617842-INS-896 |
| chr14-91628430-INS-203 | chr14-91628430-INS-205 |
| chr14-91630032-INS-396 | chr14-91630032-INS-396 |
| chr14-91638999-INS-360 | chr14-91638999-INS-360 |
| chr14-91663213-INS-341 | chr14-91663213-INS-340 |
| chr14-91702009-INS-52 | chr14-91702009-INS-54 |
| chr14-91704528-INS-240 | chr14-91704528-INS-246 |
| chr14-91730328-INS-52 | chr14-91730330-INS-51 |
| chr14-91741549-INS-283 | chr14-91741549-INS-283 |
| chr14-91765382-INS-1821 | chr14-91765383-INS-1820 |
| chr14-91811370-INS-1946 | chr14-91811371-INS-1946 |
| chr14-91829627-INS-64 | chr14-91829629-INS-56 |
| chr14-91844339-INS-469 | chr14-91844339-INS-473 |
| chr14-91847979-INS-110 | chr14-91847979-INS-110 |
| chr14-91855774-INS-4736 | chr14-91855775-INS-4738 |
| chr14-91876699-INS-6556 | chr14-91876700-INS-6556 |
| chr14-91881516-INS-452 | chr14-91881516-INS-452 |
| chr14-91893200-INS-325 | chr14-91893200-INS-325 |
| chr14-91899496-INS-88 | chr14-91899491-INS-120 |
| chr14-91953843-INS-372 | chr14-91953843-INS-372 |
| chr14-91957898-INS-6034 | chr14-91957899-INS-6036 |
| chr14-91959076-INS-7103 | chr14-91959077-INS-7103 |
| chr14-91960010-INS-351 | chr14-91960010-INS-351 |
| chr14-92020752-INS-6789 | chr14-92020753-INS-6789 |
| chr14-92036998-INS-665 | chr14-92036998-INS-665 |
| chr14-92087412-INS-6457 | chr14-92087413-INS-6457 |
| chr14-92122830-INS-165 | chr14-92122830-INS-166 |
| chr14-92141240-INS-99 | chr14-92141240-INS-99 |
| chr14-92150225-INS-76 | chr14-92150225-INS-78 |
| chr14-92188452-INS-248 | chr14-92188452-INS-248 |
| chr14-92198023-INS-193 | chr14-92198023-INS-193 |
| chr14-92205847-INS-305 | chr14-92205847-INS-305 |
| chr14-92259267-INS-193 | chr14-92259267-INS-193 |
| chr14-92269104-INS-67 | chr14-92269104-INS-67 |
| chr14-92302874-INS-501 | chr14-92302874-INS-501 |
| chr14-92322755-INS-1349 | chr14-92322756-INS-1347 |
| chr14-92367446-INS-1488 | chr14-92367447-INS-1488 |
| chr14-92376658-INS-68 | chr14-92376660-INS-88 |
| chr14-92392872-INS-146 | chr14-92392891-INS-152 |
| chr14-92400857-INS-4606 | chr14-92400858-INS-4606 |
| chr14-92416020-INS-340 | chr14-92416020-INS-340 |
| chr14-92448378-INS-526 | chr14-92448378-INS-526 |
| chr14-92500046-INS-123 | chr14-92500046-INS-123 |

| | |
|---|---|
| chr14-92546154-INS-583 | chr14-92546154-INS-583 |
| chr14-92552221-INS-7106 | chr14-92552222-INS-7106 |
| chr14-92566882-INS-361 | chr14-92566882-INS-361 |
| chr14-92614553-INS-2205 | chr14-92614554-INS-2205 |
| chr14-92617170-INS-387 | chr14-92617170-INS-387 |
| chr14-92631704-INS-72 | chr14-92631704-INS-72 |
| chr14-92638888-INS-321 | chr14-92638889-INS-321 |
| chr14-92652082-INS-187 | chr14-92652082-INS-187 |
| chr14-92692474-INS-196 | chr14-92692474-INS-196 |
| chr14-92693965-INS-156 | chr14-92693965-INS-156 |
| chr14-92748742-INS-361 | chr14-92748742-INS-361 |
| chr14-92755093-INS-83 | chr14-92755119-INS-66 |
| chr14-92764039-INS-64 | chr14-92764039-INS-62 |
| chr14-92770118-INS-670 | chr14-92770118-INS-683 |
| chr14-92795729-INS-577 | chr14-92795729-INS-577 |
| chr14-92808625-INS-56 | chr14-92808625-INS-58 |
| chr14-92855309-INS-71 | chr14-92855309-INS-71 |
| chr14-92874583-INS-239 | chr14-92874583-INS-239 |
| chr14-92878220-INS-346 | chr14-92878220-INS-338 |
| chr14-92888912-INS-55 | chr14-92888912-INS-55 |
| chr14-92901506-INS-994 | chr14-92901506-INS-993 |
| chr14-92904198-INS-291 | chr14-92904198-INS-291 |
| chr14-92918314-INS-212 | chr14-92918314-INS-213 |
| chr14-92939988-INS-2652 | chr14-92939989-INS-2645 |
| chr14-93047685-INS-108 | chr14-93047672-INS-128 |
| chr14-93048730-INS-475 | chr14-93048730-INS-475 |
| chr14-93088844-INS-441 | chr14-93088844-INS-441 |
| chr14-93093848-INS-60 | chr14-93093829-INS-82 |
| chr14-93107086-INS-533 | chr14-93107086-INS-533 |
| chr14-93110635-INS-436 | chr14-93110635-INS-436 |
| chr14-93158507-INS-64 | chr14-93158507-INS-64 |
| chr14-93186492-INS-90 | chr14-93186492-INS-90 |
| chr14-93220365-INS-360 | chr14-93220365-INS-362 |
| chr14-93224888-INS-66 | chr14-93224896-INS-80 |
| chr14-93238539-INS-6898 | chr14-93238540-INS-6897 |
| chr14-93244942-INS-270 | chr14-93244942-INS-270 |
| chr14-93263989-INS-206 | chr14-93264028-INS-196 |
| chr14-93269511-INS-104 | chr14-93269531-INS-72 |
| chr14-93297548-INS-1150 | chr14-93297562-INS-1151 |
| chr14-93301444-INS-1429 | chr14-93301445-INS-1424 |
| chr14-93302217-INS-194 | chr14-93302209-INS-182 |
| chr14-93353987-INS-486 | chr14-93353987-INS-486 |
| chr14-93368683-INS-53 | chr14-93368684-INS-53 |
| chr14-93377451-INS-58 | chr14-93377452-INS-59 |
| chr14-93393591-INS-723 | chr14-93393591-INS-724 |
| chr14-93396222-INS-82 | chr14-93396222-INS-80 |
| chr14-93403973-INS-863 | chr14-93403973-INS-865 |
| chr14-93410342-INS-106 | chr14-93410342-INS-116 |
| chr14-93496462-INS-70 | chr14-93496462-INS-68 |
| chr14-93508137-INS-230 | chr14-93508137-INS-230 |
| chr14-93509735-INS-81 | chr14-93509735-INS-83 |
| chr14-93525116-INS-505 | chr14-93525116-INS-505 |
| chr14-93533652-INS-52 | chr14-93533668-INS-78 |
| chr14-93547746-INS-446 | chr14-93547746-INS-446 |
| chr14-93578138-INS-74 | chr14-93578138-INS-72 |
| chr14-93605116-INS-84 | chr14-93605124-INS-80 |
| chr14-93656398-INS-72 | chr14-93656398-INS-74 |
| chr14-93665948-INS-934 | chr14-93665948-INS-941 |
| chr14-93667018-INS-200 | chr14-93667018-INS-200 |
| chr14-93676630-INS-342 | chr14-93676630-INS-342 |
| chr14-93776706-INS-6710 | chr14-93776707-INS-6712 |
| chr14-93812441-INS-334 | chr14-93812441-INS-334 |
| chr14-93814990-INS-69 | chr14-93814990-INS-109 |
| chr14-93820204-INS-2334 | chr14-93820205-INS-2334 |
| chr14-93833531-INS-597 | chr14-93833531-INS-597 |
| chr14-93873110-INS-121 | chr14-93873110-INS-121 |
| chr14-93883957-INS-136 | chr14-93883957-INS-144 |
| chr14-93907597-INS-3331 | chr14-93907598-INS-3332 |
| chr14-93958820-INS-78 | chr14-93958820-INS-78 |
| chr14-93987701-INS-66 | chr14-93987705-INS-68 |
| chr14-93992472-INS-162 | chr14-93992472-INS-168 |
| chr14-93994051-INS-53 | chr14-93994084-INS-50 |
| chr14-93997823-INS-683 | chr14-93997823-INS-683 |
| chr14-94040525-INS-57 | chr14-94040525-INS-57 |
| chr14-94076207-INS-863 | chr14-94076207-INS-864 |
| chr14-94077007-INS-121 | chr14-94077007-INS-120 |
| chr14-94081747-INS-52 | chr14-94081747-INS-58 |

| | |
|---|---|
| chr14-94157865-INS-1097 | chr14-94157866-INS-1097 |
| chr14-94175578-INS-134 | chr14-94175578-INS-134 |
| chr14-94177515-INS-64 | chr14-94177515-INS-64 |
| chr14-94189786-INS-51 | chr14-94189786-INS-51 |
| chr14-94193226-INS-63 | chr14-94193226-INS-63 |
| chr14-94250103-INS-230 | chr14-94250103-INS-234 |
| chr14-94276929-INS-171 | chr14-94276929-INS-172 |
| chr14-94296649-INS-400 | chr14-94296649-INS-400 |
| chr14-94303539-INS-717 | chr14-94303539-INS-717 |
| chr14-94306703-INS-694 | chr14-94306700-INS-696 |
| chr14-94308859-INS-132 | chr14-94308853-INS-144 |
| chr14-94337430-INS-72 | chr14-94337430-INS-72 |
| chr14-94343495-INS-5161 | chr14-94343496-INS-5161 |
| chr14-94364427-INS-194 | chr14-94364427-INS-194 |
| chr14-94393440-INS-72 | chr14-94393440-INS-72 |
| chr14-94404404-INS-475 | chr14-94404404-INS-475 |
| chr14-94405900-INS-66 | chr14-94405900-INS-72 |
| chr14-94408380-INS-908 | chr14-94408380-INS-908 |
| chr14-94427480-INS-60 | chr14-94427480-INS-60 |
| chr14-94436298-INS-61 | chr14-94436283-INS-62 |
| chr14-94439089-INS-676 | chr14-94439089-INS-672 |
| chr14-94448417-INS-72 | chr14-94448417-INS-76 |
| chr14-94449612-INS-549 | chr14-94449612-INS-549 |
| chr14-94483347-INS-66 | chr14-94483347-INS-78 |
| chr14-94486413-INS-6093 | chr14-94486414-INS-6093 |
| chr14-94487106-INS-434 | chr14-94487106-INS-435 |
| chr14-94514840-INS-104 | chr14-94514840-INS-104 |
| chr14-94515533-INS-1432 | chr14-94515534-INS-1421 |
| chr14-94528877-INS-596 | chr14-94528877-INS-595 |
| chr14-94533112-INS-550 | chr14-94533112-INS-548 |
| chr14-94549096-INS-136 | chr14-94549096-INS-130 |
| chr14-94554232-INS-171 | chr14-94554232-INS-159 |
| chr14-94576243-INS-868 | chr14-94576243-INS-868 |
| chr14-94626425-INS-292 | chr14-94626425-INS-292 |
| chr14-94637543-INS-156 | chr14-94637543-INS-151 |
| chr14-94639224-INS-6513 | chr14-94639225-INS-6514 |
| chr14-94645116-INS-7187 | chr14-94645117-INS-7187 |
| chr14-94730962-INS-51 | chr14-94730962-INS-51 |
| chr14-94844837-INS-1262 | chr14-94844838-INS-1262 |
| chr14-94865269-INS-121 | chr14-94865271-INS-146 |
| chr14-94906275-INS-55 | chr14-94906275-INS-70 |
| chr14-94908435-INS-74 | chr14-94908435-INS-62 |
| chr14-94909926-INS-50 | chr14-94909926-INS-50 |
| chr14-94915120-INS-59 | chr14-94915120-INS-62 |
| chr14-94936259-INS-59 | chr14-94936259-INS-59 |
| chr14-94955563-INS-62 | chr14-94955565-INS-70 |
| chr14-94969419-INS-498 | chr14-94969419-INS-498 |
| chr14-94986307-INS-65 | chr14-94986307-INS-65 |
| chr14-94993927-INS-75 | chr14-94993927-INS-68 |
| chr14-95007233-INS-65 | chr14-95007233-INS-65 |
| chr14-95011324-INS-60 | chr14-95011324-INS-60 |
| chr14-95030264-INS-53 | chr14-95030264-INS-53 |
| chr14-95053316-INS-662 | chr14-95053316-INS-662 |
| chr14-95060466-INS-66 | chr14-95060466-INS-74 |
| chr14-95067422-INS-56 | chr14-95067422-INS-56 |
| chr14-95071116-INS-254 | chr14-95071116-INS-254 |
| chr14-95076140-INS-212 | chr14-95076140-INS-212 |
| chr14-95099935-INS-7588 | chr14-95099936-INS-7588 |
| chr14-95101172-INS-79 | chr14-95101170-INS-84 |
| chr14-95103746-INS-52 | chr14-95103747-INS-78 |
| chr14-95118126-INS-218 | chr14-95118126-INS-218 |
| chr14-95131562-INS-281 | chr14-95131562-INS-281 |
| chr14-95159223-INS-168 | chr14-95159223-INS-172 |
| chr14-95163980-INS-2930 | chr14-95163981-INS-2930 |
| chr14-95175215-INS-131 | chr14-95175265-INS-126 |
| chr14-95258544-INS-172 | chr14-95258544-INS-172 |
| chr14-95265452-INS-54 | chr14-95265461-INS-52 |
| chr14-95268797-INS-624 | chr14-95268797-INS-624 |
| chr14-95273935-INS-1304 | chr14-95273936-INS-1304 |
| chr14-95280124-INS-702 | chr14-95280124-INS-701 |
| chr14-95309619-INS-170 | chr14-95309619-INS-188 |
| chr14-95317760-INS-1014 | chr14-95317761-INS-1015 |
| chr14-95331195-INS-483 | chr14-95331229-INS-439 |
| chr14-95333557-INS-151 | chr14-95333557-INS-151 |
| chr14-95362892-INS-6487 | chr14-95362893-INS-6487 |
| chr14-95375923-INS-2913 | chr14-95375924-INS-2913 |
| chr14-95384073-INS-511 | chr14-95384073-INS-511 |

| | |
|---|---|
| chr14-95391233-INS-82 | chr14-95391254-INS-65 |
| chr14-95392584-INS-76 | chr14-95392584-INS-76 |
| chr14-95408474-INS-238 | chr14-95408474-INS-239 |
| chr14-95425151-INS-489 | chr14-95425157-INS-496 |
| chr14-95445407-INS-52 | chr14-95445407-INS-52 |
| chr14-95470523-INS-297 | chr14-95470523-INS-297 |
| chr14-95474906-INS-57 | chr14-95474906-INS-61 |
| chr14-95483816-INS-2380 | chr14-95483817-INS-2380 |
| chr14-95487028-INS-1026 | chr14-95487029-INS-1026 |
| chr14-95491912-INS-86 | chr14-95491908-INS-88 |
| chr14-95502196-INS-68 | chr14-95502208-INS-50 |
| chr14-95503581-INS-56 | chr14-95503581-INS-56 |
| chr14-95515290-INS-52 | chr14-95515290-INS-52 |
| chr14-95574189-INS-364 | chr14-95574189-INS-364 |
| chr14-95596292-INS-52 | chr14-95596292-INS-52 |
| chr14-95611022-INS-1216 | chr14-95611023-INS-1216 |
| chr14-95613755-INS-60 | chr14-95613755-INS-60 |
| chr14-95622392-INS-73 | chr14-95622388-INS-61 |
| chr14-95636876-INS-54 | chr14-95636876-INS-55 |
| chr14-95638189-INS-66 | chr14-95638193-INS-110 |
| chr14-95644196-INS-136 | chr14-95644196-INS-136 |
| chr14-95658932-INS-54 | chr14-95658932-INS-53 |
| chr14-95668001-INS-67 | chr14-95668000-INS-68 |
| chr14-95682147-INS-60 | chr14-95682147-INS-60 |
| chr14-95692536-INS-1266 | chr14-95692537-INS-1265 |
| chr14-95751185-INS-740 | chr14-95751185-INS-748 |
| chr14-95771675-INS-2156 | chr14-95771676-INS-2156 |
| chr14-95773564-INS-262 | chr14-95773564-INS-262 |
| chr14-95814745-INS-64 | chr14-95814745-INS-58 |
| chr14-95839714-INS-52 | chr14-95839714-INS-54 |
| chr14-95852517-INS-70 | chr14-95852517-INS-70 |
| chr14-95875852-INS-90 | chr14-95875858-INS-90 |
| chr14-95889053-INS-54 | chr14-95889053-INS-54 |
| chr14-95890660-INS-121 | chr14-95890660-INS-120 |
| chr14-95924358-INS-166 | chr14-95924358-INS-171 |
| chr14-95927154-INS-68 | chr14-95927152-INS-64 |
| chr14-95935440-INS-57 | chr14-95935440-INS-57 |
| chr14-95953788-INS-66 | chr14-95953788-INS-68 |
| chr14-95961497-INS-137 | chr14-95961497-INS-137 |
| chr14-95966797-INS-62 | chr14-95966797-INS-62 |
| chr14-96029914-INS-6864 | chr14-96029915-INS-6864 |
| chr14-96042335-INS-76 | chr14-96042335-INS-80 |
| chr14-96043436-INS-55 | chr14-96043436-INS-59 |
| chr14-96062483-INS-389 | chr14-96062483-INS-389 |
| chr14-96071189-INS-764 | chr14-96071189-INS-764 |
| chr14-96073194-INS-204 | chr14-96073194-INS-204 |
| chr14-96124197-INS-282 | chr14-96124197-INS-282 |
| chr14-96134066-INS-315 | chr14-96134066-INS-315 |
| chr14-96137494-INS-56 | chr14-96137494-INS-56 |
| chr14-96179210-INS-117 | chr14-96179210-INS-117 |
| chr14-96194781-INS-52 | chr14-96194781-INS-60 |
| chr14-96202827-INS-199 | chr14-96202827-INS-201 |
| chr14-96216277-INS-393 | chr14-96216277-INS-393 |
| chr14-96217263-INS-1294 | chr14-96217264-INS-1294 |
| chr14-96217715-INS-162 | chr14-96217715-INS-162 |
| chr14-96221237-INS-459 | chr14-96221237-INS-459 |
| chr14-96229681-INS-59 | chr14-96229681-INS-59 |
| chr14-96241043-INS-1240 | chr14-96241044-INS-1241 |
| chr14-96263279-INS-6135 | chr14-96263280-INS-6135 |
| chr14-96284612-INS-593 | chr14-96284612-INS-593 |
| chr14-96304510-INS-54 | chr14-96304532-INS-50 |
| chr14-96330203-INS-64 | chr14-96330203-INS-64 |
| chr14-96345187-INS-97 | chr14-96345187-INS-97 |
| chr14-96357008-INS-152 | chr14-96357008-INS-152 |
| chr14-96405876-INS-76 | chr14-96405876-INS-78 |
| chr14-96411938-INS-749 | chr14-96411938-INS-749 |
| chr14-96433691-INS-440 | chr14-96433691-INS-440 |
| chr14-96436848-INS-55 | chr14-96436848-INS-55 |
| chr14-96448902-INS-1339 | chr14-96448903-INS-1339 |
| chr14-96455531-INS-351 | chr14-96455531-INS-351 |
| chr14-96468894-INS-58 | chr14-96468894-INS-60 |
| chr14-96493069-INS-176 | chr14-96493069-INS-175 |
| chr14-96499831-INS-124 | chr14-96499831-INS-124 |
| chr14-96513669-INS-911 | chr14-96513669-INS-913 |
| chr14-96514797-INS-85 | chr14-96514797-INS-89 |
| chr14-96517059-INS-80 | chr14-96517024-INS-94 |
| chr14-96538217-INS-210 | chr14-96538217-INS-210 |

| | |
|---|---|
| chr14-96570404-INS-232 | chr14-96570404-INS-232 |
| chr14-96580291-INS-55 | chr14-96580291-INS-89 |
| chr14-96607535-INS-88 | chr14-96607535-INS-88 |
| chr14-96642600-INS-201 | chr14-96642600-INS-201 |
| chr14-96677206-INS-54 | chr14-96677206-INS-63 |
| chr14-96689365-INS-183 | chr14-96689365-INS-183 |
| chr14-96691813-INS-278 | chr14-96691813-INS-278 |
| chr14-96696501-INS-786 | chr14-96696501-INS-786 |
| chr14-96713885-INS-67 | chr14-96713885-INS-67 |
| chr14-96727946-INS-96 | chr14-96727946-INS-96 |
| chr14-96729537-INS-248 | chr14-96729537-INS-248 |
| chr14-96738108-INS-461 | chr14-96738108-INS-461 |
| chr14-96758879-INS-434 | chr14-96758879-INS-434 |
| chr14-96813057-INS-62 | chr14-96813057-INS-62 |
| chr14-96815650-INS-3565 | chr14-96815651-INS-3565 |
| chr14-96815688-INS-979 | chr14-96815688-INS-979 |
| chr14-96816677-INS-51 | chr14-96816670-INS-55 |
| chr14-96823554-INS-301 | chr14-96823554-INS-301 |
| chr14-96850739-INS-136 | chr14-96850739-INS-136 |
| chr14-96856640-INS-88 | chr14-96856640-INS-96 |
| chr14-96872863-INS-65 | chr14-96872863-INS-65 |
| chr14-96894775-INS-54 | chr14-96894775-INS-54 |
| chr14-96896546-INS-246 | chr14-96896566-INS-272 |
| chr14-96930645-INS-796 | chr14-96930645-INS-795 |
| chr14-96931484-INS-139 | chr14-96931484-INS-139 |
| chr14-96944046-INS-56 | chr14-96944046-INS-56 |
| chr14-96945518-INS-1352 | chr14-96945519-INS-1377 |
| chr14-96954764-INS-256 | chr14-96954791-INS-244 |
| chr14-96966302-INS-426 | chr14-96966302-INS-426 |
| chr14-96973940-INS-365 | chr14-96973940-INS-365 |
| chr14-97011553-INS-410 | chr14-97011553-INS-410 |
| chr14-97050266-INS-728 | chr14-97050266-INS-728 |
| chr14-97051715-INS-56 | chr14-97051715-INS-64 |
| chr14-97057986-INS-108 | chr14-97057986-INS-108 |
| chr14-97066491-INS-401 | chr14-97066491-INS-401 |
| chr14-97076180-INS-57 | chr14-97076180-INS-51 |
| chr14-97078794-INS-236 | chr14-97078794-INS-236 |
| chr14-97085599-INS-51 | chr14-97085600-INS-96 |
| chr14-97112746-INS-70 | chr14-97112746-INS-74 |
| chr14-97126250-INS-156 | chr14-97126250-INS-156 |
| chr14-97143183-INS-64 | chr14-97143184-INS-64 |
| chr14-97152112-INS-61 | chr14-97152113-INS-65 |
| chr14-97174808-INS-64 | chr14-97174808-INS-64 |
| chr14-97177223-INS-65 | chr14-97177223-INS-55 |
| chr14-97180054-INS-92 | chr14-97180050-INS-72 |
| chr14-97181769-INS-52 | chr14-97181777-INS-50 |
| chr14-97196610-INS-194 | chr14-97196610-INS-194 |
| chr14-97218474-INS-5470 | chr14-97218475-INS-5470 |
| chr14-97238257-INS-56 | chr14-97238257-INS-56 |
| chr14-97239967-INS-204 | chr14-97239967-INS-204 |
| chr14-97249587-INS-458 | chr14-97249587-INS-458 |
| chr14-97253563-INS-224 | chr14-97253563-INS-224 |
| chr14-97264036-INS-68 | chr14-97264033-INS-54 |
| chr14-97326453-INS-754 | chr14-97326454-INS-755 |
| chr14-97395444-INS-2914 | chr14-97395445-INS-2914 |
| chr14-97414172-INS-869 | chr14-97414172-INS-870 |
| chr14-97448214-INS-92 | chr14-97448214-INS-92 |
| chr14-97458248-INS-58 | chr14-97458245-INS-54 |
| chr14-97470798-INS-625 | chr14-97470798-INS-625 |
| chr14-97488750-INS-89 | chr14-97488750-INS-92 |
| chr14-97498553-INS-60 | chr14-97498553-INS-60 |
| chr14-97519596-INS-52 | chr14-97519596-INS-52 |
| chr14-97526115-INS-94 | chr14-97526100-INS-98 |
| chr14-97544357-INS-63 | chr14-97544357-INS-63 |
| chr14-97589064-INS-58 | chr14-97589064-INS-64 |
| chr14-97591687-INS-73 | chr14-97591687-INS-74 |
| chr14-97594561-INS-69 | chr14-97594564-INS-69 |
| chr14-97596608-INS-60 | chr14-97596608-INS-64 |
| chr14-97609440-INS-3475 | chr14-97609441-INS-3475 |
| chr14-97619815-INS-855 | chr14-97619815-INS-867 |
| chr14-97635775-INS-60 | chr14-97635775-INS-60 |
| chr14-97657626-INS-644 | chr14-97657626-INS-644 |
| chr14-97682472-INS-6379 | chr14-97682473-INS-6379 |
| chr14-97703097-INS-137 | chr14-97703098-INS-140 |
| chr14-97764186-INS-88 | chr14-97764186-INS-92 |
| chr14-97769293-INS-624 | chr14-97769293-INS-624 |
| chr14-97784380-INS-72 | chr14-97784380-INS-57 |

| | |
|---|---|
| chr14-97787874-INS-156 | chr14-97787874-INS-156 |
| chr14-97790358-INS-100 | chr14-97790358-INS-100 |
| chr14-97794900-INS-162 | chr14-97794900-INS-162 |
| chr14-97798304-INS-62 | chr14-97798304-INS-62 |
| chr14-97844946-INS-56 | chr14-97844946-INS-52 |
| chr14-97858763-INS-59 | chr14-97858763-INS-60 |
| chr14-97867563-INS-2801 | chr14-97867564-INS-2801 |
| chr14-97874402-INS-6361 | chr14-97874403-INS-6361 |
| chr14-97880564-INS-277 | chr14-97880564-INS-277 |
| chr14-97881754-INS-435 | chr14-97881754-INS-435 |
| chr14-97885235-INS-58 | chr14-97885235-INS-58 |
| chr14-97947586-INS-148 | chr14-97947586-INS-148 |
| chr14-97986886-INS-3311 | chr14-97986887-INS-3310 |
| chr14-98028042-INS-683 | chr14-98028042-INS-683 |
| chr14-98055004-INS-227 | chr14-98055018-INS-242 |
| chr14-98061633-INS-52 | chr14-98061633-INS-51 |
| chr14-98076465-INS-355 | chr14-98076465-INS-355 |
| chr14-98091549-INS-765 | chr14-98091549-INS-766 |
| chr14-98126076-INS-714 | chr14-98126076-INS-714 |
| chr14-98237995-INS-67 | chr14-98237995-INS-67 |
| chr14-98253385-INS-573 | chr14-98253385-INS-573 |
| chr14-98292024-INS-83 | chr14-98292018-INS-99 |
| chr14-98357264-INS-137 | chr14-98357264-INS-137 |
| chr14-98430066-INS-2245 | chr14-98430067-INS-2245 |
| chr14-98454037-INS-64 | chr14-98454047-INS-52 |
| chr14-98456251-INS-228 | chr14-98456243-INS-241 |
| chr14-98490783-INS-70 | chr14-98490783-INS-70 |
| chr14-98502560-INS-135 | chr14-98502560-INS-140 |
| chr14-98540706-INS-88 | chr14-98540708-INS-88 |
| chr14-98553127-INS-86 | chr14-98553127-INS-86 |
| chr14-98554603-INS-54 | chr14-98554603-INS-54 |
| chr14-98561815-INS-52 | chr14-98561815-INS-84 |
| chr14-98575620-INS-92 | chr14-98575620-INS-92 |
| chr14-98581367-INS-2289 | chr14-98581368-INS-2289 |
| chr14-98587397-INS-66 | chr14-98587397-INS-66 |
| chr14-98598421-INS-66 | chr14-98598421-INS-64 |
| chr14-98600406-INS-235 | chr14-98600406-INS-235 |
| chr14-98607269-INS-66 | chr14-98607269-INS-66 |
| chr14-98609557-INS-128 | chr14-98609557-INS-128 |
| chr14-98610438-INS-347 | chr14-98610439-INS-347 |
| chr14-98631697-INS-76 | chr14-98631710-INS-78 |
| chr14-98640218-INS-243 | chr14-98640218-INS-235 |
| chr14-98650903-INS-429 | chr14-98650906-INS-446 |
| chr14-98664569-INS-60 | chr14-98664569-INS-60 |
| chr14-98724376-INS-701 | chr14-98724376-INS-700 |
| chr14-98749575-INS-4764 | chr14-98749576-INS-4766 |
| chr14-98762337-INS-1577 | chr14-98762338-INS-1577 |
| chr14-98766956-INS-76 | chr14-98766956-INS-84 |
| chr14-98769103-INS-117 | chr14-98769103-INS-117 |
| chr14-98792799-INS-64 | chr14-98792799-INS-64 |
| chr14-98800303-INS-246 | chr14-98800303-INS-246 |
| chr14-98816833-INS-57 | chr14-98816833-INS-57 |
| chr14-98833129-INS-385 | chr14-98833129-INS-385 |
| chr14-98852451-INS-76 | chr14-98852451-INS-112 |
| chr14-98853505-INS-52 | chr14-98853505-INS-70 |
| chr14-98855256-INS-6834 | chr14-98855257-INS-6834 |
| chr14-98896696-INS-6461 | chr14-98896697-INS-6462 |
| chr14-98908724-INS-498 | chr14-98908724-INS-498 |
| chr14-98921452-INS-6464 | chr14-98921453-INS-6466 |
| chr14-98935928-INS-66 | chr14-98935928-INS-68 |
| chr14-98956002-INS-58 | chr14-98955998-INS-62 |
| chr14-98999606-INS-148 | chr14-98999606-INS-148 |
| chr14-99007306-INS-500 | chr14-99007352-INS-498 |
| chr14-99024362-INS-72 | chr14-99024362-INS-72 |
| chr14-99041305-INS-834 | chr14-99041305-INS-834 |
| chr14-99049447-INS-1110 | chr14-99049448-INS-1112 |
| chr14-99052236-INS-54 | chr14-99052236-INS-54 |
| chr14-99106372-INS-66 | chr14-99106372-INS-66 |
| chr14-99106846-INS-279 | chr14-99106846-INS-276 |
| chr14-99108531-INS-80 | chr14-99108531-INS-80 |
| chr14-99116683-INS-94 | chr14-99116683-INS-94 |
| chr14-99119690-INS-420 | chr14-99119690-INS-423 |
| chr14-99133562-INS-81 | chr14-99133562-INS-75 |
| chr14-99153855-INS-515 | chr14-99153855-INS-515 |
| chr14-99154540-INS-52 | chr14-99154540-INS-52 |
| chr14-99174710-INS-517 | chr14-99174710-INS-518 |
| chr14-99186784-INS-80 | chr14-99186784-INS-112 |

| | |
|---|---|
| chr14-99193851-INS-370 | chr14-99193843-INS-371 |
| chr14-99210358-INS-272 | chr14-99210358-INS-272 |
| chr14-99211425-INS-6408 | chr14-99211426-INS-6408 |
| chr14-99232225-INS-53 | chr14-99232229-INS-50 |
| chr14-99233122-INS-688 | chr14-99233126-INS-680 |
| chr14-99240976-INS-60 | chr14-99240976-INS-60 |
| chr14-99263245-INS-2104 | chr14-99263246-INS-2106 |
| chr14-99263542-INS-87 | chr14-99263542-INS-87 |
| chr14-99297862-INS-504 | chr14-99297862-INS-480 |
| chr14-99335725-INS-55 | chr14-99335725-INS-55 |
| chr14-99340677-INS-81 | chr14-99340677-INS-80 |
| chr14-99360442-INS-66 | chr14-99360442-INS-69 |
| chr14-99375239-INS-353 | chr14-99375253-INS-349 |
| chr14-99391691-INS-138 | chr14-99391691-INS-138 |
| chr14-99441140-INS-1050 | chr14-99441141-INS-1050 |
| chr14-99441141-INS-185 | chr14-99441141-INS-185 |
| chr14-99449576-INS-52 | chr14-99449576-INS-52 |
| chr14-99455574-INS-134 | chr14-99455572-INS-136 |
| chr14-99458706-INS-84 | chr14-99458706-INS-84 |
| chr14-99492932-INS-201 | chr14-99492932-INS-195 |
| chr14-99502741-INS-181 | chr14-99502741-INS-201 |
| chr14-99511174-INS-483 | chr14-99511174-INS-483 |
| chr14-99527711-INS-65 | chr14-99527728-INS-66 |
| chr14-99556525-INS-191 | chr14-99556525-INS-191 |
| chr14-99558348-INS-136 | chr14-99558348-INS-164 |
| chr14-99573590-INS-160 | chr14-99573588-INS-112 |
| chr14-99585141-INS-217 | chr14-99585141-INS-217 |
| chr14-99598063-INS-67 | chr14-99598063-INS-67 |
| chr14-99599802-INS-209 | chr14-99599802-INS-208 |
| chr14-99608646-INS-66 | chr14-99608646-INS-63 |
| chr14-99621413-INS-459 | chr14-99621400-INS-466 |
| chr14-99622851-INS-217 | chr14-99622851-INS-197 |
| chr14-99628122-INS-80 | chr14-99628122-INS-74 |
| chr14-99664514-INS-205 | chr14-99664514-INS-205 |
| chr14-99674688-INS-59 | chr14-99674688-INS-62 |
| chr14-99681547-INS-63 | chr14-99681547-INS-54 |
| chr14-99690744-INS-623 | chr14-99690744-INS-623 |
| chr14-99697751-INS-52 | chr14-99697751-INS-52 |
| chr14-99716121-INS-589 | chr14-99716121-INS-589 |
| chr14-99779578-INS-241 | chr14-99779578-INS-241 |
| chr14-99784736-INS-77 | chr14-99784733-INS-60 |
| chr14-99825765-INS-186 | chr14-99825765-INS-186 |
| chr14-99886859-INS-190 | chr14-99886859-INS-190 |
| chr14-99906158-INS-2549 | chr14-99906159-INS-2549 |
| chr14-99926626-INS-220 | chr14-99926626-INS-220 |
| chr14-99933674-INS-120 | chr14-99933674-INS-136 |
| chr14-99935714-INS-189 | chr14-99935714-INS-189 |
| chr14-99937935-INS-67 | chr14-99937935-INS-67 |
| chr14-99970205-INS-185 | chr14-99970205-INS-185 |
| chr14-99988070-INS-63 | chr14-99988070-INS-63 |
| chr14-99996011-INS-88 | chr14-99996011-INS-89 |
| chr14-99999540-INS-71 | chr14-99999540-INS-71 |
| chr14-100013263-INS-67 | chr14-100013263-INS-67 |
| chr14-100015329-INS-427 | chr14-100015329-INS-427 |
| chr14-100033521-INS-257 | chr14-100033521-INS-256 |
| chr14-100047131-INS-101 | chr14-100047131-INS-99 |
| chr14-100052559-INS-95 | chr14-100052581-INS-118 |
| chr14-100059220-INS-1431 | chr14-100059221-INS-1462 |
| chr14-100071122-INS-2876 | chr14-100071123-INS-2876 |
| chr14-100078533-INS-62 | chr14-100078533-INS-62 |
| chr14-100086094-INS-164 | chr14-100086093-INS-144 |
| chr14-100099621-INS-380 | chr14-100099621-INS-380 |
| chr14-100126519-INS-116 | chr14-100126519-INS-116 |
| chr14-100142081-INS-51 | chr14-100142081-INS-60 |
| chr14-100158118-INS-2710 | chr14-100158119-INS-2710 |
| chr14-100187495-INS-626 | chr14-100187495-INS-626 |
| chr14-100196356-INS-68 | chr14-100196368-INS-56 |
| chr14-100197688-INS-216 | chr14-100197688-INS-214 |
| chr14-100230926-INS-117 | chr14-100230922-INS-117 |
| chr14-100234154-INS-188 | chr14-100234154-INS-189 |
| chr14-100248804-INS-6812 | chr14-100248805-INS-6812 |
| chr14-100249534-INS-54 | chr14-100249534-INS-54 |
| chr14-100284582-INS-90 | chr14-100284582-INS-90 |
| chr14-100326756-INS-65 | chr14-100326755-INS-80 |
| chr14-100368884-INS-60 | chr14-100368884-INS-60 |
| chr14-100389463-INS-740 | chr14-100389463-INS-765 |
| chr14-100392703-INS-87 | chr14-100392717-INS-58 |

| | |
|---|---|
| chr14-100398644-INS-1134 | chr14-100398645-INS-1135 |
| chr14-100420989-INS-210 | chr14-100420989-INS-211 |
| chr14-100431402-INS-179 | chr14-100431402-INS-179 |
| chr14-100435430-INS-54 | chr14-100435430-INS-79 |
| chr14-100442451-INS-1046 | chr14-100442452-INS-1046 |
| chr14-100469389-INS-155 | chr14-100469385-INS-158 |
| chr14-100482528-INS-132 | chr14-100482528-INS-132 |
| chr14-100551434-INS-65 | chr14-100551434-INS-67 |
| chr14-100582080-INS-6607 | chr14-100582081-INS-6607 |
| chr14-100583041-INS-477 | chr14-100583041-INS-477 |
| chr14-100583579-INS-52 | chr14-100583582-INS-93 |
| chr14-100599092-INS-84 | chr14-100599092-INS-86 |
| chr14-100628379-INS-60 | chr14-100628379-INS-62 |
| chr14-100643238-INS-60 | chr14-100643238-INS-60 |
| chr14-100643485-INS-6240 | chr14-100643486-INS-6240 |
| chr14-100650857-INS-78 | chr14-100650857-INS-76 |
| chr14-100652083-INS-58 | chr14-100652083-INS-58 |
| chr14-100654776-INS-110 | chr14-100654776-INS-90 |
| chr14-100657276-INS-66 | chr14-100657276-INS-62 |
| chr14-100676578-INS-183 | chr14-100676578-INS-183 |
| chr14-100707390-INS-100 | chr14-100707390-INS-50 |
| chr14-100709139-INS-3199 | chr14-100709140-INS-3199 |
| chr14-100733250-INS-60 | chr14-100733250-INS-60 |
| chr14-100758605-INS-6265 | chr14-100758606-INS-6265 |
| chr14-100773211-INS-203 | chr14-100773211-INS-203 |
| chr14-100781277-INS-68 | chr14-100781277-INS-60 |
| chr14-100790681-INS-68 | chr14-100790682-INS-59 |
| chr14-100807881-INS-485 | chr14-100807881-INS-485 |
| chr14-100809536-INS-80 | chr14-100809536-INS-85 |
| chr14-100842536-INS-75 | chr14-100842523-INS-80 |
| chr14-100848424-INS-107 | chr14-100848424-INS-120 |
| chr14-100858266-INS-61 | chr14-100858266-INS-61 |
| chr14-100935110-INS-650 | chr14-100935110-INS-650 |
| chr14-100945816-INS-53 | chr14-100945816-INS-53 |
| chr14-100968640-INS-477 | chr14-100968640-INS-482 |
| chr14-100972565-INS-105 | chr14-100972565-INS-105 |
| chr14-100990512-INS-81 | chr14-100990512-INS-81 |
| chr14-100995391-INS-1271 | chr14-100995392-INS-1271 |
| chr14-101002824-INS-370 | chr14-101002824-INS-370 |
| chr14-101020160-INS-749 | chr14-101020160-INS-749 |
| chr14-101028838-INS-52 | chr14-101028838-INS-52 |
| chr14-101055567-INS-267 | chr14-101055567-INS-271 |
| chr14-101073401-INS-365 | chr14-101073401-INS-365 |
| chr14-101124819-INS-5401 | chr14-101124820-INS-5401 |
| chr14-101143060-INS-6232 | chr14-101143061-INS-6232 |
| chr14-101148665-INS-55 | chr14-101148661-INS-63 |
| chr14-101165645-INS-637 | chr14-101165645-INS-637 |
| chr14-101178404-INS-279 | chr14-101178404-INS-279 |
| chr14-101180008-INS-52 | chr14-101180008-INS-60 |
| chr14-101211129-INS-60 | chr14-101211129-INS-55 |
| chr14-101221865-INS-150 | chr14-101221874-INS-126 |
| chr14-101241103-INS-76 | chr14-101241103-INS-72 |
| chr14-101248371-INS-81 | chr14-101248372-INS-81 |
| chr14-101268629-INS-2990 | chr14-101268630-INS-2986 |
| chr14-101271284-INS-542 | chr14-101271285-INS-542 |
| chr14-101272594-INS-6645 | chr14-101272595-INS-6645 |
| chr14-101274965-INS-3200 | chr14-101274966-INS-3200 |
| chr14-101281988-INS-67 | chr14-101281988-INS-67 |
| chr14-101284854-INS-58 | chr14-101284855-INS-62 |
| chr14-101286962-INS-898 | chr14-101286962-INS-898 |
| chr14-101291897-INS-81 | chr14-101291897-INS-81 |
| chr14-101304031-INS-59 | chr14-101304031-INS-59 |
| chr14-101347496-INS-197 | chr14-101347496-INS-238 |
| chr14-101365209-INS-80 | chr14-101365231-INS-78 |
| chr14-101366789-INS-405 | chr14-101366789-INS-404 |
| chr14-101370738-INS-66 | chr14-101370740-INS-68 |
| chr14-101372080-INS-57 | chr14-101372037-INS-54 |
| chr14-101380558-INS-5962 | chr14-101380559-INS-5957 |
| chr14-101384904-INS-68 | chr14-101384904-INS-70 |
| chr14-101424394-INS-69 | chr14-101424395-INS-57 |
| chr14-101429115-INS-6590 | chr14-101429116-INS-6590 |
| chr14-101431845-INS-1892 | chr14-101431846-INS-1892 |
| chr14-101441502-INS-265 | chr14-101441498-INS-312 |
| chr14-101452494-INS-56 | chr14-101452494-INS-76 |
| chr14-101458000-INS-82 | chr14-101458000-INS-82 |
| chr14-101475806-INS-361 | chr14-101475800-INS-361 |
| chr14-101481563-INS-78 | chr14-101481563-INS-79 |

| | |
|---|---|
| chr14-101482300-INS-70 | chr14-101482300-INS-70 |
| chr14-101493160-INS-54 | chr14-101493160-INS-71 |
| chr14-101494204-INS-162 | chr14-101494204-INS-160 |
| chr14-101506456-INS-62 | chr14-101506422-INS-62 |
| chr14-101515772-INS-1227 | chr14-101515773-INS-1221 |
| chr14-101516458-INS-2939 | chr14-101516459-INS-2939 |
| chr14-101531926-INS-175 | chr14-101531926-INS-175 |
| chr14-101533746-INS-59 | chr14-101533718-INS-59 |
| chr14-101535002-INS-64 | chr14-101535002-INS-68 |
| chr14-101545294-INS-423 | chr14-101545294-INS-423 |
| chr14-101561299-INS-377 | chr14-101561299-INS-377 |
| chr14-101567875-INS-551 | chr14-101567875-INS-551 |
| chr14-101573391-INS-176 | chr14-101573391-INS-176 |
| chr14-101593623-INS-220 | chr14-101593623-INS-227 |
| chr14-101600054-INS-62 | chr14-101600054-INS-52 |
| chr14-101611827-INS-55 | chr14-101611827-INS-55 |
| chr14-101615653-INS-84 | chr14-101615647-INS-88 |
| chr14-101625919-INS-453 | chr14-101625919-INS-453 |
| chr14-101629781-INS-498 | chr14-101629781-INS-498 |
| chr14-101656228-INS-326 | chr14-101656228-INS-326 |
| chr14-101659275-INS-187 | chr14-101659275-INS-188 |
| chr14-101664331-INS-69 | chr14-101664302-INS-60 |
| chr14-101670496-INS-408 | chr14-101670504-INS-376 |
| chr14-101677338-INS-529 | chr14-101677338-INS-529 |
| chr14-101680945-INS-1991 | chr14-101680946-INS-1991 |
| chr14-101687619-INS-60 | chr14-101687619-INS-70 |
| chr14-101688536-INS-164 | chr14-101688536-INS-164 |
| chr14-101691798-INS-60 | chr14-101691798-INS-66 |
| chr14-101697342-INS-8947 | chr14-101697343-INS-8947 |
| chr14-101699134-INS-599 | chr14-101699134-INS-599 |
| chr14-101701234-INS-72 | chr14-101701234-INS-72 |
| chr14-101739759-INS-68 | chr14-101739759-INS-76 |
| chr14-101746467-INS-77 | chr14-101746467-INS-77 |
| chr14-101750262-INS-105 | chr14-101750262-INS-84 |
| chr14-101763486-INS-4976 | chr14-101763487-INS-4978 |
| chr14-101789830-INS-60 | chr14-101789830-INS-80 |
| chr14-101798202-INS-51 | chr14-101798202-INS-51 |
| chr14-101804064-INS-56 | chr14-101804064-INS-56 |
| chr14-101809245-INS-200 | chr14-101809245-INS-200 |
| chr14-101833413-INS-502 | chr14-101833413-INS-502 |
| chr14-101849666-INS-95 | chr14-101849666-INS-105 |
| chr14-101851066-INS-144 | chr14-101851066-INS-108 |
| chr14-101855513-INS-160 | chr14-101855513-INS-160 |
| chr14-101899555-INS-100 | chr14-101899555-INS-99 |
| chr14-101919354-INS-2121 | chr14-101919355-INS-2091 |
| chr14-101933275-INS-157 | chr14-101933275-INS-157 |
| chr14-101939499-INS-165 | chr14-101939499-INS-175 |
| chr14-101947291-INS-55 | chr14-101947291-INS-55 |
| chr14-101965139-INS-55 | chr14-101965159-INS-57 |
| chr14-101968968-INS-64 | chr14-101968968-INS-64 |
| chr14-101972236-INS-114 | chr14-101972237-INS-141 |
| chr14-101977033-INS-1606 | chr14-101977034-INS-1606 |
| chr14-101984444-INS-66 | chr14-101984444-INS-66 |
| chr14-101991635-INS-175 | chr14-101991635-INS-175 |
| chr14-102019092-INS-153 | chr14-102019092-INS-142 |
| chr14-102038940-INS-206 | chr14-102038940-INS-206 |
| chr14-102061268-INS-56 | chr14-102061268-INS-56 |
| chr14-102115106-INS-74 | chr14-102115106-INS-74 |
| chr14-102138742-INS-76 | chr14-102138738-INS-76 |
| chr14-102149195-INS-272 | chr14-102149195-INS-273 |
| chr14-102211188-INS-67 | chr14-102211188-INS-66 |
| chr14-102230277-INS-990 | chr14-102230277-INS-990 |
| chr14-102255076-INS-54 | chr14-102255026-INS-63 |
| chr14-102259537-INS-52 | chr14-102259538-INS-72 |
| chr14-102297898-INS-471 | chr14-102297898-INS-472 |
| chr14-102309641-INS-56 | chr14-102309641-INS-56 |
| chr14-102321266-INS-98 | chr14-102321266-INS-98 |
| chr14-102322700-INS-70 | chr14-102322714-INS-79 |
| chr14-102334550-INS-485 | chr14-102334526-INS-485 |
| chr14-102345554-INS-255 | chr14-102345554-INS-255 |
| chr14-102405564-INS-61 | chr14-102405564-INS-61 |
| chr14-102406670-INS-89 | chr14-102406670-INS-89 |
| chr14-102410197-INS-354 | chr14-102410197-INS-355 |
| chr14-102428185-INS-93 | chr14-102428185-INS-93 |
| chr14-102432293-INS-5320 | chr14-102432294-INS-5320 |
| chr14-102433740-INS-52 | chr14-102433740-INS-52 |
| chr14-102484983-INS-106 | chr14-102484983-INS-97 |

| | |
|---|---|
| chr14-102492550-INS-1553 | chr14-102492551-INS-1553 |
| chr14-102505812-INS-1053 | chr14-102505813-INS-1053 |
| chr14-102505813-INS-97 | chr14-102505813-INS-97 |
| chr14-102575296-INS-306 | chr14-102575296-INS-306 |
| chr14-102578534-INS-593 | chr14-102578534-INS-593 |
| chr14-102580025-INS-108 | chr14-102580025-INS-64 |
| chr14-102604308-INS-265 | chr14-102604308-INS-227 |
| chr14-102612668-INS-404 | chr14-102612668-INS-404 |
| chr14-102614886-INS-534 | chr14-102614886-INS-534 |
| chr14-102624735-INS-6614 | chr14-102624736-INS-6614 |
| chr14-102629442-INS-277 | chr14-102629442-INS-277 |
| chr14-102668440-INS-53 | chr14-102668440-INS-51 |
| chr14-102690890-INS-50 | chr14-102690904-INS-68 |
| chr14-102706857-INS-1040 | chr14-102706858-INS-1041 |
| chr14-102709908-INS-106 | chr14-102709908-INS-89 |
| chr14-102712605-INS-86 | chr14-102712605-INS-80 |
| chr14-102756907-INS-68 | chr14-102756907-INS-93 |
| chr14-102812996-INS-823 | chr14-102812996-INS-823 |
| chr14-102850059-INS-374 | chr14-102850059-INS-374 |
| chr14-102879528-INS-87 | chr14-102879528-INS-87 |
| chr14-102884116-INS-234 | chr14-102884116-INS-238 |
| chr14-102902594-INS-340 | chr14-102902594-INS-326 |
| chr14-102920311-INS-84 | chr14-102920311-INS-84 |
| chr14-102944838-INS-64 | chr14-102944813-INS-50 |
| chr14-102966103-INS-510 | chr14-102966104-INS-510 |
| chr14-102986002-INS-170 | chr14-102986002-INS-140 |
| chr14-102987072-INS-79 | chr14-102987072-INS-79 |
| chr14-103013690-INS-3909 | chr14-103013691-INS-3909 |
| chr14-103030863-INS-400 | chr14-103030863-INS-400 |
| chr14-103036526-INS-59 | chr14-103036526-INS-59 |
| chr14-103061277-INS-58 | chr14-103061327-INS-58 |
| chr14-103069613-INS-160 | chr14-103069605-INS-190 |
| chr14-103077192-INS-76 | chr14-103077192-INS-76 |
| chr14-103111933-INS-200 | chr14-103111933-INS-200 |
| chr14-103115080-INS-160 | chr14-103115080-INS-160 |
| chr14-103158577-INS-569 | chr14-103158577-INS-572 |
| chr14-103170370-INS-68 | chr14-103170371-INS-66 |
| chr14-103173458-INS-58 | chr14-103173476-INS-50 |
| chr14-103176599-INS-56 | chr14-103176598-INS-60 |
| chr14-103184862-INS-7295 | chr14-103184863-INS-7299 |
| chr14-103201310-INS-68 | chr14-103201310-INS-72 |
| chr14-103211274-INS-60 | chr14-103211274-INS-60 |
| chr14-103228133-INS-52 | chr14-103228133-INS-50 |
| chr14-103233257-INS-1026 | chr14-103233258-INS-1026 |
| chr14-103242744-INS-173 | chr14-103242744-INS-173 |
| chr14-103248401-INS-292 | chr14-103248401-INS-277 |
| chr14-103262480-INS-136 | chr14-103262482-INS-168 |
| chr14-103264429-INS-134 | chr14-103264453-INS-110 |
| chr14-103325043-INS-73 | chr14-103325043-INS-73 |
| chr14-103325923-INS-351 | chr14-103325923-INS-351 |
| chr14-103330361-INS-69 | chr14-103330361-INS-69 |
| chr14-103348351-INS-201 | chr14-103348351-INS-201 |
| chr14-103356584-INS-215 | chr14-103356584-INS-214 |
| chr14-103380071-INS-51 | chr14-103380071-INS-51 |
| chr14-103473965-INS-268 | chr14-103473965-INS-268 |
| chr14-103478515-INS-205 | chr14-103478515-INS-205 |
| chr14-103492311-INS-186 | chr14-103492348-INS-191 |
| chr14-103510861-INS-197 | chr14-103510861-INS-197 |
| chr14-103520956-INS-6503 | chr14-103520957-INS-6504 |
| chr14-103558971-INS-6399 | chr14-103558972-INS-6398 |
| chr14-103606446-INS-56 | chr14-103606446-INS-64 |
| chr14-103720595-INS-131 | chr14-103720595-INS-134 |
| chr14-103755303-INS-55 | chr14-103755303-INS-55 |
| chr14-103780381-INS-78 | chr14-103780381-INS-123 |
| chr14-103785217-INS-70 | chr14-103785198-INS-107 |
| chr14-103792595-INS-64 | chr14-103792567-INS-79 |
| chr14-103810001-INS-648 | chr14-103810001-INS-650 |
| chr14-103824381-INS-51 | chr14-103824381-INS-51 |
| chr14-103825917-INS-56 | chr14-103825917-INS-56 |
| chr14-103834595-INS-598 | chr14-103834595-INS-598 |
| chr14-103845151-INS-132 | chr14-103845151-INS-132 |
| chr14-103897247-INS-224 | chr14-103897247-INS-224 |
| chr14-103907399-INS-73 | chr14-103907399-INS-73 |
| chr14-103908980-INS-1165 | chr14-103908981-INS-1167 |
| chr14-103913777-INS-84 | chr14-103913776-INS-84 |
| chr14-103933617-INS-88 | chr14-103933617-INS-88 |
| chr14-103958074-INS-175 | chr14-103958068-INS-184 |

| | |
|---|---|
| chr14-103984413-INS-63 | chr14-103984413-INS-63 |
| chr14-103985480-INS-110 | chr14-103985480-INS-110 |
| chr14-103989524-INS-140 | chr14-103989528-INS-112 |
| chr14-104003259-INS-1248 | chr14-104003260-INS-1248 |
| chr14-104008813-INS-3331 | chr14-104008814-INS-3331 |
| chr14-104010145-INS-84 | chr14-104010127-INS-80 |
| chr14-104039283-INS-52 | chr14-104039276-INS-62 |
| chr14-104042770-INS-460 | chr14-104042770-INS-460 |
| chr14-104103288-INS-242 | chr14-104103293-INS-250 |
| chr14-104112372-INS-64 | chr14-104112372-INS-54 |
| chr14-104245189-INS-629 | chr14-104245189-INS-629 |
| chr14-104282867-INS-70 | chr14-104282867-INS-70 |
| chr14-104370421-INS-60 | chr14-104370439-INS-66 |
| chr14-104391428-INS-62 | chr14-104391428-INS-62 |
| chr14-104399770-INS-110 | chr14-104399798-INS-62 |
| chr14-104410016-INS-152 | chr14-104410009-INS-116 |
| chr14-104426368-INS-116 | chr14-104426368-INS-116 |
| chr14-104446677-INS-103 | chr14-104446675-INS-104 |
| chr14-104450642-INS-70 | chr14-104450642-INS-70 |
| chr14-104451832-INS-2600 | chr14-104451833-INS-2600 |
| chr14-104469875-INS-3107 | chr14-104469876-INS-3107 |
| chr14-104490921-INS-6994 | chr14-104490922-INS-6995 |
| chr14-104515900-INS-63 | chr14-104515900-INS-63 |
| chr14-104517138-INS-288 | chr14-104517138-INS-291 |
| chr14-104530278-INS-1477 | chr14-104530279-INS-1477 |
| chr14-104531663-INS-232 | chr14-104531663-INS-231 |
| chr14-104543996-INS-70 | chr14-104544007-INS-71 |
| chr14-104562290-INS-7129 | chr14-104562291-INS-7130 |
| chr14-104580944-INS-198 | chr14-104580944-INS-198 |
| chr14-104591168-INS-149 | chr14-104591168-INS-196 |
| chr14-104605869-INS-433 | chr14-104605869-INS-433 |
| chr14-104608290-INS-681 | chr14-104608290-INS-681 |
| chr14-104615279-INS-75 | chr14-104615279-INS-75 |
| chr14-104645876-INS-186 | chr14-104645876-INS-186 |
| chr14-104656292-INS-64 | chr14-104656292-INS-57 |
| chr14-104671875-INS-182 | chr14-104671875-INS-182 |
| chr14-104674772-INS-54 | chr14-104674772-INS-54 |
| chr14-104678213-INS-484 | chr14-104678213-INS-484 |
| chr14-104681380-INS-83 | chr14-104681380-INS-70 |
| chr14-104710083-INS-54 | chr14-104710033-INS-89 |
| chr14-104725837-INS-58 | chr14-104725819-INS-61 |
| chr14-104760863-INS-81 | chr14-104760823-INS-72 |
| chr14-104763403-INS-63 | chr14-104763403-INS-93 |
| chr14-104808322-INS-652 | chr14-104808322-INS-652 |
| chr14-104850664-INS-463 | chr14-104850664-INS-462 |
| chr14-104867836-INS-60 | chr14-104867836-INS-60 |
| chr14-104898905-INS-59 | chr14-104898905-INS-59 |
| chr14-104952661-INS-575 | chr14-104952661-INS-554 |
| chr14-104989180-INS-61 | chr14-104989180-INS-61 |
| chr14-105031796-INS-160 | chr14-105031796-INS-162 |
| chr14-105041196-INS-213 | chr14-105041196-INS-215 |
| chr14-105042675-INS-526 | chr14-105042675-INS-526 |
| chr14-105045148-INS-59 | chr14-105045148-INS-50 |
| chr14-105057997-INS-94 | chr14-105057995-INS-108 |
| chr14-105069611-INS-789 | chr14-105069611-INS-789 |
| chr14-105089498-INS-189 | chr14-105089498-INS-192 |
| chr14-105096331-INS-939 | chr14-105096331-INS-939 |
| chr14-105106055-INS-65 | chr14-105106055-INS-65 |
| chr14-105129527-INS-73 | chr14-105129528-INS-73 |
| chr14-105157149-INS-1182 | chr14-105157150-INS-1182 |
| chr14-105177792-INS-804 | chr14-105177792-INS-804 |
| chr14-105179865-INS-61 | chr14-105179865-INS-61 |
| chr14-105207360-INS-5294 | chr14-105207361-INS-5294 |
| chr14-105213612-INS-9036 | chr14-105213613-INS-9038 |
| chr14-105219357-INS-247 | chr14-105219357-INS-247 |
| chr14-105232886-INS-441 | chr14-105232886-INS-441 |
| chr14-105275089-INS-487 | chr14-105275089-INS-487 |
| chr14-105291381-INS-123 | chr14-105291381-INS-123 |
| chr14-105300895-INS-100 | chr14-105300887-INS-72 |
| chr14-105309608-INS-210 | chr14-105309608-INS-210 |
| chr14-105356619-INS-271 | chr14-105356619-INS-272 |
| chr14-105360186-INS-74 | chr14-105360152-INS-66 |
| chr14-105411410-INS-236 | chr14-105411410-INS-236 |
| chr14-105429140-INS-205 | chr14-105429140-INS-205 |
| chr14-105431657-INS-176 | chr14-105431657-INS-176 |
| chr14-105435244-INS-187 | chr14-105435244-INS-187 |
| chr14-105478412-INS-74 | chr14-105478412-INS-74 |

| | |
|---|---|
| chr14-105480565-INS-92 | chr14-105480565-INS-100 |
| chr14-105484811-INS-88 | chr14-105484811-INS-88 |
| chr14-105499753-INS-69 | chr14-105499753-INS-77 |
| chr14-105513590-INS-66 | chr14-105513599-INS-67 |
| chr14-105533960-INS-212 | chr14-105533960-INS-210 |
| chr14-105540951-INS-443 | chr14-105540951-INS-442 |
| chr14-105550154-INS-214 | chr14-105550154-INS-214 |
| chr14-105563648-INS-70 | chr14-105563648-INS-66 |
| chr14-105598560-INS-115 | chr14-105598560-INS-119 |
| chr14-105608030-INS-202 | chr14-105608030-INS-202 |
| chr14-105620193-INS-2471 | chr14-105620194-INS-2471 |
| chr14-105650453-INS-211 | chr14-105650453-INS-211 |
| chr14-105659288-INS-4471 | chr14-105659289-INS-4471 |
| chr14-105660390-INS-74 | chr14-105660391-INS-52 |
| chr14-105685850-INS-222 | chr14-105685850-INS-222 |
| chr14-105691762-INS-6679 | chr14-105691763-INS-6679 |
| chr14-105732361-INS-78 | chr14-105732361-INS-78 |
| chr14-105742297-INS-115 | chr14-105742297-INS-115 |
| chr14-105759659-INS-102 | chr14-105759648-INS-108 |
| chr14-105763316-INS-7145 | chr14-105763317-INS-7146 |
| chr14-105764487-INS-64 | chr14-105764487-INS-64 |
| chr14-105773335-INS-69 | chr14-105773384-INS-69 |
| chr14-105773935-INS-732 | chr14-105773944-INS-731 |
| chr14-105782183-INS-189 | chr14-105782183-INS-189 |
| chr14-105820336-INS-124 | chr14-105820336-INS-132 |
| chr14-105858429-INS-80 | chr14-105858429-INS-80 |
| chr14-105859326-INS-6482 | chr14-105859327-INS-6484 |
| chr14-105865308-INS-70 | chr14-105865304-INS-54 |
| chr14-105890611-INS-288 | chr14-105890611-INS-288 |
| chr14-105917760-INS-70 | chr14-105917760-INS-60 |
| chr14-105921608-INS-174 | chr14-105921608-INS-174 |
| chr14-105944176-INS-60 | chr14-105944176-INS-60 |
| chr14-105986546-INS-147 | chr14-105986546-INS-145 |
| chr14-105989524-INS-281 | chr14-105989524-INS-268 |
| chr14-106025620-INS-52 | chr14-106025620-INS-58 |
| chr14-106043307-INS-199 | chr14-106043307-INS-199 |
| chr14-106049478-INS-56 | chr14-106049480-INS-72 |
| chr14-106085691-INS-7215 | chr14-106085692-INS-7215 |
| chr14-106098729-INS-183 | chr14-106098729-INS-183 |
| chr14-106104669-INS-63 | chr14-106104664-INS-79 |
| chr14-106128107-INS-58 | chr14-106128107-INS-58 |
| chr14-106137452-INS-66 | chr14-106137452-INS-66 |
| chr14-106148839-INS-206 | chr14-106148839-INS-206 |
| chr14-106174319-INS-209 | chr14-106174319-INS-209 |
| chr14-106220779-INS-1997 | chr14-106220780-INS-1996 |
| chr14-106221216-INS-5415 | chr14-106221217-INS-5415 |
| chr14-106225827-INS-210 | chr14-106225827-INS-211 |
| chr14-106227786-INS-399 | chr14-106227786-INS-399 |
| chr14-106272103-INS-4149 | chr14-106272104-INS-4149 |
| chr14-106279856-INS-64 | chr14-106279856-INS-64 |
| chr14-106312289-INS-140 | chr14-106312289-INS-108 |
| chr14-106327886-INS-203 | chr14-106327886-INS-203 |
| chr14-106343868-INS-79 | chr14-106343898-INS-79 |
| chr14-106351672-INS-569 | chr14-106351672-INS-569 |
| chr14-106363080-INS-191 | chr14-106363080-INS-189 |
| chr14-106390357-INS-6551 | chr14-106390358-INS-6552 |
| chr14-106395079-INS-100 | chr14-106395079-INS-100 |
| chr14-106447644-INS-331 | chr14-106447644-INS-331 |
| chr14-106470687-INS-1201 | chr14-106470688-INS-1201 |
| chr14-106485490-INS-1213 | chr14-106485491-INS-1213 |
| chr14-106544686-INS-424 | chr14-106544686-INS-424 |
| chr14-106580928-INS-203 | chr14-106580928-INS-203 |
| chr14-106588382-INS-78 | chr14-106588382-INS-78 |
| chr14-106605153-INS-53 | chr14-106605141-INS-82 |
| chr14-106776823-INS-3943 | chr14-106776824-INS-3943 |
| chr14-106794907-INS-57 | chr14-106794907-INS-65 |
| chr14-106800317-INS-806 | chr14-106800317-INS-806 |
| chr14-106806714-INS-92 | chr14-106806716-INS-52 |
| chr14-106814975-INS-1867 | chr14-106814976-INS-1867 |
| chr14-106843408-INS-53 | chr14-106843408-INS-55 |
| chr14-106864167-INS-635 | chr14-106864167-INS-635 |
| chr14-106865313-INS-70 | chr14-106865313-INS-68 |
| chr14-106868851-INS-164 | chr14-106868849-INS-180 |
| chr14-106869044-INS-4090 | chr14-106869045-INS-4080 |
| chr14-106893863-INS-60 | chr14-106893863-INS-60 |
| chr14-106924056-INS-5085 | chr14-106924057-INS-5085 |
| chr14-106927345-INS-55 | chr14-106927345-INS-55 |

| | |
|---|---|
| chr14-106931298-INS-74 | chr14-106931298-INS-76 |
| chr14-106932860-INS-53 | chr14-106932850-INS-78 |
| chr14-106944272-INS-244 | chr14-106944272-INS-244 |
| chr14-106974914-INS-65 | chr14-106974914-INS-65 |
| chr14-106977869-INS-201 | chr14-106977869-INS-198 |
| chr14-106986707-INS-219 | chr14-106986707-INS-218 |
| chr14-106992741-INS-2718 | chr14-106992742-INS-2718 |
| chr14-106994862-INS-1205 | chr14-106994863-INS-1205 |
| chr14-107008702-INS-109 | chr14-107008700-INS-112 |
| chr14-107013017-INS-120 | chr14-107013017-INS-120 |
| chr14-107026997-INS-200 | chr14-107026997-INS-200 |
| chr14-107028499-INS-190 | chr14-107028495-INS-179 |
| chr14-107060737-INS-73 | chr14-107060737-INS-64 |
| chr14-107067690-INS-1891 | chr14-107067691-INS-1891 |
| chr14-107070873-INS-1858 | chr14-107070874-INS-1858 |
| chr14-107071248-INS-343 | chr14-107071248-INS-343 |
| chr14-107103784-INS-6499 | chr14-107103785-INS-6499 |
| chr14-107103790-INS-1496 | chr14-107103791-INS-1496 |
| chr14-107104703-INS-69 | chr14-107104703-INS-69 |
| chr14-107106686-INS-1132 | chr14-107106687-INS-1132 |
| chr14-107124726-INS-119 | chr14-107124726-INS-123 |
| chr14-107138163-INS-265 | chr14-107138179-INS-254 |
| chr14-107150879-INS-796 | chr14-107150879-INS-797 |
| chr14-107153905-INS-4335 | chr14-107153906-INS-4335 |
| chr14-107160262-INS-54 | chr14-107160284-INS-54 |
| chr14-107174872-INS-82 | chr14-107174872-INS-82 |
| chr14-107186231-INS-550 | chr14-107186231-INS-578 |
| chr14-107189463-INS-408 | chr14-107189463-INS-408 |
| chr14-107191417-INS-204 | chr14-107191409-INS-200 |
| chr14-107215315-INS-488 | chr14-107215315-INS-488 |
| chr14-107243401-INS-315 | chr14-107243401-INS-315 |
| chr14-107251320-INS-60 | chr14-107251320-INS-62 |
| chr14-107259073-INS-524 | chr14-107259073-INS-524 |
| chr14-107261533-INS-97 | chr14-107261538-INS-97 |
| chr14-107268733-INS-2270 | chr14-107268734-INS-2260 |
| chr14-107276273-INS-3112 | chr14-107276274-INS-3112 |
| chr14-107299977-INS-6916 | chr14-107299978-INS-6916 |
| chr14-107311518-INS-52 | chr14-107311518-INS-52 |
| chr14-107328366-INS-1219 | chr14-107328367-INS-1219 |
| chr14-107347903-INS-54 | chr14-107347903-INS-54 |
| chr14-107353059-INS-498 | chr14-107353059-INS-500 |
| chr14-107354246-INS-54 | chr14-107354222-INS-72 |
| chr14-107367662-INS-611 | chr14-107367662-INS-611 |
| chr14-107383083-INS-188 | chr14-107383083-INS-188 |
| chr14-107415441-INS-400 | chr14-107415441-INS-400 |
| chr14-107422526-INS-278 | chr14-107422526-INS-278 |
| chr14-107461814-INS-653 | chr14-107461814-INS-663 |
| chr14-107466899-INS-345 | chr14-107466899-INS-344 |
| chr14-107483196-INS-376 | chr14-107483196-INS-382 |
| chr14-107484098-INS-523 | chr14-107484098-INS-523 |
| chr14-107484640-INS-1444 | chr14-107484641-INS-1445 |
| chr14-107491246-INS-422 | chr14-107491246-INS-422 |
| chr14-107530680-INS-4167 | chr14-107530681-INS-4167 |
| chr14-107533201-INS-343 | chr14-107533201-INS-343 |
| chr14-107545443-INS-66 | chr14-107545443-INS-58 |
| chr14-107563172-INS-377 | chr14-107563172-INS-378 |
| chr14-107573402-INS-1415 | chr14-107573403-INS-1415 |
| chr14-107581811-INS-107 | chr14-107581808-INS-109 |
| chr14-107601187-INS-52 | chr14-107601187-INS-52 |
| chr14-107615295-INS-52 | chr14-107615295-INS-54 |
| chr14-107625346-INS-367 | chr14-107625346-INS-366 |
| chr14-107674662-INS-169 | chr14-107674662-INS-170 |
| chr14-107686405-INS-54 | chr14-107686442-INS-64 |
| chr14-107698591-INS-6383 | chr14-107698592-INS-6383 |
| chr14-107706698-INS-54 | chr14-107706698-INS-52 |
| chr14-107724741-INS-57 | chr14-107724741-INS-57 |
| chr14-107727831-INS-6386 | chr14-107727832-INS-6387 |
| chr14-107744982-INS-6927 | chr14-107744983-INS-6927 |
| chr14-107759435-INS-977 | chr14-107759446-INS-977 |
| chr14-107800446-INS-80 | chr14-107800446-INS-80 |
| chr14-107806890-INS-952 | chr14-107806882-INS-970 |
| chr14-107829440-INS-131 | chr14-107829440-INS-131 |
| chr14-107856101-INS-121 | chr14-107856101-INS-121 |
| chr14-107872210-INS-77 | chr14-107872210-INS-86 |
| chr14-107884627-INS-549 | chr14-107884624-INS-550 |
| chr14-107956939-INS-4680 | chr14-107956940-INS-4680 |
| chr14-107980465-INS-67 | chr14-107980465-INS-67 |

| | |
|---|---|
| chr14-107989329-INS-168 | chr14-107989329-INS-199 |
| chr14-108079682-INS-56 | chr14-108079682-INS-64 |
| chr14-108088278-INS-74 | chr14-108088287-INS-90 |
| chr14-108107682-INS-125 | chr14-108107682-INS-125 |
| chr14-108122637-INS-852 | chr14-108122637-INS-851 |
| chr14-108147792-INS-215 | chr14-108147792-INS-215 |
| chr14-108176553-INS-92 | chr14-108176553-INS-99 |
| chr14-108191546-INS-72 | chr14-108191546-INS-66 |
| chr14-108194593-INS-115 | chr14-108194593-INS-107 |
| chr14-108198475-INS-54 | chr14-108198475-INS-52 |
| chr14-108205771-INS-213 | chr14-108205771-INS-213 |
| chr14-108251137-INS-56 | chr14-108251138-INS-58 |
| chr14-108268123-INS-60 | chr14-108268121-INS-62 |
| chr14-108282099-INS-54 | chr14-108282099-INS-53 |
| chr14-108323887-INS-2274 | chr14-108323888-INS-2276 |
| chr14-108335439-INS-85 | chr14-108335440-INS-70 |
| chr14-108350252-INS-434 | chr14-108350252-INS-434 |
| chr14-108353731-INS-5012 | chr14-108353732-INS-5012 |
| chr14-108354354-INS-248 | chr14-108354354-INS-257 |
| chr14-108371709-INS-89 | chr14-108371705-INS-96 |
| chr14-108382622-INS-56 | chr14-108382622-INS-56 |
| chr14-108451925-INS-990 | chr14-108451925-INS-990 |
| chr14-108466742-INS-103 | chr14-108466742-INS-103 |
| chr14-108517732-INS-92 | chr14-108517732-INS-92 |
| chr14-108531531-INS-246 | chr14-108531531-INS-246 |
| chr14-108542541-INS-342 | chr14-108542541-INS-342 |
| chr14-108637774-INS-51 | chr14-108637768-INS-54 |
| chr14-108651107-INS-59 | chr14-108651107-INS-59 |
| chr14-108660938-INS-75 | chr14-108660941-INS-63 |
| chr14-108675235-INS-84 | chr14-108675235-INS-76 |
| chr14-108675665-INS-862 | chr14-108675666-INS-862 |
| chr14-108676177-INS-322 | chr14-108676177-INS-323 |
| chr14-108681692-INS-505 | chr14-108681692-INS-505 |
| chr14-108693219-INS-54 | chr14-108693219-INS-54 |
| chr14-108731199-INS-383 | chr14-108731199-INS-381 |
| chr14-108740366-INS-55 | chr14-108740366-INS-55 |
| chr14-108765067-INS-190 | chr14-108765067-INS-190 |
| chr14-108772097-INS-85 | chr14-108772088-INS-90 |
| chr14-108809630-INS-56 | chr14-108809630-INS-60 |
| chr14-108812607-INS-61 | chr14-108812607-INS-61 |
| chr14-108835631-INS-237 | chr14-108835631-INS-237 |
| chr14-108837061-INS-6378 | chr14-108837062-INS-6378 |
| chr14-108837727-INS-217 | chr14-108837727-INS-215 |
| chr14-108848945-INS-955 | chr14-108848949-INS-958 |
| chr14-108851007-INS-156 | chr14-108851007-INS-171 |
| chr14-108860876-INS-821 | chr14-108860876-INS-822 |
| chr14-108876438-INS-503 | chr14-108876437-INS-510 |
| chr14-108881489-INS-251 | chr14-108881489-INS-251 |
| chr14-108890105-INS-490 | chr14-108890107-INS-489 |
| chr14-108906924-INS-418 | chr14-108906924-INS-418 |
| chr14-108953523-INS-68 | chr14-108953523-INS-68 |
| chr14-108971082-INS-54 | chr14-108971083-INS-54 |
| chr14-108976433-INS-811 | chr14-108976433-INS-812 |
| chr14-108995333-INS-612 | chr14-108995333-INS-612 |
| chr14-109016270-INS-100 | chr14-109016270-INS-60 |
| chr14-109020219-INS-149 | chr14-109020219-INS-153 |
| chr14-109063245-INS-404 | chr14-109063245-INS-404 |
| chr14-109096126-INS-176 | chr14-109096126-INS-176 |
| chr14-109127879-INS-92 | chr14-109127879-INS-92 |
| chr14-109194169-INS-6415 | chr14-109194170-INS-6415 |
| chr14-109197101-INS-855 | chr14-109197101-INS-857 |
| chr14-109231375-INS-6171 | chr14-109231376-INS-6169 |
| chr14-109245612-INS-69 | chr14-109245612-INS-70 |
| chr14-109280781-INS-1008 | chr14-109280782-INS-1012 |
| chr14-109290381-INS-128 | chr14-109290382-INS-120 |
| chr14-109300717-INS-564 | chr14-109300717-INS-564 |
| chr14-109322044-INS-56 | chr14-109322044-INS-56 |
| chr14-109327315-INS-62 | chr14-109327315-INS-62 |
| chr14-109330416-INS-2063 | chr14-109330417-INS-2059 |
| chr14-109330723-INS-319 | chr14-109330723-INS-320 |
| chr14-109332263-INS-155 | chr14-109332263-INS-155 |
| chr14-109336455-INS-566 | chr14-109336448-INS-564 |
| chr14-109355029-INS-6424 | chr14-109355030-INS-6424 |
| chr14-109368278-INS-170 | chr14-109368277-INS-171 |
| chr14-109370623-INS-76 | chr14-109370624-INS-52 |
| chr14-109387085-INS-62 | chr14-109387085-INS-62 |
| chr14-109414983-INS-96 | chr14-109414983-INS-96 |

| | |
|---|---|
| chr14-109420750-INS-887 | chr14-109420750-INS-869 |
| chr14-109431750-INS-85 | chr14-109431750-INS-85 |
| chr14-109442651-INS-112 | chr14-109442651-INS-112 |
| chr14-109444574-INS-86 | chr14-109444565-INS-93 |
| chr14-109448432-INS-60 | chr14-109448432-INS-60 |
| chr14-109451088-INS-122 | chr14-109451087-INS-123 |
| chr14-109452024-INS-672 | chr14-109452024-INS-674 |
| chr14-109469466-INS-59 | chr14-109469466-INS-59 |
| chr14-109471997-INS-98 | chr14-109472039-INS-80 |
| chr14-109473723-INS-5861 | chr14-109473724-INS-5861 |
| chr14-109484947-INS-268 | chr14-109484947-INS-268 |
| chr14-109495562-INS-2335 | chr14-109495563-INS-2338 |
| chr14-109502180-INS-3678 | chr14-109502181-INS-3641 |
| chr14-109502205-INS-237 | chr14-109502205-INS-237 |
| chr14-109507838-INS-96 | chr14-109507838-INS-64 |
| chr14-109510836-INS-91 | chr14-109510840-INS-95 |
| chr14-109511062-INS-943 | chr14-109511062-INS-943 |
| chr14-109526798-INS-138 | chr14-109526798-INS-138 |
| chr14-109529077-INS-70 | chr14-109529077-INS-120 |
| chr14-109538110-INS-342 | chr14-109538110-INS-342 |
| chr14-109561162-INS-58 | chr14-109561160-INS-96 |
| chr14-109561821-INS-73 | chr14-109561821-INS-73 |
| chr14-109565206-INS-6324 | chr14-109565207-INS-6324 |
| chr14-109571815-INS-276 | chr14-109571815-INS-291 |
| chr14-109602130-INS-727 | chr14-109602130-INS-727 |
| chr14-109622595-INS-52 | chr14-109622595-INS-52 |
| chr14-109651629-INS-179 | chr14-109651623-INS-190 |
| chr14-109653046-INS-270 | chr14-109653046-INS-270 |
| chr14-109653757-INS-51 | chr14-109653757-INS-51 |
| chr14-109669603-INS-1169 | chr14-109669601-INS-1168 |
| chr14-109681648-INS-98 | chr14-109681648-INS-98 |
| chr14-109685450-INS-55 | chr14-109685450-INS-55 |
| chr14-109686557-INS-149 | chr14-109686557-INS-149 |
| chr14-109691706-INS-3117 | chr14-109691707-INS-3117 |
| chr14-109692779-INS-60 | chr14-109692779-INS-60 |
| chr14-109697595-INS-232 | chr14-109697595-INS-232 |
| chr14-109714257-INS-181 | chr14-109714257-INS-181 |
| chr14-109723953-INS-60 | chr14-109723953-INS-60 |
| chr14-109726054-INS-143 | chr14-109726054-INS-140 |
| chr14-109731104-INS-79 | chr14-109731105-INS-83 |
| chr14-109735228-INS-201 | chr14-109735228-INS-201 |
| chr14-109752411-INS-2661 | chr14-109752412-INS-2661 |
| chr14-109764644-INS-436 | chr14-109764644-INS-436 |
| chr14-109766464-INS-762 | chr14-109766464-INS-762 |
| chr14-109829171-INS-74 | chr14-109829171-INS-52 |
| chr14-109852090-INS-867 | chr14-109852090-INS-867 |
| chr14-109898157-INS-74 | chr14-109898157-INS-75 |
| chr14-109904352-INS-56 | chr14-109904352-INS-60 |
| chr14-109908037-INS-87 | chr14-109908037-INS-87 |
| chr14-109913079-INS-64 | chr14-109913079-INS-68 |
| chr14-109925310-INS-96 | chr14-109925310-INS-102 |
| chr14-109967176-INS-438 | chr14-109967176-INS-438 |
| chr14-110024142-INS-56 | chr14-110024142-INS-56 |
| chr14-110028959-INS-150 | chr14-110028959-INS-132 |
| chr14-110052049-INS-54 | chr14-110052049-INS-54 |
| chr14-110054517-INS-323 | chr14-110054517-INS-323 |
| chr14-110074570-INS-58 | chr14-110074570-INS-58 |
| chr14-110077173-INS-91 | chr14-110077173-INS-75 |
| chr14-110086135-INS-162 | chr14-110086135-INS-164 |
| chr14-110089061-INS-148 | chr14-110089061-INS-148 |
| chr14-110191657-INS-562 | chr14-110191657-INS-562 |
| chr14-110204125-INS-509 | chr14-110204125-INS-509 |
| chr14-110248347-INS-62 | chr14-110248384-INS-58 |
| chr14-110276180-INS-60 | chr14-110276180-INS-100 |
| chr14-110305366-INS-84 | chr14-110305367-INS-74 |
| chr14-110313448-INS-54 | chr14-110313449-INS-54 |
| chr14-110314832-INS-104 | chr14-110314832-INS-100 |
| chr14-110327602-INS-52 | chr14-110327602-INS-52 |
| chr14-110350644-INS-787 | chr14-110350644-INS-787 |
| chr14-110356384-INS-2037 | chr14-110356385-INS-2037 |
| chr14-110359113-INS-56 | chr14-110359113-INS-56 |
| chr14-110370168-INS-112 | chr14-110370168-INS-104 |
| chr14-110383590-INS-149 | chr14-110383590-INS-153 |
| chr14-110388543-INS-71 | chr14-110388526-INS-55 |
| chr14-110414389-INS-2300 | chr14-110414390-INS-2301 |
| chr14-110414541-INS-62 | chr14-110414542-INS-79 |
| chr14-110427694-INS-112 | chr14-110427694-INS-112 |

| | |
|---|---|
| chr14-110432880-INS-52 | chr14-110432881-INS-52 |
| chr14-110437717-INS-573 | chr14-110437732-INS-571 |
| chr14-110451418-INS-963 | chr14-110451418-INS-963 |
| chr14-110456794-INS-88 | chr14-110456794-INS-88 |
| chr14-110516269-INS-75 | chr14-110516269-INS-75 |
| chr14-110568614-INS-80 | chr14-110568614-INS-74 |
| chr14-110572937-INS-444 | chr14-110572937-INS-444 |
| chr14-110604800-INS-52 | chr14-110604800-INS-64 |
| chr14-110612570-INS-442 | chr14-110612570-INS-444 |
| chr14-110649716-INS-66 | chr14-110649716-INS-71 |
| chr14-110661236-INS-1172 | chr14-110661237-INS-1173 |
| chr14-110664133-INS-61 | chr14-110664133-INS-71 |
| chr14-110668431-INS-76 | chr14-110668431-INS-76 |
| chr14-110669587-INS-256 | chr14-110669587-INS-256 |
| chr14-110711056-INS-64 | chr14-110711056-INS-60 |
| chr14-110711545-INS-80 | chr14-110711545-INS-80 |
| chr14-110727864-INS-919 | chr14-110727864-INS-920 |
| chr14-110766115-INS-92 | chr14-110766115-INS-96 |
| chr14-110781684-INS-69 | chr14-110781684-INS-69 |
| chr14-110795796-INS-55 | chr14-110795796-INS-55 |
| chr14-110800460-INS-363 | chr14-110800510-INS-363 |
| chr14-110809901-INS-274 | chr14-110809901-INS-274 |
| chr14-110815901-INS-78 | chr14-110815901-INS-78 |
| chr14-110827418-INS-84 | chr14-110827418-INS-84 |
| chr14-110833775-INS-1316 | chr14-110833776-INS-1316 |
| chr14-110846470-INS-723 | chr14-110846470-INS-723 |
| chr14-110914247-INS-210 | chr14-110914247-INS-210 |
| chr14-110921845-INS-84 | chr14-110921845-INS-85 |
| chr14-110985873-INS-152 | chr14-110985873-INS-156 |
| chr14-110998360-INS-52 | chr14-110998360-INS-64 |
| chr14-111003761-INS-5631 | chr14-111003762-INS-5631 |
| chr14-111005327-INS-70 | chr14-111005327-INS-72 |
| chr14-111013615-INS-281 | chr14-111013615-INS-281 |
| chr14-111021038-INS-92 | chr14-111021038-INS-114 |
| chr14-111023303-INS-155 | chr14-111023303-INS-155 |
| chr14-111027419-INS-64 | chr14-111027424-INS-56 |
| chr14-111055118-INS-55 | chr14-111055108-INS-56 |
| chr14-111097771-INS-222 | chr14-111097771-INS-198 |
| chr14-111165473-INS-54 | chr14-111165473-INS-52 |
| chr14-111175734-INS-68 | chr14-111175734-INS-70 |
| chr14-111202907-INS-71 | chr14-111202907-INS-71 |
| chr14-111215652-INS-110 | chr14-111215613-INS-124 |
| chr14-111233183-INS-114 | chr14-111233183-INS-114 |
| chr14-111242535-INS-78 | chr14-111242535-INS-54 |
| chr14-111256776-INS-622 | chr14-111256777-INS-620 |
| chr14-111265007-INS-182 | chr14-111265007-INS-182 |
| chr14-111272158-INS-117 | chr14-111272158-INS-117 |
| chr14-111279383-INS-210 | chr14-111279383-INS-211 |
| chr14-111281361-INS-53 | chr14-111281361-INS-53 |
| chr14-111294572-INS-58 | chr14-111294572-INS-58 |
| chr14-111331102-INS-232 | chr14-111331102-INS-230 |
| chr14-111335664-INS-54 | chr14-111335664-INS-54 |
| chr14-111361612-INS-151 | chr14-111361612-INS-151 |
| chr14-111362734-INS-440 | chr14-111362734-INS-440 |
| chr14-111365761-INS-60 | chr14-111365761-INS-70 |
| chr14-111368472-INS-55 | chr14-111368472-INS-55 |
| chr14-111374238-INS-481 | chr14-111374238-INS-481 |
| chr14-111383201-INS-105 | chr14-111383201-INS-100 |
| chr14-111392508-INS-57 | chr14-111392508-INS-57 |
| chr14-111423687-INS-60 | chr14-111423687-INS-60 |
| chr14-111427377-INS-882 | chr14-111427377-INS-882 |
| chr14-111437295-INS-133 | chr14-111437295-INS-133 |
| chr14-111491452-INS-2753 | chr14-111491453-INS-2753 |
| chr14-111527228-INS-497 | chr14-111527228-INS-497 |
| chr14-111534165-INS-427 | chr14-111534165-INS-427 |
| chr14-111547869-INS-6410 | chr14-111547870-INS-6411 |
| chr14-111551321-INS-117 | chr14-111551321-INS-116 |
| chr14-111556828-INS-422 | chr14-111556828-INS-416 |
| chr14-111582866-INS-203 | chr14-111582833-INS-188 |
| chr14-111587833-INS-613 | chr14-111587833-INS-622 |
| chr14-111625676-INS-72 | chr14-111625676-INS-68 |
| chr14-111641268-INS-393 | chr14-111641268-INS-400 |
| chr14-111642892-INS-887 | chr14-111642888-INS-891 |
| chr14-111690985-INS-356 | chr14-111690985-INS-381 |
| chr14-111767285-INS-474 | chr14-111767285-INS-481 |
| chr14-111780291-INS-162 | chr14-111780291-INS-159 |
| chr14-111820013-INS-97 | chr14-111820013-INS-54 |

| | |
|---|---|
| chr14-111827336-INS-871 | chr14-111827337-INS-867 |
| chr14-111830849-INS-134 | chr14-111830849-INS-135 |
| chr14-111833137-INS-52 | chr14-111833137-INS-60 |
| chr14-111841163-INS-841 | chr14-111841163-INS-840 |
| chr14-111843166-INS-244 | chr14-111843166-INS-244 |
| chr14-111879297-INS-413 | chr14-111879299-INS-443 |
| chr14-111937900-INS-144 | chr14-111937904-INS-143 |
| chr14-111986305-INS-74 | chr14-111986305-INS-74 |
| chr14-111996962-INS-90 | chr14-111996962-INS-78 |
| chr14-112026186-INS-473 | chr14-112026190-INS-448 |
| chr14-112077957-INS-102 | chr14-112077957-INS-102 |
| chr14-112081238-INS-579 | chr14-112081238-INS-579 |
| chr14-112093103-INS-483 | chr14-112093103-INS-482 |
| chr14-112097630-INS-71 | chr14-112097630-INS-83 |
| chr14-112119966-INS-68 | chr14-112119966-INS-58 |
| chr14-112121073-INS-54 | chr14-112121073-INS-54 |
| chr14-112130816-INS-637 | chr14-112130814-INS-631 |
| chr14-112138185-INS-56 | chr14-112138191-INS-52 |
| chr14-112140780-INS-212 | chr14-112140780-INS-212 |
| chr14-112149478-INS-140 | chr14-112149478-INS-140 |
| chr14-112160263-INS-526 | chr14-112160263-INS-526 |
| chr14-112171148-INS-328 | chr14-112171148-INS-328 |
| chr14-112191352-INS-62 | chr14-112191352-INS-66 |
| chr14-112211519-INS-66 | chr14-112211519-INS-64 |
| chr14-112217879-INS-930 | chr14-112217879-INS-929 |
| chr14-112241942-INS-94 | chr14-112241955-INS-50 |
| chr14-112249148-INS-80 | chr14-112249148-INS-80 |
| chr14-112271010-INS-56 | chr14-112270975-INS-92 |
| chr14-112274687-INS-69 | chr14-112274687-INS-69 |
| chr14-112304257-INS-75 | chr14-112304257-INS-71 |
| chr14-112313512-INS-134 | chr14-112313512-INS-129 |
| chr14-112324793-INS-70 | chr14-112324793-INS-72 |
| chr14-112326357-INS-429 | chr14-112326357-INS-429 |
| chr14-112330547-INS-953 | chr14-112330547-INS-953 |
| chr14-112336342-INS-907 | chr14-112336342-INS-907 |
| chr14-112336913-INS-183 | chr14-112336914-INS-181 |
| chr14-112414169-INS-55 | chr14-112414169-INS-55 |
| chr14-112416801-INS-436 | chr14-112416801-INS-436 |
| chr14-112419894-INS-389 | chr14-112419894-INS-389 |
| chr14-112421809-INS-62 | chr14-112421832-INS-52 |
| chr14-112427485-INS-86 | chr14-112427485-INS-86 |
| chr14-112436432-INS-137 | chr14-112436432-INS-140 |
| chr14-112451963-INS-7227 | chr14-112451964-INS-7227 |
| chr14-112466362-INS-52 | chr14-112466362-INS-58 |
| chr14-112481069-INS-88 | chr14-112481069-INS-88 |
| chr14-112493907-INS-56 | chr14-112493907-INS-55 |
| chr14-112519851-INS-234 | chr14-112519866-INS-249 |
| chr14-112520824-INS-76 | chr14-112520824-INS-72 |
| chr14-112566078-INS-117 | chr14-112566078-INS-117 |
| chr14-112578517-INS-76 | chr14-112578517-INS-76 |
| chr14-112582441-INS-66 | chr14-112582411-INS-66 |
| chr14-112591766-INS-54 | chr14-112591766-INS-66 |
| chr14-112595497-INS-93 | chr14-112595497-INS-95 |
| chr14-112672558-INS-101 | chr14-112672558-INS-110 |
| chr14-112687384-INS-329 | chr14-112687384-INS-329 |
| chr14-112705284-INS-348 | chr14-112705284-INS-348 |
| chr14-112714696-INS-202 | chr14-112714702-INS-202 |
| chr14-112742568-INS-81 | chr14-112742568-INS-83 |
| chr14-112752611-INS-105 | chr14-112752611-INS-103 |
| chr14-112754761-INS-106 | chr14-112754763-INS-114 |
| chr14-112761246-INS-60 | chr14-112761246-INS-60 |
| chr14-112764002-INS-54 | chr14-112764002-INS-70 |
| chr14-112769617-INS-346 | chr14-112769617-INS-346 |
| chr14-112784444-INS-60 | chr14-112784444-INS-72 |
| chr14-112881214-INS-4137 | chr14-112881215-INS-4137 |
| chr14-112970758-INS-1757 | chr14-112970759-INS-1758 |
| chr14-112998673-INS-112 | chr14-112998673-INS-94 |
| chr14-113002068-INS-6186 | chr14-113002069-INS-6187 |
| chr14-113016754-INS-69 | chr14-113016737-INS-56 |
| chr14-113081763-INS-106 | chr14-113081763-INS-84 |
| chr14-113092909-INS-5298 | chr14-113092910-INS-5298 |
| chr14-113105412-INS-385 | chr14-113105412-INS-385 |
| chr14-113107990-INS-134 | chr14-113107990-INS-127 |
| chr14-113114190-INS-306 | chr14-113114190-INS-310 |
| chr14-113133282-INS-54 | chr14-113133282-INS-66 |
| chr14-113195626-INS-76 | chr14-113195626-INS-65 |
| chr14-113231814-INS-249 | chr14-113231814-INS-249 |

| | |
|---|---|
| chr14-113248865-INS-70 | chr14-113248865-INS-50 |
| chr14-113318051-INS-1288 | chr14-113318052-INS-1287 |
| chr14-113392500-INS-698 | chr14-113392500-INS-699 |
| chr14-113393709-INS-583 | chr14-113393704-INS-588 |
| chr14-113408384-INS-201 | chr14-113408384-INS-204 |
| chr14-113457504-INS-138 | chr14-113457505-INS-128 |
| chr14-113485434-INS-100 | chr14-113485434-INS-100 |
| chr14-113574302-INS-52 | chr14-113574302-INS-52 |
| chr14-113601186-INS-69 | chr14-113601188-INS-72 |
| chr14-113697699-INS-6398 | chr14-113697700-INS-6397 |
| chr14-113727387-INS-188 | chr14-113727384-INS-150 |
| chr14-113747191-INS-1893 | chr14-113747192-INS-1920 |
| chr14-113749858-INS-80 | chr14-113749848-INS-55 |
| chr14-113755067-INS-202 | chr14-113755067-INS-202 |
| chr14-113771730-INS-546 | chr14-113771731-INS-544 |
| chr14-113779553-INS-443 | chr14-113779553-INS-444 |
| chr14-113782127-INS-990 | chr14-113782127-INS-990 |
| chr14-113827130-INS-638 | chr14-113827130-INS-637 |
| chr14-113852805-INS-102 | chr14-113852805-INS-102 |
| chr14-113856246-INS-66 | chr14-113856246-INS-66 |
| chr14-113872669-INS-1025 | chr14-113872670-INS-1025 |
| chr14-113911107-INS-206 | chr14-113911109-INS-208 |
| chr14-113920478-INS-51 | chr14-113920484-INS-57 |
| chr14-113932745-INS-131 | chr14-113932745-INS-164 |
| chr14-113938452-INS-6178 | chr14-113938453-INS-6178 |
| chr14-114008707-INS-2790 | chr14-114008708-INS-2789 |
| chr14-114060835-INS-53 | chr14-114060836-INS-54 |
| chr14-114069649-INS-84 | chr14-114069641-INS-96 |
| chr14-114077013-INS-310 | chr14-114077013-INS-320 |
| chr14-114146778-INS-1715 | chr14-114146779-INS-1715 |
| chr14-114149364-INS-925 | chr14-114149364-INS-924 |
| chr14-114154959-INS-98 | chr14-114154954-INS-98 |
| chr14-114163555-INS-150 | chr14-114163553-INS-149 |
| chr14-114180943-INS-3983 | chr14-114180944-INS-3983 |
| chr14-114194454-INS-6777 | chr14-114194455-INS-6778 |
| chr14-114195154-INS-351 | chr14-114195154-INS-351 |
| chr14-114232947-INS-245 | chr14-114232947-INS-247 |
| chr14-114266281-INS-770 | chr14-114266281-INS-770 |
| chr14-114285868-INS-116 | chr14-114285868-INS-144 |
| chr14-114327206-INS-67 | chr14-114327208-INS-100 |
| chr14-114331028-INS-580 | chr14-114331028-INS-580 |
| chr14-114390800-INS-405 | chr14-114390800-INS-405 |
| chr14-114391931-INS-2791 | chr14-114391932-INS-2798 |
| chr14-114409373-INS-112 | chr14-114409368-INS-77 |
| chr14-114449626-INS-62 | chr14-114449628-INS-58 |
| chr14-114469294-INS-1175 | chr14-114469295-INS-1178 |
| chr14-114482071-INS-60 | chr14-114482071-INS-60 |
| chr14-114488233-INS-3153 | chr14-114488234-INS-3153 |
| chr14-114494857-INS-305 | chr14-114494832-INS-286 |
| chr14-114497814-INS-82 | chr14-114497814-INS-108 |
| chr14-114504394-INS-579 | chr14-114504394-INS-595 |
| chr14-114529565-INS-845 | chr14-114529565-INS-841 |
| chr14-114535533-INS-197 | chr14-114535533-INS-196 |
| chr14-114551820-INS-401 | chr14-114551820-INS-401 |
| chr14-114553880-INS-238 | chr14-114553880-INS-239 |
| chr14-114560408-INS-77 | chr14-114560408-INS-77 |
| chr14-114572556-INS-736 | chr14-114572556-INS-735 |
| chr14-114577154-INS-210 | chr14-114577147-INS-210 |
| chr14-114580881-INS-114 | chr14-114580881-INS-114 |
| chr14-114584751-INS-506 | chr14-114584751-INS-506 |
| chr14-114603473-INS-789 | chr14-114603473-INS-790 |
| chr14-114613923-INS-89 | chr14-114613923-INS-89 |
| chr14-114627948-INS-160 | chr14-114627948-INS-160 |
| chr14-114630941-INS-727 | chr14-114630941-INS-727 |
| chr14-114637818-INS-54 | chr14-114637818-INS-54 |
| chr14-114641601-INS-72 | chr14-114641601-INS-76 |
| chr14-114645009-INS-117 | chr14-114645009-INS-101 |
| chr14-114680406-INS-112 | chr14-114680405-INS-108 |
| chr14-114685959-INS-451 | chr14-114685959-INS-451 |
| chr14-114694332-INS-871 | chr14-114694332-INS-877 |
| chr14-114702307-INS-1421 | chr14-114702308-INS-1419 |
| chr14-114761298-INS-68 | chr14-114761312-INS-80 |
| chr14-114766793-INS-105 | chr14-114766796-INS-105 |
| chr14-114808545-INS-54 | chr14-114808545-INS-52 |
| chr14-114823602-INS-6527 | chr14-114823603-INS-6528 |
| chr14-114829778-INS-100 | chr14-114829778-INS-100 |
| chr14-114844706-INS-269 | chr14-114844706-INS-269 |

| | |
|---|---|
| chr14-114862561-INS-144 | chr14-114862561-INS-144 |
| chr14-114865052-INS-1018 | chr14-114865053-INS-1017 |
| chr14-114880151-INS-88 | chr14-114880151-INS-70 |
| chr14-114888276-INS-550 | chr14-114888276-INS-555 |
| chr14-114955116-INS-5644 | chr14-114955117-INS-5640 |
| chr14-114968698-INS-363 | chr14-114968698-INS-363 |
| chr14-114986645-INS-6650 | chr14-114986646-INS-6645 |
| chr14-115007785-INS-4545 | chr14-115007786-INS-4545 |
| chr14-115020540-INS-75 | chr14-115020540-INS-88 |
| chr14-115026928-INS-135 | chr14-115026928-INS-135 |
| chr14-115030125-INS-52 | chr14-115030139-INS-51 |
| chr14-115040696-INS-5261 | chr14-115040697-INS-5261 |
| chr14-115068352-INS-143 | chr14-115068348-INS-158 |
| chr14-115074014-INS-83 | chr14-115074015-INS-84 |
| chr14-115080929-INS-99 | chr14-115080929-INS-99 |
| chr14-115082429-INS-129 | chr14-115082423-INS-136 |
| chr14-115117836-INS-74 | chr14-115117836-INS-74 |
| chr14-115130798-INS-737 | chr14-115130798-INS-731 |
| chr14-115198661-INS-60 | chr14-115198660-INS-50 |
| chr14-115200866-INS-53 | chr14-115200866-INS-61 |
| chr14-115207027-INS-261 | chr14-115207027-INS-235 |
| chr14-115216574-INS-96 | chr14-115216614-INS-72 |
| chr14-115223898-INS-61 | chr14-115223898-INS-57 |
| chr14-115265079-INS-60 | chr14-115265079-INS-60 |
| chr14-115285974-INS-344 | chr14-115285974-INS-347 |
| chr14-115313845-INS-54 | chr14-115313845-INS-66 |
| chr14-115322595-INS-279 | chr14-115322595-INS-279 |
| chr14-115345763-INS-130 | chr14-115345760-INS-143 |
| chr14-115366238-INS-76 | chr14-115366230-INS-52 |
| chr14-115372182-INS-105 | chr14-115372182-INS-104 |
| chr14-115386318-INS-62 | chr14-115386318-INS-62 |
| chr14-115402353-INS-160 | chr14-115402353-INS-163 |
| chr14-115420097-INS-113 | chr14-115420082-INS-115 |
| chr14-115485308-INS-88 | chr14-115485312-INS-84 |
| chr14-115492141-INS-985 | chr14-115492143-INS-989 |
| chr14-115506909-INS-528 | chr14-115506909-INS-527 |
| chr14-115520464-INS-84 | chr14-115520460-INS-82 |
| chr14-115531544-INS-191 | chr14-115531544-INS-191 |
| chr14-115546836-INS-144 | chr14-115546836-INS-144 |
| chr14-115561758-INS-6669 | chr14-115561759-INS-6668 |
| chr14-115570161-INS-127 | chr14-115570161-INS-127 |
| chr14-115600152-INS-54 | chr14-115600152-INS-54 |
| chr14-115603141-INS-299 | chr14-115603171-INS-270 |
| chr14-115615678-INS-2675 | chr14-115615679-INS-2678 |
| chr14-115629170-INS-214 | chr14-115629171-INS-214 |
| chr14-115677388-INS-502 | chr14-115677388-INS-503 |
| chr14-115687899-INS-690 | chr14-115687899-INS-690 |
| chr14-115702884-INS-74 | chr14-115702921-INS-58 |
| chr14-115710771-INS-98 | chr14-115710806-INS-92 |
| chr14-115714484-INS-124 | chr14-115714482-INS-163 |
| chr14-115715224-INS-865 | chr14-115715224-INS-850 |
| chr14-115741160-INS-371 | chr14-115741160-INS-370 |
| chr14-115745729-INS-935 | chr14-115745727-INS-939 |
| chr14-115752335-INS-6518 | chr14-115752336-INS-6523 |
| chr14-115753237-INS-362 | chr14-115753239-INS-362 |
| chr14-115795183-INS-168 | chr14-115795183-INS-168 |
| chr14-115807958-INS-225 | chr14-115807958-INS-229 |
| chr14-115869487-INS-89 | chr14-115869482-INS-71 |
| chr14-115887242-INS-714 | chr14-115887242-INS-717 |
| chr14-115903483-INS-138 | chr14-115903483-INS-146 |
| chr14-115904865-INS-54 | chr14-115904875-INS-77 |
| chr14-115914518-INS-106 | chr14-115914518-INS-106 |
| chr14-115921314-INS-64 | chr14-115921335-INS-89 |
| chr14-115930188-INS-2014 | chr14-115930189-INS-2014 |
| chr14-115980713-INS-185 | chr14-115980713-INS-185 |
| chr14-115991082-INS-522 | chr14-115991082-INS-522 |
| chr14-116009168-INS-2001 | chr14-116009169-INS-2001 |
| chr14-116032112-INS-135 | chr14-116032112-INS-135 |
| chr14-116055663-INS-125 | chr14-116055663-INS-125 |
| chr14-116059558-INS-82 | chr14-116059558-INS-82 |
| chr14-116091455-INS-6434 | chr14-116091456-INS-6435 |
| chr14-116091456-INS-497 | chr14-116091456-INS-497 |
| chr14-116100114-INS-473 | chr14-116100114-INS-473 |
| chr14-116110008-INS-428 | chr14-116110008-INS-428 |
| chr14-116175526-INS-1229 | chr14-116175527-INS-1229 |
| chr14-116227647-INS-56 | chr14-116227647-INS-54 |
| chr14-116238094-INS-677 | chr14-116238094-INS-677 |

| | |
|---|---|
| chr14-116240086-INS-61 | chr14-116240086-INS-63 |
| chr14-116260100-INS-58 | chr14-116260100-INS-76 |
| chr14-116284203-INS-483 | chr14-116284203-INS-482 |
| chr14-116319206-INS-400 | chr14-116319206-INS-401 |
| chr14-116369405-INS-6186 | chr14-116369406-INS-6186 |
| chr14-116398240-INS-306 | chr14-116398240-INS-306 |
| chr14-116400635-INS-7169 | chr14-116400636-INS-7169 |
| chr14-116401477-INS-78 | chr14-116401477-INS-56 |
| chr14-116421739-INS-137 | chr14-116421739-INS-137 |
| chr14-116449695-INS-2164 | chr14-116449696-INS-2165 |
| chr14-116525476-INS-135 | chr14-116525476-INS-135 |
| chr14-116527635-INS-82 | chr14-116527619-INS-52 |
| chr14-116534019-INS-58 | chr14-116534019-INS-60 |
| chr14-116553472-INS-62 | chr14-116553472-INS-62 |
| chr14-116608612-INS-147 | chr14-116608612-INS-130 |
| chr14-116610252-INS-197 | chr14-116610252-INS-202 |
| chr14-116611196-INS-484 | chr14-116611196-INS-482 |
| chr14-116618592-INS-2158 | chr14-116618593-INS-2158 |
| chr14-116621310-INS-92 | chr14-116621304-INS-89 |
| chr14-116649339-INS-186 | chr14-116649341-INS-217 |
| chr14-116664808-INS-62 | chr14-116664808-INS-62 |
| chr14-116689779-INS-639 | chr14-116689779-INS-640 |
| chr14-116703094-INS-412 | chr14-116703094-INS-412 |
| chr14-116712931-INS-320 | chr14-116712931-INS-314 |
| chr14-116738173-INS-230 | chr14-116738199-INS-238 |
| chr14-116739924-INS-61 | chr14-116739928-INS-64 |
| chr14-116799614-INS-3307 | chr14-116799615-INS-3307 |
| chr14-116854794-INS-120 | chr14-116854794-INS-123 |
| chr14-116959984-INS-64 | chr14-116959984-INS-52 |
| chr14-116990205-INS-59 | chr14-116990205-INS-54 |
| chr14-117014365-INS-96 | chr14-117014389-INS-80 |
| chr14-117035980-INS-70 | chr14-117035980-INS-98 |
| chr14-117037039-INS-139 | chr14-117037039-INS-139 |
| chr14-117091417-INS-59 | chr14-117091421-INS-53 |
| chr14-117098390-INS-96 | chr14-117098363-INS-112 |
| chr14-117151418-INS-76 | chr14-117151418-INS-82 |
| chr14-117162180-INS-70 | chr14-117162180-INS-54 |
| chr14-117173051-INS-60 | chr14-117173051-INS-66 |
| chr14-117183423-INS-79 | chr14-117183398-INS-80 |
| chr14-117251991-INS-130 | chr14-117251991-INS-130 |
| chr14-117315646-INS-267 | chr14-117315664-INS-243 |
| chr14-117359762-INS-101 | chr14-117359762-INS-89 |
| chr14-117360983-INS-526 | chr14-117360983-INS-532 |
| chr14-117428783-INS-523 | chr14-117428783-INS-523 |
| chr14-117487670-INS-1295 | chr14-117487671-INS-1295 |
| chr14-117509631-INS-352 | chr14-117509631-INS-355 |
| chr14-117522960-INS-62 | chr14-117522960-INS-66 |
| chr14-117540741-INS-101 | chr14-117540741-INS-80 |
| chr14-117555345-INS-799 | chr14-117555345-INS-822 |
| chr14-117596726-INS-139 | chr14-117596726-INS-140 |
| chr14-117656871-INS-199 | chr14-117656871-INS-199 |
| chr14-117694352-INS-52 | chr14-117694352-INS-52 |
| chr14-117698719-INS-6479 | chr14-117698720-INS-6479 |
| chr14-117725240-INS-6891 | chr14-117725241-INS-6891 |
| chr14-117751808-INS-82 | chr14-117751808-INS-60 |
| chr14-117817474-INS-124 | chr14-117817474-INS-127 |
| chr14-117821916-INS-53 | chr14-117821916-INS-53 |
| chr14-117856570-INS-60 | chr14-117856570-INS-54 |
| chr14-117880768-INS-105 | chr14-117880766-INS-143 |
| chr14-117910386-INS-168 | chr14-117910386-INS-168 |
| chr14-117918067-INS-6890 | chr14-117918068-INS-6890 |
| chr14-117921296-INS-89 | chr14-117921296-INS-89 |
| chr14-117978920-INS-180 | chr14-117978920-INS-181 |
| chr14-118007392-INS-93 | chr14-118007380-INS-114 |
| chr14-118034508-INS-56 | chr14-118034508-INS-56 |
| chr14-118040410-INS-463 | chr14-118040417-INS-461 |
| chr14-118044798-INS-68 | chr14-118044798-INS-68 |
| chr14-118051816-INS-6907 | chr14-118051817-INS-6907 |
| chr14-118059272-INS-84 | chr14-118059272-INS-84 |
| chr14-118065194-INS-62 | chr14-118065194-INS-64 |
| chr14-118077575-INS-66 | chr14-118077575-INS-66 |
| chr14-118079897-INS-90 | chr14-118079897-INS-94 |
| chr14-118084070-INS-60 | chr14-118084071-INS-60 |
| chr14-118106971-INS-1493 | chr14-118106972-INS-1493 |
| chr14-118110207-INS-108 | chr14-118110207-INS-108 |
| chr14-118147594-INS-70 | chr14-118147594-INS-74 |
| chr14-118150362-INS-90 | chr14-118150362-INS-90 |

| | |
|---|---|
| chr14-118157138-INS-65 | chr14-118157138-INS-65 |
| chr14-118168972-INS-146 | chr14-118168972-INS-148 |
| chr14-118171133-INS-419 | chr14-118171133-INS-419 |
| chr14-118176751-INS-124 | chr14-118176776-INS-126 |
| chr14-118191174-INS-120 | chr14-118191174-INS-120 |
| chr14-118199492-INS-76 | chr14-118199492-INS-76 |
| chr14-118202668-INS-56 | chr14-118202668-INS-56 |
| chr14-118209376-INS-100 | chr14-118209376-INS-100 |
| chr14-118218243-INS-735 | chr14-118218236-INS-734 |
| chr14-118275777-INS-62 | chr14-118275777-INS-93 |
| chr14-118301019-INS-148 | chr14-118301019-INS-148 |
| chr14-118335780-INS-286 | chr14-118335780-INS-286 |
| chr14-118342105-INS-102 | chr14-118342105-INS-106 |
| chr14-118347177-INS-436 | chr14-118347177-INS-436 |
| chr14-118352468-INS-72 | chr14-118352468-INS-72 |
| chr14-118361915-INS-145 | chr14-118361915-INS-153 |
| chr14-118383188-INS-946 | chr14-118383189-INS-946 |
| chr14-118391410-INS-3168 | chr14-118391411-INS-3168 |
| chr14-118409465-INS-160 | chr14-118409487-INS-174 |
| chr14-118425037-INS-120 | chr14-118425068-INS-94 |
| chr14-118450972-INS-76 | chr14-118450971-INS-79 |
| chr14-118454835-INS-109 | chr14-118454835-INS-108 |
| chr14-118506353-INS-401 | chr14-118506353-INS-395 |
| chr14-118519737-INS-564 | chr14-118519737-INS-564 |
| chr14-118527998-INS-56 | chr14-118527998-INS-58 |
| chr14-118562756-INS-120 | chr14-118562756-INS-104 |
| chr14-118563507-INS-732 | chr14-118563507-INS-732 |
| chr14-118564594-INS-62 | chr14-118564594-INS-64 |
| chr14-118566684-INS-81 | chr14-118566684-INS-72 |
| chr14-118590781-INS-75 | chr14-118590781-INS-73 |
| chr14-118620793-INS-88 | chr14-118620805-INS-102 |
| chr14-118626329-INS-341 | chr14-118626322-INS-364 |
| chr14-118633247-INS-76 | chr14-118633209-INS-92 |
| chr14-118644211-INS-333 | chr14-118644211-INS-333 |
| chr14-118644897-INS-116 | chr14-118644897-INS-116 |
| chr14-118647382-INS-54 | chr14-118647383-INS-54 |
| chr14-118654780-INS-520 | chr14-118654780-INS-520 |
| chr14-118659337-INS-87 | chr14-118659338-INS-85 |
| chr14-118661277-INS-60 | chr14-118661277-INS-60 |
| chr14-118667082-INS-129 | chr14-118667082-INS-105 |
| chr14-118669846-INS-80 | chr14-118669846-INS-62 |
| chr14-118704322-INS-52 | chr14-118704322-INS-52 |
| chr14-118760114-INS-71 | chr14-118760114-INS-66 |
| chr14-118769967-INS-491 | chr14-118769967-INS-491 |
| chr14-118784531-INS-68 | chr14-118784531-INS-68 |
| chr14-118785197-INS-138 | chr14-118785158-INS-137 |
| chr14-118791638-INS-85 | chr14-118791630-INS-118 |
| chr14-118809515-INS-114 | chr14-118809515-INS-152 |
| chr14-118820914-INS-554 | chr14-118820914-INS-554 |
| chr14-118822096-INS-97 | chr14-118822096-INS-97 |
| chr14-118831395-INS-55 | chr14-118831395-INS-55 |
| chr14-118835751-INS-88 | chr14-118835751-INS-88 |
| chr14-118837710-INS-70 | chr14-118837710-INS-70 |
| chr14-118844429-INS-82 | chr14-118844428-INS-76 |
| chr14-118851625-INS-124 | chr14-118851625-INS-128 |
| chr14-118890887-INS-64 | chr14-118890887-INS-72 |
| chr14-118921623-INS-398 | chr14-118921623-INS-400 |
| chr14-118925501-INS-231 | chr14-118925501-INS-231 |
| chr14-118927303-INS-68 | chr14-118927303-INS-56 |
| chr14-118928398-INS-142 | chr14-118928398-INS-142 |
| chr14-118969354-INS-168 | chr14-118969354-INS-183 |
| chr14-118985687-INS-88 | chr14-118985687-INS-84 |
| chr14-119018514-INS-58 | chr14-119018535-INS-54 |
| chr14-119027921-INS-111 | chr14-119027921-INS-111 |
| chr14-119043901-INS-75 | chr14-119043898-INS-75 |
| chr14-119065928-INS-192 | chr14-119065928-INS-194 |
| chr14-119070250-INS-62 | chr14-119070250-INS-62 |
| chr14-119085677-INS-126 | chr14-119085678-INS-102 |
| chr14-119128352-INS-234 | chr14-119128352-INS-234 |
| chr14-119132622-INS-66 | chr14-119132622-INS-66 |
| chr14-119136567-INS-60 | chr14-119136567-INS-56 |
| chr14-119138331-INS-3666 | chr14-119138332-INS-3666 |
| chr14-119141420-INS-103 | chr14-119141420-INS-105 |
| chr14-119169929-INS-5311 | chr14-119169930-INS-5312 |
| chr14-119170829-INS-106 | chr14-119170833-INS-70 |
| chr14-119180277-INS-73 | chr14-119180277-INS-73 |
| chr14-119192728-INS-64 | chr14-119192728-INS-64 |

| | |
|---|---|
| chr14-119220006-INS-58 | chr14-119220006-INS-58 |
| chr14-119238239-INS-88 | chr14-119238239-INS-88 |
| chr14-119241895-INS-7252 | chr14-119241896-INS-7252 |
| chr14-119254807-INS-109 | chr14-119254807-INS-91 |
| chr14-119268044-INS-56 | chr14-119268044-INS-52 |
| chr14-119272994-INS-113 | chr14-119272994-INS-113 |
| chr14-119274710-INS-152 | chr14-119274710-INS-152 |
| chr14-119304641-INS-7204 | chr14-119304642-INS-7204 |
| chr14-119321170-INS-54 | chr14-119321164-INS-50 |
| chr14-119333713-INS-1770 | chr14-119333716-INS-1778 |
| chr14-119339946-INS-96 | chr14-119339946-INS-90 |
| chr14-119347350-INS-107 | chr14-119347350-INS-107 |
| chr14-119356165-INS-6279 | chr14-119356166-INS-6277 |
| chr14-119370588-INS-2438 | chr14-119370589-INS-2438 |
| chr14-119380216-INS-511 | chr14-119380216-INS-521 |
| chr14-119392036-INS-95 | chr14-119392036-INS-85 |
| chr14-119432318-INS-722 | chr14-119432324-INS-724 |
| chr14-119450275-INS-389 | chr14-119450275-INS-389 |
| chr14-119451669-INS-172 | chr14-119451669-INS-172 |
| chr14-119454460-INS-71 | chr14-119454460-INS-63 |
| chr14-119458397-INS-324 | chr14-119458397-INS-330 |
| chr14-119472055-INS-164 | chr14-119472056-INS-164 |
| chr14-119472783-INS-82 | chr14-119472783-INS-78 |
| chr14-119481051-INS-60 | chr14-119481051-INS-60 |
| chr14-119482617-INS-222 | chr14-119482617-INS-222 |
| chr14-119489477-INS-155 | chr14-119489477-INS-155 |
| chr14-119529956-INS-342 | chr14-119529956-INS-343 |
| chr14-119539209-INS-249 | chr14-119539209-INS-249 |
| chr14-119547292-INS-62 | chr14-119547292-INS-62 |
| chr14-119550236-INS-714 | chr14-119550236-INS-714 |
| chr14-119566187-INS-7462 | chr14-119566188-INS-7462 |
| chr14-119571709-INS-102 | chr14-119571709-INS-106 |
| chr14-119599382-INS-61 | chr14-119599338-INS-55 |
| chr14-119637146-INS-84 | chr14-119637140-INS-57 |
| chr14-119638421-INS-184 | chr14-119638421-INS-184 |
| chr14-119641268-INS-87 | chr14-119641265-INS-78 |
| chr14-119661402-INS-79 | chr14-119661402-INS-79 |
| chr14-119693286-INS-87 | chr14-119693286-INS-87 |
| chr14-119699931-INS-1042 | chr14-119699932-INS-1042 |
| chr14-119724981-INS-81 | chr14-119724981-INS-81 |
| chr14-119748818-INS-63 | chr14-119748818-INS-64 |
| chr14-119750370-INS-56 | chr14-119750370-INS-62 |
| chr14-119752446-INS-295 | chr14-119752446-INS-338 |
| chr14-119780943-INS-812 | chr14-119780943-INS-811 |
| chr14-119782909-INS-114 | chr14-119782903-INS-152 |
| chr14-119791240-INS-87 | chr14-119791238-INS-57 |
| chr14-119795233-INS-1209 | chr14-119795234-INS-1209 |
| chr14-119807703-INS-137 | chr14-119807703-INS-149 |
| chr14-119856147-INS-51 | chr14-119856147-INS-51 |
| chr14-119860382-INS-556 | chr14-119860372-INS-552 |
| chr14-119879068-INS-970 | chr14-119879068-INS-972 |
| chr14-119880580-INS-496 | chr14-119880580-INS-496 |
| chr14-119881619-INS-72 | chr14-119881619-INS-74 |
| chr14-119896471-INS-165 | chr14-119896471-INS-165 |
| chr14-119902758-INS-7781 | chr14-119902759-INS-7781 |
| chr14-119936562-INS-50 | chr14-119936562-INS-54 |
| chr14-119942514-INS-63 | chr14-119942528-INS-84 |
| chr14-119948820-INS-142 | chr14-119948820-INS-142 |
| chr14-119982158-INS-58 | chr14-119982158-INS-54 |
| chr14-120005688-INS-101 | chr14-120005688-INS-101 |
| chr14-120029091-INS-58 | chr14-120029101-INS-58 |
| chr14-120035796-INS-91 | chr14-120035796-INS-89 |
| chr14-120040973-INS-62 | chr14-120040974-INS-58 |
| chr14-120052754-INS-359 | chr14-120052754-INS-359 |
| chr14-120070731-INS-100 | chr14-120070731-INS-98 |
| chr14-120075466-INS-206 | chr14-120075466-INS-210 |
| chr14-120092469-INS-55 | chr14-120092469-INS-55 |
| chr14-120101112-INS-56 | chr14-120101112-INS-56 |
| chr14-120109503-INS-98 | chr14-120109503-INS-98 |
| chr14-120116902-INS-223 | chr14-120116902-INS-223 |
| chr14-120130218-INS-112 | chr14-120130218-INS-112 |
| chr14-120137573-INS-54 | chr14-120137573-INS-58 |
| chr14-120145197-INS-451 | chr14-120145197-INS-451 |
| chr14-120166360-INS-51 | chr14-120166360-INS-51 |
| chr14-120189381-INS-92 | chr14-120189416-INS-59 |
| chr14-120207182-INS-60 | chr14-120207182-INS-60 |
| chr14-120222995-INS-124 | chr14-120222995-INS-124 |

| | |
|---|---|
| chr14-120225063-INS-132 | chr14-120225063-INS-129 |
| chr14-120226500-INS-888 | chr14-120226513-INS-866 |
| chr14-120239813-INS-81 | chr14-120239813-INS-78 |
| chr14-120255152-INS-90 | chr14-120255164-INS-70 |
| chr14-120267938-INS-115 | chr14-120267938-INS-115 |
| chr14-120287120-INS-413 | chr14-120287120-INS-413 |
| chr14-120299334-INS-212 | chr14-120299334-INS-212 |
| chr14-120312328-INS-152 | chr14-120312337-INS-161 |
| chr14-120315250-INS-54 | chr14-120315250-INS-58 |
| chr14-120322141-INS-117 | chr14-120322141-INS-157 |
| chr14-120361398-INS-60 | chr14-120361398-INS-106 |
| chr14-120370182-INS-147 | chr14-120370181-INS-186 |
| chr14-120405882-INS-355 | chr14-120405882-INS-355 |
| chr14-120409148-INS-6442 | chr14-120409149-INS-6442 |
| chr14-120430427-INS-265 | chr14-120430427-INS-265 |
| chr14-120442727-INS-82 | chr14-120442726-INS-71 |
| chr14-120455495-INS-88 | chr14-120455481-INS-86 |
| chr14-120456982-INS-95 | chr14-120456983-INS-50 |
| chr14-120457526-INS-77 | chr14-120457526-INS-77 |
| chr14-120461708-INS-56 | chr14-120461708-INS-58 |
| chr14-120472213-INS-72 | chr14-120472213-INS-72 |
| chr14-120478326-INS-673 | chr14-120478326-INS-673 |
| chr14-120481736-INS-323 | chr14-120481736-INS-323 |
| chr14-120492062-INS-56 | chr14-120492060-INS-59 |
| chr14-120499982-INS-366 | chr14-120499982-INS-354 |
| chr14-120546098-INS-62 | chr14-120546106-INS-52 |
| chr14-120569562-INS-258 | chr14-120569562-INS-258 |
| chr14-120576260-INS-79 | chr14-120576260-INS-79 |
| chr14-120591913-INS-104 | chr14-120591913-INS-106 |
| chr14-120597167-INS-54 | chr14-120597167-INS-54 |
| chr14-120609189-INS-99 | chr14-120609189-INS-99 |
| chr14-120621573-INS-102 | chr14-120621573-INS-68 |
| chr14-120624528-INS-69 | chr14-120624528-INS-69 |
| chr14-120656450-INS-73 | chr14-120656450-INS-71 |
| chr14-120669548-INS-60 | chr14-120669548-INS-60 |
| chr14-120682902-INS-5717 | chr14-120682903-INS-5713 |
| chr14-120712868-INS-124 | chr14-120712868-INS-124 |
| chr14-120718133-INS-127 | chr14-120718133-INS-127 |
| chr14-120755525-INS-6712 | chr14-120755526-INS-6712 |
| chr14-120768978-INS-150 | chr14-120768978-INS-157 |
| chr14-120773044-INS-4344 | chr14-120773045-INS-4344 |
| chr14-120776821-INS-246 | chr14-120776821-INS-246 |
| chr14-120784754-INS-446 | chr14-120784754-INS-445 |
| chr14-120802624-INS-62 | chr14-120802624-INS-62 |
| chr14-120805216-INS-330 | chr14-120805216-INS-330 |
| chr14-120808103-INS-58 | chr14-120808103-INS-58 |
| chr14-120843124-INS-400 | chr14-120843124-INS-400 |
| chr14-120849557-INS-52 | chr14-120849557-INS-50 |
| chr14-120850714-INS-498 | chr14-120850714-INS-498 |
| chr14-120853846-INS-6581 | chr14-120853847-INS-6581 |
| chr14-120868789-INS-165 | chr14-120868789-INS-165 |
| chr14-120873054-INS-154 | chr14-120873054-INS-152 |
| chr14-120895776-INS-69 | chr14-120895760-INS-67 |
| chr14-120926112-INS-60 | chr14-120926112-INS-56 |
| chr14-120937187-INS-282 | chr14-120937187-INS-285 |
| chr14-120941207-INS-211 | chr14-120941207-INS-211 |
| chr14-120942542-INS-56 | chr14-120942549-INS-54 |
| chr14-120945711-INS-963 | chr14-120945711-INS-963 |
| chr14-120964215-INS-70 | chr14-120964215-INS-50 |
| chr14-120977157-INS-273 | chr14-120977167-INS-276 |
| chr14-120988471-INS-436 | chr14-120988471-INS-436 |
| chr14-120990340-INS-76 | chr14-120990340-INS-57 |
| chr14-120998368-INS-389 | chr14-120998369-INS-371 |
| chr14-121026622-INS-506 | chr14-121026622-INS-506 |
| chr14-121028508-INS-991 | chr14-121028508-INS-988 |
| chr14-121029535-INS-60 | chr14-121029535-INS-80 |
| chr14-121063647-INS-207 | chr14-121063653-INS-208 |
| chr14-121074108-INS-133 | chr14-121074108-INS-137 |
| chr14-121084187-INS-600 | chr14-121084184-INS-591 |
| chr14-121105287-INS-208 | chr14-121105287-INS-175 |
| chr14-121121646-INS-194 | chr14-121121646-INS-194 |
| chr14-121122925-INS-63 | chr14-121122925-INS-63 |
| chr14-121131762-INS-223 | chr14-121131762-INS-223 |
| chr14-121135814-INS-118 | chr14-121135827-INS-76 |
| chr14-121138618-INS-199 | chr14-121138615-INS-196 |
| chr14-121146720-INS-68 | chr14-121146720-INS-68 |
| chr14-121180368-INS-64 | chr14-121180368-INS-64 |

| | |
|---|---|
| chr14-121190402-INS-52 | chr14-121190402-INS-54 |
| chr14-121194291-INS-4522 | chr14-121194292-INS-4522 |
| chr14-121204868-INS-234 | chr14-121204868-INS-234 |
| chr14-121206658-INS-504 | chr14-121206661-INS-504 |
| chr14-121207509-INS-56 | chr14-121207509-INS-56 |
| chr14-121216515-INS-151 | chr14-121216515-INS-151 |
| chr14-121220583-INS-407 | chr14-121220583-INS-406 |
| chr14-121221834-INS-304 | chr14-121221834-INS-304 |
| chr14-121224845-INS-350 | chr14-121224845-INS-350 |
| chr14-121227589-INS-208 | chr14-121227589-INS-208 |
| chr14-121232985-INS-148 | chr14-121232985-INS-148 |
| chr14-121235659-INS-148 | chr14-121235660-INS-188 |
| chr14-121237251-INS-276 | chr14-121237251-INS-276 |
| chr14-121241929-INS-174 | chr14-121241929-INS-174 |
| chr14-121265831-INS-210 | chr14-121265831-INS-210 |
| chr14-121266771-INS-212 | chr14-121266771-INS-212 |
| chr14-121270685-INS-203 | chr14-121270685-INS-207 |
| chr14-121279310-INS-1018 | chr14-121279311-INS-1018 |
| chr14-121280269-INS-209 | chr14-121280269-INS-212 |
| chr14-121287059-INS-96 | chr14-121287059-INS-96 |
| chr14-121296051-INS-104 | chr14-121296051-INS-104 |
| chr14-121296598-INS-7443 | chr14-121296599-INS-7443 |
| chr14-121310785-INS-178 | chr14-121310781-INS-198 |
| chr14-121322172-INS-1225 | chr14-121322173-INS-1274 |
| chr14-121342987-INS-96 | chr14-121342987-INS-97 |
| chr14-121352185-INS-90 | chr14-121352185-INS-60 |
| chr14-121368015-INS-549 | chr14-121368015-INS-550 |
| chr14-121375590-INS-64 | chr14-121375590-INS-64 |
| chr14-121380146-INS-95 | chr14-121380146-INS-109 |
| chr14-121384179-INS-166 | chr14-121384179-INS-166 |
| chr14-121396072-INS-62 | chr14-121396076-INS-51 |
| chr14-121400752-INS-140 | chr14-121400752-INS-140 |
| chr14-121402698-INS-69 | chr14-121402698-INS-69 |
| chr14-121422026-INS-216 | chr14-121422026-INS-182 |
| chr14-121433434-INS-51 | chr14-121433434-INS-51 |
| chr14-121500115-INS-170 | chr14-121500115-INS-170 |
| chr14-121503449-INS-57 | chr14-121503449-INS-57 |
| chr14-121538061-INS-168 | chr14-121538061-INS-168 |
| chr14-121542990-INS-6529 | chr14-121542991-INS-6529 |
| chr14-121545934-INS-114 | chr14-121545967-INS-126 |
| chr14-121555306-INS-80 | chr14-121555301-INS-95 |
| chr14-121572915-INS-78 | chr14-121572915-INS-78 |
| chr14-121588323-INS-1029 | chr14-121588324-INS-1029 |
| chr14-121594518-INS-181 | chr14-121594518-INS-181 |
| chr14-121601006-INS-211 | chr14-121601006-INS-211 |
| chr14-121608854-INS-63 | chr14-121608854-INS-63 |
| chr14-121622757-INS-179 | chr14-121622757-INS-189 |
| chr14-121626185-INS-67 | chr14-121626188-INS-71 |
| chr14-121630791-INS-213 | chr14-121630791-INS-213 |
| chr14-121633209-INS-71 | chr14-121633208-INS-72 |
| chr14-121637214-INS-170 | chr14-121637214-INS-169 |
| chr14-121659032-INS-200 | chr14-121659032-INS-200 |
| chr14-121674813-INS-167 | chr14-121674808-INS-173 |
| chr14-121681464-INS-84 | chr14-121681465-INS-54 |
| chr14-121685619-INS-7184 | chr14-121685620-INS-7184 |
| chr14-121686855-INS-124 | chr14-121686858-INS-124 |
| chr14-121699148-INS-51 | chr14-121699109-INS-70 |
| chr14-121704919-INS-198 | chr14-121704919-INS-198 |
| chr14-121705179-INS-588 | chr14-121705179-INS-596 |
| chr14-121707494-INS-104 | chr14-121707520-INS-112 |
| chr14-121736028-INS-71 | chr14-121736028-INS-71 |
| chr14-121755429-INS-374 | chr14-121755429-INS-374 |
| chr14-121758881-INS-4469 | chr14-121758882-INS-4469 |
| chr14-121761305-INS-53 | chr14-121761305-INS-53 |
| chr14-121765650-INS-157 | chr14-121765650-INS-157 |
| chr14-121766216-INS-1172 | chr14-121766217-INS-1188 |
| chr14-121780426-INS-148 | chr14-121780449-INS-118 |
| chr14-121786714-INS-69 | chr14-121786714-INS-60 |
| chr14-121791025-INS-219 | chr14-121791036-INS-237 |
| chr14-121793124-INS-60 | chr14-121793085-INS-52 |
| chr14-121798582-INS-301 | chr14-121798582-INS-301 |
| chr14-121831403-INS-68 | chr14-121831403-INS-51 |
| chr14-121841064-INS-58 | chr14-121841064-INS-54 |
| chr14-121860202-INS-140 | chr14-121860174-INS-144 |
| chr14-121861540-INS-105 | chr14-121861534-INS-70 |
| chr14-121868712-INS-62 | chr14-121868712-INS-62 |
| chr14-121894030-INS-144 | chr14-121894030-INS-144 |

| | |
|---|---|
| chr14-121902507-INS-386 | chr14-121902507-INS-386 |
| chr14-121922617-INS-53 | chr14-121922617-INS-53 |
| chr14-121925998-INS-6442 | chr14-121925999-INS-6442 |
| chr14-121937276-INS-125 | chr14-121937276-INS-125 |
| chr14-121941892-INS-209 | chr14-121941892-INS-210 |
| chr14-121949304-INS-114 | chr14-121949310-INS-120 |
| chr14-121956821-INS-200 | chr14-121956821-INS-200 |
| chr14-121966201-INS-74 | chr14-121966201-INS-76 |
| chr14-121969416-INS-55 | chr14-121969416-INS-55 |
| chr14-121974434-INS-126 | chr14-121974434-INS-126 |
| chr14-121978787-INS-68 | chr14-121978787-INS-68 |
| chr14-121981786-INS-942 | chr14-121981786-INS-942 |
| chr14-121983070-INS-173 | chr14-121983070-INS-174 |
| chr14-121989263-INS-108 | chr14-121989263-INS-108 |
| chr14-122008381-INS-241 | chr14-122008381-INS-236 |
| chr14-122022396-INS-54 | chr14-122022396-INS-54 |
| chr14-122030318-INS-73 | chr14-122030318-INS-77 |
| chr14-122043805-INS-217 | chr14-122043805-INS-217 |
| chr14-122052226-INS-185 | chr14-122052226-INS-187 |
| chr14-122065206-INS-76 | chr14-122065206-INS-76 |
| chr14-122075250-INS-5402 | chr14-122075251-INS-5402 |
| chr14-122076549-INS-204 | chr14-122076549-INS-204 |
| chr14-122082145-INS-602 | chr14-122082145-INS-602 |
| chr14-122087134-INS-507 | chr14-122087134-INS-507 |
| chr14-122094744-INS-61 | chr14-122094744-INS-61 |
| chr14-122153690-INS-66 | chr14-122153690-INS-66 |
| chr14-122163057-INS-81 | chr14-122163027-INS-131 |
| chr14-122171773-INS-57 | chr14-122171773-INS-57 |
| chr14-122176942-INS-196 | chr14-122176942-INS-203 |
| chr14-122181414-INS-100 | chr14-122181414-INS-91 |
| chr14-122212275-INS-91 | chr14-122212275-INS-91 |
| chr14-122217261-INS-127 | chr14-122217243-INS-128 |
| chr14-122224643-INS-117 | chr14-122224643-INS-117 |
| chr14-122228498-INS-5435 | chr14-122228499-INS-5435 |
| chr14-122234849-INS-60 | chr14-122234845-INS-57 |
| chr14-122259068-INS-110 | chr14-122259068-INS-110 |
| chr14-122263107-INS-475 | chr14-122263107-INS-475 |
| chr14-122274220-INS-195 | chr14-122274220-INS-199 |
| chr14-122276478-INS-64 | chr14-122276478-INS-64 |
| chr14-122280547-INS-724 | chr14-122280547-INS-724 |
| chr14-122298094-INS-62 | chr14-122298094-INS-62 |
| chr14-122302214-INS-348 | chr14-122302217-INS-352 |
| chr14-122303518-INS-152 | chr14-122303518-INS-152 |
| chr14-122317716-INS-242 | chr14-122317716-INS-242 |
| chr14-122336467-INS-56 | chr14-122336473-INS-55 |
| chr14-122337993-INS-56 | chr14-122337993-INS-52 |
| chr14-122358533-INS-154 | chr14-122358527-INS-151 |
| chr14-122397929-INS-78 | chr14-122397929-INS-78 |
| chr14-122415811-INS-182 | chr14-122415811-INS-167 |
| chr14-122421217-INS-76 | chr14-122421217-INS-76 |
| chr14-122423879-INS-132 | chr14-122423923-INS-86 |
| chr14-122430873-INS-158 | chr14-122430873-INS-153 |
| chr14-122459479-INS-213 | chr14-122459479-INS-213 |
| chr14-122473030-INS-104 | chr14-122473030-INS-116 |
| chr14-122474981-INS-199 | chr14-122474981-INS-202 |
| chr14-122475918-INS-54 | chr14-122475919-INS-54 |
| chr14-122479392-INS-218 | chr14-122479393-INS-219 |
| chr14-122511764-INS-67 | chr14-122511764-INS-67 |
| chr14-122524830-INS-204 | chr14-122524830-INS-205 |
| chr14-122531600-INS-199 | chr14-122531600-INS-199 |
| chr14-122540513-INS-80 | chr14-122540513-INS-80 |
| chr14-122545427-INS-7166 | chr14-122545428-INS-7166 |
| chr14-122555336-INS-216 | chr14-122555336-INS-219 |
| chr14-122648110-INS-52 | chr14-122648074-INS-80 |
| chr14-122671963-INS-8989 | chr14-122671964-INS-8989 |
| chr14-122685398-INS-72 | chr14-122685398-INS-68 |
| chr14-122767932-INS-51 | chr14-122767932-INS-52 |
| chr14-122776486-INS-74 | chr14-122776486-INS-58 |
| chr14-122790605-INS-5582 | chr14-122790606-INS-5582 |
| chr14-122807558-INS-209 | chr14-122807558-INS-209 |
| chr14-122817095-INS-118 | chr14-122817095-INS-119 |
| chr14-122818189-INS-742 | chr14-122818190-INS-784 |
| chr14-122867843-INS-90 | chr14-122867843-INS-96 |
| chr14-122890466-INS-76 | chr14-122890466-INS-76 |
| chr14-122936031-INS-60 | chr14-122936027-INS-92 |
| chr14-122956012-INS-52 | chr14-122955988-INS-52 |
| chr14-123021162-INS-62 | chr14-123021162-INS-72 |

| | |
|---|---|
| chr14-123028291-INS-57 | chr14-123028291-INS-57 |
| chr14-123029547-INS-364 | chr14-123029547-INS-364 |
| chr14-123038506-INS-214 | chr14-123038506-INS-214 |
| chr14-123050072-INS-201 | chr14-123050072-INS-201 |
| chr14-123133667-INS-109 | chr14-123133667-INS-109 |
| chr14-123168671-INS-189 | chr14-123168671-INS-168 |
| chr14-123179744-INS-569 | chr14-123179744-INS-567 |
| chr14-123205144-INS-52 | chr14-123205142-INS-80 |
| chr14-123233441-INS-251 | chr14-123233441-INS-251 |
| chr14-123241670-INS-6726 | chr14-123241671-INS-6726 |
| chr14-123247697-INS-91 | chr14-123247697-INS-91 |
| chr14-123252113-INS-482 | chr14-123252096-INS-520 |
| chr14-123286671-INS-191 | chr14-123286663-INS-199 |
| chr14-123298777-INS-279 | chr14-123298777-INS-276 |
| chr14-123323352-INS-644 | chr14-123323352-INS-645 |
| chr14-123325996-INS-55 | chr14-123325996-INS-86 |
| chr14-123331770-INS-57 | chr14-123331770-INS-63 |
| chr14-123333700-INS-1210 | chr14-123333701-INS-1210 |
| chr14-123352790-INS-2387 | chr14-123352791-INS-2384 |
| chr14-123378472-INS-7207 | chr14-123378473-INS-7210 |
| chr14-123382315-INS-6840 | chr14-123382316-INS-6841 |
| chr14-123391881-INS-260 | chr14-123391876-INS-266 |
| chr14-123399711-INS-7046 | chr14-123399712-INS-7046 |
| chr14-123463356-INS-347 | chr14-123463356-INS-347 |
| chr14-123464665-INS-469 | chr14-123464665-INS-469 |
| chr14-123487478-INS-585 | chr14-123487478-INS-585 |
| chr14-123540534-INS-62 | chr14-123540526-INS-59 |
| chr14-123592198-INS-6248 | chr14-123592199-INS-6246 |
| chr14-123596351-INS-326 | chr14-123596351-INS-321 |
| chr14-123600773-INS-243 | chr14-123600773-INS-244 |
| chr14-123609557-INS-773 | chr14-123609557-INS-775 |
| chr14-123609701-INS-52 | chr14-123609701-INS-52 |
| chr14-123644061-INS-177 | chr14-123644061-INS-188 |
| chr14-123775796-INS-147 | chr14-123775796-INS-147 |
| chr14-123800611-INS-54 | chr14-123800611-INS-60 |
| chr14-123896150-INS-203 | chr14-123896150-INS-203 |
| chr14-123910914-INS-4557 | chr14-123910915-INS-4557 |
| chr14-123936694-INS-2987 | chr14-123936695-INS-2987 |
| chr14-123948951-INS-676 | chr14-123948951-INS-677 |
| chr14-123983536-INS-1862 | chr14-123983537-INS-1862 |
| chr14-123985532-INS-2245 | chr14-123985519-INS-2246 |
| chr14-124021818-INS-80 | chr14-124021821-INS-74 |
| chr14-124023342-INS-225 | chr14-124023342-INS-225 |
| chr14-124152504-INS-700 | chr14-124152504-INS-700 |
| chr14-124169963-INS-952 | chr14-124169963-INS-952 |
| chr14-124317083-INS-54 | chr14-124317084-INS-50 |
| chr14-124318256-INS-192 | chr14-124318256-INS-192 |
| chr14-124378682-INS-73 | chr14-124378682-INS-73 |
| chr14-124411380-INS-801 | chr14-124411380-INS-795 |
| chr14-124462505-INS-396 | chr14-124462505-INS-396 |
| chr14-124520839-INS-275 | chr14-124520839-INS-280 |
| chr14-124540074-INS-912 | chr14-124540074-INS-913 |
| chr14-124625971-INS-63 | chr14-124625968-INS-59 |
| chr14-124631254-INS-668 | chr14-124631254-INS-668 |
| chr14-124643124-INS-7166 | chr14-124643125-INS-7166 |
| chr14-124693884-INS-2315 | chr14-124693885-INS-2316 |
| chr14-124760952-INS-121 | chr14-124760948-INS-87 |
| chr14-124763637-INS-1282 | chr14-124763638-INS-1283 |
| chr14-124764879-INS-60 | chr14-124764879-INS-50 |
| chr14-124790782-INS-6887 | chr14-124790783-INS-6888 |
| chr14-124814563-INS-496 | chr14-124814563-INS-496 |
| chr14-124815809-INS-57 | chr14-124815809-INS-57 |
| chr14-124817530-INS-209 | chr14-124817530-INS-210 |
| chr14-124828440-INS-62 | chr14-124828440-INS-58 |
| chr14-124834071-INS-1710 | chr14-124834072-INS-1712 |
| chr14-124836661-INS-162 | chr14-124836661-INS-162 |
| chr14-124841024-INS-408 | chr14-124841024-INS-406 |
| chr14-124864530-INS-1268 | chr14-124864531-INS-1274 |
| chr14-124878232-INS-1034 | chr14-124878233-INS-1036 |
| chr14-124908078-INS-80 | chr14-124908082-INS-112 |
| chr14-124918615-INS-207 | chr14-124918615-INS-215 |
| chr14-124940987-INS-216 | chr14-124940987-INS-216 |
| chr14-124949706-INS-598 | chr14-124949706-INS-597 |
| chr14-124954385-INS-1264 | chr14-124954386-INS-1264 |
| chr14-124974552-INS-201 | chr14-124974552-INS-201 |
| chr14-124982649-INS-4716 | chr14-124982650-INS-4716 |
| chr14-124987306-INS-96 | chr14-124987320-INS-126 |

| | |
|---|---|
| chr15-3093940-INS-116 | chr15-3093927-INS-105 |
| chr15-3186351-INS-214 | chr15-3186351-INS-214 |
| chr15-3204249-INS-52 | chr15-3204249-INS-54 |
| chr15-3210820-INS-55 | chr15-3210820-INS-55 |
| chr15-3216298-INS-8321 | chr15-3216299-INS-8321 |
| chr15-3230870-INS-6639 | chr15-3230871-INS-6639 |
| chr15-3237251-INS-6436 | chr15-3237252-INS-6442 |
| chr15-3251021-INS-892 | chr15-3251021-INS-893 |
| chr15-3282799-INS-509 | chr15-3282799-INS-508 |
| chr15-3318480-INS-396 | chr15-3318480-INS-400 |
| chr15-3334230-INS-64 | chr15-3334230-INS-56 |
| chr15-3378731-INS-173 | chr15-3378731-INS-173 |
| chr15-3423271-INS-216 | chr15-3423271-INS-216 |
| chr15-3425003-INS-61 | chr15-3425003-INS-61 |
| chr15-3472414-INS-494 | chr15-3472414-INS-474 |
| chr15-3523704-INS-100 | chr15-3523704-INS-84 |
| chr15-3538367-INS-93 | chr15-3538379-INS-63 |
| chr15-3545631-INS-499 | chr15-3545631-INS-499 |
| chr15-3570003-INS-67 | chr15-3570042-INS-64 |
| chr15-3573598-INS-76 | chr15-3573609-INS-100 |
| chr15-3622759-INS-56 | chr15-3622759-INS-58 |
| chr15-3626389-INS-64 | chr15-3626389-INS-64 |
| chr15-3640032-INS-881 | chr15-3640032-INS-881 |
| chr15-3654522-INS-501 | chr15-3654522-INS-501 |
| chr15-3709990-INS-6215 | chr15-3709991-INS-6215 |
| chr15-3715639-INS-2397 | chr15-3715640-INS-2397 |
| chr15-3751888-INS-58 | chr15-3751894-INS-99 |
| chr15-3764841-INS-643 | chr15-3764841-INS-643 |
| chr15-3770029-INS-567 | chr15-3770029-INS-580 |
| chr15-3770724-INS-211 | chr15-3770724-INS-211 |
| chr15-3798444-INS-93 | chr15-3798444-INS-55 |
| chr15-3830644-INS-54 | chr15-3830644-INS-52 |
| chr15-3856274-INS-75 | chr15-3856274-INS-71 |
| chr15-3903590-INS-56 | chr15-3903590-INS-56 |
| chr15-3930130-INS-208 | chr15-3930130-INS-191 |
| chr15-3957805-INS-84 | chr15-3957805-INS-84 |
| chr15-3960597-INS-137 | chr15-3960597-INS-144 |
| chr15-3971033-INS-357 | chr15-3971033-INS-357 |
| chr15-4036256-INS-58 | chr15-4036256-INS-58 |
| chr15-4048505-INS-609 | chr15-4048506-INS-609 |
| chr15-4111225-INS-6545 | chr15-4111226-INS-6545 |
| chr15-4125735-INS-60 | chr15-4125759-INS-56 |
| chr15-4132680-INS-212 | chr15-4132680-INS-212 |
| chr15-4133261-INS-766 | chr15-4133261-INS-771 |
| chr15-4185363-INS-56 | chr15-4185363-INS-53 |
| chr15-4198683-INS-6473 | chr15-4198684-INS-6473 |
| chr15-4205301-INS-398 | chr15-4205301-INS-399 |
| chr15-4216383-INS-92 | chr15-4216419-INS-65 |
| chr15-4217934-INS-2894 | chr15-4217935-INS-2894 |
| chr15-4231567-INS-56 | chr15-4231567-INS-58 |
| chr15-4253403-INS-85 | chr15-4253403-INS-70 |
| chr15-4287780-INS-4386 | chr15-4287781-INS-4386 |
| chr15-4292525-INS-5972 | chr15-4292526-INS-5972 |
| chr15-4301143-INS-65 | chr15-4301143-INS-65 |
| chr15-4346536-INS-76 | chr15-4346536-INS-78 |
| chr15-4370934-INS-106 | chr15-4370934-INS-59 |
| chr15-4374842-INS-506 | chr15-4374842-INS-506 |
| chr15-4384142-INS-235 | chr15-4384148-INS-244 |
| chr15-4423257-INS-54 | chr15-4423257-INS-58 |
| chr15-4464463-INS-57 | chr15-4464464-INS-52 |
| chr15-4476626-INS-381 | chr15-4476626-INS-380 |
| chr15-4478279-INS-90 | chr15-4478279-INS-90 |
| chr15-4537012-INS-134 | chr15-4537048-INS-116 |
| chr15-4556798-INS-3020 | chr15-4556799-INS-3023 |
| chr15-4579659-INS-1336 | chr15-4579660-INS-1329 |
| chr15-4600146-INS-3542 | chr15-4600147-INS-3542 |
| chr15-4635103-INS-93 | chr15-4635103-INS-84 |
| chr15-4637572-INS-64 | chr15-4637572-INS-64 |
| chr15-4690288-INS-51 | chr15-4690288-INS-52 |
| chr15-4697793-INS-54 | chr15-4697793-INS-54 |
| chr15-4701628-INS-636 | chr15-4701628-INS-636 |
| chr15-4711780-INS-202 | chr15-4711780-INS-202 |
| chr15-4734468-INS-88 | chr15-4734468-INS-68 |
| chr15-4736246-INS-87 | chr15-4736246-INS-87 |
| chr15-4765760-INS-52 | chr15-4765760-INS-50 |
| chr15-4798588-INS-143 | chr15-4798574-INS-164 |
| chr15-4808169-INS-833 | chr15-4808169-INS-834 |

| | |
|---|---|
| chr15-4823650-INS-189 | chr15-4823650-INS-189 |
| chr15-4831763-INS-1080 | chr15-4831764-INS-1079 |
| chr15-4837379-INS-477 | chr15-4837379-INS-477 |
| chr15-4846807-INS-434 | chr15-4846807-INS-434 |
| chr15-4865452-INS-6658 | chr15-4865453-INS-6658 |
| chr15-4871488-INS-52 | chr15-4871488-INS-52 |
| chr15-4879434-INS-405 | chr15-4879434-INS-405 |
| chr15-4908170-INS-382 | chr15-4908170-INS-382 |
| chr15-4935313-INS-56 | chr15-4935313-INS-58 |
| chr15-4958539-INS-1829 | chr15-4958544-INS-1828 |
| chr15-4961660-INS-318 | chr15-4961660-INS-319 |
| chr15-4963456-INS-63 | chr15-4963456-INS-63 |
| chr15-4989450-INS-1550 | chr15-4989451-INS-1536 |
| chr15-5024677-INS-92 | chr15-5024685-INS-92 |
| chr15-5043811-INS-7058 | chr15-5043812-INS-7058 |
| chr15-5057511-INS-7271 | chr15-5057512-INS-7271 |
| chr15-5102157-INS-107 | chr15-5102157-INS-143 |
| chr15-5107714-INS-213 | chr15-5107714-INS-214 |
| chr15-5111954-INS-529 | chr15-5111954-INS-525 |
| chr15-5112686-INS-157 | chr15-5112686-INS-157 |
| chr15-5151793-INS-312 | chr15-5151813-INS-332 |
| chr15-5154427-INS-226 | chr15-5154427-INS-227 |
| chr15-5162033-INS-60 | chr15-5162033-INS-60 |
| chr15-5183023-INS-206 | chr15-5183024-INS-206 |
| chr15-5186757-INS-72 | chr15-5186757-INS-64 |
| chr15-5192146-INS-59 | chr15-5192146-INS-59 |
| chr15-5204433-INS-68 | chr15-5204433-INS-68 |
| chr15-5212872-INS-58 | chr15-5212872-INS-52 |
| chr15-5218579-INS-206 | chr15-5218579-INS-205 |
| chr15-5283387-INS-103 | chr15-5283387-INS-103 |
| chr15-5285057-INS-2146 | chr15-5285058-INS-2146 |
| chr15-5285827-INS-1548 | chr15-5285828-INS-1540 |
| chr15-5324582-INS-670 | chr15-5324582-INS-674 |
| chr15-5335014-INS-98 | chr15-5335014-INS-98 |
| chr15-5344158-INS-64 | chr15-5344162-INS-50 |
| chr15-5346098-INS-88 | chr15-5346098-INS-112 |
| chr15-5372209-INS-52 | chr15-5372209-INS-52 |
| chr15-5372971-INS-134 | chr15-5372971-INS-108 |
| chr15-5378787-INS-99 | chr15-5378787-INS-99 |
| chr15-5381638-INS-2214 | chr15-5381663-INS-2210 |
| chr15-5388898-INS-109 | chr15-5388922-INS-90 |
| chr15-5419210-INS-196 | chr15-5419210-INS-191 |
| chr15-5438055-INS-303 | chr15-5438055-INS-303 |
| chr15-5520887-INS-378 | chr15-5520887-INS-378 |
| chr15-5532448-INS-154 | chr15-5532448-INS-130 |
| chr15-5534159-INS-454 | chr15-5534159-INS-454 |
| chr15-5542598-INS-54 | chr15-5542598-INS-54 |
| chr15-5651807-INS-426 | chr15-5651807-INS-426 |
| chr15-5676064-INS-121 | chr15-5676064-INS-125 |
| chr15-5680954-INS-658 | chr15-5680954-INS-657 |
| chr15-5691568-INS-76 | chr15-5691568-INS-100 |
| chr15-5705396-INS-2466 | chr15-5705397-INS-2468 |
| chr15-5743970-INS-2236 | chr15-5743971-INS-2236 |
| chr15-5747292-INS-1584 | chr15-5747293-INS-1584 |
| chr15-5776983-INS-134 | chr15-5776983-INS-134 |
| chr15-5782448-INS-59 | chr15-5782448-INS-70 |
| chr15-5828586-INS-1534 | chr15-5828587-INS-1533 |
| chr15-5852250-INS-498 | chr15-5852250-INS-498 |
| chr15-5854593-INS-436 | chr15-5854594-INS-427 |
| chr15-5858974-INS-204 | chr15-5858974-INS-204 |
| chr15-5875895-INS-2253 | chr15-5875896-INS-2253 |
| chr15-5879669-INS-493 | chr15-5879669-INS-493 |
| chr15-5891815-INS-51 | chr15-5891853-INS-55 |
| chr15-5919130-INS-63 | chr15-5919150-INS-71 |
| chr15-5922760-INS-70 | chr15-5922761-INS-58 |
| chr15-5931252-INS-481 | chr15-5931252-INS-481 |
| chr15-5935681-INS-6403 | chr15-5935682-INS-6403 |
| chr15-5936912-INS-54 | chr15-5936912-INS-52 |
| chr15-5968117-INS-95 | chr15-5968132-INS-55 |
| chr15-5989624-INS-52 | chr15-5989624-INS-52 |
| chr15-5996797-INS-134 | chr15-5996792-INS-164 |
| chr15-5998906-INS-103 | chr15-5998906-INS-103 |
| chr15-6030265-INS-6840 | chr15-6030266-INS-6840 |
| chr15-6092859-INS-115 | chr15-6092816-INS-114 |
| chr15-6099378-INS-96 | chr15-6099378-INS-96 |
| chr15-6101933-INS-64 | chr15-6101933-INS-64 |
| chr15-6122394-INS-6633 | chr15-6122395-INS-6635 |

| | |
|---|---|
| chr15-6148780-INS-52 | chr15-6148780-INS-52 |
| chr15-6187176-INS-110 | chr15-6187176-INS-110 |
| chr15-6192800-INS-1178 | chr15-6192801-INS-1178 |
| chr15-6197197-INS-151 | chr15-6197197-INS-151 |
| chr15-6205972-INS-62 | chr15-6205972-INS-105 |
| chr15-6212066-INS-345 | chr15-6212066-INS-341 |
| chr15-6238425-INS-256 | chr15-6238422-INS-236 |
| chr15-6241876-INS-67 | chr15-6241873-INS-68 |
| chr15-6245947-INS-86 | chr15-6245952-INS-70 |
| chr15-6278783-INS-95 | chr15-6278783-INS-95 |
| chr15-6320936-INS-52 | chr15-6320937-INS-78 |
| chr15-6323342-INS-62 | chr15-6323333-INS-58 |
| chr15-6335774-INS-130 | chr15-6335804-INS-110 |
| chr15-6347551-INS-218 | chr15-6347551-INS-183 |
| chr15-6348645-INS-77 | chr15-6348645-INS-77 |
| chr15-6356997-INS-3236 | chr15-6356998-INS-3237 |
| chr15-6366140-INS-80 | chr15-6366124-INS-64 |
| chr15-6368345-INS-102 | chr15-6368345-INS-103 |
| chr15-6381634-INS-105 | chr15-6381677-INS-105 |
| chr15-6439390-INS-178 | chr15-6439390-INS-170 |
| chr15-6449750-INS-276 | chr15-6449713-INS-284 |
| chr15-6513651-INS-437 | chr15-6513651-INS-437 |
| chr15-6552083-INS-105 | chr15-6552080-INS-104 |
| chr15-6602402-INS-174 | chr15-6602402-INS-174 |
| chr15-6631550-INS-349 | chr15-6631550-INS-336 |
| chr15-6632884-INS-68 | chr15-6632903-INS-66 |
| chr15-6728357-INS-242 | chr15-6728357-INS-242 |
| chr15-6752273-INS-55 | chr15-6752290-INS-60 |
| chr15-6767080-INS-400 | chr15-6767080-INS-400 |
| chr15-6771437-INS-143 | chr15-6771413-INS-152 |
| chr15-6800267-INS-319 | chr15-6800267-INS-319 |
| chr15-6830465-INS-400 | chr15-6830465-INS-402 |
| chr15-6890117-INS-120 | chr15-6890117-INS-120 |
| chr15-6890564-INS-202 | chr15-6890564-INS-202 |
| chr15-6906857-INS-245 | chr15-6906857-INS-253 |
| chr15-6910082-INS-3880 | chr15-6910083-INS-3896 |
| chr15-6912286-INS-584 | chr15-6912286-INS-546 |
| chr15-6917166-INS-75 | chr15-6917166-INS-75 |
| chr15-6929974-INS-440 | chr15-6929974-INS-440 |
| chr15-7026971-INS-208 | chr15-7026971-INS-207 |
| chr15-7056374-INS-1895 | chr15-7056375-INS-1895 |
| chr15-7060618-INS-52 | chr15-7060618-INS-50 |
| chr15-7113275-INS-6413 | chr15-7113276-INS-6413 |
| chr15-7121182-INS-1061 | chr15-7121183-INS-1014 |
| chr15-7127629-INS-56 | chr15-7127629-INS-52 |
| chr15-7135394-INS-218 | chr15-7135394-INS-218 |
| chr15-7151359-INS-56 | chr15-7151359-INS-58 |
| chr15-7188246-INS-84 | chr15-7188246-INS-86 |
| chr15-7192719-INS-205 | chr15-7192719-INS-206 |
| chr15-7248260-INS-54 | chr15-7248252-INS-74 |
| chr15-7291675-INS-74 | chr15-7291675-INS-74 |
| chr15-7324882-INS-135 | chr15-7324883-INS-134 |
| chr15-7343102-INS-56 | chr15-7343102-INS-56 |
| chr15-7359262-INS-140 | chr15-7359259-INS-113 |
| chr15-7375286-INS-51 | chr15-7375286-INS-51 |
| chr15-7382029-INS-58 | chr15-7382029-INS-58 |
| chr15-7406054-INS-86 | chr15-7406057-INS-66 |
| chr15-7410989-INS-192 | chr15-7410989-INS-192 |
| chr15-7421937-INS-52 | chr15-7421937-INS-56 |
| chr15-7441162-INS-99 | chr15-7441159-INS-134 |
| chr15-7449077-INS-62 | chr15-7449083-INS-59 |
| chr15-7470733-INS-396 | chr15-7470733-INS-396 |
| chr15-7515164-INS-124 | chr15-7515166-INS-110 |
| chr15-7539703-INS-80 | chr15-7539701-INS-66 |
| chr15-7555688-INS-74 | chr15-7555688-INS-65 |
| chr15-7583040-INS-65 | chr15-7583044-INS-102 |
| chr15-7587049-INS-70 | chr15-7587075-INS-77 |
| chr15-7588520-INS-2698 | chr15-7588521-INS-2698 |
| chr15-7594633-INS-52 | chr15-7594633-INS-58 |
| chr15-7595633-INS-415 | chr15-7595633-INS-415 |
| chr15-7598045-INS-88 | chr15-7598045-INS-93 |
| chr15-7636168-INS-435 | chr15-7636154-INS-436 |
| chr15-7658459-INS-1888 | chr15-7658460-INS-1888 |
| chr15-7684083-INS-201 | chr15-7684083-INS-201 |
| chr15-7691709-INS-108 | chr15-7691668-INS-135 |
| chr15-7709537-INS-604 | chr15-7709537-INS-604 |
| chr15-7754938-INS-60 | chr15-7754935-INS-62 |

| | |
|---|---|
| chr15-7773561-INS-61 | chr15-7773561-INS-61 |
| chr15-7797451-INS-518 | chr15-7797451-INS-522 |
| chr15-7822020-INS-1892 | chr15-7822021-INS-1892 |
| chr15-7835232-INS-113 | chr15-7835233-INS-142 |
| chr15-7903801-INS-52 | chr15-7903829-INS-74 |
| chr15-7905571-INS-651 | chr15-7905571-INS-645 |
| chr15-7966138-INS-766 | chr15-7966138-INS-766 |
| chr15-7972347-INS-254 | chr15-7972347-INS-254 |
| chr15-7987834-INS-6435 | chr15-7987835-INS-6435 |
| chr15-8000204-INS-230 | chr15-8000204-INS-231 |
| chr15-8029772-INS-4383 | chr15-8029773-INS-4383 |
| chr15-8051259-INS-348 | chr15-8051259-INS-348 |
| chr15-8053181-INS-60 | chr15-8053186-INS-63 |
| chr15-8058220-INS-765 | chr15-8058220-INS-774 |
| chr15-8065058-INS-57 | chr15-8065058-INS-57 |
| chr15-8069449-INS-214 | chr15-8069449-INS-215 |
| chr15-8072984-INS-53 | chr15-8072984-INS-53 |
| chr15-8075829-INS-61 | chr15-8075829-INS-82 |
| chr15-8118570-INS-203 | chr15-8118570-INS-203 |
| chr15-8127635-INS-310 | chr15-8127640-INS-315 |
| chr15-8145206-INS-224 | chr15-8145206-INS-223 |
| chr15-8151733-INS-59 | chr15-8151733-INS-59 |
| chr15-8158176-INS-92 | chr15-8158176-INS-109 |
| chr15-8166818-INS-126 | chr15-8166818-INS-126 |
| chr15-8168491-INS-229 | chr15-8168491-INS-229 |
| chr15-8175293-INS-283 | chr15-8175293-INS-283 |
| chr15-8181379-INS-99 | chr15-8181379-INS-99 |
| chr15-8185559-INS-64 | chr15-8185546-INS-63 |
| chr15-8199954-INS-170 | chr15-8199954-INS-170 |
| chr15-8209914-INS-123 | chr15-8209914-INS-92 |
| chr15-8219440-INS-182 | chr15-8219440-INS-173 |
| chr15-8240867-INS-84 | chr15-8240903-INS-82 |
| chr15-8249218-INS-210 | chr15-8249218-INS-205 |
| chr15-8254536-INS-56 | chr15-8254536-INS-56 |
| chr15-8255508-INS-205 | chr15-8255508-INS-205 |
| chr15-8260152-INS-100 | chr15-8260152-INS-100 |
| chr15-8289110-INS-214 | chr15-8289110-INS-214 |
| chr15-8302350-INS-747 | chr15-8302350-INS-762 |
| chr15-8309335-INS-112 | chr15-8309363-INS-84 |
| chr15-8312837-INS-62 | chr15-8312837-INS-62 |
| chr15-8313611-INS-6463 | chr15-8313612-INS-6463 |
| chr15-8318208-INS-327 | chr15-8318211-INS-328 |
| chr15-8324928-INS-83 | chr15-8324928-INS-83 |
| chr15-8365454-INS-801 | chr15-8365454-INS-804 |
| chr15-8442245-INS-380 | chr15-8442245-INS-400 |
| chr15-8461830-INS-92 | chr15-8461819-INS-116 |
| chr15-8486057-INS-124 | chr15-8486057-INS-99 |
| chr15-8487489-INS-212 | chr15-8487489-INS-211 |
| chr15-8492703-INS-68 | chr15-8492702-INS-112 |
| chr15-8501092-INS-201 | chr15-8501092-INS-201 |
| chr15-8503153-INS-170 | chr15-8503153-INS-170 |
| chr15-8523953-INS-87 | chr15-8523953-INS-129 |
| chr15-8572515-INS-377 | chr15-8572516-INS-382 |
| chr15-8573358-INS-55 | chr15-8573359-INS-50 |
| chr15-8574834-INS-120 | chr15-8574804-INS-160 |
| chr15-8597570-INS-88 | chr15-8597555-INS-74 |
| chr15-8617795-INS-133 | chr15-8617795-INS-133 |
| chr15-8618874-INS-53 | chr15-8618874-INS-52 |
| chr15-8623205-INS-65 | chr15-8623207-INS-66 |
| chr15-8644187-INS-143 | chr15-8644187-INS-127 |
| chr15-8677097-INS-62 | chr15-8677098-INS-58 |
| chr15-8681408-INS-247 | chr15-8681408-INS-211 |
| chr15-8723261-INS-78 | chr15-8723261-INS-57 |
| chr15-8735167-INS-201 | chr15-8735167-INS-201 |
| chr15-8746242-INS-98 | chr15-8746272-INS-77 |
| chr15-8749815-INS-58 | chr15-8749815-INS-58 |
| chr15-8751096-INS-362 | chr15-8751098-INS-389 |
| chr15-8762692-INS-186 | chr15-8762701-INS-168 |
| chr15-8790725-INS-183 | chr15-8790725-INS-183 |
| chr15-8822318-INS-110 | chr15-8822318-INS-110 |
| chr15-8854607-INS-80 | chr15-8854607-INS-80 |
| chr15-8861704-INS-198 | chr15-8861705-INS-199 |
| chr15-8894541-INS-84 | chr15-8894539-INS-92 |
| chr15-8917111-INS-102 | chr15-8917111-INS-102 |
| chr15-8918383-INS-358 | chr15-8918383-INS-356 |
| chr15-8939860-INS-65 | chr15-8939860-INS-65 |
| chr15-8994633-INS-144 | chr15-8994633-INS-168 |

| | |
|---|---|
| chr15-9009473-INS-400 | chr15-9009473-INS-400 |
| chr15-9014616-INS-54 | chr15-9014616-INS-54 |
| chr15-9023982-INS-95 | chr15-9023969-INS-83 |
| chr15-9028296-INS-482 | chr15-9028294-INS-486 |
| chr15-9031950-INS-330 | chr15-9031950-INS-330 |
| chr15-9093034-INS-1137 | chr15-9093035-INS-1143 |
| chr15-9103152-INS-78 | chr15-9103156-INS-78 |
| chr15-9105072-INS-504 | chr15-9105072-INS-504 |
| chr15-9115886-INS-261 | chr15-9115886-INS-261 |
| chr15-9133677-INS-115 | chr15-9133677-INS-115 |
| chr15-9154468-INS-223 | chr15-9154468-INS-224 |
| chr15-9164564-INS-60 | chr15-9164564-INS-59 |
| chr15-9195382-INS-270 | chr15-9195382-INS-270 |
| chr15-9203049-INS-222 | chr15-9203049-INS-222 |
| chr15-9237124-INS-61 | chr15-9237124-INS-72 |
| chr15-9245096-INS-2492 | chr15-9245097-INS-2492 |
| chr15-9246191-INS-156 | chr15-9246191-INS-165 |
| chr15-9273347-INS-365 | chr15-9273347-INS-362 |
| chr15-9369807-INS-56 | chr15-9369807-INS-62 |
| chr15-9484134-INS-612 | chr15-9484134-INS-611 |
| chr15-9505736-INS-2570 | chr15-9505737-INS-2569 |
| chr15-9556689-INS-52 | chr15-9556689-INS-58 |
| chr15-9563634-INS-408 | chr15-9563634-INS-408 |
| chr15-9575812-INS-295 | chr15-9575812-INS-294 |
| chr15-9595882-INS-52 | chr15-9595870-INS-68 |
| chr15-9642450-INS-66 | chr15-9642450-INS-60 |
| chr15-9657538-INS-228 | chr15-9657534-INS-227 |
| chr15-9701893-INS-6490 | chr15-9701894-INS-6490 |
| chr15-9735663-INS-154 | chr15-9735663-INS-182 |
| chr15-9735663-INS-782 | chr15-9735663-INS-784 |
| chr15-9739450-INS-223 | chr15-9739450-INS-213 |
| chr15-9743217-INS-909 | chr15-9743217-INS-909 |
| chr15-9756502-INS-242 | chr15-9756502-INS-216 |
| chr15-9758581-INS-7085 | chr15-9758576-INS-7089 |
| chr15-9764921-INS-113 | chr15-9764921-INS-113 |
| chr15-9783748-INS-62 | chr15-9783712-INS-72 |
| chr15-9793527-INS-313 | chr15-9793533-INS-319 |
| chr15-9802860-INS-64 | chr15-9802860-INS-62 |
| chr15-9829247-INS-51 | chr15-9829247-INS-72 |
| chr15-9835284-INS-1180 | chr15-9835285-INS-1181 |
| chr15-9841662-INS-163 | chr15-9841660-INS-164 |
| chr15-9844489-INS-165 | chr15-9844489-INS-165 |
| chr15-9848179-INS-150 | chr15-9848174-INS-184 |
| chr15-9863276-INS-61 | chr15-9863276-INS-61 |
| chr15-9871238-INS-58 | chr15-9871258-INS-52 |
| chr15-9914376-INS-193 | chr15-9914376-INS-193 |
| chr15-9924163-INS-199 | chr15-9924163-INS-212 |
| chr15-9940415-INS-56 | chr15-9940415-INS-52 |
| chr15-9944529-INS-67 | chr15-9944529-INS-65 |
| chr15-9945431-INS-531 | chr15-9945431-INS-531 |
| chr15-9961904-INS-82 | chr15-9961904-INS-82 |
| chr15-10076583-INS-57 | chr15-10076583-INS-57 |
| chr15-10103727-INS-52 | chr15-10103727-INS-56 |
| chr15-10121821-INS-55 | chr15-10121821-INS-55 |
| chr15-10122360-INS-53 | chr15-10122360-INS-53 |
| chr15-10141341-INS-211 | chr15-10141341-INS-211 |
| chr15-10150706-INS-248 | chr15-10150706-INS-258 |
| chr15-10232798-INS-57 | chr15-10232798-INS-56 |
| chr15-10236570-INS-500 | chr15-10236577-INS-497 |
| chr15-10258433-INS-58 | chr15-10258433-INS-57 |
| chr15-10265293-INS-199 | chr15-10265293-INS-200 |
| chr15-10267546-INS-83 | chr15-10267550-INS-82 |
| chr15-10274702-INS-61 | chr15-10274702-INS-61 |
| chr15-10290627-INS-5013 | chr15-10290628-INS-5006 |
| chr15-10291883-INS-84 | chr15-10291883-INS-84 |
| chr15-10296274-INS-199 | chr15-10296274-INS-199 |
| chr15-10302213-INS-217 | chr15-10302213-INS-217 |
| chr15-10302314-INS-7168 | chr15-10302315-INS-7168 |
| chr15-10315043-INS-59 | chr15-10315043-INS-81 |
| chr15-10322959-INS-6397 | chr15-10322960-INS-6397 |
| chr15-10359214-INS-88 | chr15-10359260-INS-64 |
| chr15-10421856-INS-2827 | chr15-10421857-INS-2827 |
| chr15-10430356-INS-5408 | chr15-10430357-INS-5408 |
| chr15-10433148-INS-90 | chr15-10433148-INS-110 |
| chr15-10439174-INS-55 | chr15-10439174-INS-55 |
| chr15-10443908-INS-838 | chr15-10443908-INS-838 |
| chr15-10452000-INS-222 | chr15-10452000-INS-222 |

| | |
|---|---|
| chr15-10458760-INS-270 | chr15-10458760-INS-280 |
| chr15-10471325-INS-179 | chr15-10471325-INS-178 |
| chr15-10494939-INS-179 | chr15-10494981-INS-146 |
| chr15-10496909-INS-514 | chr15-10496910-INS-514 |
| chr15-10541307-INS-206 | chr15-10541307-INS-218 |
| chr15-10557890-INS-516 | chr15-10557890-INS-517 |
| chr15-10584683-INS-204 | chr15-10584683-INS-205 |
| chr15-10588945-INS-84 | chr15-10588945-INS-84 |
| chr15-10593688-INS-206 | chr15-10593688-INS-206 |
| chr15-10601752-INS-86 | chr15-10601756-INS-80 |
| chr15-10627212-INS-143 | chr15-10627212-INS-143 |
| chr15-10651824-INS-408 | chr15-10651824-INS-408 |
| chr15-10665124-INS-400 | chr15-10665125-INS-400 |
| chr15-10676104-INS-72 | chr15-10676115-INS-71 |
| chr15-10678126-INS-203 | chr15-10678126-INS-203 |
| chr15-10718725-INS-946 | chr15-10718705-INS-950 |
| chr15-10726174-INS-118 | chr15-10726174-INS-118 |
| chr15-10731000-INS-56 | chr15-10731000-INS-62 |
| chr15-10743594-INS-72 | chr15-10743594-INS-62 |
| chr15-10752935-INS-273 | chr15-10752935-INS-256 |
| chr15-10765585-INS-92 | chr15-10765575-INS-67 |
| chr15-10796008-INS-180 | chr15-10795992-INS-182 |
| chr15-10798353-INS-150 | chr15-10798357-INS-150 |
| chr15-10801645-INS-76 | chr15-10801631-INS-75 |
| chr15-10809249-INS-2930 | chr15-10809250-INS-2931 |
| chr15-10812723-INS-428 | chr15-10812723-INS-428 |
| chr15-10824771-INS-118 | chr15-10824771-INS-106 |
| chr15-10825791-INS-60 | chr15-10825791-INS-60 |
| chr15-10843772-INS-62 | chr15-10843772-INS-58 |
| chr15-10869124-INS-2102 | chr15-10869125-INS-2080 |
| chr15-10870880-INS-57 | chr15-10870880-INS-56 |
| chr15-10876210-INS-54 | chr15-10876210-INS-57 |
| chr15-10890168-INS-90 | chr15-10890168-INS-98 |
| chr15-10898185-INS-78 | chr15-10898186-INS-106 |
| chr15-10903270-INS-51 | chr15-10903270-INS-51 |
| chr15-10905736-INS-102 | chr15-10905736-INS-102 |
| chr15-10922260-INS-56 | chr15-10922260-INS-56 |
| chr15-10924245-INS-95 | chr15-10924245-INS-86 |
| chr15-10974346-INS-1315 | chr15-10974347-INS-1320 |
| chr15-10988416-INS-67 | chr15-10988416-INS-67 |
| chr15-11018836-INS-52 | chr15-11018836-INS-52 |
| chr15-11045819-INS-309 | chr15-11045819-INS-311 |
| chr15-11058132-INS-502 | chr15-11058132-INS-502 |
| chr15-11062150-INS-184 | chr15-11062150-INS-184 |
| chr15-11070128-INS-52 | chr15-11070148-INS-51 |
| chr15-11086359-INS-413 | chr15-11086359-INS-406 |
| chr15-11095263-INS-178 | chr15-11095263-INS-156 |
| chr15-11154395-INS-70 | chr15-11154420-INS-50 |
| chr15-11180449-INS-202 | chr15-11180449-INS-204 |
| chr15-11213127-INS-58 | chr15-11213127-INS-58 |
| chr15-11218886-INS-152 | chr15-11218886-INS-128 |
| chr15-11261769-INS-183 | chr15-11261769-INS-181 |
| chr15-11270888-INS-3150 | chr15-11270885-INS-3158 |
| chr15-11295709-INS-129 | chr15-11295709-INS-129 |
| chr15-11314942-INS-70 | chr15-11314942-INS-70 |
| chr15-11338096-INS-55 | chr15-11338096-INS-50 |
| chr15-11342060-INS-66 | chr15-11342060-INS-71 |
| chr15-11357905-INS-944 | chr15-11357905-INS-944 |
| chr15-11362162-INS-5225 | chr15-11362163-INS-5225 |
| chr15-11376069-INS-491 | chr15-11376071-INS-492 |
| chr15-11377048-INS-403 | chr15-11377048-INS-403 |
| chr15-11406736-INS-51 | chr15-11406736-INS-63 |
| chr15-11418444-INS-103 | chr15-11418444-INS-111 |
| chr15-11439200-INS-428 | chr15-11439200-INS-428 |
| chr15-11442565-INS-58 | chr15-11442562-INS-88 |
| chr15-11450242-INS-6237 | chr15-11450243-INS-6237 |
| chr15-11480189-INS-98 | chr15-11480190-INS-98 |
| chr15-11551983-INS-202 | chr15-11551983-INS-201 |
| chr15-11559229-INS-68 | chr15-11559229-INS-68 |
| chr15-11575511-INS-2996 | chr15-11575512-INS-2974 |
| chr15-11591519-INS-519 | chr15-11591519-INS-519 |
| chr15-11607057-INS-130 | chr15-11607062-INS-131 |
| chr15-11611510-INS-221 | chr15-11611511-INS-221 |
| chr15-11633717-INS-119 | chr15-11633717-INS-119 |
| chr15-11656884-INS-568 | chr15-11656884-INS-567 |
| chr15-11663565-INS-104 | chr15-11663565-INS-71 |
| chr15-11676688-INS-208 | chr15-11676688-INS-208 |

| | |
|---|---|
| chr15-11680305-INS-4400 | chr15-11680306-INS-4401 |
| chr15-11689391-INS-58 | chr15-11689391-INS-56 |
| chr15-11706113-INS-8648 | chr15-11706106-INS-8648 |
| chr15-11718550-INS-90 | chr15-11718550-INS-90 |
| chr15-11722820-INS-203 | chr15-11722820-INS-204 |
| chr15-11739827-INS-84 | chr15-11739864-INS-62 |
| chr15-11766730-INS-374 | chr15-11766730-INS-374 |
| chr15-11819749-INS-1865 | chr15-11819750-INS-1865 |
| chr15-11832031-INS-201 | chr15-11832031-INS-202 |
| chr15-11867613-INS-7271 | chr15-11867614-INS-7271 |
| chr15-11884902-INS-1066 | chr15-11884903-INS-1066 |
| chr15-11888055-INS-94 | chr15-11888044-INS-116 |
| chr15-11891480-INS-81 | chr15-11891474-INS-96 |
| chr15-11892855-INS-91 | chr15-11892855-INS-93 |
| chr15-11900874-INS-1574 | chr15-11900875-INS-1538 |
| chr15-11947098-INS-1208 | chr15-11947099-INS-1208 |
| chr15-11947838-INS-209 | chr15-11947838-INS-209 |
| chr15-11979079-INS-68 | chr15-11979079-INS-68 |
| chr15-12015011-INS-51 | chr15-12015008-INS-69 |
| chr15-12021195-INS-64 | chr15-12021195-INS-64 |
| chr15-12029602-INS-60 | chr15-12029602-INS-60 |
| chr15-12032266-INS-297 | chr15-12032266-INS-297 |
| chr15-12034455-INS-85 | chr15-12034455-INS-85 |
| chr15-12049791-INS-55 | chr15-12049772-INS-55 |
| chr15-12075199-INS-116 | chr15-12075173-INS-114 |
| chr15-12084001-INS-6487 | chr15-12084002-INS-6487 |
| chr15-12085520-INS-341 | chr15-12085523-INS-343 |
| chr15-12100676-INS-96 | chr15-12100660-INS-72 |
| chr15-12124706-INS-203 | chr15-12124706-INS-203 |
| chr15-12126727-INS-69 | chr15-12126727-INS-69 |
| chr15-12127115-INS-210 | chr15-12127128-INS-219 |
| chr15-12169544-INS-208 | chr15-12169544-INS-208 |
| chr15-12191973-INS-102 | chr15-12191973-INS-84 |
| chr15-12199557-INS-205 | chr15-12199557-INS-206 |
| chr15-12208394-INS-146 | chr15-12208394-INS-146 |
| chr15-12250098-INS-57 | chr15-12250098-INS-57 |
| chr15-12256325-INS-206 | chr15-12256325-INS-206 |
| chr15-12274683-INS-208 | chr15-12274684-INS-183 |
| chr15-12275604-INS-65 | chr15-12275604-INS-53 |
| chr15-12296969-INS-113 | chr15-12296969-INS-107 |
| chr15-12299829-INS-498 | chr15-12299829-INS-498 |
| chr15-12312851-INS-177 | chr15-12312852-INS-171 |
| chr15-12319915-INS-6448 | chr15-12319916-INS-6448 |
| chr15-12342843-INS-96 | chr15-12342839-INS-54 |
| chr15-12355324-INS-148 | chr15-12355324-INS-143 |
| chr15-12386565-INS-6403 | chr15-12386566-INS-6402 |
| chr15-12386566-INS-498 | chr15-12386566-INS-498 |
| chr15-12396307-INS-87 | chr15-12396312-INS-92 |
| chr15-12402280-INS-54 | chr15-12402280-INS-68 |
| chr15-12414607-INS-213 | chr15-12414607-INS-212 |
| chr15-12459530-INS-79 | chr15-12459531-INS-53 |
| chr15-12486215-INS-56 | chr15-12486215-INS-58 |
| chr15-12488930-INS-191 | chr15-12488930-INS-177 |
| chr15-12525990-INS-66 | chr15-12525990-INS-66 |
| chr15-12552585-INS-164 | chr15-12552585-INS-164 |
| chr15-12562954-INS-68 | chr15-12562931-INS-76 |
| chr15-12570041-INS-118 | chr15-12570041-INS-123 |
| chr15-12571536-INS-87 | chr15-12571536-INS-87 |
| chr15-12576436-INS-106 | chr15-12576436-INS-148 |
| chr15-12580321-INS-107 | chr15-12580321-INS-111 |
| chr15-12585397-INS-96 | chr15-12585397-INS-138 |
| chr15-12646528-INS-113 | chr15-12646528-INS-113 |
| chr15-12678484-INS-140 | chr15-12678484-INS-140 |
| chr15-12704904-INS-164 | chr15-12704904-INS-165 |
| chr15-12725765-INS-77 | chr15-12725779-INS-69 |
| chr15-12745614-INS-323 | chr15-12745614-INS-311 |
| chr15-12746283-INS-204 | chr15-12746283-INS-204 |
| chr15-12748941-INS-74 | chr15-12748937-INS-62 |
| chr15-12760093-INS-64 | chr15-12760093-INS-54 |
| chr15-12766644-INS-62 | chr15-12766644-INS-62 |
| chr15-12786946-INS-1132 | chr15-12786947-INS-1132 |
| chr15-12787315-INS-198 | chr15-12787323-INS-199 |
| chr15-12789030-INS-70 | chr15-12789058-INS-58 |
| chr15-12790281-INS-637 | chr15-12790281-INS-637 |
| chr15-12801085-INS-62 | chr15-12801085-INS-66 |
| chr15-12822387-INS-254 | chr15-12822387-INS-254 |
| chr15-12831626-INS-6460 | chr15-12831627-INS-6452 |

| | |
|---|---|
| chr15-12842356-INS-83 | chr15-12842356-INS-83 |
| chr15-12879482-INS-81 | chr15-12879482-INS-81 |
| chr15-12884382-INS-60 | chr15-12884382-INS-60 |
| chr15-12893901-INS-55 | chr15-12893901-INS-55 |
| chr15-12901737-INS-54 | chr15-12901737-INS-62 |
| chr15-12913624-INS-88 | chr15-12913624-INS-88 |
| chr15-12930485-INS-62 | chr15-12930485-INS-62 |
| chr15-12953755-INS-52 | chr15-12953755-INS-54 |
| chr15-12969398-INS-298 | chr15-12969394-INS-302 |
| chr15-12992873-INS-6286 | chr15-12992874-INS-6287 |
| chr15-13000138-INS-56 | chr15-13000138-INS-56 |
| chr15-13017593-INS-83 | chr15-13017586-INS-65 |
| chr15-13059478-INS-170 | chr15-13059478-INS-181 |
| chr15-13077167-INS-61 | chr15-13077167-INS-61 |
| chr15-13085695-INS-108 | chr15-13085686-INS-113 |
| chr15-13096107-INS-8828 | chr15-13096108-INS-8830 |
| chr15-13100572-INS-66 | chr15-13100572-INS-66 |
| chr15-13116140-INS-129 | chr15-13116140-INS-129 |
| chr15-13121720-INS-60 | chr15-13121720-INS-60 |
| chr15-13131835-INS-63 | chr15-13131835-INS-63 |
| chr15-13150947-INS-75 | chr15-13150947-INS-75 |
| chr15-13157376-INS-169 | chr15-13157376-INS-161 |
| chr15-13185534-INS-87 | chr15-13185534-INS-87 |
| chr15-13192175-INS-101 | chr15-13192175-INS-100 |
| chr15-13198272-INS-58 | chr15-13198272-INS-58 |
| chr15-13207860-INS-84 | chr15-13207860-INS-79 |
| chr15-13218265-INS-236 | chr15-13218265-INS-254 |
| chr15-13233456-INS-62 | chr15-13233466-INS-65 |
| chr15-13238009-INS-88 | chr15-13238033-INS-82 |
| chr15-13241737-INS-54 | chr15-13241737-INS-54 |
| chr15-13249293-INS-92 | chr15-13249293-INS-92 |
| chr15-13290766-INS-52 | chr15-13290766-INS-52 |
| chr15-13292791-INS-96 | chr15-13292791-INS-100 |
| chr15-13310429-INS-78 | chr15-13310380-INS-78 |
| chr15-13313847-INS-340 | chr15-13313847-INS-340 |
| chr15-13316205-INS-53 | chr15-13316206-INS-53 |
| chr15-13343237-INS-60 | chr15-13343237-INS-60 |
| chr15-13370844-INS-981 | chr15-13370851-INS-998 |
| chr15-13382948-INS-67 | chr15-13382949-INS-59 |
| chr15-13421196-INS-306 | chr15-13421203-INS-318 |
| chr15-13427922-INS-5157 | chr15-13427923-INS-5157 |
| chr15-13433067-INS-80 | chr15-13433077-INS-98 |
| chr15-13464529-INS-107 | chr15-13464527-INS-76 |
| chr15-13471708-INS-348 | chr15-13471708-INS-348 |
| chr15-13497778-INS-6462 | chr15-13497779-INS-6462 |
| chr15-13502976-INS-196 | chr15-13502976-INS-149 |
| chr15-13536576-INS-51 | chr15-13536576-INS-51 |
| chr15-13561927-INS-52 | chr15-13561927-INS-55 |
| chr15-13705047-INS-6211 | chr15-13705048-INS-6212 |
| chr15-13722559-INS-137 | chr15-13722559-INS-141 |
| chr15-13738060-INS-212 | chr15-13738060-INS-209 |
| chr15-13752428-INS-157 | chr15-13752442-INS-156 |
| chr15-13757430-INS-77 | chr15-13757430-INS-71 |
| chr15-13764123-INS-66 | chr15-13764123-INS-88 |
| chr15-13769539-INS-6297 | chr15-13769540-INS-6298 |
| chr15-13774320-INS-487 | chr15-13774320-INS-487 |
| chr15-13779263-INS-390 | chr15-13779263-INS-390 |
| chr15-13795766-INS-52 | chr15-13795766-INS-52 |
| chr15-13810878-INS-1973 | chr15-13810879-INS-1973 |
| chr15-13819606-INS-79 | chr15-13819608-INS-61 |
| chr15-13874613-INS-1111 | chr15-13874611-INS-1109 |
| chr15-13887105-INS-261 | chr15-13887105-INS-265 |
| chr15-13900253-INS-230 | chr15-13900253-INS-190 |
| chr15-13934665-INS-70 | chr15-13934665-INS-104 |
| chr15-13941875-INS-983 | chr15-13941875-INS-979 |
| chr15-13960809-INS-1831 | chr15-13960810-INS-1831 |
| chr15-13967885-INS-100 | chr15-13967885-INS-102 |
| chr15-13969806-INS-196 | chr15-13969806-INS-200 |
| chr15-13973320-INS-6228 | chr15-13973321-INS-6229 |
| chr15-14007572-INS-6428 | chr15-14007573-INS-6421 |
| chr15-14011883-INS-61 | chr15-14011883-INS-61 |
| chr15-14050144-INS-8016 | chr15-14050145-INS-8016 |
| chr15-14061920-INS-956 | chr15-14061920-INS-958 |
| chr15-14080926-INS-683 | chr15-14080926-INS-683 |
| chr15-14086676-INS-1013 | chr15-14086677-INS-1013 |
| chr15-14105019-INS-2587 | chr15-14105020-INS-2587 |
| chr15-14119403-INS-72 | chr15-14119403-INS-72 |

| | |
|---|---|
| chr15-14131507-INS-3033 | chr15-14131508-INS-3017 |
| chr15-14140621-INS-69 | chr15-14140621-INS-69 |
| chr15-14150793-INS-365 | chr15-14150793-INS-376 |
| chr15-14166494-INS-78 | chr15-14166494-INS-78 |
| chr15-14176665-INS-72 | chr15-14176647-INS-78 |
| chr15-14183457-INS-195 | chr15-14183457-INS-193 |
| chr15-14199289-INS-68 | chr15-14199292-INS-52 |
| chr15-14212974-INS-1497 | chr15-14212975-INS-1494 |
| chr15-14216395-INS-547 | chr15-14216395-INS-547 |
| chr15-14218575-INS-681 | chr15-14218575-INS-681 |
| chr15-14238728-INS-80 | chr15-14238728-INS-80 |
| chr15-14243326-INS-6459 | chr15-14243327-INS-6460 |
| chr15-14246096-INS-80 | chr15-14246086-INS-64 |
| chr15-14246931-INS-183 | chr15-14246931-INS-183 |
| chr15-14247658-INS-4813 | chr15-14247659-INS-4813 |
| chr15-14260878-INS-198 | chr15-14260878-INS-197 |
| chr15-14267566-INS-2864 | chr15-14267567-INS-2864 |
| chr15-14273252-INS-469 | chr15-14273252-INS-469 |
| chr15-14308093-INS-765 | chr15-14308093-INS-765 |
| chr15-14311093-INS-101 | chr15-14311093-INS-101 |
| chr15-14326604-INS-333 | chr15-14326607-INS-318 |
| chr15-14329346-INS-65 | chr15-14329367-INS-74 |
| chr15-14349734-INS-139 | chr15-14349734-INS-139 |
| chr15-14355475-INS-2499 | chr15-14355476-INS-2494 |
| chr15-14383567-INS-377 | chr15-14383567-INS-377 |
| chr15-14399720-INS-126 | chr15-14399720-INS-130 |
| chr15-14407171-INS-594 | chr15-14407171-INS-593 |
| chr15-14418473-INS-86 | chr15-14418473-INS-86 |
| chr15-14422300-INS-66 | chr15-14422300-INS-68 |
| chr15-14424663-INS-120 | chr15-14424668-INS-164 |
| chr15-14453938-INS-57 | chr15-14453938-INS-57 |
| chr15-14454401-INS-1888 | chr15-14454402-INS-1888 |
| chr15-14457855-INS-83 | chr15-14457855-INS-83 |
| chr15-14462567-INS-175 | chr15-14462567-INS-175 |
| chr15-14466827-INS-75 | chr15-14466827-INS-75 |
| chr15-14468946-INS-53 | chr15-14468946-INS-53 |
| chr15-14469880-INS-191 | chr15-14469880-INS-191 |
| chr15-14510594-INS-117 | chr15-14510594-INS-117 |
| chr15-14512085-INS-221 | chr15-14512085-INS-221 |
| chr15-14521804-INS-150 | chr15-14521804-INS-148 |
| chr15-14531804-INS-199 | chr15-14531804-INS-199 |
| chr15-14612371-INS-235 | chr15-14612371-INS-235 |
| chr15-14617221-INS-177 | chr15-14617221-INS-177 |
| chr15-14621958-INS-542 | chr15-14621958-INS-539 |
| chr15-14628518-INS-60 | chr15-14628518-INS-70 |
| chr15-14652996-INS-203 | chr15-14652991-INS-194 |
| chr15-14691446-INS-460 | chr15-14691446-INS-460 |
| chr15-14692402-INS-6056 | chr15-14692403-INS-6055 |
| chr15-14701428-INS-386 | chr15-14701428-INS-386 |
| chr15-14720772-INS-72 | chr15-14720773-INS-74 |
| chr15-14749576-INS-114 | chr15-14749566-INS-74 |
| chr15-14770350-INS-439 | chr15-14770350-INS-439 |
| chr15-14780208-INS-678 | chr15-14780208-INS-678 |
| chr15-14783682-INS-69 | chr15-14783682-INS-69 |
| chr15-14809170-INS-106 | chr15-14809170-INS-129 |
| chr15-14825746-INS-53 | chr15-14825746-INS-53 |
| chr15-14832427-INS-193 | chr15-14832427-INS-193 |
| chr15-14861705-INS-270 | chr15-14861705-INS-265 |
| chr15-14868226-INS-463 | chr15-14868226-INS-463 |
| chr15-14885345-INS-56 | chr15-14885345-INS-56 |
| chr15-14913956-INS-1629 | chr15-14913957-INS-1621 |
| chr15-14922507-INS-85 | chr15-14922507-INS-85 |
| chr15-14928051-INS-53 | chr15-14928051-INS-53 |
| chr15-14946919-INS-598 | chr15-14946915-INS-602 |
| chr15-14975179-INS-70 | chr15-14975179-INS-60 |
| chr15-14997447-INS-53 | chr15-14997447-INS-80 |
| chr15-15023563-INS-182 | chr15-15023563-INS-182 |
| chr15-15076863-INS-509 | chr15-15076863-INS-510 |
| chr15-15093316-INS-102 | chr15-15093316-INS-102 |
| chr15-15167487-INS-81 | chr15-15167487-INS-81 |
| chr15-15182328-INS-2586 | chr15-15182329-INS-2586 |
| chr15-15184122-INS-3287 | chr15-15184123-INS-3287 |
| chr15-15220426-INS-56 | chr15-15220426-INS-54 |
| chr15-15239727-INS-114 | chr15-15239727-INS-114 |
| chr15-15242476-INS-1055 | chr15-15242477-INS-1054 |
| chr15-15304756-INS-66 | chr15-15304756-INS-60 |
| chr15-15351235-INS-68 | chr15-15351235-INS-68 |

| | |
|---|---|
| chr15-15364535-INS-72 | chr15-15364535-INS-96 |
| chr15-15381216-INS-312 | chr15-15381216-INS-308 |
| chr15-15382992-INS-109 | chr15-15382992-INS-113 |
| chr15-15391105-INS-165 | chr15-15391105-INS-165 |
| chr15-15402832-INS-58 | chr15-15402832-INS-64 |
| chr15-15423508-INS-59 | chr15-15423508-INS-59 |
| chr15-15433148-INS-124 | chr15-15433148-INS-118 |
| chr15-15607941-INS-60 | chr15-15607941-INS-60 |
| chr15-15617792-INS-71 | chr15-15617759-INS-58 |
| chr15-15625463-INS-6930 | chr15-15625464-INS-6930 |
| chr15-15632793-INS-404 | chr15-15632787-INS-437 |
| chr15-15651212-INS-498 | chr15-15651212-INS-498 |
| chr15-15659241-INS-64 | chr15-15659241-INS-64 |
| chr15-15667798-INS-6551 | chr15-15667799-INS-6551 |
| chr15-15696506-INS-119 | chr15-15696506-INS-119 |
| chr15-15705583-INS-581 | chr15-15705583-INS-581 |
| chr15-15718910-INS-619 | chr15-15718910-INS-620 |
| chr15-15775393-INS-54 | chr15-15775393-INS-54 |
| chr15-15776371-INS-51 | chr15-15776371-INS-51 |
| chr15-15777166-INS-568 | chr15-15777166-INS-568 |
| chr15-15777805-INS-68 | chr15-15777805-INS-68 |
| chr15-15792714-INS-222 | chr15-15792714-INS-182 |
| chr15-15828417-INS-52 | chr15-15828425-INS-50 |
| chr15-15835432-INS-139 | chr15-15835432-INS-139 |
| chr15-15886961-INS-1591 | chr15-15886962-INS-1590 |
| chr15-15968577-INS-238 | chr15-15968577-INS-238 |
| chr15-15983750-INS-428 | chr15-15983750-INS-428 |
| chr15-16015732-INS-72 | chr15-16015754-INS-82 |
| chr15-16030535-INS-637 | chr15-16030535-INS-637 |
| chr15-16039605-INS-52 | chr15-16039605-INS-54 |
| chr15-16047958-INS-6957 | chr15-16047959-INS-6957 |
| chr15-16114156-INS-2102 | chr15-16114157-INS-2102 |
| chr15-16230751-INS-339 | chr15-16230751-INS-339 |
| chr15-16244230-INS-929 | chr15-16244230-INS-929 |
| chr15-16268334-INS-112 | chr15-16268326-INS-150 |
| chr15-16285535-INS-56 | chr15-16285535-INS-56 |
| chr15-16291940-INS-6669 | chr15-16291941-INS-6669 |
| chr15-16302609-INS-64 | chr15-16302609-INS-62 |
| chr15-16321575-INS-149 | chr15-16321575-INS-100 |
| chr15-16322531-INS-82 | chr15-16322534-INS-126 |
| chr15-16344247-INS-56 | chr15-16344247-INS-58 |
| chr15-16349229-INS-938 | chr15-16349229-INS-938 |
| chr15-16407489-INS-499 | chr15-16407489-INS-499 |
| chr15-16421470-INS-342 | chr15-16421470-INS-343 |
| chr15-16429295-INS-434 | chr15-16429295-INS-434 |
| chr15-16436075-INS-67 | chr15-16436075-INS-67 |
| chr15-16448453-INS-128 | chr15-16448453-INS-128 |
| chr15-16475052-INS-2796 | chr15-16475053-INS-2796 |
| chr15-16507435-INS-176 | chr15-16507436-INS-136 |
| chr15-16519780-INS-745 | chr15-16519780-INS-745 |
| chr15-16584575-INS-102 | chr15-16584575-INS-102 |
| chr15-16596126-INS-1876 | chr15-16596127-INS-1876 |
| chr15-16633806-INS-383 | chr15-16633780-INS-389 |
| chr15-16639480-INS-720 | chr15-16639480-INS-720 |
| chr15-16644516-INS-3518 | chr15-16644517-INS-3518 |
| chr15-16646664-INS-104 | chr15-16646666-INS-72 |
| chr15-16648893-INS-349 | chr15-16648893-INS-331 |
| chr15-16651187-INS-76 | chr15-16651187-INS-99 |
| chr15-16655622-INS-65 | chr15-16655592-INS-66 |
| chr15-16668608-INS-66 | chr15-16668608-INS-68 |
| chr15-16674516-INS-85 | chr15-16674516-INS-87 |
| chr15-16682931-INS-82 | chr15-16682931-INS-82 |
| chr15-16700677-INS-52 | chr15-16700677-INS-73 |
| chr15-16705918-INS-6164 | chr15-16705919-INS-6164 |
| chr15-16723195-INS-151 | chr15-16723195-INS-151 |
| chr15-16739530-INS-95 | chr15-16739528-INS-89 |
| chr15-16776577-INS-4376 | chr15-16776578-INS-4377 |
| chr15-16805505-INS-89 | chr15-16805505-INS-100 |
| chr15-16909632-INS-91 | chr15-16909632-INS-79 |
| chr15-16912053-INS-53 | chr15-16912053-INS-53 |
| chr15-16966805-INS-60 | chr15-16966806-INS-60 |
| chr15-16972528-INS-170 | chr15-16972528-INS-170 |
| chr15-16975478-INS-62 | chr15-16975476-INS-64 |
| chr15-16980286-INS-2250 | chr15-16980303-INS-2227 |
| chr15-17011009-INS-2972 | chr15-17011010-INS-2972 |
| chr15-17014858-INS-2721 | chr15-17014859-INS-2721 |
| chr15-17046418-INS-1685 | chr15-17046419-INS-1684 |

| | |
|---|---|
| chr15-17112774-INS-2688 | chr15-17112775-INS-2688 |
| chr15-17122779-INS-52 | chr15-17122780-INS-52 |
| chr15-17149522-INS-117 | chr15-17149532-INS-103 |
| chr15-17211446-INS-70 | chr15-17211446-INS-70 |
| chr15-17218038-INS-212 | chr15-17218038-INS-212 |
| chr15-17228914-INS-505 | chr15-17228914-INS-505 |
| chr15-17231553-INS-96 | chr15-17231553-INS-112 |
| chr15-17232691-INS-51 | chr15-17232691-INS-51 |
| chr15-17243224-INS-163 | chr15-17243224-INS-163 |
| chr15-17246462-INS-430 | chr15-17246462-INS-430 |
| chr15-17253023-INS-4393 | chr15-17253024-INS-4394 |
| chr15-17254894-INS-51 | chr15-17254894-INS-60 |
| chr15-17287878-INS-373 | chr15-17287878-INS-373 |
| chr15-17339319-INS-133 | chr15-17339319-INS-133 |
| chr15-17350804-INS-873 | chr15-17350804-INS-874 |
| chr15-17351618-INS-495 | chr15-17351618-INS-495 |
| chr15-17361573-INS-135 | chr15-17361559-INS-143 |
| chr15-17408944-INS-132 | chr15-17408940-INS-140 |
| chr15-17437932-INS-63 | chr15-17437932-INS-63 |
| chr15-17442583-INS-975 | chr15-17442586-INS-978 |
| chr15-17489593-INS-414 | chr15-17489593-INS-414 |
| chr15-17491489-INS-133 | chr15-17491489-INS-133 |
| chr15-17492269-INS-773 | chr15-17492269-INS-773 |
| chr15-17495620-INS-6673 | chr15-17495621-INS-6673 |
| chr15-17498692-INS-424 | chr15-17498692-INS-424 |
| chr15-17503748-INS-3567 | chr15-17503749-INS-3567 |
| chr15-17509302-INS-60 | chr15-17509302-INS-62 |
| chr15-17515313-INS-66 | chr15-17515313-INS-66 |
| chr15-17527020-INS-365 | chr15-17527020-INS-366 |
| chr15-17528100-INS-4061 | chr15-17528101-INS-4061 |
| chr15-17541754-INS-61 | chr15-17541754-INS-52 |
| chr15-17546783-INS-8903 | chr15-17546784-INS-8903 |
| chr15-17548338-INS-88 | chr15-17548361-INS-53 |
| chr15-17573184-INS-75 | chr15-17573184-INS-75 |
| chr15-17576322-INS-2707 | chr15-17576323-INS-2707 |
| chr15-17597672-INS-759 | chr15-17597672-INS-759 |
| chr15-17631090-INS-415 | chr15-17631090-INS-414 |
| chr15-17641193-INS-383 | chr15-17641193-INS-383 |
| chr15-17651489-INS-1160 | chr15-17651490-INS-1161 |
| chr15-17660638-INS-88 | chr15-17660638-INS-89 |
| chr15-17662530-INS-134 | chr15-17662548-INS-102 |
| chr15-17666436-INS-86 | chr15-17666436-INS-86 |
| chr15-17680801-INS-692 | chr15-17680801-INS-692 |
| chr15-17695308-INS-99 | chr15-17695308-INS-100 |
| chr15-17697659-INS-175 | chr15-17697659-INS-175 |
| chr15-17702251-INS-80 | chr15-17702251-INS-80 |
| chr15-17704403-INS-58 | chr15-17704403-INS-58 |
| chr15-17709328-INS-636 | chr15-17709323-INS-629 |
| chr15-17716696-INS-144 | chr15-17716696-INS-144 |
| chr15-17720936-INS-156 | chr15-17720936-INS-147 |
| chr15-17723768-INS-1156 | chr15-17723772-INS-1154 |
| chr15-17724733-INS-452 | chr15-17724732-INS-451 |
| chr15-17758364-INS-97 | chr15-17758364-INS-104 |
| chr15-17795293-INS-134 | chr15-17795293-INS-134 |
| chr15-17813485-INS-55 | chr15-17813485-INS-55 |
| chr15-17826926-INS-51 | chr15-17826926-INS-51 |
| chr15-17844414-INS-65 | chr15-17844414-INS-69 |
| chr15-17859997-INS-54 | chr15-17860010-INS-61 |
| chr15-17877777-INS-758 | chr15-17877777-INS-757 |
| chr15-17879730-INS-470 | chr15-17879730-INS-470 |
| chr15-17883442-INS-434 | chr15-17883443-INS-402 |
| chr15-17911996-INS-174 | chr15-17911996-INS-176 |
| chr15-17915469-INS-162 | chr15-17915469-INS-166 |
| chr15-17927591-INS-71 | chr15-17927595-INS-55 |
| chr15-17936930-INS-171 | chr15-17936930-INS-172 |
| chr15-17953385-INS-118 | chr15-17953405-INS-100 |
| chr15-17965844-INS-52 | chr15-17965844-INS-52 |
| chr15-17978422-INS-348 | chr15-17978422-INS-348 |
| chr15-17997040-INS-64 | chr15-17997040-INS-68 |
| chr15-18005362-INS-1333 | chr15-18005363-INS-1333 |
| chr15-18007439-INS-59 | chr15-18007418-INS-75 |
| chr15-18018711-INS-60 | chr15-18018711-INS-64 |
| chr15-18039612-INS-186 | chr15-18039612-INS-186 |
| chr15-18052072-INS-84 | chr15-18052074-INS-82 |
| chr15-18095736-INS-405 | chr15-18095736-INS-405 |
| chr15-18103720-INS-134 | chr15-18103720-INS-134 |
| chr15-18119811-INS-199 | chr15-18119811-INS-199 |

| | |
|---|---|
| chr15-18132272-INS-136 | chr15-18132272-INS-140 |
| chr15-18138924-INS-74 | chr15-18138923-INS-72 |
| chr15-18139732-INS-6266 | chr15-18139733-INS-6266 |
| chr15-18142336-INS-120 | chr15-18142336-INS-120 |
| chr15-18148653-INS-330 | chr15-18148653-INS-330 |
| chr15-18184767-INS-6189 | chr15-18184768-INS-6189 |
| chr15-18188197-INS-2077 | chr15-18188198-INS-2079 |
| chr15-18197925-INS-62 | chr15-18197925-INS-62 |
| chr15-18204373-INS-503 | chr15-18204373-INS-503 |
| chr15-18279838-INS-100 | chr15-18279808-INS-104 |
| chr15-18289416-INS-52 | chr15-18289416-INS-52 |
| chr15-18291654-INS-3481 | chr15-18291655-INS-3481 |
| chr15-18302175-INS-209 | chr15-18302175-INS-209 |
| chr15-18325802-INS-53 | chr15-18325802-INS-83 |
| chr15-18364973-INS-744 | chr15-18364973-INS-750 |
| chr15-18425530-INS-228 | chr15-18425530-INS-192 |
| chr15-18450907-INS-52 | chr15-18450907-INS-56 |
| chr15-18504644-INS-769 | chr15-18504662-INS-769 |
| chr15-18584801-INS-65 | chr15-18584801-INS-65 |
| chr15-18633765-INS-133 | chr15-18633765-INS-134 |
| chr15-18644256-INS-207 | chr15-18644256-INS-207 |
| chr15-18757082-INS-6386 | chr15-18757083-INS-6386 |
| chr15-18757974-INS-72 | chr15-18757974-INS-75 |
| chr15-18821356-INS-52 | chr15-18821356-INS-52 |
| chr15-18838678-INS-84 | chr15-18838678-INS-84 |
| chr15-18908396-INS-112 | chr15-18908396-INS-113 |
| chr15-18922462-INS-60 | chr15-18922445-INS-75 |
| chr15-18927348-INS-264 | chr15-18927348-INS-264 |
| chr15-18936521-INS-499 | chr15-18936521-INS-499 |
| chr15-18937287-INS-55 | chr15-18937299-INS-78 |
| chr15-18944464-INS-76 | chr15-18944464-INS-76 |
| chr15-18954829-INS-52 | chr15-18954839-INS-54 |
| chr15-18977467-INS-84 | chr15-18977467-INS-82 |
| chr15-18982535-INS-142 | chr15-18982535-INS-142 |
| chr15-18982889-INS-3494 | chr15-18982890-INS-3492 |
| chr15-18987475-INS-279 | chr15-18987475-INS-279 |
| chr15-18999283-INS-646 | chr15-18999249-INS-663 |
| chr15-19012646-INS-434 | chr15-19012646-INS-432 |
| chr15-19089989-INS-375 | chr15-19089989-INS-375 |
| chr15-19091740-INS-364 | chr15-19091741-INS-364 |
| chr15-19147108-INS-75 | chr15-19147108-INS-75 |
| chr15-19172514-INS-72 | chr15-19172514-INS-72 |
| chr15-19200828-INS-84 | chr15-19200828-INS-54 |
| chr15-19246568-INS-1925 | chr15-19246569-INS-1924 |
| chr15-19264228-INS-4523 | chr15-19264229-INS-4530 |
| chr15-19265334-INS-728 | chr15-19265334-INS-726 |
| chr15-19276007-INS-355 | chr15-19276007-INS-355 |
| chr15-19298548-INS-54 | chr15-19298548-INS-60 |
| chr15-19322762-INS-6306 | chr15-19322763-INS-6315 |
| chr15-19332311-INS-88 | chr15-19332311-INS-88 |
| chr15-19337882-INS-125 | chr15-19337872-INS-135 |
| chr15-19353810-INS-52 | chr15-19353810-INS-50 |
| chr15-19355204-INS-62 | chr15-19355202-INS-64 |
| chr15-19356483-INS-221 | chr15-19356483-INS-221 |
| chr15-19367090-INS-585 | chr15-19367090-INS-585 |
| chr15-19375739-INS-75 | chr15-19375773-INS-66 |
| chr15-19378166-INS-219 | chr15-19378166-INS-219 |
| chr15-19382727-INS-1346 | chr15-19382728-INS-1346 |
| chr15-19413852-INS-201 | chr15-19413852-INS-201 |
| chr15-19420907-INS-229 | chr15-19420907-INS-234 |
| chr15-19433825-INS-476 | chr15-19433825-INS-477 |
| chr15-19457876-INS-1104 | chr15-19457877-INS-1100 |
| chr15-19460566-INS-459 | chr15-19460566-INS-459 |
| chr15-19465824-INS-195 | chr15-19465824-INS-196 |
| chr15-19490111-INS-71 | chr15-19490111-INS-75 |
| chr15-19491745-INS-56 | chr15-19491745-INS-60 |
| chr15-19520263-INS-555 | chr15-19520263-INS-555 |
| chr15-19545254-INS-4484 | chr15-19545255-INS-4484 |
| chr15-19571166-INS-3290 | chr15-19571167-INS-3290 |
| chr15-19576608-INS-2500 | chr15-19576609-INS-2513 |
| chr15-19596169-INS-942 | chr15-19596169-INS-945 |
| chr15-19605623-INS-1240 | chr15-19605622-INS-1216 |
| chr15-19660382-INS-55 | chr15-19660382-INS-52 |
| chr15-19698117-INS-140 | chr15-19698117-INS-146 |
| chr15-19700038-INS-656 | chr15-19700038-INS-655 |
| chr15-19745092-INS-3726 | chr15-19745093-INS-3726 |
| chr15-19746230-INS-148 | chr15-19746230-INS-148 |

| | |
|---|---|
| chr15-19750649-INS-5023 | chr15-19750650-INS-5023 |
| chr15-19762319-INS-604 | chr15-19762325-INS-606 |
| chr15-19768532-INS-78 | chr15-19768532-INS-78 |
| chr15-19770341-INS-515 | chr15-19770341-INS-515 |
| chr15-19777506-INS-225 | chr15-19777506-INS-226 |
| chr15-19777524-INS-1914 | chr15-19777525-INS-1915 |
| chr15-19786131-INS-56 | chr15-19786131-INS-56 |
| chr15-19802912-INS-150 | chr15-19802912-INS-152 |
| chr15-19820528-INS-436 | chr15-19820528-INS-436 |
| chr15-19837098-INS-1888 | chr15-19837099-INS-1888 |
| chr15-19856456-INS-112 | chr15-19856456-INS-120 |
| chr15-19874375-INS-390 | chr15-19874375-INS-390 |
| chr15-19900275-INS-6665 | chr15-19900276-INS-6665 |
| chr15-19920027-INS-198 | chr15-19920027-INS-198 |
| chr15-19935619-INS-451 | chr15-19935619-INS-498 |
| chr15-19946728-INS-399 | chr15-19946728-INS-399 |
| chr15-19962506-INS-281 | chr15-19962506-INS-281 |
| chr15-19971916-INS-75 | chr15-19971916-INS-81 |
| chr15-19994205-INS-355 | chr15-19994205-INS-355 |
| chr15-20011049-INS-174 | chr15-20011049-INS-174 |
| chr15-20031622-INS-60 | chr15-20031622-INS-60 |
| chr15-20063407-INS-6355 | chr15-20063408-INS-6358 |
| chr15-20072810-INS-62 | chr15-20072810-INS-58 |
| chr15-20083083-INS-53 | chr15-20083083-INS-54 |
| chr15-20102355-INS-6402 | chr15-20102356-INS-6402 |
| chr15-20113752-INS-1609 | chr15-20113753-INS-1616 |
| chr15-20120688-INS-1285 | chr15-20120689-INS-1286 |
| chr15-20130076-INS-837 | chr15-20130076-INS-837 |
| chr15-20181117-INS-52 | chr15-20181117-INS-52 |
| chr15-20235479-INS-6430 | chr15-20235480-INS-6430 |
| chr15-20257520-INS-6536 | chr15-20257521-INS-6537 |
| chr15-20258169-INS-165 | chr15-20258169-INS-165 |
| chr15-20261010-INS-570 | chr15-20261010-INS-570 |
| chr15-20270610-INS-1408 | chr15-20270611-INS-1408 |
| chr15-20296991-INS-6659 | chr15-20296992-INS-6663 |
| chr15-20304457-INS-532 | chr15-20304457-INS-532 |
| chr15-20311983-INS-566 | chr15-20311983-INS-566 |
| chr15-20340810-INS-170 | chr15-20340816-INS-171 |
| chr15-20349372-INS-208 | chr15-20349372-INS-208 |
| chr15-20379922-INS-6437 | chr15-20379923-INS-6437 |
| chr15-20397286-INS-608 | chr15-20397286-INS-608 |
| chr15-20397797-INS-882 | chr15-20397797-INS-882 |
| chr15-20400835-INS-573 | chr15-20400835-INS-573 |
| chr15-20441402-INS-2425 | chr15-20441403-INS-2425 |
| chr15-20447438-INS-54 | chr15-20447438-INS-52 |
| chr15-20476127-INS-1307 | chr15-20476128-INS-1307 |
| chr15-20487611-INS-2647 | chr15-20487612-INS-2647 |
| chr15-20499156-INS-208 | chr15-20499156-INS-208 |
| chr15-20521965-INS-3950 | chr15-20521966-INS-3950 |
| chr15-20538690-INS-75 | chr15-20538702-INS-59 |
| chr15-20553239-INS-414 | chr15-20553239-INS-414 |
| chr15-20553348-INS-6259 | chr15-20553350-INS-6254 |
| chr15-20562647-INS-484 | chr15-20562647-INS-484 |
| chr15-20586992-INS-479 | chr15-20586992-INS-479 |
| chr15-20631185-INS-570 | chr15-20631185-INS-570 |
| chr15-20640693-INS-276 | chr15-20640690-INS-280 |
| chr15-20649440-INS-176 | chr15-20649442-INS-174 |
| chr15-20670181-INS-172 | chr15-20670181-INS-192 |
| chr15-20681660-INS-280 | chr15-20681660-INS-280 |
| chr15-20683000-INS-7130 | chr15-20683001-INS-7130 |
| chr15-20685407-INS-534 | chr15-20685407-INS-534 |
| chr15-20686575-INS-224 | chr15-20686575-INS-224 |
| chr15-20689236-INS-382 | chr15-20689236-INS-382 |
| chr15-20695625-INS-63 | chr15-20695625-INS-66 |
| chr15-20715694-INS-84 | chr15-20715692-INS-83 |
| chr15-20725006-INS-1144 | chr15-20725007-INS-1145 |
| chr15-20740526-INS-211 | chr15-20740526-INS-211 |
| chr15-20747234-INS-56 | chr15-20747184-INS-94 |
| chr15-20758985-INS-515 | chr15-20758985-INS-515 |
| chr15-20768598-INS-74 | chr15-20768598-INS-56 |
| chr15-20783385-INS-877 | chr15-20783385-INS-877 |
| chr15-20787766-INS-184 | chr15-20787766-INS-184 |
| chr15-20801395-INS-52 | chr15-20801376-INS-72 |
| chr15-20810049-INS-52 | chr15-20810049-INS-50 |
| chr15-20815670-INS-4411 | chr15-20815671-INS-4411 |
| chr15-20819925-INS-70 | chr15-20819925-INS-70 |
| chr15-20840372-INS-85 | chr15-20840379-INS-120 |

| | |
|---|---|
| chr15-20848060-INS-57 | chr15-20848039-INS-89 |
| chr15-20930315-INS-338 | chr15-20930306-INS-345 |
| chr15-20934710-INS-571 | chr15-20934710-INS-571 |
| chr15-20951646-INS-56 | chr15-20951646-INS-62 |
| chr15-20959328-INS-51 | chr15-20959328-INS-51 |
| chr15-20964462-INS-66 | chr15-20964431-INS-88 |
| chr15-20967873-INS-4779 | chr15-20967874-INS-4779 |
| chr15-21027518-INS-51 | chr15-21027518-INS-53 |
| chr15-21096661-INS-1890 | chr15-21096662-INS-1890 |
| chr15-21098446-INS-573 | chr15-21098446-INS-573 |
| chr15-21100926-INS-80 | chr15-21100908-INS-97 |
| chr15-21165394-INS-62 | chr15-21165394-INS-60 |
| chr15-21188626-INS-168 | chr15-21188626-INS-192 |
| chr15-21226658-INS-5413 | chr15-21226659-INS-5413 |
| chr15-21277177-INS-199 | chr15-21277177-INS-199 |
| chr15-21279566-INS-246 | chr15-21279566-INS-247 |
| chr15-21381127-INS-1682 | chr15-21381128-INS-1684 |
| chr15-21388298-INS-6376 | chr15-21388299-INS-6376 |
| chr15-21411652-INS-6730 | chr15-21411653-INS-6730 |
| chr15-21444073-INS-208 | chr15-21444073-INS-208 |
| chr15-21460830-INS-218 | chr15-21460830-INS-218 |
| chr15-21499942-INS-52 | chr15-21499938-INS-52 |
| chr15-21517073-INS-63 | chr15-21517073-INS-63 |
| chr15-21583142-INS-60 | chr15-21583142-INS-52 |
| chr15-21606207-INS-286 | chr15-21606207-INS-286 |
| chr15-21624221-INS-318 | chr15-21624221-INS-318 |
| chr15-21638151-INS-80 | chr15-21638151-INS-82 |
| chr15-21642025-INS-150 | chr15-21642025-INS-150 |
| chr15-21644167-INS-1311 | chr15-21644168-INS-1311 |
| chr15-21681197-INS-94 | chr15-21681197-INS-70 |
| chr15-21709234-INS-91 | chr15-21709234-INS-91 |
| chr15-21735491-INS-610 | chr15-21735491-INS-610 |
| chr15-21739571-INS-7108 | chr15-21739572-INS-7109 |
| chr15-21754437-INS-1014 | chr15-21754438-INS-1014 |
| chr15-21768659-INS-128 | chr15-21768659-INS-128 |
| chr15-21792536-INS-51 | chr15-21792536-INS-51 |
| chr15-21844044-INS-4573 | chr15-21844045-INS-4573 |
| chr15-21915205-INS-6939 | chr15-21915206-INS-6939 |
| chr15-21934580-INS-188 | chr15-21934580-INS-188 |
| chr15-21935826-INS-2879 | chr15-21935827-INS-2879 |
| chr15-21945337-INS-5486 | chr15-21945338-INS-5486 |
| chr15-21995161-INS-510 | chr15-21995161-INS-511 |
| chr15-22054263-INS-102 | chr15-22054263-INS-104 |
| chr15-22103328-INS-902 | chr15-22103326-INS-897 |
| chr15-22107826-INS-51 | chr15-22107826-INS-72 |
| chr15-22116473-INS-6917 | chr15-22116474-INS-6916 |
| chr15-22117373-INS-155 | chr15-22117378-INS-190 |
| chr15-22141444-INS-7229 | chr15-22141445-INS-7229 |
| chr15-22182128-INS-4464 | chr15-22182129-INS-4464 |
| chr15-22195276-INS-210 | chr15-22195276-INS-211 |
| chr15-22199651-INS-58 | chr15-22199651-INS-58 |
| chr15-22214889-INS-60 | chr15-22214889-INS-72 |
| chr15-22221289-INS-4786 | chr15-22221290-INS-4786 |
| chr15-22228795-INS-4281 | chr15-22228796-INS-4276 |
| chr15-22247263-INS-120 | chr15-22247236-INS-132 |
| chr15-22253785-INS-448 | chr15-22253785-INS-449 |
| chr15-22260540-INS-332 | chr15-22260540-INS-335 |
| chr15-22320497-INS-3988 | chr15-22320498-INS-3987 |
| chr15-22323789-INS-226 | chr15-22323789-INS-226 |
| chr15-22357732-INS-51 | chr15-22357697-INS-58 |
| chr15-22362144-INS-2333 | chr15-22362145-INS-2333 |
| chr15-22378986-INS-1864 | chr15-22378987-INS-1864 |
| chr15-22387985-INS-136 | chr15-22387985-INS-124 |
| chr15-22392562-INS-6154 | chr15-22392565-INS-6140 |
| chr15-22395165-INS-53 | chr15-22395165-INS-53 |
| chr15-22397356-INS-557 | chr15-22397356-INS-557 |
| chr15-22399275-INS-2721 | chr15-22399276-INS-2721 |
| chr15-22401815-INS-1744 | chr15-22401816-INS-1744 |
| chr15-22412231-INS-78 | chr15-22412231-INS-66 |
| chr15-22447085-INS-54 | chr15-22447085-INS-52 |
| chr15-22501570-INS-6368 | chr15-22501571-INS-6368 |
| chr15-22525485-INS-166 | chr15-22525485-INS-185 |
| chr15-22536684-INS-536 | chr15-22536684-INS-556 |
| chr15-22565034-INS-168 | chr15-22565034-INS-167 |
| chr15-22586307-INS-70 | chr15-22586285-INS-63 |
| chr15-22602153-INS-7188 | chr15-22602154-INS-7188 |
| chr15-22602511-INS-65 | chr15-22602511-INS-65 |

| | |
|---|---|
| chr15-22608222-INS-58 | chr15-22608222-INS-51 |
| chr15-22635784-INS-201 | chr15-22635784-INS-201 |
| chr15-22645794-INS-55 | chr15-22645794-INS-55 |
| chr15-22656132-INS-565 | chr15-22656132-INS-566 |
| chr15-22667670-INS-55 | chr15-22667649-INS-66 |
| chr15-22674274-INS-109 | chr15-22674274-INS-134 |
| chr15-22734103-INS-6481 | chr15-22734104-INS-6481 |
| chr15-22744741-INS-181 | chr15-22744741-INS-180 |
| chr15-22748308-INS-52 | chr15-22748308-INS-52 |
| chr15-22775056-INS-78 | chr15-22775056-INS-90 |
| chr15-22782771-INS-90 | chr15-22782771-INS-80 |
| chr15-22786491-INS-50 | chr15-22786491-INS-62 |
| chr15-22791338-INS-222 | chr15-22791338-INS-222 |
| chr15-22816542-INS-356 | chr15-22816543-INS-375 |
| chr15-22848275-INS-327 | chr15-22848275-INS-329 |
| chr15-22921262-INS-254 | chr15-22921262-INS-254 |
| chr15-22932235-INS-791 | chr15-22932235-INS-791 |
| chr15-22953878-INS-63 | chr15-22953878-INS-51 |
| chr15-22966479-INS-7254 | chr15-22966480-INS-7254 |
| chr15-22974411-INS-60 | chr15-22974411-INS-80 |
| chr15-22981333-INS-1467 | chr15-22981334-INS-1473 |
| chr15-22988248-INS-118 | chr15-22988248-INS-118 |
| chr15-23021065-INS-610 | chr15-23021065-INS-610 |
| chr15-23078743-INS-59 | chr15-23078743-INS-59 |
| chr15-23103276-INS-79 | chr15-23103251-INS-81 |
| chr15-23105572-INS-697 | chr15-23105572-INS-698 |
| chr15-23124636-INS-830 | chr15-23124636-INS-830 |
| chr15-23129762-INS-1522 | chr15-23129763-INS-1522 |
| chr15-23156641-INS-1118 | chr15-23156642-INS-1118 |
| chr15-23177865-INS-689 | chr15-23177865-INS-690 |
| chr15-23199115-INS-314 | chr15-23199116-INS-278 |
| chr15-23206795-INS-349 | chr15-23206798-INS-352 |
| chr15-23250563-INS-58 | chr15-23250563-INS-56 |
| chr15-23256871-INS-146 | chr15-23256871-INS-146 |
| chr15-23261771-INS-491 | chr15-23261771-INS-491 |
| chr15-23314424-INS-70 | chr15-23314424-INS-66 |
| chr15-23318988-INS-1936 | chr15-23318989-INS-1936 |
| chr15-23332205-INS-211 | chr15-23332205-INS-211 |
| chr15-23333699-INS-344 | chr15-23333699-INS-344 |
| chr15-23364966-INS-7433 | chr15-23364967-INS-7434 |
| chr15-23381656-INS-148 | chr15-23381656-INS-166 |
| chr15-23381688-INS-6722 | chr15-23381689-INS-6722 |
| chr15-23384054-INS-345 | chr15-23384054-INS-345 |
| chr15-23431017-INS-518 | chr15-23431018-INS-517 |
| chr15-23461372-INS-90 | chr15-23461361-INS-59 |
| chr15-23479291-INS-54 | chr15-23479291-INS-52 |
| chr15-23490295-INS-463 | chr15-23490295-INS-463 |
| chr15-23505002-INS-373 | chr15-23505002-INS-373 |
| chr15-23509051-INS-201 | chr15-23509051-INS-201 |
| chr15-23514627-INS-7129 | chr15-23514628-INS-7129 |
| chr15-23541900-INS-115 | chr15-23541894-INS-113 |
| chr15-23545981-INS-3140 | chr15-23545982-INS-3140 |
| chr15-23549075-INS-1454 | chr15-23549076-INS-1455 |
| chr15-23552507-INS-1251 | chr15-23552508-INS-1251 |
| chr15-23601608-INS-603 | chr15-23601608-INS-607 |
| chr15-23626727-INS-65 | chr15-23626727-INS-65 |
| chr15-23627036-INS-3444 | chr15-23627037-INS-3444 |
| chr15-23634292-INS-368 | chr15-23634292-INS-366 |
| chr15-23644212-INS-52 | chr15-23644212-INS-50 |
| chr15-23646124-INS-4101 | chr15-23646125-INS-4102 |
| chr15-23662379-INS-7074 | chr15-23662380-INS-7074 |
| chr15-23729502-INS-208 | chr15-23729502-INS-208 |
| chr15-23796051-INS-81 | chr15-23796054-INS-81 |
| chr15-23844843-INS-153 | chr15-23844841-INS-111 |
| chr15-23847072-INS-7437 | chr15-23847073-INS-7437 |
| chr15-23858435-INS-62 | chr15-23858435-INS-62 |
| chr15-23866054-INS-130 | chr15-23866058-INS-140 |
| chr15-23882307-INS-72 | chr15-23882307-INS-74 |
| chr15-23944782-INS-68 | chr15-23944783-INS-80 |
| chr15-23996780-INS-4243 | chr15-23996781-INS-4243 |
| chr15-24012872-INS-92 | chr15-24012873-INS-92 |
| chr15-24050201-INS-145 | chr15-24050201-INS-145 |
| chr15-24077198-INS-6402 | chr15-24077199-INS-6400 |
| chr15-24126340-INS-200 | chr15-24126340-INS-200 |
| chr15-24129190-INS-68 | chr15-24129190-INS-68 |
| chr15-24154252-INS-84 | chr15-24154252-INS-84 |
| chr15-24219399-INS-267 | chr15-24219399-INS-267 |

| | |
|---|---|
| chr15-24265751-INS-94 | chr15-24265751-INS-105 |
| chr15-24268860-INS-136 | chr15-24268861-INS-146 |
| chr15-24270227-INS-137 | chr15-24270227-INS-144 |
| chr15-24326999-INS-882 | chr15-24326999-INS-882 |
| chr15-24350970-INS-133 | chr15-24350970-INS-133 |
| chr15-24356889-INS-206 | chr15-24356889-INS-207 |
| chr15-24388090-INS-183 | chr15-24388118-INS-190 |
| chr15-24393131-INS-6443 | chr15-24393132-INS-6443 |
| chr15-24399360-INS-122 | chr15-24399360-INS-122 |
| chr15-24423542-INS-72 | chr15-24423544-INS-96 |
| chr15-24451587-INS-7538 | chr15-24451588-INS-7540 |
| chr15-24473914-INS-77 | chr15-24473914-INS-77 |
| chr15-24555693-INS-4612 | chr15-24555694-INS-4612 |
| chr15-24581858-INS-7138 | chr15-24581859-INS-7138 |
| chr15-24587942-INS-108 | chr15-24587942-INS-114 |
| chr15-24595379-INS-68 | chr15-24595379-INS-68 |
| chr15-24603142-INS-52 | chr15-24603138-INS-52 |
| chr15-24626984-INS-104 | chr15-24626984-INS-103 |
| chr15-24630390-INS-960 | chr15-24630390-INS-960 |
| chr15-24631328-INS-81 | chr15-24631313-INS-75 |
| chr15-24652685-INS-2843 | chr15-24652686-INS-2843 |
| chr15-24720602-INS-2300 | chr15-24720603-INS-2300 |
| chr15-24798720-INS-1164 | chr15-24798721-INS-1163 |
| chr15-24816053-INS-292 | chr15-24816053-INS-298 |
| chr15-24824265-INS-234 | chr15-24824265-INS-234 |
| chr15-24850666-INS-549 | chr15-24850666-INS-549 |
| chr15-24852619-INS-932 | chr15-24852619-INS-932 |
| chr15-24859787-INS-62 | chr15-24859787-INS-60 |
| chr15-24874727-INS-6922 | chr15-24874728-INS-6921 |
| chr15-24889562-INS-59 | chr15-24889562-INS-65 |
| chr15-24928278-INS-86 | chr15-24928285-INS-68 |
| chr15-24932919-INS-301 | chr15-24932919-INS-301 |
| chr15-24961995-INS-428 | chr15-24961995-INS-428 |
| chr15-25036052-INS-90 | chr15-25036064-INS-91 |
| chr15-25050934-INS-52 | chr15-25050935-INS-64 |
| chr15-25057978-INS-82 | chr15-25057971-INS-82 |
| chr15-25058788-INS-2415 | chr15-25058789-INS-2415 |
| chr15-25059476-INS-203 | chr15-25059476-INS-203 |
| chr15-25074474-INS-135 | chr15-25074474-INS-137 |
| chr15-25094695-INS-883 | chr15-25094698-INS-883 |
| chr15-25146121-INS-88 | chr15-25146122-INS-59 |
| chr15-25148561-INS-66 | chr15-25148561-INS-54 |
| chr15-25163051-INS-1373 | chr15-25163053-INS-1375 |
| chr15-25171374-INS-3117 | chr15-25171375-INS-3117 |
| chr15-25171679-INS-80 | chr15-25171677-INS-70 |
| chr15-25241885-INS-98 | chr15-25241897-INS-80 |
| chr15-25258930-INS-88 | chr15-25258930-INS-88 |
| chr15-25266343-INS-51 | chr15-25266343-INS-51 |
| chr15-25266827-INS-235 | chr15-25266827-INS-237 |
| chr15-25283267-INS-177 | chr15-25283274-INS-180 |
| chr15-25290285-INS-934 | chr15-25290285-INS-938 |
| chr15-25304614-INS-453 | chr15-25304614-INS-452 |
| chr15-25306724-INS-1036 | chr15-25306725-INS-1035 |
| chr15-25324683-INS-79 | chr15-25324684-INS-73 |
| chr15-25372280-INS-56 | chr15-25372263-INS-70 |
| chr15-25375585-INS-54 | chr15-25375585-INS-54 |
| chr15-25385842-INS-58 | chr15-25385842-INS-58 |
| chr15-25396874-INS-200 | chr15-25396874-INS-200 |
| chr15-25419213-INS-82 | chr15-25419213-INS-88 |
| chr15-25438848-INS-66 | chr15-25438848-INS-74 |
| chr15-25467289-INS-58 | chr15-25467289-INS-58 |
| chr15-25470611-INS-91 | chr15-25470611-INS-91 |
| chr15-25497071-INS-61 | chr15-25497071-INS-67 |
| chr15-25520928-INS-64 | chr15-25520928-INS-76 |
| chr15-25528356-INS-91 | chr15-25528356-INS-72 |
| chr15-25537419-INS-341 | chr15-25537419-INS-341 |
| chr15-25551159-INS-68 | chr15-25551155-INS-72 |
| chr15-25557563-INS-66 | chr15-25557563-INS-66 |
| chr15-25589455-INS-93 | chr15-25589455-INS-93 |
| chr15-25609118-INS-214 | chr15-25609118-INS-214 |
| chr15-25624859-INS-60 | chr15-25624859-INS-60 |
| chr15-25692119-INS-84 | chr15-25692111-INS-130 |
| chr15-25694781-INS-6440 | chr15-25694782-INS-6440 |
| chr15-25703028-INS-210 | chr15-25703028-INS-210 |
| chr15-25707677-INS-178 | chr15-25707677-INS-179 |
| chr15-25718936-INS-65 | chr15-25718936-INS-65 |
| chr15-25745854-INS-458 | chr15-25745838-INS-479 |

| | |
|---|---|
| chr15-25747879-INS-199 | chr15-25747879-INS-199 |
| chr15-25765836-INS-197 | chr15-25765836-INS-196 |
| chr15-25767358-INS-118 | chr15-25767358-INS-119 |
| chr15-25790978-INS-182 | chr15-25790978-INS-178 |
| chr15-25797205-INS-126 | chr15-25797205-INS-124 |
| chr15-25807705-INS-78 | chr15-25807705-INS-78 |
| chr15-25818772-INS-215 | chr15-25818772-INS-215 |
| chr15-25887867-INS-72 | chr15-25887867-INS-72 |
| chr15-25889489-INS-71 | chr15-25889489-INS-71 |
| chr15-25895575-INS-240 | chr15-25895575-INS-241 |
| chr15-25898700-INS-77 | chr15-25898700-INS-77 |
| chr15-25903500-INS-68 | chr15-25903500-INS-68 |
| chr15-25922939-INS-241 | chr15-25922940-INS-242 |
| chr15-25952919-INS-179 | chr15-25952919-INS-179 |
| chr15-25955249-INS-284 | chr15-25955249-INS-284 |
| chr15-25975210-INS-4477 | chr15-25975211-INS-4479 |
| chr15-26034511-INS-138 | chr15-26034507-INS-135 |
| chr15-26043182-INS-1934 | chr15-26043183-INS-1934 |
| chr15-26082183-INS-62 | chr15-26082187-INS-51 |
| chr15-26110080-INS-1233 | chr15-26110081-INS-1240 |
| chr15-26184512-INS-639 | chr15-26184512-INS-639 |
| chr15-26201586-INS-59 | chr15-26201586-INS-59 |
| chr15-26202568-INS-205 | chr15-26202568-INS-201 |
| chr15-26206650-INS-52 | chr15-26206650-INS-52 |
| chr15-26209242-INS-185 | chr15-26209242-INS-185 |
| chr15-26210859-INS-496 | chr15-26210855-INS-505 |
| chr15-26217639-INS-58 | chr15-26217639-INS-60 |
| chr15-26218393-INS-714 | chr15-26218393-INS-715 |
| chr15-26273408-INS-98 | chr15-26273408-INS-98 |
| chr15-26305629-INS-70 | chr15-26305635-INS-55 |
| chr15-26338406-INS-4870 | chr15-26338407-INS-4870 |
| chr15-26371600-INS-68 | chr15-26371600-INS-68 |
| chr15-26383002-INS-156 | chr15-26383002-INS-156 |
| chr15-26414976-INS-72 | chr15-26414973-INS-62 |
| chr15-26416061-INS-80 | chr15-26416061-INS-82 |
| chr15-26428541-INS-51 | chr15-26428542-INS-56 |
| chr15-26439761-INS-64 | chr15-26439761-INS-60 |
| chr15-26446033-INS-56 | chr15-26446033-INS-56 |
| chr15-26448237-INS-55 | chr15-26448237-INS-55 |
| chr15-26503780-INS-136 | chr15-26503780-INS-136 |
| chr15-26515477-INS-260 | chr15-26515475-INS-270 |
| chr15-26517330-INS-67 | chr15-26517330-INS-67 |
| chr15-26537518-INS-512 | chr15-26537518-INS-512 |
| chr15-26570054-INS-74 | chr15-26570054-INS-74 |
| chr15-26571121-INS-60 | chr15-26571121-INS-60 |
| chr15-26599073-INS-373 | chr15-26599073-INS-373 |
| chr15-26669732-INS-145 | chr15-26669727-INS-120 |
| chr15-26680980-INS-212 | chr15-26680980-INS-212 |
| chr15-26694313-INS-484 | chr15-26694313-INS-484 |
| chr15-26719930-INS-207 | chr15-26719932-INS-219 |
| chr15-26774229-INS-616 | chr15-26774229-INS-616 |
| chr15-26785213-INS-123 | chr15-26785213-INS-119 |
| chr15-26798225-INS-108 | chr15-26798225-INS-108 |
| chr15-26836557-INS-124 | chr15-26836555-INS-140 |
| chr15-26840418-INS-115 | chr15-26840417-INS-160 |
| chr15-26840727-INS-1562 | chr15-26840728-INS-1561 |
| chr15-26899502-INS-81 | chr15-26899502-INS-81 |
| chr15-26938221-INS-6372 | chr15-26938222-INS-6372 |
| chr15-26946588-INS-7119 | chr15-26946589-INS-7120 |
| chr15-26949206-INS-72 | chr15-26949206-INS-78 |
| chr15-26964465-INS-843 | chr15-26964465-INS-843 |
| chr15-26975940-INS-153 | chr15-26975940-INS-156 |
| chr15-26985700-INS-61 | chr15-26985700-INS-61 |
| chr15-27003912-INS-1071 | chr15-27003913-INS-1071 |
| chr15-27056145-INS-121 | chr15-27056135-INS-170 |
| chr15-27069848-INS-827 | chr15-27069848-INS-828 |
| chr15-27124793-INS-4466 | chr15-27124794-INS-4466 |
| chr15-27144713-INS-321 | chr15-27144713-INS-321 |
| chr15-27162226-INS-350 | chr15-27162226-INS-351 |
| chr15-27206198-INS-87 | chr15-27206198-INS-87 |
| chr15-27239692-INS-69 | chr15-27239692-INS-69 |
| chr15-27245973-INS-51 | chr15-27245964-INS-76 |
| chr15-27258504-INS-5218 | chr15-27258505-INS-5218 |
| chr15-27333370-INS-178 | chr15-27333370-INS-178 |
| chr15-27342173-INS-239 | chr15-27342197-INS-258 |
| chr15-27345140-INS-146 | chr15-27345140-INS-146 |
| chr15-27347426-INS-72 | chr15-27347459-INS-84 |

| | |
|---|---|
| chr15-27351054-INS-126 | chr15-27351054-INS-126 |
| chr15-27354283-INS-619 | chr15-27354283-INS-619 |
| chr15-27372329-INS-54 | chr15-27372329-INS-52 |
| chr15-27383278-INS-53 | chr15-27383278-INS-57 |
| chr15-27420745-INS-194 | chr15-27420733-INS-195 |
| chr15-27435145-INS-72 | chr15-27435145-INS-72 |
| chr15-27437396-INS-132 | chr15-27437396-INS-132 |
| chr15-27489909-INS-127 | chr15-27489909-INS-127 |
| chr15-27546771-INS-247 | chr15-27546771-INS-243 |
| chr15-27551269-INS-67 | chr15-27551269-INS-67 |
| chr15-27564224-INS-66 | chr15-27564224-INS-66 |
| chr15-27605189-INS-56 | chr15-27605189-INS-56 |
| chr15-27653190-INS-6428 | chr15-27653191-INS-6428 |
| chr15-27708203-INS-96 | chr15-27708203-INS-84 |
| chr15-27716144-INS-197 | chr15-27716144-INS-197 |
| chr15-27793054-INS-117 | chr15-27793054-INS-117 |
| chr15-27825572-INS-72 | chr15-27825572-INS-72 |
| chr15-27855494-INS-65 | chr15-27855494-INS-65 |
| chr15-27863518-INS-220 | chr15-27863518-INS-221 |
| chr15-27885956-INS-136 | chr15-27885956-INS-136 |
| chr15-27897140-INS-66 | chr15-27897140-INS-66 |
| chr15-27900226-INS-132 | chr15-27900226-INS-132 |
| chr15-27916832-INS-78 | chr15-27916832-INS-78 |
| chr15-27940532-INS-110 | chr15-27940532-INS-110 |
| chr15-27953629-INS-71 | chr15-27953629-INS-71 |
| chr15-28001779-INS-226 | chr15-28001779-INS-220 |
| chr15-28022671-INS-104 | chr15-28022673-INS-108 |
| chr15-28039135-INS-88 | chr15-28039135-INS-88 |
| chr15-28110628-INS-68 | chr15-28110628-INS-68 |
| chr15-28127376-INS-204 | chr15-28127376-INS-204 |
| chr15-28159781-INS-76 | chr15-28159781-INS-82 |
| chr15-28167745-INS-59 | chr15-28167745-INS-102 |
| chr15-28210625-INS-56 | chr15-28210617-INS-64 |
| chr15-28211798-INS-83 | chr15-28211798-INS-83 |
| chr15-28222084-INS-68 | chr15-28222084-INS-68 |
| chr15-28240587-INS-185 | chr15-28240587-INS-185 |
| chr15-28249331-INS-202 | chr15-28249331-INS-202 |
| chr15-28250569-INS-449 | chr15-28250569-INS-449 |
| chr15-28278805-INS-282 | chr15-28278805-INS-282 |
| chr15-28288282-INS-439 | chr15-28288282-INS-439 |
| chr15-28317522-INS-76 | chr15-28317526-INS-80 |
| chr15-28339775-INS-69 | chr15-28339775-INS-75 |
| chr15-28341650-INS-914 | chr15-28341650-INS-915 |
| chr15-28365471-INS-399 | chr15-28365471-INS-399 |
| chr15-28369795-INS-107 | chr15-28369782-INS-125 |
| chr15-28375706-INS-59 | chr15-28375706-INS-64 |
| chr15-28423510-INS-146 | chr15-28423510-INS-140 |
| chr15-28444890-INS-4656 | chr15-28444891-INS-4656 |
| chr15-28468467-INS-402 | chr15-28468467-INS-402 |
| chr15-28472599-INS-252 | chr15-28472599-INS-252 |
| chr15-28524255-INS-84 | chr15-28524255-INS-54 |
| chr15-28542735-INS-2240 | chr15-28542736-INS-2240 |
| chr15-28544749-INS-84 | chr15-28544749-INS-84 |
| chr15-28554749-INS-420 | chr15-28554749-INS-425 |
| chr15-28577843-INS-6464 | chr15-28577844-INS-6464 |
| chr15-28594628-INS-578 | chr15-28594628-INS-560 |
| chr15-28611598-INS-68 | chr15-28611598-INS-70 |
| chr15-28613291-INS-90 | chr15-28613243-INS-116 |
| chr15-28619841-INS-294 | chr15-28619841-INS-294 |
| chr15-28639414-INS-157 | chr15-28639423-INS-138 |
| chr15-28653808-INS-70 | chr15-28653795-INS-82 |
| chr15-28674195-INS-878 | chr15-28674195-INS-885 |
| chr15-28678177-INS-82 | chr15-28678177-INS-88 |
| chr15-28715854-INS-2940 | chr15-28715855-INS-2940 |
| chr15-28731819-INS-59 | chr15-28731849-INS-66 |
| chr15-28759148-INS-370 | chr15-28759150-INS-370 |
| chr15-28787023-INS-272 | chr15-28787023-INS-264 |
| chr15-28799738-INS-3421 | chr15-28799739-INS-3421 |
| chr15-28825612-INS-543 | chr15-28825612-INS-543 |
| chr15-28903008-INS-469 | chr15-28903008-INS-469 |
| chr15-28904767-INS-188 | chr15-28904767-INS-188 |
| chr15-28918196-INS-5880 | chr15-28918197-INS-5881 |
| chr15-28939036-INS-62 | chr15-28939036-INS-62 |
| chr15-28968255-INS-6828 | chr15-28968256-INS-6828 |
| chr15-28984788-INS-1880 | chr15-28984789-INS-1880 |
| chr15-28987911-INS-72 | chr15-28987911-INS-72 |
| chr15-29003970-INS-56 | chr15-29003970-INS-50 |

| | |
|---|---|
| chr15-29020691-INS-533 | chr15-29020691-INS-538 |
| chr15-29021350-INS-51 | chr15-29021351-INS-52 |
| chr15-29024718-INS-3117 | chr15-29024719-INS-3118 |
| chr15-29026991-INS-497 | chr15-29026991-INS-497 |
| chr15-29029422-INS-6211 | chr15-29029423-INS-6211 |
| chr15-29047549-INS-1513 | chr15-29047550-INS-1479 |
| chr15-29065363-INS-310 | chr15-29065363-INS-310 |
| chr15-29071934-INS-265 | chr15-29071934-INS-263 |
| chr15-29082354-INS-254 | chr15-29082354-INS-254 |
| chr15-29094491-INS-6244 | chr15-29094492-INS-6244 |
| chr15-29115859-INS-62 | chr15-29115865-INS-56 |
| chr15-29144381-INS-195 | chr15-29144381-INS-195 |
| chr15-29146348-INS-427 | chr15-29146348-INS-428 |
| chr15-29176780-INS-1907 | chr15-29176781-INS-1907 |
| chr15-29178388-INS-4332 | chr15-29178389-INS-4332 |
| chr15-29195171-INS-7227 | chr15-29195172-INS-7228 |
| chr15-29199724-INS-56 | chr15-29199724-INS-56 |
| chr15-29209675-INS-52 | chr15-29209675-INS-52 |
| chr15-29210586-INS-68 | chr15-29210586-INS-68 |
| chr15-29249197-INS-87 | chr15-29249186-INS-93 |
| chr15-29270508-INS-334 | chr15-29270508-INS-334 |
| chr15-29317348-INS-6992 | chr15-29317349-INS-6992 |
| chr15-29332566-INS-3347 | chr15-29332567-INS-3347 |
| chr15-29356589-INS-58 | chr15-29356589-INS-58 |
| chr15-29411156-INS-6428 | chr15-29411157-INS-6428 |
| chr15-29419417-INS-354 | chr15-29419417-INS-354 |
| chr15-29451718-INS-342 | chr15-29451718-INS-335 |
| chr15-29499855-INS-72 | chr15-29499855-INS-72 |
| chr15-29517369-INS-6543 | chr15-29517370-INS-6543 |
| chr15-29522182-INS-158 | chr15-29522160-INS-195 |
| chr15-29526661-INS-97 | chr15-29526703-INS-60 |
| chr15-29604483-INS-56 | chr15-29604483-INS-52 |
| chr15-29605754-INS-267 | chr15-29605754-INS-243 |
| chr15-29621953-INS-51 | chr15-29621954-INS-62 |
| chr15-29626266-INS-160 | chr15-29626266-INS-160 |
| chr15-29691230-INS-201 | chr15-29691230-INS-201 |
| chr15-29741080-INS-265 | chr15-29741080-INS-265 |
| chr15-29742285-INS-862 | chr15-29742285-INS-862 |
| chr15-29743808-INS-801 | chr15-29743808-INS-801 |
| chr15-29752466-INS-268 | chr15-29752466-INS-268 |
| chr15-29830895-INS-6392 | chr15-29830896-INS-6393 |
| chr15-29873707-INS-370 | chr15-29873707-INS-369 |
| chr15-29900952-INS-404 | chr15-29900952-INS-404 |
| chr15-29924886-INS-2819 | chr15-29924887-INS-2819 |
| chr15-29931050-INS-56 | chr15-29931050-INS-64 |
| chr15-29955652-INS-132 | chr15-29955652-INS-132 |
| chr15-29959820-INS-200 | chr15-29959820-INS-202 |
| chr15-29963244-INS-67 | chr15-29963245-INS-54 |
| chr15-29978871-INS-3258 | chr15-29978872-INS-3258 |
| chr15-29980145-INS-86 | chr15-29980145-INS-104 |
| chr15-30040202-INS-494 | chr15-30040202-INS-494 |
| chr15-30079173-INS-489 | chr15-30079173-INS-489 |
| chr15-30082491-INS-81 | chr15-30082491-INS-81 |
| chr15-30106349-INS-135 | chr15-30106349-INS-135 |
| chr15-30156118-INS-226 | chr15-30156118-INS-226 |
| chr15-30160509-INS-401 | chr15-30160509-INS-401 |
| chr15-30165378-INS-208 | chr15-30165378-INS-210 |
| chr15-30214198-INS-51 | chr15-30214198-INS-51 |
| chr15-30232125-INS-88 | chr15-30232125-INS-88 |
| chr15-30236053-INS-797 | chr15-30236053-INS-797 |
| chr15-30238698-INS-278 | chr15-30238698-INS-278 |
| chr15-30246778-INS-92 | chr15-30246778-INS-92 |
| chr15-30265534-INS-68 | chr15-30265534-INS-68 |
| chr15-30269360-INS-125 | chr15-30269360-INS-119 |
| chr15-30271141-INS-7048 | chr15-30271142-INS-7048 |
| chr15-30287771-INS-97 | chr15-30287785-INS-58 |
| chr15-30311788-INS-1700 | chr15-30311789-INS-1700 |
| chr15-30342683-INS-52 | chr15-30342683-INS-52 |
| chr15-30352962-INS-60 | chr15-30352962-INS-62 |
| chr15-30357045-INS-4325 | chr15-30357046-INS-4325 |
| chr15-30374832-INS-443 | chr15-30374832-INS-443 |
| chr15-30402888-INS-442 | chr15-30402888-INS-442 |
| chr15-30406229-INS-57 | chr15-30406229-INS-57 |
| chr15-30419960-INS-52 | chr15-30419960-INS-68 |
| chr15-30434684-INS-587 | chr15-30434684-INS-587 |
| chr15-30469893-INS-461 | chr15-30469893-INS-461 |
| chr15-30485635-INS-6553 | chr15-30485636-INS-6554 |

| | |
|---|---|
| chr15-30486132-INS-142 | chr15-30486162-INS-140 |
| chr15-30488526-INS-55 | chr15-30488482-INS-60 |
| chr15-30494085-INS-134 | chr15-30494085-INS-132 |
| chr15-30504554-INS-4555 | chr15-30504555-INS-4555 |
| chr15-30514119-INS-205 | chr15-30514119-INS-206 |
| chr15-30536864-INS-139 | chr15-30536864-INS-139 |
| chr15-30545922-INS-104 | chr15-30545922-INS-116 |
| chr15-30547108-INS-56 | chr15-30547108-INS-52 |
| chr15-30549293-INS-872 | chr15-30549293-INS-872 |
| chr15-30553718-INS-54 | chr15-30553725-INS-57 |
| chr15-30608985-INS-859 | chr15-30608985-INS-859 |
| chr15-30609592-INS-53 | chr15-30609605-INS-55 |
| chr15-30620878-INS-90 | chr15-30620878-INS-96 |
| chr15-30623911-INS-51 | chr15-30623911-INS-51 |
| chr15-30640645-INS-178 | chr15-30640645-INS-177 |
| chr15-30669901-INS-298 | chr15-30669901-INS-298 |
| chr15-30674052-INS-58 | chr15-30674052-INS-58 |
| chr15-30759269-INS-209 | chr15-30759269-INS-209 |
| chr15-30787207-INS-86 | chr15-30787218-INS-65 |
| chr15-30795664-INS-52 | chr15-30795664-INS-52 |
| chr15-30804553-INS-967 | chr15-30804553-INS-968 |
| chr15-30810247-INS-64 | chr15-30810247-INS-64 |
| chr15-30813513-INS-62 | chr15-30813513-INS-62 |
| chr15-30878342-INS-56 | chr15-30878342-INS-60 |
| chr15-30881680-INS-53 | chr15-30881680-INS-51 |
| chr15-30887545-INS-104 | chr15-30887545-INS-128 |
| chr15-30908529-INS-385 | chr15-30908515-INS-405 |
| chr15-30923876-INS-160 | chr15-30923876-INS-160 |
| chr15-30946409-INS-114 | chr15-30946409-INS-118 |
| chr15-30947518-INS-84 | chr15-30947518-INS-84 |
| chr15-30971103-INS-434 | chr15-30971103-INS-434 |
| chr15-30980524-INS-226 | chr15-30980524-INS-232 |
| chr15-30986620-INS-113 | chr15-30986622-INS-107 |
| chr15-31033967-INS-256 | chr15-31033967-INS-256 |
| chr15-31039661-INS-78 | chr15-31039661-INS-78 |
| chr15-31042026-INS-56 | chr15-31041998-INS-62 |
| chr15-31056063-INS-77 | chr15-31056063-INS-95 |
| chr15-31100379-INS-904 | chr15-31100379-INS-897 |
| chr15-31101297-INS-302 | chr15-31101297-INS-309 |
| chr15-31168323-INS-225 | chr15-31168296-INS-251 |
| chr15-31172021-INS-96 | chr15-31172022-INS-96 |
| chr15-31179699-INS-284 | chr15-31179699-INS-284 |
| chr15-31184284-INS-171 | chr15-31184282-INS-170 |
| chr15-31189875-INS-370 | chr15-31189859-INS-372 |
| chr15-31197340-INS-57 | chr15-31197340-INS-58 |
| chr15-31226586-INS-255 | chr15-31226586-INS-255 |
| chr15-31231034-INS-1890 | chr15-31231035-INS-1890 |
| chr15-31283015-INS-58 | chr15-31283015-INS-58 |
| chr15-31294865-INS-53 | chr15-31294865-INS-53 |
| chr15-31302099-INS-56 | chr15-31302099-INS-58 |
| chr15-31308194-INS-57 | chr15-31308194-INS-57 |
| chr15-31314641-INS-84 | chr15-31314615-INS-86 |
| chr15-31317460-INS-85 | chr15-31317441-INS-60 |
| chr15-31327005-INS-964 | chr15-31327005-INS-946 |
| chr15-31329409-INS-61 | chr15-31329410-INS-61 |
| chr15-31331786-INS-332 | chr15-31331785-INS-333 |
| chr15-31347104-INS-75 | chr15-31347101-INS-103 |
| chr15-31360308-INS-72 | chr15-31360308-INS-72 |
| chr15-31385152-INS-206 | chr15-31385152-INS-203 |
| chr15-31386273-INS-208 | chr15-31386273-INS-208 |
| chr15-31405899-INS-100 | chr15-31405899-INS-100 |
| chr15-31408472-INS-55 | chr15-31408472-INS-55 |
| chr15-31410846-INS-233 | chr15-31410847-INS-216 |
| chr15-31427595-INS-80 | chr15-31427595-INS-62 |
| chr15-31430469-INS-213 | chr15-31430469-INS-213 |
| chr15-31432364-INS-52 | chr15-31432346-INS-73 |
| chr15-31435379-INS-79 | chr15-31435376-INS-79 |
| chr15-31450763-INS-71 | chr15-31450763-INS-71 |
| chr15-31470065-INS-212 | chr15-31470066-INS-209 |
| chr15-31492109-INS-291 | chr15-31492109-INS-291 |
| chr15-31529258-INS-178 | chr15-31529258-INS-178 |
| chr15-31538854-INS-5109 | chr15-31538855-INS-5109 |
| chr15-31540470-INS-175 | chr15-31540470-INS-176 |
| chr15-31561359-INS-171 | chr15-31561359-INS-171 |
| chr15-31563872-INS-578 | chr15-31563872-INS-578 |
| chr15-31567155-INS-63 | chr15-31567155-INS-63 |
| chr15-31577223-INS-196 | chr15-31577223-INS-196 |

| | |
|---|---|
| chr15-31579398-INS-214 | chr15-31579398-INS-214 |
| chr15-31593424-INS-52 | chr15-31593424-INS-52 |
| chr15-31610751-INS-493 | chr15-31610751-INS-493 |
| chr15-31622313-INS-1477 | chr15-31622314-INS-1477 |
| chr15-31623620-INS-241 | chr15-31623620-INS-240 |
| chr15-31633932-INS-61 | chr15-31633928-INS-62 |
| chr15-31635866-INS-395 | chr15-31635866-INS-395 |
| chr15-31639197-INS-135 | chr15-31639197-INS-135 |
| chr15-31692061-INS-498 | chr15-31692061-INS-498 |
| chr15-31696757-INS-66 | chr15-31696757-INS-66 |
| chr15-31698783-INS-518 | chr15-31698783-INS-518 |
| chr15-31721428-INS-68 | chr15-31721429-INS-68 |
| chr15-31728284-INS-177 | chr15-31728284-INS-193 |
| chr15-31744506-INS-7930 | chr15-31744507-INS-7930 |
| chr15-31747999-INS-1055 | chr15-31748000-INS-1055 |
| chr15-31767751-INS-382 | chr15-31767752-INS-383 |
| chr15-31800942-INS-78 | chr15-31800942-INS-78 |
| chr15-31804067-INS-193 | chr15-31804067-INS-193 |
| chr15-31806791-INS-64 | chr15-31806791-INS-64 |
| chr15-31810659-INS-57 | chr15-31810659-INS-57 |
| chr15-31813856-INS-162 | chr15-31813856-INS-158 |
| chr15-31824804-INS-1649 | chr15-31824805-INS-1672 |
| chr15-31860320-INS-83 | chr15-31860320-INS-95 |
| chr15-31868840-INS-77 | chr15-31868839-INS-81 |
| chr15-31889432-INS-171 | chr15-31889432-INS-171 |
| chr15-31919305-INS-160 | chr15-31919305-INS-160 |
| chr15-31924891-INS-605 | chr15-31924891-INS-605 |
| chr15-31936719-INS-179 | chr15-31936719-INS-179 |
| chr15-31975305-INS-202 | chr15-31975305-INS-202 |
| chr15-31990591-INS-58 | chr15-31990591-INS-58 |
| chr15-31991302-INS-6326 | chr15-31991303-INS-6326 |
| chr15-31994847-INS-182 | chr15-31994847-INS-173 |
| chr15-31999086-INS-209 | chr15-31999086-INS-209 |
| chr15-32013611-INS-86 | chr15-32013611-INS-88 |
| chr15-32040362-INS-70 | chr15-32040362-INS-70 |
| chr15-32054248-INS-71 | chr15-32054248-INS-52 |
| chr15-32066719-INS-736 | chr15-32066719-INS-736 |
| chr15-32077706-INS-255 | chr15-32077706-INS-256 |
| chr15-32085137-INS-152 | chr15-32085137-INS-153 |
| chr15-32086259-INS-6180 | chr15-32086260-INS-6182 |
| chr15-32136438-INS-776 | chr15-32136438-INS-777 |
| chr15-32139557-INS-2988 | chr15-32139558-INS-2988 |
| chr15-32151343-INS-390 | chr15-32151343-INS-390 |
| chr15-32195164-INS-624 | chr15-32195169-INS-628 |
| chr15-32197382-INS-6826 | chr15-32197383-INS-6826 |
| chr15-32201035-INS-85 | chr15-32201035-INS-125 |
| chr15-32228303-INS-111 | chr15-32228309-INS-118 |
| chr15-32258658-INS-58 | chr15-32258658-INS-58 |
| chr15-32268589-INS-180 | chr15-32268589-INS-192 |
| chr15-32286713-INS-1921 | chr15-32286714-INS-1922 |
| chr15-32322701-INS-179 | chr15-32322701-INS-179 |
| chr15-32331596-INS-54 | chr15-32331619-INS-53 |
| chr15-32361535-INS-221 | chr15-32361535-INS-221 |
| chr15-32364333-INS-124 | chr15-32364333-INS-124 |
| chr15-32367297-INS-1398 | chr15-32367298-INS-1442 |
| chr15-32385733-INS-439 | chr15-32385733-INS-439 |
| chr15-32386600-INS-54 | chr15-32386600-INS-54 |
| chr15-32395967-INS-212 | chr15-32395967-INS-212 |
| chr15-32398806-INS-915 | chr15-32398806-INS-915 |
| chr15-32409391-INS-685 | chr15-32409391-INS-686 |
| chr15-32429129-INS-56 | chr15-32429129-INS-56 |
| chr15-32435694-INS-426 | chr15-32435694-INS-425 |
| chr15-32469187-INS-6425 | chr15-32469188-INS-6425 |
| chr15-32488416-INS-2096 | chr15-32488417-INS-2096 |
| chr15-32490180-INS-71 | chr15-32490180-INS-71 |
| chr15-32511170-INS-79 | chr15-32511170-INS-85 |
| chr15-32521978-INS-1374 | chr15-32521991-INS-1361 |
| chr15-32522332-INS-7809 | chr15-32522333-INS-7809 |
| chr15-32536073-INS-336 | chr15-32536073-INS-336 |
| chr15-32573088-INS-79 | chr15-32573088-INS-85 |
| chr15-32585311-INS-153 | chr15-32585336-INS-162 |
| chr15-32604815-INS-68 | chr15-32604815-INS-68 |
| chr15-32611889-INS-203 | chr15-32611889-INS-200 |
| chr15-32618527-INS-74 | chr15-32618527-INS-78 |
| chr15-32651212-INS-1018 | chr15-32651213-INS-1018 |
| chr15-32653395-INS-54 | chr15-32653395-INS-54 |
| chr15-32673349-INS-3156 | chr15-32673350-INS-3156 |

| | |
|---|---|
| chr15-32706388-INS-982 | chr15-32706388-INS-982 |
| chr15-32728727-INS-73 | chr15-32728727-INS-73 |
| chr15-32735619-INS-128 | chr15-32735619-INS-128 |
| chr15-32744761-INS-1149 | chr15-32744762-INS-1149 |
| chr15-32749833-INS-94 | chr15-32749833-INS-94 |
| chr15-32752579-INS-79 | chr15-32752579-INS-79 |
| chr15-32753905-INS-459 | chr15-32753905-INS-460 |
| chr15-32766861-INS-81 | chr15-32766861-INS-81 |
| chr15-32782406-INS-792 | chr15-32782406-INS-792 |
| chr15-32795535-INS-160 | chr15-32795535-INS-160 |
| chr15-32812296-INS-225 | chr15-32812334-INS-204 |
| chr15-32828279-INS-163 | chr15-32828279-INS-163 |
| chr15-32852948-INS-1604 | chr15-32852949-INS-1604 |
| chr15-32883064-INS-69 | chr15-32883064-INS-69 |
| chr15-32894580-INS-4571 | chr15-32894581-INS-4571 |
| chr15-32898268-INS-1195 | chr15-32898269-INS-1195 |
| chr15-32903162-INS-72 | chr15-32903162-INS-72 |
| chr15-32929144-INS-52 | chr15-32929150-INS-62 |
| chr15-32934550-INS-63 | chr15-32934550-INS-62 |
| chr15-32974892-INS-80 | chr15-32974892-INS-80 |
| chr15-33029382-INS-133 | chr15-33029383-INS-134 |
| chr15-33039887-INS-345 | chr15-33039887-INS-345 |
| chr15-33057514-INS-180 | chr15-33057514-INS-180 |
| chr15-33063087-INS-53 | chr15-33063087-INS-53 |
| chr15-33071798-INS-287 | chr15-33071798-INS-287 |
| chr15-33085376-INS-54 | chr15-33085376-INS-56 |
| chr15-33101666-INS-1836 | chr15-33101667-INS-1829 |
| chr15-33104629-INS-140 | chr15-33104629-INS-140 |
| chr15-33111034-INS-213 | chr15-33111034-INS-213 |
| chr15-33118412-INS-989 | chr15-33118412-INS-989 |
| chr15-33147208-INS-1078 | chr15-33147209-INS-1078 |
| chr15-33178894-INS-55 | chr15-33178894-INS-60 |
| chr15-33208520-INS-81 | chr15-33208520-INS-87 |
| chr15-33220193-INS-100 | chr15-33220193-INS-102 |
| chr15-33225195-INS-1864 | chr15-33225196-INS-1864 |
| chr15-33230229-INS-52 | chr15-33230229-INS-54 |
| chr15-33243157-INS-104 | chr15-33243157-INS-104 |
| chr15-33258292-INS-203 | chr15-33258292-INS-203 |
| chr15-33302319-INS-4798 | chr15-33302320-INS-4798 |
| chr15-33311706-INS-669 | chr15-33311706-INS-669 |
| chr15-33319400-INS-236 | chr15-33319400-INS-230 |
| chr15-33341778-INS-93 | chr15-33341783-INS-89 |
| chr15-33342834-INS-6901 | chr15-33342834-INS-6900 |
| chr15-33344967-INS-52 | chr15-33344967-INS-52 |
| chr15-33356487-INS-54 | chr15-33356487-INS-50 |
| chr15-33358187-INS-486 | chr15-33358184-INS-486 |
| chr15-33379901-INS-76 | chr15-33379901-INS-76 |
| chr15-33382616-INS-6221 | chr15-33382617-INS-6214 |
| chr15-33386522-INS-51 | chr15-33386522-INS-51 |
| chr15-33395424-INS-5561 | chr15-33395425-INS-5561 |
| chr15-33400121-INS-1864 | chr15-33400122-INS-1864 |
| chr15-33423751-INS-76 | chr15-33423751-INS-73 |
| chr15-33424990-INS-758 | chr15-33424990-INS-758 |
| chr15-33506102-INS-143 | chr15-33506110-INS-146 |
| chr15-33508679-INS-133 | chr15-33508679-INS-133 |
| chr15-33512917-INS-2917 | chr15-33512918-INS-2917 |
| chr15-33525301-INS-82 | chr15-33525301-INS-86 |
| chr15-33551824-INS-110 | chr15-33551824-INS-78 |
| chr15-33558775-INS-562 | chr15-33558775-INS-562 |
| chr15-33567824-INS-977 | chr15-33567824-INS-977 |
| chr15-33617901-INS-133 | chr15-33617901-INS-133 |
| chr15-33623752-INS-605 | chr15-33623752-INS-605 |
| chr15-33643442-INS-857 | chr15-33643442-INS-857 |
| chr15-33653527-INS-193 | chr15-33653527-INS-196 |
| chr15-33665479-INS-201 | chr15-33665479-INS-201 |
| chr15-33672363-INS-295 | chr15-33672363-INS-295 |
| chr15-33712313-INS-81 | chr15-33712313-INS-81 |
| chr15-33761530-INS-550 | chr15-33761530-INS-550 |
| chr15-33845388-INS-3110 | chr15-33845389-INS-3110 |
| chr15-33845912-INS-201 | chr15-33845912-INS-201 |
| chr15-33847622-INS-58 | chr15-33847614-INS-58 |
| chr15-33853879-INS-56 | chr15-33853879-INS-56 |
| chr15-33897479-INS-2642 | chr15-33897480-INS-2642 |
| chr15-33901287-INS-79 | chr15-33901287-INS-79 |
| chr15-33930970-INS-55 | chr15-33930970-INS-60 |
| chr15-33942993-INS-475 | chr15-33942993-INS-475 |
| chr15-33996612-INS-208 | chr15-33996613-INS-208 |

| | |
|---|---|
| chr15-34022462-INS-986 | chr15-34022462-INS-986 |
| chr15-34035052-INS-102 | chr15-34035038-INS-101 |
| chr15-34039928-INS-229 | chr15-34039928-INS-228 |
| chr15-34063072-INS-183 | chr15-34063072-INS-183 |
| chr15-34064572-INS-131 | chr15-34064572-INS-122 |
| chr15-34067055-INS-98 | chr15-34067055-INS-94 |
| chr15-34095846-INS-65 | chr15-34095846-INS-65 |
| chr15-34097074-INS-121 | chr15-34097074-INS-121 |
| chr15-34103987-INS-80 | chr15-34103987-INS-90 |
| chr15-34130775-INS-84 | chr15-34130768-INS-98 |
| chr15-34146959-INS-86 | chr15-34146959-INS-86 |
| chr15-34191478-INS-342 | chr15-34191478-INS-342 |
| chr15-34206747-INS-209 | chr15-34206747-INS-209 |
| chr15-34208490-INS-219 | chr15-34208490-INS-219 |
| chr15-34220153-INS-73 | chr15-34220153-INS-73 |
| chr15-34226186-INS-430 | chr15-34226186-INS-436 |
| chr15-34242778-INS-171 | chr15-34242778-INS-171 |
| chr15-34258335-INS-113 | chr15-34258335-INS-115 |
| chr15-34259802-INS-2667 | chr15-34259803-INS-2667 |
| chr15-34280671-INS-171 | chr15-34280671-INS-171 |
| chr15-34317059-INS-146 | chr15-34317059-INS-135 |
| chr15-34323218-INS-431 | chr15-34323218-INS-431 |
| chr15-34325096-INS-92 | chr15-34325111-INS-99 |
| chr15-34328436-INS-210 | chr15-34328436-INS-210 |
| chr15-34354105-INS-169 | chr15-34354100-INS-170 |
| chr15-34360956-INS-3468 | chr15-34360957-INS-3469 |
| chr15-34364141-INS-820 | chr15-34364141-INS-820 |
| chr15-34371893-INS-208 | chr15-34371893-INS-208 |
| chr15-34386792-INS-112 | chr15-34386792-INS-112 |
| chr15-34398077-INS-213 | chr15-34398077-INS-213 |
| chr15-34428019-INS-182 | chr15-34428019-INS-182 |
| chr15-34430491-INS-214 | chr15-34430491-INS-214 |
| chr15-34441316-INS-105 | chr15-34441316-INS-110 |
| chr15-34444599-INS-222 | chr15-34444599-INS-223 |
| chr15-34450722-INS-67 | chr15-34450722-INS-70 |
| chr15-34452897-INS-54 | chr15-34452897-INS-54 |
| chr15-34464222-INS-223 | chr15-34464222-INS-223 |
| chr15-34472800-INS-220 | chr15-34472800-INS-220 |
| chr15-34482083-INS-113 | chr15-34482083-INS-113 |
| chr15-34544844-INS-2678 | chr15-34544845-INS-2678 |
| chr15-34566725-INS-51 | chr15-34566725-INS-51 |
| chr15-34600787-INS-74 | chr15-34600787-INS-104 |
| chr15-34614642-INS-54 | chr15-34614642-INS-56 |
| chr15-34623887-INS-135 | chr15-34623887-INS-135 |
| chr15-34646979-INS-173 | chr15-34646979-INS-170 |
| chr15-34650674-INS-96 | chr15-34650674-INS-96 |
| chr15-34659845-INS-239 | chr15-34659845-INS-245 |
| chr15-34662138-INS-2179 | chr15-34662139-INS-2179 |
| chr15-34698945-INS-256 | chr15-34698972-INS-256 |
| chr15-34722611-INS-87 | chr15-34722611-INS-100 |
| chr15-34761119-INS-697 | chr15-34761119-INS-697 |
| chr15-34773377-INS-51 | chr15-34773377-INS-51 |
| chr15-34774903-INS-72 | chr15-34774903-INS-72 |
| chr15-34784370-INS-177 | chr15-34784370-INS-177 |
| chr15-34794342-INS-57 | chr15-34794342-INS-57 |
| chr15-34802899-INS-7078 | chr15-34802899-INS-7077 |
| chr15-34804936-INS-199 | chr15-34804936-INS-189 |
| chr15-34805641-INS-56 | chr15-34805633-INS-68 |
| chr15-34820152-INS-110 | chr15-34820152-INS-110 |
| chr15-34821886-INS-91 | chr15-34821886-INS-89 |
| chr15-34827191-INS-401 | chr15-34827191-INS-401 |
| chr15-34832114-INS-144 | chr15-34832114-INS-144 |
| chr15-34862900-INS-176 | chr15-34862900-INS-176 |
| chr15-34867101-INS-114 | chr15-34867101-INS-114 |
| chr15-34934265-INS-85 | chr15-34934265-INS-92 |
| chr15-34956951-INS-92 | chr15-34956963-INS-91 |
| chr15-34960323-INS-475 | chr15-34960323-INS-475 |
| chr15-34961701-INS-214 | chr15-34961701-INS-214 |
| chr15-34994075-INS-51 | chr15-34994075-INS-51 |
| chr15-35039810-INS-156 | chr15-35039810-INS-156 |
| chr15-35043089-INS-6412 | chr15-35043090-INS-6412 |
| chr15-35057355-INS-6987 | chr15-35057343-INS-6988 |
| chr15-35079169-INS-6505 | chr15-35079170-INS-6505 |
| chr15-35096500-INS-186 | chr15-35096500-INS-186 |
| chr15-35134769-INS-446 | chr15-35134769-INS-446 |
| chr15-35151897-INS-354 | chr15-35151897-INS-354 |
| chr15-35164402-INS-208 | chr15-35164402-INS-208 |

| | |
|---|---|
| chr15-35166106-INS-60 | chr15-35166106-INS-52 |
| chr15-35166825-INS-7172 | chr15-35166826-INS-7172 |
| chr15-35172593-INS-3613 | chr15-35172594-INS-3605 |
| chr15-35189003-INS-52 | chr15-35189003-INS-52 |
| chr15-35215066-INS-71 | chr15-35215066-INS-81 |
| chr15-35224498-INS-6752 | chr15-35224499-INS-6754 |
| chr15-35239706-INS-219 | chr15-35239706-INS-220 |
| chr15-35242552-INS-1892 | chr15-35242553-INS-1885 |
| chr15-35259371-INS-193 | chr15-35259371-INS-193 |
| chr15-35267576-INS-599 | chr15-35267577-INS-599 |
| chr15-35288210-INS-92 | chr15-35288210-INS-92 |
| chr15-35330212-INS-100 | chr15-35330212-INS-100 |
| chr15-35330929-INS-472 | chr15-35330929-INS-472 |
| chr15-35352846-INS-54 | chr15-35352846-INS-56 |
| chr15-35362459-INS-219 | chr15-35362459-INS-219 |
| chr15-35463553-INS-7152 | chr15-35463554-INS-7152 |
| chr15-35515709-INS-60 | chr15-35515709-INS-60 |
| chr15-35517990-INS-6319 | chr15-35517991-INS-6319 |
| chr15-35520535-INS-401 | chr15-35520535-INS-401 |
| chr15-35578985-INS-180 | chr15-35578985-INS-180 |
| chr15-35614652-INS-60 | chr15-35614652-INS-60 |
| chr15-35617052-INS-60 | chr15-35617052-INS-54 |
| chr15-35623421-INS-3316 | chr15-35623422-INS-3316 |
| chr15-35626878-INS-328 | chr15-35626878-INS-328 |
| chr15-35629077-INS-970 | chr15-35629077-INS-972 |
| chr15-35693200-INS-2654 | chr15-35693201-INS-2654 |
| chr15-35711627-INS-470 | chr15-35711627-INS-471 |
| chr15-35715021-INS-2998 | chr15-35715022-INS-2998 |
| chr15-35733393-INS-6643 | chr15-35733394-INS-6644 |
| chr15-35790428-INS-84 | chr15-35790424-INS-88 |
| chr15-35804826-INS-112 | chr15-35804826-INS-112 |
| chr15-35835036-INS-217 | chr15-35835036-INS-217 |
| chr15-35851163-INS-54 | chr15-35851163-INS-54 |
| chr15-35854256-INS-279 | chr15-35854256-INS-279 |
| chr15-35863913-INS-65 | chr15-35863913-INS-66 |
| chr15-35894403-INS-62 | chr15-35894403-INS-62 |
| chr15-35922722-INS-2696 | chr15-35922723-INS-2696 |
| chr15-35929182-INS-56 | chr15-35929182-INS-56 |
| chr15-35946856-INS-484 | chr15-35946856-INS-484 |
| chr15-35968321-INS-318 | chr15-35968321-INS-318 |
| chr15-35979611-INS-3113 | chr15-35979612-INS-3113 |
| chr15-35984025-INS-207 | chr15-35984025-INS-207 |
| chr15-35991706-INS-200 | chr15-35991706-INS-200 |
| chr15-36002399-INS-78 | chr15-36002399-INS-60 |
| chr15-36018016-INS-217 | chr15-36018016-INS-217 |
| chr15-36032083-INS-1892 | chr15-36032084-INS-1892 |
| chr15-36045096-INS-59 | chr15-36045096-INS-59 |
| chr15-36056361-INS-153 | chr15-36056361-INS-153 |
| chr15-36076078-INS-199 | chr15-36076078-INS-199 |
| chr15-36083223-INS-295 | chr15-36083223-INS-295 |
| chr15-36107620-INS-203 | chr15-36107620-INS-203 |
| chr15-36124045-INS-241 | chr15-36124036-INS-266 |
| chr15-36130092-INS-56 | chr15-36130092-INS-56 |
| chr15-36140016-INS-89 | chr15-36140016-INS-89 |
| chr15-36169272-INS-92 | chr15-36169272-INS-92 |
| chr15-36186045-INS-89 | chr15-36186045-INS-89 |
| chr15-36194718-INS-207 | chr15-36194718-INS-207 |
| chr15-36194721-INS-818 | chr15-36194721-INS-818 |
| chr15-36198052-INS-214 | chr15-36198052-INS-215 |
| chr15-36209956-INS-196 | chr15-36209956-INS-196 |
| chr15-36217025-INS-186 | chr15-36217025-INS-186 |
| chr15-36220225-INS-60 | chr15-36220225-INS-52 |
| chr15-36227636-INS-175 | chr15-36227636-INS-175 |
| chr15-36231283-INS-121 | chr15-36231283-INS-121 |
| chr15-36237230-INS-149 | chr15-36237229-INS-149 |
| chr15-36243122-INS-61 | chr15-36243122-INS-61 |
| chr15-36243956-INS-99 | chr15-36243956-INS-99 |
| chr15-36253077-INS-81 | chr15-36253078-INS-78 |
| chr15-36285436-INS-66 | chr15-36285436-INS-66 |
| chr15-36291622-INS-79 | chr15-36291622-INS-79 |
| chr15-36298131-INS-855 | chr15-36298131-INS-849 |
| chr15-36302271-INS-51 | chr15-36302271-INS-68 |
| chr15-36310414-INS-402 | chr15-36310414-INS-406 |
| chr15-36324252-INS-188 | chr15-36324252-INS-215 |
| chr15-36325343-INS-63 | chr15-36325343-INS-75 |
| chr15-36328048-INS-115 | chr15-36328048-INS-115 |
| chr15-36333440-INS-55 | chr15-36333441-INS-55 |

| | |
|---|---|
| chr15-36353369-INS-54 | chr15-36353369-INS-54 |
| chr15-36371432-INS-64 | chr15-36371430-INS-66 |
| chr15-36383411-INS-176 | chr15-36383377-INS-189 |
| chr15-36391288-INS-66 | chr15-36391288-INS-64 |
| chr15-36431225-INS-232 | chr15-36431225-INS-232 |
| chr15-36433872-INS-62 | chr15-36433896-INS-64 |
| chr15-36446593-INS-112 | chr15-36446593-INS-112 |
| chr15-36452484-INS-54 | chr15-36452448-INS-54 |
| chr15-36453625-INS-94 | chr15-36453625-INS-94 |
| chr15-36481938-INS-215 | chr15-36481938-INS-215 |
| chr15-36484684-INS-64 | chr15-36484684-INS-64 |
| chr15-36523813-INS-65 | chr15-36523813-INS-65 |
| chr15-36553031-INS-52 | chr15-36553030-INS-99 |
| chr15-36562028-INS-82 | chr15-36562004-INS-66 |
| chr15-36567957-INS-118 | chr15-36567950-INS-126 |
| chr15-36573933-INS-983 | chr15-36573933-INS-983 |
| chr15-36578194-INS-120 | chr15-36578194-INS-98 |
| chr15-36589473-INS-182 | chr15-36589473-INS-194 |
| chr15-36619469-INS-181 | chr15-36619469-INS-183 |
| chr15-36629355-INS-84 | chr15-36629355-INS-90 |
| chr15-36630786-INS-218 | chr15-36630787-INS-205 |
| chr15-36634894-INS-57 | chr15-36634894-INS-61 |
| chr15-36637977-INS-169 | chr15-36637977-INS-128 |
| chr15-36642211-INS-536 | chr15-36642211-INS-536 |
| chr15-36649769-INS-57 | chr15-36649769-INS-59 |
| chr15-36654780-INS-51 | chr15-36654780-INS-53 |
| chr15-36659389-INS-217 | chr15-36659389-INS-217 |
| chr15-36661633-INS-194 | chr15-36661633-INS-194 |
| chr15-36669362-INS-211 | chr15-36669362-INS-211 |
| chr15-36672569-INS-80 | chr15-36672551-INS-50 |
| chr15-36698585-INS-178 | chr15-36698583-INS-162 |
| chr15-36744209-INS-440 | chr15-36744209-INS-440 |
| chr15-36757714-INS-62 | chr15-36757714-INS-64 |
| chr15-36817531-INS-76 | chr15-36817531-INS-64 |
| chr15-36842206-INS-56 | chr15-36842222-INS-56 |
| chr15-36849281-INS-111 | chr15-36849281-INS-111 |
| chr15-36906717-INS-93 | chr15-36906717-INS-69 |
| chr15-36926359-INS-52 | chr15-36926359-INS-58 |
| chr15-36944468-INS-68 | chr15-36944468-INS-68 |
| chr15-36969849-INS-290 | chr15-36969848-INS-311 |
| chr15-37006109-INS-198 | chr15-37006109-INS-197 |
| chr15-37010419-INS-58 | chr15-37010419-INS-58 |
| chr15-37013888-INS-52 | chr15-37013888-INS-52 |
| chr15-37019233-INS-74 | chr15-37019233-INS-74 |
| chr15-37032441-INS-208 | chr15-37032441-INS-208 |
| chr15-37033521-INS-468 | chr15-37033521-INS-468 |
| chr15-37043256-INS-3159 | chr15-37043257-INS-3159 |
| chr15-37046234-INS-70 | chr15-37046265-INS-70 |
| chr15-37049867-INS-458 | chr15-37049867-INS-458 |
| chr15-37086055-INS-515 | chr15-37086055-INS-515 |
| chr15-37088252-INS-54 | chr15-37088262-INS-54 |
| chr15-37113563-INS-97 | chr15-37113567-INS-61 |
| chr15-37123235-INS-308 | chr15-37123235-INS-313 |
| chr15-37139481-INS-54 | chr15-37139481-INS-56 |
| chr15-37152790-INS-60 | chr15-37152790-INS-60 |
| chr15-37158468-INS-205 | chr15-37158468-INS-200 |
| chr15-37165945-INS-58 | chr15-37165946-INS-59 |
| chr15-37170305-INS-130 | chr15-37170305-INS-130 |
| chr15-37172057-INS-54 | chr15-37172057-INS-54 |
| chr15-37174864-INS-62 | chr15-37174864-INS-61 |
| chr15-37182980-INS-201 | chr15-37182980-INS-199 |
| chr15-37187748-INS-56 | chr15-37187748-INS-56 |
| chr15-37189817-INS-69 | chr15-37189817-INS-69 |
| chr15-37200453-INS-223 | chr15-37200453-INS-223 |
| chr15-37210547-INS-59 | chr15-37210553-INS-84 |
| chr15-37229827-INS-211 | chr15-37229827-INS-211 |
| chr15-37239465-INS-96 | chr15-37239473-INS-92 |
| chr15-37245512-INS-206 | chr15-37245512-INS-206 |
| chr15-37263123-INS-384 | chr15-37263123-INS-384 |
| chr15-37265747-INS-157 | chr15-37265747-INS-177 |
| chr15-37290776-INS-157 | chr15-37290776-INS-160 |
| chr15-37297963-INS-89 | chr15-37297963-INS-89 |
| chr15-37434410-INS-75 | chr15-37434410-INS-76 |
| chr15-37442271-INS-190 | chr15-37442271-INS-190 |
| chr15-37446548-INS-52 | chr15-37446550-INS-78 |
| chr15-37461420-INS-545 | chr15-37461426-INS-547 |
| chr15-37465367-INS-197 | chr15-37465395-INS-196 |

| | |
|---|---|
| chr15-37482411-INS-6708 | chr15-37482412-INS-6708 |
| chr15-37483158-INS-407 | chr15-37483164-INS-399 |
| chr15-37573485-INS-209 | chr15-37573486-INS-209 |
| chr15-37574188-INS-178 | chr15-37574188-INS-178 |
| chr15-37607623-INS-50 | chr15-37607623-INS-62 |
| chr15-37619300-INS-374 | chr15-37619300-INS-374 |
| chr15-37629900-INS-62 | chr15-37629900-INS-62 |
| chr15-37631640-INS-64 | chr15-37631636-INS-62 |
| chr15-37632992-INS-477 | chr15-37632992-INS-477 |
| chr15-37702642-INS-84 | chr15-37702643-INS-70 |
| chr15-37715728-INS-75 | chr15-37715728-INS-75 |
| chr15-37716821-INS-122 | chr15-37716821-INS-98 |
| chr15-37755865-INS-74 | chr15-37755865-INS-74 |
| chr15-37759388-INS-51 | chr15-37759364-INS-65 |
| chr15-37763913-INS-150 | chr15-37763895-INS-144 |
| chr15-37813899-INS-1216 | chr15-37813900-INS-1216 |
| chr15-37826996-INS-147 | chr15-37826986-INS-159 |
| chr15-37834195-INS-52 | chr15-37834195-INS-82 |
| chr15-37847426-INS-334 | chr15-37847426-INS-325 |
| chr15-37856642-INS-2650 | chr15-37856643-INS-2656 |
| chr15-37860822-INS-428 | chr15-37860822-INS-428 |
| chr15-37871062-INS-173 | chr15-37871063-INS-177 |
| chr15-37909081-INS-99 | chr15-37909131-INS-62 |
| chr15-37928467-INS-54 | chr15-37928467-INS-54 |
| chr15-37950889-INS-129 | chr15-37950889-INS-86 |
| chr15-37962015-INS-83 | chr15-37962015-INS-83 |
| chr15-37963822-INS-933 | chr15-37963822-INS-933 |
| chr15-37966611-INS-60 | chr15-37966611-INS-61 |
| chr15-37971097-INS-55 | chr15-37971097-INS-56 |
| chr15-37987604-INS-179 | chr15-37987605-INS-179 |
| chr15-38011656-INS-506 | chr15-38011650-INS-509 |
| chr15-38017592-INS-52 | chr15-38017592-INS-54 |
| chr15-38039348-INS-205 | chr15-38039348-INS-205 |
| chr15-38039930-INS-205 | chr15-38039930-INS-205 |
| chr15-38044089-INS-201 | chr15-38044089-INS-201 |
| chr15-38057834-INS-58 | chr15-38057834-INS-58 |
| chr15-38059501-INS-206 | chr15-38059501-INS-206 |
| chr15-38066792-INS-211 | chr15-38066792-INS-211 |
| chr15-38083634-INS-70 | chr15-38083673-INS-74 |
| chr15-38086151-INS-216 | chr15-38086151-INS-216 |
| chr15-38091716-INS-633 | chr15-38091716-INS-633 |
| chr15-38105932-INS-105 | chr15-38105932-INS-105 |
| chr15-38107854-INS-68 | chr15-38107835-INS-70 |
| chr15-38111818-INS-214 | chr15-38111818-INS-214 |
| chr15-38112346-INS-217 | chr15-38112346-INS-217 |
| chr15-38113803-INS-53 | chr15-38113798-INS-70 |
| chr15-38118669-INS-64 | chr15-38118669-INS-68 |
| chr15-38124224-INS-358 | chr15-38124224-INS-398 |
| chr15-38149345-INS-174 | chr15-38149345-INS-174 |
| chr15-38152626-INS-466 | chr15-38152626-INS-466 |
| chr15-38158532-INS-113 | chr15-38158532-INS-132 |
| chr15-38177556-INS-88 | chr15-38177556-INS-88 |
| chr15-38183911-INS-669 | chr15-38183911-INS-669 |
| chr15-38184503-INS-55 | chr15-38184500-INS-60 |
| chr15-38196457-INS-118 | chr15-38196457-INS-118 |
| chr15-38202058-INS-184 | chr15-38202058-INS-185 |
| chr15-38232545-INS-215 | chr15-38232545-INS-215 |
| chr15-38242414-INS-75 | chr15-38242414-INS-80 |
| chr15-38254326-INS-210 | chr15-38254326-INS-210 |
| chr15-38255784-INS-4573 | chr15-38255785-INS-4575 |
| chr15-38267953-INS-72 | chr15-38267953-INS-72 |
| chr15-38277738-INS-198 | chr15-38277735-INS-199 |
| chr15-38283474-INS-69 | chr15-38283474-INS-72 |
| chr15-38288473-INS-232 | chr15-38288473-INS-224 |
| chr15-38322088-INS-111 | chr15-38322055-INS-153 |
| chr15-38343935-INS-456 | chr15-38343936-INS-448 |
| chr15-38355607-INS-211 | chr15-38355608-INS-210 |
| chr15-38373161-INS-120 | chr15-38373161-INS-120 |
| chr15-38410613-INS-54 | chr15-38410613-INS-52 |
| chr15-38425189-INS-759 | chr15-38425189-INS-759 |
| chr15-38444474-INS-173 | chr15-38444474-INS-173 |
| chr15-38447400-INS-102 | chr15-38447400-INS-54 |
| chr15-38449740-INS-528 | chr15-38449740-INS-533 |
| chr15-38474609-INS-64 | chr15-38474609-INS-64 |
| chr15-38478124-INS-130 | chr15-38478124-INS-132 |
| chr15-38497002-INS-200 | chr15-38497002-INS-200 |
| chr15-38542929-INS-58 | chr15-38542929-INS-58 |

| | |
|---|---|
| chr15-38547755-INS-120 | chr15-38547755-INS-120 |
| chr15-38554715-INS-632 | chr15-38554712-INS-633 |
| chr15-38558183-INS-235 | chr15-38558183-INS-235 |
| chr15-38560526-INS-91 | chr15-38560526-INS-91 |
| chr15-38564996-INS-79 | chr15-38565002-INS-75 |
| chr15-38575456-INS-94 | chr15-38575460-INS-82 |
| chr15-38575849-INS-201 | chr15-38575849-INS-201 |
| chr15-38579703-INS-52 | chr15-38579703-INS-52 |
| chr15-38594162-INS-59 | chr15-38594162-INS-60 |
| chr15-38597102-INS-198 | chr15-38597082-INS-192 |
| chr15-38606873-INS-201 | chr15-38606873-INS-201 |
| chr15-38618450-INS-2994 | chr15-38618451-INS-2993 |
| chr15-38633631-INS-1311 | chr15-38633637-INS-1303 |
| chr15-38649723-INS-232 | chr15-38649723-INS-232 |
| chr15-38658709-INS-124 | chr15-38658709-INS-124 |
| chr15-38677532-INS-200 | chr15-38677532-INS-200 |
| chr15-38705377-INS-74 | chr15-38705357-INS-71 |
| chr15-38714463-INS-85 | chr15-38714463-INS-85 |
| chr15-38716826-INS-210 | chr15-38716826-INS-210 |
| chr15-38720443-INS-54 | chr15-38720443-INS-54 |
| chr15-38721394-INS-77 | chr15-38721401-INS-78 |
| chr15-38729932-INS-173 | chr15-38729932-INS-175 |
| chr15-38731395-INS-359 | chr15-38731395-INS-359 |
| chr15-38735251-INS-250 | chr15-38735251-INS-250 |
| chr15-38753268-INS-348 | chr15-38753268-INS-358 |
| chr15-38773697-INS-3201 | chr15-38773698-INS-3201 |
| chr15-38777148-INS-270 | chr15-38777148-INS-260 |
| chr15-38791475-INS-56 | chr15-38791495-INS-51 |
| chr15-38811003-INS-166 | chr15-38811003-INS-124 |
| chr15-38855981-INS-63 | chr15-38855952-INS-107 |
| chr15-38868996-INS-177 | chr15-38868996-INS-181 |
| chr15-38893866-INS-59 | chr15-38893866-INS-59 |
| chr15-38902654-INS-234 | chr15-38902654-INS-235 |
| chr15-38909270-INS-61 | chr15-38909228-INS-70 |
| chr15-38913237-INS-6691 | chr15-38913238-INS-6692 |
| chr15-38934290-INS-66 | chr15-38934290-INS-66 |
| chr15-38945554-INS-66 | chr15-38945554-INS-66 |
| chr15-38992877-INS-70 | chr15-38992877-INS-70 |
| chr15-39008741-INS-511 | chr15-39008741-INS-545 |
| chr15-39017474-INS-146 | chr15-39017474-INS-142 |
| chr15-39029099-INS-532 | chr15-39029099-INS-484 |
| chr15-39065743-INS-75 | chr15-39065743-INS-50 |
| chr15-39071819-INS-79 | chr15-39071819-INS-120 |
| chr15-39094650-INS-64 | chr15-39094600-INS-72 |
| chr15-39097433-INS-461 | chr15-39097433-INS-462 |
| chr15-39114950-INS-215 | chr15-39114950-INS-215 |
| chr15-39126875-INS-6266 | chr15-39126876-INS-6266 |
| chr15-39127934-INS-208 | chr15-39127934-INS-208 |
| chr15-39137918-INS-198 | chr15-39137918-INS-198 |
| chr15-39140246-INS-207 | chr15-39140246-INS-207 |
| chr15-39141800-INS-216 | chr15-39141800-INS-216 |
| chr15-39146072-INS-60 | chr15-39146072-INS-62 |
| chr15-39147941-INS-149 | chr15-39147941-INS-149 |
| chr15-39158242-INS-6442 | chr15-39158243-INS-6442 |
| chr15-39170820-INS-80 | chr15-39170820-INS-86 |
| chr15-39183718-INS-68 | chr15-39183718-INS-58 |
| chr15-39186456-INS-179 | chr15-39186456-INS-179 |
| chr15-39195749-INS-56 | chr15-39195749-INS-56 |
| chr15-39197672-INS-66 | chr15-39197672-INS-64 |
| chr15-39198769-INS-98 | chr15-39198769-INS-92 |
| chr15-39218510-INS-7150 | chr15-39218511-INS-7149 |
| chr15-39228703-INS-58 | chr15-39228703-INS-90 |
| chr15-39230030-INS-603 | chr15-39230030-INS-612 |
| chr15-39251126-INS-464 | chr15-39251126-INS-464 |
| chr15-39267000-INS-145 | chr15-39267000-INS-154 |
| chr15-39307738-INS-53 | chr15-39307738-INS-53 |
| chr15-39327139-INS-221 | chr15-39327139-INS-221 |
| chr15-39336303-INS-544 | chr15-39336303-INS-545 |
| chr15-39338518-INS-589 | chr15-39338518-INS-590 |
| chr15-39339447-INS-112 | chr15-39339455-INS-141 |
| chr15-39376309-INS-55 | chr15-39376309-INS-55 |
| chr15-39390327-INS-90 | chr15-39390327-INS-90 |
| chr15-39425038-INS-78 | chr15-39425026-INS-79 |
| chr15-39441303-INS-379 | chr15-39441303-INS-380 |
| chr15-39467502-INS-82 | chr15-39467502-INS-86 |
| chr15-39511408-INS-81 | chr15-39511408-INS-62 |
| chr15-39533502-INS-6745 | chr15-39533503-INS-6750 |

| | |
|---|---|
| chr15-39540651-INS-66 | chr15-39540651-INS-62 |
| chr15-39548209-INS-78 | chr15-39548209-INS-78 |
| chr15-39572088-INS-285 | chr15-39572088-INS-285 |
| chr15-39577203-INS-54 | chr15-39577203-INS-54 |
| chr15-39589988-INS-207 | chr15-39589988-INS-207 |
| chr15-39613837-INS-6390 | chr15-39613838-INS-6389 |
| chr15-39623327-INS-89 | chr15-39623278-INS-98 |
| chr15-39639283-INS-72 | chr15-39639281-INS-74 |
| chr15-39644882-INS-182 | chr15-39644882-INS-189 |
| chr15-39648651-INS-158 | chr15-39648651-INS-162 |
| chr15-39664916-INS-69 | chr15-39664913-INS-69 |
| chr15-39684025-INS-72 | chr15-39684001-INS-72 |
| chr15-39684107-INS-840 | chr15-39684107-INS-840 |
| chr15-39696065-INS-73 | chr15-39696065-INS-82 |
| chr15-39711953-INS-358 | chr15-39711953-INS-358 |
| chr15-39714108-INS-55 | chr15-39714108-INS-55 |
| chr15-39716054-INS-5668 | chr15-39716055-INS-5668 |
| chr15-39723932-INS-63 | chr15-39723932-INS-60 |
| chr15-39756200-INS-750 | chr15-39756200-INS-749 |
| chr15-39759748-INS-1875 | chr15-39759749-INS-1875 |
| chr15-39865650-INS-56 | chr15-39865650-INS-64 |
| chr15-39869988-INS-70 | chr15-39870026-INS-70 |
| chr15-39903978-INS-78 | chr15-39903978-INS-78 |
| chr15-39906312-INS-498 | chr15-39906312-INS-498 |
| chr15-39910547-INS-66 | chr15-39910547-INS-68 |
| chr15-39921564-INS-53 | chr15-39921564-INS-53 |
| chr15-39927922-INS-53 | chr15-39927925-INS-51 |
| chr15-39947104-INS-63 | chr15-39947104-INS-63 |
| chr15-39956269-INS-214 | chr15-39956269-INS-214 |
| chr15-39958688-INS-187 | chr15-39958688-INS-187 |
| chr15-39964833-INS-55 | chr15-39964833-INS-55 |
| chr15-39971656-INS-60 | chr15-39971652-INS-76 |
| chr15-40007091-INS-52 | chr15-40007091-INS-64 |
| chr15-40015131-INS-89 | chr15-40015132-INS-72 |
| chr15-40018799-INS-534 | chr15-40018799-INS-534 |
| chr15-40034674-INS-72 | chr15-40034674-INS-72 |
| chr15-40051414-INS-295 | chr15-40051368-INS-315 |
| chr15-40076996-INS-402 | chr15-40076996-INS-403 |
| chr15-40099199-INS-1600 | chr15-40099200-INS-1600 |
| chr15-40120432-INS-3219 | chr15-40120433-INS-3219 |
| chr15-40127707-INS-106 | chr15-40127707-INS-106 |
| chr15-40154631-INS-1032 | chr15-40154628-INS-1025 |
| chr15-40157584-INS-6652 | chr15-40157585-INS-6652 |
| chr15-40177365-INS-73 | chr15-40177365-INS-73 |
| chr15-40189148-INS-70 | chr15-40189148-INS-70 |
| chr15-40197035-INS-152 | chr15-40197035-INS-144 |
| chr15-40202339-INS-76 | chr15-40202339-INS-75 |
| chr15-40204701-INS-609 | chr15-40204701-INS-609 |
| chr15-40217826-INS-71 | chr15-40217828-INS-69 |
| chr15-40265169-INS-133 | chr15-40265169-INS-133 |
| chr15-40266034-INS-2952 | chr15-40266035-INS-2952 |
| chr15-40286853-INS-54 | chr15-40286853-INS-50 |
| chr15-40311254-INS-201 | chr15-40311211-INS-201 |
| chr15-40312431-INS-95 | chr15-40312468-INS-78 |
| chr15-40322009-INS-565 | chr15-40322009-INS-565 |
| chr15-40328187-INS-75 | chr15-40328187-INS-75 |
| chr15-40403190-INS-340 | chr15-40403190-INS-340 |
| chr15-40419133-INS-237 | chr15-40419128-INS-276 |
| chr15-40434658-INS-6550 | chr15-40434659-INS-6537 |
| chr15-40434917-INS-56 | chr15-40434917-INS-56 |
| chr15-40470601-INS-715 | chr15-40470601-INS-715 |
| chr15-40481601-INS-54 | chr15-40481601-INS-54 |
| chr15-40491551-INS-94 | chr15-40491503-INS-95 |
| chr15-40507693-INS-54 | chr15-40507693-INS-54 |
| chr15-40562406-INS-345 | chr15-40562406-INS-345 |
| chr15-40587980-INS-431 | chr15-40587980-INS-431 |
| chr15-40601047-INS-226 | chr15-40601047-INS-226 |
| chr15-40620244-INS-58 | chr15-40620244-INS-58 |
| chr15-40621809-INS-2631 | chr15-40621810-INS-2631 |
| chr15-40623549-INS-1776 | chr15-40623550-INS-1775 |
| chr15-40629034-INS-54 | chr15-40629034-INS-56 |
| chr15-40639937-INS-65 | chr15-40639937-INS-57 |
| chr15-40643629-INS-105 | chr15-40643604-INS-108 |
| chr15-40644463-INS-6753 | chr15-40644464-INS-6754 |
| chr15-40678138-INS-88 | chr15-40678149-INS-89 |
| chr15-40680992-INS-419 | chr15-40680992-INS-419 |
| chr15-40682120-INS-51 | chr15-40682120-INS-53 |

| | |
|---|---|
| chr15-40750919-INS-68 | chr15-40750919-INS-72 |
| chr15-40784829-INS-110 | chr15-40784831-INS-108 |
| chr15-40811661-INS-113 | chr15-40811661-INS-113 |
| chr15-40816034-INS-68 | chr15-40816034-INS-60 |
| chr15-40858566-INS-742 | chr15-40858566-INS-761 |
| chr15-40900135-INS-84 | chr15-40900135-INS-92 |
| chr15-40921522-INS-161 | chr15-40921522-INS-161 |
| chr15-40925626-INS-122 | chr15-40925634-INS-126 |
| chr15-40956380-INS-69 | chr15-40956344-INS-72 |
| chr15-40969617-INS-401 | chr15-40969617-INS-401 |
| chr15-40986378-INS-52 | chr15-40986373-INS-57 |
| chr15-40989770-INS-58 | chr15-40989771-INS-61 |
| chr15-40994324-INS-259 | chr15-40994324-INS-259 |
| chr15-41014710-INS-68 | chr15-41014710-INS-68 |
| chr15-41036399-INS-193 | chr15-41036399-INS-196 |
| chr15-41055766-INS-6452 | chr15-41055767-INS-6465 |
| chr15-41056675-INS-90 | chr15-41056675-INS-90 |
| chr15-41120053-INS-405 | chr15-41120053-INS-405 |
| chr15-41121472-INS-169 | chr15-41121472-INS-169 |
| chr15-41126069-INS-467 | chr15-41126069-INS-467 |
| chr15-41160721-INS-6246 | chr15-41160722-INS-6246 |
| chr15-41163771-INS-52 | chr15-41163771-INS-52 |
| chr15-41165656-INS-427 | chr15-41165656-INS-427 |
| chr15-41167494-INS-2955 | chr15-41167495-INS-2955 |
| chr15-41171492-INS-56 | chr15-41171492-INS-56 |
| chr15-41184568-INS-90 | chr15-41184568-INS-75 |
| chr15-41188269-INS-120 | chr15-41188261-INS-100 |
| chr15-41190147-INS-64 | chr15-41190147-INS-64 |
| chr15-41211589-INS-551 | chr15-41211589-INS-551 |
| chr15-41216039-INS-97 | chr15-41216024-INS-143 |
| chr15-41218536-INS-107 | chr15-41218536-INS-107 |
| chr15-41221461-INS-108 | chr15-41221461-INS-94 |
| chr15-41254998-INS-91 | chr15-41254978-INS-114 |
| chr15-41257830-INS-1487 | chr15-41257831-INS-1487 |
| chr15-41270971-INS-52 | chr15-41270971-INS-52 |
| chr15-41284366-INS-66 | chr15-41284366-INS-68 |
| chr15-41294703-INS-56 | chr15-41294703-INS-58 |
| chr15-41304758-INS-64 | chr15-41304798-INS-52 |
| chr15-41323703-INS-177 | chr15-41323703-INS-177 |
| chr15-41337265-INS-76 | chr15-41337265-INS-68 |
| chr15-41344612-INS-54 | chr15-41344612-INS-58 |
| chr15-41349828-INS-57 | chr15-41349828-INS-57 |
| chr15-41375101-INS-157 | chr15-41375101-INS-157 |
| chr15-41378009-INS-369 | chr15-41377999-INS-384 |
| chr15-41409881-INS-171 | chr15-41409914-INS-153 |
| chr15-41413719-INS-1863 | chr15-41413720-INS-1863 |
| chr15-41413720-INS-374 | chr15-41413720-INS-374 |
| chr15-41441254-INS-778 | chr15-41441254-INS-778 |
| chr15-41460550-INS-157 | chr15-41460596-INS-150 |
| chr15-41492375-INS-368 | chr15-41492375-INS-364 |
| chr15-41498242-INS-60 | chr15-41498242-INS-56 |
| chr15-41502254-INS-410 | chr15-41502286-INS-376 |
| chr15-41513889-INS-453 | chr15-41513889-INS-452 |
| chr15-41523260-INS-56 | chr15-41523260-INS-58 |
| chr15-41545189-INS-96 | chr15-41545189-INS-96 |
| chr15-41571814-INS-5269 | chr15-41571815-INS-5269 |
| chr15-41583612-INS-232 | chr15-41583612-INS-231 |
| chr15-41594169-INS-70 | chr15-41594169-INS-70 |
| chr15-41636142-INS-179 | chr15-41636152-INS-210 |
| chr15-41672277-INS-71 | chr15-41672277-INS-71 |
| chr15-41762948-INS-53 | chr15-41762948-INS-53 |
| chr15-41763996-INS-83 | chr15-41763996-INS-83 |
| chr15-41775558-INS-54 | chr15-41775558-INS-55 |
| chr15-41796240-INS-424 | chr15-41796240-INS-424 |
| chr15-41804382-INS-2896 | chr15-41804383-INS-2916 |
| chr15-41809951-INS-66 | chr15-41809947-INS-72 |
| chr15-41852299-INS-156 | chr15-41852299-INS-156 |
| chr15-41877479-INS-115 | chr15-41877483-INS-128 |
| chr15-41907686-INS-6242 | chr15-41907687-INS-6242 |
| chr15-41941161-INS-6256 | chr15-41941162-INS-6255 |
| chr15-41953705-INS-90 | chr15-41953705-INS-50 |
| chr15-41957962-INS-1716 | chr15-41957963-INS-1717 |
| chr15-41965546-INS-65 | chr15-41965546-INS-66 |
| chr15-41982366-INS-83 | chr15-41982366-INS-73 |
| chr15-41994295-INS-92 | chr15-41994314-INS-57 |
| chr15-42008001-INS-253 | chr15-42008001-INS-253 |
| chr15-42033240-INS-54 | chr15-42033240-INS-54 |

| | |
|---|---|
| chr15-42076647-INS-2222 | chr15-42076648-INS-2222 |
| chr15-42082385-INS-6164 | chr15-42082386-INS-6165 |
| chr15-42095562-INS-1965 | chr15-42095563-INS-1965 |
| chr15-42106788-INS-6456 | chr15-42106789-INS-6456 |
| chr15-42144595-INS-2757 | chr15-42144596-INS-2757 |
| chr15-42144994-INS-500 | chr15-42144994-INS-498 |
| chr15-42151888-INS-212 | chr15-42151888-INS-213 |
| chr15-42168202-INS-107 | chr15-42168207-INS-105 |
| chr15-42169949-INS-205 | chr15-42169949-INS-209 |
| chr15-42171248-INS-75 | chr15-42171248-INS-74 |
| chr15-42177628-INS-305 | chr15-42177628-INS-305 |
| chr15-42193692-INS-104 | chr15-42193692-INS-106 |
| chr15-42227654-INS-58 | chr15-42227654-INS-58 |
| chr15-42256085-INS-56 | chr15-42256095-INS-52 |
| chr15-42278249-INS-344 | chr15-42278249-INS-344 |
| chr15-42285927-INS-4034 | chr15-42285928-INS-4034 |
| chr15-42298972-INS-249 | chr15-42298972-INS-249 |
| chr15-42353346-INS-192 | chr15-42353346-INS-192 |
| chr15-42424016-INS-59 | chr15-42424016-INS-59 |
| chr15-42442947-INS-1368 | chr15-42442948-INS-1366 |
| chr15-42444508-INS-691 | chr15-42444508-INS-691 |
| chr15-42478893-INS-77 | chr15-42478893-INS-77 |
| chr15-42502907-INS-314 | chr15-42502915-INS-316 |
| chr15-42519756-INS-118 | chr15-42519756-INS-118 |
| chr15-42523373-INS-52 | chr15-42523373-INS-51 |
| chr15-42529542-INS-70 | chr15-42529542-INS-79 |
| chr15-42531335-INS-81 | chr15-42531335-INS-81 |
| chr15-42553823-INS-82 | chr15-42553823-INS-82 |
| chr15-42575462-INS-6662 | chr15-42575463-INS-6663 |
| chr15-42580444-INS-765 | chr15-42580444-INS-766 |
| chr15-42584195-INS-96 | chr15-42584203-INS-52 |
| chr15-42585816-INS-499 | chr15-42585816-INS-499 |
| chr15-42602979-INS-2762 | chr15-42602980-INS-2762 |
| chr15-42607762-INS-7470 | chr15-42607763-INS-7470 |
| chr15-42616876-INS-416 | chr15-42616876-INS-416 |
| chr15-42618310-INS-390 | chr15-42618310-INS-390 |
| chr15-42635092-INS-108 | chr15-42635092-INS-108 |
| chr15-42642852-INS-5339 | chr15-42642853-INS-5339 |
| chr15-42673550-INS-117 | chr15-42673553-INS-112 |
| chr15-42696406-INS-56 | chr15-42696406-INS-52 |
| chr15-42712870-INS-80 | chr15-42712870-INS-100 |
| chr15-42754528-INS-206 | chr15-42754528-INS-206 |
| chr15-42755753-INS-85 | chr15-42755753-INS-114 |
| chr15-42819507-INS-944 | chr15-42819507-INS-944 |
| chr15-42833900-INS-64 | chr15-42833900-INS-64 |
| chr15-42863946-INS-56 | chr15-42863933-INS-76 |
| chr15-42871833-INS-2871 | chr15-42871834-INS-2871 |
| chr15-42889479-INS-70 | chr15-42889453-INS-72 |
| chr15-42895969-INS-55 | chr15-42895969-INS-63 |
| chr15-42932786-INS-140 | chr15-42932786-INS-146 |
| chr15-42948541-INS-82 | chr15-42948541-INS-88 |
| chr15-42950035-INS-148 | chr15-42950029-INS-151 |
| chr15-42956075-INS-60 | chr15-42956075-INS-58 |
| chr15-42972933-INS-67 | chr15-42972933-INS-67 |
| chr15-42975276-INS-111 | chr15-42975276-INS-119 |
| chr15-42987345-INS-393 | chr15-42987345-INS-408 |
| chr15-43028168-INS-82 | chr15-43028166-INS-86 |
| chr15-43030512-INS-63 | chr15-43030512-INS-91 |
| chr15-43050071-INS-178 | chr15-43050090-INS-129 |
| chr15-43100974-INS-6592 | chr15-43100975-INS-6592 |
| chr15-43108299-INS-3921 | chr15-43108300-INS-3921 |
| chr15-43130207-INS-467 | chr15-43130207-INS-467 |
| chr15-43156541-INS-114 | chr15-43156541-INS-114 |
| chr15-43158529-INS-343 | chr15-43158529-INS-343 |
| chr15-43216095-INS-1616 | chr15-43216096-INS-1616 |
| chr15-43237985-INS-6943 | chr15-43237986-INS-6944 |
| chr15-43239484-INS-697 | chr15-43239484-INS-697 |
| chr15-43246915-INS-1115 | chr15-43246916-INS-1088 |
| chr15-43268617-INS-1201 | chr15-43268618-INS-1203 |
| chr15-43275600-INS-91 | chr15-43275600-INS-91 |
| chr15-43283614-INS-6244 | chr15-43283615-INS-6242 |
| chr15-43293453-INS-67 | chr15-43293453-INS-51 |
| chr15-43304329-INS-212 | chr15-43304329-INS-212 |
| chr15-43362286-INS-574 | chr15-43362286-INS-574 |
| chr15-43367272-INS-5392 | chr15-43367273-INS-5392 |
| chr15-43437051-INS-2052 | chr15-43437052-INS-2052 |
| chr15-43449822-INS-194 | chr15-43449822-INS-194 |

| | |
|---|---|
| chr15-43470700-INS-3525 | chr15-43470701-INS-3525 |
| chr15-43506207-INS-62 | chr15-43506207-INS-66 |
| chr15-43566022-INS-5258 | chr15-43566023-INS-5258 |
| chr15-43574487-INS-116 | chr15-43574487-INS-116 |
| chr15-43592463-INS-212 | chr15-43592463-INS-212 |
| chr15-43618262-INS-6593 | chr15-43618263-INS-6593 |
| chr15-43639420-INS-317 | chr15-43639420-INS-317 |
| chr15-43650002-INS-199 | chr15-43650003-INS-202 |
| chr15-43651099-INS-560 | chr15-43651092-INS-570 |
| chr15-43740866-INS-231 | chr15-43740866-INS-231 |
| chr15-43763481-INS-545 | chr15-43763486-INS-541 |
| chr15-43763545-INS-5268 | chr15-43763546-INS-5266 |
| chr15-43769193-INS-1886 | chr15-43769194-INS-1886 |
| chr15-43775210-INS-240 | chr15-43775210-INS-241 |
| chr15-43798009-INS-175 | chr15-43798009-INS-171 |
| chr15-43800431-INS-130 | chr15-43800431-INS-118 |
| chr15-43802599-INS-495 | chr15-43802599-INS-495 |
| chr15-43807595-INS-1880 | chr15-43807596-INS-1882 |
| chr15-43854276-INS-279 | chr15-43854276-INS-282 |
| chr15-43883429-INS-1897 | chr15-43883430-INS-1897 |
| chr15-43936159-INS-241 | chr15-43936159-INS-241 |
| chr15-43936350-INS-1057 | chr15-43936351-INS-1041 |
| chr15-43937888-INS-54 | chr15-43937879-INS-52 |
| chr15-43950182-INS-199 | chr15-43950182-INS-199 |
| chr15-43957078-INS-6709 | chr15-43957079-INS-6710 |
| chr15-43964561-INS-2255 | chr15-43964562-INS-2256 |
| chr15-43982325-INS-1618 | chr15-43982326-INS-1631 |
| chr15-44011333-INS-7364 | chr15-44011334-INS-7364 |
| chr15-44020892-INS-211 | chr15-44020892-INS-211 |
| chr15-44033273-INS-1807 | chr15-44033274-INS-1808 |
| chr15-44059471-INS-61 | chr15-44059473-INS-75 |
| chr15-44065176-INS-74 | chr15-44065176-INS-80 |
| chr15-44074931-INS-55 | chr15-44074931-INS-50 |
| chr15-44096133-INS-402 | chr15-44096133-INS-402 |
| chr15-44113791-INS-199 | chr15-44113792-INS-199 |
| chr15-44147577-INS-736 | chr15-44147577-INS-736 |
| chr15-44166681-INS-199 | chr15-44166681-INS-199 |
| chr15-44198670-INS-234 | chr15-44198670-INS-235 |
| chr15-44207519-INS-69 | chr15-44207519-INS-69 |
| chr15-44211315-INS-80 | chr15-44211315-INS-50 |
| chr15-44231184-INS-3232 | chr15-44231185-INS-3232 |
| chr15-44231185-INS-514 | chr15-44231185-INS-514 |
| chr15-44249624-INS-82 | chr15-44249624-INS-82 |
| chr15-44253825-INS-7098 | chr15-44253826-INS-7098 |
| chr15-44267122-INS-206 | chr15-44267122-INS-206 |
| chr15-44269981-INS-296 | chr15-44269981-INS-296 |
| chr15-44274542-INS-101 | chr15-44274542-INS-77 |
| chr15-44309809-INS-1892 | chr15-44309810-INS-1892 |
| chr15-44321327-INS-125 | chr15-44321327-INS-121 |
| chr15-44362134-INS-513 | chr15-44362134-INS-515 |
| chr15-44385708-INS-56 | chr15-44385708-INS-56 |
| chr15-44396804-INS-247 | chr15-44396804-INS-247 |
| chr15-44419007-INS-76 | chr15-44419007-INS-60 |
| chr15-44529831-INS-194 | chr15-44529831-INS-195 |
| chr15-44639162-INS-1408 | chr15-44639163-INS-1404 |
| chr15-44660275-INS-183 | chr15-44660275-INS-183 |
| chr15-44722251-INS-512 | chr15-44722251-INS-510 |
| chr15-44817320-INS-202 | chr15-44817320-INS-203 |
| chr15-44835702-INS-398 | chr15-44835702-INS-401 |
| chr15-44846069-INS-424 | chr15-44846069-INS-424 |
| chr15-44866871-INS-208 | chr15-44866871-INS-208 |
| chr15-44879652-INS-4528 | chr15-44879653-INS-4528 |
| chr15-44902333-INS-76 | chr15-44902325-INS-98 |
| chr15-44912162-INS-54 | chr15-44912162-INS-60 |
| chr15-44930670-INS-60 | chr15-44930670-INS-60 |
| chr15-44933563-INS-77 | chr15-44933563-INS-77 |
| chr15-44934794-INS-224 | chr15-44934794-INS-224 |
| chr15-44948988-INS-60 | chr15-44948988-INS-52 |
| chr15-44964276-INS-230 | chr15-44964266-INS-237 |
| chr15-44993062-INS-57 | chr15-44993036-INS-57 |
| chr15-45001602-INS-53 | chr15-45001601-INS-57 |
| chr15-45016316-INS-718 | chr15-45016316-INS-719 |
| chr15-45043927-INS-735 | chr15-45043927-INS-737 |
| chr15-45053622-INS-126 | chr15-45053622-INS-136 |
| chr15-45057086-INS-84 | chr15-45057086-INS-80 |
| chr15-45087298-INS-63 | chr15-45087298-INS-56 |
| chr15-45098874-INS-6413 | chr15-45098875-INS-6413 |

| | |
|---|---|
| chr15-45099495-INS-83 | chr15-45099495-INS-94 |
| chr15-45114777-INS-828 | chr15-45114777-INS-828 |
| chr15-45159547-INS-3335 | chr15-45159548-INS-3338 |
| chr15-45165901-INS-68 | chr15-45165901-INS-52 |
| chr15-45172175-INS-198 | chr15-45172167-INS-198 |
| chr15-45180430-INS-87 | chr15-45180430-INS-87 |
| chr15-45240633-INS-168 | chr15-45240633-INS-195 |
| chr15-45246938-INS-6315 | chr15-45246939-INS-6315 |
| chr15-45389138-INS-66 | chr15-45389138-INS-66 |
| chr15-45403759-INS-5626 | chr15-45403760-INS-5626 |
| chr15-45407523-INS-247 | chr15-45407494-INS-247 |
| chr15-45424185-INS-83 | chr15-45424185-INS-83 |
| chr15-45426206-INS-58 | chr15-45426204-INS-58 |
| chr15-45436637-INS-126 | chr15-45436637-INS-106 |
| chr15-45445046-INS-484 | chr15-45445046-INS-484 |
| chr15-45541814-INS-67 | chr15-45541841-INS-66 |
| chr15-45587138-INS-315 | chr15-45587138-INS-315 |
| chr15-45608238-INS-452 | chr15-45608238-INS-452 |
| chr15-45661741-INS-4553 | chr15-45661742-INS-4553 |
| chr15-45662785-INS-121 | chr15-45662786-INS-121 |
| chr15-45664670-INS-76 | chr15-45664654-INS-76 |
| chr15-45674504-INS-614 | chr15-45674504-INS-617 |
| chr15-45702804-INS-151 | chr15-45702804-INS-151 |
| chr15-45711481-INS-247 | chr15-45711481-INS-247 |
| chr15-45717639-INS-772 | chr15-45717639-INS-771 |
| chr15-45726196-INS-81 | chr15-45726196-INS-93 |
| chr15-45744711-INS-7314 | chr15-45744712-INS-7316 |
| chr15-45757588-INS-8853 | chr15-45757589-INS-8854 |
| chr15-45800265-INS-58 | chr15-45800265-INS-60 |
| chr15-45824080-INS-3058 | chr15-45824081-INS-3058 |
| chr15-45825842-INS-1093 | chr15-45825843-INS-1093 |
| chr15-45839941-INS-52 | chr15-45839941-INS-60 |
| chr15-45842041-INS-51 | chr15-45842041-INS-54 |
| chr15-45849638-INS-70 | chr15-45849638-INS-70 |
| chr15-45861289-INS-7089 | chr15-45861290-INS-7089 |
| chr15-45864344-INS-395 | chr15-45864344-INS-396 |
| chr15-45865048-INS-1035 | chr15-45865049-INS-1035 |
| chr15-45877398-INS-82 | chr15-45877394-INS-85 |
| chr15-45885327-INS-962 | chr15-45885327-INS-960 |
| chr15-45908947-INS-1920 | chr15-45908948-INS-1920 |
| chr15-45926429-INS-82 | chr15-45926429-INS-85 |
| chr15-45944719-INS-117 | chr15-45944719-INS-120 |
| chr15-45955877-INS-374 | chr15-45955877-INS-378 |
| chr15-45974883-INS-105 | chr15-45974883-INS-105 |
| chr15-45980671-INS-122 | chr15-45980671-INS-122 |
| chr15-45992381-INS-71 | chr15-45992381-INS-67 |
| chr15-45996180-INS-382 | chr15-45996180-INS-382 |
| chr15-46034823-INS-58 | chr15-46034823-INS-74 |
| chr15-46038805-INS-115 | chr15-46038805-INS-117 |
| chr15-46130638-INS-1969 | chr15-46130639-INS-1969 |
| chr15-46144131-INS-56 | chr15-46144127-INS-52 |
| chr15-46167936-INS-100 | chr15-46167925-INS-102 |
| chr15-46186339-INS-63 | chr15-46186343-INS-66 |
| chr15-46198861-INS-141 | chr15-46198859-INS-145 |
| chr15-46212348-INS-177 | chr15-46212348-INS-175 |
| chr15-46234876-INS-206 | chr15-46234876-INS-205 |
| chr15-46274492-INS-86 | chr15-46274492-INS-80 |
| chr15-46282736-INS-2106 | chr15-46282737-INS-2107 |
| chr15-46283138-INS-385 | chr15-46283138-INS-385 |
| chr15-46289212-INS-120 | chr15-46289220-INS-110 |
| chr15-46315533-INS-848 | chr15-46315533-INS-850 |
| chr15-46316638-INS-64 | chr15-46316638-INS-64 |
| chr15-46319956-INS-53 | chr15-46319991-INS-51 |
| chr15-46343594-INS-2963 | chr15-46343594-INS-2974 |
| chr15-46345803-INS-6144 | chr15-46345804-INS-6144 |
| chr15-46348223-INS-1774 | chr15-46348224-INS-1774 |
| chr15-46350943-INS-3585 | chr15-46350944-INS-3585 |
| chr15-46372610-INS-4285 | chr15-46372611-INS-4285 |
| chr15-46377673-INS-575 | chr15-46377673-INS-575 |
| chr15-46388535-INS-2987 | chr15-46388536-INS-2986 |
| chr15-46437961-INS-169 | chr15-46437961-INS-169 |
| chr15-46467256-INS-6473 | chr15-46467257-INS-6477 |
| chr15-46493415-INS-1022 | chr15-46493416-INS-1020 |
| chr15-46495807-INS-475 | chr15-46495807-INS-475 |
| chr15-46496649-INS-120 | chr15-46496664-INS-92 |
| chr15-46503137-INS-416 | chr15-46503137-INS-416 |
| chr15-46506866-INS-70 | chr15-46506865-INS-70 |

| | |
|---|---|
| chr15-46508197-INS-190 | chr15-46508197-INS-189 |
| chr15-46510400-INS-5265 | chr15-46510401-INS-5265 |
| chr15-46518408-INS-228 | chr15-46518408-INS-230 |
| chr15-46534979-INS-2880 | chr15-46534980-INS-2880 |
| chr15-46536505-INS-180 | chr15-46536505-INS-180 |
| chr15-46552079-INS-125 | chr15-46552079-INS-124 |
| chr15-46553959-INS-2774 | chr15-46553960-INS-2773 |
| chr15-46576367-INS-131 | chr15-46576380-INS-87 |
| chr15-46585062-INS-1280 | chr15-46585063-INS-1262 |
| chr15-46590815-INS-162 | chr15-46590815-INS-162 |
| chr15-46619880-INS-555 | chr15-46619880-INS-555 |
| chr15-46644213-INS-694 | chr15-46644213-INS-694 |
| chr15-46661515-INS-6803 | chr15-46661516-INS-6801 |
| chr15-46672856-INS-56 | chr15-46672856-INS-64 |
| chr15-46691050-INS-166 | chr15-46691050-INS-166 |
| chr15-46694310-INS-821 | chr15-46694310-INS-824 |
| chr15-46715891-INS-507 | chr15-46715891-INS-507 |
| chr15-46718832-INS-95 | chr15-46718832-INS-83 |
| chr15-46730844-INS-219 | chr15-46730844-INS-219 |
| chr15-46780783-INS-127 | chr15-46780783-INS-127 |
| chr15-46781924-INS-96 | chr15-46781924-INS-96 |
| chr15-46789626-INS-320 | chr15-46789626-INS-320 |
| chr15-46803324-INS-3350 | chr15-46803325-INS-3370 |
| chr15-46821005-INS-172 | chr15-46821005-INS-172 |
| chr15-46845391-INS-2954 | chr15-46845392-INS-2954 |
| chr15-46849345-INS-6192 | chr15-46849346-INS-6158 |
| chr15-46858143-INS-52 | chr15-46858143-INS-54 |
| chr15-46881883-INS-84 | chr15-46881883-INS-54 |
| chr15-46886279-INS-53 | chr15-46886257-INS-53 |
| chr15-46941949-INS-366 | chr15-46941949-INS-360 |
| chr15-46953441-INS-580 | chr15-46953441-INS-562 |
| chr15-46968238-INS-77 | chr15-46968218-INS-120 |
| chr15-46995291-INS-7166 | chr15-46995292-INS-7166 |
| chr15-47034769-INS-390 | chr15-47034769-INS-390 |
| chr15-47083377-INS-96 | chr15-47083377-INS-96 |
| chr15-47095643-INS-60 | chr15-47095643-INS-68 |
| chr15-47100095-INS-64 | chr15-47100099-INS-60 |
| chr15-47142787-INS-72 | chr15-47142787-INS-58 |
| chr15-47171635-INS-6724 | chr15-47171636-INS-6727 |
| chr15-47181350-INS-61 | chr15-47181333-INS-82 |
| chr15-47196484-INS-2155 | chr15-47196485-INS-2155 |
| chr15-47248608-INS-901 | chr15-47248608-INS-901 |
| chr15-47268515-INS-455 | chr15-47268515-INS-455 |
| chr15-47338114-INS-57 | chr15-47338114-INS-57 |
| chr15-47359423-INS-2230 | chr15-47359424-INS-2224 |
| chr15-47361547-INS-506 | chr15-47361547-INS-507 |
| chr15-47454848-INS-173 | chr15-47454848-INS-136 |
| chr15-47476287-INS-76 | chr15-47476304-INS-76 |
| chr15-47492400-INS-393 | chr15-47492400-INS-393 |
| chr15-47502367-INS-651 | chr15-47502367-INS-651 |
| chr15-47528242-INS-73 | chr15-47528242-INS-71 |
| chr15-47541264-INS-60 | chr15-47541264-INS-60 |
| chr15-47549106-INS-84 | chr15-47549106-INS-78 |
| chr15-47584832-INS-5542 | chr15-47584833-INS-5542 |
| chr15-47596443-INS-1252 | chr15-47596444-INS-1256 |
| chr15-47645512-INS-52 | chr15-47645512-INS-56 |
| chr15-47720309-INS-657 | chr15-47720309-INS-669 |
| chr15-47748522-INS-51 | chr15-47748538-INS-70 |
| chr15-47751122-INS-120 | chr15-47751122-INS-121 |
| chr15-47759068-INS-78 | chr15-47759068-INS-78 |
| chr15-47808096-INS-7115 | chr15-47808097-INS-7115 |
| chr15-47840645-INS-66 | chr15-47840645-INS-84 |
| chr15-47845863-INS-70 | chr15-47845864-INS-72 |
| chr15-47851562-INS-1633 | chr15-47851563-INS-1631 |
| chr15-47852322-INS-1546 | chr15-47852328-INS-1504 |
| chr15-47858324-INS-86 | chr15-47858324-INS-86 |
| chr15-47859058-INS-6371 | chr15-47859059-INS-6371 |
| chr15-47864726-INS-391 | chr15-47864726-INS-390 |
| chr15-47939056-INS-71 | chr15-47939056-INS-71 |
| chr15-47945270-INS-55 | chr15-47945270-INS-55 |
| chr15-47966585-INS-97 | chr15-47966598-INS-55 |
| chr15-47974793-INS-60 | chr15-47974787-INS-59 |
| chr15-47999556-INS-78 | chr15-47999541-INS-92 |
| chr15-48006040-INS-581 | chr15-48006040-INS-581 |
| chr15-48033803-INS-4832 | chr15-48033804-INS-4832 |
| chr15-48066240-INS-361 | chr15-48066240-INS-360 |
| chr15-48090923-INS-4795 | chr15-48090918-INS-4801 |

| | |
|---|---|
| chr15-48095343-INS-148 | chr15-48095343-INS-140 |
| chr15-48099783-INS-54 | chr15-48099783-INS-54 |
| chr15-48100560-INS-465 | chr15-48100560-INS-465 |
| chr15-48119864-INS-199 | chr15-48119864-INS-201 |
| chr15-48144677-INS-54 | chr15-48144677-INS-54 |
| chr15-48150464-INS-192 | chr15-48150464-INS-184 |
| chr15-48162050-INS-61 | chr15-48162050-INS-62 |
| chr15-48171885-INS-295 | chr15-48171885-INS-295 |
| chr15-48174648-INS-52 | chr15-48174648-INS-72 |
| chr15-48187558-INS-67 | chr15-48187569-INS-83 |
| chr15-48191471-INS-1150 | chr15-48191472-INS-1150 |
| chr15-48270899-INS-57 | chr15-48270899-INS-69 |
| chr15-48285601-INS-750 | chr15-48285602-INS-750 |
| chr15-48338831-INS-104 | chr15-48338831-INS-104 |
| chr15-48343445-INS-68 | chr15-48343445-INS-70 |
| chr15-48347233-INS-2509 | chr15-48347234-INS-2508 |
| chr15-48369524-INS-6246 | chr15-48369525-INS-6246 |
| chr15-48394378-INS-911 | chr15-48394378-INS-911 |
| chr15-48423330-INS-365 | chr15-48423330-INS-365 |
| chr15-48445831-INS-61 | chr15-48445831-INS-69 |
| chr15-48451542-INS-200 | chr15-48451542-INS-200 |
| chr15-48455981-INS-60 | chr15-48455981-INS-60 |
| chr15-48470721-INS-128 | chr15-48470721-INS-171 |
| chr15-48489257-INS-71 | chr15-48489254-INS-59 |
| chr15-48490217-INS-106 | chr15-48490217-INS-106 |
| chr15-48495064-INS-1184 | chr15-48495065-INS-1184 |
| chr15-48557302-INS-2961 | chr15-48557303-INS-2961 |
| chr15-48557303-INS-479 | chr15-48557303-INS-479 |
| chr15-48604179-INS-87 | chr15-48604179-INS-83 |
| chr15-48643524-INS-69 | chr15-48643524-INS-51 |
| chr15-48681290-INS-318 | chr15-48681291-INS-281 |
| chr15-48682479-INS-603 | chr15-48682469-INS-601 |
| chr15-48686443-INS-51 | chr15-48686457-INS-86 |
| chr15-48690802-INS-71 | chr15-48690803-INS-50 |
| chr15-48696857-INS-67 | chr15-48696857-INS-67 |
| chr15-48708696-INS-51 | chr15-48708696-INS-51 |
| chr15-48730511-INS-6411 | chr15-48730512-INS-6412 |
| chr15-48747172-INS-171 | chr15-48747172-INS-186 |
| chr15-48751202-INS-80 | chr15-48751224-INS-64 |
| chr15-48765413-INS-104 | chr15-48765413-INS-104 |
| chr15-48776882-INS-745 | chr15-48776882-INS-745 |
| chr15-48796146-INS-2193 | chr15-48796147-INS-2194 |
| chr15-48798079-INS-102 | chr15-48798082-INS-137 |
| chr15-48804433-INS-69 | chr15-48804425-INS-83 |
| chr15-48823044-INS-70 | chr15-48823044-INS-70 |
| chr15-48850789-INS-62 | chr15-48850789-INS-66 |
| chr15-48886149-INS-1880 | chr15-48886150-INS-1880 |
| chr15-48903672-INS-418 | chr15-48903672-INS-420 |
| chr15-48920426-INS-396 | chr15-48920426-INS-396 |
| chr15-48941592-INS-762 | chr15-48941592-INS-761 |
| chr15-48983226-INS-416 | chr15-48983226-INS-416 |
| chr15-48999657-INS-125 | chr15-48999657-INS-125 |
| chr15-49019122-INS-82 | chr15-49019129-INS-57 |
| chr15-49052623-INS-53 | chr15-49052623-INS-53 |
| chr15-49075717-INS-6403 | chr15-49075718-INS-6403 |
| chr15-49082070-INS-2100 | chr15-49082073-INS-2096 |
| chr15-49099135-INS-1926 | chr15-49099136-INS-1919 |
| chr15-49155261-INS-1540 | chr15-49155262-INS-1539 |
| chr15-49168625-INS-201 | chr15-49168625-INS-201 |
| chr15-49183525-INS-454 | chr15-49183525-INS-454 |
| chr15-49249454-INS-62 | chr15-49249455-INS-68 |
| chr15-49297169-INS-73 | chr15-49297187-INS-74 |
| chr15-49299114-INS-56 | chr15-49299113-INS-54 |
| chr15-49309843-INS-52 | chr15-49309866-INS-52 |
| chr15-49312745-INS-999 | chr15-49312747-INS-1000 |
| chr15-49314295-INS-440 | chr15-49314295-INS-440 |
| chr15-49316976-INS-126 | chr15-49316976-INS-126 |
| chr15-49345048-INS-177 | chr15-49345048-INS-186 |
| chr15-49369712-INS-56 | chr15-49369708-INS-54 |
| chr15-49387663-INS-128 | chr15-49387658-INS-143 |
| chr15-49392421-INS-590 | chr15-49392421-INS-592 |
| chr15-49394943-INS-794 | chr15-49394943-INS-794 |
| chr15-49403011-INS-351 | chr15-49403011-INS-358 |
| chr15-49412340-INS-67 | chr15-49412340-INS-67 |
| chr15-49414360-INS-52 | chr15-49414360-INS-52 |
| chr15-49417463-INS-325 | chr15-49417456-INS-334 |
| chr15-49424851-INS-459 | chr15-49424851-INS-460 |

| | |
|---|---|
| chr15-49458068-INS-136 | chr15-49458095-INS-102 |
| chr15-49475325-INS-525 | chr15-49475325-INS-525 |
| chr15-49476688-INS-363 | chr15-49476688-INS-363 |
| chr15-49497902-INS-187 | chr15-49497902-INS-187 |
| chr15-49510555-INS-6876 | chr15-49510556-INS-6885 |
| chr15-49529316-INS-81 | chr15-49529316-INS-53 |
| chr15-49555885-INS-154 | chr15-49555880-INS-143 |
| chr15-49564299-INS-74 | chr15-49564316-INS-50 |
| chr15-49579703-INS-58 | chr15-49579703-INS-60 |
| chr15-49582489-INS-669 | chr15-49582489-INS-669 |
| chr15-49600504-INS-150 | chr15-49600504-INS-150 |
| chr15-49649030-INS-7109 | chr15-49649031-INS-7109 |
| chr15-49657421-INS-6445 | chr15-49657422-INS-6445 |
| chr15-49671569-INS-58 | chr15-49671569-INS-58 |
| chr15-49677616-INS-362 | chr15-49677616-INS-362 |
| chr15-49689713-INS-4522 | chr15-49689714-INS-4522 |
| chr15-49695528-INS-54 | chr15-49695528-INS-53 |
| chr15-49698709-INS-91 | chr15-49698709-INS-91 |
| chr15-49709271-INS-122 | chr15-49709263-INS-128 |
| chr15-49713230-INS-65 | chr15-49713192-INS-60 |
| chr15-49725979-INS-437 | chr15-49725979-INS-437 |
| chr15-49731613-INS-1805 | chr15-49731614-INS-1805 |
| chr15-49748815-INS-80 | chr15-49748830-INS-51 |
| chr15-49764596-INS-884 | chr15-49764596-INS-884 |
| chr15-49772311-INS-2098 | chr15-49772312-INS-2099 |
| chr15-49793950-INS-655 | chr15-49793950-INS-655 |
| chr15-49795415-INS-6448 | chr15-49795416-INS-6448 |
| chr15-49850068-INS-84 | chr15-49850068-INS-90 |
| chr15-49853660-INS-309 | chr15-49853660-INS-309 |
| chr15-49862148-INS-2776 | chr15-49862149-INS-2776 |
| chr15-49863166-INS-205 | chr15-49863166-INS-205 |
| chr15-49946140-INS-210 | chr15-49946140-INS-210 |
| chr15-49959129-INS-6685 | chr15-49959130-INS-6685 |
| chr15-49970412-INS-52 | chr15-49970412-INS-52 |
| chr15-49972993-INS-51 | chr15-49972993-INS-53 |
| chr15-49993043-INS-212 | chr15-49993043-INS-212 |
| chr15-50037946-INS-148 | chr15-50037946-INS-131 |
| chr15-50039232-INS-246 | chr15-50039232-INS-246 |
| chr15-50040058-INS-423 | chr15-50040058-INS-423 |
| chr15-50056022-INS-11403 | chr15-50056023-INS-11404 |
| chr15-50096038-INS-112 | chr15-50096034-INS-151 |
| chr15-50125787-INS-106 | chr15-50125787-INS-72 |
| chr15-50182875-INS-177 | chr15-50182875-INS-177 |
| chr15-50189863-INS-6754 | chr15-50189864-INS-6754 |
| chr15-50200886-INS-6800 | chr15-50200887-INS-6800 |
| chr15-50213790-INS-204 | chr15-50213791-INS-204 |
| chr15-50248267-INS-58 | chr15-50248270-INS-64 |
| chr15-50256493-INS-555 | chr15-50256493-INS-555 |
| chr15-50270860-INS-52 | chr15-50270860-INS-54 |
| chr15-50279443-INS-729 | chr15-50279443-INS-729 |
| chr15-50295387-INS-209 | chr15-50295387-INS-209 |
| chr15-50303390-INS-5083 | chr15-50303391-INS-5083 |
| chr15-50308409-INS-96 | chr15-50308405-INS-120 |
| chr15-50312618-INS-81 | chr15-50312626-INS-90 |
| chr15-50331816-INS-170 | chr15-50331816-INS-170 |
| chr15-50332340-INS-2408 | chr15-50332341-INS-2410 |
| chr15-50342671-INS-4336 | chr15-50342672-INS-4336 |
| chr15-50348434-INS-7107 | chr15-50348435-INS-7107 |
| chr15-50363372-INS-562 | chr15-50363373-INS-562 |
| chr15-50372566-INS-697 | chr15-50372566-INS-698 |
| chr15-50410501-INS-110 | chr15-50410501-INS-108 |
| chr15-50518413-INS-6706 | chr15-50518414-INS-6708 |
| chr15-50553796-INS-6401 | chr15-50553797-INS-6400 |
| chr15-50598641-INS-725 | chr15-50598641-INS-709 |
| chr15-50606866-INS-76 | chr15-50606866-INS-108 |
| chr15-50635923-INS-55 | chr15-50635926-INS-68 |
| chr15-50638709-INS-349 | chr15-50638709-INS-349 |
| chr15-50735787-INS-180 | chr15-50735787-INS-180 |
| chr15-50759852-INS-60 | chr15-50759819-INS-88 |
| chr15-50777942-INS-202 | chr15-50777942-INS-202 |
| chr15-50793797-INS-55 | chr15-50793797-INS-55 |
| chr15-50797348-INS-6709 | chr15-50797349-INS-6718 |
| chr15-50800000-INS-106 | chr15-50800003-INS-109 |
| chr15-50800699-INS-87 | chr15-50800698-INS-84 |
| chr15-50806360-INS-1498 | chr15-50806361-INS-1523 |
| chr15-50810940-INS-72 | chr15-50810955-INS-62 |
| chr15-50814549-INS-216 | chr15-50814549-INS-216 |

| | |
|---|---|
| chr15-50847446-INS-355 | chr15-50847446-INS-383 |
| chr15-50881220-INS-545 | chr15-50881220-INS-545 |
| chr15-50918443-INS-312 | chr15-50918443-INS-312 |
| chr15-50923410-INS-465 | chr15-50923410-INS-465 |
| chr15-50989963-INS-360 | chr15-50989963-INS-360 |
| chr15-51031561-INS-52 | chr15-51031562-INS-52 |
| chr15-51093885-INS-60 | chr15-51093885-INS-60 |
| chr15-51125051-INS-216 | chr15-51125051-INS-216 |
| chr15-51126107-INS-6681 | chr15-51126108-INS-6682 |
| chr15-51140814-INS-9074 | chr15-51140815-INS-9078 |
| chr15-51183221-INS-66 | chr15-51183221-INS-68 |
| chr15-51191999-INS-249 | chr15-51191999-INS-247 |
| chr15-51211408-INS-270 | chr15-51211408-INS-274 |
| chr15-51222613-INS-6739 | chr15-51222614-INS-6739 |
| chr15-51235951-INS-128 | chr15-51235951-INS-125 |
| chr15-51258391-INS-150 | chr15-51258391-INS-170 |
| chr15-51271359-INS-54 | chr15-51271359-INS-52 |
| chr15-51276113-INS-7113 | chr15-51276114-INS-7113 |
| chr15-51297742-INS-5228 | chr15-51297743-INS-5225 |
| chr15-51313228-INS-3898 | chr15-51313229-INS-3898 |
| chr15-51313917-INS-560 | chr15-51313917-INS-560 |
| chr15-51331517-INS-69 | chr15-51331517-INS-74 |
| chr15-51332886-INS-3237 | chr15-51332887-INS-3237 |
| chr15-51372619-INS-86 | chr15-51372619-INS-86 |
| chr15-51382111-INS-56 | chr15-51382111-INS-58 |
| chr15-51405888-INS-1925 | chr15-51405889-INS-1925 |
| chr15-51431272-INS-6487 | chr15-51431273-INS-6487 |
| chr15-51445123-INS-89 | chr15-51445123-INS-89 |
| chr15-51491959-INS-388 | chr15-51491959-INS-389 |
| chr15-51505039-INS-96 | chr15-51505060-INS-103 |
| chr15-51509505-INS-1878 | chr15-51509506-INS-1878 |
| chr15-51519015-INS-459 | chr15-51519016-INS-459 |
| chr15-51525880-INS-136 | chr15-51525880-INS-136 |
| chr15-51543770-INS-300 | chr15-51543771-INS-297 |
| chr15-51550362-INS-6376 | chr15-51550363-INS-6376 |
| chr15-51556023-INS-6506 | chr15-51556024-INS-6507 |
| chr15-51560768-INS-262 | chr15-51560765-INS-259 |
| chr15-51572272-INS-62 | chr15-51572272-INS-50 |
| chr15-51602254-INS-78 | chr15-51602254-INS-78 |
| chr15-51607875-INS-69 | chr15-51607875-INS-84 |
| chr15-51626723-INS-52 | chr15-51626701-INS-50 |
| chr15-51645751-INS-60 | chr15-51645751-INS-60 |
| chr15-51653774-INS-80 | chr15-51653774-INS-80 |
| chr15-51686743-INS-68 | chr15-51686743-INS-68 |
| chr15-51745281-INS-230 | chr15-51745281-INS-229 |
| chr15-51746533-INS-84 | chr15-51746523-INS-84 |
| chr15-51749594-INS-126 | chr15-51749594-INS-126 |
| chr15-51755518-INS-223 | chr15-51755518-INS-222 |
| chr15-51758455-INS-202 | chr15-51758455-INS-200 |
| chr15-51762341-INS-711 | chr15-51762338-INS-690 |
| chr15-51765246-INS-404 | chr15-51765246-INS-404 |
| chr15-51768064-INS-887 | chr15-51768064-INS-885 |
| chr15-51852883-INS-52 | chr15-51852883-INS-58 |
| chr15-51865964-INS-5042 | chr15-51865965-INS-5042 |
| chr15-51871492-INS-174 | chr15-51871492-INS-175 |
| chr15-51871819-INS-5547 | chr15-51871820-INS-5547 |
| chr15-51874134-INS-208 | chr15-51874134-INS-209 |
| chr15-51880283-INS-9129 | chr15-51880284-INS-9129 |
| chr15-51889022-INS-899 | chr15-51889022-INS-899 |
| chr15-51917049-INS-217 | chr15-51917049-INS-217 |
| chr15-51936082-INS-971 | chr15-51936082-INS-971 |
| chr15-51940027-INS-552 | chr15-51940027-INS-553 |
| chr15-51941213-INS-197 | chr15-51941213-INS-197 |
| chr15-51948447-INS-115 | chr15-51948443-INS-74 |
| chr15-51962135-INS-207 | chr15-51962135-INS-207 |
| chr15-51980797-INS-182 | chr15-51980797-INS-182 |
| chr15-51997569-INS-51 | chr15-51997523-INS-64 |
| chr15-52022295-INS-52 | chr15-52022295-INS-52 |
| chr15-52042614-INS-245 | chr15-52042614-INS-245 |
| chr15-52045290-INS-11531 | chr15-52045291-INS-11535 |
| chr15-52066187-INS-53 | chr15-52066187-INS-51 |
| chr15-52079915-INS-55 | chr15-52079965-INS-55 |
| chr15-52090121-INS-55 | chr15-52090121-INS-66 |
| chr15-52091358-INS-120 | chr15-52091358-INS-118 |
| chr15-52106747-INS-172 | chr15-52106750-INS-169 |
| chr15-52117031-INS-2187 | chr15-52117032-INS-2187 |
| chr15-52120963-INS-163 | chr15-52120963-INS-163 |

| | |
|---|---|
| chr15-52130989-INS-4284 | chr15-52130990-INS-4284 |
| chr15-52147419-INS-965 | chr15-52147419-INS-967 |
| chr15-52178975-INS-1213 | chr15-52178976-INS-1213 |
| chr15-52194497-INS-7111 | chr15-52194498-INS-7111 |
| chr15-52195239-INS-107 | chr15-52195239-INS-105 |
| chr15-52220541-INS-74 | chr15-52220541-INS-72 |
| chr15-52234047-INS-53 | chr15-52234058-INS-81 |
| chr15-52238500-INS-95 | chr15-52238500-INS-95 |
| chr15-52240527-INS-533 | chr15-52240527-INS-533 |
| chr15-52267158-INS-66 | chr15-52267158-INS-68 |
| chr15-52277644-INS-206 | chr15-52277644-INS-207 |
| chr15-52306288-INS-52 | chr15-52306288-INS-52 |
| chr15-52313796-INS-57 | chr15-52313796-INS-57 |
| chr15-52322714-INS-116 | chr15-52322720-INS-72 |
| chr15-52355058-INS-1881 | chr15-52355059-INS-1890 |
| chr15-52358543-INS-74 | chr15-52358520-INS-68 |
| chr15-52385342-INS-1206 | chr15-52385343-INS-1198 |
| chr15-52405355-INS-53 | chr15-52405355-INS-54 |
| chr15-52434110-INS-69 | chr15-52434110-INS-67 |
| chr15-52444879-INS-185 | chr15-52444879-INS-186 |
| chr15-52445994-INS-666 | chr15-52445994-INS-667 |
| chr15-52476490-INS-54 | chr15-52476490-INS-54 |
| chr15-52541013-INS-70 | chr15-52541013-INS-68 |
| chr15-52545858-INS-3055 | chr15-52545859-INS-3057 |
| chr15-52579829-INS-81 | chr15-52579825-INS-64 |
| chr15-52587030-INS-64 | chr15-52587030-INS-64 |
| chr15-52628163-INS-54 | chr15-52628163-INS-50 |
| chr15-52646874-INS-6797 | chr15-52646875-INS-6797 |
| chr15-52687887-INS-1649 | chr15-52687896-INS-1641 |
| chr15-52693694-INS-2343 | chr15-52693695-INS-2343 |
| chr15-52751530-INS-7134 | chr15-52751531-INS-7134 |
| chr15-52776246-INS-97 | chr15-52776246-INS-97 |
| chr15-52790755-INS-100 | chr15-52790755-INS-108 |
| chr15-52831815-INS-947 | chr15-52831815-INS-948 |
| chr15-52838600-INS-62 | chr15-52838592-INS-50 |
| chr15-52859511-INS-56 | chr15-52859511-INS-72 |
| chr15-52871508-INS-88 | chr15-52871508-INS-88 |
| chr15-52873581-INS-77 | chr15-52873581-INS-80 |
| chr15-52876868-INS-190 | chr15-52876842-INS-221 |
| chr15-52880612-INS-74 | chr15-52880612-INS-74 |
| chr15-52910471-INS-131 | chr15-52910473-INS-132 |
| chr15-52943546-INS-1889 | chr15-52943547-INS-1889 |
| chr15-52999058-INS-213 | chr15-52999058-INS-213 |
| chr15-53004539-INS-60 | chr15-53004539-INS-60 |
| chr15-53007551-INS-202 | chr15-53007551-INS-202 |
| chr15-53041489-INS-206 | chr15-53041489-INS-206 |
| chr15-53091168-INS-99 | chr15-53091168-INS-78 |
| chr15-53157233-INS-54 | chr15-53157233-INS-54 |
| chr15-53177719-INS-71 | chr15-53177719-INS-76 |
| chr15-53193679-INS-87 | chr15-53193679-INS-83 |
| chr15-53194649-INS-2473 | chr15-53194650-INS-2469 |
| chr15-53211772-INS-206 | chr15-53211772-INS-206 |
| chr15-53237441-INS-791 | chr15-53237441-INS-780 |
| chr15-53255657-INS-802 | chr15-53255657-INS-802 |
| chr15-53256936-INS-241 | chr15-53256936-INS-241 |
| chr15-53259403-INS-54 | chr15-53259403-INS-54 |
| chr15-53262431-INS-311 | chr15-53262431-INS-311 |
| chr15-53289754-INS-60 | chr15-53289754-INS-66 |
| chr15-53304528-INS-60 | chr15-53304528-INS-72 |
| chr15-53304801-INS-788 | chr15-53304801-INS-788 |
| chr15-53314585-INS-1744 | chr15-53314586-INS-1751 |
| chr15-53315970-INS-56 | chr15-53315970-INS-60 |
| chr15-53326009-INS-58 | chr15-53326009-INS-60 |
| chr15-53363401-INS-272 | chr15-53363401-INS-272 |
| chr15-53367719-INS-52 | chr15-53367719-INS-52 |
| chr15-53392500-INS-239 | chr15-53392500-INS-222 |
| chr15-53463847-INS-90 | chr15-53463847-INS-90 |
| chr15-53482337-INS-127 | chr15-53482337-INS-129 |
| chr15-53496257-INS-506 | chr15-53496257-INS-509 |
| chr15-53505630-INS-682 | chr15-53505630-INS-682 |
| chr15-53510293-INS-551 | chr15-53510293-INS-546 |
| chr15-53522109-INS-504 | chr15-53522109-INS-504 |
| chr15-53534751-INS-1890 | chr15-53534752-INS-1890 |
| chr15-53564984-INS-7118 | chr15-53564985-INS-7118 |
| chr15-53584857-INS-65 | chr15-53584857-INS-65 |
| chr15-53594783-INS-271 | chr15-53594783-INS-272 |
| chr15-53623514-INS-388 | chr15-53623514-INS-389 |

| | |
|---|---|
| chr15-53634200-INS-930 | chr15-53634200-INS-930 |
| chr15-53638064-INS-953 | chr15-53638064-INS-953 |
| chr15-53650030-INS-2269 | chr15-53650031-INS-2269 |
| chr15-53670808-INS-126 | chr15-53670808-INS-126 |
| chr15-53750062-INS-1684 | chr15-53750063-INS-1684 |
| chr15-53764442-INS-681 | chr15-53764442-INS-666 |
| chr15-53771833-INS-95 | chr15-53771833-INS-95 |
| chr15-53797511-INS-293 | chr15-53797511-INS-294 |
| chr15-53799850-INS-52 | chr15-53799850-INS-54 |
| chr15-53819546-INS-6648 | chr15-53819547-INS-6649 |
| chr15-53847113-INS-441 | chr15-53847113-INS-441 |
| chr15-53851658-INS-54 | chr15-53851672-INS-82 |
| chr15-53852655-INS-66 | chr15-53852645-INS-72 |
| chr15-53856855-INS-122 | chr15-53856855-INS-164 |
| chr15-53870349-INS-324 | chr15-53870349-INS-304 |
| chr15-53874979-INS-58 | chr15-53874979-INS-58 |
| chr15-53883388-INS-67 | chr15-53883388-INS-53 |
| chr15-53900944-INS-416 | chr15-53900944-INS-416 |
| chr15-53908211-INS-52 | chr15-53908211-INS-52 |
| chr15-53920873-INS-926 | chr15-53920873-INS-926 |
| chr15-53925210-INS-203 | chr15-53925210-INS-205 |
| chr15-53928385-INS-2737 | chr15-53928371-INS-2726 |
| chr15-53941400-INS-6538 | chr15-53941401-INS-6538 |
| chr15-53943772-INS-1889 | chr15-53943773-INS-1889 |
| chr15-53976417-INS-4178 | chr15-53976418-INS-4178 |
| chr15-53983346-INS-578 | chr15-53983346-INS-579 |
| chr15-53991040-INS-6261 | chr15-53991041-INS-6261 |
| chr15-54034067-INS-50 | chr15-54034065-INS-62 |
| chr15-54034898-INS-63 | chr15-54034898-INS-59 |
| chr15-54078911-INS-60 | chr15-54078911-INS-62 |
| chr15-54097555-INS-217 | chr15-54097555-INS-217 |
| chr15-54100942-INS-425 | chr15-54100942-INS-425 |
| chr15-54113560-INS-211 | chr15-54113560-INS-211 |
| chr15-54143700-INS-60 | chr15-54143700-INS-60 |
| chr15-54165652-INS-1500 | chr15-54165653-INS-1503 |
| chr15-54186546-INS-526 | chr15-54186546-INS-526 |
| chr15-54208098-INS-1894 | chr15-54208099-INS-1894 |
| chr15-54220329-INS-539 | chr15-54220329-INS-540 |
| chr15-54254516-INS-530 | chr15-54254516-INS-530 |
| chr15-54280359-INS-99 | chr15-54280359-INS-88 |
| chr15-54284161-INS-90 | chr15-54284162-INS-60 |
| chr15-54289119-INS-1061 | chr15-54289120-INS-1081 |
| chr15-54333225-INS-77 | chr15-54333225-INS-77 |
| chr15-54336384-INS-183 | chr15-54336384-INS-183 |
| chr15-54346688-INS-7074 | chr15-54346689-INS-7074 |
| chr15-54348478-INS-2157 | chr15-54348479-INS-2157 |
| chr15-54353313-INS-179 | chr15-54353313-INS-179 |
| chr15-54372340-INS-60 | chr15-54372340-INS-60 |
| chr15-54381560-INS-491 | chr15-54381560-INS-491 |
| chr15-54382773-INS-196 | chr15-54382773-INS-196 |
| chr15-54396552-INS-2829 | chr15-54396553-INS-2829 |
| chr15-54419964-INS-6251 | chr15-54419965-INS-6251 |
| chr15-54453234-INS-167 | chr15-54453234-INS-167 |
| chr15-54473192-INS-53 | chr15-54473193-INS-56 |
| chr15-54477431-INS-123 | chr15-54477431-INS-145 |
| chr15-54493220-INS-68 | chr15-54493208-INS-75 |
| chr15-54528117-INS-299 | chr15-54528124-INS-306 |
| chr15-54528390-INS-1750 | chr15-54528391-INS-1771 |
| chr15-54544535-INS-62 | chr15-54544535-INS-62 |
| chr15-54554750-INS-1239 | chr15-54554751-INS-1239 |
| chr15-54595972-INS-126 | chr15-54595952-INS-138 |
| chr15-54624333-INS-6112 | chr15-54624334-INS-6112 |
| chr15-54660875-INS-647 | chr15-54660875-INS-648 |
| chr15-54663911-INS-230 | chr15-54663911-INS-215 |
| chr15-54664903-INS-65 | chr15-54664903-INS-67 |
| chr15-54667047-INS-51 | chr15-54667047-INS-72 |
| chr15-54676093-INS-72 | chr15-54676089-INS-69 |
| chr15-54679378-INS-5212 | chr15-54679379-INS-5212 |
| chr15-54690022-INS-94 | chr15-54690038-INS-103 |
| chr15-54700767-INS-970 | chr15-54700767-INS-970 |
| chr15-54732581-INS-257 | chr15-54732581-INS-258 |
| chr15-54751830-INS-60 | chr15-54751830-INS-60 |
| chr15-54802495-INS-221 | chr15-54802495-INS-220 |
| chr15-54856696-INS-208 | chr15-54856696-INS-206 |
| chr15-54865144-INS-364 | chr15-54865144-INS-367 |
| chr15-54868979-INS-78 | chr15-54868975-INS-57 |
| chr15-54914430-INS-182 | chr15-54914450-INS-139 |

| | |
|---|---|
| chr15-54916046-INS-211 | chr15-54916066-INS-224 |
| chr15-54922120-INS-219 | chr15-54922120-INS-218 |
| chr15-54950303-INS-213 | chr15-54950309-INS-220 |
| chr15-54955105-INS-52 | chr15-54955105-INS-64 |
| chr15-54960673-INS-247 | chr15-54960673-INS-251 |
| chr15-54964670-INS-61 | chr15-54964670-INS-64 |
| chr15-54966579-INS-1996 | chr15-54966580-INS-1996 |
| chr15-54982821-INS-162 | chr15-54982821-INS-162 |
| chr15-54992046-INS-204 | chr15-54992039-INS-204 |
| chr15-54999675-INS-84 | chr15-54999686-INS-106 |
| chr15-55011391-INS-93 | chr15-55011391-INS-93 |
| chr15-55017068-INS-200 | chr15-55017068-INS-200 |
| chr15-55017684-INS-1890 | chr15-55017685-INS-1890 |
| chr15-55034921-INS-100 | chr15-55034925-INS-93 |
| chr15-55043679-INS-84 | chr15-55043699-INS-88 |
| chr15-55051593-INS-62 | chr15-55051594-INS-66 |
| chr15-55053187-INS-195 | chr15-55053174-INS-199 |
| chr15-55058063-INS-94 | chr15-55058063-INS-94 |
| chr15-55098807-INS-148 | chr15-55098807-INS-160 |
| chr15-55105815-INS-210 | chr15-55105815-INS-208 |
| chr15-55108370-INS-78 | chr15-55108370-INS-78 |
| chr15-55119103-INS-166 | chr15-55119103-INS-163 |
| chr15-55132011-INS-114 | chr15-55132011-INS-114 |
| chr15-55132110-INS-549 | chr15-55132110-INS-549 |
| chr15-55139025-INS-87 | chr15-55139016-INS-68 |
| chr15-55141209-INS-196 | chr15-55141209-INS-195 |
| chr15-55143270-INS-53 | chr15-55143270-INS-57 |
| chr15-55144845-INS-92 | chr15-55144845-INS-92 |
| chr15-55148606-INS-53 | chr15-55148606-INS-55 |
| chr15-55149424-INS-512 | chr15-55149424-INS-512 |
| chr15-55153488-INS-112 | chr15-55153488-INS-114 |
| chr15-55154914-INS-120 | chr15-55154914-INS-120 |
| chr15-55156977-INS-215 | chr15-55156975-INS-196 |
| chr15-55162425-INS-498 | chr15-55162425-INS-498 |
| chr15-55168228-INS-636 | chr15-55168228-INS-649 |
| chr15-55190947-INS-193 | chr15-55190947-INS-193 |
| chr15-55206301-INS-211 | chr15-55206304-INS-211 |
| chr15-55228427-INS-57 | chr15-55228427-INS-61 |
| chr15-55236475-INS-62 | chr15-55236475-INS-71 |
| chr15-55288198-INS-201 | chr15-55288198-INS-199 |
| chr15-55291463-INS-134 | chr15-55291463-INS-134 |
| chr15-55295201-INS-167 | chr15-55295201-INS-165 |
| chr15-55311289-INS-95 | chr15-55311289-INS-100 |
| chr15-55318179-INS-105 | chr15-55318179-INS-105 |
| chr15-55327863-INS-244 | chr15-55327863-INS-244 |
| chr15-55342268-INS-52 | chr15-55342272-INS-56 |
| chr15-55366951-INS-210 | chr15-55366951-INS-210 |
| chr15-55372778-INS-65 | chr15-55372778-INS-65 |
| chr15-55382395-INS-614 | chr15-55382395-INS-614 |
| chr15-55384350-INS-125 | chr15-55384350-INS-125 |
| chr15-55411451-INS-1863 | chr15-55411452-INS-1863 |
| chr15-55414046-INS-53 | chr15-55414046-INS-91 |
| chr15-55437830-INS-520 | chr15-55437830-INS-519 |
| chr15-55445079-INS-76 | chr15-55445079-INS-76 |
| chr15-55487097-INS-569 | chr15-55487097-INS-569 |
| chr15-55491253-INS-430 | chr15-55491253-INS-430 |
| chr15-55513156-INS-100 | chr15-55513176-INS-118 |
| chr15-55525152-INS-433 | chr15-55525152-INS-433 |
| chr15-55558813-INS-3546 | chr15-55558814-INS-3546 |
| chr15-55561272-INS-124 | chr15-55561272-INS-124 |
| chr15-55561426-INS-803 | chr15-55561426-INS-803 |
| chr15-55562839-INS-1522 | chr15-55562840-INS-1524 |
| chr15-55563363-INS-2727 | chr15-55563364-INS-2729 |
| chr15-55578102-INS-114 | chr15-55578093-INS-126 |
| chr15-55604273-INS-948 | chr15-55604273-INS-948 |
| chr15-55646089-INS-513 | chr15-55646089-INS-528 |
| chr15-55649380-INS-463 | chr15-55649380-INS-463 |
| chr15-55671317-INS-55 | chr15-55671305-INS-81 |
| chr15-55697800-INS-72 | chr15-55697800-INS-72 |
| chr15-55703210-INS-96 | chr15-55703210-INS-96 |
| chr15-55762097-INS-186 | chr15-55762097-INS-186 |
| chr15-55766989-INS-54 | chr15-55766989-INS-54 |
| chr15-55770756-INS-132 | chr15-55770756-INS-83 |
| chr15-55775994-INS-7135 | chr15-55775995-INS-7135 |
| chr15-55793261-INS-6591 | chr15-55793262-INS-6591 |
| chr15-55812305-INS-64 | chr15-55812305-INS-66 |
| chr15-55822781-INS-57 | chr15-55822754-INS-63 |

| | |
|---|---|
| chr15-55827961-INS-80 | chr15-55827961-INS-74 |
| chr15-55838069-INS-53 | chr15-55838053-INS-64 |
| chr15-55842903-INS-411 | chr15-55842903-INS-412 |
| chr15-55851769-INS-82 | chr15-55851769-INS-130 |
| chr15-55873607-INS-52 | chr15-55873608-INS-70 |
| chr15-55890574-INS-114 | chr15-55890574-INS-99 |
| chr15-55905871-INS-1256 | chr15-55905872-INS-1256 |
| chr15-55910172-INS-266 | chr15-55910172-INS-266 |
| chr15-55914954-INS-62 | chr15-55914910-INS-88 |
| chr15-55921062-INS-504 | chr15-55921062-INS-504 |
| chr15-55927437-INS-183 | chr15-55927437-INS-183 |
| chr15-55929788-INS-56 | chr15-55929782-INS-52 |
| chr15-55972568-INS-155 | chr15-55972568-INS-156 |
| chr15-55995790-INS-56 | chr15-55995789-INS-72 |
| chr15-56001760-INS-210 | chr15-56001760-INS-211 |
| chr15-56007961-INS-431 | chr15-56007970-INS-430 |
| chr15-56025784-INS-250 | chr15-56025784-INS-250 |
| chr15-56040397-INS-54 | chr15-56040397-INS-54 |
| chr15-56076813-INS-6401 | chr15-56076814-INS-6401 |
| chr15-56079210-INS-2021 | chr15-56079211-INS-2041 |
| chr15-56098457-INS-100 | chr15-56098457-INS-100 |
| chr15-56160016-INS-176 | chr15-56160016-INS-176 |
| chr15-56185108-INS-51 | chr15-56185108-INS-72 |
| chr15-56213369-INS-70 | chr15-56213369-INS-70 |
| chr15-56239078-INS-1046 | chr15-56239079-INS-1047 |
| chr15-56284827-INS-3044 | chr15-56284828-INS-3043 |
| chr15-56287945-INS-64 | chr15-56287945-INS-60 |
| chr15-56326579-INS-183 | chr15-56326579-INS-183 |
| chr15-56340063-INS-3137 | chr15-56340064-INS-3136 |
| chr15-56345038-INS-7086 | chr15-56345039-INS-7086 |
| chr15-56351821-INS-7140 | chr15-56351822-INS-7140 |
| chr15-56361326-INS-294 | chr15-56361326-INS-294 |
| chr15-56473731-INS-168 | chr15-56473718-INS-144 |
| chr15-56508947-INS-77 | chr15-56508982-INS-70 |
| chr15-56550848-INS-429 | chr15-56550848-INS-426 |
| chr15-56572887-INS-134 | chr15-56572887-INS-134 |
| chr15-56634049-INS-105 | chr15-56634049-INS-112 |
| chr15-56646026-INS-76 | chr15-56646026-INS-54 |
| chr15-56672696-INS-83 | chr15-56672696-INS-83 |
| chr15-56691274-INS-750 | chr15-56691274-INS-750 |
| chr15-56694155-INS-124 | chr15-56694205-INS-111 |
| chr15-56723740-INS-72 | chr15-56723725-INS-64 |
| chr15-56773221-INS-120 | chr15-56773221-INS-120 |
| chr15-56777618-INS-399 | chr15-56777618-INS-399 |
| chr15-56852763-INS-200 | chr15-56852763-INS-200 |
| chr15-56854528-INS-214 | chr15-56854528-INS-214 |
| chr15-56854719-INS-1729 | chr15-56854720-INS-1729 |
| chr15-56869090-INS-85 | chr15-56869090-INS-50 |
| chr15-56903092-INS-76 | chr15-56903092-INS-50 |
| chr15-56905543-INS-3062 | chr15-56905544-INS-3076 |
| chr15-56910827-INS-68 | chr15-56910827-INS-68 |
| chr15-56927926-INS-8457 | chr15-56927927-INS-8457 |
| chr15-56950667-INS-6474 | chr15-56950668-INS-6471 |
| chr15-56952979-INS-115 | chr15-56952979-INS-115 |
| chr15-56955780-INS-3619 | chr15-56955781-INS-3602 |
| chr15-57003191-INS-1207 | chr15-57003192-INS-1207 |
| chr15-57020427-INS-116 | chr15-57020427-INS-116 |
| chr15-57024485-INS-365 | chr15-57024485-INS-366 |
| chr15-57066252-INS-74 | chr15-57066252-INS-73 |
| chr15-57078765-INS-55 | chr15-57078765-INS-78 |
| chr15-57107027-INS-6782 | chr15-57107028-INS-6782 |
| chr15-57115333-INS-75 | chr15-57115300-INS-96 |
| chr15-57138414-INS-56 | chr15-57138414-INS-54 |
| chr15-57184095-INS-58 | chr15-57184095-INS-50 |
| chr15-57197941-INS-157 | chr15-57197941-INS-157 |
| chr15-57202724-INS-208 | chr15-57202724-INS-208 |
| chr15-57210257-INS-194 | chr15-57210257-INS-195 |
| chr15-57230655-INS-6455 | chr15-57230656-INS-6456 |
| chr15-57250210-INS-52 | chr15-57250210-INS-58 |
| chr15-57253894-INS-2749 | chr15-57253895-INS-2748 |
| chr15-57255655-INS-52 | chr15-57255655-INS-54 |
| chr15-57264356-INS-345 | chr15-57264356-INS-346 |
| chr15-57275704-INS-1476 | chr15-57275705-INS-1476 |
| chr15-57292592-INS-66 | chr15-57292592-INS-66 |
| chr15-57314949-INS-7029 | chr15-57314950-INS-7031 |
| chr15-57316408-INS-52 | chr15-57316408-INS-57 |
| chr15-57347529-INS-56 | chr15-57347529-INS-56 |

| | |
|---|---|
| chr15-57369906-INS-59 | chr15-57369904-INS-56 |
| chr15-57374783-INS-539 | chr15-57374783-INS-539 |
| chr15-57376780-INS-356 | chr15-57376780-INS-356 |
| chr15-57394170-INS-938 | chr15-57394170-INS-939 |
| chr15-57399507-INS-792 | chr15-57399507-INS-793 |
| chr15-57407063-INS-70 | chr15-57407063-INS-64 |
| chr15-57413321-INS-7159 | chr15-57413322-INS-7160 |
| chr15-57415182-INS-51 | chr15-57415182-INS-78 |
| chr15-57418753-INS-2045 | chr15-57418763-INS-2047 |
| chr15-57439810-INS-399 | chr15-57439810-INS-399 |
| chr15-57459666-INS-59 | chr15-57459666-INS-63 |
| chr15-57464592-INS-999 | chr15-57464592-INS-1000 |
| chr15-57465237-INS-54 | chr15-57465237-INS-56 |
| chr15-57502024-INS-63 | chr15-57502024-INS-72 |
| chr15-57510650-INS-153 | chr15-57510650-INS-153 |
| chr15-57515544-INS-215 | chr15-57515546-INS-216 |
| chr15-57543991-INS-114 | chr15-57543991-INS-108 |
| chr15-57545945-INS-52 | chr15-57545945-INS-52 |
| chr15-57605787-INS-6228 | chr15-57605788-INS-6228 |
| chr15-57628250-INS-74 | chr15-57628265-INS-70 |
| chr15-57671804-INS-73 | chr15-57671804-INS-79 |
| chr15-57675543-INS-132 | chr15-57675543-INS-132 |
| chr15-57683614-INS-58 | chr15-57683614-INS-58 |
| chr15-57687159-INS-5245 | chr15-57687160-INS-5245 |
| chr15-57693075-INS-60 | chr15-57693089-INS-68 |
| chr15-57694886-INS-517 | chr15-57694886-INS-517 |
| chr15-57701374-INS-66 | chr15-57701374-INS-70 |
| chr15-57718161-INS-54 | chr15-57718147-INS-71 |
| chr15-57732834-INS-53 | chr15-57732834-INS-53 |
| chr15-57753209-INS-56 | chr15-57753209-INS-56 |
| chr15-57757350-INS-96 | chr15-57757350-INS-99 |
| chr15-57763160-INS-208 | chr15-57763160-INS-210 |
| chr15-57790608-INS-360 | chr15-57790608-INS-380 |
| chr15-57798118-INS-70 | chr15-57798118-INS-68 |
| chr15-57822365-INS-88 | chr15-57822365-INS-88 |
| chr15-57831643-INS-821 | chr15-57831643-INS-821 |
| chr15-57863165-INS-108 | chr15-57863165-INS-108 |
| chr15-57865985-INS-268 | chr15-57865985-INS-288 |
| chr15-57870393-INS-6507 | chr15-57870394-INS-6507 |
| chr15-57887437-INS-218 | chr15-57887437-INS-222 |
| chr15-57891427-INS-826 | chr15-57891427-INS-826 |
| chr15-57894818-INS-76 | chr15-57894823-INS-103 |
| chr15-57897816-INS-97 | chr15-57897816-INS-97 |
| chr15-57904012-INS-55 | chr15-57904013-INS-55 |
| chr15-57905565-INS-7091 | chr15-57905566-INS-7091 |
| chr15-57926043-INS-251 | chr15-57926044-INS-252 |
| chr15-57928399-INS-202 | chr15-57928399-INS-203 |
| chr15-57929194-INS-168 | chr15-57929194-INS-168 |
| chr15-57935643-INS-236 | chr15-57935643-INS-236 |
| chr15-57944762-INS-52 | chr15-57944762-INS-52 |
| chr15-57954436-INS-92 | chr15-57954434-INS-94 |
| chr15-57962616-INS-5418 | chr15-57962617-INS-5418 |
| chr15-57980526-INS-115 | chr15-57980526-INS-116 |
| chr15-57981366-INS-199 | chr15-57981366-INS-199 |
| chr15-57987599-INS-198 | chr15-57987599-INS-198 |
| chr15-57989646-INS-3399 | chr15-57989647-INS-3399 |
| chr15-57991749-INS-211 | chr15-57991749-INS-211 |
| chr15-58007028-INS-118 | chr15-58007028-INS-118 |
| chr15-58009257-INS-74 | chr15-58009257-INS-52 |
| chr15-58012658-INS-239 | chr15-58012658-INS-206 |
| chr15-58019737-INS-422 | chr15-58019737-INS-422 |
| chr15-58034559-INS-209 | chr15-58034559-INS-209 |
| chr15-58053519-INS-183 | chr15-58053519-INS-183 |
| chr15-58057568-INS-87 | chr15-58057575-INS-108 |
| chr15-58080084-INS-230 | chr15-58080084-INS-230 |
| chr15-58085549-INS-161 | chr15-58085549-INS-161 |
| chr15-58102589-INS-73 | chr15-58102589-INS-73 |
| chr15-58122849-INS-64 | chr15-58122849-INS-52 |
| chr15-58128108-INS-963 | chr15-58128108-INS-963 |
| chr15-58133177-INS-90 | chr15-58133180-INS-91 |
| chr15-58139171-INS-92 | chr15-58139171-INS-60 |
| chr15-58145388-INS-66 | chr15-58145396-INS-51 |
| chr15-58178714-INS-56 | chr15-58178714-INS-56 |
| chr15-58180524-INS-87 | chr15-58180524-INS-87 |
| chr15-58193332-INS-64 | chr15-58193357-INS-56 |
| chr15-58207836-INS-182 | chr15-58207836-INS-164 |
| chr15-58236074-INS-372 | chr15-58236074-INS-372 |

| | |
|---|---|
| chr15-58245810-INS-204 | chr15-58245810-INS-204 |
| chr15-58249321-INS-84 | chr15-58249321-INS-84 |
| chr15-58264145-INS-87 | chr15-58264145-INS-87 |
| chr15-58397459-INS-51 | chr15-58397459-INS-60 |
| chr15-58413379-INS-1658 | chr15-58413380-INS-1636 |
| chr15-58462633-INS-58 | chr15-58462633-INS-50 |
| chr15-58517048-INS-6591 | chr15-58517049-INS-6592 |
| chr15-58527847-INS-156 | chr15-58527847-INS-162 |
| chr15-58558832-INS-99 | chr15-58558832-INS-99 |
| chr15-58580654-INS-52 | chr15-58580654-INS-54 |
| chr15-58586422-INS-129 | chr15-58586422-INS-132 |
| chr15-58607310-INS-709 | chr15-58607310-INS-709 |
| chr15-58610722-INS-97 | chr15-58610722-INS-102 |
| chr15-58621564-INS-86 | chr15-58621565-INS-86 |
| chr15-58632238-INS-74 | chr15-58632238-INS-74 |
| chr15-58637928-INS-81 | chr15-58637928-INS-81 |
| chr15-58641015-INS-2941 | chr15-58641015-INS-2936 |
| chr15-58645411-INS-90 | chr15-58645411-INS-90 |
| chr15-58671172-INS-204 | chr15-58671177-INS-208 |
| chr15-58679604-INS-72 | chr15-58679592-INS-72 |
| chr15-58691482-INS-74 | chr15-58691479-INS-75 |
| chr15-58701215-INS-1890 | chr15-58701216-INS-1890 |
| chr15-58711255-INS-1887 | chr15-58711256-INS-1887 |
| chr15-58711861-INS-432 | chr15-58711861-INS-432 |
| chr15-58718127-INS-3167 | chr15-58718128-INS-3147 |
| chr15-58723982-INS-256 | chr15-58724008-INS-257 |
| chr15-58728376-INS-1996 | chr15-58728377-INS-1996 |
| chr15-58737558-INS-216 | chr15-58737558-INS-216 |
| chr15-58751423-INS-79 | chr15-58751423-INS-79 |
| chr15-58769143-INS-1044 | chr15-58769144-INS-1044 |
| chr15-58787569-INS-78 | chr15-58787569-INS-98 |
| chr15-58829008-INS-1018 | chr15-58829009-INS-1018 |
| chr15-58830050-INS-206 | chr15-58830050-INS-206 |
| chr15-58839952-INS-166 | chr15-58839953-INS-157 |
| chr15-58848137-INS-78 | chr15-58848139-INS-86 |
| chr15-58857595-INS-64 | chr15-58857595-INS-64 |
| chr15-58879885-INS-104 | chr15-58879885-INS-104 |
| chr15-58905712-INS-547 | chr15-58905712-INS-547 |
| chr15-58916067-INS-62 | chr15-58916067-INS-88 |
| chr15-58922861-INS-266 | chr15-58922836-INS-265 |
| chr15-58923472-INS-88 | chr15-58923472-INS-64 |
| chr15-58943963-INS-112 | chr15-58943983-INS-102 |
| chr15-59008662-INS-3187 | chr15-59008663-INS-3184 |
| chr15-59011769-INS-81 | chr15-59011769-INS-81 |
| chr15-59017555-INS-52 | chr15-59017557-INS-50 |
| chr15-59020741-INS-104 | chr15-59020741-INS-104 |
| chr15-59053389-INS-1186 | chr15-59053390-INS-1174 |
| chr15-59054856-INS-63 | chr15-59054856-INS-63 |
| chr15-59056189-INS-156 | chr15-59056189-INS-132 |
| chr15-59066111-INS-217 | chr15-59066111-INS-217 |
| chr15-59087628-INS-136 | chr15-59087628-INS-136 |
| chr15-59092858-INS-52 | chr15-59092858-INS-56 |
| chr15-59099823-INS-523 | chr15-59099823-INS-522 |
| chr15-59100898-INS-200 | chr15-59100898-INS-199 |
| chr15-59115833-INS-77 | chr15-59115833-INS-50 |
| chr15-59120594-INS-167 | chr15-59120590-INS-142 |
| chr15-59150437-INS-777 | chr15-59150437-INS-777 |
| chr15-59162467-INS-427 | chr15-59162467-INS-427 |
| chr15-59167398-INS-119 | chr15-59167398-INS-119 |
| chr15-59184424-INS-242 | chr15-59184378-INS-242 |
| chr15-59202077-INS-220 | chr15-59202077-INS-224 |
| chr15-59219521-INS-165 | chr15-59219497-INS-180 |
| chr15-59234423-INS-124 | chr15-59234423-INS-124 |
| chr15-59272175-INS-106 | chr15-59272175-INS-106 |
| chr15-59287544-INS-1023 | chr15-59287547-INS-1019 |
| chr15-59327714-INS-220 | chr15-59327714-INS-220 |
| chr15-59346222-INS-198 | chr15-59346240-INS-192 |
| chr15-59354352-INS-159 | chr15-59354350-INS-159 |
| chr15-59362235-INS-162 | chr15-59362235-INS-127 |
| chr15-59366077-INS-188 | chr15-59366077-INS-188 |
| chr15-59377585-INS-6238 | chr15-59377586-INS-6239 |
| chr15-59378661-INS-51 | chr15-59378661-INS-52 |
| chr15-59395662-INS-172 | chr15-59395662-INS-172 |
| chr15-59399383-INS-244 | chr15-59399383-INS-262 |
| chr15-59411926-INS-72 | chr15-59411901-INS-67 |
| chr15-59415034-INS-68 | chr15-59415034-INS-75 |
| chr15-59421696-INS-245 | chr15-59421696-INS-245 |

| | |
|---|---|
| chr15-59430630-INS-58 | chr15-59430630-INS-58 |
| chr15-59433787-INS-58 | chr15-59433787-INS-60 |
| chr15-59440709-INS-166 | chr15-59440709-INS-166 |
| chr15-59454002-INS-400 | chr15-59454002-INS-400 |
| chr15-59477343-INS-92 | chr15-59477388-INS-92 |
| chr15-59493325-INS-79 | chr15-59493325-INS-53 |
| chr15-59494018-INS-59 | chr15-59494018-INS-90 |
| chr15-59498651-INS-228 | chr15-59498651-INS-236 |
| chr15-59503131-INS-174 | chr15-59503132-INS-167 |
| chr15-59506276-INS-5544 | chr15-59506277-INS-5545 |
| chr15-59510647-INS-202 | chr15-59510647-INS-202 |
| chr15-59528651-INS-278 | chr15-59528651-INS-280 |
| chr15-59531104-INS-72 | chr15-59531104-INS-70 |
| chr15-59548872-INS-88 | chr15-59548863-INS-95 |
| chr15-59550535-INS-64 | chr15-59550535-INS-64 |
| chr15-59575672-INS-810 | chr15-59575672-INS-810 |
| chr15-59583512-INS-72 | chr15-59583512-INS-56 |
| chr15-59590018-INS-103 | chr15-59590018-INS-103 |
| chr15-59599757-INS-66 | chr15-59599759-INS-68 |
| chr15-59629594-INS-245 | chr15-59629595-INS-280 |
| chr15-59630830-INS-6641 | chr15-59630831-INS-6641 |
| chr15-59633495-INS-106 | chr15-59633495-INS-64 |
| chr15-59640860-INS-1611 | chr15-59640869-INS-1609 |
| chr15-59648210-INS-54 | chr15-59648210-INS-51 |
| chr15-59653383-INS-54 | chr15-59653383-INS-54 |
| chr15-59681587-INS-164 | chr15-59681587-INS-164 |
| chr15-59729009-INS-83 | chr15-59729009-INS-83 |
| chr15-59747946-INS-71 | chr15-59747946-INS-62 |
| chr15-59753872-INS-197 | chr15-59753872-INS-197 |
| chr15-59758484-INS-84 | chr15-59758524-INS-62 |
| chr15-59814632-INS-437 | chr15-59814632-INS-437 |
| chr15-59820636-INS-92 | chr15-59820636-INS-92 |
| chr15-59860963-INS-64 | chr15-59860963-INS-60 |
| chr15-59866116-INS-3096 | chr15-59866117-INS-3095 |
| chr15-59867485-INS-103 | chr15-59867485-INS-54 |
| chr15-59909591-INS-1362 | chr15-59909592-INS-1369 |
| chr15-59919638-INS-109 | chr15-59919638-INS-109 |
| chr15-59936996-INS-208 | chr15-59936996-INS-208 |
| chr15-59939278-INS-194 | chr15-59939278-INS-194 |
| chr15-59972124-INS-419 | chr15-59972124-INS-403 |
| chr15-59996382-INS-58 | chr15-59996385-INS-63 |
| chr15-60006378-INS-440 | chr15-60006378-INS-440 |
| chr15-60021144-INS-606 | chr15-60021144-INS-606 |
| chr15-60031259-INS-56 | chr15-60031259-INS-60 |
| chr15-60037933-INS-895 | chr15-60037933-INS-895 |
| chr15-60044320-INS-103 | chr15-60044320-INS-103 |
| chr15-60062282-INS-6330 | chr15-60062283-INS-6330 |
| chr15-60069535-INS-88 | chr15-60069536-INS-71 |
| chr15-60091049-INS-134 | chr15-60091033-INS-119 |
| chr15-60097852-INS-6374 | chr15-60097853-INS-6376 |
| chr15-60122573-INS-59 | chr15-60122573-INS-62 |
| chr15-60131677-INS-313 | chr15-60131677-INS-313 |
| chr15-60196259-INS-1493 | chr15-60196260-INS-1493 |
| chr15-60197414-INS-53 | chr15-60197414-INS-53 |
| chr15-60213873-INS-446 | chr15-60213873-INS-426 |
| chr15-60225853-INS-212 | chr15-60225853-INS-212 |
| chr15-60230542-INS-1572 | chr15-60230547-INS-1546 |
| chr15-60236736-INS-54 | chr15-60236736-INS-78 |
| chr15-60263667-INS-62 | chr15-60263668-INS-70 |
| chr15-60304266-INS-60 | chr15-60304266-INS-60 |
| chr15-60330195-INS-218 | chr15-60330195-INS-213 |
| chr15-60370169-INS-1437 | chr15-60370170-INS-1437 |
| chr15-60433093-INS-53 | chr15-60433050-INS-66 |
| chr15-60434180-INS-1893 | chr15-60434181-INS-1893 |
| chr15-60452962-INS-480 | chr15-60452962-INS-480 |
| chr15-60476191-INS-130 | chr15-60476191-INS-144 |
| chr15-60476675-INS-6637 | chr15-60476676-INS-6637 |
| chr15-60495333-INS-53 | chr15-60495333-INS-51 |
| chr15-60498415-INS-165 | chr15-60498415-INS-164 |
| chr15-60540599-INS-1894 | chr15-60540600-INS-1894 |
| chr15-60581154-INS-214 | chr15-60581153-INS-209 |
| chr15-60587817-INS-783 | chr15-60587817-INS-793 |
| chr15-60637059-INS-227 | chr15-60637059-INS-227 |
| chr15-60641674-INS-59 | chr15-60641674-INS-55 |
| chr15-60660485-INS-76 | chr15-60660485-INS-52 |
| chr15-60690637-INS-166 | chr15-60690637-INS-166 |
| chr15-60697630-INS-57 | chr15-60697630-INS-57 |

| | |
|---|---|
| chr15-60699431-INS-140 | chr15-60699424-INS-153 |
| chr15-60733095-INS-106 | chr15-60733100-INS-106 |
| chr15-60750075-INS-1894 | chr15-60750076-INS-1894 |
| chr15-60798490-INS-193 | chr15-60798489-INS-208 |
| chr15-60802509-INS-267 | chr15-60802509-INS-266 |
| chr15-60809389-INS-83 | chr15-60809389-INS-83 |
| chr15-60853034-INS-520 | chr15-60853034-INS-535 |
| chr15-60870716-INS-237 | chr15-60870716-INS-237 |
| chr15-60878055-INS-493 | chr15-60878056-INS-495 |
| chr15-60886030-INS-1221 | chr15-60886031-INS-1221 |
| chr15-60888979-INS-54 | chr15-60888979-INS-54 |
| chr15-60890309-INS-6404 | chr15-60890310-INS-6403 |
| chr15-60898665-INS-209 | chr15-60898665-INS-208 |
| chr15-60902096-INS-3214 | chr15-60902097-INS-3214 |
| chr15-60904983-INS-2465 | chr15-60904984-INS-2465 |
| chr15-60911310-INS-635 | chr15-60911310-INS-627 |
| chr15-60945565-INS-254 | chr15-60945565-INS-254 |
| chr15-60965552-INS-526 | chr15-60965552-INS-529 |
| chr15-60993750-INS-84 | chr15-60993750-INS-83 |
| chr15-61001406-INS-928 | chr15-61001406-INS-937 |
| chr15-61004080-INS-56 | chr15-61004080-INS-58 |
| chr15-61041387-INS-390 | chr15-61041387-INS-390 |
| chr15-61106124-INS-2876 | chr15-61106125-INS-2876 |
| chr15-61123452-INS-685 | chr15-61123452-INS-685 |
| chr15-61139480-INS-66 | chr15-61139519-INS-52 |
| chr15-61149709-INS-2201 | chr15-61149710-INS-2201 |
| chr15-61168597-INS-1021 | chr15-61168598-INS-1021 |
| chr15-61186111-INS-209 | chr15-61186111-INS-200 |
| chr15-61194258-INS-1575 | chr15-61194259-INS-1575 |
| chr15-61197010-INS-400 | chr15-61197010-INS-399 |
| chr15-61215028-INS-98 | chr15-61215028-INS-98 |
| chr15-61217425-INS-6413 | chr15-61217426-INS-6412 |
| chr15-61230572-INS-770 | chr15-61230572-INS-776 |
| chr15-61233067-INS-4108 | chr15-61233068-INS-4109 |
| chr15-61263300-INS-58 | chr15-61263300-INS-60 |
| chr15-61298596-INS-817 | chr15-61298596-INS-842 |
| chr15-61308237-INS-6466 | chr15-61308238-INS-6469 |
| chr15-61351205-INS-88 | chr15-61351205-INS-88 |
| chr15-61356167-INS-193 | chr15-61356167-INS-193 |
| chr15-61381425-INS-87 | chr15-61381425-INS-66 |
| chr15-61412238-INS-243 | chr15-61412238-INS-245 |
| chr15-61420930-INS-72 | chr15-61420919-INS-118 |
| chr15-61465488-INS-511 | chr15-61465488-INS-510 |
| chr15-61477891-INS-522 | chr15-61477891-INS-523 |
| chr15-61492107-INS-101 | chr15-61492113-INS-140 |
| chr15-61516731-INS-846 | chr15-61516731-INS-846 |
| chr15-61536586-INS-7239 | chr15-61536587-INS-7239 |
| chr15-61544688-INS-401 | chr15-61544688-INS-401 |
| chr15-61559570-INS-51 | chr15-61559570-INS-51 |
| chr15-61565669-INS-766 | chr15-61565669-INS-766 |
| chr15-61572218-INS-1064 | chr15-61572219-INS-1049 |
| chr15-61585053-INS-2963 | chr15-61585054-INS-2964 |
| chr15-61619485-INS-674 | chr15-61619488-INS-684 |
| chr15-61668461-INS-6671 | chr15-61668462-INS-6673 |
| chr15-61669255-INS-268 | chr15-61669255-INS-268 |
| chr15-61676893-INS-93 | chr15-61676900-INS-126 |
| chr15-61718287-INS-97 | chr15-61718287-INS-97 |
| chr15-61725361-INS-92 | chr15-61725396-INS-64 |
| chr15-61737930-INS-60 | chr15-61737931-INS-58 |
| chr15-61738455-INS-7147 | chr15-61738456-INS-7148 |
| chr15-61813142-INS-64 | chr15-61813118-INS-104 |
| chr15-61815751-INS-90 | chr15-61815751-INS-80 |
| chr15-61840263-INS-241 | chr15-61840263-INS-242 |
| chr15-61846815-INS-64 | chr15-61846815-INS-68 |
| chr15-61868885-INS-217 | chr15-61868885-INS-217 |
| chr15-61892897-INS-66 | chr15-61892897-INS-66 |
| chr15-61926149-INS-198 | chr15-61926151-INS-202 |
| chr15-61938094-INS-174 | chr15-61938094-INS-174 |
| chr15-61946779-INS-51 | chr15-61946829-INS-51 |
| chr15-61960607-INS-228 | chr15-61960607-INS-227 |
| chr15-61983099-INS-102 | chr15-61983099-INS-102 |
| chr15-61997506-INS-80 | chr15-61997506-INS-80 |
| chr15-61998601-INS-106 | chr15-61998601-INS-117 |
| chr15-62010241-INS-78 | chr15-62010241-INS-62 |
| chr15-62046933-INS-80 | chr15-62046933-INS-70 |
| chr15-62047567-INS-316 | chr15-62047567-INS-316 |
| chr15-62059197-INS-202 | chr15-62059198-INS-202 |

| | |
|---|---|
| chr15-62100293-INS-3984 | chr15-62100294-INS-3984 |
| chr15-62106069-INS-874 | chr15-62106069-INS-874 |
| chr15-62118401-INS-108 | chr15-62118408-INS-102 |
| chr15-62140664-INS-84 | chr15-62140660-INS-84 |
| chr15-62149274-INS-7108 | chr15-62149275-INS-7108 |
| chr15-62164363-INS-101 | chr15-62164363-INS-98 |
| chr15-62206905-INS-64 | chr15-62206905-INS-64 |
| chr15-62211967-INS-802 | chr15-62211967-INS-803 |
| chr15-62214566-INS-506 | chr15-62214566-INS-506 |
| chr15-62223652-INS-68 | chr15-62223689-INS-64 |
| chr15-62255203-INS-51 | chr15-62255203-INS-63 |
| chr15-62289845-INS-3145 | chr15-62289846-INS-3145 |
| chr15-62315018-INS-52 | chr15-62315021-INS-61 |
| chr15-62335565-INS-91 | chr15-62335565-INS-105 |
| chr15-62345745-INS-106 | chr15-62345745-INS-106 |
| chr15-62349741-INS-151 | chr15-62349741-INS-152 |
| chr15-62351155-INS-1223 | chr15-62351156-INS-1223 |
| chr15-62352529-INS-246 | chr15-62352529-INS-246 |
| chr15-62361923-INS-58 | chr15-62361923-INS-61 |
| chr15-62386954-INS-822 | chr15-62386954-INS-821 |
| chr15-62417982-INS-141 | chr15-62417982-INS-141 |
| chr15-62443271-INS-765 | chr15-62443271-INS-766 |
| chr15-62451976-INS-68 | chr15-62452026-INS-68 |
| chr15-62454138-INS-259 | chr15-62454138-INS-241 |
| chr15-62459887-INS-51 | chr15-62459887-INS-51 |
| chr15-62475736-INS-88 | chr15-62475736-INS-56 |
| chr15-62487151-INS-6613 | chr15-62487152-INS-6614 |
| chr15-62513787-INS-65 | chr15-62513787-INS-60 |
| chr15-62536697-INS-671 | chr15-62536700-INS-674 |
| chr15-62542781-INS-1204 | chr15-62542782-INS-1204 |
| chr15-62542953-INS-167 | chr15-62542953-INS-167 |
| chr15-62550165-INS-341 | chr15-62550165-INS-341 |
| chr15-62554417-INS-412 | chr15-62554417-INS-412 |
| chr15-62556872-INS-1893 | chr15-62556873-INS-1893 |
| chr15-62572303-INS-967 | chr15-62572303-INS-968 |
| chr15-62580714-INS-61 | chr15-62580714-INS-61 |
| chr15-62587563-INS-2853 | chr15-62587564-INS-2855 |
| chr15-62623758-INS-1368 | chr15-62623759-INS-1369 |
| chr15-62626477-INS-58 | chr15-62626450-INS-66 |
| chr15-62630458-INS-54 | chr15-62630458-INS-54 |
| chr15-62634732-INS-891 | chr15-62634732-INS-892 |
| chr15-62649408-INS-3352 | chr15-62649409-INS-3354 |
| chr15-62709470-INS-84 | chr15-62709474-INS-66 |
| chr15-62709830-INS-7233 | chr15-62709831-INS-7233 |
| chr15-62709831-INS-373 | chr15-62709831-INS-373 |
| chr15-62713131-INS-54 | chr15-62713131-INS-102 |
| chr15-62745936-INS-436 | chr15-62745936-INS-436 |
| chr15-62764006-INS-70 | chr15-62764006-INS-68 |
| chr15-62779129-INS-1676 | chr15-62779130-INS-1677 |
| chr15-62781277-INS-70 | chr15-62781237-INS-70 |
| chr15-62787600-INS-1852 | chr15-62787601-INS-1852 |
| chr15-62791860-INS-6528 | chr15-62791857-INS-6511 |
| chr15-62822456-INS-201 | chr15-62822456-INS-201 |
| chr15-62849896-INS-239 | chr15-62849896-INS-241 |
| chr15-62873925-INS-57 | chr15-62873929-INS-62 |
| chr15-62899649-INS-204 | chr15-62899649-INS-204 |
| chr15-62917096-INS-1353 | chr15-62917097-INS-1354 |
| chr15-62988475-INS-64 | chr15-62988491-INS-54 |
| chr15-62994265-INS-513 | chr15-62994265-INS-513 |
| chr15-63014040-INS-461 | chr15-63014040-INS-460 |
| chr15-63018981-INS-405 | chr15-63018981-INS-404 |
| chr15-63065102-INS-215 | chr15-63065102-INS-215 |
| chr15-63065739-INS-82 | chr15-63065739-INS-64 |
| chr15-63110449-INS-295 | chr15-63110449-INS-298 |
| chr15-63157252-INS-105 | chr15-63157252-INS-105 |
| chr15-63164747-INS-52 | chr15-63164747-INS-52 |
| chr15-63171073-INS-473 | chr15-63171073-INS-473 |
| chr15-63180353-INS-2039 | chr15-63180354-INS-2031 |
| chr15-63219199-INS-728 | chr15-63219199-INS-728 |
| chr15-63221625-INS-2764 | chr15-63221626-INS-2761 |
| chr15-63255033-INS-483 | chr15-63255034-INS-483 |
| chr15-63270257-INS-60 | chr15-63270221-INS-72 |
| chr15-63278882-INS-54 | chr15-63278882-INS-50 |
| chr15-63317437-INS-73 | chr15-63317436-INS-74 |
| chr15-63327074-INS-6923 | chr15-63327075-INS-6923 |
| chr15-63336585-INS-682 | chr15-63336586-INS-682 |
| chr15-63344738-INS-180 | chr15-63344738-INS-132 |

| | |
|---|---|
| chr15-63355851-INS-775 | chr15-63355851-INS-775 |
| chr15-63399441-INS-75 | chr15-63399441-INS-74 |
| chr15-63408506-INS-336 | chr15-63408506-INS-336 |
| chr15-63409730-INS-606 | chr15-63409730-INS-606 |
| chr15-63432796-INS-204 | chr15-63432796-INS-204 |
| chr15-63468671-INS-6891 | chr15-63468672-INS-6892 |
| chr15-63478569-INS-623 | chr15-63478569-INS-616 |
| chr15-63486027-INS-344 | chr15-63486027-INS-344 |
| chr15-63506682-INS-118 | chr15-63506683-INS-111 |
| chr15-63571474-INS-212 | chr15-63571474-INS-212 |
| chr15-63581823-INS-60 | chr15-63581823-INS-91 |
| chr15-63590548-INS-54 | chr15-63590548-INS-53 |
| chr15-63590977-INS-880 | chr15-63590975-INS-883 |
| chr15-63608087-INS-190 | chr15-63608087-INS-190 |
| chr15-63647375-INS-117 | chr15-63647375-INS-117 |
| chr15-63826766-INS-66 | chr15-63826766-INS-66 |
| chr15-63834174-INS-66 | chr15-63834174-INS-68 |
| chr15-63886899-INS-72 | chr15-63886899-INS-64 |
| chr15-63938071-INS-70 | chr15-63938069-INS-58 |
| chr15-63945334-INS-84 | chr15-63945338-INS-80 |
| chr15-63952070-INS-58 | chr15-63952059-INS-64 |
| chr15-64039812-INS-176 | chr15-64039812-INS-176 |
| chr15-64047530-INS-743 | chr15-64047530-INS-744 |
| chr15-64071420-INS-6265 | chr15-64071421-INS-6262 |
| chr15-64080571-INS-73 | chr15-64080529-INS-69 |
| chr15-64100855-INS-71 | chr15-64100855-INS-71 |
| chr15-64117797-INS-1897 | chr15-64117798-INS-1897 |
| chr15-64134522-INS-55 | chr15-64134522-INS-55 |
| chr15-64150404-INS-4458 | chr15-64150405-INS-4458 |
| chr15-64179570-INS-66 | chr15-64179553-INS-78 |
| chr15-64201749-INS-96 | chr15-64201749-INS-90 |
| chr15-64212416-INS-54 | chr15-64212422-INS-58 |
| chr15-64233082-INS-99 | chr15-64233082-INS-72 |
| chr15-64234527-INS-7079 | chr15-64234528-INS-7079 |
| chr15-64235061-INS-64 | chr15-64235061-INS-64 |
| chr15-64238868-INS-102 | chr15-64238868-INS-102 |
| chr15-64244084-INS-632 | chr15-64244084-INS-632 |
| chr15-64260551-INS-565 | chr15-64260551-INS-565 |
| chr15-64265940-INS-60 | chr15-64265940-INS-60 |
| chr15-64267784-INS-264 | chr15-64267790-INS-268 |
| chr15-64272696-INS-60 | chr15-64272696-INS-58 |
| chr15-64275198-INS-110 | chr15-64275198-INS-104 |
| chr15-64277747-INS-60 | chr15-64277747-INS-56 |
| chr15-64302610-INS-66 | chr15-64302609-INS-54 |
| chr15-64308595-INS-1750 | chr15-64308596-INS-1750 |
| chr15-64314389-INS-59 | chr15-64314389-INS-57 |
| chr15-64321441-INS-201 | chr15-64321441-INS-201 |
| chr15-64337058-INS-54 | chr15-64337108-INS-54 |
| chr15-64371147-INS-110 | chr15-64371147-INS-110 |
| chr15-64378110-INS-188 | chr15-64378110-INS-189 |
| chr15-64402340-INS-56 | chr15-64402340-INS-54 |
| chr15-64423578-INS-114 | chr15-64423578-INS-114 |
| chr15-64428598-INS-2085 | chr15-64428599-INS-2085 |
| chr15-64436649-INS-478 | chr15-64436649-INS-478 |
| chr15-64437326-INS-82 | chr15-64437326-INS-71 |
| chr15-64449428-INS-88 | chr15-64449426-INS-90 |
| chr15-64460142-INS-281 | chr15-64460143-INS-293 |
| chr15-64474854-INS-1888 | chr15-64474855-INS-1888 |
| chr15-64496873-INS-85 | chr15-64496855-INS-102 |
| chr15-64500141-INS-55 | chr15-64500141-INS-55 |
| chr15-64504756-INS-226 | chr15-64504756-INS-230 |
| chr15-64507160-INS-51 | chr15-64507160-INS-51 |
| chr15-64517179-INS-53 | chr15-64517179-INS-50 |
| chr15-64525958-INS-666 | chr15-64525958-INS-666 |
| chr15-64569697-INS-69 | chr15-64569697-INS-69 |
| chr15-64574997-INS-324 | chr15-64574997-INS-324 |
| chr15-64579108-INS-6984 | chr15-64579109-INS-6983 |
| chr15-64582592-INS-149 | chr15-64582592-INS-149 |
| chr15-64627816-INS-68 | chr15-64627817-INS-71 |
| chr15-64650701-INS-553 | chr15-64650701-INS-554 |
| chr15-64684616-INS-231 | chr15-64684616-INS-231 |
| chr15-64700645-INS-731 | chr15-64700645-INS-731 |
| chr15-64715701-INS-65 | chr15-64715701-INS-65 |
| chr15-64746522-INS-134 | chr15-64746522-INS-134 |
| chr15-64757386-INS-248 | chr15-64757386-INS-274 |
| chr15-64759463-INS-249 | chr15-64759463-INS-249 |
| chr15-64766093-INS-147 | chr15-64766093-INS-170 |

| | |
|---|---|
| chr15-64772829-INS-50 | chr15-64772829-INS-62 |
| chr15-64777662-INS-84 | chr15-64777662-INS-96 |
| chr15-64828708-INS-100 | chr15-64828708-INS-52 |
| chr15-64854022-INS-65 | chr15-64854022-INS-65 |
| chr15-64881006-INS-73 | chr15-64881006-INS-53 |
| chr15-64883571-INS-864 | chr15-64883571-INS-864 |
| chr15-64890826-INS-166 | chr15-64890826-INS-166 |
| chr15-64928901-INS-1761 | chr15-64928902-INS-1761 |
| chr15-64950919-INS-7110 | chr15-64950920-INS-7110 |
| chr15-64954702-INS-890 | chr15-64954705-INS-888 |
| chr15-64966277-INS-119 | chr15-64966277-INS-119 |
| chr15-65012872-INS-104 | chr15-65012855-INS-81 |
| chr15-65030264-INS-214 | chr15-65030247-INS-214 |
| chr15-65070333-INS-136 | chr15-65070326-INS-145 |
| chr15-65072082-INS-935 | chr15-65072082-INS-935 |
| chr15-65075069-INS-511 | chr15-65075069-INS-511 |
| chr15-65085855-INS-391 | chr15-65085855-INS-391 |
| chr15-65102280-INS-59 | chr15-65102280-INS-59 |
| chr15-65135414-INS-727 | chr15-65135414-INS-727 |
| chr15-65154675-INS-180 | chr15-65154675-INS-180 |
| chr15-65166373-INS-6410 | chr15-65166374-INS-6411 |
| chr15-65168022-INS-58 | chr15-65168022-INS-58 |
| chr15-65211513-INS-106 | chr15-65211503-INS-149 |
| chr15-65224889-INS-279 | chr15-65224889-INS-279 |
| chr15-65266881-INS-52 | chr15-65266881-INS-52 |
| chr15-65281871-INS-164 | chr15-65281871-INS-164 |
| chr15-65297309-INS-61 | chr15-65297309-INS-69 |
| chr15-65347947-INS-190 | chr15-65347915-INS-180 |
| chr15-65352055-INS-1888 | chr15-65352056-INS-1888 |
| chr15-65355240-INS-328 | chr15-65355240-INS-329 |
| chr15-65382033-INS-2774 | chr15-65382034-INS-2774 |
| chr15-65396383-INS-110 | chr15-65396383-INS-144 |
| chr15-65412384-INS-406 | chr15-65412384-INS-405 |
| chr15-65426835-INS-68 | chr15-65426835-INS-64 |
| chr15-65430222-INS-6725 | chr15-65430223-INS-6725 |
| chr15-65443372-INS-214 | chr15-65443334-INS-214 |
| chr15-65444765-INS-2246 | chr15-65444774-INS-2261 |
| chr15-65466530-INS-259 | chr15-65466530-INS-261 |
| chr15-65470534-INS-201 | chr15-65470549-INS-201 |
| chr15-65497554-INS-6602 | chr15-65497555-INS-6602 |
| chr15-65533477-INS-389 | chr15-65533477-INS-389 |
| chr15-65551506-INS-1154 | chr15-65551507-INS-1154 |
| chr15-65557741-INS-8277 | chr15-65557742-INS-8277 |
| chr15-65560598-INS-170 | chr15-65560598-INS-170 |
| chr15-65596138-INS-64 | chr15-65596127-INS-54 |
| chr15-65618706-INS-118 | chr15-65618706-INS-118 |
| chr15-65620369-INS-672 | chr15-65620369-INS-672 |
| chr15-65623398-INS-191 | chr15-65623399-INS-191 |
| chr15-65624442-INS-824 | chr15-65624442-INS-824 |
| chr15-65640364-INS-92 | chr15-65640364-INS-92 |
| chr15-65703986-INS-137 | chr15-65703986-INS-137 |
| chr15-65706413-INS-3747 | chr15-65706414-INS-3747 |
| chr15-65740942-INS-63 | chr15-65740932-INS-54 |
| chr15-65755542-INS-528 | chr15-65755542-INS-528 |
| chr15-65762065-INS-149 | chr15-65762049-INS-154 |
| chr15-65770472-INS-57 | chr15-65770472-INS-61 |
| chr15-65778703-INS-908 | chr15-65778703-INS-908 |
| chr15-65812756-INS-5311 | chr15-65812757-INS-5311 |
| chr15-65840372-INS-167 | chr15-65840372-INS-168 |
| chr15-65855422-INS-54 | chr15-65855422-INS-54 |
| chr15-65907536-INS-93 | chr15-65907536-INS-50 |
| chr15-65914285-INS-400 | chr15-65914285-INS-400 |
| chr15-65917629-INS-81 | chr15-65917629-INS-81 |
| chr15-65948714-INS-56 | chr15-65948716-INS-56 |
| chr15-65962712-INS-66 | chr15-65962707-INS-60 |
| chr15-65964318-INS-156 | chr15-65964318-INS-167 |
| chr15-65966495-INS-384 | chr15-65966495-INS-384 |
| chr15-65994911-INS-55 | chr15-65994911-INS-55 |
| chr15-66020942-INS-98 | chr15-66020942-INS-91 |
| chr15-66026823-INS-96 | chr15-66026823-INS-52 |
| chr15-66042467-INS-159 | chr15-66042467-INS-183 |
| chr15-66076410-INS-856 | chr15-66076410-INS-856 |
| chr15-66077317-INS-1948 | chr15-66077318-INS-1948 |
| chr15-66084980-INS-221 | chr15-66084980-INS-221 |
| chr15-66099603-INS-471 | chr15-66099603-INS-470 |
| chr15-66100562-INS-79 | chr15-66100562-INS-79 |
| chr15-66102301-INS-162 | chr15-66102301-INS-162 |

| | |
|---|---|
| chr15-66141661-INS-933 | chr15-66141661-INS-936 |
| chr15-66149861-INS-632 | chr15-66149861-INS-632 |
| chr15-66175860-INS-443 | chr15-66175871-INS-416 |
| chr15-66189541-INS-86 | chr15-66189541-INS-86 |
| chr15-66215287-INS-137 | chr15-66215287-INS-137 |
| chr15-66217475-INS-75 | chr15-66217475-INS-76 |
| chr15-66219337-INS-96 | chr15-66219330-INS-97 |
| chr15-66236309-INS-417 | chr15-66236309-INS-417 |
| chr15-66242858-INS-131 | chr15-66242858-INS-132 |
| chr15-66243801-INS-59 | chr15-66243801-INS-59 |
| chr15-66264261-INS-1832 | chr15-66264262-INS-1833 |
| chr15-66318089-INS-394 | chr15-66318089-INS-394 |
| chr15-66320664-INS-1359 | chr15-66320665-INS-1360 |
| chr15-66323452-INS-74 | chr15-66323452-INS-72 |
| chr15-66344442-INS-373 | chr15-66344442-INS-373 |
| chr15-66349541-INS-78 | chr15-66349554-INS-78 |
| chr15-66397388-INS-506 | chr15-66397388-INS-506 |
| chr15-66401229-INS-159 | chr15-66401229-INS-159 |
| chr15-66403128-INS-64 | chr15-66403128-INS-56 |
| chr15-66409258-INS-62 | chr15-66409258-INS-60 |
| chr15-66413667-INS-76 | chr15-66413657-INS-88 |
| chr15-66415987-INS-83 | chr15-66415987-INS-90 |
| chr15-66448942-INS-6572 | chr15-66448943-INS-6574 |
| chr15-66463195-INS-154 | chr15-66463195-INS-153 |
| chr15-66511384-INS-479 | chr15-66511384-INS-480 |
| chr15-66521079-INS-280 | chr15-66521079-INS-280 |
| chr15-66524528-INS-99 | chr15-66524529-INS-77 |
| chr15-66525312-INS-63 | chr15-66525313-INS-59 |
| chr15-66542242-INS-83 | chr15-66542242-INS-79 |
| chr15-66577854-INS-74 | chr15-66577854-INS-74 |
| chr15-66591919-INS-92 | chr15-66591919-INS-92 |
| chr15-66603513-INS-52 | chr15-66603541-INS-62 |
| chr15-66680249-INS-452 | chr15-66680249-INS-482 |
| chr15-66695380-INS-153 | chr15-66695398-INS-126 |
| chr15-66761805-INS-86 | chr15-66761805-INS-96 |
| chr15-66792799-INS-85 | chr15-66792799-INS-85 |
| chr15-66795341-INS-94 | chr15-66795341-INS-94 |
| chr15-66819483-INS-60 | chr15-66819483-INS-64 |
| chr15-66868586-INS-198 | chr15-66868586-INS-198 |
| chr15-66877954-INS-201 | chr15-66877954-INS-201 |
| chr15-66893941-INS-205 | chr15-66893941-INS-205 |
| chr15-66897266-INS-6807 | chr15-66897267-INS-6807 |
| chr15-66902796-INS-292 | chr15-66902796-INS-292 |
| chr15-66912595-INS-1301 | chr15-66912596-INS-1302 |
| chr15-66939414-INS-125 | chr15-66939414-INS-139 |
| chr15-66963804-INS-242 | chr15-66963804-INS-242 |
| chr15-66967724-INS-60 | chr15-66967724-INS-54 |
| chr15-66975686-INS-120 | chr15-66975686-INS-120 |
| chr15-66992054-INS-54 | chr15-66992054-INS-54 |
| chr15-67017432-INS-59 | chr15-67017432-INS-88 |
| chr15-67031541-INS-154 | chr15-67031542-INS-154 |
| chr15-67065276-INS-979 | chr15-67065276-INS-980 |
| chr15-67074605-INS-503 | chr15-67074605-INS-485 |
| chr15-67077913-INS-54 | chr15-67077913-INS-66 |
| chr15-67083694-INS-246 | chr15-67083694-INS-251 |
| chr15-67127088-INS-76 | chr15-67127088-INS-104 |
| chr15-67127983-INS-235 | chr15-67127983-INS-242 |
| chr15-67133227-INS-60 | chr15-67133227-INS-60 |
| chr15-67149965-INS-564 | chr15-67149965-INS-564 |
| chr15-67193938-INS-279 | chr15-67193938-INS-279 |
| chr15-67205259-INS-522 | chr15-67205259-INS-522 |
| chr15-67216955-INS-142 | chr15-67216955-INS-143 |
| chr15-67222872-INS-242 | chr15-67222873-INS-243 |
| chr15-67284281-INS-949 | chr15-67284281-INS-944 |
| chr15-67319793-INS-133 | chr15-67319793-INS-146 |
| chr15-67331229-INS-291 | chr15-67331229-INS-284 |
| chr15-67349187-INS-2729 | chr15-67349188-INS-2729 |
| chr15-67412518-INS-74 | chr15-67412518-INS-78 |
| chr15-67416381-INS-1093 | chr15-67416382-INS-1097 |
| chr15-67428941-INS-94 | chr15-67428941-INS-94 |
| chr15-67437157-INS-474 | chr15-67437157-INS-474 |
| chr15-67455844-INS-7267 | chr15-67455845-INS-7267 |
| chr15-67468962-INS-597 | chr15-67468962-INS-597 |
| chr15-67476053-INS-6676 | chr15-67476054-INS-6678 |
| chr15-67479460-INS-55 | chr15-67479460-INS-60 |
| chr15-67482300-INS-54 | chr15-67482321-INS-57 |
| chr15-67486377-INS-55 | chr15-67486377-INS-67 |

| | |
|---|---|
| chr15-67514100-INS-4060 | chr15-67514101-INS-4060 |
| chr15-67527577-INS-149 | chr15-67527577-INS-149 |
| chr15-67538876-INS-210 | chr15-67538876-INS-210 |
| chr15-67543538-INS-67 | chr15-67543524-INS-88 |
| chr15-67545020-INS-415 | chr15-67545020-INS-416 |
| chr15-67547587-INS-978 | chr15-67547587-INS-981 |
| chr15-67577540-INS-1016 | chr15-67577541-INS-1007 |
| chr15-67578642-INS-6584 | chr15-67578643-INS-6584 |
| chr15-67593748-INS-6649 | chr15-67593749-INS-6649 |
| chr15-67597647-INS-104 | chr15-67597652-INS-140 |
| chr15-67602559-INS-244 | chr15-67602559-INS-248 |
| chr15-67603548-INS-317 | chr15-67603549-INS-317 |
| chr15-67613262-INS-65 | chr15-67613262-INS-65 |
| chr15-67624210-INS-167 | chr15-67624210-INS-192 |
| chr15-67636848-INS-64 | chr15-67636856-INS-64 |
| chr15-67641496-INS-54 | chr15-67641454-INS-58 |
| chr15-67656316-INS-339 | chr15-67656316-INS-339 |
| chr15-67676485-INS-971 | chr15-67676485-INS-971 |
| chr15-67683794-INS-125 | chr15-67683794-INS-125 |
| chr15-67711349-INS-547 | chr15-67711349-INS-548 |
| chr15-67720352-INS-59 | chr15-67720352-INS-56 |
| chr15-67721985-INS-56 | chr15-67721985-INS-74 |
| chr15-67740822-INS-1496 | chr15-67740823-INS-1496 |
| chr15-67741470-INS-364 | chr15-67741470-INS-364 |
| chr15-67758961-INS-60 | chr15-67758961-INS-54 |
| chr15-67782615-INS-894 | chr15-67782615-INS-894 |
| chr15-67797803-INS-54 | chr15-67797803-INS-54 |
| chr15-67801409-INS-256 | chr15-67801409-INS-256 |
| chr15-67815610-INS-78 | chr15-67815610-INS-78 |
| chr15-67825227-INS-326 | chr15-67825227-INS-326 |
| chr15-67826011-INS-137 | chr15-67826011-INS-167 |
| chr15-67911551-INS-290 | chr15-67911551-INS-289 |
| chr15-67928635-INS-53 | chr15-67928635-INS-53 |
| chr15-67933499-INS-52 | chr15-67933499-INS-52 |
| chr15-67934998-INS-161 | chr15-67934998-INS-163 |
| chr15-67939664-INS-81 | chr15-67939664-INS-81 |
| chr15-67961049-INS-54 | chr15-67961049-INS-58 |
| chr15-67969200-INS-62 | chr15-67969200-INS-64 |
| chr15-67975993-INS-6402 | chr15-67975994-INS-6402 |
| chr15-67999854-INS-1099 | chr15-67999855-INS-1107 |
| chr15-68024701-INS-1611 | chr15-68024702-INS-1612 |
| chr15-68028169-INS-1078 | chr15-68028170-INS-1074 |
| chr15-68034988-INS-67 | chr15-68034988-INS-52 |
| chr15-68055957-INS-89 | chr15-68055957-INS-89 |
| chr15-68083995-INS-87 | chr15-68083995-INS-87 |
| chr15-68091199-INS-135 | chr15-68091199-INS-140 |
| chr15-68099802-INS-105 | chr15-68099802-INS-60 |
| chr15-68134983-INS-90 | chr15-68134983-INS-92 |
| chr15-68171619-INS-204 | chr15-68171621-INS-234 |
| chr15-68176719-INS-4377 | chr15-68176720-INS-4377 |
| chr15-68183347-INS-744 | chr15-68183346-INS-755 |
| chr15-68215045-INS-59 | chr15-68215045-INS-59 |
| chr15-68226247-INS-74 | chr15-68226233-INS-60 |
| chr15-68240209-INS-214 | chr15-68240163-INS-214 |
| chr15-68274946-INS-1292 | chr15-68274947-INS-1292 |
| chr15-68330018-INS-259 | chr15-68330018-INS-259 |
| chr15-68376313-INS-70 | chr15-68376307-INS-84 |
| chr15-68394161-INS-54 | chr15-68394131-INS-63 |
| chr15-68402540-INS-161 | chr15-68402554-INS-176 |
| chr15-68438693-INS-846 | chr15-68438693-INS-846 |
| chr15-68440382-INS-68 | chr15-68440382-INS-68 |
| chr15-68444464-INS-686 | chr15-68444464-INS-686 |
| chr15-68446286-INS-54 | chr15-68446286-INS-54 |
| chr15-68464127-INS-684 | chr15-68464127-INS-691 |
| chr15-68466486-INS-503 | chr15-68466486-INS-503 |
| chr15-68466740-INS-7061 | chr15-68466741-INS-7061 |
| chr15-68487524-INS-62 | chr15-68487525-INS-56 |
| chr15-68497215-INS-57 | chr15-68497202-INS-60 |
| chr15-68533599-INS-54 | chr15-68533600-INS-54 |
| chr15-68537551-INS-1329 | chr15-68537552-INS-1329 |
| chr15-68539863-INS-56 | chr15-68539863-INS-54 |
| chr15-68569440-INS-56 | chr15-68569440-INS-56 |
| chr15-68613842-INS-5213 | chr15-68613843-INS-5213 |
| chr15-68640691-INS-601 | chr15-68640691-INS-601 |
| chr15-68652922-INS-318 | chr15-68652922-INS-318 |
| chr15-68655954-INS-84 | chr15-68655977-INS-52 |
| chr15-68660211-INS-2989 | chr15-68660212-INS-2989 |

| | |
|---|---|
| chr15-68676635-INS-62 | chr15-68676635-INS-62 |
| chr15-68677265-INS-585 | chr15-68677265-INS-585 |
| chr15-68691246-INS-6796 | chr15-68691247-INS-6796 |
| chr15-68707215-INS-214 | chr15-68707215-INS-214 |
| chr15-68714031-INS-59 | chr15-68714031-INS-59 |
| chr15-68715055-INS-7148 | chr15-68715056-INS-7148 |
| chr15-68715541-INS-63 | chr15-68715538-INS-68 |
| chr15-68719517-INS-6403 | chr15-68719518-INS-6403 |
| chr15-68720656-INS-190 | chr15-68720656-INS-190 |
| chr15-68748528-INS-570 | chr15-68748528-INS-570 |
| chr15-68751379-INS-57 | chr15-68751379-INS-51 |
| chr15-68783966-INS-50 | chr15-68783942-INS-72 |
| chr15-68808158-INS-63 | chr15-68808158-INS-65 |
| chr15-68830870-INS-91 | chr15-68830870-INS-95 |
| chr15-68887539-INS-695 | chr15-68887539-INS-697 |
| chr15-68903757-INS-72 | chr15-68903757-INS-74 |
| chr15-68981371-INS-98 | chr15-68981371-INS-98 |
| chr15-69024577-INS-500 | chr15-69024577-INS-500 |
| chr15-69025793-INS-78 | chr15-69025793-INS-84 |
| chr15-69031131-INS-77 | chr15-69031131-INS-111 |
| chr15-69055047-INS-58 | chr15-69055049-INS-94 |
| chr15-69115606-INS-1899 | chr15-69115607-INS-1899 |
| chr15-69160132-INS-63 | chr15-69160132-INS-51 |
| chr15-69190353-INS-850 | chr15-69190353-INS-849 |
| chr15-69199043-INS-6013 | chr15-69199044-INS-6013 |
| chr15-69213399-INS-185 | chr15-69213386-INS-158 |
| chr15-69217442-INS-6223 | chr15-69217443-INS-6223 |
| chr15-69237124-INS-179 | chr15-69237124-INS-181 |
| chr15-69270957-INS-305 | chr15-69270957-INS-305 |
| chr15-69326939-INS-78 | chr15-69326939-INS-74 |
| chr15-69373968-INS-96 | chr15-69374008-INS-56 |
| chr15-69394354-INS-910 | chr15-69394354-INS-909 |
| chr15-69433340-INS-136 | chr15-69433340-INS-136 |
| chr15-69443501-INS-893 | chr15-69443501-INS-893 |
| chr15-69450465-INS-243 | chr15-69450465-INS-243 |
| chr15-69470110-INS-887 | chr15-69470110-INS-881 |
| chr15-69498151-INS-56 | chr15-69498152-INS-60 |
| chr15-69505493-INS-108 | chr15-69505493-INS-108 |
| chr15-69541302-INS-84 | chr15-69541302-INS-74 |
| chr15-69569788-INS-889 | chr15-69569788-INS-887 |
| chr15-69571605-INS-400 | chr15-69571605-INS-400 |
| chr15-69573547-INS-212 | chr15-69573547-INS-212 |
| chr15-69580297-INS-986 | chr15-69580297-INS-985 |
| chr15-69586926-INS-92 | chr15-69586926-INS-94 |
| chr15-69588890-INS-102 | chr15-69588890-INS-80 |
| chr15-69653351-INS-84 | chr15-69653351-INS-84 |
| chr15-69657584-INS-7065 | chr15-69657585-INS-7065 |
| chr15-69659102-INS-55 | chr15-69659130-INS-65 |
| chr15-69735538-INS-649 | chr15-69735538-INS-649 |
| chr15-69750281-INS-233 | chr15-69750281-INS-233 |
| chr15-69756414-INS-417 | chr15-69756414-INS-417 |
| chr15-69758134-INS-3169 | chr15-69758135-INS-3169 |
| chr15-69777409-INS-50 | chr15-69777407-INS-94 |
| chr15-69782456-INS-185 | chr15-69782457-INS-185 |
| chr15-69904984-INS-263 | chr15-69904978-INS-279 |
| chr15-69907357-INS-387 | chr15-69907357-INS-387 |
| chr15-69916995-INS-74 | chr15-69916985-INS-77 |
| chr15-69943933-INS-453 | chr15-69943933-INS-453 |
| chr15-69967778-INS-142 | chr15-69967794-INS-116 |
| chr15-69968884-INS-87 | chr15-69968884-INS-80 |
| chr15-69979425-INS-754 | chr15-69979425-INS-753 |
| chr15-69997258-INS-52 | chr15-69997258-INS-52 |
| chr15-70017545-INS-54 | chr15-70017545-INS-57 |
| chr15-70046645-INS-82 | chr15-70046643-INS-78 |
| chr15-70050176-INS-54 | chr15-70050176-INS-56 |
| chr15-70065273-INS-60 | chr15-70065273-INS-60 |
| chr15-70097362-INS-407 | chr15-70097362-INS-407 |
| chr15-70127298-INS-1766 | chr15-70127308-INS-1759 |
| chr15-70137043-INS-71 | chr15-70137043-INS-83 |
| chr15-70198157-INS-74 | chr15-70198157-INS-76 |
| chr15-70216910-INS-52 | chr15-70216907-INS-74 |
| chr15-70223858-INS-819 | chr15-70223858-INS-819 |
| chr15-70225115-INS-389 | chr15-70225115-INS-389 |
| chr15-70261810-INS-70 | chr15-70261810-INS-60 |
| chr15-70278998-INS-56 | chr15-70278998-INS-54 |
| chr15-70281727-INS-4818 | chr15-70281728-INS-4818 |
| chr15-70281975-INS-439 | chr15-70281975-INS-439 |

| | |
|---|---|
| chr15-70315161-INS-6958 | chr15-70315162-INS-6958 |
| chr15-70330031-INS-1553 | chr15-70330032-INS-1552 |
| chr15-70355831-INS-68 | chr15-70355831-INS-72 |
| chr15-70363603-INS-944 | chr15-70363603-INS-944 |
| chr15-70389505-INS-1709 | chr15-70389506-INS-1710 |
| chr15-70403898-INS-199 | chr15-70403881-INS-199 |
| chr15-70420307-INS-82 | chr15-70420307-INS-82 |
| chr15-70428411-INS-80 | chr15-70428411-INS-80 |
| chr15-70440065-INS-55 | chr15-70440065-INS-55 |
| chr15-70445858-INS-392 | chr15-70445858-INS-392 |
| chr15-70456991-INS-1092 | chr15-70456992-INS-1092 |
| chr15-70466055-INS-882 | chr15-70466055-INS-881 |
| chr15-70466969-INS-1951 | chr15-70466970-INS-1951 |
| chr15-70521956-INS-763 | chr15-70521956-INS-757 |
| chr15-70558683-INS-51 | chr15-70558684-INS-53 |
| chr15-70591763-INS-56 | chr15-70591763-INS-56 |
| chr15-70593514-INS-72 | chr15-70593514-INS-52 |
| chr15-70602339-INS-106 | chr15-70602339-INS-106 |
| chr15-70608232-INS-67 | chr15-70608245-INS-52 |
| chr15-70629293-INS-69 | chr15-70629293-INS-71 |
| chr15-70631514-INS-2379 | chr15-70631515-INS-2379 |
| chr15-70635994-INS-4479 | chr15-70635995-INS-4479 |
| chr15-70651033-INS-460 | chr15-70651033-INS-460 |
| chr15-70681993-INS-72 | chr15-70681993-INS-64 |
| chr15-70692986-INS-228 | chr15-70692986-INS-228 |
| chr15-70715233-INS-120 | chr15-70715230-INS-123 |
| chr15-70731841-INS-617 | chr15-70731841-INS-616 |
| chr15-70779001-INS-925 | chr15-70779001-INS-925 |
| chr15-70788370-INS-94 | chr15-70788355-INS-62 |
| chr15-70807886-INS-565 | chr15-70807886-INS-579 |
| chr15-70832160-INS-1405 | chr15-70832161-INS-1405 |
| chr15-70858259-INS-79 | chr15-70858259-INS-79 |
| chr15-70875600-INS-56 | chr15-70875600-INS-56 |
| chr15-70892502-INS-383 | chr15-70892502-INS-382 |
| chr15-70895496-INS-366 | chr15-70895496-INS-368 |
| chr15-70940455-INS-62 | chr15-70940463-INS-62 |
| chr15-70951448-INS-6924 | chr15-70951449-INS-6925 |
| chr15-70951449-INS-365 | chr15-70951449-INS-365 |
| chr15-71001676-INS-158 | chr15-71001688-INS-148 |
| chr15-71002414-INS-52 | chr15-71002409-INS-74 |
| chr15-71003056-INS-6897 | chr15-71003057-INS-6897 |
| chr15-71003985-INS-132 | chr15-71003985-INS-132 |
| chr15-71006854-INS-84 | chr15-71006854-INS-84 |
| chr15-71012607-INS-669 | chr15-71012607-INS-668 |
| chr15-71023202-INS-1887 | chr15-71023203-INS-1887 |
| chr15-71044707-INS-2043 | chr15-71044708-INS-2043 |
| chr15-71046352-INS-6357 | chr15-71046353-INS-6357 |
| chr15-71047370-INS-110 | chr15-71047370-INS-111 |
| chr15-71052260-INS-777 | chr15-71052260-INS-777 |
| chr15-71057365-INS-1323 | chr15-71057366-INS-1324 |
| chr15-71095342-INS-4245 | chr15-71095343-INS-4245 |
| chr15-71122534-INS-5193 | chr15-71122535-INS-5193 |
| chr15-71149893-INS-68 | chr15-71149893-INS-74 |
| chr15-71149992-INS-1993 | chr15-71149993-INS-1994 |
| chr15-71191185-INS-674 | chr15-71191185-INS-675 |
| chr15-71210034-INS-106 | chr15-71210034-INS-144 |
| chr15-71210440-INS-1864 | chr15-71210441-INS-1864 |
| chr15-71241741-INS-453 | chr15-71241741-INS-453 |
| chr15-71268739-INS-901 | chr15-71268739-INS-901 |
| chr15-71271271-INS-53 | chr15-71271271-INS-53 |
| chr15-71425016-INS-74 | chr15-71425016-INS-74 |
| chr15-71445896-INS-70 | chr15-71445878-INS-111 |
| chr15-71450476-INS-147 | chr15-71450476-INS-139 |
| chr15-71454969-INS-56 | chr15-71454969-INS-60 |
| chr15-71478564-INS-226 | chr15-71478564-INS-221 |
| chr15-71566970-INS-130 | chr15-71566970-INS-130 |
| chr15-71576727-INS-75 | chr15-71576727-INS-71 |
| chr15-71586614-INS-214 | chr15-71586626-INS-244 |
| chr15-71590309-INS-60 | chr15-71590309-INS-60 |
| chr15-71641831-INS-75 | chr15-71641824-INS-64 |
| chr15-71668527-INS-116 | chr15-71668525-INS-118 |
| chr15-71685294-INS-320 | chr15-71685294-INS-320 |
| chr15-71690926-INS-140 | chr15-71690926-INS-140 |
| chr15-71700173-INS-329 | chr15-71700173-INS-329 |
| chr15-71707921-INS-901 | chr15-71707921-INS-898 |
| chr15-71723850-INS-269 | chr15-71723850-INS-269 |
| chr15-71728469-INS-7291 | chr15-71728470-INS-7291 |

| | |
|---|---|
| chr15-71730311-INS-62 | chr15-71730311-INS-66 |
| chr15-71763432-INS-55 | chr15-71763432-INS-55 |
| chr15-71776181-INS-56 | chr15-71776181-INS-62 |
| chr15-71777270-INS-52 | chr15-71777270-INS-52 |
| chr15-71792037-INS-60 | chr15-71792037-INS-58 |
| chr15-71795678-INS-65 | chr15-71795678-INS-65 |
| chr15-71820192-INS-53 | chr15-71820192-INS-51 |
| chr15-71828329-INS-72 | chr15-71828280-INS-88 |
| chr15-71835648-INS-230 | chr15-71835648-INS-230 |
| chr15-71837942-INS-58 | chr15-71837942-INS-78 |
| chr15-71839850-INS-139 | chr15-71839850-INS-139 |
| chr15-71844948-INS-657 | chr15-71844949-INS-651 |
| chr15-71857172-INS-1212 | chr15-71857169-INS-1209 |
| chr15-71861147-INS-301 | chr15-71861159-INS-277 |
| chr15-71879331-INS-68 | chr15-71879331-INS-76 |
| chr15-71886191-INS-1156 | chr15-71886192-INS-1156 |
| chr15-71911099-INS-6415 | chr15-71911100-INS-6415 |
| chr15-71923382-INS-87 | chr15-71923382-INS-97 |
| chr15-71938155-INS-218 | chr15-71938156-INS-227 |
| chr15-71973038-INS-106 | chr15-71973054-INS-85 |
| chr15-71991053-INS-1198 | chr15-71991054-INS-1198 |
| chr15-71996849-INS-1218 | chr15-71996850-INS-1218 |
| chr15-72000104-INS-145 | chr15-72000104-INS-145 |
| chr15-72011769-INS-7130 | chr15-72011770-INS-7130 |
| chr15-72048976-INS-3256 | chr15-72048977-INS-3256 |
| chr15-72050476-INS-6447 | chr15-72050477-INS-6447 |
| chr15-72085225-INS-176 | chr15-72085225-INS-177 |
| chr15-72103366-INS-96 | chr15-72103363-INS-72 |
| chr15-72135004-INS-556 | chr15-72135004-INS-556 |
| chr15-72165225-INS-4732 | chr15-72165226-INS-4732 |
| chr15-72166191-INS-307 | chr15-72166191-INS-307 |
| chr15-72173080-INS-60 | chr15-72173078-INS-62 |
| chr15-72179433-INS-5264 | chr15-72179434-INS-5264 |
| chr15-72181661-INS-684 | chr15-72181661-INS-684 |
| chr15-72194705-INS-1001 | chr15-72194706-INS-1001 |
| chr15-72225423-INS-6254 | chr15-72225424-INS-6255 |
| chr15-72238545-INS-6190 | chr15-72238546-INS-6191 |
| chr15-72253995-INS-52 | chr15-72253995-INS-54 |
| chr15-72255221-INS-50 | chr15-72255205-INS-54 |
| chr15-72260237-INS-81 | chr15-72260237-INS-81 |
| chr15-72267634-INS-221 | chr15-72267634-INS-224 |
| chr15-72283331-INS-494 | chr15-72283331-INS-494 |
| chr15-72328878-INS-402 | chr15-72328878-INS-402 |
| chr15-72339018-INS-6413 | chr15-72339019-INS-6413 |
| chr15-72341205-INS-64 | chr15-72341205-INS-68 |
| chr15-72345474-INS-65 | chr15-72345462-INS-75 |
| chr15-72347082-INS-55 | chr15-72347082-INS-55 |
| chr15-72396804-INS-558 | chr15-72396804-INS-558 |
| chr15-72423627-INS-52 | chr15-72423627-INS-58 |
| chr15-72425890-INS-158 | chr15-72425890-INS-158 |
| chr15-72436623-INS-342 | chr15-72436623-INS-342 |
| chr15-72439426-INS-127 | chr15-72439433-INS-168 |
| chr15-72450138-INS-1534 | chr15-72450139-INS-1518 |
| chr15-72461158-INS-2703 | chr15-72461159-INS-2703 |
| chr15-72478271-INS-56 | chr15-72478271-INS-56 |
| chr15-72526714-INS-60 | chr15-72526714-INS-60 |
| chr15-72540099-INS-6403 | chr15-72540100-INS-6403 |
| chr15-72558515-INS-427 | chr15-72558515-INS-436 |
| chr15-72604001-INS-767 | chr15-72604001-INS-760 |
| chr15-72621941-INS-70 | chr15-72621941-INS-80 |
| chr15-72642005-INS-56 | chr15-72641991-INS-67 |
| chr15-72720750-INS-68 | chr15-72720750-INS-68 |
| chr15-72763359-INS-401 | chr15-72763359-INS-402 |
| chr15-72777566-INS-58 | chr15-72777585-INS-85 |
| chr15-72797704-INS-105 | chr15-72797705-INS-65 |
| chr15-72813020-INS-52 | chr15-72813018-INS-58 |
| chr15-72820121-INS-84 | chr15-72820121-INS-84 |
| chr15-72851548-INS-194 | chr15-72851548-INS-194 |
| chr15-72856372-INS-238 | chr15-72856372-INS-238 |
| chr15-72865945-INS-56 | chr15-72865945-INS-55 |
| chr15-72897685-INS-198 | chr15-72897685-INS-198 |
| chr15-72900419-INS-58 | chr15-72900422-INS-62 |
| chr15-72947196-INS-829 | chr15-72947196-INS-827 |
| chr15-72948407-INS-76 | chr15-72948407-INS-88 |
| chr15-72949772-INS-61 | chr15-72949772-INS-61 |
| chr15-73007437-INS-358 | chr15-73007437-INS-358 |
| chr15-73014919-INS-319 | chr15-73014919-INS-319 |

| | |
|---|---|
| chr15-73035236-INS-202 | chr15-73035236-INS-218 |
| chr15-73048130-INS-50 | chr15-73048131-INS-61 |
| chr15-73058224-INS-67 | chr15-73058226-INS-53 |
| chr15-73067034-INS-505 | chr15-73067034-INS-505 |
| chr15-73088558-INS-212 | chr15-73088558-INS-211 |
| chr15-73137214-INS-108 | chr15-73137214-INS-108 |
| chr15-73159244-INS-881 | chr15-73159244-INS-880 |
| chr15-73186576-INS-1889 | chr15-73186577-INS-1889 |
| chr15-73188678-INS-209 | chr15-73188678-INS-209 |
| chr15-73205295-INS-179 | chr15-73205295-INS-177 |
| chr15-73290893-INS-108 | chr15-73290909-INS-76 |
| chr15-73314179-INS-56 | chr15-73314171-INS-76 |
| chr15-73317409-INS-203 | chr15-73317409-INS-202 |
| chr15-73329902-INS-701 | chr15-73329902-INS-702 |
| chr15-73330588-INS-56 | chr15-73330598-INS-51 |
| chr15-73333325-INS-192 | chr15-73333325-INS-192 |
| chr15-73341104-INS-163 | chr15-73341104-INS-163 |
| chr15-73346111-INS-84 | chr15-73346111-INS-84 |
| chr15-73350929-INS-206 | chr15-73350929-INS-205 |
| chr15-73356048-INS-371 | chr15-73356048-INS-366 |
| chr15-73357281-INS-84 | chr15-73357281-INS-74 |
| chr15-73357864-INS-505 | chr15-73357864-INS-521 |
| chr15-73367594-INS-109 | chr15-73367594-INS-109 |
| chr15-73381514-INS-56 | chr15-73381514-INS-65 |
| chr15-73400904-INS-210 | chr15-73400904-INS-210 |
| chr15-73403702-INS-221 | chr15-73403702-INS-221 |
| chr15-73409432-INS-52 | chr15-73409432-INS-52 |
| chr15-73432545-INS-187 | chr15-73432545-INS-187 |
| chr15-73439680-INS-64 | chr15-73439680-INS-64 |
| chr15-73447778-INS-191 | chr15-73447778-INS-191 |
| chr15-73473256-INS-148 | chr15-73473256-INS-148 |
| chr15-73503466-INS-64 | chr15-73503466-INS-58 |
| chr15-73520618-INS-198 | chr15-73520618-INS-198 |
| chr15-73528104-INS-69 | chr15-73528104-INS-69 |
| chr15-73574180-INS-72 | chr15-73574180-INS-75 |
| chr15-73577944-INS-78 | chr15-73577942-INS-94 |
| chr15-73587578-INS-67 | chr15-73587577-INS-66 |
| chr15-73609277-INS-96 | chr15-73609277-INS-93 |
| chr15-73626006-INS-68 | chr15-73626006-INS-68 |
| chr15-73639433-INS-58 | chr15-73639433-INS-74 |
| chr15-73656060-INS-295 | chr15-73656061-INS-295 |
| chr15-73677766-INS-6442 | chr15-73677767-INS-6442 |
| chr15-73705533-INS-5313 | chr15-73705534-INS-5313 |
| chr15-73722171-INS-84 | chr15-73722171-INS-84 |
| chr15-73766473-INS-441 | chr15-73766473-INS-441 |
| chr15-73782288-INS-6473 | chr15-73782289-INS-6473 |
| chr15-73783628-INS-538 | chr15-73783628-INS-538 |
| chr15-73801444-INS-67 | chr15-73801444-INS-67 |
| chr15-73821375-INS-80 | chr15-73821375-INS-80 |
| chr15-73823004-INS-747 | chr15-73823004-INS-746 |
| chr15-73850192-INS-148 | chr15-73850192-INS-160 |
| chr15-73875741-INS-144 | chr15-73875741-INS-144 |
| chr15-73881413-INS-116 | chr15-73881413-INS-120 |
| chr15-73967355-INS-142 | chr15-73967330-INS-139 |
| chr15-73997954-INS-58 | chr15-73997955-INS-50 |
| chr15-74059611-INS-59 | chr15-74059611-INS-59 |
| chr15-74064618-INS-52 | chr15-74064618-INS-54 |
| chr15-74067698-INS-64 | chr15-74067698-INS-64 |
| chr15-74078389-INS-51 | chr15-74078389-INS-57 |
| chr15-74082085-INS-69 | chr15-74082085-INS-69 |
| chr15-74103810-INS-113 | chr15-74103810-INS-113 |
| chr15-74105495-INS-920 | chr15-74105495-INS-921 |
| chr15-74130965-INS-1035 | chr15-74130966-INS-1037 |
| chr15-74152566-INS-94 | chr15-74152566-INS-94 |
| chr15-74159757-INS-89 | chr15-74159757-INS-89 |
| chr15-74162605-INS-143 | chr15-74162605-INS-150 |
| chr15-74169509-INS-216 | chr15-74169509-INS-216 |
| chr15-74188157-INS-204 | chr15-74188157-INS-204 |
| chr15-74191669-INS-81 | chr15-74191669-INS-77 |
| chr15-74245803-INS-76 | chr15-74245795-INS-54 |
| chr15-74250359-INS-69 | chr15-74250359-INS-59 |
| chr15-74267453-INS-52 | chr15-74267454-INS-54 |
| chr15-74279875-INS-206 | chr15-74279875-INS-222 |
| chr15-74286943-INS-81 | chr15-74286943-INS-107 |
| chr15-74288342-INS-56 | chr15-74288342-INS-68 |
| chr15-74360136-INS-302 | chr15-74360153-INS-268 |
| chr15-74367260-INS-86 | chr15-74367260-INS-86 |

| | |
|---|---|
| chr15-74386740-INS-75 | chr15-74386740-INS-62 |
| chr15-74395025-INS-121 | chr15-74395025-INS-121 |
| chr15-74417487-INS-286 | chr15-74417487-INS-286 |
| chr15-74445220-INS-66 | chr15-74445220-INS-66 |
| chr15-74524884-INS-210 | chr15-74524884-INS-210 |
| chr15-74525953-INS-364 | chr15-74525953-INS-364 |
| chr15-74553049-INS-206 | chr15-74553049-INS-207 |
| chr15-74560168-INS-50 | chr15-74560179-INS-71 |
| chr15-74561635-INS-1593 | chr15-74561636-INS-1593 |
| chr15-74562238-INS-160 | chr15-74562238-INS-160 |
| chr15-74566757-INS-990 | chr15-74566757-INS-990 |
| chr15-74597301-INS-65 | chr15-74597301-INS-65 |
| chr15-74603987-INS-71 | chr15-74603987-INS-71 |
| chr15-74617997-INS-472 | chr15-74617997-INS-472 |
| chr15-74619282-INS-214 | chr15-74619282-INS-213 |
| chr15-74628051-INS-175 | chr15-74628051-INS-175 |
| chr15-74640882-INS-56 | chr15-74640882-INS-54 |
| chr15-74669877-INS-67 | chr15-74669877-INS-67 |
| chr15-74673049-INS-60 | chr15-74673049-INS-60 |
| chr15-74706276-INS-129 | chr15-74706274-INS-131 |
| chr15-74737642-INS-498 | chr15-74737642-INS-498 |
| chr15-74753097-INS-66 | chr15-74753097-INS-62 |
| chr15-74759773-INS-52 | chr15-74759773-INS-52 |
| chr15-74766529-INS-144 | chr15-74766529-INS-146 |
| chr15-74788701-INS-65 | chr15-74788701-INS-65 |
| chr15-74803816-INS-535 | chr15-74803794-INS-535 |
| chr15-74808695-INS-515 | chr15-74808695-INS-515 |
| chr15-74825534-INS-536 | chr15-74825534-INS-536 |
| chr15-74829320-INS-434 | chr15-74829320-INS-434 |
| chr15-74866541-INS-134 | chr15-74866541-INS-134 |
| chr15-74932425-INS-445 | chr15-74932425-INS-445 |
| chr15-75010401-INS-55 | chr15-75010401-INS-100 |
| chr15-75020473-INS-144 | chr15-75020491-INS-126 |
| chr15-75028375-INS-60 | chr15-75028375-INS-60 |
| chr15-75062084-INS-161 | chr15-75062088-INS-184 |
| chr15-75063885-INS-180 | chr15-75063885-INS-180 |
| chr15-75086021-INS-74 | chr15-75086001-INS-79 |
| chr15-75095783-INS-53 | chr15-75095782-INS-56 |
| chr15-75116216-INS-552 | chr15-75116219-INS-547 |
| chr15-75138826-INS-65 | chr15-75138826-INS-73 |
| chr15-75139932-INS-578 | chr15-75139941-INS-590 |
| chr15-75173353-INS-205 | chr15-75173353-INS-205 |
| chr15-75184593-INS-297 | chr15-75184606-INS-291 |
| chr15-75212570-INS-475 | chr15-75212570-INS-475 |
| chr15-75217942-INS-444 | chr15-75217942-INS-444 |
| chr15-75218087-INS-2095 | chr15-75218056-INS-2118 |
| chr15-75223897-INS-737 | chr15-75223897-INS-742 |
| chr15-75227518-INS-1077 | chr15-75227519-INS-1077 |
| chr15-75230214-INS-72 | chr15-75230214-INS-72 |
| chr15-75235371-INS-212 | chr15-75235371-INS-213 |
| chr15-75244628-INS-592 | chr15-75244628-INS-592 |
| chr15-75253511-INS-199 | chr15-75253511-INS-199 |
| chr15-75280741-INS-332 | chr15-75280742-INS-332 |
| chr15-75289995-INS-4823 | chr15-75289996-INS-4823 |
| chr15-75297009-INS-1023 | chr15-75297010-INS-989 |
| chr15-75297708-INS-220 | chr15-75297706-INS-226 |
| chr15-75314459-INS-310 | chr15-75314459-INS-310 |
| chr15-75350499-INS-204 | chr15-75350499-INS-204 |
| chr15-75356671-INS-9809 | chr15-75356672-INS-9810 |
| chr15-75405789-INS-459 | chr15-75405789-INS-459 |
| chr15-75415487-INS-6476 | chr15-75415488-INS-6476 |
| chr15-75449770-INS-2833 | chr15-75449783-INS-2826 |
| chr15-75450789-INS-394 | chr15-75450790-INS-400 |
| chr15-75547631-INS-128 | chr15-75547631-INS-128 |
| chr15-75552898-INS-787 | chr15-75552898-INS-762 |
| chr15-75622265-INS-192 | chr15-75622265-INS-192 |
| chr15-75640688-INS-206 | chr15-75640688-INS-206 |
| chr15-75749751-INS-215 | chr15-75749751-INS-215 |
| chr15-75771302-INS-497 | chr15-75771302-INS-492 |
| chr15-75787845-INS-4728 | chr15-75787846-INS-4728 |
| chr15-75835215-INS-90 | chr15-75835208-INS-89 |
| chr15-75846252-INS-203 | chr15-75846252-INS-203 |
| chr15-75891298-INS-786 | chr15-75891298-INS-785 |
| chr15-75893758-INS-310 | chr15-75893758-INS-281 |
| chr15-75895680-INS-100 | chr15-75895680-INS-72 |
| chr15-75914628-INS-111 | chr15-75914628-INS-61 |
| chr15-75949408-INS-284 | chr15-75949408-INS-280 |

| | |
|---|---|
| chr15-75954079-INS-211 | chr15-75954078-INS-213 |
| chr15-76034637-INS-202 | chr15-76034637-INS-202 |
| chr15-76037303-INS-198 | chr15-76037303-INS-198 |
| chr15-76038955-INS-86 | chr15-76038953-INS-59 |
| chr15-76044111-INS-204 | chr15-76044111-INS-205 |
| chr15-76050932-INS-222 | chr15-76050932-INS-223 |
| chr15-76052938-INS-5581 | chr15-76052939-INS-5581 |
| chr15-76099228-INS-224 | chr15-76099223-INS-210 |
| chr15-76151825-INS-62 | chr15-76151825-INS-62 |
| chr15-76162151-INS-65 | chr15-76162151-INS-65 |
| chr15-76193196-INS-68 | chr15-76193196-INS-68 |
| chr15-76197458-INS-215 | chr15-76197458-INS-214 |
| chr15-76252456-INS-208 | chr15-76252456-INS-207 |
| chr15-76267133-INS-220 | chr15-76267133-INS-220 |
| chr15-76271484-INS-192 | chr15-76271484-INS-192 |
| chr15-76290822-INS-401 | chr15-76290822-INS-401 |
| chr15-76293364-INS-226 | chr15-76293364-INS-226 |
| chr15-76296935-INS-132 | chr15-76296935-INS-132 |
| chr15-76299598-INS-6439 | chr15-76299599-INS-6439 |
| chr15-76300318-INS-87 | chr15-76300318-INS-91 |
| chr15-76307620-INS-53 | chr15-76307620-INS-53 |
| chr15-76315469-INS-199 | chr15-76315469-INS-199 |
| chr15-76330242-INS-219 | chr15-76330242-INS-219 |
| chr15-76357098-INS-222 | chr15-76357098-INS-222 |
| chr15-76363599-INS-210 | chr15-76363599-INS-211 |
| chr15-76369734-INS-1418 | chr15-76369735-INS-1417 |
| chr15-76426155-INS-214 | chr15-76426155-INS-215 |
| chr15-76435177-INS-148 | chr15-76435184-INS-132 |
| chr15-76452263-INS-124 | chr15-76452263-INS-125 |
| chr15-76455897-INS-195 | chr15-76455897-INS-190 |
| chr15-76457473-INS-214 | chr15-76457473-INS-219 |
| chr15-76466594-INS-211 | chr15-76466595-INS-202 |
| chr15-76488880-INS-225 | chr15-76488880-INS-227 |
| chr15-76503661-INS-123 | chr15-76503661-INS-123 |
| chr15-76557488-INS-131 | chr15-76557488-INS-131 |
| chr15-76565813-INS-224 | chr15-76565813-INS-223 |
| chr15-76603422-INS-178 | chr15-76603433-INS-178 |
| chr15-76616263-INS-117 | chr15-76616266-INS-158 |
| chr15-76633645-INS-120 | chr15-76633645-INS-120 |
| chr15-76643351-INS-90 | chr15-76643351-INS-90 |
| chr15-76718265-INS-7238 | chr15-76718266-INS-7238 |
| chr15-76718756-INS-65 | chr15-76718756-INS-56 |
| chr15-76722810-INS-60 | chr15-76722810-INS-63 |
| chr15-76747287-INS-206 | chr15-76747287-INS-206 |
| chr15-76753333-INS-170 | chr15-76753343-INS-181 |
| chr15-76765951-INS-214 | chr15-76765951-INS-214 |
| chr15-76778444-INS-58 | chr15-76778444-INS-58 |
| chr15-76779495-INS-107 | chr15-76779496-INS-60 |
| chr15-76811298-INS-75 | chr15-76811299-INS-75 |
| chr15-76816775-INS-142 | chr15-76816775-INS-142 |
| chr15-76822046-INS-350 | chr15-76822046-INS-350 |
| chr15-76833666-INS-142 | chr15-76833662-INS-149 |
| chr15-76835096-INS-112 | chr15-76835096-INS-112 |
| chr15-76836293-INS-226 | chr15-76836293-INS-184 |
| chr15-76836838-INS-230 | chr15-76836838-INS-218 |
| chr15-76860032-INS-124 | chr15-76860033-INS-124 |
| chr15-76865333-INS-201 | chr15-76865333-INS-201 |
| chr15-76868556-INS-156 | chr15-76868556-INS-167 |
| chr15-76902032-INS-416 | chr15-76902032-INS-416 |
| chr15-76910073-INS-80 | chr15-76910064-INS-56 |
| chr15-76912507-INS-111 | chr15-76912507-INS-111 |
| chr15-76917515-INS-54 | chr15-76917545-INS-51 |
| chr15-76922749-INS-90 | chr15-76922739-INS-90 |
| chr15-76944283-INS-3049 | chr15-76944284-INS-3049 |
| chr15-76948452-INS-263 | chr15-76948452-INS-263 |
| chr15-76950176-INS-56 | chr15-76950177-INS-54 |
| chr15-76951354-INS-399 | chr15-76951354-INS-399 |
| chr15-77013953-INS-53 | chr15-77013953-INS-53 |
| chr15-77078701-INS-981 | chr15-77078701-INS-982 |
| chr15-77089781-INS-150 | chr15-77089781-INS-150 |
| chr15-77139132-INS-60 | chr15-77139132-INS-65 |
| chr15-77161782-INS-235 | chr15-77161782-INS-235 |
| chr15-77172820-INS-236 | chr15-77172820-INS-236 |
| chr15-77198415-INS-184 | chr15-77198416-INS-184 |
| chr15-77199707-INS-66 | chr15-77199704-INS-57 |
| chr15-77217464-INS-229 | chr15-77217464-INS-224 |
| chr15-77251984-INS-990 | chr15-77251982-INS-960 |

| | |
|---|---|
| chr15-77308439-INS-1547 | chr15-77308440-INS-1569 |
| chr15-77332986-INS-197 | chr15-77332985-INS-195 |
| chr15-77334880-INS-196 | chr15-77334880-INS-197 |
| chr15-77345788-INS-5634 | chr15-77345789-INS-5634 |
| chr15-77411542-INS-63 | chr15-77411542-INS-63 |
| chr15-77432224-INS-396 | chr15-77432224-INS-384 |
| chr15-77441824-INS-72 | chr15-77441824-INS-72 |
| chr15-77445339-INS-304 | chr15-77445339-INS-311 |
| chr15-77451929-INS-952 | chr15-77451929-INS-983 |
| chr15-77461390-INS-783 | chr15-77461390-INS-783 |
| chr15-77463682-INS-4609 | chr15-77463683-INS-4609 |
| chr15-77487679-INS-66 | chr15-77487679-INS-75 |
| chr15-77498320-INS-208 | chr15-77498320-INS-208 |
| chr15-77507388-INS-90 | chr15-77507388-INS-72 |
| chr15-77525689-INS-634 | chr15-77525689-INS-628 |
| chr15-77593855-INS-68 | chr15-77593855-INS-68 |
| chr15-77598080-INS-130 | chr15-77598080-INS-130 |
| chr15-77643122-INS-186 | chr15-77643122-INS-185 |
| chr15-77649826-INS-87 | chr15-77649826-INS-87 |
| chr15-77669647-INS-165 | chr15-77669647-INS-172 |
| chr15-77691225-INS-1884 | chr15-77691226-INS-1884 |
| chr15-77694010-INS-60 | chr15-77693992-INS-54 |
| chr15-77732083-INS-55 | chr15-77732083-INS-55 |
| chr15-77736730-INS-246 | chr15-77736730-INS-247 |
| chr15-77757438-INS-213 | chr15-77757438-INS-213 |
| chr15-77762143-INS-1491 | chr15-77762144-INS-1494 |
| chr15-77762512-INS-124 | chr15-77762512-INS-155 |
| chr15-77768479-INS-226 | chr15-77768479-INS-228 |
| chr15-77769025-INS-246 | chr15-77769025-INS-246 |
| chr15-77773751-INS-194 | chr15-77773751-INS-214 |
| chr15-77816137-INS-176 | chr15-77816137-INS-179 |
| chr15-77821954-INS-219 | chr15-77821954-INS-219 |
| chr15-77824032-INS-220 | chr15-77824032-INS-220 |
| chr15-77835017-INS-7104 | chr15-77835018-INS-7104 |
| chr15-77859567-INS-81 | chr15-77859567-INS-50 |
| chr15-77869541-INS-80 | chr15-77869541-INS-56 |
| chr15-77874954-INS-67 | chr15-77874954-INS-55 |
| chr15-77901372-INS-54 | chr15-77901375-INS-68 |
| chr15-77943148-INS-90 | chr15-77943136-INS-90 |
| chr15-77951091-INS-149 | chr15-77951096-INS-172 |
| chr15-78010688-INS-1015 | chr15-78010721-INS-1016 |
| chr15-78012697-INS-101 | chr15-78012697-INS-86 |
| chr15-78074031-INS-96 | chr15-78074031-INS-96 |
| chr15-78098387-INS-202 | chr15-78098388-INS-202 |
| chr15-78137416-INS-75 | chr15-78137416-INS-75 |
| chr15-78211079-INS-54 | chr15-78211051-INS-72 |
| chr15-78273593-INS-90 | chr15-78273583-INS-110 |
| chr15-78284761-INS-222 | chr15-78284761-INS-222 |
| chr15-78297551-INS-182 | chr15-78297551-INS-182 |
| chr15-78301789-INS-145 | chr15-78301785-INS-130 |
| chr15-78335341-INS-106 | chr15-78335341-INS-132 |
| chr15-78342585-INS-100 | chr15-78342619-INS-80 |
| chr15-78386565-INS-170 | chr15-78386561-INS-196 |
| chr15-78447671-INS-169 | chr15-78447671-INS-172 |
| chr15-78515292-INS-75 | chr15-78515292-INS-75 |
| chr15-78523694-INS-52 | chr15-78523694-INS-52 |
| chr15-78624649-INS-65 | chr15-78624658-INS-65 |
| chr15-78631327-INS-61 | chr15-78631328-INS-61 |
| chr15-78647080-INS-158 | chr15-78647080-INS-191 |
| chr15-78650792-INS-210 | chr15-78650792-INS-208 |
| chr15-78683018-INS-202 | chr15-78683018-INS-202 |
| chr15-78686194-INS-112 | chr15-78686194-INS-110 |
| chr15-78721069-INS-80 | chr15-78721069-INS-52 |
| chr15-78736994-INS-51 | chr15-78736994-INS-75 |
| chr15-78801367-INS-213 | chr15-78801367-INS-213 |
| chr15-78821172-INS-59 | chr15-78821172-INS-59 |
| chr15-78828899-INS-102 | chr15-78828899-INS-105 |
| chr15-78833995-INS-94 | chr15-78833993-INS-50 |
| chr15-78834630-INS-303 | chr15-78834630-INS-283 |
| chr15-78843430-INS-400 | chr15-78843430-INS-400 |
| chr15-78845551-INS-77 | chr15-78845551-INS-96 |
| chr15-78849998-INS-223 | chr15-78849998-INS-224 |
| chr15-78864648-INS-112 | chr15-78864648-INS-112 |
| chr15-78954282-INS-157 | chr15-78954282-INS-159 |
| chr15-78961888-INS-202 | chr15-78961888-INS-202 |
| chr15-78967533-INS-210 | chr15-78967533-INS-209 |
| chr15-79005671-INS-225 | chr15-79005671-INS-225 |

| | |
|---|---|
| chr15-79054388-INS-72 | chr15-79054388-INS-76 |
| chr15-79101581-INS-476 | chr15-79101581-INS-457 |
| chr15-79102944-INS-163 | chr15-79102944-INS-165 |
| chr15-79151097-INS-188 | chr15-79151097-INS-188 |
| chr15-79157642-INS-62 | chr15-79157642-INS-64 |
| chr15-79161851-INS-62 | chr15-79161851-INS-62 |
| chr15-79168509-INS-79 | chr15-79168509-INS-52 |
| chr15-79177733-INS-76 | chr15-79177748-INS-80 |
| chr15-79181096-INS-484 | chr15-79181096-INS-483 |
| chr15-79208319-INS-216 | chr15-79208319-INS-216 |
| chr15-79216715-INS-179 | chr15-79216715-INS-179 |
| chr15-79224502-INS-145 | chr15-79224502-INS-145 |
| chr15-79254217-INS-208 | chr15-79254217-INS-208 |
| chr15-79259723-INS-198 | chr15-79259723-INS-198 |
| chr15-79267685-INS-851 | chr15-79267685-INS-851 |
| chr15-79281754-INS-113 | chr15-79281754-INS-113 |
| chr15-79283360-INS-210 | chr15-79283383-INS-187 |
| chr15-79311582-INS-162 | chr15-79311582-INS-162 |
| chr15-79331295-INS-54 | chr15-79331293-INS-54 |
| chr15-79341507-INS-191 | chr15-79341507-INS-183 |
| chr15-79397468-INS-271 | chr15-79397468-INS-287 |
| chr15-79441236-INS-446 | chr15-79441236-INS-446 |
| chr15-79455994-INS-161 | chr15-79455995-INS-157 |
| chr15-79457171-INS-166 | chr15-79457171-INS-165 |
| chr15-79468038-INS-7110 | chr15-79468039-INS-7110 |
| chr15-79470321-INS-206 | chr15-79470321-INS-207 |
| chr15-79471743-INS-333 | chr15-79471743-INS-333 |
| chr15-79479484-INS-220 | chr15-79479484-INS-220 |
| chr15-79491952-INS-182 | chr15-79491952-INS-183 |
| chr15-79521584-INS-56 | chr15-79521584-INS-56 |
| chr15-79526035-INS-154 | chr15-79526035-INS-154 |
| chr15-79532334-INS-65 | chr15-79532330-INS-68 |
| chr15-79537704-INS-205 | chr15-79537704-INS-240 |
| chr15-79545028-INS-55 | chr15-79545034-INS-57 |
| chr15-79552465-INS-639 | chr15-79552465-INS-639 |
| chr15-79582493-INS-108 | chr15-79582493-INS-100 |
| chr15-79622361-INS-78 | chr15-79622365-INS-72 |
| chr15-79628082-INS-944 | chr15-79628082-INS-927 |
| chr15-79634493-INS-221 | chr15-79634493-INS-221 |
| chr15-79635418-INS-304 | chr15-79635418-INS-304 |
| chr15-79636042-INS-2869 | chr15-79636043-INS-2884 |
| chr15-79641883-INS-215 | chr15-79641883-INS-219 |
| chr15-79668275-INS-208 | chr15-79668275-INS-208 |
| chr15-79692904-INS-193 | chr15-79692904-INS-193 |
| chr15-79700503-INS-66 | chr15-79700503-INS-66 |
| chr15-79707815-INS-432 | chr15-79707815-INS-432 |
| chr15-79722805-INS-213 | chr15-79722805-INS-213 |
| chr15-79729592-INS-192 | chr15-79729592-INS-192 |
| chr15-79733944-INS-215 | chr15-79733944-INS-215 |
| chr15-79737786-INS-80 | chr15-79737786-INS-88 |
| chr15-79741389-INS-76 | chr15-79741388-INS-80 |
| chr15-79744374-INS-626 | chr15-79744374-INS-626 |
| chr15-79746315-INS-222 | chr15-79746316-INS-221 |
| chr15-79747735-INS-425 | chr15-79747737-INS-399 |
| chr15-79760762-INS-210 | chr15-79760762-INS-210 |
| chr15-79766392-INS-140 | chr15-79766392-INS-143 |
| chr15-79770497-INS-137 | chr15-79770516-INS-149 |
| chr15-79821131-INS-192 | chr15-79821131-INS-192 |
| chr15-79823222-INS-199 | chr15-79823222-INS-199 |
| chr15-79824724-INS-99 | chr15-79824724-INS-111 |
| chr15-79828406-INS-195 | chr15-79828406-INS-195 |
| chr15-79839738-INS-54 | chr15-79839738-INS-61 |
| chr15-79843013-INS-210 | chr15-79843013-INS-210 |
| chr15-79844496-INS-218 | chr15-79844496-INS-220 |
| chr15-79864817-INS-220 | chr15-79864817-INS-220 |
| chr15-79895198-INS-226 | chr15-79895198-INS-226 |
| chr15-79904970-INS-285 | chr15-79904970-INS-290 |
| chr15-79915550-INS-393 | chr15-79915548-INS-405 |
| chr15-79916253-INS-217 | chr15-79916253-INS-227 |
| chr15-79925998-INS-192 | chr15-79925998-INS-193 |
| chr15-79930908-INS-62 | chr15-79930936-INS-57 |
| chr15-79943482-INS-194 | chr15-79943482-INS-192 |
| chr15-79952547-INS-56 | chr15-79952558-INS-70 |
| chr15-79953668-INS-198 | chr15-79953668-INS-198 |
| chr15-79955288-INS-52 | chr15-79955288-INS-60 |
| chr15-79969182-INS-210 | chr15-79969182-INS-204 |
| chr15-79970540-INS-72 | chr15-79970565-INS-82 |

| | |
|---|---|
| chr15-79973167-INS-201 | chr15-79973167-INS-201 |
| chr15-80029857-INS-235 | chr15-80029857-INS-223 |
| chr15-80056799-INS-129 | chr15-80056798-INS-175 |
| chr15-80066358-INS-158 | chr15-80066358-INS-158 |
| chr15-80072369-INS-121 | chr15-80072369-INS-121 |
| chr15-80079113-INS-1002 | chr15-80079114-INS-991 |
| chr15-80083099-INS-159 | chr15-80083099-INS-159 |
| chr15-80091616-INS-72 | chr15-80091616-INS-68 |
| chr15-80094566-INS-100 | chr15-80094566-INS-104 |
| chr15-80155305-INS-141 | chr15-80155310-INS-147 |
| chr15-80164348-INS-64 | chr15-80164348-INS-72 |
| chr15-80184566-INS-77 | chr15-80184566-INS-66 |
| chr15-80192655-INS-160 | chr15-80192655-INS-156 |
| chr15-80207694-INS-74 | chr15-80207694-INS-93 |
| chr15-80222036-INS-63 | chr15-80222036-INS-59 |
| chr15-80240566-INS-75 | chr15-80240566-INS-84 |
| chr15-80254760-INS-147 | chr15-80254759-INS-146 |
| chr15-80285338-INS-72 | chr15-80285338-INS-72 |
| chr15-80296309-INS-60 | chr15-80296309-INS-60 |
| chr15-80296518-INS-6437 | chr15-80296519-INS-6438 |
| chr15-80342685-INS-75 | chr15-80342685-INS-75 |
| chr15-80343742-INS-90 | chr15-80343742-INS-90 |
| chr15-80362298-INS-65 | chr15-80362298-INS-65 |
| chr15-80381444-INS-1099 | chr15-80381445-INS-1099 |
| chr15-80390868-INS-68 | chr15-80390868-INS-67 |
| chr15-80430704-INS-58 | chr15-80430704-INS-81 |
| chr15-80441644-INS-221 | chr15-80441644-INS-222 |
| chr15-80473915-INS-631 | chr15-80473906-INS-620 |
| chr15-80503720-INS-239 | chr15-80503720-INS-239 |
| chr15-80589313-INS-1333 | chr15-80589314-INS-1325 |
| chr15-80592367-INS-189 | chr15-80592368-INS-192 |
| chr15-80593449-INS-86 | chr15-80593423-INS-84 |
| chr15-80602793-INS-93 | chr15-80602793-INS-93 |
| chr15-80625313-INS-197 | chr15-80625313-INS-197 |
| chr15-80645255-INS-284 | chr15-80645255-INS-284 |
| chr15-80754274-INS-235 | chr15-80754274-INS-238 |
| chr15-80784116-INS-171 | chr15-80784116-INS-171 |
| chr15-80785072-INS-207 | chr15-80785072-INS-206 |
| chr15-80793604-INS-272 | chr15-80793604-INS-273 |
| chr15-80831537-INS-63 | chr15-80831537-INS-84 |
| chr15-80833532-INS-126 | chr15-80833531-INS-91 |
| chr15-80838320-INS-190 | chr15-80838320-INS-190 |
| chr15-80839542-INS-5237 | chr15-80839543-INS-5237 |
| chr15-80847089-INS-293 | chr15-80847089-INS-293 |
| chr15-80854321-INS-409 | chr15-80854321-INS-404 |
| chr15-80868089-INS-54 | chr15-80868089-INS-72 |
| chr15-80886524-INS-68 | chr15-80886524-INS-68 |
| chr15-80890263-INS-212 | chr15-80890263-INS-210 |
| chr15-80917667-INS-206 | chr15-80917665-INS-210 |
| chr15-80918194-INS-205 | chr15-80918194-INS-206 |
| chr15-80924985-INS-193 | chr15-80924985-INS-193 |
| chr15-80948003-INS-56 | chr15-80948003-INS-56 |
| chr15-80954160-INS-868 | chr15-80954160-INS-868 |
| chr15-80981302-INS-224 | chr15-80981302-INS-224 |
| chr15-80999052-INS-200 | chr15-80999052-INS-200 |
| chr15-81004939-INS-223 | chr15-81004939-INS-223 |
| chr15-81007989-INS-52 | chr15-81007989-INS-52 |
| chr15-81009603-INS-222 | chr15-81009603-INS-223 |
| chr15-81012294-INS-215 | chr15-81012294-INS-215 |
| chr15-81014103-INS-197 | chr15-81014103-INS-197 |
| chr15-81020383-INS-98 | chr15-81020390-INS-61 |
| chr15-81024798-INS-65 | chr15-81024798-INS-65 |
| chr15-81037386-INS-208 | chr15-81037386-INS-208 |
| chr15-81045674-INS-98 | chr15-81045674-INS-98 |
| chr15-81070100-INS-134 | chr15-81070100-INS-134 |
| chr15-81072620-INS-213 | chr15-81072620-INS-213 |
| chr15-81084903-INS-56 | chr15-81084903-INS-56 |
| chr15-81093429-INS-235 | chr15-81093429-INS-235 |
| chr15-81098245-INS-88 | chr15-81098245-INS-87 |
| chr15-81100161-INS-148 | chr15-81100179-INS-151 |
| chr15-81100645-INS-1241 | chr15-81100652-INS-1233 |
| chr15-81102821-INS-88 | chr15-81102821-INS-60 |
| chr15-81109141-INS-72 | chr15-81109141-INS-70 |
| chr15-81128521-INS-145 | chr15-81128521-INS-145 |
| chr15-81150375-INS-147 | chr15-81150375-INS-126 |
| chr15-81165143-INS-73 | chr15-81165143-INS-73 |
| chr15-81198667-INS-208 | chr15-81198667-INS-208 |

| | |
|---|---|
| chr15-81200608-INS-61 | chr15-81200608-INS-61 |
| chr15-81203420-INS-6462 | chr15-81203421-INS-6462 |
| chr15-81209419-INS-217 | chr15-81209419-INS-216 |
| chr15-81211388-INS-365 | chr15-81211388-INS-365 |
| chr15-81222579-INS-216 | chr15-81222579-INS-216 |
| chr15-81238794-INS-234 | chr15-81238794-INS-234 |
| chr15-81252577-INS-215 | chr15-81252577-INS-214 |
| chr15-81271867-INS-201 | chr15-81271867-INS-202 |
| chr15-81277327-INS-139 | chr15-81277329-INS-142 |
| chr15-81308300-INS-188 | chr15-81308300-INS-185 |
| chr15-81315491-INS-225 | chr15-81315491-INS-226 |
| chr15-81330153-INS-183 | chr15-81330153-INS-183 |
| chr15-81333447-INS-282 | chr15-81333447-INS-282 |
| chr15-81335820-INS-56 | chr15-81335846-INS-64 |
| chr15-81343845-INS-60 | chr15-81343845-INS-60 |
| chr15-81357693-INS-240 | chr15-81357693-INS-240 |
| chr15-81388876-INS-203 | chr15-81388876-INS-203 |
| chr15-81412849-INS-56 | chr15-81412842-INS-50 |
| chr15-81415118-INS-206 | chr15-81415118-INS-201 |
| chr15-81426598-INS-181 | chr15-81426598-INS-180 |
| chr15-81431675-INS-212 | chr15-81431675-INS-212 |
| chr15-81432695-INS-70 | chr15-81432692-INS-74 |
| chr15-81450612-INS-216 | chr15-81450612-INS-211 |
| chr15-81499434-INS-203 | chr15-81499434-INS-203 |
| chr15-81510977-INS-519 | chr15-81510977-INS-528 |
| chr15-81551714-INS-211 | chr15-81551714-INS-213 |
| chr15-81563987-INS-105 | chr15-81563987-INS-105 |
| chr15-81573335-INS-95 | chr15-81573335-INS-95 |
| chr15-81585090-INS-97 | chr15-81585090-INS-97 |
| chr15-81590315-INS-74 | chr15-81590315-INS-82 |
| chr15-81608539-INS-57 | chr15-81608539-INS-57 |
| chr15-81648847-INS-355 | chr15-81648847-INS-354 |
| chr15-81670826-INS-207 | chr15-81670826-INS-207 |
| chr15-81683222-INS-197 | chr15-81683222-INS-198 |
| chr15-81704510-INS-203 | chr15-81704510-INS-203 |
| chr15-81722325-INS-188 | chr15-81722318-INS-196 |
| chr15-81725948-INS-200 | chr15-81725948-INS-200 |
| chr15-81733700-INS-67 | chr15-81733700-INS-66 |
| chr15-81736511-INS-216 | chr15-81736511-INS-216 |
| chr15-81745743-INS-54 | chr15-81745743-INS-57 |
| chr15-81748736-INS-165 | chr15-81748736-INS-165 |
| chr15-81760713-INS-121 | chr15-81760713-INS-121 |
| chr15-81773934-INS-263 | chr15-81773934-INS-263 |
| chr15-81793430-INS-105 | chr15-81793424-INS-134 |
| chr15-81799567-INS-189 | chr15-81799567-INS-190 |
| chr15-81838129-INS-53 | chr15-81838129-INS-53 |
| chr15-81848844-INS-200 | chr15-81848844-INS-200 |
| chr15-81856168-INS-203 | chr15-81856168-INS-202 |
| chr15-81871469-INS-197 | chr15-81871469-INS-197 |
| chr15-81908859-INS-221 | chr15-81908860-INS-215 |
| chr15-81912406-INS-66 | chr15-81912406-INS-66 |
| chr15-81918045-INS-412 | chr15-81918045-INS-412 |
| chr15-81937956-INS-66 | chr15-81937956-INS-66 |
| chr15-81940822-INS-184 | chr15-81940822-INS-185 |
| chr15-81959733-INS-117 | chr15-81959738-INS-72 |
| chr15-81969756-INS-56 | chr15-81969772-INS-54 |
| chr15-81972092-INS-96 | chr15-81972092-INS-96 |
| chr15-81974507-INS-69 | chr15-81974507-INS-69 |
| chr15-81978546-INS-1881 | chr15-81978547-INS-1889 |
| chr15-81979263-INS-194 | chr15-81979263-INS-194 |
| chr15-81989028-INS-211 | chr15-81989028-INS-211 |
| chr15-82006104-INS-179 | chr15-82006104-INS-183 |
| chr15-82062776-INS-1569 | chr15-82062777-INS-1579 |
| chr15-82124208-INS-158 | chr15-82124208-INS-157 |
| chr15-82126735-INS-217 | chr15-82126735-INS-217 |
| chr15-82131101-INS-9045 | chr15-82131102-INS-9045 |
| chr15-82150897-INS-86 | chr15-82150896-INS-97 |
| chr15-82165851-INS-112 | chr15-82165851-INS-105 |
| chr15-82172189-INS-169 | chr15-82172194-INS-173 |
| chr15-82204195-INS-211 | chr15-82204196-INS-206 |
| chr15-82216495-INS-88 | chr15-82216492-INS-91 |
| chr15-82304832-INS-78 | chr15-82304832-INS-60 |
| chr15-82310614-INS-203 | chr15-82310614-INS-204 |
| chr15-82417897-INS-328 | chr15-82417897-INS-328 |
| chr15-82437517-INS-8144 | chr15-82437518-INS-8145 |
| chr15-82471291-INS-400 | chr15-82471291-INS-400 |
| chr15-82472672-INS-799 | chr15-82472672-INS-799 |

| | |
|---|---|
| chr15-82531361-INS-1283 | chr15-82531362-INS-1283 |
| chr15-82532041-INS-461 | chr15-82532041-INS-461 |
| chr15-82534793-INS-58 | chr15-82534799-INS-59 |
| chr15-82541111-INS-1897 | chr15-82541112-INS-1897 |
| chr15-82549803-INS-58 | chr15-82549803-INS-58 |
| chr15-82624188-INS-1273 | chr15-82624189-INS-1271 |
| chr15-82634253-INS-511 | chr15-82634253-INS-510 |
| chr15-82637838-INS-6204 | chr15-82637839-INS-6195 |
| chr15-82753702-INS-54 | chr15-82753702-INS-54 |
| chr15-82765015-INS-244 | chr15-82765022-INS-259 |
| chr15-82770708-INS-219 | chr15-82770709-INS-211 |
| chr15-82781633-INS-461 | chr15-82781633-INS-461 |
| chr15-82785529-INS-178 | chr15-82785529-INS-179 |
| chr15-82798491-INS-60 | chr15-82798491-INS-50 |
| chr15-82816636-INS-210 | chr15-82816636-INS-210 |
| chr15-82820764-INS-60 | chr15-82820764-INS-60 |
| chr15-82840910-INS-57 | chr15-82840910-INS-57 |
| chr15-82856426-INS-212 | chr15-82856426-INS-212 |
| chr15-82891282-INS-182 | chr15-82891282-INS-181 |
| chr15-82907639-INS-6473 | chr15-82907640-INS-6473 |
| chr15-82932925-INS-56 | chr15-82932925-INS-56 |
| chr15-82943419-INS-113 | chr15-82943419-INS-88 |
| chr15-82970489-INS-76 | chr15-82970489-INS-96 |
| chr15-82972710-INS-104 | chr15-82972710-INS-76 |
| chr15-82983722-INS-233 | chr15-82983722-INS-233 |
| chr15-82992591-INS-569 | chr15-82992591-INS-569 |
| chr15-82995791-INS-5265 | chr15-82995792-INS-5265 |
| chr15-83023226-INS-157 | chr15-83023226-INS-141 |
| chr15-83038988-INS-57 | chr15-83038988-INS-57 |
| chr15-83063950-INS-201 | chr15-83063950-INS-201 |
| chr15-83065075-INS-210 | chr15-83065078-INS-214 |
| chr15-83073432-INS-184 | chr15-83073432-INS-184 |
| chr15-83086945-INS-161 | chr15-83086945-INS-161 |
| chr15-83097742-INS-181 | chr15-83097743-INS-174 |
| chr15-83125268-INS-4497 | chr15-83125269-INS-4497 |
| chr15-83146329-INS-7195 | chr15-83146330-INS-7195 |
| chr15-83203428-INS-192 | chr15-83203428-INS-192 |
| chr15-83209554-INS-2880 | chr15-83209555-INS-2880 |
| chr15-83294990-INS-188 | chr15-83294990-INS-187 |
| chr15-83336327-INS-448 | chr15-83336327-INS-448 |
| chr15-83360018-INS-82 | chr15-83360018-INS-82 |
| chr15-83385665-INS-63 | chr15-83385673-INS-59 |
| chr15-83399224-INS-1825 | chr15-83399225-INS-1825 |
| chr15-83417291-INS-172 | chr15-83417291-INS-172 |
| chr15-83423085-INS-401 | chr15-83423085-INS-401 |
| chr15-83433970-INS-200 | chr15-83433970-INS-199 |
| chr15-83445271-INS-188 | chr15-83445271-INS-189 |
| chr15-83451705-INS-134 | chr15-83451705-INS-134 |
| chr15-83467688-INS-65 | chr15-83467688-INS-65 |
| chr15-83478891-INS-246 | chr15-83478891-INS-248 |
| chr15-83497391-INS-121 | chr15-83497391-INS-82 |
| chr15-83516444-INS-89 | chr15-83516444-INS-98 |
| chr15-83526236-INS-124 | chr15-83526236-INS-122 |
| chr15-83554830-INS-76 | chr15-83554830-INS-78 |
| chr15-83591677-INS-152 | chr15-83591677-INS-152 |
| chr15-83629370-INS-96 | chr15-83629370-INS-96 |
| chr15-83652470-INS-62 | chr15-83652470-INS-63 |
| chr15-83655424-INS-1865 | chr15-83655425-INS-1865 |
| chr15-83693981-INS-66 | chr15-83693981-INS-66 |
| chr15-83725162-INS-386 | chr15-83725162-INS-385 |
| chr15-83776408-INS-83 | chr15-83776408-INS-83 |
| chr15-83843361-INS-54 | chr15-83843361-INS-54 |
| chr15-83854693-INS-153 | chr15-83854693-INS-153 |
| chr15-83860837-INS-100 | chr15-83860837-INS-100 |
| chr15-83862936-INS-56 | chr15-83862918-INS-56 |
| chr15-83940550-INS-325 | chr15-83940550-INS-325 |
| chr15-84006450-INS-54 | chr15-84006462-INS-54 |
| chr15-84015478-INS-62 | chr15-84015478-INS-62 |
| chr15-84026600-INS-201 | chr15-84026609-INS-198 |
| chr15-84039894-INS-54 | chr15-84039894-INS-54 |
| chr15-84040772-INS-255 | chr15-84040773-INS-210 |
| chr15-84048323-INS-195 | chr15-84048323-INS-197 |
| chr15-84069907-INS-183 | chr15-84069907-INS-183 |
| chr15-84071980-INS-104 | chr15-84071980-INS-104 |
| chr15-84075853-INS-151 | chr15-84075853-INS-151 |
| chr15-84079213-INS-172 | chr15-84079213-INS-175 |
| chr15-84080104-INS-498 | chr15-84080104-INS-498 |

| | |
|---|---|
| chr15-84108351-INS-227 | chr15-84108351-INS-228 |
| chr15-84109392-INS-151 | chr15-84109392-INS-153 |
| chr15-84133878-INS-467 | chr15-84133878-INS-467 |
| chr15-84160306-INS-184 | chr15-84160306-INS-188 |
| chr15-84167944-INS-179 | chr15-84167944-INS-191 |
| chr15-84172433-INS-94 | chr15-84172433-INS-62 |
| chr15-84186926-INS-77 | chr15-84186926-INS-87 |
| chr15-84198149-INS-51 | chr15-84198149-INS-51 |
| chr15-84202931-INS-619 | chr15-84202931-INS-618 |
| chr15-84215710-INS-62 | chr15-84215707-INS-78 |
| chr15-84228405-INS-113 | chr15-84228405-INS-118 |
| chr15-84239399-INS-216 | chr15-84239399-INS-216 |
| chr15-84274177-INS-55 | chr15-84274177-INS-55 |
| chr15-84292492-INS-60 | chr15-84292492-INS-60 |
| chr15-84295419-INS-152 | chr15-84295419-INS-152 |
| chr15-84296372-INS-76 | chr15-84296372-INS-76 |
| chr15-84311150-INS-354 | chr15-84311166-INS-338 |
| chr15-84311172-INS-983 | chr15-84311148-INS-1029 |
| chr15-84361969-INS-81 | chr15-84361969-INS-81 |
| chr15-84379321-INS-210 | chr15-84379321-INS-210 |
| chr15-84399233-INS-304 | chr15-84399239-INS-319 |
| chr15-84402189-INS-52 | chr15-84402148-INS-56 |
| chr15-84454159-INS-60 | chr15-84454159-INS-58 |
| chr15-84463606-INS-225 | chr15-84463606-INS-225 |
| chr15-84474759-INS-170 | chr15-84474759-INS-171 |
| chr15-84508928-INS-87 | chr15-84508936-INS-100 |
| chr15-84545039-INS-211 | chr15-84545039-INS-211 |
| chr15-84557402-INS-204 | chr15-84557402-INS-204 |
| chr15-84569251-INS-246 | chr15-84569251-INS-246 |
| chr15-84588365-INS-319 | chr15-84588365-INS-311 |
| chr15-84589351-INS-57 | chr15-84589351-INS-57 |
| chr15-84596542-INS-156 | chr15-84596542-INS-156 |
| chr15-84601307-INS-57 | chr15-84601292-INS-51 |
| chr15-84627566-INS-184 | chr15-84627565-INS-232 |
| chr15-84664695-INS-212 | chr15-84664695-INS-208 |
| chr15-84705099-INS-56 | chr15-84705099-INS-56 |
| chr15-84715319-INS-210 | chr15-84715319-INS-210 |
| chr15-84726199-INS-202 | chr15-84726199-INS-202 |
| chr15-84728847-INS-199 | chr15-84728847-INS-208 |
| chr15-84744730-INS-198 | chr15-84744730-INS-198 |
| chr15-84767675-INS-96 | chr15-84767675-INS-112 |
| chr15-84810747-INS-215 | chr15-84810747-INS-215 |
| chr15-84816441-INS-58 | chr15-84816441-INS-56 |
| chr15-84839101-INS-62 | chr15-84839101-INS-64 |
| chr15-84848081-INS-498 | chr15-84848081-INS-498 |
| chr15-84878441-INS-651 | chr15-84878441-INS-651 |
| chr15-84888899-INS-71 | chr15-84888902-INS-62 |
| chr15-84912441-INS-62 | chr15-84912441-INS-56 |
| chr15-84919052-INS-768 | chr15-84919052-INS-768 |
| chr15-84929526-INS-119 | chr15-84929526-INS-96 |
| chr15-84932809-INS-85 | chr15-84932809-INS-85 |
| chr15-84933591-INS-201 | chr15-84933591-INS-199 |
| chr15-84947632-INS-3378 | chr15-84947633-INS-3378 |
| chr15-84985909-INS-207 | chr15-84985909-INS-209 |
| chr15-85022609-INS-4137 | chr15-85022610-INS-4137 |
| chr15-85039663-INS-52 | chr15-85039663-INS-52 |
| chr15-85042811-INS-4538 | chr15-85042812-INS-4538 |
| chr15-85052220-INS-118 | chr15-85052226-INS-104 |
| chr15-85053670-INS-112 | chr15-85053671-INS-108 |
| chr15-85098390-INS-53 | chr15-85098390-INS-53 |
| chr15-85124774-INS-53 | chr15-85124775-INS-53 |
| chr15-85137057-INS-84 | chr15-85137057-INS-125 |
| chr15-85149040-INS-6440 | chr15-85149041-INS-6440 |
| chr15-85170191-INS-979 | chr15-85170191-INS-979 |
| chr15-85171150-INS-219 | chr15-85171150-INS-219 |
| chr15-85174662-INS-218 | chr15-85174662-INS-217 |
| chr15-85183228-INS-1894 | chr15-85183229-INS-1893 |
| chr15-85199490-INS-1973 | chr15-85199491-INS-1971 |
| chr15-85202175-INS-266 | chr15-85202178-INS-261 |
| chr15-85205042-INS-118 | chr15-85205086-INS-116 |
| chr15-85209233-INS-92 | chr15-85209233-INS-72 |
| chr15-85210913-INS-179 | chr15-85210913-INS-179 |
| chr15-85253654-INS-60 | chr15-85253654-INS-60 |
| chr15-85276543-INS-140 | chr15-85276526-INS-144 |
| chr15-85302758-INS-1176 | chr15-85302759-INS-1176 |
| chr15-85323925-INS-52 | chr15-85323925-INS-54 |
| chr15-85377219-INS-344 | chr15-85377240-INS-303 |

| | |
|---|---|
| chr15-85468567-INS-54 | chr15-85468571-INS-50 |
| chr15-85472888-INS-52 | chr15-85472878-INS-66 |
| chr15-85499192-INS-507 | chr15-85499192-INS-507 |
| chr15-85508887-INS-257 | chr15-85508887-INS-261 |
| chr15-85518549-INS-174 | chr15-85518549-INS-174 |
| chr15-85578285-INS-151 | chr15-85578285-INS-151 |
| chr15-85592941-INS-504 | chr15-85592941-INS-504 |
| chr15-85597285-INS-773 | chr15-85597285-INS-773 |
| chr15-85645065-INS-53 | chr15-85645056-INS-59 |
| chr15-85656928-INS-94 | chr15-85656920-INS-102 |
| chr15-85669752-INS-616 | chr15-85669707-INS-612 |
| chr15-85681334-INS-187 | chr15-85681334-INS-187 |
| chr15-85698981-INS-155 | chr15-85698981-INS-155 |
| chr15-85717265-INS-474 | chr15-85717265-INS-475 |
| chr15-85728969-INS-200 | chr15-85728969-INS-200 |
| chr15-85732220-INS-105 | chr15-85732220-INS-80 |
| chr15-85748949-INS-95 | chr15-85748947-INS-109 |
| chr15-85754771-INS-214 | chr15-85754771-INS-214 |
| chr15-85788646-INS-65 | chr15-85788646-INS-70 |
| chr15-85791297-INS-243 | chr15-85791320-INS-256 |
| chr15-85797880-INS-274 | chr15-85797876-INS-290 |
| chr15-85815571-INS-74 | chr15-85815571-INS-74 |
| chr15-85818025-INS-84 | chr15-85818025-INS-84 |
| chr15-85824695-INS-528 | chr15-85824695-INS-528 |
| chr15-85826689-INS-206 | chr15-85826689-INS-206 |
| chr15-85865645-INS-193 | chr15-85865646-INS-193 |
| chr15-85866826-INS-66 | chr15-85866826-INS-66 |
| chr15-85871908-INS-204 | chr15-85871908-INS-204 |
| chr15-85881200-INS-82 | chr15-85881200-INS-72 |
| chr15-85882151-INS-73 | chr15-85882199-INS-58 |
| chr15-85886028-INS-217 | chr15-85886030-INS-218 |
| chr15-85891354-INS-242 | chr15-85891354-INS-242 |
| chr15-85939423-INS-69 | chr15-85939423-INS-69 |
| chr15-85944668-INS-437 | chr15-85944668-INS-438 |
| chr15-85979803-INS-201 | chr15-85979804-INS-206 |
| chr15-85989972-INS-58 | chr15-85990004-INS-54 |
| chr15-85996197-INS-155 | chr15-85996197-INS-131 |
| chr15-86004107-INS-91 | chr15-86004105-INS-140 |
| chr15-86030641-INS-977 | chr15-86030641-INS-977 |
| chr15-86037017-INS-60 | chr15-86037017-INS-60 |
| chr15-86041333-INS-106 | chr15-86041315-INS-123 |
| chr15-86068723-INS-97 | chr15-86068758-INS-50 |
| chr15-86075324-INS-212 | chr15-86075324-INS-212 |
| chr15-86091954-INS-178 | chr15-86091954-INS-178 |
| chr15-86103430-INS-156 | chr15-86103430-INS-156 |
| chr15-86110244-INS-208 | chr15-86110244-INS-208 |
| chr15-86133614-INS-1588 | chr15-86133615-INS-1588 |
| chr15-86183422-INS-1449 | chr15-86183423-INS-1449 |
| chr15-86183436-INS-69 | chr15-86183436-INS-69 |
| chr15-86210666-INS-83 | chr15-86210667-INS-83 |
| chr15-86246146-INS-64 | chr15-86246146-INS-60 |
| chr15-86298862-INS-91 | chr15-86298862-INS-91 |
| chr15-86304387-INS-119 | chr15-86304387-INS-119 |
| chr15-86309054-INS-132 | chr15-86309054-INS-151 |
| chr15-86323154-INS-64 | chr15-86323154-INS-62 |
| chr15-86336460-INS-55 | chr15-86336460-INS-55 |
| chr15-86355366-INS-110 | chr15-86355366-INS-110 |
| chr15-86361113-INS-1574 | chr15-86361114-INS-1574 |
| chr15-86380923-INS-103 | chr15-86380923-INS-68 |
| chr15-86399928-INS-5055 | chr15-86399929-INS-5055 |
| chr15-86411665-INS-1425 | chr15-86411666-INS-1425 |
| chr15-86412612-INS-246 | chr15-86412612-INS-246 |
| chr15-86439480-INS-108 | chr15-86439480-INS-120 |
| chr15-86443139-INS-62 | chr15-86443139-INS-62 |
| chr15-86445036-INS-53 | chr15-86445034-INS-56 |
| chr15-86450795-INS-101 | chr15-86450795-INS-101 |
| chr15-86454567-INS-476 | chr15-86454567-INS-476 |
| chr15-86459970-INS-76 | chr15-86459970-INS-68 |
| chr15-86464258-INS-287 | chr15-86464258-INS-287 |
| chr15-86483027-INS-64 | chr15-86483027-INS-72 |
| chr15-86496810-INS-52 | chr15-86496810-INS-52 |
| chr15-86498288-INS-234 | chr15-86498288-INS-234 |
| chr15-86506978-INS-58 | chr15-86506934-INS-66 |
| chr15-86527825-INS-76 | chr15-86527825-INS-64 |
| chr15-86530024-INS-86 | chr15-86530024-INS-86 |
| chr15-86530521-INS-222 | chr15-86530521-INS-223 |
| chr15-86534647-INS-6367 | chr15-86534648-INS-6365 |

| | |
|---|---|
| chr15-86555509-INS-64 | chr15-86555509-INS-64 |
| chr15-86604993-INS-1056 | chr15-86604994-INS-1028 |
| chr15-86612734-INS-5313 | chr15-86612735-INS-5313 |
| chr15-86661150-INS-249 | chr15-86661150-INS-249 |
| chr15-86667638-INS-112 | chr15-86667638-INS-130 |
| chr15-86676490-INS-53 | chr15-86676490-INS-57 |
| chr15-86680771-INS-607 | chr15-86680771-INS-607 |
| chr15-86730590-INS-78 | chr15-86730590-INS-78 |
| chr15-86745220-INS-144 | chr15-86745220-INS-117 |
| chr15-86764878-INS-51 | chr15-86764873-INS-53 |
| chr15-86785985-INS-57 | chr15-86785985-INS-57 |
| chr15-86789411-INS-65 | chr15-86789411-INS-65 |
| chr15-86802770-INS-132 | chr15-86802770-INS-132 |
| chr15-86835398-INS-156 | chr15-86835389-INS-163 |
| chr15-86838320-INS-52 | chr15-86838321-INS-56 |
| chr15-86847874-INS-477 | chr15-86847874-INS-477 |
| chr15-86861838-INS-289 | chr15-86861838-INS-289 |
| chr15-86909337-INS-6675 | chr15-86909338-INS-6675 |
| chr15-86959695-INS-73 | chr15-86959695-INS-70 |
| chr15-87004621-INS-54 | chr15-87004621-INS-50 |
| chr15-87087658-INS-3539 | chr15-87087659-INS-3539 |
| chr15-87136460-INS-773 | chr15-87136460-INS-773 |
| chr15-87152567-INS-80 | chr15-87152567-INS-80 |
| chr15-87171692-INS-1696 | chr15-87171693-INS-1696 |
| chr15-87195548-INS-176 | chr15-87195548-INS-176 |
| chr15-87196778-INS-7260 | chr15-87196779-INS-7260 |
| chr15-87238781-INS-132 | chr15-87238781-INS-132 |
| chr15-87244388-INS-66 | chr15-87244388-INS-76 |
| chr15-87246248-INS-128 | chr15-87246265-INS-96 |
| chr15-87247997-INS-100 | chr15-87247965-INS-88 |
| chr15-87263650-INS-528 | chr15-87263650-INS-528 |
| chr15-87291186-INS-66 | chr15-87291186-INS-66 |
| chr15-87314298-INS-1021 | chr15-87314299-INS-1013 |
| chr15-87328836-INS-52 | chr15-87328836-INS-52 |
| chr15-87329889-INS-92 | chr15-87329889-INS-92 |
| chr15-87379518-INS-66 | chr15-87379539-INS-93 |
| chr15-87381261-INS-53 | chr15-87381261-INS-53 |
| chr15-87425475-INS-249 | chr15-87425475-INS-249 |
| chr15-87476829-INS-64 | chr15-87476829-INS-58 |
| chr15-87500966-INS-57 | chr15-87500966-INS-57 |
| chr15-87531684-INS-1862 | chr15-87531685-INS-1862 |
| chr15-87533467-INS-76 | chr15-87533467-INS-76 |
| chr15-87542841-INS-88 | chr15-87542841-INS-88 |
| chr15-87563262-INS-3073 | chr15-87563263-INS-3073 |
| chr15-87578664-INS-103 | chr15-87578664-INS-103 |
| chr15-87602849-INS-84 | chr15-87602849-INS-84 |
| chr15-87637234-INS-88 | chr15-87637234-INS-88 |
| chr15-87697607-INS-532 | chr15-87697607-INS-532 |
| chr15-87733371-INS-6466 | chr15-87733372-INS-6465 |
| chr15-87778773-INS-249 | chr15-87778768-INS-272 |
| chr15-87780070-INS-68 | chr15-87780065-INS-87 |
| chr15-87807121-INS-62 | chr15-87807104-INS-56 |
| chr15-87826866-INS-1123 | chr15-87826870-INS-1106 |
| chr15-87842346-INS-247 | chr15-87842346-INS-248 |
| chr15-87843698-INS-69 | chr15-87843695-INS-77 |
| chr15-87845568-INS-208 | chr15-87845568-INS-212 |
| chr15-87926316-INS-124 | chr15-87926316-INS-124 |
| chr15-87957965-INS-63 | chr15-87957965-INS-63 |
| chr15-87965079-INS-341 | chr15-87965079-INS-341 |
| chr15-87974105-INS-2999 | chr15-87974106-INS-3001 |
| chr15-87985959-INS-1600 | chr15-87985960-INS-1599 |
| chr15-87986428-INS-76 | chr15-87986428-INS-57 |
| chr15-88042602-INS-6683 | chr15-88042603-INS-6682 |
| chr15-88070435-INS-3271 | chr15-88070436-INS-3271 |
| chr15-88073796-INS-58 | chr15-88073796-INS-58 |
| chr15-88096143-INS-72 | chr15-88096143-INS-54 |
| chr15-88113485-INS-6229 | chr15-88113486-INS-6229 |
| chr15-88114923-INS-123 | chr15-88114923-INS-123 |
| chr15-88154753-INS-80 | chr15-88154753-INS-82 |
| chr15-88163520-INS-54 | chr15-88163485-INS-84 |
| chr15-88176614-INS-157 | chr15-88176614-INS-157 |
| chr15-88192217-INS-56 | chr15-88192217-INS-56 |
| chr15-88206165-INS-2883 | chr15-88206166-INS-2872 |
| chr15-88213519-INS-212 | chr15-88213517-INS-190 |
| chr15-88231537-INS-64 | chr15-88231533-INS-68 |
| chr15-88298346-INS-379 | chr15-88298346-INS-379 |
| chr15-88333537-INS-60 | chr15-88333537-INS-52 |

| | |
|---|---|
| chr15-88467632-INS-56 | chr15-88467632-INS-56 |
| chr15-88481619-INS-90 | chr15-88481619-INS-88 |
| chr15-88482653-INS-401 | chr15-88482653-INS-401 |
| chr15-88491257-INS-121 | chr15-88491257-INS-121 |
| chr15-88517555-INS-94 | chr15-88517556-INS-132 |
| chr15-88536292-INS-363 | chr15-88536292-INS-362 |
| chr15-88552903-INS-7092 | chr15-88552904-INS-7093 |
| chr15-88553567-INS-72 | chr15-88553566-INS-69 |
| chr15-88565468-INS-62 | chr15-88565468-INS-62 |
| chr15-88566681-INS-705 | chr15-88566681-INS-705 |
| chr15-88568874-INS-144 | chr15-88568878-INS-132 |
| chr15-88582234-INS-865 | chr15-88582234-INS-865 |
| chr15-88588222-INS-139 | chr15-88588222-INS-139 |
| chr15-88612632-INS-187 | chr15-88612632-INS-187 |
| chr15-88631268-INS-216 | chr15-88631268-INS-217 |
| chr15-88641714-INS-96 | chr15-88641714-INS-64 |
| chr15-88647265-INS-66 | chr15-88647265-INS-66 |
| chr15-88673376-INS-6440 | chr15-88673377-INS-6440 |
| chr15-88673930-INS-338 | chr15-88673931-INS-334 |
| chr15-88675031-INS-73 | chr15-88675031-INS-68 |
| chr15-88731414-INS-76 | chr15-88731414-INS-76 |
| chr15-88764439-INS-202 | chr15-88764436-INS-200 |
| chr15-88769540-INS-229 | chr15-88769540-INS-229 |
| chr15-88770428-INS-52 | chr15-88770424-INS-52 |
| chr15-88836726-INS-135 | chr15-88836726-INS-95 |
| chr15-88858454-INS-59 | chr15-88858454-INS-59 |
| chr15-88885949-INS-200 | chr15-88885949-INS-201 |
| chr15-88931758-INS-51 | chr15-88931751-INS-53 |
| chr15-89005640-INS-206 | chr15-89005640-INS-206 |
| chr15-89011505-INS-216 | chr15-89011505-INS-216 |
| chr15-89013747-INS-54 | chr15-89013747-INS-74 |
| chr15-89035314-INS-69 | chr15-89035330-INS-79 |
| chr15-89078555-INS-185 | chr15-89078555-INS-185 |
| chr15-89081475-INS-68 | chr15-89081476-INS-68 |
| chr15-89082310-INS-137 | chr15-89082311-INS-88 |
| chr15-89109797-INS-52 | chr15-89109800-INS-66 |
| chr15-89277872-INS-61 | chr15-89277872-INS-73 |
| chr15-89283470-INS-52 | chr15-89283470-INS-52 |
| chr15-89295299-INS-51 | chr15-89295299-INS-51 |
| chr15-89335657-INS-208 | chr15-89335657-INS-207 |
| chr15-89336577-INS-52 | chr15-89336579-INS-62 |
| chr15-89424219-INS-57 | chr15-89424219-INS-57 |
| chr15-89458923-INS-191 | chr15-89458925-INS-192 |
| chr15-89462285-INS-170 | chr15-89462290-INS-174 |
| chr15-89470796-INS-8326 | chr15-89470797-INS-8326 |
| chr15-89473332-INS-675 | chr15-89473332-INS-648 |
| chr15-89478422-INS-178 | chr15-89478420-INS-184 |
| chr15-89483965-INS-202 | chr15-89483965-INS-202 |
| chr15-89484781-INS-231 | chr15-89484781-INS-231 |
| chr15-89493606-INS-85 | chr15-89493605-INS-109 |
| chr15-89531236-INS-60 | chr15-89531250-INS-80 |
| chr15-89572183-INS-932 | chr15-89572183-INS-932 |
| chr15-89600317-INS-83 | chr15-89600298-INS-84 |
| chr15-89602150-INS-70 | chr15-89602150-INS-70 |
| chr15-89612694-INS-60 | chr15-89612694-INS-60 |
| chr15-89617310-INS-466 | chr15-89617310-INS-466 |
| chr15-89620201-INS-540 | chr15-89620201-INS-540 |
| chr15-89653216-INS-1354 | chr15-89653217-INS-1354 |
| chr15-89663878-INS-6748 | chr15-89663879-INS-6754 |
| chr15-89667942-INS-84 | chr15-89667942-INS-84 |
| chr15-89677993-INS-62 | chr15-89677993-INS-68 |
| chr15-89684989-INS-137 | chr15-89684989-INS-151 |
| chr15-89705232-INS-230 | chr15-89705242-INS-230 |
| chr15-89716509-INS-56 | chr15-89716509-INS-60 |
| chr15-89720075-INS-55 | chr15-89720075-INS-55 |
| chr15-89768896-INS-77 | chr15-89768896-INS-77 |
| chr15-89858548-INS-217 | chr15-89858532-INS-220 |
| chr15-89863394-INS-476 | chr15-89863394-INS-476 |
| chr15-89885073-INS-154 | chr15-89885073-INS-156 |
| chr15-89897172-INS-54 | chr15-89897172-INS-54 |
| chr15-89903478-INS-66 | chr15-89903478-INS-66 |
| chr15-89905273-INS-1527 | chr15-89905292-INS-1517 |
| chr15-89905323-INS-73 | chr15-89905323-INS-73 |
| chr15-89910827-INS-201 | chr15-89910827-INS-201 |
| chr15-89918227-INS-60 | chr15-89918250-INS-50 |
| chr15-89926211-INS-497 | chr15-89926211-INS-497 |
| chr15-89941658-INS-63 | chr15-89941662-INS-76 |

| | |
|---|---|
| chr15-89947299-INS-285 | chr15-89947299-INS-285 |
| chr15-89948061-INS-92 | chr15-89948061-INS-92 |
| chr15-89948583-INS-925 | chr15-89948583-INS-966 |
| chr15-89953677-INS-1433 | chr15-89953684-INS-1426 |
| chr15-89981764-INS-59 | chr15-89981764-INS-68 |
| chr15-89985621-INS-60 | chr15-89985621-INS-60 |
| chr15-89998161-INS-473 | chr15-89998161-INS-483 |
| chr15-90013215-INS-315 | chr15-90013215-INS-315 |
| chr15-90015025-INS-96 | chr15-90015025-INS-120 |
| chr15-90021007-INS-106 | chr15-90021007-INS-106 |
| chr15-90022442-INS-74 | chr15-90022442-INS-74 |
| chr15-90036931-INS-76 | chr15-90036931-INS-54 |
| chr15-90043785-INS-70 | chr15-90043785-INS-70 |
| chr15-90064198-INS-224 | chr15-90064198-INS-226 |
| chr15-90068284-INS-90 | chr15-90068284-INS-90 |
| chr15-90094900-INS-216 | chr15-90094900-INS-216 |
| chr15-90123596-INS-195 | chr15-90123596-INS-197 |
| chr15-90137902-INS-95 | chr15-90137902-INS-95 |
| chr15-90154200-INS-441 | chr15-90154200-INS-441 |
| chr15-90164690-INS-60 | chr15-90164690-INS-60 |
| chr15-90193922-INS-84 | chr15-90193918-INS-75 |
| chr15-90203404-INS-67 | chr15-90203404-INS-67 |
| chr15-90222966-INS-599 | chr15-90222966-INS-599 |
| chr15-90230015-INS-121 | chr15-90229987-INS-159 |
| chr15-90233262-INS-390 | chr15-90233262-INS-390 |
| chr15-90259042-INS-94 | chr15-90259042-INS-94 |
| chr15-90303191-INS-371 | chr15-90303191-INS-408 |
| chr15-90305168-INS-141 | chr15-90305168-INS-141 |
| chr15-90318691-INS-201 | chr15-90318691-INS-201 |
| chr15-90329129-INS-72 | chr15-90329129-INS-76 |
| chr15-90330600-INS-78 | chr15-90330600-INS-80 |
| chr15-90350163-INS-53 | chr15-90350163-INS-67 |
| chr15-90385995-INS-135 | chr15-90385999-INS-143 |
| chr15-90388440-INS-72 | chr15-90388438-INS-92 |
| chr15-90390467-INS-60 | chr15-90390467-INS-60 |
| chr15-90424533-INS-76 | chr15-90424499-INS-70 |
| chr15-90425234-INS-907 | chr15-90425234-INS-915 |
| chr15-90453606-INS-92 | chr15-90453606-INS-92 |
| chr15-90500914-INS-152 | chr15-90500914-INS-129 |
| chr15-90506107-INS-126 | chr15-90506149-INS-130 |
| chr15-90517913-INS-75 | chr15-90517913-INS-75 |
| chr15-90554374-INS-62 | chr15-90554374-INS-64 |
| chr15-90565669-INS-197 | chr15-90565669-INS-197 |
| chr15-90576540-INS-1962 | chr15-90576541-INS-1962 |
| chr15-90581344-INS-1550 | chr15-90581345-INS-1551 |
| chr15-90587089-INS-116 | chr15-90587089-INS-108 |
| chr15-90612734-INS-7122 | chr15-90612735-INS-7122 |
| chr15-90615674-INS-53 | chr15-90615681-INS-77 |
| chr15-90620467-INS-75 | chr15-90620467-INS-79 |
| chr15-90660119-INS-187 | chr15-90660119-INS-187 |
| chr15-90664611-INS-56 | chr15-90664601-INS-82 |
| chr15-90682092-INS-126 | chr15-90682092-INS-96 |
| chr15-90696249-INS-122 | chr15-90696252-INS-98 |
| chr15-90720723-INS-87 | chr15-90720688-INS-81 |
| chr15-90724314-INS-98 | chr15-90724314-INS-142 |
| chr15-90730231-INS-922 | chr15-90730231-INS-922 |
| chr15-90776872-INS-58 | chr15-90776872-INS-58 |
| chr15-90787420-INS-237 | chr15-90787420-INS-236 |
| chr15-90790182-INS-360 | chr15-90790182-INS-360 |
| chr15-90795829-INS-104 | chr15-90795829-INS-104 |
| chr15-90827782-INS-856 | chr15-90827782-INS-856 |
| chr15-90872365-INS-80 | chr15-90872365-INS-80 |
| chr15-90893298-INS-54 | chr15-90893297-INS-56 |
| chr15-90923951-INS-85 | chr15-90923951-INS-62 |
| chr15-90928777-INS-589 | chr15-90928777-INS-589 |
| chr15-90937989-INS-211 | chr15-90937989-INS-211 |
| chr15-90950213-INS-78 | chr15-90950213-INS-78 |
| chr15-90956568-INS-158 | chr15-90956568-INS-156 |
| chr15-90973141-INS-333 | chr15-90973141-INS-334 |
| chr15-90975701-INS-90 | chr15-90975702-INS-92 |
| chr15-90993243-INS-200 | chr15-90993243-INS-200 |
| chr15-91028724-INS-59 | chr15-91028724-INS-56 |
| chr15-91050648-INS-64 | chr15-91050655-INS-73 |
| chr15-91069128-INS-185 | chr15-91069128-INS-200 |
| chr15-91117904-INS-551 | chr15-91117904-INS-551 |
| chr15-91166427-INS-84 | chr15-91166427-INS-73 |
| chr15-91216742-INS-317 | chr15-91216742-INS-317 |

| | |
|---|---|
| chr15-91223712-INS-54 | chr15-91223712-INS-54 |
| chr15-91226491-INS-147 | chr15-91226491-INS-148 |
| chr15-91267459-INS-706 | chr15-91267459-INS-706 |
| chr15-91292843-INS-81 | chr15-91292843-INS-81 |
| chr15-91326663-INS-766 | chr15-91326663-INS-765 |
| chr15-91330966-INS-120 | chr15-91330966-INS-120 |
| chr15-91361330-INS-1143 | chr15-91361331-INS-1143 |
| chr15-91392495-INS-144 | chr15-91392489-INS-147 |
| chr15-91396768-INS-73 | chr15-91396768-INS-74 |
| chr15-91402943-INS-59 | chr15-91402934-INS-59 |
| chr15-91405240-INS-76 | chr15-91405249-INS-85 |
| chr15-91408054-INS-213 | chr15-91408054-INS-213 |
| chr15-91414544-INS-464 | chr15-91414544-INS-464 |
| chr15-91422167-INS-215 | chr15-91422167-INS-215 |
| chr15-91427518-INS-395 | chr15-91427518-INS-394 |
| chr15-91436483-INS-264 | chr15-91436483-INS-256 |
| chr15-91474196-INS-70 | chr15-91474187-INS-74 |
| chr15-91475856-INS-358 | chr15-91475856-INS-358 |
| chr15-91496184-INS-5543 | chr15-91496185-INS-5543 |
| chr15-91501075-INS-52 | chr15-91501073-INS-50 |
| chr15-91508501-INS-51 | chr15-91508501-INS-52 |
| chr15-91517833-INS-68 | chr15-91517833-INS-68 |
| chr15-91520883-INS-122 | chr15-91520883-INS-122 |
| chr15-91546704-INS-70 | chr15-91546723-INS-79 |
| chr15-91547422-INS-59 | chr15-91547422-INS-59 |
| chr15-91565762-INS-402 | chr15-91565762-INS-404 |
| chr15-91618553-INS-178 | chr15-91618549-INS-180 |
| chr15-91620155-INS-1894 | chr15-91620156-INS-1894 |
| chr15-91637896-INS-71 | chr15-91637896-INS-71 |
| chr15-91655553-INS-54 | chr15-91655553-INS-55 |
| chr15-91667141-INS-86 | chr15-91667141-INS-56 |
| chr15-91670239-INS-212 | chr15-91670239-INS-224 |
| chr15-91677236-INS-60 | chr15-91677246-INS-52 |
| chr15-91677969-INS-70 | chr15-91677939-INS-78 |
| chr15-91686460-INS-52 | chr15-91686460-INS-52 |
| chr15-91702276-INS-398 | chr15-91702277-INS-398 |
| chr15-91702796-INS-7137 | chr15-91702797-INS-7138 |
| chr15-91705197-INS-2726 | chr15-91705198-INS-2726 |
| chr15-91710221-INS-88 | chr15-91710221-INS-88 |
| chr15-91720114-INS-263 | chr15-91720114-INS-263 |
| chr15-91743030-INS-50 | chr15-91743047-INS-60 |
| chr15-91747531-INS-114 | chr15-91747531-INS-114 |
| chr15-91756304-INS-85 | chr15-91756294-INS-89 |
| chr15-91759937-INS-52 | chr15-91759936-INS-78 |
| chr15-91766091-INS-304 | chr15-91766091-INS-304 |
| chr15-91821038-INS-5758 | chr15-91821039-INS-5793 |
| chr15-91823287-INS-53 | chr15-91823286-INS-59 |
| chr15-91839573-INS-88 | chr15-91839573-INS-91 |
| chr15-91865543-INS-194 | chr15-91865543-INS-194 |
| chr15-91880346-INS-390 | chr15-91880346-INS-397 |
| chr15-91927062-INS-6127 | chr15-91927063-INS-6127 |
| chr15-91954170-INS-58 | chr15-91954170-INS-69 |
| chr15-91958111-INS-206 | chr15-91958111-INS-206 |
| chr15-91965300-INS-2256 | chr15-91965301-INS-2256 |
| chr15-91978188-INS-1300 | chr15-91978189-INS-1290 |
| chr15-91983091-INS-6383 | chr15-91983092-INS-6383 |
| chr15-91986135-INS-314 | chr15-91986135-INS-324 |
| chr15-91998921-INS-1448 | chr15-91998922-INS-1448 |
| chr15-91999898-INS-52 | chr15-91999899-INS-62 |
| chr15-92017224-INS-57 | chr15-92017224-INS-58 |
| chr15-92038746-INS-114 | chr15-92038746-INS-115 |
| chr15-92040664-INS-343 | chr15-92040664-INS-343 |
| chr15-92065414-INS-96 | chr15-92065435-INS-72 |
| chr15-92069634-INS-128 | chr15-92069634-INS-128 |
| chr15-92078868-INS-196 | chr15-92078868-INS-197 |
| chr15-92079098-INS-524 | chr15-92079098-INS-524 |
| chr15-92084032-INS-399 | chr15-92084032-INS-399 |
| chr15-92113415-INS-6498 | chr15-92113416-INS-6498 |
| chr15-92140176-INS-266 | chr15-92140176-INS-266 |
| chr15-92141320-INS-252 | chr15-92141320-INS-256 |
| chr15-92160028-INS-113 | chr15-92160028-INS-109 |
| chr15-92174628-INS-60 | chr15-92174628-INS-62 |
| chr15-92178856-INS-60 | chr15-92178856-INS-52 |
| chr15-92180545-INS-76 | chr15-92180545-INS-76 |
| chr15-92186552-INS-209 | chr15-92186552-INS-209 |
| chr15-92217537-INS-198 | chr15-92217537-INS-175 |
| chr15-92223387-INS-494 | chr15-92223387-INS-494 |

| | |
|---|---|
| chr15-92231719-INS-551 | chr15-92231719-INS-574 |
| chr15-92245063-INS-153 | chr15-92245063-INS-153 |
| chr15-92251034-INS-74 | chr15-92251032-INS-70 |
| chr15-92261061-INS-5258 | chr15-92261062-INS-5258 |
| chr15-92271574-INS-507 | chr15-92271575-INS-490 |
| chr15-92275171-INS-53 | chr15-92275171-INS-53 |
| chr15-92290522-INS-103 | chr15-92290522-INS-118 |
| chr15-92298341-INS-6928 | chr15-92298342-INS-6930 |
| chr15-92307401-INS-416 | chr15-92307401-INS-416 |
| chr15-92309306-INS-60 | chr15-92309275-INS-53 |
| chr15-92311205-INS-489 | chr15-92311205-INS-485 |
| chr15-92330352-INS-462 | chr15-92330352-INS-465 |
| chr15-92353422-INS-399 | chr15-92353421-INS-425 |
| chr15-92364886-INS-54 | chr15-92364886-INS-56 |
| chr15-92438999-INS-141 | chr15-92438999-INS-141 |
| chr15-92462237-INS-144 | chr15-92462232-INS-156 |
| chr15-92528882-INS-365 | chr15-92528892-INS-391 |
| chr15-92530988-INS-89 | chr15-92530988-INS-82 |
| chr15-92547327-INS-1497 | chr15-92547334-INS-1491 |
| chr15-92582343-INS-66 | chr15-92582324-INS-83 |
| chr15-92599388-INS-92 | chr15-92599388-INS-92 |
| chr15-92602482-INS-59 | chr15-92602482-INS-59 |
| chr15-92636929-INS-372 | chr15-92636929-INS-372 |
| chr15-92647298-INS-84 | chr15-92647292-INS-78 |
| chr15-92655346-INS-115 | chr15-92655361-INS-118 |
| chr15-92659769-INS-61 | chr15-92659769-INS-61 |
| chr15-92726792-INS-354 | chr15-92726792-INS-354 |
| chr15-92727655-INS-83 | chr15-92727636-INS-70 |
| chr15-92782472-INS-429 | chr15-92782472-INS-429 |
| chr15-92793744-INS-829 | chr15-92793744-INS-829 |
| chr15-92806470-INS-72 | chr15-92806470-INS-70 |
| chr15-92818280-INS-299 | chr15-92818276-INS-300 |
| chr15-92819309-INS-66 | chr15-92819309-INS-66 |
| chr15-92837997-INS-6403 | chr15-92837998-INS-6403 |
| chr15-92858306-INS-401 | chr15-92858306-INS-401 |
| chr15-92859852-INS-445 | chr15-92859856-INS-472 |
| chr15-92865210-INS-414 | chr15-92865210-INS-414 |
| chr15-92869100-INS-484 | chr15-92869100-INS-484 |
| chr15-92885115-INS-278 | chr15-92885115-INS-278 |
| chr15-92888880-INS-314 | chr15-92888880-INS-318 |
| chr15-92913627-INS-50 | chr15-92913630-INS-58 |
| chr15-92921351-INS-60 | chr15-92921347-INS-52 |
| chr15-92922133-INS-2024 | chr15-92922138-INS-2025 |
| chr15-92940991-INS-103 | chr15-92940991-INS-103 |
| chr15-92966603-INS-60 | chr15-92966603-INS-60 |
| chr15-92972451-INS-160 | chr15-92972451-INS-160 |
| chr15-93001283-INS-52 | chr15-93001283-INS-52 |
| chr15-93006522-INS-62 | chr15-93006522-INS-62 |
| chr15-93057952-INS-925 | chr15-93057952-INS-893 |
| chr15-93060858-INS-84 | chr15-93060858-INS-112 |
| chr15-93075838-INS-298 | chr15-93075838-INS-298 |
| chr15-93114251-INS-299 | chr15-93114251-INS-299 |
| chr15-93124444-INS-84 | chr15-93124444-INS-78 |
| chr15-93128161-INS-131 | chr15-93128187-INS-107 |
| chr15-93151560-INS-52 | chr15-93151560-INS-54 |
| chr15-93157253-INS-681 | chr15-93157253-INS-681 |
| chr15-93167759-INS-56 | chr15-93167759-INS-58 |
| chr15-93174851-INS-195 | chr15-93174851-INS-195 |
| chr15-93178712-INS-189 | chr15-93178712-INS-189 |
| chr15-93194547-INS-715 | chr15-93194547-INS-715 |
| chr15-93231726-INS-105 | chr15-93231726-INS-105 |
| chr15-93243238-INS-247 | chr15-93243237-INS-249 |
| chr15-93278206-INS-84 | chr15-93278206-INS-84 |
| chr15-93282456-INS-164 | chr15-93282456-INS-155 |
| chr15-93300933-INS-52 | chr15-93300933-INS-66 |
| chr15-93324600-INS-195 | chr15-93324600-INS-200 |
| chr15-93333919-INS-54 | chr15-93333921-INS-68 |
| chr15-93426108-INS-54 | chr15-93426108-INS-54 |
| chr15-93452154-INS-109 | chr15-93452154-INS-109 |
| chr15-93462317-INS-80 | chr15-93462317-INS-80 |
| chr15-93483322-INS-114 | chr15-93483322-INS-114 |
| chr15-93498686-INS-196 | chr15-93498686-INS-196 |
| chr15-93503409-INS-72 | chr15-93503387-INS-55 |
| chr15-93510607-INS-547 | chr15-93510607-INS-547 |
| chr15-93515507-INS-4373 | chr15-93515508-INS-4373 |
| chr15-93517411-INS-66 | chr15-93517411-INS-67 |
| chr15-93521847-INS-2717 | chr15-93521848-INS-2717 |

| | |
|---|---|
| chr15-93523973-INS-392 | chr15-93523973-INS-392 |
| chr15-93531693-INS-346 | chr15-93531693-INS-354 |
| chr15-93562630-INS-447 | chr15-93562630-INS-447 |
| chr15-93566098-INS-135 | chr15-93566098-INS-135 |
| chr15-93608999-INS-100 | chr15-93608999-INS-100 |
| chr15-93617911-INS-52 | chr15-93617912-INS-97 |
| chr15-93621404-INS-214 | chr15-93621404-INS-212 |
| chr15-93623418-INS-1386 | chr15-93623419-INS-1408 |
| chr15-93624985-INS-327 | chr15-93624985-INS-327 |
| chr15-93633059-INS-63 | chr15-93633056-INS-67 |
| chr15-93656113-INS-6425 | chr15-93656114-INS-6425 |
| chr15-93659102-INS-82 | chr15-93659102-INS-82 |
| chr15-93674549-INS-64 | chr15-93674595-INS-64 |
| chr15-93679731-INS-68 | chr15-93679730-INS-51 |
| chr15-93695021-INS-58 | chr15-93695021-INS-55 |
| chr15-93695833-INS-77 | chr15-93695848-INS-50 |
| chr15-93698193-INS-70 | chr15-93698193-INS-70 |
| chr15-93725705-INS-79 | chr15-93725705-INS-79 |
| chr15-93737953-INS-4759 | chr15-93737954-INS-4759 |
| chr15-93745199-INS-77 | chr15-93745199-INS-77 |
| chr15-93780002-INS-51 | chr15-93780003-INS-51 |
| chr15-93781345-INS-100 | chr15-93781345-INS-100 |
| chr15-93806269-INS-132 | chr15-93806269-INS-133 |
| chr15-93835530-INS-4162 | chr15-93835531-INS-4162 |
| chr15-93853909-INS-358 | chr15-93853909-INS-358 |
| chr15-93865237-INS-831 | chr15-93865237-INS-831 |
| chr15-93866860-INS-76 | chr15-93866860-INS-112 |
| chr15-93872504-INS-88 | chr15-93872504-INS-98 |
| chr15-93874089-INS-96 | chr15-93874089-INS-96 |
| chr15-93898450-INS-260 | chr15-93898450-INS-260 |
| chr15-93908913-INS-257 | chr15-93908913-INS-257 |
| chr15-93943015-INS-196 | chr15-93943015-INS-197 |
| chr15-93952038-INS-109 | chr15-93952038-INS-110 |
| chr15-94022001-INS-198 | chr15-94021999-INS-198 |
| chr15-94034572-INS-475 | chr15-94034572-INS-475 |
| chr15-94065237-INS-52 | chr15-94065237-INS-52 |
| chr15-94065822-INS-52 | chr15-94065822-INS-52 |
| chr15-94067867-INS-2764 | chr15-94067868-INS-2788 |
| chr15-94068499-INS-52 | chr15-94068499-INS-52 |
| chr15-94076018-INS-990 | chr15-94076018-INS-988 |
| chr15-94105051-INS-209 | chr15-94105051-INS-209 |
| chr15-94171712-INS-62 | chr15-94171712-INS-62 |
| chr15-94201023-INS-124 | chr15-94201023-INS-74 |
| chr15-94253663-INS-90 | chr15-94253663-INS-90 |
| chr15-94281281-INS-57 | chr15-94281281-INS-57 |
| chr15-94286847-INS-104 | chr15-94286848-INS-78 |
| chr15-94306905-INS-63 | chr15-94306905-INS-66 |
| chr15-94327003-INS-62 | chr15-94327015-INS-62 |
| chr15-94357405-INS-76 | chr15-94357405-INS-90 |
| chr15-94367738-INS-54 | chr15-94367738-INS-54 |
| chr15-94370447-INS-302 | chr15-94370447-INS-302 |
| chr15-94388469-INS-64 | chr15-94388468-INS-64 |
| chr15-94393723-INS-173 | chr15-94393723-INS-167 |
| chr15-94397779-INS-150 | chr15-94397779-INS-150 |
| chr15-94402074-INS-6171 | chr15-94402075-INS-6172 |
| chr15-94404710-INS-69 | chr15-94404710-INS-63 |
| chr15-94435026-INS-52 | chr15-94435026-INS-52 |
| chr15-94446424-INS-221 | chr15-94446424-INS-221 |
| chr15-94453219-INS-56 | chr15-94453219-INS-56 |
| chr15-94458251-INS-130 | chr15-94458251-INS-130 |
| chr15-94460947-INS-90 | chr15-94460947-INS-90 |
| chr15-94468272-INS-202 | chr15-94468272-INS-202 |
| chr15-94471081-INS-110 | chr15-94471082-INS-102 |
| chr15-94492720-INS-153 | chr15-94492720-INS-159 |
| chr15-94496324-INS-64 | chr15-94496324-INS-75 |
| chr15-94502550-INS-191 | chr15-94502550-INS-191 |
| chr15-94517763-INS-68 | chr15-94517763-INS-68 |
| chr15-94538298-INS-312 | chr15-94538302-INS-293 |
| chr15-94543300-INS-51 | chr15-94543339-INS-59 |
| chr15-94551173-INS-94 | chr15-94551133-INS-100 |
| chr15-94563428-INS-105 | chr15-94563421-INS-108 |
| chr15-94567542-INS-68 | chr15-94567543-INS-74 |
| chr15-94579701-INS-223 | chr15-94579701-INS-223 |
| chr15-94614747-INS-240 | chr15-94614747-INS-240 |
| chr15-94687659-INS-217 | chr15-94687659-INS-217 |
| chr15-94729442-INS-80 | chr15-94729442-INS-88 |
| chr15-94741765-INS-77 | chr15-94741750-INS-86 |

| | |
|---|---|
| chr15-94761628-INS-130 | chr15-94761628-INS-130 |
| chr15-94768277-INS-405 | chr15-94768277-INS-394 |
| chr15-94775633-INS-6395 | chr15-94775634-INS-6395 |
| chr15-94789161-INS-185 | chr15-94789161-INS-185 |
| chr15-94798548-INS-81 | chr15-94798548-INS-81 |
| chr15-94837891-INS-91 | chr15-94837891-INS-91 |
| chr15-94849025-INS-87 | chr15-94849025-INS-87 |
| chr15-94859979-INS-53 | chr15-94859980-INS-101 |
| chr15-94861878-INS-198 | chr15-94861878-INS-197 |
| chr15-94870182-INS-80 | chr15-94870205-INS-70 |
| chr15-94892990-INS-151 | chr15-94892990-INS-154 |
| chr15-94897090-INS-142 | chr15-94897133-INS-108 |
| chr15-94900455-INS-468 | chr15-94900455-INS-468 |
| chr15-95014581-INS-104 | chr15-95014581-INS-108 |
| chr15-95021623-INS-1401 | chr15-95021624-INS-1401 |
| chr15-95026613-INS-2341 | chr15-95026613-INS-2337 |
| chr15-95047324-INS-506 | chr15-95047324-INS-506 |
| chr15-95060424-INS-312 | chr15-95060425-INS-312 |
| chr15-95065175-INS-388 | chr15-95065175-INS-388 |
| chr15-95086498-INS-70 | chr15-95086478-INS-64 |
| chr15-95116678-INS-627 | chr15-95116678-INS-621 |
| chr15-95129602-INS-299 | chr15-95129604-INS-300 |
| chr15-95131571-INS-77 | chr15-95131571-INS-77 |
| chr15-95133346-INS-74 | chr15-95133346-INS-74 |
| chr15-95147033-INS-56 | chr15-95147033-INS-56 |
| chr15-95176448-INS-62 | chr15-95176448-INS-62 |
| chr15-95179357-INS-52 | chr15-95179360-INS-52 |
| chr15-95230979-INS-100 | chr15-95230982-INS-99 |
| chr15-95233351-INS-498 | chr15-95233351-INS-496 |
| chr15-95264423-INS-61 | chr15-95264423-INS-68 |
| chr15-95283624-INS-104 | chr15-95283624-INS-104 |
| chr15-95300522-INS-52 | chr15-95300524-INS-68 |
| chr15-95341793-INS-86 | chr15-95341792-INS-86 |
| chr15-95354677-INS-61 | chr15-95354677-INS-61 |
| chr15-95369445-INS-84 | chr15-95369445-INS-84 |
| chr15-95378996-INS-65 | chr15-95378996-INS-65 |
| chr15-95399305-INS-6644 | chr15-95399306-INS-6644 |
| chr15-95417124-INS-881 | chr15-95417124-INS-881 |
| chr15-95432883-INS-6895 | chr15-95432869-INS-6892 |
| chr15-95445929-INS-765 | chr15-95445929-INS-765 |
| chr15-95459917-INS-53 | chr15-95459917-INS-53 |
| chr15-95474185-INS-56 | chr15-95474185-INS-56 |
| chr15-95475945-INS-87 | chr15-95475945-INS-89 |
| chr15-95485643-INS-52 | chr15-95485643-INS-70 |
| chr15-95494632-INS-63 | chr15-95494632-INS-67 |
| chr15-95499719-INS-202 | chr15-95499711-INS-218 |
| chr15-95529581-INS-75 | chr15-95529581-INS-77 |
| chr15-95530868-INS-62 | chr15-95530868-INS-60 |
| chr15-95563591-INS-134 | chr15-95563591-INS-134 |
| chr15-95601485-INS-110 | chr15-95601485-INS-93 |
| chr15-95605716-INS-880 | chr15-95605716-INS-854 |
| chr15-95617971-INS-63 | chr15-95617971-INS-51 |
| chr15-95635042-INS-296 | chr15-95635042-INS-296 |
| chr15-95642495-INS-202 | chr15-95642495-INS-202 |
| chr15-95643877-INS-111 | chr15-95643877-INS-102 |
| chr15-95662480-INS-107 | chr15-95662481-INS-114 |
| chr15-95669379-INS-96 | chr15-95669379-INS-96 |
| chr15-95695117-INS-60 | chr15-95695117-INS-60 |
| chr15-95769361-INS-64 | chr15-95769363-INS-62 |
| chr15-95770546-INS-64 | chr15-95770546-INS-64 |
| chr15-95771336-INS-60 | chr15-95771336-INS-60 |
| chr15-95776979-INS-183 | chr15-95776979-INS-180 |
| chr15-95789747-INS-136 | chr15-95789747-INS-136 |
| chr15-95798548-INS-143 | chr15-95798546-INS-135 |
| chr15-95819658-INS-272 | chr15-95819658-INS-272 |
| chr15-95835854-INS-183 | chr15-95835854-INS-184 |
| chr15-95839309-INS-114 | chr15-95839310-INS-80 |
| chr15-95851363-INS-181 | chr15-95851363-INS-181 |
| chr15-95940197-INS-120 | chr15-95940197-INS-120 |
| chr15-95962168-INS-204 | chr15-95962168-INS-204 |
| chr15-95967088-INS-74 | chr15-95967088-INS-74 |
| chr15-95968516-INS-196 | chr15-95968516-INS-190 |
| chr15-95973585-INS-492 | chr15-95973585-INS-492 |
| chr15-95991100-INS-138 | chr15-95991100-INS-138 |
| chr15-95996971-INS-1121 | chr15-95996972-INS-1121 |
| chr15-95998757-INS-435 | chr15-95998759-INS-432 |
| chr15-96005792-INS-59 | chr15-96005777-INS-74 |

| | |
|---|---|
| chr15-96006685-INS-434 | chr15-96006685-INS-434 |
| chr15-96016130-INS-137 | chr15-96016125-INS-160 |
| chr15-96017089-INS-186 | chr15-96017089-INS-186 |
| chr15-96049613-INS-256 | chr15-96049613-INS-256 |
| chr15-96088107-INS-201 | chr15-96088107-INS-201 |
| chr15-96120264-INS-168 | chr15-96120264-INS-169 |
| chr15-96128954-INS-108 | chr15-96128998-INS-100 |
| chr15-96154697-INS-76 | chr15-96154697-INS-76 |
| chr15-96166384-INS-51 | chr15-96166384-INS-51 |
| chr15-96177236-INS-58 | chr15-96177236-INS-58 |
| chr15-96180004-INS-207 | chr15-96180004-INS-207 |
| chr15-96242591-INS-50 | chr15-96242591-INS-50 |
| chr15-96278623-INS-136 | chr15-96278611-INS-120 |
| chr15-96303800-INS-207 | chr15-96303800-INS-207 |
| chr15-96307133-INS-1410 | chr15-96307122-INS-1404 |
| chr15-96307514-INS-114 | chr15-96307514-INS-116 |
| chr15-96330854-INS-58 | chr15-96330854-INS-58 |
| chr15-96335264-INS-58 | chr15-96335264-INS-58 |
| chr15-96347059-INS-175 | chr15-96347059-INS-175 |
| chr15-96373384-INS-206 | chr15-96373384-INS-206 |
| chr15-96385454-INS-80 | chr15-96385454-INS-80 |
| chr15-96387967-INS-234 | chr15-96388006-INS-237 |
| chr15-96389308-INS-173 | chr15-96389308-INS-186 |
| chr15-96393738-INS-417 | chr15-96393738-INS-417 |
| chr15-96395926-INS-184 | chr15-96395915-INS-183 |
| chr15-96402384-INS-78 | chr15-96402411-INS-88 |
| chr15-96422015-INS-266 | chr15-96422015-INS-266 |
| chr15-96424792-INS-121 | chr15-96424792-INS-120 |
| chr15-96430058-INS-57 | chr15-96430045-INS-59 |
| chr15-96442232-INS-206 | chr15-96442232-INS-205 |
| chr15-96447055-INS-402 | chr15-96447055-INS-402 |
| chr15-96458400-INS-84 | chr15-96458400-INS-84 |
| chr15-96495396-INS-617 | chr15-96495396-INS-617 |
| chr15-96497516-INS-54 | chr15-96497516-INS-50 |
| chr15-96502829-INS-109 | chr15-96502829-INS-115 |
| chr15-96507416-INS-101 | chr15-96507416-INS-119 |
| chr15-96513926-INS-63 | chr15-96513926-INS-63 |
| chr15-96517491-INS-69 | chr15-96517491-INS-73 |
| chr15-96520621-INS-537 | chr15-96520621-INS-537 |
| chr15-96576547-INS-205 | chr15-96576547-INS-208 |
| chr15-96619158-INS-75 | chr15-96619159-INS-66 |
| chr15-96656144-INS-97 | chr15-96656130-INS-93 |
| chr15-96684484-INS-59 | chr15-96684484-INS-57 |
| chr15-96696002-INS-51 | chr15-96696002-INS-51 |
| chr15-96701708-INS-54 | chr15-96701698-INS-50 |
| chr15-96712330-INS-444 | chr15-96712330-INS-444 |
| chr15-96717569-INS-130 | chr15-96717569-INS-130 |
| chr15-96721751-INS-396 | chr15-96721751-INS-381 |
| chr15-96743883-INS-57 | chr15-96743910-INS-69 |
| chr15-96745875-INS-80 | chr15-96745875-INS-80 |
| chr15-96761330-INS-468 | chr15-96761313-INS-508 |
| chr15-96773590-INS-718 | chr15-96773591-INS-719 |
| chr15-96814005-INS-170 | chr15-96814005-INS-174 |
| chr15-96832865-INS-390 | chr15-96832865-INS-390 |
| chr15-96844479-INS-67 | chr15-96844479-INS-67 |
| chr15-96850048-INS-213 | chr15-96850048-INS-214 |
| chr15-96887561-INS-84 | chr15-96887561-INS-84 |
| chr15-96905408-INS-52 | chr15-96905408-INS-82 |
| chr15-96918121-INS-207 | chr15-96918121-INS-208 |
| chr15-96937544-INS-208 | chr15-96937544-INS-186 |
| chr15-96945932-INS-51 | chr15-96945932-INS-53 |
| chr15-97008883-INS-54 | chr15-97008874-INS-60 |
| chr15-97031221-INS-60 | chr15-97031221-INS-64 |
| chr15-97032084-INS-273 | chr15-97032084-INS-273 |
| chr15-97046677-INS-94 | chr15-97046677-INS-94 |
| chr15-97065819-INS-83 | chr15-97065819-INS-81 |
| chr15-97099549-INS-82 | chr15-97099549-INS-76 |
| chr15-97110716-INS-65 | chr15-97110716-INS-65 |
| chr15-97113048-INS-74 | chr15-97113048-INS-74 |
| chr15-97159799-INS-60 | chr15-97159799-INS-90 |
| chr15-97177462-INS-211 | chr15-97177462-INS-211 |
| chr15-97205266-INS-1443 | chr15-97205267-INS-1443 |
| chr15-97207540-INS-119 | chr15-97207577-INS-106 |
| chr15-97217827-INS-146 | chr15-97217827-INS-146 |
| chr15-97225622-INS-219 | chr15-97225622-INS-218 |
| chr15-97256142-INS-227 | chr15-97256142-INS-227 |
| chr15-97284027-INS-55 | chr15-97284027-INS-55 |

| | |
|---|---|
| chr15-97297619-INS-133 | chr15-97297619-INS-118 |
| chr15-97300846-INS-188 | chr15-97300846-INS-161 |
| chr15-97305098-INS-69 | chr15-97305098-INS-69 |
| chr15-97321606-INS-255 | chr15-97321606-INS-256 |
| chr15-97330968-INS-153 | chr15-97330968-INS-165 |
| chr15-97341856-INS-222 | chr15-97341856-INS-224 |
| chr15-97362096-INS-199 | chr15-97362096-INS-199 |
| chr15-97364597-INS-53 | chr15-97364597-INS-53 |
| chr15-97388290-INS-57 | chr15-97388290-INS-58 |
| chr15-97425455-INS-141 | chr15-97425455-INS-141 |
| chr15-97440983-INS-465 | chr15-97441004-INS-474 |
| chr15-97445728-INS-66 | chr15-97445737-INS-88 |
| chr15-97462086-INS-108 | chr15-97462086-INS-108 |
| chr15-97477230-INS-379 | chr15-97477230-INS-379 |
| chr15-97479274-INS-77 | chr15-97479273-INS-71 |
| chr15-97485846-INS-168 | chr15-97485846-INS-178 |
| chr15-97512527-INS-205 | chr15-97512527-INS-200 |
| chr15-97515010-INS-52 | chr15-97515010-INS-52 |
| chr15-97523243-INS-168 | chr15-97523237-INS-172 |
| chr15-97532340-INS-171 | chr15-97532340-INS-181 |
| chr15-97538291-INS-388 | chr15-97538291-INS-351 |
| chr15-97541867-INS-65 | chr15-97541867-INS-67 |
| chr15-97545236-INS-53 | chr15-97545236-INS-53 |
| chr15-97549272-INS-163 | chr15-97549272-INS-176 |
| chr15-97604698-INS-193 | chr15-97604698-INS-193 |
| chr15-97611082-INS-201 | chr15-97611082-INS-201 |
| chr15-97672505-INS-684 | chr15-97672504-INS-650 |
| chr15-97715728-INS-89 | chr15-97715693-INS-75 |
| chr15-97732079-INS-66 | chr15-97732083-INS-54 |
| chr15-97745738-INS-56 | chr15-97745738-INS-58 |
| chr15-97749521-INS-68 | chr15-97749521-INS-100 |
| chr15-97774243-INS-103 | chr15-97774243-INS-58 |
| chr15-97782614-INS-52 | chr15-97782614-INS-52 |
| chr15-97845207-INS-81 | chr15-97845207-INS-81 |
| chr15-97856190-INS-191 | chr15-97856198-INS-202 |
| chr15-97859340-INS-209 | chr15-97859340-INS-209 |
| chr15-97860371-INS-197 | chr15-97860371-INS-197 |
| chr15-97865410-INS-176 | chr15-97865410-INS-176 |
| chr15-97867292-INS-63 | chr15-97867292-INS-61 |
| chr15-97896893-INS-65 | chr15-97896893-INS-65 |
| chr15-97908838-INS-87 | chr15-97908838-INS-87 |
| chr15-97910169-INS-220 | chr15-97910134-INS-210 |
| chr15-97913196-INS-239 | chr15-97913196-INS-239 |
| chr15-97918758-INS-210 | chr15-97918758-INS-211 |
| chr15-97923071-INS-215 | chr15-97923085-INS-221 |
| chr15-97924232-INS-819 | chr15-97924232-INS-819 |
| chr15-97937995-INS-54 | chr15-97937995-INS-58 |
| chr15-97952622-INS-62 | chr15-97952626-INS-70 |
| chr15-97966884-INS-818 | chr15-97966884-INS-818 |
| chr15-97980837-INS-172 | chr15-97980837-INS-172 |
| chr15-98022176-INS-143 | chr15-98022176-INS-143 |
| chr15-98042430-INS-118 | chr15-98042430-INS-74 |
| chr15-98051864-INS-160 | chr15-98051864-INS-160 |
| chr15-98056766-INS-206 | chr15-98056766-INS-206 |
| chr15-98089576-INS-1892 | chr15-98089577-INS-1892 |
| chr15-98107374-INS-288 | chr15-98107374-INS-288 |
| chr15-98110728-INS-52 | chr15-98110728-INS-56 |
| chr15-98118001-INS-209 | chr15-98118001-INS-209 |
| chr15-98127734-INS-112 | chr15-98127734-INS-111 |
| chr15-98134526-INS-141 | chr15-98134526-INS-105 |
| chr15-98145209-INS-833 | chr15-98145209-INS-834 |
| chr15-98169736-INS-108 | chr15-98169736-INS-108 |
| chr15-98183237-INS-860 | chr15-98183237-INS-860 |
| chr15-98189983-INS-229 | chr15-98189983-INS-230 |
| chr15-98195392-INS-120 | chr15-98195392-INS-108 |
| chr15-98198999-INS-104 | chr15-98198999-INS-104 |
| chr15-98239339-INS-84 | chr15-98239339-INS-84 |
| chr15-98242420-INS-85 | chr15-98242420-INS-85 |
| chr15-98247684-INS-58 | chr15-98247684-INS-58 |
| chr15-98256620-INS-6440 | chr15-98256621-INS-6440 |
| chr15-98268707-INS-4782 | chr15-98268708-INS-4782 |
| chr15-98279704-INS-214 | chr15-98279704-INS-215 |
| chr15-98282666-INS-218 | chr15-98282666-INS-218 |
| chr15-98285968-INS-116 | chr15-98285968-INS-134 |
| chr15-98288501-INS-58 | chr15-98288501-INS-58 |
| chr15-98304049-INS-309 | chr15-98304049-INS-343 |
| chr15-98318611-INS-137 | chr15-98318611-INS-137 |

| | |
|---|---|
| chr15-98321243-INS-52 | chr15-98321241-INS-50 |
| chr15-98324805-INS-115 | chr15-98324805-INS-115 |
| chr15-98328445-INS-825 | chr15-98328445-INS-825 |
| chr15-98331106-INS-144 | chr15-98331100-INS-150 |
| chr15-98339670-INS-119 | chr15-98339670-INS-121 |
| chr15-98343499-INS-67 | chr15-98343499-INS-59 |
| chr15-98349705-INS-519 | chr15-98349705-INS-519 |
| chr15-98358221-INS-441 | chr15-98358221-INS-441 |
| chr15-98360671-INS-329 | chr15-98360669-INS-338 |
| chr15-98365548-INS-173 | chr15-98365548-INS-173 |
| chr15-98371811-INS-78 | chr15-98371811-INS-78 |
| chr15-98377079-INS-85 | chr15-98377075-INS-85 |
| chr15-98387105-INS-214 | chr15-98387105-INS-242 |
| chr15-98397761-INS-60 | chr15-98397761-INS-52 |
| chr15-98398774-INS-70 | chr15-98398757-INS-70 |
| chr15-98401989-INS-169 | chr15-98401989-INS-169 |
| chr15-98404824-INS-171 | chr15-98404813-INS-170 |
| chr15-98422686-INS-183 | chr15-98422686-INS-183 |
| chr15-98445908-INS-78 | chr15-98445908-INS-78 |
| chr15-98448944-INS-217 | chr15-98448944-INS-220 |
| chr15-98458655-INS-124 | chr15-98458650-INS-124 |
| chr15-98462706-INS-218 | chr15-98462706-INS-218 |
| chr15-98463750-INS-214 | chr15-98463750-INS-214 |
| chr15-98473321-INS-212 | chr15-98473321-INS-211 |
| chr15-98480379-INS-207 | chr15-98480379-INS-207 |
| chr15-98513506-INS-213 | chr15-98513506-INS-216 |
| chr15-98522477-INS-205 | chr15-98522477-INS-205 |
| chr15-98538442-INS-56 | chr15-98538442-INS-50 |
| chr15-98553180-INS-185 | chr15-98553210-INS-184 |
| chr15-98556001-INS-549 | chr15-98556001-INS-549 |
| chr15-98563389-INS-102 | chr15-98563389-INS-102 |
| chr15-98574352-INS-212 | chr15-98574352-INS-212 |
| chr15-98592929-INS-7104 | chr15-98592930-INS-7104 |
| chr15-98601411-INS-65 | chr15-98601411-INS-80 |
| chr15-98623697-INS-476 | chr15-98623697-INS-476 |
| chr15-98641818-INS-203 | chr15-98641818-INS-232 |
| chr15-98642791-INS-72 | chr15-98642791-INS-72 |
| chr15-98652379-INS-91 | chr15-98652379-INS-91 |
| chr15-98664571-INS-405 | chr15-98664571-INS-406 |
| chr15-98667760-INS-161 | chr15-98667760-INS-167 |
| chr15-98697819-INS-174 | chr15-98697819-INS-174 |
| chr15-98708836-INS-56 | chr15-98708836-INS-56 |
| chr15-98725394-INS-8985 | chr15-98725395-INS-8985 |
| chr15-98726816-INS-195 | chr15-98726816-INS-195 |
| chr15-98727886-INS-85 | chr15-98727886-INS-70 |
| chr15-98748903-INS-57 | chr15-98748903-INS-87 |
| chr15-98819350-INS-199 | chr15-98819350-INS-199 |
| chr15-98822290-INS-190 | chr15-98822287-INS-194 |
| chr15-98824687-INS-138 | chr15-98824687-INS-134 |
| chr15-98852233-INS-64 | chr15-98852233-INS-58 |
| chr15-98853651-INS-62 | chr15-98853651-INS-64 |
| chr15-98858867-INS-208 | chr15-98858867-INS-207 |
| chr15-98860312-INS-76 | chr15-98860312-INS-76 |
| chr15-98864181-INS-111 | chr15-98864181-INS-111 |
| chr15-98886960-INS-56 | chr15-98886963-INS-74 |
| chr15-98891733-INS-5928 | chr15-98891734-INS-5928 |
| chr15-98895762-INS-478 | chr15-98895762-INS-485 |
| chr15-98901271-INS-121 | chr15-98901271-INS-121 |
| chr15-98902532-INS-202 | chr15-98902532-INS-202 |
| chr15-98906915-INS-55 | chr15-98906915-INS-55 |
| chr15-98913026-INS-458 | chr15-98913026-INS-458 |
| chr15-98921144-INS-286 | chr15-98921144-INS-286 |
| chr15-98923317-INS-200 | chr15-98923317-INS-200 |
| chr15-98932496-INS-214 | chr15-98932496-INS-214 |
| chr15-98936978-INS-184 | chr15-98936978-INS-184 |
| chr15-98960442-INS-110 | chr15-98960433-INS-114 |
| chr15-98963232-INS-184 | chr15-98963232-INS-184 |
| chr15-98964371-INS-82 | chr15-98964372-INS-82 |
| chr15-98964900-INS-160 | chr15-98964900-INS-160 |
| chr15-98966504-INS-63 | chr15-98966504-INS-63 |
| chr15-98967233-INS-214 | chr15-98967233-INS-214 |
| chr15-98988001-INS-53 | chr15-98988035-INS-53 |
| chr15-99013104-INS-229 | chr15-99013104-INS-229 |
| chr15-99018087-INS-51 | chr15-99018057-INS-66 |
| chr15-99019917-INS-231 | chr15-99019917-INS-229 |
| chr15-99030790-INS-108 | chr15-99030822-INS-108 |
| chr15-99037220-INS-260 | chr15-99037206-INS-280 |

| | |
|---|---|
| chr15-99068144-INS-3185 | chr15-99068145-INS-3185 |
| chr15-99070650-INS-272 | chr15-99070650-INS-276 |
| chr15-99088006-INS-132 | chr15-99088010-INS-149 |
| chr15-99093492-INS-140 | chr15-99093492-INS-140 |
| chr15-99098833-INS-122 | chr15-99098833-INS-122 |
| chr15-99134243-INS-105 | chr15-99134243-INS-105 |
| chr15-99150898-INS-105 | chr15-99150898-INS-105 |
| chr15-99154995-INS-2820 | chr15-99154996-INS-2820 |
| chr15-99158770-INS-243 | chr15-99158770-INS-251 |
| chr15-99165297-INS-221 | chr15-99165297-INS-221 |
| chr15-99173222-INS-202 | chr15-99173222-INS-201 |
| chr15-99180778-INS-244 | chr15-99180778-INS-244 |
| chr15-99236330-INS-203 | chr15-99236312-INS-201 |
| chr15-99243607-INS-701 | chr15-99243607-INS-701 |
| chr15-99277136-INS-211 | chr15-99277136-INS-211 |
| chr15-99282439-INS-60 | chr15-99282439-INS-60 |
| chr15-99302747-INS-209 | chr15-99302747-INS-210 |
| chr15-99304292-INS-219 | chr15-99304292-INS-220 |
| chr15-99328358-INS-60 | chr15-99328358-INS-60 |
| chr15-99337731-INS-110 | chr15-99337751-INS-115 |
| chr15-99343946-INS-190 | chr15-99343946-INS-194 |
| chr15-99358741-INS-126 | chr15-99358746-INS-139 |
| chr15-99381690-INS-200 | chr15-99381690-INS-200 |
| chr15-99432191-INS-98 | chr15-99432179-INS-117 |
| chr15-99450122-INS-133 | chr15-99450122-INS-133 |
| chr15-99502346-INS-140 | chr15-99502346-INS-140 |
| chr15-99504030-INS-265 | chr15-99504030-INS-262 |
| chr15-99506849-INS-83 | chr15-99506849-INS-83 |
| chr15-99540735-INS-130 | chr15-99540735-INS-130 |
| chr15-99551537-INS-95 | chr15-99551537-INS-108 |
| chr15-99553305-INS-67 | chr15-99553305-INS-67 |
| chr15-99557275-INS-170 | chr15-99557242-INS-211 |
| chr15-99578898-INS-59 | chr15-99578898-INS-59 |
| chr15-99588449-INS-129 | chr15-99588449-INS-129 |
| chr15-99596233-INS-1896 | chr15-99596234-INS-1896 |
| chr15-99606907-INS-212 | chr15-99606907-INS-212 |
| chr15-99619575-INS-72 | chr15-99619575-INS-72 |
| chr15-99626507-INS-230 | chr15-99626508-INS-229 |
| chr15-99632778-INS-241 | chr15-99632779-INS-241 |
| chr15-99641686-INS-58 | chr15-99641696-INS-61 |
| chr15-99644476-INS-498 | chr15-99644476-INS-498 |
| chr15-99653730-INS-205 | chr15-99653730-INS-205 |
| chr15-99660345-INS-178 | chr15-99660345-INS-178 |
| chr15-99665479-INS-235 | chr15-99665479-INS-223 |
| chr15-99671807-INS-699 | chr15-99671807-INS-699 |
| chr15-99682839-INS-196 | chr15-99682839-INS-197 |
| chr15-99690681-INS-221 | chr15-99690681-INS-221 |
| chr15-99695257-INS-193 | chr15-99695257-INS-193 |
| chr15-99698935-INS-195 | chr15-99698935-INS-195 |
| chr15-99707295-INS-51 | chr15-99707295-INS-57 |
| chr15-99711888-INS-205 | chr15-99711888-INS-204 |
| chr15-99729062-INS-208 | chr15-99729062-INS-208 |
| chr15-99741140-INS-57 | chr15-99741140-INS-61 |
| chr15-99744471-INS-222 | chr15-99744471-INS-222 |
| chr15-99746878-INS-61 | chr15-99746878-INS-61 |
| chr15-99756399-INS-56 | chr15-99756399-INS-56 |
| chr15-99793131-INS-197 | chr15-99793131-INS-197 |
| chr15-99805162-INS-281 | chr15-99805162-INS-281 |
| chr15-99811239-INS-68 | chr15-99811239-INS-54 |
| chr15-99833376-INS-116 | chr15-99833376-INS-116 |
| chr15-99849854-INS-150 | chr15-99849833-INS-139 |
| chr15-99856225-INS-8986 | chr15-99856226-INS-8986 |
| chr15-99895658-INS-201 | chr15-99895658-INS-201 |
| chr15-99926392-INS-5380 | chr15-99926393-INS-5380 |
| chr15-99964567-INS-185 | chr15-99964567-INS-185 |
| chr15-100018291-INS-396 | chr15-100018291-INS-396 |
| chr15-100061015-INS-7186 | chr15-100061016-INS-7186 |
| chr15-100069920-INS-138 | chr15-100069920-INS-140 |
| chr15-100075363-INS-203 | chr15-100075363-INS-203 |
| chr15-100086852-INS-140 | chr15-100086852-INS-149 |
| chr15-100089979-INS-1890 | chr15-100089980-INS-1890 |
| chr15-100133733-INS-189 | chr15-100133733-INS-191 |
| chr15-100160965-INS-190 | chr15-100160965-INS-192 |
| chr15-100167500-INS-701 | chr15-100167500-INS-701 |
| chr15-100191161-INS-227 | chr15-100191161-INS-227 |
| chr15-100230910-INS-134 | chr15-100230910-INS-134 |
| chr15-100276678-INS-134 | chr15-100276678-INS-134 |

| | |
|---|---|
| chr15-100294096-INS-75 | chr15-100294114-INS-72 |
| chr15-100298772-INS-68 | chr15-100298772-INS-68 |
| chr15-100304154-INS-104 | chr15-100304154-INS-104 |
| chr15-100335964-INS-60 | chr15-100335972-INS-60 |
| chr15-100340601-INS-54 | chr15-100340601-INS-54 |
| chr15-100344053-INS-411 | chr15-100344053-INS-411 |
| chr15-100350452-INS-146 | chr15-100350452-INS-144 |
| chr15-100378530-INS-55 | chr15-100378530-INS-55 |
| chr15-100385631-INS-86 | chr15-100385631-INS-86 |
| chr15-100416377-INS-239 | chr15-100416377-INS-239 |
| chr15-100430955-INS-260 | chr15-100430955-INS-260 |
| chr15-100455888-INS-72 | chr15-100455888-INS-72 |
| chr15-100499227-INS-60 | chr15-100499227-INS-60 |
| chr15-100546336-INS-196 | chr15-100546337-INS-196 |
| chr15-100598829-INS-133 | chr15-100598829-INS-134 |
| chr15-100600412-INS-62 | chr15-100600404-INS-92 |
| chr15-100601918-INS-78 | chr15-100601918-INS-78 |
| chr15-100611709-INS-254 | chr15-100611709-INS-254 |
| chr15-100634763-INS-81 | chr15-100634763-INS-82 |
| chr15-100638766-INS-900 | chr15-100638766-INS-899 |
| chr15-100668123-INS-59 | chr15-100668123-INS-59 |
| chr15-100672002-INS-96 | chr15-100672002-INS-96 |
| chr15-100679683-INS-221 | chr15-100679683-INS-222 |
| chr15-100698916-INS-99 | chr15-100698916-INS-99 |
| chr15-100700503-INS-63 | chr15-100700504-INS-67 |
| chr15-100739707-INS-217 | chr15-100739707-INS-217 |
| chr15-100748054-INS-145 | chr15-100748054-INS-145 |
| chr15-100755514-INS-293 | chr15-100755514-INS-293 |
| chr15-100756052-INS-88 | chr15-100756034-INS-134 |
| chr15-100795152-INS-70 | chr15-100795152-INS-50 |
| chr15-100833654-INS-215 | chr15-100833654-INS-215 |
| chr15-100898860-INS-395 | chr15-100898860-INS-395 |
| chr15-100904231-INS-63 | chr15-100904231-INS-85 |
| chr15-100947722-INS-51 | chr15-100947722-INS-51 |
| chr15-100986919-INS-123 | chr15-100986919-INS-123 |
| chr15-101012081-INS-59 | chr15-101012083-INS-57 |
| chr15-101018726-INS-111 | chr15-101018726-INS-111 |
| chr15-101021665-INS-96 | chr15-101021665-INS-96 |
| chr15-101035749-INS-177 | chr15-101035749-INS-176 |
| chr15-101083333-INS-219 | chr15-101083333-INS-219 |
| chr15-101093833-INS-165 | chr15-101093868-INS-168 |
| chr15-101103532-INS-76 | chr15-101103532-INS-76 |
| chr15-101288716-INS-325 | chr15-101288716-INS-285 |
| chr15-101354423-INS-166 | chr15-101354423-INS-166 |
| chr15-101359956-INS-173 | chr15-101359956-INS-173 |
| chr15-101387701-INS-1891 | chr15-101387702-INS-1891 |
| chr15-101396075-INS-86 | chr15-101396075-INS-51 |
| chr15-101415324-INS-64 | chr15-101415300-INS-72 |
| chr15-101445748-INS-98 | chr15-101445748-INS-120 |
| chr15-101458299-INS-80 | chr15-101458301-INS-66 |
| chr15-101496391-INS-86 | chr15-101496391-INS-96 |
| chr15-101542892-INS-75 | chr15-101542892-INS-81 |
| chr15-101582328-INS-227 | chr15-101582329-INS-227 |
| chr15-101583295-INS-141 | chr15-101583295-INS-141 |
| chr15-101628955-INS-120 | chr15-101628955-INS-80 |
| chr15-101631598-INS-75 | chr15-101631613-INS-70 |
| chr15-101674189-INS-6920 | chr15-101674190-INS-6920 |
| chr15-101688243-INS-52 | chr15-101688235-INS-53 |
| chr15-101718288-INS-60 | chr15-101718288-INS-60 |
| chr15-101728385-INS-84 | chr15-101728379-INS-90 |
| chr15-101750681-INS-90 | chr15-101750682-INS-90 |
| chr15-101791763-INS-57 | chr15-101791763-INS-57 |
| chr15-101796285-INS-76 | chr15-101796285-INS-76 |
| chr15-101819982-INS-83 | chr15-101820002-INS-103 |
| chr15-101851374-INS-58 | chr15-101851374-INS-58 |
| chr15-101860413-INS-100 | chr15-101860413-INS-100 |
| chr15-101873293-INS-90 | chr15-101873293-INS-90 |
| chr15-101875244-INS-127 | chr15-101875243-INS-143 |
| chr15-101887828-INS-62 | chr15-101887828-INS-62 |
| chr15-101918286-INS-5552 | chr15-101918287-INS-5552 |
| chr15-101921358-INS-89 | chr15-101921358-INS-89 |
| chr15-101922761-INS-6475 | chr15-101922762-INS-6475 |
| chr15-101932983-INS-331 | chr15-101932983-INS-331 |
| chr15-101941067-INS-69 | chr15-101941067-INS-57 |
| chr15-101955324-INS-1273 | chr15-101955325-INS-1273 |
| chr15-101979239-INS-71 | chr15-101979239-INS-71 |
| chr15-102002454-INS-223 | chr15-102002454-INS-223 |

| | |
|---|---|
| chr15-102018370-INS-217 | chr15-102018370-INS-217 |
| chr15-102049795-INS-547 | chr15-102049795-INS-550 |
| chr15-102050966-INS-211 | chr15-102050966-INS-211 |
| chr15-102057198-INS-203 | chr15-102057198-INS-203 |
| chr15-102064926-INS-213 | chr15-102064926-INS-213 |
| chr15-102107732-INS-270 | chr15-102107732-INS-270 |
| chr15-102127979-INS-54 | chr15-102127979-INS-54 |
| chr15-102174327-INS-268 | chr15-102174327-INS-268 |
| chr15-102203869-INS-147 | chr15-102203869-INS-133 |
| chr15-102220286-INS-200 | chr15-102220285-INS-203 |
| chr15-102223265-INS-158 | chr15-102223265-INS-158 |
| chr15-102241618-INS-199 | chr15-102241618-INS-199 |
| chr15-102246337-INS-55 | chr15-102246337-INS-55 |
| chr15-102249913-INS-56 | chr15-102249913-INS-56 |
| chr15-102268294-INS-184 | chr15-102268294-INS-181 |
| chr15-102286977-INS-234 | chr15-102286977-INS-218 |
| chr15-102308811-INS-171 | chr15-102308802-INS-179 |
| chr15-102318185-INS-60 | chr15-102318185-INS-60 |
| chr15-102324794-INS-203 | chr15-102324794-INS-203 |
| chr15-102345862-INS-181 | chr15-102345862-INS-180 |
| chr15-102359029-INS-56 | chr15-102359029-INS-60 |
| chr15-102365346-INS-187 | chr15-102365356-INS-165 |
| chr15-102406585-INS-181 | chr15-102406585-INS-182 |
| chr15-102414158-INS-62 | chr15-102414188-INS-59 |
| chr15-102415108-INS-237 | chr15-102415108-INS-237 |
| chr15-102481893-INS-254 | chr15-102481893-INS-244 |
| chr15-102523907-INS-230 | chr15-102523907-INS-230 |
| chr15-102545283-INS-166 | chr15-102545283-INS-174 |
| chr15-102577051-INS-64 | chr15-102577051-INS-66 |
| chr15-102584585-INS-209 | chr15-102584585-INS-210 |
| chr15-102590168-INS-216 | chr15-102590169-INS-214 |
| chr15-102612793-INS-209 | chr15-102612793-INS-210 |
| chr15-102615418-INS-217 | chr15-102615418-INS-217 |
| chr15-102654678-INS-5345 | chr15-102654679-INS-5344 |
| chr15-102670831-INS-56 | chr15-102670802-INS-58 |
| chr15-102673673-INS-215 | chr15-102673676-INS-218 |
| chr15-102695142-INS-190 | chr15-102695142-INS-188 |
| chr15-102737464-INS-168 | chr15-102737464-INS-182 |
| chr15-102769778-INS-52 | chr15-102769778-INS-50 |
| chr15-102776006-INS-54 | chr15-102776006-INS-54 |
| chr15-102781905-INS-113 | chr15-102781905-INS-113 |
| chr15-102788936-INS-4225 | chr15-102788937-INS-4225 |
| chr15-102792653-INS-63 | chr15-102792653-INS-71 |
| chr15-102801329-INS-205 | chr15-102801329-INS-206 |
| chr15-102832796-INS-78 | chr15-102832796-INS-66 |
| chr15-102855500-INS-88 | chr15-102855500-INS-90 |
| chr15-102898810-INS-54 | chr15-102898808-INS-54 |
| chr15-102957419-INS-63 | chr15-102957429-INS-91 |
| chr15-102981938-INS-56 | chr15-102981938-INS-57 |
| chr15-102993185-INS-842 | chr15-102993185-INS-842 |
| chr15-103020127-INS-208 | chr15-103020127-INS-208 |
| chr15-103190914-INS-216 | chr15-103190914-INS-215 |
| chr15-103195147-INS-200 | chr15-103195158-INS-199 |
| chr15-103209914-INS-64 | chr15-103209914-INS-84 |
| chr15-103266250-INS-76 | chr15-103266250-INS-80 |
| chr15-103342390-INS-1563 | chr15-103342391-INS-1563 |
| chr15-103344345-INS-216 | chr15-103344348-INS-247 |
| chr15-103429377-INS-6040 | chr15-103429378-INS-6040 |
| chr15-103444323-INS-67 | chr15-103444323-INS-67 |
| chr15-103454550-INS-72 | chr15-103454550-INS-72 |
| chr15-103468202-INS-191 | chr15-103468202-INS-191 |
| chr15-103483178-INS-58 | chr15-103483178-INS-58 |
| chr15-103495312-INS-3731 | chr15-103495313-INS-3731 |
| chr15-103508089-INS-203 | chr15-103508089-INS-203 |
| chr15-103528641-INS-1890 | chr15-103528642-INS-1890 |
| chr15-103536419-INS-1705 | chr15-103536420-INS-1705 |
| chr15-103560052-INS-54 | chr15-103560052-INS-54 |
| chr15-103576713-INS-68 | chr15-103576713-INS-53 |
| chr15-103615759-INS-62 | chr15-103615753-INS-110 |
| chr15-103623731-INS-80 | chr15-103623731-INS-80 |
| chr15-103652125-INS-68 | chr15-103652125-INS-68 |
| chr15-103773738-INS-57 | chr15-103773738-INS-57 |
| chr15-103785143-INS-7144 | chr15-103785144-INS-7144 |
| chr15-103806113-INS-323 | chr15-103806113-INS-323 |
| chr15-103933496-INS-60 | chr15-103933496-INS-62 |
| chr16-3278620-INS-62 | chr16-3278620-INS-62 |
| chr16-3305927-INS-64 | chr16-3305928-INS-72 |

| | |
|---|---|
| chr16-3356733-INS-4831 | chr16-3356734-INS-4831 |
| chr16-3458117-INS-2907 | chr16-3458118-INS-2907 |
| chr16-3461043-INS-120 | chr16-3461043-INS-120 |
| chr16-3463784-INS-854 | chr16-3463784-INS-854 |
| chr16-3468401-INS-170 | chr16-3468401-INS-170 |
| chr16-3479182-INS-60 | chr16-3479182-INS-60 |
| chr16-3493090-INS-931 | chr16-3493090-INS-931 |
| chr16-3523525-INS-82 | chr16-3523525-INS-88 |
| chr16-3537038-INS-118 | chr16-3537034-INS-106 |
| chr16-3554152-INS-4881 | chr16-3554153-INS-4881 |
| chr16-3556546-INS-80 | chr16-3556546-INS-80 |
| chr16-3601027-INS-84 | chr16-3601027-INS-84 |
| chr16-3615266-INS-96 | chr16-3615267-INS-96 |
| chr16-3621998-INS-4712 | chr16-3621999-INS-4712 |
| chr16-3632009-INS-131 | chr16-3632035-INS-99 |
| chr16-3649102-INS-62 | chr16-3649102-INS-80 |
| chr16-3662952-INS-202 | chr16-3662952-INS-202 |
| chr16-3669503-INS-123 | chr16-3669462-INS-123 |
| chr16-3670582-INS-332 | chr16-3670597-INS-300 |
| chr16-3700401-INS-201 | chr16-3700401-INS-201 |
| chr16-3742849-INS-208 | chr16-3742849-INS-210 |
| chr16-3760961-INS-63 | chr16-3760969-INS-66 |
| chr16-3776947-INS-117 | chr16-3776947-INS-117 |
| chr16-3779139-INS-111 | chr16-3779139-INS-111 |
| chr16-3788084-INS-84 | chr16-3788084-INS-84 |
| chr16-3789740-INS-322 | chr16-3789741-INS-324 |
| chr16-3792386-INS-99 | chr16-3792364-INS-99 |
| chr16-3824992-INS-207 | chr16-3824992-INS-207 |
| chr16-3832726-INS-746 | chr16-3832726-INS-746 |
| chr16-3865698-INS-220 | chr16-3865698-INS-220 |
| chr16-3888124-INS-221 | chr16-3888124-INS-221 |
| chr16-3923953-INS-107 | chr16-3923953-INS-85 |
| chr16-3935857-INS-165 | chr16-3935857-INS-165 |
| chr16-3952744-INS-205 | chr16-3952744-INS-205 |
| chr16-3989986-INS-974 | chr16-3989981-INS-1005 |
| chr16-4041155-INS-3796 | chr16-4041156-INS-3796 |
| chr16-4056132-INS-194 | chr16-4056132-INS-196 |
| chr16-4057766-INS-140 | chr16-4057766-INS-140 |
| chr16-4092563-INS-66 | chr16-4092563-INS-102 |
| chr16-4146142-INS-164 | chr16-4146142-INS-163 |
| chr16-4160145-INS-56 | chr16-4160145-INS-62 |
| chr16-4175620-INS-1189 | chr16-4175621-INS-1189 |
| chr16-4179345-INS-200 | chr16-4179345-INS-200 |
| chr16-4243177-INS-167 | chr16-4243177-INS-171 |
| chr16-4264410-INS-75 | chr16-4264410-INS-75 |
| chr16-4292319-INS-112 | chr16-4292319-INS-112 |
| chr16-4313895-INS-498 | chr16-4313895-INS-498 |
| chr16-4334192-INS-211 | chr16-4334192-INS-211 |
| chr16-4357887-INS-165 | chr16-4357895-INS-154 |
| chr16-4358463-INS-854 | chr16-4358463-INS-855 |
| chr16-4394605-INS-199 | chr16-4394605-INS-199 |
| chr16-4398276-INS-216 | chr16-4398276-INS-217 |
| chr16-4427473-INS-88 | chr16-4427473-INS-107 |
| chr16-4484743-INS-1892 | chr16-4484744-INS-1892 |
| chr16-4497969-INS-66 | chr16-4497969-INS-59 |
| chr16-4500502-INS-137 | chr16-4500502-INS-132 |
| chr16-4518531-INS-132 | chr16-4518531-INS-132 |
| chr16-4522721-INS-56 | chr16-4522685-INS-104 |
| chr16-4560361-INS-102 | chr16-4560361-INS-102 |
| chr16-4598198-INS-4176 | chr16-4598199-INS-4176 |
| chr16-4626792-INS-724 | chr16-4626792-INS-726 |
| chr16-4644585-INS-247 | chr16-4644585-INS-247 |
| chr16-4645423-INS-248 | chr16-4645423-INS-248 |
| chr16-4645764-INS-7086 | chr16-4645765-INS-7086 |
| chr16-4656228-INS-158 | chr16-4656228-INS-158 |
| chr16-4672243-INS-60 | chr16-4672243-INS-55 |
| chr16-4674360-INS-55 | chr16-4674341-INS-54 |
| chr16-4685073-INS-92 | chr16-4685073-INS-108 |
| chr16-4690412-INS-206 | chr16-4690412-INS-205 |
| chr16-4707568-INS-165 | chr16-4707568-INS-167 |
| chr16-4723365-INS-145 | chr16-4723365-INS-145 |
| chr16-4737836-INS-72 | chr16-4737836-INS-72 |
| chr16-4750170-INS-221 | chr16-4750170-INS-222 |
| chr16-4774521-INS-200 | chr16-4774521-INS-200 |
| chr16-4817520-INS-69 | chr16-4817547-INS-90 |
| chr16-4820908-INS-4986 | chr16-4820909-INS-4986 |
| chr16-4881861-INS-129 | chr16-4881861-INS-129 |

| | |
|---|---|
| chr16-4896952-INS-54 | chr16-4896952-INS-54 |
| chr16-4947403-INS-285 | chr16-4947403-INS-291 |
| chr16-4973527-INS-294 | chr16-4973545-INS-291 |
| chr16-4985395-INS-56 | chr16-4985394-INS-50 |
| chr16-4991728-INS-52 | chr16-4991728-INS-52 |
| chr16-5000353-INS-62 | chr16-5000360-INS-56 |
| chr16-5007500-INS-85 | chr16-5007500-INS-85 |
| chr16-5010375-INS-63 | chr16-5010375-INS-57 |
| chr16-5014466-INS-56 | chr16-5014465-INS-88 |
| chr16-5026783-INS-264 | chr16-5026784-INS-264 |
| chr16-5041161-INS-116 | chr16-5041161-INS-116 |
| chr16-5050090-INS-52 | chr16-5050090-INS-52 |
| chr16-5078488-INS-506 | chr16-5078488-INS-510 |
| chr16-5081443-INS-428 | chr16-5081443-INS-431 |
| chr16-5082865-INS-658 | chr16-5082865-INS-658 |
| chr16-5088244-INS-213 | chr16-5088244-INS-213 |
| chr16-5090278-INS-476 | chr16-5090278-INS-480 |
| chr16-5094690-INS-278 | chr16-5094690-INS-278 |
| chr16-5095576-INS-167 | chr16-5095576-INS-176 |
| chr16-5102269-INS-64 | chr16-5102271-INS-64 |
| chr16-5122511-INS-76 | chr16-5122511-INS-76 |
| chr16-5156467-INS-1235 | chr16-5156468-INS-1235 |
| chr16-5165548-INS-2026 | chr16-5165549-INS-2026 |
| chr16-5169094-INS-71 | chr16-5169094-INS-67 |
| chr16-5192272-INS-80 | chr16-5192272-INS-80 |
| chr16-5213821-INS-66 | chr16-5213821-INS-77 |
| chr16-5239515-INS-6197 | chr16-5239516-INS-6197 |
| chr16-5245250-INS-63 | chr16-5245250-INS-93 |
| chr16-5260239-INS-190 | chr16-5260216-INS-174 |
| chr16-5283302-INS-540 | chr16-5283303-INS-527 |
| chr16-5356190-INS-5390 | chr16-5356191-INS-5390 |
| chr16-5465194-INS-58 | chr16-5465194-INS-58 |
| chr16-5539846-INS-1297 | chr16-5539847-INS-1297 |
| chr16-5552025-INS-149 | chr16-5552025-INS-149 |
| chr16-5553593-INS-71 | chr16-5553593-INS-71 |
| chr16-5563975-INS-135 | chr16-5563975-INS-135 |
| chr16-5603844-INS-100 | chr16-5603844-INS-100 |
| chr16-5611523-INS-54 | chr16-5611523-INS-56 |
| chr16-5621581-INS-56 | chr16-5621581-INS-60 |
| chr16-5636704-INS-357 | chr16-5636704-INS-357 |
| chr16-5678206-INS-196 | chr16-5678206-INS-196 |
| chr16-5685194-INS-58 | chr16-5685194-INS-56 |
| chr16-5709517-INS-143 | chr16-5709517-INS-143 |
| chr16-5723637-INS-57 | chr16-5723637-INS-57 |
| chr16-5749384-INS-70 | chr16-5749372-INS-54 |
| chr16-5793055-INS-388 | chr16-5793055-INS-388 |
| chr16-5827023-INS-94 | chr16-5827023-INS-95 |
| chr16-5835705-INS-53 | chr16-5835675-INS-53 |
| chr16-5854213-INS-131 | chr16-5854213-INS-131 |
| chr16-5886584-INS-66 | chr16-5886585-INS-72 |
| chr16-5894350-INS-4349 | chr16-5894351-INS-4349 |
| chr16-5928469-INS-344 | chr16-5928475-INS-356 |
| chr16-5932809-INS-96 | chr16-5932809-INS-96 |
| chr16-5947023-INS-349 | chr16-5947023-INS-333 |
| chr16-5997533-INS-241 | chr16-5997533-INS-241 |
| chr16-6013389-INS-73 | chr16-6013389-INS-73 |
| chr16-6025872-INS-64 | chr16-6025826-INS-113 |
| chr16-6031194-INS-155 | chr16-6031194-INS-155 |
| chr16-6055789-INS-54 | chr16-6055789-INS-58 |
| chr16-6058688-INS-120 | chr16-6058688-INS-120 |
| chr16-6064420-INS-718 | chr16-6064420-INS-705 |
| chr16-6077670-INS-214 | chr16-6077670-INS-213 |
| chr16-6180968-INS-56 | chr16-6180968-INS-54 |
| chr16-6204710-INS-1087 | chr16-6204711-INS-1087 |
| chr16-6207657-INS-100 | chr16-6207657-INS-100 |
| chr16-6231193-INS-58 | chr16-6231193-INS-60 |
| chr16-6232592-INS-68 | chr16-6232592-INS-68 |
| chr16-6254921-INS-158 | chr16-6254921-INS-164 |
| chr16-6301404-INS-75 | chr16-6301404-INS-75 |
| chr16-6320541-INS-275 | chr16-6320541-INS-275 |
| chr16-6336049-INS-79 | chr16-6336082-INS-51 |
| chr16-6348878-INS-128 | chr16-6348878-INS-126 |
| chr16-6361521-INS-646 | chr16-6361521-INS-646 |
| chr16-6365503-INS-52 | chr16-6365504-INS-52 |
| chr16-6380949-INS-81 | chr16-6380949-INS-81 |
| chr16-6385043-INS-60 | chr16-6385043-INS-56 |
| chr16-6397946-INS-71 | chr16-6397946-INS-72 |

| | |
|---|---|
| chr16-6400000-INS-51 | chr16-6400000-INS-57 |
| chr16-6425771-INS-145 | chr16-6425776-INS-171 |
| chr16-6435865-INS-95 | chr16-6435865-INS-100 |
| chr16-6439699-INS-155 | chr16-6439699-INS-153 |
| chr16-6453484-INS-132 | chr16-6453458-INS-142 |
| chr16-6492435-INS-70 | chr16-6492435-INS-74 |
| chr16-6494335-INS-50 | chr16-6494335-INS-50 |
| chr16-6508009-INS-52 | chr16-6508009-INS-52 |
| chr16-6535096-INS-63 | chr16-6535096-INS-63 |
| chr16-6557078-INS-54 | chr16-6557078-INS-54 |
| chr16-6614690-INS-72 | chr16-6614690-INS-72 |
| chr16-6640270-INS-325 | chr16-6640270-INS-325 |
| chr16-6643061-INS-200 | chr16-6643061-INS-203 |
| chr16-6651030-INS-55 | chr16-6651030-INS-55 |
| chr16-6728048-INS-203 | chr16-6728048-INS-204 |
| chr16-6746768-INS-57 | chr16-6746768-INS-57 |
| chr16-6751032-INS-86 | chr16-6751032-INS-88 |
| chr16-6816878-INS-58 | chr16-6816878-INS-58 |
| chr16-6818585-INS-93 | chr16-6818585-INS-56 |
| chr16-6857173-INS-57 | chr16-6857173-INS-56 |
| chr16-6870858-INS-60 | chr16-6870858-INS-62 |
| chr16-6880116-INS-406 | chr16-6880116-INS-406 |
| chr16-6913437-INS-282 | chr16-6913437-INS-282 |
| chr16-6925308-INS-53 | chr16-6925304-INS-51 |
| chr16-6926562-INS-53 | chr16-6926562-INS-53 |
| chr16-6955721-INS-6557 | chr16-6955722-INS-6557 |
| chr16-7030399-INS-52 | chr16-7030399-INS-52 |
| chr16-7050186-INS-102 | chr16-7050186-INS-102 |
| chr16-7056590-INS-124 | chr16-7056590-INS-124 |
| chr16-7070452-INS-130 | chr16-7070452-INS-135 |
| chr16-7087045-INS-228 | chr16-7087045-INS-228 |
| chr16-7158228-INS-66 | chr16-7158228-INS-68 |
| chr16-7198627-INS-56 | chr16-7198641-INS-57 |
| chr16-7200947-INS-326 | chr16-7200947-INS-326 |
| chr16-7223128-INS-55 | chr16-7223128-INS-55 |
| chr16-7248757-INS-431 | chr16-7248748-INS-434 |
| chr16-7249826-INS-1321 | chr16-7249835-INS-1309 |
| chr16-7255396-INS-54 | chr16-7255396-INS-54 |
| chr16-7268174-INS-52 | chr16-7268174-INS-52 |
| chr16-7292925-INS-52 | chr16-7292925-INS-52 |
| chr16-7314758-INS-206 | chr16-7314758-INS-206 |
| chr16-7383576-INS-133 | chr16-7383595-INS-108 |
| chr16-7390121-INS-144 | chr16-7390122-INS-147 |
| chr16-7396203-INS-98 | chr16-7396203-INS-98 |
| chr16-7417827-INS-758 | chr16-7417827-INS-758 |
| chr16-7456920-INS-92 | chr16-7456920-INS-92 |
| chr16-7470237-INS-314 | chr16-7470237-INS-314 |
| chr16-7514123-INS-365 | chr16-7514123-INS-365 |
| chr16-7524915-INS-6327 | chr16-7524916-INS-6327 |
| chr16-7531135-INS-1893 | chr16-7531136-INS-1895 |
| chr16-7554746-INS-358 | chr16-7554746-INS-358 |
| chr16-7566279-INS-179 | chr16-7566279-INS-179 |
| chr16-7580370-INS-7173 | chr16-7580371-INS-7173 |
| chr16-7616335-INS-501 | chr16-7616335-INS-501 |
| chr16-7630506-INS-63 | chr16-7630506-INS-63 |
| chr16-7657565-INS-52 | chr16-7657565-INS-52 |
| chr16-7669754-INS-52 | chr16-7669754-INS-52 |
| chr16-7716443-INS-1073 | chr16-7716444-INS-1073 |
| chr16-7788538-INS-102 | chr16-7788538-INS-108 |
| chr16-7793000-INS-60 | chr16-7793000-INS-60 |
| chr16-7803258-INS-280 | chr16-7803258-INS-281 |
| chr16-7820603-INS-131 | chr16-7820603-INS-127 |
| chr16-7850780-INS-917 | chr16-7850780-INS-917 |
| chr16-7902289-INS-358 | chr16-7902289-INS-359 |
| chr16-7919207-INS-302 | chr16-7919207-INS-302 |
| chr16-7925144-INS-51 | chr16-7925144-INS-51 |
| chr16-7952639-INS-1445 | chr16-7952640-INS-1445 |
| chr16-7953478-INS-283 | chr16-7953478-INS-283 |
| chr16-7960762-INS-3299 | chr16-7960763-INS-3301 |
| chr16-7967140-INS-208 | chr16-7967140-INS-207 |
| chr16-7967860-INS-204 | chr16-7967874-INS-204 |
| chr16-7985218-INS-120 | chr16-7985218-INS-120 |
| chr16-8017870-INS-80 | chr16-8017874-INS-86 |
| chr16-8036969-INS-138 | chr16-8036959-INS-162 |
| chr16-8093170-INS-79 | chr16-8093188-INS-72 |
| chr16-8124181-INS-115 | chr16-8124181-INS-115 |
| chr16-8131527-INS-393 | chr16-8131527-INS-394 |

| | |
|---|---|
| chr16-8138348-INS-139 | chr16-8138320-INS-175 |
| chr16-8140048-INS-95 | chr16-8140038-INS-58 |
| chr16-8141748-INS-192 | chr16-8141748-INS-191 |
| chr16-8160511-INS-4845 | chr16-8160512-INS-4845 |
| chr16-8160800-INS-207 | chr16-8160800-INS-208 |
| chr16-8165733-INS-5397 | chr16-8165734-INS-5397 |
| chr16-8207099-INS-72 | chr16-8207109-INS-74 |
| chr16-8225613-INS-55 | chr16-8225613-INS-98 |
| chr16-8240316-INS-84 | chr16-8240316-INS-84 |
| chr16-8245476-INS-54 | chr16-8245476-INS-54 |
| chr16-8252847-INS-209 | chr16-8252847-INS-209 |
| chr16-8253925-INS-651 | chr16-8253925-INS-651 |
| chr16-8277333-INS-66 | chr16-8277333-INS-66 |
| chr16-8288229-INS-543 | chr16-8288229-INS-543 |
| chr16-8295091-INS-106 | chr16-8295099-INS-94 |
| chr16-8316822-INS-100 | chr16-8316822-INS-100 |
| chr16-8318998-INS-207 | chr16-8318998-INS-207 |
| chr16-8320387-INS-54 | chr16-8320431-INS-54 |
| chr16-8324055-INS-204 | chr16-8324055-INS-204 |
| chr16-8360753-INS-244 | chr16-8360753-INS-244 |
| chr16-8362128-INS-51 | chr16-8362128-INS-51 |
| chr16-8404542-INS-210 | chr16-8404542-INS-206 |
| chr16-8424549-INS-210 | chr16-8424549-INS-210 |
| chr16-8450914-INS-128 | chr16-8450914-INS-129 |
| chr16-8471244-INS-1249 | chr16-8471245-INS-1264 |
| chr16-8483031-INS-142 | chr16-8483031-INS-142 |
| chr16-8486689-INS-202 | chr16-8486689-INS-202 |
| chr16-8545614-INS-59 | chr16-8545614-INS-58 |
| chr16-8547807-INS-54 | chr16-8547807-INS-54 |
| chr16-8550000-INS-212 | chr16-8550000-INS-212 |
| chr16-8574039-INS-194 | chr16-8574039-INS-199 |
| chr16-8575423-INS-87 | chr16-8575423-INS-88 |
| chr16-8588186-INS-65 | chr16-8588204-INS-97 |
| chr16-8598174-INS-212 | chr16-8598174-INS-212 |
| chr16-8600814-INS-56 | chr16-8600813-INS-52 |
| chr16-8609886-INS-60 | chr16-8609886-INS-66 |
| chr16-8613267-INS-189 | chr16-8613267-INS-179 |
| chr16-8616918-INS-197 | chr16-8616918-INS-197 |
| chr16-8622365-INS-56 | chr16-8622365-INS-72 |
| chr16-8625339-INS-123 | chr16-8625343-INS-107 |
| chr16-8627325-INS-165 | chr16-8627325-INS-165 |
| chr16-8633686-INS-231 | chr16-8633686-INS-231 |
| chr16-8698615-INS-187 | chr16-8698615-INS-193 |
| chr16-8710748-INS-56 | chr16-8710748-INS-51 |
| chr16-8712325-INS-393 | chr16-8712325-INS-396 |
| chr16-8714005-INS-215 | chr16-8714005-INS-215 |
| chr16-8717403-INS-964 | chr16-8717403-INS-983 |
| chr16-8732777-INS-51 | chr16-8732774-INS-54 |
| chr16-8735542-INS-4286 | chr16-8735543-INS-4286 |
| chr16-8778851-INS-450 | chr16-8778851-INS-450 |
| chr16-8844671-INS-60 | chr16-8844671-INS-60 |
| chr16-8895185-INS-152 | chr16-8895196-INS-172 |
| chr16-8907048-INS-68 | chr16-8907047-INS-72 |
| chr16-8908426-INS-952 | chr16-8908426-INS-952 |
| chr16-8920960-INS-6215 | chr16-8920961-INS-6215 |
| chr16-8944761-INS-330 | chr16-8944761-INS-330 |
| chr16-8957429-INS-136 | chr16-8957429-INS-105 |
| chr16-8964477-INS-57 | chr16-8964477-INS-57 |
| chr16-8964756-INS-356 | chr16-8964756-INS-356 |
| chr16-8977357-INS-268 | chr16-8977357-INS-268 |
| chr16-8979136-INS-152 | chr16-8979136-INS-152 |
| chr16-8982078-INS-1009 | chr16-8982079-INS-1009 |
| chr16-8995846-INS-51 | chr16-8995846-INS-55 |
| chr16-9001717-INS-117 | chr16-9001717-INS-125 |
| chr16-9022979-INS-182 | chr16-9022979-INS-176 |
| chr16-9037161-INS-70 | chr16-9037156-INS-66 |
| chr16-9038579-INS-72 | chr16-9038582-INS-64 |
| chr16-9062745-INS-98 | chr16-9062745-INS-110 |
| chr16-9067687-INS-67 | chr16-9067687-INS-67 |
| chr16-9093891-INS-7483 | chr16-9093892-INS-7483 |
| chr16-9108551-INS-6463 | chr16-9108552-INS-6464 |
| chr16-9166443-INS-67 | chr16-9166448-INS-68 |
| chr16-9170255-INS-80 | chr16-9170287-INS-58 |
| chr16-9174328-INS-6400 | chr16-9174329-INS-6400 |
| chr16-9201425-INS-779 | chr16-9201425-INS-779 |
| chr16-9214228-INS-337 | chr16-9214226-INS-340 |
| chr16-9228983-INS-6541 | chr16-9228984-INS-6545 |

| | |
|---|---|
| chr16-9231584-INS-394 | chr16-9231584-INS-394 |
| chr16-9286291-INS-54 | chr16-9286291-INS-54 |
| chr16-9297805-INS-376 | chr16-9297805-INS-376 |
| chr16-9308291-INS-69 | chr16-9308284-INS-116 |
| chr16-9335809-INS-6443 | chr16-9335810-INS-6443 |
| chr16-9342821-INS-166 | chr16-9342826-INS-178 |
| chr16-9370888-INS-189 | chr16-9370893-INS-144 |
| chr16-9372741-INS-167 | chr16-9372741-INS-171 |
| chr16-9381799-INS-68 | chr16-9381799-INS-56 |
| chr16-9447375-INS-6385 | chr16-9447376-INS-6386 |
| chr16-9452806-INS-115 | chr16-9452806-INS-113 |
| chr16-9465475-INS-53 | chr16-9465442-INS-53 |
| chr16-9492509-INS-522 | chr16-9492509-INS-523 |
| chr16-9501182-INS-93 | chr16-9501182-INS-93 |
| chr16-9526837-INS-73 | chr16-9526837-INS-72 |
| chr16-9532820-INS-117 | chr16-9532820-INS-117 |
| chr16-9535014-INS-58 | chr16-9535014-INS-58 |
| chr16-9555085-INS-79 | chr16-9555086-INS-66 |
| chr16-9561133-INS-59 | chr16-9561133-INS-59 |
| chr16-9573297-INS-71 | chr16-9573297-INS-72 |
| chr16-9594282-INS-60 | chr16-9594282-INS-62 |
| chr16-9600151-INS-54 | chr16-9600176-INS-54 |
| chr16-9622473-INS-186 | chr16-9622473-INS-187 |
| chr16-9634612-INS-110 | chr16-9634612-INS-142 |
| chr16-9660750-INS-242 | chr16-9660750-INS-243 |
| chr16-9681820-INS-55 | chr16-9681837-INS-84 |
| chr16-9707863-INS-109 | chr16-9707863-INS-109 |
| chr16-9731395-INS-62 | chr16-9731395-INS-62 |
| chr16-9740099-INS-59 | chr16-9740130-INS-51 |
| chr16-9761322-INS-439 | chr16-9761321-INS-487 |
| chr16-9762058-INS-60 | chr16-9762058-INS-60 |
| chr16-9785185-INS-176 | chr16-9785185-INS-176 |
| chr16-9798157-INS-68 | chr16-9798157-INS-68 |
| chr16-9820698-INS-98 | chr16-9820698-INS-84 |
| chr16-9848134-INS-326 | chr16-9848134-INS-326 |
| chr16-9860376-INS-308 | chr16-9860376-INS-315 |
| chr16-9864383-INS-101 | chr16-9864383-INS-101 |
| chr16-9877595-INS-112 | chr16-9877595-INS-112 |
| chr16-9884840-INS-82 | chr16-9884840-INS-82 |
| chr16-9893043-INS-52 | chr16-9893056-INS-52 |
| chr16-9895727-INS-84 | chr16-9895727-INS-84 |
| chr16-9897575-INS-6413 | chr16-9897576-INS-6413 |
| chr16-9897647-INS-238 | chr16-9897647-INS-238 |
| chr16-9901379-INS-86 | chr16-9901379-INS-62 |
| chr16-9902874-INS-184 | chr16-9902874-INS-183 |
| chr16-9905287-INS-662 | chr16-9905287-INS-662 |
| chr16-9908710-INS-53 | chr16-9908710-INS-53 |
| chr16-9916975-INS-188 | chr16-9916975-INS-195 |
| chr16-9923132-INS-60 | chr16-9923132-INS-64 |
| chr16-9937041-INS-100 | chr16-9937041-INS-100 |
| chr16-9947105-INS-176 | chr16-9947105-INS-176 |
| chr16-9954188-INS-136 | chr16-9954160-INS-124 |
| chr16-9979376-INS-6389 | chr16-9979377-INS-6389 |
| chr16-9982549-INS-68 | chr16-9982549-INS-103 |
| chr16-10035834-INS-6850 | chr16-10035835-INS-6851 |
| chr16-10037669-INS-576 | chr16-10037669-INS-571 |
| chr16-10041265-INS-699 | chr16-10041265-INS-698 |
| chr16-10074323-INS-8943 | chr16-10074324-INS-8943 |
| chr16-10074571-INS-86 | chr16-10074571-INS-86 |
| chr16-10106762-INS-90 | chr16-10106730-INS-75 |
| chr16-10127329-INS-63 | chr16-10127329-INS-62 |
| chr16-10132670-INS-275 | chr16-10132680-INS-275 |
| chr16-10151335-INS-2477 | chr16-10151336-INS-2477 |
| chr16-10164230-INS-52 | chr16-10164230-INS-52 |
| chr16-10191769-INS-68 | chr16-10191779-INS-57 |
| chr16-10195072-INS-85 | chr16-10195072-INS-77 |
| chr16-10224882-INS-154 | chr16-10224882-INS-154 |
| chr16-10233336-INS-60 | chr16-10233336-INS-76 |
| chr16-10246784-INS-178 | chr16-10246824-INS-161 |
| chr16-10251873-INS-5208 | chr16-10251874-INS-5233 |
| chr16-10253051-INS-205 | chr16-10253051-INS-202 |
| chr16-10260195-INS-90 | chr16-10260208-INS-90 |
| chr16-10266366-INS-168 | chr16-10266366-INS-168 |
| chr16-10268571-INS-68 | chr16-10268579-INS-58 |
| chr16-10271909-INS-75 | chr16-10271909-INS-75 |
| chr16-10274756-INS-70 | chr16-10274756-INS-70 |
| chr16-10283854-INS-76 | chr16-10283857-INS-72 |

| | |
|---|---|
| chr16-10285562-INS-547 | chr16-10285562-INS-547 |
| chr16-10294765-INS-819 | chr16-10294765-INS-830 |
| chr16-10310260-INS-80 | chr16-10310251-INS-85 |
| chr16-10312217-INS-179 | chr16-10312217-INS-174 |
| chr16-10382011-INS-424 | chr16-10382012-INS-424 |
| chr16-10433113-INS-208 | chr16-10433113-INS-209 |
| chr16-10448401-INS-97 | chr16-10448401-INS-103 |
| chr16-10477879-INS-59 | chr16-10477879-INS-63 |
| chr16-10500832-INS-97 | chr16-10500823-INS-94 |
| chr16-10539596-INS-234 | chr16-10539596-INS-234 |
| chr16-10555144-INS-54 | chr16-10555102-INS-62 |
| chr16-10573159-INS-87 | chr16-10573159-INS-89 |
| chr16-10579609-INS-111 | chr16-10579609-INS-111 |
| chr16-10609165-INS-56 | chr16-10609165-INS-56 |
| chr16-10611449-INS-83 | chr16-10611449-INS-83 |
| chr16-10621036-INS-54 | chr16-10621036-INS-52 |
| chr16-10640517-INS-69 | chr16-10640490-INS-81 |
| chr16-10647688-INS-66 | chr16-10647698-INS-82 |
| chr16-10665994-INS-84 | chr16-10665995-INS-84 |
| chr16-10691037-INS-71 | chr16-10691038-INS-87 |
| chr16-10698150-INS-201 | chr16-10698150-INS-199 |
| chr16-10702908-INS-103 | chr16-10702908-INS-91 |
| chr16-10705687-INS-475 | chr16-10705687-INS-475 |
| chr16-10713909-INS-246 | chr16-10713909-INS-246 |
| chr16-10735616-INS-71 | chr16-10735618-INS-74 |
| chr16-10736539-INS-72 | chr16-10736539-INS-72 |
| chr16-10744502-INS-56 | chr16-10744502-INS-63 |
| chr16-10746925-INS-3978 | chr16-10746926-INS-3978 |
| chr16-10750609-INS-1478 | chr16-10750565-INS-1476 |
| chr16-10755876-INS-124 | chr16-10755876-INS-124 |
| chr16-10795186-INS-209 | chr16-10795186-INS-209 |
| chr16-10802383-INS-122 | chr16-10802383-INS-124 |
| chr16-10804324-INS-53 | chr16-10804308-INS-53 |
| chr16-10806913-INS-144 | chr16-10806919-INS-102 |
| chr16-10833542-INS-277 | chr16-10833542-INS-277 |
| chr16-10835484-INS-86 | chr16-10835484-INS-112 |
| chr16-10853307-INS-75 | chr16-10853307-INS-74 |
| chr16-10859040-INS-135 | chr16-10859040-INS-135 |
| chr16-10880352-INS-57 | chr16-10880362-INS-91 |
| chr16-10913408-INS-76 | chr16-10913400-INS-67 |
| chr16-10920936-INS-78 | chr16-10920936-INS-66 |
| chr16-10928185-INS-216 | chr16-10928185-INS-217 |
| chr16-10931751-INS-123 | chr16-10931755-INS-75 |
| chr16-10954318-INS-907 | chr16-10954318-INS-905 |
| chr16-10984061-INS-198 | chr16-10984062-INS-198 |
| chr16-11010179-INS-192 | chr16-11010179-INS-192 |
| chr16-11017062-INS-380 | chr16-11017062-INS-380 |
| chr16-11022930-INS-477 | chr16-11022930-INS-496 |
| chr16-11069731-INS-198 | chr16-11069731-INS-207 |
| chr16-11088968-INS-1892 | chr16-11088969-INS-1892 |
| chr16-11091583-INS-92 | chr16-11091583-INS-92 |
| chr16-11096360-INS-232 | chr16-11096360-INS-232 |
| chr16-11111962-INS-388 | chr16-11111962-INS-388 |
| chr16-11118888-INS-200 | chr16-11118888-INS-201 |
| chr16-11120348-INS-56 | chr16-11120348-INS-64 |
| chr16-11121903-INS-560 | chr16-11121904-INS-560 |
| chr16-11126590-INS-164 | chr16-11126590-INS-168 |
| chr16-11150266-INS-126 | chr16-11150266-INS-112 |
| chr16-11179015-INS-1292 | chr16-11179016-INS-1244 |
| chr16-11179204-INS-429 | chr16-11179205-INS-421 |
| chr16-11184590-INS-177 | chr16-11184571-INS-197 |
| chr16-11199795-INS-714 | chr16-11199795-INS-714 |
| chr16-11212947-INS-80 | chr16-11212947-INS-56 |
| chr16-11237235-INS-1152 | chr16-11237236-INS-1152 |
| chr16-11266665-INS-80 | chr16-11266665-INS-80 |
| chr16-11298526-INS-250 | chr16-11298526-INS-249 |
| chr16-11318107-INS-1106 | chr16-11318108-INS-1106 |
| chr16-11332979-INS-162 | chr16-11332979-INS-162 |
| chr16-11334025-INS-59 | chr16-11334000-INS-60 |
| chr16-11341515-INS-115 | chr16-11341515-INS-115 |
| chr16-11362475-INS-206 | chr16-11362475-INS-207 |
| chr16-11371140-INS-147 | chr16-11371140-INS-148 |
| chr16-11379506-INS-57 | chr16-11379506-INS-57 |
| chr16-11401709-INS-643 | chr16-11401709-INS-643 |
| chr16-11415847-INS-5549 | chr16-11415848-INS-5549 |
| chr16-11422493-INS-113 | chr16-11422517-INS-131 |
| chr16-11426291-INS-63 | chr16-11426291-INS-58 |

| | |
|---|---|
| chr16-11448840-INS-269 | chr16-11448840-INS-269 |
| chr16-11463827-INS-52 | chr16-11463827-INS-52 |
| chr16-11473193-INS-451 | chr16-11473193-INS-451 |
| chr16-11488201-INS-84 | chr16-11488201-INS-84 |
| chr16-11516615-INS-63 | chr16-11516615-INS-63 |
| chr16-11523226-INS-196 | chr16-11523226-INS-196 |
| chr16-11616259-INS-65 | chr16-11616260-INS-69 |
| chr16-11636165-INS-625 | chr16-11636165-INS-624 |
| chr16-11649285-INS-377 | chr16-11649281-INS-372 |
| chr16-11699161-INS-342 | chr16-11699161-INS-342 |
| chr16-11753305-INS-110 | chr16-11753312-INS-86 |
| chr16-11875811-INS-304 | chr16-11875821-INS-304 |
| chr16-11917745-INS-55 | chr16-11917745-INS-50 |
| chr16-11961977-INS-84 | chr16-11961975-INS-84 |
| chr16-11998539-INS-85 | chr16-11998539-INS-88 |
| chr16-12094183-INS-52 | chr16-12094183-INS-52 |
| chr16-12129674-INS-129 | chr16-12129675-INS-154 |
| chr16-12139562-INS-128 | chr16-12139562-INS-128 |
| chr16-12164872-INS-64 | chr16-12164872-INS-64 |
| chr16-12188409-INS-210 | chr16-12188409-INS-210 |
| chr16-12196350-INS-69 | chr16-12196350-INS-69 |
| chr16-12197893-INS-77 | chr16-12197896-INS-89 |
| chr16-12203621-INS-853 | chr16-12203621-INS-822 |
| chr16-12205240-INS-138 | chr16-12205240-INS-138 |
| chr16-12212955-INS-59 | chr16-12212955-INS-60 |
| chr16-12232550-INS-187 | chr16-12232550-INS-180 |
| chr16-12273286-INS-52 | chr16-12273286-INS-52 |
| chr16-12288702-INS-498 | chr16-12288702-INS-498 |
| chr16-12322261-INS-51 | chr16-12322261-INS-51 |
| chr16-12328518-INS-75 | chr16-12328518-INS-79 |
| chr16-12347639-INS-66 | chr16-12347639-INS-66 |
| chr16-12348238-INS-627 | chr16-12348238-INS-625 |
| chr16-12350002-INS-222 | chr16-12350002-INS-222 |
| chr16-12356025-INS-174 | chr16-12356025-INS-174 |
| chr16-12391753-INS-829 | chr16-12391753-INS-829 |
| chr16-12395369-INS-70 | chr16-12395345-INS-79 |
| chr16-12452789-INS-3327 | chr16-12452790-INS-3328 |
| chr16-12461596-INS-78 | chr16-12461596-INS-78 |
| chr16-12472309-INS-140 | chr16-12472309-INS-112 |
| chr16-12475449-INS-76 | chr16-12475449-INS-102 |
| chr16-12477117-INS-111 | chr16-12477126-INS-120 |
| chr16-12500092-INS-81 | chr16-12500092-INS-81 |
| chr16-12521785-INS-90 | chr16-12521785-INS-90 |
| chr16-12575852-INS-1392 | chr16-12575853-INS-1406 |
| chr16-12601155-INS-56 | chr16-12601155-INS-52 |
| chr16-12609696-INS-4096 | chr16-12609697-INS-4123 |
| chr16-12623118-INS-4101 | chr16-12623119-INS-4090 |
| chr16-12641499-INS-534 | chr16-12641499-INS-534 |
| chr16-12647939-INS-3738 | chr16-12647940-INS-3738 |
| chr16-12669924-INS-164 | chr16-12669924-INS-164 |
| chr16-12675075-INS-6350 | chr16-12675076-INS-6349 |
| chr16-12681254-INS-87 | chr16-12681254-INS-91 |
| chr16-12683209-INS-65 | chr16-12683209-INS-65 |
| chr16-12716174-INS-545 | chr16-12716174-INS-545 |
| chr16-12721883-INS-68 | chr16-12721883-INS-68 |
| chr16-12723051-INS-57 | chr16-12723041-INS-57 |
| chr16-12734269-INS-268 | chr16-12734269-INS-240 |
| chr16-12761266-INS-97 | chr16-12761266-INS-114 |
| chr16-12764246-INS-55 | chr16-12764235-INS-83 |
| chr16-12779808-INS-183 | chr16-12779808-INS-183 |
| chr16-12780471-INS-1003 | chr16-12780472-INS-1013 |
| chr16-12782348-INS-120 | chr16-12782356-INS-116 |
| chr16-12823205-INS-65 | chr16-12823205-INS-66 |
| chr16-12827778-INS-95 | chr16-12827775-INS-97 |
| chr16-12844848-INS-81 | chr16-12844848-INS-82 |
| chr16-12879899-INS-94 | chr16-12879899-INS-94 |
| chr16-12940466-INS-175 | chr16-12940466-INS-175 |
| chr16-12945108-INS-68 | chr16-12945087-INS-113 |
| chr16-13003642-INS-612 | chr16-13003646-INS-615 |
| chr16-13030460-INS-219 | chr16-13030460-INS-219 |
| chr16-13062735-INS-135 | chr16-13062710-INS-137 |
| chr16-13113917-INS-7080 | chr16-13113918-INS-7080 |
| chr16-13223297-INS-97 | chr16-13223297-INS-60 |
| chr16-13239784-INS-418 | chr16-13239784-INS-419 |
| chr16-13241096-INS-1041 | chr16-13241097-INS-1074 |
| chr16-13242003-INS-872 | chr16-13242002-INS-873 |
| chr16-13247001-INS-124 | chr16-13247001-INS-124 |

| | |
|---|---|
| chr16-13253445-INS-135 | chr16-13253445-INS-122 |
| chr16-13267706-INS-71 | chr16-13267706-INS-71 |
| chr16-13286025-INS-218 | chr16-13286025-INS-219 |
| chr16-13291878-INS-56 | chr16-13291878-INS-78 |
| chr16-13316012-INS-74 | chr16-13316012-INS-50 |
| chr16-13327493-INS-677 | chr16-13327493-INS-676 |
| chr16-13329064-INS-6401 | chr16-13329065-INS-6401 |
| chr16-13341792-INS-86 | chr16-13341788-INS-96 |
| chr16-13346020-INS-70 | chr16-13346035-INS-56 |
| chr16-13348021-INS-76 | chr16-13348021-INS-76 |
| chr16-13359213-INS-129 | chr16-13359213-INS-129 |
| chr16-13392683-INS-53 | chr16-13392651-INS-68 |
| chr16-13410824-INS-541 | chr16-13410826-INS-545 |
| chr16-13422764-INS-60 | chr16-13422764-INS-80 |
| chr16-13423445-INS-225 | chr16-13423445-INS-225 |
| chr16-13434920-INS-5215 | chr16-13434921-INS-5215 |
| chr16-13451701-INS-195 | chr16-13451701-INS-195 |
| chr16-13459398-INS-183 | chr16-13459398-INS-183 |
| chr16-13459965-INS-212 | chr16-13459965-INS-212 |
| chr16-13466200-INS-780 | chr16-13466200-INS-828 |
| chr16-13473962-INS-208 | chr16-13473962-INS-208 |
| chr16-13482164-INS-203 | chr16-13482164-INS-201 |
| chr16-13491596-INS-881 | chr16-13491596-INS-882 |
| chr16-13499522-INS-160 | chr16-13499522-INS-160 |
| chr16-13508881-INS-246 | chr16-13508881-INS-234 |
| chr16-13517219-INS-218 | chr16-13517219-INS-220 |
| chr16-13540663-INS-51 | chr16-13540667-INS-93 |
| chr16-13564751-INS-9041 | chr16-13564748-INS-9045 |
| chr16-13569330-INS-105 | chr16-13569342-INS-108 |
| chr16-13578516-INS-399 | chr16-13578516-INS-399 |
| chr16-13587474-INS-232 | chr16-13587474-INS-232 |
| chr16-13597612-INS-202 | chr16-13597612-INS-208 |
| chr16-13605809-INS-91 | chr16-13605809-INS-91 |
| chr16-13609934-INS-199 | chr16-13609934-INS-206 |
| chr16-13613539-INS-102 | chr16-13613539-INS-102 |
| chr16-13615132-INS-294 | chr16-13615132-INS-294 |
| chr16-13625714-INS-89 | chr16-13625714-INS-87 |
| chr16-13648022-INS-197 | chr16-13648022-INS-197 |
| chr16-13651545-INS-128 | chr16-13651545-INS-113 |
| chr16-13653455-INS-1005 | chr16-13653456-INS-1001 |
| chr16-13663626-INS-212 | chr16-13663626-INS-213 |
| chr16-13666483-INS-206 | chr16-13666483-INS-203 |
| chr16-13694703-INS-80 | chr16-13694703-INS-100 |
| chr16-13708230-INS-201 | chr16-13708230-INS-201 |
| chr16-13710090-INS-63 | chr16-13710090-INS-82 |
| chr16-13714118-INS-1891 | chr16-13714119-INS-1891 |
| chr16-13726397-INS-53 | chr16-13726397-INS-53 |
| chr16-13734720-INS-208 | chr16-13734720-INS-205 |
| chr16-13735702-INS-154 | chr16-13735702-INS-154 |
| chr16-13778850-INS-51 | chr16-13778850-INS-53 |
| chr16-13788340-INS-203 | chr16-13788340-INS-203 |
| chr16-13790392-INS-160 | chr16-13790392-INS-158 |
| chr16-13799135-INS-8992 | chr16-13799136-INS-8992 |
| chr16-13822242-INS-85 | chr16-13822242-INS-85 |
| chr16-13832990-INS-58 | chr16-13832990-INS-58 |
| chr16-13836710-INS-63 | chr16-13836710-INS-63 |
| chr16-13855675-INS-5143 | chr16-13855676-INS-5143 |
| chr16-13907201-INS-231 | chr16-13907201-INS-233 |
| chr16-13909332-INS-385 | chr16-13909334-INS-395 |
| chr16-13910659-INS-326 | chr16-13910659-INS-326 |
| chr16-13921830-INS-2396 | chr16-13921831-INS-2396 |
| chr16-13953777-INS-218 | chr16-13953777-INS-218 |
| chr16-13992505-INS-7321 | chr16-13992506-INS-7321 |
| chr16-14015801-INS-101 | chr16-14015801-INS-138 |
| chr16-14016811-INS-1026 | chr16-14016812-INS-1022 |
| chr16-14027755-INS-1889 | chr16-14027756-INS-1889 |
| chr16-14038704-INS-330 | chr16-14038704-INS-330 |
| chr16-14040260-INS-198 | chr16-14040262-INS-183 |
| chr16-14048430-INS-229 | chr16-14048430-INS-229 |
| chr16-14057740-INS-51 | chr16-14057740-INS-51 |
| chr16-14059700-INS-52 | chr16-14059700-INS-52 |
| chr16-14061163-INS-55 | chr16-14061163-INS-55 |
| chr16-14074524-INS-62 | chr16-14074524-INS-62 |
| chr16-14083965-INS-64 | chr16-14083965-INS-64 |
| chr16-14084545-INS-96 | chr16-14084549-INS-71 |
| chr16-14106838-INS-80 | chr16-14106838-INS-80 |
| chr16-14126674-INS-210 | chr16-14126674-INS-211 |

| | |
|---|---|
| chr16-14173164-INS-183 | chr16-14173164-INS-181 |
| chr16-14174167-INS-425 | chr16-14174167-INS-438 |
| chr16-14205732-INS-51 | chr16-14205732-INS-54 |
| chr16-14219962-INS-60 | chr16-14219962-INS-56 |
| chr16-14253080-INS-6442 | chr16-14253081-INS-6442 |
| chr16-14298569-INS-52 | chr16-14298569-INS-52 |
| chr16-14334229-INS-56 | chr16-14334229-INS-50 |
| chr16-14335030-INS-89 | chr16-14335030-INS-88 |
| chr16-14364965-INS-221 | chr16-14364965-INS-213 |
| chr16-14463623-INS-160 | chr16-14463623-INS-160 |
| chr16-14464494-INS-800 | chr16-14464495-INS-800 |
| chr16-14470963-INS-298 | chr16-14470963-INS-295 |
| chr16-14536143-INS-78 | chr16-14536143-INS-52 |
| chr16-14562299-INS-124 | chr16-14562299-INS-126 |
| chr16-14588509-INS-360 | chr16-14588509-INS-360 |
| chr16-14596486-INS-6535 | chr16-14596487-INS-6535 |
| chr16-14599002-INS-56 | chr16-14599002-INS-52 |
| chr16-14648159-INS-61 | chr16-14648159-INS-71 |
| chr16-14651084-INS-55 | chr16-14651089-INS-55 |
| chr16-14659267-INS-96 | chr16-14659267-INS-88 |
| chr16-14671958-INS-223 | chr16-14671947-INS-231 |
| chr16-14719912-INS-165 | chr16-14719901-INS-165 |
| chr16-14795625-INS-569 | chr16-14795625-INS-569 |
| chr16-14813974-INS-443 | chr16-14813974-INS-443 |
| chr16-14818144-INS-630 | chr16-14818144-INS-631 |
| chr16-14827584-INS-375 | chr16-14827584-INS-377 |
| chr16-14833591-INS-6237 | chr16-14833592-INS-6237 |
| chr16-14842576-INS-66 | chr16-14842576-INS-66 |
| chr16-14897844-INS-2149 | chr16-14897845-INS-2145 |
| chr16-14907589-INS-55 | chr16-14907589-INS-55 |
| chr16-14919604-INS-54 | chr16-14919604-INS-50 |
| chr16-14928869-INS-4802 | chr16-14928870-INS-4802 |
| chr16-14991187-INS-91 | chr16-14991187-INS-91 |
| chr16-15008788-INS-834 | chr16-15008788-INS-884 |
| chr16-15009708-INS-440 | chr16-15009708-INS-440 |
| chr16-15018947-INS-73 | chr16-15018947-INS-73 |
| chr16-15023017-INS-3130 | chr16-15023018-INS-3130 |
| chr16-15029502-INS-766 | chr16-15029502-INS-773 |
| chr16-15142352-INS-54 | chr16-15142352-INS-57 |
| chr16-15171522-INS-245 | chr16-15171522-INS-229 |
| chr16-15193625-INS-1410 | chr16-15193626-INS-1409 |
| chr16-15222217-INS-208 | chr16-15222217-INS-208 |
| chr16-15288817-INS-605 | chr16-15288817-INS-605 |
| chr16-15313557-INS-225 | chr16-15313557-INS-225 |
| chr16-15331681-INS-1397 | chr16-15331682-INS-1369 |
| chr16-15338575-INS-192 | chr16-15338575-INS-192 |
| chr16-15339544-INS-59 | chr16-15339544-INS-61 |
| chr16-15360768-INS-243 | chr16-15360768-INS-238 |
| chr16-15378745-INS-448 | chr16-15378745-INS-449 |
| chr16-15405204-INS-250 | chr16-15405207-INS-259 |
| chr16-15408948-INS-55 | chr16-15408948-INS-61 |
| chr16-15431014-INS-255 | chr16-15431040-INS-236 |
| chr16-15437336-INS-132 | chr16-15437336-INS-134 |
| chr16-15453386-INS-202 | chr16-15453386-INS-202 |
| chr16-15475616-INS-102 | chr16-15475616-INS-142 |
| chr16-15478367-INS-77 | chr16-15478367-INS-61 |
| chr16-15488027-INS-407 | chr16-15488027-INS-407 |
| chr16-15495513-INS-754 | chr16-15495513-INS-754 |
| chr16-15512836-INS-762 | chr16-15512835-INS-762 |
| chr16-15536113-INS-61 | chr16-15536113-INS-61 |
| chr16-15536760-INS-210 | chr16-15536760-INS-211 |
| chr16-15544147-INS-232 | chr16-15544147-INS-232 |
| chr16-15548545-INS-225 | chr16-15548545-INS-225 |
| chr16-15555010-INS-298 | chr16-15555003-INS-298 |
| chr16-15558713-INS-212 | chr16-15558713-INS-212 |
| chr16-15571421-INS-64 | chr16-15571421-INS-64 |
| chr16-15680818-INS-54 | chr16-15680818-INS-54 |
| chr16-15684182-INS-132 | chr16-15684182-INS-136 |
| chr16-15726706-INS-62 | chr16-15726706-INS-62 |
| chr16-15779829-INS-55 | chr16-15779829-INS-55 |
| chr16-15783272-INS-119 | chr16-15783272-INS-119 |
| chr16-15812385-INS-194 | chr16-15812385-INS-194 |
| chr16-15856512-INS-214 | chr16-15856512-INS-214 |
| chr16-15861805-INS-52 | chr16-15861806-INS-60 |
| chr16-15865450-INS-110 | chr16-15865450-INS-79 |
| chr16-15878727-INS-219 | chr16-15878727-INS-224 |
| chr16-15879380-INS-400 | chr16-15879380-INS-400 |

| | |
|---|---|
| chr16-15895203-INS-168 | chr16-15895203-INS-169 |
| chr16-15917179-INS-6192 | chr16-15917180-INS-6192 |
| chr16-15922187-INS-648 | chr16-15922187-INS-605 |
| chr16-15923647-INS-743 | chr16-15923647-INS-743 |
| chr16-15933630-INS-58 | chr16-15933630-INS-54 |
| chr16-15964695-INS-74 | chr16-15964695-INS-74 |
| chr16-15966786-INS-401 | chr16-15966786-INS-401 |
| chr16-15973040-INS-3088 | chr16-15973041-INS-3088 |
| chr16-15990272-INS-199 | chr16-15990272-INS-188 |
| chr16-16015797-INS-63 | chr16-16015797-INS-63 |
| chr16-16036865-INS-52 | chr16-16036865-INS-52 |
| chr16-16054274-INS-212 | chr16-16054274-INS-213 |
| chr16-16066686-INS-106 | chr16-16066686-INS-106 |
| chr16-16094279-INS-397 | chr16-16094285-INS-398 |
| chr16-16120375-INS-163 | chr16-16120375-INS-173 |
| chr16-16128429-INS-5341 | chr16-16128430-INS-5341 |
| chr16-16158580-INS-213 | chr16-16158580-INS-210 |
| chr16-16178372-INS-238 | chr16-16178372-INS-234 |
| chr16-16180127-INS-251 | chr16-16180127-INS-251 |
| chr16-16195595-INS-86 | chr16-16195594-INS-77 |
| chr16-16209592-INS-228 | chr16-16209592-INS-228 |
| chr16-16227840-INS-356 | chr16-16227840-INS-356 |
| chr16-16256899-INS-59 | chr16-16256899-INS-63 |
| chr16-16261841-INS-90 | chr16-16261865-INS-90 |
| chr16-16270172-INS-593 | chr16-16270172-INS-608 |
| chr16-16298494-INS-2297 | chr16-16298495-INS-2289 |
| chr16-16313837-INS-89 | chr16-16313837-INS-66 |
| chr16-16314997-INS-64 | chr16-16314997-INS-52 |
| chr16-16317403-INS-288 | chr16-16317403-INS-282 |
| chr16-16338981-INS-787 | chr16-16338981-INS-784 |
| chr16-16390674-INS-6965 | chr16-16390675-INS-6965 |
| chr16-16393100-INS-120 | chr16-16393100-INS-105 |
| chr16-16401127-INS-76 | chr16-16401127-INS-76 |
| chr16-16417577-INS-59 | chr16-16417577-INS-59 |
| chr16-16424864-INS-193 | chr16-16424856-INS-203 |
| chr16-16427266-INS-215 | chr16-16427266-INS-215 |
| chr16-16449932-INS-2262 | chr16-16449933-INS-2261 |
| chr16-16459394-INS-64 | chr16-16459394-INS-66 |
| chr16-16465982-INS-6592 | chr16-16465983-INS-6592 |
| chr16-16504826-INS-200 | chr16-16504826-INS-200 |
| chr16-16538865-INS-211 | chr16-16538865-INS-211 |
| chr16-16545168-INS-667 | chr16-16545168-INS-667 |
| chr16-16549392-INS-4181 | chr16-16549393-INS-4184 |
| chr16-16551850-INS-565 | chr16-16551850-INS-565 |
| chr16-16565841-INS-282 | chr16-16565841-INS-285 |
| chr16-16584262-INS-61 | chr16-16584262-INS-61 |
| chr16-16599439-INS-78 | chr16-16599439-INS-78 |
| chr16-16635984-INS-154 | chr16-16635973-INS-154 |
| chr16-16646245-INS-719 | chr16-16646239-INS-711 |
| chr16-16646403-INS-60 | chr16-16646420-INS-79 |
| chr16-16654277-INS-462 | chr16-16654277-INS-462 |
| chr16-16666814-INS-51 | chr16-16666810-INS-51 |
| chr16-16726495-INS-206 | chr16-16726495-INS-194 |
| chr16-16764683-INS-70 | chr16-16764683-INS-70 |
| chr16-16767597-INS-123 | chr16-16767595-INS-94 |
| chr16-16770335-INS-550 | chr16-16770335-INS-554 |
| chr16-16775905-INS-397 | chr16-16775905-INS-397 |
| chr16-16776441-INS-65 | chr16-16776436-INS-68 |
| chr16-16782268-INS-545 | chr16-16782268-INS-545 |
| chr16-16793851-INS-1212 | chr16-16793852-INS-1214 |
| chr16-16837714-INS-181 | chr16-16837714-INS-188 |
| chr16-16924719-INS-165 | chr16-16924713-INS-200 |
| chr16-16933345-INS-226 | chr16-16933345-INS-226 |
| chr16-16978890-INS-211 | chr16-16978890-INS-211 |
| chr16-16979922-INS-58 | chr16-16979922-INS-58 |
| chr16-17003536-INS-78 | chr16-17003536-INS-78 |
| chr16-17007182-INS-185 | chr16-17007182-INS-186 |
| chr16-17015895-INS-54 | chr16-17015895-INS-54 |
| chr16-17023391-INS-64 | chr16-17023388-INS-69 |
| chr16-17025086-INS-254 | chr16-17025086-INS-267 |
| chr16-17034414-INS-54 | chr16-17034414-INS-54 |
| chr16-17068463-INS-200 | chr16-17068463-INS-200 |
| chr16-17111222-INS-210 | chr16-17111222-INS-210 |
| chr16-17114587-INS-444 | chr16-17114587-INS-444 |
| chr16-17144901-INS-67 | chr16-17144901-INS-62 |
| chr16-17145724-INS-589 | chr16-17145724-INS-597 |
| chr16-17159876-INS-203 | chr16-17159876-INS-208 |

| | |
|---|---|
| chr16-17188065-INS-215 | chr16-17188065-INS-215 |
| chr16-17243976-INS-211 | chr16-17243976-INS-213 |
| chr16-17251037-INS-56 | chr16-17251036-INS-80 |
| chr16-17252310-INS-206 | chr16-17252310-INS-206 |
| chr16-17272230-INS-176 | chr16-17272230-INS-176 |
| chr16-17288575-INS-52 | chr16-17288575-INS-52 |
| chr16-17296858-INS-200 | chr16-17296858-INS-200 |
| chr16-17322407-INS-60 | chr16-17322407-INS-108 |
| chr16-17374595-INS-437 | chr16-17374595-INS-416 |
| chr16-17378184-INS-64 | chr16-17378180-INS-54 |
| chr16-17382006-INS-3119 | chr16-17382007-INS-3119 |
| chr16-17382406-INS-126 | chr16-17382396-INS-150 |
| chr16-17409558-INS-2589 | chr16-17409559-INS-2591 |
| chr16-17418051-INS-434 | chr16-17418051-INS-434 |
| chr16-17420417-INS-932 | chr16-17420417-INS-932 |
| chr16-17432766-INS-73 | chr16-17432767-INS-70 |
| chr16-17467467-INS-68 | chr16-17467467-INS-68 |
| chr16-17495737-INS-112 | chr16-17495737-INS-122 |
| chr16-17506561-INS-68 | chr16-17506561-INS-68 |
| chr16-17519324-INS-69 | chr16-17519336-INS-92 |
| chr16-17530052-INS-104 | chr16-17530052-INS-68 |
| chr16-17546312-INS-7189 | chr16-17546294-INS-7208 |
| chr16-17553008-INS-342 | chr16-17553008-INS-342 |
| chr16-17599969-INS-80 | chr16-17599969-INS-54 |
| chr16-17618373-INS-125 | chr16-17618345-INS-123 |
| chr16-17696570-INS-104 | chr16-17696570-INS-104 |
| chr16-17706865-INS-189 | chr16-17706865-INS-189 |
| chr16-17723007-INS-102 | chr16-17723007-INS-102 |
| chr16-17760296-INS-205 | chr16-17760296-INS-205 |
| chr16-17815377-INS-201 | chr16-17815377-INS-203 |
| chr16-17821689-INS-67 | chr16-17821689-INS-67 |
| chr16-17826495-INS-204 | chr16-17826496-INS-217 |
| chr16-17845129-INS-204 | chr16-17845129-INS-212 |
| chr16-17860815-INS-206 | chr16-17860815-INS-210 |
| chr16-17863864-INS-96 | chr16-17863864-INS-60 |
| chr16-17868300-INS-249 | chr16-17868300-INS-244 |
| chr16-17893515-INS-57 | chr16-17893515-INS-61 |
| chr16-17909821-INS-93 | chr16-17909821-INS-94 |
| chr16-17911944-INS-194 | chr16-17911944-INS-194 |
| chr16-17930639-INS-102 | chr16-17930626-INS-84 |
| chr16-17946281-INS-76 | chr16-17946281-INS-78 |
| chr16-17961549-INS-189 | chr16-17961549-INS-199 |
| chr16-18023182-INS-341 | chr16-18023182-INS-341 |
| chr16-18028206-INS-269 | chr16-18028161-INS-270 |
| chr16-18075925-INS-80 | chr16-18075925-INS-80 |
| chr16-18090235-INS-206 | chr16-18090227-INS-218 |
| chr16-18115674-INS-392 | chr16-18115674-INS-398 |
| chr16-18133778-INS-231 | chr16-18133778-INS-232 |
| chr16-18145679-INS-166 | chr16-18145679-INS-173 |
| chr16-18149273-INS-314 | chr16-18149273-INS-314 |
| chr16-18150410-INS-643 | chr16-18150361-INS-652 |
| chr16-18170203-INS-430 | chr16-18170203-INS-430 |
| chr16-18172931-INS-214 | chr16-18172931-INS-216 |
| chr16-18253128-INS-213 | chr16-18253128-INS-213 |
| chr16-18259200-INS-228 | chr16-18259200-INS-232 |
| chr16-18303613-INS-58 | chr16-18303613-INS-58 |
| chr16-18310933-INS-241 | chr16-18310934-INS-242 |
| chr16-18313352-INS-972 | chr16-18313352-INS-974 |
| chr16-18315434-INS-211 | chr16-18315434-INS-211 |
| chr16-18366365-INS-66 | chr16-18366365-INS-66 |
| chr16-18386669-INS-58 | chr16-18386663-INS-58 |
| chr16-18454427-INS-84 | chr16-18454427-INS-84 |
| chr16-18468629-INS-114 | chr16-18468629-INS-114 |
| chr16-18476766-INS-205 | chr16-18476766-INS-206 |
| chr16-18479747-INS-204 | chr16-18479747-INS-204 |
| chr16-18501737-INS-191 | chr16-18501737-INS-195 |
| chr16-18503245-INS-213 | chr16-18503245-INS-213 |
| chr16-18508558-INS-62 | chr16-18508558-INS-64 |
| chr16-18513563-INS-453 | chr16-18513563-INS-427 |
| chr16-18521704-INS-100 | chr16-18521718-INS-92 |
| chr16-18563324-INS-6440 | chr16-18563325-INS-6440 |
| chr16-18564967-INS-898 | chr16-18564967-INS-919 |
| chr16-18569304-INS-82 | chr16-18569319-INS-78 |
| chr16-18588038-INS-254 | chr16-18588038-INS-254 |
| chr16-18616773-INS-209 | chr16-18616773-INS-209 |
| chr16-18674595-INS-99 | chr16-18674555-INS-105 |
| chr16-18689044-INS-77 | chr16-18689036-INS-54 |

| | |
|---|---|
| chr16-18692016-INS-96 | chr16-18692016-INS-96 |
| chr16-18711484-INS-54 | chr16-18711484-INS-56 |
| chr16-18725412-INS-229 | chr16-18725412-INS-225 |
| chr16-18780947-INS-93 | chr16-18780947-INS-93 |
| chr16-18783478-INS-195 | chr16-18783478-INS-197 |
| chr16-18791626-INS-3683 | chr16-18791627-INS-3683 |
| chr16-18902227-INS-195 | chr16-18902227-INS-195 |
| chr16-18922932-INS-415 | chr16-18922932-INS-415 |
| chr16-19009604-INS-72 | chr16-19009604-INS-62 |
| chr16-19077505-INS-105 | chr16-19077505-INS-105 |
| chr16-19100447-INS-2546 | chr16-19100448-INS-2546 |
| chr16-19125808-INS-401 | chr16-19125808-INS-401 |
| chr16-19211730-INS-178 | chr16-19211730-INS-166 |
| chr16-19265038-INS-575 | chr16-19265038-INS-575 |
| chr16-19283408-INS-774 | chr16-19283408-INS-772 |
| chr16-19324489-INS-60 | chr16-19324490-INS-86 |
| chr16-19328704-INS-473 | chr16-19328704-INS-473 |
| chr16-19379009-INS-402 | chr16-19379009-INS-402 |
| chr16-19399675-INS-398 | chr16-19399675-INS-398 |
| chr16-19500632-INS-198 | chr16-19500599-INS-198 |
| chr16-19669911-INS-6253 | chr16-19669912-INS-6253 |
| chr16-19673880-INS-981 | chr16-19673880-INS-981 |
| chr16-19680967-INS-221 | chr16-19680967-INS-221 |
| chr16-19688802-INS-56 | chr16-19688778-INS-53 |
| chr16-19713698-INS-686 | chr16-19713698-INS-689 |
| chr16-19724670-INS-243 | chr16-19724653-INS-239 |
| chr16-19744396-INS-680 | chr16-19744396-INS-680 |
| chr16-19766370-INS-443 | chr16-19766370-INS-443 |
| chr16-19779767-INS-64 | chr16-19779767-INS-64 |
| chr16-19817180-INS-494 | chr16-19817180-INS-494 |
| chr16-19820466-INS-116 | chr16-19820514-INS-68 |
| chr16-19828833-INS-104 | chr16-19828833-INS-67 |
| chr16-19833412-INS-100 | chr16-19833412-INS-85 |
| chr16-19852273-INS-400 | chr16-19852273-INS-400 |
| chr16-19868190-INS-325 | chr16-19868190-INS-325 |
| chr16-19870719-INS-339 | chr16-19870719-INS-338 |
| chr16-19877243-INS-6434 | chr16-19877244-INS-6435 |
| chr16-19899894-INS-214 | chr16-19899894-INS-214 |
| chr16-19915331-INS-120 | chr16-19915331-INS-120 |
| chr16-19919753-INS-197 | chr16-19919753-INS-196 |
| chr16-19949038-INS-70 | chr16-19949032-INS-69 |
| chr16-19950369-INS-200 | chr16-19950361-INS-201 |
| chr16-19952379-INS-3861 | chr16-19952380-INS-3861 |
| chr16-19996040-INS-74 | chr16-19996040-INS-74 |
| chr16-19998667-INS-172 | chr16-19998667-INS-168 |
| chr16-20015383-INS-122 | chr16-20015383-INS-122 |
| chr16-20081789-INS-185 | chr16-20081790-INS-187 |
| chr16-20084462-INS-205 | chr16-20084462-INS-205 |
| chr16-20093617-INS-137 | chr16-20093617-INS-137 |
| chr16-20105916-INS-214 | chr16-20105916-INS-210 |
| chr16-20133590-INS-126 | chr16-20133590-INS-136 |
| chr16-20148706-INS-264 | chr16-20148706-INS-266 |
| chr16-20182623-INS-161 | chr16-20182623-INS-161 |
| chr16-20183658-INS-60 | chr16-20183659-INS-57 |
| chr16-20223951-INS-54 | chr16-20223951-INS-62 |
| chr16-20254350-INS-108 | chr16-20254350-INS-106 |
| chr16-20262383-INS-984 | chr16-20262383-INS-984 |
| chr16-20270358-INS-646 | chr16-20270358-INS-646 |
| chr16-20271127-INS-70 | chr16-20271127-INS-70 |
| chr16-20273854-INS-62 | chr16-20273854-INS-62 |
| chr16-20284279-INS-202 | chr16-20284279-INS-203 |
| chr16-20286413-INS-203 | chr16-20286413-INS-203 |
| chr16-20294855-INS-186 | chr16-20294855-INS-192 |
| chr16-20331683-INS-197 | chr16-20331683-INS-198 |
| chr16-20401636-INS-179 | chr16-20401636-INS-203 |
| chr16-20402784-INS-415 | chr16-20402784-INS-415 |
| chr16-20436227-INS-55 | chr16-20436227-INS-90 |
| chr16-20475222-INS-214 | chr16-20475222-INS-214 |
| chr16-20486490-INS-79 | chr16-20486479-INS-72 |
| chr16-20503648-INS-200 | chr16-20503648-INS-200 |
| chr16-20519553-INS-239 | chr16-20519553-INS-240 |
| chr16-20567933-INS-566 | chr16-20567945-INS-571 |
| chr16-20590408-INS-194 | chr16-20590408-INS-194 |
| chr16-20605312-INS-5266 | chr16-20605313-INS-5266 |
| chr16-20615813-INS-317 | chr16-20615813-INS-327 |
| chr16-20666307-INS-290 | chr16-20666307-INS-270 |
| chr16-20702839-INS-205 | chr16-20702839-INS-202 |

| | |
|---|---|
| chr16-20712522-INS-310 | chr16-20712522-INS-310 |
| chr16-20714422-INS-280 | chr16-20714422-INS-280 |
| chr16-20791787-INS-514 | chr16-20791788-INS-514 |
| chr16-20822588-INS-238 | chr16-20822588-INS-238 |
| chr16-20845732-INS-2743 | chr16-20845733-INS-2743 |
| chr16-20860358-INS-214 | chr16-20860354-INS-217 |
| chr16-20865586-INS-92 | chr16-20865586-INS-92 |
| chr16-20890325-INS-55 | chr16-20890325-INS-55 |
| chr16-20891421-INS-153 | chr16-20891421-INS-153 |
| chr16-20896638-INS-231 | chr16-20896638-INS-231 |
| chr16-20899224-INS-924 | chr16-20899224-INS-924 |
| chr16-20900864-INS-71 | chr16-20900867-INS-101 |
| chr16-20902089-INS-221 | chr16-20902089-INS-222 |
| chr16-20916132-INS-60 | chr16-20916132-INS-60 |
| chr16-20941628-INS-425 | chr16-20941628-INS-425 |
| chr16-20944935-INS-388 | chr16-20944935-INS-388 |
| chr16-20958854-INS-8969 | chr16-20958855-INS-8969 |
| chr16-20976935-INS-475 | chr16-20976935-INS-475 |
| chr16-20982829-INS-108 | chr16-20982829-INS-84 |
| chr16-20988428-INS-475 | chr16-20988428-INS-475 |
| chr16-21010891-INS-170 | chr16-21010891-INS-179 |
| chr16-21031036-INS-151 | chr16-21031036-INS-148 |
| chr16-21099999-INS-97 | chr16-21099962-INS-97 |
| chr16-21117694-INS-187 | chr16-21117694-INS-186 |
| chr16-21130586-INS-92 | chr16-21130586-INS-92 |
| chr16-21132874-INS-313 | chr16-21132878-INS-306 |
| chr16-21134135-INS-162 | chr16-21134135-INS-173 |
| chr16-21135755-INS-110 | chr16-21135771-INS-140 |
| chr16-21160687-INS-58 | chr16-21160687-INS-58 |
| chr16-21228119-INS-324 | chr16-21228120-INS-369 |
| chr16-21263192-INS-100 | chr16-21263192-INS-100 |
| chr16-21264860-INS-140 | chr16-21264860-INS-140 |
| chr16-21274079-INS-214 | chr16-21274079-INS-214 |
| chr16-21283952-INS-94 | chr16-21283966-INS-54 |
| chr16-21289107-INS-57 | chr16-21289086-INS-59 |
| chr16-21305679-INS-59 | chr16-21305679-INS-59 |
| chr16-21312131-INS-78 | chr16-21312131-INS-84 |
| chr16-21327460-INS-893 | chr16-21327460-INS-893 |
| chr16-21357761-INS-72 | chr16-21357761-INS-64 |
| chr16-21367118-INS-61 | chr16-21367100-INS-79 |
| chr16-21368875-INS-244 | chr16-21368875-INS-245 |
| chr16-21401699-INS-1260 | chr16-21401700-INS-1260 |
| chr16-21429257-INS-120 | chr16-21429257-INS-156 |
| chr16-21430991-INS-53 | chr16-21430991-INS-53 |
| chr16-21440637-INS-175 | chr16-21440637-INS-179 |
| chr16-21442195-INS-54 | chr16-21442195-INS-52 |
| chr16-21449426-INS-144 | chr16-21449426-INS-135 |
| chr16-21510119-INS-7084 | chr16-21510120-INS-7084 |
| chr16-21528334-INS-638 | chr16-21528334-INS-638 |
| chr16-21535197-INS-514 | chr16-21535197-INS-514 |
| chr16-21547696-INS-107 | chr16-21547696-INS-111 |
| chr16-21558318-INS-85 | chr16-21558290-INS-90 |
| chr16-21602732-INS-98 | chr16-21602733-INS-98 |
| chr16-21604836-INS-207 | chr16-21604836-INS-207 |
| chr16-21616116-INS-176 | chr16-21616141-INS-178 |
| chr16-21631043-INS-1764 | chr16-21631044-INS-1764 |
| chr16-21660523-INS-758 | chr16-21660523-INS-758 |
| chr16-21669992-INS-75 | chr16-21669992-INS-75 |
| chr16-21698811-INS-176 | chr16-21698846-INS-170 |
| chr16-21724452-INS-601 | chr16-21724452-INS-602 |
| chr16-21750227-INS-199 | chr16-21750227-INS-199 |
| chr16-21767761-INS-312 | chr16-21767763-INS-313 |
| chr16-21769440-INS-85 | chr16-21769440-INS-85 |
| chr16-21785353-INS-69 | chr16-21785326-INS-70 |
| chr16-21788790-INS-316 | chr16-21788790-INS-318 |
| chr16-21789867-INS-618 | chr16-21789883-INS-599 |
| chr16-21804906-INS-51 | chr16-21804906-INS-51 |
| chr16-21838758-INS-203 | chr16-21838758-INS-203 |
| chr16-21845713-INS-70 | chr16-21845702-INS-52 |
| chr16-21846824-INS-212 | chr16-21846824-INS-212 |
| chr16-21870919-INS-193 | chr16-21870933-INS-181 |
| chr16-21873592-INS-61 | chr16-21873592-INS-63 |
| chr16-21901551-INS-204 | chr16-21901551-INS-204 |
| chr16-21906493-INS-249 | chr16-21906490-INS-255 |
| chr16-21923024-INS-213 | chr16-21923024-INS-215 |
| chr16-21942910-INS-157 | chr16-21942910-INS-146 |
| chr16-21951309-INS-222 | chr16-21951309-INS-222 |

| | |
|---|---|
| chr16-21953834-INS-80 | chr16-21953834-INS-84 |
| chr16-22009766-INS-321 | chr16-22009766-INS-321 |
| chr16-22024186-INS-51 | chr16-22024186-INS-59 |
| chr16-22059096-INS-195 | chr16-22059096-INS-195 |
| chr16-22062216-INS-221 | chr16-22062216-INS-186 |
| chr16-22090808-INS-196 | chr16-22090808-INS-196 |
| chr16-22100244-INS-5009 | chr16-22100245-INS-5010 |
| chr16-22108127-INS-330 | chr16-22108127-INS-330 |
| chr16-22109278-INS-206 | chr16-22109278-INS-204 |
| chr16-22118848-INS-634 | chr16-22118848-INS-634 |
| chr16-22120604-INS-206 | chr16-22120604-INS-206 |
| chr16-22121899-INS-201 | chr16-22121899-INS-196 |
| chr16-22125327-INS-327 | chr16-22125327-INS-327 |
| chr16-22142545-INS-206 | chr16-22142545-INS-204 |
| chr16-22178380-INS-181 | chr16-22178380-INS-181 |
| chr16-22182184-INS-211 | chr16-22182184-INS-211 |
| chr16-22187794-INS-64 | chr16-22187792-INS-65 |
| chr16-22197017-INS-147 | chr16-22197017-INS-154 |
| chr16-22243647-INS-66 | chr16-22243647-INS-66 |
| chr16-22246542-INS-72 | chr16-22246542-INS-70 |
| chr16-22249995-INS-201 | chr16-22249995-INS-200 |
| chr16-22281203-INS-142 | chr16-22281201-INS-146 |
| chr16-22289583-INS-629 | chr16-22289574-INS-671 |
| chr16-22291806-INS-197 | chr16-22291806-INS-198 |
| chr16-22292487-INS-197 | chr16-22292487-INS-197 |
| chr16-22301112-INS-220 | chr16-22301112-INS-220 |
| chr16-22317372-INS-971 | chr16-22317372-INS-971 |
| chr16-22331036-INS-216 | chr16-22331036-INS-216 |
| chr16-22331957-INS-102 | chr16-22331955-INS-85 |
| chr16-22363719-INS-124 | chr16-22363719-INS-126 |
| chr16-22389655-INS-213 | chr16-22389655-INS-213 |
| chr16-22391121-INS-66 | chr16-22391121-INS-66 |
| chr16-22392146-INS-51 | chr16-22392146-INS-54 |
| chr16-22402533-INS-125 | chr16-22402535-INS-129 |
| chr16-22429253-INS-400 | chr16-22429253-INS-400 |
| chr16-22431093-INS-95 | chr16-22431093-INS-99 |
| chr16-22434502-INS-214 | chr16-22434502-INS-214 |
| chr16-22462055-INS-7323 | chr16-22462056-INS-7323 |
| chr16-22478632-INS-64 | chr16-22478632-INS-64 |
| chr16-22493426-INS-126 | chr16-22493424-INS-127 |
| chr16-22497925-INS-136 | chr16-22497925-INS-135 |
| chr16-22500022-INS-65 | chr16-22500022-INS-65 |
| chr16-22514477-INS-3882 | chr16-22514478-INS-3882 |
| chr16-22515808-INS-7251 | chr16-22515809-INS-7251 |
| chr16-22527463-INS-126 | chr16-22527468-INS-91 |
| chr16-22555060-INS-66 | chr16-22555060-INS-66 |
| chr16-22556172-INS-392 | chr16-22556172-INS-397 |
| chr16-22580971-INS-109 | chr16-22580966-INS-124 |
| chr16-22594317-INS-208 | chr16-22594317-INS-209 |
| chr16-22604410-INS-137 | chr16-22604408-INS-149 |
| chr16-22632836-INS-166 | chr16-22632846-INS-136 |
| chr16-22648861-INS-639 | chr16-22648861-INS-639 |
| chr16-22653708-INS-74 | chr16-22653708-INS-56 |
| chr16-22698774-INS-206 | chr16-22698774-INS-209 |
| chr16-22703000-INS-209 | chr16-22703000-INS-209 |
| chr16-22703195-INS-620 | chr16-22703195-INS-605 |
| chr16-22703879-INS-51 | chr16-22703879-INS-51 |
| chr16-22707905-INS-52 | chr16-22707905-INS-52 |
| chr16-22727597-INS-216 | chr16-22727597-INS-216 |
| chr16-22728581-INS-363 | chr16-22728581-INS-363 |
| chr16-22756500-INS-116 | chr16-22756500-INS-116 |
| chr16-22763332-INS-6202 | chr16-22763333-INS-6203 |
| chr16-22769598-INS-124 | chr16-22769598-INS-133 |
| chr16-22783280-INS-722 | chr16-22783280-INS-748 |
| chr16-22793177-INS-151 | chr16-22793178-INS-151 |
| chr16-22805850-INS-163 | chr16-22805850-INS-134 |
| chr16-22838185-INS-549 | chr16-22838185-INS-549 |
| chr16-22855600-INS-5027 | chr16-22855601-INS-5027 |
| chr16-22869997-INS-202 | chr16-22869997-INS-202 |
| chr16-22882140-INS-72 | chr16-22882140-INS-90 |
| chr16-22887325-INS-207 | chr16-22887325-INS-210 |
| chr16-22949166-INS-131 | chr16-22949166-INS-131 |
| chr16-22967692-INS-724 | chr16-22967692-INS-724 |
| chr16-22992521-INS-63 | chr16-22992521-INS-63 |
| chr16-22994577-INS-203 | chr16-22994577-INS-203 |
| chr16-22996242-INS-129 | chr16-22996242-INS-80 |
| chr16-23009702-INS-120 | chr16-23009702-INS-125 |

| | |
|---|---|
| chr16-23051038-INS-95 | chr16-23051034-INS-90 |
| chr16-23070291-INS-54 | chr16-23070291-INS-52 |
| chr16-23088375-INS-86 | chr16-23088375-INS-86 |
| chr16-23116791-INS-198 | chr16-23116791-INS-201 |
| chr16-23135326-INS-1216 | chr16-23135327-INS-1216 |
| chr16-23160768-INS-96 | chr16-23160743-INS-120 |
| chr16-23161987-INS-201 | chr16-23161987-INS-202 |
| chr16-23164219-INS-60 | chr16-23164219-INS-66 |
| chr16-23209738-INS-137 | chr16-23209717-INS-129 |
| chr16-23212945-INS-62 | chr16-23212945-INS-65 |
| chr16-23225786-INS-140 | chr16-23225787-INS-140 |
| chr16-23234205-INS-1699 | chr16-23234213-INS-1693 |
| chr16-23277372-INS-84 | chr16-23277372-INS-64 |
| chr16-23361590-INS-52 | chr16-23361590-INS-52 |
| chr16-23398339-INS-159 | chr16-23398339-INS-158 |
| chr16-23407125-INS-734 | chr16-23407125-INS-736 |
| chr16-23429236-INS-73 | chr16-23429208-INS-74 |
| chr16-23476777-INS-974 | chr16-23476777-INS-975 |
| chr16-23501204-INS-743 | chr16-23501204-INS-743 |
| chr16-23523017-INS-70 | chr16-23523017-INS-72 |
| chr16-23535918-INS-4497 | chr16-23535919-INS-4497 |
| chr16-23545235-INS-1311 | chr16-23545236-INS-1310 |
| chr16-23546248-INS-55 | chr16-23546248-INS-55 |
| chr16-23559584-INS-493 | chr16-23559579-INS-491 |
| chr16-23561585-INS-62 | chr16-23561585-INS-77 |
| chr16-23565764-INS-6671 | chr16-23565765-INS-6656 |
| chr16-23577996-INS-156 | chr16-23577996-INS-156 |
| chr16-23600106-INS-399 | chr16-23600106-INS-399 |
| chr16-23617121-INS-628 | chr16-23617121-INS-628 |
| chr16-23635701-INS-75 | chr16-23635701-INS-75 |
| chr16-23680046-INS-63 | chr16-23680047-INS-54 |
| chr16-23714081-INS-539 | chr16-23714081-INS-539 |
| chr16-23719744-INS-6714 | chr16-23719745-INS-6714 |
| chr16-23721390-INS-896 | chr16-23721390-INS-896 |
| chr16-23730926-INS-442 | chr16-23730926-INS-442 |
| chr16-23760084-INS-174 | chr16-23760084-INS-176 |
| chr16-23776664-INS-69 | chr16-23776665-INS-108 |
| chr16-23778056-INS-54 | chr16-23778061-INS-69 |
| chr16-23836289-INS-53 | chr16-23836285-INS-59 |
| chr16-23877395-INS-62 | chr16-23877395-INS-60 |
| chr16-23931706-INS-176 | chr16-23931706-INS-184 |
| chr16-23936223-INS-58 | chr16-23936229-INS-100 |
| chr16-23972309-INS-153 | chr16-23972309-INS-196 |
| chr16-24006483-INS-58 | chr16-24006488-INS-60 |
| chr16-24029635-INS-56 | chr16-24029635-INS-58 |
| chr16-24031788-INS-208 | chr16-24031788-INS-208 |
| chr16-24122716-INS-90 | chr16-24122716-INS-90 |
| chr16-24139556-INS-184 | chr16-24139556-INS-204 |
| chr16-24184021-INS-85 | chr16-24184016-INS-96 |
| chr16-24191119-INS-821 | chr16-24191119-INS-821 |
| chr16-24219726-INS-71 | chr16-24219726-INS-69 |
| chr16-24244026-INS-186 | chr16-24244026-INS-179 |
| chr16-24267057-INS-55 | chr16-24267057-INS-55 |
| chr16-24276729-INS-60 | chr16-24276729-INS-60 |
| chr16-24282154-INS-63 | chr16-24282154-INS-63 |
| chr16-24286006-INS-95 | chr16-24286032-INS-95 |
| chr16-24327662-INS-92 | chr16-24327660-INS-92 |
| chr16-24346862-INS-78 | chr16-24346862-INS-79 |
| chr16-24357731-INS-119 | chr16-24357731-INS-120 |
| chr16-24417811-INS-165 | chr16-24417811-INS-165 |
| chr16-24418606-INS-462 | chr16-24418606-INS-463 |
| chr16-24422539-INS-7209 | chr16-24422540-INS-7209 |
| chr16-24424336-INS-58 | chr16-24424336-INS-58 |
| chr16-24430326-INS-143 | chr16-24430326-INS-143 |
| chr16-24529547-INS-312 | chr16-24529547-INS-312 |
| chr16-24609749-INS-58 | chr16-24609749-INS-60 |
| chr16-24630840-INS-63 | chr16-24630840-INS-66 |
| chr16-24631481-INS-181 | chr16-24631468-INS-199 |
| chr16-24694086-INS-55 | chr16-24694086-INS-60 |
| chr16-24704487-INS-371 | chr16-24704487-INS-371 |
| chr16-24743045-INS-192 | chr16-24743045-INS-192 |
| chr16-24746981-INS-117 | chr16-24746981-INS-105 |
| chr16-24750191-INS-152 | chr16-24750191-INS-152 |
| chr16-24816286-INS-146 | chr16-24816286-INS-146 |
| chr16-24830825-INS-3088 | chr16-24830826-INS-3088 |
| chr16-24858365-INS-157 | chr16-24858365-INS-157 |
| chr16-24861340-INS-57 | chr16-24861361-INS-55 |

| | |
|---|---|
| chr16-24928904-INS-540 | chr16-24928904-INS-544 |
| chr16-24930412-INS-402 | chr16-24930412-INS-402 |
| chr16-24958746-INS-73 | chr16-24958738-INS-72 |
| chr16-24994934-INS-102 | chr16-24994943-INS-90 |
| chr16-25052141-INS-1611 | chr16-25052142-INS-1612 |
| chr16-25099618-INS-59 | chr16-25099621-INS-59 |
| chr16-25115365-INS-445 | chr16-25115365-INS-445 |
| chr16-25155246-INS-2615 | chr16-25155247-INS-2614 |
| chr16-25169397-INS-144 | chr16-25169398-INS-132 |
| chr16-25177414-INS-156 | chr16-25177410-INS-171 |
| chr16-25198368-INS-76 | chr16-25198368-INS-100 |
| chr16-25203262-INS-900 | chr16-25203254-INS-938 |
| chr16-25214339-INS-85 | chr16-25214339-INS-85 |
| chr16-25239914-INS-52 | chr16-25239914-INS-54 |
| chr16-25246285-INS-250 | chr16-25246282-INS-237 |
| chr16-25253747-INS-72 | chr16-25253732-INS-60 |
| chr16-25267902-INS-63 | chr16-25267896-INS-60 |
| chr16-25275074-INS-75 | chr16-25275074-INS-70 |
| chr16-25288262-INS-7050 | chr16-25288263-INS-7051 |
| chr16-25334035-INS-152 | chr16-25334035-INS-152 |
| chr16-25355099-INS-671 | chr16-25355099-INS-631 |
| chr16-25370901-INS-6681 | chr16-25370902-INS-6681 |
| chr16-25410211-INS-74 | chr16-25410247-INS-78 |
| chr16-25413126-INS-443 | chr16-25413126-INS-443 |
| chr16-25423329-INS-381 | chr16-25423329-INS-381 |
| chr16-25446228-INS-975 | chr16-25446228-INS-969 |
| chr16-25476148-INS-96 | chr16-25476159-INS-83 |
| chr16-25490912-INS-132 | chr16-25490912-INS-138 |
| chr16-25682285-INS-576 | chr16-25682285-INS-572 |
| chr16-25716169-INS-96 | chr16-25716169-INS-62 |
| chr16-25769369-INS-118 | chr16-25769369-INS-119 |
| chr16-25820478-INS-106 | chr16-25820483-INS-110 |
| chr16-25843300-INS-947 | chr16-25843300-INS-947 |
| chr16-25872027-INS-6855 | chr16-25872028-INS-6855 |
| chr16-25934196-INS-88 | chr16-25934196-INS-88 |
| chr16-25946968-INS-212 | chr16-25946968-INS-212 |
| chr16-26057954-INS-56 | chr16-26057954-INS-64 |
| chr16-26114210-INS-62 | chr16-26114210-INS-54 |
| chr16-26168373-INS-811 | chr16-26168373-INS-813 |
| chr16-26249460-INS-6483 | chr16-26249461-INS-6484 |
| chr16-26260991-INS-233 | chr16-26260991-INS-218 |
| chr16-26307905-INS-1427 | chr16-26307906-INS-1427 |
| chr16-26340501-INS-202 | chr16-26340503-INS-206 |
| chr16-26341602-INS-218 | chr16-26341602-INS-218 |
| chr16-26343677-INS-79 | chr16-26343678-INS-86 |
| chr16-26377191-INS-97 | chr16-26377191-INS-97 |
| chr16-26415590-INS-54 | chr16-26415601-INS-91 |
| chr16-26432312-INS-62 | chr16-26432312-INS-58 |
| chr16-26433713-INS-132 | chr16-26433713-INS-132 |
| chr16-26436380-INS-67 | chr16-26436376-INS-109 |
| chr16-26458251-INS-59 | chr16-26458275-INS-52 |
| chr16-26491283-INS-440 | chr16-26491283-INS-440 |
| chr16-26563259-INS-92 | chr16-26563254-INS-63 |
| chr16-26578993-INS-142 | chr16-26578993-INS-154 |
| chr16-26621815-INS-57 | chr16-26621815-INS-67 |
| chr16-26637588-INS-569 | chr16-26637588-INS-576 |
| chr16-26676883-INS-138 | chr16-26676883-INS-139 |
| chr16-26718756-INS-104 | chr16-26718756-INS-104 |
| chr16-26730258-INS-116 | chr16-26730258-INS-116 |
| chr16-26732451-INS-92 | chr16-26732447-INS-112 |
| chr16-26751107-INS-332 | chr16-26751107-INS-332 |
| chr16-26756512-INS-564 | chr16-26756512-INS-566 |
| chr16-26789054-INS-68 | chr16-26789055-INS-68 |
| chr16-26825937-INS-58 | chr16-26825935-INS-64 |
| chr16-26832721-INS-452 | chr16-26832721-INS-482 |
| chr16-26849317-INS-87 | chr16-26849317-INS-87 |
| chr16-26872567-INS-104 | chr16-26872567-INS-106 |
| chr16-26881697-INS-741 | chr16-26881701-INS-740 |
| chr16-26888123-INS-210 | chr16-26888123-INS-209 |
| chr16-26920876-INS-585 | chr16-26920876-INS-585 |
| chr16-26945848-INS-207 | chr16-26945848-INS-208 |
| chr16-26954407-INS-184 | chr16-26954407-INS-183 |
| chr16-27009268-INS-64 | chr16-27009268-INS-88 |
| chr16-27048660-INS-109 | chr16-27048660-INS-113 |
| chr16-27074037-INS-157 | chr16-27074037-INS-157 |
| chr16-27102432-INS-119 | chr16-27102426-INS-100 |
| chr16-27124352-INS-280 | chr16-27124352-INS-277 |

| | |
|---|---|
| chr16-27149762-INS-64 | chr16-27149762-INS-64 |
| chr16-27152032-INS-454 | chr16-27152032-INS-454 |
| chr16-27171948-INS-343 | chr16-27171948-INS-321 |
| chr16-27175852-INS-137 | chr16-27175852-INS-137 |
| chr16-27192205-INS-358 | chr16-27192205-INS-358 |
| chr16-27203218-INS-166 | chr16-27203218-INS-166 |
| chr16-27273313-INS-6308 | chr16-27273314-INS-6309 |
| chr16-27274657-INS-98 | chr16-27274657-INS-98 |
| chr16-27298931-INS-686 | chr16-27298931-INS-686 |
| chr16-27317148-INS-94 | chr16-27317148-INS-94 |
| chr16-27323443-INS-100 | chr16-27323443-INS-104 |
| chr16-27328499-INS-172 | chr16-27328501-INS-189 |
| chr16-27336685-INS-878 | chr16-27336685-INS-876 |
| chr16-27363323-INS-225 | chr16-27363323-INS-224 |
| chr16-27375386-INS-546 | chr16-27375386-INS-546 |
| chr16-27375986-INS-84 | chr16-27375986-INS-82 |
| chr16-27383224-INS-450 | chr16-27383224-INS-450 |
| chr16-27391351-INS-58 | chr16-27391351-INS-58 |
| chr16-27402696-INS-4922 | chr16-27402697-INS-4929 |
| chr16-27429864-INS-56 | chr16-27429874-INS-70 |
| chr16-27455201-INS-633 | chr16-27455206-INS-634 |
| chr16-27469742-INS-216 | chr16-27469742-INS-216 |
| chr16-27486116-INS-544 | chr16-27486116-INS-544 |
| chr16-27503492-INS-105 | chr16-27503492-INS-107 |
| chr16-27510391-INS-210 | chr16-27510391-INS-210 |
| chr16-27541594-INS-6500 | chr16-27541595-INS-6501 |
| chr16-27542989-INS-54 | chr16-27542989-INS-56 |
| chr16-27553911-INS-298 | chr16-27553911-INS-298 |
| chr16-27561011-INS-60 | chr16-27561011-INS-64 |
| chr16-27567657-INS-400 | chr16-27567657-INS-400 |
| chr16-27572048-INS-57 | chr16-27572048-INS-57 |
| chr16-27590941-INS-757 | chr16-27590941-INS-758 |
| chr16-27608908-INS-438 | chr16-27608908-INS-438 |
| chr16-27640023-INS-202 | chr16-27640023-INS-202 |
| chr16-27645694-INS-57 | chr16-27645692-INS-60 |
| chr16-27665398-INS-189 | chr16-27665398-INS-189 |
| chr16-27679181-INS-102 | chr16-27679181-INS-102 |
| chr16-27750838-INS-60 | chr16-27750838-INS-60 |
| chr16-27754252-INS-7056 | chr16-27754253-INS-7056 |
| chr16-27766618-INS-56 | chr16-27766618-INS-50 |
| chr16-27783596-INS-7684 | chr16-27783597-INS-7685 |
| chr16-27827603-INS-609 | chr16-27827603-INS-612 |
| chr16-27870369-INS-102 | chr16-27870386-INS-112 |
| chr16-27893379-INS-173 | chr16-27893354-INS-182 |
| chr16-27897066-INS-507 | chr16-27897066-INS-507 |
| chr16-27910824-INS-387 | chr16-27910824-INS-387 |
| chr16-27925456-INS-60 | chr16-27925456-INS-60 |
| chr16-27953348-INS-82 | chr16-27953348-INS-92 |
| chr16-27962560-INS-1281 | chr16-27962561-INS-1268 |
| chr16-27996858-INS-245 | chr16-27996858-INS-229 |
| chr16-28005185-INS-180 | chr16-28005185-INS-174 |
| chr16-28092043-INS-52 | chr16-28092043-INS-52 |
| chr16-28145655-INS-305 | chr16-28145655-INS-308 |
| chr16-28163925-INS-442 | chr16-28163937-INS-432 |
| chr16-28184032-INS-444 | chr16-28184032-INS-444 |
| chr16-28239388-INS-124 | chr16-28239388-INS-163 |
| chr16-28333721-INS-76 | chr16-28333721-INS-80 |
| chr16-28360163-INS-68 | chr16-28360163-INS-68 |
| chr16-28422421-INS-433 | chr16-28422421-INS-437 |
| chr16-28432752-INS-54 | chr16-28432752-INS-56 |
| chr16-28433225-INS-1636 | chr16-28433226-INS-1636 |
| chr16-28474436-INS-61 | chr16-28474436-INS-62 |
| chr16-28476309-INS-72 | chr16-28476311-INS-74 |
| chr16-28508137-INS-51 | chr16-28508137-INS-51 |
| chr16-28517187-INS-527 | chr16-28517187-INS-529 |
| chr16-28524859-INS-59 | chr16-28524859-INS-60 |
| chr16-28541662-INS-335 | chr16-28541662-INS-335 |
| chr16-28544316-INS-233 | chr16-28544316-INS-234 |
| chr16-28547192-INS-300 | chr16-28547192-INS-300 |
| chr16-28550171-INS-67 | chr16-28550174-INS-89 |
| chr16-28564777-INS-307 | chr16-28564777-INS-307 |
| chr16-28573848-INS-57 | chr16-28573848-INS-57 |
| chr16-28580106-INS-88 | chr16-28580102-INS-70 |
| chr16-28590299-INS-572 | chr16-28590299-INS-572 |
| chr16-28594859-INS-1890 | chr16-28594860-INS-1890 |
| chr16-28597239-INS-648 | chr16-28597223-INS-647 |
| chr16-28604178-INS-175 | chr16-28604178-INS-153 |

| | |
|---|---|
| chr16-28614411-INS-259 | chr16-28614411-INS-260 |
| chr16-28630991-INS-185 | chr16-28630991-INS-198 |
| chr16-28634344-INS-59 | chr16-28634344-INS-59 |
| chr16-28642102-INS-85 | chr16-28642102-INS-83 |
| chr16-28669644-INS-92 | chr16-28669644-INS-92 |
| chr16-28680864-INS-62 | chr16-28680864-INS-62 |
| chr16-28685437-INS-98 | chr16-28685437-INS-92 |
| chr16-28769014-INS-72 | chr16-28769021-INS-56 |
| chr16-28772636-INS-84 | chr16-28772641-INS-95 |
| chr16-28777458-INS-134 | chr16-28777458-INS-125 |
| chr16-28786844-INS-75 | chr16-28786844-INS-75 |
| chr16-28792450-INS-153 | chr16-28792450-INS-153 |
| chr16-28810174-INS-231 | chr16-28810174-INS-231 |
| chr16-28846494-INS-147 | chr16-28846494-INS-147 |
| chr16-28857882-INS-909 | chr16-28857882-INS-895 |
| chr16-28883627-INS-110 | chr16-28883615-INS-84 |
| chr16-28910292-INS-440 | chr16-28910292-INS-440 |
| chr16-28947244-INS-76 | chr16-28947244-INS-90 |
| chr16-28953516-INS-521 | chr16-28953517-INS-521 |
| chr16-28961380-INS-57 | chr16-28961380-INS-60 |
| chr16-28985804-INS-87 | chr16-28985804-INS-88 |
| chr16-29000515-INS-303 | chr16-29000515-INS-303 |
| chr16-29013786-INS-99 | chr16-29013786-INS-99 |
| chr16-29019836-INS-7074 | chr16-29019837-INS-7074 |
| chr16-29059812-INS-52 | chr16-29059802-INS-96 |
| chr16-29061846-INS-6235 | chr16-29061833-INS-6238 |
| chr16-29065824-INS-6362 | chr16-29065825-INS-6363 |
| chr16-29070835-INS-2771 | chr16-29070836-INS-2771 |
| chr16-29085332-INS-69 | chr16-29085332-INS-77 |
| chr16-29103257-INS-67 | chr16-29103257-INS-69 |
| chr16-29119962-INS-623 | chr16-29119962-INS-623 |
| chr16-29131718-INS-69 | chr16-29131718-INS-70 |
| chr16-29209429-INS-62 | chr16-29209430-INS-58 |
| chr16-29227572-INS-105 | chr16-29227572-INS-55 |
| chr16-29270370-INS-305 | chr16-29270370-INS-309 |
| chr16-29356544-INS-155 | chr16-29356544-INS-155 |
| chr16-29364844-INS-65 | chr16-29364844-INS-65 |
| chr16-29410264-INS-199 | chr16-29410264-INS-199 |
| chr16-29427680-INS-123 | chr16-29427665-INS-133 |
| chr16-29432125-INS-198 | chr16-29432125-INS-200 |
| chr16-29459160-INS-96 | chr16-29459160-INS-96 |
| chr16-29473878-INS-206 | chr16-29473878-INS-206 |
| chr16-29477330-INS-58 | chr16-29477330-INS-108 |
| chr16-29482647-INS-418 | chr16-29482647-INS-418 |
| chr16-29488118-INS-473 | chr16-29488118-INS-473 |
| chr16-29515872-INS-126 | chr16-29515872-INS-126 |
| chr16-29520399-INS-65 | chr16-29520399-INS-65 |
| chr16-29522822-INS-303 | chr16-29522822-INS-304 |
| chr16-29539884-INS-82 | chr16-29539885-INS-69 |
| chr16-29572492-INS-380 | chr16-29572492-INS-380 |
| chr16-29598353-INS-83 | chr16-29598353-INS-89 |
| chr16-29637397-INS-115 | chr16-29637397-INS-128 |
| chr16-29703388-INS-53 | chr16-29703388-INS-53 |
| chr16-29704872-INS-79 | chr16-29704884-INS-90 |
| chr16-29709531-INS-63 | chr16-29709531-INS-63 |
| chr16-29713974-INS-62 | chr16-29713934-INS-74 |
| chr16-29720169-INS-213 | chr16-29720169-INS-212 |
| chr16-29751656-INS-512 | chr16-29751656-INS-512 |
| chr16-29840514-INS-65 | chr16-29840514-INS-81 |
| chr16-29869521-INS-62 | chr16-29869545-INS-54 |
| chr16-29908789-INS-6413 | chr16-29908790-INS-6413 |
| chr16-29910277-INS-54 | chr16-29910289-INS-55 |
| chr16-29950999-INS-668 | chr16-29950999-INS-667 |
| chr16-29965553-INS-80 | chr16-29965536-INS-85 |
| chr16-29981766-INS-216 | chr16-29981766-INS-216 |
| chr16-29985729-INS-678 | chr16-29985728-INS-676 |
| chr16-29987013-INS-4748 | chr16-29987014-INS-4748 |
| chr16-29993780-INS-168 | chr16-29993773-INS-172 |
| chr16-29994934-INS-176 | chr16-29994934-INS-148 |
| chr16-30018729-INS-52 | chr16-30018729-INS-60 |
| chr16-30026240-INS-439 | chr16-30026240-INS-439 |
| chr16-30067680-INS-204 | chr16-30067688-INS-206 |
| chr16-30071441-INS-196 | chr16-30071441-INS-198 |
| chr16-30072867-INS-214 | chr16-30072867-INS-214 |
| chr16-30093762-INS-230 | chr16-30093762-INS-230 |
| chr16-30098610-INS-228 | chr16-30098610-INS-222 |
| chr16-30109237-INS-240 | chr16-30109237-INS-240 |

| | |
|---|---|
| chr16-30170547-INS-205 | chr16-30170547-INS-208 |
| chr16-30190608-INS-212 | chr16-30190608-INS-212 |
| chr16-30202095-INS-192 | chr16-30202095-INS-188 |
| chr16-30211068-INS-204 | chr16-30211068-INS-205 |
| chr16-30214894-INS-208 | chr16-30214894-INS-208 |
| chr16-30217718-INS-183 | chr16-30217718-INS-195 |
| chr16-30228965-INS-832 | chr16-30228965-INS-832 |
| chr16-30231284-INS-126 | chr16-30231289-INS-122 |
| chr16-30233780-INS-52 | chr16-30233776-INS-52 |
| chr16-30237383-INS-201 | chr16-30237395-INS-214 |
| chr16-30238185-INS-250 | chr16-30238185-INS-251 |
| chr16-30242880-INS-71 | chr16-30242875-INS-79 |
| chr16-30247482-INS-198 | chr16-30247483-INS-196 |
| chr16-30254530-INS-505 | chr16-30254530-INS-463 |
| chr16-30265823-INS-201 | chr16-30265823-INS-200 |
| chr16-30266546-INS-217 | chr16-30266546-INS-217 |
| chr16-30272038-INS-89 | chr16-30272039-INS-79 |
| chr16-30282242-INS-80 | chr16-30282250-INS-64 |
| chr16-30288197-INS-4722 | chr16-30288198-INS-4722 |
| chr16-30298915-INS-58 | chr16-30298919-INS-54 |
| chr16-30307483-INS-201 | chr16-30307483-INS-201 |
| chr16-30311921-INS-511 | chr16-30311921-INS-510 |
| chr16-30351164-INS-56 | chr16-30351164-INS-56 |
| chr16-30354573-INS-207 | chr16-30354573-INS-211 |
| chr16-30363642-INS-70 | chr16-30363642-INS-70 |
| chr16-30377026-INS-179 | chr16-30377026-INS-191 |
| chr16-30392451-INS-176 | chr16-30392451-INS-168 |
| chr16-30404613-INS-611 | chr16-30404613-INS-611 |
| chr16-30412752-INS-111 | chr16-30412752-INS-75 |
| chr16-30417241-INS-89 | chr16-30417241-INS-89 |
| chr16-30419162-INS-128 | chr16-30419162-INS-122 |
| chr16-30424072-INS-82 | chr16-30424072-INS-84 |
| chr16-30437420-INS-857 | chr16-30437420-INS-857 |
| chr16-30444557-INS-117 | chr16-30444557-INS-117 |
| chr16-30467714-INS-331 | chr16-30467714-INS-331 |
| chr16-30503434-INS-233 | chr16-30503434-INS-238 |
| chr16-30505230-INS-390 | chr16-30505230-INS-390 |
| chr16-30510347-INS-182 | chr16-30510347-INS-181 |
| chr16-30523227-INS-132 | chr16-30523227-INS-148 |
| chr16-30530689-INS-66 | chr16-30530687-INS-50 |
| chr16-30579568-INS-159 | chr16-30579595-INS-147 |
| chr16-30663053-INS-56 | chr16-30663053-INS-58 |
| chr16-30671245-INS-53 | chr16-30671245-INS-53 |
| chr16-30672658-INS-510 | chr16-30672658-INS-542 |
| chr16-30707298-INS-54 | chr16-30707299-INS-54 |
| chr16-30717886-INS-1895 | chr16-30717887-INS-1895 |
| chr16-30720532-INS-54 | chr16-30720532-INS-66 |
| chr16-30755120-INS-6399 | chr16-30755121-INS-6399 |
| chr16-30773422-INS-198 | chr16-30773422-INS-198 |
| chr16-30788644-INS-56 | chr16-30788644-INS-56 |
| chr16-30800873-INS-56 | chr16-30800873-INS-56 |
| chr16-30809450-INS-578 | chr16-30809450-INS-532 |
| chr16-30818496-INS-653 | chr16-30818496-INS-651 |
| chr16-30829758-INS-122 | chr16-30829758-INS-122 |
| chr16-30877026-INS-7301 | chr16-30877027-INS-7301 |
| chr16-30892456-INS-80 | chr16-30892456-INS-70 |
| chr16-30893978-INS-1892 | chr16-30893979-INS-1892 |
| chr16-30959680-INS-195 | chr16-30959680-INS-195 |
| chr16-30974178-INS-146 | chr16-30974178-INS-146 |
| chr16-30975312-INS-239 | chr16-30975313-INS-239 |
| chr16-30977161-INS-105 | chr16-30977161-INS-105 |
| chr16-30980010-INS-60 | chr16-30980010-INS-60 |
| chr16-30990701-INS-75 | chr16-30990696-INS-81 |
| chr16-31014380-INS-287 | chr16-31014380-INS-287 |
| chr16-31033415-INS-55 | chr16-31033415-INS-60 |
| chr16-31035511-INS-70 | chr16-31035511-INS-72 |
| chr16-31085966-INS-77 | chr16-31085971-INS-81 |
| chr16-31146855-INS-5549 | chr16-31146856-INS-5549 |
| chr16-31166164-INS-688 | chr16-31166164-INS-687 |
| chr16-31255043-INS-114 | chr16-31255043-INS-114 |
| chr16-31264444-INS-51 | chr16-31264445-INS-69 |
| chr16-31267179-INS-121 | chr16-31267179-INS-121 |
| chr16-31281263-INS-51 | chr16-31281263-INS-51 |
| chr16-31287400-INS-431 | chr16-31287400-INS-431 |
| chr16-31337585-INS-90 | chr16-31337585-INS-90 |
| chr16-31362636-INS-60 | chr16-31362616-INS-76 |
| chr16-31363242-INS-100 | chr16-31363242-INS-100 |

| | |
|---|---|
| chr16-31368116-INS-284 | chr16-31368116-INS-282 |
| chr16-31376906-INS-5547 | chr16-31376907-INS-5547 |
| chr16-31395351-INS-72 | chr16-31395351-INS-72 |
| chr16-31456297-INS-194 | chr16-31456297-INS-194 |
| chr16-31463429-INS-894 | chr16-31463447-INS-898 |
| chr16-31487384-INS-51 | chr16-31487384-INS-51 |
| chr16-31497583-INS-241 | chr16-31497583-INS-241 |
| chr16-31519059-INS-56 | chr16-31519059-INS-63 |
| chr16-31527232-INS-60 | chr16-31527232-INS-62 |
| chr16-31537595-INS-3072 | chr16-31537596-INS-3072 |
| chr16-31557765-INS-227 | chr16-31557765-INS-227 |
| chr16-31643980-INS-1153 | chr16-31643981-INS-1153 |
| chr16-31703645-INS-80 | chr16-31703645-INS-80 |
| chr16-31707648-INS-1061 | chr16-31707649-INS-1061 |
| chr16-31726918-INS-71 | chr16-31726928-INS-50 |
| chr16-31733234-INS-168 | chr16-31733234-INS-168 |
| chr16-31759916-INS-6399 | chr16-31759917-INS-6399 |
| chr16-31770331-INS-211 | chr16-31770326-INS-191 |
| chr16-31774096-INS-120 | chr16-31774096-INS-120 |
| chr16-31788141-INS-148 | chr16-31788141-INS-148 |
| chr16-31828689-INS-54 | chr16-31828689-INS-51 |
| chr16-31848126-INS-147 | chr16-31848126-INS-147 |
| chr16-31854185-INS-189 | chr16-31854182-INS-190 |
| chr16-31858103-INS-207 | chr16-31858103-INS-207 |
| chr16-31869099-INS-210 | chr16-31869099-INS-210 |
| chr16-31919167-INS-204 | chr16-31919167-INS-204 |
| chr16-31920577-INS-4874 | chr16-31920580-INS-4862 |
| chr16-31957217-INS-101 | chr16-31957217-INS-101 |
| chr16-32025561-INS-730 | chr16-32025561-INS-730 |
| chr16-32027545-INS-56 | chr16-32027545-INS-56 |
| chr16-32039124-INS-211 | chr16-32039124-INS-211 |
| chr16-32069991-INS-212 | chr16-32069989-INS-211 |
| chr16-32076347-INS-196 | chr16-32076347-INS-196 |
| chr16-32094165-INS-72 | chr16-32094165-INS-72 |
| chr16-32101051-INS-253 | chr16-32101051-INS-254 |
| chr16-32105825-INS-181 | chr16-32105825-INS-181 |
| chr16-32107456-INS-197 | chr16-32107456-INS-196 |
| chr16-32109091-INS-167 | chr16-32109091-INS-170 |
| chr16-32116472-INS-184 | chr16-32116472-INS-185 |
| chr16-32122024-INS-208 | chr16-32122024-INS-209 |
| chr16-32125957-INS-207 | chr16-32125957-INS-207 |
| chr16-32136440-INS-64 | chr16-32136440-INS-64 |
| chr16-32137181-INS-211 | chr16-32137182-INS-210 |
| chr16-32143473-INS-67 | chr16-32143473-INS-61 |
| chr16-32152872-INS-216 | chr16-32152872-INS-217 |
| chr16-32160697-INS-215 | chr16-32160697-INS-212 |
| chr16-32178783-INS-56 | chr16-32178783-INS-56 |
| chr16-32208709-INS-1886 | chr16-32208710-INS-1886 |
| chr16-32231313-INS-200 | chr16-32231313-INS-200 |
| chr16-32245777-INS-120 | chr16-32245777-INS-120 |
| chr16-32259084-INS-211 | chr16-32259084-INS-212 |
| chr16-32301153-INS-194 | chr16-32301203-INS-176 |
| chr16-32324139-INS-202 | chr16-32324139-INS-202 |
| chr16-32333261-INS-232 | chr16-32333261-INS-230 |
| chr16-32366053-INS-1143 | chr16-32366054-INS-1131 |
| chr16-32369703-INS-56 | chr16-32369682-INS-56 |
| chr16-32373008-INS-221 | chr16-32373008-INS-221 |
| chr16-32381081-INS-397 | chr16-32381081-INS-397 |
| chr16-32394939-INS-403 | chr16-32394939-INS-404 |
| chr16-32440902-INS-211 | chr16-32440902-INS-212 |
| chr16-32455553-INS-67 | chr16-32455553-INS-115 |
| chr16-32459454-INS-120 | chr16-32459454-INS-108 |
| chr16-32472838-INS-154 | chr16-32472838-INS-150 |
| chr16-32476835-INS-56 | chr16-32476835-INS-56 |
| chr16-32496603-INS-117 | chr16-32496603-INS-117 |
| chr16-32506221-INS-238 | chr16-32506221-INS-238 |
| chr16-32516882-INS-58 | chr16-32516861-INS-54 |
| chr16-32531654-INS-205 | chr16-32531654-INS-205 |
| chr16-32537274-INS-201 | chr16-32537274-INS-201 |
| chr16-32539877-INS-150 | chr16-32539882-INS-152 |
| chr16-32543728-INS-137 | chr16-32543728-INS-137 |
| chr16-32546689-INS-59 | chr16-32546689-INS-66 |
| chr16-32568023-INS-52 | chr16-32568023-INS-56 |
| chr16-32583437-INS-88 | chr16-32583437-INS-88 |
| chr16-32592493-INS-81 | chr16-32592493-INS-81 |
| chr16-32607508-INS-334 | chr16-32607508-INS-334 |
| chr16-32616066-INS-92 | chr16-32616066-INS-96 |

| | |
|---|---|
| chr16-32624403-INS-82 | chr16-32624403-INS-82 |
| chr16-32626186-INS-134 | chr16-32626186-INS-96 |
| chr16-32639046-INS-209 | chr16-32639088-INS-187 |
| chr16-32647815-INS-54 | chr16-32647815-INS-54 |
| chr16-32672142-INS-214 | chr16-32672142-INS-215 |
| chr16-32678216-INS-99 | chr16-32678212-INS-86 |
| chr16-32690803-INS-98 | chr16-32690818-INS-100 |
| chr16-32694391-INS-4717 | chr16-32694392-INS-4717 |
| chr16-32705111-INS-52 | chr16-32705111-INS-52 |
| chr16-32710777-INS-7512 | chr16-32710778-INS-7512 |
| chr16-32712229-INS-96 | chr16-32712273-INS-64 |
| chr16-32736319-INS-196 | chr16-32736319-INS-194 |
| chr16-32750768-INS-147 | chr16-32750768-INS-155 |
| chr16-32780838-INS-207 | chr16-32780838-INS-206 |
| chr16-32781443-INS-118 | chr16-32781443-INS-118 |
| chr16-32800120-INS-196 | chr16-32800120-INS-196 |
| chr16-32825428-INS-245 | chr16-32825428-INS-245 |
| chr16-32848039-INS-92 | chr16-32848039-INS-105 |
| chr16-32863549-INS-7081 | chr16-32863550-INS-7081 |
| chr16-32868708-INS-215 | chr16-32868708-INS-215 |
| chr16-32871616-INS-939 | chr16-32871616-INS-939 |
| chr16-32900355-INS-192 | chr16-32900355-INS-192 |
| chr16-32901797-INS-206 | chr16-32901797-INS-206 |
| chr16-32905965-INS-89 | chr16-32905964-INS-114 |
| chr16-32907017-INS-64 | chr16-32907017-INS-64 |
| chr16-32918931-INS-385 | chr16-32918931-INS-371 |
| chr16-32928489-INS-203 | chr16-32928489-INS-202 |
| chr16-32929545-INS-248 | chr16-32929545-INS-279 |
| chr16-32948887-INS-200 | chr16-32948887-INS-202 |
| chr16-32961047-INS-433 | chr16-32961047-INS-433 |
| chr16-32972341-INS-203 | chr16-32972341-INS-203 |
| chr16-32974000-INS-54 | chr16-32973998-INS-54 |
| chr16-32986436-INS-196 | chr16-32986436-INS-196 |
| chr16-32989327-INS-201 | chr16-32989327-INS-201 |
| chr16-32993960-INS-63 | chr16-32993960-INS-65 |
| chr16-33000011-INS-221 | chr16-33000011-INS-221 |
| chr16-33010982-INS-517 | chr16-33010982-INS-517 |
| chr16-33016217-INS-84 | chr16-33016217-INS-85 |
| chr16-33019032-INS-210 | chr16-33019032-INS-210 |
| chr16-33027995-INS-5770 | chr16-33027996-INS-5770 |
| chr16-33096306-INS-50 | chr16-33096300-INS-98 |
| chr16-33110115-INS-214 | chr16-33110115-INS-214 |
| chr16-33118264-INS-113 | chr16-33118265-INS-102 |
| chr16-33126357-INS-166 | chr16-33126357-INS-166 |
| chr16-33129046-INS-226 | chr16-33129046-INS-227 |
| chr16-33143678-INS-56 | chr16-33143678-INS-58 |
| chr16-33144173-INS-498 | chr16-33144173-INS-498 |
| chr16-33150960-INS-120 | chr16-33150960-INS-144 |
| chr16-33156029-INS-92 | chr16-33156029-INS-52 |
| chr16-33214084-INS-198 | chr16-33214084-INS-200 |
| chr16-33214098-INS-927 | chr16-33214098-INS-927 |
| chr16-33226301-INS-4827 | chr16-33226302-INS-4827 |
| chr16-33258305-INS-83 | chr16-33258305-INS-83 |
| chr16-33265652-INS-84 | chr16-33265652-INS-89 |
| chr16-33267293-INS-59 | chr16-33267293-INS-59 |
| chr16-33269569-INS-6445 | chr16-33269570-INS-6445 |
| chr16-33272894-INS-222 | chr16-33272894-INS-224 |
| chr16-33275793-INS-67 | chr16-33275793-INS-67 |
| chr16-33301147-INS-200 | chr16-33301187-INS-165 |
| chr16-33301961-INS-193 | chr16-33301948-INS-194 |
| chr16-33305591-INS-270 | chr16-33305601-INS-310 |
| chr16-33331007-INS-211 | chr16-33331007-INS-211 |
| chr16-33351660-INS-400 | chr16-33351660-INS-400 |
| chr16-33359334-INS-69 | chr16-33359334-INS-69 |
| chr16-33362890-INS-125 | chr16-33362890-INS-125 |
| chr16-33372782-INS-172 | chr16-33372782-INS-172 |
| chr16-33374688-INS-60 | chr16-33374688-INS-52 |
| chr16-33377146-INS-95 | chr16-33377146-INS-96 |
| chr16-33387489-INS-113 | chr16-33387494-INS-119 |
| chr16-33391236-INS-109 | chr16-33391236-INS-109 |
| chr16-33412423-INS-58 | chr16-33412423-INS-76 |
| chr16-33418801-INS-870 | chr16-33418801-INS-867 |
| chr16-33483093-INS-142 | chr16-33483093-INS-142 |
| chr16-33499460-INS-92 | chr16-33499460-INS-110 |
| chr16-33502853-INS-104 | chr16-33502853-INS-104 |
| chr16-33529801-INS-184 | chr16-33529789-INS-198 |
| chr16-33543297-INS-80 | chr16-33543296-INS-128 |

| | |
|---|---|
| chr16-33567813-INS-228 | chr16-33567813-INS-229 |
| chr16-33578907-INS-200 | chr16-33578907-INS-201 |
| chr16-33592865-INS-81 | chr16-33592866-INS-60 |
| chr16-33600746-INS-95 | chr16-33600730-INS-105 |
| chr16-33604519-INS-282 | chr16-33604501-INS-294 |
| chr16-33618029-INS-65 | chr16-33618029-INS-56 |
| chr16-33631790-INS-224 | chr16-33631790-INS-225 |
| chr16-33647077-INS-160 | chr16-33647079-INS-187 |
| chr16-33649203-INS-81 | chr16-33649224-INS-84 |
| chr16-33658654-INS-606 | chr16-33658639-INS-654 |
| chr16-33659842-INS-70 | chr16-33659835-INS-64 |
| chr16-33666028-INS-525 | chr16-33666028-INS-525 |
| chr16-33720994-INS-246 | chr16-33720994-INS-248 |
| chr16-33735836-INS-648 | chr16-33735831-INS-628 |
| chr16-33769116-INS-7218 | chr16-33769117-INS-7218 |
| chr16-33778613-INS-155 | chr16-33778613-INS-155 |
| chr16-33789129-INS-355 | chr16-33789129-INS-355 |
| chr16-33810107-INS-100 | chr16-33810107-INS-136 |
| chr16-33812624-INS-329 | chr16-33812620-INS-292 |
| chr16-33816447-INS-532 | chr16-33816456-INS-539 |
| chr16-33835931-INS-54 | chr16-33835931-INS-54 |
| chr16-33853028-INS-198 | chr16-33853028-INS-198 |
| chr16-33889396-INS-77 | chr16-33889396-INS-81 |
| chr16-33899081-INS-154 | chr16-33899081-INS-156 |
| chr16-33901755-INS-284 | chr16-33901755-INS-288 |
| chr16-33932270-INS-74 | chr16-33932249-INS-56 |
| chr16-34007722-INS-85 | chr16-34007730-INS-86 |
| chr16-34013669-INS-72 | chr16-34013669-INS-60 |
| chr16-34022963-INS-95 | chr16-34022950-INS-70 |
| chr16-34043514-INS-57 | chr16-34043506-INS-59 |
| chr16-34060019-INS-156 | chr16-34059992-INS-195 |
| chr16-34084325-INS-1664 | chr16-34084326-INS-1664 |
| chr16-34151436-INS-52 | chr16-34151436-INS-52 |
| chr16-34180166-INS-66 | chr16-34180166-INS-51 |
| chr16-34180789-INS-6252 | chr16-34180790-INS-6252 |
| chr16-34189518-INS-119 | chr16-34189520-INS-93 |
| chr16-34197495-INS-225 | chr16-34197495-INS-225 |
| chr16-34242251-INS-110 | chr16-34242251-INS-110 |
| chr16-34282849-INS-167 | chr16-34282849-INS-167 |
| chr16-34293127-INS-114 | chr16-34293128-INS-110 |
| chr16-34324022-INS-76 | chr16-34324022-INS-74 |
| chr16-34358362-INS-80 | chr16-34358362-INS-120 |
| chr16-34401345-INS-78 | chr16-34401372-INS-51 |
| chr16-34420098-INS-58 | chr16-34420098-INS-54 |
| chr16-34421406-INS-159 | chr16-34421406-INS-160 |
| chr16-34423890-INS-89 | chr16-34423890-INS-89 |
| chr16-34429737-INS-139 | chr16-34429738-INS-173 |
| chr16-34554801-INS-198 | chr16-34554801-INS-198 |
| chr16-34565253-INS-58 | chr16-34565253-INS-52 |
| chr16-34572196-INS-517 | chr16-34572221-INS-513 |
| chr16-34616568-INS-70 | chr16-34616568-INS-68 |
| chr16-34641041-INS-205 | chr16-34641045-INS-203 |
| chr16-34656537-INS-54 | chr16-34656537-INS-52 |
| chr16-34659521-INS-678 | chr16-34659521-INS-666 |
| chr16-34663134-INS-70 | chr16-34663134-INS-74 |
| chr16-34665405-INS-121 | chr16-34665405-INS-121 |
| chr16-34707819-INS-56 | chr16-34707819-INS-62 |
| chr16-34714297-INS-416 | chr16-34714297-INS-416 |
| chr16-34770519-INS-62 | chr16-34770519-INS-58 |
| chr16-34771835-INS-62 | chr16-34771836-INS-62 |
| chr16-34785210-INS-191 | chr16-34785210-INS-191 |
| chr16-34796943-INS-67 | chr16-34796943-INS-67 |
| chr16-34831260-INS-201 | chr16-34831260-INS-201 |
| chr16-34841749-INS-215 | chr16-34841749-INS-213 |
| chr16-34845007-INS-56 | chr16-34845007-INS-59 |
| chr16-34857310-INS-58 | chr16-34857310-INS-58 |
| chr16-34865144-INS-7142 | chr16-34865145-INS-7142 |
| chr16-34909944-INS-209 | chr16-34909944-INS-209 |
| chr16-34983775-INS-66 | chr16-34983760-INS-107 |
| chr16-34992890-INS-75 | chr16-34992890-INS-75 |
| chr16-34994100-INS-71 | chr16-34994100-INS-71 |
| chr16-35001481-INS-142 | chr16-35001481-INS-142 |
| chr16-35017088-INS-145 | chr16-35017088-INS-145 |
| chr16-35019992-INS-149 | chr16-35019992-INS-149 |
| chr16-35057679-INS-65 | chr16-35057679-INS-66 |
| chr16-35061357-INS-140 | chr16-35061357-INS-140 |
| chr16-35066797-INS-83 | chr16-35066797-INS-83 |

| | |
|---|---|
| chr16-35114845-INS-498 | chr16-35114845-INS-498 |
| chr16-35143564-INS-536 | chr16-35143564-INS-539 |
| chr16-35152692-INS-465 | chr16-35152692-INS-465 |
| chr16-35178490-INS-60 | chr16-35178490-INS-60 |
| chr16-35180354-INS-4771 | chr16-35180355-INS-4771 |
| chr16-35188043-INS-52 | chr16-35188043-INS-52 |
| chr16-35219090-INS-95 | chr16-35219090-INS-95 |
| chr16-35239342-INS-70 | chr16-35239342-INS-56 |
| chr16-35320148-INS-73 | chr16-35320159-INS-79 |
| chr16-35332329-INS-117 | chr16-35332329-INS-117 |
| chr16-35378700-INS-69 | chr16-35378700-INS-69 |
| chr16-35390581-INS-71 | chr16-35390577-INS-83 |
| chr16-35441920-INS-115 | chr16-35441920-INS-115 |
| chr16-35481230-INS-107 | chr16-35481230-INS-108 |
| chr16-35490478-INS-52 | chr16-35490478-INS-54 |
| chr16-35504373-INS-66 | chr16-35504373-INS-54 |
| chr16-35512546-INS-464 | chr16-35512546-INS-472 |
| chr16-35512597-INS-1111 | chr16-35512601-INS-1103 |
| chr16-35554958-INS-656 | chr16-35554958-INS-654 |
| chr16-35557892-INS-53 | chr16-35557876-INS-71 |
| chr16-35560411-INS-465 | chr16-35560411-INS-465 |
| chr16-35596600-INS-216 | chr16-35596600-INS-216 |
| chr16-35597888-INS-180 | chr16-35597888-INS-180 |
| chr16-35607213-INS-210 | chr16-35607213-INS-210 |
| chr16-35642103-INS-4812 | chr16-35642104-INS-4812 |
| chr16-35667148-INS-139 | chr16-35667148-INS-139 |
| chr16-35697727-INS-97 | chr16-35697727-INS-97 |
| chr16-35699780-INS-4209 | chr16-35699781-INS-4209 |
| chr16-35701946-INS-83 | chr16-35701939-INS-86 |
| chr16-35702563-INS-169 | chr16-35702563-INS-169 |
| chr16-35705410-INS-99 | chr16-35705410-INS-101 |
| chr16-35714924-INS-596 | chr16-35714924-INS-596 |
| chr16-35726220-INS-53 | chr16-35726210-INS-78 |
| chr16-35733712-INS-62 | chr16-35733712-INS-61 |
| chr16-35761000-INS-78 | chr16-35761000-INS-69 |
| chr16-35765730-INS-72 | chr16-35765738-INS-96 |
| chr16-35772399-INS-410 | chr16-35772398-INS-413 |
| chr16-35781072-INS-202 | chr16-35781072-INS-232 |
| chr16-35811596-INS-193 | chr16-35811596-INS-194 |
| chr16-35821983-INS-110 | chr16-35821983-INS-107 |
| chr16-35826541-INS-103 | chr16-35826541-INS-103 |
| chr16-35830580-INS-289 | chr16-35830580-INS-295 |
| chr16-35846052-INS-203 | chr16-35846052-INS-203 |
| chr16-35878950-INS-56 | chr16-35878966-INS-61 |
| chr16-35920462-INS-327 | chr16-35920462-INS-363 |
| chr16-35927477-INS-7211 | chr16-35927478-INS-7209 |
| chr16-35965983-INS-1780 | chr16-35965984-INS-1780 |
| chr16-36040237-INS-1877 | chr16-36040238-INS-1891 |
| chr16-36042056-INS-67 | chr16-36042056-INS-67 |
| chr16-36052467-INS-162 | chr16-36052467-INS-162 |
| chr16-36076673-INS-57 | chr16-36076648-INS-62 |
| chr16-36160620-INS-65 | chr16-36160620-INS-58 |
| chr16-36276117-INS-373 | chr16-36276117-INS-381 |
| chr16-36284847-INS-186 | chr16-36284847-INS-187 |
| chr16-36286760-INS-53 | chr16-36286760-INS-53 |
| chr16-36331198-INS-92 | chr16-36331190-INS-59 |
| chr16-36364091-INS-165 | chr16-36364091-INS-173 |
| chr16-36475278-INS-6442 | chr16-36475279-INS-6442 |
| chr16-36476986-INS-247 | chr16-36476985-INS-242 |
| chr16-36483243-INS-7151 | chr16-36483244-INS-7151 |
| chr16-36486065-INS-51 | chr16-36486065-INS-52 |
| chr16-36507034-INS-255 | chr16-36507032-INS-269 |
| chr16-36519411-INS-162 | chr16-36519411-INS-120 |
| chr16-36533364-INS-88 | chr16-36533358-INS-81 |
| chr16-36544505-INS-60 | chr16-36544505-INS-60 |
| chr16-36550675-INS-96 | chr16-36550675-INS-92 |
| chr16-36580837-INS-169 | chr16-36580837-INS-170 |
| chr16-36604439-INS-326 | chr16-36604439-INS-326 |
| chr16-36701413-INS-123 | chr16-36701413-INS-123 |
| chr16-36772903-INS-57 | chr16-36772903-INS-78 |
| chr16-36834419-INS-166 | chr16-36834419-INS-166 |
| chr16-36838654-INS-96 | chr16-36838654-INS-51 |
| chr16-36908419-INS-6355 | chr16-36908420-INS-6356 |
| chr16-36926426-INS-66 | chr16-36926409-INS-79 |
| chr16-36949143-INS-67 | chr16-36949143-INS-78 |
| chr16-36950600-INS-86 | chr16-36950600-INS-62 |
| chr16-36989131-INS-60 | chr16-36989113-INS-90 |

| | |
|---|---|
| chr16-37036993-INS-226 | chr16-37036993-INS-226 |
| chr16-37070879-INS-104 | chr16-37070879-INS-104 |
| chr16-37071317-INS-765 | chr16-37071317-INS-765 |
| chr16-37071901-INS-52 | chr16-37071901-INS-50 |
| chr16-37097751-INS-497 | chr16-37097751-INS-497 |
| chr16-37107708-INS-70 | chr16-37107708-INS-80 |
| chr16-37150169-INS-60 | chr16-37150169-INS-50 |
| chr16-37159157-INS-7342 | chr16-37159158-INS-7344 |
| chr16-37170726-INS-345 | chr16-37170726-INS-346 |
| chr16-37194450-INS-6447 | chr16-37194451-INS-6447 |
| chr16-37240481-INS-294 | chr16-37240481-INS-294 |
| chr16-37287963-INS-76 | chr16-37287960-INS-102 |
| chr16-37300416-INS-69 | chr16-37300416-INS-50 |
| chr16-37388601-INS-68 | chr16-37388601-INS-68 |
| chr16-37466432-INS-7229 | chr16-37466433-INS-7227 |
| chr16-37488872-INS-2760 | chr16-37488873-INS-2761 |
| chr16-37490240-INS-1860 | chr16-37490241-INS-1860 |
| chr16-37501658-INS-118 | chr16-37501651-INS-150 |
| chr16-37566666-INS-1067 | chr16-37566667-INS-1069 |
| chr16-37568233-INS-6434 | chr16-37568234-INS-6434 |
| chr16-37574337-INS-79 | chr16-37574337-INS-57 |
| chr16-37587573-INS-104 | chr16-37587561-INS-104 |
| chr16-37650746-INS-52 | chr16-37650746-INS-50 |
| chr16-37656311-INS-100 | chr16-37656311-INS-100 |
| chr16-37665481-INS-9045 | chr16-37665482-INS-9045 |
| chr16-37674222-INS-489 | chr16-37674222-INS-489 |
| chr16-37677256-INS-66 | chr16-37677256-INS-59 |
| chr16-37819485-INS-62 | chr16-37819487-INS-62 |
| chr16-37840599-INS-240 | chr16-37840599-INS-240 |
| chr16-37849772-INS-60 | chr16-37849742-INS-60 |
| chr16-37855908-INS-212 | chr16-37855908-INS-212 |
| chr16-37857079-INS-211 | chr16-37857079-INS-212 |
| chr16-37866323-INS-170 | chr16-37866323-INS-170 |
| chr16-37875642-INS-57 | chr16-37875642-INS-57 |
| chr16-37995779-INS-405 | chr16-37995779-INS-405 |
| chr16-37998324-INS-54 | chr16-37998324-INS-58 |
| chr16-38006976-INS-62 | chr16-38006976-INS-62 |
| chr16-38014890-INS-84 | chr16-38014873-INS-72 |
| chr16-38083661-INS-180 | chr16-38083661-INS-181 |
| chr16-38087819-INS-78 | chr16-38087820-INS-66 |
| chr16-38116203-INS-4991 | chr16-38116204-INS-4995 |
| chr16-38142159-INS-485 | chr16-38142159-INS-437 |
| chr16-38154924-INS-1234 | chr16-38154925-INS-1235 |
| chr16-38159417-INS-226 | chr16-38159417-INS-222 |
| chr16-38184696-INS-780 | chr16-38184696-INS-804 |
| chr16-38188832-INS-81 | chr16-38188832-INS-81 |
| chr16-38278530-INS-165 | chr16-38278501-INS-163 |
| chr16-38298136-INS-52 | chr16-38298136-INS-56 |
| chr16-38331480-INS-167 | chr16-38331480-INS-167 |
| chr16-38336206-INS-214 | chr16-38336206-INS-214 |
| chr16-38359261-INS-57 | chr16-38359261-INS-57 |
| chr16-38363759-INS-143 | chr16-38363760-INS-107 |
| chr16-38372364-INS-189 | chr16-38372364-INS-189 |
| chr16-38404048-INS-119 | chr16-38404048-INS-119 |
| chr16-38410486-INS-1888 | chr16-38410487-INS-1888 |
| chr16-38414216-INS-76 | chr16-38414216-INS-76 |
| chr16-38432610-INS-72 | chr16-38432610-INS-74 |
| chr16-38442713-INS-61 | chr16-38442713-INS-61 |
| chr16-38481640-INS-111 | chr16-38481640-INS-111 |
| chr16-38505152-INS-66 | chr16-38505152-INS-66 |
| chr16-38545364-INS-76 | chr16-38545364-INS-78 |
| chr16-38554416-INS-61 | chr16-38554407-INS-60 |
| chr16-38560354-INS-2322 | chr16-38560393-INS-2284 |
| chr16-38586854-INS-700 | chr16-38586854-INS-700 |
| chr16-38592092-INS-125 | chr16-38592104-INS-126 |
| chr16-38602022-INS-159 | chr16-38602022-INS-159 |
| chr16-38604535-INS-114 | chr16-38604509-INS-115 |
| chr16-38609816-INS-111 | chr16-38609811-INS-105 |
| chr16-38627160-INS-6303 | chr16-38627161-INS-6305 |
| chr16-38715690-INS-1257 | chr16-38715691-INS-1258 |
| chr16-38727433-INS-74 | chr16-38727433-INS-74 |
| chr16-38745740-INS-7188 | chr16-38745741-INS-7188 |
| chr16-38747883-INS-72 | chr16-38747883-INS-72 |
| chr16-38749609-INS-122 | chr16-38749609-INS-124 |
| chr16-38793890-INS-236 | chr16-38793890-INS-236 |
| chr16-38794278-INS-1803 | chr16-38794279-INS-1775 |
| chr16-38807807-INS-387 | chr16-38807807-INS-392 |

| | |
|---|---|
| chr16-38928522-INS-990 | chr16-38928522-INS-990 |
| chr16-38939871-INS-95 | chr16-38939872-INS-78 |
| chr16-38963132-INS-53 | chr16-38963132-INS-56 |
| chr16-38972950-INS-6273 | chr16-38972951-INS-6285 |
| chr16-38980242-INS-71 | chr16-38980242-INS-71 |
| chr16-39008965-INS-102 | chr16-39008965-INS-102 |
| chr16-39069139-INS-6409 | chr16-39069140-INS-6415 |
| chr16-39082135-INS-3743 | chr16-39082136-INS-3738 |
| chr16-39097675-INS-400 | chr16-39097680-INS-415 |
| chr16-39115313-INS-76 | chr16-39115313-INS-78 |
| chr16-39118626-INS-400 | chr16-39118626-INS-400 |
| chr16-39142260-INS-83 | chr16-39142246-INS-92 |
| chr16-39148598-INS-861 | chr16-39148598-INS-861 |
| chr16-39157521-INS-53 | chr16-39157521-INS-50 |
| chr16-39175980-INS-1306 | chr16-39175981-INS-1306 |
| chr16-39178308-INS-342 | chr16-39178308-INS-354 |
| chr16-39183576-INS-50 | chr16-39183576-INS-50 |
| chr16-39224890-INS-1304 | chr16-39224891-INS-1304 |
| chr16-39274052-INS-72 | chr16-39274052-INS-72 |
| chr16-39283734-INS-2783 | chr16-39283735-INS-2783 |
| chr16-39347534-INS-53 | chr16-39347534-INS-53 |
| chr16-39399475-INS-99 | chr16-39399475-INS-99 |
| chr16-39480325-INS-350 | chr16-39480348-INS-352 |
| chr16-39521611-INS-2830 | chr16-39521612-INS-2830 |
| chr16-39546206-INS-65 | chr16-39546206-INS-60 |
| chr16-39592627-INS-284 | chr16-39592627-INS-284 |
| chr16-39595657-INS-764 | chr16-39595657-INS-764 |
| chr16-39596932-INS-171 | chr16-39596932-INS-171 |
| chr16-39604316-INS-55 | chr16-39604316-INS-55 |
| chr16-39644976-INS-498 | chr16-39644976-INS-498 |
| chr16-39657297-INS-3478 | chr16-39657298-INS-3454 |
| chr16-39693914-INS-769 | chr16-39693914-INS-769 |
| chr16-39703276-INS-3616 | chr16-39703277-INS-3616 |
| chr16-39719373-INS-108 | chr16-39719373-INS-116 |
| chr16-39735948-INS-85 | chr16-39735948-INS-85 |
| chr16-39752497-INS-230 | chr16-39752497-INS-203 |
| chr16-39777730-INS-216 | chr16-39777730-INS-216 |
| chr16-39790930-INS-70 | chr16-39790908-INS-52 |
| chr16-39847450-INS-151 | chr16-39847457-INS-135 |
| chr16-39856007-INS-7866 | chr16-39856008-INS-7868 |
| chr16-39862346-INS-150 | chr16-39862346-INS-151 |
| chr16-39863788-INS-67 | chr16-39863788-INS-66 |
| chr16-39874467-INS-197 | chr16-39874467-INS-197 |
| chr16-39932295-INS-69 | chr16-39932275-INS-51 |
| chr16-39935408-INS-770 | chr16-39935408-INS-770 |
| chr16-39942501-INS-51 | chr16-39942501-INS-65 |
| chr16-39970912-INS-215 | chr16-39970912-INS-236 |
| chr16-39978754-INS-7528 | chr16-39978755-INS-7534 |
| chr16-39981583-INS-113 | chr16-39981583-INS-115 |
| chr16-39992433-INS-89 | chr16-39992433-INS-91 |
| chr16-40021664-INS-118 | chr16-40021664-INS-119 |
| chr16-40035140-INS-1165 | chr16-40035141-INS-1165 |
| chr16-40067801-INS-364 | chr16-40067823-INS-317 |
| chr16-40110354-INS-451 | chr16-40110344-INS-451 |
| chr16-40117991-INS-66 | chr16-40117991-INS-72 |
| chr16-40122371-INS-134 | chr16-40122371-INS-134 |
| chr16-40179494-INS-52 | chr16-40179490-INS-56 |
| chr16-40196433-INS-105 | chr16-40196433-INS-100 |
| chr16-40221523-INS-58 | chr16-40221523-INS-68 |
| chr16-40231256-INS-624 | chr16-40231256-INS-624 |
| chr16-40242122-INS-94 | chr16-40242122-INS-88 |
| chr16-40266306-INS-102 | chr16-40266306-INS-77 |
| chr16-40276803-INS-68 | chr16-40276754-INS-72 |
| chr16-40281762-INS-196 | chr16-40281763-INS-196 |
| chr16-40313832-INS-4415 | chr16-40313833-INS-4415 |
| chr16-40320075-INS-168 | chr16-40320075-INS-168 |
| chr16-40321047-INS-220 | chr16-40321047-INS-220 |
| chr16-40364495-INS-68 | chr16-40364495-INS-100 |
| chr16-40449595-INS-78 | chr16-40449595-INS-78 |
| chr16-40450016-INS-291 | chr16-40450016-INS-291 |
| chr16-40450348-INS-6865 | chr16-40450349-INS-6865 |
| chr16-40490007-INS-418 | chr16-40490007-INS-418 |
| chr16-40491665-INS-51 | chr16-40491689-INS-51 |
| chr16-40506533-INS-52 | chr16-40506533-INS-52 |
| chr16-40523843-INS-3649 | chr16-40523844-INS-3649 |
| chr16-40525069-INS-1766 | chr16-40525070-INS-1766 |
| chr16-40554054-INS-53 | chr16-40554069-INS-64 |

| | |
|---|---|
| chr16-40587437-INS-6244 | chr16-40587438-INS-6245 |
| chr16-40598723-INS-1389 | chr16-40598724-INS-1389 |
| chr16-40600109-INS-1603 | chr16-40600110-INS-1604 |
| chr16-40601914-INS-2586 | chr16-40601915-INS-2587 |
| chr16-40603947-INS-1622 | chr16-40603948-INS-1622 |
| chr16-40610806-INS-113 | chr16-40610805-INS-114 |
| chr16-40649063-INS-289 | chr16-40649063-INS-289 |
| chr16-40657558-INS-395 | chr16-40657556-INS-367 |
| chr16-40715201-INS-7330 | chr16-40715202-INS-7330 |
| chr16-40739577-INS-115 | chr16-40739577-INS-127 |
| chr16-40867714-INS-4737 | chr16-40867715-INS-4737 |
| chr16-40873422-INS-646 | chr16-40873422-INS-646 |
| chr16-40975121-INS-624 | chr16-40975121-INS-624 |
| chr16-41025090-INS-52 | chr16-41025071-INS-98 |
| chr16-41040736-INS-107 | chr16-41040736-INS-107 |
| chr16-41044017-INS-1521 | chr16-41044018-INS-1522 |
| chr16-41074241-INS-1712 | chr16-41074242-INS-1712 |
| chr16-41085325-INS-4530 | chr16-41085326-INS-4530 |
| chr16-41100040-INS-63 | chr16-41100040-INS-63 |
| chr16-41141468-INS-62 | chr16-41141468-INS-62 |
| chr16-41164936-INS-6454 | chr16-41164937-INS-6455 |
| chr16-41207364-INS-1511 | chr16-41207365-INS-1511 |
| chr16-41211068-INS-396 | chr16-41211068-INS-396 |
| chr16-41244275-INS-82 | chr16-41244275-INS-56 |
| chr16-41261552-INS-436 | chr16-41261552-INS-419 |
| chr16-41286377-INS-64 | chr16-41286377-INS-66 |
| chr16-41335394-INS-54 | chr16-41335402-INS-58 |
| chr16-41355370-INS-6440 | chr16-41355371-INS-6440 |
| chr16-41368648-INS-391 | chr16-41368648-INS-391 |
| chr16-41374076-INS-952 | chr16-41374076-INS-952 |
| chr16-41404011-INS-225 | chr16-41404011-INS-226 |
| chr16-41450568-INS-400 | chr16-41450569-INS-399 |
| chr16-41513969-INS-114 | chr16-41513970-INS-162 |
| chr16-41579947-INS-227 | chr16-41579947-INS-228 |
| chr16-41603037-INS-187 | chr16-41603037-INS-188 |
| chr16-41664917-INS-973 | chr16-41664917-INS-965 |
| chr16-41676863-INS-1886 | chr16-41676864-INS-1891 |
| chr16-41742619-INS-794 | chr16-41742619-INS-806 |
| chr16-41787068-INS-126 | chr16-41787041-INS-137 |
| chr16-41801693-INS-1927 | chr16-41801694-INS-1917 |
| chr16-41804196-INS-4797 | chr16-41804197-INS-4797 |
| chr16-41815584-INS-882 | chr16-41815584-INS-882 |
| chr16-41833286-INS-58 | chr16-41833286-INS-50 |
| chr16-41848906-INS-2288 | chr16-41848907-INS-2289 |
| chr16-41849108-INS-395 | chr16-41849108-INS-395 |
| chr16-41910330-INS-54 | chr16-41910330-INS-54 |
| chr16-41935276-INS-66 | chr16-41935276-INS-66 |
| chr16-41939905-INS-58 | chr16-41939905-INS-58 |
| chr16-41980947-INS-58 | chr16-41980948-INS-86 |
| chr16-41985002-INS-702 | chr16-41985002-INS-702 |
| chr16-41987392-INS-64 | chr16-41987392-INS-60 |
| chr16-42022384-INS-130 | chr16-42022384-INS-130 |
| chr16-42025500-INS-94 | chr16-42025512-INS-60 |
| chr16-42025505-INS-7192 | chr16-42025506-INS-7192 |
| chr16-42040254-INS-199 | chr16-42040254-INS-199 |
| chr16-42053197-INS-236 | chr16-42053197-INS-241 |
| chr16-42069816-INS-75 | chr16-42069816-INS-122 |
| chr16-42103242-INS-76 | chr16-42103242-INS-76 |
| chr16-42116914-INS-54 | chr16-42116914-INS-54 |
| chr16-42128378-INS-253 | chr16-42128378-INS-251 |
| chr16-42146619-INS-317 | chr16-42146619-INS-317 |
| chr16-42174522-INS-83 | chr16-42174522-INS-84 |
| chr16-42178288-INS-50 | chr16-42178282-INS-58 |
| chr16-42210091-INS-292 | chr16-42210091-INS-307 |
| chr16-42220700-INS-318 | chr16-42220700-INS-318 |
| chr16-42288863-INS-4555 | chr16-42288864-INS-4555 |
| chr16-42293525-INS-81 | chr16-42293525-INS-81 |
| chr16-42304853-INS-164 | chr16-42304853-INS-169 |
| chr16-42320932-INS-6413 | chr16-42320933-INS-6411 |
| chr16-42357924-INS-161 | chr16-42357924-INS-161 |
| chr16-42369932-INS-60 | chr16-42369932-INS-54 |
| chr16-42423458-INS-203 | chr16-42423458-INS-202 |
| chr16-42456244-INS-6614 | chr16-42456245-INS-6614 |
| chr16-42481424-INS-2142 | chr16-42481425-INS-2142 |
| chr16-42494749-INS-56 | chr16-42494749-INS-57 |
| chr16-42505388-INS-88 | chr16-42505388-INS-70 |
| chr16-42506603-INS-449 | chr16-42506603-INS-449 |

| | |
|---|---|
| chr16-42546076-INS-6132 | chr16-42546077-INS-6132 |
| chr16-42557519-INS-98 | chr16-42557519-INS-99 |
| chr16-42576124-INS-155 | chr16-42576124-INS-160 |
| chr16-42619284-INS-6921 | chr16-42619285-INS-6922 |
| chr16-42765432-INS-81 | chr16-42765432-INS-70 |
| chr16-42768076-INS-278 | chr16-42768076-INS-278 |
| chr16-42795865-INS-111 | chr16-42795865-INS-111 |
| chr16-42831706-INS-118 | chr16-42831706-INS-120 |
| chr16-42835671-INS-60 | chr16-42835671-INS-78 |
| chr16-42861221-INS-206 | chr16-42861221-INS-206 |
| chr16-42953195-INS-102 | chr16-42953195-INS-102 |
| chr16-43015824-INS-64 | chr16-43015824-INS-52 |
| chr16-43057954-INS-191 | chr16-43057957-INS-202 |
| chr16-43146728-INS-2736 | chr16-43146729-INS-2736 |
| chr16-43161358-INS-56 | chr16-43161358-INS-56 |
| chr16-43231676-INS-510 | chr16-43231676-INS-510 |
| chr16-43347462-INS-109 | chr16-43347462-INS-109 |
| chr16-43376092-INS-55 | chr16-43376070-INS-56 |
| chr16-43468574-INS-54 | chr16-43468583-INS-54 |
| chr16-43551352-INS-481 | chr16-43551352-INS-481 |
| chr16-43559587-INS-106 | chr16-43559587-INS-106 |
| chr16-43562962-INS-828 | chr16-43562962-INS-828 |
| chr16-43570216-INS-271 | chr16-43570216-INS-295 |
| chr16-43628655-INS-116 | chr16-43628655-INS-116 |
| chr16-43657935-INS-213 | chr16-43657935-INS-213 |
| chr16-43680283-INS-52 | chr16-43680283-INS-52 |
| chr16-43691022-INS-422 | chr16-43691022-INS-415 |
| chr16-43743597-INS-523 | chr16-43743596-INS-514 |
| chr16-43837695-INS-94 | chr16-43837695-INS-108 |
| chr16-43886792-INS-74 | chr16-43886792-INS-77 |
| chr16-43917028-INS-220 | chr16-43917028-INS-221 |
| chr16-43937632-INS-408 | chr16-43937632-INS-406 |
| chr16-43939649-INS-70 | chr16-43939649-INS-70 |
| chr16-43941185-INS-56 | chr16-43941164-INS-53 |
| chr16-43947079-INS-176 | chr16-43947080-INS-154 |
| chr16-43975685-INS-424 | chr16-43975685-INS-426 |
| chr16-43991840-INS-66 | chr16-43991840-INS-72 |
| chr16-44000729-INS-69 | chr16-44000729-INS-70 |
| chr16-44034909-INS-104 | chr16-44034909-INS-104 |
| chr16-44051447-INS-73 | chr16-44051447-INS-73 |
| chr16-44087606-INS-104 | chr16-44087606-INS-104 |
| chr16-44101681-INS-74 | chr16-44101681-INS-76 |
| chr16-44106644-INS-4351 | chr16-44106645-INS-4351 |
| chr16-44115463-INS-57 | chr16-44115463-INS-54 |
| chr16-44121095-INS-63 | chr16-44121093-INS-69 |
| chr16-44139558-INS-195 | chr16-44139558-INS-195 |
| chr16-44141996-INS-87 | chr16-44141996-INS-87 |
| chr16-44145466-INS-54 | chr16-44145466-INS-50 |
| chr16-44148813-INS-102 | chr16-44148813-INS-150 |
| chr16-44155869-INS-6432 | chr16-44155870-INS-6432 |
| chr16-44165158-INS-51 | chr16-44165165-INS-66 |
| chr16-44170568-INS-210 | chr16-44170568-INS-210 |
| chr16-44189442-INS-4077 | chr16-44189443-INS-4077 |
| chr16-44192281-INS-186 | chr16-44192281-INS-207 |
| chr16-44203988-INS-116 | chr16-44203988-INS-117 |
| chr16-44261060-INS-57 | chr16-44261060-INS-57 |
| chr16-44283230-INS-55 | chr16-44283230-INS-64 |
| chr16-44285377-INS-169 | chr16-44285377-INS-171 |
| chr16-44286852-INS-662 | chr16-44286852-INS-662 |
| chr16-44291519-INS-688 | chr16-44291519-INS-688 |
| chr16-44294326-INS-58 | chr16-44294326-INS-58 |
| chr16-44342730-INS-574 | chr16-44342730-INS-593 |
| chr16-44357796-INS-59 | chr16-44357758-INS-74 |
| chr16-44388356-INS-126 | chr16-44388356-INS-90 |
| chr16-44402200-INS-8419 | chr16-44402201-INS-8418 |
| chr16-44414984-INS-208 | chr16-44414984-INS-204 |
| chr16-44431334-INS-322 | chr16-44431334-INS-334 |
| chr16-44444208-INS-52 | chr16-44444161-INS-58 |
| chr16-44461346-INS-126 | chr16-44461346-INS-126 |
| chr16-44497278-INS-54 | chr16-44497272-INS-84 |
| chr16-44506257-INS-106 | chr16-44506257-INS-104 |
| chr16-44517202-INS-373 | chr16-44517202-INS-377 |
| chr16-44527003-INS-68 | chr16-44527039-INS-76 |
| chr16-44575707-INS-5248 | chr16-44575708-INS-5248 |
| chr16-44593689-INS-82 | chr16-44593693-INS-96 |
| chr16-44623750-INS-5178 | chr16-44623751-INS-5178 |
| chr16-44732055-INS-738 | chr16-44732055-INS-738 |

| | |
|---|---|
| chr16-44740445-INS-107 | chr16-44740445-INS-107 |
| chr16-44757909-INS-56 | chr16-44757909-INS-75 |
| chr16-44764540-INS-90 | chr16-44764546-INS-98 |
| chr16-44785770-INS-87 | chr16-44785770-INS-97 |
| chr16-44820643-INS-64 | chr16-44820643-INS-64 |
| chr16-44827135-INS-644 | chr16-44827137-INS-658 |
| chr16-44828325-INS-205 | chr16-44828325-INS-206 |
| chr16-44847843-INS-56 | chr16-44847841-INS-56 |
| chr16-44854663-INS-429 | chr16-44854663-INS-429 |
| chr16-44901636-INS-2801 | chr16-44901637-INS-2801 |
| chr16-44921832-INS-141 | chr16-44921824-INS-102 |
| chr16-44938261-INS-1001 | chr16-44938262-INS-1001 |
| chr16-44940625-INS-93 | chr16-44940607-INS-64 |
| chr16-44950891-INS-72 | chr16-44950891-INS-72 |
| chr16-44959154-INS-199 | chr16-44959154-INS-199 |
| chr16-44977508-INS-196 | chr16-44977508-INS-195 |
| chr16-45020573-INS-125 | chr16-45020580-INS-144 |
| chr16-45026935-INS-1014 | chr16-45026936-INS-1015 |
| chr16-45035853-INS-61 | chr16-45035845-INS-77 |
| chr16-45045513-INS-169 | chr16-45045513-INS-169 |
| chr16-45050825-INS-51 | chr16-45050815-INS-72 |
| chr16-45094868-INS-6219 | chr16-45094869-INS-6218 |
| chr16-45097039-INS-78 | chr16-45097039-INS-84 |
| chr16-45115175-INS-58 | chr16-45115175-INS-58 |
| chr16-45131430-INS-2916 | chr16-45131431-INS-2917 |
| chr16-45142663-INS-749 | chr16-45142663-INS-749 |
| chr16-45160291-INS-115 | chr16-45160291-INS-115 |
| chr16-45162866-INS-56 | chr16-45162834-INS-58 |
| chr16-45164062-INS-245 | chr16-45164062-INS-245 |
| chr16-45171071-INS-62 | chr16-45171071-INS-62 |
| chr16-45180962-INS-107 | chr16-45180962-INS-107 |
| chr16-45200667-INS-286 | chr16-45200667-INS-288 |
| chr16-45202855-INS-52 | chr16-45202856-INS-64 |
| chr16-45208776-INS-112 | chr16-45208776-INS-116 |
| chr16-45249060-INS-317 | chr16-45249060-INS-317 |
| chr16-45254703-INS-88 | chr16-45254693-INS-54 |
| chr16-45255432-INS-51 | chr16-45255435-INS-52 |
| chr16-45280321-INS-425 | chr16-45280276-INS-427 |
| chr16-45291745-INS-55 | chr16-45291746-INS-104 |
| chr16-45296316-INS-396 | chr16-45296316-INS-396 |
| chr16-45312224-INS-52 | chr16-45312224-INS-51 |
| chr16-45313109-INS-4980 | chr16-45313110-INS-4980 |
| chr16-45350133-INS-203 | chr16-45350134-INS-241 |
| chr16-45361001-INS-66 | chr16-45361000-INS-66 |
| chr16-45366233-INS-54 | chr16-45366233-INS-54 |
| chr16-45436213-INS-74 | chr16-45436246-INS-81 |
| chr16-45491513-INS-176 | chr16-45491513-INS-181 |
| chr16-45493593-INS-224 | chr16-45493593-INS-224 |
| chr16-45495719-INS-795 | chr16-45495719-INS-795 |
| chr16-45503239-INS-453 | chr16-45503239-INS-454 |
| chr16-45508408-INS-60 | chr16-45508430-INS-55 |
| chr16-45521827-INS-184 | chr16-45521827-INS-193 |
| chr16-45529129-INS-92 | chr16-45529129-INS-92 |
| chr16-45532299-INS-67 | chr16-45532299-INS-50 |
| chr16-45573904-INS-52 | chr16-45573904-INS-52 |
| chr16-45599656-INS-70 | chr16-45599646-INS-76 |
| chr16-45629006-INS-100 | chr16-45629002-INS-104 |
| chr16-45635397-INS-709 | chr16-45635397-INS-709 |
| chr16-45648329-INS-190 | chr16-45648329-INS-190 |
| chr16-45652030-INS-155 | chr16-45652030-INS-155 |
| chr16-45660803-INS-176 | chr16-45660765-INS-183 |
| chr16-45683550-INS-300 | chr16-45683550-INS-300 |
| chr16-45687726-INS-111 | chr16-45687726-INS-107 |
| chr16-45692364-INS-746 | chr16-45692364-INS-746 |
| chr16-45709207-INS-133 | chr16-45709207-INS-133 |
| chr16-45718671-INS-124 | chr16-45718671-INS-119 |
| chr16-45730890-INS-1744 | chr16-45730891-INS-1742 |
| chr16-45742680-INS-168 | chr16-45742680-INS-168 |
| chr16-45756257-INS-98 | chr16-45756257-INS-139 |
| chr16-45759828-INS-6442 | chr16-45759829-INS-6442 |
| chr16-45769485-INS-52 | chr16-45769485-INS-52 |
| chr16-45771207-INS-61 | chr16-45771169-INS-50 |
| chr16-45778748-INS-119 | chr16-45778748-INS-119 |
| chr16-45786646-INS-197 | chr16-45786646-INS-197 |
| chr16-45792145-INS-132 | chr16-45792145-INS-156 |
| chr16-45805876-INS-109 | chr16-45805876-INS-109 |
| chr16-45812328-INS-370 | chr16-45812328-INS-370 |

| | |
|---|---|
| chr16-45815016-INS-276 | chr16-45815016-INS-325 |
| chr16-45837239-INS-119 | chr16-45837239-INS-119 |
| chr16-45845947-INS-122 | chr16-45845947-INS-126 |
| chr16-45862219-INS-53 | chr16-45862219-INS-53 |
| chr16-45864411-INS-60 | chr16-45864361-INS-60 |
| chr16-45864531-INS-996 | chr16-45864531-INS-994 |
| chr16-45877784-INS-52 | chr16-45877784-INS-52 |
| chr16-45891116-INS-739 | chr16-45891115-INS-753 |
| chr16-45959519-INS-89 | chr16-45959519-INS-82 |
| chr16-45979598-INS-60 | chr16-45979598-INS-60 |
| chr16-45985213-INS-117 | chr16-45985215-INS-113 |
| chr16-45996894-INS-180 | chr16-45996894-INS-180 |
| chr16-46004959-INS-400 | chr16-46004959-INS-400 |
| chr16-46006369-INS-269 | chr16-46006369-INS-270 |
| chr16-46015050-INS-4156 | chr16-46015051-INS-4156 |
| chr16-46050544-INS-8481 | chr16-46050545-INS-8481 |
| chr16-46079949-INS-3547 | chr16-46079950-INS-3544 |
| chr16-46080167-INS-1017 | chr16-46080168-INS-1017 |
| chr16-46092513-INS-66 | chr16-46092513-INS-76 |
| chr16-46107679-INS-60 | chr16-46107679-INS-62 |
| chr16-46114804-INS-78 | chr16-46114804-INS-78 |
| chr16-46124133-INS-85 | chr16-46124140-INS-52 |
| chr16-46126140-INS-103 | chr16-46126140-INS-102 |
| chr16-46132807-INS-199 | chr16-46132807-INS-198 |
| chr16-46133514-INS-339 | chr16-46133518-INS-340 |
| chr16-46135356-INS-534 | chr16-46135356-INS-534 |
| chr16-46140787-INS-148 | chr16-46140787-INS-148 |
| chr16-46150445-INS-55 | chr16-46150445-INS-50 |
| chr16-46152134-INS-204 | chr16-46152134-INS-204 |
| chr16-46159937-INS-354 | chr16-46159937-INS-354 |
| chr16-46170183-INS-504 | chr16-46170183-INS-504 |
| chr16-46183393-INS-68 | chr16-46183395-INS-69 |
| chr16-46233824-INS-345 | chr16-46233824-INS-345 |
| chr16-46243669-INS-196 | chr16-46243669-INS-197 |
| chr16-46273428-INS-244 | chr16-46273428-INS-293 |
| chr16-46283508-INS-66 | chr16-46283508-INS-66 |
| chr16-46303111-INS-247 | chr16-46303111-INS-248 |
| chr16-46314707-INS-128 | chr16-46314712-INS-125 |
| chr16-46323282-INS-52 | chr16-46323282-INS-52 |
| chr16-46326262-INS-329 | chr16-46326262-INS-329 |
| chr16-46331166-INS-198 | chr16-46331166-INS-198 |
| chr16-46340193-INS-6411 | chr16-46340194-INS-6408 |
| chr16-46372339-INS-70 | chr16-46372339-INS-70 |
| chr16-46376279-INS-790 | chr16-46376279-INS-790 |
| chr16-46388213-INS-6862 | chr16-46388225-INS-6860 |
| chr16-46403159-INS-187 | chr16-46403159-INS-188 |
| chr16-46437363-INS-58 | chr16-46437355-INS-50 |
| chr16-46450333-INS-6234 | chr16-46450334-INS-6234 |
| chr16-46496362-INS-53 | chr16-46496362-INS-52 |
| chr16-46501683-INS-70 | chr16-46501683-INS-57 |
| chr16-46502963-INS-119 | chr16-46502963-INS-115 |
| chr16-46535927-INS-146 | chr16-46535927-INS-146 |
| chr16-46572442-INS-4801 | chr16-46572443-INS-4801 |
| chr16-46609049-INS-440 | chr16-46609049-INS-440 |
| chr16-46628605-INS-54 | chr16-46628557-INS-58 |
| chr16-46702204-INS-590 | chr16-46702204-INS-590 |
| chr16-46755853-INS-102 | chr16-46755853-INS-58 |
| chr16-46757399-INS-56 | chr16-46757399-INS-56 |
| chr16-46759346-INS-358 | chr16-46759346-INS-357 |
| chr16-46774459-INS-61 | chr16-46774459-INS-52 |
| chr16-46778904-INS-6390 | chr16-46778905-INS-6390 |
| chr16-46795209-INS-50 | chr16-46795209-INS-52 |
| chr16-46814232-INS-11137 | chr16-46814233-INS-11138 |
| chr16-46822360-INS-68 | chr16-46822360-INS-72 |
| chr16-46841464-INS-67 | chr16-46841464-INS-70 |
| chr16-46851244-INS-1027 | chr16-46851245-INS-1027 |
| chr16-46853311-INS-68 | chr16-46853311-INS-104 |
| chr16-46853942-INS-503 | chr16-46853938-INS-504 |
| chr16-46854987-INS-99 | chr16-46854987-INS-99 |
| chr16-46871474-INS-60 | chr16-46871474-INS-60 |
| chr16-46875165-INS-63 | chr16-46875159-INS-71 |
| chr16-46929650-INS-63 | chr16-46929650-INS-63 |
| chr16-46930969-INS-67 | chr16-46930969-INS-87 |
| chr16-46945237-INS-80 | chr16-46945237-INS-80 |
| chr16-46947555-INS-602 | chr16-46947555-INS-598 |
| chr16-46957864-INS-212 | chr16-46957864-INS-212 |
| chr16-46964380-INS-470 | chr16-46964380-INS-470 |

| | |
|---|---|
| chr16-46969008-INS-201 | chr16-46969008-INS-201 |
| chr16-46980069-INS-900 | chr16-46980069-INS-911 |
| chr16-47034447-INS-86 | chr16-47034447-INS-81 |
| chr16-47045619-INS-133 | chr16-47045619-INS-159 |
| chr16-47055383-INS-6235 | chr16-47055384-INS-6235 |
| chr16-47059974-INS-60 | chr16-47059974-INS-56 |
| chr16-47172805-INS-586 | chr16-47172805-INS-588 |
| chr16-47178456-INS-62 | chr16-47178456-INS-62 |
| chr16-47180326-INS-258 | chr16-47180330-INS-255 |
| chr16-47184625-INS-62 | chr16-47184625-INS-62 |
| chr16-47218516-INS-6233 | chr16-47218517-INS-6233 |
| chr16-47257030-INS-55 | chr16-47257030-INS-55 |
| chr16-47269742-INS-52 | chr16-47269742-INS-52 |
| chr16-47317203-INS-175 | chr16-47317203-INS-175 |
| chr16-47323599-INS-158 | chr16-47323599-INS-158 |
| chr16-47328811-INS-56 | chr16-47328811-INS-60 |
| chr16-47330663-INS-6460 | chr16-47330664-INS-6460 |
| chr16-47341148-INS-61 | chr16-47341149-INS-61 |
| chr16-47343448-INS-4706 | chr16-47343449-INS-4706 |
| chr16-47400899-INS-125 | chr16-47400899-INS-125 |
| chr16-47413903-INS-56 | chr16-47413903-INS-60 |
| chr16-47421067-INS-1615 | chr16-47421068-INS-1614 |
| chr16-47424045-INS-534 | chr16-47424045-INS-534 |
| chr16-47428321-INS-63 | chr16-47428321-INS-63 |
| chr16-47488028-INS-51 | chr16-47488028-INS-51 |
| chr16-47496899-INS-90 | chr16-47496899-INS-90 |
| chr16-47516905-INS-158 | chr16-47516910-INS-172 |
| chr16-47596238-INS-100 | chr16-47596238-INS-108 |
| chr16-47606079-INS-84 | chr16-47606072-INS-85 |
| chr16-47627672-INS-956 | chr16-47627672-INS-955 |
| chr16-47629367-INS-58 | chr16-47629367-INS-58 |
| chr16-47671355-INS-52 | chr16-47671355-INS-50 |
| chr16-47672939-INS-54 | chr16-47672939-INS-58 |
| chr16-47699602-INS-155 | chr16-47699602-INS-109 |
| chr16-47714634-INS-177 | chr16-47714633-INS-140 |
| chr16-47748287-INS-498 | chr16-47748287-INS-498 |
| chr16-47752712-INS-791 | chr16-47752712-INS-792 |
| chr16-47768679-INS-53 | chr16-47768679-INS-50 |
| chr16-47772762-INS-961 | chr16-47772762-INS-979 |
| chr16-47779830-INS-58 | chr16-47779830-INS-74 |
| chr16-47792903-INS-74 | chr16-47792903-INS-86 |
| chr16-47795995-INS-692 | chr16-47795995-INS-693 |
| chr16-47830752-INS-60 | chr16-47830752-INS-60 |
| chr16-47864446-INS-216 | chr16-47864446-INS-216 |
| chr16-47867197-INS-55 | chr16-47867189-INS-50 |
| chr16-47867313-INS-4750 | chr16-47867314-INS-4749 |
| chr16-47875362-INS-226 | chr16-47875362-INS-226 |
| chr16-47890228-INS-151 | chr16-47890228-INS-145 |
| chr16-47893931-INS-526 | chr16-47893931-INS-526 |
| chr16-47894879-INS-54 | chr16-47894879-INS-56 |
| chr16-47949051-INS-4346 | chr16-47949052-INS-4346 |
| chr16-47955850-INS-118 | chr16-47955850-INS-133 |
| chr16-47969927-INS-382 | chr16-47969927-INS-395 |
| chr16-47973945-INS-112 | chr16-47973945-INS-115 |
| chr16-47989369-INS-1726 | chr16-47989370-INS-1726 |
| chr16-47993349-INS-6780 | chr16-47993350-INS-6783 |
| chr16-48000940-INS-463 | chr16-48000940-INS-464 |
| chr16-48001766-INS-68 | chr16-48001766-INS-70 |
| chr16-48019021-INS-56 | chr16-48019021-INS-58 |
| chr16-48020281-INS-69 | chr16-48020281-INS-69 |
| chr16-48021701-INS-56 | chr16-48021701-INS-60 |
| chr16-48032089-INS-209 | chr16-48032089-INS-209 |
| chr16-48038055-INS-238 | chr16-48038056-INS-239 |
| chr16-48042725-INS-109 | chr16-48042725-INS-109 |
| chr16-48053807-INS-697 | chr16-48053807-INS-697 |
| chr16-48055613-INS-181 | chr16-48055613-INS-172 |
| chr16-48058896-INS-369 | chr16-48058896-INS-370 |
| chr16-48065826-INS-561 | chr16-48065823-INS-565 |
| chr16-48071044-INS-114 | chr16-48071044-INS-120 |
| chr16-48082775-INS-107 | chr16-48082774-INS-82 |
| chr16-48090590-INS-202 | chr16-48090590-INS-202 |
| chr16-48125351-INS-206 | chr16-48125351-INS-203 |
| chr16-48141239-INS-70 | chr16-48141238-INS-72 |
| chr16-48149172-INS-422 | chr16-48149172-INS-424 |
| chr16-48150404-INS-86 | chr16-48150404-INS-85 |
| chr16-48151382-INS-167 | chr16-48151383-INS-167 |
| chr16-48152728-INS-87 | chr16-48152728-INS-83 |

| | |
|---|---|
| chr16-48157277-INS-108 | chr16-48157276-INS-148 |
| chr16-48159327-INS-184 | chr16-48159327-INS-184 |
| chr16-48169058-INS-79 | chr16-48169059-INS-77 |
| chr16-48191082-INS-216 | chr16-48191082-INS-216 |
| chr16-48195278-INS-109 | chr16-48195281-INS-132 |
| chr16-48208913-INS-51 | chr16-48208909-INS-59 |
| chr16-48222494-INS-204 | chr16-48222494-INS-204 |
| chr16-48253257-INS-264 | chr16-48253254-INS-267 |
| chr16-48271398-INS-56 | chr16-48271398-INS-67 |
| chr16-48272404-INS-156 | chr16-48272404-INS-155 |
| chr16-48282173-INS-4475 | chr16-48282174-INS-4474 |
| chr16-48287768-INS-381 | chr16-48287768-INS-384 |
| chr16-48330128-INS-52 | chr16-48330128-INS-52 |
| chr16-48336578-INS-53 | chr16-48336578-INS-61 |
| chr16-48352190-INS-52 | chr16-48352190-INS-52 |
| chr16-48389424-INS-454 | chr16-48389424-INS-454 |
| chr16-48400666-INS-110 | chr16-48400649-INS-146 |
| chr16-48412913-INS-69 | chr16-48412913-INS-66 |
| chr16-48418889-INS-53 | chr16-48418889-INS-55 |
| chr16-48421514-INS-79 | chr16-48421514-INS-79 |
| chr16-48424061-INS-460 | chr16-48424061-INS-467 |
| chr16-48468869-INS-95 | chr16-48468867-INS-82 |
| chr16-48484958-INS-211 | chr16-48484958-INS-175 |
| chr16-48493739-INS-210 | chr16-48493739-INS-210 |
| chr16-48497141-INS-56 | chr16-48497141-INS-56 |
| chr16-48499315-INS-565 | chr16-48499315-INS-565 |
| chr16-48515519-INS-206 | chr16-48515519-INS-207 |
| chr16-48523673-INS-235 | chr16-48523673-INS-234 |
| chr16-48535895-INS-100 | chr16-48535895-INS-101 |
| chr16-48543567-INS-51 | chr16-48543542-INS-58 |
| chr16-48546137-INS-427 | chr16-48546137-INS-426 |
| chr16-48548923-INS-203 | chr16-48548923-INS-202 |
| chr16-48572061-INS-55 | chr16-48572061-INS-55 |
| chr16-48575497-INS-55 | chr16-48575497-INS-63 |
| chr16-48590808-INS-152 | chr16-48590808-INS-153 |
| chr16-48597572-INS-83 | chr16-48597572-INS-79 |
| chr16-48600156-INS-203 | chr16-48600156-INS-203 |
| chr16-48624506-INS-83 | chr16-48624506-INS-83 |
| chr16-48642395-INS-267 | chr16-48642395-INS-274 |
| chr16-48642879-INS-655 | chr16-48642879-INS-656 |
| chr16-48658177-INS-620 | chr16-48658177-INS-618 |
| chr16-48660080-INS-8343 | chr16-48660081-INS-8343 |
| chr16-48707933-INS-83 | chr16-48707941-INS-77 |
| chr16-48736815-INS-457 | chr16-48736815-INS-457 |
| chr16-48752664-INS-78 | chr16-48752664-INS-76 |
| chr16-48761176-INS-2282 | chr16-48761177-INS-2282 |
| chr16-48811266-INS-433 | chr16-48811266-INS-433 |
| chr16-48821562-INS-59 | chr16-48821562-INS-71 |
| chr16-48832434-INS-135 | chr16-48832434-INS-135 |
| chr16-48864242-INS-183 | chr16-48864242-INS-183 |
| chr16-48876806-INS-317 | chr16-48876806-INS-316 |
| chr16-48906157-INS-510 | chr16-48906157-INS-504 |
| chr16-48927003-INS-66 | chr16-48927003-INS-68 |
| chr16-48930507-INS-4008 | chr16-48930508-INS-4007 |
| chr16-48947092-INS-60 | chr16-48947092-INS-56 |
| chr16-48952196-INS-7146 | chr16-48952197-INS-7148 |
| chr16-48958919-INS-56 | chr16-48958919-INS-62 |
| chr16-48976171-INS-360 | chr16-48976171-INS-360 |
| chr16-49010931-INS-63 | chr16-49010931-INS-63 |
| chr16-49028713-INS-98 | chr16-49028713-INS-104 |
| chr16-49032102-INS-76 | chr16-49032092-INS-57 |
| chr16-49037335-INS-588 | chr16-49037335-INS-588 |
| chr16-49046486-INS-192 | chr16-49046486-INS-192 |
| chr16-49049913-INS-103 | chr16-49049913-INS-130 |
| chr16-49052834-INS-3309 | chr16-49052835-INS-3309 |
| chr16-49074406-INS-65 | chr16-49074406-INS-60 |
| chr16-49084327-INS-56 | chr16-49084327-INS-60 |
| chr16-49088935-INS-54 | chr16-49088935-INS-54 |
| chr16-49091842-INS-209 | chr16-49091842-INS-209 |
| chr16-49104043-INS-67 | chr16-49104044-INS-82 |
| chr16-49116381-INS-96 | chr16-49116381-INS-92 |
| chr16-49118356-INS-992 | chr16-49118356-INS-984 |
| chr16-49136453-INS-126 | chr16-49136453-INS-126 |
| chr16-49145790-INS-56 | chr16-49145790-INS-56 |
| chr16-49152544-INS-204 | chr16-49152544-INS-204 |
| chr16-49153527-INS-176 | chr16-49153527-INS-176 |
| chr16-49160604-INS-384 | chr16-49160604-INS-384 |

| | |
|---|---|
| chr16-49163022-INS-832 | chr16-49163022-INS-832 |
| chr16-49204450-INS-2145 | chr16-49204451-INS-2145 |
| chr16-49204451-INS-257 | chr16-49204451-INS-257 |
| chr16-49227272-INS-2298 | chr16-49227273-INS-2298 |
| chr16-49239286-INS-99 | chr16-49239299-INS-84 |
| chr16-49242941-INS-51 | chr16-49242941-INS-50 |
| chr16-49261683-INS-432 | chr16-49261683-INS-432 |
| chr16-49264325-INS-2145 | chr16-49264326-INS-2147 |
| chr16-49279111-INS-218 | chr16-49279111-INS-218 |
| chr16-49315776-INS-107 | chr16-49315781-INS-91 |
| chr16-49327198-INS-212 | chr16-49327198-INS-212 |
| chr16-49334618-INS-949 | chr16-49334618-INS-948 |
| chr16-49379877-INS-53 | chr16-49379877-INS-50 |
| chr16-49397059-INS-751 | chr16-49397059-INS-751 |
| chr16-49402050-INS-111 | chr16-49402050-INS-105 |
| chr16-49418723-INS-59 | chr16-49418727-INS-72 |
| chr16-49425442-INS-54 | chr16-49425442-INS-52 |
| chr16-49443281-INS-325 | chr16-49443278-INS-322 |
| chr16-49475285-INS-419 | chr16-49475285-INS-419 |
| chr16-49484805-INS-316 | chr16-49484805-INS-316 |
| chr16-49506264-INS-55 | chr16-49506262-INS-50 |
| chr16-49515095-INS-213 | chr16-49515095-INS-213 |
| chr16-49531069-INS-700 | chr16-49531069-INS-701 |
| chr16-49580577-INS-162 | chr16-49580569-INS-170 |
| chr16-49592345-INS-100 | chr16-49592345-INS-100 |
| chr16-49596815-INS-56 | chr16-49596815-INS-56 |
| chr16-49628584-INS-207 | chr16-49628584-INS-207 |
| chr16-49668829-INS-100 | chr16-49668829-INS-120 |
| chr16-49753733-INS-2121 | chr16-49753734-INS-2121 |
| chr16-49776693-INS-6664 | chr16-49776694-INS-6667 |
| chr16-49803192-INS-138 | chr16-49803192-INS-162 |
| chr16-49804567-INS-76 | chr16-49804567-INS-80 |
| chr16-49855796-INS-2292 | chr16-49855797-INS-2292 |
| chr16-49869837-INS-52 | chr16-49869837-INS-54 |
| chr16-49887568-INS-58 | chr16-49887568-INS-58 |
| chr16-49894688-INS-60 | chr16-49894688-INS-60 |
| chr16-49897664-INS-61 | chr16-49897664-INS-61 |
| chr16-49903471-INS-54 | chr16-49903471-INS-50 |
| chr16-49923176-INS-199 | chr16-49923176-INS-199 |
| chr16-49924163-INS-603 | chr16-49924163-INS-603 |
| chr16-49933143-INS-129 | chr16-49933143-INS-129 |
| chr16-49944522-INS-59 | chr16-49944521-INS-59 |
| chr16-49953742-INS-199 | chr16-49953742-INS-199 |
| chr16-49957723-INS-1432 | chr16-49957724-INS-1433 |
| chr16-49959274-INS-135 | chr16-49959274-INS-135 |
| chr16-49964449-INS-71 | chr16-49964452-INS-73 |
| chr16-49979660-INS-148 | chr16-49979662-INS-179 |
| chr16-49987229-INS-412 | chr16-49987230-INS-412 |
| chr16-50007705-INS-66 | chr16-50007705-INS-69 |
| chr16-50065091-INS-60 | chr16-50065091-INS-60 |
| chr16-50070722-INS-52 | chr16-50070722-INS-50 |
| chr16-50091213-INS-288 | chr16-50091213-INS-288 |
| chr16-50133003-INS-70 | chr16-50133003-INS-70 |
| chr16-50134419-INS-1436 | chr16-50134420-INS-1436 |
| chr16-50144178-INS-1851 | chr16-50144179-INS-1834 |
| chr16-50178097-INS-99 | chr16-50178097-INS-111 |
| chr16-50181887-INS-54 | chr16-50181887-INS-54 |
| chr16-50225013-INS-55 | chr16-50225013-INS-55 |
| chr16-50246116-INS-65 | chr16-50246116-INS-67 |
| chr16-50273265-INS-64 | chr16-50273265-INS-58 |
| chr16-50280284-INS-60 | chr16-50280239-INS-55 |
| chr16-50281469-INS-202 | chr16-50281469-INS-202 |
| chr16-50319954-INS-232 | chr16-50319948-INS-251 |
| chr16-50360698-INS-64 | chr16-50360698-INS-64 |
| chr16-50364818-INS-509 | chr16-50364831-INS-522 |
| chr16-50377452-INS-60 | chr16-50377452-INS-60 |
| chr16-50380726-INS-842 | chr16-50380726-INS-842 |
| chr16-50383185-INS-52 | chr16-50383185-INS-54 |
| chr16-50403177-INS-56 | chr16-50403196-INS-62 |
| chr16-50404703-INS-116 | chr16-50404703-INS-116 |
| chr16-50431250-INS-208 | chr16-50431250-INS-212 |
| chr16-50477641-INS-460 | chr16-50477641-INS-461 |
| chr16-50482209-INS-126 | chr16-50482209-INS-116 |
| chr16-50485060-INS-172 | chr16-50485060-INS-129 |
| chr16-50507448-INS-3056 | chr16-50507449-INS-3056 |
| chr16-50512787-INS-381 | chr16-50512787-INS-381 |
| chr16-50515855-INS-176 | chr16-50515855-INS-176 |

| | |
|---|---|
| chr16-50556193-INS-54 | chr16-50556193-INS-54 |
| chr16-50562389-INS-61 | chr16-50562410-INS-73 |
| chr16-50563415-INS-59 | chr16-50563406-INS-60 |
| chr16-50583472-INS-168 | chr16-50583472-INS-140 |
| chr16-50586589-INS-56 | chr16-50586589-INS-56 |
| chr16-50596827-INS-67 | chr16-50596827-INS-84 |
| chr16-50598275-INS-324 | chr16-50598275-INS-324 |
| chr16-50617049-INS-310 | chr16-50617049-INS-309 |
| chr16-50627516-INS-173 | chr16-50627516-INS-173 |
| chr16-50636144-INS-52 | chr16-50636144-INS-54 |
| chr16-50644351-INS-68 | chr16-50644351-INS-68 |
| chr16-50707321-INS-6405 | chr16-50707322-INS-6405 |
| chr16-50720586-INS-84 | chr16-50720586-INS-68 |
| chr16-50748190-INS-719 | chr16-50748190-INS-719 |
| chr16-50750737-INS-112 | chr16-50750737-INS-112 |
| chr16-50753135-INS-6485 | chr16-50753136-INS-6486 |
| chr16-50760145-INS-177 | chr16-50760145-INS-156 |
| chr16-50765295-INS-374 | chr16-50765295-INS-379 |
| chr16-50784833-INS-242 | chr16-50784833-INS-252 |
| chr16-50798700-INS-4488 | chr16-50798701-INS-4489 |
| chr16-50807480-INS-221 | chr16-50807480-INS-220 |
| chr16-50808673-INS-209 | chr16-50808673-INS-209 |
| chr16-50810829-INS-192 | chr16-50810829-INS-192 |
| chr16-50830800-INS-60 | chr16-50830808-INS-57 |
| chr16-50837371-INS-128 | chr16-50837371-INS-122 |
| chr16-50860207-INS-323 | chr16-50860207-INS-323 |
| chr16-50879120-INS-356 | chr16-50879120-INS-356 |
| chr16-50884912-INS-1903 | chr16-50884913-INS-1904 |
| chr16-50890666-INS-50 | chr16-50890665-INS-57 |
| chr16-50895582-INS-54 | chr16-50895582-INS-56 |
| chr16-50898781-INS-4353 | chr16-50898782-INS-4353 |
| chr16-50917401-INS-53 | chr16-50917401-INS-53 |
| chr16-50928423-INS-6488 | chr16-50928424-INS-6488 |
| chr16-50940762-INS-51 | chr16-50940762-INS-51 |
| chr16-50994609-INS-108 | chr16-50994609-INS-108 |
| chr16-51014332-INS-55 | chr16-51014332-INS-55 |
| chr16-51027374-INS-72 | chr16-51027374-INS-64 |
| chr16-51048799-INS-76 | chr16-51048799-INS-52 |
| chr16-51117081-INS-399 | chr16-51117081-INS-399 |
| chr16-51187737-INS-176 | chr16-51187737-INS-177 |
| chr16-51198617-INS-1059 | chr16-51198618-INS-1059 |
| chr16-51221542-INS-761 | chr16-51221542-INS-761 |
| chr16-51238816-INS-58 | chr16-51238816-INS-58 |
| chr16-51263530-INS-939 | chr16-51263530-INS-939 |
| chr16-51291774-INS-52 | chr16-51291774-INS-52 |
| chr16-51348455-INS-103 | chr16-51348455-INS-106 |
| chr16-51367333-INS-641 | chr16-51367333-INS-641 |
| chr16-51374592-INS-2842 | chr16-51374593-INS-2879 |
| chr16-51375414-INS-5246 | chr16-51375415-INS-5247 |
| chr16-51385805-INS-61 | chr16-51385805-INS-62 |
| chr16-51387442-INS-50 | chr16-51387442-INS-52 |
| chr16-51411614-INS-136 | chr16-51411614-INS-136 |
| chr16-51453662-INS-62 | chr16-51453662-INS-62 |
| chr16-51464001-INS-6188 | chr16-51464002-INS-6188 |
| chr16-51497725-INS-69 | chr16-51497725-INS-54 |
| chr16-51498888-INS-667 | chr16-51498888-INS-666 |
| chr16-51518499-INS-328 | chr16-51518499-INS-328 |
| chr16-51520134-INS-400 | chr16-51520134-INS-400 |
| chr16-51534358-INS-735 | chr16-51534358-INS-735 |
| chr16-51561718-INS-60 | chr16-51561714-INS-78 |
| chr16-51578066-INS-677 | chr16-51578067-INS-705 |
| chr16-51591134-INS-4790 | chr16-51591135-INS-4790 |
| chr16-51683115-INS-356 | chr16-51683115-INS-358 |
| chr16-51719723-INS-6442 | chr16-51719724-INS-6442 |
| chr16-51806164-INS-116 | chr16-51806144-INS-138 |
| chr16-51812687-INS-6180 | chr16-51812688-INS-6178 |
| chr16-51813355-INS-70 | chr16-51813355-INS-58 |
| chr16-51825077-INS-68 | chr16-51825079-INS-72 |
| chr16-51832168-INS-59 | chr16-51832174-INS-56 |
| chr16-51842422-INS-80 | chr16-51842422-INS-80 |
| chr16-51844638-INS-81 | chr16-51844638-INS-105 |
| chr16-51845474-INS-166 | chr16-51845476-INS-125 |
| chr16-51978908-INS-6662 | chr16-51978909-INS-6662 |
| chr16-51989330-INS-72 | chr16-51989330-INS-73 |
| chr16-51992383-INS-250 | chr16-51992383-INS-250 |
| chr16-52060535-INS-56 | chr16-52060535-INS-60 |
| chr16-52064076-INS-226 | chr16-52064076-INS-226 |

| | |
|---|---|
| chr16-52070044-INS-56 | chr16-52069999-INS-90 |
| chr16-52260068-INS-53 | chr16-52260068-INS-53 |
| chr16-52269937-INS-59 | chr16-52269937-INS-63 |
| chr16-52280436-INS-56 | chr16-52280436-INS-74 |
| chr16-52341122-INS-848 | chr16-52341122-INS-848 |
| chr16-52343376-INS-97 | chr16-52343376-INS-97 |
| chr16-52348094-INS-852 | chr16-52348094-INS-852 |
| chr16-52372869-INS-55 | chr16-52372869-INS-78 |
| chr16-52373651-INS-364 | chr16-52373651-INS-362 |
| chr16-52389179-INS-63 | chr16-52389179-INS-63 |
| chr16-52414314-INS-70 | chr16-52414314-INS-70 |
| chr16-52439478-INS-225 | chr16-52439478-INS-225 |
| chr16-52454989-INS-133 | chr16-52454989-INS-115 |
| chr16-52456017-INS-210 | chr16-52456018-INS-210 |
| chr16-52458376-INS-54 | chr16-52458357-INS-68 |
| chr16-52460299-INS-186 | chr16-52460299-INS-172 |
| chr16-52527084-INS-3574 | chr16-52527099-INS-3580 |
| chr16-52531084-INS-11729 | chr16-52531085-INS-11737 |
| chr16-52536458-INS-102 | chr16-52536454-INS-118 |
| chr16-52545134-INS-6198 | chr16-52545135-INS-6198 |
| chr16-52563401-INS-223 | chr16-52563401-INS-223 |
| chr16-52573652-INS-431 | chr16-52573652-INS-436 |
| chr16-52585902-INS-3419 | chr16-52585903-INS-3418 |
| chr16-52594807-INS-92 | chr16-52594808-INS-62 |
| chr16-52627189-INS-60 | chr16-52627189-INS-60 |
| chr16-52627227-INS-2281 | chr16-52627228-INS-2280 |
| chr16-52636579-INS-60 | chr16-52636585-INS-51 |
| chr16-52668470-INS-920 | chr16-52668470-INS-920 |
| chr16-52675963-INS-56 | chr16-52675963-INS-58 |
| chr16-52678510-INS-179 | chr16-52678508-INS-181 |
| chr16-52689608-INS-57 | chr16-52689609-INS-90 |
| chr16-52695763-INS-56 | chr16-52695752-INS-66 |
| chr16-52717202-INS-164 | chr16-52717202-INS-166 |
| chr16-52742298-INS-67 | chr16-52742280-INS-72 |
| chr16-52745616-INS-78 | chr16-52745616-INS-66 |
| chr16-52749193-INS-8144 | chr16-52749194-INS-8162 |
| chr16-52755858-INS-3181 | chr16-52755859-INS-3181 |
| chr16-52757154-INS-64 | chr16-52757154-INS-64 |
| chr16-52761616-INS-381 | chr16-52761608-INS-387 |
| chr16-52774851-INS-102 | chr16-52774851-INS-102 |
| chr16-52817818-INS-96 | chr16-52817796-INS-114 |
| chr16-52827840-INS-56 | chr16-52827840-INS-58 |
| chr16-52853114-INS-199 | chr16-52853114-INS-200 |
| chr16-52871794-INS-6984 | chr16-52871795-INS-6985 |
| chr16-52877511-INS-76 | chr16-52877511-INS-76 |
| chr16-52881665-INS-63 | chr16-52881665-INS-68 |
| chr16-52887136-INS-890 | chr16-52887136-INS-890 |
| chr16-52893293-INS-874 | chr16-52893293-INS-874 |
| chr16-52903073-INS-140 | chr16-52903091-INS-143 |
| chr16-52979001-INS-72 | chr16-52979001-INS-72 |
| chr16-53033810-INS-3261 | chr16-53033811-INS-3261 |
| chr16-53045749-INS-64 | chr16-53045774-INS-74 |
| chr16-53051035-INS-703 | chr16-53051035-INS-700 |
| chr16-53084304-INS-4839 | chr16-53084305-INS-4845 |
| chr16-53110123-INS-86 | chr16-53110123-INS-86 |
| chr16-53111132-INS-56 | chr16-53111083-INS-64 |
| chr16-53112569-INS-445 | chr16-53112569-INS-445 |
| chr16-53117103-INS-581 | chr16-53117103-INS-600 |
| chr16-53154542-INS-137 | chr16-53154542-INS-138 |
| chr16-53163138-INS-1012 | chr16-53163139-INS-1012 |
| chr16-53207921-INS-578 | chr16-53207921-INS-578 |
| chr16-53224851-INS-198 | chr16-53224851-INS-198 |
| chr16-53293236-INS-301 | chr16-53293236-INS-301 |
| chr16-53300814-INS-128 | chr16-53300814-INS-136 |
| chr16-53317645-INS-497 | chr16-53317645-INS-497 |
| chr16-53324004-INS-242 | chr16-53324004-INS-243 |
| chr16-53340177-INS-199 | chr16-53340177-INS-199 |
| chr16-53381926-INS-98 | chr16-53381926-INS-96 |
| chr16-53417889-INS-51 | chr16-53417889-INS-51 |
| chr16-53444378-INS-202 | chr16-53444378-INS-202 |
| chr16-53479637-INS-362 | chr16-53479634-INS-362 |
| chr16-53494927-INS-94 | chr16-53494927-INS-86 |
| chr16-53508973-INS-60 | chr16-53508973-INS-68 |
| chr16-53521100-INS-62 | chr16-53521100-INS-62 |
| chr16-53543143-INS-165 | chr16-53543143-INS-165 |
| chr16-53563972-INS-375 | chr16-53563972-INS-375 |
| chr16-53585261-INS-1870 | chr16-53585262-INS-1872 |

| | |
|---|---|
| chr16-53610373-INS-1510 | chr16-53610374-INS-1508 |
| chr16-53654590-INS-690 | chr16-53654590-INS-690 |
| chr16-53714224-INS-733 | chr16-53714224-INS-733 |
| chr16-53718338-INS-421 | chr16-53718338-INS-423 |
| chr16-53738808-INS-524 | chr16-53738808-INS-524 |
| chr16-53755776-INS-1931 | chr16-53755777-INS-1935 |
| chr16-53770424-INS-427 | chr16-53770424-INS-427 |
| chr16-53800111-INS-60 | chr16-53800138-INS-56 |
| chr16-53854069-INS-2916 | chr16-53854070-INS-2913 |
| chr16-53856089-INS-280 | chr16-53856090-INS-311 |
| chr16-53864885-INS-1927 | chr16-53864886-INS-1920 |
| chr16-53864886-INS-511 | chr16-53864886-INS-511 |
| chr16-53875157-INS-64 | chr16-53875163-INS-52 |
| chr16-53914054-INS-4004 | chr16-53914055-INS-4005 |
| chr16-53936908-INS-732 | chr16-53936908-INS-725 |
| chr16-53940323-INS-418 | chr16-53940323-INS-393 |
| chr16-53976826-INS-1153 | chr16-53976827-INS-1153 |
| chr16-53992473-INS-186 | chr16-53992473-INS-186 |
| chr16-54012940-INS-776 | chr16-54012940-INS-773 |
| chr16-54027539-INS-152 | chr16-54027539-INS-152 |
| chr16-54045893-INS-781 | chr16-54045893-INS-782 |
| chr16-54075675-INS-910 | chr16-54075675-INS-933 |
| chr16-54081775-INS-81 | chr16-54081775-INS-81 |
| chr16-54137108-INS-316 | chr16-54137108-INS-316 |
| chr16-54153985-INS-69 | chr16-54153985-INS-111 |
| chr16-54156935-INS-383 | chr16-54156907-INS-374 |
| chr16-54202936-INS-1406 | chr16-54202937-INS-1403 |
| chr16-54214943-INS-1899 | chr16-54214944-INS-1899 |
| chr16-54230082-INS-70 | chr16-54230082-INS-70 |
| chr16-54273116-INS-2447 | chr16-54273117-INS-2427 |
| chr16-54320189-INS-56 | chr16-54320189-INS-56 |
| chr16-54328068-INS-203 | chr16-54328068-INS-203 |
| chr16-54348014-INS-55 | chr16-54348014-INS-55 |
| chr16-54350703-INS-1447 | chr16-54350704-INS-1447 |
| chr16-54356946-INS-146 | chr16-54356946-INS-149 |
| chr16-54372774-INS-188 | chr16-54372774-INS-156 |
| chr16-54405336-INS-4488 | chr16-54405337-INS-4488 |
| chr16-54416885-INS-4037 | chr16-54416886-INS-4038 |
| chr16-54418254-INS-930 | chr16-54418254-INS-930 |
| chr16-54423623-INS-6712 | chr16-54423624-INS-6713 |
| chr16-54442804-INS-983 | chr16-54442804-INS-982 |
| chr16-54445270-INS-4180 | chr16-54445271-INS-4179 |
| chr16-54454989-INS-677 | chr16-54454989-INS-673 |
| chr16-54491807-INS-442 | chr16-54491807-INS-442 |
| chr16-54492391-INS-6394 | chr16-54492392-INS-6394 |
| chr16-54503775-INS-60 | chr16-54503797-INS-63 |
| chr16-54521957-INS-61 | chr16-54521957-INS-61 |
| chr16-54559305-INS-6568 | chr16-54559306-INS-6568 |
| chr16-54573189-INS-323 | chr16-54573189-INS-323 |
| chr16-54574041-INS-99 | chr16-54574042-INS-99 |
| chr16-54581014-INS-56 | chr16-54581014-INS-56 |
| chr16-54598051-INS-6215 | chr16-54598052-INS-6215 |
| chr16-54642162-INS-564 | chr16-54642162-INS-564 |
| chr16-54669199-INS-54 | chr16-54669199-INS-54 |
| chr16-54727976-INS-69 | chr16-54727976-INS-70 |
| chr16-54743213-INS-71 | chr16-54743213-INS-71 |
| chr16-54745242-INS-184 | chr16-54745242-INS-152 |
| chr16-54797672-INS-454 | chr16-54797672-INS-452 |
| chr16-54807590-INS-1859 | chr16-54807591-INS-1859 |
| chr16-54815868-INS-52 | chr16-54815868-INS-78 |
| chr16-54824882-INS-146 | chr16-54824882-INS-146 |
| chr16-54894743-INS-55 | chr16-54894743-INS-55 |
| chr16-54908873-INS-507 | chr16-54908873-INS-507 |
| chr16-54952699-INS-6314 | chr16-54952700-INS-6306 |
| chr16-54964247-INS-985 | chr16-54964247-INS-985 |
| chr16-54989942-INS-110 | chr16-54989942-INS-106 |
| chr16-55014438-INS-5009 | chr16-55014439-INS-5007 |
| chr16-55021980-INS-2916 | chr16-55021981-INS-2916 |
| chr16-55031709-INS-6381 | chr16-55031710-INS-6381 |
| chr16-55037611-INS-1498 | chr16-55037612-INS-1498 |
| chr16-55111138-INS-79 | chr16-55111137-INS-78 |
| chr16-55133973-INS-79 | chr16-55133968-INS-52 |
| chr16-55150371-INS-90 | chr16-55150371-INS-90 |
| chr16-55161796-INS-6369 | chr16-55161797-INS-6369 |
| chr16-55163229-INS-228 | chr16-55163230-INS-228 |
| chr16-55199172-INS-90 | chr16-55199172-INS-90 |
| chr16-55202289-INS-81 | chr16-55202289-INS-78 |

| | |
|---|---|
| chr16-55228648-INS-52 | chr16-55228654-INS-52 |
| chr16-55240448-INS-595 | chr16-55240448-INS-583 |
| chr16-55246662-INS-266 | chr16-55246662-INS-266 |
| chr16-55290746-INS-63 | chr16-55290747-INS-105 |
| chr16-55304404-INS-2329 | chr16-55304405-INS-2328 |
| chr16-55312894-INS-322 | chr16-55312894-INS-322 |
| chr16-55324116-INS-76 | chr16-55324116-INS-76 |
| chr16-55328092-INS-910 | chr16-55328089-INS-873 |
| chr16-55337347-INS-57 | chr16-55337348-INS-50 |
| chr16-55383474-INS-6802 | chr16-55383475-INS-6802 |
| chr16-55389816-INS-145 | chr16-55389825-INS-120 |
| chr16-55398979-INS-204 | chr16-55398979-INS-204 |
| chr16-55405407-INS-64 | chr16-55405407-INS-58 |
| chr16-55424391-INS-58 | chr16-55424391-INS-74 |
| chr16-55427754-INS-328 | chr16-55427754-INS-328 |
| chr16-55436503-INS-59 | chr16-55436503-INS-59 |
| chr16-55439032-INS-606 | chr16-55439032-INS-606 |
| chr16-55483609-INS-2698 | chr16-55483610-INS-2698 |
| chr16-55485142-INS-826 | chr16-55485143-INS-809 |
| chr16-55488065-INS-590 | chr16-55488065-INS-591 |
| chr16-55524507-INS-88 | chr16-55524507-INS-90 |
| chr16-55528670-INS-401 | chr16-55528670-INS-401 |
| chr16-55535104-INS-92 | chr16-55535104-INS-91 |
| chr16-55613028-INS-456 | chr16-55613028-INS-456 |
| chr16-55626842-INS-480 | chr16-55626842-INS-480 |
| chr16-55628775-INS-7088 | chr16-55628776-INS-7088 |
| chr16-55645940-INS-202 | chr16-55645940-INS-202 |
| chr16-55660123-INS-52 | chr16-55660123-INS-52 |
| chr16-55682876-INS-998 | chr16-55682876-INS-998 |
| chr16-55685067-INS-252 | chr16-55685079-INS-235 |
| chr16-55693810-INS-105 | chr16-55693810-INS-84 |
| chr16-55700697-INS-245 | chr16-55700697-INS-248 |
| chr16-55703434-INS-962 | chr16-55703434-INS-963 |
| chr16-55709881-INS-214 | chr16-55709881-INS-216 |
| chr16-55724259-INS-313 | chr16-55724259-INS-306 |
| chr16-55727227-INS-338 | chr16-55727228-INS-364 |
| chr16-55747271-INS-107 | chr16-55747271-INS-59 |
| chr16-55759810-INS-112 | chr16-55759802-INS-112 |
| chr16-55760699-INS-287 | chr16-55760699-INS-286 |
| chr16-55765277-INS-62 | chr16-55765277-INS-63 |
| chr16-55770072-INS-56 | chr16-55770056-INS-56 |
| chr16-55812081-INS-141 | chr16-55812081-INS-141 |
| chr16-55883178-INS-187 | chr16-55883178-INS-187 |
| chr16-55887439-INS-339 | chr16-55887438-INS-349 |
| chr16-55891032-INS-195 | chr16-55891037-INS-195 |
| chr16-55892337-INS-51 | chr16-55892337-INS-56 |
| chr16-55903377-INS-73 | chr16-55903386-INS-54 |
| chr16-55934796-INS-68 | chr16-55934796-INS-68 |
| chr16-55956843-INS-184 | chr16-55956843-INS-184 |
| chr16-55973395-INS-72 | chr16-55973395-INS-68 |
| chr16-55983602-INS-207 | chr16-55983602-INS-207 |
| chr16-56027711-INS-2410 | chr16-56027712-INS-2407 |
| chr16-56038938-INS-195 | chr16-56038938-INS-193 |
| chr16-56050360-INS-4880 | chr16-56050361-INS-4880 |
| chr16-56059692-INS-211 | chr16-56059692-INS-213 |
| chr16-56063807-INS-139 | chr16-56063804-INS-127 |
| chr16-56089229-INS-439 | chr16-56089229-INS-439 |
| chr16-56095943-INS-1165 | chr16-56095944-INS-1176 |
| chr16-56128543-INS-55 | chr16-56128544-INS-62 |
| chr16-56141711-INS-226 | chr16-56141711-INS-227 |
| chr16-56176257-INS-65 | chr16-56176258-INS-55 |
| chr16-56197723-INS-105 | chr16-56197723-INS-105 |
| chr16-56199438-INS-3627 | chr16-56199439-INS-3627 |
| chr16-56213090-INS-53 | chr16-56213087-INS-70 |
| chr16-56232031-INS-197 | chr16-56232031-INS-195 |
| chr16-56296356-INS-96 | chr16-56296356-INS-127 |
| chr16-56333059-INS-54 | chr16-56333059-INS-54 |
| chr16-56334002-INS-477 | chr16-56334002-INS-479 |
| chr16-56339935-INS-60 | chr16-56339935-INS-60 |
| chr16-56390462-INS-6590 | chr16-56390463-INS-6590 |
| chr16-56411503-INS-58 | chr16-56411503-INS-90 |
| chr16-56432497-INS-56 | chr16-56432497-INS-56 |
| chr16-56442676-INS-60 | chr16-56442672-INS-52 |
| chr16-56445526-INS-63 | chr16-56445526-INS-65 |
| chr16-56475520-INS-6539 | chr16-56475521-INS-6539 |
| chr16-56492123-INS-205 | chr16-56492123-INS-207 |
| chr16-56546006-INS-732 | chr16-56546006-INS-732 |

| | |
|---|---|
| chr16-56550946-INS-800 | chr16-56550946-INS-800 |
| chr16-56552262-INS-125 | chr16-56552262-INS-125 |
| chr16-56586521-INS-54 | chr16-56586521-INS-54 |
| chr16-56600321-INS-2183 | chr16-56600322-INS-2183 |
| chr16-56606594-INS-292 | chr16-56606594-INS-292 |
| chr16-56607281-INS-3691 | chr16-56607282-INS-3692 |
| chr16-56610258-INS-85 | chr16-56610258-INS-85 |
| chr16-56635514-INS-524 | chr16-56635514-INS-525 |
| chr16-56637482-INS-289 | chr16-56637482-INS-294 |
| chr16-56654592-INS-2436 | chr16-56654593-INS-2438 |
| chr16-56655401-INS-3321 | chr16-56655402-INS-3321 |
| chr16-56659304-INS-54 | chr16-56659304-INS-54 |
| chr16-56695489-INS-3629 | chr16-56695490-INS-3629 |
| chr16-56721863-INS-2850 | chr16-56721864-INS-2850 |
| chr16-56729376-INS-74 | chr16-56729376-INS-56 |
| chr16-56742165-INS-5213 | chr16-56742166-INS-5213 |
| chr16-56754171-INS-60 | chr16-56754171-INS-62 |
| chr16-56805193-INS-830 | chr16-56805193-INS-830 |
| chr16-56810900-INS-646 | chr16-56810900-INS-646 |
| chr16-56835475-INS-2828 | chr16-56835476-INS-2828 |
| chr16-56849776-INS-6800 | chr16-56849777-INS-6800 |
| chr16-56865909-INS-63 | chr16-56865909-INS-63 |
| chr16-56887107-INS-106 | chr16-56887107-INS-107 |
| chr16-56903622-INS-97 | chr16-56903622-INS-97 |
| chr16-56954227-INS-157 | chr16-56954227-INS-157 |
| chr16-56962733-INS-159 | chr16-56962733-INS-154 |
| chr16-57017576-INS-432 | chr16-57017576-INS-432 |
| chr16-57029559-INS-87 | chr16-57029559-INS-87 |
| chr16-57031662-INS-55 | chr16-57031661-INS-52 |
| chr16-57055400-INS-1038 | chr16-57055401-INS-1048 |
| chr16-57059009-INS-64 | chr16-57059009-INS-64 |
| chr16-57084992-INS-98 | chr16-57084992-INS-104 |
| chr16-57091910-INS-408 | chr16-57091910-INS-408 |
| chr16-57093685-INS-72 | chr16-57093685-INS-82 |
| chr16-57100278-INS-117 | chr16-57100274-INS-118 |
| chr16-57128458-INS-62 | chr16-57128458-INS-66 |
| chr16-57149149-INS-65 | chr16-57149149-INS-65 |
| chr16-57154434-INS-61 | chr16-57154434-INS-61 |
| chr16-57222264-INS-2976 | chr16-57222265-INS-2976 |
| chr16-57279575-INS-52 | chr16-57279575-INS-54 |
| chr16-57305518-INS-426 | chr16-57305518-INS-423 |
| chr16-57306562-INS-60 | chr16-57306562-INS-60 |
| chr16-57360392-INS-68 | chr16-57360392-INS-67 |
| chr16-57404551-INS-75 | chr16-57404548-INS-78 |
| chr16-57420536-INS-271 | chr16-57420536-INS-259 |
| chr16-57426478-INS-100 | chr16-57426479-INS-111 |
| chr16-57456604-INS-194 | chr16-57456604-INS-197 |
| chr16-57468450-INS-63 | chr16-57468450-INS-60 |
| chr16-57523709-INS-72 | chr16-57523709-INS-78 |
| chr16-57533810-INS-159 | chr16-57533815-INS-138 |
| chr16-57541125-INS-177 | chr16-57541126-INS-181 |
| chr16-57609535-INS-2751 | chr16-57609536-INS-2751 |
| chr16-57681159-INS-105 | chr16-57681159-INS-105 |
| chr16-57691747-INS-6586 | chr16-57691748-INS-6589 |
| chr16-57726010-INS-56 | chr16-57725979-INS-62 |
| chr16-57729486-INS-377 | chr16-57729486-INS-377 |
| chr16-57756322-INS-1363 | chr16-57756323-INS-1352 |
| chr16-57767556-INS-64 | chr16-57767532-INS-104 |
| chr16-57769299-INS-174 | chr16-57769311-INS-126 |
| chr16-57783632-INS-80 | chr16-57783632-INS-80 |
| chr16-57831426-INS-663 | chr16-57831426-INS-661 |
| chr16-57840067-INS-6472 | chr16-57840068-INS-6472 |
| chr16-57842678-INS-78 | chr16-57842678-INS-86 |
| chr16-57844137-INS-242 | chr16-57844137-INS-242 |
| chr16-57854233-INS-70 | chr16-57854234-INS-70 |
| chr16-57860592-INS-163 | chr16-57860592-INS-167 |
| chr16-57873818-INS-779 | chr16-57873818-INS-779 |
| chr16-57920339-INS-167 | chr16-57920339-INS-168 |
| chr16-57946182-INS-114 | chr16-57946182-INS-102 |
| chr16-57964763-INS-64 | chr16-57964762-INS-96 |
| chr16-57999612-INS-1341 | chr16-57999613-INS-1341 |
| chr16-58045088-INS-880 | chr16-58045088-INS-881 |
| chr16-58111813-INS-220 | chr16-58111813-INS-220 |
| chr16-58147430-INS-2424 | chr16-58147431-INS-2423 |
| chr16-58180010-INS-400 | chr16-58180010-INS-400 |
| chr16-58184480-INS-208 | chr16-58184480-INS-208 |
| chr16-58208872-INS-361 | chr16-58208872-INS-361 |

| | |
|---|---|
| chr16-58212081-INS-88 | chr16-58212081-INS-116 |
| chr16-58219735-INS-7491 | chr16-58219736-INS-7491 |
| chr16-58234217-INS-64 | chr16-58234217-INS-66 |
| chr16-58237147-INS-62 | chr16-58237147-INS-58 |
| chr16-58258036-INS-113 | chr16-58258036-INS-113 |
| chr16-58264908-INS-72 | chr16-58264932-INS-72 |
| chr16-58266497-INS-99 | chr16-58266497-INS-99 |
| chr16-58291877-INS-223 | chr16-58291877-INS-223 |
| chr16-58333833-INS-70 | chr16-58333833-INS-70 |
| chr16-58337818-INS-187 | chr16-58337827-INS-186 |
| chr16-58339147-INS-1167 | chr16-58339148-INS-1172 |
| chr16-58355432-INS-51 | chr16-58355432-INS-58 |
| chr16-58356655-INS-52 | chr16-58356655-INS-52 |
| chr16-58372977-INS-123 | chr16-58372977-INS-129 |
| chr16-58395370-INS-122 | chr16-58395370-INS-122 |
| chr16-58410535-INS-4391 | chr16-58410536-INS-4391 |
| chr16-58431012-INS-6283 | chr16-58431013-INS-6284 |
| chr16-58478468-INS-62 | chr16-58478468-INS-62 |
| chr16-58498871-INS-1033 | chr16-58498872-INS-1030 |
| chr16-58562610-INS-1170 | chr16-58562611-INS-1170 |
| chr16-58565215-INS-62 | chr16-58565226-INS-68 |
| chr16-58635771-INS-7228 | chr16-58635772-INS-7230 |
| chr16-58636054-INS-112 | chr16-58636054-INS-112 |
| chr16-58643480-INS-593 | chr16-58643480-INS-593 |
| chr16-58647253-INS-146 | chr16-58647253-INS-146 |
| chr16-58654521-INS-505 | chr16-58654521-INS-504 |
| chr16-58673282-INS-4129 | chr16-58673283-INS-4129 |
| chr16-58680166-INS-624 | chr16-58680166-INS-624 |
| chr16-58728773-INS-72 | chr16-58728773-INS-72 |
| chr16-58729032-INS-432 | chr16-58729040-INS-424 |
| chr16-58740208-INS-1094 | chr16-58740219-INS-1084 |
| chr16-58809020-INS-107 | chr16-58809045-INS-110 |
| chr16-58827184-INS-252 | chr16-58827184-INS-249 |
| chr16-58833473-INS-2933 | chr16-58833474-INS-2934 |
| chr16-58848964-INS-84 | chr16-58848964-INS-96 |
| chr16-59059368-INS-1668 | chr16-59059369-INS-1667 |
| chr16-59060177-INS-64 | chr16-59060177-INS-64 |
| chr16-59070621-INS-2650 | chr16-59070622-INS-2650 |
| chr16-59106658-INS-62 | chr16-59106669-INS-61 |
| chr16-59138968-INS-171 | chr16-59138968-INS-173 |
| chr16-59140360-INS-60 | chr16-59140360-INS-68 |
| chr16-59172027-INS-284 | chr16-59172049-INS-270 |
| chr16-59232081-INS-617 | chr16-59232081-INS-617 |
| chr16-59235350-INS-78 | chr16-59235350-INS-58 |
| chr16-59244601-INS-54 | chr16-59244601-INS-54 |
| chr16-59248883-INS-4499 | chr16-59248884-INS-4499 |
| chr16-59305087-INS-52 | chr16-59305087-INS-52 |
| chr16-59329587-INS-63 | chr16-59329587-INS-63 |
| chr16-59348922-INS-53 | chr16-59348924-INS-70 |
| chr16-59370228-INS-238 | chr16-59370228-INS-236 |
| chr16-59370228-INS-924 | chr16-59370228-INS-926 |
| chr16-59429312-INS-276 | chr16-59429331-INS-297 |
| chr16-59437963-INS-202 | chr16-59437963-INS-202 |
| chr16-59445232-INS-224 | chr16-59445244-INS-212 |
| chr16-59449132-INS-51 | chr16-59449132-INS-51 |
| chr16-59484790-INS-684 | chr16-59484790-INS-700 |
| chr16-59490473-INS-52 | chr16-59490500-INS-69 |
| chr16-59529304-INS-57 | chr16-59529299-INS-106 |
| chr16-59533043-INS-174 | chr16-59533043-INS-174 |
| chr16-59535501-INS-139 | chr16-59535501-INS-137 |
| chr16-59567853-INS-1892 | chr16-59567854-INS-1878 |
| chr16-59568372-INS-166 | chr16-59568372-INS-167 |
| chr16-59573060-INS-795 | chr16-59573052-INS-807 |
| chr16-59579445-INS-108 | chr16-59579455-INS-62 |
| chr16-59583367-INS-56 | chr16-59583367-INS-54 |
| chr16-59594044-INS-56 | chr16-59594044-INS-96 |
| chr16-59605232-INS-76 | chr16-59605232-INS-76 |
| chr16-59609688-INS-60 | chr16-59609688-INS-65 |
| chr16-59628945-INS-61 | chr16-59628945-INS-63 |
| chr16-59647579-INS-6375 | chr16-59647566-INS-6376 |
| chr16-59674070-INS-821 | chr16-59674070-INS-821 |
| chr16-59726442-INS-695 | chr16-59726442-INS-695 |
| chr16-59728561-INS-147 | chr16-59728561-INS-147 |
| chr16-59730385-INS-58 | chr16-59730385-INS-64 |
| chr16-59731164-INS-546 | chr16-59731164-INS-546 |
| chr16-59746490-INS-95 | chr16-59746490-INS-95 |
| chr16-59750453-INS-6944 | chr16-59750454-INS-6946 |

| | |
|---|---|
| chr16-59767307-INS-425 | chr16-59767307-INS-425 |
| chr16-59796139-INS-125 | chr16-59796139-INS-125 |
| chr16-59797308-INS-62 | chr16-59797308-INS-110 |
| chr16-59814523-INS-6200 | chr16-59814524-INS-6200 |
| chr16-59826591-INS-150 | chr16-59826591-INS-150 |
| chr16-59862923-INS-1104 | chr16-59862924-INS-1097 |
| chr16-59875215-INS-6242 | chr16-59875216-INS-6246 |
| chr16-59887761-INS-7996 | chr16-59887762-INS-7996 |
| chr16-59899485-INS-90 | chr16-59899485-INS-90 |
| chr16-59904725-INS-143 | chr16-59904725-INS-145 |
| chr16-59951591-INS-75 | chr16-59951594-INS-50 |
| chr16-59961286-INS-71 | chr16-59961286-INS-71 |
| chr16-60013825-INS-111 | chr16-60013825-INS-123 |
| chr16-60020036-INS-201 | chr16-60020036-INS-201 |
| chr16-60025019-INS-936 | chr16-60025019-INS-936 |
| chr16-60037424-INS-81 | chr16-60037424-INS-81 |
| chr16-60059361-INS-7085 | chr16-60059362-INS-7085 |
| chr16-60081065-INS-952 | chr16-60081065-INS-952 |
| chr16-60115987-INS-1488 | chr16-60115988-INS-1488 |
| chr16-60139444-INS-149 | chr16-60139444-INS-161 |
| chr16-60156054-INS-172 | chr16-60156054-INS-172 |
| chr16-60157035-INS-3747 | chr16-60157036-INS-3747 |
| chr16-60161811-INS-184 | chr16-60161811-INS-144 |
| chr16-60167014-INS-591 | chr16-60167014-INS-595 |
| chr16-60172984-INS-129 | chr16-60172984-INS-129 |
| chr16-60186872-INS-180 | chr16-60186872-INS-180 |
| chr16-60191258-INS-2838 | chr16-60191259-INS-2838 |
| chr16-60241101-INS-220 | chr16-60241101-INS-221 |
| chr16-60242932-INS-66 | chr16-60242932-INS-66 |
| chr16-60257138-INS-214 | chr16-60257138-INS-214 |
| chr16-60275799-INS-398 | chr16-60275799-INS-400 |
| chr16-60293635-INS-519 | chr16-60293635-INS-518 |
| chr16-60294236-INS-324 | chr16-60294236-INS-324 |
| chr16-60296589-INS-838 | chr16-60296589-INS-837 |
| chr16-60353282-INS-399 | chr16-60353282-INS-399 |
| chr16-60354287-INS-102 | chr16-60354285-INS-106 |
| chr16-60366248-INS-405 | chr16-60366249-INS-403 |
| chr16-60391001-INS-60 | chr16-60391005-INS-68 |
| chr16-60400393-INS-217 | chr16-60400393-INS-220 |
| chr16-60433994-INS-305 | chr16-60433994-INS-304 |
| chr16-60442017-INS-6401 | chr16-60442018-INS-6401 |
| chr16-60446892-INS-192 | chr16-60446892-INS-192 |
| chr16-60451097-INS-62 | chr16-60451097-INS-64 |
| chr16-60454517-INS-171 | chr16-60454517-INS-170 |
| chr16-60472774-INS-1122 | chr16-60472769-INS-1109 |
| chr16-60486451-INS-123 | chr16-60486451-INS-129 |
| chr16-60497414-INS-87 | chr16-60497384-INS-135 |
| chr16-60539615-INS-118 | chr16-60539615-INS-127 |
| chr16-60586940-INS-318 | chr16-60586940-INS-318 |
| chr16-60591024-INS-94 | chr16-60591024-INS-93 |
| chr16-60605772-INS-76 | chr16-60605772-INS-76 |
| chr16-60609821-INS-438 | chr16-60609821-INS-438 |
| chr16-60628730-INS-780 | chr16-60628731-INS-781 |
| chr16-60704701-INS-2318 | chr16-60704702-INS-2318 |
| chr16-60738190-INS-63 | chr16-60738164-INS-60 |
| chr16-60810225-INS-84 | chr16-60810233-INS-69 |
| chr16-60825922-INS-57 | chr16-60825922-INS-59 |
| chr16-60863730-INS-725 | chr16-60863730-INS-725 |
| chr16-60871413-INS-651 | chr16-60871413-INS-637 |
| chr16-60872980-INS-62 | chr16-60872980-INS-62 |
| chr16-60886700-INS-7157 | chr16-60886701-INS-7157 |
| chr16-60887394-INS-156 | chr16-60887394-INS-156 |
| chr16-60901977-INS-68 | chr16-60901978-INS-52 |
| chr16-60927832-INS-82 | chr16-60927832-INS-82 |
| chr16-60934535-INS-6870 | chr16-60934536-INS-6870 |
| chr16-60951569-INS-206 | chr16-60951569-INS-207 |
| chr16-60955649-INS-135 | chr16-60955649-INS-135 |
| chr16-61063116-INS-62 | chr16-61063116-INS-62 |
| chr16-61081814-INS-352 | chr16-61081814-INS-352 |
| chr16-61088302-INS-63 | chr16-61088302-INS-81 |
| chr16-61114548-INS-592 | chr16-61114548-INS-593 |
| chr16-61145298-INS-5264 | chr16-61145299-INS-5264 |
| chr16-61162158-INS-390 | chr16-61162158-INS-390 |
| chr16-61175616-INS-2237 | chr16-61175617-INS-2238 |
| chr16-61235621-INS-560 | chr16-61235621-INS-560 |
| chr16-61260434-INS-185 | chr16-61260434-INS-185 |
| chr16-61261038-INS-828 | chr16-61261038-INS-816 |

| | |
|---|---|
| chr16-61290417-INS-1094 | chr16-61290418-INS-1094 |
| chr16-61290829-INS-67 | chr16-61290829-INS-68 |
| chr16-61319993-INS-184 | chr16-61319993-INS-192 |
| chr16-61334173-INS-6540 | chr16-61334174-INS-6541 |
| chr16-61383422-INS-62 | chr16-61383422-INS-64 |
| chr16-61386126-INS-74 | chr16-61386124-INS-74 |
| chr16-61388794-INS-52 | chr16-61388794-INS-54 |
| chr16-61392148-INS-69 | chr16-61392149-INS-69 |
| chr16-61434989-INS-110 | chr16-61434989-INS-117 |
| chr16-61436912-INS-6413 | chr16-61436913-INS-6413 |
| chr16-61441545-INS-99 | chr16-61441545-INS-100 |
| chr16-61449932-INS-74 | chr16-61449933-INS-75 |
| chr16-61480243-INS-65 | chr16-61480243-INS-65 |
| chr16-61505798-INS-195 | chr16-61505798-INS-195 |
| chr16-61516618-INS-52 | chr16-61516618-INS-52 |
| chr16-61626747-INS-207 | chr16-61626747-INS-207 |
| chr16-61628627-INS-86 | chr16-61628627-INS-81 |
| chr16-61659857-INS-168 | chr16-61659857-INS-168 |
| chr16-61663935-INS-656 | chr16-61663935-INS-656 |
| chr16-61673740-INS-212 | chr16-61673740-INS-212 |
| chr16-61689488-INS-565 | chr16-61689489-INS-566 |
| chr16-61705209-INS-328 | chr16-61705209-INS-328 |
| chr16-61712313-INS-170 | chr16-61712313-INS-168 |
| chr16-61754055-INS-702 | chr16-61754055-INS-699 |
| chr16-61800206-INS-56 | chr16-61800206-INS-58 |
| chr16-61816553-INS-72 | chr16-61816553-INS-68 |
| chr16-61821580-INS-59 | chr16-61821580-INS-59 |
| chr16-61867148-INS-68 | chr16-61867148-INS-62 |
| chr16-61876611-INS-64 | chr16-61876611-INS-64 |
| chr16-61889463-INS-70 | chr16-61889463-INS-76 |
| chr16-61894556-INS-54 | chr16-61894556-INS-59 |
| chr16-61902511-INS-64 | chr16-61902481-INS-66 |
| chr16-61920115-INS-60 | chr16-61920115-INS-64 |
| chr16-61946018-INS-812 | chr16-61946018-INS-813 |
| chr16-61947948-INS-900 | chr16-61947948-INS-900 |
| chr16-61996354-INS-153 | chr16-61996355-INS-149 |
| chr16-62012106-INS-438 | chr16-62012106-INS-438 |
| chr16-62050814-INS-2212 | chr16-62050812-INS-2212 |
| chr16-62053488-INS-86 | chr16-62053491-INS-84 |
| chr16-62062886-INS-5418 | chr16-62062887-INS-5418 |
| chr16-62068878-INS-100 | chr16-62068883-INS-118 |
| chr16-62070755-INS-1894 | chr16-62070756-INS-1894 |
| chr16-62077216-INS-415 | chr16-62077216-INS-415 |
| chr16-62089261-INS-4791 | chr16-62089262-INS-4791 |
| chr16-62094893-INS-60 | chr16-62094876-INS-54 |
| chr16-62125710-INS-396 | chr16-62125710-INS-396 |
| chr16-62128934-INS-7018 | chr16-62128935-INS-7018 |
| chr16-62144062-INS-52 | chr16-62144062-INS-52 |
| chr16-62217249-INS-741 | chr16-62217249-INS-741 |
| chr16-62251953-INS-53 | chr16-62251952-INS-54 |
| chr16-62256649-INS-1213 | chr16-62256650-INS-1214 |
| chr16-62275998-INS-81 | chr16-62275998-INS-81 |
| chr16-62297724-INS-449 | chr16-62297724-INS-448 |
| chr16-62299194-INS-156 | chr16-62299194-INS-156 |
| chr16-62434549-INS-463 | chr16-62434549-INS-473 |
| chr16-62481254-INS-78 | chr16-62481224-INS-104 |
| chr16-62518705-INS-880 | chr16-62518705-INS-881 |
| chr16-62540130-INS-790 | chr16-62540130-INS-769 |
| chr16-62553037-INS-4229 | chr16-62553038-INS-4229 |
| chr16-62567345-INS-6374 | chr16-62567346-INS-6374 |
| chr16-62569600-INS-2821 | chr16-62569601-INS-2829 |
| chr16-62570582-INS-666 | chr16-62570582-INS-661 |
| chr16-62571976-INS-71 | chr16-62571976-INS-71 |
| chr16-62610028-INS-427 | chr16-62610028-INS-427 |
| chr16-62623706-INS-61 | chr16-62623710-INS-65 |
| chr16-62629356-INS-62 | chr16-62629356-INS-62 |
| chr16-62635392-INS-90 | chr16-62635392-INS-92 |
| chr16-62639736-INS-56 | chr16-62639736-INS-52 |
| chr16-62649016-INS-74 | chr16-62649016-INS-74 |
| chr16-62653969-INS-134 | chr16-62653969-INS-134 |
| chr16-62683087-INS-529 | chr16-62683087-INS-535 |
| chr16-62702644-INS-252 | chr16-62702644-INS-253 |
| chr16-62710911-INS-64 | chr16-62710931-INS-72 |
| chr16-62713186-INS-90 | chr16-62713186-INS-90 |
| chr16-62752131-INS-55 | chr16-62752131-INS-71 |
| chr16-62774444-INS-67 | chr16-62774444-INS-95 |
| chr16-62792731-INS-196 | chr16-62792731-INS-196 |

| | |
|---|---|
| chr16-62797342-INS-254 | chr16-62797342-INS-254 |
| chr16-62833380-INS-742 | chr16-62833380-INS-742 |
| chr16-62835270-INS-114 | chr16-62835270-INS-83 |
| chr16-62851202-INS-5311 | chr16-62851203-INS-5311 |
| chr16-62894467-INS-70 | chr16-62894467-INS-70 |
| chr16-62915503-INS-178 | chr16-62915503-INS-179 |
| chr16-62931865-INS-69 | chr16-62931865-INS-54 |
| chr16-62932709-INS-131 | chr16-62932709-INS-131 |
| chr16-62940656-INS-66 | chr16-62940656-INS-62 |
| chr16-62945567-INS-549 | chr16-62945567-INS-549 |
| chr16-62959993-INS-899 | chr16-62959993-INS-899 |
| chr16-63001557-INS-88 | chr16-63001557-INS-88 |
| chr16-63029152-INS-235 | chr16-63029153-INS-266 |
| chr16-63035769-INS-113 | chr16-63035770-INS-113 |
| chr16-63048479-INS-90 | chr16-63048479-INS-90 |
| chr16-63062502-INS-1479 | chr16-63062503-INS-1480 |
| chr16-63066749-INS-54 | chr16-63066749-INS-56 |
| chr16-63091717-INS-728 | chr16-63091717-INS-728 |
| chr16-63104367-INS-136 | chr16-63104375-INS-120 |
| chr16-63120455-INS-1745 | chr16-63120456-INS-1744 |
| chr16-63125752-INS-1177 | chr16-63125753-INS-1177 |
| chr16-63136415-INS-69 | chr16-63136415-INS-72 |
| chr16-63167105-INS-2603 | chr16-63167106-INS-2602 |
| chr16-63172827-INS-54 | chr16-63172827-INS-90 |
| chr16-63182284-INS-69 | chr16-63182284-INS-105 |
| chr16-63213720-INS-3347 | chr16-63213721-INS-3347 |
| chr16-63231991-INS-409 | chr16-63231989-INS-410 |
| chr16-63243301-INS-4840 | chr16-63243302-INS-4840 |
| chr16-63269670-INS-54 | chr16-63269670-INS-60 |
| chr16-63287874-INS-107 | chr16-63287874-INS-107 |
| chr16-63310833-INS-63 | chr16-63310833-INS-63 |
| chr16-63311280-INS-6083 | chr16-63311293-INS-6085 |
| chr16-63314377-INS-231 | chr16-63314377-INS-231 |
| chr16-63317182-INS-760 | chr16-63317182-INS-764 |
| chr16-63320407-INS-941 | chr16-63320407-INS-944 |
| chr16-63347163-INS-7079 | chr16-63347164-INS-7079 |
| chr16-63353057-INS-205 | chr16-63353057-INS-207 |
| chr16-63373927-INS-132 | chr16-63373927-INS-132 |
| chr16-63376620-INS-132 | chr16-63376620-INS-140 |
| chr16-63390428-INS-451 | chr16-63390428-INS-452 |
| chr16-63425672-INS-66 | chr16-63425676-INS-58 |
| chr16-63448715-INS-130 | chr16-63448715-INS-130 |
| chr16-63471798-INS-380 | chr16-63471798-INS-380 |
| chr16-63492559-INS-78 | chr16-63492559-INS-78 |
| chr16-63494774-INS-75 | chr16-63494774-INS-64 |
| chr16-63500135-INS-63 | chr16-63500135-INS-55 |
| chr16-63504348-INS-86 | chr16-63504348-INS-78 |
| chr16-63521664-INS-66 | chr16-63521664-INS-66 |
| chr16-63552702-INS-460 | chr16-63552702-INS-460 |
| chr16-63556621-INS-64 | chr16-63556622-INS-53 |
| chr16-63576499-INS-88 | chr16-63576499-INS-70 |
| chr16-63583463-INS-141 | chr16-63583463-INS-129 |
| chr16-63591369-INS-225 | chr16-63591369-INS-225 |
| chr16-63620136-INS-52 | chr16-63620136-INS-80 |
| chr16-63622209-INS-52 | chr16-63622203-INS-58 |
| chr16-63645001-INS-120 | chr16-63645001-INS-120 |
| chr16-63651015-INS-62 | chr16-63651015-INS-62 |
| chr16-63653749-INS-780 | chr16-63653750-INS-782 |
| chr16-63658333-INS-1308 | chr16-63658334-INS-1308 |
| chr16-63676360-INS-54 | chr16-63676360-INS-54 |
| chr16-63708599-INS-842 | chr16-63708599-INS-842 |
| chr16-63718936-INS-118 | chr16-63718940-INS-114 |
| chr16-63760281-INS-112 | chr16-63760281-INS-104 |
| chr16-63765868-INS-499 | chr16-63765868-INS-499 |
| chr16-63772553-INS-737 | chr16-63772553-INS-741 |
| chr16-63797837-INS-64 | chr16-63797837-INS-64 |
| chr16-63804796-INS-52 | chr16-63804796-INS-52 |
| chr16-63807196-INS-108 | chr16-63807196-INS-111 |
| chr16-63843907-INS-103 | chr16-63843907-INS-103 |
| chr16-63848826-INS-61 | chr16-63848826-INS-72 |
| chr16-63851127-INS-52 | chr16-63851127-INS-68 |
| chr16-63863013-INS-170 | chr16-63863013-INS-170 |
| chr16-63871923-INS-56 | chr16-63871923-INS-72 |
| chr16-63893336-INS-582 | chr16-63893336-INS-582 |
| chr16-63893516-INS-4524 | chr16-63893517-INS-4524 |
| chr16-63907474-INS-1955 | chr16-63907475-INS-1955 |
| chr16-63908046-INS-5159 | chr16-63908047-INS-5159 |

| | |
|---|---|
| chr16-63923792-INS-152 | chr16-63923792-INS-150 |
| chr16-63924804-INS-178 | chr16-63924804-INS-169 |
| chr16-63978743-INS-2744 | chr16-63978744-INS-2744 |
| chr16-64004794-INS-930 | chr16-64004794-INS-930 |
| chr16-64007698-INS-390 | chr16-64007698-INS-390 |
| chr16-64010398-INS-6158 | chr16-64010399-INS-6158 |
| chr16-64016994-INS-150 | chr16-64016994-INS-150 |
| chr16-64025902-INS-54 | chr16-64025902-INS-56 |
| chr16-64067659-INS-97 | chr16-64067659-INS-120 |
| chr16-64069500-INS-82 | chr16-64069500-INS-86 |
| chr16-64069730-INS-1008 | chr16-64069742-INS-997 |
| chr16-64080567-INS-4683 | chr16-64080568-INS-4671 |
| chr16-64117307-INS-66 | chr16-64117273-INS-66 |
| chr16-64132814-INS-81 | chr16-64132819-INS-64 |
| chr16-64134864-INS-520 | chr16-64134864-INS-520 |
| chr16-64181365-INS-60 | chr16-64181372-INS-89 |
| chr16-64206504-INS-701 | chr16-64206504-INS-701 |
| chr16-64238590-INS-66 | chr16-64238590-INS-66 |
| chr16-64249364-INS-822 | chr16-64249364-INS-822 |
| chr16-64250315-INS-51 | chr16-64250315-INS-53 |
| chr16-64261037-INS-6208 | chr16-64261038-INS-6207 |
| chr16-64311761-INS-70 | chr16-64311762-INS-64 |
| chr16-64358502-INS-152 | chr16-64358502-INS-152 |
| chr16-64373263-INS-2401 | chr16-64373264-INS-2401 |
| chr16-64375833-INS-115 | chr16-64375833-INS-103 |
| chr16-64378244-INS-838 | chr16-64378244-INS-838 |
| chr16-64384265-INS-200 | chr16-64384265-INS-200 |
| chr16-64412792-INS-52 | chr16-64412792-INS-52 |
| chr16-64432788-INS-524 | chr16-64432794-INS-520 |
| chr16-64433537-INS-80 | chr16-64433537-INS-66 |
| chr16-64468233-INS-282 | chr16-64468233-INS-282 |
| chr16-64479621-INS-6659 | chr16-64479622-INS-6659 |
| chr16-64480230-INS-347 | chr16-64480230-INS-345 |
| chr16-64483115-INS-137 | chr16-64483115-INS-137 |
| chr16-64500774-INS-51 | chr16-64500774-INS-51 |
| chr16-64508721-INS-399 | chr16-64508721-INS-399 |
| chr16-64516419-INS-77 | chr16-64516419-INS-79 |
| chr16-64533295-INS-276 | chr16-64533295-INS-277 |
| chr16-64547034-INS-413 | chr16-64547034-INS-415 |
| chr16-64556300-INS-580 | chr16-64556300-INS-580 |
| chr16-64572808-INS-91 | chr16-64572791-INS-99 |
| chr16-64608383-INS-84 | chr16-64608383-INS-84 |
| chr16-64620776-INS-179 | chr16-64620776-INS-178 |
| chr16-64625138-INS-510 | chr16-64625138-INS-510 |
| chr16-64642056-INS-169 | chr16-64642060-INS-170 |
| chr16-64650142-INS-171 | chr16-64650142-INS-169 |
| chr16-64657357-INS-184 | chr16-64657357-INS-184 |
| chr16-64675311-INS-273 | chr16-64675311-INS-274 |
| chr16-64686619-INS-77 | chr16-64686619-INS-66 |
| chr16-64693144-INS-68 | chr16-64693144-INS-99 |
| chr16-64694803-INS-215 | chr16-64694803-INS-215 |
| chr16-64701166-INS-277 | chr16-64701149-INS-295 |
| chr16-64705908-INS-101 | chr16-64705908-INS-62 |
| chr16-64717483-INS-208 | chr16-64717484-INS-176 |
| chr16-64725125-INS-255 | chr16-64725125-INS-277 |
| chr16-64726463-INS-102 | chr16-64726463-INS-102 |
| chr16-64773999-INS-130 | chr16-64773999-INS-128 |
| chr16-64808666-INS-52 | chr16-64808666-INS-56 |
| chr16-64823758-INS-358 | chr16-64823761-INS-354 |
| chr16-64832927-INS-71 | chr16-64832927-INS-77 |
| chr16-64845572-INS-2476 | chr16-64845573-INS-2471 |
| chr16-64949509-INS-52 | chr16-64949509-INS-52 |
| chr16-64983809-INS-147 | chr16-64983809-INS-155 |
| chr16-64986160-INS-7112 | chr16-64986161-INS-7112 |
| chr16-65030745-INS-420 | chr16-65030745-INS-420 |
| chr16-65093281-INS-165 | chr16-65093280-INS-166 |
| chr16-65104115-INS-3363 | chr16-65104116-INS-3363 |
| chr16-65107957-INS-1988 | chr16-65107958-INS-1991 |
| chr16-65111334-INS-73 | chr16-65111334-INS-73 |
| chr16-65118829-INS-214 | chr16-65118829-INS-214 |
| chr16-65138914-INS-6795 | chr16-65138915-INS-6795 |
| chr16-65152872-INS-58 | chr16-65152872-INS-58 |
| chr16-65184301-INS-108 | chr16-65184300-INS-109 |
| chr16-65188919-INS-2757 | chr16-65188920-INS-2757 |
| chr16-65226101-INS-497 | chr16-65226101-INS-497 |
| chr16-65246820-INS-156 | chr16-65246820-INS-157 |
| chr16-65248345-INS-56 | chr16-65248343-INS-56 |

| | |
|---|---|
| chr16-65256189-INS-6733 | chr16-65256190-INS-6733 |
| chr16-65265980-INS-258 | chr16-65265980-INS-265 |
| chr16-65283959-INS-541 | chr16-65283959-INS-546 |
| chr16-65285612-INS-959 | chr16-65285600-INS-959 |
| chr16-65299388-INS-51 | chr16-65299388-INS-51 |
| chr16-65334129-INS-249 | chr16-65334129-INS-249 |
| chr16-65351085-INS-178 | chr16-65351085-INS-178 |
| chr16-65364921-INS-243 | chr16-65364921-INS-243 |
| chr16-65366051-INS-105 | chr16-65366051-INS-105 |
| chr16-65375406-INS-78 | chr16-65375405-INS-52 |
| chr16-65420061-INS-485 | chr16-65420061-INS-488 |
| chr16-65425889-INS-110 | chr16-65425889-INS-110 |
| chr16-65493109-INS-282 | chr16-65493148-INS-281 |
| chr16-65495331-INS-1492 | chr16-65495332-INS-1493 |
| chr16-65497789-INS-574 | chr16-65497789-INS-574 |
| chr16-65500863-INS-3380 | chr16-65500864-INS-3373 |
| chr16-65511331-INS-331 | chr16-65511331-INS-327 |
| chr16-65540932-INS-3588 | chr16-65540933-INS-3588 |
| chr16-65550013-INS-465 | chr16-65549998-INS-467 |
| chr16-65581830-INS-204 | chr16-65581830-INS-204 |
| chr16-65605457-INS-4336 | chr16-65605458-INS-4336 |
| chr16-65610032-INS-54 | chr16-65610032-INS-51 |
| chr16-65649232-INS-5091 | chr16-65649233-INS-5091 |
| chr16-65677533-INS-186 | chr16-65677533-INS-186 |
| chr16-65763769-INS-103 | chr16-65763769-INS-103 |
| chr16-65767970-INS-60 | chr16-65767970-INS-60 |
| chr16-65793628-INS-124 | chr16-65793613-INS-132 |
| chr16-65806738-INS-68 | chr16-65806739-INS-59 |
| chr16-65823135-INS-88 | chr16-65823135-INS-74 |
| chr16-65828781-INS-628 | chr16-65828781-INS-628 |
| chr16-65830100-INS-541 | chr16-65830100-INS-540 |
| chr16-65832648-INS-810 | chr16-65832650-INS-827 |
| chr16-65867156-INS-310 | chr16-65867156-INS-305 |
| chr16-65867942-INS-828 | chr16-65867942-INS-828 |
| chr16-65871768-INS-5252 | chr16-65871769-INS-5252 |
| chr16-65894719-INS-389 | chr16-65894719-INS-389 |
| chr16-65951133-INS-66 | chr16-65951163-INS-67 |
| chr16-65968776-INS-3129 | chr16-65968777-INS-3130 |
| chr16-65977847-INS-64 | chr16-65977847-INS-64 |
| chr16-66012466-INS-2663 | chr16-66012467-INS-2663 |
| chr16-66018237-INS-118 | chr16-66018207-INS-166 |
| chr16-66049019-INS-56 | chr16-66049019-INS-58 |
| chr16-66115932-INS-533 | chr16-66115932-INS-533 |
| chr16-66135191-INS-60 | chr16-66135191-INS-56 |
| chr16-66140498-INS-6175 | chr16-66140499-INS-6175 |
| chr16-66168591-INS-2297 | chr16-66168592-INS-2297 |
| chr16-66180899-INS-128 | chr16-66180899-INS-136 |
| chr16-66193248-INS-929 | chr16-66193248-INS-930 |
| chr16-66223453-INS-137 | chr16-66223453-INS-137 |
| chr16-66272312-INS-64 | chr16-66272312-INS-64 |
| chr16-66355291-INS-56 | chr16-66355299-INS-58 |
| chr16-66377705-INS-400 | chr16-66377705-INS-400 |
| chr16-66380105-INS-60 | chr16-66380105-INS-80 |
| chr16-66383225-INS-109 | chr16-66383225-INS-121 |
| chr16-66388031-INS-56 | chr16-66388031-INS-56 |
| chr16-66407875-INS-65 | chr16-66407850-INS-51 |
| chr16-66437555-INS-918 | chr16-66437555-INS-902 |
| chr16-66469536-INS-457 | chr16-66469536-INS-457 |
| chr16-66543876-INS-472 | chr16-66543876-INS-495 |
| chr16-66559267-INS-354 | chr16-66559267-INS-351 |
| chr16-66583285-INS-70 | chr16-66583285-INS-70 |
| chr16-66618653-INS-74 | chr16-66618653-INS-98 |
| chr16-66625528-INS-63 | chr16-66625527-INS-66 |
| chr16-66627477-INS-3456 | chr16-66627478-INS-3446 |
| chr16-66647500-INS-4478 | chr16-66647501-INS-4478 |
| chr16-66665046-INS-60 | chr16-66665046-INS-58 |
| chr16-66676488-INS-1346 | chr16-66676489-INS-1348 |
| chr16-66692013-INS-357 | chr16-66692013-INS-351 |
| chr16-66693626-INS-212 | chr16-66693625-INS-252 |
| chr16-66708316-INS-93 | chr16-66708317-INS-97 |
| chr16-66732672-INS-70 | chr16-66732672-INS-58 |
| chr16-66739114-INS-3482 | chr16-66739115-INS-3484 |
| chr16-66742129-INS-6210 | chr16-66742130-INS-6210 |
| chr16-66765770-INS-378 | chr16-66765770-INS-378 |
| chr16-66768839-INS-253 | chr16-66768839-INS-253 |
| chr16-66820955-INS-416 | chr16-66820955-INS-416 |
| chr16-66824279-INS-52 | chr16-66824279-INS-70 |

| | |
|---|---|
| chr16-66884147-INS-90 | chr16-66884147-INS-78 |
| chr16-66886548-INS-66 | chr16-66886548-INS-66 |
| chr16-66891504-INS-100 | chr16-66891504-INS-100 |
| chr16-66899951-INS-587 | chr16-66899951-INS-587 |
| chr16-66926746-INS-84 | chr16-66926746-INS-102 |
| chr16-66934932-INS-58 | chr16-66934932-INS-64 |
| chr16-66954806-INS-90 | chr16-66954806-INS-105 |
| chr16-66959674-INS-248 | chr16-66959674-INS-264 |
| chr16-66967706-INS-182 | chr16-66967706-INS-182 |
| chr16-67003797-INS-611 | chr16-67003797-INS-611 |
| chr16-67034013-INS-70 | chr16-67033999-INS-52 |
| chr16-67064852-INS-1613 | chr16-67064853-INS-1613 |
| chr16-67074333-INS-75 | chr16-67074333-INS-82 |
| chr16-67077940-INS-157 | chr16-67077941-INS-181 |
| chr16-67098379-INS-712 | chr16-67098379-INS-711 |
| chr16-67132369-INS-2121 | chr16-67132370-INS-2076 |
| chr16-67162046-INS-114 | chr16-67162046-INS-114 |
| chr16-67162642-INS-55 | chr16-67162642-INS-58 |
| chr16-67191393-INS-521 | chr16-67191403-INS-517 |
| chr16-67211421-INS-64 | chr16-67211441-INS-56 |
| chr16-67219695-INS-66 | chr16-67219721-INS-56 |
| chr16-67230735-INS-6283 | chr16-67230736-INS-6286 |
| chr16-67288440-INS-1147 | chr16-67288441-INS-1147 |
| chr16-67289021-INS-113 | chr16-67289021-INS-113 |
| chr16-67301240-INS-108 | chr16-67301240-INS-91 |
| chr16-67395104-INS-410 | chr16-67395104-INS-410 |
| chr16-67469571-INS-70 | chr16-67469571-INS-70 |
| chr16-67479552-INS-85 | chr16-67479552-INS-85 |
| chr16-67484359-INS-281 | chr16-67484359-INS-281 |
| chr16-67508242-INS-4475 | chr16-67508243-INS-4475 |
| chr16-67521107-INS-154 | chr16-67521107-INS-125 |
| chr16-67534180-INS-244 | chr16-67534180-INS-244 |
| chr16-67556188-INS-263 | chr16-67556162-INS-249 |
| chr16-67600811-INS-6101 | chr16-67600812-INS-6085 |
| chr16-67612240-INS-690 | chr16-67612240-INS-690 |
| chr16-67644123-INS-5467 | chr16-67644124-INS-5467 |
| chr16-67651149-INS-734 | chr16-67651149-INS-734 |
| chr16-67652043-INS-113 | chr16-67652043-INS-146 |
| chr16-67679988-INS-70 | chr16-67679975-INS-50 |
| chr16-67686619-INS-145 | chr16-67686619-INS-145 |
| chr16-67696644-INS-7101 | chr16-67696645-INS-7103 |
| chr16-67701291-INS-1611 | chr16-67701292-INS-1611 |
| chr16-67746476-INS-1122 | chr16-67746477-INS-1124 |
| chr16-67786579-INS-266 | chr16-67786579-INS-300 |
| chr16-67787830-INS-62 | chr16-67787830-INS-66 |
| chr16-67828597-INS-62 | chr16-67828597-INS-66 |
| chr16-67866205-INS-58 | chr16-67866195-INS-66 |
| chr16-67909320-INS-1455 | chr16-67909321-INS-1455 |
| chr16-67923874-INS-53 | chr16-67923874-INS-53 |
| chr16-67927236-INS-298 | chr16-67927236-INS-297 |
| chr16-67934715-INS-2928 | chr16-67934716-INS-2928 |
| chr16-67965315-INS-4716 | chr16-67965316-INS-4716 |
| chr16-67977608-INS-267 | chr16-67977608-INS-247 |
| chr16-67984832-INS-916 | chr16-67984858-INS-930 |
| chr16-67985764-INS-65 | chr16-67985764-INS-55 |
| chr16-67993559-INS-73 | chr16-67993559-INS-73 |
| chr16-68004001-INS-214 | chr16-68004001-INS-215 |
| chr16-68052527-INS-54 | chr16-68052527-INS-54 |
| chr16-68059435-INS-76 | chr16-68059435-INS-56 |
| chr16-68069134-INS-389 | chr16-68069134-INS-389 |
| chr16-68069955-INS-571 | chr16-68069955-INS-571 |
| chr16-68070795-INS-233 | chr16-68070795-INS-233 |
| chr16-68081837-INS-100 | chr16-68081837-INS-100 |
| chr16-68084030-INS-72 | chr16-68084032-INS-58 |
| chr16-68092867-INS-275 | chr16-68092867-INS-275 |
| chr16-68122491-INS-404 | chr16-68122491-INS-405 |
| chr16-68125030-INS-102 | chr16-68125038-INS-52 |
| chr16-68150592-INS-724 | chr16-68150592-INS-710 |
| chr16-68158524-INS-127 | chr16-68158525-INS-127 |
| chr16-68169202-INS-64 | chr16-68169202-INS-56 |
| chr16-68170681-INS-445 | chr16-68170681-INS-447 |
| chr16-68190086-INS-351 | chr16-68190086-INS-344 |
| chr16-68213558-INS-85 | chr16-68213558-INS-85 |
| chr16-68234396-INS-656 | chr16-68234396-INS-657 |
| chr16-68283902-INS-358 | chr16-68283902-INS-360 |
| chr16-68314953-INS-87 | chr16-68314953-INS-87 |
| chr16-68321764-INS-574 | chr16-68321764-INS-574 |

| | |
|---|---|
| chr16-68353983-INS-969 | chr16-68353983-INS-969 |
| chr16-68365235-INS-221 | chr16-68365235-INS-219 |
| chr16-68378289-INS-62 | chr16-68378289-INS-50 |
| chr16-68385734-INS-329 | chr16-68385734-INS-336 |
| chr16-68393437-INS-188 | chr16-68393437-INS-188 |
| chr16-68424165-INS-118 | chr16-68424165-INS-128 |
| chr16-68425946-INS-193 | chr16-68425946-INS-194 |
| chr16-68428058-INS-53 | chr16-68428058-INS-54 |
| chr16-68435752-INS-84 | chr16-68435753-INS-84 |
| chr16-68437849-INS-78 | chr16-68437849-INS-94 |
| chr16-68446279-INS-52 | chr16-68446275-INS-68 |
| chr16-68455141-INS-70 | chr16-68455141-INS-72 |
| chr16-68456473-INS-52 | chr16-68456473-INS-52 |
| chr16-68460207-INS-68 | chr16-68460185-INS-68 |
| chr16-68486404-INS-82 | chr16-68486404-INS-64 |
| chr16-68514663-INS-96 | chr16-68514663-INS-96 |
| chr16-68526193-INS-65 | chr16-68526194-INS-56 |
| chr16-68544257-INS-584 | chr16-68544257-INS-586 |
| chr16-68611959-INS-144 | chr16-68611959-INS-108 |
| chr16-68617695-INS-66 | chr16-68617695-INS-66 |
| chr16-68660332-INS-120 | chr16-68660332-INS-96 |
| chr16-68669600-INS-302 | chr16-68669600-INS-283 |
| chr16-68693938-INS-180 | chr16-68693938-INS-180 |
| chr16-68730964-INS-2178 | chr16-68730965-INS-2178 |
| chr16-68745161-INS-79 | chr16-68745161-INS-81 |
| chr16-68827309-INS-80 | chr16-68827309-INS-80 |
| chr16-68866414-INS-164 | chr16-68866412-INS-162 |
| chr16-68875134-INS-54 | chr16-68875134-INS-60 |
| chr16-68906578-INS-222 | chr16-68906569-INS-212 |
| chr16-68946451-INS-74 | chr16-68946471-INS-80 |
| chr16-68986010-INS-53 | chr16-68986010-INS-53 |
| chr16-68990823-INS-365 | chr16-68990823-INS-365 |
| chr16-68992476-INS-223 | chr16-68992476-INS-223 |
| chr16-69019020-INS-194 | chr16-69019020-INS-194 |
| chr16-69058013-INS-116 | chr16-69058013-INS-122 |
| chr16-69058843-INS-55 | chr16-69058843-INS-55 |
| chr16-69060837-INS-72 | chr16-69060841-INS-76 |
| chr16-69088458-INS-96 | chr16-69088458-INS-78 |
| chr16-69103360-INS-295 | chr16-69103361-INS-264 |
| chr16-69119370-INS-56 | chr16-69119370-INS-54 |
| chr16-69152667-INS-103 | chr16-69152667-INS-73 |
| chr16-69156028-INS-638 | chr16-69156028-INS-638 |
| chr16-69252431-INS-804 | chr16-69252431-INS-790 |
| chr16-69253470-INS-79 | chr16-69253471-INS-102 |
| chr16-69254487-INS-66 | chr16-69254487-INS-66 |
| chr16-69265700-INS-203 | chr16-69265700-INS-203 |
| chr16-69268308-INS-60 | chr16-69268308-INS-69 |
| chr16-69269918-INS-738 | chr16-69269918-INS-745 |
| chr16-69275751-INS-78 | chr16-69275755-INS-122 |
| chr16-69285064-INS-6423 | chr16-69285065-INS-6423 |
| chr16-69304159-INS-93 | chr16-69304159-INS-97 |
| chr16-69314716-INS-106 | chr16-69314716-INS-106 |
| chr16-69344590-INS-574 | chr16-69344590-INS-574 |
| chr16-69370328-INS-122 | chr16-69370328-INS-128 |
| chr16-69394630-INS-956 | chr16-69394630-INS-955 |
| chr16-69416544-INS-94 | chr16-69416532-INS-94 |
| chr16-69433810-INS-593 | chr16-69433810-INS-593 |
| chr16-69443795-INS-1265 | chr16-69443796-INS-1267 |
| chr16-69446858-INS-76 | chr16-69446858-INS-56 |
| chr16-69503698-INS-76 | chr16-69503710-INS-78 |
| chr16-69550572-INS-127 | chr16-69550546-INS-108 |
| chr16-69559955-INS-6553 | chr16-69559956-INS-6553 |
| chr16-69563287-INS-86 | chr16-69563285-INS-70 |
| chr16-69563906-INS-62 | chr16-69563906-INS-78 |
| chr16-69576318-INS-6217 | chr16-69576319-INS-6218 |
| chr16-69584666-INS-58 | chr16-69584666-INS-62 |
| chr16-69588891-INS-220 | chr16-69588891-INS-221 |
| chr16-69592775-INS-62 | chr16-69592775-INS-62 |
| chr16-69598756-INS-76 | chr16-69598756-INS-76 |
| chr16-69606981-INS-1893 | chr16-69606982-INS-1893 |
| chr16-69612563-INS-64 | chr16-69612563-INS-64 |
| chr16-69697254-INS-55 | chr16-69697256-INS-56 |
| chr16-69730478-INS-54 | chr16-69730478-INS-56 |
| chr16-69755736-INS-604 | chr16-69755736-INS-604 |
| chr16-69789925-INS-68 | chr16-69789923-INS-70 |
| chr16-69799549-INS-1108 | chr16-69799550-INS-1111 |
| chr16-69800840-INS-264 | chr16-69800840-INS-264 |

| | |
|---|---|
| chr16-69808871-INS-72 | chr16-69808871-INS-78 |
| chr16-69819517-INS-2146 | chr16-69819518-INS-2146 |
| chr16-69853670-INS-7017 | chr16-69853671-INS-7018 |
| chr16-69860222-INS-89 | chr16-69860267-INS-50 |
| chr16-69872057-INS-54 | chr16-69872057-INS-56 |
| chr16-69875095-INS-84 | chr16-69875095-INS-68 |
| chr16-69891539-INS-1466 | chr16-69891540-INS-1466 |
| chr16-69902640-INS-58 | chr16-69902614-INS-76 |
| chr16-69907902-INS-95 | chr16-69907881-INS-85 |
| chr16-69925437-INS-1785 | chr16-69925453-INS-1813 |
| chr16-69947350-INS-104 | chr16-69947350-INS-92 |
| chr16-69953936-INS-7104 | chr16-69953937-INS-7104 |
| chr16-69982911-INS-126 | chr16-69982911-INS-126 |
| chr16-69983808-INS-3008 | chr16-69983809-INS-3008 |
| chr16-69988595-INS-144 | chr16-69988595-INS-144 |
| chr16-70003997-INS-56 | chr16-70003997-INS-56 |
| chr16-70043356-INS-304 | chr16-70043356-INS-309 |
| chr16-70054634-INS-58 | chr16-70054634-INS-58 |
| chr16-70082635-INS-105 | chr16-70082635-INS-96 |
| chr16-70118526-INS-78 | chr16-70118526-INS-78 |
| chr16-70161550-INS-95 | chr16-70161550-INS-95 |
| chr16-70170196-INS-417 | chr16-70170196-INS-408 |
| chr16-70185941-INS-4748 | chr16-70185942-INS-4748 |
| chr16-70190721-INS-102 | chr16-70190720-INS-58 |
| chr16-70194978-INS-124 | chr16-70194978-INS-124 |
| chr16-70197220-INS-2863 | chr16-70197221-INS-2864 |
| chr16-70203515-INS-431 | chr16-70203515-INS-430 |
| chr16-70209140-INS-67 | chr16-70209138-INS-74 |
| chr16-70210387-INS-58 | chr16-70210387-INS-58 |
| chr16-70228695-INS-93 | chr16-70228695-INS-110 |
| chr16-70252573-INS-78 | chr16-70252573-INS-112 |
| chr16-70254955-INS-100 | chr16-70254955-INS-100 |
| chr16-70260835-INS-76 | chr16-70260835-INS-77 |
| chr16-70302202-INS-2271 | chr16-70302203-INS-2271 |
| chr16-70304227-INS-80 | chr16-70304227-INS-88 |
| chr16-70328527-INS-116 | chr16-70328527-INS-116 |
| chr16-70381644-INS-52 | chr16-70381652-INS-56 |
| chr16-70384616-INS-442 | chr16-70384616-INS-442 |
| chr16-70392023-INS-7126 | chr16-70392024-INS-7126 |
| chr16-70400737-INS-6359 | chr16-70400738-INS-6360 |
| chr16-70404762-INS-222 | chr16-70404762-INS-224 |
| chr16-70416346-INS-4382 | chr16-70416347-INS-4382 |
| chr16-70501055-INS-921 | chr16-70501055-INS-921 |
| chr16-70518829-INS-74 | chr16-70518829-INS-74 |
| chr16-70531992-INS-98 | chr16-70531995-INS-86 |
| chr16-70535770-INS-63 | chr16-70535786-INS-67 |
| chr16-70549943-INS-534 | chr16-70549943-INS-534 |
| chr16-70574872-INS-64 | chr16-70574887-INS-90 |
| chr16-70596501-INS-8138 | chr16-70596501-INS-8135 |
| chr16-70613421-INS-56 | chr16-70613421-INS-56 |
| chr16-70628001-INS-92 | chr16-70628001-INS-88 |
| chr16-70638071-INS-1894 | chr16-70638072-INS-1894 |
| chr16-70684802-INS-441 | chr16-70684802-INS-474 |
| chr16-70703798-INS-267 | chr16-70703798-INS-271 |
| chr16-70705865-INS-731 | chr16-70705864-INS-732 |
| chr16-70735244-INS-194 | chr16-70735244-INS-195 |
| chr16-70772730-INS-70 | chr16-70772730-INS-70 |
| chr16-70784332-INS-3092 | chr16-70784333-INS-3092 |
| chr16-70792816-INS-212 | chr16-70792816-INS-213 |
| chr16-70815737-INS-2377 | chr16-70815738-INS-2377 |
| chr16-70817123-INS-94 | chr16-70817123-INS-94 |
| chr16-70820757-INS-7086 | chr16-70820758-INS-7089 |
| chr16-70873624-INS-58 | chr16-70873624-INS-60 |
| chr16-70907171-INS-52 | chr16-70907171-INS-50 |
| chr16-70928959-INS-52 | chr16-70928936-INS-52 |
| chr16-70940368-INS-81 | chr16-70940381-INS-109 |
| chr16-70942476-INS-52 | chr16-70942476-INS-56 |
| chr16-70982229-INS-199 | chr16-70982229-INS-199 |
| chr16-71045117-INS-782 | chr16-71045117-INS-785 |
| chr16-71057067-INS-2535 | chr16-71057068-INS-2535 |
| chr16-71072937-INS-53 | chr16-71072910-INS-56 |
| chr16-71087209-INS-159 | chr16-71087209-INS-159 |
| chr16-71088852-INS-52 | chr16-71088834-INS-56 |
| chr16-71090899-INS-483 | chr16-71090899-INS-483 |
| chr16-71093022-INS-2042 | chr16-71093023-INS-2042 |
| chr16-71094008-INS-3344 | chr16-71094009-INS-3344 |
| chr16-71108823-INS-72 | chr16-71108823-INS-72 |

| | |
|---|---|
| chr16-71117463-INS-68 | chr16-71117424-INS-60 |
| chr16-71139286-INS-6642 | chr16-71139287-INS-6640 |
| chr16-71150195-INS-366 | chr16-71150195-INS-366 |
| chr16-71166846-INS-204 | chr16-71166846-INS-204 |
| chr16-71174414-INS-272 | chr16-71174415-INS-273 |
| chr16-71178111-INS-1137 | chr16-71178112-INS-1137 |
| chr16-71236060-INS-526 | chr16-71236060-INS-526 |
| chr16-71262675-INS-64 | chr16-71262705-INS-68 |
| chr16-71293972-INS-85 | chr16-71293970-INS-90 |
| chr16-71301639-INS-63 | chr16-71301639-INS-80 |
| chr16-71311249-INS-7089 | chr16-71311250-INS-7088 |
| chr16-71319480-INS-212 | chr16-71319480-INS-212 |
| chr16-71333170-INS-285 | chr16-71333170-INS-288 |
| chr16-71369730-INS-178 | chr16-71369730-INS-178 |
| chr16-71394013-INS-132 | chr16-71394013-INS-132 |
| chr16-71403318-INS-747 | chr16-71403318-INS-766 |
| chr16-71417462-INS-95 | chr16-71417450-INS-107 |
| chr16-71421869-INS-51 | chr16-71421860-INS-51 |
| chr16-71430703-INS-68 | chr16-71430703-INS-56 |
| chr16-71443044-INS-54 | chr16-71443049-INS-60 |
| chr16-71444054-INS-60 | chr16-71444044-INS-96 |
| chr16-71446374-INS-2026 | chr16-71446375-INS-2026 |
| chr16-71466563-INS-412 | chr16-71466564-INS-409 |
| chr16-71480026-INS-68 | chr16-71480026-INS-78 |
| chr16-71481895-INS-60 | chr16-71481895-INS-56 |
| chr16-71505795-INS-514 | chr16-71505796-INS-549 |
| chr16-71509909-INS-76 | chr16-71509908-INS-64 |
| chr16-71518559-INS-2333 | chr16-71518560-INS-2319 |
| chr16-71542067-INS-94 | chr16-71542079-INS-112 |
| chr16-71580591-INS-426 | chr16-71580591-INS-427 |
| chr16-71623455-INS-96 | chr16-71623455-INS-96 |
| chr16-71628402-INS-72 | chr16-71628402-INS-74 |
| chr16-71630054-INS-107 | chr16-71630054-INS-108 |
| chr16-71631980-INS-99 | chr16-71631980-INS-101 |
| chr16-71669494-INS-441 | chr16-71669496-INS-443 |
| chr16-71681413-INS-65 | chr16-71681413-INS-71 |
| chr16-71690493-INS-56 | chr16-71690493-INS-56 |
| chr16-71697545-INS-50 | chr16-71697545-INS-58 |
| chr16-71706797-INS-557 | chr16-71706797-INS-556 |
| chr16-71718071-INS-1122 | chr16-71718072-INS-1122 |
| chr16-71744082-INS-6888 | chr16-71744083-INS-6888 |
| chr16-71785984-INS-6317 | chr16-71785985-INS-6318 |
| chr16-71797793-INS-353 | chr16-71797793-INS-373 |
| chr16-71798966-INS-267 | chr16-71798966-INS-265 |
| chr16-71826560-INS-68 | chr16-71826560-INS-68 |
| chr16-71839869-INS-534 | chr16-71839869-INS-531 |
| chr16-71849842-INS-1138 | chr16-71849843-INS-1159 |
| chr16-71859763-INS-213 | chr16-71859763-INS-214 |
| chr16-71871955-INS-76 | chr16-71871955-INS-78 |
| chr16-71884378-INS-384 | chr16-71884378-INS-383 |
| chr16-71901083-INS-127 | chr16-71901083-INS-133 |
| chr16-71913122-INS-335 | chr16-71913122-INS-333 |
| chr16-71915438-INS-343 | chr16-71915438-INS-343 |
| chr16-71920440-INS-535 | chr16-71920440-INS-535 |
| chr16-71934745-INS-78 | chr16-71934745-INS-58 |
| chr16-71962977-INS-56 | chr16-71962977-INS-52 |
| chr16-71986883-INS-979 | chr16-71986883-INS-979 |
| chr16-72031645-INS-56 | chr16-72031645-INS-56 |
| chr16-72067650-INS-60 | chr16-72067650-INS-60 |
| chr16-72072380-INS-1360 | chr16-72072381-INS-1360 |
| chr16-72081485-INS-6437 | chr16-72081486-INS-6437 |
| chr16-72090987-INS-79 | chr16-72090987-INS-52 |
| chr16-72093027-INS-291 | chr16-72093028-INS-290 |
| chr16-72137416-INS-391 | chr16-72137416-INS-391 |
| chr16-72149808-INS-121 | chr16-72149806-INS-121 |
| chr16-72168409-INS-76 | chr16-72168409-INS-76 |
| chr16-72173030-INS-848 | chr16-72173030-INS-849 |
| chr16-72176368-INS-1515 | chr16-72176369-INS-1515 |
| chr16-72183747-INS-875 | chr16-72183747-INS-874 |
| chr16-72185461-INS-74 | chr16-72185461-INS-72 |
| chr16-72188068-INS-401 | chr16-72188068-INS-400 |
| chr16-72190228-INS-605 | chr16-72190228-INS-605 |
| chr16-72192030-INS-66 | chr16-72192030-INS-66 |
| chr16-72196419-INS-88 | chr16-72196419-INS-88 |
| chr16-72205874-INS-946 | chr16-72205878-INS-945 |
| chr16-72212648-INS-132 | chr16-72212600-INS-168 |
| chr16-72223974-INS-194 | chr16-72223974-INS-196 |

| | |
|---|---|
| chr16-72226297-INS-52 | chr16-72226297-INS-54 |
| chr16-72265895-INS-68 | chr16-72265896-INS-53 |
| chr16-72312841-INS-56 | chr16-72312841-INS-71 |
| chr16-72375021-INS-100 | chr16-72375021-INS-100 |
| chr16-72387011-INS-6445 | chr16-72387012-INS-6447 |
| chr16-72407071-INS-199 | chr16-72407071-INS-199 |
| chr16-72409602-INS-67 | chr16-72409602-INS-52 |
| chr16-72413606-INS-110 | chr16-72413606-INS-110 |
| chr16-72481641-INS-68 | chr16-72481622-INS-50 |
| chr16-72508862-INS-340 | chr16-72508862-INS-344 |
| chr16-72546824-INS-59 | chr16-72546824-INS-57 |
| chr16-72547042-INS-498 | chr16-72547042-INS-498 |
| chr16-72551070-INS-58 | chr16-72551070-INS-58 |
| chr16-72565790-INS-136 | chr16-72565790-INS-136 |
| chr16-72572872-INS-63 | chr16-72572872-INS-63 |
| chr16-72606297-INS-70 | chr16-72606297-INS-70 |
| chr16-72641221-INS-115 | chr16-72641221-INS-115 |
| chr16-72648903-INS-78 | chr16-72648903-INS-78 |
| chr16-72654628-INS-103 | chr16-72654628-INS-103 |
| chr16-72695613-INS-6445 | chr16-72695614-INS-6445 |
| chr16-72745567-INS-156 | chr16-72745567-INS-156 |
| chr16-72752452-INS-65 | chr16-72752452-INS-55 |
| chr16-72787443-INS-166 | chr16-72787443-INS-166 |
| chr16-72847270-INS-113 | chr16-72847270-INS-113 |
| chr16-72855585-INS-6695 | chr16-72855586-INS-6697 |
| chr16-72860661-INS-160 | chr16-72860647-INS-150 |
| chr16-72891685-INS-52 | chr16-72891685-INS-56 |
| chr16-72898838-INS-157 | chr16-72898838-INS-153 |
| chr16-72906710-INS-831 | chr16-72906710-INS-831 |
| chr16-72927221-INS-6252 | chr16-72927222-INS-6252 |
| chr16-72928034-INS-69 | chr16-72928034-INS-114 |
| chr16-72952625-INS-383 | chr16-72952625-INS-383 |
| chr16-72953210-INS-104 | chr16-72953203-INS-104 |
| chr16-72967383-INS-72 | chr16-72967363-INS-88 |
| chr16-72970703-INS-282 | chr16-72970697-INS-289 |
| chr16-72971380-INS-64 | chr16-72971380-INS-60 |
| chr16-73004574-INS-63 | chr16-73004575-INS-59 |
| chr16-73006680-INS-76 | chr16-73006676-INS-60 |
| chr16-73018043-INS-56 | chr16-73018043-INS-70 |
| chr16-73041328-INS-55 | chr16-73041313-INS-75 |
| chr16-73064937-INS-164 | chr16-73064937-INS-164 |
| chr16-73077934-INS-181 | chr16-73077934-INS-181 |
| chr16-73094809-INS-181 | chr16-73094857-INS-165 |
| chr16-73099705-INS-6382 | chr16-73099706-INS-6382 |
| chr16-73138005-INS-144 | chr16-73138005-INS-144 |
| chr16-73150745-INS-65 | chr16-73150745-INS-65 |
| chr16-73175420-INS-415 | chr16-73175421-INS-416 |
| chr16-73190443-INS-53 | chr16-73190443-INS-58 |
| chr16-73206923-INS-206 | chr16-73206923-INS-207 |
| chr16-73207858-INS-82 | chr16-73207858-INS-88 |
| chr16-73219036-INS-639 | chr16-73219036-INS-637 |
| chr16-73237233-INS-52 | chr16-73237233-INS-52 |
| chr16-73264308-INS-164 | chr16-73264308-INS-164 |
| chr16-73268074-INS-74 | chr16-73268074-INS-76 |
| chr16-73343081-INS-386 | chr16-73343081-INS-386 |
| chr16-73358307-INS-1921 | chr16-73358308-INS-1921 |
| chr16-73361977-INS-142 | chr16-73361977-INS-142 |
| chr16-73372197-INS-7191 | chr16-73372198-INS-7191 |
| chr16-73382593-INS-962 | chr16-73382598-INS-982 |
| chr16-73399289-INS-154 | chr16-73399289-INS-165 |
| chr16-73402039-INS-853 | chr16-73402039-INS-855 |
| chr16-73435812-INS-396 | chr16-73435812-INS-395 |
| chr16-73437255-INS-2752 | chr16-73437256-INS-2767 |
| chr16-73437411-INS-607 | chr16-73437411-INS-607 |
| chr16-73485948-INS-7113 | chr16-73485949-INS-7114 |
| chr16-73517850-INS-116 | chr16-73517838-INS-124 |
| chr16-73560689-INS-66 | chr16-73560689-INS-70 |
| chr16-73562534-INS-6806 | chr16-73562535-INS-6806 |
| chr16-73573611-INS-201 | chr16-73573611-INS-201 |
| chr16-73579924-INS-796 | chr16-73579924-INS-796 |
| chr16-73594642-INS-54 | chr16-73594642-INS-60 |
| chr16-73600107-INS-53 | chr16-73600107-INS-58 |
| chr16-73608913-INS-1792 | chr16-73608917-INS-1781 |
| chr16-73611663-INS-6670 | chr16-73611664-INS-6670 |
| chr16-73617094-INS-1711 | chr16-73617095-INS-1710 |
| chr16-73618548-INS-474 | chr16-73618548-INS-474 |
| chr16-73627781-INS-68 | chr16-73627777-INS-62 |

| | |
|---|---|
| chr16-73628530-INS-88 | chr16-73628530-INS-88 |
| chr16-73644058-INS-1883 | chr16-73644059-INS-1883 |
| chr16-73647590-INS-349 | chr16-73647590-INS-349 |
| chr16-73714684-INS-198 | chr16-73714688-INS-198 |
| chr16-73763487-INS-52 | chr16-73763487-INS-56 |
| chr16-73765618-INS-3181 | chr16-73765619-INS-3181 |
| chr16-73853545-INS-61 | chr16-73853545-INS-59 |
| chr16-73906907-INS-455 | chr16-73906907-INS-455 |
| chr16-73908713-INS-60 | chr16-73908713-INS-56 |
| chr16-73916283-INS-7075 | chr16-73916284-INS-7076 |
| chr16-73960517-INS-422 | chr16-73960517-INS-422 |
| chr16-73966361-INS-56 | chr16-73966361-INS-50 |
| chr16-73973961-INS-192 | chr16-73973961-INS-192 |
| chr16-73985800-INS-465 | chr16-73985800-INS-466 |
| chr16-73993857-INS-108 | chr16-73993859-INS-62 |
| chr16-74015553-INS-1102 | chr16-74015554-INS-1109 |
| chr16-74015984-INS-1727 | chr16-74015985-INS-1729 |
| chr16-74016861-INS-116 | chr16-74016861-INS-116 |
| chr16-74017912-INS-80 | chr16-74017912-INS-74 |
| chr16-74025788-INS-60 | chr16-74025788-INS-72 |
| chr16-74160167-INS-201 | chr16-74160167-INS-201 |
| chr16-74212081-INS-53 | chr16-74212115-INS-63 |
| chr16-74214054-INS-69 | chr16-74214056-INS-79 |
| chr16-74226908-INS-654 | chr16-74226908-INS-653 |
| chr16-74238240-INS-242 | chr16-74238240-INS-242 |
| chr16-74270958-INS-518 | chr16-74270958-INS-517 |
| chr16-74276523-INS-62 | chr16-74276523-INS-62 |
| chr16-74291205-INS-846 | chr16-74291199-INS-850 |
| chr16-74317216-INS-74 | chr16-74317216-INS-69 |
| chr16-74326408-INS-162 | chr16-74326408-INS-162 |
| chr16-74331975-INS-56 | chr16-74331975-INS-57 |
| chr16-74336637-INS-326 | chr16-74336639-INS-356 |
| chr16-74339986-INS-158 | chr16-74339986-INS-152 |
| chr16-74352693-INS-205 | chr16-74352693-INS-205 |
| chr16-74367366-INS-182 | chr16-74367366-INS-182 |
| chr16-74391689-INS-823 | chr16-74391689-INS-823 |
| chr16-74403575-INS-86 | chr16-74403549-INS-80 |
| chr16-74407070-INS-52 | chr16-74407070-INS-52 |
| chr16-74422485-INS-56 | chr16-74422485-INS-56 |
| chr16-74426824-INS-71 | chr16-74426824-INS-71 |
| chr16-74429179-INS-60 | chr16-74429179-INS-60 |
| chr16-74439607-INS-113 | chr16-74439623-INS-72 |
| chr16-74450985-INS-228 | chr16-74450985-INS-228 |
| chr16-74463933-INS-6610 | chr16-74463934-INS-6610 |
| chr16-74468968-INS-57 | chr16-74468968-INS-63 |
| chr16-74489390-INS-996 | chr16-74489390-INS-996 |
| chr16-74502675-INS-464 | chr16-74502675-INS-464 |
| chr16-74540915-INS-834 | chr16-74540915-INS-833 |
| chr16-74557340-INS-72 | chr16-74557340-INS-76 |
| chr16-74558273-INS-354 | chr16-74558273-INS-354 |
| chr16-74566403-INS-56 | chr16-74566403-INS-56 |
| chr16-74587834-INS-65 | chr16-74587834-INS-80 |
| chr16-74618813-INS-2054 | chr16-74618814-INS-2054 |
| chr16-74625382-INS-159 | chr16-74625382-INS-135 |
| chr16-74657347-INS-54 | chr16-74657359-INS-88 |
| chr16-74679920-INS-56 | chr16-74679920-INS-52 |
| chr16-74689944-INS-413 | chr16-74689944-INS-413 |
| chr16-74713143-INS-63 | chr16-74713139-INS-67 |
| chr16-74734439-INS-51 | chr16-74734439-INS-51 |
| chr16-74752844-INS-66 | chr16-74752844-INS-68 |
| chr16-74814932-INS-70 | chr16-74814932-INS-68 |
| chr16-74831248-INS-875 | chr16-74831248-INS-875 |
| chr16-74845483-INS-2495 | chr16-74845484-INS-2495 |
| chr16-74875769-INS-4054 | chr16-74875770-INS-4054 |
| chr16-74910180-INS-84 | chr16-74910180-INS-88 |
| chr16-74911466-INS-1897 | chr16-74911467-INS-1897 |
| chr16-74915888-INS-481 | chr16-74915888-INS-481 |
| chr16-74924373-INS-52 | chr16-74924373-INS-68 |
| chr16-74927802-INS-62 | chr16-74927802-INS-64 |
| chr16-74934509-INS-460 | chr16-74934509-INS-460 |
| chr16-74977420-INS-80 | chr16-74977400-INS-55 |
| chr16-74983675-INS-86 | chr16-74983691-INS-50 |
| chr16-75007633-INS-849 | chr16-75007633-INS-849 |
| chr16-75021598-INS-58 | chr16-75021611-INS-52 |
| chr16-75033773-INS-76 | chr16-75033773-INS-76 |
| chr16-75047175-INS-97 | chr16-75047175-INS-97 |
| chr16-75065242-INS-824 | chr16-75065242-INS-824 |

| | |
|---|---|
| chr16-75093261-INS-194 | chr16-75093261-INS-194 |
| chr16-75109062-INS-460 | chr16-75109062-INS-463 |
| chr16-75113136-INS-55 | chr16-75113136-INS-55 |
| chr16-75125769-INS-120 | chr16-75125769-INS-114 |
| chr16-75134592-INS-80 | chr16-75134594-INS-88 |
| chr16-75175389-INS-133 | chr16-75175389-INS-143 |
| chr16-75178177-INS-54 | chr16-75178177-INS-56 |
| chr16-75196730-INS-915 | chr16-75196730-INS-889 |
| chr16-75201956-INS-2178 | chr16-75201957-INS-2178 |
| chr16-75322349-INS-56 | chr16-75322332-INS-58 |
| chr16-75325433-INS-194 | chr16-75325433-INS-194 |
| chr16-75332272-INS-70 | chr16-75332235-INS-86 |
| chr16-75348374-INS-418 | chr16-75348374-INS-424 |
| chr16-75368120-INS-1618 | chr16-75368121-INS-1618 |
| chr16-75368121-INS-802 | chr16-75368121-INS-803 |
| chr16-75381647-INS-52 | chr16-75381647-INS-52 |
| chr16-75408021-INS-69 | chr16-75408021-INS-85 |
| chr16-75415016-INS-160 | chr16-75415002-INS-158 |
| chr16-75427271-INS-63 | chr16-75427271-INS-63 |
| chr16-75444448-INS-54 | chr16-75444448-INS-52 |
| chr16-75449592-INS-154 | chr16-75449608-INS-151 |
| chr16-75472362-INS-199 | chr16-75472362-INS-199 |
| chr16-75509956-INS-327 | chr16-75509956-INS-328 |
| chr16-75534220-INS-208 | chr16-75534220-INS-208 |
| chr16-75564805-INS-130 | chr16-75564805-INS-91 |
| chr16-75570780-INS-72 | chr16-75570780-INS-72 |
| chr16-75576037-INS-3234 | chr16-75576038-INS-3234 |
| chr16-75582478-INS-282 | chr16-75582478-INS-282 |
| chr16-75585062-INS-266 | chr16-75585062-INS-267 |
| chr16-75586679-INS-213 | chr16-75586683-INS-223 |
| chr16-75622994-INS-68 | chr16-75622994-INS-67 |
| chr16-75636609-INS-358 | chr16-75636609-INS-358 |
| chr16-75643989-INS-66 | chr16-75643989-INS-66 |
| chr16-75655570-INS-7918 | chr16-75655571-INS-7919 |
| chr16-75663055-INS-58 | chr16-75663055-INS-58 |
| chr16-75666860-INS-66 | chr16-75666860-INS-72 |
| chr16-75745494-INS-56 | chr16-75745494-INS-56 |
| chr16-75761800-INS-174 | chr16-75761800-INS-174 |
| chr16-75766945-INS-6572 | chr16-75766946-INS-6572 |
| chr16-75769936-INS-3246 | chr16-75769937-INS-3246 |
| chr16-75804415-INS-58 | chr16-75804415-INS-58 |
| chr16-75822504-INS-1134 | chr16-75822505-INS-1134 |
| chr16-75828759-INS-104 | chr16-75828759-INS-72 |
| chr16-75834633-INS-91 | chr16-75834633-INS-91 |
| chr16-75875016-INS-71 | chr16-75875016-INS-71 |
| chr16-75887547-INS-86 | chr16-75887547-INS-86 |
| chr16-75889415-INS-550 | chr16-75889415-INS-550 |
| chr16-75919868-INS-54 | chr16-75919868-INS-57 |
| chr16-75924972-INS-2816 | chr16-75924973-INS-2816 |
| chr16-75981436-INS-128 | chr16-75981436-INS-116 |
| chr16-76003713-INS-80 | chr16-76003710-INS-110 |
| chr16-76005512-INS-71 | chr16-76005516-INS-67 |
| chr16-76106744-INS-69 | chr16-76106756-INS-51 |
| chr16-76142879-INS-54 | chr16-76142879-INS-54 |
| chr16-76157034-INS-186 | chr16-76157034-INS-186 |
| chr16-76163696-INS-132 | chr16-76163699-INS-133 |
| chr16-76179772-INS-61 | chr16-76179772-INS-83 |
| chr16-76182104-INS-653 | chr16-76182104-INS-654 |
| chr16-76202596-INS-54 | chr16-76202596-INS-54 |
| chr16-76222256-INS-187 | chr16-76222256-INS-188 |
| chr16-76232705-INS-100 | chr16-76232674-INS-121 |
| chr16-76255055-INS-736 | chr16-76255055-INS-740 |
| chr16-76279868-INS-173 | chr16-76279867-INS-174 |
| chr16-76282175-INS-113 | chr16-76282187-INS-102 |
| chr16-76326470-INS-4579 | chr16-76326471-INS-4579 |
| chr16-76333050-INS-66 | chr16-76333050-INS-72 |
| chr16-76354855-INS-367 | chr16-76354855-INS-367 |
| chr16-76361649-INS-177 | chr16-76361649-INS-177 |
| chr16-76370804-INS-88 | chr16-76370814-INS-114 |
| chr16-76388423-INS-75 | chr16-76388423-INS-79 |
| chr16-76399606-INS-108 | chr16-76399606-INS-96 |
| chr16-76408901-INS-69 | chr16-76408930-INS-52 |
| chr16-76417033-INS-496 | chr16-76417033-INS-496 |
| chr16-76431793-INS-7114 | chr16-76431794-INS-7115 |
| chr16-76477976-INS-5240 | chr16-76477977-INS-5240 |
| chr16-76496027-INS-2574 | chr16-76496028-INS-2574 |
| chr16-76497176-INS-3146 | chr16-76497177-INS-3146 |

| | |
|---|---|
| chr16-76510124-INS-934 | chr16-76510124-INS-932 |
| chr16-76574316-INS-237 | chr16-76574316-INS-238 |
| chr16-76605106-INS-416 | chr16-76605106-INS-415 |
| chr16-76612752-INS-54 | chr16-76612752-INS-56 |
| chr16-76646781-INS-72 | chr16-76646781-INS-87 |
| chr16-76670329-INS-67 | chr16-76670329-INS-67 |
| chr16-76690119-INS-129 | chr16-76690119-INS-129 |
| chr16-76695285-INS-69 | chr16-76695285-INS-66 |
| chr16-76711731-INS-58 | chr16-76711731-INS-58 |
| chr16-76711978-INS-4171 | chr16-76711979-INS-4171 |
| chr16-76716041-INS-6399 | chr16-76716042-INS-6398 |
| chr16-76721137-INS-131 | chr16-76721137-INS-140 |
| chr16-76736043-INS-92 | chr16-76736043-INS-92 |
| chr16-76743124-INS-52 | chr16-76743124-INS-56 |
| chr16-76760283-INS-77 | chr16-76760283-INS-85 |
| chr16-76768784-INS-989 | chr16-76768784-INS-991 |
| chr16-76769552-INS-102 | chr16-76769552-INS-90 |
| chr16-76778491-INS-88 | chr16-76778491-INS-88 |
| chr16-76799845-INS-532 | chr16-76799845-INS-536 |
| chr16-76813201-INS-77 | chr16-76813200-INS-88 |
| chr16-76824294-INS-2171 | chr16-76824295-INS-2173 |
| chr16-76846111-INS-53 | chr16-76846111-INS-53 |
| chr16-76852476-INS-67 | chr16-76852476-INS-67 |
| chr16-76857274-INS-130 | chr16-76857315-INS-130 |
| chr16-76870565-INS-199 | chr16-76870565-INS-199 |
| chr16-76885169-INS-554 | chr16-76885169-INS-543 |
| chr16-76929082-INS-218 | chr16-76929082-INS-218 |
| chr16-76936414-INS-52 | chr16-76936415-INS-52 |
| chr16-76937436-INS-7004 | chr16-76937437-INS-7004 |
| chr16-76938695-INS-302 | chr16-76938695-INS-301 |
| chr16-76979054-INS-209 | chr16-76979054-INS-209 |
| chr16-76999050-INS-3161 | chr16-76999051-INS-3161 |
| chr16-77039766-INS-6716 | chr16-77039767-INS-6718 |
| chr16-77068636-INS-893 | chr16-77068636-INS-893 |
| chr16-77096492-INS-6236 | chr16-77096493-INS-6236 |
| chr16-77100614-INS-66 | chr16-77100592-INS-88 |
| chr16-77101813-INS-362 | chr16-77101813-INS-362 |
| chr16-77106429-INS-54 | chr16-77106425-INS-70 |
| chr16-77115513-INS-205 | chr16-77115513-INS-206 |
| chr16-77147213-INS-66 | chr16-77147213-INS-66 |
| chr16-77155290-INS-194 | chr16-77155290-INS-194 |
| chr16-77180430-INS-58 | chr16-77180430-INS-58 |
| chr16-77183587-INS-78 | chr16-77183587-INS-84 |
| chr16-77196601-INS-130 | chr16-77196610-INS-142 |
| chr16-77263422-INS-778 | chr16-77263422-INS-779 |
| chr16-77302927-INS-65 | chr16-77302927-INS-66 |
| chr16-77339106-INS-75 | chr16-77339106-INS-64 |
| chr16-77340448-INS-7130 | chr16-77340449-INS-7130 |
| chr16-77360612-INS-61 | chr16-77360612-INS-61 |
| chr16-77369121-INS-703 | chr16-77369118-INS-710 |
| chr16-77379048-INS-84 | chr16-77379048-INS-84 |
| chr16-77381976-INS-52 | chr16-77381976-INS-52 |
| chr16-77383298-INS-76 | chr16-77383298-INS-76 |
| chr16-77385963-INS-175 | chr16-77385963-INS-180 |
| chr16-77397798-INS-87 | chr16-77397801-INS-82 |
| chr16-77434111-INS-162 | chr16-77434111-INS-162 |
| chr16-77442240-INS-92 | chr16-77442240-INS-96 |
| chr16-77444534-INS-118 | chr16-77444534-INS-118 |
| chr16-77526411-INS-289 | chr16-77526411-INS-293 |
| chr16-77532128-INS-185 | chr16-77532128-INS-185 |
| chr16-77535167-INS-105 | chr16-77535152-INS-131 |
| chr16-77579246-INS-333 | chr16-77579246-INS-333 |
| chr16-77603927-INS-154 | chr16-77603935-INS-179 |
| chr16-77671437-INS-561 | chr16-77671437-INS-580 |
| chr16-77683821-INS-54 | chr16-77683821-INS-72 |
| chr16-77707034-INS-1607 | chr16-77707035-INS-1607 |
| chr16-77747120-INS-917 | chr16-77747120-INS-917 |
| chr16-77749540-INS-216 | chr16-77749540-INS-216 |
| chr16-77760364-INS-72 | chr16-77760364-INS-72 |
| chr16-77767767-INS-64 | chr16-77767767-INS-64 |
| chr16-77776710-INS-60 | chr16-77776710-INS-66 |
| chr16-77792850-INS-864 | chr16-77792850-INS-864 |
| chr16-77802472-INS-255 | chr16-77802472-INS-255 |
| chr16-77846962-INS-100 | chr16-77846963-INS-100 |
| chr16-77848764-INS-271 | chr16-77848764-INS-273 |
| chr16-77856543-INS-7231 | chr16-77856544-INS-7231 |
| chr16-77871667-INS-96 | chr16-77871667-INS-104 |

| | |
|---|---|
| chr16-77965148-INS-478 | chr16-77965148-INS-473 |
| chr16-77971082-INS-401 | chr16-77971082-INS-401 |
| chr16-77972896-INS-94 | chr16-77972896-INS-94 |
| chr16-77995144-INS-846 | chr16-77995144-INS-844 |
| chr16-78004471-INS-72 | chr16-78004486-INS-90 |
| chr16-78007981-INS-493 | chr16-78007981-INS-493 |
| chr16-78039027-INS-52 | chr16-78039027-INS-68 |
| chr16-78041152-INS-129 | chr16-78041154-INS-141 |
| chr16-78053480-INS-451 | chr16-78053480-INS-451 |
| chr16-78065018-INS-73 | chr16-78065018-INS-68 |
| chr16-78073354-INS-332 | chr16-78073354-INS-332 |
| chr16-78093058-INS-134 | chr16-78093034-INS-112 |
| chr16-78105577-INS-212 | chr16-78105577-INS-247 |
| chr16-78122315-INS-67 | chr16-78122315-INS-67 |
| chr16-78125595-INS-151 | chr16-78125595-INS-151 |
| chr16-78163559-INS-210 | chr16-78163559-INS-210 |
| chr16-78183140-INS-56 | chr16-78183140-INS-60 |
| chr16-78184691-INS-201 | chr16-78184691-INS-201 |
| chr16-78191129-INS-3262 | chr16-78191130-INS-3262 |
| chr16-78192110-INS-185 | chr16-78192110-INS-186 |
| chr16-78202913-INS-107 | chr16-78202914-INS-58 |
| chr16-78258785-INS-268 | chr16-78258785-INS-268 |
| chr16-78263028-INS-121 | chr16-78263028-INS-87 |
| chr16-78264529-INS-821 | chr16-78264529-INS-821 |
| chr16-78265348-INS-467 | chr16-78265348-INS-467 |
| chr16-78282693-INS-202 | chr16-78282693-INS-205 |
| chr16-78320205-INS-221 | chr16-78320205-INS-223 |
| chr16-78337552-INS-57 | chr16-78337552-INS-63 |
| chr16-78356322-INS-78 | chr16-78356322-INS-78 |
| chr16-78370548-INS-124 | chr16-78370548-INS-124 |
| chr16-78384045-INS-56 | chr16-78384045-INS-62 |
| chr16-78391564-INS-64 | chr16-78391564-INS-64 |
| chr16-78393231-INS-195 | chr16-78393231-INS-195 |
| chr16-78415386-INS-194 | chr16-78415386-INS-194 |
| chr16-78423258-INS-52 | chr16-78423212-INS-62 |
| chr16-78432631-INS-123 | chr16-78432631-INS-123 |
| chr16-78436468-INS-162 | chr16-78436474-INS-140 |
| chr16-78437375-INS-166 | chr16-78437375-INS-167 |
| chr16-78462223-INS-100 | chr16-78462221-INS-100 |
| chr16-78575370-INS-236 | chr16-78575370-INS-235 |
| chr16-78590863-INS-3954 | chr16-78590864-INS-3954 |
| chr16-78600249-INS-405 | chr16-78600249-INS-405 |
| chr16-78611604-INS-362 | chr16-78611604-INS-362 |
| chr16-78612677-INS-2553 | chr16-78612678-INS-2535 |
| chr16-78614652-INS-421 | chr16-78614652-INS-422 |
| chr16-78618940-INS-1216 | chr16-78618941-INS-1222 |
| chr16-78620561-INS-60 | chr16-78620561-INS-60 |
| chr16-78656992-INS-72 | chr16-78656992-INS-79 |
| chr16-78672466-INS-277 | chr16-78672466-INS-275 |
| chr16-78704967-INS-169 | chr16-78704966-INS-127 |
| chr16-78720893-INS-60 | chr16-78720898-INS-70 |
| chr16-78755316-INS-66 | chr16-78755316-INS-72 |
| chr16-78756306-INS-57 | chr16-78756306-INS-57 |
| chr16-78760699-INS-528 | chr16-78760699-INS-528 |
| chr16-78762979-INS-159 | chr16-78762979-INS-159 |
| chr16-78765246-INS-457 | chr16-78765246-INS-457 |
| chr16-78776906-INS-60 | chr16-78776906-INS-64 |
| chr16-78812258-INS-270 | chr16-78812258-INS-270 |
| chr16-78860681-INS-789 | chr16-78860681-INS-789 |
| chr16-78869471-INS-52 | chr16-78869471-INS-52 |
| chr16-78871638-INS-53 | chr16-78871638-INS-53 |
| chr16-78884446-INS-75 | chr16-78884446-INS-75 |
| chr16-78907146-INS-549 | chr16-78907154-INS-547 |
| chr16-79042363-INS-220 | chr16-79042363-INS-220 |
| chr16-79076349-INS-54 | chr16-79076349-INS-68 |
| chr16-79101282-INS-882 | chr16-79101282-INS-882 |
| chr16-79102744-INS-1213 | chr16-79102745-INS-1202 |
| chr16-79109081-INS-72 | chr16-79109081-INS-68 |
| chr16-79131078-INS-365 | chr16-79131078-INS-366 |
| chr16-79141411-INS-2858 | chr16-79141412-INS-2860 |
| chr16-79141691-INS-76 | chr16-79141699-INS-98 |
| chr16-79149946-INS-965 | chr16-79149946-INS-932 |
| chr16-79151419-INS-65 | chr16-79151419-INS-68 |
| chr16-79162805-INS-70 | chr16-79162805-INS-70 |
| chr16-79173044-INS-56 | chr16-79173044-INS-56 |
| chr16-79175015-INS-70 | chr16-79175013-INS-66 |
| chr16-79190680-INS-60 | chr16-79190680-INS-58 |

| | |
|---|---|
| chr16-79226500-INS-372 | chr16-79226500-INS-372 |
| chr16-79245226-INS-180 | chr16-79245225-INS-149 |
| chr16-79259332-INS-67 | chr16-79259332-INS-67 |
| chr16-79288447-INS-68 | chr16-79288411-INS-90 |
| chr16-79306449-INS-114 | chr16-79306447-INS-115 |
| chr16-79340053-INS-155 | chr16-79340053-INS-155 |
| chr16-79344793-INS-970 | chr16-79344793-INS-970 |
| chr16-79358552-INS-7169 | chr16-79358553-INS-7171 |
| chr16-79364445-INS-70 | chr16-79364446-INS-67 |
| chr16-79422448-INS-56 | chr16-79422448-INS-59 |
| chr16-79426897-INS-80 | chr16-79426890-INS-115 |
| chr16-79442352-INS-120 | chr16-79442352-INS-119 |
| chr16-79490963-INS-119 | chr16-79490963-INS-119 |
| chr16-79492399-INS-779 | chr16-79492399-INS-779 |
| chr16-79511872-INS-51 | chr16-79511824-INS-52 |
| chr16-79541389-INS-82 | chr16-79541389-INS-82 |
| chr16-79571161-INS-62 | chr16-79571161-INS-62 |
| chr16-79572602-INS-59 | chr16-79572602-INS-58 |
| chr16-79593569-INS-122 | chr16-79593569-INS-140 |
| chr16-79610370-INS-364 | chr16-79610370-INS-364 |
| chr16-79638422-INS-86 | chr16-79638422-INS-67 |
| chr16-79672694-INS-189 | chr16-79672694-INS-189 |
| chr16-79681377-INS-88 | chr16-79681377-INS-88 |
| chr16-79690894-INS-132 | chr16-79690894-INS-132 |
| chr16-79700867-INS-2529 | chr16-79700868-INS-2529 |
| chr16-79759076-INS-88 | chr16-79759068-INS-92 |
| chr16-79776911-INS-380 | chr16-79776911-INS-380 |
| chr16-79782123-INS-80 | chr16-79782123-INS-50 |
| chr16-79791835-INS-3343 | chr16-79791836-INS-3343 |
| chr16-79793142-INS-3009 | chr16-79793143-INS-3009 |
| chr16-79816468-INS-1845 | chr16-79816469-INS-1845 |
| chr16-79830031-INS-232 | chr16-79830031-INS-240 |
| chr16-79851101-INS-55 | chr16-79851101-INS-55 |
| chr16-79853314-INS-2084 | chr16-79853315-INS-2084 |
| chr16-79859384-INS-79 | chr16-79859384-INS-79 |
| chr16-79886165-INS-80 | chr16-79886165-INS-80 |
| chr16-79925762-INS-441 | chr16-79925762-INS-441 |
| chr16-79951118-INS-3067 | chr16-79951119-INS-3067 |
| chr16-79971188-INS-108 | chr16-79971188-INS-109 |
| chr16-79976247-INS-929 | chr16-79976247-INS-929 |
| chr16-79978167-INS-52 | chr16-79978160-INS-50 |
| chr16-79980213-INS-204 | chr16-79980213-INS-174 |
| chr16-79999052-INS-704 | chr16-79999052-INS-703 |
| chr16-80032832-INS-227 | chr16-80032832-INS-234 |
| chr16-80048943-INS-100 | chr16-80048943-INS-104 |
| chr16-80057087-INS-120 | chr16-80057087-INS-100 |
| chr16-80078626-INS-118 | chr16-80078626-INS-118 |
| chr16-80113569-INS-219 | chr16-80113569-INS-218 |
| chr16-80116781-INS-68 | chr16-80116782-INS-60 |
| chr16-80127877-INS-170 | chr16-80127866-INS-158 |
| chr16-80155272-INS-1494 | chr16-80155273-INS-1494 |
| chr16-80193259-INS-1111 | chr16-80193261-INS-1107 |
| chr16-80226513-INS-1652 | chr16-80226514-INS-1650 |
| chr16-80229203-INS-68 | chr16-80229203-INS-70 |
| chr16-80252934-INS-332 | chr16-80252934-INS-332 |
| chr16-80261185-INS-54 | chr16-80261185-INS-54 |
| chr16-80268421-INS-66 | chr16-80268421-INS-66 |
| chr16-80276343-INS-828 | chr16-80276343-INS-828 |
| chr16-80276548-INS-5701 | chr16-80276551-INS-5703 |
| chr16-80276869-INS-82 | chr16-80276869-INS-84 |
| chr16-80314215-INS-754 | chr16-80314215-INS-754 |
| chr16-80317077-INS-145 | chr16-80317077-INS-145 |
| chr16-80323844-INS-1334 | chr16-80323843-INS-1343 |
| chr16-80326099-INS-626 | chr16-80326099-INS-623 |
| chr16-80331023-INS-159 | chr16-80331023-INS-159 |
| chr16-80334504-INS-704 | chr16-80334504-INS-704 |
| chr16-80348459-INS-6633 | chr16-80348460-INS-6633 |
| chr16-80354094-INS-195 | chr16-80354094-INS-195 |
| chr16-80369005-INS-86 | chr16-80369005-INS-86 |
| chr16-80372344-INS-50 | chr16-80372344-INS-52 |
| chr16-80375920-INS-177 | chr16-80375920-INS-177 |
| chr16-80379804-INS-5353 | chr16-80379805-INS-5353 |
| chr16-80387619-INS-2213 | chr16-80387620-INS-2213 |
| chr16-80410468-INS-51 | chr16-80410468-INS-51 |
| chr16-80420304-INS-275 | chr16-80420321-INS-270 |
| chr16-80421482-INS-408 | chr16-80421482-INS-408 |
| chr16-80422377-INS-288 | chr16-80422375-INS-266 |

| | |
|---|---|
| chr16-80437490-INS-102 | chr16-80437490-INS-72 |
| chr16-80447419-INS-71 | chr16-80447378-INS-88 |
| chr16-80536418-INS-2690 | chr16-80536419-INS-2690 |
| chr16-80539298-INS-256 | chr16-80539298-INS-258 |
| chr16-80553075-INS-1213 | chr16-80553076-INS-1213 |
| chr16-80556248-INS-1158 | chr16-80556249-INS-1158 |
| chr16-80571199-INS-408 | chr16-80571199-INS-408 |
| chr16-80613678-INS-54 | chr16-80613678-INS-56 |
| chr16-80621856-INS-64 | chr16-80621856-INS-68 |
| chr16-80627297-INS-348 | chr16-80627297-INS-348 |
| chr16-80641348-INS-868 | chr16-80641348-INS-869 |
| chr16-80661803-INS-4579 | chr16-80661804-INS-4581 |
| chr16-80685712-INS-76 | chr16-80685712-INS-62 |
| chr16-80725902-INS-791 | chr16-80725902-INS-791 |
| chr16-80755536-INS-116 | chr16-80755536-INS-116 |
| chr16-80785869-INS-54 | chr16-80785869-INS-54 |
| chr16-80851144-INS-75 | chr16-80851144-INS-75 |
| chr16-80864873-INS-373 | chr16-80864873-INS-373 |
| chr16-80867905-INS-72 | chr16-80867905-INS-92 |
| chr16-80879415-INS-88 | chr16-80879415-INS-88 |
| chr16-80917504-INS-286 | chr16-80917504-INS-286 |
| chr16-80923729-INS-165 | chr16-80923729-INS-165 |
| chr16-80960193-INS-55 | chr16-80960193-INS-55 |
| chr16-80980318-INS-65 | chr16-80980330-INS-62 |
| chr16-80990267-INS-67 | chr16-80990266-INS-71 |
| chr16-80996514-INS-559 | chr16-80996514-INS-520 |
| chr16-81006574-INS-108 | chr16-81006575-INS-108 |
| chr16-81045492-INS-64 | chr16-81045492-INS-66 |
| chr16-81064514-INS-70 | chr16-81064514-INS-70 |
| chr16-81066673-INS-128 | chr16-81066673-INS-132 |
| chr16-81120653-INS-112 | chr16-81120656-INS-111 |
| chr16-81138896-INS-215 | chr16-81138896-INS-215 |
| chr16-81142553-INS-71 | chr16-81142553-INS-71 |
| chr16-81172140-INS-56 | chr16-81172140-INS-58 |
| chr16-81202464-INS-56 | chr16-81202464-INS-56 |
| chr16-81213753-INS-68 | chr16-81213753-INS-68 |
| chr16-81218832-INS-67 | chr16-81218832-INS-69 |
| chr16-81220586-INS-761 | chr16-81220592-INS-760 |
| chr16-81224446-INS-164 | chr16-81224447-INS-139 |
| chr16-81230115-INS-81 | chr16-81230092-INS-76 |
| chr16-81232910-INS-64 | chr16-81232910-INS-64 |
| chr16-81253031-INS-66 | chr16-81253031-INS-66 |
| chr16-81256794-INS-254 | chr16-81256789-INS-254 |
| chr16-81265992-INS-58 | chr16-81265992-INS-53 |
| chr16-81293598-INS-1119 | chr16-81293599-INS-1119 |
| chr16-81305151-INS-2277 | chr16-81305152-INS-2277 |
| chr16-81345660-INS-433 | chr16-81345660-INS-433 |
| chr16-81359130-INS-89 | chr16-81359130-INS-89 |
| chr16-81372589-INS-64 | chr16-81372600-INS-50 |
| chr16-81401182-INS-164 | chr16-81401182-INS-164 |
| chr16-81428558-INS-57 | chr16-81428558-INS-55 |
| chr16-81450563-INS-74 | chr16-81450563-INS-74 |
| chr16-81464500-INS-50 | chr16-81464497-INS-100 |
| chr16-81473506-INS-1888 | chr16-81473507-INS-1888 |
| chr16-81508198-INS-245 | chr16-81508199-INS-236 |
| chr16-81512368-INS-75 | chr16-81512368-INS-75 |
| chr16-81513248-INS-285 | chr16-81513248-INS-285 |
| chr16-81515049-INS-199 | chr16-81515049-INS-199 |
| chr16-81521252-INS-598 | chr16-81521252-INS-598 |
| chr16-81542878-INS-643 | chr16-81542878-INS-643 |
| chr16-81563150-INS-486 | chr16-81563150-INS-486 |
| chr16-81566972-INS-80 | chr16-81567000-INS-64 |
| chr16-81573216-INS-56 | chr16-81573216-INS-60 |
| chr16-81609344-INS-70 | chr16-81609344-INS-70 |
| chr16-81612219-INS-529 | chr16-81612219-INS-529 |
| chr16-81624882-INS-284 | chr16-81624880-INS-289 |
| chr16-81630582-INS-58 | chr16-81630582-INS-58 |
| chr16-81685614-INS-298 | chr16-81685614-INS-298 |
| chr16-81692779-INS-553 | chr16-81692779-INS-553 |
| chr16-81703934-INS-85 | chr16-81703934-INS-85 |
| chr16-81707016-INS-129 | chr16-81707016-INS-128 |
| chr16-81709902-INS-64 | chr16-81709902-INS-64 |
| chr16-81716085-INS-54 | chr16-81716085-INS-55 |
| chr16-81717847-INS-53 | chr16-81717817-INS-63 |
| chr16-81728222-INS-6567 | chr16-81728223-INS-6567 |
| chr16-81729066-INS-345 | chr16-81729066-INS-345 |
| chr16-81739324-INS-972 | chr16-81739324-INS-966 |

| | |
|---|---|
| chr16-81742135-INS-908 | chr16-81742136-INS-912 |
| chr16-81745267-INS-358 | chr16-81745267-INS-358 |
| chr16-81747497-INS-124 | chr16-81747497-INS-124 |
| chr16-81748530-INS-96 | chr16-81748539-INS-52 |
| chr16-81754507-INS-112 | chr16-81754508-INS-112 |
| chr16-81774521-INS-191 | chr16-81774521-INS-190 |
| chr16-81786557-INS-100 | chr16-81786557-INS-102 |
| chr16-81788750-INS-63 | chr16-81788750-INS-64 |
| chr16-81809525-INS-111 | chr16-81809525-INS-112 |
| chr16-81821221-INS-358 | chr16-81821221-INS-358 |
| chr16-81822250-INS-1533 | chr16-81822251-INS-1533 |
| chr16-81824004-INS-764 | chr16-81824004-INS-768 |
| chr16-81834746-INS-640 | chr16-81834746-INS-641 |
| chr16-81847831-INS-4952 | chr16-81847832-INS-4952 |
| chr16-81863476-INS-62 | chr16-81863482-INS-52 |
| chr16-81870332-INS-204 | chr16-81870333-INS-184 |
| chr16-81872302-INS-726 | chr16-81872302-INS-721 |
| chr16-81888218-INS-212 | chr16-81888218-INS-212 |
| chr16-81938756-INS-64 | chr16-81938756-INS-64 |
| chr16-81951804-INS-66 | chr16-81951804-INS-70 |
| chr16-81993920-INS-434 | chr16-81993923-INS-460 |
| chr16-81998612-INS-210 | chr16-81998624-INS-218 |
| chr16-82019080-INS-573 | chr16-82019080-INS-573 |
| chr16-82025642-INS-133 | chr16-82025631-INS-100 |
| chr16-82068224-INS-400 | chr16-82068224-INS-403 |
| chr16-82070572-INS-795 | chr16-82070572-INS-795 |
| chr16-82085124-INS-60 | chr16-82085124-INS-60 |
| chr16-82095221-INS-326 | chr16-82095221-INS-337 |
| chr16-82108783-INS-51 | chr16-82108783-INS-51 |
| chr16-82164558-INS-156 | chr16-82164558-INS-156 |
| chr16-82166739-INS-5299 | chr16-82166740-INS-5299 |
| chr16-82210871-INS-4807 | chr16-82210872-INS-4807 |
| chr16-82218257-INS-2003 | chr16-82218258-INS-2003 |
| chr16-82222512-INS-57 | chr16-82222512-INS-56 |
| chr16-82230693-INS-72 | chr16-82230695-INS-72 |
| chr16-82293067-INS-592 | chr16-82293067-INS-593 |
| chr16-82333536-INS-221 | chr16-82333558-INS-244 |
| chr16-82353030-INS-70 | chr16-82353030-INS-54 |
| chr16-82368617-INS-67 | chr16-82368592-INS-67 |
| chr16-82400339-INS-6667 | chr16-82400340-INS-6668 |
| chr16-82405509-INS-201 | chr16-82405509-INS-201 |
| chr16-82413162-INS-6423 | chr16-82413163-INS-6429 |
| chr16-82447442-INS-351 | chr16-82447442-INS-351 |
| chr16-82469072-INS-89 | chr16-82469072-INS-106 |
| chr16-82480303-INS-93 | chr16-82480303-INS-93 |
| chr16-82489434-INS-2923 | chr16-82489433-INS-2919 |
| chr16-82511457-INS-1772 | chr16-82511458-INS-1775 |
| chr16-82540706-INS-4309 | chr16-82540707-INS-4309 |
| chr16-82549454-INS-417 | chr16-82549454-INS-422 |
| chr16-82560673-INS-2306 | chr16-82560674-INS-2308 |
| chr16-82568316-INS-66 | chr16-82568316-INS-66 |
| chr16-82595098-INS-1522 | chr16-82595099-INS-1523 |
| chr16-82597510-INS-674 | chr16-82597510-INS-674 |
| chr16-82611975-INS-68 | chr16-82611975-INS-68 |
| chr16-82615336-INS-845 | chr16-82615336-INS-848 |
| chr16-82653864-INS-77 | chr16-82653870-INS-67 |
| chr16-82655766-INS-63 | chr16-82655767-INS-51 |
| chr16-82693718-INS-54 | chr16-82693718-INS-70 |
| chr16-82805112-INS-55 | chr16-82805100-INS-55 |
| chr16-82820757-INS-65 | chr16-82820759-INS-60 |
| chr16-82823936-INS-204 | chr16-82823935-INS-213 |
| chr16-82837580-INS-208 | chr16-82837580-INS-208 |
| chr16-82844820-INS-4987 | chr16-82844821-INS-4987 |
| chr16-82867127-INS-138 | chr16-82867127-INS-138 |
| chr16-82900670-INS-64 | chr16-82900670-INS-84 |
| chr16-82909510-INS-54 | chr16-82909510-INS-54 |
| chr16-82914876-INS-528 | chr16-82914876-INS-528 |
| chr16-82932985-INS-490 | chr16-82932985-INS-490 |
| chr16-82971273-INS-60 | chr16-82971273-INS-60 |
| chr16-82975002-INS-68 | chr16-82975002-INS-63 |
| chr16-82988440-INS-482 | chr16-82988440-INS-482 |
| chr16-83032796-INS-1071 | chr16-83032797-INS-1071 |
| chr16-83070923-INS-72 | chr16-83070924-INS-62 |
| chr16-83079121-INS-54 | chr16-83079121-INS-56 |
| chr16-83093944-INS-413 | chr16-83093944-INS-459 |
| chr16-83121783-INS-532 | chr16-83121783-INS-531 |
| chr16-83171796-INS-69 | chr16-83171796-INS-81 |

| | |
|---|---|
| chr16-83172264-INS-3067 | chr16-83172265-INS-3067 |
| chr16-83175499-INS-417 | chr16-83175499-INS-408 |
| chr16-83175973-INS-604 | chr16-83175973-INS-604 |
| chr16-83179868-INS-571 | chr16-83179868-INS-577 |
| chr16-83216680-INS-160 | chr16-83216680-INS-160 |
| chr16-83280945-INS-212 | chr16-83280945-INS-212 |
| chr16-83295328-INS-182 | chr16-83295328-INS-177 |
| chr16-83339731-INS-78 | chr16-83339731-INS-77 |
| chr16-83379876-INS-718 | chr16-83379876-INS-719 |
| chr16-83395731-INS-76 | chr16-83395731-INS-76 |
| chr16-83408410-INS-113 | chr16-83408410-INS-113 |
| chr16-83443777-INS-55 | chr16-83443752-INS-67 |
| chr16-83455364-INS-176 | chr16-83455364-INS-176 |
| chr16-83457034-INS-65 | chr16-83457041-INS-61 |
| chr16-83465766-INS-319 | chr16-83465766-INS-315 |
| chr16-83509107-INS-705 | chr16-83509107-INS-705 |
| chr16-83524106-INS-617 | chr16-83524096-INS-617 |
| chr16-83530794-INS-416 | chr16-83530794-INS-416 |
| chr16-83541102-INS-66 | chr16-83541085-INS-74 |
| chr16-83543474-INS-58 | chr16-83543474-INS-58 |
| chr16-83553680-INS-5295 | chr16-83553680-INS-5289 |
| chr16-83561115-INS-451 | chr16-83561115-INS-451 |
| chr16-83572386-INS-52 | chr16-83572386-INS-54 |
| chr16-83575124-INS-184 | chr16-83575112-INS-196 |
| chr16-83635666-INS-422 | chr16-83635666-INS-422 |
| chr16-83690750-INS-77 | chr16-83690750-INS-77 |
| chr16-83695820-INS-73 | chr16-83695834-INS-56 |
| chr16-83739890-INS-824 | chr16-83739890-INS-825 |
| chr16-83763087-INS-82 | chr16-83763087-INS-67 |
| chr16-83769820-INS-102 | chr16-83769820-INS-58 |
| chr16-83783929-INS-70 | chr16-83783923-INS-61 |
| chr16-83795322-INS-60 | chr16-83795322-INS-56 |
| chr16-83800233-INS-453 | chr16-83800241-INS-423 |
| chr16-83809202-INS-59 | chr16-83809202-INS-59 |
| chr16-83843838-INS-65 | chr16-83843832-INS-101 |
| chr16-83872326-INS-54 | chr16-83872326-INS-54 |
| chr16-83947254-INS-6475 | chr16-83947255-INS-6475 |
| chr16-83982111-INS-2903 | chr16-83982112-INS-2903 |
| chr16-83982473-INS-54 | chr16-83982474-INS-84 |
| chr16-83993728-INS-757 | chr16-83993728-INS-757 |
| chr16-83996086-INS-359 | chr16-83996086-INS-359 |
| chr16-84022961-INS-139 | chr16-84022961-INS-139 |
| chr16-84038966-INS-61 | chr16-84038966-INS-61 |
| chr16-84045363-INS-233 | chr16-84045363-INS-229 |
| chr16-84064576-INS-164 | chr16-84064576-INS-191 |
| chr16-84071280-INS-68 | chr16-84071282-INS-92 |
| chr16-84116309-INS-272 | chr16-84116309-INS-272 |
| chr16-84145774-INS-64 | chr16-84145774-INS-64 |
| chr16-84148117-INS-57 | chr16-84148117-INS-51 |
| chr16-84204998-INS-59 | chr16-84204998-INS-53 |
| chr16-84208205-INS-384 | chr16-84208205-INS-384 |
| chr16-84209200-INS-458 | chr16-84209200-INS-457 |
| chr16-84220678-INS-184 | chr16-84220678-INS-183 |
| chr16-84257430-INS-520 | chr16-84257430-INS-519 |
| chr16-84264735-INS-6600 | chr16-84264736-INS-6600 |
| chr16-84265733-INS-67 | chr16-84265733-INS-71 |
| chr16-84290214-INS-6441 | chr16-84290215-INS-6441 |
| chr16-84306138-INS-176 | chr16-84306138-INS-176 |
| chr16-84335355-INS-313 | chr16-84335355-INS-313 |
| chr16-84339483-INS-77 | chr16-84339483-INS-77 |
| chr16-84347641-INS-235 | chr16-84347641-INS-236 |
| chr16-84354518-INS-77 | chr16-84354518-INS-77 |
| chr16-84355369-INS-52 | chr16-84355369-INS-52 |
| chr16-84361892-INS-340 | chr16-84361892-INS-326 |
| chr16-84380716-INS-90 | chr16-84380722-INS-94 |
| chr16-84391889-INS-83 | chr16-84391889-INS-78 |
| chr16-84393704-INS-146 | chr16-84393704-INS-150 |
| chr16-84400428-INS-66 | chr16-84400428-INS-70 |
| chr16-84409754-INS-7166 | chr16-84409755-INS-7170 |
| chr16-84450228-INS-130 | chr16-84450228-INS-130 |
| chr16-84467841-INS-169 | chr16-84467841-INS-175 |
| chr16-84473372-INS-195 | chr16-84473372-INS-195 |
| chr16-84478860-INS-200 | chr16-84478860-INS-200 |
| chr16-84502509-INS-80 | chr16-84502509-INS-83 |
| chr16-84505472-INS-178 | chr16-84505472-INS-189 |
| chr16-84514601-INS-64 | chr16-84514601-INS-66 |
| chr16-84521408-INS-59 | chr16-84521407-INS-57 |

| | |
|---|---|
| chr16-84524585-INS-215 | chr16-84524573-INS-232 |
| chr16-84534700-INS-1179 | chr16-84534700-INS-1178 |
| chr16-84541340-INS-108 | chr16-84541347-INS-61 |
| chr16-84542418-INS-188 | chr16-84542409-INS-198 |
| chr16-84544924-INS-218 | chr16-84544910-INS-229 |
| chr16-84549677-INS-366 | chr16-84549677-INS-367 |
| chr16-84554281-INS-239 | chr16-84554281-INS-239 |
| chr16-84556982-INS-197 | chr16-84556982-INS-200 |
| chr16-84561001-INS-901 | chr16-84561001-INS-901 |
| chr16-84562387-INS-643 | chr16-84562391-INS-642 |
| chr16-84578650-INS-113 | chr16-84578650-INS-113 |
| chr16-84583354-INS-52 | chr16-84583354-INS-60 |
| chr16-84584488-INS-54 | chr16-84584488-INS-52 |
| chr16-84586439-INS-316 | chr16-84586439-INS-316 |
| chr16-84590086-INS-54 | chr16-84590086-INS-50 |
| chr16-84591369-INS-66 | chr16-84591366-INS-66 |
| chr16-84610313-INS-194 | chr16-84610313-INS-194 |
| chr16-84636185-INS-178 | chr16-84636185-INS-178 |
| chr16-84659872-INS-68 | chr16-84659872-INS-68 |
| chr16-84667968-INS-668 | chr16-84667968-INS-668 |
| chr16-84698463-INS-181 | chr16-84698463-INS-181 |
| chr16-84701368-INS-72 | chr16-84701368-INS-72 |
| chr16-84743655-INS-484 | chr16-84743655-INS-500 |
| chr16-84791147-INS-964 | chr16-84791147-INS-964 |
| chr16-84805001-INS-292 | chr16-84805001-INS-292 |
| chr16-84807679-INS-54 | chr16-84807679-INS-54 |
| chr16-84844342-INS-100 | chr16-84844342-INS-100 |
| chr16-84848321-INS-225 | chr16-84848321-INS-225 |
| chr16-84861811-INS-60 | chr16-84861811-INS-68 |
| chr16-84875249-INS-52 | chr16-84875249-INS-50 |
| chr16-84882140-INS-75 | chr16-84882140-INS-73 |
| chr16-84902005-INS-65 | chr16-84902005-INS-70 |
| chr16-84924860-INS-6262 | chr16-84924861-INS-6262 |
| chr16-84935724-INS-62 | chr16-84935724-INS-73 |
| chr16-84966358-INS-53 | chr16-84966372-INS-52 |
| chr16-84976875-INS-109 | chr16-84976875-INS-139 |
| chr16-84979119-INS-52 | chr16-84979127-INS-68 |
| chr16-84981758-INS-62 | chr16-84981755-INS-51 |
| chr16-85023131-INS-114 | chr16-85023131-INS-114 |
| chr16-85027789-INS-7114 | chr16-85027790-INS-7114 |
| chr16-85031279-INS-214 | chr16-85031279-INS-214 |
| chr16-85041366-INS-2465 | chr16-85041367-INS-2465 |
| chr16-85053181-INS-54 | chr16-85053183-INS-54 |
| chr16-85060379-INS-105 | chr16-85060379-INS-105 |
| chr16-85067044-INS-172 | chr16-85067044-INS-178 |
| chr16-85100166-INS-6397 | chr16-85100167-INS-6398 |
| chr16-85100292-INS-414 | chr16-85100292-INS-414 |
| chr16-85107641-INS-1152 | chr16-85107642-INS-1153 |
| chr16-85155574-INS-54 | chr16-85155574-INS-58 |
| chr16-85204848-INS-668 | chr16-85204845-INS-662 |
| chr16-85245401-INS-162 | chr16-85245401-INS-162 |
| chr16-85255057-INS-148 | chr16-85255057-INS-148 |
| chr16-85265085-INS-63 | chr16-85265085-INS-95 |
| chr16-85266425-INS-61 | chr16-85266425-INS-93 |
| chr16-85268029-INS-182 | chr16-85268029-INS-182 |
| chr16-85295770-INS-936 | chr16-85295770-INS-933 |
| chr16-85310870-INS-71 | chr16-85310871-INS-61 |
| chr16-85312682-INS-169 | chr16-85312682-INS-191 |
| chr16-85320017-INS-160 | chr16-85320017-INS-160 |
| chr16-85339240-INS-155 | chr16-85339205-INS-155 |
| chr16-85355157-INS-63 | chr16-85355157-INS-62 |
| chr16-85369119-INS-76 | chr16-85369085-INS-76 |
| chr16-85396007-INS-457 | chr16-85396007-INS-456 |
| chr16-85430971-INS-52 | chr16-85430971-INS-52 |
| chr16-85444726-INS-185 | chr16-85444726-INS-184 |
| chr16-85467192-INS-653 | chr16-85467192-INS-653 |
| chr16-85484479-INS-77 | chr16-85484479-INS-76 |
| chr16-85493967-INS-53 | chr16-85493967-INS-53 |
| chr16-85505126-INS-473 | chr16-85505126-INS-473 |
| chr16-85506402-INS-53 | chr16-85506402-INS-53 |
| chr16-85527870-INS-55 | chr16-85527870-INS-50 |
| chr16-85575501-INS-78 | chr16-85575501-INS-80 |
| chr16-85601652-INS-98 | chr16-85601652-INS-92 |
| chr16-85606274-INS-540 | chr16-85606274-INS-541 |
| chr16-85639290-INS-6579 | chr16-85639291-INS-6579 |
| chr16-85691074-INS-350 | chr16-85691076-INS-357 |
| chr16-85702095-INS-6440 | chr16-85702096-INS-6440 |

| | |
|---|---|
| chr16-85752339-INS-436 | chr16-85752339-INS-436 |
| chr16-85781735-INS-984 | chr16-85781735-INS-984 |
| chr16-85784305-INS-1186 | chr16-85784306-INS-1184 |
| chr16-85787564-INS-84 | chr16-85787564-INS-50 |
| chr16-85802565-INS-2803 | chr16-85802566-INS-2803 |
| chr16-85841338-INS-91 | chr16-85841384-INS-68 |
| chr16-85844032-INS-133 | chr16-85844025-INS-149 |
| chr16-85849863-INS-333 | chr16-85849860-INS-333 |
| chr16-85868688-INS-51 | chr16-85868688-INS-51 |
| chr16-85888971-INS-355 | chr16-85888971-INS-353 |
| chr16-85890481-INS-98 | chr16-85890480-INS-105 |
| chr16-85901664-INS-76 | chr16-85901664-INS-85 |
| chr16-85913903-INS-60 | chr16-85913908-INS-74 |
| chr16-85914316-INS-460 | chr16-85914316-INS-461 |
| chr16-85921903-INS-168 | chr16-85921903-INS-168 |
| chr16-85925985-INS-64 | chr16-85925985-INS-64 |
| chr16-85930629-INS-99 | chr16-85930623-INS-108 |
| chr16-85935010-INS-67 | chr16-85935010-INS-67 |
| chr16-85942139-INS-752 | chr16-85942139-INS-744 |
| chr16-85944618-INS-70 | chr16-85944618-INS-68 |
| chr16-85967607-INS-194 | chr16-85967607-INS-194 |
| chr16-85980267-INS-762 | chr16-85980267-INS-763 |
| chr16-85984439-INS-78 | chr16-85984439-INS-78 |
| chr16-85991801-INS-229 | chr16-85991801-INS-229 |
| chr16-86000744-INS-76 | chr16-86000742-INS-68 |
| chr16-86004767-INS-903 | chr16-86004767-INS-903 |
| chr16-86005656-INS-6588 | chr16-86005657-INS-6588 |
| chr16-86014724-INS-69 | chr16-86014715-INS-94 |
| chr16-86025653-INS-61 | chr16-86025653-INS-53 |
| chr16-86036511-INS-6287 | chr16-86036512-INS-6287 |
| chr16-86038559-INS-395 | chr16-86038559-INS-395 |
| chr16-86061345-INS-60 | chr16-86061345-INS-60 |
| chr16-86081343-INS-592 | chr16-86081343-INS-592 |
| chr16-86095257-INS-1483 | chr16-86095258-INS-1484 |
| chr16-86113888-INS-64 | chr16-86113888-INS-54 |
| chr16-86134972-INS-82 | chr16-86134972-INS-102 |
| chr16-86163979-INS-52 | chr16-86163979-INS-52 |
| chr16-86220298-INS-417 | chr16-86220298-INS-419 |
| chr16-86221553-INS-287 | chr16-86221553-INS-287 |
| chr16-86229281-INS-391 | chr16-86229281-INS-391 |
| chr16-86258078-INS-4548 | chr16-86258079-INS-4549 |
| chr16-86295943-INS-60 | chr16-86295943-INS-68 |
| chr16-86306643-INS-953 | chr16-86306643-INS-954 |
| chr16-86317396-INS-406 | chr16-86317396-INS-406 |
| chr16-86318161-INS-941 | chr16-86318161-INS-941 |
| chr16-86326797-INS-62 | chr16-86326797-INS-62 |
| chr16-86333725-INS-83 | chr16-86333725-INS-84 |
| chr16-86339291-INS-116 | chr16-86339291-INS-116 |
| chr16-86342413-INS-364 | chr16-86342413-INS-364 |
| chr16-86373412-INS-802 | chr16-86373412-INS-802 |
| chr16-86383471-INS-290 | chr16-86383471-INS-290 |
| chr16-86390908-INS-452 | chr16-86390908-INS-452 |
| chr16-86435023-INS-85 | chr16-86435023-INS-85 |
| chr16-86516509-INS-98 | chr16-86516509-INS-82 |
| chr16-86590619-INS-561 | chr16-86590619-INS-561 |
| chr16-86602361-INS-74 | chr16-86602361-INS-74 |
| chr16-86605144-INS-155 | chr16-86605142-INS-157 |
| chr16-86636640-INS-216 | chr16-86636658-INS-223 |
| chr16-86640564-INS-251 | chr16-86640564-INS-252 |
| chr16-86691614-INS-745 | chr16-86691614-INS-745 |
| chr16-86699380-INS-58 | chr16-86699380-INS-56 |
| chr16-86704700-INS-76 | chr16-86704700-INS-76 |
| chr16-86731673-INS-382 | chr16-86731703-INS-384 |
| chr16-86740569-INS-136 | chr16-86740569-INS-90 |
| chr16-86756661-INS-364 | chr16-86756665-INS-368 |
| chr16-86801188-INS-548 | chr16-86801188-INS-548 |
| chr16-86826525-INS-6618 | chr16-86826533-INS-6613 |
| chr16-86844488-INS-138 | chr16-86844491-INS-135 |
| chr16-86845235-INS-55 | chr16-86845210-INS-80 |
| chr16-86852189-INS-920 | chr16-86852189-INS-921 |
| chr16-86859339-INS-827 | chr16-86859339-INS-827 |
| chr16-86885754-INS-2774 | chr16-86885755-INS-2774 |
| chr16-86896525-INS-53 | chr16-86896525-INS-50 |
| chr16-86903563-INS-60 | chr16-86903563-INS-108 |
| chr16-86932881-INS-52 | chr16-86932881-INS-56 |
| chr16-86954834-INS-62 | chr16-86954834-INS-64 |
| chr16-86963040-INS-164 | chr16-86963040-INS-160 |

| | |
|---|---|
| chr16-86966738-INS-51 | chr16-86966738-INS-51 |
| chr16-86970222-INS-509 | chr16-86970222-INS-509 |
| chr16-86976954-INS-111 | chr16-86976954-INS-111 |
| chr16-86986882-INS-232 | chr16-86986886-INS-192 |
| chr16-87000072-INS-55 | chr16-87000072-INS-66 |
| chr16-87002253-INS-1063 | chr16-87002254-INS-1063 |
| chr16-87020444-INS-6218 | chr16-87020445-INS-6218 |
| chr16-87026536-INS-51 | chr16-87026536-INS-51 |
| chr16-87093102-INS-64 | chr16-87093102-INS-58 |
| chr16-87113264-INS-70 | chr16-87113259-INS-71 |
| chr16-87133765-INS-102 | chr16-87133765-INS-111 |
| chr16-87135756-INS-7119 | chr16-87135757-INS-7119 |
| chr16-87138862-INS-56 | chr16-87138862-INS-58 |
| chr16-87147192-INS-105 | chr16-87147192-INS-105 |
| chr16-87187658-INS-212 | chr16-87187658-INS-213 |
| chr16-87193061-INS-111 | chr16-87193061-INS-111 |
| chr16-87207046-INS-213 | chr16-87207046-INS-213 |
| chr16-87215590-INS-172 | chr16-87215590-INS-172 |
| chr16-87241601-INS-182 | chr16-87241601-INS-183 |
| chr16-87266075-INS-213 | chr16-87266075-INS-213 |
| chr16-87295858-INS-87 | chr16-87295858-INS-107 |
| chr16-87297176-INS-77 | chr16-87297176-INS-77 |
| chr16-87366453-INS-114 | chr16-87366453-INS-114 |
| chr16-87373032-INS-59 | chr16-87373032-INS-59 |
| chr16-87377560-INS-177 | chr16-87377560-INS-190 |
| chr16-87379096-INS-6696 | chr16-87379097-INS-6694 |
| chr16-87386328-INS-57 | chr16-87386328-INS-56 |
| chr16-87389788-INS-716 | chr16-87389788-INS-684 |
| chr16-87409083-INS-181 | chr16-87409091-INS-176 |
| chr16-87412005-INS-52 | chr16-87412006-INS-52 |
| chr16-87424621-INS-60 | chr16-87424622-INS-62 |
| chr16-87429682-INS-171 | chr16-87429682-INS-171 |
| chr16-87430564-INS-282 | chr16-87430564-INS-285 |
| chr16-87431526-INS-224 | chr16-87431526-INS-224 |
| chr16-87433090-INS-225 | chr16-87433090-INS-225 |
| chr16-87445951-INS-216 | chr16-87445955-INS-216 |
| chr16-87454138-INS-323 | chr16-87454138-INS-299 |
| chr16-87460305-INS-106 | chr16-87460305-INS-107 |
| chr16-87464456-INS-3238 | chr16-87464457-INS-3238 |
| chr16-87466787-INS-67 | chr16-87466787-INS-79 |
| chr16-87471104-INS-70 | chr16-87471103-INS-72 |
| chr16-87491219-INS-170 | chr16-87491219-INS-170 |
| chr16-87501343-INS-53 | chr16-87501343-INS-53 |
| chr16-87503608-INS-687 | chr16-87503608-INS-687 |
| chr16-87517782-INS-178 | chr16-87517784-INS-183 |
| chr16-87519883-INS-99 | chr16-87519883-INS-99 |
| chr16-87520842-INS-68 | chr16-87520820-INS-52 |
| chr16-87628760-INS-52 | chr16-87628741-INS-52 |
| chr16-87644041-INS-75 | chr16-87644041-INS-75 |
| chr16-87645636-INS-1259 | chr16-87645637-INS-1256 |
| chr16-87654136-INS-72 | chr16-87654136-INS-76 |
| chr16-87660187-INS-351 | chr16-87660187-INS-351 |
| chr16-87662604-INS-877 | chr16-87662604-INS-883 |
| chr16-87681261-INS-173 | chr16-87681265-INS-162 |
| chr16-87681400-INS-6849 | chr16-87681401-INS-6850 |
| chr16-87696496-INS-116 | chr16-87696496-INS-115 |
| chr16-87729531-INS-82 | chr16-87729519-INS-80 |
| chr16-87732198-INS-4417 | chr16-87732199-INS-4417 |
| chr16-87732552-INS-545 | chr16-87732552-INS-545 |
| chr16-87756631-INS-87 | chr16-87756631-INS-88 |
| chr16-87771353-INS-66 | chr16-87771353-INS-66 |
| chr16-87773715-INS-63 | chr16-87773715-INS-63 |
| chr16-87778319-INS-58 | chr16-87778320-INS-52 |
| chr16-87784592-INS-73 | chr16-87784592-INS-73 |
| chr16-87787243-INS-288 | chr16-87787255-INS-288 |
| chr16-87792111-INS-74 | chr16-87792117-INS-106 |
| chr16-87833900-INS-103 | chr16-87833900-INS-103 |
| chr16-87844606-INS-60 | chr16-87844606-INS-60 |
| chr16-87858420-INS-448 | chr16-87858420-INS-452 |
| chr16-87884856-INS-108 | chr16-87884856-INS-100 |
| chr16-87888560-INS-618 | chr16-87888560-INS-618 |
| chr16-87890144-INS-80 | chr16-87890138-INS-96 |
| chr16-87922613-INS-69 | chr16-87922613-INS-71 |
| chr16-87931568-INS-84 | chr16-87931568-INS-81 |
| chr16-87933958-INS-59 | chr16-87933958-INS-58 |
| chr16-87952768-INS-92 | chr16-87952769-INS-94 |
| chr16-87954314-INS-6683 | chr16-87954315-INS-6683 |

| | |
|---|---|
| chr16-87972066-INS-1163 | chr16-87972074-INS-1167 |
| chr16-87972970-INS-559 | chr16-87972970-INS-556 |
| chr16-87991324-INS-219 | chr16-87991324-INS-219 |
| chr16-87992734-INS-272 | chr16-87992734-INS-272 |
| chr16-87997014-INS-199 | chr16-87996998-INS-192 |
| chr16-88017927-INS-202 | chr16-88017927-INS-202 |
| chr16-88029547-INS-151 | chr16-88029527-INS-141 |
| chr16-88045286-INS-58 | chr16-88045287-INS-78 |
| chr16-88053841-INS-176 | chr16-88053841-INS-176 |
| chr16-88055004-INS-656 | chr16-88055004-INS-656 |
| chr16-88066153-INS-66 | chr16-88066153-INS-50 |
| chr16-88067190-INS-7087 | chr16-88067191-INS-7088 |
| chr16-88088989-INS-56 | chr16-88088989-INS-56 |
| chr16-88091959-INS-1537 | chr16-88091960-INS-1537 |
| chr16-88099999-INS-3679 | chr16-88100000-INS-3679 |
| chr16-88117147-INS-2837 | chr16-88117148-INS-2837 |
| chr16-88120561-INS-344 | chr16-88120561-INS-344 |
| chr16-88150664-INS-3132 | chr16-88150665-INS-3132 |
| chr16-88155246-INS-399 | chr16-88155246-INS-399 |
| chr16-88170324-INS-64 | chr16-88170324-INS-64 |
| chr16-88175208-INS-88 | chr16-88175208-INS-84 |
| chr16-88186714-INS-124 | chr16-88186714-INS-124 |
| chr16-88216509-INS-55 | chr16-88216509-INS-55 |
| chr16-88247342-INS-135 | chr16-88247342-INS-88 |
| chr16-88254342-INS-121 | chr16-88254342-INS-121 |
| chr16-88255217-INS-901 | chr16-88255217-INS-949 |
| chr16-88274006-INS-331 | chr16-88274006-INS-331 |
| chr16-88281531-INS-142 | chr16-88281531-INS-144 |
| chr16-88281531-INS-686 | chr16-88281531-INS-686 |
| chr16-88316845-INS-81 | chr16-88316845-INS-129 |
| chr16-88367680-INS-476 | chr16-88367680-INS-474 |
| chr16-88394389-INS-69 | chr16-88394389-INS-71 |
| chr16-88395245-INS-236 | chr16-88395245-INS-236 |
| chr16-88410190-INS-98 | chr16-88410167-INS-106 |
| chr16-88436736-INS-201 | chr16-88436736-INS-203 |
| chr16-88446649-INS-78 | chr16-88446649-INS-78 |
| chr16-88469217-INS-61 | chr16-88469221-INS-55 |
| chr16-88485306-INS-306 | chr16-88485306-INS-306 |
| chr16-88488573-INS-70 | chr16-88488606-INS-56 |
| chr16-88492314-INS-214 | chr16-88492314-INS-213 |
| chr16-88508262-INS-1744 | chr16-88508263-INS-1745 |
| chr16-88527889-INS-53 | chr16-88527897-INS-77 |
| chr16-88543276-INS-86 | chr16-88543276-INS-90 |
| chr16-88547180-INS-74 | chr16-88547180-INS-74 |
| chr16-88559625-INS-78 | chr16-88559607-INS-102 |
| chr16-88574947-INS-60 | chr16-88574947-INS-60 |
| chr16-88577980-INS-68 | chr16-88577980-INS-66 |
| chr16-88583388-INS-52 | chr16-88583370-INS-70 |
| chr16-88586974-INS-52 | chr16-88586974-INS-64 |
| chr16-88630081-INS-100 | chr16-88630081-INS-100 |
| chr16-88662644-INS-52 | chr16-88662635-INS-56 |
| chr16-88666138-INS-90 | chr16-88666138-INS-90 |
| chr16-88684293-INS-5565 | chr16-88684294-INS-5565 |
| chr16-88702759-INS-769 | chr16-88702751-INS-766 |
| chr16-88704716-INS-72 | chr16-88704716-INS-72 |
| chr16-88735273-INS-384 | chr16-88735273-INS-384 |
| chr16-88809491-INS-174 | chr16-88809491-INS-174 |
| chr16-88811764-INS-421 | chr16-88811764-INS-421 |
| chr16-88840996-INS-2100 | chr16-88840997-INS-2100 |
| chr16-88843487-INS-6395 | chr16-88843488-INS-6401 |
| chr16-88880351-INS-70 | chr16-88880350-INS-50 |
| chr16-88884325-INS-6079 | chr16-88884326-INS-6079 |
| chr16-88885597-INS-815 | chr16-88885597-INS-815 |
| chr16-88895486-INS-6389 | chr16-88895487-INS-6389 |
| chr16-88898156-INS-2092 | chr16-88898157-INS-2092 |
| chr16-88923296-INS-52 | chr16-88923292-INS-52 |
| chr16-88923977-INS-149 | chr16-88924024-INS-108 |
| chr16-88957302-INS-413 | chr16-88957302-INS-417 |
| chr16-88971754-INS-78 | chr16-88971754-INS-78 |
| chr16-89045174-INS-6399 | chr16-89045175-INS-6399 |
| chr16-89045187-INS-148 | chr16-89045172-INS-148 |
| chr16-89056442-INS-2990 | chr16-89056443-INS-2990 |
| chr16-89076308-INS-105 | chr16-89076308-INS-105 |
| chr16-89103404-INS-280 | chr16-89103394-INS-292 |
| chr16-89104497-INS-141 | chr16-89104497-INS-184 |
| chr16-89126405-INS-212 | chr16-89126405-INS-212 |
| chr16-89138447-INS-64 | chr16-89138448-INS-64 |

| | |
|---|---|
| chr16-89146223-INS-1460 | chr16-89146263-INS-1458 |
| chr16-89148953-INS-64 | chr16-89148953-INS-64 |
| chr16-89176879-INS-170 | chr16-89176879-INS-170 |
| chr16-89183068-INS-81 | chr16-89183068-INS-105 |
| chr16-89193763-INS-192 | chr16-89193763-INS-192 |
| chr16-89252536-INS-1198 | chr16-89252537-INS-1199 |
| chr16-89272003-INS-401 | chr16-89272003-INS-401 |
| chr16-89315415-INS-78 | chr16-89315422-INS-63 |
| chr16-89323134-INS-668 | chr16-89323174-INS-668 |
| chr16-89360877-INS-56 | chr16-89360877-INS-62 |
| chr16-89363433-INS-67 | chr16-89363433-INS-54 |
| chr16-89388492-INS-111 | chr16-89388485-INS-145 |
| chr16-89397179-INS-700 | chr16-89397179-INS-700 |
| chr16-89442239-INS-117 | chr16-89442284-INS-70 |
| chr16-89463339-INS-6454 | chr16-89463340-INS-6454 |
| chr16-89477998-INS-7115 | chr16-89477999-INS-7115 |
| chr16-89478136-INS-1116 | chr16-89478137-INS-1117 |
| chr16-89493218-INS-3075 | chr16-89493219-INS-3075 |
| chr16-89496047-INS-58 | chr16-89496047-INS-50 |
| chr16-89505792-INS-6413 | chr16-89505793-INS-6413 |
| chr16-89510695-INS-74 | chr16-89510695-INS-74 |
| chr16-89525499-INS-71 | chr16-89525499-INS-71 |
| chr16-89536287-INS-70 | chr16-89536287-INS-72 |
| chr16-89557056-INS-225 | chr16-89557056-INS-181 |
| chr16-89601810-INS-66 | chr16-89601810-INS-66 |
| chr16-89632863-INS-63 | chr16-89632863-INS-63 |
| chr16-89635076-INS-56 | chr16-89635076-INS-54 |
| chr16-89639281-INS-395 | chr16-89639281-INS-402 |
| chr16-89654633-INS-575 | chr16-89654633-INS-575 |
| chr16-89655654-INS-60 | chr16-89655654-INS-60 |
| chr16-89667376-INS-422 | chr16-89667376-INS-419 |
| chr16-89676669-INS-191 | chr16-89676669-INS-191 |
| chr16-89683824-INS-70 | chr16-89683824-INS-85 |
| chr16-89722991-INS-66 | chr16-89722991-INS-66 |
| chr16-89736755-INS-264 | chr16-89736755-INS-264 |
| chr16-89738657-INS-1278 | chr16-89738658-INS-1266 |
| chr16-89750409-INS-81 | chr16-89750409-INS-106 |
| chr16-89768906-INS-82 | chr16-89768906-INS-77 |
| chr16-89779154-INS-102 | chr16-89779154-INS-102 |
| chr16-89789442-INS-493 | chr16-89789442-INS-501 |
| chr16-89799616-INS-68 | chr16-89799616-INS-68 |
| chr16-89850787-INS-245 | chr16-89850787-INS-245 |
| chr16-89908078-INS-56 | chr16-89908050-INS-72 |
| chr16-89947574-INS-177 | chr16-89947574-INS-169 |
| chr16-89950747-INS-285 | chr16-89950747-INS-285 |
| chr16-89957438-INS-72 | chr16-89957438-INS-52 |
| chr16-89970151-INS-253 | chr16-89970151-INS-253 |
| chr16-89981562-INS-245 | chr16-89981562-INS-245 |
| chr16-89989280-INS-199 | chr16-89989280-INS-200 |
| chr16-89993408-INS-75 | chr16-89993408-INS-75 |
| chr16-89997445-INS-106 | chr16-89997445-INS-69 |
| chr16-90011784-INS-230 | chr16-90011784-INS-237 |
| chr16-90015821-INS-217 | chr16-90015821-INS-217 |
| chr16-90019373-INS-180 | chr16-90019373-INS-180 |
| chr16-90043247-INS-280 | chr16-90043247-INS-280 |
| chr16-90055411-INS-60 | chr16-90055411-INS-60 |
| chr16-90073630-INS-236 | chr16-90073630-INS-236 |
| chr16-90083090-INS-647 | chr16-90083090-INS-647 |
| chr16-90087194-INS-485 | chr16-90087196-INS-480 |
| chr16-90088102-INS-198 | chr16-90088102-INS-187 |
| chr16-90094682-INS-117 | chr16-90094682-INS-123 |
| chr16-90097377-INS-112 | chr16-90097377-INS-139 |
| chr16-90118614-INS-74 | chr16-90118614-INS-80 |
| chr16-90131919-INS-208 | chr16-90131919-INS-208 |
| chr16-90150922-INS-206 | chr16-90150922-INS-206 |
| chr16-90158626-INS-205 | chr16-90158626-INS-214 |
| chr16-90160228-INS-64 | chr16-90160228-INS-61 |
| chr16-90161690-INS-116 | chr16-90161690-INS-116 |
| chr16-90171109-INS-326 | chr16-90171109-INS-327 |
| chr16-90177062-INS-226 | chr16-90177062-INS-249 |
| chr16-90177742-INS-54 | chr16-90177742-INS-54 |
| chr16-90181442-INS-209 | chr16-90181442-INS-209 |
| chr16-90187500-INS-56 | chr16-90187500-INS-56 |
| chr16-90201648-INS-93 | chr16-90201632-INS-127 |
| chr16-90230127-INS-178 | chr16-90230127-INS-178 |
| chr16-90242041-INS-211 | chr16-90242041-INS-211 |
| chr16-90280869-INS-61 | chr16-90280869-INS-61 |

| | |
|---|---|
| chr16-90304735-INS-86 | chr16-90304735-INS-86 |
| chr16-90307017-INS-54 | chr16-90306992-INS-65 |
| chr16-90312058-INS-82 | chr16-90312058-INS-84 |
| chr16-90313174-INS-138 | chr16-90313177-INS-117 |
| chr16-90320026-INS-312 | chr16-90320028-INS-305 |
| chr16-90333072-INS-309 | chr16-90333072-INS-309 |
| chr16-90408608-INS-205 | chr16-90408606-INS-201 |
| chr16-90420326-INS-60 | chr16-90420326-INS-62 |
| chr16-90425298-INS-57 | chr16-90425264-INS-57 |
| chr16-90429963-INS-51 | chr16-90429963-INS-50 |
| chr16-90442435-INS-57 | chr16-90442435-INS-61 |
| chr16-90450236-INS-80 | chr16-90450236-INS-80 |
| chr16-90453257-INS-130 | chr16-90453257-INS-130 |
| chr16-90459304-INS-439 | chr16-90459304-INS-439 |
| chr16-90466753-INS-347 | chr16-90466753-INS-359 |
| chr16-90476598-INS-90 | chr16-90476598-INS-90 |
| chr16-90511014-INS-76 | chr16-90511021-INS-64 |
| chr16-90513256-INS-325 | chr16-90513256-INS-325 |
| chr16-90532349-INS-84 | chr16-90532353-INS-99 |
| chr16-90541233-INS-210 | chr16-90541233-INS-210 |
| chr16-90543600-INS-68 | chr16-90543600-INS-68 |
| chr16-90562896-INS-104 | chr16-90562896-INS-104 |
| chr16-90597072-INS-192 | chr16-90597072-INS-192 |
| chr16-90638067-INS-56 | chr16-90638067-INS-63 |
| chr16-90658094-INS-274 | chr16-90658088-INS-248 |
| chr16-90678248-INS-434 | chr16-90678248-INS-434 |
| chr16-90699336-INS-174 | chr16-90699336-INS-174 |
| chr16-90706295-INS-56 | chr16-90706293-INS-60 |
| chr16-90710680-INS-53 | chr16-90710676-INS-85 |
| chr16-90712673-INS-86 | chr16-90712673-INS-92 |
| chr16-90714619-INS-654 | chr16-90714619-INS-656 |
| chr16-90769671-INS-2853 | chr16-90769672-INS-2853 |
| chr16-90793051-INS-68 | chr16-90793051-INS-68 |
| chr16-90821241-INS-192 | chr16-90821241-INS-193 |
| chr16-90822338-INS-62 | chr16-90822338-INS-62 |
| chr16-90826897-INS-78 | chr16-90826897-INS-78 |
| chr16-90848966-INS-64 | chr16-90848966-INS-64 |
| chr16-90854510-INS-209 | chr16-90854510-INS-209 |
| chr16-90855948-INS-300 | chr16-90855948-INS-300 |
| chr16-90879122-INS-68 | chr16-90879111-INS-64 |
| chr16-90889348-INS-7181 | chr16-90889349-INS-7181 |
| chr16-90910036-INS-3386 | chr16-90910037-INS-3386 |
| chr16-90921575-INS-327 | chr16-90921575-INS-327 |
| chr16-90923782-INS-60 | chr16-90923782-INS-60 |
| chr16-90929274-INS-126 | chr16-90929274-INS-126 |
| chr16-90944780-INS-523 | chr16-90944780-INS-523 |
| chr16-90961971-INS-216 | chr16-90961971-INS-216 |
| chr16-90967898-INS-53 | chr16-90967898-INS-53 |
| chr16-90972181-INS-60 | chr16-90972181-INS-60 |
| chr16-91011145-INS-123 | chr16-91011145-INS-124 |
| chr16-91021378-INS-92 | chr16-91021384-INS-60 |
| chr16-91031874-INS-76 | chr16-91031874-INS-76 |
| chr16-91037714-INS-129 | chr16-91037714-INS-129 |
| chr16-91062136-INS-219 | chr16-91062136-INS-219 |
| chr16-91114591-INS-149 | chr16-91114591-INS-149 |
| chr16-91124174-INS-61 | chr16-91124174-INS-61 |
| chr16-91142741-INS-579 | chr16-91142741-INS-575 |
| chr16-91148834-INS-74 | chr16-91148828-INS-66 |
| chr16-91167179-INS-378 | chr16-91167179-INS-378 |
| chr16-91173426-INS-209 | chr16-91173426-INS-209 |
| chr16-91183101-INS-242 | chr16-91183101-INS-236 |
| chr16-91221250-INS-626 | chr16-91221250-INS-624 |
| chr16-91226079-INS-160 | chr16-91226079-INS-174 |
| chr16-91241536-INS-144 | chr16-91241585-INS-109 |
| chr16-91276535-INS-2414 | chr16-91276536-INS-2414 |
| chr16-91297469-INS-52 | chr16-91297469-INS-54 |
| chr16-91318075-INS-137 | chr16-91318076-INS-116 |
| chr16-91320492-INS-135 | chr16-91320486-INS-176 |
| chr16-91323040-INS-60 | chr16-91323040-INS-60 |
| chr16-91327246-INS-168 | chr16-91327246-INS-166 |
| chr16-91332796-INS-94 | chr16-91332790-INS-60 |
| chr16-91354103-INS-64 | chr16-91354103-INS-64 |
| chr16-91355427-INS-210 | chr16-91355427-INS-210 |
| chr16-91373529-INS-93 | chr16-91373529-INS-93 |
| chr16-91416826-INS-254 | chr16-91416826-INS-246 |
| chr16-91420813-INS-189 | chr16-91420814-INS-188 |
| chr16-91423522-INS-409 | chr16-91423522-INS-409 |

| | |
|---|---|
| chr16-91428856-INS-65 | chr16-91428856-INS-65 |
| chr16-91429264-INS-199 | chr16-91429264-INS-199 |
| chr16-91431028-INS-245 | chr16-91431028-INS-247 |
| chr16-91488249-INS-111 | chr16-91488249-INS-110 |
| chr16-91489946-INS-225 | chr16-91489946-INS-226 |
| chr16-91493622-INS-54 | chr16-91493623-INS-54 |
| chr16-91507342-INS-5512 | chr16-91507343-INS-5512 |
| chr16-91510400-INS-76 | chr16-91510400-INS-76 |
| chr16-91520614-INS-140 | chr16-91520614-INS-140 |
| chr16-91523938-INS-79 | chr16-91523938-INS-79 |
| chr16-91528517-INS-80 | chr16-91528517-INS-70 |
| chr16-91536383-INS-80 | chr16-91536383-INS-65 |
| chr16-91539286-INS-67 | chr16-91539286-INS-67 |
| chr16-91554797-INS-204 | chr16-91554797-INS-205 |
| chr16-91564354-INS-69 | chr16-91564354-INS-67 |
| chr16-91589092-INS-184 | chr16-91589092-INS-184 |
| chr16-91592339-INS-52 | chr16-91592311-INS-60 |
| chr16-91595280-INS-51 | chr16-91595270-INS-56 |
| chr16-91598165-INS-64 | chr16-91598173-INS-61 |
| chr16-91604422-INS-82 | chr16-91604422-INS-82 |
| chr16-91640224-INS-1882 | chr16-91640225-INS-1882 |
| chr16-91665209-INS-101 | chr16-91665209-INS-93 |
| chr16-91672774-INS-76 | chr16-91672774-INS-76 |
| chr16-91681124-INS-65 | chr16-91681125-INS-57 |
| chr16-91699813-INS-217 | chr16-91699813-INS-217 |
| chr16-91706366-INS-90 | chr16-91706366-INS-90 |
| chr16-91706700-INS-6403 | chr16-91706701-INS-6403 |
| chr16-91714256-INS-138 | chr16-91714256-INS-112 |
| chr16-91736081-INS-174 | chr16-91736097-INS-174 |
| chr16-91738236-INS-64 | chr16-91738236-INS-84 |
| chr16-91750045-INS-51 | chr16-91750026-INS-52 |
| chr16-91795612-INS-63 | chr16-91795612-INS-63 |
| chr16-91807961-INS-414 | chr16-91807962-INS-414 |
| chr16-91815725-INS-185 | chr16-91815725-INS-186 |
| chr16-91837473-INS-72 | chr16-91837473-INS-72 |
| chr16-91843727-INS-83 | chr16-91843727-INS-86 |
| chr16-91865232-INS-55 | chr16-91865235-INS-57 |
| chr16-91890991-INS-70 | chr16-91890976-INS-71 |
| chr16-91910404-INS-52 | chr16-91910404-INS-52 |
| chr16-91917101-INS-110 | chr16-91917105-INS-127 |
| chr16-91918254-INS-97 | chr16-91918255-INS-57 |
| chr16-91922541-INS-61 | chr16-91922541-INS-51 |
| chr16-91926203-INS-216 | chr16-91926203-INS-216 |
| chr16-91927531-INS-207 | chr16-91927531-INS-207 |
| chr16-91933244-INS-160 | chr16-91933235-INS-160 |
| chr16-91935488-INS-132 | chr16-91935488-INS-110 |
| chr16-91938216-INS-66 | chr16-91938218-INS-50 |
| chr16-91951129-INS-342 | chr16-91951129-INS-350 |
| chr16-91964567-INS-51 | chr16-91964568-INS-51 |
| chr16-91967248-INS-558 | chr16-91967248-INS-559 |
| chr16-91979060-INS-279 | chr16-91979060-INS-272 |
| chr16-91987317-INS-104 | chr16-91987317-INS-104 |
| chr16-91999035-INS-94 | chr16-91999035-INS-94 |
| chr16-92004491-INS-357 | chr16-92004491-INS-361 |
| chr16-92087232-INS-502 | chr16-92087232-INS-502 |
| chr16-92145936-INS-73 | chr16-92145936-INS-73 |
| chr16-92157344-INS-124 | chr16-92157390-INS-127 |
| chr16-92175042-INS-59 | chr16-92175067-INS-59 |
| chr16-92187766-INS-188 | chr16-92187767-INS-188 |
| chr16-92199938-INS-108 | chr16-92199938-INS-115 |
| chr16-92204806-INS-232 | chr16-92204806-INS-232 |
| chr16-92236654-INS-72 | chr16-92236654-INS-72 |
| chr16-92240204-INS-81 | chr16-92240204-INS-81 |
| chr16-92245675-INS-565 | chr16-92245709-INS-556 |
| chr16-92255318-INS-51 | chr16-92255319-INS-51 |
| chr16-92268712-INS-242 | chr16-92268712-INS-234 |
| chr16-92280769-INS-420 | chr16-92280769-INS-420 |
| chr16-92283771-INS-193 | chr16-92283772-INS-191 |
| chr16-92285332-INS-146 | chr16-92285339-INS-159 |
| chr16-92287663-INS-98 | chr16-92287663-INS-68 |
| chr16-92289747-INS-82 | chr16-92289760-INS-117 |
| chr16-92301635-INS-108 | chr16-92301635-INS-90 |
| chr16-92351555-INS-84 | chr16-92351555-INS-84 |
| chr16-92369564-INS-110 | chr16-92369564-INS-110 |
| chr16-92430923-INS-57 | chr16-92430923-INS-57 |
| chr16-92448302-INS-347 | chr16-92448302-INS-311 |
| chr16-92457628-INS-56 | chr16-92457628-INS-56 |

| | |
|---|---|
| chr16-92480361-INS-79 | chr16-92480361-INS-79 |
| chr16-92491847-INS-89 | chr16-92491847-INS-90 |
| chr16-92513948-INS-73 | chr16-92513948-INS-77 |
| chr16-92524511-INS-250 | chr16-92524511-INS-250 |
| chr16-92526689-INS-216 | chr16-92526689-INS-216 |
| chr16-92553553-INS-88 | chr16-92553555-INS-89 |
| chr16-92555920-INS-55 | chr16-92555920-INS-55 |
| chr16-92583959-INS-85 | chr16-92583951-INS-121 |
| chr16-92590728-INS-357 | chr16-92590728-INS-357 |
| chr16-92603226-INS-1888 | chr16-92603227-INS-1888 |
| chr16-92631550-INS-160 | chr16-92631549-INS-139 |
| chr16-92645809-INS-6403 | chr16-92645810-INS-6403 |
| chr16-92667759-INS-66 | chr16-92667759-INS-66 |
| chr16-92675804-INS-52 | chr16-92675804-INS-50 |
| chr16-92699388-INS-140 | chr16-92699389-INS-139 |
| chr16-92724404-INS-142 | chr16-92724404-INS-148 |
| chr16-92732706-INS-65 | chr16-92732705-INS-67 |
| chr16-92737388-INS-701 | chr16-92737388-INS-710 |
| chr16-92738130-INS-63 | chr16-92738130-INS-63 |
| chr16-92761843-INS-161 | chr16-92761843-INS-161 |
| chr16-92763011-INS-112 | chr16-92763011-INS-108 |
| chr16-92766607-INS-199 | chr16-92766607-INS-199 |
| chr16-92771564-INS-56 | chr16-92771564-INS-56 |
| chr16-92778863-INS-217 | chr16-92778863-INS-217 |
| chr16-92798315-INS-76 | chr16-92798316-INS-62 |
| chr16-92799368-INS-135 | chr16-92799368-INS-135 |
| chr16-92804365-INS-58 | chr16-92804365-INS-58 |
| chr16-92814222-INS-378 | chr16-92814222-INS-378 |
| chr16-92826854-INS-60 | chr16-92826854-INS-60 |
| chr16-92856377-INS-71 | chr16-92856377-INS-71 |
| chr16-92861735-INS-65 | chr16-92861778-INS-56 |
| chr16-92867288-INS-54 | chr16-92867288-INS-52 |
| chr16-92870594-INS-3026 | chr16-92870595-INS-3026 |
| chr16-92877735-INS-851 | chr16-92877735-INS-851 |
| chr16-92880482-INS-235 | chr16-92880462-INS-245 |
| chr16-92886183-INS-70 | chr16-92886183-INS-70 |
| chr16-92890788-INS-65 | chr16-92890772-INS-85 |
| chr16-92933138-INS-51 | chr16-92933126-INS-57 |
| chr16-92944326-INS-93 | chr16-92944326-INS-93 |
| chr16-92999814-INS-60 | chr16-92999818-INS-58 |
| chr16-93006951-INS-136 | chr16-93006939-INS-148 |
| chr16-93021802-INS-149 | chr16-93021802-INS-149 |
| chr16-93051872-INS-1892 | chr16-93051873-INS-1892 |
| chr16-93059348-INS-936 | chr16-93059348-INS-936 |
| chr16-93075581-INS-185 | chr16-93075581-INS-185 |
| chr16-93099685-INS-64 | chr16-93099651-INS-60 |
| chr16-93114201-INS-402 | chr16-93114201-INS-402 |
| chr16-93123599-INS-112 | chr16-93123599-INS-112 |
| chr16-93125390-INS-160 | chr16-93125390-INS-160 |
| chr16-93156677-INS-144 | chr16-93156677-INS-144 |
| chr16-93211581-INS-54 | chr16-93211581-INS-56 |
| chr16-93222628-INS-66 | chr16-93222628-INS-66 |
| chr16-93266864-INS-1749 | chr16-93266865-INS-1749 |
| chr16-93275608-INS-4512 | chr16-93275609-INS-4512 |
| chr16-93276525-INS-81 | chr16-93276525-INS-56 |
| chr16-93282758-INS-63 | chr16-93282758-INS-63 |
| chr16-93303998-INS-63 | chr16-93303998-INS-65 |
| chr16-93335386-INS-126 | chr16-93335386-INS-124 |
| chr16-93347140-INS-395 | chr16-93347140-INS-395 |
| chr16-93373027-INS-629 | chr16-93373027-INS-625 |
| chr16-93380787-INS-122 | chr16-93380787-INS-98 |
| chr16-93392076-INS-171 | chr16-93392076-INS-162 |
| chr16-93393126-INS-240 | chr16-93393126-INS-242 |
| chr16-93410759-INS-67 | chr16-93410760-INS-66 |
| chr16-93433892-INS-192 | chr16-93433892-INS-192 |
| chr16-93458605-INS-341 | chr16-93458605-INS-341 |
| chr16-93469977-INS-326 | chr16-93469977-INS-326 |
| chr16-93474896-INS-77 | chr16-93474896-INS-77 |
| chr16-93510436-INS-177 | chr16-93510436-INS-177 |
| chr16-93520659-INS-216 | chr16-93520659-INS-216 |
| chr16-93539775-INS-94 | chr16-93539777-INS-116 |
| chr16-93556379-INS-87 | chr16-93556379-INS-87 |
| chr16-93559670-INS-4801 | chr16-93559671-INS-4801 |
| chr16-93564734-INS-107 | chr16-93564734-INS-62 |
| chr16-93568408-INS-216 | chr16-93568408-INS-216 |
| chr16-93599934-INS-288 | chr16-93599934-INS-241 |
| chr16-93602794-INS-66 | chr16-93602794-INS-66 |

| | |
|---|---|
| chr16-93606675-INS-220 | chr16-93606675-INS-221 |
| chr16-93617717-INS-190 | chr16-93617717-INS-191 |
| chr16-93621614-INS-124 | chr16-93621614-INS-155 |
| chr16-93622588-INS-196 | chr16-93622588-INS-196 |
| chr16-93639390-INS-324 | chr16-93639390-INS-324 |
| chr16-93661159-INS-388 | chr16-93661159-INS-388 |
| chr16-93666108-INS-221 | chr16-93666108-INS-221 |
| chr16-93673547-INS-206 | chr16-93673547-INS-206 |
| chr16-93692062-INS-64 | chr16-93692062-INS-64 |
| chr16-93711697-INS-52 | chr16-93711697-INS-64 |
| chr16-93728810-INS-56 | chr16-93728810-INS-58 |
| chr16-93740415-INS-222 | chr16-93740415-INS-223 |
| chr16-93744402-INS-170 | chr16-93744402-INS-169 |
| chr16-93750284-INS-64 | chr16-93750284-INS-64 |
| chr16-93750984-INS-186 | chr16-93750984-INS-186 |
| chr16-93771626-INS-77 | chr16-93771626-INS-84 |
| chr16-93778152-INS-54 | chr16-93778152-INS-54 |
| chr16-93782806-INS-58 | chr16-93782806-INS-58 |
| chr16-93786125-INS-956 | chr16-93786125-INS-962 |
| chr16-93797614-INS-1358 | chr16-93797615-INS-1360 |
| chr16-93806962-INS-198 | chr16-93806962-INS-198 |
| chr16-93811662-INS-52 | chr16-93811662-INS-52 |
| chr16-93819424-INS-224 | chr16-93819417-INS-254 |
| chr16-93823249-INS-107 | chr16-93823249-INS-107 |
| chr16-93847688-INS-201 | chr16-93847688-INS-202 |
| chr16-93862939-INS-134 | chr16-93862938-INS-129 |
| chr16-93866215-INS-248 | chr16-93866215-INS-281 |
| chr16-93887342-INS-69 | chr16-93887342-INS-66 |
| chr16-93891586-INS-112 | chr16-93891557-INS-121 |
| chr16-93916684-INS-101 | chr16-93916686-INS-62 |
| chr16-93919386-INS-84 | chr16-93919397-INS-56 |
| chr16-93923640-INS-66 | chr16-93923640-INS-66 |
| chr16-93945629-INS-551 | chr16-93945630-INS-551 |
| chr16-93947710-INS-73 | chr16-93947710-INS-62 |
| chr16-93968737-INS-187 | chr16-93968737-INS-188 |
| chr16-93976902-INS-508 | chr16-93976902-INS-508 |
| chr16-93987511-INS-6439 | chr16-93987512-INS-6439 |
| chr16-93987882-INS-91 | chr16-93987882-INS-91 |
| chr16-93995833-INS-223 | chr16-93995833-INS-221 |
| chr16-94014287-INS-51 | chr16-94014287-INS-52 |
| chr16-94020172-INS-85 | chr16-94020172-INS-85 |
| chr16-94040726-INS-181 | chr16-94040726-INS-182 |
| chr16-94059996-INS-114 | chr16-94059996-INS-114 |
| chr16-94109325-INS-7081 | chr16-94109326-INS-7081 |
| chr16-94112696-INS-51 | chr16-94112677-INS-54 |
| chr16-94120385-INS-69 | chr16-94120385-INS-90 |
| chr16-94124258-INS-70 | chr16-94124270-INS-52 |
| chr16-94132130-INS-114 | chr16-94132147-INS-138 |
| chr16-94153782-INS-446 | chr16-94153782-INS-446 |
| chr16-94156453-INS-58 | chr16-94156453-INS-58 |
| chr16-94182889-INS-54 | chr16-94182889-INS-53 |
| chr16-94202618-INS-126 | chr16-94202618-INS-106 |
| chr16-94205036-INS-52 | chr16-94205027-INS-62 |
| chr16-94211390-INS-188 | chr16-94211390-INS-189 |
| chr16-94227994-INS-171 | chr16-94227994-INS-195 |
| chr16-94238846-INS-210 | chr16-94238846-INS-210 |
| chr16-94251362-INS-212 | chr16-94251362-INS-211 |
| chr16-94271787-INS-59 | chr16-94271787-INS-59 |
| chr16-94275017-INS-99 | chr16-94275017-INS-100 |
| chr16-94295380-INS-550 | chr16-94295380-INS-550 |
| chr16-94315005-INS-214 | chr16-94315005-INS-215 |
| chr16-94333483-INS-1893 | chr16-94333484-INS-1893 |
| chr16-94336098-INS-193 | chr16-94336098-INS-193 |
| chr16-94354641-INS-488 | chr16-94354641-INS-469 |
| chr16-94361280-INS-77 | chr16-94361280-INS-77 |
| chr16-94379334-INS-204 | chr16-94379334-INS-205 |
| chr16-94389506-INS-400 | chr16-94389506-INS-400 |
| chr16-94409223-INS-53 | chr16-94409223-INS-54 |
| chr16-94420225-INS-62 | chr16-94420225-INS-62 |
| chr16-94429734-INS-123 | chr16-94429734-INS-123 |
| chr16-94447831-INS-90 | chr16-94447851-INS-54 |
| chr16-94452856-INS-222 | chr16-94452856-INS-222 |
| chr16-94455173-INS-56 | chr16-94455173-INS-56 |
| chr16-94459274-INS-70 | chr16-94459281-INS-61 |
| chr16-94468108-INS-72 | chr16-94468108-INS-72 |
| chr16-94499899-INS-87 | chr16-94499899-INS-87 |
| chr16-94526447-INS-60 | chr16-94526447-INS-60 |

| | |
|---|---|
| chr16-94534302-INS-511 | chr16-94534302-INS-511 |
| chr16-94541482-INS-71 | chr16-94541482-INS-71 |
| chr16-94564368-INS-286 | chr16-94564368-INS-291 |
| chr16-94566587-INS-52 | chr16-94566588-INS-53 |
| chr16-94601237-INS-84 | chr16-94601237-INS-84 |
| chr16-94603267-INS-53 | chr16-94603267-INS-52 |
| chr16-94604388-INS-415 | chr16-94604388-INS-415 |
| chr16-94657743-INS-176 | chr16-94657744-INS-144 |
| chr16-94667578-INS-68 | chr16-94667578-INS-68 |
| chr16-94680262-INS-383 | chr16-94680263-INS-427 |
| chr16-94688631-INS-253 | chr16-94688631-INS-253 |
| chr16-94696013-INS-54 | chr16-94696007-INS-54 |
| chr16-94706403-INS-146 | chr16-94706403-INS-146 |
| chr16-94743149-INS-198 | chr16-94743149-INS-166 |
| chr16-94746257-INS-51 | chr16-94746257-INS-51 |
| chr16-94753553-INS-348 | chr16-94753553-INS-348 |
| chr16-94788255-INS-58 | chr16-94788255-INS-66 |
| chr16-94790433-INS-64 | chr16-94790433-INS-64 |
| chr16-94802629-INS-68 | chr16-94802629-INS-68 |
| chr16-94809476-INS-352 | chr16-94809476-INS-352 |
| chr16-94822551-INS-82 | chr16-94822551-INS-82 |
| chr16-94834802-INS-54 | chr16-94834802-INS-60 |
| chr16-94850059-INS-53 | chr16-94850055-INS-57 |
| chr16-94876374-INS-71 | chr16-94876375-INS-73 |
| chr16-94877953-INS-319 | chr16-94877953-INS-319 |
| chr16-94903851-INS-522 | chr16-94903851-INS-521 |
| chr16-94909829-INS-476 | chr16-94909829-INS-476 |
| chr16-94916815-INS-88 | chr16-94916815-INS-88 |
| chr16-94983899-INS-60 | chr16-94983899-INS-60 |
| chr16-94984745-INS-6406 | chr16-94984746-INS-6406 |
| chr16-94990268-INS-57 | chr16-94990268-INS-59 |
| chr16-94995578-INS-915 | chr16-94995578-INS-915 |
| chr16-95001665-INS-180 | chr16-95001665-INS-180 |
| chr16-95009770-INS-88 | chr16-95009770-INS-87 |
| chr16-95021011-INS-78 | chr16-95021011-INS-84 |
| chr16-95055889-INS-105 | chr16-95055890-INS-60 |
| chr16-95056398-INS-150 | chr16-95056399-INS-150 |
| chr16-95057323-INS-53 | chr16-95057323-INS-53 |
| chr16-95065947-INS-185 | chr16-95065947-INS-182 |
| chr16-95099026-INS-64 | chr16-95099027-INS-68 |
| chr16-95118520-INS-80 | chr16-95118520-INS-80 |
| chr16-95128224-INS-99 | chr16-95128224-INS-99 |
| chr16-95136606-INS-441 | chr16-95136606-INS-441 |
| chr16-95139223-INS-3083 | chr16-95139224-INS-3083 |
| chr16-95150393-INS-265 | chr16-95150393-INS-268 |
| chr16-95192732-INS-68 | chr16-95192732-INS-68 |
| chr16-95194415-INS-125 | chr16-95194415-INS-125 |
| chr16-95199119-INS-116 | chr16-95199119-INS-116 |
| chr16-95203347-INS-484 | chr16-95203347-INS-485 |
| chr16-95236245-INS-135 | chr16-95236245-INS-135 |
| chr16-95256594-INS-65 | chr16-95256592-INS-51 |
| chr16-95260621-INS-136 | chr16-95260621-INS-136 |
| chr16-95264320-INS-54 | chr16-95264320-INS-56 |
| chr16-95284580-INS-60 | chr16-95284581-INS-53 |
| chr16-95292748-INS-70 | chr16-95292748-INS-58 |
| chr16-95296541-INS-205 | chr16-95296541-INS-205 |
| chr16-95306795-INS-73 | chr16-95306791-INS-59 |
| chr16-95426148-INS-55 | chr16-95426148-INS-57 |
| chr16-95430664-INS-54 | chr16-95430664-INS-60 |
| chr16-95439231-INS-57 | chr16-95439231-INS-57 |
| chr16-95471914-INS-53 | chr16-95471914-INS-53 |
| chr16-95480888-INS-100 | chr16-95480888-INS-100 |
| chr16-95569964-INS-147 | chr16-95569958-INS-150 |
| chr16-95575851-INS-358 | chr16-95575851-INS-358 |
| chr16-95583697-INS-424 | chr16-95583697-INS-424 |
| chr16-95652707-INS-110 | chr16-95652710-INS-130 |
| chr16-95675242-INS-52 | chr16-95675222-INS-55 |
| chr16-95707190-INS-201 | chr16-95707190-INS-201 |
| chr16-95766037-INS-432 | chr16-95766037-INS-424 |
| chr16-95767450-INS-70 | chr16-95767445-INS-62 |
| chr16-95768686-INS-179 | chr16-95768686-INS-179 |
| chr16-95786152-INS-68 | chr16-95786152-INS-68 |
| chr16-95836977-INS-79 | chr16-95836968-INS-70 |
| chr16-95841040-INS-99 | chr16-95841040-INS-99 |
| chr16-95859467-INS-60 | chr16-95859467-INS-60 |
| chr16-95861777-INS-80 | chr16-95861777-INS-80 |
| chr16-95876129-INS-211 | chr16-95876129-INS-207 |

| | |
|---|---|
| chr16-95887392-INS-118 | chr16-95887390-INS-148 |
| chr16-95907461-INS-177 | chr16-95907453-INS-175 |
| chr16-95915293-INS-303 | chr16-95915293-INS-303 |
| chr16-95915434-INS-767 | chr16-95915435-INS-765 |
| chr16-95923833-INS-88 | chr16-95923833-INS-88 |
| chr16-95929929-INS-52 | chr16-95929929-INS-52 |
| chr16-95938101-INS-103 | chr16-95938106-INS-113 |
| chr16-95974012-INS-74 | chr16-95974012-INS-76 |
| chr16-95978659-INS-63 | chr16-95978659-INS-63 |
| chr16-95982640-INS-202 | chr16-95982640-INS-201 |
| chr16-96000072-INS-187 | chr16-96000072-INS-187 |
| chr16-96010053-INS-64 | chr16-96010053-INS-64 |
| chr16-96013666-INS-89 | chr16-96013666-INS-89 |
| chr16-96025381-INS-491 | chr16-96025381-INS-491 |
| chr16-96032111-INS-154 | chr16-96032111-INS-140 |
| chr16-96059488-INS-248 | chr16-96059488-INS-248 |
| chr16-96123275-INS-53 | chr16-96123275-INS-53 |
| chr16-96144344-INS-64 | chr16-96144305-INS-64 |
| chr16-96150840-INS-2318 | chr16-96150841-INS-2318 |
| chr16-96152512-INS-396 | chr16-96152512-INS-393 |
| chr16-96171905-INS-6247 | chr16-96171906-INS-6248 |
| chr16-96176130-INS-1892 | chr16-96176131-INS-1892 |
| chr16-96177225-INS-100 | chr16-96177225-INS-100 |
| chr16-96196007-INS-514 | chr16-96196007-INS-514 |
| chr16-96262820-INS-102 | chr16-96262820-INS-92 |
| chr16-96281395-INS-60 | chr16-96281419-INS-58 |
| chr16-96309520-INS-227 | chr16-96309520-INS-207 |
| chr16-96319348-INS-98 | chr16-96319348-INS-98 |
| chr16-96321070-INS-98 | chr16-96321070-INS-93 |
| chr16-96358594-INS-309 | chr16-96358594-INS-309 |
| chr16-96381543-INS-146 | chr16-96381543-INS-143 |
| chr16-96410368-INS-105 | chr16-96410367-INS-97 |
| chr16-96425906-INS-74 | chr16-96425906-INS-101 |
| chr16-96434096-INS-129 | chr16-96434096-INS-135 |
| chr16-96452641-INS-108 | chr16-96452641-INS-114 |
| chr16-96464060-INS-64 | chr16-96464028-INS-52 |
| chr16-96478569-INS-119 | chr16-96478569-INS-119 |
| chr16-96482727-INS-70 | chr16-96482727-INS-74 |
| chr16-96490637-INS-86 | chr16-96490675-INS-92 |
| chr16-96521908-INS-56 | chr16-96521908-INS-58 |
| chr16-96548376-INS-52 | chr16-96548376-INS-52 |
| chr16-96566065-INS-2012 | chr16-96566066-INS-2012 |
| chr16-96584879-INS-168 | chr16-96584879-INS-168 |
| chr16-96586430-INS-80 | chr16-96586430-INS-80 |
| chr16-96598053-INS-153 | chr16-96598037-INS-173 |
| chr16-96688738-INS-419 | chr16-96688738-INS-421 |
| chr16-96692428-INS-111 | chr16-96692415-INS-120 |
| chr16-96713219-INS-64 | chr16-96713219-INS-68 |
| chr16-96759321-INS-386 | chr16-96759321-INS-386 |
| chr16-96764502-INS-50 | chr16-96764473-INS-54 |
| chr16-96777272-INS-1952 | chr16-96777273-INS-1952 |
| chr16-96813520-INS-448 | chr16-96813520-INS-448 |
| chr16-96901101-INS-92 | chr16-96901101-INS-52 |
| chr16-96905034-INS-132 | chr16-96905034-INS-132 |
| chr16-96935405-INS-181 | chr16-96935405-INS-193 |
| chr16-96938120-INS-92 | chr16-96938120-INS-92 |
| chr16-96941092-INS-88 | chr16-96941092-INS-88 |
| chr16-96943130-INS-244 | chr16-96943130-INS-236 |
| chr16-96974105-INS-63 | chr16-96974105-INS-62 |
| chr16-97012582-INS-58 | chr16-97012568-INS-60 |
| chr16-97027209-INS-131 | chr16-97027218-INS-160 |
| chr16-97060791-INS-62 | chr16-97060791-INS-60 |
| chr16-97072908-INS-1704 | chr16-97072909-INS-1704 |
| chr16-97082117-INS-68 | chr16-97082078-INS-108 |
| chr16-97093189-INS-7066 | chr16-97093190-INS-7067 |
| chr16-97114146-INS-445 | chr16-97114146-INS-445 |
| chr16-97117051-INS-63 | chr16-97117051-INS-63 |
| chr16-97121854-INS-4700 | chr16-97121855-INS-4700 |
| chr16-97121910-INS-51 | chr16-97121910-INS-59 |
| chr16-97135994-INS-71 | chr16-97135994-INS-71 |
| chr16-97142910-INS-97 | chr16-97142912-INS-94 |
| chr16-97144982-INS-96 | chr16-97144982-INS-101 |
| chr16-97165804-INS-184 | chr16-97165804-INS-168 |
| chr16-97195714-INS-56 | chr16-97195714-INS-52 |
| chr16-97209410-INS-88 | chr16-97209410-INS-88 |
| chr16-97233466-INS-826 | chr16-97233466-INS-831 |
| chr16-97286644-INS-339 | chr16-97286645-INS-339 |

| | |
|---|---|
| chr16-97311008-INS-207 | chr16-97311008-INS-209 |
| chr16-97337365-INS-4086 | chr16-97337366-INS-4086 |
| chr16-97344629-INS-6390 | chr16-97344630-INS-6390 |
| chr16-97371048-INS-478 | chr16-97371048-INS-477 |
| chr16-97376962-INS-656 | chr16-97376962-INS-656 |
| chr16-97382608-INS-64 | chr16-97382608-INS-64 |
| chr16-97446100-INS-1240 | chr16-97446101-INS-1239 |
| chr16-97459364-INS-62 | chr16-97459364-INS-62 |
| chr16-97463353-INS-57 | chr16-97463353-INS-57 |
| chr16-97464950-INS-75 | chr16-97464950-INS-75 |
| chr16-97481083-INS-63 | chr16-97481083-INS-62 |
| chr16-97500884-INS-202 | chr16-97500884-INS-202 |
| chr16-97521706-INS-52 | chr16-97521706-INS-58 |
| chr16-97526524-INS-208 | chr16-97526524-INS-209 |
| chr16-97532630-INS-254 | chr16-97532630-INS-254 |
| chr16-97555584-INS-163 | chr16-97555584-INS-163 |
| chr16-97561878-INS-96 | chr16-97561869-INS-92 |
| chr16-97589382-INS-90 | chr16-97589382-INS-99 |
| chr16-97612710-INS-187 | chr16-97612710-INS-187 |
| chr16-97680987-INS-62 | chr16-97680977-INS-66 |
| chr16-97699430-INS-548 | chr16-97699430-INS-548 |
| chr16-97747152-INS-230 | chr16-97747152-INS-228 |
| chr16-97779236-INS-3003 | chr16-97779237-INS-3003 |
| chr16-97796863-INS-215 | chr16-97796863-INS-215 |
| chr16-97846417-INS-93 | chr16-97846417-INS-109 |
| chr17-3056299-INS-89 | chr17-3056299-INS-89 |
| chr17-3060923-INS-165 | chr17-3060923-INS-165 |
| chr17-3063733-INS-498 | chr17-3063732-INS-498 |
| chr17-3080014-INS-84 | chr17-3080014-INS-84 |
| chr17-3088050-INS-62 | chr17-3088050-INS-62 |
| chr17-3093058-INS-433 | chr17-3093058-INS-434 |
| chr17-3125709-INS-221 | chr17-3125709-INS-221 |
| chr17-3138028-INS-199 | chr17-3138028-INS-199 |
| chr17-3145973-INS-440 | chr17-3145973-INS-440 |
| chr17-3192763-INS-368 | chr17-3192763-INS-368 |
| chr17-3224704-INS-201 | chr17-3224704-INS-202 |
| chr17-3267726-INS-199 | chr17-3267726-INS-199 |
| chr17-3300385-INS-158 | chr17-3300385-INS-164 |
| chr17-3328437-INS-76 | chr17-3328433-INS-76 |
| chr17-3329080-INS-54 | chr17-3329064-INS-77 |
| chr17-3334580-INS-57 | chr17-3334580-INS-57 |
| chr17-3372594-INS-98 | chr17-3372594-INS-98 |
| chr17-3378346-INS-53 | chr17-3378346-INS-53 |
| chr17-3383048-INS-133 | chr17-3383027-INS-122 |
| chr17-3408062-INS-165 | chr17-3408062-INS-165 |
| chr17-3410947-INS-2811 | chr17-3410948-INS-2812 |
| chr17-3436781-INS-91 | chr17-3436781-INS-91 |
| chr17-3446772-INS-243 | chr17-3446772-INS-243 |
| chr17-3450438-INS-93 | chr17-3450438-INS-93 |
| chr17-3456720-INS-77 | chr17-3456720-INS-81 |
| chr17-3467959-INS-73 | chr17-3467959-INS-73 |
| chr17-3485782-INS-66 | chr17-3485782-INS-66 |
| chr17-3492112-INS-100 | chr17-3492112-INS-100 |
| chr17-3494722-INS-78 | chr17-3494722-INS-78 |
| chr17-3515102-INS-3167 | chr17-3515103-INS-3168 |
| chr17-3516608-INS-56 | chr17-3516608-INS-56 |
| chr17-3527274-INS-61 | chr17-3527274-INS-61 |
| chr17-3565991-INS-62 | chr17-3565991-INS-66 |
| chr17-3579837-INS-69 | chr17-3579837-INS-70 |
| chr17-3599255-INS-220 | chr17-3599255-INS-220 |
| chr17-3604751-INS-103 | chr17-3604751-INS-103 |
| chr17-3629720-INS-8420 | chr17-3629721-INS-8420 |
| chr17-3634625-INS-7114 | chr17-3634626-INS-7114 |
| chr17-3639306-INS-65 | chr17-3639306-INS-65 |
| chr17-3670108-INS-319 | chr17-3670108-INS-319 |
| chr17-3679040-INS-3264 | chr17-3679041-INS-3265 |
| chr17-3682059-INS-228 | chr17-3682059-INS-230 |
| chr17-3696700-INS-52 | chr17-3696700-INS-54 |
| chr17-3701365-INS-72 | chr17-3701381-INS-74 |
| chr17-3716716-INS-53 | chr17-3716680-INS-56 |
| chr17-3721618-INS-224 | chr17-3721618-INS-224 |
| chr17-3742973-INS-499 | chr17-3742973-INS-494 |
| chr17-3774973-INS-76 | chr17-3774965-INS-84 |
| chr17-3797156-INS-6483 | chr17-3797157-INS-6483 |
| chr17-3805155-INS-102 | chr17-3805155-INS-102 |
| chr17-3817763-INS-2747 | chr17-3817764-INS-2747 |
| chr17-3819474-INS-6269 | chr17-3819475-INS-6269 |

| | |
|---|---|
| chr17-3825243-INS-58 | chr17-3825243-INS-60 |
| chr17-3890993-INS-65 | chr17-3890993-INS-65 |
| chr17-3904170-INS-980 | chr17-3904172-INS-978 |
| chr17-3908044-INS-1128 | chr17-3908045-INS-1130 |
| chr17-3909658-INS-656 | chr17-3909658-INS-656 |
| chr17-3916575-INS-54 | chr17-3916573-INS-58 |
| chr17-3927640-INS-64 | chr17-3927640-INS-66 |
| chr17-3928061-INS-6418 | chr17-3928062-INS-6418 |
| chr17-3935163-INS-645 | chr17-3935163-INS-645 |
| chr17-3943061-INS-84 | chr17-3943065-INS-52 |
| chr17-3944267-INS-103 | chr17-3944267-INS-103 |
| chr17-3961256-INS-1864 | chr17-3961257-INS-1864 |
| chr17-3977515-INS-7236 | chr17-3977516-INS-7236 |
| chr17-3989776-INS-1259 | chr17-3989777-INS-1299 |
| chr17-4049642-INS-7158 | chr17-4049643-INS-7158 |
| chr17-4050213-INS-61 | chr17-4050213-INS-61 |
| chr17-4085980-INS-4572 | chr17-4085981-INS-4572 |
| chr17-4101580-INS-54 | chr17-4101580-INS-60 |
| chr17-4109360-INS-2751 | chr17-4109361-INS-2751 |
| chr17-4150217-INS-70 | chr17-4150220-INS-66 |
| chr17-4176408-INS-102 | chr17-4176408-INS-108 |
| chr17-4186088-INS-664 | chr17-4186132-INS-662 |
| chr17-4194214-INS-81 | chr17-4194214-INS-94 |
| chr17-4206588-INS-71 | chr17-4206589-INS-51 |
| chr17-4213311-INS-691 | chr17-4213311-INS-691 |
| chr17-4232678-INS-5979 | chr17-4232679-INS-5979 |
| chr17-4242209-INS-52 | chr17-4242209-INS-52 |
| chr17-4311310-INS-56 | chr17-4311310-INS-58 |
| chr17-4314485-INS-2933 | chr17-4314486-INS-2933 |
| chr17-4322665-INS-61 | chr17-4322689-INS-66 |
| chr17-4349909-INS-84 | chr17-4349908-INS-86 |
| chr17-4356579-INS-414 | chr17-4356579-INS-414 |
| chr17-4365447-INS-1894 | chr17-4365448-INS-1894 |
| chr17-4383823-INS-54 | chr17-4383835-INS-84 |
| chr17-4403044-INS-58 | chr17-4403044-INS-60 |
| chr17-4407176-INS-378 | chr17-4407176-INS-382 |
| chr17-4410514-INS-74 | chr17-4410498-INS-58 |
| chr17-4412567-INS-56 | chr17-4412567-INS-50 |
| chr17-4417261-INS-79 | chr17-4417261-INS-64 |
| chr17-4434775-INS-72 | chr17-4434775-INS-74 |
| chr17-4456844-INS-655 | chr17-4456844-INS-655 |
| chr17-4494751-INS-6772 | chr17-4494752-INS-6772 |
| chr17-4531474-INS-165 | chr17-4531474-INS-165 |
| chr17-4537016-INS-7225 | chr17-4537017-INS-7225 |
| chr17-4539835-INS-170 | chr17-4539825-INS-152 |
| chr17-4542718-INS-362 | chr17-4542718-INS-362 |
| chr17-4560499-INS-6504 | chr17-4560500-INS-6507 |
| chr17-4577545-INS-130 | chr17-4577534-INS-150 |
| chr17-4583456-INS-95 | chr17-4583454-INS-71 |
| chr17-4609278-INS-67 | chr17-4609278-INS-63 |
| chr17-4615090-INS-1213 | chr17-4615091-INS-1212 |
| chr17-4620932-INS-77 | chr17-4620932-INS-77 |
| chr17-4628989-INS-203 | chr17-4628989-INS-203 |
| chr17-4656041-INS-239 | chr17-4656041-INS-239 |
| chr17-4680836-INS-256 | chr17-4680836-INS-256 |
| chr17-4689766-INS-113 | chr17-4689766-INS-112 |
| chr17-4692895-INS-480 | chr17-4692895-INS-480 |
| chr17-4709867-INS-76 | chr17-4709867-INS-76 |
| chr17-4760729-INS-60 | chr17-4760729-INS-60 |
| chr17-4782475-INS-90 | chr17-4782475-INS-90 |
| chr17-4789014-INS-130 | chr17-4788993-INS-133 |
| chr17-4803028-INS-60 | chr17-4803028-INS-60 |
| chr17-4806516-INS-58 | chr17-4806516-INS-60 |
| chr17-4827868-INS-90 | chr17-4827868-INS-90 |
| chr17-4837320-INS-62 | chr17-4837320-INS-62 |
| chr17-4850770-INS-993 | chr17-4850770-INS-993 |
| chr17-4858745-INS-55 | chr17-4858741-INS-90 |
| chr17-4867318-INS-173 | chr17-4867318-INS-178 |
| chr17-4880522-INS-203 | chr17-4880522-INS-203 |
| chr17-4882799-INS-70 | chr17-4882799-INS-70 |
| chr17-4895423-INS-59 | chr17-4895426-INS-60 |
| chr17-4917138-INS-63 | chr17-4917138-INS-70 |
| chr17-4950088-INS-461 | chr17-4950103-INS-475 |
| chr17-4961000-INS-198 | chr17-4961000-INS-198 |
| chr17-4987284-INS-524 | chr17-4987284-INS-537 |
| chr17-4996698-INS-82 | chr17-4996698-INS-86 |
| chr17-5018739-INS-65 | chr17-5018692-INS-85 |

| | |
|---|---|
| chr17-5029194-INS-112 | chr17-5029194-INS-115 |
| chr17-5061824-INS-106 | chr17-5061824-INS-106 |
| chr17-5078166-INS-75 | chr17-5078166-INS-75 |
| chr17-5083063-INS-238 | chr17-5083063-INS-238 |
| chr17-5088607-INS-106 | chr17-5088607-INS-106 |
| chr17-5090017-INS-561 | chr17-5090017-INS-561 |
| chr17-5091279-INS-61 | chr17-5091279-INS-61 |
| chr17-5097942-INS-51 | chr17-5097942-INS-51 |
| chr17-5133526-INS-119 | chr17-5133526-INS-119 |
| chr17-5140023-INS-136 | chr17-5140023-INS-136 |
| chr17-5151484-INS-95 | chr17-5151484-INS-95 |
| chr17-5159490-INS-125 | chr17-5159490-INS-125 |
| chr17-5159778-INS-432 | chr17-5159778-INS-432 |
| chr17-5167039-INS-68 | chr17-5166995-INS-72 |
| chr17-5171976-INS-145 | chr17-5171976-INS-145 |
| chr17-5183760-INS-59 | chr17-5183775-INS-66 |
| chr17-5191765-INS-624 | chr17-5191766-INS-624 |
| chr17-5209951-INS-94 | chr17-5209940-INS-94 |
| chr17-5231542-INS-425 | chr17-5231545-INS-428 |
| chr17-5275198-INS-218 | chr17-5275198-INS-216 |
| chr17-5309085-INS-84 | chr17-5309094-INS-56 |
| chr17-5320651-INS-80 | chr17-5320651-INS-80 |
| chr17-5339024-INS-221 | chr17-5339024-INS-221 |
| chr17-5402307-INS-103 | chr17-5402307-INS-103 |
| chr17-5432999-INS-110 | chr17-5432999-INS-110 |
| chr17-5438672-INS-819 | chr17-5438672-INS-818 |
| chr17-5450390-INS-117 | chr17-5450390-INS-117 |
| chr17-5459017-INS-214 | chr17-5459017-INS-214 |
| chr17-5471226-INS-54 | chr17-5471226-INS-54 |
| chr17-5475964-INS-570 | chr17-5475971-INS-523 |
| chr17-5501323-INS-615 | chr17-5501324-INS-615 |
| chr17-5550133-INS-85 | chr17-5550133-INS-84 |
| chr17-5564299-INS-59 | chr17-5564299-INS-63 |
| chr17-5584052-INS-72 | chr17-5584050-INS-65 |
| chr17-5592894-INS-5547 | chr17-5592895-INS-5547 |
| chr17-5601104-INS-56 | chr17-5601104-INS-60 |
| chr17-5604987-INS-70 | chr17-5604987-INS-70 |
| chr17-5612230-INS-195 | chr17-5612230-INS-195 |
| chr17-5633786-INS-1150 | chr17-5633787-INS-1150 |
| chr17-5640291-INS-84 | chr17-5640291-INS-66 |
| chr17-5642622-INS-80 | chr17-5642590-INS-107 |
| chr17-5705172-INS-60 | chr17-5705151-INS-63 |
| chr17-5728883-INS-60 | chr17-5728876-INS-51 |
| chr17-5733196-INS-55 | chr17-5733196-INS-55 |
| chr17-5791335-INS-110 | chr17-5791335-INS-115 |
| chr17-5802487-INS-453 | chr17-5802487-INS-453 |
| chr17-5807706-INS-1080 | chr17-5807707-INS-1080 |
| chr17-5811474-INS-185 | chr17-5811474-INS-193 |
| chr17-5818401-INS-685 | chr17-5818401-INS-687 |
| chr17-5874435-INS-60 | chr17-5874435-INS-60 |
| chr17-5875682-INS-69 | chr17-5875682-INS-69 |
| chr17-5878188-INS-97 | chr17-5878188-INS-97 |
| chr17-5902003-INS-82 | chr17-5902015-INS-67 |
| chr17-5914828-INS-416 | chr17-5914872-INS-417 |
| chr17-5980207-INS-208 | chr17-5980207-INS-208 |
| chr17-5986129-INS-106 | chr17-5986129-INS-110 |
| chr17-5986842-INS-202 | chr17-5986842-INS-195 |
| chr17-6012786-INS-51 | chr17-6012786-INS-52 |
| chr17-6071281-INS-63 | chr17-6071281-INS-63 |
| chr17-6097115-INS-75 | chr17-6097115-INS-75 |
| chr17-6111122-INS-135 | chr17-6111122-INS-135 |
| chr17-6145074-INS-275 | chr17-6145082-INS-300 |
| chr17-6174181-INS-87 | chr17-6174181-INS-58 |
| chr17-6196119-INS-351 | chr17-6196119-INS-351 |
| chr17-6214235-INS-3340 | chr17-6214255-INS-3318 |
| chr17-6253344-INS-143 | chr17-6253344-INS-165 |
| chr17-6255379-INS-87 | chr17-6255379-INS-87 |
| chr17-6280877-INS-80 | chr17-6280877-INS-82 |
| chr17-6292344-INS-204 | chr17-6292344-INS-203 |
| chr17-6300303-INS-812 | chr17-6300303-INS-813 |
| chr17-6305142-INS-220 | chr17-6305142-INS-220 |
| chr17-6373941-INS-220 | chr17-6373941-INS-218 |
| chr17-6374092-INS-630 | chr17-6374092-INS-630 |
| chr17-6658227-INS-1300 | chr17-6658228-INS-1300 |
| chr17-6765392-INS-813 | chr17-6765392-INS-813 |
| chr17-6773384-INS-204 | chr17-6773384-INS-205 |
| chr17-6824881-INS-503 | chr17-6824882-INS-503 |

| | |
|---|---|
| chr17-6933553-INS-61 | chr17-6933553-INS-61 |
| chr17-6954296-INS-236 | chr17-6954296-INS-236 |
| chr17-6983915-INS-211 | chr17-6983915-INS-211 |
| chr17-6986103-INS-192 | chr17-6986103-INS-189 |
| chr17-6988203-INS-211 | chr17-6988203-INS-211 |
| chr17-6997782-INS-926 | chr17-6997783-INS-926 |
| chr17-7012653-INS-176 | chr17-7012616-INS-180 |
| chr17-7020663-INS-66 | chr17-7020663-INS-66 |
| chr17-7065273-INS-107 | chr17-7065273-INS-107 |
| chr17-7105943-INS-816 | chr17-7105938-INS-816 |
| chr17-7137589-INS-208 | chr17-7137589-INS-208 |
| chr17-7142254-INS-719 | chr17-7142254-INS-719 |
| chr17-7146795-INS-220 | chr17-7146795-INS-220 |
| chr17-7147300-INS-985 | chr17-7147296-INS-979 |
| chr17-7164517-INS-352 | chr17-7164517-INS-352 |
| chr17-7165234-INS-58 | chr17-7165234-INS-62 |
| chr17-7167447-INS-288 | chr17-7167482-INS-287 |
| chr17-7171590-INS-5412 | chr17-7171591-INS-5414 |
| chr17-7172288-INS-54 | chr17-7172289-INS-54 |
| chr17-7175274-INS-210 | chr17-7175274-INS-211 |
| chr17-7176150-INS-170 | chr17-7176150-INS-170 |
| chr17-7181978-INS-64 | chr17-7181978-INS-64 |
| chr17-7214985-INS-67 | chr17-7214985-INS-67 |
| chr17-7231821-INS-180 | chr17-7231821-INS-181 |
| chr17-7257799-INS-71 | chr17-7257801-INS-72 |
| chr17-7262500-INS-79 | chr17-7262500-INS-79 |
| chr17-7320804-INS-119 | chr17-7320804-INS-119 |
| chr17-7450993-INS-1392 | chr17-7450994-INS-1395 |
| chr17-7458110-INS-220 | chr17-7458110-INS-251 |
| chr17-7463473-INS-72 | chr17-7463473-INS-72 |
| chr17-7496431-INS-1888 | chr17-7496432-INS-1888 |
| chr17-7504155-INS-56 | chr17-7504155-INS-56 |
| chr17-7544555-INS-488 | chr17-7544555-INS-488 |
| chr17-7573945-INS-371 | chr17-7573945-INS-371 |
| chr17-7584684-INS-80 | chr17-7584660-INS-82 |
| chr17-7594976-INS-54 | chr17-7594976-INS-56 |
| chr17-7600100-INS-54 | chr17-7600100-INS-54 |
| chr17-7604090-INS-54 | chr17-7604090-INS-54 |
| chr17-7699147-INS-56 | chr17-7699147-INS-56 |
| chr17-7733084-INS-232 | chr17-7733084-INS-249 |
| chr17-7743036-INS-524 | chr17-7743036-INS-524 |
| chr17-7756968-INS-61 | chr17-7756968-INS-61 |
| chr17-7782609-INS-51 | chr17-7782610-INS-59 |
| chr17-7804406-INS-190 | chr17-7804406-INS-190 |
| chr17-7816017-INS-52 | chr17-7816017-INS-52 |
| chr17-7818859-INS-71 | chr17-7818859-INS-67 |
| chr17-7819964-INS-1839 | chr17-7819965-INS-1838 |
| chr17-7824362-INS-146 | chr17-7824363-INS-145 |
| chr17-7832242-INS-75 | chr17-7832243-INS-75 |
| chr17-7842148-INS-203 | chr17-7842148-INS-203 |
| chr17-7857446-INS-93 | chr17-7857447-INS-81 |
| chr17-7864339-INS-6559 | chr17-7864340-INS-6568 |
| chr17-7888339-INS-52 | chr17-7888339-INS-52 |
| chr17-7911497-INS-85 | chr17-7911497-INS-88 |
| chr17-7915845-INS-189 | chr17-7915845-INS-189 |
| chr17-7928814-INS-198 | chr17-7928814-INS-207 |
| chr17-7940933-INS-774 | chr17-7940933-INS-774 |
| chr17-7999548-INS-245 | chr17-7999548-INS-245 |
| chr17-8021724-INS-61 | chr17-8021724-INS-63 |
| chr17-8053157-INS-440 | chr17-8053157-INS-440 |
| chr17-8053983-INS-99 | chr17-8054016-INS-66 |
| chr17-8057032-INS-52 | chr17-8057032-INS-54 |
| chr17-8068275-INS-56 | chr17-8068275-INS-52 |
| chr17-8091193-INS-322 | chr17-8091193-INS-349 |
| chr17-8096663-INS-144 | chr17-8096688-INS-162 |
| chr17-8109876-INS-66 | chr17-8109868-INS-68 |
| chr17-8115706-INS-64 | chr17-8115706-INS-66 |
| chr17-8137158-INS-125 | chr17-8137158-INS-128 |
| chr17-8139594-INS-145 | chr17-8139594-INS-150 |
| chr17-8154437-INS-128 | chr17-8154437-INS-128 |
| chr17-8164849-INS-54 | chr17-8164849-INS-54 |
| chr17-8198180-INS-54 | chr17-8198180-INS-54 |
| chr17-8204020-INS-299 | chr17-8204020-INS-300 |
| chr17-8205870-INS-64 | chr17-8205870-INS-64 |
| chr17-8343220-INS-1014 | chr17-8343221-INS-1013 |
| chr17-8350446-INS-79 | chr17-8350462-INS-97 |
| chr17-8375636-INS-482 | chr17-8375636-INS-482 |

| | |
|---|---|
| chr17-8387849-INS-223 | chr17-8387849-INS-224 |
| chr17-8428119-INS-163 | chr17-8428119-INS-167 |
| chr17-8469359-INS-2320 | chr17-8469360-INS-2320 |
| chr17-8521682-INS-350 | chr17-8521682-INS-350 |
| chr17-8529212-INS-181 | chr17-8529212-INS-182 |
| chr17-8548125-INS-56 | chr17-8548125-INS-56 |
| chr17-8568465-INS-251 | chr17-8568465-INS-250 |
| chr17-8584742-INS-234 | chr17-8584742-INS-234 |
| chr17-8596182-INS-50 | chr17-8596182-INS-50 |
| chr17-8604358-INS-93 | chr17-8604358-INS-93 |
| chr17-8608733-INS-169 | chr17-8608733-INS-169 |
| chr17-8656777-INS-91 | chr17-8656744-INS-91 |
| chr17-8682430-INS-135 | chr17-8682430-INS-135 |
| chr17-8686103-INS-101 | chr17-8686103-INS-105 |
| chr17-8694837-INS-1866 | chr17-8694838-INS-1866 |
| chr17-8708951-INS-437 | chr17-8708951-INS-437 |
| chr17-8847256-INS-72 | chr17-8847257-INS-61 |
| chr17-8850590-INS-71 | chr17-8850597-INS-61 |
| chr17-8885289-INS-156 | chr17-8885289-INS-156 |
| chr17-8919051-INS-54 | chr17-8919051-INS-54 |
| chr17-8952606-INS-93 | chr17-8952608-INS-94 |
| chr17-8956807-INS-143 | chr17-8956807-INS-143 |
| chr17-8970353-INS-84 | chr17-8970353-INS-84 |
| chr17-8973069-INS-60 | chr17-8973111-INS-64 |
| chr17-9008094-INS-60 | chr17-9008094-INS-60 |
| chr17-9009338-INS-94 | chr17-9009341-INS-51 |
| chr17-9012556-INS-128 | chr17-9012561-INS-96 |
| chr17-9049474-INS-94 | chr17-9049474-INS-62 |
| chr17-9052387-INS-228 | chr17-9052387-INS-203 |
| chr17-9065188-INS-114 | chr17-9065188-INS-114 |
| chr17-9109724-INS-118 | chr17-9109730-INS-138 |
| chr17-9124993-INS-58 | chr17-9124999-INS-81 |
| chr17-9239423-INS-358 | chr17-9239423-INS-358 |
| chr17-9249633-INS-104 | chr17-9249623-INS-114 |
| chr17-9267392-INS-204 | chr17-9267392-INS-203 |
| chr17-9271005-INS-51 | chr17-9271005-INS-51 |
| chr17-9277034-INS-101 | chr17-9277034-INS-97 |
| chr17-9278395-INS-1087 | chr17-9278396-INS-1087 |
| chr17-9304132-INS-100 | chr17-9304132-INS-100 |
| chr17-9316260-INS-142 | chr17-9316260-INS-145 |
| chr17-9333496-INS-53 | chr17-9333496-INS-79 |
| chr17-9339789-INS-397 | chr17-9339789-INS-397 |
| chr17-9346450-INS-61 | chr17-9346450-INS-61 |
| chr17-9351617-INS-141 | chr17-9351617-INS-147 |
| chr17-9355156-INS-56 | chr17-9355156-INS-76 |
| chr17-9366343-INS-92 | chr17-9366343-INS-96 |
| chr17-9425833-INS-186 | chr17-9425832-INS-189 |
| chr17-9434265-INS-150 | chr17-9434265-INS-108 |
| chr17-9455560-INS-61 | chr17-9455560-INS-61 |
| chr17-9480316-INS-54 | chr17-9480318-INS-56 |
| chr17-9486100-INS-110 | chr17-9486092-INS-88 |
| chr17-9489450-INS-110 | chr17-9489467-INS-116 |
| chr17-9512964-INS-181 | chr17-9512964-INS-182 |
| chr17-9525458-INS-721 | chr17-9525458-INS-721 |
| chr17-9580827-INS-206 | chr17-9580827-INS-206 |
| chr17-9592568-INS-122 | chr17-9592568-INS-122 |
| chr17-9594032-INS-961 | chr17-9594032-INS-945 |
| chr17-9615797-INS-145 | chr17-9615794-INS-117 |
| chr17-9622318-INS-110 | chr17-9622318-INS-117 |
| chr17-9671613-INS-60 | chr17-9671613-INS-62 |
| chr17-9680136-INS-4830 | chr17-9680137-INS-4830 |
| chr17-9696455-INS-54 | chr17-9696455-INS-54 |
| chr17-9726485-INS-108 | chr17-9726485-INS-108 |
| chr17-9742646-INS-7090 | chr17-9742647-INS-7090 |
| chr17-9773636-INS-86 | chr17-9773643-INS-82 |
| chr17-9789122-INS-54 | chr17-9789122-INS-54 |
| chr17-9811355-INS-6442 | chr17-9811356-INS-6442 |
| chr17-9856933-INS-60 | chr17-9856933-INS-60 |
| chr17-9864020-INS-6899 | chr17-9864021-INS-6899 |
| chr17-9889176-INS-61 | chr17-9889176-INS-62 |
| chr17-9904009-INS-78 | chr17-9904021-INS-80 |
| chr17-9943417-INS-56 | chr17-9943415-INS-58 |
| chr17-10013457-INS-350 | chr17-10013457-INS-350 |
| chr17-10035977-INS-550 | chr17-10035977-INS-550 |
| chr17-10049748-INS-7116 | chr17-10049749-INS-7116 |
| chr17-10051865-INS-135 | chr17-10051865-INS-136 |
| chr17-10078741-INS-2099 | chr17-10078742-INS-2099 |

| | |
|---|---|
| chr17-10084328-INS-54 | chr17-10084286-INS-65 |
| chr17-10106475-INS-54 | chr17-10106475-INS-54 |
| chr17-10106795-INS-6506 | chr17-10106796-INS-6507 |
| chr17-10121612-INS-474 | chr17-10121612-INS-474 |
| chr17-10134814-INS-1895 | chr17-10134815-INS-1895 |
| chr17-10166184-INS-54 | chr17-10166184-INS-54 |
| chr17-10208105-INS-2066 | chr17-10208106-INS-2066 |
| chr17-10208287-INS-680 | chr17-10208287-INS-680 |
| chr17-10218830-INS-82 | chr17-10218830-INS-82 |
| chr17-10270977-INS-66 | chr17-10270977-INS-66 |
| chr17-10284684-INS-7172 | chr17-10284685-INS-7218 |
| chr17-10304344-INS-55 | chr17-10304344-INS-55 |
| chr17-10307194-INS-72 | chr17-10307194-INS-66 |
| chr17-10313360-INS-133 | chr17-10313360-INS-133 |
| chr17-10332588-INS-5568 | chr17-10332589-INS-5568 |
| chr17-10445840-INS-60 | chr17-10445840-INS-60 |
| chr17-10490176-INS-210 | chr17-10490176-INS-210 |
| chr17-10522953-INS-64 | chr17-10522953-INS-64 |
| chr17-10538307-INS-105 | chr17-10538307-INS-105 |
| chr17-10543071-INS-153 | chr17-10543071-INS-153 |
| chr17-10545433-INS-208 | chr17-10545433-INS-208 |
| chr17-10553072-INS-251 | chr17-10553072-INS-267 |
| chr17-10559633-INS-123 | chr17-10559633-INS-123 |
| chr17-10599124-INS-95 | chr17-10599124-INS-95 |
| chr17-10643643-INS-178 | chr17-10643643-INS-182 |
| chr17-10698614-INS-62 | chr17-10698614-INS-62 |
| chr17-10700056-INS-72 | chr17-10700056-INS-60 |
| chr17-10707893-INS-189 | chr17-10707893-INS-189 |
| chr17-10709295-INS-70 | chr17-10709295-INS-50 |
| chr17-10712018-INS-159 | chr17-10712004-INS-173 |
| chr17-10744021-INS-132 | chr17-10744045-INS-120 |
| chr17-10789771-INS-84 | chr17-10789771-INS-90 |
| chr17-10843922-INS-57 | chr17-10843922-INS-53 |
| chr17-10843932-INS-582 | chr17-10843923-INS-593 |
| chr17-10853646-INS-77 | chr17-10853675-INS-97 |
| chr17-10873045-INS-441 | chr17-10873045-INS-441 |
| chr17-10891532-INS-87 | chr17-10891532-INS-58 |
| chr17-10900515-INS-676 | chr17-10900518-INS-676 |
| chr17-10909968-INS-87 | chr17-10909969-INS-55 |
| chr17-10916344-INS-56 | chr17-10916344-INS-56 |
| chr17-10934267-INS-210 | chr17-10934267-INS-210 |
| chr17-10944983-INS-62 | chr17-10944983-INS-68 |
| chr17-10951499-INS-54 | chr17-10951499-INS-54 |
| chr17-10953340-INS-64 | chr17-10953340-INS-64 |
| chr17-10973507-INS-56 | chr17-10973507-INS-56 |
| chr17-10983355-INS-77 | chr17-10983355-INS-77 |
| chr17-10993628-INS-6635 | chr17-10993629-INS-6635 |
| chr17-11010658-INS-58 | chr17-11010658-INS-62 |
| chr17-11028310-INS-3466 | chr17-11028311-INS-3466 |
| chr17-11031677-INS-205 | chr17-11031677-INS-206 |
| chr17-11051454-INS-172 | chr17-11051454-INS-168 |
| chr17-11063694-INS-86 | chr17-11063694-INS-82 |
| chr17-11117931-INS-177 | chr17-11117931-INS-185 |
| chr17-11126596-INS-186 | chr17-11126596-INS-186 |
| chr17-11132379-INS-7143 | chr17-11132380-INS-7143 |
| chr17-11134544-INS-59 | chr17-11134544-INS-59 |
| chr17-11149804-INS-7476 | chr17-11149805-INS-7476 |
| chr17-11172295-INS-155 | chr17-11172295-INS-155 |
| chr17-11172787-INS-8480 | chr17-11172788-INS-8487 |
| chr17-11177241-INS-64 | chr17-11177241-INS-62 |
| chr17-11200993-INS-68 | chr17-11200993-INS-68 |
| chr17-11222125-INS-339 | chr17-11222125-INS-339 |
| chr17-11251443-INS-138 | chr17-11251442-INS-138 |
| chr17-11257811-INS-51 | chr17-11257811-INS-75 |
| chr17-11263180-INS-114 | chr17-11263180-INS-105 |
| chr17-11265973-INS-206 | chr17-11265973-INS-206 |
| chr17-11268050-INS-78 | chr17-11268013-INS-117 |
| chr17-11284382-INS-80 | chr17-11284383-INS-85 |
| chr17-11300116-INS-75 | chr17-11300116-INS-75 |
| chr17-11353828-INS-1286 | chr17-11353829-INS-1286 |
| chr17-11369990-INS-86 | chr17-11369990-INS-86 |
| chr17-11400857-INS-737 | chr17-11400857-INS-714 |
| chr17-11403984-INS-119 | chr17-11403984-INS-131 |
| chr17-11431085-INS-51 | chr17-11431085-INS-53 |
| chr17-11435884-INS-96 | chr17-11435849-INS-126 |
| chr17-11444300-INS-6455 | chr17-11444301-INS-6455 |
| chr17-11481094-INS-956 | chr17-11481094-INS-956 |

| | |
|---|---|
| chr17-11517615-INS-402 | chr17-11517615-INS-402 |
| chr17-11536481-INS-75 | chr17-11536484-INS-59 |
| chr17-11543314-INS-1925 | chr17-11543315-INS-1925 |
| chr17-11587290-INS-1297 | chr17-11587291-INS-1297 |
| chr17-11589186-INS-179 | chr17-11589186-INS-179 |
| chr17-11602002-INS-6251 | chr17-11602003-INS-6251 |
| chr17-11659288-INS-2236 | chr17-11659289-INS-2236 |
| chr17-11674047-INS-429 | chr17-11674047-INS-429 |
| chr17-11706915-INS-56 | chr17-11706906-INS-56 |
| chr17-11732470-INS-64 | chr17-11732471-INS-64 |
| chr17-11734476-INS-68 | chr17-11734476-INS-68 |
| chr17-11749208-INS-181 | chr17-11749208-INS-189 |
| chr17-11756999-INS-788 | chr17-11756999-INS-788 |
| chr17-11781563-INS-66 | chr17-11781563-INS-68 |
| chr17-11815951-INS-253 | chr17-11815951-INS-253 |
| chr17-11847223-INS-112 | chr17-11847203-INS-112 |
| chr17-11867026-INS-1858 | chr17-11867027-INS-1858 |
| chr17-11892180-INS-2815 | chr17-11892181-INS-2815 |
| chr17-11937240-INS-52 | chr17-11937240-INS-52 |
| chr17-11945346-INS-100 | chr17-11945346-INS-52 |
| chr17-11950552-INS-149 | chr17-11950552-INS-137 |
| chr17-11981158-INS-54 | chr17-11981159-INS-54 |
| chr17-11994912-INS-60 | chr17-11994949-INS-54 |
| chr17-11998435-INS-54 | chr17-11998435-INS-52 |
| chr17-12010833-INS-58 | chr17-12010833-INS-58 |
| chr17-12011947-INS-120 | chr17-12011947-INS-132 |
| chr17-12029057-INS-128 | chr17-12029057-INS-128 |
| chr17-12036317-INS-271 | chr17-12036317-INS-262 |
| chr17-12069498-INS-84 | chr17-12069498-INS-84 |
| chr17-12107394-INS-180 | chr17-12107394-INS-180 |
| chr17-12121193-INS-6873 | chr17-12121194-INS-6874 |
| chr17-12124651-INS-126 | chr17-12124651-INS-126 |
| chr17-12167975-INS-52 | chr17-12167975-INS-52 |
| chr17-12180505-INS-52 | chr17-12180505-INS-52 |
| chr17-12206073-INS-100 | chr17-12206073-INS-100 |
| chr17-12244768-INS-62 | chr17-12244768-INS-62 |
| chr17-12250141-INS-210 | chr17-12250141-INS-210 |
| chr17-12255189-INS-56 | chr17-12255189-INS-58 |
| chr17-12262822-INS-291 | chr17-12262822-INS-292 |
| chr17-12294369-INS-277 | chr17-12294362-INS-277 |
| chr17-12310136-INS-3671 | chr17-12310137-INS-3671 |
| chr17-12317738-INS-88 | chr17-12317755-INS-79 |
| chr17-12333184-INS-80 | chr17-12333184-INS-100 |
| chr17-12336555-INS-60 | chr17-12336555-INS-60 |
| chr17-12371643-INS-84 | chr17-12371643-INS-86 |
| chr17-12375062-INS-53 | chr17-12375062-INS-56 |
| chr17-12377373-INS-488 | chr17-12377373-INS-488 |
| chr17-12399241-INS-56 | chr17-12399241-INS-56 |
| chr17-12421976-INS-393 | chr17-12421976-INS-393 |
| chr17-12472326-INS-54 | chr17-12472326-INS-54 |
| chr17-12494473-INS-1215 | chr17-12494474-INS-1215 |
| chr17-12496321-INS-76 | chr17-12496321-INS-76 |
| chr17-12498912-INS-376 | chr17-12498912-INS-376 |
| chr17-12539821-INS-64 | chr17-12539837-INS-68 |
| chr17-12560996-INS-56 | chr17-12560996-INS-62 |
| chr17-12568824-INS-52 | chr17-12568824-INS-52 |
| chr17-12580410-INS-219 | chr17-12580410-INS-219 |
| chr17-12650330-INS-69 | chr17-12650330-INS-73 |
| chr17-12681775-INS-3318 | chr17-12681776-INS-3312 |
| chr17-12691063-INS-53 | chr17-12691063-INS-53 |
| chr17-12693676-INS-183 | chr17-12693686-INS-183 |
| chr17-12729046-INS-58 | chr17-12729046-INS-58 |
| chr17-12742084-INS-54 | chr17-12742084-INS-58 |
| chr17-12757920-INS-3945 | chr17-12757921-INS-3945 |
| chr17-12764598-INS-208 | chr17-12764614-INS-201 |
| chr17-12792550-INS-3144 | chr17-12792551-INS-3144 |
| chr17-12915071-INS-209 | chr17-12915071-INS-209 |
| chr17-12939110-INS-207 | chr17-12939110-INS-219 |
| chr17-12952318-INS-53 | chr17-12952318-INS-53 |
| chr17-12974003-INS-1868 | chr17-12974004-INS-1869 |
| chr17-13000077-INS-204 | chr17-13000077-INS-204 |
| chr17-13017709-INS-984 | chr17-13017709-INS-985 |
| chr17-13025839-INS-6270 | chr17-13025840-INS-6271 |
| chr17-13103342-INS-456 | chr17-13103342-INS-456 |
| chr17-13133672-INS-71 | chr17-13133671-INS-60 |
| chr17-13156147-INS-55 | chr17-13156147-INS-55 |
| chr17-13156729-INS-98 | chr17-13156729-INS-98 |

| | |
|---|---|
| chr17-13157384-INS-208 | chr17-13157384-INS-207 |
| chr17-13199051-INS-1450 | chr17-13199053-INS-1443 |
| chr17-13206780-INS-370 | chr17-13206780-INS-373 |
| chr17-13219794-INS-231 | chr17-13219794-INS-230 |
| chr17-13223048-INS-200 | chr17-13223048-INS-200 |
| chr17-13411734-INS-52 | chr17-13411734-INS-50 |
| chr17-13567476-INS-70 | chr17-13567460-INS-94 |
| chr17-13579088-INS-88 | chr17-13579062-INS-87 |
| chr17-13601772-INS-477 | chr17-13601772-INS-467 |
| chr17-13687387-INS-117 | chr17-13687387-INS-90 |
| chr17-13731723-INS-620 | chr17-13731723-INS-620 |
| chr17-13737128-INS-9378 | chr17-13737128-INS-9377 |
| chr17-13791971-INS-7102 | chr17-13791972-INS-7102 |
| chr17-13888342-INS-6927 | chr17-13888343-INS-6927 |
| chr17-13898026-INS-446 | chr17-13898026-INS-417 |
| chr17-13917065-INS-218 | chr17-13917021-INS-218 |
| chr17-13944011-INS-66 | chr17-13944011-INS-66 |
| chr17-13956018-INS-201 | chr17-13956018-INS-205 |
| chr17-13995602-INS-201 | chr17-13995602-INS-201 |
| chr17-14005396-INS-7137 | chr17-14005397-INS-7137 |
| chr17-14036329-INS-738 | chr17-14036329-INS-736 |
| chr17-14037552-INS-54 | chr17-14037552-INS-56 |
| chr17-14139424-INS-327 | chr17-14139424-INS-327 |
| chr17-14150751-INS-123 | chr17-14150751-INS-120 |
| chr17-14156821-INS-53 | chr17-14156821-INS-53 |
| chr17-14171054-INS-54 | chr17-14171054-INS-54 |
| chr17-14215694-INS-367 | chr17-14215694-INS-365 |
| chr17-14267713-INS-401 | chr17-14267712-INS-400 |
| chr17-14268185-INS-445 | chr17-14268185-INS-445 |
| chr17-14277396-INS-117 | chr17-14277375-INS-123 |
| chr17-14337571-INS-6270 | chr17-14337572-INS-6273 |
| chr17-14384634-INS-252 | chr17-14384636-INS-269 |
| chr17-14397215-INS-268 | chr17-14397215-INS-229 |
| chr17-14409323-INS-111 | chr17-14409323-INS-108 |
| chr17-14410221-INS-400 | chr17-14410221-INS-400 |
| chr17-14446967-INS-80 | chr17-14446967-INS-100 |
| chr17-14467186-INS-140 | chr17-14467186-INS-136 |
| chr17-14469170-INS-58 | chr17-14469170-INS-58 |
| chr17-14471631-INS-575 | chr17-14471631-INS-575 |
| chr17-14479330-INS-195 | chr17-14479330-INS-195 |
| chr17-14540091-INS-89 | chr17-14540091-INS-101 |
| chr17-14561354-INS-5374 | chr17-14561355-INS-5374 |
| chr17-14588757-INS-73 | chr17-14588757-INS-71 |
| chr17-14595143-INS-812 | chr17-14595143-INS-812 |
| chr17-14609855-INS-2272 | chr17-14609856-INS-2272 |
| chr17-14613891-INS-51 | chr17-14613891-INS-51 |
| chr17-14618076-INS-91 | chr17-14618074-INS-99 |
| chr17-14636393-INS-400 | chr17-14636393-INS-400 |
| chr17-14663720-INS-3031 | chr17-14663721-INS-3031 |
| chr17-14703471-INS-306 | chr17-14703471-INS-302 |
| chr17-14715427-INS-422 | chr17-14715427-INS-424 |
| chr17-14736147-INS-53 | chr17-14736155-INS-65 |
| chr17-14797881-INS-111 | chr17-14797917-INS-120 |
| chr17-14810844-INS-116 | chr17-14810844-INS-116 |
| chr17-14828244-INS-162 | chr17-14828244-INS-162 |
| chr17-14843488-INS-2899 | chr17-14843489-INS-2899 |
| chr17-14862064-INS-155 | chr17-14862064-INS-155 |
| chr17-14960196-INS-87 | chr17-14960196-INS-87 |
| chr17-14963610-INS-209 | chr17-14963610-INS-210 |
| chr17-14974774-INS-97 | chr17-14974774-INS-91 |
| chr17-15057530-INS-6442 | chr17-15057531-INS-6442 |
| chr17-15058016-INS-400 | chr17-15058016-INS-400 |
| chr17-15064956-INS-833 | chr17-15064956-INS-834 |
| chr17-15068037-INS-103 | chr17-15068037-INS-103 |
| chr17-15096095-INS-205 | chr17-15096095-INS-164 |
| chr17-15124534-INS-998 | chr17-15124534-INS-998 |
| chr17-15145670-INS-206 | chr17-15145670-INS-206 |
| chr17-15155981-INS-213 | chr17-15155981-INS-213 |
| chr17-15165800-INS-64 | chr17-15165800-INS-64 |
| chr17-15296927-INS-59 | chr17-15296927-INS-59 |
| chr17-15302609-INS-98 | chr17-15302609-INS-50 |
| chr17-15330774-INS-227 | chr17-15330770-INS-223 |
| chr17-15375783-INS-84 | chr17-15375783-INS-84 |
| chr17-15411374-INS-380 | chr17-15411374-INS-380 |
| chr17-15487514-INS-123 | chr17-15487511-INS-132 |
| chr17-15492980-INS-64 | chr17-15492980-INS-72 |
| chr17-15534681-INS-111 | chr17-15534681-INS-116 |

| | |
|---|---|
| chr17-15545295-INS-86 | chr17-15545295-INS-86 |
| chr17-15556498-INS-3202 | chr17-15556499-INS-3202 |
| chr17-15571451-INS-57 | chr17-15571451-INS-59 |
| chr17-15581918-INS-111 | chr17-15581918-INS-111 |
| chr17-15591827-INS-60 | chr17-15591830-INS-75 |
| chr17-15606587-INS-114 | chr17-15606626-INS-84 |
| chr17-15630126-INS-81 | chr17-15630126-INS-81 |
| chr17-15634724-INS-118 | chr17-15634722-INS-108 |
| chr17-15642589-INS-57 | chr17-15642589-INS-57 |
| chr17-15654914-INS-64 | chr17-15654908-INS-92 |
| chr17-15666381-INS-133 | chr17-15666381-INS-132 |
| chr17-15668611-INS-209 | chr17-15668611-INS-216 |
| chr17-15684084-INS-84 | chr17-15684092-INS-89 |
| chr17-15691523-INS-73 | chr17-15691523-INS-73 |
| chr17-15692813-INS-201 | chr17-15692813-INS-201 |
| chr17-15695567-INS-309 | chr17-15695567-INS-309 |
| chr17-15719822-INS-549 | chr17-15719822-INS-549 |
| chr17-15724492-INS-115 | chr17-15724492-INS-115 |
| chr17-15732945-INS-79 | chr17-15732945-INS-79 |
| chr17-15738956-INS-446 | chr17-15738952-INS-446 |
| chr17-15750756-INS-63 | chr17-15750757-INS-60 |
| chr17-15757015-INS-5006 | chr17-15757016-INS-5006 |
| chr17-15758126-INS-163 | chr17-15758126-INS-156 |
| chr17-15762395-INS-194 | chr17-15762395-INS-202 |
| chr17-15764370-INS-84 | chr17-15764370-INS-84 |
| chr17-15771092-INS-301 | chr17-15771092-INS-298 |
| chr17-15779922-INS-58 | chr17-15779922-INS-56 |
| chr17-15785695-INS-520 | chr17-15785695-INS-520 |
| chr17-15794470-INS-64 | chr17-15794472-INS-60 |
| chr17-15806863-INS-75 | chr17-15806897-INS-72 |
| chr17-15869487-INS-263 | chr17-15869487-INS-282 |
| chr17-15885664-INS-290 | chr17-15885664-INS-278 |
| chr17-15895402-INS-91 | chr17-15895402-INS-91 |
| chr17-15904319-INS-191 | chr17-15904319-INS-179 |
| chr17-15911008-INS-267 | chr17-15911008-INS-267 |
| chr17-15934363-INS-68 | chr17-15934361-INS-56 |
| chr17-15943293-INS-509 | chr17-15943293-INS-509 |
| chr17-15950313-INS-135 | chr17-15950313-INS-135 |
| chr17-15961754-INS-148 | chr17-15961754-INS-148 |
| chr17-15979986-INS-77 | chr17-15979986-INS-77 |
| chr17-16013275-INS-51 | chr17-16013270-INS-78 |
| chr17-16019330-INS-55 | chr17-16019334-INS-52 |
| chr17-16020509-INS-128 | chr17-16020509-INS-128 |
| chr17-16053208-INS-133 | chr17-16053208-INS-136 |
| chr17-16056633-INS-102 | chr17-16056629-INS-69 |
| chr17-16057925-INS-224 | chr17-16057918-INS-226 |
| chr17-16064063-INS-53 | chr17-16064064-INS-57 |
| chr17-16071274-INS-212 | chr17-16071274-INS-212 |
| chr17-16075114-INS-189 | chr17-16075114-INS-189 |
| chr17-16079635-INS-93 | chr17-16079637-INS-94 |
| chr17-16107076-INS-96 | chr17-16107071-INS-62 |
| chr17-16117027-INS-108 | chr17-16117027-INS-108 |
| chr17-16121312-INS-610 | chr17-16121312-INS-622 |
| chr17-16144925-INS-723 | chr17-16144925-INS-726 |
| chr17-16209050-INS-326 | chr17-16209050-INS-326 |
| chr17-16213750-INS-4611 | chr17-16213751-INS-4611 |
| chr17-16214999-INS-52 | chr17-16214998-INS-62 |
| chr17-16253305-INS-59 | chr17-16253306-INS-57 |
| chr17-16259681-INS-66 | chr17-16259681-INS-66 |
| chr17-16283936-INS-6398 | chr17-16283937-INS-6403 |
| chr17-16291632-INS-6751 | chr17-16291633-INS-6751 |
| chr17-16297525-INS-103 | chr17-16297525-INS-107 |
| chr17-16311451-INS-98 | chr17-16311451-INS-78 |
| chr17-16313938-INS-52 | chr17-16313971-INS-56 |
| chr17-16332683-INS-683 | chr17-16332683-INS-682 |
| chr17-16337749-INS-831 | chr17-16337749-INS-832 |
| chr17-16378531-INS-102 | chr17-16378531-INS-102 |
| chr17-16383208-INS-237 | chr17-16383208-INS-238 |
| chr17-16384023-INS-2923 | chr17-16384024-INS-2923 |
| chr17-16413971-INS-7101 | chr17-16413972-INS-7101 |
| chr17-16424780-INS-137 | chr17-16424780-INS-137 |
| chr17-16436858-INS-66 | chr17-16436858-INS-67 |
| chr17-16446161-INS-78 | chr17-16446179-INS-52 |
| chr17-16495328-INS-74 | chr17-16495328-INS-74 |
| chr17-16497592-INS-2190 | chr17-16497593-INS-2190 |
| chr17-16506991-INS-54 | chr17-16506991-INS-54 |
| chr17-16522963-INS-4572 | chr17-16522964-INS-4578 |

| | |
|---|---|
| chr17-16536894-INS-299 | chr17-16536894-INS-298 |
| chr17-16537954-INS-75 | chr17-16537954-INS-60 |
| chr17-16561679-INS-244 | chr17-16561679-INS-244 |
| chr17-16583816-INS-202 | chr17-16583819-INS-204 |
| chr17-16597149-INS-58 | chr17-16597149-INS-60 |
| chr17-16637379-INS-67 | chr17-16637379-INS-109 |
| chr17-16640823-INS-80 | chr17-16640823-INS-80 |
| chr17-16674808-INS-53 | chr17-16674808-INS-57 |
| chr17-16695907-INS-70 | chr17-16695888-INS-80 |
| chr17-16704773-INS-52 | chr17-16704773-INS-52 |
| chr17-16719903-INS-154 | chr17-16719903-INS-154 |
| chr17-16729142-INS-54 | chr17-16729142-INS-56 |
| chr17-16740733-INS-119 | chr17-16740733-INS-122 |
| chr17-16747997-INS-170 | chr17-16747997-INS-170 |
| chr17-16750128-INS-2160 | chr17-16750129-INS-2180 |
| chr17-16752929-INS-103 | chr17-16752920-INS-117 |
| chr17-16765852-INS-6371 | chr17-16765853-INS-6371 |
| chr17-16809430-INS-52 | chr17-16809430-INS-58 |
| chr17-16810093-INS-205 | chr17-16810093-INS-205 |
| chr17-16817379-INS-78 | chr17-16817379-INS-77 |
| chr17-16824087-INS-524 | chr17-16824087-INS-524 |
| chr17-16828569-INS-59 | chr17-16828553-INS-54 |
| chr17-16829518-INS-5447 | chr17-16829519-INS-5447 |
| chr17-16846178-INS-5335 | chr17-16846179-INS-5335 |
| chr17-16863222-INS-100 | chr17-16863222-INS-88 |
| chr17-16887266-INS-162 | chr17-16887266-INS-161 |
| chr17-16904779-INS-401 | chr17-16904779-INS-401 |
| chr17-16906679-INS-56 | chr17-16906679-INS-56 |
| chr17-16911249-INS-5489 | chr17-16911250-INS-5489 |
| chr17-16920419-INS-2998 | chr17-16920420-INS-2998 |
| chr17-17024427-INS-56 | chr17-17024427-INS-52 |
| chr17-17034429-INS-9803 | chr17-17034430-INS-9803 |
| chr17-17069847-INS-406 | chr17-17069847-INS-406 |
| chr17-17083686-INS-188 | chr17-17083686-INS-188 |
| chr17-17118491-INS-481 | chr17-17118491-INS-481 |
| chr17-17120245-INS-66 | chr17-17120223-INS-98 |
| chr17-17131549-INS-582 | chr17-17131549-INS-582 |
| chr17-17132690-INS-573 | chr17-17132690-INS-573 |
| chr17-17143528-INS-88 | chr17-17143528-INS-92 |
| chr17-17167124-INS-51 | chr17-17167145-INS-57 |
| chr17-17227440-INS-233 | chr17-17227440-INS-235 |
| chr17-17387923-INS-2540 | chr17-17387930-INS-2540 |
| chr17-17414752-INS-58 | chr17-17414732-INS-60 |
| chr17-17427416-INS-72 | chr17-17427416-INS-72 |
| chr17-17455766-INS-2461 | chr17-17455767-INS-2462 |
| chr17-17548760-INS-203 | chr17-17548760-INS-203 |
| chr17-17587804-INS-5798 | chr17-17587805-INS-5799 |
| chr17-17589838-INS-1061 | chr17-17589839-INS-1091 |
| chr17-17591495-INS-58 | chr17-17591493-INS-59 |
| chr17-17592654-INS-60 | chr17-17592654-INS-55 |
| chr17-17610565-INS-219 | chr17-17610565-INS-218 |
| chr17-17618755-INS-819 | chr17-17618755-INS-820 |
| chr17-17629180-INS-223 | chr17-17629180-INS-224 |
| chr17-17656406-INS-201 | chr17-17656406-INS-201 |
| chr17-17661718-INS-1335 | chr17-17661719-INS-1333 |
| chr17-17676802-INS-204 | chr17-17676802-INS-204 |
| chr17-17681396-INS-226 | chr17-17681396-INS-226 |
| chr17-17701735-INS-2629 | chr17-17701736-INS-2629 |
| chr17-17702731-INS-117 | chr17-17702727-INS-125 |
| chr17-17708787-INS-211 | chr17-17708787-INS-211 |
| chr17-17713907-INS-350 | chr17-17713907-INS-348 |
| chr17-17718076-INS-695 | chr17-17718077-INS-695 |
| chr17-17731002-INS-51 | chr17-17731002-INS-54 |
| chr17-17817441-INS-84 | chr17-17817413-INS-116 |
| chr17-17837613-INS-146 | chr17-17837637-INS-96 |
| chr17-17877830-INS-51 | chr17-17877830-INS-51 |
| chr17-17884552-INS-1892 | chr17-17884553-INS-1892 |
| chr17-17900917-INS-1010 | chr17-17900922-INS-1012 |
| chr17-17919881-INS-323 | chr17-17919881-INS-323 |
| chr17-17929274-INS-345 | chr17-17929274-INS-345 |
| chr17-17941357-INS-6722 | chr17-17941358-INS-6721 |
| chr17-17943687-INS-198 | chr17-17943687-INS-198 |
| chr17-17948302-INS-7122 | chr17-17948303-INS-7122 |
| chr17-17990482-INS-7445 | chr17-17990483-INS-7445 |
| chr17-18002098-INS-1237 | chr17-18002099-INS-1233 |
| chr17-18003119-INS-4418 | chr17-18003120-INS-4374 |
| chr17-18008290-INS-57 | chr17-18008290-INS-57 |

| | |
|---|---|
| chr17-18053682-INS-64 | chr17-18053682-INS-64 |
| chr17-18064872-INS-55 | chr17-18064872-INS-50 |
| chr17-18072899-INS-61 | chr17-18072870-INS-52 |
| chr17-18076032-INS-54 | chr17-18076032-INS-56 |
| chr17-18082179-INS-76 | chr17-18082179-INS-76 |
| chr17-18135047-INS-520 | chr17-18135047-INS-519 |
| chr17-18196539-INS-4188 | chr17-18196540-INS-4192 |
| chr17-18196540-INS-343 | chr17-18196540-INS-343 |
| chr17-18213724-INS-4741 | chr17-18213725-INS-4741 |
| chr17-18226680-INS-100 | chr17-18226680-INS-106 |
| chr17-18229674-INS-241 | chr17-18229674-INS-241 |
| chr17-18239295-INS-1010 | chr17-18239296-INS-1011 |
| chr17-18343639-INS-97 | chr17-18343639-INS-97 |
| chr17-18423345-INS-6357 | chr17-18423346-INS-6357 |
| chr17-18446041-INS-5174 | chr17-18446042-INS-5174 |
| chr17-18502002-INS-1891 | chr17-18502003-INS-1891 |
| chr17-18515810-INS-1288 | chr17-18515811-INS-1289 |
| chr17-18532314-INS-6475 | chr17-18532315-INS-6476 |
| chr17-18593963-INS-485 | chr17-18593963-INS-485 |
| chr17-18617242-INS-160 | chr17-18617242-INS-160 |
| chr17-18628641-INS-64 | chr17-18628641-INS-66 |
| chr17-18641918-INS-60 | chr17-18641918-INS-64 |
| chr17-18687435-INS-2770 | chr17-18687436-INS-2770 |
| chr17-18696397-INS-6436 | chr17-18696398-INS-6436 |
| chr17-18697695-INS-225 | chr17-18697695-INS-226 |
| chr17-18740318-INS-112 | chr17-18740318-INS-109 |
| chr17-18765526-INS-53 | chr17-18765525-INS-54 |
| chr17-18851153-INS-139 | chr17-18851153-INS-139 |
| chr17-18851971-INS-5503 | chr17-18851972-INS-5503 |
| chr17-18866161-INS-62 | chr17-18866161-INS-62 |
| chr17-18867134-INS-63 | chr17-18867134-INS-63 |
| chr17-18867568-INS-7250 | chr17-18867569-INS-7250 |
| chr17-18896148-INS-5169 | chr17-18896149-INS-5170 |
| chr17-18896344-INS-60 | chr17-18896344-INS-58 |
| chr17-18905217-INS-120 | chr17-18905217-INS-120 |
| chr17-18922969-INS-2184 | chr17-18922970-INS-2184 |
| chr17-18946581-INS-125 | chr17-18946581-INS-125 |
| chr17-18990936-INS-3048 | chr17-18990937-INS-3048 |
| chr17-19020662-INS-201 | chr17-19020662-INS-201 |
| chr17-19184116-INS-182 | chr17-19184116-INS-182 |
| chr17-19242709-INS-3932 | chr17-19242710-INS-3932 |
| chr17-19250178-INS-476 | chr17-19250178-INS-476 |
| chr17-19290206-INS-1232 | chr17-19290207-INS-1233 |
| chr17-19307119-INS-425 | chr17-19307119-INS-425 |
| chr17-19336312-INS-60 | chr17-19336312-INS-50 |
| chr17-19417690-INS-498 | chr17-19417690-INS-498 |
| chr17-19543052-INS-73 | chr17-19543052-INS-73 |
| chr17-19549362-INS-55 | chr17-19549362-INS-55 |
| chr17-19631531-INS-2841 | chr17-19631532-INS-2841 |
| chr17-19712672-INS-116 | chr17-19712672-INS-152 |
| chr17-19737494-INS-2978 | chr17-19737495-INS-2979 |
| chr17-19738686-INS-1467 | chr17-19738687-INS-1463 |
| chr17-19740251-INS-4822 | chr17-19740252-INS-4822 |
| chr17-19752938-INS-55 | chr17-19752938-INS-50 |
| chr17-19768943-INS-65 | chr17-19768939-INS-63 |
| chr17-19874985-INS-121 | chr17-19874985-INS-121 |
| chr17-19910745-INS-6962 | chr17-19910746-INS-6964 |
| chr17-19934076-INS-54 | chr17-19934078-INS-52 |
| chr17-19934928-INS-161 | chr17-19934920-INS-138 |
| chr17-19981528-INS-1301 | chr17-19981529-INS-1301 |
| chr17-20026354-INS-52 | chr17-20026354-INS-56 |
| chr17-20035990-INS-443 | chr17-20036018-INS-453 |
| chr17-20064211-INS-130 | chr17-20064211-INS-132 |
| chr17-20080479-INS-378 | chr17-20080479-INS-378 |
| chr17-20117355-INS-209 | chr17-20117355-INS-209 |
| chr17-20131909-INS-12926 | chr17-20131910-INS-12930 |
| chr17-20157313-INS-879 | chr17-20157313-INS-879 |
| chr17-20188904-INS-568 | chr17-20188904-INS-568 |
| chr17-20203619-INS-86 | chr17-20203619-INS-96 |
| chr17-20236394-INS-312 | chr17-20236394-INS-316 |
| chr17-20248575-INS-5324 | chr17-20248576-INS-5324 |
| chr17-20265848-INS-834 | chr17-20265848-INS-834 |
| chr17-20288802-INS-10195 | chr17-20288803-INS-10194 |
| chr17-20347809-INS-64 | chr17-20347810-INS-69 |
| chr17-20361348-INS-771 | chr17-20361348-INS-772 |
| chr17-20408695-INS-55 | chr17-20408696-INS-50 |
| chr17-20470385-INS-3055 | chr17-20470386-INS-3054 |

| | |
|---|---|
| chr17-20471659-INS-56 | chr17-20471654-INS-64 |
| chr17-20475601-INS-6207 | chr17-20475602-INS-6211 |
| chr17-20517570-INS-519 | chr17-20517570-INS-519 |
| chr17-20552363-INS-6897 | chr17-20552364-INS-6898 |
| chr17-20574244-INS-6555 | chr17-20574245-INS-6555 |
| chr17-20585163-INS-57 | chr17-20585139-INS-90 |
| chr17-20589933-INS-236 | chr17-20589933-INS-236 |
| chr17-20611044-INS-214 | chr17-20611044-INS-214 |
| chr17-20650684-INS-463 | chr17-20650685-INS-463 |
| chr17-20651575-INS-132 | chr17-20651575-INS-132 |
| chr17-20700460-INS-447 | chr17-20700460-INS-446 |
| chr17-20718063-INS-51 | chr17-20718063-INS-53 |
| chr17-20731068-INS-57 | chr17-20731074-INS-80 |
| chr17-20757419-INS-2891 | chr17-20757420-INS-2888 |
| chr17-20785670-INS-399 | chr17-20785670-INS-399 |
| chr17-20789725-INS-456 | chr17-20789725-INS-456 |
| chr17-20815762-INS-917 | chr17-20815762-INS-917 |
| chr17-20832774-INS-54 | chr17-20832783-INS-70 |
| chr17-21014172-INS-86 | chr17-21014172-INS-89 |
| chr17-21024010-INS-4378 | chr17-21024011-INS-4378 |
| chr17-21078274-INS-152 | chr17-21078274-INS-154 |
| chr17-21115442-INS-102 | chr17-21115442-INS-114 |
| chr17-21128226-INS-439 | chr17-21128226-INS-439 |
| chr17-21173680-INS-53 | chr17-21173680-INS-55 |
| chr17-21184395-INS-208 | chr17-21184395-INS-212 |
| chr17-21213543-INS-330 | chr17-21213546-INS-318 |
| chr17-21238247-INS-62 | chr17-21238247-INS-62 |
| chr17-21251613-INS-7310 | chr17-21251614-INS-7295 |
| chr17-21283181-INS-6603 | chr17-21283182-INS-6604 |
| chr17-21284166-INS-4699 | chr17-21284167-INS-4699 |
| chr17-21300411-INS-224 | chr17-21300411-INS-224 |
| chr17-21304894-INS-544 | chr17-21304894-INS-544 |
| chr17-21306296-INS-56 | chr17-21306296-INS-52 |
| chr17-21310375-INS-60 | chr17-21310375-INS-60 |
| chr17-21438999-INS-78 | chr17-21439038-INS-78 |
| chr17-21447126-INS-7025 | chr17-21447127-INS-7027 |
| chr17-21519053-INS-211 | chr17-21519053-INS-211 |
| chr17-21526586-INS-3069 | chr17-21526587-INS-3069 |
| chr17-21535766-INS-96 | chr17-21535766-INS-96 |
| chr17-21542173-INS-12604 | chr17-21542173-INS-12604 |
| chr17-21543488-INS-585 | chr17-21543488-INS-585 |
| chr17-21556136-INS-51 | chr17-21556134-INS-60 |
| chr17-21572609-INS-517 | chr17-21572609-INS-517 |
| chr17-21597465-INS-431 | chr17-21597465-INS-430 |
| chr17-21616379-INS-209 | chr17-21616379-INS-209 |
| chr17-21660004-INS-1844 | chr17-21660005-INS-1844 |
| chr17-21669138-INS-631 | chr17-21669138-INS-631 |
| chr17-21669954-INS-105 | chr17-21669980-INS-68 |
| chr17-21704612-INS-1167 | chr17-21704613-INS-1167 |
| chr17-21712850-INS-672 | chr17-21712850-INS-672 |
| chr17-21739796-INS-4175 | chr17-21739798-INS-4174 |
| chr17-21788995-INS-199 | chr17-21788995-INS-199 |
| chr17-21838641-INS-60 | chr17-21838641-INS-60 |
| chr17-21846893-INS-4394 | chr17-21846894-INS-4394 |
| chr17-21891365-INS-56 | chr17-21891315-INS-96 |
| chr17-21904218-INS-236 | chr17-21904218-INS-237 |
| chr17-21926178-INS-767 | chr17-21926178-INS-767 |
| chr17-21934618-INS-762 | chr17-21934618-INS-762 |
| chr17-21959983-INS-211 | chr17-21959983-INS-211 |
| chr17-21974382-INS-147 | chr17-21974382-INS-147 |
| chr17-21981872-INS-53 | chr17-21981872-INS-56 |
| chr17-21988751-INS-354 | chr17-21988751-INS-354 |
| chr17-22034304-INS-1890 | chr17-22034305-INS-1890 |
| chr17-22073727-INS-6441 | chr17-22073728-INS-6441 |
| chr17-22144676-INS-7121 | chr17-22144677-INS-7122 |
| chr17-22174530-INS-185 | chr17-22174530-INS-188 |
| chr17-22205382-INS-4385 | chr17-22205383-INS-4385 |
| chr17-22215510-INS-7342 | chr17-22215511-INS-7343 |
| chr17-22337543-INS-678 | chr17-22337543-INS-676 |
| chr17-22346292-INS-313 | chr17-22346292-INS-313 |
| chr17-22346997-INS-754 | chr17-22346997-INS-753 |
| chr17-22424062-INS-480 | chr17-22424062-INS-480 |
| chr17-22447399-INS-579 | chr17-22447399-INS-579 |
| chr17-22479793-INS-65 | chr17-22479793-INS-65 |
| chr17-22531694-INS-6662 | chr17-22531695-INS-6662 |
| chr17-22550674-INS-99 | chr17-22550670-INS-101 |
| chr17-22568309-INS-3192 | chr17-22568310-INS-3192 |

| | |
|---|---|
| chr17-22603013-INS-896 | chr17-22603009-INS-895 |
| chr17-22691150-INS-84 | chr17-22691150-INS-84 |
| chr17-22716161-INS-101 | chr17-22716161-INS-90 |
| chr17-22743699-INS-706 | chr17-22743699-INS-708 |
| chr17-22816984-INS-5510 | chr17-22816985-INS-5509 |
| chr17-22838961-INS-3542 | chr17-22838962-INS-3534 |
| chr17-22841925-INS-881 | chr17-22841926-INS-884 |
| chr17-22865241-INS-435 | chr17-22865241-INS-435 |
| chr17-22880079-INS-5217 | chr17-22880080-INS-5217 |
| chr17-22922446-INS-208 | chr17-22922446-INS-208 |
| chr17-22928942-INS-201 | chr17-22928942-INS-201 |
| chr17-22974564-INS-417 | chr17-22974564-INS-417 |
| chr17-23021769-INS-64 | chr17-23021769-INS-62 |
| chr17-23026855-INS-206 | chr17-23026855-INS-206 |
| chr17-23086527-INS-241 | chr17-23086528-INS-265 |
| chr17-23097822-INS-426 | chr17-23097822-INS-426 |
| chr17-23099884-INS-565 | chr17-23099884-INS-565 |
| chr17-23123843-INS-363 | chr17-23123843-INS-363 |
| chr17-23182479-INS-6348 | chr17-23182480-INS-6348 |
| chr17-23186544-INS-468 | chr17-23186544-INS-470 |
| chr17-23198883-INS-52 | chr17-23198883-INS-56 |
| chr17-23209017-INS-54 | chr17-23209017-INS-54 |
| chr17-23374822-INS-88 | chr17-23374822-INS-90 |
| chr17-23391633-INS-1353 | chr17-23391634-INS-1353 |
| chr17-23407720-INS-60 | chr17-23407720-INS-60 |
| chr17-23414797-INS-95 | chr17-23414787-INS-107 |
| chr17-23420994-INS-53 | chr17-23420994-INS-53 |
| chr17-23432041-INS-6291 | chr17-23432042-INS-6291 |
| chr17-23447489-INS-205 | chr17-23447489-INS-205 |
| chr17-23456849-INS-132 | chr17-23456849-INS-132 |
| chr17-23480221-INS-63 | chr17-23480222-INS-63 |
| chr17-23482138-INS-571 | chr17-23482138-INS-571 |
| chr17-23489199-INS-608 | chr17-23489199-INS-608 |
| chr17-23494897-INS-976 | chr17-23494897-INS-976 |
| chr17-23497987-INS-73 | chr17-23497987-INS-79 |
| chr17-23524107-INS-3120 | chr17-23524108-INS-3120 |
| chr17-23553565-INS-235 | chr17-23553565-INS-235 |
| chr17-23556735-INS-70 | chr17-23556735-INS-70 |
| chr17-23574189-INS-8989 | chr17-23574189-INS-8974 |
| chr17-23605221-INS-66 | chr17-23605221-INS-66 |
| chr17-23622023-INS-185 | chr17-23622023-INS-183 |
| chr17-23640726-INS-331 | chr17-23640722-INS-331 |
| chr17-23651084-INS-499 | chr17-23651084-INS-499 |
| chr17-23695946-INS-793 | chr17-23695946-INS-793 |
| chr17-23700005-INS-1488 | chr17-23700006-INS-1488 |
| chr17-23704921-INS-883 | chr17-23704921-INS-883 |
| chr17-23744333-INS-499 | chr17-23744333-INS-499 |
| chr17-23746984-INS-53 | chr17-23746984-INS-54 |
| chr17-23752419-INS-5246 | chr17-23752420-INS-5246 |
| chr17-23808317-INS-202 | chr17-23808318-INS-202 |
| chr17-23816591-INS-1285 | chr17-23816592-INS-1285 |
| chr17-23816612-INS-178 | chr17-23816612-INS-179 |
| chr17-23826700-INS-116 | chr17-23826700-INS-116 |
| chr17-23832650-INS-58 | chr17-23832650-INS-59 |
| chr17-23843320-INS-127 | chr17-23843339-INS-101 |
| chr17-23846190-INS-166 | chr17-23846190-INS-166 |
| chr17-23853186-INS-55 | chr17-23853186-INS-55 |
| chr17-23856208-INS-76 | chr17-23856208-INS-72 |
| chr17-23857177-INS-212 | chr17-23857177-INS-212 |
| chr17-23859530-INS-201 | chr17-23859530-INS-201 |
| chr17-23864806-INS-212 | chr17-23864806-INS-213 |
| chr17-23874460-INS-220 | chr17-23874471-INS-214 |
| chr17-23897709-INS-62 | chr17-23897709-INS-57 |
| chr17-23904300-INS-204 | chr17-23904300-INS-202 |
| chr17-23942347-INS-214 | chr17-23942347-INS-214 |
| chr17-23949267-INS-54 | chr17-23949264-INS-57 |
| chr17-23957004-INS-55 | chr17-23957004-INS-55 |
| chr17-23986270-INS-225 | chr17-23986270-INS-226 |
| chr17-23991171-INS-205 | chr17-23991171-INS-204 |
| chr17-24014676-INS-209 | chr17-24014676-INS-209 |
| chr17-24018131-INS-497 | chr17-24018131-INS-497 |
| chr17-24019929-INS-199 | chr17-24019929-INS-199 |
| chr17-24027761-INS-220 | chr17-24027761-INS-219 |
| chr17-24084008-INS-175 | chr17-24084008-INS-175 |
| chr17-24085158-INS-81 | chr17-24085158-INS-81 |
| chr17-24102783-INS-206 | chr17-24102783-INS-205 |
| chr17-24103796-INS-81 | chr17-24103796-INS-81 |

| | |
|---|---|
| chr17-24109870-INS-125 | chr17-24109852-INS-150 |
| chr17-24145933-INS-58 | chr17-24145933-INS-58 |
| chr17-24169818-INS-94 | chr17-24169818-INS-96 |
| chr17-24171488-INS-84 | chr17-24171488-INS-84 |
| chr17-24177732-INS-51 | chr17-24177732-INS-51 |
| chr17-24180011-INS-206 | chr17-24180011-INS-206 |
| chr17-24186570-INS-255 | chr17-24186570-INS-257 |
| chr17-24187799-INS-88 | chr17-24187799-INS-88 |
| chr17-24201559-INS-207 | chr17-24201559-INS-207 |
| chr17-24207445-INS-206 | chr17-24207445-INS-207 |
| chr17-24213420-INS-57 | chr17-24213420-INS-57 |
| chr17-24231421-INS-124 | chr17-24231432-INS-88 |
| chr17-24245648-INS-72 | chr17-24245673-INS-63 |
| chr17-24247524-INS-207 | chr17-24247524-INS-206 |
| chr17-24254600-INS-54 | chr17-24254578-INS-72 |
| chr17-24269034-INS-61 | chr17-24269034-INS-61 |
| chr17-24341800-INS-217 | chr17-24341800-INS-194 |
| chr17-24362789-INS-266 | chr17-24362789-INS-267 |
| chr17-24363853-INS-51 | chr17-24363855-INS-50 |
| chr17-24367776-INS-220 | chr17-24367776-INS-219 |
| chr17-24394457-INS-209 | chr17-24394457-INS-208 |
| chr17-24447882-INS-56 | chr17-24447882-INS-58 |
| chr17-24468417-INS-215 | chr17-24468417-INS-214 |
| chr17-24490216-INS-203 | chr17-24490216-INS-203 |
| chr17-24493209-INS-75 | chr17-24493209-INS-75 |
| chr17-24513908-INS-64 | chr17-24513908-INS-76 |
| chr17-24530629-INS-106 | chr17-24530629-INS-106 |
| chr17-24538101-INS-242 | chr17-24538101-INS-238 |
| chr17-24555791-INS-2628 | chr17-24555792-INS-2628 |
| chr17-24557657-INS-158 | chr17-24557657-INS-165 |
| chr17-24562010-INS-2543 | chr17-24562011-INS-2543 |
| chr17-24600469-INS-215 | chr17-24600469-INS-213 |
| chr17-24614511-INS-68 | chr17-24614512-INS-82 |
| chr17-24629765-INS-104 | chr17-24629765-INS-104 |
| chr17-24650057-INS-1257 | chr17-24650058-INS-1257 |
| chr17-24650866-INS-297 | chr17-24650866-INS-297 |
| chr17-24655680-INS-196 | chr17-24655680-INS-196 |
| chr17-24659265-INS-209 | chr17-24659265-INS-209 |
| chr17-24671650-INS-209 | chr17-24671650-INS-204 |
| chr17-24676233-INS-54 | chr17-24676237-INS-58 |
| chr17-24743527-INS-178 | chr17-24743527-INS-177 |
| chr17-24747883-INS-205 | chr17-24747883-INS-205 |
| chr17-24762570-INS-155 | chr17-24762589-INS-154 |
| chr17-24781180-INS-422 | chr17-24781180-INS-422 |
| chr17-24801075-INS-201 | chr17-24801075-INS-202 |
| chr17-24809249-INS-54 | chr17-24809248-INS-58 |
| chr17-24828700-INS-181 | chr17-24828700-INS-183 |
| chr17-24846566-INS-419 | chr17-24846566-INS-419 |
| chr17-24859711-INS-54 | chr17-24859711-INS-54 |
| chr17-24872600-INS-58 | chr17-24872600-INS-54 |
| chr17-24881216-INS-53 | chr17-24881247-INS-61 |
| chr17-24894094-INS-214 | chr17-24894094-INS-215 |
| chr17-24896700-INS-98 | chr17-24896700-INS-98 |
| chr17-24902753-INS-60 | chr17-24902753-INS-56 |
| chr17-24903575-INS-839 | chr17-24903575-INS-839 |
| chr17-24906772-INS-216 | chr17-24906772-INS-214 |
| chr17-24907682-INS-211 | chr17-24907682-INS-212 |
| chr17-24916476-INS-270 | chr17-24916476-INS-271 |
| chr17-24926811-INS-67 | chr17-24926812-INS-60 |
| chr17-24927884-INS-203 | chr17-24927884-INS-203 |
| chr17-24931119-INS-72 | chr17-24931119-INS-72 |
| chr17-24949371-INS-232 | chr17-24949371-INS-232 |
| chr17-24974671-INS-210 | chr17-24974671-INS-210 |
| chr17-24988439-INS-88 | chr17-24988471-INS-56 |
| chr17-24999085-INS-422 | chr17-24999085-INS-422 |
| chr17-25008491-INS-232 | chr17-25008491-INS-234 |
| chr17-25017336-INS-125 | chr17-25017335-INS-133 |
| chr17-25028694-INS-68 | chr17-25028694-INS-70 |
| chr17-25051172-INS-62 | chr17-25051172-INS-72 |
| chr17-25056858-INS-228 | chr17-25056858-INS-228 |
| chr17-25059685-INS-182 | chr17-25059685-INS-180 |
| chr17-25123459-INS-256 | chr17-25123459-INS-256 |
| chr17-25134124-INS-79 | chr17-25134124-INS-79 |
| chr17-25140558-INS-1032 | chr17-25140559-INS-1032 |
| chr17-25147655-INS-57 | chr17-25147655-INS-53 |
| chr17-25156242-INS-203 | chr17-25156242-INS-200 |
| chr17-25157466-INS-205 | chr17-25157466-INS-206 |

| | |
|---|---|
| chr17-25168341-INS-403 | chr17-25168341-INS-403 |
| chr17-25173552-INS-205 | chr17-25173552-INS-205 |
| chr17-25198677-INS-207 | chr17-25198677-INS-207 |
| chr17-25199317-INS-51 | chr17-25199317-INS-52 |
| chr17-25217626-INS-228 | chr17-25217626-INS-224 |
| chr17-25223726-INS-60 | chr17-25223726-INS-60 |
| chr17-25265110-INS-139 | chr17-25265110-INS-129 |
| chr17-25291064-INS-64 | chr17-25291064-INS-64 |
| chr17-25314802-INS-54 | chr17-25314802-INS-54 |
| chr17-25324871-INS-217 | chr17-25324871-INS-216 |
| chr17-25330349-INS-178 | chr17-25330349-INS-178 |
| chr17-25360659-INS-73 | chr17-25360659-INS-58 |
| chr17-25411001-INS-196 | chr17-25411001-INS-198 |
| chr17-25452109-INS-240 | chr17-25452109-INS-240 |
| chr17-25455145-INS-194 | chr17-25455145-INS-194 |
| chr17-25460941-INS-60 | chr17-25460941-INS-68 |
| chr17-25479180-INS-109 | chr17-25479180-INS-109 |
| chr17-25506820-INS-244 | chr17-25506820-INS-237 |
| chr17-25510707-INS-210 | chr17-25510707-INS-209 |
| chr17-25523877-INS-670 | chr17-25523878-INS-677 |
| chr17-25542646-INS-58 | chr17-25542651-INS-60 |
| chr17-25555921-INS-57 | chr17-25555921-INS-57 |
| chr17-25561708-INS-84 | chr17-25561695-INS-112 |
| chr17-25566705-INS-123 | chr17-25566705-INS-123 |
| chr17-25572050-INS-237 | chr17-25572050-INS-243 |
| chr17-25575897-INS-185 | chr17-25575897-INS-185 |
| chr17-25576816-INS-89 | chr17-25576816-INS-111 |
| chr17-25619342-INS-213 | chr17-25619342-INS-214 |
| chr17-25656978-INS-206 | chr17-25656979-INS-185 |
| chr17-25660941-INS-196 | chr17-25660941-INS-201 |
| chr17-25670814-INS-116 | chr17-25670826-INS-92 |
| chr17-25688411-INS-236 | chr17-25688411-INS-236 |
| chr17-25708077-INS-505 | chr17-25708077-INS-505 |
| chr17-25726104-INS-888 | chr17-25726104-INS-868 |
| chr17-25735668-INS-54 | chr17-25735668-INS-57 |
| chr17-25781970-INS-74 | chr17-25781933-INS-67 |
| chr17-25835626-INS-1149 | chr17-25835627-INS-1149 |
| chr17-25836110-INS-72 | chr17-25836094-INS-72 |
| chr17-25839809-INS-323 | chr17-25839814-INS-347 |
| chr17-25846544-INS-51 | chr17-25846544-INS-51 |
| chr17-25868661-INS-94 | chr17-25868661-INS-106 |
| chr17-25884416-INS-206 | chr17-25884416-INS-207 |
| chr17-25888402-INS-190 | chr17-25888402-INS-184 |
| chr17-25892224-INS-195 | chr17-25892224-INS-195 |
| chr17-25894887-INS-62 | chr17-25894887-INS-62 |
| chr17-25902661-INS-222 | chr17-25902661-INS-222 |
| chr17-25926332-INS-184 | chr17-25926332-INS-185 |
| chr17-25931996-INS-60 | chr17-25931996-INS-64 |
| chr17-26016979-INS-194 | chr17-26016979-INS-194 |
| chr17-26070065-INS-211 | chr17-26070065-INS-212 |
| chr17-26089242-INS-212 | chr17-26089242-INS-212 |
| chr17-26096413-INS-121 | chr17-26096363-INS-125 |
| chr17-26104392-INS-222 | chr17-26104392-INS-224 |
| chr17-26108224-INS-74 | chr17-26108224-INS-84 |
| chr17-26210785-INS-66 | chr17-26210786-INS-66 |
| chr17-26212413-INS-205 | chr17-26212413-INS-204 |
| chr17-26233608-INS-202 | chr17-26233608-INS-200 |
| chr17-26248276-INS-246 | chr17-26248276-INS-246 |
| chr17-26264161-INS-101 | chr17-26264161-INS-97 |
| chr17-26269769-INS-186 | chr17-26269769-INS-195 |
| chr17-26276501-INS-60 | chr17-26276501-INS-60 |
| chr17-26284968-INS-288 | chr17-26284966-INS-296 |
| chr17-26297071-INS-198 | chr17-26297071-INS-198 |
| chr17-26306529-INS-76 | chr17-26306529-INS-76 |
| chr17-26308160-INS-205 | chr17-26308160-INS-205 |
| chr17-26321395-INS-165 | chr17-26321395-INS-165 |
| chr17-26328603-INS-217 | chr17-26328603-INS-218 |
| chr17-26406549-INS-212 | chr17-26406549-INS-209 |
| chr17-26413842-INS-55 | chr17-26413842-INS-55 |
| chr17-26433897-INS-300 | chr17-26433897-INS-320 |
| chr17-26434708-INS-209 | chr17-26434708-INS-208 |
| chr17-26444063-INS-193 | chr17-26444063-INS-193 |
| chr17-26447781-INS-96 | chr17-26447781-INS-97 |
| chr17-26449006-INS-203 | chr17-26449006-INS-203 |
| chr17-26449607-INS-255 | chr17-26449607-INS-255 |
| chr17-26450968-INS-199 | chr17-26450964-INS-195 |
| chr17-26453630-INS-1063 | chr17-26453631-INS-1059 |

| | |
|---|---|
| chr17-26471065-INS-189 | chr17-26471065-INS-182 |
| chr17-26474690-INS-119 | chr17-26474690-INS-119 |
| chr17-26480692-INS-239 | chr17-26480692-INS-237 |
| chr17-26490701-INS-78 | chr17-26490701-INS-78 |
| chr17-26520831-INS-654 | chr17-26520831-INS-654 |
| chr17-26521539-INS-107 | chr17-26521539-INS-107 |
| chr17-26554904-INS-299 | chr17-26554904-INS-299 |
| chr17-26563422-INS-223 | chr17-26563422-INS-223 |
| chr17-26577765-INS-2901 | chr17-26577766-INS-2901 |
| chr17-26580808-INS-82 | chr17-26580796-INS-72 |
| chr17-26585955-INS-59 | chr17-26585955-INS-60 |
| chr17-26614706-INS-100 | chr17-26614706-INS-50 |
| chr17-26690086-INS-747 | chr17-26690086-INS-747 |
| chr17-26696488-INS-212 | chr17-26696488-INS-212 |
| chr17-26701298-INS-699 | chr17-26701298-INS-696 |
| chr17-26757883-INS-246 | chr17-26757889-INS-273 |
| chr17-26758782-INS-212 | chr17-26758767-INS-227 |
| chr17-26784938-INS-57 | chr17-26784938-INS-57 |
| chr17-26786144-INS-292 | chr17-26786144-INS-289 |
| chr17-26807374-INS-68 | chr17-26807374-INS-68 |
| chr17-26809267-INS-56 | chr17-26809267-INS-62 |
| chr17-26814536-INS-56 | chr17-26814536-INS-64 |
| chr17-26828486-INS-168 | chr17-26828486-INS-167 |
| chr17-26867703-INS-60 | chr17-26867703-INS-52 |
| chr17-26888252-INS-211 | chr17-26888252-INS-211 |
| chr17-26891767-INS-53 | chr17-26891767-INS-53 |
| chr17-26907053-INS-105 | chr17-26907054-INS-127 |
| chr17-26908516-INS-85 | chr17-26908516-INS-85 |
| chr17-26911606-INS-210 | chr17-26911606-INS-210 |
| chr17-26945476-INS-539 | chr17-26945476-INS-539 |
| chr17-26958273-INS-366 | chr17-26958273-INS-372 |
| chr17-26984399-INS-56 | chr17-26984376-INS-56 |
| chr17-26986548-INS-72 | chr17-26986548-INS-72 |
| chr17-27000745-INS-218 | chr17-27000745-INS-218 |
| chr17-27023088-INS-207 | chr17-27023088-INS-207 |
| chr17-27050679-INS-700 | chr17-27050680-INS-696 |
| chr17-27059003-INS-68 | chr17-27059003-INS-68 |
| chr17-27095901-INS-60 | chr17-27095901-INS-62 |
| chr17-27098768-INS-211 | chr17-27098768-INS-212 |
| chr17-27100989-INS-68 | chr17-27100989-INS-64 |
| chr17-27110774-INS-108 | chr17-27110774-INS-108 |
| chr17-27121545-INS-5355 | chr17-27121546-INS-5355 |
| chr17-27205874-INS-105 | chr17-27205874-INS-85 |
| chr17-27211628-INS-201 | chr17-27211628-INS-201 |
| chr17-27216080-INS-120 | chr17-27216080-INS-120 |
| chr17-27253894-INS-82 | chr17-27253894-INS-82 |
| chr17-27273623-INS-201 | chr17-27273625-INS-196 |
| chr17-27280945-INS-203 | chr17-27280945-INS-203 |
| chr17-27298147-INS-71 | chr17-27298170-INS-55 |
| chr17-27315423-INS-75 | chr17-27315423-INS-74 |
| chr17-27354315-INS-341 | chr17-27354315-INS-341 |
| chr17-27442134-INS-201 | chr17-27442134-INS-201 |
| chr17-27452220-INS-230 | chr17-27452220-INS-233 |
| chr17-27452873-INS-440 | chr17-27452873-INS-440 |
| chr17-27471235-INS-7173 | chr17-27471236-INS-7173 |
| chr17-27539975-INS-3475 | chr17-27539976-INS-3477 |
| chr17-27637491-INS-84 | chr17-27637491-INS-86 |
| chr17-27659405-INS-56 | chr17-27659406-INS-56 |
| chr17-27706109-INS-62 | chr17-27706109-INS-64 |
| chr17-27711736-INS-94 | chr17-27711736-INS-94 |
| chr17-27751510-INS-3849 | chr17-27751511-INS-3849 |
| chr17-27753214-INS-67 | chr17-27753214-INS-67 |
| chr17-27756102-INS-207 | chr17-27756102-INS-207 |
| chr17-27781206-INS-70 | chr17-27781206-INS-80 |
| chr17-27790105-INS-624 | chr17-27790105-INS-625 |
| chr17-27797034-INS-194 | chr17-27797034-INS-198 |
| chr17-27808708-INS-91 | chr17-27808687-INS-91 |
| chr17-27812588-INS-92 | chr17-27812596-INS-85 |
| chr17-27817020-INS-223 | chr17-27817020-INS-226 |
| chr17-27820637-INS-6443 | chr17-27820638-INS-6443 |
| chr17-27821424-INS-220 | chr17-27821424-INS-220 |
| chr17-27835913-INS-144 | chr17-27835913-INS-144 |
| chr17-27837606-INS-220 | chr17-27837606-INS-210 |
| chr17-27844662-INS-207 | chr17-27844662-INS-207 |
| chr17-27847400-INS-623 | chr17-27847399-INS-642 |
| chr17-27856744-INS-170 | chr17-27856744-INS-170 |
| chr17-27873722-INS-190 | chr17-27873722-INS-187 |

| | |
|---|---|
| chr17-27876520-INS-225 | chr17-27876520-INS-224 |
| chr17-27895853-INS-218 | chr17-27895853-INS-218 |
| chr17-27918255-INS-203 | chr17-27918255-INS-206 |
| chr17-27922545-INS-80 | chr17-27922548-INS-68 |
| chr17-27924348-INS-211 | chr17-27924348-INS-211 |
| chr17-27946868-INS-88 | chr17-27946868-INS-90 |
| chr17-27948725-INS-212 | chr17-27948725-INS-212 |
| chr17-27954538-INS-333 | chr17-27954538-INS-334 |
| chr17-27999494-INS-5017 | chr17-27999495-INS-5017 |
| chr17-28029224-INS-229 | chr17-28029224-INS-240 |
| chr17-28042869-INS-211 | chr17-28042869-INS-207 |
| chr17-28050754-INS-526 | chr17-28050754-INS-528 |
| chr17-28055946-INS-202 | chr17-28055946-INS-202 |
| chr17-28067539-INS-244 | chr17-28067539-INS-244 |
| chr17-28071123-INS-150 | chr17-28071131-INS-157 |
| chr17-28075201-INS-197 | chr17-28075201-INS-198 |
| chr17-28084439-INS-198 | chr17-28084439-INS-200 |
| chr17-28103336-INS-56 | chr17-28103336-INS-56 |
| chr17-28104643-INS-82 | chr17-28104643-INS-82 |
| chr17-28109658-INS-65 | chr17-28109658-INS-75 |
| chr17-28115605-INS-72 | chr17-28115605-INS-54 |
| chr17-28153215-INS-202 | chr17-28153215-INS-202 |
| chr17-28162259-INS-207 | chr17-28162259-INS-208 |
| chr17-28206938-INS-212 | chr17-28206938-INS-210 |
| chr17-28215302-INS-209 | chr17-28215302-INS-209 |
| chr17-28218832-INS-98 | chr17-28218832-INS-98 |
| chr17-28247940-INS-216 | chr17-28247940-INS-216 |
| chr17-28260461-INS-100 | chr17-28260455-INS-70 |
| chr17-28263026-INS-149 | chr17-28263026-INS-149 |
| chr17-28283008-INS-204 | chr17-28283008-INS-204 |
| chr17-28302842-INS-79 | chr17-28302842-INS-52 |
| chr17-28311508-INS-203 | chr17-28311508-INS-203 |
| chr17-28317019-INS-245 | chr17-28317019-INS-246 |
| chr17-28318415-INS-52 | chr17-28318415-INS-54 |
| chr17-28330212-INS-160 | chr17-28330212-INS-160 |
| chr17-28347075-INS-53 | chr17-28347076-INS-57 |
| chr17-28352683-INS-72 | chr17-28352683-INS-78 |
| chr17-28407845-INS-707 | chr17-28407845-INS-707 |
| chr17-28413793-INS-207 | chr17-28413793-INS-207 |
| chr17-28429164-INS-51 | chr17-28429164-INS-88 |
| chr17-28494507-INS-2022 | chr17-28494508-INS-2005 |
| chr17-28516978-INS-99 | chr17-28516978-INS-99 |
| chr17-28580431-INS-119 | chr17-28580431-INS-140 |
| chr17-28588776-INS-100 | chr17-28588784-INS-58 |
| chr17-28596410-INS-357 | chr17-28596410-INS-354 |
| chr17-28601159-INS-197 | chr17-28601160-INS-205 |
| chr17-28604496-INS-145 | chr17-28604496-INS-119 |
| chr17-28607248-INS-118 | chr17-28607248-INS-118 |
| chr17-28624822-INS-375 | chr17-28624822-INS-375 |
| chr17-28637130-INS-222 | chr17-28637130-INS-223 |
| chr17-28639010-INS-169 | chr17-28639010-INS-169 |
| chr17-28663865-INS-199 | chr17-28663865-INS-199 |
| chr17-28684424-INS-247 | chr17-28684408-INS-251 |
| chr17-28688941-INS-51 | chr17-28688941-INS-62 |
| chr17-28693498-INS-235 | chr17-28693498-INS-236 |
| chr17-28699763-INS-216 | chr17-28699763-INS-221 |
| chr17-28721043-INS-96 | chr17-28721053-INS-80 |
| chr17-28722998-INS-63 | chr17-28722998-INS-57 |
| chr17-28752088-INS-262 | chr17-28752088-INS-266 |
| chr17-28762852-INS-435 | chr17-28762852-INS-435 |
| chr17-28763972-INS-507 | chr17-28763972-INS-507 |
| chr17-28769451-INS-84 | chr17-28769451-INS-88 |
| chr17-28783853-INS-206 | chr17-28783854-INS-206 |
| chr17-28789497-INS-526 | chr17-28789497-INS-526 |
| chr17-28792501-INS-259 | chr17-28792501-INS-259 |
| chr17-28823218-INS-126 | chr17-28823218-INS-126 |
| chr17-28842889-INS-213 | chr17-28842889-INS-206 |
| chr17-28848820-INS-212 | chr17-28848820-INS-213 |
| chr17-28860226-INS-60 | chr17-28860226-INS-60 |
| chr17-28866797-INS-68 | chr17-28866797-INS-52 |
| chr17-28872132-INS-110 | chr17-28872140-INS-108 |
| chr17-28885676-INS-207 | chr17-28885676-INS-210 |
| chr17-28894888-INS-967 | chr17-28894888-INS-978 |
| chr17-28897181-INS-200 | chr17-28897181-INS-200 |
| chr17-28901242-INS-236 | chr17-28901242-INS-215 |
| chr17-28914309-INS-90 | chr17-28914309-INS-92 |
| chr17-28920740-INS-61 | chr17-28920740-INS-61 |

| | |
|---|---|
| chr17-28922689-INS-225 | chr17-28922689-INS-225 |
| chr17-28973032-INS-196 | chr17-28973032-INS-196 |
| chr17-28973998-INS-56 | chr17-28973998-INS-73 |
| chr17-28980166-INS-212 | chr17-28980166-INS-212 |
| chr17-29007192-INS-200 | chr17-29007192-INS-200 |
| chr17-29013644-INS-198 | chr17-29013650-INS-210 |
| chr17-29016874-INS-179 | chr17-29016874-INS-179 |
| chr17-29022019-INS-190 | chr17-29022019-INS-190 |
| chr17-29058890-INS-66 | chr17-29058890-INS-66 |
| chr17-29062409-INS-221 | chr17-29062409-INS-223 |
| chr17-29064308-INS-203 | chr17-29064308-INS-203 |
| chr17-29090055-INS-1893 | chr17-29090056-INS-1893 |
| chr17-29122702-INS-211 | chr17-29122702-INS-210 |
| chr17-29124061-INS-684 | chr17-29124061-INS-684 |
| chr17-29128017-INS-60 | chr17-29128004-INS-84 |
| chr17-29130616-INS-198 | chr17-29130616-INS-198 |
| chr17-29132617-INS-88 | chr17-29132617-INS-88 |
| chr17-29158718-INS-7065 | chr17-29158719-INS-7066 |
| chr17-29183687-INS-56 | chr17-29183687-INS-58 |
| chr17-29212542-INS-213 | chr17-29212542-INS-214 |
| chr17-29234118-INS-80 | chr17-29234124-INS-72 |
| chr17-29244968-INS-449 | chr17-29244968-INS-449 |
| chr17-29266491-INS-146 | chr17-29266491-INS-154 |
| chr17-29282729-INS-440 | chr17-29282729-INS-440 |
| chr17-29290573-INS-105 | chr17-29290573-INS-106 |
| chr17-29297820-INS-59 | chr17-29297833-INS-71 |
| chr17-29305864-INS-210 | chr17-29305864-INS-210 |
| chr17-29309437-INS-237 | chr17-29309437-INS-238 |
| chr17-29340831-INS-187 | chr17-29340831-INS-187 |
| chr17-29343365-INS-400 | chr17-29343365-INS-399 |
| chr17-29344161-INS-65 | chr17-29344161-INS-65 |
| chr17-29412504-INS-64 | chr17-29412523-INS-78 |
| chr17-29435576-INS-531 | chr17-29435576-INS-531 |
| chr17-29438488-INS-220 | chr17-29438488-INS-220 |
| chr17-29476534-INS-144 | chr17-29476534-INS-144 |
| chr17-29477888-INS-232 | chr17-29477888-INS-232 |
| chr17-29484524-INS-59 | chr17-29484524-INS-58 |
| chr17-29504928-INS-3144 | chr17-29504929-INS-3144 |
| chr17-29517229-INS-202 | chr17-29517229-INS-202 |
| chr17-29525994-INS-74 | chr17-29525994-INS-71 |
| chr17-29533076-INS-217 | chr17-29533076-INS-217 |
| chr17-29536222-INS-220 | chr17-29536222-INS-220 |
| chr17-29537235-INS-221 | chr17-29537235-INS-222 |
| chr17-29547263-INS-114 | chr17-29547263-INS-114 |
| chr17-29550031-INS-107 | chr17-29550031-INS-107 |
| chr17-29562610-INS-228 | chr17-29562610-INS-225 |
| chr17-29569096-INS-204 | chr17-29569096-INS-204 |
| chr17-29571741-INS-218 | chr17-29571735-INS-207 |
| chr17-29573457-INS-51 | chr17-29573457-INS-51 |
| chr17-29594243-INS-388 | chr17-29594243-INS-398 |
| chr17-29595319-INS-114 | chr17-29595319-INS-110 |
| chr17-29598689-INS-462 | chr17-29598689-INS-462 |
| chr17-29602159-INS-172 | chr17-29602159-INS-173 |
| chr17-29613972-INS-203 | chr17-29613972-INS-204 |
| chr17-29634124-INS-205 | chr17-29634124-INS-205 |
| chr17-29652178-INS-192 | chr17-29652178-INS-192 |
| chr17-29662672-INS-91 | chr17-29662664-INS-102 |
| chr17-29663937-INS-72 | chr17-29663937-INS-72 |
| chr17-29666696-INS-164 | chr17-29666696-INS-164 |
| chr17-29671199-INS-96 | chr17-29671214-INS-100 |
| chr17-29691786-INS-66 | chr17-29691784-INS-63 |
| chr17-29697879-INS-59 | chr17-29697879-INS-75 |
| chr17-29716541-INS-99 | chr17-29716539-INS-72 |
| chr17-29742771-INS-482 | chr17-29742771-INS-482 |
| chr17-29746177-INS-5150 | chr17-29746178-INS-5150 |
| chr17-29748400-INS-76 | chr17-29748400-INS-68 |
| chr17-29754504-INS-70 | chr17-29754502-INS-72 |
| chr17-29756390-INS-192 | chr17-29756390-INS-191 |
| chr17-29758793-INS-223 | chr17-29758793-INS-224 |
| chr17-29777627-INS-112 | chr17-29777627-INS-112 |
| chr17-29780666-INS-114 | chr17-29780666-INS-114 |
| chr17-29783842-INS-208 | chr17-29783842-INS-208 |
| chr17-29785549-INS-52 | chr17-29785549-INS-52 |
| chr17-29789875-INS-207 | chr17-29789875-INS-207 |
| chr17-29792769-INS-216 | chr17-29792769-INS-216 |
| chr17-29794392-INS-219 | chr17-29794392-INS-220 |
| chr17-29797172-INS-193 | chr17-29797172-INS-151 |

| | |
|---|---|
| chr17-29820719-INS-117 | chr17-29820719-INS-117 |
| chr17-29829285-INS-56 | chr17-29829287-INS-73 |
| chr17-29838354-INS-71 | chr17-29838354-INS-71 |
| chr17-29842646-INS-207 | chr17-29842646-INS-207 |
| chr17-29850952-INS-229 | chr17-29850952-INS-230 |
| chr17-29857263-INS-92 | chr17-29857263-INS-92 |
| chr17-29869022-INS-233 | chr17-29869022-INS-235 |
| chr17-29871925-INS-114 | chr17-29871912-INS-115 |
| chr17-29878709-INS-270 | chr17-29878709-INS-269 |
| chr17-29879219-INS-208 | chr17-29879220-INS-203 |
| chr17-29881039-INS-212 | chr17-29881039-INS-212 |
| chr17-29910482-INS-57 | chr17-29910482-INS-57 |
| chr17-29941819-INS-229 | chr17-29941819-INS-222 |
| chr17-29944593-INS-54 | chr17-29944550-INS-60 |
| chr17-29954089-INS-84 | chr17-29954089-INS-84 |
| chr17-29956426-INS-53 | chr17-29956426-INS-55 |
| chr17-29960240-INS-172 | chr17-29960240-INS-172 |
| chr17-29967148-INS-62 | chr17-29967148-INS-68 |
| chr17-29968598-INS-217 | chr17-29968599-INS-171 |
| chr17-29972462-INS-8985 | chr17-29972463-INS-8985 |
| chr17-29977854-INS-5567 | chr17-29977855-INS-5567 |
| chr17-29978426-INS-161 | chr17-29978426-INS-161 |
| chr17-29991448-INS-66 | chr17-29991435-INS-68 |
| chr17-30007108-INS-206 | chr17-30007108-INS-206 |
| chr17-30016527-INS-164 | chr17-30016527-INS-164 |
| chr17-30041776-INS-165 | chr17-30041776-INS-165 |
| chr17-30046826-INS-63 | chr17-30046795-INS-60 |
| chr17-30077861-INS-52 | chr17-30077861-INS-52 |
| chr17-30137032-INS-54 | chr17-30137033-INS-54 |
| chr17-30140861-INS-82 | chr17-30140859-INS-50 |
| chr17-30181696-INS-58 | chr17-30181701-INS-66 |
| chr17-30205212-INS-82 | chr17-30205212-INS-82 |
| chr17-30213887-INS-200 | chr17-30213887-INS-206 |
| chr17-30255421-INS-62 | chr17-30255421-INS-62 |
| chr17-30258740-INS-227 | chr17-30258740-INS-228 |
| chr17-30264222-INS-7109 | chr17-30264223-INS-7109 |
| chr17-30293601-INS-207 | chr17-30293601-INS-207 |
| chr17-30331685-INS-55 | chr17-30331685-INS-55 |
| chr17-30508260-INS-54 | chr17-30508274-INS-54 |
| chr17-30550921-INS-72 | chr17-30550921-INS-72 |
| chr17-30555448-INS-150 | chr17-30555448-INS-157 |
| chr17-30616143-INS-189 | chr17-30616143-INS-189 |
| chr17-30640589-INS-211 | chr17-30640589-INS-210 |
| chr17-30646329-INS-65 | chr17-30646329-INS-65 |
| chr17-30685172-INS-187 | chr17-30685172-INS-187 |
| chr17-30695659-INS-96 | chr17-30695660-INS-64 |
| chr17-30716094-INS-1894 | chr17-30716095-INS-1894 |
| chr17-30750150-INS-3117 | chr17-30750151-INS-3117 |
| chr17-30758745-INS-206 | chr17-30758745-INS-206 |
| chr17-30770528-INS-51 | chr17-30770528-INS-55 |
| chr17-30810875-INS-236 | chr17-30810875-INS-236 |
| chr17-30815340-INS-551 | chr17-30815340-INS-548 |
| chr17-30817001-INS-78 | chr17-30817001-INS-78 |
| chr17-30828034-INS-1206 | chr17-30828035-INS-1204 |
| chr17-30843742-INS-58 | chr17-30843742-INS-50 |
| chr17-30850140-INS-104 | chr17-30850140-INS-97 |
| chr17-30872434-INS-214 | chr17-30872434-INS-214 |
| chr17-30874432-INS-204 | chr17-30874432-INS-203 |
| chr17-30885117-INS-208 | chr17-30885117-INS-209 |
| chr17-30899295-INS-209 | chr17-30899295-INS-209 |
| chr17-30905836-INS-96 | chr17-30905836-INS-104 |
| chr17-30921484-INS-72 | chr17-30921484-INS-72 |
| chr17-30944646-INS-339 | chr17-30944646-INS-339 |
| chr17-30948481-INS-56 | chr17-30948481-INS-56 |
| chr17-30974272-INS-59 | chr17-30974272-INS-62 |
| chr17-30978468-INS-64 | chr17-30978436-INS-50 |
| chr17-30979842-INS-53 | chr17-30979842-INS-53 |
| chr17-30985259-INS-221 | chr17-30985259-INS-209 |
| chr17-30993025-INS-111 | chr17-30993017-INS-109 |
| chr17-30993144-INS-2998 | chr17-30993145-INS-2999 |
| chr17-30995843-INS-207 | chr17-30995843-INS-207 |
| chr17-31053335-INS-617 | chr17-31053335-INS-617 |
| chr17-31070494-INS-123 | chr17-31070494-INS-132 |
| chr17-31080474-INS-145 | chr17-31080505-INS-132 |
| chr17-31097593-INS-121 | chr17-31097593-INS-121 |
| chr17-31104956-INS-403 | chr17-31104956-INS-403 |
| chr17-31111541-INS-52 | chr17-31111545-INS-55 |

| | |
|---|---|
| chr17-31112882-INS-52 | chr17-31112882-INS-52 |
| chr17-31114969-INS-240 | chr17-31114969-INS-240 |
| chr17-31136903-INS-67 | chr17-31136903-INS-67 |
| chr17-31138998-INS-462 | chr17-31138998-INS-462 |
| chr17-31175614-INS-93 | chr17-31175606-INS-115 |
| chr17-31203835-INS-279 | chr17-31203835-INS-277 |
| chr17-31209222-INS-180 | chr17-31209222-INS-175 |
| chr17-31222706-INS-126 | chr17-31222706-INS-126 |
| chr17-31227388-INS-52 | chr17-31227391-INS-51 |
| chr17-31230857-INS-722 | chr17-31230857-INS-724 |
| chr17-31252886-INS-82 | chr17-31252886-INS-82 |
| chr17-31254586-INS-271 | chr17-31254586-INS-271 |
| chr17-31256238-INS-176 | chr17-31256238-INS-176 |
| chr17-31281958-INS-90 | chr17-31281958-INS-135 |
| chr17-31284158-INS-69 | chr17-31284158-INS-71 |
| chr17-31287479-INS-60 | chr17-31287479-INS-60 |
| chr17-31292091-INS-70 | chr17-31292091-INS-74 |
| chr17-31295771-INS-120 | chr17-31295771-INS-120 |
| chr17-31307567-INS-210 | chr17-31307567-INS-210 |
| chr17-31320346-INS-70 | chr17-31320333-INS-50 |
| chr17-31345684-INS-52 | chr17-31345684-INS-52 |
| chr17-31350180-INS-215 | chr17-31350180-INS-215 |
| chr17-31354898-INS-82 | chr17-31354898-INS-84 |
| chr17-31369620-INS-82 | chr17-31369620-INS-88 |
| chr17-31376207-INS-65 | chr17-31376207-INS-65 |
| chr17-31409630-INS-77 | chr17-31409634-INS-88 |
| chr17-31426183-INS-70 | chr17-31426183-INS-70 |
| chr17-31428988-INS-859 | chr17-31428988-INS-858 |
| chr17-31465694-INS-203 | chr17-31465694-INS-204 |
| chr17-31470014-INS-493 | chr17-31470014-INS-501 |
| chr17-31479158-INS-60 | chr17-31479197-INS-51 |
| chr17-31481333-INS-54 | chr17-31481337-INS-91 |
| chr17-31486408-INS-210 | chr17-31486408-INS-210 |
| chr17-31494916-INS-313 | chr17-31494916-INS-313 |
| chr17-31524736-INS-60 | chr17-31524726-INS-64 |
| chr17-31546908-INS-54 | chr17-31546908-INS-54 |
| chr17-31554206-INS-176 | chr17-31554206-INS-180 |
| chr17-31563143-INS-97 | chr17-31563143-INS-97 |
| chr17-31565704-INS-205 | chr17-31565704-INS-205 |
| chr17-31569432-INS-5534 | chr17-31569433-INS-5534 |
| chr17-31569995-INS-338 | chr17-31570006-INS-339 |
| chr17-31576716-INS-54 | chr17-31576716-INS-54 |
| chr17-31578334-INS-60 | chr17-31578334-INS-60 |
| chr17-31586583-INS-445 | chr17-31586583-INS-444 |
| chr17-31601604-INS-201 | chr17-31601604-INS-201 |
| chr17-31616393-INS-52 | chr17-31616375-INS-57 |
| chr17-31627658-INS-209 | chr17-31627658-INS-209 |
| chr17-31641154-INS-61 | chr17-31641154-INS-61 |
| chr17-31651103-INS-184 | chr17-31651103-INS-178 |
| chr17-31657252-INS-87 | chr17-31657258-INS-85 |
| chr17-31663627-INS-183 | chr17-31663627-INS-183 |
| chr17-31673333-INS-69 | chr17-31673333-INS-69 |
| chr17-31682436-INS-556 | chr17-31682436-INS-556 |
| chr17-31695812-INS-70 | chr17-31695812-INS-74 |
| chr17-31702537-INS-102 | chr17-31702537-INS-102 |
| chr17-31712527-INS-113 | chr17-31712565-INS-80 |
| chr17-31724727-INS-206 | chr17-31724727-INS-206 |
| chr17-31735942-INS-126 | chr17-31735942-INS-131 |
| chr17-31737853-INS-195 | chr17-31737853-INS-195 |
| chr17-31743379-INS-59 | chr17-31743379-INS-62 |
| chr17-31745173-INS-682 | chr17-31745173-INS-690 |
| chr17-31748209-INS-216 | chr17-31748209-INS-216 |
| chr17-31788302-INS-177 | chr17-31788302-INS-178 |
| chr17-31796797-INS-66 | chr17-31796797-INS-62 |
| chr17-31806475-INS-209 | chr17-31806475-INS-209 |
| chr17-31823486-INS-91 | chr17-31823486-INS-97 |
| chr17-31867576-INS-142 | chr17-31867576-INS-142 |
| chr17-31884447-INS-65 | chr17-31884447-INS-70 |
| chr17-31906177-INS-176 | chr17-31906177-INS-180 |
| chr17-31910766-INS-230 | chr17-31910766-INS-229 |
| chr17-31923145-INS-78 | chr17-31923133-INS-65 |
| chr17-31936040-INS-2005 | chr17-31936041-INS-2005 |
| chr17-31936749-INS-74 | chr17-31936749-INS-74 |
| chr17-31941564-INS-603 | chr17-31941564-INS-605 |
| chr17-31958524-INS-63 | chr17-31958523-INS-61 |
| chr17-32008928-INS-109 | chr17-32008928-INS-109 |
| chr17-32020523-INS-65 | chr17-32020521-INS-63 |

| | |
|---|---|
| chr17-32034869-INS-91 | chr17-32034879-INS-73 |
| chr17-32076910-INS-190 | chr17-32076885-INS-201 |
| chr17-32079811-INS-3961 | chr17-32079812-INS-3961 |
| chr17-32090093-INS-7134 | chr17-32090094-INS-7134 |
| chr17-32095237-INS-2635 | chr17-32095238-INS-2635 |
| chr17-32095385-INS-158 | chr17-32095385-INS-156 |
| chr17-32107502-INS-66 | chr17-32107502-INS-60 |
| chr17-32110093-INS-56 | chr17-32110093-INS-56 |
| chr17-32123707-INS-71 | chr17-32123707-INS-59 |
| chr17-32129186-INS-116 | chr17-32129186-INS-116 |
| chr17-32179885-INS-6440 | chr17-32179886-INS-6440 |
| chr17-32180984-INS-203 | chr17-32180984-INS-201 |
| chr17-32193520-INS-497 | chr17-32193520-INS-497 |
| chr17-32209759-INS-128 | chr17-32209759-INS-129 |
| chr17-32211571-INS-242 | chr17-32211571-INS-234 |
| chr17-32263669-INS-164 | chr17-32263669-INS-164 |
| chr17-32286892-INS-227 | chr17-32286892-INS-227 |
| chr17-32289828-INS-230 | chr17-32289828-INS-229 |
| chr17-32296600-INS-68 | chr17-32296600-INS-70 |
| chr17-32304507-INS-95 | chr17-32304507-INS-96 |
| chr17-32386817-INS-219 | chr17-32386817-INS-222 |
| chr17-32388352-INS-208 | chr17-32388352-INS-206 |
| chr17-32390365-INS-211 | chr17-32390378-INS-211 |
| chr17-32433276-INS-56 | chr17-32433276-INS-50 |
| chr17-32434333-INS-104 | chr17-32434332-INS-150 |
| chr17-32505935-INS-97 | chr17-32505935-INS-109 |
| chr17-32507103-INS-160 | chr17-32507103-INS-160 |
| chr17-32509929-INS-219 | chr17-32509929-INS-218 |
| chr17-32514736-INS-1922 | chr17-32514737-INS-1927 |
| chr17-32549181-INS-163 | chr17-32549181-INS-163 |
| chr17-32615060-INS-79 | chr17-32615060-INS-76 |
| chr17-32620700-INS-208 | chr17-32620700-INS-209 |
| chr17-32671064-INS-76 | chr17-32671068-INS-84 |
| chr17-32674028-INS-410 | chr17-32674028-INS-411 |
| chr17-32679330-INS-872 | chr17-32679330-INS-874 |
| chr17-32695270-INS-72 | chr17-32695270-INS-92 |
| chr17-32726566-INS-56 | chr17-32726566-INS-52 |
| chr17-32742711-INS-57 | chr17-32742711-INS-57 |
| chr17-32754358-INS-1011 | chr17-32754359-INS-1011 |
| chr17-32755416-INS-4897 | chr17-32755417-INS-4897 |
| chr17-32757485-INS-76 | chr17-32757485-INS-79 |
| chr17-32888705-INS-216 | chr17-32888711-INS-216 |
| chr17-32891503-INS-205 | chr17-32891503-INS-207 |
| chr17-32904853-INS-75 | chr17-32904853-INS-75 |
| chr17-32911607-INS-162 | chr17-32911607-INS-162 |
| chr17-32928557-INS-511 | chr17-32928555-INS-511 |
| chr17-32941184-INS-62 | chr17-32941180-INS-81 |
| chr17-32959481-INS-103 | chr17-32959481-INS-103 |
| chr17-33018017-INS-452 | chr17-33018017-INS-452 |
| chr17-33019707-INS-217 | chr17-33019707-INS-218 |
| chr17-33021383-INS-343 | chr17-33021383-INS-343 |
| chr17-33025598-INS-204 | chr17-33025598-INS-204 |
| chr17-33026796-INS-105 | chr17-33026796-INS-105 |
| chr17-33029328-INS-68 | chr17-33029333-INS-61 |
| chr17-33044948-INS-93 | chr17-33044948-INS-107 |
| chr17-33074284-INS-70 | chr17-33074284-INS-70 |
| chr17-33088376-INS-2688 | chr17-33088377-INS-2688 |
| chr17-33090674-INS-65 | chr17-33090674-INS-65 |
| chr17-33112968-INS-53 | chr17-33112968-INS-53 |
| chr17-33117548-INS-505 | chr17-33117561-INS-475 |
| chr17-33133330-INS-182 | chr17-33133337-INS-188 |
| chr17-33139454-INS-247 | chr17-33139454-INS-247 |
| chr17-33151190-INS-69 | chr17-33151190-INS-74 |
| chr17-33170512-INS-89 | chr17-33170512-INS-89 |
| chr17-33173032-INS-61 | chr17-33173032-INS-62 |
| chr17-33188236-INS-55 | chr17-33188236-INS-55 |
| chr17-33204372-INS-4583 | chr17-33204373-INS-4584 |
| chr17-33210589-INS-535 | chr17-33210589-INS-535 |
| chr17-33217098-INS-232 | chr17-33217098-INS-231 |
| chr17-33218968-INS-749 | chr17-33218968-INS-749 |
| chr17-33230582-INS-191 | chr17-33230578-INS-192 |
| chr17-33278723-INS-344 | chr17-33278723-INS-348 |
| chr17-33289642-INS-343 | chr17-33289642-INS-343 |
| chr17-33292718-INS-5102 | chr17-33292719-INS-5103 |
| chr17-33313723-INS-84 | chr17-33313723-INS-55 |
| chr17-33331280-INS-2685 | chr17-33331281-INS-2685 |
| chr17-33393519-INS-57 | chr17-33393519-INS-57 |

| | |
|---|---|
| chr17-33401622-INS-174 | chr17-33401622-INS-174 |
| chr17-33425903-INS-6889 | chr17-33425904-INS-6889 |
| chr17-33433730-INS-65 | chr17-33433730-INS-59 |
| chr17-33446339-INS-2999 | chr17-33446340-INS-2996 |
| chr17-33460066-INS-2249 | chr17-33460067-INS-2249 |
| chr17-33527065-INS-1071 | chr17-33527066-INS-1070 |
| chr17-33534596-INS-209 | chr17-33534596-INS-209 |
| chr17-33545268-INS-194 | chr17-33545268-INS-195 |
| chr17-33546216-INS-2841 | chr17-33546217-INS-2841 |
| chr17-33556719-INS-74 | chr17-33556724-INS-99 |
| chr17-33607599-INS-72 | chr17-33607598-INS-72 |
| chr17-33611652-INS-56 | chr17-33611652-INS-58 |
| chr17-33615679-INS-98 | chr17-33615679-INS-98 |
| chr17-33646307-INS-7050 | chr17-33646308-INS-7051 |
| chr17-33655062-INS-279 | chr17-33655062-INS-279 |
| chr17-33682690-INS-197 | chr17-33682690-INS-197 |
| chr17-33725483-INS-208 | chr17-33725483-INS-208 |
| chr17-33727295-INS-52 | chr17-33727295-INS-52 |
| chr17-33730398-INS-84 | chr17-33730386-INS-132 |
| chr17-33731105-INS-52 | chr17-33731106-INS-52 |
| chr17-33744481-INS-114 | chr17-33744479-INS-118 |
| chr17-33760414-INS-290 | chr17-33760414-INS-290 |
| chr17-33777752-INS-88 | chr17-33777752-INS-88 |
| chr17-33779496-INS-92 | chr17-33779496-INS-92 |
| chr17-33807386-INS-70 | chr17-33807387-INS-60 |
| chr17-33824118-INS-429 | chr17-33824139-INS-389 |
| chr17-33916764-INS-58 | chr17-33916764-INS-60 |
| chr17-33920072-INS-88 | chr17-33920064-INS-64 |
| chr17-33928291-INS-151 | chr17-33928291-INS-151 |
| chr17-33945192-INS-175 | chr17-33945192-INS-175 |
| chr17-33951879-INS-183 | chr17-33951879-INS-183 |
| chr17-33982041-INS-674 | chr17-33982041-INS-674 |
| chr17-33985390-INS-174 | chr17-33985390-INS-174 |
| chr17-33990982-INS-58 | chr17-33990982-INS-60 |
| chr17-34010449-INS-179 | chr17-34010449-INS-187 |
| chr17-34024367-INS-68 | chr17-34024367-INS-68 |
| chr17-34048507-INS-205 | chr17-34048507-INS-205 |
| chr17-34051742-INS-217 | chr17-34051749-INS-238 |
| chr17-34053487-INS-227 | chr17-34053487-INS-227 |
| chr17-34068697-INS-273 | chr17-34068697-INS-269 |
| chr17-34073749-INS-211 | chr17-34073749-INS-212 |
| chr17-34078089-INS-211 | chr17-34078089-INS-209 |
| chr17-34081876-INS-213 | chr17-34081876-INS-212 |
| chr17-34084368-INS-71 | chr17-34084368-INS-71 |
| chr17-34093447-INS-601 | chr17-34093443-INS-614 |
| chr17-34095350-INS-231 | chr17-34095350-INS-232 |
| chr17-34105662-INS-198 | chr17-34105662-INS-198 |
| chr17-34112678-INS-244 | chr17-34112678-INS-199 |
| chr17-34113687-INS-168 | chr17-34113687-INS-168 |
| chr17-34123329-INS-191 | chr17-34123329-INS-191 |
| chr17-34144138-INS-82 | chr17-34144138-INS-82 |
| chr17-34158273-INS-2909 | chr17-34158274-INS-2909 |
| chr17-34163437-INS-207 | chr17-34163437-INS-207 |
| chr17-34176263-INS-76 | chr17-34176263-INS-78 |
| chr17-34183726-INS-179 | chr17-34183726-INS-179 |
| chr17-34194120-INS-356 | chr17-34194120-INS-356 |
| chr17-34196111-INS-197 | chr17-34196111-INS-197 |
| chr17-34202263-INS-51 | chr17-34202264-INS-52 |
| chr17-34204473-INS-98 | chr17-34204473-INS-98 |
| chr17-34205454-INS-199 | chr17-34205454-INS-199 |
| chr17-34209829-INS-63 | chr17-34209829-INS-61 |
| chr17-34222335-INS-196 | chr17-34222335-INS-196 |
| chr17-34226806-INS-401 | chr17-34226806-INS-401 |
| chr17-34234958-INS-212 | chr17-34234958-INS-212 |
| chr17-34237209-INS-425 | chr17-34237209-INS-425 |
| chr17-34253826-INS-181 | chr17-34253826-INS-181 |
| chr17-34275680-INS-74 | chr17-34275680-INS-75 |
| chr17-34286777-INS-51 | chr17-34286778-INS-51 |
| chr17-34294582-INS-68 | chr17-34294582-INS-64 |
| chr17-34307208-INS-54 | chr17-34307208-INS-54 |
| chr17-34325905-INS-520 | chr17-34325906-INS-520 |
| chr17-34345154-INS-186 | chr17-34345154-INS-186 |
| chr17-34346887-INS-196 | chr17-34346887-INS-196 |
| chr17-34353769-INS-203 | chr17-34353769-INS-203 |
| chr17-34360822-INS-54 | chr17-34360822-INS-54 |
| chr17-34362669-INS-64 | chr17-34362667-INS-64 |
| chr17-34371179-INS-336 | chr17-34371179-INS-336 |

| | |
|---|---|
| chr17-34374045-INS-175 | chr17-34374045-INS-175 |
| chr17-34380050-INS-73 | chr17-34380050-INS-73 |
| chr17-34388381-INS-118 | chr17-34388381-INS-108 |
| chr17-34392755-INS-130 | chr17-34392755-INS-130 |
| chr17-34397161-INS-242 | chr17-34397161-INS-251 |
| chr17-34398723-INS-57 | chr17-34398723-INS-57 |
| chr17-34412374-INS-94 | chr17-34412374-INS-96 |
| chr17-34422149-INS-290 | chr17-34422149-INS-290 |
| chr17-34432223-INS-174 | chr17-34432223-INS-166 |
| chr17-34453418-INS-538 | chr17-34453418-INS-533 |
| chr17-34463785-INS-76 | chr17-34463783-INS-77 |
| chr17-34470120-INS-498 | chr17-34470120-INS-498 |
| chr17-34471194-INS-55 | chr17-34471194-INS-55 |
| chr17-34475114-INS-54 | chr17-34475101-INS-54 |
| chr17-34489631-INS-428 | chr17-34489631-INS-428 |
| chr17-34490778-INS-394 | chr17-34490778-INS-394 |
| chr17-34498537-INS-74 | chr17-34498538-INS-74 |
| chr17-34499752-INS-70 | chr17-34499753-INS-57 |
| chr17-34500752-INS-442 | chr17-34500752-INS-442 |
| chr17-34510975-INS-242 | chr17-34510975-INS-242 |
| chr17-34522356-INS-126 | chr17-34522356-INS-130 |
| chr17-34527583-INS-52 | chr17-34527583-INS-52 |
| chr17-34559749-INS-127 | chr17-34559744-INS-127 |
| chr17-34565711-INS-630 | chr17-34565711-INS-630 |
| chr17-34579848-INS-58 | chr17-34579852-INS-74 |
| chr17-34606490-INS-359 | chr17-34606490-INS-380 |
| chr17-34633624-INS-59 | chr17-34633582-INS-66 |
| chr17-34639087-INS-66 | chr17-34639087-INS-66 |
| chr17-34729269-INS-76 | chr17-34729269-INS-62 |
| chr17-34748922-INS-89 | chr17-34748922-INS-90 |
| chr17-34751154-INS-68 | chr17-34751154-INS-80 |
| chr17-34756343-INS-188 | chr17-34756343-INS-188 |
| chr17-34759156-INS-224 | chr17-34759156-INS-224 |
| chr17-34777212-INS-249 | chr17-34777211-INS-250 |
| chr17-34853870-INS-56 | chr17-34853870-INS-56 |
| chr17-34855550-INS-76 | chr17-34855551-INS-76 |
| chr17-34856442-INS-54 | chr17-34856442-INS-54 |
| chr17-34882672-INS-174 | chr17-34882672-INS-174 |
| chr17-34888654-INS-57 | chr17-34888654-INS-57 |
| chr17-34940964-INS-136 | chr17-34940964-INS-136 |
| chr17-34946405-INS-65 | chr17-34946405-INS-65 |
| chr17-34958402-INS-213 | chr17-34958402-INS-213 |
| chr17-34987156-INS-204 | chr17-34987156-INS-204 |
| chr17-34991029-INS-315 | chr17-34991032-INS-323 |
| chr17-35012460-INS-78 | chr17-35012460-INS-78 |
| chr17-35097300-INS-223 | chr17-35097300-INS-222 |
| chr17-35120933-INS-922 | chr17-35120933-INS-924 |
| chr17-35139074-INS-206 | chr17-35139074-INS-210 |
| chr17-35172162-INS-69 | chr17-35172162-INS-75 |
| chr17-35248065-INS-52 | chr17-35248065-INS-51 |
| chr17-35256272-INS-203 | chr17-35256272-INS-203 |
| chr17-35264605-INS-209 | chr17-35264605-INS-207 |
| chr17-35301333-INS-242 | chr17-35301333-INS-244 |
| chr17-35304287-INS-205 | chr17-35304287-INS-205 |
| chr17-35305482-INS-1888 | chr17-35305483-INS-1888 |
| chr17-35317082-INS-215 | chr17-35317082-INS-215 |
| chr17-35321382-INS-62 | chr17-35321382-INS-62 |
| chr17-35324791-INS-97 | chr17-35324791-INS-97 |
| chr17-35329179-INS-158 | chr17-35329179-INS-170 |
| chr17-35353609-INS-221 | chr17-35353609-INS-220 |
| chr17-35396878-INS-212 | chr17-35396878-INS-208 |
| chr17-35401622-INS-1399 | chr17-35401623-INS-1423 |
| chr17-35416790-INS-55 | chr17-35416795-INS-50 |
| chr17-35446529-INS-209 | chr17-35446529-INS-209 |
| chr17-35508274-INS-322 | chr17-35508274-INS-322 |
| chr17-35524852-INS-6403 | chr17-35524853-INS-6403 |
| chr17-35526592-INS-330 | chr17-35526592-INS-330 |
| chr17-35532827-INS-194 | chr17-35532822-INS-194 |
| chr17-35544917-INS-232 | chr17-35544917-INS-232 |
| chr17-35564987-INS-312 | chr17-35564987-INS-313 |
| chr17-35605010-INS-165 | chr17-35605032-INS-120 |
| chr17-35611030-INS-83 | chr17-35611031-INS-83 |
| chr17-35616626-INS-201 | chr17-35616613-INS-205 |
| chr17-35650013-INS-56 | chr17-35650014-INS-56 |
| chr17-35666525-INS-89 | chr17-35666525-INS-90 |
| chr17-35742230-INS-325 | chr17-35742230-INS-323 |
| chr17-35773806-INS-236 | chr17-35773794-INS-213 |

| | |
|---|---|
| chr17-35786619-INS-223 | chr17-35786619-INS-222 |
| chr17-35787896-INS-60 | chr17-35787896-INS-60 |
| chr17-35803261-INS-52 | chr17-35803260-INS-52 |
| chr17-35811270-INS-172 | chr17-35811270-INS-172 |
| chr17-35833153-INS-202 | chr17-35833150-INS-195 |
| chr17-35836582-INS-190 | chr17-35836582-INS-190 |
| chr17-35845567-INS-402 | chr17-35845567-INS-402 |
| chr17-35858675-INS-63 | chr17-35858673-INS-111 |
| chr17-35863664-INS-171 | chr17-35863665-INS-171 |
| chr17-35865365-INS-228 | chr17-35865365-INS-229 |
| chr17-35872201-INS-211 | chr17-35872201-INS-211 |
| chr17-35874572-INS-530 | chr17-35874572-INS-530 |
| chr17-35907033-INS-455 | chr17-35907033-INS-455 |
| chr17-35910023-INS-237 | chr17-35910023-INS-237 |
| chr17-35923656-INS-68 | chr17-35923632-INS-88 |
| chr17-35933305-INS-90 | chr17-35933305-INS-91 |
| chr17-35936021-INS-184 | chr17-35936021-INS-191 |
| chr17-35937871-INS-7483 | chr17-35937872-INS-7483 |
| chr17-35947124-INS-70 | chr17-35947124-INS-76 |
| chr17-35955552-INS-52 | chr17-35955532-INS-53 |
| chr17-35958126-INS-212 | chr17-35958126-INS-212 |
| chr17-35960047-INS-263 | chr17-35960047-INS-263 |
| chr17-35964640-INS-54 | chr17-35964640-INS-54 |
| chr17-35995715-INS-228 | chr17-35995715-INS-227 |
| chr17-36001539-INS-223 | chr17-36001539-INS-223 |
| chr17-36007382-INS-64 | chr17-36007383-INS-52 |
| chr17-36017566-INS-73 | chr17-36017543-INS-81 |
| chr17-36041788-INS-54 | chr17-36041788-INS-54 |
| chr17-36046472-INS-207 | chr17-36046472-INS-207 |
| chr17-36061835-INS-62 | chr17-36061835-INS-62 |
| chr17-36086715-INS-220 | chr17-36086715-INS-220 |
| chr17-36127520-INS-214 | chr17-36127520-INS-215 |
| chr17-36182149-INS-170 | chr17-36182149-INS-170 |
| chr17-36187424-INS-210 | chr17-36187424-INS-207 |
| chr17-36243870-INS-173 | chr17-36243870-INS-173 |
| chr17-36245552-INS-201 | chr17-36245552-INS-201 |
| chr17-36256654-INS-93 | chr17-36256654-INS-93 |
| chr17-36259049-INS-216 | chr17-36259049-INS-217 |
| chr17-36273282-INS-51 | chr17-36273282-INS-51 |
| chr17-36275020-INS-286 | chr17-36275020-INS-286 |
| chr17-36277069-INS-60 | chr17-36277069-INS-60 |
| chr17-36282292-INS-204 | chr17-36282292-INS-204 |
| chr17-36292669-INS-335 | chr17-36292669-INS-335 |
| chr17-36301209-INS-276 | chr17-36301210-INS-291 |
| chr17-36306589-INS-200 | chr17-36306589-INS-200 |
| chr17-36309726-INS-465 | chr17-36309728-INS-481 |
| chr17-36331776-INS-55 | chr17-36331776-INS-55 |
| chr17-36335739-INS-461 | chr17-36335739-INS-462 |
| chr17-36337820-INS-390 | chr17-36337820-INS-390 |
| chr17-36338556-INS-196 | chr17-36338556-INS-196 |
| chr17-36434606-INS-92 | chr17-36434606-INS-92 |
| chr17-36435730-INS-175 | chr17-36435730-INS-175 |
| chr17-36438845-INS-528 | chr17-36438845-INS-528 |
| chr17-36503630-INS-75 | chr17-36503630-INS-50 |
| chr17-36506950-INS-81 | chr17-36506950-INS-81 |
| chr17-36544588-INS-1913 | chr17-36544589-INS-1913 |
| chr17-36553834-INS-55 | chr17-36553834-INS-55 |
| chr17-36613302-INS-205 | chr17-36613302-INS-205 |
| chr17-36614308-INS-1898 | chr17-36614309-INS-1898 |
| chr17-36616107-INS-118 | chr17-36616107-INS-102 |
| chr17-36643054-INS-72 | chr17-36643068-INS-60 |
| chr17-36650676-INS-139 | chr17-36650680-INS-141 |
| chr17-36667495-INS-102 | chr17-36667495-INS-102 |
| chr17-36693798-INS-60 | chr17-36693800-INS-61 |
| chr17-36733402-INS-7797 | chr17-36733403-INS-7797 |
| chr17-36738720-INS-494 | chr17-36738720-INS-494 |
| chr17-36739298-INS-498 | chr17-36739298-INS-498 |
| chr17-36744899-INS-69 | chr17-36744899-INS-69 |
| chr17-36754212-INS-6883 | chr17-36754213-INS-6885 |
| chr17-36808592-INS-54 | chr17-36808592-INS-53 |
| chr17-36811249-INS-6435 | chr17-36811250-INS-6435 |
| chr17-36817881-INS-427 | chr17-36817881-INS-427 |
| chr17-36839624-INS-211 | chr17-36839624-INS-209 |
| chr17-36862150-INS-1862 | chr17-36862151-INS-1862 |
| chr17-36866260-INS-74 | chr17-36866252-INS-74 |
| chr17-36868728-INS-498 | chr17-36868728-INS-498 |
| chr17-36892357-INS-72 | chr17-36892357-INS-72 |

| | |
|---|---|
| chr17-36910526-INS-58 | chr17-36910526-INS-71 |
| chr17-36916893-INS-1034 | chr17-36916894-INS-998 |
| chr17-36920981-INS-208 | chr17-36920981-INS-208 |
| chr17-36939211-INS-75 | chr17-36939211-INS-75 |
| chr17-36973128-INS-202 | chr17-36973128-INS-202 |
| chr17-36999988-INS-1185 | chr17-36999989-INS-1185 |
| chr17-37045814-INS-601 | chr17-37045814-INS-601 |
| chr17-37050973-INS-229 | chr17-37050973-INS-228 |
| chr17-37089161-INS-71 | chr17-37089143-INS-83 |
| chr17-37107918-INS-5090 | chr17-37107919-INS-5090 |
| chr17-37113316-INS-69 | chr17-37113316-INS-81 |
| chr17-37117165-INS-214 | chr17-37117165-INS-214 |
| chr17-37118241-INS-577 | chr17-37118241-INS-577 |
| chr17-37119029-INS-134 | chr17-37119029-INS-130 |
| chr17-37133594-INS-212 | chr17-37133594-INS-212 |
| chr17-37135411-INS-514 | chr17-37135411-INS-514 |
| chr17-37139796-INS-55 | chr17-37139796-INS-67 |
| chr17-37141742-INS-63 | chr17-37141742-INS-63 |
| chr17-37144732-INS-486 | chr17-37144732-INS-486 |
| chr17-37154250-INS-213 | chr17-37154250-INS-213 |
| chr17-37176144-INS-212 | chr17-37176144-INS-212 |
| chr17-37236655-INS-171 | chr17-37236655-INS-171 |
| chr17-37238175-INS-215 | chr17-37238175-INS-215 |
| chr17-37290129-INS-320 | chr17-37290129-INS-320 |
| chr17-37336167-INS-208 | chr17-37336167-INS-208 |
| chr17-37342380-INS-339 | chr17-37342380-INS-339 |
| chr17-37342970-INS-5284 | chr17-37342971-INS-5284 |
| chr17-37345030-INS-94 | chr17-37345030-INS-84 |
| chr17-37379850-INS-79 | chr17-37379850-INS-79 |
| chr17-37389069-INS-53 | chr17-37389069-INS-53 |
| chr17-37407619-INS-69 | chr17-37407607-INS-70 |
| chr17-37415090-INS-419 | chr17-37415090-INS-419 |
| chr17-37424717-INS-118 | chr17-37424717-INS-109 |
| chr17-37433466-INS-58 | chr17-37433466-INS-58 |
| chr17-37458212-INS-6407 | chr17-37458213-INS-6407 |
| chr17-37464159-INS-1103 | chr17-37464160-INS-1103 |
| chr17-37476074-INS-75 | chr17-37476074-INS-95 |
| chr17-37483461-INS-80 | chr17-37483456-INS-59 |
| chr17-37509788-INS-74 | chr17-37509788-INS-74 |
| chr17-37527071-INS-1752 | chr17-37527072-INS-1751 |
| chr17-37569340-INS-110 | chr17-37569340-INS-85 |
| chr17-37608078-INS-204 | chr17-37608078-INS-204 |
| chr17-37619593-INS-56 | chr17-37619593-INS-56 |
| chr17-37657921-INS-203 | chr17-37657921-INS-204 |
| chr17-37658894-INS-868 | chr17-37658894-INS-866 |
| chr17-37666469-INS-198 | chr17-37666469-INS-198 |
| chr17-37699459-INS-103 | chr17-37699459-INS-57 |
| chr17-37707337-INS-467 | chr17-37707337-INS-467 |
| chr17-37748094-INS-78 | chr17-37748094-INS-78 |
| chr17-37766339-INS-731 | chr17-37766339-INS-731 |
| chr17-37776054-INS-2864 | chr17-37776055-INS-2865 |
| chr17-37794857-INS-2024 | chr17-37794858-INS-2023 |
| chr17-37832404-INS-52 | chr17-37832404-INS-52 |
| chr17-37846589-INS-59 | chr17-37846589-INS-59 |
| chr17-37876435-INS-4338 | chr17-37876436-INS-4338 |
| chr17-37896846-INS-6707 | chr17-37896847-INS-6708 |
| chr17-37918052-INS-853 | chr17-37918052-INS-853 |
| chr17-37937655-INS-2703 | chr17-37937656-INS-2703 |
| chr17-37940766-INS-64 | chr17-37940766-INS-66 |
| chr17-37964767-INS-3333 | chr17-37964751-INS-3333 |
| chr17-37975815-INS-277 | chr17-37975815-INS-277 |
| chr17-37984248-INS-6401 | chr17-37984249-INS-6401 |
| chr17-38030540-INS-52 | chr17-38030522-INS-84 |
| chr17-38045222-INS-7189 | chr17-38045223-INS-7188 |
| chr17-38051735-INS-8327 | chr17-38051736-INS-8327 |
| chr17-38096451-INS-6837 | chr17-38096452-INS-6837 |
| chr17-38105954-INS-6446 | chr17-38105955-INS-6446 |
| chr17-38157319-INS-57 | chr17-38157319-INS-58 |
| chr17-38164412-INS-208 | chr17-38164412-INS-208 |
| chr17-38255498-INS-333 | chr17-38255498-INS-333 |
| chr17-38264595-INS-179 | chr17-38264595-INS-179 |
| chr17-38292087-INS-7411 | chr17-38292088-INS-7412 |
| chr17-38621722-INS-4176 | chr17-38621723-INS-4176 |
| chr17-38667610-INS-6442 | chr17-38667611-INS-6442 |
| chr17-38791549-INS-5142 | chr17-38791550-INS-5141 |
| chr17-38835135-INS-2596 | chr17-38835136-INS-2599 |
| chr17-38858720-INS-2345 | chr17-38858721-INS-2345 |

| | |
|---|---|
| chr17-39006515-INS-6872 | chr17-39006516-INS-6873 |
| chr17-39015966-INS-174 | chr17-39015966-INS-170 |
| chr17-39030274-INS-544 | chr17-39030274-INS-544 |
| chr17-39053414-INS-120 | chr17-39053414-INS-120 |
| chr17-39059164-INS-56 | chr17-39059164-INS-56 |
| chr17-39082449-INS-3665 | chr17-39082450-INS-3658 |
| chr17-39103858-INS-4916 | chr17-39103859-INS-4916 |
| chr17-39109063-INS-6859 | chr17-39109064-INS-6860 |
| chr17-39113346-INS-443 | chr17-39113346-INS-443 |
| chr17-39181560-INS-64 | chr17-39181560-INS-64 |
| chr17-39246676-INS-563 | chr17-39246676-INS-560 |
| chr17-39273734-INS-6481 | chr17-39273735-INS-6481 |
| chr17-39295946-INS-4968 | chr17-39295947-INS-4969 |
| chr17-39321627-INS-7084 | chr17-39321628-INS-7084 |
| chr17-39423501-INS-160 | chr17-39423501-INS-160 |
| chr17-39423723-INS-5237 | chr17-39423724-INS-5237 |
| chr17-39458482-INS-6401 | chr17-39458483-INS-6401 |
| chr17-39562712-INS-5351 | chr17-39562713-INS-5351 |
| chr17-39813052-INS-4720 | chr17-39813053-INS-4720 |
| chr17-39841070-INS-222 | chr17-39841070-INS-222 |
| chr17-39895508-INS-5728 | chr17-39895509-INS-5728 |
| chr17-39988434-INS-73 | chr17-39988417-INS-69 |
| chr17-39997692-INS-54 | chr17-39997654-INS-54 |
| chr17-40013268-INS-81 | chr17-40013268-INS-66 |
| chr17-40017593-INS-52 | chr17-40017586-INS-68 |
| chr17-40079580-INS-214 | chr17-40079580-INS-214 |
| chr17-40085420-INS-51 | chr17-40085420-INS-51 |
| chr17-40108553-INS-1672 | chr17-40108554-INS-1669 |
| chr17-40126664-INS-593 | chr17-40126665-INS-587 |
| chr17-40144468-INS-52 | chr17-40144468-INS-52 |
| chr17-40160096-INS-1192 | chr17-40160097-INS-1194 |
| chr17-40168915-INS-1812 | chr17-40168916-INS-1812 |
| chr17-40185209-INS-3573 | chr17-40185210-INS-3575 |
| chr17-40212075-INS-3326 | chr17-40212076-INS-3326 |
| chr17-40232426-INS-2562 | chr17-40232427-INS-2548 |
| chr17-40285799-INS-6410 | chr17-40285847-INS-6410 |
| chr17-40437898-INS-208 | chr17-40437898-INS-208 |
| chr17-40534848-INS-156 | chr17-40534848-INS-156 |
| chr17-40655857-INS-53 | chr17-40655857-INS-53 |
| chr17-40693505-INS-183 | chr17-40693505-INS-183 |
| chr17-40710588-INS-87 | chr17-40710588-INS-92 |
| chr17-40725971-INS-6208 | chr17-40725972-INS-6208 |
| chr17-40846359-INS-456 | chr17-40846359-INS-456 |
| chr17-40852197-INS-212 | chr17-40852197-INS-212 |
| chr17-40892484-INS-6452 | chr17-40892485-INS-6452 |
| chr17-40916688-INS-60 | chr17-40916688-INS-60 |
| chr17-40944252-INS-7192 | chr17-40944253-INS-7192 |
| chr17-40944253-INS-362 | chr17-40944253-INS-362 |
| chr17-40945241-INS-361 | chr17-40945241-INS-361 |
| chr17-40965580-INS-67 | chr17-40965580-INS-109 |
| chr17-40970306-INS-609 | chr17-40970306-INS-609 |
| chr17-41020400-INS-140 | chr17-41020401-INS-127 |
| chr17-41038998-INS-6403 | chr17-41038999-INS-6403 |
| chr17-41103505-INS-311 | chr17-41103505-INS-311 |
| chr17-41194088-INS-454 | chr17-41194088-INS-454 |
| chr17-41200567-INS-52 | chr17-41200567-INS-52 |
| chr17-41218440-INS-528 | chr17-41218441-INS-528 |
| chr17-41220615-INS-156 | chr17-41220615-INS-156 |
| chr17-41221508-INS-1954 | chr17-41221509-INS-1954 |
| chr17-41222608-INS-59 | chr17-41222608-INS-59 |
| chr17-41243925-INS-287 | chr17-41243925-INS-285 |
| chr17-41259272-INS-80 | chr17-41259272-INS-80 |
| chr17-41265323-INS-90 | chr17-41265323-INS-92 |
| chr17-41270911-INS-570 | chr17-41270914-INS-571 |
| chr17-41381708-INS-1143 | chr17-41381709-INS-1137 |
| chr17-41404112-INS-714 | chr17-41404112-INS-715 |
| chr17-41411018-INS-1828 | chr17-41411019-INS-1829 |
| chr17-41419489-INS-140 | chr17-41419489-INS-140 |
| chr17-41428121-INS-110 | chr17-41428121-INS-110 |
| chr17-41436336-INS-58 | chr17-41436336-INS-58 |
| chr17-41438306-INS-84 | chr17-41438306-INS-84 |
| chr17-41446754-INS-50 | chr17-41446754-INS-50 |
| chr17-41465714-INS-58 | chr17-41465714-INS-58 |
| chr17-41482938-INS-332 | chr17-41482938-INS-332 |
| chr17-41508288-INS-70 | chr17-41508288-INS-70 |
| chr17-41509587-INS-56 | chr17-41509587-INS-52 |
| chr17-41558798-INS-60 | chr17-41558798-INS-60 |

| | |
|---|---|
| chr17-41566859-INS-134 | chr17-41566862-INS-126 |
| chr17-41567671-INS-396 | chr17-41567671-INS-396 |
| chr17-41569047-INS-202 | chr17-41569047-INS-202 |
| chr17-41574352-INS-549 | chr17-41574352-INS-549 |
| chr17-41590317-INS-94 | chr17-41590333-INS-52 |
| chr17-41593504-INS-6439 | chr17-41593505-INS-6440 |
| chr17-41629423-INS-2559 | chr17-41629424-INS-2559 |
| chr17-41677410-INS-2512 | chr17-41677411-INS-2513 |
| chr17-41688789-INS-109 | chr17-41688789-INS-113 |
| chr17-41689780-INS-6461 | chr17-41689781-INS-6461 |
| chr17-41697138-INS-3195 | chr17-41697139-INS-3195 |
| chr17-41701061-INS-214 | chr17-41701061-INS-214 |
| chr17-41704567-INS-5548 | chr17-41704568-INS-5548 |
| chr17-41713905-INS-5454 | chr17-41713906-INS-5454 |
| chr17-41716500-INS-6217 | chr17-41716501-INS-6217 |
| chr17-41734734-INS-1374 | chr17-41734735-INS-1375 |
| chr17-41736005-INS-4133 | chr17-41736006-INS-4133 |
| chr17-41738484-INS-122 | chr17-41738484-INS-122 |
| chr17-41746825-INS-4347 | chr17-41746826-INS-4347 |
| chr17-41756588-INS-209 | chr17-41756588-INS-209 |
| chr17-41759894-INS-394 | chr17-41759894-INS-394 |
| chr17-41769630-INS-457 | chr17-41769630-INS-457 |
| chr17-41773187-INS-57 | chr17-41773175-INS-78 |
| chr17-41794879-INS-85 | chr17-41794879-INS-84 |
| chr17-41804677-INS-6930 | chr17-41804678-INS-6930 |
| chr17-41833852-INS-198 | chr17-41833852-INS-198 |
| chr17-41848219-INS-201 | chr17-41848219-INS-201 |
| chr17-41859857-INS-741 | chr17-41859857-INS-741 |
| chr17-41866488-INS-6452 | chr17-41866489-INS-6452 |
| chr17-41914858-INS-56 | chr17-41914858-INS-56 |
| chr17-41920913-INS-672 | chr17-41920918-INS-679 |
| chr17-41923930-INS-75 | chr17-41923959-INS-54 |
| chr17-41929415-INS-6228 | chr17-41929416-INS-6227 |
| chr17-41944488-INS-111 | chr17-41944488-INS-111 |
| chr17-41949410-INS-85 | chr17-41949410-INS-85 |
| chr17-42058259-INS-54 | chr17-42058259-INS-58 |
| chr17-42116628-INS-182 | chr17-42116634-INS-186 |
| chr17-42122945-INS-7085 | chr17-42122946-INS-7085 |
| chr17-42159446-INS-358 | chr17-42159446-INS-358 |
| chr17-42164151-INS-54 | chr17-42164151-INS-60 |
| chr17-42209241-INS-375 | chr17-42209241-INS-376 |
| chr17-42210318-INS-156 | chr17-42210318-INS-158 |
| chr17-42225602-INS-201 | chr17-42225602-INS-201 |
| chr17-42244712-INS-497 | chr17-42244712-INS-497 |
| chr17-42247875-INS-60 | chr17-42247875-INS-60 |
| chr17-42287458-INS-70 | chr17-42287458-INS-66 |
| chr17-42296946-INS-83 | chr17-42296946-INS-83 |
| chr17-42345361-INS-310 | chr17-42345361-INS-310 |
| chr17-42347275-INS-399 | chr17-42347275-INS-399 |
| chr17-42364695-INS-6465 | chr17-42364696-INS-6466 |
| chr17-42373361-INS-1405 | chr17-42373362-INS-1406 |
| chr17-42427557-INS-67 | chr17-42427557-INS-95 |
| chr17-42433847-INS-7221 | chr17-42433848-INS-7221 |
| chr17-42434149-INS-2228 | chr17-42434150-INS-2228 |
| chr17-42443830-INS-88 | chr17-42443830-INS-88 |
| chr17-42446825-INS-485 | chr17-42446825-INS-501 |
| chr17-42449097-INS-1940 | chr17-42449098-INS-1940 |
| chr17-42457988-INS-6907 | chr17-42457989-INS-6907 |
| chr17-42520944-INS-471 | chr17-42520944-INS-471 |
| chr17-42527048-INS-1294 | chr17-42527049-INS-1294 |
| chr17-42527682-INS-420 | chr17-42527689-INS-426 |
| chr17-42540070-INS-127 | chr17-42540070-INS-127 |
| chr17-42556810-INS-150 | chr17-42556810-INS-153 |
| chr17-42596138-INS-110 | chr17-42596143-INS-84 |
| chr17-42611056-INS-5309 | chr17-42611057-INS-5309 |
| chr17-42648209-INS-56 | chr17-42648209-INS-56 |
| chr17-42657822-INS-884 | chr17-42657822-INS-885 |
| chr17-42671541-INS-91 | chr17-42671541-INS-64 |
| chr17-42677070-INS-244 | chr17-42677070-INS-244 |
| chr17-42714578-INS-52 | chr17-42714578-INS-52 |
| chr17-42716406-INS-187 | chr17-42716406-INS-187 |
| chr17-42746089-INS-457 | chr17-42746089-INS-457 |
| chr17-42746496-INS-765 | chr17-42746496-INS-762 |
| chr17-42755368-INS-128 | chr17-42755368-INS-128 |
| chr17-42762113-INS-72 | chr17-42762113-INS-72 |
| chr17-42774740-INS-7186 | chr17-42774741-INS-7186 |
| chr17-42775179-INS-576 | chr17-42775179-INS-576 |

| | |
|---|---|
| chr17-42780201-INS-52 | chr17-42780201-INS-52 |
| chr17-42801280-INS-108 | chr17-42801280-INS-116 |
| chr17-42824801-INS-178 | chr17-42824801-INS-178 |
| chr17-42837974-INS-6332 | chr17-42837975-INS-6327 |
| chr17-42843992-INS-400 | chr17-42843992-INS-400 |
| chr17-42871025-INS-1248 | chr17-42871025-INS-1241 |
| chr17-42890511-INS-508 | chr17-42890512-INS-508 |
| chr17-42894294-INS-56 | chr17-42894294-INS-56 |
| chr17-42902282-INS-7223 | chr17-42902283-INS-7223 |
| chr17-42941766-INS-741 | chr17-42941766-INS-741 |
| chr17-42943460-INS-4808 | chr17-42943461-INS-4841 |
| chr17-42966302-INS-480 | chr17-42966302-INS-480 |
| chr17-42970639-INS-66 | chr17-42970639-INS-66 |
| chr17-42988545-INS-136 | chr17-42988545-INS-138 |
| chr17-42993265-INS-104 | chr17-42993265-INS-120 |
| chr17-43004444-INS-59 | chr17-43004444-INS-60 |
| chr17-43043611-INS-95 | chr17-43043611-INS-94 |
| chr17-43067725-INS-575 | chr17-43067725-INS-575 |
| chr17-43081540-INS-437 | chr17-43081540-INS-437 |
| chr17-43086156-INS-3319 | chr17-43086157-INS-3319 |
| chr17-43096352-INS-102 | chr17-43096362-INS-107 |
| chr17-43113431-INS-72 | chr17-43113431-INS-74 |
| chr17-43132226-INS-111 | chr17-43132226-INS-66 |
| chr17-43135526-INS-70 | chr17-43135526-INS-88 |
| chr17-43167324-INS-757 | chr17-43167324-INS-759 |
| chr17-43174720-INS-206 | chr17-43174720-INS-206 |
| chr17-43181392-INS-53 | chr17-43181392-INS-55 |
| chr17-43209167-INS-63 | chr17-43209167-INS-63 |
| chr17-43214619-INS-212 | chr17-43214619-INS-226 |
| chr17-43243627-INS-65 | chr17-43243627-INS-79 |
| chr17-43247816-INS-92 | chr17-43247816-INS-92 |
| chr17-43259374-INS-90 | chr17-43259376-INS-113 |
| chr17-43260379-INS-64 | chr17-43260379-INS-64 |
| chr17-43268589-INS-109 | chr17-43268589-INS-78 |
| chr17-43271597-INS-208 | chr17-43271597-INS-212 |
| chr17-43326562-INS-110 | chr17-43326562-INS-70 |
| chr17-43341867-INS-55 | chr17-43341883-INS-50 |
| chr17-43343372-INS-575 | chr17-43343372-INS-584 |
| chr17-43354030-INS-302 | chr17-43354030-INS-302 |
| chr17-43379668-INS-160 | chr17-43379696-INS-152 |
| chr17-43399052-INS-56 | chr17-43399052-INS-76 |
| chr17-43424025-INS-74 | chr17-43424025-INS-74 |
| chr17-43429464-INS-53 | chr17-43429464-INS-74 |
| chr17-43431453-INS-66 | chr17-43431437-INS-50 |
| chr17-43459432-INS-304 | chr17-43459432-INS-304 |
| chr17-43459994-INS-78 | chr17-43459994-INS-78 |
| chr17-43488845-INS-1568 | chr17-43488846-INS-1568 |
| chr17-43534084-INS-255 | chr17-43534085-INS-255 |
| chr17-43536617-INS-108 | chr17-43536617-INS-110 |
| chr17-43539317-INS-187 | chr17-43539317-INS-186 |
| chr17-43541618-INS-65 | chr17-43541610-INS-77 |
| chr17-43549425-INS-404 | chr17-43549425-INS-404 |
| chr17-43552779-INS-98 | chr17-43552779-INS-82 |
| chr17-43556996-INS-81 | chr17-43556998-INS-74 |
| chr17-43585387-INS-62 | chr17-43585355-INS-87 |
| chr17-43595929-INS-64 | chr17-43595929-INS-64 |
| chr17-43601253-INS-5008 | chr17-43601254-INS-5008 |
| chr17-43602506-INS-64 | chr17-43602506-INS-56 |
| chr17-43616962-INS-60 | chr17-43616962-INS-65 |
| chr17-43625796-INS-456 | chr17-43625796-INS-466 |
| chr17-43630150-INS-52 | chr17-43630150-INS-52 |
| chr17-43631761-INS-74 | chr17-43631761-INS-72 |
| chr17-43632679-INS-158 | chr17-43632679-INS-158 |
| chr17-43647659-INS-52 | chr17-43647659-INS-54 |
| chr17-43651665-INS-264 | chr17-43651665-INS-255 |
| chr17-43653092-INS-891 | chr17-43653092-INS-891 |
| chr17-43653727-INS-52 | chr17-43653727-INS-52 |
| chr17-43698426-INS-85 | chr17-43698426-INS-50 |
| chr17-43707288-INS-63 | chr17-43707288-INS-63 |
| chr17-43709094-INS-58 | chr17-43709074-INS-76 |
| chr17-43724861-INS-244 | chr17-43724861-INS-256 |
| chr17-43726686-INS-68 | chr17-43726686-INS-68 |
| chr17-43729325-INS-63 | chr17-43729294-INS-69 |
| chr17-43757326-INS-483 | chr17-43757326-INS-483 |
| chr17-43772903-INS-53 | chr17-43772903-INS-56 |
| chr17-43779800-INS-649 | chr17-43779800-INS-649 |
| chr17-43804789-INS-84 | chr17-43804779-INS-78 |

| | |
|---|---|
| chr17-43831560-INS-94 | chr17-43831560-INS-111 |
| chr17-43833700-INS-121 | chr17-43833700-INS-142 |
| chr17-43834637-INS-124 | chr17-43834637-INS-126 |
| chr17-43837643-INS-340 | chr17-43837644-INS-340 |
| chr17-43847834-INS-295 | chr17-43847834-INS-295 |
| chr17-43866424-INS-84 | chr17-43866424-INS-84 |
| chr17-43871718-INS-195 | chr17-43871718-INS-212 |
| chr17-43873673-INS-316 | chr17-43873673-INS-316 |
| chr17-43878331-INS-246 | chr17-43878331-INS-231 |
| chr17-43894756-INS-87 | chr17-43894756-INS-86 |
| chr17-43908009-INS-66 | chr17-43908009-INS-66 |
| chr17-43909353-INS-62 | chr17-43909353-INS-62 |
| chr17-43921745-INS-65 | chr17-43921745-INS-75 |
| chr17-43928927-INS-55 | chr17-43928927-INS-55 |
| chr17-43942833-INS-283 | chr17-43942833-INS-269 |
| chr17-43957265-INS-108 | chr17-43957273-INS-108 |
| chr17-43964411-INS-156 | chr17-43964411-INS-156 |
| chr17-43968044-INS-365 | chr17-43968043-INS-410 |
| chr17-43974326-INS-220 | chr17-43974326-INS-220 |
| chr17-43981194-INS-62 | chr17-43981194-INS-62 |
| chr17-43988543-INS-64 | chr17-43988543-INS-74 |
| chr17-43997124-INS-60 | chr17-43997124-INS-60 |
| chr17-44005479-INS-313 | chr17-44005479-INS-313 |
| chr17-44011717-INS-6213 | chr17-44011718-INS-6213 |
| chr17-44013188-INS-153 | chr17-44013229-INS-126 |
| chr17-44028671-INS-128 | chr17-44028671-INS-116 |
| chr17-44033437-INS-124 | chr17-44033437-INS-124 |
| chr17-44044619-INS-63 | chr17-44044619-INS-63 |
| chr17-44049188-INS-74 | chr17-44049188-INS-74 |
| chr17-44074967-INS-113 | chr17-44074967-INS-105 |
| chr17-44081064-INS-139 | chr17-44081064-INS-139 |
| chr17-44087281-INS-118 | chr17-44087281-INS-122 |
| chr17-44094319-INS-75 | chr17-44094319-INS-55 |
| chr17-44106797-INS-670 | chr17-44106797-INS-670 |
| chr17-44111568-INS-67 | chr17-44111568-INS-67 |
| chr17-44123607-INS-95 | chr17-44123607-INS-96 |
| chr17-44127098-INS-6914 | chr17-44127099-INS-6915 |
| chr17-44136311-INS-251 | chr17-44136311-INS-255 |
| chr17-44154430-INS-184 | chr17-44154430-INS-184 |
| chr17-44155834-INS-6939 | chr17-44155835-INS-6939 |
| chr17-44158921-INS-62 | chr17-44158921-INS-70 |
| chr17-44164336-INS-148 | chr17-44164336-INS-148 |
| chr17-44192986-INS-90 | chr17-44192986-INS-90 |
| chr17-44217661-INS-169 | chr17-44217661-INS-153 |
| chr17-44230989-INS-95 | chr17-44230992-INS-85 |
| chr17-44286381-INS-146 | chr17-44286381-INS-149 |
| chr17-44324276-INS-178 | chr17-44324276-INS-178 |
| chr17-44346969-INS-64 | chr17-44346969-INS-64 |
| chr17-44349909-INS-71 | chr17-44349909-INS-50 |
| chr17-44368838-INS-879 | chr17-44368838-INS-886 |
| chr17-44372031-INS-103 | chr17-44372031-INS-103 |
| chr17-44381180-INS-343 | chr17-44381180-INS-343 |
| chr17-44383946-INS-58 | chr17-44383946-INS-58 |
| chr17-44419323-INS-59 | chr17-44419324-INS-51 |
| chr17-44433995-INS-98 | chr17-44433969-INS-98 |
| chr17-44444267-INS-387 | chr17-44444267-INS-385 |
| chr17-44447960-INS-72 | chr17-44447960-INS-73 |
| chr17-44466897-INS-54 | chr17-44466897-INS-64 |
| chr17-44498496-INS-72 | chr17-44498514-INS-70 |
| chr17-44521356-INS-345 | chr17-44521356-INS-345 |
| chr17-44527537-INS-60 | chr17-44527537-INS-62 |
| chr17-44537050-INS-82 | chr17-44537050-INS-50 |
| chr17-44540351-INS-952 | chr17-44540351-INS-952 |
| chr17-44552515-INS-106 | chr17-44552515-INS-106 |
| chr17-44593743-INS-56 | chr17-44593743-INS-56 |
| chr17-44597720-INS-6882 | chr17-44597721-INS-6878 |
| chr17-44599649-INS-518 | chr17-44599649-INS-519 |
| chr17-44601376-INS-64 | chr17-44601376-INS-64 |
| chr17-44612850-INS-170 | chr17-44612855-INS-168 |
| chr17-44622566-INS-51 | chr17-44622566-INS-52 |
| chr17-44635703-INS-187 | chr17-44635703-INS-188 |
| chr17-44667896-INS-65 | chr17-44667896-INS-65 |
| chr17-44683724-INS-52 | chr17-44683724-INS-58 |
| chr17-44696807-INS-53 | chr17-44696839-INS-51 |
| chr17-44698332-INS-88 | chr17-44698328-INS-92 |
| chr17-44711354-INS-52 | chr17-44711354-INS-52 |
| chr17-44732663-INS-74 | chr17-44732663-INS-56 |

| | |
|---|---|
| chr17-44757301-INS-185 | chr17-44757301-INS-185 |
| chr17-44765635-INS-67 | chr17-44765635-INS-68 |
| chr17-44780263-INS-899 | chr17-44780259-INS-900 |
| chr17-44799550-INS-566 | chr17-44799551-INS-566 |
| chr17-44800066-INS-61 | chr17-44800024-INS-72 |
| chr17-44803016-INS-53 | chr17-44803016-INS-55 |
| chr17-44832307-INS-75 | chr17-44832307-INS-74 |
| chr17-44855459-INS-90 | chr17-44855459-INS-130 |
| chr17-44861086-INS-198 | chr17-44861086-INS-204 |
| chr17-44861795-INS-109 | chr17-44861792-INS-85 |
| chr17-44897662-INS-76 | chr17-44897662-INS-58 |
| chr17-44909711-INS-96 | chr17-44909711-INS-96 |
| chr17-44916090-INS-81 | chr17-44916090-INS-84 |
| chr17-44931289-INS-166 | chr17-44931272-INS-175 |
| chr17-44952414-INS-58 | chr17-44952415-INS-50 |
| chr17-44960634-INS-77 | chr17-44960634-INS-75 |
| chr17-44966332-INS-122 | chr17-44966332-INS-122 |
| chr17-45028408-INS-54 | chr17-45028408-INS-54 |
| chr17-45076659-INS-52 | chr17-45076660-INS-60 |
| chr17-45078260-INS-1629 | chr17-45078261-INS-1629 |
| chr17-45091505-INS-322 | chr17-45091505-INS-322 |
| chr17-45119301-INS-75 | chr17-45119301-INS-74 |
| chr17-45139487-INS-71 | chr17-45139487-INS-71 |
| chr17-45142451-INS-124 | chr17-45142451-INS-124 |
| chr17-45146497-INS-85 | chr17-45146497-INS-85 |
| chr17-45149294-INS-208 | chr17-45149294-INS-207 |
| chr17-45173413-INS-358 | chr17-45173413-INS-359 |
| chr17-45204088-INS-195 | chr17-45204082-INS-187 |
| chr17-45212674-INS-482 | chr17-45212674-INS-483 |
| chr17-45237912-INS-8508 | chr17-45237913-INS-8507 |
| chr17-45291560-INS-80 | chr17-45291560-INS-65 |
| chr17-45295127-INS-2103 | chr17-45295128-INS-2102 |
| chr17-45322844-INS-80 | chr17-45322844-INS-80 |
| chr17-45339243-INS-121 | chr17-45339243-INS-121 |
| chr17-45351683-INS-1862 | chr17-45351684-INS-1862 |
| chr17-45352378-INS-54 | chr17-45352343-INS-82 |
| chr17-45388667-INS-1088 | chr17-45388668-INS-1089 |
| chr17-45412760-INS-2198 | chr17-45412761-INS-2198 |
| chr17-45428250-INS-77 | chr17-45428250-INS-77 |
| chr17-45436764-INS-2234 | chr17-45436765-INS-2234 |
| chr17-45461748-INS-514 | chr17-45461748-INS-516 |
| chr17-45469902-INS-60 | chr17-45469902-INS-60 |
| chr17-45473624-INS-52 | chr17-45473594-INS-52 |
| chr17-45476687-INS-65 | chr17-45476687-INS-65 |
| chr17-45496860-INS-74 | chr17-45496860-INS-56 |
| chr17-45501213-INS-570 | chr17-45501213-INS-570 |
| chr17-45503108-INS-63 | chr17-45503108-INS-75 |
| chr17-45512480-INS-478 | chr17-45512482-INS-502 |
| chr17-45524668-INS-126 | chr17-45524668-INS-85 |
| chr17-45525786-INS-234 | chr17-45525786-INS-234 |
| chr17-45548809-INS-72 | chr17-45548809-INS-72 |
| chr17-45556553-INS-224 | chr17-45556553-INS-224 |
| chr17-45576053-INS-204 | chr17-45576053-INS-204 |
| chr17-45624484-INS-310 | chr17-45624484-INS-310 |
| chr17-45627749-INS-64 | chr17-45627741-INS-52 |
| chr17-45634309-INS-65 | chr17-45634309-INS-66 |
| chr17-45645432-INS-90 | chr17-45645427-INS-54 |
| chr17-45653384-INS-5175 | chr17-45653385-INS-5175 |
| chr17-45671807-INS-204 | chr17-45671807-INS-156 |
| chr17-45691178-INS-230 | chr17-45691178-INS-230 |
| chr17-45694027-INS-400 | chr17-45694027-INS-400 |
| chr17-45697281-INS-88 | chr17-45697281-INS-88 |
| chr17-45706321-INS-128 | chr17-45706321-INS-128 |
| chr17-45719764-INS-174 | chr17-45719764-INS-174 |
| chr17-45728247-INS-205 | chr17-45728247-INS-209 |
| chr17-45736105-INS-217 | chr17-45736105-INS-222 |
| chr17-45737985-INS-108 | chr17-45737985-INS-108 |
| chr17-45761047-INS-178 | chr17-45761047-INS-178 |
| chr17-45774265-INS-72 | chr17-45774265-INS-57 |
| chr17-45782655-INS-155 | chr17-45782655-INS-167 |
| chr17-45783915-INS-198 | chr17-45783915-INS-197 |
| chr17-45788489-INS-6402 | chr17-45788490-INS-6402 |
| chr17-45796340-INS-90 | chr17-45796336-INS-113 |
| chr17-45800586-INS-55 | chr17-45800586-INS-51 |
| chr17-45813699-INS-274 | chr17-45813699-INS-273 |
| chr17-45816021-INS-161 | chr17-45816012-INS-147 |
| chr17-45831404-INS-211 | chr17-45831404-INS-211 |

| | |
|---|---|
| chr17-45846970-INS-113 | chr17-45846972-INS-107 |
| chr17-45870791-INS-53 | chr17-45870791-INS-53 |
| chr17-45889882-INS-2831 | chr17-45889883-INS-2822 |
| chr17-45891569-INS-233 | chr17-45891569-INS-233 |
| chr17-45925276-INS-388 | chr17-45925276-INS-388 |
| chr17-45929250-INS-210 | chr17-45929250-INS-210 |
| chr17-45938591-INS-434 | chr17-45938591-INS-434 |
| chr17-45962548-INS-204 | chr17-45962548-INS-202 |
| chr17-45994264-INS-1864 | chr17-45994265-INS-1864 |
| chr17-46000680-INS-76 | chr17-46000664-INS-110 |
| chr17-46017456-INS-61 | chr17-46017456-INS-63 |
| chr17-46019503-INS-56 | chr17-46019503-INS-68 |
| chr17-46057453-INS-75 | chr17-46057453-INS-75 |
| chr17-46060702-INS-80 | chr17-46060700-INS-92 |
| chr17-46085861-INS-119 | chr17-46085861-INS-168 |
| chr17-46087616-INS-304 | chr17-46087616-INS-296 |
| chr17-46093933-INS-220 | chr17-46093933-INS-220 |
| chr17-46135202-INS-102 | chr17-46135202-INS-108 |
| chr17-46141398-INS-344 | chr17-46141398-INS-344 |
| chr17-46148635-INS-115 | chr17-46148635-INS-106 |
| chr17-46151535-INS-279 | chr17-46151535-INS-279 |
| chr17-46179803-INS-54 | chr17-46179772-INS-56 |
| chr17-46210688-INS-208 | chr17-46210688-INS-208 |
| chr17-46213443-INS-55 | chr17-46213443-INS-55 |
| chr17-46220349-INS-76 | chr17-46220349-INS-76 |
| chr17-46236130-INS-100 | chr17-46236130-INS-100 |
| chr17-46241834-INS-56 | chr17-46241834-INS-56 |
| chr17-46251869-INS-92 | chr17-46251869-INS-91 |
| chr17-46253545-INS-109 | chr17-46253545-INS-110 |
| chr17-46267018-INS-56 | chr17-46267018-INS-58 |
| chr17-46273898-INS-106 | chr17-46273898-INS-104 |
| chr17-46289757-INS-65 | chr17-46289757-INS-65 |
| chr17-46371861-INS-52 | chr17-46371861-INS-52 |
| chr17-46380429-INS-56 | chr17-46380429-INS-56 |
| chr17-46398170-INS-63 | chr17-46398170-INS-63 |
| chr17-46418857-INS-55 | chr17-46418881-INS-69 |
| chr17-46421632-INS-145 | chr17-46421632-INS-125 |
| chr17-46469919-INS-66 | chr17-46469915-INS-64 |
| chr17-46484308-INS-252 | chr17-46484308-INS-253 |
| chr17-46501905-INS-204 | chr17-46501905-INS-208 |
| chr17-46508981-INS-285 | chr17-46508981-INS-284 |
| chr17-46517706-INS-179 | chr17-46517706-INS-179 |
| chr17-46561299-INS-57 | chr17-46561299-INS-57 |
| chr17-46566736-INS-92 | chr17-46566732-INS-97 |
| chr17-46570366-INS-117 | chr17-46570366-INS-117 |
| chr17-46572880-INS-61 | chr17-46572880-INS-61 |
| chr17-46586609-INS-360 | chr17-46586609-INS-360 |
| chr17-46603200-INS-207 | chr17-46603200-INS-207 |
| chr17-46604820-INS-68 | chr17-46604820-INS-76 |
| chr17-46607182-INS-93 | chr17-46607182-INS-93 |
| chr17-46633183-INS-99 | chr17-46633227-INS-50 |
| chr17-46645267-INS-1186 | chr17-46645268-INS-1186 |
| chr17-46646419-INS-86 | chr17-46646419-INS-86 |
| chr17-46650949-INS-6537 | chr17-46650950-INS-6537 |
| chr17-46653395-INS-108 | chr17-46653395-INS-122 |
| chr17-46661900-INS-211 | chr17-46661900-INS-211 |
| chr17-46685543-INS-216 | chr17-46685543-INS-217 |
| chr17-46689966-INS-3682 | chr17-46689967-INS-3682 |
| chr17-46693596-INS-203 | chr17-46693596-INS-203 |
| chr17-46723441-INS-51 | chr17-46723441-INS-72 |
| chr17-46728315-INS-90 | chr17-46728315-INS-86 |
| chr17-46742804-INS-281 | chr17-46742804-INS-281 |
| chr17-46758899-INS-90 | chr17-46758899-INS-90 |
| chr17-46772650-INS-55 | chr17-46772650-INS-55 |
| chr17-46776394-INS-200 | chr17-46776394-INS-198 |
| chr17-46831897-INS-220 | chr17-46831897-INS-220 |
| chr17-46840330-INS-201 | chr17-46840319-INS-187 |
| chr17-46870542-INS-61 | chr17-46870542-INS-61 |
| chr17-46878413-INS-212 | chr17-46878413-INS-212 |
| chr17-46904886-INS-144 | chr17-46904886-INS-144 |
| chr17-46910465-INS-213 | chr17-46910465-INS-213 |
| chr17-46917412-INS-871 | chr17-46917412-INS-871 |
| chr17-46945211-INS-67 | chr17-46945211-INS-67 |
| chr17-46949207-INS-189 | chr17-46949207-INS-190 |
| chr17-47009339-INS-212 | chr17-47009339-INS-204 |
| chr17-47048471-INS-56 | chr17-47048471-INS-56 |
| chr17-47056430-INS-52 | chr17-47056430-INS-57 |

| | |
|---|---|
| chr17-47061551-INS-81 | chr17-47061556-INS-71 |
| chr17-47075150-INS-159 | chr17-47075150-INS-159 |
| chr17-47082389-INS-193 | chr17-47082389-INS-187 |
| chr17-47087628-INS-54 | chr17-47087628-INS-54 |
| chr17-47092276-INS-206 | chr17-47092276-INS-207 |
| chr17-47104005-INS-193 | chr17-47104006-INS-192 |
| chr17-47114566-INS-198 | chr17-47114566-INS-198 |
| chr17-47119088-INS-84 | chr17-47119088-INS-84 |
| chr17-47126682-INS-233 | chr17-47126682-INS-235 |
| chr17-47130910-INS-55 | chr17-47130910-INS-55 |
| chr17-47145716-INS-68 | chr17-47145716-INS-60 |
| chr17-47150271-INS-217 | chr17-47150271-INS-217 |
| chr17-47155628-INS-403 | chr17-47155629-INS-397 |
| chr17-47164924-INS-204 | chr17-47164924-INS-204 |
| chr17-47179835-INS-157 | chr17-47179835-INS-157 |
| chr17-47183323-INS-141 | chr17-47183323-INS-141 |
| chr17-47211873-INS-209 | chr17-47211873-INS-209 |
| chr17-47218930-INS-164 | chr17-47218930-INS-164 |
| chr17-47225686-INS-56 | chr17-47225712-INS-68 |
| chr17-47228123-INS-61 | chr17-47228123-INS-61 |
| chr17-47247887-INS-251 | chr17-47247887-INS-253 |
| chr17-47274164-INS-294 | chr17-47274164-INS-294 |
| chr17-47294487-INS-225 | chr17-47294487-INS-252 |
| chr17-47308576-INS-200 | chr17-47308569-INS-204 |
| chr17-47329339-INS-196 | chr17-47329339-INS-192 |
| chr17-47349263-INS-505 | chr17-47349263-INS-505 |
| chr17-47360662-INS-200 | chr17-47360662-INS-209 |
| chr17-47373979-INS-181 | chr17-47373979-INS-181 |
| chr17-47378535-INS-208 | chr17-47378531-INS-209 |
| chr17-47404806-INS-54 | chr17-47404806-INS-90 |
| chr17-47407244-INS-111 | chr17-47407271-INS-111 |
| chr17-47424289-INS-66 | chr17-47424289-INS-66 |
| chr17-47428356-INS-53 | chr17-47428356-INS-62 |
| chr17-47443996-INS-60 | chr17-47443996-INS-60 |
| chr17-47476335-INS-53 | chr17-47476335-INS-53 |
| chr17-47482603-INS-164 | chr17-47482603-INS-164 |
| chr17-47548769-INS-72 | chr17-47548769-INS-72 |
| chr17-47555483-INS-218 | chr17-47555474-INS-255 |
| chr17-47559044-INS-70 | chr17-47559045-INS-70 |
| chr17-47559594-INS-59 | chr17-47559594-INS-60 |
| chr17-47592043-INS-231 | chr17-47592044-INS-232 |
| chr17-47592992-INS-52 | chr17-47592992-INS-52 |
| chr17-47596989-INS-78 | chr17-47596989-INS-78 |
| chr17-47626791-INS-59 | chr17-47626788-INS-74 |
| chr17-47640948-INS-164 | chr17-47640951-INS-166 |
| chr17-47649829-INS-302 | chr17-47649834-INS-294 |
| chr17-47668323-INS-50 | chr17-47668283-INS-83 |
| chr17-47674696-INS-67 | chr17-47674696-INS-67 |
| chr17-47679039-INS-219 | chr17-47679039-INS-219 |
| chr17-47695493-INS-1418 | chr17-47695480-INS-1401 |
| chr17-47695676-INS-204 | chr17-47695676-INS-205 |
| chr17-47695716-INS-4399 | chr17-47695717-INS-4399 |
| chr17-47707699-INS-102 | chr17-47707699-INS-102 |
| chr17-47708496-INS-205 | chr17-47708496-INS-205 |
| chr17-47713691-INS-3258 | chr17-47713692-INS-3258 |
| chr17-47714253-INS-114 | chr17-47714253-INS-67 |
| chr17-47715230-INS-53 | chr17-47715230-INS-55 |
| chr17-47718556-INS-214 | chr17-47718556-INS-214 |
| chr17-47741107-INS-86 | chr17-47741108-INS-105 |
| chr17-47746936-INS-203 | chr17-47746936-INS-203 |
| chr17-47752071-INS-56 | chr17-47752071-INS-56 |
| chr17-47758866-INS-209 | chr17-47758866-INS-209 |
| chr17-47766154-INS-66 | chr17-47766154-INS-66 |
| chr17-47793647-INS-78 | chr17-47793647-INS-78 |
| chr17-47795274-INS-224 | chr17-47795274-INS-224 |
| chr17-47803466-INS-185 | chr17-47803466-INS-185 |
| chr17-47814727-INS-234 | chr17-47814727-INS-220 |
| chr17-47817613-INS-218 | chr17-47817613-INS-218 |
| chr17-47823484-INS-164 | chr17-47823484-INS-164 |
| chr17-47826356-INS-52 | chr17-47826356-INS-52 |
| chr17-47828458-INS-261 | chr17-47828458-INS-262 |
| chr17-47831828-INS-273 | chr17-47831826-INS-269 |
| chr17-47845991-INS-52 | chr17-47845991-INS-51 |
| chr17-47868907-INS-100 | chr17-47868907-INS-100 |
| chr17-47872170-INS-247 | chr17-47872170-INS-247 |
| chr17-47873217-INS-205 | chr17-47873217-INS-205 |
| chr17-47878934-INS-6442 | chr17-47878935-INS-6442 |

| | |
|---|---|
| chr17-47884836-INS-65 | chr17-47884836-INS-115 |
| chr17-47887044-INS-178 | chr17-47887044-INS-178 |
| chr17-47887664-INS-199 | chr17-47887664-INS-199 |
| chr17-47893235-INS-96 | chr17-47893235-INS-96 |
| chr17-47894162-INS-198 | chr17-47894162-INS-198 |
| chr17-47913826-INS-858 | chr17-47913828-INS-846 |
| chr17-47937337-INS-2837 | chr17-47937338-INS-2853 |
| chr17-47939556-INS-128 | chr17-47939556-INS-128 |
| chr17-47945122-INS-797 | chr17-47945122-INS-799 |
| chr17-47968960-INS-54 | chr17-47968960-INS-54 |
| chr17-48008393-INS-181 | chr17-48008393-INS-181 |
| chr17-48020860-INS-192 | chr17-48020855-INS-218 |
| chr17-48021810-INS-277 | chr17-48021810-INS-277 |
| chr17-48028563-INS-102 | chr17-48028563-INS-104 |
| chr17-48032988-INS-194 | chr17-48032988-INS-196 |
| chr17-48035089-INS-197 | chr17-48035089-INS-197 |
| chr17-48092973-INS-51 | chr17-48092974-INS-51 |
| chr17-48100140-INS-117 | chr17-48100101-INS-117 |
| chr17-48116904-INS-234 | chr17-48116915-INS-205 |
| chr17-48118608-INS-217 | chr17-48118608-INS-215 |
| chr17-48120266-INS-181 | chr17-48120266-INS-181 |
| chr17-48137113-INS-207 | chr17-48137113-INS-207 |
| chr17-48138975-INS-289 | chr17-48138975-INS-292 |
| chr17-48160650-INS-4490 | chr17-48160651-INS-4494 |
| chr17-48164157-INS-99 | chr17-48164157-INS-99 |
| chr17-48205582-INS-204 | chr17-48205582-INS-205 |
| chr17-48207140-INS-99 | chr17-48207140-INS-99 |
| chr17-48259664-INS-51 | chr17-48259664-INS-63 |
| chr17-48261093-INS-214 | chr17-48261093-INS-213 |
| chr17-48268021-INS-93 | chr17-48268022-INS-75 |
| chr17-48291143-INS-58 | chr17-48291143-INS-58 |
| chr17-48295651-INS-215 | chr17-48295651-INS-214 |
| chr17-48302028-INS-129 | chr17-48302017-INS-141 |
| chr17-48322894-INS-610 | chr17-48322894-INS-580 |
| chr17-48323615-INS-151 | chr17-48323615-INS-151 |
| chr17-48328346-INS-96 | chr17-48328350-INS-71 |
| chr17-48333519-INS-673 | chr17-48333519-INS-675 |
| chr17-48346434-INS-149 | chr17-48346434-INS-149 |
| chr17-48378258-INS-378 | chr17-48378250-INS-388 |
| chr17-48446294-INS-3947 | chr17-48446295-INS-3947 |
| chr17-48514113-INS-96 | chr17-48514113-INS-96 |
| chr17-48520318-INS-492 | chr17-48520318-INS-492 |
| chr17-48522722-INS-59 | chr17-48522721-INS-59 |
| chr17-48527842-INS-129 | chr17-48527842-INS-129 |
| chr17-48557452-INS-62 | chr17-48557452-INS-62 |
| chr17-48585858-INS-213 | chr17-48585862-INS-232 |
| chr17-48592372-INS-118 | chr17-48592371-INS-92 |
| chr17-48606689-INS-61 | chr17-48606666-INS-92 |
| chr17-48628990-INS-107 | chr17-48628990-INS-107 |
| chr17-48634444-INS-157 | chr17-48634444-INS-152 |
| chr17-48647929-INS-62 | chr17-48647900-INS-62 |
| chr17-48651601-INS-108 | chr17-48651601-INS-112 |
| chr17-48674710-INS-208 | chr17-48674710-INS-208 |
| chr17-48684732-INS-216 | chr17-48684737-INS-229 |
| chr17-48693092-INS-6430 | chr17-48693093-INS-6388 |
| chr17-48700668-INS-56 | chr17-48700668-INS-56 |
| chr17-48732665-INS-76 | chr17-48732665-INS-76 |
| chr17-48745628-INS-96 | chr17-48745624-INS-106 |
| chr17-48746663-INS-78 | chr17-48746685-INS-56 |
| chr17-48749117-INS-151 | chr17-48749117-INS-151 |
| chr17-48753895-INS-430 | chr17-48753895-INS-429 |
| chr17-48756497-INS-61 | chr17-48756501-INS-53 |
| chr17-48769840-INS-155 | chr17-48769838-INS-155 |
| chr17-48781972-INS-57 | chr17-48781972-INS-59 |
| chr17-48784406-INS-420 | chr17-48784406-INS-420 |
| chr17-48792450-INS-55 | chr17-48792450-INS-54 |
| chr17-48807276-INS-213 | chr17-48807276-INS-213 |
| chr17-48809369-INS-91 | chr17-48809369-INS-73 |
| chr17-48817332-INS-85 | chr17-48817332-INS-54 |
| chr17-48842601-INS-58 | chr17-48842571-INS-56 |
| chr17-48862346-INS-52 | chr17-48862346-INS-62 |
| chr17-48883860-INS-70 | chr17-48883850-INS-72 |
| chr17-48888594-INS-240 | chr17-48888594-INS-240 |
| chr17-48892716-INS-63 | chr17-48892714-INS-72 |
| chr17-48893848-INS-74 | chr17-48893849-INS-74 |
| chr17-48894763-INS-130 | chr17-48894764-INS-95 |
| chr17-48899517-INS-80 | chr17-48899517-INS-80 |

| | |
|---|---|
| chr17-48923973-INS-81 | chr17-48923970-INS-86 |
| chr17-48948491-INS-60 | chr17-48948487-INS-66 |
| chr17-48957378-INS-62 | chr17-48957378-INS-62 |
| chr17-48970547-INS-880 | chr17-48970547-INS-880 |
| chr17-48975545-INS-182 | chr17-48975545-INS-182 |
| chr17-49007329-INS-536 | chr17-49007329-INS-536 |
| chr17-49020277-INS-57 | chr17-49020277-INS-57 |
| chr17-49026617-INS-118 | chr17-49026617-INS-118 |
| chr17-49031134-INS-71 | chr17-49031141-INS-62 |
| chr17-49036344-INS-497 | chr17-49036344-INS-497 |
| chr17-49063033-INS-85 | chr17-49063033-INS-85 |
| chr17-49071278-INS-52 | chr17-49071278-INS-50 |
| chr17-49097357-INS-80 | chr17-49097357-INS-80 |
| chr17-49101152-INS-107 | chr17-49101152-INS-107 |
| chr17-49114255-INS-386 | chr17-49114255-INS-386 |
| chr17-49116413-INS-7284 | chr17-49116414-INS-7284 |
| chr17-49169006-INS-414 | chr17-49168990-INS-430 |
| chr17-49189696-INS-810 | chr17-49189696-INS-810 |
| chr17-49203394-INS-57 | chr17-49203394-INS-65 |
| chr17-49210518-INS-68 | chr17-49210518-INS-70 |
| chr17-49215684-INS-79 | chr17-49215676-INS-76 |
| chr17-49232962-INS-98 | chr17-49232962-INS-50 |
| chr17-49260691-INS-54 | chr17-49260691-INS-54 |
| chr17-49298152-INS-62 | chr17-49298152-INS-62 |
| chr17-49300261-INS-87 | chr17-49300261-INS-81 |
| chr17-49306205-INS-434 | chr17-49306205-INS-434 |
| chr17-49308687-INS-54 | chr17-49308687-INS-64 |
| chr17-49310064-INS-82 | chr17-49310064-INS-82 |
| chr17-49313605-INS-90 | chr17-49313605-INS-96 |
| chr17-49320437-INS-78 | chr17-49320437-INS-72 |
| chr17-49376283-INS-60 | chr17-49376283-INS-60 |
| chr17-49379275-INS-4844 | chr17-49379276-INS-4844 |
| chr17-49382018-INS-143 | chr17-49382018-INS-101 |
| chr17-49413023-INS-66 | chr17-49413052-INS-56 |
| chr17-49415521-INS-2737 | chr17-49415522-INS-2737 |
| chr17-49424812-INS-162 | chr17-49424812-INS-162 |
| chr17-49435742-INS-60 | chr17-49435742-INS-54 |
| chr17-49445837-INS-203 | chr17-49445837-INS-205 |
| chr17-49461678-INS-1893 | chr17-49461679-INS-1893 |
| chr17-49463033-INS-410 | chr17-49463033-INS-410 |
| chr17-49464147-INS-101 | chr17-49464147-INS-93 |
| chr17-49468497-INS-53 | chr17-49468497-INS-52 |
| chr17-49480149-INS-84 | chr17-49480149-INS-78 |
| chr17-49486138-INS-80 | chr17-49486138-INS-78 |
| chr17-49504590-INS-52 | chr17-49504590-INS-52 |
| chr17-49516037-INS-54 | chr17-49516037-INS-54 |
| chr17-49526528-INS-92 | chr17-49526528-INS-68 |
| chr17-49528360-INS-297 | chr17-49528360-INS-297 |
| chr17-49534948-INS-56 | chr17-49534948-INS-52 |
| chr17-49568327-INS-2816 | chr17-49568328-INS-2801 |
| chr17-49570474-INS-60 | chr17-49570474-INS-52 |
| chr17-49615939-INS-188 | chr17-49615933-INS-181 |
| chr17-49644691-INS-228 | chr17-49644691-INS-216 |
| chr17-49660711-INS-105 | chr17-49660711-INS-102 |
| chr17-49672772-INS-1344 | chr17-49672773-INS-1344 |
| chr17-49692275-INS-56 | chr17-49692249-INS-52 |
| chr17-49746042-INS-54 | chr17-49746042-INS-54 |
| chr17-49839697-INS-1898 | chr17-49839698-INS-1898 |
| chr17-49916859-INS-57 | chr17-49916818-INS-60 |
| chr17-49942706-INS-5598 | chr17-49942707-INS-5598 |
| chr17-49949103-INS-953 | chr17-49949103-INS-954 |
| chr17-49959066-INS-51 | chr17-49959067-INS-65 |
| chr17-49983080-INS-70 | chr17-49983066-INS-86 |
| chr17-49986457-INS-102 | chr17-49986457-INS-102 |
| chr17-50094219-INS-80 | chr17-50094219-INS-52 |
| chr17-50094353-INS-3841 | chr17-50094354-INS-3841 |
| chr17-50119939-INS-70 | chr17-50119977-INS-70 |
| chr17-50130200-INS-206 | chr17-50130200-INS-205 |
| chr17-50141107-INS-52 | chr17-50141119-INS-50 |
| chr17-50162713-INS-89 | chr17-50162713-INS-87 |
| chr17-50164718-INS-333 | chr17-50164719-INS-337 |
| chr17-50173663-INS-139 | chr17-50173654-INS-147 |
| chr17-50178414-INS-64 | chr17-50178414-INS-64 |
| chr17-50195729-INS-498 | chr17-50195729-INS-498 |
| chr17-50198000-INS-202 | chr17-50198000-INS-202 |
| chr17-50254736-INS-51 | chr17-50254761-INS-58 |
| chr17-50255955-INS-69 | chr17-50255955-INS-69 |

| | |
|---|---|
| chr17-50258316-INS-484 | chr17-50258316-INS-484 |
| chr17-50264953-INS-78 | chr17-50264953-INS-105 |
| chr17-50268888-INS-178 | chr17-50268888-INS-176 |
| chr17-50297400-INS-7132 | chr17-50297401-INS-7132 |
| chr17-50303526-INS-5845 | chr17-50303527-INS-5846 |
| chr17-50333174-INS-488 | chr17-50333174-INS-502 |
| chr17-50350908-INS-412 | chr17-50350909-INS-401 |
| chr17-50359529-INS-51 | chr17-50359529-INS-51 |
| chr17-50364939-INS-74 | chr17-50364939-INS-74 |
| chr17-50393349-INS-164 | chr17-50393349-INS-158 |
| chr17-50412679-INS-2552 | chr17-50412680-INS-2553 |
| chr17-50423975-INS-368 | chr17-50423975-INS-368 |
| chr17-50439941-INS-415 | chr17-50439941-INS-415 |
| chr17-50465492-INS-78 | chr17-50465492-INS-88 |
| chr17-50477628-INS-53 | chr17-50477628-INS-53 |
| chr17-50499069-INS-7097 | chr17-50499065-INS-7098 |
| chr17-50524913-INS-96 | chr17-50524913-INS-96 |
| chr17-50531020-INS-93 | chr17-50531020-INS-93 |
| chr17-50554497-INS-60 | chr17-50554497-INS-52 |
| chr17-50575557-INS-4623 | chr17-50575558-INS-4623 |
| chr17-50576545-INS-2507 | chr17-50576546-INS-2505 |
| chr17-50618528-INS-999 | chr17-50618528-INS-999 |
| chr17-50622500-INS-77 | chr17-50622500-INS-50 |
| chr17-50629146-INS-410 | chr17-50629146-INS-410 |
| chr17-50633446-INS-196 | chr17-50633446-INS-196 |
| chr17-50652908-INS-368 | chr17-50652883-INS-366 |
| chr17-50661211-INS-69 | chr17-50661217-INS-53 |
| chr17-50670535-INS-74 | chr17-50670535-INS-104 |
| chr17-50673691-INS-250 | chr17-50673691-INS-258 |
| chr17-50699319-INS-59 | chr17-50699319-INS-59 |
| chr17-50707967-INS-152 | chr17-50707966-INS-151 |
| chr17-50721033-INS-3789 | chr17-50721034-INS-3790 |
| chr17-50746198-INS-107 | chr17-50746198-INS-106 |
| chr17-50762046-INS-70 | chr17-50762046-INS-66 |
| chr17-50768858-INS-100 | chr17-50768875-INS-101 |
| chr17-50803308-INS-220 | chr17-50803308-INS-220 |
| chr17-50820816-INS-133 | chr17-50820816-INS-133 |
| chr17-50823385-INS-58 | chr17-50823385-INS-58 |
| chr17-50836561-INS-387 | chr17-50836561-INS-367 |
| chr17-50882755-INS-266 | chr17-50882755-INS-266 |
| chr17-50890551-INS-1117 | chr17-50890552-INS-1117 |
| chr17-50892473-INS-71 | chr17-50892468-INS-65 |
| chr17-50912518-INS-7115 | chr17-50912519-INS-7115 |
| chr17-50921282-INS-54 | chr17-50921282-INS-66 |
| chr17-50923269-INS-131 | chr17-50923269-INS-131 |
| chr17-50988518-INS-715 | chr17-50988518-INS-718 |
| chr17-51011972-INS-2707 | chr17-51011973-INS-2707 |
| chr17-51020124-INS-53 | chr17-51020103-INS-52 |
| chr17-51036157-INS-226 | chr17-51036157-INS-226 |
| chr17-51045154-INS-3562 | chr17-51045155-INS-3562 |
| chr17-51061112-INS-6241 | chr17-51061113-INS-6241 |
| chr17-51110501-INS-108 | chr17-51110501-INS-108 |
| chr17-51134370-INS-93 | chr17-51134371-INS-97 |
| chr17-51144994-INS-7328 | chr17-51144995-INS-7331 |
| chr17-51155430-INS-64 | chr17-51155430-INS-64 |
| chr17-51167546-INS-64 | chr17-51167546-INS-66 |
| chr17-51176957-INS-6446 | chr17-51176958-INS-6446 |
| chr17-51234024-INS-212 | chr17-51234024-INS-212 |
| chr17-51267268-INS-237 | chr17-51267269-INS-237 |
| chr17-51322061-INS-50 | chr17-51322061-INS-60 |
| chr17-51360193-INS-447 | chr17-51360193-INS-447 |
| chr17-51393068-INS-1846 | chr17-51393069-INS-1848 |
| chr17-51451816-INS-52 | chr17-51451816-INS-52 |
| chr17-51491166-INS-86 | chr17-51491164-INS-67 |
| chr17-51513540-INS-108 | chr17-51513541-INS-110 |
| chr17-51522646-INS-406 | chr17-51522646-INS-422 |
| chr17-51529091-INS-58 | chr17-51529091-INS-58 |
| chr17-51548283-INS-205 | chr17-51548283-INS-205 |
| chr17-51564488-INS-326 | chr17-51564488-INS-326 |
| chr17-51598888-INS-58 | chr17-51598888-INS-58 |
| chr17-51636732-INS-6628 | chr17-51636733-INS-6631 |
| chr17-51637979-INS-105 | chr17-51637979-INS-60 |
| chr17-51684136-INS-217 | chr17-51684125-INS-220 |
| chr17-51716925-INS-58 | chr17-51716905-INS-68 |
| chr17-51723642-INS-64 | chr17-51723642-INS-80 |
| chr17-51746170-INS-595 | chr17-51746170-INS-595 |
| chr17-51748294-INS-1104 | chr17-51748295-INS-1104 |

| | |
|---|---|
| chr17-51772368-INS-911 | chr17-51772368-INS-911 |
| chr17-51775394-INS-398 | chr17-51775394-INS-400 |
| chr17-51781312-INS-1569 | chr17-51781313-INS-1563 |
| chr17-51859802-INS-62 | chr17-51859802-INS-62 |
| chr17-51904047-INS-398 | chr17-51904047-INS-398 |
| chr17-51904922-INS-52 | chr17-51904922-INS-76 |
| chr17-51926596-INS-472 | chr17-51926596-INS-498 |
| chr17-52000357-INS-78 | chr17-52000357-INS-78 |
| chr17-52003486-INS-436 | chr17-52003486-INS-436 |
| chr17-52008544-INS-231 | chr17-52008544-INS-238 |
| chr17-52021304-INS-219 | chr17-52021302-INS-219 |
| chr17-52051377-INS-143 | chr17-52051377-INS-100 |
| chr17-52062996-INS-332 | chr17-52062996-INS-352 |
| chr17-52145046-INS-57 | chr17-52145046-INS-57 |
| chr17-52162715-INS-76 | chr17-52162715-INS-78 |
| chr17-52168655-INS-62 | chr17-52168655-INS-68 |
| chr17-52195256-INS-56 | chr17-52195256-INS-56 |
| chr17-52196283-INS-250 | chr17-52196283-INS-250 |
| chr17-52196463-INS-1805 | chr17-52196481-INS-1788 |
| chr17-52202922-INS-56 | chr17-52202926-INS-50 |
| chr17-52204654-INS-85 | chr17-52204654-INS-85 |
| chr17-52246358-INS-79 | chr17-52246358-INS-79 |
| chr17-52250910-INS-7103 | chr17-52250911-INS-7104 |
| chr17-52264683-INS-588 | chr17-52264683-INS-588 |
| chr17-52269309-INS-65 | chr17-52269287-INS-74 |
| chr17-52269702-INS-2433 | chr17-52269703-INS-2433 |
| chr17-52288221-INS-653 | chr17-52288221-INS-653 |
| chr17-52309263-INS-1270 | chr17-52309264-INS-1270 |
| chr17-52315592-INS-63 | chr17-52315593-INS-60 |
| chr17-52324553-INS-6449 | chr17-52324554-INS-6449 |
| chr17-52336310-INS-156 | chr17-52336297-INS-193 |
| chr17-52339915-INS-225 | chr17-52339957-INS-197 |
| chr17-52356513-INS-653 | chr17-52356513-INS-653 |
| chr17-52388403-INS-2164 | chr17-52388404-INS-2170 |
| chr17-52394557-INS-102 | chr17-52394565-INS-52 |
| chr17-52409431-INS-143 | chr17-52409431-INS-143 |
| chr17-52419709-INS-120 | chr17-52419697-INS-123 |
| chr17-52450616-INS-903 | chr17-52450616-INS-903 |
| chr17-52470309-INS-432 | chr17-52470309-INS-432 |
| chr17-52513098-INS-442 | chr17-52513098-INS-442 |
| chr17-52546650-INS-59 | chr17-52546647-INS-74 |
| chr17-52547762-INS-2813 | chr17-52547763-INS-2813 |
| chr17-52554700-INS-80 | chr17-52554700-INS-80 |
| chr17-52604779-INS-1001 | chr17-52604780-INS-1001 |
| chr17-52607473-INS-345 | chr17-52607473-INS-345 |
| chr17-52608881-INS-232 | chr17-52608881-INS-232 |
| chr17-52609326-INS-6706 | chr17-52609327-INS-6706 |
| chr17-52642789-INS-7043 | chr17-52642791-INS-7041 |
| chr17-52657551-INS-203 | chr17-52657538-INS-243 |
| chr17-52674639-INS-184 | chr17-52674639-INS-196 |
| chr17-52677888-INS-1669 | chr17-52677889-INS-1669 |
| chr17-52678795-INS-366 | chr17-52678795-INS-327 |
| chr17-52701401-INS-210 | chr17-52701401-INS-210 |
| chr17-52752229-INS-63 | chr17-52752250-INS-60 |
| chr17-52774050-INS-187 | chr17-52774051-INS-187 |
| chr17-52808348-INS-54 | chr17-52808348-INS-56 |
| chr17-52809976-INS-95 | chr17-52809976-INS-95 |
| chr17-52827333-INS-63 | chr17-52827333-INS-66 |
| chr17-52847189-INS-198 | chr17-52847189-INS-198 |
| chr17-52849425-INS-60 | chr17-52849425-INS-60 |
| chr17-52857875-INS-64 | chr17-52857875-INS-65 |
| chr17-52860352-INS-896 | chr17-52860352-INS-896 |
| chr17-52862917-INS-6380 | chr17-52862918-INS-6381 |
| chr17-52863135-INS-106 | chr17-52863135-INS-107 |
| chr17-52866333-INS-69 | chr17-52866349-INS-67 |
| chr17-52872412-INS-106 | chr17-52872412-INS-105 |
| chr17-52874067-INS-2083 | chr17-52874068-INS-2083 |
| chr17-52902917-INS-107 | chr17-52902916-INS-131 |
| chr17-52979218-INS-1676 | chr17-52979219-INS-1676 |
| chr17-52982861-INS-184 | chr17-52982861-INS-184 |
| chr17-52997626-INS-109 | chr17-52997622-INS-112 |
| chr17-52998304-INS-2283 | chr17-52998306-INS-2284 |
| chr17-53034168-INS-1394 | chr17-53034169-INS-1398 |
| chr17-53038798-INS-238 | chr17-53038798-INS-256 |
| chr17-53088024-INS-618 | chr17-53088024-INS-617 |
| chr17-53094032-INS-97 | chr17-53094032-INS-104 |
| chr17-53098860-INS-201 | chr17-53098846-INS-210 |

| | |
|---|---|
| chr17-53117256-INS-94 | chr17-53117256-INS-94 |
| chr17-53119277-INS-113 | chr17-53119277-INS-113 |
| chr17-53121783-INS-102 | chr17-53121783-INS-102 |
| chr17-53122621-INS-53 | chr17-53122621-INS-53 |
| chr17-53125919-INS-6765 | chr17-53125920-INS-6765 |
| chr17-53127504-INS-51 | chr17-53127508-INS-61 |
| chr17-53130827-INS-53 | chr17-53130827-INS-53 |
| chr17-53167975-INS-80 | chr17-53167973-INS-84 |
| chr17-53176613-INS-228 | chr17-53176613-INS-227 |
| chr17-53206551-INS-157 | chr17-53206551-INS-157 |
| chr17-53219456-INS-74 | chr17-53219452-INS-95 |
| chr17-53224640-INS-745 | chr17-53224640-INS-745 |
| chr17-53247310-INS-196 | chr17-53247310-INS-196 |
| chr17-53251992-INS-111 | chr17-53251992-INS-111 |
| chr17-53259859-INS-170 | chr17-53259859-INS-170 |
| chr17-53266068-INS-729 | chr17-53266068-INS-727 |
| chr17-53287145-INS-113 | chr17-53287145-INS-85 |
| chr17-53288243-INS-92 | chr17-53288243-INS-92 |
| chr17-53306909-INS-502 | chr17-53306910-INS-502 |
| chr17-53307496-INS-441 | chr17-53307497-INS-441 |
| chr17-53324203-INS-1530 | chr17-53324204-INS-1531 |
| chr17-53349100-INS-957 | chr17-53349100-INS-957 |
| chr17-53352922-INS-460 | chr17-53352932-INS-468 |
| chr17-53356025-INS-445 | chr17-53356025-INS-445 |
| chr17-53361124-INS-394 | chr17-53361124-INS-394 |
| chr17-53386982-INS-759 | chr17-53386982-INS-808 |
| chr17-53404506-INS-53 | chr17-53404506-INS-53 |
| chr17-53438863-INS-52 | chr17-53438863-INS-52 |
| chr17-53459577-INS-66 | chr17-53459577-INS-66 |
| chr17-53468343-INS-128 | chr17-53468343-INS-79 |
| chr17-53475031-INS-606 | chr17-53475031-INS-606 |
| chr17-53476917-INS-2507 | chr17-53476918-INS-2507 |
| chr17-53502029-INS-204 | chr17-53502029-INS-201 |
| chr17-53506564-INS-613 | chr17-53506564-INS-615 |
| chr17-53508348-INS-62 | chr17-53508353-INS-63 |
| chr17-53508535-INS-6370 | chr17-53508536-INS-6371 |
| chr17-53579745-INS-62 | chr17-53579745-INS-62 |
| chr17-53609436-INS-71 | chr17-53609436-INS-75 |
| chr17-53621140-INS-6285 | chr17-53621141-INS-6285 |
| chr17-53632169-INS-162 | chr17-53632171-INS-166 |
| chr17-53636146-INS-180 | chr17-53636163-INS-157 |
| chr17-53638375-INS-352 | chr17-53638375-INS-352 |
| chr17-53644303-INS-75 | chr17-53644318-INS-79 |
| chr17-53668660-INS-2789 | chr17-53668661-INS-2789 |
| chr17-53699098-INS-823 | chr17-53699098-INS-824 |
| chr17-53704935-INS-108 | chr17-53704935-INS-108 |
| chr17-53716587-INS-193 | chr17-53716587-INS-194 |
| chr17-53723858-INS-575 | chr17-53723858-INS-575 |
| chr17-53726052-INS-1171 | chr17-53726053-INS-1195 |
| chr17-53728471-INS-168 | chr17-53728471-INS-168 |
| chr17-53735207-INS-84 | chr17-53735207-INS-62 |
| chr17-53745501-INS-67 | chr17-53745501-INS-67 |
| chr17-53751516-INS-76 | chr17-53751519-INS-64 |
| chr17-53762489-INS-63 | chr17-53762489-INS-63 |
| chr17-53767060-INS-464 | chr17-53767060-INS-464 |
| chr17-53771330-INS-64 | chr17-53771330-INS-64 |
| chr17-53781450-INS-65 | chr17-53781458-INS-86 |
| chr17-53807118-INS-109 | chr17-53807118-INS-130 |
| chr17-53821832-INS-195 | chr17-53821832-INS-195 |
| chr17-53851878-INS-54 | chr17-53851873-INS-54 |
| chr17-53868801-INS-107 | chr17-53868801-INS-107 |
| chr17-53883084-INS-231 | chr17-53883084-INS-231 |
| chr17-53887846-INS-199 | chr17-53887847-INS-198 |
| chr17-53889246-INS-343 | chr17-53889246-INS-343 |
| chr17-53905929-INS-143 | chr17-53905929-INS-143 |
| chr17-53906481-INS-363 | chr17-53906481-INS-362 |
| chr17-53921505-INS-113 | chr17-53921505-INS-108 |
| chr17-53924031-INS-70 | chr17-53924031-INS-70 |
| chr17-53933518-INS-136 | chr17-53933518-INS-136 |
| chr17-53964460-INS-241 | chr17-53964460-INS-241 |
| chr17-53968847-INS-281 | chr17-53968847-INS-283 |
| chr17-53991923-INS-195 | chr17-53991923-INS-195 |
| chr17-53999465-INS-108 | chr17-53999465-INS-108 |
| chr17-54000724-INS-200 | chr17-54000724-INS-200 |
| chr17-54007498-INS-1238 | chr17-54007499-INS-1239 |
| chr17-54008674-INS-56 | chr17-54008674-INS-61 |
| chr17-54012341-INS-572 | chr17-54012341-INS-572 |

| | |
|---|---|
| chr17-54014049-INS-132 | chr17-54014018-INS-131 |
| chr17-54020305-INS-57 | chr17-54020301-INS-74 |
| chr17-54021568-INS-100 | chr17-54021568-INS-100 |
| chr17-54025375-INS-70 | chr17-54025375-INS-70 |
| chr17-54037470-INS-51 | chr17-54037470-INS-53 |
| chr17-54093326-INS-68 | chr17-54093326-INS-68 |
| chr17-54130213-INS-80 | chr17-54130211-INS-83 |
| chr17-54131605-INS-58 | chr17-54131605-INS-58 |
| chr17-54132849-INS-140 | chr17-54132849-INS-140 |
| chr17-54151864-INS-453 | chr17-54151864-INS-456 |
| chr17-54167326-INS-74 | chr17-54167326-INS-74 |
| chr17-54169280-INS-69 | chr17-54169280-INS-69 |
| chr17-54184064-INS-785 | chr17-54184064-INS-786 |
| chr17-54196169-INS-271 | chr17-54196169-INS-271 |
| chr17-54196559-INS-6468 | chr17-54196560-INS-6468 |
| chr17-54203396-INS-334 | chr17-54203396-INS-334 |
| chr17-54218096-INS-832 | chr17-54218096-INS-830 |
| chr17-54228628-INS-765 | chr17-54228628-INS-765 |
| chr17-54244265-INS-198 | chr17-54244265-INS-198 |
| chr17-54271301-INS-982 | chr17-54271301-INS-992 |
| chr17-54277597-INS-366 | chr17-54277597-INS-366 |
| chr17-54279974-INS-137 | chr17-54279974-INS-136 |
| chr17-54286984-INS-527 | chr17-54286984-INS-528 |
| chr17-54305091-INS-6249 | chr17-54305092-INS-6249 |
| chr17-54326023-INS-75 | chr17-54326023-INS-75 |
| chr17-54332220-INS-79 | chr17-54332218-INS-88 |
| chr17-54336148-INS-105 | chr17-54336148-INS-66 |
| chr17-54337865-INS-73 | chr17-54337865-INS-73 |
| chr17-54368571-INS-946 | chr17-54368571-INS-942 |
| chr17-54377381-INS-6184 | chr17-54377382-INS-6183 |
| chr17-54378660-INS-56 | chr17-54378672-INS-52 |
| chr17-54391407-INS-58 | chr17-54391407-INS-58 |
| chr17-54400059-INS-51 | chr17-54400059-INS-55 |
| chr17-54450027-INS-113 | chr17-54450009-INS-104 |
| chr17-54451622-INS-481 | chr17-54451622-INS-485 |
| chr17-54453084-INS-263 | chr17-54453084-INS-263 |
| chr17-54454873-INS-125 | chr17-54454905-INS-114 |
| chr17-54470540-INS-60 | chr17-54470540-INS-60 |
| chr17-54474051-INS-6590 | chr17-54474052-INS-6590 |
| chr17-54477027-INS-51 | chr17-54477027-INS-51 |
| chr17-54481547-INS-60 | chr17-54481547-INS-76 |
| chr17-54493012-INS-840 | chr17-54493012-INS-840 |
| chr17-54514069-INS-80 | chr17-54514069-INS-101 |
| chr17-54531080-INS-60 | chr17-54531080-INS-61 |
| chr17-54535737-INS-66 | chr17-54535737-INS-58 |
| chr17-54598463-INS-122 | chr17-54598496-INS-96 |
| chr17-54618503-INS-462 | chr17-54618503-INS-462 |
| chr17-54627592-INS-903 | chr17-54627598-INS-902 |
| chr17-54644827-INS-728 | chr17-54644827-INS-727 |
| chr17-54667035-INS-210 | chr17-54667035-INS-210 |
| chr17-54668822-INS-75 | chr17-54668828-INS-55 |
| chr17-54692585-INS-5033 | chr17-54692586-INS-5033 |
| chr17-54695901-INS-62 | chr17-54695907-INS-54 |
| chr17-54699409-INS-347 | chr17-54699409-INS-347 |
| chr17-54705606-INS-787 | chr17-54705606-INS-784 |
| chr17-54737366-INS-200 | chr17-54737366-INS-200 |
| chr17-54757640-INS-7083 | chr17-54757641-INS-7068 |
| chr17-54757749-INS-318 | chr17-54757749-INS-318 |
| chr17-54766012-INS-6689 | chr17-54766013-INS-6689 |
| chr17-54778032-INS-196 | chr17-54778032-INS-196 |
| chr17-54792669-INS-1510 | chr17-54792670-INS-1515 |
| chr17-54795702-INS-220 | chr17-54795702-INS-220 |
| chr17-54808032-INS-60 | chr17-54808032-INS-60 |
| chr17-54815354-INS-431 | chr17-54815354-INS-431 |
| chr17-54845912-INS-6440 | chr17-54845913-INS-6440 |
| chr17-54886033-INS-204 | chr17-54886033-INS-204 |
| chr17-54892726-INS-75 | chr17-54892726-INS-75 |
| chr17-54905148-INS-7076 | chr17-54905149-INS-7076 |
| chr17-54906458-INS-272 | chr17-54906458-INS-272 |
| chr17-54911528-INS-283 | chr17-54911528-INS-283 |
| chr17-54932142-INS-74 | chr17-54932109-INS-85 |
| chr17-54957905-INS-7175 | chr17-54957906-INS-7175 |
| chr17-55003608-INS-52 | chr17-55003608-INS-56 |
| chr17-55011321-INS-61 | chr17-55011321-INS-61 |
| chr17-55015246-INS-166 | chr17-55015246-INS-166 |
| chr17-55038311-INS-58 | chr17-55038310-INS-73 |
| chr17-55042635-INS-381 | chr17-55042635-INS-381 |

| | |
|---|---|
| chr17-55044800-INS-572 | chr17-55044800-INS-574 |
| chr17-55049791-INS-523 | chr17-55049791-INS-522 |
| chr17-55101493-INS-281 | chr17-55101493-INS-259 |
| chr17-55115925-INS-289 | chr17-55115925-INS-287 |
| chr17-55124275-INS-86 | chr17-55124273-INS-70 |
| chr17-55141964-INS-58 | chr17-55141928-INS-91 |
| chr17-55144786-INS-81 | chr17-55144786-INS-81 |
| chr17-55201558-INS-145 | chr17-55201560-INS-154 |
| chr17-55247200-INS-70 | chr17-55247200-INS-60 |
| chr17-55269416-INS-136 | chr17-55269416-INS-163 |
| chr17-55287643-INS-148 | chr17-55287643-INS-136 |
| chr17-55297536-INS-430 | chr17-55297536-INS-430 |
| chr17-55343348-INS-7169 | chr17-55343349-INS-7170 |
| chr17-55373177-INS-199 | chr17-55373177-INS-200 |
| chr17-55400191-INS-158 | chr17-55400191-INS-158 |
| chr17-55403691-INS-3632 | chr17-55403692-INS-3635 |
| chr17-55452498-INS-387 | chr17-55452498-INS-387 |
| chr17-55517796-INS-125 | chr17-55517796-INS-125 |
| chr17-55518072-INS-535 | chr17-55518072-INS-535 |
| chr17-55543480-INS-329 | chr17-55543480-INS-329 |
| chr17-55544875-INS-490 | chr17-55544875-INS-490 |
| chr17-55546710-INS-619 | chr17-55546711-INS-618 |
| chr17-55547503-INS-4511 | chr17-55547504-INS-4511 |
| chr17-55568564-INS-204 | chr17-55568564-INS-204 |
| chr17-55597733-INS-76 | chr17-55597733-INS-50 |
| chr17-55600781-INS-59 | chr17-55600781-INS-59 |
| chr17-55625190-INS-311 | chr17-55625160-INS-324 |
| chr17-55630091-INS-64 | chr17-55630091-INS-68 |
| chr17-55633710-INS-1803 | chr17-55633711-INS-1803 |
| chr17-55640727-INS-56 | chr17-55640727-INS-50 |
| chr17-55651642-INS-53 | chr17-55651626-INS-80 |
| chr17-55664268-INS-806 | chr17-55664268-INS-806 |
| chr17-55668035-INS-610 | chr17-55668035-INS-610 |
| chr17-55672258-INS-88 | chr17-55672259-INS-78 |
| chr17-55677690-INS-890 | chr17-55677690-INS-890 |
| chr17-55735748-INS-61 | chr17-55735748-INS-61 |
| chr17-55781686-INS-375 | chr17-55781693-INS-371 |
| chr17-55784508-INS-2362 | chr17-55784509-INS-2363 |
| chr17-55788546-INS-2340 | chr17-55788547-INS-2340 |
| chr17-55791036-INS-450 | chr17-55791036-INS-450 |
| chr17-55793366-INS-54 | chr17-55793366-INS-57 |
| chr17-55796504-INS-60 | chr17-55796487-INS-78 |
| chr17-55813504-INS-366 | chr17-55813504-INS-366 |
| chr17-55822678-INS-193 | chr17-55822678-INS-197 |
| chr17-55834588-INS-1102 | chr17-55834589-INS-1107 |
| chr17-55835404-INS-58 | chr17-55835404-INS-58 |
| chr17-55895255-INS-1321 | chr17-55895256-INS-1321 |
| chr17-55940651-INS-81 | chr17-55940651-INS-81 |
| chr17-55979658-INS-354 | chr17-55979658-INS-354 |
| chr17-56022238-INS-5338 | chr17-56022239-INS-5338 |
| chr17-56023372-INS-353 | chr17-56023372-INS-353 |
| chr17-56090045-INS-413 | chr17-56090045-INS-412 |
| chr17-56100748-INS-558 | chr17-56100748-INS-556 |
| chr17-56101480-INS-329 | chr17-56101480-INS-329 |
| chr17-56153558-INS-662 | chr17-56153558-INS-662 |
| chr17-56195761-INS-298 | chr17-56195764-INS-297 |
| chr17-56209812-INS-1548 | chr17-56209813-INS-1554 |
| chr17-56223864-INS-216 | chr17-56223864-INS-216 |
| chr17-56230107-INS-207 | chr17-56230107-INS-207 |
| chr17-56233477-INS-923 | chr17-56233477-INS-923 |
| chr17-56243226-INS-5352 | chr17-56243227-INS-5352 |
| chr17-56245362-INS-66 | chr17-56245362-INS-66 |
| chr17-56247793-INS-209 | chr17-56247793-INS-207 |
| chr17-56272649-INS-96 | chr17-56272650-INS-89 |
| chr17-56282161-INS-100 | chr17-56282162-INS-72 |
| chr17-56306353-INS-221 | chr17-56306353-INS-230 |
| chr17-56339674-INS-100 | chr17-56339674-INS-100 |
| chr17-56346225-INS-413 | chr17-56346225-INS-413 |
| chr17-56417641-INS-70 | chr17-56417625-INS-66 |
| chr17-56457728-INS-105 | chr17-56457728-INS-101 |
| chr17-56491860-INS-199 | chr17-56491860-INS-199 |
| chr17-56500558-INS-104 | chr17-56500557-INS-118 |
| chr17-56514896-INS-164 | chr17-56514896-INS-164 |
| chr17-56520671-INS-71 | chr17-56520672-INS-57 |
| chr17-56543279-INS-70 | chr17-56543259-INS-109 |
| chr17-56554140-INS-198 | chr17-56554140-INS-195 |
| chr17-56582349-INS-211 | chr17-56582349-INS-211 |

| | |
|---|---|
| chr17-56585671-INS-182 | chr17-56585671-INS-181 |
| chr17-56596253-INS-226 | chr17-56596237-INS-214 |
| chr17-56602574-INS-80 | chr17-56602574-INS-80 |
| chr17-56636737-INS-1187 | chr17-56636738-INS-1200 |
| chr17-56785813-INS-67 | chr17-56785813-INS-52 |
| chr17-56791414-INS-60 | chr17-56791414-INS-60 |
| chr17-56798573-INS-82 | chr17-56798587-INS-84 |
| chr17-56836119-INS-1113 | chr17-56836105-INS-1122 |
| chr17-56838717-INS-200 | chr17-56838717-INS-200 |
| chr17-56840192-INS-484 | chr17-56840192-INS-484 |
| chr17-56841077-INS-277 | chr17-56841077-INS-277 |
| chr17-56846742-INS-199 | chr17-56846742-INS-199 |
| chr17-56867771-INS-72 | chr17-56867771-INS-72 |
| chr17-56952928-INS-204 | chr17-56952929-INS-204 |
| chr17-56956722-INS-62 | chr17-56956722-INS-62 |
| chr17-56962230-INS-88 | chr17-56962230-INS-94 |
| chr17-57025396-INS-205 | chr17-57025396-INS-221 |
| chr17-57084688-INS-82 | chr17-57084688-INS-82 |
| chr17-57173105-INS-186 | chr17-57173105-INS-182 |
| chr17-57182318-INS-211 | chr17-57182318-INS-211 |
| chr17-57185299-INS-343 | chr17-57185299-INS-343 |
| chr17-57271957-INS-264 | chr17-57271957-INS-260 |
| chr17-57289743-INS-199 | chr17-57289743-INS-199 |
| chr17-57294132-INS-152 | chr17-57294132-INS-155 |
| chr17-57301086-INS-142 | chr17-57301086-INS-134 |
| chr17-57308128-INS-211 | chr17-57308128-INS-211 |
| chr17-57311889-INS-97 | chr17-57311889-INS-97 |
| chr17-57314606-INS-56 | chr17-57314606-INS-56 |
| chr17-57323997-INS-199 | chr17-57323997-INS-198 |
| chr17-57342140-INS-208 | chr17-57342140-INS-208 |
| chr17-57349881-INS-83 | chr17-57349883-INS-67 |
| chr17-57357456-INS-61 | chr17-57357456-INS-61 |
| chr17-57380283-INS-64 | chr17-57380283-INS-60 |
| chr17-57391804-INS-139 | chr17-57391804-INS-139 |
| chr17-57396619-INS-643 | chr17-57396619-INS-643 |
| chr17-57425873-INS-52 | chr17-57425873-INS-52 |
| chr17-57437597-INS-210 | chr17-57437597-INS-210 |
| chr17-57439277-INS-60 | chr17-57439277-INS-60 |
| chr17-57468090-INS-177 | chr17-57468090-INS-177 |
| chr17-57470339-INS-73 | chr17-57470339-INS-73 |
| chr17-57487240-INS-95 | chr17-57487240-INS-52 |
| chr17-57523020-INS-79 | chr17-57523015-INS-79 |
| chr17-57527785-INS-75 | chr17-57527785-INS-75 |
| chr17-57550896-INS-105 | chr17-57550896-INS-115 |
| chr17-57601199-INS-371 | chr17-57601182-INS-377 |
| chr17-57607651-INS-64 | chr17-57607651-INS-64 |
| chr17-57642227-INS-52 | chr17-57642227-INS-56 |
| chr17-57662100-INS-56 | chr17-57662100-INS-56 |
| chr17-57668219-INS-70 | chr17-57668219-INS-116 |
| chr17-57710534-INS-573 | chr17-57710534-INS-573 |
| chr17-57793615-INS-106 | chr17-57793615-INS-114 |
| chr17-57807407-INS-6225 | chr17-57807408-INS-6225 |
| chr17-57808808-INS-503 | chr17-57808808-INS-492 |
| chr17-57828165-INS-54 | chr17-57828165-INS-56 |
| chr17-57932402-INS-63 | chr17-57932402-INS-65 |
| chr17-57974765-INS-76 | chr17-57974765-INS-76 |
| chr17-57978203-INS-963 | chr17-57978188-INS-964 |
| chr17-57979829-INS-80 | chr17-57979829-INS-82 |
| chr17-58015316-INS-215 | chr17-58015316-INS-215 |
| chr17-58033642-INS-57 | chr17-58033642-INS-59 |
| chr17-58047903-INS-366 | chr17-58047903-INS-366 |
| chr17-58070955-INS-879 | chr17-58070955-INS-877 |
| chr17-58073876-INS-66 | chr17-58073876-INS-62 |
| chr17-58109154-INS-80 | chr17-58109131-INS-63 |
| chr17-58112748-INS-51 | chr17-58112748-INS-51 |
| chr17-58117225-INS-179 | chr17-58117225-INS-177 |
| chr17-58158480-INS-202 | chr17-58158480-INS-202 |
| chr17-58165761-INS-350 | chr17-58165761-INS-351 |
| chr17-58169928-INS-72 | chr17-58169928-INS-73 |
| chr17-58197681-INS-130 | chr17-58197681-INS-130 |
| chr17-58234137-INS-72 | chr17-58234137-INS-78 |
| chr17-58235708-INS-404 | chr17-58235713-INS-405 |
| chr17-58244830-INS-96 | chr17-58244830-INS-102 |
| chr17-58250830-INS-4886 | chr17-58250831-INS-4886 |
| chr17-58335223-INS-99 | chr17-58335223-INS-99 |
| chr17-58343786-INS-392 | chr17-58343786-INS-392 |
| chr17-58346876-INS-55 | chr17-58346885-INS-94 |

| | |
|---|---|
| chr17-58353253-INS-945 | chr17-58353253-INS-945 |
| chr17-58369523-INS-62 | chr17-58369523-INS-70 |
| chr17-58370773-INS-60 | chr17-58370783-INS-51 |
| chr17-58384463-INS-6895 | chr17-58384464-INS-6897 |
| chr17-58386857-INS-114 | chr17-58386857-INS-114 |
| chr17-58437016-INS-57 | chr17-58437016-INS-57 |
| chr17-58440078-INS-57 | chr17-58440078-INS-57 |
| chr17-58442628-INS-54 | chr17-58442628-INS-54 |
| chr17-58459385-INS-766 | chr17-58459385-INS-733 |
| chr17-58463288-INS-67 | chr17-58463288-INS-71 |
| chr17-58475178-INS-114 | chr17-58475178-INS-123 |
| chr17-58501993-INS-77 | chr17-58501993-INS-77 |
| chr17-58508499-INS-52 | chr17-58508499-INS-52 |
| chr17-58550416-INS-301 | chr17-58550416-INS-301 |
| chr17-58580217-INS-528 | chr17-58580230-INS-545 |
| chr17-58644487-INS-59 | chr17-58644487-INS-59 |
| chr17-58679462-INS-203 | chr17-58679462-INS-203 |
| chr17-58683506-INS-99 | chr17-58683506-INS-99 |
| chr17-58686594-INS-214 | chr17-58686594-INS-192 |
| chr17-58704635-INS-6336 | chr17-58704636-INS-6336 |
| chr17-58730496-INS-74 | chr17-58730496-INS-74 |
| chr17-58732261-INS-82 | chr17-58732291-INS-76 |
| chr17-58736393-INS-535 | chr17-58736393-INS-545 |
| chr17-58741681-INS-858 | chr17-58741681-INS-858 |
| chr17-58758674-INS-69 | chr17-58758686-INS-63 |
| chr17-58776623-INS-76 | chr17-58776623-INS-81 |
| chr17-58788070-INS-122 | chr17-58788070-INS-122 |
| chr17-58789166-INS-143 | chr17-58789166-INS-136 |
| chr17-58826478-INS-3557 | chr17-58826479-INS-3554 |
| chr17-58865708-INS-7950 | chr17-58865709-INS-7952 |
| chr17-58873454-INS-407 | chr17-58873454-INS-407 |
| chr17-58874955-INS-65 | chr17-58874955-INS-66 |
| chr17-58875414-INS-776 | chr17-58875414-INS-776 |
| chr17-58876624-INS-76 | chr17-58876624-INS-76 |
| chr17-58887679-INS-169 | chr17-58887679-INS-167 |
| chr17-58902977-INS-66 | chr17-58902977-INS-66 |
| chr17-58905334-INS-54 | chr17-58905334-INS-56 |
| chr17-58918394-INS-190 | chr17-58918394-INS-190 |
| chr17-58920399-INS-63 | chr17-58920399-INS-51 |
| chr17-58936397-INS-67 | chr17-58936392-INS-67 |
| chr17-58938130-INS-68 | chr17-58938129-INS-52 |
| chr17-58949596-INS-60 | chr17-58949596-INS-63 |
| chr17-58960596-INS-221 | chr17-58960608-INS-225 |
| chr17-58964436-INS-692 | chr17-58964436-INS-692 |
| chr17-58973642-INS-818 | chr17-58973642-INS-818 |
| chr17-58988877-INS-525 | chr17-58988877-INS-525 |
| chr17-59001272-INS-492 | chr17-59001272-INS-492 |
| chr17-59006282-INS-948 | chr17-59006269-INS-948 |
| chr17-59020688-INS-60 | chr17-59020688-INS-60 |
| chr17-59024507-INS-201 | chr17-59024507-INS-201 |
| chr17-59043864-INS-63 | chr17-59043864-INS-63 |
| chr17-59047898-INS-55 | chr17-59047898-INS-66 |
| chr17-59057513-INS-521 | chr17-59057513-INS-521 |
| chr17-59059045-INS-68 | chr17-59059043-INS-50 |
| chr17-59079610-INS-100 | chr17-59079610-INS-52 |
| chr17-59083570-INS-548 | chr17-59083570-INS-549 |
| chr17-59087167-INS-181 | chr17-59087167-INS-196 |
| chr17-59109644-INS-556 | chr17-59109644-INS-551 |
| chr17-59119953-INS-52 | chr17-59119953-INS-64 |
| chr17-59125134-INS-89 | chr17-59125134-INS-95 |
| chr17-59127614-INS-6479 | chr17-59127615-INS-6480 |
| chr17-59128453-INS-62 | chr17-59128453-INS-64 |
| chr17-59135682-INS-51 | chr17-59135697-INS-60 |
| chr17-59162814-INS-619 | chr17-59162814-INS-619 |
| chr17-59168555-INS-977 | chr17-59168555-INS-977 |
| chr17-59215009-INS-954 | chr17-59215025-INS-938 |
| chr17-59216508-INS-70 | chr17-59216508-INS-70 |
| chr17-59221697-INS-103 | chr17-59221697-INS-103 |
| chr17-59224307-INS-62 | chr17-59224307-INS-63 |
| chr17-59285210-INS-438 | chr17-59285210-INS-438 |
| chr17-59303216-INS-527 | chr17-59303216-INS-527 |
| chr17-59306934-INS-98 | chr17-59306934-INS-102 |
| chr17-59316424-INS-95 | chr17-59316447-INS-115 |
| chr17-59374155-INS-6567 | chr17-59374156-INS-6569 |
| chr17-59382907-INS-101 | chr17-59382943-INS-62 |
| chr17-59413956-INS-56 | chr17-59413944-INS-66 |
| chr17-59437389-INS-214 | chr17-59437389-INS-214 |

| | |
|---|---|
| chr17-59445140-INS-99 | chr17-59445140-INS-102 |
| chr17-59485993-INS-1279 | chr17-59485994-INS-1279 |
| chr17-59499392-INS-6382 | chr17-59499393-INS-6384 |
| chr17-59526184-INS-425 | chr17-59526184-INS-433 |
| chr17-59527725-INS-165 | chr17-59527725-INS-165 |
| chr17-59531469-INS-99 | chr17-59531469-INS-100 |
| chr17-59571511-INS-64 | chr17-59571511-INS-68 |
| chr17-59573667-INS-6555 | chr17-59573668-INS-6555 |
| chr17-59646272-INS-105 | chr17-59646272-INS-105 |
| chr17-59701071-INS-78 | chr17-59701071-INS-66 |
| chr17-59702656-INS-90 | chr17-59702656-INS-96 |
| chr17-59702983-INS-2035 | chr17-59702984-INS-2035 |
| chr17-59706871-INS-6369 | chr17-59706872-INS-6369 |
| chr17-59714579-INS-64 | chr17-59714579-INS-64 |
| chr17-59719416-INS-128 | chr17-59719416-INS-120 |
| chr17-59745039-INS-299 | chr17-59745039-INS-297 |
| chr17-59752444-INS-505 | chr17-59752444-INS-500 |
| chr17-59801820-INS-177 | chr17-59801820-INS-177 |
| chr17-59803651-INS-303 | chr17-59803651-INS-304 |
| chr17-59817057-INS-412 | chr17-59817057-INS-412 |
| chr17-59832063-INS-106 | chr17-59832063-INS-132 |
| chr17-59854042-INS-285 | chr17-59854042-INS-285 |
| chr17-59874773-INS-214 | chr17-59874773-INS-214 |
| chr17-59910044-INS-5931 | chr17-59910045-INS-5929 |
| chr17-59913078-INS-134 | chr17-59913078-INS-134 |
| chr17-59956129-INS-64 | chr17-59956127-INS-52 |
| chr17-59963273-INS-54 | chr17-59963273-INS-66 |
| chr17-59964991-INS-336 | chr17-59964992-INS-336 |
| chr17-59975684-INS-497 | chr17-59975685-INS-497 |
| chr17-59988548-INS-90 | chr17-59988548-INS-90 |
| chr17-59997836-INS-915 | chr17-59997840-INS-911 |
| chr17-60020258-INS-7230 | chr17-60020259-INS-7230 |
| chr17-60025962-INS-649 | chr17-60025963-INS-644 |
| chr17-60027574-INS-455 | chr17-60027574-INS-455 |
| chr17-60028857-INS-180 | chr17-60028857-INS-173 |
| chr17-60075457-INS-91 | chr17-60075487-INS-69 |
| chr17-60098054-INS-61 | chr17-60098009-INS-60 |
| chr17-60104468-INS-55 | chr17-60104468-INS-53 |
| chr17-60113688-INS-51 | chr17-60113688-INS-51 |
| chr17-60126440-INS-63 | chr17-60126436-INS-70 |
| chr17-60180357-INS-354 | chr17-60180362-INS-351 |
| chr17-60199093-INS-6163 | chr17-60199094-INS-6164 |
| chr17-60228172-INS-108 | chr17-60228172-INS-108 |
| chr17-60233127-INS-1995 | chr17-60233128-INS-1995 |
| chr17-60236257-INS-4122 | chr17-60236258-INS-4100 |
| chr17-60254345-INS-169 | chr17-60254345-INS-169 |
| chr17-60259976-INS-609 | chr17-60259976-INS-609 |
| chr17-60265830-INS-216 | chr17-60265830-INS-178 |
| chr17-60268510-INS-165 | chr17-60268503-INS-131 |
| chr17-60269247-INS-2846 | chr17-60269248-INS-2847 |
| chr17-60277914-INS-74 | chr17-60277908-INS-60 |
| chr17-60282458-INS-6257 | chr17-60282459-INS-6258 |
| chr17-60302502-INS-1751 | chr17-60302503-INS-1751 |
| chr17-60311678-INS-547 | chr17-60311678-INS-547 |
| chr17-60322152-INS-80 | chr17-60322152-INS-60 |
| chr17-60331470-INS-957 | chr17-60331470-INS-953 |
| chr17-60334449-INS-65 | chr17-60334449-INS-65 |
| chr17-60367817-INS-859 | chr17-60367817-INS-869 |
| chr17-60369223-INS-83 | chr17-60369223-INS-71 |
| chr17-60391324-INS-201 | chr17-60391324-INS-201 |
| chr17-60425251-INS-215 | chr17-60425251-INS-215 |
| chr17-60439616-INS-96 | chr17-60439616-INS-85 |
| chr17-60447784-INS-769 | chr17-60447784-INS-766 |
| chr17-60458484-INS-7119 | chr17-60458485-INS-7119 |
| chr17-60469682-INS-3212 | chr17-60469683-INS-3212 |
| chr17-60479543-INS-3176 | chr17-60479544-INS-3177 |
| chr17-60505914-INS-6452 | chr17-60505915-INS-6452 |
| chr17-60506228-INS-216 | chr17-60506228-INS-216 |
| chr17-60507350-INS-91 | chr17-60507350-INS-95 |
| chr17-60513690-INS-125 | chr17-60513691-INS-100 |
| chr17-60520799-INS-164 | chr17-60520799-INS-164 |
| chr17-60525103-INS-151 | chr17-60525105-INS-154 |
| chr17-60543564-INS-58 | chr17-60543564-INS-58 |
| chr17-60587521-INS-113 | chr17-60587521-INS-113 |
| chr17-60637024-INS-182 | chr17-60637024-INS-182 |
| chr17-60644378-INS-51 | chr17-60644378-INS-57 |
| chr17-60649022-INS-601 | chr17-60649022-INS-601 |

| | |
|---|---|
| chr17-60656423-INS-498 | chr17-60656423-INS-498 |
| chr17-60664065-INS-82 | chr17-60664065-INS-122 |
| chr17-60685006-INS-422 | chr17-60685006-INS-422 |
| chr17-60779241-INS-120 | chr17-60779241-INS-132 |
| chr17-60796040-INS-461 | chr17-60796040-INS-462 |
| chr17-60811031-INS-585 | chr17-60811031-INS-586 |
| chr17-60818085-INS-127 | chr17-60818085-INS-127 |
| chr17-60825224-INS-927 | chr17-60825224-INS-927 |
| chr17-60847162-INS-1138 | chr17-60847163-INS-1120 |
| chr17-60849321-INS-6985 | chr17-60849322-INS-7003 |
| chr17-60924155-INS-132 | chr17-60924155-INS-134 |
| chr17-60952837-INS-52 | chr17-60952808-INS-52 |
| chr17-60979762-INS-769 | chr17-60979762-INS-765 |
| chr17-60981179-INS-58 | chr17-60981191-INS-64 |
| chr17-60990948-INS-5043 | chr17-60990949-INS-5038 |
| chr17-61029478-INS-390 | chr17-61029478-INS-390 |
| chr17-61052664-INS-6416 | chr17-61052665-INS-6415 |
| chr17-61071480-INS-58 | chr17-61071480-INS-68 |
| chr17-61086440-INS-3301 | chr17-61086441-INS-3300 |
| chr17-61135952-INS-512 | chr17-61135952-INS-527 |
| chr17-61149100-INS-83 | chr17-61149100-INS-81 |
| chr17-61152258-INS-992 | chr17-61152258-INS-992 |
| chr17-61248284-INS-82 | chr17-61248285-INS-84 |
| chr17-61281127-INS-245 | chr17-61281127-INS-245 |
| chr17-61294611-INS-325 | chr17-61294611-INS-325 |
| chr17-61299232-INS-625 | chr17-61299232-INS-625 |
| chr17-61336311-INS-472 | chr17-61336311-INS-473 |
| chr17-61420179-INS-1365 | chr17-61420180-INS-1363 |
| chr17-61445941-INS-148 | chr17-61445941-INS-112 |
| chr17-61456578-INS-89 | chr17-61456578-INS-64 |
| chr17-61538310-INS-58 | chr17-61538310-INS-64 |
| chr17-61552070-INS-3188 | chr17-61552071-INS-3189 |
| chr17-61554311-INS-196 | chr17-61554311-INS-196 |
| chr17-61558400-INS-72 | chr17-61558400-INS-52 |
| chr17-61588383-INS-60 | chr17-61588383-INS-55 |
| chr17-61624798-INS-52 | chr17-61624798-INS-94 |
| chr17-61653301-INS-582 | chr17-61653301-INS-582 |
| chr17-61664946-INS-521 | chr17-61664946-INS-521 |
| chr17-61672540-INS-365 | chr17-61672540-INS-374 |
| chr17-61702831-INS-60 | chr17-61702831-INS-60 |
| chr17-61723158-INS-140 | chr17-61723201-INS-100 |
| chr17-61724783-INS-625 | chr17-61724783-INS-625 |
| chr17-61731655-INS-114 | chr17-61731655-INS-105 |
| chr17-61735361-INS-71 | chr17-61735361-INS-65 |
| chr17-61756491-INS-87 | chr17-61756491-INS-88 |
| chr17-61765181-INS-482 | chr17-61765181-INS-483 |
| chr17-61773947-INS-968 | chr17-61773947-INS-973 |
| chr17-61786011-INS-102 | chr17-61786011-INS-102 |
| chr17-61795105-INS-144 | chr17-61795105-INS-150 |
| chr17-61799815-INS-98 | chr17-61799815-INS-98 |
| chr17-61848851-INS-66 | chr17-61848851-INS-67 |
| chr17-61863566-INS-303 | chr17-61863566-INS-303 |
| chr17-61923844-INS-55 | chr17-61923844-INS-55 |
| chr17-61952385-INS-100 | chr17-61952385-INS-120 |
| chr17-61963223-INS-966 | chr17-61963223-INS-966 |
| chr17-61977046-INS-72 | chr17-61977046-INS-70 |
| chr17-61981676-INS-200 | chr17-61981676-INS-200 |
| chr17-62011649-INS-72 | chr17-62011649-INS-94 |
| chr17-62059896-INS-121 | chr17-62059896-INS-121 |
| chr17-62070508-INS-64 | chr17-62070508-INS-56 |
| chr17-62129254-INS-685 | chr17-62129254-INS-671 |
| chr17-62134671-INS-63 | chr17-62134696-INS-60 |
| chr17-62178122-INS-89 | chr17-62178122-INS-96 |
| chr17-62190304-INS-3306 | chr17-62190305-INS-3306 |
| chr17-62195118-INS-4822 | chr17-62195119-INS-4822 |
| chr17-62195685-INS-70 | chr17-62195685-INS-74 |
| chr17-62204324-INS-74 | chr17-62204324-INS-74 |
| chr17-62226220-INS-1277 | chr17-62226221-INS-1281 |
| chr17-62249748-INS-2953 | chr17-62249747-INS-2997 |
| chr17-62250914-INS-383 | chr17-62250914-INS-382 |
| chr17-62265730-INS-243 | chr17-62265735-INS-243 |
| chr17-62274302-INS-76 | chr17-62274302-INS-58 |
| chr17-62305462-INS-279 | chr17-62305467-INS-261 |
| chr17-62323158-INS-4087 | chr17-62323159-INS-4087 |
| chr17-62334051-INS-412 | chr17-62334068-INS-426 |
| chr17-62343444-INS-63 | chr17-62343469-INS-60 |
| chr17-62349750-INS-104 | chr17-62349754-INS-60 |

| | |
|---|---|
| chr17-62388890-INS-67 | chr17-62388890-INS-68 |
| chr17-62406039-INS-7081 | chr17-62406040-INS-7081 |
| chr17-62426790-INS-389 | chr17-62426790-INS-390 |
| chr17-62468941-INS-896 | chr17-62468948-INS-914 |
| chr17-62508859-INS-162 | chr17-62508859-INS-168 |
| chr17-62594141-INS-85 | chr17-62594141-INS-57 |
| chr17-62611637-INS-535 | chr17-62611637-INS-537 |
| chr17-62614476-INS-93 | chr17-62614476-INS-93 |
| chr17-62662994-INS-6769 | chr17-62662995-INS-6769 |
| chr17-62664005-INS-64 | chr17-62664005-INS-64 |
| chr17-62735282-INS-105 | chr17-62735282-INS-110 |
| chr17-62758526-INS-602 | chr17-62758526-INS-602 |
| chr17-62764749-INS-6483 | chr17-62764750-INS-6482 |
| chr17-62792131-INS-133 | chr17-62792132-INS-141 |
| chr17-62806499-INS-96 | chr17-62806503-INS-124 |
| chr17-62817059-INS-79 | chr17-62817059-INS-79 |
| chr17-62824546-INS-58 | chr17-62824550-INS-68 |
| chr17-62840228-INS-103 | chr17-62840228-INS-103 |
| chr17-62849660-INS-97 | chr17-62849660-INS-78 |
| chr17-62862957-INS-5831 | chr17-62862958-INS-5831 |
| chr17-62868993-INS-2134 | chr17-62868994-INS-2134 |
| chr17-62871489-INS-81 | chr17-62871489-INS-81 |
| chr17-62879761-INS-169 | chr17-62879761-INS-169 |
| chr17-62917843-INS-77 | chr17-62917852-INS-82 |
| chr17-62937815-INS-373 | chr17-62937815-INS-382 |
| chr17-62955088-INS-60 | chr17-62955088-INS-60 |
| chr17-62991384-INS-254 | chr17-62991384-INS-247 |
| chr17-62992962-INS-362 | chr17-62992962-INS-362 |
| chr17-63014200-INS-174 | chr17-63014203-INS-175 |
| chr17-63057167-INS-65 | chr17-63057149-INS-50 |
| chr17-63075360-INS-208 | chr17-63075360-INS-209 |
| chr17-63090509-INS-69 | chr17-63090509-INS-69 |
| chr17-63099177-INS-89 | chr17-63099175-INS-88 |
| chr17-63118536-INS-89 | chr17-63118536-INS-74 |
| chr17-63150684-INS-58 | chr17-63150693-INS-74 |
| chr17-63206460-INS-151 | chr17-63206460-INS-160 |
| chr17-63248455-INS-348 | chr17-63248455-INS-345 |
| chr17-63251338-INS-118 | chr17-63251358-INS-136 |
| chr17-63254331-INS-968 | chr17-63254331-INS-968 |
| chr17-63285993-INS-336 | chr17-63285993-INS-336 |
| chr17-63300065-INS-58 | chr17-63300065-INS-69 |
| chr17-63309571-INS-56 | chr17-63309575-INS-78 |
| chr17-63317868-INS-404 | chr17-63317868-INS-404 |
| chr17-63325046-INS-3231 | chr17-63325047-INS-3231 |
| chr17-63327684-INS-84 | chr17-63327645-INS-99 |
| chr17-63396427-INS-506 | chr17-63396427-INS-506 |
| chr17-63416909-INS-60 | chr17-63416909-INS-60 |
| chr17-63426907-INS-87 | chr17-63426907-INS-87 |
| chr17-63458012-INS-6000 | chr17-63458013-INS-6000 |
| chr17-63486206-INS-63 | chr17-63486206-INS-63 |
| chr17-63548137-INS-3976 | chr17-63548131-INS-3977 |
| chr17-63581883-INS-73 | chr17-63581837-INS-74 |
| chr17-63611121-INS-1688 | chr17-63611122-INS-1689 |
| chr17-63641367-INS-74 | chr17-63641394-INS-86 |
| chr17-63642974-INS-80 | chr17-63642974-INS-80 |
| chr17-63690546-INS-66 | chr17-63690546-INS-66 |
| chr17-63719693-INS-465 | chr17-63719693-INS-465 |
| chr17-63720657-INS-296 | chr17-63720608-INS-302 |
| chr17-63728877-INS-1890 | chr17-63728878-INS-1890 |
| chr17-63744707-INS-54 | chr17-63744707-INS-56 |
| chr17-63786048-INS-211 | chr17-63786048-INS-211 |
| chr17-63818891-INS-73 | chr17-63818885-INS-91 |
| chr17-63819621-INS-332 | chr17-63819622-INS-310 |
| chr17-63829790-INS-200 | chr17-63829790-INS-200 |
| chr17-63844208-INS-188 | chr17-63844208-INS-188 |
| chr17-63853433-INS-209 | chr17-63853433-INS-208 |
| chr17-63866153-INS-204 | chr17-63866153-INS-204 |
| chr17-63868481-INS-135 | chr17-63868481-INS-136 |
| chr17-63881962-INS-173 | chr17-63881962-INS-173 |
| chr17-63902958-INS-219 | chr17-63902958-INS-219 |
| chr17-63928079-INS-162 | chr17-63928079-INS-166 |
| chr17-63954641-INS-136 | chr17-63954641-INS-144 |
| chr17-63971576-INS-56 | chr17-63971576-INS-78 |
| chr17-63986840-INS-57 | chr17-63986840-INS-66 |
| chr17-63990408-INS-74 | chr17-63990408-INS-74 |
| chr17-63995674-INS-3171 | chr17-63995678-INS-3170 |
| chr17-64012957-INS-2809 | chr17-64012958-INS-2809 |

| | |
|---|---|
| chr17-64028505-INS-7080 | chr17-64028506-INS-7080 |
| chr17-64033228-INS-2759 | chr17-64033229-INS-2760 |
| chr17-64041056-INS-105 | chr17-64041075-INS-103 |
| chr17-64042055-INS-1380 | chr17-64042056-INS-1380 |
| chr17-64046221-INS-106 | chr17-64046225-INS-114 |
| chr17-64083288-INS-69 | chr17-64083288-INS-69 |
| chr17-64101933-INS-60 | chr17-64101926-INS-68 |
| chr17-64121785-INS-82 | chr17-64121785-INS-81 |
| chr17-64151952-INS-82 | chr17-64151948-INS-84 |
| chr17-64182226-INS-73 | chr17-64182226-INS-73 |
| chr17-64214021-INS-52 | chr17-64214021-INS-52 |
| chr17-64249731-INS-83 | chr17-64249750-INS-88 |
| chr17-64261109-INS-241 | chr17-64261109-INS-236 |
| chr17-64264796-INS-52 | chr17-64264796-INS-56 |
| chr17-64282423-INS-143 | chr17-64282419-INS-126 |
| chr17-64283196-INS-90 | chr17-64283196-INS-90 |
| chr17-64302327-INS-52 | chr17-64302327-INS-52 |
| chr17-64306927-INS-90 | chr17-64306927-INS-90 |
| chr17-64320791-INS-6339 | chr17-64320792-INS-6339 |
| chr17-64364790-INS-1890 | chr17-64364791-INS-1890 |
| chr17-64386329-INS-87 | chr17-64386300-INS-104 |
| chr17-64389003-INS-218 | chr17-64389003-INS-218 |
| chr17-64431227-INS-61 | chr17-64431236-INS-51 |
| chr17-64434787-INS-2951 | chr17-64434788-INS-2955 |
| chr17-64436094-INS-61 | chr17-64436094-INS-61 |
| chr17-64452182-INS-105 | chr17-64452182-INS-103 |
| chr17-64462870-INS-1735 | chr17-64462871-INS-1735 |
| chr17-64480727-INS-64 | chr17-64480727-INS-64 |
| chr17-64487477-INS-113 | chr17-64487477-INS-113 |
| chr17-64497161-INS-54 | chr17-64497161-INS-54 |
| chr17-64516726-INS-64 | chr17-64516726-INS-64 |
| chr17-64540146-INS-68 | chr17-64540146-INS-72 |
| chr17-64545988-INS-399 | chr17-64545988-INS-399 |
| chr17-64550665-INS-6403 | chr17-64550666-INS-6403 |
| chr17-64552743-INS-247 | chr17-64552743-INS-244 |
| chr17-64556266-INS-939 | chr17-64556266-INS-939 |
| chr17-64573686-INS-313 | chr17-64573686-INS-314 |
| chr17-64576746-INS-154 | chr17-64576746-INS-154 |
| chr17-64584077-INS-1222 | chr17-64584078-INS-1220 |
| chr17-64593227-INS-103 | chr17-64593227-INS-103 |
| chr17-64595883-INS-496 | chr17-64595886-INS-504 |
| chr17-64636989-INS-62 | chr17-64636989-INS-54 |
| chr17-64645929-INS-56 | chr17-64645929-INS-56 |
| chr17-64686390-INS-1029 | chr17-64686391-INS-1029 |
| chr17-64741901-INS-53 | chr17-64741901-INS-53 |
| chr17-64742444-INS-519 | chr17-64742444-INS-519 |
| chr17-64769864-INS-445 | chr17-64769864-INS-445 |
| chr17-64772672-INS-166 | chr17-64772672-INS-140 |
| chr17-64811510-INS-52 | chr17-64811510-INS-53 |
| chr17-64823552-INS-65 | chr17-64823594-INS-51 |
| chr17-64840839-INS-436 | chr17-64840839-INS-435 |
| chr17-64861224-INS-68 | chr17-64861224-INS-80 |
| chr17-64866162-INS-51 | chr17-64866162-INS-60 |
| chr17-64867426-INS-117 | chr17-64867426-INS-78 |
| chr17-64887828-INS-58 | chr17-64887828-INS-52 |
| chr17-64915773-INS-280 | chr17-64915766-INS-255 |
| chr17-65053815-INS-124 | chr17-65053815-INS-124 |
| chr17-65055065-INS-190 | chr17-65055065-INS-190 |
| chr17-65070382-INS-8164 | chr17-65070382-INS-8166 |
| chr17-65083450-INS-64 | chr17-65083450-INS-62 |
| chr17-65098954-INS-115 | chr17-65098954-INS-79 |
| chr17-65105249-INS-59 | chr17-65105249-INS-59 |
| chr17-65120605-INS-390 | chr17-65120605-INS-390 |
| chr17-65149394-INS-703 | chr17-65149394-INS-704 |
| chr17-65150430-INS-107 | chr17-65150462-INS-82 |
| chr17-65158487-INS-74 | chr17-65158487-INS-50 |
| chr17-65167656-INS-208 | chr17-65167665-INS-229 |
| chr17-65176918-INS-6879 | chr17-65176918-INS-6879 |
| chr17-65233064-INS-92 | chr17-65233096-INS-84 |
| chr17-65247190-INS-69 | chr17-65247191-INS-70 |
| chr17-65271377-INS-1238 | chr17-65271378-INS-1238 |
| chr17-65273350-INS-516 | chr17-65273308-INS-538 |
| chr17-65276712-INS-5113 | chr17-65276713-INS-5113 |
| chr17-65292191-INS-52 | chr17-65292191-INS-54 |
| chr17-65316586-INS-136 | chr17-65316586-INS-136 |
| chr17-65380244-INS-184 | chr17-65380242-INS-178 |
| chr17-65386294-INS-101 | chr17-65386294-INS-101 |

| | |
|---|---|
| chr17-65406655-INS-54 | chr17-65406655-INS-60 |
| chr17-65407908-INS-133 | chr17-65407908-INS-133 |
| chr17-65425675-INS-125 | chr17-65425662-INS-90 |
| chr17-65454047-INS-203 | chr17-65454047-INS-201 |
| chr17-65457319-INS-204 | chr17-65457319-INS-204 |
| chr17-65474644-INS-75 | chr17-65474644-INS-75 |
| chr17-65475947-INS-108 | chr17-65475947-INS-108 |
| chr17-65485793-INS-96 | chr17-65485793-INS-96 |
| chr17-65505226-INS-202 | chr17-65505226-INS-204 |
| chr17-65575208-INS-64 | chr17-65575208-INS-54 |
| chr17-65578235-INS-68 | chr17-65578231-INS-80 |
| chr17-65585688-INS-430 | chr17-65585688-INS-404 |
| chr17-65614651-INS-128 | chr17-65614651-INS-128 |
| chr17-65647504-INS-140 | chr17-65647512-INS-144 |
| chr17-65656833-INS-59 | chr17-65656833-INS-59 |
| chr17-65714066-INS-52 | chr17-65714066-INS-54 |
| chr17-65725256-INS-105 | chr17-65725256-INS-105 |
| chr17-65741654-INS-190 | chr17-65741654-INS-190 |
| chr17-65749030-INS-330 | chr17-65749061-INS-309 |
| chr17-65801097-INS-794 | chr17-65801097-INS-800 |
| chr17-65801657-INS-57 | chr17-65801691-INS-66 |
| chr17-65808808-INS-552 | chr17-65808808-INS-552 |
| chr17-65856932-INS-278 | chr17-65856927-INS-260 |
| chr17-65859176-INS-63 | chr17-65859176-INS-63 |
| chr17-65896294-INS-52 | chr17-65896294-INS-52 |
| chr17-65903558-INS-140 | chr17-65903558-INS-140 |
| chr17-65915541-INS-208 | chr17-65915541-INS-209 |
| chr17-65971420-INS-163 | chr17-65971420-INS-163 |
| chr17-65976940-INS-127 | chr17-65976940-INS-127 |
| chr17-65979609-INS-456 | chr17-65979609-INS-452 |
| chr17-65997006-INS-66 | chr17-65997006-INS-66 |
| chr17-66006288-INS-512 | chr17-66006288-INS-512 |
| chr17-66045664-INS-226 | chr17-66045664-INS-226 |
| chr17-66061685-INS-225 | chr17-66061685-INS-225 |
| chr17-66114436-INS-56 | chr17-66114436-INS-56 |
| chr17-66115897-INS-234 | chr17-66115897-INS-234 |
| chr17-66120081-INS-57 | chr17-66120081-INS-60 |
| chr17-66129770-INS-219 | chr17-66129770-INS-219 |
| chr17-66141729-INS-205 | chr17-66141729-INS-205 |
| chr17-66176170-INS-195 | chr17-66176170-INS-195 |
| chr17-66181313-INS-200 | chr17-66181313-INS-200 |
| chr17-66189373-INS-219 | chr17-66189373-INS-217 |
| chr17-66197301-INS-5548 | chr17-66197302-INS-5548 |
| chr17-66201684-INS-5302 | chr17-66201685-INS-5302 |
| chr17-66203409-INS-151 | chr17-66203405-INS-158 |
| chr17-66211549-INS-197 | chr17-66211549-INS-197 |
| chr17-66227791-INS-588 | chr17-66227791-INS-589 |
| chr17-66229189-INS-7111 | chr17-66229190-INS-7111 |
| chr17-66229294-INS-190 | chr17-66229294-INS-199 |
| chr17-66241661-INS-210 | chr17-66241661-INS-210 |
| chr17-66259401-INS-86 | chr17-66259378-INS-124 |
| chr17-66268615-INS-146 | chr17-66268615-INS-146 |
| chr17-66304227-INS-96 | chr17-66304227-INS-96 |
| chr17-66320958-INS-187 | chr17-66320958-INS-187 |
| chr17-66363813-INS-223 | chr17-66363813-INS-223 |
| chr17-66371754-INS-56 | chr17-66371770-INS-56 |
| chr17-66377575-INS-192 | chr17-66377571-INS-197 |
| chr17-66381559-INS-213 | chr17-66381559-INS-213 |
| chr17-66414481-INS-55 | chr17-66414482-INS-52 |
| chr17-66419917-INS-102 | chr17-66419917-INS-104 |
| chr17-66420655-INS-215 | chr17-66420655-INS-223 |
| chr17-66434999-INS-203 | chr17-66434999-INS-228 |
| chr17-66437246-INS-205 | chr17-66437246-INS-206 |
| chr17-66440613-INS-128 | chr17-66440613-INS-112 |
| chr17-66471336-INS-599 | chr17-66471336-INS-599 |
| chr17-66484450-INS-6488 | chr17-66484451-INS-6488 |
| chr17-66491767-INS-119 | chr17-66491767-INS-119 |
| chr17-66584639-INS-209 | chr17-66584639-INS-209 |
| chr17-66623297-INS-51 | chr17-66623267-INS-60 |
| chr17-66624568-INS-229 | chr17-66624568-INS-231 |
| chr17-66628024-INS-108 | chr17-66628024-INS-108 |
| chr17-66636746-INS-72 | chr17-66636746-INS-72 |
| chr17-66661082-INS-216 | chr17-66661082-INS-216 |
| chr17-66677420-INS-271 | chr17-66677420-INS-271 |
| chr17-66693915-INS-457 | chr17-66693915-INS-457 |
| chr17-66696909-INS-65 | chr17-66696909-INS-65 |
| chr17-66708222-INS-570 | chr17-66708222-INS-570 |

| | |
|---|---|
| chr17-66721070-INS-390 | chr17-66721070-INS-390 |
| chr17-66735757-INS-266 | chr17-66735757-INS-267 |
| chr17-66736743-INS-101 | chr17-66736743-INS-101 |
| chr17-66750342-INS-734 | chr17-66750335-INS-740 |
| chr17-66758802-INS-558 | chr17-66758802-INS-568 |
| chr17-66781932-INS-510 | chr17-66781932-INS-510 |
| chr17-66790542-INS-68 | chr17-66790542-INS-68 |
| chr17-66803176-INS-214 | chr17-66803176-INS-212 |
| chr17-66816413-INS-313 | chr17-66816413-INS-315 |
| chr17-66840109-INS-249 | chr17-66840096-INS-290 |
| chr17-66845658-INS-201 | chr17-66845658-INS-203 |
| chr17-66869124-INS-200 | chr17-66869124-INS-200 |
| chr17-66888157-INS-89 | chr17-66888157-INS-90 |
| chr17-66888608-INS-1694 | chr17-66888609-INS-1693 |
| chr17-66892383-INS-102 | chr17-66892383-INS-102 |
| chr17-66908526-INS-200 | chr17-66908526-INS-200 |
| chr17-66917564-INS-63 | chr17-66917564-INS-65 |
| chr17-66925434-INS-441 | chr17-66925434-INS-441 |
| chr17-66952460-INS-54 | chr17-66952460-INS-54 |
| chr17-66969183-INS-150 | chr17-66969180-INS-156 |
| chr17-67030663-INS-853 | chr17-67030663-INS-853 |
| chr17-67051319-INS-68 | chr17-67051319-INS-68 |
| chr17-67063933-INS-68 | chr17-67063971-INS-62 |
| chr17-67076912-INS-400 | chr17-67076912-INS-400 |
| chr17-67080734-INS-89 | chr17-67080734-INS-62 |
| chr17-67081976-INS-57 | chr17-67081976-INS-53 |
| chr17-67122447-INS-339 | chr17-67122447-INS-339 |
| chr17-67154931-INS-150 | chr17-67154927-INS-144 |
| chr17-67171255-INS-72 | chr17-67171255-INS-60 |
| chr17-67195231-INS-54 | chr17-67195231-INS-56 |
| chr17-67200188-INS-2028 | chr17-67200189-INS-2028 |
| chr17-67225293-INS-70 | chr17-67225293-INS-80 |
| chr17-67284940-INS-94 | chr17-67284940-INS-114 |
| chr17-67304518-INS-557 | chr17-67304545-INS-560 |
| chr17-67367413-INS-72 | chr17-67367378-INS-96 |
| chr17-67369659-INS-79 | chr17-67369659-INS-84 |
| chr17-67381414-INS-59 | chr17-67381414-INS-59 |
| chr17-67385867-INS-311 | chr17-67385867-INS-311 |
| chr17-67396111-INS-114 | chr17-67396083-INS-100 |
| chr17-67399984-INS-57 | chr17-67399984-INS-57 |
| chr17-67404269-INS-75 | chr17-67404232-INS-93 |
| chr17-67409588-INS-62 | chr17-67409588-INS-74 |
| chr17-67431156-INS-116 | chr17-67431121-INS-142 |
| chr17-67441933-INS-140 | chr17-67441933-INS-140 |
| chr17-67442979-INS-98 | chr17-67442979-INS-98 |
| chr17-67451304-INS-76 | chr17-67451304-INS-76 |
| chr17-67473796-INS-264 | chr17-67473796-INS-264 |
| chr17-67488483-INS-1032 | chr17-67488484-INS-1032 |
| chr17-67492237-INS-103 | chr17-67492237-INS-103 |
| chr17-67509008-INS-64 | chr17-67508985-INS-88 |
| chr17-67512534-INS-204 | chr17-67512531-INS-210 |
| chr17-67549021-INS-54 | chr17-67549021-INS-52 |
| chr17-67556558-INS-65 | chr17-67556558-INS-65 |
| chr17-67570268-INS-96 | chr17-67570268-INS-96 |
| chr17-67606946-INS-92 | chr17-67606946-INS-92 |
| chr17-67618429-INS-81 | chr17-67618429-INS-82 |
| chr17-67622387-INS-100 | chr17-67622387-INS-100 |
| chr17-67636294-INS-6506 | chr17-67636295-INS-6506 |
| chr17-67637111-INS-61 | chr17-67637111-INS-61 |
| chr17-67637516-INS-210 | chr17-67637516-INS-209 |
| chr17-67642407-INS-76 | chr17-67642409-INS-74 |
| chr17-67677032-INS-68 | chr17-67677032-INS-79 |
| chr17-67682640-INS-201 | chr17-67682640-INS-205 |
| chr17-67686835-INS-111 | chr17-67686836-INS-109 |
| chr17-67694790-INS-106 | chr17-67694789-INS-110 |
| chr17-67699011-INS-8016 | chr17-67699012-INS-8001 |
| chr17-67726834-INS-315 | chr17-67726834-INS-304 |
| chr17-67739854-INS-100 | chr17-67739862-INS-125 |
| chr17-67741653-INS-254 | chr17-67741653-INS-254 |
| chr17-67746134-INS-4500 | chr17-67746135-INS-4500 |
| chr17-67760495-INS-870 | chr17-67760495-INS-871 |
| chr17-67764584-INS-130 | chr17-67764584-INS-130 |
| chr17-67769679-INS-463 | chr17-67769679-INS-463 |
| chr17-67787733-INS-440 | chr17-67787733-INS-440 |
| chr17-67791280-INS-138 | chr17-67791280-INS-138 |
| chr17-67806655-INS-200 | chr17-67806655-INS-200 |
| chr17-67832620-INS-80 | chr17-67832600-INS-91 |

| | |
|---|---|
| chr17-67838080-INS-70 | chr17-67838083-INS-70 |
| chr17-67863601-INS-615 | chr17-67863601-INS-582 |
| chr17-67875523-INS-68 | chr17-67875554-INS-70 |
| chr17-67876244-INS-542 | chr17-67876254-INS-566 |
| chr17-67906037-INS-457 | chr17-67906039-INS-457 |
| chr17-67910709-INS-61 | chr17-67910709-INS-61 |
| chr17-67950902-INS-281 | chr17-67950902-INS-280 |
| chr17-67966622-INS-121 | chr17-67966587-INS-154 |
| chr17-67974845-INS-110 | chr17-67974833-INS-122 |
| chr17-67990149-INS-396 | chr17-67990149-INS-394 |
| chr17-67993864-INS-211 | chr17-67993864-INS-212 |
| chr17-67999962-INS-156 | chr17-67999962-INS-156 |
| chr17-68057275-INS-147 | chr17-68057315-INS-119 |
| chr17-68061112-INS-89 | chr17-68061112-INS-89 |
| chr17-68079138-INS-86 | chr17-68079138-INS-86 |
| chr17-68090250-INS-91 | chr17-68090250-INS-91 |
| chr17-68101375-INS-542 | chr17-68101375-INS-542 |
| chr17-68163539-INS-122 | chr17-68163539-INS-118 |
| chr17-68176811-INS-228 | chr17-68176811-INS-228 |
| chr17-68183662-INS-216 | chr17-68183657-INS-220 |
| chr17-68197210-INS-493 | chr17-68197210-INS-498 |
| chr17-68228104-INS-1543 | chr17-68228105-INS-1543 |
| chr17-68236895-INS-431 | chr17-68236890-INS-436 |
| chr17-68238139-INS-358 | chr17-68238139-INS-358 |
| chr17-68241042-INS-257 | chr17-68241042-INS-257 |
| chr17-68264960-INS-54 | chr17-68264960-INS-54 |
| chr17-68272788-INS-3871 | chr17-68272789-INS-3871 |
| chr17-68275392-INS-294 | chr17-68275410-INS-322 |
| chr17-68289348-INS-86 | chr17-68289348-INS-86 |
| chr17-68299962-INS-207 | chr17-68299962-INS-207 |
| chr17-68312780-INS-54 | chr17-68312780-INS-54 |
| chr17-68330302-INS-52 | chr17-68330302-INS-52 |
| chr17-68330814-INS-84 | chr17-68330814-INS-68 |
| chr17-68341312-INS-72 | chr17-68341312-INS-72 |
| chr17-68346086-INS-108 | chr17-68346086-INS-110 |
| chr17-68349671-INS-908 | chr17-68349671-INS-911 |
| chr17-68354570-INS-110 | chr17-68354548-INS-96 |
| chr17-68369345-INS-1203 | chr17-68369346-INS-1203 |
| chr17-68371159-INS-277 | chr17-68371159-INS-277 |
| chr17-68380394-INS-58 | chr17-68380394-INS-56 |
| chr17-68406633-INS-213 | chr17-68406633-INS-178 |
| chr17-68412416-INS-81 | chr17-68412465-INS-75 |
| chr17-68431932-INS-137 | chr17-68431932-INS-125 |
| chr17-68436536-INS-91 | chr17-68436536-INS-88 |
| chr17-68439623-INS-60 | chr17-68439623-INS-60 |
| chr17-68448222-INS-63 | chr17-68448222-INS-63 |
| chr17-68449482-INS-125 | chr17-68449486-INS-135 |
| chr17-68474227-INS-171 | chr17-68474227-INS-171 |
| chr17-68485378-INS-71 | chr17-68485414-INS-75 |
| chr17-68487644-INS-248 | chr17-68487644-INS-250 |
| chr17-68499096-INS-881 | chr17-68499096-INS-881 |
| chr17-68510911-INS-84 | chr17-68510911-INS-94 |
| chr17-68526810-INS-95 | chr17-68526810-INS-85 |
| chr17-68550854-INS-139 | chr17-68550855-INS-139 |
| chr17-68578614-INS-59 | chr17-68578614-INS-59 |
| chr17-68592910-INS-60 | chr17-68592938-INS-82 |
| chr17-68642096-INS-84 | chr17-68642096-INS-84 |
| chr17-68650260-INS-225 | chr17-68650260-INS-225 |
| chr17-68667247-INS-69 | chr17-68667231-INS-55 |
| chr17-68672920-INS-58 | chr17-68672920-INS-63 |
| chr17-68727558-INS-7284 | chr17-68727559-INS-7287 |
| chr17-68735922-INS-102 | chr17-68735922-INS-102 |
| chr17-68740283-INS-116 | chr17-68740283-INS-127 |
| chr17-68756892-INS-1107 | chr17-68756893-INS-1107 |
| chr17-68795029-INS-92 | chr17-68795029-INS-92 |
| chr17-68819538-INS-175 | chr17-68819538-INS-198 |
| chr17-68823487-INS-146 | chr17-68823490-INS-168 |
| chr17-68828635-INS-70 | chr17-68828633-INS-118 |
| chr17-68833788-INS-615 | chr17-68833788-INS-615 |
| chr17-68852992-INS-52 | chr17-68852992-INS-78 |
| chr17-68878434-INS-56 | chr17-68878434-INS-56 |
| chr17-68905256-INS-74 | chr17-68905256-INS-96 |
| chr17-68915750-INS-271 | chr17-68915750-INS-270 |
| chr17-68920957-INS-79 | chr17-68920957-INS-79 |
| chr17-68938336-INS-56 | chr17-68938336-INS-50 |
| chr17-68940879-INS-129 | chr17-68940879-INS-126 |
| chr17-68976190-INS-58 | chr17-68976190-INS-58 |

| | |
|---|---|
| chr17-68982966-INS-64 | chr17-68982966-INS-66 |
| chr17-68990935-INS-57 | chr17-68990935-INS-57 |
| chr17-69034783-INS-199 | chr17-69034783-INS-199 |
| chr17-69040258-INS-68 | chr17-69040258-INS-56 |
| chr17-69042677-INS-220 | chr17-69042677-INS-219 |
| chr17-69043209-INS-62 | chr17-69043209-INS-62 |
| chr17-69080172-INS-166 | chr17-69080172-INS-170 |
| chr17-69086673-INS-106 | chr17-69086673-INS-106 |
| chr17-69098206-INS-75 | chr17-69098206-INS-78 |
| chr17-69108808-INS-707 | chr17-69108808-INS-707 |
| chr17-69111386-INS-959 | chr17-69111386-INS-956 |
| chr17-69122502-INS-161 | chr17-69122505-INS-207 |
| chr17-69162385-INS-88 | chr17-69162398-INS-71 |
| chr17-69180685-INS-556 | chr17-69180685-INS-583 |
| chr17-69191192-INS-62 | chr17-69191212-INS-52 |
| chr17-69207779-INS-84 | chr17-69207779-INS-70 |
| chr17-69296668-INS-52 | chr17-69296668-INS-52 |
| chr17-69301128-INS-99 | chr17-69301128-INS-99 |
| chr17-69319406-INS-699 | chr17-69319406-INS-699 |
| chr17-69322709-INS-56 | chr17-69322700-INS-54 |
| chr17-69325349-INS-54 | chr17-69325349-INS-54 |
| chr17-69342773-INS-64 | chr17-69342773-INS-64 |
| chr17-69371549-INS-351 | chr17-69371549-INS-352 |
| chr17-69389764-INS-4915 | chr17-69389765-INS-4915 |
| chr17-69415003-INS-52 | chr17-69414991-INS-50 |
| chr17-69417794-INS-132 | chr17-69417835-INS-93 |
| chr17-69435728-INS-927 | chr17-69435728-INS-923 |
| chr17-69440311-INS-820 | chr17-69440309-INS-821 |
| chr17-69457718-INS-133 | chr17-69457729-INS-95 |
| chr17-69461860-INS-88 | chr17-69461860-INS-76 |
| chr17-69463189-INS-67 | chr17-69463190-INS-57 |
| chr17-69472022-INS-52 | chr17-69472022-INS-54 |
| chr17-69498322-INS-61 | chr17-69498322-INS-61 |
| chr17-69548190-INS-145 | chr17-69548180-INS-173 |
| chr17-69597605-INS-96 | chr17-69597603-INS-77 |
| chr17-69606062-INS-84 | chr17-69606090-INS-70 |
| chr17-69609869-INS-5240 | chr17-69609870-INS-5240 |
| chr17-69664843-INS-793 | chr17-69664844-INS-785 |
| chr17-69665263-INS-159 | chr17-69665271-INS-130 |
| chr17-69698571-INS-220 | chr17-69698571-INS-220 |
| chr17-69732410-INS-157 | chr17-69732410-INS-122 |
| chr17-69738556-INS-193 | chr17-69738556-INS-193 |
| chr17-69747395-INS-102 | chr17-69747395-INS-102 |
| chr17-69764293-INS-1382 | chr17-69764294-INS-1384 |
| chr17-69783867-INS-197 | chr17-69783868-INS-192 |
| chr17-69791331-INS-53 | chr17-69791331-INS-78 |
| chr17-69802543-INS-89 | chr17-69802543-INS-82 |
| chr17-69868040-INS-94 | chr17-69868040-INS-97 |
| chr17-69882125-INS-128 | chr17-69882125-INS-144 |
| chr17-69884430-INS-84 | chr17-69884418-INS-114 |
| chr17-69887659-INS-56 | chr17-69887659-INS-60 |
| chr17-69893859-INS-785 | chr17-69893850-INS-780 |
| chr17-69908744-INS-70 | chr17-69908739-INS-69 |
| chr17-69938242-INS-60 | chr17-69938246-INS-56 |
| chr17-69940468-INS-2543 | chr17-69940469-INS-2550 |
| chr17-69941315-INS-68 | chr17-69941348-INS-75 |
| chr17-69951154-INS-142 | chr17-69951142-INS-126 |
| chr17-69964306-INS-78 | chr17-69964306-INS-78 |
| chr17-69983683-INS-72 | chr17-69983683-INS-72 |
| chr17-70113552-INS-57 | chr17-70113552-INS-53 |
| chr17-70127573-INS-199 | chr17-70127573-INS-199 |
| chr17-70130701-INS-5013 | chr17-70130702-INS-5013 |
| chr17-70133345-INS-706 | chr17-70133345-INS-705 |
| chr17-70134874-INS-526 | chr17-70134874-INS-542 |
| chr17-70152706-INS-706 | chr17-70152706-INS-706 |
| chr17-70186939-INS-168 | chr17-70186939-INS-168 |
| chr17-70192317-INS-553 | chr17-70192317-INS-549 |
| chr17-70207001-INS-2013 | chr17-70207002-INS-2013 |
| chr17-70213341-INS-54 | chr17-70213314-INS-56 |
| chr17-70218471-INS-82 | chr17-70218471-INS-82 |
| chr17-70271882-INS-90 | chr17-70271882-INS-82 |
| chr17-70295479-INS-85 | chr17-70295479-INS-85 |
| chr17-70330224-INS-84 | chr17-70330224-INS-84 |
| chr17-70338148-INS-79 | chr17-70338132-INS-72 |
| chr17-70338669-INS-1234 | chr17-70338670-INS-1236 |
| chr17-70402975-INS-189 | chr17-70402975-INS-195 |
| chr17-70417045-INS-182 | chr17-70417045-INS-191 |

| | |
|---|---|
| chr17-70417747-INS-208 | chr17-70417747-INS-208 |
| chr17-70438665-INS-160 | chr17-70438665-INS-162 |
| chr17-70478405-INS-82 | chr17-70478405-INS-94 |
| chr17-70488932-INS-500 | chr17-70488932-INS-525 |
| chr17-70493718-INS-6401 | chr17-70493719-INS-6401 |
| chr17-70520510-INS-884 | chr17-70520510-INS-884 |
| chr17-70549890-INS-121 | chr17-70549890-INS-121 |
| chr17-70639470-INS-191 | chr17-70639470-INS-190 |
| chr17-70647383-INS-315 | chr17-70647383-INS-316 |
| chr17-70661429-INS-387 | chr17-70661429-INS-387 |
| chr17-70666350-INS-931 | chr17-70666350-INS-931 |
| chr17-70668483-INS-1172 | chr17-70668484-INS-1167 |
| chr17-70671164-INS-198 | chr17-70671165-INS-198 |
| chr17-70688213-INS-94 | chr17-70688213-INS-94 |
| chr17-70704179-INS-52 | chr17-70704179-INS-52 |
| chr17-70704709-INS-67 | chr17-70704709-INS-67 |
| chr17-70710741-INS-92 | chr17-70710741-INS-94 |
| chr17-70723149-INS-1987 | chr17-70723150-INS-1987 |
| chr17-70726708-INS-207 | chr17-70726708-INS-207 |
| chr17-70744879-INS-212 | chr17-70744900-INS-212 |
| chr17-70760908-INS-62 | chr17-70760908-INS-62 |
| chr17-70796911-INS-6681 | chr17-70796912-INS-6682 |
| chr17-70818189-INS-616 | chr17-70818189-INS-616 |
| chr17-70821626-INS-309 | chr17-70821626-INS-309 |
| chr17-70831740-INS-50 | chr17-70831737-INS-98 |
| chr17-70849848-INS-55 | chr17-70849849-INS-52 |
| chr17-70852580-INS-76 | chr17-70852581-INS-83 |
| chr17-70865135-INS-2629 | chr17-70865136-INS-2630 |
| chr17-70865503-INS-74 | chr17-70865543-INS-71 |
| chr17-70865813-INS-315 | chr17-70865813-INS-314 |
| chr17-70880499-INS-819 | chr17-70880499-INS-819 |
| chr17-70901584-INS-52 | chr17-70901584-INS-52 |
| chr17-70910887-INS-548 | chr17-70910887-INS-547 |
| chr17-70977990-INS-76 | chr17-70977990-INS-72 |
| chr17-70983937-INS-826 | chr17-70983937-INS-812 |
| chr17-70986478-INS-6445 | chr17-70986479-INS-6445 |
| chr17-70986622-INS-209 | chr17-70986622-INS-209 |
| chr17-71003664-INS-62 | chr17-71003671-INS-55 |
| chr17-71005560-INS-163 | chr17-71005560-INS-182 |
| chr17-71009658-INS-90 | chr17-71009658-INS-90 |
| chr17-71011820-INS-53 | chr17-71011821-INS-52 |
| chr17-71016218-INS-441 | chr17-71016218-INS-441 |
| chr17-71017221-INS-113 | chr17-71017221-INS-113 |
| chr17-71024902-INS-261 | chr17-71024902-INS-264 |
| chr17-71025924-INS-68 | chr17-71025932-INS-57 |
| chr17-71031461-INS-92 | chr17-71031452-INS-96 |
| chr17-71040795-INS-85 | chr17-71040784-INS-110 |
| chr17-71050201-INS-52 | chr17-71050201-INS-54 |
| chr17-71056810-INS-205 | chr17-71056810-INS-205 |
| chr17-71066103-INS-218 | chr17-71066103-INS-218 |
| chr17-71071544-INS-174 | chr17-71071544-INS-183 |
| chr17-71073307-INS-184 | chr17-71073307-INS-183 |
| chr17-71075055-INS-203 | chr17-71075055-INS-200 |
| chr17-71083435-INS-207 | chr17-71083435-INS-207 |
| chr17-71085187-INS-219 | chr17-71085187-INS-225 |
| chr17-71086455-INS-203 | chr17-71086455-INS-204 |
| chr17-71087920-INS-126 | chr17-71087915-INS-130 |
| chr17-71100771-INS-209 | chr17-71100771-INS-209 |
| chr17-71106737-INS-206 | chr17-71106747-INS-206 |
| chr17-71108713-INS-68 | chr17-71108721-INS-56 |
| chr17-71130074-INS-70 | chr17-71130066-INS-90 |
| chr17-71139084-INS-440 | chr17-71139084-INS-440 |
| chr17-71146839-INS-207 | chr17-71146839-INS-205 |
| chr17-71149055-INS-128 | chr17-71149055-INS-144 |
| chr17-71160871-INS-71 | chr17-71160871-INS-87 |
| chr17-71161887-INS-214 | chr17-71161887-INS-214 |
| chr17-71163506-INS-271 | chr17-71163506-INS-277 |
| chr17-71179722-INS-7111 | chr17-71179723-INS-7111 |
| chr17-71181376-INS-58 | chr17-71181383-INS-62 |
| chr17-71198238-INS-143 | chr17-71198238-INS-143 |
| chr17-71200800-INS-208 | chr17-71200800-INS-208 |
| chr17-71202964-INS-219 | chr17-71202964-INS-219 |
| chr17-71208055-INS-687 | chr17-71208055-INS-693 |
| chr17-71212529-INS-113 | chr17-71212529-INS-113 |
| chr17-71234876-INS-216 | chr17-71234876-INS-215 |
| chr17-71239958-INS-58 | chr17-71239958-INS-58 |
| chr17-71249374-INS-64 | chr17-71249374-INS-69 |

| | |
|---|---|
| chr17-71261128-INS-108 | chr17-71261128-INS-58 |
| chr17-71262221-INS-102 | chr17-71262221-INS-102 |
| chr17-71266637-INS-56 | chr17-71266638-INS-52 |
| chr17-71280039-INS-442 | chr17-71280053-INS-445 |
| chr17-71318924-INS-542 | chr17-71318924-INS-542 |
| chr17-71327047-INS-52 | chr17-71327047-INS-54 |
| chr17-71337271-INS-202 | chr17-71337271-INS-203 |
| chr17-71340285-INS-602 | chr17-71340286-INS-606 |
| chr17-71341986-INS-135 | chr17-71341986-INS-135 |
| chr17-71346397-INS-265 | chr17-71346397-INS-265 |
| chr17-71350008-INS-203 | chr17-71350008-INS-203 |
| chr17-71371224-INS-69 | chr17-71371174-INS-69 |
| chr17-71383293-INS-118 | chr17-71383293-INS-118 |
| chr17-71386926-INS-71 | chr17-71386917-INS-77 |
| chr17-71391583-INS-93 | chr17-71391583-INS-93 |
| chr17-71392780-INS-346 | chr17-71392780-INS-346 |
| chr17-71420790-INS-70 | chr17-71420790-INS-110 |
| chr17-71453742-INS-97 | chr17-71453735-INS-91 |
| chr17-71454470-INS-540 | chr17-71454470-INS-540 |
| chr17-71468897-INS-207 | chr17-71468897-INS-207 |
| chr17-71500838-INS-154 | chr17-71500838-INS-154 |
| chr17-71529624-INS-64 | chr17-71529624-INS-64 |
| chr17-71535137-INS-170 | chr17-71535137-INS-170 |
| chr17-71574061-INS-60 | chr17-71574061-INS-60 |
| chr17-71586979-INS-198 | chr17-71586979-INS-198 |
| chr17-71611716-INS-75 | chr17-71611716-INS-75 |
| chr17-71616871-INS-74 | chr17-71616871-INS-76 |
| chr17-71637358-INS-224 | chr17-71637358-INS-224 |
| chr17-71642413-INS-327 | chr17-71642413-INS-327 |
| chr17-71644641-INS-532 | chr17-71644641-INS-532 |
| chr17-71702927-INS-4040 | chr17-71702928-INS-4040 |
| chr17-71704897-INS-1894 | chr17-71704898-INS-1894 |
| chr17-71710297-INS-109 | chr17-71710297-INS-109 |
| chr17-71720451-INS-210 | chr17-71720451-INS-210 |
| chr17-71723652-INS-216 | chr17-71723652-INS-217 |
| chr17-71784441-INS-386 | chr17-71784441-INS-386 |
| chr17-71786341-INS-219 | chr17-71786341-INS-222 |
| chr17-71791639-INS-226 | chr17-71791653-INS-227 |
| chr17-71795598-INS-220 | chr17-71795598-INS-222 |
| chr17-71800745-INS-145 | chr17-71800715-INS-180 |
| chr17-71802934-INS-185 | chr17-71802934-INS-185 |
| chr17-71820594-INS-65 | chr17-71820594-INS-65 |
| chr17-71824893-INS-231 | chr17-71824893-INS-231 |
| chr17-71833325-INS-1895 | chr17-71833326-INS-1895 |
| chr17-71844886-INS-209 | chr17-71844886-INS-209 |
| chr17-71848001-INS-135 | chr17-71848001-INS-135 |
| chr17-71851028-INS-216 | chr17-71851028-INS-216 |
| chr17-71853755-INS-1337 | chr17-71853756-INS-1335 |
| chr17-71861792-INS-193 | chr17-71861792-INS-195 |
| chr17-71866885-INS-204 | chr17-71866885-INS-204 |
| chr17-71882114-INS-213 | chr17-71882114-INS-217 |
| chr17-71897493-INS-219 | chr17-71897493-INS-219 |
| chr17-71899714-INS-210 | chr17-71899714-INS-210 |
| chr17-71914170-INS-56 | chr17-71914170-INS-56 |
| chr17-71916753-INS-5542 | chr17-71916754-INS-5542 |
| chr17-71917148-INS-1596 | chr17-71917149-INS-1596 |
| chr17-71922769-INS-58 | chr17-71922769-INS-58 |
| chr17-71931024-INS-174 | chr17-71931024-INS-174 |
| chr17-71941861-INS-198 | chr17-71941861-INS-198 |
| chr17-71945059-INS-190 | chr17-71945059-INS-182 |
| chr17-71948770-INS-63 | chr17-71948770-INS-65 |
| chr17-71958652-INS-199 | chr17-71958652-INS-199 |
| chr17-71967221-INS-957 | chr17-71967227-INS-926 |
| chr17-71971466-INS-56 | chr17-71971477-INS-66 |
| chr17-71976692-INS-557 | chr17-71976692-INS-572 |
| chr17-71986851-INS-104 | chr17-71986871-INS-100 |
| chr17-72024648-INS-144 | chr17-72024640-INS-144 |
| chr17-72031105-INS-213 | chr17-72031105-INS-213 |
| chr17-72038764-INS-58 | chr17-72038759-INS-74 |
| chr17-72064512-INS-80 | chr17-72064527-INS-82 |
| chr17-72093431-INS-750 | chr17-72093431-INS-750 |
| chr17-72094568-INS-145 | chr17-72094568-INS-145 |
| chr17-72102300-INS-902 | chr17-72102300-INS-901 |
| chr17-72108438-INS-146 | chr17-72108438-INS-146 |
| chr17-72131952-INS-497 | chr17-72131952-INS-497 |
| chr17-72169854-INS-324 | chr17-72169854-INS-325 |
| chr17-72178033-INS-197 | chr17-72178033-INS-193 |

| | |
|---|---|
| chr17-72186600-INS-230 | chr17-72186600-INS-247 |
| chr17-72200407-INS-99 | chr17-72200407-INS-51 |
| chr17-72201873-INS-72 | chr17-72201873-INS-76 |
| chr17-72205828-INS-178 | chr17-72205828-INS-178 |
| chr17-72229709-INS-96 | chr17-72229709-INS-93 |
| chr17-72236860-INS-99 | chr17-72236860-INS-93 |
| chr17-72239811-INS-314 | chr17-72239825-INS-309 |
| chr17-72244128-INS-76 | chr17-72244128-INS-80 |
| chr17-72253489-INS-6403 | chr17-72253490-INS-6403 |
| chr17-72256584-INS-557 | chr17-72256584-INS-559 |
| chr17-72267825-INS-1952 | chr17-72267826-INS-1950 |
| chr17-72281124-INS-7272 | chr17-72281125-INS-7273 |
| chr17-72284657-INS-126 | chr17-72284657-INS-129 |
| chr17-72299917-INS-53 | chr17-72299917-INS-53 |
| chr17-72329778-INS-80 | chr17-72329778-INS-82 |
| chr17-72332810-INS-67 | chr17-72332811-INS-52 |
| chr17-72349054-INS-113 | chr17-72349042-INS-113 |
| chr17-72351364-INS-100 | chr17-72351329-INS-89 |
| chr17-72357193-INS-65 | chr17-72357193-INS-67 |
| chr17-72366772-INS-3150 | chr17-72366773-INS-3150 |
| chr17-72368452-INS-52 | chr17-72368444-INS-60 |
| chr17-72373020-INS-99 | chr17-72373024-INS-91 |
| chr17-72384240-INS-113 | chr17-72384276-INS-112 |
| chr17-72390708-INS-52 | chr17-72390708-INS-52 |
| chr17-72403457-INS-103 | chr17-72403457-INS-102 |
| chr17-72428510-INS-486 | chr17-72428510-INS-486 |
| chr17-72429329-INS-301 | chr17-72429329-INS-300 |
| chr17-72442273-INS-100 | chr17-72442273-INS-101 |
| chr17-72446459-INS-72 | chr17-72446459-INS-74 |
| chr17-72448362-INS-78 | chr17-72448362-INS-88 |
| chr17-72453179-INS-62 | chr17-72453179-INS-62 |
| chr17-72488180-INS-529 | chr17-72488180-INS-529 |
| chr17-72490928-INS-132 | chr17-72490929-INS-138 |
| chr17-72491899-INS-51 | chr17-72491855-INS-84 |
| chr17-72496415-INS-66 | chr17-72496415-INS-78 |
| chr17-72560167-INS-72 | chr17-72560214-INS-68 |
| chr17-72565785-INS-498 | chr17-72565785-INS-498 |
| chr17-72579082-INS-182 | chr17-72579082-INS-182 |
| chr17-72588558-INS-6754 | chr17-72588559-INS-6755 |
| chr17-72612459-INS-572 | chr17-72612459-INS-572 |
| chr17-72627197-INS-100 | chr17-72627195-INS-78 |
| chr17-72638778-INS-54 | chr17-72638748-INS-56 |
| chr17-72690348-INS-279 | chr17-72690348-INS-279 |
| chr17-72697292-INS-56 | chr17-72697296-INS-52 |
| chr17-72702614-INS-401 | chr17-72702614-INS-401 |
| chr17-72712256-INS-62 | chr17-72712256-INS-50 |
| chr17-72734670-INS-1933 | chr17-72734671-INS-1928 |
| chr17-72772218-INS-258 | chr17-72772206-INS-282 |
| chr17-72786870-INS-52 | chr17-72786870-INS-52 |
| chr17-72813202-INS-203 | chr17-72813210-INS-206 |
| chr17-72845478-INS-486 | chr17-72845478-INS-492 |
| chr17-72863346-INS-1687 | chr17-72863347-INS-1687 |
| chr17-72872862-INS-52 | chr17-72872854-INS-94 |
| chr17-72890850-INS-70 | chr17-72890850-INS-70 |
| chr17-72892067-INS-83 | chr17-72892067-INS-74 |
| chr17-72920333-INS-52 | chr17-72920333-INS-52 |
| chr17-72941950-INS-212 | chr17-72941950-INS-212 |
| chr17-72972340-INS-62 | chr17-72972345-INS-52 |
| chr17-72986640-INS-192 | chr17-72986640-INS-168 |
| chr17-73012127-INS-87 | chr17-73012127-INS-87 |
| chr17-73025749-INS-115 | chr17-73025710-INS-115 |
| chr17-73060078-INS-376 | chr17-73060078-INS-376 |
| chr17-73064990-INS-497 | chr17-73064995-INS-498 |
| chr17-73085612-INS-100 | chr17-73085612-INS-58 |
| chr17-73090159-INS-498 | chr17-73090159-INS-498 |
| chr17-73091652-INS-210 | chr17-73091652-INS-205 |
| chr17-73097694-INS-158 | chr17-73097694-INS-122 |
| chr17-73119863-INS-201 | chr17-73119863-INS-201 |
| chr17-73122610-INS-395 | chr17-73122610-INS-395 |
| chr17-73128425-INS-78 | chr17-73128450-INS-90 |
| chr17-73130804-INS-159 | chr17-73130804-INS-158 |
| chr17-73134243-INS-4323 | chr17-73134244-INS-4323 |
| chr17-73135138-INS-223 | chr17-73135138-INS-219 |
| chr17-73173199-INS-121 | chr17-73173199-INS-121 |
| chr17-73176022-INS-75 | chr17-73176022-INS-75 |
| chr17-73179482-INS-285 | chr17-73179482-INS-256 |
| chr17-73184285-INS-273 | chr17-73184285-INS-273 |

| | |
|---|---|
| chr17-73216913-INS-4386 | chr17-73216914-INS-4386 |
| chr17-73298530-INS-124 | chr17-73298530-INS-128 |
| chr17-73305136-INS-1016 | chr17-73305137-INS-1016 |
| chr17-73307663-INS-62 | chr17-73307663-INS-62 |
| chr17-73331436-INS-545 | chr17-73331436-INS-540 |
| chr17-73334960-INS-139 | chr17-73334961-INS-179 |
| chr17-73337735-INS-366 | chr17-73337735-INS-366 |
| chr17-73355975-INS-199 | chr17-73355975-INS-200 |
| chr17-73366642-INS-932 | chr17-73366642-INS-932 |
| chr17-73380183-INS-59 | chr17-73380183-INS-59 |
| chr17-73384504-INS-52 | chr17-73384504-INS-52 |
| chr17-73387221-INS-715 | chr17-73387217-INS-719 |
| chr17-73391813-INS-80 | chr17-73391813-INS-80 |
| chr17-73395951-INS-176 | chr17-73395965-INS-172 |
| chr17-73396505-INS-200 | chr17-73396505-INS-200 |
| chr17-73421729-INS-207 | chr17-73421729-INS-207 |
| chr17-73437022-INS-561 | chr17-73437022-INS-554 |
| chr17-73467404-INS-192 | chr17-73467404-INS-193 |
| chr17-73484895-INS-60 | chr17-73484895-INS-60 |
| chr17-73490189-INS-692 | chr17-73490189-INS-692 |
| chr17-73506867-INS-53 | chr17-73506867-INS-53 |
| chr17-73515424-INS-64 | chr17-73515424-INS-66 |
| chr17-73525763-INS-56 | chr17-73525763-INS-60 |
| chr17-73525766-INS-5584 | chr17-73525767-INS-5584 |
| chr17-73568264-INS-145 | chr17-73568264-INS-145 |
| chr17-73569666-INS-66 | chr17-73569666-INS-66 |
| chr17-73574329-INS-71 | chr17-73574319-INS-97 |
| chr17-73576648-INS-530 | chr17-73576648-INS-530 |
| chr17-73616085-INS-78 | chr17-73616085-INS-78 |
| chr17-73618674-INS-103 | chr17-73618674-INS-103 |
| chr17-73619429-INS-226 | chr17-73619429-INS-227 |
| chr17-73662804-INS-65 | chr17-73662804-INS-60 |
| chr17-73666090-INS-97 | chr17-73666088-INS-98 |
| chr17-73695367-INS-1801 | chr17-73695368-INS-1801 |
| chr17-73724865-INS-6737 | chr17-73724865-INS-6757 |
| chr17-73724866-INS-555 | chr17-73724865-INS-574 |
| chr17-73800373-INS-62 | chr17-73800369-INS-74 |
| chr17-73817970-INS-65 | chr17-73817970-INS-65 |
| chr17-73838380-INS-54 | chr17-73838380-INS-56 |
| chr17-73847395-INS-277 | chr17-73847393-INS-277 |
| chr17-73855516-INS-126 | chr17-73855516-INS-126 |
| chr17-73864908-INS-64 | chr17-73864908-INS-64 |
| chr17-73878736-INS-62 | chr17-73878736-INS-62 |
| chr17-73885945-INS-171 | chr17-73885945-INS-171 |
| chr17-73900489-INS-117 | chr17-73900489-INS-117 |
| chr17-73903705-INS-56 | chr17-73903683-INS-50 |
| chr17-73904746-INS-52 | chr17-73904754-INS-56 |
| chr17-73911821-INS-87 | chr17-73911821-INS-74 |
| chr17-73913855-INS-66 | chr17-73913855-INS-66 |
| chr17-73953606-INS-58 | chr17-73953596-INS-56 |
| chr17-73970346-INS-2759 | chr17-73970347-INS-2759 |
| chr17-73974926-INS-735 | chr17-73974926-INS-735 |
| chr17-73987035-INS-505 | chr17-73987035-INS-505 |
| chr17-74004234-INS-141 | chr17-74004234-INS-141 |
| chr17-74013492-INS-520 | chr17-74013492-INS-524 |
| chr17-74053299-INS-81 | chr17-74053299-INS-81 |
| chr17-74085401-INS-130 | chr17-74085395-INS-140 |
| chr17-74100576-INS-5240 | chr17-74100577-INS-5240 |
| chr17-74124452-INS-164 | chr17-74124452-INS-164 |
| chr17-74133974-INS-58 | chr17-74133974-INS-72 |
| chr17-74142529-INS-6299 | chr17-74142530-INS-6299 |
| chr17-74160568-INS-79 | chr17-74160568-INS-79 |
| chr17-74171852-INS-353 | chr17-74171852-INS-343 |
| chr17-74176597-INS-123 | chr17-74176597-INS-123 |
| chr17-74181339-INS-209 | chr17-74181339-INS-209 |
| chr17-74210668-INS-54 | chr17-74210668-INS-50 |
| chr17-74212555-INS-117 | chr17-74212555-INS-118 |
| chr17-74224161-INS-128 | chr17-74224161-INS-116 |
| chr17-74231675-INS-785 | chr17-74231675-INS-787 |
| chr17-74254847-INS-88 | chr17-74254847-INS-80 |
| chr17-74295737-INS-7080 | chr17-74295738-INS-7080 |
| chr17-74321080-INS-2965 | chr17-74321081-INS-2920 |
| chr17-74335729-INS-100 | chr17-74335725-INS-104 |
| chr17-74366266-INS-216 | chr17-74366266-INS-216 |
| chr17-74379098-INS-103 | chr17-74379098-INS-71 |
| chr17-74388793-INS-194 | chr17-74388793-INS-194 |
| chr17-74392329-INS-1221 | chr17-74392330-INS-1236 |

| | |
|---|---|
| chr17-74392715-INS-761 | chr17-74392718-INS-749 |
| chr17-74399155-INS-1181 | chr17-74399156-INS-1181 |
| chr17-74420978-INS-422 | chr17-74420978-INS-422 |
| chr17-74441529-INS-78 | chr17-74441529-INS-78 |
| chr17-74443687-INS-65 | chr17-74443687-INS-64 |
| chr17-74444217-INS-335 | chr17-74444217-INS-347 |
| chr17-74452673-INS-58 | chr17-74452627-INS-58 |
| chr17-74472782-INS-213 | chr17-74472807-INS-199 |
| chr17-74491827-INS-226 | chr17-74491827-INS-226 |
| chr17-74516156-INS-175 | chr17-74516156-INS-176 |
| chr17-74519562-INS-51 | chr17-74519562-INS-51 |
| chr17-74558766-INS-208 | chr17-74558766-INS-207 |
| chr17-74583541-INS-141 | chr17-74583541-INS-142 |
| chr17-74590011-INS-124 | chr17-74590012-INS-122 |
| chr17-74592593-INS-202 | chr17-74592593-INS-202 |
| chr17-74596587-INS-205 | chr17-74596587-INS-205 |
| chr17-74605345-INS-61 | chr17-74605345-INS-61 |
| chr17-74607769-INS-221 | chr17-74607769-INS-221 |
| chr17-74617853-INS-79 | chr17-74617875-INS-81 |
| chr17-74622386-INS-153 | chr17-74622386-INS-153 |
| chr17-74624668-INS-51 | chr17-74624668-INS-51 |
| chr17-74632501-INS-180 | chr17-74632501-INS-180 |
| chr17-74634613-INS-104 | chr17-74634612-INS-108 |
| chr17-74637653-INS-259 | chr17-74637653-INS-261 |
| chr17-74640491-INS-181 | chr17-74640491-INS-188 |
| chr17-74655743-INS-248 | chr17-74655743-INS-248 |
| chr17-74661357-INS-205 | chr17-74661357-INS-206 |
| chr17-74670466-INS-210 | chr17-74670466-INS-210 |
| chr17-74677983-INS-201 | chr17-74677983-INS-201 |
| chr17-74685037-INS-201 | chr17-74685037-INS-201 |
| chr17-74686424-INS-825 | chr17-74686424-INS-825 |
| chr17-74687713-INS-116 | chr17-74687713-INS-112 |
| chr17-74701241-INS-192 | chr17-74701241-INS-192 |
| chr17-74702248-INS-199 | chr17-74702248-INS-199 |
| chr17-74709471-INS-51 | chr17-74709471-INS-51 |
| chr17-74714111-INS-129 | chr17-74714111-INS-131 |
| chr17-74722050-INS-186 | chr17-74722051-INS-186 |
| chr17-74741204-INS-84 | chr17-74741204-INS-84 |
| chr17-74743132-INS-183 | chr17-74743132-INS-185 |
| chr17-74760985-INS-161 | chr17-74760986-INS-161 |
| chr17-74768514-INS-290 | chr17-74768514-INS-290 |
| chr17-74777995-INS-512 | chr17-74777995-INS-512 |
| chr17-74779884-INS-168 | chr17-74779884-INS-168 |
| chr17-74786708-INS-59 | chr17-74786695-INS-53 |
| chr17-74792756-INS-64 | chr17-74792729-INS-68 |
| chr17-74793639-INS-472 | chr17-74793639-INS-472 |
| chr17-74801457-INS-129 | chr17-74801457-INS-129 |
| chr17-74807516-INS-348 | chr17-74807516-INS-348 |
| chr17-74809678-INS-53 | chr17-74809678-INS-58 |
| chr17-74811221-INS-194 | chr17-74811220-INS-196 |
| chr17-74814446-INS-179 | chr17-74814446-INS-179 |
| chr17-74816259-INS-359 | chr17-74816260-INS-359 |
| chr17-74818962-INS-184 | chr17-74818962-INS-183 |
| chr17-74824549-INS-232 | chr17-74824549-INS-232 |
| chr17-74865305-INS-408 | chr17-74865300-INS-417 |
| chr17-74871505-INS-206 | chr17-74871505-INS-206 |
| chr17-74879471-INS-53 | chr17-74879472-INS-53 |
| chr17-74882190-INS-284 | chr17-74882190-INS-284 |
| chr17-74898637-INS-64 | chr17-74898642-INS-65 |
| chr17-74919541-INS-215 | chr17-74919541-INS-215 |
| chr17-74973859-INS-353 | chr17-74973859-INS-353 |
| chr17-74990383-INS-205 | chr17-74990383-INS-205 |
| chr17-74997678-INS-113 | chr17-74997678-INS-115 |
| chr17-75000289-INS-196 | chr17-75000289-INS-196 |
| chr17-75008688-INS-529 | chr17-75008689-INS-530 |
| chr17-75013475-INS-190 | chr17-75013475-INS-190 |
| chr17-75021427-INS-5274 | chr17-75021428-INS-5274 |
| chr17-75025186-INS-218 | chr17-75025186-INS-219 |
| chr17-75027857-INS-228 | chr17-75027857-INS-229 |
| chr17-75050143-INS-439 | chr17-75050143-INS-439 |
| chr17-75074426-INS-117 | chr17-75074428-INS-121 |
| chr17-75087696-INS-201 | chr17-75087696-INS-199 |
| chr17-75089140-INS-3120 | chr17-75089141-INS-3087 |
| chr17-75102621-INS-54 | chr17-75102621-INS-54 |
| chr17-75109864-INS-205 | chr17-75109864-INS-205 |
| chr17-75114419-INS-60 | chr17-75114419-INS-60 |
| chr17-75133846-INS-6443 | chr17-75133847-INS-6443 |

| | |
|---|---|
| chr17-75137552-INS-362 | chr17-75137552-INS-362 |
| chr17-75138629-INS-226 | chr17-75138584-INS-271 |
| chr17-75184707-INS-207 | chr17-75184707-INS-207 |
| chr17-75205464-INS-208 | chr17-75205464-INS-208 |
| chr17-75222656-INS-65 | chr17-75222656-INS-65 |
| chr17-75223896-INS-234 | chr17-75223896-INS-234 |
| chr17-75235392-INS-52 | chr17-75235392-INS-52 |
| chr17-75245604-INS-60 | chr17-75245604-INS-60 |
| chr17-75253364-INS-88 | chr17-75253364-INS-80 |
| chr17-75254336-INS-464 | chr17-75254336-INS-464 |
| chr17-75259480-INS-533 | chr17-75259480-INS-533 |
| chr17-75260843-INS-144 | chr17-75260843-INS-144 |
| chr17-75280049-INS-182 | chr17-75280049-INS-183 |
| chr17-75284358-INS-82 | chr17-75284377-INS-110 |
| chr17-75291090-INS-6447 | chr17-75291091-INS-6447 |
| chr17-75294119-INS-69 | chr17-75294119-INS-66 |
| chr17-75311882-INS-78 | chr17-75311882-INS-78 |
| chr17-75315785-INS-224 | chr17-75315785-INS-225 |
| chr17-75317584-INS-90 | chr17-75317579-INS-98 |
| chr17-75327498-INS-152 | chr17-75327498-INS-152 |
| chr17-75365515-INS-53 | chr17-75365515-INS-53 |
| chr17-75369793-INS-53 | chr17-75369794-INS-50 |
| chr17-75376568-INS-821 | chr17-75376568-INS-822 |
| chr17-75379794-INS-7010 | chr17-75379795-INS-7010 |
| chr17-75391972-INS-1460 | chr17-75391973-INS-1460 |
| chr17-75408001-INS-5549 | chr17-75408002-INS-5549 |
| chr17-75420684-INS-60 | chr17-75420684-INS-50 |
| chr17-75441665-INS-963 | chr17-75441665-INS-963 |
| chr17-75442324-INS-330 | chr17-75442306-INS-344 |
| chr17-75447136-INS-231 | chr17-75447136-INS-231 |
| chr17-75450578-INS-200 | chr17-75450578-INS-200 |
| chr17-75452764-INS-86 | chr17-75452766-INS-79 |
| chr17-75456475-INS-1736 | chr17-75456476-INS-1736 |
| chr17-75469878-INS-193 | chr17-75469878-INS-193 |
| chr17-75472856-INS-64 | chr17-75472856-INS-64 |
| chr17-75476457-INS-69 | chr17-75476457-INS-69 |
| chr17-75480316-INS-65 | chr17-75480316-INS-65 |
| chr17-75481536-INS-62 | chr17-75481536-INS-65 |
| chr17-75495486-INS-62 | chr17-75495488-INS-54 |
| chr17-75519998-INS-101 | chr17-75519998-INS-101 |
| chr17-75520894-INS-210 | chr17-75520908-INS-240 |
| chr17-75522579-INS-95 | chr17-75522579-INS-110 |
| chr17-75527456-INS-64 | chr17-75527456-INS-62 |
| chr17-75533750-INS-170 | chr17-75533750-INS-170 |
| chr17-75562144-INS-98 | chr17-75562144-INS-98 |
| chr17-75575865-INS-107 | chr17-75575866-INS-66 |
| chr17-75582123-INS-67 | chr17-75582123-INS-67 |
| chr17-75604418-INS-392 | chr17-75604418-INS-407 |
| chr17-75608625-INS-51 | chr17-75608637-INS-52 |
| chr17-75642476-INS-56 | chr17-75642476-INS-60 |
| chr17-75673354-INS-73 | chr17-75673328-INS-74 |
| chr17-75703558-INS-215 | chr17-75703558-INS-215 |
| chr17-75718471-INS-65 | chr17-75718470-INS-77 |
| chr17-75721844-INS-414 | chr17-75721844-INS-414 |
| chr17-75728659-INS-115 | chr17-75728659-INS-123 |
| chr17-75742211-INS-212 | chr17-75742211-INS-230 |
| chr17-75752670-INS-198 | chr17-75752670-INS-198 |
| chr17-75758902-INS-212 | chr17-75758902-INS-212 |
| chr17-75760615-INS-80 | chr17-75760597-INS-77 |
| chr17-75768141-INS-532 | chr17-75768141-INS-497 |
| chr17-75770507-INS-197 | chr17-75770507-INS-200 |
| chr17-75786687-INS-93 | chr17-75786646-INS-94 |
| chr17-75820124-INS-201 | chr17-75820124-INS-201 |
| chr17-75855723-INS-56 | chr17-75855723-INS-52 |
| chr17-75862642-INS-2179 | chr17-75862643-INS-2179 |
| chr17-75868100-INS-686 | chr17-75868100-INS-686 |
| chr17-75868900-INS-381 | chr17-75868900-INS-347 |
| chr17-75892815-INS-72 | chr17-75892819-INS-64 |
| chr17-75894384-INS-445 | chr17-75894384-INS-445 |
| chr17-75896992-INS-240 | chr17-75896990-INS-244 |
| chr17-75915344-INS-68 | chr17-75915344-INS-68 |
| chr17-75915392-INS-3816 | chr17-75915393-INS-3816 |
| chr17-75916828-INS-52 | chr17-75916828-INS-52 |
| chr17-75923651-INS-6384 | chr17-75923652-INS-6385 |
| chr17-75924138-INS-56 | chr17-75924142-INS-56 |
| chr17-75932866-INS-2118 | chr17-75932867-INS-2118 |
| chr17-75943403-INS-54 | chr17-75943403-INS-54 |

| | |
|---|---|
| chr17-75957669-INS-169 | chr17-75957666-INS-172 |
| chr17-75976829-INS-83 | chr17-75976829-INS-83 |
| chr17-75978926-INS-6415 | chr17-75978927-INS-6415 |
| chr17-75978968-INS-430 | chr17-75978968-INS-430 |
| chr17-75997621-INS-6473 | chr17-75997622-INS-6473 |
| chr17-76015960-INS-338 | chr17-76015960-INS-338 |
| chr17-76026148-INS-422 | chr17-76026149-INS-397 |
| chr17-76032509-INS-865 | chr17-76032509-INS-865 |
| chr17-76039719-INS-504 | chr17-76039719-INS-504 |
| chr17-76042030-INS-1226 | chr17-76042031-INS-1226 |
| chr17-76066411-INS-4527 | chr17-76066412-INS-4527 |
| chr17-76075440-INS-377 | chr17-76075438-INS-379 |
| chr17-76077573-INS-114 | chr17-76077573-INS-110 |
| chr17-76082206-INS-408 | chr17-76082198-INS-416 |
| chr17-76082968-INS-121 | chr17-76082968-INS-121 |
| chr17-76089139-INS-569 | chr17-76089139-INS-569 |
| chr17-76089273-INS-7485 | chr17-76089274-INS-7488 |
| chr17-76097525-INS-389 | chr17-76097525-INS-389 |
| chr17-76104536-INS-670 | chr17-76104536-INS-669 |
| chr17-76125908-INS-97 | chr17-76125908-INS-97 |
| chr17-76133030-INS-53 | chr17-76133030-INS-52 |
| chr17-76144067-INS-117 | chr17-76144067-INS-117 |
| chr17-76154929-INS-189 | chr17-76154930-INS-186 |
| chr17-76194892-INS-133 | chr17-76194865-INS-140 |
| chr17-76226020-INS-121 | chr17-76226020-INS-101 |
| chr17-76252711-INS-73 | chr17-76252711-INS-70 |
| chr17-76305086-INS-68 | chr17-76305086-INS-68 |
| chr17-76308491-INS-3324 | chr17-76308492-INS-3324 |
| chr17-76309563-INS-52 | chr17-76309561-INS-60 |
| chr17-76333300-INS-79 | chr17-76333300-INS-79 |
| chr17-76335491-INS-140 | chr17-76335491-INS-151 |
| chr17-76338004-INS-256 | chr17-76338004-INS-234 |
| chr17-76361388-INS-4012 | chr17-76361389-INS-4012 |
| chr17-76373930-INS-54 | chr17-76373930-INS-54 |
| chr17-76381647-INS-70 | chr17-76381647-INS-62 |
| chr17-76446595-INS-151 | chr17-76446595-INS-159 |
| chr17-76473910-INS-1144 | chr17-76473911-INS-1144 |
| chr17-76474815-INS-1239 | chr17-76474816-INS-1240 |
| chr17-76481455-INS-1892 | chr17-76481456-INS-1892 |
| chr17-76512068-INS-60 | chr17-76512087-INS-63 |
| chr17-76513057-INS-53 | chr17-76513058-INS-50 |
| chr17-76545886-INS-56 | chr17-76545886-INS-62 |
| chr17-76560230-INS-334 | chr17-76560230-INS-334 |
| chr17-76561437-INS-64 | chr17-76561437-INS-64 |
| chr17-76568249-INS-148 | chr17-76568249-INS-148 |
| chr17-76581828-INS-390 | chr17-76581828-INS-390 |
| chr17-76590551-INS-6206 | chr17-76590552-INS-6206 |
| chr17-76592084-INS-52 | chr17-76592084-INS-52 |
| chr17-76607774-INS-57 | chr17-76607774-INS-57 |
| chr17-76626487-INS-189 | chr17-76626487-INS-189 |
| chr17-76639086-INS-391 | chr17-76639086-INS-391 |
| chr17-76656395-INS-454 | chr17-76656395-INS-438 |
| chr17-76675309-INS-63 | chr17-76675309-INS-51 |
| chr17-76684801-INS-54 | chr17-76684801-INS-54 |
| chr17-76711171-INS-50 | chr17-76711169-INS-64 |
| chr17-76719011-INS-468 | chr17-76719011-INS-468 |
| chr17-76727714-INS-179 | chr17-76727714-INS-149 |
| chr17-76731093-INS-56 | chr17-76731093-INS-56 |
| chr17-76774595-INS-56 | chr17-76774596-INS-60 |
| chr17-76775062-INS-124 | chr17-76775062-INS-124 |
| chr17-76777340-INS-659 | chr17-76777340-INS-659 |
| chr17-76806040-INS-80 | chr17-76806040-INS-80 |
| chr17-76817265-INS-906 | chr17-76817265-INS-903 |
| chr17-76860896-INS-89 | chr17-76860896-INS-87 |
| chr17-76869759-INS-153 | chr17-76869759-INS-153 |
| chr17-76913201-INS-58 | chr17-76913201-INS-58 |
| chr17-76941950-INS-92 | chr17-76941950-INS-75 |
| chr17-76969278-INS-181 | chr17-76969267-INS-185 |
| chr17-76974032-INS-291 | chr17-76974032-INS-263 |
| chr17-76977983-INS-4704 | chr17-76977984-INS-4704 |
| chr17-76986220-INS-58 | chr17-76986220-INS-52 |
| chr17-77004408-INS-527 | chr17-77004408-INS-527 |
| chr17-77067043-INS-73 | chr17-77067043-INS-90 |
| chr17-77095166-INS-423 | chr17-77095166-INS-423 |
| chr17-77109739-INS-802 | chr17-77109739-INS-803 |
| chr17-77127353-INS-2696 | chr17-77127354-INS-2696 |
| chr17-77143704-INS-56 | chr17-77143704-INS-60 |

| | |
|---|---|
| chr17-77175557-INS-66 | chr17-77175557-INS-66 |
| chr17-77186303-INS-63 | chr17-77186303-INS-91 |
| chr17-77188557-INS-77 | chr17-77188557-INS-65 |
| chr17-77191466-INS-432 | chr17-77191466-INS-432 |
| chr17-77225927-INS-87 | chr17-77225927-INS-87 |
| chr17-77226252-INS-676 | chr17-77226252-INS-676 |
| chr17-77228394-INS-614 | chr17-77228394-INS-614 |
| chr17-77237722-INS-124 | chr17-77237714-INS-104 |
| chr17-77239361-INS-52 | chr17-77239361-INS-56 |
| chr17-77246018-INS-319 | chr17-77246018-INS-319 |
| chr17-77267839-INS-728 | chr17-77267839-INS-729 |
| chr17-77270279-INS-683 | chr17-77270279-INS-683 |
| chr17-77275601-INS-750 | chr17-77275601-INS-750 |
| chr17-77281217-INS-4757 | chr17-77281218-INS-4757 |
| chr17-77326613-INS-6652 | chr17-77326614-INS-6652 |
| chr17-77327912-INS-279 | chr17-77327912-INS-279 |
| chr17-77336872-INS-63 | chr17-77336872-INS-57 |
| chr17-77346232-INS-727 | chr17-77346232-INS-727 |
| chr17-77355655-INS-64 | chr17-77355656-INS-94 |
| chr17-77390298-INS-521 | chr17-77390298-INS-521 |
| chr17-77399869-INS-765 | chr17-77399869-INS-766 |
| chr17-77423692-INS-62 | chr17-77423692-INS-62 |
| chr17-77450248-INS-167 | chr17-77450248-INS-167 |
| chr17-77454187-INS-520 | chr17-77454187-INS-520 |
| chr17-77479288-INS-6609 | chr17-77479289-INS-6615 |
| chr17-77500778-INS-96 | chr17-77500778-INS-90 |
| chr17-77540005-INS-212 | chr17-77540005-INS-212 |
| chr17-77552031-INS-114 | chr17-77552031-INS-114 |
| chr17-77564986-INS-3176 | chr17-77564987-INS-3179 |
| chr17-77569805-INS-1939 | chr17-77569806-INS-1940 |
| chr17-77584145-INS-192 | chr17-77584145-INS-222 |
| chr17-77619245-INS-132 | chr17-77619271-INS-114 |
| chr17-77641856-INS-4754 | chr17-77641857-INS-4754 |
| chr17-77672669-INS-158 | chr17-77672669-INS-158 |
| chr17-77686894-INS-646 | chr17-77686894-INS-646 |
| chr17-77700552-INS-58 | chr17-77700552-INS-58 |
| chr17-77700684-INS-7212 | chr17-77700685-INS-7212 |
| chr17-77706852-INS-86 | chr17-77706852-INS-86 |
| chr17-77724651-INS-72 | chr17-77724651-INS-72 |
| chr17-77742604-INS-72 | chr17-77742589-INS-96 |
| chr17-77749864-INS-68 | chr17-77749864-INS-72 |
| chr17-77750729-INS-1225 | chr17-77750730-INS-1225 |
| chr17-77756752-INS-405 | chr17-77756752-INS-405 |
| chr17-77779060-INS-100 | chr17-77779060-INS-102 |
| chr17-77819723-INS-159 | chr17-77819720-INS-161 |
| chr17-77827822-INS-55 | chr17-77827822-INS-55 |
| chr17-77838927-INS-310 | chr17-77838927-INS-311 |
| chr17-77902567-INS-214 | chr17-77902567-INS-214 |
| chr17-77904981-INS-5643 | chr17-77904982-INS-5691 |
| chr17-77925662-INS-94 | chr17-77925662-INS-94 |
| chr17-77930312-INS-122 | chr17-77930312-INS-122 |
| chr17-77963849-INS-84 | chr17-77963849-INS-88 |
| chr17-77992967-INS-121 | chr17-77992967-INS-121 |
| chr17-78034975-INS-443 | chr17-78034975-INS-443 |
| chr17-78038360-INS-76 | chr17-78038360-INS-72 |
| chr17-78049842-INS-794 | chr17-78049842-INS-794 |
| chr17-78060348-INS-60 | chr17-78060348-INS-98 |
| chr17-78101007-INS-218 | chr17-78101007-INS-218 |
| chr17-78114119-INS-972 | chr17-78114119-INS-971 |
| chr17-78115498-INS-937 | chr17-78115498-INS-937 |
| chr17-78156948-INS-193 | chr17-78156948-INS-193 |
| chr17-78161933-INS-82 | chr17-78161933-INS-50 |
| chr17-78196016-INS-4671 | chr17-78196017-INS-4671 |
| chr17-78215113-INS-1716 | chr17-78215114-INS-1719 |
| chr17-78222061-INS-173 | chr17-78222061-INS-173 |
| chr17-78240077-INS-76 | chr17-78240077-INS-86 |
| chr17-78247106-INS-69 | chr17-78247106-INS-69 |
| chr17-78283641-INS-112 | chr17-78283650-INS-74 |
| chr17-78284690-INS-85 | chr17-78284690-INS-85 |
| chr17-78306962-INS-4821 | chr17-78306972-INS-4809 |
| chr17-78327410-INS-63 | chr17-78327410-INS-63 |
| chr17-78328265-INS-401 | chr17-78328265-INS-401 |
| chr17-78351388-INS-125 | chr17-78351388-INS-123 |
| chr17-78367569-INS-96 | chr17-78367569-INS-96 |
| chr17-78411761-INS-293 | chr17-78411761-INS-308 |
| chr17-78431925-INS-116 | chr17-78431925-INS-116 |
| chr17-78443171-INS-67 | chr17-78443220-INS-50 |

| | |
|---|---|
| chr17-78486794-INS-109 | chr17-78486814-INS-123 |
| chr17-78502737-INS-289 | chr17-78502737-INS-289 |
| chr17-78551978-INS-78 | chr17-78551978-INS-80 |
| chr17-78553805-INS-100 | chr17-78553805-INS-100 |
| chr17-78561357-INS-73 | chr17-78561357-INS-72 |
| chr17-78607752-INS-84 | chr17-78607752-INS-84 |
| chr17-78618710-INS-56 | chr17-78618710-INS-56 |
| chr17-78638669-INS-56 | chr17-78638669-INS-58 |
| chr17-78642430-INS-151 | chr17-78642430-INS-159 |
| chr17-78697599-INS-76 | chr17-78697585-INS-78 |
| chr17-78714599-INS-101 | chr17-78714599-INS-126 |
| chr17-78727823-INS-130 | chr17-78727823-INS-122 |
| chr17-78733593-INS-64 | chr17-78733593-INS-62 |
| chr17-78737253-INS-156 | chr17-78737253-INS-141 |
| chr17-78740081-INS-57 | chr17-78740081-INS-57 |
| chr17-78749416-INS-357 | chr17-78749416-INS-360 |
| chr17-78765145-INS-234 | chr17-78765104-INS-232 |
| chr17-78785462-INS-111 | chr17-78785462-INS-111 |
| chr17-78805047-INS-58 | chr17-78805057-INS-51 |
| chr17-78820697-INS-3331 | chr17-78820698-INS-3340 |
| chr17-78826242-INS-621 | chr17-78826242-INS-622 |
| chr17-78834473-INS-55 | chr17-78834475-INS-103 |
| chr17-78836449-INS-54 | chr17-78836449-INS-99 |
| chr17-78846453-INS-52 | chr17-78846469-INS-52 |
| chr17-78851734-INS-6881 | chr17-78851735-INS-6881 |
| chr17-78865571-INS-190 | chr17-78865571-INS-190 |
| chr17-78866944-INS-68 | chr17-78866944-INS-68 |
| chr17-78905167-INS-140 | chr17-78905167-INS-140 |
| chr17-78926619-INS-199 | chr17-78926619-INS-199 |
| chr17-78933090-INS-162 | chr17-78933072-INS-149 |
| chr17-78935602-INS-52 | chr17-78935602-INS-58 |
| chr17-78936926-INS-977 | chr17-78936926-INS-977 |
| chr17-78969017-INS-778 | chr17-78969032-INS-790 |
| chr17-78989443-INS-64 | chr17-78989443-INS-110 |
| chr17-79026818-INS-1742 | chr17-79026819-INS-1742 |
| chr17-79039374-INS-153 | chr17-79039374-INS-153 |
| chr17-79068741-INS-132 | chr17-79068741-INS-132 |
| chr17-79074872-INS-166 | chr17-79074872-INS-166 |
| chr17-79081390-INS-112 | chr17-79081390-INS-88 |
| chr17-79081988-INS-977 | chr17-79081988-INS-977 |
| chr17-79084870-INS-78 | chr17-79084870-INS-78 |
| chr17-79117024-INS-172 | chr17-79117024-INS-172 |
| chr17-79117142-INS-602 | chr17-79117142-INS-602 |
| chr17-79127997-INS-56 | chr17-79127997-INS-56 |
| chr17-79141845-INS-57 | chr17-79141845-INS-59 |
| chr17-79159430-INS-52 | chr17-79159430-INS-52 |
| chr17-79186246-INS-197 | chr17-79186246-INS-197 |
| chr17-79186936-INS-171 | chr17-79186936-INS-169 |
| chr17-79199087-INS-231 | chr17-79199087-INS-231 |
| chr17-79222943-INS-235 | chr17-79222943-INS-235 |
| chr17-79230226-INS-207 | chr17-79230226-INS-207 |
| chr17-79247754-INS-139 | chr17-79247754-INS-139 |
| chr17-79274255-INS-53 | chr17-79274253-INS-59 |
| chr17-79285618-INS-205 | chr17-79285618-INS-205 |
| chr17-79326736-INS-157 | chr17-79326736-INS-157 |
| chr17-79334855-INS-66 | chr17-79334873-INS-68 |
| chr17-79340620-INS-51 | chr17-79340620-INS-51 |
| chr17-79368415-INS-112 | chr17-79368415-INS-114 |
| chr17-79374621-INS-71 | chr17-79374619-INS-77 |
| chr17-79390494-INS-147 | chr17-79390494-INS-147 |
| chr17-79394048-INS-111 | chr17-79394048-INS-111 |
| chr17-79408473-INS-57 | chr17-79408463-INS-96 |
| chr17-79431882-INS-231 | chr17-79431882-INS-231 |
| chr17-79438752-INS-205 | chr17-79438752-INS-205 |
| chr17-79447627-INS-162 | chr17-79447627-INS-162 |
| chr17-79453600-INS-91 | chr17-79453600-INS-91 |
| chr17-79480697-INS-71 | chr17-79480659-INS-64 |
| chr17-79499161-INS-133 | chr17-79499161-INS-129 |
| chr17-79504795-INS-542 | chr17-79504795-INS-542 |
| chr17-79516602-INS-128 | chr17-79516602-INS-128 |
| chr17-79530375-INS-57 | chr17-79530371-INS-52 |
| chr17-79536103-INS-79 | chr17-79536081-INS-85 |
| chr17-79550690-INS-51 | chr17-79550679-INS-52 |
| chr17-79599933-INS-116 | chr17-79599917-INS-113 |
| chr17-79635428-INS-58 | chr17-79635428-INS-63 |
| chr17-79641144-INS-70 | chr17-79641144-INS-70 |
| chr17-79646900-INS-72 | chr17-79646900-INS-72 |

| | |
|---|---|
| chr17-79659599-INS-62 | chr17-79659599-INS-62 |
| chr17-79693515-INS-58 | chr17-79693515-INS-58 |
| chr17-79698577-INS-5107 | chr17-79698578-INS-5125 |
| chr17-79729993-INS-54 | chr17-79729993-INS-52 |
| chr17-79755195-INS-317 | chr17-79755195-INS-317 |
| chr17-79772984-INS-84 | chr17-79772984-INS-84 |
| chr17-79807426-INS-194 | chr17-79807426-INS-206 |
| chr17-79809274-INS-1502 | chr17-79809275-INS-1503 |
| chr17-79811236-INS-169 | chr17-79811236-INS-172 |
| chr17-79816085-INS-180 | chr17-79816086-INS-180 |
| chr17-79816565-INS-409 | chr17-79816565-INS-409 |
| chr17-79818303-INS-64 | chr17-79818303-INS-64 |
| chr17-79823123-INS-195 | chr17-79823123-INS-195 |
| chr17-79826956-INS-66 | chr17-79826956-INS-66 |
| chr17-79836829-INS-66 | chr17-79836827-INS-68 |
| chr17-79862481-INS-77 | chr17-79862489-INS-62 |
| chr17-79874590-INS-960 | chr17-79874590-INS-960 |
| chr17-79880909-INS-104 | chr17-79880909-INS-104 |
| chr17-79898594-INS-432 | chr17-79898594-INS-434 |
| chr17-79940652-INS-209 | chr17-79940652-INS-213 |
| chr17-79961484-INS-56 | chr17-79961484-INS-52 |
| chr17-79966297-INS-174 | chr17-79966297-INS-174 |
| chr17-79990979-INS-716 | chr17-79990979-INS-717 |
| chr17-79998871-INS-56 | chr17-79998875-INS-54 |
| chr17-80018168-INS-73 | chr17-80018166-INS-62 |
| chr17-80053639-INS-52 | chr17-80053639-INS-52 |
| chr17-80054927-INS-691 | chr17-80054927-INS-691 |
| chr17-80058553-INS-60 | chr17-80058564-INS-57 |
| chr17-80073167-INS-3029 | chr17-80073168-INS-3029 |
| chr17-80088365-INS-68 | chr17-80088365-INS-68 |
| chr17-80101547-INS-128 | chr17-80101547-INS-122 |
| chr17-80106882-INS-201 | chr17-80106882-INS-211 |
| chr17-80120001-INS-1210 | chr17-80120002-INS-1210 |
| chr17-80131291-INS-363 | chr17-80131291-INS-368 |
| chr17-80135089-INS-58 | chr17-80135087-INS-66 |
| chr17-80140280-INS-291 | chr17-80140280-INS-291 |
| chr17-80142433-INS-84 | chr17-80142433-INS-84 |
| chr17-80173294-INS-139 | chr17-80173289-INS-102 |
| chr17-80188096-INS-99 | chr17-80188097-INS-91 |
| chr17-80189694-INS-64 | chr17-80189694-INS-64 |
| chr17-80192179-INS-477 | chr17-80192179-INS-477 |
| chr17-80201294-INS-111 | chr17-80201294-INS-111 |
| chr17-80212509-INS-151 | chr17-80212511-INS-148 |
| chr17-80219081-INS-124 | chr17-80219074-INS-120 |
| chr17-80233089-INS-72 | chr17-80233089-INS-66 |
| chr17-80242605-INS-189 | chr17-80242605-INS-189 |
| chr17-80243641-INS-305 | chr17-80243641-INS-305 |
| chr17-80272717-INS-76 | chr17-80272717-INS-79 |
| chr17-80275841-INS-519 | chr17-80275841-INS-519 |
| chr17-80282054-INS-55 | chr17-80282054-INS-52 |
| chr17-80286452-INS-168 | chr17-80286455-INS-168 |
| chr17-80288235-INS-60 | chr17-80288235-INS-64 |
| chr17-80289189-INS-167 | chr17-80289189-INS-162 |
| chr17-80290333-INS-1354 | chr17-80290334-INS-1353 |
| chr17-80318388-INS-530 | chr17-80318388-INS-530 |
| chr17-80319471-INS-56 | chr17-80319471-INS-58 |
| chr17-80327157-INS-3941 | chr17-80327158-INS-3941 |
| chr17-80330513-INS-200 | chr17-80330513-INS-200 |
| chr17-80351115-INS-238 | chr17-80351115-INS-238 |
| chr17-80361197-INS-67 | chr17-80361197-INS-68 |
| chr17-80362272-INS-71 | chr17-80362272-INS-76 |
| chr17-80365426-INS-69 | chr17-80365426-INS-69 |
| chr17-80366666-INS-87 | chr17-80366666-INS-87 |
| chr17-80375051-INS-81 | chr17-80375051-INS-82 |
| chr17-80378841-INS-304 | chr17-80378841-INS-304 |
| chr17-80388191-INS-129 | chr17-80388191-INS-129 |
| chr17-80392989-INS-4788 | chr17-80392990-INS-4788 |
| chr17-80396341-INS-108 | chr17-80396355-INS-137 |
| chr17-80411391-INS-64 | chr17-80411384-INS-68 |
| chr17-80419775-INS-88 | chr17-80419775-INS-119 |
| chr17-80431116-INS-64 | chr17-80431116-INS-64 |
| chr17-80447104-INS-196 | chr17-80447104-INS-199 |
| chr17-80454169-INS-268 | chr17-80454169-INS-268 |
| chr17-80457439-INS-351 | chr17-80457439-INS-351 |
| chr17-80471017-INS-183 | chr17-80471017-INS-184 |
| chr17-80493883-INS-173 | chr17-80493883-INS-179 |
| chr17-80500340-INS-688 | chr17-80500340-INS-688 |

| | |
|---|---|
| chr17-80501530-INS-323 | chr17-80501531-INS-323 |
| chr17-80504142-INS-456 | chr17-80504142-INS-456 |
| chr17-80517702-INS-200 | chr17-80517702-INS-200 |
| chr17-80529367-INS-63 | chr17-80529383-INS-61 |
| chr17-80531712-INS-177 | chr17-80531712-INS-179 |
| chr17-80537272-INS-52 | chr17-80537272-INS-54 |
| chr17-80538008-INS-239 | chr17-80538008-INS-245 |
| chr17-80539010-INS-420 | chr17-80539010-INS-420 |
| chr17-80542905-INS-60 | chr17-80542905-INS-60 |
| chr17-80547119-INS-147 | chr17-80547119-INS-147 |
| chr17-80555201-INS-197 | chr17-80555201-INS-197 |
| chr17-80570926-INS-360 | chr17-80570919-INS-337 |
| chr17-80575397-INS-54 | chr17-80575397-INS-54 |
| chr17-80589165-INS-203 | chr17-80589165-INS-203 |
| chr17-80591616-INS-343 | chr17-80591616-INS-343 |
| chr17-80602342-INS-216 | chr17-80602342-INS-216 |
| chr17-80608420-INS-70 | chr17-80608420-INS-78 |
| chr17-80616509-INS-54 | chr17-80616509-INS-54 |
| chr17-80617619-INS-96 | chr17-80617619-INS-92 |
| chr17-80650787-INS-66 | chr17-80650787-INS-66 |
| chr17-80664539-INS-1381 | chr17-80664540-INS-1382 |
| chr17-80665294-INS-91 | chr17-80665294-INS-91 |
| chr17-80666836-INS-329 | chr17-80666836-INS-337 |
| chr17-80671545-INS-286 | chr17-80671545-INS-286 |
| chr17-80680123-INS-80 | chr17-80680123-INS-72 |
| chr17-80693582-INS-138 | chr17-80693582-INS-139 |
| chr17-80709219-INS-91 | chr17-80709219-INS-91 |
| chr17-80722613-INS-463 | chr17-80722613-INS-463 |
| chr17-80743594-INS-150 | chr17-80743589-INS-151 |
| chr17-80761677-INS-118 | chr17-80761677-INS-118 |
| chr17-80772870-INS-147 | chr17-80772870-INS-147 |
| chr17-80805574-INS-55 | chr17-80805617-INS-52 |
| chr17-80829575-INS-63 | chr17-80829575-INS-66 |
| chr17-80852045-INS-169 | chr17-80852045-INS-175 |
| chr17-80859281-INS-231 | chr17-80859281-INS-219 |
| chr17-80864913-INS-373 | chr17-80864913-INS-373 |
| chr17-80870553-INS-288 | chr17-80870553-INS-289 |
| chr17-80891947-INS-164 | chr17-80891947-INS-164 |
| chr17-80905522-INS-135 | chr17-80905522-INS-135 |
| chr17-80908010-INS-54 | chr17-80908010-INS-54 |
| chr17-80941859-INS-1490 | chr17-80941860-INS-1490 |
| chr17-80956328-INS-171 | chr17-80956339-INS-190 |
| chr17-80975441-INS-181 | chr17-80975430-INS-221 |
| chr17-81000985-INS-205 | chr17-81000985-INS-205 |
| chr17-81005158-INS-71 | chr17-81005188-INS-71 |
| chr17-81009010-INS-176 | chr17-81009010-INS-177 |
| chr17-81047856-INS-558 | chr17-81047855-INS-568 |
| chr17-81067075-INS-77 | chr17-81067075-INS-77 |
| chr17-81078023-INS-892 | chr17-81078023-INS-891 |
| chr17-81079429-INS-1475 | chr17-81079430-INS-1475 |
| chr17-81086627-INS-62 | chr17-81086627-INS-68 |
| chr17-81101260-INS-177 | chr17-81101260-INS-178 |
| chr17-81121445-INS-121 | chr17-81121445-INS-111 |
| chr17-81126900-INS-266 | chr17-81126900-INS-246 |
| chr17-81129286-INS-52 | chr17-81129286-INS-54 |
| chr17-81169947-INS-370 | chr17-81169947-INS-369 |
| chr17-81176452-INS-129 | chr17-81176452-INS-129 |
| chr17-81193456-INS-92 | chr17-81193456-INS-72 |
| chr17-81202277-INS-6486 | chr17-81202278-INS-6486 |
| chr17-81204758-INS-54 | chr17-81204758-INS-54 |
| chr17-81215002-INS-70 | chr17-81215002-INS-75 |
| chr17-81224170-INS-689 | chr17-81224170-INS-689 |
| chr17-81255692-INS-68 | chr17-81255708-INS-72 |
| chr17-81274049-INS-52 | chr17-81274049-INS-58 |
| chr17-81281946-INS-155 | chr17-81281946-INS-155 |
| chr17-81303786-INS-170 | chr17-81303786-INS-170 |
| chr17-81307147-INS-529 | chr17-81307147-INS-530 |
| chr17-81317194-INS-62 | chr17-81317194-INS-62 |
| chr17-81330828-INS-52 | chr17-81330828-INS-55 |
| chr17-81338327-INS-97 | chr17-81338327-INS-130 |
| chr17-81344197-INS-85 | chr17-81344197-INS-132 |
| chr17-81354864-INS-195 | chr17-81354864-INS-195 |
| chr17-81405554-INS-104 | chr17-81405554-INS-112 |
| chr17-81411334-INS-82 | chr17-81411342-INS-58 |
| chr17-81449158-INS-75 | chr17-81449159-INS-58 |
| chr17-81450306-INS-206 | chr17-81450306-INS-204 |
| chr17-81451509-INS-50 | chr17-81451508-INS-57 |

| | |
|---|---|
| chr17-81475293-INS-59 | chr17-81475293-INS-59 |
| chr17-81505728-INS-212 | chr17-81505728-INS-215 |
| chr17-81510481-INS-328 | chr17-81510481-INS-328 |
| chr17-81522882-INS-86 | chr17-81522881-INS-104 |
| chr17-81624785-INS-153 | chr17-81624777-INS-189 |
| chr17-81651279-INS-113 | chr17-81651279-INS-113 |
| chr17-81743497-INS-51 | chr17-81743497-INS-51 |
| chr17-81758034-INS-68 | chr17-81758037-INS-52 |
| chr17-81777473-INS-57 | chr17-81777473-INS-57 |
| chr17-81804576-INS-56 | chr17-81804576-INS-56 |
| chr17-81812483-INS-208 | chr17-81812483-INS-208 |
| chr17-81858518-INS-183 | chr17-81858518-INS-183 |
| chr17-81893773-INS-89 | chr17-81893761-INS-102 |
| chr17-81905441-INS-54 | chr17-81905441-INS-55 |
| chr17-81947280-INS-105 | chr17-81947309-INS-84 |
| chr17-81969970-INS-159 | chr17-81969970-INS-112 |
| chr17-82003944-INS-197 | chr17-82003944-INS-197 |
| chr17-82047173-INS-684 | chr17-82047185-INS-684 |
| chr17-82057923-INS-7038 | chr17-82057924-INS-7041 |
| chr17-82063269-INS-52 | chr17-82063269-INS-56 |
| chr17-82091261-INS-51 | chr17-82091261-INS-51 |
| chr17-82109985-INS-1860 | chr17-82109986-INS-1860 |
| chr17-82113180-INS-4394 | chr17-82113181-INS-4394 |
| chr17-82149032-INS-93 | chr17-82149032-INS-80 |
| chr17-82149843-INS-7047 | chr17-82149844-INS-7049 |
| chr17-82256817-INS-261 | chr17-82256817-INS-261 |
| chr17-82259307-INS-867 | chr17-82259307-INS-868 |
| chr17-82272316-INS-350 | chr17-82272316-INS-350 |
| chr17-82286011-INS-416 | chr17-82286014-INS-416 |
| chr17-82289166-INS-3910 | chr17-82289167-INS-3902 |
| chr17-82329046-INS-6789 | chr17-82329047-INS-6792 |
| chr17-82361613-INS-95 | chr17-82361613-INS-95 |
| chr17-82373614-INS-162 | chr17-82373614-INS-162 |
| chr17-82441489-INS-57 | chr17-82441489-INS-57 |
| chr17-82444558-INS-60 | chr17-82444558-INS-56 |
| chr17-82461486-INS-79 | chr17-82461497-INS-60 |
| chr17-82477662-INS-178 | chr17-82477662-INS-178 |
| chr17-82505396-INS-154 | chr17-82505396-INS-176 |
| chr17-82520434-INS-438 | chr17-82520434-INS-454 |
| chr17-82539538-INS-934 | chr17-82539538-INS-928 |
| chr17-82560383-INS-439 | chr17-82560383-INS-439 |
| chr17-82565637-INS-5550 | chr17-82565638-INS-5550 |
| chr17-82573290-INS-131 | chr17-82573290-INS-131 |
| chr17-82578947-INS-120 | chr17-82578947-INS-120 |
| chr17-82593079-INS-231 | chr17-82593076-INS-234 |
| chr17-82599501-INS-84 | chr17-82599485-INS-60 |
| chr17-82620818-INS-322 | chr17-82620818-INS-322 |
| chr17-82710077-INS-442 | chr17-82710077-INS-442 |
| chr17-82754347-INS-75 | chr17-82754347-INS-75 |
| chr17-82799400-INS-6826 | chr17-82799401-INS-6826 |
| chr17-82853330-INS-1082 | chr17-82853331-INS-1120 |
| chr17-82856486-INS-2264 | chr17-82856487-INS-2249 |
| chr17-82922630-INS-3874 | chr17-82922631-INS-3874 |
| chr17-82928380-INS-70 | chr17-82928380-INS-72 |
| chr17-82940336-INS-2244 | chr17-82940333-INS-2245 |
| chr17-82948043-INS-136 | chr17-82948043-INS-136 |
| chr17-82956812-INS-94 | chr17-82956812-INS-89 |
| chr17-82984204-INS-5509 | chr17-82984205-INS-5509 |
| chr17-82990581-INS-199 | chr17-82990581-INS-199 |
| chr17-82994726-INS-237 | chr17-82994726-INS-237 |
| chr17-83030019-INS-61 | chr17-83030030-INS-64 |
| chr17-83042725-INS-58 | chr17-83042725-INS-58 |
| chr17-83068360-INS-334 | chr17-83068360-INS-334 |
| chr17-83076759-INS-165 | chr17-83076759-INS-165 |
| chr17-83113948-INS-56 | chr17-83113948-INS-60 |
| chr17-83115836-INS-522 | chr17-83115836-INS-517 |
| chr17-83128474-INS-52 | chr17-83128450-INS-60 |
| chr17-83148159-INS-124 | chr17-83148160-INS-77 |
| chr17-83153828-INS-488 | chr17-83153828-INS-488 |
| chr17-83170602-INS-544 | chr17-83170603-INS-552 |
| chr17-83178090-INS-95 | chr17-83178098-INS-108 |
| chr17-83180379-INS-66 | chr17-83180379-INS-54 |
| chr17-83187357-INS-237 | chr17-83187357-INS-237 |
| chr17-83216013-INS-61 | chr17-83216014-INS-61 |
| chr17-83232381-INS-69 | chr17-83232381-INS-73 |
| chr17-83233072-INS-3503 | chr17-83233069-INS-3491 |
| chr17-83248588-INS-6577 | chr17-83248589-INS-6577 |

| | |
|---|---|
| chr17-83251562-INS-67 | chr17-83251562-INS-79 |
| chr17-83256876-INS-111 | chr17-83256870-INS-106 |
| chr17-83265261-INS-76 | chr17-83265215-INS-118 |
| chr17-83269001-INS-2854 | chr17-83269002-INS-2853 |
| chr17-83279307-INS-58 | chr17-83279307-INS-86 |
| chr17-83302763-INS-498 | chr17-83302763-INS-498 |
| chr17-83321559-INS-7079 | chr17-83321560-INS-7079 |
| chr17-83354680-INS-94 | chr17-83354695-INS-129 |
| chr17-83363294-INS-66 | chr17-83363294-INS-66 |
| chr17-83368755-INS-422 | chr17-83368755-INS-422 |
| chr17-83385771-INS-60 | chr17-83385769-INS-86 |
| chr17-83413959-INS-189 | chr17-83413947-INS-192 |
| chr17-83433590-INS-7166 | chr17-83433591-INS-7166 |
| chr17-83439832-INS-203 | chr17-83439832-INS-209 |
| chr17-83459679-INS-225 | chr17-83459679-INS-225 |
| chr17-83504403-INS-52 | chr17-83504403-INS-52 |
| chr17-83510733-INS-144 | chr17-83510732-INS-148 |
| chr17-83519223-INS-202 | chr17-83519223-INS-201 |
| chr17-83551536-INS-66 | chr17-83551536-INS-66 |
| chr17-83559883-INS-80 | chr17-83559883-INS-80 |
| chr17-83571962-INS-86 | chr17-83571946-INS-102 |
| chr17-83590434-INS-62 | chr17-83590434-INS-62 |
| chr17-83597694-INS-68 | chr17-83597694-INS-68 |
| chr17-83602494-INS-68 | chr17-83602492-INS-54 |
| chr17-83623575-INS-60 | chr17-83623575-INS-50 |
| chr17-83626077-INS-90 | chr17-83626078-INS-54 |
| chr17-83634709-INS-198 | chr17-83634709-INS-198 |
| chr17-83637519-INS-67 | chr17-83637519-INS-66 |
| chr17-83639245-INS-223 | chr17-83639245-INS-224 |
| chr17-83640621-INS-63 | chr17-83640621-INS-63 |
| chr17-83650350-INS-237 | chr17-83650350-INS-219 |
| chr17-83664248-INS-560 | chr17-83664248-INS-603 |
| chr17-83674623-INS-7180 | chr17-83674624-INS-7180 |
| chr17-83682426-INS-190 | chr17-83682426-INS-191 |
| chr17-83696892-INS-136 | chr17-83696892-INS-136 |
| chr17-83702565-INS-80 | chr17-83702565-INS-52 |
| chr17-83712791-INS-396 | chr17-83712791-INS-396 |
| chr17-83724094-INS-213 | chr17-83724094-INS-212 |
| chr17-83728126-INS-205 | chr17-83728126-INS-206 |
| chr17-83749479-INS-211 | chr17-83749479-INS-235 |
| chr17-83754104-INS-2950 | chr17-83754105-INS-2950 |
| chr17-83754852-INS-82 | chr17-83754852-INS-82 |
| chr17-83760898-INS-155 | chr17-83760898-INS-155 |
| chr17-83762781-INS-53 | chr17-83762781-INS-76 |
| chr17-83779994-INS-84 | chr17-83779994-INS-84 |
| chr17-83782740-INS-60 | chr17-83782740-INS-60 |
| chr17-83816385-INS-174 | chr17-83816385-INS-174 |
| chr17-83818673-INS-196 | chr17-83818673-INS-171 |
| chr17-83825932-INS-67 | chr17-83825932-INS-68 |
| chr17-83842703-INS-727 | chr17-83842703-INS-727 |
| chr17-83856844-INS-75 | chr17-83856844-INS-75 |
| chr17-83863135-INS-130 | chr17-83863135-INS-130 |
| chr17-83872514-INS-68 | chr17-83872514-INS-96 |
| chr17-83873911-INS-199 | chr17-83873911-INS-199 |
| chr17-83886380-INS-54 | chr17-83886380-INS-54 |
| chr17-83890538-INS-200 | chr17-83890538-INS-200 |
| chr17-83898304-INS-62 | chr17-83898304-INS-62 |
| chr17-83906451-INS-209 | chr17-83906441-INS-206 |
| chr17-83910065-INS-68 | chr17-83910065-INS-68 |
| chr17-83915050-INS-553 | chr17-83915022-INS-536 |
| chr17-83916749-INS-199 | chr17-83916750-INS-198 |
| chr17-83925963-INS-155 | chr17-83925963-INS-155 |
| chr17-83956455-INS-69 | chr17-83956452-INS-78 |
| chr17-83964572-INS-58 | chr17-83964572-INS-58 |
| chr17-83965862-INS-207 | chr17-83965862-INS-207 |
| chr17-83972138-INS-75 | chr17-83972138-INS-65 |
| chr17-83973944-INS-168 | chr17-83973945-INS-176 |
| chr17-83977608-INS-60 | chr17-83977608-INS-50 |
| chr17-83982258-INS-162 | chr17-83982258-INS-162 |
| chr17-83994391-INS-134 | chr17-83994391-INS-135 |
| chr17-84005207-INS-78 | chr17-84005186-INS-68 |
| chr17-84006875-INS-75 | chr17-84006849-INS-84 |
| chr17-84022229-INS-52 | chr17-84022229-INS-56 |
| chr17-84059984-INS-124 | chr17-84059966-INS-140 |
| chr17-84065042-INS-200 | chr17-84065042-INS-200 |
| chr17-84073646-INS-372 | chr17-84073646-INS-380 |
| chr17-84125155-INS-194 | chr17-84125155-INS-194 |

| | |
|---|---|
| chr17-84143398-INS-177 | chr17-84143398-INS-177 |
| chr17-84144779-INS-200 | chr17-84144777-INS-200 |
| chr17-84147300-INS-61 | chr17-84147300-INS-66 |
| chr17-84152339-INS-178 | chr17-84152339-INS-178 |
| chr17-84156185-INS-214 | chr17-84156185-INS-212 |
| chr17-84159765-INS-182 | chr17-84159765-INS-183 |
| chr17-84164875-INS-52 | chr17-84164898-INS-78 |
| chr17-84181741-INS-84 | chr17-84181745-INS-80 |
| chr17-84196069-INS-55 | chr17-84196069-INS-54 |
| chr17-84205790-INS-1091 | chr17-84205791-INS-1085 |
| chr17-84232980-INS-116 | chr17-84232948-INS-104 |
| chr17-84237061-INS-88 | chr17-84237061-INS-88 |
| chr17-84238978-INS-122 | chr17-84238978-INS-122 |
| chr17-84248178-INS-405 | chr17-84248178-INS-405 |
| chr17-84280667-INS-51 | chr17-84280667-INS-51 |
| chr17-84282609-INS-254 | chr17-84282609-INS-254 |
| chr17-84296269-INS-60 | chr17-84296269-INS-60 |
| chr17-84300185-INS-274 | chr17-84300187-INS-272 |
| chr17-84300614-INS-6453 | chr17-84300615-INS-6453 |
| chr17-84348578-INS-7112 | chr17-84348579-INS-7112 |
| chr17-84357188-INS-146 | chr17-84357188-INS-126 |
| chr17-84360322-INS-204 | chr17-84360322-INS-203 |
| chr17-84385056-INS-52 | chr17-84385054-INS-52 |
| chr17-84401606-INS-185 | chr17-84401598-INS-187 |
| chr17-84408541-INS-59 | chr17-84408541-INS-75 |
| chr17-84409457-INS-101 | chr17-84409457-INS-101 |
| chr17-84413647-INS-58 | chr17-84413647-INS-60 |
| chr17-84429192-INS-123 | chr17-84429192-INS-124 |
| chr17-84430571-INS-244 | chr17-84430572-INS-222 |
| chr17-84434588-INS-66 | chr17-84434588-INS-66 |
| chr17-84444275-INS-248 | chr17-84444275-INS-248 |
| chr17-84457636-INS-191 | chr17-84457636-INS-191 |
| chr17-84498780-INS-211 | chr17-84498780-INS-213 |
| chr17-84514953-INS-219 | chr17-84514953-INS-221 |
| chr17-84533309-INS-216 | chr17-84533309-INS-216 |
| chr17-84554566-INS-54 | chr17-84554566-INS-96 |
| chr17-84564204-INS-162 | chr17-84564198-INS-170 |
| chr17-84568143-INS-72 | chr17-84568143-INS-77 |
| chr17-84583505-INS-72 | chr17-84583505-INS-96 |
| chr17-84585540-INS-58 | chr17-84585540-INS-62 |
| chr17-84593327-INS-128 | chr17-84593327-INS-128 |
| chr17-84615122-INS-56 | chr17-84615122-INS-68 |
| chr17-84618694-INS-7651 | chr17-84618695-INS-7651 |
| chr17-84623476-INS-78 | chr17-84623522-INS-64 |
| chr17-84636200-INS-88 | chr17-84636200-INS-88 |
| chr17-84645253-INS-74 | chr17-84645252-INS-84 |
| chr17-84652845-INS-487 | chr17-84652845-INS-487 |
| chr17-84660803-INS-111 | chr17-84660813-INS-99 |
| chr17-84688495-INS-181 | chr17-84688495-INS-217 |
| chr17-84702704-INS-202 | chr17-84702704-INS-202 |
| chr17-84707492-INS-1109 | chr17-84707493-INS-1109 |
| chr17-84714316-INS-201 | chr17-84714316-INS-202 |
| chr17-84731210-INS-156 | chr17-84731213-INS-149 |
| chr17-84735670-INS-203 | chr17-84735670-INS-205 |
| chr17-84774632-INS-53 | chr17-84774632-INS-53 |
| chr17-84808557-INS-214 | chr17-84808557-INS-218 |
| chr17-84810770-INS-219 | chr17-84810770-INS-220 |
| chr17-84854066-INS-395 | chr17-84854066-INS-395 |
| chr17-84854962-INS-195 | chr17-84854962-INS-195 |
| chr17-84894514-INS-884 | chr17-84894514-INS-885 |
| chr17-84895399-INS-279 | chr17-84895399-INS-259 |
| chr17-84908409-INS-84 | chr17-84908409-INS-73 |
| chr17-84928028-INS-173 | chr17-84928028-INS-173 |
| chr17-84933431-INS-125 | chr17-84933431-INS-113 |
| chr17-84940202-INS-196 | chr17-84940202-INS-196 |
| chr17-84942214-INS-60 | chr17-84942215-INS-60 |
| chr17-84955893-INS-106 | chr17-84955893-INS-106 |
| chr17-84975326-INS-78 | chr17-84975325-INS-73 |
| chr17-84981958-INS-63 | chr17-84981979-INS-50 |
| chr17-84984498-INS-54 | chr17-84984498-INS-52 |
| chr17-84987793-INS-66 | chr17-84987793-INS-52 |
| chr17-84995186-INS-69 | chr17-84995186-INS-69 |
| chr17-85002867-INS-70 | chr17-85002892-INS-54 |
| chr17-85005546-INS-74 | chr17-85005546-INS-74 |
| chr17-85038850-INS-155 | chr17-85038850-INS-155 |
| chr17-85044556-INS-92 | chr17-85044556-INS-98 |
| chr17-85049148-INS-56 | chr17-85049148-INS-60 |

| | |
|---|---|
| chr17-85055493-INS-54 | chr17-85055493-INS-54 |
| chr17-85066955-INS-125 | chr17-85066955-INS-119 |
| chr17-85072767-INS-199 | chr17-85072767-INS-200 |
| chr17-85079803-INS-59 | chr17-85079803-INS-59 |
| chr17-85090689-INS-273 | chr17-85090689-INS-273 |
| chr17-85110191-INS-56 | chr17-85110191-INS-56 |
| chr17-85118595-INS-397 | chr17-85118595-INS-397 |
| chr17-85136333-INS-113 | chr17-85136333-INS-108 |
| chr17-85159695-INS-60 | chr17-85159695-INS-60 |
| chr17-85175334-INS-139 | chr17-85175334-INS-143 |
| chr17-85188244-INS-353 | chr17-85188244-INS-333 |
| chr17-85197806-INS-142 | chr17-85197806-INS-94 |
| chr17-85204953-INS-59 | chr17-85204953-INS-59 |
| chr17-85207942-INS-75 | chr17-85207905-INS-67 |
| chr17-85217077-INS-359 | chr17-85217077-INS-363 |
| chr17-85222462-INS-96 | chr17-85222463-INS-62 |
| chr17-85230196-INS-172 | chr17-85230196-INS-174 |
| chr17-85231830-INS-63 | chr17-85231830-INS-65 |
| chr17-85236649-INS-333 | chr17-85236658-INS-320 |
| chr17-85244904-INS-73 | chr17-85244904-INS-73 |
| chr17-85255340-INS-1285 | chr17-85255341-INS-1285 |
| chr17-85347318-INS-124 | chr17-85347319-INS-152 |
| chr17-85360796-INS-120 | chr17-85360801-INS-100 |
| chr17-85361913-INS-60 | chr17-85361914-INS-64 |
| chr17-85363055-INS-139 | chr17-85363055-INS-92 |
| chr17-85389199-INS-102 | chr17-85389199-INS-102 |
| chr17-85404394-INS-52 | chr17-85404394-INS-52 |
| chr17-85471038-INS-70 | chr17-85471039-INS-57 |
| chr17-85498709-INS-56 | chr17-85498709-INS-56 |
| chr17-85518060-INS-58 | chr17-85518060-INS-62 |
| chr17-85551188-INS-56 | chr17-85551187-INS-64 |
| chr17-85560783-INS-180 | chr17-85560783-INS-180 |
| chr17-85593657-INS-58 | chr17-85593690-INS-61 |
| chr17-85609586-INS-210 | chr17-85609586-INS-210 |
| chr17-85624306-INS-210 | chr17-85624306-INS-211 |
| chr17-85635331-INS-63 | chr17-85635331-INS-57 |
| chr17-85639537-INS-73 | chr17-85639537-INS-77 |
| chr17-85642525-INS-216 | chr17-85642523-INS-198 |
| chr17-85645908-INS-249 | chr17-85645908-INS-237 |
| chr17-85706693-INS-60 | chr17-85706688-INS-87 |
| chr17-85731071-INS-76 | chr17-85731071-INS-73 |
| chr17-85743050-INS-63 | chr17-85743050-INS-63 |
| chr17-85746094-INS-351 | chr17-85746094-INS-354 |
| chr17-85828757-INS-132 | chr17-85828757-INS-117 |
| chr17-85829413-INS-178 | chr17-85829413-INS-178 |
| chr17-85840966-INS-375 | chr17-85840967-INS-392 |
| chr17-85867786-INS-96 | chr17-85867786-INS-144 |
| chr17-85872241-INS-52 | chr17-85872241-INS-58 |
| chr17-85885344-INS-102 | chr17-85885344-INS-102 |
| chr17-85900705-INS-136 | chr17-85900705-INS-128 |
| chr17-85937694-INS-91 | chr17-85937694-INS-95 |
| chr17-85940868-INS-64 | chr17-85940868-INS-62 |
| chr17-85954845-INS-108 | chr17-85954845-INS-108 |
| chr17-85974937-INS-96 | chr17-85974937-INS-80 |
| chr17-85993512-INS-73 | chr17-85993512-INS-77 |
| chr17-86009897-INS-103 | chr17-86009897-INS-96 |
| chr17-86014884-INS-156 | chr17-86014884-INS-169 |
| chr17-86023313-INS-492 | chr17-86023313-INS-492 |
| chr17-86045408-INS-497 | chr17-86045408-INS-497 |
| chr17-86046673-INS-306 | chr17-86046673-INS-256 |
| chr17-86052137-INS-75 | chr17-86052137-INS-75 |
| chr17-86064716-INS-245 | chr17-86064716-INS-257 |
| chr17-86066657-INS-207 | chr17-86066657-INS-207 |
| chr17-86072262-INS-56 | chr17-86072262-INS-56 |
| chr17-86132500-INS-52 | chr17-86132500-INS-52 |
| chr17-86137694-INS-934 | chr17-86137694-INS-931 |
| chr17-86163004-INS-103 | chr17-86163004-INS-103 |
| chr17-86171310-INS-410 | chr17-86171310-INS-416 |
| chr17-86191893-INS-60 | chr17-86191893-INS-60 |
| chr17-86195832-INS-56 | chr17-86195788-INS-88 |
| chr17-86216429-INS-1486 | chr17-86216430-INS-1486 |
| chr17-86249648-INS-60 | chr17-86249648-INS-60 |
| chr17-86262327-INS-92 | chr17-86262327-INS-92 |
| chr17-86267115-INS-52 | chr17-86267115-INS-56 |
| chr17-86271058-INS-4557 | chr17-86271059-INS-4558 |
| chr17-86308413-INS-524 | chr17-86308401-INS-546 |
| chr17-86310634-INS-65 | chr17-86310634-INS-60 |

| | |
|---|---|
| chr17-86380729-INS-516 | chr17-86380729-INS-516 |
| chr17-86391862-INS-142 | chr17-86391862-INS-142 |
| chr17-86400407-INS-92 | chr17-86400407-INS-94 |
| chr17-86418124-INS-60 | chr17-86418124-INS-55 |
| chr17-86431261-INS-87 | chr17-86431262-INS-87 |
| chr17-86465081-INS-56 | chr17-86465081-INS-56 |
| chr17-86468654-INS-96 | chr17-86468654-INS-96 |
| chr17-86470693-INS-6487 | chr17-86470694-INS-6487 |
| chr17-86497146-INS-65 | chr17-86497146-INS-65 |
| chr17-86514999-INS-210 | chr17-86514999-INS-206 |
| chr17-86521655-INS-150 | chr17-86521655-INS-142 |
| chr17-86534782-INS-445 | chr17-86534782-INS-438 |
| chr17-86536237-INS-84 | chr17-86536237-INS-84 |
| chr17-86540563-INS-76 | chr17-86540563-INS-76 |
| chr17-86549573-INS-236 | chr17-86549573-INS-236 |
| chr17-86559649-INS-61 | chr17-86559649-INS-61 |
| chr17-86562922-INS-59 | chr17-86562922-INS-55 |
| chr17-86591996-INS-52 | chr17-86591997-INS-70 |
| chr17-86648813-INS-62 | chr17-86648813-INS-60 |
| chr17-86664875-INS-825 | chr17-86664850-INS-800 |
| chr17-86704822-INS-81 | chr17-86704822-INS-84 |
| chr17-86713804-INS-6720 | chr17-86713805-INS-6722 |
| chr17-86724931-INS-206 | chr17-86724931-INS-243 |
| chr17-86731778-INS-66 | chr17-86731778-INS-66 |
| chr17-86747527-INS-56 | chr17-86747527-INS-58 |
| chr17-86749344-INS-74 | chr17-86749344-INS-74 |
| chr17-86753676-INS-372 | chr17-86753676-INS-372 |
| chr17-86761285-INS-60 | chr17-86761285-INS-60 |
| chr17-86805331-INS-108 | chr17-86805331-INS-108 |
| chr17-86820198-INS-54 | chr17-86820198-INS-54 |
| chr17-86823299-INS-134 | chr17-86823299-INS-134 |
| chr17-86845726-INS-61 | chr17-86845694-INS-61 |
| chr17-86871936-INS-52 | chr17-86871936-INS-52 |
| chr17-86896768-INS-55 | chr17-86896768-INS-55 |
| chr17-86907268-INS-208 | chr17-86907268-INS-208 |
| chr17-86910734-INS-273 | chr17-86910734-INS-273 |
| chr17-86935002-INS-210 | chr17-86934995-INS-220 |
| chr17-86962873-INS-75 | chr17-86962873-INS-72 |
| chr17-86984839-INS-52 | chr17-86984839-INS-52 |
| chr17-86984866-INS-1985 | chr17-86984867-INS-1985 |
| chr17-86986394-INS-145 | chr17-86986394-INS-145 |
| chr17-86996774-INS-547 | chr17-86996774-INS-511 |
| chr17-87000993-INS-155 | chr17-87000989-INS-183 |
| chr17-87004757-INS-123 | chr17-87004757-INS-123 |
| chr17-87011042-INS-84 | chr17-87011042-INS-84 |
| chr17-87013591-INS-96 | chr17-87013622-INS-113 |
| chr17-87038121-INS-94 | chr17-87038121-INS-94 |
| chr17-87043439-INS-165 | chr17-87043439-INS-165 |
| chr17-87054851-INS-195 | chr17-87054851-INS-194 |
| chr17-87091252-INS-230 | chr17-87091252-INS-230 |
| chr17-87095263-INS-251 | chr17-87095263-INS-251 |
| chr17-87105109-INS-200 | chr17-87105109-INS-200 |
| chr17-87110973-INS-724 | chr17-87110973-INS-724 |
| chr17-87117803-INS-671 | chr17-87117803-INS-659 |
| chr17-87127099-INS-54 | chr17-87127099-INS-56 |
| chr17-87139125-INS-86 | chr17-87139110-INS-100 |
| chr17-87144493-INS-96 | chr17-87144482-INS-101 |
| chr17-87153730-INS-59 | chr17-87153730-INS-77 |
| chr17-87169424-INS-157 | chr17-87169424-INS-157 |
| chr17-87173993-INS-54 | chr17-87173993-INS-54 |
| chr17-87196099-INS-103 | chr17-87196093-INS-107 |
| chr17-87202249-INS-230 | chr17-87202249-INS-230 |
| chr17-87209416-INS-392 | chr17-87209416-INS-392 |
| chr17-87223358-INS-96 | chr17-87223358-INS-99 |
| chr17-87235163-INS-466 | chr17-87235163-INS-466 |
| chr17-87247981-INS-188 | chr17-87247981-INS-188 |
| chr17-87315616-INS-70 | chr17-87315630-INS-73 |
| chr17-87317776-INS-64 | chr17-87317776-INS-68 |
| chr17-87325176-INS-398 | chr17-87325176-INS-398 |
| chr17-87338996-INS-58 | chr17-87338996-INS-58 |
| chr17-87349431-INS-52 | chr17-87349431-INS-52 |
| chr17-87355500-INS-81 | chr17-87355500-INS-108 |
| chr17-87360064-INS-95 | chr17-87360065-INS-81 |
| chr17-87360917-INS-110 | chr17-87360917-INS-110 |
| chr17-87388629-INS-69 | chr17-87388629-INS-69 |
| chr17-87397286-INS-87 | chr17-87397290-INS-126 |
| chr17-87433908-INS-5386 | chr17-87433909-INS-5386 |

| | |
|---|---|
| chr17-87456918-INS-78 | chr17-87456918-INS-77 |
| chr17-87459267-INS-160 | chr17-87459267-INS-160 |
| chr17-87472745-INS-5372 | chr17-87472746-INS-5372 |
| chr17-87493946-INS-2161 | chr17-87493947-INS-2143 |
| chr17-87509504-INS-88 | chr17-87509500-INS-56 |
| chr17-87514119-INS-81 | chr17-87514115-INS-82 |
| chr17-87517949-INS-250 | chr17-87517953-INS-248 |
| chr17-87518726-INS-126 | chr17-87518678-INS-156 |
| chr17-87522502-INS-126 | chr17-87522502-INS-126 |
| chr17-87524868-INS-174 | chr17-87524868-INS-202 |
| chr17-87529660-INS-63 | chr17-87529660-INS-63 |
| chr17-87535893-INS-210 | chr17-87535893-INS-210 |
| chr17-87545912-INS-55 | chr17-87545912-INS-55 |
| chr17-87555935-INS-117 | chr17-87555935-INS-157 |
| chr17-87565897-INS-304 | chr17-87565897-INS-316 |
| chr17-87566932-INS-242 | chr17-87566932-INS-242 |
| chr17-87602265-INS-74 | chr17-87602277-INS-58 |
| chr17-87631257-INS-580 | chr17-87631257-INS-580 |
| chr17-87632245-INS-125 | chr17-87632245-INS-125 |
| chr17-87640986-INS-7080 | chr17-87640987-INS-7080 |
| chr17-87673628-INS-64 | chr17-87673628-INS-60 |
| chr17-87682176-INS-55 | chr17-87682176-INS-55 |
| chr17-87695734-INS-373 | chr17-87695734-INS-417 |
| chr17-87714135-INS-368 | chr17-87714086-INS-395 |
| chr17-87720635-INS-378 | chr17-87720635-INS-378 |
| chr17-87722584-INS-84 | chr17-87722584-INS-84 |
| chr17-87778976-INS-359 | chr17-87779019-INS-360 |
| chr17-87784854-INS-1251 | chr17-87784855-INS-1251 |
| chr17-87812735-INS-2333 | chr17-87812736-INS-2329 |
| chr17-87818261-INS-220 | chr17-87818261-INS-220 |
| chr17-87827047-INS-125 | chr17-87827047-INS-127 |
| chr17-87829222-INS-216 | chr17-87829222-INS-216 |
| chr17-87844583-INS-76 | chr17-87844583-INS-68 |
| chr17-87864030-INS-164 | chr17-87864030-INS-164 |
| chr17-87885779-INS-344 | chr17-87885779-INS-344 |
| chr17-87892689-INS-209 | chr17-87892689-INS-209 |
| chr17-87900571-INS-167 | chr17-87900571-INS-167 |
| chr17-87903352-INS-183 | chr17-87903352-INS-183 |
| chr17-87914794-INS-69 | chr17-87914795-INS-78 |
| chr17-87923080-INS-88 | chr17-87923080-INS-90 |
| chr17-87941111-INS-223 | chr17-87941111-INS-224 |
| chr17-87954276-INS-1969 | chr17-87954277-INS-1969 |
| chr17-87987599-INS-198 | chr17-87987599-INS-198 |
| chr17-87989766-INS-107 | chr17-87989766-INS-146 |
| chr17-88006057-INS-207 | chr17-88006057-INS-205 |
| chr17-88010271-INS-144 | chr17-88010271-INS-96 |
| chr17-88019881-INS-212 | chr17-88019881-INS-238 |
| chr17-88031165-INS-196 | chr17-88031165-INS-196 |
| chr17-88037913-INS-55 | chr17-88037914-INS-55 |
| chr17-88045529-INS-145 | chr17-88045529-INS-145 |
| chr17-88063601-INS-63 | chr17-88063603-INS-66 |
| chr17-88082836-INS-192 | chr17-88082836-INS-192 |
| chr17-88086901-INS-61 | chr17-88086902-INS-66 |
| chr17-88108584-INS-60 | chr17-88108586-INS-52 |
| chr17-88109596-INS-311 | chr17-88109603-INS-323 |
| chr17-88112206-INS-7172 | chr17-88112207-INS-7173 |
| chr17-88126596-INS-247 | chr17-88126608-INS-253 |
| chr17-88129268-INS-99 | chr17-88129268-INS-105 |
| chr17-88133485-INS-62 | chr17-88133451-INS-50 |
| chr17-88135033-INS-197 | chr17-88135031-INS-214 |
| chr17-88141111-INS-69 | chr17-88141093-INS-66 |
| chr17-88153775-INS-57 | chr17-88153775-INS-57 |
| chr17-88155421-INS-360 | chr17-88155427-INS-363 |
| chr17-88160321-INS-79 | chr17-88160322-INS-80 |
| chr17-88168165-INS-138 | chr17-88168211-INS-135 |
| chr17-88202209-INS-62 | chr17-88202209-INS-64 |
| chr17-88222698-INS-184 | chr17-88222698-INS-186 |
| chr17-88248505-INS-182 | chr17-88248505-INS-182 |
| chr17-88272630-INS-76 | chr17-88272630-INS-76 |
| chr17-88275237-INS-124 | chr17-88275237-INS-130 |
| chr17-88317809-INS-238 | chr17-88317809-INS-238 |
| chr17-88350692-INS-206 | chr17-88350692-INS-206 |
| chr17-88391251-INS-52 | chr17-88391252-INS-68 |
| chr17-88394355-INS-212 | chr17-88394354-INS-229 |
| chr17-88397827-INS-86 | chr17-88397810-INS-54 |
| chr17-88408991-INS-143 | chr17-88408991-INS-144 |
| chr17-88441157-INS-58 | chr17-88441157-INS-87 |

| | |
|---|---|
| chr17-88480657-INS-90 | chr17-88480657-INS-88 |
| chr17-88486840-INS-439 | chr17-88486840-INS-439 |
| chr17-88496686-INS-189 | chr17-88496686-INS-189 |
| chr17-88498068-INS-64 | chr17-88498068-INS-64 |
| chr17-88501960-INS-67 | chr17-88501960-INS-67 |
| chr17-88504337-INS-242 | chr17-88504337-INS-260 |
| chr17-88508096-INS-178 | chr17-88508097-INS-178 |
| chr17-88514694-INS-60 | chr17-88514712-INS-54 |
| chr17-88530913-INS-89 | chr17-88530913-INS-108 |
| chr17-88540532-INS-279 | chr17-88540577-INS-257 |
| chr17-88543711-INS-4841 | chr17-88543712-INS-4841 |
| chr17-88577026-INS-343 | chr17-88577026-INS-343 |
| chr17-88595103-INS-8328 | chr17-88595104-INS-8327 |
| chr17-88599413-INS-176 | chr17-88599413-INS-170 |
| chr17-88601245-INS-219 | chr17-88601245-INS-219 |
| chr17-88606228-INS-115 | chr17-88606228-INS-113 |
| chr17-88610770-INS-204 | chr17-88610770-INS-207 |
| chr17-88612349-INS-181 | chr17-88612349-INS-186 |
| chr17-88628062-INS-261 | chr17-88628062-INS-260 |
| chr17-88630845-INS-868 | chr17-88630845-INS-887 |
| chr17-88631808-INS-405 | chr17-88631794-INS-384 |
| chr17-88642961-INS-550 | chr17-88642961-INS-550 |
| chr17-88664026-INS-92 | chr17-88664026-INS-79 |
| chr17-88666455-INS-197 | chr17-88666455-INS-197 |
| chr17-88681492-INS-78 | chr17-88681492-INS-78 |
| chr17-88682765-INS-55 | chr17-88682771-INS-54 |
| chr17-88696659-INS-64 | chr17-88696659-INS-66 |
| chr17-88714449-INS-65 | chr17-88714449-INS-65 |
| chr17-88719713-INS-206 | chr17-88719713-INS-205 |
| chr17-88720740-INS-204 | chr17-88720740-INS-204 |
| chr17-88726980-INS-191 | chr17-88726980-INS-188 |
| chr17-88730656-INS-496 | chr17-88730656-INS-502 |
| chr17-88738494-INS-110 | chr17-88738494-INS-110 |
| chr17-88742089-INS-110 | chr17-88742088-INS-105 |
| chr17-88746158-INS-137 | chr17-88746158-INS-138 |
| chr17-88782537-INS-108 | chr17-88782537-INS-108 |
| chr17-88802731-INS-189 | chr17-88802731-INS-147 |
| chr17-88817487-INS-57 | chr17-88817487-INS-57 |
| chr17-88833526-INS-2986 | chr17-88833527-INS-2987 |
| chr17-88834029-INS-51 | chr17-88834028-INS-86 |
| chr17-88839760-INS-82 | chr17-88839760-INS-88 |
| chr17-88861190-INS-70 | chr17-88861190-INS-65 |
| chr17-88862123-INS-54 | chr17-88862123-INS-54 |
| chr17-88870919-INS-5305 | chr17-88870920-INS-5305 |
| chr17-88898454-INS-135 | chr17-88898454-INS-135 |
| chr17-88916932-INS-54 | chr17-88916932-INS-54 |
| chr17-88932306-INS-99 | chr17-88932306-INS-99 |
| chr17-88937123-INS-141 | chr17-88937123-INS-141 |
| chr17-88947686-INS-176 | chr17-88947686-INS-172 |
| chr17-88964212-INS-807 | chr17-88964212-INS-807 |
| chr17-88972413-INS-72 | chr17-88972413-INS-72 |
| chr17-88977909-INS-56 | chr17-88977919-INS-78 |
| chr17-88980291-INS-102 | chr17-88980300-INS-136 |
| chr17-88982423-INS-78 | chr17-88982423-INS-80 |
| chr17-88999036-INS-55 | chr17-88999036-INS-55 |
| chr17-89003103-INS-400 | chr17-89003103-INS-400 |
| chr17-89008484-INS-71 | chr17-89008504-INS-71 |
| chr17-89011046-INS-56 | chr17-89011046-INS-56 |
| chr17-89015394-INS-72 | chr17-89015394-INS-72 |
| chr17-89039727-INS-754 | chr17-89039727-INS-753 |
| chr17-89044251-INS-295 | chr17-89044249-INS-297 |
| chr17-89053922-INS-68 | chr17-89053922-INS-68 |
| chr17-89058992-INS-52 | chr17-89058992-INS-64 |
| chr17-89062828-INS-342 | chr17-89062828-INS-342 |
| chr17-89074531-INS-56 | chr17-89074531-INS-50 |
| chr17-89085989-INS-296 | chr17-89085989-INS-296 |
| chr17-89102799-INS-217 | chr17-89102799-INS-217 |
| chr17-89115306-INS-60 | chr17-89115306-INS-60 |
| chr17-89120500-INS-96 | chr17-89120500-INS-96 |
| chr17-89122806-INS-128 | chr17-89122806-INS-128 |
| chr17-89136937-INS-54 | chr17-89136937-INS-54 |
| chr17-89140171-INS-51 | chr17-89140171-INS-51 |
| chr17-89150787-INS-159 | chr17-89150787-INS-159 |
| chr17-89153330-INS-382 | chr17-89153295-INS-373 |
| chr17-89172862-INS-134 | chr17-89172863-INS-100 |
| chr17-89179438-INS-56 | chr17-89179438-INS-56 |
| chr17-89186019-INS-346 | chr17-89186019-INS-347 |

| | |
|---|---|
| chr17-89196363-INS-77 | chr17-89196363-INS-82 |
| chr17-89205713-INS-458 | chr17-89205713-INS-458 |
| chr17-89238081-INS-136 | chr17-89238095-INS-118 |
| chr17-89239111-INS-233 | chr17-89239111-INS-233 |
| chr17-89265246-INS-99 | chr17-89265230-INS-132 |
| chr17-89299349-INS-65 | chr17-89299350-INS-97 |
| chr17-89317802-INS-291 | chr17-89317802-INS-300 |
| chr17-89330151-INS-484 | chr17-89330151-INS-484 |
| chr17-89374911-INS-76 | chr17-89374911-INS-76 |
| chr17-89391890-INS-472 | chr17-89391890-INS-473 |
| chr17-89404172-INS-6360 | chr17-89404173-INS-6360 |
| chr17-89431934-INS-299 | chr17-89431907-INS-289 |
| chr17-89467268-INS-326 | chr17-89467268-INS-326 |
| chr17-89470291-INS-60 | chr17-89470291-INS-60 |
| chr17-89498156-INS-375 | chr17-89498156-INS-375 |
| chr17-89524960-INS-56 | chr17-89524960-INS-56 |
| chr17-89527805-INS-437 | chr17-89527805-INS-437 |
| chr17-89532660-INS-125 | chr17-89532660-INS-125 |
| chr17-89556653-INS-160 | chr17-89556654-INS-139 |
| chr17-89562325-INS-69 | chr17-89562325-INS-71 |
| chr17-89562975-INS-516 | chr17-89562975-INS-515 |
| chr17-89583898-INS-68 | chr17-89583898-INS-85 |
| chr17-89593163-INS-318 | chr17-89593160-INS-317 |
| chr17-89602993-INS-4725 | chr17-89602994-INS-4725 |
| chr17-89603772-INS-56 | chr17-89603772-INS-50 |
| chr17-89617645-INS-52 | chr17-89617645-INS-52 |
| chr17-89629283-INS-80 | chr17-89629283-INS-80 |
| chr17-89643807-INS-4527 | chr17-89643808-INS-4527 |
| chr17-89651614-INS-114 | chr17-89651632-INS-92 |
| chr17-89657603-INS-73 | chr17-89657603-INS-65 |
| chr17-89659682-INS-985 | chr17-89659683-INS-982 |
| chr17-89690147-INS-98 | chr17-89690147-INS-100 |
| chr17-89719082-INS-52 | chr17-89719082-INS-98 |
| chr17-89734199-INS-704 | chr17-89734199-INS-704 |
| chr17-89737864-INS-2104 | chr17-89737865-INS-2104 |
| chr17-89746794-INS-165 | chr17-89746794-INS-169 |
| chr17-89770485-INS-79 | chr17-89770485-INS-86 |
| chr17-89772836-INS-574 | chr17-89772836-INS-574 |
| chr17-89776967-INS-274 | chr17-89776956-INS-238 |
| chr17-89803117-INS-202 | chr17-89803118-INS-205 |
| chr17-89821599-INS-56 | chr17-89821588-INS-58 |
| chr17-89840203-INS-91 | chr17-89840208-INS-93 |
| chr17-89864967-INS-817 | chr17-89864967-INS-817 |
| chr17-89885170-INS-137 | chr17-89885170-INS-137 |
| chr17-89936024-INS-124 | chr17-89936029-INS-84 |
| chr17-89937522-INS-70 | chr17-89937522-INS-70 |
| chr17-89939741-INS-305 | chr17-89939741-INS-287 |
| chr17-89951902-INS-73 | chr17-89951942-INS-64 |
| chr17-89978621-INS-683 | chr17-89978621-INS-684 |
| chr17-89979985-INS-94 | chr17-89979985-INS-72 |
| chr17-90019723-INS-352 | chr17-90019723-INS-352 |
| chr17-90024126-INS-7632 | chr17-90024137-INS-7613 |
| chr17-90047801-INS-70 | chr17-90047801-INS-70 |
| chr17-90057585-INS-354 | chr17-90057585-INS-354 |
| chr17-90067697-INS-6917 | chr17-90067698-INS-6923 |
| chr17-90069533-INS-67 | chr17-90069533-INS-67 |
| chr17-90075306-INS-141 | chr17-90075328-INS-105 |
| chr17-90099271-INS-75 | chr17-90099271-INS-79 |
| chr17-90164180-INS-210 | chr17-90164180-INS-212 |
| chr17-90179932-INS-84 | chr17-90179968-INS-80 |
| chr17-90180314-INS-363 | chr17-90180314-INS-363 |
| chr17-90214939-INS-113 | chr17-90214939-INS-113 |
| chr17-90228548-INS-136 | chr17-90228551-INS-164 |
| chr17-90229972-INS-110 | chr17-90229974-INS-122 |
| chr17-90250836-INS-205 | chr17-90250836-INS-205 |
| chr17-90282906-INS-310 | chr17-90282900-INS-310 |
| chr17-90288503-INS-67 | chr17-90288494-INS-56 |
| chr17-90292579-INS-56 | chr17-90292581-INS-56 |
| chr17-90320025-INS-1209 | chr17-90320026-INS-1208 |
| chr17-90351413-INS-260 | chr17-90351413-INS-260 |
| chr17-90357107-INS-51 | chr17-90357107-INS-51 |
| chr17-90360959-INS-56 | chr17-90360959-INS-56 |
| chr17-90409063-INS-123 | chr17-90409063-INS-90 |
| chr17-90423403-INS-226 | chr17-90423403-INS-225 |
| chr17-90427077-INS-59 | chr17-90427077-INS-59 |
| chr17-90433241-INS-1894 | chr17-90433242-INS-1894 |
| chr17-90444879-INS-1885 | chr17-90444880-INS-1885 |

| | |
|---|---|
| chr17-90448605-INS-66 | chr17-90448605-INS-66 |
| chr17-90450161-INS-65 | chr17-90450184-INS-87 |
| chr17-90452912-INS-365 | chr17-90452912-INS-365 |
| chr17-90491822-INS-182 | chr17-90491822-INS-149 |
| chr17-90499599-INS-56 | chr17-90499599-INS-50 |
| chr17-90519253-INS-91 | chr17-90519253-INS-91 |
| chr17-90535084-INS-354 | chr17-90535084-INS-354 |
| chr17-90546089-INS-57 | chr17-90546089-INS-57 |
| chr17-90555979-INS-7952 | chr17-90555980-INS-7952 |
| chr17-90572423-INS-201 | chr17-90572423-INS-200 |
| chr17-90579003-INS-211 | chr17-90579003-INS-211 |
| chr17-90585112-INS-66 | chr17-90585117-INS-70 |
| chr17-90587135-INS-177 | chr17-90587135-INS-177 |
| chr17-90611460-INS-1732 | chr17-90611465-INS-1729 |
| chr17-90614465-INS-6673 | chr17-90614466-INS-6673 |
| chr17-90654053-INS-2291 | chr17-90654054-INS-2291 |
| chr17-90666539-INS-6945 | chr17-90666540-INS-6945 |
| chr17-90714375-INS-74 | chr17-90714375-INS-74 |
| chr17-90780441-INS-115 | chr17-90780441-INS-116 |
| chr17-90798937-INS-3420 | chr17-90798938-INS-3418 |
| chr17-90806438-INS-522 | chr17-90806438-INS-522 |
| chr17-90870734-INS-99 | chr17-90870734-INS-99 |
| chr17-90875596-INS-60 | chr17-90875596-INS-60 |
| chr17-90909462-INS-90 | chr17-90909470-INS-86 |
| chr17-90940085-INS-52 | chr17-90940086-INS-57 |
| chr17-90958226-INS-64 | chr17-90958210-INS-85 |
| chr17-90985770-INS-764 | chr17-90985770-INS-756 |
| chr17-90997667-INS-58 | chr17-90997667-INS-54 |
| chr17-91015674-INS-56 | chr17-91015674-INS-52 |
| chr17-91016353-INS-1880 | chr17-91016354-INS-1881 |
| chr17-91051443-INS-351 | chr17-91051443-INS-351 |
| chr17-91062752-INS-235 | chr17-91062752-INS-235 |
| chr17-91074879-INS-1320 | chr17-91074880-INS-1327 |
| chr17-91112966-INS-156 | chr17-91112964-INS-158 |
| chr17-91122513-INS-790 | chr17-91122513-INS-790 |
| chr17-91127026-INS-54 | chr17-91127026-INS-50 |
| chr17-91138044-INS-2421 | chr17-91138045-INS-2421 |
| chr17-91144120-INS-903 | chr17-91144120-INS-903 |
| chr17-91159766-INS-4147 | chr17-91159767-INS-4147 |
| chr17-91172873-INS-102 | chr17-91172873-INS-102 |
| chr17-91177480-INS-1091 | chr17-91177481-INS-1091 |
| chr17-91210296-INS-1321 | chr17-91210297-INS-1321 |
| chr17-91219268-INS-6758 | chr17-91219269-INS-6758 |
| chr17-91256794-INS-807 | chr17-91256794-INS-809 |
| chr17-91262355-INS-5235 | chr17-91262356-INS-5235 |
| chr17-91262991-INS-210 | chr17-91262991-INS-210 |
| chr17-91300453-INS-212 | chr17-91300453-INS-212 |
| chr17-91305959-INS-628 | chr17-91305959-INS-630 |
| chr17-91315415-INS-54 | chr17-91315426-INS-59 |
| chr17-91318747-INS-59 | chr17-91318747-INS-59 |
| chr17-91321873-INS-4387 | chr17-91321874-INS-4387 |
| chr17-91330155-INS-57 | chr17-91330155-INS-57 |
| chr17-91344779-INS-7191 | chr17-91344780-INS-7191 |
| chr17-91392346-INS-64 | chr17-91392344-INS-64 |
| chr17-91411403-INS-68 | chr17-91411403-INS-70 |
| chr17-91420710-INS-122 | chr17-91420710-INS-118 |
| chr17-91438851-INS-442 | chr17-91438851-INS-442 |
| chr17-91444730-INS-534 | chr17-91444749-INS-545 |
| chr17-91461443-INS-53 | chr17-91461443-INS-55 |
| chr17-91475540-INS-2731 | chr17-91475541-INS-2731 |
| chr17-91490585-INS-415 | chr17-91490585-INS-416 |
| chr17-91491806-INS-4074 | chr17-91491807-INS-4074 |
| chr17-91499769-INS-238 | chr17-91499767-INS-242 |
| chr17-91508945-INS-243 | chr17-91508945-INS-243 |
| chr17-91511374-INS-134 | chr17-91511374-INS-134 |
| chr17-91517358-INS-134 | chr17-91517358-INS-135 |
| chr17-91527464-INS-1364 | chr17-91527465-INS-1364 |
| chr17-91578849-INS-497 | chr17-91578849-INS-497 |
| chr17-91592149-INS-345 | chr17-91592149-INS-345 |
| chr17-91597277-INS-60 | chr17-91597277-INS-60 |
| chr17-91599044-INS-65 | chr17-91599044-INS-69 |
| chr17-91604258-INS-74 | chr17-91604247-INS-50 |
| chr17-91634441-INS-63 | chr17-91634441-INS-63 |
| chr17-91700648-INS-53 | chr17-91700636-INS-53 |
| chr17-91711082-INS-59 | chr17-91711082-INS-60 |
| chr17-91748353-INS-236 | chr17-91748353-INS-236 |
| chr17-91758722-INS-57 | chr17-91758722-INS-57 |

| | |
|---|---|
| chr17-91770174-INS-880 | chr17-91770174-INS-882 |
| chr17-91785329-INS-481 | chr17-91785329-INS-481 |
| chr17-91788493-INS-399 | chr17-91788493-INS-399 |
| chr17-91816752-INS-1602 | chr17-91816753-INS-1603 |
| chr17-91838153-INS-208 | chr17-91838153-INS-208 |
| chr17-91865591-INS-577 | chr17-91865591-INS-577 |
| chr17-91982129-INS-135 | chr17-91982109-INS-114 |
| chr17-92005146-INS-69 | chr17-92005146-INS-69 |
| chr17-92019065-INS-143 | chr17-92019055-INS-158 |
| chr17-92029248-INS-62 | chr17-92029248-INS-66 |
| chr17-92038464-INS-214 | chr17-92038464-INS-214 |
| chr17-92063358-INS-78 | chr17-92063358-INS-78 |
| chr17-92066345-INS-51 | chr17-92066345-INS-59 |
| chr17-92070179-INS-132 | chr17-92070179-INS-132 |
| chr17-92072808-INS-196 | chr17-92072808-INS-196 |
| chr17-92076813-INS-594 | chr17-92076813-INS-605 |
| chr17-92104740-INS-181 | chr17-92104740-INS-182 |
| chr17-92119170-INS-52 | chr17-92119170-INS-52 |
| chr17-92123206-INS-6739 | chr17-92123207-INS-6739 |
| chr17-92131874-INS-435 | chr17-92131874-INS-435 |
| chr17-92157066-INS-7100 | chr17-92157067-INS-7100 |
| chr17-92163945-INS-108 | chr17-92163945-INS-104 |
| chr17-92198168-INS-62 | chr17-92198168-INS-64 |
| chr17-92237568-INS-863 | chr17-92237568-INS-862 |
| chr17-92240593-INS-53 | chr17-92240593-INS-59 |
| chr17-92263816-INS-7102 | chr17-92263817-INS-7102 |
| chr17-92268576-INS-644 | chr17-92268576-INS-644 |
| chr17-92271134-INS-90 | chr17-92271134-INS-87 |
| chr17-92272063-INS-377 | chr17-92272063-INS-377 |
| chr17-92328456-INS-128 | chr17-92328462-INS-108 |
| chr17-92361784-INS-51 | chr17-92361784-INS-53 |
| chr17-92369687-INS-3330 | chr17-92369688-INS-3331 |
| chr17-92369688-INS-483 | chr17-92369688-INS-483 |
| chr17-92384828-INS-220 | chr17-92384828-INS-219 |
| chr17-92400872-INS-479 | chr17-92400873-INS-472 |
| chr17-92402152-INS-876 | chr17-92402152-INS-874 |
| chr17-92407884-INS-137 | chr17-92407884-INS-137 |
| chr17-92440820-INS-7144 | chr17-92440821-INS-7144 |
| chr17-92472946-INS-1789 | chr17-92472947-INS-1801 |
| chr17-92475083-INS-50 | chr17-92475083-INS-56 |
| chr17-92477654-INS-194 | chr17-92477654-INS-219 |
| chr17-92508850-INS-4388 | chr17-92508851-INS-4384 |
| chr17-92520466-INS-7284 | chr17-92520467-INS-7284 |
| chr17-92527454-INS-72 | chr17-92527454-INS-72 |
| chr17-92542933-INS-4056 | chr17-92542934-INS-4056 |
| chr17-92597353-INS-948 | chr17-92597353-INS-948 |
| chr17-92634695-INS-62 | chr17-92634695-INS-68 |
| chr17-92638997-INS-101 | chr17-92638997-INS-101 |
| chr17-92640955-INS-389 | chr17-92640955-INS-389 |
| chr17-92664288-INS-1092 | chr17-92664289-INS-1093 |
| chr17-92667673-INS-196 | chr17-92667674-INS-200 |
| chr17-92668450-INS-6691 | chr17-92668451-INS-6693 |
| chr17-92673691-INS-280 | chr17-92673691-INS-279 |
| chr17-92675567-INS-78 | chr17-92675533-INS-92 |
| chr17-92684925-INS-117 | chr17-92684925-INS-117 |
| chr17-92692061-INS-56 | chr17-92692057-INS-58 |
| chr17-92722492-INS-613 | chr17-92722492-INS-613 |
| chr17-92757452-INS-133 | chr17-92757452-INS-133 |
| chr17-92773538-INS-843 | chr17-92773538-INS-843 |
| chr17-92791685-INS-1671 | chr17-92791686-INS-1663 |
| chr17-92796113-INS-52 | chr17-92796110-INS-50 |
| chr17-92806160-INS-111 | chr17-92806160-INS-92 |
| chr17-92808873-INS-76 | chr17-92808873-INS-70 |
| chr17-92812467-INS-118 | chr17-92812467-INS-118 |
| chr17-92820131-INS-54 | chr17-92820145-INS-60 |
| chr17-92848515-INS-1556 | chr17-92848516-INS-1556 |
| chr17-92856048-INS-3251 | chr17-92856047-INS-3249 |
| chr17-92862333-INS-274 | chr17-92862327-INS-276 |
| chr17-92909863-INS-477 | chr17-92909863-INS-477 |
| chr17-92917553-INS-7984 | chr17-92917553-INS-7969 |
| chr17-92941087-INS-6700 | chr17-92941088-INS-6700 |
| chr17-92950530-INS-2933 | chr17-92950531-INS-2933 |
| chr17-92951885-INS-196 | chr17-92951885-INS-196 |
| chr17-92958399-INS-3215 | chr17-92958402-INS-3193 |
| chr17-92972471-INS-5547 | chr17-92972472-INS-5547 |
| chr17-92978023-INS-159 | chr17-92978023-INS-157 |
| chr17-92980505-INS-429 | chr17-92980505-INS-422 |

| | |
|---|---|
| chr17-92993630-INS-390 | chr17-92993630-INS-390 |
| chr17-93002483-INS-216 | chr17-93002483-INS-216 |
| chr17-93018583-INS-3799 | chr17-93018584-INS-3799 |
| chr17-93028270-INS-7086 | chr17-93028271-INS-7086 |
| chr17-93032820-INS-87 | chr17-93032820-INS-74 |
| chr17-93040622-INS-166 | chr17-93040622-INS-166 |
| chr17-93044970-INS-190 | chr17-93044970-INS-190 |
| chr17-93059008-INS-235 | chr17-93059009-INS-235 |
| chr17-93089312-INS-128 | chr17-93089313-INS-129 |
| chr17-93100170-INS-790 | chr17-93100170-INS-777 |
| chr17-93120079-INS-52 | chr17-93120079-INS-58 |
| chr17-93138812-INS-218 | chr17-93138811-INS-238 |
| chr17-93163913-INS-1219 | chr17-93163914-INS-1230 |
| chr17-93164513-INS-6714 | chr17-93164506-INS-6702 |
| chr17-93175527-INS-220 | chr17-93175527-INS-220 |
| chr17-93177454-INS-54 | chr17-93177454-INS-50 |
| chr17-93193608-INS-1316 | chr17-93193609-INS-1316 |
| chr17-93225876-INS-137 | chr17-93225876-INS-109 |
| chr17-93247847-INS-64 | chr17-93247872-INS-74 |
| chr17-93257103-INS-1894 | chr17-93257104-INS-1894 |
| chr17-93266877-INS-415 | chr17-93266868-INS-420 |
| chr17-93287330-INS-498 | chr17-93287330-INS-498 |
| chr17-93288210-INS-2963 | chr17-93288211-INS-2963 |
| chr17-93311890-INS-104 | chr17-93311892-INS-82 |
| chr17-93313107-INS-825 | chr17-93313107-INS-825 |
| chr17-93357738-INS-6458 | chr17-93357739-INS-6458 |
| chr17-93383795-INS-173 | chr17-93383795-INS-175 |
| chr17-93403963-INS-354 | chr17-93403963-INS-356 |
| chr17-93489652-INS-383 | chr17-93489641-INS-410 |
| chr17-93517998-INS-56 | chr17-93518042-INS-60 |
| chr17-93529363-INS-159 | chr17-93529364-INS-134 |
| chr17-93574496-INS-198 | chr17-93574496-INS-198 |
| chr17-93578099-INS-481 | chr17-93578099-INS-481 |
| chr17-93612190-INS-255 | chr17-93612190-INS-254 |
| chr17-93632103-INS-585 | chr17-93632103-INS-585 |
| chr17-93665870-INS-4038 | chr17-93665871-INS-4038 |
| chr17-93686838-INS-423 | chr17-93686838-INS-423 |
| chr17-93732095-INS-197 | chr17-93732095-INS-197 |
| chr17-93736403-INS-7214 | chr17-93736404-INS-7215 |
| chr17-93741939-INS-212 | chr17-93741939-INS-209 |
| chr17-93743777-INS-7089 | chr17-93743778-INS-7089 |
| chr17-93750255-INS-6173 | chr17-93750256-INS-6174 |
| chr17-93766877-INS-209 | chr17-93766877-INS-210 |
| chr17-93774655-INS-231 | chr17-93774655-INS-232 |
| chr17-93783389-INS-481 | chr17-93783389-INS-481 |
| chr17-93786319-INS-243 | chr17-93786319-INS-243 |
| chr17-93794804-INS-160 | chr17-93794804-INS-160 |
| chr17-93808709-INS-5030 | chr17-93808710-INS-5030 |
| chr17-93831598-INS-328 | chr17-93831598-INS-328 |
| chr17-93851480-INS-358 | chr17-93851480-INS-374 |
| chr17-93856442-INS-496 | chr17-93856442-INS-497 |
| chr17-93898322-INS-293 | chr17-93898322-INS-293 |
| chr17-93912755-INS-339 | chr17-93912755-INS-340 |
| chr17-93974700-INS-2349 | chr17-93974701-INS-2349 |
| chr17-94011500-INS-382 | chr17-94011500-INS-382 |
| chr17-94012193-INS-3298 | chr17-94012194-INS-3312 |
| chr17-94024856-INS-253 | chr17-94024856-INS-253 |
| chr17-94042766-INS-5235 | chr17-94042767-INS-5235 |
| chr17-94074455-INS-383 | chr17-94074455-INS-383 |
| chr17-94093497-INS-321 | chr17-94093497-INS-321 |
| chr17-94111005-INS-3235 | chr17-94111006-INS-3233 |
| chr17-94126386-INS-13374 | chr17-94126387-INS-13376 |
| chr17-94136321-INS-1287 | chr17-94136322-INS-1288 |
| chr17-94143415-INS-137 | chr17-94143415-INS-137 |
| chr17-94147418-INS-1703 | chr17-94147419-INS-1658 |
| chr17-94169407-INS-5480 | chr17-94169408-INS-5480 |
| chr17-94178942-INS-1250 | chr17-94178943-INS-1250 |
| chr17-94203234-INS-433 | chr17-94203234-INS-433 |
| chr17-94205392-INS-4943 | chr17-94205393-INS-4943 |
| chr17-94230694-INS-86 | chr17-94230697-INS-111 |
| chr17-94233724-INS-342 | chr17-94233724-INS-342 |
| chr17-94247596-INS-441 | chr17-94247596-INS-441 |
| chr17-94261871-INS-937 | chr17-94261871-INS-937 |
| chr17-94268920-INS-58 | chr17-94268920-INS-56 |
| chr17-94272939-INS-1900 | chr17-94272940-INS-1900 |
| chr17-94273882-INS-5698 | chr17-94273883-INS-5698 |
| chr17-94277047-INS-54 | chr17-94277052-INS-69 |

| | |
|---|---|
| chr17-94307134-INS-1593 | chr17-94307135-INS-1593 |
| chr17-94312380-INS-618 | chr17-94312380-INS-619 |
| chr17-94342804-INS-201 | chr17-94342804-INS-202 |
| chr17-94350317-INS-54 | chr17-94350317-INS-54 |
| chr17-94354585-INS-2395 | chr17-94354586-INS-2395 |
| chr17-94368571-INS-64 | chr17-94368528-INS-76 |
| chr17-94375239-INS-292 | chr17-94375239-INS-293 |
| chr17-94378258-INS-461 | chr17-94378258-INS-461 |
| chr17-94384725-INS-502 | chr17-94384756-INS-510 |
| chr17-94399677-INS-1169 | chr17-94399678-INS-1170 |
| chr17-94460092-INS-6577 | chr17-94460093-INS-6582 |
| chr17-94504358-INS-52 | chr17-94504384-INS-68 |
| chr17-94524027-INS-6875 | chr17-94524028-INS-6875 |
| chr17-94569904-INS-110 | chr17-94569904-INS-92 |
| chr17-94627479-INS-405 | chr17-94627479-INS-405 |
| chr17-94646306-INS-4373 | chr17-94646307-INS-4373 |
| chr17-94662648-INS-80 | chr17-94662621-INS-75 |
| chr17-94678075-INS-6295 | chr17-94678076-INS-6295 |
| chr17-94688244-INS-70 | chr17-94688244-INS-74 |
| chr17-94722446-INS-963 | chr17-94722446-INS-963 |
| chr17-94731388-INS-1534 | chr17-94731389-INS-1534 |
| chr17-94733470-INS-431 | chr17-94733470-INS-434 |
| chr17-94736389-INS-7366 | chr17-94736390-INS-7366 |
| chr17-94755880-INS-58 | chr17-94755880-INS-62 |
| chr17-94806252-INS-378 | chr17-94806252-INS-377 |
| chr17-94810904-INS-6729 | chr17-94810905-INS-6726 |
| chr17-94815880-INS-425 | chr17-94815880-INS-425 |
| chr17-94853461-INS-2685 | chr17-94853462-INS-2685 |
| chr17-94893685-INS-211 | chr17-94893685-INS-211 |
| chr17-94899179-INS-227 | chr17-94899179-INS-228 |
| chr17-94905329-INS-122 | chr17-94905345-INS-153 |
| chr17-94924348-INS-62 | chr17-94924345-INS-64 |
| chr17-94934416-INS-1319 | chr17-94934417-INS-1319 |
| chr17-94942061-INS-52 | chr17-94942061-INS-52 |
| chr17-94943100-INS-98 | chr17-94943100-INS-110 |
| chr17-94946978-INS-100 | chr17-94946978-INS-60 |
| chr17-94951151-INS-199 | chr17-94951151-INS-199 |
| chr17-95007825-INS-499 | chr17-95007825-INS-499 |
| chr17-95013815-INS-7130 | chr17-95013816-INS-7130 |
| chr17-95016246-INS-228 | chr17-95016246-INS-228 |
| chr17-95023497-INS-144 | chr17-95023497-INS-144 |
| chr17-95045545-INS-153 | chr17-95045545-INS-153 |
| chr17-95058362-INS-51 | chr17-95058362-INS-53 |
| chr17-95064087-INS-723 | chr17-95064087-INS-723 |
| chr17-95107299-INS-5232 | chr17-95107300-INS-5232 |
| chr17-95132352-INS-6533 | chr17-95132353-INS-6534 |
| chr17-95132843-INS-90 | chr17-95132843-INS-120 |
| chr17-95145088-INS-60 | chr17-95145085-INS-84 |
| chr17-95187575-INS-222 | chr17-95187575-INS-225 |
| chr18-3046544-INS-508 | chr18-3046544-INS-513 |
| chr18-3084264-INS-6440 | chr18-3084265-INS-6440 |
| chr18-3116691-INS-15173 | chr18-3116692-INS-15169 |
| chr18-3164598-INS-72 | chr18-3164598-INS-72 |
| chr18-3171443-INS-59 | chr18-3171443-INS-71 |
| chr18-3199529-INS-80 | chr18-3199527-INS-75 |
| chr18-3253356-INS-385 | chr18-3253356-INS-384 |
| chr18-3335083-INS-205 | chr18-3335083-INS-205 |
| chr18-3463101-INS-102 | chr18-3463101-INS-104 |
| chr18-3485797-INS-179 | chr18-3485797-INS-169 |
| chr18-3527260-INS-116 | chr18-3527260-INS-120 |
| chr18-3545779-INS-66 | chr18-3545774-INS-55 |
| chr18-3560801-INS-109 | chr18-3560803-INS-72 |
| chr18-3658391-INS-304 | chr18-3658391-INS-305 |
| chr18-3689851-INS-68 | chr18-3689879-INS-64 |
| chr18-3696196-INS-283 | chr18-3696196-INS-283 |
| chr18-3697533-INS-81 | chr18-3697569-INS-56 |
| chr18-3724425-INS-54 | chr18-3724425-INS-54 |
| chr18-3766499-INS-54 | chr18-3766466-INS-61 |
| chr18-3790957-INS-242 | chr18-3790957-INS-244 |
| chr18-3804621-INS-71 | chr18-3804621-INS-72 |
| chr18-3841040-INS-56 | chr18-3841040-INS-58 |
| chr18-3857721-INS-62 | chr18-3857721-INS-62 |
| chr18-3908559-INS-486 | chr18-3908559-INS-487 |
| chr18-3913208-INS-58 | chr18-3913208-INS-50 |
| chr18-3954129-INS-56 | chr18-3954129-INS-56 |
| chr18-3955220-INS-106 | chr18-3955220-INS-108 |
| chr18-3961013-INS-83 | chr18-3961038-INS-96 |

| | |
|---|---|
| chr18-3962943-INS-59 | chr18-3962943-INS-59 |
| chr18-3967542-INS-55 | chr18-3967542-INS-55 |
| chr18-3975382-INS-82 | chr18-3975377-INS-83 |
| chr18-4029543-INS-59 | chr18-4029543-INS-59 |
| chr18-4045204-INS-54 | chr18-4045204-INS-54 |
| chr18-4172552-INS-66 | chr18-4172535-INS-63 |
| chr18-4180167-INS-156 | chr18-4180167-INS-158 |
| chr18-4184806-INS-450 | chr18-4184806-INS-445 |
| chr18-4200340-INS-56 | chr18-4200340-INS-54 |
| chr18-4277547-INS-278 | chr18-4277547-INS-269 |
| chr18-4292423-INS-212 | chr18-4292423-INS-212 |
| chr18-4320868-INS-198 | chr18-4320868-INS-196 |
| chr18-4330541-INS-792 | chr18-4330541-INS-822 |
| chr18-4377589-INS-273 | chr18-4377589-INS-273 |
| chr18-4381201-INS-214 | chr18-4381201-INS-214 |
| chr18-4395285-INS-197 | chr18-4395285-INS-198 |
| chr18-4436230-INS-59 | chr18-4436230-INS-59 |
| chr18-4437705-INS-81 | chr18-4437681-INS-82 |
| chr18-4439388-INS-1224 | chr18-4439389-INS-1225 |
| chr18-4444172-INS-326 | chr18-4444172-INS-326 |
| chr18-4446245-INS-2441 | chr18-4446246-INS-2441 |
| chr18-4485813-INS-235 | chr18-4485813-INS-235 |
| chr18-4487894-INS-51 | chr18-4487894-INS-51 |
| chr18-4498673-INS-125 | chr18-4498673-INS-125 |
| chr18-4502527-INS-52 | chr18-4502527-INS-52 |
| chr18-4527696-INS-299 | chr18-4527696-INS-299 |
| chr18-4529317-INS-110 | chr18-4529317-INS-110 |
| chr18-4546543-INS-68 | chr18-4546543-INS-70 |
| chr18-4550773-INS-448 | chr18-4550773-INS-456 |
| chr18-4553566-INS-118 | chr18-4553566-INS-76 |
| chr18-4556570-INS-355 | chr18-4556570-INS-355 |
| chr18-4559245-INS-96 | chr18-4559245-INS-96 |
| chr18-4576462-INS-453 | chr18-4576462-INS-454 |
| chr18-4595782-INS-203 | chr18-4595782-INS-206 |
| chr18-4596156-INS-688 | chr18-4596156-INS-688 |
| chr18-4655104-INS-78 | chr18-4655104-INS-80 |
| chr18-4752520-INS-1023 | chr18-4752521-INS-1019 |
| chr18-4762483-INS-6051 | chr18-4762484-INS-6051 |
| chr18-4765849-INS-115 | chr18-4765849-INS-114 |
| chr18-4767829-INS-99 | chr18-4767845-INS-105 |
| chr18-4785081-INS-343 | chr18-4785081-INS-343 |
| chr18-4791611-INS-56 | chr18-4791611-INS-56 |
| chr18-4801350-INS-255 | chr18-4801350-INS-255 |
| chr18-4806310-INS-274 | chr18-4806310-INS-275 |
| chr18-4809274-INS-977 | chr18-4809274-INS-977 |
| chr18-4812331-INS-103 | chr18-4812331-INS-103 |
| chr18-4813481-INS-62 | chr18-4813481-INS-54 |
| chr18-4829831-INS-450 | chr18-4829831-INS-450 |
| chr18-4852623-INS-89 | chr18-4852623-INS-89 |
| chr18-4856244-INS-94 | chr18-4856262-INS-54 |
| chr18-4857456-INS-104 | chr18-4857456-INS-104 |
| chr18-4862942-INS-6491 | chr18-4862943-INS-6494 |
| chr18-4873888-INS-96 | chr18-4873928-INS-96 |
| chr18-4891233-INS-54 | chr18-4891233-INS-52 |
| chr18-4902199-INS-3143 | chr18-4902200-INS-3143 |
| chr18-4910660-INS-91 | chr18-4910670-INS-98 |
| chr18-4912219-INS-218 | chr18-4912219-INS-219 |
| chr18-4945017-INS-165 | chr18-4945016-INS-161 |
| chr18-4950029-INS-246 | chr18-4950029-INS-244 |
| chr18-4950327-INS-1909 | chr18-4950328-INS-1911 |
| chr18-4982077-INS-143 | chr18-4982078-INS-127 |
| chr18-4989384-INS-83 | chr18-4989384-INS-83 |
| chr18-5001530-INS-98 | chr18-5001530-INS-56 |
| chr18-5009703-INS-983 | chr18-5009703-INS-982 |
| chr18-5016549-INS-146 | chr18-5016548-INS-136 |
| chr18-5079533-INS-406 | chr18-5079533-INS-406 |
| chr18-5093105-INS-63 | chr18-5093105-INS-63 |
| chr18-5132657-INS-54 | chr18-5132657-INS-60 |
| chr18-5151246-INS-416 | chr18-5151246-INS-416 |
| chr18-5154200-INS-64 | chr18-5154200-INS-64 |
| chr18-5164984-INS-62 | chr18-5164984-INS-62 |
| chr18-5197424-INS-395 | chr18-5197424-INS-395 |
| chr18-5242633-INS-3280 | chr18-5242628-INS-3280 |
| chr18-5252036-INS-66 | chr18-5252036-INS-60 |
| chr18-5256271-INS-520 | chr18-5256271-INS-520 |
| chr18-5283813-INS-52 | chr18-5283813-INS-54 |
| chr18-5316454-INS-832 | chr18-5316454-INS-832 |

| | |
|---|---|
| chr18-5328527-INS-1892 | chr18-5328528-INS-1892 |
| chr18-5338386-INS-1226 | chr18-5338387-INS-1227 |
| chr18-5348053-INS-6442 | chr18-5348054-INS-6442 |
| chr18-5363209-INS-294 | chr18-5363209-INS-262 |
| chr18-5369965-INS-52 | chr18-5369965-INS-52 |
| chr18-5419981-INS-315 | chr18-5419981-INS-321 |
| chr18-5431485-INS-64 | chr18-5431485-INS-64 |
| chr18-5484461-INS-88 | chr18-5484461-INS-132 |
| chr18-5495577-INS-100 | chr18-5495577-INS-100 |
| chr18-5537613-INS-64 | chr18-5537613-INS-56 |
| chr18-5539478-INS-2764 | chr18-5539479-INS-2764 |
| chr18-5543632-INS-84 | chr18-5543614-INS-60 |
| chr18-5584545-INS-6340 | chr18-5584546-INS-6341 |
| chr18-5612127-INS-287 | chr18-5612127-INS-287 |
| chr18-5620253-INS-154 | chr18-5620253-INS-154 |
| chr18-5640867-INS-324 | chr18-5640873-INS-340 |
| chr18-5675209-INS-10174 | chr18-5675202-INS-10165 |
| chr18-5738507-INS-70 | chr18-5738507-INS-68 |
| chr18-5765179-INS-56 | chr18-5765179-INS-56 |
| chr18-5778161-INS-100 | chr18-5778166-INS-80 |
| chr18-5779750-INS-52 | chr18-5779750-INS-52 |
| chr18-5804888-INS-864 | chr18-5804908-INS-877 |
| chr18-5830784-INS-308 | chr18-5830784-INS-298 |
| chr18-5832292-INS-182 | chr18-5832292-INS-182 |
| chr18-5859652-INS-1050 | chr18-5859653-INS-1050 |
| chr18-5909606-INS-201 | chr18-5909606-INS-201 |
| chr18-5928905-INS-7178 | chr18-5928906-INS-7178 |
| chr18-5932910-INS-51 | chr18-5932910-INS-76 |
| chr18-5940871-INS-52 | chr18-5940868-INS-50 |
| chr18-5993566-INS-2669 | chr18-5993567-INS-2669 |
| chr18-6009190-INS-205 | chr18-6009190-INS-204 |
| chr18-6011128-INS-69 | chr18-6011128-INS-60 |
| chr18-6031318-INS-104 | chr18-6031318-INS-110 |
| chr18-6046974-INS-81 | chr18-6046974-INS-81 |
| chr18-6099245-INS-504 | chr18-6099245-INS-503 |
| chr18-6110744-INS-210 | chr18-6110743-INS-213 |
| chr18-6112753-INS-206 | chr18-6112753-INS-206 |
| chr18-6122877-INS-212 | chr18-6122877-INS-212 |
| chr18-6144659-INS-69 | chr18-6144699-INS-76 |
| chr18-6148739-INS-59 | chr18-6148739-INS-59 |
| chr18-6155943-INS-81 | chr18-6155943-INS-57 |
| chr18-6164870-INS-123 | chr18-6164887-INS-82 |
| chr18-6170412-INS-199 | chr18-6170412-INS-199 |
| chr18-6178293-INS-104 | chr18-6178289-INS-90 |
| chr18-6183196-INS-196 | chr18-6183196-INS-196 |
| chr18-6210832-INS-684 | chr18-6210832-INS-684 |
| chr18-6231912-INS-176 | chr18-6231912-INS-178 |
| chr18-6235028-INS-66 | chr18-6235028-INS-66 |
| chr18-6252693-INS-472 | chr18-6252693-INS-472 |
| chr18-6268418-INS-64 | chr18-6268415-INS-66 |
| chr18-6292788-INS-628 | chr18-6292788-INS-629 |
| chr18-6309720-INS-72 | chr18-6309720-INS-70 |
| chr18-6317118-INS-6651 | chr18-6317119-INS-6651 |
| chr18-6318332-INS-1405 | chr18-6318333-INS-1405 |
| chr18-6371816-INS-84 | chr18-6371816-INS-84 |
| chr18-6389502-INS-208 | chr18-6389505-INS-209 |
| chr18-6408034-INS-710 | chr18-6408026-INS-747 |
| chr18-6423272-INS-224 | chr18-6423272-INS-224 |
| chr18-6444818-INS-64 | chr18-6444818-INS-64 |
| chr18-6466778-INS-259 | chr18-6466778-INS-260 |
| chr18-6520254-INS-68 | chr18-6520254-INS-68 |
| chr18-6522433-INS-357 | chr18-6522433-INS-358 |
| chr18-6526510-INS-390 | chr18-6526510-INS-390 |
| chr18-6530498-INS-2453 | chr18-6530499-INS-2445 |
| chr18-6538953-INS-1891 | chr18-6538954-INS-1891 |
| chr18-6554471-INS-6402 | chr18-6554472-INS-6402 |
| chr18-6565795-INS-804 | chr18-6565795-INS-805 |
| chr18-6581051-INS-68 | chr18-6581051-INS-68 |
| chr18-6597838-INS-323 | chr18-6597838-INS-323 |
| chr18-6600568-INS-327 | chr18-6600568-INS-327 |
| chr18-6614520-INS-175 | chr18-6614520-INS-182 |
| chr18-6615246-INS-459 | chr18-6615246-INS-459 |
| chr18-6677602-INS-408 | chr18-6677602-INS-408 |
| chr18-6703442-INS-169 | chr18-6703443-INS-168 |
| chr18-6732192-INS-458 | chr18-6732192-INS-458 |
| chr18-6754135-INS-291 | chr18-6754135-INS-291 |
| chr18-6755695-INS-70 | chr18-6755695-INS-80 |

| | |
|---|---|
| chr18-6758995-INS-5446 | chr18-6758996-INS-5445 |
| chr18-6771263-INS-218 | chr18-6771263-INS-218 |
| chr18-6775528-INS-187 | chr18-6775528-INS-188 |
| chr18-6811137-INS-120 | chr18-6811137-INS-118 |
| chr18-6828538-INS-280 | chr18-6828538-INS-279 |
| chr18-6836906-INS-82 | chr18-6836901-INS-74 |
| chr18-6837792-INS-134 | chr18-6837818-INS-123 |
| chr18-6882329-INS-60 | chr18-6882329-INS-60 |
| chr18-6883118-INS-382 | chr18-6883118-INS-382 |
| chr18-6985241-INS-212 | chr18-6985241-INS-212 |
| chr18-6990725-INS-59 | chr18-6990725-INS-59 |
| chr18-7051787-INS-7030 | chr18-7051788-INS-7030 |
| chr18-7082801-INS-431 | chr18-7082801-INS-431 |
| chr18-7083496-INS-52 | chr18-7083496-INS-56 |
| chr18-7113652-INS-68 | chr18-7113647-INS-81 |
| chr18-7116162-INS-302 | chr18-7116162-INS-333 |
| chr18-7131529-INS-64 | chr18-7131529-INS-82 |
| chr18-7132901-INS-391 | chr18-7132901-INS-392 |
| chr18-7150018-INS-939 | chr18-7150018-INS-940 |
| chr18-7196130-INS-75 | chr18-7196130-INS-75 |
| chr18-7198446-INS-164 | chr18-7198446-INS-164 |
| chr18-7215225-INS-130 | chr18-7215225-INS-156 |
| chr18-7216641-INS-291 | chr18-7216641-INS-309 |
| chr18-7220179-INS-67 | chr18-7220179-INS-67 |
| chr18-7341549-INS-6447 | chr18-7341550-INS-6450 |
| chr18-7369497-INS-369 | chr18-7369497-INS-371 |
| chr18-7398268-INS-629 | chr18-7398268-INS-628 |
| chr18-7437230-INS-177 | chr18-7437230-INS-177 |
| chr18-7451249-INS-187 | chr18-7451249-INS-187 |
| chr18-7465507-INS-3193 | chr18-7465508-INS-3193 |
| chr18-7472824-INS-239 | chr18-7472824-INS-239 |
| chr18-7490652-INS-439 | chr18-7490652-INS-465 |
| chr18-7494115-INS-351 | chr18-7494082-INS-351 |
| chr18-7519366-INS-428 | chr18-7519400-INS-425 |
| chr18-7539051-INS-2312 | chr18-7539052-INS-2317 |
| chr18-7547497-INS-88 | chr18-7547514-INS-84 |
| chr18-7574127-INS-74 | chr18-7574127-INS-74 |
| chr18-7583228-INS-59 | chr18-7583228-INS-59 |
| chr18-7589084-INS-385 | chr18-7589084-INS-385 |
| chr18-7612130-INS-72 | chr18-7612130-INS-108 |
| chr18-7630586-INS-54 | chr18-7630586-INS-54 |
| chr18-7632935-INS-104 | chr18-7632935-INS-122 |
| chr18-7641505-INS-60 | chr18-7641505-INS-56 |
| chr18-7648905-INS-106 | chr18-7648909-INS-60 |
| chr18-7658659-INS-6412 | chr18-7658660-INS-6412 |
| chr18-7665510-INS-224 | chr18-7665510-INS-224 |
| chr18-7682808-INS-675 | chr18-7682808-INS-675 |
| chr18-7706464-INS-178 | chr18-7706464-INS-175 |
| chr18-7745274-INS-7119 | chr18-7745275-INS-7119 |
| chr18-7751113-INS-99 | chr18-7751147-INS-100 |
| chr18-7753463-INS-3639 | chr18-7753464-INS-3637 |
| chr18-7800949-INS-328 | chr18-7800949-INS-328 |
| chr18-7832696-INS-680 | chr18-7832696-INS-681 |
| chr18-7837490-INS-197 | chr18-7837490-INS-177 |
| chr18-7838901-INS-170 | chr18-7838901-INS-171 |
| chr18-7858722-INS-66 | chr18-7858731-INS-67 |
| chr18-7875941-INS-220 | chr18-7875941-INS-220 |
| chr18-7889128-INS-680 | chr18-7889128-INS-680 |
| chr18-7917695-INS-72 | chr18-7917695-INS-110 |
| chr18-7946875-INS-367 | chr18-7946875-INS-349 |
| chr18-8008098-INS-1310 | chr18-8008099-INS-1310 |
| chr18-8046773-INS-286 | chr18-8046774-INS-286 |
| chr18-8059568-INS-222 | chr18-8059568-INS-222 |
| chr18-8078781-INS-53 | chr18-8078781-INS-53 |
| chr18-8083959-INS-62 | chr18-8083959-INS-63 |
| chr18-8089910-INS-123 | chr18-8089910-INS-123 |
| chr18-8096155-INS-183 | chr18-8096155-INS-190 |
| chr18-8106550-INS-5504 | chr18-8106551-INS-5504 |
| chr18-8147663-INS-54 | chr18-8147659-INS-58 |
| chr18-8179036-INS-56 | chr18-8179036-INS-76 |
| chr18-8189790-INS-2077 | chr18-8189791-INS-2077 |
| chr18-8193567-INS-711 | chr18-8193567-INS-711 |
| chr18-8199273-INS-56 | chr18-8199273-INS-58 |
| chr18-8202057-INS-137 | chr18-8202057-INS-139 |
| chr18-8210557-INS-5664 | chr18-8210558-INS-5664 |
| chr18-8225746-INS-3816 | chr18-8225747-INS-3816 |
| chr18-8238790-INS-6380 | chr18-8238791-INS-6380 |

| | |
|---|---|
| chr18-8264971-INS-512 | chr18-8264971-INS-512 |
| chr18-8288440-INS-84 | chr18-8288440-INS-114 |
| chr18-8324364-INS-7259 | chr18-8324365-INS-7260 |
| chr18-8327528-INS-52 | chr18-8327528-INS-52 |
| chr18-8349842-INS-1257 | chr18-8349843-INS-1257 |
| chr18-8412369-INS-348 | chr18-8412368-INS-348 |
| chr18-8414604-INS-398 | chr18-8414605-INS-398 |
| chr18-8415111-INS-382 | chr18-8415111-INS-382 |
| chr18-8458299-INS-58 | chr18-8458286-INS-64 |
| chr18-8475139-INS-52 | chr18-8475139-INS-50 |
| chr18-8494871-INS-6258 | chr18-8494872-INS-6259 |
| chr18-8522044-INS-65 | chr18-8522070-INS-66 |
| chr18-8539353-INS-110 | chr18-8539353-INS-110 |
| chr18-8551255-INS-60 | chr18-8551255-INS-54 |
| chr18-8562860-INS-52 | chr18-8562870-INS-60 |
| chr18-8603922-INS-58 | chr18-8603922-INS-71 |
| chr18-8605266-INS-74 | chr18-8605266-INS-71 |
| chr18-8630478-INS-850 | chr18-8630478-INS-850 |
| chr18-8631463-INS-100 | chr18-8631461-INS-63 |
| chr18-8651275-INS-447 | chr18-8651275-INS-447 |
| chr18-8657226-INS-6388 | chr18-8657227-INS-6388 |
| chr18-8663860-INS-51 | chr18-8663860-INS-51 |
| chr18-8697713-INS-180 | chr18-8697713-INS-180 |
| chr18-8719164-INS-10793 | chr18-8719165-INS-10794 |
| chr18-8734058-INS-381 | chr18-8734058-INS-381 |
| chr18-8751502-INS-55 | chr18-8751502-INS-67 |
| chr18-8801148-INS-6455 | chr18-8801149-INS-6456 |
| chr18-8809609-INS-159 | chr18-8809609-INS-159 |
| chr18-8825723-INS-60 | chr18-8825723-INS-63 |
| chr18-8837902-INS-77 | chr18-8837926-INS-64 |
| chr18-8865791-INS-90 | chr18-8865791-INS-55 |
| chr18-8872111-INS-68 | chr18-8872111-INS-56 |
| chr18-8874433-INS-54 | chr18-8874433-INS-54 |
| chr18-8876501-INS-84 | chr18-8876501-INS-84 |
| chr18-8882976-INS-191 | chr18-8882976-INS-191 |
| chr18-8889010-INS-120 | chr18-8889047-INS-98 |
| chr18-8906268-INS-52 | chr18-8906268-INS-52 |
| chr18-8948714-INS-58 | chr18-8948714-INS-58 |
| chr18-8952105-INS-66 | chr18-8952105-INS-84 |
| chr18-8953301-INS-68 | chr18-8953291-INS-106 |
| chr18-8975589-INS-80 | chr18-8975589-INS-76 |
| chr18-8990084-INS-203 | chr18-8990084-INS-203 |
| chr18-9010164-INS-104 | chr18-9010158-INS-107 |
| chr18-9013410-INS-56 | chr18-9013410-INS-56 |
| chr18-9014102-INS-182 | chr18-9014102-INS-182 |
| chr18-9029813-INS-90 | chr18-9029813-INS-72 |
| chr18-9039805-INS-230 | chr18-9039805-INS-230 |
| chr18-9041255-INS-635 | chr18-9041207-INS-669 |
| chr18-9042597-INS-138 | chr18-9042597-INS-141 |
| chr18-9044737-INS-200 | chr18-9044737-INS-200 |
| chr18-9059075-INS-1865 | chr18-9059076-INS-1865 |
| chr18-9070031-INS-196 | chr18-9070031-INS-196 |
| chr18-9098037-INS-6703 | chr18-9098038-INS-6703 |
| chr18-9117813-INS-214 | chr18-9117813-INS-214 |
| chr18-9135281-INS-74 | chr18-9135280-INS-90 |
| chr18-9140569-INS-62 | chr18-9140569-INS-72 |
| chr18-9147089-INS-104 | chr18-9147089-INS-120 |
| chr18-9157514-INS-7281 | chr18-9157515-INS-7281 |
| chr18-9167754-INS-4080 | chr18-9167755-INS-4080 |
| chr18-9176127-INS-93 | chr18-9176127-INS-93 |
| chr18-9186657-INS-7122 | chr18-9186657-INS-7122 |
| chr18-9242849-INS-143 | chr18-9242849-INS-156 |
| chr18-9270243-INS-60 | chr18-9270252-INS-84 |
| chr18-9272246-INS-217 | chr18-9272246-INS-217 |
| chr18-9272721-INS-2676 | chr18-9272722-INS-2676 |
| chr18-9297895-INS-225 | chr18-9297895-INS-225 |
| chr18-9299123-INS-75 | chr18-9299123-INS-71 |
| chr18-9320790-INS-2367 | chr18-9320783-INS-2334 |
| chr18-9336635-INS-1229 | chr18-9336636-INS-1229 |
| chr18-9373687-INS-108 | chr18-9373687-INS-108 |
| chr18-9388950-INS-64 | chr18-9388950-INS-64 |
| chr18-9413903-INS-215 | chr18-9413904-INS-215 |
| chr18-9437404-INS-249 | chr18-9437404-INS-250 |
| chr18-9455243-INS-7234 | chr18-9455244-INS-7234 |
| chr18-9457540-INS-480 | chr18-9457540-INS-480 |
| chr18-9464382-INS-1483 | chr18-9464383-INS-1483 |
| chr18-9465483-INS-56 | chr18-9465483-INS-56 |

| | |
|---|---|
| chr18-9468929-INS-206 | chr18-9468929-INS-206 |
| chr18-9494484-INS-71 | chr18-9494484-INS-71 |
| chr18-9496993-INS-179 | chr18-9496993-INS-179 |
| chr18-9502166-INS-54 | chr18-9502166-INS-54 |
| chr18-9559739-INS-8852 | chr18-9559740-INS-8852 |
| chr18-9563163-INS-435 | chr18-9563163-INS-435 |
| chr18-9608643-INS-113 | chr18-9608643-INS-113 |
| chr18-9620218-INS-1440 | chr18-9620219-INS-1440 |
| chr18-9620644-INS-106 | chr18-9620644-INS-122 |
| chr18-9660155-INS-285 | chr18-9660149-INS-278 |
| chr18-9678281-INS-63 | chr18-9678302-INS-60 |
| chr18-9691347-INS-1895 | chr18-9691348-INS-1895 |
| chr18-9691554-INS-52 | chr18-9691554-INS-52 |
| chr18-9712553-INS-83 | chr18-9712553-INS-87 |
| chr18-9712569-INS-721 | chr18-9712565-INS-732 |
| chr18-9716608-INS-69 | chr18-9716608-INS-78 |
| chr18-9719326-INS-6219 | chr18-9719327-INS-6219 |
| chr18-9752636-INS-111 | chr18-9752636-INS-157 |
| chr18-9795509-INS-75 | chr18-9795509-INS-50 |
| chr18-9797268-INS-122 | chr18-9797268-INS-122 |
| chr18-9802255-INS-876 | chr18-9802255-INS-876 |
| chr18-9833012-INS-144 | chr18-9833012-INS-121 |
| chr18-9881189-INS-457 | chr18-9881189-INS-457 |
| chr18-9940275-INS-66 | chr18-9940275-INS-66 |
| chr18-9941956-INS-880 | chr18-9941956-INS-880 |
| chr18-9964082-INS-157 | chr18-9964082-INS-156 |
| chr18-9967835-INS-173 | chr18-9967835-INS-175 |
| chr18-9985977-INS-133 | chr18-9985977-INS-133 |
| chr18-10015494-INS-172 | chr18-10015494-INS-172 |
| chr18-10033280-INS-2463 | chr18-10033281-INS-2462 |
| chr18-10033873-INS-200 | chr18-10033873-INS-200 |
| chr18-10037147-INS-80 | chr18-10037147-INS-80 |
| chr18-10038487-INS-203 | chr18-10038487-INS-203 |
| chr18-10039383-INS-1912 | chr18-10039384-INS-1928 |
| chr18-10045960-INS-115 | chr18-10045960-INS-115 |
| chr18-10057933-INS-64 | chr18-10057933-INS-66 |
| chr18-10076086-INS-60 | chr18-10076086-INS-60 |
| chr18-10079609-INS-200 | chr18-10079609-INS-200 |
| chr18-10112517-INS-52 | chr18-10112517-INS-52 |
| chr18-10113821-INS-2013 | chr18-10113822-INS-2013 |
| chr18-10145042-INS-402 | chr18-10145042-INS-402 |
| chr18-10165246-INS-74 | chr18-10165246-INS-74 |
| chr18-10174793-INS-2851 | chr18-10174794-INS-2852 |
| chr18-10191483-INS-1892 | chr18-10191484-INS-1892 |
| chr18-10242774-INS-204 | chr18-10242774-INS-203 |
| chr18-10248975-INS-169 | chr18-10248975-INS-169 |
| chr18-10285688-INS-495 | chr18-10285688-INS-495 |
| chr18-10317272-INS-333 | chr18-10317272-INS-333 |
| chr18-10321731-INS-72 | chr18-10321731-INS-72 |
| chr18-10329543-INS-219 | chr18-10329543-INS-234 |
| chr18-10337638-INS-129 | chr18-10337639-INS-149 |
| chr18-10352883-INS-72 | chr18-10352851-INS-88 |
| chr18-10361113-INS-100 | chr18-10361113-INS-100 |
| chr18-10397689-INS-64 | chr18-10397689-INS-64 |
| chr18-10409971-INS-6899 | chr18-10409972-INS-6900 |
| chr18-10414922-INS-165 | chr18-10414922-INS-165 |
| chr18-10475408-INS-58 | chr18-10475408-INS-58 |
| chr18-10497421-INS-66 | chr18-10497421-INS-58 |
| chr18-10522666-INS-144 | chr18-10522666-INS-144 |
| chr18-10536029-INS-206 | chr18-10536029-INS-206 |
| chr18-10538710-INS-79 | chr18-10538759-INS-50 |
| chr18-10546428-INS-212 | chr18-10546428-INS-213 |
| chr18-10565802-INS-54 | chr18-10565802-INS-52 |
| chr18-10572562-INS-81 | chr18-10572562-INS-81 |
| chr18-10574295-INS-52 | chr18-10574295-INS-52 |
| chr18-10595058-INS-57 | chr18-10595058-INS-57 |
| chr18-10612889-INS-74 | chr18-10612889-INS-111 |
| chr18-10623528-INS-120 | chr18-10623528-INS-120 |
| chr18-10633720-INS-54 | chr18-10633724-INS-55 |
| chr18-10663999-INS-115 | chr18-10663999-INS-137 |
| chr18-10735085-INS-182 | chr18-10735085-INS-182 |
| chr18-10739230-INS-76 | chr18-10739230-INS-76 |
| chr18-10754368-INS-182 | chr18-10754368-INS-181 |
| chr18-10772992-INS-483 | chr18-10772992-INS-483 |
| chr18-10795885-INS-108 | chr18-10795885-INS-108 |
| chr18-10802583-INS-56 | chr18-10802583-INS-55 |
| chr18-10806658-INS-377 | chr18-10806658-INS-415 |

| | |
|---|---|
| chr18-10824276-INS-52 | chr18-10824276-INS-53 |
| chr18-10826874-INS-717 | chr18-10826874-INS-718 |
| chr18-10861153-INS-201 | chr18-10861153-INS-201 |
| chr18-10864530-INS-54 | chr18-10864530-INS-54 |
| chr18-10865819-INS-186 | chr18-10865819-INS-186 |
| chr18-10903532-INS-62 | chr18-10903532-INS-62 |
| chr18-10928634-INS-80 | chr18-10928634-INS-80 |
| chr18-10944403-INS-414 | chr18-10944403-INS-414 |
| chr18-10946272-INS-88 | chr18-10946268-INS-52 |
| chr18-10961946-INS-58 | chr18-10961946-INS-54 |
| chr18-10993934-INS-153 | chr18-10993934-INS-156 |
| chr18-11011535-INS-70 | chr18-11011536-INS-75 |
| chr18-11019656-INS-65 | chr18-11019656-INS-60 |
| chr18-11026427-INS-136 | chr18-11026427-INS-179 |
| chr18-11034289-INS-54 | chr18-11034289-INS-54 |
| chr18-11042166-INS-52 | chr18-11042144-INS-51 |
| chr18-11062032-INS-205 | chr18-11062032-INS-205 |
| chr18-11084001-INS-263 | chr18-11084001-INS-263 |
| chr18-11090501-INS-69 | chr18-11090501-INS-71 |
| chr18-11093547-INS-1862 | chr18-11093548-INS-1862 |
| chr18-11103241-INS-53 | chr18-11103241-INS-54 |
| chr18-11110023-INS-416 | chr18-11110015-INS-436 |
| chr18-11110974-INS-70 | chr18-11110974-INS-72 |
| chr18-11130513-INS-244 | chr18-11130513-INS-244 |
| chr18-11141864-INS-50 | chr18-11141864-INS-52 |
| chr18-11142978-INS-118 | chr18-11142978-INS-85 |
| chr18-11153433-INS-158 | chr18-11153433-INS-158 |
| chr18-11183194-INS-201 | chr18-11183194-INS-201 |
| chr18-11193561-INS-289 | chr18-11193517-INS-306 |
| chr18-11216705-INS-154 | chr18-11216705-INS-154 |
| chr18-11219982-INS-101 | chr18-11219982-INS-77 |
| chr18-11223743-INS-109 | chr18-11223743-INS-155 |
| chr18-11228895-INS-96 | chr18-11228916-INS-117 |
| chr18-11231489-INS-70 | chr18-11231489-INS-82 |
| chr18-11238866-INS-148 | chr18-11238869-INS-148 |
| chr18-11248165-INS-7186 | chr18-11248166-INS-7164 |
| chr18-11288621-INS-542 | chr18-11288595-INS-543 |
| chr18-11289882-INS-194 | chr18-11289882-INS-198 |
| chr18-11318904-INS-109 | chr18-11318952-INS-109 |
| chr18-11329506-INS-88 | chr18-11329506-INS-88 |
| chr18-11334116-INS-327 | chr18-11334117-INS-327 |
| chr18-11334341-INS-871 | chr18-11334341-INS-871 |
| chr18-11352029-INS-178 | chr18-11352029-INS-178 |
| chr18-11387181-INS-201 | chr18-11387181-INS-201 |
| chr18-11414495-INS-227 | chr18-11414495-INS-227 |
| chr18-11415207-INS-75 | chr18-11415208-INS-75 |
| chr18-11415832-INS-60 | chr18-11415832-INS-60 |
| chr18-11416841-INS-84 | chr18-11416841-INS-84 |
| chr18-11462368-INS-74 | chr18-11462368-INS-74 |
| chr18-11477972-INS-386 | chr18-11477972-INS-386 |
| chr18-11504469-INS-53 | chr18-11504469-INS-54 |
| chr18-11516484-INS-123 | chr18-11516484-INS-123 |
| chr18-11519805-INS-67 | chr18-11519805-INS-67 |
| chr18-11544592-INS-93 | chr18-11544592-INS-93 |
| chr18-11556691-INS-69 | chr18-11556691-INS-69 |
| chr18-11558611-INS-82 | chr18-11558611-INS-84 |
| chr18-11559157-INS-64 | chr18-11559157-INS-64 |
| chr18-11580858-INS-130 | chr18-11580858-INS-130 |
| chr18-11598399-INS-140 | chr18-11598399-INS-140 |
| chr18-11600972-INS-464 | chr18-11600972-INS-464 |
| chr18-11618607-INS-55 | chr18-11618607-INS-55 |
| chr18-11621343-INS-51 | chr18-11621343-INS-51 |
| chr18-11631241-INS-204 | chr18-11631241-INS-204 |
| chr18-11633490-INS-82 | chr18-11633490-INS-82 |
| chr18-11633731-INS-981 | chr18-11633731-INS-973 |
| chr18-11659544-INS-66 | chr18-11659544-INS-66 |
| chr18-11665113-INS-232 | chr18-11665113-INS-225 |
| chr18-11674291-INS-110 | chr18-11674291-INS-110 |
| chr18-11679039-INS-168 | chr18-11679039-INS-120 |
| chr18-11702997-INS-6381 | chr18-11702998-INS-6382 |
| chr18-11707054-INS-7114 | chr18-11707054-INS-7114 |
| chr18-11721379-INS-73 | chr18-11721379-INS-73 |
| chr18-11723425-INS-64 | chr18-11723425-INS-60 |
| chr18-11729132-INS-328 | chr18-11729132-INS-328 |
| chr18-11731627-INS-90 | chr18-11731651-INS-88 |
| chr18-11733381-INS-546 | chr18-11733381-INS-546 |
| chr18-11746014-INS-91 | chr18-11746014-INS-91 |

| | |
|---|---|
| chr18-11768318-INS-199 | chr18-11768318-INS-199 |
| chr18-11784371-INS-80 | chr18-11784371-INS-92 |
| chr18-11792249-INS-242 | chr18-11792249-INS-243 |
| chr18-11807089-INS-204 | chr18-11807089-INS-204 |
| chr18-11856775-INS-107 | chr18-11856775-INS-105 |
| chr18-11859632-INS-213 | chr18-11859632-INS-213 |
| chr18-11891383-INS-529 | chr18-11891383-INS-529 |
| chr18-11901797-INS-213 | chr18-11901797-INS-213 |
| chr18-11904149-INS-55 | chr18-11904158-INS-65 |
| chr18-11913354-INS-84 | chr18-11913354-INS-64 |
| chr18-11916505-INS-377 | chr18-11916505-INS-377 |
| chr18-11917876-INS-189 | chr18-11917857-INS-189 |
| chr18-11924824-INS-76 | chr18-11924824-INS-71 |
| chr18-11937192-INS-214 | chr18-11937192-INS-215 |
| chr18-11960563-INS-180 | chr18-11960563-INS-180 |
| chr18-11987151-INS-743 | chr18-11987151-INS-743 |
| chr18-11996474-INS-79 | chr18-11996474-INS-79 |
| chr18-12001040-INS-59 | chr18-12001040-INS-59 |
| chr18-12005622-INS-60 | chr18-12005626-INS-67 |
| chr18-12014609-INS-189 | chr18-12014603-INS-192 |
| chr18-12050135-INS-57 | chr18-12050135-INS-57 |
| chr18-12055369-INS-76 | chr18-12055369-INS-62 |
| chr18-12067403-INS-56 | chr18-12067399-INS-58 |
| chr18-12083597-INS-203 | chr18-12083597-INS-203 |
| chr18-12120528-INS-56 | chr18-12120529-INS-56 |
| chr18-12122573-INS-59 | chr18-12122573-INS-59 |
| chr18-12128808-INS-994 | chr18-12128808-INS-1003 |
| chr18-12133563-INS-54 | chr18-12133563-INS-60 |
| chr18-12141143-INS-86 | chr18-12141143-INS-86 |
| chr18-12141793-INS-69 | chr18-12141793-INS-69 |
| chr18-12152772-INS-196 | chr18-12152772-INS-196 |
| chr18-12156054-INS-164 | chr18-12156054-INS-164 |
| chr18-12161370-INS-59 | chr18-12161370-INS-63 |
| chr18-12166706-INS-194 | chr18-12166696-INS-195 |
| chr18-12169321-INS-115 | chr18-12169321-INS-119 |
| chr18-12185091-INS-219 | chr18-12185091-INS-219 |
| chr18-12191529-INS-380 | chr18-12191529-INS-380 |
| chr18-12205051-INS-201 | chr18-12205051-INS-201 |
| chr18-12221838-INS-195 | chr18-12221838-INS-196 |
| chr18-12227757-INS-59 | chr18-12227757-INS-59 |
| chr18-12231084-INS-226 | chr18-12231085-INS-212 |
| chr18-12234859-INS-215 | chr18-12234859-INS-215 |
| chr18-12238207-INS-57 | chr18-12238207-INS-57 |
| chr18-12243159-INS-54 | chr18-12243159-INS-54 |
| chr18-12250378-INS-200 | chr18-12250378-INS-200 |
| chr18-12253568-INS-60 | chr18-12253568-INS-60 |
| chr18-12279143-INS-205 | chr18-12279143-INS-205 |
| chr18-12287710-INS-354 | chr18-12287710-INS-354 |
| chr18-12303040-INS-232 | chr18-12303040-INS-232 |
| chr18-12306255-INS-121 | chr18-12306255-INS-121 |
| chr18-12308323-INS-207 | chr18-12308323-INS-207 |
| chr18-12313185-INS-213 | chr18-12313176-INS-226 |
| chr18-12314915-INS-208 | chr18-12314915-INS-208 |
| chr18-12319521-INS-74 | chr18-12319525-INS-75 |
| chr18-12332870-INS-797 | chr18-12332870-INS-797 |
| chr18-12342107-INS-493 | chr18-12342107-INS-493 |
| chr18-12342771-INS-76 | chr18-12342771-INS-76 |
| chr18-12348670-INS-94 | chr18-12348665-INS-98 |
| chr18-12350716-INS-51 | chr18-12350716-INS-51 |
| chr18-12356647-INS-60 | chr18-12356648-INS-57 |
| chr18-12360544-INS-70 | chr18-12360545-INS-70 |
| chr18-12361054-INS-101 | chr18-12361055-INS-101 |
| chr18-12365651-INS-66 | chr18-12365651-INS-66 |
| chr18-12372628-INS-173 | chr18-12372628-INS-175 |
| chr18-12378647-INS-319 | chr18-12378647-INS-319 |
| chr18-12379549-INS-217 | chr18-12379549-INS-217 |
| chr18-12406044-INS-583 | chr18-12406044-INS-583 |
| chr18-12410850-INS-406 | chr18-12410850-INS-406 |
| chr18-12432955-INS-103 | chr18-12432955-INS-103 |
| chr18-12444912-INS-62 | chr18-12444912-INS-65 |
| chr18-12484152-INS-72 | chr18-12484152-INS-72 |
| chr18-12491714-INS-88 | chr18-12491714-INS-92 |
| chr18-12499379-INS-56 | chr18-12499379-INS-56 |
| chr18-12511865-INS-69 | chr18-12511865-INS-69 |
| chr18-12513935-INS-72 | chr18-12513936-INS-71 |
| chr18-12515599-INS-300 | chr18-12515599-INS-300 |
| chr18-12523717-INS-207 | chr18-12523717-INS-207 |

| | |
|---|---|
| chr18-12542430-INS-145 | chr18-12542430-INS-145 |
| chr18-12544315-INS-59 | chr18-12544315-INS-59 |
| chr18-12547426-INS-53 | chr18-12547426-INS-53 |
| chr18-12552369-INS-67 | chr18-12552369-INS-67 |
| chr18-12553476-INS-96 | chr18-12553476-INS-96 |
| chr18-12585478-INS-204 | chr18-12585479-INS-204 |
| chr18-12592972-INS-54 | chr18-12592946-INS-54 |
| chr18-12609123-INS-967 | chr18-12609123-INS-965 |
| chr18-12623089-INS-82 | chr18-12623089-INS-82 |
| chr18-12629170-INS-67 | chr18-12629170-INS-67 |
| chr18-12640301-INS-1255 | chr18-12640302-INS-1243 |
| chr18-12640724-INS-56 | chr18-12640724-INS-54 |
| chr18-12647141-INS-124 | chr18-12647133-INS-137 |
| chr18-12648385-INS-215 | chr18-12648385-INS-215 |
| chr18-12668215-INS-153 | chr18-12668215-INS-153 |
| chr18-12676070-INS-150 | chr18-12676070-INS-150 |
| chr18-12681724-INS-479 | chr18-12681724-INS-479 |
| chr18-12708508-INS-165 | chr18-12708507-INS-169 |
| chr18-12711187-INS-322 | chr18-12711187-INS-322 |
| chr18-12721541-INS-84 | chr18-12721541-INS-85 |
| chr18-12740977-INS-69 | chr18-12740977-INS-69 |
| chr18-12742166-INS-73 | chr18-12742166-INS-73 |
| chr18-12770641-INS-56 | chr18-12770641-INS-56 |
| chr18-12800994-INS-4878 | chr18-12800995-INS-4878 |
| chr18-12809813-INS-160 | chr18-12809813-INS-160 |
| chr18-12811276-INS-184 | chr18-12811276-INS-184 |
| chr18-12817530-INS-68 | chr18-12817530-INS-68 |
| chr18-12818520-INS-80 | chr18-12818520-INS-80 |
| chr18-12827970-INS-170 | chr18-12827970-INS-170 |
| chr18-12833322-INS-77 | chr18-12833322-INS-77 |
| chr18-12847216-INS-93 | chr18-12847208-INS-62 |
| chr18-12907781-INS-176 | chr18-12907781-INS-176 |
| chr18-12908328-INS-188 | chr18-12908328-INS-188 |
| chr18-12919395-INS-357 | chr18-12919395-INS-355 |
| chr18-12923349-INS-207 | chr18-12923349-INS-207 |
| chr18-12959841-INS-1190 | chr18-12959842-INS-1190 |
| chr18-12969858-INS-7112 | chr18-12969859-INS-7112 |
| chr18-12973725-INS-64 | chr18-12973725-INS-58 |
| chr18-12974752-INS-58 | chr18-12974752-INS-58 |
| chr18-12983788-INS-186 | chr18-12983788-INS-186 |
| chr18-12987980-INS-87 | chr18-12987980-INS-87 |
| chr18-13015029-INS-98 | chr18-13015029-INS-98 |
| chr18-13018402-INS-80 | chr18-13018406-INS-94 |
| chr18-13020484-INS-171 | chr18-13020484-INS-171 |
| chr18-13033497-INS-75 | chr18-13033497-INS-61 |
| chr18-13037293-INS-95 | chr18-13037293-INS-95 |
| chr18-13044837-INS-96 | chr18-13044837-INS-96 |
| chr18-13059120-INS-136 | chr18-13059120-INS-90 |
| chr18-13061635-INS-58 | chr18-13061635-INS-58 |
| chr18-13068590-INS-192 | chr18-13068590-INS-192 |
| chr18-13079715-INS-203 | chr18-13079715-INS-203 |
| chr18-13084083-INS-52 | chr18-13084083-INS-52 |
| chr18-13092452-INS-215 | chr18-13092452-INS-216 |
| chr18-13098029-INS-81 | chr18-13098029-INS-81 |
| chr18-13114221-INS-209 | chr18-13114221-INS-208 |
| chr18-13127638-INS-66 | chr18-13127638-INS-66 |
| chr18-13134230-INS-1243 | chr18-13134231-INS-1242 |
| chr18-13139421-INS-246 | chr18-13139421-INS-248 |
| chr18-13150353-INS-51 | chr18-13150373-INS-66 |
| chr18-13153437-INS-61 | chr18-13153437-INS-63 |
| chr18-13156108-INS-135 | chr18-13156108-INS-135 |
| chr18-13183240-INS-69 | chr18-13183242-INS-87 |
| chr18-13187809-INS-138 | chr18-13187809-INS-134 |
| chr18-13199959-INS-2914 | chr18-13199960-INS-2914 |
| chr18-13208764-INS-54 | chr18-13208764-INS-52 |
| chr18-13217149-INS-2036 | chr18-13217150-INS-2036 |
| chr18-13217238-INS-8274 | chr18-13217229-INS-8274 |
| chr18-13263521-INS-215 | chr18-13263521-INS-179 |
| chr18-13279256-INS-420 | chr18-13279256-INS-420 |
| chr18-13329475-INS-63 | chr18-13329475-INS-63 |
| chr18-13402935-INS-62 | chr18-13402935-INS-62 |
| chr18-13441953-INS-400 | chr18-13441953-INS-400 |
| chr18-13448011-INS-52 | chr18-13447999-INS-68 |
| chr18-13469847-INS-106 | chr18-13469847-INS-106 |
| chr18-13493416-INS-7121 | chr18-13493417-INS-7121 |
| chr18-13498768-INS-68 | chr18-13498775-INS-88 |
| chr18-13517046-INS-389 | chr18-13517046-INS-389 |

| | |
|---|---|
| chr18-13526409-INS-409 | chr18-13526409-INS-408 |
| chr18-13529290-INS-90 | chr18-13529282-INS-71 |
| chr18-13550361-INS-166 | chr18-13550361-INS-166 |
| chr18-13551499-INS-735 | chr18-13551499-INS-735 |
| chr18-13552331-INS-66 | chr18-13552331-INS-66 |
| chr18-13597216-INS-56 | chr18-13597216-INS-56 |
| chr18-13611619-INS-70 | chr18-13611619-INS-120 |
| chr18-13622692-INS-106 | chr18-13622711-INS-114 |
| chr18-13636499-INS-108 | chr18-13636513-INS-92 |
| chr18-13643187-INS-73 | chr18-13643187-INS-73 |
| chr18-13673465-INS-101 | chr18-13673500-INS-80 |
| chr18-13673498-INS-1187 | chr18-13673499-INS-1187 |
| chr18-13697837-INS-52 | chr18-13697833-INS-50 |
| chr18-13698939-INS-64 | chr18-13698939-INS-64 |
| chr18-13715187-INS-212 | chr18-13715187-INS-212 |
| chr18-13730948-INS-72 | chr18-13730948-INS-72 |
| chr18-13744444-INS-281 | chr18-13744444-INS-282 |
| chr18-13758917-INS-58 | chr18-13758917-INS-52 |
| chr18-13762745-INS-52 | chr18-13762742-INS-60 |
| chr18-13801950-INS-286 | chr18-13801950-INS-286 |
| chr18-13807650-INS-75 | chr18-13807650-INS-75 |
| chr18-13859799-INS-54 | chr18-13859799-INS-69 |
| chr18-13908595-INS-56 | chr18-13908594-INS-58 |
| chr18-13955689-INS-3884 | chr18-13955690-INS-3886 |
| chr18-14025473-INS-58 | chr18-14025473-INS-58 |
| chr18-14079295-INS-102 | chr18-14079295-INS-115 |
| chr18-14096880-INS-260 | chr18-14096880-INS-280 |
| chr18-14132653-INS-101 | chr18-14132653-INS-101 |
| chr18-14149342-INS-440 | chr18-14149342-INS-440 |
| chr18-14157677-INS-205 | chr18-14157677-INS-205 |
| chr18-14159909-INS-95 | chr18-14159893-INS-78 |
| chr18-14207680-INS-2733 | chr18-14207681-INS-2734 |
| chr18-14219606-INS-222 | chr18-14219606-INS-222 |
| chr18-14226146-INS-234 | chr18-14226146-INS-203 |
| chr18-14227134-INS-167 | chr18-14227134-INS-173 |
| chr18-14228896-INS-59 | chr18-14228896-INS-63 |
| chr18-14252212-INS-62 | chr18-14252201-INS-90 |
| chr18-14259964-INS-89 | chr18-14259978-INS-69 |
| chr18-14281471-INS-6442 | chr18-14281472-INS-6442 |
| chr18-14282026-INS-70 | chr18-14282026-INS-70 |
| chr18-14288861-INS-2935 | chr18-14288862-INS-2935 |
| chr18-14351929-INS-576 | chr18-14351926-INS-589 |
| chr18-14400641-INS-56 | chr18-14400645-INS-72 |
| chr18-14402532-INS-168 | chr18-14402576-INS-137 |
| chr18-14442718-INS-54 | chr18-14442685-INS-64 |
| chr18-14458917-INS-174 | chr18-14458913-INS-176 |
| chr18-14465416-INS-54 | chr18-14465416-INS-52 |
| chr18-14500624-INS-87 | chr18-14500624-INS-64 |
| chr18-14511975-INS-71 | chr18-14511991-INS-56 |
| chr18-14533845-INS-81 | chr18-14533845-INS-54 |
| chr18-14537639-INS-70 | chr18-14537639-INS-70 |
| chr18-14540738-INS-4899 | chr18-14540739-INS-4899 |
| chr18-14560541-INS-122 | chr18-14560584-INS-78 |
| chr18-14564653-INS-142 | chr18-14564653-INS-142 |
| chr18-14566157-INS-1892 | chr18-14566158-INS-1892 |
| chr18-14596588-INS-54 | chr18-14596591-INS-80 |
| chr18-14600394-INS-61 | chr18-14600421-INS-57 |
| chr18-14605461-INS-112 | chr18-14605461-INS-155 |
| chr18-14624871-INS-75 | chr18-14624871-INS-110 |
| chr18-14675290-INS-6216 | chr18-14675291-INS-6216 |
| chr18-14727153-INS-587 | chr18-14727154-INS-599 |
| chr18-14748988-INS-92 | chr18-14748988-INS-92 |
| chr18-14790899-INS-1896 | chr18-14790900-INS-1896 |
| chr18-14794833-INS-343 | chr18-14794833-INS-343 |
| chr18-14799029-INS-216 | chr18-14799029-INS-216 |
| chr18-14822590-INS-203 | chr18-14822590-INS-203 |
| chr18-14877719-INS-4382 | chr18-14877720-INS-4382 |
| chr18-14896524-INS-6758 | chr18-14896525-INS-6763 |
| chr18-14913091-INS-196 | chr18-14913091-INS-197 |
| chr18-14922265-INS-199 | chr18-14922265-INS-199 |
| chr18-14950195-INS-222 | chr18-14950195-INS-221 |
| chr18-14976144-INS-2033 | chr18-14976145-INS-2034 |
| chr18-15008460-INS-248 | chr18-15008460-INS-248 |
| chr18-15043952-INS-940 | chr18-15043952-INS-940 |
| chr18-15047184-INS-54 | chr18-15047184-INS-56 |
| chr18-15051236-INS-56 | chr18-15051236-INS-56 |
| chr18-15054121-INS-122 | chr18-15054121-INS-98 |

| | |
|---|---|
| chr18-15058127-INS-727 | chr18-15058127-INS-723 |
| chr18-15062591-INS-75 | chr18-15062593-INS-73 |
| chr18-15068015-INS-179 | chr18-15068015-INS-210 |
| chr18-15096978-INS-80 | chr18-15096978-INS-80 |
| chr18-15100763-INS-74 | chr18-15100763-INS-74 |
| chr18-15102333-INS-54 | chr18-15102333-INS-56 |
| chr18-15110584-INS-6104 | chr18-15110585-INS-6103 |
| chr18-15111507-INS-232 | chr18-15111507-INS-190 |
| chr18-15131902-INS-80 | chr18-15131912-INS-86 |
| chr18-15137868-INS-201 | chr18-15137868-INS-201 |
| chr18-15140195-INS-56 | chr18-15140195-INS-70 |
| chr18-15143314-INS-57 | chr18-15143314-INS-54 |
| chr18-15153460-INS-231 | chr18-15153460-INS-229 |
| chr18-15181376-INS-93 | chr18-15181376-INS-104 |
| chr18-15225381-INS-205 | chr18-15225381-INS-205 |
| chr18-15241368-INS-291 | chr18-15241368-INS-273 |
| chr18-15292378-INS-774 | chr18-15292378-INS-775 |
| chr18-15301327-INS-3853 | chr18-15301328-INS-3859 |
| chr18-15347769-INS-66 | chr18-15347769-INS-54 |
| chr18-15354799-INS-122 | chr18-15354788-INS-159 |
| chr18-15378878-INS-188 | chr18-15378878-INS-174 |
| chr18-15381486-INS-105 | chr18-15381486-INS-82 |
| chr18-15411857-INS-556 | chr18-15411857-INS-573 |
| chr18-15432267-INS-174 | chr18-15432267-INS-169 |
| chr18-15439849-INS-52 | chr18-15439842-INS-85 |
| chr18-15452953-INS-64 | chr18-15452953-INS-64 |
| chr18-15477775-INS-129 | chr18-15477775-INS-129 |
| chr18-15540540-INS-402 | chr18-15540519-INS-401 |
| chr18-15547062-INS-344 | chr18-15547062-INS-344 |
| chr18-15554130-INS-94 | chr18-15554130-INS-92 |
| chr18-15571735-INS-89 | chr18-15571735-INS-89 |
| chr18-15572610-INS-348 | chr18-15572610-INS-348 |
| chr18-15574184-INS-229 | chr18-15574184-INS-274 |
| chr18-15580518-INS-90 | chr18-15580532-INS-90 |
| chr18-15646778-INS-90 | chr18-15646778-INS-90 |
| chr18-15691394-INS-991 | chr18-15691394-INS-991 |
| chr18-15698789-INS-56 | chr18-15698789-INS-74 |
| chr18-15791772-INS-144 | chr18-15791772-INS-144 |
| chr18-15803773-INS-139 | chr18-15803773-INS-147 |
| chr18-15812917-INS-96 | chr18-15812917-INS-78 |
| chr18-15902283-INS-68 | chr18-15902283-INS-50 |
| chr18-15914077-INS-60 | chr18-15914060-INS-60 |
| chr18-15918004-INS-399 | chr18-15918004-INS-399 |
| chr18-15924465-INS-1156 | chr18-15924461-INS-1157 |
| chr18-15941228-INS-4486 | chr18-15941229-INS-4486 |
| chr18-15949589-INS-74 | chr18-15949590-INS-70 |
| chr18-15957997-INS-419 | chr18-15957997-INS-416 |
| chr18-16003463-INS-96 | chr18-16003463-INS-144 |
| chr18-16023053-INS-54 | chr18-16023053-INS-52 |
| chr18-16070724-INS-267 | chr18-16070684-INS-272 |
| chr18-16104786-INS-61 | chr18-16104786-INS-89 |
| chr18-16126224-INS-72 | chr18-16126224-INS-72 |
| chr18-16132670-INS-550 | chr18-16132670-INS-548 |
| chr18-16135491-INS-4701 | chr18-16135492-INS-4701 |
| chr18-16174357-INS-175 | chr18-16174357-INS-180 |
| chr18-16177017-INS-86 | chr18-16177017-INS-130 |
| chr18-16179525-INS-102 | chr18-16179525-INS-102 |
| chr18-16181369-INS-6805 | chr18-16181370-INS-6807 |
| chr18-16192414-INS-231 | chr18-16192414-INS-237 |
| chr18-16198523-INS-184 | chr18-16198523-INS-184 |
| chr18-16227877-INS-82 | chr18-16227877-INS-82 |
| chr18-16231470-INS-67 | chr18-16231470-INS-67 |
| chr18-16241065-INS-100 | chr18-16241065-INS-100 |
| chr18-16334281-INS-84 | chr18-16334268-INS-58 |
| chr18-16355610-INS-214 | chr18-16355612-INS-228 |
| chr18-16355752-INS-1804 | chr18-16355753-INS-1804 |
| chr18-16367135-INS-64 | chr18-16367135-INS-64 |
| chr18-16394988-INS-7068 | chr18-16394989-INS-7068 |
| chr18-16399388-INS-2768 | chr18-16399389-INS-2768 |
| chr18-16413705-INS-164 | chr18-16413705-INS-136 |
| chr18-16461409-INS-646 | chr18-16461409-INS-646 |
| chr18-16465146-INS-332 | chr18-16465146-INS-332 |
| chr18-16477267-INS-440 | chr18-16477267-INS-440 |
| chr18-16525885-INS-5121 | chr18-16525886-INS-5122 |
| chr18-16551334-INS-70 | chr18-16551334-INS-66 |
| chr18-16562049-INS-6595 | chr18-16562050-INS-6598 |
| chr18-16569870-INS-1889 | chr18-16569871-INS-1889 |

| | |
|---|---|
| chr18-16571332-INS-89 | chr18-16571337-INS-112 |
| chr18-16586466-INS-235 | chr18-16586466-INS-235 |
| chr18-16593400-INS-372 | chr18-16593400-INS-372 |
| chr18-16667188-INS-2594 | chr18-16667189-INS-2594 |
| chr18-16687328-INS-67 | chr18-16687328-INS-72 |
| chr18-16690227-INS-1872 | chr18-16690228-INS-1872 |
| chr18-16705174-INS-120 | chr18-16705174-INS-120 |
| chr18-16720029-INS-234 | chr18-16720029-INS-234 |
| chr18-16727209-INS-6446 | chr18-16727210-INS-6446 |
| chr18-16775379-INS-65 | chr18-16775379-INS-65 |
| chr18-16776230-INS-51 | chr18-16776210-INS-50 |
| chr18-16795362-INS-94 | chr18-16795362-INS-99 |
| chr18-16800765-INS-84 | chr18-16800765-INS-74 |
| chr18-16805445-INS-60 | chr18-16805445-INS-60 |
| chr18-16874373-INS-68 | chr18-16874373-INS-61 |
| chr18-16896442-INS-456 | chr18-16896442-INS-456 |
| chr18-16956837-INS-68 | chr18-16956837-INS-76 |
| chr18-16993598-INS-70 | chr18-16993609-INS-50 |
| chr18-17001875-INS-1348 | chr18-17001876-INS-1348 |
| chr18-17018832-INS-538 | chr18-17018832-INS-538 |
| chr18-17055895-INS-4484 | chr18-17055896-INS-4484 |
| chr18-17058143-INS-910 | chr18-17058143-INS-905 |
| chr18-17102689-INS-236 | chr18-17102689-INS-238 |
| chr18-17157553-INS-2005 | chr18-17157554-INS-2005 |
| chr18-17192050-INS-324 | chr18-17192050-INS-324 |
| chr18-17196884-INS-442 | chr18-17196884-INS-442 |
| chr18-17209702-INS-3181 | chr18-17209703-INS-3181 |
| chr18-17212127-INS-7025 | chr18-17212128-INS-7025 |
| chr18-17216668-INS-295 | chr18-17216668-INS-297 |
| chr18-17239694-INS-6525 | chr18-17239695-INS-6526 |
| chr18-17242659-INS-120 | chr18-17242659-INS-120 |
| chr18-17257328-INS-498 | chr18-17257328-INS-498 |
| chr18-17266076-INS-55 | chr18-17266107-INS-50 |
| chr18-17270465-INS-3968 | chr18-17270466-INS-3969 |
| chr18-17275712-INS-330 | chr18-17275712-INS-330 |
| chr18-17278165-INS-362 | chr18-17278165-INS-360 |
| chr18-17280219-INS-60 | chr18-17280217-INS-50 |
| chr18-17301195-INS-52 | chr18-17301195-INS-52 |
| chr18-17312318-INS-2836 | chr18-17312319-INS-2836 |
| chr18-17314401-INS-68 | chr18-17314402-INS-76 |
| chr18-17349684-INS-987 | chr18-17349684-INS-987 |
| chr18-17373350-INS-6436 | chr18-17373351-INS-6436 |
| chr18-17382238-INS-5545 | chr18-17382239-INS-5545 |
| chr18-17434353-INS-1524 | chr18-17434354-INS-1524 |
| chr18-17466511-INS-1037 | chr18-17466512-INS-1041 |
| chr18-17470048-INS-1892 | chr18-17470049-INS-1892 |
| chr18-17470049-INS-400 | chr18-17470049-INS-400 |
| chr18-17472016-INS-3130 | chr18-17472017-INS-3130 |
| chr18-17474029-INS-88 | chr18-17474029-INS-88 |
| chr18-17505438-INS-5834 | chr18-17505439-INS-5834 |
| chr18-17506264-INS-64 | chr18-17506264-INS-64 |
| chr18-17514465-INS-445 | chr18-17514465-INS-444 |
| chr18-17544522-INS-88 | chr18-17544522-INS-88 |
| chr18-17548366-INS-2767 | chr18-17548367-INS-2767 |
| chr18-17556947-INS-58 | chr18-17556947-INS-60 |
| chr18-17577375-INS-215 | chr18-17577375-INS-207 |
| chr18-17587155-INS-214 | chr18-17587155-INS-214 |
| chr18-17638210-INS-76 | chr18-17638210-INS-76 |
| chr18-17686934-INS-56 | chr18-17686934-INS-53 |
| chr18-17741690-INS-169 | chr18-17741690-INS-169 |
| chr18-17766863-INS-2067 | chr18-17766864-INS-2067 |
| chr18-17785445-INS-498 | chr18-17785445-INS-498 |
| chr18-17793376-INS-86 | chr18-17793387-INS-54 |
| chr18-17810632-INS-3964 | chr18-17810633-INS-3964 |
| chr18-17835011-INS-1224 | chr18-17835012-INS-1224 |
| chr18-17844798-INS-268 | chr18-17844798-INS-268 |
| chr18-17847434-INS-212 | chr18-17847420-INS-211 |
| chr18-17876033-INS-3405 | chr18-17876034-INS-3406 |
| chr18-17877046-INS-64 | chr18-17877046-INS-64 |
| chr18-17884280-INS-2779 | chr18-17884281-INS-2777 |
| chr18-17885061-INS-74 | chr18-17885061-INS-76 |
| chr18-17897260-INS-67 | chr18-17897260-INS-100 |
| chr18-17913578-INS-298 | chr18-17913578-INS-298 |
| chr18-17928324-INS-146 | chr18-17928350-INS-136 |
| chr18-17941051-INS-53 | chr18-17941051-INS-53 |
| chr18-17942058-INS-74 | chr18-17942058-INS-78 |
| chr18-17952316-INS-363 | chr18-17952316-INS-364 |

| | |
|---|---|
| chr18-17961264-INS-206 | chr18-17961267-INS-205 |
| chr18-18036126-INS-6498 | chr18-18036127-INS-6498 |
| chr18-18049822-INS-7082 | chr18-18049823-INS-7082 |
| chr18-18050438-INS-59 | chr18-18050438-INS-55 |
| chr18-18052170-INS-532 | chr18-18052170-INS-536 |
| chr18-18060009-INS-231 | chr18-18060009-INS-231 |
| chr18-18094384-INS-52 | chr18-18094384-INS-56 |
| chr18-18110930-INS-6805 | chr18-18110931-INS-6805 |
| chr18-18119031-INS-56 | chr18-18119031-INS-56 |
| chr18-18137895-INS-394 | chr18-18137895-INS-393 |
| chr18-18147081-INS-66 | chr18-18147081-INS-54 |
| chr18-18160534-INS-119 | chr18-18160536-INS-139 |
| chr18-18169579-INS-778 | chr18-18169579-INS-778 |
| chr18-18189707-INS-70 | chr18-18189707-INS-76 |
| chr18-18193760-INS-196 | chr18-18193760-INS-196 |
| chr18-18204749-INS-1864 | chr18-18204750-INS-1864 |
| chr18-18234051-INS-374 | chr18-18234051-INS-375 |
| chr18-18252722-INS-3912 | chr18-18252723-INS-3912 |
| chr18-18271121-INS-73 | chr18-18271121-INS-75 |
| chr18-18300621-INS-101 | chr18-18300621-INS-101 |
| chr18-18312975-INS-123 | chr18-18312975-INS-73 |
| chr18-18313939-INS-68 | chr18-18313939-INS-68 |
| chr18-18316090-INS-70 | chr18-18316095-INS-53 |
| chr18-18347037-INS-6582 | chr18-18347038-INS-6583 |
| chr18-18368993-INS-6244 | chr18-18368994-INS-6244 |
| chr18-18379525-INS-194 | chr18-18379525-INS-194 |
| chr18-18426958-INS-231 | chr18-18426958-INS-231 |
| chr18-18448242-INS-70 | chr18-18448242-INS-70 |
| chr18-18483084-INS-6861 | chr18-18483085-INS-6862 |
| chr18-18489316-INS-147 | chr18-18489318-INS-147 |
| chr18-18519418-INS-2598 | chr18-18519419-INS-2598 |
| chr18-18519419-INS-437 | chr18-18519419-INS-437 |
| chr18-18537412-INS-126 | chr18-18537412-INS-132 |
| chr18-18537458-INS-4286 | chr18-18537459-INS-4287 |
| chr18-18545578-INS-349 | chr18-18545578-INS-350 |
| chr18-18590809-INS-4234 | chr18-18590810-INS-4240 |
| chr18-18640708-INS-1318 | chr18-18640722-INS-1315 |
| chr18-18655788-INS-225 | chr18-18655788-INS-225 |
| chr18-18706269-INS-1487 | chr18-18706270-INS-1487 |
| chr18-18718130-INS-210 | chr18-18718136-INS-222 |
| chr18-18751458-INS-411 | chr18-18751458-INS-416 |
| chr18-18816778-INS-174 | chr18-18816778-INS-174 |
| chr18-18828345-INS-1214 | chr18-18828346-INS-1214 |
| chr18-18845863-INS-2841 | chr18-18845864-INS-2845 |
| chr18-18907790-INS-6634 | chr18-18907791-INS-6634 |
| chr18-18912483-INS-60 | chr18-18912483-INS-72 |
| chr18-18923426-INS-626 | chr18-18923426-INS-590 |
| chr18-18938463-INS-112 | chr18-18938463-INS-112 |
| chr18-19004873-INS-6566 | chr18-19004874-INS-6566 |
| chr18-19029084-INS-5954 | chr18-19029085-INS-5954 |
| chr18-19034726-INS-1893 | chr18-19034727-INS-1893 |
| chr18-19097408-INS-56 | chr18-19097425-INS-88 |
| chr18-19107148-INS-57 | chr18-19107148-INS-57 |
| chr18-19127288-INS-98 | chr18-19127288-INS-96 |
| chr18-19144604-INS-7482 | chr18-19144605-INS-7482 |
| chr18-19159262-INS-188 | chr18-19159262-INS-188 |
| chr18-19166318-INS-487 | chr18-19166318-INS-487 |
| chr18-19168914-INS-54 | chr18-19168914-INS-54 |
| chr18-19259486-INS-6396 | chr18-19259487-INS-6396 |
| chr18-19299996-INS-655 | chr18-19299996-INS-664 |
| chr18-19308361-INS-106 | chr18-19308361-INS-106 |
| chr18-19330461-INS-6256 | chr18-19330462-INS-6256 |
| chr18-19333208-INS-182 | chr18-19333208-INS-182 |
| chr18-19351856-INS-135 | chr18-19351860-INS-151 |
| chr18-19354261-INS-336 | chr18-19354261-INS-331 |
| chr18-19372291-INS-65 | chr18-19372291-INS-65 |
| chr18-19391713-INS-126 | chr18-19391731-INS-104 |
| chr18-19443378-INS-477 | chr18-19443378-INS-477 |
| chr18-19445287-INS-224 | chr18-19445287-INS-224 |
| chr18-19450995-INS-249 | chr18-19450995-INS-244 |
| chr18-19466206-INS-2247 | chr18-19466207-INS-2247 |
| chr18-19470706-INS-742 | chr18-19470706-INS-745 |
| chr18-19496597-INS-52 | chr18-19496597-INS-52 |
| chr18-19508952-INS-97 | chr18-19508952-INS-97 |
| chr18-19525333-INS-79 | chr18-19525333-INS-79 |
| chr18-19547420-INS-3866 | chr18-19547421-INS-3866 |
| chr18-19547962-INS-106 | chr18-19547962-INS-106 |

| | |
|---|---|
| chr18-19554244-INS-569 | chr18-19554244-INS-567 |
| chr18-19587963-INS-306 | chr18-19587959-INS-307 |
| chr18-19596778-INS-501 | chr18-19596776-INS-499 |
| chr18-19619572-INS-62 | chr18-19619572-INS-62 |
| chr18-19632869-INS-323 | chr18-19632869-INS-323 |
| chr18-19633654-INS-1872 | chr18-19633655-INS-1872 |
| chr18-19645146-INS-249 | chr18-19645146-INS-252 |
| chr18-19655808-INS-80 | chr18-19655814-INS-64 |
| chr18-19694461-INS-422 | chr18-19694461-INS-422 |
| chr18-19718321-INS-6173 | chr18-19718322-INS-6174 |
| chr18-19768842-INS-434 | chr18-19768842-INS-468 |
| chr18-19769924-INS-782 | chr18-19769924-INS-782 |
| chr18-19785237-INS-705 | chr18-19785237-INS-705 |
| chr18-19787383-INS-260 | chr18-19787383-INS-252 |
| chr18-19791228-INS-52 | chr18-19791228-INS-52 |
| chr18-19817949-INS-51 | chr18-19817938-INS-50 |
| chr18-19838251-INS-1109 | chr18-19838252-INS-1109 |
| chr18-19863056-INS-469 | chr18-19863056-INS-458 |
| chr18-19878644-INS-674 | chr18-19878644-INS-674 |
| chr18-19879411-INS-53 | chr18-19879385-INS-100 |
| chr18-19884512-INS-2683 | chr18-19884513-INS-2683 |
| chr18-19886953-INS-6377 | chr18-19886954-INS-6376 |
| chr18-19901430-INS-174 | chr18-19901421-INS-156 |
| chr18-19908367-INS-400 | chr18-19908367-INS-400 |
| chr18-19909041-INS-1229 | chr18-19909042-INS-1229 |
| chr18-19911963-INS-1332 | chr18-19911964-INS-1329 |
| chr18-19923944-INS-56 | chr18-19923946-INS-57 |
| chr18-19926142-INS-102 | chr18-19926142-INS-102 |
| chr18-19938056-INS-3823 | chr18-19938057-INS-3823 |
| chr18-19938057-INS-448 | chr18-19938057-INS-448 |
| chr18-19945149-INS-201 | chr18-19945149-INS-201 |
| chr18-19961763-INS-358 | chr18-19961763-INS-359 |
| chr18-19969031-INS-186 | chr18-19969031-INS-186 |
| chr18-19975860-INS-4399 | chr18-19975861-INS-4399 |
| chr18-19978762-INS-177 | chr18-19978762-INS-177 |
| chr18-19983433-INS-68 | chr18-19983433-INS-68 |
| chr18-19995753-INS-212 | chr18-19995753-INS-212 |
| chr18-20026408-INS-184 | chr18-20026408-INS-184 |
| chr18-20052142-INS-211 | chr18-20052142-INS-211 |
| chr18-20062026-INS-385 | chr18-20062026-INS-385 |
| chr18-20079995-INS-3019 | chr18-20079999-INS-3010 |
| chr18-20087502-INS-464 | chr18-20087502-INS-471 |
| chr18-20092407-INS-958 | chr18-20092407-INS-968 |
| chr18-20098447-INS-149 | chr18-20098447-INS-149 |
| chr18-20111690-INS-241 | chr18-20111693-INS-229 |
| chr18-20130067-INS-107 | chr18-20130067-INS-99 |
| chr18-20133024-INS-60 | chr18-20133040-INS-84 |
| chr18-20148454-INS-54 | chr18-20148454-INS-56 |
| chr18-20157635-INS-61 | chr18-20157631-INS-61 |
| chr18-20171026-INS-76 | chr18-20171026-INS-76 |
| chr18-20177146-INS-139 | chr18-20177146-INS-132 |
| chr18-20208769-INS-852 | chr18-20208769-INS-852 |
| chr18-20210813-INS-647 | chr18-20210813-INS-647 |
| chr18-20235779-INS-56 | chr18-20235779-INS-56 |
| chr18-20238199-INS-868 | chr18-20238199-INS-868 |
| chr18-20264842-INS-149 | chr18-20264842-INS-149 |
| chr18-20269767-INS-197 | chr18-20269767-INS-197 |
| chr18-20288257-INS-91 | chr18-20288257-INS-91 |
| chr18-20308388-INS-58 | chr18-20308389-INS-66 |
| chr18-20360933-INS-631 | chr18-20360933-INS-625 |
| chr18-20371464-INS-464 | chr18-20371464-INS-465 |
| chr18-20385538-INS-278 | chr18-20385538-INS-276 |
| chr18-20407291-INS-434 | chr18-20407291-INS-434 |
| chr18-20475735-INS-52 | chr18-20475735-INS-52 |
| chr18-20524485-INS-96 | chr18-20524451-INS-96 |
| chr18-20558644-INS-77 | chr18-20558616-INS-92 |
| chr18-20567286-INS-76 | chr18-20567286-INS-76 |
| chr18-20568746-INS-68 | chr18-20568746-INS-60 |
| chr18-20598725-INS-65 | chr18-20598725-INS-65 |
| chr18-20636892-INS-360 | chr18-20636892-INS-360 |
| chr18-20661870-INS-173 | chr18-20661870-INS-142 |
| chr18-20664178-INS-223 | chr18-20664178-INS-223 |
| chr18-20667752-INS-560 | chr18-20667752-INS-560 |
| chr18-20696765-INS-58 | chr18-20696769-INS-53 |
| chr18-20726729-INS-168 | chr18-20726729-INS-168 |
| chr18-20728421-INS-706 | chr18-20728421-INS-716 |
| chr18-20764793-INS-199 | chr18-20764793-INS-198 |

| | |
|---|---|
| chr18-20777664-INS-110 | chr18-20777664-INS-74 |
| chr18-20786239-INS-155 | chr18-20786239-INS-155 |
| chr18-20786363-INS-1100 | chr18-20786364-INS-1100 |
| chr18-20818354-INS-197 | chr18-20818354-INS-197 |
| chr18-20827692-INS-169 | chr18-20827692-INS-183 |
| chr18-20829034-INS-793 | chr18-20829034-INS-793 |
| chr18-20833632-INS-104 | chr18-20833632-INS-104 |
| chr18-20836690-INS-205 | chr18-20836690-INS-205 |
| chr18-20844685-INS-240 | chr18-20844685-INS-240 |
| chr18-20854322-INS-53 | chr18-20854322-INS-54 |
| chr18-20865266-INS-106 | chr18-20865266-INS-105 |
| chr18-20870685-INS-92 | chr18-20870685-INS-92 |
| chr18-20886740-INS-256 | chr18-20886740-INS-233 |
| chr18-20895475-INS-538 | chr18-20895475-INS-538 |
| chr18-20932728-INS-169 | chr18-20932728-INS-169 |
| chr18-21009646-INS-201 | chr18-21009646-INS-201 |
| chr18-21070587-INS-202 | chr18-21070587-INS-203 |
| chr18-21078629-INS-204 | chr18-21078629-INS-204 |
| chr18-21087078-INS-70 | chr18-21087071-INS-76 |
| chr18-21090730-INS-183 | chr18-21090730-INS-183 |
| chr18-21106701-INS-64 | chr18-21106701-INS-52 |
| chr18-21124714-INS-434 | chr18-21124714-INS-433 |
| chr18-21126389-INS-194 | chr18-21126389-INS-195 |
| chr18-21128390-INS-365 | chr18-21128390-INS-365 |
| chr18-21130465-INS-122 | chr18-21130466-INS-123 |
| chr18-21138057-INS-205 | chr18-21138057-INS-205 |
| chr18-21141568-INS-203 | chr18-21141568-INS-203 |
| chr18-21151199-INS-191 | chr18-21151200-INS-175 |
| chr18-21167085-INS-629 | chr18-21167085-INS-629 |
| chr18-21178739-INS-61 | chr18-21178739-INS-61 |
| chr18-21220924-INS-82 | chr18-21220924-INS-82 |
| chr18-21223524-INS-72 | chr18-21223524-INS-98 |
| chr18-21245181-INS-4765 | chr18-21245182-INS-4765 |
| chr18-21264277-INS-224 | chr18-21264277-INS-224 |
| chr18-21279832-INS-146 | chr18-21279833-INS-133 |
| chr18-21321071-INS-1491 | chr18-21321072-INS-1481 |
| chr18-21364274-INS-170 | chr18-21364274-INS-176 |
| chr18-21369503-INS-522 | chr18-21369503-INS-524 |
| chr18-21442278-INS-458 | chr18-21442278-INS-458 |
| chr18-21445251-INS-230 | chr18-21445246-INS-266 |
| chr18-21455268-INS-54 | chr18-21455268-INS-54 |
| chr18-21461335-INS-202 | chr18-21461335-INS-202 |
| chr18-21476647-INS-196 | chr18-21476647-INS-196 |
| chr18-21516099-INS-104 | chr18-21516099-INS-104 |
| chr18-21524624-INS-204 | chr18-21524624-INS-204 |
| chr18-21570056-INS-153 | chr18-21570056-INS-153 |
| chr18-21571201-INS-1557 | chr18-21571202-INS-1557 |
| chr18-21578647-INS-400 | chr18-21578647-INS-400 |
| chr18-21596856-INS-2304 | chr18-21596857-INS-2304 |
| chr18-21606531-INS-66 | chr18-21606505-INS-52 |
| chr18-21611289-INS-58 | chr18-21611239-INS-82 |
| chr18-21624119-INS-129 | chr18-21624119-INS-129 |
| chr18-21631588-INS-104 | chr18-21631625-INS-55 |
| chr18-21657830-INS-151 | chr18-21657825-INS-152 |
| chr18-21664851-INS-150 | chr18-21664851-INS-150 |
| chr18-21666809-INS-100 | chr18-21666808-INS-59 |
| chr18-21699714-INS-410 | chr18-21699714-INS-410 |
| chr18-21746055-INS-371 | chr18-21746055-INS-371 |
| chr18-21761807-INS-54 | chr18-21761807-INS-60 |
| chr18-21842285-INS-8413 | chr18-21842286-INS-8413 |
| chr18-21853033-INS-67 | chr18-21853033-INS-67 |
| chr18-21855874-INS-88 | chr18-21855874-INS-68 |
| chr18-21861511-INS-51 | chr18-21861511-INS-51 |
| chr18-21882796-INS-162 | chr18-21882796-INS-164 |
| chr18-21898837-INS-60 | chr18-21898837-INS-56 |
| chr18-21956798-INS-54 | chr18-21956798-INS-54 |
| chr18-21971368-INS-67 | chr18-21971326-INS-67 |
| chr18-22011765-INS-77 | chr18-22011765-INS-93 |
| chr18-22034308-INS-52 | chr18-22034308-INS-52 |
| chr18-22050566-INS-6869 | chr18-22050567-INS-6869 |
| chr18-22103622-INS-59 | chr18-22103622-INS-59 |
| chr18-22216405-INS-78 | chr18-22216405-INS-76 |
| chr18-22218335-INS-306 | chr18-22218335-INS-306 |
| chr18-22227779-INS-1502 | chr18-22227780-INS-1502 |
| chr18-22236981-INS-5517 | chr18-22236982-INS-5549 |
| chr18-22258871-INS-1058 | chr18-22258872-INS-1058 |
| chr18-22270195-INS-52 | chr18-22270196-INS-68 |

| | |
|---|---|
| chr18-22323417-INS-399 | chr18-22323417-INS-399 |
| chr18-22341541-INS-51 | chr18-22341580-INS-60 |
| chr18-22394965-INS-400 | chr18-22394965-INS-400 |
| chr18-22398198-INS-60 | chr18-22398198-INS-62 |
| chr18-22406617-INS-197 | chr18-22406617-INS-197 |
| chr18-22410943-INS-175 | chr18-22410944-INS-175 |
| chr18-22416061-INS-460 | chr18-22416061-INS-460 |
| chr18-22424858-INS-70 | chr18-22424898-INS-60 |
| chr18-22428887-INS-702 | chr18-22428887-INS-703 |
| chr18-22448443-INS-105 | chr18-22448443-INS-89 |
| chr18-22486740-INS-2336 | chr18-22486741-INS-2337 |
| chr18-22547291-INS-1997 | chr18-22547292-INS-2000 |
| chr18-22569667-INS-94 | chr18-22569668-INS-59 |
| chr18-22572420-INS-617 | chr18-22572420-INS-617 |
| chr18-22670373-INS-580 | chr18-22670373-INS-580 |
| chr18-22672869-INS-364 | chr18-22672869-INS-365 |
| chr18-22684185-INS-322 | chr18-22684185-INS-322 |
| chr18-22721919-INS-829 | chr18-22721919-INS-829 |
| chr18-22748747-INS-76 | chr18-22748747-INS-60 |
| chr18-22755898-INS-123 | chr18-22755898-INS-123 |
| chr18-22756579-INS-176 | chr18-22756584-INS-175 |
| chr18-22764772-INS-192 | chr18-22764755-INS-192 |
| chr18-22778786-INS-201 | chr18-22778786-INS-202 |
| chr18-22782140-INS-396 | chr18-22782140-INS-397 |
| chr18-22791473-INS-76 | chr18-22791473-INS-78 |
| chr18-22811276-INS-82 | chr18-22811276-INS-82 |
| chr18-22830802-INS-173 | chr18-22830802-INS-200 |
| chr18-22859097-INS-67 | chr18-22859097-INS-67 |
| chr18-22893933-INS-587 | chr18-22893933-INS-586 |
| chr18-22896923-INS-702 | chr18-22896923-INS-702 |
| chr18-22917519-INS-79 | chr18-22917519-INS-82 |
| chr18-22932284-INS-129 | chr18-22932284-INS-129 |
| chr18-22951951-INS-62 | chr18-22951951-INS-62 |
| chr18-22990732-INS-58 | chr18-22990730-INS-54 |
| chr18-23018661-INS-52 | chr18-23018661-INS-52 |
| chr18-23033956-INS-79 | chr18-23033956-INS-99 |
| chr18-23060186-INS-120 | chr18-23060186-INS-120 |
| chr18-23075012-INS-120 | chr18-23075012-INS-120 |
| chr18-23085184-INS-58 | chr18-23085184-INS-60 |
| chr18-23182281-INS-60 | chr18-23182287-INS-80 |
| chr18-23204069-INS-66 | chr18-23204069-INS-82 |
| chr18-23217237-INS-110 | chr18-23217247-INS-100 |
| chr18-23238516-INS-133 | chr18-23238516-INS-131 |
| chr18-23274979-INS-72 | chr18-23274979-INS-84 |
| chr18-23324862-INS-133 | chr18-23324862-INS-133 |
| chr18-23328121-INS-52 | chr18-23328121-INS-60 |
| chr18-23330706-INS-182 | chr18-23330706-INS-182 |
| chr18-23346063-INS-258 | chr18-23346063-INS-258 |
| chr18-23387744-INS-75 | chr18-23387744-INS-55 |
| chr18-23427153-INS-115 | chr18-23427163-INS-148 |
| chr18-23486898-INS-77 | chr18-23486890-INS-52 |
| chr18-23498138-INS-203 | chr18-23498138-INS-213 |
| chr18-23520114-INS-53 | chr18-23520114-INS-62 |
| chr18-23521379-INS-743 | chr18-23521379-INS-744 |
| chr18-23530109-INS-164 | chr18-23530109-INS-166 |
| chr18-23531742-INS-224 | chr18-23531741-INS-227 |
| chr18-23575264-INS-71 | chr18-23575264-INS-72 |
| chr18-23592256-INS-1127 | chr18-23592257-INS-1129 |
| chr18-23675417-INS-54 | chr18-23675417-INS-54 |
| chr18-23727092-INS-120 | chr18-23727092-INS-120 |
| chr18-23731533-INS-76 | chr18-23731533-INS-76 |
| chr18-23746693-INS-109 | chr18-23746693-INS-109 |
| chr18-23782399-INS-51 | chr18-23782399-INS-51 |
| chr18-23794822-INS-239 | chr18-23794822-INS-239 |
| chr18-23796073-INS-65 | chr18-23796073-INS-90 |
| chr18-23818767-INS-159 | chr18-23818767-INS-159 |
| chr18-23838161-INS-830 | chr18-23838161-INS-830 |
| chr18-23859098-INS-64 | chr18-23859098-INS-80 |
| chr18-23871975-INS-99 | chr18-23871995-INS-99 |
| chr18-23876476-INS-619 | chr18-23876476-INS-622 |
| chr18-23890772-INS-215 | chr18-23890772-INS-216 |
| chr18-23902213-INS-126 | chr18-23902216-INS-130 |
| chr18-23908146-INS-79 | chr18-23908146-INS-79 |
| chr18-23931355-INS-72 | chr18-23931355-INS-76 |
| chr18-23967366-INS-154 | chr18-23967366-INS-156 |
| chr18-23998087-INS-58 | chr18-23998087-INS-58 |
| chr18-24018995-INS-58 | chr18-24018995-INS-58 |

| | |
|---|---|
| chr18-24051422-INS-72 | chr18-24051422-INS-68 |
| chr18-24074810-INS-133 | chr18-24074810-INS-163 |
| chr18-24099890-INS-172 | chr18-24099890-INS-172 |
| chr18-24111437-INS-60 | chr18-24111399-INS-66 |
| chr18-24135482-INS-1161 | chr18-24135483-INS-1161 |
| chr18-24141675-INS-94 | chr18-24141675-INS-114 |
| chr18-24181711-INS-194 | chr18-24181711-INS-194 |
| chr18-24204225-INS-549 | chr18-24204225-INS-549 |
| chr18-24206790-INS-796 | chr18-24206790-INS-813 |
| chr18-24222724-INS-7296 | chr18-24222725-INS-7296 |
| chr18-24231882-INS-372 | chr18-24231878-INS-378 |
| chr18-24252895-INS-203 | chr18-24252895-INS-203 |
| chr18-24278609-INS-7135 | chr18-24278610-INS-7135 |
| chr18-24283879-INS-438 | chr18-24283914-INS-438 |
| chr18-24289018-INS-122 | chr18-24289018-INS-122 |
| chr18-24302971-INS-101 | chr18-24302971-INS-101 |
| chr18-24324255-INS-78 | chr18-24324255-INS-78 |
| chr18-24345059-INS-105 | chr18-24345059-INS-107 |
| chr18-24347792-INS-225 | chr18-24347792-INS-225 |
| chr18-24368154-INS-214 | chr18-24368154-INS-214 |
| chr18-24369554-INS-1046 | chr18-24369555-INS-1045 |
| chr18-24403865-INS-108 | chr18-24403865-INS-108 |
| chr18-24410340-INS-399 | chr18-24410340-INS-399 |
| chr18-24413368-INS-124 | chr18-24413368-INS-124 |
| chr18-24433329-INS-124 | chr18-24433329-INS-105 |
| chr18-24446930-INS-55 | chr18-24446930-INS-55 |
| chr18-24476434-INS-54 | chr18-24476428-INS-72 |
| chr18-24482865-INS-192 | chr18-24482865-INS-180 |
| chr18-24496040-INS-1375 | chr18-24496041-INS-1375 |
| chr18-24499616-INS-252 | chr18-24499614-INS-252 |
| chr18-24511975-INS-767 | chr18-24511975-INS-767 |
| chr18-24515148-INS-7088 | chr18-24515149-INS-7088 |
| chr18-24516730-INS-73 | chr18-24516730-INS-76 |
| chr18-24518866-INS-520 | chr18-24518866-INS-520 |
| chr18-24541649-INS-183 | chr18-24541649-INS-183 |
| chr18-24542521-INS-1007 | chr18-24542521-INS-1008 |
| chr18-24560651-INS-146 | chr18-24560651-INS-132 |
| chr18-24582311-INS-104 | chr18-24582311-INS-104 |
| chr18-24592456-INS-692 | chr18-24592456-INS-692 |
| chr18-24614903-INS-54 | chr18-24614903-INS-56 |
| chr18-24629042-INS-60 | chr18-24629042-INS-60 |
| chr18-24649261-INS-451 | chr18-24649261-INS-462 |
| chr18-24663878-INS-104 | chr18-24663879-INS-106 |
| chr18-24664555-INS-206 | chr18-24664555-INS-207 |
| chr18-24672053-INS-53 | chr18-24672053-INS-53 |
| chr18-24757229-INS-69 | chr18-24757234-INS-63 |
| chr18-24761436-INS-204 | chr18-24761436-INS-205 |
| chr18-24775947-INS-194 | chr18-24775947-INS-195 |
| chr18-24777225-INS-241 | chr18-24777225-INS-217 |
| chr18-24810510-INS-70 | chr18-24810510-INS-70 |
| chr18-24825872-INS-56 | chr18-24825872-INS-56 |
| chr18-24836075-INS-328 | chr18-24836075-INS-328 |
| chr18-24856570-INS-85 | chr18-24856570-INS-85 |
| chr18-24894663-INS-266 | chr18-24894663-INS-294 |
| chr18-24909130-INS-156 | chr18-24909130-INS-156 |
| chr18-24920033-INS-74 | chr18-24920033-INS-74 |
| chr18-24947600-INS-80 | chr18-24947627-INS-81 |
| chr18-24955250-INS-1071 | chr18-24955251-INS-1071 |
| chr18-24973200-INS-53 | chr18-24973181-INS-56 |
| chr18-24976621-INS-6644 | chr18-24976622-INS-6644 |
| chr18-24996761-INS-210 | chr18-24996761-INS-210 |
| chr18-25000070-INS-191 | chr18-25000065-INS-201 |
| chr18-25003776-INS-213 | chr18-25003776-INS-213 |
| chr18-25006112-INS-1029 | chr18-25006113-INS-1028 |
| chr18-25017919-INS-56 | chr18-25017919-INS-56 |
| chr18-25024489-INS-2228 | chr18-25024490-INS-2229 |
| chr18-25032249-INS-205 | chr18-25032249-INS-248 |
| chr18-25083111-INS-73 | chr18-25083111-INS-76 |
| chr18-25116225-INS-401 | chr18-25116225-INS-401 |
| chr18-25133622-INS-56 | chr18-25133622-INS-54 |
| chr18-25144606-INS-105 | chr18-25144606-INS-81 |
| chr18-25149315-INS-78 | chr18-25149316-INS-67 |
| chr18-25151789-INS-1717 | chr18-25151790-INS-1712 |
| chr18-25154817-INS-68 | chr18-25154825-INS-66 |
| chr18-25161694-INS-251 | chr18-25161694-INS-251 |
| chr18-25182890-INS-216 | chr18-25182890-INS-216 |
| chr18-25182897-INS-876 | chr18-25182897-INS-868 |

| | |
|---|---|
| chr18-25200299-INS-5552 | chr18-25200300-INS-5552 |
| chr18-25216750-INS-116 | chr18-25216750-INS-122 |
| chr18-25242358-INS-82 | chr18-25242357-INS-94 |
| chr18-25244045-INS-381 | chr18-25244045-INS-382 |
| chr18-25259247-INS-100 | chr18-25259250-INS-122 |
| chr18-25292829-INS-271 | chr18-25292829-INS-271 |
| chr18-25314482-INS-55 | chr18-25314482-INS-83 |
| chr18-25366948-INS-1476 | chr18-25366912-INS-1471 |
| chr18-25378952-INS-107 | chr18-25378956-INS-111 |
| chr18-25381710-INS-147 | chr18-25381710-INS-146 |
| chr18-25383870-INS-95 | chr18-25383870-INS-95 |
| chr18-25396613-INS-157 | chr18-25396613-INS-157 |
| chr18-25410299-INS-1078 | chr18-25410302-INS-1080 |
| chr18-25413171-INS-864 | chr18-25413163-INS-868 |
| chr18-25416927-INS-223 | chr18-25416927-INS-223 |
| chr18-25441169-INS-280 | chr18-25441169-INS-280 |
| chr18-25455709-INS-54 | chr18-25455709-INS-76 |
| chr18-25460991-INS-144 | chr18-25460991-INS-144 |
| chr18-25469280-INS-109 | chr18-25469280-INS-109 |
| chr18-25473567-INS-104 | chr18-25473567-INS-104 |
| chr18-25474582-INS-644 | chr18-25474582-INS-644 |
| chr18-25479531-INS-565 | chr18-25479531-INS-565 |
| chr18-25486234-INS-1511 | chr18-25486235-INS-1511 |
| chr18-25507652-INS-82 | chr18-25507702-INS-58 |
| chr18-25532869-INS-53 | chr18-25532869-INS-54 |
| chr18-25540135-INS-1891 | chr18-25540136-INS-1891 |
| chr18-25545407-INS-100 | chr18-25545407-INS-60 |
| chr18-25574839-INS-3410 | chr18-25574840-INS-3410 |
| chr18-25577076-INS-161 | chr18-25577076-INS-161 |
| chr18-25624794-INS-3043 | chr18-25624795-INS-3043 |
| chr18-25669554-INS-565 | chr18-25669554-INS-565 |
| chr18-25715100-INS-65 | chr18-25715100-INS-53 |
| chr18-25722419-INS-74 | chr18-25722419-INS-80 |
| chr18-25727926-INS-70 | chr18-25727926-INS-72 |
| chr18-25740877-INS-100 | chr18-25740885-INS-63 |
| chr18-25778709-INS-148 | chr18-25778675-INS-144 |
| chr18-25794090-INS-67 | chr18-25794125-INS-66 |
| chr18-25823944-INS-67 | chr18-25823930-INS-72 |
| chr18-25835042-INS-136 | chr18-25835046-INS-104 |
| chr18-25869392-INS-76 | chr18-25869392-INS-74 |
| chr18-25900067-INS-203 | chr18-25900067-INS-196 |
| chr18-25937331-INS-76 | chr18-25937331-INS-76 |
| chr18-25998825-INS-1234 | chr18-25998826-INS-1234 |
| chr18-26001521-INS-88 | chr18-26001521-INS-88 |
| chr18-26026583-INS-267 | chr18-26026603-INS-270 |
| chr18-26028486-INS-246 | chr18-26028486-INS-246 |
| chr18-26072693-INS-89 | chr18-26072691-INS-89 |
| chr18-26087951-INS-102 | chr18-26087939-INS-83 |
| chr18-26090519-INS-229 | chr18-26090519-INS-229 |
| chr18-26144876-INS-52 | chr18-26144876-INS-52 |
| chr18-26146341-INS-462 | chr18-26146341-INS-463 |
| chr18-26153812-INS-89 | chr18-26153842-INS-54 |
| chr18-26160631-INS-135 | chr18-26160631-INS-138 |
| chr18-26164268-INS-97 | chr18-26164268-INS-97 |
| chr18-26170901-INS-331 | chr18-26170901-INS-331 |
| chr18-26191926-INS-1560 | chr18-26191927-INS-1560 |
| chr18-26194766-INS-119 | chr18-26194766-INS-141 |
| chr18-26198861-INS-646 | chr18-26198861-INS-646 |
| chr18-26232573-INS-65 | chr18-26232573-INS-65 |
| chr18-26266730-INS-5544 | chr18-26266731-INS-5544 |
| chr18-26289608-INS-529 | chr18-26289608-INS-529 |
| chr18-26296403-INS-55 | chr18-26296403-INS-55 |
| chr18-26348684-INS-120 | chr18-26348684-INS-120 |
| chr18-26351855-INS-745 | chr18-26351855-INS-746 |
| chr18-26389954-INS-777 | chr18-26389954-INS-777 |
| chr18-26409408-INS-988 | chr18-26409408-INS-988 |
| chr18-26420061-INS-401 | chr18-26420061-INS-401 |
| chr18-26443833-INS-54 | chr18-26443833-INS-54 |
| chr18-26496356-INS-399 | chr18-26496356-INS-399 |
| chr18-26535310-INS-6442 | chr18-26535311-INS-6445 |
| chr18-26567559-INS-104 | chr18-26567559-INS-104 |
| chr18-26633509-INS-518 | chr18-26633509-INS-518 |
| chr18-26669306-INS-792 | chr18-26669306-INS-802 |
| chr18-26683934-INS-57 | chr18-26683908-INS-90 |
| chr18-26685185-INS-7078 | chr18-26685186-INS-7078 |
| chr18-26699433-INS-612 | chr18-26699433-INS-613 |
| chr18-26749860-INS-60 | chr18-26749860-INS-68 |

| | |
|---|---|
| chr18-26835180-INS-2313 | chr18-26835181-INS-2313 |
| chr18-26847405-INS-335 | chr18-26847405-INS-335 |
| chr18-26858328-INS-6231 | chr18-26858329-INS-6232 |
| chr18-26949460-INS-72 | chr18-26949460-INS-80 |
| chr18-26993251-INS-190 | chr18-26993251-INS-190 |
| chr18-27008685-INS-508 | chr18-27008685-INS-508 |
| chr18-27027108-INS-61 | chr18-27027108-INS-61 |
| chr18-27040067-INS-1495 | chr18-27040068-INS-1492 |
| chr18-27042440-INS-54 | chr18-27042440-INS-60 |
| chr18-27089867-INS-161 | chr18-27089867-INS-160 |
| chr18-27145384-INS-914 | chr18-27145384-INS-914 |
| chr18-27148225-INS-52 | chr18-27148225-INS-64 |
| chr18-27209621-INS-184 | chr18-27209617-INS-196 |
| chr18-27244124-INS-551 | chr18-27244124-INS-550 |
| chr18-27258560-INS-680 | chr18-27258560-INS-680 |
| chr18-27301716-INS-68 | chr18-27301716-INS-68 |
| chr18-27319790-INS-7110 | chr18-27319791-INS-7111 |
| chr18-27326171-INS-333 | chr18-27326171-INS-333 |
| chr18-27356886-INS-277 | chr18-27356886-INS-277 |
| chr18-27389844-INS-52 | chr18-27389844-INS-54 |
| chr18-27398336-INS-719 | chr18-27398336-INS-719 |
| chr18-27399335-INS-514 | chr18-27399335-INS-514 |
| chr18-27406974-INS-2384 | chr18-27406975-INS-2387 |
| chr18-27451558-INS-52 | chr18-27451558-INS-52 |
| chr18-27453997-INS-2460 | chr18-27453998-INS-2461 |
| chr18-27462713-INS-62 | chr18-27462713-INS-62 |
| chr18-27470742-INS-1587 | chr18-27470743-INS-1587 |
| chr18-27492387-INS-561 | chr18-27492405-INS-572 |
| chr18-27519206-INS-192 | chr18-27519206-INS-192 |
| chr18-27564694-INS-108 | chr18-27564694-INS-108 |
| chr18-27571175-INS-126 | chr18-27571175-INS-126 |
| chr18-27583405-INS-419 | chr18-27583405-INS-421 |
| chr18-27626845-INS-553 | chr18-27626845-INS-553 |
| chr18-27643566-INS-69 | chr18-27643566-INS-110 |
| chr18-27651531-INS-1775 | chr18-27651532-INS-1776 |
| chr18-27674636-INS-58 | chr18-27674636-INS-61 |
| chr18-27693919-INS-1047 | chr18-27693920-INS-1047 |
| chr18-27702012-INS-59 | chr18-27702013-INS-58 |
| chr18-27711904-INS-58 | chr18-27711904-INS-52 |
| chr18-27716459-INS-66 | chr18-27716459-INS-69 |
| chr18-27726597-INS-6738 | chr18-27726598-INS-6738 |
| chr18-27740568-INS-128 | chr18-27740568-INS-128 |
| chr18-27744618-INS-4231 | chr18-27744619-INS-4233 |
| chr18-27777997-INS-66 | chr18-27777997-INS-68 |
| chr18-27788967-INS-56 | chr18-27788967-INS-60 |
| chr18-27792570-INS-62 | chr18-27792566-INS-66 |
| chr18-27807266-INS-70 | chr18-27807266-INS-70 |
| chr18-27809953-INS-88 | chr18-27809951-INS-90 |
| chr18-27818105-INS-92 | chr18-27818105-INS-92 |
| chr18-27862345-INS-2861 | chr18-27862346-INS-2861 |
| chr18-27881540-INS-4522 | chr18-27881541-INS-4522 |
| chr18-27889034-INS-432 | chr18-27889034-INS-428 |
| chr18-27906885-INS-54 | chr18-27906885-INS-60 |
| chr18-27918740-INS-556 | chr18-27918740-INS-556 |
| chr18-27920732-INS-88 | chr18-27920725-INS-54 |
| chr18-27966142-INS-3856 | chr18-27966143-INS-3856 |
| chr18-27973753-INS-106 | chr18-27973748-INS-104 |
| chr18-27995883-INS-1974 | chr18-27995884-INS-1950 |
| chr18-28015912-INS-54 | chr18-28015912-INS-56 |
| chr18-28057675-INS-318 | chr18-28057663-INS-322 |
| chr18-28093687-INS-498 | chr18-28093687-INS-498 |
| chr18-28110278-INS-1894 | chr18-28110279-INS-1894 |
| chr18-28141723-INS-6431 | chr18-28141724-INS-6431 |
| chr18-28182546-INS-60 | chr18-28182517-INS-54 |
| chr18-28203009-INS-101 | chr18-28203009-INS-110 |
| chr18-28207719-INS-166 | chr18-28207719-INS-166 |
| chr18-28212232-INS-108 | chr18-28212234-INS-62 |
| chr18-28223909-INS-136 | chr18-28223909-INS-136 |
| chr18-28246853-INS-61 | chr18-28246849-INS-77 |
| chr18-28321881-INS-54 | chr18-28321881-INS-54 |
| chr18-28335728-INS-91 | chr18-28335731-INS-87 |
| chr18-28336466-INS-515 | chr18-28336466-INS-513 |
| chr18-28349951-INS-202 | chr18-28349951-INS-251 |
| chr18-28395836-INS-96 | chr18-28395836-INS-96 |
| chr18-28409365-INS-99 | chr18-28409354-INS-99 |
| chr18-28416836-INS-163 | chr18-28416836-INS-114 |
| chr18-28455884-INS-7173 | chr18-28455885-INS-7173 |

| | |
|---|---|
| chr18-28496719-INS-111 | chr18-28496689-INS-156 |
| chr18-28498792-INS-85 | chr18-28498823-INS-90 |
| chr18-28549622-INS-75 | chr18-28549622-INS-75 |
| chr18-28564488-INS-63 | chr18-28564452-INS-54 |
| chr18-28590316-INS-204 | chr18-28590316-INS-207 |
| chr18-28621212-INS-88 | chr18-28621213-INS-85 |
| chr18-28635695-INS-136 | chr18-28635695-INS-136 |
| chr18-28645166-INS-179 | chr18-28645176-INS-153 |
| chr18-28661058-INS-189 | chr18-28661058-INS-190 |
| chr18-28662597-INS-651 | chr18-28662597-INS-650 |
| chr18-28678678-INS-545 | chr18-28678678-INS-544 |
| chr18-28681117-INS-61 | chr18-28681117-INS-63 |
| chr18-28686663-INS-898 | chr18-28686626-INS-924 |
| chr18-28699861-INS-102 | chr18-28699861-INS-122 |
| chr18-28704846-INS-681 | chr18-28704846-INS-681 |
| chr18-28718663-INS-411 | chr18-28718663-INS-413 |
| chr18-28733110-INS-2341 | chr18-28733111-INS-2341 |
| chr18-28807731-INS-52 | chr18-28807710-INS-52 |
| chr18-28815760-INS-108 | chr18-28815760-INS-108 |
| chr18-28851116-INS-82 | chr18-28851123-INS-100 |
| chr18-28861669-INS-7270 | chr18-28861670-INS-7270 |
| chr18-28872296-INS-64 | chr18-28872294-INS-90 |
| chr18-28874005-INS-448 | chr18-28874005-INS-448 |
| chr18-28900579-INS-8728 | chr18-28900580-INS-8728 |
| chr18-28920643-INS-120 | chr18-28920643-INS-106 |
| chr18-28926933-INS-970 | chr18-28926933-INS-970 |
| chr18-28930303-INS-157 | chr18-28930303-INS-157 |
| chr18-28940784-INS-526 | chr18-28940784-INS-526 |
| chr18-28990162-INS-55 | chr18-28990162-INS-61 |
| chr18-29026107-INS-50 | chr18-29026107-INS-54 |
| chr18-29035478-INS-168 | chr18-29035478-INS-168 |
| chr18-29056531-INS-65 | chr18-29056550-INS-89 |
| chr18-29061794-INS-1895 | chr18-29061795-INS-1895 |
| chr18-29061892-INS-168 | chr18-29061892-INS-168 |
| chr18-29086935-INS-613 | chr18-29086935-INS-613 |
| chr18-29088693-INS-1888 | chr18-29088694-INS-1888 |
| chr18-29103927-INS-400 | chr18-29103927-INS-400 |
| chr18-29106297-INS-525 | chr18-29106297-INS-539 |
| chr18-29107367-INS-50 | chr18-29107336-INS-60 |
| chr18-29163073-INS-1349 | chr18-29163074-INS-1369 |
| chr18-29170963-INS-55 | chr18-29170963-INS-79 |
| chr18-29176730-INS-299 | chr18-29176730-INS-299 |
| chr18-29194715-INS-76 | chr18-29194739-INS-66 |
| chr18-29223826-INS-185 | chr18-29223826-INS-185 |
| chr18-29225291-INS-670 | chr18-29225291-INS-671 |
| chr18-29229881-INS-90 | chr18-29229913-INS-97 |
| chr18-29242636-INS-54 | chr18-29242637-INS-54 |
| chr18-29254197-INS-296 | chr18-29254197-INS-296 |
| chr18-29311034-INS-63 | chr18-29311036-INS-61 |
| chr18-29317021-INS-336 | chr18-29317021-INS-340 |
| chr18-29320157-INS-612 | chr18-29320157-INS-612 |
| chr18-29356227-INS-67 | chr18-29356227-INS-68 |
| chr18-29363413-INS-55 | chr18-29363413-INS-55 |
| chr18-29383151-INS-70 | chr18-29383151-INS-70 |
| chr18-29393062-INS-90 | chr18-29393062-INS-90 |
| chr18-29440023-INS-510 | chr18-29440023-INS-510 |
| chr18-29441757-INS-156 | chr18-29441757-INS-156 |
| chr18-29452975-INS-52 | chr18-29452975-INS-52 |
| chr18-29495772-INS-112 | chr18-29495780-INS-114 |
| chr18-29499227-INS-90 | chr18-29499227-INS-52 |
| chr18-29501851-INS-201 | chr18-29501851-INS-203 |
| chr18-29516289-INS-82 | chr18-29516289-INS-52 |
| chr18-29546644-INS-310 | chr18-29546644-INS-310 |
| chr18-29552498-INS-6524 | chr18-29552499-INS-6528 |
| chr18-29580002-INS-450 | chr18-29580002-INS-450 |
| chr18-29665854-INS-3151 | chr18-29665855-INS-3153 |
| chr18-29666756-INS-6437 | chr18-29666757-INS-6437 |
| chr18-29669847-INS-223 | chr18-29669847-INS-222 |
| chr18-29698457-INS-56 | chr18-29698457-INS-60 |
| chr18-29714602-INS-5405 | chr18-29714603-INS-5405 |
| chr18-29755975-INS-73 | chr18-29755975-INS-73 |
| chr18-29763760-INS-52 | chr18-29763760-INS-51 |
| chr18-29782861-INS-61 | chr18-29782861-INS-62 |
| chr18-29831699-INS-1419 | chr18-29831700-INS-1419 |
| chr18-29859639-INS-93 | chr18-29859639-INS-75 |
| chr18-29860818-INS-98 | chr18-29860818-INS-132 |
| chr18-29883046-INS-51 | chr18-29883046-INS-51 |

| | |
|---|---|
| chr18-29903271-INS-791 | chr18-29903271-INS-792 |
| chr18-29931830-INS-52 | chr18-29931828-INS-53 |
| chr18-29973412-INS-1952 | chr18-29973413-INS-1953 |
| chr18-30032599-INS-58 | chr18-30032599-INS-64 |
| chr18-30044246-INS-205 | chr18-30044246-INS-205 |
| chr18-30098860-INS-152 | chr18-30098860-INS-152 |
| chr18-30110383-INS-204 | chr18-30110383-INS-204 |
| chr18-30125507-INS-52 | chr18-30125507-INS-52 |
| chr18-30126878-INS-52 | chr18-30126878-INS-78 |
| chr18-30134511-INS-6222 | chr18-30134512-INS-6222 |
| chr18-30175806-INS-88 | chr18-30175791-INS-102 |
| chr18-30178321-INS-63 | chr18-30178321-INS-83 |
| chr18-30230671-INS-344 | chr18-30230671-INS-344 |
| chr18-30233737-INS-4856 | chr18-30233738-INS-4856 |
| chr18-30246694-INS-59 | chr18-30246694-INS-54 |
| chr18-30273892-INS-7085 | chr18-30273893-INS-7085 |
| chr18-30276356-INS-60 | chr18-30276359-INS-78 |
| chr18-30279470-INS-53 | chr18-30279487-INS-50 |
| chr18-30283476-INS-937 | chr18-30283476-INS-938 |
| chr18-30306891-INS-108 | chr18-30306893-INS-124 |
| chr18-30315833-INS-177 | chr18-30315833-INS-177 |
| chr18-30337217-INS-290 | chr18-30337217-INS-290 |
| chr18-30363428-INS-52 | chr18-30363412-INS-64 |
| chr18-30377568-INS-6456 | chr18-30377569-INS-6458 |
| chr18-30378699-INS-550 | chr18-30378699-INS-552 |
| chr18-30387625-INS-199 | chr18-30387625-INS-199 |
| chr18-30419606-INS-79 | chr18-30419606-INS-79 |
| chr18-30424910-INS-82 | chr18-30424910-INS-82 |
| chr18-30433245-INS-76 | chr18-30433225-INS-96 |
| chr18-30439213-INS-6401 | chr18-30439214-INS-6401 |
| chr18-30463861-INS-674 | chr18-30463861-INS-674 |
| chr18-30464760-INS-164 | chr18-30464760-INS-164 |
| chr18-30494303-INS-72 | chr18-30494303-INS-72 |
| chr18-30496967-INS-195 | chr18-30496967-INS-195 |
| chr18-30510027-INS-87 | chr18-30510027-INS-87 |
| chr18-30521905-INS-51 | chr18-30521905-INS-51 |
| chr18-30527108-INS-399 | chr18-30527108-INS-399 |
| chr18-30529623-INS-78 | chr18-30529614-INS-52 |
| chr18-30535611-INS-60 | chr18-30535611-INS-60 |
| chr18-30548281-INS-418 | chr18-30548281-INS-418 |
| chr18-30555498-INS-497 | chr18-30555498-INS-497 |
| chr18-30560192-INS-94 | chr18-30560192-INS-100 |
| chr18-30562309-INS-84 | chr18-30562309-INS-78 |
| chr18-30572785-INS-51 | chr18-30572785-INS-53 |
| chr18-30577030-INS-60 | chr18-30577030-INS-56 |
| chr18-30589305-INS-54 | chr18-30589305-INS-54 |
| chr18-30593891-INS-145 | chr18-30593891-INS-145 |
| chr18-30613978-INS-462 | chr18-30613978-INS-463 |
| chr18-30629946-INS-112 | chr18-30629946-INS-92 |
| chr18-30635714-INS-56 | chr18-30635714-INS-50 |
| chr18-30664170-INS-52 | chr18-30664170-INS-50 |
| chr18-30672863-INS-6150 | chr18-30672864-INS-6150 |
| chr18-30693927-INS-428 | chr18-30693927-INS-428 |
| chr18-30694590-INS-112 | chr18-30694590-INS-106 |
| chr18-30698455-INS-77 | chr18-30698458-INS-60 |
| chr18-30707877-INS-883 | chr18-30707877-INS-885 |
| chr18-30725621-INS-55 | chr18-30725623-INS-51 |
| chr18-30726816-INS-103 | chr18-30726816-INS-103 |
| chr18-30760936-INS-59 | chr18-30760936-INS-55 |
| chr18-30763491-INS-67 | chr18-30763499-INS-73 |
| chr18-30780916-INS-1156 | chr18-30780917-INS-1156 |
| chr18-30782378-INS-58 | chr18-30782378-INS-58 |
| chr18-30808514-INS-6389 | chr18-30808515-INS-6375 |
| chr18-30819560-INS-102 | chr18-30819560-INS-100 |
| chr18-30820145-INS-6168 | chr18-30820146-INS-6171 |
| chr18-30826656-INS-55 | chr18-30826656-INS-67 |
| chr18-30840441-INS-57 | chr18-30840441-INS-57 |
| chr18-30875766-INS-537 | chr18-30875766-INS-536 |
| chr18-30898661-INS-1327 | chr18-30898662-INS-1328 |
| chr18-30899277-INS-51 | chr18-30899277-INS-51 |
| chr18-30951544-INS-130 | chr18-30951544-INS-136 |
| chr18-30955261-INS-306 | chr18-30955261-INS-327 |
| chr18-30956748-INS-171 | chr18-30956748-INS-171 |
| chr18-30964321-INS-380 | chr18-30964321-INS-380 |
| chr18-31004704-INS-363 | chr18-31004704-INS-362 |
| chr18-31009531-INS-80 | chr18-31009535-INS-84 |
| chr18-31028456-INS-171 | chr18-31028457-INS-171 |

| | |
|---|---|
| chr18-31031160-INS-124 | chr18-31031160-INS-124 |
| chr18-31054969-INS-249 | chr18-31054969-INS-268 |
| chr18-31059343-INS-82 | chr18-31059343-INS-84 |
| chr18-31117590-INS-75 | chr18-31117590-INS-65 |
| chr18-31133157-INS-668 | chr18-31133157-INS-668 |
| chr18-31139279-INS-148 | chr18-31139279-INS-148 |
| chr18-31185448-INS-1472 | chr18-31185449-INS-1476 |
| chr18-31241160-INS-2591 | chr18-31241161-INS-2591 |
| chr18-31249014-INS-6943 | chr18-31249015-INS-6944 |
| chr18-31254121-INS-6900 | chr18-31254122-INS-6902 |
| chr18-31263742-INS-428 | chr18-31263742-INS-428 |
| chr18-31264751-INS-56 | chr18-31264724-INS-55 |
| chr18-31273653-INS-746 | chr18-31273653-INS-747 |
| chr18-31276477-INS-6208 | chr18-31276478-INS-6205 |
| chr18-31276740-INS-205 | chr18-31276740-INS-205 |
| chr18-31282827-INS-74 | chr18-31282827-INS-74 |
| chr18-31300529-INS-189 | chr18-31300529-INS-189 |
| chr18-31309318-INS-779 | chr18-31309318-INS-779 |
| chr18-31330580-INS-78 | chr18-31330580-INS-70 |
| chr18-31335146-INS-55 | chr18-31335161-INS-50 |
| chr18-31348260-INS-700 | chr18-31348260-INS-701 |
| chr18-31356011-INS-221 | chr18-31356011-INS-221 |
| chr18-31394976-INS-83 | chr18-31394978-INS-83 |
| chr18-31434545-INS-332 | chr18-31434568-INS-344 |
| chr18-31499644-INS-86 | chr18-31499647-INS-86 |
| chr18-31517104-INS-400 | chr18-31517104-INS-400 |
| chr18-31521280-INS-75 | chr18-31521277-INS-76 |
| chr18-31522703-INS-118 | chr18-31522703-INS-118 |
| chr18-31535116-INS-152 | chr18-31535116-INS-152 |
| chr18-31540626-INS-453 | chr18-31540626-INS-453 |
| chr18-31564041-INS-721 | chr18-31564041-INS-721 |
| chr18-31587249-INS-195 | chr18-31587249-INS-185 |
| chr18-31596433-INS-6103 | chr18-31596431-INS-6099 |
| chr18-31631225-INS-64 | chr18-31631223-INS-68 |
| chr18-31635429-INS-965 | chr18-31635429-INS-965 |
| chr18-31664343-INS-1137 | chr18-31664344-INS-1137 |
| chr18-31672825-INS-295 | chr18-31672825-INS-295 |
| chr18-31676845-INS-1892 | chr18-31676846-INS-1892 |
| chr18-31700253-INS-301 | chr18-31700253-INS-312 |
| chr18-31702164-INS-66 | chr18-31702164-INS-84 |
| chr18-31711490-INS-4843 | chr18-31711491-INS-4843 |
| chr18-31726314-INS-60 | chr18-31726314-INS-61 |
| chr18-31745512-INS-185 | chr18-31745513-INS-197 |
| chr18-31748471-INS-345 | chr18-31748471-INS-345 |
| chr18-31751167-INS-208 | chr18-31751167-INS-208 |
| chr18-31759033-INS-7498 | chr18-31759034-INS-7541 |
| chr18-31792836-INS-361 | chr18-31792836-INS-361 |
| chr18-31835753-INS-98 | chr18-31835753-INS-93 |
| chr18-31841038-INS-215 | chr18-31841038-INS-215 |
| chr18-31860479-INS-126 | chr18-31860479-INS-142 |
| chr18-31865851-INS-803 | chr18-31865851-INS-811 |
| chr18-31866870-INS-169 | chr18-31866870-INS-170 |
| chr18-31871087-INS-165 | chr18-31871088-INS-166 |
| chr18-31879795-INS-82 | chr18-31879795-INS-85 |
| chr18-31883403-INS-144 | chr18-31883403-INS-120 |
| chr18-31919732-INS-151 | chr18-31919732-INS-113 |
| chr18-31932203-INS-226 | chr18-31932203-INS-226 |
| chr18-31936772-INS-60 | chr18-31936772-INS-60 |
| chr18-31939483-INS-95 | chr18-31939483-INS-95 |
| chr18-31952529-INS-161 | chr18-31952529-INS-161 |
| chr18-31961169-INS-209 | chr18-31961169-INS-209 |
| chr18-31975444-INS-199 | chr18-31975444-INS-199 |
| chr18-31986465-INS-71 | chr18-31986465-INS-71 |
| chr18-32000641-INS-219 | chr18-32000641-INS-221 |
| chr18-32015162-INS-60 | chr18-32015162-INS-89 |
| chr18-32067640-INS-64 | chr18-32067608-INS-68 |
| chr18-32072766-INS-153 | chr18-32072769-INS-139 |
| chr18-32123916-INS-106 | chr18-32123916-INS-94 |
| chr18-32124944-INS-91 | chr18-32124940-INS-85 |
| chr18-32126409-INS-200 | chr18-32126409-INS-199 |
| chr18-32130622-INS-498 | chr18-32130622-INS-498 |
| chr18-32140526-INS-141 | chr18-32140526-INS-141 |
| chr18-32157293-INS-87 | chr18-32157293-INS-87 |
| chr18-32166328-INS-198 | chr18-32166328-INS-198 |
| chr18-32170403-INS-207 | chr18-32170403-INS-207 |
| chr18-32194745-INS-92 | chr18-32194745-INS-76 |
| chr18-32195580-INS-200 | chr18-32195580-INS-200 |

| | |
|---|---|
| chr18-32236955-INS-407 | chr18-32236955-INS-407 |
| chr18-32261539-INS-189 | chr18-32261539-INS-188 |
| chr18-32264751-INS-56 | chr18-32264759-INS-52 |
| chr18-32291393-INS-734 | chr18-32291393-INS-736 |
| chr18-32313536-INS-135 | chr18-32313536-INS-136 |
| chr18-32323912-INS-103 | chr18-32323908-INS-90 |
| chr18-32325811-INS-76 | chr18-32325811-INS-76 |
| chr18-32333944-INS-135 | chr18-32333944-INS-135 |
| chr18-32351311-INS-84 | chr18-32351311-INS-84 |
| chr18-32376526-INS-2040 | chr18-32376527-INS-2040 |
| chr18-32406238-INS-84 | chr18-32406236-INS-82 |
| chr18-32415237-INS-108 | chr18-32415237-INS-112 |
| chr18-32445694-INS-164 | chr18-32445682-INS-191 |
| chr18-32458222-INS-590 | chr18-32458222-INS-590 |
| chr18-32500254-INS-4830 | chr18-32500255-INS-4830 |
| chr18-32515629-INS-62 | chr18-32515629-INS-62 |
| chr18-32518646-INS-54 | chr18-32518646-INS-52 |
| chr18-32573706-INS-64 | chr18-32573706-INS-64 |
| chr18-32576657-INS-54 | chr18-32576657-INS-54 |
| chr18-32585132-INS-225 | chr18-32585132-INS-226 |
| chr18-32610126-INS-433 | chr18-32610126-INS-433 |
| chr18-32615819-INS-6401 | chr18-32615820-INS-6401 |
| chr18-32619826-INS-440 | chr18-32619826-INS-440 |
| chr18-32628521-INS-68 | chr18-32628520-INS-60 |
| chr18-32637352-INS-129 | chr18-32637367-INS-95 |
| chr18-32638898-INS-57 | chr18-32638907-INS-67 |
| chr18-32657218-INS-6449 | chr18-32657219-INS-6451 |
| chr18-32681813-INS-54 | chr18-32681813-INS-55 |
| chr18-32719292-INS-1618 | chr18-32719293-INS-1640 |
| chr18-32731743-INS-154 | chr18-32731743-INS-150 |
| chr18-32750444-INS-115 | chr18-32750444-INS-115 |
| chr18-32754281-INS-217 | chr18-32754281-INS-217 |
| chr18-32759059-INS-1768 | chr18-32759060-INS-1767 |
| chr18-32763438-INS-107 | chr18-32763464-INS-72 |
| chr18-32767053-INS-2806 | chr18-32767054-INS-2797 |
| chr18-32781797-INS-100 | chr18-32781797-INS-102 |
| chr18-32815878-INS-214 | chr18-32815878-INS-214 |
| chr18-32824993-INS-57 | chr18-32824993-INS-57 |
| chr18-32843341-INS-112 | chr18-32843347-INS-86 |
| chr18-32854028-INS-62 | chr18-32854028-INS-54 |
| chr18-32859501-INS-71 | chr18-32859480-INS-80 |
| chr18-32866099-INS-179 | chr18-32866099-INS-179 |
| chr18-32920524-INS-70 | chr18-32920524-INS-70 |
| chr18-32932732-INS-62 | chr18-32932732-INS-62 |
| chr18-32933087-INS-217 | chr18-32933087-INS-216 |
| chr18-32934679-INS-186 | chr18-32934679-INS-186 |
| chr18-32994729-INS-150 | chr18-32994729-INS-150 |
| chr18-33025931-INS-244 | chr18-33025931-INS-216 |
| chr18-33054209-INS-97 | chr18-33054209-INS-97 |
| chr18-33054775-INS-3680 | chr18-33054778-INS-3658 |
| chr18-33077390-INS-57 | chr18-33077382-INS-97 |
| chr18-33117341-INS-55 | chr18-33117341-INS-55 |
| chr18-33134156-INS-399 | chr18-33134156-INS-386 |
| chr18-33141323-INS-132 | chr18-33141349-INS-128 |
| chr18-33191867-INS-56 | chr18-33191867-INS-56 |
| chr18-33198091-INS-213 | chr18-33198091-INS-213 |
| chr18-33218213-INS-523 | chr18-33218213-INS-523 |
| chr18-33223550-INS-82 | chr18-33223550-INS-84 |
| chr18-33240455-INS-504 | chr18-33240455-INS-504 |
| chr18-33255559-INS-204 | chr18-33255559-INS-204 |
| chr18-33260693-INS-185 | chr18-33260693-INS-183 |
| chr18-33276519-INS-3087 | chr18-33276520-INS-3083 |
| chr18-33281062-INS-6858 | chr18-33281063-INS-6860 |
| chr18-33294713-INS-80 | chr18-33294713-INS-80 |
| chr18-33301259-INS-6751 | chr18-33301260-INS-6753 |
| chr18-33322041-INS-5476 | chr18-33322042-INS-5478 |
| chr18-33337494-INS-222 | chr18-33337494-INS-222 |
| chr18-33367554-INS-68 | chr18-33367554-INS-74 |
| chr18-33383022-INS-104 | chr18-33383022-INS-96 |
| chr18-33388908-INS-243 | chr18-33388908-INS-242 |
| chr18-33395550-INS-1166 | chr18-33395551-INS-1166 |
| chr18-33404795-INS-84 | chr18-33404802-INS-59 |
| chr18-33415374-INS-155 | chr18-33415359-INS-153 |
| chr18-33434134-INS-464 | chr18-33434134-INS-464 |
| chr18-33498385-INS-144 | chr18-33498385-INS-143 |
| chr18-33502100-INS-227 | chr18-33502100-INS-227 |
| chr18-33513779-INS-56 | chr18-33513779-INS-56 |

| | |
|---|---|
| chr18-33554622-INS-2776 | chr18-33554623-INS-2776 |
| chr18-33555162-INS-6191 | chr18-33555163-INS-6191 |
| chr18-33596726-INS-66 | chr18-33596726-INS-64 |
| chr18-33598864-INS-68 | chr18-33598864-INS-66 |
| chr18-33610498-INS-57 | chr18-33610498-INS-57 |
| chr18-33624295-INS-63 | chr18-33624287-INS-76 |
| chr18-33643761-INS-6170 | chr18-33643762-INS-6170 |
| chr18-33664614-INS-53 | chr18-33664578-INS-63 |
| chr18-33667475-INS-2062 | chr18-33667476-INS-2062 |
| chr18-33690008-INS-61 | chr18-33689980-INS-100 |
| chr18-33690728-INS-6653 | chr18-33690729-INS-6654 |
| chr18-33696708-INS-752 | chr18-33696708-INS-753 |
| chr18-33724128-INS-356 | chr18-33724128-INS-356 |
| chr18-33736127-INS-51 | chr18-33736127-INS-54 |
| chr18-33744122-INS-60 | chr18-33744116-INS-66 |
| chr18-33767600-INS-60 | chr18-33767600-INS-70 |
| chr18-33790655-INS-72 | chr18-33790609-INS-69 |
| chr18-33793818-INS-186 | chr18-33793819-INS-200 |
| chr18-33798672-INS-393 | chr18-33798672-INS-393 |
| chr18-33823297-INS-1246 | chr18-33823298-INS-1248 |
| chr18-33846896-INS-206 | chr18-33846896-INS-206 |
| chr18-33852729-INS-537 | chr18-33852729-INS-538 |
| chr18-33857620-INS-72 | chr18-33857624-INS-78 |
| chr18-33864443-INS-6888 | chr18-33864444-INS-6888 |
| chr18-33868091-INS-107 | chr18-33868091-INS-107 |
| chr18-33872816-INS-215 | chr18-33872816-INS-215 |
| chr18-33874576-INS-136 | chr18-33874577-INS-124 |
| chr18-33890399-INS-443 | chr18-33890378-INS-447 |
| chr18-33901011-INS-693 | chr18-33901011-INS-693 |
| chr18-33912105-INS-90 | chr18-33912105-INS-89 |
| chr18-33926507-INS-84 | chr18-33926509-INS-86 |
| chr18-33935308-INS-58 | chr18-33935308-INS-58 |
| chr18-33958987-INS-209 | chr18-33958987-INS-209 |
| chr18-34002396-INS-93 | chr18-34002396-INS-93 |
| chr18-34016242-INS-186 | chr18-34016242-INS-189 |
| chr18-34031865-INS-106 | chr18-34031865-INS-107 |
| chr18-34052623-INS-209 | chr18-34052623-INS-209 |
| chr18-34053147-INS-1885 | chr18-34053148-INS-1885 |
| chr18-34061651-INS-208 | chr18-34061651-INS-208 |
| chr18-34062172-INS-168 | chr18-34062172-INS-168 |
| chr18-34078887-INS-268 | chr18-34078887-INS-268 |
| chr18-34079917-INS-312 | chr18-34079917-INS-279 |
| chr18-34092397-INS-238 | chr18-34092397-INS-256 |
| chr18-34094842-INS-280 | chr18-34094840-INS-291 |
| chr18-34124203-INS-200 | chr18-34124203-INS-200 |
| chr18-34125449-INS-189 | chr18-34125449-INS-189 |
| chr18-34129372-INS-669 | chr18-34129372-INS-666 |
| chr18-34138246-INS-130 | chr18-34138246-INS-150 |
| chr18-34150699-INS-203 | chr18-34150699-INS-203 |
| chr18-34153844-INS-56 | chr18-34153844-INS-56 |
| chr18-34167245-INS-60 | chr18-34167254-INS-84 |
| chr18-34171052-INS-397 | chr18-34171052-INS-398 |
| chr18-34178127-INS-194 | chr18-34178127-INS-194 |
| chr18-34180318-INS-520 | chr18-34180319-INS-520 |
| chr18-34184841-INS-168 | chr18-34184841-INS-168 |
| chr18-34201030-INS-58 | chr18-34201030-INS-58 |
| chr18-34216509-INS-51 | chr18-34216509-INS-51 |
| chr18-34240845-INS-702 | chr18-34240845-INS-702 |
| chr18-34245264-INS-1864 | chr18-34245265-INS-1864 |
| chr18-34247387-INS-119 | chr18-34247387-INS-83 |
| chr18-34255615-INS-60 | chr18-34255615-INS-60 |
| chr18-34260757-INS-71 | chr18-34260757-INS-71 |
| chr18-34268235-INS-77 | chr18-34268235-INS-77 |
| chr18-34272661-INS-71 | chr18-34272661-INS-97 |
| chr18-34284132-INS-343 | chr18-34284132-INS-342 |
| chr18-34290802-INS-190 | chr18-34290802-INS-160 |
| chr18-34293952-INS-209 | chr18-34293952-INS-209 |
| chr18-34297880-INS-120 | chr18-34297880-INS-120 |
| chr18-34303422-INS-7214 | chr18-34303423-INS-7214 |
| chr18-34304964-INS-1859 | chr18-34304965-INS-1859 |
| chr18-34306097-INS-82 | chr18-34306097-INS-82 |
| chr18-34318079-INS-272 | chr18-34318074-INS-275 |
| chr18-34345583-INS-221 | chr18-34345570-INS-208 |
| chr18-34377474-INS-214 | chr18-34377474-INS-214 |
| chr18-34402396-INS-176 | chr18-34402396-INS-176 |
| chr18-34432807-INS-1892 | chr18-34432808-INS-1892 |
| chr18-34440308-INS-65 | chr18-34440308-INS-65 |

| | |
|---|---|
| chr18-34520441-INS-234 | chr18-34520441-INS-233 |
| chr18-34522485-INS-158 | chr18-34522482-INS-159 |
| chr18-34529858-INS-201 | chr18-34529858-INS-201 |
| chr18-34566733-INS-61 | chr18-34566734-INS-65 |
| chr18-34579189-INS-90 | chr18-34579189-INS-90 |
| chr18-34599837-INS-132 | chr18-34599837-INS-132 |
| chr18-34620314-INS-136 | chr18-34620314-INS-136 |
| chr18-34645085-INS-86 | chr18-34645085-INS-90 |
| chr18-34650573-INS-1220 | chr18-34650574-INS-1220 |
| chr18-34663895-INS-69 | chr18-34663895-INS-59 |
| chr18-34685481-INS-82 | chr18-34685481-INS-82 |
| chr18-34694394-INS-65 | chr18-34694394-INS-66 |
| chr18-34701863-INS-102 | chr18-34701863-INS-102 |
| chr18-34703851-INS-102 | chr18-34703851-INS-68 |
| chr18-34711229-INS-221 | chr18-34711229-INS-220 |
| chr18-34722186-INS-109 | chr18-34722186-INS-109 |
| chr18-34727397-INS-114 | chr18-34727398-INS-80 |
| chr18-34778822-INS-174 | chr18-34778822-INS-174 |
| chr18-34790253-INS-119 | chr18-34790253-INS-119 |
| chr18-34791934-INS-64 | chr18-34791934-INS-64 |
| chr18-34807518-INS-207 | chr18-34807518-INS-208 |
| chr18-34822241-INS-207 | chr18-34822241-INS-210 |
| chr18-34829744-INS-52 | chr18-34829744-INS-52 |
| chr18-34833746-INS-186 | chr18-34833746-INS-179 |
| chr18-34851140-INS-199 | chr18-34851140-INS-200 |
| chr18-34874401-INS-236 | chr18-34874401-INS-235 |
| chr18-34878614-INS-731 | chr18-34878614-INS-730 |
| chr18-34886831-INS-55 | chr18-34886831-INS-55 |
| chr18-34922089-INS-212 | chr18-34922089-INS-232 |
| chr18-34924169-INS-276 | chr18-34924162-INS-286 |
| chr18-34986168-INS-52 | chr18-34986144-INS-54 |
| chr18-34987395-INS-156 | chr18-34987395-INS-168 |
| chr18-34990514-INS-186 | chr18-34990514-INS-186 |
| chr18-35007162-INS-86 | chr18-35007162-INS-84 |
| chr18-35012646-INS-206 | chr18-35012646-INS-206 |
| chr18-35019698-INS-517 | chr18-35019698-INS-517 |
| chr18-35028759-INS-59 | chr18-35028772-INS-50 |
| chr18-35044409-INS-203 | chr18-35044409-INS-203 |
| chr18-35045472-INS-180 | chr18-35045472-INS-180 |
| chr18-35068960-INS-216 | chr18-35068960-INS-216 |
| chr18-35074221-INS-544 | chr18-35074221-INS-544 |
| chr18-35097135-INS-84 | chr18-35097105-INS-117 |
| chr18-35107855-INS-213 | chr18-35107855-INS-213 |
| chr18-35124532-INS-149 | chr18-35124532-INS-181 |
| chr18-35133640-INS-74 | chr18-35133640-INS-74 |
| chr18-35137568-INS-235 | chr18-35137568-INS-235 |
| chr18-35139422-INS-210 | chr18-35139422-INS-210 |
| chr18-35176039-INS-230 | chr18-35176039-INS-231 |
| chr18-35194334-INS-66 | chr18-35194334-INS-68 |
| chr18-35199896-INS-74 | chr18-35199896-INS-74 |
| chr18-35210388-INS-6413 | chr18-35210389-INS-6413 |
| chr18-35220967-INS-52 | chr18-35220943-INS-66 |
| chr18-35225269-INS-229 | chr18-35225269-INS-228 |
| chr18-35228789-INS-6334 | chr18-35228790-INS-6334 |
| chr18-35233180-INS-325 | chr18-35233180-INS-325 |
| chr18-35243529-INS-182 | chr18-35243529-INS-183 |
| chr18-35254827-INS-187 | chr18-35254827-INS-187 |
| chr18-35261527-INS-177 | chr18-35261527-INS-177 |
| chr18-35302113-INS-123 | chr18-35302113-INS-123 |
| chr18-35315111-INS-223 | chr18-35315111-INS-225 |
| chr18-35410270-INS-195 | chr18-35410270-INS-195 |
| chr18-35454346-INS-142 | chr18-35454351-INS-149 |
| chr18-35477911-INS-170 | chr18-35477911-INS-170 |
| chr18-35496838-INS-572 | chr18-35496838-INS-572 |
| chr18-35498886-INS-796 | chr18-35498886-INS-798 |
| chr18-35510671-INS-62 | chr18-35510657-INS-54 |
| chr18-35526223-INS-225 | chr18-35526223-INS-226 |
| chr18-35553254-INS-55 | chr18-35553254-INS-90 |
| chr18-35560427-INS-108 | chr18-35560427-INS-114 |
| chr18-35569788-INS-70 | chr18-35569789-INS-72 |
| chr18-35578563-INS-85 | chr18-35578578-INS-87 |
| chr18-35587931-INS-206 | chr18-35587932-INS-165 |
| chr18-35590933-INS-52 | chr18-35590949-INS-61 |
| chr18-35593432-INS-184 | chr18-35593433-INS-185 |
| chr18-35673741-INS-71 | chr18-35673741-INS-71 |
| chr18-35675215-INS-256 | chr18-35675215-INS-256 |
| chr18-35676832-INS-435 | chr18-35676832-INS-435 |

| | |
|---|---|
| chr18-35683170-INS-76 | chr18-35683163-INS-60 |
| chr18-35691075-INS-1453 | chr18-35691077-INS-1458 |
| chr18-35736674-INS-576 | chr18-35736674-INS-576 |
| chr18-35769314-INS-171 | chr18-35769321-INS-185 |
| chr18-35771664-INS-226 | chr18-35771664-INS-226 |
| chr18-35792610-INS-215 | chr18-35792611-INS-216 |
| chr18-35804943-INS-70 | chr18-35804943-INS-64 |
| chr18-35851614-INS-70 | chr18-35851607-INS-70 |
| chr18-35852437-INS-57 | chr18-35852437-INS-57 |
| chr18-35857466-INS-67 | chr18-35857466-INS-69 |
| chr18-35859003-INS-194 | chr18-35859003-INS-194 |
| chr18-35869990-INS-84 | chr18-35869990-INS-84 |
| chr18-35881808-INS-622 | chr18-35881808-INS-622 |
| chr18-35899038-INS-102 | chr18-35899038-INS-102 |
| chr18-35912558-INS-198 | chr18-35912558-INS-198 |
| chr18-35934537-INS-192 | chr18-35934537-INS-193 |
| chr18-35980163-INS-206 | chr18-35980163-INS-206 |
| chr18-35986068-INS-425 | chr18-35986068-INS-424 |
| chr18-36032975-INS-224 | chr18-36032975-INS-224 |
| chr18-36036251-INS-210 | chr18-36036251-INS-210 |
| chr18-36043107-INS-179 | chr18-36043107-INS-179 |
| chr18-36049494-INS-199 | chr18-36049494-INS-200 |
| chr18-36056583-INS-55 | chr18-36056606-INS-59 |
| chr18-36068284-INS-58 | chr18-36068284-INS-58 |
| chr18-36075286-INS-60 | chr18-36075271-INS-52 |
| chr18-36082526-INS-111 | chr18-36082526-INS-112 |
| chr18-36088410-INS-87 | chr18-36088410-INS-90 |
| chr18-36089023-INS-102 | chr18-36089023-INS-108 |
| chr18-36101482-INS-64 | chr18-36101482-INS-64 |
| chr18-36134648-INS-60 | chr18-36134648-INS-60 |
| chr18-36214228-INS-122 | chr18-36214229-INS-74 |
| chr18-36237098-INS-149 | chr18-36237099-INS-166 |
| chr18-36245503-INS-73 | chr18-36245553-INS-73 |
| chr18-36255358-INS-95 | chr18-36255372-INS-78 |
| chr18-36264273-INS-305 | chr18-36264273-INS-304 |
| chr18-36275470-INS-82 | chr18-36275470-INS-54 |
| chr18-36317081-INS-206 | chr18-36317081-INS-205 |
| chr18-36356393-INS-122 | chr18-36356393-INS-165 |
| chr18-36374435-INS-157 | chr18-36374425-INS-133 |
| chr18-36380417-INS-52 | chr18-36380417-INS-55 |
| chr18-36384367-INS-128 | chr18-36384364-INS-130 |
| chr18-36386157-INS-329 | chr18-36386157-INS-330 |
| chr18-36392007-INS-222 | chr18-36392007-INS-222 |
| chr18-36392165-INS-1583 | chr18-36392166-INS-1582 |
| chr18-36408109-INS-90 | chr18-36408109-INS-90 |
| chr18-36479848-INS-52 | chr18-36479865-INS-74 |
| chr18-36492677-INS-107 | chr18-36492677-INS-107 |
| chr18-36508733-INS-171 | chr18-36508733-INS-175 |
| chr18-36511709-INS-131 | chr18-36511709-INS-132 |
| chr18-36564996-INS-171 | chr18-36564996-INS-172 |
| chr18-36578506-INS-201 | chr18-36578506-INS-201 |
| chr18-36648385-INS-5411 | chr18-36648386-INS-5411 |
| chr18-36679021-INS-209 | chr18-36679021-INS-212 |
| chr18-36680807-INS-203 | chr18-36680808-INS-203 |
| chr18-36686482-INS-74 | chr18-36686482-INS-74 |
| chr18-36707289-INS-161 | chr18-36707289-INS-161 |
| chr18-36722071-INS-182 | chr18-36722071-INS-182 |
| chr18-36735648-INS-211 | chr18-36735648-INS-211 |
| chr18-36747042-INS-84 | chr18-36747042-INS-84 |
| chr18-36783306-INS-58 | chr18-36783306-INS-58 |
| chr18-36823629-INS-174 | chr18-36823629-INS-174 |
| chr18-36838236-INS-194 | chr18-36838236-INS-195 |
| chr18-36842666-INS-68 | chr18-36842638-INS-114 |
| chr18-36876148-INS-60 | chr18-36876103-INS-66 |
| chr18-36881021-INS-55 | chr18-36881021-INS-55 |
| chr18-36903745-INS-107 | chr18-36903745-INS-111 |
| chr18-36917760-INS-124 | chr18-36917759-INS-92 |
| chr18-36934249-INS-240 | chr18-36934249-INS-239 |
| chr18-36957771-INS-504 | chr18-36957771-INS-504 |
| chr18-36959607-INS-7172 | chr18-36959608-INS-7172 |
| chr18-36974232-INS-6401 | chr18-36974233-INS-6401 |
| chr18-36985315-INS-1124 | chr18-36985316-INS-1124 |
| chr18-37045772-INS-140 | chr18-37045772-INS-95 |
| chr18-37051586-INS-73 | chr18-37051586-INS-73 |
| chr18-37057358-INS-53 | chr18-37057358-INS-53 |
| chr18-37060916-INS-84 | chr18-37060916-INS-72 |
| chr18-37067101-INS-55 | chr18-37067102-INS-57 |

| | |
|---|---|
| chr18-37070771-INS-59 | chr18-37070771-INS-75 |
| chr18-37090823-INS-103 | chr18-37090823-INS-64 |
| chr18-37123374-INS-84 | chr18-37123374-INS-52 |
| chr18-37134899-INS-391 | chr18-37134899-INS-394 |
| chr18-37161955-INS-79 | chr18-37161955-INS-71 |
| chr18-37162692-INS-64 | chr18-37162657-INS-84 |
| chr18-37176033-INS-66 | chr18-37176033-INS-66 |
| chr18-37208049-INS-197 | chr18-37208005-INS-197 |
| chr18-37244485-INS-56 | chr18-37244479-INS-76 |
| chr18-37260710-INS-505 | chr18-37260710-INS-505 |
| chr18-37295843-INS-1441 | chr18-37295844-INS-1441 |
| chr18-37338343-INS-55 | chr18-37338343-INS-55 |
| chr18-37344983-INS-410 | chr18-37344983-INS-410 |
| chr18-37375292-INS-223 | chr18-37375292-INS-223 |
| chr18-37392859-INS-97 | chr18-37392858-INS-143 |
| chr18-37406771-INS-676 | chr18-37406771-INS-676 |
| chr18-37425560-INS-432 | chr18-37425554-INS-438 |
| chr18-37479463-INS-199 | chr18-37479463-INS-199 |
| chr18-37485299-INS-179 | chr18-37485299-INS-179 |
| chr18-37548222-INS-166 | chr18-37548222-INS-166 |
| chr18-37553627-INS-129 | chr18-37553627-INS-165 |
| chr18-37606031-INS-180 | chr18-37606031-INS-172 |
| chr18-37672967-INS-5695 | chr18-37672968-INS-5695 |
| chr18-37693066-INS-430 | chr18-37693066-INS-430 |
| chr18-37698313-INS-199 | chr18-37698309-INS-203 |
| chr18-37752329-INS-358 | chr18-37752329-INS-313 |
| chr18-37758754-INS-55 | chr18-37758754-INS-55 |
| chr18-37786970-INS-571 | chr18-37786970-INS-571 |
| chr18-37805058-INS-106 | chr18-37805057-INS-142 |
| chr18-37850906-INS-57 | chr18-37850906-INS-57 |
| chr18-37897227-INS-228 | chr18-37897222-INS-233 |
| chr18-37897781-INS-66 | chr18-37897781-INS-68 |
| chr18-37902289-INS-52 | chr18-37902289-INS-56 |
| chr18-37907305-INS-191 | chr18-37907305-INS-187 |
| chr18-37911656-INS-68 | chr18-37911656-INS-66 |
| chr18-37992225-INS-60 | chr18-37992225-INS-62 |
| chr18-37995439-INS-216 | chr18-37995439-INS-218 |
| chr18-38004179-INS-202 | chr18-38004179-INS-203 |
| chr18-38006496-INS-138 | chr18-38006496-INS-145 |
| chr18-38032973-INS-166 | chr18-38033002-INS-120 |
| chr18-38044146-INS-91 | chr18-38044146-INS-91 |
| chr18-38050391-INS-184 | chr18-38050391-INS-183 |
| chr18-38061559-INS-130 | chr18-38061563-INS-106 |
| chr18-38170404-INS-400 | chr18-38170404-INS-400 |
| chr18-38171805-INS-58 | chr18-38171805-INS-60 |
| chr18-38185113-INS-72 | chr18-38185098-INS-78 |
| chr18-38203488-INS-217 | chr18-38203488-INS-217 |
| chr18-38246429-INS-217 | chr18-38246429-INS-218 |
| chr18-38257225-INS-106 | chr18-38257225-INS-104 |
| chr18-38283713-INS-114 | chr18-38283713-INS-118 |
| chr18-38292029-INS-71 | chr18-38292029-INS-73 |
| chr18-38302842-INS-134 | chr18-38302842-INS-134 |
| chr18-38306234-INS-166 | chr18-38306263-INS-168 |
| chr18-38317006-INS-170 | chr18-38317006-INS-170 |
| chr18-38363836-INS-52 | chr18-38363836-INS-62 |
| chr18-38377661-INS-231 | chr18-38377661-INS-231 |
| chr18-38406636-INS-172 | chr18-38406636-INS-172 |
| chr18-38429184-INS-232 | chr18-38429184-INS-232 |
| chr18-38431334-INS-413 | chr18-38431334-INS-413 |
| chr18-38434077-INS-55 | chr18-38434082-INS-99 |
| chr18-38445132-INS-129 | chr18-38445132-INS-129 |
| chr18-38456068-INS-160 | chr18-38456068-INS-160 |
| chr18-38458130-INS-85 | chr18-38458130-INS-85 |
| chr18-38480393-INS-191 | chr18-38480343-INS-205 |
| chr18-38494440-INS-112 | chr18-38494440-INS-75 |
| chr18-38495511-INS-406 | chr18-38495511-INS-406 |
| chr18-38509870-INS-96 | chr18-38509870-INS-72 |
| chr18-38515272-INS-72 | chr18-38515272-INS-72 |
| chr18-38525112-INS-72 | chr18-38525112-INS-69 |
| chr18-38528316-INS-825 | chr18-38528316-INS-825 |
| chr18-38528446-INS-102 | chr18-38528446-INS-102 |
| chr18-38563495-INS-221 | chr18-38563495-INS-221 |
| chr18-38574344-INS-194 | chr18-38574328-INS-206 |
| chr18-38603080-INS-210 | chr18-38603080-INS-211 |
| chr18-38622831-INS-59 | chr18-38622879-INS-52 |
| chr18-38625051-INS-169 | chr18-38625051-INS-168 |
| chr18-38637147-INS-178 | chr18-38637147-INS-178 |

| | |
|---|---|
| chr18-38645186-INS-145 | chr18-38645186-INS-195 |
| chr18-38666458-INS-53 | chr18-38666415-INS-53 |
| chr18-38669098-INS-170 | chr18-38669098-INS-182 |
| chr18-38691673-INS-170 | chr18-38691670-INS-200 |
| chr18-38701002-INS-169 | chr18-38701002-INS-169 |
| chr18-38705041-INS-228 | chr18-38705041-INS-227 |
| chr18-38744713-INS-68 | chr18-38744677-INS-58 |
| chr18-38748347-INS-1892 | chr18-38748348-INS-1892 |
| chr18-38771731-INS-205 | chr18-38771731-INS-205 |
| chr18-38775196-INS-396 | chr18-38775196-INS-396 |
| chr18-38783929-INS-198 | chr18-38783929-INS-198 |
| chr18-38792392-INS-76 | chr18-38792392-INS-85 |
| chr18-38795123-INS-85 | chr18-38795094-INS-117 |
| chr18-38806762-INS-198 | chr18-38806762-INS-197 |
| chr18-38807812-INS-51 | chr18-38807812-INS-85 |
| chr18-38812266-INS-59 | chr18-38812262-INS-72 |
| chr18-38815994-INS-188 | chr18-38815993-INS-195 |
| chr18-38820668-INS-155 | chr18-38820668-INS-155 |
| chr18-38840149-INS-224 | chr18-38840149-INS-223 |
| chr18-38847820-INS-53 | chr18-38847820-INS-53 |
| chr18-38853644-INS-513 | chr18-38853633-INS-513 |
| chr18-38887112-INS-477 | chr18-38887112-INS-477 |
| chr18-38890878-INS-63 | chr18-38890878-INS-63 |
| chr18-38905785-INS-113 | chr18-38905763-INS-86 |
| chr18-38910476-INS-51 | chr18-38910476-INS-51 |
| chr18-38922551-INS-64 | chr18-38922552-INS-64 |
| chr18-38938533-INS-68 | chr18-38938541-INS-67 |
| chr18-38942868-INS-1893 | chr18-38942869-INS-1893 |
| chr18-38947589-INS-80 | chr18-38947585-INS-60 |
| chr18-38957738-INS-71 | chr18-38957738-INS-71 |
| chr18-38975646-INS-209 | chr18-38975646-INS-209 |
| chr18-39066565-INS-86 | chr18-39066565-INS-86 |
| chr18-39080988-INS-72 | chr18-39080988-INS-72 |
| chr18-39083708-INS-498 | chr18-39083708-INS-498 |
| chr18-39084357-INS-636 | chr18-39084357-INS-636 |
| chr18-39087636-INS-1287 | chr18-39087637-INS-1287 |
| chr18-39089593-INS-5141 | chr18-39089594-INS-5141 |
| chr18-39093475-INS-76 | chr18-39093475-INS-76 |
| chr18-39110982-INS-102 | chr18-39110982-INS-102 |
| chr18-39170726-INS-544 | chr18-39170727-INS-495 |
| chr18-39209599-INS-68 | chr18-39209599-INS-76 |
| chr18-39241927-INS-113 | chr18-39241927-INS-113 |
| chr18-39252427-INS-61 | chr18-39252427-INS-61 |
| chr18-39256987-INS-170 | chr18-39256987-INS-169 |
| chr18-39291798-INS-54 | chr18-39291798-INS-54 |
| chr18-39346021-INS-71 | chr18-39346050-INS-50 |
| chr18-39354813-INS-54 | chr18-39354813-INS-54 |
| chr18-39383215-INS-52 | chr18-39383215-INS-50 |
| chr18-39388361-INS-116 | chr18-39388365-INS-74 |
| chr18-39405971-INS-143 | chr18-39405959-INS-124 |
| chr18-39423217-INS-77 | chr18-39423217-INS-77 |
| chr18-39442887-INS-7086 | chr18-39442888-INS-7086 |
| chr18-39451690-INS-71 | chr18-39451690-INS-71 |
| chr18-39462522-INS-894 | chr18-39462522-INS-905 |
| chr18-39464106-INS-338 | chr18-39464106-INS-338 |
| chr18-39470700-INS-55 | chr18-39470700-INS-55 |
| chr18-39484493-INS-60 | chr18-39484493-INS-50 |
| chr18-39517579-INS-70 | chr18-39517579-INS-72 |
| chr18-39519727-INS-58 | chr18-39519705-INS-62 |
| chr18-39525610-INS-259 | chr18-39525610-INS-259 |
| chr18-39570389-INS-144 | chr18-39570389-INS-126 |
| chr18-39574784-INS-102 | chr18-39574784-INS-102 |
| chr18-39594662-INS-60 | chr18-39594662-INS-60 |
| chr18-39633573-INS-109 | chr18-39633573-INS-109 |
| chr18-39661811-INS-3661 | chr18-39661812-INS-3662 |
| chr18-39684741-INS-415 | chr18-39684741-INS-415 |
| chr18-39701201-INS-64 | chr18-39701184-INS-82 |
| chr18-39702450-INS-57 | chr18-39702450-INS-57 |
| chr18-39712877-INS-61 | chr18-39712877-INS-61 |
| chr18-39744095-INS-103 | chr18-39744095-INS-127 |
| chr18-39757707-INS-79 | chr18-39757707-INS-62 |
| chr18-39788278-INS-463 | chr18-39788279-INS-463 |
| chr18-39788782-INS-371 | chr18-39788782-INS-371 |
| chr18-39792867-INS-66 | chr18-39792867-INS-90 |
| chr18-39795730-INS-404 | chr18-39795730-INS-403 |
| chr18-39808202-INS-377 | chr18-39808202-INS-377 |
| chr18-39866804-INS-943 | chr18-39866804-INS-943 |

| | |
|---|---|
| chr18-39871461-INS-115 | chr18-39871461-INS-115 |
| chr18-39874662-INS-62 | chr18-39874662-INS-80 |
| chr18-39886656-INS-60 | chr18-39886656-INS-104 |
| chr18-39899897-INS-183 | chr18-39899897-INS-183 |
| chr18-39913203-INS-1014 | chr18-39913204-INS-1014 |
| chr18-39936003-INS-708 | chr18-39936003-INS-708 |
| chr18-39954513-INS-76 | chr18-39954509-INS-56 |
| chr18-39968212-INS-102 | chr18-39968206-INS-100 |
| chr18-39970142-INS-58 | chr18-39970178-INS-56 |
| chr18-39984544-INS-175 | chr18-39984544-INS-132 |
| chr18-40001542-INS-82 | chr18-40001542-INS-82 |
| chr18-40013178-INS-136 | chr18-40013183-INS-137 |
| chr18-40018786-INS-57 | chr18-40018786-INS-51 |
| chr18-40027951-INS-700 | chr18-40027951-INS-700 |
| chr18-40036003-INS-130 | chr18-40036004-INS-136 |
| chr18-40040113-INS-165 | chr18-40040113-INS-163 |
| chr18-40067331-INS-52 | chr18-40067331-INS-52 |
| chr18-40089873-INS-203 | chr18-40089873-INS-203 |
| chr18-40148520-INS-97 | chr18-40148518-INS-101 |
| chr18-40172585-INS-1091 | chr18-40172584-INS-1125 |
| chr18-40180054-INS-109 | chr18-40180054-INS-109 |
| chr18-40203500-INS-90 | chr18-40203500-INS-90 |
| chr18-40212226-INS-72 | chr18-40212226-INS-76 |
| chr18-40212909-INS-498 | chr18-40212909-INS-498 |
| chr18-40269252-INS-383 | chr18-40269252-INS-383 |
| chr18-40271399-INS-961 | chr18-40271399-INS-961 |
| chr18-40294942-INS-264 | chr18-40294942-INS-256 |
| chr18-40309001-INS-53 | chr18-40309001-INS-64 |
| chr18-40344031-INS-203 | chr18-40344031-INS-202 |
| chr18-40346134-INS-217 | chr18-40346134-INS-218 |
| chr18-40370947-INS-104 | chr18-40370947-INS-104 |
| chr18-40375684-INS-51 | chr18-40375684-INS-55 |
| chr18-40413173-INS-63 | chr18-40413173-INS-87 |
| chr18-40416389-INS-568 | chr18-40416389-INS-568 |
| chr18-40470492-INS-148 | chr18-40470496-INS-124 |
| chr18-40478556-INS-82 | chr18-40478556-INS-82 |
| chr18-40494641-INS-77 | chr18-40494641-INS-65 |
| chr18-40500660-INS-6345 | chr18-40500661-INS-6345 |
| chr18-40506222-INS-72 | chr18-40506222-INS-70 |
| chr18-40511973-INS-350 | chr18-40511973-INS-351 |
| chr18-40517789-INS-401 | chr18-40517789-INS-401 |
| chr18-40541154-INS-127 | chr18-40541154-INS-127 |
| chr18-40553264-INS-282 | chr18-40553264-INS-292 |
| chr18-40560189-INS-62 | chr18-40560189-INS-62 |
| chr18-40566449-INS-846 | chr18-40566449-INS-846 |
| chr18-40575522-INS-102 | chr18-40575522-INS-102 |
| chr18-40581541-INS-112 | chr18-40581541-INS-112 |
| chr18-40634527-INS-66 | chr18-40634527-INS-56 |
| chr18-40653953-INS-222 | chr18-40653953-INS-213 |
| chr18-40686299-INS-116 | chr18-40686299-INS-116 |
| chr18-40692701-INS-125 | chr18-40692701-INS-89 |
| chr18-40699249-INS-406 | chr18-40699249-INS-403 |
| chr18-40702599-INS-306 | chr18-40702599-INS-266 |
| chr18-40704732-INS-284 | chr18-40704732-INS-284 |
| chr18-40712915-INS-6406 | chr18-40712916-INS-6406 |
| chr18-40745212-INS-798 | chr18-40745212-INS-798 |
| chr18-40750350-INS-64 | chr18-40750350-INS-64 |
| chr18-40765643-INS-61 | chr18-40765643-INS-61 |
| chr18-40767144-INS-737 | chr18-40767144-INS-739 |
| chr18-40798717-INS-62 | chr18-40798717-INS-65 |
| chr18-40805952-INS-52 | chr18-40805952-INS-76 |
| chr18-40825248-INS-64 | chr18-40825225-INS-96 |
| chr18-40854825-INS-2802 | chr18-40854826-INS-2801 |
| chr18-40909624-INS-1001 | chr18-40909625-INS-1001 |
| chr18-40923935-INS-119 | chr18-40923941-INS-113 |
| chr18-40942646-INS-110 | chr18-40942646-INS-110 |
| chr18-40949969-INS-52 | chr18-40950002-INS-60 |
| chr18-40953990-INS-6275 | chr18-40953991-INS-6279 |
| chr18-40991225-INS-73 | chr18-40991235-INS-54 |
| chr18-41000276-INS-65 | chr18-41000276-INS-52 |
| chr18-41001476-INS-335 | chr18-41001476-INS-335 |
| chr18-41050757-INS-88 | chr18-41050757-INS-88 |
| chr18-41075920-INS-234 | chr18-41075920-INS-234 |
| chr18-41077408-INS-80 | chr18-41077412-INS-75 |
| chr18-41096940-INS-531 | chr18-41096940-INS-530 |
| chr18-41101030-INS-345 | chr18-41101030-INS-345 |
| chr18-41219004-INS-106 | chr18-41219004-INS-106 |

| | |
|---|---|
| chr18-41259025-INS-8979 | chr18-41259026-INS-8982 |
| chr18-41261411-INS-3256 | chr18-41261412-INS-3256 |
| chr18-41274705-INS-132 | chr18-41274723-INS-135 |
| chr18-41312641-INS-636 | chr18-41312640-INS-636 |
| chr18-41328713-INS-88 | chr18-41328719-INS-56 |
| chr18-41353652-INS-516 | chr18-41353652-INS-516 |
| chr18-41360313-INS-126 | chr18-41360314-INS-110 |
| chr18-41369061-INS-56 | chr18-41369061-INS-60 |
| chr18-41377245-INS-64 | chr18-41377266-INS-64 |
| chr18-41383893-INS-58 | chr18-41383905-INS-70 |
| chr18-41386427-INS-895 | chr18-41386427-INS-896 |
| chr18-41400045-INS-441 | chr18-41400045-INS-441 |
| chr18-41430498-INS-2832 | chr18-41430499-INS-2832 |
| chr18-41438970-INS-1565 | chr18-41438971-INS-1565 |
| chr18-41452168-INS-483 | chr18-41452168-INS-483 |
| chr18-41476077-INS-56 | chr18-41476077-INS-56 |
| chr18-41509647-INS-76 | chr18-41509619-INS-65 |
| chr18-41530406-INS-1303 | chr18-41530407-INS-1300 |
| chr18-41546147-INS-2950 | chr18-41546148-INS-2950 |
| chr18-41548413-INS-138 | chr18-41548413-INS-147 |
| chr18-41577065-INS-870 | chr18-41577062-INS-879 |
| chr18-41585407-INS-66 | chr18-41585410-INS-72 |
| chr18-41612009-INS-60 | chr18-41612010-INS-55 |
| chr18-41620305-INS-54 | chr18-41620305-INS-54 |
| chr18-41640318-INS-424 | chr18-41640318-INS-423 |
| chr18-41684486-INS-212 | chr18-41684486-INS-212 |
| chr18-41710830-INS-101 | chr18-41710830-INS-101 |
| chr18-41725575-INS-58 | chr18-41725572-INS-55 |
| chr18-41747521-INS-629 | chr18-41747521-INS-651 |
| chr18-41753756-INS-108 | chr18-41753759-INS-115 |
| chr18-41772047-INS-4580 | chr18-41772048-INS-4580 |
| chr18-41772588-INS-123 | chr18-41772588-INS-123 |
| chr18-41775618-INS-136 | chr18-41775618-INS-127 |
| chr18-41776437-INS-324 | chr18-41776437-INS-325 |
| chr18-41847234-INS-146 | chr18-41847235-INS-145 |
| chr18-41899603-INS-232 | chr18-41899603-INS-239 |
| chr18-41911293-INS-1503 | chr18-41911294-INS-1503 |
| chr18-41926301-INS-462 | chr18-41926301-INS-462 |
| chr18-41932665-INS-410 | chr18-41932666-INS-410 |
| chr18-41941686-INS-588 | chr18-41941686-INS-588 |
| chr18-41943455-INS-2371 | chr18-41943450-INS-2373 |
| chr18-41949694-INS-4978 | chr18-41949695-INS-4980 |
| chr18-41959390-INS-214 | chr18-41959390-INS-214 |
| chr18-41965613-INS-151 | chr18-41965613-INS-130 |
| chr18-41967299-INS-554 | chr18-41967299-INS-576 |
| chr18-41968645-INS-119 | chr18-41968645-INS-132 |
| chr18-41971946-INS-6385 | chr18-41971947-INS-6385 |
| chr18-41977441-INS-126 | chr18-41977441-INS-127 |
| chr18-41984211-INS-85 | chr18-41984211-INS-70 |
| chr18-41998544-INS-208 | chr18-41998544-INS-208 |
| chr18-42015481-INS-245 | chr18-42015481-INS-245 |
| chr18-42016212-INS-368 | chr18-42016212-INS-368 |
| chr18-42030315-INS-68 | chr18-42030315-INS-76 |
| chr18-42078539-INS-57 | chr18-42078539-INS-57 |
| chr18-42107829-INS-2753 | chr18-42107830-INS-2753 |
| chr18-42128625-INS-138 | chr18-42128625-INS-180 |
| chr18-42129450-INS-404 | chr18-42129450-INS-407 |
| chr18-42146668-INS-4519 | chr18-42146669-INS-4519 |
| chr18-42175174-INS-5278 | chr18-42175175-INS-5278 |
| chr18-42176648-INS-113 | chr18-42176648-INS-111 |
| chr18-42194149-INS-533 | chr18-42194149-INS-534 |
| chr18-42206530-INS-210 | chr18-42206556-INS-197 |
| chr18-42249642-INS-57 | chr18-42249643-INS-69 |
| chr18-42252185-INS-55 | chr18-42252195-INS-63 |
| chr18-42265663-INS-90 | chr18-42265663-INS-81 |
| chr18-42282549-INS-71 | chr18-42282548-INS-75 |
| chr18-42301640-INS-206 | chr18-42301640-INS-206 |
| chr18-42304383-INS-61 | chr18-42304384-INS-62 |
| chr18-42309868-INS-52 | chr18-42309868-INS-52 |
| chr18-42310296-INS-130 | chr18-42310296-INS-130 |
| chr18-42312472-INS-116 | chr18-42312472-INS-116 |
| chr18-42320945-INS-180 | chr18-42320945-INS-180 |
| chr18-42335031-INS-135 | chr18-42335031-INS-91 |
| chr18-42363869-INS-84 | chr18-42363869-INS-69 |
| chr18-42379459-INS-174 | chr18-42379459-INS-175 |
| chr18-42380733-INS-197 | chr18-42380733-INS-197 |
| chr18-42381764-INS-179 | chr18-42381764-INS-179 |

| | |
|---|---|
| chr18-42395633-INS-373 | chr18-42395633-INS-375 |
| chr18-42400456-INS-198 | chr18-42400456-INS-197 |
| chr18-42404162-INS-233 | chr18-42404162-INS-232 |
| chr18-42414548-INS-208 | chr18-42414548-INS-211 |
| chr18-42423265-INS-212 | chr18-42423265-INS-212 |
| chr18-42496005-INS-479 | chr18-42496005-INS-479 |
| chr18-42498896-INS-73 | chr18-42498896-INS-73 |
| chr18-42505827-INS-204 | chr18-42505828-INS-206 |
| chr18-42524813-INS-312 | chr18-42524813-INS-312 |
| chr18-42525513-INS-655 | chr18-42525513-INS-655 |
| chr18-42535613-INS-55 | chr18-42535613-INS-75 |
| chr18-42537647-INS-413 | chr18-42537647-INS-414 |
| chr18-42542985-INS-64 | chr18-42542985-INS-63 |
| chr18-42548503-INS-54 | chr18-42548509-INS-51 |
| chr18-42550357-INS-249 | chr18-42550377-INS-233 |
| chr18-42553051-INS-52 | chr18-42553055-INS-56 |
| chr18-42567998-INS-1890 | chr18-42567999-INS-1890 |
| chr18-42579353-INS-176 | chr18-42579353-INS-176 |
| chr18-42585974-INS-80 | chr18-42585974-INS-104 |
| chr18-42589265-INS-200 | chr18-42589265-INS-202 |
| chr18-42593119-INS-52 | chr18-42593119-INS-52 |
| chr18-42599883-INS-162 | chr18-42599883-INS-162 |
| chr18-42622383-INS-229 | chr18-42622384-INS-225 |
| chr18-42627315-INS-504 | chr18-42627315-INS-504 |
| chr18-42634520-INS-142 | chr18-42634520-INS-140 |
| chr18-42639340-INS-110 | chr18-42639325-INS-113 |
| chr18-42640209-INS-179 | chr18-42640209-INS-180 |
| chr18-42653990-INS-211 | chr18-42653990-INS-211 |
| chr18-42665750-INS-82 | chr18-42665730-INS-72 |
| chr18-42671860-INS-192 | chr18-42671860-INS-187 |
| chr18-42679198-INS-2648 | chr18-42679208-INS-2648 |
| chr18-42719765-INS-312 | chr18-42719765-INS-294 |
| chr18-42791612-INS-90 | chr18-42791612-INS-90 |
| chr18-42796842-INS-212 | chr18-42796842-INS-221 |
| chr18-42809068-INS-55 | chr18-42809068-INS-55 |
| chr18-42811205-INS-168 | chr18-42811205-INS-168 |
| chr18-42822081-INS-251 | chr18-42822081-INS-251 |
| chr18-42841041-INS-76 | chr18-42841041-INS-80 |
| chr18-42894592-INS-392 | chr18-42894592-INS-392 |
| chr18-42925292-INS-57 | chr18-42925292-INS-57 |
| chr18-42960188-INS-55 | chr18-42960188-INS-55 |
| chr18-42963914-INS-95 | chr18-42963914-INS-85 |
| chr18-42970519-INS-53 | chr18-42970523-INS-74 |
| chr18-42972242-INS-66 | chr18-42972242-INS-66 |
| chr18-42992312-INS-170 | chr18-42992312-INS-170 |
| chr18-43009480-INS-92 | chr18-43009480-INS-92 |
| chr18-43038788-INS-90 | chr18-43038788-INS-78 |
| chr18-43040323-INS-84 | chr18-43040321-INS-84 |
| chr18-43049266-INS-124 | chr18-43049267-INS-124 |
| chr18-43059795-INS-70 | chr18-43059804-INS-50 |
| chr18-43112110-INS-102 | chr18-43112110-INS-102 |
| chr18-43176078-INS-75 | chr18-43176078-INS-68 |
| chr18-43197045-INS-7389 | chr18-43197046-INS-7390 |
| chr18-43223707-INS-163 | chr18-43223707-INS-207 |
| chr18-43230751-INS-75 | chr18-43230751-INS-75 |
| chr18-43234205-INS-64 | chr18-43234205-INS-91 |
| chr18-43253045-INS-306 | chr18-43253075-INS-276 |
| chr18-43257217-INS-6374 | chr18-43257218-INS-6374 |
| chr18-43269861-INS-422 | chr18-43269861-INS-422 |
| chr18-43274054-INS-104 | chr18-43274054-INS-104 |
| chr18-43312397-INS-6626 | chr18-43312398-INS-6621 |
| chr18-43323428-INS-6374 | chr18-43323429-INS-6372 |
| chr18-43339586-INS-56 | chr18-43339586-INS-56 |
| chr18-43347884-INS-62 | chr18-43347884-INS-90 |
| chr18-43359991-INS-81 | chr18-43359992-INS-69 |
| chr18-43378273-INS-206 | chr18-43378273-INS-206 |
| chr18-43391340-INS-76 | chr18-43391340-INS-56 |
| chr18-43428548-INS-204 | chr18-43428548-INS-204 |
| chr18-43434814-INS-482 | chr18-43434814-INS-482 |
| chr18-43441402-INS-51 | chr18-43441402-INS-54 |
| chr18-43446909-INS-97 | chr18-43446909-INS-102 |
| chr18-43508324-INS-68 | chr18-43508325-INS-52 |
| chr18-43514588-INS-251 | chr18-43514588-INS-251 |
| chr18-43540805-INS-153 | chr18-43540805-INS-153 |
| chr18-43549117-INS-138 | chr18-43549117-INS-163 |
| chr18-43551371-INS-90 | chr18-43551371-INS-100 |
| chr18-43553214-INS-574 | chr18-43553214-INS-574 |

| | |
|---|---|
| chr18-43572875-INS-209 | chr18-43572875-INS-215 |
| chr18-43582881-INS-60 | chr18-43582881-INS-50 |
| chr18-43586308-INS-58 | chr18-43586319-INS-51 |
| chr18-43586693-INS-891 | chr18-43586693-INS-891 |
| chr18-43593945-INS-501 | chr18-43593945-INS-502 |
| chr18-43599461-INS-282 | chr18-43599461-INS-281 |
| chr18-43605312-INS-164 | chr18-43605312-INS-172 |
| chr18-43613671-INS-54 | chr18-43613671-INS-54 |
| chr18-43618149-INS-90 | chr18-43618149-INS-76 |
| chr18-43656846-INS-76 | chr18-43656846-INS-76 |
| chr18-43681009-INS-58 | chr18-43680995-INS-54 |
| chr18-43741690-INS-90 | chr18-43741696-INS-83 |
| chr18-43767686-INS-199 | chr18-43767686-INS-199 |
| chr18-43773010-INS-72 | chr18-43773010-INS-72 |
| chr18-43858491-INS-229 | chr18-43858491-INS-229 |
| chr18-43861829-INS-6620 | chr18-43861830-INS-6620 |
| chr18-43867446-INS-236 | chr18-43867446-INS-238 |
| chr18-43885219-INS-6281 | chr18-43885220-INS-6281 |
| chr18-43896113-INS-56 | chr18-43896125-INS-53 |
| chr18-43904813-INS-810 | chr18-43904813-INS-810 |
| chr18-43910273-INS-474 | chr18-43910273-INS-472 |
| chr18-43914587-INS-777 | chr18-43914587-INS-777 |
| chr18-43921138-INS-66 | chr18-43921138-INS-66 |
| chr18-43931497-INS-182 | chr18-43931497-INS-185 |
| chr18-43935723-INS-578 | chr18-43935723-INS-548 |
| chr18-43938566-INS-268 | chr18-43938565-INS-277 |
| chr18-43953115-INS-495 | chr18-43953115-INS-495 |
| chr18-43953996-INS-194 | chr18-43953988-INS-202 |
| chr18-43960287-INS-73 | chr18-43960287-INS-73 |
| chr18-43996236-INS-55 | chr18-43996236-INS-56 |
| chr18-44015477-INS-54 | chr18-44015477-INS-54 |
| chr18-44049408-INS-66 | chr18-44049408-INS-75 |
| chr18-44054843-INS-2162 | chr18-44054829-INS-2155 |
| chr18-44054882-INS-149 | chr18-44054882-INS-146 |
| chr18-44055815-INS-158 | chr18-44055815-INS-158 |
| chr18-44064705-INS-6901 | chr18-44064706-INS-6903 |
| chr18-44066504-INS-50 | chr18-44066504-INS-56 |
| chr18-44079741-INS-169 | chr18-44079741-INS-169 |
| chr18-44089871-INS-249 | chr18-44089871-INS-249 |
| chr18-44090762-INS-448 | chr18-44090763-INS-448 |
| chr18-44139643-INS-367 | chr18-44139643-INS-367 |
| chr18-44158706-INS-796 | chr18-44158706-INS-796 |
| chr18-44210292-INS-100 | chr18-44210288-INS-106 |
| chr18-44218374-INS-66 | chr18-44218373-INS-96 |
| chr18-44221585-INS-114 | chr18-44221585-INS-121 |
| chr18-44231598-INS-88 | chr18-44231598-INS-78 |
| chr18-44234990-INS-66 | chr18-44234990-INS-65 |
| chr18-44247011-INS-58 | chr18-44247011-INS-61 |
| chr18-44281626-INS-6652 | chr18-44281627-INS-6652 |
| chr18-44292603-INS-68 | chr18-44292639-INS-52 |
| chr18-44316461-INS-7083 | chr18-44316462-INS-7083 |
| chr18-44323061-INS-88 | chr18-44323041-INS-58 |
| chr18-44332976-INS-1339 | chr18-44332977-INS-1339 |
| chr18-44386375-INS-67 | chr18-44386375-INS-67 |
| chr18-44387576-INS-352 | chr18-44387576-INS-352 |
| chr18-44399128-INS-1065 | chr18-44399129-INS-1065 |
| chr18-44414870-INS-66 | chr18-44414872-INS-68 |
| chr18-44433850-INS-434 | chr18-44433850-INS-432 |
| chr18-44441340-INS-76 | chr18-44441340-INS-74 |
| chr18-44442337-INS-52 | chr18-44442312-INS-74 |
| chr18-44444899-INS-234 | chr18-44444899-INS-234 |
| chr18-44450981-INS-66 | chr18-44450981-INS-76 |
| chr18-44451375-INS-2737 | chr18-44451376-INS-2737 |
| chr18-44493590-INS-116 | chr18-44493590-INS-132 |
| chr18-44499942-INS-195 | chr18-44499943-INS-194 |
| chr18-44545491-INS-90 | chr18-44545491-INS-90 |
| chr18-44564846-INS-86 | chr18-44564846-INS-86 |
| chr18-44596261-INS-480 | chr18-44596261-INS-480 |
| chr18-44598292-INS-76 | chr18-44598292-INS-82 |
| chr18-44612204-INS-234 | chr18-44612204-INS-222 |
| chr18-44630948-INS-209 | chr18-44630948-INS-209 |
| chr18-44666608-INS-94 | chr18-44666608-INS-94 |
| chr18-44670063-INS-673 | chr18-44670063-INS-678 |
| chr18-44685556-INS-261 | chr18-44685556-INS-279 |
| chr18-44687710-INS-212 | chr18-44687710-INS-212 |
| chr18-44698734-INS-88 | chr18-44698742-INS-66 |
| chr18-44767150-INS-2400 | chr18-44767151-INS-2400 |

| | |
|---|---|
| chr18-44769317-INS-87 | chr18-44769306-INS-62 |
| chr18-44783929-INS-215 | chr18-44783926-INS-188 |
| chr18-44813194-INS-250 | chr18-44813194-INS-249 |
| chr18-44820070-INS-6659 | chr18-44820071-INS-6670 |
| chr18-44823335-INS-751 | chr18-44823334-INS-763 |
| chr18-44843248-INS-101 | chr18-44843247-INS-100 |
| chr18-44852668-INS-156 | chr18-44852668-INS-156 |
| chr18-44867127-INS-54 | chr18-44867127-INS-55 |
| chr18-44885808-INS-366 | chr18-44885808-INS-366 |
| chr18-44890926-INS-1154 | chr18-44890927-INS-1151 |
| chr18-44906019-INS-54 | chr18-44906019-INS-56 |
| chr18-44916906-INS-6285 | chr18-44916907-INS-6304 |
| chr18-44930111-INS-999 | chr18-44930111-INS-995 |
| chr18-44947928-INS-6800 | chr18-44947929-INS-6800 |
| chr18-44955829-INS-72 | chr18-44955829-INS-72 |
| chr18-44964734-INS-176 | chr18-44964736-INS-178 |
| chr18-44971844-INS-6666 | chr18-44971845-INS-6666 |
| chr18-44977942-INS-200 | chr18-44977942-INS-200 |
| chr18-45001092-INS-145 | chr18-45001092-INS-145 |
| chr18-45013220-INS-338 | chr18-45013220-INS-338 |
| chr18-45039089-INS-73 | chr18-45039089-INS-73 |
| chr18-45042949-INS-440 | chr18-45042949-INS-440 |
| chr18-45052555-INS-200 | chr18-45052555-INS-200 |
| chr18-45058160-INS-68 | chr18-45058160-INS-68 |
| chr18-45084576-INS-413 | chr18-45084576-INS-413 |
| chr18-45102379-INS-7114 | chr18-45102380-INS-7114 |
| chr18-45111076-INS-498 | chr18-45111076-INS-498 |
| chr18-45173711-INS-60 | chr18-45173711-INS-60 |
| chr18-45192443-INS-276 | chr18-45192443-INS-276 |
| chr18-45195368-INS-912 | chr18-45195368-INS-912 |
| chr18-45204735-INS-338 | chr18-45204735-INS-338 |
| chr18-45214692-INS-5400 | chr18-45214693-INS-5400 |
| chr18-45249242-INS-222 | chr18-45249271-INS-186 |
| chr18-45261664-INS-930 | chr18-45261681-INS-940 |
| chr18-45300411-INS-145 | chr18-45300411-INS-146 |
| chr18-45331306-INS-55 | chr18-45331306-INS-57 |
| chr18-45336454-INS-94 | chr18-45336465-INS-86 |
| chr18-45381206-INS-705 | chr18-45381206-INS-704 |
| chr18-45382264-INS-6447 | chr18-45382265-INS-6447 |
| chr18-45389820-INS-89 | chr18-45389820-INS-97 |
| chr18-45408704-INS-56 | chr18-45408704-INS-56 |
| chr18-45431619-INS-72 | chr18-45431617-INS-74 |
| chr18-45438294-INS-1592 | chr18-45438295-INS-1593 |
| chr18-45438725-INS-103 | chr18-45438725-INS-103 |
| chr18-45461810-INS-513 | chr18-45461810-INS-475 |
| chr18-45466158-INS-543 | chr18-45466160-INS-553 |
| chr18-45471507-INS-62 | chr18-45471507-INS-60 |
| chr18-45479096-INS-652 | chr18-45479096-INS-652 |
| chr18-45512950-INS-172 | chr18-45512950-INS-174 |
| chr18-45524685-INS-217 | chr18-45524685-INS-217 |
| chr18-45530958-INS-64 | chr18-45530958-INS-62 |
| chr18-45566986-INS-186 | chr18-45566986-INS-186 |
| chr18-45574570-INS-325 | chr18-45574570-INS-325 |
| chr18-45575519-INS-79 | chr18-45575519-INS-63 |
| chr18-45577273-INS-1138 | chr18-45577274-INS-1138 |
| chr18-45579618-INS-1315 | chr18-45579619-INS-1315 |
| chr18-45586347-INS-54 | chr18-45586356-INS-54 |
| chr18-45603229-INS-3697 | chr18-45603230-INS-3721 |
| chr18-45651993-INS-62 | chr18-45651997-INS-68 |
| chr18-45655342-INS-156 | chr18-45655342-INS-156 |
| chr18-45701996-INS-98 | chr18-45701996-INS-98 |
| chr18-45709219-INS-200 | chr18-45709219-INS-200 |
| chr18-45780200-INS-79 | chr18-45780200-INS-99 |
| chr18-45860305-INS-254 | chr18-45860305-INS-256 |
| chr18-45909283-INS-51 | chr18-45909283-INS-71 |
| chr18-45919479-INS-6523 | chr18-45919480-INS-6523 |
| chr18-45919993-INS-460 | chr18-45919993-INS-460 |
| chr18-45934257-INS-185 | chr18-45934257-INS-187 |
| chr18-45972068-INS-761 | chr18-45972068-INS-762 |
| chr18-45990689-INS-205 | chr18-45990691-INS-202 |
| chr18-46003130-INS-232 | chr18-46003124-INS-226 |
| chr18-46018345-INS-122 | chr18-46018345-INS-98 |
| chr18-46027642-INS-155 | chr18-46027642-INS-155 |
| chr18-46053204-INS-66 | chr18-46053221-INS-94 |
| chr18-46087384-INS-6806 | chr18-46087385-INS-6806 |
| chr18-46112307-INS-6638 | chr18-46112308-INS-6637 |
| chr18-46117889-INS-56 | chr18-46117881-INS-52 |

| | |
|---|---|
| chr18-46118869-INS-98 | chr18-46118869-INS-98 |
| chr18-46144390-INS-59 | chr18-46144388-INS-50 |
| chr18-46171113-INS-373 | chr18-46171113-INS-373 |
| chr18-46204873-INS-6827 | chr18-46204874-INS-6833 |
| chr18-46233729-INS-6804 | chr18-46233730-INS-6804 |
| chr18-46235024-INS-88 | chr18-46235024-INS-74 |
| chr18-46268076-INS-222 | chr18-46268076-INS-185 |
| chr18-46273159-INS-220 | chr18-46273159-INS-221 |
| chr18-46289163-INS-186 | chr18-46289164-INS-190 |
| chr18-46290807-INS-215 | chr18-46290807-INS-194 |
| chr18-46337799-INS-56 | chr18-46337799-INS-59 |
| chr18-46349804-INS-64 | chr18-46349804-INS-64 |
| chr18-46352393-INS-7054 | chr18-46352394-INS-7055 |
| chr18-46355158-INS-75 | chr18-46355158-INS-121 |
| chr18-46370482-INS-63 | chr18-46370475-INS-86 |
| chr18-46406070-INS-68 | chr18-46406070-INS-70 |
| chr18-46409629-INS-696 | chr18-46409630-INS-694 |
| chr18-46418444-INS-52 | chr18-46418444-INS-52 |
| chr18-46424943-INS-197 | chr18-46424943-INS-198 |
| chr18-46432125-INS-172 | chr18-46432125-INS-172 |
| chr18-46432993-INS-882 | chr18-46432993-INS-882 |
| chr18-46455072-INS-187 | chr18-46455072-INS-189 |
| chr18-46471222-INS-255 | chr18-46471222-INS-254 |
| chr18-46483381-INS-55 | chr18-46483381-INS-55 |
| chr18-46485133-INS-153 | chr18-46485147-INS-158 |
| chr18-46496881-INS-2124 | chr18-46496881-INS-2118 |
| chr18-46497136-INS-201 | chr18-46497136-INS-201 |
| chr18-46500409-INS-56 | chr18-46500409-INS-53 |
| chr18-46513638-INS-68 | chr18-46513635-INS-71 |
| chr18-46517006-INS-1411 | chr18-46517007-INS-1450 |
| chr18-46518333-INS-375 | chr18-46518334-INS-375 |
| chr18-46535138-INS-4444 | chr18-46535139-INS-4444 |
| chr18-46536590-INS-191 | chr18-46536590-INS-185 |
| chr18-46553018-INS-393 | chr18-46553018-INS-391 |
| chr18-46555825-INS-58 | chr18-46555825-INS-53 |
| chr18-46567415-INS-311 | chr18-46567415-INS-311 |
| chr18-46586363-INS-165 | chr18-46586364-INS-175 |
| chr18-46594534-INS-257 | chr18-46594534-INS-219 |
| chr18-46603819-INS-60 | chr18-46603771-INS-68 |
| chr18-46628523-INS-209 | chr18-46628523-INS-210 |
| chr18-46629675-INS-108 | chr18-46629675-INS-108 |
| chr18-46668959-INS-221 | chr18-46668959-INS-223 |
| chr18-46679423-INS-62 | chr18-46679423-INS-54 |
| chr18-46685320-INS-102 | chr18-46685280-INS-102 |
| chr18-46690624-INS-188 | chr18-46690624-INS-188 |
| chr18-46713702-INS-204 | chr18-46713706-INS-203 |
| chr18-46716685-INS-6399 | chr18-46716686-INS-6399 |
| chr18-46717841-INS-73 | chr18-46717841-INS-73 |
| chr18-46758342-INS-114 | chr18-46758337-INS-72 |
| chr18-46763706-INS-206 | chr18-46763706-INS-207 |
| chr18-46764821-INS-200 | chr18-46764801-INS-206 |
| chr18-46778358-INS-436 | chr18-46778358-INS-438 |
| chr18-46782874-INS-351 | chr18-46782874-INS-351 |
| chr18-46854074-INS-91 | chr18-46854074-INS-101 |
| chr18-46877033-INS-1886 | chr18-46877034-INS-1886 |
| chr18-46897635-INS-188 | chr18-46897635-INS-172 |
| chr18-46901537-INS-63 | chr18-46901521-INS-76 |
| chr18-46932838-INS-84 | chr18-46932838-INS-86 |
| chr18-46935245-INS-99 | chr18-46935245-INS-59 |
| chr18-47003719-INS-195 | chr18-47003719-INS-198 |
| chr18-47025650-INS-65 | chr18-47025650-INS-55 |
| chr18-47028570-INS-853 | chr18-47028570-INS-853 |
| chr18-47035723-INS-169 | chr18-47035723-INS-169 |
| chr18-47038430-INS-56 | chr18-47038430-INS-56 |
| chr18-47068703-INS-198 | chr18-47068703-INS-198 |
| chr18-47071161-INS-122 | chr18-47071161-INS-122 |
| chr18-47076812-INS-1344 | chr18-47076813-INS-1344 |
| chr18-47079740-INS-54 | chr18-47079740-INS-57 |
| chr18-47080676-INS-225 | chr18-47080676-INS-225 |
| chr18-47095526-INS-217 | chr18-47095526-INS-217 |
| chr18-47104744-INS-76 | chr18-47104744-INS-74 |
| chr18-47112011-INS-232 | chr18-47112011-INS-233 |
| chr18-47131129-INS-1089 | chr18-47131130-INS-1089 |
| chr18-47204814-INS-52 | chr18-47204814-INS-50 |
| chr18-47208153-INS-6287 | chr18-47208154-INS-6288 |
| chr18-47238413-INS-222 | chr18-47238413-INS-222 |
| chr18-47261739-INS-53 | chr18-47261738-INS-54 |

| | |
|---|---|
| chr18-47287284-INS-140 | chr18-47287282-INS-139 |
| chr18-47372590-INS-186 | chr18-47372590-INS-186 |
| chr18-47393710-INS-78 | chr18-47393710-INS-78 |
| chr18-47418841-INS-208 | chr18-47418841-INS-208 |
| chr18-47421153-INS-409 | chr18-47421153-INS-409 |
| chr18-47422818-INS-65 | chr18-47422818-INS-65 |
| chr18-47496466-INS-193 | chr18-47496466-INS-192 |
| chr18-47519224-INS-75 | chr18-47519224-INS-75 |
| chr18-47530060-INS-60 | chr18-47530060-INS-52 |
| chr18-47563556-INS-7246 | chr18-47563557-INS-7246 |
| chr18-47566907-INS-66 | chr18-47566907-INS-66 |
| chr18-47573188-INS-63 | chr18-47573188-INS-61 |
| chr18-47587797-INS-276 | chr18-47587797-INS-276 |
| chr18-47612841-INS-66 | chr18-47612841-INS-66 |
| chr18-47631057-INS-109 | chr18-47631057-INS-109 |
| chr18-47654222-INS-60 | chr18-47654222-INS-55 |
| chr18-47671991-INS-5548 | chr18-47671992-INS-5548 |
| chr18-47706376-INS-71 | chr18-47706376-INS-63 |
| chr18-47712262-INS-70 | chr18-47712262-INS-70 |
| chr18-47728961-INS-201 | chr18-47728961-INS-201 |
| chr18-47742948-INS-127 | chr18-47742948-INS-79 |
| chr18-47746723-INS-78 | chr18-47746723-INS-78 |
| chr18-47763871-INS-227 | chr18-47763871-INS-223 |
| chr18-47772607-INS-76 | chr18-47772607-INS-82 |
| chr18-47807125-INS-682 | chr18-47807125-INS-681 |
| chr18-47819649-INS-649 | chr18-47819649-INS-649 |
| chr18-47835682-INS-186 | chr18-47835682-INS-186 |
| chr18-47855230-INS-66 | chr18-47855230-INS-66 |
| chr18-47882342-INS-172 | chr18-47882358-INS-156 |
| chr18-47884895-INS-82 | chr18-47884895-INS-86 |
| chr18-47933819-INS-1319 | chr18-47933820-INS-1319 |
| chr18-47946811-INS-297 | chr18-47946811-INS-297 |
| chr18-47971669-INS-188 | chr18-47971669-INS-200 |
| chr18-47974701-INS-335 | chr18-47974701-INS-338 |
| chr18-47983661-INS-6415 | chr18-47983662-INS-6416 |
| chr18-47987093-INS-55 | chr18-47987093-INS-55 |
| chr18-48012032-INS-477 | chr18-48012032-INS-477 |
| chr18-48065033-INS-1703 | chr18-48065034-INS-1703 |
| chr18-48072684-INS-64 | chr18-48072686-INS-66 |
| chr18-48073517-INS-328 | chr18-48073517-INS-328 |
| chr18-48076918-INS-6747 | chr18-48076919-INS-6747 |
| chr18-48092706-INS-7919 | chr18-48092707-INS-7919 |
| chr18-48119958-INS-166 | chr18-48119958-INS-166 |
| chr18-48120170-INS-1808 | chr18-48120171-INS-1806 |
| chr18-48142540-INS-52 | chr18-48142540-INS-56 |
| chr18-48146030-INS-2552 | chr18-48146031-INS-2552 |
| chr18-48161503-INS-986 | chr18-48161503-INS-986 |
| chr18-48171710-INS-6663 | chr18-48171711-INS-6664 |
| chr18-48185393-INS-68 | chr18-48185393-INS-60 |
| chr18-48186551-INS-90 | chr18-48186556-INS-128 |
| chr18-48195127-INS-413 | chr18-48195127-INS-413 |
| chr18-48206201-INS-349 | chr18-48206201-INS-302 |
| chr18-48222765-INS-128 | chr18-48222765-INS-124 |
| chr18-48250196-INS-117 | chr18-48250196-INS-117 |
| chr18-48257071-INS-52 | chr18-48257071-INS-56 |
| chr18-48260443-INS-283 | chr18-48260443-INS-278 |
| chr18-48267008-INS-140 | chr18-48267008-INS-140 |
| chr18-48270043-INS-60 | chr18-48270043-INS-64 |
| chr18-48275295-INS-52 | chr18-48275295-INS-52 |
| chr18-48288470-INS-79 | chr18-48288470-INS-79 |
| chr18-48292073-INS-90 | chr18-48292073-INS-90 |
| chr18-48306945-INS-80 | chr18-48306945-INS-104 |
| chr18-48312513-INS-399 | chr18-48312513-INS-399 |
| chr18-48324701-INS-70 | chr18-48324701-INS-68 |
| chr18-48359983-INS-53 | chr18-48359978-INS-63 |
| chr18-48405051-INS-344 | chr18-48405051-INS-344 |
| chr18-48405132-INS-6904 | chr18-48405133-INS-6904 |
| chr18-48411880-INS-76 | chr18-48411880-INS-114 |
| chr18-48420972-INS-85 | chr18-48420972-INS-85 |
| chr18-48470650-INS-1390 | chr18-48470651-INS-1390 |
| chr18-48493487-INS-855 | chr18-48493487-INS-858 |
| chr18-48504935-INS-51 | chr18-48504935-INS-51 |
| chr18-48513177-INS-889 | chr18-48513177-INS-893 |
| chr18-48518106-INS-60 | chr18-48518106-INS-60 |
| chr18-48530149-INS-76 | chr18-48530146-INS-93 |
| chr18-48544300-INS-51 | chr18-48544300-INS-50 |
| chr18-48554624-INS-56 | chr18-48554624-INS-56 |

| | |
|---|---|
| chr18-48568782-INS-84 | chr18-48568782-INS-84 |
| chr18-48570110-INS-82 | chr18-48570110-INS-91 |
| chr18-48582978-INS-121 | chr18-48582978-INS-121 |
| chr18-48585016-INS-56 | chr18-48585013-INS-105 |
| chr18-48587889-INS-680 | chr18-48587889-INS-680 |
| chr18-48596819-INS-59 | chr18-48596819-INS-74 |
| chr18-48602187-INS-785 | chr18-48602187-INS-786 |
| chr18-48618190-INS-582 | chr18-48618190-INS-582 |
| chr18-48624775-INS-434 | chr18-48624775-INS-434 |
| chr18-48634295-INS-659 | chr18-48634295-INS-659 |
| chr18-48680565-INS-6250 | chr18-48680566-INS-6253 |
| chr18-48693205-INS-522 | chr18-48693205-INS-522 |
| chr18-48698130-INS-837 | chr18-48698130-INS-838 |
| chr18-48734487-INS-65 | chr18-48734467-INS-61 |
| chr18-48739285-INS-523 | chr18-48739285-INS-523 |
| chr18-48744490-INS-51 | chr18-48744490-INS-51 |
| chr18-48746329-INS-87 | chr18-48746330-INS-87 |
| chr18-48781830-INS-582 | chr18-48781830-INS-583 |
| chr18-48798437-INS-92 | chr18-48798437-INS-54 |
| chr18-48909961-INS-109 | chr18-48909961-INS-144 |
| chr18-48911487-INS-216 | chr18-48911487-INS-217 |
| chr18-48917215-INS-6197 | chr18-48917216-INS-6198 |
| chr18-48919879-INS-319 | chr18-48919875-INS-322 |
| chr18-48950771-INS-131 | chr18-48950774-INS-92 |
| chr18-48951419-INS-200 | chr18-48951420-INS-200 |
| chr18-48976059-INS-572 | chr18-48976059-INS-564 |
| chr18-49067081-INS-136 | chr18-49067075-INS-139 |
| chr18-49098645-INS-169 | chr18-49098650-INS-170 |
| chr18-49103885-INS-2683 | chr18-49103886-INS-2698 |
| chr18-49105077-INS-616 | chr18-49105077-INS-616 |
| chr18-49106241-INS-649 | chr18-49106241-INS-657 |
| chr18-49107873-INS-1232 | chr18-49107874-INS-1232 |
| chr18-49117105-INS-97 | chr18-49117105-INS-96 |
| chr18-49120625-INS-70 | chr18-49120627-INS-56 |
| chr18-49146835-INS-64 | chr18-49146835-INS-64 |
| chr18-49175496-INS-6485 | chr18-49175497-INS-6485 |
| chr18-49192518-INS-2537 | chr18-49192519-INS-2537 |
| chr18-49209405-INS-54 | chr18-49209405-INS-72 |
| chr18-49215296-INS-59 | chr18-49215296-INS-57 |
| chr18-49215855-INS-5642 | chr18-49215856-INS-5642 |
| chr18-49244227-INS-2172 | chr18-49244228-INS-2172 |
| chr18-49271477-INS-7341 | chr18-49271478-INS-7342 |
| chr18-49277051-INS-55 | chr18-49277051-INS-63 |
| chr18-49293407-INS-54 | chr18-49293403-INS-62 |
| chr18-49305704-INS-85 | chr18-49305707-INS-80 |
| chr18-49309229-INS-98 | chr18-49309229-INS-104 |
| chr18-49316088-INS-204 | chr18-49316080-INS-164 |
| chr18-49319026-INS-1010 | chr18-49319046-INS-992 |
| chr18-49326375-INS-627 | chr18-49326375-INS-627 |
| chr18-49327253-INS-548 | chr18-49327253-INS-548 |
| chr18-49347862-INS-98 | chr18-49347862-INS-108 |
| chr18-49398575-INS-102 | chr18-49398575-INS-103 |
| chr18-49403142-INS-7268 | chr18-49403143-INS-7268 |
| chr18-49413562-INS-2439 | chr18-49413571-INS-2435 |
| chr18-49428367-INS-6223 | chr18-49428368-INS-6223 |
| chr18-49430333-INS-52 | chr18-49430332-INS-72 |
| chr18-49431803-INS-2725 | chr18-49431804-INS-2728 |
| chr18-49432555-INS-363 | chr18-49432555-INS-363 |
| chr18-49434640-INS-52 | chr18-49434640-INS-89 |
| chr18-49445712-INS-1287 | chr18-49445713-INS-1287 |
| chr18-49458901-INS-57 | chr18-49458890-INS-66 |
| chr18-49462406-INS-536 | chr18-49462406-INS-556 |
| chr18-49472072-INS-545 | chr18-49472072-INS-545 |
| chr18-49493924-INS-55 | chr18-49493921-INS-73 |
| chr18-49503807-INS-6621 | chr18-49503808-INS-6620 |
| chr18-49518075-INS-117 | chr18-49518075-INS-127 |
| chr18-49564309-INS-697 | chr18-49564309-INS-697 |
| chr18-49568866-INS-59 | chr18-49568866-INS-59 |
| chr18-49584156-INS-274 | chr18-49584157-INS-274 |
| chr18-49613838-INS-103 | chr18-49613838-INS-103 |
| chr18-49616232-INS-56 | chr18-49616232-INS-56 |
| chr18-49618892-INS-217 | chr18-49618892-INS-217 |
| chr18-49627724-INS-625 | chr18-49627724-INS-631 |
| chr18-49641204-INS-151 | chr18-49641204-INS-151 |
| chr18-49653138-INS-196 | chr18-49653138-INS-196 |
| chr18-49672523-INS-6375 | chr18-49672524-INS-6375 |
| chr18-49770797-INS-790 | chr18-49770797-INS-790 |

| | |
|---|---|
| chr18-49775973-INS-79 | chr18-49775964-INS-53 |
| chr18-49785880-INS-466 | chr18-49785880-INS-466 |
| chr18-49788258-INS-6919 | chr18-49788259-INS-6918 |
| chr18-49876724-INS-259 | chr18-49876724-INS-272 |
| chr18-49884313-INS-60 | chr18-49884313-INS-58 |
| chr18-49893893-INS-202 | chr18-49893893-INS-202 |
| chr18-49916730-INS-181 | chr18-49916730-INS-174 |
| chr18-49925095-INS-201 | chr18-49925095-INS-201 |
| chr18-49927494-INS-109 | chr18-49927494-INS-107 |
| chr18-49940969-INS-81 | chr18-49940981-INS-61 |
| chr18-49962478-INS-66 | chr18-49962478-INS-70 |
| chr18-50050520-INS-206 | chr18-50050520-INS-206 |
| chr18-50088552-INS-480 | chr18-50088553-INS-480 |
| chr18-50106650-INS-11266 | chr18-50106651-INS-11266 |
| chr18-50106651-INS-440 | chr18-50106651-INS-440 |
| chr18-50112073-INS-168 | chr18-50112073-INS-168 |
| chr18-50149788-INS-198 | chr18-50149788-INS-198 |
| chr18-50157647-INS-276 | chr18-50157647-INS-276 |
| chr18-50199668-INS-2854 | chr18-50199669-INS-2854 |
| chr18-50208277-INS-773 | chr18-50208277-INS-773 |
| chr18-50229206-INS-72 | chr18-50229206-INS-55 |
| chr18-50232054-INS-189 | chr18-50232054-INS-187 |
| chr18-50255367-INS-52 | chr18-50255347-INS-98 |
| chr18-50268121-INS-218 | chr18-50268121-INS-218 |
| chr18-50273961-INS-81 | chr18-50273969-INS-82 |
| chr18-50305795-INS-205 | chr18-50305795-INS-205 |
| chr18-50314690-INS-240 | chr18-50314690-INS-240 |
| chr18-50316578-INS-529 | chr18-50316578-INS-533 |
| chr18-50333306-INS-6449 | chr18-50333307-INS-6450 |
| chr18-50339779-INS-4936 | chr18-50339780-INS-4937 |
| chr18-50353852-INS-365 | chr18-50353852-INS-365 |
| chr18-50373242-INS-407 | chr18-50373242-INS-407 |
| chr18-50385346-INS-1390 | chr18-50385347-INS-1390 |
| chr18-50386506-INS-208 | chr18-50386511-INS-208 |
| chr18-50408314-INS-222 | chr18-50408314-INS-222 |
| chr18-50433117-INS-519 | chr18-50433117-INS-519 |
| chr18-50434383-INS-615 | chr18-50434354-INS-614 |
| chr18-50477419-INS-205 | chr18-50477419-INS-205 |
| chr18-50510183-INS-472 | chr18-50510183-INS-472 |
| chr18-50515730-INS-6626 | chr18-50515731-INS-6629 |
| chr18-50516409-INS-399 | chr18-50516410-INS-399 |
| chr18-50520866-INS-534 | chr18-50520866-INS-534 |
| chr18-50543417-INS-670 | chr18-50543417-INS-672 |
| chr18-50611544-INS-576 | chr18-50611544-INS-576 |
| chr18-50613040-INS-2651 | chr18-50613041-INS-2612 |
| chr18-50624386-INS-52 | chr18-50624377-INS-50 |
| chr18-50627981-INS-6654 | chr18-50627982-INS-6655 |
| chr18-50641631-INS-6713 | chr18-50641632-INS-6713 |
| chr18-50660114-INS-202 | chr18-50660114-INS-203 |
| chr18-50680024-INS-113 | chr18-50680025-INS-113 |
| chr18-50686245-INS-124 | chr18-50686232-INS-100 |
| chr18-50691187-INS-65 | chr18-50691209-INS-50 |
| chr18-50716194-INS-7098 | chr18-50716195-INS-7098 |
| chr18-50747507-INS-6432 | chr18-50747508-INS-6433 |
| chr18-50752412-INS-782 | chr18-50752412-INS-782 |
| chr18-50802068-INS-108 | chr18-50802068-INS-110 |
| chr18-50809516-INS-5188 | chr18-50809517-INS-5188 |
| chr18-50817397-INS-98 | chr18-50817437-INS-60 |
| chr18-50848872-INS-6886 | chr18-50848873-INS-6887 |
| chr18-50851168-INS-918 | chr18-50851168-INS-919 |
| chr18-50861008-INS-101 | chr18-50861008-INS-121 |
| chr18-50888806-INS-419 | chr18-50888806-INS-419 |
| chr18-50961969-INS-772 | chr18-50961969-INS-772 |
| chr18-50998780-INS-63 | chr18-50998781-INS-63 |
| chr18-51003699-INS-154 | chr18-51003699-INS-154 |
| chr18-51018562-INS-266 | chr18-51018562-INS-266 |
| chr18-51033735-INS-1389 | chr18-51033736-INS-1389 |
| chr18-51072062-INS-64 | chr18-51072062-INS-72 |
| chr18-51074172-INS-495 | chr18-51074172-INS-495 |
| chr18-51078459-INS-59 | chr18-51078455-INS-68 |
| chr18-51189122-INS-86 | chr18-51189122-INS-88 |
| chr18-51203654-INS-66 | chr18-51203654-INS-66 |
| chr18-51221076-INS-101 | chr18-51221074-INS-107 |
| chr18-51292523-INS-60 | chr18-51292534-INS-73 |
| chr18-51303940-INS-672 | chr18-51303940-INS-677 |
| chr18-51306161-INS-54 | chr18-51306162-INS-68 |
| chr18-51308349-INS-102 | chr18-51308349-INS-123 |

| | |
|---|---|
| chr18-51329124-INS-273 | chr18-51329124-INS-273 |
| chr18-51337012-INS-164 | chr18-51337012-INS-119 |
| chr18-51355746-INS-56 | chr18-51355746-INS-56 |
| chr18-51375496-INS-55 | chr18-51375475-INS-72 |
| chr18-51376449-INS-160 | chr18-51376464-INS-111 |
| chr18-51394376-INS-68 | chr18-51394376-INS-68 |
| chr18-51409128-INS-390 | chr18-51409128-INS-390 |
| chr18-51425401-INS-71 | chr18-51425401-INS-75 |
| chr18-51497132-INS-58 | chr18-51497125-INS-68 |
| chr18-51517240-INS-390 | chr18-51517240-INS-390 |
| chr18-51539667-INS-278 | chr18-51539667-INS-276 |
| chr18-51544285-INS-514 | chr18-51544285-INS-539 |
| chr18-51548049-INS-54 | chr18-51548049-INS-54 |
| chr18-51549361-INS-63 | chr18-51549361-INS-63 |
| chr18-51587682-INS-767 | chr18-51587682-INS-766 |
| chr18-51599124-INS-184 | chr18-51599120-INS-152 |
| chr18-51601823-INS-2397 | chr18-51601824-INS-2397 |
| chr18-51611461-INS-429 | chr18-51611461-INS-429 |
| chr18-51615648-INS-3843 | chr18-51615649-INS-3843 |
| chr18-51624611-INS-714 | chr18-51624611-INS-714 |
| chr18-51641876-INS-66 | chr18-51641876-INS-66 |
| chr18-51642207-INS-6196 | chr18-51642208-INS-6201 |
| chr18-51668435-INS-80 | chr18-51668435-INS-74 |
| chr18-51691885-INS-121 | chr18-51691885-INS-121 |
| chr18-51706597-INS-99 | chr18-51706597-INS-72 |
| chr18-51709050-INS-834 | chr18-51709050-INS-834 |
| chr18-51732783-INS-215 | chr18-51732784-INS-215 |
| chr18-51737667-INS-52 | chr18-51737666-INS-51 |
| chr18-51743534-INS-56 | chr18-51743534-INS-58 |
| chr18-51751280-INS-279 | chr18-51751280-INS-279 |
| chr18-51752421-INS-1888 | chr18-51752422-INS-1888 |
| chr18-51755521-INS-666 | chr18-51755521-INS-666 |
| chr18-51757141-INS-1886 | chr18-51757142-INS-1886 |
| chr18-51802187-INS-78 | chr18-51802187-INS-80 |
| chr18-51811383-INS-68 | chr18-51811383-INS-56 |
| chr18-51814890-INS-8021 | chr18-51814891-INS-8022 |
| chr18-51843980-INS-214 | chr18-51843980-INS-210 |
| chr18-51850217-INS-1893 | chr18-51850218-INS-1893 |
| chr18-51858389-INS-1958 | chr18-51858390-INS-1958 |
| chr18-51861790-INS-53 | chr18-51861790-INS-56 |
| chr18-51889393-INS-146 | chr18-51889393-INS-118 |
| chr18-51894437-INS-54 | chr18-51894437-INS-56 |
| chr18-51913277-INS-90 | chr18-51913277-INS-86 |
| chr18-51922003-INS-69 | chr18-51922025-INS-77 |
| chr18-51925073-INS-6368 | chr18-51925074-INS-6368 |
| chr18-51959113-INS-884 | chr18-51959113-INS-884 |
| chr18-51966845-INS-4248 | chr18-51966846-INS-4249 |
| chr18-51973722-INS-66 | chr18-51973722-INS-66 |
| chr18-51975200-INS-269 | chr18-51975200-INS-268 |
| chr18-51989631-INS-2886 | chr18-51989632-INS-2886 |
| chr18-51993806-INS-120 | chr18-51993807-INS-125 |
| chr18-51997105-INS-6618 | chr18-51997106-INS-6618 |
| chr18-52026030-INS-189 | chr18-52026030-INS-198 |
| chr18-52048558-INS-4708 | chr18-52048559-INS-4708 |
| chr18-52059096-INS-609 | chr18-52059096-INS-609 |
| chr18-52092366-INS-2488 | chr18-52092367-INS-2490 |
| chr18-52095385-INS-204 | chr18-52095385-INS-231 |
| chr18-52103451-INS-565 | chr18-52103451-INS-555 |
| chr18-52131286-INS-113 | chr18-52131287-INS-113 |
| chr18-52133155-INS-55 | chr18-52133155-INS-59 |
| chr18-52151240-INS-60 | chr18-52151240-INS-68 |
| chr18-52165992-INS-71 | chr18-52165992-INS-65 |
| chr18-52188731-INS-983 | chr18-52188744-INS-1003 |
| chr18-52242093-INS-129 | chr18-52242093-INS-132 |
| chr18-52254668-INS-1486 | chr18-52254669-INS-1486 |
| chr18-52255705-INS-5185 | chr18-52255706-INS-5185 |
| chr18-52273082-INS-97 | chr18-52273072-INS-129 |
| chr18-52280371-INS-52 | chr18-52280371-INS-52 |
| chr18-52282904-INS-55 | chr18-52282904-INS-55 |
| chr18-52288694-INS-123 | chr18-52288691-INS-119 |
| chr18-52289803-INS-3003 | chr18-52289804-INS-3003 |
| chr18-52302693-INS-637 | chr18-52302693-INS-637 |
| chr18-52305394-INS-4049 | chr18-52305395-INS-4049 |
| chr18-52306182-INS-52 | chr18-52306180-INS-60 |
| chr18-52307362-INS-2386 | chr18-52307363-INS-2386 |
| chr18-52310266-INS-184 | chr18-52310266-INS-184 |
| chr18-52311066-INS-594 | chr18-52311064-INS-593 |

| | |
|---|---|
| chr18-52326934-INS-2946 | chr18-52326935-INS-2946 |
| chr18-52362833-INS-438 | chr18-52362833-INS-438 |
| chr18-52384583-INS-522 | chr18-52384583-INS-522 |
| chr18-52412925-INS-231 | chr18-52412925-INS-231 |
| chr18-52418096-INS-67 | chr18-52418096-INS-67 |
| chr18-52427923-INS-1459 | chr18-52427924-INS-1459 |
| chr18-52440380-INS-564 | chr18-52440380-INS-563 |
| chr18-52452140-INS-114 | chr18-52452140-INS-116 |
| chr18-52462438-INS-388 | chr18-52462434-INS-367 |
| chr18-52492069-INS-2506 | chr18-52492070-INS-2507 |
| chr18-52499162-INS-64 | chr18-52499162-INS-64 |
| chr18-52543727-INS-105 | chr18-52543727-INS-105 |
| chr18-52572755-INS-349 | chr18-52572755-INS-349 |
| chr18-52590780-INS-56 | chr18-52590780-INS-58 |
| chr18-52591195-INS-186 | chr18-52591195-INS-186 |
| chr18-52596055-INS-551 | chr18-52596055-INS-554 |
| chr18-52625231-INS-74 | chr18-52625231-INS-76 |
| chr18-52630137-INS-216 | chr18-52630137-INS-231 |
| chr18-52632647-INS-61 | chr18-52632647-INS-61 |
| chr18-52642009-INS-237 | chr18-52642009-INS-240 |
| chr18-52642785-INS-379 | chr18-52642785-INS-378 |
| chr18-52660693-INS-76 | chr18-52660682-INS-116 |
| chr18-52689791-INS-7081 | chr18-52689792-INS-7081 |
| chr18-52705850-INS-211 | chr18-52705833-INS-196 |
| chr18-52709274-INS-581 | chr18-52709274-INS-581 |
| chr18-52723038-INS-89 | chr18-52723006-INS-86 |
| chr18-52729956-INS-179 | chr18-52729956-INS-179 |
| chr18-52739210-INS-246 | chr18-52739219-INS-256 |
| chr18-52745427-INS-66 | chr18-52745427-INS-66 |
| chr18-52765633-INS-72 | chr18-52765633-INS-88 |
| chr18-52811435-INS-418 | chr18-52811437-INS-419 |
| chr18-52820593-INS-86 | chr18-52820593-INS-86 |
| chr18-52856022-INS-112 | chr18-52856020-INS-120 |
| chr18-52856442-INS-700 | chr18-52856442-INS-700 |
| chr18-52864549-INS-77 | chr18-52864551-INS-54 |
| chr18-52922710-INS-151 | chr18-52922710-INS-195 |
| chr18-52950367-INS-70 | chr18-52950367-INS-90 |
| chr18-52953703-INS-73 | chr18-52953703-INS-72 |
| chr18-52955714-INS-52 | chr18-52955714-INS-50 |
| chr18-52966956-INS-249 | chr18-52966956-INS-249 |
| chr18-53025848-INS-2639 | chr18-53025849-INS-2645 |
| chr18-53052183-INS-52 | chr18-53052183-INS-74 |
| chr18-53059726-INS-268 | chr18-53059727-INS-232 |
| chr18-53069754-INS-126 | chr18-53069754-INS-126 |
| chr18-53074596-INS-54 | chr18-53074596-INS-54 |
| chr18-53077571-INS-60 | chr18-53077571-INS-62 |
| chr18-53081621-INS-635 | chr18-53081621-INS-635 |
| chr18-53092998-INS-58 | chr18-53092998-INS-58 |
| chr18-53114332-INS-345 | chr18-53114334-INS-360 |
| chr18-53136582-INS-953 | chr18-53136582-INS-955 |
| chr18-53189508-INS-359 | chr18-53189523-INS-360 |
| chr18-53218014-INS-58 | chr18-53218060-INS-55 |
| chr18-53224375-INS-56 | chr18-53224373-INS-56 |
| chr18-53226873-INS-58 | chr18-53226873-INS-60 |
| chr18-53235150-INS-6354 | chr18-53235151-INS-6355 |
| chr18-53251155-INS-72 | chr18-53251155-INS-75 |
| chr18-53252764-INS-52 | chr18-53252764-INS-50 |
| chr18-53276544-INS-823 | chr18-53276544-INS-822 |
| chr18-53279743-INS-55 | chr18-53279743-INS-55 |
| chr18-53284260-INS-70 | chr18-53284260-INS-54 |
| chr18-53298508-INS-2889 | chr18-53298509-INS-2889 |
| chr18-53312601-INS-59 | chr18-53312601-INS-59 |
| chr18-53318869-INS-205 | chr18-53318869-INS-206 |
| chr18-53332485-INS-72 | chr18-53332486-INS-68 |
| chr18-53375688-INS-80 | chr18-53375688-INS-50 |
| chr18-53381943-INS-180 | chr18-53381943-INS-180 |
| chr18-53385107-INS-415 | chr18-53385107-INS-415 |
| chr18-53429428-INS-54 | chr18-53429410-INS-50 |
| chr18-53435860-INS-401 | chr18-53435860-INS-401 |
| chr18-53451714-INS-122 | chr18-53451714-INS-120 |
| chr18-53506237-INS-608 | chr18-53506237-INS-608 |
| chr18-53522829-INS-396 | chr18-53522829-INS-396 |
| chr18-53536060-INS-6412 | chr18-53536061-INS-6412 |
| chr18-53554781-INS-75 | chr18-53554781-INS-54 |
| chr18-53583446-INS-54 | chr18-53583446-INS-60 |
| chr18-53588118-INS-151 | chr18-53588118-INS-151 |
| chr18-53596744-INS-52 | chr18-53596744-INS-54 |

| | |
|---|---|
| chr18-53618328-INS-139 | chr18-53618328-INS-139 |
| chr18-53620615-INS-229 | chr18-53620615-INS-208 |
| chr18-53645163-INS-176 | chr18-53645163-INS-185 |
| chr18-53656493-INS-66 | chr18-53656521-INS-54 |
| chr18-53663496-INS-59 | chr18-53663492-INS-93 |
| chr18-53667633-INS-4604 | chr18-53667634-INS-4604 |
| chr18-53674368-INS-110 | chr18-53674368-INS-110 |
| chr18-53676758-INS-60 | chr18-53676740-INS-54 |
| chr18-53684514-INS-67 | chr18-53684521-INS-68 |
| chr18-53700490-INS-54 | chr18-53700496-INS-51 |
| chr18-53732733-INS-109 | chr18-53732733-INS-109 |
| chr18-53735989-INS-180 | chr18-53735989-INS-165 |
| chr18-53747881-INS-942 | chr18-53747880-INS-947 |
| chr18-53780272-INS-191 | chr18-53780275-INS-204 |
| chr18-53785631-INS-73 | chr18-53785582-INS-102 |
| chr18-53795439-INS-57 | chr18-53795439-INS-70 |
| chr18-53796946-INS-68 | chr18-53796946-INS-71 |
| chr18-53797865-INS-2724 | chr18-53797866-INS-2724 |
| chr18-53810336-INS-100 | chr18-53810337-INS-78 |
| chr18-53813846-INS-452 | chr18-53813846-INS-451 |
| chr18-53848862-INS-206 | chr18-53848862-INS-206 |
| chr18-53856120-INS-92 | chr18-53856120-INS-92 |
| chr18-53858621-INS-74 | chr18-53858621-INS-72 |
| chr18-53868272-INS-216 | chr18-53868272-INS-216 |
| chr18-53879347-INS-54 | chr18-53879331-INS-50 |
| chr18-53881848-INS-683 | chr18-53881848-INS-683 |
| chr18-53885271-INS-54 | chr18-53885271-INS-54 |
| chr18-53901852-INS-184 | chr18-53901852-INS-184 |
| chr18-53926518-INS-74 | chr18-53926517-INS-92 |
| chr18-53941531-INS-57 | chr18-53941531-INS-57 |
| chr18-53943168-INS-58 | chr18-53943168-INS-58 |
| chr18-54022315-INS-105 | chr18-54022315-INS-60 |
| chr18-54044604-INS-6469 | chr18-54044605-INS-6469 |
| chr18-54050300-INS-100 | chr18-54050347-INS-60 |
| chr18-54058455-INS-52 | chr18-54058455-INS-66 |
| chr18-54091394-INS-183 | chr18-54091394-INS-181 |
| chr18-54098660-INS-56 | chr18-54098660-INS-56 |
| chr18-54102812-INS-284 | chr18-54102812-INS-284 |
| chr18-54104869-INS-374 | chr18-54104869-INS-374 |
| chr18-54109031-INS-178 | chr18-54109031-INS-178 |
| chr18-54110856-INS-5313 | chr18-54110857-INS-5313 |
| chr18-54122775-INS-52 | chr18-54122775-INS-52 |
| chr18-54170182-INS-60 | chr18-54170182-INS-62 |
| chr18-54178743-INS-149 | chr18-54178769-INS-157 |
| chr18-54202538-INS-2800 | chr18-54202539-INS-2800 |
| chr18-54214384-INS-169 | chr18-54214408-INS-151 |
| chr18-54215944-INS-66 | chr18-54215944-INS-67 |
| chr18-54267432-INS-105 | chr18-54267432-INS-105 |
| chr18-54282319-INS-54 | chr18-54282319-INS-54 |
| chr18-54320946-INS-6987 | chr18-54320947-INS-6987 |
| chr18-54354824-INS-200 | chr18-54354824-INS-200 |
| chr18-54382523-INS-68 | chr18-54382523-INS-68 |
| chr18-54456342-INS-150 | chr18-54456342-INS-150 |
| chr18-54480019-INS-328 | chr18-54480019-INS-328 |
| chr18-54512943-INS-3059 | chr18-54512944-INS-3061 |
| chr18-54528381-INS-416 | chr18-54528381-INS-416 |
| chr18-54531075-INS-272 | chr18-54531075-INS-272 |
| chr18-54533938-INS-1011 | chr18-54533939-INS-1009 |
| chr18-54541985-INS-51 | chr18-54541985-INS-51 |
| chr18-54550346-INS-2260 | chr18-54550347-INS-2260 |
| chr18-54555169-INS-3073 | chr18-54555170-INS-3073 |
| chr18-54568590-INS-6628 | chr18-54568591-INS-6628 |
| chr18-54592196-INS-56 | chr18-54592196-INS-62 |
| chr18-54616306-INS-449 | chr18-54616306-INS-449 |
| chr18-54622507-INS-85 | chr18-54622507-INS-75 |
| chr18-54660396-INS-519 | chr18-54660396-INS-519 |
| chr18-54700112-INS-290 | chr18-54700113-INS-288 |
| chr18-54747780-INS-2977 | chr18-54747781-INS-2977 |
| chr18-54751279-INS-54 | chr18-54751279-INS-50 |
| chr18-54752693-INS-654 | chr18-54752693-INS-682 |
| chr18-54797809-INS-498 | chr18-54797809-INS-498 |
| chr18-54800018-INS-100 | chr18-54800018-INS-80 |
| chr18-54818508-INS-78 | chr18-54818508-INS-78 |
| chr18-54819918-INS-68 | chr18-54819929-INS-52 |
| chr18-54851043-INS-6870 | chr18-54851044-INS-6871 |
| chr18-54886138-INS-66 | chr18-54886138-INS-66 |
| chr18-54898927-INS-169 | chr18-54898929-INS-189 |

| | |
|---|---|
| chr18-54902539-INS-303 | chr18-54902539-INS-298 |
| chr18-54910924-INS-102 | chr18-54910947-INS-60 |
| chr18-54919897-INS-55 | chr18-54919897-INS-50 |
| chr18-54924620-INS-2847 | chr18-54924621-INS-2847 |
| chr18-54981308-INS-255 | chr18-54981308-INS-256 |
| chr18-54984081-INS-282 | chr18-54984071-INS-323 |
| chr18-54994372-INS-50 | chr18-54994355-INS-52 |
| chr18-55045871-INS-68 | chr18-55045862-INS-92 |
| chr18-55063931-INS-70 | chr18-55063931-INS-60 |
| chr18-55073056-INS-78 | chr18-55073056-INS-64 |
| chr18-55076711-INS-76 | chr18-55076711-INS-60 |
| chr18-55115471-INS-72 | chr18-55115471-INS-68 |
| chr18-55143995-INS-105 | chr18-55143995-INS-85 |
| chr18-55152821-INS-1090 | chr18-55152822-INS-1090 |
| chr18-55168275-INS-203 | chr18-55168275-INS-203 |
| chr18-55185256-INS-447 | chr18-55185256-INS-447 |
| chr18-55185948-INS-180 | chr18-55185945-INS-168 |
| chr18-55194031-INS-111 | chr18-55194073-INS-70 |
| chr18-55204498-INS-65 | chr18-55204498-INS-53 |
| chr18-55264101-INS-62 | chr18-55264103-INS-66 |
| chr18-55264753-INS-87 | chr18-55264753-INS-96 |
| chr18-55291422-INS-70 | chr18-55291426-INS-58 |
| chr18-55308740-INS-295 | chr18-55308740-INS-299 |
| chr18-55312392-INS-362 | chr18-55312392-INS-352 |
| chr18-55318500-INS-80 | chr18-55318500-INS-80 |
| chr18-55325528-INS-59 | chr18-55325528-INS-51 |
| chr18-55348180-INS-328 | chr18-55348180-INS-328 |
| chr18-55364736-INS-117 | chr18-55364761-INS-129 |
| chr18-55366561-INS-50 | chr18-55366563-INS-56 |
| chr18-55373256-INS-474 | chr18-55373256-INS-486 |
| chr18-55388667-INS-92 | chr18-55388667-INS-76 |
| chr18-55397628-INS-106 | chr18-55397628-INS-110 |
| chr18-55401011-INS-135 | chr18-55401011-INS-141 |
| chr18-55409896-INS-132 | chr18-55409896-INS-135 |
| chr18-55430005-INS-331 | chr18-55430005-INS-331 |
| chr18-55439270-INS-75 | chr18-55439270-INS-75 |
| chr18-55442204-INS-141 | chr18-55442197-INS-125 |
| chr18-55487446-INS-119 | chr18-55487419-INS-124 |
| chr18-55490210-INS-83 | chr18-55490207-INS-85 |
| chr18-55496127-INS-51 | chr18-55496127-INS-70 |
| chr18-55517578-INS-58 | chr18-55517578-INS-58 |
| chr18-55533035-INS-119 | chr18-55533035-INS-119 |
| chr18-55569975-INS-59 | chr18-55569944-INS-53 |
| chr18-55571412-INS-90 | chr18-55571412-INS-126 |
| chr18-55572808-INS-87 | chr18-55572808-INS-88 |
| chr18-55593528-INS-630 | chr18-55593528-INS-630 |
| chr18-55601197-INS-428 | chr18-55601183-INS-417 |
| chr18-55609438-INS-64 | chr18-55609438-INS-71 |
| chr18-55638189-INS-113 | chr18-55638189-INS-97 |
| chr18-55643566-INS-76 | chr18-55643566-INS-72 |
| chr18-55656145-INS-6389 | chr18-55656146-INS-6385 |
| chr18-55729751-INS-1018 | chr18-55729752-INS-1019 |
| chr18-55760572-INS-52 | chr18-55760572-INS-54 |
| chr18-55766358-INS-62 | chr18-55766358-INS-70 |
| chr18-55768039-INS-322 | chr18-55768039-INS-322 |
| chr18-55806706-INS-192 | chr18-55806706-INS-192 |
| chr18-55821165-INS-3217 | chr18-55821166-INS-3216 |
| chr18-55829906-INS-105 | chr18-55829906-INS-105 |
| chr18-55835327-INS-82 | chr18-55835327-INS-82 |
| chr18-55842239-INS-6430 | chr18-55842240-INS-6430 |
| chr18-55848498-INS-6441 | chr18-55848499-INS-6441 |
| chr18-55851111-INS-775 | chr18-55851111-INS-776 |
| chr18-55856998-INS-110 | chr18-55857005-INS-110 |
| chr18-55897472-INS-7139 | chr18-55897473-INS-7139 |
| chr18-55931090-INS-2172 | chr18-55931090-INS-2165 |
| chr18-55944415-INS-85 | chr18-55944415-INS-100 |
| chr18-55952454-INS-216 | chr18-55952452-INS-256 |
| chr18-55976854-INS-52 | chr18-55976854-INS-52 |
| chr18-55988232-INS-56 | chr18-55988228-INS-50 |
| chr18-55999545-INS-106 | chr18-55999537-INS-76 |
| chr18-56012757-INS-6436 | chr18-56012758-INS-6435 |
| chr18-56014256-INS-662 | chr18-56014256-INS-662 |
| chr18-56021101-INS-70 | chr18-56021099-INS-72 |
| chr18-56026664-INS-504 | chr18-56026664-INS-516 |
| chr18-56030158-INS-400 | chr18-56030158-INS-400 |
| chr18-56038844-INS-417 | chr18-56038844-INS-418 |
| chr18-56111185-INS-68 | chr18-56111185-INS-76 |

| | |
|---|---|
| chr18-56130690-INS-5321 | chr18-56130691-INS-5320 |
| chr18-56140441-INS-118 | chr18-56140441-INS-119 |
| chr18-56161925-INS-318 | chr18-56161925-INS-318 |
| chr18-56171522-INS-1868 | chr18-56171523-INS-1868 |
| chr18-56186556-INS-6633 | chr18-56186557-INS-6633 |
| chr18-56190061-INS-321 | chr18-56190061-INS-321 |
| chr18-56214015-INS-2470 | chr18-56214016-INS-2470 |
| chr18-56224642-INS-1906 | chr18-56224643-INS-1907 |
| chr18-56233890-INS-4087 | chr18-56233891-INS-4087 |
| chr18-56245645-INS-117 | chr18-56245645-INS-118 |
| chr18-56259548-INS-117 | chr18-56259548-INS-133 |
| chr18-56266341-INS-74 | chr18-56266341-INS-50 |
| chr18-56288513-INS-305 | chr18-56288521-INS-313 |
| chr18-56294136-INS-51 | chr18-56294103-INS-51 |
| chr18-56305180-INS-8695 | chr18-56305181-INS-8695 |
| chr18-56309296-INS-149 | chr18-56309296-INS-149 |
| chr18-56312140-INS-1176 | chr18-56312141-INS-1178 |
| chr18-56349059-INS-92 | chr18-56349040-INS-64 |
| chr18-56380085-INS-166 | chr18-56380086-INS-154 |
| chr18-56385499-INS-64 | chr18-56385499-INS-52 |
| chr18-56415821-INS-53 | chr18-56415821-INS-56 |
| chr18-56433233-INS-96 | chr18-56433233-INS-96 |
| chr18-56446436-INS-66 | chr18-56446434-INS-76 |
| chr18-56448808-INS-92 | chr18-56448808-INS-96 |
| chr18-56450073-INS-264 | chr18-56450073-INS-264 |
| chr18-56489899-INS-208 | chr18-56489899-INS-207 |
| chr18-56500416-INS-431 | chr18-56500416-INS-431 |
| chr18-56503154-INS-59 | chr18-56503154-INS-59 |
| chr18-56509389-INS-60 | chr18-56509389-INS-54 |
| chr18-56518689-INS-507 | chr18-56518689-INS-507 |
| chr18-56525281-INS-1886 | chr18-56525282-INS-1886 |
| chr18-56551316-INS-65 | chr18-56551316-INS-51 |
| chr18-56553483-INS-534 | chr18-56553483-INS-528 |
| chr18-56571347-INS-1303 | chr18-56571348-INS-1303 |
| chr18-56571348-INS-243 | chr18-56571348-INS-243 |
| chr18-56580699-INS-229 | chr18-56580699-INS-227 |
| chr18-56582273-INS-52 | chr18-56582273-INS-53 |
| chr18-56587934-INS-2444 | chr18-56587935-INS-2444 |
| chr18-56610724-INS-204 | chr18-56610724-INS-203 |
| chr18-56619639-INS-171 | chr18-56619639-INS-172 |
| chr18-56628212-INS-67 | chr18-56628212-INS-67 |
| chr18-56629757-INS-223 | chr18-56629757-INS-223 |
| chr18-56645765-INS-493 | chr18-56645765-INS-498 |
| chr18-56653763-INS-341 | chr18-56653763-INS-329 |
| chr18-56654868-INS-114 | chr18-56654868-INS-114 |
| chr18-56657708-INS-220 | chr18-56657708-INS-220 |
| chr18-56681018-INS-70 | chr18-56681020-INS-102 |
| chr18-56712452-INS-208 | chr18-56712452-INS-208 |
| chr18-56714597-INS-288 | chr18-56714597-INS-288 |
| chr18-56726808-INS-88 | chr18-56726808-INS-94 |
| chr18-56743312-INS-1174 | chr18-56743313-INS-1174 |
| chr18-56744654-INS-205 | chr18-56744654-INS-204 |
| chr18-56760785-INS-69 | chr18-56760785-INS-69 |
| chr18-56774321-INS-207 | chr18-56774321-INS-207 |
| chr18-56779609-INS-61 | chr18-56779609-INS-61 |
| chr18-56784827-INS-51 | chr18-56784827-INS-56 |
| chr18-56785086-INS-681 | chr18-56785086-INS-682 |
| chr18-56805542-INS-202 | chr18-56805542-INS-203 |
| chr18-56806323-INS-215 | chr18-56806323-INS-215 |
| chr18-56807342-INS-224 | chr18-56807342-INS-223 |
| chr18-56815838-INS-220 | chr18-56815838-INS-221 |
| chr18-56830510-INS-218 | chr18-56830510-INS-218 |
| chr18-56839237-INS-52 | chr18-56839237-INS-52 |
| chr18-56865104-INS-207 | chr18-56865104-INS-212 |
| chr18-56904006-INS-216 | chr18-56904006-INS-215 |
| chr18-56937781-INS-105 | chr18-56937781-INS-105 |
| chr18-56941444-INS-325 | chr18-56941444-INS-325 |
| chr18-56950028-INS-231 | chr18-56950028-INS-231 |
| chr18-56986951-INS-101 | chr18-56986945-INS-72 |
| chr18-56994266-INS-51 | chr18-56994266-INS-55 |
| chr18-57036106-INS-55 | chr18-57036106-INS-62 |
| chr18-57066976-INS-139 | chr18-57066976-INS-139 |
| chr18-57070660-INS-210 | chr18-57070660-INS-210 |
| chr18-57100150-INS-362 | chr18-57100150-INS-362 |
| chr18-57118430-INS-1390 | chr18-57118431-INS-1346 |
| chr18-57127359-INS-91 | chr18-57127359-INS-96 |
| chr18-57127557-INS-828 | chr18-57127557-INS-828 |

| | |
|---|---|
| chr18-57136051-INS-55 | chr18-57136051-INS-55 |
| chr18-57157081-INS-843 | chr18-57157081-INS-843 |
| chr18-57164157-INS-3070 | chr18-57164158-INS-3069 |
| chr18-57168532-INS-63 | chr18-57168532-INS-63 |
| chr18-57171336-INS-3359 | chr18-57171337-INS-3357 |
| chr18-57172268-INS-90 | chr18-57172268-INS-90 |
| chr18-57180924-INS-68 | chr18-57180912-INS-52 |
| chr18-57183425-INS-846 | chr18-57183425-INS-853 |
| chr18-57208537-INS-713 | chr18-57208537-INS-713 |
| chr18-57220267-INS-554 | chr18-57220267-INS-554 |
| chr18-57256163-INS-87 | chr18-57256161-INS-55 |
| chr18-57309665-INS-74 | chr18-57309683-INS-58 |
| chr18-57316111-INS-143 | chr18-57316111-INS-143 |
| chr18-57342092-INS-108 | chr18-57342108-INS-70 |
| chr18-57346681-INS-72 | chr18-57346683-INS-62 |
| chr18-57347638-INS-200 | chr18-57347638-INS-200 |
| chr18-57355737-INS-138 | chr18-57355742-INS-99 |
| chr18-57357271-INS-106 | chr18-57357271-INS-110 |
| chr18-57364807-INS-801 | chr18-57364807-INS-849 |
| chr18-57364807-INS-90 | chr18-57364807-INS-93 |
| chr18-57370885-INS-55 | chr18-57370885-INS-50 |
| chr18-57387599-INS-137 | chr18-57387598-INS-106 |
| chr18-57400321-INS-78 | chr18-57400319-INS-52 |
| chr18-57434059-INS-180 | chr18-57434059-INS-180 |
| chr18-57455434-INS-102 | chr18-57455442-INS-52 |
| chr18-57489061-INS-215 | chr18-57489046-INS-196 |
| chr18-57494732-INS-72 | chr18-57494733-INS-76 |
| chr18-57499626-INS-66 | chr18-57499626-INS-66 |
| chr18-57512469-INS-114 | chr18-57512515-INS-65 |
| chr18-57530540-INS-53 | chr18-57530540-INS-57 |
| chr18-57551633-INS-52 | chr18-57551633-INS-52 |
| chr18-57557052-INS-559 | chr18-57557052-INS-558 |
| chr18-57575042-INS-52 | chr18-57575042-INS-52 |
| chr18-57587961-INS-190 | chr18-57587961-INS-190 |
| chr18-57657545-INS-90 | chr18-57657515-INS-96 |
| chr18-57672452-INS-69 | chr18-57672452-INS-69 |
| chr18-57692406-INS-131 | chr18-57692406-INS-84 |
| chr18-57732256-INS-145 | chr18-57732236-INS-182 |
| chr18-57817323-INS-171 | chr18-57817323-INS-171 |
| chr18-57827468-INS-90 | chr18-57827468-INS-90 |
| chr18-57843375-INS-54 | chr18-57843375-INS-52 |
| chr18-57849513-INS-118 | chr18-57849511-INS-120 |
| chr18-57872287-INS-53 | chr18-57872287-INS-57 |
| chr18-57891961-INS-70 | chr18-57891961-INS-50 |
| chr18-57912289-INS-93 | chr18-57912289-INS-93 |
| chr18-57916045-INS-171 | chr18-57916046-INS-171 |
| chr18-57940804-INS-199 | chr18-57940804-INS-198 |
| chr18-57947193-INS-337 | chr18-57947193-INS-337 |
| chr18-57951827-INS-143 | chr18-57951827-INS-143 |
| chr18-57960550-INS-2000 | chr18-57960551-INS-2000 |
| chr18-57963082-INS-118 | chr18-57963083-INS-126 |
| chr18-57970947-INS-2096 | chr18-57970948-INS-2069 |
| chr18-57971691-INS-5174 | chr18-57971692-INS-5174 |
| chr18-57989007-INS-73 | chr18-57989007-INS-73 |
| chr18-57995686-INS-6365 | chr18-57995687-INS-6367 |
| chr18-57999854-INS-327 | chr18-57999854-INS-328 |
| chr18-58000495-INS-75 | chr18-58000495-INS-77 |
| chr18-58004933-INS-95 | chr18-58004934-INS-71 |
| chr18-58060655-INS-202 | chr18-58060655-INS-201 |
| chr18-58064643-INS-1151 | chr18-58064644-INS-1147 |
| chr18-58109119-INS-65 | chr18-58109088-INS-60 |
| chr18-58111989-INS-120 | chr18-58111989-INS-124 |
| chr18-58113140-INS-117 | chr18-58113146-INS-76 |
| chr18-58163173-INS-256 | chr18-58163189-INS-224 |
| chr18-58164298-INS-166 | chr18-58164298-INS-166 |
| chr18-58175387-INS-122 | chr18-58175343-INS-127 |
| chr18-58205318-INS-178 | chr18-58205318-INS-178 |
| chr18-58214703-INS-754 | chr18-58214703-INS-754 |
| chr18-58230800-INS-276 | chr18-58230802-INS-304 |
| chr18-58237872-INS-4474 | chr18-58237873-INS-4474 |
| chr18-58253319-INS-6838 | chr18-58253320-INS-6838 |
| chr18-58262486-INS-175 | chr18-58262486-INS-174 |
| chr18-58263805-INS-894 | chr18-58263805-INS-895 |
| chr18-58306370-INS-73 | chr18-58306386-INS-63 |
| chr18-58332035-INS-78 | chr18-58332035-INS-80 |
| chr18-58334938-INS-1889 | chr18-58334939-INS-1889 |
| chr18-58366449-INS-95 | chr18-58366449-INS-95 |

| | |
|---|---|
| chr18-58383070-INS-114 | chr18-58383070-INS-108 |
| chr18-58404412-INS-74 | chr18-58404412-INS-76 |
| chr18-58414297-INS-62 | chr18-58414297-INS-64 |
| chr18-58445105-INS-510 | chr18-58445105-INS-510 |
| chr18-58473278-INS-75 | chr18-58473278-INS-75 |
| chr18-58504549-INS-208 | chr18-58504549-INS-208 |
| chr18-58526861-INS-439 | chr18-58526861-INS-439 |
| chr18-58531158-INS-206 | chr18-58531158-INS-200 |
| chr18-58537355-INS-407 | chr18-58537355-INS-407 |
| chr18-58543157-INS-60 | chr18-58543157-INS-60 |
| chr18-58574590-INS-925 | chr18-58574590-INS-925 |
| chr18-58582279-INS-57 | chr18-58582291-INS-62 |
| chr18-58583615-INS-2338 | chr18-58583616-INS-2338 |
| chr18-58610358-INS-51 | chr18-58610351-INS-69 |
| chr18-58649968-INS-117 | chr18-58649945-INS-91 |
| chr18-58656384-INS-130 | chr18-58656384-INS-130 |
| chr18-58677390-INS-62 | chr18-58677390-INS-62 |
| chr18-58694069-INS-118 | chr18-58694067-INS-79 |
| chr18-58701863-INS-196 | chr18-58701863-INS-196 |
| chr18-58713835-INS-60 | chr18-58713835-INS-60 |
| chr18-58721442-INS-201 | chr18-58721442-INS-201 |
| chr18-58736275-INS-54 | chr18-58736275-INS-54 |
| chr18-58738791-INS-64 | chr18-58738791-INS-64 |
| chr18-58752227-INS-942 | chr18-58752227-INS-942 |
| chr18-58757812-INS-241 | chr18-58757812-INS-262 |
| chr18-58760697-INS-199 | chr18-58760697-INS-199 |
| chr18-58766101-INS-56 | chr18-58766059-INS-54 |
| chr18-58770428-INS-218 | chr18-58770429-INS-266 |
| chr18-58803461-INS-145 | chr18-58803461-INS-145 |
| chr18-58808986-INS-68 | chr18-58808986-INS-68 |
| chr18-58827289-INS-6217 | chr18-58827290-INS-6217 |
| chr18-58833492-INS-498 | chr18-58833492-INS-498 |
| chr18-58870105-INS-88 | chr18-58870105-INS-76 |
| chr18-58903338-INS-468 | chr18-58903338-INS-459 |
| chr18-58905188-INS-6204 | chr18-58905189-INS-6205 |
| chr18-58911661-INS-6907 | chr18-58911662-INS-6907 |
| chr18-58914418-INS-82 | chr18-58914418-INS-82 |
| chr18-58930356-INS-79 | chr18-58930356-INS-79 |
| chr18-58950023-INS-239 | chr18-58950023-INS-241 |
| chr18-58962562-INS-564 | chr18-58962562-INS-564 |
| chr18-58965182-INS-5512 | chr18-58965183-INS-5512 |
| chr18-58986247-INS-90 | chr18-58986246-INS-96 |
| chr18-59008973-INS-76 | chr18-59008973-INS-82 |
| chr18-59041232-INS-75 | chr18-59041232-INS-91 |
| chr18-59042373-INS-141 | chr18-59042373-INS-141 |
| chr18-59057225-INS-272 | chr18-59057225-INS-272 |
| chr18-59058238-INS-56 | chr18-59058238-INS-56 |
| chr18-59074845-INS-927 | chr18-59074845-INS-923 |
| chr18-59093106-INS-211 | chr18-59093106-INS-211 |
| chr18-59099023-INS-77 | chr18-59099023-INS-77 |
| chr18-59112638-INS-268 | chr18-59112638-INS-268 |
| chr18-59123466-INS-155 | chr18-59123466-INS-155 |
| chr18-59133067-INS-62 | chr18-59133067-INS-75 |
| chr18-59135997-INS-1890 | chr18-59135998-INS-1890 |
| chr18-59147365-INS-60 | chr18-59147365-INS-50 |
| chr18-59160403-INS-52 | chr18-59160403-INS-54 |
| chr18-59171688-INS-6451 | chr18-59171689-INS-6451 |
| chr18-59181792-INS-64 | chr18-59181792-INS-66 |
| chr18-59210142-INS-478 | chr18-59210142-INS-479 |
| chr18-59213655-INS-63 | chr18-59213655-INS-63 |
| chr18-59230360-INS-54 | chr18-59230360-INS-50 |
| chr18-59247132-INS-937 | chr18-59247132-INS-937 |
| chr18-59262269-INS-139 | chr18-59262269-INS-144 |
| chr18-59281254-INS-1706 | chr18-59281255-INS-1706 |
| chr18-59298719-INS-67 | chr18-59298719-INS-84 |
| chr18-59300674-INS-257 | chr18-59300674-INS-267 |
| chr18-59307361-INS-60 | chr18-59307361-INS-65 |
| chr18-59328782-INS-73 | chr18-59328782-INS-73 |
| chr18-59371728-INS-87 | chr18-59371728-INS-87 |
| chr18-59372689-INS-589 | chr18-59372690-INS-589 |
| chr18-59374191-INS-74 | chr18-59374190-INS-74 |
| chr18-59378979-INS-3454 | chr18-59378980-INS-3460 |
| chr18-59381850-INS-125 | chr18-59381852-INS-94 |
| chr18-59385656-INS-60 | chr18-59385646-INS-62 |
| chr18-59408898-INS-420 | chr18-59408898-INS-420 |
| chr18-59410694-INS-55 | chr18-59410691-INS-61 |
| chr18-59430972-INS-1240 | chr18-59430973-INS-1240 |

| | |
|---|---|
| chr18-59438696-INS-72 | chr18-59438719-INS-62 |
| chr18-59497776-INS-218 | chr18-59497776-INS-218 |
| chr18-59498772-INS-110 | chr18-59498790-INS-94 |
| chr18-59524832-INS-169 | chr18-59524815-INS-144 |
| chr18-59536666-INS-114 | chr18-59536666-INS-112 |
| chr18-59549747-INS-63 | chr18-59549727-INS-67 |
| chr18-59566734-INS-127 | chr18-59566734-INS-131 |
| chr18-59574575-INS-94 | chr18-59574575-INS-94 |
| chr18-59577958-INS-80 | chr18-59577964-INS-87 |
| chr18-59578379-INS-328 | chr18-59578379-INS-328 |
| chr18-59585381-INS-58 | chr18-59585381-INS-62 |
| chr18-59650331-INS-169 | chr18-59650331-INS-169 |
| chr18-59660474-INS-187 | chr18-59660474-INS-190 |
| chr18-59662352-INS-208 | chr18-59662352-INS-208 |
| chr18-59671629-INS-6237 | chr18-59671630-INS-6237 |
| chr18-59672052-INS-660 | chr18-59672053-INS-660 |
| chr18-59704496-INS-401 | chr18-59704496-INS-401 |
| chr18-59713805-INS-74 | chr18-59713805-INS-76 |
| chr18-59738082-INS-403 | chr18-59738082-INS-403 |
| chr18-59757371-INS-239 | chr18-59757371-INS-237 |
| chr18-59761498-INS-3944 | chr18-59761499-INS-3944 |
| chr18-59771875-INS-3280 | chr18-59771876-INS-3280 |
| chr18-59781070-INS-569 | chr18-59781070-INS-569 |
| chr18-59789143-INS-328 | chr18-59789143-INS-328 |
| chr18-59794281-INS-64 | chr18-59794281-INS-68 |
| chr18-59817366-INS-54 | chr18-59817366-INS-84 |
| chr18-59819640-INS-7299 | chr18-59819641-INS-7300 |
| chr18-59824548-INS-358 | chr18-59824548-INS-358 |
| chr18-59852882-INS-641 | chr18-59852882-INS-641 |
| chr18-59857649-INS-754 | chr18-59857649-INS-754 |
| chr18-59862288-INS-55 | chr18-59862288-INS-58 |
| chr18-59871045-INS-104 | chr18-59871045-INS-104 |
| chr18-59890744-INS-58 | chr18-59890744-INS-64 |
| chr18-59896093-INS-393 | chr18-59896093-INS-390 |
| chr18-59915404-INS-90 | chr18-59915404-INS-90 |
| chr18-59948209-INS-2842 | chr18-59948210-INS-2844 |
| chr18-59952192-INS-88 | chr18-59952192-INS-84 |
| chr18-59975782-INS-382 | chr18-59975782-INS-382 |
| chr18-60019168-INS-93 | chr18-60019168-INS-93 |
| chr18-60023213-INS-386 | chr18-60023213-INS-386 |
| chr18-60025114-INS-324 | chr18-60025114-INS-324 |
| chr18-60026511-INS-292 | chr18-60026511-INS-292 |
| chr18-60027122-INS-669 | chr18-60027122-INS-669 |
| chr18-60062810-INS-59 | chr18-60062810-INS-71 |
| chr18-60069009-INS-64 | chr18-60069009-INS-64 |
| chr18-60106230-INS-56 | chr18-60106200-INS-62 |
| chr18-60117101-INS-54 | chr18-60117101-INS-54 |
| chr18-60118295-INS-1397 | chr18-60118296-INS-1408 |
| chr18-60129772-INS-93 | chr18-60129772-INS-93 |
| chr18-60135825-INS-52 | chr18-60135825-INS-52 |
| chr18-60144784-INS-71 | chr18-60144784-INS-93 |
| chr18-60166288-INS-162 | chr18-60166288-INS-157 |
| chr18-60170869-INS-106 | chr18-60170869-INS-106 |
| chr18-60187549-INS-56 | chr18-60187553-INS-52 |
| chr18-60190254-INS-76 | chr18-60190254-INS-70 |
| chr18-60194479-INS-105 | chr18-60194479-INS-115 |
| chr18-60222435-INS-6889 | chr18-60222436-INS-6887 |
| chr18-60228067-INS-3009 | chr18-60228068-INS-3009 |
| chr18-60236047-INS-163 | chr18-60236047-INS-163 |
| chr18-60238289-INS-1449 | chr18-60238290-INS-1450 |
| chr18-60239511-INS-6191 | chr18-60239512-INS-6200 |
| chr18-60243300-INS-6670 | chr18-60243301-INS-6670 |
| chr18-60249750-INS-1743 | chr18-60249751-INS-1743 |
| chr18-60268047-INS-351 | chr18-60268047-INS-373 |
| chr18-60272626-INS-170 | chr18-60272626-INS-170 |
| chr18-60276694-INS-98 | chr18-60276694-INS-106 |
| chr18-60283730-INS-57 | chr18-60283730-INS-57 |
| chr18-60284889-INS-330 | chr18-60284889-INS-330 |
| chr18-60288710-INS-383 | chr18-60288710-INS-387 |
| chr18-60307716-INS-111 | chr18-60307716-INS-114 |
| chr18-60309626-INS-738 | chr18-60309626-INS-737 |
| chr18-60312616-INS-66 | chr18-60312616-INS-54 |
| chr18-60314090-INS-437 | chr18-60314090-INS-437 |
| chr18-60324520-INS-62 | chr18-60324533-INS-50 |
| chr18-60327154-INS-855 | chr18-60327154-INS-856 |
| chr18-60335126-INS-460 | chr18-60335126-INS-464 |
| chr18-60349651-INS-448 | chr18-60349651-INS-450 |

| | |
|---|---|
| chr18-60463152-INS-215 | chr18-60463152-INS-215 |
| chr18-60474543-INS-4800 | chr18-60474544-INS-4800 |
| chr18-60496590-INS-188 | chr18-60496590-INS-193 |
| chr18-60503453-INS-92 | chr18-60503453-INS-92 |
| chr18-60509947-INS-486 | chr18-60509947-INS-486 |
| chr18-60594021-INS-100 | chr18-60594037-INS-56 |
| chr18-60599725-INS-62 | chr18-60599725-INS-64 |
| chr18-60611864-INS-88 | chr18-60611864-INS-76 |
| chr18-60617461-INS-239 | chr18-60617461-INS-238 |
| chr18-60637810-INS-58 | chr18-60637810-INS-59 |
| chr18-60643591-INS-1568 | chr18-60643592-INS-1568 |
| chr18-60650384-INS-257 | chr18-60650384-INS-257 |
| chr18-60668256-INS-511 | chr18-60668256-INS-511 |
| chr18-60709286-INS-96 | chr18-60709270-INS-116 |
| chr18-60711847-INS-56 | chr18-60711847-INS-58 |
| chr18-60717438-INS-59 | chr18-60717433-INS-73 |
| chr18-60742088-INS-108 | chr18-60742088-INS-112 |
| chr18-60747378-INS-53 | chr18-60747396-INS-52 |
| chr18-60754839-INS-53 | chr18-60754839-INS-53 |
| chr18-60770826-INS-57 | chr18-60770826-INS-57 |
| chr18-60774246-INS-53 | chr18-60774233-INS-88 |
| chr18-60806416-INS-64 | chr18-60806416-INS-64 |
| chr18-60809290-INS-699 | chr18-60809290-INS-699 |
| chr18-60826076-INS-56 | chr18-60826076-INS-50 |
| chr18-60834758-INS-406 | chr18-60834758-INS-406 |
| chr18-60853835-INS-702 | chr18-60853835-INS-702 |
| chr18-60858291-INS-208 | chr18-60858289-INS-170 |
| chr18-60859015-INS-175 | chr18-60859015-INS-175 |
| chr18-60871462-INS-196 | chr18-60871462-INS-196 |
| chr18-60902583-INS-194 | chr18-60902583-INS-194 |
| chr18-60908514-INS-123 | chr18-60908514-INS-125 |
| chr18-60915607-INS-216 | chr18-60915607-INS-218 |
| chr18-60917787-INS-248 | chr18-60917787-INS-248 |
| chr18-60919415-INS-60 | chr18-60919408-INS-65 |
| chr18-60920673-INS-641 | chr18-60920690-INS-637 |
| chr18-60928854-INS-228 | chr18-60928854-INS-228 |
| chr18-60929504-INS-71 | chr18-60929504-INS-57 |
| chr18-60932793-INS-210 | chr18-60932793-INS-210 |
| chr18-60933296-INS-3939 | chr18-60933297-INS-3939 |
| chr18-60966370-INS-152 | chr18-60966364-INS-119 |
| chr18-60984185-INS-124 | chr18-60984185-INS-124 |
| chr18-60986957-INS-4543 | chr18-60986958-INS-4543 |
| chr18-60991276-INS-82 | chr18-60991278-INS-78 |
| chr18-61005749-INS-74 | chr18-61005768-INS-96 |
| chr18-61007135-INS-56 | chr18-61007136-INS-56 |
| chr18-61017818-INS-60 | chr18-61017818-INS-60 |
| chr18-61063920-INS-52 | chr18-61063920-INS-52 |
| chr18-61072774-INS-87 | chr18-61072774-INS-87 |
| chr18-61074621-INS-270 | chr18-61074621-INS-266 |
| chr18-61080816-INS-84 | chr18-61080816-INS-84 |
| chr18-61082983-INS-56 | chr18-61082983-INS-58 |
| chr18-61084078-INS-84 | chr18-61084078-INS-88 |
| chr18-61089702-INS-86 | chr18-61089735-INS-86 |
| chr18-61115384-INS-79 | chr18-61115384-INS-79 |
| chr18-61123328-INS-56 | chr18-61123328-INS-56 |
| chr18-61132345-INS-112 | chr18-61132345-INS-66 |
| chr18-61145918-INS-114 | chr18-61145918-INS-124 |
| chr18-61165338-INS-1892 | chr18-61165339-INS-1892 |
| chr18-61167259-INS-160 | chr18-61167259-INS-172 |
| chr18-61233229-INS-72 | chr18-61233229-INS-74 |
| chr18-61238764-INS-71 | chr18-61238764-INS-73 |
| chr18-61271157-INS-141 | chr18-61271157-INS-141 |
| chr18-61293189-INS-58 | chr18-61293189-INS-54 |
| chr18-61298524-INS-757 | chr18-61298524-INS-757 |
| chr18-61328154-INS-164 | chr18-61328154-INS-164 |
| chr18-61342329-INS-259 | chr18-61342329-INS-259 |
| chr18-61343396-INS-2820 | chr18-61343397-INS-2821 |
| chr18-61366048-INS-60 | chr18-61366048-INS-60 |
| chr18-61367595-INS-126 | chr18-61367595-INS-93 |
| chr18-61402558-INS-2962 | chr18-61402559-INS-2962 |
| chr18-61410225-INS-218 | chr18-61410225-INS-219 |
| chr18-61435497-INS-52 | chr18-61435497-INS-52 |
| chr18-61472591-INS-92 | chr18-61472591-INS-92 |
| chr18-61482261-INS-373 | chr18-61482261-INS-371 |
| chr18-61486364-INS-68 | chr18-61486384-INS-76 |
| chr18-61504741-INS-68 | chr18-61504741-INS-72 |
| chr18-61536640-INS-205 | chr18-61536640-INS-249 |

| | |
|---|---|
| chr18-61545397-INS-174 | chr18-61545397-INS-174 |
| chr18-61555884-INS-204 | chr18-61555880-INS-225 |
| chr18-61558316-INS-100 | chr18-61558316-INS-75 |
| chr18-61576623-INS-160 | chr18-61576649-INS-155 |
| chr18-61585317-INS-84 | chr18-61585322-INS-86 |
| chr18-61592381-INS-219 | chr18-61592395-INS-237 |
| chr18-61596862-INS-90 | chr18-61596862-INS-90 |
| chr18-61607933-INS-85 | chr18-61607932-INS-64 |
| chr18-61608365-INS-166 | chr18-61608333-INS-180 |
| chr18-61611887-INS-214 | chr18-61611887-INS-218 |
| chr18-61641468-INS-66 | chr18-61641468-INS-57 |
| chr18-61650617-INS-233 | chr18-61650617-INS-232 |
| chr18-61656199-INS-1888 | chr18-61656200-INS-1888 |
| chr18-61666968-INS-64 | chr18-61666968-INS-67 |
| chr18-61700039-INS-202 | chr18-61700039-INS-202 |
| chr18-61737046-INS-221 | chr18-61737046-INS-221 |
| chr18-61765562-INS-148 | chr18-61765562-INS-148 |
| chr18-61775259-INS-72 | chr18-61775242-INS-71 |
| chr18-61803399-INS-74 | chr18-61803399-INS-70 |
| chr18-61813688-INS-696 | chr18-61813688-INS-696 |
| chr18-61817955-INS-5559 | chr18-61817956-INS-5559 |
| chr18-61839365-INS-87 | chr18-61839365-INS-87 |
| chr18-61855432-INS-122 | chr18-61855432-INS-85 |
| chr18-61878938-INS-194 | chr18-61878924-INS-193 |
| chr18-61921280-INS-52 | chr18-61921267-INS-60 |
| chr18-61924232-INS-81 | chr18-61924232-INS-75 |
| chr18-61929982-INS-308 | chr18-61929982-INS-312 |
| chr18-61931455-INS-101 | chr18-61931455-INS-101 |
| chr18-61981821-INS-535 | chr18-61981827-INS-510 |
| chr18-62007054-INS-72 | chr18-62007048-INS-56 |
| chr18-62029611-INS-122 | chr18-62029611-INS-116 |
| chr18-62031501-INS-65 | chr18-62031496-INS-100 |
| chr18-62110123-INS-168 | chr18-62110123-INS-166 |
| chr18-62117736-INS-72 | chr18-62117736-INS-75 |
| chr18-62142844-INS-82 | chr18-62142844-INS-52 |
| chr18-62164094-INS-80 | chr18-62164094-INS-80 |
| chr18-62191899-INS-124 | chr18-62191899-INS-120 |
| chr18-62238376-INS-54 | chr18-62238377-INS-50 |
| chr18-62277921-INS-126 | chr18-62277921-INS-129 |
| chr18-62303237-INS-71 | chr18-62303236-INS-93 |
| chr18-62362387-INS-128 | chr18-62362390-INS-132 |
| chr18-62369471-INS-3610 | chr18-62369472-INS-3566 |
| chr18-62370361-INS-339 | chr18-62370361-INS-340 |
| chr18-62405423-INS-78 | chr18-62405421-INS-54 |
| chr18-62414842-INS-74 | chr18-62414842-INS-76 |
| chr18-62445149-INS-7084 | chr18-62445150-INS-7084 |
| chr18-62473390-INS-58 | chr18-62473390-INS-62 |
| chr18-62484048-INS-338 | chr18-62484058-INS-335 |
| chr18-62489022-INS-99 | chr18-62488977-INS-100 |
| chr18-62542328-INS-191 | chr18-62542319-INS-186 |
| chr18-62548930-INS-52 | chr18-62548930-INS-52 |
| chr18-62560712-INS-212 | chr18-62560712-INS-212 |
| chr18-62577464-INS-62 | chr18-62577464-INS-62 |
| chr18-62620417-INS-69 | chr18-62620417-INS-70 |
| chr18-62674789-INS-977 | chr18-62674789-INS-977 |
| chr18-62696798-INS-58 | chr18-62696791-INS-54 |
| chr18-62719716-INS-54 | chr18-62719716-INS-54 |
| chr18-62729746-INS-846 | chr18-62729746-INS-846 |
| chr18-62734653-INS-1485 | chr18-62734654-INS-1486 |
| chr18-62739023-INS-1207 | chr18-62739024-INS-1204 |
| chr18-62743654-INS-629 | chr18-62743654-INS-628 |
| chr18-62751323-INS-800 | chr18-62751323-INS-799 |
| chr18-62778103-INS-94 | chr18-62778103-INS-94 |
| chr18-62810348-INS-61 | chr18-62810348-INS-64 |
| chr18-62832258-INS-2605 | chr18-62832259-INS-2605 |
| chr18-62840324-INS-70 | chr18-62840324-INS-69 |
| chr18-62856787-INS-2735 | chr18-62856788-INS-2736 |
| chr18-62876725-INS-63 | chr18-62876725-INS-62 |
| chr18-62886564-INS-764 | chr18-62886564-INS-766 |
| chr18-62906732-INS-78 | chr18-62906732-INS-63 |
| chr18-62936666-INS-167 | chr18-62936666-INS-167 |
| chr18-62951549-INS-438 | chr18-62951549-INS-439 |
| chr18-62959497-INS-126 | chr18-62959501-INS-118 |
| chr18-62979443-INS-69 | chr18-62979443-INS-67 |
| chr18-62985787-INS-407 | chr18-62985787-INS-407 |
| chr18-62994717-INS-590 | chr18-62994717-INS-593 |
| chr18-63003575-INS-128 | chr18-63003575-INS-128 |

| | |
|---|---|
| chr18-63065282-INS-106 | chr18-63065283-INS-100 |
| chr18-63091134-INS-73 | chr18-63091132-INS-77 |
| chr18-63119704-INS-166 | chr18-63119704-INS-166 |
| chr18-63122402-INS-281 | chr18-63122402-INS-281 |
| chr18-63147047-INS-6442 | chr18-63147048-INS-6442 |
| chr18-63167452-INS-212 | chr18-63167452-INS-212 |
| chr18-63205276-INS-67 | chr18-63205239-INS-67 |
| chr18-63240491-INS-102 | chr18-63240491-INS-70 |
| chr18-63264381-INS-185 | chr18-63264381-INS-182 |
| chr18-63286554-INS-68 | chr18-63286554-INS-68 |
| chr18-63293300-INS-78 | chr18-63293300-INS-70 |
| chr18-63306461-INS-100 | chr18-63306461-INS-100 |
| chr18-63314589-INS-72 | chr18-63314589-INS-72 |
| chr18-63373106-INS-187 | chr18-63373106-INS-187 |
| chr18-63443428-INS-97 | chr18-63443428-INS-97 |
| chr18-63444591-INS-65 | chr18-63444591-INS-70 |
| chr18-63449335-INS-880 | chr18-63449335-INS-880 |
| chr18-63467324-INS-104 | chr18-63467317-INS-104 |
| chr18-63477662-INS-77 | chr18-63477662-INS-77 |
| chr18-63490822-INS-58 | chr18-63490822-INS-85 |
| chr18-63495459-INS-3020 | chr18-63495460-INS-3018 |
| chr18-63502965-INS-62 | chr18-63502965-INS-50 |
| chr18-63530426-INS-104 | chr18-63530426-INS-104 |
| chr18-63562673-INS-4801 | chr18-63562674-INS-4801 |
| chr18-63575809-INS-127 | chr18-63575809-INS-81 |
| chr18-63582221-INS-212 | chr18-63582221-INS-212 |
| chr18-63608213-INS-604 | chr18-63608213-INS-604 |
| chr18-63622190-INS-112 | chr18-63622190-INS-112 |
| chr18-63632498-INS-348 | chr18-63632498-INS-328 |
| chr18-63640330-INS-525 | chr18-63640330-INS-525 |
| chr18-63641579-INS-62 | chr18-63641580-INS-54 |
| chr18-63644801-INS-74 | chr18-63644801-INS-74 |
| chr18-63660960-INS-60 | chr18-63660960-INS-60 |
| chr18-63685109-INS-383 | chr18-63685109-INS-383 |
| chr18-63687424-INS-411 | chr18-63687424-INS-411 |
| chr18-63696670-INS-6491 | chr18-63696671-INS-6489 |
| chr18-63697314-INS-385 | chr18-63697314-INS-386 |
| chr18-63699346-INS-72 | chr18-63699346-INS-80 |
| chr18-63714960-INS-56 | chr18-63714960-INS-52 |
| chr18-63718495-INS-553 | chr18-63718495-INS-535 |
| chr18-63731006-INS-4517 | chr18-63731007-INS-4517 |
| chr18-63737627-INS-783 | chr18-63737627-INS-783 |
| chr18-63744741-INS-55 | chr18-63744741-INS-70 |
| chr18-63748409-INS-94 | chr18-63748409-INS-82 |
| chr18-63752077-INS-490 | chr18-63752077-INS-490 |
| chr18-63768107-INS-7156 | chr18-63768108-INS-7156 |
| chr18-63772497-INS-2163 | chr18-63772498-INS-2171 |
| chr18-63780899-INS-229 | chr18-63780899-INS-229 |
| chr18-63789384-INS-97 | chr18-63789383-INS-97 |
| chr18-63800524-INS-54 | chr18-63800524-INS-54 |
| chr18-63808108-INS-56 | chr18-63808108-INS-60 |
| chr18-63835964-INS-211 | chr18-63835964-INS-211 |
| chr18-63842170-INS-60 | chr18-63842174-INS-56 |
| chr18-63867712-INS-56 | chr18-63867712-INS-52 |
| chr18-63896175-INS-419 | chr18-63896175-INS-419 |
| chr18-63948890-INS-57 | chr18-63948890-INS-82 |
| chr18-63954517-INS-216 | chr18-63954517-INS-216 |
| chr18-63955699-INS-2801 | chr18-63955700-INS-2802 |
| chr18-63978857-INS-559 | chr18-63978857-INS-559 |
| chr18-64014799-INS-115 | chr18-64014799-INS-115 |
| chr18-64038630-INS-256 | chr18-64038630-INS-256 |
| chr18-64071008-INS-55 | chr18-64071008-INS-55 |
| chr18-64073221-INS-58 | chr18-64073259-INS-62 |
| chr18-64113802-INS-108 | chr18-64113802-INS-72 |
| chr18-64182432-INS-201 | chr18-64182432-INS-201 |
| chr18-64203158-INS-236 | chr18-64203158-INS-236 |
| chr18-64204235-INS-720 | chr18-64204235-INS-716 |
| chr18-64206063-INS-221 | chr18-64206063-INS-221 |
| chr18-64215611-INS-65 | chr18-64215611-INS-70 |
| chr18-64283033-INS-116 | chr18-64283033-INS-118 |
| chr18-64289322-INS-397 | chr18-64289322-INS-384 |
| chr18-64291346-INS-238 | chr18-64291345-INS-254 |
| chr18-64320814-INS-70 | chr18-64320814-INS-73 |
| chr18-64342999-INS-74 | chr18-64342999-INS-54 |
| chr18-64346883-INS-843 | chr18-64346883-INS-844 |
| chr18-64358149-INS-194 | chr18-64358149-INS-194 |
| chr18-64369412-INS-498 | chr18-64369412-INS-498 |

| | |
|---|---|
| chr18-64380382-INS-138 | chr18-64380382-INS-138 |
| chr18-64385014-INS-62 | chr18-64385002-INS-51 |
| chr18-64409590-INS-644 | chr18-64409590-INS-644 |
| chr18-64422920-INS-57 | chr18-64422920-INS-60 |
| chr18-64424067-INS-199 | chr18-64424067-INS-210 |
| chr18-64431311-INS-56 | chr18-64431311-INS-56 |
| chr18-64463098-INS-199 | chr18-64463098-INS-199 |
| chr18-64503135-INS-177 | chr18-64503135-INS-173 |
| chr18-64550245-INS-92 | chr18-64550245-INS-92 |
| chr18-64556975-INS-99 | chr18-64557024-INS-54 |
| chr18-64566423-INS-201 | chr18-64566443-INS-172 |
| chr18-64571691-INS-205 | chr18-64571691-INS-207 |
| chr18-64580564-INS-167 | chr18-64580564-INS-166 |
| chr18-64602032-INS-57 | chr18-64602032-INS-57 |
| chr18-64607334-INS-263 | chr18-64607334-INS-265 |
| chr18-64628871-INS-214 | chr18-64628871-INS-214 |
| chr18-64644314-INS-68 | chr18-64644314-INS-56 |
| chr18-64679398-INS-52 | chr18-64679398-INS-50 |
| chr18-64702323-INS-222 | chr18-64702323-INS-222 |
| chr18-64748430-INS-134 | chr18-64748436-INS-105 |
| chr18-64753313-INS-61 | chr18-64753313-INS-50 |
| chr18-64762977-INS-1053 | chr18-64762978-INS-1053 |
| chr18-64781904-INS-5543 | chr18-64781905-INS-5543 |
| chr18-64818211-INS-58 | chr18-64818211-INS-70 |
| chr18-64819640-INS-336 | chr18-64819640-INS-336 |
| chr18-64821405-INS-206 | chr18-64821405-INS-206 |
| chr18-64824058-INS-69 | chr18-64824058-INS-69 |
| chr18-64843177-INS-376 | chr18-64843176-INS-370 |
| chr18-64844013-INS-132 | chr18-64843989-INS-173 |
| chr18-64851734-INS-400 | chr18-64851734-INS-400 |
| chr18-64866690-INS-173 | chr18-64866690-INS-173 |
| chr18-64873236-INS-116 | chr18-64873236-INS-84 |
| chr18-64877268-INS-60 | chr18-64877260-INS-80 |
| chr18-64897145-INS-65 | chr18-64897145-INS-92 |
| chr18-64901604-INS-115 | chr18-64901604-INS-124 |
| chr18-64926436-INS-2818 | chr18-64926437-INS-2814 |
| chr18-64927451-INS-6632 | chr18-64927452-INS-6632 |
| chr18-64928752-INS-128 | chr18-64928752-INS-152 |
| chr18-64958052-INS-52 | chr18-64958052-INS-54 |
| chr18-64960849-INS-514 | chr18-64960850-INS-514 |
| chr18-64962591-INS-74 | chr18-64962591-INS-76 |
| chr18-64964957-INS-441 | chr18-64964957-INS-441 |
| chr18-64972417-INS-458 | chr18-64972417-INS-458 |
| chr18-64981356-INS-182 | chr18-64981357-INS-176 |
| chr18-64997817-INS-72 | chr18-64997815-INS-104 |
| chr18-65005409-INS-52 | chr18-65005409-INS-52 |
| chr18-65008473-INS-202 | chr18-65008473-INS-191 |
| chr18-65011204-INS-233 | chr18-65011204-INS-254 |
| chr18-65042099-INS-233 | chr18-65042099-INS-233 |
| chr18-65045241-INS-77 | chr18-65045241-INS-71 |
| chr18-65060569-INS-58 | chr18-65060590-INS-66 |
| chr18-65062072-INS-9045 | chr18-65062073-INS-9045 |
| chr18-65073997-INS-52 | chr18-65073979-INS-60 |
| chr18-65108123-INS-275 | chr18-65108123-INS-275 |
| chr18-65110263-INS-58 | chr18-65110263-INS-58 |
| chr18-65128409-INS-447 | chr18-65128409-INS-440 |
| chr18-65148660-INS-67 | chr18-65148682-INS-67 |
| chr18-65189258-INS-55 | chr18-65189258-INS-55 |
| chr18-65200290-INS-129 | chr18-65200290-INS-129 |
| chr18-65237473-INS-312 | chr18-65237473-INS-312 |
| chr18-65246537-INS-52 | chr18-65246537-INS-54 |
| chr18-65250009-INS-57 | chr18-65250009-INS-57 |
| chr18-65260139-INS-61 | chr18-65260139-INS-61 |
| chr18-65273131-INS-401 | chr18-65273131-INS-401 |
| chr18-65279156-INS-90 | chr18-65279138-INS-76 |
| chr18-65285936-INS-124 | chr18-65285937-INS-124 |
| chr18-65292066-INS-111 | chr18-65292066-INS-111 |
| chr18-65310971-INS-192 | chr18-65310954-INS-203 |
| chr18-65329807-INS-96 | chr18-65329807-INS-96 |
| chr18-65340216-INS-74 | chr18-65340216-INS-74 |
| chr18-65396000-INS-218 | chr18-65396000-INS-219 |
| chr18-65397639-INS-56 | chr18-65397611-INS-53 |
| chr18-65409061-INS-66 | chr18-65409061-INS-78 |
| chr18-65413216-INS-52 | chr18-65413224-INS-53 |
| chr18-65418267-INS-422 | chr18-65418267-INS-422 |
| chr18-65423871-INS-7162 | chr18-65423872-INS-7162 |
| chr18-65434410-INS-277 | chr18-65434404-INS-234 |

| | |
|---|---|
| chr18-65446245-INS-84 | chr18-65446245-INS-84 |
| chr18-65459453-INS-54 | chr18-65459456-INS-76 |
| chr18-65469064-INS-146 | chr18-65469064-INS-146 |
| chr18-65473549-INS-399 | chr18-65473549-INS-399 |
| chr18-65474893-INS-6437 | chr18-65474894-INS-6437 |
| chr18-65490354-INS-62 | chr18-65490360-INS-59 |
| chr18-65494622-INS-60 | chr18-65494622-INS-63 |
| chr18-65496791-INS-53 | chr18-65496791-INS-53 |
| chr18-65504648-INS-56 | chr18-65504646-INS-56 |
| chr18-65506855-INS-65 | chr18-65506855-INS-79 |
| chr18-65510669-INS-72 | chr18-65510669-INS-72 |
| chr18-65519340-INS-291 | chr18-65519340-INS-291 |
| chr18-65546568-INS-80 | chr18-65546568-INS-80 |
| chr18-65553198-INS-55 | chr18-65553198-INS-55 |
| chr18-65554551-INS-142 | chr18-65554551-INS-142 |
| chr18-65586389-INS-58 | chr18-65586389-INS-60 |
| chr18-65599934-INS-52 | chr18-65599934-INS-52 |
| chr18-65641041-INS-172 | chr18-65641044-INS-194 |
| chr18-65643126-INS-93 | chr18-65643107-INS-102 |
| chr18-65662176-INS-176 | chr18-65662177-INS-176 |
| chr18-65666334-INS-103 | chr18-65666334-INS-103 |
| chr18-65670437-INS-509 | chr18-65670437-INS-509 |
| chr18-65673284-INS-238 | chr18-65673284-INS-240 |
| chr18-65675494-INS-211 | chr18-65675494-INS-211 |
| chr18-65689577-INS-369 | chr18-65689577-INS-369 |
| chr18-65696473-INS-141 | chr18-65696473-INS-142 |
| chr18-65721983-INS-207 | chr18-65721983-INS-222 |
| chr18-65771142-INS-173 | chr18-65771142-INS-185 |
| chr18-65804942-INS-64 | chr18-65804942-INS-64 |
| chr18-65825283-INS-5277 | chr18-65825284-INS-5276 |
| chr18-65837762-INS-55 | chr18-65837761-INS-66 |
| chr18-65842555-INS-145 | chr18-65842555-INS-148 |
| chr18-65843350-INS-110 | chr18-65843350-INS-110 |
| chr18-65844942-INS-113 | chr18-65844942-INS-113 |
| chr18-65848479-INS-56 | chr18-65848479-INS-60 |
| chr18-65865042-INS-112 | chr18-65865042-INS-84 |
| chr18-65880221-INS-135 | chr18-65880221-INS-135 |
| chr18-65881926-INS-207 | chr18-65881926-INS-207 |
| chr18-65884730-INS-216 | chr18-65884730-INS-216 |
| chr18-65886493-INS-226 | chr18-65886493-INS-226 |
| chr18-65888747-INS-99 | chr18-65888747-INS-93 |
| chr18-65908790-INS-242 | chr18-65908790-INS-242 |
| chr18-65914089-INS-73 | chr18-65914086-INS-55 |
| chr18-65916127-INS-744 | chr18-65916127-INS-744 |
| chr18-65938125-INS-54 | chr18-65938125-INS-54 |
| chr18-65994248-INS-911 | chr18-65994248-INS-911 |
| chr18-66000724-INS-228 | chr18-66000724-INS-227 |
| chr18-66004444-INS-204 | chr18-66004444-INS-204 |
| chr18-66022122-INS-60 | chr18-66022122-INS-60 |
| chr18-66030772-INS-328 | chr18-66030779-INS-330 |
| chr18-66076669-INS-109 | chr18-66076669-INS-109 |
| chr18-66082299-INS-241 | chr18-66082299-INS-241 |
| chr18-66090588-INS-154 | chr18-66090586-INS-147 |
| chr18-66099086-INS-78 | chr18-66099086-INS-54 |
| chr18-66111624-INS-69 | chr18-66111624-INS-68 |
| chr18-66112401-INS-6958 | chr18-66112402-INS-6958 |
| chr18-66155048-INS-152 | chr18-66155048-INS-152 |
| chr18-66163091-INS-55 | chr18-66163091-INS-55 |
| chr18-66186417-INS-61 | chr18-66186417-INS-93 |
| chr18-66196704-INS-371 | chr18-66196716-INS-375 |
| chr18-66263951-INS-113 | chr18-66263951-INS-113 |
| chr18-66307286-INS-691 | chr18-66307286-INS-691 |
| chr18-66314763-INS-326 | chr18-66314763-INS-328 |
| chr18-66315459-INS-63 | chr18-66315459-INS-66 |
| chr18-66316785-INS-504 | chr18-66316785-INS-505 |
| chr18-66318952-INS-67 | chr18-66318952-INS-67 |
| chr18-66333437-INS-65 | chr18-66333459-INS-73 |
| chr18-66334582-INS-58 | chr18-66334582-INS-58 |
| chr18-66338921-INS-397 | chr18-66338921-INS-403 |
| chr18-66354490-INS-150 | chr18-66354490-INS-150 |
| chr18-66360698-INS-659 | chr18-66360698-INS-659 |
| chr18-66369562-INS-64 | chr18-66369562-INS-96 |
| chr18-66378879-INS-182 | chr18-66378879-INS-182 |
| chr18-66393979-INS-74 | chr18-66393980-INS-50 |
| chr18-66407880-INS-54 | chr18-66407880-INS-54 |
| chr18-66413451-INS-60 | chr18-66413451-INS-62 |
| chr18-66415700-INS-144 | chr18-66415700-INS-102 |

| | |
|---|---|
| chr18-66417102-INS-53 | chr18-66417102-INS-69 |
| chr18-66438532-INS-6441 | chr18-66438533-INS-6441 |
| chr18-66438533-INS-497 | chr18-66438533-INS-497 |
| chr18-66470548-INS-63 | chr18-66470548-INS-95 |
| chr18-66475946-INS-134 | chr18-66475946-INS-136 |
| chr18-66491112-INS-67 | chr18-66491114-INS-90 |
| chr18-66494679-INS-410 | chr18-66494678-INS-412 |
| chr18-66507620-INS-227 | chr18-66507620-INS-178 |
| chr18-66524040-INS-101 | chr18-66524040-INS-104 |
| chr18-66528019-INS-168 | chr18-66528019-INS-168 |
| chr18-66548901-INS-68 | chr18-66548903-INS-94 |
| chr18-66549492-INS-211 | chr18-66549492-INS-211 |
| chr18-66558726-INS-60 | chr18-66558728-INS-65 |
| chr18-66565347-INS-7002 | chr18-66565348-INS-7007 |
| chr18-66576546-INS-51 | chr18-66576546-INS-66 |
| chr18-66606666-INS-88 | chr18-66606647-INS-110 |
| chr18-66608397-INS-93 | chr18-66608397-INS-93 |
| chr18-66610338-INS-56 | chr18-66610347-INS-63 |
| chr18-66611880-INS-262 | chr18-66611880-INS-265 |
| chr18-66627119-INS-112 | chr18-66627119-INS-112 |
| chr18-66651473-INS-70 | chr18-66651433-INS-118 |
| chr18-66656093-INS-58 | chr18-66656093-INS-58 |
| chr18-66665706-INS-118 | chr18-66665706-INS-118 |
| chr18-66669714-INS-553 | chr18-66669714-INS-553 |
| chr18-66685303-INS-332 | chr18-66685303-INS-332 |
| chr18-66687086-INS-213 | chr18-66687086-INS-179 |
| chr18-66712031-INS-279 | chr18-66712031-INS-279 |
| chr18-66725087-INS-711 | chr18-66725087-INS-711 |
| chr18-66727423-INS-93 | chr18-66727423-INS-93 |
| chr18-66757104-INS-188 | chr18-66757104-INS-188 |
| chr18-66774031-INS-55 | chr18-66774032-INS-55 |
| chr18-66775741-INS-115 | chr18-66775741-INS-116 |
| chr18-66781674-INS-79 | chr18-66781674-INS-79 |
| chr18-66818913-INS-51 | chr18-66818910-INS-54 |
| chr18-66837427-INS-94 | chr18-66837427-INS-94 |
| chr18-66841532-INS-384 | chr18-66841532-INS-384 |
| chr18-66848310-INS-7114 | chr18-66848311-INS-7114 |
| chr18-66856100-INS-62 | chr18-66856100-INS-62 |
| chr18-66877671-INS-447 | chr18-66877671-INS-447 |
| chr18-66879252-INS-2659 | chr18-66879253-INS-2659 |
| chr18-66903367-INS-60 | chr18-66903366-INS-94 |
| chr18-66905525-INS-91 | chr18-66905526-INS-74 |
| chr18-66914067-INS-538 | chr18-66914067-INS-552 |
| chr18-66932189-INS-217 | chr18-66932189-INS-217 |
| chr18-66936446-INS-73 | chr18-66936446-INS-71 |
| chr18-66942159-INS-514 | chr18-66942159-INS-514 |
| chr18-66948536-INS-95 | chr18-66948536-INS-95 |
| chr18-66951786-INS-363 | chr18-66951786-INS-363 |
| chr18-66952340-INS-52 | chr18-66952340-INS-52 |
| chr18-67025559-INS-328 | chr18-67025559-INS-328 |
| chr18-67051793-INS-220 | chr18-67051795-INS-202 |
| chr18-67058848-INS-75 | chr18-67058848-INS-85 |
| chr18-67063161-INS-69 | chr18-67063161-INS-71 |
| chr18-67070384-INS-70 | chr18-67070384-INS-70 |
| chr18-67071656-INS-53 | chr18-67071656-INS-55 |
| chr18-67087437-INS-201 | chr18-67087437-INS-201 |
| chr18-67113512-INS-55 | chr18-67113510-INS-56 |
| chr18-67129947-INS-195 | chr18-67129947-INS-195 |
| chr18-67152531-INS-52 | chr18-67152531-INS-52 |
| chr18-67158717-INS-84 | chr18-67158722-INS-85 |
| chr18-67161476-INS-117 | chr18-67161476-INS-117 |
| chr18-67250175-INS-60 | chr18-67250175-INS-70 |
| chr18-67251679-INS-118 | chr18-67251679-INS-92 |
| chr18-67252674-INS-76 | chr18-67252674-INS-76 |
| chr18-67255697-INS-144 | chr18-67255697-INS-134 |
| chr18-67256813-INS-60 | chr18-67256813-INS-60 |
| chr18-67265044-INS-51 | chr18-67265045-INS-57 |
| chr18-67293889-INS-56 | chr18-67293889-INS-57 |
| chr18-67319706-INS-821 | chr18-67319706-INS-780 |
| chr18-67332014-INS-935 | chr18-67332014-INS-903 |
| chr18-67342396-INS-184 | chr18-67342396-INS-184 |
| chr18-67367273-INS-117 | chr18-67367267-INS-89 |
| chr18-67382342-INS-6223 | chr18-67382343-INS-6223 |
| chr18-67388969-INS-54 | chr18-67388967-INS-56 |
| chr18-67399876-INS-63 | chr18-67399884-INS-84 |
| chr18-67409076-INS-85 | chr18-67409076-INS-131 |
| chr18-67440118-INS-485 | chr18-67440118-INS-485 |

| | |
|---|---|
| chr18-67446268-INS-53 | chr18-67446268-INS-53 |
| chr18-67447554-INS-340 | chr18-67447554-INS-340 |
| chr18-67475315-INS-55 | chr18-67475315-INS-55 |
| chr18-67480980-INS-6442 | chr18-67480981-INS-6442 |
| chr18-67481307-INS-92 | chr18-67481307-INS-92 |
| chr18-67490142-INS-217 | chr18-67490142-INS-218 |
| chr18-67511564-INS-136 | chr18-67511564-INS-136 |
| chr18-67512725-INS-79 | chr18-67512725-INS-89 |
| chr18-67532739-INS-208 | chr18-67532739-INS-208 |
| chr18-67552595-INS-97 | chr18-67552595-INS-97 |
| chr18-67554495-INS-1240 | chr18-67554496-INS-1222 |
| chr18-67569095-INS-216 | chr18-67569073-INS-200 |
| chr18-67589073-INS-210 | chr18-67589073-INS-210 |
| chr18-67600961-INS-58 | chr18-67600961-INS-58 |
| chr18-67616394-INS-182 | chr18-67616394-INS-182 |
| chr18-67617273-INS-191 | chr18-67617273-INS-194 |
| chr18-67627679-INS-225 | chr18-67627679-INS-224 |
| chr18-67636760-INS-52 | chr18-67636760-INS-52 |
| chr18-67647955-INS-58 | chr18-67647955-INS-60 |
| chr18-67657995-INS-200 | chr18-67657996-INS-192 |
| chr18-67677334-INS-95 | chr18-67677334-INS-95 |
| chr18-67680581-INS-52 | chr18-67680582-INS-84 |
| chr18-67692410-INS-52 | chr18-67692410-INS-52 |
| chr18-67702295-INS-242 | chr18-67702336-INS-198 |
| chr18-67722447-INS-214 | chr18-67722447-INS-214 |
| chr18-67726493-INS-133 | chr18-67726493-INS-133 |
| chr18-67740050-INS-51 | chr18-67740044-INS-57 |
| chr18-67746846-INS-141 | chr18-67746846-INS-128 |
| chr18-67753456-INS-61 | chr18-67753447-INS-73 |
| chr18-67766114-INS-373 | chr18-67766114-INS-373 |
| chr18-67771033-INS-212 | chr18-67771033-INS-212 |
| chr18-67776316-INS-212 | chr18-67776316-INS-211 |
| chr18-67784467-INS-206 | chr18-67784467-INS-208 |
| chr18-67788028-INS-127 | chr18-67788028-INS-125 |
| chr18-67789717-INS-57 | chr18-67789717-INS-50 |
| chr18-67792472-INS-215 | chr18-67792472-INS-215 |
| chr18-67798415-INS-188 | chr18-67798415-INS-187 |
| chr18-67820537-INS-190 | chr18-67820537-INS-190 |
| chr18-67821880-INS-133 | chr18-67821880-INS-160 |
| chr18-67830482-INS-712 | chr18-67830482-INS-712 |
| chr18-67843646-INS-216 | chr18-67843646-INS-217 |
| chr18-67853913-INS-3291 | chr18-67853914-INS-3291 |
| chr18-67855211-INS-220 | chr18-67855211-INS-220 |
| chr18-67868563-INS-249 | chr18-67868564-INS-225 |
| chr18-67889923-INS-63 | chr18-67889920-INS-84 |
| chr18-67893362-INS-54 | chr18-67893362-INS-54 |
| chr18-67897373-INS-216 | chr18-67897373-INS-216 |
| chr18-67909697-INS-57 | chr18-67909697-INS-57 |
| chr18-67915717-INS-121 | chr18-67915722-INS-128 |
| chr18-67918344-INS-195 | chr18-67918344-INS-195 |
| chr18-67920024-INS-218 | chr18-67920024-INS-218 |
| chr18-67920973-INS-225 | chr18-67920973-INS-216 |
| chr18-67924349-INS-79 | chr18-67924351-INS-92 |
| chr18-67926767-INS-58 | chr18-67926767-INS-87 |
| chr18-67947173-INS-205 | chr18-67947173-INS-205 |
| chr18-67957871-INS-226 | chr18-67957871-INS-226 |
| chr18-67970692-INS-62 | chr18-67970693-INS-62 |
| chr18-67993253-INS-6452 | chr18-67993254-INS-6452 |
| chr18-67994956-INS-60 | chr18-67994956-INS-60 |
| chr18-67995739-INS-201 | chr18-67995739-INS-201 |
| chr18-67998258-INS-213 | chr18-67998258-INS-213 |
| chr18-68002134-INS-80 | chr18-68002132-INS-91 |
| chr18-68020047-INS-522 | chr18-68020047-INS-512 |
| chr18-68061446-INS-135 | chr18-68061470-INS-161 |
| chr18-68077973-INS-128 | chr18-68077973-INS-128 |
| chr18-68138288-INS-180 | chr18-68138288-INS-180 |
| chr18-68154466-INS-84 | chr18-68154466-INS-84 |
| chr18-68167966-INS-1579 | chr18-68167967-INS-1580 |
| chr18-68189162-INS-75 | chr18-68189162-INS-68 |
| chr18-68217056-INS-70 | chr18-68217056-INS-70 |
| chr18-68224918-INS-240 | chr18-68224918-INS-240 |
| chr18-68225307-INS-5183 | chr18-68225308-INS-5183 |
| chr18-68246262-INS-217 | chr18-68246262-INS-214 |
| chr18-68247634-INS-220 | chr18-68247634-INS-220 |
| chr18-68327109-INS-7083 | chr18-68327110-INS-7083 |
| chr18-68339098-INS-441 | chr18-68339098-INS-441 |
| chr18-68339623-INS-59 | chr18-68339625-INS-58 |

| | |
|---|---|
| chr18-68341807-INS-386 | chr18-68341807-INS-386 |
| chr18-68343010-INS-5553 | chr18-68343011-INS-5553 |
| chr18-68346958-INS-51 | chr18-68346959-INS-51 |
| chr18-68356550-INS-242 | chr18-68356550-INS-242 |
| chr18-68380318-INS-624 | chr18-68380318-INS-624 |
| chr18-68402261-INS-77 | chr18-68402261-INS-77 |
| chr18-68410399-INS-64 | chr18-68410399-INS-64 |
| chr18-68442734-INS-212 | chr18-68442734-INS-213 |
| chr18-68449993-INS-221 | chr18-68449993-INS-221 |
| chr18-68478851-INS-54 | chr18-68478851-INS-66 |
| chr18-68488372-INS-812 | chr18-68488372-INS-812 |
| chr18-68543463-INS-52 | chr18-68543465-INS-84 |
| chr18-68581964-INS-182 | chr18-68581964-INS-169 |
| chr18-68591967-INS-55 | chr18-68591967-INS-55 |
| chr18-68604468-INS-91 | chr18-68604470-INS-91 |
| chr18-68605612-INS-81 | chr18-68605612-INS-81 |
| chr18-68607972-INS-445 | chr18-68607972-INS-446 |
| chr18-68634210-INS-85 | chr18-68634214-INS-76 |
| chr18-68691639-INS-569 | chr18-68691639-INS-567 |
| chr18-68734848-INS-52 | chr18-68734848-INS-76 |
| chr18-68736634-INS-71 | chr18-68736634-INS-71 |
| chr18-68822876-INS-51 | chr18-68822876-INS-51 |
| chr18-68883085-INS-127 | chr18-68883085-INS-127 |
| chr18-68886933-INS-66 | chr18-68886933-INS-68 |
| chr18-68887946-INS-108 | chr18-68887946-INS-123 |
| chr18-68890014-INS-61 | chr18-68890014-INS-63 |
| chr18-68891706-INS-61 | chr18-68891706-INS-61 |
| chr18-68911638-INS-84 | chr18-68911638-INS-84 |
| chr18-68925620-INS-237 | chr18-68925620-INS-236 |
| chr18-68928360-INS-115 | chr18-68928360-INS-98 |
| chr18-68934449-INS-1072 | chr18-68934450-INS-1066 |
| chr18-68947195-INS-53 | chr18-68947195-INS-53 |
| chr18-68958848-INS-58 | chr18-68958834-INS-58 |
| chr18-68968123-INS-396 | chr18-68968123-INS-425 |
| chr18-68981531-INS-52 | chr18-68981548-INS-72 |
| chr18-68998747-INS-72 | chr18-68998760-INS-64 |
| chr18-69031601-INS-55 | chr18-69031602-INS-68 |
| chr18-69032016-INS-856 | chr18-69032016-INS-880 |
| chr18-69059914-INS-63 | chr18-69059914-INS-65 |
| chr18-69067444-INS-66 | chr18-69067444-INS-55 |
| chr18-69075188-INS-135 | chr18-69075209-INS-161 |
| chr18-69082468-INS-244 | chr18-69082468-INS-247 |
| chr18-69086515-INS-284 | chr18-69086515-INS-284 |
| chr18-69089447-INS-60 | chr18-69089422-INS-50 |
| chr18-69099523-INS-6255 | chr18-69099524-INS-6256 |
| chr18-69113696-INS-154 | chr18-69113695-INS-150 |
| chr18-69130129-INS-410 | chr18-69130129-INS-408 |
| chr18-69135527-INS-6261 | chr18-69135528-INS-6261 |
| chr18-69139394-INS-54 | chr18-69139371-INS-78 |
| chr18-69155556-INS-105 | chr18-69155572-INS-56 |
| chr18-69161026-INS-348 | chr18-69161026-INS-348 |
| chr18-69163177-INS-56 | chr18-69163176-INS-64 |
| chr18-69166701-INS-58 | chr18-69166662-INS-73 |
| chr18-69174461-INS-92 | chr18-69174461-INS-98 |
| chr18-69177102-INS-55 | chr18-69177095-INS-51 |
| chr18-69178179-INS-95 | chr18-69178179-INS-95 |
| chr18-69182978-INS-98 | chr18-69182967-INS-108 |
| chr18-69212498-INS-226 | chr18-69212498-INS-228 |
| chr18-69240364-INS-621 | chr18-69240364-INS-621 |
| chr18-69281602-INS-381 | chr18-69281602-INS-381 |
| chr18-69356415-INS-70 | chr18-69356415-INS-70 |
| chr18-69370352-INS-185 | chr18-69370351-INS-157 |
| chr18-69381639-INS-586 | chr18-69381631-INS-552 |
| chr18-69405785-INS-68 | chr18-69405785-INS-103 |
| chr18-69415360-INS-271 | chr18-69415360-INS-224 |
| chr18-69459439-INS-6316 | chr18-69459440-INS-6316 |
| chr18-69465826-INS-70 | chr18-69465826-INS-60 |
| chr18-69545488-INS-288 | chr18-69545488-INS-288 |
| chr18-69552692-INS-145 | chr18-69552692-INS-145 |
| chr18-69574762-INS-1357 | chr18-69574763-INS-1357 |
| chr18-69602548-INS-663 | chr18-69602548-INS-663 |
| chr18-69618279-INS-117 | chr18-69618279-INS-117 |
| chr18-69666903-INS-125 | chr18-69666903-INS-131 |
| chr18-69676325-INS-139 | chr18-69676325-INS-109 |
| chr18-69749347-INS-96 | chr18-69749347-INS-95 |
| chr18-69756190-INS-580 | chr18-69756190-INS-581 |
| chr18-69800109-INS-71 | chr18-69800109-INS-76 |

| | |
|---|---|
| chr18-69828009-INS-421 | chr18-69828009-INS-376 |
| chr18-69836028-INS-174 | chr18-69836032-INS-136 |
| chr18-69844350-INS-59 | chr18-69844350-INS-59 |
| chr18-69873032-INS-191 | chr18-69873032-INS-191 |
| chr18-69895180-INS-140 | chr18-69895174-INS-156 |
| chr18-69905831-INS-214 | chr18-69905831-INS-213 |
| chr18-69920592-INS-125 | chr18-69920593-INS-128 |
| chr18-69928981-INS-90 | chr18-69928981-INS-90 |
| chr18-69943982-INS-64 | chr18-69943996-INS-63 |
| chr18-69960596-INS-68 | chr18-69960596-INS-68 |
| chr18-69963019-INS-55 | chr18-69963020-INS-55 |
| chr18-69989999-INS-76 | chr18-69989999-INS-52 |
| chr18-69993679-INS-5795 | chr18-69993680-INS-5795 |
| chr18-70009361-INS-504 | chr18-70009361-INS-504 |
| chr18-70024283-INS-198 | chr18-70024300-INS-209 |
| chr18-70036146-INS-114 | chr18-70036170-INS-116 |
| chr18-70037835-INS-7182 | chr18-70037836-INS-7181 |
| chr18-70045022-INS-52 | chr18-70045022-INS-52 |
| chr18-70064047-INS-829 | chr18-70064047-INS-828 |
| chr18-70071332-INS-127 | chr18-70071333-INS-95 |
| chr18-70130304-INS-74 | chr18-70130304-INS-66 |
| chr18-70131376-INS-65 | chr18-70131376-INS-75 |
| chr18-70135789-INS-580 | chr18-70135789-INS-593 |
| chr18-70151740-INS-102 | chr18-70151740-INS-114 |
| chr18-70155537-INS-88 | chr18-70155537-INS-102 |
| chr18-70161431-INS-198 | chr18-70161431-INS-198 |
| chr18-70162864-INS-78 | chr18-70162864-INS-78 |
| chr18-70169212-INS-387 | chr18-70169213-INS-387 |
| chr18-70178401-INS-60 | chr18-70178402-INS-72 |
| chr18-70193846-INS-51 | chr18-70193846-INS-51 |
| chr18-70213097-INS-103 | chr18-70213097-INS-103 |
| chr18-70269134-INS-82 | chr18-70269119-INS-120 |
| chr18-70275605-INS-692 | chr18-70275605-INS-692 |
| chr18-70284020-INS-1707 | chr18-70284021-INS-1707 |
| chr18-70329344-INS-64 | chr18-70329344-INS-50 |
| chr18-70331456-INS-56 | chr18-70331455-INS-57 |
| chr18-70337577-INS-4818 | chr18-70337578-INS-4818 |
| chr18-70350917-INS-60 | chr18-70350917-INS-60 |
| chr18-70356292-INS-150 | chr18-70356292-INS-150 |
| chr18-70380284-INS-978 | chr18-70380284-INS-978 |
| chr18-70393575-INS-98 | chr18-70393575-INS-98 |
| chr18-70393866-INS-1888 | chr18-70393867-INS-1888 |
| chr18-70399114-INS-514 | chr18-70399114-INS-507 |
| chr18-70407806-INS-52 | chr18-70407806-INS-52 |
| chr18-70423669-INS-168 | chr18-70423679-INS-148 |
| chr18-70435050-INS-69 | chr18-70435050-INS-84 |
| chr18-70451074-INS-779 | chr18-70451074-INS-778 |
| chr18-70474146-INS-125 | chr18-70474146-INS-128 |
| chr18-70491573-INS-193 | chr18-70491573-INS-193 |
| chr18-70493820-INS-205 | chr18-70493820-INS-205 |
| chr18-70508236-INS-182 | chr18-70508258-INS-135 |
| chr18-70539821-INS-53 | chr18-70539821-INS-65 |
| chr18-70546560-INS-64 | chr18-70546560-INS-64 |
| chr18-70555174-INS-592 | chr18-70555187-INS-591 |
| chr18-70564362-INS-54 | chr18-70564360-INS-75 |
| chr18-70572325-INS-100 | chr18-70572325-INS-100 |
| chr18-70606856-INS-75 | chr18-70606856-INS-73 |
| chr18-70617782-INS-103 | chr18-70617782-INS-103 |
| chr18-70629594-INS-282 | chr18-70629598-INS-300 |
| chr18-70640996-INS-65 | chr18-70640996-INS-65 |
| chr18-70676225-INS-5295 | chr18-70676226-INS-5295 |
| chr18-70676812-INS-54 | chr18-70676812-INS-61 |
| chr18-70684060-INS-211 | chr18-70684068-INS-187 |
| chr18-70691857-INS-1823 | chr18-70691858-INS-1823 |
| chr18-70706388-INS-51 | chr18-70706388-INS-51 |
| chr18-70743811-INS-154 | chr18-70743811-INS-154 |
| chr18-70745539-INS-153 | chr18-70745539-INS-155 |
| chr18-70747385-INS-100 | chr18-70747385-INS-100 |
| chr18-70750682-INS-186 | chr18-70750682-INS-186 |
| chr18-70756152-INS-605 | chr18-70756152-INS-605 |
| chr18-70760396-INS-6442 | chr18-70760397-INS-6442 |
| chr18-70767102-INS-55 | chr18-70767102-INS-55 |
| chr18-70791479-INS-56 | chr18-70791479-INS-56 |
| chr18-70794822-INS-427 | chr18-70794822-INS-427 |
| chr18-70798745-INS-74 | chr18-70798743-INS-85 |
| chr18-70801590-INS-66 | chr18-70801590-INS-50 |
| chr18-70810625-INS-194 | chr18-70810625-INS-194 |

| | |
|---|---|
| chr18-70820749-INS-111 | chr18-70820749-INS-75 |
| chr18-70822122-INS-92 | chr18-70822122-INS-92 |
| chr18-70839175-INS-212 | chr18-70839175-INS-212 |
| chr18-70939347-INS-1951 | chr18-70939348-INS-1951 |
| chr18-70968444-INS-188 | chr18-70968444-INS-188 |
| chr18-70984587-INS-149 | chr18-70984587-INS-149 |
| chr18-71004485-INS-78 | chr18-71004484-INS-80 |
| chr18-71014724-INS-72 | chr18-71014724-INS-72 |
| chr18-71023697-INS-2911 | chr18-71023698-INS-2911 |
| chr18-71044631-INS-3297 | chr18-71044632-INS-3297 |
| chr18-71052875-INS-66 | chr18-71052841-INS-78 |
| chr18-71073792-INS-108 | chr18-71073794-INS-76 |
| chr18-71093753-INS-52 | chr18-71093753-INS-52 |
| chr18-71105029-INS-92 | chr18-71105028-INS-78 |
| chr18-71109477-INS-54 | chr18-71109477-INS-54 |
| chr18-71174012-INS-62 | chr18-71174012-INS-72 |
| chr18-71178315-INS-83 | chr18-71178315-INS-83 |
| chr18-71194542-INS-69 | chr18-71194538-INS-97 |
| chr18-71212859-INS-111 | chr18-71212859-INS-119 |
| chr18-71223830-INS-1133 | chr18-71223831-INS-1133 |
| chr18-71245769-INS-76 | chr18-71245769-INS-80 |
| chr18-71271978-INS-403 | chr18-71271978-INS-403 |
| chr18-71313033-INS-993 | chr18-71313033-INS-994 |
| chr18-71327073-INS-75 | chr18-71327073-INS-103 |
| chr18-71346673-INS-73 | chr18-71346673-INS-73 |
| chr18-71361958-INS-115 | chr18-71361958-INS-103 |
| chr18-71375419-INS-52 | chr18-71375419-INS-74 |
| chr18-71382447-INS-60 | chr18-71382447-INS-60 |
| chr18-71452460-INS-987 | chr18-71452460-INS-986 |
| chr18-71484965-INS-3233 | chr18-71484966-INS-3250 |
| chr18-71495502-INS-56 | chr18-71495502-INS-56 |
| chr18-71497585-INS-105 | chr18-71497585-INS-105 |
| chr18-71525532-INS-1505 | chr18-71525533-INS-1504 |
| chr18-71527813-INS-61 | chr18-71527813-INS-62 |
| chr18-71552908-INS-104 | chr18-71552908-INS-104 |
| chr18-71570840-INS-70 | chr18-71570867-INS-57 |
| chr18-71574399-INS-78 | chr18-71574405-INS-81 |
| chr18-71587699-INS-68 | chr18-71587699-INS-72 |
| chr18-71604516-INS-54 | chr18-71604516-INS-54 |
| chr18-71615002-INS-187 | chr18-71615002-INS-186 |
| chr18-71642376-INS-287 | chr18-71642376-INS-285 |
| chr18-71673300-INS-79 | chr18-71673300-INS-79 |
| chr18-71727673-INS-154 | chr18-71727667-INS-155 |
| chr18-71732543-INS-54 | chr18-71732522-INS-61 |
| chr18-71749687-INS-52 | chr18-71749682-INS-91 |
| chr18-71780327-INS-214 | chr18-71780327-INS-213 |
| chr18-71781064-INS-1854 | chr18-71781065-INS-1854 |
| chr18-71796854-INS-224 | chr18-71796854-INS-238 |
| chr18-71810956-INS-1399 | chr18-71810957-INS-1400 |
| chr18-71810957-INS-498 | chr18-71810957-INS-498 |
| chr18-71851789-INS-58 | chr18-71851789-INS-58 |
| chr18-71852662-INS-200 | chr18-71852662-INS-201 |
| chr18-71868186-INS-826 | chr18-71868186-INS-826 |
| chr18-71894869-INS-728 | chr18-71894869-INS-728 |
| chr18-71911400-INS-84 | chr18-71911400-INS-86 |
| chr18-71955827-INS-391 | chr18-71955827-INS-392 |
| chr18-71970611-INS-6450 | chr18-71970612-INS-6452 |
| chr18-72013837-INS-108 | chr18-72013815-INS-127 |
| chr18-72036857-INS-72 | chr18-72036857-INS-72 |
| chr18-72061832-INS-187 | chr18-72061832-INS-203 |
| chr18-72095770-INS-60 | chr18-72095770-INS-52 |
| chr18-72121380-INS-68 | chr18-72121384-INS-68 |
| chr18-72146465-INS-78 | chr18-72146465-INS-81 |
| chr18-72177147-INS-471 | chr18-72177147-INS-471 |
| chr18-72180019-INS-60 | chr18-72180019-INS-60 |
| chr18-72193904-INS-88 | chr18-72193904-INS-90 |
| chr18-72208263-INS-56 | chr18-72208227-INS-50 |
| chr18-72269527-INS-213 | chr18-72269527-INS-213 |
| chr18-72270653-INS-497 | chr18-72270653-INS-500 |
| chr18-72281497-INS-250 | chr18-72281497-INS-240 |
| chr18-72352965-INS-96 | chr18-72352989-INS-108 |
| chr18-72360837-INS-5252 | chr18-72360838-INS-5252 |
| chr18-72364494-INS-86 | chr18-72364494-INS-87 |
| chr18-72384800-INS-164 | chr18-72384800-INS-164 |
| chr18-72386309-INS-126 | chr18-72386309-INS-126 |
| chr18-72403704-INS-894 | chr18-72403704-INS-894 |
| chr18-72440228-INS-740 | chr18-72440228-INS-740 |

| | |
|---|---|
| chr18-72440420-INS-6904 | chr18-72440421-INS-6904 |
| chr18-72449105-INS-234 | chr18-72449087-INS-242 |
| chr18-72452398-INS-86 | chr18-72452404-INS-82 |
| chr18-72454465-INS-181 | chr18-72454465-INS-183 |
| chr18-72482069-INS-187 | chr18-72482067-INS-191 |
| chr18-72502827-INS-69 | chr18-72502811-INS-72 |
| chr18-72510437-INS-156 | chr18-72510403-INS-204 |
| chr18-72513219-INS-154 | chr18-72513171-INS-158 |
| chr18-72515010-INS-220 | chr18-72515010-INS-220 |
| chr18-72516084-INS-777 | chr18-72516084-INS-777 |
| chr18-72527104-INS-524 | chr18-72527104-INS-524 |
| chr18-72545145-INS-181 | chr18-72545145-INS-158 |
| chr18-72546239-INS-6438 | chr18-72546240-INS-6440 |
| chr18-72558771-INS-51 | chr18-72558771-INS-59 |
| chr18-72562757-INS-1654 | chr18-72562758-INS-1654 |
| chr18-72568242-INS-6399 | chr18-72568243-INS-6399 |
| chr18-72574569-INS-58 | chr18-72574584-INS-54 |
| chr18-72597546-INS-281 | chr18-72597546-INS-281 |
| chr18-72609635-INS-57 | chr18-72609635-INS-57 |
| chr18-72654958-INS-203 | chr18-72654958-INS-203 |
| chr18-72661715-INS-57 | chr18-72661715-INS-52 |
| chr18-72663140-INS-828 | chr18-72663140-INS-828 |
| chr18-72663675-INS-298 | chr18-72663675-INS-298 |
| chr18-72678987-INS-54 | chr18-72678989-INS-64 |
| chr18-72681817-INS-101 | chr18-72681817-INS-101 |
| chr18-72694071-INS-52 | chr18-72694071-INS-52 |
| chr18-72696002-INS-67 | chr18-72696002-INS-73 |
| chr18-72727531-INS-387 | chr18-72727531-INS-385 |
| chr18-72786837-INS-51 | chr18-72786837-INS-59 |
| chr18-72797773-INS-75 | chr18-72797773-INS-85 |
| chr18-72823563-INS-76 | chr18-72823562-INS-80 |
| chr18-72828857-INS-6347 | chr18-72828858-INS-6347 |
| chr18-72845448-INS-78 | chr18-72845448-INS-76 |
| chr18-72877384-INS-83 | chr18-72877393-INS-87 |
| chr18-72905504-INS-441 | chr18-72905504-INS-480 |
| chr18-72936772-INS-1393 | chr18-72936773-INS-1394 |
| chr18-72945795-INS-806 | chr18-72945795-INS-806 |
| chr18-72960790-INS-87 | chr18-72960790-INS-111 |
| chr18-72974043-INS-52 | chr18-72974058-INS-53 |
| chr18-72998753-INS-436 | chr18-72998753-INS-433 |
| chr18-73010678-INS-201 | chr18-73010678-INS-201 |
| chr18-73014728-INS-685 | chr18-73014728-INS-686 |
| chr18-73036839-INS-6212 | chr18-73036840-INS-6210 |
| chr18-73056549-INS-78 | chr18-73056549-INS-64 |
| chr18-73061836-INS-6202 | chr18-73061837-INS-6202 |
| chr18-73065891-INS-72 | chr18-73065891-INS-60 |
| chr18-73075579-INS-68 | chr18-73075569-INS-62 |
| chr18-73119671-INS-61 | chr18-73119671-INS-61 |
| chr18-73122179-INS-564 | chr18-73122179-INS-564 |
| chr18-73131939-INS-552 | chr18-73131939-INS-555 |
| chr18-73134149-INS-751 | chr18-73134149-INS-770 |
| chr18-73136099-INS-53 | chr18-73136099-INS-65 |
| chr18-73150591-INS-327 | chr18-73150591-INS-327 |
| chr18-73160316-INS-337 | chr18-73160316-INS-337 |
| chr18-73173845-INS-54 | chr18-73173845-INS-54 |
| chr18-73191879-INS-87 | chr18-73191839-INS-117 |
| chr18-73288429-INS-7403 | chr18-73288430-INS-7405 |
| chr18-73291826-INS-101 | chr18-73291826-INS-101 |
| chr18-73298131-INS-72 | chr18-73298166-INS-53 |
| chr18-73305668-INS-364 | chr18-73305668-INS-364 |
| chr18-73313631-INS-400 | chr18-73313631-INS-400 |
| chr18-73321666-INS-395 | chr18-73321666-INS-395 |
| chr18-73329019-INS-64 | chr18-73329036-INS-68 |
| chr18-73373644-INS-7210 | chr18-73373645-INS-7210 |
| chr18-73377647-INS-60 | chr18-73377647-INS-54 |
| chr18-73401698-INS-52 | chr18-73401698-INS-52 |
| chr18-73423222-INS-401 | chr18-73423222-INS-401 |
| chr18-73448965-INS-135 | chr18-73449006-INS-98 |
| chr18-73457515-INS-83 | chr18-73457515-INS-85 |
| chr18-73464185-INS-415 | chr18-73464179-INS-400 |
| chr18-73487778-INS-266 | chr18-73487792-INS-262 |
| chr18-73491501-INS-68 | chr18-73491501-INS-68 |
| chr18-73498406-INS-2317 | chr18-73498407-INS-2347 |
| chr18-73506029-INS-514 | chr18-73506029-INS-514 |
| chr18-73511533-INS-91 | chr18-73511533-INS-91 |
| chr18-73559024-INS-60 | chr18-73559024-INS-60 |
| chr18-73573584-INS-80 | chr18-73573584-INS-118 |

| | |
|---|---|
| chr18-73575270-INS-227 | chr18-73575270-INS-227 |
| chr18-73580642-INS-52 | chr18-73580642-INS-50 |
| chr18-73583333-INS-337 | chr18-73583333-INS-342 |
| chr18-73587127-INS-5445 | chr18-73587128-INS-5445 |
| chr18-73594732-INS-60 | chr18-73594732-INS-58 |
| chr18-73610510-INS-168 | chr18-73610510-INS-168 |
| chr18-73625403-INS-209 | chr18-73625403-INS-209 |
| chr18-73628948-INS-53 | chr18-73628947-INS-51 |
| chr18-73631751-INS-51 | chr18-73631751-INS-51 |
| chr18-73643169-INS-212 | chr18-73643169-INS-212 |
| chr18-73649488-INS-1890 | chr18-73649489-INS-1890 |
| chr18-73665869-INS-52 | chr18-73665859-INS-55 |
| chr18-73667801-INS-71 | chr18-73667802-INS-53 |
| chr18-73676239-INS-1933 | chr18-73676240-INS-1934 |
| chr18-73677423-INS-75 | chr18-73677411-INS-84 |
| chr18-73681402-INS-200 | chr18-73681402-INS-200 |
| chr18-73691137-INS-60 | chr18-73691137-INS-60 |
| chr18-73694788-INS-176 | chr18-73694788-INS-176 |
| chr18-73707911-INS-56 | chr18-73707911-INS-52 |
| chr18-73716191-INS-6434 | chr18-73716192-INS-6434 |
| chr18-73738490-INS-3966 | chr18-73738491-INS-3970 |
| chr18-73741400-INS-168 | chr18-73741400-INS-168 |
| chr18-73745965-INS-356 | chr18-73745965-INS-356 |
| chr18-73774275-INS-54 | chr18-73774228-INS-54 |
| chr18-73782408-INS-170 | chr18-73782408-INS-149 |
| chr18-73785681-INS-340 | chr18-73785681-INS-298 |
| chr18-73794203-INS-110 | chr18-73794203-INS-112 |
| chr18-73838897-INS-209 | chr18-73838897-INS-208 |
| chr18-73848595-INS-54 | chr18-73848595-INS-54 |
| chr18-73861018-INS-346 | chr18-73861018-INS-346 |
| chr18-73900727-INS-54 | chr18-73900731-INS-60 |
| chr18-73908926-INS-52 | chr18-73908926-INS-52 |
| chr18-73909421-INS-205 | chr18-73909421-INS-205 |
| chr18-73911935-INS-52 | chr18-73911935-INS-56 |
| chr18-73915679-INS-315 | chr18-73915679-INS-315 |
| chr18-73933164-INS-116 | chr18-73933164-INS-116 |
| chr18-73937075-INS-94 | chr18-73937075-INS-94 |
| chr18-73941666-INS-493 | chr18-73941666-INS-493 |
| chr18-73978219-INS-90 | chr18-73978219-INS-90 |
| chr18-73996971-INS-52 | chr18-73996971-INS-55 |
| chr18-74008221-INS-141 | chr18-74008222-INS-141 |
| chr18-74012017-INS-50 | chr18-74012017-INS-54 |
| chr18-74033907-INS-60 | chr18-74033908-INS-69 |
| chr18-74043547-INS-56 | chr18-74043547-INS-58 |
| chr18-74053191-INS-74 | chr18-74053191-INS-74 |
| chr18-74057404-INS-360 | chr18-74057404-INS-360 |
| chr18-74070587-INS-594 | chr18-74070588-INS-599 |
| chr18-74115472-INS-90 | chr18-74115475-INS-110 |
| chr18-74126713-INS-78 | chr18-74126713-INS-84 |
| chr18-74134235-INS-176 | chr18-74134236-INS-176 |
| chr18-74137653-INS-55 | chr18-74137653-INS-60 |
| chr18-74141578-INS-52 | chr18-74141578-INS-52 |
| chr18-74143840-INS-67 | chr18-74143840-INS-67 |
| chr18-74161749-INS-545 | chr18-74161749-INS-547 |
| chr18-74182846-INS-162 | chr18-74182850-INS-151 |
| chr18-74184126-INS-116 | chr18-74184126-INS-112 |
| chr18-74192395-INS-182 | chr18-74192395-INS-182 |
| chr18-74248602-INS-209 | chr18-74248597-INS-238 |
| chr18-74264811-INS-5208 | chr18-74264812-INS-5208 |
| chr18-74267690-INS-940 | chr18-74267690-INS-940 |
| chr18-74281867-INS-152 | chr18-74281867-INS-152 |
| chr18-74297322-INS-112 | chr18-74297322-INS-112 |
| chr18-74300368-INS-1586 | chr18-74300369-INS-1586 |
| chr18-74307589-INS-51 | chr18-74307589-INS-51 |
| chr18-74310120-INS-76 | chr18-74310135-INS-53 |
| chr18-74334363-INS-101 | chr18-74334363-INS-101 |
| chr18-74336226-INS-205 | chr18-74336227-INS-172 |
| chr18-74347583-INS-64 | chr18-74347583-INS-76 |
| chr18-74364632-INS-110 | chr18-74364632-INS-111 |
| chr18-74410767-INS-58 | chr18-74410779-INS-68 |
| chr18-74438982-INS-52 | chr18-74438982-INS-52 |
| chr18-74462822-INS-510 | chr18-74462822-INS-509 |
| chr18-74469098-INS-86 | chr18-74469098-INS-92 |
| chr18-74473740-INS-57 | chr18-74473740-INS-96 |
| chr18-74524777-INS-90 | chr18-74524777-INS-97 |
| chr18-74533794-INS-217 | chr18-74533794-INS-218 |
| chr18-74550752-INS-6772 | chr18-74550753-INS-6772 |

| | |
|---|---|
| chr18-74553567-INS-70 | chr18-74553567-INS-70 |
| chr18-74562083-INS-6488 | chr18-74562084-INS-6488 |
| chr18-74606938-INS-445 | chr18-74606938-INS-445 |
| chr18-74615043-INS-1050 | chr18-74615043-INS-1040 |
| chr18-74616772-INS-95 | chr18-74616772-INS-92 |
| chr18-74623117-INS-78 | chr18-74623117-INS-52 |
| chr18-74633584-INS-65 | chr18-74633584-INS-61 |
| chr18-74662279-INS-351 | chr18-74662279-INS-345 |
| chr18-74689394-INS-162 | chr18-74689435-INS-151 |
| chr18-74693939-INS-182 | chr18-74693939-INS-174 |
| chr18-74694521-INS-497 | chr18-74694521-INS-497 |
| chr18-74720621-INS-89 | chr18-74720608-INS-90 |
| chr18-74746340-INS-78 | chr18-74746340-INS-87 |
| chr18-74812788-INS-95 | chr18-74812825-INS-96 |
| chr18-74817909-INS-56 | chr18-74817906-INS-50 |
| chr18-74821835-INS-67 | chr18-74821823-INS-79 |
| chr18-74847096-INS-84 | chr18-74847096-INS-84 |
| chr18-74903138-INS-218 | chr18-74903138-INS-218 |
| chr18-74922210-INS-72 | chr18-74922168-INS-98 |
| chr18-74928020-INS-213 | chr18-74928020-INS-212 |
| chr18-74929487-INS-148 | chr18-74929487-INS-148 |
| chr18-74943661-INS-2776 | chr18-74943662-INS-2776 |
| chr18-74966652-INS-436 | chr18-74966655-INS-448 |
| chr18-74968729-INS-110 | chr18-74968730-INS-110 |
| chr18-74980393-INS-78 | chr18-74980393-INS-78 |
| chr18-74985981-INS-309 | chr18-74985981-INS-309 |
| chr18-74989735-INS-910 | chr18-74989735-INS-915 |
| chr18-74992834-INS-208 | chr18-74992834-INS-208 |
| chr18-74997146-INS-446 | chr18-74997146-INS-444 |
| chr18-75002382-INS-84 | chr18-75002382-INS-84 |
| chr18-75025686-INS-105 | chr18-75025686-INS-83 |
| chr18-75034107-INS-396 | chr18-75034107-INS-398 |
| chr18-75070354-INS-51 | chr18-75070354-INS-51 |
| chr18-75076725-INS-126 | chr18-75076718-INS-168 |
| chr18-75099056-INS-52 | chr18-75099056-INS-56 |
| chr18-75129836-INS-132 | chr18-75129836-INS-132 |
| chr18-75145594-INS-101 | chr18-75145594-INS-101 |
| chr18-75149726-INS-201 | chr18-75149726-INS-205 |
| chr18-75158790-INS-64 | chr18-75158779-INS-67 |
| chr18-75167450-INS-212 | chr18-75167450-INS-212 |
| chr18-75173221-INS-54 | chr18-75173221-INS-54 |
| chr18-75180020-INS-153 | chr18-75180020-INS-145 |
| chr18-75190237-INS-170 | chr18-75190237-INS-170 |
| chr18-75191162-INS-760 | chr18-75191162-INS-774 |
| chr18-75192980-INS-112 | chr18-75192980-INS-147 |
| chr18-75204234-INS-943 | chr18-75204234-INS-944 |
| chr18-75206596-INS-96 | chr18-75206596-INS-102 |
| chr18-75218775-INS-66 | chr18-75218775-INS-66 |
| chr18-75221706-INS-90 | chr18-75221706-INS-90 |
| chr18-75228253-INS-216 | chr18-75228253-INS-216 |
| chr18-75230716-INS-56 | chr18-75230716-INS-56 |
| chr18-75235261-INS-397 | chr18-75235261-INS-397 |
| chr18-75270536-INS-114 | chr18-75270526-INS-76 |
| chr18-75272950-INS-69 | chr18-75272950-INS-69 |
| chr18-75286956-INS-202 | chr18-75286956-INS-202 |
| chr18-75303998-INS-490 | chr18-75303998-INS-490 |
| chr18-75308262-INS-264 | chr18-75308262-INS-264 |
| chr18-75334770-INS-666 | chr18-75334770-INS-662 |
| chr18-75339863-INS-120 | chr18-75339863-INS-80 |
| chr18-75348931-INS-156 | chr18-75348931-INS-156 |
| chr18-75363532-INS-58 | chr18-75363532-INS-58 |
| chr18-75381835-INS-213 | chr18-75381835-INS-213 |
| chr18-75385534-INS-199 | chr18-75385534-INS-199 |
| chr18-75401287-INS-212 | chr18-75401287-INS-210 |
| chr18-75402835-INS-225 | chr18-75402835-INS-226 |
| chr18-75420563-INS-220 | chr18-75420563-INS-222 |
| chr18-75425705-INS-183 | chr18-75425695-INS-168 |
| chr18-75433490-INS-87 | chr18-75433477-INS-58 |
| chr18-75443174-INS-126 | chr18-75443174-INS-126 |
| chr18-75460579-INS-52 | chr18-75460579-INS-52 |
| chr18-75464346-INS-86 | chr18-75464346-INS-91 |
| chr18-75470793-INS-65 | chr18-75470793-INS-66 |
| chr18-75479115-INS-466 | chr18-75479115-INS-466 |
| chr18-75480631-INS-207 | chr18-75480631-INS-209 |
| chr18-75481928-INS-81 | chr18-75481937-INS-87 |
| chr18-75484208-INS-51 | chr18-75484208-INS-53 |
| chr18-75487545-INS-211 | chr18-75487545-INS-211 |

| | |
|---|---|
| chr18-75518645-INS-223 | chr18-75518645-INS-223 |
| chr18-75537121-INS-193 | chr18-75537129-INS-165 |
| chr18-75539808-INS-123 | chr18-75539808-INS-123 |
| chr18-75546687-INS-198 | chr18-75546687-INS-197 |
| chr18-75561292-INS-99 | chr18-75561292-INS-100 |
| chr18-75569088-INS-82 | chr18-75569088-INS-102 |
| chr18-75606273-INS-206 | chr18-75606273-INS-206 |
| chr18-75613582-INS-70 | chr18-75613582-INS-70 |
| chr18-75615156-INS-152 | chr18-75615156-INS-152 |
| chr18-75616736-INS-488 | chr18-75616736-INS-492 |
| chr18-75620131-INS-338 | chr18-75620131-INS-339 |
| chr18-75621434-INS-201 | chr18-75621434-INS-202 |
| chr18-75623729-INS-219 | chr18-75623729-INS-219 |
| chr18-75629708-INS-236 | chr18-75629708-INS-236 |
| chr18-75643177-INS-205 | chr18-75643177-INS-205 |
| chr18-75655487-INS-92 | chr18-75655487-INS-92 |
| chr18-75665364-INS-377 | chr18-75665364-INS-387 |
| chr18-75666916-INS-80 | chr18-75666916-INS-80 |
| chr18-75670444-INS-54 | chr18-75670444-INS-54 |
| chr18-75695096-INS-4981 | chr18-75695097-INS-4981 |
| chr18-75702479-INS-205 | chr18-75702479-INS-205 |
| chr18-75703828-INS-178 | chr18-75703828-INS-178 |
| chr18-75707738-INS-393 | chr18-75707738-INS-392 |
| chr18-75757618-INS-2215 | chr18-75757619-INS-2215 |
| chr18-75761759-INS-69 | chr18-75761759-INS-106 |
| chr18-75764493-INS-54 | chr18-75764493-INS-54 |
| chr18-75765797-INS-104 | chr18-75765797-INS-102 |
| chr18-75782693-INS-112 | chr18-75782693-INS-110 |
| chr18-75804844-INS-55 | chr18-75804876-INS-71 |
| chr18-75885267-INS-878 | chr18-75885267-INS-878 |
| chr18-75888091-INS-233 | chr18-75888092-INS-233 |
| chr18-75904827-INS-101 | chr18-75904827-INS-101 |
| chr18-75911120-INS-809 | chr18-75911120-INS-809 |
| chr18-75913742-INS-54 | chr18-75913742-INS-54 |
| chr18-75923672-INS-205 | chr18-75923672-INS-203 |
| chr18-75929557-INS-195 | chr18-75929557-INS-195 |
| chr18-75949508-INS-78 | chr18-75949508-INS-78 |
| chr18-75951726-INS-60 | chr18-75951726-INS-60 |
| chr18-75976498-INS-52 | chr18-75976498-INS-50 |
| chr18-75983418-INS-653 | chr18-75983418-INS-652 |
| chr18-75991331-INS-57 | chr18-75991360-INS-54 |
| chr18-75993141-INS-54 | chr18-75993140-INS-54 |
| chr18-76013627-INS-60 | chr18-76013627-INS-90 |
| chr18-76052676-INS-52 | chr18-76052676-INS-58 |
| chr18-76074326-INS-338 | chr18-76074326-INS-338 |
| chr18-76083196-INS-71 | chr18-76083196-INS-71 |
| chr18-76086718-INS-71 | chr18-76086718-INS-71 |
| chr18-76091807-INS-63 | chr18-76091808-INS-62 |
| chr18-76101753-INS-52 | chr18-76101753-INS-54 |
| chr18-76107647-INS-55 | chr18-76107647-INS-55 |
| chr18-76124288-INS-114 | chr18-76124288-INS-114 |
| chr18-76143400-INS-59 | chr18-76143400-INS-59 |
| chr18-76146074-INS-54 | chr18-76146074-INS-52 |
| chr18-76187426-INS-55 | chr18-76187426-INS-50 |
| chr18-76203564-INS-74 | chr18-76203564-INS-74 |
| chr18-76211175-INS-60 | chr18-76211175-INS-60 |
| chr18-76220162-INS-252 | chr18-76220162-INS-252 |
| chr18-76284640-INS-74 | chr18-76284640-INS-82 |
| chr18-76310612-INS-68 | chr18-76310564-INS-96 |
| chr18-76339799-INS-248 | chr18-76339799-INS-255 |
| chr18-76349636-INS-51 | chr18-76349636-INS-71 |
| chr18-76393268-INS-1891 | chr18-76393269-INS-1891 |
| chr18-76403747-INS-440 | chr18-76403747-INS-440 |
| chr18-76403981-INS-3036 | chr18-76403982-INS-3034 |
| chr18-76457012-INS-90 | chr18-76457034-INS-54 |
| chr18-76460283-INS-314 | chr18-76460283-INS-316 |
| chr18-76474289-INS-80 | chr18-76474289-INS-80 |
| chr18-76487836-INS-51 | chr18-76487836-INS-51 |
| chr18-76489946-INS-1576 | chr18-76489947-INS-1573 |
| chr18-76546903-INS-130 | chr18-76546903-INS-130 |
| chr18-76566616-INS-115 | chr18-76566616-INS-115 |
| chr18-76598817-INS-343 | chr18-76598817-INS-343 |
| chr18-76611270-INS-180 | chr18-76611232-INS-197 |
| chr18-76615130-INS-789 | chr18-76615130-INS-825 |
| chr18-76625540-INS-54 | chr18-76625540-INS-57 |
| chr18-76638487-INS-141 | chr18-76638479-INS-140 |
| chr18-76646045-INS-87 | chr18-76646045-INS-89 |

| | |
|---|---|
| chr18-76691275-INS-93 | chr18-76691278-INS-72 |
| chr18-76693326-INS-54 | chr18-76693326-INS-56 |
| chr18-76718958-INS-66 | chr18-76718958-INS-62 |
| chr18-76722393-INS-111 | chr18-76722393-INS-95 |
| chr18-76744591-INS-197 | chr18-76744591-INS-199 |
| chr18-76745621-INS-87 | chr18-76745621-INS-87 |
| chr18-76776076-INS-58 | chr18-76776076-INS-58 |
| chr18-76788415-INS-75 | chr18-76788415-INS-75 |
| chr18-76798432-INS-66 | chr18-76798432-INS-78 |
| chr18-76800311-INS-70 | chr18-76800314-INS-68 |
| chr18-76808010-INS-60 | chr18-76808010-INS-60 |
| chr18-76814826-INS-204 | chr18-76814826-INS-206 |
| chr18-76837381-INS-74 | chr18-76837381-INS-74 |
| chr18-76844897-INS-93 | chr18-76844916-INS-105 |
| chr18-76882104-INS-70 | chr18-76882101-INS-101 |
| chr18-76913954-INS-140 | chr18-76913954-INS-140 |
| chr18-76919392-INS-61 | chr18-76919392-INS-61 |
| chr18-76926467-INS-87 | chr18-76926467-INS-104 |
| chr18-76928943-INS-273 | chr18-76928928-INS-304 |
| chr18-76932854-INS-80 | chr18-76932853-INS-56 |
| chr18-76945769-INS-65 | chr18-76945769-INS-57 |
| chr18-76986621-INS-138 | chr18-76986621-INS-90 |
| chr18-77012103-INS-58 | chr18-77012103-INS-58 |
| chr18-77013580-INS-99 | chr18-77013586-INS-121 |
| chr18-77032480-INS-207 | chr18-77032480-INS-209 |
| chr18-77038270-INS-56 | chr18-77038274-INS-68 |
| chr18-77067280-INS-52 | chr18-77067280-INS-86 |
| chr18-77076895-INS-94 | chr18-77076895-INS-141 |
| chr18-77113459-INS-199 | chr18-77113460-INS-198 |
| chr18-77120866-INS-548 | chr18-77120864-INS-560 |
| chr18-77124768-INS-1864 | chr18-77124769-INS-1864 |
| chr18-77131385-INS-88 | chr18-77131385-INS-79 |
| chr18-77139987-INS-241 | chr18-77139987-INS-241 |
| chr18-77142333-INS-53 | chr18-77142333-INS-53 |
| chr18-77172924-INS-80 | chr18-77172929-INS-77 |
| chr18-77187412-INS-226 | chr18-77187412-INS-226 |
| chr18-77242886-INS-58 | chr18-77242886-INS-58 |
| chr18-77257096-INS-163 | chr18-77257096-INS-196 |
| chr18-77258755-INS-142 | chr18-77258755-INS-142 |
| chr18-77260682-INS-56 | chr18-77260682-INS-56 |
| chr18-77283504-INS-64 | chr18-77283520-INS-68 |
| chr18-77291299-INS-58 | chr18-77291339-INS-52 |
| chr18-77292754-INS-165 | chr18-77292781-INS-157 |
| chr18-77318643-INS-205 | chr18-77318643-INS-205 |
| chr18-77324585-INS-96 | chr18-77324588-INS-96 |
| chr18-77330688-INS-65 | chr18-77330694-INS-55 |
| chr18-77345665-INS-155 | chr18-77345693-INS-155 |
| chr18-77349829-INS-144 | chr18-77349835-INS-140 |
| chr18-77359397-INS-618 | chr18-77359397-INS-619 |
| chr18-77375361-INS-872 | chr18-77375361-INS-872 |
| chr18-77386210-INS-6442 | chr18-77386211-INS-6442 |
| chr18-77420854-INS-426 | chr18-77420889-INS-397 |
| chr18-77422361-INS-66 | chr18-77422361-INS-50 |
| chr18-77430384-INS-94 | chr18-77430384-INS-63 |
| chr18-77437777-INS-98 | chr18-77437777-INS-98 |
| chr18-77440136-INS-142 | chr18-77440136-INS-145 |
| chr18-77471090-INS-86 | chr18-77471091-INS-72 |
| chr18-77472981-INS-64 | chr18-77472981-INS-66 |
| chr18-77474776-INS-213 | chr18-77474776-INS-213 |
| chr18-77498652-INS-433 | chr18-77498652-INS-433 |
| chr18-77519853-INS-54 | chr18-77519854-INS-78 |
| chr18-77526331-INS-54 | chr18-77526301-INS-87 |
| chr18-77543431-INS-90 | chr18-77543431-INS-92 |
| chr18-77552208-INS-222 | chr18-77552208-INS-222 |
| chr18-77552742-INS-879 | chr18-77552742-INS-879 |
| chr18-77591643-INS-82 | chr18-77591643-INS-78 |
| chr18-77603158-INS-52 | chr18-77603158-INS-52 |
| chr18-77622459-INS-200 | chr18-77622459-INS-200 |
| chr18-77655038-INS-77 | chr18-77655038-INS-99 |
| chr18-77664744-INS-202 | chr18-77664744-INS-202 |
| chr18-77666244-INS-159 | chr18-77666244-INS-158 |
| chr18-77668312-INS-205 | chr18-77668315-INS-199 |
| chr18-77677155-INS-989 | chr18-77677155-INS-989 |
| chr18-77678327-INS-198 | chr18-77678334-INS-199 |
| chr18-77689443-INS-107 | chr18-77689443-INS-107 |
| chr18-77699977-INS-220 | chr18-77699977-INS-220 |
| chr18-77704400-INS-117 | chr18-77704400-INS-117 |

| | |
|---|---|
| chr18-77707091-INS-53 | chr18-77707092-INS-58 |
| chr18-77712078-INS-199 | chr18-77712078-INS-199 |
| chr18-77731491-INS-216 | chr18-77731491-INS-215 |
| chr18-77734477-INS-205 | chr18-77734477-INS-205 |
| chr18-77738454-INS-201 | chr18-77738454-INS-201 |
| chr18-77746532-INS-176 | chr18-77746532-INS-176 |
| chr18-77748013-INS-230 | chr18-77748013-INS-230 |
| chr18-77754344-INS-72 | chr18-77754344-INS-76 |
| chr18-77789697-INS-52 | chr18-77789697-INS-52 |
| chr18-77818430-INS-663 | chr18-77818434-INS-706 |
| chr18-77829112-INS-76 | chr18-77829099-INS-80 |
| chr18-77834537-INS-201 | chr18-77834537-INS-201 |
| chr18-77842553-INS-219 | chr18-77842553-INS-219 |
| chr18-77843747-INS-207 | chr18-77843743-INS-200 |
| chr18-77847713-INS-195 | chr18-77847713-INS-195 |
| chr18-77850445-INS-200 | chr18-77850445-INS-201 |
| chr18-77854401-INS-245 | chr18-77854399-INS-259 |
| chr18-77860012-INS-208 | chr18-77860012-INS-208 |
| chr18-77887090-INS-171 | chr18-77887090-INS-172 |
| chr18-77889876-INS-62 | chr18-77889856-INS-90 |
| chr18-77899466-INS-77 | chr18-77899466-INS-67 |
| chr18-77911403-INS-227 | chr18-77911403-INS-227 |
| chr18-77919600-INS-162 | chr18-77919594-INS-177 |
| chr18-77923515-INS-230 | chr18-77923551-INS-238 |
| chr18-77934727-INS-192 | chr18-77934727-INS-193 |
| chr18-77938614-INS-181 | chr18-77938614-INS-181 |
| chr18-77944844-INS-198 | chr18-77944844-INS-198 |
| chr18-77946085-INS-196 | chr18-77946093-INS-198 |
| chr18-77947502-INS-651 | chr18-77947503-INS-647 |
| chr18-77949465-INS-58 | chr18-77949465-INS-58 |
| chr18-77956481-INS-219 | chr18-77956481-INS-219 |
| chr18-77963248-INS-143 | chr18-77963248-INS-139 |
| chr18-77968913-INS-60 | chr18-77968913-INS-60 |
| chr18-77973340-INS-186 | chr18-77973385-INS-184 |
| chr18-78006453-INS-75 | chr18-78006411-INS-75 |
| chr18-78027628-INS-485 | chr18-78027628-INS-485 |
| chr18-78057639-INS-120 | chr18-78057639-INS-134 |
| chr18-78074324-INS-140 | chr18-78074324-INS-140 |
| chr18-78076990-INS-200 | chr18-78076990-INS-200 |
| chr18-78094174-INS-151 | chr18-78094187-INS-162 |
| chr18-78103446-INS-114 | chr18-78103470-INS-102 |
| chr18-78133900-INS-3117 | chr18-78133901-INS-3117 |
| chr18-78171435-INS-3190 | chr18-78171436-INS-3186 |
| chr18-78174606-INS-168 | chr18-78174630-INS-119 |
| chr18-78177523-INS-123 | chr18-78177523-INS-123 |
| chr18-78194447-INS-149 | chr18-78194411-INS-197 |
| chr18-78288175-INS-85 | chr18-78288175-INS-85 |
| chr18-78357901-INS-78 | chr18-78357901-INS-60 |
| chr18-78360798-INS-61 | chr18-78360798-INS-61 |
| chr18-78364120-INS-96 | chr18-78364120-INS-96 |
| chr18-78384842-INS-86 | chr18-78384842-INS-86 |
| chr18-78388960-INS-606 | chr18-78388958-INS-610 |
| chr18-78397612-INS-80 | chr18-78397651-INS-77 |
| chr18-78404626-INS-96 | chr18-78404626-INS-103 |
| chr18-78412270-INS-157 | chr18-78412270-INS-157 |
| chr18-78450691-INS-64 | chr18-78450691-INS-88 |
| chr18-78460581-INS-232 | chr18-78460581-INS-232 |
| chr18-78489581-INS-155 | chr18-78489583-INS-155 |
| chr18-78571511-INS-56 | chr18-78571511-INS-52 |
| chr18-78628532-INS-78 | chr18-78628532-INS-78 |
| chr18-78650137-INS-66 | chr18-78650137-INS-66 |
| chr18-78665222-INS-118 | chr18-78665222-INS-118 |
| chr18-78672766-INS-95 | chr18-78672766-INS-100 |
| chr18-78697120-INS-876 | chr18-78697120-INS-903 |
| chr18-78739642-INS-1068 | chr18-78739643-INS-1068 |
| chr18-78780945-INS-1893 | chr18-78780946-INS-1893 |
| chr18-78792448-INS-256 | chr18-78792448-INS-256 |
| chr18-78861363-INS-93 | chr18-78861367-INS-115 |
| chr18-78867227-INS-58 | chr18-78867227-INS-58 |
| chr18-78882759-INS-56 | chr18-78882760-INS-62 |
| chr18-78887447-INS-106 | chr18-78887447-INS-106 |
| chr18-78970201-INS-326 | chr18-78970201-INS-326 |
| chr18-78978913-INS-69 | chr18-78978922-INS-75 |
| chr18-79054861-INS-70 | chr18-79054861-INS-70 |
| chr18-79058578-INS-233 | chr18-79058569-INS-201 |
| chr18-79072128-INS-516 | chr18-79072128-INS-537 |
| chr18-79103919-INS-93 | chr18-79103921-INS-93 |

| | |
|---|---|
| chr18-79122646-INS-57 | chr18-79122646-INS-57 |
| chr18-79159799-INS-78 | chr18-79159799-INS-78 |
| chr18-79170721-INS-64 | chr18-79170721-INS-70 |
| chr18-79174894-INS-67 | chr18-79174894-INS-63 |
| chr18-79184828-INS-129 | chr18-79184838-INS-134 |
| chr18-79193723-INS-464 | chr18-79193723-INS-464 |
| chr18-79211220-INS-6195 | chr18-79211221-INS-6195 |
| chr18-79221098-INS-480 | chr18-79221098-INS-480 |
| chr18-79227904-INS-54 | chr18-79227904-INS-54 |
| chr18-79238607-INS-65 | chr18-79238607-INS-65 |
| chr18-79240495-INS-887 | chr18-79240495-INS-887 |
| chr18-79243136-INS-1930 | chr18-79243137-INS-1932 |
| chr18-79252623-INS-98 | chr18-79252623-INS-83 |
| chr18-79274771-INS-81 | chr18-79274761-INS-116 |
| chr18-79285555-INS-76 | chr18-79285553-INS-56 |
| chr18-79312339-INS-6428 | chr18-79312340-INS-6428 |
| chr18-79358696-INS-284 | chr18-79358697-INS-284 |
| chr18-79376097-INS-68 | chr18-79376097-INS-68 |
| chr18-79388115-INS-705 | chr18-79388115-INS-700 |
| chr18-79409054-INS-199 | chr18-79409054-INS-199 |
| chr18-79484750-INS-472 | chr18-79484751-INS-472 |
| chr18-79515342-INS-454 | chr18-79515342-INS-454 |
| chr18-79532418-INS-64 | chr18-79532418-INS-64 |
| chr18-79548955-INS-104 | chr18-79548955-INS-110 |
| chr18-79606001-INS-148 | chr18-79606001-INS-147 |
| chr18-79637169-INS-169 | chr18-79637169-INS-169 |
| chr18-79669981-INS-68 | chr18-79669981-INS-68 |
| chr18-79696843-INS-1619 | chr18-79696844-INS-1607 |
| chr18-79697622-INS-584 | chr18-79697622-INS-584 |
| chr18-79713755-INS-929 | chr18-79713755-INS-929 |
| chr18-79722860-INS-326 | chr18-79722860-INS-326 |
| chr18-79744145-INS-118 | chr18-79744142-INS-74 |
| chr18-79792518-INS-214 | chr18-79792518-INS-214 |
| chr18-79794475-INS-68 | chr18-79794475-INS-60 |
| chr18-79797091-INS-75 | chr18-79797091-INS-81 |
| chr18-79806852-INS-837 | chr18-79806852-INS-837 |
| chr18-79822374-INS-208 | chr18-79822374-INS-208 |
| chr18-79824805-INS-290 | chr18-79824805-INS-290 |
| chr18-79831041-INS-6436 | chr18-79831042-INS-6436 |
| chr18-79831720-INS-67 | chr18-79831717-INS-83 |
| chr18-79862900-INS-354 | chr18-79862900-INS-354 |
| chr18-79906586-INS-553 | chr18-79906586-INS-545 |
| chr18-79907921-INS-206 | chr18-79907921-INS-205 |
| chr18-79910677-INS-54 | chr18-79910687-INS-64 |
| chr18-79922458-INS-66 | chr18-79922458-INS-66 |
| chr18-79925983-INS-105 | chr18-79925996-INS-106 |
| chr18-79943787-INS-74 | chr18-79943767-INS-67 |
| chr18-79960234-INS-436 | chr18-79960234-INS-436 |
| chr18-79976612-INS-399 | chr18-79976612-INS-399 |
| chr18-79983077-INS-6440 | chr18-79983078-INS-6440 |
| chr18-80001578-INS-126 | chr18-80001579-INS-133 |
| chr18-80010849-INS-84 | chr18-80010849-INS-84 |
| chr18-80012408-INS-281 | chr18-80012408-INS-282 |
| chr18-80025614-INS-62 | chr18-80025614-INS-68 |
| chr18-80026871-INS-4562 | chr18-80026872-INS-4562 |
| chr18-80032122-INS-57 | chr18-80032127-INS-50 |
| chr18-80038119-INS-51 | chr18-80038119-INS-55 |
| chr18-80097493-INS-66 | chr18-80097493-INS-66 |
| chr18-80141609-INS-199 | chr18-80141609-INS-199 |
| chr18-80152714-INS-215 | chr18-80152714-INS-214 |
| chr18-80175376-INS-54 | chr18-80175376-INS-54 |
| chr18-80179837-INS-209 | chr18-80179837-INS-210 |
| chr18-80186241-INS-62 | chr18-80186241-INS-62 |
| chr18-80208389-INS-134 | chr18-80208383-INS-168 |
| chr18-80218716-INS-164 | chr18-80218716-INS-186 |
| chr18-80231914-INS-206 | chr18-80231914-INS-206 |
| chr18-80245169-INS-462 | chr18-80245169-INS-465 |
| chr18-80248809-INS-398 | chr18-80248810-INS-398 |
| chr18-80252723-INS-207 | chr18-80252723-INS-207 |
| chr18-80256223-INS-204 | chr18-80256238-INS-207 |
| chr18-80262835-INS-162 | chr18-80262835-INS-162 |
| chr18-80271198-INS-299 | chr18-80271198-INS-299 |
| chr18-80289167-INS-193 | chr18-80289167-INS-193 |
| chr18-80289988-INS-186 | chr18-80289988-INS-186 |
| chr18-80290604-INS-207 | chr18-80290604-INS-207 |
| chr18-80293322-INS-202 | chr18-80293322-INS-202 |
| chr18-80310101-INS-72 | chr18-80310101-INS-72 |

| | |
|---|---|
| chr18-80318192-INS-214 | chr18-80318192-INS-214 |
| chr18-80335070-INS-57 | chr18-80335070-INS-57 |
| chr18-80349768-INS-106 | chr18-80349768-INS-106 |
| chr18-80369541-INS-111 | chr18-80369541-INS-113 |
| chr18-80376314-INS-170 | chr18-80376317-INS-166 |
| chr18-80378254-INS-117 | chr18-80378265-INS-132 |
| chr18-80400423-INS-129 | chr18-80400427-INS-104 |
| chr18-80417987-INS-229 | chr18-80417987-INS-229 |
| chr18-80434574-INS-101 | chr18-80434574-INS-101 |
| chr18-80449973-INS-680 | chr18-80449973-INS-681 |
| chr18-80455131-INS-56 | chr18-80455131-INS-56 |
| chr18-80476608-INS-64 | chr18-80476608-INS-64 |
| chr18-80522405-INS-339 | chr18-80522405-INS-339 |
| chr18-80523686-INS-89 | chr18-80523684-INS-89 |
| chr18-80537132-INS-82 | chr18-80537121-INS-78 |
| chr18-80570159-INS-360 | chr18-80570159-INS-358 |
| chr18-80604370-INS-76 | chr18-80604370-INS-76 |
| chr18-80607215-INS-90 | chr18-80607215-INS-90 |
| chr18-80627646-INS-68 | chr18-80627646-INS-68 |
| chr18-80644515-INS-74 | chr18-80644515-INS-50 |
| chr18-80647816-INS-62 | chr18-80647816-INS-62 |
| chr18-80667941-INS-57 | chr18-80667941-INS-61 |
| chr18-80669704-INS-197 | chr18-80669704-INS-197 |
| chr18-80670554-INS-123 | chr18-80670554-INS-124 |
| chr18-80694537-INS-161 | chr18-80694537-INS-161 |
| chr18-80726970-INS-170 | chr18-80726970-INS-172 |
| chr18-80762341-INS-52 | chr18-80762341-INS-57 |
| chr18-80771446-INS-75 | chr18-80771446-INS-75 |
| chr18-80822795-INS-1564 | chr18-80822796-INS-1564 |
| chr18-80835969-INS-89 | chr18-80835969-INS-101 |
| chr18-80844539-INS-185 | chr18-80844539-INS-186 |
| chr18-80851071-INS-200 | chr18-80851074-INS-164 |
| chr18-80853127-INS-69 | chr18-80853127-INS-69 |
| chr18-80866012-INS-489 | chr18-80866012-INS-489 |
| chr18-80868796-INS-52 | chr18-80868796-INS-54 |
| chr18-80886309-INS-60 | chr18-80886310-INS-60 |
| chr18-80888834-INS-227 | chr18-80888837-INS-191 |
| chr18-80932680-INS-687 | chr18-80932680-INS-687 |
| chr18-80938617-INS-131 | chr18-80938617-INS-131 |
| chr18-80940438-INS-345 | chr18-80940438-INS-345 |
| chr18-80941586-INS-72 | chr18-80941586-INS-72 |
| chr18-80986864-INS-119 | chr18-80986864-INS-119 |
| chr18-80992950-INS-53 | chr18-80992980-INS-50 |
| chr18-80999427-INS-96 | chr18-80999403-INS-128 |
| chr18-81020814-INS-64 | chr18-81020807-INS-72 |
| chr18-81043657-INS-118 | chr18-81043657-INS-118 |
| chr18-81055595-INS-52 | chr18-81055570-INS-98 |
| chr18-81059388-INS-73 | chr18-81059386-INS-73 |
| chr18-81074861-INS-84 | chr18-81074865-INS-88 |
| chr18-81090206-INS-7250 | chr18-81090207-INS-7250 |
| chr18-81106860-INS-60 | chr18-81106861-INS-60 |
| chr18-81114125-INS-81 | chr18-81114125-INS-81 |
| chr18-81116760-INS-280 | chr18-81116760-INS-280 |
| chr18-81117686-INS-64 | chr18-81117686-INS-87 |
| chr18-81137106-INS-182 | chr18-81137106-INS-194 |
| chr18-81143722-INS-102 | chr18-81143722-INS-102 |
| chr18-81144127-INS-216 | chr18-81144132-INS-217 |
| chr18-81149655-INS-1892 | chr18-81149656-INS-1892 |
| chr18-81181738-INS-6403 | chr18-81181739-INS-6403 |
| chr18-81190659-INS-640 | chr18-81190659-INS-640 |
| chr18-81212443-INS-509 | chr18-81212443-INS-509 |
| chr18-81226660-INS-6455 | chr18-81226661-INS-6456 |
| chr18-81242739-INS-57 | chr18-81242739-INS-57 |
| chr18-81246246-INS-209 | chr18-81246246-INS-209 |
| chr18-81251694-INS-1895 | chr18-81251695-INS-1895 |
| chr18-81282416-INS-61 | chr18-81282416-INS-63 |
| chr18-81301712-INS-105 | chr18-81301712-INS-95 |
| chr18-81310453-INS-122 | chr18-81310481-INS-80 |
| chr18-81312377-INS-93 | chr18-81312371-INS-63 |
| chr18-81355611-INS-54 | chr18-81355611-INS-72 |
| chr18-81364494-INS-497 | chr18-81364494-INS-500 |
| chr18-81372782-INS-58 | chr18-81372782-INS-58 |
| chr18-81396114-INS-303 | chr18-81396119-INS-337 |
| chr18-81405578-INS-62 | chr18-81405564-INS-64 |
| chr18-81426445-INS-152 | chr18-81426445-INS-152 |
| chr18-81449105-INS-78 | chr18-81449105-INS-79 |
| chr18-81486524-INS-82 | chr18-81486520-INS-62 |

| | |
|---|---|
| chr18-81527660-INS-155 | chr18-81527660-INS-152 |
| chr18-81530286-INS-74 | chr18-81530286-INS-72 |
| chr18-81577669-INS-58 | chr18-81577698-INS-54 |
| chr18-81589398-INS-276 | chr18-81589398-INS-276 |
| chr18-81596617-INS-309 | chr18-81596617-INS-301 |
| chr18-81598567-INS-78 | chr18-81598567-INS-78 |
| chr18-81602651-INS-269 | chr18-81602652-INS-270 |
| chr18-81611810-INS-439 | chr18-81611810-INS-438 |
| chr18-81613921-INS-66 | chr18-81613921-INS-72 |
| chr18-81650200-INS-71 | chr18-81650202-INS-61 |
| chr18-81674534-INS-76 | chr18-81674534-INS-76 |
| chr18-81697314-INS-73 | chr18-81697314-INS-73 |
| chr18-81701152-INS-6244 | chr18-81701153-INS-6245 |
| chr18-81729514-INS-487 | chr18-81729514-INS-495 |
| chr18-81732663-INS-221 | chr18-81732663-INS-226 |
| chr18-81740264-INS-238 | chr18-81740269-INS-272 |
| chr18-81769344-INS-56 | chr18-81769344-INS-56 |
| chr18-81774483-INS-263 | chr18-81774483-INS-269 |
| chr18-81782926-INS-210 | chr18-81782943-INS-198 |
| chr18-81805336-INS-205 | chr18-81805336-INS-205 |
| chr18-81810310-INS-223 | chr18-81810310-INS-223 |
| chr18-81819744-INS-400 | chr18-81819744-INS-396 |
| chr18-81825034-INS-864 | chr18-81825034-INS-864 |
| chr18-81868741-INS-2577 | chr18-81868742-INS-2577 |
| chr18-81881329-INS-703 | chr18-81881329-INS-707 |
| chr18-81940043-INS-76 | chr18-81940043-INS-120 |
| chr18-81949633-INS-131 | chr18-81949633-INS-153 |
| chr18-81959100-INS-648 | chr18-81959111-INS-651 |
| chr18-81962233-INS-1614 | chr18-81962234-INS-1618 |
| chr18-81974105-INS-87 | chr18-81974116-INS-86 |
| chr18-81977963-INS-53 | chr18-81977963-INS-53 |
| chr18-81982853-INS-422 | chr18-81982846-INS-422 |
| chr18-82034244-INS-52 | chr18-82034244-INS-54 |
| chr18-82036388-INS-176 | chr18-82036408-INS-195 |
| chr18-82037845-INS-380 | chr18-82037845-INS-380 |
| chr18-82039566-INS-58 | chr18-82039566-INS-90 |
| chr18-82042446-INS-121 | chr18-82042446-INS-121 |
| chr18-82052139-INS-162 | chr18-82052142-INS-163 |
| chr18-82063791-INS-588 | chr18-82063791-INS-588 |
| chr18-82069876-INS-343 | chr18-82069876-INS-343 |
| chr18-82109809-INS-383 | chr18-82109809-INS-386 |
| chr18-82148335-INS-336 | chr18-82148347-INS-348 |
| chr18-82252613-INS-54 | chr18-82252613-INS-54 |
| chr18-82263471-INS-219 | chr18-82263471-INS-219 |
| chr18-82267142-INS-5637 | chr18-82267143-INS-5637 |
| chr18-82270435-INS-959 | chr18-82270435-INS-958 |
| chr18-82298169-INS-103 | chr18-82298169-INS-104 |
| chr18-82304483-INS-54 | chr18-82304483-INS-57 |
| chr18-82308618-INS-219 | chr18-82308618-INS-218 |
| chr18-82320094-INS-170 | chr18-82320094-INS-170 |
| chr18-82325345-INS-7076 | chr18-82325346-INS-7076 |
| chr18-82336631-INS-53 | chr18-82336616-INS-66 |
| chr18-82364884-INS-76 | chr18-82364864-INS-69 |
| chr18-82368735-INS-163 | chr18-82368735-INS-178 |
| chr18-82377352-INS-62 | chr18-82377352-INS-58 |
| chr18-82384849-INS-7089 | chr18-82384850-INS-7089 |
| chr18-82387032-INS-112 | chr18-82387032-INS-112 |
| chr18-82399850-INS-122 | chr18-82399850-INS-121 |
| chr18-82411112-INS-94 | chr18-82411112-INS-94 |
| chr18-82440145-INS-160 | chr18-82440145-INS-160 |
| chr18-82443370-INS-7081 | chr18-82443371-INS-7081 |
| chr18-82454204-INS-70 | chr18-82454204-INS-66 |
| chr18-82476051-INS-58 | chr18-82476045-INS-74 |
| chr18-82480515-INS-60 | chr18-82480515-INS-60 |
| chr18-82528361-INS-108 | chr18-82528361-INS-112 |
| chr18-82535472-INS-163 | chr18-82535472-INS-163 |
| chr18-82547824-INS-3632 | chr18-82547825-INS-3632 |
| chr18-82548149-INS-71 | chr18-82548149-INS-76 |
| chr18-82596480-INS-107 | chr18-82596480-INS-107 |
| chr18-82606187-INS-70 | chr18-82606187-INS-78 |
| chr18-82636385-INS-5026 | chr18-82636385-INS-5026 |
| chr18-82657059-INS-76 | chr18-82657059-INS-64 |
| chr18-82662978-INS-235 | chr18-82662978-INS-235 |
| chr18-82678719-INS-58 | chr18-82678719-INS-60 |
| chr18-82681618-INS-98 | chr18-82681618-INS-98 |
| chr18-82683260-INS-211 | chr18-82683260-INS-211 |
| chr18-82689009-INS-64 | chr18-82689009-INS-64 |

| | |
|---|---|
| chr18-82704346-INS-70 | chr18-82704336-INS-101 |
| chr18-82727713-INS-213 | chr18-82727713-INS-213 |
| chr18-82731406-INS-141 | chr18-82731406-INS-136 |
| chr18-82738870-INS-64 | chr18-82738870-INS-74 |
| chr18-82742049-INS-75 | chr18-82742049-INS-59 |
| chr18-82749390-INS-202 | chr18-82749390-INS-202 |
| chr18-82750090-INS-202 | chr18-82750090-INS-205 |
| chr18-82752907-INS-94 | chr18-82752907-INS-96 |
| chr18-82757057-INS-364 | chr18-82757057-INS-364 |
| chr18-82758591-INS-53 | chr18-82758591-INS-53 |
| chr18-82771747-INS-64 | chr18-82771731-INS-75 |
| chr18-82775585-INS-179 | chr18-82775585-INS-179 |
| chr18-82787392-INS-60 | chr18-82787392-INS-64 |
| chr18-82804599-INS-84 | chr18-82804599-INS-100 |
| chr18-82837588-INS-91 | chr18-82837588-INS-91 |
| chr18-82848017-INS-96 | chr18-82848017-INS-98 |
| chr18-82853996-INS-415 | chr18-82853996-INS-415 |
| chr18-82859527-INS-53 | chr18-82859528-INS-53 |
| chr18-82948768-INS-54 | chr18-82948768-INS-54 |
| chr18-82962066-INS-95 | chr18-82962066-INS-95 |
| chr18-82978759-INS-378 | chr18-82978759-INS-378 |
| chr18-83000771-INS-51 | chr18-83000761-INS-55 |
| chr18-83011619-INS-159 | chr18-83011619-INS-159 |
| chr18-83031461-INS-614 | chr18-83031459-INS-626 |
| chr18-83042199-INS-182 | chr18-83042172-INS-180 |
| chr18-83049614-INS-6180 | chr18-83049615-INS-6180 |
| chr18-83058730-INS-56 | chr18-83058730-INS-56 |
| chr18-83072700-INS-60 | chr18-83072689-INS-102 |
| chr18-83081443-INS-1894 | chr18-83081444-INS-1894 |
| chr18-83137212-INS-79 | chr18-83137208-INS-75 |
| chr18-83150767-INS-86 | chr18-83150767-INS-86 |
| chr18-83154089-INS-97 | chr18-83154084-INS-104 |
| chr18-83175841-INS-51 | chr18-83175841-INS-51 |
| chr18-83214299-INS-1117 | chr18-83214300-INS-1117 |
| chr18-83222293-INS-80 | chr18-83222303-INS-100 |
| chr18-83248702-INS-6207 | chr18-83248703-INS-6209 |
| chr18-83248930-INS-60 | chr18-83248929-INS-62 |
| chr18-83253576-INS-1173 | chr18-83253577-INS-1173 |
| chr18-83257109-INS-84 | chr18-83257097-INS-124 |
| chr18-83271272-INS-66 | chr18-83271271-INS-66 |
| chr18-83307897-INS-53 | chr18-83307884-INS-78 |
| chr18-83311793-INS-136 | chr18-83311793-INS-136 |
| chr18-83312670-INS-4125 | chr18-83312671-INS-4125 |
| chr18-83317111-INS-281 | chr18-83317111-INS-281 |
| chr18-83331028-INS-63 | chr18-83331040-INS-99 |
| chr18-83347099-INS-6423 | chr18-83347100-INS-6423 |
| chr18-83348581-INS-70 | chr18-83348586-INS-79 |
| chr18-83351657-INS-128 | chr18-83351657-INS-128 |
| chr18-83352570-INS-54 | chr18-83352570-INS-96 |
| chr18-83361602-INS-262 | chr18-83361602-INS-262 |
| chr18-83399377-INS-5204 | chr18-83399378-INS-5204 |
| chr18-83486323-INS-60 | chr18-83486323-INS-54 |
| chr18-83503107-INS-84 | chr18-83503108-INS-64 |
| chr18-83530605-INS-54 | chr18-83530605-INS-56 |
| chr18-83555265-INS-1276 | chr18-83555261-INS-1284 |
| chr18-83586794-INS-54 | chr18-83586794-INS-81 |
| chr18-83594685-INS-113 | chr18-83594685-INS-113 |
| chr18-83602714-INS-571 | chr18-83602714-INS-571 |
| chr18-83607283-INS-96 | chr18-83607289-INS-58 |
| chr18-83615365-INS-58 | chr18-83615363-INS-54 |
| chr18-83623078-INS-60 | chr18-83623078-INS-60 |
| chr18-83624071-INS-68 | chr18-83624054-INS-72 |
| chr18-83627913-INS-58 | chr18-83627878-INS-90 |
| chr18-83641120-INS-470 | chr18-83641122-INS-472 |
| chr18-83665075-INS-398 | chr18-83665075-INS-398 |
| chr18-83765693-INS-52 | chr18-83765693-INS-52 |
| chr18-83803987-INS-124 | chr18-83803995-INS-124 |
| chr18-83820180-INS-671 | chr18-83820180-INS-671 |
| chr18-83837266-INS-71 | chr18-83837266-INS-71 |
| chr18-83842202-INS-133 | chr18-83842199-INS-142 |
| chr18-83854720-INS-56 | chr18-83854720-INS-56 |
| chr18-83862332-INS-266 | chr18-83862332-INS-267 |
| chr18-83876012-INS-494 | chr18-83876008-INS-512 |
| chr18-83889371-INS-80 | chr18-83889359-INS-86 |
| chr18-83892479-INS-6351 | chr18-83892480-INS-6354 |
| chr18-83895184-INS-64 | chr18-83895184-INS-64 |
| chr18-83907681-INS-158 | chr18-83907681-INS-153 |

| | |
|---|---|
| chr18-83939318-INS-4852 | chr18-83939319-INS-4852 |
| chr18-83939846-INS-343 | chr18-83939849-INS-344 |
| chr18-83945216-INS-105 | chr18-83945216-INS-105 |
| chr18-83950373-INS-210 | chr18-83950380-INS-222 |
| chr18-83958923-INS-291 | chr18-83958923-INS-291 |
| chr18-83961089-INS-85 | chr18-83961089-INS-85 |
| chr18-83969263-INS-1869 | chr18-83969264-INS-1869 |
| chr18-83992957-INS-60 | chr18-83992957-INS-60 |
| chr18-84005662-INS-246 | chr18-84005662-INS-249 |
| chr18-84010279-INS-69 | chr18-84010279-INS-69 |
| chr18-84019993-INS-142 | chr18-84019993-INS-142 |
| chr18-84035186-INS-50 | chr18-84035198-INS-70 |
| chr18-84086139-INS-54 | chr18-84086139-INS-54 |
| chr18-84100359-INS-59 | chr18-84100359-INS-64 |
| chr18-84126872-INS-81 | chr18-84126872-INS-81 |
| chr18-84131500-INS-51 | chr18-84131478-INS-77 |
| chr18-84173575-INS-234 | chr18-84173575-INS-235 |
| chr18-84176162-INS-52 | chr18-84176175-INS-56 |
| chr18-84190516-INS-65 | chr18-84190516-INS-65 |
| chr18-84221290-INS-63 | chr18-84221290-INS-90 |
| chr18-84247762-INS-62 | chr18-84247762-INS-62 |
| chr18-84258953-INS-76 | chr18-84258953-INS-76 |
| chr18-84291076-INS-89 | chr18-84291061-INS-68 |
| chr18-84316640-INS-545 | chr18-84316640-INS-546 |
| chr18-84319066-INS-52 | chr18-84319066-INS-52 |
| chr18-84337806-INS-60 | chr18-84337757-INS-71 |
| chr18-84338947-INS-135 | chr18-84338947-INS-144 |
| chr18-84376663-INS-68 | chr18-84376633-INS-78 |
| chr18-84407940-INS-108 | chr18-84407940-INS-108 |
| chr18-84423176-INS-61 | chr18-84423175-INS-61 |
| chr18-84471365-INS-96 | chr18-84471365-INS-64 |
| chr18-84488465-INS-56 | chr18-84488465-INS-56 |
| chr18-84490544-INS-53 | chr18-84490544-INS-53 |
| chr18-84493565-INS-1862 | chr18-84493566-INS-1862 |
| chr18-84493769-INS-228 | chr18-84493769-INS-229 |
| chr18-84497644-INS-93 | chr18-84497644-INS-94 |
| chr18-84515642-INS-116 | chr18-84515643-INS-115 |
| chr18-84522116-INS-276 | chr18-84522116-INS-278 |
| chr18-84541882-INS-92 | chr18-84541893-INS-85 |
| chr18-84547222-INS-6466 | chr18-84547223-INS-6467 |
| chr18-84609365-INS-87 | chr18-84609365-INS-86 |
| chr18-84638260-INS-232 | chr18-84638260-INS-232 |
| chr18-84640679-INS-407 | chr18-84640680-INS-408 |
| chr18-84645154-INS-479 | chr18-84645154-INS-479 |
| chr18-84653497-INS-201 | chr18-84653497-INS-201 |
| chr18-84669428-INS-817 | chr18-84669428-INS-815 |
| chr18-84675890-INS-58 | chr18-84675890-INS-60 |
| chr18-84680868-INS-173 | chr18-84680868-INS-173 |
| chr18-84707723-INS-54 | chr18-84707723-INS-54 |
| chr18-84723000-INS-181 | chr18-84723000-INS-181 |
| chr18-84723950-INS-62 | chr18-84723950-INS-62 |
| chr18-84743161-INS-880 | chr18-84743161-INS-880 |
| chr18-84743167-INS-60 | chr18-84743167-INS-60 |
| chr18-84750293-INS-630 | chr18-84750336-INS-633 |
| chr18-84754042-INS-75 | chr18-84754042-INS-75 |
| chr18-84759454-INS-148 | chr18-84759454-INS-148 |
| chr18-84768127-INS-1565 | chr18-84768128-INS-1566 |
| chr18-84787989-INS-272 | chr18-84788026-INS-272 |
| chr18-84789118-INS-638 | chr18-84789118-INS-643 |
| chr18-84798531-INS-231 | chr18-84798531-INS-244 |
| chr18-84799076-INS-6386 | chr18-84799077-INS-6386 |
| chr18-84805151-INS-205 | chr18-84805151-INS-207 |
| chr18-84813845-INS-685 | chr18-84813845-INS-691 |
| chr18-84818414-INS-511 | chr18-84818415-INS-511 |
| chr18-84825734-INS-150 | chr18-84825734-INS-150 |
| chr18-84857953-INS-4871 | chr18-84857954-INS-4871 |
| chr18-84864239-INS-5617 | chr18-84864240-INS-5615 |
| chr18-84870176-INS-60 | chr18-84870154-INS-100 |
| chr18-84875778-INS-417 | chr18-84875778-INS-422 |
| chr18-84881246-INS-120 | chr18-84881246-INS-120 |
| chr18-84900462-INS-166 | chr18-84900462-INS-166 |
| chr18-84904646-INS-176 | chr18-84904646-INS-176 |
| chr18-84905692-INS-60 | chr18-84905692-INS-60 |
| chr18-84920779-INS-125 | chr18-84920780-INS-78 |
| chr18-84925933-INS-52 | chr18-84925933-INS-91 |
| chr18-84954599-INS-1236 | chr18-84954600-INS-1236 |
| chr18-84960146-INS-195 | chr18-84960146-INS-195 |

| | |
|---|---|
| chr18-84973315-INS-58 | chr18-84973318-INS-58 |
| chr18-84978715-INS-179 | chr18-84978715-INS-178 |
| chr18-84989198-INS-70 | chr18-84989198-INS-70 |
| chr18-84995624-INS-202 | chr18-84995624-INS-205 |
| chr18-85013890-INS-498 | chr18-85013890-INS-522 |
| chr18-85029275-INS-7160 | chr18-85029276-INS-7160 |
| chr18-85038249-INS-203 | chr18-85038249-INS-203 |
| chr18-85071969-INS-399 | chr18-85071969-INS-399 |
| chr18-85081103-INS-207 | chr18-85081103-INS-207 |
| chr18-85103005-INS-91 | chr18-85103041-INS-60 |
| chr18-85113559-INS-193 | chr18-85113559-INS-194 |
| chr18-85117500-INS-143 | chr18-85117501-INS-146 |
| chr18-85133910-INS-52 | chr18-85133903-INS-56 |
| chr18-85137979-INS-2343 | chr18-85137980-INS-2342 |
| chr18-85140940-INS-6663 | chr18-85140941-INS-6662 |
| chr18-85142221-INS-77 | chr18-85142221-INS-78 |
| chr18-85150257-INS-88 | chr18-85150257-INS-88 |
| chr18-85155571-INS-96 | chr18-85155571-INS-96 |
| chr18-85168960-INS-96 | chr18-85168960-INS-96 |
| chr18-85181343-INS-51 | chr18-85181343-INS-51 |
| chr18-85203607-INS-112 | chr18-85203607-INS-112 |
| chr18-85219716-INS-70 | chr18-85219716-INS-76 |
| chr18-85221750-INS-722 | chr18-85221750-INS-722 |
| chr18-85235214-INS-74 | chr18-85235173-INS-75 |
| chr18-85240028-INS-51 | chr18-85240024-INS-63 |
| chr18-85250796-INS-6236 | chr18-85250797-INS-6237 |
| chr18-85266558-INS-156 | chr18-85266548-INS-152 |
| chr18-85278562-INS-73 | chr18-85278560-INS-94 |
| chr18-85286339-INS-64 | chr18-85286339-INS-64 |
| chr18-85290584-INS-54 | chr18-85290562-INS-60 |
| chr18-85306051-INS-468 | chr18-85306051-INS-468 |
| chr18-85306644-INS-54 | chr18-85306678-INS-60 |
| chr18-85309655-INS-124 | chr18-85309656-INS-124 |
| chr18-85321059-INS-53 | chr18-85321062-INS-59 |
| chr18-85343253-INS-212 | chr18-85343253-INS-212 |
| chr18-85367430-INS-1886 | chr18-85367431-INS-1886 |
| chr18-85373457-INS-1458 | chr18-85373458-INS-1459 |
| chr18-85373849-INS-54 | chr18-85373842-INS-58 |
| chr18-85384011-INS-481 | chr18-85384011-INS-483 |
| chr18-85386484-INS-55 | chr18-85386484-INS-63 |
| chr18-85424477-INS-276 | chr18-85424477-INS-276 |
| chr18-85425557-INS-387 | chr18-85425557-INS-387 |
| chr18-85452257-INS-836 | chr18-85452259-INS-855 |
| chr18-85505497-INS-2203 | chr18-85505498-INS-2203 |
| chr18-85510602-INS-6388 | chr18-85510603-INS-6388 |
| chr18-85513097-INS-2803 | chr18-85513098-INS-2800 |
| chr18-85515612-INS-276 | chr18-85515612-INS-246 |
| chr18-85517534-INS-222 | chr18-85517534-INS-222 |
| chr18-85518734-INS-53 | chr18-85518737-INS-50 |
| chr18-85620101-INS-171 | chr18-85620101-INS-171 |
| chr18-85653112-INS-6415 | chr18-85653113-INS-6415 |
| chr18-85653210-INS-445 | chr18-85653210-INS-446 |
| chr18-85666447-INS-6496 | chr18-85666448-INS-6496 |
| chr18-85668739-INS-97 | chr18-85668739-INS-97 |
| chr18-85703936-INS-80 | chr18-85703936-INS-54 |
| chr18-85709063-INS-86 | chr18-85709063-INS-86 |
| chr18-85721949-INS-157 | chr18-85721947-INS-157 |
| chr18-85726069-INS-162 | chr18-85726069-INS-162 |
| chr18-85744603-INS-7143 | chr18-85744604-INS-7144 |
| chr18-85771938-INS-58 | chr18-85771938-INS-94 |
| chr18-85810748-INS-392 | chr18-85810748-INS-392 |
| chr18-85845089-INS-201 | chr18-85845089-INS-201 |
| chr18-85863598-INS-259 | chr18-85863598-INS-271 |
| chr18-85889177-INS-516 | chr18-85889177-INS-516 |
| chr18-85897502-INS-461 | chr18-85897502-INS-450 |
| chr18-85898888-INS-51 | chr18-85898892-INS-54 |
| chr18-85945387-INS-660 | chr18-85945387-INS-661 |
| chr18-85956007-INS-809 | chr18-85956007-INS-809 |
| chr18-85973951-INS-3707 | chr18-85973952-INS-3687 |
| chr18-85983556-INS-4826 | chr18-85983557-INS-4826 |
| chr18-85987432-INS-52 | chr18-85987432-INS-57 |
| chr18-86023908-INS-213 | chr18-86023908-INS-213 |
| chr18-86052874-INS-52 | chr18-86052846-INS-64 |
| chr18-86084382-INS-124 | chr18-86084382-INS-124 |
| chr18-86091155-INS-128 | chr18-86091155-INS-113 |
| chr18-86143668-INS-76 | chr18-86143668-INS-82 |
| chr18-86246607-INS-62 | chr18-86246606-INS-64 |

| | |
|---|---|
| chr18-86293372-INS-119 | chr18-86293372-INS-80 |
| chr18-86305506-INS-55 | chr18-86305506-INS-67 |
| chr18-86371230-INS-1889 | chr18-86371231-INS-1889 |
| chr18-86378842-INS-612 | chr18-86378842-INS-612 |
| chr18-86399352-INS-6332 | chr18-86399353-INS-6334 |
| chr18-86420216-INS-186 | chr18-86420216-INS-185 |
| chr18-86465667-INS-66 | chr18-86465667-INS-66 |
| chr18-86471488-INS-71 | chr18-86471489-INS-59 |
| chr18-86495312-INS-57 | chr18-86495312-INS-57 |
| chr18-86510637-INS-75 | chr18-86510637-INS-75 |
| chr18-86562500-INS-3032 | chr18-86562501-INS-3032 |
| chr18-86565715-INS-3176 | chr18-86565716-INS-3155 |
| chr18-86570108-INS-121 | chr18-86570108-INS-123 |
| chr18-86599094-INS-64 | chr18-86599094-INS-64 |
| chr18-86615566-INS-744 | chr18-86615566-INS-743 |
| chr18-86618678-INS-123 | chr18-86618678-INS-106 |
| chr18-86625886-INS-181 | chr18-86625886-INS-182 |
| chr18-86653298-INS-210 | chr18-86653298-INS-210 |
| chr18-86657440-INS-2000 | chr18-86657441-INS-2000 |
| chr18-86658210-INS-115 | chr18-86658210-INS-115 |
| chr18-86668970-INS-1325 | chr18-86668976-INS-1323 |
| chr18-86678080-INS-3274 | chr18-86678081-INS-3268 |
| chr18-86678081-INS-517 | chr18-86678081-INS-517 |
| chr18-86684524-INS-100 | chr18-86684518-INS-136 |
| chr18-86713685-INS-74 | chr18-86713659-INS-60 |
| chr18-86718901-INS-125 | chr18-86718901-INS-135 |
| chr18-86738811-INS-79 | chr18-86738811-INS-79 |
| chr18-86749513-INS-122 | chr18-86749541-INS-81 |
| chr18-86805724-INS-113 | chr18-86805724-INS-113 |
| chr18-86828723-INS-67 | chr18-86828723-INS-103 |
| chr18-86852393-INS-199 | chr18-86852393-INS-199 |
| chr18-86935411-INS-98 | chr18-86935411-INS-98 |
| chr18-86951664-INS-193 | chr18-86951664-INS-193 |
| chr18-86970054-INS-201 | chr18-86970054-INS-201 |
| chr18-86999712-INS-535 | chr18-86999713-INS-535 |
| chr18-87011043-INS-76 | chr18-87011044-INS-100 |
| chr18-87038812-INS-390 | chr18-87038812-INS-390 |
| chr18-87059303-INS-68 | chr18-87059340-INS-66 |
| chr18-87088609-INS-51 | chr18-87088609-INS-51 |
| chr18-87091251-INS-674 | chr18-87091251-INS-674 |
| chr18-87095310-INS-1516 | chr18-87095311-INS-1508 |
| chr18-87134014-INS-101 | chr18-87134009-INS-101 |
| chr18-87192628-INS-76 | chr18-87192628-INS-81 |
| chr18-87195045-INS-706 | chr18-87195045-INS-706 |
| chr18-87198371-INS-385 | chr18-87198377-INS-390 |
| chr18-87201609-INS-6398 | chr18-87201610-INS-6397 |
| chr18-87218245-INS-571 | chr18-87218245-INS-606 |
| chr18-87244336-INS-196 | chr18-87244336-INS-197 |
| chr18-87257280-INS-193 | chr18-87257280-INS-154 |
| chr18-87259247-INS-58 | chr18-87259247-INS-60 |
| chr18-87261905-INS-853 | chr18-87261905-INS-853 |
| chr18-87340588-INS-6679 | chr18-87340589-INS-6681 |
| chr18-87363479-INS-416 | chr18-87363479-INS-416 |
| chr18-87404813-INS-105 | chr18-87404802-INS-137 |
| chr18-87409869-INS-106 | chr18-87409870-INS-77 |
| chr18-87460506-INS-343 | chr18-87460506-INS-343 |
| chr18-87464803-INS-441 | chr18-87464803-INS-441 |
| chr18-87479017-INS-92 | chr18-87479017-INS-92 |
| chr18-87497056-INS-206 | chr18-87497056-INS-206 |
| chr18-87499092-INS-93 | chr18-87499092-INS-93 |
| chr18-87503513-INS-118 | chr18-87503540-INS-111 |
| chr18-87507855-INS-69 | chr18-87507855-INS-57 |
| chr18-87525323-INS-6458 | chr18-87525324-INS-6459 |
| chr18-87528860-INS-58 | chr18-87528860-INS-60 |
| chr18-87529381-INS-6881 | chr18-87529382-INS-6882 |
| chr18-87539906-INS-70 | chr18-87539906-INS-70 |
| chr18-87554932-INS-1635 | chr18-87554933-INS-1629 |
| chr18-87567804-INS-501 | chr18-87567804-INS-501 |
| chr18-87569012-INS-65 | chr18-87569011-INS-51 |
| chr18-87601714-INS-391 | chr18-87601714-INS-391 |
| chr18-87624805-INS-69 | chr18-87624766-INS-69 |
| chr18-87643163-INS-52 | chr18-87643163-INS-50 |
| chr18-87658781-INS-380 | chr18-87658781-INS-380 |
| chr18-87659341-INS-133 | chr18-87659372-INS-112 |
| chr18-87665790-INS-1288 | chr18-87665791-INS-1289 |
| chr18-87669133-INS-9972 | chr18-87669134-INS-9971 |
| chr18-87684488-INS-352 | chr18-87684488-INS-383 |

| | |
|---|---|
| chr18-87732090-INS-70 | chr18-87732090-INS-70 |
| chr18-87766484-INS-451 | chr18-87766484-INS-451 |
| chr18-87770175-INS-690 | chr18-87770176-INS-691 |
| chr18-87781750-INS-92 | chr18-87781750-INS-100 |
| chr18-87784553-INS-549 | chr18-87784553-INS-549 |
| chr18-87870534-INS-63 | chr18-87870534-INS-63 |
| chr18-87870850-INS-6259 | chr18-87870850-INS-6261 |
| chr18-87877218-INS-225 | chr18-87877218-INS-225 |
| chr18-87880133-INS-58 | chr18-87880133-INS-58 |
| chr18-87919181-INS-68 | chr18-87919181-INS-70 |
| chr18-87922804-INS-903 | chr18-87922804-INS-903 |
| chr18-87980806-INS-72 | chr18-87980806-INS-64 |
| chr18-88044436-INS-58 | chr18-88044436-INS-62 |
| chr18-88090363-INS-685 | chr18-88090363-INS-689 |
| chr18-88166384-INS-175 | chr18-88166384-INS-175 |
| chr18-88174136-INS-108 | chr18-88174165-INS-94 |
| chr18-88194714-INS-104 | chr18-88194714-INS-54 |
| chr18-88208059-INS-118 | chr18-88208059-INS-119 |
| chr18-88209904-INS-6756 | chr18-88209905-INS-6756 |
| chr18-88222608-INS-4548 | chr18-88222609-INS-4541 |
| chr18-88222609-INS-390 | chr18-88222609-INS-390 |
| chr18-88229887-INS-59 | chr18-88229879-INS-55 |
| chr18-88245263-INS-6651 | chr18-88245264-INS-6652 |
| chr18-88277335-INS-127 | chr18-88277335-INS-127 |
| chr18-88301828-INS-6576 | chr18-88301829-INS-6576 |
| chr18-88314780-INS-58 | chr18-88314780-INS-58 |
| chr18-88321808-INS-208 | chr18-88321808-INS-208 |
| chr18-88359631-INS-706 | chr18-88359630-INS-711 |
| chr18-88398137-INS-3806 | chr18-88398138-INS-3806 |
| chr18-88417440-INS-536 | chr18-88417440-INS-535 |
| chr18-88447279-INS-720 | chr18-88447277-INS-718 |
| chr18-88468029-INS-5414 | chr18-88468030-INS-5414 |
| chr18-88472810-INS-1453 | chr18-88472811-INS-1455 |
| chr18-88474890-INS-985 | chr18-88474890-INS-984 |
| chr18-88504914-INS-1643 | chr18-88504915-INS-1646 |
| chr18-88507937-INS-88 | chr18-88507937-INS-90 |
| chr18-88523343-INS-104 | chr18-88523343-INS-104 |
| chr18-88530207-INS-1918 | chr18-88530208-INS-1918 |
| chr18-88534054-INS-813 | chr18-88534054-INS-813 |
| chr18-88546099-INS-87 | chr18-88546097-INS-90 |
| chr18-88546811-INS-88 | chr18-88546811-INS-94 |
| chr18-88570828-INS-74 | chr18-88570828-INS-74 |
| chr18-88585196-INS-53 | chr18-88585196-INS-77 |
| chr18-88594485-INS-100 | chr18-88594485-INS-103 |
| chr18-88602298-INS-312 | chr18-88602298-INS-312 |
| chr18-88661214-INS-237 | chr18-88661214-INS-237 |
| chr18-88669099-INS-824 | chr18-88669099-INS-824 |
| chr18-88717709-INS-68 | chr18-88717709-INS-77 |
| chr18-88727527-INS-405 | chr18-88727527-INS-405 |
| chr18-88736561-INS-82 | chr18-88736561-INS-54 |
| chr18-88757386-INS-6947 | chr18-88757387-INS-6947 |
| chr18-88766108-INS-216 | chr18-88766108-INS-217 |
| chr18-88769090-INS-1425 | chr18-88769091-INS-1425 |
| chr18-88776657-INS-67 | chr18-88776657-INS-65 |
| chr18-88796329-INS-830 | chr18-88796329-INS-861 |
| chr18-88804637-INS-596 | chr18-88804637-INS-632 |
| chr18-88825768-INS-179 | chr18-88825771-INS-181 |
| chr18-88834145-INS-1863 | chr18-88834150-INS-1859 |
| chr18-88839119-INS-328 | chr18-88839119-INS-328 |
| chr18-88851960-INS-327 | chr18-88851960-INS-327 |
| chr18-88903338-INS-2425 | chr18-88903339-INS-2426 |
| chr18-88921138-INS-537 | chr18-88921138-INS-537 |
| chr18-88945545-INS-54 | chr18-88945545-INS-54 |
| chr18-88946335-INS-708 | chr18-88946335-INS-708 |
| chr18-88974033-INS-56 | chr18-88974033-INS-56 |
| chr18-89042022-INS-3119 | chr18-89042023-INS-3119 |
| chr18-89078826-INS-351 | chr18-89078826-INS-351 |
| chr18-89099105-INS-155 | chr18-89099105-INS-155 |
| chr18-89116811-INS-390 | chr18-89116811-INS-390 |
| chr18-89172152-INS-2606 | chr18-89172153-INS-2606 |
| chr18-89178273-INS-123 | chr18-89178273-INS-123 |
| chr18-89268423-INS-62 | chr18-89268423-INS-56 |
| chr18-89292373-INS-865 | chr18-89292373-INS-866 |
| chr18-89301932-INS-6581 | chr18-89301933-INS-6581 |
| chr18-89336685-INS-329 | chr18-89336685-INS-329 |
| chr18-89338583-INS-62 | chr18-89338583-INS-66 |
| chr18-89354258-INS-554 | chr18-89354258-INS-554 |

| | |
|---|---|
| chr18-89360512-INS-1833 | chr18-89360513-INS-1833 |
| chr18-89377951-INS-60 | chr18-89377951-INS-60 |
| chr18-89407357-INS-658 | chr18-89407357-INS-658 |
| chr18-89411928-INS-72 | chr18-89411928-INS-72 |
| chr18-89438891-INS-5328 | chr18-89438892-INS-5326 |
| chr18-89446593-INS-72 | chr18-89446593-INS-72 |
| chr18-89474386-INS-6297 | chr18-89474387-INS-6297 |
| chr18-89499749-INS-7323 | chr18-89499750-INS-7323 |
| chr18-89505197-INS-3698 | chr18-89505198-INS-3707 |
| chr18-89512829-INS-66 | chr18-89512829-INS-78 |
| chr18-89532182-INS-3251 | chr18-89532183-INS-3252 |
| chr18-89537997-INS-57 | chr18-89537997-INS-58 |
| chr18-89538873-INS-1187 | chr18-89538862-INS-1181 |
| chr18-89548310-INS-1304 | chr18-89548311-INS-1304 |
| chr18-89561159-INS-6306 | chr18-89561160-INS-6305 |
| chr18-89562655-INS-1823 | chr18-89562656-INS-1823 |
| chr18-89563329-INS-60 | chr18-89563329-INS-50 |
| chr18-89583808-INS-394 | chr18-89583808-INS-394 |
| chr18-89618603-INS-978 | chr18-89618603-INS-978 |
| chr18-89639116-INS-52 | chr18-89639116-INS-52 |
| chr18-89677460-INS-1435 | chr18-89677461-INS-1435 |
| chr18-89698803-INS-86 | chr18-89698777-INS-102 |
| chr18-89721414-INS-6191 | chr18-89721415-INS-6191 |
| chr18-89723459-INS-455 | chr18-89723459-INS-455 |
| chr18-89758129-INS-220 | chr18-89758129-INS-220 |
| chr18-89819483-INS-1641 | chr18-89819484-INS-1639 |
| chr18-89829014-INS-6649 | chr18-89829015-INS-6650 |
| chr18-89862059-INS-668 | chr18-89862059-INS-668 |
| chr18-89864071-INS-347 | chr18-89864071-INS-347 |
| chr18-89866648-INS-1038 | chr18-89866649-INS-1037 |
| chr18-89885047-INS-872 | chr18-89885047-INS-872 |
| chr18-89906337-INS-422 | chr18-89906337-INS-422 |
| chr18-89909624-INS-206 | chr18-89909624-INS-209 |
| chr18-89924643-INS-61 | chr18-89924643-INS-76 |
| chr18-89926569-INS-769 | chr18-89926569-INS-769 |
| chr18-90015097-INS-171 | chr18-90015097-INS-171 |
| chr18-90031210-INS-498 | chr18-90031210-INS-498 |
| chr18-90032661-INS-94 | chr18-90032640-INS-111 |
| chr18-90034227-INS-56 | chr18-90034227-INS-52 |
| chr18-90050923-INS-94 | chr18-90050923-INS-94 |
| chr18-90071055-INS-393 | chr18-90071056-INS-393 |
| chr18-90072083-INS-573 | chr18-90072083-INS-573 |
| chr18-90080090-INS-64 | chr18-90080090-INS-67 |
| chr18-90084345-INS-55 | chr18-90084345-INS-55 |
| chr18-90110152-INS-204 | chr18-90110152-INS-204 |
| chr18-90119947-INS-64 | chr18-90119947-INS-64 |
| chr18-90140594-INS-693 | chr18-90140594-INS-695 |
| chr18-90168862-INS-7795 | chr18-90168863-INS-7795 |
| chr18-90188270-INS-241 | chr18-90188270-INS-241 |
| chr18-90189080-INS-6498 | chr18-90189081-INS-6497 |
| chr18-90214350-INS-3071 | chr18-90214351-INS-3071 |
| chr18-90215053-INS-2327 | chr18-90215054-INS-2326 |
| chr18-90303689-INS-108 | chr18-90303689-INS-132 |
| chr18-90309026-INS-56 | chr18-90309026-INS-56 |
| chr18-90339747-INS-9704 | chr18-90339748-INS-9705 |
| chr18-90421769-INS-4676 | chr18-90421770-INS-4675 |
| chr18-90423236-INS-207 | chr18-90423236-INS-208 |
| chr18-90563895-INS-1702 | chr18-90563896-INS-1705 |
| chr18-90610533-INS-232 | chr18-90610533-INS-214 |
| chr19-3096802-INS-55 | chr19-3096802-INS-51 |
| chr19-3112612-INS-1099 | chr19-3112613-INS-1099 |
| chr19-3255655-INS-160 | chr19-3255655-INS-168 |
| chr19-3262746-INS-6360 | chr19-3262747-INS-6360 |
| chr19-3277853-INS-76 | chr19-3277853-INS-76 |
| chr19-3297091-INS-201 | chr19-3297071-INS-201 |
| chr19-3327673-INS-210 | chr19-3327673-INS-210 |
| chr19-3348275-INS-99 | chr19-3348275-INS-99 |
| chr19-3368142-INS-1064 | chr19-3368143-INS-1095 |
| chr19-3403200-INS-55 | chr19-3403200-INS-55 |
| chr19-3404293-INS-81 | chr19-3404280-INS-71 |
| chr19-3437560-INS-6454 | chr19-3437561-INS-6454 |
| chr19-3471002-INS-52 | chr19-3471002-INS-52 |
| chr19-3499153-INS-238 | chr19-3499153-INS-241 |
| chr19-3500816-INS-65 | chr19-3500816-INS-65 |
| chr19-3535466-INS-449 | chr19-3535466-INS-449 |
| chr19-3552391-INS-108 | chr19-3552391-INS-110 |
| chr19-3560265-INS-67 | chr19-3560266-INS-67 |

| | |
|---|---|
| chr19-3562641-INS-154 | chr19-3562641-INS-154 |
| chr19-3578443-INS-3520 | chr19-3578444-INS-3523 |
| chr19-3591188-INS-70 | chr19-3591188-INS-67 |
| chr19-3617107-INS-201 | chr19-3617107-INS-203 |
| chr19-3620070-INS-210 | chr19-3620070-INS-209 |
| chr19-3622163-INS-96 | chr19-3622163-INS-88 |
| chr19-3640414-INS-558 | chr19-3640415-INS-555 |
| chr19-3642819-INS-206 | chr19-3642819-INS-206 |
| chr19-3647658-INS-111 | chr19-3647658-INS-111 |
| chr19-3648673-INS-223 | chr19-3648673-INS-224 |
| chr19-3653655-INS-160 | chr19-3653655-INS-128 |
| chr19-3656675-INS-103 | chr19-3656675-INS-103 |
| chr19-3676007-INS-72 | chr19-3676023-INS-73 |
| chr19-3684527-INS-132 | chr19-3684527-INS-132 |
| chr19-3702794-INS-237 | chr19-3702794-INS-240 |
| chr19-3718450-INS-200 | chr19-3718450-INS-200 |
| chr19-3743553-INS-209 | chr19-3743553-INS-215 |
| chr19-3766332-INS-108 | chr19-3766332-INS-108 |
| chr19-3774723-INS-214 | chr19-3774723-INS-214 |
| chr19-3783917-INS-207 | chr19-3783917-INS-213 |
| chr19-3800316-INS-53 | chr19-3800316-INS-53 |
| chr19-3877945-INS-311 | chr19-3877945-INS-313 |
| chr19-3878885-INS-214 | chr19-3878885-INS-214 |
| chr19-3884314-INS-63 | chr19-3884314-INS-63 |
| chr19-3894159-INS-229 | chr19-3894159-INS-228 |
| chr19-3905364-INS-202 | chr19-3905364-INS-201 |
| chr19-3915858-INS-128 | chr19-3915858-INS-122 |
| chr19-3930853-INS-97 | chr19-3930853-INS-97 |
| chr19-3958026-INS-62 | chr19-3958026-INS-62 |
| chr19-3983784-INS-435 | chr19-3983784-INS-435 |
| chr19-4000806-INS-87 | chr19-4000806-INS-111 |
| chr19-4041248-INS-112 | chr19-4041248-INS-112 |
| chr19-4074504-INS-75 | chr19-4074519-INS-50 |
| chr19-4079737-INS-61 | chr19-4079749-INS-64 |
| chr19-4280902-INS-161 | chr19-4280894-INS-162 |
| chr19-4338239-INS-168 | chr19-4338239-INS-145 |
| chr19-4352970-INS-64 | chr19-4352970-INS-63 |
| chr19-4381449-INS-216 | chr19-4381449-INS-217 |
| chr19-4387657-INS-74 | chr19-4387657-INS-74 |
| chr19-4388098-INS-212 | chr19-4388098-INS-212 |
| chr19-4414347-INS-220 | chr19-4414347-INS-220 |
| chr19-4416049-INS-235 | chr19-4416049-INS-236 |
| chr19-4440858-INS-221 | chr19-4440858-INS-221 |
| chr19-4450824-INS-6415 | chr19-4450825-INS-6415 |
| chr19-4463276-INS-192 | chr19-4463276-INS-193 |
| chr19-4490471-INS-82 | chr19-4490471-INS-82 |
| chr19-4493711-INS-210 | chr19-4493711-INS-210 |
| chr19-4509407-INS-473 | chr19-4509407-INS-473 |
| chr19-4518441-INS-130 | chr19-4518441-INS-118 |
| chr19-4521531-INS-54 | chr19-4521531-INS-54 |
| chr19-4532274-INS-82 | chr19-4532274-INS-88 |
| chr19-4543616-INS-90 | chr19-4543616-INS-93 |
| chr19-4544294-INS-61 | chr19-4544279-INS-74 |
| chr19-4550105-INS-75 | chr19-4550105-INS-78 |
| chr19-4554344-INS-88 | chr19-4554344-INS-52 |
| chr19-4588885-INS-207 | chr19-4588891-INS-239 |
| chr19-4599225-INS-133 | chr19-4599225-INS-133 |
| chr19-4607210-INS-76 | chr19-4607211-INS-73 |
| chr19-4608154-INS-280 | chr19-4608154-INS-280 |
| chr19-4615251-INS-79 | chr19-4615251-INS-79 |
| chr19-4618908-INS-229 | chr19-4618908-INS-229 |
| chr19-4630110-INS-69 | chr19-4630111-INS-66 |
| chr19-4632781-INS-199 | chr19-4632781-INS-199 |
| chr19-4635334-INS-65 | chr19-4635334-INS-65 |
| chr19-4641007-INS-756 | chr19-4641007-INS-756 |
| chr19-4645133-INS-96 | chr19-4645134-INS-96 |
| chr19-4677970-INS-367 | chr19-4677970-INS-368 |
| chr19-4696074-INS-60 | chr19-4696074-INS-60 |
| chr19-4710578-INS-128 | chr19-4710579-INS-173 |
| chr19-4720353-INS-200 | chr19-4720353-INS-201 |
| chr19-4726631-INS-63 | chr19-4726633-INS-63 |
| chr19-4759150-INS-86 | chr19-4759150-INS-86 |
| chr19-4804362-INS-212 | chr19-4804362-INS-212 |
| chr19-4810359-INS-123 | chr19-4810359-INS-123 |
| chr19-4810901-INS-80 | chr19-4810901-INS-80 |
| chr19-4823894-INS-171 | chr19-4823894-INS-184 |
| chr19-4849905-INS-64 | chr19-4849905-INS-62 |

| | |
|---|---|
| chr19-4856846-INS-360 | chr19-4856846-INS-348 |
| chr19-4866876-INS-387 | chr19-4866876-INS-385 |
| chr19-4884870-INS-74 | chr19-4884879-INS-75 |
| chr19-4915927-INS-78 | chr19-4915927-INS-78 |
| chr19-4919223-INS-188 | chr19-4919223-INS-188 |
| chr19-4923286-INS-55 | chr19-4923293-INS-55 |
| chr19-4935893-INS-208 | chr19-4935893-INS-208 |
| chr19-4993253-INS-115 | chr19-4993253-INS-125 |
| chr19-5005159-INS-83 | chr19-5005159-INS-83 |
| chr19-5013897-INS-216 | chr19-5013897-INS-216 |
| chr19-5020758-INS-479 | chr19-5020759-INS-479 |
| chr19-5022210-INS-51 | chr19-5022211-INS-55 |
| chr19-5024210-INS-213 | chr19-5024210-INS-212 |
| chr19-5033238-INS-116 | chr19-5033238-INS-124 |
| chr19-5061378-INS-796 | chr19-5061379-INS-792 |
| chr19-5069796-INS-58 | chr19-5069796-INS-50 |
| chr19-5152646-INS-213 | chr19-5152646-INS-214 |
| chr19-5175813-INS-233 | chr19-5175813-INS-243 |
| chr19-5180969-INS-1890 | chr19-5180970-INS-1890 |
| chr19-5197252-INS-51 | chr19-5197252-INS-51 |
| chr19-5199080-INS-1455 | chr19-5199086-INS-1446 |
| chr19-5199725-INS-218 | chr19-5199725-INS-218 |
| chr19-5202161-INS-71 | chr19-5202161-INS-75 |
| chr19-5215649-INS-215 | chr19-5215665-INS-226 |
| chr19-5219803-INS-198 | chr19-5219806-INS-199 |
| chr19-5222613-INS-207 | chr19-5222613-INS-211 |
| chr19-5247092-INS-498 | chr19-5247092-INS-498 |
| chr19-5256041-INS-231 | chr19-5256041-INS-232 |
| chr19-5262486-INS-56 | chr19-5262464-INS-105 |
| chr19-5271762-INS-217 | chr19-5271762-INS-217 |
| chr19-5285234-INS-176 | chr19-5285234-INS-180 |
| chr19-5286457-INS-5527 | chr19-5286458-INS-5527 |
| chr19-5308504-INS-60 | chr19-5308504-INS-60 |
| chr19-5327983-INS-266 | chr19-5327983-INS-266 |
| chr19-5331449-INS-204 | chr19-5331449-INS-204 |
| chr19-5339535-INS-60 | chr19-5339535-INS-60 |
| chr19-5374361-INS-391 | chr19-5374361-INS-391 |
| chr19-5401349-INS-52 | chr19-5401349-INS-52 |
| chr19-5406609-INS-405 | chr19-5406611-INS-402 |
| chr19-5425095-INS-343 | chr19-5425095-INS-343 |
| chr19-5445982-INS-184 | chr19-5445982-INS-184 |
| chr19-5449868-INS-112 | chr19-5449868-INS-138 |
| chr19-5457434-INS-228 | chr19-5457434-INS-266 |
| chr19-5459934-INS-224 | chr19-5459934-INS-207 |
| chr19-5479857-INS-372 | chr19-5479857-INS-374 |
| chr19-5492499-INS-51 | chr19-5492499-INS-51 |
| chr19-5541890-INS-209 | chr19-5541890-INS-207 |
| chr19-5551478-INS-195 | chr19-5551478-INS-195 |
| chr19-5554783-INS-150 | chr19-5554783-INS-150 |
| chr19-5561991-INS-84 | chr19-5561991-INS-84 |
| chr19-5563448-INS-57 | chr19-5563452-INS-53 |
| chr19-5568574-INS-54 | chr19-5568574-INS-54 |
| chr19-5575215-INS-209 | chr19-5575215-INS-209 |
| chr19-5591807-INS-1328 | chr19-5591814-INS-1312 |
| chr19-5623516-INS-328 | chr19-5623516-INS-324 |
| chr19-5625230-INS-123 | chr19-5625230-INS-123 |
| chr19-5630631-INS-57 | chr19-5630631-INS-57 |
| chr19-5636458-INS-107 | chr19-5636458-INS-107 |
| chr19-5641750-INS-211 | chr19-5641750-INS-211 |
| chr19-5643110-INS-403 | chr19-5643110-INS-403 |
| chr19-5673172-INS-202 | chr19-5673172-INS-202 |
| chr19-5679018-INS-75 | chr19-5679020-INS-67 |
| chr19-5685060-INS-107 | chr19-5685060-INS-142 |
| chr19-5761384-INS-64 | chr19-5761384-INS-64 |
| chr19-5788625-INS-203 | chr19-5788625-INS-203 |
| chr19-5859356-INS-86 | chr19-5859356-INS-70 |
| chr19-5864902-INS-72 | chr19-5864903-INS-58 |
| chr19-5869964-INS-443 | chr19-5869964-INS-441 |
| chr19-5873259-INS-211 | chr19-5873259-INS-211 |
| chr19-5889056-INS-5379 | chr19-5889057-INS-5379 |
| chr19-5899943-INS-173 | chr19-5899943-INS-173 |
| chr19-5911788-INS-94 | chr19-5911788-INS-144 |
| chr19-5919991-INS-94 | chr19-5919991-INS-141 |
| chr19-5946258-INS-206 | chr19-5946257-INS-222 |
| chr19-5983685-INS-58 | chr19-5983688-INS-62 |
| chr19-6000007-INS-158 | chr19-6000007-INS-158 |
| chr19-6009503-INS-139 | chr19-6009503-INS-119 |

| | |
|---|---|
| chr19-6029903-INS-134 | chr19-6029903-INS-134 |
| chr19-6035343-INS-66 | chr19-6035295-INS-67 |
| chr19-6059822-INS-178 | chr19-6059822-INS-179 |
| chr19-6074970-INS-150 | chr19-6074970-INS-150 |
| chr19-6077412-INS-174 | chr19-6077412-INS-174 |
| chr19-6078915-INS-102 | chr19-6078915-INS-90 |
| chr19-6085276-INS-107 | chr19-6085276-INS-107 |
| chr19-6087195-INS-204 | chr19-6087195-INS-204 |
| chr19-6090552-INS-120 | chr19-6090552-INS-120 |
| chr19-6094175-INS-82 | chr19-6094156-INS-132 |
| chr19-6106755-INS-108 | chr19-6106755-INS-107 |
| chr19-6109605-INS-1017 | chr19-6109606-INS-1046 |
| chr19-6138076-INS-646 | chr19-6138076-INS-649 |
| chr19-6143351-INS-7151 | chr19-6143352-INS-7151 |
| chr19-6148029-INS-126 | chr19-6148029-INS-126 |
| chr19-6180982-INS-171 | chr19-6180982-INS-175 |
| chr19-6183735-INS-68 | chr19-6183735-INS-68 |
| chr19-6209823-INS-210 | chr19-6209823-INS-241 |
| chr19-6210407-INS-55 | chr19-6210407-INS-53 |
| chr19-6210844-INS-163 | chr19-6210844-INS-163 |
| chr19-6219012-INS-78 | chr19-6219012-INS-78 |
| chr19-6230436-INS-82 | chr19-6230419-INS-82 |
| chr19-6231503-INS-90 | chr19-6231503-INS-106 |
| chr19-6244034-INS-127 | chr19-6244034-INS-148 |
| chr19-6247333-INS-210 | chr19-6247333-INS-212 |
| chr19-6287410-INS-55 | chr19-6287410-INS-55 |
| chr19-6288592-INS-329 | chr19-6288592-INS-324 |
| chr19-6295038-INS-64 | chr19-6295038-INS-63 |
| chr19-6297127-INS-209 | chr19-6297124-INS-200 |
| chr19-6344043-INS-228 | chr19-6344043-INS-228 |
| chr19-6352029-INS-168 | chr19-6352029-INS-126 |
| chr19-6366712-INS-217 | chr19-6366712-INS-216 |
| chr19-6373303-INS-53 | chr19-6373277-INS-53 |
| chr19-6376608-INS-72 | chr19-6376596-INS-68 |
| chr19-6397385-INS-55 | chr19-6397385-INS-55 |
| chr19-6421472-INS-208 | chr19-6421472-INS-208 |
| chr19-6526433-INS-209 | chr19-6526433-INS-209 |
| chr19-6586075-INS-63 | chr19-6586075-INS-71 |
| chr19-6601277-INS-221 | chr19-6601277-INS-224 |
| chr19-6606611-INS-52 | chr19-6606611-INS-52 |
| chr19-6617692-INS-74 | chr19-6617692-INS-74 |
| chr19-6618779-INS-488 | chr19-6618785-INS-483 |
| chr19-6649323-INS-104 | chr19-6649323-INS-104 |
| chr19-6685930-INS-56 | chr19-6685931-INS-54 |
| chr19-6704112-INS-231 | chr19-6704112-INS-208 |
| chr19-6723489-INS-537 | chr19-6723489-INS-537 |
| chr19-6818983-INS-171 | chr19-6818983-INS-171 |
| chr19-6869514-INS-220 | chr19-6869514-INS-220 |
| chr19-6895261-INS-103 | chr19-6895261-INS-55 |
| chr19-6989570-INS-200 | chr19-6989570-INS-202 |
| chr19-7002135-INS-201 | chr19-7002135-INS-201 |
| chr19-7029253-INS-87 | chr19-7029253-INS-87 |
| chr19-7039857-INS-217 | chr19-7039857-INS-217 |
| chr19-7050818-INS-91 | chr19-7050818-INS-91 |
| chr19-7167620-INS-90 | chr19-7167608-INS-60 |
| chr19-7188416-INS-389 | chr19-7188416-INS-386 |
| chr19-7237583-INS-60 | chr19-7237583-INS-54 |
| chr19-7277374-INS-67 | chr19-7277374-INS-67 |
| chr19-7321804-INS-206 | chr19-7321804-INS-205 |
| chr19-7325622-INS-150 | chr19-7325642-INS-131 |
| chr19-7342471-INS-53 | chr19-7342471-INS-57 |
| chr19-7344882-INS-204 | chr19-7344882-INS-202 |
| chr19-7347766-INS-64 | chr19-7347766-INS-64 |
| chr19-7350407-INS-114 | chr19-7350407-INS-114 |
| chr19-7362330-INS-126 | chr19-7362331-INS-124 |
| chr19-7367642-INS-155 | chr19-7367642-INS-155 |
| chr19-7373791-INS-172 | chr19-7373791-INS-172 |
| chr19-7390466-INS-229 | chr19-7390466-INS-229 |
| chr19-7417822-INS-207 | chr19-7417822-INS-208 |
| chr19-7436346-INS-1210 | chr19-7436347-INS-1210 |
| chr19-7484211-INS-54 | chr19-7484197-INS-67 |
| chr19-7510481-INS-234 | chr19-7510480-INS-205 |
| chr19-7521381-INS-202 | chr19-7521381-INS-204 |
| chr19-7533864-INS-4842 | chr19-7533865-INS-4842 |
| chr19-7537915-INS-56 | chr19-7537915-INS-76 |
| chr19-7549833-INS-76 | chr19-7549824-INS-67 |
| chr19-7577312-INS-209 | chr19-7577312-INS-209 |

| | |
|---|---|
| chr19-7586183-INS-203 | chr19-7586198-INS-209 |
| chr19-7714134-INS-94 | chr19-7714149-INS-105 |
| chr19-7722339-INS-67 | chr19-7722339-INS-67 |
| chr19-7871177-INS-119 | chr19-7871177-INS-120 |
| chr19-7900726-INS-358 | chr19-7900733-INS-364 |
| chr19-7903341-INS-89 | chr19-7903341-INS-89 |
| chr19-8032340-INS-80 | chr19-8032351-INS-79 |
| chr19-8047725-INS-54 | chr19-8047727-INS-55 |
| chr19-8057127-INS-497 | chr19-8057127-INS-497 |
| chr19-8076969-INS-1279 | chr19-8076970-INS-1279 |
| chr19-8231919-INS-172 | chr19-8231919-INS-174 |
| chr19-8271788-INS-213 | chr19-8271788-INS-213 |
| chr19-8276979-INS-2652 | chr19-8276980-INS-2651 |
| chr19-8283128-INS-52 | chr19-8283128-INS-53 |
| chr19-8406257-INS-64 | chr19-8406257-INS-88 |
| chr19-8411828-INS-4715 | chr19-8411829-INS-4715 |
| chr19-8460328-INS-6500 | chr19-8460329-INS-6500 |
| chr19-8496427-INS-400 | chr19-8496427-INS-400 |
| chr19-8522732-INS-2375 | chr19-8522733-INS-2375 |
| chr19-8525858-INS-7432 | chr19-8525859-INS-7432 |
| chr19-8546621-INS-73 | chr19-8546621-INS-73 |
| chr19-8557648-INS-92 | chr19-8557648-INS-92 |
| chr19-8566576-INS-59 | chr19-8566576-INS-61 |
| chr19-8624518-INS-117 | chr19-8624518-INS-120 |
| chr19-8629907-INS-141 | chr19-8629907-INS-124 |
| chr19-8651373-INS-421 | chr19-8651393-INS-398 |
| chr19-8665993-INS-239 | chr19-8665993-INS-241 |
| chr19-8667952-INS-218 | chr19-8667963-INS-209 |
| chr19-8679107-INS-280 | chr19-8679107-INS-280 |
| chr19-8706559-INS-188 | chr19-8706559-INS-187 |
| chr19-8711355-INS-211 | chr19-8711355-INS-211 |
| chr19-8725148-INS-188 | chr19-8725178-INS-189 |
| chr19-8749109-INS-57 | chr19-8749109-INS-57 |
| chr19-8754930-INS-52 | chr19-8754930-INS-52 |
| chr19-8758871-INS-85 | chr19-8758871-INS-85 |
| chr19-8762137-INS-168 | chr19-8762137-INS-172 |
| chr19-8765867-INS-201 | chr19-8765867-INS-201 |
| chr19-8772908-INS-190 | chr19-8772908-INS-190 |
| chr19-8840742-INS-76 | chr19-8840718-INS-119 |
| chr19-8845147-INS-135 | chr19-8845128-INS-135 |
| chr19-8846424-INS-255 | chr19-8846454-INS-210 |
| chr19-8855354-INS-212 | chr19-8855354-INS-211 |
| chr19-8858770-INS-53 | chr19-8858770-INS-56 |
| chr19-8864004-INS-207 | chr19-8864004-INS-226 |
| chr19-8897511-INS-207 | chr19-8897511-INS-207 |
| chr19-8909646-INS-81 | chr19-8909647-INS-54 |
| chr19-8932435-INS-211 | chr19-8932435-INS-210 |
| chr19-8934071-INS-196 | chr19-8934071-INS-196 |
| chr19-8947024-INS-87 | chr19-8947032-INS-87 |
| chr19-8969234-INS-60 | chr19-8969234-INS-60 |
| chr19-9006176-INS-1193 | chr19-9006177-INS-1195 |
| chr19-9041562-INS-203 | chr19-9041555-INS-242 |
| chr19-9042717-INS-128 | chr19-9042718-INS-109 |
| chr19-9067530-INS-649 | chr19-9067530-INS-653 |
| chr19-9076459-INS-61 | chr19-9076459-INS-65 |
| chr19-9085895-INS-333 | chr19-9085891-INS-337 |
| chr19-9115852-INS-415 | chr19-9115852-INS-415 |
| chr19-9186511-INS-88 | chr19-9186511-INS-88 |
| chr19-9211090-INS-783 | chr19-9211090-INS-816 |
| chr19-9365132-INS-103 | chr19-9365133-INS-102 |
| chr19-9447678-INS-786 | chr19-9447678-INS-786 |
| chr19-9449790-INS-55 | chr19-9449790-INS-54 |
| chr19-9454496-INS-61 | chr19-9454497-INS-73 |
| chr19-9494498-INS-400 | chr19-9494498-INS-400 |
| chr19-9505539-INS-3597 | chr19-9505531-INS-3593 |
| chr19-9562120-INS-213 | chr19-9562120-INS-207 |
| chr19-9577068-INS-2997 | chr19-9577069-INS-2997 |
| chr19-9583828-INS-103 | chr19-9583828-INS-103 |
| chr19-9605169-INS-1006 | chr19-9605170-INS-1004 |
| chr19-9609927-INS-6424 | chr19-9609928-INS-6424 |
| chr19-9616723-INS-146 | chr19-9616723-INS-146 |
| chr19-9633427-INS-2679 | chr19-9633428-INS-2667 |
| chr19-9638091-INS-81 | chr19-9638091-INS-81 |
| chr19-9644036-INS-77 | chr19-9644036-INS-77 |
| chr19-9834474-INS-53 | chr19-9834454-INS-69 |
| chr19-9882427-INS-201 | chr19-9882427-INS-201 |
| chr19-9892394-INS-66 | chr19-9892394-INS-66 |

| | |
|---|---|
| chr19-9905526-INS-93 | chr19-9905526-INS-80 |
| chr19-9919146-INS-59 | chr19-9919146-INS-59 |
| chr19-9943379-INS-187 | chr19-9943379-INS-188 |
| chr19-9970358-INS-230 | chr19-9970358-INS-232 |
| chr19-9972131-INS-96 | chr19-9972113-INS-96 |
| chr19-9973915-INS-1119 | chr19-9973916-INS-1119 |
| chr19-9974086-INS-224 | chr19-9974078-INS-228 |
| chr19-9982379-INS-198 | chr19-9982377-INS-198 |
| chr19-10000434-INS-197 | chr19-10000434-INS-193 |
| chr19-10028946-INS-100 | chr19-10028946-INS-50 |
| chr19-10051765-INS-77 | chr19-10051765-INS-61 |
| chr19-10061935-INS-174 | chr19-10061935-INS-174 |
| chr19-10083287-INS-55 | chr19-10083287-INS-55 |
| chr19-10088661-INS-400 | chr19-10088661-INS-400 |
| chr19-10098127-INS-844 | chr19-10098127-INS-844 |
| chr19-10098871-INS-218 | chr19-10098871-INS-218 |
| chr19-10118817-INS-85 | chr19-10118818-INS-84 |
| chr19-10148081-INS-206 | chr19-10148064-INS-207 |
| chr19-10153834-INS-202 | chr19-10153834-INS-201 |
| chr19-10194974-INS-55 | chr19-10194974-INS-55 |
| chr19-10204323-INS-89 | chr19-10204323-INS-89 |
| chr19-10284543-INS-80 | chr19-10284543-INS-86 |
| chr19-10290468-INS-160 | chr19-10290468-INS-174 |
| chr19-10323543-INS-107 | chr19-10323543-INS-112 |
| chr19-10340255-INS-130 | chr19-10340255-INS-130 |
| chr19-10358417-INS-444 | chr19-10358413-INS-464 |
| chr19-10382934-INS-86 | chr19-10382934-INS-92 |
| chr19-10431571-INS-64 | chr19-10431571-INS-62 |
| chr19-10481035-INS-52 | chr19-10481035-INS-58 |
| chr19-10490217-INS-106 | chr19-10490217-INS-110 |
| chr19-10497848-INS-213 | chr19-10497848-INS-214 |
| chr19-10512968-INS-66 | chr19-10512972-INS-74 |
| chr19-10530159-INS-599 | chr19-10530159-INS-599 |
| chr19-10538882-INS-243 | chr19-10538883-INS-242 |
| chr19-10567375-INS-197 | chr19-10567375-INS-197 |
| chr19-10575025-INS-52 | chr19-10575025-INS-54 |
| chr19-10581489-INS-102 | chr19-10581489-INS-102 |
| chr19-10620845-INS-203 | chr19-10620845-INS-203 |
| chr19-10624870-INS-105 | chr19-10624870-INS-68 |
| chr19-10626736-INS-208 | chr19-10626736-INS-208 |
| chr19-10644882-INS-211 | chr19-10644882-INS-211 |
| chr19-10656928-INS-59 | chr19-10656928-INS-60 |
| chr19-10671905-INS-5286 | chr19-10671906-INS-5286 |
| chr19-10676767-INS-82 | chr19-10676767-INS-82 |
| chr19-10685170-INS-67 | chr19-10685170-INS-67 |
| chr19-10706589-INS-165 | chr19-10706589-INS-171 |
| chr19-10714712-INS-62 | chr19-10714712-INS-60 |
| chr19-10722003-INS-171 | chr19-10722004-INS-181 |
| chr19-10728898-INS-59 | chr19-10728898-INS-60 |
| chr19-10749987-INS-59 | chr19-10749987-INS-53 |
| chr19-10789391-INS-200 | chr19-10789391-INS-200 |
| chr19-10794735-INS-102 | chr19-10794735-INS-102 |
| chr19-10800873-INS-62 | chr19-10800875-INS-52 |
| chr19-10814004-INS-200 | chr19-10814004-INS-200 |
| chr19-10830050-INS-57 | chr19-10830050-INS-58 |
| chr19-10871349-INS-58 | chr19-10871349-INS-62 |
| chr19-10887339-INS-79 | chr19-10887339-INS-79 |
| chr19-10906000-INS-637 | chr19-10905991-INS-633 |
| chr19-10907091-INS-60 | chr19-10907079-INS-52 |
| chr19-10909369-INS-529 | chr19-10909369-INS-523 |
| chr19-10944028-INS-489 | chr19-10944028-INS-489 |
| chr19-11006609-INS-439 | chr19-11006609-INS-439 |
| chr19-11008746-INS-82 | chr19-11008746-INS-80 |
| chr19-11020885-INS-744 | chr19-11020888-INS-744 |
| chr19-11041062-INS-199 | chr19-11041062-INS-199 |
| chr19-11088262-INS-52 | chr19-11088262-INS-52 |
| chr19-11122726-INS-277 | chr19-11122726-INS-277 |
| chr19-11160363-INS-94 | chr19-11160378-INS-52 |
| chr19-11162632-INS-401 | chr19-11162632-INS-401 |
| chr19-11220350-INS-225 | chr19-11220350-INS-227 |
| chr19-11223942-INS-189 | chr19-11223942-INS-182 |
| chr19-11230689-INS-125 | chr19-11230689-INS-125 |
| chr19-11252011-INS-594 | chr19-11252010-INS-564 |
| chr19-11297771-INS-239 | chr19-11297771-INS-243 |
| chr19-11340906-INS-81 | chr19-11340906-INS-81 |
| chr19-11353742-INS-814 | chr19-11353742-INS-808 |
| chr19-11382203-INS-62 | chr19-11382203-INS-62 |

| | |
|---|---|
| chr19-11390106-INS-200 | chr19-11390106-INS-200 |
| chr19-11410105-INS-64 | chr19-11410105-INS-64 |
| chr19-11473647-INS-201 | chr19-11473647-INS-201 |
| chr19-11486251-INS-328 | chr19-11486251-INS-328 |
| chr19-11489276-INS-419 | chr19-11489276-INS-419 |
| chr19-11514617-INS-837 | chr19-11514617-INS-853 |
| chr19-11516224-INS-51 | chr19-11516224-INS-51 |
| chr19-11530126-INS-539 | chr19-11530126-INS-539 |
| chr19-11530641-INS-6462 | chr19-11530642-INS-6463 |
| chr19-11560068-INS-920 | chr19-11560068-INS-920 |
| chr19-11601994-INS-60 | chr19-11601994-INS-50 |
| chr19-11606452-INS-165 | chr19-11606452-INS-165 |
| chr19-11612306-INS-566 | chr19-11612306-INS-566 |
| chr19-11615181-INS-54 | chr19-11615181-INS-50 |
| chr19-11662223-INS-3845 | chr19-11662226-INS-3840 |
| chr19-11702756-INS-86 | chr19-11702748-INS-96 |
| chr19-11711471-INS-73 | chr19-11711471-INS-73 |
| chr19-11712729-INS-181 | chr19-11712729-INS-181 |
| chr19-11720096-INS-488 | chr19-11720096-INS-488 |
| chr19-11752806-INS-103 | chr19-11752806-INS-103 |
| chr19-11768279-INS-549 | chr19-11768279-INS-549 |
| chr19-11777757-INS-91 | chr19-11777713-INS-110 |
| chr19-11781241-INS-56 | chr19-11781241-INS-56 |
| chr19-11796641-INS-87 | chr19-11796641-INS-89 |
| chr19-11801130-INS-57 | chr19-11801130-INS-57 |
| chr19-11803257-INS-177 | chr19-11803257-INS-170 |
| chr19-11827971-INS-55 | chr19-11827971-INS-55 |
| chr19-11840276-INS-61 | chr19-11840276-INS-61 |
| chr19-11844506-INS-71 | chr19-11844508-INS-51 |
| chr19-11865551-INS-212 | chr19-11865551-INS-212 |
| chr19-11888092-INS-223 | chr19-11888092-INS-224 |
| chr19-11900953-INS-213 | chr19-11900953-INS-213 |
| chr19-11933634-INS-56 | chr19-11933634-INS-56 |
| chr19-11934557-INS-204 | chr19-11934557-INS-204 |
| chr19-11935120-INS-212 | chr19-11935120-INS-212 |
| chr19-11938820-INS-89 | chr19-11938820-INS-89 |
| chr19-11956558-INS-80 | chr19-11956558-INS-80 |
| chr19-11972875-INS-71 | chr19-11972875-INS-71 |
| chr19-11998950-INS-160 | chr19-11998950-INS-160 |
| chr19-12019138-INS-56 | chr19-12019138-INS-53 |
| chr19-12022320-INS-70 | chr19-12022320-INS-73 |
| chr19-12025999-INS-405 | chr19-12025999-INS-405 |
| chr19-12031509-INS-72 | chr19-12031509-INS-68 |
| chr19-12043658-INS-569 | chr19-12043658-INS-569 |
| chr19-12089382-INS-76 | chr19-12089382-INS-76 |
| chr19-12120111-INS-3422 | chr19-12120112-INS-3422 |
| chr19-12124674-INS-1962 | chr19-12124675-INS-1962 |
| chr19-12186444-INS-6712 | chr19-12186445-INS-6712 |
| chr19-12191512-INS-102 | chr19-12191512-INS-102 |
| chr19-12236846-INS-649 | chr19-12236846-INS-649 |
| chr19-12248856-INS-4034 | chr19-12248857-INS-4034 |
| chr19-12258101-INS-67 | chr19-12258101-INS-67 |
| chr19-12292863-INS-120 | chr19-12292863-INS-141 |
| chr19-12369877-INS-209 | chr19-12369877-INS-209 |
| chr19-12383080-INS-2720 | chr19-12383081-INS-2720 |
| chr19-12393127-INS-65 | chr19-12393127-INS-66 |
| chr19-12394252-INS-140 | chr19-12394204-INS-178 |
| chr19-12399061-INS-77 | chr19-12399040-INS-55 |
| chr19-12419057-INS-1218 | chr19-12419058-INS-1218 |
| chr19-12462525-INS-84 | chr19-12462525-INS-84 |
| chr19-12490112-INS-60 | chr19-12490112-INS-66 |
| chr19-12498791-INS-155 | chr19-12498791-INS-155 |
| chr19-12515183-INS-6401 | chr19-12515184-INS-6401 |
| chr19-12538903-INS-917 | chr19-12538898-INS-930 |
| chr19-12577399-INS-126 | chr19-12577399-INS-126 |
| chr19-12593778-INS-686 | chr19-12593778-INS-681 |
| chr19-12594440-INS-6203 | chr19-12594441-INS-6202 |
| chr19-12621225-INS-4020 | chr19-12621226-INS-4020 |
| chr19-12630365-INS-60 | chr19-12630365-INS-60 |
| chr19-12633996-INS-499 | chr19-12633996-INS-499 |
| chr19-12637989-INS-159 | chr19-12637989-INS-161 |
| chr19-12640079-INS-97 | chr19-12640079-INS-94 |
| chr19-12676183-INS-187 | chr19-12676187-INS-194 |
| chr19-12715960-INS-757 | chr19-12715960-INS-774 |
| chr19-12717996-INS-116 | chr19-12717997-INS-73 |
| chr19-12720042-INS-166 | chr19-12720042-INS-166 |
| chr19-12720661-INS-126 | chr19-12720661-INS-126 |

| | |
|---|---|
| chr19-12723240-INS-74 | chr19-12723240-INS-76 |
| chr19-12730515-INS-92 | chr19-12730523-INS-60 |
| chr19-12766059-INS-2005 | chr19-12766060-INS-2005 |
| chr19-12766664-INS-135 | chr19-12766664-INS-135 |
| chr19-12802057-INS-60 | chr19-12802057-INS-60 |
| chr19-12813135-INS-3298 | chr19-12813136-INS-3298 |
| chr19-12842475-INS-6390 | chr19-12842476-INS-6390 |
| chr19-12869262-INS-646 | chr19-12869262-INS-645 |
| chr19-12888748-INS-502 | chr19-12888748-INS-502 |
| chr19-12894781-INS-6017 | chr19-12894782-INS-6017 |
| chr19-12908254-INS-151 | chr19-12908254-INS-149 |
| chr19-12915468-INS-225 | chr19-12915468-INS-225 |
| chr19-12932204-INS-78 | chr19-12932204-INS-78 |
| chr19-13006102-INS-64 | chr19-13006102-INS-64 |
| chr19-13009450-INS-72 | chr19-13009453-INS-83 |
| chr19-13012131-INS-547 | chr19-13012131-INS-545 |
| chr19-13013326-INS-902 | chr19-13013326-INS-902 |
| chr19-13075910-INS-69 | chr19-13075943-INS-57 |
| chr19-13087606-INS-57 | chr19-13087606-INS-57 |
| chr19-13135169-INS-83 | chr19-13135169-INS-83 |
| chr19-13157586-INS-851 | chr19-13157586-INS-851 |
| chr19-13177627-INS-197 | chr19-13177668-INS-197 |
| chr19-13187560-INS-832 | chr19-13187560-INS-832 |
| chr19-13206585-INS-54 | chr19-13206585-INS-51 |
| chr19-13240088-INS-1201 | chr19-13240089-INS-1202 |
| chr19-13241636-INS-56 | chr19-13241593-INS-52 |
| chr19-13268394-INS-1443 | chr19-13268396-INS-1428 |
| chr19-13313445-INS-999 | chr19-13313445-INS-996 |
| chr19-13358855-INS-1280 | chr19-13358856-INS-1280 |
| chr19-13365147-INS-7155 | chr19-13365148-INS-7156 |
| chr19-13380343-INS-3113 | chr19-13380344-INS-3112 |
| chr19-13381278-INS-54 | chr19-13381278-INS-65 |
| chr19-13443376-INS-108 | chr19-13443376-INS-108 |
| chr19-13454118-INS-306 | chr19-13454118-INS-306 |
| chr19-13573422-INS-652 | chr19-13573422-INS-652 |
| chr19-13579843-INS-58 | chr19-13579834-INS-62 |
| chr19-13606540-INS-298 | chr19-13606540-INS-283 |
| chr19-13612322-INS-5553 | chr19-13612323-INS-5553 |
| chr19-13646391-INS-1576 | chr19-13646392-INS-1576 |
| chr19-13683797-INS-58 | chr19-13683789-INS-62 |
| chr19-13761273-INS-185 | chr19-13761274-INS-187 |
| chr19-13776788-INS-3319 | chr19-13776833-INS-3309 |
| chr19-13813100-INS-198 | chr19-13813100-INS-197 |
| chr19-13821588-INS-51 | chr19-13821574-INS-97 |
| chr19-13837871-INS-80 | chr19-13837871-INS-68 |
| chr19-13843280-INS-173 | chr19-13843280-INS-173 |
| chr19-13844798-INS-58 | chr19-13844798-INS-69 |
| chr19-13872434-INS-212 | chr19-13872434-INS-212 |
| chr19-13931967-INS-526 | chr19-13931967-INS-526 |
| chr19-13934411-INS-160 | chr19-13934411-INS-164 |
| chr19-13944045-INS-312 | chr19-13944045-INS-312 |
| chr19-13945850-INS-56 | chr19-13945842-INS-54 |
| chr19-13957403-INS-54 | chr19-13957403-INS-54 |
| chr19-13975004-INS-414 | chr19-13975004-INS-418 |
| chr19-13976101-INS-204 | chr19-13976101-INS-204 |
| chr19-13997508-INS-72 | chr19-13997536-INS-72 |
| chr19-14023672-INS-168 | chr19-14023696-INS-168 |
| chr19-14026952-INS-59 | chr19-14026952-INS-68 |
| chr19-14043053-INS-57 | chr19-14043053-INS-57 |
| chr19-14065044-INS-58 | chr19-14065044-INS-58 |
| chr19-14072697-INS-513 | chr19-14072697-INS-513 |
| chr19-14081683-INS-81 | chr19-14081683-INS-65 |
| chr19-14129625-INS-82 | chr19-14129633-INS-66 |
| chr19-14152958-INS-907 | chr19-14152958-INS-906 |
| chr19-14159701-INS-52 | chr19-14159701-INS-52 |
| chr19-14164699-INS-81 | chr19-14164699-INS-117 |
| chr19-14165466-INS-6672 | chr19-14165452-INS-6661 |
| chr19-14188112-INS-50 | chr19-14188095-INS-77 |
| chr19-14219431-INS-223 | chr19-14219431-INS-223 |
| chr19-14224008-INS-134 | chr19-14224019-INS-152 |
| chr19-14273540-INS-78 | chr19-14273540-INS-78 |
| chr19-14283874-INS-645 | chr19-14283874-INS-645 |
| chr19-14288657-INS-319 | chr19-14288657-INS-315 |
| chr19-14292297-INS-120 | chr19-14292297-INS-120 |
| chr19-14312599-INS-317 | chr19-14312598-INS-322 |
| chr19-14330428-INS-6432 | chr19-14330429-INS-6432 |
| chr19-14332572-INS-209 | chr19-14332572-INS-210 |

| | |
|---|---|
| chr19-14353910-INS-117 | chr19-14353910-INS-114 |
| chr19-14370721-INS-60 | chr19-14370721-INS-86 |
| chr19-14376784-INS-197 | chr19-14376784-INS-197 |
| chr19-14383079-INS-66 | chr19-14383079-INS-60 |
| chr19-14397741-INS-2075 | chr19-14397742-INS-2075 |
| chr19-14408777-INS-64 | chr19-14408777-INS-104 |
| chr19-14432859-INS-405 | chr19-14432859-INS-408 |
| chr19-14440350-INS-169 | chr19-14440325-INS-173 |
| chr19-14494241-INS-56 | chr19-14494245-INS-101 |
| chr19-14505758-INS-111 | chr19-14505782-INS-80 |
| chr19-14519880-INS-114 | chr19-14519876-INS-128 |
| chr19-14538133-INS-127 | chr19-14538133-INS-127 |
| chr19-14547546-INS-227 | chr19-14547546-INS-228 |
| chr19-14596319-INS-202 | chr19-14596319-INS-203 |
| chr19-14597598-INS-152 | chr19-14597603-INS-122 |
| chr19-14606278-INS-298 | chr19-14606278-INS-306 |
| chr19-14626752-INS-488 | chr19-14626752-INS-488 |
| chr19-14630531-INS-58 | chr19-14630535-INS-68 |
| chr19-14661225-INS-55 | chr19-14661225-INS-55 |
| chr19-14666547-INS-5001 | chr19-14666548-INS-5004 |
| chr19-14668645-INS-201 | chr19-14668646-INS-201 |
| chr19-14690595-INS-5552 | chr19-14690596-INS-5552 |
| chr19-14702522-INS-98 | chr19-14702511-INS-59 |
| chr19-14715009-INS-170 | chr19-14715009-INS-170 |
| chr19-14726296-INS-51 | chr19-14726296-INS-51 |
| chr19-14747447-INS-64 | chr19-14747411-INS-60 |
| chr19-14751233-INS-400 | chr19-14751233-INS-376 |
| chr19-14796803-INS-78 | chr19-14796803-INS-64 |
| chr19-14801586-INS-861 | chr19-14801586-INS-861 |
| chr19-14808582-INS-313 | chr19-14808579-INS-313 |
| chr19-14846408-INS-352 | chr19-14846408-INS-352 |
| chr19-14893766-INS-213 | chr19-14893766-INS-213 |
| chr19-14907431-INS-3142 | chr19-14907432-INS-3142 |
| chr19-14952416-INS-54 | chr19-14952401-INS-54 |
| chr19-14966546-INS-710 | chr19-14966546-INS-710 |
| chr19-14991242-INS-7008 | chr19-14991243-INS-7008 |
| chr19-15000562-INS-2302 | chr19-15000563-INS-2312 |
| chr19-15002537-INS-190 | chr19-15002537-INS-190 |
| chr19-15015221-INS-203 | chr19-15015221-INS-203 |
| chr19-15017798-INS-76 | chr19-15017798-INS-85 |
| chr19-15022229-INS-274 | chr19-15022229-INS-274 |
| chr19-15050337-INS-284 | chr19-15050375-INS-291 |
| chr19-15106785-INS-59 | chr19-15106785-INS-56 |
| chr19-15120291-INS-82 | chr19-15120289-INS-82 |
| chr19-15125369-INS-100 | chr19-15125367-INS-110 |
| chr19-15125510-INS-6881 | chr19-15125511-INS-6881 |
| chr19-15159531-INS-8770 | chr19-15159532-INS-8770 |
| chr19-15168606-INS-2483 | chr19-15168607-INS-2483 |
| chr19-15226594-INS-119 | chr19-15226590-INS-126 |
| chr19-15246223-INS-58 | chr19-15246223-INS-70 |
| chr19-15285075-INS-54 | chr19-15285075-INS-54 |
| chr19-15314269-INS-74 | chr19-15314269-INS-76 |
| chr19-15353008-INS-498 | chr19-15353008-INS-498 |
| chr19-15383819-INS-320 | chr19-15383819-INS-320 |
| chr19-15396795-INS-195 | chr19-15396794-INS-194 |
| chr19-15445814-INS-120 | chr19-15445814-INS-84 |
| chr19-15459472-INS-72 | chr19-15459472-INS-76 |
| chr19-15460648-INS-246 | chr19-15460648-INS-246 |
| chr19-15495154-INS-56 | chr19-15495154-INS-58 |
| chr19-15502056-INS-75 | chr19-15502056-INS-51 |
| chr19-15503611-INS-92 | chr19-15503634-INS-66 |
| chr19-15537528-INS-144 | chr19-15537528-INS-144 |
| chr19-15540015-INS-93 | chr19-15540015-INS-93 |
| chr19-15550053-INS-112 | chr19-15550047-INS-116 |
| chr19-15557079-INS-884 | chr19-15557079-INS-884 |
| chr19-15559849-INS-107 | chr19-15559849-INS-107 |
| chr19-15603166-INS-732 | chr19-15603169-INS-743 |
| chr19-15603196-INS-155 | chr19-15603177-INS-152 |
| chr19-15629479-INS-328 | chr19-15629479-INS-328 |
| chr19-15638932-INS-136 | chr19-15638932-INS-136 |
| chr19-15644980-INS-115 | chr19-15644980-INS-136 |
| chr19-15650465-INS-2201 | chr19-15650466-INS-2201 |
| chr19-15652933-INS-110 | chr19-15652933-INS-90 |
| chr19-15690986-INS-208 | chr19-15690986-INS-208 |
| chr19-15710868-INS-901 | chr19-15710868-INS-901 |
| chr19-15756625-INS-57 | chr19-15756625-INS-57 |
| chr19-15792133-INS-207 | chr19-15792133-INS-207 |

| | |
|---|---|
| chr19-15793569-INS-138 | chr19-15793593-INS-146 |
| chr19-15904219-INS-55 | chr19-15904219-INS-55 |
| chr19-15917183-INS-74 | chr19-15917177-INS-73 |
| chr19-15921588-INS-209 | chr19-15921588-INS-210 |
| chr19-15929921-INS-610 | chr19-15929921-INS-609 |
| chr19-15940735-INS-106 | chr19-15940735-INS-106 |
| chr19-15948762-INS-61 | chr19-15948762-INS-61 |
| chr19-15954098-INS-7199 | chr19-15954099-INS-7199 |
| chr19-15973909-INS-109 | chr19-15973909-INS-110 |
| chr19-15979916-INS-118 | chr19-15979916-INS-134 |
| chr19-15982050-INS-76 | chr19-15982050-INS-79 |
| chr19-15988263-INS-210 | chr19-15988263-INS-210 |
| chr19-15996185-INS-197 | chr19-15996185-INS-197 |
| chr19-16002365-INS-54 | chr19-16002365-INS-56 |
| chr19-16003848-INS-115 | chr19-16003848-INS-115 |
| chr19-16014186-INS-98 | chr19-16014169-INS-98 |
| chr19-16014412-INS-1660 | chr19-16014413-INS-1660 |
| chr19-16102519-INS-130 | chr19-16102519-INS-148 |
| chr19-16104081-INS-91 | chr19-16104081-INS-67 |
| chr19-16108343-INS-327 | chr19-16108343-INS-327 |
| chr19-16152527-INS-527 | chr19-16152527-INS-519 |
| chr19-16197240-INS-99 | chr19-16197240-INS-54 |
| chr19-16203147-INS-116 | chr19-16203147-INS-116 |
| chr19-16230050-INS-94 | chr19-16230047-INS-92 |
| chr19-16247218-INS-154 | chr19-16247218-INS-154 |
| chr19-16268410-INS-64 | chr19-16268410-INS-52 |
| chr19-16310418-INS-85 | chr19-16310418-INS-85 |
| chr19-16429817-INS-166 | chr19-16429803-INS-196 |
| chr19-16467351-INS-6329 | chr19-16467358-INS-6330 |
| chr19-16490435-INS-64 | chr19-16490435-INS-65 |
| chr19-16565695-INS-74 | chr19-16565670-INS-52 |
| chr19-16570504-INS-64 | chr19-16570504-INS-64 |
| chr19-16623397-INS-132 | chr19-16623397-INS-132 |
| chr19-16637228-INS-296 | chr19-16637223-INS-315 |
| chr19-16646249-INS-84 | chr19-16646249-INS-63 |
| chr19-16689927-INS-198 | chr19-16689927-INS-196 |
| chr19-16703899-INS-236 | chr19-16703899-INS-236 |
| chr19-16705322-INS-552 | chr19-16705322-INS-552 |
| chr19-16724593-INS-307 | chr19-16724593-INS-307 |
| chr19-16748346-INS-76 | chr19-16748331-INS-93 |
| chr19-16761282-INS-138 | chr19-16761282-INS-138 |
| chr19-16763180-INS-4485 | chr19-16763181-INS-4486 |
| chr19-16772223-INS-60 | chr19-16772223-INS-50 |
| chr19-16779338-INS-76 | chr19-16779338-INS-54 |
| chr19-16782683-INS-90 | chr19-16782655-INS-71 |
| chr19-16795746-INS-52 | chr19-16795746-INS-52 |
| chr19-16807926-INS-52 | chr19-16807926-INS-52 |
| chr19-16824953-INS-3044 | chr19-16824962-INS-3034 |
| chr19-16834075-INS-55 | chr19-16834076-INS-54 |
| chr19-16843269-INS-177 | chr19-16843269-INS-177 |
| chr19-16860911-INS-482 | chr19-16860911-INS-482 |
| chr19-16864122-INS-155 | chr19-16864113-INS-161 |
| chr19-16884312-INS-108 | chr19-16884312-INS-98 |
| chr19-16888744-INS-196 | chr19-16888744-INS-196 |
| chr19-16889732-INS-1705 | chr19-16889733-INS-1705 |
| chr19-16892253-INS-107 | chr19-16892253-INS-107 |
| chr19-16902344-INS-2820 | chr19-16902345-INS-2821 |
| chr19-16922333-INS-361 | chr19-16922333-INS-360 |
| chr19-16929335-INS-76 | chr19-16929335-INS-82 |
| chr19-16940596-INS-56 | chr19-16940596-INS-66 |
| chr19-16951230-INS-62 | chr19-16951230-INS-62 |
| chr19-16983234-INS-286 | chr19-16983240-INS-279 |
| chr19-17000364-INS-166 | chr19-17000364-INS-168 |
| chr19-17053175-INS-83 | chr19-17053175-INS-83 |
| chr19-17075891-INS-502 | chr19-17075891-INS-502 |
| chr19-17087425-INS-154 | chr19-17087425-INS-154 |
| chr19-17092345-INS-747 | chr19-17092345-INS-747 |
| chr19-17112501-INS-50 | chr19-17112501-INS-56 |
| chr19-17117353-INS-110 | chr19-17117353-INS-111 |
| chr19-17145543-INS-52 | chr19-17145543-INS-52 |
| chr19-17148259-INS-103 | chr19-17148259-INS-101 |
| chr19-17153987-INS-56 | chr19-17153987-INS-56 |
| chr19-17168821-INS-52 | chr19-17168779-INS-56 |
| chr19-17180662-INS-961 | chr19-17180662-INS-961 |
| chr19-17202732-INS-162 | chr19-17202732-INS-179 |
| chr19-17204541-INS-181 | chr19-17204539-INS-173 |
| chr19-17208523-INS-56 | chr19-17208523-INS-54 |

| | |
|---|---|
| chr19-17219976-INS-52 | chr19-17219976-INS-57 |
| chr19-17221621-INS-234 | chr19-17221621-INS-234 |
| chr19-17228349-INS-96 | chr19-17228349-INS-96 |
| chr19-17242048-INS-191 | chr19-17242048-INS-195 |
| chr19-17243083-INS-60 | chr19-17243083-INS-57 |
| chr19-17255388-INS-151 | chr19-17255388-INS-151 |
| chr19-17267692-INS-73 | chr19-17267692-INS-73 |
| chr19-17270066-INS-52 | chr19-17270066-INS-52 |
| chr19-17275871-INS-110 | chr19-17275885-INS-90 |
| chr19-17291117-INS-79 | chr19-17291117-INS-79 |
| chr19-17293069-INS-77 | chr19-17293069-INS-61 |
| chr19-17295802-INS-109 | chr19-17295802-INS-130 |
| chr19-17297052-INS-55 | chr19-17297052-INS-51 |
| chr19-17311824-INS-725 | chr19-17311824-INS-720 |
| chr19-17335416-INS-165 | chr19-17335416-INS-165 |
| chr19-17353116-INS-97 | chr19-17353116-INS-137 |
| chr19-17354236-INS-606 | chr19-17354237-INS-595 |
| chr19-17354781-INS-156 | chr19-17354781-INS-156 |
| chr19-17356477-INS-90 | chr19-17356478-INS-95 |
| chr19-17368240-INS-686 | chr19-17368204-INS-699 |
| chr19-17384643-INS-216 | chr19-17384630-INS-215 |
| chr19-17400480-INS-2988 | chr19-17400481-INS-2988 |
| chr19-17406401-INS-76 | chr19-17406397-INS-80 |
| chr19-17425679-INS-65 | chr19-17425679-INS-55 |
| chr19-17431766-INS-99 | chr19-17431766-INS-99 |
| chr19-17459262-INS-132 | chr19-17459262-INS-132 |
| chr19-17483980-INS-7008 | chr19-17483981-INS-7008 |
| chr19-17490865-INS-400 | chr19-17490865-INS-400 |
| chr19-17503837-INS-78 | chr19-17503837-INS-54 |
| chr19-17511568-INS-65 | chr19-17511568-INS-100 |
| chr19-17514688-INS-315 | chr19-17514688-INS-315 |
| chr19-17548741-INS-72 | chr19-17548741-INS-72 |
| chr19-17563357-INS-399 | chr19-17563357-INS-399 |
| chr19-17569683-INS-290 | chr19-17569683-INS-296 |
| chr19-17582322-INS-94 | chr19-17582322-INS-94 |
| chr19-17585421-INS-78 | chr19-17585421-INS-64 |
| chr19-17612700-INS-160 | chr19-17612700-INS-163 |
| chr19-17648437-INS-94 | chr19-17648434-INS-70 |
| chr19-17668957-INS-438 | chr19-17668957-INS-436 |
| chr19-17677353-INS-61 | chr19-17677353-INS-61 |
| chr19-17681492-INS-86 | chr19-17681492-INS-94 |
| chr19-17687810-INS-6695 | chr19-17687811-INS-6695 |
| chr19-17703402-INS-6885 | chr19-17703403-INS-6885 |
| chr19-17708509-INS-72 | chr19-17708509-INS-72 |
| chr19-17721085-INS-84 | chr19-17721085-INS-72 |
| chr19-17729280-INS-108 | chr19-17729284-INS-108 |
| chr19-17731881-INS-79 | chr19-17731881-INS-79 |
| chr19-17767855-INS-82 | chr19-17767851-INS-102 |
| chr19-17779920-INS-140 | chr19-17779920-INS-140 |
| chr19-17780952-INS-66 | chr19-17780952-INS-66 |
| chr19-17784040-INS-358 | chr19-17784040-INS-358 |
| chr19-17803136-INS-60 | chr19-17803136-INS-60 |
| chr19-17809832-INS-52 | chr19-17809832-INS-50 |
| chr19-17826357-INS-66 | chr19-17826314-INS-70 |
| chr19-17833331-INS-170 | chr19-17833331-INS-175 |
| chr19-17846882-INS-216 | chr19-17846882-INS-216 |
| chr19-17860903-INS-564 | chr19-17860903-INS-564 |
| chr19-17867721-INS-1828 | chr19-17867722-INS-1828 |
| chr19-17898457-INS-60 | chr19-17898457-INS-55 |
| chr19-17900345-INS-6310 | chr19-17900346-INS-6310 |
| chr19-17907084-INS-60 | chr19-17907084-INS-60 |
| chr19-17907675-INS-389 | chr19-17907675-INS-389 |
| chr19-17908603-INS-114 | chr19-17908608-INS-74 |
| chr19-17925405-INS-64 | chr19-17925405-INS-60 |
| chr19-17973361-INS-51 | chr19-17973358-INS-52 |
| chr19-17982593-INS-1196 | chr19-17982594-INS-1193 |
| chr19-17995736-INS-72 | chr19-17995739-INS-92 |
| chr19-18004021-INS-86 | chr19-18004035-INS-58 |
| chr19-18021422-INS-4379 | chr19-18021423-INS-4379 |
| chr19-18026647-INS-80 | chr19-18026647-INS-80 |
| chr19-18032377-INS-218 | chr19-18032377-INS-217 |
| chr19-18057594-INS-97 | chr19-18057594-INS-117 |
| chr19-18090873-INS-52 | chr19-18090873-INS-52 |
| chr19-18094594-INS-54 | chr19-18094561-INS-54 |
| chr19-18100237-INS-57 | chr19-18100237-INS-71 |
| chr19-18128645-INS-152 | chr19-18128645-INS-156 |
| chr19-18150421-INS-52 | chr19-18150422-INS-56 |

| | |
|---|---|
| chr19-18151421-INS-527 | chr19-18151435-INS-542 |
| chr19-18155410-INS-6466 | chr19-18155411-INS-6466 |
| chr19-18171832-INS-496 | chr19-18171833-INS-496 |
| chr19-18189099-INS-121 | chr19-18189099-INS-125 |
| chr19-18191632-INS-117 | chr19-18191632-INS-131 |
| chr19-18201535-INS-1647 | chr19-18201536-INS-1649 |
| chr19-18237913-INS-358 | chr19-18237913-INS-358 |
| chr19-18257882-INS-68 | chr19-18257882-INS-70 |
| chr19-18284638-INS-193 | chr19-18284650-INS-207 |
| chr19-18295990-INS-4062 | chr19-18295991-INS-4062 |
| chr19-18335158-INS-173 | chr19-18335158-INS-173 |
| chr19-18363036-INS-56 | chr19-18363038-INS-72 |
| chr19-18370648-INS-56 | chr19-18370648-INS-53 |
| chr19-18371204-INS-510 | chr19-18371204-INS-510 |
| chr19-18383368-INS-6473 | chr19-18383369-INS-6473 |
| chr19-18383584-INS-56 | chr19-18383584-INS-58 |
| chr19-18409910-INS-143 | chr19-18409910-INS-143 |
| chr19-18419729-INS-124 | chr19-18419729-INS-124 |
| chr19-18435905-INS-7152 | chr19-18435906-INS-7154 |
| chr19-18451536-INS-51 | chr19-18451514-INS-78 |
| chr19-18468952-INS-169 | chr19-18468953-INS-168 |
| chr19-18496174-INS-445 | chr19-18496174-INS-445 |
| chr19-18498007-INS-60 | chr19-18498036-INS-60 |
| chr19-18504766-INS-193 | chr19-18504766-INS-194 |
| chr19-18517117-INS-76 | chr19-18517105-INS-56 |
| chr19-18517762-INS-590 | chr19-18517762-INS-590 |
| chr19-18525873-INS-107 | chr19-18525874-INS-107 |
| chr19-18536611-INS-1889 | chr19-18536603-INS-1880 |
| chr19-18559646-INS-165 | chr19-18559642-INS-171 |
| chr19-18593161-INS-192 | chr19-18593161-INS-192 |
| chr19-18595418-INS-570 | chr19-18595418-INS-569 |
| chr19-18603448-INS-74 | chr19-18603448-INS-74 |
| chr19-18612531-INS-58 | chr19-18612531-INS-58 |
| chr19-18619727-INS-212 | chr19-18619727-INS-212 |
| chr19-18625426-INS-54 | chr19-18625416-INS-50 |
| chr19-18631598-INS-589 | chr19-18631598-INS-589 |
| chr19-18646158-INS-210 | chr19-18646158-INS-210 |
| chr19-18650984-INS-51 | chr19-18650984-INS-51 |
| chr19-18660083-INS-899 | chr19-18660083-INS-906 |
| chr19-18660142-INS-52 | chr19-18660164-INS-58 |
| chr19-18680354-INS-65 | chr19-18680354-INS-65 |
| chr19-18683752-INS-57 | chr19-18683752-INS-57 |
| chr19-18685973-INS-55 | chr19-18685973-INS-55 |
| chr19-18687217-INS-142 | chr19-18687188-INS-151 |
| chr19-18699515-INS-90 | chr19-18699535-INS-107 |
| chr19-18715140-INS-302 | chr19-18715147-INS-309 |
| chr19-18722763-INS-3227 | chr19-18722764-INS-3227 |
| chr19-18742916-INS-1344 | chr19-18742917-INS-1344 |
| chr19-18763047-INS-60 | chr19-18763041-INS-73 |
| chr19-18807123-INS-152 | chr19-18807123-INS-152 |
| chr19-18809797-INS-58 | chr19-18809797-INS-50 |
| chr19-18815944-INS-57 | chr19-18815945-INS-57 |
| chr19-18825993-INS-199 | chr19-18825993-INS-199 |
| chr19-18840325-INS-5246 | chr19-18840326-INS-5246 |
| chr19-18856078-INS-75 | chr19-18856078-INS-90 |
| chr19-18870911-INS-420 | chr19-18870911-INS-421 |
| chr19-18872981-INS-808 | chr19-18872972-INS-818 |
| chr19-18875434-INS-56 | chr19-18875434-INS-51 |
| chr19-18883407-INS-121 | chr19-18883407-INS-129 |
| chr19-18888848-INS-60 | chr19-18888850-INS-56 |
| chr19-18921599-INS-53 | chr19-18921618-INS-55 |
| chr19-18934900-INS-656 | chr19-18934900-INS-656 |
| chr19-18980715-INS-56 | chr19-18980672-INS-80 |
| chr19-19009085-INS-69 | chr19-19009085-INS-71 |
| chr19-19040170-INS-124 | chr19-19040170-INS-124 |
| chr19-19043265-INS-80 | chr19-19043265-INS-80 |
| chr19-19044192-INS-452 | chr19-19044192-INS-452 |
| chr19-19066324-INS-66 | chr19-19066334-INS-68 |
| chr19-19071222-INS-6897 | chr19-19071223-INS-6897 |
| chr19-19113728-INS-225 | chr19-19113728-INS-224 |
| chr19-19142569-INS-60 | chr19-19142568-INS-56 |
| chr19-19147627-INS-54 | chr19-19147611-INS-64 |
| chr19-19160251-INS-65 | chr19-19160251-INS-65 |
| chr19-19186276-INS-58 | chr19-19186276-INS-54 |
| chr19-19192043-INS-51 | chr19-19192044-INS-51 |
| chr19-19205107-INS-58 | chr19-19205107-INS-58 |
| chr19-19213598-INS-1303 | chr19-19213599-INS-1303 |

| | |
|---|---|
| chr19-19218814-INS-214 | chr19-19218814-INS-214 |
| chr19-19229558-INS-76 | chr19-19229558-INS-74 |
| chr19-19239833-INS-340 | chr19-19239833-INS-340 |
| chr19-19240964-INS-224 | chr19-19240964-INS-225 |
| chr19-19268429-INS-53 | chr19-19268429-INS-55 |
| chr19-19274333-INS-53 | chr19-19274334-INS-53 |
| chr19-19292792-INS-383 | chr19-19292793-INS-385 |
| chr19-19297221-INS-511 | chr19-19297217-INS-514 |
| chr19-19298702-INS-51 | chr19-19298702-INS-95 |
| chr19-19316237-INS-146 | chr19-19316237-INS-161 |
| chr19-19332734-INS-89 | chr19-19332759-INS-96 |
| chr19-19398178-INS-594 | chr19-19398178-INS-594 |
| chr19-19401156-INS-72 | chr19-19401156-INS-72 |
| chr19-19417137-INS-76 | chr19-19417146-INS-64 |
| chr19-19419253-INS-7335 | chr19-19419254-INS-7336 |
| chr19-19425282-INS-69 | chr19-19425282-INS-65 |
| chr19-19429305-INS-203 | chr19-19429305-INS-203 |
| chr19-19450008-INS-131 | chr19-19450008-INS-131 |
| chr19-19461948-INS-685 | chr19-19461948-INS-636 |
| chr19-19471910-INS-113 | chr19-19471910-INS-114 |
| chr19-19500526-INS-52 | chr19-19500524-INS-74 |
| chr19-19517348-INS-2910 | chr19-19517349-INS-2910 |
| chr19-19546993-INS-145 | chr19-19546993-INS-145 |
| chr19-19547599-INS-3643 | chr19-19547600-INS-3643 |
| chr19-19563388-INS-1633 | chr19-19563389-INS-1634 |
| chr19-19573718-INS-62 | chr19-19573718-INS-62 |
| chr19-19576173-INS-213 | chr19-19576173-INS-223 |
| chr19-19591439-INS-417 | chr19-19591439-INS-417 |
| chr19-19603218-INS-1152 | chr19-19603219-INS-1151 |
| chr19-19607710-INS-56 | chr19-19607710-INS-56 |
| chr19-19609447-INS-70 | chr19-19609439-INS-78 |
| chr19-19619508-INS-200 | chr19-19619508-INS-200 |
| chr19-19628630-INS-64 | chr19-19628630-INS-72 |
| chr19-19660193-INS-52 | chr19-19660193-INS-52 |
| chr19-19662389-INS-92 | chr19-19662389-INS-80 |
| chr19-19699698-INS-217 | chr19-19699698-INS-221 |
| chr19-19707278-INS-400 | chr19-19707278-INS-400 |
| chr19-19710508-INS-360 | chr19-19710508-INS-360 |
| chr19-19720938-INS-592 | chr19-19720938-INS-600 |
| chr19-19732071-INS-199 | chr19-19732071-INS-199 |
| chr19-19807843-INS-60 | chr19-19807843-INS-60 |
| chr19-19821053-INS-2765 | chr19-19821054-INS-2765 |
| chr19-19830240-INS-91 | chr19-19830240-INS-91 |
| chr19-19832216-INS-76 | chr19-19832216-INS-76 |
| chr19-19887651-INS-646 | chr19-19887651-INS-668 |
| chr19-19907061-INS-57 | chr19-19907061-INS-53 |
| chr19-19918183-INS-6807 | chr19-19918184-INS-6807 |
| chr19-19933762-INS-151 | chr19-19933762-INS-163 |
| chr19-19943393-INS-482 | chr19-19943405-INS-472 |
| chr19-19944335-INS-6472 | chr19-19944336-INS-6473 |
| chr19-19945261-INS-925 | chr19-19945261-INS-925 |
| chr19-19960289-INS-89 | chr19-19960289-INS-89 |
| chr19-19965519-INS-186 | chr19-19965520-INS-186 |
| chr19-19976092-INS-71 | chr19-19976092-INS-75 |
| chr19-20002052-INS-132 | chr19-20002052-INS-132 |
| chr19-20011905-INS-67 | chr19-20011895-INS-111 |
| chr19-20017769-INS-294 | chr19-20017769-INS-294 |
| chr19-20020260-INS-519 | chr19-20020260-INS-522 |
| chr19-20030549-INS-3910 | chr19-20030550-INS-3910 |
| chr19-20042921-INS-54 | chr19-20042935-INS-84 |
| chr19-20049742-INS-902 | chr19-20049742-INS-902 |
| chr19-20054723-INS-249 | chr19-20054723-INS-261 |
| chr19-20074812-INS-171 | chr19-20074812-INS-171 |
| chr19-20098471-INS-96 | chr19-20098471-INS-109 |
| chr19-20114348-INS-76 | chr19-20114334-INS-60 |
| chr19-20131301-INS-905 | chr19-20131301-INS-905 |
| chr19-20138488-INS-200 | chr19-20138488-INS-200 |
| chr19-20145872-INS-452 | chr19-20145872-INS-452 |
| chr19-20186632-INS-200 | chr19-20186632-INS-200 |
| chr19-20189582-INS-186 | chr19-20189582-INS-189 |
| chr19-20225628-INS-148 | chr19-20225628-INS-151 |
| chr19-20227887-INS-283 | chr19-20227887-INS-291 |
| chr19-20232144-INS-460 | chr19-20232144-INS-460 |
| chr19-20244900-INS-112 | chr19-20244900-INS-116 |
| chr19-20322566-INS-1133 | chr19-20322567-INS-1129 |
| chr19-20325638-INS-427 | chr19-20325638-INS-427 |
| chr19-20332693-INS-1891 | chr19-20332694-INS-1891 |

| | |
|---|---|
| chr19-20373770-INS-54 | chr19-20373770-INS-54 |
| chr19-20404202-INS-60 | chr19-20404202-INS-62 |
| chr19-20407903-INS-106 | chr19-20407934-INS-105 |
| chr19-20424480-INS-63 | chr19-20424518-INS-70 |
| chr19-20442308-INS-56 | chr19-20442308-INS-54 |
| chr19-20444298-INS-179 | chr19-20444298-INS-173 |
| chr19-20446616-INS-312 | chr19-20446616-INS-312 |
| chr19-20467375-INS-65 | chr19-20467375-INS-57 |
| chr19-20477880-INS-76 | chr19-20477875-INS-58 |
| chr19-20481940-INS-473 | chr19-20481940-INS-473 |
| chr19-20529069-INS-6215 | chr19-20529070-INS-6215 |
| chr19-20563172-INS-94 | chr19-20563156-INS-82 |
| chr19-20589574-INS-76 | chr19-20589596-INS-50 |
| chr19-20594784-INS-196 | chr19-20594784-INS-196 |
| chr19-20602426-INS-179 | chr19-20602426-INS-179 |
| chr19-20604990-INS-817 | chr19-20604990-INS-817 |
| chr19-20605805-INS-70 | chr19-20605805-INS-70 |
| chr19-20607559-INS-2664 | chr19-20607560-INS-2663 |
| chr19-20627362-INS-211 | chr19-20627362-INS-211 |
| chr19-20632823-INS-95 | chr19-20632823-INS-51 |
| chr19-20633910-INS-110 | chr19-20633910-INS-110 |
| chr19-20634756-INS-4166 | chr19-20634757-INS-4166 |
| chr19-20637699-INS-510 | chr19-20637699-INS-510 |
| chr19-20639061-INS-109 | chr19-20639061-INS-109 |
| chr19-20642073-INS-221 | chr19-20642073-INS-220 |
| chr19-20656946-INS-138 | chr19-20656946-INS-138 |
| chr19-20678958-INS-6565 | chr19-20678959-INS-6565 |
| chr19-20683544-INS-82 | chr19-20683554-INS-92 |
| chr19-20695651-INS-1355 | chr19-20695652-INS-1355 |
| chr19-20708576-INS-64 | chr19-20708576-INS-68 |
| chr19-20714511-INS-201 | chr19-20714511-INS-201 |
| chr19-20717902-INS-607 | chr19-20717902-INS-607 |
| chr19-20718699-INS-61 | chr19-20718699-INS-50 |
| chr19-20737363-INS-104 | chr19-20737363-INS-104 |
| chr19-20767630-INS-1251 | chr19-20767631-INS-1252 |
| chr19-20781025-INS-90 | chr19-20781007-INS-93 |
| chr19-20833079-INS-210 | chr19-20833079-INS-210 |
| chr19-20844080-INS-749 | chr19-20844080-INS-749 |
| chr19-20845267-INS-51 | chr19-20845267-INS-51 |
| chr19-20851562-INS-52 | chr19-20851562-INS-52 |
| chr19-20869405-INS-211 | chr19-20869405-INS-212 |
| chr19-20881323-INS-425 | chr19-20881323-INS-425 |
| chr19-20885914-INS-4944 | chr19-20885915-INS-4944 |
| chr19-20896402-INS-440 | chr19-20896402-INS-440 |
| chr19-20899881-INS-327 | chr19-20899885-INS-323 |
| chr19-20903336-INS-74 | chr19-20903336-INS-78 |
| chr19-20912256-INS-51 | chr19-20912259-INS-59 |
| chr19-20922879-INS-251 | chr19-20922879-INS-267 |
| chr19-20924470-INS-85 | chr19-20924470-INS-85 |
| chr19-20954619-INS-298 | chr19-20954619-INS-298 |
| chr19-20957508-INS-536 | chr19-20957508-INS-536 |
| chr19-20987655-INS-3791 | chr19-20987656-INS-3791 |
| chr19-20993744-INS-62 | chr19-20993744-INS-70 |
| chr19-21016761-INS-68 | chr19-21016761-INS-68 |
| chr19-21047636-INS-70 | chr19-21047636-INS-70 |
| chr19-21065787-INS-54 | chr19-21065787-INS-54 |
| chr19-21071116-INS-386 | chr19-21071116-INS-389 |
| chr19-21076405-INS-2112 | chr19-21076406-INS-2112 |
| chr19-21111255-INS-60 | chr19-21111255-INS-75 |
| chr19-21119118-INS-131 | chr19-21119112-INS-132 |
| chr19-21121595-INS-56 | chr19-21121595-INS-56 |
| chr19-21134198-INS-93 | chr19-21134196-INS-98 |
| chr19-21151146-INS-52 | chr19-21151154-INS-56 |
| chr19-21156419-INS-149 | chr19-21156419-INS-152 |
| chr19-21180704-INS-211 | chr19-21180689-INS-198 |
| chr19-21186559-INS-805 | chr19-21186559-INS-804 |
| chr19-21191084-INS-66 | chr19-21191084-INS-66 |
| chr19-21192824-INS-70 | chr19-21192824-INS-66 |
| chr19-21195684-INS-212 | chr19-21195684-INS-212 |
| chr19-21210019-INS-196 | chr19-21210019-INS-196 |
| chr19-21214122-INS-301 | chr19-21214122-INS-300 |
| chr19-21266945-INS-190 | chr19-21266946-INS-187 |
| chr19-21267237-INS-861 | chr19-21267238-INS-864 |
| chr19-21273160-INS-56 | chr19-21273160-INS-56 |
| chr19-21280332-INS-76 | chr19-21280302-INS-58 |
| chr19-21284472-INS-51 | chr19-21284472-INS-51 |
| chr19-21290147-INS-74 | chr19-21290145-INS-85 |

| | |
|---|---|
| chr19-21291247-INS-962 | chr19-21291247-INS-921 |
| chr19-21307622-INS-74 | chr19-21307622-INS-80 |
| chr19-21319132-INS-60 | chr19-21319113-INS-87 |
| chr19-21330694-INS-175 | chr19-21330694-INS-175 |
| chr19-21345969-INS-72 | chr19-21345969-INS-80 |
| chr19-21354729-INS-570 | chr19-21354730-INS-575 |
| chr19-21362369-INS-87 | chr19-21362369-INS-98 |
| chr19-21365263-INS-215 | chr19-21365263-INS-215 |
| chr19-21379595-INS-215 | chr19-21379595-INS-207 |
| chr19-21384661-INS-123 | chr19-21384661-INS-126 |
| chr19-21401548-INS-106 | chr19-21401555-INS-83 |
| chr19-21409680-INS-52 | chr19-21409680-INS-52 |
| chr19-21428849-INS-54 | chr19-21428849-INS-60 |
| chr19-21435953-INS-610 | chr19-21435953-INS-609 |
| chr19-21445799-INS-52 | chr19-21445759-INS-52 |
| chr19-21452923-INS-175 | chr19-21452923-INS-175 |
| chr19-21469347-INS-55 | chr19-21469347-INS-55 |
| chr19-21478068-INS-109 | chr19-21478068-INS-109 |
| chr19-21480752-INS-56 | chr19-21480752-INS-56 |
| chr19-21481309-INS-188 | chr19-21481310-INS-188 |
| chr19-21484120-INS-64 | chr19-21484120-INS-64 |
| chr19-21496176-INS-97 | chr19-21496176-INS-100 |
| chr19-21530058-INS-347 | chr19-21530093-INS-358 |
| chr19-21531922-INS-69 | chr19-21531922-INS-73 |
| chr19-21540726-INS-344 | chr19-21540726-INS-344 |
| chr19-21542496-INS-395 | chr19-21542501-INS-399 |
| chr19-21566667-INS-100 | chr19-21566667-INS-100 |
| chr19-21581680-INS-424 | chr19-21581680-INS-424 |
| chr19-21583746-INS-86 | chr19-21583746-INS-86 |
| chr19-21594478-INS-475 | chr19-21594478-INS-474 |
| chr19-21605816-INS-100 | chr19-21605816-INS-100 |
| chr19-21610551-INS-7203 | chr19-21610552-INS-7203 |
| chr19-21612360-INS-126 | chr19-21612360-INS-126 |
| chr19-21651797-INS-155 | chr19-21651797-INS-156 |
| chr19-21666586-INS-74 | chr19-21666586-INS-74 |
| chr19-21673423-INS-212 | chr19-21673423-INS-212 |
| chr19-21675554-INS-208 | chr19-21675554-INS-207 |
| chr19-21677706-INS-965 | chr19-21677706-INS-965 |
| chr19-21688808-INS-206 | chr19-21688808-INS-206 |
| chr19-21697345-INS-56 | chr19-21697345-INS-56 |
| chr19-21713360-INS-62 | chr19-21713360-INS-62 |
| chr19-21717360-INS-60 | chr19-21717361-INS-56 |
| chr19-21725255-INS-81 | chr19-21725255-INS-85 |
| chr19-21733210-INS-163 | chr19-21733211-INS-169 |
| chr19-21734457-INS-552 | chr19-21734457-INS-552 |
| chr19-21745597-INS-54 | chr19-21745597-INS-54 |
| chr19-21762230-INS-51 | chr19-21762230-INS-51 |
| chr19-21773124-INS-186 | chr19-21773124-INS-187 |
| chr19-21774083-INS-422 | chr19-21774083-INS-432 |
| chr19-21786077-INS-103 | chr19-21786077-INS-103 |
| chr19-21787067-INS-56 | chr19-21787067-INS-56 |
| chr19-21789440-INS-160 | chr19-21789440-INS-160 |
| chr19-21798553-INS-76 | chr19-21798554-INS-76 |
| chr19-21827084-INS-1526 | chr19-21827085-INS-1527 |
| chr19-21828985-INS-214 | chr19-21828985-INS-214 |
| chr19-21845177-INS-53 | chr19-21845177-INS-53 |
| chr19-21877705-INS-72 | chr19-21877694-INS-58 |
| chr19-21918576-INS-80 | chr19-21918576-INS-80 |
| chr19-21926365-INS-509 | chr19-21926365-INS-509 |
| chr19-21941729-INS-72 | chr19-21941729-INS-72 |
| chr19-21971091-INS-887 | chr19-21971091-INS-886 |
| chr19-21975326-INS-98 | chr19-21975326-INS-92 |
| chr19-21995460-INS-116 | chr19-21995460-INS-116 |
| chr19-22031058-INS-171 | chr19-22031034-INS-190 |
| chr19-22037266-INS-101 | chr19-22037272-INS-99 |
| chr19-22038026-INS-60 | chr19-22038026-INS-65 |
| chr19-22051845-INS-410 | chr19-22051845-INS-410 |
| chr19-22056145-INS-171 | chr19-22056145-INS-171 |
| chr19-22077264-INS-106 | chr19-22077264-INS-97 |
| chr19-22084168-INS-6785 | chr19-22084169-INS-6785 |
| chr19-22109140-INS-74 | chr19-22109094-INS-96 |
| chr19-22142058-INS-60 | chr19-22142058-INS-68 |
| chr19-22183008-INS-74 | chr19-22183008-INS-72 |
| chr19-22212500-INS-55 | chr19-22212500-INS-55 |
| chr19-22230809-INS-564 | chr19-22230809-INS-564 |
| chr19-22232093-INS-56 | chr19-22232086-INS-57 |
| chr19-22236076-INS-154 | chr19-22236076-INS-154 |

| | |
|---|---|
| chr19-22273609-INS-62 | chr19-22273609-INS-62 |
| chr19-22275298-INS-71 | chr19-22275298-INS-70 |
| chr19-22287861-INS-787 | chr19-22287861-INS-787 |
| chr19-22299330-INS-120 | chr19-22299330-INS-120 |
| chr19-22304157-INS-68 | chr19-22304162-INS-64 |
| chr19-22306162-INS-222 | chr19-22306162-INS-222 |
| chr19-22323590-INS-195 | chr19-22323590-INS-195 |
| chr19-22415598-INS-285 | chr19-22415598-INS-287 |
| chr19-22472632-INS-143 | chr19-22472632-INS-141 |
| chr19-22476228-INS-242 | chr19-22476248-INS-266 |
| chr19-22480321-INS-182 | chr19-22480321-INS-181 |
| chr19-22490970-INS-336 | chr19-22490970-INS-336 |
| chr19-22499536-INS-670 | chr19-22499536-INS-682 |
| chr19-22514810-INS-52 | chr19-22514810-INS-52 |
| chr19-22519037-INS-56 | chr19-22519016-INS-55 |
| chr19-22528289-INS-84 | chr19-22528289-INS-84 |
| chr19-22529658-INS-157 | chr19-22529658-INS-157 |
| chr19-22530425-INS-60 | chr19-22530425-INS-60 |
| chr19-22541990-INS-52 | chr19-22541990-INS-52 |
| chr19-22543067-INS-68 | chr19-22543067-INS-68 |
| chr19-22562038-INS-138 | chr19-22562038-INS-138 |
| chr19-22565922-INS-83 | chr19-22565920-INS-71 |
| chr19-22585392-INS-62 | chr19-22585392-INS-62 |
| chr19-22593075-INS-497 | chr19-22593075-INS-497 |
| chr19-22605317-INS-147 | chr19-22605317-INS-147 |
| chr19-22612192-INS-270 | chr19-22612192-INS-270 |
| chr19-22630907-INS-213 | chr19-22630905-INS-216 |
| chr19-22639336-INS-52 | chr19-22639336-INS-52 |
| chr19-22643890-INS-59 | chr19-22643890-INS-56 |
| chr19-22654010-INS-77 | chr19-22654003-INS-75 |
| chr19-22666077-INS-68 | chr19-22666077-INS-68 |
| chr19-22717409-INS-6458 | chr19-22717410-INS-6459 |
| chr19-22755626-INS-107 | chr19-22755626-INS-107 |
| chr19-22798987-INS-82 | chr19-22798987-INS-51 |
| chr19-22817509-INS-602 | chr19-22817509-INS-603 |
| chr19-22840936-INS-86 | chr19-22840924-INS-111 |
| chr19-22870776-INS-72 | chr19-22870776-INS-68 |
| chr19-22873935-INS-109 | chr19-22873935-INS-117 |
| chr19-22885967-INS-62 | chr19-22885968-INS-62 |
| chr19-22899220-INS-100 | chr19-22899222-INS-96 |
| chr19-22925428-INS-498 | chr19-22925428-INS-498 |
| chr19-22930173-INS-57 | chr19-22930173-INS-57 |
| chr19-22938894-INS-50 | chr19-22938894-INS-58 |
| chr19-22972975-INS-210 | chr19-22972975-INS-209 |
| chr19-22982143-INS-465 | chr19-22982143-INS-456 |
| chr19-22991289-INS-78 | chr19-22991326-INS-58 |
| chr19-22999107-INS-208 | chr19-22999107-INS-208 |
| chr19-23005506-INS-213 | chr19-23005506-INS-213 |
| chr19-23006327-INS-75 | chr19-23006327-INS-71 |
| chr19-23011091-INS-138 | chr19-23011091-INS-139 |
| chr19-23020605-INS-174 | chr19-23020605-INS-209 |
| chr19-23027063-INS-211 | chr19-23027063-INS-212 |
| chr19-23027802-INS-194 | chr19-23027802-INS-194 |
| chr19-23033046-INS-56 | chr19-23033046-INS-56 |
| chr19-23042840-INS-195 | chr19-23042840-INS-195 |
| chr19-23045285-INS-219 | chr19-23045285-INS-219 |
| chr19-23048857-INS-203 | chr19-23048857-INS-203 |
| chr19-23054052-INS-78 | chr19-23054052-INS-68 |
| chr19-23059439-INS-178 | chr19-23059440-INS-176 |
| chr19-23073503-INS-786 | chr19-23073547-INS-785 |
| chr19-23079106-INS-75 | chr19-23079106-INS-75 |
| chr19-23090284-INS-201 | chr19-23090284-INS-201 |
| chr19-23094960-INS-200 | chr19-23094960-INS-200 |
| chr19-23098844-INS-196 | chr19-23098844-INS-198 |
| chr19-23100573-INS-92 | chr19-23100573-INS-92 |
| chr19-23105743-INS-315 | chr19-23105743-INS-315 |
| chr19-23134214-INS-52 | chr19-23134214-INS-52 |
| chr19-23148808-INS-204 | chr19-23148808-INS-204 |
| chr19-23151584-INS-55 | chr19-23151584-INS-59 |
| chr19-23154332-INS-211 | chr19-23154333-INS-196 |
| chr19-23154870-INS-377 | chr19-23154871-INS-377 |
| chr19-23164018-INS-246 | chr19-23164018-INS-246 |
| chr19-23165940-INS-386 | chr19-23165940-INS-387 |
| chr19-23167063-INS-220 | chr19-23167063-INS-219 |
| chr19-23171829-INS-50 | chr19-23171829-INS-50 |
| chr19-23178158-INS-54 | chr19-23178158-INS-56 |
| chr19-23188472-INS-114 | chr19-23188472-INS-114 |

| | |
|---|---|
| chr19-23189402-INS-427 | chr19-23189402-INS-429 |
| chr19-23194824-INS-58 | chr19-23194824-INS-58 |
| chr19-23229893-INS-89 | chr19-23229897-INS-72 |
| chr19-23231438-INS-608 | chr19-23231438-INS-608 |
| chr19-23237406-INS-75 | chr19-23237396-INS-68 |
| chr19-23239129-INS-65 | chr19-23239129-INS-65 |
| chr19-23242087-INS-81 | chr19-23242087-INS-81 |
| chr19-23256219-INS-118 | chr19-23256220-INS-79 |
| chr19-23261475-INS-5314 | chr19-23261476-INS-5316 |
| chr19-23264833-INS-215 | chr19-23264833-INS-218 |
| chr19-23289103-INS-364 | chr19-23289103-INS-364 |
| chr19-23314478-INS-1635 | chr19-23314479-INS-1635 |
| chr19-23315330-INS-399 | chr19-23315331-INS-399 |
| chr19-23351658-INS-61 | chr19-23351658-INS-61 |
| chr19-23376281-INS-56 | chr19-23376281-INS-58 |
| chr19-23383000-INS-1641 | chr19-23383001-INS-1648 |
| chr19-23383636-INS-65 | chr19-23383636-INS-65 |
| chr19-23386151-INS-237 | chr19-23386151-INS-235 |
| chr19-23387596-INS-3672 | chr19-23387597-INS-3664 |
| chr19-23392651-INS-66 | chr19-23392651-INS-60 |
| chr19-23449217-INS-203 | chr19-23449217-INS-203 |
| chr19-23458898-INS-59 | chr19-23458898-INS-59 |
| chr19-23470813-INS-54 | chr19-23470813-INS-60 |
| chr19-23501467-INS-456 | chr19-23501467-INS-456 |
| chr19-23536093-INS-401 | chr19-23536093-INS-401 |
| chr19-23546511-INS-64 | chr19-23546511-INS-52 |
| chr19-23549119-INS-123 | chr19-23549125-INS-128 |
| chr19-23556300-INS-52 | chr19-23556301-INS-60 |
| chr19-23567767-INS-66 | chr19-23567764-INS-63 |
| chr19-23583470-INS-89 | chr19-23583471-INS-62 |
| chr19-23592502-INS-175 | chr19-23592502-INS-175 |
| chr19-23604681-INS-67 | chr19-23604681-INS-67 |
| chr19-23606781-INS-7104 | chr19-23606782-INS-7104 |
| chr19-23626867-INS-106 | chr19-23626867-INS-106 |
| chr19-23629110-INS-92 | chr19-23629122-INS-52 |
| chr19-23654905-INS-55 | chr19-23654913-INS-71 |
| chr19-23656618-INS-555 | chr19-23656618-INS-555 |
| chr19-23691748-INS-99 | chr19-23691748-INS-99 |
| chr19-23710671-INS-253 | chr19-23710671-INS-253 |
| chr19-23736702-INS-54 | chr19-23736702-INS-52 |
| chr19-23749284-INS-58 | chr19-23749260-INS-66 |
| chr19-23753052-INS-110 | chr19-23753055-INS-123 |
| chr19-23760018-INS-212 | chr19-23760018-INS-216 |
| chr19-23775979-INS-399 | chr19-23775979-INS-399 |
| chr19-23790734-INS-95 | chr19-23790734-INS-95 |
| chr19-23823173-INS-61 | chr19-23823174-INS-61 |
| chr19-23847055-INS-52 | chr19-23847055-INS-79 |
| chr19-23852784-INS-67 | chr19-23852784-INS-77 |
| chr19-23859586-INS-114 | chr19-23859586-INS-102 |
| chr19-23906342-INS-1888 | chr19-23906343-INS-1888 |
| chr19-23937405-INS-137 | chr19-23937405-INS-137 |
| chr19-23941274-INS-390 | chr19-23941274-INS-390 |
| chr19-23965408-INS-400 | chr19-23965408-INS-400 |
| chr19-23971129-INS-94 | chr19-23971130-INS-74 |
| chr19-23973388-INS-58 | chr19-23973388-INS-60 |
| chr19-23974051-INS-202 | chr19-23974051-INS-202 |
| chr19-23990061-INS-56 | chr19-23990061-INS-56 |
| chr19-23991992-INS-136 | chr19-23991992-INS-136 |
| chr19-24012541-INS-52 | chr19-24012541-INS-52 |
| chr19-24016463-INS-62 | chr19-24016463-INS-62 |
| chr19-24030660-INS-71 | chr19-24030660-INS-63 |
| chr19-24073783-INS-177 | chr19-24073783-INS-177 |
| chr19-24125025-INS-204 | chr19-24125025-INS-198 |
| chr19-24168499-INS-86 | chr19-24168498-INS-106 |
| chr19-24169862-INS-117 | chr19-24169862-INS-140 |
| chr19-24177165-INS-847 | chr19-24177166-INS-800 |
| chr19-24177166-INS-273 | chr19-24177166-INS-303 |
| chr19-24203284-INS-196 | chr19-24203284-INS-206 |
| chr19-24223604-INS-208 | chr19-24223604-INS-208 |
| chr19-24228276-INS-203 | chr19-24228276-INS-201 |
| chr19-24229727-INS-208 | chr19-24229727-INS-208 |
| chr19-24231955-INS-65 | chr19-24231955-INS-65 |
| chr19-24251174-INS-174 | chr19-24251174-INS-174 |
| chr19-24268601-INS-103 | chr19-24268601-INS-123 |
| chr19-24269906-INS-56 | chr19-24269906-INS-56 |
| chr19-24271859-INS-214 | chr19-24271860-INS-200 |
| chr19-24277918-INS-267 | chr19-24277918-INS-265 |

| | |
|---|---|
| chr19-24292574-INS-202 | chr19-24292574-INS-202 |
| chr19-24295114-INS-206 | chr19-24295114-INS-206 |
| chr19-24314506-INS-225 | chr19-24314506-INS-224 |
| chr19-24316891-INS-226 | chr19-24316891-INS-226 |
| chr19-24324178-INS-125 | chr19-24324179-INS-125 |
| chr19-24327043-INS-60 | chr19-24327043-INS-60 |
| chr19-24357076-INS-232 | chr19-24357076-INS-222 |
| chr19-24406198-INS-62 | chr19-24406152-INS-72 |
| chr19-24424156-INS-7517 | chr19-24424157-INS-7521 |
| chr19-24438822-INS-54 | chr19-24438822-INS-54 |
| chr19-24446641-INS-54 | chr19-24446641-INS-75 |
| chr19-24473182-INS-58 | chr19-24473182-INS-58 |
| chr19-24481181-INS-193 | chr19-24481181-INS-197 |
| chr19-24495308-INS-252 | chr19-24495308-INS-252 |
| chr19-24511652-INS-100 | chr19-24511668-INS-88 |
| chr19-24516995-INS-118 | chr19-24516995-INS-118 |
| chr19-24518592-INS-210 | chr19-24518592-INS-210 |
| chr19-24542612-INS-68 | chr19-24542613-INS-66 |
| chr19-24546846-INS-58 | chr19-24546846-INS-52 |
| chr19-24567595-INS-136 | chr19-24567595-INS-136 |
| chr19-24568364-INS-685 | chr19-24568364-INS-685 |
| chr19-24600958-INS-150 | chr19-24600958-INS-150 |
| chr19-24603367-INS-352 | chr19-24603367-INS-308 |
| chr19-24624712-INS-51 | chr19-24624696-INS-64 |
| chr19-24648710-INS-60 | chr19-24648710-INS-60 |
| chr19-24713300-INS-61 | chr19-24713300-INS-61 |
| chr19-24723614-INS-86 | chr19-24723614-INS-86 |
| chr19-24733058-INS-53 | chr19-24733058-INS-53 |
| chr19-24737681-INS-193 | chr19-24737681-INS-183 |
| chr19-24744738-INS-183 | chr19-24744738-INS-173 |
| chr19-24750832-INS-64 | chr19-24750832-INS-64 |
| chr19-24756830-INS-260 | chr19-24756830-INS-260 |
| chr19-24774304-INS-96 | chr19-24774304-INS-96 |
| chr19-24776140-INS-68 | chr19-24776140-INS-68 |
| chr19-24797246-INS-141 | chr19-24797246-INS-141 |
| chr19-24818721-INS-322 | chr19-24818722-INS-322 |
| chr19-24842831-INS-199 | chr19-24842831-INS-195 |
| chr19-24844987-INS-88 | chr19-24844987-INS-88 |
| chr19-24851035-INS-60 | chr19-24851035-INS-60 |
| chr19-24858501-INS-113 | chr19-24858496-INS-121 |
| chr19-24865529-INS-197 | chr19-24865529-INS-197 |
| chr19-24898771-INS-179 | chr19-24898771-INS-181 |
| chr19-24902585-INS-64 | chr19-24902585-INS-64 |
| chr19-24913083-INS-113 | chr19-24913083-INS-79 |
| chr19-24916094-INS-124 | chr19-24916094-INS-120 |
| chr19-24949424-INS-355 | chr19-24949424-INS-355 |
| chr19-24949943-INS-83 | chr19-24949943-INS-83 |
| chr19-24973002-INS-3009 | chr19-24973017-INS-3003 |
| chr19-24983468-INS-604 | chr19-24983468-INS-610 |
| chr19-24984660-INS-227 | chr19-24984659-INS-228 |
| chr19-24992424-INS-60 | chr19-24992424-INS-66 |
| chr19-24998424-INS-90 | chr19-24998424-INS-90 |
| chr19-25011828-INS-6811 | chr19-25011829-INS-6812 |
| chr19-25013700-INS-93 | chr19-25013700-INS-89 |
| chr19-25017211-INS-71 | chr19-25017211-INS-71 |
| chr19-25023984-INS-439 | chr19-25023984-INS-439 |
| chr19-25059106-INS-128 | chr19-25059106-INS-132 |
| chr19-25066127-INS-983 | chr19-25066127-INS-983 |
| chr19-25071410-INS-54 | chr19-25071410-INS-54 |
| chr19-25074270-INS-3052 | chr19-25074271-INS-3052 |
| chr19-25087546-INS-661 | chr19-25087546-INS-661 |
| chr19-25109688-INS-206 | chr19-25109688-INS-206 |
| chr19-25119014-INS-56 | chr19-25119014-INS-56 |
| chr19-25135223-INS-58 | chr19-25135223-INS-58 |
| chr19-25157230-INS-148 | chr19-25157230-INS-148 |
| chr19-25191461-INS-170 | chr19-25191461-INS-170 |
| chr19-25196115-INS-99 | chr19-25196115-INS-99 |
| chr19-25196920-INS-6442 | chr19-25196921-INS-6442 |
| chr19-25209411-INS-85 | chr19-25209411-INS-51 |
| chr19-25213301-INS-88 | chr19-25213292-INS-90 |
| chr19-25225655-INS-152 | chr19-25225655-INS-152 |
| chr19-25232083-INS-463 | chr19-25232083-INS-463 |
| chr19-25237549-INS-390 | chr19-25237549-INS-390 |
| chr19-25238447-INS-81 | chr19-25238447-INS-81 |
| chr19-25240134-INS-62 | chr19-25240134-INS-62 |
| chr19-25263131-INS-202 | chr19-25263131-INS-204 |
| chr19-25279110-INS-77 | chr19-25279110-INS-77 |

| | |
|---|---|
| chr19-25301764-INS-169 | chr19-25301764-INS-169 |
| chr19-25324937-INS-51 | chr19-25324940-INS-50 |
| chr19-25340338-INS-80 | chr19-25340338-INS-80 |
| chr19-25350983-INS-207 | chr19-25350983-INS-207 |
| chr19-25352998-INS-838 | chr19-25352972-INS-842 |
| chr19-25358720-INS-497 | chr19-25358720-INS-497 |
| chr19-25367548-INS-90 | chr19-25367548-INS-138 |
| chr19-25371499-INS-70 | chr19-25371499-INS-72 |
| chr19-25380172-INS-245 | chr19-25380172-INS-245 |
| chr19-25389445-INS-95 | chr19-25389445-INS-80 |
| chr19-25416184-INS-2662 | chr19-25416185-INS-2662 |
| chr19-25433835-INS-298 | chr19-25433835-INS-299 |
| chr19-25437398-INS-83 | chr19-25437398-INS-83 |
| chr19-25456159-INS-382 | chr19-25456159-INS-382 |
| chr19-25482229-INS-93 | chr19-25482229-INS-89 |
| chr19-25526353-INS-51 | chr19-25526351-INS-65 |
| chr19-25530871-INS-52 | chr19-25530872-INS-75 |
| chr19-25560599-INS-315 | chr19-25560599-INS-315 |
| chr19-25570751-INS-501 | chr19-25570751-INS-496 |
| chr19-25572630-INS-94 | chr19-25572630-INS-55 |
| chr19-25575783-INS-126 | chr19-25575783-INS-126 |
| chr19-25595254-INS-63 | chr19-25595254-INS-63 |
| chr19-25609192-INS-62 | chr19-25609192-INS-109 |
| chr19-25613532-INS-73 | chr19-25613532-INS-73 |
| chr19-25614623-INS-530 | chr19-25614623-INS-530 |
| chr19-25615403-INS-157 | chr19-25615403-INS-157 |
| chr19-25618038-INS-82 | chr19-25618038-INS-82 |
| chr19-25619875-INS-73 | chr19-25619875-INS-90 |
| chr19-25622246-INS-432 | chr19-25622246-INS-432 |
| chr19-25636174-INS-352 | chr19-25636174-INS-351 |
| chr19-25680761-INS-79 | chr19-25680809-INS-56 |
| chr19-25684538-INS-168 | chr19-25684537-INS-198 |
| chr19-25695266-INS-110 | chr19-25695266-INS-109 |
| chr19-25706203-INS-134 | chr19-25706203-INS-138 |
| chr19-25728045-INS-65 | chr19-25728045-INS-65 |
| chr19-25731063-INS-199 | chr19-25731063-INS-199 |
| chr19-25737400-INS-2727 | chr19-25737401-INS-2727 |
| chr19-25766059-INS-6367 | chr19-25766060-INS-6367 |
| chr19-25768019-INS-81 | chr19-25768049-INS-56 |
| chr19-25785134-INS-315 | chr19-25785134-INS-317 |
| chr19-25789143-INS-58 | chr19-25789137-INS-64 |
| chr19-25790135-INS-123 | chr19-25790135-INS-123 |
| chr19-25802065-INS-728 | chr19-25802065-INS-729 |
| chr19-25831380-INS-102 | chr19-25831370-INS-122 |
| chr19-25849895-INS-87 | chr19-25849895-INS-87 |
| chr19-25901722-INS-68 | chr19-25901719-INS-73 |
| chr19-25946347-INS-714 | chr19-25946347-INS-715 |
| chr19-25954740-INS-75 | chr19-25954747-INS-78 |
| chr19-25963068-INS-82 | chr19-25963068-INS-92 |
| chr19-25966046-INS-52 | chr19-25966046-INS-52 |
| chr19-25967752-INS-303 | chr19-25967746-INS-311 |
| chr19-25971610-INS-422 | chr19-25971610-INS-422 |
| chr19-25985628-INS-51 | chr19-25985601-INS-84 |
| chr19-25987101-INS-540 | chr19-25987101-INS-540 |
| chr19-25996474-INS-138 | chr19-25996474-INS-138 |
| chr19-26016685-INS-7212 | chr19-26016686-INS-7212 |
| chr19-26052009-INS-1367 | chr19-26052010-INS-1367 |
| chr19-26052427-INS-60 | chr19-26052427-INS-72 |
| chr19-26076864-INS-63 | chr19-26076864-INS-67 |
| chr19-26088987-INS-52 | chr19-26089003-INS-64 |
| chr19-26114028-INS-95 | chr19-26114028-INS-95 |
| chr19-26121998-INS-60 | chr19-26121998-INS-60 |
| chr19-26170753-INS-167 | chr19-26170753-INS-166 |
| chr19-26182694-INS-400 | chr19-26182694-INS-400 |
| chr19-26238855-INS-58 | chr19-26238855-INS-58 |
| chr19-26240436-INS-64 | chr19-26240436-INS-64 |
| chr19-26244715-INS-65 | chr19-26244715-INS-69 |
| chr19-26300642-INS-64 | chr19-26300642-INS-66 |
| chr19-26336904-INS-4667 | chr19-26336905-INS-4667 |
| chr19-26379226-INS-76 | chr19-26379214-INS-87 |
| chr19-26405808-INS-450 | chr19-26405808-INS-449 |
| chr19-26451098-INS-54 | chr19-26451098-INS-54 |
| chr19-26533460-INS-54 | chr19-26533460-INS-54 |
| chr19-26572718-INS-59 | chr19-26572715-INS-87 |
| chr19-26597961-INS-173 | chr19-26597961-INS-155 |
| chr19-26617434-INS-300 | chr19-26617434-INS-300 |
| chr19-26622451-INS-78 | chr19-26622443-INS-82 |

| | |
|---|---|
| chr19-26631473-INS-54 | chr19-26631473-INS-54 |
| chr19-26661582-INS-62 | chr19-26661582-INS-58 |
| chr19-26681542-INS-232 | chr19-26681542-INS-232 |
| chr19-26685359-INS-60 | chr19-26685359-INS-60 |
| chr19-26697171-INS-75 | chr19-26697171-INS-75 |
| chr19-26784187-INS-103 | chr19-26784188-INS-100 |
| chr19-26786100-INS-54 | chr19-26786100-INS-54 |
| chr19-26799072-INS-235 | chr19-26799051-INS-273 |
| chr19-26815641-INS-536 | chr19-26815641-INS-537 |
| chr19-26823672-INS-3268 | chr19-26823673-INS-3268 |
| chr19-26829288-INS-65 | chr19-26829288-INS-65 |
| chr19-26832502-INS-111 | chr19-26832502-INS-114 |
| chr19-26835346-INS-401 | chr19-26835347-INS-402 |
| chr19-26901198-INS-81 | chr19-26901198-INS-80 |
| chr19-26903887-INS-60 | chr19-26903887-INS-60 |
| chr19-26952695-INS-6748 | chr19-26952696-INS-6748 |
| chr19-26960003-INS-128 | chr19-26960003-INS-128 |
| chr19-26962736-INS-72 | chr19-26962736-INS-72 |
| chr19-26967047-INS-90 | chr19-26967047-INS-90 |
| chr19-26971267-INS-68 | chr19-26971267-INS-68 |
| chr19-26978678-INS-103 | chr19-26978698-INS-80 |
| chr19-27023541-INS-3460 | chr19-27023542-INS-3461 |
| chr19-27026408-INS-2921 | chr19-27026409-INS-2922 |
| chr19-27036145-INS-100 | chr19-27036145-INS-106 |
| chr19-27067892-INS-1113 | chr19-27067893-INS-1113 |
| chr19-27067893-INS-582 | chr19-27067893-INS-534 |
| chr19-27084099-INS-556 | chr19-27084099-INS-562 |
| chr19-27119449-INS-69 | chr19-27119449-INS-56 |
| chr19-27172516-INS-84 | chr19-27172516-INS-78 |
| chr19-27210620-INS-194 | chr19-27210620-INS-195 |
| chr19-27243728-INS-56 | chr19-27243728-INS-74 |
| chr19-27255484-INS-203 | chr19-27255484-INS-203 |
| chr19-27263778-INS-512 | chr19-27263778-INS-505 |
| chr19-27279577-INS-4339 | chr19-27279578-INS-4339 |
| chr19-27294713-INS-72 | chr19-27294713-INS-72 |
| chr19-27343380-INS-209 | chr19-27343380-INS-209 |
| chr19-27369500-INS-219 | chr19-27369500-INS-219 |
| chr19-27429584-INS-117 | chr19-27429585-INS-69 |
| chr19-27460005-INS-1434 | chr19-27460006-INS-1434 |
| chr19-27465482-INS-100 | chr19-27465482-INS-72 |
| chr19-27482140-INS-207 | chr19-27482140-INS-207 |
| chr19-27484158-INS-233 | chr19-27484158-INS-229 |
| chr19-27494148-INS-802 | chr19-27494148-INS-802 |
| chr19-27522212-INS-54 | chr19-27522211-INS-54 |
| chr19-27531386-INS-133 | chr19-27531386-INS-99 |
| chr19-27575641-INS-231 | chr19-27575641-INS-235 |
| chr19-27613298-INS-535 | chr19-27613298-INS-535 |
| chr19-27614640-INS-208 | chr19-27614642-INS-209 |
| chr19-27631023-INS-52 | chr19-27631018-INS-51 |
| chr19-27644970-INS-54 | chr19-27644970-INS-54 |
| chr19-27652752-INS-6635 | chr19-27652753-INS-6637 |
| chr19-27663570-INS-106 | chr19-27663570-INS-106 |
| chr19-27701606-INS-190 | chr19-27701606-INS-182 |
| chr19-27705529-INS-57 | chr19-27705529-INS-57 |
| chr19-27717067-INS-68 | chr19-27717069-INS-72 |
| chr19-27719839-INS-93 | chr19-27719839-INS-93 |
| chr19-27824103-INS-80 | chr19-27824103-INS-78 |
| chr19-27863931-INS-460 | chr19-27863931-INS-460 |
| chr19-27951649-INS-246 | chr19-27951669-INS-222 |
| chr19-27956194-INS-143 | chr19-27956231-INS-151 |
| chr19-28003483-INS-66 | chr19-28003483-INS-66 |
| chr19-28016190-INS-72 | chr19-28016190-INS-72 |
| chr19-28091278-INS-70 | chr19-28091278-INS-70 |
| chr19-28100564-INS-76 | chr19-28100564-INS-76 |
| chr19-28140126-INS-210 | chr19-28140120-INS-210 |
| chr19-28150347-INS-594 | chr19-28150347-INS-594 |
| chr19-28151581-INS-62 | chr19-28151535-INS-80 |
| chr19-28167030-INS-70 | chr19-28167031-INS-65 |
| chr19-28202202-INS-196 | chr19-28202202-INS-196 |
| chr19-28219599-INS-255 | chr19-28219599-INS-249 |
| chr19-28232448-INS-99 | chr19-28232448-INS-94 |
| chr19-28244909-INS-265 | chr19-28244909-INS-262 |
| chr19-28246316-INS-87 | chr19-28246316-INS-97 |
| chr19-28263330-INS-2131 | chr19-28263331-INS-2131 |
| chr19-28282013-INS-86 | chr19-28282013-INS-86 |
| chr19-28284873-INS-188 | chr19-28284873-INS-193 |
| chr19-28333777-INS-542 | chr19-28333777-INS-542 |

| | |
|---|---|
| chr19-28416303-INS-68 | chr19-28416303-INS-70 |
| chr19-28422221-INS-224 | chr19-28422221-INS-224 |
| chr19-28447202-INS-99 | chr19-28447202-INS-103 |
| chr19-28448728-INS-64 | chr19-28448761-INS-50 |
| chr19-28471266-INS-330 | chr19-28471266-INS-322 |
| chr19-28484629-INS-412 | chr19-28484630-INS-420 |
| chr19-28489012-INS-230 | chr19-28489012-INS-229 |
| chr19-28526959-INS-63 | chr19-28526959-INS-63 |
| chr19-28535084-INS-120 | chr19-28535084-INS-76 |
| chr19-28542468-INS-378 | chr19-28542468-INS-378 |
| chr19-28546257-INS-128 | chr19-28546270-INS-130 |
| chr19-28607276-INS-92 | chr19-28607316-INS-68 |
| chr19-28609552-INS-77 | chr19-28609534-INS-52 |
| chr19-28692625-INS-54 | chr19-28692625-INS-58 |
| chr19-28695231-INS-62 | chr19-28695231-INS-64 |
| chr19-28701520-INS-57 | chr19-28701520-INS-61 |
| chr19-28706559-INS-78 | chr19-28706559-INS-62 |
| chr19-28714395-INS-262 | chr19-28714395-INS-263 |
| chr19-28719325-INS-499 | chr19-28719325-INS-499 |
| chr19-28726322-INS-213 | chr19-28726322-INS-213 |
| chr19-28766856-INS-197 | chr19-28766856-INS-197 |
| chr19-28768543-INS-61 | chr19-28768543-INS-81 |
| chr19-28775956-INS-82 | chr19-28775956-INS-84 |
| chr19-28778532-INS-52 | chr19-28778532-INS-52 |
| chr19-28781963-INS-301 | chr19-28781963-INS-316 |
| chr19-28817353-INS-54 | chr19-28817353-INS-56 |
| chr19-28819492-INS-59 | chr19-28819492-INS-59 |
| chr19-28825913-INS-106 | chr19-28825913-INS-102 |
| chr19-28848274-INS-381 | chr19-28848274-INS-385 |
| chr19-28858144-INS-56 | chr19-28858144-INS-58 |
| chr19-28875572-INS-197 | chr19-28875572-INS-205 |
| chr19-28923503-INS-55 | chr19-28923503-INS-60 |
| chr19-28932617-INS-51 | chr19-28932617-INS-51 |
| chr19-28933850-INS-106 | chr19-28933850-INS-106 |
| chr19-28948336-INS-290 | chr19-28948336-INS-290 |
| chr19-28949851-INS-221 | chr19-28949851-INS-221 |
| chr19-28977272-INS-351 | chr19-28977272-INS-351 |
| chr19-28986722-INS-304 | chr19-28986722-INS-292 |
| chr19-29015895-INS-116 | chr19-29015895-INS-119 |
| chr19-29036608-INS-76 | chr19-29036608-INS-76 |
| chr19-29051171-INS-104 | chr19-29051171-INS-100 |
| chr19-29064347-INS-135 | chr19-29064347-INS-143 |
| chr19-29068919-INS-73 | chr19-29068919-INS-75 |
| chr19-29073097-INS-182 | chr19-29073098-INS-181 |
| chr19-29108261-INS-205 | chr19-29108261-INS-205 |
| chr19-29113980-INS-60 | chr19-29113980-INS-60 |
| chr19-29116058-INS-210 | chr19-29116058-INS-211 |
| chr19-29136753-INS-54 | chr19-29136756-INS-75 |
| chr19-29147526-INS-56 | chr19-29147526-INS-58 |
| chr19-29150071-INS-8511 | chr19-29150071-INS-8554 |
| chr19-29156442-INS-62 | chr19-29156442-INS-62 |
| chr19-29170073-INS-210 | chr19-29170073-INS-210 |
| chr19-29178342-INS-1166 | chr19-29178343-INS-1167 |
| chr19-29194166-INS-80 | chr19-29194166-INS-80 |
| chr19-29202365-INS-339 | chr19-29202365-INS-339 |
| chr19-29214909-INS-80 | chr19-29214909-INS-80 |
| chr19-29216530-INS-96 | chr19-29216530-INS-96 |
| chr19-29223301-INS-213 | chr19-29223301-INS-218 |
| chr19-29288945-INS-193 | chr19-29288945-INS-193 |
| chr19-29298023-INS-186 | chr19-29297992-INS-186 |
| chr19-29299246-INS-195 | chr19-29299246-INS-195 |
| chr19-29322394-INS-81 | chr19-29322394-INS-81 |
| chr19-29362755-INS-86 | chr19-29362755-INS-86 |
| chr19-29364518-INS-78 | chr19-29364518-INS-112 |
| chr19-29376796-INS-57 | chr19-29376796-INS-81 |
| chr19-29392324-INS-108 | chr19-29392315-INS-90 |
| chr19-29446764-INS-199 | chr19-29446764-INS-199 |
| chr19-29453747-INS-223 | chr19-29453747-INS-223 |
| chr19-29457071-INS-208 | chr19-29457071-INS-209 |
| chr19-29506200-INS-135 | chr19-29506200-INS-134 |
| chr19-29558993-INS-119 | chr19-29558993-INS-119 |
| chr19-29602072-INS-975 | chr19-29602072-INS-976 |
| chr19-29658298-INS-116 | chr19-29658298-INS-116 |
| chr19-29676508-INS-68 | chr19-29676514-INS-61 |
| chr19-29687361-INS-58 | chr19-29687361-INS-70 |
| chr19-29722352-INS-639 | chr19-29722352-INS-638 |
| chr19-29804644-INS-167 | chr19-29804644-INS-167 |

| | |
|---|---|
| chr19-29822609-INS-566 | chr19-29822609-INS-566 |
| chr19-29828970-INS-7167 | chr19-29828971-INS-7167 |
| chr19-29831001-INS-126 | chr19-29831005-INS-127 |
| chr19-29846505-INS-3388 | chr19-29846506-INS-3388 |
| chr19-29860481-INS-80 | chr19-29860485-INS-52 |
| chr19-29864790-INS-76 | chr19-29864790-INS-92 |
| chr19-29879465-INS-479 | chr19-29879465-INS-479 |
| chr19-29921224-INS-67 | chr19-29921224-INS-65 |
| chr19-29927364-INS-471 | chr19-29927364-INS-471 |
| chr19-29928466-INS-6597 | chr19-29928467-INS-6597 |
| chr19-29933420-INS-53 | chr19-29933420-INS-97 |
| chr19-29963454-INS-347 | chr19-29963454-INS-347 |
| chr19-29964688-INS-2781 | chr19-29964689-INS-2781 |
| chr19-29965998-INS-1221 | chr19-29965999-INS-1221 |
| chr19-29975404-INS-60 | chr19-29975411-INS-86 |
| chr19-29988651-INS-297 | chr19-29988653-INS-306 |
| chr19-30001935-INS-199 | chr19-30001951-INS-199 |
| chr19-30053408-INS-81 | chr19-30053408-INS-81 |
| chr19-30058969-INS-373 | chr19-30058969-INS-373 |
| chr19-30086208-INS-224 | chr19-30086208-INS-225 |
| chr19-30089995-INS-218 | chr19-30089995-INS-218 |
| chr19-30113757-INS-146 | chr19-30113757-INS-146 |
| chr19-30114984-INS-140 | chr19-30114984-INS-140 |
| chr19-30116445-INS-51 | chr19-30116445-INS-51 |
| chr19-30131014-INS-56 | chr19-30131014-INS-56 |
| chr19-30133343-INS-545 | chr19-30133343-INS-545 |
| chr19-30139319-INS-74 | chr19-30139338-INS-80 |
| chr19-30159126-INS-219 | chr19-30159127-INS-219 |
| chr19-30170412-INS-197 | chr19-30170412-INS-197 |
| chr19-30174649-INS-60 | chr19-30174649-INS-60 |
| chr19-30180533-INS-270 | chr19-30180580-INS-264 |
| chr19-30188856-INS-53 | chr19-30188856-INS-53 |
| chr19-30191700-INS-73 | chr19-30191700-INS-73 |
| chr19-30209773-INS-242 | chr19-30209773-INS-242 |
| chr19-30224662-INS-180 | chr19-30224662-INS-180 |
| chr19-30226980-INS-172 | chr19-30226979-INS-159 |
| chr19-30234858-INS-51 | chr19-30234858-INS-56 |
| chr19-30252012-INS-6566 | chr19-30252013-INS-6565 |
| chr19-30262971-INS-78 | chr19-30262957-INS-78 |
| chr19-30267383-INS-175 | chr19-30267383-INS-145 |
| chr19-30311577-INS-99 | chr19-30311577-INS-99 |
| chr19-30318375-INS-51 | chr19-30318375-INS-55 |
| chr19-30334395-INS-96 | chr19-30334395-INS-94 |
| chr19-30340411-INS-450 | chr19-30340411-INS-450 |
| chr19-30387780-INS-78 | chr19-30387780-INS-78 |
| chr19-30394222-INS-68 | chr19-30394222-INS-68 |
| chr19-30397012-INS-214 | chr19-30397012-INS-214 |
| chr19-30414132-INS-95 | chr19-30414132-INS-89 |
| chr19-30441897-INS-94 | chr19-30441897-INS-94 |
| chr19-30453708-INS-111 | chr19-30453708-INS-111 |
| chr19-30460411-INS-6866 | chr19-30460412-INS-6866 |
| chr19-30460532-INS-150 | chr19-30460532-INS-132 |
| chr19-30469314-INS-162 | chr19-30469314-INS-162 |
| chr19-30490917-INS-419 | chr19-30490917-INS-420 |
| chr19-30491574-INS-76 | chr19-30491574-INS-76 |
| chr19-30510754-INS-180 | chr19-30510754-INS-173 |
| chr19-30555825-INS-102 | chr19-30555870-INS-106 |
| chr19-30557676-INS-216 | chr19-30557676-INS-216 |
| chr19-30560616-INS-1982 | chr19-30560617-INS-1982 |
| chr19-30571626-INS-88 | chr19-30571578-INS-123 |
| chr19-30595199-INS-261 | chr19-30595199-INS-261 |
| chr19-30630493-INS-63 | chr19-30630493-INS-63 |
| chr19-30675551-INS-82 | chr19-30675551-INS-82 |
| chr19-30690297-INS-307 | chr19-30690297-INS-317 |
| chr19-30696206-INS-391 | chr19-30696206-INS-391 |
| chr19-30698176-INS-54 | chr19-30698176-INS-58 |
| chr19-30715426-INS-430 | chr19-30715426-INS-424 |
| chr19-30767800-INS-6187 | chr19-30767801-INS-6188 |
| chr19-30772417-INS-761 | chr19-30772417-INS-763 |
| chr19-30778237-INS-144 | chr19-30778237-INS-124 |
| chr19-30800720-INS-1087 | chr19-30800723-INS-1088 |
| chr19-30804130-INS-152 | chr19-30804130-INS-152 |
| chr19-30812084-INS-70 | chr19-30812084-INS-76 |
| chr19-30817732-INS-54 | chr19-30817732-INS-54 |
| chr19-30821024-INS-101 | chr19-30820998-INS-80 |
| chr19-30831978-INS-57 | chr19-30831978-INS-57 |
| chr19-30835017-INS-6706 | chr19-30835018-INS-6706 |

| | |
|---|---|
| chr19-30835698-INS-241 | chr19-30835698-INS-241 |
| chr19-30850552-INS-79 | chr19-30850548-INS-83 |
| chr19-30890776-INS-129 | chr19-30890776-INS-129 |
| chr19-30896248-INS-80 | chr19-30896248-INS-80 |
| chr19-30903887-INS-325 | chr19-30903887-INS-325 |
| chr19-30923160-INS-58 | chr19-30923160-INS-58 |
| chr19-30957281-INS-66 | chr19-30957281-INS-68 |
| chr19-30979268-INS-77 | chr19-30979268-INS-77 |
| chr19-30980249-INS-506 | chr19-30980249-INS-506 |
| chr19-31006858-INS-148 | chr19-31006902-INS-143 |
| chr19-31023714-INS-60 | chr19-31023714-INS-60 |
| chr19-31041259-INS-520 | chr19-31041259-INS-520 |
| chr19-31079697-INS-163 | chr19-31079697-INS-163 |
| chr19-31089248-INS-205 | chr19-31089248-INS-208 |
| chr19-31097747-INS-69 | chr19-31097751-INS-66 |
| chr19-31100849-INS-153 | chr19-31100849-INS-153 |
| chr19-31133621-INS-882 | chr19-31133621-INS-882 |
| chr19-31142389-INS-385 | chr19-31142389-INS-389 |
| chr19-31192826-INS-2005 | chr19-31192827-INS-2002 |
| chr19-31209415-INS-279 | chr19-31209415-INS-239 |
| chr19-31225966-INS-919 | chr19-31225966-INS-918 |
| chr19-31273649-INS-133 | chr19-31273649-INS-127 |
| chr19-31286996-INS-921 | chr19-31286996-INS-923 |
| chr19-31288908-INS-84 | chr19-31288908-INS-84 |
| chr19-31386906-INS-70 | chr19-31386891-INS-82 |
| chr19-31388727-INS-1202 | chr19-31388712-INS-1201 |
| chr19-31408070-INS-100 | chr19-31408070-INS-101 |
| chr19-31433871-INS-432 | chr19-31433871-INS-432 |
| chr19-31434964-INS-1428 | chr19-31434965-INS-1428 |
| chr19-31441699-INS-135 | chr19-31441679-INS-119 |
| chr19-31482429-INS-6748 | chr19-31482430-INS-6748 |
| chr19-31488692-INS-114 | chr19-31488692-INS-113 |
| chr19-31517220-INS-863 | chr19-31517220-INS-863 |
| chr19-31529575-INS-72 | chr19-31529573-INS-80 |
| chr19-31530057-INS-1267 | chr19-31530058-INS-1267 |
| chr19-31581868-INS-276 | chr19-31581868-INS-276 |
| chr19-31609599-INS-68 | chr19-31609599-INS-102 |
| chr19-31635472-INS-83 | chr19-31635493-INS-105 |
| chr19-31732926-INS-53 | chr19-31732927-INS-53 |
| chr19-31765247-INS-122 | chr19-31765247-INS-120 |
| chr19-31794320-INS-6372 | chr19-31794321-INS-6373 |
| chr19-31797950-INS-52 | chr19-31797950-INS-60 |
| chr19-31864576-INS-197 | chr19-31864575-INS-197 |
| chr19-31939233-INS-392 | chr19-31939233-INS-392 |
| chr19-31957240-INS-913 | chr19-31957240-INS-913 |
| chr19-31959765-INS-639 | chr19-31959766-INS-686 |
| chr19-31974290-INS-64 | chr19-31974290-INS-64 |
| chr19-31998851-INS-285 | chr19-31998851-INS-285 |
| chr19-32022964-INS-355 | chr19-32022964-INS-355 |
| chr19-32038177-INS-82 | chr19-32038176-INS-68 |
| chr19-32057658-INS-360 | chr19-32057658-INS-360 |
| chr19-32061105-INS-519 | chr19-32061105-INS-518 |
| chr19-32070698-INS-191 | chr19-32070698-INS-191 |
| chr19-32073941-INS-84 | chr19-32073941-INS-76 |
| chr19-32189160-INS-178 | chr19-32189160-INS-177 |
| chr19-32234326-INS-401 | chr19-32234326-INS-417 |
| chr19-32255181-INS-52 | chr19-32255181-INS-60 |
| chr19-32354918-INS-66 | chr19-32354918-INS-72 |
| chr19-32407877-INS-106 | chr19-32407877-INS-106 |
| chr19-32435168-INS-58 | chr19-32435200-INS-59 |
| chr19-32486878-INS-62 | chr19-32486878-INS-72 |
| chr19-32489961-INS-57 | chr19-32489961-INS-57 |
| chr19-32523944-INS-7252 | chr19-32523947-INS-7255 |
| chr19-32536901-INS-110 | chr19-32536871-INS-138 |
| chr19-32559180-INS-631 | chr19-32559180-INS-633 |
| chr19-32583205-INS-75 | chr19-32583216-INS-111 |
| chr19-32592163-INS-203 | chr19-32592163-INS-207 |
| chr19-32609179-INS-116 | chr19-32609213-INS-104 |
| chr19-32611459-INS-116 | chr19-32611459-INS-131 |
| chr19-32613634-INS-52 | chr19-32613634-INS-54 |
| chr19-32633736-INS-435 | chr19-32633736-INS-435 |
| chr19-32663404-INS-117 | chr19-32663405-INS-102 |
| chr19-32679381-INS-228 | chr19-32679381-INS-229 |
| chr19-32683488-INS-70 | chr19-32683488-INS-60 |
| chr19-32699367-INS-59 | chr19-32699320-INS-74 |
| chr19-32701463-INS-213 | chr19-32701463-INS-213 |
| chr19-32707112-INS-191 | chr19-32707112-INS-192 |

| | |
|---|---|
| chr19-32709761-INS-71 | chr19-32709761-INS-84 |
| chr19-32723959-INS-134 | chr19-32723959-INS-137 |
| chr19-32724283-INS-752 | chr19-32724283-INS-754 |
| chr19-32726257-INS-208 | chr19-32726248-INS-202 |
| chr19-32739994-INS-207 | chr19-32739994-INS-204 |
| chr19-32763490-INS-116 | chr19-32763504-INS-144 |
| chr19-32767667-INS-244 | chr19-32767667-INS-244 |
| chr19-32779060-INS-204 | chr19-32779060-INS-203 |
| chr19-32803692-INS-616 | chr19-32803692-INS-616 |
| chr19-32805355-INS-358 | chr19-32805355-INS-358 |
| chr19-32816854-INS-131 | chr19-32816852-INS-161 |
| chr19-32822640-INS-70 | chr19-32822640-INS-73 |
| chr19-32827510-INS-62 | chr19-32827510-INS-62 |
| chr19-32829089-INS-86 | chr19-32829101-INS-94 |
| chr19-32830759-INS-171 | chr19-32830759-INS-189 |
| chr19-32840965-INS-147 | chr19-32840965-INS-149 |
| chr19-32857245-INS-198 | chr19-32857246-INS-198 |
| chr19-32871140-INS-98 | chr19-32871141-INS-99 |
| chr19-32909515-INS-71 | chr19-32909513-INS-80 |
| chr19-32942965-INS-186 | chr19-32942965-INS-182 |
| chr19-32953949-INS-260 | chr19-32953949-INS-260 |
| chr19-32959121-INS-60 | chr19-32959147-INS-50 |
| chr19-32976657-INS-1892 | chr19-32976658-INS-1892 |
| chr19-32982461-INS-213 | chr19-32982461-INS-214 |
| chr19-33003417-INS-67 | chr19-33003417-INS-73 |
| chr19-33017775-INS-59 | chr19-33017764-INS-72 |
| chr19-33030560-INS-105 | chr19-33030560-INS-109 |
| chr19-33034843-INS-126 | chr19-33034847-INS-156 |
| chr19-33051742-INS-90 | chr19-33051759-INS-68 |
| chr19-33084867-INS-55 | chr19-33084867-INS-55 |
| chr19-33104222-INS-1709 | chr19-33104223-INS-1709 |
| chr19-33140732-INS-357 | chr19-33140732-INS-357 |
| chr19-33177176-INS-52 | chr19-33177176-INS-52 |
| chr19-33192780-INS-87 | chr19-33192780-INS-63 |
| chr19-33225158-INS-7024 | chr19-33225159-INS-7024 |
| chr19-33255174-INS-198 | chr19-33255174-INS-198 |
| chr19-33287388-INS-129 | chr19-33287388-INS-129 |
| chr19-33328923-INS-203 | chr19-33328923-INS-203 |
| chr19-33354133-INS-654 | chr19-33354133-INS-654 |
| chr19-33375331-INS-541 | chr19-33375331-INS-541 |
| chr19-33407570-INS-548 | chr19-33407570-INS-548 |
| chr19-33429337-INS-2159 | chr19-33429338-INS-2171 |
| chr19-33489910-INS-56 | chr19-33489910-INS-56 |
| chr19-33508295-INS-53 | chr19-33508295-INS-55 |
| chr19-33509492-INS-4895 | chr19-33509494-INS-4890 |
| chr19-33521080-INS-710 | chr19-33521080-INS-710 |
| chr19-33527886-INS-7084 | chr19-33527887-INS-7084 |
| chr19-33620608-INS-214 | chr19-33620608-INS-214 |
| chr19-33646127-INS-110 | chr19-33646127-INS-110 |
| chr19-33765330-INS-234 | chr19-33765330-INS-234 |
| chr19-33799569-INS-489 | chr19-33799569-INS-489 |
| chr19-33839725-INS-68 | chr19-33839725-INS-70 |
| chr19-33846008-INS-380 | chr19-33846008-INS-380 |
| chr19-33864283-INS-62 | chr19-33864283-INS-62 |
| chr19-33866252-INS-59 | chr19-33866252-INS-59 |
| chr19-33893022-INS-225 | chr19-33893022-INS-225 |
| chr19-33926148-INS-971 | chr19-33926148-INS-971 |
| chr19-33984538-INS-107 | chr19-33984538-INS-148 |
| chr19-33997165-INS-58 | chr19-33997165-INS-58 |
| chr19-34033874-INS-9676 | chr19-34033875-INS-9676 |
| chr19-34057134-INS-72 | chr19-34057134-INS-72 |
| chr19-34063116-INS-76 | chr19-34063120-INS-56 |
| chr19-34105395-INS-52 | chr19-34105395-INS-56 |
| chr19-34123832-INS-332 | chr19-34123832-INS-332 |
| chr19-34164963-INS-225 | chr19-34164963-INS-225 |
| chr19-34179130-INS-52 | chr19-34179130-INS-52 |
| chr19-34179767-INS-69 | chr19-34179767-INS-69 |
| chr19-34230322-INS-117 | chr19-34230322-INS-119 |
| chr19-34234376-INS-208 | chr19-34234368-INS-211 |
| chr19-34247232-INS-400 | chr19-34247232-INS-395 |
| chr19-34252584-INS-72 | chr19-34252584-INS-72 |
| chr19-34261505-INS-88 | chr19-34261505-INS-89 |
| chr19-34288881-INS-56 | chr19-34288881-INS-64 |
| chr19-34292518-INS-62 | chr19-34292515-INS-70 |
| chr19-34306172-INS-51 | chr19-34306172-INS-55 |
| chr19-34318463-INS-6281 | chr19-34318464-INS-6281 |
| chr19-34329486-INS-255 | chr19-34329487-INS-293 |

| | |
|---|---|
| chr19-34394652-INS-128 | chr19-34394652-INS-130 |
| chr19-34408261-INS-73 | chr19-34408269-INS-51 |
| chr19-34411249-INS-359 | chr19-34411249-INS-355 |
| chr19-34428978-INS-393 | chr19-34428978-INS-393 |
| chr19-34429841-INS-626 | chr19-34429843-INS-641 |
| chr19-34457988-INS-167 | chr19-34457988-INS-170 |
| chr19-34481586-INS-97 | chr19-34481586-INS-97 |
| chr19-34484961-INS-215 | chr19-34484942-INS-206 |
| chr19-34520774-INS-556 | chr19-34520774-INS-556 |
| chr19-34557251-INS-52 | chr19-34557251-INS-52 |
| chr19-34592972-INS-54 | chr19-34592972-INS-54 |
| chr19-34618944-INS-115 | chr19-34618944-INS-106 |
| chr19-34645731-INS-3015 | chr19-34645732-INS-3015 |
| chr19-34680052-INS-127 | chr19-34680052-INS-127 |
| chr19-34733010-INS-1912 | chr19-34733011-INS-1912 |
| chr19-34738937-INS-216 | chr19-34738937-INS-216 |
| chr19-34765219-INS-434 | chr19-34765219-INS-434 |
| chr19-34767038-INS-506 | chr19-34767038-INS-506 |
| chr19-34772742-INS-92 | chr19-34772720-INS-121 |
| chr19-34776643-INS-62 | chr19-34776644-INS-58 |
| chr19-34783631-INS-93 | chr19-34783631-INS-78 |
| chr19-34808797-INS-54 | chr19-34808785-INS-58 |
| chr19-34822919-INS-251 | chr19-34822919-INS-254 |
| chr19-34840970-INS-115 | chr19-34840970-INS-107 |
| chr19-34880681-INS-207 | chr19-34880681-INS-207 |
| chr19-34888592-INS-122 | chr19-34888592-INS-122 |
| chr19-34892643-INS-284 | chr19-34892650-INS-299 |
| chr19-34916578-INS-66 | chr19-34916578-INS-66 |
| chr19-34940439-INS-568 | chr19-34940439-INS-568 |
| chr19-34960914-INS-601 | chr19-34960914-INS-601 |
| chr19-35017786-INS-5087 | chr19-35017787-INS-5087 |
| chr19-35022204-INS-277 | chr19-35022204-INS-281 |
| chr19-35068949-INS-280 | chr19-35068949-INS-281 |
| chr19-35073123-INS-62 | chr19-35073123-INS-62 |
| chr19-35087870-INS-96 | chr19-35087870-INS-96 |
| chr19-35099168-INS-379 | chr19-35099168-INS-379 |
| chr19-35132919-INS-72 | chr19-35132952-INS-83 |
| chr19-35175188-INS-1116 | chr19-35175189-INS-1116 |
| chr19-35183288-INS-81 | chr19-35183288-INS-81 |
| chr19-35190436-INS-64 | chr19-35190436-INS-64 |
| chr19-35199801-INS-62 | chr19-35199801-INS-66 |
| chr19-35247903-INS-213 | chr19-35247898-INS-219 |
| chr19-35274088-INS-80 | chr19-35274088-INS-56 |
| chr19-35294011-INS-585 | chr19-35294011-INS-586 |
| chr19-35324489-INS-6346 | chr19-35324490-INS-6347 |
| chr19-35339939-INS-7283 | chr19-35339940-INS-7285 |
| chr19-35368870-INS-1051 | chr19-35368871-INS-1051 |
| chr19-35377212-INS-52 | chr19-35377212-INS-54 |
| chr19-35404086-INS-2146 | chr19-35404087-INS-2146 |
| chr19-35413846-INS-595 | chr19-35413846-INS-615 |
| chr19-35433448-INS-197 | chr19-35433448-INS-197 |
| chr19-35461003-INS-216 | chr19-35461003-INS-216 |
| chr19-35482413-INS-368 | chr19-35482413-INS-368 |
| chr19-35484378-INS-334 | chr19-35484378-INS-334 |
| chr19-35485887-INS-7543 | chr19-35485888-INS-7544 |
| chr19-35488386-INS-89 | chr19-35488384-INS-93 |
| chr19-35495321-INS-5784 | chr19-35495322-INS-5784 |
| chr19-35532651-INS-440 | chr19-35532651-INS-440 |
| chr19-35550819-INS-72 | chr19-35550819-INS-72 |
| chr19-35564454-INS-113 | chr19-35564454-INS-113 |
| chr19-35579714-INS-79 | chr19-35579714-INS-78 |
| chr19-35580588-INS-56 | chr19-35580588-INS-52 |
| chr19-35587257-INS-309 | chr19-35587257-INS-309 |
| chr19-35600260-INS-183 | chr19-35600260-INS-183 |
| chr19-35611769-INS-358 | chr19-35611769-INS-358 |
| chr19-35656010-INS-51 | chr19-35656010-INS-51 |
| chr19-35707602-INS-70 | chr19-35707602-INS-65 |
| chr19-35719951-INS-96 | chr19-35719951-INS-92 |
| chr19-35724717-INS-432 | chr19-35724717-INS-432 |
| chr19-35733515-INS-422 | chr19-35733515-INS-423 |
| chr19-35734294-INS-657 | chr19-35734294-INS-658 |
| chr19-35769080-INS-6104 | chr19-35769081-INS-6104 |
| chr19-35789724-INS-109 | chr19-35789724-INS-81 |
| chr19-35809100-INS-928 | chr19-35809100-INS-927 |
| chr19-35814038-INS-953 | chr19-35814038-INS-952 |
| chr19-35818175-INS-65 | chr19-35818175-INS-77 |
| chr19-35866639-INS-238 | chr19-35866639-INS-232 |

| | |
|---|---|
| chr19-35878539-INS-322 | chr19-35878539-INS-322 |
| chr19-35897634-INS-212 | chr19-35897634-INS-212 |
| chr19-35903149-INS-83 | chr19-35903149-INS-83 |
| chr19-35926382-INS-70 | chr19-35926382-INS-51 |
| chr19-35951952-INS-447 | chr19-35951956-INS-445 |
| chr19-35958776-INS-188 | chr19-35958767-INS-191 |
| chr19-35964687-INS-175 | chr19-35964690-INS-127 |
| chr19-35967330-INS-115 | chr19-35967330-INS-115 |
| chr19-36020932-INS-54 | chr19-36020930-INS-104 |
| chr19-36038772-INS-127 | chr19-36038772-INS-123 |
| chr19-36086004-INS-472 | chr19-36086004-INS-465 |
| chr19-36086789-INS-78 | chr19-36086799-INS-68 |
| chr19-36096560-INS-72 | chr19-36096577-INS-72 |
| chr19-36127075-INS-200 | chr19-36127083-INS-172 |
| chr19-36158735-INS-173 | chr19-36158735-INS-176 |
| chr19-36181800-INS-114 | chr19-36181796-INS-116 |
| chr19-36185769-INS-93 | chr19-36185793-INS-66 |
| chr19-36198310-INS-205 | chr19-36198310-INS-215 |
| chr19-36210336-INS-90 | chr19-36210336-INS-74 |
| chr19-36230818-INS-78 | chr19-36230818-INS-80 |
| chr19-36267275-INS-52 | chr19-36267264-INS-54 |
| chr19-36275004-INS-52 | chr19-36275004-INS-52 |
| chr19-36302346-INS-187 | chr19-36302346-INS-164 |
| chr19-36315779-INS-73 | chr19-36315779-INS-73 |
| chr19-36321707-INS-151 | chr19-36321707-INS-153 |
| chr19-36331876-INS-204 | chr19-36331876-INS-202 |
| chr19-36374419-INS-68 | chr19-36374419-INS-70 |
| chr19-36439414-INS-53 | chr19-36439414-INS-53 |
| chr19-36448894-INS-271 | chr19-36448894-INS-271 |
| chr19-36466354-INS-70 | chr19-36466354-INS-90 |
| chr19-36485590-INS-527 | chr19-36485590-INS-527 |
| chr19-36505313-INS-64 | chr19-36505313-INS-66 |
| chr19-36507779-INS-1079 | chr19-36507780-INS-1079 |
| chr19-36515831-INS-867 | chr19-36515831-INS-876 |
| chr19-36521381-INS-52 | chr19-36521381-INS-52 |
| chr19-36552597-INS-175 | chr19-36552597-INS-176 |
| chr19-36558807-INS-198 | chr19-36558807-INS-198 |
| chr19-36561237-INS-6286 | chr19-36561238-INS-6283 |
| chr19-36562618-INS-148 | chr19-36562619-INS-185 |
| chr19-36604199-INS-139 | chr19-36604215-INS-137 |
| chr19-36628473-INS-77 | chr19-36628473-INS-69 |
| chr19-36643842-INS-209 | chr19-36643842-INS-209 |
| chr19-36697384-INS-74 | chr19-36697384-INS-90 |
| chr19-36701864-INS-188 | chr19-36701864-INS-177 |
| chr19-36713955-INS-61 | chr19-36713955-INS-61 |
| chr19-36744716-INS-6711 | chr19-36744717-INS-6712 |
| chr19-36747923-INS-206 | chr19-36747923-INS-206 |
| chr19-36749096-INS-56 | chr19-36749096-INS-58 |
| chr19-36759030-INS-215 | chr19-36759030-INS-214 |
| chr19-36765373-INS-129 | chr19-36765373-INS-129 |
| chr19-36770375-INS-54 | chr19-36770375-INS-66 |
| chr19-36785823-INS-209 | chr19-36785823-INS-209 |
| chr19-36797812-INS-80 | chr19-36797812-INS-84 |
| chr19-36799445-INS-200 | chr19-36799445-INS-200 |
| chr19-36845362-INS-104 | chr19-36845362-INS-104 |
| chr19-36848162-INS-212 | chr19-36848162-INS-212 |
| chr19-36878415-INS-172 | chr19-36878415-INS-190 |
| chr19-36902049-INS-66 | chr19-36902049-INS-66 |
| chr19-36905947-INS-208 | chr19-36905955-INS-237 |
| chr19-36908878-INS-115 | chr19-36908878-INS-115 |
| chr19-36917314-INS-197 | chr19-36917314-INS-197 |
| chr19-36932526-INS-55 | chr19-36932527-INS-60 |
| chr19-36943265-INS-204 | chr19-36943265-INS-204 |
| chr19-36945553-INS-244 | chr19-36945553-INS-245 |
| chr19-36952992-INS-185 | chr19-36952992-INS-181 |
| chr19-36978066-INS-177 | chr19-36978066-INS-177 |
| chr19-36984635-INS-202 | chr19-36984635-INS-202 |
| chr19-37019658-INS-203 | chr19-37019658-INS-203 |
| chr19-37036646-INS-199 | chr19-37036646-INS-199 |
| chr19-37040226-INS-191 | chr19-37040232-INS-197 |
| chr19-37073028-INS-88 | chr19-37073031-INS-88 |
| chr19-37099904-INS-206 | chr19-37099904-INS-207 |
| chr19-37109992-INS-112 | chr19-37109992-INS-112 |
| chr19-37112338-INS-197 | chr19-37112338-INS-197 |
| chr19-37146284-INS-142 | chr19-37146273-INS-149 |
| chr19-37168098-INS-72 | chr19-37168098-INS-72 |
| chr19-37195773-INS-142 | chr19-37195773-INS-142 |

| | |
|---|---|
| chr19-37200589-INS-199 | chr19-37200589-INS-199 |
| chr19-37201531-INS-135 | chr19-37201535-INS-163 |
| chr19-37213167-INS-210 | chr19-37213167-INS-210 |
| chr19-37225293-INS-129 | chr19-37225293-INS-132 |
| chr19-37243730-INS-118 | chr19-37243730-INS-118 |
| chr19-37253176-INS-234 | chr19-37253176-INS-234 |
| chr19-37266346-INS-199 | chr19-37266346-INS-199 |
| chr19-37295904-INS-176 | chr19-37295904-INS-176 |
| chr19-37301754-INS-132 | chr19-37301754-INS-132 |
| chr19-37315118-INS-56 | chr19-37315143-INS-56 |
| chr19-37324995-INS-170 | chr19-37324995-INS-170 |
| chr19-37330378-INS-207 | chr19-37330378-INS-208 |
| chr19-37371484-INS-94 | chr19-37371484-INS-94 |
| chr19-37389745-INS-82 | chr19-37389745-INS-121 |
| chr19-37396284-INS-350 | chr19-37396284-INS-364 |
| chr19-37412421-INS-403 | chr19-37412421-INS-403 |
| chr19-37413808-INS-78 | chr19-37413808-INS-75 |
| chr19-37420596-INS-213 | chr19-37420596-INS-212 |
| chr19-37437649-INS-7106 | chr19-37437650-INS-7106 |
| chr19-37439501-INS-214 | chr19-37439501-INS-213 |
| chr19-37464431-INS-483 | chr19-37464431-INS-499 |
| chr19-37465510-INS-97 | chr19-37465510-INS-97 |
| chr19-37482456-INS-110 | chr19-37482456-INS-104 |
| chr19-37488689-INS-63 | chr19-37488710-INS-74 |
| chr19-37493616-INS-108 | chr19-37493617-INS-124 |
| chr19-37518521-INS-133 | chr19-37518521-INS-133 |
| chr19-37532697-INS-87 | chr19-37532697-INS-92 |
| chr19-37533853-INS-78 | chr19-37533853-INS-78 |
| chr19-37540844-INS-475 | chr19-37540844-INS-485 |
| chr19-37545145-INS-60 | chr19-37545150-INS-50 |
| chr19-37555464-INS-84 | chr19-37555464-INS-84 |
| chr19-37567515-INS-112 | chr19-37567515-INS-112 |
| chr19-37570882-INS-653 | chr19-37570882-INS-653 |
| chr19-37578007-INS-199 | chr19-37578007-INS-199 |
| chr19-37583107-INS-179 | chr19-37583107-INS-179 |
| chr19-37587613-INS-207 | chr19-37587613-INS-207 |
| chr19-37603137-INS-78 | chr19-37603137-INS-78 |
| chr19-37616482-INS-71 | chr19-37616482-INS-71 |
| chr19-37623463-INS-213 | chr19-37623463-INS-213 |
| chr19-37634631-INS-223 | chr19-37634631-INS-221 |
| chr19-37646005-INS-207 | chr19-37646005-INS-207 |
| chr19-37678940-INS-387 | chr19-37678940-INS-387 |
| chr19-37688692-INS-101 | chr19-37688727-INS-97 |
| chr19-37736330-INS-4585 | chr19-37736331-INS-4585 |
| chr19-37749115-INS-65 | chr19-37749115-INS-50 |
| chr19-37787906-INS-154 | chr19-37787906-INS-154 |
| chr19-37809029-INS-64 | chr19-37809029-INS-60 |
| chr19-37828227-INS-1338 | chr19-37828228-INS-1337 |
| chr19-37838970-INS-82 | chr19-37838970-INS-82 |
| chr19-37846232-INS-520 | chr19-37846232-INS-569 |
| chr19-37860459-INS-109 | chr19-37860459-INS-74 |
| chr19-37868125-INS-75 | chr19-37868122-INS-82 |
| chr19-37872031-INS-398 | chr19-37872030-INS-399 |
| chr19-37897417-INS-116 | chr19-37897376-INS-147 |
| chr19-37901531-INS-71 | chr19-37901524-INS-62 |
| chr19-37903567-INS-95 | chr19-37903567-INS-75 |
| chr19-37906576-INS-74 | chr19-37906577-INS-75 |
| chr19-37909033-INS-260 | chr19-37909033-INS-260 |
| chr19-37926001-INS-350 | chr19-37926001-INS-350 |
| chr19-37971864-INS-376 | chr19-37971864-INS-377 |
| chr19-37982059-INS-69 | chr19-37982059-INS-69 |
| chr19-38006934-INS-60 | chr19-38006934-INS-60 |
| chr19-38008300-INS-187 | chr19-38008300-INS-187 |
| chr19-38016666-INS-214 | chr19-38016666-INS-224 |
| chr19-38017534-INS-119 | chr19-38017547-INS-123 |
| chr19-38018975-INS-198 | chr19-38018975-INS-202 |
| chr19-38027251-INS-175 | chr19-38027251-INS-179 |
| chr19-38028515-INS-84 | chr19-38028515-INS-84 |
| chr19-38035031-INS-401 | chr19-38035031-INS-401 |
| chr19-38036364-INS-205 | chr19-38036364-INS-205 |
| chr19-38052161-INS-88 | chr19-38052161-INS-88 |
| chr19-38088086-INS-66 | chr19-38088086-INS-54 |
| chr19-38097855-INS-700 | chr19-38097855-INS-700 |
| chr19-38103363-INS-87 | chr19-38103363-INS-88 |
| chr19-38119117-INS-80 | chr19-38119117-INS-78 |
| chr19-38127711-INS-78 | chr19-38127711-INS-82 |
| chr19-38131025-INS-248 | chr19-38131025-INS-248 |

| | |
|---|---|
| chr19-38134721-INS-1532 | chr19-38134722-INS-1520 |
| chr19-38141279-INS-177 | chr19-38141279-INS-177 |
| chr19-38167954-INS-67 | chr19-38167954-INS-71 |
| chr19-38192556-INS-190 | chr19-38192556-INS-190 |
| chr19-38236593-INS-210 | chr19-38236593-INS-210 |
| chr19-38264499-INS-101 | chr19-38264454-INS-110 |
| chr19-38275967-INS-61 | chr19-38275950-INS-52 |
| chr19-38277336-INS-481 | chr19-38277336-INS-481 |
| chr19-38284130-INS-456 | chr19-38284130-INS-456 |
| chr19-38327349-INS-311 | chr19-38327349-INS-311 |
| chr19-38336192-INS-112 | chr19-38336192-INS-114 |
| chr19-38344263-INS-55 | chr19-38344263-INS-55 |
| chr19-38395509-INS-269 | chr19-38395509-INS-269 |
| chr19-38411201-INS-183 | chr19-38411201-INS-183 |
| chr19-38428217-INS-217 | chr19-38428217-INS-217 |
| chr19-38435125-INS-170 | chr19-38435125-INS-170 |
| chr19-38438285-INS-75 | chr19-38438285-INS-75 |
| chr19-38444534-INS-345 | chr19-38444534-INS-345 |
| chr19-38445852-INS-241 | chr19-38445852-INS-229 |
| chr19-38459531-INS-167 | chr19-38459531-INS-145 |
| chr19-38461868-INS-207 | chr19-38461868-INS-207 |
| chr19-38474436-INS-132 | chr19-38474436-INS-132 |
| chr19-38478061-INS-87 | chr19-38478058-INS-69 |
| chr19-38486035-INS-198 | chr19-38486035-INS-203 |
| chr19-38511904-INS-124 | chr19-38511886-INS-160 |
| chr19-38518639-INS-343 | chr19-38518641-INS-338 |
| chr19-38521080-INS-52 | chr19-38521080-INS-50 |
| chr19-38526142-INS-195 | chr19-38526142-INS-195 |
| chr19-38530120-INS-480 | chr19-38530120-INS-480 |
| chr19-38542585-INS-204 | chr19-38542585-INS-204 |
| chr19-38549980-INS-222 | chr19-38549980-INS-222 |
| chr19-38570058-INS-77 | chr19-38570058-INS-77 |
| chr19-38575159-INS-52 | chr19-38575159-INS-52 |
| chr19-38598354-INS-154 | chr19-38598353-INS-195 |
| chr19-38624844-INS-56 | chr19-38624844-INS-58 |
| chr19-38630846-INS-194 | chr19-38630846-INS-194 |
| chr19-38650711-INS-65 | chr19-38650711-INS-65 |
| chr19-38654985-INS-212 | chr19-38654985-INS-212 |
| chr19-38692090-INS-495 | chr19-38692090-INS-495 |
| chr19-38720012-INS-189 | chr19-38720012-INS-189 |
| chr19-38731555-INS-118 | chr19-38731555-INS-108 |
| chr19-38739537-INS-333 | chr19-38739537-INS-292 |
| chr19-38745939-INS-181 | chr19-38745939-INS-182 |
| chr19-38756948-INS-203 | chr19-38756948-INS-201 |
| chr19-38760537-INS-87 | chr19-38760537-INS-87 |
| chr19-38765693-INS-90 | chr19-38765693-INS-90 |
| chr19-38773648-INS-52 | chr19-38773648-INS-52 |
| chr19-38776697-INS-208 | chr19-38776697-INS-208 |
| chr19-38795706-INS-75 | chr19-38795706-INS-84 |
| chr19-38800192-INS-198 | chr19-38800192-INS-201 |
| chr19-38804598-INS-221 | chr19-38804598-INS-221 |
| chr19-38810472-INS-195 | chr19-38810473-INS-195 |
| chr19-38812053-INS-230 | chr19-38812053-INS-230 |
| chr19-38818988-INS-56 | chr19-38818991-INS-61 |
| chr19-38823940-INS-149 | chr19-38823940-INS-149 |
| chr19-38837259-INS-1110 | chr19-38837262-INS-1109 |
| chr19-38837934-INS-501 | chr19-38837934-INS-535 |
| chr19-38838979-INS-70 | chr19-38838979-INS-70 |
| chr19-38853341-INS-68 | chr19-38853325-INS-58 |
| chr19-38855610-INS-1031 | chr19-38855611-INS-1033 |
| chr19-38890026-INS-104 | chr19-38890043-INS-89 |
| chr19-38930284-INS-73 | chr19-38930327-INS-68 |
| chr19-38963017-INS-430 | chr19-38963017-INS-420 |
| chr19-38979196-INS-1578 | chr19-38979197-INS-1618 |
| chr19-38979197-INS-134 | chr19-38979193-INS-132 |
| chr19-39013291-INS-5179 | chr19-39013292-INS-5179 |
| chr19-39025318-INS-52 | chr19-39025318-INS-52 |
| chr19-39029222-INS-107 | chr19-39029222-INS-113 |
| chr19-39100872-INS-52 | chr19-39100873-INS-52 |
| chr19-39206207-INS-2485 | chr19-39206230-INS-2504 |
| chr19-39275794-INS-451 | chr19-39275794-INS-451 |
| chr19-39303319-INS-6573 | chr19-39303320-INS-6573 |
| chr19-39312848-INS-50 | chr19-39312848-INS-60 |
| chr19-39319349-INS-270 | chr19-39319349-INS-272 |
| chr19-39358028-INS-3375 | chr19-39358029-INS-3374 |
| chr19-39387984-INS-58 | chr19-39387994-INS-59 |
| chr19-39416603-INS-587 | chr19-39416603-INS-587 |

| | |
|---|---|
| chr19-39477522-INS-399 | chr19-39477522-INS-399 |
| chr19-39530321-INS-200 | chr19-39530321-INS-199 |
| chr19-39535857-INS-9523 | chr19-39535856-INS-9509 |
| chr19-39546070-INS-52 | chr19-39546071-INS-77 |
| chr19-39562899-INS-6589 | chr19-39562900-INS-6589 |
| chr19-39574444-INS-54 | chr19-39574444-INS-52 |
| chr19-39638982-INS-59 | chr19-39638996-INS-79 |
| chr19-39645718-INS-64 | chr19-39645718-INS-64 |
| chr19-39662297-INS-151 | chr19-39662297-INS-151 |
| chr19-39734684-INS-80 | chr19-39734687-INS-97 |
| chr19-39770610-INS-247 | chr19-39770610-INS-247 |
| chr19-39800313-INS-57 | chr19-39800288-INS-57 |
| chr19-39849442-INS-6440 | chr19-39849443-INS-6440 |
| chr19-39995324-INS-557 | chr19-39995324-INS-557 |
| chr19-40045247-INS-2950 | chr19-40045248-INS-2950 |
| chr19-40070296-INS-810 | chr19-40070296-INS-810 |
| chr19-40088034-INS-4761 | chr19-40088035-INS-4761 |
| chr19-40113608-INS-581 | chr19-40113609-INS-581 |
| chr19-40144502-INS-121 | chr19-40144502-INS-120 |
| chr19-40168202-INS-70 | chr19-40168202-INS-68 |
| chr19-40175149-INS-232 | chr19-40175149-INS-228 |
| chr19-40184268-INS-208 | chr19-40184268-INS-208 |
| chr19-40199939-INS-622 | chr19-40199939-INS-622 |
| chr19-40241774-INS-490 | chr19-40241774-INS-491 |
| chr19-40244933-INS-173 | chr19-40244933-INS-173 |
| chr19-40251083-INS-50 | chr19-40251071-INS-88 |
| chr19-40257833-INS-69 | chr19-40257850-INS-67 |
| chr19-40261556-INS-321 | chr19-40261556-INS-320 |
| chr19-40264289-INS-97 | chr19-40264289-INS-102 |
| chr19-40268395-INS-74 | chr19-40268397-INS-76 |
| chr19-40273273-INS-172 | chr19-40273273-INS-172 |
| chr19-40292875-INS-106 | chr19-40292875-INS-106 |
| chr19-40316839-INS-65 | chr19-40316839-INS-67 |
| chr19-40393392-INS-52 | chr19-40393392-INS-55 |
| chr19-40421456-INS-332 | chr19-40421456-INS-333 |
| chr19-40432158-INS-54 | chr19-40432158-INS-54 |
| chr19-40485326-INS-735 | chr19-40485326-INS-735 |
| chr19-40492827-INS-214 | chr19-40492827-INS-215 |
| chr19-40507164-INS-201 | chr19-40507170-INS-166 |
| chr19-40517837-INS-195 | chr19-40517837-INS-195 |
| chr19-40519310-INS-201 | chr19-40519314-INS-210 |
| chr19-40522685-INS-183 | chr19-40522685-INS-182 |
| chr19-40524528-INS-413 | chr19-40524528-INS-413 |
| chr19-40531828-INS-4800 | chr19-40531829-INS-4800 |
| chr19-40535452-INS-703 | chr19-40535452-INS-676 |
| chr19-40542335-INS-69 | chr19-40542335-INS-69 |
| chr19-40558465-INS-52 | chr19-40558465-INS-52 |
| chr19-40567302-INS-202 | chr19-40567302-INS-202 |
| chr19-40574222-INS-147 | chr19-40574222-INS-147 |
| chr19-40595626-INS-60 | chr19-40595626-INS-64 |
| chr19-40602697-INS-131 | chr19-40602697-INS-130 |
| chr19-40613836-INS-60 | chr19-40613833-INS-60 |
| chr19-40619860-INS-451 | chr19-40619860-INS-451 |
| chr19-40623201-INS-211 | chr19-40623201-INS-206 |
| chr19-40626876-INS-88 | chr19-40626876-INS-88 |
| chr19-40639117-INS-636 | chr19-40639117-INS-636 |
| chr19-40659015-INS-80 | chr19-40659015-INS-80 |
| chr19-40662344-INS-77 | chr19-40662344-INS-77 |
| chr19-40675345-INS-52 | chr19-40675345-INS-52 |
| chr19-40706972-INS-56 | chr19-40706972-INS-56 |
| chr19-40710309-INS-168 | chr19-40710305-INS-160 |
| chr19-40720349-INS-51 | chr19-40720359-INS-52 |
| chr19-40733647-INS-105 | chr19-40733647-INS-105 |
| chr19-40744339-INS-191 | chr19-40744345-INS-173 |
| chr19-40761902-INS-123 | chr19-40761902-INS-123 |
| chr19-40802557-INS-83 | chr19-40802557-INS-83 |
| chr19-40839172-INS-425 | chr19-40839172-INS-426 |
| chr19-40843192-INS-54 | chr19-40843189-INS-63 |
| chr19-40846630-INS-105 | chr19-40846630-INS-105 |
| chr19-40861367-INS-52 | chr19-40861368-INS-64 |
| chr19-40891446-INS-540 | chr19-40891446-INS-540 |
| chr19-40909281-INS-107 | chr19-40909281-INS-107 |
| chr19-40915134-INS-66 | chr19-40915134-INS-73 |
| chr19-40918564-INS-94 | chr19-40918564-INS-94 |
| chr19-40919861-INS-56 | chr19-40919862-INS-56 |
| chr19-40933500-INS-166 | chr19-40933520-INS-164 |
| chr19-40940129-INS-72 | chr19-40940130-INS-56 |

| | |
|---|---|
| chr19-40943915-INS-72 | chr19-40943915-INS-86 |
| chr19-40953358-INS-116 | chr19-40953358-INS-112 |
| chr19-40972717-INS-70 | chr19-40972717-INS-75 |
| chr19-41001852-INS-62 | chr19-41001852-INS-64 |
| chr19-41003268-INS-87 | chr19-41003268-INS-88 |
| chr19-41048685-INS-211 | chr19-41048685-INS-211 |
| chr19-41086158-INS-69 | chr19-41086158-INS-69 |
| chr19-41111414-INS-462 | chr19-41111414-INS-462 |
| chr19-41115793-INS-336 | chr19-41115793-INS-336 |
| chr19-41160796-INS-230 | chr19-41160796-INS-230 |
| chr19-41168801-INS-78 | chr19-41168801-INS-78 |
| chr19-41266556-INS-6440 | chr19-41266557-INS-6440 |
| chr19-41298670-INS-95 | chr19-41298670-INS-76 |
| chr19-41302592-INS-228 | chr19-41302592-INS-228 |
| chr19-41308076-INS-94 | chr19-41308076-INS-94 |
| chr19-41357436-INS-170 | chr19-41357436-INS-167 |
| chr19-41378064-INS-75 | chr19-41378064-INS-75 |
| chr19-41380934-INS-101 | chr19-41380934-INS-101 |
| chr19-41383064-INS-244 | chr19-41383064-INS-205 |
| chr19-41391488-INS-70 | chr19-41391488-INS-72 |
| chr19-41403188-INS-211 | chr19-41403189-INS-209 |
| chr19-41405343-INS-82 | chr19-41405343-INS-68 |
| chr19-41423646-INS-153 | chr19-41423646-INS-152 |
| chr19-41431531-INS-213 | chr19-41431531-INS-216 |
| chr19-41439013-INS-197 | chr19-41439013-INS-194 |
| chr19-41478909-INS-512 | chr19-41478909-INS-512 |
| chr19-41521080-INS-57 | chr19-41521080-INS-57 |
| chr19-41529123-INS-204 | chr19-41529123-INS-204 |
| chr19-41530425-INS-455 | chr19-41530425-INS-455 |
| chr19-41541844-INS-211 | chr19-41541844-INS-211 |
| chr19-41626656-INS-111 | chr19-41626656-INS-105 |
| chr19-41630088-INS-5646 | chr19-41630089-INS-5646 |
| chr19-41643213-INS-200 | chr19-41643213-INS-200 |
| chr19-41644892-INS-120 | chr19-41644892-INS-120 |
| chr19-41670022-INS-70 | chr19-41670022-INS-70 |
| chr19-41699371-INS-82 | chr19-41699365-INS-58 |
| chr19-41702085-INS-56 | chr19-41702085-INS-52 |
| chr19-41724890-INS-328 | chr19-41724890-INS-328 |
| chr19-41748548-INS-292 | chr19-41748548-INS-292 |
| chr19-41749381-INS-70 | chr19-41749358-INS-114 |
| chr19-41775295-INS-198 | chr19-41775295-INS-198 |
| chr19-41804012-INS-206 | chr19-41804012-INS-206 |
| chr19-41808104-INS-219 | chr19-41808104-INS-219 |
| chr19-41808821-INS-55 | chr19-41808821-INS-58 |
| chr19-41812891-INS-86 | chr19-41812891-INS-86 |
| chr19-41815402-INS-68 | chr19-41815402-INS-60 |
| chr19-41824865-INS-213 | chr19-41824889-INS-174 |
| chr19-41843845-INS-215 | chr19-41843845-INS-213 |
| chr19-41852546-INS-101 | chr19-41852546-INS-106 |
| chr19-41864706-INS-203 | chr19-41864706-INS-201 |
| chr19-41891260-INS-203 | chr19-41891260-INS-203 |
| chr19-41894599-INS-213 | chr19-41894599-INS-213 |
| chr19-41898469-INS-198 | chr19-41898489-INS-211 |
| chr19-41907387-INS-206 | chr19-41907387-INS-206 |
| chr19-41908563-INS-203 | chr19-41908563-INS-204 |
| chr19-41925066-INS-154 | chr19-41925066-INS-154 |
| chr19-41930615-INS-86 | chr19-41930615-INS-86 |
| chr19-41931852-INS-196 | chr19-41931852-INS-195 |
| chr19-41983205-INS-52 | chr19-41983171-INS-78 |
| chr19-41996080-INS-2680 | chr19-41996081-INS-2680 |
| chr19-42002699-INS-250 | chr19-42002699-INS-231 |
| chr19-42031309-INS-199 | chr19-42031309-INS-197 |
| chr19-42046728-INS-206 | chr19-42046728-INS-206 |
| chr19-42052724-INS-177 | chr19-42052724-INS-177 |
| chr19-42065621-INS-211 | chr19-42065621-INS-210 |
| chr19-42066713-INS-155 | chr19-42066713-INS-155 |
| chr19-42076300-INS-220 | chr19-42076300-INS-220 |
| chr19-42083644-INS-928 | chr19-42083644-INS-930 |
| chr19-42094674-INS-193 | chr19-42094674-INS-195 |
| chr19-42100252-INS-177 | chr19-42100253-INS-168 |
| chr19-42101423-INS-844 | chr19-42101423-INS-844 |
| chr19-42135929-INS-85 | chr19-42135929-INS-85 |
| chr19-42153465-INS-999 | chr19-42153465-INS-979 |
| chr19-42157189-INS-363 | chr19-42157189-INS-354 |
| chr19-42160761-INS-257 | chr19-42160761-INS-257 |
| chr19-42173337-INS-193 | chr19-42173337-INS-236 |
| chr19-42188488-INS-201 | chr19-42188488-INS-200 |

| | |
|---|---|
| chr19-42194292-INS-207 | chr19-42194292-INS-209 |
| chr19-42210802-INS-57 | chr19-42210802-INS-57 |
| chr19-42220700-INS-101 | chr19-42220700-INS-101 |
| chr19-42241330-INS-51 | chr19-42241330-INS-51 |
| chr19-42249074-INS-102 | chr19-42249074-INS-106 |
| chr19-42254035-INS-80 | chr19-42254035-INS-80 |
| chr19-42293681-INS-207 | chr19-42293681-INS-207 |
| chr19-42298380-INS-60 | chr19-42298350-INS-74 |
| chr19-42326393-INS-62 | chr19-42326391-INS-74 |
| chr19-42348634-INS-62 | chr19-42348634-INS-62 |
| chr19-42353867-INS-57 | chr19-42353869-INS-58 |
| chr19-42354887-INS-54 | chr19-42354887-INS-56 |
| chr19-42358582-INS-129 | chr19-42358582-INS-130 |
| chr19-42365396-INS-100 | chr19-42365396-INS-148 |
| chr19-42383072-INS-70 | chr19-42383072-INS-50 |
| chr19-42383526-INS-379 | chr19-42383526-INS-379 |
| chr19-42391896-INS-77 | chr19-42391900-INS-73 |
| chr19-42413806-INS-55 | chr19-42413806-INS-55 |
| chr19-42442009-INS-220 | chr19-42442009-INS-171 |
| chr19-42461511-INS-321 | chr19-42461511-INS-323 |
| chr19-42576120-INS-68 | chr19-42576120-INS-108 |
| chr19-42597713-INS-127 | chr19-42597713-INS-133 |
| chr19-42609881-INS-3006 | chr19-42609882-INS-3006 |
| chr19-42617484-INS-157 | chr19-42617486-INS-168 |
| chr19-42628530-INS-133 | chr19-42628530-INS-145 |
| chr19-42665176-INS-229 | chr19-42665176-INS-229 |
| chr19-42684511-INS-112 | chr19-42684511-INS-112 |
| chr19-42705984-INS-581 | chr19-42705984-INS-580 |
| chr19-42717656-INS-71 | chr19-42717656-INS-71 |
| chr19-42726154-INS-55 | chr19-42726154-INS-55 |
| chr19-42733702-INS-152 | chr19-42733699-INS-128 |
| chr19-42735126-INS-523 | chr19-42735126-INS-523 |
| chr19-42772183-INS-70 | chr19-42772183-INS-82 |
| chr19-42802226-INS-97 | chr19-42802226-INS-97 |
| chr19-42808915-INS-103 | chr19-42808915-INS-103 |
| chr19-42820535-INS-244 | chr19-42820535-INS-196 |
| chr19-42836761-INS-200 | chr19-42836761-INS-200 |
| chr19-42865872-INS-82 | chr19-42865873-INS-124 |
| chr19-42872676-INS-586 | chr19-42872676-INS-596 |
| chr19-42877165-INS-129 | chr19-42877165-INS-147 |
| chr19-42877774-INS-235 | chr19-42877774-INS-239 |
| chr19-42890962-INS-226 | chr19-42890962-INS-226 |
| chr19-42905315-INS-56 | chr19-42905319-INS-72 |
| chr19-42960524-INS-1784 | chr19-42960527-INS-1780 |
| chr19-42964437-INS-165 | chr19-42964437-INS-165 |
| chr19-42973009-INS-6691 | chr19-42973010-INS-6692 |
| chr19-43020386-INS-1535 | chr19-43020387-INS-1534 |
| chr19-43033215-INS-60 | chr19-43033215-INS-60 |
| chr19-43033922-INS-8318 | chr19-43033923-INS-8317 |
| chr19-43054688-INS-52 | chr19-43054688-INS-60 |
| chr19-43056874-INS-56 | chr19-43056874-INS-58 |
| chr19-43074195-INS-247 | chr19-43074195-INS-247 |
| chr19-43081151-INS-107 | chr19-43081151-INS-107 |
| chr19-43084060-INS-57 | chr19-43084060-INS-60 |
| chr19-43109810-INS-1912 | chr19-43109811-INS-1912 |
| chr19-43142563-INS-226 | chr19-43142563-INS-228 |
| chr19-43151457-INS-130 | chr19-43151457-INS-130 |
| chr19-43173137-INS-581 | chr19-43173137-INS-583 |
| chr19-43178050-INS-1468 | chr19-43178051-INS-1468 |
| chr19-43187183-INS-197 | chr19-43187183-INS-197 |
| chr19-43191462-INS-64 | chr19-43191462-INS-64 |
| chr19-43199643-INS-102 | chr19-43199643-INS-107 |
| chr19-43203914-INS-646 | chr19-43203914-INS-646 |
| chr19-43217636-INS-56 | chr19-43217636-INS-60 |
| chr19-43251127-INS-661 | chr19-43251127-INS-659 |
| chr19-43265767-INS-141 | chr19-43265767-INS-141 |
| chr19-43284907-INS-57 | chr19-43284907-INS-54 |
| chr19-43341143-INS-94 | chr19-43341143-INS-94 |
| chr19-43356972-INS-804 | chr19-43356972-INS-804 |
| chr19-43476236-INS-206 | chr19-43476236-INS-206 |
| chr19-43495564-INS-54 | chr19-43495545-INS-54 |
| chr19-43502850-INS-200 | chr19-43502850-INS-200 |
| chr19-43506648-INS-204 | chr19-43506648-INS-204 |
| chr19-43520087-INS-209 | chr19-43520087-INS-217 |
| chr19-43524381-INS-2041 | chr19-43524382-INS-2061 |
| chr19-43543557-INS-59 | chr19-43543557-INS-65 |
| chr19-43561435-INS-140 | chr19-43561429-INS-135 |

| | |
|---|---|
| chr19-43574023-INS-55 | chr19-43574023-INS-55 |
| chr19-43609319-INS-201 | chr19-43609319-INS-209 |
| chr19-43621785-INS-66 | chr19-43621785-INS-70 |
| chr19-43626139-INS-184 | chr19-43626139-INS-184 |
| chr19-43629841-INS-80 | chr19-43629841-INS-80 |
| chr19-43646316-INS-84 | chr19-43646316-INS-58 |
| chr19-43650784-INS-209 | chr19-43650784-INS-208 |
| chr19-43662242-INS-497 | chr19-43662242-INS-497 |
| chr19-43696219-INS-350 | chr19-43696220-INS-350 |
| chr19-43703930-INS-255 | chr19-43703930-INS-255 |
| chr19-43704893-INS-249 | chr19-43704879-INS-293 |
| chr19-43744400-INS-94 | chr19-43744396-INS-95 |
| chr19-43746323-INS-200 | chr19-43746323-INS-208 |
| chr19-43752477-INS-80 | chr19-43752477-INS-80 |
| chr19-43753026-INS-188 | chr19-43753026-INS-184 |
| chr19-43765888-INS-6452 | chr19-43765889-INS-6452 |
| chr19-43775656-INS-94 | chr19-43775656-INS-96 |
| chr19-43783554-INS-96 | chr19-43783553-INS-103 |
| chr19-43785539-INS-320 | chr19-43785552-INS-327 |
| chr19-43791711-INS-98 | chr19-43791711-INS-128 |
| chr19-43794601-INS-5599 | chr19-43794602-INS-5614 |
| chr19-43811998-INS-56 | chr19-43811998-INS-62 |
| chr19-43836982-INS-199 | chr19-43836982-INS-199 |
| chr19-43854393-INS-165 | chr19-43854393-INS-165 |
| chr19-43859413-INS-93 | chr19-43859413-INS-93 |
| chr19-43864418-INS-206 | chr19-43864418-INS-206 |
| chr19-43875774-INS-183 | chr19-43875774-INS-184 |
| chr19-43896683-INS-83 | chr19-43896683-INS-83 |
| chr19-43901382-INS-62 | chr19-43901382-INS-66 |
| chr19-43903291-INS-151 | chr19-43903291-INS-152 |
| chr19-43907591-INS-51 | chr19-43907591-INS-51 |
| chr19-43933996-INS-183 | chr19-43933997-INS-144 |
| chr19-43947063-INS-52 | chr19-43947062-INS-60 |
| chr19-43986495-INS-62 | chr19-43986495-INS-62 |
| chr19-43988044-INS-1452 | chr19-43988045-INS-1452 |
| chr19-43990903-INS-146 | chr19-43990917-INS-148 |
| chr19-44019034-INS-199 | chr19-44019034-INS-198 |
| chr19-44028299-INS-222 | chr19-44028299-INS-214 |
| chr19-44036545-INS-66 | chr19-44036545-INS-66 |
| chr19-44048126-INS-87 | chr19-44048126-INS-87 |
| chr19-44055016-INS-193 | chr19-44055016-INS-193 |
| chr19-44078623-INS-62 | chr19-44078623-INS-62 |
| chr19-44087735-INS-181 | chr19-44087735-INS-181 |
| chr19-44108172-INS-228 | chr19-44108155-INS-255 |
| chr19-44116717-INS-200 | chr19-44116717-INS-200 |
| chr19-44128929-INS-207 | chr19-44128929-INS-207 |
| chr19-44143402-INS-52 | chr19-44143402-INS-58 |
| chr19-44146771-INS-72 | chr19-44146771-INS-72 |
| chr19-44147658-INS-142 | chr19-44147658-INS-161 |
| chr19-44176435-INS-235 | chr19-44176435-INS-243 |
| chr19-44189990-INS-132 | chr19-44189990-INS-132 |
| chr19-44193406-INS-172 | chr19-44193406-INS-180 |
| chr19-44196505-INS-578 | chr19-44196506-INS-577 |
| chr19-44225632-INS-210 | chr19-44225632-INS-210 |
| chr19-44227730-INS-87 | chr19-44227739-INS-78 |
| chr19-44250857-INS-202 | chr19-44250857-INS-202 |
| chr19-44252231-INS-204 | chr19-44252231-INS-204 |
| chr19-44278295-INS-202 | chr19-44278295-INS-202 |
| chr19-44305878-INS-85 | chr19-44305878-INS-84 |
| chr19-44330900-INS-58 | chr19-44330900-INS-58 |
| chr19-44343836-INS-52 | chr19-44343836-INS-52 |
| chr19-44348849-INS-350 | chr19-44348849-INS-350 |
| chr19-44353165-INS-72 | chr19-44353165-INS-72 |
| chr19-44353843-INS-353 | chr19-44353843-INS-401 |
| chr19-44361340-INS-205 | chr19-44361340-INS-205 |
| chr19-44392887-INS-184 | chr19-44392887-INS-184 |
| chr19-44419381-INS-452 | chr19-44419381-INS-452 |
| chr19-44426287-INS-600 | chr19-44426287-INS-600 |
| chr19-44450005-INS-211 | chr19-44450005-INS-211 |
| chr19-44479276-INS-97 | chr19-44479276-INS-95 |
| chr19-44496002-INS-236 | chr19-44496002-INS-270 |
| chr19-44515364-INS-70 | chr19-44515403-INS-51 |
| chr19-44535731-INS-109 | chr19-44535731-INS-113 |
| chr19-44549414-INS-195 | chr19-44549414-INS-194 |
| chr19-44602426-INS-357 | chr19-44602426-INS-357 |
| chr19-44688867-INS-206 | chr19-44688867-INS-211 |
| chr19-44702334-INS-837 | chr19-44702330-INS-840 |

| | |
|---|---|
| chr19-44706924-INS-74 | chr19-44706924-INS-74 |
| chr19-44709992-INS-5549 | chr19-44709993-INS-5549 |
| chr19-44711066-INS-184 | chr19-44711066-INS-186 |
| chr19-44727364-INS-212 | chr19-44727364-INS-212 |
| chr19-44757018-INS-195 | chr19-44757018-INS-195 |
| chr19-44764566-INS-91 | chr19-44764566-INS-91 |
| chr19-44770384-INS-207 | chr19-44770384-INS-207 |
| chr19-44776110-INS-117 | chr19-44776110-INS-117 |
| chr19-44804264-INS-94 | chr19-44804264-INS-94 |
| chr19-44833709-INS-59 | chr19-44833709-INS-52 |
| chr19-44860008-INS-525 | chr19-44859988-INS-495 |
| chr19-44868331-INS-54 | chr19-44868332-INS-54 |
| chr19-44870459-INS-58 | chr19-44870423-INS-82 |
| chr19-44901704-INS-162 | chr19-44901704-INS-162 |
| chr19-44902118-INS-120 | chr19-44902118-INS-120 |
| chr19-44914776-INS-1890 | chr19-44914777-INS-1890 |
| chr19-44928208-INS-67 | chr19-44928208-INS-67 |
| chr19-44935885-INS-227 | chr19-44935885-INS-227 |
| chr19-44938230-INS-77 | chr19-44938230-INS-86 |
| chr19-44968107-INS-283 | chr19-44968107-INS-283 |
| chr19-44976567-INS-339 | chr19-44976567-INS-339 |
| chr19-44982264-INS-184 | chr19-44982264-INS-184 |
| chr19-44984462-INS-52 | chr19-44984478-INS-74 |
| chr19-45039969-INS-201 | chr19-45039969-INS-201 |
| chr19-45048889-INS-214 | chr19-45048889-INS-208 |
| chr19-45050519-INS-198 | chr19-45050519-INS-197 |
| chr19-45052263-INS-218 | chr19-45052263-INS-220 |
| chr19-45059244-INS-112 | chr19-45059244-INS-112 |
| chr19-45071830-INS-1893 | chr19-45071831-INS-1893 |
| chr19-45090165-INS-220 | chr19-45090165-INS-220 |
| chr19-45099487-INS-219 | chr19-45099487-INS-220 |
| chr19-45113010-INS-89 | chr19-45113010-INS-79 |
| chr19-45116680-INS-171 | chr19-45116680-INS-171 |
| chr19-45123810-INS-673 | chr19-45123810-INS-673 |
| chr19-45124836-INS-51 | chr19-45124836-INS-56 |
| chr19-45134855-INS-219 | chr19-45134855-INS-219 |
| chr19-45155224-INS-56 | chr19-45155224-INS-68 |
| chr19-45157694-INS-50 | chr19-45157694-INS-54 |
| chr19-45164318-INS-52 | chr19-45164318-INS-52 |
| chr19-45182823-INS-108 | chr19-45182823-INS-99 |
| chr19-45184465-INS-596 | chr19-45184465-INS-596 |
| chr19-45205298-INS-212 | chr19-45205298-INS-214 |
| chr19-45209444-INS-69 | chr19-45209423-INS-50 |
| chr19-45244592-INS-62 | chr19-45244592-INS-53 |
| chr19-45270131-INS-181 | chr19-45270131-INS-178 |
| chr19-45271585-INS-195 | chr19-45271585-INS-203 |
| chr19-45274419-INS-547 | chr19-45274419-INS-547 |
| chr19-45282814-INS-75 | chr19-45282842-INS-90 |
| chr19-45288085-INS-211 | chr19-45288085-INS-212 |
| chr19-45297061-INS-76 | chr19-45297070-INS-56 |
| chr19-45314483-INS-76 | chr19-45314484-INS-54 |
| chr19-45329375-INS-212 | chr19-45329375-INS-211 |
| chr19-45332154-INS-118 | chr19-45332154-INS-118 |
| chr19-45351265-INS-54 | chr19-45351263-INS-56 |
| chr19-45359963-INS-128 | chr19-45359974-INS-138 |
| chr19-45364967-INS-203 | chr19-45364967-INS-203 |
| chr19-45399174-INS-146 | chr19-45399174-INS-128 |
| chr19-45418735-INS-239 | chr19-45418735-INS-241 |
| chr19-45493244-INS-68 | chr19-45493240-INS-62 |
| chr19-45520235-INS-62 | chr19-45520235-INS-86 |
| chr19-45557146-INS-230 | chr19-45557146-INS-241 |
| chr19-45573198-INS-102 | chr19-45573208-INS-76 |
| chr19-45582075-INS-52 | chr19-45582075-INS-84 |
| chr19-45594458-INS-115 | chr19-45594458-INS-115 |
| chr19-45607218-INS-225 | chr19-45607218-INS-225 |
| chr19-45631493-INS-200 | chr19-45631493-INS-202 |
| chr19-45641224-INS-170 | chr19-45641224-INS-166 |
| chr19-45645999-INS-201 | chr19-45645999-INS-199 |
| chr19-45653718-INS-158 | chr19-45653718-INS-158 |
| chr19-45660161-INS-197 | chr19-45660171-INS-206 |
| chr19-45665768-INS-95 | chr19-45665748-INS-133 |
| chr19-45672615-INS-209 | chr19-45672615-INS-211 |
| chr19-45674365-INS-267 | chr19-45674365-INS-267 |
| chr19-45675255-INS-214 | chr19-45675255-INS-215 |
| chr19-45682616-INS-81 | chr19-45682616-INS-66 |
| chr19-45706789-INS-222 | chr19-45706789-INS-222 |
| chr19-45709410-INS-207 | chr19-45709410-INS-206 |

| | |
|---|---|
| chr19-45725025-INS-58 | chr19-45725025-INS-58 |
| chr19-45754527-INS-212 | chr19-45754527-INS-212 |
| chr19-45760961-INS-207 | chr19-45760961-INS-207 |
| chr19-45761511-INS-162 | chr19-45761511-INS-162 |
| chr19-45785027-INS-210 | chr19-45785027-INS-210 |
| chr19-45785966-INS-74 | chr19-45785966-INS-50 |
| chr19-45788314-INS-215 | chr19-45788314-INS-215 |
| chr19-45808256-INS-213 | chr19-45808256-INS-213 |
| chr19-45812132-INS-198 | chr19-45812132-INS-198 |
| chr19-45821042-INS-248 | chr19-45821042-INS-222 |
| chr19-45827158-INS-175 | chr19-45827159-INS-180 |
| chr19-45829054-INS-203 | chr19-45829054-INS-202 |
| chr19-45835173-INS-161 | chr19-45835173-INS-151 |
| chr19-45839874-INS-844 | chr19-45839874-INS-845 |
| chr19-45849203-INS-209 | chr19-45849203-INS-209 |
| chr19-45852231-INS-209 | chr19-45852231-INS-209 |
| chr19-45865106-INS-232 | chr19-45865106-INS-232 |
| chr19-45882950-INS-54 | chr19-45882950-INS-55 |
| chr19-45885642-INS-54 | chr19-45885595-INS-100 |
| chr19-45893233-INS-1890 | chr19-45893234-INS-1890 |
| chr19-45897378-INS-640 | chr19-45897378-INS-624 |
| chr19-45906691-INS-198 | chr19-45906691-INS-196 |
| chr19-45912134-INS-352 | chr19-45912121-INS-352 |
| chr19-45930067-INS-237 | chr19-45930067-INS-237 |
| chr19-45945034-INS-694 | chr19-45945034-INS-694 |
| chr19-45947951-INS-207 | chr19-45947951-INS-207 |
| chr19-45949828-INS-211 | chr19-45949828-INS-212 |
| chr19-45952069-INS-212 | chr19-45952069-INS-212 |
| chr19-45954463-INS-389 | chr19-45954463-INS-389 |
| chr19-45962653-INS-182 | chr19-45962650-INS-174 |
| chr19-45970082-INS-224 | chr19-45970082-INS-226 |
| chr19-45975429-INS-201 | chr19-45975429-INS-200 |
| chr19-45983335-INS-213 | chr19-45983335-INS-213 |
| chr19-45984781-INS-170 | chr19-45984781-INS-171 |
| chr19-45989877-INS-68 | chr19-45989871-INS-70 |
| chr19-46001013-INS-218 | chr19-46001013-INS-219 |
| chr19-46002339-INS-174 | chr19-46002339-INS-180 |
| chr19-46007371-INS-198 | chr19-46007371-INS-198 |
| chr19-46009728-INS-287 | chr19-46009728-INS-287 |
| chr19-46038272-INS-185 | chr19-46038273-INS-175 |
| chr19-46076300-INS-219 | chr19-46076300-INS-219 |
| chr19-46079531-INS-219 | chr19-46079531-INS-219 |
| chr19-46088921-INS-201 | chr19-46088921-INS-201 |
| chr19-46108593-INS-176 | chr19-46108593-INS-176 |
| chr19-46115572-INS-61 | chr19-46115525-INS-59 |
| chr19-46118449-INS-298 | chr19-46118449-INS-298 |
| chr19-46120103-INS-53 | chr19-46120101-INS-54 |
| chr19-46120757-INS-184 | chr19-46120757-INS-185 |
| chr19-46134555-INS-71 | chr19-46134555-INS-67 |
| chr19-46147828-INS-235 | chr19-46147828-INS-237 |
| chr19-46157209-INS-58 | chr19-46157204-INS-72 |
| chr19-46177576-INS-535 | chr19-46177576-INS-535 |
| chr19-46194636-INS-103 | chr19-46194656-INS-65 |
| chr19-46202922-INS-64 | chr19-46202922-INS-64 |
| chr19-46203460-INS-414 | chr19-46203460-INS-413 |
| chr19-46221215-INS-101 | chr19-46221223-INS-127 |
| chr19-46226226-INS-212 | chr19-46226226-INS-212 |
| chr19-46265369-INS-250 | chr19-46265369-INS-250 |
| chr19-46284013-INS-2058 | chr19-46284014-INS-2058 |
| chr19-46286436-INS-264 | chr19-46286436-INS-263 |
| chr19-46291322-INS-191 | chr19-46291323-INS-193 |
| chr19-46296205-INS-85 | chr19-46296205-INS-86 |
| chr19-46324846-INS-77 | chr19-46324846-INS-76 |
| chr19-46328738-INS-229 | chr19-46328738-INS-225 |
| chr19-46331093-INS-50 | chr19-46331094-INS-57 |
| chr19-46333111-INS-59 | chr19-46333111-INS-59 |
| chr19-46334399-INS-204 | chr19-46334399-INS-204 |
| chr19-46343414-INS-225 | chr19-46343414-INS-228 |
| chr19-46350078-INS-222 | chr19-46350078-INS-220 |
| chr19-46361244-INS-139 | chr19-46361244-INS-139 |
| chr19-46372881-INS-86 | chr19-46372881-INS-86 |
| chr19-46374481-INS-52 | chr19-46374485-INS-64 |
| chr19-46379885-INS-204 | chr19-46379849-INS-215 |
| chr19-46382163-INS-505 | chr19-46382148-INS-518 |
| chr19-46392926-INS-511 | chr19-46392930-INS-513 |
| chr19-46395874-INS-201 | chr19-46395874-INS-201 |
| chr19-46400836-INS-392 | chr19-46400836-INS-395 |

| | |
|---|---|
| chr19-46405731-INS-221 | chr19-46405731-INS-219 |
| chr19-46416807-INS-224 | chr19-46416807-INS-224 |
| chr19-46418756-INS-56 | chr19-46418756-INS-56 |
| chr19-46419538-INS-213 | chr19-46419538-INS-213 |
| chr19-46451586-INS-56 | chr19-46451538-INS-80 |
| chr19-46453427-INS-216 | chr19-46453427-INS-216 |
| chr19-46499557-INS-69 | chr19-46499557-INS-62 |
| chr19-46523025-INS-69 | chr19-46523002-INS-74 |
| chr19-46535258-INS-161 | chr19-46535275-INS-162 |
| chr19-46539162-INS-213 | chr19-46539162-INS-213 |
| chr19-46539708-INS-217 | chr19-46539708-INS-218 |
| chr19-46548150-INS-201 | chr19-46548150-INS-200 |
| chr19-46552035-INS-322 | chr19-46552035-INS-322 |
| chr19-46553454-INS-188 | chr19-46553454-INS-188 |
| chr19-46554734-INS-206 | chr19-46554734-INS-227 |
| chr19-46565615-INS-208 | chr19-46565615-INS-200 |
| chr19-46567774-INS-146 | chr19-46567784-INS-154 |
| chr19-46568702-INS-352 | chr19-46568702-INS-352 |
| chr19-46577485-INS-205 | chr19-46577485-INS-205 |
| chr19-46597349-INS-160 | chr19-46597349-INS-160 |
| chr19-46617537-INS-592 | chr19-46617537-INS-593 |
| chr19-46626947-INS-58 | chr19-46626947-INS-60 |
| chr19-46628705-INS-203 | chr19-46628706-INS-203 |
| chr19-46646176-INS-211 | chr19-46646175-INS-212 |
| chr19-46674884-INS-2089 | chr19-46674885-INS-2091 |
| chr19-46680254-INS-108 | chr19-46680254-INS-105 |
| chr19-46701184-INS-70 | chr19-46701184-INS-59 |
| chr19-46704951-INS-201 | chr19-46704951-INS-202 |
| chr19-46712776-INS-94 | chr19-46712776-INS-94 |
| chr19-46719451-INS-69 | chr19-46719451-INS-69 |
| chr19-46722556-INS-204 | chr19-46722556-INS-204 |
| chr19-46735422-INS-72 | chr19-46735422-INS-72 |
| chr19-46769126-INS-51 | chr19-46769126-INS-51 |
| chr19-46793721-INS-229 | chr19-46793721-INS-229 |
| chr19-46798096-INS-379 | chr19-46798096-INS-379 |
| chr19-46799781-INS-135 | chr19-46799781-INS-140 |
| chr19-46804374-INS-199 | chr19-46804373-INS-203 |
| chr19-46805338-INS-53 | chr19-46805331-INS-72 |
| chr19-46821353-INS-71 | chr19-46821353-INS-71 |
| chr19-46823441-INS-195 | chr19-46823441-INS-195 |
| chr19-46824407-INS-496 | chr19-46824407-INS-496 |
| chr19-46837726-INS-237 | chr19-46837726-INS-237 |
| chr19-46856240-INS-60 | chr19-46856240-INS-60 |
| chr19-46899801-INS-217 | chr19-46899801-INS-217 |
| chr19-46907821-INS-207 | chr19-46907821-INS-207 |
| chr19-46914593-INS-2758 | chr19-46914594-INS-2758 |
| chr19-46942034-INS-172 | chr19-46942034-INS-173 |
| chr19-46954723-INS-202 | chr19-46954723-INS-202 |
| chr19-46962754-INS-200 | chr19-46962754-INS-200 |
| chr19-46980636-INS-174 | chr19-46980636-INS-186 |
| chr19-46983005-INS-238 | chr19-46983005-INS-240 |
| chr19-46991433-INS-109 | chr19-46991448-INS-105 |
| chr19-47004977-INS-70 | chr19-47004977-INS-70 |
| chr19-47009970-INS-164 | chr19-47009954-INS-178 |
| chr19-47010532-INS-210 | chr19-47010539-INS-214 |
| chr19-47033861-INS-215 | chr19-47033861-INS-215 |
| chr19-47035716-INS-156 | chr19-47035716-INS-156 |
| chr19-47036688-INS-472 | chr19-47036688-INS-477 |
| chr19-47040989-INS-128 | chr19-47040989-INS-124 |
| chr19-47045747-INS-239 | chr19-47045747-INS-239 |
| chr19-47051366-INS-187 | chr19-47051366-INS-188 |
| chr19-47059265-INS-201 | chr19-47059265-INS-201 |
| chr19-47061884-INS-54 | chr19-47061885-INS-64 |
| chr19-47088115-INS-197 | chr19-47088115-INS-197 |
| chr19-47092775-INS-208 | chr19-47092775-INS-207 |
| chr19-47101259-INS-210 | chr19-47101259-INS-210 |
| chr19-47135625-INS-197 | chr19-47135642-INS-202 |
| chr19-47137766-INS-202 | chr19-47137766-INS-202 |
| chr19-47141295-INS-78 | chr19-47141295-INS-104 |
| chr19-47149179-INS-94 | chr19-47149182-INS-102 |
| chr19-47150556-INS-286 | chr19-47150556-INS-282 |
| chr19-47151346-INS-191 | chr19-47151347-INS-180 |
| chr19-47152734-INS-209 | chr19-47152734-INS-214 |
| chr19-47154805-INS-217 | chr19-47154805-INS-215 |
| chr19-47155585-INS-203 | chr19-47155585-INS-202 |
| chr19-47162657-INS-852 | chr19-47162673-INS-866 |
| chr19-47162784-INS-57 | chr19-47162759-INS-95 |

| | |
|---|---|
| chr19-47171047-INS-200 | chr19-47171047-INS-200 |
| chr19-47172240-INS-69 | chr19-47172240-INS-56 |
| chr19-47174682-INS-130 | chr19-47174678-INS-158 |
| chr19-47181610-INS-212 | chr19-47181610-INS-212 |
| chr19-47202881-INS-190 | chr19-47202881-INS-190 |
| chr19-47226276-INS-104 | chr19-47226282-INS-81 |
| chr19-47234465-INS-61 | chr19-47234465-INS-57 |
| chr19-47315753-INS-54 | chr19-47315753-INS-54 |
| chr19-47333601-INS-213 | chr19-47333601-INS-213 |
| chr19-47336969-INS-213 | chr19-47336969-INS-213 |
| chr19-47342203-INS-176 | chr19-47342203-INS-177 |
| chr19-47357863-INS-54 | chr19-47357863-INS-54 |
| chr19-47368947-INS-61 | chr19-47368965-INS-64 |
| chr19-47382445-INS-199 | chr19-47382445-INS-199 |
| chr19-47396911-INS-63 | chr19-47396911-INS-63 |
| chr19-47421521-INS-117 | chr19-47421523-INS-102 |
| chr19-47451085-INS-78 | chr19-47451067-INS-84 |
| chr19-47461010-INS-198 | chr19-47461010-INS-209 |
| chr19-47474020-INS-68 | chr19-47474023-INS-55 |
| chr19-47486502-INS-208 | chr19-47486502-INS-209 |
| chr19-47494975-INS-134 | chr19-47495019-INS-104 |
| chr19-47498417-INS-97 | chr19-47498417-INS-88 |
| chr19-47500298-INS-64 | chr19-47500298-INS-58 |
| chr19-47507068-INS-201 | chr19-47507068-INS-201 |
| chr19-47518981-INS-481 | chr19-47518981-INS-481 |
| chr19-47525794-INS-123 | chr19-47525794-INS-123 |
| chr19-47527862-INS-135 | chr19-47527863-INS-132 |
| chr19-47534188-INS-66 | chr19-47534188-INS-66 |
| chr19-47550170-INS-215 | chr19-47550170-INS-215 |
| chr19-47553572-INS-4549 | chr19-47553573-INS-4549 |
| chr19-47555190-INS-281 | chr19-47555190-INS-281 |
| chr19-47558293-INS-201 | chr19-47558293-INS-201 |
| chr19-47577146-INS-62 | chr19-47577146-INS-64 |
| chr19-47584257-INS-203 | chr19-47584257-INS-202 |
| chr19-47604204-INS-233 | chr19-47604204-INS-231 |
| chr19-47612078-INS-202 | chr19-47612078-INS-207 |
| chr19-47623030-INS-108 | chr19-47623030-INS-108 |
| chr19-47643518-INS-428 | chr19-47643515-INS-440 |
| chr19-47660733-INS-56 | chr19-47660733-INS-56 |
| chr19-47669314-INS-140 | chr19-47669314-INS-130 |
| chr19-47688996-INS-157 | chr19-47688996-INS-146 |
| chr19-47695768-INS-59 | chr19-47695768-INS-59 |
| chr19-47720991-INS-193 | chr19-47720991-INS-194 |
| chr19-47728965-INS-181 | chr19-47728965-INS-181 |
| chr19-47741927-INS-206 | chr19-47741927-INS-206 |
| chr19-47749442-INS-193 | chr19-47749443-INS-193 |
| chr19-47752960-INS-644 | chr19-47752962-INS-627 |
| chr19-47758322-INS-234 | chr19-47758322-INS-238 |
| chr19-47760570-INS-343 | chr19-47760571-INS-316 |
| chr19-47782737-INS-60 | chr19-47782740-INS-55 |
| chr19-47807844-INS-206 | chr19-47807844-INS-206 |
| chr19-47837838-INS-411 | chr19-47837838-INS-411 |
| chr19-47849770-INS-53 | chr19-47849771-INS-53 |
| chr19-47882139-INS-2871 | chr19-47882140-INS-2872 |
| chr19-47910670-INS-62 | chr19-47910654-INS-74 |
| chr19-47919792-INS-206 | chr19-47919792-INS-207 |
| chr19-47968364-INS-63 | chr19-47968365-INS-58 |
| chr19-47979004-INS-92 | chr19-47979004-INS-97 |
| chr19-47990014-INS-172 | chr19-47990014-INS-171 |
| chr19-47993811-INS-72 | chr19-47993811-INS-56 |
| chr19-47999911-INS-185 | chr19-47999941-INS-146 |
| chr19-48003337-INS-60 | chr19-48003333-INS-52 |
| chr19-48006802-INS-52 | chr19-48006801-INS-60 |
| chr19-48011910-INS-414 | chr19-48011910-INS-462 |
| chr19-48021916-INS-228 | chr19-48021916-INS-228 |
| chr19-48034001-INS-81 | chr19-48034001-INS-81 |
| chr19-48040032-INS-96 | chr19-48040062-INS-96 |
| chr19-48084583-INS-379 | chr19-48084584-INS-379 |
| chr19-48099461-INS-68 | chr19-48099440-INS-64 |
| chr19-48099974-INS-293 | chr19-48099974-INS-297 |
| chr19-48105651-INS-94 | chr19-48105651-INS-134 |
| chr19-48107074-INS-1316 | chr19-48107078-INS-1320 |
| chr19-48122168-INS-1296 | chr19-48122169-INS-1296 |
| chr19-48122229-INS-85 | chr19-48122229-INS-83 |
| chr19-48137859-INS-191 | chr19-48137859-INS-191 |
| chr19-48150253-INS-64 | chr19-48150253-INS-64 |
| chr19-48166448-INS-68 | chr19-48166448-INS-68 |

| | |
|---|---|
| chr19-48208563-INS-63 | chr19-48208563-INS-69 |
| chr19-48212327-INS-77 | chr19-48212335-INS-81 |
| chr19-48213702-INS-76 | chr19-48213700-INS-78 |
| chr19-48242333-INS-123 | chr19-48242333-INS-123 |
| chr19-48260108-INS-62 | chr19-48260101-INS-69 |
| chr19-48263718-INS-94 | chr19-48263709-INS-75 |
| chr19-48267058-INS-279 | chr19-48267058-INS-275 |
| chr19-48278621-INS-156 | chr19-48278625-INS-157 |
| chr19-48300172-INS-834 | chr19-48300172-INS-834 |
| chr19-48301298-INS-185 | chr19-48301298-INS-186 |
| chr19-48335670-INS-146 | chr19-48335670-INS-146 |
| chr19-48338101-INS-162 | chr19-48338102-INS-165 |
| chr19-48416491-INS-119 | chr19-48416491-INS-114 |
| chr19-48420028-INS-80 | chr19-48420023-INS-72 |
| chr19-48424094-INS-912 | chr19-48424094-INS-915 |
| chr19-48427377-INS-56 | chr19-48427377-INS-50 |
| chr19-48431485-INS-1216 | chr19-48431486-INS-1217 |
| chr19-48465446-INS-81 | chr19-48465446-INS-83 |
| chr19-48468262-INS-119 | chr19-48468262-INS-119 |
| chr19-48507552-INS-66 | chr19-48507555-INS-61 |
| chr19-48543185-INS-58 | chr19-48543185-INS-58 |
| chr19-48546153-INS-113 | chr19-48546153-INS-113 |
| chr19-48611977-INS-579 | chr19-48611977-INS-580 |
| chr19-48618831-INS-56 | chr19-48618831-INS-56 |
| chr19-48682977-INS-88 | chr19-48682983-INS-56 |
| chr19-48692193-INS-6600 | chr19-48692194-INS-6602 |
| chr19-48722816-INS-66 | chr19-48722799-INS-67 |
| chr19-48737142-INS-144 | chr19-48737142-INS-144 |
| chr19-48752358-INS-78 | chr19-48752358-INS-54 |
| chr19-48877201-INS-70 | chr19-48877201-INS-80 |
| chr19-48939276-INS-5484 | chr19-48939277-INS-5484 |
| chr19-48955286-INS-3219 | chr19-48955287-INS-3227 |
| chr19-48986152-INS-116 | chr19-48986152-INS-98 |
| chr19-48992611-INS-60 | chr19-48992611-INS-70 |
| chr19-48998896-INS-147 | chr19-48998896-INS-147 |
| chr19-49015895-INS-71 | chr19-49015904-INS-66 |
| chr19-49018855-INS-65 | chr19-49018855-INS-71 |
| chr19-49037114-INS-245 | chr19-49037114-INS-246 |
| chr19-49072258-INS-105 | chr19-49072253-INS-68 |
| chr19-49079883-INS-74 | chr19-49079883-INS-74 |
| chr19-49172647-INS-605 | chr19-49172647-INS-607 |
| chr19-49176914-INS-112 | chr19-49176914-INS-110 |
| chr19-49197906-INS-86 | chr19-49197908-INS-90 |
| chr19-49226079-INS-62 | chr19-49226079-INS-72 |
| chr19-49255386-INS-2957 | chr19-49255387-INS-2957 |
| chr19-49256104-INS-104 | chr19-49256104-INS-84 |
| chr19-49277347-INS-66 | chr19-49277347-INS-68 |
| chr19-49294453-INS-68 | chr19-49294453-INS-68 |
| chr19-49296759-INS-66 | chr19-49296759-INS-71 |
| chr19-49302986-INS-307 | chr19-49302986-INS-306 |
| chr19-49309196-INS-78 | chr19-49309196-INS-78 |
| chr19-49313002-INS-106 | chr19-49313002-INS-106 |
| chr19-49339034-INS-327 | chr19-49339034-INS-327 |
| chr19-49409615-INS-708 | chr19-49409602-INS-713 |
| chr19-49416189-INS-52 | chr19-49416189-INS-56 |
| chr19-49466150-INS-70 | chr19-49466150-INS-112 |
| chr19-49472959-INS-313 | chr19-49472956-INS-316 |
| chr19-49545355-INS-73 | chr19-49545355-INS-73 |
| chr19-49548258-INS-431 | chr19-49548256-INS-429 |
| chr19-49550469-INS-54 | chr19-49550469-INS-56 |
| chr19-49617311-INS-63 | chr19-49617311-INS-83 |
| chr19-49621660-INS-97 | chr19-49621660-INS-97 |
| chr19-49637258-INS-529 | chr19-49637258-INS-529 |
| chr19-49661703-INS-6335 | chr19-49661704-INS-6338 |
| chr19-49670622-INS-967 | chr19-49670622-INS-967 |
| chr19-49676304-INS-440 | chr19-49676304-INS-440 |
| chr19-49680573-INS-172 | chr19-49680573-INS-172 |
| chr19-49690109-INS-84 | chr19-49690107-INS-51 |
| chr19-49708775-INS-205 | chr19-49708775-INS-220 |
| chr19-49731327-INS-91 | chr19-49731327-INS-91 |
| chr19-49778043-INS-169 | chr19-49778044-INS-169 |
| chr19-49794810-INS-114 | chr19-49794810-INS-84 |
| chr19-49799925-INS-1866 | chr19-49799926-INS-1865 |
| chr19-49816556-INS-1183 | chr19-49816557-INS-1220 |
| chr19-49816577-INS-168 | chr19-49816577-INS-173 |
| chr19-49829468-INS-52 | chr19-49829468-INS-50 |
| chr19-49848051-INS-6150 | chr19-49848052-INS-6144 |

| | |
|---|---|
| chr19-49849169-INS-1357 | chr19-49849170-INS-1359 |
| chr19-49859314-INS-2187 | chr19-49859315-INS-2187 |
| chr19-49871572-INS-261 | chr19-49871572-INS-261 |
| chr19-49873823-INS-4693 | chr19-49873824-INS-4693 |
| chr19-49887556-INS-85 | chr19-49887556-INS-81 |
| chr19-49927065-INS-53 | chr19-49927071-INS-88 |
| chr19-49940402-INS-115 | chr19-49940402-INS-115 |
| chr19-49946206-INS-82 | chr19-49946206-INS-102 |
| chr19-49967444-INS-6557 | chr19-49967445-INS-6557 |
| chr19-50005028-INS-143 | chr19-50005028-INS-143 |
| chr19-50007031-INS-2884 | chr19-50007032-INS-2886 |
| chr19-50010204-INS-219 | chr19-50010204-INS-219 |
| chr19-50042917-INS-55 | chr19-50042902-INS-58 |
| chr19-50051985-INS-4640 | chr19-50051986-INS-4637 |
| chr19-50077172-INS-55 | chr19-50077172-INS-55 |
| chr19-50080412-INS-52 | chr19-50080412-INS-51 |
| chr19-50110528-INS-51 | chr19-50110528-INS-51 |
| chr19-50113036-INS-406 | chr19-50113036-INS-406 |
| chr19-50171705-INS-165 | chr19-50171705-INS-171 |
| chr19-50179835-INS-105 | chr19-50179835-INS-124 |
| chr19-50209764-INS-2529 | chr19-50209765-INS-2529 |
| chr19-50221322-INS-64 | chr19-50221322-INS-66 |
| chr19-50231234-INS-481 | chr19-50231234-INS-481 |
| chr19-50237087-INS-6606 | chr19-50237088-INS-6606 |
| chr19-50268794-INS-6457 | chr19-50268795-INS-6457 |
| chr19-50307595-INS-607 | chr19-50307610-INS-558 |
| chr19-50323108-INS-64 | chr19-50323114-INS-100 |
| chr19-50335492-INS-70 | chr19-50335504-INS-51 |
| chr19-50349415-INS-54 | chr19-50349415-INS-52 |
| chr19-50357602-INS-56 | chr19-50357600-INS-56 |
| chr19-50386769-INS-3664 | chr19-50386770-INS-3665 |
| chr19-50402161-INS-791 | chr19-50402161-INS-791 |
| chr19-50414901-INS-212 | chr19-50414901-INS-212 |
| chr19-50448036-INS-113 | chr19-50448071-INS-112 |
| chr19-50469512-INS-237 | chr19-50469512-INS-285 |
| chr19-50511117-INS-56 | chr19-50511117-INS-56 |
| chr19-50555850-INS-52 | chr19-50555850-INS-50 |
| chr19-50595789-INS-368 | chr19-50595789-INS-376 |
| chr19-50598098-INS-69 | chr19-50598098-INS-70 |
| chr19-50656568-INS-463 | chr19-50656568-INS-463 |
| chr19-50658791-INS-207 | chr19-50658804-INS-188 |
| chr19-50692947-INS-3113 | chr19-50692948-INS-3113 |
| chr19-50716474-INS-66 | chr19-50716474-INS-84 |
| chr19-50718624-INS-56 | chr19-50718624-INS-58 |
| chr19-50735637-INS-74 | chr19-50735635-INS-75 |
| chr19-50750075-INS-116 | chr19-50750090-INS-68 |
| chr19-50772040-INS-70 | chr19-50772038-INS-56 |
| chr19-50780027-INS-91 | chr19-50780027-INS-78 |
| chr19-50797705-INS-69 | chr19-50797705-INS-69 |
| chr19-50804743-INS-270 | chr19-50804744-INS-271 |
| chr19-50806952-INS-69 | chr19-50806952-INS-68 |
| chr19-50811789-INS-128 | chr19-50811789-INS-131 |
| chr19-50816116-INS-339 | chr19-50816116-INS-339 |
| chr19-50833059-INS-116 | chr19-50833085-INS-72 |
| chr19-50881326-INS-60 | chr19-50881326-INS-70 |
| chr19-50895239-INS-97 | chr19-50895230-INS-108 |
| chr19-50900059-INS-540 | chr19-50900070-INS-551 |
| chr19-50913291-INS-200 | chr19-50913291-INS-200 |
| chr19-50937113-INS-623 | chr19-50937114-INS-631 |
| chr19-50940741-INS-52 | chr19-50940741-INS-74 |
| chr19-50947118-INS-99 | chr19-50947114-INS-63 |
| chr19-50957012-INS-1380 | chr19-50957013-INS-1379 |
| chr19-50962194-INS-66 | chr19-50962194-INS-66 |
| chr19-50971438-INS-52 | chr19-50971438-INS-50 |
| chr19-50981681-INS-105 | chr19-50981681-INS-105 |
| chr19-51001802-INS-90 | chr19-51001788-INS-80 |
| chr19-51009130-INS-113 | chr19-51009130-INS-113 |
| chr19-51011538-INS-89 | chr19-51011538-INS-80 |
| chr19-51029690-INS-810 | chr19-51029690-INS-811 |
| chr19-51031433-INS-126 | chr19-51031433-INS-126 |
| chr19-51036859-INS-94 | chr19-51036859-INS-70 |
| chr19-51063848-INS-79 | chr19-51063848-INS-81 |
| chr19-51072859-INS-184 | chr19-51072859-INS-184 |
| chr19-51088955-INS-73 | chr19-51088955-INS-74 |
| chr19-51120533-INS-78 | chr19-51120535-INS-58 |
| chr19-51137378-INS-75 | chr19-51137378-INS-75 |
| chr19-51160657-INS-168 | chr19-51160657-INS-168 |

| | |
|---|---|
| chr19-51194100-INS-52 | chr19-51194100-INS-54 |
| chr19-51201850-INS-2772 | chr19-51201851-INS-2770 |
| chr19-51211240-INS-98 | chr19-51211240-INS-110 |
| chr19-51221450-INS-121 | chr19-51221450-INS-77 |
| chr19-51241386-INS-486 | chr19-51241386-INS-486 |
| chr19-51242106-INS-6400 | chr19-51242107-INS-6400 |
| chr19-51254862-INS-171 | chr19-51254900-INS-164 |
| chr19-51293375-INS-212 | chr19-51293375-INS-214 |
| chr19-51349272-INS-63 | chr19-51349272-INS-51 |
| chr19-51376329-INS-382 | chr19-51376329-INS-382 |
| chr19-51392621-INS-684 | chr19-51392621-INS-687 |
| chr19-51430196-INS-276 | chr19-51430196-INS-286 |
| chr19-51467840-INS-60 | chr19-51467840-INS-108 |
| chr19-51512260-INS-1541 | chr19-51512261-INS-1541 |
| chr19-51521810-INS-191 | chr19-51521810-INS-190 |
| chr19-51522998-INS-63 | chr19-51522979-INS-50 |
| chr19-51548656-INS-976 | chr19-51548656-INS-976 |
| chr19-51561242-INS-116 | chr19-51561242-INS-116 |
| chr19-51562655-INS-64 | chr19-51562664-INS-54 |
| chr19-51567164-INS-76 | chr19-51567164-INS-62 |
| chr19-51575045-INS-124 | chr19-51575045-INS-124 |
| chr19-51593783-INS-58 | chr19-51593783-INS-60 |
| chr19-51600669-INS-68 | chr19-51600673-INS-50 |
| chr19-51610279-INS-111 | chr19-51610284-INS-71 |
| chr19-51617072-INS-73 | chr19-51617056-INS-66 |
| chr19-51635696-INS-96 | chr19-51635696-INS-118 |
| chr19-51653146-INS-1681 | chr19-51653147-INS-1680 |
| chr19-51682117-INS-72 | chr19-51682085-INS-94 |
| chr19-51692201-INS-584 | chr19-51692201-INS-584 |
| chr19-51707341-INS-79 | chr19-51707341-INS-118 |
| chr19-51719095-INS-58 | chr19-51719095-INS-58 |
| chr19-51723508-INS-401 | chr19-51723508-INS-401 |
| chr19-51735565-INS-466 | chr19-51735565-INS-467 |
| chr19-51736438-INS-92 | chr19-51736438-INS-94 |
| chr19-51745079-INS-52 | chr19-51745079-INS-54 |
| chr19-51760040-INS-873 | chr19-51760040-INS-872 |
| chr19-51772882-INS-52 | chr19-51772927-INS-52 |
| chr19-51777241-INS-66 | chr19-51777260-INS-69 |
| chr19-51781075-INS-61 | chr19-51781075-INS-61 |
| chr19-51790701-INS-109 | chr19-51790707-INS-69 |
| chr19-51821550-INS-183 | chr19-51821550-INS-183 |
| chr19-51828652-INS-7105 | chr19-51828653-INS-7105 |
| chr19-51838810-INS-59 | chr19-51838810-INS-61 |
| chr19-51840860-INS-170 | chr19-51840873-INS-166 |
| chr19-51860210-INS-6175 | chr19-51860211-INS-6175 |
| chr19-51865202-INS-64 | chr19-51865203-INS-64 |
| chr19-51868812-INS-358 | chr19-51868812-INS-358 |
| chr19-51900520-INS-69 | chr19-51900498-INS-64 |
| chr19-51912446-INS-346 | chr19-51912446-INS-347 |
| chr19-51945082-INS-160 | chr19-51945082-INS-160 |
| chr19-51984259-INS-438 | chr19-51984259-INS-438 |
| chr19-52015501-INS-64 | chr19-52015501-INS-64 |
| chr19-52028379-INS-92 | chr19-52028379-INS-92 |
| chr19-52034134-INS-55 | chr19-52034134-INS-55 |
| chr19-52038836-INS-81 | chr19-52038851-INS-69 |
| chr19-52043121-INS-506 | chr19-52043121-INS-491 |
| chr19-52046313-INS-63 | chr19-52046313-INS-63 |
| chr19-52066136-INS-51 | chr19-52066153-INS-62 |
| chr19-52076938-INS-54 | chr19-52076938-INS-56 |
| chr19-52086877-INS-230 | chr19-52086883-INS-237 |
| chr19-52152376-INS-76 | chr19-52152388-INS-72 |
| chr19-52155205-INS-195 | chr19-52155205-INS-195 |
| chr19-52197740-INS-407 | chr19-52197740-INS-406 |
| chr19-52202221-INS-637 | chr19-52202221-INS-637 |
| chr19-52206232-INS-93 | chr19-52206232-INS-93 |
| chr19-52223448-INS-187 | chr19-52223448-INS-187 |
| chr19-52246030-INS-1330 | chr19-52246031-INS-1330 |
| chr19-52270744-INS-58 | chr19-52270744-INS-60 |
| chr19-52309222-INS-114 | chr19-52309222-INS-94 |
| chr19-52315748-INS-421 | chr19-52315748-INS-385 |
| chr19-52367448-INS-60 | chr19-52367446-INS-64 |
| chr19-52411666-INS-58 | chr19-52411666-INS-81 |
| chr19-52447183-INS-145 | chr19-52447183-INS-147 |
| chr19-52451055-INS-106 | chr19-52451055-INS-106 |
| chr19-52462285-INS-70 | chr19-52462285-INS-70 |
| chr19-52470387-INS-499 | chr19-52470387-INS-499 |
| chr19-52496215-INS-162 | chr19-52496215-INS-162 |

| | |
|---|---|
| chr19-52502275-INS-91 | chr19-52502278-INS-92 |
| chr19-52503270-INS-394 | chr19-52503270-INS-394 |
| chr19-52505133-INS-308 | chr19-52505134-INS-333 |
| chr19-52513590-INS-61 | chr19-52513552-INS-71 |
| chr19-52518912-INS-637 | chr19-52518912-INS-637 |
| chr19-52539396-INS-73 | chr19-52539395-INS-74 |
| chr19-52541076-INS-94 | chr19-52541076-INS-94 |
| chr19-52549917-INS-53 | chr19-52549917-INS-53 |
| chr19-52572288-INS-2913 | chr19-52572289-INS-2913 |
| chr19-52585172-INS-1955 | chr19-52585173-INS-1955 |
| chr19-52623025-INS-441 | chr19-52623025-INS-441 |
| chr19-52677710-INS-124 | chr19-52677710-INS-128 |
| chr19-52680707-INS-80 | chr19-52680707-INS-80 |
| chr19-52685791-INS-51 | chr19-52685791-INS-51 |
| chr19-52694764-INS-175 | chr19-52694778-INS-195 |
| chr19-52705728-INS-309 | chr19-52705728-INS-309 |
| chr19-52748177-INS-429 | chr19-52748177-INS-430 |
| chr19-52758600-INS-143 | chr19-52758600-INS-143 |
| chr19-52761037-INS-2784 | chr19-52761038-INS-2784 |
| chr19-52767055-INS-202 | chr19-52767055-INS-202 |
| chr19-52781561-INS-173 | chr19-52781561-INS-173 |
| chr19-52823375-INS-91 | chr19-52823379-INS-87 |
| chr19-52866317-INS-53 | chr19-52866287-INS-53 |
| chr19-52878049-INS-4353 | chr19-52878050-INS-4353 |
| chr19-52883188-INS-63 | chr19-52883188-INS-63 |
| chr19-52893427-INS-183 | chr19-52893410-INS-205 |
| chr19-52899423-INS-325 | chr19-52899423-INS-325 |
| chr19-52909103-INS-76 | chr19-52909106-INS-85 |
| chr19-52920575-INS-1315 | chr19-52920576-INS-1315 |
| chr19-52943226-INS-160 | chr19-52943226-INS-160 |
| chr19-52946412-INS-4473 | chr19-52946413-INS-4473 |
| chr19-52972764-INS-586 | chr19-52972764-INS-570 |
| chr19-53012430-INS-180 | chr19-53012430-INS-180 |
| chr19-53057243-INS-412 | chr19-53057243-INS-412 |
| chr19-53065638-INS-60 | chr19-53065638-INS-60 |
| chr19-53082034-INS-72 | chr19-53082034-INS-72 |
| chr19-53085220-INS-1891 | chr19-53085221-INS-1891 |
| chr19-53094569-INS-177 | chr19-53094556-INS-173 |
| chr19-53115428-INS-236 | chr19-53115430-INS-228 |
| chr19-53127845-INS-208 | chr19-53127845-INS-206 |
| chr19-53139894-INS-93 | chr19-53139894-INS-93 |
| chr19-53155232-INS-228 | chr19-53155232-INS-228 |
| chr19-53165314-INS-196 | chr19-53165315-INS-186 |
| chr19-53171097-INS-476 | chr19-53171097-INS-477 |
| chr19-53185023-INS-60 | chr19-53185023-INS-60 |
| chr19-53196837-INS-52 | chr19-53196837-INS-52 |
| chr19-53216987-INS-100 | chr19-53216987-INS-100 |
| chr19-53239265-INS-56 | chr19-53239265-INS-54 |
| chr19-53254758-INS-60 | chr19-53254748-INS-70 |
| chr19-53256662-INS-109 | chr19-53256654-INS-134 |
| chr19-53269620-INS-104 | chr19-53269620-INS-105 |
| chr19-53280895-INS-6238 | chr19-53280896-INS-6239 |
| chr19-53295120-INS-66 | chr19-53295119-INS-79 |
| chr19-53324860-INS-218 | chr19-53324860-INS-212 |
| chr19-53350901-INS-51 | chr19-53350882-INS-57 |
| chr19-53359356-INS-58 | chr19-53359356-INS-54 |
| chr19-53366785-INS-7090 | chr19-53366786-INS-7090 |
| chr19-53387385-INS-52 | chr19-53387384-INS-52 |
| chr19-53414551-INS-104 | chr19-53414543-INS-105 |
| chr19-53433982-INS-7162 | chr19-53433983-INS-7162 |
| chr19-53455848-INS-4793 | chr19-53455849-INS-4793 |
| chr19-53521944-INS-205 | chr19-53521944-INS-207 |
| chr19-53535288-INS-90 | chr19-53535267-INS-120 |
| chr19-53538572-INS-106 | chr19-53538572-INS-84 |
| chr19-53545081-INS-65 | chr19-53545081-INS-65 |
| chr19-53555582-INS-212 | chr19-53555582-INS-208 |
| chr19-53561088-INS-188 | chr19-53561089-INS-185 |
| chr19-53564396-INS-69 | chr19-53564395-INS-69 |
| chr19-53570317-INS-214 | chr19-53570317-INS-219 |
| chr19-53571551-INS-154 | chr19-53571551-INS-167 |
| chr19-53585068-INS-76 | chr19-53585069-INS-72 |
| chr19-53599990-INS-209 | chr19-53599990-INS-210 |
| chr19-53639183-INS-70 | chr19-53639183-INS-61 |
| chr19-53640198-INS-81 | chr19-53640199-INS-108 |
| chr19-53640810-INS-3781 | chr19-53640811-INS-3782 |
| chr19-53645820-INS-92 | chr19-53645828-INS-112 |
| chr19-53656158-INS-352 | chr19-53656159-INS-370 |

| | |
|---|---|
| chr19-53656981-INS-69 | chr19-53656981-INS-69 |
| chr19-53672973-INS-213 | chr19-53672974-INS-213 |
| chr19-53675690-INS-6612 | chr19-53675691-INS-6614 |
| chr19-53687903-INS-90 | chr19-53687903-INS-90 |
| chr19-53726539-INS-535 | chr19-53726539-INS-537 |
| chr19-53771946-INS-54 | chr19-53771946-INS-66 |
| chr19-53795131-INS-80 | chr19-53795131-INS-80 |
| chr19-53874958-INS-72 | chr19-53874958-INS-72 |
| chr19-53881803-INS-54 | chr19-53881803-INS-54 |
| chr19-53895136-INS-697 | chr19-53895136-INS-702 |
| chr19-53910609-INS-89 | chr19-53910609-INS-87 |
| chr19-53975420-INS-54 | chr19-53975423-INS-54 |
| chr19-54018424-INS-54 | chr19-54018424-INS-58 |
| chr19-54026907-INS-214 | chr19-54026909-INS-213 |
| chr19-54064883-INS-64 | chr19-54064883-INS-64 |
| chr19-54092166-INS-176 | chr19-54092166-INS-183 |
| chr19-54093687-INS-82 | chr19-54093683-INS-96 |
| chr19-54095438-INS-442 | chr19-54095438-INS-434 |
| chr19-54104630-INS-346 | chr19-54104630-INS-346 |
| chr19-54167403-INS-290 | chr19-54167403-INS-290 |
| chr19-54178333-INS-66 | chr19-54178333-INS-66 |
| chr19-54182646-INS-60 | chr19-54182646-INS-60 |
| chr19-54186642-INS-81 | chr19-54186646-INS-89 |
| chr19-54192127-INS-654 | chr19-54192127-INS-653 |
| chr19-54197404-INS-242 | chr19-54197404-INS-227 |
| chr19-54201209-INS-78 | chr19-54201209-INS-105 |
| chr19-54212134-INS-671 | chr19-54212134-INS-669 |
| chr19-54215859-INS-70 | chr19-54215859-INS-72 |
| chr19-54222189-INS-196 | chr19-54222189-INS-196 |
| chr19-54241864-INS-58 | chr19-54241864-INS-58 |
| chr19-54249733-INS-55 | chr19-54249733-INS-55 |
| chr19-54252593-INS-420 | chr19-54252593-INS-420 |
| chr19-54283413-INS-64 | chr19-54283413-INS-62 |
| chr19-54285852-INS-65 | chr19-54285852-INS-83 |
| chr19-54288586-INS-80 | chr19-54288586-INS-80 |
| chr19-54312654-INS-66 | chr19-54312654-INS-60 |
| chr19-54320658-INS-120 | chr19-54320678-INS-134 |
| chr19-54359320-INS-151 | chr19-54359320-INS-151 |
| chr19-54366258-INS-469 | chr19-54366258-INS-469 |
| chr19-54391788-INS-96 | chr19-54391788-INS-96 |
| chr19-54429497-INS-87 | chr19-54429497-INS-87 |
| chr19-54437209-INS-7097 | chr19-54437210-INS-7097 |
| chr19-54443824-INS-191 | chr19-54443824-INS-191 |
| chr19-54449628-INS-2295 | chr19-54449629-INS-2296 |
| chr19-54452068-INS-498 | chr19-54452068-INS-498 |
| chr19-54457655-INS-56 | chr19-54457655-INS-56 |
| chr19-54468735-INS-54 | chr19-54468735-INS-56 |
| chr19-54509317-INS-85 | chr19-54509317-INS-69 |
| chr19-54585224-INS-90 | chr19-54585224-INS-90 |
| chr19-54663009-INS-1203 | chr19-54663010-INS-1203 |
| chr19-54685473-INS-630 | chr19-54685473-INS-625 |
| chr19-54802888-INS-146 | chr19-54802888-INS-101 |
| chr19-54806774-INS-915 | chr19-54806791-INS-920 |
| chr19-54809128-INS-1864 | chr19-54809129-INS-1864 |
| chr19-54835237-INS-55 | chr19-54835237-INS-55 |
| chr19-54849364-INS-54 | chr19-54849364-INS-66 |
| chr19-54889166-INS-847 | chr19-54889178-INS-861 |
| chr19-54892042-INS-90 | chr19-54892042-INS-90 |
| chr19-54895343-INS-601 | chr19-54895343-INS-597 |
| chr19-54916995-INS-403 | chr19-54916995-INS-403 |
| chr19-54923496-INS-615 | chr19-54923496-INS-615 |
| chr19-54933757-INS-70 | chr19-54933757-INS-61 |
| chr19-54939499-INS-225 | chr19-54939499-INS-223 |
| chr19-54944001-INS-132 | chr19-54944001-INS-132 |
| chr19-54945568-INS-7062 | chr19-54945569-INS-7063 |
| chr19-54946558-INS-94 | chr19-54946549-INS-116 |
| chr19-54948633-INS-56 | chr19-54948633-INS-56 |
| chr19-54951042-INS-1103 | chr19-54951040-INS-1104 |
| chr19-54956968-INS-296 | chr19-54956968-INS-292 |
| chr19-54959855-INS-78 | chr19-54959858-INS-57 |
| chr19-54970007-INS-73 | chr19-54970007-INS-73 |
| chr19-55017323-INS-106 | chr19-55017317-INS-90 |
| chr19-55029437-INS-87 | chr19-55029422-INS-94 |
| chr19-55031303-INS-105 | chr19-55031303-INS-105 |
| chr19-55050327-INS-101 | chr19-55050326-INS-149 |
| chr19-55053376-INS-631 | chr19-55053376-INS-638 |
| chr19-55076681-INS-101 | chr19-55076681-INS-107 |

| | |
|---|---|
| chr19-55083420-INS-901 | chr19-55083420-INS-901 |
| chr19-55100141-INS-83 | chr19-55100141-INS-82 |
| chr19-55101648-INS-322 | chr19-55101648-INS-322 |
| chr19-55158091-INS-592 | chr19-55158091-INS-592 |
| chr19-55230129-INS-854 | chr19-55230129-INS-855 |
| chr19-55252122-INS-51 | chr19-55252122-INS-55 |
| chr19-55266430-INS-64 | chr19-55266454-INS-64 |
| chr19-55267505-INS-200 | chr19-55267505-INS-200 |
| chr19-55313257-INS-182 | chr19-55313257-INS-182 |
| chr19-55430534-INS-196 | chr19-55430534-INS-195 |
| chr19-55484099-INS-82 | chr19-55484099-INS-82 |
| chr19-55516892-INS-56 | chr19-55516892-INS-56 |
| chr19-55521477-INS-53 | chr19-55521447-INS-64 |
| chr19-55540574-INS-102 | chr19-55540574-INS-102 |
| chr19-55553806-INS-346 | chr19-55553806-INS-347 |
| chr19-55596144-INS-52 | chr19-55596144-INS-60 |
| chr19-55602055-INS-89 | chr19-55602082-INS-59 |
| chr19-55612088-INS-96 | chr19-55612130-INS-54 |
| chr19-55650261-INS-647 | chr19-55650261-INS-602 |
| chr19-55658380-INS-297 | chr19-55658374-INS-258 |
| chr19-55685154-INS-56 | chr19-55685154-INS-68 |
| chr19-55704484-INS-58 | chr19-55704484-INS-62 |
| chr19-55714539-INS-331 | chr19-55714539-INS-331 |
| chr19-55715973-INS-64 | chr19-55715973-INS-52 |
| chr19-55719882-INS-134 | chr19-55719882-INS-134 |
| chr19-55739834-INS-56 | chr19-55739834-INS-56 |
| chr19-55749123-INS-197 | chr19-55749123-INS-197 |
| chr19-55759023-INS-288 | chr19-55759071-INS-240 |
| chr19-55759440-INS-431 | chr19-55759441-INS-431 |
| chr19-55765880-INS-70 | chr19-55765880-INS-55 |
| chr19-55799844-INS-122 | chr19-55799850-INS-140 |
| chr19-55811584-INS-53 | chr19-55811584-INS-55 |
| chr19-55820002-INS-160 | chr19-55820002-INS-173 |
| chr19-55822962-INS-184 | chr19-55822962-INS-134 |
| chr19-55824375-INS-313 | chr19-55824375-INS-333 |
| chr19-55825581-INS-68 | chr19-55825581-INS-68 |
| chr19-55847489-INS-181 | chr19-55847489-INS-175 |
| chr19-55922401-INS-482 | chr19-55922401-INS-482 |
| chr19-55949878-INS-447 | chr19-55949868-INS-427 |
| chr19-55953162-INS-168 | chr19-55953162-INS-168 |
| chr19-55972434-INS-56 | chr19-55972434-INS-56 |
| chr19-56014225-INS-79 | chr19-56014224-INS-78 |
| chr19-56034646-INS-108 | chr19-56034656-INS-138 |
| chr19-56038115-INS-366 | chr19-56038115-INS-365 |
| chr19-56039782-INS-258 | chr19-56039783-INS-231 |
| chr19-56056438-INS-60 | chr19-56056438-INS-60 |
| chr19-56094564-INS-62 | chr19-56094561-INS-62 |
| chr19-56100918-INS-66 | chr19-56100918-INS-62 |
| chr19-56102843-INS-57 | chr19-56102848-INS-74 |
| chr19-56126321-INS-101 | chr19-56126321-INS-101 |
| chr19-56141764-INS-51 | chr19-56141746-INS-55 |
| chr19-56184014-INS-83 | chr19-56184014-INS-83 |
| chr19-56201266-INS-454 | chr19-56201266-INS-454 |
| chr19-56235235-INS-57 | chr19-56235239-INS-55 |
| chr19-56247589-INS-55 | chr19-56247589-INS-57 |
| chr19-56265639-INS-175 | chr19-56265638-INS-174 |
| chr19-56278948-INS-52 | chr19-56278948-INS-52 |
| chr19-56281412-INS-340 | chr19-56281412-INS-340 |
| chr19-56317323-INS-775 | chr19-56317323-INS-779 |
| chr19-56351686-INS-69 | chr19-56351686-INS-81 |
| chr19-56353598-INS-182 | chr19-56353598-INS-186 |
| chr19-56363207-INS-51 | chr19-56363207-INS-50 |
| chr19-56367956-INS-144 | chr19-56367956-INS-120 |
| chr19-56372686-INS-83 | chr19-56372686-INS-83 |
| chr19-56374912-INS-99 | chr19-56374912-INS-97 |
| chr19-56378858-INS-65 | chr19-56378857-INS-71 |
| chr19-56382920-INS-55 | chr19-56382920-INS-55 |
| chr19-56412970-INS-811 | chr19-56412970-INS-785 |
| chr19-56415923-INS-56 | chr19-56415923-INS-56 |
| chr19-56417615-INS-202 | chr19-56417615-INS-202 |
| chr19-56429249-INS-517 | chr19-56429274-INS-518 |
| chr19-56431159-INS-120 | chr19-56431159-INS-120 |
| chr19-56452516-INS-829 | chr19-56452517-INS-834 |
| chr19-56453599-INS-201 | chr19-56453603-INS-205 |
| chr19-56464382-INS-62 | chr19-56464382-INS-62 |
| chr19-56480063-INS-375 | chr19-56480063-INS-329 |
| chr19-56487086-INS-1185 | chr19-56487087-INS-1185 |

| | |
|---|---|
| chr19-56490609-INS-438 | chr19-56490610-INS-438 |
| chr19-56499760-INS-5552 | chr19-56499761-INS-5552 |
| chr19-56501611-INS-117 | chr19-56501611-INS-117 |
| chr19-56512464-INS-625 | chr19-56512464-INS-626 |
| chr19-56554463-INS-200 | chr19-56554463-INS-200 |
| chr19-56561285-INS-106 | chr19-56561285-INS-72 |
| chr19-56573124-INS-68 | chr19-56573124-INS-70 |
| chr19-56603807-INS-108 | chr19-56603807-INS-108 |
| chr19-56609259-INS-69 | chr19-56609259-INS-69 |
| chr19-56615773-INS-64 | chr19-56615773-INS-64 |
| chr19-56622529-INS-51 | chr19-56622529-INS-51 |
| chr19-56648973-INS-57 | chr19-56648973-INS-57 |
| chr19-56654782-INS-84 | chr19-56654782-INS-84 |
| chr19-56684308-INS-83 | chr19-56684327-INS-83 |
| chr19-56734752-INS-77 | chr19-56734752-INS-83 |
| chr19-56755267-INS-8454 | chr19-56755268-INS-8454 |
| chr19-56771975-INS-272 | chr19-56771975-INS-272 |
| chr19-56791433-INS-162 | chr19-56791433-INS-162 |
| chr19-56805103-INS-86 | chr19-56805103-INS-86 |
| chr19-56836168-INS-1852 | chr19-56836169-INS-1852 |
| chr19-56845465-INS-60 | chr19-56845465-INS-60 |
| chr19-56864872-INS-71 | chr19-56864872-INS-71 |
| chr19-56892498-INS-63 | chr19-56892498-INS-51 |
| chr19-56945383-INS-57 | chr19-56945383-INS-57 |
| chr19-56980104-INS-74 | chr19-56980104-INS-74 |
| chr19-56987096-INS-62 | chr19-56987096-INS-78 |
| chr19-57016256-INS-1891 | chr19-57016257-INS-1891 |
| chr19-57072756-INS-118 | chr19-57072756-INS-118 |
| chr19-57114180-INS-103 | chr19-57114180-INS-103 |
| chr19-57163194-INS-53 | chr19-57163194-INS-53 |
| chr19-57181632-INS-68 | chr19-57181632-INS-68 |
| chr19-57193294-INS-86 | chr19-57193294-INS-86 |
| chr19-57210866-INS-120 | chr19-57210866-INS-120 |
| chr19-57211782-INS-817 | chr19-57211782-INS-811 |
| chr19-57216726-INS-54 | chr19-57216726-INS-56 |
| chr19-57223683-INS-65 | chr19-57223683-INS-89 |
| chr19-57232362-INS-103 | chr19-57232363-INS-103 |
| chr19-57239681-INS-365 | chr19-57239703-INS-369 |
| chr19-57259395-INS-90 | chr19-57259395-INS-90 |
| chr19-57259926-INS-175 | chr19-57259926-INS-174 |
| chr19-57313473-INS-54 | chr19-57313474-INS-54 |
| chr19-57313725-INS-1202 | chr19-57313726-INS-1202 |
| chr19-57313977-INS-198 | chr19-57313977-INS-198 |
| chr19-57320308-INS-714 | chr19-57320308-INS-714 |
| chr19-57332912-INS-88 | chr19-57332881-INS-78 |
| chr19-57335594-INS-54 | chr19-57335594-INS-54 |
| chr19-57338711-INS-199 | chr19-57338711-INS-199 |
| chr19-57339874-INS-60 | chr19-57339874-INS-60 |
| chr19-57342555-INS-225 | chr19-57342555-INS-225 |
| chr19-57345237-INS-50 | chr19-57345217-INS-68 |
| chr19-57382839-INS-196 | chr19-57382839-INS-195 |
| chr19-57394515-INS-134 | chr19-57394521-INS-136 |
| chr19-57402660-INS-1892 | chr19-57402661-INS-1892 |
| chr19-57416773-INS-1890 | chr19-57416774-INS-1891 |
| chr19-57418504-INS-54 | chr19-57418503-INS-83 |
| chr19-57439087-INS-580 | chr19-57439087-INS-580 |
| chr19-57442463-INS-7137 | chr19-57442464-INS-7137 |
| chr19-57444157-INS-52 | chr19-57444157-INS-52 |
| chr19-57465639-INS-1339 | chr19-57465640-INS-1339 |
| chr19-57473047-INS-201 | chr19-57473047-INS-201 |
| chr19-57474127-INS-67 | chr19-57474127-INS-67 |
| chr19-57483199-INS-243 | chr19-57483199-INS-243 |
| chr19-57486578-INS-86 | chr19-57486578-INS-88 |
| chr19-57487237-INS-606 | chr19-57487237-INS-606 |
| chr19-57489828-INS-481 | chr19-57489828-INS-481 |
| chr19-57518010-INS-336 | chr19-57518010-INS-333 |
| chr19-57522132-INS-82 | chr19-57522130-INS-98 |
| chr19-57538909-INS-103 | chr19-57538910-INS-103 |
| chr19-57540921-INS-111 | chr19-57540921-INS-113 |
| chr19-57570907-INS-95 | chr19-57570907-INS-95 |
| chr19-57584880-INS-106 | chr19-57584880-INS-106 |
| chr19-57609466-INS-131 | chr19-57609466-INS-131 |
| chr19-57631186-INS-54 | chr19-57631186-INS-54 |
| chr19-57639536-INS-104 | chr19-57639536-INS-104 |
| chr19-57647179-INS-110 | chr19-57647179-INS-108 |
| chr19-57653317-INS-59 | chr19-57653317-INS-53 |
| chr19-57654979-INS-3659 | chr19-57654980-INS-3659 |

| | |
|---|---|
| chr19-57660761-INS-80 | chr19-57660761-INS-80 |
| chr19-57664136-INS-142 | chr19-57664136-INS-142 |
| chr19-57667927-INS-62 | chr19-57667927-INS-62 |
| chr19-57727312-INS-143 | chr19-57727335-INS-146 |
| chr19-57767046-INS-64 | chr19-57767046-INS-64 |
| chr19-57769427-INS-264 | chr19-57769465-INS-271 |
| chr19-57776932-INS-96 | chr19-57776932-INS-126 |
| chr19-57802450-INS-104 | chr19-57802450-INS-104 |
| chr19-57868701-INS-52 | chr19-57868701-INS-52 |
| chr19-57880469-INS-83 | chr19-57880470-INS-71 |
| chr19-57886264-INS-144 | chr19-57886264-INS-144 |
| chr19-57915053-INS-2782 | chr19-57915054-INS-2783 |
| chr19-57926812-INS-108 | chr19-57926812-INS-108 |
| chr19-58009288-INS-431 | chr19-58009288-INS-431 |
| chr19-58033905-INS-109 | chr19-58033944-INS-109 |
| chr19-58046083-INS-56 | chr19-58046083-INS-56 |
| chr19-58047429-INS-134 | chr19-58047429-INS-134 |
| chr19-58061123-INS-57 | chr19-58061123-INS-55 |
| chr19-58069230-INS-102 | chr19-58069230-INS-95 |
| chr19-58092439-INS-73 | chr19-58092439-INS-66 |
| chr19-58097194-INS-93 | chr19-58097192-INS-101 |
| chr19-58109552-INS-80 | chr19-58109552-INS-80 |
| chr19-58162425-INS-6469 | chr19-58162426-INS-6469 |
| chr19-58193319-INS-168 | chr19-58193319-INS-168 |
| chr19-58220921-INS-128 | chr19-58220921-INS-151 |
| chr19-58251431-INS-105 | chr19-58251431-INS-105 |
| chr19-58253019-INS-455 | chr19-58253019-INS-455 |
| chr19-58263543-INS-62 | chr19-58263527-INS-64 |
| chr19-58266301-INS-52 | chr19-58266290-INS-52 |
| chr19-58275694-INS-467 | chr19-58275701-INS-477 |
| chr19-58280501-INS-101 | chr19-58280501-INS-143 |
| chr19-58304521-INS-64 | chr19-58304521-INS-67 |
| chr19-58307077-INS-110 | chr19-58307028-INS-146 |
| chr19-58394082-INS-231 | chr19-58394082-INS-229 |
| chr19-58401582-INS-77 | chr19-58401582-INS-85 |
| chr19-58402890-INS-164 | chr19-58402890-INS-180 |
| chr19-58435989-INS-645 | chr19-58435989-INS-642 |
| chr19-58455868-INS-56 | chr19-58455866-INS-56 |
| chr19-58470356-INS-224 | chr19-58470356-INS-224 |
| chr19-58555465-INS-589 | chr19-58555465-INS-589 |
| chr19-58574627-INS-94 | chr19-58574627-INS-78 |
| chr19-58595839-INS-270 | chr19-58595839-INS-270 |
| chr19-58599238-INS-76 | chr19-58599204-INS-84 |
| chr19-58600422-INS-229 | chr19-58600422-INS-229 |
| chr19-58673275-INS-6121 | chr19-58673276-INS-6124 |
| chr19-58699354-INS-322 | chr19-58699354-INS-322 |
| chr19-58733151-INS-108 | chr19-58733151-INS-108 |
| chr19-58757875-INS-78 | chr19-58757875-INS-78 |
| chr19-58816628-INS-58 | chr19-58816628-INS-58 |
| chr19-58823715-INS-73 | chr19-58823715-INS-73 |
| chr19-58828788-INS-108 | chr19-58828788-INS-108 |
| chr19-58836992-INS-494 | chr19-58836992-INS-494 |
| chr19-58840347-INS-89 | chr19-58840347-INS-89 |
| chr19-58866713-INS-66 | chr19-58866713-INS-72 |
| chr19-58901654-INS-57 | chr19-58901654-INS-57 |
| chr19-58905315-INS-1347 | chr19-58905316-INS-1348 |
| chr19-58915992-INS-92 | chr19-58915992-INS-92 |
| chr19-58940519-INS-93 | chr19-58940519-INS-93 |
| chr19-59076918-INS-146 | chr19-59076918-INS-146 |
| chr19-59080602-INS-58 | chr19-59080602-INS-54 |
| chr19-59087371-INS-100 | chr19-59087371-INS-84 |
| chr19-59092030-INS-287 | chr19-59092030-INS-287 |
| chr19-59103965-INS-3155 | chr19-59103966-INS-3155 |
| chr19-59105581-INS-52 | chr19-59105581-INS-52 |
| chr19-59106539-INS-248 | chr19-59106539-INS-248 |
| chr19-59150889-INS-80 | chr19-59150889-INS-80 |
| chr19-59164935-INS-68 | chr19-59164935-INS-69 |
| chr19-59167587-INS-208 | chr19-59167587-INS-208 |
| chr19-59174885-INS-217 | chr19-59174886-INS-170 |
| chr19-59195277-INS-184 | chr19-59195277-INS-184 |
| chr19-59198178-INS-72 | chr19-59198178-INS-74 |
| chr19-59216716-INS-389 | chr19-59216717-INS-388 |
| chr19-59220523-INS-1220 | chr19-59220524-INS-1221 |
| chr19-59239528-INS-94 | chr19-59239528-INS-94 |
| chr19-59268605-INS-168 | chr19-59268605-INS-168 |
| chr19-59281624-INS-100 | chr19-59281624-INS-100 |
| chr19-59328515-INS-120 | chr19-59328515-INS-117 |

| | |
|---|---|
| chr19-59369287-INS-129 | chr19-59369287-INS-133 |
| chr19-59371611-INS-220 | chr19-59371611-INS-220 |
| chr19-59381868-INS-78 | chr19-59381868-INS-105 |
| chr19-59421710-INS-65 | chr19-59421710-INS-58 |
| chr19-59425699-INS-66 | chr19-59425699-INS-66 |
| chr19-59463826-INS-1791 | chr19-59463827-INS-1791 |
| chr19-59475229-INS-51 | chr19-59475260-INS-59 |
| chr19-59487133-INS-86 | chr19-59487134-INS-99 |
| chr19-59490811-INS-642 | chr19-59490811-INS-642 |
| chr19-59495102-INS-413 | chr19-59495102-INS-413 |
| chr19-59523024-INS-197 | chr19-59523024-INS-198 |
| chr19-59530570-INS-51 | chr19-59530571-INS-55 |
| chr19-59541410-INS-160 | chr19-59541407-INS-163 |
| chr19-59622528-INS-193 | chr19-59622494-INS-232 |
| chr19-59654426-INS-157 | chr19-59654426-INS-152 |
| chr19-59659801-INS-335 | chr19-59659801-INS-335 |
| chr19-59672265-INS-790 | chr19-59672265-INS-790 |
| chr19-59691709-INS-197 | chr19-59691709-INS-197 |
| chr19-59701550-INS-54 | chr19-59701574-INS-56 |
| chr19-59720476-INS-148 | chr19-59720476-INS-148 |
| chr19-59753157-INS-441 | chr19-59753157-INS-441 |
| chr19-59762317-INS-201 | chr19-59762317-INS-201 |
| chr19-59777067-INS-80 | chr19-59777067-INS-86 |
| chr19-59795461-INS-785 | chr19-59795461-INS-785 |
| chr19-59858584-INS-64 | chr19-59858584-INS-64 |
| chr19-59866788-INS-1756 | chr19-59866789-INS-1756 |
| chr19-59882558-INS-52 | chr19-59882558-INS-50 |
| chr19-59883947-INS-65 | chr19-59883941-INS-82 |
| chr19-59906167-INS-80 | chr19-59906167-INS-80 |
| chr19-59939083-INS-107 | chr19-59939083-INS-107 |
| chr19-59947799-INS-492 | chr19-59947799-INS-492 |
| chr19-59976000-INS-987 | chr19-59976000-INS-1023 |
| chr19-60026679-INS-6443 | chr19-60026680-INS-6443 |
| chr19-60028209-INS-52 | chr19-60028209-INS-56 |
| chr19-60055487-INS-182 | chr19-60055487-INS-182 |
| chr19-60067776-INS-1199 | chr19-60067777-INS-1199 |
| chr19-60081095-INS-652 | chr19-60081096-INS-652 |
| chr19-60087040-INS-400 | chr19-60087040-INS-400 |
| chr19-60127698-INS-52 | chr19-60127698-INS-51 |
| chr19-60180239-INS-63 | chr19-60180239-INS-63 |
| chr19-60237638-INS-68 | chr19-60237638-INS-68 |
| chr19-60303874-INS-427 | chr19-60303874-INS-426 |
| chr19-60367071-INS-79 | chr19-60367071-INS-79 |
| chr19-60391606-INS-118 | chr19-60391620-INS-118 |
| chr19-60404355-INS-498 | chr19-60404355-INS-498 |
| chr19-60411906-INS-316 | chr19-60411911-INS-317 |
| chr19-60413971-INS-746 | chr19-60413971-INS-747 |
| chr19-60459156-INS-4330 | chr19-60459157-INS-4330 |
| chr19-60505528-INS-2415 | chr19-60505529-INS-2415 |
| chr19-60530492-INS-225 | chr19-60530492-INS-225 |
| chr19-60538610-INS-278 | chr19-60538611-INS-278 |
| chr19-60539165-INS-6414 | chr19-60539166-INS-6415 |
| chr19-60544151-INS-215 | chr19-60544151-INS-215 |
| chr19-60552476-INS-54 | chr19-60552480-INS-52 |
| chr19-60571183-INS-6206 | chr19-60571184-INS-6206 |
| chr19-60584383-INS-56 | chr19-60584383-INS-50 |
| chr19-60611537-INS-212 | chr19-60611537-INS-219 |
| chr19-60617667-INS-61 | chr19-60617667-INS-61 |
| chr19-60665233-INS-214 | chr19-60665233-INS-213 |
| chr19-60670666-INS-62 | chr19-60670668-INS-60 |
| chr19-60678182-INS-203 | chr19-60678182-INS-203 |
| chr19-60685307-INS-184 | chr19-60685307-INS-184 |
| chr19-60686226-INS-342 | chr19-60686226-INS-342 |
| chr19-60693780-INS-80 | chr19-60693781-INS-60 |
| chr19-60712907-INS-241 | chr19-60712903-INS-245 |
| chr19-60724289-INS-213 | chr19-60724289-INS-213 |
| chr19-60741097-INS-60 | chr19-60741097-INS-66 |
| chr19-60786262-INS-52 | chr19-60786262-INS-54 |
| chr19-60794390-INS-97 | chr19-60794390-INS-97 |
| chr19-60818721-INS-219 | chr19-60818721-INS-219 |
| chr19-60825210-INS-375 | chr19-60825210-INS-376 |
| chr19-60828743-INS-98 | chr19-60828743-INS-98 |
| chr19-60835701-INS-172 | chr19-60835701-INS-172 |
| chr19-60837032-INS-187 | chr19-60837032-INS-161 |
| chr19-60840022-INS-422 | chr19-60840022-INS-421 |
| chr19-60840598-INS-315 | chr19-60840598-INS-336 |
| chr19-60852013-INS-198 | chr19-60852013-INS-198 |

| | |
|---|---|
| chr19-60864325-INS-239 | chr19-60864325-INS-239 |
| chr19-60874437-INS-398 | chr19-60874437-INS-398 |
| chr19-60894747-INS-201 | chr19-60894747-INS-201 |
| chr19-60927621-INS-181 | chr19-60927621-INS-183 |
| chr19-60992719-INS-457 | chr19-60992719-INS-457 |
| chr19-61088688-INS-6401 | chr19-61088689-INS-6401 |
| chr19-61088689-INS-497 | chr19-61088689-INS-497 |
| chr19-61089718-INS-56 | chr19-61089718-INS-56 |
| chr19-61113578-INS-238 | chr19-61113578-INS-239 |
| chr19-61121801-INS-61 | chr19-61121801-INS-61 |
| chr19-61123757-INS-196 | chr19-61123757-INS-230 |
| chr19-61139585-INS-876 | chr19-61139586-INS-878 |
| chr19-61148494-INS-210 | chr19-61148494-INS-210 |
| chr19-61166725-INS-559 | chr19-61166725-INS-559 |
| chr19-61174656-INS-1714 | chr19-61174657-INS-1713 |
| chr19-61273468-INS-1076 | chr19-61273469-INS-1082 |
| chr19-61310885-INS-214 | chr19-61310885-INS-214 |
| chr2-3071884-INS-807 | chr2-3071884-INS-807 |
| chr2-3092338-INS-188 | chr2-3092338-INS-198 |
| chr2-3135794-INS-698 | chr2-3135794-INS-698 |
| chr2-3151786-INS-1646 | chr2-3151787-INS-1646 |
| chr2-3161482-INS-1870 | chr2-3161483-INS-1870 |
| chr2-3217706-INS-52 | chr2-3217706-INS-52 |
| chr2-3218792-INS-122 | chr2-3218792-INS-122 |
| chr2-3229534-INS-6991 | chr2-3229535-INS-6991 |
| chr2-3243027-INS-6437 | chr2-3243028-INS-6437 |
| chr2-3271793-INS-174 | chr2-3271826-INS-151 |
| chr2-3354590-INS-113 | chr2-3354590-INS-120 |
| chr2-3382517-INS-672 | chr2-3382517-INS-672 |
| chr2-3426224-INS-205 | chr2-3426224-INS-205 |
| chr2-3436391-INS-56 | chr2-3436391-INS-56 |
| chr2-3458321-INS-172 | chr2-3458321-INS-172 |
| chr2-3499021-INS-217 | chr2-3499021-INS-217 |
| chr2-3504593-INS-346 | chr2-3504593-INS-346 |
| chr2-3567456-INS-84 | chr2-3567450-INS-87 |
| chr2-3568833-INS-4521 | chr2-3568834-INS-4521 |
| chr2-3573342-INS-850 | chr2-3573342-INS-850 |
| chr2-3574005-INS-2339 | chr2-3574006-INS-2339 |
| chr2-3577179-INS-90 | chr2-3577179-INS-90 |
| chr2-3581130-INS-340 | chr2-3581130-INS-340 |
| chr2-3607487-INS-595 | chr2-3607487-INS-595 |
| chr2-3638784-INS-97 | chr2-3638804-INS-97 |
| chr2-3664470-INS-95 | chr2-3664470-INS-114 |
| chr2-3666618-INS-198 | chr2-3666618-INS-196 |
| chr2-3680731-INS-53 | chr2-3680768-INS-62 |
| chr2-3692483-INS-92 | chr2-3692482-INS-93 |
| chr2-3706581-INS-66 | chr2-3706581-INS-74 |
| chr2-3723026-INS-605 | chr2-3723026-INS-605 |
| chr2-3734502-INS-120 | chr2-3734481-INS-128 |
| chr2-3743151-INS-200 | chr2-3743151-INS-200 |
| chr2-3775562-INS-52 | chr2-3775562-INS-52 |
| chr2-3795926-INS-4097 | chr2-3795927-INS-4097 |
| chr2-3804735-INS-452 | chr2-3804735-INS-452 |
| chr2-3880786-INS-105 | chr2-3880787-INS-95 |
| chr2-3886713-INS-60 | chr2-3886713-INS-60 |
| chr2-3903002-INS-499 | chr2-3903002-INS-499 |
| chr2-3940563-INS-89 | chr2-3940561-INS-54 |
| chr2-3942690-INS-187 | chr2-3942690-INS-187 |
| chr2-3955030-INS-92 | chr2-3955030-INS-92 |
| chr2-4043660-INS-60 | chr2-4043660-INS-60 |
| chr2-4061367-INS-85 | chr2-4061367-INS-84 |
| chr2-4066967-INS-5550 | chr2-4066968-INS-5550 |
| chr2-4079502-INS-56 | chr2-4079502-INS-58 |
| chr2-4093820-INS-51 | chr2-4093821-INS-56 |
| chr2-4107770-INS-420 | chr2-4107770-INS-420 |
| chr2-4121996-INS-52 | chr2-4122002-INS-70 |
| chr2-4125752-INS-95 | chr2-4125752-INS-55 |
| chr2-4141740-INS-134 | chr2-4141786-INS-122 |
| chr2-4178576-INS-84 | chr2-4178577-INS-56 |
| chr2-4186890-INS-68 | chr2-4186876-INS-76 |
| chr2-4210546-INS-88 | chr2-4210546-INS-92 |
| chr2-4212353-INS-822 | chr2-4212353-INS-822 |
| chr2-4234676-INS-214 | chr2-4234676-INS-214 |
| chr2-4235090-INS-438 | chr2-4235090-INS-438 |
| chr2-4244061-INS-85 | chr2-4244061-INS-85 |
| chr2-4246425-INS-52 | chr2-4246424-INS-64 |
| chr2-4272639-INS-114 | chr2-4272639-INS-114 |

| | |
|---|---|
| chr2-4286003-INS-100 | chr2-4286003-INS-100 |
| chr2-4334844-INS-257 | chr2-4334844-INS-239 |
| chr2-4348816-INS-403 | chr2-4348817-INS-401 |
| chr2-4351723-INS-51 | chr2-4351723-INS-51 |
| chr2-4366312-INS-57 | chr2-4366312-INS-55 |
| chr2-4474404-INS-816 | chr2-4474404-INS-816 |
| chr2-4499002-INS-284 | chr2-4499002-INS-284 |
| chr2-4516759-INS-206 | chr2-4516759-INS-206 |
| chr2-4518389-INS-206 | chr2-4518389-INS-206 |
| chr2-4523398-INS-835 | chr2-4523398-INS-834 |
| chr2-4535438-INS-57 | chr2-4535438-INS-57 |
| chr2-4537393-INS-134 | chr2-4537393-INS-134 |
| chr2-4623679-INS-264 | chr2-4623679-INS-264 |
| chr2-4641145-INS-88 | chr2-4641145-INS-124 |
| chr2-4644319-INS-217 | chr2-4644319-INS-216 |
| chr2-4657148-INS-61 | chr2-4657148-INS-61 |
| chr2-4659372-INS-496 | chr2-4659372-INS-496 |
| chr2-4668818-INS-241 | chr2-4668818-INS-241 |
| chr2-4669623-INS-5220 | chr2-4669624-INS-5218 |
| chr2-4691380-INS-51 | chr2-4691380-INS-51 |
| chr2-4721368-INS-199 | chr2-4721368-INS-200 |
| chr2-4783005-INS-168 | chr2-4783005-INS-168 |
| chr2-4790646-INS-3211 | chr2-4790647-INS-3211 |
| chr2-4812099-INS-108 | chr2-4812099-INS-108 |
| chr2-4831484-INS-122 | chr2-4831484-INS-122 |
| chr2-4880167-INS-206 | chr2-4880167-INS-206 |
| chr2-4928964-INS-7200 | chr2-4928965-INS-7200 |
| chr2-4952364-INS-786 | chr2-4952364-INS-787 |
| chr2-4985274-INS-105 | chr2-4985274-INS-60 |
| chr2-5002763-INS-124 | chr2-5002742-INS-149 |
| chr2-5007338-INS-94 | chr2-5007324-INS-85 |
| chr2-5008608-INS-62 | chr2-5008608-INS-62 |
| chr2-5074462-INS-172 | chr2-5074462-INS-172 |
| chr2-5079603-INS-108 | chr2-5079603-INS-108 |
| chr2-5084068-INS-78 | chr2-5084068-INS-72 |
| chr2-5087180-INS-72 | chr2-5087181-INS-63 |
| chr2-5090397-INS-120 | chr2-5090373-INS-166 |
| chr2-5096630-INS-205 | chr2-5096630-INS-211 |
| chr2-5100809-INS-58 | chr2-5100809-INS-66 |
| chr2-5103354-INS-174 | chr2-5103354-INS-174 |
| chr2-5114255-INS-109 | chr2-5114255-INS-109 |
| chr2-5116726-INS-56 | chr2-5116726-INS-56 |
| chr2-5118771-INS-432 | chr2-5118777-INS-439 |
| chr2-5124022-INS-363 | chr2-5124023-INS-363 |
| chr2-5130643-INS-72 | chr2-5130643-INS-79 |
| chr2-5139166-INS-70 | chr2-5139166-INS-70 |
| chr2-5148837-INS-55 | chr2-5148837-INS-55 |
| chr2-5163506-INS-143 | chr2-5163506-INS-117 |
| chr2-5166559-INS-116 | chr2-5166560-INS-74 |
| chr2-5188063-INS-180 | chr2-5188063-INS-181 |
| chr2-5193176-INS-58 | chr2-5193203-INS-64 |
| chr2-5199586-INS-1768 | chr2-5199587-INS-1769 |
| chr2-5211614-INS-6606 | chr2-5211615-INS-6606 |
| chr2-5226721-INS-597 | chr2-5226721-INS-597 |
| chr2-5239490-INS-5554 | chr2-5239491-INS-5555 |
| chr2-5240645-INS-295 | chr2-5240645-INS-295 |
| chr2-5244897-INS-169 | chr2-5244879-INS-214 |
| chr2-5247344-INS-98 | chr2-5247345-INS-98 |
| chr2-5260433-INS-2769 | chr2-5260434-INS-2765 |
| chr2-5290288-INS-486 | chr2-5290284-INS-488 |
| chr2-5320287-INS-4537 | chr2-5320288-INS-4537 |
| chr2-5328324-INS-105 | chr2-5328324-INS-105 |
| chr2-5342161-INS-78 | chr2-5342161-INS-78 |
| chr2-5348453-INS-74 | chr2-5348453-INS-74 |
| chr2-5392365-INS-52 | chr2-5392365-INS-54 |
| chr2-5407139-INS-336 | chr2-5407139-INS-336 |
| chr2-5468664-INS-227 | chr2-5468664-INS-227 |
| chr2-5482466-INS-57 | chr2-5482468-INS-57 |
| chr2-5493860-INS-695 | chr2-5493860-INS-695 |
| chr2-5499982-INS-59 | chr2-5499982-INS-59 |
| chr2-5509011-INS-1873 | chr2-5509012-INS-1873 |
| chr2-5511497-INS-178 | chr2-5511497-INS-178 |
| chr2-5522271-INS-119 | chr2-5522271-INS-121 |
| chr2-5531082-INS-71 | chr2-5531092-INS-84 |
| chr2-5539759-INS-52 | chr2-5539759-INS-50 |
| chr2-5541331-INS-108 | chr2-5541339-INS-138 |
| chr2-5544480-INS-987 | chr2-5544480-INS-990 |

| | |
|---|---|
| chr2-5550689-INS-681 | chr2-5550689-INS-681 |
| chr2-5625335-INS-4080 | chr2-5625336-INS-4080 |
| chr2-5626033-INS-53 | chr2-5626033-INS-53 |
| chr2-5642947-INS-3429 | chr2-5642948-INS-3434 |
| chr2-5651347-INS-98 | chr2-5651347-INS-110 |
| chr2-5658365-INS-58 | chr2-5658363-INS-56 |
| chr2-5662616-INS-208 | chr2-5662616-INS-208 |
| chr2-5666518-INS-53 | chr2-5666523-INS-56 |
| chr2-5687830-INS-155 | chr2-5687830-INS-155 |
| chr2-5707311-INS-141 | chr2-5707311-INS-144 |
| chr2-5713655-INS-68 | chr2-5713655-INS-68 |
| chr2-5716017-INS-75 | chr2-5716017-INS-76 |
| chr2-5726475-INS-53 | chr2-5726455-INS-100 |
| chr2-5738858-INS-129 | chr2-5738858-INS-129 |
| chr2-5742865-INS-90 | chr2-5742878-INS-107 |
| chr2-5774158-INS-170 | chr2-5774158-INS-169 |
| chr2-5774860-INS-170 | chr2-5774863-INS-171 |
| chr2-5787395-INS-421 | chr2-5787395-INS-421 |
| chr2-5787402-INS-2965 | chr2-5787403-INS-2965 |
| chr2-5811688-INS-5367 | chr2-5811689-INS-5367 |
| chr2-5835759-INS-6473 | chr2-5835760-INS-6472 |
| chr2-5856438-INS-98 | chr2-5856438-INS-98 |
| chr2-5874425-INS-77 | chr2-5874425-INS-77 |
| chr2-5914688-INS-414 | chr2-5914688-INS-416 |
| chr2-5948695-INS-5376 | chr2-5948696-INS-5376 |
| chr2-5948908-INS-768 | chr2-5948908-INS-768 |
| chr2-5949706-INS-55 | chr2-5949706-INS-59 |
| chr2-5979163-INS-52 | chr2-5979148-INS-62 |
| chr2-5988798-INS-186 | chr2-5988798-INS-187 |
| chr2-5997774-INS-107 | chr2-5997774-INS-57 |
| chr2-6063708-INS-1397 | chr2-6063709-INS-1390 |
| chr2-6066491-INS-460 | chr2-6066491-INS-460 |
| chr2-6083700-INS-551 | chr2-6083700-INS-551 |
| chr2-6095321-INS-117 | chr2-6095296-INS-117 |
| chr2-6119240-INS-132 | chr2-6119240-INS-145 |
| chr2-6144727-INS-67 | chr2-6144729-INS-73 |
| chr2-6163689-INS-290 | chr2-6163689-INS-244 |
| chr2-6178580-INS-522 | chr2-6178580-INS-522 |
| chr2-6184406-INS-82 | chr2-6184407-INS-85 |
| chr2-6187064-INS-143 | chr2-6187064-INS-147 |
| chr2-6200065-INS-54 | chr2-6200065-INS-56 |
| chr2-6218384-INS-555 | chr2-6218384-INS-555 |
| chr2-6240931-INS-190 | chr2-6240932-INS-208 |
| chr2-6250406-INS-502 | chr2-6250406-INS-502 |
| chr2-6281799-INS-60 | chr2-6281799-INS-64 |
| chr2-6370432-INS-899 | chr2-6370433-INS-899 |
| chr2-6376818-INS-196 | chr2-6376818-INS-196 |
| chr2-6406887-INS-147 | chr2-6406887-INS-147 |
| chr2-6408932-INS-4950 | chr2-6408933-INS-4950 |
| chr2-6411564-INS-937 | chr2-6411564-INS-939 |
| chr2-6460455-INS-132 | chr2-6460489-INS-108 |
| chr2-6464264-INS-707 | chr2-6464264-INS-707 |
| chr2-6477632-INS-82 | chr2-6477632-INS-82 |
| chr2-6484956-INS-60 | chr2-6484956-INS-60 |
| chr2-6487697-INS-76 | chr2-6487697-INS-76 |
| chr2-6501481-INS-65 | chr2-6501481-INS-65 |
| chr2-6525639-INS-450 | chr2-6525639-INS-449 |
| chr2-6527663-INS-217 | chr2-6527663-INS-215 |
| chr2-6534870-INS-82 | chr2-6534870-INS-82 |
| chr2-6536029-INS-126 | chr2-6536029-INS-117 |
| chr2-6538930-INS-61 | chr2-6538924-INS-61 |
| chr2-6564956-INS-72 | chr2-6564953-INS-109 |
| chr2-6577077-INS-441 | chr2-6577077-INS-441 |
| chr2-6645003-INS-70 | chr2-6644973-INS-54 |
| chr2-6665596-INS-190 | chr2-6665596-INS-195 |
| chr2-6670598-INS-107 | chr2-6670598-INS-67 |
| chr2-6679166-INS-59 | chr2-6679124-INS-55 |
| chr2-6748563-INS-53 | chr2-6748563-INS-53 |
| chr2-6749439-INS-198 | chr2-6749439-INS-198 |
| chr2-6773343-INS-194 | chr2-6773343-INS-194 |
| chr2-6783405-INS-56 | chr2-6783380-INS-56 |
| chr2-6844922-INS-212 | chr2-6844922-INS-212 |
| chr2-6953162-INS-222 | chr2-6953162-INS-222 |
| chr2-6958279-INS-92 | chr2-6958273-INS-96 |
| chr2-6979485-INS-293 | chr2-6979485-INS-293 |
| chr2-6992533-INS-125 | chr2-6992523-INS-156 |
| chr2-7002832-INS-326 | chr2-7002827-INS-329 |

| | |
|---|---|
| chr2-7045643-INS-68 | chr2-7045643-INS-88 |
| chr2-7056642-INS-119 | chr2-7056654-INS-100 |
| chr2-7063514-INS-62 | chr2-7063514-INS-62 |
| chr2-7082057-INS-114 | chr2-7082057-INS-116 |
| chr2-7092971-INS-80 | chr2-7092971-INS-80 |
| chr2-7114277-INS-52 | chr2-7114277-INS-56 |
| chr2-7116421-INS-200 | chr2-7116421-INS-200 |
| chr2-7205668-INS-567 | chr2-7205669-INS-566 |
| chr2-7215140-INS-263 | chr2-7215140-INS-286 |
| chr2-7243904-INS-54 | chr2-7243904-INS-52 |
| chr2-7245067-INS-54 | chr2-7245067-INS-62 |
| chr2-7289068-INS-90 | chr2-7289068-INS-59 |
| chr2-7291457-INS-4747 | chr2-7291458-INS-4729 |
| chr2-7306732-INS-78 | chr2-7306733-INS-81 |
| chr2-7337750-INS-78 | chr2-7337750-INS-80 |
| chr2-7402762-INS-62 | chr2-7402761-INS-86 |
| chr2-7444292-INS-54 | chr2-7444292-INS-56 |
| chr2-7472995-INS-343 | chr2-7472995-INS-343 |
| chr2-7478380-INS-123 | chr2-7478410-INS-126 |
| chr2-7485688-INS-73 | chr2-7485683-INS-68 |
| chr2-7487327-INS-60 | chr2-7487327-INS-60 |
| chr2-7490719-INS-64 | chr2-7490713-INS-102 |
| chr2-7506705-INS-97 | chr2-7506705-INS-96 |
| chr2-7568813-INS-1474 | chr2-7568814-INS-1474 |
| chr2-7649061-INS-197 | chr2-7649061-INS-197 |
| chr2-7666974-INS-5430 | chr2-7666975-INS-5430 |
| chr2-7667878-INS-271 | chr2-7667879-INS-267 |
| chr2-7707751-INS-53 | chr2-7707751-INS-53 |
| chr2-7722612-INS-254 | chr2-7722613-INS-296 |
| chr2-7731696-INS-82 | chr2-7731696-INS-82 |
| chr2-7748942-INS-108 | chr2-7748988-INS-61 |
| chr2-7756686-INS-371 | chr2-7756683-INS-370 |
| chr2-7815370-INS-68 | chr2-7815370-INS-70 |
| chr2-7822744-INS-392 | chr2-7822744-INS-399 |
| chr2-7825581-INS-541 | chr2-7825581-INS-541 |
| chr2-7846950-INS-169 | chr2-7846950-INS-169 |
| chr2-7880590-INS-108 | chr2-7880591-INS-112 |
| chr2-7919842-INS-179 | chr2-7919842-INS-177 |
| chr2-7988695-INS-3721 | chr2-7988696-INS-3721 |
| chr2-8019471-INS-201 | chr2-8019471-INS-201 |
| chr2-8032977-INS-162 | chr2-8032973-INS-168 |
| chr2-8042625-INS-212 | chr2-8042625-INS-212 |
| chr2-8055035-INS-96 | chr2-8055035-INS-96 |
| chr2-8068175-INS-1272 | chr2-8068176-INS-1272 |
| chr2-8101836-INS-107 | chr2-8101836-INS-107 |
| chr2-8145263-INS-56 | chr2-8145263-INS-58 |
| chr2-8159153-INS-323 | chr2-8159171-INS-343 |
| chr2-8179913-INS-650 | chr2-8179913-INS-650 |
| chr2-8222902-INS-6844 | chr2-8222903-INS-6844 |
| chr2-8235874-INS-107 | chr2-8235874-INS-107 |
| chr2-8263080-INS-53 | chr2-8263080-INS-53 |
| chr2-8300527-INS-60 | chr2-8300531-INS-72 |
| chr2-8325540-INS-1909 | chr2-8325541-INS-1903 |
| chr2-8343214-INS-430 | chr2-8343214-INS-431 |
| chr2-8381224-INS-150 | chr2-8381239-INS-150 |
| chr2-8409050-INS-6504 | chr2-8409051-INS-6503 |
| chr2-8414298-INS-7097 | chr2-8414299-INS-7097 |
| chr2-8438210-INS-66 | chr2-8438249-INS-60 |
| chr2-8443971-INS-65 | chr2-8443971-INS-65 |
| chr2-8464934-INS-79 | chr2-8464938-INS-64 |
| chr2-8475903-INS-94 | chr2-8475903-INS-91 |
| chr2-8531642-INS-111 | chr2-8531642-INS-111 |
| chr2-8552434-INS-6160 | chr2-8552435-INS-6161 |
| chr2-8553730-INS-2345 | chr2-8553731-INS-2345 |
| chr2-8565753-INS-101 | chr2-8565719-INS-102 |
| chr2-8580607-INS-66 | chr2-8580607-INS-66 |
| chr2-8582319-INS-261 | chr2-8582319-INS-267 |
| chr2-8589603-INS-3716 | chr2-8589604-INS-3716 |
| chr2-8606269-INS-6242 | chr2-8606270-INS-6241 |
| chr2-8644406-INS-79 | chr2-8644393-INS-52 |
| chr2-8664761-INS-52 | chr2-8664751-INS-62 |
| chr2-8678417-INS-1030 | chr2-8678418-INS-1030 |
| chr2-8688910-INS-152 | chr2-8688910-INS-152 |
| chr2-8716412-INS-755 | chr2-8716411-INS-749 |
| chr2-8745758-INS-4743 | chr2-8745759-INS-4743 |
| chr2-8755944-INS-52 | chr2-8755944-INS-50 |
| chr2-8775019-INS-361 | chr2-8775019-INS-361 |

| | |
|---|---|
| chr2-8801546-INS-74 | chr2-8801546-INS-66 |
| chr2-8820643-INS-254 | chr2-8820643-INS-254 |
| chr2-8821353-INS-68 | chr2-8821353-INS-70 |
| chr2-8830097-INS-967 | chr2-8830097-INS-967 |
| chr2-8889982-INS-6190 | chr2-8889983-INS-6190 |
| chr2-8915620-INS-70 | chr2-8915620-INS-70 |
| chr2-8919675-INS-1006 | chr2-8919676-INS-998 |
| chr2-8941401-INS-3631 | chr2-8941402-INS-3621 |
| chr2-8948277-INS-178 | chr2-8948275-INS-173 |
| chr2-8951475-INS-217 | chr2-8951475-INS-217 |
| chr2-8957336-INS-285 | chr2-8957336-INS-287 |
| chr2-8970583-INS-60 | chr2-8970583-INS-64 |
| chr2-8990598-INS-165 | chr2-8990596-INS-163 |
| chr2-9011114-INS-80 | chr2-9011114-INS-80 |
| chr2-9054191-INS-7053 | chr2-9054192-INS-7053 |
| chr2-9110511-INS-58 | chr2-9110511-INS-54 |
| chr2-9117184-INS-644 | chr2-9117184-INS-652 |
| chr2-9119247-INS-462 | chr2-9119247-INS-462 |
| chr2-9128018-INS-690 | chr2-9128018-INS-690 |
| chr2-9154550-INS-709 | chr2-9154550-INS-709 |
| chr2-9164801-INS-6497 | chr2-9164802-INS-6542 |
| chr2-9168060-INS-87 | chr2-9168050-INS-91 |
| chr2-9217560-INS-424 | chr2-9217560-INS-424 |
| chr2-9236595-INS-464 | chr2-9236596-INS-464 |
| chr2-9237401-INS-90 | chr2-9237363-INS-110 |
| chr2-9252526-INS-6205 | chr2-9252527-INS-6205 |
| chr2-9281012-INS-891 | chr2-9281012-INS-894 |
| chr2-9303274-INS-52 | chr2-9303274-INS-56 |
| chr2-9326727-INS-399 | chr2-9326727-INS-399 |
| chr2-9354795-INS-154 | chr2-9354795-INS-156 |
| chr2-9367173-INS-54 | chr2-9367207-INS-64 |
| chr2-9374895-INS-638 | chr2-9374895-INS-638 |
| chr2-9375781-INS-51 | chr2-9375781-INS-51 |
| chr2-9378922-INS-85 | chr2-9378922-INS-89 |
| chr2-9413073-INS-196 | chr2-9413073-INS-196 |
| chr2-9430437-INS-183 | chr2-9430437-INS-183 |
| chr2-9441905-INS-374 | chr2-9441905-INS-374 |
| chr2-9463958-INS-78 | chr2-9463958-INS-84 |
| chr2-9465045-INS-53 | chr2-9465045-INS-53 |
| chr2-9489565-INS-52 | chr2-9489565-INS-51 |
| chr2-9501993-INS-60 | chr2-9501993-INS-70 |
| chr2-9506428-INS-90 | chr2-9506378-INS-97 |
| chr2-9523221-INS-57 | chr2-9523221-INS-57 |
| chr2-9558600-INS-57 | chr2-9558600-INS-57 |
| chr2-9597336-INS-659 | chr2-9597336-INS-659 |
| chr2-9597867-INS-54 | chr2-9597868-INS-54 |
| chr2-9605389-INS-265 | chr2-9605389-INS-273 |
| chr2-9606718-INS-625 | chr2-9606718-INS-623 |
| chr2-9632683-INS-95 | chr2-9632649-INS-120 |
| chr2-9639923-INS-696 | chr2-9639923-INS-687 |
| chr2-9640755-INS-825 | chr2-9640755-INS-825 |
| chr2-9641859-INS-54 | chr2-9641859-INS-54 |
| chr2-9648502-INS-6468 | chr2-9648503-INS-6468 |
| chr2-9659375-INS-92 | chr2-9659375-INS-94 |
| chr2-9688056-INS-68 | chr2-9688056-INS-68 |
| chr2-9689261-INS-68 | chr2-9689261-INS-70 |
| chr2-9694877-INS-68 | chr2-9694885-INS-74 |
| chr2-9699973-INS-56 | chr2-9699973-INS-60 |
| chr2-9708198-INS-90 | chr2-9708200-INS-86 |
| chr2-9731845-INS-87 | chr2-9731844-INS-100 |
| chr2-9762955-INS-94 | chr2-9762997-INS-56 |
| chr2-9766808-INS-72 | chr2-9766808-INS-56 |
| chr2-9796426-INS-60 | chr2-9796426-INS-52 |
| chr2-9901396-INS-258 | chr2-9901396-INS-258 |
| chr2-9973366-INS-56 | chr2-9973374-INS-94 |
| chr2-9979498-INS-207 | chr2-9979498-INS-206 |
| chr2-10010914-INS-54 | chr2-10010914-INS-54 |
| chr2-10021434-INS-6442 | chr2-10021435-INS-6442 |
| chr2-10024371-INS-418 | chr2-10024371-INS-417 |
| chr2-10040292-INS-58 | chr2-10040292-INS-54 |
| chr2-10058943-INS-193 | chr2-10058943-INS-192 |
| chr2-10062017-INS-96 | chr2-10062017-INS-96 |
| chr2-10089983-INS-409 | chr2-10089983-INS-419 |
| chr2-10095389-INS-179 | chr2-10095389-INS-179 |
| chr2-10127940-INS-108 | chr2-10127940-INS-108 |
| chr2-10139353-INS-64 | chr2-10139317-INS-107 |
| chr2-10149931-INS-75 | chr2-10149931-INS-74 |

| | |
|---|---|
| chr2-10152745-INS-80 | chr2-10152745-INS-64 |
| chr2-10164282-INS-57 | chr2-10164280-INS-85 |
| chr2-10174226-INS-158 | chr2-10174226-INS-159 |
| chr2-10235159-INS-171 | chr2-10235159-INS-147 |
| chr2-10308881-INS-245 | chr2-10308881-INS-245 |
| chr2-10312801-INS-70 | chr2-10312801-INS-70 |
| chr2-10314348-INS-130 | chr2-10314348-INS-130 |
| chr2-10319130-INS-5218 | chr2-10319131-INS-5218 |
| chr2-10332261-INS-448 | chr2-10332261-INS-464 |
| chr2-10345730-INS-169 | chr2-10345730-INS-170 |
| chr2-10355160-INS-673 | chr2-10355160-INS-674 |
| chr2-10362473-INS-5345 | chr2-10362474-INS-5345 |
| chr2-10387650-INS-173 | chr2-10387682-INS-145 |
| chr2-10406521-INS-56 | chr2-10406521-INS-72 |
| chr2-10417865-INS-90 | chr2-10417873-INS-54 |
| chr2-10426261-INS-439 | chr2-10426261-INS-447 |
| chr2-10429050-INS-119 | chr2-10429050-INS-140 |
| chr2-10437364-INS-81 | chr2-10437364-INS-89 |
| chr2-10439519-INS-76 | chr2-10439524-INS-94 |
| chr2-10462951-INS-80 | chr2-10462955-INS-76 |
| chr2-10471212-INS-72 | chr2-10471212-INS-72 |
| chr2-10528283-INS-328 | chr2-10528260-INS-308 |
| chr2-10563461-INS-280 | chr2-10563459-INS-278 |
| chr2-10573744-INS-1414 | chr2-10573745-INS-1439 |
| chr2-10575166-INS-55 | chr2-10575166-INS-57 |
| chr2-10613914-INS-1012 | chr2-10613915-INS-1012 |
| chr2-10680140-INS-60 | chr2-10680140-INS-60 |
| chr2-10681563-INS-52 | chr2-10681563-INS-56 |
| chr2-10693343-INS-1199 | chr2-10693344-INS-1199 |
| chr2-10714982-INS-6392 | chr2-10714983-INS-6392 |
| chr2-10724665-INS-196 | chr2-10724665-INS-198 |
| chr2-10733637-INS-922 | chr2-10733637-INS-922 |
| chr2-10739143-INS-83 | chr2-10739143-INS-56 |
| chr2-10756831-INS-60 | chr2-10756817-INS-80 |
| chr2-10758041-INS-60 | chr2-10758041-INS-60 |
| chr2-10783054-INS-130 | chr2-10783054-INS-128 |
| chr2-10841884-INS-76 | chr2-10841868-INS-50 |
| chr2-10858854-INS-56 | chr2-10858850-INS-80 |
| chr2-10863269-INS-253 | chr2-10863269-INS-253 |
| chr2-10864710-INS-178 | chr2-10864710-INS-162 |
| chr2-10865696-INS-337 | chr2-10865696-INS-337 |
| chr2-10869024-INS-4923 | chr2-10869025-INS-4923 |
| chr2-10886586-INS-62 | chr2-10886586-INS-68 |
| chr2-10888512-INS-164 | chr2-10888513-INS-164 |
| chr2-10919555-INS-397 | chr2-10919555-INS-397 |
| chr2-10927291-INS-69 | chr2-10927291-INS-95 |
| chr2-10954111-INS-7500 | chr2-10954112-INS-7498 |
| chr2-10965017-INS-113 | chr2-10965017-INS-98 |
| chr2-10969159-INS-127 | chr2-10969159-INS-97 |
| chr2-10973638-INS-56 | chr2-10973638-INS-56 |
| chr2-10976170-INS-71 | chr2-10976172-INS-65 |
| chr2-11006919-INS-56 | chr2-11006919-INS-61 |
| chr2-11008232-INS-124 | chr2-11008232-INS-124 |
| chr2-11031473-INS-96 | chr2-11031473-INS-90 |
| chr2-11038529-INS-61 | chr2-11038529-INS-61 |
| chr2-11042302-INS-113 | chr2-11042314-INS-118 |
| chr2-11056096-INS-52 | chr2-11056096-INS-52 |
| chr2-11059318-INS-63 | chr2-11059321-INS-55 |
| chr2-11086689-INS-503 | chr2-11086689-INS-503 |
| chr2-11089704-INS-60 | chr2-11089704-INS-60 |
| chr2-11094155-INS-115 | chr2-11094155-INS-66 |
| chr2-11097486-INS-52 | chr2-11097507-INS-56 |
| chr2-11106903-INS-571 | chr2-11106903-INS-582 |
| chr2-11108088-INS-263 | chr2-11108088-INS-285 |
| chr2-11144090-INS-117 | chr2-11144090-INS-117 |
| chr2-11145819-INS-306 | chr2-11145819-INS-306 |
| chr2-11159569-INS-60 | chr2-11159566-INS-63 |
| chr2-11167280-INS-174 | chr2-11167280-INS-134 |
| chr2-11172657-INS-212 | chr2-11172657-INS-212 |
| chr2-11218725-INS-52 | chr2-11218725-INS-54 |
| chr2-11225198-INS-84 | chr2-11225198-INS-66 |
| chr2-11285899-INS-79 | chr2-11285899-INS-67 |
| chr2-11287823-INS-193 | chr2-11287823-INS-193 |
| chr2-11297351-INS-71 | chr2-11297350-INS-75 |
| chr2-11313897-INS-572 | chr2-11313897-INS-577 |
| chr2-11317279-INS-71 | chr2-11317279-INS-102 |
| chr2-11318356-INS-84 | chr2-11318356-INS-84 |

| | |
|---|---|
| chr2-11335842-INS-297 | chr2-11335842-INS-306 |
| chr2-11350378-INS-87 | chr2-11350378-INS-88 |
| chr2-11353327-INS-144 | chr2-11353327-INS-116 |
| chr2-11373786-INS-6129 | chr2-11373787-INS-6129 |
| chr2-11402438-INS-60 | chr2-11402438-INS-65 |
| chr2-11409421-INS-196 | chr2-11409421-INS-196 |
| chr2-11422187-INS-54 | chr2-11422156-INS-58 |
| chr2-11445210-INS-70 | chr2-11445210-INS-70 |
| chr2-11455195-INS-193 | chr2-11455195-INS-193 |
| chr2-11465771-INS-108 | chr2-11465787-INS-63 |
| chr2-11475716-INS-192 | chr2-11475716-INS-192 |
| chr2-11514209-INS-199 | chr2-11514209-INS-200 |
| chr2-11522633-INS-211 | chr2-11522633-INS-211 |
| chr2-11533641-INS-53 | chr2-11533661-INS-64 |
| chr2-11543522-INS-272 | chr2-11543522-INS-273 |
| chr2-11554379-INS-866 | chr2-11554379-INS-859 |
| chr2-11554399-INS-57 | chr2-11554399-INS-57 |
| chr2-11571680-INS-161 | chr2-11571682-INS-174 |
| chr2-11575000-INS-74 | chr2-11575000-INS-78 |
| chr2-11578244-INS-53 | chr2-11578245-INS-51 |
| chr2-11602517-INS-100 | chr2-11602517-INS-50 |
| chr2-11606595-INS-183 | chr2-11606595-INS-187 |
| chr2-11616442-INS-6453 | chr2-11616443-INS-6453 |
| chr2-11636519-INS-252 | chr2-11636519-INS-270 |
| chr2-11638225-INS-55 | chr2-11638225-INS-55 |
| chr2-11644369-INS-182 | chr2-11644369-INS-182 |
| chr2-11671651-INS-2769 | chr2-11671652-INS-2769 |
| chr2-11687085-INS-431 | chr2-11687085-INS-430 |
| chr2-11703237-INS-204 | chr2-11703237-INS-205 |
| chr2-11706886-INS-519 | chr2-11706886-INS-532 |
| chr2-11725490-INS-55 | chr2-11725490-INS-55 |
| chr2-11731442-INS-82 | chr2-11731442-INS-82 |
| chr2-11736223-INS-52 | chr2-11736223-INS-52 |
| chr2-11741414-INS-6453 | chr2-11741415-INS-6453 |
| chr2-11741851-INS-715 | chr2-11741851-INS-715 |
| chr2-11786089-INS-168 | chr2-11786089-INS-168 |
| chr2-11796108-INS-53 | chr2-11796108-INS-53 |
| chr2-11799951-INS-701 | chr2-11799951-INS-700 |
| chr2-11829778-INS-7855 | chr2-11829779-INS-7857 |
| chr2-11873450-INS-78 | chr2-11873450-INS-78 |
| chr2-11890148-INS-80 | chr2-11890152-INS-100 |
| chr2-11899130-INS-108 | chr2-11899130-INS-108 |
| chr2-11907678-INS-418 | chr2-11907678-INS-418 |
| chr2-11928127-INS-157 | chr2-11928127-INS-149 |
| chr2-11983454-INS-66 | chr2-11983454-INS-66 |
| chr2-11993373-INS-422 | chr2-11993373-INS-430 |
| chr2-12018030-INS-54 | chr2-12018030-INS-54 |
| chr2-12043833-INS-534 | chr2-12043833-INS-534 |
| chr2-12069630-INS-130 | chr2-12069630-INS-146 |
| chr2-12092785-INS-64 | chr2-12092785-INS-64 |
| chr2-12099600-INS-398 | chr2-12099600-INS-398 |
| chr2-12116444-INS-125 | chr2-12116444-INS-129 |
| chr2-12126972-INS-402 | chr2-12126972-INS-399 |
| chr2-12133234-INS-341 | chr2-12133234-INS-341 |
| chr2-12170401-INS-58 | chr2-12170401-INS-60 |
| chr2-12173163-INS-72 | chr2-12173156-INS-66 |
| chr2-12208957-INS-132 | chr2-12208957-INS-132 |
| chr2-12214919-INS-6454 | chr2-12214920-INS-6454 |
| chr2-12216131-INS-192 | chr2-12216131-INS-192 |
| chr2-12225197-INS-73 | chr2-12225197-INS-73 |
| chr2-12235318-INS-56 | chr2-12235318-INS-58 |
| chr2-12239410-INS-61 | chr2-12239410-INS-65 |
| chr2-12242149-INS-86 | chr2-12242149-INS-88 |
| chr2-12252445-INS-115 | chr2-12252445-INS-117 |
| chr2-12255619-INS-722 | chr2-12255619-INS-722 |
| chr2-12259331-INS-134 | chr2-12259350-INS-134 |
| chr2-12286458-INS-712 | chr2-12286458-INS-710 |
| chr2-12349536-INS-55 | chr2-12349536-INS-55 |
| chr2-12372726-INS-56 | chr2-12372726-INS-56 |
| chr2-12437377-INS-135 | chr2-12437377-INS-136 |
| chr2-12442089-INS-103 | chr2-12442089-INS-103 |
| chr2-12447542-INS-214 | chr2-12447542-INS-213 |
| chr2-12466121-INS-60 | chr2-12466121-INS-56 |
| chr2-12467861-INS-90 | chr2-12467861-INS-90 |
| chr2-12531348-INS-209 | chr2-12531348-INS-207 |
| chr2-12536287-INS-389 | chr2-12536287-INS-389 |
| chr2-12543374-INS-203 | chr2-12543374-INS-203 |

| | |
|---|---|
| chr2-12555581-INS-730 | chr2-12555581-INS-748 |
| chr2-12595810-INS-427 | chr2-12595810-INS-428 |
| chr2-12619928-INS-201 | chr2-12619928-INS-201 |
| chr2-12634658-INS-62 | chr2-12634658-INS-64 |
| chr2-12675377-INS-6405 | chr2-12675378-INS-6405 |
| chr2-12682343-INS-361 | chr2-12682334-INS-367 |
| chr2-12684951-INS-60 | chr2-12684943-INS-100 |
| chr2-12696408-INS-55 | chr2-12696408-INS-52 |
| chr2-12711009-INS-124 | chr2-12711009-INS-124 |
| chr2-12798105-INS-96 | chr2-12798105-INS-65 |
| chr2-12828358-INS-104 | chr2-12828358-INS-104 |
| chr2-12832115-INS-843 | chr2-12832115-INS-845 |
| chr2-12855386-INS-740 | chr2-12855386-INS-741 |
| chr2-12857343-INS-74 | chr2-12857343-INS-74 |
| chr2-12902651-INS-247 | chr2-12902651-INS-235 |
| chr2-12911426-INS-154 | chr2-12911426-INS-154 |
| chr2-12913367-INS-2322 | chr2-12913368-INS-2322 |
| chr2-12931654-INS-86 | chr2-12931656-INS-94 |
| chr2-12933627-INS-62 | chr2-12933636-INS-54 |
| chr2-12972481-INS-234 | chr2-12972481-INS-234 |
| chr2-12976208-INS-68 | chr2-12976223-INS-64 |
| chr2-12994537-INS-80 | chr2-12994537-INS-60 |
| chr2-13036346-INS-72 | chr2-13036349-INS-58 |
| chr2-13042186-INS-100 | chr2-13042187-INS-81 |
| chr2-13052625-INS-3015 | chr2-13052626-INS-3015 |
| chr2-13139882-INS-68 | chr2-13139882-INS-68 |
| chr2-13172231-INS-71 | chr2-13172231-INS-77 |
| chr2-13193465-INS-188 | chr2-13193465-INS-188 |
| chr2-13262597-INS-1890 | chr2-13262598-INS-1890 |
| chr2-13272004-INS-67 | chr2-13272004-INS-67 |
| chr2-13286475-INS-54 | chr2-13286475-INS-54 |
| chr2-13348504-INS-77 | chr2-13348483-INS-81 |
| chr2-13348746-INS-518 | chr2-13348746-INS-518 |
| chr2-13495495-INS-63 | chr2-13495496-INS-89 |
| chr2-13520709-INS-59 | chr2-13520709-INS-59 |
| chr2-13567211-INS-6508 | chr2-13567212-INS-6509 |
| chr2-13594305-INS-612 | chr2-13594305-INS-612 |
| chr2-13595890-INS-394 | chr2-13595890-INS-444 |
| chr2-13608484-INS-459 | chr2-13608484-INS-459 |
| chr2-13615693-INS-122 | chr2-13615693-INS-122 |
| chr2-13622333-INS-416 | chr2-13622334-INS-416 |
| chr2-13630513-INS-84 | chr2-13630513-INS-84 |
| chr2-13631975-INS-110 | chr2-13631975-INS-91 |
| chr2-13697777-INS-60 | chr2-13697777-INS-60 |
| chr2-13706121-INS-381 | chr2-13706121-INS-380 |
| chr2-13708630-INS-238 | chr2-13708630-INS-240 |
| chr2-13729435-INS-91 | chr2-13729435-INS-92 |
| chr2-13748517-INS-63 | chr2-13748532-INS-52 |
| chr2-13752270-INS-2674 | chr2-13752271-INS-2674 |
| chr2-13767406-INS-249 | chr2-13767401-INS-282 |
| chr2-13797447-INS-81 | chr2-13797447-INS-80 |
| chr2-13839009-INS-341 | chr2-13839009-INS-341 |
| chr2-13866211-INS-146 | chr2-13866211-INS-146 |
| chr2-13875116-INS-201 | chr2-13875116-INS-201 |
| chr2-13893428-INS-60 | chr2-13893428-INS-60 |
| chr2-13896279-INS-181 | chr2-13896279-INS-182 |
| chr2-13943429-INS-226 | chr2-13943429-INS-221 |
| chr2-13965905-INS-967 | chr2-13965905-INS-967 |
| chr2-14008159-INS-480 | chr2-14008159-INS-480 |
| chr2-14033541-INS-212 | chr2-14033541-INS-212 |
| chr2-14036516-INS-5229 | chr2-14036517-INS-5229 |
| chr2-14040359-INS-174 | chr2-14040359-INS-129 |
| chr2-14084327-INS-236 | chr2-14084329-INS-247 |
| chr2-14104373-INS-90 | chr2-14104373-INS-90 |
| chr2-14129876-INS-207 | chr2-14129876-INS-207 |
| chr2-14142691-INS-74 | chr2-14142691-INS-67 |
| chr2-14168707-INS-56 | chr2-14168707-INS-73 |
| chr2-14180424-INS-61 | chr2-14180424-INS-72 |
| chr2-14201385-INS-104 | chr2-14201385-INS-104 |
| chr2-14205487-INS-171 | chr2-14205463-INS-204 |
| chr2-14221241-INS-54 | chr2-14221266-INS-55 |
| chr2-14231037-INS-5827 | chr2-14231023-INS-5822 |
| chr2-14268497-INS-141 | chr2-14268497-INS-141 |
| chr2-14329401-INS-60 | chr2-14329401-INS-60 |
| chr2-14331705-INS-100 | chr2-14331705-INS-102 |
| chr2-14339352-INS-294 | chr2-14339353-INS-294 |
| chr2-14347741-INS-70 | chr2-14347735-INS-78 |

| | |
|---|---|
| chr2-14404345-INS-86 | chr2-14404345-INS-92 |
| chr2-14423432-INS-232 | chr2-14423434-INS-233 |
| chr2-14432302-INS-695 | chr2-14432302-INS-695 |
| chr2-14462291-INS-528 | chr2-14462291-INS-528 |
| chr2-14481119-INS-51 | chr2-14481119-INS-51 |
| chr2-14498136-INS-121 | chr2-14498136-INS-121 |
| chr2-14502423-INS-4614 | chr2-14502424-INS-4614 |
| chr2-14525494-INS-62 | chr2-14525494-INS-86 |
| chr2-14534545-INS-6587 | chr2-14534546-INS-6587 |
| chr2-14537317-INS-213 | chr2-14537317-INS-213 |
| chr2-14551538-INS-81 | chr2-14551538-INS-108 |
| chr2-14591152-INS-485 | chr2-14591152-INS-485 |
| chr2-14599929-INS-486 | chr2-14599929-INS-484 |
| chr2-14602968-INS-57 | chr2-14602968-INS-57 |
| chr2-14663500-INS-6619 | chr2-14663501-INS-6620 |
| chr2-14671237-INS-81 | chr2-14671233-INS-77 |
| chr2-14693335-INS-811 | chr2-14693335-INS-812 |
| chr2-14693464-INS-52 | chr2-14693462-INS-84 |
| chr2-14713259-INS-96 | chr2-14713250-INS-76 |
| chr2-14728687-INS-54 | chr2-14728687-INS-54 |
| chr2-14770732-INS-539 | chr2-14770732-INS-552 |
| chr2-14778223-INS-6985 | chr2-14778224-INS-6985 |
| chr2-14792540-INS-497 | chr2-14792541-INS-497 |
| chr2-14814597-INS-70 | chr2-14814598-INS-66 |
| chr2-14826426-INS-489 | chr2-14826426-INS-490 |
| chr2-14829633-INS-79 | chr2-14829633-INS-79 |
| chr2-14846819-INS-89 | chr2-14846819-INS-81 |
| chr2-14874021-INS-59 | chr2-14873998-INS-59 |
| chr2-14892467-INS-86 | chr2-14892440-INS-85 |
| chr2-14914116-INS-71 | chr2-14914116-INS-97 |
| chr2-14920272-INS-63 | chr2-14920272-INS-63 |
| chr2-14929793-INS-118 | chr2-14929793-INS-118 |
| chr2-14936553-INS-144 | chr2-14936553-INS-143 |
| chr2-14978242-INS-867 | chr2-14978242-INS-867 |
| chr2-14980699-INS-60 | chr2-14980699-INS-60 |
| chr2-15011800-INS-66 | chr2-15011800-INS-82 |
| chr2-15016516-INS-3049 | chr2-15016517-INS-3049 |
| chr2-15021504-INS-245 | chr2-15021504-INS-254 |
| chr2-15043323-INS-822 | chr2-15043323-INS-822 |
| chr2-15047297-INS-124 | chr2-15047297-INS-124 |
| chr2-15052915-INS-54 | chr2-15052924-INS-54 |
| chr2-15098546-INS-7626 | chr2-15098535-INS-7627 |
| chr2-15119055-INS-651 | chr2-15119055-INS-651 |
| chr2-15146311-INS-158 | chr2-15146311-INS-158 |
| chr2-15160362-INS-1720 | chr2-15160363-INS-1721 |
| chr2-15172367-INS-64 | chr2-15172367-INS-64 |
| chr2-15174971-INS-423 | chr2-15174971-INS-389 |
| chr2-15176562-INS-85 | chr2-15176561-INS-80 |
| chr2-15191937-INS-58 | chr2-15191935-INS-54 |
| chr2-15214340-INS-4156 | chr2-15214341-INS-4156 |
| chr2-15233091-INS-354 | chr2-15233091-INS-354 |
| chr2-15245552-INS-75 | chr2-15245552-INS-75 |
| chr2-15270408-INS-56 | chr2-15270432-INS-50 |
| chr2-15280163-INS-3502 | chr2-15280164-INS-3502 |
| chr2-15297813-INS-960 | chr2-15297813-INS-938 |
| chr2-15298672-INS-513 | chr2-15298672-INS-513 |
| chr2-15299848-INS-519 | chr2-15299848-INS-519 |
| chr2-15305304-INS-88 | chr2-15305299-INS-53 |
| chr2-15314730-INS-65 | chr2-15314730-INS-65 |
| chr2-15323342-INS-4338 | chr2-15323343-INS-4338 |
| chr2-15335135-INS-11648 | chr2-15335136-INS-11675 |
| chr2-15341328-INS-393 | chr2-15341328-INS-394 |
| chr2-15378633-INS-177 | chr2-15378633-INS-178 |
| chr2-15389970-INS-97 | chr2-15389970-INS-97 |
| chr2-15396152-INS-138 | chr2-15396152-INS-138 |
| chr2-15403097-INS-1599 | chr2-15403098-INS-1597 |
| chr2-15404072-INS-6571 | chr2-15404088-INS-6600 |
| chr2-15475343-INS-428 | chr2-15475343-INS-428 |
| chr2-15486190-INS-6821 | chr2-15486191-INS-6822 |
| chr2-15502243-INS-160 | chr2-15502243-INS-161 |
| chr2-15528702-INS-93 | chr2-15528702-INS-93 |
| chr2-15589949-INS-55 | chr2-15589949-INS-55 |
| chr2-15595562-INS-67 | chr2-15595562-INS-70 |
| chr2-15618731-INS-390 | chr2-15618731-INS-390 |
| chr2-15626042-INS-67 | chr2-15626042-INS-71 |
| chr2-15643158-INS-177 | chr2-15643165-INS-178 |
| chr2-15659252-INS-86 | chr2-15659248-INS-67 |

| | |
|---|---|
| chr2-15675631-INS-54 | chr2-15675634-INS-58 |
| chr2-15678091-INS-54 | chr2-15678115-INS-71 |
| chr2-15683735-INS-6404 | chr2-15683736-INS-6404 |
| chr2-15703270-INS-2737 | chr2-15703271-INS-2737 |
| chr2-15706683-INS-766 | chr2-15706683-INS-767 |
| chr2-15715290-INS-3234 | chr2-15715325-INS-3227 |
| chr2-15781794-INS-60 | chr2-15781794-INS-55 |
| chr2-15790941-INS-162 | chr2-15790941-INS-128 |
| chr2-15811573-INS-51 | chr2-15811573-INS-51 |
| chr2-15836518-INS-6449 | chr2-15836519-INS-6446 |
| chr2-15838352-INS-242 | chr2-15838352-INS-242 |
| chr2-15849580-INS-86 | chr2-15849580-INS-88 |
| chr2-15851121-INS-200 | chr2-15851121-INS-200 |
| chr2-15852789-INS-198 | chr2-15852789-INS-198 |
| chr2-15854726-INS-669 | chr2-15854726-INS-669 |
| chr2-15867560-INS-72 | chr2-15867560-INS-72 |
| chr2-15882422-INS-350 | chr2-15882422-INS-350 |
| chr2-15907695-INS-64 | chr2-15907665-INS-72 |
| chr2-15943618-INS-227 | chr2-15943618-INS-226 |
| chr2-15947827-INS-175 | chr2-15947827-INS-181 |
| chr2-15951022-INS-664 | chr2-15951022-INS-648 |
| chr2-15955192-INS-611 | chr2-15955192-INS-609 |
| chr2-15960984-INS-198 | chr2-15960958-INS-236 |
| chr2-15976514-INS-7195 | chr2-15976515-INS-7195 |
| chr2-16038255-INS-121 | chr2-16038255-INS-127 |
| chr2-16072056-INS-61 | chr2-16072056-INS-62 |
| chr2-16075947-INS-96 | chr2-16075947-INS-96 |
| chr2-16132297-INS-62 | chr2-16132296-INS-66 |
| chr2-16172168-INS-423 | chr2-16172168-INS-423 |
| chr2-16218167-INS-1334 | chr2-16218168-INS-1334 |
| chr2-16222107-INS-354 | chr2-16222107-INS-354 |
| chr2-16228506-INS-51 | chr2-16228506-INS-51 |
| chr2-16229794-INS-287 | chr2-16229794-INS-287 |
| chr2-16249252-INS-4388 | chr2-16249253-INS-4388 |
| chr2-16255036-INS-110 | chr2-16255036-INS-142 |
| chr2-16255879-INS-242 | chr2-16255879-INS-228 |
| chr2-16278518-INS-72 | chr2-16278519-INS-72 |
| chr2-16285442-INS-57 | chr2-16285439-INS-50 |
| chr2-16333644-INS-3019 | chr2-16333645-INS-3019 |
| chr2-16342068-INS-60 | chr2-16342068-INS-60 |
| chr2-16359497-INS-504 | chr2-16359497-INS-504 |
| chr2-16407077-INS-88 | chr2-16407077-INS-93 |
| chr2-16426015-INS-181 | chr2-16426015-INS-181 |
| chr2-16427777-INS-58 | chr2-16427772-INS-96 |
| chr2-16443216-INS-74 | chr2-16443216-INS-74 |
| chr2-16445095-INS-280 | chr2-16445095-INS-280 |
| chr2-16450231-INS-57 | chr2-16450232-INS-61 |
| chr2-16459686-INS-53 | chr2-16459686-INS-53 |
| chr2-16477020-INS-198 | chr2-16477022-INS-214 |
| chr2-16478283-INS-121 | chr2-16478285-INS-100 |
| chr2-16479081-INS-732 | chr2-16479081-INS-730 |
| chr2-16480833-INS-157 | chr2-16480833-INS-157 |
| chr2-16495486-INS-783 | chr2-16495486-INS-783 |
| chr2-16504795-INS-198 | chr2-16504795-INS-198 |
| chr2-16508904-INS-208 | chr2-16508904-INS-208 |
| chr2-16516259-INS-70 | chr2-16516233-INS-99 |
| chr2-16532664-INS-366 | chr2-16532664-INS-365 |
| chr2-16612203-INS-66 | chr2-16612203-INS-60 |
| chr2-16617060-INS-855 | chr2-16617060-INS-855 |
| chr2-16640064-INS-457 | chr2-16640064-INS-457 |
| chr2-16640684-INS-74 | chr2-16640684-INS-70 |
| chr2-16733685-INS-60 | chr2-16733685-INS-56 |
| chr2-16816163-INS-57 | chr2-16816163-INS-57 |
| chr2-16827499-INS-6150 | chr2-16827508-INS-6148 |
| chr2-16836485-INS-204 | chr2-16836485-INS-206 |
| chr2-16842860-INS-404 | chr2-16842860-INS-404 |
| chr2-16863339-INS-980 | chr2-16863339-INS-980 |
| chr2-16863692-INS-107 | chr2-16863692-INS-107 |
| chr2-16882188-INS-536 | chr2-16882188-INS-536 |
| chr2-16965822-INS-108 | chr2-16965822-INS-115 |
| chr2-17094744-INS-5181 | chr2-17094745-INS-5181 |
| chr2-17139323-INS-61 | chr2-17139323-INS-61 |
| chr2-17171205-INS-6343 | chr2-17171206-INS-6343 |
| chr2-17178058-INS-1052 | chr2-17178059-INS-1052 |
| chr2-17182163-INS-177 | chr2-17182163-INS-177 |
| chr2-17235460-INS-155 | chr2-17235460-INS-155 |
| chr2-17252073-INS-604 | chr2-17252072-INS-606 |

| | |
|---|---|
| chr2-17287987-INS-4050 | chr2-17287988-INS-4050 |
| chr2-17306030-INS-82 | chr2-17306030-INS-86 |
| chr2-17309060-INS-177 | chr2-17309060-INS-177 |
| chr2-17339627-INS-272 | chr2-17339627-INS-276 |
| chr2-17365732-INS-74 | chr2-17365732-INS-74 |
| chr2-17412351-INS-144 | chr2-17412351-INS-160 |
| chr2-17433045-INS-68 | chr2-17433033-INS-52 |
| chr2-17456411-INS-67 | chr2-17456411-INS-67 |
| chr2-17561597-INS-249 | chr2-17561597-INS-250 |
| chr2-17568316-INS-1890 | chr2-17568317-INS-1890 |
| chr2-17625189-INS-902 | chr2-17625189-INS-910 |
| chr2-17630633-INS-1768 | chr2-17630634-INS-1772 |
| chr2-17681448-INS-683 | chr2-17681448-INS-683 |
| chr2-17684349-INS-52 | chr2-17684349-INS-52 |
| chr2-17719875-INS-82 | chr2-17719886-INS-52 |
| chr2-17778140-INS-113 | chr2-17778140-INS-81 |
| chr2-17785517-INS-75 | chr2-17785517-INS-75 |
| chr2-17793904-INS-60 | chr2-17793884-INS-76 |
| chr2-17817942-INS-208 | chr2-17817942-INS-208 |
| chr2-17838024-INS-99 | chr2-17838024-INS-99 |
| chr2-17839823-INS-66 | chr2-17839823-INS-66 |
| chr2-17854119-INS-149 | chr2-17854119-INS-149 |
| chr2-17867906-INS-58 | chr2-17867906-INS-62 |
| chr2-17879810-INS-72 | chr2-17879810-INS-100 |
| chr2-17880873-INS-924 | chr2-17880873-INS-924 |
| chr2-17888866-INS-524 | chr2-17888866-INS-524 |
| chr2-17891501-INS-125 | chr2-17891501-INS-125 |
| chr2-17903477-INS-124 | chr2-17903477-INS-93 |
| chr2-17906753-INS-201 | chr2-17906753-INS-203 |
| chr2-17910066-INS-277 | chr2-17910066-INS-278 |
| chr2-17912390-INS-62 | chr2-17912355-INS-60 |
| chr2-17931797-INS-72 | chr2-17931793-INS-52 |
| chr2-17946816-INS-262 | chr2-17946816-INS-262 |
| chr2-17948391-INS-209 | chr2-17948391-INS-209 |
| chr2-17955212-INS-84 | chr2-17955212-INS-100 |
| chr2-17960819-INS-217 | chr2-17960819-INS-218 |
| chr2-17963860-INS-497 | chr2-17963860-INS-497 |
| chr2-17964907-INS-6454 | chr2-17964908-INS-6454 |
| chr2-17967136-INS-778 | chr2-17967136-INS-776 |
| chr2-17973876-INS-80 | chr2-17973876-INS-90 |
| chr2-17975935-INS-66 | chr2-17975935-INS-66 |
| chr2-17977470-INS-313 | chr2-17977470-INS-320 |
| chr2-17980053-INS-199 | chr2-17980053-INS-199 |
| chr2-17981591-INS-481 | chr2-17981591-INS-481 |
| chr2-17992638-INS-260 | chr2-17992643-INS-215 |
| chr2-18006220-INS-214 | chr2-18006220-INS-209 |
| chr2-18012580-INS-202 | chr2-18012580-INS-202 |
| chr2-18019796-INS-196 | chr2-18019796-INS-196 |
| chr2-18094630-INS-219 | chr2-18094635-INS-232 |
| chr2-18118658-INS-56 | chr2-18118658-INS-56 |
| chr2-18134482-INS-221 | chr2-18134482-INS-222 |
| chr2-18135301-INS-989 | chr2-18135305-INS-952 |
| chr2-18142646-INS-215 | chr2-18142646-INS-214 |
| chr2-18149236-INS-301 | chr2-18149226-INS-343 |
| chr2-18150342-INS-219 | chr2-18150342-INS-215 |
| chr2-18160935-INS-211 | chr2-18160935-INS-211 |
| chr2-18176352-INS-74 | chr2-18176352-INS-85 |
| chr2-18191771-INS-140 | chr2-18191771-INS-140 |
| chr2-18194595-INS-52 | chr2-18194597-INS-50 |
| chr2-18217914-INS-776 | chr2-18217914-INS-783 |
| chr2-18254056-INS-575 | chr2-18254056-INS-575 |
| chr2-18279893-INS-239 | chr2-18279877-INS-243 |
| chr2-18285919-INS-821 | chr2-18285919-INS-814 |
| chr2-18304596-INS-203 | chr2-18304596-INS-203 |
| chr2-18306705-INS-367 | chr2-18306705-INS-369 |
| chr2-18337191-INS-951 | chr2-18337191-INS-968 |
| chr2-18341781-INS-98 | chr2-18341781-INS-62 |
| chr2-18355873-INS-84 | chr2-18355873-INS-104 |
| chr2-18367090-INS-88 | chr2-18367090-INS-88 |
| chr2-18373365-INS-62 | chr2-18373365-INS-63 |
| chr2-18374998-INS-1901 | chr2-18374999-INS-1888 |
| chr2-18399865-INS-416 | chr2-18399865-INS-416 |
| chr2-18413860-INS-172 | chr2-18413860-INS-172 |
| chr2-18420689-INS-111 | chr2-18420690-INS-112 |
| chr2-18422270-INS-503 | chr2-18422270-INS-503 |
| chr2-18433372-INS-69 | chr2-18433374-INS-55 |
| chr2-18434298-INS-4555 | chr2-18434299-INS-4555 |

| | |
|---|---|
| chr2-18440530-INS-236 | chr2-18440530-INS-236 |
| chr2-18451651-INS-372 | chr2-18451651-INS-372 |
| chr2-18456938-INS-5040 | chr2-18456939-INS-5040 |
| chr2-18468038-INS-194 | chr2-18468038-INS-195 |
| chr2-18471442-INS-145 | chr2-18471442-INS-145 |
| chr2-18497393-INS-431 | chr2-18497393-INS-431 |
| chr2-18520214-INS-401 | chr2-18520214-INS-401 |
| chr2-18521832-INS-5308 | chr2-18521833-INS-5308 |
| chr2-18552202-INS-71 | chr2-18552200-INS-55 |
| chr2-18553367-INS-74 | chr2-18553374-INS-85 |
| chr2-18573701-INS-118 | chr2-18573699-INS-119 |
| chr2-18575092-INS-51 | chr2-18575092-INS-51 |
| chr2-18579642-INS-187 | chr2-18579642-INS-187 |
| chr2-18605624-INS-94 | chr2-18605610-INS-115 |
| chr2-18617543-INS-207 | chr2-18617543-INS-207 |
| chr2-18634234-INS-104 | chr2-18634234-INS-104 |
| chr2-18648057-INS-416 | chr2-18648057-INS-416 |
| chr2-18653735-INS-114 | chr2-18653730-INS-114 |
| chr2-18673139-INS-122 | chr2-18673139-INS-122 |
| chr2-18678226-INS-52 | chr2-18678247-INS-61 |
| chr2-18707678-INS-417 | chr2-18707678-INS-416 |
| chr2-18730070-INS-64 | chr2-18730070-INS-64 |
| chr2-18734094-INS-3490 | chr2-18734095-INS-3490 |
| chr2-18740840-INS-103 | chr2-18740837-INS-87 |
| chr2-18743398-INS-111 | chr2-18743398-INS-111 |
| chr2-18772813-INS-159 | chr2-18772832-INS-176 |
| chr2-18811801-INS-133 | chr2-18811801-INS-168 |
| chr2-18817276-INS-60 | chr2-18817276-INS-56 |
| chr2-18828721-INS-85 | chr2-18828721-INS-85 |
| chr2-18868908-INS-72 | chr2-18868908-INS-72 |
| chr2-18899029-INS-53 | chr2-18899029-INS-55 |
| chr2-18905715-INS-773 | chr2-18905715-INS-773 |
| chr2-18910796-INS-295 | chr2-18910803-INS-294 |
| chr2-18927120-INS-120 | chr2-18927120-INS-120 |
| chr2-18939639-INS-76 | chr2-18939639-INS-76 |
| chr2-18972142-INS-1173 | chr2-18972143-INS-1173 |
| chr2-18976231-INS-431 | chr2-18976231-INS-431 |
| chr2-18980669-INS-96 | chr2-18980669-INS-96 |
| chr2-18992435-INS-211 | chr2-18992435-INS-211 |
| chr2-19017229-INS-6599 | chr2-19017230-INS-6599 |
| chr2-19038327-INS-182 | chr2-19038327-INS-186 |
| chr2-19059794-INS-76 | chr2-19059794-INS-50 |
| chr2-19070267-INS-66 | chr2-19070267-INS-88 |
| chr2-19082953-INS-1043 | chr2-19082954-INS-1039 |
| chr2-19126461-INS-88 | chr2-19126504-INS-54 |
| chr2-19127402-INS-210 | chr2-19127402-INS-210 |
| chr2-19136596-INS-65 | chr2-19136596-INS-59 |
| chr2-19146045-INS-50 | chr2-19146045-INS-62 |
| chr2-19157477-INS-161 | chr2-19157477-INS-181 |
| chr2-19163685-INS-6544 | chr2-19163686-INS-6545 |
| chr2-19177244-INS-84 | chr2-19177244-INS-85 |
| chr2-19197408-INS-209 | chr2-19197408-INS-209 |
| chr2-19199638-INS-113 | chr2-19199638-INS-126 |
| chr2-19236276-INS-177 | chr2-19236276-INS-177 |
| chr2-19240140-INS-2096 | chr2-19240141-INS-2096 |
| chr2-19252388-INS-52 | chr2-19252388-INS-54 |
| chr2-19266891-INS-58 | chr2-19266891-INS-78 |
| chr2-19277181-INS-89 | chr2-19277181-INS-78 |
| chr2-19319295-INS-605 | chr2-19319295-INS-607 |
| chr2-19322625-INS-102 | chr2-19322626-INS-126 |
| chr2-19332869-INS-1613 | chr2-19332870-INS-1617 |
| chr2-19335985-INS-66 | chr2-19335985-INS-70 |
| chr2-19348299-INS-224 | chr2-19348299-INS-224 |
| chr2-19374526-INS-51 | chr2-19374526-INS-57 |
| chr2-19377303-INS-53 | chr2-19377303-INS-53 |
| chr2-19383716-INS-174 | chr2-19383704-INS-187 |
| chr2-19385646-INS-66 | chr2-19385646-INS-66 |
| chr2-19387092-INS-154 | chr2-19387092-INS-164 |
| chr2-19424551-INS-94 | chr2-19424551-INS-94 |
| chr2-19428869-INS-92 | chr2-19428893-INS-93 |
| chr2-19474792-INS-111 | chr2-19474799-INS-125 |
| chr2-19541426-INS-123 | chr2-19541426-INS-123 |
| chr2-19560014-INS-310 | chr2-19560014-INS-310 |
| chr2-19576265-INS-203 | chr2-19576265-INS-203 |
| chr2-19594200-INS-223 | chr2-19594200-INS-223 |
| chr2-19603017-INS-146 | chr2-19603017-INS-151 |
| chr2-19650555-INS-416 | chr2-19650555-INS-416 |

| | |
|---|---|
| chr2-19701345-INS-55 | chr2-19701345-INS-55 |
| chr2-19729859-INS-57 | chr2-19729859-INS-57 |
| chr2-19732823-INS-54 | chr2-19732823-INS-58 |
| chr2-19750924-INS-7104 | chr2-19750925-INS-7104 |
| chr2-19757251-INS-168 | chr2-19757252-INS-158 |
| chr2-19773117-INS-446 | chr2-19773117-INS-446 |
| chr2-19834116-INS-6730 | chr2-19834117-INS-6730 |
| chr2-19846547-INS-256 | chr2-19846547-INS-256 |
| chr2-19853772-INS-276 | chr2-19853772-INS-276 |
| chr2-19860759-INS-282 | chr2-19860759-INS-282 |
| chr2-19890810-INS-480 | chr2-19890810-INS-480 |
| chr2-19930202-INS-6535 | chr2-19930203-INS-6535 |
| chr2-19936721-INS-84 | chr2-19936721-INS-81 |
| chr2-19960328-INS-611 | chr2-19960328-INS-611 |
| chr2-19983185-INS-63 | chr2-19983189-INS-64 |
| chr2-19984351-INS-66 | chr2-19984351-INS-68 |
| chr2-20055944-INS-60 | chr2-20055944-INS-60 |
| chr2-20059179-INS-153 | chr2-20059179-INS-153 |
| chr2-20074255-INS-85 | chr2-20074255-INS-85 |
| chr2-20083645-INS-84 | chr2-20083645-INS-60 |
| chr2-20088640-INS-323 | chr2-20088640-INS-323 |
| chr2-20121233-INS-2996 | chr2-20121234-INS-2996 |
| chr2-20134141-INS-922 | chr2-20134141-INS-922 |
| chr2-20166365-INS-119 | chr2-20166359-INS-110 |
| chr2-20188502-INS-802 | chr2-20188502-INS-802 |
| chr2-20202258-INS-272 | chr2-20202258-INS-272 |
| chr2-20214135-INS-158 | chr2-20214135-INS-158 |
| chr2-20290861-INS-2090 | chr2-20290862-INS-2090 |
| chr2-20298005-INS-210 | chr2-20298005-INS-211 |
| chr2-20301964-INS-91 | chr2-20301963-INS-95 |
| chr2-20314079-INS-60 | chr2-20314079-INS-60 |
| chr2-20318377-INS-456 | chr2-20318377-INS-469 |
| chr2-20360551-INS-91 | chr2-20360551-INS-91 |
| chr2-20365665-INS-72 | chr2-20365665-INS-73 |
| chr2-20368765-INS-6415 | chr2-20368766-INS-6415 |
| chr2-20373745-INS-297 | chr2-20373745-INS-297 |
| chr2-20412454-INS-66 | chr2-20412408-INS-81 |
| chr2-20443719-INS-174 | chr2-20443719-INS-190 |
| chr2-20444567-INS-182 | chr2-20444567-INS-182 |
| chr2-20478457-INS-64 | chr2-20478455-INS-70 |
| chr2-20570279-INS-76 | chr2-20570279-INS-76 |
| chr2-20582518-INS-192 | chr2-20582518-INS-180 |
| chr2-20590301-INS-234 | chr2-20590301-INS-234 |
| chr2-20605068-INS-86 | chr2-20605068-INS-97 |
| chr2-20630968-INS-77 | chr2-20630968-INS-77 |
| chr2-20636439-INS-204 | chr2-20636439-INS-204 |
| chr2-20637382-INS-67 | chr2-20637382-INS-67 |
| chr2-20639134-INS-208 | chr2-20639141-INS-205 |
| chr2-20648257-INS-214 | chr2-20648257-INS-214 |
| chr2-20652257-INS-96 | chr2-20652297-INS-62 |
| chr2-20656665-INS-515 | chr2-20656665-INS-515 |
| chr2-20664263-INS-98 | chr2-20664263-INS-72 |
| chr2-20669775-INS-823 | chr2-20669775-INS-823 |
| chr2-20677579-INS-210 | chr2-20677536-INS-204 |
| chr2-20681461-INS-186 | chr2-20681461-INS-186 |
| chr2-20683725-INS-5313 | chr2-20683726-INS-5313 |
| chr2-20684152-INS-132 | chr2-20684152-INS-132 |
| chr2-20706242-INS-181 | chr2-20706242-INS-193 |
| chr2-20717297-INS-110 | chr2-20717297-INS-110 |
| chr2-20768873-INS-67 | chr2-20768873-INS-66 |
| chr2-20775281-INS-208 | chr2-20775281-INS-208 |
| chr2-20792284-INS-400 | chr2-20792284-INS-400 |
| chr2-20793683-INS-66 | chr2-20793683-INS-66 |
| chr2-20828439-INS-226 | chr2-20828439-INS-227 |
| chr2-20836222-INS-60 | chr2-20836222-INS-60 |
| chr2-20844115-INS-823 | chr2-20844115-INS-823 |
| chr2-20845606-INS-68 | chr2-20845606-INS-68 |
| chr2-20847504-INS-66 | chr2-20847502-INS-70 |
| chr2-20855935-INS-218 | chr2-20855935-INS-218 |
| chr2-20879586-INS-62 | chr2-20879586-INS-62 |
| chr2-20884681-INS-224 | chr2-20884681-INS-224 |
| chr2-20886799-INS-60 | chr2-20886799-INS-64 |
| chr2-20906405-INS-205 | chr2-20906405-INS-205 |
| chr2-20930350-INS-90 | chr2-20930350-INS-95 |
| chr2-20933637-INS-78 | chr2-20933637-INS-78 |
| chr2-20958186-INS-216 | chr2-20958186-INS-216 |
| chr2-20993212-INS-108 | chr2-20993212-INS-108 |

| | |
|---|---|
| chr2-21031037-INS-565 | chr2-21031037-INS-565 |
| chr2-21049620-INS-7182 | chr2-21049621-INS-7182 |
| chr2-21050467-INS-65 | chr2-21050467-INS-65 |
| chr2-21067661-INS-546 | chr2-21067661-INS-546 |
| chr2-21097393-INS-56 | chr2-21097393-INS-60 |
| chr2-21118731-INS-2321 | chr2-21118720-INS-2305 |
| chr2-21123133-INS-140 | chr2-21123165-INS-150 |
| chr2-21135032-INS-53 | chr2-21135032-INS-54 |
| chr2-21137642-INS-798 | chr2-21137642-INS-803 |
| chr2-21212290-INS-51 | chr2-21212290-INS-51 |
| chr2-21222023-INS-69 | chr2-21222023-INS-69 |
| chr2-21224504-INS-75 | chr2-21224504-INS-75 |
| chr2-21239484-INS-68 | chr2-21239484-INS-62 |
| chr2-21250817-INS-362 | chr2-21250818-INS-362 |
| chr2-21260725-INS-613 | chr2-21260725-INS-625 |
| chr2-21268858-INS-199 | chr2-21268858-INS-199 |
| chr2-21275783-INS-74 | chr2-21275795-INS-76 |
| chr2-21282604-INS-57 | chr2-21282603-INS-66 |
| chr2-21286537-INS-132 | chr2-21286537-INS-132 |
| chr2-21301878-INS-305 | chr2-21301878-INS-305 |
| chr2-21319195-INS-375 | chr2-21319195-INS-383 |
| chr2-21326158-INS-84 | chr2-21326158-INS-84 |
| chr2-21329462-INS-3366 | chr2-21329463-INS-3366 |
| chr2-21342391-INS-102 | chr2-21342391-INS-102 |
| chr2-21343352-INS-154 | chr2-21343352-INS-115 |
| chr2-21388079-INS-1321 | chr2-21388080-INS-1317 |
| chr2-21405577-INS-369 | chr2-21405577-INS-369 |
| chr2-21420799-INS-52 | chr2-21420783-INS-84 |
| chr2-21424225-INS-2535 | chr2-21424226-INS-2535 |
| chr2-21456217-INS-1025 | chr2-21456218-INS-1023 |
| chr2-21457960-INS-257 | chr2-21457960-INS-258 |
| chr2-21481245-INS-66 | chr2-21481245-INS-66 |
| chr2-21516154-INS-51 | chr2-21516154-INS-52 |
| chr2-21539424-INS-1655 | chr2-21539425-INS-1655 |
| chr2-21549259-INS-6038 | chr2-21549260-INS-6038 |
| chr2-21557571-INS-201 | chr2-21557571-INS-201 |
| chr2-21560519-INS-131 | chr2-21560519-INS-131 |
| chr2-21563714-INS-54 | chr2-21563714-INS-54 |
| chr2-21566086-INS-196 | chr2-21566096-INS-208 |
| chr2-21568301-INS-51 | chr2-21568301-INS-51 |
| chr2-21647417-INS-193 | chr2-21647417-INS-193 |
| chr2-21653364-INS-207 | chr2-21653364-INS-207 |
| chr2-21655443-INS-227 | chr2-21655443-INS-239 |
| chr2-21661357-INS-197 | chr2-21661357-INS-197 |
| chr2-21663490-INS-218 | chr2-21663490-INS-215 |
| chr2-21692809-INS-411 | chr2-21692809-INS-410 |
| chr2-21761618-INS-214 | chr2-21761618-INS-214 |
| chr2-21774665-INS-56 | chr2-21774665-INS-56 |
| chr2-21782883-INS-56 | chr2-21782883-INS-50 |
| chr2-21807290-INS-66 | chr2-21807290-INS-66 |
| chr2-21845115-INS-64 | chr2-21845115-INS-68 |
| chr2-21859410-INS-304 | chr2-21859410-INS-305 |
| chr2-21894067-INS-141 | chr2-21894074-INS-145 |
| chr2-21897577-INS-130 | chr2-21897577-INS-131 |
| chr2-21965443-INS-195 | chr2-21965443-INS-196 |
| chr2-22017992-INS-66 | chr2-22017992-INS-68 |
| chr2-22027930-INS-106 | chr2-22027930-INS-106 |
| chr2-22054509-INS-212 | chr2-22054496-INS-227 |
| chr2-22060483-INS-200 | chr2-22060483-INS-200 |
| chr2-22132094-INS-64 | chr2-22132094-INS-72 |
| chr2-22151050-INS-72 | chr2-22151050-INS-72 |
| chr2-22160570-INS-55 | chr2-22160544-INS-60 |
| chr2-22174413-INS-76 | chr2-22174413-INS-74 |
| chr2-22210139-INS-214 | chr2-22210162-INS-214 |
| chr2-22247081-INS-56 | chr2-22247077-INS-60 |
| chr2-22254880-INS-74 | chr2-22254880-INS-74 |
| chr2-22256452-INS-5116 | chr2-22256453-INS-5116 |
| chr2-22261830-INS-1371 | chr2-22261831-INS-1373 |
| chr2-22267249-INS-124 | chr2-22267249-INS-126 |
| chr2-22300984-INS-60 | chr2-22300997-INS-72 |
| chr2-22320486-INS-539 | chr2-22320486-INS-539 |
| chr2-22329911-INS-141 | chr2-22329911-INS-143 |
| chr2-22367160-INS-76 | chr2-22367160-INS-76 |
| chr2-22389534-INS-51 | chr2-22389534-INS-55 |
| chr2-22411704-INS-203 | chr2-22411704-INS-203 |
| chr2-22460911-INS-73 | chr2-22460911-INS-73 |
| chr2-22464371-INS-135 | chr2-22464371-INS-129 |

| | |
|---|---|
| chr2-22485600-INS-404 | chr2-22485560-INS-434 |
| chr2-22499523-INS-409 | chr2-22499523-INS-408 |
| chr2-22511606-INS-90 | chr2-22511606-INS-90 |
| chr2-22532340-INS-204 | chr2-22532340-INS-184 |
| chr2-22537210-INS-51 | chr2-22537210-INS-51 |
| chr2-22546322-INS-231 | chr2-22546322-INS-236 |
| chr2-22577190-INS-91 | chr2-22577190-INS-91 |
| chr2-22579783-INS-860 | chr2-22579783-INS-887 |
| chr2-22587838-INS-81 | chr2-22587838-INS-81 |
| chr2-22593505-INS-198 | chr2-22593505-INS-198 |
| chr2-22604018-INS-57 | chr2-22604018-INS-69 |
| chr2-22629909-INS-72 | chr2-22629909-INS-72 |
| chr2-22654418-INS-56 | chr2-22654418-INS-52 |
| chr2-22656893-INS-94 | chr2-22656893-INS-94 |
| chr2-22659066-INS-768 | chr2-22659066-INS-770 |
| chr2-22679008-INS-372 | chr2-22679007-INS-401 |
| chr2-22682483-INS-82 | chr2-22682483-INS-82 |
| chr2-22745893-INS-209 | chr2-22745893-INS-209 |
| chr2-22781698-INS-215 | chr2-22781698-INS-216 |
| chr2-22795623-INS-412 | chr2-22795623-INS-411 |
| chr2-22821461-INS-247 | chr2-22821461-INS-247 |
| chr2-22864090-INS-99 | chr2-22864090-INS-88 |
| chr2-22887732-INS-265 | chr2-22887732-INS-263 |
| chr2-22897537-INS-167 | chr2-22897537-INS-167 |
| chr2-22933264-INS-89 | chr2-22933264-INS-89 |
| chr2-22938448-INS-78 | chr2-22938411-INS-91 |
| chr2-22963577-INS-380 | chr2-22963577-INS-380 |
| chr2-23012579-INS-161 | chr2-23012580-INS-166 |
| chr2-23018907-INS-128 | chr2-23018907-INS-128 |
| chr2-23051642-INS-201 | chr2-23051642-INS-201 |
| chr2-23054745-INS-196 | chr2-23054745-INS-196 |
| chr2-23075697-INS-88 | chr2-23075697-INS-88 |
| chr2-23077166-INS-210 | chr2-23077166-INS-210 |
| chr2-23112615-INS-57 | chr2-23112615-INS-59 |
| chr2-23167902-INS-399 | chr2-23167903-INS-398 |
| chr2-23208803-INS-72 | chr2-23208803-INS-72 |
| chr2-23226424-INS-457 | chr2-23226424-INS-448 |
| chr2-23316875-INS-636 | chr2-23316876-INS-636 |
| chr2-23325191-INS-116 | chr2-23325191-INS-121 |
| chr2-23443737-INS-4985 | chr2-23443738-INS-4985 |
| chr2-23512724-INS-2478 | chr2-23512725-INS-2478 |
| chr2-23522371-INS-678 | chr2-23522372-INS-682 |
| chr2-23529136-INS-131 | chr2-23529143-INS-131 |
| chr2-23567037-INS-739 | chr2-23567038-INS-739 |
| chr2-23573488-INS-78 | chr2-23573488-INS-78 |
| chr2-23629736-INS-7547 | chr2-23629737-INS-7549 |
| chr2-23714312-INS-72 | chr2-23714308-INS-52 |
| chr2-23724070-INS-70 | chr2-23724070-INS-90 |
| chr2-23761521-INS-426 | chr2-23761521-INS-427 |
| chr2-23765646-INS-58 | chr2-23765646-INS-58 |
| chr2-23859876-INS-345 | chr2-23859877-INS-345 |
| chr2-23910287-INS-236 | chr2-23910287-INS-231 |
| chr2-23924534-INS-77 | chr2-23924534-INS-77 |
| chr2-23962879-INS-110 | chr2-23962879-INS-110 |
| chr2-23992375-INS-78 | chr2-23992378-INS-51 |
| chr2-24064920-INS-655 | chr2-24064920-INS-655 |
| chr2-24072468-INS-133 | chr2-24072468-INS-152 |
| chr2-24075597-INS-209 | chr2-24075597-INS-209 |
| chr2-24076660-INS-96 | chr2-24076660-INS-96 |
| chr2-24089424-INS-61 | chr2-24089424-INS-61 |
| chr2-24094597-INS-421 | chr2-24094597-INS-468 |
| chr2-24103058-INS-556 | chr2-24103067-INS-556 |
| chr2-24123853-INS-1645 | chr2-24123854-INS-1609 |
| chr2-24212112-INS-51 | chr2-24212112-INS-51 |
| chr2-24241920-INS-107 | chr2-24241925-INS-68 |
| chr2-24281184-INS-451 | chr2-24281184-INS-451 |
| chr2-24322031-INS-51 | chr2-24322032-INS-50 |
| chr2-24338768-INS-168 | chr2-24338803-INS-157 |
| chr2-24353921-INS-58 | chr2-24353921-INS-62 |
| chr2-24375216-INS-67 | chr2-24375206-INS-102 |
| chr2-24381562-INS-357 | chr2-24381562-INS-357 |
| chr2-24382680-INS-134 | chr2-24382702-INS-84 |
| chr2-24388445-INS-55 | chr2-24388445-INS-55 |
| chr2-24395876-INS-64 | chr2-24395876-INS-64 |
| chr2-24397905-INS-84 | chr2-24397905-INS-84 |
| chr2-24452582-INS-63 | chr2-24452620-INS-63 |
| chr2-24461919-INS-55 | chr2-24461963-INS-57 |

| | |
|---|---|
| chr2-24463393-INS-60 | chr2-24463393-INS-68 |
| chr2-24473810-INS-408 | chr2-24473810-INS-412 |
| chr2-24509125-INS-82 | chr2-24509125-INS-84 |
| chr2-24519877-INS-105 | chr2-24519877-INS-105 |
| chr2-24524605-INS-398 | chr2-24524605-INS-398 |
| chr2-24554379-INS-2098 | chr2-24554380-INS-2098 |
| chr2-24604057-INS-1365 | chr2-24604058-INS-1365 |
| chr2-24619218-INS-180 | chr2-24619218-INS-202 |
| chr2-24628124-INS-61 | chr2-24628124-INS-60 |
| chr2-24657320-INS-62 | chr2-24657270-INS-109 |
| chr2-24660302-INS-480 | chr2-24660302-INS-433 |
| chr2-24670650-INS-169 | chr2-24670650-INS-169 |
| chr2-24672561-INS-476 | chr2-24672561-INS-476 |
| chr2-24686469-INS-174 | chr2-24686469-INS-180 |
| chr2-24750957-INS-206 | chr2-24750967-INS-207 |
| chr2-24752122-INS-83 | chr2-24752126-INS-58 |
| chr2-24756859-INS-230 | chr2-24756859-INS-230 |
| chr2-24762383-INS-174 | chr2-24762383-INS-179 |
| chr2-24786548-INS-200 | chr2-24786548-INS-201 |
| chr2-24788491-INS-4975 | chr2-24788492-INS-4966 |
| chr2-24799203-INS-202 | chr2-24799203-INS-202 |
| chr2-24814663-INS-346 | chr2-24814653-INS-346 |
| chr2-24819068-INS-98 | chr2-24819068-INS-96 |
| chr2-24838794-INS-118 | chr2-24838794-INS-118 |
| chr2-24846117-INS-198 | chr2-24846117-INS-198 |
| chr2-24854420-INS-398 | chr2-24854420-INS-398 |
| chr2-24860847-INS-108 | chr2-24860847-INS-60 |
| chr2-24862157-INS-205 | chr2-24862157-INS-210 |
| chr2-24866719-INS-198 | chr2-24866733-INS-211 |
| chr2-24872831-INS-203 | chr2-24872831-INS-199 |
| chr2-24893293-INS-539 | chr2-24893293-INS-531 |
| chr2-24902178-INS-123 | chr2-24902178-INS-114 |
| chr2-24905672-INS-88 | chr2-24905680-INS-94 |
| chr2-24915880-INS-65 | chr2-24915877-INS-84 |
| chr2-24958340-INS-109 | chr2-24958340-INS-104 |
| chr2-24962218-INS-506 | chr2-24962218-INS-508 |
| chr2-24997031-INS-408 | chr2-24997031-INS-408 |
| chr2-24999810-INS-1362 | chr2-24999811-INS-1362 |
| chr2-25006585-INS-170 | chr2-25006585-INS-144 |
| chr2-25020147-INS-84 | chr2-25020147-INS-126 |
| chr2-25039539-INS-510 | chr2-25039539-INS-510 |
| chr2-25215116-INS-82 | chr2-25215116-INS-82 |
| chr2-25232270-INS-66 | chr2-25232270-INS-65 |
| chr2-25258373-INS-732 | chr2-25258373-INS-732 |
| chr2-25270331-INS-177 | chr2-25270331-INS-177 |
| chr2-25275979-INS-472 | chr2-25275980-INS-472 |
| chr2-25306194-INS-205 | chr2-25306194-INS-206 |
| chr2-25364684-INS-107 | chr2-25364673-INS-120 |
| chr2-25404917-INS-257 | chr2-25404915-INS-262 |
| chr2-25416066-INS-69 | chr2-25416066-INS-69 |
| chr2-25423893-INS-124 | chr2-25423893-INS-93 |
| chr2-25464520-INS-196 | chr2-25464520-INS-196 |
| chr2-25465701-INS-88 | chr2-25465742-INS-85 |
| chr2-25580899-INS-177 | chr2-25580896-INS-181 |
| chr2-25620360-INS-249 | chr2-25620360-INS-249 |
| chr2-25676293-INS-56 | chr2-25676293-INS-56 |
| chr2-25680269-INS-201 | chr2-25680269-INS-202 |
| chr2-25720430-INS-338 | chr2-25720430-INS-384 |
| chr2-25723927-INS-2395 | chr2-25723928-INS-2395 |
| chr2-25750457-INS-92 | chr2-25750457-INS-92 |
| chr2-25755939-INS-55 | chr2-25755939-INS-75 |
| chr2-25757398-INS-225 | chr2-25757394-INS-228 |
| chr2-25776834-INS-172 | chr2-25776834-INS-172 |
| chr2-25851041-INS-56 | chr2-25851041-INS-56 |
| chr2-25859836-INS-7461 | chr2-25859837-INS-7462 |
| chr2-25870990-INS-197 | chr2-25870990-INS-197 |
| chr2-25881555-INS-201 | chr2-25881555-INS-201 |
| chr2-25898297-INS-204 | chr2-25898297-INS-205 |
| chr2-25910718-INS-208 | chr2-25910718-INS-209 |
| chr2-25912369-INS-110 | chr2-25912375-INS-104 |
| chr2-25924109-INS-77 | chr2-25924109-INS-77 |
| chr2-25960250-INS-5205 | chr2-25960251-INS-5205 |
| chr2-25969819-INS-180 | chr2-25969819-INS-180 |
| chr2-25991118-INS-5311 | chr2-25991119-INS-5311 |
| chr2-26023779-INS-74 | chr2-26023779-INS-60 |
| chr2-26032697-INS-104 | chr2-26032683-INS-101 |
| chr2-26041976-INS-221 | chr2-26041976-INS-221 |

| | |
|---|---|
| chr2-26044543-INS-91 | chr2-26044543-INS-106 |
| chr2-26045643-INS-105 | chr2-26045643-INS-115 |
| chr2-26055363-INS-57 | chr2-26055363-INS-50 |
| chr2-26063852-INS-217 | chr2-26063852-INS-217 |
| chr2-26079035-INS-146 | chr2-26079035-INS-146 |
| chr2-26102481-INS-378 | chr2-26102481-INS-344 |
| chr2-26108163-INS-52 | chr2-26108163-INS-52 |
| chr2-26112834-INS-6130 | chr2-26112835-INS-6130 |
| chr2-26131893-INS-221 | chr2-26131893-INS-225 |
| chr2-26133196-INS-82 | chr2-26133196-INS-82 |
| chr2-26143590-INS-358 | chr2-26143590-INS-358 |
| chr2-26154371-INS-217 | chr2-26154371-INS-221 |
| chr2-26178522-INS-147 | chr2-26178523-INS-101 |
| chr2-26180275-INS-75 | chr2-26180275-INS-74 |
| chr2-26200446-INS-210 | chr2-26200446-INS-210 |
| chr2-26219328-INS-206 | chr2-26219328-INS-208 |
| chr2-26221352-INS-209 | chr2-26221352-INS-209 |
| chr2-26248348-INS-67 | chr2-26248348-INS-67 |
| chr2-26261155-INS-146 | chr2-26261155-INS-147 |
| chr2-26269389-INS-231 | chr2-26269389-INS-233 |
| chr2-26297117-INS-74 | chr2-26297117-INS-74 |
| chr2-26340095-INS-206 | chr2-26340095-INS-208 |
| chr2-26389220-INS-564 | chr2-26389220-INS-565 |
| chr2-26444538-INS-201 | chr2-26444537-INS-216 |
| chr2-26456783-INS-206 | chr2-26456783-INS-206 |
| chr2-26474850-INS-118 | chr2-26474850-INS-118 |
| chr2-26507345-INS-199 | chr2-26507345-INS-202 |
| chr2-26511200-INS-211 | chr2-26511200-INS-227 |
| chr2-26543699-INS-65 | chr2-26543699-INS-65 |
| chr2-26547188-INS-206 | chr2-26547188-INS-206 |
| chr2-26554873-INS-196 | chr2-26554873-INS-196 |
| chr2-26562353-INS-52 | chr2-26562353-INS-52 |
| chr2-26563143-INS-1037 | chr2-26563144-INS-1071 |
| chr2-26563386-INS-259 | chr2-26563387-INS-257 |
| chr2-26569059-INS-6312 | chr2-26569060-INS-6311 |
| chr2-26572231-INS-84 | chr2-26572231-INS-85 |
| chr2-26579573-INS-187 | chr2-26579573-INS-187 |
| chr2-26620364-INS-120 | chr2-26620364-INS-144 |
| chr2-26629262-INS-66 | chr2-26629262-INS-66 |
| chr2-26639255-INS-1072 | chr2-26639256-INS-1072 |
| chr2-26659763-INS-167 | chr2-26659763-INS-167 |
| chr2-26660529-INS-2037 | chr2-26660530-INS-2037 |
| chr2-26674517-INS-84 | chr2-26674517-INS-54 |
| chr2-26679146-INS-77 | chr2-26679146-INS-77 |
| chr2-26700174-INS-200 | chr2-26700174-INS-200 |
| chr2-26706039-INS-78 | chr2-26706039-INS-58 |
| chr2-26715794-INS-3223 | chr2-26715795-INS-3233 |
| chr2-26723053-INS-198 | chr2-26723053-INS-198 |
| chr2-26728333-INS-216 | chr2-26728333-INS-216 |
| chr2-26765600-INS-2896 | chr2-26765601-INS-2896 |
| chr2-26825791-INS-108 | chr2-26825797-INS-111 |
| chr2-26827840-INS-78 | chr2-26827840-INS-69 |
| chr2-26847429-INS-6455 | chr2-26847430-INS-6455 |
| chr2-26858337-INS-57 | chr2-26858337-INS-58 |
| chr2-26875997-INS-181 | chr2-26875997-INS-178 |
| chr2-26882873-INS-149 | chr2-26882873-INS-149 |
| chr2-26902742-INS-66 | chr2-26902742-INS-88 |
| chr2-26903579-INS-200 | chr2-26903579-INS-200 |
| chr2-26909673-INS-233 | chr2-26909673-INS-233 |
| chr2-26921061-INS-141 | chr2-26921061-INS-155 |
| chr2-26933035-INS-204 | chr2-26933035-INS-204 |
| chr2-26998572-INS-58 | chr2-26998572-INS-58 |
| chr2-27045706-INS-99 | chr2-27045706-INS-99 |
| chr2-27080349-INS-134 | chr2-27080350-INS-99 |
| chr2-27106654-INS-103 | chr2-27106652-INS-109 |
| chr2-27140501-INS-442 | chr2-27140496-INS-451 |
| chr2-27229857-INS-68 | chr2-27229857-INS-68 |
| chr2-27246726-INS-87 | chr2-27246726-INS-89 |
| chr2-27255061-INS-60 | chr2-27255061-INS-60 |
| chr2-27266889-INS-54 | chr2-27266889-INS-56 |
| chr2-27272222-INS-60 | chr2-27272222-INS-60 |
| chr2-27296748-INS-60 | chr2-27296748-INS-76 |
| chr2-27300474-INS-86 | chr2-27300474-INS-86 |
| chr2-27315015-INS-133 | chr2-27314997-INS-119 |
| chr2-27331610-INS-137 | chr2-27331610-INS-171 |
| chr2-27376691-INS-231 | chr2-27376691-INS-231 |
| chr2-27377939-INS-230 | chr2-27377939-INS-225 |

| | |
|---|---|
| chr2-27385231-INS-377 | chr2-27385229-INS-344 |
| chr2-27392277-INS-489 | chr2-27392277-INS-489 |
| chr2-27400441-INS-240 | chr2-27400441-INS-240 |
| chr2-27406261-INS-61 | chr2-27406261-INS-60 |
| chr2-27416810-INS-109 | chr2-27416810-INS-111 |
| chr2-27428952-INS-62 | chr2-27428952-INS-64 |
| chr2-27530542-INS-104 | chr2-27530561-INS-70 |
| chr2-27557477-INS-6433 | chr2-27557478-INS-6433 |
| chr2-27574417-INS-52 | chr2-27574417-INS-52 |
| chr2-27658085-INS-497 | chr2-27658085-INS-480 |
| chr2-27660414-INS-56 | chr2-27660414-INS-58 |
| chr2-27685564-INS-138 | chr2-27685564-INS-110 |
| chr2-27707485-INS-351 | chr2-27707485-INS-351 |
| chr2-27731854-INS-54 | chr2-27731854-INS-54 |
| chr2-27761863-INS-78 | chr2-27761863-INS-78 |
| chr2-27775313-INS-100 | chr2-27775313-INS-100 |
| chr2-27804398-INS-73 | chr2-27804398-INS-69 |
| chr2-27806023-INS-4478 | chr2-27806024-INS-4478 |
| chr2-27807571-INS-75 | chr2-27807571-INS-67 |
| chr2-27838686-INS-101 | chr2-27838687-INS-101 |
| chr2-27841099-INS-94 | chr2-27841099-INS-98 |
| chr2-27851394-INS-124 | chr2-27851394-INS-124 |
| chr2-27863543-INS-84 | chr2-27863543-INS-84 |
| chr2-27894428-INS-81 | chr2-27894428-INS-81 |
| chr2-27956417-INS-54 | chr2-27956417-INS-54 |
| chr2-27964976-INS-80 | chr2-27964976-INS-80 |
| chr2-27996475-INS-936 | chr2-27996475-INS-936 |
| chr2-27996477-INS-98 | chr2-27996477-INS-98 |
| chr2-27998257-INS-549 | chr2-27998257-INS-549 |
| chr2-28045356-INS-129 | chr2-28045356-INS-129 |
| chr2-28055048-INS-1883 | chr2-28055049-INS-1883 |
| chr2-28056153-INS-52 | chr2-28056128-INS-83 |
| chr2-28065935-INS-119 | chr2-28065935-INS-119 |
| chr2-28092502-INS-220 | chr2-28092502-INS-220 |
| chr2-28110595-INS-68 | chr2-28110595-INS-68 |
| chr2-28124476-INS-157 | chr2-28124476-INS-157 |
| chr2-28158750-INS-130 | chr2-28158751-INS-128 |
| chr2-28172394-INS-746 | chr2-28172394-INS-746 |
| chr2-28206389-INS-639 | chr2-28206389-INS-640 |
| chr2-28209797-INS-225 | chr2-28209797-INS-227 |
| chr2-28243942-INS-67 | chr2-28243942-INS-68 |
| chr2-28301663-INS-57 | chr2-28301663-INS-57 |
| chr2-28307155-INS-60 | chr2-28307151-INS-56 |
| chr2-28318321-INS-111 | chr2-28318340-INS-106 |
| chr2-28319664-INS-5529 | chr2-28319665-INS-5529 |
| chr2-28323373-INS-120 | chr2-28323373-INS-138 |
| chr2-28346543-INS-62 | chr2-28346543-INS-62 |
| chr2-28350901-INS-174 | chr2-28350901-INS-174 |
| chr2-28366477-INS-204 | chr2-28366477-INS-204 |
| chr2-28390006-INS-4533 | chr2-28390007-INS-4533 |
| chr2-28431161-INS-1892 | chr2-28431162-INS-1892 |
| chr2-28454494-INS-113 | chr2-28454494-INS-121 |
| chr2-28483119-INS-65 | chr2-28483119-INS-50 |
| chr2-28486759-INS-56 | chr2-28486759-INS-56 |
| chr2-28490810-INS-498 | chr2-28490810-INS-498 |
| chr2-28493117-INS-234 | chr2-28493117-INS-228 |
| chr2-28500570-INS-53 | chr2-28500570-INS-53 |
| chr2-28516683-INS-206 | chr2-28516683-INS-215 |
| chr2-28528683-INS-102 | chr2-28528683-INS-102 |
| chr2-28560654-INS-219 | chr2-28560654-INS-219 |
| chr2-28608629-INS-761 | chr2-28608652-INS-752 |
| chr2-28609564-INS-148 | chr2-28609564-INS-150 |
| chr2-28611904-INS-496 | chr2-28611904-INS-467 |
| chr2-28623630-INS-59 | chr2-28623630-INS-59 |
| chr2-28674323-INS-204 | chr2-28674323-INS-230 |
| chr2-28691792-INS-223 | chr2-28691792-INS-255 |
| chr2-28700270-INS-76 | chr2-28700270-INS-76 |
| chr2-28739018-INS-208 | chr2-28739018-INS-208 |
| chr2-28743818-INS-129 | chr2-28743817-INS-80 |
| chr2-28754847-INS-176 | chr2-28754847-INS-178 |
| chr2-28762201-INS-63 | chr2-28762201-INS-63 |
| chr2-28828430-INS-168 | chr2-28828430-INS-177 |
| chr2-28841029-INS-58 | chr2-28841029-INS-62 |
| chr2-28850163-INS-52 | chr2-28850163-INS-80 |
| chr2-28857666-INS-199 | chr2-28857666-INS-198 |
| chr2-28882938-INS-688 | chr2-28882938-INS-688 |
| chr2-28896072-INS-204 | chr2-28896072-INS-204 |

| | |
|---|---|
| chr2-28927203-INS-96 | chr2-28927203-INS-96 |
| chr2-28946073-INS-60 | chr2-28946073-INS-60 |
| chr2-28957298-INS-126 | chr2-28957298-INS-126 |
| chr2-28957781-INS-209 | chr2-28957781-INS-214 |
| chr2-28961991-INS-332 | chr2-28961991-INS-332 |
| chr2-28990471-INS-197 | chr2-28990471-INS-205 |
| chr2-29021260-INS-192 | chr2-29021260-INS-192 |
| chr2-29028841-INS-107 | chr2-29028841-INS-105 |
| chr2-29051533-INS-104 | chr2-29051533-INS-104 |
| chr2-29057834-INS-202 | chr2-29057834-INS-202 |
| chr2-29060637-INS-222 | chr2-29060637-INS-222 |
| chr2-29061398-INS-54 | chr2-29061398-INS-54 |
| chr2-29081620-INS-82 | chr2-29081620-INS-82 |
| chr2-29173782-INS-76 | chr2-29173782-INS-57 |
| chr2-29175751-INS-55 | chr2-29175751-INS-55 |
| chr2-29184220-INS-77 | chr2-29184220-INS-77 |
| chr2-29197973-INS-75 | chr2-29197973-INS-75 |
| chr2-29216764-INS-88 | chr2-29216764-INS-50 |
| chr2-29244184-INS-210 | chr2-29244184-INS-210 |
| chr2-29297249-INS-52 | chr2-29297249-INS-92 |
| chr2-29304293-INS-88 | chr2-29304294-INS-64 |
| chr2-29325133-INS-145 | chr2-29325133-INS-145 |
| chr2-29345429-INS-1387 | chr2-29345430-INS-1387 |
| chr2-29375042-INS-515 | chr2-29375042-INS-515 |
| chr2-29429751-INS-369 | chr2-29429770-INS-368 |
| chr2-29437988-INS-64 | chr2-29437979-INS-52 |
| chr2-29439671-INS-96 | chr2-29439671-INS-96 |
| chr2-29461728-INS-56 | chr2-29461738-INS-62 |
| chr2-29479024-INS-108 | chr2-29479025-INS-108 |
| chr2-29497398-INS-202 | chr2-29497398-INS-202 |
| chr2-29519135-INS-208 | chr2-29519135-INS-208 |
| chr2-29521873-INS-128 | chr2-29521856-INS-102 |
| chr2-29527178-INS-57 | chr2-29527178-INS-72 |
| chr2-29537134-INS-362 | chr2-29537134-INS-362 |
| chr2-29640080-INS-210 | chr2-29640080-INS-210 |
| chr2-29642537-INS-193 | chr2-29642537-INS-194 |
| chr2-29654651-INS-72 | chr2-29654651-INS-72 |
| chr2-29656133-INS-199 | chr2-29656133-INS-199 |
| chr2-29660793-INS-97 | chr2-29660793-INS-97 |
| chr2-29668584-INS-198 | chr2-29668584-INS-208 |
| chr2-29682283-INS-207 | chr2-29682283-INS-207 |
| chr2-29688275-INS-83 | chr2-29688289-INS-108 |
| chr2-29709962-INS-56 | chr2-29709962-INS-58 |
| chr2-29715164-INS-209 | chr2-29715164-INS-209 |
| chr2-29755666-INS-217 | chr2-29755666-INS-216 |
| chr2-29811726-INS-213 | chr2-29811726-INS-217 |
| chr2-29813228-INS-83 | chr2-29813228-INS-83 |
| chr2-29822694-INS-163 | chr2-29822694-INS-167 |
| chr2-29864396-INS-207 | chr2-29864396-INS-210 |
| chr2-29885704-INS-866 | chr2-29885704-INS-821 |
| chr2-29902326-INS-480 | chr2-29902326-INS-467 |
| chr2-29933271-INS-229 | chr2-29933271-INS-221 |
| chr2-29949324-INS-191 | chr2-29949319-INS-197 |
| chr2-29963015-INS-203 | chr2-29963015-INS-205 |
| chr2-29963686-INS-219 | chr2-29963686-INS-219 |
| chr2-29978235-INS-105 | chr2-29978235-INS-105 |
| chr2-29979571-INS-204 | chr2-29979571-INS-204 |
| chr2-29989908-INS-56 | chr2-29989908-INS-58 |
| chr2-30005904-INS-177 | chr2-30005904-INS-178 |
| chr2-30010167-INS-71 | chr2-30010169-INS-75 |
| chr2-30017171-INS-67 | chr2-30017190-INS-58 |
| chr2-30019368-INS-52 | chr2-30019368-INS-60 |
| chr2-30027934-INS-169 | chr2-30027934-INS-169 |
| chr2-30037317-INS-234 | chr2-30037317-INS-236 |
| chr2-30052195-INS-234 | chr2-30052215-INS-188 |
| chr2-30094643-INS-209 | chr2-30094644-INS-209 |
| chr2-30101086-INS-55 | chr2-30101086-INS-55 |
| chr2-30118605-INS-202 | chr2-30118605-INS-202 |
| chr2-30134103-INS-249 | chr2-30134104-INS-243 |
| chr2-30136979-INS-176 | chr2-30136979-INS-177 |
| chr2-30151242-INS-209 | chr2-30151242-INS-208 |
| chr2-30154332-INS-141 | chr2-30154363-INS-105 |
| chr2-30157500-INS-376 | chr2-30157500-INS-376 |
| chr2-30163985-INS-182 | chr2-30163985-INS-182 |
| chr2-30165894-INS-96 | chr2-30165894-INS-96 |
| chr2-30168166-INS-121 | chr2-30168166-INS-113 |
| chr2-30182579-INS-265 | chr2-30182579-INS-265 |

| | |
|---|---|
| chr2-30183964-INS-259 | chr2-30183964-INS-254 |
| chr2-30252505-INS-202 | chr2-30252505-INS-203 |
| chr2-30279284-INS-237 | chr2-30279284-INS-237 |
| chr2-30307189-INS-199 | chr2-30307189-INS-199 |
| chr2-30317852-INS-106 | chr2-30317852-INS-106 |
| chr2-30321342-INS-206 | chr2-30321342-INS-217 |
| chr2-30375700-INS-168 | chr2-30375693-INS-203 |
| chr2-30379013-INS-287 | chr2-30379013-INS-289 |
| chr2-30389847-INS-416 | chr2-30389847-INS-416 |
| chr2-30401255-INS-65 | chr2-30401255-INS-65 |
| chr2-30445414-INS-88 | chr2-30445414-INS-88 |
| chr2-30452850-INS-58 | chr2-30452851-INS-59 |
| chr2-30461080-INS-166 | chr2-30461079-INS-166 |
| chr2-30480852-INS-221 | chr2-30480852-INS-223 |
| chr2-30492390-INS-198 | chr2-30492388-INS-200 |
| chr2-30497756-INS-52 | chr2-30497756-INS-70 |
| chr2-30500520-INS-4802 | chr2-30500521-INS-4802 |
| chr2-30579493-INS-54 | chr2-30579493-INS-54 |
| chr2-30617953-INS-71 | chr2-30617949-INS-79 |
| chr2-30622621-INS-183 | chr2-30622621-INS-186 |
| chr2-30656070-INS-162 | chr2-30656070-INS-162 |
| chr2-30683743-INS-204 | chr2-30683743-INS-204 |
| chr2-30709451-INS-132 | chr2-30709451-INS-135 |
| chr2-30710924-INS-232 | chr2-30710924-INS-233 |
| chr2-30720473-INS-583 | chr2-30720473-INS-583 |
| chr2-30727067-INS-549 | chr2-30727067-INS-549 |
| chr2-30735908-INS-58 | chr2-30735908-INS-64 |
| chr2-30763536-INS-225 | chr2-30763536-INS-225 |
| chr2-30777843-INS-150 | chr2-30777843-INS-150 |
| chr2-30779474-INS-173 | chr2-30779474-INS-173 |
| chr2-30797935-INS-169 | chr2-30797935-INS-169 |
| chr2-30821979-INS-212 | chr2-30821979-INS-212 |
| chr2-30827910-INS-64 | chr2-30827910-INS-64 |
| chr2-30832030-INS-174 | chr2-30832030-INS-182 |
| chr2-30839585-INS-149 | chr2-30839588-INS-193 |
| chr2-30898120-INS-82 | chr2-30898120-INS-61 |
| chr2-30905129-INS-206 | chr2-30905129-INS-207 |
| chr2-30920068-INS-89 | chr2-30920068-INS-56 |
| chr2-30961122-INS-204 | chr2-30961122-INS-202 |
| chr2-30978038-INS-452 | chr2-30978044-INS-458 |
| chr2-30978912-INS-81 | chr2-30978912-INS-81 |
| chr2-30989885-INS-62 | chr2-30989886-INS-63 |
| chr2-31006012-INS-775 | chr2-31006012-INS-773 |
| chr2-31019795-INS-92 | chr2-31019770-INS-99 |
| chr2-31023870-INS-198 | chr2-31023870-INS-201 |
| chr2-31027833-INS-197 | chr2-31027833-INS-197 |
| chr2-31057295-INS-206 | chr2-31057295-INS-206 |
| chr2-31059924-INS-219 | chr2-31059924-INS-219 |
| chr2-31067502-INS-220 | chr2-31067502-INS-216 |
| chr2-31068820-INS-170 | chr2-31068820-INS-170 |
| chr2-31086703-INS-57 | chr2-31086660-INS-57 |
| chr2-31124935-INS-1862 | chr2-31124936-INS-1862 |
| chr2-31126504-INS-210 | chr2-31126490-INS-208 |
| chr2-31127037-INS-242 | chr2-31127037-INS-245 |
| chr2-31128651-INS-225 | chr2-31128651-INS-225 |
| chr2-31158531-INS-1427 | chr2-31158532-INS-1427 |
| chr2-31159741-INS-375 | chr2-31159741-INS-375 |
| chr2-31199316-INS-53 | chr2-31199317-INS-69 |
| chr2-31234882-INS-86 | chr2-31234882-INS-88 |
| chr2-31249664-INS-193 | chr2-31249664-INS-173 |
| chr2-31255998-INS-82 | chr2-31255998-INS-82 |
| chr2-31304846-INS-56 | chr2-31304846-INS-58 |
| chr2-31326688-INS-60 | chr2-31326688-INS-60 |
| chr2-31328576-INS-60 | chr2-31328576-INS-60 |
| chr2-31350904-INS-80 | chr2-31350904-INS-85 |
| chr2-31369221-INS-160 | chr2-31369221-INS-160 |
| chr2-31427972-INS-220 | chr2-31427972-INS-221 |
| chr2-31435136-INS-54 | chr2-31435104-INS-50 |
| chr2-31446228-INS-84 | chr2-31446227-INS-96 |
| chr2-31522518-INS-206 | chr2-31522495-INS-251 |
| chr2-31543124-INS-97 | chr2-31543124-INS-73 |
| chr2-31597292-INS-1011 | chr2-31597293-INS-1012 |
| chr2-31616548-INS-218 | chr2-31616533-INS-215 |
| chr2-31621723-INS-5485 | chr2-31621724-INS-5485 |
| chr2-31624552-INS-242 | chr2-31624548-INS-244 |
| chr2-31633483-INS-204 | chr2-31633483-INS-205 |
| chr2-31656462-INS-145 | chr2-31656462-INS-187 |

| | |
|---|---|
| chr2-31663079-INS-7120 | chr2-31663080-INS-7120 |
| chr2-31686278-INS-207 | chr2-31686278-INS-206 |
| chr2-31738841-INS-183 | chr2-31738841-INS-189 |
| chr2-31746131-INS-51 | chr2-31746131-INS-51 |
| chr2-31767801-INS-75 | chr2-31767801-INS-75 |
| chr2-31770128-INS-221 | chr2-31770128-INS-222 |
| chr2-31871608-INS-51 | chr2-31871608-INS-66 |
| chr2-31911158-INS-2893 | chr2-31911159-INS-2895 |
| chr2-31932270-INS-59 | chr2-31932268-INS-59 |
| chr2-31941345-INS-61 | chr2-31941345-INS-61 |
| chr2-31945316-INS-67 | chr2-31945316-INS-67 |
| chr2-31947796-INS-1892 | chr2-31947797-INS-1892 |
| chr2-31970154-INS-211 | chr2-31970154-INS-209 |
| chr2-31997743-INS-64 | chr2-31997743-INS-68 |
| chr2-32010369-INS-70 | chr2-32010369-INS-70 |
| chr2-32015259-INS-74 | chr2-32015235-INS-103 |
| chr2-32021966-INS-197 | chr2-32021966-INS-196 |
| chr2-32036132-INS-192 | chr2-32036179-INS-194 |
| chr2-32068449-INS-198 | chr2-32068449-INS-198 |
| chr2-32071166-INS-62 | chr2-32071166-INS-78 |
| chr2-32084137-INS-103 | chr2-32084137-INS-103 |
| chr2-32134010-INS-168 | chr2-32134010-INS-170 |
| chr2-32157631-INS-56 | chr2-32157631-INS-85 |
| chr2-32158469-INS-202 | chr2-32158469-INS-202 |
| chr2-32160540-INS-130 | chr2-32160540-INS-158 |
| chr2-32167426-INS-215 | chr2-32167426-INS-215 |
| chr2-32248982-INS-406 | chr2-32248982-INS-406 |
| chr2-32262906-INS-60 | chr2-32262906-INS-62 |
| chr2-32281891-INS-232 | chr2-32281891-INS-232 |
| chr2-32303956-INS-245 | chr2-32303956-INS-245 |
| chr2-32351822-INS-203 | chr2-32351822-INS-205 |
| chr2-32358434-INS-100 | chr2-32358434-INS-84 |
| chr2-32450231-INS-217 | chr2-32450231-INS-216 |
| chr2-32489325-INS-825 | chr2-32489325-INS-825 |
| chr2-32507441-INS-62 | chr2-32507441-INS-62 |
| chr2-32540065-INS-68 | chr2-32540071-INS-56 |
| chr2-32557372-INS-104 | chr2-32557372-INS-104 |
| chr2-32558685-INS-105 | chr2-32558685-INS-105 |
| chr2-32565484-INS-140 | chr2-32565478-INS-165 |
| chr2-32578660-INS-64 | chr2-32578660-INS-64 |
| chr2-32592575-INS-210 | chr2-32592575-INS-210 |
| chr2-32618317-INS-355 | chr2-32618317-INS-355 |
| chr2-32663258-INS-171 | chr2-32663258-INS-171 |
| chr2-32670280-INS-206 | chr2-32670280-INS-206 |
| chr2-32683661-INS-207 | chr2-32683657-INS-207 |
| chr2-32688806-INS-1523 | chr2-32688807-INS-1523 |
| chr2-32727157-INS-226 | chr2-32727157-INS-226 |
| chr2-32758343-INS-325 | chr2-32758343-INS-325 |
| chr2-32783170-INS-204 | chr2-32783170-INS-204 |
| chr2-32796245-INS-135 | chr2-32796245-INS-135 |
| chr2-32804201-INS-68 | chr2-32804201-INS-68 |
| chr2-32845391-INS-186 | chr2-32845391-INS-185 |
| chr2-32849714-INS-191 | chr2-32849714-INS-191 |
| chr2-32915096-INS-64 | chr2-32915096-INS-56 |
| chr2-32950854-INS-228 | chr2-32950854-INS-228 |
| chr2-33065909-INS-199 | chr2-33065909-INS-199 |
| chr2-33229262-INS-384 | chr2-33229233-INS-381 |
| chr2-33315845-INS-327 | chr2-33315845-INS-327 |
| chr2-33317436-INS-208 | chr2-33317436-INS-208 |
| chr2-33323227-INS-318 | chr2-33323227-INS-302 |
| chr2-33368100-INS-66 | chr2-33368100-INS-66 |
| chr2-33385625-INS-208 | chr2-33385625-INS-203 |
| chr2-33409375-INS-180 | chr2-33409375-INS-180 |
| chr2-33425120-INS-52 | chr2-33425120-INS-52 |
| chr2-33443651-INS-177 | chr2-33443651-INS-177 |
| chr2-33448319-INS-762 | chr2-33448319-INS-765 |
| chr2-33482234-INS-64 | chr2-33482234-INS-64 |
| chr2-33504208-INS-360 | chr2-33504208-INS-330 |
| chr2-33587766-INS-187 | chr2-33587766-INS-187 |
| chr2-33685465-INS-131 | chr2-33685465-INS-131 |
| chr2-33761985-INS-84 | chr2-33761985-INS-84 |
| chr2-33795590-INS-54 | chr2-33795590-INS-56 |
| chr2-33838849-INS-80 | chr2-33838849-INS-80 |
| chr2-33857714-INS-53 | chr2-33857714-INS-55 |
| chr2-33860634-INS-178 | chr2-33860634-INS-179 |
| chr2-33876017-INS-103 | chr2-33876017-INS-111 |
| chr2-33877786-INS-344 | chr2-33877786-INS-344 |

| | |
|---|---|
| chr2-33878537-INS-361 | chr2-33878537-INS-362 |
| chr2-33908781-INS-58 | chr2-33908787-INS-50 |
| chr2-33919303-INS-80 | chr2-33919303-INS-80 |
| chr2-33966107-INS-565 | chr2-33966096-INS-568 |
| chr2-34003184-INS-376 | chr2-34003184-INS-376 |
| chr2-34015745-INS-60 | chr2-34015745-INS-60 |
| chr2-34054309-INS-7367 | chr2-34054310-INS-7367 |
| chr2-34069855-INS-56 | chr2-34069855-INS-56 |
| chr2-34190745-INS-56 | chr2-34190745-INS-56 |
| chr2-34196583-INS-124 | chr2-34196583-INS-124 |
| chr2-34223126-INS-72 | chr2-34223126-INS-84 |
| chr2-34278034-INS-70 | chr2-34278034-INS-70 |
| chr2-34307809-INS-58 | chr2-34307809-INS-62 |
| chr2-34360873-INS-205 | chr2-34360873-INS-205 |
| chr2-34376025-INS-4332 | chr2-34376026-INS-4330 |
| chr2-34376222-INS-51 | chr2-34376222-INS-51 |
| chr2-34379164-INS-58 | chr2-34379164-INS-58 |
| chr2-34444637-INS-84 | chr2-34444637-INS-84 |
| chr2-34536564-INS-71 | chr2-34536564-INS-71 |
| chr2-34554280-INS-62 | chr2-34554280-INS-62 |
| chr2-34577174-INS-105 | chr2-34577154-INS-96 |
| chr2-34611344-INS-58 | chr2-34611344-INS-58 |
| chr2-34636795-INS-3266 | chr2-34636796-INS-3266 |
| chr2-34646171-INS-77 | chr2-34646171-INS-77 |
| chr2-34654721-INS-282 | chr2-34654721-INS-282 |
| chr2-34659681-INS-207 | chr2-34659681-INS-207 |
| chr2-34666682-INS-192 | chr2-34666682-INS-192 |
| chr2-34676085-INS-186 | chr2-34676085-INS-186 |
| chr2-34691563-INS-191 | chr2-34691563-INS-191 |
| chr2-34714098-INS-1779 | chr2-34714099-INS-1798 |
| chr2-34722993-INS-563 | chr2-34722993-INS-563 |
| chr2-34724313-INS-209 | chr2-34724313-INS-209 |
| chr2-34725068-INS-207 | chr2-34725068-INS-206 |
| chr2-34730901-INS-5312 | chr2-34730902-INS-5312 |
| chr2-34732974-INS-77 | chr2-34732974-INS-77 |
| chr2-34734116-INS-115 | chr2-34734127-INS-119 |
| chr2-34735735-INS-207 | chr2-34735735-INS-207 |
| chr2-34751426-INS-80 | chr2-34751426-INS-116 |
| chr2-34775397-INS-57 | chr2-34775397-INS-58 |
| chr2-34782202-INS-56 | chr2-34782202-INS-56 |
| chr2-34783685-INS-148 | chr2-34783685-INS-148 |
| chr2-34807742-INS-228 | chr2-34807742-INS-228 |
| chr2-34836281-INS-137 | chr2-34836281-INS-133 |
| chr2-34860870-INS-6473 | chr2-34860871-INS-6473 |
| chr2-34887013-INS-88 | chr2-34887013-INS-88 |
| chr2-34889524-INS-62 | chr2-34889524-INS-64 |
| chr2-34909809-INS-55 | chr2-34909809-INS-55 |
| chr2-34923184-INS-77 | chr2-34923185-INS-77 |
| chr2-34956879-INS-756 | chr2-34956879-INS-756 |
| chr2-35003896-INS-100 | chr2-35003889-INS-64 |
| chr2-35026413-INS-129 | chr2-35026413-INS-131 |
| chr2-35053644-INS-389 | chr2-35053644-INS-389 |
| chr2-35085971-INS-58 | chr2-35085971-INS-60 |
| chr2-35097710-INS-505 | chr2-35097710-INS-505 |
| chr2-35114372-INS-444 | chr2-35114372-INS-444 |
| chr2-35125734-INS-194 | chr2-35125734-INS-194 |
| chr2-35139438-INS-347 | chr2-35139438-INS-348 |
| chr2-35143282-INS-100 | chr2-35143282-INS-101 |
| chr2-35239539-INS-381 | chr2-35239539-INS-381 |
| chr2-35260482-INS-519 | chr2-35260482-INS-505 |
| chr2-35272195-INS-6216 | chr2-35272196-INS-6216 |
| chr2-35316776-INS-131 | chr2-35316776-INS-130 |
| chr2-35319876-INS-58 | chr2-35319876-INS-70 |
| chr2-35349323-INS-5699 | chr2-35349324-INS-5699 |
| chr2-35423343-INS-57 | chr2-35423343-INS-60 |
| chr2-35434078-INS-89 | chr2-35434078-INS-89 |
| chr2-35452274-INS-1890 | chr2-35452275-INS-1890 |
| chr2-35466637-INS-195 | chr2-35466637-INS-195 |
| chr2-35532581-INS-187 | chr2-35532596-INS-137 |
| chr2-35585791-INS-415 | chr2-35585791-INS-423 |
| chr2-35632706-INS-174 | chr2-35632706-INS-174 |
| chr2-35685127-INS-56 | chr2-35685125-INS-84 |
| chr2-35705892-INS-56 | chr2-35705892-INS-56 |
| chr2-35789611-INS-74 | chr2-35789591-INS-78 |
| chr2-35798464-INS-125 | chr2-35798434-INS-125 |
| chr2-35824205-INS-64 | chr2-35824205-INS-64 |
| chr2-35828705-INS-51 | chr2-35828702-INS-54 |

| | |
|---|---|
| chr2-35831625-INS-163 | chr2-35831629-INS-180 |
| chr2-35840641-INS-5522 | chr2-35840642-INS-5522 |
| chr2-36055422-INS-952 | chr2-36055422-INS-952 |
| chr2-36060180-INS-1470 | chr2-36060181-INS-1471 |
| chr2-36083896-INS-169 | chr2-36083899-INS-174 |
| chr2-36090102-INS-88 | chr2-36090102-INS-88 |
| chr2-36159095-INS-65 | chr2-36159099-INS-64 |
| chr2-36172832-INS-5311 | chr2-36172833-INS-5311 |
| chr2-36191241-INS-1840 | chr2-36191242-INS-1824 |
| chr2-36205778-INS-7966 | chr2-36205779-INS-7966 |
| chr2-36375284-INS-84 | chr2-36375302-INS-56 |
| chr2-36383985-INS-72 | chr2-36383985-INS-72 |
| chr2-36421061-INS-370 | chr2-36421061-INS-370 |
| chr2-36439083-INS-624 | chr2-36439084-INS-624 |
| chr2-36461901-INS-2484 | chr2-36461902-INS-2484 |
| chr2-36477579-INS-179 | chr2-36477579-INS-179 |
| chr2-36501933-INS-504 | chr2-36501933-INS-504 |
| chr2-36528721-INS-176 | chr2-36528720-INS-184 |
| chr2-36603813-INS-54 | chr2-36603813-INS-52 |
| chr2-36630495-INS-2244 | chr2-36630496-INS-2244 |
| chr2-36681679-INS-55 | chr2-36681704-INS-55 |
| chr2-36727365-INS-4551 | chr2-36727366-INS-4551 |
| chr2-36727716-INS-88 | chr2-36727716-INS-88 |
| chr2-36777291-INS-432 | chr2-36777291-INS-432 |
| chr2-36844356-INS-472 | chr2-36844356-INS-472 |
| chr2-36867892-INS-275 | chr2-36867892-INS-275 |
| chr2-36952245-INS-68 | chr2-36952200-INS-79 |
| chr2-36956125-INS-60 | chr2-36956125-INS-63 |
| chr2-36969317-INS-542 | chr2-36969317-INS-542 |
| chr2-37012032-INS-963 | chr2-37012032-INS-963 |
| chr2-37015435-INS-84 | chr2-37015461-INS-60 |
| chr2-37023086-INS-113 | chr2-37023087-INS-113 |
| chr2-37025079-INS-56 | chr2-37025079-INS-58 |
| chr2-37037656-INS-155 | chr2-37037656-INS-155 |
| chr2-37060020-INS-199 | chr2-37059993-INS-199 |
| chr2-37098081-INS-407 | chr2-37098081-INS-407 |
| chr2-37134835-INS-1347 | chr2-37134836-INS-1347 |
| chr2-37240011-INS-3661 | chr2-37240012-INS-3664 |
| chr2-37286737-INS-61 | chr2-37286737-INS-61 |
| chr2-37307355-INS-160 | chr2-37307355-INS-161 |
| chr2-37320278-INS-77 | chr2-37320278-INS-77 |
| chr2-37344798-INS-220 | chr2-37344798-INS-221 |
| chr2-37351660-INS-1890 | chr2-37351661-INS-1890 |
| chr2-37361988-INS-597 | chr2-37361988-INS-597 |
| chr2-37409688-INS-98 | chr2-37409689-INS-89 |
| chr2-37416410-INS-231 | chr2-37416410-INS-232 |
| chr2-37431986-INS-72 | chr2-37432004-INS-60 |
| chr2-37520406-INS-92 | chr2-37520406-INS-92 |
| chr2-37543494-INS-69 | chr2-37543494-INS-65 |
| chr2-37544576-INS-87 | chr2-37544576-INS-90 |
| chr2-37600428-INS-268 | chr2-37600428-INS-268 |
| chr2-37615271-INS-67 | chr2-37615271-INS-67 |
| chr2-37653390-INS-199 | chr2-37653390-INS-199 |
| chr2-37662315-INS-212 | chr2-37662315-INS-212 |
| chr2-37664454-INS-80 | chr2-37664454-INS-99 |
| chr2-37676911-INS-175 | chr2-37676911-INS-182 |
| chr2-37750719-INS-204 | chr2-37750719-INS-204 |
| chr2-37761067-INS-202 | chr2-37761067-INS-202 |
| chr2-37766932-INS-76 | chr2-37766932-INS-76 |
| chr2-37770180-INS-54 | chr2-37770180-INS-54 |
| chr2-37873918-INS-88 | chr2-37873918-INS-60 |
| chr2-37894323-INS-57 | chr2-37894323-INS-57 |
| chr2-37910710-INS-6476 | chr2-37910711-INS-6476 |
| chr2-37969111-INS-52 | chr2-37969149-INS-54 |
| chr2-38015762-INS-96 | chr2-38015762-INS-92 |
| chr2-38044138-INS-211 | chr2-38044138-INS-212 |
| chr2-38082005-INS-75 | chr2-38082005-INS-75 |
| chr2-38199308-INS-6707 | chr2-38199309-INS-6707 |
| chr2-38206668-INS-426 | chr2-38206667-INS-426 |
| chr2-38260194-INS-52 | chr2-38260192-INS-73 |
| chr2-38263411-INS-183 | chr2-38263423-INS-198 |
| chr2-38302001-INS-212 | chr2-38302001-INS-212 |
| chr2-38302736-INS-54 | chr2-38302744-INS-56 |
| chr2-38378964-INS-192 | chr2-38378964-INS-192 |
| chr2-38422196-INS-53 | chr2-38422196-INS-67 |
| chr2-38431216-INS-64 | chr2-38431216-INS-57 |
| chr2-38486096-INS-75 | chr2-38486081-INS-83 |

| | |
|---|---|
| chr2-38498600-INS-98 | chr2-38498600-INS-98 |
| chr2-38530270-INS-200 | chr2-38530270-INS-200 |
| chr2-38552239-INS-53 | chr2-38552239-INS-53 |
| chr2-38624785-INS-86 | chr2-38624785-INS-88 |
| chr2-38639951-INS-951 | chr2-38639951-INS-951 |
| chr2-38699042-INS-67 | chr2-38698992-INS-85 |
| chr2-38732164-INS-216 | chr2-38732164-INS-216 |
| chr2-38735279-INS-498 | chr2-38735279-INS-498 |
| chr2-38746308-INS-54 | chr2-38746308-INS-54 |
| chr2-38775545-INS-215 | chr2-38775545-INS-215 |
| chr2-38782517-INS-445 | chr2-38782517-INS-445 |
| chr2-38798564-INS-85 | chr2-38798564-INS-90 |
| chr2-38799841-INS-215 | chr2-38799841-INS-214 |
| chr2-38817420-INS-84 | chr2-38817452-INS-62 |
| chr2-38821361-INS-132 | chr2-38821361-INS-136 |
| chr2-38843136-INS-65 | chr2-38843136-INS-65 |
| chr2-38845661-INS-7149 | chr2-38845662-INS-7149 |
| chr2-38848763-INS-204 | chr2-38848749-INS-227 |
| chr2-38884456-INS-186 | chr2-38884456-INS-186 |
| chr2-38952090-INS-66 | chr2-38952090-INS-68 |
| chr2-39014823-INS-5549 | chr2-39014824-INS-5549 |
| chr2-39022671-INS-136 | chr2-39022671-INS-136 |
| chr2-39136675-INS-212 | chr2-39136675-INS-212 |
| chr2-39185235-INS-592 | chr2-39185235-INS-592 |
| chr2-39265669-INS-274 | chr2-39265669-INS-275 |
| chr2-39279615-INS-6385 | chr2-39279616-INS-6386 |
| chr2-39398777-INS-52 | chr2-39398777-INS-56 |
| chr2-39407800-INS-204 | chr2-39407800-INS-204 |
| chr2-39444012-INS-3178 | chr2-39444013-INS-3178 |
| chr2-39452257-INS-190 | chr2-39452257-INS-188 |
| chr2-39459924-INS-6401 | chr2-39459925-INS-6401 |
| chr2-39495543-INS-102 | chr2-39495543-INS-105 |
| chr2-39541350-INS-57 | chr2-39541350-INS-66 |
| chr2-39555192-INS-1563 | chr2-39555193-INS-1562 |
| chr2-39594996-INS-7007 | chr2-39594997-INS-7007 |
| chr2-39618223-INS-52 | chr2-39618223-INS-72 |
| chr2-39682563-INS-310 | chr2-39682563-INS-310 |
| chr2-39686636-INS-550 | chr2-39686636-INS-550 |
| chr2-39748805-INS-2786 | chr2-39748806-INS-2786 |
| chr2-39760146-INS-327 | chr2-39760146-INS-327 |
| chr2-39767832-INS-7233 | chr2-39767833-INS-7234 |
| chr2-39781790-INS-417 | chr2-39781790-INS-429 |
| chr2-39782820-INS-201 | chr2-39782820-INS-201 |
| chr2-39816246-INS-84 | chr2-39816221-INS-94 |
| chr2-39882692-INS-84 | chr2-39882692-INS-84 |
| chr2-39897329-INS-361 | chr2-39897329-INS-332 |
| chr2-39918656-INS-60 | chr2-39918656-INS-54 |
| chr2-39943668-INS-6698 | chr2-39943669-INS-6697 |
| chr2-40060022-INS-201 | chr2-40060022-INS-201 |
| chr2-40070413-INS-4338 | chr2-40070414-INS-4338 |
| chr2-40084745-INS-6416 | chr2-40084746-INS-6420 |
| chr2-40102680-INS-172 | chr2-40102680-INS-173 |
| chr2-40145347-INS-73 | chr2-40145322-INS-120 |
| chr2-40148411-INS-352 | chr2-40148411-INS-352 |
| chr2-40165930-INS-88 | chr2-40165930-INS-98 |
| chr2-40184736-INS-56 | chr2-40184733-INS-55 |
| chr2-40198201-INS-209 | chr2-40198201-INS-207 |
| chr2-40200482-INS-438 | chr2-40200482-INS-438 |
| chr2-40212608-INS-54 | chr2-40212608-INS-58 |
| chr2-40229446-INS-642 | chr2-40229446-INS-642 |
| chr2-40267155-INS-50 | chr2-40267155-INS-52 |
| chr2-40271745-INS-66 | chr2-40271745-INS-82 |
| chr2-40315239-INS-218 | chr2-40315239-INS-218 |
| chr2-40377854-INS-592 | chr2-40377854-INS-592 |
| chr2-40383893-INS-56 | chr2-40383893-INS-50 |
| chr2-40392280-INS-6260 | chr2-40392281-INS-6259 |
| chr2-40422242-INS-245 | chr2-40422242-INS-245 |
| chr2-40425107-INS-92 | chr2-40425107-INS-92 |
| chr2-40484049-INS-88 | chr2-40484049-INS-112 |
| chr2-40495759-INS-506 | chr2-40495759-INS-506 |
| chr2-40498603-INS-4846 | chr2-40498604-INS-4846 |
| chr2-40501194-INS-146 | chr2-40501194-INS-146 |
| chr2-40564771-INS-441 | chr2-40564771-INS-441 |
| chr2-40565177-INS-848 | chr2-40565177-INS-853 |
| chr2-40580245-INS-85 | chr2-40580245-INS-85 |
| chr2-40590784-INS-659 | chr2-40590784-INS-659 |
| chr2-40599646-INS-69 | chr2-40599643-INS-57 |

| | |
|---|---|
| chr2-40610954-INS-6265 | chr2-40610955-INS-6263 |
| chr2-40616474-INS-50 | chr2-40616474-INS-54 |
| chr2-40625195-INS-76 | chr2-40625185-INS-68 |
| chr2-40634646-INS-52 | chr2-40634646-INS-64 |
| chr2-40653884-INS-201 | chr2-40653884-INS-201 |
| chr2-40666344-INS-64 | chr2-40666355-INS-81 |
| chr2-40677940-INS-214 | chr2-40677940-INS-214 |
| chr2-40707278-INS-137 | chr2-40707278-INS-142 |
| chr2-40729186-INS-1410 | chr2-40729187-INS-1410 |
| chr2-40734299-INS-1412 | chr2-40734300-INS-1413 |
| chr2-40751140-INS-220 | chr2-40751140-INS-220 |
| chr2-40768005-INS-169 | chr2-40768005-INS-157 |
| chr2-40792981-INS-303 | chr2-40792981-INS-293 |
| chr2-40839987-INS-56 | chr2-40839987-INS-56 |
| chr2-40868266-INS-110 | chr2-40868266-INS-109 |
| chr2-40890060-INS-128 | chr2-40890060-INS-128 |
| chr2-40899513-INS-1083 | chr2-40899514-INS-1083 |
| chr2-40916889-INS-643 | chr2-40916889-INS-643 |
| chr2-40932207-INS-1217 | chr2-40932208-INS-1217 |
| chr2-40963223-INS-74 | chr2-40963223-INS-74 |
| chr2-40977522-INS-66 | chr2-40977522-INS-66 |
| chr2-40978754-INS-121 | chr2-40978745-INS-169 |
| chr2-41018171-INS-75 | chr2-41018171-INS-81 |
| chr2-41034869-INS-66 | chr2-41034869-INS-66 |
| chr2-41037128-INS-629 | chr2-41037128-INS-629 |
| chr2-41038971-INS-365 | chr2-41038972-INS-377 |
| chr2-41067389-INS-2270 | chr2-41067390-INS-2270 |
| chr2-41083020-INS-324 | chr2-41083020-INS-324 |
| chr2-41113405-INS-163 | chr2-41113405-INS-163 |
| chr2-41115158-INS-65 | chr2-41115156-INS-81 |
| chr2-41130548-INS-89 | chr2-41130548-INS-89 |
| chr2-41134470-INS-209 | chr2-41134470-INS-209 |
| chr2-41178071-INS-109 | chr2-41178059-INS-136 |
| chr2-41180658-INS-78 | chr2-41180658-INS-78 |
| chr2-41196190-INS-453 | chr2-41196190-INS-452 |
| chr2-41196725-INS-6801 | chr2-41196726-INS-6802 |
| chr2-41203809-INS-113 | chr2-41203809-INS-113 |
| chr2-41215507-INS-1486 | chr2-41215508-INS-1486 |
| chr2-41240624-INS-6293 | chr2-41240633-INS-6280 |
| chr2-41256426-INS-62 | chr2-41256426-INS-62 |
| chr2-41264456-INS-855 | chr2-41264456-INS-856 |
| chr2-41326845-INS-991 | chr2-41326845-INS-991 |
| chr2-41329185-INS-483 | chr2-41329185-INS-485 |
| chr2-41377376-INS-137 | chr2-41377376-INS-142 |
| chr2-41399594-INS-133 | chr2-41399594-INS-133 |
| chr2-41408118-INS-84 | chr2-41408118-INS-84 |
| chr2-41425562-INS-122 | chr2-41425562-INS-122 |
| chr2-41426236-INS-1242 | chr2-41426237-INS-1243 |
| chr2-41472615-INS-464 | chr2-41472615-INS-460 |
| chr2-41474783-INS-425 | chr2-41474753-INS-419 |
| chr2-41525699-INS-64 | chr2-41525699-INS-65 |
| chr2-41529088-INS-56 | chr2-41529088-INS-64 |
| chr2-41559781-INS-6592 | chr2-41559782-INS-6596 |
| chr2-41609801-INS-3380 | chr2-41609808-INS-3379 |
| chr2-41677040-INS-51 | chr2-41677040-INS-51 |
| chr2-41692562-INS-453 | chr2-41692562-INS-465 |
| chr2-41750963-INS-60 | chr2-41750963-INS-62 |
| chr2-41797503-INS-1194 | chr2-41797504-INS-1194 |
| chr2-41872796-INS-70 | chr2-41872796-INS-70 |
| chr2-41893888-INS-60 | chr2-41893888-INS-72 |
| chr2-41957211-INS-163 | chr2-41957211-INS-163 |
| chr2-42053467-INS-342 | chr2-42053455-INS-375 |
| chr2-42071752-INS-98 | chr2-42071755-INS-86 |
| chr2-42076061-INS-426 | chr2-42076061-INS-426 |
| chr2-42133480-INS-52 | chr2-42133476-INS-56 |
| chr2-42150509-INS-96 | chr2-42150510-INS-95 |
| chr2-42177771-INS-54 | chr2-42177771-INS-60 |
| chr2-42190265-INS-122 | chr2-42190265-INS-122 |
| chr2-42224511-INS-97 | chr2-42224511-INS-111 |
| chr2-42238693-INS-440 | chr2-42238693-INS-440 |
| chr2-42258606-INS-267 | chr2-42258606-INS-267 |
| chr2-42261977-INS-478 | chr2-42261973-INS-481 |
| chr2-42296612-INS-1016 | chr2-42296616-INS-1008 |
| chr2-42303253-INS-183 | chr2-42303253-INS-183 |
| chr2-42317253-INS-55 | chr2-42317253-INS-75 |
| chr2-42326386-INS-62 | chr2-42326386-INS-62 |
| chr2-42372611-INS-86 | chr2-42372611-INS-86 |

| | |
|---|---|
| chr2-42374285-INS-160 | chr2-42374285-INS-159 |
| chr2-42412387-INS-62 | chr2-42412401-INS-50 |
| chr2-42434233-INS-52 | chr2-42434237-INS-94 |
| chr2-42435062-INS-6162 | chr2-42435063-INS-6162 |
| chr2-42475950-INS-50 | chr2-42475950-INS-50 |
| chr2-42516784-INS-56 | chr2-42516784-INS-52 |
| chr2-42546167-INS-242 | chr2-42546167-INS-242 |
| chr2-42559427-INS-104 | chr2-42559426-INS-62 |
| chr2-42566806-INS-74 | chr2-42566806-INS-74 |
| chr2-42575192-INS-4159 | chr2-42575193-INS-4151 |
| chr2-42589229-INS-6894 | chr2-42589230-INS-6894 |
| chr2-42656575-INS-7195 | chr2-42656576-INS-7195 |
| chr2-42662494-INS-1722 | chr2-42662495-INS-1721 |
| chr2-42687845-INS-246 | chr2-42687845-INS-246 |
| chr2-42690931-INS-916 | chr2-42690931-INS-942 |
| chr2-42738350-INS-342 | chr2-42738348-INS-344 |
| chr2-42774399-INS-112 | chr2-42774399-INS-100 |
| chr2-42823330-INS-193 | chr2-42823330-INS-193 |
| chr2-42839214-INS-171 | chr2-42839214-INS-168 |
| chr2-42847534-INS-432 | chr2-42847534-INS-432 |
| chr2-42863264-INS-507 | chr2-42863265-INS-507 |
| chr2-42871270-INS-100 | chr2-42871270-INS-100 |
| chr2-42909937-INS-81 | chr2-42909937-INS-81 |
| chr2-42912648-INS-51 | chr2-42912648-INS-55 |
| chr2-42916275-INS-50 | chr2-42916275-INS-50 |
| chr2-42930986-INS-939 | chr2-42930986-INS-944 |
| chr2-42960923-INS-213 | chr2-42960923-INS-213 |
| chr2-42976898-INS-173 | chr2-42976898-INS-173 |
| chr2-42981314-INS-778 | chr2-42981314-INS-779 |
| chr2-42993392-INS-1660 | chr2-42993393-INS-1661 |
| chr2-42995536-INS-52 | chr2-42995536-INS-52 |
| chr2-43002628-INS-54 | chr2-43002628-INS-56 |
| chr2-43003120-INS-4618 | chr2-43003121-INS-4619 |
| chr2-43032635-INS-416 | chr2-43032636-INS-416 |
| chr2-43033435-INS-50 | chr2-43033435-INS-60 |
| chr2-43091287-INS-67 | chr2-43091287-INS-67 |
| chr2-43105103-INS-7156 | chr2-43105104-INS-7157 |
| chr2-43149628-INS-66 | chr2-43149628-INS-66 |
| chr2-43153518-INS-581 | chr2-43153518-INS-582 |
| chr2-43156589-INS-289 | chr2-43156589-INS-289 |
| chr2-43169534-INS-6381 | chr2-43169530-INS-6380 |
| chr2-43188247-INS-204 | chr2-43188247-INS-205 |
| chr2-43197798-INS-79 | chr2-43197798-INS-79 |
| chr2-43235317-INS-173 | chr2-43235321-INS-180 |
| chr2-43240160-INS-179 | chr2-43240160-INS-179 |
| chr2-43266758-INS-200 | chr2-43266758-INS-202 |
| chr2-43345093-INS-98 | chr2-43345093-INS-98 |
| chr2-43351893-INS-76 | chr2-43351893-INS-56 |
| chr2-43378732-INS-133 | chr2-43378732-INS-133 |
| chr2-43415434-INS-153 | chr2-43415466-INS-117 |
| chr2-43431669-INS-52 | chr2-43431671-INS-58 |
| chr2-43463560-INS-724 | chr2-43463560-INS-690 |
| chr2-43464195-INS-62 | chr2-43464195-INS-63 |
| chr2-43479335-INS-66 | chr2-43479335-INS-50 |
| chr2-43525676-INS-190 | chr2-43525678-INS-190 |
| chr2-43597952-INS-110 | chr2-43597964-INS-123 |
| chr2-43607526-INS-94 | chr2-43607525-INS-68 |
| chr2-43623188-INS-364 | chr2-43623188-INS-365 |
| chr2-43627506-INS-80 | chr2-43627506-INS-78 |
| chr2-43634964-INS-65 | chr2-43634964-INS-65 |
| chr2-43643853-INS-99 | chr2-43643853-INS-105 |
| chr2-43645491-INS-401 | chr2-43645491-INS-402 |
| chr2-43668887-INS-103 | chr2-43668877-INS-134 |
| chr2-43680715-INS-64 | chr2-43680715-INS-66 |
| chr2-43769251-INS-800 | chr2-43769251-INS-803 |
| chr2-43773465-INS-200 | chr2-43773465-INS-201 |
| chr2-43788855-INS-66 | chr2-43788851-INS-70 |
| chr2-43790601-INS-718 | chr2-43790596-INS-714 |
| chr2-43800967-INS-161 | chr2-43800967-INS-161 |
| chr2-43811167-INS-6739 | chr2-43811168-INS-6741 |
| chr2-43850729-INS-76 | chr2-43850729-INS-62 |
| chr2-43904662-INS-63 | chr2-43904662-INS-63 |
| chr2-43931644-INS-72 | chr2-43931662-INS-86 |
| chr2-43963032-INS-4678 | chr2-43963033-INS-4679 |
| chr2-44009928-INS-56 | chr2-44009949-INS-59 |
| chr2-44068125-INS-52 | chr2-44068125-INS-53 |
| chr2-44114608-INS-108 | chr2-44114608-INS-108 |

| | |
|---|---|
| chr2-44121697-INS-1544 | chr2-44121698-INS-1543 |
| chr2-44305007-INS-1460 | chr2-44305008-INS-1461 |
| chr2-44307165-INS-6437 | chr2-44307166-INS-6437 |
| chr2-44311281-INS-580 | chr2-44311281-INS-577 |
| chr2-44314101-INS-247 | chr2-44314098-INS-282 |
| chr2-44326091-INS-1810 | chr2-44326092-INS-1811 |
| chr2-44342794-INS-76 | chr2-44342794-INS-80 |
| chr2-44367786-INS-351 | chr2-44367786-INS-312 |
| chr2-44371774-INS-4816 | chr2-44371775-INS-4816 |
| chr2-44378261-INS-690 | chr2-44378262-INS-688 |
| chr2-44414733-INS-60 | chr2-44414733-INS-50 |
| chr2-44422732-INS-7135 | chr2-44422733-INS-7135 |
| chr2-44471720-INS-148 | chr2-44471712-INS-153 |
| chr2-44575031-INS-55 | chr2-44575031-INS-50 |
| chr2-44702953-INS-88 | chr2-44702941-INS-88 |
| chr2-44742923-INS-6801 | chr2-44742924-INS-6801 |
| chr2-44770565-INS-56 | chr2-44770556-INS-50 |
| chr2-44797902-INS-3713 | chr2-44797903-INS-3712 |
| chr2-44798166-INS-404 | chr2-44798166-INS-432 |
| chr2-44935643-INS-112 | chr2-44935643-INS-114 |
| chr2-44988577-INS-62 | chr2-44988580-INS-54 |
| chr2-45012166-INS-65 | chr2-45012166-INS-50 |
| chr2-45027787-INS-82 | chr2-45027787-INS-81 |
| chr2-45029840-INS-60 | chr2-45029840-INS-60 |
| chr2-45070015-INS-75 | chr2-45070033-INS-54 |
| chr2-45102508-INS-180 | chr2-45102525-INS-190 |
| chr2-45183321-INS-75 | chr2-45183321-INS-75 |
| chr2-45205729-INS-76 | chr2-45205698-INS-60 |
| chr2-45221053-INS-58 | chr2-45221053-INS-61 |
| chr2-45227945-INS-92 | chr2-45227945-INS-90 |
| chr2-45330681-INS-65 | chr2-45330681-INS-65 |
| chr2-45336868-INS-58 | chr2-45336866-INS-80 |
| chr2-45342427-INS-787 | chr2-45342427-INS-788 |
| chr2-45347759-INS-208 | chr2-45347759-INS-208 |
| chr2-45358220-INS-78 | chr2-45358243-INS-74 |
| chr2-45361778-INS-80 | chr2-45361778-INS-89 |
| chr2-45419548-INS-6655 | chr2-45419549-INS-6656 |
| chr2-45452965-INS-1213 | chr2-45452953-INS-1216 |
| chr2-45475927-INS-513 | chr2-45475927-INS-514 |
| chr2-45479967-INS-1969 | chr2-45479968-INS-1969 |
| chr2-45502532-INS-63 | chr2-45502532-INS-99 |
| chr2-45504929-INS-1595 | chr2-45504930-INS-1604 |
| chr2-45515638-INS-212 | chr2-45515636-INS-179 |
| chr2-45537636-INS-6264 | chr2-45537637-INS-6272 |
| chr2-45555895-INS-60 | chr2-45555895-INS-56 |
| chr2-45618302-INS-6599 | chr2-45618303-INS-6600 |
| chr2-45705264-INS-2425 | chr2-45705265-INS-2425 |
| chr2-45765202-INS-249 | chr2-45765202-INS-249 |
| chr2-45876377-INS-56 | chr2-45876392-INS-85 |
| chr2-45927140-INS-1324 | chr2-45927139-INS-1322 |
| chr2-45959840-INS-459 | chr2-45959840-INS-458 |
| chr2-45960992-INS-215 | chr2-45960991-INS-215 |
| chr2-45964332-INS-902 | chr2-45964332-INS-902 |
| chr2-45978291-INS-70 | chr2-45978291-INS-82 |
| chr2-46001389-INS-65 | chr2-46001389-INS-65 |
| chr2-46047023-INS-177 | chr2-46047023-INS-174 |
| chr2-46053394-INS-74 | chr2-46053394-INS-88 |
| chr2-46055387-INS-194 | chr2-46055387-INS-194 |
| chr2-46057636-INS-244 | chr2-46057636-INS-245 |
| chr2-46066259-INS-308 | chr2-46066259-INS-309 |
| chr2-46087423-INS-1941 | chr2-46087420-INS-1945 |
| chr2-46096664-INS-185 | chr2-46096664-INS-191 |
| chr2-46123358-INS-82 | chr2-46123358-INS-82 |
| chr2-46124911-INS-1936 | chr2-46124912-INS-1935 |
| chr2-46132871-INS-72 | chr2-46132871-INS-72 |
| chr2-46141542-INS-1805 | chr2-46141543-INS-1810 |
| chr2-46142594-INS-2233 | chr2-46142595-INS-2233 |
| chr2-46179000-INS-84 | chr2-46178994-INS-50 |
| chr2-46226965-INS-137 | chr2-46226965-INS-128 |
| chr2-46242091-INS-54 | chr2-46242091-INS-54 |
| chr2-46257771-INS-53 | chr2-46257771-INS-53 |
| chr2-46292949-INS-179 | chr2-46292949-INS-178 |
| chr2-46307081-INS-61 | chr2-46307111-INS-51 |
| chr2-46310567-INS-86 | chr2-46310555-INS-112 |
| chr2-46335982-INS-215 | chr2-46335975-INS-247 |
| chr2-46354166-INS-56 | chr2-46354166-INS-56 |
| chr2-46396788-INS-59 | chr2-46396788-INS-59 |

| | |
|---|---|
| chr2-46400621-INS-629 | chr2-46400621-INS-629 |
| chr2-46513512-INS-70 | chr2-46513512-INS-70 |
| chr2-46513709-INS-6141 | chr2-46513710-INS-6141 |
| chr2-46539124-INS-277 | chr2-46539124-INS-277 |
| chr2-46546731-INS-1889 | chr2-46546732-INS-1889 |
| chr2-46551960-INS-1977 | chr2-46551961-INS-1977 |
| chr2-46604614-INS-827 | chr2-46604614-INS-827 |
| chr2-46651664-INS-6516 | chr2-46651665-INS-6516 |
| chr2-46661276-INS-365 | chr2-46661248-INS-366 |
| chr2-46665071-INS-52 | chr2-46665071-INS-64 |
| chr2-46667761-INS-3895 | chr2-46667762-INS-3895 |
| chr2-46726506-INS-81 | chr2-46726506-INS-81 |
| chr2-46740633-INS-857 | chr2-46740633-INS-857 |
| chr2-46753168-INS-1178 | chr2-46753169-INS-1177 |
| chr2-46781100-INS-2916 | chr2-46781101-INS-2919 |
| chr2-46781210-INS-6463 | chr2-46781211-INS-6463 |
| chr2-46783709-INS-197 | chr2-46783709-INS-199 |
| chr2-46817117-INS-60 | chr2-46817117-INS-68 |
| chr2-46819122-INS-399 | chr2-46819122-INS-399 |
| chr2-46826701-INS-55 | chr2-46826701-INS-55 |
| chr2-46841871-INS-71 | chr2-46841871-INS-67 |
| chr2-46864296-INS-955 | chr2-46864296-INS-955 |
| chr2-46865668-INS-189 | chr2-46865669-INS-192 |
| chr2-46876355-INS-4530 | chr2-46876356-INS-4530 |
| chr2-46885494-INS-304 | chr2-46885490-INS-305 |
| chr2-46890535-INS-4384 | chr2-46890536-INS-4384 |
| chr2-46898802-INS-623 | chr2-46898803-INS-623 |
| chr2-46923024-INS-643 | chr2-46923024-INS-645 |
| chr2-46931795-INS-1003 | chr2-46931796-INS-1003 |
| chr2-46932438-INS-6879 | chr2-46932439-INS-6879 |
| chr2-46948112-INS-67 | chr2-46948112-INS-67 |
| chr2-46950332-INS-53 | chr2-46950326-INS-54 |
| chr2-47041266-INS-67 | chr2-47041266-INS-106 |
| chr2-47043606-INS-853 | chr2-47043606-INS-853 |
| chr2-47062387-INS-84 | chr2-47062387-INS-94 |
| chr2-47082903-INS-54 | chr2-47082947-INS-54 |
| chr2-47084235-INS-136 | chr2-47084251-INS-104 |
| chr2-47098980-INS-1625 | chr2-47098981-INS-1626 |
| chr2-47110269-INS-721 | chr2-47110269-INS-721 |
| chr2-47163920-INS-703 | chr2-47163920-INS-703 |
| chr2-47168179-INS-106 | chr2-47168212-INS-111 |
| chr2-47198374-INS-521 | chr2-47198374-INS-521 |
| chr2-47208621-INS-146 | chr2-47208642-INS-171 |
| chr2-47310587-INS-10567 | chr2-47310588-INS-10564 |
| chr2-47321664-INS-3351 | chr2-47321665-INS-3351 |
| chr2-47349309-INS-2495 | chr2-47349310-INS-2479 |
| chr2-47349551-INS-50 | chr2-47349553-INS-60 |
| chr2-47363816-INS-54 | chr2-47363816-INS-54 |
| chr2-47383710-INS-58 | chr2-47383710-INS-58 |
| chr2-47392842-INS-157 | chr2-47392842-INS-157 |
| chr2-47409004-INS-854 | chr2-47409004-INS-855 |
| chr2-47413761-INS-56 | chr2-47413761-INS-58 |
| chr2-47417806-INS-207 | chr2-47417806-INS-208 |
| chr2-47444317-INS-464 | chr2-47444317-INS-464 |
| chr2-47465418-INS-92 | chr2-47465419-INS-76 |
| chr2-47466385-INS-72 | chr2-47466386-INS-102 |
| chr2-47470962-INS-70 | chr2-47470962-INS-60 |
| chr2-47484446-INS-7428 | chr2-47484447-INS-7428 |
| chr2-47487020-INS-263 | chr2-47487020-INS-263 |
| chr2-47512644-INS-869 | chr2-47512644-INS-868 |
| chr2-47559632-INS-215 | chr2-47559587-INS-212 |
| chr2-47572964-INS-64 | chr2-47572964-INS-64 |
| chr2-47621310-INS-1587 | chr2-47621311-INS-1587 |
| chr2-47672520-INS-356 | chr2-47672520-INS-356 |
| chr2-47678839-INS-2109 | chr2-47678840-INS-2109 |
| chr2-47700951-INS-2263 | chr2-47700952-INS-2263 |
| chr2-47704569-INS-148 | chr2-47704571-INS-104 |
| chr2-47716553-INS-55 | chr2-47716553-INS-57 |
| chr2-47738322-INS-114 | chr2-47738322-INS-114 |
| chr2-47755865-INS-56 | chr2-47755865-INS-56 |
| chr2-47772270-INS-72 | chr2-47772270-INS-90 |
| chr2-47773562-INS-98 | chr2-47773591-INS-96 |
| chr2-47782829-INS-56 | chr2-47782829-INS-58 |
| chr2-47799708-INS-142 | chr2-47799708-INS-142 |
| chr2-47806196-INS-3871 | chr2-47806197-INS-3885 |
| chr2-47837333-INS-51 | chr2-47837320-INS-56 |
| chr2-47869173-INS-1318 | chr2-47869174-INS-1318 |

| | |
|---|---|
| chr2-47885005-INS-2813 | chr2-47885006-INS-2813 |
| chr2-47891105-INS-61 | chr2-47891105-INS-61 |
| chr2-47896789-INS-90 | chr2-47896789-INS-90 |
| chr2-47906921-INS-675 | chr2-47906921-INS-675 |
| chr2-47920973-INS-362 | chr2-47920973-INS-362 |
| chr2-47996163-INS-99 | chr2-47996163-INS-101 |
| chr2-48056900-INS-2245 | chr2-48056901-INS-2245 |
| chr2-48086550-INS-154 | chr2-48086539-INS-162 |
| chr2-48087716-INS-197 | chr2-48087716-INS-197 |
| chr2-48156987-INS-465 | chr2-48156987-INS-465 |
| chr2-48159767-INS-6448 | chr2-48159768-INS-6448 |
| chr2-48175307-INS-58 | chr2-48175307-INS-66 |
| chr2-48195141-INS-232 | chr2-48195141-INS-230 |
| chr2-48195504-INS-7251 | chr2-48195505-INS-7252 |
| chr2-48244234-INS-116 | chr2-48244234-INS-116 |
| chr2-48303611-INS-345 | chr2-48303611-INS-345 |
| chr2-48306948-INS-148 | chr2-48306948-INS-113 |
| chr2-48317604-INS-643 | chr2-48317604-INS-644 |
| chr2-48327165-INS-246 | chr2-48327179-INS-269 |
| chr2-48329226-INS-52 | chr2-48329226-INS-50 |
| chr2-48341763-INS-819 | chr2-48341763-INS-819 |
| chr2-48342568-INS-1331 | chr2-48342566-INS-1297 |
| chr2-48402032-INS-70 | chr2-48402032-INS-70 |
| chr2-48412383-INS-1226 | chr2-48412384-INS-1226 |
| chr2-48416811-INS-193 | chr2-48416809-INS-198 |
| chr2-48430203-INS-7099 | chr2-48430204-INS-7100 |
| chr2-48430296-INS-96 | chr2-48430296-INS-96 |
| chr2-48432280-INS-245 | chr2-48432280-INS-245 |
| chr2-48443892-INS-60 | chr2-48443935-INS-62 |
| chr2-48445452-INS-103 | chr2-48445452-INS-103 |
| chr2-48453903-INS-156 | chr2-48453903-INS-156 |
| chr2-48491792-INS-266 | chr2-48491792-INS-266 |
| chr2-48505060-INS-7054 | chr2-48505061-INS-7055 |
| chr2-48518284-INS-54 | chr2-48518284-INS-54 |
| chr2-48520149-INS-1149 | chr2-48520150-INS-1152 |
| chr2-48555207-INS-52 | chr2-48555207-INS-54 |
| chr2-48561482-INS-52 | chr2-48561482-INS-56 |
| chr2-48562798-INS-142 | chr2-48562801-INS-133 |
| chr2-48564218-INS-400 | chr2-48564218-INS-400 |
| chr2-48587198-INS-73 | chr2-48587198-INS-77 |
| chr2-48612985-INS-54 | chr2-48612985-INS-52 |
| chr2-48619357-INS-402 | chr2-48619357-INS-402 |
| chr2-48627191-INS-54 | chr2-48627194-INS-64 |
| chr2-48698427-INS-58 | chr2-48698427-INS-58 |
| chr2-48727529-INS-396 | chr2-48727529-INS-397 |
| chr2-48812205-INS-156 | chr2-48812207-INS-147 |
| chr2-48825344-INS-151 | chr2-48825344-INS-151 |
| chr2-48869960-INS-56 | chr2-48869960-INS-60 |
| chr2-48962442-INS-190 | chr2-48962442-INS-195 |
| chr2-48974279-INS-504 | chr2-48974279-INS-504 |
| chr2-48979365-INS-438 | chr2-48979365-INS-439 |
| chr2-48994919-INS-6459 | chr2-48994920-INS-6461 |
| chr2-48998453-INS-6684 | chr2-48998454-INS-6685 |
| chr2-49063266-INS-6702 | chr2-49063267-INS-6703 |
| chr2-49092455-INS-54 | chr2-49092455-INS-52 |
| chr2-49122726-INS-75 | chr2-49122726-INS-75 |
| chr2-49174432-INS-384 | chr2-49174433-INS-384 |
| chr2-49181321-INS-399 | chr2-49181321-INS-399 |
| chr2-49226965-INS-55 | chr2-49226965-INS-55 |
| chr2-49249907-INS-54 | chr2-49249907-INS-54 |
| chr2-49309410-INS-307 | chr2-49309410-INS-306 |
| chr2-49313787-INS-53 | chr2-49313787-INS-53 |
| chr2-49331745-INS-209 | chr2-49331745-INS-209 |
| chr2-49466397-INS-206 | chr2-49466397-INS-206 |
| chr2-49468331-INS-85 | chr2-49468331-INS-97 |
| chr2-49483938-INS-205 | chr2-49483938-INS-203 |
| chr2-49491509-INS-401 | chr2-49491509-INS-401 |
| chr2-49498519-INS-326 | chr2-49498519-INS-326 |
| chr2-49500734-INS-204 | chr2-49500734-INS-204 |
| chr2-49505365-INS-355 | chr2-49505365-INS-355 |
| chr2-49512103-INS-219 | chr2-49512103-INS-219 |
| chr2-49531145-INS-68 | chr2-49531145-INS-68 |
| chr2-49532294-INS-203 | chr2-49532287-INS-207 |
| chr2-49553301-INS-400 | chr2-49553301-INS-400 |
| chr2-49555675-INS-200 | chr2-49555675-INS-200 |
| chr2-49561500-INS-119 | chr2-49561500-INS-119 |
| chr2-49572438-INS-77 | chr2-49572477-INS-80 |

| | |
|---|---|
| chr2-49602859-INS-162 | chr2-49602859-INS-162 |
| chr2-49655461-INS-221 | chr2-49655462-INS-209 |
| chr2-49658073-INS-143 | chr2-49658073-INS-143 |
| chr2-49683509-INS-60 | chr2-49683510-INS-60 |
| chr2-49692281-INS-80 | chr2-49692281-INS-80 |
| chr2-49744143-INS-64 | chr2-49744101-INS-98 |
| chr2-49829758-INS-94 | chr2-49829758-INS-94 |
| chr2-49852564-INS-55 | chr2-49852564-INS-51 |
| chr2-49858238-INS-146 | chr2-49858238-INS-149 |
| chr2-49872784-INS-506 | chr2-49872784-INS-504 |
| chr2-49879864-INS-84 | chr2-49879864-INS-60 |
| chr2-50016221-INS-54 | chr2-50016174-INS-77 |
| chr2-50030412-INS-882 | chr2-50030412-INS-882 |
| chr2-50039171-INS-57 | chr2-50039171-INS-57 |
| chr2-50092374-INS-156 | chr2-50092374-INS-150 |
| chr2-50102437-INS-68 | chr2-50102437-INS-74 |
| chr2-50114529-INS-66 | chr2-50114529-INS-68 |
| chr2-50125262-INS-116 | chr2-50125262-INS-107 |
| chr2-50153140-INS-130 | chr2-50153140-INS-130 |
| chr2-50166133-INS-6820 | chr2-50166134-INS-6821 |
| chr2-50209165-INS-617 | chr2-50209165-INS-596 |
| chr2-50218647-INS-54 | chr2-50218647-INS-54 |
| chr2-50220787-INS-400 | chr2-50220787-INS-399 |
| chr2-50238141-INS-183 | chr2-50238141-INS-183 |
| chr2-50253019-INS-99 | chr2-50253020-INS-70 |
| chr2-50262851-INS-52 | chr2-50262892-INS-52 |
| chr2-50268502-INS-379 | chr2-50268502-INS-346 |
| chr2-50282450-INS-119 | chr2-50282450-INS-119 |
| chr2-50297070-INS-86 | chr2-50297070-INS-136 |
| chr2-50298362-INS-171 | chr2-50298362-INS-171 |
| chr2-50317686-INS-1007 | chr2-50317687-INS-1007 |
| chr2-50332774-INS-256 | chr2-50332775-INS-256 |
| chr2-50336702-INS-65 | chr2-50336702-INS-55 |
| chr2-50356215-INS-6517 | chr2-50356216-INS-6518 |
| chr2-50381985-INS-66 | chr2-50381985-INS-70 |
| chr2-50383610-INS-457 | chr2-50383610-INS-457 |
| chr2-50409753-INS-82 | chr2-50409753-INS-84 |
| chr2-50415182-INS-490 | chr2-50415182-INS-490 |
| chr2-50437093-INS-542 | chr2-50437093-INS-542 |
| chr2-50447176-INS-418 | chr2-50447176-INS-429 |
| chr2-50475123-INS-102 | chr2-50475123-INS-103 |
| chr2-50480823-INS-74 | chr2-50480823-INS-76 |
| chr2-50591858-INS-60 | chr2-50591858-INS-66 |
| chr2-50592758-INS-950 | chr2-50592758-INS-988 |
| chr2-50609550-INS-400 | chr2-50609550-INS-400 |
| chr2-50651908-INS-467 | chr2-50651908-INS-467 |
| chr2-50664182-INS-70 | chr2-50664182-INS-70 |
| chr2-50687590-INS-2728 | chr2-50687591-INS-2758 |
| chr2-50698244-INS-470 | chr2-50698244-INS-470 |
| chr2-50704682-INS-144 | chr2-50704682-INS-144 |
| chr2-50710706-INS-86 | chr2-50710706-INS-86 |
| chr2-50721295-INS-82 | chr2-50721295-INS-82 |
| chr2-50769096-INS-1094 | chr2-50769097-INS-1094 |
| chr2-50811594-INS-105 | chr2-50811636-INS-108 |
| chr2-50847204-INS-203 | chr2-50847204-INS-203 |
| chr2-50878149-INS-198 | chr2-50878149-INS-198 |
| chr2-50907188-INS-201 | chr2-50907188-INS-201 |
| chr2-50911396-INS-211 | chr2-50911396-INS-211 |
| chr2-50924264-INS-53 | chr2-50924264-INS-53 |
| chr2-50975743-INS-59 | chr2-50975743-INS-59 |
| chr2-50982971-INS-233 | chr2-50982971-INS-233 |
| chr2-50989023-INS-104 | chr2-50989060-INS-83 |
| chr2-50994039-INS-462 | chr2-50994037-INS-475 |
| chr2-51008401-INS-138 | chr2-51008401-INS-138 |
| chr2-51015799-INS-94 | chr2-51015799-INS-94 |
| chr2-51024040-INS-56 | chr2-51024040-INS-56 |
| chr2-51049994-INS-258 | chr2-51049994-INS-258 |
| chr2-51105286-INS-91 | chr2-51105286-INS-95 |
| chr2-51109279-INS-218 | chr2-51109279-INS-218 |
| chr2-51151012-INS-61 | chr2-51151012-INS-66 |
| chr2-51193414-INS-91 | chr2-51193414-INS-91 |
| chr2-51209491-INS-118 | chr2-51209491-INS-118 |
| chr2-51223682-INS-331 | chr2-51223682-INS-331 |
| chr2-51234132-INS-99 | chr2-51234132-INS-97 |
| chr2-51262645-INS-598 | chr2-51262645-INS-597 |
| chr2-51265388-INS-434 | chr2-51265388-INS-434 |
| chr2-51272340-INS-232 | chr2-51272340-INS-232 |

| | |
|---|---|
| chr2-51297633-INS-56 | chr2-51297633-INS-56 |
| chr2-51300864-INS-53 | chr2-51300864-INS-53 |
| chr2-51305183-INS-56 | chr2-51305183-INS-58 |
| chr2-51315140-INS-381 | chr2-51315140-INS-384 |
| chr2-51321094-INS-164 | chr2-51321094-INS-165 |
| chr2-51324493-INS-161 | chr2-51324493-INS-161 |
| chr2-51326870-INS-65 | chr2-51326870-INS-67 |
| chr2-51333463-INS-63 | chr2-51333463-INS-67 |
| chr2-51339695-INS-124 | chr2-51339734-INS-121 |
| chr2-51347459-INS-154 | chr2-51347459-INS-154 |
| chr2-51368364-INS-55 | chr2-51368364-INS-55 |
| chr2-51369870-INS-80 | chr2-51369870-INS-72 |
| chr2-51382176-INS-211 | chr2-51382176-INS-211 |
| chr2-51384473-INS-57 | chr2-51384471-INS-61 |
| chr2-51385551-INS-86 | chr2-51385551-INS-86 |
| chr2-51390859-INS-52 | chr2-51390860-INS-56 |
| chr2-51393892-INS-896 | chr2-51393892-INS-896 |
| chr2-51400394-INS-120 | chr2-51400394-INS-156 |
| chr2-51456529-INS-717 | chr2-51456529-INS-718 |
| chr2-51478031-INS-139 | chr2-51478031-INS-139 |
| chr2-51479352-INS-506 | chr2-51479352-INS-505 |
| chr2-51514354-INS-80 | chr2-51514354-INS-80 |
| chr2-51523473-INS-1082 | chr2-51523474-INS-1084 |
| chr2-51524850-INS-236 | chr2-51524850-INS-243 |
| chr2-51530793-INS-53 | chr2-51530778-INS-74 |
| chr2-51538504-INS-156 | chr2-51538488-INS-162 |
| chr2-51549132-INS-4372 | chr2-51549137-INS-4367 |
| chr2-51557701-INS-196 | chr2-51557701-INS-196 |
| chr2-51589641-INS-342 | chr2-51589641-INS-342 |
| chr2-51592386-INS-58 | chr2-51592354-INS-70 |
| chr2-51596626-INS-94 | chr2-51596627-INS-79 |
| chr2-51600777-INS-53 | chr2-51600777-INS-57 |
| chr2-51608846-INS-6081 | chr2-51608847-INS-6081 |
| chr2-51613844-INS-328 | chr2-51613844-INS-328 |
| chr2-51615528-INS-67 | chr2-51615528-INS-67 |
| chr2-51619284-INS-52 | chr2-51619284-INS-62 |
| chr2-51622780-INS-119 | chr2-51622780-INS-101 |
| chr2-51624204-INS-54 | chr2-51624234-INS-58 |
| chr2-51630376-INS-431 | chr2-51630376-INS-431 |
| chr2-51633760-INS-79 | chr2-51633760-INS-79 |
| chr2-51635756-INS-205 | chr2-51635756-INS-201 |
| chr2-51663319-INS-6484 | chr2-51663320-INS-6487 |
| chr2-51663893-INS-61 | chr2-51663893-INS-61 |
| chr2-51672675-INS-56 | chr2-51672675-INS-56 |
| chr2-51686867-INS-6134 | chr2-51686868-INS-6134 |
| chr2-51693003-INS-89 | chr2-51693003-INS-89 |
| chr2-51705149-INS-66 | chr2-51705149-INS-66 |
| chr2-51705553-INS-1891 | chr2-51705554-INS-1891 |
| chr2-51714748-INS-3088 | chr2-51714749-INS-3088 |
| chr2-51730164-INS-205 | chr2-51730164-INS-210 |
| chr2-51753992-INS-476 | chr2-51753994-INS-480 |
| chr2-51760688-INS-80 | chr2-51760682-INS-86 |
| chr2-51763343-INS-54 | chr2-51763343-INS-54 |
| chr2-51791281-INS-60 | chr2-51791242-INS-60 |
| chr2-51843636-INS-101 | chr2-51843636-INS-101 |
| chr2-51847882-INS-52 | chr2-51847882-INS-52 |
| chr2-51851165-INS-129 | chr2-51851163-INS-131 |
| chr2-51855601-INS-86 | chr2-51855601-INS-86 |
| chr2-51903838-INS-60 | chr2-51903838-INS-62 |
| chr2-51907251-INS-660 | chr2-51907252-INS-652 |
| chr2-51919151-INS-314 | chr2-51919151-INS-314 |
| chr2-51922090-INS-96 | chr2-51922087-INS-114 |
| chr2-51925848-INS-119 | chr2-51925873-INS-96 |
| chr2-51930472-INS-249 | chr2-51930472-INS-249 |
| chr2-51931891-INS-71 | chr2-51931880-INS-54 |
| chr2-52025547-INS-282 | chr2-52025547-INS-282 |
| chr2-52055775-INS-51 | chr2-52055775-INS-51 |
| chr2-52099084-INS-253 | chr2-52099084-INS-255 |
| chr2-52102064-INS-60 | chr2-52102064-INS-68 |
| chr2-52103751-INS-169 | chr2-52103751-INS-163 |
| chr2-52129719-INS-66 | chr2-52129719-INS-66 |
| chr2-52147182-INS-310 | chr2-52147182-INS-310 |
| chr2-52151789-INS-98 | chr2-52151789-INS-98 |
| chr2-52159074-INS-52 | chr2-52159074-INS-52 |
| chr2-52188467-INS-77 | chr2-52188467-INS-77 |
| chr2-52216920-INS-294 | chr2-52216920-INS-297 |
| chr2-52237543-INS-6358 | chr2-52237544-INS-6360 |

| | |
|---|---|
| chr2-52242753-INS-62 | chr2-52242753-INS-68 |
| chr2-52251892-INS-568 | chr2-52251892-INS-569 |
| chr2-52256967-INS-70 | chr2-52256967-INS-70 |
| chr2-52280375-INS-686 | chr2-52280375-INS-686 |
| chr2-52297193-INS-97 | chr2-52297193-INS-97 |
| chr2-52306089-INS-138 | chr2-52306089-INS-138 |
| chr2-52345213-INS-184 | chr2-52345213-INS-159 |
| chr2-52353217-INS-57 | chr2-52353217-INS-57 |
| chr2-52359665-INS-91 | chr2-52359683-INS-50 |
| chr2-52370110-INS-216 | chr2-52370110-INS-215 |
| chr2-52391917-INS-121 | chr2-52391921-INS-80 |
| chr2-52399683-INS-624 | chr2-52399683-INS-624 |
| chr2-52412892-INS-95 | chr2-52412892-INS-60 |
| chr2-52417558-INS-90 | chr2-52417558-INS-99 |
| chr2-52418932-INS-57 | chr2-52418932-INS-57 |
| chr2-52425515-INS-216 | chr2-52425516-INS-225 |
| chr2-52451053-INS-371 | chr2-52451053-INS-371 |
| chr2-52454203-INS-60 | chr2-52454203-INS-60 |
| chr2-52523850-INS-61 | chr2-52523850-INS-61 |
| chr2-52532682-INS-569 | chr2-52532682-INS-570 |
| chr2-52535092-INS-52 | chr2-52535092-INS-52 |
| chr2-52538885-INS-90 | chr2-52538885-INS-96 |
| chr2-52540742-INS-98 | chr2-52540742-INS-98 |
| chr2-52542243-INS-51 | chr2-52542243-INS-51 |
| chr2-52550908-INS-207 | chr2-52550908-INS-207 |
| chr2-52560647-INS-62 | chr2-52560653-INS-64 |
| chr2-52562191-INS-114 | chr2-52562216-INS-114 |
| chr2-52616078-INS-228 | chr2-52616078-INS-228 |
| chr2-52628367-INS-75 | chr2-52628367-INS-75 |
| chr2-52659169-INS-85 | chr2-52659157-INS-69 |
| chr2-52661742-INS-82 | chr2-52661740-INS-55 |
| chr2-52665224-INS-73 | chr2-52665210-INS-94 |
| chr2-52667115-INS-74 | chr2-52667112-INS-83 |
| chr2-52667425-INS-72 | chr2-52667425-INS-68 |
| chr2-52671842-INS-66 | chr2-52671846-INS-51 |
| chr2-52684802-INS-543 | chr2-52684802-INS-550 |
| chr2-52701677-INS-7795 | chr2-52701678-INS-7788 |
| chr2-52702997-INS-203 | chr2-52702997-INS-207 |
| chr2-52732553-INS-200 | chr2-52732553-INS-200 |
| chr2-52742066-INS-122 | chr2-52742066-INS-103 |
| chr2-52744896-INS-195 | chr2-52744897-INS-195 |
| chr2-52762327-INS-162 | chr2-52762330-INS-135 |
| chr2-52839109-INS-216 | chr2-52839109-INS-216 |
| chr2-52883737-INS-193 | chr2-52883737-INS-194 |
| chr2-52908778-INS-62 | chr2-52908778-INS-62 |
| chr2-52926982-INS-60 | chr2-52926982-INS-60 |
| chr2-52930638-INS-201 | chr2-52930646-INS-205 |
| chr2-52934177-INS-133 | chr2-52934176-INS-133 |
| chr2-52947019-INS-204 | chr2-52947019-INS-204 |
| chr2-52952806-INS-426 | chr2-52952806-INS-400 |
| chr2-52975343-INS-204 | chr2-52975343-INS-203 |
| chr2-53014308-INS-78 | chr2-53014345-INS-66 |
| chr2-53021329-INS-206 | chr2-53021329-INS-206 |
| chr2-53085061-INS-503 | chr2-53085061-INS-503 |
| chr2-53125740-INS-243 | chr2-53125740-INS-243 |
| chr2-53150779-INS-114 | chr2-53150755-INS-155 |
| chr2-53155008-INS-78 | chr2-53155000-INS-98 |
| chr2-53155702-INS-548 | chr2-53155702-INS-549 |
| chr2-53183869-INS-4709 | chr2-53183870-INS-4709 |
| chr2-53219536-INS-4400 | chr2-53219540-INS-4395 |
| chr2-53220221-INS-2782 | chr2-53220222-INS-2782 |
| chr2-53223865-INS-501 | chr2-53223865-INS-501 |
| chr2-53233043-INS-1113 | chr2-53233044-INS-1113 |
| chr2-53243944-INS-416 | chr2-53243944-INS-416 |
| chr2-53306802-INS-58 | chr2-53306802-INS-56 |
| chr2-53354732-INS-113 | chr2-53354743-INS-82 |
| chr2-53362823-INS-55 | chr2-53362806-INS-56 |
| chr2-53366980-INS-1899 | chr2-53366996-INS-1895 |
| chr2-53401532-INS-410 | chr2-53401532-INS-410 |
| chr2-53402870-INS-78 | chr2-53402870-INS-78 |
| chr2-53406756-INS-676 | chr2-53406756-INS-681 |
| chr2-53436813-INS-489 | chr2-53436822-INS-491 |
| chr2-53439734-INS-52 | chr2-53439722-INS-55 |
| chr2-53450398-INS-102 | chr2-53450398-INS-111 |
| chr2-53454939-INS-116 | chr2-53454939-INS-165 |
| chr2-53457091-INS-1774 | chr2-53457092-INS-1770 |
| chr2-53473748-INS-54 | chr2-53473748-INS-56 |

| | |
|---|---|
| chr2-53475302-INS-72 | chr2-53475302-INS-73 |
| chr2-53489213-INS-943 | chr2-53489213-INS-943 |
| chr2-53497067-INS-6207 | chr2-53497068-INS-6209 |
| chr2-53507748-INS-198 | chr2-53507748-INS-197 |
| chr2-53522055-INS-344 | chr2-53522055-INS-344 |
| chr2-53534230-INS-199 | chr2-53534230-INS-199 |
| chr2-53555593-INS-432 | chr2-53555593-INS-432 |
| chr2-53581596-INS-167 | chr2-53581596-INS-163 |
| chr2-53604941-INS-501 | chr2-53604941-INS-499 |
| chr2-53621279-INS-342 | chr2-53621279-INS-339 |
| chr2-53645038-INS-51 | chr2-53645038-INS-51 |
| chr2-53702235-INS-8725 | chr2-53702236-INS-8723 |
| chr2-53715586-INS-677 | chr2-53715586-INS-669 |
| chr2-53721533-INS-143 | chr2-53721533-INS-152 |
| chr2-53726485-INS-6493 | chr2-53726486-INS-6493 |
| chr2-53730325-INS-1800 | chr2-53730326-INS-1794 |
| chr2-53763554-INS-1400 | chr2-53763555-INS-1402 |
| chr2-53765698-INS-7230 | chr2-53765699-INS-7230 |
| chr2-53798752-INS-91 | chr2-53798750-INS-94 |
| chr2-53815818-INS-212 | chr2-53815818-INS-212 |
| chr2-53840600-INS-164 | chr2-53840600-INS-162 |
| chr2-53845069-INS-5935 | chr2-53845069-INS-5935 |
| chr2-53851055-INS-1699 | chr2-53851056-INS-1683 |
| chr2-53859241-INS-3362 | chr2-53859243-INS-3363 |
| chr2-53906066-INS-54 | chr2-53906066-INS-54 |
| chr2-53920342-INS-62 | chr2-53920342-INS-58 |
| chr2-53939852-INS-77 | chr2-53939852-INS-77 |
| chr2-53948108-INS-3897 | chr2-53948109-INS-3897 |
| chr2-53969099-INS-58 | chr2-53969099-INS-58 |
| chr2-53981911-INS-56 | chr2-53981911-INS-56 |
| chr2-53992384-INS-62 | chr2-53992384-INS-64 |
| chr2-54015416-INS-197 | chr2-54015412-INS-208 |
| chr2-54024574-INS-344 | chr2-54024574-INS-344 |
| chr2-54034294-INS-7397 | chr2-54034295-INS-7397 |
| chr2-54042349-INS-61 | chr2-54042355-INS-69 |
| chr2-54049717-INS-933 | chr2-54049717-INS-933 |
| chr2-54061082-INS-503 | chr2-54061082-INS-503 |
| chr2-54079704-INS-7310 | chr2-54079705-INS-7310 |
| chr2-54115958-INS-150 | chr2-54115958-INS-150 |
| chr2-54162453-INS-217 | chr2-54162453-INS-216 |
| chr2-54188932-INS-1814 | chr2-54188933-INS-1813 |
| chr2-54205224-INS-75 | chr2-54205224-INS-75 |
| chr2-54230593-INS-7156 | chr2-54230594-INS-7158 |
| chr2-54240726-INS-160 | chr2-54240726-INS-160 |
| chr2-54254273-INS-213 | chr2-54254273-INS-213 |
| chr2-54271061-INS-52 | chr2-54271061-INS-52 |
| chr2-54284066-INS-132 | chr2-54284066-INS-132 |
| chr2-54305486-INS-265 | chr2-54305486-INS-265 |
| chr2-54312142-INS-591 | chr2-54312142-INS-591 |
| chr2-54319501-INS-64 | chr2-54319501-INS-74 |
| chr2-54342172-INS-66 | chr2-54342172-INS-66 |
| chr2-54349906-INS-103 | chr2-54349912-INS-64 |
| chr2-54352049-INS-400 | chr2-54352049-INS-400 |
| chr2-54367197-INS-87 | chr2-54367197-INS-87 |
| chr2-54376214-INS-6386 | chr2-54376215-INS-6386 |
| chr2-54398592-INS-7094 | chr2-54398593-INS-7093 |
| chr2-54430366-INS-94 | chr2-54430386-INS-95 |
| chr2-54432715-INS-6126 | chr2-54432716-INS-6126 |
| chr2-54476111-INS-818 | chr2-54476111-INS-818 |
| chr2-54510345-INS-159 | chr2-54510345-INS-170 |
| chr2-54526314-INS-576 | chr2-54526314-INS-579 |
| chr2-54565876-INS-518 | chr2-54565876-INS-518 |
| chr2-54567600-INS-615 | chr2-54567601-INS-612 |
| chr2-54618252-INS-2994 | chr2-54618253-INS-2994 |
| chr2-54631756-INS-212 | chr2-54631756-INS-212 |
| chr2-54656004-INS-783 | chr2-54656004-INS-784 |
| chr2-54678842-INS-62 | chr2-54678840-INS-66 |
| chr2-54702892-INS-873 | chr2-54702892-INS-873 |
| chr2-54733830-INS-100 | chr2-54733830-INS-100 |
| chr2-54801039-INS-353 | chr2-54801039-INS-354 |
| chr2-54818735-INS-4348 | chr2-54818736-INS-4348 |
| chr2-54836833-INS-116 | chr2-54836833-INS-116 |
| chr2-54863835-INS-127 | chr2-54863835-INS-127 |
| chr2-54867358-INS-139 | chr2-54867358-INS-139 |
| chr2-54892020-INS-62 | chr2-54892020-INS-62 |
| chr2-54898024-INS-2238 | chr2-54898028-INS-2234 |
| chr2-54905628-INS-52 | chr2-54905628-INS-60 |

| | |
|---|---|
| chr2-54937031-INS-671 | chr2-54937031-INS-671 |
| chr2-54938601-INS-226 | chr2-54938601-INS-226 |
| chr2-54970550-INS-310 | chr2-54970551-INS-322 |
| chr2-54976849-INS-81 | chr2-54976828-INS-87 |
| chr2-54992193-INS-124 | chr2-54992193-INS-88 |
| chr2-55015686-INS-401 | chr2-55015686-INS-402 |
| chr2-55016815-INS-6250 | chr2-55016816-INS-6250 |
| chr2-55043960-INS-3093 | chr2-55043961-INS-3095 |
| chr2-55065803-INS-6453 | chr2-55065804-INS-6449 |
| chr2-55067816-INS-694 | chr2-55067816-INS-696 |
| chr2-55154349-INS-110 | chr2-55154352-INS-109 |
| chr2-55161710-INS-56 | chr2-55161710-INS-56 |
| chr2-55214194-INS-214 | chr2-55214194-INS-214 |
| chr2-55248386-INS-225 | chr2-55248386-INS-228 |
| chr2-55266374-INS-6443 | chr2-55266375-INS-6444 |
| chr2-55271466-INS-68 | chr2-55271466-INS-84 |
| chr2-55275079-INS-84 | chr2-55275079-INS-82 |
| chr2-55358616-INS-84 | chr2-55358616-INS-81 |
| chr2-55363938-INS-425 | chr2-55363938-INS-426 |
| chr2-55372816-INS-702 | chr2-55372816-INS-702 |
| chr2-55379477-INS-477 | chr2-55379477-INS-477 |
| chr2-55413338-INS-804 | chr2-55413343-INS-808 |
| chr2-55443187-INS-1485 | chr2-55443188-INS-1485 |
| chr2-55444916-INS-58 | chr2-55444910-INS-51 |
| chr2-55460275-INS-74 | chr2-55460287-INS-92 |
| chr2-55476033-INS-279 | chr2-55476033-INS-279 |
| chr2-55480391-INS-58 | chr2-55480380-INS-98 |
| chr2-55511948-INS-5218 | chr2-55511949-INS-5218 |
| chr2-55557160-INS-58 | chr2-55557160-INS-56 |
| chr2-55592922-INS-61 | chr2-55592922-INS-61 |
| chr2-55593245-INS-5035 | chr2-55593246-INS-5035 |
| chr2-55616959-INS-2324 | chr2-55616960-INS-2324 |
| chr2-55636193-INS-6353 | chr2-55636194-INS-6353 |
| chr2-55700086-INS-215 | chr2-55700086-INS-215 |
| chr2-55745417-INS-1704 | chr2-55745418-INS-1704 |
| chr2-55770765-INS-75 | chr2-55770762-INS-78 |
| chr2-55774515-INS-4957 | chr2-55774516-INS-4957 |
| chr2-55774983-INS-68 | chr2-55774983-INS-68 |
| chr2-55798337-INS-59 | chr2-55798338-INS-54 |
| chr2-55798910-INS-538 | chr2-55798910-INS-539 |
| chr2-55801745-INS-494 | chr2-55801745-INS-495 |
| chr2-55812600-INS-74 | chr2-55812600-INS-76 |
| chr2-55819629-INS-74 | chr2-55819629-INS-69 |
| chr2-55860992-INS-78 | chr2-55860992-INS-78 |
| chr2-55872523-INS-132 | chr2-55872523-INS-132 |
| chr2-55902879-INS-6582 | chr2-55902880-INS-6582 |
| chr2-55917645-INS-133 | chr2-55917645-INS-139 |
| chr2-55929596-INS-2002 | chr2-55929597-INS-1998 |
| chr2-55930307-INS-65 | chr2-55930308-INS-65 |
| chr2-55967477-INS-214 | chr2-55967477-INS-218 |
| chr2-55974087-INS-1743 | chr2-55974088-INS-1743 |
| chr2-56019122-INS-76 | chr2-56019122-INS-60 |
| chr2-56026994-INS-424 | chr2-56026994-INS-424 |
| chr2-56039045-INS-205 | chr2-56039045-INS-205 |
| chr2-56051142-INS-325 | chr2-56051142-INS-325 |
| chr2-56055449-INS-441 | chr2-56055449-INS-443 |
| chr2-56101787-INS-168 | chr2-56101787-INS-168 |
| chr2-56146980-INS-54 | chr2-56146980-INS-50 |
| chr2-56160830-INS-4171 | chr2-56160831-INS-4171 |
| chr2-56162669-INS-1617 | chr2-56162670-INS-1617 |
| chr2-56164169-INS-5564 | chr2-56164170-INS-5564 |
| chr2-56172229-INS-1892 | chr2-56172230-INS-1892 |
| chr2-56179423-INS-280 | chr2-56179423-INS-284 |
| chr2-56199694-INS-160 | chr2-56199694-INS-160 |
| chr2-56222183-INS-60 | chr2-56222183-INS-78 |
| chr2-56232430-INS-6564 | chr2-56232431-INS-6564 |
| chr2-56236688-INS-2601 | chr2-56236689-INS-2603 |
| chr2-56246169-INS-1209 | chr2-56246170-INS-1169 |
| chr2-56260392-INS-184 | chr2-56260392-INS-184 |
| chr2-56279413-INS-2096 | chr2-56279414-INS-2098 |
| chr2-56279753-INS-596 | chr2-56279753-INS-596 |
| chr2-56290950-INS-65 | chr2-56290950-INS-65 |
| chr2-56298734-INS-1862 | chr2-56298735-INS-1862 |
| chr2-56343012-INS-52 | chr2-56343012-INS-54 |
| chr2-56387076-INS-6174 | chr2-56387077-INS-6174 |
| chr2-56425368-INS-331 | chr2-56425368-INS-331 |
| chr2-56472692-INS-7102 | chr2-56472693-INS-7102 |

| | |
|---|---|
| chr2-56521194-INS-402 | chr2-56521194-INS-402 |
| chr2-56526992-INS-3099 | chr2-56526995-INS-3093 |
| chr2-56555823-INS-68 | chr2-56555823-INS-68 |
| chr2-56579739-INS-373 | chr2-56579739-INS-373 |
| chr2-56598034-INS-3619 | chr2-56598035-INS-3618 |
| chr2-56600771-INS-60 | chr2-56600771-INS-72 |
| chr2-56617677-INS-52 | chr2-56617719-INS-56 |
| chr2-56619483-INS-113 | chr2-56619483-INS-119 |
| chr2-56666746-INS-123 | chr2-56666746-INS-123 |
| chr2-56709938-INS-466 | chr2-56709938-INS-466 |
| chr2-56714171-INS-67 | chr2-56714156-INS-55 |
| chr2-56753728-INS-56 | chr2-56753728-INS-50 |
| chr2-56768946-INS-208 | chr2-56768946-INS-208 |
| chr2-56782331-INS-56 | chr2-56782331-INS-50 |
| chr2-56806934-INS-77 | chr2-56806934-INS-77 |
| chr2-56861645-INS-6737 | chr2-56861646-INS-6735 |
| chr2-56868360-INS-107 | chr2-56868392-INS-114 |
| chr2-56927000-INS-3911 | chr2-56927001-INS-3911 |
| chr2-56935891-INS-6454 | chr2-56935892-INS-6453 |
| chr2-56943946-INS-96 | chr2-56943933-INS-92 |
| chr2-56959868-INS-4165 | chr2-56959869-INS-4165 |
| chr2-56966114-INS-2854 | chr2-56966115-INS-2854 |
| chr2-56968033-INS-164 | chr2-56968033-INS-179 |
| chr2-56968623-INS-1278 | chr2-56968624-INS-1278 |
| chr2-57019376-INS-270 | chr2-57019376-INS-270 |
| chr2-57032456-INS-626 | chr2-57032456-INS-631 |
| chr2-57046311-INS-124 | chr2-57046310-INS-128 |
| chr2-57070290-INS-58 | chr2-57070290-INS-68 |
| chr2-57088216-INS-202 | chr2-57088219-INS-200 |
| chr2-57097420-INS-76 | chr2-57097408-INS-92 |
| chr2-57123457-INS-162 | chr2-57123457-INS-162 |
| chr2-57147313-INS-187 | chr2-57147313-INS-187 |
| chr2-57149655-INS-80 | chr2-57149654-INS-56 |
| chr2-57168628-INS-64 | chr2-57168631-INS-56 |
| chr2-57178953-INS-145 | chr2-57178953-INS-170 |
| chr2-57185514-INS-114 | chr2-57185514-INS-114 |
| chr2-57207747-INS-523 | chr2-57207744-INS-526 |
| chr2-57212760-INS-450 | chr2-57212760-INS-450 |
| chr2-57273387-INS-6562 | chr2-57273388-INS-6564 |
| chr2-57273851-INS-58 | chr2-57273867-INS-61 |
| chr2-57284672-INS-138 | chr2-57284672-INS-138 |
| chr2-57315620-INS-1591 | chr2-57315621-INS-1587 |
| chr2-57511656-INS-7327 | chr2-57511657-INS-7330 |
| chr2-57603521-INS-69 | chr2-57603521-INS-87 |
| chr2-57605928-INS-66 | chr2-57605928-INS-66 |
| chr2-57610433-INS-285 | chr2-57610433-INS-315 |
| chr2-57615702-INS-7248 | chr2-57615703-INS-7248 |
| chr2-57622121-INS-918 | chr2-57622121-INS-918 |
| chr2-57640712-INS-5176 | chr2-57640713-INS-5176 |
| chr2-57641043-INS-134 | chr2-57641043-INS-119 |
| chr2-57679444-INS-6507 | chr2-57679445-INS-6507 |
| chr2-57686275-INS-1625 | chr2-57686276-INS-1626 |
| chr2-57688369-INS-192 | chr2-57688369-INS-184 |
| chr2-57701912-INS-238 | chr2-57701912-INS-239 |
| chr2-57706534-INS-144 | chr2-57706534-INS-160 |
| chr2-57713186-INS-175 | chr2-57713202-INS-175 |
| chr2-57714440-INS-98 | chr2-57714440-INS-98 |
| chr2-57759274-INS-1488 | chr2-57759275-INS-1490 |
| chr2-57777492-INS-52 | chr2-57777490-INS-58 |
| chr2-57801915-INS-56 | chr2-57801911-INS-73 |
| chr2-57804713-INS-339 | chr2-57804713-INS-339 |
| chr2-57805080-INS-999 | chr2-57805080-INS-998 |
| chr2-57831116-INS-412 | chr2-57831116-INS-412 |
| chr2-57835334-INS-119 | chr2-57835334-INS-123 |
| chr2-57844020-INS-1889 | chr2-57844021-INS-1889 |
| chr2-57857110-INS-68 | chr2-57857110-INS-68 |
| chr2-57883780-INS-69 | chr2-57883780-INS-69 |
| chr2-57886271-INS-78 | chr2-57886271-INS-85 |
| chr2-57958880-INS-5941 | chr2-57958881-INS-5942 |
| chr2-58020800-INS-60 | chr2-58020800-INS-60 |
| chr2-58022231-INS-69 | chr2-58022227-INS-71 |
| chr2-58047015-INS-52 | chr2-58047015-INS-54 |
| chr2-58052486-INS-52 | chr2-58052486-INS-60 |
| chr2-58058035-INS-6894 | chr2-58058036-INS-6894 |
| chr2-58072919-INS-6326 | chr2-58072920-INS-6327 |
| chr2-58073266-INS-82 | chr2-58073266-INS-60 |
| chr2-58084097-INS-82 | chr2-58084097-INS-80 |

| | |
|---|---|
| chr2-58090917-INS-1736 | chr2-58090918-INS-1735 |
| chr2-58106269-INS-56 | chr2-58106269-INS-56 |
| chr2-58140202-INS-62 | chr2-58140202-INS-84 |
| chr2-58143132-INS-53 | chr2-58143132-INS-53 |
| chr2-58170124-INS-89 | chr2-58170121-INS-90 |
| chr2-58222047-INS-75 | chr2-58222047-INS-61 |
| chr2-58229017-INS-54 | chr2-58229019-INS-50 |
| chr2-58268335-INS-257 | chr2-58268335-INS-257 |
| chr2-58269302-INS-399 | chr2-58269302-INS-399 |
| chr2-58277274-INS-72 | chr2-58277274-INS-72 |
| chr2-58278197-INS-52 | chr2-58278197-INS-52 |
| chr2-58285467-INS-6324 | chr2-58285468-INS-6322 |
| chr2-58303479-INS-704 | chr2-58303479-INS-704 |
| chr2-58308845-INS-68 | chr2-58308845-INS-68 |
| chr2-58311514-INS-504 | chr2-58311514-INS-504 |
| chr2-58333931-INS-51 | chr2-58333931-INS-61 |
| chr2-58339886-INS-3144 | chr2-58339887-INS-3144 |
| chr2-58388010-INS-56 | chr2-58388010-INS-70 |
| chr2-58441488-INS-52 | chr2-58441488-INS-88 |
| chr2-58455775-INS-80 | chr2-58455775-INS-68 |
| chr2-58461282-INS-537 | chr2-58461282-INS-537 |
| chr2-58463407-INS-80 | chr2-58463407-INS-58 |
| chr2-58481092-INS-64 | chr2-58481092-INS-64 |
| chr2-58494638-INS-251 | chr2-58494638-INS-251 |
| chr2-58500431-INS-191 | chr2-58500432-INS-191 |
| chr2-58518172-INS-51 | chr2-58518191-INS-55 |
| chr2-58523030-INS-443 | chr2-58523030-INS-443 |
| chr2-58530305-INS-70 | chr2-58530305-INS-70 |
| chr2-58534872-INS-399 | chr2-58534872-INS-398 |
| chr2-58557077-INS-1866 | chr2-58557078-INS-1866 |
| chr2-58584882-INS-97 | chr2-58584890-INS-93 |
| chr2-58595618-INS-53 | chr2-58595618-INS-52 |
| chr2-58606286-INS-206 | chr2-58606286-INS-206 |
| chr2-58621788-INS-134 | chr2-58621788-INS-135 |
| chr2-58630526-INS-119 | chr2-58630532-INS-79 |
| chr2-58651888-INS-62 | chr2-58651888-INS-63 |
| chr2-58684663-INS-118 | chr2-58684662-INS-144 |
| chr2-58685630-INS-51 | chr2-58685630-INS-51 |
| chr2-58686650-INS-108 | chr2-58686649-INS-72 |
| chr2-58692716-INS-51 | chr2-58692716-INS-56 |
| chr2-58709386-INS-80 | chr2-58709386-INS-80 |
| chr2-58735460-INS-120 | chr2-58735460-INS-120 |
| chr2-58752897-INS-206 | chr2-58752897-INS-204 |
| chr2-58785019-INS-7121 | chr2-58785020-INS-7121 |
| chr2-58851793-INS-110 | chr2-58851793-INS-98 |
| chr2-58865520-INS-222 | chr2-58865520-INS-222 |
| chr2-58867650-INS-67 | chr2-58867639-INS-94 |
| chr2-58881401-INS-66 | chr2-58881401-INS-66 |
| chr2-58910525-INS-187 | chr2-58910525-INS-187 |
| chr2-58920055-INS-56 | chr2-58920031-INS-94 |
| chr2-58921300-INS-6411 | chr2-58921301-INS-6376 |
| chr2-58931373-INS-67 | chr2-58931373-INS-64 |
| chr2-58939039-INS-85 | chr2-58939039-INS-91 |
| chr2-58976145-INS-382 | chr2-58976141-INS-387 |
| chr2-58986435-INS-523 | chr2-58986435-INS-527 |
| chr2-58993113-INS-56 | chr2-58993080-INS-100 |
| chr2-59003422-INS-56 | chr2-59003422-INS-56 |
| chr2-59010069-INS-64 | chr2-59010069-INS-64 |
| chr2-59053754-INS-392 | chr2-59053754-INS-391 |
| chr2-59064388-INS-57 | chr2-59064393-INS-57 |
| chr2-59074098-INS-72 | chr2-59074098-INS-72 |
| chr2-59074861-INS-1207 | chr2-59074862-INS-1207 |
| chr2-59116013-INS-288 | chr2-59116013-INS-288 |
| chr2-59121881-INS-54 | chr2-59121881-INS-56 |
| chr2-59125897-INS-820 | chr2-59125897-INS-796 |
| chr2-59133682-INS-188 | chr2-59133682-INS-188 |
| chr2-59178972-INS-209 | chr2-59178972-INS-205 |
| chr2-59200420-INS-98 | chr2-59200420-INS-86 |
| chr2-59210138-INS-127 | chr2-59210147-INS-103 |
| chr2-59219138-INS-51 | chr2-59219138-INS-51 |
| chr2-59224857-INS-74 | chr2-59224857-INS-74 |
| chr2-59231163-INS-208 | chr2-59231163-INS-208 |
| chr2-59252220-INS-468 | chr2-59252220-INS-468 |
| chr2-59260038-INS-173 | chr2-59260038-INS-173 |
| chr2-59272788-INS-198 | chr2-59272788-INS-182 |
| chr2-59280877-INS-84 | chr2-59280901-INS-50 |
| chr2-59299041-INS-52 | chr2-59299041-INS-64 |

| | |
|---|---|
| chr2-59307530-INS-108 | chr2-59307546-INS-113 |
| chr2-59347736-INS-146 | chr2-59347736-INS-146 |
| chr2-59348399-INS-492 | chr2-59348399-INS-495 |
| chr2-59351907-INS-59 | chr2-59351907-INS-59 |
| chr2-59356845-INS-401 | chr2-59356843-INS-393 |
| chr2-59360683-INS-397 | chr2-59360683-INS-397 |
| chr2-59388683-INS-214 | chr2-59388683-INS-214 |
| chr2-59432562-INS-243 | chr2-59432562-INS-270 |
| chr2-59469807-INS-79 | chr2-59469807-INS-79 |
| chr2-59478061-INS-56 | chr2-59478061-INS-56 |
| chr2-59488855-INS-220 | chr2-59488855-INS-221 |
| chr2-59493265-INS-70 | chr2-59493290-INS-75 |
| chr2-59505003-INS-218 | chr2-59505003-INS-219 |
| chr2-59534961-INS-413 | chr2-59534950-INS-417 |
| chr2-59560425-INS-54 | chr2-59560425-INS-55 |
| chr2-59569678-INS-74 | chr2-59569679-INS-57 |
| chr2-59576974-INS-192 | chr2-59576974-INS-192 |
| chr2-59587070-INS-182 | chr2-59587070-INS-182 |
| chr2-59588653-INS-320 | chr2-59588653-INS-320 |
| chr2-59617632-INS-169 | chr2-59617632-INS-169 |
| chr2-59626541-INS-209 | chr2-59626541-INS-209 |
| chr2-59641315-INS-88 | chr2-59641315-INS-88 |
| chr2-59671994-INS-88 | chr2-59672037-INS-70 |
| chr2-59677637-INS-386 | chr2-59677637-INS-386 |
| chr2-59685258-INS-213 | chr2-59685258-INS-213 |
| chr2-59689577-INS-58 | chr2-59689577-INS-58 |
| chr2-59698421-INS-3430 | chr2-59698422-INS-3430 |
| chr2-59787363-INS-749 | chr2-59787363-INS-749 |
| chr2-59788703-INS-64 | chr2-59788703-INS-64 |
| chr2-59807614-INS-202 | chr2-59807614-INS-202 |
| chr2-59839537-INS-950 | chr2-59839537-INS-950 |
| chr2-59842722-INS-56 | chr2-59842722-INS-60 |
| chr2-59888644-INS-214 | chr2-59888644-INS-215 |
| chr2-59900785-INS-58 | chr2-59900817-INS-60 |
| chr2-59901469-INS-204 | chr2-59901469-INS-208 |
| chr2-59973290-INS-220 | chr2-59973290-INS-224 |
| chr2-59988526-INS-92 | chr2-59988526-INS-92 |
| chr2-59990455-INS-190 | chr2-59990455-INS-190 |
| chr2-60058491-INS-200 | chr2-60058491-INS-204 |
| chr2-60100747-INS-122 | chr2-60100747-INS-90 |
| chr2-60106154-INS-182 | chr2-60106154-INS-182 |
| chr2-60106947-INS-189 | chr2-60106947-INS-187 |
| chr2-60120320-INS-1313 | chr2-60120321-INS-1313 |
| chr2-60129173-INS-331 | chr2-60129173-INS-331 |
| chr2-60135290-INS-74 | chr2-60135290-INS-74 |
| chr2-60137547-INS-183 | chr2-60137549-INS-186 |
| chr2-60160538-INS-208 | chr2-60160538-INS-208 |
| chr2-60166042-INS-674 | chr2-60166035-INS-658 |
| chr2-60261587-INS-156 | chr2-60261587-INS-156 |
| chr2-60287541-INS-60 | chr2-60287539-INS-60 |
| chr2-60299744-INS-134 | chr2-60299744-INS-134 |
| chr2-60305299-INS-60 | chr2-60305299-INS-66 |
| chr2-60305655-INS-210 | chr2-60305655-INS-215 |
| chr2-60327879-INS-1020 | chr2-60327880-INS-1021 |
| chr2-60328566-INS-60 | chr2-60328566-INS-85 |
| chr2-60340432-INS-484 | chr2-60340432-INS-484 |
| chr2-60360086-INS-88 | chr2-60360086-INS-88 |
| chr2-60364872-INS-457 | chr2-60364872-INS-457 |
| chr2-60386221-INS-237 | chr2-60386221-INS-240 |
| chr2-60459233-INS-100 | chr2-60459233-INS-98 |
| chr2-60477049-INS-51 | chr2-60477049-INS-51 |
| chr2-60491231-INS-333 | chr2-60491231-INS-335 |
| chr2-60506602-INS-363 | chr2-60506602-INS-363 |
| chr2-60531180-INS-516 | chr2-60531180-INS-513 |
| chr2-60558750-INS-1777 | chr2-60558751-INS-1777 |
| chr2-60560990-INS-54 | chr2-60560990-INS-54 |
| chr2-60573328-INS-748 | chr2-60573328-INS-746 |
| chr2-60593092-INS-480 | chr2-60593092-INS-480 |
| chr2-60637459-INS-60 | chr2-60637434-INS-57 |
| chr2-60649291-INS-344 | chr2-60649291-INS-344 |
| chr2-60669827-INS-138 | chr2-60669827-INS-138 |
| chr2-60728833-INS-92 | chr2-60728833-INS-92 |
| chr2-60730191-INS-82 | chr2-60730191-INS-82 |
| chr2-60746040-INS-196 | chr2-60746040-INS-194 |
| chr2-60754001-INS-68 | chr2-60753997-INS-108 |
| chr2-60764223-INS-93 | chr2-60764227-INS-71 |
| chr2-60771451-INS-122 | chr2-60771443-INS-116 |

| | |
|---|---|
| chr2-60774715-INS-1868 | chr2-60774716-INS-1868 |
| chr2-60781416-INS-52 | chr2-60781416-INS-96 |
| chr2-60805963-INS-6687 | chr2-60805964-INS-6688 |
| chr2-60814549-INS-79 | chr2-60814549-INS-72 |
| chr2-60818219-INS-267 | chr2-60818219-INS-267 |
| chr2-60867748-INS-3070 | chr2-60867749-INS-3070 |
| chr2-60882010-INS-51 | chr2-60882011-INS-51 |
| chr2-60901814-INS-92 | chr2-60901814-INS-92 |
| chr2-60907249-INS-333 | chr2-60907249-INS-333 |
| chr2-60947598-INS-108 | chr2-60947598-INS-72 |
| chr2-60960329-INS-76 | chr2-60960329-INS-76 |
| chr2-60968553-INS-462 | chr2-60968542-INS-479 |
| chr2-60979079-INS-60 | chr2-60979079-INS-60 |
| chr2-60994645-INS-57 | chr2-60994610-INS-69 |
| chr2-61000207-INS-74 | chr2-61000207-INS-74 |
| chr2-61031511-INS-77 | chr2-61031511-INS-77 |
| chr2-61036735-INS-3527 | chr2-61036736-INS-3527 |
| chr2-61041640-INS-91 | chr2-61041650-INS-71 |
| chr2-61048818-INS-401 | chr2-61048818-INS-401 |
| chr2-61057409-INS-75 | chr2-61057409-INS-63 |
| chr2-61078402-INS-6979 | chr2-61078403-INS-6979 |
| chr2-61081015-INS-215 | chr2-61081015-INS-215 |
| chr2-61131919-INS-476 | chr2-61131919-INS-476 |
| chr2-61151002-INS-6271 | chr2-61151003-INS-6271 |
| chr2-61209288-INS-82 | chr2-61209288-INS-82 |
| chr2-61212146-INS-6482 | chr2-61212147-INS-6485 |
| chr2-61227606-INS-92 | chr2-61227563-INS-125 |
| chr2-61255285-INS-177 | chr2-61255285-INS-167 |
| chr2-61275143-INS-201 | chr2-61275143-INS-201 |
| chr2-61282491-INS-78 | chr2-61282491-INS-78 |
| chr2-61296537-INS-124 | chr2-61296537-INS-124 |
| chr2-61327046-INS-118 | chr2-61327046-INS-118 |
| chr2-61372554-INS-498 | chr2-61372554-INS-498 |
| chr2-61403766-INS-665 | chr2-61403767-INS-658 |
| chr2-61407189-INS-443 | chr2-61407189-INS-437 |
| chr2-61446414-INS-1934 | chr2-61446415-INS-1948 |
| chr2-61448407-INS-769 | chr2-61448408-INS-760 |
| chr2-61497434-INS-7966 | chr2-61497435-INS-7963 |
| chr2-61511407-INS-170 | chr2-61511407-INS-195 |
| chr2-61522653-INS-3354 | chr2-61522654-INS-3354 |
| chr2-61524202-INS-68 | chr2-61524202-INS-68 |
| chr2-61533203-INS-910 | chr2-61533203-INS-910 |
| chr2-61533683-INS-53 | chr2-61533683-INS-53 |
| chr2-61621339-INS-87 | chr2-61621339-INS-93 |
| chr2-61626884-INS-54 | chr2-61626884-INS-54 |
| chr2-61655559-INS-94 | chr2-61655559-INS-98 |
| chr2-61692124-INS-216 | chr2-61692124-INS-217 |
| chr2-61710704-INS-60 | chr2-61710704-INS-60 |
| chr2-61721851-INS-171 | chr2-61721851-INS-173 |
| chr2-61723148-INS-6494 | chr2-61723149-INS-6496 |
| chr2-61728826-INS-2677 | chr2-61728827-INS-2677 |
| chr2-61740937-INS-6669 | chr2-61740938-INS-6671 |
| chr2-61826805-INS-452 | chr2-61826805-INS-454 |
| chr2-61832186-INS-7546 | chr2-61832187-INS-7547 |
| chr2-61840153-INS-634 | chr2-61840153-INS-635 |
| chr2-61857256-INS-96 | chr2-61857262-INS-66 |
| chr2-61859153-INS-2150 | chr2-61859154-INS-2150 |
| chr2-61872149-INS-1854 | chr2-61872150-INS-1854 |
| chr2-61897846-INS-199 | chr2-61897847-INS-200 |
| chr2-61898770-INS-6459 | chr2-61898771-INS-6461 |
| chr2-61899809-INS-207 | chr2-61899809-INS-207 |
| chr2-61929534-INS-6419 | chr2-61929535-INS-6420 |
| chr2-61970265-INS-122 | chr2-61970265-INS-122 |
| chr2-61972021-INS-960 | chr2-61972021-INS-960 |
| chr2-61973940-INS-60 | chr2-61973940-INS-50 |
| chr2-61985568-INS-188 | chr2-61985568-INS-188 |
| chr2-62005721-INS-204 | chr2-62005721-INS-204 |
| chr2-62021495-INS-2082 | chr2-62021496-INS-2082 |
| chr2-62022806-INS-391 | chr2-62022806-INS-391 |
| chr2-62034488-INS-2785 | chr2-62034489-INS-2772 |
| chr2-62049517-INS-88 | chr2-62049517-INS-90 |
| chr2-62089888-INS-831 | chr2-62089888-INS-831 |
| chr2-62108530-INS-6442 | chr2-62108531-INS-6442 |
| chr2-62111391-INS-1134 | chr2-62111392-INS-1131 |
| chr2-62272257-INS-150 | chr2-62272259-INS-154 |
| chr2-62294027-INS-63 | chr2-62294027-INS-62 |
| chr2-62303238-INS-100 | chr2-62303238-INS-124 |

| | |
|---|---|
| chr2-62323132-INS-261 | chr2-62323132-INS-267 |
| chr2-62324554-INS-7523 | chr2-62324555-INS-7523 |
| chr2-62334442-INS-106 | chr2-62334442-INS-110 |
| chr2-62400319-INS-74 | chr2-62400320-INS-84 |
| chr2-62421778-INS-277 | chr2-62421778-INS-277 |
| chr2-62424089-INS-219 | chr2-62424089-INS-219 |
| chr2-62438534-INS-61 | chr2-62438529-INS-62 |
| chr2-62466749-INS-94 | chr2-62466749-INS-94 |
| chr2-62472177-INS-212 | chr2-62472177-INS-212 |
| chr2-62498047-INS-1238 | chr2-62498048-INS-1237 |
| chr2-62517255-INS-90 | chr2-62517255-INS-94 |
| chr2-62539929-INS-1890 | chr2-62539930-INS-1890 |
| chr2-62557428-INS-1518 | chr2-62557429-INS-1526 |
| chr2-62562406-INS-797 | chr2-62562406-INS-797 |
| chr2-62579718-INS-64 | chr2-62579718-INS-64 |
| chr2-62589603-INS-80 | chr2-62589603-INS-88 |
| chr2-62597710-INS-54 | chr2-62597710-INS-50 |
| chr2-62602134-INS-1692 | chr2-62602135-INS-1692 |
| chr2-62604543-INS-57 | chr2-62604543-INS-57 |
| chr2-62610887-INS-146 | chr2-62610887-INS-147 |
| chr2-62640807-INS-83 | chr2-62640807-INS-81 |
| chr2-62673265-INS-80 | chr2-62673265-INS-88 |
| chr2-62686414-INS-82 | chr2-62686417-INS-69 |
| chr2-62714570-INS-185 | chr2-62714570-INS-185 |
| chr2-62721710-INS-137 | chr2-62721707-INS-133 |
| chr2-62722072-INS-740 | chr2-62722072-INS-740 |
| chr2-62733497-INS-6222 | chr2-62733498-INS-6223 |
| chr2-62766779-INS-214 | chr2-62766774-INS-212 |
| chr2-62777706-INS-159 | chr2-62777709-INS-151 |
| chr2-62837519-INS-350 | chr2-62837519-INS-350 |
| chr2-62856007-INS-451 | chr2-62856007-INS-451 |
| chr2-62858248-INS-206 | chr2-62858248-INS-200 |
| chr2-62865186-INS-296 | chr2-62865186-INS-296 |
| chr2-62882773-INS-52 | chr2-62882773-INS-54 |
| chr2-62885829-INS-105 | chr2-62885829-INS-103 |
| chr2-62911664-INS-82 | chr2-62911664-INS-82 |
| chr2-62915774-INS-216 | chr2-62915787-INS-215 |
| chr2-62921967-INS-101 | chr2-62921967-INS-101 |
| chr2-62932324-INS-52 | chr2-62932318-INS-58 |
| chr2-62937213-INS-2567 | chr2-62937214-INS-2563 |
| chr2-63008205-INS-66 | chr2-63008205-INS-68 |
| chr2-63059539-INS-142 | chr2-63059540-INS-146 |
| chr2-63074409-INS-104 | chr2-63074409-INS-104 |
| chr2-63112448-INS-7126 | chr2-63112449-INS-7126 |
| chr2-63113619-INS-1529 | chr2-63113620-INS-1530 |
| chr2-63125352-INS-540 | chr2-63125352-INS-540 |
| chr2-63150477-INS-6451 | chr2-63150478-INS-6452 |
| chr2-63163441-INS-915 | chr2-63163441-INS-915 |
| chr2-63169170-INS-807 | chr2-63169170-INS-803 |
| chr2-63182514-INS-181 | chr2-63182514-INS-184 |
| chr2-63204511-INS-212 | chr2-63204511-INS-212 |
| chr2-63240818-INS-183 | chr2-63240818-INS-184 |
| chr2-63253890-INS-528 | chr2-63253890-INS-530 |
| chr2-63273049-INS-68 | chr2-63273049-INS-52 |
| chr2-63296962-INS-220 | chr2-63296962-INS-220 |
| chr2-63358657-INS-64 | chr2-63358657-INS-64 |
| chr2-63380419-INS-223 | chr2-63380429-INS-232 |
| chr2-63397660-INS-1322 | chr2-63397655-INS-1307 |
| chr2-63449161-INS-410 | chr2-63449161-INS-411 |
| chr2-63490768-INS-940 | chr2-63490768-INS-940 |
| chr2-63520612-INS-100 | chr2-63520612-INS-56 |
| chr2-63525168-INS-1177 | chr2-63525169-INS-1178 |
| chr2-63544401-INS-56 | chr2-63544401-INS-88 |
| chr2-63591045-INS-64 | chr2-63591045-INS-60 |
| chr2-63599605-INS-276 | chr2-63599605-INS-276 |
| chr2-63609225-INS-110 | chr2-63609225-INS-111 |
| chr2-63621911-INS-66 | chr2-63621911-INS-58 |
| chr2-63661431-INS-236 | chr2-63661431-INS-250 |
| chr2-63667738-INS-125 | chr2-63667739-INS-125 |
| chr2-63684465-INS-96 | chr2-63684425-INS-96 |
| chr2-63696478-INS-52 | chr2-63696478-INS-52 |
| chr2-63703507-INS-121 | chr2-63703507-INS-121 |
| chr2-63737177-INS-5887 | chr2-63737178-INS-5890 |
| chr2-63781337-INS-53 | chr2-63781337-INS-53 |
| chr2-63785586-INS-97 | chr2-63785582-INS-82 |
| chr2-63796204-INS-62 | chr2-63796204-INS-58 |
| chr2-63796801-INS-86 | chr2-63796783-INS-55 |

| | |
|---|---|
| chr2-63801159-INS-72 | chr2-63801139-INS-64 |
| chr2-63827977-INS-54 | chr2-63827977-INS-56 |
| chr2-63831031-INS-53 | chr2-63831031-INS-53 |
| chr2-63842026-INS-74 | chr2-63842026-INS-74 |
| chr2-63878165-INS-92 | chr2-63878165-INS-94 |
| chr2-63930646-INS-110 | chr2-63930632-INS-88 |
| chr2-63948649-INS-82 | chr2-63948649-INS-84 |
| chr2-63991685-INS-196 | chr2-63991685-INS-195 |
| chr2-64001173-INS-144 | chr2-64001173-INS-144 |
| chr2-64043717-INS-485 | chr2-64043717-INS-485 |
| chr2-64058778-INS-199 | chr2-64058778-INS-199 |
| chr2-64095381-INS-77 | chr2-64095381-INS-77 |
| chr2-64116820-INS-1726 | chr2-64116821-INS-1726 |
| chr2-64184312-INS-54 | chr2-64184312-INS-54 |
| chr2-64233683-INS-4953 | chr2-64233684-INS-4954 |
| chr2-64339938-INS-502 | chr2-64339938-INS-503 |
| chr2-64410262-INS-54 | chr2-64410262-INS-54 |
| chr2-64431673-INS-65 | chr2-64431673-INS-69 |
| chr2-64446190-INS-77 | chr2-64446190-INS-77 |
| chr2-64490740-INS-533 | chr2-64490740-INS-533 |
| chr2-64545832-INS-322 | chr2-64545832-INS-322 |
| chr2-64585708-INS-56 | chr2-64585708-INS-52 |
| chr2-64596944-INS-60 | chr2-64596944-INS-106 |
| chr2-64613395-INS-130 | chr2-64613395-INS-94 |
| chr2-64632148-INS-77 | chr2-64632148-INS-67 |
| chr2-64697591-INS-72 | chr2-64697615-INS-80 |
| chr2-64753868-INS-53 | chr2-64753868-INS-53 |
| chr2-64761561-INS-56 | chr2-64761601-INS-58 |
| chr2-64787023-INS-80 | chr2-64787023-INS-68 |
| chr2-64827127-INS-396 | chr2-64827127-INS-396 |
| chr2-64832547-INS-5547 | chr2-64832548-INS-5547 |
| chr2-64850714-INS-988 | chr2-64850714-INS-988 |
| chr2-64862625-INS-85 | chr2-64862625-INS-85 |
| chr2-64867937-INS-58 | chr2-64867937-INS-52 |
| chr2-64879679-INS-3701 | chr2-64879680-INS-3701 |
| chr2-64890913-INS-358 | chr2-64890913-INS-358 |
| chr2-64893687-INS-105 | chr2-64893687-INS-105 |
| chr2-64898550-INS-1598 | chr2-64898538-INS-1603 |
| chr2-64935430-INS-201 | chr2-64935431-INS-200 |
| chr2-64945158-INS-58 | chr2-64945158-INS-60 |
| chr2-64967021-INS-161 | chr2-64967021-INS-161 |
| chr2-64992082-INS-56 | chr2-64992082-INS-56 |
| chr2-65010819-INS-52 | chr2-65010820-INS-52 |
| chr2-65031772-INS-482 | chr2-65031772-INS-482 |
| chr2-65040711-INS-295 | chr2-65040711-INS-295 |
| chr2-65087982-INS-64 | chr2-65087982-INS-64 |
| chr2-65115031-INS-155 | chr2-65115031-INS-155 |
| chr2-65128919-INS-68 | chr2-65128922-INS-64 |
| chr2-65130586-INS-152 | chr2-65130586-INS-152 |
| chr2-65141895-INS-1888 | chr2-65141896-INS-1888 |
| chr2-65142769-INS-59 | chr2-65142789-INS-62 |
| chr2-65165182-INS-420 | chr2-65165182-INS-425 |
| chr2-65172140-INS-212 | chr2-65172140-INS-212 |
| chr2-65175026-INS-91 | chr2-65175026-INS-91 |
| chr2-65199666-INS-162 | chr2-65199667-INS-162 |
| chr2-65202262-INS-64 | chr2-65202262-INS-64 |
| chr2-65232230-INS-117 | chr2-65232230-INS-117 |
| chr2-65235834-INS-171 | chr2-65235834-INS-171 |
| chr2-65242797-INS-77 | chr2-65242797-INS-85 |
| chr2-65261377-INS-92 | chr2-65261377-INS-92 |
| chr2-65280435-INS-51 | chr2-65280436-INS-53 |
| chr2-65306429-INS-191 | chr2-65306429-INS-219 |
| chr2-65311893-INS-64 | chr2-65311893-INS-50 |
| chr2-65317940-INS-200 | chr2-65317940-INS-223 |
| chr2-65322197-INS-192 | chr2-65322197-INS-191 |
| chr2-65358986-INS-6421 | chr2-65358987-INS-6421 |
| chr2-65398948-INS-95 | chr2-65398952-INS-98 |
| chr2-65406286-INS-223 | chr2-65406286-INS-223 |
| chr2-65441983-INS-95 | chr2-65441983-INS-100 |
| chr2-65447916-INS-52 | chr2-65447916-INS-52 |
| chr2-65517658-INS-218 | chr2-65517658-INS-218 |
| chr2-65527192-INS-7289 | chr2-65527193-INS-7289 |
| chr2-65545780-INS-6232 | chr2-65545781-INS-6232 |
| chr2-65564100-INS-214 | chr2-65564100-INS-215 |
| chr2-65567944-INS-282 | chr2-65567944-INS-282 |
| chr2-65602090-INS-238 | chr2-65602090-INS-238 |
| chr2-65612586-INS-136 | chr2-65612586-INS-138 |

| | |
|---|---|
| chr2-65613980-INS-132 | chr2-65613980-INS-132 |
| chr2-65614936-INS-91 | chr2-65614937-INS-50 |
| chr2-65632284-INS-95 | chr2-65632234-INS-144 |
| chr2-65634871-INS-536 | chr2-65634871-INS-536 |
| chr2-65643519-INS-64 | chr2-65643519-INS-80 |
| chr2-65654494-INS-57 | chr2-65654494-INS-57 |
| chr2-65659122-INS-314 | chr2-65659122-INS-313 |
| chr2-65697758-INS-68 | chr2-65697758-INS-68 |
| chr2-65732930-INS-85 | chr2-65732930-INS-85 |
| chr2-65754889-INS-70 | chr2-65754928-INS-81 |
| chr2-65766337-INS-4074 | chr2-65766338-INS-4074 |
| chr2-65790604-INS-2157 | chr2-65790605-INS-2157 |
| chr2-65807504-INS-6260 | chr2-65807505-INS-6256 |
| chr2-65843067-INS-390 | chr2-65843068-INS-390 |
| chr2-65903764-INS-68 | chr2-65903740-INS-62 |
| chr2-65905372-INS-429 | chr2-65905372-INS-429 |
| chr2-65945609-INS-64 | chr2-65945609-INS-60 |
| chr2-65960966-INS-59 | chr2-65960966-INS-97 |
| chr2-65976227-INS-55 | chr2-65976226-INS-58 |
| chr2-65982405-INS-6196 | chr2-65982406-INS-6197 |
| chr2-66005765-INS-84 | chr2-66005765-INS-92 |
| chr2-66058533-INS-56 | chr2-66058533-INS-56 |
| chr2-66064629-INS-568 | chr2-66064629-INS-534 |
| chr2-66077931-INS-133 | chr2-66077931-INS-133 |
| chr2-66084199-INS-4461 | chr2-66084200-INS-4461 |
| chr2-66128599-INS-214 | chr2-66128596-INS-210 |
| chr2-66131992-INS-112 | chr2-66131987-INS-136 |
| chr2-66135663-INS-108 | chr2-66135664-INS-67 |
| chr2-66145290-INS-247 | chr2-66145272-INS-282 |
| chr2-66190560-INS-70 | chr2-66190560-INS-70 |
| chr2-66194070-INS-67 | chr2-66194070-INS-95 |
| chr2-66207877-INS-175 | chr2-66207877-INS-163 |
| chr2-66224418-INS-94 | chr2-66224417-INS-104 |
| chr2-66244674-INS-255 | chr2-66244674-INS-255 |
| chr2-66268818-INS-573 | chr2-66268799-INS-575 |
| chr2-66270080-INS-52 | chr2-66270080-INS-52 |
| chr2-66283343-INS-406 | chr2-66283343-INS-406 |
| chr2-66305236-INS-150 | chr2-66305236-INS-150 |
| chr2-66335665-INS-52 | chr2-66335666-INS-52 |
| chr2-66343481-INS-914 | chr2-66343481-INS-914 |
| chr2-66402544-INS-646 | chr2-66402544-INS-652 |
| chr2-66410017-INS-56 | chr2-66410017-INS-56 |
| chr2-66416251-INS-90 | chr2-66416251-INS-86 |
| chr2-66483600-INS-2101 | chr2-66483601-INS-2104 |
| chr2-66512450-INS-64 | chr2-66512450-INS-68 |
| chr2-66534327-INS-807 | chr2-66534327-INS-817 |
| chr2-66536155-INS-94 | chr2-66536155-INS-104 |
| chr2-66600656-INS-151 | chr2-66600657-INS-117 |
| chr2-66662780-INS-72 | chr2-66662772-INS-52 |
| chr2-66664165-INS-72 | chr2-66664165-INS-95 |
| chr2-66664817-INS-558 | chr2-66664817-INS-533 |
| chr2-66685228-INS-451 | chr2-66685228-INS-451 |
| chr2-66701663-INS-396 | chr2-66701663-INS-396 |
| chr2-66740075-INS-6413 | chr2-66740076-INS-6413 |
| chr2-66752257-INS-86 | chr2-66752257-INS-86 |
| chr2-66754767-INS-4348 | chr2-66754768-INS-4347 |
| chr2-66758501-INS-180 | chr2-66758517-INS-209 |
| chr2-66768089-INS-7102 | chr2-66768090-INS-7102 |
| chr2-66778428-INS-444 | chr2-66778428-INS-444 |
| chr2-66790489-INS-90 | chr2-66790496-INS-90 |
| chr2-66793218-INS-72 | chr2-66793218-INS-56 |
| chr2-66829696-INS-208 | chr2-66829696-INS-207 |
| chr2-66859546-INS-122 | chr2-66859546-INS-155 |
| chr2-66868530-INS-2142 | chr2-66868531-INS-2146 |
| chr2-66874813-INS-2603 | chr2-66874814-INS-2603 |
| chr2-66876968-INS-629 | chr2-66876968-INS-629 |
| chr2-66882668-INS-73 | chr2-66882668-INS-73 |
| chr2-66900271-INS-601 | chr2-66900271-INS-598 |
| chr2-66907305-INS-761 | chr2-66907305-INS-761 |
| chr2-66942119-INS-6676 | chr2-66942120-INS-6676 |
| chr2-66942837-INS-1016 | chr2-66942838-INS-1016 |
| chr2-66951285-INS-7485 | chr2-66951286-INS-7485 |
| chr2-66955069-INS-100 | chr2-66955069-INS-80 |
| chr2-66975429-INS-2806 | chr2-66975430-INS-2806 |
| chr2-66977688-INS-64 | chr2-66977688-INS-64 |
| chr2-66993161-INS-291 | chr2-66993161-INS-290 |
| chr2-66996751-INS-572 | chr2-66996751-INS-563 |

| | |
|---|---|
| chr2-66997939-INS-1100 | chr2-66997940-INS-1106 |
| chr2-67012064-INS-6895 | chr2-67012065-INS-6894 |
| chr2-67020435-INS-125 | chr2-67020407-INS-131 |
| chr2-67023943-INS-440 | chr2-67023943-INS-440 |
| chr2-67026829-INS-52 | chr2-67026829-INS-52 |
| chr2-67029290-INS-6329 | chr2-67029291-INS-6329 |
| chr2-67033013-INS-634 | chr2-67033013-INS-634 |
| chr2-67038824-INS-1985 | chr2-67038825-INS-1985 |
| chr2-67043998-INS-219 | chr2-67043998-INS-219 |
| chr2-67049481-INS-5977 | chr2-67049482-INS-5977 |
| chr2-67060547-INS-80 | chr2-67060547-INS-106 |
| chr2-67064663-INS-58 | chr2-67064663-INS-56 |
| chr2-67066973-INS-52 | chr2-67066973-INS-54 |
| chr2-67096881-INS-52 | chr2-67096881-INS-64 |
| chr2-67105535-INS-1456 | chr2-67105536-INS-1456 |
| chr2-67124458-INS-54 | chr2-67124458-INS-56 |
| chr2-67160318-INS-76 | chr2-67160318-INS-76 |
| chr2-67179592-INS-359 | chr2-67179592-INS-359 |
| chr2-67204224-INS-56 | chr2-67204224-INS-84 |
| chr2-67210505-INS-81 | chr2-67210505-INS-82 |
| chr2-67218029-INS-71 | chr2-67218035-INS-85 |
| chr2-67233997-INS-166 | chr2-67234000-INS-170 |
| chr2-67236771-INS-403 | chr2-67236771-INS-405 |
| chr2-67259198-INS-228 | chr2-67259198-INS-227 |
| chr2-67275423-INS-57 | chr2-67275423-INS-50 |
| chr2-67328987-INS-50 | chr2-67328987-INS-60 |
| chr2-67363695-INS-51 | chr2-67363649-INS-64 |
| chr2-67368559-INS-72 | chr2-67368559-INS-56 |
| chr2-67381659-INS-264 | chr2-67381659-INS-264 |
| chr2-67400533-INS-6450 | chr2-67400534-INS-6452 |
| chr2-67404627-INS-52 | chr2-67404627-INS-52 |
| chr2-67432815-INS-72 | chr2-67432815-INS-72 |
| chr2-67446419-INS-79 | chr2-67446419-INS-79 |
| chr2-67455888-INS-6927 | chr2-67455889-INS-6922 |
| chr2-67457256-INS-6593 | chr2-67457257-INS-6607 |
| chr2-67462790-INS-825 | chr2-67462790-INS-825 |
| chr2-67475010-INS-60 | chr2-67475010-INS-60 |
| chr2-67489577-INS-84 | chr2-67489577-INS-89 |
| chr2-67527865-INS-1051 | chr2-67527866-INS-1051 |
| chr2-67532365-INS-53 | chr2-67532363-INS-55 |
| chr2-67537202-INS-51 | chr2-67537196-INS-94 |
| chr2-67560318-INS-1384 | chr2-67560319-INS-1384 |
| chr2-67572978-INS-107 | chr2-67572978-INS-93 |
| chr2-67577432-INS-58 | chr2-67577433-INS-70 |
| chr2-67600797-INS-1354 | chr2-67600798-INS-1354 |
| chr2-67649709-INS-57 | chr2-67649709-INS-70 |
| chr2-67661634-INS-6524 | chr2-67661635-INS-6524 |
| chr2-67663730-INS-122 | chr2-67663724-INS-99 |
| chr2-67683855-INS-1962 | chr2-67683856-INS-1964 |
| chr2-67690372-INS-70 | chr2-67690372-INS-54 |
| chr2-67696231-INS-64 | chr2-67696231-INS-66 |
| chr2-67709097-INS-186 | chr2-67709120-INS-165 |
| chr2-67727880-INS-6935 | chr2-67727881-INS-6937 |
| chr2-67798239-INS-158 | chr2-67798239-INS-206 |
| chr2-67799140-INS-1426 | chr2-67799141-INS-1415 |
| chr2-67800271-INS-78 | chr2-67800271-INS-80 |
| chr2-67801369-INS-385 | chr2-67801369-INS-385 |
| chr2-67815768-INS-50 | chr2-67815768-INS-68 |
| chr2-67824720-INS-64 | chr2-67824720-INS-60 |
| chr2-67827499-INS-160 | chr2-67827499-INS-160 |
| chr2-67837200-INS-66 | chr2-67837201-INS-81 |
| chr2-67840765-INS-800 | chr2-67840765-INS-766 |
| chr2-67884625-INS-6588 | chr2-67884626-INS-6588 |
| chr2-67897346-INS-54 | chr2-67897346-INS-61 |
| chr2-67909410-INS-193 | chr2-67909410-INS-173 |
| chr2-67925563-INS-2303 | chr2-67925564-INS-2304 |
| chr2-67930112-INS-50 | chr2-67930113-INS-56 |
| chr2-67940372-INS-1452 | chr2-67940373-INS-1452 |
| chr2-67941563-INS-52 | chr2-67941563-INS-55 |
| chr2-67956904-INS-62 | chr2-67956903-INS-61 |
| chr2-67958236-INS-87 | chr2-67958216-INS-112 |
| chr2-67976997-INS-83 | chr2-67976997-INS-95 |
| chr2-67993442-INS-54 | chr2-67993442-INS-54 |
| chr2-68032021-INS-72 | chr2-68032021-INS-86 |
| chr2-68037126-INS-53 | chr2-68037126-INS-53 |
| chr2-68082983-INS-1590 | chr2-68082984-INS-1604 |
| chr2-68083202-INS-124 | chr2-68083202-INS-124 |

| | |
|---|---|
| chr2-68085183-INS-1890 | chr2-68085184-INS-1890 |
| chr2-68098812-INS-72 | chr2-68098805-INS-96 |
| chr2-68120792-INS-4544 | chr2-68120793-INS-4544 |
| chr2-68124570-INS-440 | chr2-68124570-INS-440 |
| chr2-68130402-INS-66 | chr2-68130401-INS-66 |
| chr2-68135720-INS-182 | chr2-68135720-INS-178 |
| chr2-68142422-INS-52 | chr2-68142422-INS-52 |
| chr2-68161359-INS-725 | chr2-68161359-INS-726 |
| chr2-68166074-INS-92 | chr2-68166098-INS-102 |
| chr2-68173204-INS-55 | chr2-68173204-INS-60 |
| chr2-68212301-INS-74 | chr2-68212301-INS-74 |
| chr2-68217779-INS-73 | chr2-68217788-INS-79 |
| chr2-68226159-INS-172 | chr2-68226159-INS-172 |
| chr2-68273897-INS-60 | chr2-68273897-INS-60 |
| chr2-68312649-INS-6357 | chr2-68312650-INS-6357 |
| chr2-68324005-INS-778 | chr2-68324005-INS-754 |
| chr2-68324738-INS-56 | chr2-68324738-INS-56 |
| chr2-68339906-INS-61 | chr2-68339906-INS-59 |
| chr2-68396831-INS-78 | chr2-68396831-INS-78 |
| chr2-68410953-INS-1065 | chr2-68410954-INS-1065 |
| chr2-68440121-INS-105 | chr2-68440121-INS-125 |
| chr2-68461037-INS-720 | chr2-68461037-INS-706 |
| chr2-68467186-INS-359 | chr2-68467186-INS-359 |
| chr2-68478252-INS-6539 | chr2-68478253-INS-6541 |
| chr2-68494923-INS-384 | chr2-68494926-INS-386 |
| chr2-68516716-INS-143 | chr2-68516716-INS-143 |
| chr2-68519047-INS-63 | chr2-68519047-INS-63 |
| chr2-68545040-INS-52 | chr2-68545040-INS-52 |
| chr2-68565718-INS-6225 | chr2-68565719-INS-6225 |
| chr2-68577166-INS-195 | chr2-68577166-INS-194 |
| chr2-68588455-INS-319 | chr2-68588438-INS-319 |
| chr2-68607702-INS-149 | chr2-68607702-INS-150 |
| chr2-68612299-INS-119 | chr2-68612299-INS-120 |
| chr2-68640411-INS-1371 | chr2-68640412-INS-1371 |
| chr2-68642038-INS-364 | chr2-68642038-INS-364 |
| chr2-68655009-INS-64 | chr2-68655007-INS-78 |
| chr2-68662269-INS-466 | chr2-68662269-INS-466 |
| chr2-68682720-INS-980 | chr2-68682720-INS-980 |
| chr2-68685497-INS-62 | chr2-68685497-INS-66 |
| chr2-68699234-INS-88 | chr2-68699234-INS-88 |
| chr2-68742675-INS-143 | chr2-68742675-INS-143 |
| chr2-68769032-INS-89 | chr2-68769032-INS-95 |
| chr2-68787930-INS-1471 | chr2-68787931-INS-1470 |
| chr2-68798599-INS-82 | chr2-68798599-INS-82 |
| chr2-68801469-INS-52 | chr2-68801469-INS-64 |
| chr2-68849207-INS-105 | chr2-68849207-INS-105 |
| chr2-68885845-INS-307 | chr2-68885845-INS-307 |
| chr2-68891371-INS-97 | chr2-68891371-INS-98 |
| chr2-68919953-INS-5141 | chr2-68919954-INS-5141 |
| chr2-68930321-INS-75 | chr2-68930321-INS-75 |
| chr2-68945067-INS-69 | chr2-68945067-INS-69 |
| chr2-68977926-INS-491 | chr2-68977926-INS-491 |
| chr2-68994050-INS-211 | chr2-68994050-INS-211 |
| chr2-69007551-INS-3054 | chr2-69007552-INS-3054 |
| chr2-69012947-INS-859 | chr2-69012947-INS-859 |
| chr2-69028975-INS-187 | chr2-69028975-INS-187 |
| chr2-69030693-INS-470 | chr2-69030693-INS-463 |
| chr2-69031834-INS-140 | chr2-69031834-INS-112 |
| chr2-69041669-INS-68 | chr2-69041663-INS-80 |
| chr2-69056614-INS-96 | chr2-69056614-INS-96 |
| chr2-69067551-INS-60 | chr2-69067551-INS-60 |
| chr2-69079044-INS-53 | chr2-69079044-INS-53 |
| chr2-69085466-INS-60 | chr2-69085466-INS-60 |
| chr2-69086415-INS-6256 | chr2-69086416-INS-6256 |
| chr2-69086797-INS-70 | chr2-69086797-INS-72 |
| chr2-69096912-INS-76 | chr2-69096912-INS-110 |
| chr2-69110926-INS-145 | chr2-69110926-INS-144 |
| chr2-69122691-INS-544 | chr2-69122691-INS-547 |
| chr2-69169153-INS-77 | chr2-69169153-INS-77 |
| chr2-69190690-INS-267 | chr2-69190690-INS-270 |
| chr2-69195695-INS-82 | chr2-69195714-INS-54 |
| chr2-69197743-INS-72 | chr2-69197743-INS-70 |
| chr2-69200498-INS-669 | chr2-69200498-INS-669 |
| chr2-69204695-INS-270 | chr2-69204682-INS-270 |
| chr2-69221327-INS-262 | chr2-69221342-INS-288 |
| chr2-69229452-INS-84 | chr2-69229448-INS-54 |
| chr2-69242676-INS-153 | chr2-69242676-INS-151 |

| | |
|---|---|
| chr2-69250263-INS-260 | chr2-69250264-INS-264 |
| chr2-69274439-INS-208 | chr2-69274439-INS-208 |
| chr2-69323625-INS-441 | chr2-69323625-INS-471 |
| chr2-69328134-INS-196 | chr2-69328133-INS-200 |
| chr2-69363769-INS-222 | chr2-69363771-INS-227 |
| chr2-69372474-INS-114 | chr2-69372490-INS-140 |
| chr2-69383726-INS-138 | chr2-69383726-INS-138 |
| chr2-69404960-INS-62 | chr2-69404961-INS-58 |
| chr2-69405713-INS-52 | chr2-69405714-INS-52 |
| chr2-69409214-INS-60 | chr2-69409211-INS-53 |
| chr2-69412057-INS-228 | chr2-69412057-INS-230 |
| chr2-69423308-INS-197 | chr2-69423308-INS-200 |
| chr2-69463546-INS-5165 | chr2-69463547-INS-5165 |
| chr2-69467021-INS-6641 | chr2-69467022-INS-6641 |
| chr2-69554803-INS-58 | chr2-69554803-INS-50 |
| chr2-69570234-INS-206 | chr2-69570234-INS-206 |
| chr2-69589906-INS-125 | chr2-69589889-INS-99 |
| chr2-69598448-INS-209 | chr2-69598448-INS-209 |
| chr2-69615702-INS-560 | chr2-69615702-INS-560 |
| chr2-69617835-INS-196 | chr2-69617835-INS-192 |
| chr2-69618799-INS-1491 | chr2-69618800-INS-1491 |
| chr2-69625743-INS-226 | chr2-69625743-INS-227 |
| chr2-69628872-INS-56 | chr2-69628875-INS-56 |
| chr2-69631389-INS-59 | chr2-69631389-INS-59 |
| chr2-69632220-INS-204 | chr2-69632220-INS-205 |
| chr2-69634761-INS-423 | chr2-69634758-INS-424 |
| chr2-69636843-INS-213 | chr2-69636843-INS-213 |
| chr2-69668470-INS-132 | chr2-69668470-INS-133 |
| chr2-69673680-INS-196 | chr2-69673680-INS-197 |
| chr2-69689422-INS-214 | chr2-69689422-INS-214 |
| chr2-69713937-INS-521 | chr2-69713937-INS-521 |
| chr2-69714551-INS-219 | chr2-69714508-INS-213 |
| chr2-69731828-INS-54 | chr2-69731828-INS-54 |
| chr2-69734855-INS-209 | chr2-69734855-INS-209 |
| chr2-69739015-INS-209 | chr2-69739015-INS-210 |
| chr2-69740579-INS-82 | chr2-69740579-INS-88 |
| chr2-69742082-INS-777 | chr2-69742082-INS-777 |
| chr2-69754263-INS-212 | chr2-69754263-INS-215 |
| chr2-69756085-INS-53 | chr2-69756082-INS-58 |
| chr2-69764958-INS-212 | chr2-69764958-INS-212 |
| chr2-69789753-INS-210 | chr2-69789753-INS-210 |
| chr2-69793688-INS-64 | chr2-69793688-INS-64 |
| chr2-69804182-INS-66 | chr2-69804182-INS-72 |
| chr2-69808818-INS-206 | chr2-69808818-INS-209 |
| chr2-69821046-INS-58 | chr2-69821046-INS-52 |
| chr2-69844882-INS-58 | chr2-69844882-INS-58 |
| chr2-69883909-INS-215 | chr2-69883909-INS-214 |
| chr2-69896592-INS-130 | chr2-69896588-INS-94 |
| chr2-69985173-INS-51 | chr2-69985173-INS-51 |
| chr2-69997465-INS-2738 | chr2-69997466-INS-2738 |
| chr2-70011341-INS-835 | chr2-70011341-INS-838 |
| chr2-70047963-INS-63 | chr2-70047963-INS-63 |
| chr2-70053433-INS-52 | chr2-70053433-INS-54 |
| chr2-70057467-INS-59 | chr2-70057467-INS-55 |
| chr2-70089132-INS-62 | chr2-70089132-INS-50 |
| chr2-70109203-INS-111 | chr2-70109203-INS-74 |
| chr2-70116722-INS-181 | chr2-70116722-INS-181 |
| chr2-70150130-INS-212 | chr2-70150130-INS-212 |
| chr2-70178619-INS-146 | chr2-70178619-INS-164 |
| chr2-70181324-INS-52 | chr2-70181304-INS-63 |
| chr2-70204290-INS-58 | chr2-70204290-INS-58 |
| chr2-70217069-INS-116 | chr2-70217069-INS-164 |
| chr2-70240345-INS-130 | chr2-70240346-INS-140 |
| chr2-70250742-INS-95 | chr2-70250742-INS-95 |
| chr2-70275226-INS-198 | chr2-70275230-INS-204 |
| chr2-70276706-INS-143 | chr2-70276706-INS-145 |
| chr2-70284069-INS-150 | chr2-70284069-INS-150 |
| chr2-70287013-INS-1094 | chr2-70287014-INS-1093 |
| chr2-70294723-INS-61 | chr2-70294723-INS-61 |
| chr2-70298532-INS-251 | chr2-70298533-INS-209 |
| chr2-70309431-INS-69 | chr2-70309431-INS-50 |
| chr2-70323551-INS-264 | chr2-70323551-INS-264 |
| chr2-70330040-INS-58 | chr2-70330040-INS-58 |
| chr2-70330672-INS-438 | chr2-70330684-INS-448 |
| chr2-70354610-INS-64 | chr2-70354610-INS-64 |
| chr2-70372359-INS-116 | chr2-70372375-INS-69 |
| chr2-70408946-INS-4447 | chr2-70408947-INS-4447 |

| | |
|---|---|
| chr2-70448235-INS-195 | chr2-70448235-INS-195 |
| chr2-70457181-INS-282 | chr2-70457181-INS-281 |
| chr2-70474362-INS-127 | chr2-70474362-INS-129 |
| chr2-70489960-INS-4802 | chr2-70489961-INS-4802 |
| chr2-70494744-INS-187 | chr2-70494744-INS-188 |
| chr2-70502600-INS-60 | chr2-70502600-INS-60 |
| chr2-70536321-INS-78 | chr2-70536321-INS-80 |
| chr2-70563647-INS-52 | chr2-70563647-INS-78 |
| chr2-70584295-INS-102 | chr2-70584295-INS-108 |
| chr2-70613097-INS-146 | chr2-70613097-INS-144 |
| chr2-70656460-INS-64 | chr2-70656468-INS-86 |
| chr2-70692859-INS-156 | chr2-70692859-INS-116 |
| chr2-70697482-INS-58 | chr2-70697482-INS-70 |
| chr2-70699130-INS-203 | chr2-70699130-INS-203 |
| chr2-70740107-INS-52 | chr2-70740107-INS-52 |
| chr2-70767679-INS-327 | chr2-70767679-INS-327 |
| chr2-70778548-INS-56 | chr2-70778548-INS-64 |
| chr2-70788659-INS-399 | chr2-70788659-INS-399 |
| chr2-70800457-INS-60 | chr2-70800457-INS-58 |
| chr2-70807951-INS-226 | chr2-70807951-INS-226 |
| chr2-70833044-INS-266 | chr2-70833044-INS-266 |
| chr2-70835009-INS-52 | chr2-70835009-INS-54 |
| chr2-70840378-INS-54 | chr2-70840361-INS-57 |
| chr2-70851493-INS-70 | chr2-70851493-INS-70 |
| chr2-70866069-INS-120 | chr2-70866068-INS-129 |
| chr2-70894463-INS-263 | chr2-70894463-INS-263 |
| chr2-70915924-INS-88 | chr2-70915924-INS-92 |
| chr2-70949125-INS-80 | chr2-70949125-INS-82 |
| chr2-70951519-INS-184 | chr2-70951519-INS-184 |
| chr2-70966558-INS-190 | chr2-70966556-INS-190 |
| chr2-70967684-INS-167 | chr2-70967684-INS-167 |
| chr2-70969337-INS-60 | chr2-70969337-INS-60 |
| chr2-70973305-INS-64 | chr2-70973305-INS-62 |
| chr2-70980630-INS-60 | chr2-70980630-INS-60 |
| chr2-70995608-INS-491 | chr2-70995608-INS-491 |
| chr2-71008206-INS-106 | chr2-71008206-INS-106 |
| chr2-71018259-INS-485 | chr2-71018260-INS-485 |
| chr2-71052456-INS-310 | chr2-71052456-INS-318 |
| chr2-71065219-INS-60 | chr2-71065219-INS-60 |
| chr2-71083681-INS-57 | chr2-71083679-INS-57 |
| chr2-71089703-INS-138 | chr2-71089703-INS-138 |
| chr2-71095091-INS-150 | chr2-71095091-INS-150 |
| chr2-71119273-INS-56 | chr2-71119273-INS-60 |
| chr2-71120210-INS-62 | chr2-71120210-INS-54 |
| chr2-71127540-INS-341 | chr2-71127540-INS-341 |
| chr2-71147560-INS-181 | chr2-71147560-INS-180 |
| chr2-71166561-INS-70 | chr2-71166561-INS-70 |
| chr2-71170322-INS-99 | chr2-71170324-INS-100 |
| chr2-71172307-INS-194 | chr2-71172307-INS-194 |
| chr2-71203278-INS-211 | chr2-71203278-INS-211 |
| chr2-71208717-INS-201 | chr2-71208718-INS-154 |
| chr2-71212252-INS-952 | chr2-71212252-INS-950 |
| chr2-71213017-INS-188 | chr2-71213017-INS-188 |
| chr2-71224829-INS-90 | chr2-71224829-INS-90 |
| chr2-71256247-INS-333 | chr2-71256247-INS-333 |
| chr2-71257315-INS-206 | chr2-71257315-INS-206 |
| chr2-71264621-INS-355 | chr2-71264621-INS-355 |
| chr2-71281873-INS-80 | chr2-71281863-INS-102 |
| chr2-71302790-INS-52 | chr2-71302791-INS-56 |
| chr2-71318112-INS-84 | chr2-71318113-INS-85 |
| chr2-71322443-INS-157 | chr2-71322434-INS-175 |
| chr2-71329702-INS-73 | chr2-71329702-INS-73 |
| chr2-71354996-INS-56 | chr2-71354996-INS-50 |
| chr2-71358305-INS-76 | chr2-71358305-INS-64 |
| chr2-71378139-INS-62 | chr2-71378140-INS-66 |
| chr2-71386391-INS-160 | chr2-71386391-INS-162 |
| chr2-71387935-INS-128 | chr2-71387936-INS-128 |
| chr2-71392015-INS-254 | chr2-71392015-INS-257 |
| chr2-71393028-INS-239 | chr2-71393028-INS-218 |
| chr2-71410047-INS-64 | chr2-71410047-INS-54 |
| chr2-71430142-INS-213 | chr2-71430142-INS-213 |
| chr2-71451797-INS-52 | chr2-71451797-INS-54 |
| chr2-71461230-INS-120 | chr2-71461230-INS-70 |
| chr2-71468169-INS-57 | chr2-71468159-INS-64 |
| chr2-71469843-INS-2359 | chr2-71469838-INS-2355 |
| chr2-71477273-INS-296 | chr2-71477273-INS-306 |
| chr2-71505503-INS-52 | chr2-71505503-INS-52 |

| | |
|---|---|
| chr2-71507608-INS-428 | chr2-71507608-INS-427 |
| chr2-71523587-INS-198 | chr2-71523587-INS-198 |
| chr2-71539066-INS-58 | chr2-71539067-INS-58 |
| chr2-71545686-INS-196 | chr2-71545686-INS-196 |
| chr2-71547925-INS-56 | chr2-71547925-INS-56 |
| chr2-71585448-INS-68 | chr2-71585448-INS-68 |
| chr2-71660829-INS-202 | chr2-71660829-INS-202 |
| chr2-71665132-INS-54 | chr2-71665132-INS-54 |
| chr2-71685418-INS-213 | chr2-71685418-INS-213 |
| chr2-71693506-INS-899 | chr2-71693506-INS-899 |
| chr2-71696959-INS-1154 | chr2-71696960-INS-1142 |
| chr2-71704609-INS-1888 | chr2-71704610-INS-1888 |
| chr2-71706177-INS-615 | chr2-71706177-INS-615 |
| chr2-71710068-INS-189 | chr2-71710068-INS-189 |
| chr2-71761677-INS-3102 | chr2-71761678-INS-3102 |
| chr2-71763506-INS-478 | chr2-71763506-INS-478 |
| chr2-71764780-INS-426 | chr2-71764780-INS-426 |
| chr2-71769043-INS-106 | chr2-71769043-INS-106 |
| chr2-71780082-INS-94 | chr2-71780082-INS-100 |
| chr2-71783804-INS-545 | chr2-71783804-INS-545 |
| chr2-71790020-INS-98 | chr2-71790021-INS-98 |
| chr2-71799516-INS-207 | chr2-71799516-INS-214 |
| chr2-71830147-INS-74 | chr2-71830147-INS-76 |
| chr2-71861103-INS-52 | chr2-71861103-INS-54 |
| chr2-71865356-INS-3144 | chr2-71865357-INS-3144 |
| chr2-71868609-INS-752 | chr2-71868609-INS-752 |
| chr2-71911585-INS-69 | chr2-71911585-INS-73 |
| chr2-71916848-INS-66 | chr2-71916848-INS-66 |
| chr2-71931957-INS-87 | chr2-71931957-INS-87 |
| chr2-71958768-INS-515 | chr2-71958768-INS-515 |
| chr2-71970123-INS-64 | chr2-71970123-INS-64 |
| chr2-71988994-INS-110 | chr2-71988994-INS-110 |
| chr2-71996810-INS-76 | chr2-71996825-INS-75 |
| chr2-72008413-INS-71 | chr2-72008415-INS-55 |
| chr2-72010922-INS-56 | chr2-72010891-INS-74 |
| chr2-72018210-INS-176 | chr2-72018210-INS-176 |
| chr2-72144657-INS-303 | chr2-72144657-INS-303 |
| chr2-72146968-INS-175 | chr2-72146968-INS-175 |
| chr2-72149068-INS-136 | chr2-72149068-INS-136 |
| chr2-72216977-INS-117 | chr2-72216977-INS-153 |
| chr2-72273474-INS-142 | chr2-72273474-INS-142 |
| chr2-72280049-INS-57 | chr2-72280049-INS-57 |
| chr2-72285332-INS-72 | chr2-72285331-INS-56 |
| chr2-72293920-INS-96 | chr2-72293920-INS-96 |
| chr2-72305812-INS-92 | chr2-72305810-INS-101 |
| chr2-72321090-INS-137 | chr2-72321090-INS-156 |
| chr2-72325966-INS-55 | chr2-72325966-INS-63 |
| chr2-72343824-INS-51 | chr2-72343848-INS-61 |
| chr2-72363089-INS-395 | chr2-72363089-INS-395 |
| chr2-72375114-INS-208 | chr2-72375114-INS-208 |
| chr2-72379243-INS-508 | chr2-72379243-INS-508 |
| chr2-72410910-INS-140 | chr2-72410914-INS-136 |
| chr2-72421168-INS-79 | chr2-72421168-INS-79 |
| chr2-72437703-INS-443 | chr2-72437703-INS-443 |
| chr2-72466904-INS-64 | chr2-72466899-INS-58 |
| chr2-72473269-INS-57 | chr2-72473269-INS-57 |
| chr2-72504681-INS-100 | chr2-72504672-INS-130 |
| chr2-72509637-INS-64 | chr2-72509651-INS-74 |
| chr2-72515560-INS-1506 | chr2-72515561-INS-1506 |
| chr2-72528689-INS-279 | chr2-72528689-INS-279 |
| chr2-72532040-INS-285 | chr2-72532041-INS-287 |
| chr2-72534384-INS-54 | chr2-72534384-INS-60 |
| chr2-72536196-INS-492 | chr2-72536196-INS-492 |
| chr2-72541173-INS-71 | chr2-72541173-INS-51 |
| chr2-72558096-INS-156 | chr2-72558096-INS-152 |
| chr2-72575355-INS-74 | chr2-72575355-INS-74 |
| chr2-72603041-INS-63 | chr2-72603041-INS-57 |
| chr2-72634879-INS-99 | chr2-72634869-INS-94 |
| chr2-72649044-INS-56 | chr2-72649044-INS-56 |
| chr2-72670003-INS-80 | chr2-72670003-INS-80 |
| chr2-72673161-INS-100 | chr2-72673189-INS-54 |
| chr2-72691949-INS-72 | chr2-72691949-INS-72 |
| chr2-72731955-INS-83 | chr2-72731955-INS-83 |
| chr2-72757847-INS-292 | chr2-72757842-INS-329 |
| chr2-72758573-INS-3379 | chr2-72758574-INS-3380 |
| chr2-72797295-INS-607 | chr2-72797296-INS-608 |
| chr2-72819413-INS-218 | chr2-72819413-INS-218 |

| | |
|---|---|
| chr2-72853756-INS-124 | chr2-72853756-INS-144 |
| chr2-72858912-INS-986 | chr2-72858912-INS-988 |
| chr2-72873189-INS-59 | chr2-72873189-INS-59 |
| chr2-72885853-INS-358 | chr2-72885853-INS-358 |
| chr2-72891428-INS-58 | chr2-72891428-INS-58 |
| chr2-72896063-INS-215 | chr2-72896063-INS-215 |
| chr2-72899560-INS-135 | chr2-72899560-INS-135 |
| chr2-72918585-INS-118 | chr2-72918585-INS-118 |
| chr2-72921035-INS-50 | chr2-72921035-INS-88 |
| chr2-72931517-INS-101 | chr2-72931517-INS-101 |
| chr2-72959714-INS-157 | chr2-72959714-INS-157 |
| chr2-72983087-INS-207 | chr2-72983087-INS-206 |
| chr2-73008469-INS-331 | chr2-73008469-INS-331 |
| chr2-73015604-INS-214 | chr2-73015604-INS-214 |
| chr2-73018369-INS-75 | chr2-73018369-INS-55 |
| chr2-73025379-INS-1004 | chr2-73025380-INS-1004 |
| chr2-73026978-INS-54 | chr2-73026978-INS-54 |
| chr2-73126970-INS-309 | chr2-73126970-INS-309 |
| chr2-73132718-INS-227 | chr2-73132718-INS-226 |
| chr2-73180696-INS-91 | chr2-73180696-INS-89 |
| chr2-73207040-INS-73 | chr2-73207040-INS-73 |
| chr2-73226731-INS-341 | chr2-73226731-INS-341 |
| chr2-73262517-INS-76 | chr2-73262517-INS-76 |
| chr2-73292858-INS-55 | chr2-73292906-INS-57 |
| chr2-73367326-INS-66 | chr2-73367329-INS-64 |
| chr2-73379191-INS-216 | chr2-73379191-INS-216 |
| chr2-73381956-INS-77 | chr2-73381956-INS-77 |
| chr2-73383231-INS-54 | chr2-73383210-INS-66 |
| chr2-73386395-INS-52 | chr2-73386395-INS-56 |
| chr2-73388742-INS-62 | chr2-73388742-INS-62 |
| chr2-73392058-INS-434 | chr2-73392058-INS-434 |
| chr2-73398113-INS-62 | chr2-73398113-INS-68 |
| chr2-73425951-INS-58 | chr2-73425933-INS-53 |
| chr2-73451088-INS-2426 | chr2-73451089-INS-2444 |
| chr2-73471985-INS-406 | chr2-73471985-INS-406 |
| chr2-73480774-INS-62 | chr2-73480774-INS-62 |
| chr2-73568195-INS-58 | chr2-73568195-INS-58 |
| chr2-73575553-INS-207 | chr2-73575553-INS-207 |
| chr2-73600088-INS-50 | chr2-73600088-INS-74 |
| chr2-73631669-INS-54 | chr2-73631669-INS-54 |
| chr2-73657123-INS-397 | chr2-73657123-INS-397 |
| chr2-73658723-INS-55 | chr2-73658723-INS-50 |
| chr2-73660058-INS-301 | chr2-73660058-INS-278 |
| chr2-73726917-INS-467 | chr2-73726917-INS-467 |
| chr2-73734012-INS-64 | chr2-73734012-INS-72 |
| chr2-73736570-INS-127 | chr2-73736541-INS-171 |
| chr2-73776087-INS-223 | chr2-73776087-INS-230 |
| chr2-73786283-INS-52 | chr2-73786294-INS-58 |
| chr2-73828483-INS-66 | chr2-73828480-INS-69 |
| chr2-73840820-INS-75 | chr2-73840820-INS-75 |
| chr2-73869193-INS-53 | chr2-73869193-INS-53 |
| chr2-73888186-INS-299 | chr2-73888225-INS-300 |
| chr2-73900629-INS-134 | chr2-73900627-INS-110 |
| chr2-73920599-INS-64 | chr2-73920599-INS-68 |
| chr2-73928573-INS-124 | chr2-73928574-INS-84 |
| chr2-73940249-INS-667 | chr2-73940249-INS-667 |
| chr2-73945676-INS-78 | chr2-73945676-INS-90 |
| chr2-73992417-INS-78 | chr2-73992417-INS-78 |
| chr2-74003338-INS-507 | chr2-74003338-INS-507 |
| chr2-74006865-INS-60 | chr2-74006865-INS-60 |
| chr2-74016166-INS-62 | chr2-74016166-INS-62 |
| chr2-74046366-INS-99 | chr2-74046366-INS-99 |
| chr2-74103334-INS-62 | chr2-74103334-INS-82 |
| chr2-74107544-INS-70 | chr2-74107544-INS-72 |
| chr2-74144789-INS-51 | chr2-74144765-INS-54 |
| chr2-74146119-INS-4810 | chr2-74146120-INS-4815 |
| chr2-74157936-INS-511 | chr2-74157936-INS-511 |
| chr2-74183878-INS-59 | chr2-74183851-INS-69 |
| chr2-74226415-INS-71 | chr2-74226415-INS-88 |
| chr2-74259773-INS-698 | chr2-74259773-INS-702 |
| chr2-74278004-INS-55 | chr2-74278004-INS-56 |
| chr2-74287302-INS-78 | chr2-74287307-INS-116 |
| chr2-74315511-INS-198 | chr2-74315511-INS-198 |
| chr2-74329295-INS-6476 | chr2-74329296-INS-6476 |
| chr2-74332824-INS-391 | chr2-74332825-INS-391 |
| chr2-74339620-INS-90 | chr2-74339620-INS-90 |
| chr2-74392333-INS-76 | chr2-74392321-INS-72 |

| | |
|---|---|
| chr2-74427754-INS-667 | chr2-74427754-INS-667 |
| chr2-74436542-INS-1510 | chr2-74436543-INS-1510 |
| chr2-74451589-INS-147 | chr2-74451590-INS-150 |
| chr2-74454026-INS-78 | chr2-74454026-INS-72 |
| chr2-74459491-INS-121 | chr2-74459451-INS-134 |
| chr2-74475270-INS-189 | chr2-74475270-INS-189 |
| chr2-74493424-INS-56 | chr2-74493424-INS-55 |
| chr2-74621696-INS-138 | chr2-74621696-INS-138 |
| chr2-74623234-INS-67 | chr2-74623234-INS-67 |
| chr2-74625443-INS-113 | chr2-74625443-INS-93 |
| chr2-74636645-INS-109 | chr2-74636645-INS-109 |
| chr2-74638513-INS-172 | chr2-74638513-INS-172 |
| chr2-74648609-INS-255 | chr2-74648609-INS-219 |
| chr2-74692688-INS-54 | chr2-74692688-INS-54 |
| chr2-74739631-INS-55 | chr2-74739631-INS-71 |
| chr2-74742788-INS-55 | chr2-74742788-INS-60 |
| chr2-74754667-INS-51 | chr2-74754667-INS-50 |
| chr2-74761677-INS-135 | chr2-74761677-INS-170 |
| chr2-74778072-INS-56 | chr2-74778082-INS-72 |
| chr2-74802609-INS-118 | chr2-74802609-INS-149 |
| chr2-74807683-INS-60 | chr2-74807700-INS-51 |
| chr2-74813837-INS-176 | chr2-74813838-INS-176 |
| chr2-74816640-INS-113 | chr2-74816639-INS-114 |
| chr2-74823827-INS-104 | chr2-74823827-INS-56 |
| chr2-74853856-INS-153 | chr2-74853856-INS-153 |
| chr2-74874519-INS-54 | chr2-74874513-INS-60 |
| chr2-74881642-INS-85 | chr2-74881642-INS-85 |
| chr2-74899929-INS-111 | chr2-74899944-INS-74 |
| chr2-74907520-INS-70 | chr2-74907520-INS-60 |
| chr2-74908222-INS-827 | chr2-74908222-INS-827 |
| chr2-74908803-INS-90 | chr2-74908803-INS-59 |
| chr2-74913644-INS-115 | chr2-74913644-INS-129 |
| chr2-74923176-INS-383 | chr2-74923176-INS-383 |
| chr2-74929902-INS-70 | chr2-74929902-INS-70 |
| chr2-74949639-INS-280 | chr2-74949639-INS-280 |
| chr2-74951222-INS-52 | chr2-74951222-INS-50 |
| chr2-74979779-INS-101 | chr2-74979779-INS-114 |
| chr2-75008987-INS-3111 | chr2-75008988-INS-3111 |
| chr2-75030391-INS-78 | chr2-75030391-INS-89 |
| chr2-75103607-INS-128 | chr2-75103607-INS-120 |
| chr2-75128939-INS-593 | chr2-75128939-INS-589 |
| chr2-75191448-INS-60 | chr2-75191448-INS-60 |
| chr2-75287449-INS-72 | chr2-75287449-INS-72 |
| chr2-75296002-INS-797 | chr2-75296002-INS-816 |
| chr2-75340301-INS-52 | chr2-75340301-INS-52 |
| chr2-75342584-INS-447 | chr2-75342584-INS-447 |
| chr2-75348962-INS-136 | chr2-75348962-INS-136 |
| chr2-75353133-INS-52 | chr2-75353133-INS-52 |
| chr2-75358568-INS-1894 | chr2-75358569-INS-1894 |
| chr2-75367638-INS-99 | chr2-75367638-INS-102 |
| chr2-75401959-INS-204 | chr2-75401959-INS-204 |
| chr2-75405651-INS-66 | chr2-75405601-INS-94 |
| chr2-75412136-INS-98 | chr2-75412136-INS-98 |
| chr2-75412962-INS-202 | chr2-75412962-INS-202 |
| chr2-75421266-INS-51 | chr2-75421266-INS-51 |
| chr2-75431534-INS-200 | chr2-75431534-INS-201 |
| chr2-75452481-INS-257 | chr2-75452481-INS-259 |
| chr2-75465686-INS-196 | chr2-75465687-INS-189 |
| chr2-75466884-INS-497 | chr2-75466884-INS-486 |
| chr2-75474991-INS-193 | chr2-75474991-INS-190 |
| chr2-75504382-INS-214 | chr2-75504382-INS-214 |
| chr2-75530139-INS-202 | chr2-75530139-INS-201 |
| chr2-75553047-INS-90 | chr2-75553037-INS-104 |
| chr2-75557605-INS-84 | chr2-75557605-INS-84 |
| chr2-75571374-INS-209 | chr2-75571374-INS-211 |
| chr2-75581264-INS-206 | chr2-75581264-INS-205 |
| chr2-75603796-INS-188 | chr2-75603797-INS-191 |
| chr2-75606517-INS-102 | chr2-75606517-INS-102 |
| chr2-75618043-INS-91 | chr2-75618043-INS-91 |
| chr2-75619353-INS-66 | chr2-75619353-INS-66 |
| chr2-75633236-INS-56 | chr2-75633211-INS-80 |
| chr2-75671785-INS-212 | chr2-75671785-INS-216 |
| chr2-75711503-INS-207 | chr2-75711503-INS-207 |
| chr2-75797368-INS-3136 | chr2-75797369-INS-3136 |
| chr2-75803531-INS-96 | chr2-75803539-INS-118 |
| chr2-75831491-INS-67 | chr2-75831491-INS-56 |
| chr2-75837653-INS-72 | chr2-75837653-INS-72 |

| | |
|---|---|
| chr2-75873864-INS-141 | chr2-75873864-INS-141 |
| chr2-75885917-INS-632 | chr2-75885920-INS-633 |
| chr2-75920161-INS-607 | chr2-75920161-INS-611 |
| chr2-75926474-INS-54 | chr2-75926474-INS-54 |
| chr2-75936093-INS-71 | chr2-75936093-INS-71 |
| chr2-75943226-INS-56 | chr2-75943226-INS-56 |
| chr2-75967667-INS-724 | chr2-75967667-INS-724 |
| chr2-75975401-INS-76 | chr2-75975401-INS-81 |
| chr2-76011636-INS-74 | chr2-76011636-INS-52 |
| chr2-76013070-INS-97 | chr2-76013070-INS-97 |
| chr2-76017613-INS-52 | chr2-76017613-INS-52 |
| chr2-76023248-INS-70 | chr2-76023248-INS-70 |
| chr2-76041989-INS-92 | chr2-76041990-INS-72 |
| chr2-76088358-INS-81 | chr2-76088358-INS-81 |
| chr2-76103095-INS-199 | chr2-76103095-INS-175 |
| chr2-76113671-INS-180 | chr2-76113671-INS-180 |
| chr2-76120338-INS-229 | chr2-76120338-INS-229 |
| chr2-76145421-INS-63 | chr2-76145421-INS-63 |
| chr2-76177332-INS-60 | chr2-76177332-INS-60 |
| chr2-76178808-INS-307 | chr2-76178808-INS-307 |
| chr2-76180598-INS-52 | chr2-76180598-INS-54 |
| chr2-76186874-INS-100 | chr2-76186874-INS-100 |
| chr2-76196690-INS-65 | chr2-76196690-INS-58 |
| chr2-76197352-INS-69 | chr2-76197353-INS-69 |
| chr2-76199428-INS-99 | chr2-76199428-INS-99 |
| chr2-76220755-INS-53 | chr2-76220755-INS-53 |
| chr2-76226911-INS-126 | chr2-76226911-INS-126 |
| chr2-76242129-INS-153 | chr2-76242129-INS-181 |
| chr2-76243610-INS-52 | chr2-76243610-INS-54 |
| chr2-76257056-INS-234 | chr2-76257056-INS-234 |
| chr2-76296054-INS-604 | chr2-76296054-INS-604 |
| chr2-76313751-INS-5288 | chr2-76313752-INS-5288 |
| chr2-76337661-INS-196 | chr2-76337661-INS-196 |
| chr2-76404026-INS-108 | chr2-76404026-INS-112 |
| chr2-76437957-INS-2637 | chr2-76437958-INS-2630 |
| chr2-76441658-INS-153 | chr2-76441658-INS-149 |
| chr2-76443082-INS-1688 | chr2-76443083-INS-1688 |
| chr2-76443930-INS-4590 | chr2-76443931-INS-4590 |
| chr2-76454102-INS-502 | chr2-76454102-INS-501 |
| chr2-76490653-INS-417 | chr2-76490653-INS-417 |
| chr2-76500600-INS-560 | chr2-76500600-INS-559 |
| chr2-76590955-INS-75 | chr2-76590955-INS-75 |
| chr2-76644333-INS-66 | chr2-76644333-INS-54 |
| chr2-76668712-INS-198 | chr2-76668712-INS-198 |
| chr2-76699255-INS-53 | chr2-76699255-INS-51 |
| chr2-76824402-INS-71 | chr2-76824402-INS-71 |
| chr2-76825832-INS-188 | chr2-76825832-INS-188 |
| chr2-76827963-INS-100 | chr2-76827963-INS-100 |
| chr2-76838235-INS-209 | chr2-76838235-INS-209 |
| chr2-76842952-INS-91 | chr2-76842952-INS-91 |
| chr2-76869450-INS-61 | chr2-76869450-INS-61 |
| chr2-76876822-INS-6975 | chr2-76876823-INS-6975 |
| chr2-76886070-INS-323 | chr2-76886070-INS-322 |
| chr2-76890879-INS-60 | chr2-76890900-INS-67 |
| chr2-76902363-INS-967 | chr2-76902363-INS-961 |
| chr2-76911174-INS-54 | chr2-76911174-INS-54 |
| chr2-76911756-INS-217 | chr2-76911756-INS-218 |
| chr2-76913009-INS-212 | chr2-76913009-INS-212 |
| chr2-76929634-INS-203 | chr2-76929634-INS-203 |
| chr2-76937102-INS-1289 | chr2-76937103-INS-1289 |
| chr2-76941100-INS-93 | chr2-76941128-INS-111 |
| chr2-76943197-INS-129 | chr2-76943197-INS-129 |
| chr2-76946797-INS-203 | chr2-76946797-INS-204 |
| chr2-76980793-INS-78 | chr2-76980793-INS-78 |
| chr2-77001378-INS-58 | chr2-77001378-INS-50 |
| chr2-77044150-INS-921 | chr2-77044150-INS-921 |
| chr2-77082251-INS-100 | chr2-77082251-INS-100 |
| chr2-77088147-INS-5624 | chr2-77088148-INS-5616 |
| chr2-77133386-INS-82 | chr2-77133386-INS-81 |
| chr2-77141418-INS-51 | chr2-77141397-INS-54 |
| chr2-77144064-INS-52 | chr2-77144064-INS-52 |
| chr2-77144389-INS-2829 | chr2-77144390-INS-2830 |
| chr2-77149466-INS-147 | chr2-77149468-INS-190 |
| chr2-77164415-INS-72 | chr2-77164415-INS-76 |
| chr2-77169010-INS-195 | chr2-77169010-INS-151 |
| chr2-77185568-INS-50 | chr2-77185572-INS-54 |
| chr2-77212866-INS-149 | chr2-77212866-INS-149 |

| | |
|---|---|
| chr2-77265469-INS-90 | chr2-77265470-INS-90 |
| chr2-77271240-INS-176 | chr2-77271250-INS-178 |
| chr2-77428941-INS-160 | chr2-77428941-INS-160 |
| chr2-77435918-INS-214 | chr2-77435918-INS-214 |
| chr2-77450539-INS-60 | chr2-77450539-INS-60 |
| chr2-77461476-INS-7870 | chr2-77461477-INS-7870 |
| chr2-77480162-INS-327 | chr2-77480162-INS-327 |
| chr2-77494348-INS-716 | chr2-77494348-INS-717 |
| chr2-77499733-INS-180 | chr2-77499733-INS-181 |
| chr2-77523225-INS-68 | chr2-77523225-INS-68 |
| chr2-77536971-INS-232 | chr2-77536971-INS-187 |
| chr2-77541184-INS-248 | chr2-77541184-INS-212 |
| chr2-77542423-INS-72 | chr2-77542423-INS-80 |
| chr2-77547541-INS-126 | chr2-77547541-INS-126 |
| chr2-77549154-INS-104 | chr2-77549154-INS-69 |
| chr2-77557299-INS-879 | chr2-77557299-INS-879 |
| chr2-77576325-INS-64 | chr2-77576325-INS-64 |
| chr2-77576736-INS-1004 | chr2-77576737-INS-1004 |
| chr2-77579317-INS-54 | chr2-77579317-INS-64 |
| chr2-77584329-INS-119 | chr2-77584329-INS-119 |
| chr2-77590662-INS-66 | chr2-77590662-INS-66 |
| chr2-77622653-INS-130 | chr2-77622653-INS-130 |
| chr2-77676994-INS-430 | chr2-77676994-INS-430 |
| chr2-77686181-INS-70 | chr2-77686181-INS-70 |
| chr2-77857642-INS-190 | chr2-77857642-INS-225 |
| chr2-77867358-INS-356 | chr2-77867358-INS-356 |
| chr2-77868256-INS-82 | chr2-77868244-INS-110 |
| chr2-77876280-INS-134 | chr2-77876280-INS-141 |
| chr2-77921978-INS-1290 | chr2-77921979-INS-1290 |
| chr2-77942692-INS-88 | chr2-77942692-INS-87 |
| chr2-77943890-INS-484 | chr2-77943890-INS-484 |
| chr2-77992619-INS-64 | chr2-77992619-INS-64 |
| chr2-78011821-INS-55 | chr2-78011821-INS-55 |
| chr2-78016731-INS-227 | chr2-78016731-INS-232 |
| chr2-78030119-INS-104 | chr2-78030119-INS-106 |
| chr2-78040170-INS-6134 | chr2-78040171-INS-6132 |
| chr2-78067075-INS-6491 | chr2-78067076-INS-6491 |
| chr2-78088618-INS-6263 | chr2-78088619-INS-6263 |
| chr2-78097806-INS-54 | chr2-78097806-INS-56 |
| chr2-78102989-INS-624 | chr2-78102990-INS-621 |
| chr2-78119110-INS-84 | chr2-78119106-INS-87 |
| chr2-78122585-INS-6484 | chr2-78122586-INS-6484 |
| chr2-78143911-INS-323 | chr2-78143911-INS-323 |
| chr2-78155436-INS-6238 | chr2-78155437-INS-6238 |
| chr2-78188369-INS-342 | chr2-78188369-INS-342 |
| chr2-78241690-INS-7099 | chr2-78241691-INS-7091 |
| chr2-78245176-INS-1084 | chr2-78245177-INS-1131 |
| chr2-78251424-INS-201 | chr2-78251424-INS-204 |
| chr2-78266124-INS-65 | chr2-78266148-INS-65 |
| chr2-78334204-INS-308 | chr2-78334204-INS-304 |
| chr2-78362646-INS-125 | chr2-78362646-INS-129 |
| chr2-78382187-INS-62 | chr2-78382187-INS-68 |
| chr2-78402705-INS-57 | chr2-78402696-INS-85 |
| chr2-78435058-INS-211 | chr2-78435058-INS-211 |
| chr2-78454334-INS-92 | chr2-78454325-INS-70 |
| chr2-78471817-INS-386 | chr2-78471817-INS-337 |
| chr2-78492698-INS-208 | chr2-78492698-INS-208 |
| chr2-78503868-INS-68 | chr2-78503868-INS-70 |
| chr2-78517906-INS-76 | chr2-78517906-INS-78 |
| chr2-78543041-INS-6871 | chr2-78543042-INS-6871 |
| chr2-78564656-INS-643 | chr2-78564656-INS-643 |
| chr2-78581691-INS-1268 | chr2-78581692-INS-1268 |
| chr2-78647535-INS-116 | chr2-78647535-INS-116 |
| chr2-78674570-INS-52 | chr2-78674570-INS-52 |
| chr2-78704867-INS-54 | chr2-78704867-INS-56 |
| chr2-78712389-INS-72 | chr2-78712389-INS-68 |
| chr2-78755966-INS-55 | chr2-78755966-INS-56 |
| chr2-78770472-INS-124 | chr2-78770472-INS-120 |
| chr2-78796735-INS-255 | chr2-78796735-INS-255 |
| chr2-78826303-INS-52 | chr2-78826303-INS-74 |
| chr2-78835942-INS-406 | chr2-78835942-INS-407 |
| chr2-78885170-INS-319 | chr2-78885170-INS-318 |
| chr2-78907155-INS-164 | chr2-78907155-INS-118 |
| chr2-78922378-INS-51 | chr2-78922378-INS-51 |
| chr2-78939197-INS-154 | chr2-78939197-INS-156 |
| chr2-78974299-INS-1888 | chr2-78974300-INS-1888 |
| chr2-79013071-INS-62 | chr2-79013071-INS-66 |

| | |
|---|---|
| chr2-79024882-INS-4400 | chr2-79024883-INS-4400 |
| chr2-79044078-INS-168 | chr2-79044078-INS-168 |
| chr2-79108947-INS-100 | chr2-79108947-INS-72 |
| chr2-79135689-INS-2209 | chr2-79135690-INS-2209 |
| chr2-79186976-INS-498 | chr2-79186976-INS-498 |
| chr2-79189988-INS-729 | chr2-79189988-INS-730 |
| chr2-79190649-INS-125 | chr2-79190651-INS-125 |
| chr2-79195325-INS-542 | chr2-79195325-INS-542 |
| chr2-79228467-INS-59 | chr2-79228463-INS-72 |
| chr2-79231278-INS-111 | chr2-79231278-INS-111 |
| chr2-79239371-INS-828 | chr2-79239371-INS-828 |
| chr2-79266353-INS-78 | chr2-79266353-INS-78 |
| chr2-79273725-INS-2013 | chr2-79273726-INS-2014 |
| chr2-79354555-INS-76 | chr2-79354555-INS-80 |
| chr2-79357124-INS-52 | chr2-79357124-INS-50 |
| chr2-79363610-INS-54 | chr2-79363610-INS-54 |
| chr2-79389852-INS-2931 | chr2-79389853-INS-2931 |
| chr2-79399120-INS-7100 | chr2-79399121-INS-7100 |
| chr2-79425519-INS-115 | chr2-79425519-INS-114 |
| chr2-79440571-INS-75 | chr2-79440570-INS-76 |
| chr2-79453937-INS-243 | chr2-79453937-INS-243 |
| chr2-79498429-INS-231 | chr2-79498429-INS-235 |
| chr2-79536361-INS-56 | chr2-79536361-INS-68 |
| chr2-79596297-INS-498 | chr2-79596297-INS-498 |
| chr2-79601342-INS-61 | chr2-79601342-INS-61 |
| chr2-79604784-INS-60 | chr2-79604784-INS-60 |
| chr2-79619915-INS-60 | chr2-79619915-INS-55 |
| chr2-79644622-INS-72 | chr2-79644622-INS-72 |
| chr2-79650916-INS-593 | chr2-79650917-INS-592 |
| chr2-79652473-INS-632 | chr2-79652473-INS-623 |
| chr2-79654323-INS-99 | chr2-79654323-INS-81 |
| chr2-79691031-INS-963 | chr2-79691031-INS-963 |
| chr2-79717427-INS-68 | chr2-79717427-INS-68 |
| chr2-79724581-INS-96 | chr2-79724581-INS-112 |
| chr2-79752136-INS-61 | chr2-79752121-INS-72 |
| chr2-79752919-INS-1057 | chr2-79752920-INS-1055 |
| chr2-79767036-INS-212 | chr2-79767036-INS-212 |
| chr2-79781407-INS-493 | chr2-79781407-INS-493 |
| chr2-79782179-INS-903 | chr2-79782179-INS-903 |
| chr2-79794604-INS-996 | chr2-79794604-INS-996 |
| chr2-79794895-INS-58 | chr2-79794895-INS-62 |
| chr2-79835712-INS-434 | chr2-79835712-INS-416 |
| chr2-79849362-INS-4518 | chr2-79849363-INS-4518 |
| chr2-79878753-INS-51 | chr2-79878770-INS-59 |
| chr2-79889202-INS-237 | chr2-79889200-INS-237 |
| chr2-79890975-INS-63 | chr2-79890971-INS-73 |
| chr2-79916664-INS-136 | chr2-79916664-INS-135 |
| chr2-79928167-INS-52 | chr2-79928167-INS-52 |
| chr2-79986152-INS-158 | chr2-79986154-INS-148 |
| chr2-79987229-INS-2857 | chr2-79987230-INS-2857 |
| chr2-79992407-INS-7222 | chr2-79992408-INS-7223 |
| chr2-80029461-INS-6701 | chr2-80029462-INS-6703 |
| chr2-80046703-INS-656 | chr2-80046703-INS-656 |
| chr2-80061686-INS-73 | chr2-80061686-INS-73 |
| chr2-80079831-INS-120 | chr2-80079831-INS-88 |
| chr2-80094597-INS-152 | chr2-80094597-INS-156 |
| chr2-80101065-INS-56 | chr2-80101065-INS-52 |
| chr2-80150613-INS-5190 | chr2-80150614-INS-5190 |
| chr2-80154205-INS-225 | chr2-80154205-INS-224 |
| chr2-80162762-INS-208 | chr2-80162762-INS-207 |
| chr2-80174216-INS-178 | chr2-80174216-INS-178 |
| chr2-80193720-INS-359 | chr2-80193722-INS-377 |
| chr2-80201737-INS-82 | chr2-80201737-INS-82 |
| chr2-80203502-INS-430 | chr2-80203502-INS-450 |
| chr2-80215431-INS-170 | chr2-80215431-INS-169 |
| chr2-80229032-INS-831 | chr2-80229032-INS-828 |
| chr2-80241015-INS-200 | chr2-80241002-INS-201 |
| chr2-80359011-INS-73 | chr2-80359011-INS-73 |
| chr2-80360202-INS-130 | chr2-80360202-INS-130 |
| chr2-80365749-INS-528 | chr2-80365749-INS-528 |
| chr2-80380822-INS-192 | chr2-80380822-INS-195 |
| chr2-80416776-INS-4350 | chr2-80416777-INS-4350 |
| chr2-80467164-INS-80 | chr2-80467164-INS-66 |
| chr2-80477515-INS-94 | chr2-80477485-INS-96 |
| chr2-80493466-INS-341 | chr2-80493466-INS-341 |
| chr2-80526016-INS-472 | chr2-80526016-INS-472 |
| chr2-80551620-INS-221 | chr2-80551620-INS-224 |

| | |
|---|---|
| chr2-80586332-INS-2795 | chr2-80586333-INS-2793 |
| chr2-80695698-INS-579 | chr2-80695698-INS-611 |
| chr2-80706856-INS-52 | chr2-80706856-INS-52 |
| chr2-80708603-INS-7064 | chr2-80708604-INS-7065 |
| chr2-80723300-INS-500 | chr2-80723300-INS-500 |
| chr2-80756574-INS-204 | chr2-80756574-INS-204 |
| chr2-80767524-INS-280 | chr2-80767530-INS-285 |
| chr2-80780406-INS-170 | chr2-80780406-INS-170 |
| chr2-80824091-INS-82 | chr2-80824091-INS-82 |
| chr2-80860654-INS-663 | chr2-80860654-INS-663 |
| chr2-80862905-INS-7259 | chr2-80862906-INS-7259 |
| chr2-80888680-INS-246 | chr2-80888680-INS-246 |
| chr2-80917983-INS-93 | chr2-80917983-INS-93 |
| chr2-80932586-INS-72 | chr2-80932566-INS-54 |
| chr2-80959367-INS-70 | chr2-80959367-INS-70 |
| chr2-80973378-INS-55 | chr2-80973374-INS-62 |
| chr2-81062793-INS-3282 | chr2-81062794-INS-3282 |
| chr2-81068646-INS-1890 | chr2-81068647-INS-1890 |
| chr2-81082720-INS-101 | chr2-81082720-INS-101 |
| chr2-81100507-INS-908 | chr2-81100507-INS-908 |
| chr2-81106114-INS-114 | chr2-81106114-INS-122 |
| chr2-81129497-INS-295 | chr2-81129497-INS-295 |
| chr2-81130101-INS-69 | chr2-81130101-INS-52 |
| chr2-81144863-INS-57 | chr2-81144863-INS-57 |
| chr2-81145990-INS-336 | chr2-81145990-INS-341 |
| chr2-81151406-INS-284 | chr2-81151406-INS-284 |
| chr2-81152272-INS-702 | chr2-81152272-INS-702 |
| chr2-81173342-INS-52 | chr2-81173342-INS-65 |
| chr2-81220427-INS-7026 | chr2-81220428-INS-7026 |
| chr2-81243849-INS-86 | chr2-81243849-INS-98 |
| chr2-81309910-INS-182 | chr2-81309946-INS-182 |
| chr2-81315417-INS-86 | chr2-81315417-INS-88 |
| chr2-81367250-INS-119 | chr2-81367252-INS-118 |
| chr2-81381365-INS-944 | chr2-81381365-INS-944 |
| chr2-81453795-INS-58 | chr2-81453795-INS-62 |
| chr2-81484806-INS-58 | chr2-81484800-INS-86 |
| chr2-81581010-INS-201 | chr2-81581010-INS-201 |
| chr2-81640194-INS-79 | chr2-81640194-INS-84 |
| chr2-81662129-INS-6939 | chr2-81662130-INS-6939 |
| chr2-81690863-INS-54 | chr2-81690859-INS-50 |
| chr2-81705228-INS-201 | chr2-81705228-INS-201 |
| chr2-81721203-INS-623 | chr2-81721184-INS-613 |
| chr2-81745601-INS-543 | chr2-81745601-INS-545 |
| chr2-81750013-INS-181 | chr2-81750013-INS-177 |
| chr2-81765416-INS-894 | chr2-81765416-INS-896 |
| chr2-81777468-INS-58 | chr2-81777468-INS-58 |
| chr2-81816713-INS-70 | chr2-81816713-INS-69 |
| chr2-81952014-INS-80 | chr2-81952046-INS-56 |
| chr2-82047105-INS-224 | chr2-82047104-INS-243 |
| chr2-82085004-INS-1295 | chr2-82085005-INS-1287 |
| chr2-82107831-INS-338 | chr2-82107831-INS-350 |
| chr2-82159671-INS-56 | chr2-82159671-INS-78 |
| chr2-82227473-INS-1452 | chr2-82227474-INS-1451 |
| chr2-82230804-INS-88 | chr2-82230818-INS-96 |
| chr2-82236461-INS-90 | chr2-82236461-INS-134 |
| chr2-82261242-INS-208 | chr2-82261242-INS-208 |
| chr2-82300415-INS-2438 | chr2-82300416-INS-2438 |
| chr2-82344534-INS-6422 | chr2-82344535-INS-6421 |
| chr2-82408319-INS-6450 | chr2-82408316-INS-6449 |
| chr2-82447081-INS-7312 | chr2-82447082-INS-7312 |
| chr2-82454247-INS-59 | chr2-82454247-INS-56 |
| chr2-82478859-INS-331 | chr2-82478859-INS-335 |
| chr2-82479650-INS-2487 | chr2-82479651-INS-2469 |
| chr2-82484564-INS-387 | chr2-82484564-INS-387 |
| chr2-82493594-INS-54 | chr2-82493594-INS-60 |
| chr2-82531669-INS-585 | chr2-82531669-INS-587 |
| chr2-82543771-INS-963 | chr2-82543771-INS-963 |
| chr2-82560982-INS-202 | chr2-82560982-INS-202 |
| chr2-82589490-INS-72 | chr2-82589490-INS-64 |
| chr2-82599347-INS-60 | chr2-82599347-INS-68 |
| chr2-82640146-INS-66 | chr2-82640164-INS-66 |
| chr2-82664483-INS-209 | chr2-82664483-INS-209 |
| chr2-82676859-INS-468 | chr2-82676859-INS-477 |
| chr2-82791526-INS-86 | chr2-82791526-INS-78 |
| chr2-82793910-INS-6452 | chr2-82793911-INS-6453 |
| chr2-82842628-INS-66 | chr2-82842628-INS-66 |
| chr2-82851559-INS-697 | chr2-82851538-INS-702 |

| | |
|---|---|
| chr2-82854414-INS-55 | chr2-82854422-INS-67 |
| chr2-82876067-INS-778 | chr2-82876061-INS-788 |
| chr2-82902788-INS-214 | chr2-82902788-INS-214 |
| chr2-82968246-INS-3024 | chr2-82968247-INS-3028 |
| chr2-82984272-INS-2908 | chr2-82984273-INS-2925 |
| chr2-82995804-INS-1699 | chr2-82995805-INS-1700 |
| chr2-82998068-INS-52 | chr2-82998048-INS-62 |
| chr2-83047450-INS-6176 | chr2-83047451-INS-6176 |
| chr2-83076303-INS-7702 | chr2-83076304-INS-7703 |
| chr2-83086315-INS-7088 | chr2-83086316-INS-7088 |
| chr2-83104329-INS-4794 | chr2-83104330-INS-4795 |
| chr2-83142287-INS-177 | chr2-83142287-INS-177 |
| chr2-83166946-INS-529 | chr2-83166946-INS-529 |
| chr2-83170068-INS-72 | chr2-83170068-INS-72 |
| chr2-83180702-INS-80 | chr2-83180702-INS-80 |
| chr2-83216326-INS-1169 | chr2-83216327-INS-1169 |
| chr2-83225796-INS-407 | chr2-83225785-INS-423 |
| chr2-83276278-INS-762 | chr2-83276278-INS-762 |
| chr2-83286121-INS-386 | chr2-83286121-INS-386 |
| chr2-83291819-INS-6839 | chr2-83291820-INS-6839 |
| chr2-83300748-INS-6418 | chr2-83300749-INS-6418 |
| chr2-83332152-INS-266 | chr2-83332152-INS-266 |
| chr2-83365698-INS-52 | chr2-83365698-INS-54 |
| chr2-83387047-INS-212 | chr2-83387047-INS-212 |
| chr2-83463165-INS-97 | chr2-83463161-INS-80 |
| chr2-83465392-INS-1888 | chr2-83465393-INS-1888 |
| chr2-83507208-INS-7087 | chr2-83507209-INS-7087 |
| chr2-83512102-INS-206 | chr2-83512102-INS-206 |
| chr2-83513024-INS-448 | chr2-83513024-INS-448 |
| chr2-83520534-INS-194 | chr2-83520534-INS-194 |
| chr2-83522212-INS-470 | chr2-83522212-INS-468 |
| chr2-83528514-INS-75 | chr2-83528514-INS-62 |
| chr2-83536033-INS-5406 | chr2-83536034-INS-5406 |
| chr2-83568523-INS-171 | chr2-83568560-INS-167 |
| chr2-83582299-INS-6315 | chr2-83582300-INS-6316 |
| chr2-83590070-INS-498 | chr2-83590070-INS-498 |
| chr2-83627888-INS-51 | chr2-83627888-INS-51 |
| chr2-83639093-INS-131 | chr2-83639091-INS-132 |
| chr2-83656388-INS-108 | chr2-83656388-INS-110 |
| chr2-83666610-INS-119 | chr2-83666610-INS-120 |
| chr2-83677841-INS-180 | chr2-83677841-INS-180 |
| chr2-83685542-INS-136 | chr2-83685541-INS-137 |
| chr2-83704446-INS-69 | chr2-83704442-INS-73 |
| chr2-83895147-INS-1988 | chr2-83895148-INS-1999 |
| chr2-83931901-INS-69 | chr2-83931901-INS-65 |
| chr2-83942248-INS-52 | chr2-83942248-INS-54 |
| chr2-84002422-INS-391 | chr2-84002422-INS-392 |
| chr2-84006399-INS-78 | chr2-84006399-INS-50 |
| chr2-84012745-INS-205 | chr2-84012745-INS-205 |
| chr2-84015923-INS-116 | chr2-84015923-INS-116 |
| chr2-84059064-INS-5552 | chr2-84059065-INS-5552 |
| chr2-84069299-INS-1520 | chr2-84069300-INS-1520 |
| chr2-84076043-INS-204 | chr2-84076043-INS-204 |
| chr2-84089990-INS-1893 | chr2-84089991-INS-1893 |
| chr2-84094992-INS-1237 | chr2-84094993-INS-1237 |
| chr2-84094993-INS-122 | chr2-84094993-INS-122 |
| chr2-84097408-INS-1384 | chr2-84097409-INS-1384 |
| chr2-84118387-INS-927 | chr2-84118387-INS-936 |
| chr2-84142887-INS-75 | chr2-84142887-INS-75 |
| chr2-84187960-INS-59 | chr2-84187958-INS-67 |
| chr2-84192359-INS-57 | chr2-84192358-INS-53 |
| chr2-84210368-INS-280 | chr2-84210368-INS-280 |
| chr2-84229575-INS-198 | chr2-84229574-INS-201 |
| chr2-84235549-INS-105 | chr2-84235568-INS-90 |
| chr2-84243952-INS-6797 | chr2-84243953-INS-6796 |
| chr2-84251607-INS-124 | chr2-84251607-INS-134 |
| chr2-84259528-INS-6682 | chr2-84259529-INS-6682 |
| chr2-84259696-INS-517 | chr2-84259696-INS-517 |
| chr2-84333004-INS-206 | chr2-84333004-INS-210 |
| chr2-84342994-INS-198 | chr2-84342994-INS-197 |
| chr2-84350923-INS-87 | chr2-84350923-INS-87 |
| chr2-84354440-INS-181 | chr2-84354440-INS-172 |
| chr2-84356931-INS-198 | chr2-84356931-INS-198 |
| chr2-84390119-INS-58 | chr2-84390119-INS-58 |
| chr2-84403816-INS-68 | chr2-84403816-INS-54 |
| chr2-84407025-INS-132 | chr2-84407025-INS-132 |
| chr2-84425107-INS-180 | chr2-84425107-INS-175 |

| | |
|---|---|
| chr2-84530602-INS-183 | chr2-84530602-INS-183 |
| chr2-84548953-INS-85 | chr2-84548996-INS-85 |
| chr2-84552246-INS-51 | chr2-84552246-INS-51 |
| chr2-84601535-INS-56 | chr2-84601527-INS-64 |
| chr2-84610832-INS-60 | chr2-84610832-INS-60 |
| chr2-84616433-INS-71 | chr2-84616417-INS-58 |
| chr2-84643780-INS-194 | chr2-84643779-INS-194 |
| chr2-84778247-INS-101 | chr2-84778247-INS-101 |
| chr2-84783470-INS-395 | chr2-84783470-INS-430 |
| chr2-84787147-INS-145 | chr2-84787147-INS-146 |
| chr2-84789582-INS-259 | chr2-84789582-INS-260 |
| chr2-84791411-INS-78 | chr2-84791411-INS-95 |
| chr2-84816185-INS-1441 | chr2-84816186-INS-1441 |
| chr2-84821723-INS-341 | chr2-84821723-INS-342 |
| chr2-84837816-INS-252 | chr2-84837817-INS-241 |
| chr2-84926703-INS-54 | chr2-84926664-INS-50 |
| chr2-84931247-INS-380 | chr2-84931247-INS-380 |
| chr2-84950711-INS-52 | chr2-84950711-INS-52 |
| chr2-85011694-INS-84 | chr2-85011694-INS-86 |
| chr2-85030289-INS-112 | chr2-85030289-INS-114 |
| chr2-85036681-INS-53 | chr2-85036691-INS-71 |
| chr2-85039619-INS-158 | chr2-85039617-INS-160 |
| chr2-85057694-INS-4658 | chr2-85057695-INS-4658 |
| chr2-85072369-INS-60 | chr2-85072357-INS-68 |
| chr2-85080199-INS-104 | chr2-85080199-INS-104 |
| chr2-85099494-INS-544 | chr2-85099494-INS-520 |
| chr2-85131428-INS-176 | chr2-85131428-INS-176 |
| chr2-85133377-INS-87 | chr2-85133377-INS-91 |
| chr2-85152668-INS-102 | chr2-85152668-INS-102 |
| chr2-85172143-INS-2686 | chr2-85172144-INS-2686 |
| chr2-85172547-INS-140 | chr2-85172543-INS-115 |
| chr2-85181460-INS-154 | chr2-85181460-INS-155 |
| chr2-85242045-INS-176 | chr2-85242046-INS-176 |
| chr2-85259440-INS-900 | chr2-85259442-INS-901 |
| chr2-85284585-INS-6420 | chr2-85284586-INS-6420 |
| chr2-85288184-INS-95 | chr2-85288181-INS-68 |
| chr2-85289994-INS-3070 | chr2-85289995-INS-3033 |
| chr2-85292244-INS-2910 | chr2-85292245-INS-2910 |
| chr2-85338263-INS-78 | chr2-85338263-INS-78 |
| chr2-85340041-INS-738 | chr2-85340041-INS-738 |
| chr2-85354762-INS-219 | chr2-85354754-INS-233 |
| chr2-85371801-INS-474 | chr2-85371801-INS-474 |
| chr2-85389397-INS-144 | chr2-85389397-INS-144 |
| chr2-85389849-INS-2709 | chr2-85389850-INS-2718 |
| chr2-85406819-INS-51 | chr2-85406819-INS-51 |
| chr2-85437422-INS-54 | chr2-85437422-INS-56 |
| chr2-85441806-INS-130 | chr2-85441806-INS-130 |
| chr2-85452398-INS-587 | chr2-85452398-INS-587 |
| chr2-85482022-INS-87 | chr2-85482022-INS-74 |
| chr2-85492415-INS-6467 | chr2-85492416-INS-6467 |
| chr2-85494965-INS-58 | chr2-85494959-INS-68 |
| chr2-85562471-INS-78 | chr2-85562471-INS-78 |
| chr2-85617082-INS-215 | chr2-85617082-INS-215 |
| chr2-85647496-INS-75 | chr2-85647496-INS-75 |
| chr2-85650570-INS-402 | chr2-85650570-INS-402 |
| chr2-85730150-INS-671 | chr2-85730150-INS-671 |
| chr2-85780570-INS-88 | chr2-85780562-INS-64 |
| chr2-85842962-INS-569 | chr2-85842963-INS-569 |
| chr2-85856474-INS-104 | chr2-85856474-INS-104 |
| chr2-85868152-INS-2213 | chr2-85868153-INS-2213 |
| chr2-85905403-INS-5197 | chr2-85905404-INS-5153 |
| chr2-85921864-INS-3764 | chr2-85921865-INS-3765 |
| chr2-85944392-INS-852 | chr2-85944392-INS-852 |
| chr2-85983380-INS-485 | chr2-85983380-INS-516 |
| chr2-85985905-INS-946 | chr2-85985905-INS-946 |
| chr2-85992126-INS-4312 | chr2-85992127-INS-4302 |
| chr2-86004051-INS-100 | chr2-86004047-INS-56 |
| chr2-86029646-INS-86 | chr2-86029646-INS-86 |
| chr2-86061198-INS-5315 | chr2-86061199-INS-5315 |
| chr2-86151261-INS-6710 | chr2-86151262-INS-6710 |
| chr2-86219628-INS-55 | chr2-86219628-INS-55 |
| chr2-86232144-INS-637 | chr2-86232144-INS-637 |
| chr2-86291994-INS-869 | chr2-86291994-INS-869 |
| chr2-86314955-INS-363 | chr2-86314955-INS-363 |
| chr2-86327414-INS-61 | chr2-86327414-INS-61 |
| chr2-86380017-INS-492 | chr2-86380017-INS-492 |
| chr2-86399308-INS-52 | chr2-86399308-INS-56 |

| | |
|---|---|
| chr2-86404441-INS-7100 | chr2-86404442-INS-7102 |
| chr2-86405406-INS-126 | chr2-86405405-INS-133 |
| chr2-86430646-INS-1848 | chr2-86430647-INS-1848 |
| chr2-86440674-INS-58 | chr2-86440674-INS-58 |
| chr2-86442560-INS-53 | chr2-86442560-INS-53 |
| chr2-86448526-INS-53 | chr2-86448526-INS-53 |
| chr2-86481058-INS-6790 | chr2-86481059-INS-6790 |
| chr2-86493162-INS-1799 | chr2-86493163-INS-1799 |
| chr2-86504484-INS-6451 | chr2-86504485-INS-6454 |
| chr2-86505023-INS-600 | chr2-86505023-INS-600 |
| chr2-86545206-INS-857 | chr2-86545206-INS-857 |
| chr2-86560863-INS-2215 | chr2-86560864-INS-2215 |
| chr2-86570680-INS-80 | chr2-86570680-INS-82 |
| chr2-86615182-INS-261 | chr2-86615182-INS-261 |
| chr2-86618363-INS-57 | chr2-86618363-INS-57 |
| chr2-86673202-INS-208 | chr2-86673202-INS-208 |
| chr2-86726742-INS-88 | chr2-86726742-INS-84 |
| chr2-86744166-INS-8270 | chr2-86744167-INS-8270 |
| chr2-86744902-INS-54 | chr2-86744902-INS-50 |
| chr2-86811524-INS-118 | chr2-86811524-INS-118 |
| chr2-86837657-INS-60 | chr2-86837657-INS-64 |
| chr2-86839378-INS-56 | chr2-86839378-INS-60 |
| chr2-86887901-INS-4428 | chr2-86887902-INS-4428 |
| chr2-86891238-INS-1493 | chr2-86891239-INS-1493 |
| chr2-86898035-INS-6631 | chr2-86898036-INS-6632 |
| chr2-86934336-INS-864 | chr2-86934336-INS-865 |
| chr2-87024435-INS-1677 | chr2-87024436-INS-1679 |
| chr2-87040670-INS-5143 | chr2-87040671-INS-5143 |
| chr2-87045694-INS-54 | chr2-87045701-INS-94 |
| chr2-87172596-INS-73 | chr2-87172605-INS-58 |
| chr2-87193645-INS-156 | chr2-87193645-INS-156 |
| chr2-87207756-INS-64 | chr2-87207756-INS-64 |
| chr2-87225360-INS-100 | chr2-87225360-INS-100 |
| chr2-87226710-INS-2794 | chr2-87226711-INS-2779 |
| chr2-87228542-INS-86 | chr2-87228540-INS-64 |
| chr2-87232275-INS-69 | chr2-87232275-INS-69 |
| chr2-87243339-INS-82 | chr2-87243339-INS-82 |
| chr2-87246704-INS-54 | chr2-87246654-INS-70 |
| chr2-87251332-INS-226 | chr2-87251332-INS-226 |
| chr2-87285884-INS-201 | chr2-87285894-INS-199 |
| chr2-87323940-INS-165 | chr2-87323940-INS-165 |
| chr2-87329093-INS-253 | chr2-87329093-INS-253 |
| chr2-87342925-INS-63 | chr2-87342925-INS-60 |
| chr2-87359787-INS-331 | chr2-87359788-INS-331 |
| chr2-87382209-INS-411 | chr2-87382209-INS-411 |
| chr2-87388734-INS-222 | chr2-87388734-INS-222 |
| chr2-87409947-INS-1550 | chr2-87409936-INS-1550 |
| chr2-87411946-INS-197 | chr2-87411948-INS-197 |
| chr2-87458106-INS-66 | chr2-87458106-INS-72 |
| chr2-87458114-INS-6190 | chr2-87458115-INS-6190 |
| chr2-87471486-INS-1393 | chr2-87471487-INS-1393 |
| chr2-87490597-INS-681 | chr2-87490597-INS-681 |
| chr2-87591325-INS-6797 | chr2-87591326-INS-6797 |
| chr2-87621054-INS-201 | chr2-87621054-INS-201 |
| chr2-87631415-INS-52 | chr2-87631417-INS-52 |
| chr2-87645556-INS-271 | chr2-87645556-INS-271 |
| chr2-87646574-INS-759 | chr2-87646574-INS-759 |
| chr2-87676249-INS-703 | chr2-87676247-INS-702 |
| chr2-87691392-INS-350 | chr2-87691392-INS-350 |
| chr2-87782157-INS-74 | chr2-87782150-INS-68 |
| chr2-87784041-INS-110 | chr2-87784041-INS-110 |
| chr2-87789889-INS-160 | chr2-87789889-INS-160 |
| chr2-87795540-INS-63 | chr2-87795540-INS-63 |
| chr2-87799517-INS-4610 | chr2-87799518-INS-4610 |
| chr2-87807520-INS-149 | chr2-87807520-INS-149 |
| chr2-87845423-INS-84 | chr2-87845423-INS-84 |
| chr2-87902443-INS-204 | chr2-87902443-INS-204 |
| chr2-87904157-INS-4507 | chr2-87904158-INS-4507 |
| chr2-87942963-INS-446 | chr2-87942963-INS-446 |
| chr2-87997373-INS-1905 | chr2-87997374-INS-1908 |
| chr2-88064022-INS-828 | chr2-88064022-INS-828 |
| chr2-88071781-INS-212 | chr2-88071781-INS-212 |
| chr2-88108547-INS-2125 | chr2-88108548-INS-2125 |
| chr2-88126944-INS-7340 | chr2-88126945-INS-7341 |
| chr2-88150536-INS-6440 | chr2-88150537-INS-6440 |
| chr2-88157366-INS-202 | chr2-88157366-INS-202 |
| chr2-88158949-INS-61 | chr2-88158981-INS-55 |

| | |
|---|---|
| chr2-88226404-INS-6433 | chr2-88226405-INS-6433 |
| chr2-88227534-INS-84 | chr2-88227535-INS-72 |
| chr2-88233469-INS-61 | chr2-88233455-INS-75 |
| chr2-88326526-INS-7121 | chr2-88326527-INS-7121 |
| chr2-88335324-INS-552 | chr2-88335315-INS-552 |
| chr2-88336788-INS-59 | chr2-88336788-INS-59 |
| chr2-88358396-INS-587 | chr2-88358396-INS-589 |
| chr2-88367117-INS-52 | chr2-88367117-INS-56 |
| chr2-88386688-INS-58 | chr2-88386688-INS-54 |
| chr2-88465892-INS-503 | chr2-88465892-INS-503 |
| chr2-88469123-INS-89 | chr2-88469123-INS-93 |
| chr2-88493086-INS-440 | chr2-88493087-INS-426 |
| chr2-88574843-INS-1490 | chr2-88574844-INS-1490 |
| chr2-88601851-INS-216 | chr2-88601851-INS-216 |
| chr2-88612368-INS-117 | chr2-88612368-INS-117 |
| chr2-88673207-INS-60 | chr2-88673208-INS-60 |
| chr2-88682219-INS-7049 | chr2-88682220-INS-7049 |
| chr2-88708955-INS-88 | chr2-88708955-INS-96 |
| chr2-88710260-INS-128 | chr2-88710260-INS-130 |
| chr2-88716005-INS-50 | chr2-88716013-INS-64 |
| chr2-88729049-INS-75 | chr2-88729050-INS-75 |
| chr2-88735187-INS-1605 | chr2-88735188-INS-1604 |
| chr2-88738457-INS-613 | chr2-88738457-INS-613 |
| chr2-88750091-INS-2058 | chr2-88750092-INS-2058 |
| chr2-88750690-INS-7926 | chr2-88750691-INS-7926 |
| chr2-88757583-INS-502 | chr2-88757583-INS-502 |
| chr2-88770672-INS-84 | chr2-88770672-INS-84 |
| chr2-88851708-INS-1070 | chr2-88851709-INS-1070 |
| chr2-88856178-INS-70 | chr2-88856178-INS-60 |
| chr2-88917943-INS-6648 | chr2-88917944-INS-6647 |
| chr2-88923478-INS-1156 | chr2-88923479-INS-1147 |
| chr2-88982996-INS-75 | chr2-88982958-INS-72 |
| chr2-88985225-INS-99 | chr2-88985225-INS-100 |
| chr2-88994213-INS-5240 | chr2-88994214-INS-5243 |
| chr2-89025504-INS-296 | chr2-89025504-INS-297 |
| chr2-89034542-INS-1856 | chr2-89034545-INS-1853 |
| chr2-89080475-INS-201 | chr2-89080470-INS-202 |
| chr2-89144360-INS-91 | chr2-89144359-INS-59 |
| chr2-89160696-INS-5273 | chr2-89160697-INS-5273 |
| chr2-89202369-INS-107 | chr2-89202388-INS-109 |
| chr2-89219759-INS-632 | chr2-89219759-INS-632 |
| chr2-89235718-INS-6732 | chr2-89235719-INS-6734 |
| chr2-89306010-INS-181 | chr2-89306008-INS-184 |
| chr2-89332802-INS-5344 | chr2-89332803-INS-5344 |
| chr2-89371267-INS-4107 | chr2-89371268-INS-4107 |
| chr2-89376481-INS-155 | chr2-89376481-INS-155 |
| chr2-89398635-INS-415 | chr2-89398635-INS-415 |
| chr2-89453391-INS-52 | chr2-89453377-INS-54 |
| chr2-89456071-INS-360 | chr2-89456066-INS-363 |
| chr2-89470206-INS-7123 | chr2-89470207-INS-7124 |
| chr2-89481798-INS-390 | chr2-89481798-INS-390 |
| chr2-89536625-INS-4337 | chr2-89536626-INS-4337 |
| chr2-89557372-INS-2806 | chr2-89557373-INS-2806 |
| chr2-89626361-INS-184 | chr2-89626361-INS-184 |
| chr2-89669988-INS-78 | chr2-89669988-INS-56 |
| chr2-89690724-INS-56 | chr2-89690724-INS-56 |
| chr2-89695415-INS-372 | chr2-89695415-INS-372 |
| chr2-89714295-INS-221 | chr2-89714295-INS-221 |
| chr2-89747076-INS-506 | chr2-89747076-INS-506 |
| chr2-89771112-INS-462 | chr2-89771112-INS-462 |
| chr2-89821232-INS-87 | chr2-89821232-INS-93 |
| chr2-89835610-INS-465 | chr2-89835610-INS-465 |
| chr2-89838838-INS-324 | chr2-89838838-INS-324 |
| chr2-89878334-INS-105 | chr2-89878334-INS-112 |
| chr2-89917318-INS-783 | chr2-89917314-INS-774 |
| chr2-89917853-INS-212 | chr2-89917853-INS-212 |
| chr2-90142350-INS-653 | chr2-90142350-INS-635 |
| chr2-90227609-INS-129 | chr2-90227609-INS-137 |
| chr2-90287338-INS-419 | chr2-90287338-INS-420 |
| chr2-90334242-INS-57 | chr2-90334242-INS-65 |
| chr2-90344404-INS-176 | chr2-90344408-INS-170 |
| chr2-90346175-INS-55 | chr2-90346186-INS-66 |
| chr2-90365525-INS-4841 | chr2-90365526-INS-4841 |
| chr2-90382292-INS-2903 | chr2-90382293-INS-2903 |
| chr2-90435313-INS-220 | chr2-90435313-INS-220 |
| chr2-90440772-INS-168 | chr2-90440772-INS-196 |
| chr2-90458920-INS-108 | chr2-90458920-INS-136 |

| | |
|---|---|
| chr2-90479482-INS-498 | chr2-90479482-INS-498 |
| chr2-90485747-INS-204 | chr2-90485744-INS-207 |
| chr2-90502891-INS-162 | chr2-90502891-INS-180 |
| chr2-90570781-INS-328 | chr2-90570781-INS-328 |
| chr2-90586582-INS-214 | chr2-90586582-INS-214 |
| chr2-90603058-INS-177 | chr2-90603058-INS-167 |
| chr2-90647602-INS-5219 | chr2-90647603-INS-5219 |
| chr2-90658030-INS-98 | chr2-90658077-INS-71 |
| chr2-90663678-INS-211 | chr2-90663678-INS-211 |
| chr2-90675409-INS-64 | chr2-90675409-INS-64 |
| chr2-90687883-INS-197 | chr2-90687883-INS-198 |
| chr2-90750131-INS-155 | chr2-90750131-INS-155 |
| chr2-90757885-INS-220 | chr2-90757885-INS-220 |
| chr2-90773522-INS-212 | chr2-90773522-INS-211 |
| chr2-90802298-INS-112 | chr2-90802298-INS-126 |
| chr2-90813945-INS-84 | chr2-90813945-INS-84 |
| chr2-90841451-INS-538 | chr2-90841451-INS-538 |
| chr2-90860474-INS-215 | chr2-90860474-INS-215 |
| chr2-90870195-INS-211 | chr2-90870195-INS-211 |
| chr2-90879000-INS-199 | chr2-90879000-INS-198 |
| chr2-90883939-INS-74 | chr2-90883929-INS-58 |
| chr2-90895917-INS-90 | chr2-90895917-INS-90 |
| chr2-90941479-INS-245 | chr2-90941479-INS-245 |
| chr2-91018437-INS-56 | chr2-91018449-INS-51 |
| chr2-91026340-INS-235 | chr2-91026340-INS-221 |
| chr2-91062224-INS-2990 | chr2-91062225-INS-2990 |
| chr2-91069226-INS-120 | chr2-91069226-INS-120 |
| chr2-91082762-INS-78 | chr2-91082762-INS-78 |
| chr2-91084454-INS-219 | chr2-91084454-INS-218 |
| chr2-91103062-INS-60 | chr2-91103062-INS-85 |
| chr2-91125455-INS-206 | chr2-91125455-INS-207 |
| chr2-91134117-INS-171 | chr2-91134118-INS-189 |
| chr2-91134663-INS-304 | chr2-91134663-INS-304 |
| chr2-91168142-INS-227 | chr2-91168142-INS-229 |
| chr2-91172173-INS-469 | chr2-91172173-INS-469 |
| chr2-91179713-INS-61 | chr2-91179713-INS-61 |
| chr2-91183426-INS-54 | chr2-91183426-INS-56 |
| chr2-91195720-INS-228 | chr2-91195720-INS-227 |
| chr2-91206991-INS-107 | chr2-91206987-INS-130 |
| chr2-91214956-INS-4390 | chr2-91214957-INS-4390 |
| chr2-91220685-INS-72 | chr2-91220681-INS-54 |
| chr2-91249981-INS-222 | chr2-91249981-INS-223 |
| chr2-91302042-INS-218 | chr2-91302042-INS-219 |
| chr2-91316647-INS-262 | chr2-91316647-INS-263 |
| chr2-91338136-INS-98 | chr2-91338150-INS-50 |
| chr2-91352242-INS-68 | chr2-91352238-INS-62 |
| chr2-91353521-INS-80 | chr2-91353521-INS-84 |
| chr2-91366376-INS-186 | chr2-91366376-INS-191 |
| chr2-91369992-INS-228 | chr2-91369992-INS-228 |
| chr2-91371723-INS-6399 | chr2-91371724-INS-6399 |
| chr2-91398093-INS-74 | chr2-91398093-INS-77 |
| chr2-91401340-INS-62 | chr2-91401340-INS-56 |
| chr2-91404100-INS-85 | chr2-91404100-INS-93 |
| chr2-91417216-INS-212 | chr2-91417216-INS-212 |
| chr2-91459877-INS-52 | chr2-91459906-INS-64 |
| chr2-91467715-INS-98 | chr2-91467715-INS-98 |
| chr2-91474547-INS-56 | chr2-91474547-INS-56 |
| chr2-91488253-INS-219 | chr2-91488253-INS-219 |
| chr2-91490079-INS-207 | chr2-91490079-INS-207 |
| chr2-91492960-INS-208 | chr2-91492960-INS-225 |
| chr2-91514276-INS-218 | chr2-91514276-INS-218 |
| chr2-91515974-INS-56 | chr2-91515970-INS-56 |
| chr2-91532290-INS-2079 | chr2-91532291-INS-2079 |
| chr2-91537516-INS-207 | chr2-91537516-INS-207 |
| chr2-91546140-INS-75 | chr2-91546140-INS-76 |
| chr2-91548258-INS-212 | chr2-91548258-INS-213 |
| chr2-91567107-INS-262 | chr2-91567107-INS-263 |
| chr2-91569132-INS-200 | chr2-91569132-INS-200 |
| chr2-91631811-INS-94 | chr2-91631811-INS-94 |
| chr2-91634275-INS-78 | chr2-91634275-INS-78 |
| chr2-91653491-INS-78 | chr2-91653491-INS-78 |
| chr2-91656528-INS-64 | chr2-91656528-INS-53 |
| chr2-91666779-INS-216 | chr2-91666779-INS-216 |
| chr2-91693646-INS-97 | chr2-91693645-INS-126 |
| chr2-91697271-INS-177 | chr2-91697271-INS-174 |
| chr2-91700411-INS-197 | chr2-91700411-INS-197 |
| chr2-91712353-INS-58 | chr2-91712353-INS-52 |

| | |
|---|---|
| chr2-91757348-INS-104 | chr2-91757348-INS-139 |
| chr2-91759182-INS-1864 | chr2-91759183-INS-1864 |
| chr2-91798952-INS-186 | chr2-91798952-INS-186 |
| chr2-91800902-INS-215 | chr2-91800902-INS-214 |
| chr2-91810104-INS-74 | chr2-91810104-INS-78 |
| chr2-91811421-INS-214 | chr2-91811421-INS-214 |
| chr2-91816000-INS-451 | chr2-91816000-INS-457 |
| chr2-91823212-INS-143 | chr2-91823212-INS-187 |
| chr2-91825925-INS-211 | chr2-91825925-INS-223 |
| chr2-91829171-INS-86 | chr2-91829216-INS-58 |
| chr2-91833717-INS-68 | chr2-91833717-INS-60 |
| chr2-91888681-INS-217 | chr2-91888681-INS-217 |
| chr2-91907404-INS-315 | chr2-91907404-INS-315 |
| chr2-91926817-INS-206 | chr2-91926817-INS-206 |
| chr2-91945156-INS-356 | chr2-91945156-INS-363 |
| chr2-91975509-INS-57 | chr2-91975509-INS-57 |
| chr2-91980298-INS-277 | chr2-91980298-INS-277 |
| chr2-91981426-INS-205 | chr2-91981426-INS-206 |
| chr2-91983227-INS-58 | chr2-91983214-INS-57 |
| chr2-92022549-INS-211 | chr2-92022549-INS-211 |
| chr2-92034105-INS-210 | chr2-92034105-INS-210 |
| chr2-92036292-INS-75 | chr2-92036292-INS-77 |
| chr2-92079650-INS-214 | chr2-92079643-INS-217 |
| chr2-92121448-INS-55 | chr2-92121448-INS-100 |
| chr2-92170911-INS-204 | chr2-92170911-INS-204 |
| chr2-92191726-INS-68 | chr2-92191726-INS-56 |
| chr2-92204558-INS-211 | chr2-92204558-INS-210 |
| chr2-92238128-INS-5038 | chr2-92238129-INS-5038 |
| chr2-92248082-INS-51 | chr2-92248082-INS-51 |
| chr2-92266178-INS-84 | chr2-92266178-INS-66 |
| chr2-92276154-INS-56 | chr2-92276154-INS-58 |
| chr2-92299864-INS-149 | chr2-92299864-INS-155 |
| chr2-92317066-INS-342 | chr2-92317066-INS-342 |
| chr2-92345000-INS-78 | chr2-92345000-INS-90 |
| chr2-92374508-INS-101 | chr2-92374508-INS-130 |
| chr2-92459160-INS-84 | chr2-92459160-INS-121 |
| chr2-92490703-INS-57 | chr2-92490705-INS-59 |
| chr2-92539135-INS-72 | chr2-92539135-INS-72 |
| chr2-92546053-INS-65 | chr2-92546045-INS-72 |
| chr2-92550217-INS-52 | chr2-92550217-INS-52 |
| chr2-92582792-INS-52 | chr2-92582792-INS-52 |
| chr2-92624345-INS-101 | chr2-92624345-INS-101 |
| chr2-92628339-INS-136 | chr2-92628351-INS-116 |
| chr2-92636087-INS-1889 | chr2-92636088-INS-1889 |
| chr2-92684496-INS-498 | chr2-92684488-INS-490 |
| chr2-92695930-INS-66 | chr2-92695930-INS-90 |
| chr2-92712351-INS-60 | chr2-92712351-INS-60 |
| chr2-92722283-INS-701 | chr2-92722283-INS-717 |
| chr2-92748675-INS-178 | chr2-92748675-INS-179 |
| chr2-92774691-INS-96 | chr2-92774679-INS-80 |
| chr2-92777329-INS-885 | chr2-92777329-INS-885 |
| chr2-92788505-INS-203 | chr2-92788505-INS-201 |
| chr2-92791426-INS-64 | chr2-92791427-INS-64 |
| chr2-92853970-INS-62 | chr2-92853972-INS-52 |
| chr2-92892205-INS-1496 | chr2-92892206-INS-1494 |
| chr2-92898824-INS-114 | chr2-92898824-INS-75 |
| chr2-92900939-INS-59 | chr2-92900966-INS-61 |
| chr2-92951413-INS-64 | chr2-92951413-INS-64 |
| chr2-92959445-INS-60 | chr2-92959445-INS-60 |
| chr2-92961066-INS-216 | chr2-92961059-INS-226 |
| chr2-92986058-INS-114 | chr2-92986058-INS-114 |
| chr2-92993517-INS-58 | chr2-92993517-INS-58 |
| chr2-93011335-INS-55 | chr2-93011335-INS-63 |
| chr2-93094609-INS-6969 | chr2-93094610-INS-6969 |
| chr2-93141484-INS-92 | chr2-93141485-INS-91 |
| chr2-93177824-INS-66 | chr2-93177824-INS-66 |
| chr2-93190143-INS-68 | chr2-93190143-INS-68 |
| chr2-93192590-INS-702 | chr2-93192590-INS-702 |
| chr2-93206565-INS-52 | chr2-93206565-INS-52 |
| chr2-93209590-INS-82 | chr2-93209567-INS-110 |
| chr2-93245037-INS-197 | chr2-93245037-INS-197 |
| chr2-93308338-INS-164 | chr2-93308338-INS-164 |
| chr2-93330587-INS-61 | chr2-93330587-INS-54 |
| chr2-93334748-INS-51 | chr2-93334748-INS-67 |
| chr2-93342195-INS-186 | chr2-93342195-INS-186 |
| chr2-93359145-INS-463 | chr2-93359145-INS-463 |
| chr2-93382433-INS-286 | chr2-93382433-INS-286 |

| | |
|---|---|
| chr2-93390521-INS-73 | chr2-93390521-INS-73 |
| chr2-93408722-INS-92 | chr2-93408722-INS-92 |
| chr2-93462932-INS-95 | chr2-93462932-INS-98 |
| chr2-93499514-INS-58 | chr2-93499514-INS-58 |
| chr2-93505196-INS-69 | chr2-93505160-INS-69 |
| chr2-93513755-INS-76 | chr2-93513795-INS-60 |
| chr2-93514904-INS-108 | chr2-93514904-INS-68 |
| chr2-93562440-INS-206 | chr2-93562440-INS-206 |
| chr2-93594635-INS-52 | chr2-93594635-INS-52 |
| chr2-93611592-INS-90 | chr2-93611592-INS-90 |
| chr2-93620235-INS-56 | chr2-93620236-INS-58 |
| chr2-93624328-INS-56 | chr2-93624327-INS-56 |
| chr2-93646661-INS-174 | chr2-93646661-INS-174 |
| chr2-93669711-INS-892 | chr2-93669711-INS-895 |
| chr2-93675956-INS-144 | chr2-93675956-INS-144 |
| chr2-93693607-INS-693 | chr2-93693607-INS-693 |
| chr2-93700640-INS-69 | chr2-93700641-INS-51 |
| chr2-93705371-INS-60 | chr2-93705371-INS-52 |
| chr2-93706804-INS-58 | chr2-93706804-INS-68 |
| chr2-93729226-INS-415 | chr2-93729226-INS-418 |
| chr2-93735571-INS-80 | chr2-93735570-INS-100 |
| chr2-93750098-INS-74 | chr2-93750130-INS-50 |
| chr2-93762547-INS-90 | chr2-93762547-INS-90 |
| chr2-93778461-INS-85 | chr2-93778461-INS-119 |
| chr2-93789602-INS-195 | chr2-93789602-INS-195 |
| chr2-93817543-INS-510 | chr2-93817543-INS-510 |
| chr2-93866900-INS-58 | chr2-93866900-INS-58 |
| chr2-93869927-INS-60 | chr2-93869955-INS-61 |
| chr2-93888132-INS-6902 | chr2-93888133-INS-6902 |
| chr2-93894513-INS-81 | chr2-93894513-INS-97 |
| chr2-93898035-INS-125 | chr2-93898035-INS-125 |
| chr2-93905916-INS-65 | chr2-93905931-INS-52 |
| chr2-93926570-INS-116 | chr2-93926543-INS-110 |
| chr2-93981168-INS-1907 | chr2-93981169-INS-1907 |
| chr2-93990804-INS-66 | chr2-93990804-INS-68 |
| chr2-94002432-INS-165 | chr2-94002433-INS-171 |
| chr2-94048870-INS-61 | chr2-94048858-INS-89 |
| chr2-94061862-INS-7076 | chr2-94061863-INS-7076 |
| chr2-94061893-INS-214 | chr2-94061893-INS-214 |
| chr2-94080197-INS-354 | chr2-94080191-INS-360 |
| chr2-94114216-INS-90 | chr2-94114195-INS-90 |
| chr2-94148055-INS-217 | chr2-94148055-INS-232 |
| chr2-94155288-INS-61 | chr2-94155288-INS-61 |
| chr2-94166105-INS-176 | chr2-94166105-INS-176 |
| chr2-94168428-INS-390 | chr2-94168428-INS-390 |
| chr2-94174949-INS-55 | chr2-94174949-INS-55 |
| chr2-94177033-INS-442 | chr2-94177033-INS-445 |
| chr2-94211643-INS-188 | chr2-94211643-INS-190 |
| chr2-94222049-INS-66 | chr2-94222049-INS-66 |
| chr2-94223720-INS-7100 | chr2-94223721-INS-7100 |
| chr2-94263883-INS-9053 | chr2-94263884-INS-9053 |
| chr2-94270875-INS-66 | chr2-94270875-INS-66 |
| chr2-94291714-INS-66 | chr2-94291728-INS-66 |
| chr2-94300188-INS-354 | chr2-94300188-INS-354 |
| chr2-94307538-INS-225 | chr2-94307544-INS-247 |
| chr2-94310802-INS-182 | chr2-94310802-INS-185 |
| chr2-94317504-INS-59 | chr2-94317504-INS-59 |
| chr2-94324616-INS-76 | chr2-94324616-INS-76 |
| chr2-94335892-INS-116 | chr2-94335893-INS-85 |
| chr2-94343358-INS-55 | chr2-94343341-INS-52 |
| chr2-94363097-INS-62 | chr2-94363097-INS-76 |
| chr2-94383916-INS-89 | chr2-94383912-INS-106 |
| chr2-94416396-INS-86 | chr2-94416396-INS-88 |
| chr2-94510833-INS-6323 | chr2-94510834-INS-6325 |
| chr2-94519225-INS-392 | chr2-94519225-INS-392 |
| chr2-94527258-INS-56 | chr2-94527258-INS-52 |
| chr2-94528879-INS-122 | chr2-94528863-INS-136 |
| chr2-94538140-INS-3261 | chr2-94538141-INS-3261 |
| chr2-94549367-INS-76 | chr2-94549372-INS-68 |
| chr2-94580202-INS-58 | chr2-94580202-INS-62 |
| chr2-94591211-INS-168 | chr2-94591211-INS-168 |
| chr2-94598869-INS-55 | chr2-94598869-INS-65 |
| chr2-94601797-INS-1883 | chr2-94601798-INS-1883 |
| chr2-94604027-INS-60 | chr2-94604027-INS-58 |
| chr2-94611031-INS-1961 | chr2-94611032-INS-1962 |
| chr2-94615121-INS-7113 | chr2-94615122-INS-7113 |
| chr2-94629273-INS-54 | chr2-94629273-INS-50 |

| | |
|---|---|
| chr2-94652287-INS-699 | chr2-94652287-INS-699 |
| chr2-94655366-INS-259 | chr2-94655366-INS-252 |
| chr2-94664550-INS-6811 | chr2-94664551-INS-6811 |
| chr2-94675305-INS-132 | chr2-94675305-INS-136 |
| chr2-94678872-INS-6802 | chr2-94678873-INS-6802 |
| chr2-94679697-INS-110 | chr2-94679698-INS-87 |
| chr2-94692740-INS-722 | chr2-94692740-INS-722 |
| chr2-94709555-INS-60 | chr2-94709555-INS-58 |
| chr2-94740929-INS-192 | chr2-94740929-INS-192 |
| chr2-94762155-INS-54 | chr2-94762161-INS-54 |
| chr2-94766939-INS-7290 | chr2-94766940-INS-7290 |
| chr2-94776780-INS-95 | chr2-94776772-INS-119 |
| chr2-94783747-INS-2900 | chr2-94783748-INS-2900 |
| chr2-94785062-INS-2075 | chr2-94785063-INS-2075 |
| chr2-94789408-INS-346 | chr2-94789408-INS-344 |
| chr2-94836619-INS-3764 | chr2-94836620-INS-3758 |
| chr2-94902027-INS-390 | chr2-94902027-INS-390 |
| chr2-94915677-INS-312 | chr2-94915677-INS-312 |
| chr2-94969070-INS-76 | chr2-94969069-INS-78 |
| chr2-95010262-INS-191 | chr2-95010262-INS-203 |
| chr2-95021682-INS-1517 | chr2-95021683-INS-1517 |
| chr2-95067438-INS-102 | chr2-95067438-INS-96 |
| chr2-95097608-INS-56 | chr2-95097608-INS-56 |
| chr2-95151398-INS-55 | chr2-95151398-INS-54 |
| chr2-95185511-INS-9811 | chr2-95185512-INS-9826 |
| chr2-95216730-INS-64 | chr2-95216726-INS-72 |
| chr2-95244611-INS-636 | chr2-95244612-INS-636 |
| chr2-95263017-INS-73 | chr2-95263017-INS-73 |
| chr2-95285480-INS-56 | chr2-95285512-INS-68 |
| chr2-95350925-INS-7343 | chr2-95350926-INS-7346 |
| chr2-95371195-INS-60 | chr2-95371195-INS-60 |
| chr2-95383925-INS-225 | chr2-95383925-INS-225 |
| chr2-95386604-INS-204 | chr2-95386604-INS-204 |
| chr2-95434257-INS-328 | chr2-95434257-INS-328 |
| chr2-95448446-INS-99 | chr2-95448446-INS-99 |
| chr2-95458327-INS-65 | chr2-95458328-INS-75 |
| chr2-95476296-INS-498 | chr2-95476296-INS-498 |
| chr2-95486209-INS-580 | chr2-95486209-INS-580 |
| chr2-95493291-INS-69 | chr2-95493289-INS-69 |
| chr2-95505438-INS-6591 | chr2-95505439-INS-6589 |
| chr2-95554075-INS-4320 | chr2-95554076-INS-4320 |
| chr2-95555059-INS-88 | chr2-95555059-INS-64 |
| chr2-95561955-INS-52 | chr2-95561955-INS-50 |
| chr2-95583924-INS-51 | chr2-95583924-INS-51 |
| chr2-95590047-INS-77 | chr2-95590047-INS-77 |
| chr2-95592208-INS-52 | chr2-95592207-INS-58 |
| chr2-95603559-INS-61 | chr2-95603559-INS-61 |
| chr2-95715202-INS-538 | chr2-95715202-INS-537 |
| chr2-95718069-INS-54 | chr2-95718108-INS-64 |
| chr2-95720764-INS-530 | chr2-95720764-INS-530 |
| chr2-95736829-INS-56 | chr2-95736829-INS-52 |
| chr2-95741503-INS-225 | chr2-95741503-INS-224 |
| chr2-95766657-INS-6644 | chr2-95766658-INS-6644 |
| chr2-95775480-INS-1335 | chr2-95775481-INS-1335 |
| chr2-95783412-INS-92 | chr2-95783412-INS-92 |
| chr2-95799594-INS-6452 | chr2-95799595-INS-6452 |
| chr2-95805416-INS-62 | chr2-95805416-INS-64 |
| chr2-95832952-INS-274 | chr2-95832952-INS-274 |
| chr2-95833855-INS-1871 | chr2-95833856-INS-1871 |
| chr2-95836382-INS-1829 | chr2-95836383-INS-1829 |
| chr2-95857154-INS-425 | chr2-95857154-INS-426 |
| chr2-95863457-INS-74 | chr2-95863457-INS-50 |
| chr2-95867224-INS-59 | chr2-95867224-INS-72 |
| chr2-95872423-INS-69 | chr2-95872423-INS-69 |
| chr2-95917417-INS-788 | chr2-95917417-INS-788 |
| chr2-95937211-INS-54 | chr2-95937211-INS-62 |
| chr2-95951439-INS-67 | chr2-95951439-INS-67 |
| chr2-95961972-INS-7312 | chr2-95961973-INS-7312 |
| chr2-95969654-INS-202 | chr2-95969654-INS-202 |
| chr2-95978660-INS-2509 | chr2-95978661-INS-2509 |
| chr2-95979000-INS-332 | chr2-95979000-INS-332 |
| chr2-96033522-INS-502 | chr2-96033522-INS-502 |
| chr2-96086931-INS-173 | chr2-96086931-INS-173 |
| chr2-96101290-INS-1756 | chr2-96101291-INS-1755 |
| chr2-96112626-INS-7485 | chr2-96112627-INS-7488 |
| chr2-96138335-INS-59 | chr2-96138308-INS-69 |
| chr2-96186430-INS-941 | chr2-96186430-INS-923 |

| | |
|---|---|
| chr2-96220690-INS-107 | chr2-96220690-INS-107 |
| chr2-96241859-INS-65 | chr2-96241859-INS-70 |
| chr2-96246655-INS-1837 | chr2-96246656-INS-1839 |
| chr2-96250580-INS-97 | chr2-96250568-INS-104 |
| chr2-96270830-INS-942 | chr2-96270830-INS-942 |
| chr2-96271062-INS-75 | chr2-96271062-INS-75 |
| chr2-96279371-INS-519 | chr2-96279371-INS-505 |
| chr2-96289645-INS-812 | chr2-96289645-INS-816 |
| chr2-96303992-INS-51 | chr2-96303992-INS-61 |
| chr2-96345865-INS-280 | chr2-96345865-INS-281 |
| chr2-96387077-INS-494 | chr2-96387077-INS-493 |
| chr2-96393429-INS-2628 | chr2-96393430-INS-2629 |
| chr2-96406892-INS-1384 | chr2-96406893-INS-1384 |
| chr2-96474933-INS-52 | chr2-96474933-INS-60 |
| chr2-96492242-INS-4379 | chr2-96492243-INS-4379 |
| chr2-96529010-INS-55 | chr2-96529010-INS-65 |
| chr2-96538035-INS-312 | chr2-96538035-INS-312 |
| chr2-96553150-INS-68 | chr2-96553150-INS-68 |
| chr2-96564713-INS-227 | chr2-96564713-INS-227 |
| chr2-96567651-INS-549 | chr2-96567651-INS-549 |
| chr2-96571091-INS-65 | chr2-96571091-INS-65 |
| chr2-96574349-INS-50 | chr2-96574349-INS-50 |
| chr2-96577517-INS-94 | chr2-96577517-INS-133 |
| chr2-96599659-INS-430 | chr2-96599659-INS-430 |
| chr2-96642061-INS-50 | chr2-96642061-INS-54 |
| chr2-96648669-INS-216 | chr2-96648669-INS-216 |
| chr2-96654542-INS-6210 | chr2-96654543-INS-6210 |
| chr2-96657502-INS-240 | chr2-96657502-INS-240 |
| chr2-96675199-INS-59 | chr2-96675199-INS-89 |
| chr2-96687136-INS-6691 | chr2-96687137-INS-6691 |
| chr2-96709378-INS-79 | chr2-96709379-INS-127 |
| chr2-96717312-INS-456 | chr2-96717312-INS-456 |
| chr2-96731831-INS-195 | chr2-96731831-INS-192 |
| chr2-96761766-INS-60 | chr2-96761766-INS-52 |
| chr2-96775758-INS-1603 | chr2-96775759-INS-1634 |
| chr2-96783863-INS-524 | chr2-96783863-INS-525 |
| chr2-96823283-INS-58 | chr2-96823283-INS-58 |
| chr2-96824289-INS-1035 | chr2-96824290-INS-1028 |
| chr2-96844370-INS-6373 | chr2-96844371-INS-6374 |
| chr2-96850143-INS-213 | chr2-96850143-INS-213 |
| chr2-96916892-INS-204 | chr2-96916892-INS-216 |
| chr2-96935868-INS-6450 | chr2-96935869-INS-6451 |
| chr2-96966182-INS-90 | chr2-96966174-INS-78 |
| chr2-96967270-INS-92 | chr2-96967270-INS-54 |
| chr2-96977960-INS-76 | chr2-96977960-INS-88 |
| chr2-97000844-INS-2912 | chr2-97000845-INS-2912 |
| chr2-97022574-INS-327 | chr2-97022574-INS-330 |
| chr2-97033229-INS-89 | chr2-97033229-INS-95 |
| chr2-97057647-INS-3188 | chr2-97057648-INS-3188 |
| chr2-97058674-INS-873 | chr2-97058674-INS-873 |
| chr2-97064276-INS-2969 | chr2-97064277-INS-2970 |
| chr2-97080845-INS-74 | chr2-97080845-INS-80 |
| chr2-97085944-INS-122 | chr2-97085944-INS-121 |
| chr2-97113439-INS-625 | chr2-97113439-INS-625 |
| chr2-97149793-INS-52 | chr2-97149793-INS-52 |
| chr2-97159532-INS-1639 | chr2-97159533-INS-1639 |
| chr2-97162861-INS-1479 | chr2-97162862-INS-1479 |
| chr2-97192672-INS-88 | chr2-97192672-INS-88 |
| chr2-97194295-INS-191 | chr2-97194295-INS-191 |
| chr2-97218385-INS-54 | chr2-97218385-INS-60 |
| chr2-97257689-INS-56 | chr2-97257689-INS-56 |
| chr2-97270222-INS-164 | chr2-97270222-INS-164 |
| chr2-97288252-INS-1867 | chr2-97288253-INS-1867 |
| chr2-97292679-INS-81 | chr2-97292679-INS-81 |
| chr2-97304728-INS-57 | chr2-97304728-INS-57 |
| chr2-97351435-INS-450 | chr2-97351435-INS-450 |
| chr2-97357002-INS-123 | chr2-97357002-INS-123 |
| chr2-97367058-INS-81 | chr2-97367058-INS-81 |
| chr2-97371010-INS-85 | chr2-97371010-INS-93 |
| chr2-97373310-INS-1045 | chr2-97373311-INS-1050 |
| chr2-97424927-INS-56 | chr2-97424927-INS-58 |
| chr2-97426439-INS-213 | chr2-97426439-INS-212 |
| chr2-97427619-INS-59 | chr2-97427619-INS-59 |
| chr2-97450650-INS-903 | chr2-97450650-INS-906 |
| chr2-97488221-INS-52 | chr2-97488221-INS-52 |
| chr2-97618123-INS-647 | chr2-97618123-INS-647 |
| chr2-97625575-INS-3290 | chr2-97625576-INS-3288 |

| | |
|---|---|
| chr2-97645842-INS-6378 | chr2-97645843-INS-6378 |
| chr2-97652792-INS-147 | chr2-97652792-INS-144 |
| chr2-97681564-INS-2280 | chr2-97681565-INS-2280 |
| chr2-97692154-INS-4438 | chr2-97692129-INS-4442 |
| chr2-97696507-INS-6494 | chr2-97696508-INS-6495 |
| chr2-97699572-INS-1394 | chr2-97699573-INS-1394 |
| chr2-97723196-INS-52 | chr2-97723196-INS-52 |
| chr2-97724129-INS-52 | chr2-97724130-INS-54 |
| chr2-97747112-INS-122 | chr2-97747141-INS-107 |
| chr2-97757400-INS-63 | chr2-97757401-INS-66 |
| chr2-97766154-INS-223 | chr2-97766154-INS-228 |
| chr2-97824878-INS-173 | chr2-97824878-INS-173 |
| chr2-97849112-INS-7162 | chr2-97849113-INS-7162 |
| chr2-97876373-INS-1860 | chr2-97876374-INS-1860 |
| chr2-97904047-INS-355 | chr2-97904047-INS-363 |
| chr2-97912450-INS-60 | chr2-97912453-INS-63 |
| chr2-97939843-INS-74 | chr2-97939844-INS-54 |
| chr2-97940317-INS-116 | chr2-97940304-INS-87 |
| chr2-97969711-INS-989 | chr2-97969711-INS-963 |
| chr2-97984893-INS-878 | chr2-97984897-INS-881 |
| chr2-98042537-INS-1747 | chr2-98042538-INS-1748 |
| chr2-98050394-INS-92 | chr2-98050394-INS-90 |
| chr2-98051370-INS-541 | chr2-98051370-INS-541 |
| chr2-98059700-INS-708 | chr2-98059700-INS-689 |
| chr2-98063366-INS-1299 | chr2-98063369-INS-1295 |
| chr2-98071185-INS-54 | chr2-98071185-INS-55 |
| chr2-98098094-INS-174 | chr2-98098084-INS-194 |
| chr2-98102092-INS-74 | chr2-98102092-INS-74 |
| chr2-98104141-INS-3671 | chr2-98104142-INS-3671 |
| chr2-98122286-INS-6217 | chr2-98122287-INS-6217 |
| chr2-98183527-INS-72 | chr2-98183527-INS-80 |
| chr2-98196748-INS-60 | chr2-98196748-INS-54 |
| chr2-98297176-INS-71 | chr2-98297176-INS-84 |
| chr2-98299773-INS-218 | chr2-98299773-INS-218 |
| chr2-98307643-INS-71 | chr2-98307643-INS-103 |
| chr2-98314984-INS-108 | chr2-98314984-INS-63 |
| chr2-98336702-INS-3706 | chr2-98336703-INS-3706 |
| chr2-98405285-INS-817 | chr2-98405285-INS-802 |
| chr2-98426778-INS-748 | chr2-98426778-INS-746 |
| chr2-98472973-INS-113 | chr2-98472974-INS-113 |
| chr2-98518880-INS-68 | chr2-98518880-INS-88 |
| chr2-98524982-INS-429 | chr2-98524982-INS-428 |
| chr2-98529962-INS-6913 | chr2-98529963-INS-6912 |
| chr2-98549600-INS-56 | chr2-98549589-INS-68 |
| chr2-98567550-INS-72 | chr2-98567580-INS-88 |
| chr2-98578270-INS-110 | chr2-98578274-INS-106 |
| chr2-98613855-INS-212 | chr2-98613855-INS-212 |
| chr2-98652809-INS-128 | chr2-98652809-INS-128 |
| chr2-98666826-INS-64 | chr2-98666804-INS-57 |
| chr2-98683614-INS-54 | chr2-98683653-INS-50 |
| chr2-98701393-INS-65 | chr2-98701393-INS-65 |
| chr2-98809041-INS-7144 | chr2-98809042-INS-7144 |
| chr2-98819352-INS-287 | chr2-98819352-INS-287 |
| chr2-98829462-INS-54 | chr2-98829462-INS-66 |
| chr2-98857743-INS-118 | chr2-98857743-INS-118 |
| chr2-98905780-INS-79 | chr2-98905780-INS-79 |
| chr2-98915391-INS-139 | chr2-98915390-INS-143 |
| chr2-98927287-INS-5335 | chr2-98927288-INS-5335 |
| chr2-99022555-INS-6974 | chr2-99022556-INS-6972 |
| chr2-99038797-INS-1893 | chr2-99038798-INS-1893 |
| chr2-99098460-INS-119 | chr2-99098465-INS-130 |
| chr2-99142257-INS-59 | chr2-99142257-INS-59 |
| chr2-99161634-INS-341 | chr2-99161634-INS-341 |
| chr2-99175699-INS-7095 | chr2-99175700-INS-7095 |
| chr2-99243648-INS-84 | chr2-99243648-INS-78 |
| chr2-99273422-INS-340 | chr2-99273422-INS-340 |
| chr2-99316269-INS-82 | chr2-99316269-INS-74 |
| chr2-99362770-INS-52 | chr2-99362748-INS-72 |
| chr2-99410369-INS-52 | chr2-99410369-INS-50 |
| chr2-99428749-INS-162 | chr2-99428749-INS-176 |
| chr2-99439921-INS-58 | chr2-99439921-INS-60 |
| chr2-99455294-INS-2322 | chr2-99455295-INS-2322 |
| chr2-99462453-INS-498 | chr2-99462453-INS-498 |
| chr2-99464726-INS-7109 | chr2-99464727-INS-7110 |
| chr2-99473261-INS-407 | chr2-99473261-INS-407 |
| chr2-99481147-INS-223 | chr2-99481147-INS-207 |
| chr2-99487687-INS-75 | chr2-99487687-INS-96 |

| | |
|---|---|
| chr2-99506807-INS-6775 | chr2-99506808-INS-6775 |
| chr2-99527342-INS-451 | chr2-99527337-INS-454 |
| chr2-99529743-INS-87 | chr2-99529729-INS-83 |
| chr2-99547224-INS-66 | chr2-99547224-INS-72 |
| chr2-99557755-INS-376 | chr2-99557751-INS-375 |
| chr2-99580294-INS-5983 | chr2-99580294-INS-5978 |
| chr2-99594265-INS-138 | chr2-99594265-INS-164 |
| chr2-99603072-INS-596 | chr2-99603072-INS-596 |
| chr2-99659733-INS-2203 | chr2-99659734-INS-2223 |
| chr2-99660260-INS-70 | chr2-99660260-INS-70 |
| chr2-99698196-INS-70 | chr2-99698196-INS-70 |
| chr2-99710365-INS-50 | chr2-99710365-INS-62 |
| chr2-99720871-INS-224 | chr2-99720859-INS-230 |
| chr2-99729177-INS-221 | chr2-99729177-INS-221 |
| chr2-99754724-INS-54 | chr2-99754719-INS-60 |
| chr2-99767781-INS-74 | chr2-99767783-INS-72 |
| chr2-99781686-INS-57 | chr2-99781686-INS-57 |
| chr2-99783725-INS-64 | chr2-99783725-INS-58 |
| chr2-99784938-INS-72 | chr2-99784938-INS-64 |
| chr2-99801892-INS-2757 | chr2-99801893-INS-2757 |
| chr2-99801962-INS-1231 | chr2-99801963-INS-1231 |
| chr2-99817441-INS-176 | chr2-99817441-INS-178 |
| chr2-99823077-INS-98 | chr2-99823077-INS-98 |
| chr2-99834961-INS-374 | chr2-99834961-INS-374 |
| chr2-99836522-INS-324 | chr2-99836522-INS-324 |
| chr2-99838478-INS-52 | chr2-99838478-INS-52 |
| chr2-99841944-INS-375 | chr2-99841944-INS-375 |
| chr2-99883100-INS-194 | chr2-99883100-INS-194 |
| chr2-99911566-INS-2928 | chr2-99911567-INS-2928 |
| chr2-99925461-INS-60 | chr2-99925421-INS-84 |
| chr2-99935347-INS-6208 | chr2-99935348-INS-6208 |
| chr2-99936149-INS-311 | chr2-99936149-INS-311 |
| chr2-99941367-INS-78 | chr2-99941367-INS-77 |
| chr2-99948532-INS-54 | chr2-99948532-INS-53 |
| chr2-99955836-INS-50 | chr2-99955836-INS-50 |
| chr2-99972723-INS-58 | chr2-99972723-INS-63 |
| chr2-99973284-INS-634 | chr2-99973284-INS-636 |
| chr2-100007558-INS-428 | chr2-100007558-INS-428 |
| chr2-100011627-INS-556 | chr2-100011627-INS-556 |
| chr2-100033862-INS-903 | chr2-100033862-INS-903 |
| chr2-100035260-INS-278 | chr2-100035264-INS-276 |
| chr2-100048873-INS-6170 | chr2-100048874-INS-6169 |
| chr2-100057991-INS-54 | chr2-100057991-INS-58 |
| chr2-100068863-INS-249 | chr2-100068863-INS-231 |
| chr2-100073757-INS-77 | chr2-100073757-INS-73 |
| chr2-100086480-INS-614 | chr2-100086480-INS-614 |
| chr2-100123371-INS-3377 | chr2-100123372-INS-3377 |
| chr2-100142585-INS-6457 | chr2-100142586-INS-6458 |
| chr2-100198609-INS-66 | chr2-100198609-INS-70 |
| chr2-100222309-INS-183 | chr2-100222309-INS-183 |
| chr2-100243221-INS-64 | chr2-100243221-INS-72 |
| chr2-100247863-INS-81 | chr2-100247893-INS-68 |
| chr2-100381668-INS-242 | chr2-100381670-INS-241 |
| chr2-100624210-INS-456 | chr2-100624211-INS-456 |
| chr2-100651256-INS-196 | chr2-100651256-INS-200 |
| chr2-100675719-INS-79 | chr2-100675719-INS-79 |
| chr2-100676090-INS-5981 | chr2-100676091-INS-5984 |
| chr2-100684496-INS-1084 | chr2-100684497-INS-1084 |
| chr2-100702263-INS-156 | chr2-100702263-INS-156 |
| chr2-100703766-INS-74 | chr2-100703772-INS-89 |
| chr2-100707004-INS-90 | chr2-100706962-INS-89 |
| chr2-100753480-INS-255 | chr2-100753480-INS-255 |
| chr2-100805041-INS-287 | chr2-100805041-INS-287 |
| chr2-100805266-INS-7057 | chr2-100805267-INS-7059 |
| chr2-100826330-INS-5479 | chr2-100826330-INS-5481 |
| chr2-100886596-INS-130 | chr2-100886596-INS-128 |
| chr2-100889498-INS-212 | chr2-100889498-INS-212 |
| chr2-100898998-INS-6546 | chr2-100898999-INS-6546 |
| chr2-100922021-INS-145 | chr2-100921999-INS-164 |
| chr2-100927922-INS-695 | chr2-100927922-INS-695 |
| chr2-100930456-INS-1995 | chr2-100930457-INS-1995 |
| chr2-100931239-INS-96 | chr2-100931239-INS-100 |
| chr2-100965835-INS-617 | chr2-100965835-INS-616 |
| chr2-100968918-INS-1879 | chr2-100968919-INS-1879 |
| chr2-100988632-INS-495 | chr2-100988632-INS-495 |
| chr2-100990586-INS-58 | chr2-100990586-INS-50 |
| chr2-101011740-INS-353 | chr2-101011740-INS-353 |

| | |
|---|---|
| chr2-101019357-INS-6841 | chr2-101019358-INS-6841 |
| chr2-101022406-INS-177 | chr2-101022406-INS-193 |
| chr2-101043784-INS-118 | chr2-101043784-INS-124 |
| chr2-101055538-INS-172 | chr2-101055538-INS-172 |
| chr2-101075182-INS-385 | chr2-101075182-INS-385 |
| chr2-101088361-INS-58 | chr2-101088361-INS-58 |
| chr2-101091165-INS-209 | chr2-101091165-INS-209 |
| chr2-101098824-INS-995 | chr2-101098824-INS-995 |
| chr2-101108132-INS-7162 | chr2-101108133-INS-7161 |
| chr2-101116559-INS-70 | chr2-101116559-INS-72 |
| chr2-101172304-INS-58 | chr2-101172304-INS-60 |
| chr2-101196330-INS-144 | chr2-101196330-INS-144 |
| chr2-101212510-INS-483 | chr2-101212510-INS-483 |
| chr2-101224256-INS-160 | chr2-101224234-INS-160 |
| chr2-101224893-INS-53 | chr2-101224893-INS-69 |
| chr2-101229616-INS-1648 | chr2-101229617-INS-1649 |
| chr2-101240314-INS-1277 | chr2-101240315-INS-1278 |
| chr2-101253194-INS-2983 | chr2-101253195-INS-2983 |
| chr2-101253729-INS-102 | chr2-101253729-INS-102 |
| chr2-101256300-INS-6367 | chr2-101256301-INS-6367 |
| chr2-101261754-INS-6653 | chr2-101261755-INS-6653 |
| chr2-101267421-INS-2471 | chr2-101267422-INS-2470 |
| chr2-101275605-INS-563 | chr2-101275605-INS-563 |
| chr2-101337254-INS-216 | chr2-101337254-INS-216 |
| chr2-101338272-INS-54 | chr2-101338272-INS-54 |
| chr2-101354599-INS-70 | chr2-101354599-INS-70 |
| chr2-101371797-INS-60 | chr2-101371797-INS-60 |
| chr2-101379106-INS-348 | chr2-101379106-INS-338 |
| chr2-101402277-INS-105 | chr2-101402278-INS-66 |
| chr2-101406920-INS-60 | chr2-101406920-INS-62 |
| chr2-101417087-INS-6083 | chr2-101417088-INS-6084 |
| chr2-101426003-INS-217 | chr2-101426003-INS-222 |
| chr2-101436210-INS-6568 | chr2-101436209-INS-6556 |
| chr2-101441293-INS-53 | chr2-101441293-INS-53 |
| chr2-101450719-INS-791 | chr2-101450719-INS-752 |
| chr2-101544104-INS-67 | chr2-101544110-INS-59 |
| chr2-101569820-INS-62 | chr2-101569793-INS-65 |
| chr2-101578068-INS-402 | chr2-101578068-INS-402 |
| chr2-101580626-INS-63 | chr2-101580626-INS-63 |
| chr2-101587115-INS-65 | chr2-101587115-INS-64 |
| chr2-101590439-INS-1488 | chr2-101590440-INS-1488 |
| chr2-101594543-INS-612 | chr2-101594543-INS-611 |
| chr2-101611418-INS-218 | chr2-101611418-INS-218 |
| chr2-101615544-INS-215 | chr2-101615544-INS-216 |
| chr2-101631014-INS-175 | chr2-101631014-INS-175 |
| chr2-101663269-INS-72 | chr2-101663241-INS-80 |
| chr2-101665266-INS-377 | chr2-101665266-INS-378 |
| chr2-101667906-INS-208 | chr2-101667910-INS-196 |
| chr2-101705114-INS-320 | chr2-101705114-INS-320 |
| chr2-101722837-INS-755 | chr2-101722837-INS-757 |
| chr2-101750321-INS-51 | chr2-101750321-INS-51 |
| chr2-101771671-INS-53 | chr2-101771671-INS-53 |
| chr2-101772706-INS-78 | chr2-101772706-INS-76 |
| chr2-101780282-INS-118 | chr2-101780282-INS-118 |
| chr2-101796818-INS-79 | chr2-101796818-INS-79 |
| chr2-101821543-INS-68 | chr2-101821543-INS-68 |
| chr2-101823830-INS-52 | chr2-101823830-INS-50 |
| chr2-101824710-INS-173 | chr2-101824710-INS-175 |
| chr2-101846311-INS-52 | chr2-101846311-INS-50 |
| chr2-101854104-INS-60 | chr2-101854103-INS-90 |
| chr2-101886415-INS-6792 | chr2-101886416-INS-6795 |
| chr2-101911282-INS-70 | chr2-101911282-INS-117 |
| chr2-101913137-INS-274 | chr2-101913134-INS-316 |
| chr2-101917168-INS-132 | chr2-101917154-INS-116 |
| chr2-101943466-INS-374 | chr2-101943466-INS-376 |
| chr2-101948908-INS-80 | chr2-101948908-INS-80 |
| chr2-101956789-INS-89 | chr2-101956789-INS-89 |
| chr2-101962561-INS-111 | chr2-101962561-INS-111 |
| chr2-101970751-INS-1090 | chr2-101970752-INS-1090 |
| chr2-102021977-INS-68 | chr2-102021979-INS-68 |
| chr2-102040836-INS-63 | chr2-102040836-INS-61 |
| chr2-102049381-INS-141 | chr2-102049385-INS-159 |
| chr2-102058700-INS-190 | chr2-102058700-INS-170 |
| chr2-102061688-INS-55 | chr2-102061688-INS-55 |
| chr2-102073134-INS-206 | chr2-102073134-INS-206 |
| chr2-102078571-INS-60 | chr2-102078571-INS-56 |
| chr2-102082357-INS-103 | chr2-102082357-INS-103 |

| | |
|---|---|
| chr2-102083336-INS-499 | chr2-102083336-INS-499 |
| chr2-102089569-INS-543 | chr2-102089569-INS-543 |
| chr2-102094433-INS-199 | chr2-102094433-INS-199 |
| chr2-102110077-INS-92 | chr2-102110077-INS-94 |
| chr2-102114404-INS-1875 | chr2-102114420-INS-1869 |
| chr2-102114432-INS-6589 | chr2-102114433-INS-6589 |
| chr2-102155183-INS-1019 | chr2-102155184-INS-1019 |
| chr2-102192150-INS-224 | chr2-102192150-INS-226 |
| chr2-102195917-INS-206 | chr2-102195917-INS-212 |
| chr2-102198302-INS-216 | chr2-102198302-INS-217 |
| chr2-102202922-INS-62 | chr2-102202922-INS-63 |
| chr2-102241963-INS-178 | chr2-102241963-INS-178 |
| chr2-102259464-INS-66 | chr2-102259464-INS-76 |
| chr2-102272154-INS-76 | chr2-102272154-INS-76 |
| chr2-102285598-INS-79 | chr2-102285598-INS-80 |
| chr2-102312343-INS-56 | chr2-102312343-INS-60 |
| chr2-102315970-INS-102 | chr2-102315970-INS-102 |
| chr2-102336032-INS-406 | chr2-102336032-INS-406 |
| chr2-102348959-INS-335 | chr2-102348969-INS-326 |
| chr2-102373745-INS-6384 | chr2-102373746-INS-6384 |
| chr2-102374300-INS-54 | chr2-102374300-INS-54 |
| chr2-102375351-INS-71 | chr2-102375351-INS-56 |
| chr2-102384140-INS-56 | chr2-102384140-INS-52 |
| chr2-102392414-INS-2539 | chr2-102392415-INS-2539 |
| chr2-102469073-INS-78 | chr2-102469073-INS-83 |
| chr2-102471619-INS-76 | chr2-102471619-INS-76 |
| chr2-102482007-INS-78 | chr2-102482007-INS-80 |
| chr2-102487063-INS-382 | chr2-102487063-INS-382 |
| chr2-102497627-INS-309 | chr2-102497627-INS-309 |
| chr2-102528145-INS-64 | chr2-102528145-INS-58 |
| chr2-102538434-INS-1776 | chr2-102538435-INS-1776 |
| chr2-102540528-INS-1739 | chr2-102540529-INS-1739 |
| chr2-102586852-INS-80 | chr2-102586852-INS-79 |
| chr2-102602238-INS-51 | chr2-102602238-INS-51 |
| chr2-102605838-INS-91 | chr2-102605838-INS-76 |
| chr2-102605882-INS-861 | chr2-102605872-INS-853 |
| chr2-102626923-INS-2792 | chr2-102626924-INS-2812 |
| chr2-102635152-INS-820 | chr2-102635152-INS-820 |
| chr2-102636327-INS-197 | chr2-102636327-INS-197 |
| chr2-102653644-INS-194 | chr2-102653644-INS-194 |
| chr2-102701338-INS-2005 | chr2-102701339-INS-2006 |
| chr2-102732310-INS-97 | chr2-102732310-INS-104 |
| chr2-102733937-INS-1170 | chr2-102733938-INS-1208 |
| chr2-102740988-INS-215 | chr2-102740988-INS-227 |
| chr2-102761786-INS-6559 | chr2-102761787-INS-6559 |
| chr2-102766919-INS-55 | chr2-102766919-INS-60 |
| chr2-102802237-INS-412 | chr2-102802237-INS-412 |
| chr2-102822594-INS-123 | chr2-102822594-INS-150 |
| chr2-102827341-INS-427 | chr2-102827341-INS-427 |
| chr2-102855019-INS-86 | chr2-102855019-INS-86 |
| chr2-102867332-INS-186 | chr2-102867332-INS-186 |
| chr2-102892784-INS-936 | chr2-102892784-INS-943 |
| chr2-102898811-INS-741 | chr2-102898807-INS-743 |
| chr2-102915404-INS-108 | chr2-102915404-INS-108 |
| chr2-102953396-INS-74 | chr2-102953396-INS-78 |
| chr2-102981081-INS-77 | chr2-102981083-INS-75 |
| chr2-102984266-INS-61 | chr2-102984309-INS-62 |
| chr2-103012614-INS-313 | chr2-103012614-INS-313 |
| chr2-103028425-INS-150 | chr2-103028424-INS-150 |
| chr2-103059146-INS-5252 | chr2-103059147-INS-5252 |
| chr2-103091657-INS-83 | chr2-103091656-INS-92 |
| chr2-103100030-INS-52 | chr2-103100030-INS-66 |
| chr2-103148194-INS-270 | chr2-103148194-INS-270 |
| chr2-103200661-INS-71 | chr2-103200638-INS-54 |
| chr2-103264516-INS-102 | chr2-103264516-INS-122 |
| chr2-103274823-INS-68 | chr2-103274823-INS-68 |
| chr2-103294831-INS-182 | chr2-103294831-INS-182 |
| chr2-103305888-INS-62 | chr2-103305888-INS-62 |
| chr2-103322608-INS-200 | chr2-103322608-INS-200 |
| chr2-103330730-INS-175 | chr2-103330730-INS-176 |
| chr2-103350403-INS-54 | chr2-103350376-INS-58 |
| chr2-103353080-INS-73 | chr2-103353080-INS-83 |
| chr2-103366083-INS-64 | chr2-103366081-INS-66 |
| chr2-103420169-INS-112 | chr2-103420185-INS-110 |
| chr2-103452320-INS-54 | chr2-103452320-INS-54 |
| chr2-103458865-INS-1537 | chr2-103458866-INS-1537 |
| chr2-103459120-INS-51 | chr2-103459120-INS-51 |

| | |
|---|---|
| chr2-103484401-INS-225 | chr2-103484401-INS-225 |
| chr2-103493005-INS-52 | chr2-103493027-INS-52 |
| chr2-103501394-INS-91 | chr2-103501394-INS-91 |
| chr2-103502448-INS-207 | chr2-103502448-INS-207 |
| chr2-103519819-INS-133 | chr2-103519819-INS-133 |
| chr2-103585979-INS-77 | chr2-103585979-INS-93 |
| chr2-103601525-INS-223 | chr2-103601525-INS-223 |
| chr2-103619544-INS-200 | chr2-103619544-INS-200 |
| chr2-103637083-INS-56 | chr2-103637109-INS-50 |
| chr2-103643173-INS-182 | chr2-103643173-INS-210 |
| chr2-103656684-INS-3386 | chr2-103656685-INS-3386 |
| chr2-103660653-INS-96 | chr2-103660653-INS-96 |
| chr2-103673228-INS-194 | chr2-103673228-INS-172 |
| chr2-103697186-INS-62 | chr2-103697186-INS-62 |
| chr2-103699917-INS-54 | chr2-103699917-INS-56 |
| chr2-103716743-INS-66 | chr2-103716743-INS-66 |
| chr2-103720549-INS-561 | chr2-103720549-INS-564 |
| chr2-103728010-INS-84 | chr2-103728039-INS-102 |
| chr2-103742863-INS-217 | chr2-103742863-INS-218 |
| chr2-103746973-INS-221 | chr2-103746973-INS-217 |
| chr2-103747564-INS-185 | chr2-103747564-INS-185 |
| chr2-103774778-INS-347 | chr2-103774778-INS-350 |
| chr2-103808621-INS-98 | chr2-103808661-INS-98 |
| chr2-103829802-INS-109 | chr2-103829830-INS-65 |
| chr2-103832546-INS-679 | chr2-103832546-INS-709 |
| chr2-103835218-INS-430 | chr2-103835218-INS-430 |
| chr2-103908088-INS-78 | chr2-103908059-INS-90 |
| chr2-103913893-INS-99 | chr2-103913893-INS-99 |
| chr2-103917918-INS-69 | chr2-103917918-INS-67 |
| chr2-103925384-INS-132 | chr2-103925384-INS-132 |
| chr2-103944230-INS-465 | chr2-103944230-INS-465 |
| chr2-103962980-INS-81 | chr2-103962980-INS-67 |
| chr2-104004182-INS-507 | chr2-104004182-INS-505 |
| chr2-104006773-INS-101 | chr2-104006773-INS-99 |
| chr2-104043755-INS-60 | chr2-104043755-INS-60 |
| chr2-104048263-INS-101 | chr2-104048263-INS-101 |
| chr2-104050594-INS-56 | chr2-104050594-INS-56 |
| chr2-104065715-INS-90 | chr2-104065722-INS-95 |
| chr2-104073912-INS-5375 | chr2-104073913-INS-5375 |
| chr2-104075101-INS-517 | chr2-104075101-INS-517 |
| chr2-104080177-INS-238 | chr2-104080177-INS-230 |
| chr2-104081646-INS-66 | chr2-104081646-INS-66 |
| chr2-104082918-INS-8252 | chr2-104082919-INS-8254 |
| chr2-104089351-INS-225 | chr2-104089351-INS-225 |
| chr2-104091775-INS-561 | chr2-104091775-INS-563 |
| chr2-104104137-INS-126 | chr2-104104137-INS-126 |
| chr2-104106275-INS-5447 | chr2-104106276-INS-5447 |
| chr2-104107410-INS-54 | chr2-104107410-INS-54 |
| chr2-104108117-INS-52 | chr2-104108119-INS-50 |
| chr2-104147424-INS-322 | chr2-104147424-INS-322 |
| chr2-104166153-INS-2932 | chr2-104166154-INS-2932 |
| chr2-104188456-INS-85 | chr2-104188456-INS-93 |
| chr2-104197025-INS-68 | chr2-104197025-INS-68 |
| chr2-104205053-INS-87 | chr2-104205053-INS-89 |
| chr2-104218406-INS-72 | chr2-104218412-INS-68 |
| chr2-104221108-INS-359 | chr2-104221108-INS-359 |
| chr2-104223195-INS-52 | chr2-104223195-INS-52 |
| chr2-104254629-INS-217 | chr2-104254629-INS-217 |
| chr2-104263209-INS-52 | chr2-104263226-INS-56 |
| chr2-104270073-INS-94 | chr2-104270052-INS-68 |
| chr2-104295863-INS-100 | chr2-104295863-INS-130 |
| chr2-104307737-INS-219 | chr2-104307737-INS-219 |
| chr2-104317858-INS-141 | chr2-104317858-INS-105 |
| chr2-104330171-INS-205 | chr2-104330171-INS-205 |
| chr2-104353880-INS-56 | chr2-104353898-INS-66 |
| chr2-104357076-INS-344 | chr2-104357076-INS-344 |
| chr2-104374457-INS-118 | chr2-104374457-INS-118 |
| chr2-104375670-INS-57 | chr2-104375670-INS-63 |
| chr2-104377768-INS-81 | chr2-104377781-INS-54 |
| chr2-104381358-INS-498 | chr2-104381358-INS-498 |
| chr2-104410848-INS-206 | chr2-104410848-INS-206 |
| chr2-104417723-INS-201 | chr2-104417723-INS-199 |
| chr2-104430747-INS-82 | chr2-104430747-INS-82 |
| chr2-104438194-INS-114 | chr2-104438194-INS-118 |
| chr2-104459482-INS-199 | chr2-104459482-INS-200 |
| chr2-104482985-INS-76 | chr2-104482967-INS-74 |
| chr2-104501673-INS-78 | chr2-104501673-INS-52 |

| | |
|---|---|
| chr2-104508259-INS-209 | chr2-104508259-INS-209 |
| chr2-104518628-INS-139 | chr2-104518628-INS-139 |
| chr2-104521938-INS-218 | chr2-104521938-INS-224 |
| chr2-104522739-INS-139 | chr2-104522739-INS-139 |
| chr2-104531228-INS-52 | chr2-104531228-INS-52 |
| chr2-104540281-INS-184 | chr2-104540282-INS-191 |
| chr2-104555288-INS-205 | chr2-104555288-INS-205 |
| chr2-104556819-INS-240 | chr2-104556819-INS-241 |
| chr2-104559340-INS-191 | chr2-104559340-INS-191 |
| chr2-104563432-INS-210 | chr2-104563432-INS-210 |
| chr2-104574296-INS-51 | chr2-104574296-INS-51 |
| chr2-104589041-INS-54 | chr2-104589041-INS-54 |
| chr2-104598276-INS-202 | chr2-104598276-INS-202 |
| chr2-104604361-INS-408 | chr2-104604361-INS-415 |
| chr2-104625065-INS-260 | chr2-104625065-INS-217 |
| chr2-104666233-INS-56 | chr2-104666233-INS-56 |
| chr2-104669409-INS-231 | chr2-104669409-INS-231 |
| chr2-104687994-INS-64 | chr2-104687994-INS-61 |
| chr2-104726841-INS-6380 | chr2-104726842-INS-6381 |
| chr2-104734000-INS-59 | chr2-104734000-INS-96 |
| chr2-104764263-INS-102 | chr2-104764263-INS-102 |
| chr2-104790275-INS-204 | chr2-104790275-INS-211 |
| chr2-104821214-INS-552 | chr2-104821214-INS-552 |
| chr2-104834734-INS-92 | chr2-104834734-INS-92 |
| chr2-104838538-INS-466 | chr2-104838538-INS-466 |
| chr2-104851556-INS-88 | chr2-104851556-INS-88 |
| chr2-104860324-INS-68 | chr2-104860332-INS-58 |
| chr2-104882615-INS-92 | chr2-104882565-INS-110 |
| chr2-104886440-INS-53 | chr2-104886457-INS-54 |
| chr2-104911033-INS-60 | chr2-104911033-INS-60 |
| chr2-104923540-INS-171 | chr2-104923541-INS-142 |
| chr2-104976623-INS-198 | chr2-104976623-INS-198 |
| chr2-104993118-INS-52 | chr2-104993114-INS-55 |
| chr2-105000637-INS-51 | chr2-105000637-INS-51 |
| chr2-105005431-INS-50 | chr2-105005431-INS-65 |
| chr2-105015074-INS-1110 | chr2-105015075-INS-1112 |
| chr2-105025883-INS-1584 | chr2-105025884-INS-1584 |
| chr2-105029934-INS-2767 | chr2-105029935-INS-2767 |
| chr2-105031983-INS-309 | chr2-105031983-INS-309 |
| chr2-105050210-INS-52 | chr2-105050210-INS-53 |
| chr2-105083378-INS-1776 | chr2-105083379-INS-1776 |
| chr2-105085177-INS-114 | chr2-105085177-INS-114 |
| chr2-105123605-INS-335 | chr2-105123605-INS-335 |
| chr2-105143926-INS-5393 | chr2-105143927-INS-5393 |
| chr2-105166359-INS-54 | chr2-105166359-INS-54 |
| chr2-105181873-INS-72 | chr2-105181873-INS-72 |
| chr2-105223369-INS-169 | chr2-105223369-INS-176 |
| chr2-105236250-INS-57 | chr2-105236250-INS-59 |
| chr2-105242749-INS-58 | chr2-105242749-INS-58 |
| chr2-105255427-INS-342 | chr2-105255427-INS-342 |
| chr2-105299402-INS-1067 | chr2-105299403-INS-1067 |
| chr2-105316152-INS-64 | chr2-105316177-INS-76 |
| chr2-105318540-INS-132 | chr2-105318540-INS-104 |
| chr2-105330948-INS-171 | chr2-105330948-INS-171 |
| chr2-105364557-INS-400 | chr2-105364557-INS-400 |
| chr2-105418856-INS-202 | chr2-105418856-INS-202 |
| chr2-105445286-INS-206 | chr2-105445286-INS-206 |
| chr2-105453776-INS-212 | chr2-105453776-INS-212 |
| chr2-105469139-INS-88 | chr2-105469139-INS-88 |
| chr2-105482243-INS-199 | chr2-105482243-INS-198 |
| chr2-105492410-INS-65 | chr2-105492442-INS-54 |
| chr2-105536082-INS-81 | chr2-105536082-INS-81 |
| chr2-105563681-INS-60 | chr2-105563681-INS-56 |
| chr2-105576865-INS-158 | chr2-105576865-INS-158 |
| chr2-105591671-INS-6889 | chr2-105591672-INS-6892 |
| chr2-105603447-INS-206 | chr2-105603447-INS-206 |
| chr2-105624151-INS-98 | chr2-105624151-INS-98 |
| chr2-105673587-INS-8872 | chr2-105673588-INS-8872 |
| chr2-105677230-INS-348 | chr2-105677227-INS-354 |
| chr2-105717759-INS-196 | chr2-105717759-INS-196 |
| chr2-105719650-INS-6329 | chr2-105719651-INS-6329 |
| chr2-105722905-INS-101 | chr2-105722905-INS-101 |
| chr2-105725901-INS-54 | chr2-105725901-INS-54 |
| chr2-105825557-INS-6566 | chr2-105825558-INS-6567 |
| chr2-105828339-INS-2804 | chr2-105828340-INS-2804 |
| chr2-105859491-INS-1776 | chr2-105859492-INS-1776 |
| chr2-105859594-INS-200 | chr2-105859594-INS-200 |

| | |
|---|---|
| chr2-105872202-INS-421 | chr2-105872202-INS-422 |
| chr2-105896565-INS-58 | chr2-105896544-INS-60 |
| chr2-105898813-INS-1769 | chr2-105898814-INS-1770 |
| chr2-105899391-INS-5303 | chr2-105899392-INS-5303 |
| chr2-105900226-INS-68 | chr2-105900226-INS-60 |
| chr2-105918520-INS-417 | chr2-105918520-INS-417 |
| chr2-105928486-INS-295 | chr2-105928486-INS-338 |
| chr2-105934895-INS-742 | chr2-105934895-INS-754 |
| chr2-105954163-INS-54 | chr2-105954163-INS-58 |
| chr2-105955911-INS-711 | chr2-105955911-INS-711 |
| chr2-105981052-INS-6154 | chr2-105981053-INS-6154 |
| chr2-105987860-INS-70 | chr2-105987860-INS-66 |
| chr2-105991386-INS-56 | chr2-105991390-INS-52 |
| chr2-105996370-INS-6413 | chr2-105996371-INS-6413 |
| chr2-105996889-INS-2884 | chr2-105996897-INS-2877 |
| chr2-106020698-INS-403 | chr2-106020698-INS-386 |
| chr2-106032861-INS-412 | chr2-106032861-INS-413 |
| chr2-106034140-INS-146 | chr2-106034121-INS-163 |
| chr2-106042110-INS-6363 | chr2-106042111-INS-6350 |
| chr2-106063819-INS-1889 | chr2-106063820-INS-1889 |
| chr2-106067662-INS-277 | chr2-106067662-INS-257 |
| chr2-106069968-INS-52 | chr2-106069968-INS-54 |
| chr2-106085968-INS-477 | chr2-106085968-INS-478 |
| chr2-106092643-INS-882 | chr2-106092643-INS-883 |
| chr2-106095266-INS-224 | chr2-106095266-INS-224 |
| chr2-106098505-INS-888 | chr2-106098505-INS-913 |
| chr2-106132183-INS-183 | chr2-106132177-INS-160 |
| chr2-106152380-INS-526 | chr2-106152380-INS-526 |
| chr2-106154472-INS-64 | chr2-106154472-INS-66 |
| chr2-106167813-INS-133 | chr2-106167830-INS-162 |
| chr2-106185916-INS-288 | chr2-106185916-INS-286 |
| chr2-106187092-INS-6899 | chr2-106187093-INS-6899 |
| chr2-106189573-INS-169 | chr2-106189573-INS-169 |
| chr2-106218270-INS-171 | chr2-106218270-INS-171 |
| chr2-106290384-INS-80 | chr2-106290384-INS-56 |
| chr2-106312492-INS-67 | chr2-106312492-INS-63 |
| chr2-106318506-INS-328 | chr2-106318506-INS-328 |
| chr2-106325971-INS-239 | chr2-106325972-INS-242 |
| chr2-106334513-INS-443 | chr2-106334513-INS-443 |
| chr2-106352047-INS-69 | chr2-106352048-INS-69 |
| chr2-106357227-INS-327 | chr2-106357227-INS-328 |
| chr2-106360597-INS-64 | chr2-106360597-INS-64 |
| chr2-106361294-INS-69 | chr2-106361294-INS-69 |
| chr2-106386115-INS-463 | chr2-106386116-INS-463 |
| chr2-106390206-INS-108 | chr2-106390202-INS-110 |
| chr2-106394098-INS-136 | chr2-106394098-INS-116 |
| chr2-106412585-INS-78 | chr2-106412585-INS-68 |
| chr2-106418354-INS-66 | chr2-106418311-INS-76 |
| chr2-106436138-INS-75 | chr2-106436138-INS-56 |
| chr2-106437456-INS-108 | chr2-106437456-INS-153 |
| chr2-106451003-INS-6590 | chr2-106451004-INS-6590 |
| chr2-106464303-INS-63 | chr2-106464303-INS-60 |
| chr2-106490032-INS-183 | chr2-106490031-INS-184 |
| chr2-106502147-INS-177 | chr2-106502147-INS-176 |
| chr2-106519641-INS-209 | chr2-106519641-INS-209 |
| chr2-106521055-INS-1675 | chr2-106521056-INS-1675 |
| chr2-106539494-INS-596 | chr2-106539495-INS-592 |
| chr2-106542948-INS-327 | chr2-106542948-INS-327 |
| chr2-106563579-INS-490 | chr2-106563579-INS-490 |
| chr2-106567520-INS-64 | chr2-106567524-INS-55 |
| chr2-106593198-INS-208 | chr2-106593198-INS-208 |
| chr2-106600587-INS-112 | chr2-106600587-INS-112 |
| chr2-106618974-INS-65 | chr2-106618974-INS-65 |
| chr2-106633390-INS-180 | chr2-106633390-INS-180 |
| chr2-106654578-INS-134 | chr2-106654578-INS-132 |
| chr2-106693394-INS-174 | chr2-106693394-INS-174 |
| chr2-106711154-INS-81 | chr2-106711154-INS-81 |
| chr2-106714811-INS-56 | chr2-106714811-INS-52 |
| chr2-106721454-INS-266 | chr2-106721454-INS-266 |
| chr2-106733331-INS-54 | chr2-106733331-INS-54 |
| chr2-106734204-INS-2930 | chr2-106734205-INS-2930 |
| chr2-106738270-INS-187 | chr2-106738270-INS-187 |
| chr2-106744920-INS-130 | chr2-106744957-INS-117 |
| chr2-106758519-INS-181 | chr2-106758520-INS-198 |
| chr2-106803122-INS-79 | chr2-106803122-INS-116 |
| chr2-106817118-INS-549 | chr2-106817118-INS-549 |
| chr2-106828640-INS-6213 | chr2-106828641-INS-6213 |

| | |
|---|---|
| chr2-106841970-INS-176 | chr2-106841970-INS-176 |
| chr2-106851298-INS-54 | chr2-106851298-INS-54 |
| chr2-106881273-INS-211 | chr2-106881273-INS-211 |
| chr2-106916980-INS-60 | chr2-106916983-INS-90 |
| chr2-106918988-INS-1131 | chr2-106918989-INS-1131 |
| chr2-106956065-INS-108 | chr2-106956080-INS-109 |
| chr2-106960157-INS-88 | chr2-106960167-INS-58 |
| chr2-106975681-INS-52 | chr2-106975681-INS-52 |
| chr2-106983870-INS-59 | chr2-106983870-INS-59 |
| chr2-107004395-INS-116 | chr2-107004419-INS-66 |
| chr2-107030245-INS-142 | chr2-107030245-INS-142 |
| chr2-107038670-INS-6358 | chr2-107038671-INS-6358 |
| chr2-107047283-INS-61 | chr2-107047283-INS-60 |
| chr2-107047968-INS-54 | chr2-107047968-INS-54 |
| chr2-107067723-INS-96 | chr2-107067721-INS-72 |
| chr2-107085605-INS-1316 | chr2-107085606-INS-1316 |
| chr2-107088666-INS-88 | chr2-107088666-INS-76 |
| chr2-107099176-INS-6590 | chr2-107099177-INS-6590 |
| chr2-107117858-INS-3418 | chr2-107117859-INS-3418 |
| chr2-107122298-INS-102 | chr2-107122298-INS-52 |
| chr2-107141022-INS-78 | chr2-107141022-INS-82 |
| chr2-107167925-INS-66 | chr2-107167923-INS-68 |
| chr2-107177482-INS-54 | chr2-107177482-INS-56 |
| chr2-107184233-INS-1220 | chr2-107184234-INS-1221 |
| chr2-107190391-INS-56 | chr2-107190391-INS-60 |
| chr2-107198523-INS-53 | chr2-107198523-INS-53 |
| chr2-107199929-INS-108 | chr2-107199929-INS-124 |
| chr2-107200082-INS-6686 | chr2-107200083-INS-6686 |
| chr2-107202831-INS-507 | chr2-107202831-INS-510 |
| chr2-107235600-INS-454 | chr2-107235600-INS-457 |
| chr2-107239403-INS-96 | chr2-107239404-INS-94 |
| chr2-107240039-INS-74 | chr2-107240039-INS-50 |
| chr2-107257647-INS-96 | chr2-107257609-INS-96 |
| chr2-107268810-INS-84 | chr2-107268810-INS-76 |
| chr2-107272642-INS-532 | chr2-107272642-INS-533 |
| chr2-107285983-INS-318 | chr2-107285983-INS-318 |
| chr2-107344831-INS-507 | chr2-107344831-INS-507 |
| chr2-107359272-INS-65 | chr2-107359272-INS-60 |
| chr2-107361184-INS-102 | chr2-107361180-INS-102 |
| chr2-107389790-INS-90 | chr2-107389790-INS-91 |
| chr2-107433433-INS-62 | chr2-107433433-INS-62 |
| chr2-107481184-INS-1793 | chr2-107481185-INS-1793 |
| chr2-107485004-INS-302 | chr2-107485004-INS-302 |
| chr2-107485941-INS-59 | chr2-107485941-INS-68 |
| chr2-107489194-INS-448 | chr2-107489194-INS-448 |
| chr2-107499651-INS-58 | chr2-107499651-INS-91 |
| chr2-107501852-INS-52 | chr2-107501852-INS-52 |
| chr2-107507108-INS-68 | chr2-107507108-INS-76 |
| chr2-107521223-INS-5579 | chr2-107521224-INS-5579 |
| chr2-107527508-INS-70 | chr2-107527508-INS-66 |
| chr2-107545376-INS-62 | chr2-107545376-INS-62 |
| chr2-107582092-INS-967 | chr2-107582092-INS-968 |
| chr2-107611158-INS-85 | chr2-107611158-INS-85 |
| chr2-107625735-INS-6884 | chr2-107625743-INS-6881 |
| chr2-107636953-INS-1271 | chr2-107636954-INS-1271 |
| chr2-107641404-INS-74 | chr2-107641394-INS-92 |
| chr2-107655518-INS-101 | chr2-107655518-INS-101 |
| chr2-107664248-INS-90 | chr2-107664248-INS-93 |
| chr2-107673238-INS-173 | chr2-107673238-INS-173 |
| chr2-107680622-INS-646 | chr2-107680622-INS-645 |
| chr2-107692147-INS-78 | chr2-107692147-INS-98 |
| chr2-107693043-INS-549 | chr2-107693043-INS-549 |
| chr2-107697126-INS-701 | chr2-107697126-INS-701 |
| chr2-107705977-INS-5389 | chr2-107705978-INS-5389 |
| chr2-107706348-INS-1648 | chr2-107706349-INS-1648 |
| chr2-107720192-INS-1536 | chr2-107720193-INS-1536 |
| chr2-107726339-INS-400 | chr2-107726339-INS-400 |
| chr2-107733509-INS-1831 | chr2-107733510-INS-1831 |
| chr2-107754759-INS-7289 | chr2-107754768-INS-7292 |
| chr2-107778881-INS-66 | chr2-107778873-INS-80 |
| chr2-107794071-INS-68 | chr2-107794071-INS-68 |
| chr2-107838000-INS-955 | chr2-107838000-INS-955 |
| chr2-107849721-INS-60 | chr2-107849700-INS-74 |
| chr2-107862017-INS-6645 | chr2-107862018-INS-6645 |
| chr2-107897877-INS-73 | chr2-107897887-INS-85 |
| chr2-107902153-INS-780 | chr2-107902153-INS-781 |
| chr2-107910145-INS-6653 | chr2-107910146-INS-6652 |

| | |
|---|---|
| chr2-107930250-INS-13281 | chr2-107930251-INS-13282 |
| chr2-107949381-INS-210 | chr2-107949381-INS-210 |
| chr2-107973224-INS-116 | chr2-107973223-INS-101 |
| chr2-107976069-INS-52 | chr2-107976069-INS-52 |
| chr2-107981857-INS-105 | chr2-107981866-INS-65 |
| chr2-107982770-INS-4604 | chr2-107982771-INS-4600 |
| chr2-107985221-INS-52 | chr2-107985221-INS-60 |
| chr2-107990450-INS-289 | chr2-107990450-INS-289 |
| chr2-108002934-INS-390 | chr2-108002934-INS-390 |
| chr2-108010250-INS-242 | chr2-108010250-INS-243 |
| chr2-108016358-INS-52 | chr2-108016359-INS-52 |
| chr2-108060298-INS-54 | chr2-108060298-INS-50 |
| chr2-108087678-INS-567 | chr2-108087678-INS-567 |
| chr2-108090000-INS-52 | chr2-108090000-INS-56 |
| chr2-108096960-INS-62 | chr2-108096960-INS-66 |
| chr2-108098623-INS-580 | chr2-108098623-INS-581 |
| chr2-108099483-INS-1109 | chr2-108099484-INS-1109 |
| chr2-108127379-INS-2111 | chr2-108127380-INS-2111 |
| chr2-108136742-INS-51 | chr2-108136742-INS-51 |
| chr2-108140718-INS-56 | chr2-108140718-INS-54 |
| chr2-108157005-INS-65 | chr2-108157006-INS-52 |
| chr2-108165510-INS-457 | chr2-108165510-INS-444 |
| chr2-108169036-INS-110 | chr2-108169036-INS-109 |
| chr2-108186388-INS-499 | chr2-108186388-INS-499 |
| chr2-108186983-INS-5556 | chr2-108186984-INS-5556 |
| chr2-108188277-INS-68 | chr2-108188277-INS-68 |
| chr2-108215236-INS-1748 | chr2-108215237-INS-1749 |
| chr2-108221320-INS-201 | chr2-108221320-INS-200 |
| chr2-108225892-INS-5542 | chr2-108225893-INS-5542 |
| chr2-108252750-INS-1233 | chr2-108252751-INS-1235 |
| chr2-108278631-INS-252 | chr2-108278631-INS-252 |
| chr2-108282519-INS-182 | chr2-108282519-INS-143 |
| chr2-108295440-INS-6624 | chr2-108295441-INS-6624 |
| chr2-108300664-INS-73 | chr2-108300685-INS-75 |
| chr2-108324696-INS-56 | chr2-108324696-INS-54 |
| chr2-108325291-INS-4825 | chr2-108325292-INS-4826 |
| chr2-108348716-INS-976 | chr2-108348716-INS-975 |
| chr2-108368509-INS-93 | chr2-108368509-INS-93 |
| chr2-108376790-INS-66 | chr2-108376790-INS-50 |
| chr2-108414404-INS-63 | chr2-108414404-INS-63 |
| chr2-108431545-INS-6459 | chr2-108431546-INS-6462 |
| chr2-108433969-INS-96 | chr2-108433946-INS-90 |
| chr2-108438401-INS-755 | chr2-108438401-INS-758 |
| chr2-108458680-INS-96 | chr2-108458680-INS-96 |
| chr2-108469281-INS-151 | chr2-108469281-INS-130 |
| chr2-108477742-INS-63 | chr2-108477730-INS-70 |
| chr2-108485958-INS-6410 | chr2-108485959-INS-6410 |
| chr2-108500419-INS-142 | chr2-108500419-INS-182 |
| chr2-108538250-INS-54 | chr2-108538252-INS-50 |
| chr2-108690644-INS-117 | chr2-108690644-INS-117 |
| chr2-108692954-INS-1323 | chr2-108692955-INS-1323 |
| chr2-108695992-INS-72 | chr2-108695992-INS-76 |
| chr2-108706276-INS-1333 | chr2-108706277-INS-1333 |
| chr2-108722146-INS-5491 | chr2-108722147-INS-5493 |
| chr2-108774431-INS-56 | chr2-108774431-INS-50 |
| chr2-108786437-INS-1705 | chr2-108786438-INS-1706 |
| chr2-108797543-INS-68 | chr2-108797543-INS-70 |
| chr2-108835999-INS-6527 | chr2-108836007-INS-6534 |
| chr2-108848569-INS-838 | chr2-108848569-INS-838 |
| chr2-108890593-INS-66 | chr2-108890596-INS-74 |
| chr2-108925414-INS-122 | chr2-108925414-INS-122 |
| chr2-108989326-INS-171 | chr2-108989326-INS-171 |
| chr2-108994998-INS-6693 | chr2-108994999-INS-6694 |
| chr2-109055045-INS-101 | chr2-109055045-INS-52 |
| chr2-109106280-INS-170 | chr2-109106280-INS-171 |
| chr2-109179750-INS-200 | chr2-109179748-INS-203 |
| chr2-109218675-INS-169 | chr2-109218675-INS-169 |
| chr2-109317896-INS-70 | chr2-109317896-INS-100 |
| chr2-109321113-INS-214 | chr2-109321113-INS-198 |
| chr2-109325591-INS-6764 | chr2-109325592-INS-6764 |
| chr2-109346585-INS-333 | chr2-109346585-INS-331 |
| chr2-109364478-INS-529 | chr2-109364478-INS-529 |
| chr2-109378979-INS-3044 | chr2-109378980-INS-3044 |
| chr2-109405262-INS-60 | chr2-109405276-INS-50 |
| chr2-109585484-INS-124 | chr2-109585484-INS-124 |
| chr2-109588065-INS-448 | chr2-109588065-INS-448 |
| chr2-109594396-INS-6341 | chr2-109594397-INS-6344 |

| | |
|---|---|
| chr2-109596349-INS-69 | chr2-109596345-INS-74 |
| chr2-109598510-INS-7243 | chr2-109598511-INS-7245 |
| chr2-109609476-INS-64 | chr2-109609476-INS-58 |
| chr2-109616634-INS-169 | chr2-109616634-INS-169 |
| chr2-109621609-INS-59 | chr2-109621613-INS-54 |
| chr2-109627915-INS-218 | chr2-109627915-INS-218 |
| chr2-109634969-INS-215 | chr2-109634969-INS-215 |
| chr2-109657556-INS-6863 | chr2-109657557-INS-6864 |
| chr2-109674039-INS-155 | chr2-109674039-INS-155 |
| chr2-109677478-INS-56 | chr2-109677478-INS-56 |
| chr2-109678395-INS-55 | chr2-109678395-INS-62 |
| chr2-109681692-INS-194 | chr2-109681692-INS-184 |
| chr2-109684784-INS-6304 | chr2-109684785-INS-6304 |
| chr2-109695706-INS-337 | chr2-109695706-INS-337 |
| chr2-109699566-INS-248 | chr2-109699566-INS-248 |
| chr2-109706823-INS-68 | chr2-109706823-INS-60 |
| chr2-109717294-INS-222 | chr2-109717294-INS-222 |
| chr2-109735859-INS-55 | chr2-109735859-INS-55 |
| chr2-109748790-INS-52 | chr2-109748790-INS-54 |
| chr2-109753057-INS-226 | chr2-109753057-INS-234 |
| chr2-109758665-INS-213 | chr2-109758665-INS-213 |
| chr2-109760749-INS-184 | chr2-109760749-INS-184 |
| chr2-109766463-INS-56 | chr2-109766463-INS-56 |
| chr2-109781785-INS-52 | chr2-109781785-INS-52 |
| chr2-109787707-INS-214 | chr2-109787707-INS-214 |
| chr2-109788076-INS-428 | chr2-109788076-INS-423 |
| chr2-109789009-INS-5171 | chr2-109789010-INS-5171 |
| chr2-109797226-INS-100 | chr2-109797226-INS-89 |
| chr2-109840805-INS-121 | chr2-109840805-INS-118 |
| chr2-109850774-INS-59 | chr2-109850774-INS-59 |
| chr2-109875764-INS-498 | chr2-109875764-INS-498 |
| chr2-109935019-INS-201 | chr2-109935019-INS-201 |
| chr2-110029577-INS-220 | chr2-110029577-INS-221 |
| chr2-110047702-INS-66 | chr2-110047690-INS-70 |
| chr2-110067905-INS-424 | chr2-110067905-INS-424 |
| chr2-110098924-INS-3349 | chr2-110098925-INS-3334 |
| chr2-110106138-INS-1197 | chr2-110106139-INS-1197 |
| chr2-110112256-INS-6160 | chr2-110112257-INS-6159 |
| chr2-110136510-INS-3171 | chr2-110136511-INS-3171 |
| chr2-110140888-INS-3214 | chr2-110140897-INS-3213 |
| chr2-110153309-INS-56 | chr2-110153309-INS-50 |
| chr2-110160210-INS-107 | chr2-110160213-INS-65 |
| chr2-110202855-INS-206 | chr2-110202855-INS-203 |
| chr2-110216211-INS-232 | chr2-110216211-INS-232 |
| chr2-110227044-INS-109 | chr2-110227044-INS-109 |
| chr2-110229899-INS-2299 | chr2-110229900-INS-2299 |
| chr2-110232945-INS-6680 | chr2-110232946-INS-6680 |
| chr2-110241433-INS-76 | chr2-110241433-INS-72 |
| chr2-110245038-INS-128 | chr2-110245038-INS-128 |
| chr2-110265946-INS-212 | chr2-110265946-INS-212 |
| chr2-110284885-INS-55 | chr2-110284876-INS-64 |
| chr2-110325273-INS-337 | chr2-110325273-INS-337 |
| chr2-110330798-INS-476 | chr2-110330798-INS-476 |
| chr2-110347278-INS-56 | chr2-110347283-INS-58 |
| chr2-110438182-INS-7125 | chr2-110438183-INS-7125 |
| chr2-110444750-INS-137 | chr2-110444750-INS-137 |
| chr2-110472717-INS-137 | chr2-110472717-INS-137 |
| chr2-110546089-INS-6386 | chr2-110546090-INS-6386 |
| chr2-110566807-INS-517 | chr2-110566807-INS-517 |
| chr2-110576797-INS-70 | chr2-110576797-INS-72 |
| chr2-110586341-INS-77 | chr2-110586339-INS-86 |
| chr2-110631965-INS-4338 | chr2-110631966-INS-4338 |
| chr2-110634599-INS-1379 | chr2-110634600-INS-1379 |
| chr2-110637750-INS-490 | chr2-110637750-INS-490 |
| chr2-110645892-INS-5284 | chr2-110645893-INS-5284 |
| chr2-110655718-INS-86 | chr2-110655718-INS-86 |
| chr2-110656482-INS-625 | chr2-110656482-INS-627 |
| chr2-110681362-INS-118 | chr2-110681373-INS-147 |
| chr2-110700848-INS-532 | chr2-110700848-INS-531 |
| chr2-110725243-INS-108 | chr2-110725255-INS-93 |
| chr2-110869713-INS-64 | chr2-110869713-INS-50 |
| chr2-111063205-INS-755 | chr2-111063205-INS-754 |
| chr2-111069127-INS-684 | chr2-111069127-INS-684 |
| chr2-111167395-INS-68 | chr2-111167411-INS-52 |
| chr2-111189120-INS-72 | chr2-111189120-INS-76 |
| chr2-111191413-INS-553 | chr2-111191413-INS-553 |
| chr2-111194515-INS-165 | chr2-111194515-INS-166 |

| | |
|---|---|
| chr2-111238740-INS-7110 | chr2-111238741-INS-7112 |
| chr2-111248366-INS-96 | chr2-111248367-INS-96 |
| chr2-111250381-INS-141 | chr2-111250400-INS-142 |
| chr2-111297634-INS-59 | chr2-111297634-INS-59 |
| chr2-111332178-INS-133 | chr2-111332178-INS-133 |
| chr2-111346738-INS-5657 | chr2-111346739-INS-5657 |
| chr2-111402757-INS-738 | chr2-111402757-INS-739 |
| chr2-111403835-INS-1490 | chr2-111403836-INS-1492 |
| chr2-111428907-INS-6719 | chr2-111428907-INS-6720 |
| chr2-111435619-INS-560 | chr2-111435617-INS-559 |
| chr2-111441056-INS-509 | chr2-111441056-INS-509 |
| chr2-111447524-INS-76 | chr2-111447531-INS-96 |
| chr2-111498846-INS-6847 | chr2-111498847-INS-6853 |
| chr2-111505201-INS-4733 | chr2-111505202-INS-4733 |
| chr2-111529709-INS-6203 | chr2-111529710-INS-6205 |
| chr2-111534617-INS-199 | chr2-111534617-INS-199 |
| chr2-111535034-INS-6053 | chr2-111535035-INS-6053 |
| chr2-111566248-INS-504 | chr2-111566248-INS-504 |
| chr2-111571990-INS-603 | chr2-111571990-INS-603 |
| chr2-111600275-INS-557 | chr2-111600275-INS-557 |
| chr2-111612043-INS-200 | chr2-111612043-INS-200 |
| chr2-111627709-INS-289 | chr2-111627715-INS-274 |
| chr2-111635824-INS-90 | chr2-111635824-INS-90 |
| chr2-111657317-INS-479 | chr2-111657317-INS-480 |
| chr2-111697095-INS-85 | chr2-111697095-INS-56 |
| chr2-111734572-INS-2258 | chr2-111734573-INS-2258 |
| chr2-111805984-INS-6446 | chr2-111805985-INS-6446 |
| chr2-111836264-INS-52 | chr2-111836264-INS-52 |
| chr2-111846789-INS-60 | chr2-111846808-INS-54 |
| chr2-111855686-INS-53 | chr2-111855686-INS-53 |
| chr2-111859877-INS-68 | chr2-111859877-INS-60 |
| chr2-111866072-INS-1351 | chr2-111866073-INS-1351 |
| chr2-111874084-INS-120 | chr2-111874084-INS-90 |
| chr2-111933191-INS-62 | chr2-111933191-INS-62 |
| chr2-111942081-INS-77 | chr2-111942081-INS-81 |
| chr2-111953565-INS-229 | chr2-111953565-INS-229 |
| chr2-111991179-INS-64 | chr2-111991179-INS-64 |
| chr2-111994555-INS-3271 | chr2-111994547-INS-3261 |
| chr2-112030473-INS-62 | chr2-112030465-INS-64 |
| chr2-112037287-INS-880 | chr2-112037287-INS-880 |
| chr2-112040436-INS-5030 | chr2-112040437-INS-5030 |
| chr2-112053939-INS-62 | chr2-112053933-INS-50 |
| chr2-112090463-INS-106 | chr2-112090463-INS-106 |
| chr2-112099659-INS-168 | chr2-112099659-INS-165 |
| chr2-112113555-INS-1363 | chr2-112113556-INS-1356 |
| chr2-112186126-INS-337 | chr2-112186126-INS-337 |
| chr2-112197075-INS-211 | chr2-112197075-INS-212 |
| chr2-112206969-INS-69 | chr2-112206969-INS-70 |
| chr2-112212980-INS-172 | chr2-112212981-INS-173 |
| chr2-112226999-INS-112 | chr2-112226999-INS-112 |
| chr2-112245850-INS-60 | chr2-112245850-INS-60 |
| chr2-112260092-INS-189 | chr2-112260092-INS-188 |
| chr2-112264397-INS-64 | chr2-112264391-INS-88 |
| chr2-112267040-INS-273 | chr2-112267040-INS-275 |
| chr2-112268386-INS-198 | chr2-112268386-INS-198 |
| chr2-112281647-INS-439 | chr2-112281647-INS-439 |
| chr2-112283341-INS-209 | chr2-112283341-INS-209 |
| chr2-112322697-INS-214 | chr2-112322709-INS-198 |
| chr2-112331369-INS-144 | chr2-112331370-INS-115 |
| chr2-112344221-INS-147 | chr2-112344221-INS-148 |
| chr2-112434263-INS-2015 | chr2-112434264-INS-2015 |
| chr2-112448538-INS-593 | chr2-112448538-INS-593 |
| chr2-112453700-INS-80 | chr2-112453700-INS-70 |
| chr2-112458927-INS-174 | chr2-112458927-INS-174 |
| chr2-112490255-INS-195 | chr2-112490255-INS-195 |
| chr2-112496133-INS-84 | chr2-112496134-INS-112 |
| chr2-112496993-INS-55 | chr2-112496981-INS-55 |
| chr2-112506449-INS-53 | chr2-112506449-INS-56 |
| chr2-112507129-INS-805 | chr2-112507129-INS-804 |
| chr2-112549287-INS-82 | chr2-112549287-INS-82 |
| chr2-112596489-INS-238 | chr2-112596489-INS-244 |
| chr2-112609341-INS-68 | chr2-112609341-INS-58 |
| chr2-112614197-INS-7160 | chr2-112614198-INS-7160 |
| chr2-112634461-INS-55 | chr2-112634461-INS-61 |
| chr2-112637078-INS-54 | chr2-112637078-INS-58 |
| chr2-112643314-INS-1437 | chr2-112643315-INS-1437 |
| chr2-112655881-INS-336 | chr2-112655881-INS-364 |

| | |
|---|---|
| chr2-112663682-INS-7401 | chr2-112663683-INS-7401 |
| chr2-112686942-INS-3039 | chr2-112686943-INS-3039 |
| chr2-112704580-INS-95 | chr2-112704580-INS-99 |
| chr2-112724406-INS-66 | chr2-112724406-INS-66 |
| chr2-112807101-INS-66 | chr2-112807101-INS-50 |
| chr2-112827375-INS-62 | chr2-112827375-INS-62 |
| chr2-112844249-INS-52 | chr2-112844249-INS-54 |
| chr2-112900042-INS-119 | chr2-112900042-INS-119 |
| chr2-112909493-INS-60 | chr2-112909493-INS-60 |
| chr2-112910799-INS-78 | chr2-112910799-INS-78 |
| chr2-112952579-INS-110 | chr2-112952579-INS-95 |
| chr2-112978394-INS-362 | chr2-112978394-INS-362 |
| chr2-112979099-INS-61 | chr2-112979099-INS-61 |
| chr2-113011671-INS-6727 | chr2-113011672-INS-6727 |
| chr2-113012402-INS-88 | chr2-113012400-INS-100 |
| chr2-113047915-INS-78 | chr2-113047915-INS-70 |
| chr2-113067799-INS-6321 | chr2-113067800-INS-6323 |
| chr2-113086264-INS-6448 | chr2-113086265-INS-6448 |
| chr2-113096289-INS-85 | chr2-113096289-INS-85 |
| chr2-113102917-INS-5333 | chr2-113102918-INS-5334 |
| chr2-113119021-INS-106 | chr2-113119021-INS-106 |
| chr2-113133023-INS-286 | chr2-113133023-INS-285 |
| chr2-113150601-INS-52 | chr2-113150601-INS-52 |
| chr2-113157604-INS-210 | chr2-113157604-INS-195 |
| chr2-113160073-INS-97 | chr2-113160073-INS-91 |
| chr2-113173578-INS-6249 | chr2-113173579-INS-6250 |
| chr2-113203632-INS-978 | chr2-113203632-INS-978 |
| chr2-113208567-INS-4397 | chr2-113208568-INS-4395 |
| chr2-113218339-INS-60 | chr2-113218339-INS-60 |
| chr2-113241210-INS-62 | chr2-113241210-INS-62 |
| chr2-113265303-INS-56 | chr2-113265303-INS-102 |
| chr2-113298614-INS-198 | chr2-113298614-INS-200 |
| chr2-113322171-INS-52 | chr2-113322171-INS-52 |
| chr2-113367294-INS-62 | chr2-113367294-INS-62 |
| chr2-113385519-INS-6180 | chr2-113385520-INS-6180 |
| chr2-113391854-INS-6251 | chr2-113391855-INS-6251 |
| chr2-113396467-INS-1422 | chr2-113396468-INS-1422 |
| chr2-113416944-INS-57 | chr2-113416944-INS-57 |
| chr2-113440090-INS-7468 | chr2-113440091-INS-7468 |
| chr2-113468849-INS-53 | chr2-113468849-INS-58 |
| chr2-113493709-INS-62 | chr2-113493709-INS-52 |
| chr2-113497873-INS-70 | chr2-113497873-INS-58 |
| chr2-113500026-INS-331 | chr2-113500026-INS-332 |
| chr2-113503656-INS-143 | chr2-113503657-INS-142 |
| chr2-113512774-INS-54 | chr2-113512774-INS-52 |
| chr2-113571925-INS-220 | chr2-113571925-INS-220 |
| chr2-113578622-INS-65 | chr2-113578622-INS-74 |
| chr2-113592273-INS-52 | chr2-113592273-INS-54 |
| chr2-113626094-INS-439 | chr2-113626094-INS-446 |
| chr2-113649793-INS-54 | chr2-113649754-INS-63 |
| chr2-113659748-INS-56 | chr2-113659748-INS-56 |
| chr2-113693189-INS-328 | chr2-113693189-INS-328 |
| chr2-113694814-INS-78 | chr2-113694812-INS-62 |
| chr2-113696753-INS-108 | chr2-113696753-INS-108 |
| chr2-113725028-INS-1676 | chr2-113725029-INS-1676 |
| chr2-113743644-INS-149 | chr2-113743644-INS-149 |
| chr2-113783532-INS-52 | chr2-113783532-INS-52 |
| chr2-113789212-INS-560 | chr2-113789212-INS-560 |
| chr2-113795050-INS-78 | chr2-113795050-INS-87 |
| chr2-113803907-INS-497 | chr2-113803907-INS-497 |
| chr2-113812420-INS-537 | chr2-113812420-INS-536 |
| chr2-113829479-INS-476 | chr2-113829479-INS-477 |
| chr2-113835343-INS-643 | chr2-113835343-INS-645 |
| chr2-113866942-INS-144 | chr2-113866942-INS-132 |
| chr2-113895316-INS-147 | chr2-113895316-INS-147 |
| chr2-113902880-INS-88 | chr2-113902882-INS-92 |
| chr2-113934611-INS-72 | chr2-113934611-INS-74 |
| chr2-114003704-INS-201 | chr2-114003704-INS-201 |
| chr2-114010161-INS-209 | chr2-114010161-INS-209 |
| chr2-114037872-INS-69 | chr2-114037875-INS-56 |
| chr2-114051906-INS-1100 | chr2-114051907-INS-1100 |
| chr2-114054370-INS-62 | chr2-114054370-INS-66 |
| chr2-114112035-INS-57 | chr2-114112035-INS-56 |
| chr2-114131639-INS-236 | chr2-114131639-INS-237 |
| chr2-114141443-INS-207 | chr2-114141443-INS-207 |
| chr2-114185994-INS-102 | chr2-114185994-INS-96 |
| chr2-114189758-INS-1874 | chr2-114189759-INS-1875 |

| | |
|---|---|
| chr2-114240373-INS-62 | chr2-114240373-INS-60 |
| chr2-114244089-INS-56 | chr2-114244089-INS-63 |
| chr2-114251674-INS-60 | chr2-114251661-INS-90 |
| chr2-114277267-INS-6449 | chr2-114277268-INS-6449 |
| chr2-114285161-INS-82 | chr2-114285161-INS-82 |
| chr2-114291845-INS-6240 | chr2-114291846-INS-6240 |
| chr2-114324212-INS-62 | chr2-114324176-INS-63 |
| chr2-114348410-INS-66 | chr2-114348406-INS-78 |
| chr2-114351909-INS-76 | chr2-114351909-INS-124 |
| chr2-114352769-INS-532 | chr2-114352769-INS-532 |
| chr2-114417396-INS-624 | chr2-114417397-INS-627 |
| chr2-114418178-INS-91 | chr2-114418178-INS-107 |
| chr2-114423308-INS-138 | chr2-114423306-INS-154 |
| chr2-114445222-INS-66 | chr2-114445222-INS-57 |
| chr2-114493763-INS-78 | chr2-114493757-INS-72 |
| chr2-114496435-INS-246 | chr2-114496435-INS-246 |
| chr2-114503278-INS-252 | chr2-114503278-INS-254 |
| chr2-114540485-INS-62 | chr2-114540499-INS-57 |
| chr2-114544719-INS-605 | chr2-114544719-INS-602 |
| chr2-114556855-INS-1831 | chr2-114556856-INS-1831 |
| chr2-114565046-INS-65 | chr2-114565046-INS-65 |
| chr2-114584957-INS-54 | chr2-114584957-INS-54 |
| chr2-114601936-INS-1890 | chr2-114601937-INS-1890 |
| chr2-114624941-INS-58 | chr2-114624939-INS-68 |
| chr2-114634903-INS-261 | chr2-114634900-INS-267 |
| chr2-114654313-INS-351 | chr2-114654313-INS-351 |
| chr2-114656016-INS-864 | chr2-114656016-INS-864 |
| chr2-114690276-INS-750 | chr2-114690276-INS-766 |
| chr2-114710983-INS-52 | chr2-114710989-INS-56 |
| chr2-114724399-INS-7039 | chr2-114724400-INS-7056 |
| chr2-114761218-INS-673 | chr2-114761218-INS-673 |
| chr2-114803882-INS-5184 | chr2-114803883-INS-5187 |
| chr2-114814435-INS-63 | chr2-114814435-INS-63 |
| chr2-114826311-INS-2717 | chr2-114826312-INS-2717 |
| chr2-114827716-INS-77 | chr2-114827716-INS-75 |
| chr2-114849981-INS-175 | chr2-114849981-INS-168 |
| chr2-114858276-INS-7050 | chr2-114858277-INS-7050 |
| chr2-114864793-INS-62 | chr2-114864793-INS-62 |
| chr2-114866356-INS-133 | chr2-114866356-INS-133 |
| chr2-114875069-INS-1256 | chr2-114875070-INS-1256 |
| chr2-114885287-INS-89 | chr2-114885287-INS-89 |
| chr2-114929533-INS-56 | chr2-114929533-INS-56 |
| chr2-114972733-INS-957 | chr2-114972733-INS-957 |
| chr2-114974705-INS-62 | chr2-114974701-INS-82 |
| chr2-115045603-INS-1087 | chr2-115045604-INS-1087 |
| chr2-115046030-INS-477 | chr2-115046030-INS-477 |
| chr2-115081472-INS-208 | chr2-115081468-INS-214 |
| chr2-115088424-INS-6432 | chr2-115088425-INS-6432 |
| chr2-115130454-INS-96 | chr2-115130454-INS-90 |
| chr2-115149644-INS-204 | chr2-115149644-INS-204 |
| chr2-115154768-INS-52 | chr2-115154768-INS-52 |
| chr2-115165015-INS-260 | chr2-115165015-INS-257 |
| chr2-115195113-INS-6612 | chr2-115195114-INS-6612 |
| chr2-115197534-INS-58 | chr2-115197534-INS-56 |
| chr2-115204746-INS-436 | chr2-115204749-INS-455 |
| chr2-115226538-INS-55 | chr2-115226538-INS-55 |
| chr2-115231949-INS-62 | chr2-115231949-INS-62 |
| chr2-115233472-INS-6374 | chr2-115233473-INS-6374 |
| chr2-115258034-INS-67 | chr2-115258034-INS-67 |
| chr2-115294535-INS-58 | chr2-115294535-INS-60 |
| chr2-115326969-INS-97 | chr2-115326969-INS-97 |
| chr2-115337807-INS-6703 | chr2-115337808-INS-6706 |
| chr2-115350274-INS-95 | chr2-115350274-INS-85 |
| chr2-115359815-INS-1279 | chr2-115359816-INS-1236 |
| chr2-115370845-INS-200 | chr2-115370845-INS-201 |
| chr2-115399637-INS-213 | chr2-115399637-INS-213 |
| chr2-115413840-INS-66 | chr2-115413840-INS-66 |
| chr2-115436926-INS-61 | chr2-115436893-INS-79 |
| chr2-115445513-INS-202 | chr2-115445513-INS-202 |
| chr2-115458301-INS-504 | chr2-115458301-INS-504 |
| chr2-115475716-INS-52 | chr2-115475716-INS-54 |
| chr2-115485456-INS-68 | chr2-115485457-INS-52 |
| chr2-115492266-INS-59 | chr2-115492272-INS-57 |
| chr2-115496655-INS-4812 | chr2-115496656-INS-4812 |
| chr2-115511734-INS-286 | chr2-115511734-INS-286 |
| chr2-115542596-INS-225 | chr2-115542596-INS-213 |
| chr2-115594433-INS-60 | chr2-115594433-INS-50 |

| | |
|---|---|
| chr2-115609538-INS-504 | chr2-115609538-INS-504 |
| chr2-115649838-INS-54 | chr2-115649838-INS-58 |
| chr2-115651392-INS-57 | chr2-115651392-INS-51 |
| chr2-115659105-INS-214 | chr2-115659105-INS-214 |
| chr2-115708470-INS-248 | chr2-115708470-INS-248 |
| chr2-115712306-INS-58 | chr2-115712306-INS-58 |
| chr2-115725124-INS-55 | chr2-115725124-INS-55 |
| chr2-115760733-INS-82 | chr2-115760733-INS-82 |
| chr2-115763382-INS-193 | chr2-115763382-INS-200 |
| chr2-115768057-INS-76 | chr2-115768057-INS-78 |
| chr2-115770975-INS-52 | chr2-115770975-INS-52 |
| chr2-115777661-INS-145 | chr2-115777661-INS-137 |
| chr2-115789547-INS-209 | chr2-115789547-INS-209 |
| chr2-115823777-INS-60 | chr2-115823777-INS-72 |
| chr2-115835100-INS-101 | chr2-115835100-INS-74 |
| chr2-115865781-INS-102 | chr2-115865781-INS-102 |
| chr2-115890495-INS-75 | chr2-115890495-INS-60 |
| chr2-115921062-INS-70 | chr2-115921043-INS-78 |
| chr2-115964943-INS-56 | chr2-115964943-INS-52 |
| chr2-115976217-INS-692 | chr2-115976217-INS-692 |
| chr2-115978697-INS-6178 | chr2-115978698-INS-6179 |
| chr2-115982707-INS-108 | chr2-115982707-INS-108 |
| chr2-116035446-INS-53 | chr2-116035446-INS-75 |
| chr2-116090166-INS-6741 | chr2-116090167-INS-6741 |
| chr2-116095429-INS-52 | chr2-116095429-INS-52 |
| chr2-116111870-INS-56 | chr2-116111845-INS-72 |
| chr2-116164443-INS-56 | chr2-116164443-INS-68 |
| chr2-116190357-INS-52 | chr2-116190350-INS-76 |
| chr2-116192996-INS-56 | chr2-116192996-INS-72 |
| chr2-116211556-INS-55 | chr2-116211556-INS-53 |
| chr2-116218283-INS-841 | chr2-116218283-INS-849 |
| chr2-116222527-INS-6608 | chr2-116222528-INS-6608 |
| chr2-116229262-INS-76 | chr2-116229262-INS-94 |
| chr2-116231759-INS-78 | chr2-116231759-INS-76 |
| chr2-116247818-INS-65 | chr2-116247818-INS-50 |
| chr2-116250898-INS-71 | chr2-116250898-INS-71 |
| chr2-116257129-INS-6587 | chr2-116257130-INS-6587 |
| chr2-116363337-INS-67 | chr2-116363337-INS-67 |
| chr2-116380858-INS-116 | chr2-116380858-INS-116 |
| chr2-116386631-INS-150 | chr2-116386632-INS-150 |
| chr2-116391943-INS-58 | chr2-116391944-INS-58 |
| chr2-116413460-INS-1601 | chr2-116413461-INS-1601 |
| chr2-116415437-INS-894 | chr2-116415437-INS-896 |
| chr2-116451330-INS-3140 | chr2-116451331-INS-3140 |
| chr2-116452445-INS-65 | chr2-116452445-INS-65 |
| chr2-116458030-INS-5388 | chr2-116458031-INS-5388 |
| chr2-116460391-INS-6471 | chr2-116460392-INS-6471 |
| chr2-116472289-INS-462 | chr2-116472289-INS-462 |
| chr2-116487314-INS-234 | chr2-116487314-INS-234 |
| chr2-116506008-INS-110 | chr2-116506008-INS-72 |
| chr2-116545920-INS-52 | chr2-116545920-INS-50 |
| chr2-116560191-INS-261 | chr2-116560191-INS-262 |
| chr2-116610939-INS-132 | chr2-116610942-INS-128 |
| chr2-116611886-INS-238 | chr2-116611887-INS-238 |
| chr2-116616867-INS-68 | chr2-116616867-INS-68 |
| chr2-116620445-INS-83 | chr2-116620445-INS-82 |
| chr2-116636385-INS-96 | chr2-116636385-INS-96 |
| chr2-116642671-INS-66 | chr2-116642671-INS-66 |
| chr2-116654387-INS-84 | chr2-116654389-INS-84 |
| chr2-116656252-INS-644 | chr2-116656252-INS-644 |
| chr2-116685674-INS-109 | chr2-116685674-INS-104 |
| chr2-116703348-INS-401 | chr2-116703348-INS-401 |
| chr2-116723472-INS-7190 | chr2-116723473-INS-7190 |
| chr2-116731813-INS-7488 | chr2-116731814-INS-7488 |
| chr2-116757572-INS-7126 | chr2-116757573-INS-7126 |
| chr2-116764779-INS-166 | chr2-116764779-INS-160 |
| chr2-116769344-INS-204 | chr2-116769344-INS-204 |
| chr2-116785095-INS-299 | chr2-116785095-INS-299 |
| chr2-116786218-INS-110 | chr2-116786226-INS-132 |
| chr2-116801614-INS-81 | chr2-116801614-INS-81 |
| chr2-116804412-INS-106 | chr2-116804405-INS-91 |
| chr2-116843152-INS-60 | chr2-116843152-INS-60 |
| chr2-116885470-INS-95 | chr2-116885471-INS-111 |
| chr2-116895057-INS-62 | chr2-116895057-INS-62 |
| chr2-116947676-INS-5146 | chr2-116947677-INS-5144 |
| chr2-116958870-INS-52 | chr2-116958870-INS-67 |
| chr2-116967019-INS-1064 | chr2-116967020-INS-1065 |

| | |
|---|---|
| chr2-116980873-INS-60 | chr2-116980881-INS-52 |
| chr2-117000750-INS-68 | chr2-117000750-INS-68 |
| chr2-117033614-INS-76 | chr2-117033587-INS-54 |
| chr2-117037212-INS-73 | chr2-117037203-INS-112 |
| chr2-117042172-INS-179 | chr2-117042172-INS-179 |
| chr2-117056479-INS-428 | chr2-117056479-INS-428 |
| chr2-117057595-INS-825 | chr2-117057595-INS-825 |
| chr2-117066091-INS-92 | chr2-117066082-INS-126 |
| chr2-117067847-INS-489 | chr2-117067847-INS-495 |
| chr2-117071286-INS-52 | chr2-117071286-INS-71 |
| chr2-117132059-INS-171 | chr2-117132059-INS-171 |
| chr2-117161586-INS-174 | chr2-117161586-INS-175 |
| chr2-117178280-INS-91 | chr2-117178280-INS-78 |
| chr2-117190621-INS-446 | chr2-117190621-INS-446 |
| chr2-117194266-INS-65 | chr2-117194270-INS-51 |
| chr2-117198396-INS-138 | chr2-117198396-INS-138 |
| chr2-117209324-INS-79 | chr2-117209320-INS-51 |
| chr2-117215547-INS-593 | chr2-117215552-INS-592 |
| chr2-117224985-INS-52 | chr2-117224985-INS-52 |
| chr2-117247118-INS-60 | chr2-117247118-INS-55 |
| chr2-117266018-INS-137 | chr2-117266018-INS-137 |
| chr2-117267850-INS-52 | chr2-117267850-INS-52 |
| chr2-117267999-INS-6398 | chr2-117268000-INS-6398 |
| chr2-117272509-INS-435 | chr2-117272509-INS-435 |
| chr2-117273170-INS-54 | chr2-117273170-INS-58 |
| chr2-117299571-INS-50 | chr2-117299533-INS-83 |
| chr2-117317736-INS-624 | chr2-117317736-INS-624 |
| chr2-117321403-INS-2893 | chr2-117321404-INS-2893 |
| chr2-117356226-INS-1958 | chr2-117356227-INS-1958 |
| chr2-117360380-INS-161 | chr2-117360381-INS-165 |
| chr2-117360443-INS-2631 | chr2-117360444-INS-2624 |
| chr2-117391523-INS-876 | chr2-117391523-INS-876 |
| chr2-117394898-INS-620 | chr2-117394897-INS-621 |
| chr2-117404906-INS-58 | chr2-117404906-INS-74 |
| chr2-117424298-INS-1151 | chr2-117424299-INS-1151 |
| chr2-117449381-INS-7240 | chr2-117449382-INS-7240 |
| chr2-117450007-INS-62 | chr2-117450007-INS-66 |
| chr2-117467532-INS-52 | chr2-117467532-INS-50 |
| chr2-117474160-INS-531 | chr2-117474160-INS-531 |
| chr2-117508716-INS-7124 | chr2-117508716-INS-7124 |
| chr2-117508749-INS-65 | chr2-117508717-INS-65 |
| chr2-117523417-INS-5098 | chr2-117523418-INS-5102 |
| chr2-117526663-INS-52 | chr2-117526663-INS-52 |
| chr2-117528755-INS-114 | chr2-117528755-INS-115 |
| chr2-117529010-INS-7124 | chr2-117529011-INS-7124 |
| chr2-117540045-INS-756 | chr2-117540045-INS-756 |
| chr2-117554945-INS-591 | chr2-117554943-INS-598 |
| chr2-117559131-INS-6665 | chr2-117559132-INS-6665 |
| chr2-117564391-INS-498 | chr2-117564391-INS-498 |
| chr2-117565315-INS-63 | chr2-117565288-INS-93 |
| chr2-117571875-INS-593 | chr2-117571875-INS-594 |
| chr2-117596279-INS-56 | chr2-117596279-INS-60 |
| chr2-117605818-INS-78 | chr2-117605818-INS-78 |
| chr2-117625132-INS-64 | chr2-117625132-INS-50 |
| chr2-117637255-INS-476 | chr2-117637259-INS-477 |
| chr2-117690457-INS-3100 | chr2-117690458-INS-3100 |
| chr2-117707664-INS-98 | chr2-117707664-INS-130 |
| chr2-117709346-INS-233 | chr2-117709346-INS-233 |
| chr2-117721862-INS-305 | chr2-117721865-INS-313 |
| chr2-117748976-INS-199 | chr2-117748976-INS-198 |
| chr2-117754527-INS-118 | chr2-117754527-INS-118 |
| chr2-117795252-INS-90 | chr2-117795241-INS-93 |
| chr2-117804400-INS-97 | chr2-117804400-INS-72 |
| chr2-117812241-INS-212 | chr2-117812241-INS-212 |
| chr2-117825476-INS-54 | chr2-117825476-INS-54 |
| chr2-117858387-INS-72 | chr2-117858433-INS-59 |
| chr2-117861814-INS-146 | chr2-117861817-INS-147 |
| chr2-117862620-INS-163 | chr2-117862590-INS-170 |
| chr2-117871830-INS-6681 | chr2-117871831-INS-6681 |
| chr2-117913532-INS-58 | chr2-117913532-INS-56 |
| chr2-117939967-INS-198 | chr2-117939967-INS-198 |
| chr2-117943228-INS-56 | chr2-117943228-INS-52 |
| chr2-117968706-INS-92 | chr2-117968724-INS-88 |
| chr2-117996964-INS-98 | chr2-117996964-INS-88 |
| chr2-118006444-INS-52 | chr2-118006444-INS-52 |
| chr2-118027758-INS-6466 | chr2-118027759-INS-6466 |
| chr2-118037591-INS-84 | chr2-118037591-INS-84 |

| | |
|---|---|
| chr2-118071500-INS-176 | chr2-118071500-INS-176 |
| chr2-118073943-INS-56 | chr2-118073943-INS-61 |
| chr2-118077924-INS-399 | chr2-118077924-INS-398 |
| chr2-118079454-INS-3120 | chr2-118079455-INS-3121 |
| chr2-118094901-INS-99 | chr2-118094901-INS-99 |
| chr2-118131111-INS-70 | chr2-118131111-INS-70 |
| chr2-118181616-INS-413 | chr2-118181616-INS-429 |
| chr2-118196965-INS-52 | chr2-118196966-INS-52 |
| chr2-118197918-INS-746 | chr2-118197918-INS-746 |
| chr2-118209085-INS-100 | chr2-118209085-INS-100 |
| chr2-118214148-INS-204 | chr2-118214148-INS-204 |
| chr2-118246720-INS-208 | chr2-118246707-INS-205 |
| chr2-118254019-INS-71 | chr2-118254037-INS-72 |
| chr2-118257461-INS-320 | chr2-118257462-INS-324 |
| chr2-118384702-INS-562 | chr2-118384703-INS-564 |
| chr2-118421205-INS-3122 | chr2-118421206-INS-3122 |
| chr2-118421206-INS-437 | chr2-118421206-INS-437 |
| chr2-118422571-INS-6475 | chr2-118422572-INS-6463 |
| chr2-118450724-INS-72 | chr2-118450724-INS-72 |
| chr2-118460091-INS-798 | chr2-118460091-INS-806 |
| chr2-118465340-INS-3507 | chr2-118465341-INS-3508 |
| chr2-118485809-INS-216 | chr2-118485809-INS-216 |
| chr2-118492625-INS-232 | chr2-118492625-INS-233 |
| chr2-118525923-INS-70 | chr2-118525923-INS-68 |
| chr2-118570715-INS-715 | chr2-118570715-INS-714 |
| chr2-118572328-INS-207 | chr2-118572328-INS-207 |
| chr2-118630169-INS-214 | chr2-118630169-INS-213 |
| chr2-118652751-INS-70 | chr2-118652751-INS-70 |
| chr2-118656270-INS-174 | chr2-118656270-INS-176 |
| chr2-118712816-INS-51 | chr2-118712816-INS-51 |
| chr2-118730266-INS-121 | chr2-118730276-INS-105 |
| chr2-118780136-INS-56 | chr2-118780108-INS-85 |
| chr2-118790072-INS-58 | chr2-118790077-INS-50 |
| chr2-118810728-INS-55 | chr2-118810730-INS-78 |
| chr2-118835466-INS-444 | chr2-118835466-INS-444 |
| chr2-118854692-INS-205 | chr2-118854692-INS-205 |
| chr2-118855168-INS-1712 | chr2-118855169-INS-1705 |
| chr2-118875675-INS-182 | chr2-118875675-INS-167 |
| chr2-118883215-INS-92 | chr2-118883215-INS-92 |
| chr2-118885488-INS-202 | chr2-118885488-INS-202 |
| chr2-118903426-INS-1380 | chr2-118903427-INS-1391 |
| chr2-118910425-INS-223 | chr2-118910426-INS-223 |
| chr2-118924934-INS-120 | chr2-118924925-INS-89 |
| chr2-118929523-INS-106 | chr2-118929523-INS-106 |
| chr2-118973640-INS-439 | chr2-118973640-INS-440 |
| chr2-118987696-INS-206 | chr2-118987696-INS-207 |
| chr2-119021229-INS-233 | chr2-119021229-INS-258 |
| chr2-119033862-INS-211 | chr2-119033862-INS-203 |
| chr2-119042516-INS-103 | chr2-119042516-INS-103 |
| chr2-119074678-INS-172 | chr2-119074678-INS-175 |
| chr2-119075451-INS-618 | chr2-119075451-INS-628 |
| chr2-119092273-INS-54 | chr2-119092274-INS-72 |
| chr2-119093145-INS-220 | chr2-119093153-INS-201 |
| chr2-119106954-INS-53 | chr2-119106954-INS-53 |
| chr2-119121898-INS-106 | chr2-119121898-INS-106 |
| chr2-119160418-INS-145 | chr2-119160407-INS-140 |
| chr2-119170585-INS-188 | chr2-119170585-INS-194 |
| chr2-119171796-INS-786 | chr2-119171796-INS-786 |
| chr2-119174805-INS-233 | chr2-119174805-INS-233 |
| chr2-119188132-INS-54 | chr2-119188136-INS-64 |
| chr2-119195145-INS-314 | chr2-119195145-INS-321 |
| chr2-119197299-INS-409 | chr2-119197299-INS-409 |
| chr2-119218558-INS-204 | chr2-119218558-INS-195 |
| chr2-119220858-INS-205 | chr2-119220858-INS-205 |
| chr2-119233190-INS-463 | chr2-119233190-INS-463 |
| chr2-119248746-INS-220 | chr2-119248746-INS-220 |
| chr2-119256909-INS-160 | chr2-119256861-INS-207 |
| chr2-119319799-INS-54 | chr2-119319799-INS-60 |
| chr2-119353173-INS-153 | chr2-119353175-INS-152 |
| chr2-119355229-INS-228 | chr2-119355229-INS-226 |
| chr2-119359984-INS-196 | chr2-119359984-INS-195 |
| chr2-119368955-INS-214 | chr2-119368955-INS-214 |
| chr2-119373094-INS-3171 | chr2-119373095-INS-3171 |
| chr2-119373766-INS-74 | chr2-119373766-INS-74 |
| chr2-119388021-INS-62 | chr2-119388013-INS-70 |
| chr2-119388967-INS-200 | chr2-119388967-INS-199 |
| chr2-119407014-INS-199 | chr2-119407014-INS-199 |

| | |
|---|---|
| chr2-119408069-INS-214 | chr2-119408069-INS-214 |
| chr2-119409662-INS-442 | chr2-119409662-INS-444 |
| chr2-119410520-INS-218 | chr2-119410520-INS-219 |
| chr2-119412936-INS-95 | chr2-119412936-INS-95 |
| chr2-119420110-INS-708 | chr2-119420110-INS-689 |
| chr2-119426593-INS-65 | chr2-119426593-INS-65 |
| chr2-119465264-INS-218 | chr2-119465264-INS-218 |
| chr2-119475930-INS-182 | chr2-119475908-INS-162 |
| chr2-119481204-INS-195 | chr2-119481204-INS-188 |
| chr2-119514192-INS-180 | chr2-119514209-INS-192 |
| chr2-119543987-INS-150 | chr2-119543975-INS-120 |
| chr2-119546188-INS-52 | chr2-119546188-INS-52 |
| chr2-119566384-INS-232 | chr2-119566384-INS-234 |
| chr2-119591555-INS-208 | chr2-119591555-INS-208 |
| chr2-119597340-INS-171 | chr2-119597340-INS-178 |
| chr2-119599313-INS-141 | chr2-119599306-INS-116 |
| chr2-119622603-INS-205 | chr2-119622603-INS-205 |
| chr2-119650704-INS-87 | chr2-119650704-INS-63 |
| chr2-119653345-INS-54 | chr2-119653345-INS-54 |
| chr2-119665044-INS-118 | chr2-119665046-INS-117 |
| chr2-119667432-INS-67 | chr2-119667432-INS-65 |
| chr2-119670038-INS-244 | chr2-119670038-INS-244 |
| chr2-119683381-INS-223 | chr2-119683381-INS-224 |
| chr2-119685485-INS-210 | chr2-119685485-INS-210 |
| chr2-119710279-INS-53 | chr2-119710279-INS-56 |
| chr2-119712335-INS-86 | chr2-119712326-INS-109 |
| chr2-119712620-INS-403 | chr2-119712620-INS-403 |
| chr2-119722687-INS-216 | chr2-119722687-INS-209 |
| chr2-119769375-INS-208 | chr2-119769375-INS-208 |
| chr2-119798755-INS-221 | chr2-119798755-INS-221 |
| chr2-119815442-INS-230 | chr2-119815442-INS-229 |
| chr2-119815475-INS-6466 | chr2-119815476-INS-6466 |
| chr2-119849812-INS-154 | chr2-119849812-INS-154 |
| chr2-119887476-INS-86 | chr2-119887476-INS-86 |
| chr2-119930214-INS-226 | chr2-119930214-INS-227 |
| chr2-119931771-INS-491 | chr2-119931771-INS-489 |
| chr2-119963730-INS-64 | chr2-119963730-INS-60 |
| chr2-119978230-INS-212 | chr2-119978230-INS-216 |
| chr2-119983679-INS-230 | chr2-119983679-INS-231 |
| chr2-120005507-INS-216 | chr2-120005507-INS-217 |
| chr2-120009010-INS-208 | chr2-120009010-INS-208 |
| chr2-120019521-INS-209 | chr2-120019521-INS-209 |
| chr2-120022739-INS-1892 | chr2-120022740-INS-1892 |
| chr2-120023286-INS-73 | chr2-120023286-INS-73 |
| chr2-120072939-INS-211 | chr2-120072952-INS-217 |
| chr2-120112790-INS-181 | chr2-120112790-INS-181 |
| chr2-120116460-INS-50 | chr2-120116441-INS-64 |
| chr2-120120073-INS-52 | chr2-120120073-INS-52 |
| chr2-120127069-INS-56 | chr2-120127070-INS-56 |
| chr2-120136681-INS-101 | chr2-120136681-INS-57 |
| chr2-120159201-INS-522 | chr2-120159201-INS-521 |
| chr2-120201255-INS-75 | chr2-120201255-INS-75 |
| chr2-120203672-INS-223 | chr2-120203672-INS-223 |
| chr2-120224676-INS-210 | chr2-120224676-INS-207 |
| chr2-120232241-INS-83 | chr2-120232241-INS-83 |
| chr2-120237976-INS-826 | chr2-120237976-INS-832 |
| chr2-120256918-INS-115 | chr2-120256968-INS-68 |
| chr2-120258443-INS-51 | chr2-120258443-INS-51 |
| chr2-120260690-INS-210 | chr2-120260691-INS-210 |
| chr2-120267970-INS-201 | chr2-120267970-INS-201 |
| chr2-120298323-INS-63 | chr2-120298323-INS-62 |
| chr2-120309168-INS-186 | chr2-120309168-INS-186 |
| chr2-120314368-INS-965 | chr2-120314368-INS-965 |
| chr2-120321020-INS-56 | chr2-120321020-INS-56 |
| chr2-120350269-INS-87 | chr2-120350269-INS-87 |
| chr2-120366220-INS-67 | chr2-120366220-INS-67 |
| chr2-120376941-INS-121 | chr2-120376941-INS-121 |
| chr2-120392184-INS-167 | chr2-120392184-INS-167 |
| chr2-120407756-INS-201 | chr2-120407756-INS-201 |
| chr2-120426789-INS-6179 | chr2-120426790-INS-6179 |
| chr2-120442266-INS-172 | chr2-120442266-INS-172 |
| chr2-120444791-INS-243 | chr2-120444792-INS-242 |
| chr2-120455279-INS-206 | chr2-120455279-INS-206 |
| chr2-120473227-INS-174 | chr2-120473250-INS-189 |
| chr2-120478397-INS-216 | chr2-120478397-INS-216 |
| chr2-120484447-INS-141 | chr2-120484454-INS-137 |
| chr2-120486689-INS-342 | chr2-120486689-INS-342 |

| | |
|---|---|
| chr2-120487906-INS-186 | chr2-120487906-INS-184 |
| chr2-120489978-INS-206 | chr2-120489979-INS-159 |
| chr2-120492566-INS-182 | chr2-120492566-INS-182 |
| chr2-120503804-INS-208 | chr2-120503804-INS-207 |
| chr2-120508956-INS-207 | chr2-120508956-INS-211 |
| chr2-120515004-INS-443 | chr2-120515004-INS-444 |
| chr2-120539339-INS-2272 | chr2-120539348-INS-2256 |
| chr2-120577011-INS-804 | chr2-120577011-INS-804 |
| chr2-120667190-INS-55 | chr2-120667190-INS-60 |
| chr2-120697159-INS-96 | chr2-120697159-INS-96 |
| chr2-120708206-INS-289 | chr2-120708206-INS-286 |
| chr2-120709544-INS-212 | chr2-120709544-INS-210 |
| chr2-120741756-INS-424 | chr2-120741756-INS-424 |
| chr2-120745745-INS-180 | chr2-120745745-INS-181 |
| chr2-120746682-INS-62 | chr2-120746680-INS-65 |
| chr2-120756846-INS-212 | chr2-120756846-INS-212 |
| chr2-120768596-INS-221 | chr2-120768596-INS-222 |
| chr2-120781099-INS-64 | chr2-120781099-INS-69 |
| chr2-120847564-INS-1414 | chr2-120847565-INS-1414 |
| chr2-120855758-INS-226 | chr2-120855758-INS-226 |
| chr2-120865574-INS-238 | chr2-120865574-INS-238 |
| chr2-120875188-INS-72 | chr2-120875188-INS-72 |
| chr2-120907616-INS-58 | chr2-120907616-INS-58 |
| chr2-120909448-INS-96 | chr2-120909448-INS-96 |
| chr2-120914709-INS-158 | chr2-120914709-INS-158 |
| chr2-120935162-INS-89 | chr2-120935162-INS-89 |
| chr2-120964601-INS-547 | chr2-120964601-INS-547 |
| chr2-120976636-INS-531 | chr2-120976637-INS-531 |
| chr2-120980273-INS-54 | chr2-120980273-INS-60 |
| chr2-121049466-INS-73 | chr2-121049466-INS-73 |
| chr2-121093362-INS-205 | chr2-121093362-INS-205 |
| chr2-121136770-INS-60 | chr2-121136770-INS-72 |
| chr2-121149931-INS-140 | chr2-121149931-INS-92 |
| chr2-121186897-INS-142 | chr2-121186897-INS-143 |
| chr2-121222289-INS-216 | chr2-121222289-INS-216 |
| chr2-121229408-INS-64 | chr2-121229408-INS-64 |
| chr2-121254849-INS-225 | chr2-121254849-INS-225 |
| chr2-121257588-INS-195 | chr2-121257588-INS-195 |
| chr2-121276503-INS-67 | chr2-121276503-INS-63 |
| chr2-121291564-INS-209 | chr2-121291564-INS-209 |
| chr2-121295716-INS-745 | chr2-121295717-INS-746 |
| chr2-121296572-INS-206 | chr2-121296572-INS-206 |
| chr2-121312365-INS-1863 | chr2-121312366-INS-1863 |
| chr2-121328266-INS-745 | chr2-121328267-INS-745 |
| chr2-121329122-INS-205 | chr2-121329122-INS-205 |
| chr2-121346639-INS-52 | chr2-121346641-INS-52 |
| chr2-121353549-INS-212 | chr2-121353549-INS-214 |
| chr2-121355282-INS-185 | chr2-121355282-INS-185 |
| chr2-121358794-INS-178 | chr2-121358794-INS-178 |
| chr2-121359976-INS-55 | chr2-121359976-INS-59 |
| chr2-121366634-INS-201 | chr2-121366634-INS-204 |
| chr2-121382634-INS-88 | chr2-121382634-INS-88 |
| chr2-121384630-INS-98 | chr2-121384630-INS-98 |
| chr2-121385468-INS-177 | chr2-121385468-INS-175 |
| chr2-121386784-INS-126 | chr2-121386782-INS-87 |
| chr2-121402839-INS-165 | chr2-121402839-INS-198 |
| chr2-121417832-INS-177 | chr2-121417832-INS-177 |
| chr2-121427164-INS-1891 | chr2-121427165-INS-1891 |
| chr2-121431734-INS-260 | chr2-121431734-INS-260 |
| chr2-121434693-INS-58 | chr2-121434693-INS-88 |
| chr2-121477422-INS-314 | chr2-121477422-INS-314 |
| chr2-121488086-INS-205 | chr2-121488086-INS-205 |
| chr2-121492410-INS-182 | chr2-121492410-INS-182 |
| chr2-121504743-INS-202 | chr2-121504736-INS-217 |
| chr2-121521873-INS-86 | chr2-121521873-INS-86 |
| chr2-121552142-INS-88 | chr2-121552142-INS-50 |
| chr2-121572347-INS-658 | chr2-121572347-INS-636 |
| chr2-121586907-INS-244 | chr2-121586907-INS-274 |
| chr2-121590528-INS-435 | chr2-121590529-INS-411 |
| chr2-121595074-INS-157 | chr2-121595074-INS-157 |
| chr2-121605526-INS-363 | chr2-121605526-INS-363 |
| chr2-121608763-INS-59 | chr2-121608763-INS-59 |
| chr2-121617304-INS-54 | chr2-121617304-INS-54 |
| chr2-121656215-INS-207 | chr2-121656215-INS-222 |
| chr2-121692766-INS-226 | chr2-121692766-INS-226 |
| chr2-121698695-INS-162 | chr2-121698695-INS-162 |
| chr2-121720858-INS-176 | chr2-121720844-INS-184 |

| | |
|---|---|
| chr2-121740715-INS-429 | chr2-121740715-INS-429 |
| chr2-121807549-INS-91 | chr2-121807549-INS-95 |
| chr2-121813513-INS-180 | chr2-121813513-INS-187 |
| chr2-121823511-INS-562 | chr2-121823511-INS-572 |
| chr2-121832914-INS-74 | chr2-121832914-INS-74 |
| chr2-121850815-INS-207 | chr2-121850815-INS-206 |
| chr2-121855254-INS-281 | chr2-121855254-INS-282 |
| chr2-121925718-INS-64 | chr2-121925718-INS-64 |
| chr2-121928431-INS-186 | chr2-121928431-INS-186 |
| chr2-121932910-INS-69 | chr2-121932910-INS-69 |
| chr2-121945449-INS-54 | chr2-121945449-INS-54 |
| chr2-121966561-INS-6475 | chr2-121966562-INS-6475 |
| chr2-121979840-INS-52 | chr2-121979840-INS-52 |
| chr2-121986031-INS-205 | chr2-121986031-INS-205 |
| chr2-121999068-INS-90 | chr2-121999068-INS-90 |
| chr2-122002968-INS-65 | chr2-122002984-INS-67 |
| chr2-122031567-INS-63 | chr2-122031567-INS-63 |
| chr2-122039237-INS-54 | chr2-122039237-INS-52 |
| chr2-122042229-INS-86 | chr2-122042229-INS-104 |
| chr2-122052752-INS-201 | chr2-122052752-INS-200 |
| chr2-122056529-INS-106 | chr2-122056529-INS-106 |
| chr2-122066611-INS-935 | chr2-122066611-INS-934 |
| chr2-122074794-INS-231 | chr2-122074794-INS-231 |
| chr2-122080674-INS-78 | chr2-122080674-INS-78 |
| chr2-122081598-INS-246 | chr2-122081598-INS-246 |
| chr2-122095395-INS-317 | chr2-122095395-INS-317 |
| chr2-122194095-INS-102 | chr2-122194095-INS-102 |
| chr2-122202753-INS-163 | chr2-122202753-INS-163 |
| chr2-122216237-INS-452 | chr2-122216229-INS-476 |
| chr2-122219578-INS-80 | chr2-122219578-INS-80 |
| chr2-122232769-INS-137 | chr2-122232770-INS-137 |
| chr2-122248277-INS-82 | chr2-122248277-INS-74 |
| chr2-122292693-INS-168 | chr2-122292693-INS-168 |
| chr2-122304760-INS-52 | chr2-122304760-INS-52 |
| chr2-122330656-INS-637 | chr2-122330655-INS-644 |
| chr2-122335127-INS-136 | chr2-122335127-INS-136 |
| chr2-122413018-INS-220 | chr2-122413018-INS-220 |
| chr2-122417575-INS-81 | chr2-122417575-INS-81 |
| chr2-122433587-INS-100 | chr2-122433587-INS-104 |
| chr2-122442164-INS-201 | chr2-122442164-INS-201 |
| chr2-122445040-INS-235 | chr2-122445040-INS-235 |
| chr2-122450554-INS-198 | chr2-122450554-INS-198 |
| chr2-122463385-INS-59 | chr2-122463385-INS-53 |
| chr2-122501631-INS-456 | chr2-122501631-INS-456 |
| chr2-122517387-INS-52 | chr2-122517387-INS-52 |
| chr2-122522763-INS-408 | chr2-122522763-INS-408 |
| chr2-122531762-INS-4330 | chr2-122531763-INS-4330 |
| chr2-122606170-INS-2617 | chr2-122606171-INS-2617 |
| chr2-122612172-INS-79 | chr2-122612172-INS-85 |
| chr2-122617388-INS-616 | chr2-122617388-INS-616 |
| chr2-122628349-INS-349 | chr2-122628349-INS-349 |
| chr2-122637897-INS-2203 | chr2-122637898-INS-2203 |
| chr2-122679409-INS-6549 | chr2-122679410-INS-6549 |
| chr2-122685527-INS-71 | chr2-122685527-INS-75 |
| chr2-122691468-INS-115 | chr2-122691466-INS-134 |
| chr2-122717158-INS-70 | chr2-122717158-INS-72 |
| chr2-122734756-INS-496 | chr2-122734756-INS-496 |
| chr2-122799031-INS-82 | chr2-122799052-INS-72 |
| chr2-122854227-INS-1313 | chr2-122854228-INS-1313 |
| chr2-122856786-INS-54 | chr2-122856786-INS-54 |
| chr2-122858376-INS-1888 | chr2-122858377-INS-1888 |
| chr2-122871175-INS-7725 | chr2-122871176-INS-7724 |
| chr2-122878242-INS-1193 | chr2-122878243-INS-1193 |
| chr2-122926151-INS-88 | chr2-122926151-INS-92 |
| chr2-122946195-INS-83 | chr2-122946195-INS-83 |
| chr2-122961837-INS-52 | chr2-122961837-INS-52 |
| chr2-122994644-INS-429 | chr2-122994644-INS-429 |
| chr2-123016316-INS-52 | chr2-123016316-INS-50 |
| chr2-123018210-INS-211 | chr2-123018238-INS-229 |
| chr2-123037007-INS-68 | chr2-123037007-INS-68 |
| chr2-123067697-INS-75 | chr2-123067737-INS-75 |
| chr2-123079019-INS-299 | chr2-123079019-INS-299 |
| chr2-123082397-INS-132 | chr2-123082397-INS-130 |
| chr2-123082399-INS-638 | chr2-123082399-INS-642 |
| chr2-123102081-INS-218 | chr2-123102081-INS-222 |
| chr2-123106854-INS-712 | chr2-123106854-INS-708 |
| chr2-123126845-INS-319 | chr2-123126845-INS-320 |

| | |
|---|---|
| chr2-123130086-INS-53 | chr2-123130086-INS-53 |
| chr2-123152354-INS-2599 | chr2-123152355-INS-2599 |
| chr2-123164272-INS-1780 | chr2-123164273-INS-1782 |
| chr2-123217566-INS-842 | chr2-123217569-INS-848 |
| chr2-123222003-INS-127 | chr2-123222004-INS-120 |
| chr2-123243104-INS-57 | chr2-123243104-INS-57 |
| chr2-123249955-INS-4204 | chr2-123249956-INS-4205 |
| chr2-123254181-INS-88 | chr2-123254181-INS-88 |
| chr2-123284933-INS-6603 | chr2-123284934-INS-6603 |
| chr2-123349949-INS-52 | chr2-123349947-INS-84 |
| chr2-123370053-INS-83 | chr2-123370049-INS-94 |
| chr2-123379404-INS-248 | chr2-123379404-INS-248 |
| chr2-123402387-INS-1936 | chr2-123402388-INS-1936 |
| chr2-123415440-INS-60 | chr2-123415431-INS-58 |
| chr2-123433271-INS-7376 | chr2-123433272-INS-7379 |
| chr2-123468402-INS-239 | chr2-123468402-INS-239 |
| chr2-123526445-INS-6487 | chr2-123526446-INS-6487 |
| chr2-123536083-INS-1350 | chr2-123536084-INS-1356 |
| chr2-123542960-INS-332 | chr2-123542960-INS-332 |
| chr2-123547414-INS-70 | chr2-123547414-INS-70 |
| chr2-123584974-INS-58 | chr2-123584974-INS-58 |
| chr2-123608860-INS-6875 | chr2-123608861-INS-6877 |
| chr2-123617380-INS-1665 | chr2-123617381-INS-1665 |
| chr2-123622154-INS-58 | chr2-123622154-INS-58 |
| chr2-123642637-INS-84 | chr2-123642637-INS-81 |
| chr2-123676066-INS-1069 | chr2-123676067-INS-1070 |
| chr2-123685728-INS-65 | chr2-123685728-INS-87 |
| chr2-123687033-INS-851 | chr2-123687032-INS-851 |
| chr2-123692751-INS-52 | chr2-123692751-INS-52 |
| chr2-123695891-INS-60 | chr2-123695891-INS-60 |
| chr2-123713350-INS-4223 | chr2-123713351-INS-4221 |
| chr2-123763397-INS-143 | chr2-123763397-INS-141 |
| chr2-123839841-INS-6244 | chr2-123839841-INS-6221 |
| chr2-123842807-INS-6647 | chr2-123842808-INS-6647 |
| chr2-123843708-INS-58 | chr2-123843708-INS-51 |
| chr2-123857892-INS-488 | chr2-123857892-INS-488 |
| chr2-123878569-INS-60 | chr2-123878565-INS-80 |
| chr2-123881426-INS-361 | chr2-123881426-INS-361 |
| chr2-123892779-INS-419 | chr2-123892782-INS-423 |
| chr2-123898572-INS-1179 | chr2-123898573-INS-1179 |
| chr2-123988628-INS-80 | chr2-123988628-INS-78 |
| chr2-123995195-INS-89 | chr2-123995195-INS-89 |
| chr2-124012552-INS-140 | chr2-124012573-INS-159 |
| chr2-124059468-INS-79 | chr2-124059468-INS-108 |
| chr2-124085628-INS-66 | chr2-124085584-INS-66 |
| chr2-124108507-INS-135 | chr2-124108507-INS-135 |
| chr2-124182798-INS-58 | chr2-124182798-INS-68 |
| chr2-124216608-INS-200 | chr2-124216608-INS-205 |
| chr2-124235652-INS-6209 | chr2-124235653-INS-6210 |
| chr2-124280214-INS-182 | chr2-124280214-INS-182 |
| chr2-124298495-INS-353 | chr2-124298496-INS-354 |
| chr2-124318114-INS-201 | chr2-124318114-INS-201 |
| chr2-124322495-INS-517 | chr2-124322495-INS-517 |
| chr2-124359944-INS-125 | chr2-124359944-INS-125 |
| chr2-124390168-INS-145 | chr2-124390208-INS-114 |
| chr2-124395742-INS-225 | chr2-124395742-INS-227 |
| chr2-124409189-INS-166 | chr2-124409189-INS-166 |
| chr2-124464733-INS-70 | chr2-124464733-INS-77 |
| chr2-124482455-INS-400 | chr2-124482455-INS-400 |
| chr2-124490479-INS-86 | chr2-124490479-INS-108 |
| chr2-124499782-INS-235 | chr2-124499782-INS-235 |
| chr2-124528703-INS-60 | chr2-124528703-INS-60 |
| chr2-124532807-INS-119 | chr2-124532807-INS-119 |
| chr2-124570905-INS-51 | chr2-124570905-INS-51 |
| chr2-124576544-INS-65 | chr2-124576544-INS-66 |
| chr2-124599423-INS-52 | chr2-124599423-INS-52 |
| chr2-124601882-INS-52 | chr2-124601882-INS-52 |
| chr2-124616996-INS-6936 | chr2-124616997-INS-6936 |
| chr2-124654976-INS-162 | chr2-124654976-INS-162 |
| chr2-124656582-INS-88 | chr2-124656586-INS-72 |
| chr2-124705259-INS-57 | chr2-124705234-INS-76 |
| chr2-124709405-INS-280 | chr2-124709405-INS-280 |
| chr2-124710141-INS-6164 | chr2-124710142-INS-6168 |
| chr2-124817010-INS-5315 | chr2-124817011-INS-5315 |
| chr2-124821045-INS-56 | chr2-124821045-INS-56 |
| chr2-124838770-INS-594 | chr2-124838770-INS-594 |
| chr2-124843684-INS-60 | chr2-124843684-INS-62 |

| | |
|---|---|
| chr2-124844191-INS-202 | chr2-124844191-INS-202 |
| chr2-124847522-INS-51 | chr2-124847522-INS-51 |
| chr2-124859098-INS-158 | chr2-124859098-INS-160 |
| chr2-124876090-INS-2849 | chr2-124876091-INS-2849 |
| chr2-124887116-INS-483 | chr2-124887116-INS-483 |
| chr2-124895256-INS-100 | chr2-124895304-INS-68 |
| chr2-124902779-INS-67 | chr2-124902779-INS-67 |
| chr2-124904864-INS-5743 | chr2-124904865-INS-5743 |
| chr2-124908220-INS-1500 | chr2-124908216-INS-1507 |
| chr2-124911641-INS-56 | chr2-124911641-INS-64 |
| chr2-124916284-INS-118 | chr2-124916284-INS-122 |
| chr2-124917376-INS-762 | chr2-124917376-INS-764 |
| chr2-124935039-INS-199 | chr2-124935047-INS-198 |
| chr2-124944775-INS-400 | chr2-124944775-INS-402 |
| chr2-124948708-INS-208 | chr2-124948708-INS-208 |
| chr2-124967670-INS-214 | chr2-124967670-INS-216 |
| chr2-124973396-INS-234 | chr2-124973396-INS-234 |
| chr2-124983317-INS-217 | chr2-124983317-INS-217 |
| chr2-124985868-INS-202 | chr2-124985868-INS-201 |
| chr2-124988920-INS-124 | chr2-124988920-INS-124 |
| chr2-125025215-INS-71 | chr2-125025215-INS-71 |
| chr2-125060436-INS-64 | chr2-125060436-INS-76 |
| chr2-125060958-INS-52 | chr2-125060956-INS-56 |
| chr2-125074171-INS-2976 | chr2-125074172-INS-2976 |
| chr2-125086042-INS-537 | chr2-125086042-INS-537 |
| chr2-125112422-INS-137 | chr2-125112422-INS-137 |
| chr2-125118038-INS-6476 | chr2-125118039-INS-6476 |
| chr2-125148035-INS-88 | chr2-125148033-INS-90 |
| chr2-125186455-INS-239 | chr2-125186455-INS-243 |
| chr2-125193369-INS-56 | chr2-125193369-INS-50 |
| chr2-125228386-INS-54 | chr2-125228386-INS-54 |
| chr2-125232188-INS-113 | chr2-125232188-INS-119 |
| chr2-125258305-INS-373 | chr2-125258305-INS-373 |
| chr2-125289890-INS-177 | chr2-125289890-INS-177 |
| chr2-125305724-INS-60 | chr2-125305724-INS-60 |
| chr2-125313395-INS-143 | chr2-125313395-INS-142 |
| chr2-125332781-INS-476 | chr2-125332781-INS-476 |
| chr2-125335970-INS-71 | chr2-125335971-INS-71 |
| chr2-125344553-INS-56 | chr2-125344553-INS-56 |
| chr2-125364166-INS-6310 | chr2-125364167-INS-6310 |
| chr2-125380240-INS-54 | chr2-125380240-INS-52 |
| chr2-125396034-INS-94 | chr2-125396034-INS-96 |
| chr2-125403883-INS-211 | chr2-125403883-INS-210 |
| chr2-125411351-INS-73 | chr2-125411351-INS-73 |
| chr2-125430027-INS-98 | chr2-125430021-INS-118 |
| chr2-125437524-INS-52 | chr2-125437524-INS-52 |
| chr2-125458321-INS-52 | chr2-125458321-INS-52 |
| chr2-125524311-INS-196 | chr2-125524311-INS-196 |
| chr2-125537597-INS-74 | chr2-125537597-INS-84 |
| chr2-125570973-INS-53 | chr2-125570973-INS-52 |
| chr2-125573424-INS-89 | chr2-125573424-INS-89 |
| chr2-125585433-INS-206 | chr2-125585433-INS-206 |
| chr2-125588740-INS-109 | chr2-125588738-INS-110 |
| chr2-125608134-INS-1376 | chr2-125608135-INS-1376 |
| chr2-125616527-INS-174 | chr2-125616527-INS-175 |
| chr2-125621102-INS-65 | chr2-125621102-INS-65 |
| chr2-125634147-INS-52 | chr2-125634147-INS-60 |
| chr2-125644801-INS-180 | chr2-125644801-INS-182 |
| chr2-125656502-INS-1877 | chr2-125656503-INS-1877 |
| chr2-125665515-INS-203 | chr2-125665515-INS-206 |
| chr2-125708401-INS-76 | chr2-125708401-INS-76 |
| chr2-125727666-INS-8818 | chr2-125727667-INS-8818 |
| chr2-125734495-INS-80 | chr2-125734495-INS-84 |
| chr2-125742010-INS-176 | chr2-125742010-INS-176 |
| chr2-125763870-INS-592 | chr2-125763870-INS-592 |
| chr2-125769770-INS-52 | chr2-125769770-INS-88 |
| chr2-125791630-INS-80 | chr2-125791631-INS-59 |
| chr2-125806785-INS-58 | chr2-125806785-INS-58 |
| chr2-125822941-INS-188 | chr2-125822938-INS-200 |
| chr2-125826676-INS-86 | chr2-125826676-INS-86 |
| chr2-125828818-INS-53 | chr2-125828821-INS-65 |
| chr2-125831208-INS-64 | chr2-125831208-INS-50 |
| chr2-125836037-INS-148 | chr2-125836019-INS-147 |
| chr2-125866506-INS-61 | chr2-125866506-INS-65 |
| chr2-125870879-INS-63 | chr2-125870879-INS-63 |
| chr2-125875573-INS-60 | chr2-125875573-INS-62 |
| chr2-125888046-INS-622 | chr2-125888046-INS-625 |

| | |
|---|---|
| chr2-125891826-INS-475 | chr2-125891826-INS-475 |
| chr2-125893488-INS-185 | chr2-125893488-INS-185 |
| chr2-125913359-INS-542 | chr2-125913359-INS-543 |
| chr2-125916799-INS-387 | chr2-125916799-INS-387 |
| chr2-125918384-INS-6453 | chr2-125918385-INS-6453 |
| chr2-125953531-INS-102 | chr2-125953531-INS-102 |
| chr2-125955223-INS-7271 | chr2-125955224-INS-7277 |
| chr2-125972616-INS-68 | chr2-125972617-INS-68 |
| chr2-125974763-INS-3153 | chr2-125974764-INS-3153 |
| chr2-125999976-INS-78 | chr2-125999976-INS-120 |
| chr2-126001244-INS-114 | chr2-126001244-INS-114 |
| chr2-126037365-INS-65 | chr2-126037365-INS-69 |
| chr2-126038384-INS-324 | chr2-126038384-INS-324 |
| chr2-126046411-INS-498 | chr2-126046411-INS-498 |
| chr2-126055431-INS-198 | chr2-126055431-INS-198 |
| chr2-126123963-INS-83 | chr2-126124001-INS-54 |
| chr2-126126573-INS-60 | chr2-126126573-INS-96 |
| chr2-126140367-INS-2563 | chr2-126140368-INS-2563 |
| chr2-126153402-INS-216 | chr2-126153402-INS-216 |
| chr2-126177440-INS-616 | chr2-126177440-INS-616 |
| chr2-126180988-INS-205 | chr2-126180988-INS-206 |
| chr2-126204086-INS-287 | chr2-126204086-INS-287 |
| chr2-126212447-INS-129 | chr2-126212447-INS-129 |
| chr2-126221407-INS-187 | chr2-126221407-INS-187 |
| chr2-126226829-INS-108 | chr2-126226829-INS-92 |
| chr2-126236496-INS-61 | chr2-126236492-INS-69 |
| chr2-126257294-INS-188 | chr2-126257294-INS-191 |
| chr2-126265625-INS-160 | chr2-126265625-INS-185 |
| chr2-126280041-INS-57 | chr2-126280041-INS-57 |
| chr2-126303543-INS-66 | chr2-126303543-INS-72 |
| chr2-126309395-INS-65 | chr2-126309395-INS-65 |
| chr2-126321032-INS-260 | chr2-126321032-INS-261 |
| chr2-126325997-INS-602 | chr2-126325996-INS-604 |
| chr2-126335654-INS-102 | chr2-126335654-INS-62 |
| chr2-126351348-INS-177 | chr2-126351348-INS-179 |
| chr2-126361994-INS-66 | chr2-126361994-INS-99 |
| chr2-126366926-INS-477 | chr2-126366926-INS-477 |
| chr2-126372266-INS-1208 | chr2-126372267-INS-1208 |
| chr2-126380798-INS-82 | chr2-126380823-INS-90 |
| chr2-126392994-INS-1889 | chr2-126392995-INS-1889 |
| chr2-126428987-INS-6439 | chr2-126428988-INS-6439 |
| chr2-126434663-INS-6495 | chr2-126434664-INS-6520 |
| chr2-126477627-INS-60 | chr2-126477627-INS-75 |
| chr2-126519858-INS-204 | chr2-126519858-INS-205 |
| chr2-126522400-INS-180 | chr2-126522400-INS-180 |
| chr2-126526240-INS-201 | chr2-126526240-INS-202 |
| chr2-126526521-INS-3370 | chr2-126526522-INS-3371 |
| chr2-126527815-INS-1748 | chr2-126527816-INS-1748 |
| chr2-126533145-INS-401 | chr2-126533145-INS-401 |
| chr2-126535810-INS-144 | chr2-126535852-INS-96 |
| chr2-126591279-INS-199 | chr2-126591279-INS-199 |
| chr2-126594553-INS-54 | chr2-126594551-INS-66 |
| chr2-126595340-INS-113 | chr2-126595340-INS-115 |
| chr2-126598898-INS-183 | chr2-126598898-INS-183 |
| chr2-126609225-INS-62 | chr2-126609225-INS-62 |
| chr2-126619400-INS-205 | chr2-126619400-INS-205 |
| chr2-126635390-INS-134 | chr2-126635390-INS-134 |
| chr2-126650469-INS-210 | chr2-126650469-INS-224 |
| chr2-126651486-INS-577 | chr2-126651486-INS-577 |
| chr2-126660253-INS-89 | chr2-126660250-INS-102 |
| chr2-126681794-INS-225 | chr2-126681794-INS-236 |
| chr2-126685694-INS-210 | chr2-126685694-INS-211 |
| chr2-126690283-INS-205 | chr2-126690283-INS-203 |
| chr2-126721924-INS-58 | chr2-126721924-INS-62 |
| chr2-126726733-INS-52 | chr2-126726733-INS-52 |
| chr2-126727738-INS-227 | chr2-126727738-INS-227 |
| chr2-126754740-INS-78 | chr2-126754740-INS-84 |
| chr2-126780259-INS-324 | chr2-126780259-INS-324 |
| chr2-126784417-INS-228 | chr2-126784417-INS-229 |
| chr2-126786223-INS-53 | chr2-126786223-INS-53 |
| chr2-126787357-INS-212 | chr2-126787357-INS-212 |
| chr2-126789956-INS-73 | chr2-126789956-INS-76 |
| chr2-126791588-INS-625 | chr2-126791590-INS-605 |
| chr2-126794122-INS-9047 | chr2-126794123-INS-9047 |
| chr2-126798107-INS-83 | chr2-126798107-INS-83 |
| chr2-126805058-INS-204 | chr2-126805058-INS-209 |
| chr2-126814912-INS-165 | chr2-126814912-INS-160 |

| | |
|---|---|
| chr2-126815554-INS-155 | chr2-126815530-INS-156 |
| chr2-126816566-INS-67 | chr2-126816566-INS-70 |
| chr2-126820477-INS-260 | chr2-126820477-INS-260 |
| chr2-126837502-INS-62 | chr2-126837502-INS-62 |
| chr2-126843177-INS-83 | chr2-126843177-INS-83 |
| chr2-126855904-INS-197 | chr2-126855905-INS-197 |
| chr2-126856898-INS-185 | chr2-126856898-INS-190 |
| chr2-126859790-INS-56 | chr2-126859790-INS-58 |
| chr2-126869382-INS-202 | chr2-126869382-INS-202 |
| chr2-126871016-INS-67 | chr2-126871016-INS-67 |
| chr2-126873683-INS-86 | chr2-126873676-INS-134 |
| chr2-126879468-INS-85 | chr2-126879468-INS-85 |
| chr2-126880917-INS-80 | chr2-126880917-INS-80 |
| chr2-126912427-INS-524 | chr2-126912427-INS-524 |
| chr2-126920818-INS-67 | chr2-126920818-INS-79 |
| chr2-126932402-INS-76 | chr2-126932402-INS-76 |
| chr2-126957684-INS-213 | chr2-126957684-INS-209 |
| chr2-126972778-INS-108 | chr2-126972778-INS-110 |
| chr2-127008441-INS-203 | chr2-127008441-INS-233 |
| chr2-127012382-INS-176 | chr2-127012382-INS-181 |
| chr2-127034171-INS-60 | chr2-127034138-INS-65 |
| chr2-127039852-INS-223 | chr2-127039852-INS-223 |
| chr2-127052142-INS-203 | chr2-127052142-INS-206 |
| chr2-127062002-INS-54 | chr2-127061986-INS-59 |
| chr2-127065022-INS-179 | chr2-127065022-INS-179 |
| chr2-127067170-INS-130 | chr2-127067170-INS-130 |
| chr2-127072813-INS-200 | chr2-127072813-INS-204 |
| chr2-127122864-INS-52 | chr2-127122864-INS-56 |
| chr2-127124098-INS-170 | chr2-127124098-INS-171 |
| chr2-127130690-INS-52 | chr2-127130690-INS-52 |
| chr2-127145703-INS-119 | chr2-127145703-INS-119 |
| chr2-127148338-INS-223 | chr2-127148338-INS-225 |
| chr2-127157135-INS-253 | chr2-127157135-INS-253 |
| chr2-127159861-INS-57 | chr2-127159861-INS-57 |
| chr2-127162361-INS-151 | chr2-127162361-INS-159 |
| chr2-127168141-INS-116 | chr2-127168141-INS-116 |
| chr2-127224517-INS-508 | chr2-127224517-INS-508 |
| chr2-127234786-INS-484 | chr2-127234786-INS-484 |
| chr2-127247501-INS-105 | chr2-127247501-INS-105 |
| chr2-127249945-INS-396 | chr2-127249945-INS-396 |
| chr2-127270255-INS-95 | chr2-127270255-INS-95 |
| chr2-127296287-INS-286 | chr2-127296287-INS-286 |
| chr2-127333381-INS-473 | chr2-127333381-INS-473 |
| chr2-127336107-INS-85 | chr2-127336107-INS-82 |
| chr2-127375962-INS-68 | chr2-127375957-INS-68 |
| chr2-127400583-INS-91 | chr2-127400583-INS-63 |
| chr2-127462756-INS-109 | chr2-127462755-INS-130 |
| chr2-127463372-INS-133 | chr2-127463373-INS-139 |
| chr2-127474933-INS-70 | chr2-127474933-INS-70 |
| chr2-127504154-INS-72 | chr2-127504154-INS-72 |
| chr2-127519785-INS-148 | chr2-127519785-INS-160 |
| chr2-127523662-INS-80 | chr2-127523662-INS-56 |
| chr2-127528863-INS-112 | chr2-127528886-INS-105 |
| chr2-127538149-INS-171 | chr2-127538149-INS-171 |
| chr2-127555620-INS-123 | chr2-127555621-INS-128 |
| chr2-127556586-INS-182 | chr2-127556587-INS-186 |
| chr2-127557479-INS-59 | chr2-127557477-INS-52 |
| chr2-127559784-INS-160 | chr2-127559773-INS-156 |
| chr2-127583415-INS-57 | chr2-127583414-INS-52 |
| chr2-127584050-INS-63 | chr2-127584029-INS-60 |
| chr2-127588419-INS-61 | chr2-127588419-INS-64 |
| chr2-127590721-INS-319 | chr2-127590721-INS-319 |
| chr2-127596553-INS-88 | chr2-127596553-INS-72 |
| chr2-127605584-INS-51 | chr2-127605584-INS-56 |
| chr2-127610511-INS-187 | chr2-127610511-INS-187 |
| chr2-127613283-INS-407 | chr2-127613283-INS-404 |
| chr2-127647950-INS-212 | chr2-127647950-INS-214 |
| chr2-127651958-INS-594 | chr2-127651958-INS-593 |
| chr2-127664078-INS-214 | chr2-127664078-INS-214 |
| chr2-127676783-INS-93 | chr2-127676783-INS-81 |
| chr2-127680251-INS-120 | chr2-127680251-INS-108 |
| chr2-127687490-INS-208 | chr2-127687490-INS-208 |
| chr2-127693096-INS-54 | chr2-127693096-INS-52 |
| chr2-127698309-INS-6385 | chr2-127698310-INS-6385 |
| chr2-127709681-INS-56 | chr2-127709682-INS-52 |
| chr2-127713463-INS-56 | chr2-127713463-INS-56 |
| chr2-127720239-INS-131 | chr2-127720239-INS-131 |

| | |
|---|---|
| chr2-127729076-INS-213 | chr2-127729076-INS-213 |
| chr2-127743357-INS-60 | chr2-127743357-INS-62 |
| chr2-127793127-INS-57 | chr2-127793127-INS-57 |
| chr2-127819538-INS-206 | chr2-127819538-INS-205 |
| chr2-127825275-INS-56 | chr2-127825275-INS-56 |
| chr2-127828782-INS-7117 | chr2-127828783-INS-7114 |
| chr2-127855300-INS-1207 | chr2-127855301-INS-1207 |
| chr2-127870247-INS-76 | chr2-127870247-INS-71 |
| chr2-127871448-INS-519 | chr2-127871448-INS-551 |
| chr2-127888363-INS-52 | chr2-127888390-INS-52 |
| chr2-127889909-INS-156 | chr2-127889909-INS-159 |
| chr2-127900668-INS-297 | chr2-127900668-INS-297 |
| chr2-127906578-INS-5550 | chr2-127906579-INS-5550 |
| chr2-127937020-INS-56 | chr2-127937020-INS-56 |
| chr2-127985440-INS-55 | chr2-127985440-INS-55 |
| chr2-128010605-INS-52 | chr2-128010605-INS-52 |
| chr2-128047488-INS-60 | chr2-128047488-INS-60 |
| chr2-128065862-INS-70 | chr2-128065862-INS-54 |
| chr2-128075325-INS-406 | chr2-128075325-INS-406 |
| chr2-128076114-INS-2320 | chr2-128076115-INS-2320 |
| chr2-128078365-INS-69 | chr2-128078361-INS-72 |
| chr2-128089780-INS-325 | chr2-128089780-INS-325 |
| chr2-128106656-INS-182 | chr2-128106656-INS-182 |
| chr2-128118098-INS-67 | chr2-128118098-INS-59 |
| chr2-128149106-INS-1601 | chr2-128149107-INS-1579 |
| chr2-128161830-INS-86 | chr2-128161830-INS-86 |
| chr2-128161846-INS-7128 | chr2-128161847-INS-7128 |
| chr2-128173799-INS-76 | chr2-128173802-INS-72 |
| chr2-128183050-INS-329 | chr2-128183050-INS-329 |
| chr2-128192072-INS-54 | chr2-128192072-INS-54 |
| chr2-128195317-INS-54 | chr2-128195317-INS-54 |
| chr2-128272790-INS-268 | chr2-128272790-INS-269 |
| chr2-128279902-INS-191 | chr2-128279902-INS-191 |
| chr2-128287354-INS-4843 | chr2-128287355-INS-4843 |
| chr2-128288925-INS-1079 | chr2-128288926-INS-1081 |
| chr2-128291963-INS-79 | chr2-128291963-INS-82 |
| chr2-128295685-INS-52 | chr2-128295685-INS-52 |
| chr2-128299128-INS-1889 | chr2-128299129-INS-1889 |
| chr2-128302773-INS-196 | chr2-128302773-INS-196 |
| chr2-128316905-INS-288 | chr2-128316905-INS-290 |
| chr2-128375411-INS-257 | chr2-128375411-INS-258 |
| chr2-128376495-INS-72 | chr2-128376487-INS-54 |
| chr2-128406860-INS-6419 | chr2-128406861-INS-6418 |
| chr2-128418429-INS-64 | chr2-128418434-INS-64 |
| chr2-128440883-INS-80 | chr2-128440877-INS-68 |
| chr2-128441248-INS-5167 | chr2-128441249-INS-5167 |
| chr2-128464972-INS-209 | chr2-128464972-INS-209 |
| chr2-128475114-INS-240 | chr2-128475114-INS-240 |
| chr2-128490159-INS-205 | chr2-128490159-INS-205 |
| chr2-128515187-INS-679 | chr2-128515187-INS-679 |
| chr2-128518871-INS-418 | chr2-128518871-INS-418 |
| chr2-128522581-INS-200 | chr2-128522581-INS-204 |
| chr2-128528226-INS-210 | chr2-128528226-INS-210 |
| chr2-128529809-INS-2052 | chr2-128529810-INS-2052 |
| chr2-128557451-INS-81 | chr2-128557451-INS-81 |
| chr2-128564986-INS-201 | chr2-128564986-INS-201 |
| chr2-128567855-INS-125 | chr2-128567855-INS-126 |
| chr2-128593646-INS-57 | chr2-128593646-INS-57 |
| chr2-128600351-INS-225 | chr2-128600351-INS-225 |
| chr2-128606496-INS-544 | chr2-128606496-INS-540 |
| chr2-128607216-INS-80 | chr2-128607218-INS-55 |
| chr2-128622294-INS-258 | chr2-128622294-INS-258 |
| chr2-128625518-INS-210 | chr2-128625504-INS-208 |
| chr2-128628087-INS-181 | chr2-128628087-INS-180 |
| chr2-128648495-INS-6442 | chr2-128648496-INS-6442 |
| chr2-128652570-INS-186 | chr2-128652570-INS-186 |
| chr2-128655626-INS-86 | chr2-128655626-INS-86 |
| chr2-128672604-INS-69 | chr2-128672604-INS-69 |
| chr2-128704072-INS-295 | chr2-128704072-INS-294 |
| chr2-128722628-INS-108 | chr2-128722628-INS-108 |
| chr2-128729326-INS-77 | chr2-128729326-INS-83 |
| chr2-128730678-INS-70 | chr2-128730678-INS-72 |
| chr2-128733368-INS-231 | chr2-128733368-INS-232 |
| chr2-128774200-INS-242 | chr2-128774200-INS-242 |
| chr2-128778431-INS-207 | chr2-128778431-INS-207 |
| chr2-128808724-INS-841 | chr2-128808724-INS-838 |
| chr2-128817677-INS-65 | chr2-128817677-INS-65 |

| | |
|---|---|
| chr2-128828103-INS-202 | chr2-128828103-INS-202 |
| chr2-128831003-INS-116 | chr2-128831003-INS-116 |
| chr2-128831745-INS-120 | chr2-128831745-INS-120 |
| chr2-128839913-INS-320 | chr2-128839913-INS-320 |
| chr2-128844254-INS-498 | chr2-128844254-INS-498 |
| chr2-128876240-INS-176 | chr2-128876240-INS-179 |
| chr2-128884289-INS-51 | chr2-128884289-INS-85 |
| chr2-128889243-INS-273 | chr2-128889243-INS-274 |
| chr2-128895280-INS-51 | chr2-128895263-INS-66 |
| chr2-128906469-INS-132 | chr2-128906469-INS-124 |
| chr2-128921191-INS-171 | chr2-128921191-INS-172 |
| chr2-128922836-INS-6619 | chr2-128922837-INS-6619 |
| chr2-128946623-INS-96 | chr2-128946623-INS-96 |
| chr2-128961261-INS-207 | chr2-128961261-INS-207 |
| chr2-128989942-INS-76 | chr2-128989942-INS-76 |
| chr2-128994386-INS-52 | chr2-128994386-INS-54 |
| chr2-128997691-INS-175 | chr2-128997691-INS-183 |
| chr2-129002885-INS-192 | chr2-129002869-INS-192 |
| chr2-129004588-INS-207 | chr2-129004588-INS-207 |
| chr2-129006851-INS-199 | chr2-129006851-INS-199 |
| chr2-129010488-INS-241 | chr2-129010488-INS-241 |
| chr2-129017381-INS-76 | chr2-129017407-INS-90 |
| chr2-129022825-INS-76 | chr2-129022825-INS-80 |
| chr2-129035402-INS-130 | chr2-129035402-INS-124 |
| chr2-129062369-INS-170 | chr2-129062369-INS-170 |
| chr2-129077564-INS-6460 | chr2-129077565-INS-6460 |
| chr2-129079530-INS-83 | chr2-129079530-INS-79 |
| chr2-129083422-INS-61 | chr2-129083422-INS-61 |
| chr2-129084018-INS-166 | chr2-129084029-INS-185 |
| chr2-129096081-INS-92 | chr2-129096078-INS-93 |
| chr2-129117725-INS-189 | chr2-129117725-INS-189 |
| chr2-129119577-INS-72 | chr2-129119577-INS-72 |
| chr2-129124254-INS-260 | chr2-129124254-INS-260 |
| chr2-129130138-INS-181 | chr2-129130123-INS-176 |
| chr2-129141467-INS-193 | chr2-129141467-INS-193 |
| chr2-129173951-INS-166 | chr2-129173945-INS-136 |
| chr2-129186480-INS-4799 | chr2-129186481-INS-4800 |
| chr2-129204563-INS-70 | chr2-129204569-INS-61 |
| chr2-129205504-INS-3721 | chr2-129205505-INS-3721 |
| chr2-129236482-INS-103 | chr2-129236474-INS-98 |
| chr2-129237495-INS-1341 | chr2-129237496-INS-1341 |
| chr2-129241868-INS-68 | chr2-129241868-INS-56 |
| chr2-129257099-INS-254 | chr2-129257099-INS-253 |
| chr2-129266453-INS-54 | chr2-129266453-INS-50 |
| chr2-129271468-INS-96 | chr2-129271477-INS-106 |
| chr2-129273546-INS-6714 | chr2-129273547-INS-6715 |
| chr2-129312979-INS-159 | chr2-129312979-INS-159 |
| chr2-129325685-INS-52 | chr2-129325685-INS-52 |
| chr2-129353047-INS-205 | chr2-129353049-INS-204 |
| chr2-129432087-INS-81 | chr2-129432088-INS-81 |
| chr2-129437079-INS-233 | chr2-129437079-INS-234 |
| chr2-129486894-INS-60 | chr2-129486894-INS-60 |
| chr2-129491624-INS-267 | chr2-129491624-INS-267 |
| chr2-129500131-INS-204 | chr2-129500131-INS-204 |
| chr2-129502370-INS-75 | chr2-129502370-INS-75 |
| chr2-129546676-INS-258 | chr2-129546676-INS-258 |
| chr2-129550945-INS-60 | chr2-129550945-INS-62 |
| chr2-129551840-INS-459 | chr2-129551840-INS-457 |
| chr2-129586248-INS-54 | chr2-129586248-INS-54 |
| chr2-129589277-INS-723 | chr2-129589277-INS-724 |
| chr2-129591217-INS-71 | chr2-129591217-INS-80 |
| chr2-129622039-INS-60 | chr2-129622059-INS-65 |
| chr2-129625490-INS-989 | chr2-129625490-INS-989 |
| chr2-129647567-INS-52 | chr2-129647567-INS-52 |
| chr2-129657247-INS-816 | chr2-129657247-INS-816 |
| chr2-129657247-INS-96 | chr2-129657247-INS-96 |
| chr2-129674423-INS-449 | chr2-129674423-INS-449 |
| chr2-129686197-INS-161 | chr2-129686197-INS-161 |
| chr2-129700021-INS-55 | chr2-129700028-INS-56 |
| chr2-129739432-INS-64 | chr2-129739432-INS-66 |
| chr2-129740464-INS-361 | chr2-129740464-INS-361 |
| chr2-129746587-INS-64 | chr2-129746587-INS-64 |
| chr2-129752849-INS-65 | chr2-129752849-INS-61 |
| chr2-129758512-INS-82 | chr2-129758520-INS-58 |
| chr2-129805547-INS-541 | chr2-129805547-INS-541 |
| chr2-129809608-INS-63 | chr2-129809608-INS-63 |
| chr2-129853254-INS-100 | chr2-129853254-INS-80 |

| | |
|---|---|
| chr2-129863604-INS-75 | chr2-129863614-INS-50 |
| chr2-129888081-INS-129 | chr2-129888081-INS-129 |
| chr2-129897201-INS-84 | chr2-129897201-INS-84 |
| chr2-129908664-INS-5546 | chr2-129908667-INS-5547 |
| chr2-129925135-INS-60 | chr2-129925141-INS-78 |
| chr2-129927280-INS-4958 | chr2-129927281-INS-4958 |
| chr2-129961767-INS-153 | chr2-129961794-INS-150 |
| chr2-129971358-INS-156 | chr2-129971360-INS-192 |
| chr2-129975693-INS-80 | chr2-129975693-INS-100 |
| chr2-129991835-INS-81 | chr2-129991835-INS-81 |
| chr2-130000609-INS-458 | chr2-130000607-INS-448 |
| chr2-130034575-INS-437 | chr2-130034575-INS-437 |
| chr2-130059284-INS-67 | chr2-130059284-INS-67 |
| chr2-130067546-INS-54 | chr2-130067560-INS-56 |
| chr2-130069663-INS-72 | chr2-130069663-INS-64 |
| chr2-130088211-INS-224 | chr2-130088211-INS-223 |
| chr2-130112289-INS-6471 | chr2-130112290-INS-6471 |
| chr2-130113494-INS-207 | chr2-130113494-INS-207 |
| chr2-130129957-INS-179 | chr2-130129957-INS-194 |
| chr2-130185504-INS-80 | chr2-130185504-INS-70 |
| chr2-130187474-INS-66 | chr2-130187474-INS-66 |
| chr2-130213487-INS-58 | chr2-130213487-INS-60 |
| chr2-130218578-INS-721 | chr2-130218578-INS-695 |
| chr2-130229244-INS-88 | chr2-130229259-INS-75 |
| chr2-130246279-INS-54 | chr2-130246274-INS-79 |
| chr2-130259695-INS-213 | chr2-130259695-INS-212 |
| chr2-130312395-INS-1889 | chr2-130312396-INS-1889 |
| chr2-130315278-INS-1619 | chr2-130315279-INS-1619 |
| chr2-130315957-INS-66 | chr2-130315957-INS-66 |
| chr2-130367842-INS-78 | chr2-130367842-INS-81 |
| chr2-130371396-INS-268 | chr2-130371396-INS-268 |
| chr2-130380069-INS-72 | chr2-130380069-INS-74 |
| chr2-130418923-INS-58 | chr2-130418923-INS-63 |
| chr2-130447271-INS-89 | chr2-130447271-INS-52 |
| chr2-130481989-INS-88 | chr2-130481989-INS-60 |
| chr2-130500153-INS-193 | chr2-130500153-INS-193 |
| chr2-130512404-INS-79 | chr2-130512409-INS-81 |
| chr2-130530731-INS-110 | chr2-130530738-INS-135 |
| chr2-130540025-INS-68 | chr2-130540025-INS-60 |
| chr2-130567834-INS-207 | chr2-130567834-INS-205 |
| chr2-130628140-INS-206 | chr2-130628140-INS-206 |
| chr2-130638491-INS-96 | chr2-130638488-INS-118 |
| chr2-130641587-INS-58 | chr2-130641587-INS-52 |
| chr2-130655922-INS-189 | chr2-130655922-INS-189 |
| chr2-130662082-INS-58 | chr2-130662082-INS-60 |
| chr2-130673839-INS-61 | chr2-130673839-INS-61 |
| chr2-130692667-INS-202 | chr2-130692667-INS-202 |
| chr2-130699918-INS-400 | chr2-130699918-INS-400 |
| chr2-130720667-INS-215 | chr2-130720667-INS-215 |
| chr2-130736439-INS-4168 | chr2-130736440-INS-4168 |
| chr2-130750363-INS-206 | chr2-130750363-INS-206 |
| chr2-130798850-INS-917 | chr2-130798850-INS-918 |
| chr2-130809338-INS-183 | chr2-130809338-INS-184 |
| chr2-130828363-INS-193 | chr2-130828363-INS-193 |
| chr2-130836542-INS-75 | chr2-130836539-INS-60 |
| chr2-130846908-INS-65 | chr2-130846908-INS-65 |
| chr2-130854512-INS-190 | chr2-130854521-INS-194 |
| chr2-130922084-INS-308 | chr2-130922084-INS-312 |
| chr2-130943509-INS-59 | chr2-130943509-INS-55 |
| chr2-130943550-INS-6499 | chr2-130943551-INS-6499 |
| chr2-130953067-INS-92 | chr2-130953067-INS-92 |
| chr2-130976259-INS-216 | chr2-130976224-INS-224 |
| chr2-131051015-INS-91 | chr2-131051011-INS-56 |
| chr2-131160874-INS-81 | chr2-131160874-INS-80 |
| chr2-131165539-INS-986 | chr2-131165539-INS-956 |
| chr2-131175366-INS-278 | chr2-131175366-INS-280 |
| chr2-131200342-INS-124 | chr2-131200342-INS-128 |
| chr2-131220504-INS-1469 | chr2-131220505-INS-1469 |
| chr2-131239896-INS-96 | chr2-131239897-INS-78 |
| chr2-131267664-INS-93 | chr2-131267664-INS-93 |
| chr2-131271210-INS-57 | chr2-131271210-INS-57 |
| chr2-131274762-INS-60 | chr2-131274762-INS-60 |
| chr2-131277782-INS-210 | chr2-131277782-INS-210 |
| chr2-131281507-INS-92 | chr2-131281498-INS-57 |
| chr2-131291440-INS-390 | chr2-131291440-INS-389 |
| chr2-131297517-INS-206 | chr2-131297523-INS-205 |
| chr2-131305746-INS-228 | chr2-131305746-INS-272 |

| | |
|---|---|
| chr2-131338709-INS-53 | chr2-131338709-INS-68 |
| chr2-131355114-INS-69 | chr2-131355114-INS-69 |
| chr2-131363112-INS-66 | chr2-131363112-INS-66 |
| chr2-131381771-INS-967 | chr2-131381771-INS-974 |
| chr2-131400674-INS-153 | chr2-131400674-INS-153 |
| chr2-131408924-INS-4121 | chr2-131408925-INS-4121 |
| chr2-131437773-INS-92 | chr2-131437819-INS-60 |
| chr2-131451973-INS-672 | chr2-131451973-INS-672 |
| chr2-131459679-INS-459 | chr2-131459679-INS-459 |
| chr2-131469491-INS-2844 | chr2-131469492-INS-2844 |
| chr2-131469605-INS-82 | chr2-131469605-INS-82 |
| chr2-131473825-INS-528 | chr2-131473825-INS-528 |
| chr2-131491180-INS-124 | chr2-131491180-INS-170 |
| chr2-131497296-INS-62 | chr2-131497302-INS-78 |
| chr2-131521963-INS-117 | chr2-131521963-INS-117 |
| chr2-131526190-INS-62 | chr2-131526188-INS-56 |
| chr2-131550139-INS-176 | chr2-131550139-INS-134 |
| chr2-131551885-INS-2791 | chr2-131551886-INS-2791 |
| chr2-131555487-INS-72 | chr2-131555487-INS-72 |
| chr2-131562533-INS-61 | chr2-131562508-INS-61 |
| chr2-131571677-INS-52 | chr2-131571677-INS-52 |
| chr2-131580297-INS-73 | chr2-131580297-INS-73 |
| chr2-131581469-INS-72 | chr2-131581469-INS-72 |
| chr2-131596498-INS-64 | chr2-131596458-INS-67 |
| chr2-131601056-INS-55 | chr2-131601057-INS-57 |
| chr2-131611560-INS-161 | chr2-131611560-INS-161 |
| chr2-131620035-INS-83 | chr2-131620035-INS-93 |
| chr2-131625800-INS-182 | chr2-131625800-INS-182 |
| chr2-131635549-INS-92 | chr2-131635549-INS-92 |
| chr2-131643219-INS-174 | chr2-131643219-INS-174 |
| chr2-131657657-INS-2774 | chr2-131657658-INS-2774 |
| chr2-131661502-INS-395 | chr2-131661502-INS-395 |
| chr2-131666746-INS-7107 | chr2-131666747-INS-7107 |
| chr2-131673824-INS-67 | chr2-131673824-INS-67 |
| chr2-131693474-INS-62 | chr2-131693474-INS-62 |
| chr2-131696881-INS-75 | chr2-131696881-INS-76 |
| chr2-131704619-INS-69 | chr2-131704619-INS-71 |
| chr2-131715256-INS-179 | chr2-131715256-INS-179 |
| chr2-131720919-INS-212 | chr2-131720919-INS-212 |
| chr2-131732506-INS-68 | chr2-131732506-INS-68 |
| chr2-131781364-INS-182 | chr2-131781364-INS-182 |
| chr2-131784726-INS-220 | chr2-131784726-INS-220 |
| chr2-131791726-INS-422 | chr2-131791726-INS-425 |
| chr2-131807749-INS-392 | chr2-131807749-INS-392 |
| chr2-131818313-INS-215 | chr2-131818313-INS-215 |
| chr2-131831332-INS-607 | chr2-131831332-INS-607 |
| chr2-131840857-INS-91 | chr2-131840886-INS-60 |
| chr2-131856736-INS-228 | chr2-131856736-INS-227 |
| chr2-131865424-INS-204 | chr2-131865424-INS-201 |
| chr2-131872930-INS-97 | chr2-131872930-INS-97 |
| chr2-131881912-INS-240 | chr2-131881912-INS-255 |
| chr2-131909145-INS-133 | chr2-131909145-INS-137 |
| chr2-131912167-INS-139 | chr2-131912167-INS-141 |
| chr2-131943794-INS-247 | chr2-131943794-INS-247 |
| chr2-131969020-INS-268 | chr2-131969061-INS-248 |
| chr2-131995563-INS-358 | chr2-131995563-INS-357 |
| chr2-132015780-INS-194 | chr2-132015780-INS-215 |
| chr2-132055034-INS-51 | chr2-132055031-INS-59 |
| chr2-132060162-INS-205 | chr2-132060162-INS-205 |
| chr2-132069394-INS-513 | chr2-132069394-INS-510 |
| chr2-132078864-INS-178 | chr2-132078864-INS-179 |
| chr2-132079537-INS-238 | chr2-132079537-INS-238 |
| chr2-132091098-INS-188 | chr2-132091098-INS-186 |
| chr2-132099058-INS-263 | chr2-132099058-INS-263 |
| chr2-132117961-INS-138 | chr2-132117961-INS-138 |
| chr2-132125510-INS-208 | chr2-132125510-INS-208 |
| chr2-132163840-INS-60 | chr2-132163840-INS-60 |
| chr2-132165910-INS-114 | chr2-132165910-INS-114 |
| chr2-132179107-INS-1523 | chr2-132179118-INS-1518 |
| chr2-132182020-INS-477 | chr2-132182021-INS-477 |
| chr2-132187003-INS-4438 | chr2-132187004-INS-4438 |
| chr2-132191216-INS-151 | chr2-132191216-INS-160 |
| chr2-132236160-INS-498 | chr2-132236160-INS-498 |
| chr2-132237084-INS-214 | chr2-132237088-INS-208 |
| chr2-132244926-INS-920 | chr2-132244926-INS-920 |
| chr2-132249941-INS-540 | chr2-132249941-INS-540 |
| chr2-132267079-INS-6564 | chr2-132267080-INS-6567 |

| | |
|---|---|
| chr2-132272644-INS-1445 | chr2-132272645-INS-1445 |
| chr2-132280713-INS-100 | chr2-132280713-INS-88 |
| chr2-132289179-INS-60 | chr2-132289179-INS-60 |
| chr2-132294549-INS-72 | chr2-132294549-INS-76 |
| chr2-132296191-INS-1603 | chr2-132296192-INS-1603 |
| chr2-132300123-INS-358 | chr2-132300123-INS-358 |
| chr2-132308729-INS-71 | chr2-132308733-INS-75 |
| chr2-132313689-INS-227 | chr2-132313689-INS-245 |
| chr2-132330993-INS-81 | chr2-132330993-INS-81 |
| chr2-132346763-INS-363 | chr2-132346763-INS-365 |
| chr2-132361674-INS-2883 | chr2-132361675-INS-2883 |
| chr2-132364652-INS-84 | chr2-132364657-INS-92 |
| chr2-132385471-INS-229 | chr2-132385471-INS-224 |
| chr2-132428908-INS-273 | chr2-132428908-INS-272 |
| chr2-132436329-INS-199 | chr2-132436329-INS-197 |
| chr2-132447276-INS-61 | chr2-132447280-INS-65 |
| chr2-132453219-INS-166 | chr2-132453219-INS-166 |
| chr2-132480841-INS-67 | chr2-132480834-INS-70 |
| chr2-132482141-INS-58 | chr2-132482142-INS-61 |
| chr2-132486090-INS-208 | chr2-132486090-INS-208 |
| chr2-132497354-INS-158 | chr2-132497354-INS-159 |
| chr2-132503024-INS-185 | chr2-132503024-INS-185 |
| chr2-132515275-INS-68 | chr2-132515275-INS-76 |
| chr2-132524169-INS-157 | chr2-132524169-INS-159 |
| chr2-132524525-INS-2121 | chr2-132524526-INS-2139 |
| chr2-132547614-INS-212 | chr2-132547614-INS-212 |
| chr2-132561748-INS-233 | chr2-132561748-INS-248 |
| chr2-132568803-INS-54 | chr2-132568805-INS-53 |
| chr2-132589164-INS-325 | chr2-132589164-INS-325 |
| chr2-132600472-INS-103 | chr2-132600472-INS-103 |
| chr2-132606567-INS-222 | chr2-132606567-INS-222 |
| chr2-132612235-INS-182 | chr2-132612257-INS-139 |
| chr2-132640521-INS-60 | chr2-132640521-INS-60 |
| chr2-132655660-INS-218 | chr2-132655660-INS-218 |
| chr2-132684974-INS-217 | chr2-132684974-INS-217 |
| chr2-132695775-INS-82 | chr2-132695774-INS-71 |
| chr2-132721896-INS-56 | chr2-132721896-INS-66 |
| chr2-132723825-INS-404 | chr2-132723825-INS-404 |
| chr2-132748988-INS-71 | chr2-132748988-INS-71 |
| chr2-132749406-INS-1550 | chr2-132749407-INS-1550 |
| chr2-132768036-INS-3622 | chr2-132768040-INS-3611 |
| chr2-132793405-INS-80 | chr2-132793405-INS-81 |
| chr2-132841849-INS-125 | chr2-132841849-INS-125 |
| chr2-132862250-INS-5519 | chr2-132862251-INS-5519 |
| chr2-132871651-INS-1497 | chr2-132871652-INS-1497 |
| chr2-132880919-INS-72 | chr2-132880919-INS-72 |
| chr2-132890315-INS-6245 | chr2-132890316-INS-6245 |
| chr2-132898371-INS-184 | chr2-132898371-INS-184 |
| chr2-132919065-INS-4217 | chr2-132919066-INS-4217 |
| chr2-132920455-INS-99 | chr2-132920453-INS-64 |
| chr2-132921461-INS-75 | chr2-132921468-INS-51 |
| chr2-132931506-INS-81 | chr2-132931492-INS-78 |
| chr2-132938832-INS-50 | chr2-132938833-INS-67 |
| chr2-132948283-INS-54 | chr2-132948283-INS-54 |
| chr2-132969791-INS-730 | chr2-132969776-INS-752 |
| chr2-133002622-INS-60 | chr2-133002640-INS-70 |
| chr2-133016576-INS-209 | chr2-133016576-INS-209 |
| chr2-133038479-INS-6403 | chr2-133038480-INS-6403 |
| chr2-133047117-INS-2063 | chr2-133047118-INS-2063 |
| chr2-133070449-INS-6406 | chr2-133070450-INS-6406 |
| chr2-133073386-INS-51 | chr2-133073386-INS-60 |
| chr2-133074520-INS-68 | chr2-133074516-INS-104 |
| chr2-133075607-INS-63 | chr2-133075607-INS-60 |
| chr2-133076366-INS-154 | chr2-133076366-INS-161 |
| chr2-133078091-INS-94 | chr2-133078091-INS-94 |
| chr2-133106717-INS-52 | chr2-133106718-INS-52 |
| chr2-133123929-INS-108 | chr2-133123929-INS-136 |
| chr2-133144575-INS-339 | chr2-133144577-INS-342 |
| chr2-133148882-INS-374 | chr2-133148882-INS-374 |
| chr2-133151959-INS-169 | chr2-133151959-INS-169 |
| chr2-133152853-INS-255 | chr2-133152853-INS-255 |
| chr2-133162024-INS-155 | chr2-133162058-INS-128 |
| chr2-133199142-INS-122 | chr2-133199142-INS-122 |
| chr2-133213887-INS-386 | chr2-133213887-INS-386 |
| chr2-133214957-INS-52 | chr2-133214957-INS-52 |
| chr2-133280229-INS-90 | chr2-133280229-INS-95 |
| chr2-133283252-INS-60 | chr2-133283252-INS-64 |

| | |
|---|---|
| chr2-133292741-INS-745 | chr2-133292741-INS-746 |
| chr2-133298634-INS-104 | chr2-133298634-INS-104 |
| chr2-133302422-INS-457 | chr2-133302422-INS-458 |
| chr2-133319788-INS-64 | chr2-133319788-INS-64 |
| chr2-133325402-INS-56 | chr2-133325392-INS-74 |
| chr2-133351768-INS-362 | chr2-133351768-INS-363 |
| chr2-133368700-INS-75 | chr2-133368680-INS-80 |
| chr2-133382752-INS-54 | chr2-133382752-INS-62 |
| chr2-133400234-INS-52 | chr2-133400234-INS-52 |
| chr2-133407655-INS-64 | chr2-133407655-INS-64 |
| chr2-133415810-INS-133 | chr2-133415810-INS-158 |
| chr2-133419343-INS-77 | chr2-133419343-INS-77 |
| chr2-133431649-INS-56 | chr2-133431649-INS-54 |
| chr2-133439802-INS-76 | chr2-133439805-INS-75 |
| chr2-133444138-INS-1890 | chr2-133444139-INS-1890 |
| chr2-133500740-INS-325 | chr2-133500740-INS-325 |
| chr2-133536251-INS-480 | chr2-133536251-INS-484 |
| chr2-133550177-INS-56 | chr2-133550177-INS-64 |
| chr2-133567087-INS-54 | chr2-133567087-INS-58 |
| chr2-133574178-INS-93 | chr2-133574184-INS-62 |
| chr2-133594165-INS-200 | chr2-133594165-INS-200 |
| chr2-133606671-INS-58 | chr2-133606671-INS-58 |
| chr2-133614038-INS-1262 | chr2-133614039-INS-1263 |
| chr2-133693506-INS-76 | chr2-133693506-INS-77 |
| chr2-133749639-INS-1531 | chr2-133749640-INS-1531 |
| chr2-133756274-INS-506 | chr2-133756274-INS-506 |
| chr2-133793346-INS-116 | chr2-133793346-INS-100 |
| chr2-133795310-INS-91 | chr2-133795310-INS-91 |
| chr2-133818451-INS-837 | chr2-133818447-INS-834 |
| chr2-133859758-INS-242 | chr2-133859758-INS-242 |
| chr2-133873452-INS-268 | chr2-133873452-INS-263 |
| chr2-133874590-INS-177 | chr2-133874589-INS-166 |
| chr2-133879815-INS-90 | chr2-133879815-INS-86 |
| chr2-133885553-INS-72 | chr2-133885553-INS-72 |
| chr2-133889552-INS-7178 | chr2-133889553-INS-7178 |
| chr2-133904690-INS-3350 | chr2-133904691-INS-3350 |
| chr2-133927420-INS-312 | chr2-133927424-INS-312 |
| chr2-133960494-INS-56 | chr2-133960494-INS-58 |
| chr2-133979076-INS-54 | chr2-133979076-INS-58 |
| chr2-133979626-INS-54 | chr2-133979626-INS-56 |
| chr2-133984446-INS-78 | chr2-133984446-INS-78 |
| chr2-134000219-INS-54 | chr2-134000219-INS-50 |
| chr2-134001226-INS-176 | chr2-134001226-INS-170 |
| chr2-134013882-INS-85 | chr2-134013873-INS-78 |
| chr2-134018156-INS-68 | chr2-134018156-INS-68 |
| chr2-134020409-INS-95 | chr2-134020411-INS-106 |
| chr2-134035357-INS-1885 | chr2-134035358-INS-1885 |
| chr2-134038410-INS-93 | chr2-134038410-INS-93 |
| chr2-134043101-INS-205 | chr2-134043101-INS-203 |
| chr2-134074887-INS-4919 | chr2-134074888-INS-4919 |
| chr2-134086004-INS-56 | chr2-134086004-INS-54 |
| chr2-134129626-INS-224 | chr2-134129669-INS-208 |
| chr2-134151980-INS-362 | chr2-134151980-INS-362 |
| chr2-134165340-INS-276 | chr2-134165345-INS-277 |
| chr2-134169889-INS-888 | chr2-134169880-INS-873 |
| chr2-134170687-INS-184 | chr2-134170687-INS-183 |
| chr2-134172970-INS-51 | chr2-134172971-INS-60 |
| chr2-134175679-INS-60 | chr2-134175679-INS-64 |
| chr2-134210543-INS-60 | chr2-134210511-INS-68 |
| chr2-134237883-INS-782 | chr2-134237856-INS-791 |
| chr2-134267029-INS-208 | chr2-134267029-INS-208 |
| chr2-134384340-INS-121 | chr2-134384345-INS-117 |
| chr2-134395396-INS-117 | chr2-134395430-INS-67 |
| chr2-134399408-INS-158 | chr2-134399441-INS-112 |
| chr2-134491895-INS-1978 | chr2-134491896-INS-1978 |
| chr2-134493196-INS-120 | chr2-134493196-INS-120 |
| chr2-134496078-INS-1873 | chr2-134496079-INS-1875 |
| chr2-134496636-INS-813 | chr2-134496636-INS-813 |
| chr2-134518043-INS-507 | chr2-134518043-INS-507 |
| chr2-134527350-INS-52 | chr2-134527328-INS-61 |
| chr2-134619294-INS-462 | chr2-134619294-INS-462 |
| chr2-134626621-INS-94 | chr2-134626621-INS-88 |
| chr2-134631540-INS-60 | chr2-134631540-INS-56 |
| chr2-134714760-INS-69 | chr2-134714760-INS-69 |
| chr2-134729831-INS-7508 | chr2-134729832-INS-7508 |
| chr2-134744065-INS-210 | chr2-134744065-INS-210 |
| chr2-134766837-INS-89 | chr2-134766837-INS-71 |

| | |
|---|---|
| chr2-134819397-INS-90 | chr2-134819397-INS-90 |
| chr2-134825905-INS-7160 | chr2-134825906-INS-7160 |
| chr2-134829369-INS-54 | chr2-134829372-INS-58 |
| chr2-134860902-INS-4154 | chr2-134860903-INS-4156 |
| chr2-134878574-INS-6834 | chr2-134878575-INS-6834 |
| chr2-134895537-INS-72 | chr2-134895537-INS-72 |
| chr2-134923266-INS-76 | chr2-134923266-INS-50 |
| chr2-134932524-INS-71 | chr2-134932524-INS-71 |
| chr2-134942555-INS-54 | chr2-134942555-INS-50 |
| chr2-134984571-INS-54 | chr2-134984571-INS-54 |
| chr2-135029065-INS-80 | chr2-135029105-INS-52 |
| chr2-135031607-INS-63 | chr2-135031607-INS-63 |
| chr2-135057500-INS-53 | chr2-135057500-INS-66 |
| chr2-135060102-INS-408 | chr2-135060103-INS-378 |
| chr2-135077836-INS-102 | chr2-135077836-INS-54 |
| chr2-135084550-INS-57 | chr2-135084550-INS-59 |
| chr2-135118189-INS-488 | chr2-135118189-INS-488 |
| chr2-135135789-INS-368 | chr2-135135778-INS-366 |
| chr2-135168581-INS-53 | chr2-135168589-INS-88 |
| chr2-135183429-INS-6798 | chr2-135183430-INS-6798 |
| chr2-135190807-INS-112 | chr2-135190807-INS-104 |
| chr2-135196176-INS-540 | chr2-135196176-INS-540 |
| chr2-135221520-INS-99 | chr2-135221520-INS-99 |
| chr2-135236464-INS-204 | chr2-135236464-INS-205 |
| chr2-135252068-INS-190 | chr2-135252068-INS-202 |
| chr2-135258480-INS-87 | chr2-135258487-INS-86 |
| chr2-135302361-INS-58 | chr2-135302361-INS-58 |
| chr2-135306492-INS-54 | chr2-135306492-INS-54 |
| chr2-135308431-INS-106 | chr2-135308476-INS-111 |
| chr2-135390577-INS-59 | chr2-135390577-INS-58 |
| chr2-135410059-INS-157 | chr2-135410059-INS-189 |
| chr2-135483160-INS-71 | chr2-135483160-INS-71 |
| chr2-135524217-INS-54 | chr2-135524217-INS-58 |
| chr2-135527291-INS-503 | chr2-135527293-INS-478 |
| chr2-135537491-INS-1191 | chr2-135537492-INS-1191 |
| chr2-135539154-INS-6501 | chr2-135539155-INS-6501 |
| chr2-135542744-INS-1306 | chr2-135542745-INS-1306 |
| chr2-135546147-INS-1321 | chr2-135546148-INS-1321 |
| chr2-135557326-INS-8781 | chr2-135557327-INS-8783 |
| chr2-135603938-INS-6661 | chr2-135603939-INS-6661 |
| chr2-135635350-INS-207 | chr2-135635350-INS-207 |
| chr2-135767385-INS-98 | chr2-135767384-INS-62 |
| chr2-135795316-INS-6466 | chr2-135795317-INS-6466 |
| chr2-135828254-INS-57 | chr2-135828254-INS-53 |
| chr2-135846638-INS-217 | chr2-135846638-INS-218 |
| chr2-135847668-INS-84 | chr2-135847668-INS-84 |
| chr2-135869328-INS-853 | chr2-135869328-INS-854 |
| chr2-135961029-INS-77 | chr2-135961029-INS-77 |
| chr2-135966179-INS-72 | chr2-135966183-INS-56 |
| chr2-135984793-INS-1788 | chr2-135984794-INS-1789 |
| chr2-135994272-INS-80 | chr2-135994272-INS-80 |
| chr2-136000687-INS-170 | chr2-136000687-INS-169 |
| chr2-136030347-INS-72 | chr2-136030347-INS-72 |
| chr2-136032173-INS-173 | chr2-136032173-INS-215 |
| chr2-136159733-INS-1002 | chr2-136159734-INS-1002 |
| chr2-136161238-INS-360 | chr2-136161238-INS-360 |
| chr2-136168117-INS-58 | chr2-136168117-INS-58 |
| chr2-136172540-INS-514 | chr2-136172544-INS-515 |
| chr2-136181420-INS-53 | chr2-136181420-INS-53 |
| chr2-136189507-INS-79 | chr2-136189507-INS-63 |
| chr2-136193058-INS-64 | chr2-136193058-INS-76 |
| chr2-136219413-INS-65 | chr2-136219413-INS-75 |
| chr2-136225328-INS-114 | chr2-136225328-INS-115 |
| chr2-136268068-INS-90 | chr2-136268068-INS-99 |
| chr2-136277846-INS-71 | chr2-136277846-INS-71 |
| chr2-136293049-INS-548 | chr2-136293049-INS-548 |
| chr2-136315873-INS-66 | chr2-136315903-INS-75 |
| chr2-136326751-INS-191 | chr2-136326751-INS-191 |
| chr2-136332137-INS-247 | chr2-136332137-INS-201 |
| chr2-136347358-INS-646 | chr2-136347358-INS-646 |
| chr2-136351198-INS-2457 | chr2-136351199-INS-2431 |
| chr2-136361933-INS-3884 | chr2-136361934-INS-3896 |
| chr2-136390740-INS-111 | chr2-136390740-INS-126 |
| chr2-136392878-INS-435 | chr2-136392878-INS-435 |
| chr2-136413380-INS-160 | chr2-136413380-INS-160 |
| chr2-136439718-INS-491 | chr2-136439718-INS-492 |
| chr2-136454919-INS-66 | chr2-136454919-INS-66 |

| | |
|---|---|
| chr2-136468508-INS-5506 | chr2-136468509-INS-5506 |
| chr2-136487077-INS-68 | chr2-136487077-INS-60 |
| chr2-136530033-INS-128 | chr2-136530033-INS-128 |
| chr2-136561560-INS-66 | chr2-136561560-INS-66 |
| chr2-136566618-INS-286 | chr2-136566618-INS-286 |
| chr2-136574438-INS-53 | chr2-136574438-INS-53 |
| chr2-136639078-INS-104 | chr2-136639076-INS-84 |
| chr2-136655071-INS-54 | chr2-136655071-INS-54 |
| chr2-136660739-INS-80 | chr2-136660739-INS-88 |
| chr2-136698916-INS-475 | chr2-136698916-INS-475 |
| chr2-136724385-INS-529 | chr2-136724385-INS-529 |
| chr2-136743554-INS-165 | chr2-136743554-INS-165 |
| chr2-136747848-INS-284 | chr2-136747848-INS-284 |
| chr2-136834493-INS-554 | chr2-136834493-INS-554 |
| chr2-136842968-INS-1186 | chr2-136842969-INS-1188 |
| chr2-136939554-INS-261 | chr2-136939554-INS-261 |
| chr2-136973447-INS-143 | chr2-136973447-INS-149 |
| chr2-136978637-INS-69 | chr2-136978637-INS-81 |
| chr2-136995640-INS-58 | chr2-136995640-INS-56 |
| chr2-136997414-INS-104 | chr2-136997409-INS-102 |
| chr2-137006271-INS-173 | chr2-137006271-INS-173 |
| chr2-137021150-INS-2809 | chr2-137021151-INS-2809 |
| chr2-137024516-INS-98 | chr2-137024516-INS-90 |
| chr2-137026554-INS-60 | chr2-137026559-INS-70 |
| chr2-137029197-INS-442 | chr2-137029197-INS-442 |
| chr2-137048482-INS-76 | chr2-137048482-INS-84 |
| chr2-137061009-INS-127 | chr2-137061029-INS-90 |
| chr2-137063766-INS-170 | chr2-137063766-INS-170 |
| chr2-137064487-INS-6379 | chr2-137064488-INS-6379 |
| chr2-137065321-INS-52 | chr2-137065321-INS-52 |
| chr2-137078137-INS-189 | chr2-137078136-INS-186 |
| chr2-137092798-INS-109 | chr2-137092798-INS-109 |
| chr2-137095103-INS-71 | chr2-137095097-INS-50 |
| chr2-137097840-INS-50 | chr2-137097840-INS-50 |
| chr2-137215279-INS-76 | chr2-137215279-INS-60 |
| chr2-137227865-INS-1341 | chr2-137227866-INS-1348 |
| chr2-137239348-INS-80 | chr2-137239349-INS-80 |
| chr2-137277807-INS-54 | chr2-137277807-INS-58 |
| chr2-137285088-INS-76 | chr2-137285088-INS-78 |
| chr2-137302324-INS-235 | chr2-137302371-INS-199 |
| chr2-137323219-INS-789 | chr2-137323219-INS-789 |
| chr2-137330311-INS-63 | chr2-137330311-INS-65 |
| chr2-137356805-INS-845 | chr2-137356805-INS-843 |
| chr2-137359862-INS-59 | chr2-137359838-INS-101 |
| chr2-137391981-INS-511 | chr2-137391981-INS-510 |
| chr2-137395324-INS-68 | chr2-137395324-INS-70 |
| chr2-137407523-INS-63 | chr2-137407520-INS-80 |
| chr2-137420889-INS-234 | chr2-137420889-INS-234 |
| chr2-137423281-INS-79 | chr2-137423281-INS-79 |
| chr2-137426352-INS-1099 | chr2-137426360-INS-1089 |
| chr2-137431118-INS-483 | chr2-137431118-INS-484 |
| chr2-137442221-INS-214 | chr2-137442221-INS-214 |
| chr2-137461495-INS-94 | chr2-137461495-INS-108 |
| chr2-137476139-INS-169 | chr2-137476142-INS-169 |
| chr2-137484521-INS-2939 | chr2-137484522-INS-2939 |
| chr2-137484522-INS-247 | chr2-137484522-INS-247 |
| chr2-137487357-INS-67 | chr2-137487357-INS-71 |
| chr2-137495382-INS-58 | chr2-137495382-INS-58 |
| chr2-137500542-INS-74 | chr2-137500542-INS-72 |
| chr2-137504589-INS-249 | chr2-137504577-INS-246 |
| chr2-137517531-INS-142 | chr2-137517531-INS-128 |
| chr2-137537141-INS-52 | chr2-137537141-INS-52 |
| chr2-137583695-INS-68 | chr2-137583695-INS-68 |
| chr2-137597643-INS-67 | chr2-137597643-INS-67 |
| chr2-137607562-INS-1572 | chr2-137607563-INS-1572 |
| chr2-137612389-INS-1309 | chr2-137612390-INS-1306 |
| chr2-137675634-INS-408 | chr2-137675634-INS-408 |
| chr2-137692673-INS-133 | chr2-137692673-INS-129 |
| chr2-137740642-INS-1126 | chr2-137740643-INS-1126 |
| chr2-137741776-INS-52 | chr2-137741821-INS-50 |
| chr2-137746041-INS-188 | chr2-137746041-INS-187 |
| chr2-137785299-INS-76 | chr2-137785299-INS-76 |
| chr2-137801008-INS-73 | chr2-137801008-INS-73 |
| chr2-137824509-INS-61 | chr2-137824494-INS-67 |
| chr2-137844495-INS-72 | chr2-137844495-INS-76 |
| chr2-137884462-INS-249 | chr2-137884462-INS-237 |
| chr2-137907398-INS-124 | chr2-137907398-INS-123 |

| | |
|---|---|
| chr2-137931876-INS-96 | chr2-137931876-INS-108 |
| chr2-137942806-INS-204 | chr2-137942806-INS-204 |
| chr2-137946748-INS-6440 | chr2-137946749-INS-6441 |
| chr2-137950144-INS-497 | chr2-137950144-INS-497 |
| chr2-137963435-INS-215 | chr2-137963462-INS-189 |
| chr2-137966714-INS-66 | chr2-137966714-INS-66 |
| chr2-137969160-INS-587 | chr2-137969160-INS-587 |
| chr2-137985659-INS-406 | chr2-137985659-INS-406 |
| chr2-137991089-INS-436 | chr2-137991089-INS-436 |
| chr2-137991917-INS-113 | chr2-137991888-INS-96 |
| chr2-137995027-INS-241 | chr2-137995027-INS-241 |
| chr2-138016605-INS-199 | chr2-138016562-INS-199 |
| chr2-138022510-INS-307 | chr2-138022510-INS-305 |
| chr2-138030164-INS-62 | chr2-138030164-INS-62 |
| chr2-138030995-INS-337 | chr2-138030995-INS-337 |
| chr2-138051951-INS-141 | chr2-138051964-INS-149 |
| chr2-138096781-INS-52 | chr2-138096781-INS-91 |
| chr2-138103782-INS-830 | chr2-138103782-INS-830 |
| chr2-138144051-INS-2371 | chr2-138144052-INS-2372 |
| chr2-138148494-INS-6463 | chr2-138148495-INS-6464 |
| chr2-138149792-INS-590 | chr2-138149792-INS-597 |
| chr2-138154636-INS-896 | chr2-138154636-INS-902 |
| chr2-138203904-INS-52 | chr2-138203904-INS-52 |
| chr2-138207001-INS-71 | chr2-138207001-INS-71 |
| chr2-138209914-INS-790 | chr2-138209915-INS-793 |
| chr2-138220433-INS-57 | chr2-138220433-INS-83 |
| chr2-138225035-INS-57 | chr2-138225035-INS-57 |
| chr2-138238680-INS-56 | chr2-138238723-INS-52 |
| chr2-138316450-INS-71 | chr2-138316450-INS-81 |
| chr2-138340960-INS-791 | chr2-138340960-INS-808 |
| chr2-138386891-INS-66 | chr2-138386891-INS-64 |
| chr2-138388291-INS-469 | chr2-138388291-INS-469 |
| chr2-138456247-INS-6408 | chr2-138456248-INS-6408 |
| chr2-138474877-INS-216 | chr2-138474877-INS-218 |
| chr2-138498584-INS-646 | chr2-138498584-INS-641 |
| chr2-138506847-INS-7178 | chr2-138506848-INS-7178 |
| chr2-138506848-INS-369 | chr2-138506848-INS-369 |
| chr2-138562727-INS-6298 | chr2-138562728-INS-6299 |
| chr2-138602017-INS-53 | chr2-138602017-INS-53 |
| chr2-138604577-INS-103 | chr2-138604577-INS-123 |
| chr2-138609424-INS-1331 | chr2-138609425-INS-1331 |
| chr2-138612156-INS-246 | chr2-138612156-INS-246 |
| chr2-138621383-INS-72 | chr2-138621383-INS-82 |
| chr2-138625041-INS-64 | chr2-138625041-INS-84 |
| chr2-138636670-INS-168 | chr2-138636693-INS-132 |
| chr2-138638986-INS-218 | chr2-138638981-INS-220 |
| chr2-138685525-INS-72 | chr2-138685525-INS-64 |
| chr2-138712895-INS-257 | chr2-138712895-INS-252 |
| chr2-138774791-INS-51 | chr2-138774791-INS-55 |
| chr2-138786505-INS-83 | chr2-138786508-INS-113 |
| chr2-138791710-INS-68 | chr2-138791710-INS-70 |
| chr2-138818138-INS-126 | chr2-138818123-INS-144 |
| chr2-138840206-INS-179 | chr2-138840206-INS-179 |
| chr2-138856032-INS-1172 | chr2-138856033-INS-1167 |
| chr2-138863961-INS-76 | chr2-138863961-INS-76 |
| chr2-138887576-INS-246 | chr2-138887577-INS-246 |
| chr2-138902804-INS-310 | chr2-138902804-INS-310 |
| chr2-138905600-INS-94 | chr2-138905600-INS-92 |
| chr2-138947896-INS-73 | chr2-138947868-INS-69 |
| chr2-138948713-INS-1055 | chr2-138948714-INS-1056 |
| chr2-138952411-INS-108 | chr2-138952380-INS-144 |
| chr2-138964691-INS-63 | chr2-138964675-INS-71 |
| chr2-139007987-INS-6546 | chr2-139007988-INS-6547 |
| chr2-139009234-INS-52 | chr2-139009234-INS-60 |
| chr2-139014328-INS-264 | chr2-139014328-INS-263 |
| chr2-139018424-INS-545 | chr2-139018424-INS-549 |
| chr2-139026253-INS-6346 | chr2-139026254-INS-6345 |
| chr2-139043951-INS-350 | chr2-139043951-INS-350 |
| chr2-139046470-INS-1184 | chr2-139046471-INS-1184 |
| chr2-139047465-INS-131 | chr2-139047466-INS-110 |
| chr2-139050732-INS-4974 | chr2-139050733-INS-4974 |
| chr2-139101939-INS-71 | chr2-139101939-INS-65 |
| chr2-139168186-INS-56 | chr2-139168186-INS-62 |
| chr2-139182559-INS-69 | chr2-139182559-INS-69 |
| chr2-139225973-INS-405 | chr2-139225973-INS-405 |
| chr2-139237627-INS-688 | chr2-139237627-INS-688 |
| chr2-139261143-INS-2114 | chr2-139261144-INS-2114 |

| | |
|---|---|
| chr2-139277509-INS-478 | chr2-139277509-INS-478 |
| chr2-139321109-INS-1140 | chr2-139321100-INS-1123 |
| chr2-139345202-INS-962 | chr2-139345202-INS-965 |
| chr2-139348055-INS-2020 | chr2-139348056-INS-2018 |
| chr2-139354093-INS-54 | chr2-139354093-INS-52 |
| chr2-139361977-INS-173 | chr2-139361992-INS-148 |
| chr2-139403508-INS-3895 | chr2-139403509-INS-3895 |
| chr2-139411453-INS-86 | chr2-139411453-INS-80 |
| chr2-139424489-INS-73 | chr2-139424489-INS-73 |
| chr2-139443006-INS-203 | chr2-139443006-INS-204 |
| chr2-139477504-INS-51 | chr2-139477504-INS-67 |
| chr2-139541752-INS-435 | chr2-139541752-INS-435 |
| chr2-139592457-INS-264 | chr2-139592445-INS-246 |
| chr2-139629441-INS-59 | chr2-139629442-INS-74 |
| chr2-139635505-INS-189 | chr2-139635505-INS-189 |
| chr2-139640421-INS-86 | chr2-139640394-INS-85 |
| chr2-139665276-INS-68 | chr2-139665290-INS-53 |
| chr2-139684057-INS-63 | chr2-139684057-INS-63 |
| chr2-139697230-INS-6376 | chr2-139697231-INS-6376 |
| chr2-139716830-INS-82 | chr2-139716830-INS-82 |
| chr2-139772275-INS-57 | chr2-139772276-INS-81 |
| chr2-139790922-INS-844 | chr2-139790922-INS-844 |
| chr2-139793527-INS-51 | chr2-139793527-INS-51 |
| chr2-139794640-INS-211 | chr2-139794640-INS-211 |
| chr2-139802048-INS-67 | chr2-139802048-INS-67 |
| chr2-139803354-INS-106 | chr2-139803354-INS-106 |
| chr2-139824284-INS-140 | chr2-139824284-INS-140 |
| chr2-139828405-INS-326 | chr2-139828405-INS-327 |
| chr2-139834613-INS-221 | chr2-139834613-INS-221 |
| chr2-139841291-INS-166 | chr2-139841291-INS-167 |
| chr2-139842163-INS-329 | chr2-139842163-INS-330 |
| chr2-139853648-INS-176 | chr2-139853648-INS-176 |
| chr2-139856831-INS-608 | chr2-139856831-INS-608 |
| chr2-139863392-INS-2935 | chr2-139863389-INS-2941 |
| chr2-139865105-INS-62 | chr2-139865105-INS-62 |
| chr2-139887310-INS-200 | chr2-139887310-INS-199 |
| chr2-139894824-INS-372 | chr2-139894824-INS-372 |
| chr2-139919438-INS-268 | chr2-139919438-INS-260 |
| chr2-139931546-INS-213 | chr2-139931546-INS-213 |
| chr2-139941624-INS-528 | chr2-139941624-INS-528 |
| chr2-139950587-INS-52 | chr2-139950587-INS-52 |
| chr2-139956898-INS-108 | chr2-139956898-INS-108 |
| chr2-139965979-INS-212 | chr2-139965979-INS-212 |
| chr2-139986265-INS-102 | chr2-139986265-INS-102 |
| chr2-139997013-INS-60 | chr2-139997013-INS-60 |
| chr2-140003418-INS-74 | chr2-140003418-INS-111 |
| chr2-140009650-INS-72 | chr2-140009651-INS-72 |
| chr2-140024278-INS-72 | chr2-140024256-INS-92 |
| chr2-140025607-INS-179 | chr2-140025616-INS-177 |
| chr2-140034685-INS-244 | chr2-140034685-INS-252 |
| chr2-140043835-INS-58 | chr2-140043815-INS-62 |
| chr2-140046538-INS-70 | chr2-140046538-INS-65 |
| chr2-140051065-INS-98 | chr2-140051065-INS-98 |
| chr2-140058862-INS-115 | chr2-140058862-INS-115 |
| chr2-140062693-INS-96 | chr2-140062693-INS-140 |
| chr2-140073697-INS-7188 | chr2-140073698-INS-7188 |
| chr2-140077316-INS-58 | chr2-140077316-INS-58 |
| chr2-140110892-INS-52 | chr2-140110892-INS-50 |
| chr2-140113553-INS-7008 | chr2-140113554-INS-7008 |
| chr2-140145706-INS-970 | chr2-140145706-INS-970 |
| chr2-140147722-INS-4751 | chr2-140147723-INS-4751 |
| chr2-140153448-INS-88 | chr2-140153491-INS-58 |
| chr2-140181639-INS-727 | chr2-140181639-INS-727 |
| chr2-140195438-INS-3176 | chr2-140195439-INS-3177 |
| chr2-140214779-INS-5836 | chr2-140214780-INS-5836 |
| chr2-140219520-INS-4966 | chr2-140219521-INS-4966 |
| chr2-140259144-INS-79 | chr2-140259144-INS-79 |
| chr2-140272932-INS-843 | chr2-140272932-INS-845 |
| chr2-140285429-INS-3279 | chr2-140285430-INS-3279 |
| chr2-140298087-INS-2760 | chr2-140298088-INS-2760 |
| chr2-140300513-INS-574 | chr2-140300515-INS-575 |
| chr2-140305608-INS-2797 | chr2-140305609-INS-2797 |
| chr2-140305862-INS-204 | chr2-140305862-INS-204 |
| chr2-140307732-INS-7080 | chr2-140307733-INS-7080 |
| chr2-140323391-INS-345 | chr2-140323391-INS-345 |
| chr2-140331088-INS-63 | chr2-140331113-INS-56 |
| chr2-140350754-INS-6137 | chr2-140350755-INS-6137 |

| | |
|---|---|
| chr2-140393603-INS-6911 | chr2-140393604-INS-6911 |
| chr2-140397889-INS-221 | chr2-140397889-INS-221 |
| chr2-140405337-INS-130 | chr2-140405337-INS-130 |
| chr2-140426831-INS-168 | chr2-140426831-INS-166 |
| chr2-140446275-INS-171 | chr2-140446275-INS-175 |
| chr2-140446290-INS-675 | chr2-140446290-INS-680 |
| chr2-140451784-INS-103 | chr2-140451779-INS-63 |
| chr2-140476064-INS-54 | chr2-140476053-INS-94 |
| chr2-140523649-INS-427 | chr2-140523649-INS-427 |
| chr2-140564961-INS-60 | chr2-140564961-INS-60 |
| chr2-140621303-INS-958 | chr2-140621303-INS-959 |
| chr2-140636260-INS-165 | chr2-140636244-INS-165 |
| chr2-140659816-INS-51 | chr2-140659816-INS-54 |
| chr2-140660165-INS-2873 | chr2-140660166-INS-2873 |
| chr2-140666074-INS-183 | chr2-140666074-INS-183 |
| chr2-140701631-INS-72 | chr2-140701631-INS-84 |
| chr2-140717894-INS-104 | chr2-140717888-INS-88 |
| chr2-140745161-INS-51 | chr2-140745161-INS-51 |
| chr2-140750710-INS-7063 | chr2-140750711-INS-7063 |
| chr2-140761823-INS-3933 | chr2-140761833-INS-3935 |
| chr2-140817632-INS-6372 | chr2-140817633-INS-6371 |
| chr2-140849366-INS-2736 | chr2-140849367-INS-2733 |
| chr2-140851482-INS-90 | chr2-140851482-INS-61 |
| chr2-140859523-INS-61 | chr2-140859532-INS-79 |
| chr2-140875142-INS-327 | chr2-140875142-INS-327 |
| chr2-140876508-INS-154 | chr2-140876508-INS-159 |
| chr2-140888213-INS-266 | chr2-140888213-INS-241 |
| chr2-140897487-INS-66 | chr2-140897488-INS-66 |
| chr2-140907341-INS-6622 | chr2-140907342-INS-6623 |
| chr2-140923817-INS-468 | chr2-140923817-INS-468 |
| chr2-140940705-INS-101 | chr2-140940705-INS-106 |
| chr2-140940707-INS-685 | chr2-140940707-INS-676 |
| chr2-140965187-INS-52 | chr2-140965187-INS-52 |
| chr2-140998286-INS-105 | chr2-140998288-INS-105 |
| chr2-141078156-INS-68 | chr2-141078156-INS-64 |
| chr2-141086329-INS-51 | chr2-141086329-INS-63 |
| chr2-141161672-INS-555 | chr2-141161672-INS-552 |
| chr2-141167090-INS-52 | chr2-141167090-INS-52 |
| chr2-141171903-INS-146 | chr2-141171903-INS-131 |
| chr2-141190134-INS-82 | chr2-141190144-INS-54 |
| chr2-141201996-INS-54 | chr2-141201996-INS-54 |
| chr2-141226674-INS-61 | chr2-141226674-INS-61 |
| chr2-141245529-INS-1287 | chr2-141245530-INS-1295 |
| chr2-141253792-INS-1160 | chr2-141253793-INS-1159 |
| chr2-141263967-INS-76 | chr2-141263967-INS-80 |
| chr2-141304106-INS-176 | chr2-141304106-INS-176 |
| chr2-141409174-INS-141 | chr2-141409154-INS-132 |
| chr2-141424356-INS-278 | chr2-141424356-INS-278 |
| chr2-141452834-INS-54 | chr2-141452840-INS-57 |
| chr2-141457221-INS-372 | chr2-141457221-INS-372 |
| chr2-141495123-INS-159 | chr2-141495147-INS-168 |
| chr2-141498192-INS-6083 | chr2-141498193-INS-6083 |
| chr2-141506383-INS-68 | chr2-141506383-INS-78 |
| chr2-141551442-INS-60 | chr2-141551442-INS-60 |
| chr2-141573995-INS-84 | chr2-141574018-INS-76 |
| chr2-141625665-INS-54 | chr2-141625666-INS-54 |
| chr2-141641007-INS-57 | chr2-141641007-INS-57 |
| chr2-141642743-INS-92 | chr2-141642743-INS-92 |
| chr2-141659001-INS-83 | chr2-141659004-INS-77 |
| chr2-141663660-INS-54 | chr2-141663660-INS-58 |
| chr2-141681061-INS-117 | chr2-141681061-INS-99 |
| chr2-141682832-INS-494 | chr2-141682832-INS-494 |
| chr2-141686048-INS-58 | chr2-141686048-INS-56 |
| chr2-141687361-INS-5964 | chr2-141687362-INS-5964 |
| chr2-141690887-INS-2293 | chr2-141690888-INS-2295 |
| chr2-141694373-INS-1098 | chr2-141694374-INS-1099 |
| chr2-141701604-INS-3022 | chr2-141701605-INS-3022 |
| chr2-141711210-INS-60 | chr2-141711210-INS-60 |
| chr2-141712685-INS-1809 | chr2-141712686-INS-1809 |
| chr2-141860282-INS-931 | chr2-141860282-INS-932 |
| chr2-141880332-INS-4306 | chr2-141880333-INS-4306 |
| chr2-141894807-INS-695 | chr2-141894807-INS-693 |
| chr2-141906980-INS-66 | chr2-141906980-INS-70 |
| chr2-141939599-INS-426 | chr2-141939599-INS-426 |
| chr2-141961597-INS-195 | chr2-141961597-INS-196 |
| chr2-141964161-INS-84 | chr2-141964161-INS-86 |
| chr2-141983568-INS-83 | chr2-141983568-INS-81 |

| | |
|---|---|
| chr2-142001816-INS-56 | chr2-142001816-INS-56 |
| chr2-142011680-INS-598 | chr2-142011680-INS-598 |
| chr2-142053945-INS-72 | chr2-142053945-INS-70 |
| chr2-142062642-INS-52 | chr2-142062642-INS-52 |
| chr2-142121992-INS-58 | chr2-142121992-INS-60 |
| chr2-142184011-INS-84 | chr2-142184011-INS-78 |
| chr2-142200972-INS-2985 | chr2-142200973-INS-2985 |
| chr2-142206527-INS-201 | chr2-142206527-INS-201 |
| chr2-142215802-INS-200 | chr2-142215802-INS-203 |
| chr2-142228328-INS-741 | chr2-142228328-INS-738 |
| chr2-142253489-INS-93 | chr2-142253489-INS-119 |
| chr2-142258740-INS-1287 | chr2-142258741-INS-1287 |
| chr2-142291705-INS-178 | chr2-142291705-INS-178 |
| chr2-142341818-INS-60 | chr2-142341818-INS-60 |
| chr2-142349967-INS-5430 | chr2-142349968-INS-5422 |
| chr2-142372358-INS-54 | chr2-142372358-INS-50 |
| chr2-142392365-INS-52 | chr2-142392365-INS-54 |
| chr2-142413096-INS-60 | chr2-142413096-INS-55 |
| chr2-142422296-INS-64 | chr2-142422296-INS-64 |
| chr2-142447339-INS-54 | chr2-142447339-INS-59 |
| chr2-142451011-INS-60 | chr2-142451011-INS-64 |
| chr2-142479522-INS-940 | chr2-142479518-INS-943 |
| chr2-142492514-INS-141 | chr2-142492519-INS-141 |
| chr2-142515730-INS-55 | chr2-142515730-INS-57 |
| chr2-142530508-INS-233 | chr2-142530508-INS-241 |
| chr2-142547885-INS-72 | chr2-142547885-INS-76 |
| chr2-142586897-INS-134 | chr2-142586897-INS-165 |
| chr2-142606695-INS-1808 | chr2-142606696-INS-1808 |
| chr2-142623489-INS-60 | chr2-142623489-INS-60 |
| chr2-142641831-INS-2119 | chr2-142641832-INS-2119 |
| chr2-142676266-INS-71 | chr2-142676266-INS-71 |
| chr2-142689794-INS-159 | chr2-142689794-INS-159 |
| chr2-142691931-INS-211 | chr2-142691931-INS-215 |
| chr2-142706083-INS-7220 | chr2-142706084-INS-7220 |
| chr2-142715332-INS-102 | chr2-142715332-INS-102 |
| chr2-142762561-INS-338 | chr2-142762557-INS-338 |
| chr2-142786612-INS-80 | chr2-142786612-INS-80 |
| chr2-142789764-INS-2929 | chr2-142789765-INS-2929 |
| chr2-142801224-INS-6475 | chr2-142801229-INS-6476 |
| chr2-142807302-INS-199 | chr2-142807302-INS-198 |
| chr2-142818312-INS-237 | chr2-142818312-INS-233 |
| chr2-142819866-INS-86 | chr2-142819866-INS-86 |
| chr2-142840627-INS-226 | chr2-142840627-INS-227 |
| chr2-142841585-INS-6596 | chr2-142841586-INS-6598 |
| chr2-142852524-INS-5070 | chr2-142852525-INS-5070 |
| chr2-142866707-INS-200 | chr2-142866707-INS-200 |
| chr2-142873013-INS-326 | chr2-142873013-INS-326 |
| chr2-142895763-INS-8335 | chr2-142895764-INS-8335 |
| chr2-142927278-INS-67 | chr2-142927278-INS-52 |
| chr2-142930860-INS-662 | chr2-142930860-INS-661 |
| chr2-142941225-INS-126 | chr2-142941225-INS-132 |
| chr2-142963361-INS-76 | chr2-142963362-INS-76 |
| chr2-142971681-INS-247 | chr2-142971681-INS-247 |
| chr2-142999708-INS-527 | chr2-142999708-INS-527 |
| chr2-143003986-INS-186 | chr2-143003986-INS-186 |
| chr2-143022373-INS-199 | chr2-143022363-INS-208 |
| chr2-143038631-INS-2136 | chr2-143038632-INS-2138 |
| chr2-143054370-INS-312 | chr2-143054370-INS-338 |
| chr2-143070959-INS-299 | chr2-143070959-INS-302 |
| chr2-143074081-INS-76 | chr2-143074081-INS-76 |
| chr2-143075641-INS-154 | chr2-143075641-INS-154 |
| chr2-143077139-INS-429 | chr2-143077111-INS-457 |
| chr2-143081528-INS-82 | chr2-143081528-INS-82 |
| chr2-143092004-INS-4705 | chr2-143092005-INS-4706 |
| chr2-143096206-INS-152 | chr2-143096206-INS-152 |
| chr2-143116517-INS-68 | chr2-143116517-INS-68 |
| chr2-143120942-INS-6624 | chr2-143120943-INS-6626 |
| chr2-143192127-INS-206 | chr2-143192127-INS-206 |
| chr2-143200090-INS-68 | chr2-143200088-INS-66 |
| chr2-143222918-INS-1147 | chr2-143222919-INS-1147 |
| chr2-143226486-INS-68 | chr2-143226484-INS-104 |
| chr2-143228038-INS-6676 | chr2-143228039-INS-6676 |
| chr2-143233965-INS-7083 | chr2-143233966-INS-7083 |
| chr2-143248125-INS-72 | chr2-143248121-INS-80 |
| chr2-143260449-INS-313 | chr2-143260449-INS-303 |
| chr2-143263644-INS-206 | chr2-143263646-INS-196 |
| chr2-143301678-INS-60 | chr2-143301678-INS-54 |

| | |
|---|---|
| chr2-143304350-INS-1153 | chr2-143304351-INS-1153 |
| chr2-143325845-INS-6908 | chr2-143325846-INS-6908 |
| chr2-143333756-INS-10546 | chr2-143333757-INS-10546 |
| chr2-143356039-INS-92 | chr2-143356031-INS-95 |
| chr2-143368725-INS-113 | chr2-143368725-INS-121 |
| chr2-143373021-INS-340 | chr2-143373021-INS-340 |
| chr2-143378714-INS-64 | chr2-143378714-INS-60 |
| chr2-143397242-INS-225 | chr2-143397242-INS-226 |
| chr2-143414837-INS-58 | chr2-143414809-INS-53 |
| chr2-143423418-INS-6255 | chr2-143423419-INS-6211 |
| chr2-143428319-INS-51 | chr2-143428319-INS-51 |
| chr2-143429966-INS-88 | chr2-143430002-INS-64 |
| chr2-143435127-INS-202 | chr2-143435127-INS-206 |
| chr2-143442586-INS-56 | chr2-143442586-INS-56 |
| chr2-143456170-INS-76 | chr2-143456170-INS-77 |
| chr2-143473499-INS-697 | chr2-143473499-INS-697 |
| chr2-143474682-INS-124 | chr2-143474678-INS-143 |
| chr2-143481878-INS-105 | chr2-143481881-INS-84 |
| chr2-143483944-INS-156 | chr2-143483941-INS-116 |
| chr2-143486145-INS-1247 | chr2-143486146-INS-1247 |
| chr2-143509701-INS-55 | chr2-143509701-INS-57 |
| chr2-143527347-INS-53 | chr2-143527347-INS-53 |
| chr2-143537108-INS-383 | chr2-143537108-INS-385 |
| chr2-143539764-INS-52 | chr2-143539764-INS-50 |
| chr2-143545196-INS-80 | chr2-143545228-INS-87 |
| chr2-143556260-INS-92 | chr2-143556260-INS-92 |
| chr2-143557058-INS-1025 | chr2-143557065-INS-1015 |
| chr2-143588628-INS-103 | chr2-143588628-INS-94 |
| chr2-143614677-INS-207 | chr2-143614675-INS-213 |
| chr2-143618915-INS-79 | chr2-143618915-INS-69 |
| chr2-143621014-INS-74 | chr2-143621007-INS-66 |
| chr2-143631859-INS-4805 | chr2-143631860-INS-4805 |
| chr2-143638487-INS-52 | chr2-143638487-INS-52 |
| chr2-143646648-INS-114 | chr2-143646635-INS-78 |
| chr2-143651669-INS-55 | chr2-143651669-INS-60 |
| chr2-143659995-INS-52 | chr2-143659995-INS-52 |
| chr2-143666813-INS-102 | chr2-143666813-INS-108 |
| chr2-143679891-INS-51 | chr2-143679891-INS-51 |
| chr2-143683211-INS-71 | chr2-143683211-INS-79 |
| chr2-143691264-INS-135 | chr2-143691262-INS-137 |
| chr2-143694524-INS-588 | chr2-143694524-INS-588 |
| chr2-143698789-INS-523 | chr2-143698789-INS-523 |
| chr2-143710856-INS-60 | chr2-143710843-INS-84 |
| chr2-143727661-INS-335 | chr2-143727661-INS-335 |
| chr2-143738666-INS-579 | chr2-143738666-INS-578 |
| chr2-143749907-INS-415 | chr2-143749907-INS-415 |
| chr2-143753227-INS-52 | chr2-143753227-INS-56 |
| chr2-143770940-INS-52 | chr2-143770941-INS-52 |
| chr2-143777076-INS-172 | chr2-143777076-INS-172 |
| chr2-143785315-INS-83 | chr2-143785315-INS-83 |
| chr2-143845165-INS-56 | chr2-143845165-INS-56 |
| chr2-143861979-INS-117 | chr2-143861979-INS-117 |
| chr2-143865484-INS-214 | chr2-143865484-INS-214 |
| chr2-143897624-INS-638 | chr2-143897657-INS-642 |
| chr2-143945840-INS-468 | chr2-143945840-INS-468 |
| chr2-143981181-INS-203 | chr2-143981181-INS-200 |
| chr2-143981815-INS-207 | chr2-143981815-INS-207 |
| chr2-144048167-INS-84 | chr2-144048167-INS-76 |
| chr2-144060324-INS-401 | chr2-144060324-INS-401 |
| chr2-144077868-INS-674 | chr2-144077868-INS-674 |
| chr2-144082056-INS-55 | chr2-144082056-INS-55 |
| chr2-144083316-INS-194 | chr2-144083316-INS-194 |
| chr2-144100469-INS-194 | chr2-144100469-INS-194 |
| chr2-144123634-INS-205 | chr2-144123634-INS-204 |
| chr2-144137305-INS-212 | chr2-144137305-INS-211 |
| chr2-144149432-INS-56 | chr2-144149432-INS-56 |
| chr2-144158312-INS-197 | chr2-144158312-INS-197 |
| chr2-144190819-INS-68 | chr2-144190819-INS-56 |
| chr2-144206771-INS-60 | chr2-144206771-INS-60 |
| chr2-144218658-INS-202 | chr2-144218658-INS-202 |
| chr2-144234875-INS-61 | chr2-144234875-INS-61 |
| chr2-144244897-INS-116 | chr2-144244897-INS-116 |
| chr2-144247454-INS-81 | chr2-144247454-INS-81 |
| chr2-144265972-INS-102 | chr2-144265972-INS-102 |
| chr2-144282978-INS-426 | chr2-144282978-INS-426 |
| chr2-144285602-INS-80 | chr2-144285602-INS-80 |
| chr2-144289497-INS-195 | chr2-144289497-INS-196 |

| | |
|---|---|
| chr2-144295383-INS-203 | chr2-144295383-INS-203 |
| chr2-144296778-INS-680 | chr2-144296778-INS-681 |
| chr2-144335121-INS-101 | chr2-144335121-INS-101 |
| chr2-144341377-INS-207 | chr2-144341377-INS-206 |
| chr2-144345458-INS-52 | chr2-144345458-INS-60 |
| chr2-144348367-INS-237 | chr2-144348367-INS-237 |
| chr2-144360299-INS-210 | chr2-144360299-INS-210 |
| chr2-144394544-INS-205 | chr2-144394544-INS-205 |
| chr2-144395323-INS-3063 | chr2-144395324-INS-3063 |
| chr2-144415777-INS-185 | chr2-144415777-INS-185 |
| chr2-144425925-INS-414 | chr2-144425925-INS-414 |
| chr2-144442961-INS-104 | chr2-144442961-INS-130 |
| chr2-144451320-INS-82 | chr2-144451320-INS-94 |
| chr2-144460282-INS-112 | chr2-144460282-INS-112 |
| chr2-144461922-INS-305 | chr2-144461922-INS-305 |
| chr2-144510055-INS-52 | chr2-144510055-INS-52 |
| chr2-144519570-INS-822 | chr2-144519570-INS-823 |
| chr2-144556744-INS-268 | chr2-144556744-INS-262 |
| chr2-144564187-INS-182 | chr2-144564177-INS-185 |
| chr2-144573027-INS-61 | chr2-144573027-INS-66 |
| chr2-144584183-INS-66 | chr2-144584183-INS-66 |
| chr2-144621305-INS-133 | chr2-144621305-INS-133 |
| chr2-144651288-INS-202 | chr2-144651288-INS-202 |
| chr2-144676056-INS-456 | chr2-144676056-INS-456 |
| chr2-144683532-INS-78 | chr2-144683509-INS-61 |
| chr2-144688633-INS-131 | chr2-144688633-INS-131 |
| chr2-144690102-INS-196 | chr2-144690073-INS-229 |
| chr2-144711241-INS-816 | chr2-144711241-INS-817 |
| chr2-144713607-INS-82 | chr2-144713607-INS-82 |
| chr2-144715874-INS-530 | chr2-144715874-INS-530 |
| chr2-144716957-INS-54 | chr2-144716957-INS-52 |
| chr2-144750861-INS-214 | chr2-144750861-INS-204 |
| chr2-144758256-INS-75 | chr2-144758256-INS-69 |
| chr2-144762556-INS-108 | chr2-144762556-INS-108 |
| chr2-144771269-INS-52 | chr2-144771269-INS-52 |
| chr2-144793433-INS-893 | chr2-144793433-INS-893 |
| chr2-144822794-INS-6478 | chr2-144822795-INS-6478 |
| chr2-144861208-INS-779 | chr2-144861208-INS-779 |
| chr2-144861951-INS-449 | chr2-144861951-INS-449 |
| chr2-144865184-INS-1572 | chr2-144865185-INS-1580 |
| chr2-144902996-INS-4701 | chr2-144903003-INS-4693 |
| chr2-144909769-INS-2763 | chr2-144909770-INS-2763 |
| chr2-144937771-INS-57 | chr2-144937771-INS-60 |
| chr2-144972484-INS-198 | chr2-144972484-INS-198 |
| chr2-144996019-INS-75 | chr2-144996019-INS-75 |
| chr2-145000318-INS-778 | chr2-145000318-INS-778 |
| chr2-145030240-INS-180 | chr2-145030240-INS-181 |
| chr2-145156637-INS-711 | chr2-145156637-INS-711 |
| chr2-145165495-INS-86 | chr2-145165495-INS-86 |
| chr2-145183413-INS-102 | chr2-145183414-INS-102 |
| chr2-145236063-INS-56 | chr2-145236025-INS-56 |
| chr2-145241496-INS-58 | chr2-145241496-INS-56 |
| chr2-145267435-INS-133 | chr2-145267435-INS-133 |
| chr2-145292581-INS-1221 | chr2-145292582-INS-1221 |
| chr2-145318143-INS-3258 | chr2-145318144-INS-3258 |
| chr2-145378290-INS-1391 | chr2-145378291-INS-1392 |
| chr2-145390990-INS-5844 | chr2-145390991-INS-5844 |
| chr2-145438407-INS-85 | chr2-145438407-INS-108 |
| chr2-145474439-INS-2955 | chr2-145474440-INS-2953 |
| chr2-145480961-INS-459 | chr2-145480961-INS-459 |
| chr2-145496534-INS-597 | chr2-145496535-INS-605 |
| chr2-145502092-INS-54 | chr2-145502092-INS-54 |
| chr2-145507937-INS-6507 | chr2-145507938-INS-6507 |
| chr2-145521792-INS-1892 | chr2-145521793-INS-1892 |
| chr2-145535851-INS-342 | chr2-145535851-INS-342 |
| chr2-145550134-INS-341 | chr2-145550134-INS-341 |
| chr2-145552865-INS-409 | chr2-145552865-INS-408 |
| chr2-145553880-INS-609 | chr2-145553880-INS-609 |
| chr2-145556092-INS-77 | chr2-145556092-INS-77 |
| chr2-145561283-INS-346 | chr2-145561283-INS-346 |
| chr2-145571147-INS-572 | chr2-145571147-INS-572 |
| chr2-145596035-INS-956 | chr2-145596035-INS-957 |
| chr2-145609466-INS-219 | chr2-145609466-INS-220 |
| chr2-145642794-INS-114 | chr2-145642794-INS-157 |
| chr2-145648229-INS-924 | chr2-145648229-INS-924 |
| chr2-145667343-INS-98 | chr2-145667343-INS-100 |
| chr2-145701274-INS-74 | chr2-145701274-INS-78 |

| | |
|---|---|
| chr2-145723054-INS-209 | chr2-145723054-INS-209 |
| chr2-145780807-INS-96 | chr2-145780814-INS-105 |
| chr2-145783168-INS-85 | chr2-145783168-INS-85 |
| chr2-145799788-INS-195 | chr2-145799788-INS-195 |
| chr2-145828195-INS-54 | chr2-145828195-INS-58 |
| chr2-145854595-INS-54 | chr2-145854595-INS-54 |
| chr2-145867923-INS-85 | chr2-145867923-INS-80 |
| chr2-145876061-INS-111 | chr2-145876061-INS-111 |
| chr2-145893639-INS-64 | chr2-145893639-INS-64 |
| chr2-146006205-INS-215 | chr2-146006205-INS-215 |
| chr2-146032653-INS-60 | chr2-146032653-INS-60 |
| chr2-146052259-INS-104 | chr2-146052264-INS-105 |
| chr2-146059144-INS-53 | chr2-146059144-INS-52 |
| chr2-146078485-INS-6238 | chr2-146078486-INS-6238 |
| chr2-146137413-INS-322 | chr2-146137413-INS-322 |
| chr2-146158236-INS-292 | chr2-146158236-INS-318 |
| chr2-146193438-INS-826 | chr2-146193438-INS-826 |
| chr2-146211775-INS-63 | chr2-146211776-INS-82 |
| chr2-146226400-INS-54 | chr2-146226400-INS-52 |
| chr2-146306902-INS-56 | chr2-146306902-INS-56 |
| chr2-146308166-INS-211 | chr2-146308166-INS-211 |
| chr2-146332635-INS-6432 | chr2-146332636-INS-6432 |
| chr2-146364349-INS-62 | chr2-146364354-INS-51 |
| chr2-146366133-INS-6652 | chr2-146366134-INS-6652 |
| chr2-146371733-INS-105 | chr2-146371733-INS-105 |
| chr2-146380865-INS-7074 | chr2-146380866-INS-7074 |
| chr2-146386900-INS-72 | chr2-146386900-INS-72 |
| chr2-146424055-INS-146 | chr2-146424055-INS-146 |
| chr2-146450354-INS-74 | chr2-146450362-INS-81 |
| chr2-146451874-INS-70 | chr2-146451878-INS-69 |
| chr2-146471511-INS-213 | chr2-146471511-INS-187 |
| chr2-146497976-INS-79 | chr2-146497976-INS-79 |
| chr2-146508768-INS-109 | chr2-146508768-INS-110 |
| chr2-146513263-INS-214 | chr2-146513263-INS-239 |
| chr2-146525092-INS-51 | chr2-146525111-INS-51 |
| chr2-146527180-INS-51 | chr2-146527180-INS-51 |
| chr2-146543773-INS-92 | chr2-146543773-INS-94 |
| chr2-146548606-INS-175 | chr2-146548606-INS-175 |
| chr2-146610420-INS-73 | chr2-146610390-INS-83 |
| chr2-146616009-INS-53 | chr2-146616009-INS-53 |
| chr2-146646836-INS-122 | chr2-146646836-INS-122 |
| chr2-146685593-INS-475 | chr2-146685594-INS-473 |
| chr2-146689965-INS-3863 | chr2-146689966-INS-3877 |
| chr2-146694822-INS-185 | chr2-146694822-INS-185 |
| chr2-146742041-INS-198 | chr2-146742041-INS-198 |
| chr2-146765844-INS-79 | chr2-146765840-INS-56 |
| chr2-146809545-INS-1348 | chr2-146809546-INS-1348 |
| chr2-146815763-INS-2340 | chr2-146815764-INS-2342 |
| chr2-146821042-INS-2324 | chr2-146821043-INS-2342 |
| chr2-146821512-INS-83 | chr2-146821512-INS-83 |
| chr2-146849082-INS-70 | chr2-146849082-INS-70 |
| chr2-146849547-INS-401 | chr2-146849547-INS-401 |
| chr2-146858212-INS-352 | chr2-146858212-INS-353 |
| chr2-146863598-INS-195 | chr2-146863598-INS-195 |
| chr2-146903853-INS-213 | chr2-146903853-INS-212 |
| chr2-146941108-INS-210 | chr2-146941108-INS-210 |
| chr2-146999424-INS-222 | chr2-146999424-INS-222 |
| chr2-147056086-INS-338 | chr2-147056086-INS-338 |
| chr2-147058381-INS-542 | chr2-147058381-INS-542 |
| chr2-147060256-INS-54 | chr2-147060256-INS-52 |
| chr2-147063300-INS-52 | chr2-147063300-INS-52 |
| chr2-147100391-INS-1254 | chr2-147100392-INS-1254 |
| chr2-147112869-INS-969 | chr2-147112869-INS-969 |
| chr2-147162041-INS-54 | chr2-147162041-INS-54 |
| chr2-147171254-INS-53 | chr2-147171254-INS-59 |
| chr2-147195878-INS-116 | chr2-147195878-INS-116 |
| chr2-147224840-INS-54 | chr2-147224840-INS-66 |
| chr2-147225653-INS-188 | chr2-147225653-INS-187 |
| chr2-147287059-INS-78 | chr2-147287039-INS-124 |
| chr2-147300297-INS-1686 | chr2-147300298-INS-1686 |
| chr2-147325537-INS-54 | chr2-147325537-INS-54 |
| chr2-147416346-INS-175 | chr2-147416346-INS-196 |
| chr2-147437125-INS-139 | chr2-147437125-INS-139 |
| chr2-147439754-INS-66 | chr2-147439754-INS-66 |
| chr2-147444335-INS-116 | chr2-147444343-INS-124 |
| chr2-147496421-INS-388 | chr2-147496421-INS-395 |
| chr2-147501611-INS-1340 | chr2-147501612-INS-1340 |

| | |
|---|---|
| chr2-147504034-INS-1007 | chr2-147504035-INS-1004 |
| chr2-147505604-INS-600 | chr2-147505604-INS-600 |
| chr2-147508110-INS-60 | chr2-147508111-INS-68 |
| chr2-147509541-INS-1388 | chr2-147509542-INS-1389 |
| chr2-147526861-INS-109 | chr2-147526845-INS-105 |
| chr2-147528729-INS-68 | chr2-147528763-INS-70 |
| chr2-147570171-INS-90 | chr2-147570169-INS-94 |
| chr2-147574538-INS-73 | chr2-147574538-INS-73 |
| chr2-147578597-INS-52 | chr2-147578597-INS-50 |
| chr2-147586028-INS-70 | chr2-147586028-INS-70 |
| chr2-147615989-INS-876 | chr2-147615989-INS-876 |
| chr2-147629543-INS-74 | chr2-147629543-INS-74 |
| chr2-147630141-INS-513 | chr2-147630141-INS-512 |
| chr2-147641867-INS-184 | chr2-147641867-INS-184 |
| chr2-147666780-INS-137 | chr2-147666780-INS-137 |
| chr2-147673578-INS-90 | chr2-147673572-INS-89 |
| chr2-147676959-INS-698 | chr2-147676959-INS-699 |
| chr2-147684129-INS-426 | chr2-147684129-INS-426 |
| chr2-147717886-INS-54 | chr2-147717883-INS-50 |
| chr2-147732109-INS-68 | chr2-147732109-INS-72 |
| chr2-147732936-INS-242 | chr2-147732936-INS-241 |
| chr2-147743415-INS-79 | chr2-147743415-INS-92 |
| chr2-147756026-INS-536 | chr2-147756026-INS-536 |
| chr2-147776476-INS-57 | chr2-147776476-INS-57 |
| chr2-147780778-INS-200 | chr2-147780778-INS-200 |
| chr2-147792367-INS-58 | chr2-147792368-INS-51 |
| chr2-147797406-INS-89 | chr2-147797406-INS-70 |
| chr2-147799540-INS-200 | chr2-147799541-INS-199 |
| chr2-147801103-INS-84 | chr2-147801146-INS-55 |
| chr2-147811929-INS-53 | chr2-147811929-INS-53 |
| chr2-147914026-INS-162 | chr2-147914026-INS-162 |
| chr2-147926615-INS-62 | chr2-147926615-INS-62 |
| chr2-147935563-INS-497 | chr2-147935563-INS-497 |
| chr2-147949410-INS-5293 | chr2-147949411-INS-5293 |
| chr2-147956388-INS-6489 | chr2-147956390-INS-6488 |
| chr2-147956541-INS-58 | chr2-147956541-INS-64 |
| chr2-147964421-INS-168 | chr2-147964436-INS-155 |
| chr2-147968828-INS-510 | chr2-147968828-INS-510 |
| chr2-147974824-INS-784 | chr2-147974824-INS-784 |
| chr2-147994474-INS-414 | chr2-147994474-INS-414 |
| chr2-148014194-INS-314 | chr2-148014195-INS-314 |
| chr2-148049121-INS-54 | chr2-148049121-INS-60 |
| chr2-148061361-INS-7120 | chr2-148061362-INS-7120 |
| chr2-148069710-INS-304 | chr2-148069710-INS-304 |
| chr2-148097403-INS-74 | chr2-148097403-INS-74 |
| chr2-148108955-INS-146 | chr2-148108973-INS-138 |
| chr2-148140156-INS-3163 | chr2-148140157-INS-3164 |
| chr2-148149887-INS-218 | chr2-148149887-INS-218 |
| chr2-148152929-INS-3231 | chr2-148152930-INS-3213 |
| chr2-148158623-INS-114 | chr2-148158623-INS-114 |
| chr2-148222693-INS-1107 | chr2-148222694-INS-1137 |
| chr2-148241741-INS-74 | chr2-148241741-INS-74 |
| chr2-148243559-INS-72 | chr2-148243559-INS-72 |
| chr2-148260085-INS-59 | chr2-148260085-INS-59 |
| chr2-148296591-INS-54 | chr2-148296591-INS-72 |
| chr2-148302570-INS-424 | chr2-148302570-INS-424 |
| chr2-148382938-INS-138 | chr2-148382938-INS-148 |
| chr2-148388050-INS-323 | chr2-148388050-INS-323 |
| chr2-148389125-INS-352 | chr2-148389125-INS-352 |
| chr2-148395065-INS-126 | chr2-148395065-INS-124 |
| chr2-148396987-INS-136 | chr2-148396989-INS-137 |
| chr2-148399962-INS-81 | chr2-148399962-INS-81 |
| chr2-148411266-INS-60 | chr2-148411266-INS-60 |
| chr2-148444217-INS-138 | chr2-148444219-INS-138 |
| chr2-148449036-INS-400 | chr2-148449036-INS-400 |
| chr2-148482410-INS-66 | chr2-148482410-INS-66 |
| chr2-148504995-INS-56 | chr2-148504995-INS-54 |
| chr2-148513187-INS-58 | chr2-148513187-INS-59 |
| chr2-148527522-INS-341 | chr2-148527522-INS-343 |
| chr2-148557437-INS-202 | chr2-148557437-INS-202 |
| chr2-148559644-INS-58 | chr2-148559644-INS-58 |
| chr2-148583785-INS-70 | chr2-148583745-INS-85 |
| chr2-148586412-INS-101 | chr2-148586412-INS-52 |
| chr2-148617882-INS-52 | chr2-148617882-INS-56 |
| chr2-148619317-INS-68 | chr2-148619330-INS-80 |
| chr2-148621347-INS-173 | chr2-148621347-INS-173 |
| chr2-148633997-INS-1003 | chr2-148633998-INS-1003 |

| | |
|---|---|
| chr2-148648150-INS-165 | chr2-148648150-INS-165 |
| chr2-148653964-INS-6371 | chr2-148653965-INS-6371 |
| chr2-148659692-INS-54 | chr2-148659693-INS-71 |
| chr2-148663871-INS-107 | chr2-148663909-INS-64 |
| chr2-148666956-INS-54 | chr2-148666956-INS-56 |
| chr2-148667428-INS-86 | chr2-148667428-INS-88 |
| chr2-148673380-INS-77 | chr2-148673371-INS-61 |
| chr2-148679967-INS-6413 | chr2-148679968-INS-6413 |
| chr2-148682776-INS-203 | chr2-148682776-INS-203 |
| chr2-148697185-INS-76 | chr2-148697185-INS-76 |
| chr2-148699190-INS-52 | chr2-148699190-INS-58 |
| chr2-148738830-INS-487 | chr2-148738830-INS-487 |
| chr2-148739625-INS-943 | chr2-148739625-INS-943 |
| chr2-148739848-INS-7110 | chr2-148739849-INS-7110 |
| chr2-148747194-INS-61 | chr2-148747194-INS-63 |
| chr2-148757520-INS-8462 | chr2-148757521-INS-8462 |
| chr2-148777334-INS-2148 | chr2-148777335-INS-2148 |
| chr2-148788245-INS-6475 | chr2-148788246-INS-6475 |
| chr2-148860248-INS-1069 | chr2-148860249-INS-1069 |
| chr2-148865003-INS-282 | chr2-148864985-INS-303 |
| chr2-148873751-INS-853 | chr2-148873751-INS-853 |
| chr2-148902400-INS-311 | chr2-148902400-INS-318 |
| chr2-148904049-INS-712 | chr2-148904044-INS-722 |
| chr2-148911446-INS-59 | chr2-148911436-INS-70 |
| chr2-148926620-INS-77 | chr2-148926620-INS-77 |
| chr2-148932918-INS-69 | chr2-148932919-INS-65 |
| chr2-149048662-INS-104 | chr2-149048662-INS-104 |
| chr2-149179247-INS-460 | chr2-149179248-INS-460 |
| chr2-149193816-INS-553 | chr2-149193816-INS-553 |
| chr2-149207095-INS-118 | chr2-149207076-INS-95 |
| chr2-149208384-INS-57 | chr2-149208384-INS-57 |
| chr2-149240028-INS-60 | chr2-149240028-INS-60 |
| chr2-149244546-INS-6094 | chr2-149244547-INS-6100 |
| chr2-149274466-INS-5175 | chr2-149274467-INS-5175 |
| chr2-149275313-INS-133 | chr2-149275313-INS-133 |
| chr2-149285297-INS-147 | chr2-149285297-INS-165 |
| chr2-149304788-INS-6421 | chr2-149304789-INS-6421 |
| chr2-149426171-INS-4224 | chr2-149426172-INS-4224 |
| chr2-149463406-INS-60 | chr2-149463406-INS-68 |
| chr2-149467998-INS-115 | chr2-149467998-INS-115 |
| chr2-149491299-INS-590 | chr2-149491299-INS-590 |
| chr2-149512131-INS-112 | chr2-149512131-INS-114 |
| chr2-149519739-INS-56 | chr2-149519739-INS-58 |
| chr2-149533123-INS-165 | chr2-149533133-INS-159 |
| chr2-149569687-INS-304 | chr2-149569687-INS-305 |
| chr2-149573364-INS-140 | chr2-149573364-INS-140 |
| chr2-149575300-INS-344 | chr2-149575300-INS-344 |
| chr2-149598991-INS-159 | chr2-149598991-INS-159 |
| chr2-149618379-INS-148 | chr2-149618379-INS-168 |
| chr2-149626978-INS-78 | chr2-149626978-INS-80 |
| chr2-149628655-INS-84 | chr2-149628703-INS-55 |
| chr2-149651343-INS-938 | chr2-149651343-INS-938 |
| chr2-149694703-INS-733 | chr2-149694703-INS-734 |
| chr2-149699107-INS-122 | chr2-149699107-INS-122 |
| chr2-149717116-INS-180 | chr2-149717116-INS-180 |
| chr2-149725973-INS-52 | chr2-149725973-INS-52 |
| chr2-149739912-INS-186 | chr2-149739912-INS-185 |
| chr2-149749046-INS-66 | chr2-149749046-INS-66 |
| chr2-149758222-INS-99 | chr2-149758221-INS-81 |
| chr2-149774698-INS-1887 | chr2-149774699-INS-1887 |
| chr2-149775845-INS-205 | chr2-149775845-INS-205 |
| chr2-149805092-INS-722 | chr2-149805092-INS-722 |
| chr2-149854457-INS-62 | chr2-149854438-INS-50 |
| chr2-149937123-INS-267 | chr2-149937123-INS-267 |
| chr2-149938646-INS-51 | chr2-149938647-INS-53 |
| chr2-149947428-INS-197 | chr2-149947428-INS-197 |
| chr2-149948450-INS-265 | chr2-149948451-INS-266 |
| chr2-149983711-INS-828 | chr2-149983711-INS-828 |
| chr2-149989733-INS-984 | chr2-149989733-INS-984 |
| chr2-149998862-INS-212 | chr2-149998862-INS-212 |
| chr2-150007264-INS-1828 | chr2-150007265-INS-1828 |
| chr2-150034492-INS-252 | chr2-150034492-INS-252 |
| chr2-150035793-INS-120 | chr2-150035793-INS-120 |
| chr2-150089503-INS-6434 | chr2-150089504-INS-6439 |
| chr2-150127804-INS-341 | chr2-150127804-INS-341 |
| chr2-150132993-INS-174 | chr2-150132993-INS-175 |
| chr2-150138269-INS-52 | chr2-150138269-INS-50 |

| | |
|---|---|
| chr2-150144621-INS-56 | chr2-150144621-INS-60 |
| chr2-150150533-INS-55 | chr2-150150533-INS-55 |
| chr2-150164689-INS-65 | chr2-150164689-INS-65 |
| chr2-150215391-INS-50 | chr2-150215391-INS-56 |
| chr2-150246546-INS-2895 | chr2-150246547-INS-2895 |
| chr2-150357964-INS-60 | chr2-150357964-INS-60 |
| chr2-150382824-INS-2603 | chr2-150382825-INS-2609 |
| chr2-150483152-INS-188 | chr2-150483132-INS-162 |
| chr2-150511666-INS-501 | chr2-150511666-INS-501 |
| chr2-150521479-INS-53 | chr2-150521484-INS-58 |
| chr2-150534896-INS-177 | chr2-150534896-INS-176 |
| chr2-150536668-INS-773 | chr2-150536668-INS-765 |
| chr2-150602256-INS-146 | chr2-150602256-INS-146 |
| chr2-150616729-INS-6401 | chr2-150616730-INS-6401 |
| chr2-150619314-INS-52 | chr2-150619314-INS-52 |
| chr2-150633581-INS-79 | chr2-150633581-INS-79 |
| chr2-150635582-INS-118 | chr2-150635582-INS-118 |
| chr2-150647346-INS-67 | chr2-150647346-INS-67 |
| chr2-150654465-INS-210 | chr2-150654465-INS-210 |
| chr2-150686385-INS-583 | chr2-150686385-INS-583 |
| chr2-150693712-INS-58 | chr2-150693712-INS-64 |
| chr2-150722429-INS-216 | chr2-150722429-INS-216 |
| chr2-150728719-INS-80 | chr2-150728718-INS-96 |
| chr2-150745293-INS-217 | chr2-150745293-INS-217 |
| chr2-150748419-INS-224 | chr2-150748419-INS-224 |
| chr2-150766168-INS-52 | chr2-150766168-INS-82 |
| chr2-150778281-INS-352 | chr2-150778281-INS-352 |
| chr2-150787192-INS-280 | chr2-150787192-INS-252 |
| chr2-150803056-INS-211 | chr2-150803056-INS-211 |
| chr2-150805258-INS-3131 | chr2-150805259-INS-3131 |
| chr2-150858939-INS-6374 | chr2-150858941-INS-6376 |
| chr2-150861128-INS-287 | chr2-150861128-INS-287 |
| chr2-150881779-INS-518 | chr2-150881779-INS-518 |
| chr2-150884324-INS-54 | chr2-150884324-INS-54 |
| chr2-150902676-INS-160 | chr2-150902676-INS-161 |
| chr2-150918258-INS-449 | chr2-150918258-INS-449 |
| chr2-150957985-INS-437 | chr2-150957985-INS-437 |
| chr2-150978343-INS-199 | chr2-150978350-INS-199 |
| chr2-151035443-INS-723 | chr2-151035396-INS-725 |
| chr2-151082377-INS-2660 | chr2-151082378-INS-2660 |
| chr2-151240912-INS-1341 | chr2-151240913-INS-1341 |
| chr2-151378214-INS-54 | chr2-151378214-INS-54 |
| chr2-151407838-INS-129 | chr2-151407838-INS-127 |
| chr2-151454236-INS-177 | chr2-151454236-INS-177 |
| chr2-151457952-INS-60 | chr2-151457952-INS-60 |
| chr2-151466082-INS-503 | chr2-151466082-INS-503 |
| chr2-151467345-INS-68 | chr2-151467345-INS-68 |
| chr2-151507406-INS-196 | chr2-151507406-INS-196 |
| chr2-151517370-INS-4792 | chr2-151517371-INS-4792 |
| chr2-151527822-INS-86 | chr2-151527822-INS-92 |
| chr2-151552505-INS-147 | chr2-151552505-INS-147 |
| chr2-151569024-INS-958 | chr2-151569024-INS-959 |
| chr2-151569964-INS-142 | chr2-151569964-INS-164 |
| chr2-151596131-INS-112 | chr2-151596131-INS-113 |
| chr2-151601099-INS-113 | chr2-151601099-INS-113 |
| chr2-151605309-INS-1841 | chr2-151605310-INS-1841 |
| chr2-151605538-INS-148 | chr2-151605558-INS-172 |
| chr2-151608130-INS-555 | chr2-151608130-INS-555 |
| chr2-151611661-INS-245 | chr2-151611661-INS-245 |
| chr2-151662845-INS-60 | chr2-151662845-INS-56 |
| chr2-151666891-INS-134 | chr2-151666890-INS-135 |
| chr2-151697628-INS-60 | chr2-151697629-INS-56 |
| chr2-151704389-INS-82 | chr2-151704389-INS-82 |
| chr2-151709636-INS-282 | chr2-151709642-INS-290 |
| chr2-151730900-INS-2067 | chr2-151730901-INS-2067 |
| chr2-151756824-INS-82 | chr2-151756824-INS-82 |
| chr2-151788880-INS-230 | chr2-151788881-INS-231 |
| chr2-151869487-INS-86 | chr2-151869487-INS-88 |
| chr2-151870323-INS-96 | chr2-151870370-INS-50 |
| chr2-151879643-INS-102 | chr2-151879643-INS-102 |
| chr2-151884929-INS-112 | chr2-151884930-INS-74 |
| chr2-151896850-INS-5315 | chr2-151896851-INS-5315 |
| chr2-151901979-INS-74 | chr2-151901979-INS-87 |
| chr2-151904682-INS-175 | chr2-151904664-INS-175 |
| chr2-151907704-INS-75 | chr2-151907704-INS-73 |
| chr2-151913037-INS-185 | chr2-151913017-INS-200 |
| chr2-151964222-INS-87 | chr2-151964216-INS-95 |

| | |
|---|---|
| chr2-151975875-INS-163 | chr2-151975873-INS-159 |
| chr2-151994978-INS-52 | chr2-151994978-INS-60 |
| chr2-152004242-INS-72 | chr2-152004242-INS-84 |
| chr2-152030297-INS-72 | chr2-152030309-INS-84 |
| chr2-152039632-INS-86 | chr2-152039630-INS-55 |
| chr2-152054192-INS-52 | chr2-152054192-INS-52 |
| chr2-152066484-INS-52 | chr2-152066484-INS-52 |
| chr2-152071893-INS-165 | chr2-152071893-INS-170 |
| chr2-152086574-INS-170 | chr2-152086574-INS-169 |
| chr2-152088312-INS-551 | chr2-152088312-INS-551 |
| chr2-152141752-INS-227 | chr2-152141752-INS-227 |
| chr2-152168908-INS-208 | chr2-152168908-INS-209 |
| chr2-152175194-INS-64 | chr2-152175194-INS-71 |
| chr2-152197284-INS-382 | chr2-152197284-INS-382 |
| chr2-152200315-INS-59 | chr2-152200315-INS-95 |
| chr2-152209765-INS-82 | chr2-152209737-INS-128 |
| chr2-152214866-INS-52 | chr2-152214866-INS-56 |
| chr2-152219420-INS-205 | chr2-152219420-INS-205 |
| chr2-152228386-INS-7451 | chr2-152228387-INS-7451 |
| chr2-152233560-INS-97 | chr2-152233560-INS-101 |
| chr2-152236730-INS-124 | chr2-152236730-INS-124 |
| chr2-152242681-INS-107 | chr2-152242681-INS-106 |
| chr2-152268972-INS-68 | chr2-152268972-INS-64 |
| chr2-152274036-INS-540 | chr2-152274036-INS-540 |
| chr2-152292927-INS-76 | chr2-152292927-INS-75 |
| chr2-152319993-INS-57 | chr2-152319993-INS-58 |
| chr2-152354629-INS-109 | chr2-152354629-INS-109 |
| chr2-152374806-INS-82 | chr2-152374806-INS-82 |
| chr2-152386530-INS-443 | chr2-152386530-INS-443 |
| chr2-152392093-INS-196 | chr2-152392093-INS-196 |
| chr2-152398080-INS-236 | chr2-152398080-INS-218 |
| chr2-152458457-INS-154 | chr2-152458457-INS-154 |
| chr2-152482842-INS-111 | chr2-152482842-INS-111 |
| chr2-152488463-INS-5008 | chr2-152488464-INS-5008 |
| chr2-152491533-INS-4612 | chr2-152491535-INS-4600 |
| chr2-152563803-INS-200 | chr2-152563803-INS-199 |
| chr2-152570206-INS-81 | chr2-152570206-INS-81 |
| chr2-152574815-INS-61 | chr2-152574815-INS-59 |
| chr2-152592672-INS-530 | chr2-152592672-INS-530 |
| chr2-152636884-INS-208 | chr2-152636884-INS-207 |
| chr2-152674293-INS-108 | chr2-152674293-INS-112 |
| chr2-152681040-INS-201 | chr2-152681040-INS-201 |
| chr2-152685145-INS-208 | chr2-152685145-INS-207 |
| chr2-152688911-INS-74 | chr2-152688911-INS-74 |
| chr2-152698628-INS-70 | chr2-152698628-INS-84 |
| chr2-152723456-INS-277 | chr2-152723456-INS-277 |
| chr2-152740534-INS-245 | chr2-152740534-INS-245 |
| chr2-152751024-INS-56 | chr2-152751024-INS-78 |
| chr2-152771511-INS-339 | chr2-152771538-INS-319 |
| chr2-152773154-INS-394 | chr2-152773154-INS-394 |
| chr2-152776826-INS-193 | chr2-152776826-INS-193 |
| chr2-152781575-INS-64 | chr2-152781575-INS-65 |
| chr2-152792961-INS-56 | chr2-152792961-INS-62 |
| chr2-152805900-INS-57 | chr2-152805871-INS-54 |
| chr2-152810359-INS-54 | chr2-152810359-INS-72 |
| chr2-152813533-INS-103 | chr2-152813533-INS-103 |
| chr2-152818835-INS-84 | chr2-152818827-INS-89 |
| chr2-152822451-INS-6442 | chr2-152822452-INS-6442 |
| chr2-152826469-INS-76 | chr2-152826471-INS-78 |
| chr2-152827450-INS-94 | chr2-152827451-INS-57 |
| chr2-152840014-INS-111 | chr2-152840014-INS-68 |
| chr2-152843095-INS-54 | chr2-152843095-INS-54 |
| chr2-152846745-INS-117 | chr2-152846749-INS-113 |
| chr2-152849312-INS-54 | chr2-152849312-INS-54 |
| chr2-152856374-INS-8417 | chr2-152856375-INS-8417 |
| chr2-152865495-INS-98 | chr2-152865471-INS-114 |
| chr2-152866730-INS-1884 | chr2-152866731-INS-1884 |
| chr2-152871450-INS-55 | chr2-152871450-INS-55 |
| chr2-152880932-INS-124 | chr2-152880932-INS-124 |
| chr2-152883459-INS-154 | chr2-152883459-INS-154 |
| chr2-152917885-INS-809 | chr2-152917885-INS-859 |
| chr2-152919862-INS-215 | chr2-152919862-INS-216 |
| chr2-152946858-INS-108 | chr2-152946858-INS-110 |
| chr2-152959618-INS-59 | chr2-152959618-INS-51 |
| chr2-152964919-INS-56 | chr2-152964919-INS-56 |
| chr2-152980006-INS-114 | chr2-152980006-INS-114 |
| chr2-152991259-INS-70 | chr2-152991259-INS-70 |

| | |
|---|---|
| chr2-153050162-INS-400 | chr2-153050164-INS-401 |
| chr2-153061789-INS-88 | chr2-153061789-INS-88 |
| chr2-153086311-INS-174 | chr2-153086311-INS-174 |
| chr2-153142353-INS-207 | chr2-153142353-INS-207 |
| chr2-153146250-INS-71 | chr2-153146250-INS-71 |
| chr2-153151381-INS-446 | chr2-153151381-INS-446 |
| chr2-153163496-INS-60 | chr2-153163517-INS-54 |
| chr2-153201334-INS-154 | chr2-153201334-INS-154 |
| chr2-153202412-INS-210 | chr2-153202412-INS-210 |
| chr2-153212808-INS-2300 | chr2-153212809-INS-2300 |
| chr2-153219871-INS-223 | chr2-153219871-INS-223 |
| chr2-153229431-INS-195 | chr2-153229431-INS-195 |
| chr2-153261007-INS-218 | chr2-153261007-INS-219 |
| chr2-153314355-INS-245 | chr2-153314355-INS-245 |
| chr2-153336097-INS-70 | chr2-153336097-INS-70 |
| chr2-153349043-INS-214 | chr2-153349043-INS-214 |
| chr2-153360948-INS-116 | chr2-153360948-INS-116 |
| chr2-153375320-INS-225 | chr2-153375325-INS-226 |
| chr2-153385587-INS-204 | chr2-153385587-INS-204 |
| chr2-153394870-INS-201 | chr2-153394870-INS-201 |
| chr2-153395668-INS-64 | chr2-153395668-INS-64 |
| chr2-153415493-INS-214 | chr2-153415493-INS-214 |
| chr2-153429353-INS-213 | chr2-153429354-INS-203 |
| chr2-153429763-INS-78 | chr2-153429763-INS-78 |
| chr2-153431573-INS-164 | chr2-153431573-INS-164 |
| chr2-153432532-INS-220 | chr2-153432532-INS-220 |
| chr2-153447645-INS-64 | chr2-153447645-INS-64 |
| chr2-153467082-INS-655 | chr2-153467083-INS-655 |
| chr2-153469211-INS-177 | chr2-153469211-INS-177 |
| chr2-153475726-INS-196 | chr2-153475726-INS-196 |
| chr2-153478555-INS-200 | chr2-153478555-INS-200 |
| chr2-153479565-INS-103 | chr2-153479565-INS-103 |
| chr2-153489847-INS-154 | chr2-153489847-INS-154 |
| chr2-153503750-INS-202 | chr2-153503750-INS-212 |
| chr2-153518424-INS-468 | chr2-153518424-INS-468 |
| chr2-153530346-INS-204 | chr2-153530346-INS-204 |
| chr2-153531731-INS-199 | chr2-153531731-INS-198 |
| chr2-153532291-INS-64 | chr2-153532291-INS-58 |
| chr2-153545975-INS-218 | chr2-153545975-INS-218 |
| chr2-153547530-INS-215 | chr2-153547530-INS-215 |
| chr2-153549749-INS-193 | chr2-153549753-INS-182 |
| chr2-153552002-INS-220 | chr2-153552002-INS-221 |
| chr2-153554394-INS-250 | chr2-153554394-INS-253 |
| chr2-153587923-INS-209 | chr2-153587923-INS-209 |
| chr2-153603856-INS-211 | chr2-153603856-INS-211 |
| chr2-153641123-INS-54 | chr2-153641123-INS-54 |
| chr2-153656857-INS-759 | chr2-153656857-INS-759 |
| chr2-153667870-INS-114 | chr2-153667870-INS-114 |
| chr2-153669924-INS-202 | chr2-153669924-INS-202 |
| chr2-153710766-INS-54 | chr2-153710766-INS-54 |
| chr2-153755789-INS-55 | chr2-153755789-INS-53 |
| chr2-153766064-INS-159 | chr2-153766064-INS-160 |
| chr2-153779444-INS-67 | chr2-153779444-INS-67 |
| chr2-153782548-INS-84 | chr2-153782548-INS-84 |
| chr2-153798385-INS-486 | chr2-153798385-INS-486 |
| chr2-153837086-INS-78 | chr2-153837086-INS-80 |
| chr2-153838134-INS-261 | chr2-153838134-INS-253 |
| chr2-153839614-INS-75 | chr2-153839614-INS-85 |
| chr2-153912475-INS-412 | chr2-153912475-INS-412 |
| chr2-153913627-INS-474 | chr2-153913627-INS-474 |
| chr2-153918709-INS-106 | chr2-153918709-INS-112 |
| chr2-153940142-INS-489 | chr2-153940142-INS-489 |
| chr2-153944637-INS-190 | chr2-153944637-INS-191 |
| chr2-153945947-INS-78 | chr2-153945947-INS-72 |
| chr2-153973201-INS-75 | chr2-153973201-INS-75 |
| chr2-153977894-INS-67 | chr2-153977894-INS-67 |
| chr2-153983057-INS-1101 | chr2-153983058-INS-1101 |
| chr2-153995421-INS-1144 | chr2-153995422-INS-1142 |
| chr2-153999790-INS-106 | chr2-153999790-INS-106 |
| chr2-154003242-INS-60 | chr2-154003242-INS-60 |
| chr2-154028219-INS-70 | chr2-154028220-INS-64 |
| chr2-154030229-INS-54 | chr2-154030229-INS-50 |
| chr2-154034062-INS-86 | chr2-154034062-INS-80 |
| chr2-154051105-INS-1143 | chr2-154051110-INS-1133 |
| chr2-154064726-INS-189 | chr2-154064726-INS-189 |
| chr2-154089042-INS-350 | chr2-154089042-INS-350 |
| chr2-154138627-INS-5562 | chr2-154138628-INS-5562 |

| | |
|---|---|
| chr2-154153049-INS-214 | chr2-154153049-INS-214 |
| chr2-154163464-INS-74 | chr2-154163426-INS-110 |
| chr2-154176303-INS-879 | chr2-154176303-INS-879 |
| chr2-154178237-INS-171 | chr2-154178237-INS-171 |
| chr2-154225327-INS-192 | chr2-154225327-INS-196 |
| chr2-154226298-INS-101 | chr2-154226298-INS-101 |
| chr2-154235529-INS-189 | chr2-154235530-INS-186 |
| chr2-154236236-INS-992 | chr2-154236236-INS-988 |
| chr2-154241008-INS-2345 | chr2-154241013-INS-2320 |
| chr2-154254047-INS-440 | chr2-154254047-INS-443 |
| chr2-154254941-INS-240 | chr2-154254941-INS-240 |
| chr2-154260435-INS-216 | chr2-154260436-INS-209 |
| chr2-154260992-INS-438 | chr2-154260992-INS-438 |
| chr2-154266059-INS-89 | chr2-154266059-INS-89 |
| chr2-154276795-INS-58 | chr2-154276795-INS-58 |
| chr2-154281684-INS-204 | chr2-154281684-INS-204 |
| chr2-154303897-INS-233 | chr2-154303897-INS-233 |
| chr2-154310647-INS-164 | chr2-154310647-INS-164 |
| chr2-154317042-INS-210 | chr2-154317038-INS-219 |
| chr2-154325461-INS-195 | chr2-154325461-INS-195 |
| chr2-154327427-INS-207 | chr2-154327427-INS-207 |
| chr2-154330212-INS-222 | chr2-154330212-INS-221 |
| chr2-154334801-INS-200 | chr2-154334801-INS-200 |
| chr2-154392202-INS-100 | chr2-154392202-INS-100 |
| chr2-154403995-INS-208 | chr2-154403995-INS-209 |
| chr2-154416058-INS-205 | chr2-154416058-INS-205 |
| chr2-154427118-INS-208 | chr2-154427118-INS-208 |
| chr2-154429139-INS-203 | chr2-154429139-INS-203 |
| chr2-154432001-INS-187 | chr2-154432001-INS-187 |
| chr2-154433680-INS-66 | chr2-154433680-INS-66 |
| chr2-154435984-INS-373 | chr2-154435984-INS-373 |
| chr2-154437807-INS-65 | chr2-154437807-INS-65 |
| chr2-154447368-INS-200 | chr2-154447368-INS-200 |
| chr2-154457336-INS-80 | chr2-154457336-INS-72 |
| chr2-154464999-INS-214 | chr2-154464999-INS-214 |
| chr2-154469091-INS-203 | chr2-154469091-INS-203 |
| chr2-154489371-INS-191 | chr2-154489371-INS-191 |
| chr2-154494301-INS-120 | chr2-154494348-INS-121 |
| chr2-154498008-INS-189 | chr2-154498008-INS-190 |
| chr2-154505499-INS-213 | chr2-154505499-INS-214 |
| chr2-154506036-INS-209 | chr2-154506037-INS-209 |
| chr2-154525676-INS-72 | chr2-154525676-INS-64 |
| chr2-154535439-INS-123 | chr2-154535439-INS-123 |
| chr2-154539805-INS-207 | chr2-154539805-INS-200 |
| chr2-154542124-INS-54 | chr2-154542124-INS-54 |
| chr2-154561823-INS-183 | chr2-154561823-INS-180 |
| chr2-154562808-INS-61 | chr2-154562808-INS-61 |
| chr2-154573818-INS-59 | chr2-154573818-INS-67 |
| chr2-154582168-INS-105 | chr2-154582176-INS-112 |
| chr2-154596210-INS-208 | chr2-154596210-INS-208 |
| chr2-154601751-INS-82 | chr2-154601750-INS-51 |
| chr2-154606704-INS-60 | chr2-154606704-INS-60 |
| chr2-154612293-INS-359 | chr2-154612293-INS-364 |
| chr2-154623583-INS-132 | chr2-154623596-INS-130 |
| chr2-154638839-INS-90 | chr2-154638839-INS-90 |
| chr2-154663203-INS-220 | chr2-154663203-INS-219 |
| chr2-154666117-INS-208 | chr2-154666117-INS-208 |
| chr2-154669006-INS-207 | chr2-154669006-INS-204 |
| chr2-154690565-INS-75 | chr2-154690565-INS-75 |
| chr2-154706522-INS-134 | chr2-154706522-INS-131 |
| chr2-154722090-INS-217 | chr2-154722090-INS-217 |
| chr2-154726155-INS-216 | chr2-154726155-INS-217 |
| chr2-154736654-INS-188 | chr2-154736654-INS-187 |
| chr2-154743398-INS-745 | chr2-154743398-INS-745 |
| chr2-154746068-INS-3956 | chr2-154746069-INS-3957 |
| chr2-154748794-INS-208 | chr2-154748794-INS-208 |
| chr2-154749306-INS-8753 | chr2-154749307-INS-8750 |
| chr2-154751851-INS-214 | chr2-154751851-INS-214 |
| chr2-154756945-INS-171 | chr2-154756945-INS-173 |
| chr2-154757601-INS-56 | chr2-154757601-INS-58 |
| chr2-154765554-INS-264 | chr2-154765554-INS-264 |
| chr2-154791023-INS-69 | chr2-154791004-INS-66 |
| chr2-154798589-INS-177 | chr2-154798589-INS-177 |
| chr2-154807024-INS-4720 | chr2-154807025-INS-4720 |
| chr2-154812795-INS-57 | chr2-154812795-INS-57 |
| chr2-154813881-INS-273 | chr2-154813881-INS-274 |
| chr2-154815141-INS-1891 | chr2-154815142-INS-1891 |

| | |
|---|---|
| chr2-154816145-INS-199 | chr2-154816145-INS-199 |
| chr2-154824323-INS-5561 | chr2-154824324-INS-5561 |
| chr2-154836689-INS-345 | chr2-154836689-INS-345 |
| chr2-154845696-INS-144 | chr2-154845696-INS-150 |
| chr2-154923874-INS-200 | chr2-154923874-INS-206 |
| chr2-154928533-INS-208 | chr2-154928533-INS-208 |
| chr2-154935634-INS-114 | chr2-154935638-INS-100 |
| chr2-154943441-INS-754 | chr2-154943441-INS-753 |
| chr2-154946154-INS-71 | chr2-154946154-INS-71 |
| chr2-154947033-INS-230 | chr2-154947033-INS-230 |
| chr2-154959344-INS-174 | chr2-154959344-INS-175 |
| chr2-154973024-INS-212 | chr2-154973024-INS-212 |
| chr2-154990254-INS-569 | chr2-154990254-INS-569 |
| chr2-154996003-INS-205 | chr2-154996003-INS-208 |
| chr2-154997158-INS-73 | chr2-154997158-INS-73 |
| chr2-154999335-INS-103 | chr2-154999375-INS-81 |
| chr2-155004321-INS-202 | chr2-155004321-INS-202 |
| chr2-155008902-INS-65 | chr2-155008902-INS-65 |
| chr2-155016952-INS-239 | chr2-155016951-INS-242 |
| chr2-155031105-INS-52 | chr2-155031105-INS-52 |
| chr2-155046659-INS-77 | chr2-155046659-INS-86 |
| chr2-155053819-INS-209 | chr2-155053834-INS-207 |
| chr2-155056699-INS-301 | chr2-155056698-INS-304 |
| chr2-155074871-INS-248 | chr2-155074871-INS-248 |
| chr2-155076273-INS-72 | chr2-155076273-INS-72 |
| chr2-155091588-INS-83 | chr2-155091588-INS-83 |
| chr2-155130139-INS-144 | chr2-155130139-INS-144 |
| chr2-155157733-INS-141 | chr2-155157733-INS-141 |
| chr2-155160737-INS-430 | chr2-155160737-INS-431 |
| chr2-155169803-INS-177 | chr2-155169803-INS-177 |
| chr2-155171995-INS-114 | chr2-155171995-INS-114 |
| chr2-155181851-INS-60 | chr2-155181851-INS-70 |
| chr2-155186748-INS-163 | chr2-155186748-INS-163 |
| chr2-155194135-INS-51 | chr2-155194104-INS-56 |
| chr2-155197968-INS-92 | chr2-155197968-INS-92 |
| chr2-155210729-INS-299 | chr2-155210729-INS-299 |
| chr2-155252544-INS-210 | chr2-155252544-INS-210 |
| chr2-155289757-INS-56 | chr2-155289757-INS-58 |
| chr2-155293114-INS-209 | chr2-155293114-INS-209 |
| chr2-155336637-INS-74 | chr2-155336637-INS-74 |
| chr2-155339296-INS-156 | chr2-155339296-INS-160 |
| chr2-155358610-INS-5312 | chr2-155358611-INS-5312 |
| chr2-155389366-INS-52 | chr2-155389368-INS-50 |
| chr2-155399871-INS-201 | chr2-155399871-INS-201 |
| chr2-155401460-INS-60 | chr2-155401460-INS-60 |
| chr2-155423038-INS-295 | chr2-155423038-INS-295 |
| chr2-155428017-INS-161 | chr2-155428017-INS-155 |
| chr2-155431199-INS-65 | chr2-155431199-INS-70 |
| chr2-155453217-INS-60 | chr2-155453213-INS-81 |
| chr2-155494734-INS-249 | chr2-155494734-INS-291 |
| chr2-155539727-INS-160 | chr2-155539727-INS-160 |
| chr2-155554515-INS-442 | chr2-155554515-INS-443 |
| chr2-155569364-INS-132 | chr2-155569364-INS-133 |
| chr2-155578485-INS-215 | chr2-155578485-INS-215 |
| chr2-155580913-INS-5405 | chr2-155580914-INS-5405 |
| chr2-155587028-INS-204 | chr2-155587028-INS-201 |
| chr2-155599342-INS-174 | chr2-155599342-INS-176 |
| chr2-155609622-INS-206 | chr2-155609613-INS-247 |
| chr2-155639144-INS-203 | chr2-155639144-INS-204 |
| chr2-155646800-INS-601 | chr2-155646800-INS-601 |
| chr2-155650845-INS-65 | chr2-155650841-INS-77 |
| chr2-155681823-INS-216 | chr2-155681823-INS-219 |
| chr2-155682456-INS-173 | chr2-155682457-INS-173 |
| chr2-155700554-INS-62 | chr2-155700554-INS-66 |
| chr2-155712151-INS-67 | chr2-155712151-INS-62 |
| chr2-155715952-INS-56 | chr2-155715952-INS-62 |
| chr2-155720010-INS-204 | chr2-155720010-INS-204 |
| chr2-155722916-INS-55 | chr2-155722916-INS-81 |
| chr2-155733122-INS-73 | chr2-155733122-INS-63 |
| chr2-155743138-INS-56 | chr2-155743111-INS-52 |
| chr2-155743620-INS-99 | chr2-155743620-INS-100 |
| chr2-155757814-INS-78 | chr2-155757814-INS-78 |
| chr2-155758670-INS-120 | chr2-155758670-INS-124 |
| chr2-155764499-INS-174 | chr2-155764499-INS-174 |
| chr2-155765741-INS-189 | chr2-155765741-INS-192 |
| chr2-155773904-INS-317 | chr2-155773904-INS-325 |
| chr2-155808315-INS-222 | chr2-155808315-INS-222 |

| | |
|---|---|
| chr2-155823793-INS-210 | chr2-155823793-INS-210 |
| chr2-155860386-INS-226 | chr2-155860386-INS-226 |
| chr2-155897760-INS-225 | chr2-155897760-INS-228 |
| chr2-155899883-INS-75 | chr2-155899884-INS-72 |
| chr2-155903770-INS-53 | chr2-155903770-INS-53 |
| chr2-155905967-INS-170 | chr2-155905967-INS-170 |
| chr2-155946676-INS-216 | chr2-155946676-INS-217 |
| chr2-155977216-INS-224 | chr2-155977216-INS-224 |
| chr2-155983989-INS-88 | chr2-155983989-INS-88 |
| chr2-155992806-INS-69 | chr2-155992806-INS-69 |
| chr2-156002776-INS-162 | chr2-156002776-INS-162 |
| chr2-156007785-INS-198 | chr2-156007785-INS-197 |
| chr2-156028698-INS-232 | chr2-156028698-INS-239 |
| chr2-156037713-INS-57 | chr2-156037713-INS-57 |
| chr2-156048512-INS-213 | chr2-156048512-INS-213 |
| chr2-156050273-INS-193 | chr2-156050273-INS-192 |
| chr2-156105634-INS-178 | chr2-156105634-INS-177 |
| chr2-156139645-INS-219 | chr2-156139645-INS-219 |
| chr2-156143451-INS-120 | chr2-156143451-INS-108 |
| chr2-156153744-INS-177 | chr2-156153744-INS-179 |
| chr2-156155622-INS-204 | chr2-156155622-INS-204 |
| chr2-156157903-INS-200 | chr2-156157903-INS-201 |
| chr2-156158784-INS-1894 | chr2-156158785-INS-1894 |
| chr2-156189214-INS-282 | chr2-156189214-INS-278 |
| chr2-156195826-INS-173 | chr2-156195826-INS-173 |
| chr2-156237954-INS-223 | chr2-156237954-INS-222 |
| chr2-156239837-INS-8669 | chr2-156239838-INS-8668 |
| chr2-156244298-INS-204 | chr2-156244298-INS-204 |
| chr2-156279398-INS-168 | chr2-156279403-INS-171 |
| chr2-156295174-INS-212 | chr2-156295174-INS-211 |
| chr2-156307125-INS-212 | chr2-156307125-INS-212 |
| chr2-156324030-INS-55 | chr2-156324030-INS-57 |
| chr2-156335393-INS-72 | chr2-156335393-INS-58 |
| chr2-156358829-INS-821 | chr2-156358829-INS-821 |
| chr2-156362287-INS-53 | chr2-156362287-INS-53 |
| chr2-156394261-INS-55 | chr2-156394265-INS-50 |
| chr2-156414639-INS-6441 | chr2-156414640-INS-6441 |
| chr2-156431991-INS-254 | chr2-156431991-INS-254 |
| chr2-156432674-INS-188 | chr2-156432674-INS-188 |
| chr2-156513567-INS-223 | chr2-156513567-INS-223 |
| chr2-156518383-INS-260 | chr2-156518383-INS-270 |
| chr2-156569491-INS-503 | chr2-156569488-INS-508 |
| chr2-156571096-INS-218 | chr2-156571096-INS-219 |
| chr2-156580869-INS-186 | chr2-156580889-INS-214 |
| chr2-156593279-INS-152 | chr2-156593279-INS-154 |
| chr2-156596978-INS-200 | chr2-156596978-INS-200 |
| chr2-156619864-INS-294 | chr2-156619864-INS-294 |
| chr2-156632793-INS-201 | chr2-156632793-INS-201 |
| chr2-156634698-INS-188 | chr2-156634698-INS-188 |
| chr2-156643522-INS-128 | chr2-156643522-INS-128 |
| chr2-156647669-INS-199 | chr2-156647669-INS-200 |
| chr2-156650235-INS-421 | chr2-156650235-INS-421 |
| chr2-156733343-INS-110 | chr2-156733341-INS-91 |
| chr2-156757613-INS-72 | chr2-156757613-INS-72 |
| chr2-156758736-INS-76 | chr2-156758736-INS-76 |
| chr2-156761067-INS-65 | chr2-156761067-INS-73 |
| chr2-156762061-INS-200 | chr2-156762061-INS-198 |
| chr2-156765454-INS-53 | chr2-156765454-INS-53 |
| chr2-156772757-INS-244 | chr2-156772757-INS-243 |
| chr2-156786483-INS-599 | chr2-156786483-INS-599 |
| chr2-156788599-INS-52 | chr2-156788599-INS-51 |
| chr2-156798774-INS-53 | chr2-156798774-INS-57 |
| chr2-156811831-INS-185 | chr2-156811831-INS-200 |
| chr2-156819264-INS-57 | chr2-156819264-INS-75 |
| chr2-156823044-INS-80 | chr2-156823044-INS-80 |
| chr2-156829626-INS-168 | chr2-156829626-INS-169 |
| chr2-156830603-INS-208 | chr2-156830603-INS-212 |
| chr2-156843225-INS-153 | chr2-156843225-INS-153 |
| chr2-156867963-INS-437 | chr2-156867963-INS-437 |
| chr2-156884233-INS-238 | chr2-156884238-INS-234 |
| chr2-156889838-INS-435 | chr2-156889838-INS-432 |
| chr2-156912191-INS-61 | chr2-156912191-INS-50 |
| chr2-156924740-INS-218 | chr2-156924740-INS-219 |
| chr2-156998313-INS-228 | chr2-156998314-INS-228 |
| chr2-157007154-INS-97 | chr2-157007154-INS-102 |
| chr2-157031872-INS-72 | chr2-157031872-INS-50 |
| chr2-157039563-INS-211 | chr2-157039563-INS-208 |

| | |
|---|---|
| chr2-157063782-INS-52 | chr2-157063783-INS-50 |
| chr2-157074525-INS-178 | chr2-157074525-INS-190 |
| chr2-157075472-INS-65 | chr2-157075472-INS-55 |
| chr2-157079347-INS-204 | chr2-157079347-INS-205 |
| chr2-157080950-INS-201 | chr2-157080950-INS-201 |
| chr2-157086031-INS-990 | chr2-157086048-INS-1021 |
| chr2-157088161-INS-213 | chr2-157088161-INS-213 |
| chr2-157099043-INS-207 | chr2-157099043-INS-208 |
| chr2-157102569-INS-495 | chr2-157102569-INS-492 |
| chr2-157116123-INS-663 | chr2-157116123-INS-663 |
| chr2-157117425-INS-407 | chr2-157117425-INS-407 |
| chr2-157118937-INS-175 | chr2-157118937-INS-175 |
| chr2-157119942-INS-438 | chr2-157119942-INS-438 |
| chr2-157123664-INS-146 | chr2-157123664-INS-162 |
| chr2-157125845-INS-131 | chr2-157125845-INS-131 |
| chr2-157151905-INS-54 | chr2-157151905-INS-54 |
| chr2-157175974-INS-123 | chr2-157175974-INS-123 |
| chr2-157181370-INS-52 | chr2-157181370-INS-52 |
| chr2-157193633-INS-68 | chr2-157193598-INS-56 |
| chr2-157198177-INS-283 | chr2-157198177-INS-283 |
| chr2-157206772-INS-190 | chr2-157206772-INS-190 |
| chr2-157208540-INS-5241 | chr2-157208541-INS-5241 |
| chr2-157218388-INS-434 | chr2-157218388-INS-434 |
| chr2-157221329-INS-1141 | chr2-157221325-INS-1141 |
| chr2-157225385-INS-190 | chr2-157225386-INS-190 |
| chr2-157233224-INS-201 | chr2-157233224-INS-201 |
| chr2-157237565-INS-51 | chr2-157237566-INS-65 |
| chr2-157241448-INS-53 | chr2-157241449-INS-53 |
| chr2-157244180-INS-100 | chr2-157244183-INS-64 |
| chr2-157258820-INS-502 | chr2-157258820-INS-502 |
| chr2-157276287-INS-201 | chr2-157276287-INS-201 |
| chr2-157291793-INS-205 | chr2-157291793-INS-207 |
| chr2-157317291-INS-56 | chr2-157317285-INS-62 |
| chr2-157410634-INS-76 | chr2-157410634-INS-76 |
| chr2-157483002-INS-105 | chr2-157483002-INS-63 |
| chr2-157489569-INS-397 | chr2-157489602-INS-403 |
| chr2-157507124-INS-930 | chr2-157507124-INS-930 |
| chr2-157519009-INS-478 | chr2-157519009-INS-494 |
| chr2-157531183-INS-59 | chr2-157531160-INS-52 |
| chr2-157535390-INS-68 | chr2-157535394-INS-56 |
| chr2-157538186-INS-4727 | chr2-157538187-INS-4727 |
| chr2-157546116-INS-84 | chr2-157546116-INS-75 |
| chr2-157552478-INS-208 | chr2-157552478-INS-208 |
| chr2-157559539-INS-220 | chr2-157559539-INS-220 |
| chr2-157562318-INS-77 | chr2-157562288-INS-60 |
| chr2-157578281-INS-54 | chr2-157578281-INS-72 |
| chr2-157591849-INS-70 | chr2-157591849-INS-70 |
| chr2-157614556-INS-613 | chr2-157614556-INS-613 |
| chr2-157638851-INS-257 | chr2-157638851-INS-257 |
| chr2-157655167-INS-523 | chr2-157655167-INS-523 |
| chr2-157665250-INS-175 | chr2-157665250-INS-179 |
| chr2-157709184-INS-152 | chr2-157709184-INS-152 |
| chr2-157742003-INS-220 | chr2-157742003-INS-228 |
| chr2-157742828-INS-7145 | chr2-157742829-INS-7145 |
| chr2-157746384-INS-204 | chr2-157746384-INS-204 |
| chr2-157773387-INS-214 | chr2-157773381-INS-205 |
| chr2-157775664-INS-56 | chr2-157775657-INS-80 |
| chr2-157796934-INS-210 | chr2-157796934-INS-210 |
| chr2-157798731-INS-221 | chr2-157798731-INS-221 |
| chr2-157802085-INS-204 | chr2-157802085-INS-203 |
| chr2-157806328-INS-221 | chr2-157806328-INS-222 |
| chr2-157808855-INS-88 | chr2-157808856-INS-88 |
| chr2-157841246-INS-75 | chr2-157841246-INS-75 |
| chr2-157853672-INS-164 | chr2-157853672-INS-164 |
| chr2-157858791-INS-1886 | chr2-157858792-INS-1886 |
| chr2-157860314-INS-207 | chr2-157860314-INS-207 |
| chr2-157876162-INS-206 | chr2-157876162-INS-206 |
| chr2-157892648-INS-101 | chr2-157892648-INS-150 |
| chr2-157898276-INS-203 | chr2-157898276-INS-203 |
| chr2-157909816-INS-50 | chr2-157909816-INS-63 |
| chr2-157931250-INS-633 | chr2-157931250-INS-636 |
| chr2-157942992-INS-81 | chr2-157942992-INS-58 |
| chr2-157956479-INS-62 | chr2-157956479-INS-74 |
| chr2-157966991-INS-178 | chr2-157966991-INS-179 |
| chr2-157998781-INS-212 | chr2-157998781-INS-212 |
| chr2-158048880-INS-130 | chr2-158048881-INS-112 |
| chr2-158054622-INS-56 | chr2-158054622-INS-54 |

| | |
|---|---|
| chr2-158066932-INS-127 | chr2-158066932-INS-142 |
| chr2-158089589-INS-81 | chr2-158089582-INS-108 |
| chr2-158098047-INS-85 | chr2-158098047-INS-85 |
| chr2-158114362-INS-79 | chr2-158114387-INS-94 |
| chr2-158143244-INS-55 | chr2-158143244-INS-61 |
| chr2-158153492-INS-202 | chr2-158153492-INS-202 |
| chr2-158154703-INS-153 | chr2-158154706-INS-154 |
| chr2-158157364-INS-213 | chr2-158157364-INS-216 |
| chr2-158158983-INS-231 | chr2-158158983-INS-230 |
| chr2-158175461-INS-60 | chr2-158175461-INS-54 |
| chr2-158179487-INS-57 | chr2-158179487-INS-57 |
| chr2-158182975-INS-206 | chr2-158182975-INS-206 |
| chr2-158187076-INS-221 | chr2-158187076-INS-221 |
| chr2-158204319-INS-56 | chr2-158204323-INS-60 |
| chr2-158205664-INS-62 | chr2-158205664-INS-62 |
| chr2-158214079-INS-95 | chr2-158214079-INS-95 |
| chr2-158234962-INS-54 | chr2-158234958-INS-74 |
| chr2-158236577-INS-169 | chr2-158236577-INS-135 |
| chr2-158243351-INS-159 | chr2-158243350-INS-190 |
| chr2-158271046-INS-211 | chr2-158271047-INS-192 |
| chr2-158332575-INS-66 | chr2-158332579-INS-57 |
| chr2-158336877-INS-78 | chr2-158336875-INS-56 |
| chr2-158350253-INS-330 | chr2-158350253-INS-316 |
| chr2-158353085-INS-310 | chr2-158353084-INS-306 |
| chr2-158367751-INS-54 | chr2-158367751-INS-56 |
| chr2-158377004-INS-72 | chr2-158377004-INS-74 |
| chr2-158384809-INS-118 | chr2-158384795-INS-83 |
| chr2-158388120-INS-364 | chr2-158388129-INS-369 |
| chr2-158436060-INS-51 | chr2-158436060-INS-51 |
| chr2-158436063-INS-636 | chr2-158436030-INS-671 |
| chr2-158462457-INS-481 | chr2-158462457-INS-481 |
| chr2-158463541-INS-58 | chr2-158463541-INS-58 |
| chr2-158482665-INS-86 | chr2-158482665-INS-86 |
| chr2-158484375-INS-210 | chr2-158484375-INS-208 |
| chr2-158485276-INS-222 | chr2-158485276-INS-222 |
| chr2-158495312-INS-64 | chr2-158495295-INS-55 |
| chr2-158536165-INS-270 | chr2-158536174-INS-275 |
| chr2-158555295-INS-124 | chr2-158555295-INS-124 |
| chr2-158574046-INS-727 | chr2-158574046-INS-727 |
| chr2-158608449-INS-7193 | chr2-158608450-INS-7193 |
| chr2-158615930-INS-208 | chr2-158615930-INS-208 |
| chr2-158622082-INS-82 | chr2-158622082-INS-82 |
| chr2-158634543-INS-5399 | chr2-158634544-INS-5399 |
| chr2-158635015-INS-366 | chr2-158635015-INS-366 |
| chr2-158638841-INS-149 | chr2-158638841-INS-149 |
| chr2-158643730-INS-214 | chr2-158643730-INS-214 |
| chr2-158649629-INS-74 | chr2-158649629-INS-74 |
| chr2-158663360-INS-153 | chr2-158663360-INS-153 |
| chr2-158695899-INS-174 | chr2-158695850-INS-182 |
| chr2-158702920-INS-1887 | chr2-158702921-INS-1887 |
| chr2-158712584-INS-73 | chr2-158712584-INS-73 |
| chr2-158731139-INS-123 | chr2-158731139-INS-74 |
| chr2-158742570-INS-65 | chr2-158742570-INS-65 |
| chr2-158765576-INS-805 | chr2-158765576-INS-801 |
| chr2-158775151-INS-104 | chr2-158775151-INS-104 |
| chr2-158776421-INS-5587 | chr2-158776422-INS-5588 |
| chr2-158781704-INS-50 | chr2-158781700-INS-82 |
| chr2-158788826-INS-72 | chr2-158788835-INS-74 |
| chr2-158815974-INS-194 | chr2-158815983-INS-158 |
| chr2-158820236-INS-70 | chr2-158820236-INS-64 |
| chr2-158835792-INS-224 | chr2-158835792-INS-232 |
| chr2-158872669-INS-77 | chr2-158872669-INS-77 |
| chr2-158884484-INS-134 | chr2-158884476-INS-116 |
| chr2-158886112-INS-400 | chr2-158886112-INS-400 |
| chr2-158918689-INS-6150 | chr2-158918678-INS-6149 |
| chr2-158920853-INS-65 | chr2-158920856-INS-54 |
| chr2-158930020-INS-116 | chr2-158930005-INS-124 |
| chr2-158939327-INS-491 | chr2-158939327-INS-491 |
| chr2-158944610-INS-152 | chr2-158944610-INS-156 |
| chr2-158954226-INS-862 | chr2-158954226-INS-861 |
| chr2-158966797-INS-68 | chr2-158966749-INS-84 |
| chr2-158996251-INS-122 | chr2-158996249-INS-146 |
| chr2-159019153-INS-499 | chr2-159019153-INS-499 |
| chr2-159020750-INS-605 | chr2-159020750-INS-605 |
| chr2-159027774-INS-96 | chr2-159027774-INS-80 |
| chr2-159039782-INS-68 | chr2-159039774-INS-62 |
| chr2-159048090-INS-2496 | chr2-159048091-INS-2469 |

| | |
|---|---|
| chr2-159077608-INS-133 | chr2-159077608-INS-158 |
| chr2-159081119-INS-4793 | chr2-159081120-INS-4793 |
| chr2-159119913-INS-52 | chr2-159119913-INS-52 |
| chr2-159132919-INS-100 | chr2-159132919-INS-100 |
| chr2-159157539-INS-97 | chr2-159157539-INS-92 |
| chr2-159158693-INS-65 | chr2-159158693-INS-54 |
| chr2-159199418-INS-537 | chr2-159199418-INS-537 |
| chr2-159221423-INS-142 | chr2-159221423-INS-144 |
| chr2-159223719-INS-106 | chr2-159223719-INS-106 |
| chr2-159229201-INS-202 | chr2-159229201-INS-202 |
| chr2-159262445-INS-64 | chr2-159262429-INS-68 |
| chr2-159267344-INS-84 | chr2-159267344-INS-92 |
| chr2-159282051-INS-124 | chr2-159282053-INS-112 |
| chr2-159282691-INS-51 | chr2-159282691-INS-54 |
| chr2-159308819-INS-667 | chr2-159308819-INS-667 |
| chr2-159315745-INS-176 | chr2-159315745-INS-176 |
| chr2-159321622-INS-557 | chr2-159321623-INS-557 |
| chr2-159330170-INS-56 | chr2-159330170-INS-56 |
| chr2-159345995-INS-72 | chr2-159345993-INS-62 |
| chr2-159382811-INS-816 | chr2-159382811-INS-816 |
| chr2-159387490-INS-63 | chr2-159387490-INS-60 |
| chr2-159391149-INS-90 | chr2-159391149-INS-86 |
| chr2-159393204-INS-567 | chr2-159393204-INS-567 |
| chr2-159394825-INS-6498 | chr2-159394826-INS-6498 |
| chr2-159411622-INS-432 | chr2-159411622-INS-451 |
| chr2-159453856-INS-400 | chr2-159453856-INS-400 |
| chr2-159454381-INS-60 | chr2-159454381-INS-60 |
| chr2-159461677-INS-1450 | chr2-159461678-INS-1451 |
| chr2-159510725-INS-248 | chr2-159510725-INS-248 |
| chr2-159536614-INS-52 | chr2-159536614-INS-52 |
| chr2-159552789-INS-2161 | chr2-159552790-INS-2152 |
| chr2-159558502-INS-1011 | chr2-159558503-INS-1009 |
| chr2-159563270-INS-1664 | chr2-159563271-INS-1671 |
| chr2-159580497-INS-223 | chr2-159580496-INS-183 |
| chr2-159615680-INS-346 | chr2-159615680-INS-346 |
| chr2-159627537-INS-70 | chr2-159627570-INS-70 |
| chr2-159630606-INS-208 | chr2-159630606-INS-208 |
| chr2-159638684-INS-80 | chr2-159638684-INS-80 |
| chr2-159645943-INS-434 | chr2-159645943-INS-431 |
| chr2-159648999-INS-120 | chr2-159648999-INS-76 |
| chr2-159655464-INS-70 | chr2-159655464-INS-82 |
| chr2-159667546-INS-58 | chr2-159667546-INS-86 |
| chr2-159685805-INS-178 | chr2-159685805-INS-180 |
| chr2-159721171-INS-58 | chr2-159721172-INS-62 |
| chr2-159750034-INS-52 | chr2-159750034-INS-62 |
| chr2-159755069-INS-151 | chr2-159755070-INS-151 |
| chr2-159763647-INS-55 | chr2-159763647-INS-66 |
| chr2-159797186-INS-172 | chr2-159797186-INS-172 |
| chr2-159810569-INS-107 | chr2-159810569-INS-107 |
| chr2-159864834-INS-126 | chr2-159864831-INS-129 |
| chr2-159881513-INS-152 | chr2-159881513-INS-141 |
| chr2-159975817-INS-367 | chr2-159975790-INS-369 |
| chr2-159998854-INS-74 | chr2-159998854-INS-76 |
| chr2-160017150-INS-71 | chr2-160017150-INS-64 |
| chr2-160061076-INS-132 | chr2-160061072-INS-90 |
| chr2-160061808-INS-72 | chr2-160061808-INS-74 |
| chr2-160063722-INS-52 | chr2-160063722-INS-52 |
| chr2-160160046-INS-930 | chr2-160160046-INS-930 |
| chr2-160192690-INS-520 | chr2-160192690-INS-526 |
| chr2-160222088-INS-186 | chr2-160222088-INS-179 |
| chr2-160237067-INS-58 | chr2-160237067-INS-60 |
| chr2-160253184-INS-180 | chr2-160253184-INS-175 |
| chr2-160274925-INS-498 | chr2-160274925-INS-498 |
| chr2-160275841-INS-120 | chr2-160275841-INS-124 |
| chr2-160286665-INS-110 | chr2-160286665-INS-122 |
| chr2-160288226-INS-56 | chr2-160288226-INS-54 |
| chr2-160291652-INS-58 | chr2-160291657-INS-54 |
| chr2-160329395-INS-76 | chr2-160329395-INS-76 |
| chr2-160342774-INS-125 | chr2-160342774-INS-125 |
| chr2-160376424-INS-54 | chr2-160376422-INS-70 |
| chr2-160407921-INS-211 | chr2-160407921-INS-211 |
| chr2-160411211-INS-160 | chr2-160411211-INS-160 |
| chr2-160414140-INS-2487 | chr2-160414141-INS-2487 |
| chr2-160431952-INS-102 | chr2-160431952-INS-71 |
| chr2-160437021-INS-68 | chr2-160437024-INS-86 |
| chr2-160439328-INS-84 | chr2-160439320-INS-84 |
| chr2-160441344-INS-417 | chr2-160441344-INS-417 |

| | |
|---|---|
| chr2-160449343-INS-53 | chr2-160449344-INS-56 |
| chr2-160455249-INS-133 | chr2-160455249-INS-133 |
| chr2-160478681-INS-6442 | chr2-160478682-INS-6442 |
| chr2-160484625-INS-116 | chr2-160484597-INS-130 |
| chr2-160492891-INS-217 | chr2-160492891-INS-215 |
| chr2-160498736-INS-225 | chr2-160498736-INS-225 |
| chr2-160502870-INS-175 | chr2-160502870-INS-176 |
| chr2-160512505-INS-297 | chr2-160512505-INS-291 |
| chr2-160545907-INS-1103 | chr2-160545908-INS-1101 |
| chr2-160555343-INS-230 | chr2-160555343-INS-230 |
| chr2-160557372-INS-218 | chr2-160557372-INS-218 |
| chr2-160585192-INS-919 | chr2-160585197-INS-926 |
| chr2-160646522-INS-579 | chr2-160646528-INS-581 |
| chr2-160696275-INS-4665 | chr2-160696276-INS-4665 |
| chr2-160709455-INS-165 | chr2-160709455-INS-165 |
| chr2-160768010-INS-260 | chr2-160768010-INS-228 |
| chr2-160780218-INS-124 | chr2-160780190-INS-129 |
| chr2-160802557-INS-111 | chr2-160802557-INS-111 |
| chr2-160827962-INS-69 | chr2-160827918-INS-70 |
| chr2-160880413-INS-7360 | chr2-160880414-INS-7361 |
| chr2-160901469-INS-161 | chr2-160901469-INS-161 |
| chr2-160925298-INS-513 | chr2-160925298-INS-513 |
| chr2-160961463-INS-4113 | chr2-160961464-INS-4113 |
| chr2-160976454-INS-108 | chr2-160976434-INS-96 |
| chr2-161016314-INS-132 | chr2-161016314-INS-132 |
| chr2-161017300-INS-5485 | chr2-161017301-INS-5485 |
| chr2-161026667-INS-111 | chr2-161026667-INS-111 |
| chr2-161030641-INS-55 | chr2-161030641-INS-55 |
| chr2-161032393-INS-232 | chr2-161032393-INS-260 |
| chr2-161049285-INS-58 | chr2-161049285-INS-60 |
| chr2-161050349-INS-2073 | chr2-161050350-INS-2072 |
| chr2-161051846-INS-82 | chr2-161051834-INS-108 |
| chr2-161062120-INS-5098 | chr2-161062121-INS-5098 |
| chr2-161063071-INS-194 | chr2-161063071-INS-195 |
| chr2-161103802-INS-55 | chr2-161103802-INS-55 |
| chr2-161146676-INS-373 | chr2-161146676-INS-373 |
| chr2-161146676-INS-912 | chr2-161146676-INS-902 |
| chr2-161151573-INS-185 | chr2-161151573-INS-185 |
| chr2-161153217-INS-532 | chr2-161153217-INS-532 |
| chr2-161173637-INS-1439 | chr2-161173638-INS-1439 |
| chr2-161226174-INS-51 | chr2-161226174-INS-51 |
| chr2-161237613-INS-59 | chr2-161237613-INS-59 |
| chr2-161243030-INS-1248 | chr2-161243031-INS-1248 |
| chr2-161243377-INS-137 | chr2-161243377-INS-137 |
| chr2-161292140-INS-54 | chr2-161292123-INS-58 |
| chr2-161299537-INS-104 | chr2-161299537-INS-104 |
| chr2-161320078-INS-273 | chr2-161320077-INS-245 |
| chr2-161338040-INS-4171 | chr2-161338041-INS-4171 |
| chr2-161344796-INS-505 | chr2-161344796-INS-504 |
| chr2-161457895-INS-52 | chr2-161457895-INS-52 |
| chr2-161485072-INS-110 | chr2-161485072-INS-133 |
| chr2-161503677-INS-194 | chr2-161503677-INS-194 |
| chr2-161505605-INS-96 | chr2-161505605-INS-96 |
| chr2-161507584-INS-87 | chr2-161507615-INS-99 |
| chr2-161523350-INS-72 | chr2-161523350-INS-72 |
| chr2-161538571-INS-204 | chr2-161538568-INS-178 |
| chr2-161559423-INS-80 | chr2-161559423-INS-80 |
| chr2-161587488-INS-106 | chr2-161587488-INS-98 |
| chr2-161629776-INS-82 | chr2-161629776-INS-82 |
| chr2-161653602-INS-83 | chr2-161653600-INS-82 |
| chr2-161657272-INS-115 | chr2-161657272-INS-115 |
| chr2-161675463-INS-51 | chr2-161675463-INS-50 |
| chr2-161678490-INS-447 | chr2-161678490-INS-446 |
| chr2-161689822-INS-158 | chr2-161689822-INS-158 |
| chr2-161695179-INS-142 | chr2-161695179-INS-142 |
| chr2-161700369-INS-65 | chr2-161700369-INS-65 |
| chr2-161712288-INS-2201 | chr2-161712289-INS-2201 |
| chr2-161716342-INS-72 | chr2-161716336-INS-56 |
| chr2-161729057-INS-6395 | chr2-161729058-INS-6395 |
| chr2-161751196-INS-153 | chr2-161751196-INS-154 |
| chr2-161793574-INS-126 | chr2-161793574-INS-77 |
| chr2-161801608-INS-223 | chr2-161801608-INS-223 |
| chr2-161810825-INS-3553 | chr2-161810826-INS-3553 |
| chr2-161816400-INS-6238 | chr2-161816401-INS-6238 |
| chr2-161837836-INS-116 | chr2-161837836-INS-116 |
| chr2-161888087-INS-951 | chr2-161888087-INS-952 |
| chr2-161891323-INS-432 | chr2-161891323-INS-432 |

| | |
|---|---|
| chr2-161911248-INS-70 | chr2-161911248-INS-70 |
| chr2-161961772-INS-114 | chr2-161961772-INS-108 |
| chr2-161983013-INS-537 | chr2-161983013-INS-537 |
| chr2-162056821-INS-52 | chr2-162056821-INS-54 |
| chr2-162096004-INS-972 | chr2-162096004-INS-972 |
| chr2-162111227-INS-5569 | chr2-162111228-INS-5569 |
| chr2-162111798-INS-766 | chr2-162111798-INS-766 |
| chr2-162112951-INS-56 | chr2-162112951-INS-57 |
| chr2-162133784-INS-1295 | chr2-162133771-INS-1283 |
| chr2-162145706-INS-89 | chr2-162145706-INS-89 |
| chr2-162157733-INS-493 | chr2-162157733-INS-493 |
| chr2-162200117-INS-119 | chr2-162200117-INS-119 |
| chr2-162227860-INS-213 | chr2-162227860-INS-212 |
| chr2-162237513-INS-224 | chr2-162237513-INS-224 |
| chr2-162246377-INS-86 | chr2-162246377-INS-56 |
| chr2-162264225-INS-105 | chr2-162264225-INS-105 |
| chr2-162269243-INS-56 | chr2-162269243-INS-56 |
| chr2-162276447-INS-245 | chr2-162276447-INS-245 |
| chr2-162302085-INS-110 | chr2-162302088-INS-123 |
| chr2-162308605-INS-61 | chr2-162308605-INS-61 |
| chr2-162378433-INS-132 | chr2-162378433-INS-126 |
| chr2-162420584-INS-6360 | chr2-162420585-INS-6370 |
| chr2-162437663-INS-201 | chr2-162437663-INS-201 |
| chr2-162474173-INS-52 | chr2-162474173-INS-60 |
| chr2-162481604-INS-56 | chr2-162481604-INS-56 |
| chr2-162485130-INS-85 | chr2-162485152-INS-90 |
| chr2-162488750-INS-57 | chr2-162488747-INS-60 |
| chr2-162492322-INS-101 | chr2-162492310-INS-101 |
| chr2-162513814-INS-102 | chr2-162513814-INS-102 |
| chr2-162518202-INS-315 | chr2-162518202-INS-315 |
| chr2-162523122-INS-96 | chr2-162523122-INS-96 |
| chr2-162541981-INS-389 | chr2-162541981-INS-388 |
| chr2-162565884-INS-6424 | chr2-162565885-INS-6424 |
| chr2-162590185-INS-63 | chr2-162590153-INS-54 |
| chr2-162607120-INS-166 | chr2-162607120-INS-166 |
| chr2-162608161-INS-493 | chr2-162608161-INS-493 |
| chr2-162622157-INS-90 | chr2-162622158-INS-90 |
| chr2-162649349-INS-54 | chr2-162649349-INS-54 |
| chr2-162650525-INS-117 | chr2-162650521-INS-73 |
| chr2-162660568-INS-116 | chr2-162660568-INS-116 |
| chr2-162680761-INS-1444 | chr2-162680762-INS-1444 |
| chr2-162684378-INS-203 | chr2-162684378-INS-203 |
| chr2-162686742-INS-167 | chr2-162686742-INS-165 |
| chr2-162707073-INS-217 | chr2-162707056-INS-220 |
| chr2-162712198-INS-52 | chr2-162712198-INS-54 |
| chr2-162721112-INS-208 | chr2-162721112-INS-208 |
| chr2-162725183-INS-240 | chr2-162725184-INS-195 |
| chr2-162737537-INS-104 | chr2-162737537-INS-78 |
| chr2-162745698-INS-140 | chr2-162745698-INS-140 |
| chr2-162754730-INS-54 | chr2-162754730-INS-54 |
| chr2-162765467-INS-79 | chr2-162765467-INS-79 |
| chr2-162782113-INS-810 | chr2-162782113-INS-810 |
| chr2-162797397-INS-289 | chr2-162797397-INS-289 |
| chr2-162812680-INS-111 | chr2-162812680-INS-96 |
| chr2-162822211-INS-56 | chr2-162822217-INS-60 |
| chr2-162825292-INS-54 | chr2-162825309-INS-56 |
| chr2-162830594-INS-62 | chr2-162830593-INS-109 |
| chr2-162835120-INS-206 | chr2-162835120-INS-206 |
| chr2-162854080-INS-68 | chr2-162854080-INS-68 |
| chr2-162857019-INS-55 | chr2-162857019-INS-55 |
| chr2-162874545-INS-62 | chr2-162874545-INS-62 |
| chr2-162878103-INS-173 | chr2-162878103-INS-173 |
| chr2-162898403-INS-403 | chr2-162898403-INS-403 |
| chr2-162903996-INS-232 | chr2-162903996-INS-232 |
| chr2-162908196-INS-217 | chr2-162908205-INS-215 |
| chr2-162909719-INS-860 | chr2-162909727-INS-853 |
| chr2-162926797-INS-113 | chr2-162926797-INS-137 |
| chr2-162934330-INS-892 | chr2-162934330-INS-893 |
| chr2-162940902-INS-98 | chr2-162940902-INS-97 |
| chr2-162963720-INS-58 | chr2-162963720-INS-58 |
| chr2-162988386-INS-56 | chr2-162988386-INS-56 |
| chr2-163006985-INS-198 | chr2-163006985-INS-199 |
| chr2-163022528-INS-206 | chr2-163022528-INS-206 |
| chr2-163028326-INS-98 | chr2-163028327-INS-94 |
| chr2-163040489-INS-90 | chr2-163040487-INS-93 |
| chr2-163068557-INS-153 | chr2-163068557-INS-153 |
| chr2-163100169-INS-66 | chr2-163100169-INS-50 |

| | |
|---|---|
| chr2-163104870-INS-134 | chr2-163104870-INS-134 |
| chr2-163134740-INS-171 | chr2-163134722-INS-207 |
| chr2-163134749-INS-705 | chr2-163134770-INS-681 |
| chr2-163141637-INS-552 | chr2-163141637-INS-552 |
| chr2-163169083-INS-2213 | chr2-163169084-INS-2215 |
| chr2-163181031-INS-81 | chr2-163181031-INS-81 |
| chr2-163197006-INS-165 | chr2-163197006-INS-165 |
| chr2-163236475-INS-52 | chr2-163236497-INS-64 |
| chr2-163271994-INS-55 | chr2-163271993-INS-91 |
| chr2-163274184-INS-60 | chr2-163274184-INS-60 |
| chr2-163275400-INS-190 | chr2-163275428-INS-195 |
| chr2-163305670-INS-73 | chr2-163305670-INS-73 |
| chr2-163317232-INS-180 | chr2-163317232-INS-167 |
| chr2-163319843-INS-765 | chr2-163319843-INS-765 |
| chr2-163321127-INS-62 | chr2-163321134-INS-60 |
| chr2-163326127-INS-208 | chr2-163326133-INS-224 |
| chr2-163328762-INS-643 | chr2-163328762-INS-643 |
| chr2-163339451-INS-78 | chr2-163339451-INS-60 |
| chr2-163341024-INS-132 | chr2-163341024-INS-120 |
| chr2-163348604-INS-5671 | chr2-163348605-INS-5671 |
| chr2-163357385-INS-66 | chr2-163357385-INS-64 |
| chr2-163419592-INS-52 | chr2-163419592-INS-92 |
| chr2-163421350-INS-97 | chr2-163421350-INS-97 |
| chr2-163432061-INS-52 | chr2-163432061-INS-58 |
| chr2-163436995-INS-496 | chr2-163436995-INS-512 |
| chr2-163438941-INS-62 | chr2-163438939-INS-50 |
| chr2-163448946-INS-79 | chr2-163448934-INS-67 |
| chr2-163452192-INS-64 | chr2-163452192-INS-76 |
| chr2-163474195-INS-165 | chr2-163474195-INS-175 |
| chr2-163480760-INS-289 | chr2-163480761-INS-289 |
| chr2-163487393-INS-201 | chr2-163487393-INS-201 |
| chr2-163490861-INS-7099 | chr2-163490862-INS-7099 |
| chr2-163499200-INS-286 | chr2-163499200-INS-286 |
| chr2-163520609-INS-373 | chr2-163520610-INS-344 |
| chr2-163524519-INS-99 | chr2-163524524-INS-50 |
| chr2-163525769-INS-213 | chr2-163525769-INS-187 |
| chr2-163543786-INS-98 | chr2-163543786-INS-98 |
| chr2-163555215-INS-205 | chr2-163555215-INS-205 |
| chr2-163563884-INS-157 | chr2-163563884-INS-157 |
| chr2-163574367-INS-111 | chr2-163574367-INS-111 |
| chr2-163611497-INS-61 | chr2-163611497-INS-61 |
| chr2-163627006-INS-5116 | chr2-163627007-INS-5116 |
| chr2-163644225-INS-470 | chr2-163644225-INS-470 |
| chr2-163667776-INS-144 | chr2-163667776-INS-167 |
| chr2-163721666-INS-115 | chr2-163721666-INS-116 |
| chr2-163724087-INS-80 | chr2-163724087-INS-80 |
| chr2-163746349-INS-162 | chr2-163746349-INS-160 |
| chr2-163769813-INS-66 | chr2-163769813-INS-84 |
| chr2-163784142-INS-77 | chr2-163784130-INS-58 |
| chr2-163809790-INS-56 | chr2-163809789-INS-60 |
| chr2-163814032-INS-64 | chr2-163814032-INS-64 |
| chr2-163816567-INS-84 | chr2-163816587-INS-83 |
| chr2-163827684-INS-221 | chr2-163827685-INS-213 |
| chr2-163871497-INS-898 | chr2-163871497-INS-898 |
| chr2-163875749-INS-99 | chr2-163875749-INS-101 |
| chr2-163878110-INS-93 | chr2-163878110-INS-91 |
| chr2-163883391-INS-60 | chr2-163883429-INS-66 |
| chr2-163887991-INS-58 | chr2-163887973-INS-64 |
| chr2-163903201-INS-140 | chr2-163903201-INS-136 |
| chr2-163918777-INS-115 | chr2-163918777-INS-115 |
| chr2-163921866-INS-741 | chr2-163921866-INS-741 |
| chr2-163932218-INS-200 | chr2-163932218-INS-196 |
| chr2-163945573-INS-206 | chr2-163945573-INS-205 |
| chr2-163962206-INS-72 | chr2-163962190-INS-58 |
| chr2-163982065-INS-174 | chr2-163982065-INS-174 |
| chr2-163982781-INS-78 | chr2-163982803-INS-60 |
| chr2-163984651-INS-74 | chr2-163984651-INS-74 |
| chr2-164013798-INS-1290 | chr2-164013799-INS-1290 |
| chr2-164030564-INS-86 | chr2-164030564-INS-86 |
| chr2-164032497-INS-207 | chr2-164032535-INS-181 |
| chr2-164037665-INS-203 | chr2-164037665-INS-202 |
| chr2-164043626-INS-193 | chr2-164043626-INS-192 |
| chr2-164051697-INS-245 | chr2-164051697-INS-245 |
| chr2-164066872-INS-214 | chr2-164066872-INS-191 |
| chr2-164070894-INS-58 | chr2-164070894-INS-58 |
| chr2-164082362-INS-61 | chr2-164082362-INS-61 |
| chr2-164084272-INS-479 | chr2-164084275-INS-480 |

| | |
|---|---|
| chr2-164094157-INS-74 | chr2-164094157-INS-78 |
| chr2-164109619-INS-1277 | chr2-164109620-INS-1277 |
| chr2-164133983-INS-55 | chr2-164133983-INS-55 |
| chr2-164140175-INS-1570 | chr2-164140178-INS-1570 |
| chr2-164164883-INS-80 | chr2-164164883-INS-86 |
| chr2-164166159-INS-201 | chr2-164166159-INS-179 |
| chr2-164177850-INS-66 | chr2-164177850-INS-66 |
| chr2-164191279-INS-179 | chr2-164191279-INS-179 |
| chr2-164203481-INS-5424 | chr2-164203482-INS-5424 |
| chr2-164220524-INS-141 | chr2-164220524-INS-93 |
| chr2-164255513-INS-54 | chr2-164255511-INS-54 |
| chr2-164259338-INS-190 | chr2-164259350-INS-191 |
| chr2-164288834-INS-202 | chr2-164288834-INS-202 |
| chr2-164299645-INS-64 | chr2-164299645-INS-64 |
| chr2-164336944-INS-191 | chr2-164336939-INS-193 |
| chr2-164344131-INS-80 | chr2-164344131-INS-79 |
| chr2-164346787-INS-65 | chr2-164346787-INS-65 |
| chr2-164359210-INS-78 | chr2-164359211-INS-91 |
| chr2-164359662-INS-201 | chr2-164359662-INS-201 |
| chr2-164362461-INS-163 | chr2-164362461-INS-167 |
| chr2-164379990-INS-306 | chr2-164379990-INS-341 |
| chr2-164398562-INS-76 | chr2-164398562-INS-102 |
| chr2-164409228-INS-194 | chr2-164409228-INS-194 |
| chr2-164424559-INS-348 | chr2-164424559-INS-348 |
| chr2-164475619-INS-215 | chr2-164475619-INS-210 |
| chr2-164490825-INS-108 | chr2-164490825-INS-108 |
| chr2-164502452-INS-2676 | chr2-164502453-INS-2676 |
| chr2-164514694-INS-54 | chr2-164514694-INS-70 |
| chr2-164528176-INS-369 | chr2-164528176-INS-369 |
| chr2-164529201-INS-199 | chr2-164529202-INS-183 |
| chr2-164529785-INS-216 | chr2-164529784-INS-217 |
| chr2-164531160-INS-378 | chr2-164531160-INS-374 |
| chr2-164533960-INS-52 | chr2-164533959-INS-50 |
| chr2-164556073-INS-648 | chr2-164556073-INS-648 |
| chr2-164591462-INS-216 | chr2-164591462-INS-216 |
| chr2-164671412-INS-120 | chr2-164671414-INS-98 |
| chr2-164672439-INS-184 | chr2-164672439-INS-185 |
| chr2-164686171-INS-101 | chr2-164686171-INS-101 |
| chr2-164689672-INS-110 | chr2-164689629-INS-132 |
| chr2-164705170-INS-216 | chr2-164705170-INS-219 |
| chr2-164712699-INS-209 | chr2-164712699-INS-210 |
| chr2-164723536-INS-201 | chr2-164723536-INS-201 |
| chr2-164745377-INS-194 | chr2-164745377-INS-194 |
| chr2-164745900-INS-67 | chr2-164745900-INS-67 |
| chr2-164759264-INS-185 | chr2-164759264-INS-194 |
| chr2-164778527-INS-83 | chr2-164778527-INS-83 |
| chr2-164797073-INS-5543 | chr2-164797074-INS-5543 |
| chr2-164852800-INS-266 | chr2-164852800-INS-266 |
| chr2-164906963-INS-82 | chr2-164906963-INS-98 |
| chr2-164917690-INS-92 | chr2-164917644-INS-137 |
| chr2-164941525-INS-54 | chr2-164941525-INS-54 |
| chr2-164951869-INS-84 | chr2-164951827-INS-105 |
| chr2-164959753-INS-111 | chr2-164959753-INS-111 |
| chr2-164988107-INS-91 | chr2-164988107-INS-91 |
| chr2-165015942-INS-56 | chr2-165015937-INS-59 |
| chr2-165062906-INS-95 | chr2-165062906-INS-95 |
| chr2-165064994-INS-218 | chr2-165064994-INS-218 |
| chr2-165088724-INS-477 | chr2-165088724-INS-477 |
| chr2-165092628-INS-54 | chr2-165092628-INS-54 |
| chr2-165093290-INS-197 | chr2-165093290-INS-197 |
| chr2-165114426-INS-401 | chr2-165114426-INS-401 |
| chr2-165129260-INS-187 | chr2-165129260-INS-192 |
| chr2-165132875-INS-114 | chr2-165132876-INS-112 |
| chr2-165133922-INS-192 | chr2-165133922-INS-192 |
| chr2-165162268-INS-72 | chr2-165162240-INS-70 |
| chr2-165178903-INS-104 | chr2-165178909-INS-58 |
| chr2-165200203-INS-200 | chr2-165200173-INS-200 |
| chr2-165200770-INS-285 | chr2-165200770-INS-280 |
| chr2-165204218-INS-72 | chr2-165204218-INS-72 |
| chr2-165218530-INS-54 | chr2-165218530-INS-56 |
| chr2-165226497-INS-120 | chr2-165226497-INS-125 |
| chr2-165265549-INS-133 | chr2-165265549-INS-133 |
| chr2-165277152-INS-156 | chr2-165277146-INS-174 |
| chr2-165281521-INS-57 | chr2-165281521-INS-57 |
| chr2-165291339-INS-86 | chr2-165291339-INS-86 |
| chr2-165300101-INS-100 | chr2-165300101-INS-100 |
| chr2-165310863-INS-78 | chr2-165310863-INS-80 |

| | |
|---|---|
| chr2-165316634-INS-52 | chr2-165316634-INS-52 |
| chr2-165335478-INS-108 | chr2-165335479-INS-111 |
| chr2-165353335-INS-88 | chr2-165353340-INS-66 |
| chr2-165383856-INS-379 | chr2-165383856-INS-383 |
| chr2-165419727-INS-177 | chr2-165419719-INS-220 |
| chr2-165423939-INS-205 | chr2-165423939-INS-205 |
| chr2-165429387-INS-69 | chr2-165429387-INS-69 |
| chr2-165439896-INS-88 | chr2-165439898-INS-50 |
| chr2-165450145-INS-281 | chr2-165450146-INS-282 |
| chr2-165452746-INS-72 | chr2-165452746-INS-72 |
| chr2-165458959-INS-60 | chr2-165458959-INS-60 |
| chr2-165469479-INS-175 | chr2-165469479-INS-175 |
| chr2-165502280-INS-88 | chr2-165502277-INS-88 |
| chr2-165522209-INS-63 | chr2-165522209-INS-63 |
| chr2-165525733-INS-184 | chr2-165525733-INS-183 |
| chr2-165533873-INS-54 | chr2-165533873-INS-50 |
| chr2-165552900-INS-196 | chr2-165552900-INS-196 |
| chr2-165559043-INS-177 | chr2-165559043-INS-177 |
| chr2-165562073-INS-60 | chr2-165562073-INS-60 |
| chr2-165585169-INS-187 | chr2-165585169-INS-188 |
| chr2-165589087-INS-101 | chr2-165589087-INS-117 |
| chr2-165603703-INS-56 | chr2-165603666-INS-50 |
| chr2-165607928-INS-60 | chr2-165607928-INS-50 |
| chr2-165609843-INS-175 | chr2-165609843-INS-178 |
| chr2-165610762-INS-218 | chr2-165610762-INS-217 |
| chr2-165612719-INS-148 | chr2-165612719-INS-152 |
| chr2-165617068-INS-184 | chr2-165617068-INS-184 |
| chr2-165621851-INS-53 | chr2-165621851-INS-54 |
| chr2-165626756-INS-82 | chr2-165626744-INS-82 |
| chr2-165635612-INS-226 | chr2-165635612-INS-226 |
| chr2-165637187-INS-76 | chr2-165637187-INS-69 |
| chr2-165638591-INS-368 | chr2-165638582-INS-389 |
| chr2-165664619-INS-209 | chr2-165664619-INS-209 |
| chr2-165678832-INS-88 | chr2-165678832-INS-100 |
| chr2-165680394-INS-203 | chr2-165680394-INS-220 |
| chr2-165731762-INS-257 | chr2-165731762-INS-221 |
| chr2-165734002-INS-70 | chr2-165734020-INS-90 |
| chr2-165740253-INS-413 | chr2-165740253-INS-414 |
| chr2-165749415-INS-162 | chr2-165749415-INS-162 |
| chr2-165753947-INS-71 | chr2-165753948-INS-75 |
| chr2-165758705-INS-201 | chr2-165758705-INS-201 |
| chr2-165761240-INS-76 | chr2-165761240-INS-88 |
| chr2-165766003-INS-219 | chr2-165766003-INS-220 |
| chr2-165766472-INS-201 | chr2-165766473-INS-199 |
| chr2-165768260-INS-132 | chr2-165768260-INS-132 |
| chr2-165782426-INS-195 | chr2-165782426-INS-195 |
| chr2-165796013-INS-523 | chr2-165796013-INS-524 |
| chr2-165797709-INS-58 | chr2-165797694-INS-52 |
| chr2-165802755-INS-218 | chr2-165802755-INS-218 |
| chr2-165805640-INS-194 | chr2-165805640-INS-194 |
| chr2-165807481-INS-69 | chr2-165807449-INS-70 |
| chr2-165822116-INS-75 | chr2-165822116-INS-75 |
| chr2-165824313-INS-207 | chr2-165824313-INS-208 |
| chr2-165826745-INS-57 | chr2-165826745-INS-57 |
| chr2-165858977-INS-98 | chr2-165858977-INS-64 |
| chr2-165883669-INS-69 | chr2-165883654-INS-85 |
| chr2-165885235-INS-209 | chr2-165885235-INS-209 |
| chr2-165895994-INS-74 | chr2-165895994-INS-86 |
| chr2-165924936-INS-52 | chr2-165924936-INS-52 |
| chr2-165930712-INS-63 | chr2-165930712-INS-65 |
| chr2-165949254-INS-58 | chr2-165949254-INS-58 |
| chr2-165953238-INS-333 | chr2-165953238-INS-333 |
| chr2-165980352-INS-153 | chr2-165980352-INS-150 |
| chr2-166002677-INS-152 | chr2-166002678-INS-146 |
| chr2-166026780-INS-65 | chr2-166026780-INS-65 |
| chr2-166045341-INS-66 | chr2-166045341-INS-66 |
| chr2-166053440-INS-507 | chr2-166053440-INS-507 |
| chr2-166063573-INS-194 | chr2-166063573-INS-190 |
| chr2-166068608-INS-53 | chr2-166068608-INS-55 |
| chr2-166071387-INS-57 | chr2-166071392-INS-76 |
| chr2-166093999-INS-76 | chr2-166093999-INS-76 |
| chr2-166101345-INS-153 | chr2-166101345-INS-153 |
| chr2-166126040-INS-54 | chr2-166126040-INS-50 |
| chr2-166127882-INS-128 | chr2-166127895-INS-102 |
| chr2-166139692-INS-56 | chr2-166139692-INS-50 |
| chr2-166149436-INS-6440 | chr2-166149437-INS-6440 |
| chr2-166149437-INS-497 | chr2-166149437-INS-497 |

| | |
|---|---|
| chr2-166184979-INS-105 | chr2-166184979-INS-70 |
| chr2-166192063-INS-60 | chr2-166192063-INS-60 |
| chr2-166197270-INS-6440 | chr2-166197271-INS-6440 |
| chr2-166198905-INS-60 | chr2-166198905-INS-60 |
| chr2-166231665-INS-78 | chr2-166231665-INS-78 |
| chr2-166240630-INS-87 | chr2-166240630-INS-87 |
| chr2-166257826-INS-93 | chr2-166257826-INS-90 |
| chr2-166269718-INS-400 | chr2-166269718-INS-400 |
| chr2-166277771-INS-6440 | chr2-166277772-INS-6440 |
| chr2-166289498-INS-144 | chr2-166289498-INS-96 |
| chr2-166299239-INS-51 | chr2-166299237-INS-53 |
| chr2-166328387-INS-125 | chr2-166328387-INS-125 |
| chr2-166335024-INS-130 | chr2-166335025-INS-130 |
| chr2-166354515-INS-700 | chr2-166354515-INS-700 |
| chr2-166364602-INS-88 | chr2-166364623-INS-96 |
| chr2-166381551-INS-323 | chr2-166381551-INS-323 |
| chr2-166390511-INS-100 | chr2-166390512-INS-60 |
| chr2-166405088-INS-183 | chr2-166405088-INS-183 |
| chr2-166426472-INS-75 | chr2-166426472-INS-75 |
| chr2-166462075-INS-207 | chr2-166462075-INS-206 |
| chr2-166476872-INS-132 | chr2-166476872-INS-132 |
| chr2-166491114-INS-55 | chr2-166491118-INS-62 |
| chr2-166492028-INS-72 | chr2-166492028-INS-54 |
| chr2-166511992-INS-5355 | chr2-166511993-INS-5354 |
| chr2-166548343-INS-209 | chr2-166548343-INS-209 |
| chr2-166577310-INS-55 | chr2-166577294-INS-76 |
| chr2-166584790-INS-163 | chr2-166584790-INS-163 |
| chr2-166607651-INS-183 | chr2-166607651-INS-173 |
| chr2-166609703-INS-51 | chr2-166609680-INS-51 |
| chr2-166625116-INS-198 | chr2-166625116-INS-198 |
| chr2-166633143-INS-184 | chr2-166633139-INS-164 |
| chr2-166635482-INS-64 | chr2-166635482-INS-64 |
| chr2-166673849-INS-58 | chr2-166673849-INS-68 |
| chr2-166682977-INS-208 | chr2-166682977-INS-209 |
| chr2-166704456-INS-67 | chr2-166704456-INS-67 |
| chr2-166713057-INS-204 | chr2-166713057-INS-204 |
| chr2-166721694-INS-131 | chr2-166721695-INS-141 |
| chr2-166730580-INS-195 | chr2-166730580-INS-196 |
| chr2-166785818-INS-111 | chr2-166785818-INS-74 |
| chr2-166813349-INS-215 | chr2-166813349-INS-215 |
| chr2-166815544-INS-61 | chr2-166815544-INS-62 |
| chr2-166817556-INS-253 | chr2-166817556-INS-230 |
| chr2-166824969-INS-1258 | chr2-166824970-INS-1258 |
| chr2-166844247-INS-117 | chr2-166844262-INS-126 |
| chr2-166856399-INS-729 | chr2-166856399-INS-730 |
| chr2-166864922-INS-57 | chr2-166864922-INS-57 |
| chr2-166875069-INS-143 | chr2-166875088-INS-105 |
| chr2-166895453-INS-210 | chr2-166895453-INS-211 |
| chr2-166907989-INS-197 | chr2-166907989-INS-197 |
| chr2-166908780-INS-869 | chr2-166908780-INS-879 |
| chr2-166910683-INS-148 | chr2-166910677-INS-154 |
| chr2-166950537-INS-123 | chr2-166950537-INS-123 |
| chr2-166951336-INS-116 | chr2-166951336-INS-116 |
| chr2-166952402-INS-820 | chr2-166952399-INS-806 |
| chr2-166963472-INS-4839 | chr2-166963473-INS-4839 |
| chr2-166997053-INS-88 | chr2-166997053-INS-88 |
| chr2-167039170-INS-5392 | chr2-167039171-INS-5392 |
| chr2-167090029-INS-226 | chr2-167090029-INS-226 |
| chr2-167116557-INS-54 | chr2-167116559-INS-86 |
| chr2-167118563-INS-56 | chr2-167118568-INS-76 |
| chr2-167147277-INS-137 | chr2-167147299-INS-111 |
| chr2-167150354-INS-60 | chr2-167150354-INS-57 |
| chr2-167153652-INS-192 | chr2-167153652-INS-192 |
| chr2-167154777-INS-138 | chr2-167154777-INS-158 |
| chr2-167158227-INS-150 | chr2-167158227-INS-169 |
| chr2-167163057-INS-206 | chr2-167163069-INS-207 |
| chr2-167181433-INS-212 | chr2-167181433-INS-214 |
| chr2-167199749-INS-784 | chr2-167199749-INS-766 |
| chr2-167212991-INS-244 | chr2-167212991-INS-216 |
| chr2-167278128-INS-208 | chr2-167278128-INS-206 |
| chr2-167280901-INS-102 | chr2-167280901-INS-102 |
| chr2-167286070-INS-874 | chr2-167286070-INS-874 |
| chr2-167290652-INS-60 | chr2-167290652-INS-60 |
| chr2-167311068-INS-213 | chr2-167311087-INS-241 |
| chr2-167350735-INS-208 | chr2-167350735-INS-224 |
| chr2-167370753-INS-118 | chr2-167370749-INS-95 |
| chr2-167373144-INS-54 | chr2-167373144-INS-54 |

| | |
|---|---|
| chr2-167394348-INS-222 | chr2-167394348-INS-224 |
| chr2-167418036-INS-74 | chr2-167418036-INS-74 |
| chr2-167432996-INS-63 | chr2-167433032-INS-75 |
| chr2-167458422-INS-53 | chr2-167458422-INS-53 |
| chr2-167474656-INS-77 | chr2-167474656-INS-77 |
| chr2-167478770-INS-65 | chr2-167478770-INS-69 |
| chr2-167490570-INS-6387 | chr2-167490571-INS-6387 |
| chr2-167524441-INS-67 | chr2-167524426-INS-76 |
| chr2-167545083-INS-296 | chr2-167545083-INS-296 |
| chr2-167586641-INS-60 | chr2-167586651-INS-64 |
| chr2-167655319-INS-77 | chr2-167655319-INS-77 |
| chr2-167656797-INS-252 | chr2-167656797-INS-256 |
| chr2-167658266-INS-78 | chr2-167658258-INS-80 |
| chr2-167662012-INS-61 | chr2-167662012-INS-61 |
| chr2-167671988-INS-54 | chr2-167671991-INS-50 |
| chr2-167718665-INS-1710 | chr2-167718666-INS-1710 |
| chr2-167718681-INS-228 | chr2-167718681-INS-228 |
| chr2-167770933-INS-7189 | chr2-167770934-INS-7189 |
| chr2-167820390-INS-196 | chr2-167820390-INS-196 |
| chr2-167826934-INS-60 | chr2-167826923-INS-62 |
| chr2-167856161-INS-337 | chr2-167856161-INS-361 |
| chr2-167883908-INS-589 | chr2-167883908-INS-589 |
| chr2-167904684-INS-262 | chr2-167904652-INS-254 |
| chr2-167912077-INS-199 | chr2-167912077-INS-199 |
| chr2-167916509-INS-185 | chr2-167916509-INS-185 |
| chr2-167919344-INS-68 | chr2-167919344-INS-68 |
| chr2-167924675-INS-200 | chr2-167924675-INS-200 |
| chr2-167935268-INS-206 | chr2-167935268-INS-220 |
| chr2-167961674-INS-107 | chr2-167961681-INS-82 |
| chr2-167980764-INS-406 | chr2-167980764-INS-406 |
| chr2-168005095-INS-115 | chr2-168005095-INS-117 |
| chr2-168012675-INS-105 | chr2-168012676-INS-56 |
| chr2-168033963-INS-211 | chr2-168033963-INS-211 |
| chr2-168061633-INS-360 | chr2-168061634-INS-360 |
| chr2-168109726-INS-52 | chr2-168109726-INS-95 |
| chr2-168146330-INS-599 | chr2-168146331-INS-607 |
| chr2-168148320-INS-100 | chr2-168148320-INS-100 |
| chr2-168150125-INS-54 | chr2-168150109-INS-56 |
| chr2-168165429-INS-120 | chr2-168165429-INS-120 |
| chr2-168193794-INS-260 | chr2-168193794-INS-243 |
| chr2-168239215-INS-80 | chr2-168239215-INS-80 |
| chr2-168244882-INS-57 | chr2-168244882-INS-57 |
| chr2-168249520-INS-689 | chr2-168249522-INS-650 |
| chr2-168253818-INS-163 | chr2-168253839-INS-167 |
| chr2-168257929-INS-102 | chr2-168257925-INS-63 |
| chr2-168275675-INS-170 | chr2-168275675-INS-175 |
| chr2-168288926-INS-563 | chr2-168288934-INS-575 |
| chr2-168291647-INS-65 | chr2-168291647-INS-65 |
| chr2-168302762-INS-72 | chr2-168302762-INS-72 |
| chr2-168307496-INS-91 | chr2-168307496-INS-89 |
| chr2-168311639-INS-114 | chr2-168311610-INS-101 |
| chr2-168366259-INS-103 | chr2-168366305-INS-96 |
| chr2-168394668-INS-420 | chr2-168394670-INS-373 |
| chr2-168403641-INS-64 | chr2-168403641-INS-64 |
| chr2-168410084-INS-61 | chr2-168410084-INS-61 |
| chr2-168429547-INS-62 | chr2-168429547-INS-62 |
| chr2-168438285-INS-66 | chr2-168438243-INS-110 |
| chr2-168473266-INS-182 | chr2-168473258-INS-186 |
| chr2-168508182-INS-772 | chr2-168508182-INS-773 |
| chr2-168518367-INS-68 | chr2-168518367-INS-78 |
| chr2-168545875-INS-132 | chr2-168545875-INS-132 |
| chr2-168572381-INS-53 | chr2-168572381-INS-53 |
| chr2-168576521-INS-215 | chr2-168576521-INS-216 |
| chr2-168579646-INS-58 | chr2-168579633-INS-69 |
| chr2-168601540-INS-234 | chr2-168601540-INS-234 |
| chr2-168602818-INS-51 | chr2-168602824-INS-51 |
| chr2-168612478-INS-498 | chr2-168612478-INS-498 |
| chr2-168643326-INS-238 | chr2-168643327-INS-204 |
| chr2-168653753-INS-243 | chr2-168653753-INS-243 |
| chr2-168664483-INS-169 | chr2-168664483-INS-167 |
| chr2-168666508-INS-492 | chr2-168666508-INS-497 |
| chr2-168685768-INS-72 | chr2-168685768-INS-69 |
| chr2-168694275-INS-76 | chr2-168694275-INS-76 |
| chr2-168700933-INS-198 | chr2-168700933-INS-197 |
| chr2-168703790-INS-116 | chr2-168703798-INS-112 |
| chr2-168706833-INS-175 | chr2-168706833-INS-164 |
| chr2-168721820-INS-189 | chr2-168721820-INS-189 |

| | |
|---|---|
| chr2-168732678-INS-726 | chr2-168732678-INS-726 |
| chr2-168739051-INS-237 | chr2-168739051-INS-238 |
| chr2-168742890-INS-53 | chr2-168742890-INS-57 |
| chr2-168755939-INS-70 | chr2-168755939-INS-63 |
| chr2-168778365-INS-51 | chr2-168778366-INS-53 |
| chr2-168782939-INS-100 | chr2-168782940-INS-105 |
| chr2-168790341-INS-84 | chr2-168790325-INS-100 |
| chr2-168803853-INS-60 | chr2-168803852-INS-67 |
| chr2-168807100-INS-211 | chr2-168807100-INS-211 |
| chr2-168808573-INS-51 | chr2-168808572-INS-52 |
| chr2-168828209-INS-69 | chr2-168828232-INS-69 |
| chr2-168830773-INS-71 | chr2-168830790-INS-86 |
| chr2-168837477-INS-73 | chr2-168837466-INS-113 |
| chr2-168840228-INS-114 | chr2-168840228-INS-120 |
| chr2-168841557-INS-327 | chr2-168841557-INS-327 |
| chr2-168848986-INS-6441 | chr2-168848987-INS-6441 |
| chr2-168870758-INS-75 | chr2-168870758-INS-75 |
| chr2-168883724-INS-2553 | chr2-168883725-INS-2527 |
| chr2-168889601-INS-70 | chr2-168889611-INS-58 |
| chr2-168898530-INS-59 | chr2-168898530-INS-58 |
| chr2-168923916-INS-51 | chr2-168923916-INS-50 |
| chr2-168929836-INS-100 | chr2-168929836-INS-100 |
| chr2-168955224-INS-1516 | chr2-168955225-INS-1516 |
| chr2-168981696-INS-786 | chr2-168981696-INS-786 |
| chr2-168988390-INS-55 | chr2-168988390-INS-55 |
| chr2-169032416-INS-480 | chr2-169032416-INS-487 |
| chr2-169054666-INS-52 | chr2-169054666-INS-52 |
| chr2-169072153-INS-1087 | chr2-169072154-INS-1087 |
| chr2-169076094-INS-519 | chr2-169076094-INS-519 |
| chr2-169112188-INS-158 | chr2-169112188-INS-156 |
| chr2-169125810-INS-62 | chr2-169125830-INS-64 |
| chr2-169136911-INS-64 | chr2-169136902-INS-76 |
| chr2-169157488-INS-130 | chr2-169157488-INS-114 |
| chr2-169161118-INS-216 | chr2-169161118-INS-217 |
| chr2-169185025-INS-399 | chr2-169185025-INS-399 |
| chr2-169187612-INS-1885 | chr2-169187612-INS-1887 |
| chr2-169241113-INS-802 | chr2-169241113-INS-779 |
| chr2-169272510-INS-81 | chr2-169272510-INS-81 |
| chr2-169276133-INS-803 | chr2-169276132-INS-802 |
| chr2-169300494-INS-52 | chr2-169300494-INS-54 |
| chr2-169317055-INS-135 | chr2-169317055-INS-85 |
| chr2-169327416-INS-130 | chr2-169327407-INS-139 |
| chr2-169333004-INS-498 | chr2-169333004-INS-498 |
| chr2-169336912-INS-65 | chr2-169336902-INS-71 |
| chr2-169339551-INS-80 | chr2-169339551-INS-82 |
| chr2-169344565-INS-63 | chr2-169344565-INS-63 |
| chr2-169350220-INS-56 | chr2-169350227-INS-60 |
| chr2-169353250-INS-338 | chr2-169353250-INS-338 |
| chr2-169358679-INS-70 | chr2-169358679-INS-50 |
| chr2-169364271-INS-313 | chr2-169364289-INS-317 |
| chr2-169385702-INS-198 | chr2-169385702-INS-198 |
| chr2-169473395-INS-75 | chr2-169473395-INS-75 |
| chr2-169485726-INS-65 | chr2-169485726-INS-65 |
| chr2-169492238-INS-52 | chr2-169492238-INS-52 |
| chr2-169501987-INS-397 | chr2-169501987-INS-397 |
| chr2-169506934-INS-52 | chr2-169506934-INS-52 |
| chr2-169515303-INS-112 | chr2-169515303-INS-115 |
| chr2-169525042-INS-56 | chr2-169525042-INS-56 |
| chr2-169545325-INS-74 | chr2-169545295-INS-102 |
| chr2-169547993-INS-77 | chr2-169547992-INS-55 |
| chr2-169551059-INS-72 | chr2-169551059-INS-66 |
| chr2-169555831-INS-208 | chr2-169555831-INS-204 |
| chr2-169568964-INS-213 | chr2-169568964-INS-204 |
| chr2-169570556-INS-190 | chr2-169570556-INS-190 |
| chr2-169580597-INS-63 | chr2-169580599-INS-55 |
| chr2-169598759-INS-592 | chr2-169598759-INS-594 |
| chr2-169603349-INS-67 | chr2-169603349-INS-63 |
| chr2-169606571-INS-59 | chr2-169606571-INS-59 |
| chr2-169630942-INS-64 | chr2-169630942-INS-64 |
| chr2-169652483-INS-68 | chr2-169652483-INS-68 |
| chr2-169666059-INS-64 | chr2-169666059-INS-64 |
| chr2-169680757-INS-77 | chr2-169680757-INS-77 |
| chr2-169688001-INS-83 | chr2-169688001-INS-83 |
| chr2-169694696-INS-67 | chr2-169694696-INS-67 |
| chr2-169708475-INS-92 | chr2-169708475-INS-76 |
| chr2-169709393-INS-53 | chr2-169709427-INS-59 |
| chr2-169765949-INS-104 | chr2-169765949-INS-100 |

| | |
|---|---|
| chr2-169773379-INS-387 | chr2-169773379-INS-387 |
| chr2-169781668-INS-162 | chr2-169781667-INS-155 |
| chr2-169797869-INS-53 | chr2-169797869-INS-68 |
| chr2-169843658-INS-52 | chr2-169843658-INS-52 |
| chr2-169847679-INS-69 | chr2-169847679-INS-69 |
| chr2-169852938-INS-171 | chr2-169852938-INS-171 |
| chr2-169887742-INS-328 | chr2-169887742-INS-346 |
| chr2-169891704-INS-76 | chr2-169891704-INS-76 |
| chr2-169894331-INS-51 | chr2-169894331-INS-58 |
| chr2-169896968-INS-115 | chr2-169896968-INS-78 |
| chr2-169899624-INS-87 | chr2-169899624-INS-58 |
| chr2-169922320-INS-207 | chr2-169922320-INS-214 |
| chr2-169965500-INS-93 | chr2-169965501-INS-93 |
| chr2-169987395-INS-83 | chr2-169987376-INS-70 |
| chr2-169998031-INS-51 | chr2-169998031-INS-51 |
| chr2-170009172-INS-164 | chr2-170009172-INS-164 |
| chr2-170010372-INS-715 | chr2-170010372-INS-757 |
| chr2-170020616-INS-177 | chr2-170020616-INS-176 |
| chr2-170029798-INS-185 | chr2-170029798-INS-186 |
| chr2-170031063-INS-585 | chr2-170031063-INS-593 |
| chr2-170032809-INS-208 | chr2-170032809-INS-208 |
| chr2-170033935-INS-159 | chr2-170033935-INS-159 |
| chr2-170052780-INS-204 | chr2-170052780-INS-210 |
| chr2-170079274-INS-199 | chr2-170079274-INS-198 |
| chr2-170083267-INS-70 | chr2-170083281-INS-60 |
| chr2-170089562-INS-114 | chr2-170089562-INS-117 |
| chr2-170093256-INS-760 | chr2-170093256-INS-760 |
| chr2-170106254-INS-245 | chr2-170106255-INS-239 |
| chr2-170120512-INS-100 | chr2-170120506-INS-106 |
| chr2-170129875-INS-120 | chr2-170129878-INS-116 |
| chr2-170151753-INS-171 | chr2-170151752-INS-217 |
| chr2-170157589-INS-63 | chr2-170157589-INS-84 |
| chr2-170178994-INS-112 | chr2-170178978-INS-108 |
| chr2-170190540-INS-400 | chr2-170190540-INS-400 |
| chr2-170192468-INS-113 | chr2-170192468-INS-91 |
| chr2-170218785-INS-60 | chr2-170218785-INS-52 |
| chr2-170221405-INS-60 | chr2-170221405-INS-60 |
| chr2-170244579-INS-72 | chr2-170244579-INS-70 |
| chr2-170249496-INS-226 | chr2-170249496-INS-226 |
| chr2-170253253-INS-95 | chr2-170253253-INS-95 |
| chr2-170297137-INS-914 | chr2-170297137-INS-914 |
| chr2-170310444-INS-86 | chr2-170310444-INS-86 |
| chr2-170343902-INS-50 | chr2-170343873-INS-68 |
| chr2-170347073-INS-1300 | chr2-170347074-INS-1260 |
| chr2-170354009-INS-220 | chr2-170354009-INS-212 |
| chr2-170355422-INS-920 | chr2-170355422-INS-920 |
| chr2-170374746-INS-232 | chr2-170374746-INS-233 |
| chr2-170379897-INS-63 | chr2-170379897-INS-63 |
| chr2-170384696-INS-392 | chr2-170384696-INS-392 |
| chr2-170393376-INS-90 | chr2-170393376-INS-140 |
| chr2-170418928-INS-385 | chr2-170418928-INS-385 |
| chr2-170419768-INS-160 | chr2-170419768-INS-160 |
| chr2-170425747-INS-181 | chr2-170425743-INS-181 |
| chr2-170438966-INS-89 | chr2-170438965-INS-132 |
| chr2-170457024-INS-107 | chr2-170457024-INS-107 |
| chr2-170459651-INS-56 | chr2-170459651-INS-58 |
| chr2-170474126-INS-432 | chr2-170474126-INS-430 |
| chr2-170486569-INS-115 | chr2-170486569-INS-69 |
| chr2-170488791-INS-253 | chr2-170488792-INS-244 |
| chr2-170515529-INS-54 | chr2-170515529-INS-60 |
| chr2-170522832-INS-197 | chr2-170522825-INS-207 |
| chr2-170526457-INS-7105 | chr2-170526458-INS-7105 |
| chr2-170528245-INS-146 | chr2-170528234-INS-164 |
| chr2-170529150-INS-440 | chr2-170529150-INS-440 |
| chr2-170534347-INS-76 | chr2-170534347-INS-76 |
| chr2-170553013-INS-60 | chr2-170553013-INS-60 |
| chr2-170560378-INS-58 | chr2-170560378-INS-58 |
| chr2-170568216-INS-329 | chr2-170568216-INS-329 |
| chr2-170605090-INS-103 | chr2-170605091-INS-72 |
| chr2-170613868-INS-73 | chr2-170613860-INS-115 |
| chr2-170630656-INS-483 | chr2-170630656-INS-492 |
| chr2-170638646-INS-156 | chr2-170638646-INS-156 |
| chr2-170656492-INS-215 | chr2-170656496-INS-207 |
| chr2-170664724-INS-72 | chr2-170664724-INS-72 |
| chr2-170688743-INS-549 | chr2-170688743-INS-549 |
| chr2-170690428-INS-137 | chr2-170690420-INS-105 |
| chr2-170697503-INS-209 | chr2-170697503-INS-237 |

| | |
|---|---|
| chr2-170702809-INS-142 | chr2-170702809-INS-136 |
| chr2-170716823-INS-826 | chr2-170716823-INS-826 |
| chr2-170721356-INS-68 | chr2-170721356-INS-68 |
| chr2-170739975-INS-64 | chr2-170739975-INS-64 |
| chr2-170740501-INS-54 | chr2-170740499-INS-101 |
| chr2-170741776-INS-58 | chr2-170741776-INS-58 |
| chr2-170781167-INS-51 | chr2-170781167-INS-93 |
| chr2-170800472-INS-485 | chr2-170800472-INS-489 |
| chr2-170808780-INS-64 | chr2-170808780-INS-66 |
| chr2-170833885-INS-128 | chr2-170833886-INS-131 |
| chr2-170860985-INS-87 | chr2-170860985-INS-93 |
| chr2-170874313-INS-1318 | chr2-170874314-INS-1318 |
| chr2-170881441-INS-58 | chr2-170881441-INS-62 |
| chr2-170907590-INS-122 | chr2-170907590-INS-126 |
| chr2-170920504-INS-83 | chr2-170920524-INS-85 |
| chr2-170960471-INS-614 | chr2-170960471-INS-614 |
| chr2-171005301-INS-916 | chr2-171005302-INS-916 |
| chr2-171023300-INS-60 | chr2-171023300-INS-60 |
| chr2-171164623-INS-61 | chr2-171164623-INS-61 |
| chr2-171165903-INS-311 | chr2-171165903-INS-327 |
| chr2-171168695-INS-65 | chr2-171168709-INS-72 |
| chr2-171193404-INS-106 | chr2-171193404-INS-106 |
| chr2-171212245-INS-109 | chr2-171212245-INS-111 |
| chr2-171217032-INS-113 | chr2-171217032-INS-113 |
| chr2-171240879-INS-150 | chr2-171240878-INS-154 |
| chr2-171243407-INS-454 | chr2-171243407-INS-454 |
| chr2-171251933-INS-51 | chr2-171251933-INS-51 |
| chr2-171257433-INS-76 | chr2-171257433-INS-76 |
| chr2-171271110-INS-56 | chr2-171271110-INS-56 |
| chr2-171282418-INS-126 | chr2-171282418-INS-126 |
| chr2-171321360-INS-340 | chr2-171321360-INS-342 |
| chr2-171324904-INS-169 | chr2-171324904-INS-191 |
| chr2-171354900-INS-490 | chr2-171354900-INS-493 |
| chr2-171358180-INS-148 | chr2-171358176-INS-133 |
| chr2-171359951-INS-54 | chr2-171359951-INS-54 |
| chr2-171397571-INS-169 | chr2-171397571-INS-169 |
| chr2-171401006-INS-62 | chr2-171401006-INS-62 |
| chr2-171402356-INS-350 | chr2-171402356-INS-369 |
| chr2-171451581-INS-52 | chr2-171451609-INS-53 |
| chr2-171455710-INS-74 | chr2-171455710-INS-74 |
| chr2-171469466-INS-214 | chr2-171469466-INS-215 |
| chr2-171490707-INS-276 | chr2-171490707-INS-276 |
| chr2-171503214-INS-391 | chr2-171503214-INS-391 |
| chr2-171559441-INS-53 | chr2-171559441-INS-53 |
| chr2-171579041-INS-54 | chr2-171579002-INS-88 |
| chr2-171604810-INS-300 | chr2-171604810-INS-300 |
| chr2-171606322-INS-66 | chr2-171606322-INS-68 |
| chr2-171661985-INS-54 | chr2-171661985-INS-54 |
| chr2-171669633-INS-64 | chr2-171669614-INS-80 |
| chr2-171696421-INS-68 | chr2-171696405-INS-74 |
| chr2-171718312-INS-173 | chr2-171718312-INS-173 |
| chr2-171735771-INS-56 | chr2-171735771-INS-56 |
| chr2-171752528-INS-161 | chr2-171752528-INS-161 |
| chr2-171816469-INS-328 | chr2-171816469-INS-328 |
| chr2-171823481-INS-52 | chr2-171823506-INS-50 |
| chr2-171841200-INS-247 | chr2-171841247-INS-229 |
| chr2-171886747-INS-80 | chr2-171886760-INS-84 |
| chr2-171890181-INS-246 | chr2-171890181-INS-246 |
| chr2-171894981-INS-189 | chr2-171894993-INS-208 |
| chr2-171915771-INS-72 | chr2-171915771-INS-72 |
| chr2-171933575-INS-52 | chr2-171933620-INS-57 |
| chr2-171935100-INS-404 | chr2-171935100-INS-404 |
| chr2-171943262-INS-60 | chr2-171943262-INS-50 |
| chr2-171945273-INS-244 | chr2-171945273-INS-244 |
| chr2-171948302-INS-305 | chr2-171948302-INS-305 |
| chr2-171966891-INS-120 | chr2-171966891-INS-118 |
| chr2-171977295-INS-55 | chr2-171977295-INS-57 |
| chr2-171989058-INS-85 | chr2-171989034-INS-69 |
| chr2-171991373-INS-52 | chr2-171991374-INS-54 |
| chr2-172004838-INS-212 | chr2-172004838-INS-213 |
| chr2-172009594-INS-61 | chr2-172009594-INS-64 |
| chr2-172020110-INS-1304 | chr2-172020112-INS-1300 |
| chr2-172021175-INS-97 | chr2-172021175-INS-97 |
| chr2-172023009-INS-580 | chr2-172023009-INS-581 |
| chr2-172064503-INS-201 | chr2-172064503-INS-201 |
| chr2-172074761-INS-105 | chr2-172074783-INS-120 |
| chr2-172085483-INS-54 | chr2-172085483-INS-54 |

| | |
|---|---|
| chr2-172136399-INS-137 | chr2-172136394-INS-151 |
| chr2-172140451-INS-70 | chr2-172140451-INS-70 |
| chr2-172143221-INS-169 | chr2-172143221-INS-165 |
| chr2-172155370-INS-78 | chr2-172155368-INS-92 |
| chr2-172185198-INS-235 | chr2-172185198-INS-236 |
| chr2-172204807-INS-102 | chr2-172204807-INS-102 |
| chr2-172206068-INS-62 | chr2-172206068-INS-62 |
| chr2-172235072-INS-60 | chr2-172235061-INS-60 |
| chr2-172242713-INS-51 | chr2-172242711-INS-60 |
| chr2-172247920-INS-64 | chr2-172247920-INS-62 |
| chr2-172267407-INS-192 | chr2-172267407-INS-180 |
| chr2-172286291-INS-112 | chr2-172286291-INS-112 |
| chr2-172317445-INS-194 | chr2-172317445-INS-194 |
| chr2-172324304-INS-345 | chr2-172324305-INS-345 |
| chr2-172327388-INS-65 | chr2-172327388-INS-69 |
| chr2-172332728-INS-208 | chr2-172332728-INS-208 |
| chr2-172333681-INS-63 | chr2-172333681-INS-63 |
| chr2-172340325-INS-93 | chr2-172340325-INS-93 |
| chr2-172352199-INS-70 | chr2-172352199-INS-70 |
| chr2-172357748-INS-58 | chr2-172357748-INS-60 |
| chr2-172359718-INS-181 | chr2-172359718-INS-181 |
| chr2-172364157-INS-225 | chr2-172364157-INS-226 |
| chr2-172381065-INS-142 | chr2-172381065-INS-144 |
| chr2-172448373-INS-58 | chr2-172448373-INS-58 |
| chr2-172450393-INS-277 | chr2-172450393-INS-278 |
| chr2-172468263-INS-143 | chr2-172468263-INS-149 |
| chr2-172473072-INS-78 | chr2-172473072-INS-79 |
| chr2-172475490-INS-61 | chr2-172475489-INS-62 |
| chr2-172480765-INS-170 | chr2-172480765-INS-157 |
| chr2-172487357-INS-198 | chr2-172487357-INS-198 |
| chr2-172488567-INS-190 | chr2-172488567-INS-190 |
| chr2-172500675-INS-142 | chr2-172500675-INS-142 |
| chr2-172518412-INS-184 | chr2-172518408-INS-164 |
| chr2-172525530-INS-72 | chr2-172525530-INS-72 |
| chr2-172564158-INS-204 | chr2-172564158-INS-204 |
| chr2-172565146-INS-159 | chr2-172565147-INS-160 |
| chr2-172585309-INS-61 | chr2-172585308-INS-71 |
| chr2-172615114-INS-181 | chr2-172615115-INS-182 |
| chr2-172627637-INS-306 | chr2-172627637-INS-306 |
| chr2-172644428-INS-81 | chr2-172644428-INS-81 |
| chr2-172647097-INS-336 | chr2-172647097-INS-335 |
| chr2-172653031-INS-527 | chr2-172653031-INS-527 |
| chr2-172666805-INS-8334 | chr2-172666806-INS-8334 |
| chr2-172695797-INS-66 | chr2-172695797-INS-66 |
| chr2-172716437-INS-254 | chr2-172716463-INS-242 |
| chr2-172777943-INS-271 | chr2-172777943-INS-270 |
| chr2-172787905-INS-64 | chr2-172787905-INS-96 |
| chr2-172790149-INS-672 | chr2-172790125-INS-700 |
| chr2-172795539-INS-80 | chr2-172795539-INS-80 |
| chr2-172836030-INS-60 | chr2-172836030-INS-73 |
| chr2-172847678-INS-59 | chr2-172847678-INS-59 |
| chr2-172854155-INS-52 | chr2-172854155-INS-52 |
| chr2-172873869-INS-102 | chr2-172873869-INS-111 |
| chr2-172878785-INS-60 | chr2-172878785-INS-68 |
| chr2-172895884-INS-102 | chr2-172895884-INS-102 |
| chr2-172906424-INS-52 | chr2-172906424-INS-52 |
| chr2-172907671-INS-66 | chr2-172907671-INS-66 |
| chr2-172924156-INS-52 | chr2-172924156-INS-52 |
| chr2-172925813-INS-52 | chr2-172925813-INS-52 |
| chr2-172933634-INS-63 | chr2-172933634-INS-63 |
| chr2-172945249-INS-92 | chr2-172945270-INS-106 |
| chr2-172947565-INS-99 | chr2-172947565-INS-132 |
| chr2-172950909-INS-104 | chr2-172950909-INS-104 |
| chr2-172962135-INS-105 | chr2-172962135-INS-77 |
| chr2-172971487-INS-395 | chr2-172971492-INS-400 |
| chr2-173004757-INS-91 | chr2-173004757-INS-114 |
| chr2-173040361-INS-66 | chr2-173040361-INS-84 |
| chr2-173043546-INS-683 | chr2-173043546-INS-682 |
| chr2-173044199-INS-2623 | chr2-173044200-INS-2623 |
| chr2-173059851-INS-230 | chr2-173059851-INS-230 |
| chr2-173079314-INS-214 | chr2-173079314-INS-213 |
| chr2-173084465-INS-220 | chr2-173084465-INS-220 |
| chr2-173111930-INS-62 | chr2-173111930-INS-62 |
| chr2-173140527-INS-434 | chr2-173140527-INS-434 |
| chr2-173165169-INS-65 | chr2-173165169-INS-65 |
| chr2-173170481-INS-88 | chr2-173170481-INS-88 |
| chr2-173173769-INS-79 | chr2-173173769-INS-79 |

| | |
|---|---|
| chr2-173178552-INS-164 | chr2-173178554-INS-161 |
| chr2-173179972-INS-200 | chr2-173179972-INS-206 |
| chr2-173187680-INS-152 | chr2-173187684-INS-160 |
| chr2-173192642-INS-101 | chr2-173192642-INS-114 |
| chr2-173193176-INS-321 | chr2-173193189-INS-285 |
| chr2-173197214-INS-72 | chr2-173197214-INS-78 |
| chr2-173204455-INS-157 | chr2-173204452-INS-116 |
| chr2-173209358-INS-119 | chr2-173209353-INS-95 |
| chr2-173211414-INS-513 | chr2-173211414-INS-512 |
| chr2-173220615-INS-115 | chr2-173220615-INS-121 |
| chr2-173223220-INS-61 | chr2-173223220-INS-61 |
| chr2-173229049-INS-76 | chr2-173229049-INS-76 |
| chr2-173236012-INS-94 | chr2-173236004-INS-76 |
| chr2-173258115-INS-70 | chr2-173258115-INS-69 |
| chr2-173276093-INS-52 | chr2-173276089-INS-64 |
| chr2-173280049-INS-75 | chr2-173280049-INS-75 |
| chr2-173292398-INS-150 | chr2-173292398-INS-145 |
| chr2-173304698-INS-119 | chr2-173304698-INS-119 |
| chr2-173305817-INS-134 | chr2-173305817-INS-136 |
| chr2-173311175-INS-2801 | chr2-173311176-INS-2801 |
| chr2-173311176-INS-498 | chr2-173311176-INS-498 |
| chr2-173337923-INS-498 | chr2-173337923-INS-498 |
| chr2-173351321-INS-59 | chr2-173351321-INS-59 |
| chr2-173352947-INS-1388 | chr2-173352948-INS-1383 |
| chr2-173357623-INS-208 | chr2-173357623-INS-208 |
| chr2-173363384-INS-1889 | chr2-173363385-INS-1889 |
| chr2-173366503-INS-81 | chr2-173366503-INS-82 |
| chr2-173371462-INS-70 | chr2-173371462-INS-62 |
| chr2-173385153-INS-208 | chr2-173385178-INS-228 |
| chr2-173430903-INS-192 | chr2-173430903-INS-192 |
| chr2-173458954-INS-161 | chr2-173458954-INS-161 |
| chr2-173473278-INS-74 | chr2-173473278-INS-74 |
| chr2-173475433-INS-62 | chr2-173475433-INS-62 |
| chr2-173478336-INS-85 | chr2-173478337-INS-58 |
| chr2-173478793-INS-148 | chr2-173478794-INS-148 |
| chr2-173565446-INS-176 | chr2-173565452-INS-184 |
| chr2-173587528-INS-52 | chr2-173587528-INS-52 |
| chr2-173593811-INS-183 | chr2-173593811-INS-187 |
| chr2-173595669-INS-101 | chr2-173595669-INS-101 |
| chr2-173596261-INS-1655 | chr2-173596262-INS-1655 |
| chr2-173650235-INS-653 | chr2-173650235-INS-651 |
| chr2-173658364-INS-56 | chr2-173658364-INS-56 |
| chr2-173663476-INS-615 | chr2-173663476-INS-615 |
| chr2-173679881-INS-102 | chr2-173679881-INS-102 |
| chr2-173698717-INS-317 | chr2-173698717-INS-319 |
| chr2-173704696-INS-225 | chr2-173704696-INS-225 |
| chr2-173727941-INS-64 | chr2-173727941-INS-67 |
| chr2-173795079-INS-100 | chr2-173795080-INS-118 |
| chr2-173801803-INS-553 | chr2-173801803-INS-546 |
| chr2-173916586-INS-53 | chr2-173916586-INS-57 |
| chr2-173973312-INS-110 | chr2-173973306-INS-102 |
| chr2-174005238-INS-54 | chr2-174005238-INS-57 |
| chr2-174089996-INS-3346 | chr2-174090004-INS-3338 |
| chr2-174146328-INS-709 | chr2-174146328-INS-710 |
| chr2-174162209-INS-63 | chr2-174162209-INS-66 |
| chr2-174206466-INS-78 | chr2-174206466-INS-78 |
| chr2-174229017-INS-483 | chr2-174229017-INS-483 |
| chr2-174236293-INS-54 | chr2-174236294-INS-54 |
| chr2-174244487-INS-56 | chr2-174244487-INS-56 |
| chr2-174263247-INS-94 | chr2-174263247-INS-94 |
| chr2-174268493-INS-72 | chr2-174268493-INS-72 |
| chr2-174272656-INS-368 | chr2-174272658-INS-366 |
| chr2-174282636-INS-96 | chr2-174282636-INS-102 |
| chr2-174286458-INS-140 | chr2-174286458-INS-140 |
| chr2-174289720-INS-98 | chr2-174289720-INS-98 |
| chr2-174309101-INS-56 | chr2-174309101-INS-56 |
| chr2-174314229-INS-68 | chr2-174314229-INS-68 |
| chr2-174316035-INS-68 | chr2-174316035-INS-74 |
| chr2-174326408-INS-86 | chr2-174326408-INS-86 |
| chr2-174329822-INS-123 | chr2-174329822-INS-123 |
| chr2-174331452-INS-462 | chr2-174331452-INS-462 |
| chr2-174335793-INS-78 | chr2-174335793-INS-80 |
| chr2-174340771-INS-65 | chr2-174340771-INS-60 |
| chr2-174357873-INS-54 | chr2-174357873-INS-54 |
| chr2-174362485-INS-6475 | chr2-174362486-INS-6475 |
| chr2-174363062-INS-358 | chr2-174363063-INS-358 |
| chr2-174370391-INS-73 | chr2-174370391-INS-73 |

| | |
|---|---|
| chr2-174404611-INS-97 | chr2-174404611-INS-97 |
| chr2-174416684-INS-60 | chr2-174416684-INS-60 |
| chr2-174417864-INS-78 | chr2-174417841-INS-104 |
| chr2-174422729-INS-60 | chr2-174422729-INS-60 |
| chr2-174425622-INS-65 | chr2-174425623-INS-60 |
| chr2-174438800-INS-64 | chr2-174438800-INS-52 |
| chr2-174475043-INS-74 | chr2-174475043-INS-111 |
| chr2-174491074-INS-140 | chr2-174491052-INS-157 |
| chr2-174522287-INS-86 | chr2-174522288-INS-66 |
| chr2-174522870-INS-81 | chr2-174522870-INS-108 |
| chr2-174532733-INS-90 | chr2-174532733-INS-100 |
| chr2-174567216-INS-61 | chr2-174567216-INS-61 |
| chr2-174581322-INS-220 | chr2-174581322-INS-220 |
| chr2-174591206-INS-408 | chr2-174591206-INS-409 |
| chr2-174637661-INS-58 | chr2-174637661-INS-58 |
| chr2-174648961-INS-7104 | chr2-174648962-INS-7104 |
| chr2-174654604-INS-306 | chr2-174654604-INS-306 |
| chr2-174659473-INS-64 | chr2-174659473-INS-62 |
| chr2-174671788-INS-53 | chr2-174671788-INS-54 |
| chr2-174688828-INS-1901 | chr2-174688829-INS-1901 |
| chr2-174722414-INS-74 | chr2-174722414-INS-74 |
| chr2-174744893-INS-56 | chr2-174744893-INS-64 |
| chr2-174753156-INS-7083 | chr2-174753157-INS-7083 |
| chr2-174783752-INS-52 | chr2-174783757-INS-88 |
| chr2-174785974-INS-1896 | chr2-174785975-INS-1896 |
| chr2-175054084-INS-476 | chr2-175054088-INS-484 |
| chr2-177756325-INS-3276 | chr2-177756326-INS-3279 |
| chr2-177768634-INS-448 | chr2-177768634-INS-448 |
| chr2-177774832-INS-1710 | chr2-177774833-INS-1715 |
| chr2-177817814-INS-90 | chr2-177817820-INS-96 |
| chr2-177820066-INS-450 | chr2-177820066-INS-450 |
| chr2-177821128-INS-60 | chr2-177821128-INS-60 |
| chr2-177846173-INS-51 | chr2-177846173-INS-51 |
| chr2-177847290-INS-3201 | chr2-177847291-INS-3201 |
| chr2-177857094-INS-53 | chr2-177857080-INS-84 |
| chr2-177870432-INS-58 | chr2-177870432-INS-66 |
| chr2-177880328-INS-123 | chr2-177880332-INS-123 |
| chr2-177890008-INS-92 | chr2-177890008-INS-92 |
| chr2-177935568-INS-293 | chr2-177935568-INS-293 |
| chr2-177984618-INS-2588 | chr2-177984619-INS-2588 |
| chr2-178014598-INS-4145 | chr2-178014599-INS-4146 |
| chr2-178028240-INS-646 | chr2-178028240-INS-646 |
| chr2-178034614-INS-177 | chr2-178034614-INS-179 |
| chr2-178070902-INS-561 | chr2-178070902-INS-561 |
| chr2-178095220-INS-264 | chr2-178095220-INS-270 |
| chr2-178112559-INS-79 | chr2-178112559-INS-79 |
| chr2-178131622-INS-54 | chr2-178131622-INS-56 |
| chr2-178136251-INS-748 | chr2-178136251-INS-748 |
| chr2-178166710-INS-203 | chr2-178166710-INS-159 |
| chr2-178169152-INS-3581 | chr2-178169153-INS-3581 |
| chr2-178173045-INS-66 | chr2-178173045-INS-99 |
| chr2-178175237-INS-65 | chr2-178175200-INS-103 |
| chr2-178208414-INS-63 | chr2-178208414-INS-84 |
| chr2-178219061-INS-114 | chr2-178219061-INS-92 |
| chr2-178225544-INS-120 | chr2-178225544-INS-120 |
| chr2-178241074-INS-510 | chr2-178241074-INS-509 |
| chr2-178254295-INS-52 | chr2-178254315-INS-76 |
| chr2-178259207-INS-64 | chr2-178259207-INS-64 |
| chr2-178282533-INS-162 | chr2-178282533-INS-162 |
| chr2-178283526-INS-209 | chr2-178283526-INS-209 |
| chr2-178335262-INS-55 | chr2-178335264-INS-56 |
| chr2-178338320-INS-80 | chr2-178338320-INS-80 |
| chr2-178343078-INS-165 | chr2-178343078-INS-165 |
| chr2-178391390-INS-68 | chr2-178391390-INS-68 |
| chr2-178394631-INS-399 | chr2-178394631-INS-399 |
| chr2-178398388-INS-964 | chr2-178398417-INS-975 |
| chr2-178412034-INS-87 | chr2-178412044-INS-71 |
| chr2-178422778-INS-74 | chr2-178422778-INS-74 |
| chr2-178438163-INS-60 | chr2-178438163-INS-60 |
| chr2-178469334-INS-831 | chr2-178469334-INS-831 |
| chr2-178535481-INS-66 | chr2-178535527-INS-54 |
| chr2-178541130-INS-110 | chr2-178541130-INS-110 |
| chr2-178584565-INS-78 | chr2-178584565-INS-78 |
| chr2-178591461-INS-102 | chr2-178591475-INS-103 |
| chr2-178618734-INS-61 | chr2-178618734-INS-61 |
| chr2-178626047-INS-345 | chr2-178626047-INS-345 |
| chr2-178631274-INS-63 | chr2-178631274-INS-63 |

| | |
|---|---|
| chr2-178637676-INS-60 | chr2-178637676-INS-60 |
| chr2-178642197-INS-134 | chr2-178642197-INS-134 |
| chr2-178647484-INS-88 | chr2-178647484-INS-88 |
| chr2-178648628-INS-120 | chr2-178648628-INS-128 |
| chr2-178676775-INS-198 | chr2-178676775-INS-198 |
| chr2-178689177-INS-54 | chr2-178689177-INS-68 |
| chr2-178698545-INS-113 | chr2-178698530-INS-123 |
| chr2-178703088-INS-105 | chr2-178703088-INS-105 |
| chr2-178704936-INS-52 | chr2-178704936-INS-52 |
| chr2-178771350-INS-382 | chr2-178771361-INS-389 |
| chr2-178773309-INS-56 | chr2-178773309-INS-56 |
| chr2-178807125-INS-52 | chr2-178807086-INS-55 |
| chr2-178825215-INS-101 | chr2-178825205-INS-113 |
| chr2-178845238-INS-440 | chr2-178845238-INS-438 |
| chr2-178888873-INS-60 | chr2-178888907-INS-60 |
| chr2-178888873-INS-906 | chr2-178888873-INS-906 |
| chr2-178895955-INS-1328 | chr2-178895956-INS-1328 |
| chr2-178900277-INS-81 | chr2-178900277-INS-86 |
| chr2-178968846-INS-66 | chr2-178968845-INS-77 |
| chr2-178980802-INS-84 | chr2-178980802-INS-84 |
| chr2-178992796-INS-83 | chr2-178992794-INS-84 |
| chr2-178996696-INS-56 | chr2-178996696-INS-56 |
| chr2-179002428-INS-93 | chr2-179002428-INS-93 |
| chr2-179030212-INS-58 | chr2-179030212-INS-58 |
| chr2-179031750-INS-55 | chr2-179031740-INS-67 |
| chr2-179068289-INS-140 | chr2-179068289-INS-120 |
| chr2-179068327-INS-7279 | chr2-179068328-INS-7279 |
| chr2-179069663-INS-801 | chr2-179069663-INS-806 |
| chr2-179090639-INS-64 | chr2-179090639-INS-64 |
| chr2-179092364-INS-60 | chr2-179092364-INS-64 |
| chr2-179112780-INS-56 | chr2-179112772-INS-92 |
| chr2-179122224-INS-54 | chr2-179122224-INS-54 |
| chr2-179153367-INS-104 | chr2-179153377-INS-112 |
| chr2-179155648-INS-152 | chr2-179155648-INS-152 |
| chr2-179180254-INS-70 | chr2-179180254-INS-70 |
| chr2-179213431-INS-75 | chr2-179213431-INS-85 |
| chr2-179237409-INS-475 | chr2-179237409-INS-436 |
| chr2-179242951-INS-102 | chr2-179242948-INS-105 |
| chr2-179253882-INS-100 | chr2-179253871-INS-80 |
| chr2-179265366-INS-377 | chr2-179265366-INS-377 |
| chr2-179267677-INS-700 | chr2-179267677-INS-690 |
| chr2-179280600-INS-68 | chr2-179280600-INS-68 |
| chr2-179296933-INS-60 | chr2-179296933-INS-60 |
| chr2-179301763-INS-485 | chr2-179301763-INS-485 |
| chr2-179303098-INS-136 | chr2-179303098-INS-134 |
| chr2-179330488-INS-108 | chr2-179330488-INS-113 |
| chr2-179345010-INS-87 | chr2-179345010-INS-87 |
| chr2-179358319-INS-108 | chr2-179358319-INS-108 |
| chr2-179369024-INS-76 | chr2-179368996-INS-102 |
| chr2-179419125-INS-88 | chr2-179419122-INS-116 |
| chr2-179424442-INS-202 | chr2-179424442-INS-202 |
| chr2-179441540-INS-70 | chr2-179441542-INS-66 |
| chr2-179444021-INS-66 | chr2-179444003-INS-60 |
| chr2-179449278-INS-64 | chr2-179449276-INS-78 |
| chr2-179455607-INS-96 | chr2-179455607-INS-96 |
| chr2-179459740-INS-64 | chr2-179459740-INS-64 |
| chr2-179489959-INS-140 | chr2-179489960-INS-138 |
| chr2-179496396-INS-204 | chr2-179496396-INS-205 |
| chr2-179503596-INS-105 | chr2-179503596-INS-65 |
| chr2-179520818-INS-1443 | chr2-179520819-INS-1443 |
| chr2-179526989-INS-62 | chr2-179526985-INS-66 |
| chr2-179545589-INS-51 | chr2-179545589-INS-51 |
| chr2-179593690-INS-52 | chr2-179593690-INS-52 |
| chr2-179600396-INS-99 | chr2-179600425-INS-118 |
| chr2-179604980-INS-211 | chr2-179604980-INS-211 |
| chr2-179610995-INS-57 | chr2-179610997-INS-53 |
| chr2-179631969-INS-52 | chr2-179631964-INS-55 |
| chr2-179634787-INS-194 | chr2-179634787-INS-193 |
| chr2-179637872-INS-227 | chr2-179637872-INS-227 |
| chr2-179647226-INS-889 | chr2-179647226-INS-895 |
| chr2-179649877-INS-260 | chr2-179649877-INS-260 |
| chr2-179654926-INS-400 | chr2-179654926-INS-400 |
| chr2-179664887-INS-60 | chr2-179664887-INS-60 |
| chr2-179692843-INS-58 | chr2-179692843-INS-50 |
| chr2-179722516-INS-62 | chr2-179722516-INS-62 |
| chr2-179724717-INS-60 | chr2-179724718-INS-64 |
| chr2-179765396-INS-96 | chr2-179765396-INS-96 |

| | |
|---|---|
| chr2-179798017-INS-154 | chr2-179798026-INS-115 |
| chr2-179841463-INS-69 | chr2-179841450-INS-72 |
| chr2-179847044-INS-498 | chr2-179847044-INS-498 |
| chr2-179853563-INS-100 | chr2-179853563-INS-100 |
| chr2-179874163-INS-71 | chr2-179874199-INS-84 |
| chr2-179884333-INS-72 | chr2-179884329-INS-76 |
| chr2-179954252-INS-418 | chr2-179954252-INS-418 |
| chr2-179962098-INS-191 | chr2-179962095-INS-214 |
| chr2-179980864-INS-192 | chr2-179980864-INS-192 |
| chr2-179982068-INS-54 | chr2-179982068-INS-56 |
| chr2-180167913-INS-131 | chr2-180167913-INS-133 |
| chr2-180173325-INS-246 | chr2-180173304-INS-266 |
| chr2-180203884-INS-2944 | chr2-180203885-INS-2944 |
| chr2-180212954-INS-186 | chr2-180212954-INS-202 |
| chr2-180259694-INS-248 | chr2-180259694-INS-257 |
| chr2-180269630-INS-113 | chr2-180269628-INS-86 |
| chr2-180336925-INS-222 | chr2-180336925-INS-220 |
| chr2-180349438-INS-175 | chr2-180349438-INS-176 |
| chr2-180385302-INS-202 | chr2-180385302-INS-202 |
| chr2-180386564-INS-70 | chr2-180386564-INS-72 |
| chr2-180391848-INS-56 | chr2-180391848-INS-56 |
| chr2-180426955-INS-684 | chr2-180426955-INS-684 |
| chr2-180469196-INS-225 | chr2-180469196-INS-226 |
| chr2-180472302-INS-54 | chr2-180472302-INS-54 |
| chr2-180510455-INS-56 | chr2-180510455-INS-56 |
| chr2-180511507-INS-165 | chr2-180511505-INS-172 |
| chr2-180550044-INS-205 | chr2-180550044-INS-205 |
| chr2-180569925-INS-165 | chr2-180569925-INS-165 |
| chr2-180582534-INS-96 | chr2-180582537-INS-116 |
| chr2-180770710-INS-196 | chr2-180770707-INS-211 |
| chr2-180784435-INS-68 | chr2-180784454-INS-66 |
| chr2-180808245-INS-211 | chr2-180808260-INS-208 |
| chr2-180832024-INS-226 | chr2-180832024-INS-226 |
| chr2-180839676-INS-124 | chr2-180839676-INS-124 |
| chr2-180847293-INS-58 | chr2-180847293-INS-53 |
| chr2-180890344-INS-78 | chr2-180890344-INS-50 |
| chr2-180913715-INS-191 | chr2-180913715-INS-191 |
| chr2-180977571-INS-578 | chr2-180977571-INS-578 |
| chr2-180978585-INS-173 | chr2-180978584-INS-175 |
| chr2-181010774-INS-59 | chr2-181010777-INS-55 |
| chr2-181012602-INS-211 | chr2-181012602-INS-211 |
| chr2-181013883-INS-283 | chr2-181013891-INS-275 |
| chr2-181045868-INS-66 | chr2-181045868-INS-67 |
| chr2-181085568-INS-216 | chr2-181085568-INS-216 |
| chr2-181099504-INS-1860 | chr2-181099505-INS-1860 |
| chr2-181169829-INS-920 | chr2-181169829-INS-922 |
| chr2-181182950-INS-53 | chr2-181182940-INS-56 |
| chr2-181185404-INS-325 | chr2-181185404-INS-309 |
| chr2-181204403-INS-206 | chr2-181204403-INS-206 |
| chr2-181222311-INS-238 | chr2-181222311-INS-239 |
| chr2-181271889-INS-6454 | chr2-181271890-INS-6454 |
| chr2-181333184-INS-58 | chr2-181333196-INS-58 |
| chr2-181351156-INS-561 | chr2-181351156-INS-563 |
| chr2-181373256-INS-115 | chr2-181373256-INS-68 |
| chr2-181419306-INS-192 | chr2-181419306-INS-192 |
| chr2-181434051-INS-63 | chr2-181434051-INS-63 |
| chr2-181455285-INS-202 | chr2-181455285-INS-202 |
| chr2-181472167-INS-4198 | chr2-181472168-INS-4198 |
| chr2-181507879-INS-531 | chr2-181507879-INS-531 |
| chr2-181535039-INS-181 | chr2-181535039-INS-181 |
| chr2-181536737-INS-74 | chr2-181536738-INS-80 |
| chr2-181558695-INS-59 | chr2-181558695-INS-56 |
| chr2-181564940-INS-7175 | chr2-181564941-INS-7175 |
| chr2-181574319-INS-58 | chr2-181574316-INS-50 |
| chr2-181576486-INS-416 | chr2-181576486-INS-416 |
| chr3-3010414-INS-223 | chr3-3010402-INS-224 |
| chr3-3016594-INS-443 | chr3-3016594-INS-443 |
| chr3-3024944-INS-269 | chr3-3024944-INS-269 |
| chr3-3063678-INS-313 | chr3-3063678-INS-313 |
| chr3-3080756-INS-63 | chr3-3080756-INS-63 |
| chr3-3205809-INS-60 | chr3-3205844-INS-55 |
| chr3-3211840-INS-2600 | chr3-3211841-INS-2600 |
| chr3-3297351-INS-4011 | chr3-3297352-INS-4011 |
| chr3-3384941-INS-208 | chr3-3384941-INS-208 |
| chr3-3444701-INS-471 | chr3-3444699-INS-474 |
| chr3-3545306-INS-70 | chr3-3545306-INS-86 |
| chr3-3707532-INS-928 | chr3-3707532-INS-926 |

| | |
|---|---|
| chr3-3713618-INS-97 | chr3-3713618-INS-97 |
| chr3-3729225-INS-123 | chr3-3729225-INS-147 |
| chr3-3771079-INS-2690 | chr3-3771090-INS-2692 |
| chr3-3812193-INS-69 | chr3-3812193-INS-69 |
| chr3-3849341-INS-2444 | chr3-3849342-INS-2444 |
| chr3-3852127-INS-95 | chr3-3852127-INS-63 |
| chr3-3918482-INS-7470 | chr3-3918483-INS-7470 |
| chr3-3924320-INS-4475 | chr3-3924321-INS-4475 |
| chr3-3936879-INS-776 | chr3-3936879-INS-776 |
| chr3-3957956-INS-7609 | chr3-3957957-INS-7609 |
| chr3-4013848-INS-2686 | chr3-4013849-INS-2688 |
| chr3-4075008-INS-104 | chr3-4075008-INS-104 |
| chr3-4126778-INS-85 | chr3-4126778-INS-85 |
| chr3-4127517-INS-6369 | chr3-4127518-INS-6369 |
| chr3-4201810-INS-3739 | chr3-4201811-INS-3739 |
| chr3-4271788-INS-517 | chr3-4271788-INS-517 |
| chr3-4280878-INS-62 | chr3-4280878-INS-62 |
| chr3-4283161-INS-1491 | chr3-4283162-INS-1490 |
| chr3-4308375-INS-399 | chr3-4308375-INS-399 |
| chr3-4368785-INS-54 | chr3-4368785-INS-58 |
| chr3-4445258-INS-68 | chr3-4445258-INS-84 |
| chr3-4475408-INS-615 | chr3-4475408-INS-617 |
| chr3-4523208-INS-6455 | chr3-4523209-INS-6455 |
| chr3-4547244-INS-80 | chr3-4547262-INS-80 |
| chr3-4555797-INS-3361 | chr3-4555798-INS-3364 |
| chr3-4572979-INS-6215 | chr3-4572980-INS-6214 |
| chr3-4606747-INS-2241 | chr3-4606748-INS-2239 |
| chr3-4632581-INS-6562 | chr3-4632582-INS-6562 |
| chr3-4633596-INS-218 | chr3-4633597-INS-218 |
| chr3-4675539-INS-59 | chr3-4675539-INS-59 |
| chr3-4676206-INS-1488 | chr3-4676206-INS-1489 |
| chr3-4685607-INS-105 | chr3-4685607-INS-105 |
| chr3-4731985-INS-1043 | chr3-4731986-INS-1043 |
| chr3-4746978-INS-65 | chr3-4746978-INS-65 |
| chr3-4748844-INS-4707 | chr3-4748845-INS-4707 |
| chr3-4775615-INS-86 | chr3-4775615-INS-86 |
| chr3-4856904-INS-68 | chr3-4856904-INS-68 |
| chr3-4911561-INS-55 | chr3-4911511-INS-65 |
| chr3-4918680-INS-58 | chr3-4918676-INS-57 |
| chr3-4946975-INS-4427 | chr3-4946976-INS-4427 |
| chr3-4994170-INS-670 | chr3-4994170-INS-675 |
| chr3-4996293-INS-494 | chr3-4996293-INS-494 |
| chr3-5012629-INS-2343 | chr3-5012630-INS-2340 |
| chr3-5019142-INS-1588 | chr3-5019143-INS-1589 |
| chr3-5025865-INS-498 | chr3-5025865-INS-498 |
| chr3-5044269-INS-7473 | chr3-5044270-INS-7473 |
| chr3-5064769-INS-246 | chr3-5064787-INS-247 |
| chr3-5080355-INS-245 | chr3-5080355-INS-245 |
| chr3-5096776-INS-224 | chr3-5096776-INS-224 |
| chr3-5146642-INS-75 | chr3-5146642-INS-75 |
| chr3-5147586-INS-971 | chr3-5147586-INS-973 |
| chr3-5156142-INS-92 | chr3-5156142-INS-92 |
| chr3-5194953-INS-60 | chr3-5194991-INS-54 |
| chr3-5208699-INS-64 | chr3-5208741-INS-72 |
| chr3-5215641-INS-203 | chr3-5215641-INS-203 |
| chr3-5226936-INS-414 | chr3-5226921-INS-412 |
| chr3-5237039-INS-6431 | chr3-5237040-INS-6432 |
| chr3-5240514-INS-63 | chr3-5240514-INS-66 |
| chr3-5261949-INS-6732 | chr3-5261950-INS-6730 |
| chr3-5262651-INS-86 | chr3-5262651-INS-86 |
| chr3-5264716-INS-457 | chr3-5264716-INS-457 |
| chr3-5306356-INS-6724 | chr3-5306357-INS-6728 |
| chr3-5320863-INS-596 | chr3-5320863-INS-599 |
| chr3-5382160-INS-99 | chr3-5382160-INS-99 |
| chr3-5416175-INS-5549 | chr3-5416176-INS-5549 |
| chr3-5449638-INS-73 | chr3-5449638-INS-73 |
| chr3-5462649-INS-70 | chr3-5462651-INS-74 |
| chr3-5482963-INS-177 | chr3-5482963-INS-177 |
| chr3-5607564-INS-724 | chr3-5607564-INS-724 |
| chr3-5612956-INS-342 | chr3-5612956-INS-342 |
| chr3-5618826-INS-66 | chr3-5618827-INS-58 |
| chr3-5646383-INS-6190 | chr3-5646383-INS-6190 |
| chr3-5648698-INS-1024 | chr3-5648699-INS-1024 |
| chr3-5679194-INS-124 | chr3-5679225-INS-101 |
| chr3-5708034-INS-278 | chr3-5708034-INS-278 |
| chr3-5759826-INS-517 | chr3-5759826-INS-517 |
| chr3-5792723-INS-1015 | chr3-5792724-INS-1015 |

| | |
|---|---|
| chr3-5814488-INS-3006 | chr3-5814489-INS-3006 |
| chr3-5820944-INS-294 | chr3-5820944-INS-295 |
| chr3-5833708-INS-6200 | chr3-5833709-INS-6200 |
| chr3-5847765-INS-1886 | chr3-5847766-INS-1887 |
| chr3-5859644-INS-3996 | chr3-5859645-INS-3996 |
| chr3-5874503-INS-285 | chr3-5874503-INS-285 |
| chr3-5883864-INS-6547 | chr3-5883865-INS-6547 |
| chr3-5897171-INS-489 | chr3-5897171-INS-490 |
| chr3-5920591-INS-343 | chr3-5920591-INS-343 |
| chr3-5921949-INS-72 | chr3-5921949-INS-74 |
| chr3-5929891-INS-84 | chr3-5929894-INS-68 |
| chr3-5960035-INS-59 | chr3-5960035-INS-59 |
| chr3-5969134-INS-522 | chr3-5969133-INS-503 |
| chr3-5992749-INS-320 | chr3-5992749-INS-320 |
| chr3-5998836-INS-173 | chr3-5998836-INS-178 |
| chr3-6045024-INS-170 | chr3-6045024-INS-194 |
| chr3-6051452-INS-475 | chr3-6051452-INS-475 |
| chr3-6059462-INS-549 | chr3-6059462-INS-549 |
| chr3-6069777-INS-372 | chr3-6069777-INS-348 |
| chr3-6082145-INS-2068 | chr3-6082146-INS-2068 |
| chr3-6087189-INS-2551 | chr3-6087190-INS-2552 |
| chr3-6090713-INS-6635 | chr3-6090714-INS-6635 |
| chr3-6091589-INS-4521 | chr3-6091590-INS-4521 |
| chr3-6092161-INS-330 | chr3-6092161-INS-327 |
| chr3-6094438-INS-52 | chr3-6094438-INS-52 |
| chr3-6113136-INS-82 | chr3-6113136-INS-82 |
| chr3-6115569-INS-66 | chr3-6115569-INS-66 |
| chr3-6137395-INS-56 | chr3-6137399-INS-62 |
| chr3-6188849-INS-188 | chr3-6188849-INS-174 |
| chr3-6230712-INS-942 | chr3-6230712-INS-931 |
| chr3-6233069-INS-182 | chr3-6233069-INS-185 |
| chr3-6302857-INS-1491 | chr3-6302858-INS-1490 |
| chr3-6313988-INS-6443 | chr3-6313989-INS-6443 |
| chr3-6393043-INS-1886 | chr3-6393044-INS-1886 |
| chr3-6419431-INS-126 | chr3-6419431-INS-126 |
| chr3-6458873-INS-420 | chr3-6458873-INS-420 |
| chr3-6534538-INS-213 | chr3-6534538-INS-213 |
| chr3-6543623-INS-208 | chr3-6543623-INS-208 |
| chr3-6547953-INS-4541 | chr3-6547954-INS-4542 |
| chr3-6559015-INS-78 | chr3-6559045-INS-91 |
| chr3-6574165-INS-6472 | chr3-6574166-INS-6472 |
| chr3-6589545-INS-326 | chr3-6589545-INS-310 |
| chr3-6591030-INS-67 | chr3-6591030-INS-67 |
| chr3-6600514-INS-203 | chr3-6600486-INS-203 |
| chr3-6610320-INS-206 | chr3-6610320-INS-206 |
| chr3-6634649-INS-6403 | chr3-6634650-INS-6403 |
| chr3-6647626-INS-64 | chr3-6647626-INS-52 |
| chr3-6650682-INS-54 | chr3-6650656-INS-52 |
| chr3-6663773-INS-56 | chr3-6663773-INS-80 |
| chr3-6692457-INS-408 | chr3-6692457-INS-408 |
| chr3-6700702-INS-98 | chr3-6700702-INS-98 |
| chr3-6701717-INS-6871 | chr3-6701718-INS-6871 |
| chr3-6754082-INS-2685 | chr3-6754083-INS-2685 |
| chr3-6758476-INS-60 | chr3-6758476-INS-69 |
| chr3-6761511-INS-64 | chr3-6761511-INS-64 |
| chr3-6781092-INS-6674 | chr3-6781093-INS-6676 |
| chr3-6787752-INS-101 | chr3-6787752-INS-99 |
| chr3-6794874-INS-78 | chr3-6794875-INS-78 |
| chr3-6817715-INS-206 | chr3-6817715-INS-207 |
| chr3-6848496-INS-439 | chr3-6848497-INS-439 |
| chr3-6854067-INS-6343 | chr3-6854068-INS-6344 |
| chr3-6865045-INS-102 | chr3-6865045-INS-146 |
| chr3-6870396-INS-105 | chr3-6870391-INS-108 |
| chr3-6881226-INS-2249 | chr3-6881230-INS-2246 |
| chr3-6882280-INS-122 | chr3-6882281-INS-123 |
| chr3-6899561-INS-6249 | chr3-6899562-INS-6249 |
| chr3-6910842-INS-1867 | chr3-6910843-INS-1867 |
| chr3-6910843-INS-387 | chr3-6910843-INS-387 |
| chr3-6915313-INS-212 | chr3-6915313-INS-212 |
| chr3-6925430-INS-51 | chr3-6925411-INS-80 |
| chr3-6943283-INS-5816 | chr3-6943284-INS-5816 |
| chr3-6987685-INS-4558 | chr3-6987686-INS-4558 |
| chr3-7001791-INS-6399 | chr3-7001792-INS-6400 |
| chr3-7011992-INS-64 | chr3-7011992-INS-64 |
| chr3-7012670-INS-617 | chr3-7012670-INS-617 |
| chr3-7093244-INS-286 | chr3-7093240-INS-278 |
| chr3-7117749-INS-81 | chr3-7117749-INS-81 |

| | |
|---|---|
| chr3-7220512-INS-66 | chr3-7220512-INS-66 |
| chr3-7224538-INS-62 | chr3-7224538-INS-56 |
| chr3-7226005-INS-6148 | chr3-7226006-INS-6148 |
| chr3-7279077-INS-857 | chr3-7279077-INS-869 |
| chr3-7294429-INS-84 | chr3-7294433-INS-96 |
| chr3-7302680-INS-75 | chr3-7302680-INS-75 |
| chr3-7304428-INS-549 | chr3-7304428-INS-549 |
| chr3-7342201-INS-774 | chr3-7342201-INS-774 |
| chr3-7358416-INS-331 | chr3-7358416-INS-330 |
| chr3-7394209-INS-313 | chr3-7394210-INS-316 |
| chr3-7394958-INS-72 | chr3-7394958-INS-72 |
| chr3-7399897-INS-132 | chr3-7399897-INS-128 |
| chr3-7421275-INS-406 | chr3-7421276-INS-406 |
| chr3-7422310-INS-395 | chr3-7422310-INS-395 |
| chr3-7424268-INS-84 | chr3-7424266-INS-84 |
| chr3-7427411-INS-80 | chr3-7427407-INS-64 |
| chr3-7436833-INS-485 | chr3-7436833-INS-485 |
| chr3-7444283-INS-461 | chr3-7444283-INS-459 |
| chr3-7445483-INS-80 | chr3-7445474-INS-92 |
| chr3-7486706-INS-930 | chr3-7486706-INS-931 |
| chr3-7493077-INS-60 | chr3-7493060-INS-67 |
| chr3-7522396-INS-74 | chr3-7522392-INS-78 |
| chr3-7533572-INS-55 | chr3-7533557-INS-75 |
| chr3-7539702-INS-1957 | chr3-7539703-INS-1957 |
| chr3-7546151-INS-3514 | chr3-7546152-INS-3514 |
| chr3-7558319-INS-52 | chr3-7558307-INS-57 |
| chr3-7563094-INS-91 | chr3-7563094-INS-91 |
| chr3-7573387-INS-200 | chr3-7573387-INS-168 |
| chr3-7584615-INS-2988 | chr3-7584616-INS-2989 |
| chr3-7604572-INS-52 | chr3-7604572-INS-52 |
| chr3-7620910-INS-67 | chr3-7620888-INS-65 |
| chr3-7646146-INS-78 | chr3-7646142-INS-112 |
| chr3-7658721-INS-188 | chr3-7658721-INS-188 |
| chr3-7661439-INS-173 | chr3-7661439-INS-174 |
| chr3-7662271-INS-6651 | chr3-7662272-INS-6651 |
| chr3-7667417-INS-346 | chr3-7667417-INS-346 |
| chr3-7692543-INS-203 | chr3-7692543-INS-203 |
| chr3-7717840-INS-56 | chr3-7717840-INS-56 |
| chr3-7718639-INS-427 | chr3-7718639-INS-427 |
| chr3-7735656-INS-448 | chr3-7735656-INS-448 |
| chr3-7752078-INS-141 | chr3-7752082-INS-156 |
| chr3-7755995-INS-7440 | chr3-7755996-INS-7440 |
| chr3-7768513-INS-68 | chr3-7768514-INS-68 |
| chr3-7784531-INS-7130 | chr3-7784532-INS-7130 |
| chr3-7785942-INS-6598 | chr3-7785943-INS-6598 |
| chr3-7792175-INS-161 | chr3-7792176-INS-161 |
| chr3-7797874-INS-201 | chr3-7797874-INS-201 |
| chr3-7821466-INS-246 | chr3-7821467-INS-196 |
| chr3-7844017-INS-55 | chr3-7843981-INS-70 |
| chr3-7849421-INS-219 | chr3-7849421-INS-219 |
| chr3-7882027-INS-62 | chr3-7882027-INS-62 |
| chr3-7898160-INS-129 | chr3-7898160-INS-129 |
| chr3-7907131-INS-126 | chr3-7907112-INS-164 |
| chr3-7909742-INS-410 | chr3-7909742-INS-405 |
| chr3-7916925-INS-1666 | chr3-7916926-INS-1666 |
| chr3-7936579-INS-905 | chr3-7936579-INS-905 |
| chr3-7953196-INS-61 | chr3-7953196-INS-63 |
| chr3-7955714-INS-3383 | chr3-7955715-INS-3418 |
| chr3-7968967-INS-84 | chr3-7968967-INS-90 |
| chr3-7975444-INS-424 | chr3-7975444-INS-424 |
| chr3-7983682-INS-104 | chr3-7983682-INS-104 |
| chr3-7990463-INS-60 | chr3-7990463-INS-60 |
| chr3-8004200-INS-2417 | chr3-8004201-INS-2415 |
| chr3-8004424-INS-6473 | chr3-8004425-INS-6473 |
| chr3-8014774-INS-95 | chr3-8014774-INS-95 |
| chr3-8017250-INS-82 | chr3-8017250-INS-80 |
| chr3-8018175-INS-2064 | chr3-8018176-INS-2064 |
| chr3-8020374-INS-214 | chr3-8020374-INS-214 |
| chr3-8024635-INS-99 | chr3-8024635-INS-99 |
| chr3-8031544-INS-150 | chr3-8031544-INS-175 |
| chr3-8039351-INS-527 | chr3-8039351-INS-527 |
| chr3-8059471-INS-892 | chr3-8059471-INS-892 |
| chr3-8074045-INS-315 | chr3-8074024-INS-336 |
| chr3-8076002-INS-67 | chr3-8076002-INS-67 |
| chr3-8091171-INS-108 | chr3-8091171-INS-106 |
| chr3-8100193-INS-111 | chr3-8100193-INS-111 |
| chr3-8123759-INS-54 | chr3-8123760-INS-57 |

| | |
|---|---|
| chr3-8156847-INS-436 | chr3-8156847-INS-483 |
| chr3-8171167-INS-59 | chr3-8171167-INS-81 |
| chr3-8212451-INS-448 | chr3-8212451-INS-437 |
| chr3-8268761-INS-6459 | chr3-8268762-INS-6460 |
| chr3-8309492-INS-59 | chr3-8309530-INS-54 |
| chr3-8311291-INS-59 | chr3-8311291-INS-59 |
| chr3-8321328-INS-184 | chr3-8321328-INS-184 |
| chr3-8330869-INS-119 | chr3-8330869-INS-119 |
| chr3-8348996-INS-665 | chr3-8348996-INS-666 |
| chr3-8353190-INS-333 | chr3-8353190-INS-335 |
| chr3-8361112-INS-59 | chr3-8361112-INS-59 |
| chr3-8380290-INS-54 | chr3-8380290-INS-54 |
| chr3-8382496-INS-223 | chr3-8382506-INS-200 |
| chr3-8403432-INS-198 | chr3-8403432-INS-198 |
| chr3-8407088-INS-615 | chr3-8407088-INS-615 |
| chr3-8407794-INS-199 | chr3-8407794-INS-199 |
| chr3-8420392-INS-5328 | chr3-8420393-INS-5328 |
| chr3-8475443-INS-6256 | chr3-8475444-INS-6256 |
| chr3-8516397-INS-212 | chr3-8516397-INS-212 |
| chr3-8529062-INS-1135 | chr3-8529063-INS-1135 |
| chr3-8538710-INS-275 | chr3-8538710-INS-276 |
| chr3-8539756-INS-177 | chr3-8539756-INS-193 |
| chr3-8570180-INS-702 | chr3-8570180-INS-688 |
| chr3-8587934-INS-316 | chr3-8587934-INS-316 |
| chr3-8589584-INS-199 | chr3-8589584-INS-198 |
| chr3-8591541-INS-492 | chr3-8591541-INS-496 |
| chr3-8595036-INS-87 | chr3-8595036-INS-87 |
| chr3-8600052-INS-200 | chr3-8600052-INS-176 |
| chr3-8617510-INS-67 | chr3-8617510-INS-67 |
| chr3-8628027-INS-60 | chr3-8628027-INS-65 |
| chr3-8631443-INS-68 | chr3-8631443-INS-68 |
| chr3-8691951-INS-52 | chr3-8691951-INS-56 |
| chr3-8701448-INS-70 | chr3-8701449-INS-72 |
| chr3-8741102-INS-52 | chr3-8741116-INS-62 |
| chr3-8753964-INS-55 | chr3-8753964-INS-51 |
| chr3-8760879-INS-72 | chr3-8760879-INS-72 |
| chr3-8835959-INS-473 | chr3-8835963-INS-502 |
| chr3-8844213-INS-132 | chr3-8844239-INS-92 |
| chr3-8864616-INS-53 | chr3-8864616-INS-51 |
| chr3-8922052-INS-120 | chr3-8922052-INS-120 |
| chr3-8925955-INS-626 | chr3-8925955-INS-640 |
| chr3-8928879-INS-196 | chr3-8928883-INS-190 |
| chr3-8954720-INS-280 | chr3-8954720-INS-280 |
| chr3-8971742-INS-199 | chr3-8971785-INS-199 |
| chr3-8980837-INS-116 | chr3-8980837-INS-116 |
| chr3-8986698-INS-96 | chr3-8986698-INS-96 |
| chr3-9000885-INS-5547 | chr3-9000886-INS-5547 |
| chr3-9005029-INS-69 | chr3-9005032-INS-52 |
| chr3-9021903-INS-70 | chr3-9021903-INS-70 |
| chr3-9029815-INS-61 | chr3-9029815-INS-58 |
| chr3-9036665-INS-122 | chr3-9036665-INS-94 |
| chr3-9038764-INS-249 | chr3-9038762-INS-270 |
| chr3-9043415-INS-80 | chr3-9043382-INS-96 |
| chr3-9055132-INS-383 | chr3-9055132-INS-383 |
| chr3-9058615-INS-179 | chr3-9058615-INS-179 |
| chr3-9077473-INS-50 | chr3-9077498-INS-64 |
| chr3-9084533-INS-6458 | chr3-9084534-INS-6459 |
| chr3-9107128-INS-208 | chr3-9107128-INS-208 |
| chr3-9115826-INS-51 | chr3-9115826-INS-51 |
| chr3-9124710-INS-278 | chr3-9124712-INS-277 |
| chr3-9132261-INS-220 | chr3-9132253-INS-214 |
| chr3-9135078-INS-70 | chr3-9135098-INS-70 |
| chr3-9137974-INS-226 | chr3-9137974-INS-218 |
| chr3-9147911-INS-55 | chr3-9147911-INS-55 |
| chr3-9153271-INS-134 | chr3-9153271-INS-142 |
| chr3-9171898-INS-1888 | chr3-9171899-INS-1888 |
| chr3-9206262-INS-211 | chr3-9206262-INS-213 |
| chr3-9207684-INS-213 | chr3-9207684-INS-210 |
| chr3-9209964-INS-77 | chr3-9209964-INS-77 |
| chr3-9228090-INS-201 | chr3-9228090-INS-198 |
| chr3-9260033-INS-76 | chr3-9260033-INS-76 |
| chr3-9264324-INS-86 | chr3-9264324-INS-86 |
| chr3-9274648-INS-178 | chr3-9274648-INS-179 |
| chr3-9275732-INS-52 | chr3-9275765-INS-52 |
| chr3-9303479-INS-73 | chr3-9303479-INS-73 |
| chr3-9323547-INS-100 | chr3-9323547-INS-98 |
| chr3-9353198-INS-108 | chr3-9353198-INS-108 |

| | |
|---|---|
| chr3-9356463-INS-189 | chr3-9356464-INS-186 |
| chr3-9358291-INS-74 | chr3-9358291-INS-74 |
| chr3-9375851-INS-51 | chr3-9375851-INS-51 |
| chr3-9381757-INS-53 | chr3-9381757-INS-53 |
| chr3-9396837-INS-96 | chr3-9396837-INS-96 |
| chr3-9419154-INS-56 | chr3-9419154-INS-62 |
| chr3-9427325-INS-52 | chr3-9427325-INS-56 |
| chr3-9477363-INS-128 | chr3-9477363-INS-128 |
| chr3-9478850-INS-220 | chr3-9478850-INS-220 |
| chr3-9480903-INS-64 | chr3-9480896-INS-96 |
| chr3-9508540-INS-80 | chr3-9508540-INS-74 |
| chr3-9513997-INS-67 | chr3-9513997-INS-67 |
| chr3-9522389-INS-198 | chr3-9522389-INS-198 |
| chr3-9528412-INS-96 | chr3-9528412-INS-98 |
| chr3-9532379-INS-121 | chr3-9532379-INS-128 |
| chr3-9555115-INS-341 | chr3-9555115-INS-341 |
| chr3-9571304-INS-211 | chr3-9571304-INS-213 |
| chr3-9577777-INS-126 | chr3-9577777-INS-126 |
| chr3-9600930-INS-201 | chr3-9600930-INS-201 |
| chr3-9648347-INS-560 | chr3-9648347-INS-556 |
| chr3-9655631-INS-55 | chr3-9655631-INS-50 |
| chr3-9663556-INS-198 | chr3-9663556-INS-199 |
| chr3-9667297-INS-75 | chr3-9667297-INS-75 |
| chr3-9670059-INS-60 | chr3-9670059-INS-60 |
| chr3-9678742-INS-62 | chr3-9678745-INS-58 |
| chr3-9682430-INS-132 | chr3-9682430-INS-132 |
| chr3-9719024-INS-56 | chr3-9719024-INS-74 |
| chr3-9723562-INS-75 | chr3-9723562-INS-80 |
| chr3-9733156-INS-158 | chr3-9733156-INS-158 |
| chr3-9736982-INS-127 | chr3-9736982-INS-127 |
| chr3-9748082-INS-180 | chr3-9748082-INS-180 |
| chr3-9776948-INS-87 | chr3-9776948-INS-87 |
| chr3-9787907-INS-62 | chr3-9787937-INS-66 |
| chr3-9803969-INS-164 | chr3-9803969-INS-164 |
| chr3-9820735-INS-904 | chr3-9820735-INS-904 |
| chr3-9830157-INS-73 | chr3-9830165-INS-73 |
| chr3-9841889-INS-95 | chr3-9841889-INS-95 |
| chr3-9858506-INS-469 | chr3-9858506-INS-469 |
| chr3-9859603-INS-217 | chr3-9859603-INS-213 |
| chr3-9864018-INS-54 | chr3-9864018-INS-55 |
| chr3-9882922-INS-442 | chr3-9882922-INS-447 |
| chr3-9887298-INS-180 | chr3-9887298-INS-180 |
| chr3-9890236-INS-83 | chr3-9890236-INS-83 |
| chr3-9902741-INS-60 | chr3-9902741-INS-58 |
| chr3-9911039-INS-398 | chr3-9911039-INS-398 |
| chr3-9915114-INS-1415 | chr3-9915115-INS-1436 |
| chr3-9922931-INS-117 | chr3-9922908-INS-120 |
| chr3-9926806-INS-86 | chr3-9926829-INS-86 |
| chr3-9929107-INS-447 | chr3-9929099-INS-460 |
| chr3-9947474-INS-109 | chr3-9947474-INS-113 |
| chr3-9950391-INS-60 | chr3-9950391-INS-64 |
| chr3-9953888-INS-54 | chr3-9953888-INS-54 |
| chr3-9957286-INS-96 | chr3-9957286-INS-96 |
| chr3-9960908-INS-161 | chr3-9960908-INS-161 |
| chr3-9992678-INS-1897 | chr3-9992679-INS-1897 |
| chr3-9994609-INS-56 | chr3-9994609-INS-60 |
| chr3-10005834-INS-55 | chr3-10005835-INS-55 |
| chr3-10028140-INS-58 | chr3-10028140-INS-58 |
| chr3-10037931-INS-70 | chr3-10037931-INS-70 |
| chr3-10039878-INS-92 | chr3-10039878-INS-110 |
| chr3-10050175-INS-5160 | chr3-10050176-INS-5160 |
| chr3-10067184-INS-79 | chr3-10067184-INS-79 |
| chr3-10070895-INS-213 | chr3-10070895-INS-213 |
| chr3-10100705-INS-193 | chr3-10100705-INS-193 |
| chr3-10108372-INS-57 | chr3-10108402-INS-60 |
| chr3-10110035-INS-158 | chr3-10110035-INS-158 |
| chr3-10115434-INS-101 | chr3-10115434-INS-59 |
| chr3-10116409-INS-299 | chr3-10116409-INS-340 |
| chr3-10119555-INS-5474 | chr3-10119556-INS-5474 |
| chr3-10132837-INS-183 | chr3-10132837-INS-183 |
| chr3-10142860-INS-54 | chr3-10142860-INS-60 |
| chr3-10165452-INS-54 | chr3-10165452-INS-54 |
| chr3-10174486-INS-140 | chr3-10174480-INS-123 |
| chr3-10182744-INS-435 | chr3-10182744-INS-435 |
| chr3-10209129-INS-120 | chr3-10209129-INS-120 |
| chr3-10225590-INS-678 | chr3-10225591-INS-660 |
| chr3-10226763-INS-199 | chr3-10226763-INS-199 |

| | |
|---|---|
| chr3-10227205-INS-7383 | chr3-10227206-INS-7383 |
| chr3-10233407-INS-6598 | chr3-10233408-INS-6598 |
| chr3-10240927-INS-230 | chr3-10240927-INS-207 |
| chr3-10276547-INS-123 | chr3-10276547-INS-123 |
| chr3-10308861-INS-102 | chr3-10308861-INS-102 |
| chr3-10320181-INS-401 | chr3-10320181-INS-401 |
| chr3-10328011-INS-390 | chr3-10328011-INS-362 |
| chr3-10341490-INS-121 | chr3-10341490-INS-121 |
| chr3-10366631-INS-201 | chr3-10366631-INS-196 |
| chr3-10378403-INS-71 | chr3-10378403-INS-71 |
| chr3-10387257-INS-196 | chr3-10387298-INS-171 |
| chr3-10390258-INS-63 | chr3-10390258-INS-63 |
| chr3-10407249-INS-187 | chr3-10407249-INS-195 |
| chr3-10416630-INS-188 | chr3-10416630-INS-189 |
| chr3-10421404-INS-215 | chr3-10421404-INS-215 |
| chr3-10427157-INS-203 | chr3-10427157-INS-203 |
| chr3-10428916-INS-6432 | chr3-10428917-INS-6433 |
| chr3-10436210-INS-76 | chr3-10436210-INS-76 |
| chr3-10451511-INS-165 | chr3-10451511-INS-165 |
| chr3-10455705-INS-63 | chr3-10455719-INS-91 |
| chr3-10471681-INS-128 | chr3-10471681-INS-128 |
| chr3-10480970-INS-221 | chr3-10480970-INS-222 |
| chr3-10506236-INS-171 | chr3-10506236-INS-171 |
| chr3-10509382-INS-178 | chr3-10509382-INS-178 |
| chr3-10515750-INS-874 | chr3-10515750-INS-875 |
| chr3-10516786-INS-222 | chr3-10516786-INS-218 |
| chr3-10519422-INS-565 | chr3-10519422-INS-565 |
| chr3-10526045-INS-126 | chr3-10526045-INS-126 |
| chr3-10529611-INS-136 | chr3-10529611-INS-136 |
| chr3-10539039-INS-195 | chr3-10539039-INS-195 |
| chr3-10543260-INS-57 | chr3-10543218-INS-51 |
| chr3-10557578-INS-2060 | chr3-10557579-INS-2061 |
| chr3-10573560-INS-6703 | chr3-10573561-INS-6705 |
| chr3-10577851-INS-198 | chr3-10577852-INS-245 |
| chr3-10581497-INS-8303 | chr3-10581498-INS-8310 |
| chr3-10592794-INS-90 | chr3-10592794-INS-90 |
| chr3-10606981-INS-528 | chr3-10606981-INS-528 |
| chr3-10609185-INS-453 | chr3-10609185-INS-453 |
| chr3-10618646-INS-409 | chr3-10618646-INS-409 |
| chr3-10660229-INS-125 | chr3-10660229-INS-133 |
| chr3-10663132-INS-918 | chr3-10663132-INS-919 |
| chr3-10668111-INS-208 | chr3-10668112-INS-203 |
| chr3-10694225-INS-1884 | chr3-10694226-INS-1884 |
| chr3-10719383-INS-56 | chr3-10719385-INS-56 |
| chr3-10721805-INS-81 | chr3-10721805-INS-88 |
| chr3-10728700-INS-73 | chr3-10728700-INS-73 |
| chr3-10773975-INS-54 | chr3-10773975-INS-66 |
| chr3-10844896-INS-208 | chr3-10844896-INS-214 |
| chr3-10886910-INS-347 | chr3-10886910-INS-347 |
| chr3-10888815-INS-234 | chr3-10888819-INS-252 |
| chr3-10911688-INS-201 | chr3-10911688-INS-201 |
| chr3-10917812-INS-83 | chr3-10917812-INS-70 |
| chr3-10920430-INS-107 | chr3-10920449-INS-107 |
| chr3-10941170-INS-76 | chr3-10941170-INS-76 |
| chr3-10963414-INS-132 | chr3-10963414-INS-132 |
| chr3-10979616-INS-7457 | chr3-10979617-INS-7457 |
| chr3-11020429-INS-52 | chr3-11020429-INS-88 |
| chr3-11034691-INS-56 | chr3-11034683-INS-52 |
| chr3-11043863-INS-58 | chr3-11043863-INS-58 |
| chr3-11046959-INS-160 | chr3-11046959-INS-157 |
| chr3-11049197-INS-3771 | chr3-11049198-INS-3762 |
| chr3-11082740-INS-6553 | chr3-11082741-INS-6554 |
| chr3-11108909-INS-60 | chr3-11108909-INS-62 |
| chr3-11158908-INS-342 | chr3-11158908-INS-342 |
| chr3-11175553-INS-219 | chr3-11175553-INS-219 |
| chr3-11183676-INS-51 | chr3-11183673-INS-51 |
| chr3-11248452-INS-168 | chr3-11248452-INS-141 |
| chr3-11251216-INS-205 | chr3-11251216-INS-205 |
| chr3-11253121-INS-65 | chr3-11253121-INS-68 |
| chr3-11255973-INS-198 | chr3-11255973-INS-207 |
| chr3-11264481-INS-482 | chr3-11264481-INS-498 |
| chr3-11268265-INS-465 | chr3-11268265-INS-465 |
| chr3-11304861-INS-58 | chr3-11304861-INS-58 |
| chr3-11434254-INS-559 | chr3-11434254-INS-559 |
| chr3-11472393-INS-8723 | chr3-11472394-INS-8723 |
| chr3-11475150-INS-175 | chr3-11475150-INS-175 |
| chr3-11477871-INS-121 | chr3-11477871-INS-123 |

| | |
|---|---|
| chr3-11494061-INS-73 | chr3-11494061-INS-73 |
| chr3-11500979-INS-2797 | chr3-11500980-INS-2797 |
| chr3-11504485-INS-83 | chr3-11504485-INS-63 |
| chr3-11514538-INS-83 | chr3-11514538-INS-87 |
| chr3-11526797-INS-68 | chr3-11526765-INS-88 |
| chr3-11531466-INS-3226 | chr3-11531467-INS-3227 |
| chr3-11575924-INS-65 | chr3-11575924-INS-58 |
| chr3-11578212-INS-134 | chr3-11578212-INS-146 |
| chr3-11589770-INS-214 | chr3-11589770-INS-214 |
| chr3-11600568-INS-390 | chr3-11600568-INS-390 |
| chr3-11613530-INS-878 | chr3-11613530-INS-878 |
| chr3-11625797-INS-57 | chr3-11625797-INS-57 |
| chr3-11628860-INS-204 | chr3-11628860-INS-209 |
| chr3-11633868-INS-85 | chr3-11633868-INS-73 |
| chr3-11668877-INS-227 | chr3-11668877-INS-236 |
| chr3-11680171-INS-74 | chr3-11680163-INS-68 |
| chr3-11704667-INS-6638 | chr3-11704668-INS-6632 |
| chr3-11716084-INS-641 | chr3-11716084-INS-621 |
| chr3-11722044-INS-267 | chr3-11722044-INS-268 |
| chr3-11745301-INS-931 | chr3-11745302-INS-930 |
| chr3-11778811-INS-179 | chr3-11778811-INS-179 |
| chr3-11806360-INS-540 | chr3-11806360-INS-540 |
| chr3-11828637-INS-61 | chr3-11828665-INS-56 |
| chr3-11835069-INS-1063 | chr3-11835070-INS-1063 |
| chr3-11846075-INS-54 | chr3-11846075-INS-52 |
| chr3-11882581-INS-69 | chr3-11882581-INS-63 |
| chr3-11892039-INS-101 | chr3-11892039-INS-101 |
| chr3-11902391-INS-310 | chr3-11902391-INS-310 |
| chr3-11912366-INS-59 | chr3-11912366-INS-53 |
| chr3-11935794-INS-66 | chr3-11935817-INS-82 |
| chr3-11944722-INS-106 | chr3-11944722-INS-112 |
| chr3-11975193-INS-948 | chr3-11975193-INS-948 |
| chr3-11980005-INS-85 | chr3-11980005-INS-85 |
| chr3-11982871-INS-60 | chr3-11982875-INS-54 |
| chr3-12049714-INS-290 | chr3-12049714-INS-290 |
| chr3-12068224-INS-165 | chr3-12068224-INS-165 |
| chr3-12088518-INS-78 | chr3-12088518-INS-78 |
| chr3-12141686-INS-74 | chr3-12141700-INS-66 |
| chr3-12143103-INS-518 | chr3-12143103-INS-518 |
| chr3-12144068-INS-58 | chr3-12144068-INS-58 |
| chr3-12146080-INS-97 | chr3-12146080-INS-77 |
| chr3-12190306-INS-64 | chr3-12190326-INS-64 |
| chr3-12208398-INS-2797 | chr3-12208399-INS-2797 |
| chr3-12231024-INS-989 | chr3-12231024-INS-989 |
| chr3-12265765-INS-538 | chr3-12265765-INS-538 |
| chr3-12276898-INS-433 | chr3-12276910-INS-423 |
| chr3-12298609-INS-532 | chr3-12298609-INS-532 |
| chr3-12330225-INS-197 | chr3-12330225-INS-196 |
| chr3-12446948-INS-78 | chr3-12446964-INS-64 |
| chr3-12450722-INS-68 | chr3-12450722-INS-68 |
| chr3-12453165-INS-68 | chr3-12453169-INS-52 |
| chr3-12454609-INS-592 | chr3-12454609-INS-590 |
| chr3-12456416-INS-4180 | chr3-12456417-INS-4180 |
| chr3-12474127-INS-64 | chr3-12474127-INS-64 |
| chr3-12477065-INS-75 | chr3-12477065-INS-75 |
| chr3-12495282-INS-199 | chr3-12495282-INS-199 |
| chr3-12521286-INS-193 | chr3-12521286-INS-193 |
| chr3-12523613-INS-327 | chr3-12523613-INS-327 |
| chr3-12568634-INS-75 | chr3-12568632-INS-68 |
| chr3-12570448-INS-65 | chr3-12570448-INS-65 |
| chr3-12594935-INS-6655 | chr3-12594936-INS-6654 |
| chr3-12660898-INS-3437 | chr3-12660902-INS-3434 |
| chr3-12661253-INS-115 | chr3-12661253-INS-115 |
| chr3-12670871-INS-92 | chr3-12670871-INS-92 |
| chr3-12700558-INS-56 | chr3-12700558-INS-56 |
| chr3-12741100-INS-4445 | chr3-12741101-INS-4445 |
| chr3-12750337-INS-81 | chr3-12750335-INS-72 |
| chr3-12765338-INS-55 | chr3-12765338-INS-55 |
| chr3-12798215-INS-78 | chr3-12798215-INS-78 |
| chr3-12806679-INS-2331 | chr3-12806680-INS-2331 |
| chr3-12810364-INS-713 | chr3-12810364-INS-712 |
| chr3-12814980-INS-63 | chr3-12814984-INS-61 |
| chr3-12823170-INS-1312 | chr3-12823171-INS-1312 |
| chr3-12836970-INS-59 | chr3-12836971-INS-58 |
| chr3-12838386-INS-64 | chr3-12838386-INS-66 |
| chr3-12842522-INS-336 | chr3-12842522-INS-336 |
| chr3-12849091-INS-349 | chr3-12849042-INS-358 |

| | |
|---|---|
| chr3-12849721-INS-51 | chr3-12849713-INS-52 |
| chr3-12862040-INS-653 | chr3-12862040-INS-656 |
| chr3-12875115-INS-59 | chr3-12875115-INS-93 |
| chr3-12884671-INS-452 | chr3-12884708-INS-447 |
| chr3-12908632-INS-88 | chr3-12908632-INS-88 |
| chr3-12940339-INS-119 | chr3-12940339-INS-121 |
| chr3-12962800-INS-89 | chr3-12962800-INS-89 |
| chr3-13007651-INS-542 | chr3-13007651-INS-543 |
| chr3-13028194-INS-54 | chr3-13028194-INS-95 |
| chr3-13089329-INS-64 | chr3-13089358-INS-50 |
| chr3-13094425-INS-203 | chr3-13094425-INS-203 |
| chr3-13154724-INS-189 | chr3-13154724-INS-189 |
| chr3-13178504-INS-6461 | chr3-13178505-INS-6462 |
| chr3-13196061-INS-142 | chr3-13196061-INS-158 |
| chr3-13201249-INS-5177 | chr3-13201250-INS-5177 |
| chr3-13216440-INS-149 | chr3-13216440-INS-145 |
| chr3-13217054-INS-80 | chr3-13217024-INS-112 |
| chr3-13225221-INS-363 | chr3-13225221-INS-363 |
| chr3-13294057-INS-63 | chr3-13294035-INS-63 |
| chr3-13296414-INS-390 | chr3-13296414-INS-390 |
| chr3-13317803-INS-423 | chr3-13317803-INS-423 |
| chr3-13321795-INS-380 | chr3-13321795-INS-380 |
| chr3-13324936-INS-522 | chr3-13324936-INS-522 |
| chr3-13331411-INS-731 | chr3-13331411-INS-732 |
| chr3-13342999-INS-2348 | chr3-13343000-INS-2348 |
| chr3-13369188-INS-140 | chr3-13369188-INS-144 |
| chr3-13377801-INS-444 | chr3-13377801-INS-455 |
| chr3-13382818-INS-157 | chr3-13382819-INS-157 |
| chr3-13383225-INS-562 | chr3-13383226-INS-556 |
| chr3-13402088-INS-224 | chr3-13402088-INS-224 |
| chr3-13412777-INS-6986 | chr3-13412778-INS-6989 |
| chr3-13415532-INS-162 | chr3-13415532-INS-162 |
| chr3-13436880-INS-270 | chr3-13436880-INS-312 |
| chr3-13450139-INS-70 | chr3-13450133-INS-50 |
| chr3-13452498-INS-52 | chr3-13452496-INS-58 |
| chr3-13484305-INS-799 | chr3-13484305-INS-796 |
| chr3-13509935-INS-52 | chr3-13509935-INS-50 |
| chr3-13510824-INS-6453 | chr3-13510825-INS-6452 |
| chr3-13516920-INS-118 | chr3-13516924-INS-86 |
| chr3-13530105-INS-53 | chr3-13530105-INS-53 |
| chr3-13546169-INS-434 | chr3-13546169-INS-434 |
| chr3-13558458-INS-131 | chr3-13558458-INS-127 |
| chr3-13579550-INS-265 | chr3-13579550-INS-265 |
| chr3-13598641-INS-114 | chr3-13598626-INS-129 |
| chr3-13622000-INS-54 | chr3-13621997-INS-100 |
| chr3-13631772-INS-317 | chr3-13631773-INS-300 |
| chr3-13682115-INS-201 | chr3-13682115-INS-201 |
| chr3-13686344-INS-763 | chr3-13686344-INS-764 |
| chr3-13695087-INS-279 | chr3-13695087-INS-279 |
| chr3-13704071-INS-52 | chr3-13704071-INS-84 |
| chr3-13705811-INS-3977 | chr3-13705812-INS-3978 |
| chr3-13740182-INS-457 | chr3-13740182-INS-457 |
| chr3-13740997-INS-811 | chr3-13740997-INS-811 |
| chr3-13748442-INS-67 | chr3-13748442-INS-69 |
| chr3-13774804-INS-400 | chr3-13774804-INS-400 |
| chr3-13790960-INS-106 | chr3-13790960-INS-103 |
| chr3-13827184-INS-6627 | chr3-13827185-INS-6627 |
| chr3-13828305-INS-265 | chr3-13828305-INS-265 |
| chr3-13870820-INS-641 | chr3-13870820-INS-645 |
| chr3-13883763-INS-78 | chr3-13883764-INS-80 |
| chr3-13891712-INS-52 | chr3-13891712-INS-52 |
| chr3-13925506-INS-725 | chr3-13925506-INS-725 |
| chr3-13958325-INS-75 | chr3-13958325-INS-75 |
| chr3-14018924-INS-64 | chr3-14018924-INS-64 |
| chr3-14035598-INS-6878 | chr3-14035599-INS-6877 |
| chr3-14061631-INS-132 | chr3-14061631-INS-132 |
| chr3-14090782-INS-774 | chr3-14090782-INS-774 |
| chr3-14099428-INS-72 | chr3-14099428-INS-72 |
| chr3-14136449-INS-60 | chr3-14136449-INS-59 |
| chr3-14154651-INS-102 | chr3-14154651-INS-106 |
| chr3-14191113-INS-195 | chr3-14191113-INS-192 |
| chr3-14191508-INS-891 | chr3-14191508-INS-890 |
| chr3-14215991-INS-80 | chr3-14215991-INS-50 |
| chr3-14230436-INS-5219 | chr3-14230437-INS-5219 |
| chr3-14240823-INS-581 | chr3-14240823-INS-582 |
| chr3-14248196-INS-1423 | chr3-14248197-INS-1423 |
| chr3-14252179-INS-62 | chr3-14252189-INS-60 |

| | |
|---|---|
| chr3-14266651-INS-6549 | chr3-14266652-INS-6549 |
| chr3-14275282-INS-866 | chr3-14275282-INS-880 |
| chr3-14292867-INS-827 | chr3-14292867-INS-827 |
| chr3-14316546-INS-412 | chr3-14316546-INS-411 |
| chr3-14334385-INS-62 | chr3-14334381-INS-50 |
| chr3-14350879-INS-115 | chr3-14350879-INS-115 |
| chr3-14422700-INS-945 | chr3-14422700-INS-945 |
| chr3-14429548-INS-341 | chr3-14429548-INS-344 |
| chr3-14477942-INS-7144 | chr3-14477943-INS-7144 |
| chr3-14502653-INS-557 | chr3-14502653-INS-557 |
| chr3-14571238-INS-216 | chr3-14571238-INS-216 |
| chr3-14577699-INS-7195 | chr3-14577700-INS-7195 |
| chr3-14602843-INS-92 | chr3-14602875-INS-69 |
| chr3-14626824-INS-68 | chr3-14626824-INS-68 |
| chr3-14668001-INS-211 | chr3-14668001-INS-212 |
| chr3-14700798-INS-58 | chr3-14700798-INS-56 |
| chr3-14713056-INS-117 | chr3-14713056-INS-121 |
| chr3-14717811-INS-4315 | chr3-14717812-INS-4315 |
| chr3-14740770-INS-69 | chr3-14740770-INS-71 |
| chr3-14741942-INS-169 | chr3-14741942-INS-169 |
| chr3-14749852-INS-320 | chr3-14749852-INS-320 |
| chr3-14753394-INS-112 | chr3-14753348-INS-120 |
| chr3-14800650-INS-7328 | chr3-14800651-INS-7328 |
| chr3-14832827-INS-6879 | chr3-14832828-INS-6881 |
| chr3-14865819-INS-6237 | chr3-14865820-INS-6238 |
| chr3-14876356-INS-135 | chr3-14876356-INS-135 |
| chr3-14890345-INS-57 | chr3-14890320-INS-64 |
| chr3-14891951-INS-71 | chr3-14891951-INS-71 |
| chr3-14895359-INS-81 | chr3-14895378-INS-72 |
| chr3-14934215-INS-1859 | chr3-14934216-INS-1859 |
| chr3-14970816-INS-713 | chr3-14970816-INS-713 |
| chr3-14975817-INS-251 | chr3-14975817-INS-251 |
| chr3-15035899-INS-5342 | chr3-15035900-INS-5342 |
| chr3-15064020-INS-90 | chr3-15064020-INS-90 |
| chr3-15071360-INS-167 | chr3-15071356-INS-175 |
| chr3-15075978-INS-7101 | chr3-15075979-INS-7101 |
| chr3-15083001-INS-86 | chr3-15083001-INS-86 |
| chr3-15093329-INS-66 | chr3-15093329-INS-68 |
| chr3-15103201-INS-4452 | chr3-15103202-INS-4452 |
| chr3-15118102-INS-600 | chr3-15118102-INS-598 |
| chr3-15119873-INS-198 | chr3-15119873-INS-198 |
| chr3-15120186-INS-7140 | chr3-15120187-INS-7140 |
| chr3-15131918-INS-80 | chr3-15131887-INS-80 |
| chr3-15140102-INS-63 | chr3-15140124-INS-63 |
| chr3-15164652-INS-8775 | chr3-15164653-INS-8775 |
| chr3-15165986-INS-473 | chr3-15165986-INS-473 |
| chr3-15180853-INS-63 | chr3-15180853-INS-63 |
| chr3-15194788-INS-2306 | chr3-15194789-INS-2306 |
| chr3-15204377-INS-491 | chr3-15204377-INS-491 |
| chr3-15209394-INS-105 | chr3-15209394-INS-105 |
| chr3-15212413-INS-56 | chr3-15212413-INS-56 |
| chr3-15214149-INS-101 | chr3-15214149-INS-105 |
| chr3-15245237-INS-73 | chr3-15245237-INS-73 |
| chr3-15267243-INS-116 | chr3-15267243-INS-116 |
| chr3-15286530-INS-74 | chr3-15286524-INS-84 |
| chr3-15293849-INS-324 | chr3-15293849-INS-328 |
| chr3-15296648-INS-150 | chr3-15296648-INS-150 |
| chr3-15307129-INS-1084 | chr3-15307130-INS-1083 |
| chr3-15342020-INS-59 | chr3-15342020-INS-59 |
| chr3-15355543-INS-55 | chr3-15355543-INS-76 |
| chr3-15415082-INS-567 | chr3-15415082-INS-567 |
| chr3-15473247-INS-52 | chr3-15473247-INS-54 |
| chr3-15474700-INS-64 | chr3-15474700-INS-64 |
| chr3-15578105-INS-400 | chr3-15578109-INS-400 |
| chr3-15608809-INS-108 | chr3-15608809-INS-108 |
| chr3-15635846-INS-54 | chr3-15635840-INS-54 |
| chr3-15649130-INS-224 | chr3-15649130-INS-224 |
| chr3-15705619-INS-82 | chr3-15705601-INS-61 |
| chr3-15827492-INS-567 | chr3-15827492-INS-567 |
| chr3-15941095-INS-59 | chr3-15941095-INS-59 |
| chr3-15944924-INS-66 | chr3-15944924-INS-66 |
| chr3-15963259-INS-6417 | chr3-15963260-INS-6417 |
| chr3-15968670-INS-146 | chr3-15968670-INS-127 |
| chr3-15997072-INS-2927 | chr3-15997069-INS-2923 |
| chr3-16002696-INS-6724 | chr3-16002697-INS-6724 |
| chr3-16007065-INS-64 | chr3-16007066-INS-64 |
| chr3-16058017-INS-399 | chr3-16058017-INS-399 |

| | |
|---|---|
| chr3-16079344-INS-7982 | chr3-16079344-INS-7982 |
| chr3-16094965-INS-3096 | chr3-16094966-INS-3096 |
| chr3-16105050-INS-54 | chr3-16105052-INS-79 |
| chr3-16106115-INS-196 | chr3-16106115-INS-196 |
| chr3-16108689-INS-7182 | chr3-16108690-INS-7182 |
| chr3-16121286-INS-588 | chr3-16121286-INS-585 |
| chr3-16150768-INS-60 | chr3-16150759-INS-67 |
| chr3-16152219-INS-7471 | chr3-16152220-INS-7471 |
| chr3-16159815-INS-65 | chr3-16159815-INS-65 |
| chr3-16199984-INS-1892 | chr3-16199985-INS-1892 |
| chr3-16206296-INS-196 | chr3-16206296-INS-196 |
| chr3-16265237-INS-79 | chr3-16265239-INS-79 |
| chr3-16321695-INS-396 | chr3-16321695-INS-396 |
| chr3-16322240-INS-1878 | chr3-16322252-INS-1869 |
| chr3-16331441-INS-422 | chr3-16331441-INS-422 |
| chr3-16335104-INS-125 | chr3-16335104-INS-125 |
| chr3-16351721-INS-120 | chr3-16351721-INS-106 |
| chr3-16359266-INS-90 | chr3-16359267-INS-85 |
| chr3-16363052-INS-401 | chr3-16363052-INS-401 |
| chr3-16414982-INS-133 | chr3-16414982-INS-133 |
| chr3-16437913-INS-60 | chr3-16437913-INS-52 |
| chr3-16442879-INS-453 | chr3-16442879-INS-459 |
| chr3-16452963-INS-2809 | chr3-16452964-INS-2809 |
| chr3-16475132-INS-126 | chr3-16475132-INS-126 |
| chr3-16489964-INS-12444 | chr3-16489964-INS-12445 |
| chr3-16564546-INS-75 | chr3-16564546-INS-72 |
| chr3-16575444-INS-55 | chr3-16575444-INS-69 |
| chr3-16588553-INS-56 | chr3-16588553-INS-52 |
| chr3-16644989-INS-63 | chr3-16644981-INS-92 |
| chr3-16671665-INS-536 | chr3-16671665-INS-538 |
| chr3-16697526-INS-458 | chr3-16697526-INS-458 |
| chr3-16727493-INS-329 | chr3-16727493-INS-329 |
| chr3-16733063-INS-152 | chr3-16733090-INS-149 |
| chr3-16745767-INS-82 | chr3-16745767-INS-87 |
| chr3-16770235-INS-60 | chr3-16770235-INS-52 |
| chr3-16800932-INS-6231 | chr3-16800933-INS-6230 |
| chr3-16821511-INS-409 | chr3-16821511-INS-409 |
| chr3-16845436-INS-54 | chr3-16845436-INS-54 |
| chr3-16855519-INS-54 | chr3-16855483-INS-58 |
| chr3-16856367-INS-2856 | chr3-16856368-INS-2856 |
| chr3-16856368-INS-464 | chr3-16856368-INS-464 |
| chr3-16858239-INS-61 | chr3-16858239-INS-53 |
| chr3-16895470-INS-6631 | chr3-16895471-INS-6631 |
| chr3-16924838-INS-4549 | chr3-16924839-INS-4549 |
| chr3-16936811-INS-624 | chr3-16936827-INS-622 |
| chr3-16985451-INS-468 | chr3-16985451-INS-468 |
| chr3-16988008-INS-568 | chr3-16988008-INS-545 |
| chr3-17016540-INS-79 | chr3-17016540-INS-79 |
| chr3-17078264-INS-2702 | chr3-17078265-INS-2702 |
| chr3-17138681-INS-180 | chr3-17138681-INS-179 |
| chr3-17197243-INS-231 | chr3-17197243-INS-231 |
| chr3-17209769-INS-139 | chr3-17209755-INS-126 |
| chr3-17215429-INS-416 | chr3-17215429-INS-416 |
| chr3-17235887-INS-60 | chr3-17235887-INS-75 |
| chr3-17255848-INS-277 | chr3-17255848-INS-277 |
| chr3-17261996-INS-75 | chr3-17261996-INS-80 |
| chr3-17321991-INS-390 | chr3-17321991-INS-390 |
| chr3-17330274-INS-128 | chr3-17330274-INS-109 |
| chr3-17345910-INS-557 | chr3-17345910-INS-557 |
| chr3-17357992-INS-115 | chr3-17357992-INS-115 |
| chr3-17364454-INS-63 | chr3-17364454-INS-61 |
| chr3-17396853-INS-83 | chr3-17396864-INS-74 |
| chr3-17403754-INS-57 | chr3-17403720-INS-55 |
| chr3-17406968-INS-5660 | chr3-17406969-INS-5660 |
| chr3-17414832-INS-2245 | chr3-17414833-INS-2267 |
| chr3-17416485-INS-6477 | chr3-17416486-INS-6478 |
| chr3-17420474-INS-1950 | chr3-17420475-INS-1950 |
| chr3-17453320-INS-55 | chr3-17453320-INS-55 |
| chr3-17532444-INS-197 | chr3-17532444-INS-197 |
| chr3-17533933-INS-70 | chr3-17533933-INS-70 |
| chr3-17566076-INS-97 | chr3-17566076-INS-98 |
| chr3-17601469-INS-6126 | chr3-17601470-INS-6127 |
| chr3-17608844-INS-6969 | chr3-17608845-INS-6969 |
| chr3-17614426-INS-196 | chr3-17614426-INS-196 |
| chr3-17616591-INS-1203 | chr3-17616592-INS-1204 |
| chr3-17680001-INS-80 | chr3-17680001-INS-82 |
| chr3-17695841-INS-5236 | chr3-17695842-INS-5236 |

| | |
|---|---|
| chr3-17743578-INS-130 | chr3-17743578-INS-130 |
| chr3-17768098-INS-767 | chr3-17768098-INS-767 |
| chr3-17769604-INS-1030 | chr3-17769605-INS-1031 |
| chr3-17803038-INS-119 | chr3-17803038-INS-119 |
| chr3-17831018-INS-87 | chr3-17831017-INS-85 |
| chr3-17897904-INS-344 | chr3-17897904-INS-344 |
| chr3-17903785-INS-2849 | chr3-17903786-INS-2849 |
| chr3-17910419-INS-98 | chr3-17910419-INS-78 |
| chr3-17922808-INS-70 | chr3-17922808-INS-82 |
| chr3-17934178-INS-885 | chr3-17934178-INS-898 |
| chr3-17936702-INS-66 | chr3-17936702-INS-66 |
| chr3-17944297-INS-165 | chr3-17944297-INS-165 |
| chr3-17956026-INS-208 | chr3-17956026-INS-209 |
| chr3-17956643-INS-64 | chr3-17956643-INS-64 |
| chr3-17980810-INS-57 | chr3-17980810-INS-65 |
| chr3-17982605-INS-2469 | chr3-17982606-INS-2469 |
| chr3-17998836-INS-696 | chr3-17998836-INS-714 |
| chr3-18017379-INS-224 | chr3-18017379-INS-224 |
| chr3-18035215-INS-212 | chr3-18035215-INS-212 |
| chr3-18053639-INS-88 | chr3-18053646-INS-52 |
| chr3-18092861-INS-241 | chr3-18092862-INS-243 |
| chr3-18133276-INS-62 | chr3-18133276-INS-57 |
| chr3-18186594-INS-63 | chr3-18186591-INS-74 |
| chr3-18190255-INS-1456 | chr3-18190256-INS-1459 |
| chr3-18216378-INS-6928 | chr3-18216379-INS-6931 |
| chr3-18245419-INS-54 | chr3-18245419-INS-54 |
| chr3-18257721-INS-7496 | chr3-18257722-INS-7498 |
| chr3-18269233-INS-231 | chr3-18269233-INS-232 |
| chr3-18273829-INS-498 | chr3-18273829-INS-498 |
| chr3-18276580-INS-7079 | chr3-18276581-INS-7079 |
| chr3-18313241-INS-52 | chr3-18313241-INS-50 |
| chr3-18314990-INS-455 | chr3-18314990-INS-456 |
| chr3-18316265-INS-80 | chr3-18316265-INS-77 |
| chr3-18358404-INS-6894 | chr3-18358405-INS-6895 |
| chr3-18362436-INS-6382 | chr3-18362437-INS-6382 |
| chr3-18362437-INS-504 | chr3-18362437-INS-504 |
| chr3-18364438-INS-635 | chr3-18364438-INS-636 |
| chr3-18365466-INS-1733 | chr3-18365467-INS-1733 |
| chr3-18394339-INS-65 | chr3-18394339-INS-65 |
| chr3-18420569-INS-338 | chr3-18420569-INS-338 |
| chr3-18438661-INS-206 | chr3-18438662-INS-207 |
| chr3-18441529-INS-6592 | chr3-18441530-INS-6604 |
| chr3-18442096-INS-82 | chr3-18442097-INS-83 |
| chr3-18536568-INS-224 | chr3-18536568-INS-225 |
| chr3-18545719-INS-6483 | chr3-18545720-INS-6483 |
| chr3-18547258-INS-54 | chr3-18547258-INS-54 |
| chr3-18563461-INS-62 | chr3-18563461-INS-62 |
| chr3-18571972-INS-361 | chr3-18571972-INS-361 |
| chr3-18589175-INS-7113 | chr3-18589176-INS-7114 |
| chr3-18626493-INS-328 | chr3-18626493-INS-328 |
| chr3-18635314-INS-7433 | chr3-18635315-INS-7436 |
| chr3-18637213-INS-761 | chr3-18637213-INS-752 |
| chr3-18648871-INS-1718 | chr3-18648872-INS-1719 |
| chr3-18674314-INS-54 | chr3-18674314-INS-58 |
| chr3-18701584-INS-72 | chr3-18701584-INS-72 |
| chr3-18718462-INS-78 | chr3-18718460-INS-79 |
| chr3-18729309-INS-438 | chr3-18729309-INS-431 |
| chr3-18795003-INS-172 | chr3-18795003-INS-175 |
| chr3-18831111-INS-235 | chr3-18831090-INS-282 |
| chr3-18837193-INS-153 | chr3-18837193-INS-153 |
| chr3-18892747-INS-401 | chr3-18892747-INS-401 |
| chr3-18905326-INS-102 | chr3-18905327-INS-104 |
| chr3-18915268-INS-2878 | chr3-18915266-INS-2880 |
| chr3-18921281-INS-2282 | chr3-18921282-INS-2282 |
| chr3-18923160-INS-3053 | chr3-18923161-INS-3053 |
| chr3-18940487-INS-6761 | chr3-18940488-INS-6763 |
| chr3-18943468-INS-128 | chr3-18943468-INS-128 |
| chr3-18966635-INS-83 | chr3-18966635-INS-83 |
| chr3-18971051-INS-797 | chr3-18971051-INS-797 |
| chr3-18991502-INS-1658 | chr3-18991503-INS-1662 |
| chr3-19017304-INS-192 | chr3-19017304-INS-189 |
| chr3-19043718-INS-56 | chr3-19043700-INS-75 |
| chr3-19057783-INS-80 | chr3-19057783-INS-80 |
| chr3-19067976-INS-6435 | chr3-19067977-INS-6435 |
| chr3-19097716-INS-53 | chr3-19097716-INS-53 |
| chr3-19100041-INS-379 | chr3-19100042-INS-379 |
| chr3-19106719-INS-351 | chr3-19106719-INS-351 |

| | |
|---|---|
| chr3-19127416-INS-885 | chr3-19127416-INS-885 |
| chr3-19141587-INS-7089 | chr3-19141588-INS-7089 |
| chr3-19171704-INS-625 | chr3-19171704-INS-625 |
| chr3-19186037-INS-64 | chr3-19186037-INS-70 |
| chr3-19200175-INS-64 | chr3-19200215-INS-65 |
| chr3-19202756-INS-62 | chr3-19202756-INS-64 |
| chr3-19224063-INS-51 | chr3-19224064-INS-53 |
| chr3-19227052-INS-863 | chr3-19227052-INS-875 |
| chr3-19235292-INS-7089 | chr3-19235293-INS-7089 |
| chr3-19253242-INS-167 | chr3-19253244-INS-167 |
| chr3-19300055-INS-184 | chr3-19300055-INS-154 |
| chr3-19307567-INS-1120 | chr3-19307568-INS-1120 |
| chr3-19320424-INS-306 | chr3-19320414-INS-344 |
| chr3-19350749-INS-114 | chr3-19350773-INS-87 |
| chr3-19359430-INS-1892 | chr3-19359431-INS-1892 |
| chr3-19372257-INS-188 | chr3-19372257-INS-188 |
| chr3-19383290-INS-55 | chr3-19383290-INS-55 |
| chr3-19398208-INS-56 | chr3-19398208-INS-56 |
| chr3-19402448-INS-61 | chr3-19402448-INS-61 |
| chr3-19404601-INS-2759 | chr3-19404602-INS-2759 |
| chr3-19434509-INS-2871 | chr3-19434510-INS-2870 |
| chr3-19447136-INS-600 | chr3-19447136-INS-602 |
| chr3-19449214-INS-180 | chr3-19449214-INS-188 |
| chr3-19492001-INS-53 | chr3-19492009-INS-53 |
| chr3-19506042-INS-328 | chr3-19506042-INS-327 |
| chr3-19513924-INS-64 | chr3-19513924-INS-68 |
| chr3-19516238-INS-173 | chr3-19516238-INS-173 |
| chr3-19524039-INS-934 | chr3-19524039-INS-934 |
| chr3-19526261-INS-100 | chr3-19526249-INS-138 |
| chr3-19527910-INS-136 | chr3-19527910-INS-136 |
| chr3-19535977-INS-162 | chr3-19535977-INS-179 |
| chr3-19538810-INS-68 | chr3-19538810-INS-91 |
| chr3-19584884-INS-428 | chr3-19584884-INS-429 |
| chr3-19590798-INS-65 | chr3-19590798-INS-65 |
| chr3-19592213-INS-207 | chr3-19592213-INS-207 |
| chr3-19592216-INS-2647 | chr3-19592217-INS-2647 |
| chr3-19616686-INS-136 | chr3-19616686-INS-158 |
| chr3-19659280-INS-556 | chr3-19659280-INS-544 |
| chr3-19660396-INS-291 | chr3-19660396-INS-291 |
| chr3-19661773-INS-51 | chr3-19661762-INS-52 |
| chr3-19662824-INS-217 | chr3-19662824-INS-217 |
| chr3-19679194-INS-152 | chr3-19679194-INS-151 |
| chr3-19693600-INS-4753 | chr3-19693601-INS-4752 |
| chr3-19708504-INS-84 | chr3-19708504-INS-84 |
| chr3-19733933-INS-157 | chr3-19733936-INS-154 |
| chr3-19762001-INS-6784 | chr3-19762002-INS-6785 |
| chr3-19769997-INS-595 | chr3-19769997-INS-594 |
| chr3-19771283-INS-182 | chr3-19771283-INS-209 |
| chr3-19774566-INS-2725 | chr3-19774567-INS-2725 |
| chr3-19782720-INS-62 | chr3-19782698-INS-52 |
| chr3-19811017-INS-1497 | chr3-19811018-INS-1497 |
| chr3-19844475-INS-55 | chr3-19844475-INS-55 |
| chr3-19849756-INS-58 | chr3-19849756-INS-50 |
| chr3-19886393-INS-184 | chr3-19886393-INS-184 |
| chr3-19912072-INS-121 | chr3-19912072-INS-121 |
| chr3-19915499-INS-515 | chr3-19915499-INS-523 |
| chr3-19917956-INS-249 | chr3-19917956-INS-247 |
| chr3-19922419-INS-971 | chr3-19922419-INS-967 |
| chr3-19927774-INS-326 | chr3-19927774-INS-326 |
| chr3-19961405-INS-83 | chr3-19961406-INS-83 |
| chr3-19985451-INS-350 | chr3-19985451-INS-350 |
| chr3-20031885-INS-253 | chr3-20031885-INS-256 |
| chr3-20044835-INS-108 | chr3-20044832-INS-102 |
| chr3-20066151-INS-98 | chr3-20066151-INS-148 |
| chr3-20069539-INS-613 | chr3-20069539-INS-613 |
| chr3-20076683-INS-225 | chr3-20076683-INS-225 |
| chr3-20078485-INS-70 | chr3-20078485-INS-70 |
| chr3-20079544-INS-51 | chr3-20079544-INS-51 |
| chr3-20087651-INS-92 | chr3-20087651-INS-92 |
| chr3-20102360-INS-539 | chr3-20102360-INS-539 |
| chr3-20161674-INS-236 | chr3-20161674-INS-236 |
| chr3-20164976-INS-201 | chr3-20164976-INS-206 |
| chr3-20169272-INS-50 | chr3-20169283-INS-85 |
| chr3-20192683-INS-85 | chr3-20192683-INS-85 |
| chr3-20214063-INS-52 | chr3-20214063-INS-52 |
| chr3-20229969-INS-343 | chr3-20229969-INS-343 |
| chr3-20244458-INS-54 | chr3-20244450-INS-62 |

| | |
|---|---|
| chr3-20252992-INS-374 | chr3-20252992-INS-375 |
| chr3-20257889-INS-310 | chr3-20257889-INS-310 |
| chr3-20280486-INS-70 | chr3-20280488-INS-62 |
| chr3-20281722-INS-108 | chr3-20281722-INS-108 |
| chr3-20285256-INS-54 | chr3-20285256-INS-54 |
| chr3-20296569-INS-982 | chr3-20296572-INS-986 |
| chr3-20302379-INS-262 | chr3-20302379-INS-254 |
| chr3-20305344-INS-399 | chr3-20305344-INS-399 |
| chr3-20309374-INS-85 | chr3-20309357-INS-78 |
| chr3-20342617-INS-484 | chr3-20342617-INS-484 |
| chr3-20350884-INS-92 | chr3-20350884-INS-92 |
| chr3-20371918-INS-72 | chr3-20371908-INS-84 |
| chr3-20386711-INS-439 | chr3-20386711-INS-440 |
| chr3-20388733-INS-658 | chr3-20388733-INS-659 |
| chr3-20389425-INS-156 | chr3-20389425-INS-147 |
| chr3-20409492-INS-66 | chr3-20409492-INS-70 |
| chr3-20428974-INS-73 | chr3-20428974-INS-73 |
| chr3-20450676-INS-57 | chr3-20450676-INS-57 |
| chr3-20480981-INS-53 | chr3-20480981-INS-53 |
| chr3-20481797-INS-772 | chr3-20481797-INS-760 |
| chr3-20515267-INS-747 | chr3-20515267-INS-747 |
| chr3-20530518-INS-67 | chr3-20530518-INS-69 |
| chr3-20558538-INS-57 | chr3-20558587-INS-52 |
| chr3-20585374-INS-56 | chr3-20585374-INS-60 |
| chr3-20593296-INS-95 | chr3-20593296-INS-95 |
| chr3-20600258-INS-117 | chr3-20600258-INS-119 |
| chr3-20606738-INS-850 | chr3-20606738-INS-850 |
| chr3-20613832-INS-57 | chr3-20613832-INS-57 |
| chr3-20614702-INS-56 | chr3-20614702-INS-54 |
| chr3-20619239-INS-186 | chr3-20619239-INS-186 |
| chr3-20635872-INS-825 | chr3-20635872-INS-825 |
| chr3-20651227-INS-6463 | chr3-20651228-INS-6463 |
| chr3-20662384-INS-209 | chr3-20662384-INS-209 |
| chr3-20686781-INS-7034 | chr3-20686781-INS-7034 |
| chr3-20702099-INS-411 | chr3-20702099-INS-412 |
| chr3-20723699-INS-6127 | chr3-20723700-INS-6119 |
| chr3-20724779-INS-1878 | chr3-20724780-INS-1868 |
| chr3-20762821-INS-58 | chr3-20762822-INS-60 |
| chr3-20767555-INS-58 | chr3-20767556-INS-52 |
| chr3-20880371-INS-55 | chr3-20880371-INS-55 |
| chr3-20886011-INS-981 | chr3-20886011-INS-982 |
| chr3-20958763-INS-71 | chr3-20958763-INS-71 |
| chr3-21034452-INS-74 | chr3-21034452-INS-74 |
| chr3-21035004-INS-6378 | chr3-21035005-INS-6378 |
| chr3-21094880-INS-921 | chr3-21094880-INS-921 |
| chr3-21122279-INS-179 | chr3-21122279-INS-179 |
| chr3-21145757-INS-3040 | chr3-21145751-INS-3036 |
| chr3-21168441-INS-3423 | chr3-21168442-INS-3422 |
| chr3-21184045-INS-143 | chr3-21184045-INS-143 |
| chr3-21185307-INS-90 | chr3-21185307-INS-93 |
| chr3-21210359-INS-156 | chr3-21210359-INS-156 |
| chr3-21215360-INS-1888 | chr3-21215361-INS-1888 |
| chr3-21216935-INS-3752 | chr3-21216936-INS-3752 |
| chr3-21234528-INS-1867 | chr3-21234529-INS-1867 |
| chr3-21240907-INS-482 | chr3-21240907-INS-482 |
| chr3-21295078-INS-78 | chr3-21295078-INS-78 |
| chr3-21321690-INS-110 | chr3-21321706-INS-60 |
| chr3-21333445-INS-4693 | chr3-21333446-INS-4693 |
| chr3-21403473-INS-54 | chr3-21403504-INS-71 |
| chr3-21426029-INS-90 | chr3-21426029-INS-90 |
| chr3-21449912-INS-62 | chr3-21449912-INS-64 |
| chr3-21449948-INS-7073 | chr3-21449949-INS-7073 |
| chr3-21491456-INS-2637 | chr3-21491457-INS-2637 |
| chr3-21494344-INS-6360 | chr3-21494345-INS-6360 |
| chr3-21511393-INS-6440 | chr3-21511394-INS-6440 |
| chr3-21513525-INS-69 | chr3-21513513-INS-75 |
| chr3-21529621-INS-281 | chr3-21529621-INS-284 |
| chr3-21537633-INS-1342 | chr3-21537634-INS-1342 |
| chr3-21556150-INS-2425 | chr3-21556151-INS-2425 |
| chr3-21601957-INS-117 | chr3-21601957-INS-114 |
| chr3-21626839-INS-529 | chr3-21626839-INS-542 |
| chr3-21643543-INS-134 | chr3-21643543-INS-137 |
| chr3-21655745-INS-58 | chr3-21655745-INS-57 |
| chr3-21656627-INS-1437 | chr3-21656628-INS-1428 |
| chr3-21689256-INS-130 | chr3-21689256-INS-130 |
| chr3-21698520-INS-440 | chr3-21698520-INS-440 |
| chr3-21702524-INS-1587 | chr3-21702532-INS-1585 |

| | |
|---|---|
| chr3-21724858-INS-148 | chr3-21724858-INS-148 |
| chr3-21729236-INS-246 | chr3-21729236-INS-208 |
| chr3-21730617-INS-205 | chr3-21730617-INS-216 |
| chr3-21736896-INS-62 | chr3-21736896-INS-68 |
| chr3-21786727-INS-945 | chr3-21786727-INS-900 |
| chr3-21790126-INS-4604 | chr3-21790113-INS-4592 |
| chr3-21792483-INS-3687 | chr3-21792484-INS-3687 |
| chr3-21794329-INS-1867 | chr3-21794330-INS-1867 |
| chr3-21808146-INS-500 | chr3-21808146-INS-500 |
| chr3-21817507-INS-270 | chr3-21817487-INS-271 |
| chr3-21828295-INS-57 | chr3-21828296-INS-67 |
| chr3-21829241-INS-320 | chr3-21829241-INS-320 |
| chr3-21832143-INS-758 | chr3-21832143-INS-758 |
| chr3-21832880-INS-740 | chr3-21832880-INS-740 |
| chr3-21834582-INS-198 | chr3-21834582-INS-199 |
| chr3-21843117-INS-181 | chr3-21843119-INS-182 |
| chr3-21846375-INS-61 | chr3-21846375-INS-61 |
| chr3-21859697-INS-1136 | chr3-21859698-INS-1138 |
| chr3-21902384-INS-139 | chr3-21902384-INS-148 |
| chr3-21906848-INS-59 | chr3-21906848-INS-59 |
| chr3-21911956-INS-990 | chr3-21911956-INS-990 |
| chr3-21913500-INS-217 | chr3-21913500-INS-217 |
| chr3-21923570-INS-948 | chr3-21923576-INS-978 |
| chr3-21958056-INS-6439 | chr3-21958057-INS-6437 |
| chr3-21976232-INS-218 | chr3-21976232-INS-219 |
| chr3-21987041-INS-256 | chr3-21987041-INS-256 |
| chr3-21988576-INS-177 | chr3-21988576-INS-177 |
| chr3-21998512-INS-111 | chr3-21998508-INS-102 |
| chr3-22033303-INS-211 | chr3-22033287-INS-195 |
| chr3-22050765-INS-84 | chr3-22050765-INS-68 |
| chr3-22052759-INS-56 | chr3-22052759-INS-56 |
| chr3-22055346-INS-114 | chr3-22055346-INS-92 |
| chr3-22059922-INS-284 | chr3-22059922-INS-285 |
| chr3-22063174-INS-53 | chr3-22063175-INS-51 |
| chr3-22074603-INS-170 | chr3-22074603-INS-170 |
| chr3-22081952-INS-54 | chr3-22081952-INS-54 |
| chr3-22085410-INS-52 | chr3-22085410-INS-56 |
| chr3-22098009-INS-214 | chr3-22098009-INS-216 |
| chr3-22108262-INS-126 | chr3-22108263-INS-129 |
| chr3-22123429-INS-302 | chr3-22123429-INS-302 |
| chr3-22141840-INS-80 | chr3-22141840-INS-68 |
| chr3-22163837-INS-169 | chr3-22163837-INS-147 |
| chr3-22171170-INS-65 | chr3-22171170-INS-65 |
| chr3-22186940-INS-87 | chr3-22186928-INS-99 |
| chr3-22251785-INS-126 | chr3-22251785-INS-142 |
| chr3-22256062-INS-5549 | chr3-22256063-INS-5549 |
| chr3-22262056-INS-77 | chr3-22262056-INS-77 |
| chr3-22278972-INS-80 | chr3-22278998-INS-56 |
| chr3-22281496-INS-211 | chr3-22281496-INS-212 |
| chr3-22341789-INS-1554 | chr3-22341790-INS-1521 |
| chr3-22343427-INS-58 | chr3-22343427-INS-52 |
| chr3-22351548-INS-75 | chr3-22351548-INS-75 |
| chr3-22378734-INS-512 | chr3-22378734-INS-511 |
| chr3-22384761-INS-98 | chr3-22384761-INS-77 |
| chr3-22386086-INS-271 | chr3-22386086-INS-274 |
| chr3-22454515-INS-80 | chr3-22454477-INS-122 |
| chr3-22493247-INS-136 | chr3-22493279-INS-90 |
| chr3-22533737-INS-308 | chr3-22533737-INS-281 |
| chr3-22549487-INS-197 | chr3-22549487-INS-197 |
| chr3-22552197-INS-327 | chr3-22552197-INS-327 |
| chr3-22575024-INS-199 | chr3-22575024-INS-199 |
| chr3-22596444-INS-465 | chr3-22596445-INS-465 |
| chr3-22730036-INS-201 | chr3-22730036-INS-201 |
| chr3-22792579-INS-54 | chr3-22792579-INS-56 |
| chr3-22794270-INS-68 | chr3-22794270-INS-70 |
| chr3-22800951-INS-167 | chr3-22800951-INS-167 |
| chr3-22807243-INS-4497 | chr3-22807235-INS-4488 |
| chr3-22824013-INS-208 | chr3-22824013-INS-208 |
| chr3-22853903-INS-2263 | chr3-22853913-INS-2254 |
| chr3-22857750-INS-6247 | chr3-22857751-INS-6251 |
| chr3-22864818-INS-545 | chr3-22864818-INS-546 |
| chr3-22886475-INS-6213 | chr3-22886476-INS-6213 |
| chr3-22894439-INS-1893 | chr3-22894440-INS-1893 |
| chr3-22906271-INS-463 | chr3-22906271-INS-463 |
| chr3-22975201-INS-1886 | chr3-22975202-INS-1886 |
| chr3-23055873-INS-97 | chr3-23055873-INS-95 |
| chr3-23105169-INS-75 | chr3-23105169-INS-90 |

| | |
|---|---|
| chr3-23154398-INS-55 | chr3-23154400-INS-68 |
| chr3-23166293-INS-3792 | chr3-23166294-INS-3791 |
| chr3-23169591-INS-852 | chr3-23169591-INS-855 |
| chr3-23200316-INS-72 | chr3-23200316-INS-78 |
| chr3-23212780-INS-136 | chr3-23212780-INS-137 |
| chr3-23252175-INS-6379 | chr3-23252176-INS-6379 |
| chr3-23253806-INS-382 | chr3-23253808-INS-381 |
| chr3-23265079-INS-94 | chr3-23265079-INS-136 |
| chr3-23295409-INS-1483 | chr3-23295410-INS-1483 |
| chr3-23298599-INS-2131 | chr3-23298600-INS-2133 |
| chr3-23302861-INS-203 | chr3-23302861-INS-203 |
| chr3-23349153-INS-2795 | chr3-23349154-INS-2795 |
| chr3-23388521-INS-742 | chr3-23388521-INS-743 |
| chr3-23396994-INS-120 | chr3-23396981-INS-111 |
| chr3-23403560-INS-506 | chr3-23403560-INS-503 |
| chr3-23419681-INS-715 | chr3-23419681-INS-715 |
| chr3-23464751-INS-946 | chr3-23464751-INS-948 |
| chr3-23468903-INS-317 | chr3-23468894-INS-356 |
| chr3-23533610-INS-92 | chr3-23533610-INS-92 |
| chr3-23540562-INS-1619 | chr3-23540563-INS-1619 |
| chr3-23549066-INS-2071 | chr3-23549067-INS-2071 |
| chr3-23555317-INS-208 | chr3-23555317-INS-208 |
| chr3-23563422-INS-7200 | chr3-23563423-INS-7200 |
| chr3-23564027-INS-214 | chr3-23564027-INS-214 |
| chr3-23602764-INS-214 | chr3-23602764-INS-215 |
| chr3-23611879-INS-125 | chr3-23611879-INS-126 |
| chr3-23615925-INS-5464 | chr3-23615926-INS-5464 |
| chr3-23654832-INS-7616 | chr3-23654833-INS-7617 |
| chr3-23661072-INS-84 | chr3-23661072-INS-84 |
| chr3-23675829-INS-161 | chr3-23675829-INS-161 |
| chr3-23677670-INS-78 | chr3-23677670-INS-78 |
| chr3-23680686-INS-202 | chr3-23680686-INS-202 |
| chr3-23689369-INS-53 | chr3-23689369-INS-53 |
| chr3-23694335-INS-62 | chr3-23694335-INS-62 |
| chr3-23702438-INS-131 | chr3-23702406-INS-136 |
| chr3-23710101-INS-6298 | chr3-23710102-INS-6298 |
| chr3-23752174-INS-293 | chr3-23752174-INS-293 |
| chr3-23770525-INS-1373 | chr3-23770526-INS-1373 |
| chr3-23772600-INS-100 | chr3-23772600-INS-105 |
| chr3-23790042-INS-683 | chr3-23790042-INS-683 |
| chr3-23793101-INS-6411 | chr3-23793102-INS-6411 |
| chr3-23802325-INS-572 | chr3-23802325-INS-572 |
| chr3-23809203-INS-137 | chr3-23809203-INS-137 |
| chr3-23846383-INS-812 | chr3-23846383-INS-811 |
| chr3-23847819-INS-229 | chr3-23847819-INS-228 |
| chr3-23854063-INS-168 | chr3-23854062-INS-160 |
| chr3-23856230-INS-196 | chr3-23856230-INS-196 |
| chr3-23871554-INS-432 | chr3-23871558-INS-433 |
| chr3-23874135-INS-248 | chr3-23874135-INS-243 |
| chr3-23918306-INS-371 | chr3-23918311-INS-387 |
| chr3-23938066-INS-3081 | chr3-23938067-INS-3081 |
| chr3-23944945-INS-301 | chr3-23944945-INS-288 |
| chr3-23991949-INS-6563 | chr3-23991950-INS-6563 |
| chr3-23999538-INS-1595 | chr3-23999539-INS-1595 |
| chr3-24011215-INS-6366 | chr3-24011216-INS-6366 |
| chr3-24053310-INS-2263 | chr3-24053311-INS-2257 |
| chr3-24067846-INS-83 | chr3-24067846-INS-59 |
| chr3-24069343-INS-379 | chr3-24069343-INS-379 |
| chr3-24099953-INS-510 | chr3-24099953-INS-509 |
| chr3-24108164-INS-6884 | chr3-24108165-INS-6884 |
| chr3-24113960-INS-323 | chr3-24113960-INS-323 |
| chr3-24125882-INS-105 | chr3-24125907-INS-55 |
| chr3-24127382-INS-1615 | chr3-24127383-INS-1615 |
| chr3-24188364-INS-87 | chr3-24188364-INS-69 |
| chr3-24213770-INS-1338 | chr3-24213771-INS-1333 |
| chr3-24256439-INS-68 | chr3-24256441-INS-66 |
| chr3-24257074-INS-370 | chr3-24257074-INS-384 |
| chr3-24301527-INS-193 | chr3-24301515-INS-169 |
| chr3-24310238-INS-319 | chr3-24310238-INS-330 |
| chr3-24377301-INS-466 | chr3-24377301-INS-458 |
| chr3-24390761-INS-495 | chr3-24390761-INS-495 |
| chr3-24393987-INS-189 | chr3-24394002-INS-189 |
| chr3-24420268-INS-63 | chr3-24420268-INS-70 |
| chr3-24435872-INS-81 | chr3-24435876-INS-84 |
| chr3-24443730-INS-66 | chr3-24443730-INS-66 |
| chr3-24466628-INS-74 | chr3-24466628-INS-72 |
| chr3-24481008-INS-1944 | chr3-24481009-INS-1945 |

| | |
|---|---|
| chr3-24505751-INS-357 | chr3-24505751-INS-357 |
| chr3-24514175-INS-7177 | chr3-24514176-INS-7178 |
| chr3-24520867-INS-6652 | chr3-24520868-INS-6652 |
| chr3-24528885-INS-1043 | chr3-24528886-INS-1042 |
| chr3-24534467-INS-488 | chr3-24534467-INS-490 |
| chr3-24544667-INS-54 | chr3-24544667-INS-54 |
| chr3-24577956-INS-6170 | chr3-24577957-INS-6170 |
| chr3-24599004-INS-332 | chr3-24599004-INS-332 |
| chr3-24626461-INS-60 | chr3-24626461-INS-62 |
| chr3-24677066-INS-207 | chr3-24677066-INS-207 |
| chr3-24715988-INS-7504 | chr3-24715989-INS-7504 |
| chr3-24726739-INS-6183 | chr3-24726740-INS-6183 |
| chr3-24751283-INS-6394 | chr3-24751283-INS-6399 |
| chr3-24789543-INS-3570 | chr3-24789544-INS-3571 |
| chr3-24799674-INS-7330 | chr3-24799675-INS-7330 |
| chr3-24821346-INS-67 | chr3-24821346-INS-67 |
| chr3-24829092-INS-68 | chr3-24829092-INS-69 |
| chr3-24845765-INS-69 | chr3-24845765-INS-69 |
| chr3-24862889-INS-98 | chr3-24862889-INS-71 |
| chr3-24924769-INS-6240 | chr3-24924770-INS-6240 |
| chr3-24935807-INS-96 | chr3-24935807-INS-96 |
| chr3-24969801-INS-52 | chr3-24969801-INS-52 |
| chr3-25068925-INS-153 | chr3-25068925-INS-153 |
| chr3-25111729-INS-1195 | chr3-25111730-INS-1195 |
| chr3-25119025-INS-784 | chr3-25119025-INS-784 |
| chr3-25144503-INS-374 | chr3-25144503-INS-374 |
| chr3-25165453-INS-687 | chr3-25165453-INS-709 |
| chr3-25221942-INS-155 | chr3-25221942-INS-141 |
| chr3-25256239-INS-253 | chr3-25256239-INS-253 |
| chr3-25276603-INS-3139 | chr3-25276604-INS-3139 |
| chr3-25289944-INS-6516 | chr3-25289945-INS-6518 |
| chr3-25376013-INS-60 | chr3-25376013-INS-60 |
| chr3-25385218-INS-465 | chr3-25385218-INS-465 |
| chr3-25449931-INS-4978 | chr3-25449932-INS-4978 |
| chr3-25459972-INS-456 | chr3-25459972-INS-456 |
| chr3-25510026-INS-6944 | chr3-25510027-INS-6944 |
| chr3-25553063-INS-7256 | chr3-25553064-INS-7252 |
| chr3-25567276-INS-1297 | chr3-25567277-INS-1298 |
| chr3-25614791-INS-1888 | chr3-25614792-INS-1888 |
| chr3-25643890-INS-80 | chr3-25643912-INS-97 |
| chr3-25656571-INS-3213 | chr3-25656572-INS-3213 |
| chr3-25690124-INS-949 | chr3-25690124-INS-948 |
| chr3-25699782-INS-78 | chr3-25699782-INS-78 |
| chr3-25706809-INS-57 | chr3-25706809-INS-61 |
| chr3-25751845-INS-64 | chr3-25751845-INS-61 |
| chr3-25802357-INS-56 | chr3-25802357-INS-56 |
| chr3-25827274-INS-531 | chr3-25827284-INS-506 |
| chr3-25834665-INS-1887 | chr3-25834666-INS-1887 |
| chr3-25881679-INS-1216 | chr3-25881680-INS-1219 |
| chr3-25886956-INS-69 | chr3-25886989-INS-70 |
| chr3-25913207-INS-701 | chr3-25913207-INS-698 |
| chr3-25947489-INS-69 | chr3-25947489-INS-62 |
| chr3-25984122-INS-506 | chr3-25984122-INS-506 |
| chr3-25989618-INS-177 | chr3-25989619-INS-144 |
| chr3-25992538-INS-191 | chr3-25992538-INS-191 |
| chr3-25995828-INS-155 | chr3-25995821-INS-115 |
| chr3-25998079-INS-52 | chr3-25998079-INS-74 |
| chr3-26007419-INS-225 | chr3-26007419-INS-225 |
| chr3-26016380-INS-551 | chr3-26016380-INS-551 |
| chr3-26058950-INS-303 | chr3-26058950-INS-303 |
| chr3-26076851-INS-93 | chr3-26076851-INS-93 |
| chr3-26113879-INS-107 | chr3-26113879-INS-67 |
| chr3-26120013-INS-1787 | chr3-26120014-INS-1787 |
| chr3-26125013-INS-77 | chr3-26125013-INS-77 |
| chr3-26233369-INS-2078 | chr3-26233370-INS-2078 |
| chr3-26247806-INS-935 | chr3-26247806-INS-935 |
| chr3-26303586-INS-1999 | chr3-26303587-INS-1999 |
| chr3-26331972-INS-88 | chr3-26331965-INS-124 |
| chr3-26337848-INS-9793 | chr3-26337849-INS-9797 |
| chr3-26348337-INS-62 | chr3-26348337-INS-62 |
| chr3-26351032-INS-69 | chr3-26351032-INS-69 |
| chr3-26360349-INS-71 | chr3-26360348-INS-74 |
| chr3-26368670-INS-3009 | chr3-26368671-INS-3009 |
| chr3-26418341-INS-2122 | chr3-26418342-INS-2122 |
| chr3-26465798-INS-6254 | chr3-26465799-INS-6256 |
| chr3-26485949-INS-3114 | chr3-26485950-INS-3114 |
| chr3-26496011-INS-89 | chr3-26496011-INS-89 |

| | |
|---|---|
| chr3-26508829-INS-286 | chr3-26508829-INS-258 |
| chr3-26514525-INS-141 | chr3-26514523-INS-137 |
| chr3-26529225-INS-57 | chr3-26529225-INS-63 |
| chr3-26537788-INS-194 | chr3-26537788-INS-188 |
| chr3-26539156-INS-64 | chr3-26539156-INS-64 |
| chr3-26552362-INS-398 | chr3-26552362-INS-394 |
| chr3-26590881-INS-56 | chr3-26590881-INS-56 |
| chr3-26644261-INS-58 | chr3-26644261-INS-58 |
| chr3-26645172-INS-2294 | chr3-26645173-INS-2272 |
| chr3-26650915-INS-282 | chr3-26650915-INS-282 |
| chr3-26659679-INS-96 | chr3-26659679-INS-76 |
| chr3-26661240-INS-825 | chr3-26661240-INS-825 |
| chr3-26669385-INS-6502 | chr3-26669386-INS-6503 |
| chr3-26675926-INS-1320 | chr3-26675927-INS-1320 |
| chr3-26715550-INS-1872 | chr3-26715551-INS-1873 |
| chr3-26725649-INS-54 | chr3-26725649-INS-54 |
| chr3-26735092-INS-78 | chr3-26735092-INS-80 |
| chr3-26740054-INS-120 | chr3-26740054-INS-120 |
| chr3-26744829-INS-1441 | chr3-26744830-INS-1441 |
| chr3-26752110-INS-6221 | chr3-26752111-INS-6225 |
| chr3-26755453-INS-6404 | chr3-26755454-INS-6404 |
| chr3-26772545-INS-358 | chr3-26772545-INS-358 |
| chr3-26789881-INS-658 | chr3-26789881-INS-658 |
| chr3-26812595-INS-4552 | chr3-26812596-INS-4552 |
| chr3-26833715-INS-88 | chr3-26833710-INS-80 |
| chr3-26838359-INS-214 | chr3-26838359-INS-214 |
| chr3-26839714-INS-176 | chr3-26839714-INS-176 |
| chr3-26857315-INS-6506 | chr3-26857316-INS-6506 |
| chr3-26896001-INS-86 | chr3-26896001-INS-86 |
| chr3-26922300-INS-69 | chr3-26922300-INS-69 |
| chr3-26937402-INS-3166 | chr3-26937403-INS-3167 |
| chr3-26947256-INS-76 | chr3-26947257-INS-56 |
| chr3-26979695-INS-6980 | chr3-26979696-INS-6981 |
| chr3-26984206-INS-92 | chr3-26984206-INS-56 |
| chr3-26987908-INS-284 | chr3-26987908-INS-285 |
| chr3-27088743-INS-3263 | chr3-27088744-INS-3263 |
| chr3-27089002-INS-116 | chr3-27089002-INS-120 |
| chr3-27093064-INS-203 | chr3-27093064-INS-203 |
| chr3-27104841-INS-64 | chr3-27104822-INS-71 |
| chr3-27109495-INS-148 | chr3-27109533-INS-153 |
| chr3-27115986-INS-76 | chr3-27116029-INS-66 |
| chr3-27127781-INS-228 | chr3-27127781-INS-228 |
| chr3-27136700-INS-229 | chr3-27136749-INS-230 |
| chr3-27214099-INS-57 | chr3-27214099-INS-57 |
| chr3-27217119-INS-946 | chr3-27217119-INS-946 |
| chr3-27218302-INS-326 | chr3-27218302-INS-326 |
| chr3-27223787-INS-204 | chr3-27223787-INS-202 |
| chr3-27277313-INS-54 | chr3-27277313-INS-78 |
| chr3-27278246-INS-199 | chr3-27278246-INS-199 |
| chr3-27321401-INS-87 | chr3-27321401-INS-87 |
| chr3-27341653-INS-57 | chr3-27341653-INS-57 |
| chr3-27345419-INS-589 | chr3-27345430-INS-585 |
| chr3-27346215-INS-227 | chr3-27346215-INS-230 |
| chr3-27354290-INS-138 | chr3-27354290-INS-138 |
| chr3-27358719-INS-6403 | chr3-27358720-INS-6403 |
| chr3-27374441-INS-56 | chr3-27374443-INS-52 |
| chr3-27393036-INS-408 | chr3-27393036-INS-415 |
| chr3-27534868-INS-177 | chr3-27534868-INS-177 |
| chr3-27536370-INS-71 | chr3-27536348-INS-54 |
| chr3-27564436-INS-1810 | chr3-27564437-INS-1810 |
| chr3-27606524-INS-58 | chr3-27606524-INS-60 |
| chr3-27620576-INS-85 | chr3-27620576-INS-85 |
| chr3-27635742-INS-60 | chr3-27635701-INS-75 |
| chr3-27638237-INS-180 | chr3-27638237-INS-180 |
| chr3-27650352-INS-117 | chr3-27650352-INS-72 |
| chr3-27651278-INS-149 | chr3-27651252-INS-182 |
| chr3-27663800-INS-70 | chr3-27663800-INS-72 |
| chr3-27700136-INS-1880 | chr3-27700137-INS-1880 |
| chr3-27709515-INS-56 | chr3-27709515-INS-50 |
| chr3-27759261-INS-56 | chr3-27759261-INS-56 |
| chr3-27768710-INS-54 | chr3-27768711-INS-57 |
| chr3-27788700-INS-62 | chr3-27788703-INS-66 |
| chr3-27806569-INS-80 | chr3-27806569-INS-80 |
| chr3-27827250-INS-124 | chr3-27827250-INS-124 |
| chr3-27852121-INS-649 | chr3-27852121-INS-647 |
| chr3-27883945-INS-132 | chr3-27883945-INS-132 |
| chr3-27891763-INS-66 | chr3-27891763-INS-63 |

| | |
|---|---|
| chr3-27892192-INS-476 | chr3-27892192-INS-477 |
| chr3-27894079-INS-491 | chr3-27894095-INS-443 |
| chr3-27904973-INS-2682 | chr3-27904974-INS-2682 |
| chr3-27928767-INS-67 | chr3-27928767-INS-67 |
| chr3-27973718-INS-90 | chr3-27973717-INS-66 |
| chr3-27982594-INS-191 | chr3-27982594-INS-193 |
| chr3-27985543-INS-88 | chr3-27985531-INS-82 |
| chr3-27992259-INS-76 | chr3-27992259-INS-54 |
| chr3-27999008-INS-254 | chr3-27999003-INS-260 |
| chr3-28007093-INS-196 | chr3-28007093-INS-196 |
| chr3-28011951-INS-209 | chr3-28011951-INS-210 |
| chr3-28065823-INS-153 | chr3-28065823-INS-153 |
| chr3-28135679-INS-302 | chr3-28135679-INS-299 |
| chr3-28175210-INS-1966 | chr3-28175211-INS-1978 |
| chr3-28184944-INS-386 | chr3-28184944-INS-386 |
| chr3-28190122-INS-1210 | chr3-28190123-INS-1209 |
| chr3-28204923-INS-6708 | chr3-28204924-INS-6708 |
| chr3-28206570-INS-192 | chr3-28206570-INS-192 |
| chr3-28214889-INS-66 | chr3-28214875-INS-78 |
| chr3-28216394-INS-98 | chr3-28216442-INS-99 |
| chr3-28219507-INS-67 | chr3-28219511-INS-55 |
| chr3-28292482-INS-93 | chr3-28292487-INS-57 |
| chr3-28297582-INS-60 | chr3-28297582-INS-88 |
| chr3-28336878-INS-96 | chr3-28336878-INS-96 |
| chr3-28396112-INS-133 | chr3-28396095-INS-139 |
| chr3-28426642-INS-59 | chr3-28426642-INS-59 |
| chr3-28432519-INS-616 | chr3-28432519-INS-616 |
| chr3-28437555-INS-856 | chr3-28437555-INS-856 |
| chr3-28443248-INS-248 | chr3-28443248-INS-248 |
| chr3-28445515-INS-189 | chr3-28445515-INS-211 |
| chr3-28450607-INS-59 | chr3-28450607-INS-59 |
| chr3-28463511-INS-248 | chr3-28463511-INS-248 |
| chr3-28472215-INS-64 | chr3-28472216-INS-57 |
| chr3-28537787-INS-351 | chr3-28537787-INS-351 |
| chr3-28546893-INS-99 | chr3-28546891-INS-104 |
| chr3-28562866-INS-99 | chr3-28562866-INS-99 |
| chr3-28592402-INS-108 | chr3-28592389-INS-142 |
| chr3-28601462-INS-112 | chr3-28601462-INS-112 |
| chr3-28643724-INS-53 | chr3-28643706-INS-64 |
| chr3-28674205-INS-80 | chr3-28674205-INS-78 |
| chr3-28678993-INS-80 | chr3-28678993-INS-90 |
| chr3-28698328-INS-434 | chr3-28698328-INS-429 |
| chr3-28767320-INS-611 | chr3-28767305-INS-594 |
| chr3-28774387-INS-189 | chr3-28774387-INS-189 |
| chr3-28789307-INS-52 | chr3-28789307-INS-52 |
| chr3-28795159-INS-101 | chr3-28795159-INS-101 |
| chr3-28802487-INS-1546 | chr3-28802488-INS-1546 |
| chr3-28804535-INS-469 | chr3-28804535-INS-482 |
| chr3-28821427-INS-65 | chr3-28821427-INS-65 |
| chr3-28822560-INS-484 | chr3-28822560-INS-485 |
| chr3-28841224-INS-185 | chr3-28841224-INS-185 |
| chr3-28855385-INS-401 | chr3-28855385-INS-402 |
| chr3-28867627-INS-89 | chr3-28867627-INS-89 |
| chr3-28894677-INS-482 | chr3-28894672-INS-481 |
| chr3-28896706-INS-62 | chr3-28896706-INS-61 |
| chr3-28899644-INS-176 | chr3-28899644-INS-176 |
| chr3-28902781-INS-76 | chr3-28902745-INS-118 |
| chr3-28913649-INS-132 | chr3-28913649-INS-132 |
| chr3-28967143-INS-2532 | chr3-28967144-INS-2533 |
| chr3-28985606-INS-105 | chr3-28985606-INS-105 |
| chr3-28992835-INS-54 | chr3-28992835-INS-54 |
| chr3-29021137-INS-84 | chr3-29021137-INS-84 |
| chr3-29033929-INS-102 | chr3-29033929-INS-102 |
| chr3-29055313-INS-80 | chr3-29055309-INS-72 |
| chr3-29096267-INS-75 | chr3-29096267-INS-75 |
| chr3-29109428-INS-440 | chr3-29109428-INS-440 |
| chr3-29111577-INS-79 | chr3-29111588-INS-68 |
| chr3-29117697-INS-66 | chr3-29117697-INS-66 |
| chr3-29150661-INS-84 | chr3-29150661-INS-80 |
| chr3-29163972-INS-338 | chr3-29163972-INS-340 |
| chr3-29173595-INS-64 | chr3-29173595-INS-64 |
| chr3-29176422-INS-597 | chr3-29176422-INS-597 |
| chr3-29186158-INS-65 | chr3-29186158-INS-65 |
| chr3-29213566-INS-1155 | chr3-29213567-INS-1157 |
| chr3-29218251-INS-363 | chr3-29218251-INS-363 |
| chr3-29233134-INS-549 | chr3-29233134-INS-548 |
| chr3-29257929-INS-52 | chr3-29257929-INS-52 |

| | |
|---|---|
| chr3-29291108-INS-95 | chr3-29291108-INS-95 |
| chr3-29297021-INS-99 | chr3-29297021-INS-84 |
| chr3-29303051-INS-60 | chr3-29303051-INS-63 |
| chr3-29313213-INS-60 | chr3-29313213-INS-50 |
| chr3-29323417-INS-131 | chr3-29323417-INS-131 |
| chr3-29331946-INS-92 | chr3-29331942-INS-80 |
| chr3-29344667-INS-52 | chr3-29344667-INS-54 |
| chr3-29378157-INS-7307 | chr3-29378158-INS-7307 |
| chr3-29381658-INS-200 | chr3-29381658-INS-204 |
| chr3-29392653-INS-198 | chr3-29392653-INS-198 |
| chr3-29431389-INS-1004 | chr3-29431390-INS-1003 |
| chr3-29432524-INS-81 | chr3-29432492-INS-97 |
| chr3-29433962-INS-61 | chr3-29433962-INS-61 |
| chr3-29502076-INS-6896 | chr3-29502077-INS-6897 |
| chr3-29504217-INS-52 | chr3-29504217-INS-66 |
| chr3-29564443-INS-316 | chr3-29564476-INS-296 |
| chr3-29566917-INS-57 | chr3-29566917-INS-57 |
| chr3-29572714-INS-344 | chr3-29572715-INS-346 |
| chr3-29604196-INS-165 | chr3-29604196-INS-165 |
| chr3-29651727-INS-1242 | chr3-29651726-INS-1245 |
| chr3-29688167-INS-445 | chr3-29688167-INS-444 |
| chr3-29708998-INS-7085 | chr3-29708999-INS-7085 |
| chr3-29760767-INS-113 | chr3-29760767-INS-113 |
| chr3-29770644-INS-651 | chr3-29770644-INS-651 |
| chr3-29773836-INS-200 | chr3-29773836-INS-200 |
| chr3-29781741-INS-59 | chr3-29781739-INS-62 |
| chr3-29795543-INS-197 | chr3-29795543-INS-197 |
| chr3-29808395-INS-2890 | chr3-29808396-INS-2890 |
| chr3-29809972-INS-109 | chr3-29809972-INS-109 |
| chr3-29818865-INS-6475 | chr3-29818866-INS-6475 |
| chr3-29821804-INS-68 | chr3-29821804-INS-66 |
| chr3-29827901-INS-696 | chr3-29827900-INS-681 |
| chr3-29839240-INS-4827 | chr3-29839247-INS-4812 |
| chr3-29839893-INS-52 | chr3-29839893-INS-61 |
| chr3-29842775-INS-868 | chr3-29842775-INS-874 |
| chr3-29859678-INS-6403 | chr3-29859679-INS-6403 |
| chr3-29861181-INS-256 | chr3-29861181-INS-245 |
| chr3-29940738-INS-88 | chr3-29940738-INS-87 |
| chr3-29951567-INS-65 | chr3-29951567-INS-53 |
| chr3-29978705-INS-80 | chr3-29978705-INS-84 |
| chr3-29979047-INS-503 | chr3-29979047-INS-503 |
| chr3-29993612-INS-78 | chr3-29993604-INS-56 |
| chr3-30002778-INS-53 | chr3-30002778-INS-53 |
| chr3-30038113-INS-56 | chr3-30038113-INS-56 |
| chr3-30040484-INS-181 | chr3-30040484-INS-177 |
| chr3-30048026-INS-626 | chr3-30047996-INS-664 |
| chr3-30070118-INS-55 | chr3-30070118-INS-53 |
| chr3-30128313-INS-220 | chr3-30128313-INS-220 |
| chr3-30130107-INS-84 | chr3-30130111-INS-72 |
| chr3-30132772-INS-52 | chr3-30132772-INS-50 |
| chr3-30174609-INS-93 | chr3-30174602-INS-100 |
| chr3-30222573-INS-54 | chr3-30222573-INS-50 |
| chr3-30312205-INS-54 | chr3-30312205-INS-78 |
| chr3-30334433-INS-182 | chr3-30334433-INS-151 |
| chr3-30422276-INS-99 | chr3-30422276-INS-95 |
| chr3-30439468-INS-897 | chr3-30439468-INS-894 |
| chr3-30509445-INS-215 | chr3-30509445-INS-215 |
| chr3-30533770-INS-599 | chr3-30533770-INS-599 |
| chr3-30579331-INS-77 | chr3-30579331-INS-77 |
| chr3-30580273-INS-445 | chr3-30580273-INS-445 |
| chr3-30585579-INS-357 | chr3-30585579-INS-357 |
| chr3-30603987-INS-647 | chr3-30603987-INS-647 |
| chr3-30615517-INS-381 | chr3-30615517-INS-381 |
| chr3-30624471-INS-663 | chr3-30624471-INS-663 |
| chr3-30628731-INS-584 | chr3-30628742-INS-594 |
| chr3-30642429-INS-79 | chr3-30642429-INS-90 |
| chr3-30658805-INS-56 | chr3-30658781-INS-68 |
| chr3-30677864-INS-211 | chr3-30677864-INS-199 |
| chr3-30688231-INS-191 | chr3-30688217-INS-190 |
| chr3-30724626-INS-7209 | chr3-30724627-INS-7209 |
| chr3-30758560-INS-85 | chr3-30758560-INS-90 |
| chr3-30760051-INS-1012 | chr3-30760052-INS-1012 |
| chr3-30764001-INS-60 | chr3-30764001-INS-60 |
| chr3-30767499-INS-6164 | chr3-30767500-INS-6129 |
| chr3-30774579-INS-68 | chr3-30774579-INS-84 |
| chr3-30781914-INS-146 | chr3-30781914-INS-169 |
| chr3-30796773-INS-256 | chr3-30796773-INS-257 |

| | |
|---|---|
| chr3-30812404-INS-473 | chr3-30812404-INS-473 |
| chr3-30829852-INS-203 | chr3-30829852-INS-203 |
| chr3-30830959-INS-52 | chr3-30830959-INS-52 |
| chr3-30832327-INS-179 | chr3-30832327-INS-172 |
| chr3-30840247-INS-102 | chr3-30840207-INS-104 |
| chr3-30852377-INS-102 | chr3-30852377-INS-102 |
| chr3-30870150-INS-199 | chr3-30870150-INS-199 |
| chr3-30888397-INS-1526 | chr3-30888398-INS-1503 |
| chr3-30900656-INS-114 | chr3-30900656-INS-114 |
| chr3-30901831-INS-1266 | chr3-30901832-INS-1261 |
| chr3-30905403-INS-53 | chr3-30905403-INS-60 |
| chr3-30910625-INS-56 | chr3-30910625-INS-56 |
| chr3-30936535-INS-223 | chr3-30936535-INS-223 |
| chr3-30938053-INS-636 | chr3-30938054-INS-636 |
| chr3-30940873-INS-97 | chr3-30940873-INS-68 |
| chr3-30984654-INS-200 | chr3-30984654-INS-160 |
| chr3-31019752-INS-79 | chr3-31019752-INS-79 |
| chr3-31024648-INS-84 | chr3-31024648-INS-60 |
| chr3-31042631-INS-220 | chr3-31042631-INS-220 |
| chr3-31055212-INS-95 | chr3-31055212-INS-95 |
| chr3-31075816-INS-330 | chr3-31075816-INS-379 |
| chr3-31078237-INS-155 | chr3-31078235-INS-163 |
| chr3-31094573-INS-203 | chr3-31094573-INS-203 |
| chr3-31096188-INS-60 | chr3-31096188-INS-88 |
| chr3-31103809-INS-603 | chr3-31103808-INS-608 |
| chr3-31124236-INS-383 | chr3-31124236-INS-390 |
| chr3-31130537-INS-230 | chr3-31130537-INS-231 |
| chr3-31137458-INS-88 | chr3-31137458-INS-88 |
| chr3-31141220-INS-64 | chr3-31141220-INS-68 |
| chr3-31181689-INS-289 | chr3-31181689-INS-329 |
| chr3-31184650-INS-74 | chr3-31184650-INS-74 |
| chr3-31187422-INS-70 | chr3-31187422-INS-72 |
| chr3-31195098-INS-172 | chr3-31195098-INS-172 |
| chr3-31218703-INS-70 | chr3-31218703-INS-74 |
| chr3-31229107-INS-62 | chr3-31229119-INS-93 |
| chr3-31235204-INS-90 | chr3-31235204-INS-90 |
| chr3-31245871-INS-198 | chr3-31245871-INS-198 |
| chr3-31254043-INS-296 | chr3-31254043-INS-265 |
| chr3-31263964-INS-52 | chr3-31263964-INS-52 |
| chr3-31267494-INS-63 | chr3-31267494-INS-63 |
| chr3-31278949-INS-79 | chr3-31278950-INS-104 |
| chr3-31301962-INS-626 | chr3-31301962-INS-626 |
| chr3-31312999-INS-214 | chr3-31312999-INS-170 |
| chr3-31338343-INS-122 | chr3-31338346-INS-109 |
| chr3-31340361-INS-365 | chr3-31340359-INS-356 |
| chr3-31341617-INS-58 | chr3-31341617-INS-58 |
| chr3-31343159-INS-186 | chr3-31343159-INS-186 |
| chr3-31365279-INS-435 | chr3-31365279-INS-435 |
| chr3-31402348-INS-535 | chr3-31402348-INS-535 |
| chr3-31417461-INS-66 | chr3-31417461-INS-64 |
| chr3-31420521-INS-201 | chr3-31420521-INS-201 |
| chr3-31435207-INS-60 | chr3-31435207-INS-60 |
| chr3-31461010-INS-148 | chr3-31461010-INS-144 |
| chr3-31469171-INS-330 | chr3-31469172-INS-344 |
| chr3-31486835-INS-830 | chr3-31486836-INS-831 |
| chr3-31494938-INS-60 | chr3-31494938-INS-72 |
| chr3-31496361-INS-602 | chr3-31496361-INS-602 |
| chr3-31501404-INS-456 | chr3-31501404-INS-456 |
| chr3-31505542-INS-59 | chr3-31505542-INS-59 |
| chr3-31507989-INS-1128 | chr3-31507990-INS-1128 |
| chr3-31508850-INS-6343 | chr3-31508851-INS-6345 |
| chr3-31512058-INS-55 | chr3-31512058-INS-51 |
| chr3-31557410-INS-2683 | chr3-31557411-INS-2685 |
| chr3-31567749-INS-707 | chr3-31567749-INS-709 |
| chr3-31576615-INS-1890 | chr3-31576616-INS-1890 |
| chr3-31640608-INS-104 | chr3-31640608-INS-104 |
| chr3-31669998-INS-152 | chr3-31669998-INS-152 |
| chr3-31673046-INS-1846 | chr3-31673047-INS-1846 |
| chr3-31704697-INS-207 | chr3-31704697-INS-203 |
| chr3-31743612-INS-240 | chr3-31743612-INS-240 |
| chr3-31745122-INS-83 | chr3-31745124-INS-122 |
| chr3-31747668-INS-84 | chr3-31747668-INS-76 |
| chr3-31768034-INS-556 | chr3-31768034-INS-558 |
| chr3-31774617-INS-1908 | chr3-31774623-INS-1906 |
| chr3-31788248-INS-153 | chr3-31788248-INS-153 |
| chr3-31812951-INS-210 | chr3-31812951-INS-212 |
| chr3-31823729-INS-90 | chr3-31823729-INS-90 |

| | |
|---|---|
| chr3-31825906-INS-2970 | chr3-31825907-INS-2955 |
| chr3-31836552-INS-281 | chr3-31836552-INS-289 |
| chr3-31856740-INS-64 | chr3-31856740-INS-80 |
| chr3-31857040-INS-1034 | chr3-31857041-INS-1034 |
| chr3-31880401-INS-2790 | chr3-31880402-INS-2789 |
| chr3-31930891-INS-5286 | chr3-31930892-INS-5287 |
| chr3-31947999-INS-68 | chr3-31947999-INS-68 |
| chr3-31954163-INS-160 | chr3-31954167-INS-192 |
| chr3-31986710-INS-378 | chr3-31986710-INS-378 |
| chr3-32012066-INS-57 | chr3-32012066-INS-65 |
| chr3-32013576-INS-431 | chr3-32013576-INS-431 |
| chr3-32020331-INS-53 | chr3-32020298-INS-80 |
| chr3-32039108-INS-675 | chr3-32039107-INS-678 |
| chr3-32050074-INS-394 | chr3-32050074-INS-394 |
| chr3-32071603-INS-81 | chr3-32071603-INS-81 |
| chr3-32077405-INS-384 | chr3-32077405-INS-400 |
| chr3-32080521-INS-56 | chr3-32080521-INS-56 |
| chr3-32081757-INS-52 | chr3-32081728-INS-52 |
| chr3-32083308-INS-55 | chr3-32083308-INS-55 |
| chr3-32088589-INS-516 | chr3-32088589-INS-507 |
| chr3-32106041-INS-90 | chr3-32106041-INS-90 |
| chr3-32108337-INS-6440 | chr3-32108338-INS-6438 |
| chr3-32152974-INS-193 | chr3-32152974-INS-193 |
| chr3-32205538-INS-52 | chr3-32205538-INS-88 |
| chr3-32243209-INS-92 | chr3-32243209-INS-96 |
| chr3-32245315-INS-896 | chr3-32245315-INS-897 |
| chr3-32247749-INS-110 | chr3-32247749-INS-112 |
| chr3-32250712-INS-329 | chr3-32250712-INS-328 |
| chr3-32260783-INS-197 | chr3-32260783-INS-196 |
| chr3-32275418-INS-58 | chr3-32275418-INS-58 |
| chr3-32276502-INS-385 | chr3-32276502-INS-385 |
| chr3-32285857-INS-98 | chr3-32285883-INS-72 |
| chr3-32287925-INS-55 | chr3-32287925-INS-55 |
| chr3-32294964-INS-201 | chr3-32294964-INS-201 |
| chr3-32296067-INS-7147 | chr3-32296068-INS-7163 |
| chr3-32322079-INS-130 | chr3-32322086-INS-90 |
| chr3-32342838-INS-53 | chr3-32342838-INS-53 |
| chr3-32397222-INS-214 | chr3-32397222-INS-214 |
| chr3-32408618-INS-554 | chr3-32408619-INS-554 |
| chr3-32414170-INS-224 | chr3-32414170-INS-224 |
| chr3-32423371-INS-237 | chr3-32423371-INS-237 |
| chr3-32430426-INS-51 | chr3-32430426-INS-53 |
| chr3-32445319-INS-83 | chr3-32445319-INS-89 |
| chr3-32469057-INS-217 | chr3-32469057-INS-219 |
| chr3-32520798-INS-82 | chr3-32520798-INS-123 |
| chr3-32529727-INS-471 | chr3-32529727-INS-471 |
| chr3-32532777-INS-4491 | chr3-32532778-INS-4491 |
| chr3-32535281-INS-85 | chr3-32535281-INS-89 |
| chr3-32552022-INS-129 | chr3-32552022-INS-129 |
| chr3-32569573-INS-209 | chr3-32569573-INS-210 |
| chr3-32587167-INS-207 | chr3-32587167-INS-207 |
| chr3-32627056-INS-296 | chr3-32627056-INS-313 |
| chr3-32655864-INS-127 | chr3-32655864-INS-127 |
| chr3-32693263-INS-1866 | chr3-32693264-INS-1866 |
| chr3-32696776-INS-52 | chr3-32696780-INS-64 |
| chr3-32704273-INS-248 | chr3-32704273-INS-248 |
| chr3-32714349-INS-59 | chr3-32714349-INS-59 |
| chr3-32719851-INS-52 | chr3-32719851-INS-52 |
| chr3-32736629-INS-80 | chr3-32736629-INS-56 |
| chr3-32750903-INS-195 | chr3-32750903-INS-187 |
| chr3-32759918-INS-210 | chr3-32759918-INS-211 |
| chr3-32796172-INS-236 | chr3-32796172-INS-236 |
| chr3-32799863-INS-164 | chr3-32799863-INS-164 |
| chr3-32832185-INS-381 | chr3-32832180-INS-382 |
| chr3-32840595-INS-127 | chr3-32840595-INS-128 |
| chr3-32851637-INS-92 | chr3-32851637-INS-92 |
| chr3-32860969-INS-202 | chr3-32860969-INS-202 |
| chr3-32863961-INS-107 | chr3-32863942-INS-96 |
| chr3-32895721-INS-89 | chr3-32895721-INS-89 |
| chr3-32913221-INS-84 | chr3-32913221-INS-84 |
| chr3-32922999-INS-216 | chr3-32922999-INS-216 |
| chr3-32941726-INS-66 | chr3-32941726-INS-66 |
| chr3-32944047-INS-140 | chr3-32944051-INS-96 |
| chr3-32945306-INS-87 | chr3-32945303-INS-72 |
| chr3-32963999-INS-474 | chr3-32963998-INS-474 |
| chr3-32991174-INS-428 | chr3-32991174-INS-427 |
| chr3-32992708-INS-52 | chr3-32992725-INS-60 |

| | |
|---|---|
| chr3-33005773-INS-94 | chr3-33005731-INS-119 |
| chr3-33056317-INS-2552 | chr3-33056318-INS-2552 |
| chr3-33093444-INS-60 | chr3-33093442-INS-62 |
| chr3-33095039-INS-1681 | chr3-33095040-INS-1655 |
| chr3-33102762-INS-78 | chr3-33102762-INS-66 |
| chr3-33104924-INS-111 | chr3-33104924-INS-102 |
| chr3-33107803-INS-115 | chr3-33107800-INS-121 |
| chr3-33133274-INS-162 | chr3-33133274-INS-162 |
| chr3-33149193-INS-67 | chr3-33149238-INS-68 |
| chr3-33157710-INS-403 | chr3-33157710-INS-408 |
| chr3-33161710-INS-103 | chr3-33161686-INS-113 |
| chr3-33205101-INS-1392 | chr3-33205102-INS-1381 |
| chr3-33212764-INS-65 | chr3-33212777-INS-70 |
| chr3-33218479-INS-399 | chr3-33218479-INS-399 |
| chr3-33231566-INS-56 | chr3-33231566-INS-56 |
| chr3-33275036-INS-6505 | chr3-33275037-INS-6505 |
| chr3-33305957-INS-528 | chr3-33305957-INS-528 |
| chr3-33324057-INS-200 | chr3-33324057-INS-200 |
| chr3-33352725-INS-92 | chr3-33352728-INS-70 |
| chr3-33392253-INS-74 | chr3-33392253-INS-85 |
| chr3-33420312-INS-210 | chr3-33420315-INS-214 |
| chr3-33424927-INS-58 | chr3-33424927-INS-52 |
| chr3-33426695-INS-59 | chr3-33426695-INS-59 |
| chr3-33442732-INS-91 | chr3-33442745-INS-99 |
| chr3-33445392-INS-437 | chr3-33445392-INS-437 |
| chr3-33456788-INS-62 | chr3-33456788-INS-60 |
| chr3-33469905-INS-144 | chr3-33469905-INS-134 |
| chr3-33476111-INS-343 | chr3-33476111-INS-347 |
| chr3-33479001-INS-172 | chr3-33479001-INS-173 |
| chr3-33508175-INS-1863 | chr3-33508176-INS-1863 |
| chr3-33508434-INS-56 | chr3-33508422-INS-56 |
| chr3-33511269-INS-51 | chr3-33511229-INS-80 |
| chr3-33546362-INS-7337 | chr3-33546363-INS-7310 |
| chr3-33568644-INS-2772 | chr3-33568645-INS-2755 |
| chr3-33596454-INS-71 | chr3-33596454-INS-63 |
| chr3-33607266-INS-516 | chr3-33607266-INS-482 |
| chr3-33619523-INS-56 | chr3-33619523-INS-56 |
| chr3-33638116-INS-416 | chr3-33638116-INS-416 |
| chr3-33653890-INS-215 | chr3-33653890-INS-215 |
| chr3-33721629-INS-54 | chr3-33721629-INS-50 |
| chr3-33726128-INS-464 | chr3-33726128-INS-465 |
| chr3-33747421-INS-72 | chr3-33747421-INS-72 |
| chr3-33756911-INS-689 | chr3-33756911-INS-691 |
| chr3-33761949-INS-175 | chr3-33761949-INS-178 |
| chr3-33790364-INS-627 | chr3-33790364-INS-629 |
| chr3-33797938-INS-1554 | chr3-33797939-INS-1553 |
| chr3-33800318-INS-6215 | chr3-33800319-INS-6215 |
| chr3-33806242-INS-400 | chr3-33806242-INS-400 |
| chr3-33808488-INS-505 | chr3-33808488-INS-505 |
| chr3-33814393-INS-551 | chr3-33814393-INS-566 |
| chr3-33840890-INS-199 | chr3-33840890-INS-205 |
| chr3-33856378-INS-220 | chr3-33856378-INS-222 |
| chr3-33865114-INS-380 | chr3-33865114-INS-380 |
| chr3-33879119-INS-6431 | chr3-33879120-INS-6431 |
| chr3-33896068-INS-360 | chr3-33896068-INS-360 |
| chr3-33920465-INS-60 | chr3-33920465-INS-63 |
| chr3-33966445-INS-226 | chr3-33966445-INS-226 |
| chr3-33976275-INS-4041 | chr3-33976276-INS-4041 |
| chr3-34007257-INS-205 | chr3-34007257-INS-203 |
| chr3-34016965-INS-302 | chr3-34016965-INS-302 |
| chr3-34034281-INS-70 | chr3-34034281-INS-70 |
| chr3-34045886-INS-171 | chr3-34045886-INS-170 |
| chr3-34048892-INS-457 | chr3-34048892-INS-409 |
| chr3-34051239-INS-60 | chr3-34051239-INS-62 |
| chr3-34080466-INS-72 | chr3-34080466-INS-70 |
| chr3-34087699-INS-68 | chr3-34087697-INS-78 |
| chr3-34111197-INS-642 | chr3-34111197-INS-644 |
| chr3-34150516-INS-172 | chr3-34150516-INS-172 |
| chr3-34156310-INS-197 | chr3-34156310-INS-197 |
| chr3-34171493-INS-189 | chr3-34171493-INS-189 |
| chr3-34179079-INS-336 | chr3-34179079-INS-336 |
| chr3-34221591-INS-194 | chr3-34221591-INS-194 |
| chr3-34229451-INS-69 | chr3-34229451-INS-69 |
| chr3-34243588-INS-623 | chr3-34243588-INS-625 |
| chr3-34247781-INS-164 | chr3-34247781-INS-160 |
| chr3-34274434-INS-66 | chr3-34274434-INS-66 |
| chr3-34282536-INS-94 | chr3-34282536-INS-94 |

| | |
|---|---|
| chr3-34291661-INS-187 | chr3-34291661-INS-187 |
| chr3-34302960-INS-971 | chr3-34302960-INS-972 |
| chr3-34317383-INS-131 | chr3-34317383-INS-96 |
| chr3-34326782-INS-55 | chr3-34326782-INS-51 |
| chr3-34355855-INS-66 | chr3-34355819-INS-54 |
| chr3-34408006-INS-52 | chr3-34408006-INS-52 |
| chr3-34424257-INS-95 | chr3-34424257-INS-122 |
| chr3-34440034-INS-209 | chr3-34440034-INS-209 |
| chr3-34446120-INS-58 | chr3-34446120-INS-64 |
| chr3-34474551-INS-142 | chr3-34474546-INS-139 |
| chr3-34513707-INS-57 | chr3-34513707-INS-57 |
| chr3-34515304-INS-988 | chr3-34515304-INS-965 |
| chr3-34518037-INS-70 | chr3-34518037-INS-70 |
| chr3-34534057-INS-322 | chr3-34534057-INS-322 |
| chr3-34561613-INS-67 | chr3-34561613-INS-70 |
| chr3-34588031-INS-180 | chr3-34588031-INS-180 |
| chr3-34596658-INS-78 | chr3-34596659-INS-50 |
| chr3-34607525-INS-760 | chr3-34607525-INS-760 |
| chr3-34620346-INS-81 | chr3-34620346-INS-81 |
| chr3-34633652-INS-60 | chr3-34633652-INS-60 |
| chr3-34636536-INS-213 | chr3-34636536-INS-213 |
| chr3-34649981-INS-313 | chr3-34649981-INS-312 |
| chr3-34678613-INS-65 | chr3-34678623-INS-65 |
| chr3-34681041-INS-2806 | chr3-34681042-INS-2796 |
| chr3-34682123-INS-110 | chr3-34682123-INS-112 |
| chr3-34689717-INS-67 | chr3-34689713-INS-74 |
| chr3-34719646-INS-61 | chr3-34719646-INS-66 |
| chr3-34739308-INS-230 | chr3-34739308-INS-235 |
| chr3-34747569-INS-135 | chr3-34747569-INS-135 |
| chr3-34748448-INS-60 | chr3-34748448-INS-60 |
| chr3-34752505-INS-70 | chr3-34752458-INS-75 |
| chr3-34765851-INS-203 | chr3-34765851-INS-203 |
| chr3-34780123-INS-269 | chr3-34780123-INS-234 |
| chr3-34782981-INS-202 | chr3-34782981-INS-202 |
| chr3-34798075-INS-236 | chr3-34798075-INS-257 |
| chr3-34817099-INS-60 | chr3-34817099-INS-62 |
| chr3-34825152-INS-60 | chr3-34825152-INS-60 |
| chr3-34827456-INS-68 | chr3-34827453-INS-68 |
| chr3-34833334-INS-222 | chr3-34833334-INS-222 |
| chr3-34838037-INS-62 | chr3-34838080-INS-54 |
| chr3-34857847-INS-76 | chr3-34857847-INS-75 |
| chr3-34858805-INS-70 | chr3-34858805-INS-70 |
| chr3-34874970-INS-86 | chr3-34874970-INS-104 |
| chr3-34908970-INS-77 | chr3-34908970-INS-69 |
| chr3-34912680-INS-51 | chr3-34912652-INS-50 |
| chr3-34948748-INS-220 | chr3-34948770-INS-220 |
| chr3-34952965-INS-81 | chr3-34952965-INS-81 |
| chr3-34996894-INS-64 | chr3-34996894-INS-66 |
| chr3-35010074-INS-347 | chr3-35010074-INS-347 |
| chr3-35024834-INS-4486 | chr3-35024835-INS-4486 |
| chr3-35044114-INS-751 | chr3-35044114-INS-746 |
| chr3-35055101-INS-80 | chr3-35055101-INS-80 |
| chr3-35072432-INS-64 | chr3-35072432-INS-64 |
| chr3-35087117-INS-81 | chr3-35087117-INS-81 |
| chr3-35088222-INS-134 | chr3-35088222-INS-134 |
| chr3-35102939-INS-1865 | chr3-35102940-INS-1865 |
| chr3-35119965-INS-72 | chr3-35119965-INS-74 |
| chr3-35150961-INS-214 | chr3-35150961-INS-215 |
| chr3-35216542-INS-60 | chr3-35216542-INS-68 |
| chr3-35218632-INS-403 | chr3-35218632-INS-403 |
| chr3-35279180-INS-1449 | chr3-35279181-INS-1449 |
| chr3-35315611-INS-6670 | chr3-35315612-INS-6670 |
| chr3-35325969-INS-80 | chr3-35325969-INS-80 |
| chr3-35330301-INS-138 | chr3-35330295-INS-170 |
| chr3-35337015-INS-53 | chr3-35337015-INS-53 |
| chr3-35339648-INS-4351 | chr3-35339649-INS-4351 |
| chr3-35392177-INS-825 | chr3-35392177-INS-825 |
| chr3-35441187-INS-65 | chr3-35441180-INS-50 |
| chr3-35442066-INS-1531 | chr3-35442067-INS-1531 |
| chr3-35493581-INS-2795 | chr3-35493582-INS-2795 |
| chr3-35532972-INS-166 | chr3-35532971-INS-166 |
| chr3-35535131-INS-71 | chr3-35535131-INS-57 |
| chr3-35546086-INS-434 | chr3-35546086-INS-432 |
| chr3-35552497-INS-64 | chr3-35552497-INS-66 |
| chr3-35556590-INS-426 | chr3-35556590-INS-426 |
| chr3-35573799-INS-56 | chr3-35573799-INS-56 |
| chr3-35595341-INS-67 | chr3-35595345-INS-67 |

| | |
|---|---|
| chr3-35621077-INS-272 | chr3-35621077-INS-272 |
| chr3-35625203-INS-487 | chr3-35625203-INS-487 |
| chr3-35633093-INS-269 | chr3-35633093-INS-270 |
| chr3-35638104-INS-164 | chr3-35638104-INS-164 |
| chr3-35642016-INS-80 | chr3-35642016-INS-80 |
| chr3-35651858-INS-553 | chr3-35651858-INS-553 |
| chr3-35654372-INS-327 | chr3-35654372-INS-332 |
| chr3-35656121-INS-113 | chr3-35656121-INS-114 |
| chr3-35657294-INS-138 | chr3-35657294-INS-137 |
| chr3-35671890-INS-6217 | chr3-35671890-INS-6218 |
| chr3-35673392-INS-208 | chr3-35673392-INS-208 |
| chr3-35675525-INS-469 | chr3-35675525-INS-466 |
| chr3-35678882-INS-1493 | chr3-35678883-INS-1493 |
| chr3-35688724-INS-507 | chr3-35688724-INS-507 |
| chr3-35715052-INS-65 | chr3-35715068-INS-53 |
| chr3-35722183-INS-819 | chr3-35722183-INS-819 |
| chr3-35728491-INS-190 | chr3-35728491-INS-190 |
| chr3-35730673-INS-325 | chr3-35730673-INS-324 |
| chr3-35737666-INS-3996 | chr3-35737667-INS-3996 |
| chr3-35739901-INS-131 | chr3-35739901-INS-131 |
| chr3-35745946-INS-151 | chr3-35745946-INS-159 |
| chr3-35750576-INS-152 | chr3-35750576-INS-152 |
| chr3-35756582-INS-840 | chr3-35756582-INS-840 |
| chr3-35772766-INS-63 | chr3-35772766-INS-53 |
| chr3-35783987-INS-473 | chr3-35783987-INS-473 |
| chr3-35797459-INS-52 | chr3-35797495-INS-53 |
| chr3-35820147-INS-266 | chr3-35820147-INS-266 |
| chr3-35880712-INS-269 | chr3-35880705-INS-265 |
| chr3-35882175-INS-257 | chr3-35882175-INS-220 |
| chr3-35887025-INS-127 | chr3-35887025-INS-127 |
| chr3-35915853-INS-665 | chr3-35915853-INS-665 |
| chr3-35921259-INS-317 | chr3-35921259-INS-317 |
| chr3-35937159-INS-54 | chr3-35937159-INS-54 |
| chr3-35973921-INS-405 | chr3-35973921-INS-405 |
| chr3-36021712-INS-55 | chr3-36021712-INS-55 |
| chr3-36030692-INS-64 | chr3-36030692-INS-64 |
| chr3-36041200-INS-56 | chr3-36041200-INS-56 |
| chr3-36045962-INS-192 | chr3-36045962-INS-197 |
| chr3-36050321-INS-58 | chr3-36050321-INS-58 |
| chr3-36058321-INS-104 | chr3-36058321-INS-104 |
| chr3-36060876-INS-225 | chr3-36060876-INS-235 |
| chr3-36063405-INS-56 | chr3-36063385-INS-64 |
| chr3-36071575-INS-6540 | chr3-36071576-INS-6541 |
| chr3-36080497-INS-200 | chr3-36080497-INS-200 |
| chr3-36084851-INS-136 | chr3-36084852-INS-136 |
| chr3-36088008-INS-341 | chr3-36088008-INS-341 |
| chr3-36092611-INS-360 | chr3-36092611-INS-376 |
| chr3-36099145-INS-60 | chr3-36099140-INS-54 |
| chr3-36112045-INS-56 | chr3-36112045-INS-66 |
| chr3-36150262-INS-699 | chr3-36150262-INS-698 |
| chr3-36167493-INS-70 | chr3-36167502-INS-71 |
| chr3-36168621-INS-695 | chr3-36168621-INS-695 |
| chr3-36170795-INS-1751 | chr3-36170796-INS-1751 |
| chr3-36186851-INS-390 | chr3-36186851-INS-390 |
| chr3-36207850-INS-205 | chr3-36207850-INS-205 |
| chr3-36228833-INS-209 | chr3-36228833-INS-259 |
| chr3-36238000-INS-400 | chr3-36238000-INS-400 |
| chr3-36256359-INS-6182 | chr3-36256360-INS-6182 |
| chr3-36257494-INS-103 | chr3-36257516-INS-101 |
| chr3-36272744-INS-117 | chr3-36272744-INS-123 |
| chr3-36281298-INS-125 | chr3-36281298-INS-131 |
| chr3-36293716-INS-81 | chr3-36293716-INS-81 |
| chr3-36302535-INS-685 | chr3-36302531-INS-721 |
| chr3-36318862-INS-96 | chr3-36318862-INS-72 |
| chr3-36339240-INS-408 | chr3-36339240-INS-408 |
| chr3-36348575-INS-69 | chr3-36348575-INS-72 |
| chr3-36360411-INS-178 | chr3-36360411-INS-168 |
| chr3-36371897-INS-66 | chr3-36371897-INS-66 |
| chr3-36372304-INS-200 | chr3-36372304-INS-200 |
| chr3-36394116-INS-205 | chr3-36394116-INS-205 |
| chr3-36397300-INS-2029 | chr3-36397301-INS-2029 |
| chr3-36399385-INS-72 | chr3-36399385-INS-76 |
| chr3-36424376-INS-52 | chr3-36424376-INS-52 |
| chr3-36425806-INS-118 | chr3-36425806-INS-118 |
| chr3-36515623-INS-58 | chr3-36515623-INS-58 |
| chr3-36517865-INS-6440 | chr3-36517866-INS-6440 |
| chr3-36535126-INS-74 | chr3-36535126-INS-74 |

| | |
|---|---|
| chr3-36548529-INS-166 | chr3-36548525-INS-130 |
| chr3-36554445-INS-186 | chr3-36554445-INS-155 |
| chr3-36557499-INS-195 | chr3-36557499-INS-195 |
| chr3-36563843-INS-497 | chr3-36563843-INS-497 |
| chr3-36564839-INS-7031 | chr3-36564840-INS-7030 |
| chr3-36573486-INS-198 | chr3-36573486-INS-189 |
| chr3-36588028-INS-70 | chr3-36588047-INS-62 |
| chr3-36614542-INS-64 | chr3-36614542-INS-50 |
| chr3-36615752-INS-95 | chr3-36615752-INS-95 |
| chr3-36619691-INS-54 | chr3-36619691-INS-54 |
| chr3-36634691-INS-367 | chr3-36634670-INS-394 |
| chr3-36649802-INS-52 | chr3-36649802-INS-70 |
| chr3-36672085-INS-66 | chr3-36672085-INS-66 |
| chr3-36683844-INS-63 | chr3-36683844-INS-63 |
| chr3-36686432-INS-84 | chr3-36686432-INS-84 |
| chr3-36687815-INS-165 | chr3-36687815-INS-165 |
| chr3-36690000-INS-96 | chr3-36690000-INS-80 |
| chr3-36717991-INS-56 | chr3-36717991-INS-62 |
| chr3-36726044-INS-53 | chr3-36726044-INS-56 |
| chr3-36745257-INS-188 | chr3-36745257-INS-190 |
| chr3-36748788-INS-58 | chr3-36748788-INS-52 |
| chr3-36780424-INS-84 | chr3-36780424-INS-84 |
| chr3-36801242-INS-85 | chr3-36801242-INS-85 |
| chr3-36815439-INS-209 | chr3-36815413-INS-252 |
| chr3-36827926-INS-1013 | chr3-36827927-INS-1013 |
| chr3-36852727-INS-301 | chr3-36852727-INS-307 |
| chr3-36864264-INS-58 | chr3-36864260-INS-68 |
| chr3-36871870-INS-109 | chr3-36871870-INS-98 |
| chr3-36880877-INS-354 | chr3-36880877-INS-354 |
| chr3-36912841-INS-506 | chr3-36912841-INS-506 |
| chr3-36937519-INS-58 | chr3-36937510-INS-60 |
| chr3-36942486-INS-485 | chr3-36942486-INS-485 |
| chr3-36951674-INS-56 | chr3-36951674-INS-52 |
| chr3-36962526-INS-823 | chr3-36962528-INS-823 |
| chr3-37003471-INS-98 | chr3-37003438-INS-133 |
| chr3-37039339-INS-6558 | chr3-37039340-INS-6559 |
| chr3-37078741-INS-58 | chr3-37078741-INS-67 |
| chr3-37091192-INS-7389 | chr3-37091193-INS-7390 |
| chr3-37095730-INS-393 | chr3-37095730-INS-393 |
| chr3-37100275-INS-728 | chr3-37100275-INS-730 |
| chr3-37101842-INS-229 | chr3-37101842-INS-229 |
| chr3-37106082-INS-127 | chr3-37106080-INS-110 |
| chr3-37109367-INS-492 | chr3-37109367-INS-492 |
| chr3-37114115-INS-6755 | chr3-37114116-INS-6755 |
| chr3-37115920-INS-98 | chr3-37115920-INS-52 |
| chr3-37159634-INS-63 | chr3-37159634-INS-63 |
| chr3-37170588-INS-62 | chr3-37170588-INS-62 |
| chr3-37184396-INS-70 | chr3-37184396-INS-108 |
| chr3-37204155-INS-227 | chr3-37204155-INS-221 |
| chr3-37211852-INS-461 | chr3-37211852-INS-462 |
| chr3-37242121-INS-400 | chr3-37242121-INS-400 |
| chr3-37304955-INS-78 | chr3-37304960-INS-70 |
| chr3-37309190-INS-151 | chr3-37309190-INS-151 |
| chr3-37328624-INS-234 | chr3-37328624-INS-234 |
| chr3-37329187-INS-106 | chr3-37329161-INS-120 |
| chr3-37333393-INS-410 | chr3-37333393-INS-410 |
| chr3-37350478-INS-334 | chr3-37350478-INS-334 |
| chr3-37398914-INS-202 | chr3-37398914-INS-202 |
| chr3-37404189-INS-55 | chr3-37404189-INS-55 |
| chr3-37476943-INS-209 | chr3-37476943-INS-209 |
| chr3-37482675-INS-52 | chr3-37482675-INS-52 |
| chr3-37488120-INS-157 | chr3-37488120-INS-157 |
| chr3-37501109-INS-161 | chr3-37501109-INS-143 |
| chr3-37521111-INS-70 | chr3-37521111-INS-70 |
| chr3-37526745-INS-7195 | chr3-37526746-INS-7197 |
| chr3-37573156-INS-912 | chr3-37573154-INS-915 |
| chr3-37583027-INS-207 | chr3-37583027-INS-207 |
| chr3-37625709-INS-438 | chr3-37625726-INS-439 |
| chr3-37653412-INS-388 | chr3-37653412-INS-388 |
| chr3-37685759-INS-90 | chr3-37685759-INS-90 |
| chr3-37744381-INS-207 | chr3-37744381-INS-207 |
| chr3-37756770-INS-50 | chr3-37756763-INS-54 |
| chr3-37758468-INS-431 | chr3-37758468-INS-431 |
| chr3-37765378-INS-60 | chr3-37765384-INS-54 |
| chr3-37772485-INS-108 | chr3-37772485-INS-108 |
| chr3-37781792-INS-622 | chr3-37781792-INS-622 |
| chr3-37805555-INS-51 | chr3-37805555-INS-51 |

| | |
|---|---|
| chr3-37827162-INS-76 | chr3-37827162-INS-76 |
| chr3-37868323-INS-52 | chr3-37868323-INS-50 |
| chr3-37874087-INS-572 | chr3-37874087-INS-572 |
| chr3-37878554-INS-1152 | chr3-37878555-INS-1152 |
| chr3-37887678-INS-140 | chr3-37887678-INS-140 |
| chr3-37907098-INS-4454 | chr3-37907099-INS-4454 |
| chr3-37912752-INS-195 | chr3-37912752-INS-195 |
| chr3-37915959-INS-109 | chr3-37915959-INS-109 |
| chr3-37928356-INS-118 | chr3-37928356-INS-90 |
| chr3-37934445-INS-66 | chr3-37934445-INS-70 |
| chr3-37952696-INS-59 | chr3-37952696-INS-68 |
| chr3-37970942-INS-51 | chr3-37970950-INS-58 |
| chr3-38085531-INS-7119 | chr3-38085532-INS-7119 |
| chr3-38096061-INS-64 | chr3-38096061-INS-74 |
| chr3-38102600-INS-56 | chr3-38102612-INS-54 |
| chr3-38104218-INS-2803 | chr3-38104219-INS-2807 |
| chr3-38124823-INS-60 | chr3-38124823-INS-108 |
| chr3-38132641-INS-670 | chr3-38132641-INS-669 |
| chr3-38193353-INS-167 | chr3-38193353-INS-167 |
| chr3-38219876-INS-83 | chr3-38219876-INS-83 |
| chr3-38249689-INS-167 | chr3-38249689-INS-167 |
| chr3-38262079-INS-508 | chr3-38262079-INS-508 |
| chr3-38262721-INS-1131 | chr3-38262722-INS-1131 |
| chr3-38278336-INS-78 | chr3-38278336-INS-83 |
| chr3-38321735-INS-561 | chr3-38321735-INS-561 |
| chr3-38336406-INS-61 | chr3-38336406-INS-61 |
| chr3-38340657-INS-184 | chr3-38340657-INS-184 |
| chr3-38343330-INS-60 | chr3-38343330-INS-60 |
| chr3-38348088-INS-194 | chr3-38348068-INS-175 |
| chr3-38350504-INS-475 | chr3-38350504-INS-475 |
| chr3-38352693-INS-108 | chr3-38352693-INS-108 |
| chr3-38355059-INS-52 | chr3-38355059-INS-56 |
| chr3-38360021-INS-499 | chr3-38360021-INS-499 |
| chr3-38363297-INS-54 | chr3-38363297-INS-54 |
| chr3-38375158-INS-282 | chr3-38375158-INS-285 |
| chr3-38404875-INS-58 | chr3-38404875-INS-54 |
| chr3-38406957-INS-168 | chr3-38406957-INS-168 |
| chr3-38413809-INS-52 | chr3-38413807-INS-62 |
| chr3-38419577-INS-394 | chr3-38419577-INS-394 |
| chr3-38420829-INS-2765 | chr3-38420830-INS-2768 |
| chr3-38431880-INS-9494 | chr3-38431881-INS-9494 |
| chr3-38432144-INS-72 | chr3-38432144-INS-72 |
| chr3-38435779-INS-66 | chr3-38435764-INS-68 |
| chr3-38437570-INS-84 | chr3-38437570-INS-54 |
| chr3-38449684-INS-118 | chr3-38449684-INS-94 |
| chr3-38451856-INS-54 | chr3-38451856-INS-66 |
| chr3-38462651-INS-229 | chr3-38462651-INS-229 |
| chr3-38464349-INS-77 | chr3-38464349-INS-105 |
| chr3-38501977-INS-60 | chr3-38501977-INS-60 |
| chr3-38513807-INS-136 | chr3-38513807-INS-130 |
| chr3-38520436-INS-66 | chr3-38520436-INS-99 |
| chr3-38578245-INS-7571 | chr3-38578246-INS-7572 |
| chr3-38612036-INS-84 | chr3-38612036-INS-102 |
| chr3-38633427-INS-330 | chr3-38633428-INS-282 |
| chr3-38646299-INS-53 | chr3-38646299-INS-53 |
| chr3-38656830-INS-783 | chr3-38656830-INS-786 |
| chr3-38665359-INS-60 | chr3-38665359-INS-62 |
| chr3-38673875-INS-274 | chr3-38673918-INS-268 |
| chr3-38689346-INS-240 | chr3-38689346-INS-240 |
| chr3-38722658-INS-54 | chr3-38722658-INS-54 |
| chr3-38724976-INS-82 | chr3-38724976-INS-102 |
| chr3-38762399-INS-7413 | chr3-38762400-INS-7413 |
| chr3-38787987-INS-64 | chr3-38787949-INS-68 |
| chr3-38794590-INS-70 | chr3-38794590-INS-70 |
| chr3-38834511-INS-54 | chr3-38834511-INS-62 |
| chr3-38838672-INS-94 | chr3-38838691-INS-105 |
| chr3-38841035-INS-504 | chr3-38841035-INS-504 |
| chr3-38856468-INS-51 | chr3-38856425-INS-99 |
| chr3-38864718-INS-208 | chr3-38864718-INS-208 |
| chr3-38866039-INS-113 | chr3-38866039-INS-113 |
| chr3-38868783-INS-178 | chr3-38868783-INS-179 |
| chr3-38892189-INS-89 | chr3-38892189-INS-97 |
| chr3-38961231-INS-63 | chr3-38961231-INS-63 |
| chr3-38980687-INS-59 | chr3-38980687-INS-55 |
| chr3-39027195-INS-2291 | chr3-39027196-INS-2291 |
| chr3-39027195-INS-664 | chr3-39027196-INS-682 |
| chr3-39047866-INS-63 | chr3-39047866-INS-63 |

| | |
|---|---|
| chr3-39080107-INS-204 | chr3-39080107-INS-204 |
| chr3-39087779-INS-960 | chr3-39087779-INS-961 |
| chr3-39092756-INS-66 | chr3-39092750-INS-68 |
| chr3-39101259-INS-94 | chr3-39101259-INS-96 |
| chr3-39107992-INS-58 | chr3-39107992-INS-58 |
| chr3-39115825-INS-148 | chr3-39115817-INS-161 |
| chr3-39169369-INS-96 | chr3-39169370-INS-74 |
| chr3-39186793-INS-92 | chr3-39186794-INS-61 |
| chr3-39187720-INS-57 | chr3-39187721-INS-56 |
| chr3-39195486-INS-73 | chr3-39195486-INS-87 |
| chr3-39199797-INS-5555 | chr3-39199798-INS-5555 |
| chr3-39249438-INS-87 | chr3-39249438-INS-87 |
| chr3-39250319-INS-2921 | chr3-39250320-INS-2921 |
| chr3-39258557-INS-87 | chr3-39258544-INS-99 |
| chr3-39268638-INS-91 | chr3-39268638-INS-95 |
| chr3-39310009-INS-1429 | chr3-39310010-INS-1429 |
| chr3-39310201-INS-54 | chr3-39310201-INS-50 |
| chr3-39322001-INS-190 | chr3-39322010-INS-210 |
| chr3-39338888-INS-1291 | chr3-39338889-INS-1291 |
| chr3-39347763-INS-80 | chr3-39347763-INS-76 |
| chr3-39356994-INS-1892 | chr3-39356995-INS-1892 |
| chr3-39368250-INS-74 | chr3-39368250-INS-76 |
| chr3-39382642-INS-129 | chr3-39382643-INS-133 |
| chr3-39402981-INS-148 | chr3-39402981-INS-148 |
| chr3-39406993-INS-142 | chr3-39406993-INS-142 |
| chr3-39408669-INS-74 | chr3-39408669-INS-78 |
| chr3-39431626-INS-911 | chr3-39431626-INS-908 |
| chr3-39439771-INS-13044 | chr3-39439772-INS-13049 |
| chr3-39468281-INS-6346 | chr3-39468282-INS-6347 |
| chr3-39472649-INS-883 | chr3-39472649-INS-882 |
| chr3-39483498-INS-298 | chr3-39483498-INS-300 |
| chr3-39497916-INS-56 | chr3-39497920-INS-77 |
| chr3-39508623-INS-576 | chr3-39508614-INS-563 |
| chr3-39567625-INS-1845 | chr3-39567626-INS-1845 |
| chr3-39594218-INS-649 | chr3-39594218-INS-650 |
| chr3-39597431-INS-64 | chr3-39597433-INS-50 |
| chr3-39623773-INS-104 | chr3-39623773-INS-106 |
| chr3-39640199-INS-123 | chr3-39640199-INS-123 |
| chr3-39644810-INS-97 | chr3-39644797-INS-85 |
| chr3-39677194-INS-333 | chr3-39677194-INS-333 |
| chr3-39690420-INS-60 | chr3-39690420-INS-60 |
| chr3-39691632-INS-214 | chr3-39691632-INS-208 |
| chr3-39701291-INS-70 | chr3-39701291-INS-70 |
| chr3-39734056-INS-6601 | chr3-39734057-INS-6601 |
| chr3-39742979-INS-4838 | chr3-39742980-INS-4838 |
| chr3-39748566-INS-52 | chr3-39748566-INS-52 |
| chr3-39751101-INS-6376 | chr3-39751102-INS-6376 |
| chr3-39761561-INS-60 | chr3-39761561-INS-52 |
| chr3-39776145-INS-114 | chr3-39776145-INS-114 |
| chr3-39782959-INS-250 | chr3-39782959-INS-250 |
| chr3-39793737-INS-6376 | chr3-39793738-INS-6376 |
| chr3-39805880-INS-54 | chr3-39805880-INS-58 |
| chr3-39816236-INS-73 | chr3-39816201-INS-95 |
| chr3-39836009-INS-6847 | chr3-39836010-INS-6847 |
| chr3-39844373-INS-144 | chr3-39844373-INS-144 |
| chr3-39866583-INS-73 | chr3-39866583-INS-94 |
| chr3-39867436-INS-172 | chr3-39867436-INS-172 |
| chr3-39868288-INS-1890 | chr3-39868289-INS-1890 |
| chr3-39895856-INS-2516 | chr3-39895857-INS-2516 |
| chr3-39897505-INS-256 | chr3-39897505-INS-262 |
| chr3-39902099-INS-66 | chr3-39902099-INS-66 |
| chr3-39907378-INS-54 | chr3-39907378-INS-52 |
| chr3-39994368-INS-130 | chr3-39994368-INS-177 |
| chr3-40026027-INS-63 | chr3-40026027-INS-63 |
| chr3-40089764-INS-57 | chr3-40089764-INS-65 |
| chr3-40118903-INS-200 | chr3-40118903-INS-212 |
| chr3-40122054-INS-237 | chr3-40122054-INS-219 |
| chr3-40148590-INS-114 | chr3-40148590-INS-78 |
| chr3-40151999-INS-333 | chr3-40151999-INS-333 |
| chr3-40158617-INS-78 | chr3-40158617-INS-78 |
| chr3-40167239-INS-104 | chr3-40167229-INS-110 |
| chr3-40175271-INS-104 | chr3-40175271-INS-104 |
| chr3-40204701-INS-52 | chr3-40204701-INS-52 |
| chr3-40213437-INS-1627 | chr3-40213438-INS-1627 |
| chr3-40227307-INS-54 | chr3-40227291-INS-66 |
| chr3-40231441-INS-202 | chr3-40231441-INS-202 |
| chr3-40238064-INS-102 | chr3-40238064-INS-66 |

| | |
|---|---|
| chr3-40252577-INS-61 | chr3-40252577-INS-61 |
| chr3-40269333-INS-8203 | chr3-40269334-INS-8201 |
| chr3-40274746-INS-731 | chr3-40274746-INS-731 |
| chr3-40282046-INS-62 | chr3-40282046-INS-60 |
| chr3-40294133-INS-552 | chr3-40294133-INS-558 |
| chr3-40309605-INS-153 | chr3-40309605-INS-153 |
| chr3-40357581-INS-618 | chr3-40357581-INS-618 |
| chr3-40360453-INS-52 | chr3-40360453-INS-52 |
| chr3-40367536-INS-1487 | chr3-40367537-INS-1484 |
| chr3-40426986-INS-62 | chr3-40426986-INS-72 |
| chr3-40431049-INS-6219 | chr3-40431050-INS-6219 |
| chr3-40465875-INS-415 | chr3-40465874-INS-421 |
| chr3-40471064-INS-2150 | chr3-40471065-INS-2148 |
| chr3-40577974-INS-72 | chr3-40577978-INS-56 |
| chr3-40620197-INS-60 | chr3-40620197-INS-60 |
| chr3-40636065-INS-56 | chr3-40636082-INS-56 |
| chr3-40651301-INS-123 | chr3-40651301-INS-123 |
| chr3-40677763-INS-721 | chr3-40677763-INS-721 |
| chr3-40725404-INS-84 | chr3-40725412-INS-102 |
| chr3-40815921-INS-392 | chr3-40815921-INS-392 |
| chr3-40850868-INS-64 | chr3-40850863-INS-58 |
| chr3-40861451-INS-62 | chr3-40861451-INS-66 |
| chr3-40918907-INS-762 | chr3-40918907-INS-761 |
| chr3-40938532-INS-223 | chr3-40938532-INS-223 |
| chr3-40977805-INS-57 | chr3-40977805-INS-57 |
| chr3-41001070-INS-266 | chr3-41001070-INS-267 |
| chr3-41008758-INS-63 | chr3-41008758-INS-65 |
| chr3-41105763-INS-417 | chr3-41105763-INS-418 |
| chr3-41124897-INS-275 | chr3-41124871-INS-298 |
| chr3-41126240-INS-909 | chr3-41126240-INS-908 |
| chr3-41130108-INS-820 | chr3-41130108-INS-820 |
| chr3-41131654-INS-92 | chr3-41131654-INS-92 |
| chr3-41142783-INS-528 | chr3-41142783-INS-528 |
| chr3-41155134-INS-6837 | chr3-41155135-INS-6837 |
| chr3-41184695-INS-82 | chr3-41184695-INS-86 |
| chr3-41199839-INS-61 | chr3-41199839-INS-50 |
| chr3-41209334-INS-6864 | chr3-41209335-INS-6865 |
| chr3-41248401-INS-244 | chr3-41248402-INS-246 |
| chr3-41248458-INS-53 | chr3-41248423-INS-75 |
| chr3-41268955-INS-210 | chr3-41268955-INS-210 |
| chr3-41326799-INS-63 | chr3-41326783-INS-63 |
| chr3-41330020-INS-138 | chr3-41330046-INS-150 |
| chr3-41335590-INS-164 | chr3-41335590-INS-164 |
| chr3-41350272-INS-160 | chr3-41350272-INS-160 |
| chr3-41368912-INS-97 | chr3-41368912-INS-97 |
| chr3-41399394-INS-70 | chr3-41399395-INS-56 |
| chr3-41424621-INS-78 | chr3-41424627-INS-90 |
| chr3-41456181-INS-100 | chr3-41456181-INS-100 |
| chr3-41463119-INS-349 | chr3-41463119-INS-349 |
| chr3-41473546-INS-56 | chr3-41473546-INS-56 |
| chr3-41478910-INS-62 | chr3-41478910-INS-62 |
| chr3-41493933-INS-85 | chr3-41493933-INS-85 |
| chr3-41515124-INS-52 | chr3-41515124-INS-53 |
| chr3-41530918-INS-208 | chr3-41530918-INS-208 |
| chr3-41532442-INS-161 | chr3-41532442-INS-161 |
| chr3-41556821-INS-105 | chr3-41556821-INS-81 |
| chr3-41579485-INS-7777 | chr3-41579486-INS-7777 |
| chr3-41589558-INS-631 | chr3-41589558-INS-634 |
| chr3-41606086-INS-1290 | chr3-41606087-INS-1290 |
| chr3-41608937-INS-748 | chr3-41608937-INS-748 |
| chr3-41624931-INS-143 | chr3-41624931-INS-144 |
| chr3-41636443-INS-241 | chr3-41636443-INS-241 |
| chr3-41654106-INS-89 | chr3-41654106-INS-89 |
| chr3-41685568-INS-6624 | chr3-41685569-INS-6624 |
| chr3-41738607-INS-217 | chr3-41738607-INS-217 |
| chr3-41752394-INS-221 | chr3-41752394-INS-219 |
| chr3-41784840-INS-58 | chr3-41784840-INS-58 |
| chr3-41786106-INS-2898 | chr3-41786107-INS-2894 |
| chr3-41817459-INS-115 | chr3-41817492-INS-72 |
| chr3-41854614-INS-6494 | chr3-41854615-INS-6494 |
| chr3-41879948-INS-112 | chr3-41879948-INS-112 |
| chr3-41886993-INS-226 | chr3-41886993-INS-227 |
| chr3-41904661-INS-969 | chr3-41904661-INS-968 |
| chr3-41922633-INS-387 | chr3-41922633-INS-390 |
| chr3-41929333-INS-303 | chr3-41929339-INS-320 |
| chr3-41939247-INS-55 | chr3-41939245-INS-82 |
| chr3-41939580-INS-6653 | chr3-41939581-INS-6653 |

| | |
|---|---|
| chr3-41941480-INS-94 | chr3-41941480-INS-94 |
| chr3-42001311-INS-535 | chr3-42001311-INS-535 |
| chr3-42023089-INS-96 | chr3-42023089-INS-95 |
| chr3-42047281-INS-2554 | chr3-42047282-INS-2554 |
| chr3-42051074-INS-83 | chr3-42051070-INS-59 |
| chr3-42057065-INS-813 | chr3-42057065-INS-815 |
| chr3-42060325-INS-5446 | chr3-42060326-INS-5446 |
| chr3-42097033-INS-6251 | chr3-42097034-INS-6251 |
| chr3-42098483-INS-780 | chr3-42098483-INS-780 |
| chr3-42101852-INS-721 | chr3-42101852-INS-720 |
| chr3-42157657-INS-51 | chr3-42157614-INS-57 |
| chr3-42164084-INS-80 | chr3-42164084-INS-86 |
| chr3-42174029-INS-64 | chr3-42174029-INS-68 |
| chr3-42197033-INS-208 | chr3-42197033-INS-208 |
| chr3-42197826-INS-962 | chr3-42197826-INS-962 |
| chr3-42198820-INS-138 | chr3-42198820-INS-138 |
| chr3-42215293-INS-641 | chr3-42215293-INS-641 |
| chr3-42217145-INS-78 | chr3-42217151-INS-66 |
| chr3-42220338-INS-158 | chr3-42220338-INS-158 |
| chr3-42220705-INS-378 | chr3-42220705-INS-379 |
| chr3-42232683-INS-128 | chr3-42232683-INS-132 |
| chr3-42235069-INS-591 | chr3-42235069-INS-591 |
| chr3-42239431-INS-61 | chr3-42239430-INS-63 |
| chr3-42262059-INS-108 | chr3-42262065-INS-72 |
| chr3-42305152-INS-64 | chr3-42305153-INS-56 |
| chr3-42312429-INS-57 | chr3-42312429-INS-57 |
| chr3-42327162-INS-56 | chr3-42327162-INS-54 |
| chr3-42337112-INS-52 | chr3-42337113-INS-56 |
| chr3-42368383-INS-81 | chr3-42368383-INS-111 |
| chr3-42386737-INS-351 | chr3-42386737-INS-351 |
| chr3-42393213-INS-72 | chr3-42393213-INS-66 |
| chr3-42426073-INS-183 | chr3-42426070-INS-175 |
| chr3-42436274-INS-923 | chr3-42436274-INS-923 |
| chr3-42446390-INS-54 | chr3-42446390-INS-54 |
| chr3-42473874-INS-322 | chr3-42473874-INS-322 |
| chr3-42483068-INS-67 | chr3-42483067-INS-53 |
| chr3-42494053-INS-601 | chr3-42494053-INS-602 |
| chr3-42526208-INS-102 | chr3-42526208-INS-102 |
| chr3-42535339-INS-641 | chr3-42535340-INS-665 |
| chr3-42537224-INS-6167 | chr3-42537225-INS-6167 |
| chr3-42549605-INS-6159 | chr3-42549606-INS-6159 |
| chr3-42568031-INS-1309 | chr3-42568032-INS-1310 |
| chr3-42577702-INS-198 | chr3-42577702-INS-198 |
| chr3-42649745-INS-380 | chr3-42649745-INS-380 |
| chr3-42714412-INS-6387 | chr3-42714413-INS-6388 |
| chr3-42733984-INS-6456 | chr3-42733985-INS-6456 |
| chr3-42735482-INS-211 | chr3-42735477-INS-211 |
| chr3-42736168-INS-543 | chr3-42736168-INS-543 |
| chr3-42762229-INS-241 | chr3-42762229-INS-239 |
| chr3-42772704-INS-326 | chr3-42772704-INS-326 |
| chr3-42784323-INS-1693 | chr3-42784328-INS-1688 |
| chr3-42786990-INS-369 | chr3-42786990-INS-369 |
| chr3-42797277-INS-81 | chr3-42797277-INS-81 |
| chr3-42801977-INS-1328 | chr3-42801978-INS-1328 |
| chr3-42804532-INS-86 | chr3-42804508-INS-70 |
| chr3-42837617-INS-52 | chr3-42837605-INS-58 |
| chr3-42838128-INS-6447 | chr3-42838129-INS-6447 |
| chr3-42841253-INS-88 | chr3-42841253-INS-89 |
| chr3-42880581-INS-118 | chr3-42880581-INS-120 |
| chr3-42901763-INS-540 | chr3-42901734-INS-528 |
| chr3-42905796-INS-788 | chr3-42905796-INS-791 |
| chr3-42921787-INS-6309 | chr3-42921788-INS-6307 |
| chr3-42929502-INS-586 | chr3-42929502-INS-586 |
| chr3-42931218-INS-113 | chr3-42931218-INS-113 |
| chr3-42945061-INS-268 | chr3-42945061-INS-268 |
| chr3-42975577-INS-2215 | chr3-42975578-INS-2216 |
| chr3-42983128-INS-54 | chr3-42983128-INS-54 |
| chr3-43000426-INS-2819 | chr3-43000427-INS-2819 |
| chr3-43003636-INS-69 | chr3-43003638-INS-70 |
| chr3-43036080-INS-183 | chr3-43036080-INS-183 |
| chr3-43105214-INS-102 | chr3-43105214-INS-88 |
| chr3-43109437-INS-52 | chr3-43109437-INS-52 |
| chr3-43109989-INS-1044 | chr3-43109980-INS-1032 |
| chr3-43123716-INS-5466 | chr3-43123717-INS-5466 |
| chr3-43128645-INS-52 | chr3-43128645-INS-60 |
| chr3-43174052-INS-498 | chr3-43174052-INS-498 |
| chr3-43175202-INS-1490 | chr3-43175203-INS-1490 |

| | |
|---|---|
| chr3-43206382-INS-184 | chr3-43206382-INS-184 |
| chr3-43228115-INS-917 | chr3-43228115-INS-918 |
| chr3-43229042-INS-73 | chr3-43229034-INS-88 |
| chr3-43236155-INS-4883 | chr3-43236156-INS-4883 |
| chr3-43278152-INS-1481 | chr3-43278153-INS-1481 |
| chr3-43290466-INS-112 | chr3-43290466-INS-98 |
| chr3-43366779-INS-6652 | chr3-43366780-INS-6652 |
| chr3-43369566-INS-2354 | chr3-43369567-INS-2354 |
| chr3-43399371-INS-299 | chr3-43399371-INS-300 |
| chr3-43405522-INS-56 | chr3-43405522-INS-50 |
| chr3-43422302-INS-397 | chr3-43422302-INS-397 |
| chr3-43483999-INS-112 | chr3-43483996-INS-124 |
| chr3-43497993-INS-180 | chr3-43497993-INS-180 |
| chr3-43541667-INS-7126 | chr3-43541668-INS-7126 |
| chr3-43550354-INS-2470 | chr3-43550355-INS-2470 |
| chr3-43558152-INS-53 | chr3-43558152-INS-53 |
| chr3-43613298-INS-222 | chr3-43613298-INS-222 |
| chr3-43618721-INS-52 | chr3-43618721-INS-52 |
| chr3-43631789-INS-4817 | chr3-43631790-INS-4817 |
| chr3-43722818-INS-68 | chr3-43722818-INS-68 |
| chr3-43733920-INS-96 | chr3-43733920-INS-92 |
| chr3-43748903-INS-4603 | chr3-43748904-INS-4603 |
| chr3-43762386-INS-54 | chr3-43762386-INS-60 |
| chr3-43785341-INS-56 | chr3-43785341-INS-56 |
| chr3-43786550-INS-200 | chr3-43786550-INS-200 |
| chr3-43836823-INS-69 | chr3-43836823-INS-69 |
| chr3-43879664-INS-96 | chr3-43879664-INS-96 |
| chr3-43897401-INS-215 | chr3-43897401-INS-215 |
| chr3-43933830-INS-85 | chr3-43933855-INS-63 |
| chr3-43936988-INS-89 | chr3-43936988-INS-81 |
| chr3-43940920-INS-66 | chr3-43940918-INS-51 |
| chr3-43944946-INS-3913 | chr3-43944947-INS-3911 |
| chr3-44026837-INS-57 | chr3-44026839-INS-89 |
| chr3-44095362-INS-171 | chr3-44095363-INS-125 |
| chr3-44118504-INS-6284 | chr3-44118505-INS-6284 |
| chr3-44159952-INS-86 | chr3-44159952-INS-92 |
| chr3-44171304-INS-83 | chr3-44171304-INS-81 |
| chr3-44243217-INS-7229 | chr3-44243218-INS-7229 |
| chr3-44252417-INS-7153 | chr3-44252418-INS-7153 |
| chr3-44260844-INS-76 | chr3-44260844-INS-76 |
| chr3-44273506-INS-64 | chr3-44273506-INS-56 |
| chr3-44285909-INS-1830 | chr3-44285910-INS-1829 |
| chr3-44287839-INS-112 | chr3-44287839-INS-64 |
| chr3-44290937-INS-55 | chr3-44290942-INS-80 |
| chr3-44295649-INS-1837 | chr3-44295650-INS-1838 |
| chr3-44327249-INS-61 | chr3-44327224-INS-85 |
| chr3-44339592-INS-2350 | chr3-44339593-INS-2349 |
| chr3-44354793-INS-375 | chr3-44354793-INS-373 |
| chr3-44376390-INS-471 | chr3-44376390-INS-471 |
| chr3-44390845-INS-4155 | chr3-44390846-INS-4155 |
| chr3-44396487-INS-6256 | chr3-44396488-INS-6256 |
| chr3-44399419-INS-52 | chr3-44399419-INS-50 |
| chr3-44434341-INS-81 | chr3-44434341-INS-60 |
| chr3-44438001-INS-60 | chr3-44438003-INS-54 |
| chr3-44469566-INS-77 | chr3-44469566-INS-77 |
| chr3-44487460-INS-87 | chr3-44487460-INS-51 |
| chr3-44491748-INS-1106 | chr3-44491749-INS-1106 |
| chr3-44506193-INS-52 | chr3-44506193-INS-52 |
| chr3-44532951-INS-1943 | chr3-44532952-INS-1932 |
| chr3-44554381-INS-560 | chr3-44554381-INS-559 |
| chr3-44566542-INS-2499 | chr3-44566543-INS-2498 |
| chr3-44575706-INS-1102 | chr3-44575707-INS-1102 |
| chr3-44592968-INS-760 | chr3-44592968-INS-760 |
| chr3-44632682-INS-60 | chr3-44632682-INS-60 |
| chr3-44668267-INS-60 | chr3-44668291-INS-56 |
| chr3-44670358-INS-72 | chr3-44670353-INS-80 |
| chr3-44678960-INS-4761 | chr3-44678961-INS-4771 |
| chr3-44697619-INS-52 | chr3-44697619-INS-64 |
| chr3-44753174-INS-60 | chr3-44753194-INS-77 |
| chr3-44762810-INS-390 | chr3-44762810-INS-390 |
| chr3-44774444-INS-68 | chr3-44774428-INS-52 |
| chr3-44778723-INS-6476 | chr3-44778724-INS-6476 |
| chr3-44824192-INS-52 | chr3-44824192-INS-92 |
| chr3-44848366-INS-79 | chr3-44848366-INS-83 |
| chr3-44866787-INS-457 | chr3-44866787-INS-457 |
| chr3-44873038-INS-62 | chr3-44873058-INS-74 |
| chr3-44892120-INS-998 | chr3-44892120-INS-974 |

| | |
|---|---|
| chr3-44926993-INS-2251 | chr3-44926995-INS-2257 |
| chr3-44938915-INS-58 | chr3-44938915-INS-58 |
| chr3-44953884-INS-378 | chr3-44953884-INS-393 |
| chr3-44958351-INS-548 | chr3-44958351-INS-552 |
| chr3-44980431-INS-133 | chr3-44980431-INS-134 |
| chr3-44982488-INS-4136 | chr3-44982489-INS-4131 |
| chr3-45003712-INS-54 | chr3-45003712-INS-54 |
| chr3-45038916-INS-6747 | chr3-45038917-INS-6747 |
| chr3-45092453-INS-58 | chr3-45092453-INS-58 |
| chr3-45110662-INS-1888 | chr3-45110663-INS-1888 |
| chr3-45129101-INS-1580 | chr3-45129102-INS-1565 |
| chr3-45135618-INS-54 | chr3-45135636-INS-51 |
| chr3-45147461-INS-464 | chr3-45147461-INS-464 |
| chr3-45176720-INS-65 | chr3-45176720-INS-65 |
| chr3-45177014-INS-4585 | chr3-45177015-INS-4585 |
| chr3-45191468-INS-6098 | chr3-45191469-INS-6102 |
| chr3-45210033-INS-503 | chr3-45210033-INS-495 |
| chr3-45233591-INS-129 | chr3-45233566-INS-132 |
| chr3-45237663-INS-3066 | chr3-45237664-INS-3067 |
| chr3-45271284-INS-583 | chr3-45271284-INS-583 |
| chr3-45278069-INS-141 | chr3-45278069-INS-141 |
| chr3-45319971-INS-1567 | chr3-45319972-INS-1567 |
| chr3-45350170-INS-381 | chr3-45350170-INS-381 |
| chr3-45381209-INS-81 | chr3-45381209-INS-53 |
| chr3-45400269-INS-357 | chr3-45400269-INS-357 |
| chr3-45422630-INS-65 | chr3-45422630-INS-65 |
| chr3-45475909-INS-57 | chr3-45475909-INS-61 |
| chr3-45486595-INS-6223 | chr3-45486596-INS-6223 |
| chr3-45506643-INS-451 | chr3-45506643-INS-444 |
| chr3-45545452-INS-784 | chr3-45545452-INS-784 |
| chr3-45563630-INS-6513 | chr3-45563631-INS-6517 |
| chr3-45564175-INS-386 | chr3-45564175-INS-386 |
| chr3-45584469-INS-6593 | chr3-45584470-INS-6593 |
| chr3-45586029-INS-159 | chr3-45586029-INS-158 |
| chr3-45598961-INS-1065 | chr3-45598962-INS-1062 |
| chr3-45599470-INS-76 | chr3-45599470-INS-50 |
| chr3-45614082-INS-156 | chr3-45614080-INS-186 |
| chr3-45616539-INS-71 | chr3-45616539-INS-75 |
| chr3-45616905-INS-1296 | chr3-45616906-INS-1296 |
| chr3-45621472-INS-2333 | chr3-45621473-INS-2333 |
| chr3-45626667-INS-5405 | chr3-45626668-INS-5405 |
| chr3-45627721-INS-450 | chr3-45627721-INS-450 |
| chr3-45632018-INS-64 | chr3-45632024-INS-71 |
| chr3-45654509-INS-56 | chr3-45654509-INS-56 |
| chr3-45673426-INS-59 | chr3-45673426-INS-51 |
| chr3-45675549-INS-293 | chr3-45675549-INS-293 |
| chr3-45695299-INS-628 | chr3-45695299-INS-628 |
| chr3-45711909-INS-170 | chr3-45711909-INS-171 |
| chr3-45714073-INS-632 | chr3-45714073-INS-632 |
| chr3-45721960-INS-70 | chr3-45721960-INS-74 |
| chr3-45726686-INS-54 | chr3-45726686-INS-50 |
| chr3-45730613-INS-389 | chr3-45730613-INS-391 |
| chr3-45742402-INS-53 | chr3-45742406-INS-68 |
| chr3-45753174-INS-79 | chr3-45753174-INS-79 |
| chr3-45772024-INS-6388 | chr3-45772025-INS-6386 |
| chr3-45796153-INS-758 | chr3-45796153-INS-758 |
| chr3-45810553-INS-6158 | chr3-45810554-INS-6158 |
| chr3-45824551-INS-2269 | chr3-45824552-INS-2269 |
| chr3-45826074-INS-60 | chr3-45826074-INS-70 |
| chr3-45828138-INS-129 | chr3-45828138-INS-129 |
| chr3-45836124-INS-266 | chr3-45836124-INS-277 |
| chr3-45843702-INS-270 | chr3-45843711-INS-263 |
| chr3-45844819-INS-104 | chr3-45844802-INS-82 |
| chr3-45877580-INS-475 | chr3-45877580-INS-475 |
| chr3-45910562-INS-3754 | chr3-45910563-INS-3754 |
| chr3-45911855-INS-1025 | chr3-45911856-INS-1025 |
| chr3-45913278-INS-348 | chr3-45913278-INS-348 |
| chr3-45916753-INS-60 | chr3-45916753-INS-60 |
| chr3-45951751-INS-115 | chr3-45951751-INS-115 |
| chr3-45954189-INS-6662 | chr3-45954190-INS-6663 |
| chr3-45975906-INS-296 | chr3-45975904-INS-289 |
| chr3-45976952-INS-803 | chr3-45976952-INS-805 |
| chr3-45983024-INS-56 | chr3-45983024-INS-56 |
| chr3-46002455-INS-541 | chr3-46002455-INS-541 |
| chr3-46009844-INS-366 | chr3-46009844-INS-366 |
| chr3-46011759-INS-58 | chr3-46011759-INS-60 |
| chr3-46092818-INS-1380 | chr3-46092819-INS-1380 |

| | |
|---|---|
| chr3-46118623-INS-194 | chr3-46118623-INS-194 |
| chr3-46130093-INS-7085 | chr3-46130094-INS-7085 |
| chr3-46135227-INS-52 | chr3-46135220-INS-52 |
| chr3-46168288-INS-2939 | chr3-46168289-INS-2939 |
| chr3-46182681-INS-492 | chr3-46182681-INS-492 |
| chr3-46202981-INS-225 | chr3-46202982-INS-232 |
| chr3-46221169-INS-374 | chr3-46221169-INS-374 |
| chr3-46235272-INS-99 | chr3-46235272-INS-91 |
| chr3-46287114-INS-313 | chr3-46287070-INS-313 |
| chr3-46299635-INS-381 | chr3-46299635-INS-381 |
| chr3-46317728-INS-57 | chr3-46317728-INS-88 |
| chr3-46322483-INS-6465 | chr3-46322484-INS-6465 |
| chr3-46325401-INS-877 | chr3-46325401-INS-877 |
| chr3-46335755-INS-439 | chr3-46335755-INS-439 |
| chr3-46356049-INS-448 | chr3-46356049-INS-448 |
| chr3-46380757-INS-107 | chr3-46380757-INS-104 |
| chr3-46392204-INS-90 | chr3-46392204-INS-62 |
| chr3-46432044-INS-58 | chr3-46432032-INS-59 |
| chr3-46435459-INS-64 | chr3-46435459-INS-84 |
| chr3-46461113-INS-6330 | chr3-46461114-INS-6331 |
| chr3-46479055-INS-54 | chr3-46479055-INS-54 |
| chr3-46481069-INS-60 | chr3-46481070-INS-54 |
| chr3-46497984-INS-715 | chr3-46497984-INS-716 |
| chr3-46505158-INS-1195 | chr3-46505159-INS-1195 |
| chr3-46531638-INS-366 | chr3-46531638-INS-366 |
| chr3-46549458-INS-69 | chr3-46549458-INS-55 |
| chr3-46562642-INS-480 | chr3-46562642-INS-484 |
| chr3-46596796-INS-83 | chr3-46596796-INS-78 |
| chr3-46631544-INS-359 | chr3-46631544-INS-358 |
| chr3-46638097-INS-184 | chr3-46638083-INS-204 |
| chr3-46641220-INS-591 | chr3-46641220-INS-591 |
| chr3-46668997-INS-1494 | chr3-46668998-INS-1493 |
| chr3-46672553-INS-56 | chr3-46672553-INS-72 |
| chr3-46708766-INS-52 | chr3-46708766-INS-52 |
| chr3-46711287-INS-71 | chr3-46711331-INS-59 |
| chr3-46716857-INS-52 | chr3-46716857-INS-54 |
| chr3-46717571-INS-158 | chr3-46717571-INS-158 |
| chr3-46724783-INS-970 | chr3-46724783-INS-968 |
| chr3-46727033-INS-120 | chr3-46727033-INS-75 |
| chr3-46821076-INS-90 | chr3-46821077-INS-93 |
| chr3-46836319-INS-1231 | chr3-46836307-INS-1219 |
| chr3-46869526-INS-5957 | chr3-46869527-INS-5957 |
| chr3-46879474-INS-2619 | chr3-46879475-INS-2616 |
| chr3-46895699-INS-2951 | chr3-46895700-INS-2951 |
| chr3-46905576-INS-80 | chr3-46905576-INS-82 |
| chr3-46923135-INS-2764 | chr3-46923136-INS-2764 |
| chr3-46939697-INS-132 | chr3-46939696-INS-140 |
| chr3-46950996-INS-474 | chr3-46950996-INS-474 |
| chr3-46957148-INS-82 | chr3-46957148-INS-82 |
| chr3-46982112-INS-80 | chr3-46982092-INS-63 |
| chr3-47003170-INS-76 | chr3-47003170-INS-52 |
| chr3-47009767-INS-119 | chr3-47009767-INS-119 |
| chr3-47017582-INS-222 | chr3-47017579-INS-223 |
| chr3-47034206-INS-62 | chr3-47034206-INS-62 |
| chr3-47035533-INS-57 | chr3-47035533-INS-57 |
| chr3-47038376-INS-343 | chr3-47038376-INS-343 |
| chr3-47047161-INS-611 | chr3-47047161-INS-612 |
| chr3-47049620-INS-216 | chr3-47049619-INS-215 |
| chr3-47053496-INS-70 | chr3-47053496-INS-70 |
| chr3-47067851-INS-62 | chr3-47067851-INS-64 |
| chr3-47072767-INS-341 | chr3-47072767-INS-336 |
| chr3-47074513-INS-113 | chr3-47074513-INS-113 |
| chr3-47090049-INS-73 | chr3-47090049-INS-73 |
| chr3-47121203-INS-59 | chr3-47121210-INS-83 |
| chr3-47143454-INS-56 | chr3-47143429-INS-51 |
| chr3-47154306-INS-191 | chr3-47154306-INS-190 |
| chr3-47164377-INS-293 | chr3-47164377-INS-293 |
| chr3-47176593-INS-1368 | chr3-47176594-INS-1369 |
| chr3-47189502-INS-224 | chr3-47189502-INS-224 |
| chr3-47199129-INS-142 | chr3-47199129-INS-142 |
| chr3-47224733-INS-66 | chr3-47224731-INS-68 |
| chr3-47234283-INS-75 | chr3-47234292-INS-94 |
| chr3-47251136-INS-104 | chr3-47251136-INS-82 |
| chr3-47254398-INS-80 | chr3-47254398-INS-82 |
| chr3-47257752-INS-480 | chr3-47257752-INS-480 |
| chr3-47278881-INS-712 | chr3-47278881-INS-712 |
| chr3-47281390-INS-214 | chr3-47281391-INS-214 |

| | |
|---|---|
| chr3-47319739-INS-52 | chr3-47319739-INS-52 |
| chr3-47334261-INS-84 | chr3-47334261-INS-88 |
| chr3-47349825-INS-627 | chr3-47349825-INS-626 |
| chr3-47430629-INS-92 | chr3-47430629-INS-83 |
| chr3-47434967-INS-164 | chr3-47434967-INS-164 |
| chr3-47440432-INS-74 | chr3-47440432-INS-78 |
| chr3-47455587-INS-70 | chr3-47455587-INS-54 |
| chr3-47468387-INS-63 | chr3-47468387-INS-69 |
| chr3-47473514-INS-6894 | chr3-47473515-INS-6894 |
| chr3-47515678-INS-60 | chr3-47515678-INS-60 |
| chr3-47610646-INS-464 | chr3-47610646-INS-464 |
| chr3-47616213-INS-56 | chr3-47616211-INS-54 |
| chr3-47630984-INS-555 | chr3-47630984-INS-555 |
| chr3-47635641-INS-409 | chr3-47635641-INS-409 |
| chr3-47662099-INS-81 | chr3-47662099-INS-81 |
| chr3-47663852-INS-84 | chr3-47663852-INS-84 |
| chr3-47666268-INS-107 | chr3-47666268-INS-107 |
| chr3-47673837-INS-6888 | chr3-47673838-INS-6888 |
| chr3-47681823-INS-60 | chr3-47681821-INS-66 |
| chr3-47688818-INS-358 | chr3-47688818-INS-357 |
| chr3-47697000-INS-52 | chr3-47697000-INS-50 |
| chr3-47702731-INS-56 | chr3-47702731-INS-56 |
| chr3-47715134-INS-578 | chr3-47715134-INS-578 |
| chr3-47733148-INS-70 | chr3-47733148-INS-62 |
| chr3-47788748-INS-270 | chr3-47788748-INS-270 |
| chr3-47793013-INS-52 | chr3-47793013-INS-54 |
| chr3-47819112-INS-60 | chr3-47819112-INS-54 |
| chr3-47844862-INS-106 | chr3-47844860-INS-120 |
| chr3-47845785-INS-56 | chr3-47845785-INS-57 |
| chr3-47846816-INS-338 | chr3-47846816-INS-338 |
| chr3-47849061-INS-374 | chr3-47849061-INS-390 |
| chr3-47869326-INS-127 | chr3-47869317-INS-123 |
| chr3-47898961-INS-53 | chr3-47898939-INS-63 |
| chr3-47908827-INS-52 | chr3-47908827-INS-52 |
| chr3-47914437-INS-79 | chr3-47914453-INS-84 |
| chr3-47927680-INS-212 | chr3-47927680-INS-212 |
| chr3-47941398-INS-64 | chr3-47941398-INS-56 |
| chr3-47958977-INS-201 | chr3-47958951-INS-201 |
| chr3-47967057-INS-222 | chr3-47967073-INS-215 |
| chr3-47979994-INS-3586 | chr3-47979995-INS-3586 |
| chr3-47994333-INS-1095 | chr3-47994334-INS-1095 |
| chr3-48030565-INS-220 | chr3-48030565-INS-220 |
| chr3-48045539-INS-6162 | chr3-48045540-INS-6163 |
| chr3-48062088-INS-198 | chr3-48062088-INS-198 |
| chr3-48065233-INS-66 | chr3-48065233-INS-64 |
| chr3-48070180-INS-302 | chr3-48070180-INS-305 |
| chr3-48073058-INS-105 | chr3-48073058-INS-81 |
| chr3-48078529-INS-8376 | chr3-48078530-INS-8376 |
| chr3-48082442-INS-54 | chr3-48082442-INS-55 |
| chr3-48093608-INS-6437 | chr3-48093609-INS-6437 |
| chr3-48098825-INS-59 | chr3-48098834-INS-55 |
| chr3-48107322-INS-52 | chr3-48107322-INS-52 |
| chr3-48109216-INS-54 | chr3-48109216-INS-54 |
| chr3-48117166-INS-175 | chr3-48117150-INS-181 |
| chr3-48119274-INS-82 | chr3-48119274-INS-86 |
| chr3-48131859-INS-401 | chr3-48131859-INS-401 |
| chr3-48139227-INS-85 | chr3-48139227-INS-83 |
| chr3-48154373-INS-122 | chr3-48154373-INS-122 |
| chr3-48167381-INS-148 | chr3-48167381-INS-148 |
| chr3-48188200-INS-1200 | chr3-48188201-INS-1201 |
| chr3-48196702-INS-6589 | chr3-48196703-INS-6589 |
| chr3-48200991-INS-1667 | chr3-48200992-INS-1667 |
| chr3-48224267-INS-2631 | chr3-48224268-INS-2631 |
| chr3-48252061-INS-58 | chr3-48252061-INS-58 |
| chr3-48284997-INS-52 | chr3-48284998-INS-52 |
| chr3-48308059-INS-60 | chr3-48308059-INS-60 |
| chr3-48339134-INS-52 | chr3-48339134-INS-52 |
| chr3-48372696-INS-827 | chr3-48372696-INS-827 |
| chr3-48407279-INS-246 | chr3-48407279-INS-246 |
| chr3-48417560-INS-1273 | chr3-48417561-INS-1274 |
| chr3-48446190-INS-140 | chr3-48446190-INS-138 |
| chr3-48465720-INS-52 | chr3-48465677-INS-64 |
| chr3-48477704-INS-84 | chr3-48477704-INS-84 |
| chr3-48481041-INS-305 | chr3-48481041-INS-326 |
| chr3-48483190-INS-70 | chr3-48483193-INS-70 |
| chr3-48492965-INS-6442 | chr3-48492966-INS-6442 |
| chr3-48498467-INS-2337 | chr3-48498468-INS-2338 |

| | |
|---|---|
| chr3-48517861-INS-3913 | chr3-48517862-INS-3915 |
| chr3-48528644-INS-772 | chr3-48528644-INS-773 |
| chr3-48531200-INS-456 | chr3-48531201-INS-456 |
| chr3-48537005-INS-69 | chr3-48537005-INS-69 |
| chr3-48549605-INS-149 | chr3-48549606-INS-149 |
| chr3-48551058-INS-160 | chr3-48551058-INS-160 |
| chr3-48556529-INS-58 | chr3-48556529-INS-58 |
| chr3-48587937-INS-84 | chr3-48587937-INS-84 |
| chr3-48593400-INS-193 | chr3-48593400-INS-193 |
| chr3-48610245-INS-160 | chr3-48610277-INS-129 |
| chr3-48632999-INS-205 | chr3-48632999-INS-205 |
| chr3-48655942-INS-752 | chr3-48655942-INS-752 |
| chr3-48673082-INS-106 | chr3-48673070-INS-139 |
| chr3-48723409-INS-84 | chr3-48723409-INS-85 |
| chr3-48733315-INS-62 | chr3-48733314-INS-84 |
| chr3-48769085-INS-80 | chr3-48769085-INS-68 |
| chr3-48798148-INS-223 | chr3-48798148-INS-223 |
| chr3-48814548-INS-200 | chr3-48814548-INS-196 |
| chr3-48854749-INS-64 | chr3-48854749-INS-64 |
| chr3-48876981-INS-55 | chr3-48876979-INS-63 |
| chr3-48908500-INS-166 | chr3-48908500-INS-166 |
| chr3-48909557-INS-90 | chr3-48909579-INS-78 |
| chr3-48910277-INS-56 | chr3-48910278-INS-52 |
| chr3-48947820-INS-390 | chr3-48947820-INS-390 |
| chr3-48950796-INS-142 | chr3-48950796-INS-142 |
| chr3-48966222-INS-68 | chr3-48966222-INS-68 |
| chr3-48995666-INS-99 | chr3-48995666-INS-141 |
| chr3-49014465-INS-150 | chr3-49014461-INS-153 |
| chr3-49032826-INS-105 | chr3-49032822-INS-99 |
| chr3-49036495-INS-59 | chr3-49036472-INS-51 |
| chr3-49044928-INS-735 | chr3-49044928-INS-735 |
| chr3-49083122-INS-183 | chr3-49083122-INS-183 |
| chr3-49101878-INS-219 | chr3-49101878-INS-220 |
| chr3-49109823-INS-64 | chr3-49109824-INS-64 |
| chr3-49178431-INS-54 | chr3-49178431-INS-50 |
| chr3-49190149-INS-58 | chr3-49190149-INS-56 |
| chr3-49212553-INS-75 | chr3-49212553-INS-61 |
| chr3-49245415-INS-823 | chr3-49245415-INS-824 |
| chr3-49259620-INS-180 | chr3-49259620-INS-180 |
| chr3-49265048-INS-286 | chr3-49265048-INS-286 |
| chr3-49268746-INS-142 | chr3-49268746-INS-140 |
| chr3-49277897-INS-81 | chr3-49277897-INS-65 |
| chr3-49279176-INS-215 | chr3-49279176-INS-215 |
| chr3-49286546-INS-108 | chr3-49286546-INS-94 |
| chr3-49289182-INS-70 | chr3-49289182-INS-66 |
| chr3-49299096-INS-88 | chr3-49299096-INS-92 |
| chr3-49305605-INS-58 | chr3-49305605-INS-60 |
| chr3-49351035-INS-795 | chr3-49351035-INS-802 |
| chr3-49389631-INS-117 | chr3-49389631-INS-117 |
| chr3-49405521-INS-100 | chr3-49405521-INS-84 |
| chr3-49443929-INS-119 | chr3-49443927-INS-154 |
| chr3-49454736-INS-197 | chr3-49454736-INS-197 |
| chr3-49456313-INS-109 | chr3-49456313-INS-109 |
| chr3-49464204-INS-55 | chr3-49464204-INS-55 |
| chr3-49469839-INS-56 | chr3-49469839-INS-50 |
| chr3-49476817-INS-352 | chr3-49476817-INS-353 |
| chr3-49481410-INS-71 | chr3-49481410-INS-71 |
| chr3-49486346-INS-70 | chr3-49486346-INS-68 |
| chr3-49494223-INS-60 | chr3-49494223-INS-74 |
| chr3-49514914-INS-1600 | chr3-49514915-INS-1601 |
| chr3-49517599-INS-141 | chr3-49517599-INS-141 |
| chr3-49562916-INS-68 | chr3-49562892-INS-51 |
| chr3-49579044-INS-52 | chr3-49579048-INS-52 |
| chr3-49617081-INS-6691 | chr3-49617082-INS-6693 |
| chr3-49624274-INS-201 | chr3-49624274-INS-202 |
| chr3-49629666-INS-1893 | chr3-49629667-INS-1893 |
| chr3-49643425-INS-1233 | chr3-49643426-INS-1222 |
| chr3-49647497-INS-181 | chr3-49647530-INS-195 |
| chr3-49657098-INS-7155 | chr3-49657099-INS-7155 |
| chr3-49659966-INS-79 | chr3-49659954-INS-57 |
| chr3-49749220-INS-54 | chr3-49749219-INS-50 |
| chr3-49765542-INS-52 | chr3-49765542-INS-54 |
| chr3-49771811-INS-529 | chr3-49771811-INS-529 |
| chr3-49783985-INS-85 | chr3-49783985-INS-85 |
| chr3-49806364-INS-2815 | chr3-49806366-INS-2816 |
| chr3-49826121-INS-108 | chr3-49826121-INS-106 |
| chr3-49867917-INS-1933 | chr3-49867918-INS-1933 |

| | |
|---|---|
| chr3-49868330-INS-54 | chr3-49868330-INS-54 |
| chr3-49871182-INS-3482 | chr3-49871183-INS-3486 |
| chr3-49873787-INS-377 | chr3-49873788-INS-379 |
| chr3-49877441-INS-60 | chr3-49877441-INS-60 |
| chr3-49878412-INS-91 | chr3-49878412-INS-91 |
| chr3-49884387-INS-95 | chr3-49884387-INS-95 |
| chr3-49906438-INS-139 | chr3-49906438-INS-139 |
| chr3-49907967-INS-593 | chr3-49907967-INS-593 |
| chr3-49914158-INS-300 | chr3-49914158-INS-300 |
| chr3-49923701-INS-66 | chr3-49923701-INS-66 |
| chr3-49925779-INS-60 | chr3-49925779-INS-60 |
| chr3-49941875-INS-201 | chr3-49941875-INS-201 |
| chr3-49943351-INS-284 | chr3-49943351-INS-284 |
| chr3-49971410-INS-65 | chr3-49971410-INS-72 |
| chr3-49984088-INS-529 | chr3-49984088-INS-529 |
| chr3-50007213-INS-70 | chr3-50007213-INS-60 |
| chr3-50019820-INS-53 | chr3-50019820-INS-55 |
| chr3-50022044-INS-183 | chr3-50022044-INS-183 |
| chr3-50056195-INS-175 | chr3-50056195-INS-175 |
| chr3-50065146-INS-6196 | chr3-50065150-INS-6191 |
| chr3-50065933-INS-60 | chr3-50065933-INS-60 |
| chr3-50075061-INS-446 | chr3-50075061-INS-447 |
| chr3-50077091-INS-52 | chr3-50077061-INS-76 |
| chr3-50082555-INS-203 | chr3-50082556-INS-156 |
| chr3-50096977-INS-424 | chr3-50096977-INS-424 |
| chr3-50098164-INS-78 | chr3-50098164-INS-78 |
| chr3-50099235-INS-56 | chr3-50099238-INS-50 |
| chr3-50107677-INS-498 | chr3-50107677-INS-498 |
| chr3-50138463-INS-605 | chr3-50138463-INS-606 |
| chr3-50159169-INS-50 | chr3-50159169-INS-50 |
| chr3-50165405-INS-3455 | chr3-50165406-INS-3455 |
| chr3-50196280-INS-506 | chr3-50196280-INS-506 |
| chr3-50204035-INS-64 | chr3-50204035-INS-66 |
| chr3-50211466-INS-78 | chr3-50211461-INS-88 |
| chr3-50214919-INS-490 | chr3-50214919-INS-492 |
| chr3-50227995-INS-217 | chr3-50227995-INS-212 |
| chr3-50269190-INS-6441 | chr3-50269191-INS-6441 |
| chr3-50269191-INS-498 | chr3-50269191-INS-498 |
| chr3-50274468-INS-224 | chr3-50274468-INS-225 |
| chr3-50301072-INS-75 | chr3-50301072-INS-102 |
| chr3-50304326-INS-919 | chr3-50304326-INS-919 |
| chr3-50305992-INS-2009 | chr3-50305993-INS-2006 |
| chr3-50313175-INS-2696 | chr3-50313176-INS-2698 |
| chr3-50369458-INS-54 | chr3-50369458-INS-54 |
| chr3-50439325-INS-176 | chr3-50439312-INS-170 |
| chr3-50469868-INS-55 | chr3-50469868-INS-55 |
| chr3-50490455-INS-51 | chr3-50490449-INS-51 |
| chr3-50527904-INS-205 | chr3-50527904-INS-205 |
| chr3-50529460-INS-6599 | chr3-50529461-INS-6600 |
| chr3-50569050-INS-56 | chr3-50569062-INS-78 |
| chr3-50591068-INS-191 | chr3-50591068-INS-199 |
| chr3-50593990-INS-199 | chr3-50593998-INS-170 |
| chr3-50597473-INS-60 | chr3-50597473-INS-56 |
| chr3-50599845-INS-156 | chr3-50599845-INS-156 |
| chr3-50620463-INS-1109 | chr3-50620464-INS-1109 |
| chr3-50662062-INS-107 | chr3-50662062-INS-107 |
| chr3-50689524-INS-7079 | chr3-50689525-INS-7079 |
| chr3-50692548-INS-85 | chr3-50692553-INS-86 |
| chr3-50721022-INS-212 | chr3-50721022-INS-212 |
| chr3-50727915-INS-147 | chr3-50727915-INS-147 |
| chr3-50742303-INS-927 | chr3-50742303-INS-927 |
| chr3-50746257-INS-199 | chr3-50746257-INS-199 |
| chr3-50760842-INS-200 | chr3-50760877-INS-150 |
| chr3-50811759-INS-56 | chr3-50811759-INS-60 |
| chr3-50846843-INS-2056 | chr3-50846844-INS-2056 |
| chr3-50886752-INS-433 | chr3-50886752-INS-449 |
| chr3-50975000-INS-68 | chr3-50975000-INS-68 |
| chr3-50993634-INS-60 | chr3-50993634-INS-72 |
| chr3-51006904-INS-53 | chr3-51006910-INS-52 |
| chr3-51033010-INS-60 | chr3-51033010-INS-64 |
| chr3-51035362-INS-102 | chr3-51035369-INS-106 |
| chr3-51050574-INS-108 | chr3-51050574-INS-112 |
| chr3-51070699-INS-195 | chr3-51070699-INS-195 |
| chr3-51072462-INS-1554 | chr3-51072463-INS-1557 |
| chr3-51085340-INS-79 | chr3-51085315-INS-85 |
| chr3-51088512-INS-152 | chr3-51088512-INS-153 |
| chr3-51100766-INS-125 | chr3-51100751-INS-90 |

| | |
|---|---|
| chr3-51115200-INS-209 | chr3-51115200-INS-213 |
| chr3-51121844-INS-208 | chr3-51121844-INS-209 |
| chr3-51156632-INS-218 | chr3-51156632-INS-236 |
| chr3-51216879-INS-532 | chr3-51216879-INS-532 |
| chr3-51238100-INS-205 | chr3-51238100-INS-207 |
| chr3-51252102-INS-209 | chr3-51252103-INS-209 |
| chr3-51252587-INS-237 | chr3-51252587-INS-238 |
| chr3-51255088-INS-71 | chr3-51255095-INS-63 |
| chr3-51262411-INS-52 | chr3-51262411-INS-52 |
| chr3-51283832-INS-358 | chr3-51283833-INS-358 |
| chr3-51300208-INS-179 | chr3-51300208-INS-180 |
| chr3-51308609-INS-216 | chr3-51308609-INS-216 |
| chr3-51311915-INS-141 | chr3-51311915-INS-141 |
| chr3-51330886-INS-83 | chr3-51330886-INS-102 |
| chr3-51340152-INS-99 | chr3-51340152-INS-97 |
| chr3-51343356-INS-3576 | chr3-51343357-INS-3572 |
| chr3-51360957-INS-200 | chr3-51360957-INS-200 |
| chr3-51385951-INS-577 | chr3-51385983-INS-592 |
| chr3-51387400-INS-96 | chr3-51387400-INS-96 |
| chr3-51393693-INS-69 | chr3-51393692-INS-85 |
| chr3-51403454-INS-136 | chr3-51403454-INS-136 |
| chr3-51407674-INS-54 | chr3-51407674-INS-70 |
| chr3-51410991-INS-55 | chr3-51410991-INS-103 |
| chr3-51436667-INS-72 | chr3-51436671-INS-68 |
| chr3-51475271-INS-151 | chr3-51475251-INS-175 |
| chr3-51478324-INS-201 | chr3-51478324-INS-201 |
| chr3-51494400-INS-57 | chr3-51494405-INS-85 |
| chr3-51495728-INS-184 | chr3-51495724-INS-162 |
| chr3-51497141-INS-78 | chr3-51497141-INS-78 |
| chr3-51499941-INS-56 | chr3-51499934-INS-52 |
| chr3-51505689-INS-209 | chr3-51505689-INS-213 |
| chr3-51542369-INS-194 | chr3-51542370-INS-202 |
| chr3-51543803-INS-91 | chr3-51543823-INS-120 |
| chr3-51545958-INS-134 | chr3-51545958-INS-134 |
| chr3-51546464-INS-5687 | chr3-51546465-INS-5687 |
| chr3-51556592-INS-199 | chr3-51556593-INS-200 |
| chr3-51568310-INS-51 | chr3-51568310-INS-51 |
| chr3-51576842-INS-57 | chr3-51576842-INS-59 |
| chr3-51584234-INS-326 | chr3-51584234-INS-326 |
| chr3-51587619-INS-54 | chr3-51587631-INS-68 |
| chr3-51608424-INS-52 | chr3-51608427-INS-62 |
| chr3-51619647-INS-65 | chr3-51619647-INS-61 |
| chr3-51647432-INS-88 | chr3-51647457-INS-56 |
| chr3-51657751-INS-54 | chr3-51657751-INS-54 |
| chr3-51678733-INS-192 | chr3-51678733-INS-192 |
| chr3-51730446-INS-221 | chr3-51730434-INS-205 |
| chr3-51747387-INS-114 | chr3-51747387-INS-108 |
| chr3-51803305-INS-94 | chr3-51803305-INS-90 |
| chr3-51810352-INS-78 | chr3-51810352-INS-78 |
| chr3-51830985-INS-181 | chr3-51830985-INS-182 |
| chr3-51846992-INS-56 | chr3-51846992-INS-80 |
| chr3-51868852-INS-274 | chr3-51868864-INS-291 |
| chr3-51878882-INS-195 | chr3-51878873-INS-221 |
| chr3-51881578-INS-60 | chr3-51881578-INS-64 |
| chr3-51887011-INS-56 | chr3-51886999-INS-72 |
| chr3-51890170-INS-5494 | chr3-51890171-INS-5494 |
| chr3-51898126-INS-64 | chr3-51898126-INS-64 |
| chr3-51946053-INS-78 | chr3-51946053-INS-63 |
| chr3-52003068-INS-83 | chr3-52003068-INS-83 |
| chr3-52017506-INS-56 | chr3-52017506-INS-56 |
| chr3-52023893-INS-68 | chr3-52023893-INS-68 |
| chr3-52026354-INS-415 | chr3-52026354-INS-415 |
| chr3-52036046-INS-318 | chr3-52036046-INS-318 |
| chr3-52047254-INS-210 | chr3-52047254-INS-210 |
| chr3-52053461-INS-460 | chr3-52053461-INS-460 |
| chr3-52070482-INS-90 | chr3-52070482-INS-92 |
| chr3-52070951-INS-146 | chr3-52070951-INS-140 |
| chr3-52074292-INS-51 | chr3-52074293-INS-51 |
| chr3-52114308-INS-221 | chr3-52114308-INS-221 |
| chr3-52131245-INS-504 | chr3-52131245-INS-504 |
| chr3-52132871-INS-95 | chr3-52132871-INS-95 |
| chr3-52143055-INS-201 | chr3-52143055-INS-201 |
| chr3-52158172-INS-105 | chr3-52158179-INS-77 |
| chr3-52160422-INS-132 | chr3-52160422-INS-132 |
| chr3-52204989-INS-83 | chr3-52204989-INS-64 |
| chr3-52227172-INS-52 | chr3-52227170-INS-95 |
| chr3-52228357-INS-160 | chr3-52228357-INS-160 |

| | |
|---|---|
| chr3-52284186-INS-59 | chr3-52284186-INS-59 |
| chr3-52292485-INS-225 | chr3-52292496-INS-198 |
| chr3-52294457-INS-242 | chr3-52294481-INS-244 |
| chr3-52313966-INS-188 | chr3-52313966-INS-189 |
| chr3-52318296-INS-282 | chr3-52318296-INS-279 |
| chr3-52319766-INS-59 | chr3-52319766-INS-68 |
| chr3-52327110-INS-88 | chr3-52327110-INS-88 |
| chr3-52337465-INS-58 | chr3-52337471-INS-88 |
| chr3-52352844-INS-554 | chr3-52352844-INS-558 |
| chr3-52356373-INS-127 | chr3-52356373-INS-126 |
| chr3-52361750-INS-54 | chr3-52361750-INS-54 |
| chr3-52371715-INS-145 | chr3-52371715-INS-152 |
| chr3-52384823-INS-553 | chr3-52384823-INS-553 |
| chr3-52404775-INS-79 | chr3-52404775-INS-80 |
| chr3-52491369-INS-52 | chr3-52491367-INS-54 |
| chr3-52524396-INS-130 | chr3-52524396-INS-130 |
| chr3-52573980-INS-112 | chr3-52573980-INS-71 |
| chr3-52591583-INS-328 | chr3-52591583-INS-327 |
| chr3-52620939-INS-222 | chr3-52620939-INS-222 |
| chr3-52628060-INS-1654 | chr3-52628061-INS-1654 |
| chr3-52631492-INS-1864 | chr3-52631493-INS-1864 |
| chr3-52644085-INS-151 | chr3-52644085-INS-160 |
| chr3-52669613-INS-182 | chr3-52669613-INS-184 |
| chr3-52719740-INS-119 | chr3-52719740-INS-119 |
| chr3-52724451-INS-2038 | chr3-52724452-INS-2038 |
| chr3-52728575-INS-353 | chr3-52728547-INS-356 |
| chr3-52730980-INS-80 | chr3-52730980-INS-80 |
| chr3-52738746-INS-114 | chr3-52738747-INS-86 |
| chr3-52777944-INS-328 | chr3-52777944-INS-328 |
| chr3-52830395-INS-214 | chr3-52830395-INS-215 |
| chr3-52845638-INS-51 | chr3-52845638-INS-51 |
| chr3-52856707-INS-63 | chr3-52856707-INS-55 |
| chr3-52876614-INS-62 | chr3-52876614-INS-62 |
| chr3-52895675-INS-429 | chr3-52895675-INS-429 |
| chr3-52902752-INS-56 | chr3-52902752-INS-56 |
| chr3-52926008-INS-651 | chr3-52926008-INS-651 |
| chr3-52931424-INS-114 | chr3-52931424-INS-114 |
| chr3-52959016-INS-166 | chr3-52959016-INS-166 |
| chr3-52982003-INS-1418 | chr3-52982004-INS-1418 |
| chr3-53020533-INS-54 | chr3-53020533-INS-54 |
| chr3-53059164-INS-128 | chr3-53059168-INS-96 |
| chr3-53112985-INS-73 | chr3-53112985-INS-73 |
| chr3-53119667-INS-4231 | chr3-53119668-INS-4231 |
| chr3-53130864-INS-63 | chr3-53130864-INS-63 |
| chr3-53141135-INS-72 | chr3-53141112-INS-72 |
| chr3-53158044-INS-79 | chr3-53158044-INS-79 |
| chr3-53181802-INS-143 | chr3-53181802-INS-141 |
| chr3-53185138-INS-214 | chr3-53185138-INS-214 |
| chr3-53204321-INS-152 | chr3-53204321-INS-152 |
| chr3-53213002-INS-339 | chr3-53213002-INS-339 |
| chr3-53232889-INS-85 | chr3-53232889-INS-79 |
| chr3-53235482-INS-6506 | chr3-53235483-INS-6506 |
| chr3-53244570-INS-54 | chr3-53244542-INS-67 |
| chr3-53257682-INS-338 | chr3-53257682-INS-338 |
| chr3-53294244-INS-3359 | chr3-53294245-INS-3310 |
| chr3-53294245-INS-356 | chr3-53294245-INS-356 |
| chr3-53301804-INS-438 | chr3-53301804-INS-438 |
| chr3-53328253-INS-106 | chr3-53328253-INS-106 |
| chr3-53339781-INS-333 | chr3-53339781-INS-333 |
| chr3-53346423-INS-362 | chr3-53346439-INS-377 |
| chr3-53350663-INS-312 | chr3-53350663-INS-322 |
| chr3-53355287-INS-230 | chr3-53355287-INS-231 |
| chr3-53356740-INS-479 | chr3-53356729-INS-497 |
| chr3-53409849-INS-264 | chr3-53409847-INS-306 |
| chr3-53438062-INS-51 | chr3-53438062-INS-51 |
| chr3-53450098-INS-282 | chr3-53450098-INS-280 |
| chr3-53473613-INS-7231 | chr3-53473614-INS-7231 |
| chr3-53474675-INS-163 | chr3-53474675-INS-157 |
| chr3-53494674-INS-54 | chr3-53494682-INS-69 |
| chr3-53499625-INS-72 | chr3-53499625-INS-72 |
| chr3-53527509-INS-474 | chr3-53527509-INS-483 |
| chr3-53531082-INS-64 | chr3-53531082-INS-64 |
| chr3-53539259-INS-72 | chr3-53539259-INS-78 |
| chr3-53552544-INS-5549 | chr3-53552545-INS-5549 |
| chr3-53554720-INS-1960 | chr3-53554721-INS-1960 |
| chr3-53566622-INS-5549 | chr3-53566623-INS-5549 |
| chr3-53567332-INS-390 | chr3-53567332-INS-390 |

| | |
|---|---|
| chr3-53570278-INS-400 | chr3-53570278-INS-400 |
| chr3-53582290-INS-248 | chr3-53582290-INS-248 |
| chr3-53582820-INS-2030 | chr3-53582821-INS-2028 |
| chr3-53600940-INS-68 | chr3-53600940-INS-65 |
| chr3-53630498-INS-420 | chr3-53630498-INS-421 |
| chr3-53630498-INS-66 | chr3-53630498-INS-66 |
| chr3-53633499-INS-84 | chr3-53633500-INS-84 |
| chr3-53668913-INS-264 | chr3-53668913-INS-267 |
| chr3-53675051-INS-994 | chr3-53675051-INS-993 |
| chr3-53680235-INS-66 | chr3-53680235-INS-66 |
| chr3-53696435-INS-541 | chr3-53696435-INS-528 |
| chr3-53710941-INS-66 | chr3-53710941-INS-66 |
| chr3-53724723-INS-52 | chr3-53724723-INS-66 |
| chr3-53736949-INS-122 | chr3-53736949-INS-122 |
| chr3-53737585-INS-56 | chr3-53737579-INS-54 |
| chr3-53738434-INS-56 | chr3-53738430-INS-58 |
| chr3-53742157-INS-6617 | chr3-53742158-INS-6617 |
| chr3-53752700-INS-77 | chr3-53752700-INS-75 |
| chr3-53780826-INS-51 | chr3-53780826-INS-51 |
| chr3-53785602-INS-64 | chr3-53785602-INS-64 |
| chr3-53792774-INS-61 | chr3-53792759-INS-60 |
| chr3-53794959-INS-55 | chr3-53794959-INS-55 |
| chr3-53800877-INS-189 | chr3-53800878-INS-189 |
| chr3-53816222-INS-55 | chr3-53816222-INS-57 |
| chr3-53819722-INS-157 | chr3-53819734-INS-159 |
| chr3-53861565-INS-170 | chr3-53861565-INS-170 |
| chr3-53864578-INS-6187 | chr3-53864579-INS-6187 |
| chr3-53868407-INS-53 | chr3-53868424-INS-62 |
| chr3-53880535-INS-1481 | chr3-53880536-INS-1481 |
| chr3-53903404-INS-74 | chr3-53903404-INS-74 |
| chr3-53930968-INS-160 | chr3-53930968-INS-209 |
| chr3-53933218-INS-86 | chr3-53933218-INS-86 |
| chr3-53941508-INS-69 | chr3-53941508-INS-70 |
| chr3-53954607-INS-89 | chr3-53954607-INS-92 |
| chr3-53994590-INS-61 | chr3-53994547-INS-63 |
| chr3-54022507-INS-90 | chr3-54022494-INS-54 |
| chr3-54025229-INS-85 | chr3-54025229-INS-85 |
| chr3-54093510-INS-87 | chr3-54093510-INS-63 |
| chr3-54113362-INS-160 | chr3-54113362-INS-157 |
| chr3-54145727-INS-6542 | chr3-54145728-INS-6542 |
| chr3-54166754-INS-236 | chr3-54166754-INS-239 |
| chr3-54178651-INS-67 | chr3-54178651-INS-67 |
| chr3-54203606-INS-53 | chr3-54203606-INS-53 |
| chr3-54213774-INS-186 | chr3-54213774-INS-187 |
| chr3-54217384-INS-73 | chr3-54217384-INS-82 |
| chr3-54224441-INS-195 | chr3-54224441-INS-195 |
| chr3-54234577-INS-214 | chr3-54234577-INS-214 |
| chr3-54235658-INS-601 | chr3-54235658-INS-601 |
| chr3-54243523-INS-75 | chr3-54243523-INS-75 |
| chr3-54305854-INS-111 | chr3-54305854-INS-111 |
| chr3-54321470-INS-2094 | chr3-54321486-INS-2082 |
| chr3-54338571-INS-99 | chr3-54338560-INS-141 |
| chr3-54368650-INS-84 | chr3-54368650-INS-90 |
| chr3-54375776-INS-94 | chr3-54375776-INS-87 |
| chr3-54376930-INS-886 | chr3-54376930-INS-886 |
| chr3-54378399-INS-329 | chr3-54378399-INS-329 |
| chr3-54380043-INS-180 | chr3-54380043-INS-166 |
| chr3-54385965-INS-214 | chr3-54385965-INS-215 |
| chr3-54426921-INS-1893 | chr3-54426922-INS-1893 |
| chr3-54435733-INS-6401 | chr3-54435734-INS-6401 |
| chr3-54458026-INS-104 | chr3-54458026-INS-104 |
| chr3-54479631-INS-116 | chr3-54479631-INS-116 |
| chr3-54513215-INS-72 | chr3-54513215-INS-72 |
| chr3-54566120-INS-66 | chr3-54566120-INS-62 |
| chr3-54581516-INS-61 | chr3-54581516-INS-59 |
| chr3-54586286-INS-231 | chr3-54586293-INS-210 |
| chr3-54587343-INS-66 | chr3-54587344-INS-54 |
| chr3-54616347-INS-62 | chr3-54616347-INS-58 |
| chr3-54618758-INS-81 | chr3-54618758-INS-81 |
| chr3-54651577-INS-74 | chr3-54651576-INS-83 |
| chr3-54660818-INS-272 | chr3-54660818-INS-276 |
| chr3-54678777-INS-74 | chr3-54678777-INS-74 |
| chr3-54696911-INS-99 | chr3-54696911-INS-99 |
| chr3-54699838-INS-162 | chr3-54699844-INS-178 |
| chr3-54702152-INS-423 | chr3-54702152-INS-423 |
| chr3-54721357-INS-1893 | chr3-54721358-INS-1893 |
| chr3-54724594-INS-52 | chr3-54724594-INS-50 |

| | |
|---|---|
| chr3-54740564-INS-256 | chr3-54740564-INS-256 |
| chr3-54764614-INS-51 | chr3-54764622-INS-57 |
| chr3-54770178-INS-60 | chr3-54770179-INS-59 |
| chr3-54770930-INS-95 | chr3-54770932-INS-106 |
| chr3-54784496-INS-64 | chr3-54784519-INS-52 |
| chr3-54798414-INS-1890 | chr3-54798415-INS-1890 |
| chr3-54824664-INS-1888 | chr3-54824665-INS-1888 |
| chr3-54832084-INS-94 | chr3-54832084-INS-94 |
| chr3-54848715-INS-149 | chr3-54848715-INS-151 |
| chr3-54855072-INS-160 | chr3-54855072-INS-156 |
| chr3-54869663-INS-643 | chr3-54869663-INS-639 |
| chr3-54884608-INS-379 | chr3-54884608-INS-379 |
| chr3-54894341-INS-938 | chr3-54894341-INS-938 |
| chr3-54898569-INS-341 | chr3-54898569-INS-354 |
| chr3-54937554-INS-86 | chr3-54937564-INS-90 |
| chr3-54968231-INS-168 | chr3-54968231-INS-126 |
| chr3-54981008-INS-212 | chr3-54981008-INS-206 |
| chr3-54983869-INS-215 | chr3-54983869-INS-215 |
| chr3-54988886-INS-228 | chr3-54988883-INS-186 |
| chr3-54994092-INS-130 | chr3-54994051-INS-130 |
| chr3-55060458-INS-187 | chr3-55060458-INS-187 |
| chr3-55121469-INS-1210 | chr3-55121470-INS-1210 |
| chr3-55155309-INS-107 | chr3-55155309-INS-109 |
| chr3-55162919-INS-1890 | chr3-55162920-INS-1890 |
| chr3-55180877-INS-468 | chr3-55180877-INS-468 |
| chr3-55191198-INS-326 | chr3-55191198-INS-326 |
| chr3-55201600-INS-6454 | chr3-55201601-INS-6454 |
| chr3-55211923-INS-619 | chr3-55211923-INS-619 |
| chr3-55229964-INS-70 | chr3-55229964-INS-64 |
| chr3-55243205-INS-1322 | chr3-55243206-INS-1329 |
| chr3-55266263-INS-174 | chr3-55266263-INS-175 |
| chr3-55275542-INS-78 | chr3-55275526-INS-81 |
| chr3-55277095-INS-70 | chr3-55277095-INS-70 |
| chr3-55279858-INS-329 | chr3-55279861-INS-370 |
| chr3-55282222-INS-98 | chr3-55282222-INS-98 |
| chr3-55286857-INS-70 | chr3-55286857-INS-65 |
| chr3-55326707-INS-51 | chr3-55326707-INS-57 |
| chr3-55332360-INS-52 | chr3-55332360-INS-52 |
| chr3-55342948-INS-117 | chr3-55342916-INS-141 |
| chr3-55354239-INS-76 | chr3-55354211-INS-76 |
| chr3-55362547-INS-7187 | chr3-55362548-INS-7187 |
| chr3-55397534-INS-154 | chr3-55397527-INS-121 |
| chr3-55408956-INS-54 | chr3-55408956-INS-54 |
| chr3-55475337-INS-73 | chr3-55475337-INS-76 |
| chr3-55556953-INS-56 | chr3-55556953-INS-56 |
| chr3-55580893-INS-68 | chr3-55580893-INS-68 |
| chr3-55585191-INS-77 | chr3-55585191-INS-77 |
| chr3-55594510-INS-136 | chr3-55594510-INS-136 |
| chr3-55606509-INS-215 | chr3-55606509-INS-215 |
| chr3-55620754-INS-74 | chr3-55620754-INS-74 |
| chr3-55644043-INS-68 | chr3-55644010-INS-73 |
| chr3-55650862-INS-1007 | chr3-55650863-INS-1007 |
| chr3-55714953-INS-159 | chr3-55714953-INS-159 |
| chr3-55721754-INS-70 | chr3-55721759-INS-70 |
| chr3-55747474-INS-6056 | chr3-55747475-INS-6055 |
| chr3-55751540-INS-143 | chr3-55751540-INS-143 |
| chr3-55768741-INS-58 | chr3-55768741-INS-69 |
| chr3-55789000-INS-200 | chr3-55789000-INS-200 |
| chr3-55802116-INS-57 | chr3-55802116-INS-57 |
| chr3-55818124-INS-227 | chr3-55818124-INS-227 |
| chr3-55829910-INS-243 | chr3-55829910-INS-243 |
| chr3-55844897-INS-134 | chr3-55844897-INS-133 |
| chr3-55928518-INS-324 | chr3-55928518-INS-325 |
| chr3-55960302-INS-58 | chr3-55960337-INS-50 |
| chr3-55966625-INS-87 | chr3-55966625-INS-87 |
| chr3-55971122-INS-201 | chr3-55971122-INS-201 |
| chr3-55973162-INS-198 | chr3-55973162-INS-198 |
| chr3-55997223-INS-265 | chr3-55997223-INS-265 |
| chr3-56011906-INS-392 | chr3-56011906-INS-395 |
| chr3-56014271-INS-6190 | chr3-56014272-INS-6191 |
| chr3-56021338-INS-76 | chr3-56021350-INS-88 |
| chr3-56055657-INS-51 | chr3-56055657-INS-51 |
| chr3-56058759-INS-61 | chr3-56058728-INS-62 |
| chr3-56082166-INS-60 | chr3-56082166-INS-60 |
| chr3-56084485-INS-344 | chr3-56084484-INS-348 |
| chr3-56094383-INS-4128 | chr3-56094384-INS-4128 |
| chr3-56112825-INS-861 | chr3-56112825-INS-860 |

| | |
|---|---|
| chr3-56114696-INS-7183 | chr3-56114697-INS-7185 |
| chr3-56149503-INS-183 | chr3-56149503-INS-189 |
| chr3-56174003-INS-430 | chr3-56174003-INS-421 |
| chr3-56200045-INS-2085 | chr3-56200046-INS-2085 |
| chr3-56201067-INS-184 | chr3-56201067-INS-183 |
| chr3-56213322-INS-54 | chr3-56213322-INS-78 |
| chr3-56251465-INS-372 | chr3-56251465-INS-372 |
| chr3-56257273-INS-121 | chr3-56257273-INS-106 |
| chr3-56257894-INS-731 | chr3-56257894-INS-731 |
| chr3-56260202-INS-73 | chr3-56260202-INS-54 |
| chr3-56265265-INS-536 | chr3-56265265-INS-536 |
| chr3-56272152-INS-1494 | chr3-56272153-INS-1494 |
| chr3-56299326-INS-2765 | chr3-56299327-INS-2775 |
| chr3-56331242-INS-109 | chr3-56331242-INS-134 |
| chr3-56363781-INS-127 | chr3-56363781-INS-128 |
| chr3-56380038-INS-60 | chr3-56380038-INS-60 |
| chr3-56397825-INS-3050 | chr3-56397826-INS-3051 |
| chr3-56441981-INS-984 | chr3-56441981-INS-983 |
| chr3-56478328-INS-65 | chr3-56478315-INS-114 |
| chr3-56519903-INS-1033 | chr3-56519904-INS-1032 |
| chr3-56531587-INS-64 | chr3-56531587-INS-82 |
| chr3-56539499-INS-1025 | chr3-56539500-INS-1026 |
| chr3-56580874-INS-754 | chr3-56580874-INS-754 |
| chr3-56584620-INS-64 | chr3-56584620-INS-77 |
| chr3-56601316-INS-52 | chr3-56601316-INS-52 |
| chr3-56604406-INS-7056 | chr3-56604407-INS-7056 |
| chr3-56607439-INS-73 | chr3-56607427-INS-63 |
| chr3-56621315-INS-185 | chr3-56621315-INS-184 |
| chr3-56628264-INS-71 | chr3-56628271-INS-70 |
| chr3-56639495-INS-60 | chr3-56639495-INS-60 |
| chr3-56665874-INS-65 | chr3-56665872-INS-72 |
| chr3-56692991-INS-2816 | chr3-56692992-INS-2816 |
| chr3-56693969-INS-82 | chr3-56693969-INS-84 |
| chr3-56712087-INS-6624 | chr3-56712088-INS-6625 |
| chr3-56733402-INS-52 | chr3-56733402-INS-52 |
| chr3-56793388-INS-56 | chr3-56793388-INS-92 |
| chr3-56801414-INS-87 | chr3-56801414-INS-99 |
| chr3-56805894-INS-2279 | chr3-56805895-INS-2280 |
| chr3-56852925-INS-680 | chr3-56852925-INS-681 |
| chr3-56868142-INS-100 | chr3-56868142-INS-52 |
| chr3-56874347-INS-107 | chr3-56874347-INS-115 |
| chr3-56904041-INS-410 | chr3-56904041-INS-409 |
| chr3-56907494-INS-303 | chr3-56907494-INS-304 |
| chr3-56910447-INS-381 | chr3-56910447-INS-381 |
| chr3-56914927-INS-123 | chr3-56914927-INS-125 |
| chr3-56955492-INS-112 | chr3-56955492-INS-122 |
| chr3-57023178-INS-782 | chr3-57023178-INS-782 |
| chr3-57027771-INS-653 | chr3-57027771-INS-653 |
| chr3-57038309-INS-59 | chr3-57038309-INS-59 |
| chr3-57050770-INS-97 | chr3-57050760-INS-107 |
| chr3-57061433-INS-140 | chr3-57061433-INS-174 |
| chr3-57086643-INS-54 | chr3-57086629-INS-74 |
| chr3-57093283-INS-1893 | chr3-57093284-INS-1893 |
| chr3-57107417-INS-1770 | chr3-57107418-INS-1771 |
| chr3-57115191-INS-509 | chr3-57115191-INS-505 |
| chr3-57150930-INS-543 | chr3-57150930-INS-542 |
| chr3-57155108-INS-2428 | chr3-57155109-INS-2428 |
| chr3-57188059-INS-161 | chr3-57188072-INS-162 |
| chr3-57217137-INS-486 | chr3-57217137-INS-486 |
| chr3-57218570-INS-496 | chr3-57218570-INS-496 |
| chr3-57244202-INS-491 | chr3-57244202-INS-491 |
| chr3-57248788-INS-100 | chr3-57248792-INS-94 |
| chr3-57294160-INS-100 | chr3-57294156-INS-101 |
| chr3-57296941-INS-179 | chr3-57296941-INS-179 |
| chr3-57311751-INS-1157 | chr3-57311752-INS-1157 |
| chr3-57313180-INS-66 | chr3-57313180-INS-70 |
| chr3-57323371-INS-155 | chr3-57323371-INS-155 |
| chr3-57338783-INS-346 | chr3-57338783-INS-346 |
| chr3-57346644-INS-66 | chr3-57346609-INS-60 |
| chr3-57370531-INS-548 | chr3-57370531-INS-566 |
| chr3-57383282-INS-1314 | chr3-57383283-INS-1313 |
| chr3-57386405-INS-158 | chr3-57386387-INS-152 |
| chr3-57393204-INS-293 | chr3-57393204-INS-292 |
| chr3-57428477-INS-80 | chr3-57428477-INS-56 |
| chr3-57453814-INS-182 | chr3-57453814-INS-182 |
| chr3-57462708-INS-178 | chr3-57462706-INS-162 |
| chr3-57483914-INS-56 | chr3-57483914-INS-52 |

| | |
|---|---|
| chr3-57499696-INS-208 | chr3-57499696-INS-208 |
| chr3-57521331-INS-56 | chr3-57521331-INS-58 |
| chr3-57523099-INS-207 | chr3-57523100-INS-211 |
| chr3-57524923-INS-78 | chr3-57524923-INS-74 |
| chr3-57527183-INS-204 | chr3-57527189-INS-206 |
| chr3-57535794-INS-320 | chr3-57535834-INS-308 |
| chr3-57558335-INS-246 | chr3-57558335-INS-246 |
| chr3-57559553-INS-118 | chr3-57559553-INS-118 |
| chr3-57567224-INS-883 | chr3-57567224-INS-882 |
| chr3-57597241-INS-129 | chr3-57597241-INS-129 |
| chr3-57625162-INS-102 | chr3-57625152-INS-81 |
| chr3-57630787-INS-192 | chr3-57630787-INS-170 |
| chr3-57640702-INS-154 | chr3-57640719-INS-122 |
| chr3-57659009-INS-172 | chr3-57659009-INS-173 |
| chr3-57667610-INS-100 | chr3-57667610-INS-100 |
| chr3-57684042-INS-441 | chr3-57684042-INS-441 |
| chr3-57731547-INS-76 | chr3-57731547-INS-76 |
| chr3-57757639-INS-356 | chr3-57757639-INS-355 |
| chr3-57762054-INS-128 | chr3-57762054-INS-128 |
| chr3-57786041-INS-84 | chr3-57786041-INS-68 |
| chr3-57787290-INS-226 | chr3-57787290-INS-226 |
| chr3-57789918-INS-175 | chr3-57789918-INS-175 |
| chr3-57796103-INS-89 | chr3-57796103-INS-89 |
| chr3-57834715-INS-160 | chr3-57834715-INS-175 |
| chr3-57838646-INS-80 | chr3-57838649-INS-70 |
| chr3-57841300-INS-378 | chr3-57841300-INS-379 |
| chr3-57841917-INS-56 | chr3-57841916-INS-68 |
| chr3-57844532-INS-212 | chr3-57844532-INS-212 |
| chr3-57851814-INS-106 | chr3-57851814-INS-106 |
| chr3-57864699-INS-58 | chr3-57864699-INS-60 |
| chr3-57867999-INS-60 | chr3-57867999-INS-60 |
| chr3-57880757-INS-1618 | chr3-57880758-INS-1618 |
| chr3-57884963-INS-6379 | chr3-57884964-INS-6379 |
| chr3-57893738-INS-70 | chr3-57893707-INS-104 |
| chr3-57899813-INS-538 | chr3-57899813-INS-538 |
| chr3-57905465-INS-70 | chr3-57905490-INS-89 |
| chr3-57912168-INS-107 | chr3-57912168-INS-97 |
| chr3-57945422-INS-717 | chr3-57945422-INS-718 |
| chr3-57946565-INS-368 | chr3-57946561-INS-351 |
| chr3-57947435-INS-74 | chr3-57947435-INS-76 |
| chr3-57947929-INS-140 | chr3-57947929-INS-142 |
| chr3-57954024-INS-51 | chr3-57954024-INS-54 |
| chr3-57970022-INS-63 | chr3-57970022-INS-63 |
| chr3-57980033-INS-72 | chr3-57980056-INS-60 |
| chr3-57994087-INS-154 | chr3-57994087-INS-154 |
| chr3-58006380-INS-7189 | chr3-58006381-INS-7189 |
| chr3-58035467-INS-7009 | chr3-58035468-INS-7009 |
| chr3-58050234-INS-169 | chr3-58050234-INS-169 |
| chr3-58069819-INS-3174 | chr3-58069820-INS-3174 |
| chr3-58087872-INS-146 | chr3-58087831-INS-154 |
| chr3-58094987-INS-369 | chr3-58094987-INS-376 |
| chr3-58107011-INS-1271 | chr3-58107012-INS-1271 |
| chr3-58142148-INS-56 | chr3-58142148-INS-54 |
| chr3-58155083-INS-631 | chr3-58155083-INS-637 |
| chr3-58176290-INS-133 | chr3-58176290-INS-134 |
| chr3-58211023-INS-79 | chr3-58211014-INS-62 |
| chr3-58379563-INS-153 | chr3-58379563-INS-159 |
| chr3-58382214-INS-111 | chr3-58382214-INS-111 |
| chr3-58382902-INS-196 | chr3-58382902-INS-197 |
| chr3-58398146-INS-59 | chr3-58398146-INS-59 |
| chr3-58400659-INS-209 | chr3-58400659-INS-209 |
| chr3-58451060-INS-262 | chr3-58451060-INS-262 |
| chr3-58469511-INS-181 | chr3-58469511-INS-181 |
| chr3-58475669-INS-176 | chr3-58475669-INS-173 |
| chr3-58479257-INS-182 | chr3-58479257-INS-182 |
| chr3-58481053-INS-72 | chr3-58481053-INS-72 |
| chr3-58509788-INS-241 | chr3-58509788-INS-246 |
| chr3-58524965-INS-212 | chr3-58524965-INS-212 |
| chr3-58529181-INS-175 | chr3-58529161-INS-202 |
| chr3-58546144-INS-6932 | chr3-58546145-INS-6933 |
| chr3-58575548-INS-73 | chr3-58575548-INS-73 |
| chr3-58607538-INS-72 | chr3-58607538-INS-72 |
| chr3-58613731-INS-6662 | chr3-58613732-INS-6664 |
| chr3-58700745-INS-974 | chr3-58700745-INS-972 |
| chr3-58708572-INS-255 | chr3-58708572-INS-255 |
| chr3-58760149-INS-208 | chr3-58760149-INS-209 |
| chr3-58785998-INS-593 | chr3-58785995-INS-597 |

| | |
|---|---|
| chr3-58791783-INS-55 | chr3-58791801-INS-66 |
| chr3-58802910-INS-99 | chr3-58802910-INS-99 |
| chr3-58818006-INS-7090 | chr3-58818007-INS-7088 |
| chr3-58819572-INS-194 | chr3-58819572-INS-189 |
| chr3-58823569-INS-199 | chr3-58823569-INS-199 |
| chr3-58828339-INS-6457 | chr3-58828340-INS-6457 |
| chr3-58836582-INS-66 | chr3-58836582-INS-66 |
| chr3-58852236-INS-7669 | chr3-58852232-INS-7688 |
| chr3-58852659-INS-58 | chr3-58852633-INS-70 |
| chr3-58854498-INS-70 | chr3-58854491-INS-78 |
| chr3-58865751-INS-205 | chr3-58865751-INS-205 |
| chr3-58878788-INS-255 | chr3-58878788-INS-255 |
| chr3-58887467-INS-76 | chr3-58887467-INS-69 |
| chr3-58894512-INS-110 | chr3-58894506-INS-110 |
| chr3-58958052-INS-119 | chr3-58958052-INS-81 |
| chr3-58959370-INS-218 | chr3-58959370-INS-220 |
| chr3-59029254-INS-96 | chr3-59029254-INS-96 |
| chr3-59079617-INS-5959 | chr3-59079618-INS-5959 |
| chr3-59096994-INS-159 | chr3-59096995-INS-159 |
| chr3-59207611-INS-710 | chr3-59207611-INS-690 |
| chr3-59353570-INS-52 | chr3-59353570-INS-52 |
| chr3-59357635-INS-4958 | chr3-59357636-INS-4958 |
| chr3-59361195-INS-246 | chr3-59361195-INS-246 |
| chr3-59397969-INS-213 | chr3-59397969-INS-211 |
| chr3-59398789-INS-941 | chr3-59398789-INS-942 |
| chr3-59405758-INS-262 | chr3-59405758-INS-260 |
| chr3-59472248-INS-93 | chr3-59472248-INS-95 |
| chr3-59500971-INS-6578 | chr3-59500972-INS-6577 |
| chr3-59582192-INS-64 | chr3-59582192-INS-64 |
| chr3-59590253-INS-84 | chr3-59590253-INS-60 |
| chr3-59592026-INS-140 | chr3-59592026-INS-140 |
| chr3-59627610-INS-5022 | chr3-59627611-INS-5040 |
| chr3-59635393-INS-72 | chr3-59635393-INS-72 |
| chr3-59682394-INS-5336 | chr3-59682395-INS-5336 |
| chr3-59690274-INS-304 | chr3-59690274-INS-304 |
| chr3-59693030-INS-335 | chr3-59693030-INS-335 |
| chr3-59751431-INS-2885 | chr3-59751432-INS-2885 |
| chr3-59770470-INS-153 | chr3-59770466-INS-157 |
| chr3-59837862-INS-128 | chr3-59837862-INS-129 |
| chr3-59852350-INS-202 | chr3-59852350-INS-202 |
| chr3-59883914-INS-80 | chr3-59883914-INS-70 |
| chr3-59884876-INS-6930 | chr3-59884877-INS-6930 |
| chr3-59889305-INS-328 | chr3-59889305-INS-328 |
| chr3-59899941-INS-147 | chr3-59899941-INS-147 |
| chr3-59925058-INS-281 | chr3-59925038-INS-281 |
| chr3-59933477-INS-52 | chr3-59933477-INS-54 |
| chr3-59943966-INS-6400 | chr3-59943967-INS-6400 |
| chr3-59979094-INS-68 | chr3-59979092-INS-54 |
| chr3-60036310-INS-6283 | chr3-60036311-INS-6283 |
| chr3-60054936-INS-327 | chr3-60054936-INS-327 |
| chr3-60064873-INS-7145 | chr3-60064874-INS-7145 |
| chr3-60076526-INS-68 | chr3-60076528-INS-52 |
| chr3-60124720-INS-191 | chr3-60124720-INS-188 |
| chr3-60214353-INS-5327 | chr3-60214354-INS-5325 |
| chr3-60250547-INS-145 | chr3-60250547-INS-145 |
| chr3-60261712-INS-1860 | chr3-60261713-INS-1862 |
| chr3-60267000-INS-551 | chr3-60267000-INS-551 |
| chr3-60300639-INS-54 | chr3-60300640-INS-60 |
| chr3-60314650-INS-147 | chr3-60314650-INS-138 |
| chr3-60333169-INS-94 | chr3-60333169-INS-51 |
| chr3-60344516-INS-208 | chr3-60344516-INS-208 |
| chr3-60350375-INS-108 | chr3-60350348-INS-111 |
| chr3-60370984-INS-233 | chr3-60370984-INS-218 |
| chr3-60512947-INS-129 | chr3-60512955-INS-103 |
| chr3-60544348-INS-102 | chr3-60544348-INS-102 |
| chr3-60583012-INS-55 | chr3-60583012-INS-65 |
| chr3-60597686-INS-7084 | chr3-60597687-INS-7084 |
| chr3-60638228-INS-6921 | chr3-60638229-INS-6923 |
| chr3-60643722-INS-72 | chr3-60643722-INS-72 |
| chr3-60656453-INS-669 | chr3-60656453-INS-669 |
| chr3-60657272-INS-160 | chr3-60657269-INS-164 |
| chr3-60658675-INS-399 | chr3-60658675-INS-399 |
| chr3-60659363-INS-1086 | chr3-60659364-INS-1082 |
| chr3-60671251-INS-250 | chr3-60671251-INS-250 |
| chr3-60685694-INS-7130 | chr3-60685695-INS-7130 |
| chr3-60715324-INS-7001 | chr3-60715325-INS-7000 |
| chr3-60722118-INS-6568 | chr3-60722119-INS-6568 |

| | |
|---|---|
| chr3-60731768-INS-1666 | chr3-60731769-INS-1665 |
| chr3-60743259-INS-3421 | chr3-60743260-INS-3408 |
| chr3-60746188-INS-1145 | chr3-60746186-INS-1136 |
| chr3-60776486-INS-863 | chr3-60776486-INS-863 |
| chr3-60779393-INS-150 | chr3-60779393-INS-150 |
| chr3-60801642-INS-370 | chr3-60801642-INS-370 |
| chr3-60803200-INS-76 | chr3-60803200-INS-52 |
| chr3-60809970-INS-57 | chr3-60809970-INS-57 |
| chr3-60813506-INS-285 | chr3-60813509-INS-292 |
| chr3-60840645-INS-52 | chr3-60840645-INS-52 |
| chr3-60862875-INS-233 | chr3-60862875-INS-234 |
| chr3-60962263-INS-52 | chr3-60962265-INS-50 |
| chr3-60968797-INS-80 | chr3-60968797-INS-58 |
| chr3-60975582-INS-2225 | chr3-60975583-INS-2234 |
| chr3-61025353-INS-6464 | chr3-61025354-INS-6463 |
| chr3-61030541-INS-2102 | chr3-61030542-INS-2109 |
| chr3-61076415-INS-60 | chr3-61076415-INS-54 |
| chr3-61107973-INS-1027 | chr3-61107974-INS-1027 |
| chr3-61110222-INS-3731 | chr3-61110223-INS-3730 |
| chr3-61118294-INS-6906 | chr3-61118295-INS-6905 |
| chr3-61168162-INS-204 | chr3-61168162-INS-204 |
| chr3-61201119-INS-518 | chr3-61201119-INS-514 |
| chr3-61213813-INS-89 | chr3-61213813-INS-89 |
| chr3-61235285-INS-60 | chr3-61235246-INS-84 |
| chr3-61236986-INS-84 | chr3-61236986-INS-56 |
| chr3-61298899-INS-985 | chr3-61298899-INS-985 |
| chr3-61298914-INS-2741 | chr3-61298915-INS-2742 |
| chr3-61300498-INS-6413 | chr3-61300499-INS-6410 |
| chr3-61320020-INS-390 | chr3-61320020-INS-390 |
| chr3-61382272-INS-87 | chr3-61382272-INS-87 |
| chr3-61392123-INS-1890 | chr3-61392124-INS-1890 |
| chr3-61395563-INS-63 | chr3-61395563-INS-63 |
| chr3-61458736-INS-5358 | chr3-61458737-INS-5358 |
| chr3-61458737-INS-404 | chr3-61458737-INS-404 |
| chr3-61480352-INS-497 | chr3-61480352-INS-497 |
| chr3-61501069-INS-408 | chr3-61501069-INS-408 |
| chr3-61515405-INS-584 | chr3-61515405-INS-584 |
| chr3-61528386-INS-667 | chr3-61528386-INS-667 |
| chr3-61563547-INS-2059 | chr3-61563548-INS-2059 |
| chr3-61566211-INS-348 | chr3-61566209-INS-350 |
| chr3-61583273-INS-2401 | chr3-61583274-INS-2401 |
| chr3-61586408-INS-131 | chr3-61586407-INS-148 |
| chr3-61634315-INS-61 | chr3-61634315-INS-63 |
| chr3-61635584-INS-54 | chr3-61635584-INS-50 |
| chr3-61654957-INS-5228 | chr3-61654958-INS-5228 |
| chr3-61673757-INS-1836 | chr3-61673758-INS-1836 |
| chr3-61678948-INS-230 | chr3-61678948-INS-230 |
| chr3-61722341-INS-6222 | chr3-61722342-INS-6223 |
| chr3-61726507-INS-569 | chr3-61726507-INS-570 |
| chr3-61731110-INS-468 | chr3-61731110-INS-466 |
| chr3-61794457-INS-60 | chr3-61794451-INS-66 |
| chr3-61808458-INS-176 | chr3-61808459-INS-187 |
| chr3-61852011-INS-99 | chr3-61852011-INS-88 |
| chr3-61853944-INS-55 | chr3-61853931-INS-72 |
| chr3-61867293-INS-508 | chr3-61867293-INS-489 |
| chr3-61883684-INS-2903 | chr3-61883685-INS-2903 |
| chr3-61886446-INS-414 | chr3-61886446-INS-412 |
| chr3-61918179-INS-1966 | chr3-61918180-INS-1971 |
| chr3-61925219-INS-490 | chr3-61925220-INS-486 |
| chr3-61932393-INS-807 | chr3-61932393-INS-805 |
| chr3-61934608-INS-961 | chr3-61934608-INS-958 |
| chr3-61953505-INS-69 | chr3-61953505-INS-69 |
| chr3-62067834-INS-68 | chr3-62067834-INS-68 |
| chr3-62079661-INS-85 | chr3-62079647-INS-84 |
| chr3-62107985-INS-54 | chr3-62107985-INS-54 |
| chr3-62127873-INS-116 | chr3-62127873-INS-80 |
| chr3-62154982-INS-534 | chr3-62154982-INS-534 |
| chr3-62157623-INS-6795 | chr3-62157624-INS-6797 |
| chr3-62160960-INS-64 | chr3-62160948-INS-52 |
| chr3-62163775-INS-446 | chr3-62163775-INS-446 |
| chr3-62169130-INS-188 | chr3-62169121-INS-204 |
| chr3-62173929-INS-6471 | chr3-62173930-INS-6471 |
| chr3-62195860-INS-87 | chr3-62195860-INS-87 |
| chr3-62240297-INS-1894 | chr3-62240298-INS-1894 |
| chr3-62240368-INS-72 | chr3-62240368-INS-72 |
| chr3-62306146-INS-405 | chr3-62306152-INS-397 |
| chr3-62314457-INS-82 | chr3-62314453-INS-81 |

| | |
|---|---|
| chr3-62315167-INS-499 | chr3-62315168-INS-499 |
| chr3-62316818-INS-1112 | chr3-62316819-INS-1112 |
| chr3-62395862-INS-195 | chr3-62395862-INS-199 |
| chr3-62443492-INS-2027 | chr3-62443493-INS-2025 |
| chr3-62463592-INS-690 | chr3-62463592-INS-690 |
| chr3-62480879-INS-215 | chr3-62480879-INS-215 |
| chr3-62518576-INS-465 | chr3-62518576-INS-465 |
| chr3-62547381-INS-373 | chr3-62547382-INS-373 |
| chr3-62550398-INS-627 | chr3-62550398-INS-638 |
| chr3-62557471-INS-1887 | chr3-62557472-INS-1887 |
| chr3-62562994-INS-56 | chr3-62562994-INS-52 |
| chr3-62567028-INS-439 | chr3-62567028-INS-439 |
| chr3-62582260-INS-6608 | chr3-62582261-INS-6609 |
| chr3-62603902-INS-1117 | chr3-62603903-INS-1117 |
| chr3-62628096-INS-72 | chr3-62628096-INS-72 |
| chr3-62631899-INS-4154 | chr3-62631900-INS-4154 |
| chr3-62693250-INS-180 | chr3-62693253-INS-190 |
| chr3-62693439-INS-6583 | chr3-62693440-INS-6584 |
| chr3-62751309-INS-6794 | chr3-62751310-INS-6795 |
| chr3-62788533-INS-275 | chr3-62788533-INS-275 |
| chr3-62819847-INS-6498 | chr3-62819848-INS-6492 |
| chr3-62826911-INS-117 | chr3-62826939-INS-68 |
| chr3-62835805-INS-3011 | chr3-62835806-INS-3012 |
| chr3-62885877-INS-498 | chr3-62885877-INS-498 |
| chr3-62901168-INS-54 | chr3-62901168-INS-54 |
| chr3-62909639-INS-524 | chr3-62909638-INS-533 |
| chr3-62929567-INS-688 | chr3-62929567-INS-688 |
| chr3-62933656-INS-121 | chr3-62933656-INS-121 |
| chr3-62950462-INS-84 | chr3-62950462-INS-98 |
| chr3-62957858-INS-4361 | chr3-62957859-INS-4361 |
| chr3-62990449-INS-186 | chr3-62990449-INS-186 |
| chr3-62993246-INS-1057 | chr3-62993247-INS-1062 |
| chr3-63013538-INS-1054 | chr3-63013539-INS-1054 |
| chr3-63048999-INS-7238 | chr3-63049000-INS-7238 |
| chr3-63057191-INS-1225 | chr3-63057192-INS-1225 |
| chr3-63067690-INS-276 | chr3-63067690-INS-277 |
| chr3-63070336-INS-394 | chr3-63070336-INS-402 |
| chr3-63088869-INS-52 | chr3-63088869-INS-52 |
| chr3-63098727-INS-324 | chr3-63098727-INS-324 |
| chr3-63102119-INS-6525 | chr3-63102120-INS-6526 |
| chr3-63121066-INS-268 | chr3-63121066-INS-268 |
| chr3-63128417-INS-54 | chr3-63128417-INS-62 |
| chr3-63135871-INS-169 | chr3-63135871-INS-168 |
| chr3-63138611-INS-65 | chr3-63138611-INS-58 |
| chr3-63155739-INS-51 | chr3-63155739-INS-81 |
| chr3-63168248-INS-203 | chr3-63168248-INS-203 |
| chr3-63192743-INS-56 | chr3-63192743-INS-56 |
| chr3-63297871-INS-114 | chr3-63297871-INS-118 |
| chr3-63317249-INS-52 | chr3-63317247-INS-53 |
| chr3-63342905-INS-6293 | chr3-63342906-INS-6292 |
| chr3-63363995-INS-76 | chr3-63363995-INS-76 |
| chr3-63411803-INS-143 | chr3-63411803-INS-137 |
| chr3-63416500-INS-126 | chr3-63416490-INS-120 |
| chr3-63446528-INS-479 | chr3-63446528-INS-480 |
| chr3-63486367-INS-113 | chr3-63486367-INS-113 |
| chr3-63504317-INS-417 | chr3-63504317-INS-433 |
| chr3-63506090-INS-386 | chr3-63506102-INS-393 |
| chr3-63515733-INS-52 | chr3-63515733-INS-52 |
| chr3-63603423-INS-198 | chr3-63603423-INS-199 |
| chr3-63611882-INS-566 | chr3-63611882-INS-565 |
| chr3-63622194-INS-203 | chr3-63622194-INS-200 |
| chr3-63643525-INS-5091 | chr3-63643526-INS-5091 |
| chr3-63689907-INS-63 | chr3-63689907-INS-63 |
| chr3-63698171-INS-810 | chr3-63698171-INS-810 |
| chr3-63698465-INS-3379 | chr3-63698466-INS-3379 |
| chr3-63727432-INS-1209 | chr3-63727433-INS-1211 |
| chr3-63748514-INS-203 | chr3-63748514-INS-204 |
| chr3-63753187-INS-202 | chr3-63753187-INS-198 |
| chr3-63754443-INS-72 | chr3-63754443-INS-72 |
| chr3-63756700-INS-178 | chr3-63756700-INS-178 |
| chr3-63759335-INS-54 | chr3-63759335-INS-54 |
| chr3-63780979-INS-781 | chr3-63780979-INS-778 |
| chr3-63781819-INS-289 | chr3-63781819-INS-286 |
| chr3-63795142-INS-217 | chr3-63795142-INS-218 |
| chr3-63797280-INS-56 | chr3-63797280-INS-56 |
| chr3-63815301-INS-105 | chr3-63815301-INS-123 |
| chr3-63817619-INS-117 | chr3-63817619-INS-117 |

| | |
|---|---|
| chr3-63820998-INS-75 | chr3-63820998-INS-75 |
| chr3-63889173-INS-60 | chr3-63889173-INS-66 |
| chr3-63896359-INS-7301 | chr3-63896360-INS-7301 |
| chr3-63949321-INS-1066 | chr3-63949322-INS-1066 |
| chr3-63961330-INS-5380 | chr3-63961331-INS-5380 |
| chr3-63969654-INS-6440 | chr3-63969655-INS-6440 |
| chr3-63980227-INS-6586 | chr3-63980228-INS-6586 |
| chr3-64007869-INS-74 | chr3-64007869-INS-52 |
| chr3-64026998-INS-8417 | chr3-64026999-INS-8417 |
| chr3-64085085-INS-6471 | chr3-64085086-INS-6471 |
| chr3-64090535-INS-212 | chr3-64090535-INS-212 |
| chr3-64128309-INS-7207 | chr3-64128310-INS-7207 |
| chr3-64132656-INS-7084 | chr3-64132657-INS-7084 |
| chr3-64215244-INS-244 | chr3-64215244-INS-276 |
| chr3-64291917-INS-7465 | chr3-64291918-INS-7466 |
| chr3-64672821-INS-901 | chr3-64672821-INS-901 |
| chr3-64703221-INS-6957 | chr3-64703222-INS-6959 |
| chr3-64715467-INS-6716 | chr3-64715468-INS-6719 |
| chr3-64798713-INS-251 | chr3-64798713-INS-247 |
| chr3-64839319-INS-227 | chr3-64839319-INS-229 |
| chr3-64872079-INS-59 | chr3-64872079-INS-59 |
| chr3-64880111-INS-198 | chr3-64880111-INS-194 |
| chr3-64905237-INS-1932 | chr3-64905238-INS-1933 |
| chr3-64925057-INS-53 | chr3-64925056-INS-60 |
| chr3-64943259-INS-52 | chr3-64943251-INS-72 |
| chr3-64945088-INS-753 | chr3-64945088-INS-752 |
| chr3-64948626-INS-261 | chr3-64948626-INS-261 |
| chr3-64951196-INS-165 | chr3-64951196-INS-164 |
| chr3-64964905-INS-241 | chr3-64964905-INS-241 |
| chr3-64977533-INS-56 | chr3-64977533-INS-54 |
| chr3-65007543-INS-54 | chr3-65007543-INS-102 |
| chr3-65024107-INS-137 | chr3-65024107-INS-137 |
| chr3-65031072-INS-4438 | chr3-65031073-INS-4438 |
| chr3-65068373-INS-365 | chr3-65068373-INS-375 |
| chr3-65072753-INS-6232 | chr3-65072754-INS-6232 |
| chr3-65088879-INS-316 | chr3-65088879-INS-316 |
| chr3-65124383-INS-392 | chr3-65124383-INS-392 |
| chr3-65138446-INS-203 | chr3-65138446-INS-221 |
| chr3-65147585-INS-51 | chr3-65147585-INS-51 |
| chr3-65150563-INS-80 | chr3-65150563-INS-92 |
| chr3-65151930-INS-2255 | chr3-65151931-INS-2255 |
| chr3-65168307-INS-157 | chr3-65168306-INS-171 |
| chr3-65175610-INS-64 | chr3-65175610-INS-68 |
| chr3-65215000-INS-58 | chr3-65215000-INS-58 |
| chr3-65217718-INS-945 | chr3-65217719-INS-945 |
| chr3-65224558-INS-624 | chr3-65224558-INS-630 |
| chr3-65227583-INS-211 | chr3-65227583-INS-215 |
| chr3-65255888-INS-74 | chr3-65255888-INS-104 |
| chr3-65259499-INS-60 | chr3-65259499-INS-60 |
| chr3-65276671-INS-51 | chr3-65276671-INS-51 |
| chr3-65341618-INS-204 | chr3-65341618-INS-206 |
| chr3-65350779-INS-87 | chr3-65350779-INS-72 |
| chr3-65354350-INS-445 | chr3-65354350-INS-445 |
| chr3-65369703-INS-72 | chr3-65369703-INS-72 |
| chr3-65418543-INS-3081 | chr3-65418544-INS-3067 |
| chr3-65442398-INS-204 | chr3-65442398-INS-200 |
| chr3-65447523-INS-174 | chr3-65447523-INS-173 |
| chr3-65465769-INS-1456 | chr3-65465770-INS-1456 |
| chr3-65469374-INS-4562 | chr3-65469375-INS-4562 |
| chr3-65480122-INS-7098 | chr3-65480123-INS-7098 |
| chr3-65480748-INS-201 | chr3-65480748-INS-201 |
| chr3-65521392-INS-1237 | chr3-65521393-INS-1237 |
| chr3-65539794-INS-490 | chr3-65539794-INS-490 |
| chr3-65542599-INS-52 | chr3-65542599-INS-52 |
| chr3-65545677-INS-1301 | chr3-65545678-INS-1301 |
| chr3-65546308-INS-498 | chr3-65546308-INS-498 |
| chr3-65550639-INS-359 | chr3-65550639-INS-359 |
| chr3-65554005-INS-3292 | chr3-65554006-INS-3307 |
| chr3-65568839-INS-4525 | chr3-65568846-INS-4519 |
| chr3-65578427-INS-800 | chr3-65578427-INS-814 |
| chr3-65578486-INS-142 | chr3-65578520-INS-142 |
| chr3-65584383-INS-57 | chr3-65584383-INS-57 |
| chr3-65608861-INS-4462 | chr3-65608862-INS-4462 |
| chr3-65619349-INS-1047 | chr3-65619350-INS-1050 |
| chr3-65631482-INS-484 | chr3-65631482-INS-484 |
| chr3-65633097-INS-64 | chr3-65633097-INS-66 |
| chr3-65653819-INS-863 | chr3-65653819-INS-864 |

| | |
|---|---|
| chr3-65670463-INS-66 | chr3-65670463-INS-75 |
| chr3-65679037-INS-1196 | chr3-65679038-INS-1196 |
| chr3-65684005-INS-74 | chr3-65684019-INS-54 |
| chr3-65685211-INS-117 | chr3-65685202-INS-99 |
| chr3-65714548-INS-80 | chr3-65714549-INS-59 |
| chr3-65718560-INS-316 | chr3-65718560-INS-316 |
| chr3-65719277-INS-60 | chr3-65719277-INS-60 |
| chr3-65723092-INS-6210 | chr3-65723093-INS-6210 |
| chr3-65723589-INS-68 | chr3-65723589-INS-68 |
| chr3-65776755-INS-212 | chr3-65776772-INS-213 |
| chr3-65784849-INS-5866 | chr3-65784850-INS-5867 |
| chr3-65793093-INS-84 | chr3-65793093-INS-101 |
| chr3-65802263-INS-94 | chr3-65802254-INS-55 |
| chr3-65832063-INS-752 | chr3-65832064-INS-748 |
| chr3-65852496-INS-124 | chr3-65852496-INS-124 |
| chr3-65868092-INS-203 | chr3-65868099-INS-211 |
| chr3-65895775-INS-84 | chr3-65895767-INS-132 |
| chr3-65903924-INS-382 | chr3-65903924-INS-398 |
| chr3-65921614-INS-211 | chr3-65921614-INS-211 |
| chr3-65930568-INS-206 | chr3-65930569-INS-190 |
| chr3-65941650-INS-579 | chr3-65941650-INS-583 |
| chr3-65946994-INS-461 | chr3-65946994-INS-461 |
| chr3-65951711-INS-62 | chr3-65951697-INS-62 |
| chr3-66008384-INS-80 | chr3-66008385-INS-70 |
| chr3-66015155-INS-2616 | chr3-66015156-INS-2615 |
| chr3-66024185-INS-88 | chr3-66024185-INS-114 |
| chr3-66060944-INS-641 | chr3-66060944-INS-655 |
| chr3-66073527-INS-72 | chr3-66073527-INS-74 |
| chr3-66091995-INS-2201 | chr3-66091996-INS-2201 |
| chr3-66093657-INS-2988 | chr3-66093658-INS-2988 |
| chr3-66094462-INS-2523 | chr3-66094463-INS-2516 |
| chr3-66111651-INS-52 | chr3-66111652-INS-72 |
| chr3-66142275-INS-146 | chr3-66142275-INS-146 |
| chr3-66148756-INS-91 | chr3-66148756-INS-91 |
| chr3-66152981-INS-6649 | chr3-66152982-INS-6649 |
| chr3-66153312-INS-1363 | chr3-66153313-INS-1363 |
| chr3-66153882-INS-327 | chr3-66153882-INS-327 |
| chr3-66157054-INS-58 | chr3-66157050-INS-72 |
| chr3-66164467-INS-603 | chr3-66164467-INS-601 |
| chr3-66187875-INS-58 | chr3-66187875-INS-58 |
| chr3-66188760-INS-486 | chr3-66188760-INS-491 |
| chr3-66193418-INS-62 | chr3-66193462-INS-50 |
| chr3-66226077-INS-67 | chr3-66226080-INS-52 |
| chr3-66237420-INS-60 | chr3-66237420-INS-60 |
| chr3-66249686-INS-52 | chr3-66249686-INS-52 |
| chr3-66253965-INS-1976 | chr3-66253966-INS-1973 |
| chr3-66293439-INS-81 | chr3-66293439-INS-115 |
| chr3-66307782-INS-68 | chr3-66307782-INS-72 |
| chr3-66321220-INS-823 | chr3-66321220-INS-795 |
| chr3-66336404-INS-52 | chr3-66336404-INS-52 |
| chr3-66436251-INS-56 | chr3-66436253-INS-55 |
| chr3-66458374-INS-86 | chr3-66458359-INS-86 |
| chr3-66466410-INS-422 | chr3-66466410-INS-420 |
| chr3-66479848-INS-104 | chr3-66479848-INS-72 |
| chr3-66494087-INS-74 | chr3-66494109-INS-96 |
| chr3-66497564-INS-95 | chr3-66497564-INS-95 |
| chr3-66499480-INS-652 | chr3-66499480-INS-659 |
| chr3-66507426-INS-670 | chr3-66507426-INS-670 |
| chr3-66528891-INS-95 | chr3-66528889-INS-63 |
| chr3-66574140-INS-1632 | chr3-66574141-INS-1632 |
| chr3-66625934-INS-147 | chr3-66625934-INS-108 |
| chr3-66634443-INS-559 | chr3-66634443-INS-547 |
| chr3-66637822-INS-217 | chr3-66637822-INS-217 |
| chr3-66650536-INS-95 | chr3-66650536-INS-95 |
| chr3-66658279-INS-62 | chr3-66658279-INS-62 |
| chr3-66658677-INS-6464 | chr3-66658678-INS-6464 |
| chr3-66682604-INS-60 | chr3-66682604-INS-60 |
| chr3-66707061-INS-70 | chr3-66707061-INS-66 |
| chr3-66714870-INS-382 | chr3-66714870-INS-380 |
| chr3-66728923-INS-6467 | chr3-66728924-INS-6468 |
| chr3-66745673-INS-7295 | chr3-66745674-INS-7296 |
| chr3-66774117-INS-70 | chr3-66774117-INS-70 |
| chr3-66775878-INS-1072 | chr3-66775879-INS-1083 |
| chr3-66851946-INS-1841 | chr3-66851947-INS-1839 |
| chr3-66889430-INS-52 | chr3-66889427-INS-58 |
| chr3-66914129-INS-1627 | chr3-66914130-INS-1626 |
| chr3-66954704-INS-232 | chr3-66954704-INS-242 |

| | |
|---|---|
| chr3-66959964-INS-7501 | chr3-66959965-INS-7502 |
| chr3-66966083-INS-5371 | chr3-66966084-INS-5371 |
| chr3-66967686-INS-105 | chr3-66967686-INS-143 |
| chr3-66973725-INS-89 | chr3-66973722-INS-120 |
| chr3-67027456-INS-72 | chr3-67027456-INS-72 |
| chr3-67031635-INS-71 | chr3-67031633-INS-53 |
| chr3-67036409-INS-248 | chr3-67036409-INS-248 |
| chr3-67056610-INS-5197 | chr3-67056611-INS-5197 |
| chr3-67075981-INS-85 | chr3-67075981-INS-85 |
| chr3-67128265-INS-227 | chr3-67128265-INS-227 |
| chr3-67135236-INS-60 | chr3-67135236-INS-60 |
| chr3-67153780-INS-90 | chr3-67153751-INS-74 |
| chr3-67165949-INS-6359 | chr3-67165950-INS-6359 |
| chr3-67210754-INS-918 | chr3-67210754-INS-917 |
| chr3-67211468-INS-1508 | chr3-67211475-INS-1501 |
| chr3-67240746-INS-61 | chr3-67240714-INS-66 |
| chr3-67317182-INS-157 | chr3-67317225-INS-150 |
| chr3-67352893-INS-6501 | chr3-67352894-INS-6500 |
| chr3-67362142-INS-171 | chr3-67362142-INS-171 |
| chr3-67384199-INS-218 | chr3-67384199-INS-219 |
| chr3-67386628-INS-128 | chr3-67386628-INS-128 |
| chr3-67410028-INS-204 | chr3-67410028-INS-204 |
| chr3-67438520-INS-196 | chr3-67438520-INS-195 |
| chr3-67459574-INS-67 | chr3-67459572-INS-68 |
| chr3-67478779-INS-212 | chr3-67478779-INS-213 |
| chr3-67483419-INS-126 | chr3-67483419-INS-120 |
| chr3-67489489-INS-55 | chr3-67489489-INS-55 |
| chr3-67503795-INS-161 | chr3-67503795-INS-160 |
| chr3-67535006-INS-228 | chr3-67535012-INS-261 |
| chr3-67542375-INS-298 | chr3-67542371-INS-281 |
| chr3-67549179-INS-486 | chr3-67549179-INS-486 |
| chr3-67578241-INS-55 | chr3-67578253-INS-68 |
| chr3-67602300-INS-7091 | chr3-67602301-INS-7091 |
| chr3-67608227-INS-180 | chr3-67608227-INS-180 |
| chr3-67647387-INS-95 | chr3-67647363-INS-127 |
| chr3-67688857-INS-58 | chr3-67688857-INS-76 |
| chr3-67709338-INS-80 | chr3-67709356-INS-64 |
| chr3-67849704-INS-202 | chr3-67849704-INS-188 |
| chr3-67850580-INS-620 | chr3-67850580-INS-599 |
| chr3-67857719-INS-190 | chr3-67857714-INS-189 |
| chr3-67861539-INS-54 | chr3-67861529-INS-66 |
| chr3-67864108-INS-6234 | chr3-67864109-INS-6234 |
| chr3-67869278-INS-66 | chr3-67869278-INS-66 |
| chr3-67875206-INS-191 | chr3-67875246-INS-157 |
| chr3-67885257-INS-754 | chr3-67885257-INS-750 |
| chr3-67910772-INS-52 | chr3-67910731-INS-68 |
| chr3-67939184-INS-66 | chr3-67939184-INS-66 |
| chr3-67950845-INS-52 | chr3-67950849-INS-52 |
| chr3-67961588-INS-961 | chr3-67961588-INS-979 |
| chr3-67992277-INS-94 | chr3-67992277-INS-94 |
| chr3-67995059-INS-108 | chr3-67995059-INS-108 |
| chr3-68002564-INS-6864 | chr3-68002565-INS-6864 |
| chr3-68040536-INS-280 | chr3-68040536-INS-280 |
| chr3-68042002-INS-112 | chr3-68042002-INS-116 |
| chr3-68076919-INS-85 | chr3-68076919-INS-50 |
| chr3-68077338-INS-2224 | chr3-68077339-INS-2223 |
| chr3-68099037-INS-56 | chr3-68099037-INS-56 |
| chr3-68123492-INS-156 | chr3-68123492-INS-156 |
| chr3-68126247-INS-100 | chr3-68126247-INS-100 |
| chr3-68132630-INS-7277 | chr3-68132631-INS-7277 |
| chr3-68132755-INS-76 | chr3-68132761-INS-89 |
| chr3-68140541-INS-54 | chr3-68140541-INS-50 |
| chr3-68146545-INS-149 | chr3-68146503-INS-149 |
| chr3-68163281-INS-50 | chr3-68163281-INS-56 |
| chr3-68174893-INS-6203 | chr3-68174894-INS-6203 |
| chr3-68186276-INS-200 | chr3-68186276-INS-200 |
| chr3-68193257-INS-215 | chr3-68193257-INS-215 |
| chr3-68208659-INS-52 | chr3-68208659-INS-64 |
| chr3-68225276-INS-52 | chr3-68225272-INS-56 |
| chr3-68226326-INS-112 | chr3-68226326-INS-112 |
| chr3-68237137-INS-6277 | chr3-68237138-INS-6277 |
| chr3-68278518-INS-387 | chr3-68278518-INS-387 |
| chr3-68321883-INS-1463 | chr3-68321884-INS-1463 |
| chr3-68323482-INS-1891 | chr3-68323483-INS-1891 |
| chr3-68356431-INS-6388 | chr3-68356432-INS-6388 |
| chr3-68380156-INS-53 | chr3-68380108-INS-57 |
| chr3-68423514-INS-400 | chr3-68423514-INS-400 |

| | |
|---|---|
| chr3-68425769-INS-54 | chr3-68425769-INS-54 |
| chr3-68496319-INS-892 | chr3-68496319-INS-896 |
| chr3-68516940-INS-771 | chr3-68516940-INS-771 |
| chr3-68542717-INS-208 | chr3-68542717-INS-208 |
| chr3-68547105-INS-2234 | chr3-68547106-INS-2212 |
| chr3-68548579-INS-3272 | chr3-68548580-INS-3278 |
| chr3-68562555-INS-54 | chr3-68562594-INS-50 |
| chr3-68580714-INS-200 | chr3-68580668-INS-212 |
| chr3-68596718-INS-81 | chr3-68596718-INS-81 |
| chr3-68606737-INS-497 | chr3-68606737-INS-497 |
| chr3-68631830-INS-79 | chr3-68631830-INS-79 |
| chr3-68668197-INS-52 | chr3-68668201-INS-74 |
| chr3-68750335-INS-71 | chr3-68750343-INS-79 |
| chr3-68793636-INS-6475 | chr3-68793637-INS-6475 |
| chr3-68806717-INS-83 | chr3-68806709-INS-71 |
| chr3-68830904-INS-1351 | chr3-68830905-INS-1354 |
| chr3-68842669-INS-403 | chr3-68842669-INS-403 |
| chr3-68855784-INS-52 | chr3-68855784-INS-52 |
| chr3-68897831-INS-196 | chr3-68897831-INS-198 |
| chr3-68921545-INS-207 | chr3-68921545-INS-207 |
| chr3-68928367-INS-205 | chr3-68928367-INS-205 |
| chr3-68935261-INS-97 | chr3-68935262-INS-98 |
| chr3-68937962-INS-139 | chr3-68937962-INS-139 |
| chr3-68945634-INS-208 | chr3-68945634-INS-208 |
| chr3-68947360-INS-1932 | chr3-68947361-INS-1932 |
| chr3-68951947-INS-54 | chr3-68951947-INS-54 |
| chr3-68959980-INS-60 | chr3-68959980-INS-56 |
| chr3-68961560-INS-126 | chr3-68961560-INS-126 |
| chr3-68982529-INS-82 | chr3-68982529-INS-66 |
| chr3-69015431-INS-207 | chr3-69015431-INS-209 |
| chr3-69064669-INS-103 | chr3-69064669-INS-98 |
| chr3-69076009-INS-88 | chr3-69076009-INS-96 |
| chr3-69080865-INS-269 | chr3-69080865-INS-270 |
| chr3-69084094-INS-235 | chr3-69084094-INS-236 |
| chr3-69110032-INS-3425 | chr3-69110033-INS-3425 |
| chr3-69118582-INS-246 | chr3-69118582-INS-247 |
| chr3-69122849-INS-202 | chr3-69122849-INS-200 |
| chr3-69128210-INS-54 | chr3-69128209-INS-58 |
| chr3-69135689-INS-54 | chr3-69135689-INS-54 |
| chr3-69138499-INS-56 | chr3-69138499-INS-58 |
| chr3-69144329-INS-52 | chr3-69144329-INS-68 |
| chr3-69157482-INS-64 | chr3-69157500-INS-85 |
| chr3-69160375-INS-85 | chr3-69160375-INS-88 |
| chr3-69178435-INS-368 | chr3-69178435-INS-368 |
| chr3-69188176-INS-1170 | chr3-69188177-INS-1171 |
| chr3-69191551-INS-62 | chr3-69191551-INS-62 |
| chr3-69244467-INS-186 | chr3-69244467-INS-186 |
| chr3-69248421-INS-67 | chr3-69248421-INS-77 |
| chr3-69254490-INS-81 | chr3-69254490-INS-81 |
| chr3-69270587-INS-445 | chr3-69270587-INS-446 |
| chr3-69290385-INS-226 | chr3-69290385-INS-226 |
| chr3-69295477-INS-914 | chr3-69295477-INS-915 |
| chr3-69309639-INS-90 | chr3-69309639-INS-90 |
| chr3-69329892-INS-56 | chr3-69329892-INS-54 |
| chr3-69332440-INS-64 | chr3-69332440-INS-80 |
| chr3-69352357-INS-151 | chr3-69352357-INS-151 |
| chr3-69360905-INS-82 | chr3-69360905-INS-82 |
| chr3-69399391-INS-54 | chr3-69399391-INS-54 |
| chr3-69410438-INS-108 | chr3-69410438-INS-108 |
| chr3-69439798-INS-114 | chr3-69439798-INS-118 |
| chr3-69441278-INS-492 | chr3-69441278-INS-492 |
| chr3-69446626-INS-82 | chr3-69446626-INS-82 |
| chr3-69471333-INS-275 | chr3-69471333-INS-267 |
| chr3-69496421-INS-54 | chr3-69496410-INS-60 |
| chr3-69503840-INS-195 | chr3-69503840-INS-195 |
| chr3-69514485-INS-88 | chr3-69514485-INS-92 |
| chr3-69535762-INS-176 | chr3-69535754-INS-178 |
| chr3-69545270-INS-297 | chr3-69545270-INS-297 |
| chr3-69562672-INS-323 | chr3-69562669-INS-321 |
| chr3-69566942-INS-51 | chr3-69566943-INS-51 |
| chr3-69567635-INS-51 | chr3-69567636-INS-51 |
| chr3-69612608-INS-460 | chr3-69612608-INS-460 |
| chr3-69617058-INS-92 | chr3-69617058-INS-92 |
| chr3-69626839-INS-222 | chr3-69626839-INS-214 |
| chr3-69658344-INS-147 | chr3-69658344-INS-147 |
| chr3-69671280-INS-6413 | chr3-69671281-INS-6413 |
| chr3-69679308-INS-315 | chr3-69679308-INS-315 |

| | |
|---|---|
| chr3-69722950-INS-338 | chr3-69722950-INS-338 |
| chr3-69752181-INS-455 | chr3-69752181-INS-455 |
| chr3-69789097-INS-6745 | chr3-69789098-INS-6745 |
| chr3-69829117-INS-382 | chr3-69829117-INS-382 |
| chr3-69830186-INS-827 | chr3-69830186-INS-827 |
| chr3-69865510-INS-93 | chr3-69865537-INS-57 |
| chr3-69865869-INS-298 | chr3-69865869-INS-298 |
| chr3-69867774-INS-55 | chr3-69867774-INS-55 |
| chr3-69873024-INS-132 | chr3-69873024-INS-132 |
| chr3-69911676-INS-98 | chr3-69911677-INS-124 |
| chr3-69920574-INS-70 | chr3-69920574-INS-70 |
| chr3-69923383-INS-6705 | chr3-69923384-INS-6706 |
| chr3-69923838-INS-678 | chr3-69923838-INS-678 |
| chr3-69927884-INS-615 | chr3-69927884-INS-613 |
| chr3-69941753-INS-191 | chr3-69941753-INS-191 |
| chr3-69948481-INS-127 | chr3-69948481-INS-87 |
| chr3-69987669-INS-174 | chr3-69987669-INS-174 |
| chr3-69993533-INS-1699 | chr3-69993534-INS-1684 |
| chr3-70020595-INS-455 | chr3-70020595-INS-455 |
| chr3-70029501-INS-63 | chr3-70029501-INS-68 |
| chr3-70053315-INS-2166 | chr3-70053320-INS-2158 |
| chr3-70056598-INS-6408 | chr3-70056599-INS-6406 |
| chr3-70096825-INS-851 | chr3-70096826-INS-850 |
| chr3-70126961-INS-706 | chr3-70126961-INS-707 |
| chr3-70161353-INS-73 | chr3-70161353-INS-73 |
| chr3-70186769-INS-76 | chr3-70186769-INS-76 |
| chr3-70193375-INS-485 | chr3-70193375-INS-485 |
| chr3-70196783-INS-98 | chr3-70196756-INS-103 |
| chr3-70218187-INS-82 | chr3-70218183-INS-82 |
| chr3-70228187-INS-507 | chr3-70228187-INS-507 |
| chr3-70230470-INS-95 | chr3-70230470-INS-95 |
| chr3-70237869-INS-486 | chr3-70237869-INS-487 |
| chr3-70250222-INS-6208 | chr3-70250223-INS-6208 |
| chr3-70317101-INS-65 | chr3-70317115-INS-62 |
| chr3-70360013-INS-54 | chr3-70360013-INS-56 |
| chr3-70369929-INS-296 | chr3-70369929-INS-296 |
| chr3-70377135-INS-108 | chr3-70377135-INS-108 |
| chr3-70394376-INS-107 | chr3-70394366-INS-115 |
| chr3-70404941-INS-7913 | chr3-70404942-INS-7916 |
| chr3-70442993-INS-312 | chr3-70442993-INS-312 |
| chr3-70473560-INS-1110 | chr3-70473561-INS-1110 |
| chr3-70508298-INS-1922 | chr3-70508299-INS-1967 |
| chr3-70543266-INS-342 | chr3-70543266-INS-342 |
| chr3-70656327-INS-105 | chr3-70656327-INS-75 |
| chr3-70704271-INS-54 | chr3-70704271-INS-54 |
| chr3-70787099-INS-639 | chr3-70787099-INS-641 |
| chr3-70838808-INS-62 | chr3-70838808-INS-62 |
| chr3-70883417-INS-930 | chr3-70883417-INS-937 |
| chr3-70892435-INS-92 | chr3-70892420-INS-100 |
| chr3-70923182-INS-52 | chr3-70923182-INS-54 |
| chr3-70927727-INS-6412 | chr3-70927728-INS-6412 |
| chr3-70980155-INS-212 | chr3-70980155-INS-212 |
| chr3-71000725-INS-677 | chr3-71000725-INS-676 |
| chr3-71010206-INS-2709 | chr3-71010207-INS-2709 |
| chr3-71033451-INS-64 | chr3-71033413-INS-58 |
| chr3-71047544-INS-52 | chr3-71047546-INS-53 |
| chr3-71050465-INS-661 | chr3-71050465-INS-661 |
| chr3-71072297-INS-306 | chr3-71072297-INS-306 |
| chr3-71073451-INS-114 | chr3-71073451-INS-122 |
| chr3-71079211-INS-404 | chr3-71079214-INS-401 |
| chr3-71088394-INS-50 | chr3-71088394-INS-66 |
| chr3-71094991-INS-2719 | chr3-71094992-INS-2719 |
| chr3-71105432-INS-650 | chr3-71105432-INS-654 |
| chr3-71107700-INS-183 | chr3-71107700-INS-183 |
| chr3-71111284-INS-6274 | chr3-71111285-INS-6283 |
| chr3-71116773-INS-688 | chr3-71116773-INS-691 |
| chr3-71120219-INS-55 | chr3-71120219-INS-55 |
| chr3-71138661-INS-1145 | chr3-71138662-INS-1148 |
| chr3-71184626-INS-288 | chr3-71184626-INS-315 |
| chr3-71187676-INS-58 | chr3-71187690-INS-52 |
| chr3-71193829-INS-54 | chr3-71193816-INS-68 |
| chr3-71196749-INS-194 | chr3-71196749-INS-196 |
| chr3-71199626-INS-141 | chr3-71199626-INS-141 |
| chr3-71218430-INS-78 | chr3-71218430-INS-60 |
| chr3-71222339-INS-657 | chr3-71222339-INS-657 |
| chr3-71235512-INS-2764 | chr3-71235513-INS-2763 |
| chr3-71265239-INS-68 | chr3-71265215-INS-83 |

| | |
|---|---|
| chr3-71329198-INS-308 | chr3-71329198-INS-308 |
| chr3-71348376-INS-2848 | chr3-71348377-INS-2848 |
| chr3-71354727-INS-402 | chr3-71354728-INS-402 |
| chr3-71383085-INS-334 | chr3-71383085-INS-334 |
| chr3-71384845-INS-6377 | chr3-71384846-INS-6377 |
| chr3-71392411-INS-298 | chr3-71392411-INS-297 |
| chr3-71405537-INS-1761 | chr3-71405538-INS-1796 |
| chr3-71406661-INS-6420 | chr3-71406662-INS-6439 |
| chr3-71417777-INS-2795 | chr3-71417783-INS-2785 |
| chr3-71430715-INS-52 | chr3-71430717-INS-75 |
| chr3-71448378-INS-187 | chr3-71448378-INS-188 |
| chr3-71502360-INS-1078 | chr3-71502361-INS-1078 |
| chr3-71548159-INS-3272 | chr3-71548160-INS-3272 |
| chr3-71569454-INS-6703 | chr3-71569455-INS-6703 |
| chr3-71598962-INS-175 | chr3-71598964-INS-180 |
| chr3-71620083-INS-1593 | chr3-71620084-INS-1593 |
| chr3-71627093-INS-221 | chr3-71627093-INS-228 |
| chr3-71646663-INS-103 | chr3-71646662-INS-101 |
| chr3-71652056-INS-120 | chr3-71652056-INS-75 |
| chr3-71671952-INS-58 | chr3-71671950-INS-58 |
| chr3-71677009-INS-463 | chr3-71677011-INS-460 |
| chr3-71686077-INS-6515 | chr3-71686078-INS-6516 |
| chr3-71690417-INS-337 | chr3-71690417-INS-337 |
| chr3-71715488-INS-2674 | chr3-71715489-INS-2672 |
| chr3-71715906-INS-94 | chr3-71715906-INS-94 |
| chr3-71718564-INS-447 | chr3-71718564-INS-447 |
| chr3-71725898-INS-72 | chr3-71725920-INS-56 |
| chr3-71729946-INS-52 | chr3-71729966-INS-56 |
| chr3-71741582-INS-911 | chr3-71741582-INS-911 |
| chr3-71765584-INS-63 | chr3-71765584-INS-63 |
| chr3-71767829-INS-91 | chr3-71767829-INS-91 |
| chr3-71780587-INS-78 | chr3-71780587-INS-80 |
| chr3-71852811-INS-66 | chr3-71852811-INS-66 |
| chr3-71855893-INS-52 | chr3-71855893-INS-56 |
| chr3-71858234-INS-105 | chr3-71858234-INS-105 |
| chr3-71867644-INS-76 | chr3-71867644-INS-76 |
| chr3-71903433-INS-52 | chr3-71903433-INS-52 |
| chr3-71909478-INS-4822 | chr3-71909479-INS-4822 |
| chr3-71919118-INS-7336 | chr3-71919119-INS-7336 |
| chr3-71924263-INS-88 | chr3-71924263-INS-86 |
| chr3-71929833-INS-302 | chr3-71929837-INS-296 |
| chr3-71945046-INS-72 | chr3-71945046-INS-72 |
| chr3-71947679-INS-52 | chr3-71947679-INS-52 |
| chr3-71963899-INS-6886 | chr3-71963900-INS-6886 |
| chr3-71969335-INS-194 | chr3-71969334-INS-197 |
| chr3-72001837-INS-5708 | chr3-72001838-INS-5708 |
| chr3-72002578-INS-79 | chr3-72002578-INS-79 |
| chr3-72003612-INS-275 | chr3-72003612-INS-275 |
| chr3-72005863-INS-6867 | chr3-72005864-INS-6867 |
| chr3-72009557-INS-1734 | chr3-72009558-INS-1735 |
| chr3-72027084-INS-488 | chr3-72027084-INS-510 |
| chr3-72053605-INS-4812 | chr3-72053606-INS-4814 |
| chr3-72065777-INS-120 | chr3-72065777-INS-121 |
| chr3-72068103-INS-1610 | chr3-72068116-INS-1598 |
| chr3-72072287-INS-2650 | chr3-72072288-INS-2650 |
| chr3-72074368-INS-437 | chr3-72074368-INS-437 |
| chr3-72098756-INS-6303 | chr3-72098757-INS-6303 |
| chr3-72108154-INS-62 | chr3-72108154-INS-88 |
| chr3-72118906-INS-52 | chr3-72118906-INS-52 |
| chr3-72143346-INS-104 | chr3-72143346-INS-104 |
| chr3-72145023-INS-62 | chr3-72145025-INS-88 |
| chr3-72146945-INS-578 | chr3-72146945-INS-579 |
| chr3-72149432-INS-80 | chr3-72149454-INS-60 |
| chr3-72173003-INS-1647 | chr3-72173004-INS-1648 |
| chr3-72279879-INS-140 | chr3-72279879-INS-140 |
| chr3-72295071-INS-89 | chr3-72295071-INS-60 |
| chr3-72303355-INS-100 | chr3-72303360-INS-64 |
| chr3-72305886-INS-72 | chr3-72305869-INS-50 |
| chr3-72324623-INS-460 | chr3-72324623-INS-460 |
| chr3-72326849-INS-1472 | chr3-72326850-INS-1481 |
| chr3-72347458-INS-513 | chr3-72347458-INS-513 |
| chr3-72349140-INS-753 | chr3-72349140-INS-753 |
| chr3-72361254-INS-297 | chr3-72361242-INS-320 |
| chr3-72375492-INS-6710 | chr3-72375493-INS-6704 |
| chr3-72380158-INS-110 | chr3-72380158-INS-115 |
| chr3-72410717-INS-1778 | chr3-72410718-INS-1818 |
| chr3-72461455-INS-108 | chr3-72461455-INS-108 |

| | |
|---|---|
| chr3-72490266-INS-338 | chr3-72490266-INS-338 |
| chr3-72563455-INS-6657 | chr3-72563456-INS-6660 |
| chr3-72590511-INS-117 | chr3-72590511-INS-117 |
| chr3-72597948-INS-1890 | chr3-72597949-INS-1890 |
| chr3-72609773-INS-6823 | chr3-72609774-INS-6824 |
| chr3-72637131-INS-220 | chr3-72637131-INS-238 |
| chr3-72660403-INS-498 | chr3-72660403-INS-498 |
| chr3-72663645-INS-1701 | chr3-72663646-INS-1701 |
| chr3-72663937-INS-7265 | chr3-72663938-INS-7265 |
| chr3-72682125-INS-56 | chr3-72682125-INS-52 |
| chr3-72695126-INS-2831 | chr3-72695127-INS-2804 |
| chr3-72717325-INS-2821 | chr3-72717326-INS-2821 |
| chr3-72722623-INS-6379 | chr3-72722624-INS-6378 |
| chr3-72725801-INS-2205 | chr3-72725802-INS-2205 |
| chr3-72742020-INS-8663 | chr3-72742021-INS-8662 |
| chr3-72747777-INS-2253 | chr3-72747778-INS-2253 |
| chr3-72762850-INS-7413 | chr3-72762851-INS-7413 |
| chr3-72796706-INS-52 | chr3-72796706-INS-53 |
| chr3-72817362-INS-2105 | chr3-72817363-INS-2105 |
| chr3-72832576-INS-1594 | chr3-72832579-INS-1580 |
| chr3-72847783-INS-580 | chr3-72847783-INS-579 |
| chr3-72887303-INS-395 | chr3-72887303-INS-395 |
| chr3-72909565-INS-549 | chr3-72909565-INS-549 |
| chr3-72918650-INS-1939 | chr3-72918651-INS-1943 |
| chr3-72944710-INS-92 | chr3-72944710-INS-108 |
| chr3-72986362-INS-65 | chr3-72986362-INS-65 |
| chr3-73001611-INS-204 | chr3-73001611-INS-204 |
| chr3-73003495-INS-6240 | chr3-73003496-INS-6241 |
| chr3-73012404-INS-4344 | chr3-73012405-INS-4344 |
| chr3-73029388-INS-255 | chr3-73029388-INS-236 |
| chr3-73031161-INS-6509 | chr3-73031162-INS-6509 |
| chr3-73036486-INS-733 | chr3-73036486-INS-733 |
| chr3-73059468-INS-197 | chr3-73059468-INS-193 |
| chr3-73092177-INS-197 | chr3-73092177-INS-197 |
| chr3-73101619-INS-671 | chr3-73101619-INS-672 |
| chr3-73103723-INS-96 | chr3-73103723-INS-80 |
| chr3-73130112-INS-56 | chr3-73130118-INS-52 |
| chr3-73155413-INS-1286 | chr3-73155414-INS-1287 |
| chr3-73178039-INS-120 | chr3-73178039-INS-120 |
| chr3-73181591-INS-657 | chr3-73181591-INS-656 |
| chr3-73215684-INS-400 | chr3-73215684-INS-400 |
| chr3-73239168-INS-1645 | chr3-73239169-INS-1646 |
| chr3-73263939-INS-1822 | chr3-73263940-INS-1823 |
| chr3-73268751-INS-1891 | chr3-73268752-INS-1891 |
| chr3-73273269-INS-82 | chr3-73273269-INS-76 |
| chr3-73285808-INS-79 | chr3-73285816-INS-52 |
| chr3-73287793-INS-105 | chr3-73287791-INS-106 |
| chr3-73304428-INS-1787 | chr3-73304429-INS-1788 |
| chr3-73307067-INS-120 | chr3-73307067-INS-120 |
| chr3-73331471-INS-467 | chr3-73331471-INS-467 |
| chr3-73335466-INS-191 | chr3-73335466-INS-191 |
| chr3-73348965-INS-7118 | chr3-73348966-INS-7118 |
| chr3-73349056-INS-60 | chr3-73349056-INS-68 |
| chr3-73351004-INS-89 | chr3-73351004-INS-89 |
| chr3-73363899-INS-204 | chr3-73363899-INS-204 |
| chr3-73401040-INS-52 | chr3-73401040-INS-52 |
| chr3-73415213-INS-602 | chr3-73415213-INS-602 |
| chr3-73432796-INS-170 | chr3-73432796-INS-171 |
| chr3-73443053-INS-1994 | chr3-73443058-INS-2031 |
| chr3-73444825-INS-330 | chr3-73444825-INS-330 |
| chr3-73457326-INS-779 | chr3-73457326-INS-776 |
| chr3-73478812-INS-102 | chr3-73478812-INS-96 |
| chr3-73492222-INS-236 | chr3-73492226-INS-237 |
| chr3-73497019-INS-133 | chr3-73497019-INS-133 |
| chr3-73500863-INS-1899 | chr3-73500864-INS-1899 |
| chr3-73524389-INS-122 | chr3-73524389-INS-125 |
| chr3-73543114-INS-896 | chr3-73543114-INS-896 |
| chr3-73553842-INS-56 | chr3-73553842-INS-56 |
| chr3-73572431-INS-157 | chr3-73572431-INS-164 |
| chr3-73577717-INS-117 | chr3-73577734-INS-133 |
| chr3-73578582-INS-6625 | chr3-73578582-INS-6628 |
| chr3-73582541-INS-240 | chr3-73582541-INS-240 |
| chr3-73602390-INS-611 | chr3-73602390-INS-611 |
| chr3-73621508-INS-56 | chr3-73621508-INS-56 |
| chr3-73627544-INS-81 | chr3-73627544-INS-91 |
| chr3-73651981-INS-62 | chr3-73651981-INS-62 |
| chr3-73659170-INS-197 | chr3-73659170-INS-197 |

| | |
|---|---|
| chr3-73661903-INS-78 | chr3-73661899-INS-82 |
| chr3-73735860-INS-60 | chr3-73735860-INS-65 |
| chr3-73742126-INS-80 | chr3-73742151-INS-96 |
| chr3-73744069-INS-166 | chr3-73744069-INS-167 |
| chr3-73755015-INS-2454 | chr3-73755016-INS-2435 |
| chr3-73759847-INS-229 | chr3-73759847-INS-229 |
| chr3-73768760-INS-6401 | chr3-73768761-INS-6401 |
| chr3-73792805-INS-111 | chr3-73792805-INS-153 |
| chr3-73795506-INS-2392 | chr3-73795507-INS-2399 |
| chr3-73834948-INS-72 | chr3-73834948-INS-72 |
| chr3-73861784-INS-431 | chr3-73861784-INS-433 |
| chr3-73866134-INS-1180 | chr3-73866135-INS-1183 |
| chr3-73872713-INS-111 | chr3-73872713-INS-111 |
| chr3-73887312-INS-145 | chr3-73887312-INS-146 |
| chr3-73899384-INS-61 | chr3-73899401-INS-79 |
| chr3-73906284-INS-419 | chr3-73906281-INS-417 |
| chr3-73913739-INS-81 | chr3-73913739-INS-81 |
| chr3-73950701-INS-528 | chr3-73950701-INS-535 |
| chr3-73985468-INS-250 | chr3-73985468-INS-252 |
| chr3-74027398-INS-537 | chr3-74027398-INS-537 |
| chr3-74052859-INS-54 | chr3-74052859-INS-58 |
| chr3-74057437-INS-769 | chr3-74057437-INS-775 |
| chr3-74065129-INS-78 | chr3-74065129-INS-62 |
| chr3-74069260-INS-138 | chr3-74069260-INS-138 |
| chr3-74111809-INS-52 | chr3-74111809-INS-52 |
| chr3-74166462-INS-64 | chr3-74166462-INS-70 |
| chr3-74201485-INS-54 | chr3-74201485-INS-56 |
| chr3-74214398-INS-647 | chr3-74214398-INS-647 |
| chr3-74215633-INS-2525 | chr3-74215634-INS-2526 |
| chr3-74270446-INS-52 | chr3-74270446-INS-52 |
| chr3-74294403-INS-1746 | chr3-74294404-INS-1746 |
| chr3-74297643-INS-969 | chr3-74297643-INS-972 |
| chr3-74312218-INS-65 | chr3-74312218-INS-66 |
| chr3-74330606-INS-355 | chr3-74330606-INS-356 |
| chr3-74336546-INS-54 | chr3-74336546-INS-54 |
| chr3-74348238-INS-301 | chr3-74348238-INS-301 |
| chr3-74368568-INS-142 | chr3-74368568-INS-118 |
| chr3-74395526-INS-152 | chr3-74395526-INS-152 |
| chr3-74406478-INS-465 | chr3-74406479-INS-465 |
| chr3-74421411-INS-78 | chr3-74421411-INS-100 |
| chr3-74462809-INS-199 | chr3-74462809-INS-199 |
| chr3-74463986-INS-2046 | chr3-74463987-INS-2046 |
| chr3-74479227-INS-84 | chr3-74479235-INS-92 |
| chr3-74597372-INS-57 | chr3-74597372-INS-60 |
| chr3-74624053-INS-6632 | chr3-74624054-INS-6633 |
| chr3-74703822-INS-2744 | chr3-74703823-INS-2744 |
| chr3-74714864-INS-58 | chr3-74714896-INS-59 |
| chr3-74723647-INS-52 | chr3-74723615-INS-78 |
| chr3-74772367-INS-201 | chr3-74772367-INS-201 |
| chr3-74809307-INS-60 | chr3-74809307-INS-67 |
| chr3-74839913-INS-72 | chr3-74839913-INS-52 |
| chr3-74851894-INS-7104 | chr3-74851895-INS-7104 |
| chr3-74860079-INS-587 | chr3-74860079-INS-587 |
| chr3-74875800-INS-521 | chr3-74875800-INS-521 |
| chr3-74898354-INS-4939 | chr3-74898355-INS-4938 |
| chr3-74918401-INS-230 | chr3-74918401-INS-229 |
| chr3-74940182-INS-6387 | chr3-74940183-INS-6387 |
| chr3-74997023-INS-5167 | chr3-74997024-INS-5167 |
| chr3-75005570-INS-1893 | chr3-75005571-INS-1893 |
| chr3-75040679-INS-138 | chr3-75040679-INS-110 |
| chr3-75109481-INS-1862 | chr3-75109482-INS-1862 |
| chr3-75111124-INS-6932 | chr3-75111125-INS-6932 |
| chr3-75134565-INS-1879 | chr3-75134566-INS-1897 |
| chr3-75142547-INS-6462 | chr3-75142548-INS-6448 |
| chr3-75172295-INS-58 | chr3-75172295-INS-50 |
| chr3-75185120-INS-180 | chr3-75185124-INS-198 |
| chr3-75194860-INS-7104 | chr3-75194861-INS-7104 |
| chr3-75219574-INS-58 | chr3-75219574-INS-58 |
| chr3-75243622-INS-69 | chr3-75243601-INS-77 |
| chr3-75266333-INS-382 | chr3-75266333-INS-384 |
| chr3-75291155-INS-237 | chr3-75291155-INS-237 |
| chr3-75298624-INS-503 | chr3-75298624-INS-503 |
| chr3-75311571-INS-785 | chr3-75311571-INS-785 |
| chr3-75312403-INS-58 | chr3-75312403-INS-58 |
| chr3-75314034-INS-621 | chr3-75314034-INS-621 |
| chr3-75322575-INS-212 | chr3-75322575-INS-212 |
| chr3-75358057-INS-68 | chr3-75358057-INS-68 |

| | |
|---|---|
| chr3-75360647-INS-2197 | chr3-75360648-INS-2197 |
| chr3-75373542-INS-117 | chr3-75373542-INS-117 |
| chr3-75382917-INS-3163 | chr3-75382918-INS-3163 |
| chr3-75390852-INS-499 | chr3-75390852-INS-499 |
| chr3-75396324-INS-560 | chr3-75396324-INS-561 |
| chr3-75405478-INS-58 | chr3-75405478-INS-96 |
| chr3-75408284-INS-54 | chr3-75408256-INS-104 |
| chr3-75453044-INS-83 | chr3-75453056-INS-85 |
| chr3-75454887-INS-904 | chr3-75454887-INS-894 |
| chr3-75526397-INS-5526 | chr3-75526398-INS-5526 |
| chr3-75564841-INS-52 | chr3-75564841-INS-52 |
| chr3-75567294-INS-606 | chr3-75567294-INS-606 |
| chr3-75596110-INS-503 | chr3-75596110-INS-503 |
| chr3-75601946-INS-180 | chr3-75601946-INS-180 |
| chr3-75605336-INS-57 | chr3-75605336-INS-57 |
| chr3-75607038-INS-88 | chr3-75607038-INS-88 |
| chr3-75609588-INS-91 | chr3-75609588-INS-91 |
| chr3-75614354-INS-84 | chr3-75614354-INS-79 |
| chr3-75660375-INS-74 | chr3-75660375-INS-100 |
| chr3-75662100-INS-63 | chr3-75662100-INS-53 |
| chr3-75674692-INS-482 | chr3-75674692-INS-481 |
| chr3-75706882-INS-5635 | chr3-75706883-INS-5635 |
| chr3-75710197-INS-1070 | chr3-75710198-INS-1070 |
| chr3-75740648-INS-6937 | chr3-75740649-INS-6938 |
| chr3-75744518-INS-975 | chr3-75744518-INS-975 |
| chr3-75768667-INS-469 | chr3-75768667-INS-469 |
| chr3-75831146-INS-5235 | chr3-75831147-INS-5235 |
| chr3-75853851-INS-87 | chr3-75853851-INS-123 |
| chr3-75867926-INS-70 | chr3-75867927-INS-51 |
| chr3-75873669-INS-153 | chr3-75873669-INS-142 |
| chr3-75877704-INS-81 | chr3-75877704-INS-81 |
| chr3-75905308-INS-74 | chr3-75905308-INS-76 |
| chr3-75909981-INS-618 | chr3-75909981-INS-620 |
| chr3-75911215-INS-55 | chr3-75911215-INS-55 |
| chr3-75918513-INS-390 | chr3-75918513-INS-390 |
| chr3-75918513-INS-898 | chr3-75918513-INS-900 |
| chr3-75942811-INS-3847 | chr3-75942812-INS-3847 |
| chr3-75943625-INS-1077 | chr3-75943626-INS-1078 |
| chr3-75955252-INS-1730 | chr3-75955254-INS-1725 |
| chr3-75968724-INS-291 | chr3-75968724-INS-290 |
| chr3-75994218-INS-245 | chr3-75994218-INS-226 |
| chr3-76058876-INS-6946 | chr3-76058877-INS-6947 |
| chr3-76080017-INS-2410 | chr3-76080018-INS-2410 |
| chr3-76088212-INS-632 | chr3-76088212-INS-632 |
| chr3-76091495-INS-1864 | chr3-76091496-INS-1864 |
| chr3-76093637-INS-6607 | chr3-76093638-INS-6607 |
| chr3-76123984-INS-93 | chr3-76123984-INS-53 |
| chr3-76155111-INS-555 | chr3-76155111-INS-545 |
| chr3-76157077-INS-78 | chr3-76157077-INS-78 |
| chr3-76157562-INS-98 | chr3-76157576-INS-50 |
| chr3-76174296-INS-400 | chr3-76174296-INS-400 |
| chr3-76203356-INS-70 | chr3-76203356-INS-70 |
| chr3-76212544-INS-1733 | chr3-76212545-INS-1733 |
| chr3-76216467-INS-7047 | chr3-76216468-INS-7047 |
| chr3-76224815-INS-62 | chr3-76224815-INS-54 |
| chr3-76234180-INS-65 | chr3-76234181-INS-70 |
| chr3-76238874-INS-506 | chr3-76238888-INS-477 |
| chr3-76240226-INS-6167 | chr3-76240227-INS-6167 |
| chr3-76315790-INS-6475 | chr3-76315791-INS-6475 |
| chr3-76315791-INS-503 | chr3-76315791-INS-503 |
| chr3-76325487-INS-384 | chr3-76325487-INS-384 |
| chr3-76340475-INS-7200 | chr3-76340476-INS-7199 |
| chr3-76352947-INS-6669 | chr3-76352948-INS-6669 |
| chr3-76386567-INS-11630 | chr3-76386568-INS-11622 |
| chr3-76392125-INS-55 | chr3-76392125-INS-55 |
| chr3-76417992-INS-64 | chr3-76418023-INS-64 |
| chr3-76439469-INS-7589 | chr3-76439470-INS-7592 |
| chr3-76445209-INS-161 | chr3-76445217-INS-159 |
| chr3-76493594-INS-356 | chr3-76493594-INS-346 |
| chr3-76514687-INS-4046 | chr3-76514688-INS-4046 |
| chr3-76517960-INS-194 | chr3-76517966-INS-197 |
| chr3-76558272-INS-1642 | chr3-76558273-INS-1639 |
| chr3-76558729-INS-293 | chr3-76558729-INS-293 |
| chr3-76560912-INS-167 | chr3-76560912-INS-167 |
| chr3-76571551-INS-8783 | chr3-76571552-INS-8783 |
| chr3-76579005-INS-57 | chr3-76579005-INS-60 |
| chr3-76579888-INS-6375 | chr3-76579889-INS-6375 |

| | |
|---|---|
| chr3-76654941-INS-132 | chr3-76654941-INS-132 |
| chr3-76658982-INS-176 | chr3-76658982-INS-176 |
| chr3-76662167-INS-151 | chr3-76662168-INS-151 |
| chr3-76669834-INS-199 | chr3-76669834-INS-199 |
| chr3-76683195-INS-1994 | chr3-76683196-INS-1995 |
| chr3-76709693-INS-7358 | chr3-76709694-INS-7358 |
| chr3-76710506-INS-262 | chr3-76710506-INS-262 |
| chr3-76718405-INS-388 | chr3-76718409-INS-389 |
| chr3-76730023-INS-299 | chr3-76730023-INS-299 |
| chr3-76730547-INS-284 | chr3-76730547-INS-240 |
| chr3-76781986-INS-143 | chr3-76781986-INS-143 |
| chr3-76836472-INS-79 | chr3-76836472-INS-102 |
| chr3-76864635-INS-6475 | chr3-76864636-INS-6475 |
| chr3-76887436-INS-2066 | chr3-76887437-INS-2066 |
| chr3-76918973-INS-60 | chr3-76918973-INS-65 |
| chr3-76943797-INS-102 | chr3-76943798-INS-102 |
| chr3-76950259-INS-423 | chr3-76950259-INS-423 |
| chr3-76954031-INS-63 | chr3-76954031-INS-65 |
| chr3-76959209-INS-198 | chr3-76959209-INS-198 |
| chr3-76965914-INS-152 | chr3-76965916-INS-135 |
| chr3-76970446-INS-51 | chr3-76970446-INS-51 |
| chr3-76984673-INS-768 | chr3-76984673-INS-768 |
| chr3-76993571-INS-62 | chr3-76993571-INS-68 |
| chr3-77000241-INS-60 | chr3-77000241-INS-55 |
| chr3-77027457-INS-248 | chr3-77027457-INS-249 |
| chr3-77033148-INS-2415 | chr3-77033148-INS-2420 |
| chr3-77042908-INS-371 | chr3-77042908-INS-373 |
| chr3-77083421-INS-207 | chr3-77083421-INS-211 |
| chr3-77154660-INS-82 | chr3-77154664-INS-50 |
| chr3-77155395-INS-344 | chr3-77155395-INS-344 |
| chr3-77177714-INS-53 | chr3-77177714-INS-51 |
| chr3-77194329-INS-84 | chr3-77194329-INS-80 |
| chr3-77236568-INS-88 | chr3-77236568-INS-108 |
| chr3-77241379-INS-1730 | chr3-77241380-INS-1730 |
| chr3-77287230-INS-139 | chr3-77287230-INS-139 |
| chr3-77323252-INS-725 | chr3-77323252-INS-725 |
| chr3-77326833-INS-63 | chr3-77326833-INS-63 |
| chr3-77335629-INS-126 | chr3-77335629-INS-108 |
| chr3-77350497-INS-870 | chr3-77350497-INS-870 |
| chr3-77356946-INS-80 | chr3-77356946-INS-84 |
| chr3-77363442-INS-510 | chr3-77363442-INS-504 |
| chr3-77378864-INS-2292 | chr3-77378865-INS-2290 |
| chr3-77423610-INS-2946 | chr3-77423611-INS-2952 |
| chr3-77448012-INS-498 | chr3-77448012-INS-498 |
| chr3-77452982-INS-80 | chr3-77453016-INS-80 |
| chr3-77461855-INS-4255 | chr3-77461856-INS-4255 |
| chr3-77462190-INS-54 | chr3-77462190-INS-50 |
| chr3-77464503-INS-153 | chr3-77464535-INS-122 |
| chr3-77465041-INS-4362 | chr3-77465042-INS-4363 |
| chr3-77466395-INS-3390 | chr3-77466396-INS-3390 |
| chr3-77472040-INS-333 | chr3-77472040-INS-333 |
| chr3-77490640-INS-7153 | chr3-77490641-INS-7153 |
| chr3-77496049-INS-143 | chr3-77496049-INS-143 |
| chr3-77528618-INS-549 | chr3-77528618-INS-549 |
| chr3-77533898-INS-74 | chr3-77533896-INS-67 |
| chr3-77552267-INS-62 | chr3-77552267-INS-52 |
| chr3-77572380-INS-4223 | chr3-77572381-INS-4223 |
| chr3-77612536-INS-214 | chr3-77612536-INS-214 |
| chr3-77625102-INS-6898 | chr3-77625103-INS-6901 |
| chr3-77639627-INS-155 | chr3-77639627-INS-155 |
| chr3-77643030-INS-4263 | chr3-77643031-INS-4267 |
| chr3-77645026-INS-74 | chr3-77645026-INS-75 |
| chr3-77680408-INS-5088 | chr3-77680409-INS-5088 |
| chr3-77682106-INS-343 | chr3-77682106-INS-343 |
| chr3-77691084-INS-6528 | chr3-77691085-INS-6529 |
| chr3-77691534-INS-431 | chr3-77691534-INS-431 |
| chr3-77692208-INS-306 | chr3-77692208-INS-304 |
| chr3-77697469-INS-1573 | chr3-77697470-INS-1573 |
| chr3-77729134-INS-167 | chr3-77729134-INS-139 |
| chr3-77742223-INS-611 | chr3-77742223-INS-610 |
| chr3-77750404-INS-97 | chr3-77750404-INS-97 |
| chr3-77763542-INS-6301 | chr3-77763543-INS-6304 |
| chr3-77774348-INS-7717 | chr3-77774349-INS-7718 |
| chr3-77779706-INS-58 | chr3-77779696-INS-78 |
| chr3-77814403-INS-398 | chr3-77814403-INS-398 |
| chr3-77818780-INS-441 | chr3-77818780-INS-441 |
| chr3-77821847-INS-209 | chr3-77821847-INS-208 |

| | |
|---|---|
| chr3-77882867-INS-516 | chr3-77882867-INS-516 |
| chr3-77944419-INS-480 | chr3-77944419-INS-480 |
| chr3-77952289-INS-60 | chr3-77952289-INS-60 |
| chr3-77958860-INS-53 | chr3-77958860-INS-53 |
| chr3-77962783-INS-5029 | chr3-77962784-INS-5030 |
| chr3-77968571-INS-52 | chr3-77968571-INS-50 |
| chr3-77972147-INS-506 | chr3-77972147-INS-506 |
| chr3-77978988-INS-434 | chr3-77978988-INS-434 |
| chr3-78041430-INS-54 | chr3-78041424-INS-96 |
| chr3-78042627-INS-60 | chr3-78042627-INS-60 |
| chr3-78054015-INS-553 | chr3-78054015-INS-556 |
| chr3-78068179-INS-558 | chr3-78068179-INS-558 |
| chr3-78092405-INS-5374 | chr3-78092406-INS-5374 |
| chr3-78102514-INS-56 | chr3-78102514-INS-56 |
| chr3-78126919-INS-163 | chr3-78126919-INS-164 |
| chr3-78132251-INS-312 | chr3-78132251-INS-312 |
| chr3-78157276-INS-401 | chr3-78157276-INS-401 |
| chr3-78166942-INS-1902 | chr3-78166943-INS-1902 |
| chr3-78196285-INS-130 | chr3-78196285-INS-178 |
| chr3-78201675-INS-52 | chr3-78201656-INS-54 |
| chr3-78215257-INS-3776 | chr3-78215258-INS-3770 |
| chr3-78223425-INS-1008 | chr3-78223426-INS-1008 |
| chr3-78232433-INS-84 | chr3-78232433-INS-84 |
| chr3-78255344-INS-58 | chr3-78255344-INS-56 |
| chr3-78264743-INS-180 | chr3-78264743-INS-168 |
| chr3-78273152-INS-1550 | chr3-78273153-INS-1550 |
| chr3-78295778-INS-539 | chr3-78295778-INS-539 |
| chr3-78319385-INS-63 | chr3-78319385-INS-66 |
| chr3-78322886-INS-685 | chr3-78322886-INS-685 |
| chr3-78338363-INS-6427 | chr3-78338364-INS-6427 |
| chr3-78370863-INS-1580 | chr3-78370864-INS-1587 |
| chr3-78371807-INS-176 | chr3-78371808-INS-171 |
| chr3-78376085-INS-285 | chr3-78376085-INS-285 |
| chr3-78377007-INS-58 | chr3-78377007-INS-97 |
| chr3-78424392-INS-207 | chr3-78424392-INS-207 |
| chr3-78483866-INS-6429 | chr3-78483867-INS-6429 |
| chr3-78567407-INS-2866 | chr3-78567408-INS-2866 |
| chr3-78570287-INS-6206 | chr3-78570288-INS-6206 |
| chr3-78576640-INS-6867 | chr3-78576641-INS-6867 |
| chr3-78584067-INS-1248 | chr3-78584068-INS-1249 |
| chr3-78603123-INS-586 | chr3-78603120-INS-585 |
| chr3-78606373-INS-592 | chr3-78606373-INS-588 |
| chr3-78610518-INS-339 | chr3-78610518-INS-339 |
| chr3-78639399-INS-54 | chr3-78639399-INS-56 |
| chr3-78655209-INS-72 | chr3-78655209-INS-71 |
| chr3-78669042-INS-118 | chr3-78669042-INS-120 |
| chr3-78732925-INS-106 | chr3-78732925-INS-106 |
| chr3-78742794-INS-235 | chr3-78742794-INS-237 |
| chr3-78745719-INS-67 | chr3-78745719-INS-67 |
| chr3-78755542-INS-62 | chr3-78755542-INS-90 |
| chr3-78802415-INS-189 | chr3-78802412-INS-190 |
| chr3-78818847-INS-147 | chr3-78818847-INS-148 |
| chr3-78831842-INS-79 | chr3-78831841-INS-79 |
| chr3-78834439-INS-136 | chr3-78834439-INS-136 |
| chr3-78846870-INS-614 | chr3-78846870-INS-613 |
| chr3-78864530-INS-180 | chr3-78864530-INS-174 |
| chr3-78869609-INS-62 | chr3-78869620-INS-54 |
| chr3-78870384-INS-76 | chr3-78870384-INS-76 |
| chr3-78898671-INS-96 | chr3-78898677-INS-78 |
| chr3-78900582-INS-158 | chr3-78900582-INS-175 |
| chr3-78903627-INS-344 | chr3-78903626-INS-352 |
| chr3-78908552-INS-667 | chr3-78908552-INS-667 |
| chr3-78943018-INS-65 | chr3-78943018-INS-50 |
| chr3-78943673-INS-69 | chr3-78943673-INS-69 |
| chr3-78952326-INS-53 | chr3-78952326-INS-53 |
| chr3-78960263-INS-54 | chr3-78960263-INS-56 |
| chr3-78964399-INS-1592 | chr3-78964400-INS-1591 |
| chr3-79005884-INS-59 | chr3-79005884-INS-57 |
| chr3-79007831-INS-282 | chr3-79007831-INS-282 |
| chr3-79018117-INS-213 | chr3-79018117-INS-213 |
| chr3-79020220-INS-90 | chr3-79020208-INS-60 |
| chr3-79022583-INS-77 | chr3-79022583-INS-77 |
| chr3-79022834-INS-1988 | chr3-79022835-INS-1988 |
| chr3-79023647-INS-828 | chr3-79023647-INS-828 |
| chr3-79028233-INS-78 | chr3-79028233-INS-90 |
| chr3-79066340-INS-309 | chr3-79066340-INS-309 |
| chr3-79085572-INS-380 | chr3-79085572-INS-380 |

| | |
|---|---|
| chr3-79109545-INS-410 | chr3-79109545-INS-410 |
| chr3-79121535-INS-64 | chr3-79121535-INS-64 |
| chr3-79132380-INS-54 | chr3-79132380-INS-54 |
| chr3-79137573-INS-174 | chr3-79137573-INS-176 |
| chr3-79148390-INS-709 | chr3-79148390-INS-699 |
| chr3-79150161-INS-74 | chr3-79150162-INS-74 |
| chr3-79158990-INS-56 | chr3-79158990-INS-59 |
| chr3-79173135-INS-209 | chr3-79173135-INS-208 |
| chr3-79174650-INS-221 | chr3-79174650-INS-221 |
| chr3-79179916-INS-144 | chr3-79179916-INS-180 |
| chr3-79182332-INS-69 | chr3-79182332-INS-67 |
| chr3-79208739-INS-833 | chr3-79208739-INS-836 |
| chr3-79234665-INS-56 | chr3-79234665-INS-60 |
| chr3-79236126-INS-3062 | chr3-79236127-INS-3062 |
| chr3-79267082-INS-927 | chr3-79267082-INS-927 |
| chr3-79304091-INS-95 | chr3-79304091-INS-132 |
| chr3-79315700-INS-150 | chr3-79315692-INS-172 |
| chr3-79351417-INS-80 | chr3-79351410-INS-98 |
| chr3-79378056-INS-80 | chr3-79378056-INS-80 |
| chr3-79392966-INS-62 | chr3-79392966-INS-66 |
| chr3-79404180-INS-64 | chr3-79404180-INS-64 |
| chr3-79412590-INS-56 | chr3-79412590-INS-66 |
| chr3-79414271-INS-202 | chr3-79414271-INS-201 |
| chr3-79416314-INS-170 | chr3-79416314-INS-170 |
| chr3-79417624-INS-201 | chr3-79417624-INS-201 |
| chr3-79432773-INS-752 | chr3-79432773-INS-752 |
| chr3-79436767-INS-60 | chr3-79436769-INS-56 |
| chr3-79467893-INS-206 | chr3-79467893-INS-206 |
| chr3-79477394-INS-56 | chr3-79477394-INS-56 |
| chr3-79489589-INS-216 | chr3-79489590-INS-262 |
| chr3-79495157-INS-706 | chr3-79495157-INS-706 |
| chr3-79509038-INS-222 | chr3-79509038-INS-222 |
| chr3-79512151-INS-392 | chr3-79512151-INS-402 |
| chr3-79512894-INS-54 | chr3-79512894-INS-54 |
| chr3-79548793-INS-242 | chr3-79548793-INS-244 |
| chr3-79550684-INS-194 | chr3-79550684-INS-195 |
| chr3-79551698-INS-66 | chr3-79551698-INS-66 |
| chr3-79570138-INS-84 | chr3-79570184-INS-58 |
| chr3-79571223-INS-93 | chr3-79571223-INS-93 |
| chr3-79592374-INS-68 | chr3-79592374-INS-56 |
| chr3-79593621-INS-159 | chr3-79593621-INS-119 |
| chr3-79680448-INS-100 | chr3-79680448-INS-98 |
| chr3-79681806-INS-60 | chr3-79681804-INS-66 |
| chr3-79698925-INS-290 | chr3-79698935-INS-302 |
| chr3-79708714-INS-66 | chr3-79708714-INS-66 |
| chr3-79758567-INS-53 | chr3-79758565-INS-64 |
| chr3-79766217-INS-272 | chr3-79766217-INS-282 |
| chr3-79767759-INS-226 | chr3-79767759-INS-228 |
| chr3-79778866-INS-819 | chr3-79778869-INS-817 |
| chr3-79789135-INS-74 | chr3-79789135-INS-74 |
| chr3-79797075-INS-75 | chr3-79797081-INS-62 |
| chr3-79806649-INS-221 | chr3-79806649-INS-217 |
| chr3-79809809-INS-642 | chr3-79809809-INS-678 |
| chr3-79838048-INS-147 | chr3-79838040-INS-124 |
| chr3-79838148-INS-6220 | chr3-79838149-INS-6229 |
| chr3-79882889-INS-53 | chr3-79882889-INS-53 |
| chr3-79901917-INS-184 | chr3-79901917-INS-184 |
| chr3-79902703-INS-116 | chr3-79902703-INS-116 |
| chr3-79906658-INS-294 | chr3-79906658-INS-294 |
| chr3-79907361-INS-97 | chr3-79907362-INS-83 |
| chr3-79908925-INS-294 | chr3-79908925-INS-294 |
| chr3-79927975-INS-64 | chr3-79927975-INS-64 |
| chr3-79930425-INS-600 | chr3-79930445-INS-607 |
| chr3-79934788-INS-328 | chr3-79934788-INS-328 |
| chr3-79956378-INS-182 | chr3-79956378-INS-182 |
| chr3-79961492-INS-445 | chr3-79961492-INS-445 |
| chr3-79968785-INS-55 | chr3-79968803-INS-55 |
| chr3-79969631-INS-838 | chr3-79969631-INS-837 |
| chr3-79974538-INS-7452 | chr3-79974539-INS-7452 |
| chr3-80012262-INS-163 | chr3-80012262-INS-181 |
| chr3-80013229-INS-104 | chr3-80013229-INS-104 |
| chr3-80021912-INS-504 | chr3-80021912-INS-504 |
| chr3-80059772-INS-78 | chr3-80059772-INS-74 |
| chr3-80100966-INS-7410 | chr3-80100967-INS-7410 |
| chr3-80108617-INS-52 | chr3-80108618-INS-56 |
| chr3-80123222-INS-681 | chr3-80123222-INS-681 |
| chr3-80138676-INS-127 | chr3-80138676-INS-127 |

| | |
|---|---|
| chr3-80153768-INS-52 | chr3-80153766-INS-69 |
| chr3-80155083-INS-114 | chr3-80155084-INS-115 |
| chr3-80165430-INS-6361 | chr3-80165430-INS-6361 |
| chr3-80166580-INS-283 | chr3-80166580-INS-288 |
| chr3-80169319-INS-6574 | chr3-80169320-INS-6575 |
| chr3-80173212-INS-72 | chr3-80173212-INS-72 |
| chr3-80176360-INS-52 | chr3-80176360-INS-52 |
| chr3-80177279-INS-66 | chr3-80177279-INS-96 |
| chr3-80188021-INS-152 | chr3-80188024-INS-154 |
| chr3-80189391-INS-642 | chr3-80189391-INS-642 |
| chr3-80202236-INS-3357 | chr3-80202237-INS-3357 |
| chr3-80237775-INS-416 | chr3-80237775-INS-416 |
| chr3-80265345-INS-59 | chr3-80265345-INS-59 |
| chr3-80306962-INS-120 | chr3-80306962-INS-120 |
| chr3-80310215-INS-70 | chr3-80310215-INS-76 |
| chr3-80317025-INS-1102 | chr3-80317026-INS-1102 |
| chr3-80318471-INS-343 | chr3-80318471-INS-343 |
| chr3-80320366-INS-243 | chr3-80320366-INS-243 |
| chr3-80340765-INS-90 | chr3-80340760-INS-90 |
| chr3-80353630-INS-97 | chr3-80353630-INS-55 |
| chr3-80363700-INS-115 | chr3-80363700-INS-111 |
| chr3-80368852-INS-82 | chr3-80368852-INS-88 |
| chr3-80392166-INS-118 | chr3-80392166-INS-144 |
| chr3-80397373-INS-87 | chr3-80397402-INS-80 |
| chr3-80411708-INS-1258 | chr3-80411709-INS-1258 |
| chr3-80419614-INS-1133 | chr3-80419615-INS-1134 |
| chr3-80433360-INS-61 | chr3-80433360-INS-60 |
| chr3-80448488-INS-246 | chr3-80448488-INS-246 |
| chr3-80460897-INS-56 | chr3-80460897-INS-56 |
| chr3-80461141-INS-6256 | chr3-80461142-INS-6258 |
| chr3-80463290-INS-213 | chr3-80463290-INS-213 |
| chr3-80476499-INS-6685 | chr3-80476500-INS-6685 |
| chr3-80486905-INS-212 | chr3-80486905-INS-213 |
| chr3-80488197-INS-230 | chr3-80488197-INS-230 |
| chr3-80500922-INS-1833 | chr3-80500923-INS-1835 |
| chr3-80520196-INS-7321 | chr3-80520197-INS-7321 |
| chr3-80521402-INS-95 | chr3-80521402-INS-77 |
| chr3-80525129-INS-716 | chr3-80525129-INS-716 |
| chr3-80555365-INS-156 | chr3-80555365-INS-156 |
| chr3-80561373-INS-469 | chr3-80561373-INS-469 |
| chr3-80565442-INS-66 | chr3-80565422-INS-87 |
| chr3-80567604-INS-57 | chr3-80567611-INS-63 |
| chr3-80589804-INS-60 | chr3-80589804-INS-78 |
| chr3-80647028-INS-161 | chr3-80647028-INS-161 |
| chr3-80725509-INS-253 | chr3-80725509-INS-240 |
| chr3-80771849-INS-145 | chr3-80771849-INS-146 |
| chr3-80835892-INS-6023 | chr3-80835893-INS-6023 |
| chr3-80843580-INS-75 | chr3-80843580-INS-75 |
| chr3-80847753-INS-211 | chr3-80847753-INS-212 |
| chr3-80855679-INS-60 | chr3-80855679-INS-54 |
| chr3-80889997-INS-508 | chr3-80889991-INS-506 |
| chr3-80896900-INS-7772 | chr3-80896904-INS-7765 |
| chr3-80898285-INS-403 | chr3-80898285-INS-412 |
| chr3-80900350-INS-475 | chr3-80900350-INS-472 |
| chr3-80902166-INS-199 | chr3-80902166-INS-199 |
| chr3-80905229-INS-73 | chr3-80905229-INS-64 |
| chr3-80907839-INS-65 | chr3-80907839-INS-65 |
| chr3-80920516-INS-52 | chr3-80920516-INS-52 |
| chr3-80933595-INS-6412 | chr3-80933596-INS-6413 |
| chr3-80956999-INS-1680 | chr3-80957000-INS-1681 |
| chr3-80976524-INS-187 | chr3-80976518-INS-195 |
| chr3-80992321-INS-84 | chr3-80992321-INS-72 |
| chr3-81015317-INS-56 | chr3-81015317-INS-56 |
| chr3-81025605-INS-159 | chr3-81025605-INS-155 |
| chr3-81032078-INS-132 | chr3-81032067-INS-120 |
| chr3-81042871-INS-478 | chr3-81042871-INS-478 |
| chr3-81051289-INS-55 | chr3-81051320-INS-56 |
| chr3-81057325-INS-54 | chr3-81057325-INS-54 |
| chr3-81083216-INS-127 | chr3-81083216-INS-127 |
| chr3-81084795-INS-92 | chr3-81084795-INS-92 |
| chr3-81097748-INS-193 | chr3-81097748-INS-195 |
| chr3-81098367-INS-52 | chr3-81098367-INS-54 |
| chr3-81100251-INS-836 | chr3-81100251-INS-841 |
| chr3-81115515-INS-4138 | chr3-81115516-INS-4138 |
| chr3-81125072-INS-452 | chr3-81125072-INS-452 |
| chr3-81126146-INS-12019 | chr3-81126147-INS-12025 |
| chr3-81141968-INS-276 | chr3-81141951-INS-281 |

| | |
|---|---|
| chr3-81154973-INS-66 | chr3-81154973-INS-70 |
| chr3-81170331-INS-60 | chr3-81170331-INS-60 |
| chr3-81234834-INS-6601 | chr3-81234835-INS-6604 |
| chr3-81238380-INS-69 | chr3-81238380-INS-71 |
| chr3-81241687-INS-82 | chr3-81241688-INS-84 |
| chr3-81262627-INS-887 | chr3-81262627-INS-890 |
| chr3-81287749-INS-60 | chr3-81287749-INS-64 |
| chr3-81297659-INS-1458 | chr3-81297660-INS-1471 |
| chr3-81314808-INS-52 | chr3-81314808-INS-52 |
| chr3-81316976-INS-757 | chr3-81316976-INS-756 |
| chr3-81319369-INS-1063 | chr3-81319362-INS-1057 |
| chr3-81376270-INS-77 | chr3-81376263-INS-84 |
| chr3-81386898-INS-192 | chr3-81386898-INS-192 |
| chr3-81392530-INS-373 | chr3-81392530-INS-373 |
| chr3-81417403-INS-490 | chr3-81417403-INS-490 |
| chr3-81418393-INS-100 | chr3-81418393-INS-100 |
| chr3-81418976-INS-78 | chr3-81418946-INS-81 |
| chr3-81434250-INS-53 | chr3-81434250-INS-53 |
| chr3-81438586-INS-51 | chr3-81438586-INS-51 |
| chr3-81458280-INS-6349 | chr3-81458281-INS-6349 |
| chr3-81475363-INS-129 | chr3-81475369-INS-96 |
| chr3-81477254-INS-495 | chr3-81477254-INS-495 |
| chr3-81478184-INS-76 | chr3-81478176-INS-78 |
| chr3-81494218-INS-3226 | chr3-81494219-INS-3225 |
| chr3-81496170-INS-1261 | chr3-81496171-INS-1262 |
| chr3-81523736-INS-1069 | chr3-81523737-INS-1069 |
| chr3-81568430-INS-58 | chr3-81568414-INS-50 |
| chr3-81590246-INS-4874 | chr3-81590247-INS-4874 |
| chr3-81596519-INS-77 | chr3-81596519-INS-77 |
| chr3-81662988-INS-55 | chr3-81662988-INS-55 |
| chr3-81678602-INS-2264 | chr3-81678603-INS-2262 |
| chr3-81712924-INS-322 | chr3-81712924-INS-296 |
| chr3-81724544-INS-4521 | chr3-81724545-INS-4521 |
| chr3-81726234-INS-487 | chr3-81726234-INS-485 |
| chr3-81731770-INS-276 | chr3-81731770-INS-277 |
| chr3-81754623-INS-879 | chr3-81754623-INS-879 |
| chr3-81772407-INS-922 | chr3-81772407-INS-923 |
| chr3-81782276-INS-292 | chr3-81782276-INS-292 |
| chr3-81789727-INS-74 | chr3-81789699-INS-54 |
| chr3-81791990-INS-6195 | chr3-81791991-INS-6194 |
| chr3-81823040-INS-1995 | chr3-81823041-INS-1995 |
| chr3-81853273-INS-51 | chr3-81853273-INS-51 |
| chr3-81861090-INS-6426 | chr3-81861091-INS-6426 |
| chr3-81864031-INS-70 | chr3-81864031-INS-55 |
| chr3-81867497-INS-92 | chr3-81867497-INS-92 |
| chr3-81893132-INS-195 | chr3-81893132-INS-195 |
| chr3-81903742-INS-189 | chr3-81903742-INS-189 |
| chr3-81920096-INS-60 | chr3-81920096-INS-110 |
| chr3-81927604-INS-5774 | chr3-81927605-INS-5774 |
| chr3-81936569-INS-51 | chr3-81936569-INS-51 |
| chr3-81954377-INS-141 | chr3-81954377-INS-152 |
| chr3-82043912-INS-72 | chr3-82043908-INS-74 |
| chr3-82044820-INS-53 | chr3-82044822-INS-57 |
| chr3-82053665-INS-74 | chr3-82053665-INS-74 |
| chr3-82070396-INS-5112 | chr3-82070397-INS-5112 |
| chr3-82109853-INS-151 | chr3-82109888-INS-110 |
| chr3-82167214-INS-71 | chr3-82167214-INS-97 |
| chr3-82179207-INS-69 | chr3-82179207-INS-78 |
| chr3-82207526-INS-215 | chr3-82207526-INS-215 |
| chr3-82251062-INS-6156 | chr3-82251063-INS-6156 |
| chr3-82257559-INS-54 | chr3-82257559-INS-54 |
| chr3-82263783-INS-2043 | chr3-82263784-INS-2027 |
| chr3-82287657-INS-224 | chr3-82287657-INS-224 |
| chr3-82296139-INS-134 | chr3-82296137-INS-122 |
| chr3-82303828-INS-440 | chr3-82303828-INS-440 |
| chr3-82318436-INS-53 | chr3-82318436-INS-53 |
| chr3-82323243-INS-52 | chr3-82323240-INS-52 |
| chr3-82327177-INS-139 | chr3-82327177-INS-139 |
| chr3-82333613-INS-69 | chr3-82333613-INS-69 |
| chr3-82336625-INS-371 | chr3-82336625-INS-377 |
| chr3-82338832-INS-62 | chr3-82338832-INS-62 |
| chr3-82356406-INS-2121 | chr3-82356407-INS-2109 |
| chr3-82384767-INS-89 | chr3-82384765-INS-62 |
| chr3-82411645-INS-56 | chr3-82411645-INS-90 |
| chr3-82419073-INS-412 | chr3-82419073-INS-410 |
| chr3-82430256-INS-6279 | chr3-82430257-INS-6279 |
| chr3-82459828-INS-451 | chr3-82459823-INS-458 |

| | |
|---|---|
| chr3-82461236-INS-110 | chr3-82461237-INS-118 |
| chr3-82476068-INS-58 | chr3-82476068-INS-58 |
| chr3-82494679-INS-1248 | chr3-82494680-INS-1247 |
| chr3-82495185-INS-68 | chr3-82495173-INS-64 |
| chr3-82505701-INS-950 | chr3-82505701-INS-951 |
| chr3-82511463-INS-510 | chr3-82511463-INS-510 |
| chr3-82514475-INS-210 | chr3-82514454-INS-225 |
| chr3-82533306-INS-60 | chr3-82533306-INS-60 |
| chr3-82533689-INS-210 | chr3-82533689-INS-210 |
| chr3-82544130-INS-278 | chr3-82544130-INS-278 |
| chr3-82562372-INS-53 | chr3-82562372-INS-53 |
| chr3-82567351-INS-171 | chr3-82567342-INS-176 |
| chr3-82575517-INS-640 | chr3-82575517-INS-640 |
| chr3-82611208-INS-1231 | chr3-82611218-INS-1251 |
| chr3-82621033-INS-866 | chr3-82621033-INS-866 |
| chr3-82624022-INS-3024 | chr3-82624023-INS-3024 |
| chr3-82631780-INS-2348 | chr3-82631781-INS-2328 |
| chr3-82675339-INS-138 | chr3-82675339-INS-139 |
| chr3-82675893-INS-863 | chr3-82675893-INS-863 |
| chr3-82694959-INS-9571 | chr3-82694960-INS-9572 |
| chr3-82698353-INS-195 | chr3-82698351-INS-204 |
| chr3-82700290-INS-57 | chr3-82700326-INS-66 |
| chr3-82722359-INS-171 | chr3-82722359-INS-171 |
| chr3-82752997-INS-429 | chr3-82752997-INS-429 |
| chr3-82798868-INS-103 | chr3-82798868-INS-103 |
| chr3-82802722-INS-51 | chr3-82802722-INS-51 |
| chr3-82812220-INS-217 | chr3-82812220-INS-216 |
| chr3-82814589-INS-156 | chr3-82814589-INS-156 |
| chr3-82840217-INS-200 | chr3-82840217-INS-200 |
| chr3-82844358-INS-54 | chr3-82844358-INS-52 |
| chr3-82856668-INS-548 | chr3-82856668-INS-548 |
| chr3-82859305-INS-61 | chr3-82859305-INS-61 |
| chr3-82871397-INS-644 | chr3-82871397-INS-644 |
| chr3-82898449-INS-185 | chr3-82898449-INS-183 |
| chr3-82903183-INS-125 | chr3-82903183-INS-94 |
| chr3-82927340-INS-6442 | chr3-82927341-INS-6442 |
| chr3-82984310-INS-56 | chr3-82984310-INS-52 |
| chr3-83002418-INS-8162 | chr3-83002419-INS-8162 |
| chr3-83013018-INS-97 | chr3-83013018-INS-97 |
| chr3-83023278-INS-97 | chr3-83023278-INS-103 |
| chr3-83057121-INS-52 | chr3-83057121-INS-66 |
| chr3-83062143-INS-6119 | chr3-83062144-INS-6119 |
| chr3-83106405-INS-6209 | chr3-83106406-INS-6209 |
| chr3-83115820-INS-1077 | chr3-83115821-INS-1076 |
| chr3-83121643-INS-153 | chr3-83121643-INS-153 |
| chr3-83125170-INS-826 | chr3-83125170-INS-826 |
| chr3-83144502-INS-68 | chr3-83144502-INS-66 |
| chr3-83187982-INS-6239 | chr3-83187983-INS-6239 |
| chr3-83194932-INS-68 | chr3-83194933-INS-52 |
| chr3-83210139-INS-217 | chr3-83210139-INS-217 |
| chr3-83211431-INS-122 | chr3-83211431-INS-122 |
| chr3-83215321-INS-69 | chr3-83215321-INS-69 |
| chr3-83234231-INS-137 | chr3-83234249-INS-145 |
| chr3-83251319-INS-52 | chr3-83251273-INS-56 |
| chr3-83260555-INS-66 | chr3-83260555-INS-66 |
| chr3-83273759-INS-203 | chr3-83273759-INS-202 |
| chr3-83338513-INS-131 | chr3-83338504-INS-155 |
| chr3-83355887-INS-6596 | chr3-83355888-INS-6597 |
| chr3-83366485-INS-64 | chr3-83366496-INS-52 |
| chr3-83366955-INS-3022 | chr3-83366956-INS-3022 |
| chr3-83378588-INS-529 | chr3-83378592-INS-544 |
| chr3-83405531-INS-7093 | chr3-83405532-INS-7093 |
| chr3-83409407-INS-67 | chr3-83409407-INS-68 |
| chr3-83415125-INS-6475 | chr3-83415126-INS-6475 |
| chr3-83416461-INS-198 | chr3-83416460-INS-198 |
| chr3-83429296-INS-102 | chr3-83429344-INS-92 |
| chr3-83429917-INS-91 | chr3-83429917-INS-90 |
| chr3-83477209-INS-98 | chr3-83477207-INS-143 |
| chr3-83480788-INS-380 | chr3-83480790-INS-379 |
| chr3-83501683-INS-5347 | chr3-83501684-INS-5348 |
| chr3-83514877-INS-6437 | chr3-83514878-INS-6438 |
| chr3-83546092-INS-502 | chr3-83546092-INS-502 |
| chr3-83551117-INS-184 | chr3-83551117-INS-184 |
| chr3-83554242-INS-6390 | chr3-83554243-INS-6390 |
| chr3-83566228-INS-6991 | chr3-83566229-INS-6991 |
| chr3-83566613-INS-452 | chr3-83566613-INS-452 |
| chr3-83594303-INS-287 | chr3-83594303-INS-286 |

| | |
|---|---|
| chr3-83620666-INS-215 | chr3-83620666-INS-215 |
| chr3-83626599-INS-81 | chr3-83626601-INS-67 |
| chr3-83632987-INS-1162 | chr3-83632988-INS-1162 |
| chr3-83710879-INS-2910 | chr3-83710880-INS-2910 |
| chr3-83716555-INS-116 | chr3-83716555-INS-116 |
| chr3-83722923-INS-77 | chr3-83722923-INS-77 |
| chr3-83734225-INS-91 | chr3-83734225-INS-130 |
| chr3-83742283-INS-114 | chr3-83742283-INS-114 |
| chr3-83746176-INS-76 | chr3-83746192-INS-80 |
| chr3-83758243-INS-62 | chr3-83758243-INS-62 |
| chr3-83762896-INS-201 | chr3-83762896-INS-201 |
| chr3-83770839-INS-59 | chr3-83770839-INS-73 |
| chr3-83778815-INS-191 | chr3-83778815-INS-177 |
| chr3-83781975-INS-69 | chr3-83781975-INS-62 |
| chr3-83785636-INS-113 | chr3-83785636-INS-113 |
| chr3-83792598-INS-5352 | chr3-83792599-INS-5352 |
| chr3-83807161-INS-495 | chr3-83807161-INS-495 |
| chr3-83828617-INS-60 | chr3-83828617-INS-60 |
| chr3-83848483-INS-56 | chr3-83848515-INS-54 |
| chr3-83853099-INS-183 | chr3-83853099-INS-179 |
| chr3-83865500-INS-56 | chr3-83865503-INS-56 |
| chr3-83867814-INS-368 | chr3-83867814-INS-368 |
| chr3-83870235-INS-102 | chr3-83870235-INS-102 |
| chr3-83874994-INS-6447 | chr3-83874995-INS-6447 |
| chr3-83883284-INS-64 | chr3-83883284-INS-68 |
| chr3-83892667-INS-202 | chr3-83892667-INS-202 |
| chr3-83903747-INS-87 | chr3-83903705-INS-84 |
| chr3-83920309-INS-276 | chr3-83920309-INS-276 |
| chr3-83961843-INS-54 | chr3-83961843-INS-54 |
| chr3-83974104-INS-56 | chr3-83974112-INS-68 |
| chr3-83986304-INS-1439 | chr3-83986305-INS-1439 |
| chr3-84013855-INS-7138 | chr3-84013856-INS-7138 |
| chr3-84032471-INS-3144 | chr3-84032472-INS-3144 |
| chr3-84042262-INS-197 | chr3-84042262-INS-198 |
| chr3-84090080-INS-580 | chr3-84090080-INS-590 |
| chr3-84092919-INS-73 | chr3-84092919-INS-73 |
| chr3-84114302-INS-224 | chr3-84114302-INS-224 |
| chr3-84118027-INS-211 | chr3-84118027-INS-211 |
| chr3-84148247-INS-72 | chr3-84148247-INS-86 |
| chr3-84170555-INS-1893 | chr3-84170556-INS-1893 |
| chr3-84182793-INS-58 | chr3-84182793-INS-61 |
| chr3-84198671-INS-58 | chr3-84198671-INS-80 |
| chr3-84201231-INS-96 | chr3-84201232-INS-96 |
| chr3-84227183-INS-1331 | chr3-84227184-INS-1331 |
| chr3-84241986-INS-216 | chr3-84241986-INS-216 |
| chr3-84281282-INS-60 | chr3-84281282-INS-60 |
| chr3-84288681-INS-102 | chr3-84288681-INS-102 |
| chr3-84295541-INS-148 | chr3-84295541-INS-164 |
| chr3-84331374-INS-5153 | chr3-84331375-INS-5153 |
| chr3-84331823-INS-54 | chr3-84331840-INS-66 |
| chr3-84452810-INS-548 | chr3-84452810-INS-547 |
| chr3-84464417-INS-88 | chr3-84464417-INS-88 |
| chr3-84474245-INS-59 | chr3-84474245-INS-59 |
| chr3-84479287-INS-54 | chr3-84479293-INS-72 |
| chr3-84484790-INS-209 | chr3-84484790-INS-209 |
| chr3-84487792-INS-289 | chr3-84487792-INS-290 |
| chr3-84550081-INS-85 | chr3-84550081-INS-80 |
| chr3-84554923-INS-492 | chr3-84554923-INS-492 |
| chr3-84577663-INS-375 | chr3-84577663-INS-375 |
| chr3-84586900-INS-90 | chr3-84586900-INS-60 |
| chr3-84592660-INS-113 | chr3-84592658-INS-119 |
| chr3-84593628-INS-1283 | chr3-84593629-INS-1283 |
| chr3-84600220-INS-226 | chr3-84600220-INS-226 |
| chr3-84616808-INS-1873 | chr3-84616809-INS-1873 |
| chr3-84642762-INS-1905 | chr3-84642763-INS-1899 |
| chr3-84642762-INS-414 | chr3-84642762-INS-414 |
| chr3-84695511-INS-86 | chr3-84695535-INS-57 |
| chr3-84708054-INS-441 | chr3-84708054-INS-441 |
| chr3-84746670-INS-90 | chr3-84746669-INS-57 |
| chr3-84770747-INS-183 | chr3-84770747-INS-184 |
| chr3-84791558-INS-83 | chr3-84791558-INS-83 |
| chr3-84820911-INS-222 | chr3-84820911-INS-224 |
| chr3-84864477-INS-74 | chr3-84864509-INS-62 |
| chr3-84870567-INS-198 | chr3-84870567-INS-189 |
| chr3-84879984-INS-6561 | chr3-84879985-INS-6561 |
| chr3-84989771-INS-195 | chr3-84989771-INS-225 |
| chr3-84998954-INS-63 | chr3-84998967-INS-75 |

| | |
|---|---|
| chr3-85033117-INS-56 | chr3-85033117-INS-50 |
| chr3-85038054-INS-97 | chr3-85038054-INS-123 |
| chr3-85048746-INS-64 | chr3-85048746-INS-80 |
| chr3-85079515-INS-370 | chr3-85079517-INS-375 |
| chr3-85081453-INS-64 | chr3-85081454-INS-58 |
| chr3-85099358-INS-90 | chr3-85099358-INS-60 |
| chr3-85140780-INS-60 | chr3-85140780-INS-96 |
| chr3-85148757-INS-146 | chr3-85148756-INS-146 |
| chr3-85149774-INS-68 | chr3-85149774-INS-96 |
| chr3-85155942-INS-53 | chr3-85155942-INS-51 |
| chr3-85176797-INS-154 | chr3-85176797-INS-154 |
| chr3-85217537-INS-60 | chr3-85217533-INS-63 |
| chr3-85227335-INS-7113 | chr3-85227336-INS-7113 |
| chr3-85241955-INS-186 | chr3-85241949-INS-186 |
| chr3-85255911-INS-364 | chr3-85255911-INS-364 |
| chr3-85278694-INS-76 | chr3-85278694-INS-50 |
| chr3-85319772-INS-53 | chr3-85319772-INS-74 |
| chr3-85322343-INS-56 | chr3-85322344-INS-56 |
| chr3-85325122-INS-7379 | chr3-85325123-INS-7378 |
| chr3-85368842-INS-60 | chr3-85368794-INS-109 |
| chr3-85371737-INS-55 | chr3-85371737-INS-59 |
| chr3-85374697-INS-52 | chr3-85374697-INS-52 |
| chr3-85385158-INS-52 | chr3-85385158-INS-52 |
| chr3-85399934-INS-272 | chr3-85399934-INS-291 |
| chr3-85407654-INS-208 | chr3-85407654-INS-208 |
| chr3-85408896-INS-62 | chr3-85408894-INS-88 |
| chr3-85413669-INS-118 | chr3-85413669-INS-118 |
| chr3-85415862-INS-7394 | chr3-85415863-INS-7394 |
| chr3-85446194-INS-7030 | chr3-85446195-INS-7032 |
| chr3-85517369-INS-76 | chr3-85517369-INS-76 |
| chr3-85599163-INS-60 | chr3-85599134-INS-90 |
| chr3-85653091-INS-340 | chr3-85653091-INS-340 |
| chr3-85694333-INS-335 | chr3-85694333-INS-335 |
| chr3-85700466-INS-538 | chr3-85700466-INS-538 |
| chr3-85710356-INS-128 | chr3-85710356-INS-128 |
| chr3-85722909-INS-55 | chr3-85722909-INS-55 |
| chr3-85761362-INS-56 | chr3-85761362-INS-52 |
| chr3-85780488-INS-591 | chr3-85780488-INS-591 |
| chr3-85789534-INS-203 | chr3-85789534-INS-203 |
| chr3-85791997-INS-108 | chr3-85791997-INS-144 |
| chr3-85810213-INS-139 | chr3-85810213-INS-139 |
| chr3-85813006-INS-287 | chr3-85812992-INS-259 |
| chr3-85816978-INS-102 | chr3-85816978-INS-90 |
| chr3-85819020-INS-1161 | chr3-85819021-INS-1161 |
| chr3-85845058-INS-57 | chr3-85845062-INS-72 |
| chr3-85865179-INS-155 | chr3-85865179-INS-155 |
| chr3-85913956-INS-66 | chr3-85913956-INS-66 |
| chr3-85927165-INS-3136 | chr3-85927166-INS-3134 |
| chr3-85929293-INS-217 | chr3-85929293-INS-220 |
| chr3-85932740-INS-668 | chr3-85932740-INS-668 |
| chr3-85936960-INS-107 | chr3-85936960-INS-108 |
| chr3-85937659-INS-195 | chr3-85937659-INS-194 |
| chr3-85956676-INS-145 | chr3-85956676-INS-145 |
| chr3-85959810-INS-1836 | chr3-85959811-INS-1836 |
| chr3-85966094-INS-245 | chr3-85966094-INS-245 |
| chr3-85976339-INS-183 | chr3-85976339-INS-183 |
| chr3-85994795-INS-90 | chr3-85994795-INS-90 |
| chr3-85998740-INS-188 | chr3-85998740-INS-188 |
| chr3-86006459-INS-88 | chr3-86006459-INS-88 |
| chr3-86030113-INS-198 | chr3-86030113-INS-198 |
| chr3-86031127-INS-151 | chr3-86031125-INS-151 |
| chr3-86038582-INS-229 | chr3-86038583-INS-232 |
| chr3-86056307-INS-206 | chr3-86056307-INS-206 |
| chr3-86056917-INS-7165 | chr3-86056918-INS-7165 |
| chr3-86066619-INS-208 | chr3-86066619-INS-208 |
| chr3-86075608-INS-231 | chr3-86075608-INS-231 |
| chr3-86091842-INS-6473 | chr3-86091843-INS-6473 |
| chr3-86098358-INS-60 | chr3-86098358-INS-52 |
| chr3-86102953-INS-52 | chr3-86102951-INS-54 |
| chr3-86105637-INS-210 | chr3-86105637-INS-213 |
| chr3-86123241-INS-54 | chr3-86123239-INS-58 |
| chr3-86155606-INS-7346 | chr3-86155607-INS-7346 |
| chr3-86169278-INS-214 | chr3-86169278-INS-214 |
| chr3-86177743-INS-235 | chr3-86177743-INS-234 |
| chr3-86198601-INS-6592 | chr3-86198602-INS-6589 |
| chr3-86210921-INS-125 | chr3-86210921-INS-126 |
| chr3-86214699-INS-66 | chr3-86214699-INS-66 |

| | |
|---|---|
| chr3-86298985-INS-109 | chr3-86298985-INS-109 |
| chr3-86312316-INS-95 | chr3-86312316-INS-95 |
| chr3-86315633-INS-7457 | chr3-86315634-INS-7457 |
| chr3-86335714-INS-510 | chr3-86335714-INS-512 |
| chr3-86357820-INS-261 | chr3-86357820-INS-213 |
| chr3-86361415-INS-1304 | chr3-86361416-INS-1304 |
| chr3-86362218-INS-222 | chr3-86362218-INS-229 |
| chr3-86362447-INS-2246 | chr3-86362448-INS-2246 |
| chr3-86364768-INS-1293 | chr3-86364769-INS-1293 |
| chr3-86416691-INS-82 | chr3-86416699-INS-54 |
| chr3-86484821-INS-60 | chr3-86484821-INS-58 |
| chr3-86487408-INS-174 | chr3-86487408-INS-174 |
| chr3-86507072-INS-279 | chr3-86507072-INS-280 |
| chr3-86517281-INS-369 | chr3-86517281-INS-368 |
| chr3-86534399-INS-6613 | chr3-86534400-INS-6612 |
| chr3-86540720-INS-83 | chr3-86540720-INS-83 |
| chr3-86562242-INS-1686 | chr3-86562243-INS-1690 |
| chr3-86562881-INS-56 | chr3-86562881-INS-56 |
| chr3-86564780-INS-72 | chr3-86564780-INS-72 |
| chr3-86598435-INS-198 | chr3-86598435-INS-199 |
| chr3-86607916-INS-62 | chr3-86607912-INS-67 |
| chr3-86610118-INS-99 | chr3-86610118-INS-75 |
| chr3-86628038-INS-1841 | chr3-86628039-INS-1840 |
| chr3-86682090-INS-75 | chr3-86682090-INS-75 |
| chr3-86708054-INS-7134 | chr3-86708055-INS-7134 |
| chr3-86709646-INS-201 | chr3-86709646-INS-201 |
| chr3-86717765-INS-126 | chr3-86717765-INS-126 |
| chr3-86732646-INS-152 | chr3-86732646-INS-152 |
| chr3-86741098-INS-233 | chr3-86741098-INS-233 |
| chr3-86742091-INS-222 | chr3-86742091-INS-222 |
| chr3-86746049-INS-498 | chr3-86746049-INS-498 |
| chr3-86748856-INS-3188 | chr3-86748857-INS-3188 |
| chr3-86765267-INS-136 | chr3-86765264-INS-133 |
| chr3-86784731-INS-178 | chr3-86784731-INS-178 |
| chr3-86807031-INS-181 | chr3-86807031-INS-181 |
| chr3-86811606-INS-54 | chr3-86811622-INS-52 |
| chr3-86836564-INS-5201 | chr3-86836565-INS-5201 |
| chr3-86883098-INS-1892 | chr3-86883099-INS-1892 |
| chr3-86892577-INS-67 | chr3-86892575-INS-75 |
| chr3-86955215-INS-413 | chr3-86955215-INS-413 |
| chr3-86959141-INS-146 | chr3-86959121-INS-194 |
| chr3-86978940-INS-229 | chr3-86978940-INS-230 |
| chr3-87017406-INS-94 | chr3-87017401-INS-133 |
| chr3-87023003-INS-69 | chr3-87023003-INS-69 |
| chr3-87027082-INS-213 | chr3-87027082-INS-213 |
| chr3-87032400-INS-99 | chr3-87032400-INS-57 |
| chr3-87042394-INS-58 | chr3-87042390-INS-50 |
| chr3-87046534-INS-56 | chr3-87046534-INS-61 |
| chr3-87112962-INS-3377 | chr3-87112963-INS-3377 |
| chr3-87116472-INS-2877 | chr3-87116473-INS-2877 |
| chr3-87123591-INS-197 | chr3-87123591-INS-197 |
| chr3-87185380-INS-219 | chr3-87185380-INS-219 |
| chr3-87193109-INS-52 | chr3-87193109-INS-52 |
| chr3-87237796-INS-64 | chr3-87237792-INS-60 |
| chr3-87241803-INS-62 | chr3-87241803-INS-72 |
| chr3-87249443-INS-485 | chr3-87249443-INS-485 |
| chr3-87312806-INS-56 | chr3-87312806-INS-56 |
| chr3-87337127-INS-1025 | chr3-87337128-INS-1025 |
| chr3-87340383-INS-1894 | chr3-87340384-INS-1894 |
| chr3-87347309-INS-171 | chr3-87347309-INS-171 |
| chr3-87375156-INS-1130 | chr3-87375157-INS-1130 |
| chr3-87375505-INS-439 | chr3-87375505-INS-438 |
| chr3-87381894-INS-726 | chr3-87381894-INS-726 |
| chr3-87401210-INS-62 | chr3-87401210-INS-55 |
| chr3-87411990-INS-4107 | chr3-87411991-INS-4107 |
| chr3-87417715-INS-6454 | chr3-87417716-INS-6454 |
| chr3-87420716-INS-62 | chr3-87420716-INS-56 |
| chr3-87465090-INS-478 | chr3-87465090-INS-482 |
| chr3-87496030-INS-182 | chr3-87496030-INS-182 |
| chr3-87501596-INS-255 | chr3-87501596-INS-255 |
| chr3-87507600-INS-6476 | chr3-87507601-INS-6476 |
| chr3-87517059-INS-57 | chr3-87517059-INS-57 |
| chr3-87530462-INS-164 | chr3-87530462-INS-164 |
| chr3-87530835-INS-162 | chr3-87530835-INS-162 |
| chr3-87531695-INS-79 | chr3-87531685-INS-87 |
| chr3-87556252-INS-259 | chr3-87556252-INS-259 |
| chr3-87607017-INS-70 | chr3-87607011-INS-68 |

| | |
|---|---|
| chr3-87658104-INS-94 | chr3-87658104-INS-94 |
| chr3-87667917-INS-264 | chr3-87667917-INS-220 |
| chr3-87719427-INS-84 | chr3-87719426-INS-93 |
| chr3-87731627-INS-71 | chr3-87731627-INS-71 |
| chr3-87805858-INS-54 | chr3-87805830-INS-93 |
| chr3-87865568-INS-5568 | chr3-87865569-INS-5568 |
| chr3-87905872-INS-85 | chr3-87905872-INS-85 |
| chr3-87910481-INS-151 | chr3-87910481-INS-151 |
| chr3-87914483-INS-1891 | chr3-87914484-INS-1891 |
| chr3-87951953-INS-254 | chr3-87951953-INS-252 |
| chr3-87957194-INS-56 | chr3-87957194-INS-58 |
| chr3-87974470-INS-66 | chr3-87974470-INS-66 |
| chr3-87976701-INS-219 | chr3-87976701-INS-219 |
| chr3-87983836-INS-134 | chr3-87983808-INS-182 |
| chr3-87987451-INS-204 | chr3-87987451-INS-204 |
| chr3-87988010-INS-204 | chr3-87988010-INS-200 |
| chr3-87992870-INS-205 | chr3-87992870-INS-209 |
| chr3-88000197-INS-210 | chr3-88000197-INS-210 |
| chr3-88003740-INS-63 | chr3-88003740-INS-63 |
| chr3-88020123-INS-230 | chr3-88020123-INS-230 |
| chr3-88032261-INS-76 | chr3-88032261-INS-76 |
| chr3-88080269-INS-5311 | chr3-88080270-INS-5311 |
| chr3-88156730-INS-400 | chr3-88156730-INS-399 |
| chr3-88179184-INS-171 | chr3-88179202-INS-182 |
| chr3-88184268-INS-6964 | chr3-88184269-INS-6964 |
| chr3-88186047-INS-203 | chr3-88186047-INS-202 |
| chr3-88216878-INS-52 | chr3-88216878-INS-52 |
| chr3-88226988-INS-210 | chr3-88226988-INS-210 |
| chr3-88300557-INS-175 | chr3-88300557-INS-176 |
| chr3-88310783-INS-60 | chr3-88310783-INS-60 |
| chr3-88338597-INS-4525 | chr3-88338598-INS-4490 |
| chr3-88361469-INS-54 | chr3-88361469-INS-54 |
| chr3-88379236-INS-746 | chr3-88379236-INS-746 |
| chr3-88384657-INS-5409 | chr3-88384658-INS-5409 |
| chr3-88404434-INS-244 | chr3-88404434-INS-243 |
| chr3-88416597-INS-64 | chr3-88416597-INS-82 |
| chr3-88424962-INS-203 | chr3-88424962-INS-203 |
| chr3-88429779-INS-84 | chr3-88429778-INS-58 |
| chr3-88435953-INS-84 | chr3-88435971-INS-96 |
| chr3-88467379-INS-111 | chr3-88467379-INS-111 |
| chr3-88491266-INS-301 | chr3-88491266-INS-301 |
| chr3-88506002-INS-196 | chr3-88506002-INS-206 |
| chr3-88518466-INS-118 | chr3-88518466-INS-118 |
| chr3-88557930-INS-151 | chr3-88557930-INS-161 |
| chr3-88565098-INS-231 | chr3-88565098-INS-243 |
| chr3-88566701-INS-290 | chr3-88566701-INS-279 |
| chr3-88567588-INS-69 | chr3-88567588-INS-65 |
| chr3-88568267-INS-223 | chr3-88568267-INS-219 |
| chr3-88571453-INS-4953 | chr3-88571454-INS-4953 |
| chr3-88582700-INS-198 | chr3-88582700-INS-198 |
| chr3-88584564-INS-222 | chr3-88584564-INS-222 |
| chr3-88600287-INS-113 | chr3-88600285-INS-113 |
| chr3-88602004-INS-208 | chr3-88602004-INS-208 |
| chr3-88632039-INS-123 | chr3-88632038-INS-113 |
| chr3-88635359-INS-194 | chr3-88635359-INS-196 |
| chr3-88641301-INS-1548 | chr3-88641302-INS-1548 |
| chr3-88644646-INS-54 | chr3-88644640-INS-53 |
| chr3-88647000-INS-787 | chr3-88647000-INS-796 |
| chr3-88649495-INS-252 | chr3-88649495-INS-255 |
| chr3-88661596-INS-229 | chr3-88661596-INS-228 |
| chr3-88695326-INS-54 | chr3-88695326-INS-56 |
| chr3-88701476-INS-220 | chr3-88701476-INS-212 |
| chr3-88735944-INS-199 | chr3-88735938-INS-199 |
| chr3-88748318-INS-58 | chr3-88748322-INS-54 |
| chr3-88759538-INS-78 | chr3-88759538-INS-128 |
| chr3-88789437-INS-209 | chr3-88789437-INS-210 |
| chr3-88805076-INS-190 | chr3-88805076-INS-191 |
| chr3-88826767-INS-4363 | chr3-88826768-INS-4363 |
| chr3-88861511-INS-231 | chr3-88861511-INS-230 |
| chr3-88881224-INS-185 | chr3-88881224-INS-185 |
| chr3-88941094-INS-97 | chr3-88941094-INS-93 |
| chr3-88950219-INS-54 | chr3-88950206-INS-70 |
| chr3-88956905-INS-80 | chr3-88956905-INS-80 |
| chr3-88960590-INS-168 | chr3-88960590-INS-176 |
| chr3-89024106-INS-326 | chr3-89024106-INS-326 |
| chr3-89033828-INS-385 | chr3-89033828-INS-388 |
| chr3-89035508-INS-594 | chr3-89035508-INS-594 |

| | |
|---|---|
| chr3-89043041-INS-63 | chr3-89043027-INS-53 |
| chr3-89045550-INS-726 | chr3-89045550-INS-727 |
| chr3-89067927-INS-208 | chr3-89067927-INS-177 |
| chr3-89088398-INS-73 | chr3-89088398-INS-73 |
| chr3-89088866-INS-214 | chr3-89088866-INS-182 |
| chr3-89100689-INS-89 | chr3-89100689-INS-89 |
| chr3-89102888-INS-360 | chr3-89102888-INS-360 |
| chr3-89104248-INS-215 | chr3-89104248-INS-216 |
| chr3-89129920-INS-54 | chr3-89129920-INS-50 |
| chr3-89147306-INS-211 | chr3-89147306-INS-210 |
| chr3-89155431-INS-215 | chr3-89155431-INS-213 |
| chr3-89203731-INS-212 | chr3-89203731-INS-211 |
| chr3-89211547-INS-250 | chr3-89211547-INS-255 |
| chr3-89215718-INS-70 | chr3-89215718-INS-53 |
| chr3-89271903-INS-206 | chr3-89271903-INS-206 |
| chr3-89379465-INS-6440 | chr3-89379466-INS-6440 |
| chr3-89382027-INS-84 | chr3-89382023-INS-107 |
| chr3-89438140-INS-232 | chr3-89438140-INS-231 |
| chr3-89439396-INS-737 | chr3-89439396-INS-737 |
| chr3-89440933-INS-83 | chr3-89440952-INS-94 |
| chr3-89453042-INS-64 | chr3-89453042-INS-57 |
| chr3-89462270-INS-696 | chr3-89462266-INS-690 |
| chr3-89492660-INS-58 | chr3-89492660-INS-66 |
| chr3-89511672-INS-123 | chr3-89511672-INS-122 |
| chr3-89560864-INS-66 | chr3-89560864-INS-66 |
| chr3-89655278-INS-54 | chr3-89655278-INS-53 |
| chr3-89698068-INS-84 | chr3-89698068-INS-84 |
| chr3-89710558-INS-287 | chr3-89710558-INS-287 |
| chr3-89717089-INS-5482 | chr3-89717090-INS-5482 |
| chr3-89725085-INS-415 | chr3-89725085-INS-415 |
| chr3-89744898-INS-83 | chr3-89744898-INS-85 |
| chr3-89746202-INS-229 | chr3-89746202-INS-228 |
| chr3-89754079-INS-873 | chr3-89754079-INS-912 |
| chr3-89768847-INS-57 | chr3-89768847-INS-57 |
| chr3-89773373-INS-189 | chr3-89773373-INS-201 |
| chr3-89792915-INS-52 | chr3-89792915-INS-58 |
| chr3-89798532-INS-59 | chr3-89798482-INS-77 |
| chr3-89809387-INS-212 | chr3-89809388-INS-214 |
| chr3-89821549-INS-133 | chr3-89821545-INS-105 |
| chr3-89840913-INS-199 | chr3-89840913-INS-200 |
| chr3-89862337-INS-146 | chr3-89862337-INS-152 |
| chr3-89875298-INS-55 | chr3-89875298-INS-55 |
| chr3-89891875-INS-466 | chr3-89891875-INS-472 |
| chr3-89939484-INS-209 | chr3-89939484-INS-209 |
| chr3-89954082-INS-85 | chr3-89954082-INS-89 |
| chr3-89988203-INS-97 | chr3-89988203-INS-90 |
| chr3-89989845-INS-184 | chr3-89989845-INS-177 |
| chr3-90003080-INS-219 | chr3-90003080-INS-225 |
| chr3-90004129-INS-200 | chr3-90004129-INS-200 |
| chr3-90013371-INS-57 | chr3-90013368-INS-62 |
| chr3-90036714-INS-52 | chr3-90036714-INS-64 |
| chr3-90039969-INS-7115 | chr3-90039970-INS-7115 |
| chr3-90045901-INS-78 | chr3-90045901-INS-80 |
| chr3-90051988-INS-725 | chr3-90051988-INS-725 |
| chr3-90055959-INS-164 | chr3-90055959-INS-166 |
| chr3-90066283-INS-382 | chr3-90066284-INS-381 |
| chr3-90081240-INS-210 | chr3-90081239-INS-210 |
| chr3-90082981-INS-386 | chr3-90082981-INS-387 |
| chr3-90090695-INS-80 | chr3-90090691-INS-80 |
| chr3-90105977-INS-232 | chr3-90105977-INS-231 |
| chr3-90113799-INS-234 | chr3-90113800-INS-226 |
| chr3-90137562-INS-222 | chr3-90137563-INS-222 |
| chr3-90140641-INS-211 | chr3-90140641-INS-211 |
| chr3-90142055-INS-82 | chr3-90142055-INS-84 |
| chr3-90189366-INS-197 | chr3-90189366-INS-189 |
| chr3-90198613-INS-665 | chr3-90198613-INS-646 |
| chr3-90209297-INS-216 | chr3-90209297-INS-210 |
| chr3-90220015-INS-194 | chr3-90220015-INS-193 |
| chr3-90232781-INS-72 | chr3-90232781-INS-84 |
| chr3-90241455-INS-367 | chr3-90241455-INS-366 |
| chr3-90248312-INS-184 | chr3-90248312-INS-184 |
| chr3-90274339-INS-172 | chr3-90274339-INS-188 |
| chr3-90277751-INS-201 | chr3-90277751-INS-199 |
| chr3-90313153-INS-56 | chr3-90313153-INS-56 |
| chr3-90319503-INS-204 | chr3-90319503-INS-204 |
| chr3-90326776-INS-199 | chr3-90326776-INS-199 |
| chr3-90329228-INS-157 | chr3-90329228-INS-145 |

| | |
|---|---|
| chr3-90346719-INS-57 | chr3-90346705-INS-59 |
| chr3-90349352-INS-178 | chr3-90349352-INS-181 |
| chr3-90360648-INS-127 | chr3-90360648-INS-131 |
| chr3-90362738-INS-162 | chr3-90362738-INS-159 |
| chr3-90363999-INS-141 | chr3-90364000-INS-94 |
| chr3-90382860-INS-202 | chr3-90382860-INS-204 |
| chr3-90392298-INS-70 | chr3-90392298-INS-70 |
| chr3-90398626-INS-119 | chr3-90398583-INS-117 |
| chr3-90400600-INS-1413 | chr3-90400601-INS-1413 |
| chr3-90400895-INS-237 | chr3-90400892-INS-242 |
| chr3-90426919-INS-75 | chr3-90426879-INS-92 |
| chr3-90428947-INS-735 | chr3-90428950-INS-710 |
| chr3-90452220-INS-136 | chr3-90452220-INS-136 |
| chr3-90465580-INS-878 | chr3-90465580-INS-878 |
| chr3-90467014-INS-7238 | chr3-90467015-INS-7238 |
| chr3-90471627-INS-114 | chr3-90471627-INS-120 |
| chr3-90500047-INS-211 | chr3-90500047-INS-212 |
| chr3-90509055-INS-175 | chr3-90509055-INS-153 |
| chr3-90544580-INS-117 | chr3-90544580-INS-93 |
| chr3-90545778-INS-418 | chr3-90545778-INS-419 |
| chr3-90552860-INS-1358 | chr3-90552861-INS-1358 |
| chr3-90560692-INS-55 | chr3-90560692-INS-58 |
| chr3-90566657-INS-386 | chr3-90566657-INS-386 |
| chr3-90570384-INS-52 | chr3-90570384-INS-52 |
| chr3-90571167-INS-542 | chr3-90571167-INS-553 |
| chr3-90574328-INS-199 | chr3-90574328-INS-199 |
| chr3-90583365-INS-265 | chr3-90583365-INS-265 |
| chr3-90589393-INS-6394 | chr3-90589394-INS-6396 |
| chr3-90593520-INS-782 | chr3-90593520-INS-783 |
| chr3-90620943-INS-6621 | chr3-90620944-INS-6621 |
| chr3-90622777-INS-60 | chr3-90622777-INS-60 |
| chr3-90625976-INS-837 | chr3-90625976-INS-836 |
| chr3-90646606-INS-203 | chr3-90646606-INS-203 |
| chr3-90662010-INS-6627 | chr3-90662011-INS-6631 |
| chr3-90773001-INS-6164 | chr3-90773002-INS-6164 |
| chr3-90835018-INS-376 | chr3-90835019-INS-376 |
| chr3-90850200-INS-6440 | chr3-90850201-INS-6440 |
| chr3-90936812-INS-3581 | chr3-90936813-INS-3581 |
| chr3-90965235-INS-2727 | chr3-90965236-INS-2732 |
| chr3-91048093-INS-6385 | chr3-91048094-INS-6385 |
| chr3-91076368-INS-438 | chr3-91076369-INS-438 |
| chr3-91277191-INS-219 | chr3-91277192-INS-213 |
| chr3-91284379-INS-62 | chr3-91284379-INS-52 |
| chr3-91293620-INS-586 | chr3-91293627-INS-570 |
| chr3-91388492-INS-2013 | chr3-91388493-INS-2024 |
| chr3-91439798-INS-136 | chr3-91439819-INS-132 |
| chr3-91450964-INS-70 | chr3-91450965-INS-74 |
| chr3-91517435-INS-107 | chr3-91517435-INS-107 |
| chr3-91537556-INS-5522 | chr3-91537557-INS-5522 |
| chr3-91609343-INS-428 | chr3-91609343-INS-428 |
| chr3-91670409-INS-823 | chr3-91670409-INS-823 |
| chr3-91781628-INS-97 | chr3-91781628-INS-97 |
| chr3-91808759-INS-487 | chr3-91808759-INS-487 |
| chr3-91820548-INS-5282 | chr3-91820549-INS-5282 |
| chr3-91831020-INS-132 | chr3-91831020-INS-136 |
| chr3-91838983-INS-369 | chr3-91838983-INS-369 |
| chr3-91852352-INS-6530 | chr3-91852353-INS-6530 |
| chr3-91897880-INS-228 | chr3-91897880-INS-230 |
| chr3-91901681-INS-92 | chr3-91901650-INS-84 |
| chr3-91903895-INS-5466 | chr3-91903896-INS-5466 |
| chr3-91906700-INS-80 | chr3-91906687-INS-88 |
| chr3-91912003-INS-131 | chr3-91912002-INS-137 |
| chr3-91923216-INS-97 | chr3-91923215-INS-97 |
| chr3-91927907-INS-70 | chr3-91927907-INS-70 |
| chr3-91929027-INS-194 | chr3-91929027-INS-194 |
| chr3-91931729-INS-669 | chr3-91931729-INS-669 |
| chr3-91932359-INS-1184 | chr3-91932350-INS-1162 |
| chr3-91957707-INS-1569 | chr3-91957709-INS-1563 |
| chr3-91967027-INS-202 | chr3-91967027-INS-202 |
| chr3-91969651-INS-1875 | chr3-91969652-INS-1877 |
| chr3-92017723-INS-52 | chr3-92017723-INS-54 |
| chr3-92038828-INS-444 | chr3-92038828-INS-444 |
| chr3-92055141-INS-52 | chr3-92055141-INS-58 |
| chr3-92120511-INS-126 | chr3-92120511-INS-110 |
| chr3-92190009-INS-119 | chr3-92190009-INS-137 |
| chr3-92190894-INS-82 | chr3-92190892-INS-85 |
| chr3-92200629-INS-763 | chr3-92200629-INS-769 |

| | |
|---|---|
| chr3-92221788-INS-74 | chr3-92221788-INS-74 |
| chr3-92231169-INS-72 | chr3-92231169-INS-72 |
| chr3-92277143-INS-282 | chr3-92277143-INS-285 |
| chr3-92285653-INS-6418 | chr3-92285654-INS-6418 |
| chr3-92294858-INS-3182 | chr3-92294859-INS-3194 |
| chr3-92315339-INS-7728 | chr3-92315340-INS-7728 |
| chr3-92319698-INS-467 | chr3-92319698-INS-467 |
| chr3-92373730-INS-780 | chr3-92373730-INS-782 |
| chr3-92377746-INS-308 | chr3-92377746-INS-308 |
| chr3-92389138-INS-69 | chr3-92389141-INS-62 |
| chr3-92396776-INS-71 | chr3-92396775-INS-68 |
| chr3-92405300-INS-1270 | chr3-92405301-INS-1271 |
| chr3-92422509-INS-1899 | chr3-92422510-INS-1899 |
| chr3-92434250-INS-408 | chr3-92434250-INS-409 |
| chr3-92440574-INS-76 | chr3-92440573-INS-78 |
| chr3-92444204-INS-86 | chr3-92444204-INS-64 |
| chr3-92532116-INS-171 | chr3-92532116-INS-170 |
| chr3-92571292-INS-96 | chr3-92571292-INS-96 |
| chr3-92593092-INS-60 | chr3-92593092-INS-60 |
| chr3-92630960-INS-65 | chr3-92630960-INS-65 |
| chr3-92654699-INS-328 | chr3-92654699-INS-324 |
| chr3-92679465-INS-7307 | chr3-92679466-INS-7307 |
| chr3-92718418-INS-52 | chr3-92718418-INS-52 |
| chr3-92720401-INS-504 | chr3-92720401-INS-505 |
| chr3-92739767-INS-188 | chr3-92739768-INS-187 |
| chr3-92772394-INS-87 | chr3-92772394-INS-96 |
| chr3-92796463-INS-80 | chr3-92796435-INS-88 |
| chr3-92829359-INS-6241 | chr3-92829360-INS-6244 |
| chr3-92830346-INS-52 | chr3-92830346-INS-56 |
| chr3-92846204-INS-155 | chr3-92846205-INS-105 |
| chr3-92857540-INS-60 | chr3-92857540-INS-60 |
| chr3-92865525-INS-16010 | chr3-92865497-INS-16012 |
| chr3-92873080-INS-142 | chr3-92873080-INS-136 |
| chr3-92882535-INS-1576 | chr3-92882536-INS-1578 |
| chr3-92895368-INS-65 | chr3-92895368-INS-65 |
| chr3-92907017-INS-52 | chr3-92907017-INS-60 |
| chr3-92910520-INS-185 | chr3-92910520-INS-185 |
| chr3-92918629-INS-51 | chr3-92918629-INS-51 |
| chr3-92930251-INS-62 | chr3-92930251-INS-62 |
| chr3-92970203-INS-825 | chr3-92970203-INS-825 |
| chr3-92974036-INS-643 | chr3-92974036-INS-643 |
| chr3-92976671-INS-76 | chr3-92976677-INS-60 |
| chr3-92980510-INS-3113 | chr3-92980511-INS-3113 |
| chr3-93004678-INS-657 | chr3-93004678-INS-657 |
| chr3-93006750-INS-94 | chr3-93006750-INS-93 |
| chr3-93037052-INS-564 | chr3-93037052-INS-564 |
| chr3-93063272-INS-10005 | chr3-93063273-INS-10005 |
| chr3-93069288-INS-84 | chr3-93069288-INS-84 |
| chr3-93105792-INS-52 | chr3-93105792-INS-56 |
| chr3-93110593-INS-78 | chr3-93110593-INS-78 |
| chr3-93167633-INS-2371 | chr3-93167634-INS-2371 |
| chr3-93207655-INS-444 | chr3-93207655-INS-444 |
| chr3-93262116-INS-58 | chr3-93262116-INS-58 |
| chr3-93272649-INS-2283 | chr3-93272650-INS-2276 |
| chr3-93284153-INS-219 | chr3-93284153-INS-183 |
| chr3-93304211-INS-108 | chr3-93304211-INS-108 |
| chr3-93309235-INS-136 | chr3-93309235-INS-140 |
| chr3-93319031-INS-8419 | chr3-93319032-INS-8419 |
| chr3-93324582-INS-1569 | chr3-93324583-INS-1568 |
| chr3-93326182-INS-116 | chr3-93326182-INS-116 |
| chr3-93333485-INS-322 | chr3-93333485-INS-322 |
| chr3-93334581-INS-178 | chr3-93334588-INS-179 |
| chr3-93372988-INS-4887 | chr3-93372989-INS-4887 |
| chr3-93374318-INS-528 | chr3-93374318-INS-528 |
| chr3-93391267-INS-1251 | chr3-93391268-INS-1247 |
| chr3-93398788-INS-8343 | chr3-93398789-INS-8344 |
| chr3-93403681-INS-6280 | chr3-93403682-INS-6285 |
| chr3-93404852-INS-105 | chr3-93404853-INS-105 |
| chr3-93407147-INS-1010 | chr3-93407148-INS-1017 |
| chr3-93425068-INS-153 | chr3-93425068-INS-153 |
| chr3-93429688-INS-116 | chr3-93429688-INS-104 |
| chr3-93445495-INS-6677 | chr3-93445496-INS-6677 |
| chr3-93471469-INS-1892 | chr3-93471470-INS-1892 |
| chr3-93479029-INS-94 | chr3-93479058-INS-61 |
| chr3-93481250-INS-345 | chr3-93481250-INS-345 |
| chr3-93496762-INS-428 | chr3-93496762-INS-428 |
| chr3-93513551-INS-110 | chr3-93513576-INS-60 |

| | |
|---|---|
| chr3-93515956-INS-216 | chr3-93515956-INS-215 |
| chr3-93532292-INS-7377 | chr3-93532293-INS-7378 |
| chr3-93541906-INS-66 | chr3-93541906-INS-72 |
| chr3-93545998-INS-72 | chr3-93545994-INS-90 |
| chr3-93555998-INS-1198 | chr3-93556004-INS-1194 |
| chr3-93572006-INS-328 | chr3-93572006-INS-328 |
| chr3-93589566-INS-212 | chr3-93589566-INS-211 |
| chr3-93592288-INS-207 | chr3-93592288-INS-207 |
| chr3-93612479-INS-4962 | chr3-93612480-INS-4963 |
| chr3-93739690-INS-332 | chr3-93739690-INS-332 |
| chr3-93830090-INS-416 | chr3-93830090-INS-416 |
| chr3-93954045-INS-700 | chr3-93954045-INS-706 |
| chr3-93985171-INS-526 | chr3-93985171-INS-526 |
| chr3-94000841-INS-209 | chr3-94000841-INS-209 |
| chr3-94024310-INS-1430 | chr3-94024311-INS-1431 |
| chr3-94099526-INS-2898 | chr3-94099527-INS-2899 |
| chr3-94110622-INS-62 | chr3-94110622-INS-62 |
| chr3-94141831-INS-98 | chr3-94141831-INS-92 |
| chr3-94142958-INS-7243 | chr3-94142959-INS-7243 |
| chr3-94171631-INS-130 | chr3-94171631-INS-130 |
| chr3-94220823-INS-644 | chr3-94220823-INS-641 |
| chr3-94238703-INS-66 | chr3-94238703-INS-68 |
| chr3-94301578-INS-238 | chr3-94301578-INS-236 |
| chr3-94370011-INS-54 | chr3-94370011-INS-54 |
| chr3-94375277-INS-1191 | chr3-94375278-INS-1160 |
| chr3-94400628-INS-225 | chr3-94400628-INS-225 |
| chr3-94446706-INS-218 | chr3-94446706-INS-219 |
| chr3-94471113-INS-200 | chr3-94471113-INS-199 |
| chr3-94499014-INS-7107 | chr3-94499015-INS-7107 |
| chr3-94515280-INS-188 | chr3-94515280-INS-189 |
| chr3-94533660-INS-132 | chr3-94533658-INS-134 |
| chr3-94573523-INS-3237 | chr3-94573524-INS-3237 |
| chr3-94578656-INS-208 | chr3-94578656-INS-208 |
| chr3-94597618-INS-197 | chr3-94597618-INS-197 |
| chr3-94643911-INS-1888 | chr3-94643912-INS-1888 |
| chr3-94645041-INS-4892 | chr3-94645042-INS-4892 |
| chr3-94658574-INS-521 | chr3-94658574-INS-521 |
| chr3-94659364-INS-58 | chr3-94659364-INS-70 |
| chr3-94663318-INS-109 | chr3-94663319-INS-78 |
| chr3-94712827-INS-58 | chr3-94712827-INS-70 |
| chr3-94717868-INS-55 | chr3-94717874-INS-50 |
| chr3-94722050-INS-65 | chr3-94722050-INS-51 |
| chr3-94724038-INS-250 | chr3-94724038-INS-259 |
| chr3-94733070-INS-108 | chr3-94733070-INS-108 |
| chr3-94773632-INS-61 | chr3-94773635-INS-62 |
| chr3-94781010-INS-71 | chr3-94781010-INS-71 |
| chr3-94797780-INS-7213 | chr3-94797781-INS-7213 |
| chr3-94802735-INS-88 | chr3-94802735-INS-100 |
| chr3-94831416-INS-539 | chr3-94831416-INS-537 |
| chr3-94832458-INS-76 | chr3-94832490-INS-88 |
| chr3-94836708-INS-189 | chr3-94836710-INS-173 |
| chr3-94837572-INS-202 | chr3-94837573-INS-203 |
| chr3-94857448-INS-220 | chr3-94857448-INS-215 |
| chr3-94909555-INS-232 | chr3-94909555-INS-232 |
| chr3-94924341-INS-1895 | chr3-94924342-INS-1895 |
| chr3-94946868-INS-308 | chr3-94946837-INS-309 |
| chr3-94959105-INS-3181 | chr3-94959106-INS-3181 |
| chr3-94965359-INS-100 | chr3-94965359-INS-100 |
| chr3-94975992-INS-231 | chr3-94976030-INS-220 |
| chr3-94989876-INS-201 | chr3-94989869-INS-218 |
| chr3-94991453-INS-223 | chr3-94991453-INS-223 |
| chr3-94995483-INS-225 | chr3-94995483-INS-225 |
| chr3-94998452-INS-74 | chr3-94998452-INS-76 |
| chr3-95000122-INS-209 | chr3-95000122-INS-209 |
| chr3-95016369-INS-62 | chr3-95016338-INS-62 |
| chr3-95024012-INS-171 | chr3-95024012-INS-171 |
| chr3-95050832-INS-60 | chr3-95050794-INS-94 |
| chr3-95066312-INS-255 | chr3-95066312-INS-256 |
| chr3-95104300-INS-170 | chr3-95104300-INS-170 |
| chr3-95106050-INS-92 | chr3-95106058-INS-88 |
| chr3-95113402-INS-516 | chr3-95113402-INS-516 |
| chr3-95132372-INS-173 | chr3-95132372-INS-173 |
| chr3-95144552-INS-221 | chr3-95144552-INS-221 |
| chr3-95146851-INS-51 | chr3-95146851-INS-51 |
| chr3-95152168-INS-179 | chr3-95152168-INS-187 |
| chr3-95166781-INS-199 | chr3-95166781-INS-199 |
| chr3-95179375-INS-63 | chr3-95179373-INS-64 |

| | |
|---|---|
| chr3-95181640-INS-224 | chr3-95181640-INS-226 |
| chr3-95186292-INS-199 | chr3-95186292-INS-201 |
| chr3-95189542-INS-176 | chr3-95189542-INS-176 |
| chr3-95193851-INS-312 | chr3-95193851-INS-312 |
| chr3-95212192-INS-318 | chr3-95212192-INS-318 |
| chr3-95216751-INS-7097 | chr3-95216752-INS-7097 |
| chr3-95220129-INS-222 | chr3-95220129-INS-221 |
| chr3-95242559-INS-216 | chr3-95242559-INS-216 |
| chr3-95251110-INS-82 | chr3-95251112-INS-82 |
| chr3-95268918-INS-209 | chr3-95268918-INS-210 |
| chr3-95278756-INS-62 | chr3-95278756-INS-66 |
| chr3-95292878-INS-60 | chr3-95292882-INS-70 |
| chr3-95293446-INS-594 | chr3-95293441-INS-597 |
| chr3-95308880-INS-52 | chr3-95308880-INS-52 |
| chr3-95312495-INS-475 | chr3-95312495-INS-476 |
| chr3-95320167-INS-211 | chr3-95320167-INS-211 |
| chr3-95322784-INS-206 | chr3-95322784-INS-206 |
| chr3-95350594-INS-112 | chr3-95350567-INS-137 |
| chr3-95352400-INS-101 | chr3-95352399-INS-55 |
| chr3-95374578-INS-58 | chr3-95374578-INS-58 |
| chr3-95376344-INS-74 | chr3-95376340-INS-66 |
| chr3-95405786-INS-7125 | chr3-95405787-INS-7125 |
| chr3-95409277-INS-177 | chr3-95409277-INS-172 |
| chr3-95418775-INS-90 | chr3-95418775-INS-90 |
| chr3-95428001-INS-930 | chr3-95428001-INS-930 |
| chr3-95429478-INS-640 | chr3-95429478-INS-646 |
| chr3-95462940-INS-272 | chr3-95462940-INS-272 |
| chr3-95464027-INS-2856 | chr3-95464028-INS-2839 |
| chr3-95468630-INS-249 | chr3-95468630-INS-249 |
| chr3-95483375-INS-109 | chr3-95483375-INS-116 |
| chr3-95484298-INS-206 | chr3-95484298-INS-206 |
| chr3-95489498-INS-591 | chr3-95489498-INS-592 |
| chr3-95514435-INS-94 | chr3-95514435-INS-94 |
| chr3-95530346-INS-90 | chr3-95530340-INS-99 |
| chr3-95547239-INS-208 | chr3-95547239-INS-208 |
| chr3-95557331-INS-206 | chr3-95557331-INS-206 |
| chr3-95574858-INS-314 | chr3-95574858-INS-315 |
| chr3-95576006-INS-70 | chr3-95576006-INS-70 |
| chr3-95579426-INS-58 | chr3-95579426-INS-57 |
| chr3-95585633-INS-218 | chr3-95585633-INS-218 |
| chr3-95599473-INS-55 | chr3-95599489-INS-54 |
| chr3-95602712-INS-202 | chr3-95602712-INS-202 |
| chr3-95610527-INS-398 | chr3-95610527-INS-398 |
| chr3-95611932-INS-7097 | chr3-95611933-INS-7097 |
| chr3-95618767-INS-225 | chr3-95618767-INS-225 |
| chr3-95622536-INS-201 | chr3-95622536-INS-202 |
| chr3-95625493-INS-83 | chr3-95625493-INS-83 |
| chr3-95626927-INS-207 | chr3-95626927-INS-206 |
| chr3-95628062-INS-218 | chr3-95628062-INS-218 |
| chr3-95629978-INS-183 | chr3-95629978-INS-173 |
| chr3-95633429-INS-168 | chr3-95633429-INS-169 |
| chr3-95634509-INS-61 | chr3-95634509-INS-66 |
| chr3-95637750-INS-197 | chr3-95637750-INS-230 |
| chr3-95648192-INS-265 | chr3-95648192-INS-265 |
| chr3-95665175-INS-207 | chr3-95665175-INS-207 |
| chr3-95678279-INS-216 | chr3-95678279-INS-222 |
| chr3-95687631-INS-92 | chr3-95687631-INS-96 |
| chr3-95695888-INS-198 | chr3-95695888-INS-198 |
| chr3-95708161-INS-204 | chr3-95708161-INS-204 |
| chr3-95734546-INS-398 | chr3-95734546-INS-399 |
| chr3-95740480-INS-202 | chr3-95740480-INS-204 |
| chr3-95793401-INS-51 | chr3-95793401-INS-83 |
| chr3-95815951-INS-215 | chr3-95815951-INS-214 |
| chr3-95823454-INS-381 | chr3-95823460-INS-339 |
| chr3-95833330-INS-220 | chr3-95833330-INS-220 |
| chr3-95856328-INS-176 | chr3-95856323-INS-203 |
| chr3-95868872-INS-55 | chr3-95868872-INS-55 |
| chr3-95870155-INS-56 | chr3-95870155-INS-58 |
| chr3-95877367-INS-131 | chr3-95877368-INS-131 |
| chr3-95879174-INS-219 | chr3-95879174-INS-221 |
| chr3-95881925-INS-58 | chr3-95881925-INS-62 |
| chr3-95889989-INS-217 | chr3-95889989-INS-226 |
| chr3-95923440-INS-63 | chr3-95923445-INS-100 |
| chr3-95937193-INS-66 | chr3-95937193-INS-52 |
| chr3-95943312-INS-225 | chr3-95943312-INS-225 |
| chr3-95984034-INS-86 | chr3-95984004-INS-114 |
| chr3-95989965-INS-210 | chr3-95989965-INS-210 |

| | |
|---|---|
| chr3-95998918-INS-209 | chr3-95998918-INS-209 |
| chr3-96022264-INS-221 | chr3-96022264-INS-221 |
| chr3-96022829-INS-76 | chr3-96022802-INS-109 |
| chr3-96117835-INS-68 | chr3-96117835-INS-68 |
| chr3-96190213-INS-200 | chr3-96190213-INS-201 |
| chr3-96191999-INS-60 | chr3-96191999-INS-60 |
| chr3-96204208-INS-182 | chr3-96204208-INS-181 |
| chr3-96217899-INS-2259 | chr3-96217900-INS-2259 |
| chr3-96219313-INS-240 | chr3-96219313-INS-241 |
| chr3-96231468-INS-390 | chr3-96231468-INS-382 |
| chr3-96240655-INS-199 | chr3-96240655-INS-203 |
| chr3-96251401-INS-354 | chr3-96251401-INS-354 |
| chr3-96331239-INS-165 | chr3-96331240-INS-161 |
| chr3-96349574-INS-206 | chr3-96349574-INS-204 |
| chr3-96376640-INS-325 | chr3-96376640-INS-325 |
| chr3-96380497-INS-96 | chr3-96380502-INS-101 |
| chr3-96384324-INS-6873 | chr3-96384324-INS-6881 |
| chr3-96384577-INS-124 | chr3-96384577-INS-124 |
| chr3-96403344-INS-211 | chr3-96403344-INS-212 |
| chr3-96424086-INS-85 | chr3-96424064-INS-95 |
| chr3-96428556-INS-181 | chr3-96428556-INS-181 |
| chr3-96456760-INS-272 | chr3-96456760-INS-270 |
| chr3-96460624-INS-188 | chr3-96460624-INS-189 |
| chr3-96462332-INS-217 | chr3-96462332-INS-217 |
| chr3-96475903-INS-160 | chr3-96475903-INS-159 |
| chr3-96535345-INS-201 | chr3-96535345-INS-199 |
| chr3-96541096-INS-207 | chr3-96541096-INS-207 |
| chr3-96550149-INS-201 | chr3-96550149-INS-201 |
| chr3-96566586-INS-102 | chr3-96566586-INS-114 |
| chr3-96589866-INS-61 | chr3-96589864-INS-75 |
| chr3-96595039-INS-67 | chr3-96595043-INS-59 |
| chr3-96598695-INS-400 | chr3-96598695-INS-400 |
| chr3-96653370-INS-184 | chr3-96653370-INS-184 |
| chr3-96680466-INS-214 | chr3-96680466-INS-215 |
| chr3-96720077-INS-339 | chr3-96720077-INS-347 |
| chr3-96724861-INS-234 | chr3-96724861-INS-195 |
| chr3-96733003-INS-66 | chr3-96733003-INS-68 |
| chr3-96734050-INS-204 | chr3-96734050-INS-205 |
| chr3-96760295-INS-189 | chr3-96760295-INS-190 |
| chr3-96797337-INS-419 | chr3-96797337-INS-419 |
| chr3-96800141-INS-234 | chr3-96800145-INS-252 |
| chr3-96833874-INS-180 | chr3-96833874-INS-180 |
| chr3-96849116-INS-103 | chr3-96849116-INS-97 |
| chr3-96906444-INS-5040 | chr3-96906445-INS-5040 |
| chr3-96913284-INS-2919 | chr3-96913285-INS-2919 |
| chr3-96929593-INS-212 | chr3-96929593-INS-212 |
| chr3-96974675-INS-6240 | chr3-96974676-INS-6240 |
| chr3-96979181-INS-232 | chr3-96979181-INS-232 |
| chr3-96982875-INS-7024 | chr3-96982876-INS-7024 |
| chr3-97006869-INS-206 | chr3-97006869-INS-205 |
| chr3-97014579-INS-208 | chr3-97014579-INS-207 |
| chr3-97031932-INS-98 | chr3-97031926-INS-62 |
| chr3-97034008-INS-7326 | chr3-97034009-INS-7326 |
| chr3-97038064-INS-220 | chr3-97038064-INS-220 |
| chr3-97064748-INS-146 | chr3-97064748-INS-148 |
| chr3-97067871-INS-84 | chr3-97067871-INS-105 |
| chr3-97110110-INS-72 | chr3-97110110-INS-72 |
| chr3-97145658-INS-75 | chr3-97145664-INS-100 |
| chr3-97190502-INS-56 | chr3-97190502-INS-65 |
| chr3-97211233-INS-186 | chr3-97211233-INS-194 |
| chr3-97227035-INS-84 | chr3-97227035-INS-84 |
| chr3-97229413-INS-201 | chr3-97229413-INS-201 |
| chr3-97239843-INS-476 | chr3-97239843-INS-476 |
| chr3-97250800-INS-56 | chr3-97250800-INS-56 |
| chr3-97296478-INS-7067 | chr3-97296479-INS-7067 |
| chr3-97379487-INS-81 | chr3-97379495-INS-114 |
| chr3-97391032-INS-446 | chr3-97391032-INS-452 |
| chr3-97392883-INS-140 | chr3-97392883-INS-101 |
| chr3-97435073-INS-96 | chr3-97435073-INS-96 |
| chr3-97449502-INS-1328 | chr3-97449503-INS-1330 |
| chr3-97468754-INS-154 | chr3-97468754-INS-154 |
| chr3-97471578-INS-74 | chr3-97471578-INS-56 |
| chr3-97478484-INS-51 | chr3-97478484-INS-51 |
| chr3-97519158-INS-332 | chr3-97519158-INS-334 |
| chr3-97543856-INS-64 | chr3-97543856-INS-63 |
| chr3-97561481-INS-224 | chr3-97561481-INS-243 |
| chr3-97561729-INS-7063 | chr3-97561730-INS-7063 |

| | |
|---|---|
| chr3-97592397-INS-55 | chr3-97592389-INS-53 |
| chr3-97593060-INS-74 | chr3-97593060-INS-76 |
| chr3-97626235-INS-71 | chr3-97626214-INS-59 |
| chr3-97629422-INS-6899 | chr3-97629423-INS-6902 |
| chr3-97644933-INS-126 | chr3-97644933-INS-126 |
| chr3-97654584-INS-176 | chr3-97654584-INS-174 |
| chr3-97665901-INS-69 | chr3-97665901-INS-71 |
| chr3-97692799-INS-87 | chr3-97692800-INS-86 |
| chr3-97719289-INS-71 | chr3-97719289-INS-75 |
| chr3-97721951-INS-88 | chr3-97721951-INS-126 |
| chr3-97725021-INS-84 | chr3-97725021-INS-84 |
| chr3-97745808-INS-60 | chr3-97745807-INS-76 |
| chr3-97758791-INS-101 | chr3-97758830-INS-55 |
| chr3-97792826-INS-597 | chr3-97792822-INS-608 |
| chr3-97798400-INS-395 | chr3-97798400-INS-395 |
| chr3-97810805-INS-103 | chr3-97810805-INS-103 |
| chr3-97817045-INS-591 | chr3-97817045-INS-591 |
| chr3-97858212-INS-60 | chr3-97858199-INS-61 |
| chr3-97859265-INS-78 | chr3-97859265-INS-72 |
| chr3-97864232-INS-111 | chr3-97864232-INS-123 |
| chr3-97868469-INS-73 | chr3-97868469-INS-82 |
| chr3-97885660-INS-62 | chr3-97885660-INS-62 |
| chr3-97890506-INS-342 | chr3-97890506-INS-342 |
| chr3-97929072-INS-207 | chr3-97929072-INS-206 |
| chr3-97939535-INS-108 | chr3-97939535-INS-98 |
| chr3-97969870-INS-68 | chr3-97969870-INS-52 |
| chr3-97977636-INS-54 | chr3-97977636-INS-52 |
| chr3-97988653-INS-100 | chr3-97988607-INS-120 |
| chr3-98010650-INS-582 | chr3-98010650-INS-582 |
| chr3-98017931-INS-56 | chr3-98017931-INS-56 |
| chr3-98023612-INS-78 | chr3-98023612-INS-78 |
| chr3-98048368-INS-67 | chr3-98048368-INS-63 |
| chr3-98056712-INS-56 | chr3-98056712-INS-56 |
| chr3-98072219-INS-1505 | chr3-98072220-INS-1505 |
| chr3-98089627-INS-6673 | chr3-98089628-INS-6673 |
| chr3-98102083-INS-100 | chr3-98102083-INS-100 |
| chr3-98109540-INS-66 | chr3-98109540-INS-66 |
| chr3-98113313-INS-108 | chr3-98113313-INS-108 |
| chr3-98121997-INS-120 | chr3-98121997-INS-117 |
| chr3-98123337-INS-54 | chr3-98123337-INS-54 |
| chr3-98124336-INS-3068 | chr3-98124337-INS-3068 |
| chr3-98126026-INS-66 | chr3-98126026-INS-66 |
| chr3-98135707-INS-52 | chr3-98135707-INS-52 |
| chr3-98163399-INS-255 | chr3-98163399-INS-255 |
| chr3-98173675-INS-4635 | chr3-98173676-INS-4635 |
| chr3-98179187-INS-52 | chr3-98179187-INS-52 |
| chr3-98190862-INS-1002 | chr3-98190863-INS-1002 |
| chr3-98201239-INS-69 | chr3-98201239-INS-69 |
| chr3-98218204-INS-60 | chr3-98218204-INS-60 |
| chr3-98237179-INS-60 | chr3-98237179-INS-56 |
| chr3-98269542-INS-495 | chr3-98269542-INS-495 |
| chr3-98283863-INS-72 | chr3-98283863-INS-78 |
| chr3-98296588-INS-144 | chr3-98296585-INS-192 |
| chr3-98300477-INS-132 | chr3-98300477-INS-132 |
| chr3-98303514-INS-206 | chr3-98303514-INS-207 |
| chr3-98359664-INS-65 | chr3-98359664-INS-65 |
| chr3-98371461-INS-435 | chr3-98371477-INS-443 |
| chr3-98385280-INS-1303 | chr3-98385281-INS-1303 |
| chr3-98518504-INS-2484 | chr3-98518505-INS-2483 |
| chr3-98540331-INS-66 | chr3-98540337-INS-64 |
| chr3-98575034-INS-218 | chr3-98575034-INS-218 |
| chr3-98577333-INS-207 | chr3-98577333-INS-207 |
| chr3-98578632-INS-215 | chr3-98578632-INS-215 |
| chr3-98584444-INS-122 | chr3-98584441-INS-84 |
| chr3-98597203-INS-2093 | chr3-98597204-INS-2094 |
| chr3-98613372-INS-51 | chr3-98613368-INS-58 |
| chr3-98641384-INS-3138 | chr3-98641385-INS-3138 |
| chr3-98647792-INS-112 | chr3-98647792-INS-113 |
| chr3-98661681-INS-6722 | chr3-98661682-INS-6722 |
| chr3-98663461-INS-57 | chr3-98663461-INS-81 |
| chr3-98672219-INS-1979 | chr3-98672206-INS-1967 |
| chr3-98691726-INS-2539 | chr3-98691727-INS-2539 |
| chr3-98707350-INS-659 | chr3-98707350-INS-656 |
| chr3-98710071-INS-1371 | chr3-98710072-INS-1370 |
| chr3-98711364-INS-1624 | chr3-98711365-INS-1624 |
| chr3-98719927-INS-55 | chr3-98719936-INS-56 |
| chr3-98722719-INS-275 | chr3-98722719-INS-275 |

| | |
|---|---|
| chr3-98742500-INS-7014 | chr3-98742501-INS-7058 |
| chr3-98778701-INS-51 | chr3-98778701-INS-72 |
| chr3-98785554-INS-85 | chr3-98785585-INS-96 |
| chr3-98878299-INS-1878 | chr3-98878300-INS-1878 |
| chr3-98892547-INS-2044 | chr3-98892548-INS-2044 |
| chr3-98899467-INS-6975 | chr3-98899468-INS-6976 |
| chr3-98907954-INS-7151 | chr3-98907955-INS-7151 |
| chr3-98909088-INS-58 | chr3-98909100-INS-68 |
| chr3-98927640-INS-73 | chr3-98927640-INS-73 |
| chr3-98931823-INS-69 | chr3-98931823-INS-52 |
| chr3-98934820-INS-67 | chr3-98934820-INS-68 |
| chr3-98973937-INS-251 | chr3-98973937-INS-251 |
| chr3-99016974-INS-170 | chr3-99016974-INS-169 |
| chr3-99037671-INS-141 | chr3-99037671-INS-141 |
| chr3-99055801-INS-6883 | chr3-99055802-INS-6884 |
| chr3-99060359-INS-263 | chr3-99060359-INS-263 |
| chr3-99063414-INS-55 | chr3-99063414-INS-55 |
| chr3-99067038-INS-683 | chr3-99067038-INS-683 |
| chr3-99071752-INS-620 | chr3-99071752-INS-621 |
| chr3-99079360-INS-222 | chr3-99079360-INS-216 |
| chr3-99092496-INS-201 | chr3-99092496-INS-201 |
| chr3-99093374-INS-57 | chr3-99093374-INS-57 |
| chr3-99102892-INS-111 | chr3-99102892-INS-111 |
| chr3-99102892-INS-814 | chr3-99102892-INS-814 |
| chr3-99134578-INS-107 | chr3-99134566-INS-103 |
| chr3-99137500-INS-242 | chr3-99137500-INS-242 |
| chr3-99171271-INS-66 | chr3-99171271-INS-51 |
| chr3-99173392-INS-76 | chr3-99173392-INS-76 |
| chr3-99178128-INS-69 | chr3-99178128-INS-69 |
| chr3-99180287-INS-640 | chr3-99180287-INS-640 |
| chr3-99181151-INS-151 | chr3-99181155-INS-143 |
| chr3-99205812-INS-801 | chr3-99205812-INS-801 |
| chr3-99226863-INS-762 | chr3-99226863-INS-763 |
| chr3-99227582-INS-7994 | chr3-99227583-INS-7996 |
| chr3-99231191-INS-190 | chr3-99231191-INS-189 |
| chr3-99248642-INS-84 | chr3-99248638-INS-68 |
| chr3-99264506-INS-52 | chr3-99264507-INS-50 |
| chr3-99264685-INS-841 | chr3-99264685-INS-841 |
| chr3-99298087-INS-94 | chr3-99298087-INS-94 |
| chr3-99300756-INS-722 | chr3-99300757-INS-726 |
| chr3-99312770-INS-214 | chr3-99312771-INS-213 |
| chr3-99313979-INS-584 | chr3-99313979-INS-584 |
| chr3-99363228-INS-1030 | chr3-99363229-INS-1030 |
| chr3-99364281-INS-71 | chr3-99364281-INS-75 |
| chr3-99369658-INS-198 | chr3-99369661-INS-198 |
| chr3-99386206-INS-198 | chr3-99386207-INS-198 |
| chr3-99399121-INS-2386 | chr3-99399122-INS-2386 |
| chr3-99417160-INS-74 | chr3-99417160-INS-74 |
| chr3-99432643-INS-4548 | chr3-99432644-INS-4548 |
| chr3-99437427-INS-140 | chr3-99437427-INS-140 |
| chr3-99457285-INS-605 | chr3-99457285-INS-605 |
| chr3-99465468-INS-70 | chr3-99465463-INS-78 |
| chr3-99494680-INS-64 | chr3-99494723-INS-52 |
| chr3-99501998-INS-892 | chr3-99501998-INS-929 |
| chr3-99513981-INS-3511 | chr3-99513982-INS-3473 |
| chr3-99519175-INS-53 | chr3-99519175-INS-51 |
| chr3-99539399-INS-1060 | chr3-99539400-INS-1058 |
| chr3-99569616-INS-105 | chr3-99569616-INS-81 |
| chr3-99585734-INS-128 | chr3-99585734-INS-130 |
| chr3-99586407-INS-8673 | chr3-99586408-INS-8671 |
| chr3-99596788-INS-52 | chr3-99596788-INS-54 |
| chr3-99606972-INS-53 | chr3-99606972-INS-53 |
| chr3-99624290-INS-110 | chr3-99624252-INS-120 |
| chr3-99648652-INS-425 | chr3-99648652-INS-425 |
| chr3-99656594-INS-51 | chr3-99656596-INS-56 |
| chr3-99671067-INS-87 | chr3-99671067-INS-87 |
| chr3-99712429-INS-64 | chr3-99712422-INS-88 |
| chr3-99753244-INS-67 | chr3-99753244-INS-60 |
| chr3-99815141-INS-212 | chr3-99815141-INS-212 |
| chr3-99826403-INS-92 | chr3-99826403-INS-90 |
| chr3-99851376-INS-3183 | chr3-99851377-INS-3183 |
| chr3-99861995-INS-803 | chr3-99861995-INS-803 |
| chr3-99913332-INS-64 | chr3-99913332-INS-64 |
| chr3-99916163-INS-83 | chr3-99916163-INS-88 |
| chr3-99921677-INS-2500 | chr3-99921678-INS-2499 |
| chr3-99923377-INS-6253 | chr3-99923378-INS-6253 |
| chr3-99935279-INS-116 | chr3-99935279-INS-116 |

| | |
|---|---|
| chr3-99954703-INS-2313 | chr3-99954704-INS-2352 |
| chr3-99971918-INS-60 | chr3-99971918-INS-73 |
| chr3-99997841-INS-2210 | chr3-99997842-INS-2210 |
| chr3-100016143-INS-3268 | chr3-100016144-INS-3268 |
| chr3-100079730-INS-62 | chr3-100079730-INS-54 |
| chr3-100104544-INS-201 | chr3-100104544-INS-200 |
| chr3-100169981-INS-385 | chr3-100169981-INS-384 |
| chr3-100198940-INS-1257 | chr3-100198941-INS-1257 |
| chr3-100205489-INS-967 | chr3-100205489-INS-961 |
| chr3-100223768-INS-100 | chr3-100223789-INS-113 |
| chr3-100236703-INS-6780 | chr3-100236704-INS-6781 |
| chr3-100261342-INS-57 | chr3-100261342-INS-65 |
| chr3-100282631-INS-58 | chr3-100282631-INS-96 |
| chr3-100319160-INS-118 | chr3-100319160-INS-118 |
| chr3-100362197-INS-165 | chr3-100362197-INS-165 |
| chr3-100365393-INS-4218 | chr3-100365394-INS-4218 |
| chr3-100385061-INS-100 | chr3-100385061-INS-100 |
| chr3-100397288-INS-52 | chr3-100397288-INS-52 |
| chr3-100419932-INS-70 | chr3-100419932-INS-72 |
| chr3-100422392-INS-186 | chr3-100422392-INS-225 |
| chr3-100429114-INS-238 | chr3-100429114-INS-239 |
| chr3-100439662-INS-6735 | chr3-100439663-INS-6700 |
| chr3-100445562-INS-6401 | chr3-100445563-INS-6401 |
| chr3-100448159-INS-140 | chr3-100448159-INS-140 |
| chr3-100456550-INS-5545 | chr3-100456551-INS-5545 |
| chr3-100464017-INS-56 | chr3-100464017-INS-58 |
| chr3-100466625-INS-52 | chr3-100466625-INS-52 |
| chr3-100504647-INS-226 | chr3-100504647-INS-226 |
| chr3-100508144-INS-414 | chr3-100508144-INS-416 |
| chr3-100519342-INS-121 | chr3-100519330-INS-117 |
| chr3-100567840-INS-63 | chr3-100567840-INS-57 |
| chr3-100585642-INS-527 | chr3-100585642-INS-527 |
| chr3-100603275-INS-5317 | chr3-100603276-INS-5318 |
| chr3-100625727-INS-101 | chr3-100625727-INS-99 |
| chr3-100632863-INS-126 | chr3-100632857-INS-138 |
| chr3-100638323-INS-143 | chr3-100638323-INS-144 |
| chr3-100651512-INS-56 | chr3-100651512-INS-62 |
| chr3-100657192-INS-151 | chr3-100657192-INS-119 |
| chr3-100664016-INS-61 | chr3-100664016-INS-73 |
| chr3-100673464-INS-399 | chr3-100673464-INS-399 |
| chr3-100691295-INS-2802 | chr3-100691296-INS-2802 |
| chr3-100694006-INS-105 | chr3-100694006-INS-105 |
| chr3-100694808-INS-826 | chr3-100694808-INS-826 |
| chr3-100709744-INS-7146 | chr3-100709745-INS-7146 |
| chr3-100722320-INS-160 | chr3-100722320-INS-160 |
| chr3-100743361-INS-314 | chr3-100743361-INS-306 |
| chr3-100756011-INS-72 | chr3-100756011-INS-62 |
| chr3-100781411-INS-79 | chr3-100781412-INS-112 |
| chr3-100791569-INS-92 | chr3-100791569-INS-92 |
| chr3-100814744-INS-68 | chr3-100814700-INS-78 |
| chr3-100825069-INS-384 | chr3-100825069-INS-384 |
| chr3-100826265-INS-56 | chr3-100826265-INS-59 |
| chr3-100826281-INS-589 | chr3-100826281-INS-593 |
| chr3-100847809-INS-66 | chr3-100847809-INS-66 |
| chr3-100850891-INS-247 | chr3-100850891-INS-247 |
| chr3-100869262-INS-194 | chr3-100869262-INS-195 |
| chr3-100879040-INS-176 | chr3-100879040-INS-206 |
| chr3-100912105-INS-191 | chr3-100912102-INS-171 |
| chr3-100915723-INS-201 | chr3-100915723-INS-201 |
| chr3-100923231-INS-344 | chr3-100923231-INS-344 |
| chr3-100931387-INS-53 | chr3-100931387-INS-55 |
| chr3-100965788-INS-75 | chr3-100965788-INS-75 |
| chr3-101007566-INS-51 | chr3-101007566-INS-51 |
| chr3-101018374-INS-61 | chr3-101018374-INS-76 |
| chr3-101029253-INS-334 | chr3-101029253-INS-335 |
| chr3-101035896-INS-745 | chr3-101035896-INS-745 |
| chr3-101046353-INS-251 | chr3-101046353-INS-254 |
| chr3-101059000-INS-143 | chr3-101059000-INS-143 |
| chr3-101084517-INS-591 | chr3-101084517-INS-591 |
| chr3-101092115-INS-54 | chr3-101092115-INS-55 |
| chr3-101115739-INS-124 | chr3-101115739-INS-124 |
| chr3-101118737-INS-201 | chr3-101118737-INS-201 |
| chr3-101123104-INS-112 | chr3-101123070-INS-126 |
| chr3-101123647-INS-202 | chr3-101123647-INS-202 |
| chr3-101129214-INS-229 | chr3-101129214-INS-229 |
| chr3-101138492-INS-125 | chr3-101138492-INS-132 |
| chr3-101175861-INS-57 | chr3-101175861-INS-58 |

| | |
|---|---|
| chr3-101185148-INS-234 | chr3-101185148-INS-233 |
| chr3-101218121-INS-84 | chr3-101218122-INS-84 |
| chr3-101228718-INS-111 | chr3-101228718-INS-123 |
| chr3-101243436-INS-84 | chr3-101243436-INS-84 |
| chr3-101343408-INS-74 | chr3-101343408-INS-76 |
| chr3-101347293-INS-56 | chr3-101347293-INS-56 |
| chr3-101376804-INS-72 | chr3-101376804-INS-60 |
| chr3-101382246-INS-109 | chr3-101382246-INS-109 |
| chr3-101398500-INS-53 | chr3-101398500-INS-63 |
| chr3-101425353-INS-226 | chr3-101425353-INS-218 |
| chr3-101429164-INS-203 | chr3-101429164-INS-206 |
| chr3-101435036-INS-56 | chr3-101435036-INS-75 |
| chr3-101445243-INS-78 | chr3-101445243-INS-78 |
| chr3-101455803-INS-555 | chr3-101455803-INS-558 |
| chr3-101503512-INS-125 | chr3-101503512-INS-125 |
| chr3-101529295-INS-379 | chr3-101529295-INS-379 |
| chr3-101533547-INS-54 | chr3-101533547-INS-62 |
| chr3-101538398-INS-53 | chr3-101538360-INS-59 |
| chr3-101539392-INS-5365 | chr3-101539393-INS-5365 |
| chr3-101562965-INS-55 | chr3-101562965-INS-69 |
| chr3-101569514-INS-532 | chr3-101569514-INS-521 |
| chr3-101579730-INS-62 | chr3-101579720-INS-75 |
| chr3-101584175-INS-60 | chr3-101584176-INS-60 |
| chr3-101595658-INS-96 | chr3-101595658-INS-66 |
| chr3-101600431-INS-54 | chr3-101600427-INS-66 |
| chr3-101615446-INS-98 | chr3-101615446-INS-70 |
| chr3-101626990-INS-68 | chr3-101626990-INS-68 |
| chr3-101658324-INS-84 | chr3-101658324-INS-84 |
| chr3-101690786-INS-54 | chr3-101690786-INS-54 |
| chr3-101706216-INS-81 | chr3-101706216-INS-81 |
| chr3-101718223-INS-57 | chr3-101718223-INS-58 |
| chr3-101721087-INS-52 | chr3-101721123-INS-52 |
| chr3-101726468-INS-111 | chr3-101726468-INS-111 |
| chr3-101738773-INS-76 | chr3-101738817-INS-56 |
| chr3-101740144-INS-90 | chr3-101740144-INS-96 |
| chr3-101743129-INS-60 | chr3-101743120-INS-60 |
| chr3-101755911-INS-310 | chr3-101755914-INS-267 |
| chr3-101765094-INS-164 | chr3-101765094-INS-194 |
| chr3-101778468-INS-115 | chr3-101778468-INS-138 |
| chr3-101793515-INS-72 | chr3-101793515-INS-72 |
| chr3-101835424-INS-1055 | chr3-101835425-INS-1055 |
| chr3-101840177-INS-67 | chr3-101840177-INS-85 |
| chr3-101848131-INS-207 | chr3-101848131-INS-207 |
| chr3-101852051-INS-1120 | chr3-101852047-INS-1115 |
| chr3-101853196-INS-345 | chr3-101853196-INS-336 |
| chr3-101870488-INS-1928 | chr3-101870489-INS-1928 |
| chr3-101904088-INS-214 | chr3-101904088-INS-212 |
| chr3-101910851-INS-75 | chr3-101910851-INS-72 |
| chr3-101913369-INS-80 | chr3-101913369-INS-80 |
| chr3-101944748-INS-1086 | chr3-101944748-INS-1080 |
| chr3-101950044-INS-52 | chr3-101950044-INS-52 |
| chr3-101964360-INS-123 | chr3-101964360-INS-123 |
| chr3-101978037-INS-455 | chr3-101978037-INS-455 |
| chr3-102013001-INS-189 | chr3-102013001-INS-189 |
| chr3-102079747-INS-70 | chr3-102079747-INS-70 |
| chr3-102083078-INS-64 | chr3-102083074-INS-52 |
| chr3-102093112-INS-3598 | chr3-102093113-INS-3598 |
| chr3-102118989-INS-97 | chr3-102119031-INS-66 |
| chr3-102122975-INS-152 | chr3-102122975-INS-150 |
| chr3-102154050-INS-68 | chr3-102154050-INS-83 |
| chr3-102163018-INS-220 | chr3-102162998-INS-200 |
| chr3-102183961-INS-66 | chr3-102183961-INS-60 |
| chr3-102198796-INS-376 | chr3-102198796-INS-376 |
| chr3-102222636-INS-6457 | chr3-102222637-INS-6457 |
| chr3-102247224-INS-112 | chr3-102247224-INS-135 |
| chr3-102256900-INS-66 | chr3-102256900-INS-66 |
| chr3-102259585-INS-3423 | chr3-102259586-INS-3420 |
| chr3-102278361-INS-60 | chr3-102278360-INS-68 |
| chr3-102312437-INS-4798 | chr3-102312438-INS-4798 |
| chr3-102318871-INS-248 | chr3-102318871-INS-248 |
| chr3-102327390-INS-66 | chr3-102327390-INS-54 |
| chr3-102393515-INS-115 | chr3-102393515-INS-115 |
| chr3-102404882-INS-86 | chr3-102404882-INS-86 |
| chr3-102406798-INS-56 | chr3-102406798-INS-56 |
| chr3-102408266-INS-103 | chr3-102408267-INS-96 |
| chr3-102446259-INS-56 | chr3-102446259-INS-100 |
| chr3-102450491-INS-64 | chr3-102450491-INS-66 |

| | |
|---|---|
| chr3-102458683-INS-86 | chr3-102458683-INS-84 |
| chr3-102479236-INS-595 | chr3-102479229-INS-595 |
| chr3-102489440-INS-356 | chr3-102489440-INS-356 |
| chr3-102500123-INS-161 | chr3-102500123-INS-134 |
| chr3-102509447-INS-76 | chr3-102509447-INS-76 |
| chr3-102532396-INS-56 | chr3-102532396-INS-56 |
| chr3-102577083-INS-132 | chr3-102577083-INS-120 |
| chr3-102601216-INS-117 | chr3-102601216-INS-117 |
| chr3-102637591-INS-52 | chr3-102637591-INS-54 |
| chr3-102708217-INS-6371 | chr3-102708218-INS-6371 |
| chr3-102735999-INS-2955 | chr3-102736000-INS-2955 |
| chr3-102749325-INS-84 | chr3-102749325-INS-64 |
| chr3-102754948-INS-327 | chr3-102754949-INS-289 |
| chr3-102759654-INS-84 | chr3-102759654-INS-84 |
| chr3-102773679-INS-214 | chr3-102773679-INS-214 |
| chr3-102815146-INS-174 | chr3-102815146-INS-185 |
| chr3-102821931-INS-62 | chr3-102821931-INS-62 |
| chr3-102824627-INS-548 | chr3-102824627-INS-549 |
| chr3-102834506-INS-195 | chr3-102834506-INS-196 |
| chr3-102835780-INS-4606 | chr3-102835781-INS-4606 |
| chr3-102855012-INS-54 | chr3-102855012-INS-58 |
| chr3-102870339-INS-69 | chr3-102870336-INS-104 |
| chr3-102875260-INS-369 | chr3-102875260-INS-362 |
| chr3-102877690-INS-53 | chr3-102877690-INS-53 |
| chr3-102883172-INS-198 | chr3-102883172-INS-198 |
| chr3-102884670-INS-56 | chr3-102884670-INS-56 |
| chr3-102940170-INS-214 | chr3-102940170-INS-214 |
| chr3-102941567-INS-510 | chr3-102941567-INS-508 |
| chr3-102980069-INS-4454 | chr3-102980070-INS-4454 |
| chr3-102997700-INS-317 | chr3-102997710-INS-311 |
| chr3-103003239-INS-54 | chr3-103003239-INS-69 |
| chr3-103014836-INS-254 | chr3-103014836-INS-254 |
| chr3-103017182-INS-199 | chr3-103017182-INS-198 |
| chr3-103029308-INS-159 | chr3-103029331-INS-120 |
| chr3-103040753-INS-214 | chr3-103040753-INS-210 |
| chr3-103070861-INS-207 | chr3-103070861-INS-207 |
| chr3-103088137-INS-7187 | chr3-103088138-INS-7194 |
| chr3-103108099-INS-184 | chr3-103108099-INS-185 |
| chr3-103121480-INS-4560 | chr3-103121481-INS-4560 |
| chr3-103132208-INS-205 | chr3-103132208-INS-205 |
| chr3-103134198-INS-56 | chr3-103134198-INS-56 |
| chr3-103135171-INS-204 | chr3-103135171-INS-206 |
| chr3-103138276-INS-68 | chr3-103138323-INS-68 |
| chr3-103140385-INS-125 | chr3-103140385-INS-124 |
| chr3-103150677-INS-213 | chr3-103150677-INS-213 |
| chr3-103156421-INS-7135 | chr3-103156422-INS-7135 |
| chr3-103156839-INS-91 | chr3-103156839-INS-52 |
| chr3-103157147-INS-1888 | chr3-103157148-INS-1888 |
| chr3-103178002-INS-98 | chr3-103178002-INS-98 |
| chr3-103179920-INS-2847 | chr3-103179921-INS-2871 |
| chr3-103180013-INS-118 | chr3-103180018-INS-95 |
| chr3-103282539-INS-545 | chr3-103282539-INS-545 |
| chr3-103299811-INS-496 | chr3-103299811-INS-499 |
| chr3-103316954-INS-90 | chr3-103316954-INS-90 |
| chr3-103318970-INS-192 | chr3-103318970-INS-192 |
| chr3-103349604-INS-1076 | chr3-103349605-INS-1076 |
| chr3-103351887-INS-221 | chr3-103351927-INS-227 |
| chr3-103354824-INS-79 | chr3-103354824-INS-88 |
| chr3-103368391-INS-54 | chr3-103368391-INS-54 |
| chr3-103395945-INS-5433 | chr3-103395946-INS-5433 |
| chr3-103406704-INS-55 | chr3-103406705-INS-50 |
| chr3-103409210-INS-216 | chr3-103409210-INS-218 |
| chr3-103451111-INS-53 | chr3-103451111-INS-50 |
| chr3-103467697-INS-64 | chr3-103467697-INS-64 |
| chr3-103469777-INS-2947 | chr3-103469778-INS-2947 |
| chr3-103506442-INS-7079 | chr3-103506443-INS-7079 |
| chr3-103525503-INS-93 | chr3-103525503-INS-93 |
| chr3-103591443-INS-198 | chr3-103591443-INS-198 |
| chr3-103601519-INS-94 | chr3-103601519-INS-94 |
| chr3-103602889-INS-58 | chr3-103602889-INS-50 |
| chr3-103614168-INS-64 | chr3-103614168-INS-64 |
| chr3-103627341-INS-57 | chr3-103627341-INS-64 |
| chr3-103670917-INS-58 | chr3-103670917-INS-58 |
| chr3-103678716-INS-96 | chr3-103678716-INS-50 |
| chr3-103706841-INS-210 | chr3-103706841-INS-207 |
| chr3-103756194-INS-145 | chr3-103756194-INS-180 |
| chr3-103780630-INS-88 | chr3-103780666-INS-76 |

| | |
|---|---|
| chr3-103795772-INS-7174 | chr3-103795773-INS-7174 |
| chr3-103796864-INS-77 | chr3-103796864-INS-77 |
| chr3-103832700-INS-97 | chr3-103832705-INS-66 |
| chr3-103863159-INS-250 | chr3-103863159-INS-250 |
| chr3-103916017-INS-68 | chr3-103916017-INS-72 |
| chr3-103944098-INS-373 | chr3-103944098-INS-371 |
| chr3-103970557-INS-3292 | chr3-103970558-INS-3290 |
| chr3-103989824-INS-210 | chr3-103989824-INS-214 |
| chr3-104030245-INS-214 | chr3-104030245-INS-214 |
| chr3-104139633-INS-746 | chr3-104139633-INS-746 |
| chr3-104149445-INS-4392 | chr3-104149446-INS-4392 |
| chr3-104159894-INS-72 | chr3-104159894-INS-72 |
| chr3-104208083-INS-64 | chr3-104208083-INS-64 |
| chr3-104211362-INS-56 | chr3-104211362-INS-56 |
| chr3-104215897-INS-4933 | chr3-104215898-INS-4933 |
| chr3-104236228-INS-100 | chr3-104236228-INS-100 |
| chr3-104258325-INS-6502 | chr3-104258326-INS-6510 |
| chr3-104313797-INS-53 | chr3-104313797-INS-53 |
| chr3-104319239-INS-1888 | chr3-104319240-INS-1888 |
| chr3-104325473-INS-3454 | chr3-104325474-INS-3454 |
| chr3-104340556-INS-52 | chr3-104340556-INS-54 |
| chr3-104357168-INS-214 | chr3-104357168-INS-214 |
| chr3-104369175-INS-1748 | chr3-104369176-INS-1747 |
| chr3-104393761-INS-76 | chr3-104393761-INS-68 |
| chr3-104445918-INS-101 | chr3-104445940-INS-66 |
| chr3-104449988-INS-198 | chr3-104449988-INS-198 |
| chr3-104471807-INS-58 | chr3-104471807-INS-86 |
| chr3-104482715-INS-144 | chr3-104482715-INS-144 |
| chr3-104488621-INS-72 | chr3-104488621-INS-72 |
| chr3-104569647-INS-214 | chr3-104569647-INS-214 |
| chr3-104606631-INS-215 | chr3-104606615-INS-190 |
| chr3-104613322-INS-6452 | chr3-104613323-INS-6452 |
| chr3-104619293-INS-205 | chr3-104619293-INS-205 |
| chr3-104624594-INS-180 | chr3-104624594-INS-180 |
| chr3-104633029-INS-123 | chr3-104633029-INS-125 |
| chr3-104634041-INS-51 | chr3-104634041-INS-61 |
| chr3-104643892-INS-71 | chr3-104643909-INS-59 |
| chr3-104649707-INS-229 | chr3-104649707-INS-231 |
| chr3-104652396-INS-216 | chr3-104652370-INS-217 |
| chr3-104656374-INS-64 | chr3-104656374-INS-64 |
| chr3-104657178-INS-572 | chr3-104657178-INS-572 |
| chr3-104661511-INS-66 | chr3-104661511-INS-70 |
| chr3-104664671-INS-62 | chr3-104664671-INS-66 |
| chr3-104668177-INS-584 | chr3-104668177-INS-584 |
| chr3-104670828-INS-97 | chr3-104670845-INS-80 |
| chr3-104758718-INS-60 | chr3-104758718-INS-60 |
| chr3-104764410-INS-345 | chr3-104764410-INS-345 |
| chr3-104773902-INS-293 | chr3-104773902-INS-293 |
| chr3-104792804-INS-216 | chr3-104792804-INS-216 |
| chr3-104809555-INS-93 | chr3-104809555-INS-93 |
| chr3-104812876-INS-259 | chr3-104812876-INS-259 |
| chr3-104823701-INS-179 | chr3-104823701-INS-179 |
| chr3-104830154-INS-810 | chr3-104830154-INS-810 |
| chr3-104858271-INS-204 | chr3-104858271-INS-204 |
| chr3-104860638-INS-147 | chr3-104860630-INS-112 |
| chr3-104877012-INS-1893 | chr3-104877013-INS-1893 |
| chr3-104888504-INS-82 | chr3-104888504-INS-80 |
| chr3-104900047-INS-506 | chr3-104900047-INS-506 |
| chr3-104922028-INS-1490 | chr3-104922029-INS-1490 |
| chr3-104955773-INS-104 | chr3-104955773-INS-106 |
| chr3-104960721-INS-58 | chr3-104960721-INS-58 |
| chr3-104962523-INS-72 | chr3-104962523-INS-72 |
| chr3-104963632-INS-67 | chr3-104963632-INS-67 |
| chr3-104967881-INS-135 | chr3-104967881-INS-135 |
| chr3-104988258-INS-207 | chr3-104988258-INS-207 |
| chr3-104991099-INS-174 | chr3-104991099-INS-143 |
| chr3-104993847-INS-118 | chr3-104993847-INS-118 |
| chr3-105030009-INS-110 | chr3-105030022-INS-100 |
| chr3-105033321-INS-1432 | chr3-105033322-INS-1432 |
| chr3-105035974-INS-604 | chr3-105035974-INS-607 |
| chr3-105075703-INS-1894 | chr3-105075704-INS-1894 |
| chr3-105076821-INS-54 | chr3-105076821-INS-56 |
| chr3-105097314-INS-52 | chr3-105097314-INS-58 |
| chr3-105100858-INS-59 | chr3-105100858-INS-61 |
| chr3-105108405-INS-57 | chr3-105108405-INS-55 |
| chr3-105114230-INS-72 | chr3-105114230-INS-72 |
| chr3-105116027-INS-810 | chr3-105116027-INS-808 |

| | |
|---|---|
| chr3-105136082-INS-236 | chr3-105136082-INS-236 |
| chr3-105166054-INS-53 | chr3-105166052-INS-52 |
| chr3-105178533-INS-98 | chr3-105178525-INS-96 |
| chr3-105213306-INS-55 | chr3-105213306-INS-56 |
| chr3-105215558-INS-7447 | chr3-105215559-INS-7447 |
| chr3-105218596-INS-105 | chr3-105218596-INS-58 |
| chr3-105260146-INS-1170 | chr3-105260147-INS-1170 |
| chr3-105298474-INS-51 | chr3-105298474-INS-51 |
| chr3-105305683-INS-89 | chr3-105305683-INS-93 |
| chr3-105318037-INS-77 | chr3-105318037-INS-72 |
| chr3-105330130-INS-81 | chr3-105330102-INS-66 |
| chr3-105348367-INS-52 | chr3-105348367-INS-52 |
| chr3-105352302-INS-525 | chr3-105352302-INS-526 |
| chr3-105382438-INS-53 | chr3-105382438-INS-53 |
| chr3-105388816-INS-69 | chr3-105388771-INS-72 |
| chr3-105393302-INS-6494 | chr3-105393303-INS-6495 |
| chr3-105417333-INS-693 | chr3-105417332-INS-728 |
| chr3-105420908-INS-182 | chr3-105420908-INS-187 |
| chr3-105427601-INS-612 | chr3-105427589-INS-630 |
| chr3-105434874-INS-97 | chr3-105434836-INS-88 |
| chr3-105466044-INS-104 | chr3-105466044-INS-107 |
| chr3-105469386-INS-80 | chr3-105469386-INS-69 |
| chr3-105473188-INS-54 | chr3-105473188-INS-96 |
| chr3-105477572-INS-65 | chr3-105477572-INS-77 |
| chr3-105479345-INS-54 | chr3-105479345-INS-54 |
| chr3-105531783-INS-61 | chr3-105531783-INS-63 |
| chr3-105537621-INS-758 | chr3-105537621-INS-758 |
| chr3-105549429-INS-3719 | chr3-105549430-INS-3719 |
| chr3-105553623-INS-78 | chr3-105553623-INS-78 |
| chr3-105587047-INS-255 | chr3-105587047-INS-236 |
| chr3-105622206-INS-61 | chr3-105622176-INS-50 |
| chr3-105630148-INS-300 | chr3-105630148-INS-302 |
| chr3-105723770-INS-80 | chr3-105723778-INS-88 |
| chr3-105766758-INS-140 | chr3-105766758-INS-147 |
| chr3-105771238-INS-111 | chr3-105771238-INS-114 |
| chr3-105798826-INS-171 | chr3-105798826-INS-172 |
| chr3-105818889-INS-68 | chr3-105818890-INS-68 |
| chr3-105827867-INS-1120 | chr3-105827868-INS-1120 |
| chr3-105828185-INS-63 | chr3-105828185-INS-67 |
| chr3-105838560-INS-2750 | chr3-105838561-INS-2750 |
| chr3-105843653-INS-352 | chr3-105843653-INS-357 |
| chr3-105845254-INS-74 | chr3-105845254-INS-74 |
| chr3-105846570-INS-175 | chr3-105846570-INS-175 |
| chr3-105857515-INS-224 | chr3-105857515-INS-224 |
| chr3-105886370-INS-88 | chr3-105886370-INS-88 |
| chr3-105891936-INS-57 | chr3-105891936-INS-60 |
| chr3-105892768-INS-55 | chr3-105892769-INS-53 |
| chr3-105895279-INS-64 | chr3-105895279-INS-64 |
| chr3-105920517-INS-96 | chr3-105920517-INS-100 |
| chr3-105942889-INS-750 | chr3-105942871-INS-743 |
| chr3-105951525-INS-6240 | chr3-105951526-INS-6240 |
| chr3-105967642-INS-369 | chr3-105967642-INS-369 |
| chr3-105987977-INS-3043 | chr3-105987978-INS-3043 |
| chr3-106079035-INS-220 | chr3-106079035-INS-220 |
| chr3-106135509-INS-7082 | chr3-106135510-INS-7082 |
| chr3-106224231-INS-972 | chr3-106224231-INS-974 |
| chr3-106231970-INS-6838 | chr3-106231970-INS-6838 |
| chr3-106249552-INS-5396 | chr3-106249553-INS-5397 |
| chr3-106282291-INS-1980 | chr3-106282290-INS-1979 |
| chr3-106289972-INS-3184 | chr3-106289973-INS-3192 |
| chr3-106290800-INS-56 | chr3-106290800-INS-56 |
| chr3-106305327-INS-3979 | chr3-106305328-INS-3979 |
| chr3-106309738-INS-72 | chr3-106309738-INS-72 |
| chr3-106359472-INS-65 | chr3-106359472-INS-65 |
| chr3-106378939-INS-52 | chr3-106378939-INS-52 |
| chr3-106384458-INS-97 | chr3-106384456-INS-105 |
| chr3-106413527-INS-71 | chr3-106413527-INS-79 |
| chr3-106434352-INS-1127 | chr3-106434371-INS-1104 |
| chr3-106488411-INS-742 | chr3-106488412-INS-757 |
| chr3-106489632-INS-4983 | chr3-106489633-INS-4983 |
| chr3-106517857-INS-199 | chr3-106517857-INS-199 |
| chr3-106532403-INS-909 | chr3-106532403-INS-910 |
| chr3-106550722-INS-453 | chr3-106550722-INS-444 |
| chr3-106561545-INS-5389 | chr3-106561546-INS-5389 |
| chr3-106564229-INS-80 | chr3-106564229-INS-80 |
| chr3-106588646-INS-90 | chr3-106588646-INS-90 |
| chr3-106677255-INS-187 | chr3-106677255-INS-187 |

| | |
|---|---|
| chr3-106679713-INS-68 | chr3-106679713-INS-68 |
| chr3-106687839-INS-76 | chr3-106687839-INS-76 |
| chr3-106696232-INS-368 | chr3-106696232-INS-368 |
| chr3-106817831-INS-65 | chr3-106817831-INS-65 |
| chr3-106820779-INS-80 | chr3-106820779-INS-80 |
| chr3-106852027-INS-132 | chr3-106852027-INS-132 |
| chr3-106861262-INS-94 | chr3-106861262-INS-94 |
| chr3-106868902-INS-5317 | chr3-106868903-INS-5317 |
| chr3-106881216-INS-528 | chr3-106881216-INS-529 |
| chr3-106882751-INS-79 | chr3-106882729-INS-63 |
| chr3-106889288-INS-59 | chr3-106889286-INS-64 |
| chr3-106957855-INS-223 | chr3-106957855-INS-223 |
| chr3-106968096-INS-96 | chr3-106968096-INS-90 |
| chr3-106974567-INS-189 | chr3-106974567-INS-189 |
| chr3-106987574-INS-161 | chr3-106987593-INS-152 |
| chr3-107017328-INS-102 | chr3-107017328-INS-102 |
| chr3-107032828-INS-475 | chr3-107032828-INS-475 |
| chr3-107038485-INS-3132 | chr3-107038486-INS-3132 |
| chr3-107039448-INS-104 | chr3-107039448-INS-100 |
| chr3-107041470-INS-85 | chr3-107041470-INS-91 |
| chr3-107045959-INS-81 | chr3-107045959-INS-81 |
| chr3-107048291-INS-2799 | chr3-107048292-INS-2799 |
| chr3-107052328-INS-60 | chr3-107052328-INS-60 |
| chr3-107079648-INS-295 | chr3-107079648-INS-295 |
| chr3-107081151-INS-54 | chr3-107081151-INS-80 |
| chr3-107092905-INS-71 | chr3-107092905-INS-71 |
| chr3-107099251-INS-171 | chr3-107099251-INS-171 |
| chr3-107105130-INS-364 | chr3-107105130-INS-368 |
| chr3-107106565-INS-72 | chr3-107106562-INS-92 |
| chr3-107111845-INS-3735 | chr3-107111846-INS-3755 |
| chr3-107123452-INS-112 | chr3-107123459-INS-91 |
| chr3-107137865-INS-108 | chr3-107137865-INS-108 |
| chr3-107155758-INS-74 | chr3-107155758-INS-78 |
| chr3-107173487-INS-358 | chr3-107173487-INS-358 |
| chr3-107175141-INS-98 | chr3-107175141-INS-105 |
| chr3-107186596-INS-53 | chr3-107186596-INS-73 |
| chr3-107225780-INS-61 | chr3-107225780-INS-64 |
| chr3-107241756-INS-6478 | chr3-107241757-INS-6478 |
| chr3-107251402-INS-52 | chr3-107251402-INS-52 |
| chr3-107262964-INS-135 | chr3-107262998-INS-133 |
| chr3-107274135-INS-245 | chr3-107274115-INS-258 |
| chr3-107276542-INS-275 | chr3-107276542-INS-271 |
| chr3-107287406-INS-117 | chr3-107287406-INS-115 |
| chr3-107291453-INS-236 | chr3-107291453-INS-258 |
| chr3-107297678-INS-308 | chr3-107297679-INS-292 |
| chr3-107333566-INS-1889 | chr3-107333567-INS-1889 |
| chr3-107339224-INS-133 | chr3-107339224-INS-136 |
| chr3-107358356-INS-55 | chr3-107358358-INS-82 |
| chr3-107434632-INS-52 | chr3-107434633-INS-58 |
| chr3-107445132-INS-284 | chr3-107445132-INS-285 |
| chr3-107465710-INS-60 | chr3-107465710-INS-60 |
| chr3-107473489-INS-92 | chr3-107473489-INS-90 |
| chr3-107526832-INS-200 | chr3-107526832-INS-200 |
| chr3-107531205-INS-725 | chr3-107531205-INS-735 |
| chr3-107540968-INS-62 | chr3-107540968-INS-64 |
| chr3-107546234-INS-149 | chr3-107546234-INS-149 |
| chr3-107600071-INS-954 | chr3-107600071-INS-954 |
| chr3-107626657-INS-629 | chr3-107626657-INS-631 |
| chr3-107642671-INS-481 | chr3-107642671-INS-499 |
| chr3-107661562-INS-61 | chr3-107661542-INS-52 |
| chr3-107666772-INS-64 | chr3-107666772-INS-68 |
| chr3-107755262-INS-1434 | chr3-107755263-INS-1434 |
| chr3-107768259-INS-55 | chr3-107768259-INS-53 |
| chr3-107802973-INS-126 | chr3-107802973-INS-121 |
| chr3-107809804-INS-185 | chr3-107809804-INS-186 |
| chr3-107842082-INS-62 | chr3-107842082-INS-60 |
| chr3-107852826-INS-174 | chr3-107852826-INS-174 |
| chr3-107867138-INS-204 | chr3-107867138-INS-204 |
| chr3-107895003-INS-201 | chr3-107895003-INS-201 |
| chr3-107904635-INS-7849 | chr3-107904636-INS-7849 |
| chr3-107914843-INS-80 | chr3-107914843-INS-78 |
| chr3-107917290-INS-91 | chr3-107917289-INS-97 |
| chr3-107918489-INS-328 | chr3-107918489-INS-328 |
| chr3-107931275-INS-221 | chr3-107931275-INS-221 |
| chr3-107934046-INS-141 | chr3-107934046-INS-141 |
| chr3-107944408-INS-504 | chr3-107944408-INS-504 |
| chr3-107955635-INS-71 | chr3-107955635-INS-69 |

| | |
|---|---|
| chr3-107974110-INS-53 | chr3-107974110-INS-53 |
| chr3-107978733-INS-252 | chr3-107978733-INS-252 |
| chr3-108069751-INS-394 | chr3-108069751-INS-394 |
| chr3-108091289-INS-156 | chr3-108091289-INS-156 |
| chr3-108163287-INS-203 | chr3-108163287-INS-203 |
| chr3-108169973-INS-211 | chr3-108169973-INS-209 |
| chr3-108201809-INS-202 | chr3-108201809-INS-202 |
| chr3-108209615-INS-82 | chr3-108209615-INS-82 |
| chr3-108236664-INS-358 | chr3-108236664-INS-353 |
| chr3-108244175-INS-1891 | chr3-108244176-INS-1891 |
| chr3-108269062-INS-60 | chr3-108269062-INS-60 |
| chr3-108323712-INS-54 | chr3-108323712-INS-56 |
| chr3-108328936-INS-122 | chr3-108328942-INS-75 |
| chr3-108391464-INS-95 | chr3-108391464-INS-95 |
| chr3-108412856-INS-177 | chr3-108412856-INS-178 |
| chr3-108414438-INS-77 | chr3-108414438-INS-77 |
| chr3-108420172-INS-59 | chr3-108420172-INS-59 |
| chr3-108421365-INS-56 | chr3-108421365-INS-56 |
| chr3-108444527-INS-54 | chr3-108444509-INS-58 |
| chr3-108464895-INS-397 | chr3-108464895-INS-395 |
| chr3-108490235-INS-202 | chr3-108490235-INS-202 |
| chr3-108503965-INS-2189 | chr3-108503980-INS-2173 |
| chr3-108507924-INS-62 | chr3-108507924-INS-92 |
| chr3-108520519-INS-498 | chr3-108520519-INS-498 |
| chr3-108523499-INS-414 | chr3-108523499-INS-456 |
| chr3-108530161-INS-179 | chr3-108530161-INS-179 |
| chr3-108539480-INS-210 | chr3-108539479-INS-213 |
| chr3-108568291-INS-58 | chr3-108568291-INS-58 |
| chr3-108569537-INS-162 | chr3-108569537-INS-162 |
| chr3-108583786-INS-92 | chr3-108583786-INS-92 |
| chr3-108588321-INS-211 | chr3-108588321-INS-211 |
| chr3-108636948-INS-72 | chr3-108636948-INS-108 |
| chr3-108657420-INS-100 | chr3-108657401-INS-107 |
| chr3-108658935-INS-6881 | chr3-108658936-INS-6883 |
| chr3-108673355-INS-149 | chr3-108673355-INS-149 |
| chr3-108673596-INS-5166 | chr3-108673597-INS-5165 |
| chr3-108674332-INS-315 | chr3-108674333-INS-315 |
| chr3-108691833-INS-402 | chr3-108691833-INS-402 |
| chr3-108695362-INS-80 | chr3-108695362-INS-80 |
| chr3-108702538-INS-212 | chr3-108702538-INS-212 |
| chr3-108714962-INS-198 | chr3-108714963-INS-226 |
| chr3-108733812-INS-98 | chr3-108733819-INS-88 |
| chr3-108740497-INS-205 | chr3-108740497-INS-205 |
| chr3-108756851-INS-104 | chr3-108756851-INS-72 |
| chr3-108769449-INS-165 | chr3-108769449-INS-165 |
| chr3-108774771-INS-305 | chr3-108774771-INS-306 |
| chr3-108780758-INS-66 | chr3-108780758-INS-66 |
| chr3-108799540-INS-264 | chr3-108799540-INS-264 |
| chr3-108814888-INS-210 | chr3-108814888-INS-210 |
| chr3-108816315-INS-215 | chr3-108816315-INS-215 |
| chr3-108820793-INS-181 | chr3-108820793-INS-182 |
| chr3-108824138-INS-54 | chr3-108824118-INS-56 |
| chr3-108832749-INS-343 | chr3-108832749-INS-352 |
| chr3-108835813-INS-82 | chr3-108835792-INS-96 |
| chr3-108860941-INS-61 | chr3-108860941-INS-65 |
| chr3-108862278-INS-131 | chr3-108862278-INS-132 |
| chr3-108864680-INS-152 | chr3-108864680-INS-128 |
| chr3-108866923-INS-76 | chr3-108866923-INS-76 |
| chr3-108888939-INS-367 | chr3-108888947-INS-364 |
| chr3-108906979-INS-1344 | chr3-108906980-INS-1344 |
| chr3-108915746-INS-204 | chr3-108915746-INS-204 |
| chr3-108916190-INS-1655 | chr3-108916191-INS-1656 |
| chr3-108929346-INS-56 | chr3-108929346-INS-64 |
| chr3-108966421-INS-202 | chr3-108966421-INS-202 |
| chr3-108977997-INS-603 | chr3-108977997-INS-603 |
| chr3-109012812-INS-68 | chr3-109012813-INS-100 |
| chr3-109014011-INS-6334 | chr3-109014012-INS-6334 |
| chr3-109027117-INS-168 | chr3-109027117-INS-168 |
| chr3-109065519-INS-1883 | chr3-109065520-INS-1887 |
| chr3-109094153-INS-64 | chr3-109094187-INS-50 |
| chr3-109116924-INS-59 | chr3-109116924-INS-57 |
| chr3-109126982-INS-872 | chr3-109126982-INS-872 |
| chr3-109129474-INS-401 | chr3-109129474-INS-401 |
| chr3-109132655-INS-52 | chr3-109132652-INS-60 |
| chr3-109138588-INS-67 | chr3-109138589-INS-68 |
| chr3-109145121-INS-80 | chr3-109145121-INS-80 |
| chr3-109165651-INS-204 | chr3-109165651-INS-204 |

| | |
|---|---|
| chr3-109181588-INS-7240 | chr3-109181589-INS-7241 |
| chr3-109209208-INS-1169 | chr3-109209209-INS-1169 |
| chr3-109211891-INS-377 | chr3-109211891-INS-377 |
| chr3-109226000-INS-2403 | chr3-109226001-INS-2404 |
| chr3-109229500-INS-139 | chr3-109229500-INS-139 |
| chr3-109229863-INS-593 | chr3-109229863-INS-593 |
| chr3-109231074-INS-168 | chr3-109231074-INS-168 |
| chr3-109267268-INS-176 | chr3-109267268-INS-176 |
| chr3-109278925-INS-150 | chr3-109278925-INS-105 |
| chr3-109281890-INS-174 | chr3-109281890-INS-175 |
| chr3-109290063-INS-84 | chr3-109290063-INS-84 |
| chr3-109302668-INS-61 | chr3-109302666-INS-61 |
| chr3-109309945-INS-288 | chr3-109309945-INS-288 |
| chr3-109314503-INS-56 | chr3-109314503-INS-58 |
| chr3-109365584-INS-195 | chr3-109365584-INS-195 |
| chr3-109379438-INS-218 | chr3-109379438-INS-220 |
| chr3-109393329-INS-60 | chr3-109393329-INS-54 |
| chr3-109417935-INS-56 | chr3-109417941-INS-62 |
| chr3-109433495-INS-221 | chr3-109433495-INS-219 |
| chr3-109450091-INS-56 | chr3-109450091-INS-74 |
| chr3-109460350-INS-142 | chr3-109460350-INS-98 |
| chr3-109463532-INS-53 | chr3-109463527-INS-53 |
| chr3-109497034-INS-6303 | chr3-109497035-INS-6302 |
| chr3-109502437-INS-89 | chr3-109502438-INS-89 |
| chr3-109514094-INS-1560 | chr3-109514095-INS-1564 |
| chr3-109546232-INS-67 | chr3-109546232-INS-67 |
| chr3-109561687-INS-63 | chr3-109561687-INS-63 |
| chr3-109567537-INS-74 | chr3-109567531-INS-80 |
| chr3-109576159-INS-62 | chr3-109576159-INS-62 |
| chr3-109626887-INS-62 | chr3-109626887-INS-62 |
| chr3-109631359-INS-437 | chr3-109631359-INS-480 |
| chr3-109640457-INS-62 | chr3-109640457-INS-62 |
| chr3-109653317-INS-168 | chr3-109653317-INS-169 |
| chr3-109678334-INS-100 | chr3-109678334-INS-132 |
| chr3-109705302-INS-123 | chr3-109705302-INS-123 |
| chr3-109716385-INS-74 | chr3-109716369-INS-68 |
| chr3-109732089-INS-1783 | chr3-109732090-INS-1781 |
| chr3-109748327-INS-52 | chr3-109748327-INS-54 |
| chr3-109768397-INS-62 | chr3-109768397-INS-72 |
| chr3-109776538-INS-98 | chr3-109776538-INS-98 |
| chr3-109795880-INS-307 | chr3-109795880-INS-307 |
| chr3-109818037-INS-70 | chr3-109818037-INS-70 |
| chr3-109892712-INS-82 | chr3-109892704-INS-66 |
| chr3-109904265-INS-6684 | chr3-109904266-INS-6686 |
| chr3-109912883-INS-52 | chr3-109912883-INS-54 |
| chr3-109931722-INS-57 | chr3-109931722-INS-52 |
| chr3-109952777-INS-70 | chr3-109952778-INS-72 |
| chr3-109979951-INS-52 | chr3-109979949-INS-54 |
| chr3-110003282-INS-452 | chr3-110003281-INS-459 |
| chr3-110033539-INS-58 | chr3-110033545-INS-64 |
| chr3-110062130-INS-84 | chr3-110062130-INS-84 |
| chr3-110082907-INS-194 | chr3-110082907-INS-209 |
| chr3-110096781-INS-1037 | chr3-110096782-INS-1006 |
| chr3-110101193-INS-4749 | chr3-110101194-INS-4749 |
| chr3-110116640-INS-208 | chr3-110116640-INS-208 |
| chr3-110133594-INS-90 | chr3-110133594-INS-90 |
| chr3-110197922-INS-388 | chr3-110197922-INS-389 |
| chr3-110218801-INS-126 | chr3-110218801-INS-126 |
| chr3-110223317-INS-70 | chr3-110223310-INS-55 |
| chr3-110260742-INS-7198 | chr3-110260743-INS-7198 |
| chr3-110332823-INS-1472 | chr3-110332824-INS-1493 |
| chr3-110360149-INS-78 | chr3-110360149-INS-78 |
| chr3-110363442-INS-4829 | chr3-110363443-INS-4829 |
| chr3-110374866-INS-6157 | chr3-110374867-INS-6155 |
| chr3-110428709-INS-600 | chr3-110428709-INS-600 |
| chr3-110430954-INS-54 | chr3-110430954-INS-58 |
| chr3-110449619-INS-145 | chr3-110449619-INS-145 |
| chr3-110451359-INS-61 | chr3-110451359-INS-93 |
| chr3-110455399-INS-54 | chr3-110455398-INS-67 |
| chr3-110457305-INS-829 | chr3-110457305-INS-845 |
| chr3-110473583-INS-1578 | chr3-110473584-INS-1578 |
| chr3-110477089-INS-65 | chr3-110477089-INS-65 |
| chr3-110478451-INS-479 | chr3-110478451-INS-480 |
| chr3-110496354-INS-79 | chr3-110496354-INS-83 |
| chr3-110497392-INS-6883 | chr3-110497393-INS-6900 |
| chr3-110519522-INS-2109 | chr3-110519530-INS-2107 |
| chr3-110524622-INS-167 | chr3-110524622-INS-167 |

| | |
|---|---|
| chr3-110534484-INS-103 | chr3-110534483-INS-95 |
| chr3-110545205-INS-120 | chr3-110545207-INS-94 |
| chr3-110559320-INS-64 | chr3-110559322-INS-72 |
| chr3-110570843-INS-61 | chr3-110570872-INS-60 |
| chr3-110626878-INS-54 | chr3-110626878-INS-60 |
| chr3-110629728-INS-140 | chr3-110629679-INS-159 |
| chr3-110632031-INS-5160 | chr3-110632032-INS-5160 |
| chr3-110632032-INS-417 | chr3-110632032-INS-421 |
| chr3-110634909-INS-70 | chr3-110634902-INS-70 |
| chr3-110704504-INS-709 | chr3-110704504-INS-711 |
| chr3-110723217-INS-1072 | chr3-110723218-INS-1074 |
| chr3-110724479-INS-259 | chr3-110724479-INS-259 |
| chr3-110751145-INS-3041 | chr3-110751146-INS-3041 |
| chr3-110755437-INS-2398 | chr3-110755438-INS-2398 |
| chr3-110758473-INS-230 | chr3-110758473-INS-230 |
| chr3-110786481-INS-54 | chr3-110786481-INS-102 |
| chr3-110818854-INS-52 | chr3-110818852-INS-54 |
| chr3-110828943-INS-196 | chr3-110828943-INS-196 |
| chr3-110873644-INS-388 | chr3-110873644-INS-388 |
| chr3-110888053-INS-217 | chr3-110888053-INS-218 |
| chr3-110921043-INS-69 | chr3-110921043-INS-64 |
| chr3-110934270-INS-687 | chr3-110934270-INS-687 |
| chr3-110943861-INS-204 | chr3-110943861-INS-205 |
| chr3-110945446-INS-714 | chr3-110945446-INS-714 |
| chr3-110969531-INS-457 | chr3-110969526-INS-457 |
| chr3-110995807-INS-479 | chr3-110995807-INS-479 |
| chr3-110996205-INS-3073 | chr3-110996206-INS-3073 |
| chr3-111018349-INS-107 | chr3-111018349-INS-111 |
| chr3-111022142-INS-68 | chr3-111022138-INS-65 |
| chr3-111053800-INS-399 | chr3-111053800-INS-399 |
| chr3-111058078-INS-56 | chr3-111058078-INS-56 |
| chr3-111085705-INS-99 | chr3-111085735-INS-51 |
| chr3-111095326-INS-52 | chr3-111095316-INS-66 |
| chr3-111117086-INS-997 | chr3-111117074-INS-1008 |
| chr3-111124824-INS-536 | chr3-111124824-INS-536 |
| chr3-111151335-INS-787 | chr3-111151335-INS-787 |
| chr3-111154046-INS-144 | chr3-111154046-INS-144 |
| chr3-111180269-INS-133 | chr3-111180303-INS-145 |
| chr3-111210357-INS-6402 | chr3-111210358-INS-6402 |
| chr3-111225034-INS-100 | chr3-111225040-INS-83 |
| chr3-111266213-INS-75 | chr3-111266213-INS-59 |
| chr3-111266486-INS-7136 | chr3-111266487-INS-7138 |
| chr3-111276531-INS-112 | chr3-111276531-INS-146 |
| chr3-111288807-INS-53 | chr3-111288807-INS-53 |
| chr3-111291602-INS-1573 | chr3-111291603-INS-1573 |
| chr3-111311012-INS-282 | chr3-111311012-INS-282 |
| chr3-111312222-INS-5783 | chr3-111312223-INS-5783 |
| chr3-111345213-INS-62 | chr3-111345209-INS-50 |
| chr3-111349849-INS-68 | chr3-111349849-INS-68 |
| chr3-111368643-INS-55 | chr3-111368643-INS-55 |
| chr3-111410332-INS-148 | chr3-111410332-INS-148 |
| chr3-111435535-INS-55 | chr3-111435535-INS-55 |
| chr3-111446304-INS-310 | chr3-111446304-INS-310 |
| chr3-111468444-INS-233 | chr3-111468444-INS-233 |
| chr3-111520546-INS-63 | chr3-111520546-INS-63 |
| chr3-111525874-INS-55 | chr3-111525874-INS-55 |
| chr3-111528546-INS-4347 | chr3-111528547-INS-4347 |
| chr3-111545514-INS-156 | chr3-111545514-INS-121 |
| chr3-111563672-INS-72 | chr3-111563672-INS-54 |
| chr3-111564856-INS-1607 | chr3-111564857-INS-1607 |
| chr3-111581429-INS-197 | chr3-111581429-INS-197 |
| chr3-111604666-INS-52 | chr3-111604646-INS-50 |
| chr3-111643906-INS-299 | chr3-111643906-INS-300 |
| chr3-111663497-INS-657 | chr3-111663495-INS-694 |
| chr3-111684910-INS-111 | chr3-111684910-INS-115 |
| chr3-111691936-INS-110 | chr3-111691936-INS-110 |
| chr3-111711121-INS-196 | chr3-111711121-INS-196 |
| chr3-111728847-INS-100 | chr3-111728847-INS-80 |
| chr3-111733243-INS-6440 | chr3-111733244-INS-6440 |
| chr3-111735260-INS-94 | chr3-111735260-INS-94 |
| chr3-111770426-INS-1304 | chr3-111770427-INS-1304 |
| chr3-111779693-INS-392 | chr3-111779693-INS-392 |
| chr3-111781843-INS-7236 | chr3-111781844-INS-7240 |
| chr3-111799641-INS-52 | chr3-111799624-INS-58 |
| chr3-111811867-INS-560 | chr3-111811867-INS-560 |
| chr3-111815439-INS-199 | chr3-111815416-INS-199 |
| chr3-111818008-INS-429 | chr3-111818008-INS-429 |

| | |
|---|---|
| chr3-111860660-INS-1285 | chr3-111860661-INS-1284 |
| chr3-111893811-INS-132 | chr3-111893815-INS-135 |
| chr3-111990097-INS-188 | chr3-111990097-INS-188 |
| chr3-111997926-INS-71 | chr3-111997926-INS-71 |
| chr3-112009943-INS-72 | chr3-112009943-INS-74 |
| chr3-112019200-INS-797 | chr3-112019200-INS-797 |
| chr3-112042253-INS-322 | chr3-112042253-INS-322 |
| chr3-112104250-INS-108 | chr3-112104253-INS-109 |
| chr3-112106357-INS-367 | chr3-112106357-INS-367 |
| chr3-112137110-INS-6384 | chr3-112137111-INS-6384 |
| chr3-112141623-INS-198 | chr3-112141623-INS-198 |
| chr3-112153860-INS-172 | chr3-112153860-INS-172 |
| chr3-112156747-INS-457 | chr3-112156747-INS-457 |
| chr3-112170261-INS-357 | chr3-112170261-INS-357 |
| chr3-112199418-INS-64 | chr3-112199418-INS-66 |
| chr3-112205605-INS-52 | chr3-112205605-INS-54 |
| chr3-112214742-INS-925 | chr3-112214742-INS-924 |
| chr3-112259222-INS-988 | chr3-112259222-INS-989 |
| chr3-112262741-INS-445 | chr3-112262741-INS-445 |
| chr3-112388950-INS-1202 | chr3-112388951-INS-1202 |
| chr3-112395366-INS-86 | chr3-112395366-INS-86 |
| chr3-112403068-INS-53 | chr3-112403068-INS-53 |
| chr3-112411902-INS-7050 | chr3-112411903-INS-7050 |
| chr3-112433843-INS-52 | chr3-112433832-INS-52 |
| chr3-112440668-INS-139 | chr3-112440668-INS-139 |
| chr3-112462967-INS-415 | chr3-112462964-INS-418 |
| chr3-112479661-INS-90 | chr3-112479661-INS-90 |
| chr3-112488311-INS-110 | chr3-112488311-INS-110 |
| chr3-112508903-INS-605 | chr3-112508903-INS-599 |
| chr3-112509475-INS-54 | chr3-112509475-INS-54 |
| chr3-112519860-INS-65 | chr3-112519860-INS-65 |
| chr3-112539591-INS-54 | chr3-112539591-INS-54 |
| chr3-112552876-INS-52 | chr3-112552876-INS-50 |
| chr3-112591561-INS-53 | chr3-112591561-INS-78 |
| chr3-112619538-INS-3514 | chr3-112619539-INS-3514 |
| chr3-112624214-INS-4938 | chr3-112624215-INS-4938 |
| chr3-112635659-INS-2884 | chr3-112635660-INS-2884 |
| chr3-112655525-INS-3836 | chr3-112655526-INS-3836 |
| chr3-112664529-INS-50 | chr3-112664529-INS-72 |
| chr3-112699917-INS-326 | chr3-112699917-INS-326 |
| chr3-112708626-INS-6573 | chr3-112708627-INS-6573 |
| chr3-112721868-INS-54 | chr3-112721868-INS-54 |
| chr3-112741009-INS-84 | chr3-112741012-INS-85 |
| chr3-112775619-INS-724 | chr3-112775579-INS-724 |
| chr3-112807648-INS-66 | chr3-112807648-INS-66 |
| chr3-112827305-INS-53 | chr3-112827305-INS-53 |
| chr3-112831290-INS-1888 | chr3-112831291-INS-1888 |
| chr3-112844831-INS-89 | chr3-112844831-INS-89 |
| chr3-112862901-INS-5229 | chr3-112862902-INS-5229 |
| chr3-112903435-INS-56 | chr3-112903436-INS-68 |
| chr3-112905259-INS-396 | chr3-112905259-INS-398 |
| chr3-112923805-INS-54 | chr3-112923805-INS-54 |
| chr3-112929801-INS-720 | chr3-112929799-INS-723 |
| chr3-112940560-INS-227 | chr3-112940560-INS-226 |
| chr3-112953659-INS-53 | chr3-112953627-INS-85 |
| chr3-112978136-INS-1828 | chr3-112978137-INS-1832 |
| chr3-112997747-INS-3702 | chr3-112997748-INS-3702 |
| chr3-113011071-INS-6364 | chr3-113011072-INS-6364 |
| chr3-113030253-INS-80 | chr3-113030253-INS-72 |
| chr3-113036253-INS-1285 | chr3-113036254-INS-1286 |
| chr3-113037821-INS-218 | chr3-113037810-INS-216 |
| chr3-113047088-INS-89 | chr3-113047088-INS-97 |
| chr3-113053382-INS-64 | chr3-113053382-INS-66 |
| chr3-113062716-INS-208 | chr3-113062716-INS-208 |
| chr3-113075985-INS-88 | chr3-113075985-INS-93 |
| chr3-113349981-INS-124 | chr3-113349981-INS-124 |
| chr3-113361945-INS-2027 | chr3-113361946-INS-2027 |
| chr3-113383661-INS-137 | chr3-113383661-INS-138 |
| chr3-113471600-INS-328 | chr3-113471600-INS-313 |
| chr3-113473913-INS-53 | chr3-113473913-INS-53 |
| chr3-113491320-INS-811 | chr3-113491320-INS-812 |
| chr3-113506534-INS-643 | chr3-113506552-INS-666 |
| chr3-113531723-INS-207 | chr3-113531723-INS-207 |
| chr3-113558838-INS-98 | chr3-113558838-INS-98 |
| chr3-113570326-INS-351 | chr3-113570326-INS-342 |
| chr3-113576433-INS-95 | chr3-113576433-INS-72 |
| chr3-113609298-INS-373 | chr3-113609298-INS-373 |

| | |
|---|---|
| chr3-113609908-INS-61 | chr3-113609909-INS-61 |
| chr3-113610337-INS-1446 | chr3-113610338-INS-1446 |
| chr3-113611122-INS-1889 | chr3-113611123-INS-1889 |
| chr3-113640151-INS-640 | chr3-113640151-INS-640 |
| chr3-113641297-INS-66 | chr3-113641297-INS-66 |
| chr3-113650849-INS-301 | chr3-113650849-INS-301 |
| chr3-113675357-INS-72 | chr3-113675355-INS-72 |
| chr3-113691877-INS-215 | chr3-113691877-INS-212 |
| chr3-113723115-INS-1277 | chr3-113723116-INS-1277 |
| chr3-113732999-INS-53 | chr3-113732999-INS-53 |
| chr3-113738916-INS-319 | chr3-113738916-INS-320 |
| chr3-113750188-INS-55 | chr3-113750188-INS-60 |
| chr3-113751906-INS-2639 | chr3-113751907-INS-2639 |
| chr3-113751907-INS-411 | chr3-113751907-INS-411 |
| chr3-113806576-INS-487 | chr3-113806576-INS-487 |
| chr3-113813913-INS-161 | chr3-113813913-INS-161 |
| chr3-113837006-INS-161 | chr3-113836994-INS-152 |
| chr3-113839195-INS-64 | chr3-113839195-INS-64 |
| chr3-113849462-INS-262 | chr3-113849461-INS-256 |
| chr3-113880004-INS-109 | chr3-113880004-INS-110 |
| chr3-113882181-INS-172 | chr3-113882181-INS-172 |
| chr3-113884376-INS-204 | chr3-113884376-INS-204 |
| chr3-113887024-INS-54 | chr3-113887024-INS-52 |
| chr3-113946928-INS-56 | chr3-113946928-INS-54 |
| chr3-114001064-INS-187 | chr3-114001064-INS-187 |
| chr3-114004416-INS-274 | chr3-114004398-INS-276 |
| chr3-114025416-INS-72 | chr3-114025416-INS-66 |
| chr3-114029588-INS-65 | chr3-114029601-INS-76 |
| chr3-114045026-INS-60 | chr3-114045026-INS-61 |
| chr3-114047063-INS-158 | chr3-114047063-INS-158 |
| chr3-114076599-INS-78 | chr3-114076592-INS-72 |
| chr3-114085282-INS-246 | chr3-114085282-INS-249 |
| chr3-114088148-INS-60 | chr3-114088166-INS-80 |
| chr3-114117027-INS-62 | chr3-114117027-INS-54 |
| chr3-114135570-INS-146 | chr3-114135570-INS-146 |
| chr3-114174312-INS-262 | chr3-114174312-INS-258 |
| chr3-114176208-INS-88 | chr3-114176208-INS-92 |
| chr3-114187036-INS-338 | chr3-114187036-INS-338 |
| chr3-114197547-INS-110 | chr3-114197547-INS-111 |
| chr3-114208832-INS-94 | chr3-114208832-INS-96 |
| chr3-114230667-INS-656 | chr3-114230667-INS-654 |
| chr3-114240078-INS-418 | chr3-114240078-INS-418 |
| chr3-114250676-INS-6646 | chr3-114250677-INS-6646 |
| chr3-114280486-INS-234 | chr3-114280486-INS-234 |
| chr3-114306379-INS-146 | chr3-114306379-INS-126 |
| chr3-114308507-INS-442 | chr3-114308507-INS-442 |
| chr3-114315567-INS-129 | chr3-114315593-INS-112 |
| chr3-114319868-INS-128 | chr3-114319868-INS-128 |
| chr3-114341140-INS-1077 | chr3-114341141-INS-1091 |
| chr3-114341476-INS-121 | chr3-114341465-INS-143 |
| chr3-114355947-INS-221 | chr3-114355947-INS-222 |
| chr3-114371454-INS-568 | chr3-114371454-INS-568 |
| chr3-114394597-INS-98 | chr3-114394597-INS-98 |
| chr3-114399145-INS-826 | chr3-114399145-INS-826 |
| chr3-114414642-INS-173 | chr3-114414642-INS-173 |
| chr3-114434827-INS-6746 | chr3-114434828-INS-6747 |
| chr3-114438212-INS-445 | chr3-114438212-INS-445 |
| chr3-114448549-INS-310 | chr3-114448549-INS-310 |
| chr3-114456275-INS-211 | chr3-114456275-INS-211 |
| chr3-114475222-INS-657 | chr3-114475222-INS-638 |
| chr3-114585151-INS-6565 | chr3-114585152-INS-6565 |
| chr3-114590798-INS-58 | chr3-114590798-INS-60 |
| chr3-114604009-INS-57 | chr3-114604009-INS-57 |
| chr3-114668179-INS-58 | chr3-114668179-INS-62 |
| chr3-114718186-INS-715 | chr3-114718186-INS-717 |
| chr3-114743340-INS-56 | chr3-114743340-INS-50 |
| chr3-114763024-INS-6715 | chr3-114763025-INS-6715 |
| chr3-114786220-INS-71 | chr3-114786209-INS-85 |
| chr3-114805733-INS-270 | chr3-114805733-INS-270 |
| chr3-114826134-INS-4572 | chr3-114826135-INS-4572 |
| chr3-114827510-INS-258 | chr3-114827510-INS-275 |
| chr3-114833114-INS-306 | chr3-114833114-INS-304 |
| chr3-114843552-INS-56 | chr3-114843552-INS-56 |
| chr3-114867626-INS-1532 | chr3-114867627-INS-1531 |
| chr3-114870460-INS-221 | chr3-114870460-INS-221 |
| chr3-114871067-INS-52 | chr3-114871067-INS-54 |
| chr3-114875325-INS-161 | chr3-114875325-INS-162 |

| | |
|---|---|
| chr3-114897020-INS-6065 | chr3-114897021-INS-6061 |
| chr3-114950940-INS-1335 | chr3-114950941-INS-1336 |
| chr3-114968475-INS-480 | chr3-114968475-INS-483 |
| chr3-114978901-INS-220 | chr3-114978901-INS-220 |
| chr3-115012819-INS-612 | chr3-115012819-INS-612 |
| chr3-115019435-INS-197 | chr3-115019435-INS-197 |
| chr3-115022487-INS-6431 | chr3-115022488-INS-6431 |
| chr3-115034009-INS-2464 | chr3-115034010-INS-2464 |
| chr3-115050385-INS-6799 | chr3-115050386-INS-6799 |
| chr3-115057202-INS-786 | chr3-115057202-INS-785 |
| chr3-115062179-INS-421 | chr3-115062179-INS-421 |
| chr3-115100635-INS-79 | chr3-115100635-INS-97 |
| chr3-115109921-INS-234 | chr3-115109921-INS-234 |
| chr3-115121693-INS-122 | chr3-115121688-INS-128 |
| chr3-115134935-INS-98 | chr3-115134944-INS-56 |
| chr3-115169531-INS-271 | chr3-115169531-INS-271 |
| chr3-115194466-INS-136 | chr3-115194466-INS-112 |
| chr3-115195562-INS-181 | chr3-115195562-INS-180 |
| chr3-115197939-INS-413 | chr3-115197939-INS-413 |
| chr3-115207335-INS-3661 | chr3-115207336-INS-3663 |
| chr3-115226905-INS-72 | chr3-115226905-INS-91 |
| chr3-115264128-INS-58 | chr3-115264108-INS-68 |
| chr3-115272142-INS-140 | chr3-115272142-INS-140 |
| chr3-115276064-INS-827 | chr3-115276064-INS-827 |
| chr3-115287331-INS-919 | chr3-115287329-INS-933 |
| chr3-115305679-INS-60 | chr3-115305679-INS-60 |
| chr3-115312437-INS-416 | chr3-115312437-INS-416 |
| chr3-115328089-INS-271 | chr3-115328089-INS-271 |
| chr3-115333000-INS-88 | chr3-115333015-INS-84 |
| chr3-115346709-INS-57 | chr3-115346709-INS-57 |
| chr3-115356674-INS-93 | chr3-115356675-INS-72 |
| chr3-115365791-INS-4953 | chr3-115365796-INS-4951 |
| chr3-115391321-INS-840 | chr3-115391321-INS-843 |
| chr3-115409245-INS-6421 | chr3-115409246-INS-6423 |
| chr3-115436929-INS-170 | chr3-115436929-INS-170 |
| chr3-115492089-INS-52 | chr3-115492089-INS-68 |
| chr3-115512346-INS-85 | chr3-115512346-INS-85 |
| chr3-115523685-INS-110 | chr3-115523683-INS-112 |
| chr3-115530863-INS-51 | chr3-115530864-INS-84 |
| chr3-115535000-INS-344 | chr3-115535000-INS-344 |
| chr3-115550502-INS-433 | chr3-115550502-INS-430 |
| chr3-115552399-INS-322 | chr3-115552399-INS-322 |
| chr3-115581961-INS-74 | chr3-115581943-INS-79 |
| chr3-115611739-INS-60 | chr3-115611739-INS-62 |
| chr3-115626564-INS-64 | chr3-115626564-INS-60 |
| chr3-115635904-INS-60 | chr3-115635904-INS-62 |
| chr3-115644307-INS-6758 | chr3-115644308-INS-6758 |
| chr3-115647071-INS-207 | chr3-115647071-INS-207 |
| chr3-115746864-INS-117 | chr3-115746876-INS-115 |
| chr3-115748953-INS-180 | chr3-115748953-INS-180 |
| chr3-115780811-INS-68 | chr3-115780811-INS-68 |
| chr3-115787332-INS-54 | chr3-115787332-INS-54 |
| chr3-115818283-INS-151 | chr3-115818282-INS-151 |
| chr3-115862331-INS-400 | chr3-115862331-INS-399 |
| chr3-115874538-INS-447 | chr3-115874538-INS-447 |
| chr3-115891911-INS-65 | chr3-115891911-INS-69 |
| chr3-115895443-INS-399 | chr3-115895443-INS-399 |
| chr3-115931292-INS-100 | chr3-115931292-INS-98 |
| chr3-115946661-INS-2657 | chr3-115946662-INS-2705 |
| chr3-115949657-INS-2678 | chr3-115949658-INS-2678 |
| chr3-115953728-INS-69 | chr3-115953728-INS-69 |
| chr3-115957892-INS-1892 | chr3-115957893-INS-1892 |
| chr3-115968732-INS-6968 | chr3-115968733-INS-6975 |
| chr3-116023901-INS-1557 | chr3-116023887-INS-1553 |
| chr3-116069787-INS-57 | chr3-116069818-INS-65 |
| chr3-116081528-INS-238 | chr3-116081528-INS-238 |
| chr3-116099094-INS-278 | chr3-116099098-INS-278 |
| chr3-116102859-INS-121 | chr3-116102859-INS-121 |
| chr3-116110087-INS-118 | chr3-116110087-INS-119 |
| chr3-116110903-INS-197 | chr3-116110903-INS-197 |
| chr3-116135698-INS-187 | chr3-116135698-INS-187 |
| chr3-116159283-INS-671 | chr3-116159283-INS-671 |
| chr3-116214107-INS-92 | chr3-116214107-INS-108 |
| chr3-116214608-INS-147 | chr3-116214608-INS-147 |
| chr3-116227326-INS-60 | chr3-116227326-INS-60 |
| chr3-116230158-INS-342 | chr3-116230153-INS-348 |
| chr3-116244812-INS-208 | chr3-116244812-INS-208 |

| | |
|---|---|
| chr3-116246264-INS-202 | chr3-116246264-INS-203 |
| chr3-116274829-INS-52 | chr3-116274829-INS-52 |
| chr3-116275837-INS-7232 | chr3-116275838-INS-7232 |
| chr3-116275954-INS-188 | chr3-116275954-INS-188 |
| chr3-116285065-INS-192 | chr3-116285065-INS-192 |
| chr3-116298568-INS-84 | chr3-116298568-INS-84 |
| chr3-116317794-INS-197 | chr3-116317794-INS-195 |
| chr3-116324344-INS-52 | chr3-116324344-INS-52 |
| chr3-116332542-INS-76 | chr3-116332542-INS-78 |
| chr3-116357698-INS-238 | chr3-116357698-INS-243 |
| chr3-116365894-INS-181 | chr3-116365894-INS-181 |
| chr3-116373100-INS-413 | chr3-116373100-INS-414 |
| chr3-116377661-INS-200 | chr3-116377661-INS-200 |
| chr3-116382918-INS-451 | chr3-116382918-INS-452 |
| chr3-116403435-INS-172 | chr3-116403442-INS-177 |
| chr3-116416272-INS-182 | chr3-116416272-INS-176 |
| chr3-116418365-INS-97 | chr3-116418375-INS-134 |
| chr3-116436677-INS-196 | chr3-116436677-INS-196 |
| chr3-116466633-INS-54 | chr3-116466637-INS-70 |
| chr3-116471922-INS-169 | chr3-116471922-INS-169 |
| chr3-116475473-INS-334 | chr3-116475473-INS-335 |
| chr3-116479928-INS-407 | chr3-116479928-INS-405 |
| chr3-116486863-INS-126 | chr3-116486863-INS-126 |
| chr3-116495103-INS-197 | chr3-116495103-INS-197 |
| chr3-116502755-INS-90 | chr3-116502755-INS-96 |
| chr3-116515694-INS-211 | chr3-116515694-INS-212 |
| chr3-116516528-INS-56 | chr3-116516528-INS-68 |
| chr3-116520314-INS-72 | chr3-116520314-INS-66 |
| chr3-116521651-INS-171 | chr3-116521651-INS-189 |
| chr3-116543764-INS-65 | chr3-116543759-INS-53 |
| chr3-116581863-INS-214 | chr3-116581863-INS-214 |
| chr3-116618225-INS-149 | chr3-116618225-INS-149 |
| chr3-116627055-INS-56 | chr3-116627048-INS-56 |
| chr3-116643916-INS-65 | chr3-116643924-INS-69 |
| chr3-116663198-INS-72 | chr3-116663199-INS-72 |
| chr3-116667055-INS-196 | chr3-116667055-INS-196 |
| chr3-116674080-INS-52 | chr3-116674080-INS-53 |
| chr3-116684903-INS-523 | chr3-116684903-INS-523 |
| chr3-116693722-INS-430 | chr3-116693722-INS-427 |
| chr3-116695833-INS-151 | chr3-116695833-INS-151 |
| chr3-116707203-INS-66 | chr3-116707195-INS-74 |
| chr3-116714774-INS-96 | chr3-116714774-INS-96 |
| chr3-116724776-INS-138 | chr3-116724782-INS-126 |
| chr3-116733917-INS-98 | chr3-116733938-INS-110 |
| chr3-116738271-INS-101 | chr3-116738271-INS-101 |
| chr3-116753006-INS-274 | chr3-116753006-INS-275 |
| chr3-116764991-INS-586 | chr3-116764991-INS-561 |
| chr3-116772168-INS-4353 | chr3-116772169-INS-4353 |
| chr3-116772679-INS-76 | chr3-116772681-INS-72 |
| chr3-116784431-INS-64 | chr3-116784431-INS-62 |
| chr3-116786454-INS-90 | chr3-116786451-INS-83 |
| chr3-116787677-INS-85 | chr3-116787677-INS-64 |
| chr3-116791935-INS-179 | chr3-116791942-INS-180 |
| chr3-116798880-INS-377 | chr3-116798880-INS-378 |
| chr3-116820251-INS-554 | chr3-116820251-INS-554 |
| chr3-116865641-INS-657 | chr3-116865641-INS-655 |
| chr3-116873099-INS-6392 | chr3-116873100-INS-6392 |
| chr3-116874908-INS-75 | chr3-116874908-INS-78 |
| chr3-116906279-INS-6228 | chr3-116906280-INS-6228 |
| chr3-116908491-INS-409 | chr3-116908491-INS-409 |
| chr3-116911798-INS-6916 | chr3-116911799-INS-6918 |
| chr3-116926591-INS-116 | chr3-116926592-INS-140 |
| chr3-116927515-INS-114 | chr3-116927515-INS-114 |
| chr3-116936318-INS-3211 | chr3-116936319-INS-3211 |
| chr3-116953643-INS-248 | chr3-116953643-INS-250 |
| chr3-116954588-INS-1138 | chr3-116954589-INS-1138 |
| chr3-116964221-INS-78 | chr3-116964221-INS-119 |
| chr3-116975335-INS-151 | chr3-116975335-INS-151 |
| chr3-116979489-INS-6595 | chr3-116979490-INS-6595 |
| chr3-116986270-INS-51 | chr3-116986277-INS-78 |
| chr3-117041004-INS-126 | chr3-117041004-INS-125 |
| chr3-117046382-INS-60 | chr3-117046382-INS-60 |
| chr3-117051852-INS-199 | chr3-117051852-INS-199 |
| chr3-117057000-INS-99 | chr3-117057000-INS-78 |
| chr3-117093644-INS-54 | chr3-117093644-INS-60 |
| chr3-117109264-INS-51 | chr3-117109266-INS-59 |
| chr3-117117560-INS-952 | chr3-117117560-INS-930 |

| | |
|---|---|
| chr3-117131317-INS-107 | chr3-117131317-INS-107 |
| chr3-117139306-INS-552 | chr3-117139306-INS-552 |
| chr3-117154446-INS-102 | chr3-117154446-INS-99 |
| chr3-117171515-INS-58 | chr3-117171515-INS-58 |
| chr3-117187558-INS-54 | chr3-117187558-INS-54 |
| chr3-117204832-INS-136 | chr3-117204843-INS-134 |
| chr3-117213675-INS-300 | chr3-117213675-INS-300 |
| chr3-117233613-INS-239 | chr3-117233613-INS-239 |
| chr3-117234557-INS-155 | chr3-117234557-INS-110 |
| chr3-117252425-INS-76 | chr3-117252420-INS-96 |
| chr3-117272205-INS-6487 | chr3-117272206-INS-6487 |
| chr3-117283461-INS-92 | chr3-117283461-INS-92 |
| chr3-117292033-INS-94 | chr3-117292033-INS-104 |
| chr3-117293283-INS-90 | chr3-117293289-INS-90 |
| chr3-117310696-INS-169 | chr3-117310696-INS-171 |
| chr3-117338855-INS-666 | chr3-117338855-INS-666 |
| chr3-117358556-INS-140 | chr3-117358556-INS-140 |
| chr3-117374330-INS-102 | chr3-117374330-INS-107 |
| chr3-117402101-INS-327 | chr3-117402101-INS-367 |
| chr3-117404110-INS-150 | chr3-117404110-INS-164 |
| chr3-117489049-INS-614 | chr3-117489049-INS-615 |
| chr3-117527420-INS-69 | chr3-117527420-INS-69 |
| chr3-117541804-INS-70 | chr3-117541789-INS-61 |
| chr3-117554353-INS-100 | chr3-117554346-INS-64 |
| chr3-117592381-INS-111 | chr3-117592382-INS-111 |
| chr3-117678552-INS-416 | chr3-117678552-INS-416 |
| chr3-117690209-INS-3311 | chr3-117690210-INS-3311 |
| chr3-117707818-INS-125 | chr3-117707818-INS-125 |
| chr3-117716598-INS-655 | chr3-117716598-INS-655 |
| chr3-117730579-INS-104 | chr3-117730579-INS-104 |
| chr3-117743428-INS-6180 | chr3-117743429-INS-6180 |
| chr3-117780158-INS-7097 | chr3-117780159-INS-7097 |
| chr3-117806194-INS-135 | chr3-117806194-INS-177 |
| chr3-117819160-INS-206 | chr3-117819160-INS-208 |
| chr3-117928527-INS-66 | chr3-117928527-INS-77 |
| chr3-117932725-INS-2094 | chr3-117932726-INS-2094 |
| chr3-117951935-INS-60 | chr3-117951935-INS-55 |
| chr3-117957361-INS-77 | chr3-117957361-INS-82 |
| chr3-117969283-INS-58 | chr3-117969283-INS-58 |
| chr3-117980875-INS-106 | chr3-117980875-INS-107 |
| chr3-118013271-INS-60 | chr3-118013271-INS-69 |
| chr3-118020984-INS-6440 | chr3-118020985-INS-6440 |
| chr3-118021163-INS-56 | chr3-118021163-INS-56 |
| chr3-118021651-INS-56 | chr3-118021641-INS-104 |
| chr3-118025988-INS-2623 | chr3-118025989-INS-2623 |
| chr3-118025989-INS-642 | chr3-118025989-INS-644 |
| chr3-118036514-INS-6877 | chr3-118036515-INS-6862 |
| chr3-118061837-INS-2843 | chr3-118061838-INS-2843 |
| chr3-118091292-INS-381 | chr3-118091292-INS-381 |
| chr3-118122739-INS-56 | chr3-118122724-INS-52 |
| chr3-118125353-INS-80 | chr3-118125353-INS-80 |
| chr3-118128414-INS-56 | chr3-118128448-INS-64 |
| chr3-118159258-INS-5113 | chr3-118159259-INS-5113 |
| chr3-118197681-INS-287 | chr3-118197681-INS-289 |
| chr3-118212446-INS-51 | chr3-118212446-INS-52 |
| chr3-118257771-INS-60 | chr3-118257771-INS-52 |
| chr3-118289508-INS-64 | chr3-118289508-INS-64 |
| chr3-118307975-INS-59 | chr3-118307975-INS-59 |
| chr3-118330954-INS-166 | chr3-118330954-INS-160 |
| chr3-118351010-INS-68 | chr3-118350984-INS-80 |
| chr3-118362882-INS-74 | chr3-118362886-INS-52 |
| chr3-118371654-INS-207 | chr3-118371656-INS-208 |
| chr3-118379603-INS-51 | chr3-118379593-INS-58 |
| chr3-118393549-INS-82 | chr3-118393549-INS-95 |
| chr3-118426183-INS-127 | chr3-118426183-INS-133 |
| chr3-118452140-INS-615 | chr3-118452140-INS-615 |
| chr3-118456685-INS-56 | chr3-118456685-INS-56 |
| chr3-118465183-INS-51 | chr3-118465183-INS-51 |
| chr3-118467656-INS-146 | chr3-118467656-INS-147 |
| chr3-118527246-INS-100 | chr3-118527246-INS-96 |
| chr3-118528291-INS-71 | chr3-118528293-INS-70 |
| chr3-118540692-INS-230 | chr3-118540692-INS-230 |
| chr3-118552266-INS-88 | chr3-118552289-INS-100 |
| chr3-118585122-INS-71 | chr3-118585122-INS-72 |
| chr3-118663053-INS-66 | chr3-118663053-INS-66 |
| chr3-118668836-INS-84 | chr3-118668836-INS-86 |
| chr3-118699061-INS-72 | chr3-118699061-INS-72 |

| | |
|---|---|
| chr3-118708131-INS-717 | chr3-118708131-INS-717 |
| chr3-118708752-INS-214 | chr3-118708752-INS-213 |
| chr3-118710360-INS-660 | chr3-118710360-INS-661 |
| chr3-118733677-INS-90 | chr3-118733677-INS-88 |
| chr3-118794214-INS-632 | chr3-118794214-INS-630 |
| chr3-118800945-INS-527 | chr3-118800945-INS-527 |
| chr3-118802052-INS-128 | chr3-118802052-INS-126 |
| chr3-118818194-INS-59 | chr3-118818194-INS-69 |
| chr3-118828893-INS-96 | chr3-118828893-INS-96 |
| chr3-118831428-INS-351 | chr3-118831428-INS-351 |
| chr3-118838328-INS-52 | chr3-118838328-INS-54 |
| chr3-118863862-INS-4844 | chr3-118863863-INS-4844 |
| chr3-118874261-INS-110 | chr3-118874266-INS-110 |
| chr3-118892909-INS-428 | chr3-118892909-INS-428 |
| chr3-118930644-INS-2701 | chr3-118930645-INS-2692 |
| chr3-118937588-INS-114 | chr3-118937588-INS-66 |
| chr3-118942966-INS-55 | chr3-118942966-INS-65 |
| chr3-118964631-INS-67 | chr3-118964631-INS-67 |
| chr3-118969142-INS-66 | chr3-118969142-INS-70 |
| chr3-118973928-INS-70 | chr3-118973928-INS-50 |
| chr3-118975676-INS-99 | chr3-118975692-INS-62 |
| chr3-118980861-INS-59 | chr3-118980861-INS-59 |
| chr3-118989200-INS-431 | chr3-118989200-INS-435 |
| chr3-119014009-INS-64 | chr3-119014009-INS-62 |
| chr3-119032069-INS-69 | chr3-119032069-INS-69 |
| chr3-119049582-INS-70 | chr3-119049582-INS-70 |
| chr3-119083426-INS-72 | chr3-119083426-INS-56 |
| chr3-119141329-INS-1017 | chr3-119141330-INS-1018 |
| chr3-119146329-INS-59 | chr3-119146329-INS-57 |
| chr3-119161720-INS-980 | chr3-119161720-INS-981 |
| chr3-119182791-INS-772 | chr3-119182791-INS-772 |
| chr3-119209333-INS-6503 | chr3-119209334-INS-6504 |
| chr3-119209423-INS-108 | chr3-119209420-INS-110 |
| chr3-119218528-INS-551 | chr3-119218528-INS-551 |
| chr3-119223617-INS-923 | chr3-119223617-INS-923 |
| chr3-119236681-INS-56 | chr3-119236681-INS-56 |
| chr3-119244824-INS-98 | chr3-119244824-INS-98 |
| chr3-119257107-INS-66 | chr3-119257110-INS-110 |
| chr3-119290639-INS-181 | chr3-119290639-INS-181 |
| chr3-119296212-INS-76 | chr3-119296214-INS-72 |
| chr3-119324926-INS-205 | chr3-119324926-INS-205 |
| chr3-119335578-INS-692 | chr3-119335578-INS-708 |
| chr3-119349779-INS-115 | chr3-119349779-INS-115 |
| chr3-119406006-INS-56 | chr3-119406006-INS-52 |
| chr3-119410228-INS-902 | chr3-119410223-INS-922 |
| chr3-119410238-INS-6430 | chr3-119410239-INS-6430 |
| chr3-119473190-INS-1410 | chr3-119473191-INS-1410 |
| chr3-119523480-INS-817 | chr3-119523480-INS-816 |
| chr3-119544870-INS-318 | chr3-119544870-INS-319 |
| chr3-119551130-INS-2211 | chr3-119551131-INS-2212 |
| chr3-119559841-INS-114 | chr3-119559835-INS-74 |
| chr3-119590490-INS-84 | chr3-119590490-INS-84 |
| chr3-119599757-INS-60 | chr3-119599753-INS-52 |
| chr3-119600442-INS-7080 | chr3-119600443-INS-7083 |
| chr3-119610476-INS-56 | chr3-119610476-INS-67 |
| chr3-119636977-INS-410 | chr3-119636977-INS-410 |
| chr3-119638340-INS-58 | chr3-119638340-INS-60 |
| chr3-119650915-INS-90 | chr3-119650915-INS-90 |
| chr3-119660336-INS-57 | chr3-119660336-INS-57 |
| chr3-119665560-INS-87 | chr3-119665560-INS-51 |
| chr3-119718087-INS-102 | chr3-119718087-INS-102 |
| chr3-119728479-INS-56 | chr3-119728479-INS-66 |
| chr3-119737195-INS-64 | chr3-119737195-INS-64 |
| chr3-119765110-INS-66 | chr3-119765110-INS-69 |
| chr3-119789077-INS-181 | chr3-119789077-INS-183 |
| chr3-119799172-INS-6444 | chr3-119799173-INS-6444 |
| chr3-119805231-INS-110 | chr3-119805231-INS-110 |
| chr3-119810264-INS-1850 | chr3-119810265-INS-1850 |
| chr3-119811347-INS-738 | chr3-119811347-INS-739 |
| chr3-119823977-INS-220 | chr3-119823977-INS-220 |
| chr3-119830475-INS-60 | chr3-119830475-INS-63 |
| chr3-119836912-INS-214 | chr3-119836912-INS-215 |
| chr3-119858607-INS-6649 | chr3-119858608-INS-6649 |
| chr3-119884158-INS-72 | chr3-119884158-INS-70 |
| chr3-119898109-INS-686 | chr3-119898109-INS-689 |
| chr3-119905414-INS-74 | chr3-119905414-INS-74 |
| chr3-119916045-INS-87 | chr3-119916045-INS-51 |

| | |
|---|---|
| chr3-119918457-INS-56 | chr3-119918457-INS-50 |
| chr3-119926453-INS-1271 | chr3-119926454-INS-1272 |
| chr3-119936176-INS-73 | chr3-119936152-INS-84 |
| chr3-119940527-INS-202 | chr3-119940527-INS-202 |
| chr3-119943133-INS-153 | chr3-119943133-INS-153 |
| chr3-119951667-INS-1291 | chr3-119951668-INS-1291 |
| chr3-119966145-INS-211 | chr3-119966145-INS-211 |
| chr3-119977025-INS-5550 | chr3-119977026-INS-5550 |
| chr3-119985161-INS-206 | chr3-119985161-INS-206 |
| chr3-119989664-INS-180 | chr3-119989664-INS-180 |
| chr3-120015876-INS-59 | chr3-120015876-INS-59 |
| chr3-120041751-INS-196 | chr3-120041751-INS-196 |
| chr3-120051461-INS-90 | chr3-120051437-INS-110 |
| chr3-120061343-INS-4338 | chr3-120061344-INS-4343 |
| chr3-120065712-INS-87 | chr3-120065712-INS-105 |
| chr3-120113672-INS-1860 | chr3-120113673-INS-1860 |
| chr3-120120278-INS-6651 | chr3-120120279-INS-6651 |
| chr3-120127173-INS-610 | chr3-120127173-INS-610 |
| chr3-120177842-INS-59 | chr3-120177842-INS-63 |
| chr3-120247466-INS-2614 | chr3-120247467-INS-2583 |
| chr3-120274842-INS-63 | chr3-120274842-INS-63 |
| chr3-120276462-INS-51 | chr3-120276462-INS-51 |
| chr3-120289950-INS-62 | chr3-120289950-INS-66 |
| chr3-120322948-INS-234 | chr3-120322948-INS-192 |
| chr3-120338595-INS-59 | chr3-120338595-INS-59 |
| chr3-120349217-INS-52 | chr3-120349193-INS-56 |
| chr3-120400442-INS-2472 | chr3-120400443-INS-2472 |
| chr3-120408052-INS-76 | chr3-120408052-INS-84 |
| chr3-120411123-INS-1556 | chr3-120411124-INS-1557 |
| chr3-120422672-INS-1132 | chr3-120422673-INS-1132 |
| chr3-120429822-INS-237 | chr3-120429811-INS-225 |
| chr3-120432557-INS-624 | chr3-120432557-INS-640 |
| chr3-120435394-INS-1164 | chr3-120435395-INS-1164 |
| chr3-120439556-INS-54 | chr3-120439556-INS-54 |
| chr3-120469314-INS-78 | chr3-120469323-INS-95 |
| chr3-120499850-INS-155 | chr3-120499850-INS-120 |
| chr3-120504539-INS-65 | chr3-120504539-INS-65 |
| chr3-120545832-INS-509 | chr3-120545832-INS-509 |
| chr3-120552837-INS-59 | chr3-120552837-INS-59 |
| chr3-120570296-INS-685 | chr3-120570296-INS-685 |
| chr3-120589313-INS-402 | chr3-120589313-INS-402 |
| chr3-120595005-INS-3448 | chr3-120595006-INS-3452 |
| chr3-120598446-INS-148 | chr3-120598445-INS-149 |
| chr3-120654216-INS-72 | chr3-120654216-INS-72 |
| chr3-120661206-INS-51 | chr3-120661206-INS-53 |
| chr3-120669919-INS-51 | chr3-120669887-INS-82 |
| chr3-120670938-INS-65 | chr3-120670931-INS-50 |
| chr3-120705765-INS-66 | chr3-120705811-INS-51 |
| chr3-120707640-INS-144 | chr3-120707640-INS-160 |
| chr3-120712450-INS-55 | chr3-120712450-INS-55 |
| chr3-120765738-INS-1579 | chr3-120765739-INS-1593 |
| chr3-120782268-INS-82 | chr3-120782268-INS-82 |
| chr3-120801316-INS-60 | chr3-120801317-INS-60 |
| chr3-120817406-INS-60 | chr3-120817406-INS-60 |
| chr3-120824688-INS-95 | chr3-120824688-INS-95 |
| chr3-120831664-INS-54 | chr3-120831664-INS-54 |
| chr3-120852119-INS-203 | chr3-120852119-INS-203 |
| chr3-120863628-INS-52 | chr3-120863628-INS-55 |
| chr3-120873578-INS-7009 | chr3-120873579-INS-7010 |
| chr3-120873701-INS-54 | chr3-120873701-INS-52 |
| chr3-120901840-INS-342 | chr3-120901840-INS-342 |
| chr3-120916719-INS-64 | chr3-120916707-INS-64 |
| chr3-120940156-INS-383 | chr3-120940156-INS-384 |
| chr3-120951986-INS-56 | chr3-120951986-INS-60 |
| chr3-120987619-INS-240 | chr3-120987619-INS-242 |
| chr3-121016328-INS-54 | chr3-121016328-INS-54 |
| chr3-121027190-INS-209 | chr3-121027190-INS-209 |
| chr3-121029884-INS-144 | chr3-121029884-INS-148 |
| chr3-121045451-INS-273 | chr3-121045451-INS-273 |
| chr3-121047224-INS-91 | chr3-121047250-INS-84 |
| chr3-121080052-INS-195 | chr3-121080052-INS-195 |
| chr3-121084270-INS-1700 | chr3-121084271-INS-1700 |
| chr3-121088477-INS-142 | chr3-121088477-INS-136 |
| chr3-121091999-INS-183 | chr3-121092021-INS-178 |
| chr3-121096646-INS-6234 | chr3-121096647-INS-6234 |
| chr3-121102611-INS-211 | chr3-121102611-INS-213 |
| chr3-121104900-INS-64 | chr3-121104900-INS-55 |

| | |
|---|---|
| chr3-121115615-INS-375 | chr3-121115615-INS-375 |
| chr3-121132368-INS-60 | chr3-121132368-INS-74 |
| chr3-121135816-INS-95 | chr3-121135816-INS-114 |
| chr3-121147693-INS-1895 | chr3-121147694-INS-1895 |
| chr3-121149236-INS-57 | chr3-121149236-INS-57 |
| chr3-121151174-INS-104 | chr3-121151174-INS-108 |
| chr3-121152202-INS-130 | chr3-121152220-INS-140 |
| chr3-121173902-INS-197 | chr3-121173902-INS-197 |
| chr3-121200373-INS-76 | chr3-121200373-INS-76 |
| chr3-121206556-INS-304 | chr3-121206556-INS-304 |
| chr3-121209626-INS-52 | chr3-121209626-INS-60 |
| chr3-121210920-INS-200 | chr3-121210920-INS-200 |
| chr3-121226613-INS-68 | chr3-121226645-INS-52 |
| chr3-121234768-INS-150 | chr3-121234768-INS-150 |
| chr3-121267516-INS-58 | chr3-121267516-INS-52 |
| chr3-121289997-INS-54 | chr3-121289997-INS-58 |
| chr3-121296389-INS-69 | chr3-121296389-INS-69 |
| chr3-121299443-INS-201 | chr3-121299443-INS-201 |
| chr3-121311086-INS-204 | chr3-121311086-INS-214 |
| chr3-121317034-INS-52 | chr3-121317034-INS-52 |
| chr3-121333302-INS-196 | chr3-121333302-INS-198 |
| chr3-121341218-INS-57 | chr3-121341218-INS-61 |
| chr3-121348871-INS-134 | chr3-121348873-INS-135 |
| chr3-121368585-INS-84 | chr3-121368585-INS-112 |
| chr3-121370628-INS-90 | chr3-121370628-INS-90 |
| chr3-121378164-INS-954 | chr3-121378164-INS-951 |
| chr3-121399008-INS-430 | chr3-121399008-INS-433 |
| chr3-121407833-INS-214 | chr3-121407833-INS-214 |
| chr3-121427352-INS-52 | chr3-121427333-INS-55 |
| chr3-121438933-INS-145 | chr3-121438933-INS-151 |
| chr3-121442484-INS-174 | chr3-121442484-INS-174 |
| chr3-121456086-INS-1322 | chr3-121456087-INS-1322 |
| chr3-121477729-INS-64 | chr3-121477729-INS-52 |
| chr3-121485630-INS-385 | chr3-121485630-INS-385 |
| chr3-121490219-INS-79 | chr3-121490219-INS-79 |
| chr3-121531240-INS-445 | chr3-121531235-INS-450 |
| chr3-121551026-INS-170 | chr3-121551026-INS-170 |
| chr3-121562181-INS-193 | chr3-121562181-INS-193 |
| chr3-121566452-INS-52 | chr3-121566452-INS-52 |
| chr3-121574556-INS-74 | chr3-121574556-INS-50 |
| chr3-121587625-INS-66 | chr3-121587627-INS-78 |
| chr3-121596678-INS-387 | chr3-121596678-INS-385 |
| chr3-121603155-INS-192 | chr3-121603155-INS-193 |
| chr3-121641335-INS-175 | chr3-121641343-INS-131 |
| chr3-121646940-INS-97 | chr3-121646940-INS-97 |
| chr3-121649156-INS-59 | chr3-121649157-INS-71 |
| chr3-121654527-INS-106 | chr3-121654537-INS-101 |
| chr3-121659797-INS-222 | chr3-121659827-INS-207 |
| chr3-121672204-INS-261 | chr3-121672236-INS-271 |
| chr3-121674050-INS-204 | chr3-121674050-INS-204 |
| chr3-121679802-INS-75 | chr3-121679798-INS-63 |
| chr3-121683490-INS-71 | chr3-121683492-INS-80 |
| chr3-121700930-INS-241 | chr3-121700930-INS-241 |
| chr3-121703988-INS-207 | chr3-121703988-INS-207 |
| chr3-121720333-INS-86 | chr3-121720333-INS-86 |
| chr3-121729869-INS-7294 | chr3-121729870-INS-7294 |
| chr3-121734536-INS-124 | chr3-121734536-INS-99 |
| chr3-121751229-INS-7358 | chr3-121751230-INS-7358 |
| chr3-121756026-INS-753 | chr3-121756026-INS-754 |
| chr3-121806876-INS-55 | chr3-121806876-INS-55 |
| chr3-121821460-INS-214 | chr3-121821460-INS-214 |
| chr3-121825076-INS-207 | chr3-121825076-INS-166 |
| chr3-121828492-INS-86 | chr3-121828492-INS-113 |
| chr3-121835391-INS-63 | chr3-121835391-INS-69 |
| chr3-121852927-INS-65 | chr3-121852927-INS-67 |
| chr3-121884514-INS-177 | chr3-121884514-INS-175 |
| chr3-121902274-INS-54 | chr3-121902274-INS-54 |
| chr3-121938311-INS-84 | chr3-121938311-INS-100 |
| chr3-121940175-INS-442 | chr3-121940137-INS-492 |
| chr3-121952394-INS-4266 | chr3-121952395-INS-4267 |
| chr3-121955698-INS-107 | chr3-121955698-INS-107 |
| chr3-121966627-INS-142 | chr3-121966627-INS-142 |
| chr3-121969211-INS-110 | chr3-121969211-INS-111 |
| chr3-121978799-INS-68 | chr3-121978800-INS-68 |
| chr3-121983269-INS-85 | chr3-121983269-INS-79 |
| chr3-121986328-INS-111 | chr3-121986330-INS-159 |
| chr3-121996205-INS-143 | chr3-121996205-INS-143 |

| | |
|---|---|
| chr3-121998322-INS-54 | chr3-121998322-INS-54 |
| chr3-122013397-INS-426 | chr3-122013397-INS-426 |
| chr3-122016216-INS-4515 | chr3-122016217-INS-4515 |
| chr3-122019345-INS-2844 | chr3-122019346-INS-2845 |
| chr3-122036683-INS-114 | chr3-122036683-INS-114 |
| chr3-122091747-INS-105 | chr3-122091747-INS-90 |
| chr3-122113967-INS-102 | chr3-122113967-INS-102 |
| chr3-122116494-INS-172 | chr3-122116494-INS-188 |
| chr3-122118511-INS-114 | chr3-122118511-INS-76 |
| chr3-122124106-INS-71 | chr3-122124106-INS-89 |
| chr3-122124863-INS-234 | chr3-122124863-INS-234 |
| chr3-122130251-INS-60 | chr3-122130251-INS-62 |
| chr3-122139029-INS-195 | chr3-122139029-INS-195 |
| chr3-122143709-INS-208 | chr3-122143709-INS-208 |
| chr3-122152440-INS-205 | chr3-122152440-INS-205 |
| chr3-122159336-INS-83 | chr3-122159336-INS-83 |
| chr3-122165038-INS-62 | chr3-122165038-INS-61 |
| chr3-122167032-INS-75 | chr3-122167032-INS-75 |
| chr3-122171402-INS-69 | chr3-122171402-INS-69 |
| chr3-122192870-INS-114 | chr3-122192842-INS-118 |
| chr3-122204383-INS-189 | chr3-122204383-INS-190 |
| chr3-122230272-INS-74 | chr3-122230272-INS-74 |
| chr3-122251820-INS-202 | chr3-122251820-INS-202 |
| chr3-122253901-INS-90 | chr3-122253900-INS-113 |
| chr3-122262763-INS-54 | chr3-122262763-INS-54 |
| chr3-122264261-INS-173 | chr3-122264261-INS-173 |
| chr3-122273556-INS-148 | chr3-122273556-INS-148 |
| chr3-122312094-INS-51 | chr3-122312127-INS-53 |
| chr3-122314337-INS-66 | chr3-122314351-INS-66 |
| chr3-122339070-INS-203 | chr3-122339083-INS-208 |
| chr3-122382374-INS-118 | chr3-122382374-INS-119 |
| chr3-122384718-INS-157 | chr3-122384718-INS-161 |
| chr3-122398395-INS-7119 | chr3-122398395-INS-7119 |
| chr3-122400557-INS-114 | chr3-122400557-INS-114 |
| chr3-122417750-INS-720 | chr3-122417750-INS-719 |
| chr3-122428321-INS-149 | chr3-122428321-INS-124 |
| chr3-122431558-INS-88 | chr3-122431558-INS-132 |
| chr3-122433858-INS-224 | chr3-122433858-INS-224 |
| chr3-122435665-INS-89 | chr3-122435665-INS-88 |
| chr3-122444906-INS-50 | chr3-122444906-INS-55 |
| chr3-122452423-INS-199 | chr3-122452423-INS-199 |
| chr3-122459161-INS-52 | chr3-122459161-INS-68 |
| chr3-122461267-INS-71 | chr3-122461249-INS-84 |
| chr3-122514820-INS-223 | chr3-122514843-INS-234 |
| chr3-122516093-INS-77 | chr3-122516093-INS-77 |
| chr3-122542795-INS-82 | chr3-122542795-INS-104 |
| chr3-122543874-INS-60 | chr3-122543874-INS-60 |
| chr3-122545274-INS-162 | chr3-122545274-INS-162 |
| chr3-122559073-INS-104 | chr3-122559073-INS-104 |
| chr3-122560371-INS-384 | chr3-122560371-INS-383 |
| chr3-122564307-INS-165 | chr3-122564303-INS-201 |
| chr3-122578282-INS-568 | chr3-122578282-INS-568 |
| chr3-122599028-INS-2688 | chr3-122599029-INS-2688 |
| chr3-122599196-INS-560 | chr3-122599197-INS-560 |
| chr3-122606087-INS-60 | chr3-122606087-INS-76 |
| chr3-122640661-INS-114 | chr3-122640661-INS-115 |
| chr3-122664210-INS-369 | chr3-122664210-INS-386 |
| chr3-122682770-INS-60 | chr3-122682770-INS-58 |
| chr3-122686427-INS-210 | chr3-122686427-INS-210 |
| chr3-122687449-INS-73 | chr3-122687449-INS-63 |
| chr3-122719386-INS-90 | chr3-122719386-INS-91 |
| chr3-122749802-INS-977 | chr3-122749802-INS-979 |
| chr3-122799333-INS-282 | chr3-122799337-INS-243 |
| chr3-122801188-INS-55 | chr3-122801188-INS-55 |
| chr3-122849395-INS-56 | chr3-122849400-INS-79 |
| chr3-122851524-INS-76 | chr3-122851524-INS-76 |
| chr3-122863324-INS-132 | chr3-122863324-INS-135 |
| chr3-122868158-INS-136 | chr3-122868158-INS-136 |
| chr3-122869471-INS-1856 | chr3-122869472-INS-1856 |
| chr3-122903772-INS-76 | chr3-122903772-INS-76 |
| chr3-122919214-INS-159 | chr3-122919215-INS-140 |
| chr3-122925314-INS-1525 | chr3-122925315-INS-1525 |
| chr3-122936443-INS-60 | chr3-122936443-INS-60 |
| chr3-122954192-INS-86 | chr3-122954178-INS-96 |
| chr3-122959880-INS-225 | chr3-122959880-INS-226 |
| chr3-122971113-INS-108 | chr3-122971119-INS-88 |
| chr3-122980145-INS-60 | chr3-122980145-INS-56 |

| | |
|---|---|
| chr3-122983593-INS-115 | chr3-122983593-INS-115 |
| chr3-123008286-INS-52 | chr3-123008286-INS-50 |
| chr3-123046267-INS-70 | chr3-123046267-INS-74 |
| chr3-123078891-INS-504 | chr3-123078891-INS-504 |
| chr3-123088617-INS-66 | chr3-123088617-INS-70 |
| chr3-123093882-INS-87 | chr3-123093885-INS-93 |
| chr3-123100574-INS-165 | chr3-123100574-INS-165 |
| chr3-123108190-INS-436 | chr3-123108190-INS-436 |
| chr3-123121072-INS-202 | chr3-123121073-INS-202 |
| chr3-123132829-INS-52 | chr3-123132829-INS-52 |
| chr3-123133738-INS-97 | chr3-123133738-INS-98 |
| chr3-123149544-INS-159 | chr3-123149544-INS-159 |
| chr3-123165690-INS-344 | chr3-123165690-INS-344 |
| chr3-123180089-INS-123 | chr3-123180103-INS-155 |
| chr3-123218486-INS-115 | chr3-123218486-INS-115 |
| chr3-123250983-INS-55 | chr3-123250983-INS-55 |
| chr3-123257804-INS-107 | chr3-123257804-INS-108 |
| chr3-123262281-INS-68 | chr3-123262281-INS-68 |
| chr3-123303212-INS-456 | chr3-123303212-INS-456 |
| chr3-123329791-INS-62 | chr3-123329791-INS-62 |
| chr3-123338921-INS-56 | chr3-123338899-INS-59 |
| chr3-123343577-INS-68 | chr3-123343551-INS-54 |
| chr3-123345939-INS-52 | chr3-123345939-INS-52 |
| chr3-123363562-INS-71 | chr3-123363559-INS-69 |
| chr3-123381525-INS-64 | chr3-123381525-INS-62 |
| chr3-123390869-INS-6873 | chr3-123390870-INS-6874 |
| chr3-123411456-INS-6389 | chr3-123411457-INS-6389 |
| chr3-123434786-INS-100 | chr3-123434786-INS-64 |
| chr3-123438089-INS-124 | chr3-123438089-INS-136 |
| chr3-123445334-INS-60 | chr3-123445334-INS-52 |
| chr3-123473778-INS-3328 | chr3-123473779-INS-3341 |
| chr3-123507383-INS-95 | chr3-123507383-INS-95 |
| chr3-123522058-INS-606 | chr3-123522058-INS-606 |
| chr3-123529319-INS-580 | chr3-123529319-INS-581 |
| chr3-123581263-INS-418 | chr3-123581263-INS-418 |
| chr3-123619459-INS-69 | chr3-123619459-INS-69 |
| chr3-123643072-INS-114 | chr3-123643072-INS-133 |
| chr3-123657533-INS-536 | chr3-123657533-INS-538 |
| chr3-123662256-INS-80 | chr3-123662256-INS-60 |
| chr3-123679978-INS-67 | chr3-123679978-INS-67 |
| chr3-123687171-INS-66 | chr3-123687185-INS-68 |
| chr3-123769671-INS-546 | chr3-123769671-INS-545 |
| chr3-123773134-INS-453 | chr3-123773134-INS-453 |
| chr3-123783279-INS-167 | chr3-123783279-INS-167 |
| chr3-123791425-INS-71 | chr3-123791456-INS-53 |
| chr3-123811795-INS-2754 | chr3-123811796-INS-2754 |
| chr3-123824254-INS-63 | chr3-123824250-INS-68 |
| chr3-123862257-INS-88 | chr3-123862257-INS-88 |
| chr3-123864485-INS-52 | chr3-123864485-INS-52 |
| chr3-123878547-INS-128 | chr3-123878547-INS-128 |
| chr3-123887266-INS-102 | chr3-123887267-INS-122 |
| chr3-123891877-INS-408 | chr3-123891877-INS-408 |
| chr3-123895883-INS-505 | chr3-123895883-INS-505 |
| chr3-123906259-INS-94 | chr3-123906259-INS-102 |
| chr3-123914740-INS-327 | chr3-123914740-INS-327 |
| chr3-123924085-INS-68 | chr3-123924085-INS-70 |
| chr3-123933713-INS-312 | chr3-123933713-INS-312 |
| chr3-123971705-INS-5894 | chr3-123971706-INS-5894 |
| chr3-123995998-INS-56 | chr3-123995998-INS-60 |
| chr3-124072167-INS-704 | chr3-124072167-INS-708 |
| chr3-124082700-INS-864 | chr3-124082700-INS-864 |
| chr3-124096664-INS-160 | chr3-124096677-INS-134 |
| chr3-124107639-INS-571 | chr3-124107639-INS-571 |
| chr3-124134430-INS-56 | chr3-124134430-INS-62 |
| chr3-124135844-INS-515 | chr3-124135844-INS-515 |
| chr3-124138310-INS-277 | chr3-124138310-INS-238 |
| chr3-124144351-INS-60 | chr3-124144351-INS-60 |
| chr3-124169730-INS-64 | chr3-124169732-INS-62 |
| chr3-124178994-INS-96 | chr3-124178994-INS-96 |
| chr3-124184450-INS-80 | chr3-124184450-INS-80 |
| chr3-124201083-INS-226 | chr3-124201083-INS-226 |
| chr3-124208926-INS-57 | chr3-124208926-INS-57 |
| chr3-124215341-INS-52 | chr3-124215341-INS-52 |
| chr3-124219540-INS-158 | chr3-124219540-INS-158 |
| chr3-124233695-INS-235 | chr3-124233695-INS-235 |
| chr3-124252520-INS-950 | chr3-124252520-INS-950 |
| chr3-124263569-INS-2333 | chr3-124263570-INS-2334 |

| | |
|---|---|
| chr3-124301142-INS-895 | chr3-124301142-INS-895 |
| chr3-124311052-INS-93 | chr3-124311052-INS-93 |
| chr3-124311244-INS-6189 | chr3-124311245-INS-6189 |
| chr3-124328298-INS-125 | chr3-124328298-INS-113 |
| chr3-124339683-INS-1725 | chr3-124339684-INS-1725 |
| chr3-124352017-INS-121 | chr3-124352017-INS-122 |
| chr3-124385882-INS-81 | chr3-124385883-INS-60 |
| chr3-124402102-INS-171 | chr3-124402102-INS-171 |
| chr3-124425489-INS-126 | chr3-124425487-INS-80 |
| chr3-124426158-INS-6297 | chr3-124426159-INS-6297 |
| chr3-124436633-INS-233 | chr3-124436632-INS-243 |
| chr3-124465033-INS-54 | chr3-124465033-INS-54 |
| chr3-124475819-INS-69 | chr3-124475819-INS-69 |
| chr3-124487202-INS-332 | chr3-124487202-INS-334 |
| chr3-124508178-INS-51 | chr3-124508178-INS-51 |
| chr3-124509319-INS-150 | chr3-124509319-INS-150 |
| chr3-124531172-INS-6436 | chr3-124531173-INS-6436 |
| chr3-124541891-INS-61 | chr3-124541891-INS-51 |
| chr3-124561026-INS-56 | chr3-124561026-INS-56 |
| chr3-124591519-INS-179 | chr3-124591519-INS-179 |
| chr3-124601845-INS-52 | chr3-124601845-INS-76 |
| chr3-124604302-INS-182 | chr3-124604302-INS-182 |
| chr3-124694124-INS-606 | chr3-124694124-INS-606 |
| chr3-124699748-INS-127 | chr3-124699748-INS-127 |
| chr3-124702567-INS-214 | chr3-124702567-INS-214 |
| chr3-124722996-INS-7097 | chr3-124722997-INS-7097 |
| chr3-124724451-INS-240 | chr3-124724451-INS-241 |
| chr3-124730819-INS-400 | chr3-124730820-INS-400 |
| chr3-124734456-INS-324 | chr3-124734456-INS-324 |
| chr3-124735811-INS-86 | chr3-124735811-INS-86 |
| chr3-124760311-INS-4478 | chr3-124760312-INS-4478 |
| chr3-124766875-INS-76 | chr3-124766875-INS-76 |
| chr3-124777074-INS-651 | chr3-124777074-INS-651 |
| chr3-124786821-INS-201 | chr3-124786821-INS-201 |
| chr3-124804737-INS-95 | chr3-124804737-INS-129 |
| chr3-124818202-INS-3241 | chr3-124818203-INS-3241 |
| chr3-124823473-INS-210 | chr3-124823473-INS-210 |
| chr3-124847163-INS-145 | chr3-124847163-INS-129 |
| chr3-124849206-INS-60 | chr3-124849206-INS-64 |
| chr3-124855430-INS-271 | chr3-124855430-INS-271 |
| chr3-124904212-INS-68 | chr3-124904212-INS-60 |
| chr3-124906862-INS-587 | chr3-124906862-INS-587 |
| chr3-124916720-INS-83 | chr3-124916720-INS-71 |
| chr3-124921055-INS-56 | chr3-124921055-INS-68 |
| chr3-124931656-INS-720 | chr3-124931656-INS-720 |
| chr3-124943492-INS-483 | chr3-124943492-INS-483 |
| chr3-124957116-INS-50 | chr3-124957116-INS-50 |
| chr3-125013147-INS-58 | chr3-125013147-INS-58 |
| chr3-125030218-INS-62 | chr3-125030218-INS-56 |
| chr3-125066353-INS-378 | chr3-125066353-INS-379 |
| chr3-125087154-INS-53 | chr3-125087150-INS-83 |
| chr3-125106691-INS-566 | chr3-125106691-INS-566 |
| chr3-125139086-INS-177 | chr3-125139086-INS-175 |
| chr3-125155571-INS-70 | chr3-125155571-INS-72 |
| chr3-125159583-INS-545 | chr3-125159583-INS-553 |
| chr3-125183514-INS-119 | chr3-125183514-INS-88 |
| chr3-125183689-INS-485 | chr3-125183689-INS-485 |
| chr3-125188862-INS-68 | chr3-125188862-INS-68 |
| chr3-125212694-INS-355 | chr3-125212694-INS-354 |
| chr3-125223934-INS-83 | chr3-125223934-INS-85 |
| chr3-125236636-INS-57 | chr3-125236636-INS-57 |
| chr3-125239941-INS-137 | chr3-125239941-INS-138 |
| chr3-125250222-INS-181 | chr3-125250222-INS-181 |
| chr3-125255253-INS-513 | chr3-125255253-INS-511 |
| chr3-125274760-INS-1067 | chr3-125274761-INS-1067 |
| chr3-125277782-INS-124 | chr3-125277782-INS-124 |
| chr3-125299725-INS-230 | chr3-125299725-INS-229 |
| chr3-125302752-INS-6440 | chr3-125302753-INS-6440 |
| chr3-125310773-INS-60 | chr3-125310818-INS-62 |
| chr3-125325774-INS-82 | chr3-125325774-INS-86 |
| chr3-125402290-INS-64 | chr3-125402290-INS-68 |
| chr3-125411424-INS-58 | chr3-125411424-INS-58 |
| chr3-125423710-INS-221 | chr3-125423716-INS-223 |
| chr3-125456670-INS-174 | chr3-125456670-INS-174 |
| chr3-125460410-INS-268 | chr3-125460410-INS-268 |
| chr3-125464063-INS-195 | chr3-125464055-INS-194 |
| chr3-125468741-INS-435 | chr3-125468741-INS-435 |

| | |
|---|---|
| chr3-125474440-INS-449 | chr3-125474440-INS-448 |
| chr3-125488164-INS-52 | chr3-125488164-INS-51 |
| chr3-125491150-INS-215 | chr3-125491150-INS-215 |
| chr3-125494646-INS-56 | chr3-125494647-INS-68 |
| chr3-125496890-INS-3404 | chr3-125496891-INS-3404 |
| chr3-125499059-INS-513 | chr3-125499059-INS-513 |
| chr3-125515132-INS-114 | chr3-125515132-INS-120 |
| chr3-125544633-INS-4148 | chr3-125544635-INS-4148 |
| chr3-125566140-INS-3277 | chr3-125566141-INS-3277 |
| chr3-125573151-INS-173 | chr3-125573151-INS-173 |
| chr3-125588772-INS-6209 | chr3-125588773-INS-6210 |
| chr3-125618446-INS-6579 | chr3-125618447-INS-6579 |
| chr3-125618828-INS-168 | chr3-125618828-INS-168 |
| chr3-125635546-INS-103 | chr3-125635546-INS-103 |
| chr3-125680027-INS-679 | chr3-125680027-INS-679 |
| chr3-125682354-INS-104 | chr3-125682354-INS-110 |
| chr3-125715950-INS-64 | chr3-125715950-INS-60 |
| chr3-125736764-INS-54 | chr3-125736764-INS-50 |
| chr3-125769398-INS-1488 | chr3-125769399-INS-1488 |
| chr3-125797220-INS-6261 | chr3-125797221-INS-6261 |
| chr3-125808921-INS-97 | chr3-125808921-INS-77 |
| chr3-125810813-INS-296 | chr3-125810813-INS-296 |
| chr3-125836887-INS-2534 | chr3-125836888-INS-2519 |
| chr3-125860377-INS-201 | chr3-125860377-INS-201 |
| chr3-125886816-INS-60 | chr3-125886816-INS-60 |
| chr3-125901714-INS-52 | chr3-125901714-INS-52 |
| chr3-125913854-INS-198 | chr3-125913854-INS-198 |
| chr3-125916535-INS-228 | chr3-125916535-INS-228 |
| chr3-125919105-INS-54 | chr3-125919105-INS-66 |
| chr3-125941291-INS-477 | chr3-125941291-INS-477 |
| chr3-125947181-INS-239 | chr3-125947181-INS-239 |
| chr3-125973264-INS-513 | chr3-125973256-INS-527 |
| chr3-125986024-INS-1619 | chr3-125986025-INS-1615 |
| chr3-125998864-INS-113 | chr3-125998853-INS-129 |
| chr3-126001236-INS-5137 | chr3-126001237-INS-5137 |
| chr3-126002631-INS-72 | chr3-126002631-INS-72 |
| chr3-126010379-INS-3027 | chr3-126010380-INS-3027 |
| chr3-126023994-INS-107 | chr3-126023995-INS-107 |
| chr3-126028745-INS-55 | chr3-126028745-INS-55 |
| chr3-126037271-INS-77 | chr3-126037271-INS-77 |
| chr3-126039448-INS-234 | chr3-126039448-INS-241 |
| chr3-126041833-INS-72 | chr3-126041833-INS-56 |
| chr3-126050318-INS-104 | chr3-126050319-INS-104 |
| chr3-126066984-INS-92 | chr3-126066984-INS-92 |
| chr3-126081528-INS-214 | chr3-126081528-INS-214 |
| chr3-126081944-INS-7270 | chr3-126081945-INS-7269 |
| chr3-126144173-INS-328 | chr3-126144173-INS-328 |
| chr3-126146566-INS-498 | chr3-126146566-INS-498 |
| chr3-126153080-INS-113 | chr3-126153084-INS-87 |
| chr3-126156465-INS-52 | chr3-126156465-INS-54 |
| chr3-126174756-INS-126 | chr3-126174753-INS-135 |
| chr3-126212335-INS-209 | chr3-126212335-INS-209 |
| chr3-126229044-INS-4597 | chr3-126229045-INS-4597 |
| chr3-126236565-INS-744 | chr3-126236565-INS-744 |
| chr3-126245809-INS-321 | chr3-126245809-INS-321 |
| chr3-126248933-INS-56 | chr3-126248933-INS-50 |
| chr3-126269608-INS-70 | chr3-126269608-INS-70 |
| chr3-126295772-INS-210 | chr3-126295772-INS-210 |
| chr3-126299027-INS-491 | chr3-126299027-INS-491 |
| chr3-126318087-INS-104 | chr3-126318087-INS-110 |
| chr3-126334750-INS-113 | chr3-126334750-INS-116 |
| chr3-126336612-INS-684 | chr3-126336612-INS-684 |
| chr3-126358152-INS-55 | chr3-126358152-INS-55 |
| chr3-126379656-INS-66 | chr3-126379656-INS-72 |
| chr3-126393227-INS-364 | chr3-126393204-INS-338 |
| chr3-126394593-INS-67 | chr3-126394593-INS-65 |
| chr3-126400958-INS-206 | chr3-126400958-INS-209 |
| chr3-126451198-INS-64 | chr3-126451198-INS-60 |
| chr3-126452941-INS-140 | chr3-126452941-INS-140 |
| chr3-126466727-INS-62 | chr3-126466727-INS-56 |
| chr3-126467819-INS-77 | chr3-126467819-INS-77 |
| chr3-126474954-INS-57 | chr3-126474954-INS-57 |
| chr3-126483417-INS-722 | chr3-126483417-INS-722 |
| chr3-126484635-INS-64 | chr3-126484635-INS-66 |
| chr3-126501712-INS-56 | chr3-126501712-INS-60 |
| chr3-126503276-INS-142 | chr3-126503276-INS-142 |
| chr3-126506073-INS-72 | chr3-126506073-INS-60 |

| | |
|---|---|
| chr3-126516536-INS-577 | chr3-126516536-INS-577 |
| chr3-126540686-INS-51 | chr3-126540655-INS-76 |
| chr3-126626230-INS-56 | chr3-126626204-INS-84 |
| chr3-126630221-INS-156 | chr3-126630237-INS-145 |
| chr3-126671649-INS-73 | chr3-126671679-INS-86 |
| chr3-126680264-INS-104 | chr3-126680250-INS-95 |
| chr3-126696657-INS-93 | chr3-126696657-INS-93 |
| chr3-126706404-INS-416 | chr3-126706404-INS-416 |
| chr3-126711592-INS-55 | chr3-126711593-INS-55 |
| chr3-126714717-INS-6538 | chr3-126714718-INS-6539 |
| chr3-126718102-INS-303 | chr3-126718102-INS-302 |
| chr3-126719132-INS-51 | chr3-126719129-INS-54 |
| chr3-126767862-INS-60 | chr3-126767862-INS-60 |
| chr3-126792460-INS-58 | chr3-126792460-INS-75 |
| chr3-126800212-INS-733 | chr3-126800212-INS-733 |
| chr3-126817927-INS-160 | chr3-126817927-INS-160 |
| chr3-126833667-INS-74 | chr3-126833667-INS-76 |
| chr3-126844336-INS-96 | chr3-126844336-INS-96 |
| chr3-126884653-INS-677 | chr3-126884653-INS-677 |
| chr3-126899055-INS-156 | chr3-126899071-INS-166 |
| chr3-126914712-INS-160 | chr3-126914712-INS-160 |
| chr3-126934666-INS-417 | chr3-126934666-INS-417 |
| chr3-126972072-INS-164 | chr3-126972072-INS-182 |
| chr3-126981378-INS-52 | chr3-126981378-INS-52 |
| chr3-126988168-INS-58 | chr3-126988128-INS-58 |
| chr3-126999581-INS-319 | chr3-126999581-INS-319 |
| chr3-127027113-INS-89 | chr3-127027113-INS-89 |
| chr3-127029227-INS-76 | chr3-127029235-INS-60 |
| chr3-127033212-INS-241 | chr3-127033212-INS-240 |
| chr3-127060433-INS-107 | chr3-127060433-INS-123 |
| chr3-127084484-INS-91 | chr3-127084484-INS-91 |
| chr3-127092723-INS-80 | chr3-127092723-INS-80 |
| chr3-127123319-INS-1275 | chr3-127123320-INS-1275 |
| chr3-127149751-INS-265 | chr3-127149751-INS-265 |
| chr3-127151593-INS-114 | chr3-127151594-INS-112 |
| chr3-127154018-INS-60 | chr3-127154018-INS-54 |
| chr3-127161093-INS-68 | chr3-127161093-INS-74 |
| chr3-127163985-INS-1930 | chr3-127163986-INS-1931 |
| chr3-127168442-INS-286 | chr3-127168403-INS-283 |
| chr3-127176208-INS-226 | chr3-127176208-INS-226 |
| chr3-127177563-INS-100 | chr3-127177569-INS-104 |
| chr3-127193488-INS-174 | chr3-127193488-INS-174 |
| chr3-127232569-INS-77 | chr3-127232569-INS-65 |
| chr3-127236035-INS-92 | chr3-127236035-INS-92 |
| chr3-127240494-INS-207 | chr3-127240519-INS-203 |
| chr3-127241918-INS-238 | chr3-127241918-INS-234 |
| chr3-127247761-INS-79 | chr3-127247761-INS-87 |
| chr3-127248588-INS-182 | chr3-127248588-INS-181 |
| chr3-127249309-INS-55 | chr3-127249309-INS-55 |
| chr3-127255535-INS-85 | chr3-127255507-INS-77 |
| chr3-127261112-INS-94 | chr3-127261112-INS-94 |
| chr3-127266508-INS-129 | chr3-127266507-INS-122 |
| chr3-127272608-INS-217 | chr3-127272608-INS-217 |
| chr3-127282287-INS-6928 | chr3-127282288-INS-6928 |
| chr3-127284413-INS-104 | chr3-127284413-INS-104 |
| chr3-127295395-INS-60 | chr3-127295395-INS-60 |
| chr3-127325705-INS-485 | chr3-127325708-INS-489 |
| chr3-127330977-INS-198 | chr3-127330977-INS-198 |
| chr3-127358496-INS-2727 | chr3-127358497-INS-2727 |
| chr3-127359040-INS-207 | chr3-127359041-INS-201 |
| chr3-127375930-INS-179 | chr3-127375930-INS-178 |
| chr3-127378878-INS-179 | chr3-127378878-INS-179 |
| chr3-127391017-INS-5297 | chr3-127391018-INS-5296 |
| chr3-127391255-INS-60 | chr3-127391255-INS-60 |
| chr3-127397581-INS-2424 | chr3-127397584-INS-2422 |
| chr3-127399773-INS-55 | chr3-127399773-INS-55 |
| chr3-127408534-INS-800 | chr3-127408534-INS-800 |
| chr3-127416739-INS-60 | chr3-127416739-INS-60 |
| chr3-127432348-INS-296 | chr3-127432345-INS-303 |
| chr3-127435642-INS-207 | chr3-127435642-INS-207 |
| chr3-127441239-INS-65 | chr3-127441239-INS-66 |
| chr3-127446490-INS-52 | chr3-127446475-INS-66 |
| chr3-127450795-INS-399 | chr3-127450795-INS-399 |
| chr3-127458498-INS-64 | chr3-127458498-INS-64 |
| chr3-127483597-INS-123 | chr3-127483600-INS-119 |
| chr3-127502355-INS-194 | chr3-127502355-INS-194 |
| chr3-127527662-INS-129 | chr3-127527631-INS-177 |

| | |
|---|---|
| chr3-127529499-INS-244 | chr3-127529499-INS-254 |
| chr3-127552833-INS-54 | chr3-127552834-INS-58 |
| chr3-127562650-INS-78 | chr3-127562670-INS-64 |
| chr3-127574718-INS-7117 | chr3-127574719-INS-7117 |
| chr3-127575177-INS-199 | chr3-127575177-INS-199 |
| chr3-127639816-INS-925 | chr3-127639813-INS-924 |
| chr3-127645928-INS-65 | chr3-127645928-INS-65 |
| chr3-127648010-INS-54 | chr3-127648010-INS-54 |
| chr3-127659581-INS-128 | chr3-127659580-INS-128 |
| chr3-127660040-INS-8318 | chr3-127660041-INS-8318 |
| chr3-127669080-INS-58 | chr3-127669080-INS-58 |
| chr3-127671751-INS-234 | chr3-127671751-INS-234 |
| chr3-127675682-INS-66 | chr3-127675682-INS-66 |
| chr3-127694324-INS-180 | chr3-127694330-INS-155 |
| chr3-127718132-INS-66 | chr3-127718132-INS-68 |
| chr3-127720812-INS-209 | chr3-127720812-INS-208 |
| chr3-127735287-INS-75 | chr3-127735287-INS-84 |
| chr3-127737658-INS-301 | chr3-127737658-INS-301 |
| chr3-127742799-INS-60 | chr3-127742799-INS-60 |
| chr3-127761663-INS-60 | chr3-127761655-INS-50 |
| chr3-127771196-INS-927 | chr3-127771196-INS-916 |
| chr3-127774995-INS-205 | chr3-127774995-INS-206 |
| chr3-127799530-INS-86 | chr3-127799530-INS-86 |
| chr3-127808287-INS-56 | chr3-127808287-INS-94 |
| chr3-127830853-INS-77 | chr3-127830853-INS-77 |
| chr3-127854448-INS-55 | chr3-127854448-INS-64 |
| chr3-127858762-INS-67 | chr3-127858762-INS-67 |
| chr3-127864673-INS-1889 | chr3-127864674-INS-1889 |
| chr3-127872126-INS-6232 | chr3-127872127-INS-6232 |
| chr3-127879326-INS-92 | chr3-127879326-INS-88 |
| chr3-127916764-INS-1160 | chr3-127916765-INS-1161 |
| chr3-127919186-INS-77 | chr3-127919186-INS-77 |
| chr3-127930374-INS-55 | chr3-127930365-INS-52 |
| chr3-127956926-INS-702 | chr3-127956926-INS-705 |
| chr3-127958027-INS-58 | chr3-127958026-INS-57 |
| chr3-127970157-INS-1377 | chr3-127970158-INS-1376 |
| chr3-127990994-INS-57 | chr3-127990994-INS-57 |
| chr3-128001222-INS-54 | chr3-128001222-INS-54 |
| chr3-128003709-INS-54 | chr3-128003709-INS-54 |
| chr3-128010930-INS-309 | chr3-128010941-INS-310 |
| chr3-128075586-INS-511 | chr3-128075586-INS-511 |
| chr3-128092849-INS-54 | chr3-128092849-INS-54 |
| chr3-128106877-INS-61 | chr3-128106886-INS-67 |
| chr3-128227673-INS-93 | chr3-128227673-INS-93 |
| chr3-128284168-INS-6548 | chr3-128284169-INS-6548 |
| chr3-128284343-INS-336 | chr3-128284343-INS-337 |
| chr3-128315048-INS-2494 | chr3-128315049-INS-2494 |
| chr3-128341792-INS-121 | chr3-128341792-INS-104 |
| chr3-128375678-INS-1929 | chr3-128375679-INS-1929 |
| chr3-128377020-INS-178 | chr3-128377020-INS-178 |
| chr3-128382066-INS-804 | chr3-128382066-INS-793 |
| chr3-128385182-INS-94 | chr3-128385182-INS-102 |
| chr3-128392759-INS-90 | chr3-128392756-INS-90 |
| chr3-128406332-INS-69 | chr3-128406332-INS-63 |
| chr3-128424601-INS-76 | chr3-128424601-INS-76 |
| chr3-128443961-INS-6767 | chr3-128443962-INS-6767 |
| chr3-128505862-INS-195 | chr3-128505862-INS-195 |
| chr3-128536077-INS-750 | chr3-128536077-INS-750 |
| chr3-128539293-INS-54 | chr3-128539261-INS-84 |
| chr3-128540934-INS-8055 | chr3-128540935-INS-8054 |
| chr3-128606398-INS-608 | chr3-128606398-INS-613 |
| chr3-128609309-INS-262 | chr3-128609293-INS-281 |
| chr3-128681830-INS-218 | chr3-128681830-INS-223 |
| chr3-128682302-INS-3623 | chr3-128682303-INS-3595 |
| chr3-128725815-INS-93 | chr3-128725830-INS-63 |
| chr3-128731756-INS-126 | chr3-128731756-INS-152 |
| chr3-128763613-INS-54 | chr3-128763613-INS-60 |
| chr3-128777720-INS-111 | chr3-128777757-INS-106 |
| chr3-128802542-INS-114 | chr3-128802542-INS-114 |
| chr3-128814738-INS-66 | chr3-128814738-INS-66 |
| chr3-128818734-INS-214 | chr3-128818734-INS-214 |
| chr3-128837334-INS-120 | chr3-128837334-INS-122 |
| chr3-128841349-INS-140 | chr3-128841349-INS-140 |
| chr3-128865132-INS-148 | chr3-128865132-INS-165 |
| chr3-128873271-INS-173 | chr3-128873271-INS-173 |
| chr3-128884670-INS-73 | chr3-128884670-INS-69 |
| chr3-128886099-INS-54 | chr3-128886092-INS-60 |

| | |
|---|---|
| chr3-128896180-INS-186 | chr3-128896182-INS-187 |
| chr3-128912258-INS-5264 | chr3-128912259-INS-5264 |
| chr3-128922947-INS-60 | chr3-128922947-INS-76 |
| chr3-128926880-INS-103 | chr3-128926880-INS-103 |
| chr3-129052361-INS-296 | chr3-129052361-INS-296 |
| chr3-129061291-INS-140 | chr3-129061269-INS-180 |
| chr3-129066488-INS-112 | chr3-129066491-INS-97 |
| chr3-129076059-INS-204 | chr3-129076059-INS-204 |
| chr3-129098855-INS-195 | chr3-129098855-INS-195 |
| chr3-129109700-INS-428 | chr3-129109700-INS-427 |
| chr3-129134637-INS-59 | chr3-129134637-INS-59 |
| chr3-129136149-INS-187 | chr3-129136149-INS-197 |
| chr3-129142561-INS-64 | chr3-129142557-INS-50 |
| chr3-129176539-INS-1173 | chr3-129176530-INS-1203 |
| chr3-129179983-INS-85 | chr3-129179983-INS-90 |
| chr3-129205736-INS-116 | chr3-129205736-INS-81 |
| chr3-129225618-INS-459 | chr3-129225614-INS-489 |
| chr3-129229310-INS-91 | chr3-129229311-INS-91 |
| chr3-129241215-INS-199 | chr3-129241215-INS-200 |
| chr3-129253222-INS-65 | chr3-129253222-INS-65 |
| chr3-129259507-INS-774 | chr3-129259507-INS-773 |
| chr3-129266305-INS-169 | chr3-129266305-INS-171 |
| chr3-129276600-INS-196 | chr3-129276601-INS-199 |
| chr3-129278648-INS-184 | chr3-129278648-INS-184 |
| chr3-129287672-INS-117 | chr3-129287675-INS-120 |
| chr3-129288858-INS-476 | chr3-129288858-INS-476 |
| chr3-129292349-INS-89 | chr3-129292379-INS-64 |
| chr3-129297310-INS-195 | chr3-129297310-INS-195 |
| chr3-129309145-INS-76 | chr3-129309145-INS-76 |
| chr3-129342980-INS-7158 | chr3-129342981-INS-7159 |
| chr3-129344486-INS-57 | chr3-129344486-INS-59 |
| chr3-129357133-INS-681 | chr3-129357133-INS-683 |
| chr3-129378275-INS-53 | chr3-129378275-INS-53 |
| chr3-129380707-INS-95 | chr3-129380707-INS-95 |
| chr3-129390334-INS-277 | chr3-129390334-INS-277 |
| chr3-129396606-INS-84 | chr3-129396606-INS-60 |
| chr3-129413169-INS-96 | chr3-129413174-INS-70 |
| chr3-129419897-INS-201 | chr3-129419897-INS-202 |
| chr3-129435069-INS-169 | chr3-129435065-INS-170 |
| chr3-129447714-INS-205 | chr3-129447714-INS-205 |
| chr3-129472868-INS-69 | chr3-129472868-INS-77 |
| chr3-129480916-INS-199 | chr3-129480916-INS-199 |
| chr3-129488974-INS-95 | chr3-129488973-INS-78 |
| chr3-129493078-INS-56 | chr3-129493049-INS-55 |
| chr3-129500456-INS-97 | chr3-129500488-INS-110 |
| chr3-129521071-INS-68 | chr3-129521071-INS-68 |
| chr3-129525143-INS-101 | chr3-129525143-INS-101 |
| chr3-129526232-INS-114 | chr3-129526232-INS-114 |
| chr3-129538855-INS-427 | chr3-129538855-INS-427 |
| chr3-129541251-INS-54 | chr3-129541233-INS-90 |
| chr3-129569972-INS-60 | chr3-129569972-INS-60 |
| chr3-129572432-INS-126 | chr3-129572432-INS-123 |
| chr3-129577183-INS-4406 | chr3-129577184-INS-4406 |
| chr3-129588896-INS-468 | chr3-129588896-INS-468 |
| chr3-129602877-INS-198 | chr3-129602877-INS-198 |
| chr3-129604602-INS-137 | chr3-129604602-INS-137 |
| chr3-129621982-INS-54 | chr3-129621982-INS-54 |
| chr3-129625791-INS-241 | chr3-129625791-INS-241 |
| chr3-129627705-INS-170 | chr3-129627705-INS-151 |
| chr3-129639012-INS-69 | chr3-129639012-INS-57 |
| chr3-129645474-INS-683 | chr3-129645474-INS-683 |
| chr3-129648458-INS-196 | chr3-129648453-INS-203 |
| chr3-129653205-INS-216 | chr3-129653205-INS-216 |
| chr3-129660444-INS-212 | chr3-129660444-INS-212 |
| chr3-129666470-INS-223 | chr3-129666470-INS-223 |
| chr3-129672811-INS-71 | chr3-129672811-INS-70 |
| chr3-129683345-INS-326 | chr3-129683345-INS-326 |
| chr3-129713335-INS-203 | chr3-129713335-INS-203 |
| chr3-129720820-INS-70 | chr3-129720835-INS-83 |
| chr3-129723645-INS-105 | chr3-129723645-INS-105 |
| chr3-129725539-INS-134 | chr3-129725539-INS-134 |
| chr3-129727069-INS-98 | chr3-129727069-INS-98 |
| chr3-129728074-INS-889 | chr3-129728074-INS-890 |
| chr3-129728736-INS-224 | chr3-129728736-INS-226 |
| chr3-129737025-INS-204 | chr3-129737026-INS-206 |
| chr3-129762395-INS-253 | chr3-129762393-INS-253 |
| chr3-129765461-INS-3933 | chr3-129765462-INS-3943 |

| | |
|---|---|
| chr3-129771844-INS-56 | chr3-129771844-INS-60 |
| chr3-129804454-INS-200 | chr3-129804454-INS-200 |
| chr3-129808754-INS-51 | chr3-129808754-INS-51 |
| chr3-129816036-INS-52 | chr3-129816036-INS-54 |
| chr3-129858230-INS-57 | chr3-129858230-INS-63 |
| chr3-129859606-INS-96 | chr3-129859609-INS-107 |
| chr3-129886417-INS-716 | chr3-129886417-INS-716 |
| chr3-129906178-INS-118 | chr3-129906172-INS-112 |
| chr3-129932209-INS-579 | chr3-129932231-INS-582 |
| chr3-129936926-INS-4601 | chr3-129936927-INS-4601 |
| chr3-129937434-INS-98 | chr3-129937434-INS-96 |
| chr3-129945425-INS-3174 | chr3-129945426-INS-3174 |
| chr3-129946793-INS-51 | chr3-129946787-INS-63 |
| chr3-129964977-INS-187 | chr3-129964977-INS-187 |
| chr3-129983952-INS-779 | chr3-129983952-INS-779 |
| chr3-130027443-INS-56 | chr3-130027443-INS-56 |
| chr3-130031937-INS-204 | chr3-130031937-INS-204 |
| chr3-130038109-INS-70 | chr3-130038109-INS-70 |
| chr3-130065606-INS-107 | chr3-130065606-INS-107 |
| chr3-130112558-INS-149 | chr3-130112558-INS-150 |
| chr3-130114073-INS-547 | chr3-130114073-INS-547 |
| chr3-130122102-INS-72 | chr3-130122102-INS-72 |
| chr3-130129550-INS-122 | chr3-130129550-INS-132 |
| chr3-130132734-INS-66 | chr3-130132734-INS-58 |
| chr3-130134322-INS-89 | chr3-130134322-INS-89 |
| chr3-130166065-INS-158 | chr3-130166071-INS-126 |
| chr3-130189190-INS-196 | chr3-130189190-INS-196 |
| chr3-130200798-INS-54 | chr3-130200798-INS-56 |
| chr3-130203916-INS-142 | chr3-130203916-INS-142 |
| chr3-130205624-INS-6901 | chr3-130205625-INS-6902 |
| chr3-130226044-INS-55 | chr3-130226044-INS-59 |
| chr3-130227382-INS-84 | chr3-130227382-INS-92 |
| chr3-130239493-INS-268 | chr3-130239493-INS-268 |
| chr3-130259762-INS-6696 | chr3-130259763-INS-6697 |
| chr3-130265773-INS-117 | chr3-130265773-INS-117 |
| chr3-130274504-INS-111 | chr3-130274521-INS-133 |
| chr3-130279382-INS-93 | chr3-130279382-INS-98 |
| chr3-130290615-INS-58 | chr3-130290615-INS-60 |
| chr3-130296357-INS-63 | chr3-130296357-INS-63 |
| chr3-130310006-INS-227 | chr3-130310006-INS-227 |
| chr3-130325226-INS-111 | chr3-130325226-INS-111 |
| chr3-130332324-INS-181 | chr3-130332324-INS-185 |
| chr3-130335844-INS-102 | chr3-130335844-INS-102 |
| chr3-130338190-INS-63 | chr3-130338190-INS-56 |
| chr3-130352351-INS-202 | chr3-130352351-INS-201 |
| chr3-130354020-INS-95 | chr3-130354020-INS-95 |
| chr3-130356083-INS-1071 | chr3-130356084-INS-1073 |
| chr3-130356084-INS-233 | chr3-130356084-INS-233 |
| chr3-130372896-INS-62 | chr3-130372897-INS-83 |
| chr3-130405217-INS-2142 | chr3-130405218-INS-2143 |
| chr3-130425230-INS-60 | chr3-130425182-INS-59 |
| chr3-130435414-INS-51 | chr3-130435414-INS-51 |
| chr3-130445591-INS-191 | chr3-130445591-INS-190 |
| chr3-130455321-INS-56 | chr3-130455317-INS-52 |
| chr3-130474685-INS-52 | chr3-130474685-INS-56 |
| chr3-130481936-INS-100 | chr3-130481936-INS-100 |
| chr3-130488479-INS-129 | chr3-130488479-INS-129 |
| chr3-130527157-INS-194 | chr3-130527157-INS-194 |
| chr3-130528105-INS-5412 | chr3-130528106-INS-5408 |
| chr3-130534149-INS-611 | chr3-130534149-INS-582 |
| chr3-130578721-INS-202 | chr3-130578721-INS-202 |
| chr3-130581282-INS-1753 | chr3-130581283-INS-1753 |
| chr3-130583221-INS-178 | chr3-130583221-INS-178 |
| chr3-130584086-INS-881 | chr3-130584086-INS-881 |
| chr3-130588621-INS-60 | chr3-130588621-INS-60 |
| chr3-130589569-INS-205 | chr3-130589569-INS-204 |
| chr3-130646493-INS-57 | chr3-130646494-INS-63 |
| chr3-130673637-INS-58 | chr3-130673637-INS-56 |
| chr3-130682983-INS-625 | chr3-130682983-INS-627 |
| chr3-130692075-INS-211 | chr3-130692075-INS-211 |
| chr3-130698790-INS-56 | chr3-130698790-INS-65 |
| chr3-130731956-INS-81 | chr3-130731956-INS-86 |
| chr3-130740194-INS-141 | chr3-130740188-INS-132 |
| chr3-130747593-INS-7586 | chr3-130747594-INS-7586 |
| chr3-130757010-INS-98 | chr3-130757010-INS-98 |
| chr3-130791845-INS-82 | chr3-130791845-INS-82 |
| chr3-130797208-INS-623 | chr3-130797209-INS-626 |

| | |
|---|---|
| chr3-130827184-INS-92 | chr3-130827184-INS-96 |
| chr3-130842065-INS-152 | chr3-130842065-INS-152 |
| chr3-130869128-INS-101 | chr3-130869128-INS-101 |
| chr3-130874056-INS-54 | chr3-130874056-INS-60 |
| chr3-130874893-INS-68 | chr3-130874893-INS-68 |
| chr3-130923922-INS-204 | chr3-130923923-INS-204 |
| chr3-130927431-INS-339 | chr3-130927431-INS-339 |
| chr3-130941297-INS-217 | chr3-130941297-INS-223 |
| chr3-130945587-INS-214 | chr3-130945587-INS-218 |
| chr3-130953519-INS-61 | chr3-130953519-INS-59 |
| chr3-130956080-INS-976 | chr3-130956048-INS-979 |
| chr3-130960794-INS-172 | chr3-130960794-INS-172 |
| chr3-130964804-INS-82 | chr3-130964804-INS-82 |
| chr3-131000147-INS-74 | chr3-131000128-INS-84 |
| chr3-131005636-INS-71 | chr3-131005636-INS-68 |
| chr3-131017292-INS-67 | chr3-131017292-INS-67 |
| chr3-131028236-INS-209 | chr3-131028236-INS-209 |
| chr3-131051502-INS-443 | chr3-131051502-INS-451 |
| chr3-131061346-INS-202 | chr3-131061346-INS-220 |
| chr3-131075589-INS-217 | chr3-131075589-INS-217 |
| chr3-131078474-INS-6401 | chr3-131078475-INS-6401 |
| chr3-131111517-INS-243 | chr3-131111517-INS-255 |
| chr3-131169156-INS-85 | chr3-131169156-INS-69 |
| chr3-131175008-INS-2766 | chr3-131175009-INS-2767 |
| chr3-131198507-INS-54 | chr3-131198507-INS-66 |
| chr3-131202098-INS-369 | chr3-131202098-INS-370 |
| chr3-131247863-INS-329 | chr3-131247863-INS-327 |
| chr3-131309440-INS-225 | chr3-131309440-INS-225 |
| chr3-131314390-INS-56 | chr3-131314390-INS-62 |
| chr3-131317026-INS-56 | chr3-131317026-INS-56 |
| chr3-131329390-INS-2898 | chr3-131329391-INS-2898 |
| chr3-131358416-INS-53 | chr3-131358420-INS-53 |
| chr3-131375449-INS-6160 | chr3-131375450-INS-6160 |
| chr3-131394448-INS-816 | chr3-131394448-INS-816 |
| chr3-131438101-INS-122 | chr3-131438101-INS-123 |
| chr3-131489079-INS-912 | chr3-131489075-INS-908 |
| chr3-131503597-INS-102 | chr3-131503597-INS-102 |
| chr3-131523582-INS-755 | chr3-131523582-INS-755 |
| chr3-131531434-INS-126 | chr3-131531432-INS-132 |
| chr3-131532605-INS-82 | chr3-131532605-INS-84 |
| chr3-131535548-INS-86 | chr3-131535548-INS-86 |
| chr3-131552697-INS-57 | chr3-131552697-INS-57 |
| chr3-131564199-INS-2076 | chr3-131564200-INS-2076 |
| chr3-131575904-INS-6784 | chr3-131575905-INS-6784 |
| chr3-131595126-INS-548 | chr3-131595126-INS-549 |
| chr3-131626371-INS-88 | chr3-131626371-INS-96 |
| chr3-131630795-INS-192 | chr3-131630795-INS-208 |
| chr3-131631726-INS-59 | chr3-131631720-INS-57 |
| chr3-131665059-INS-72 | chr3-131665059-INS-78 |
| chr3-131693769-INS-58 | chr3-131693769-INS-58 |
| chr3-131707698-INS-1848 | chr3-131707699-INS-1849 |
| chr3-131707784-INS-52 | chr3-131707784-INS-52 |
| chr3-131714727-INS-1936 | chr3-131714728-INS-1936 |
| chr3-131722763-INS-6386 | chr3-131722764-INS-6388 |
| chr3-131737638-INS-389 | chr3-131737638-INS-391 |
| chr3-131762985-INS-151 | chr3-131762986-INS-151 |
| chr3-131783775-INS-4841 | chr3-131783776-INS-4841 |
| chr3-131788448-INS-67 | chr3-131788448-INS-75 |
| chr3-131816627-INS-231 | chr3-131816627-INS-231 |
| chr3-131825771-INS-317 | chr3-131825771-INS-317 |
| chr3-131871213-INS-178 | chr3-131871213-INS-178 |
| chr3-131877962-INS-129 | chr3-131877989-INS-92 |
| chr3-131910233-INS-152 | chr3-131910227-INS-132 |
| chr3-131916678-INS-62 | chr3-131916678-INS-62 |
| chr3-131941366-INS-746 | chr3-131941363-INS-755 |
| chr3-131941493-INS-63 | chr3-131941493-INS-63 |
| chr3-131954450-INS-904 | chr3-131954450-INS-904 |
| chr3-131963887-INS-52 | chr3-131963887-INS-52 |
| chr3-132014524-INS-439 | chr3-132014524-INS-439 |
| chr3-132016521-INS-498 | chr3-132016521-INS-498 |
| chr3-132033977-INS-494 | chr3-132033977-INS-494 |
| chr3-132038157-INS-527 | chr3-132038157-INS-516 |
| chr3-132040451-INS-144 | chr3-132040451-INS-144 |
| chr3-132050251-INS-208 | chr3-132050248-INS-213 |
| chr3-132067773-INS-53 | chr3-132067773-INS-53 |
| chr3-132072747-INS-76 | chr3-132072747-INS-76 |
| chr3-132079062-INS-66 | chr3-132079063-INS-56 |

| | |
|---|---|
| chr3-132087040-INS-123 | chr3-132087040-INS-84 |
| chr3-132094661-INS-172 | chr3-132094661-INS-167 |
| chr3-132101886-INS-115 | chr3-132101886-INS-115 |
| chr3-132148402-INS-826 | chr3-132148396-INS-828 |
| chr3-132171587-INS-911 | chr3-132171587-INS-911 |
| chr3-132177796-INS-135 | chr3-132177793-INS-152 |
| chr3-132185957-INS-54 | chr3-132185957-INS-51 |
| chr3-132195032-INS-58 | chr3-132195032-INS-58 |
| chr3-132207730-INS-165 | chr3-132207730-INS-165 |
| chr3-132210597-INS-695 | chr3-132210597-INS-695 |
| chr3-132230500-INS-137 | chr3-132230500-INS-137 |
| chr3-132243901-INS-103 | chr3-132243901-INS-103 |
| chr3-132247027-INS-655 | chr3-132247027-INS-651 |
| chr3-132255403-INS-73 | chr3-132255403-INS-91 |
| chr3-132257322-INS-407 | chr3-132257322-INS-408 |
| chr3-132259184-INS-460 | chr3-132259184-INS-460 |
| chr3-132292340-INS-376 | chr3-132292352-INS-358 |
| chr3-132323498-INS-899 | chr3-132323498-INS-899 |
| chr3-132345302-INS-60 | chr3-132345302-INS-58 |
| chr3-132347547-INS-55 | chr3-132347547-INS-55 |
| chr3-132348841-INS-272 | chr3-132348841-INS-272 |
| chr3-132352372-INS-137 | chr3-132352373-INS-96 |
| chr3-132360876-INS-65 | chr3-132360876-INS-65 |
| chr3-132408776-INS-171 | chr3-132408776-INS-171 |
| chr3-132413259-INS-111 | chr3-132413259-INS-111 |
| chr3-132441707-INS-80 | chr3-132441707-INS-80 |
| chr3-132456419-INS-118 | chr3-132456419-INS-118 |
| chr3-132462282-INS-6548 | chr3-132462283-INS-6548 |
| chr3-132476357-INS-170 | chr3-132476357-INS-172 |
| chr3-132499650-INS-80 | chr3-132499650-INS-80 |
| chr3-132515369-INS-69 | chr3-132515369-INS-73 |
| chr3-132536065-INS-76 | chr3-132536065-INS-52 |
| chr3-132576918-INS-4362 | chr3-132576919-INS-4362 |
| chr3-132586931-INS-466 | chr3-132586931-INS-466 |
| chr3-132598263-INS-152 | chr3-132598256-INS-152 |
| chr3-132599373-INS-68 | chr3-132599373-INS-68 |
| chr3-132601155-INS-130 | chr3-132601155-INS-130 |
| chr3-132646594-INS-73 | chr3-132646594-INS-73 |
| chr3-132666312-INS-1183 | chr3-132666313-INS-1182 |
| chr3-132681058-INS-446 | chr3-132681058-INS-446 |
| chr3-132710772-INS-52 | chr3-132710776-INS-52 |
| chr3-132741382-INS-11707 | chr3-132741383-INS-11707 |
| chr3-132757112-INS-57 | chr3-132757110-INS-61 |
| chr3-132759560-INS-180 | chr3-132759560-INS-196 |
| chr3-132769215-INS-624 | chr3-132769215-INS-624 |
| chr3-132810614-INS-74 | chr3-132810614-INS-74 |
| chr3-132823937-INS-685 | chr3-132823937-INS-685 |
| chr3-132837568-INS-56 | chr3-132837568-INS-56 |
| chr3-132863897-INS-4364 | chr3-132863898-INS-4364 |
| chr3-132908091-INS-72 | chr3-132908091-INS-73 |
| chr3-132908610-INS-794 | chr3-132908605-INS-801 |
| chr3-132931356-INS-51 | chr3-132931336-INS-70 |
| chr3-132942106-INS-103 | chr3-132942106-INS-103 |
| chr3-132959341-INS-202 | chr3-132959341-INS-202 |
| chr3-132977620-INS-292 | chr3-132977620-INS-292 |
| chr3-132986551-INS-106 | chr3-132986527-INS-138 |
| chr3-132991586-INS-60 | chr3-132991596-INS-85 |
| chr3-133001173-INS-55 | chr3-133001173-INS-55 |
| chr3-133008765-INS-165 | chr3-133008765-INS-165 |
| chr3-133010963-INS-63 | chr3-133010963-INS-63 |
| chr3-133026099-INS-203 | chr3-133026099-INS-203 |
| chr3-133033103-INS-149 | chr3-133033103-INS-147 |
| chr3-133034304-INS-198 | chr3-133034311-INS-201 |
| chr3-133062912-INS-120 | chr3-133062912-INS-120 |
| chr3-133070040-INS-6485 | chr3-133070041-INS-6485 |
| chr3-133086531-INS-5147 | chr3-133086532-INS-5147 |
| chr3-133118383-INS-1865 | chr3-133118384-INS-1865 |
| chr3-133132259-INS-761 | chr3-133132259-INS-761 |
| chr3-133141127-INS-129 | chr3-133141127-INS-129 |
| chr3-133145869-INS-481 | chr3-133145869-INS-479 |
| chr3-133151882-INS-244 | chr3-133151882-INS-243 |
| chr3-133157292-INS-197 | chr3-133157292-INS-199 |
| chr3-133180634-INS-98 | chr3-133180624-INS-113 |
| chr3-133202518-INS-206 | chr3-133202518-INS-205 |
| chr3-133216049-INS-179 | chr3-133216049-INS-179 |
| chr3-133218966-INS-54 | chr3-133218966-INS-54 |
| chr3-133229080-INS-211 | chr3-133229080-INS-210 |

| | |
|---|---|
| chr3-133254197-INS-209 | chr3-133254187-INS-227 |
| chr3-133257114-INS-195 | chr3-133257114-INS-195 |
| chr3-133261055-INS-51 | chr3-133261055-INS-53 |
| chr3-133288112-INS-215 | chr3-133288112-INS-214 |
| chr3-133290102-INS-84 | chr3-133290106-INS-74 |
| chr3-133336270-INS-192 | chr3-133336270-INS-192 |
| chr3-133348495-INS-424 | chr3-133348495-INS-424 |
| chr3-133351298-INS-213 | chr3-133351298-INS-215 |
| chr3-133358412-INS-60 | chr3-133358412-INS-68 |
| chr3-133361656-INS-363 | chr3-133361656-INS-363 |
| chr3-133363692-INS-84 | chr3-133363692-INS-84 |
| chr3-133374536-INS-64 | chr3-133374536-INS-64 |
| chr3-133379466-INS-94 | chr3-133379466-INS-114 |
| chr3-133393295-INS-54 | chr3-133393254-INS-103 |
| chr3-133395165-INS-54 | chr3-133395165-INS-56 |
| chr3-133405202-INS-6272 | chr3-133405202-INS-6279 |
| chr3-133417775-INS-73 | chr3-133417775-INS-72 |
| chr3-133419631-INS-6287 | chr3-133419632-INS-6287 |
| chr3-133422594-INS-60 | chr3-133422594-INS-60 |
| chr3-133437935-INS-212 | chr3-133437935-INS-218 |
| chr3-133473333-INS-169 | chr3-133473335-INS-175 |
| chr3-133479201-INS-1860 | chr3-133479202-INS-1860 |
| chr3-133519831-INS-89 | chr3-133519831-INS-89 |
| chr3-133526187-INS-218 | chr3-133526189-INS-215 |
| chr3-133559274-INS-56 | chr3-133559275-INS-56 |
| chr3-133578858-INS-7116 | chr3-133578859-INS-7116 |
| chr3-133604600-INS-73 | chr3-133604595-INS-51 |
| chr3-133629404-INS-158 | chr3-133629404-INS-118 |
| chr3-133645992-INS-96 | chr3-133645992-INS-96 |
| chr3-133666451-INS-1236 | chr3-133666452-INS-1236 |
| chr3-133742901-INS-6376 | chr3-133742902-INS-6377 |
| chr3-133746775-INS-640 | chr3-133746775-INS-640 |
| chr3-133763292-INS-7208 | chr3-133763293-INS-7208 |
| chr3-133818426-INS-6403 | chr3-133818427-INS-6403 |
| chr3-133833148-INS-1942 | chr3-133833149-INS-1940 |
| chr3-133835812-INS-79 | chr3-133835837-INS-83 |
| chr3-133837930-INS-2597 | chr3-133837931-INS-2600 |
| chr3-133850061-INS-63 | chr3-133850061-INS-63 |
| chr3-133853454-INS-1203 | chr3-133853455-INS-1203 |
| chr3-133858343-INS-6595 | chr3-133858344-INS-6595 |
| chr3-133882256-INS-52 | chr3-133882250-INS-57 |
| chr3-133915953-INS-102 | chr3-133915961-INS-67 |
| chr3-133957985-INS-76 | chr3-133957985-INS-76 |
| chr3-133972665-INS-219 | chr3-133972665-INS-220 |
| chr3-133978303-INS-190 | chr3-133978284-INS-198 |
| chr3-134055074-INS-347 | chr3-134055074-INS-347 |
| chr3-134078104-INS-755 | chr3-134078104-INS-755 |
| chr3-134097975-INS-84 | chr3-134097975-INS-65 |
| chr3-134146136-INS-52 | chr3-134146136-INS-52 |
| chr3-134191068-INS-61 | chr3-134191093-INS-55 |
| chr3-134194809-INS-197 | chr3-134194809-INS-197 |
| chr3-134216968-INS-60 | chr3-134216968-INS-60 |
| chr3-134228461-INS-102 | chr3-134228461-INS-90 |
| chr3-134269177-INS-232 | chr3-134269181-INS-252 |
| chr3-134287826-INS-109 | chr3-134287826-INS-109 |
| chr3-134305340-INS-59 | chr3-134305340-INS-59 |
| chr3-134312270-INS-497 | chr3-134312270-INS-497 |
| chr3-134320791-INS-63 | chr3-134320791-INS-63 |
| chr3-134338161-INS-74 | chr3-134338161-INS-74 |
| chr3-134376177-INS-869 | chr3-134376177-INS-873 |
| chr3-134405561-INS-4570 | chr3-134405562-INS-4570 |
| chr3-134408310-INS-206 | chr3-134408310-INS-206 |
| chr3-134410276-INS-89 | chr3-134410276-INS-89 |
| chr3-134434472-INS-3464 | chr3-134434473-INS-3465 |
| chr3-134441733-INS-377 | chr3-134441733-INS-377 |
| chr3-134443123-INS-451 | chr3-134443123-INS-451 |
| chr3-134451297-INS-6372 | chr3-134451298-INS-6372 |
| chr3-134490500-INS-56 | chr3-134490485-INS-52 |
| chr3-134513757-INS-75 | chr3-134513757-INS-75 |
| chr3-134515060-INS-60 | chr3-134515078-INS-70 |
| chr3-134551500-INS-6464 | chr3-134551501-INS-6463 |
| chr3-134565190-INS-332 | chr3-134565190-INS-332 |
| chr3-134578660-INS-90 | chr3-134578675-INS-60 |
| chr3-134627189-INS-217 | chr3-134627189-INS-221 |
| chr3-134656996-INS-432 | chr3-134656996-INS-432 |
| chr3-134671827-INS-493 | chr3-134671827-INS-493 |
| chr3-134690339-INS-405 | chr3-134690339-INS-405 |

| | |
|---|---|
| chr3-134696032-INS-6089 | chr3-134696033-INS-6089 |
| chr3-134722312-INS-52 | chr3-134722312-INS-50 |
| chr3-134735965-INS-209 | chr3-134735965-INS-213 |
| chr3-134743914-INS-192 | chr3-134743914-INS-194 |
| chr3-134745194-INS-966 | chr3-134745194-INS-966 |
| chr3-134746887-INS-358 | chr3-134746887-INS-358 |
| chr3-134761483-INS-6161 | chr3-134761484-INS-6161 |
| chr3-134776838-INS-5098 | chr3-134776839-INS-5098 |
| chr3-134781190-INS-1622 | chr3-134781191-INS-1622 |
| chr3-134782717-INS-74 | chr3-134782717-INS-70 |
| chr3-134788811-INS-459 | chr3-134788811-INS-459 |
| chr3-134809063-INS-7168 | chr3-134809064-INS-7168 |
| chr3-134823320-INS-64 | chr3-134823320-INS-76 |
| chr3-134829935-INS-106 | chr3-134829935-INS-57 |
| chr3-134832383-INS-437 | chr3-134832384-INS-437 |
| chr3-134837958-INS-108 | chr3-134837958-INS-112 |
| chr3-134841132-INS-4081 | chr3-134841133-INS-4081 |
| chr3-134879152-INS-403 | chr3-134879152-INS-403 |
| chr3-134879658-INS-200 | chr3-134879658-INS-201 |
| chr3-134885158-INS-167 | chr3-134885158-INS-167 |
| chr3-134886156-INS-162 | chr3-134886156-INS-162 |
| chr3-134892973-INS-78 | chr3-134892973-INS-79 |
| chr3-134908919-INS-1072 | chr3-134908920-INS-1071 |
| chr3-134910286-INS-687 | chr3-134910286-INS-697 |
| chr3-134916228-INS-72 | chr3-134916228-INS-72 |
| chr3-134925042-INS-2179 | chr3-134925043-INS-2185 |
| chr3-134925558-INS-156 | chr3-134925558-INS-156 |
| chr3-134951113-INS-537 | chr3-134951113-INS-537 |
| chr3-134955040-INS-985 | chr3-134955040-INS-985 |
| chr3-134955080-INS-389 | chr3-134955040-INS-434 |
| chr3-134956504-INS-222 | chr3-134956529-INS-185 |
| chr3-134958363-INS-96 | chr3-134958365-INS-86 |
| chr3-134968875-INS-6360 | chr3-134968876-INS-6360 |
| chr3-134971874-INS-84 | chr3-134971888-INS-60 |
| chr3-134995655-INS-176 | chr3-134995655-INS-178 |
| chr3-135001158-INS-106 | chr3-135001158-INS-120 |
| chr3-135005232-INS-60 | chr3-135005232-INS-55 |
| chr3-135011277-INS-1890 | chr3-135011278-INS-1890 |
| chr3-135015104-INS-256 | chr3-135015104-INS-261 |
| chr3-135024310-INS-185 | chr3-135024310-INS-185 |
| chr3-135067729-INS-187 | chr3-135067729-INS-174 |
| chr3-135070650-INS-76 | chr3-135070650-INS-77 |
| chr3-135078740-INS-66 | chr3-135078740-INS-66 |
| chr3-135102894-INS-75 | chr3-135102894-INS-75 |
| chr3-135115715-INS-52 | chr3-135115715-INS-52 |
| chr3-135133825-INS-52 | chr3-135133825-INS-52 |
| chr3-135142136-INS-1890 | chr3-135142137-INS-1890 |
| chr3-135181415-INS-52 | chr3-135181423-INS-68 |
| chr3-135188411-INS-1888 | chr3-135188412-INS-1888 |
| chr3-135189030-INS-6394 | chr3-135189031-INS-6394 |
| chr3-135192115-INS-80 | chr3-135192115-INS-64 |
| chr3-135195830-INS-6609 | chr3-135195831-INS-6609 |
| chr3-135203903-INS-1538 | chr3-135203904-INS-1538 |
| chr3-135206880-INS-194 | chr3-135206880-INS-194 |
| chr3-135214645-INS-1864 | chr3-135214646-INS-1864 |
| chr3-135217137-INS-255 | chr3-135217137-INS-255 |
| chr3-135232726-INS-182 | chr3-135232678-INS-204 |
| chr3-135238679-INS-56 | chr3-135238679-INS-56 |
| chr3-135247383-INS-434 | chr3-135247383-INS-434 |
| chr3-135250773-INS-344 | chr3-135250773-INS-341 |
| chr3-135256126-INS-223 | chr3-135256127-INS-222 |
| chr3-135257863-INS-155 | chr3-135257863-INS-155 |
| chr3-135258461-INS-165 | chr3-135258461-INS-165 |
| chr3-135288436-INS-240 | chr3-135288436-INS-241 |
| chr3-135291527-INS-188 | chr3-135291527-INS-188 |
| chr3-135342695-INS-210 | chr3-135342695-INS-209 |
| chr3-135354509-INS-64 | chr3-135354509-INS-64 |
| chr3-135363124-INS-193 | chr3-135363125-INS-173 |
| chr3-135410621-INS-55 | chr3-135410621-INS-55 |
| chr3-135413155-INS-88 | chr3-135413155-INS-88 |
| chr3-135420101-INS-190 | chr3-135420101-INS-190 |
| chr3-135423574-INS-1233 | chr3-135423575-INS-1233 |
| chr3-135428897-INS-154 | chr3-135428897-INS-154 |
| chr3-135434532-INS-53 | chr3-135434542-INS-63 |
| chr3-135443024-INS-371 | chr3-135443024-INS-371 |
| chr3-135483501-INS-7073 | chr3-135483502-INS-7073 |
| chr3-135485787-INS-1102 | chr3-135485788-INS-1101 |

| | |
|---|---|
| chr3-135518116-INS-8623 | chr3-135518117-INS-8636 |
| chr3-135531445-INS-62 | chr3-135531453-INS-58 |
| chr3-135542412-INS-4098 | chr3-135542413-INS-4089 |
| chr3-135546788-INS-76 | chr3-135546788-INS-68 |
| chr3-135550012-INS-216 | chr3-135550012-INS-222 |
| chr3-135551040-INS-4531 | chr3-135551041-INS-4531 |
| chr3-135555906-INS-68 | chr3-135555906-INS-64 |
| chr3-135558862-INS-94 | chr3-135558862-INS-96 |
| chr3-135578542-INS-131 | chr3-135578589-INS-111 |
| chr3-135581412-INS-452 | chr3-135581412-INS-456 |
| chr3-135582091-INS-56 | chr3-135582091-INS-56 |
| chr3-135584250-INS-929 | chr3-135584250-INS-930 |
| chr3-135653757-INS-7477 | chr3-135653758-INS-7477 |
| chr3-135671094-INS-78 | chr3-135671094-INS-76 |
| chr3-135697583-INS-80 | chr3-135697583-INS-92 |
| chr3-135704544-INS-8754 | chr3-135704545-INS-8754 |
| chr3-135720034-INS-165 | chr3-135720034-INS-174 |
| chr3-135804673-INS-6392 | chr3-135804677-INS-6393 |
| chr3-135816237-INS-516 | chr3-135816237-INS-515 |
| chr3-135819564-INS-744 | chr3-135819564-INS-741 |
| chr3-135848827-INS-69 | chr3-135848817-INS-53 |
| chr3-135850050-INS-502 | chr3-135850050-INS-500 |
| chr3-135851713-INS-778 | chr3-135851713-INS-779 |
| chr3-135852769-INS-135 | chr3-135852769-INS-135 |
| chr3-135855705-INS-6653 | chr3-135855706-INS-6653 |
| chr3-135858847-INS-1110 | chr3-135858848-INS-1110 |
| chr3-135866533-INS-171 | chr3-135866533-INS-171 |
| chr3-135878580-INS-54 | chr3-135878580-INS-58 |
| chr3-135891025-INS-126 | chr3-135891025-INS-126 |
| chr3-135929616-INS-428 | chr3-135929616-INS-428 |
| chr3-135947886-INS-175 | chr3-135947886-INS-175 |
| chr3-135949057-INS-63 | chr3-135949057-INS-63 |
| chr3-136057418-INS-101 | chr3-136057387-INS-95 |
| chr3-136059088-INS-66 | chr3-136059088-INS-70 |
| chr3-136075051-INS-812 | chr3-136075050-INS-814 |
| chr3-136079734-INS-217 | chr3-136079734-INS-217 |
| chr3-136092052-INS-206 | chr3-136092052-INS-206 |
| chr3-136110091-INS-315 | chr3-136110091-INS-315 |
| chr3-136133389-INS-6322 | chr3-136133390-INS-6329 |
| chr3-136166925-INS-439 | chr3-136166926-INS-439 |
| chr3-136188372-INS-378 | chr3-136188372-INS-380 |
| chr3-136222526-INS-204 | chr3-136222526-INS-204 |
| chr3-136232985-INS-64 | chr3-136232985-INS-64 |
| chr3-136244263-INS-218 | chr3-136244264-INS-217 |
| chr3-136254286-INS-52 | chr3-136254289-INS-50 |
| chr3-136256935-INS-216 | chr3-136256922-INS-240 |
| chr3-136258926-INS-210 | chr3-136258926-INS-204 |
| chr3-136267336-INS-363 | chr3-136267336-INS-363 |
| chr3-136270736-INS-70 | chr3-136270737-INS-61 |
| chr3-136273774-INS-257 | chr3-136273774-INS-272 |
| chr3-136276768-INS-6789 | chr3-136276769-INS-6789 |
| chr3-136287033-INS-63 | chr3-136287033-INS-67 |
| chr3-136288278-INS-57 | chr3-136288278-INS-78 |
| chr3-136298875-INS-422 | chr3-136298875-INS-418 |
| chr3-136318346-INS-446 | chr3-136318347-INS-446 |
| chr3-136350575-INS-7029 | chr3-136350576-INS-7029 |
| chr3-136419747-INS-83 | chr3-136419747-INS-75 |
| chr3-136427451-INS-350 | chr3-136427438-INS-358 |
| chr3-136430381-INS-51 | chr3-136430381-INS-51 |
| chr3-136438171-INS-65 | chr3-136438153-INS-102 |
| chr3-136478934-INS-1508 | chr3-136478935-INS-1504 |
| chr3-136498825-INS-192 | chr3-136498825-INS-191 |
| chr3-136505618-INS-55 | chr3-136505633-INS-63 |
| chr3-136508044-INS-1543 | chr3-136508045-INS-1542 |
| chr3-136510297-INS-51 | chr3-136510298-INS-55 |
| chr3-136600266-INS-1891 | chr3-136600267-INS-1896 |
| chr3-136628349-INS-1888 | chr3-136628350-INS-1888 |
| chr3-136633152-INS-58 | chr3-136633152-INS-58 |
| chr3-136648302-INS-996 | chr3-136648302-INS-996 |
| chr3-136653297-INS-6723 | chr3-136653298-INS-6767 |
| chr3-136658338-INS-59 | chr3-136658338-INS-59 |
| chr3-136728607-INS-66 | chr3-136728607-INS-54 |
| chr3-136732585-INS-652 | chr3-136732585-INS-652 |
| chr3-136743293-INS-82 | chr3-136743293-INS-80 |
| chr3-136752243-INS-7245 | chr3-136752244-INS-7245 |
| chr3-136761355-INS-10993 | chr3-136761356-INS-10993 |
| chr3-136825492-INS-68 | chr3-136825492-INS-72 |

| | |
|---|---|
| chr3-136828218-INS-492 | chr3-136828218-INS-492 |
| chr3-136831497-INS-718 | chr3-136831497-INS-718 |
| chr3-136849161-INS-6193 | chr3-136849162-INS-6193 |
| chr3-136849939-INS-177 | chr3-136849939-INS-177 |
| chr3-136853766-INS-1885 | chr3-136853767-INS-1885 |
| chr3-136859938-INS-124 | chr3-136859938-INS-123 |
| chr3-136861807-INS-73 | chr3-136861804-INS-81 |
| chr3-136872632-INS-115 | chr3-136872632-INS-116 |
| chr3-136890811-INS-363 | chr3-136890811-INS-363 |
| chr3-136933936-INS-112 | chr3-136933936-INS-116 |
| chr3-136944106-INS-112 | chr3-136944106-INS-113 |
| chr3-136950451-INS-95 | chr3-136950460-INS-70 |
| chr3-136952029-INS-54 | chr3-136952029-INS-54 |
| chr3-136979124-INS-188 | chr3-136979124-INS-188 |
| chr3-136981101-INS-523 | chr3-136981101-INS-523 |
| chr3-137029676-INS-640 | chr3-137029676-INS-631 |
| chr3-137037729-INS-51 | chr3-137037753-INS-54 |
| chr3-137049111-INS-76 | chr3-137049111-INS-76 |
| chr3-137088033-INS-636 | chr3-137088033-INS-637 |
| chr3-137100521-INS-2093 | chr3-137100522-INS-2081 |
| chr3-137106375-INS-157 | chr3-137106375-INS-157 |
| chr3-137120943-INS-52 | chr3-137120943-INS-52 |
| chr3-137138633-INS-136 | chr3-137138633-INS-136 |
| chr3-137141023-INS-7239 | chr3-137141024-INS-7239 |
| chr3-137170190-INS-96 | chr3-137170190-INS-90 |
| chr3-137182113-INS-84 | chr3-137182113-INS-60 |
| chr3-137184501-INS-7128 | chr3-137184502-INS-7128 |
| chr3-137213141-INS-2780 | chr3-137213142-INS-2780 |
| chr3-137215863-INS-149 | chr3-137215871-INS-174 |
| chr3-137221850-INS-1594 | chr3-137221851-INS-1594 |
| chr3-137237082-INS-6436 | chr3-137237083-INS-6436 |
| chr3-137254603-INS-88 | chr3-137254583-INS-92 |
| chr3-137255847-INS-2622 | chr3-137255848-INS-2622 |
| chr3-137275098-INS-58 | chr3-137275098-INS-66 |
| chr3-137276630-INS-60 | chr3-137276630-INS-60 |
| chr3-137290209-INS-1538 | chr3-137290210-INS-1540 |
| chr3-137302365-INS-522 | chr3-137302365-INS-522 |
| chr3-137355720-INS-124 | chr3-137355720-INS-92 |
| chr3-137356655-INS-701 | chr3-137356655-INS-705 |
| chr3-137358216-INS-1027 | chr3-137358217-INS-1028 |
| chr3-137363477-INS-62 | chr3-137363477-INS-64 |
| chr3-137385524-INS-70 | chr3-137385522-INS-70 |
| chr3-137390925-INS-252 | chr3-137390925-INS-244 |
| chr3-137401613-INS-108 | chr3-137401613-INS-72 |
| chr3-137406177-INS-6446 | chr3-137406178-INS-6446 |
| chr3-137408010-INS-71 | chr3-137408010-INS-63 |
| chr3-137431534-INS-341 | chr3-137431534-INS-341 |
| chr3-137485372-INS-70 | chr3-137485372-INS-72 |
| chr3-137488454-INS-52 | chr3-137488454-INS-64 |
| chr3-137491774-INS-53 | chr3-137491774-INS-55 |
| chr3-137504212-INS-191 | chr3-137504212-INS-191 |
| chr3-137505102-INS-70 | chr3-137505103-INS-93 |
| chr3-137516322-INS-1332 | chr3-137516323-INS-1333 |
| chr3-137528591-INS-164 | chr3-137528592-INS-164 |
| chr3-137531353-INS-196 | chr3-137531353-INS-195 |
| chr3-137540100-INS-81 | chr3-137540100-INS-81 |
| chr3-137542719-INS-101 | chr3-137542718-INS-139 |
| chr3-137556740-INS-216 | chr3-137556742-INS-209 |
| chr3-137562826-INS-80 | chr3-137562830-INS-79 |
| chr3-137564858-INS-74 | chr3-137564848-INS-102 |
| chr3-137577894-INS-208 | chr3-137577894-INS-208 |
| chr3-137583509-INS-56 | chr3-137583509-INS-56 |
| chr3-137585125-INS-94 | chr3-137585125-INS-60 |
| chr3-137597361-INS-124 | chr3-137597361-INS-134 |
| chr3-137665094-INS-75 | chr3-137665094-INS-81 |
| chr3-137698698-INS-70 | chr3-137698698-INS-72 |
| chr3-137702790-INS-58 | chr3-137702790-INS-60 |
| chr3-137705732-INS-80 | chr3-137705732-INS-76 |
| chr3-137727078-INS-190 | chr3-137727078-INS-190 |
| chr3-137742113-INS-2217 | chr3-137742114-INS-2217 |
| chr3-137751054-INS-56 | chr3-137751043-INS-66 |
| chr3-137760562-INS-54 | chr3-137760562-INS-64 |
| chr3-137765339-INS-66 | chr3-137765339-INS-64 |
| chr3-137768391-INS-58 | chr3-137768362-INS-58 |
| chr3-137792822-INS-2523 | chr3-137792823-INS-2523 |
| chr3-137807878-INS-56 | chr3-137807878-INS-56 |
| chr3-137816855-INS-59 | chr3-137816855-INS-59 |

| | |
|---|---|
| chr3-137836945-INS-55 | chr3-137836943-INS-60 |
| chr3-137840760-INS-65 | chr3-137840760-INS-69 |
| chr3-137845326-INS-56 | chr3-137845326-INS-56 |
| chr3-137855991-INS-51 | chr3-137855991-INS-51 |
| chr3-137868919-INS-293 | chr3-137868919-INS-274 |
| chr3-137890948-INS-498 | chr3-137890948-INS-498 |
| chr3-137904956-INS-8841 | chr3-137904957-INS-8842 |
| chr3-137913894-INS-384 | chr3-137913894-INS-384 |
| chr3-137936866-INS-56 | chr3-137936866-INS-58 |
| chr3-137949519-INS-2282 | chr3-137949520-INS-2281 |
| chr3-138002468-INS-88 | chr3-138002468-INS-116 |
| chr3-138035995-INS-171 | chr3-138035995-INS-184 |
| chr3-138038373-INS-120 | chr3-138038373-INS-120 |
| chr3-138039410-INS-178 | chr3-138039410-INS-183 |
| chr3-138041992-INS-255 | chr3-138041990-INS-248 |
| chr3-138048930-INS-88 | chr3-138048930-INS-88 |
| chr3-138050471-INS-75 | chr3-138050472-INS-68 |
| chr3-138060263-INS-51 | chr3-138060263-INS-51 |
| chr3-138087833-INS-100 | chr3-138087833-INS-102 |
| chr3-138119861-INS-62 | chr3-138119861-INS-62 |
| chr3-138126744-INS-779 | chr3-138126746-INS-817 |
| chr3-138127419-INS-1488 | chr3-138127420-INS-1488 |
| chr3-138129301-INS-97 | chr3-138129301-INS-97 |
| chr3-138155881-INS-209 | chr3-138155881-INS-213 |
| chr3-138174020-INS-76 | chr3-138174014-INS-58 |
| chr3-138191073-INS-207 | chr3-138191042-INS-211 |
| chr3-138192380-INS-214 | chr3-138192380-INS-215 |
| chr3-138210111-INS-87 | chr3-138210123-INS-90 |
| chr3-138214599-INS-198 | chr3-138214599-INS-199 |
| chr3-138217606-INS-64 | chr3-138217608-INS-68 |
| chr3-138278085-INS-189 | chr3-138278085-INS-189 |
| chr3-138279722-INS-216 | chr3-138279722-INS-215 |
| chr3-138288967-INS-1532 | chr3-138288968-INS-1532 |
| chr3-138289318-INS-182 | chr3-138289318-INS-182 |
| chr3-138300822-INS-86 | chr3-138300822-INS-86 |
| chr3-138302875-INS-66 | chr3-138302875-INS-66 |
| chr3-138317234-INS-145 | chr3-138317234-INS-145 |
| chr3-138326014-INS-204 | chr3-138326014-INS-204 |
| chr3-138327376-INS-75 | chr3-138327376-INS-75 |
| chr3-138340015-INS-143 | chr3-138340015-INS-143 |
| chr3-138371880-INS-200 | chr3-138371880-INS-200 |
| chr3-138375298-INS-164 | chr3-138375298-INS-153 |
| chr3-138385969-INS-157 | chr3-138385971-INS-139 |
| chr3-138405686-INS-77 | chr3-138405690-INS-54 |
| chr3-138425483-INS-214 | chr3-138425483-INS-214 |
| chr3-138426326-INS-2753 | chr3-138426327-INS-2753 |
| chr3-138430777-INS-7156 | chr3-138430778-INS-7156 |
| chr3-138437943-INS-84 | chr3-138437943-INS-84 |
| chr3-138592763-INS-137 | chr3-138592763-INS-137 |
| chr3-138615253-INS-327 | chr3-138615253-INS-327 |
| chr3-138627659-INS-382 | chr3-138627659-INS-382 |
| chr3-138629019-INS-450 | chr3-138629020-INS-452 |
| chr3-138656988-INS-90 | chr3-138656988-INS-90 |
| chr3-138658791-INS-155 | chr3-138658791-INS-155 |
| chr3-138660637-INS-1860 | chr3-138660638-INS-1860 |
| chr3-138698812-INS-55 | chr3-138698812-INS-55 |
| chr3-138699683-INS-3972 | chr3-138699684-INS-3972 |
| chr3-138722913-INS-279 | chr3-138722913-INS-277 |
| chr3-138728179-INS-845 | chr3-138728179-INS-839 |
| chr3-138746134-INS-63 | chr3-138746136-INS-60 |
| chr3-138760096-INS-88 | chr3-138760096-INS-88 |
| chr3-138774919-INS-74 | chr3-138774919-INS-78 |
| chr3-138789204-INS-63 | chr3-138789204-INS-63 |
| chr3-138850901-INS-59 | chr3-138850901-INS-61 |
| chr3-138874219-INS-186 | chr3-138874219-INS-184 |
| chr3-138877957-INS-3986 | chr3-138877958-INS-3986 |
| chr3-138882472-INS-166 | chr3-138882472-INS-124 |
| chr3-138887163-INS-1488 | chr3-138887164-INS-1488 |
| chr3-138902639-INS-1998 | chr3-138902640-INS-1998 |
| chr3-138917564-INS-58 | chr3-138917565-INS-68 |
| chr3-138928562-INS-1187 | chr3-138928563-INS-1178 |
| chr3-138950687-INS-6437 | chr3-138950688-INS-6438 |
| chr3-138979414-INS-76 | chr3-138979404-INS-54 |
| chr3-138983017-INS-84 | chr3-138983017-INS-84 |
| chr3-138988297-INS-56 | chr3-138988297-INS-52 |
| chr3-138991047-INS-84 | chr3-138991008-INS-96 |
| chr3-139047717-INS-108 | chr3-139047691-INS-97 |

| | |
|---|---|
| chr3-139064113-INS-91 | chr3-139064113-INS-91 |
| chr3-139089231-INS-81 | chr3-139089231-INS-81 |
| chr3-139115119-INS-524 | chr3-139115119-INS-524 |
| chr3-139129919-INS-52 | chr3-139129919-INS-52 |
| chr3-139139529-INS-107 | chr3-139139529-INS-107 |
| chr3-139170323-INS-925 | chr3-139170323-INS-948 |
| chr3-139192644-INS-57 | chr3-139192645-INS-69 |
| chr3-139195554-INS-970 | chr3-139195554-INS-970 |
| chr3-139225533-INS-1347 | chr3-139225534-INS-1347 |
| chr3-139244821-INS-493 | chr3-139244821-INS-493 |
| chr3-139326487-INS-68 | chr3-139326491-INS-64 |
| chr3-139330549-INS-52 | chr3-139330549-INS-74 |
| chr3-139363395-INS-5258 | chr3-139363396-INS-5258 |
| chr3-139370808-INS-1389 | chr3-139370809-INS-1390 |
| chr3-139378614-INS-694 | chr3-139378614-INS-694 |
| chr3-139420202-INS-98 | chr3-139420188-INS-78 |
| chr3-139420533-INS-1169 | chr3-139420534-INS-1169 |
| chr3-139425445-INS-1422 | chr3-139425446-INS-1422 |
| chr3-139454248-INS-7336 | chr3-139454249-INS-7336 |
| chr3-139459570-INS-64 | chr3-139459570-INS-64 |
| chr3-139464918-INS-72 | chr3-139464921-INS-68 |
| chr3-139506342-INS-141 | chr3-139506342-INS-141 |
| chr3-139516629-INS-948 | chr3-139516629-INS-963 |
| chr3-139518255-INS-84 | chr3-139518270-INS-87 |
| chr3-139540667-INS-3107 | chr3-139540668-INS-3107 |
| chr3-139557119-INS-267 | chr3-139557119-INS-224 |
| chr3-139585163-INS-586 | chr3-139585163-INS-586 |
| chr3-139586957-INS-828 | chr3-139586957-INS-827 |
| chr3-139599212-INS-162 | chr3-139599212-INS-159 |
| chr3-139601087-INS-1888 | chr3-139601088-INS-1888 |
| chr3-139613721-INS-64 | chr3-139613721-INS-66 |
| chr3-139626783-INS-104 | chr3-139626783-INS-108 |
| chr3-139634849-INS-667 | chr3-139634849-INS-667 |
| chr3-139635953-INS-110 | chr3-139635954-INS-67 |
| chr3-139640197-INS-59 | chr3-139640198-INS-59 |
| chr3-139681175-INS-875 | chr3-139681175-INS-871 |
| chr3-139704273-INS-1346 | chr3-139704274-INS-1346 |
| chr3-139708407-INS-325 | chr3-139708407-INS-324 |
| chr3-139786095-INS-52 | chr3-139786095-INS-54 |
| chr3-139794170-INS-100 | chr3-139794125-INS-96 |
| chr3-139797507-INS-2620 | chr3-139797508-INS-2620 |
| chr3-139806359-INS-1998 | chr3-139806360-INS-1998 |
| chr3-139808702-INS-6563 | chr3-139808703-INS-6563 |
| chr3-139819759-INS-169 | chr3-139819759-INS-169 |
| chr3-139875967-INS-712 | chr3-139875967-INS-712 |
| chr3-139876514-INS-56 | chr3-139876514-INS-56 |
| chr3-139883047-INS-253 | chr3-139883047-INS-253 |
| chr3-139902438-INS-99 | chr3-139902438-INS-99 |
| chr3-139911411-INS-52 | chr3-139911411-INS-63 |
| chr3-139918799-INS-864 | chr3-139918799-INS-863 |
| chr3-139961527-INS-769 | chr3-139961527-INS-769 |
| chr3-139970468-INS-230 | chr3-139970468-INS-230 |
| chr3-139979658-INS-296 | chr3-139979658-INS-298 |
| chr3-140015083-INS-84 | chr3-140015083-INS-80 |
| chr3-140041807-INS-68 | chr3-140041807-INS-68 |
| chr3-140061082-INS-56 | chr3-140061082-INS-68 |
| chr3-140067551-INS-208 | chr3-140067551-INS-208 |
| chr3-140067913-INS-971 | chr3-140067913-INS-971 |
| chr3-140071262-INS-1080 | chr3-140071263-INS-1080 |
| chr3-140080811-INS-94 | chr3-140080811-INS-94 |
| chr3-140100039-INS-64 | chr3-140100039-INS-66 |
| chr3-140102145-INS-766 | chr3-140102145-INS-766 |
| chr3-140118895-INS-54 | chr3-140118895-INS-54 |
| chr3-140128616-INS-6862 | chr3-140128617-INS-6863 |
| chr3-140130614-INS-139 | chr3-140130614-INS-123 |
| chr3-140144575-INS-3689 | chr3-140144576-INS-3690 |
| chr3-140174515-INS-1532 | chr3-140174516-INS-1533 |
| chr3-140187407-INS-7144 | chr3-140187408-INS-7144 |
| chr3-140203792-INS-3340 | chr3-140203793-INS-3300 |
| chr3-140213840-INS-70 | chr3-140213840-INS-74 |
| chr3-140232264-INS-62 | chr3-140232267-INS-92 |
| chr3-140242790-INS-54 | chr3-140242790-INS-52 |
| chr3-140271353-INS-364 | chr3-140271353-INS-364 |
| chr3-140273528-INS-129 | chr3-140273528-INS-130 |
| chr3-140287289-INS-54 | chr3-140287289-INS-56 |
| chr3-140289459-INS-2421 | chr3-140289460-INS-2422 |
| chr3-140289941-INS-52 | chr3-140289941-INS-50 |

| | |
|---|---|
| chr3-140316900-INS-227 | chr3-140316900-INS-226 |
| chr3-140318981-INS-185 | chr3-140318969-INS-171 |
| chr3-140319524-INS-3749 | chr3-140319525-INS-3749 |
| chr3-140337568-INS-1028 | chr3-140337569-INS-992 |
| chr3-140343505-INS-52 | chr3-140343505-INS-52 |
| chr3-140360284-INS-208 | chr3-140360284-INS-208 |
| chr3-140392906-INS-56 | chr3-140392906-INS-56 |
| chr3-140394825-INS-91 | chr3-140394868-INS-86 |
| chr3-140410364-INS-88 | chr3-140410364-INS-97 |
| chr3-140418016-INS-56 | chr3-140418026-INS-51 |
| chr3-140426731-INS-4888 | chr3-140426737-INS-4876 |
| chr3-140432378-INS-117 | chr3-140432378-INS-117 |
| chr3-140440350-INS-518 | chr3-140440350-INS-518 |
| chr3-140472691-INS-1905 | chr3-140472692-INS-1905 |
| chr3-140482014-INS-561 | chr3-140482014-INS-561 |
| chr3-140505535-INS-2073 | chr3-140505536-INS-2060 |
| chr3-140510534-INS-202 | chr3-140510534-INS-202 |
| chr3-140516666-INS-60 | chr3-140516666-INS-55 |
| chr3-140544920-INS-51 | chr3-140544964-INS-51 |
| chr3-140558074-INS-65 | chr3-140558091-INS-60 |
| chr3-140569886-INS-749 | chr3-140569886-INS-749 |
| chr3-140579884-INS-5256 | chr3-140579885-INS-5256 |
| chr3-140582307-INS-307 | chr3-140582307-INS-307 |
| chr3-140624149-INS-578 | chr3-140624149-INS-578 |
| chr3-140629240-INS-247 | chr3-140629240-INS-245 |
| chr3-140640480-INS-118 | chr3-140640475-INS-84 |
| chr3-140643073-INS-183 | chr3-140643073-INS-184 |
| chr3-140656777-INS-72 | chr3-140656783-INS-52 |
| chr3-140673731-INS-132 | chr3-140673731-INS-132 |
| chr3-140688260-INS-687 | chr3-140688260-INS-689 |
| chr3-140690125-INS-88 | chr3-140690125-INS-72 |
| chr3-140691038-INS-95 | chr3-140691038-INS-95 |
| chr3-140693210-INS-68 | chr3-140693210-INS-72 |
| chr3-140694777-INS-398 | chr3-140694777-INS-398 |
| chr3-140717476-INS-69 | chr3-140717476-INS-81 |
| chr3-140726942-INS-64 | chr3-140726942-INS-52 |
| chr3-140741177-INS-66 | chr3-140741175-INS-68 |
| chr3-140744924-INS-290 | chr3-140744924-INS-290 |
| chr3-140745774-INS-60 | chr3-140745766-INS-52 |
| chr3-140749506-INS-122 | chr3-140749506-INS-122 |
| chr3-140765354-INS-72 | chr3-140765362-INS-60 |
| chr3-140779083-INS-58 | chr3-140779083-INS-81 |
| chr3-140782041-INS-84 | chr3-140782041-INS-88 |
| chr3-140783177-INS-68 | chr3-140783155-INS-56 |
| chr3-140803214-INS-111 | chr3-140803205-INS-117 |
| chr3-140815873-INS-84 | chr3-140815873-INS-87 |
| chr3-140822247-INS-626 | chr3-140822247-INS-626 |
| chr3-140826896-INS-481 | chr3-140826896-INS-481 |
| chr3-140838294-INS-1087 | chr3-140838295-INS-1088 |
| chr3-140870275-INS-131 | chr3-140870275-INS-168 |
| chr3-140876818-INS-815 | chr3-140876818-INS-815 |
| chr3-140877760-INS-52 | chr3-140877760-INS-56 |
| chr3-140904300-INS-219 | chr3-140904300-INS-220 |
| chr3-140908995-INS-830 | chr3-140908995-INS-831 |
| chr3-140910544-INS-192 | chr3-140910544-INS-192 |
| chr3-140912900-INS-2996 | chr3-140912901-INS-2996 |
| chr3-140929034-INS-673 | chr3-140929034-INS-673 |
| chr3-141010790-INS-475 | chr3-141010790-INS-475 |
| chr3-141013518-INS-6465 | chr3-141013519-INS-6466 |
| chr3-141040936-INS-92 | chr3-141040911-INS-128 |
| chr3-141047400-INS-116 | chr3-141047400-INS-124 |
| chr3-141055615-INS-104 | chr3-141055615-INS-110 |
| chr3-141056430-INS-494 | chr3-141056430-INS-494 |
| chr3-141057447-INS-510 | chr3-141057447-INS-512 |
| chr3-141061503-INS-261 | chr3-141061503-INS-278 |
| chr3-141067978-INS-68 | chr3-141067978-INS-108 |
| chr3-141073767-INS-571 | chr3-141073772-INS-575 |
| chr3-141130485-INS-173 | chr3-141130485-INS-173 |
| chr3-141147666-INS-138 | chr3-141147666-INS-132 |
| chr3-141197275-INS-54 | chr3-141197276-INS-57 |
| chr3-141270715-INS-66 | chr3-141270715-INS-60 |
| chr3-141271598-INS-1276 | chr3-141271599-INS-1275 |
| chr3-141304603-INS-50 | chr3-141304603-INS-50 |
| chr3-141329415-INS-2231 | chr3-141329416-INS-2230 |
| chr3-141336402-INS-52 | chr3-141336402-INS-52 |
| chr3-141338341-INS-72 | chr3-141338335-INS-84 |
| chr3-141342024-INS-59 | chr3-141342024-INS-59 |

| | |
|---|---|
| chr3-141386229-INS-639 | chr3-141386229-INS-639 |
| chr3-141422628-INS-290 | chr3-141422628-INS-290 |
| chr3-141430668-INS-899 | chr3-141430668-INS-897 |
| chr3-141454037-INS-76 | chr3-141454037-INS-76 |
| chr3-141507907-INS-62 | chr3-141507907-INS-62 |
| chr3-141542706-INS-2438 | chr3-141542707-INS-2438 |
| chr3-141560225-INS-64 | chr3-141560225-INS-63 |
| chr3-141561465-INS-1242 | chr3-141561466-INS-1242 |
| chr3-141562564-INS-92 | chr3-141562559-INS-98 |
| chr3-141564335-INS-6698 | chr3-141564336-INS-6698 |
| chr3-141565261-INS-258 | chr3-141565261-INS-258 |
| chr3-141595766-INS-188 | chr3-141595766-INS-155 |
| chr3-141598898-INS-105 | chr3-141598893-INS-63 |
| chr3-141609097-INS-217 | chr3-141609097-INS-217 |
| chr3-141619668-INS-478 | chr3-141619668-INS-510 |
| chr3-141636499-INS-5209 | chr3-141636500-INS-5209 |
| chr3-141640988-INS-266 | chr3-141640996-INS-306 |
| chr3-141651456-INS-6422 | chr3-141651457-INS-6422 |
| chr3-141662580-INS-3817 | chr3-141662581-INS-3818 |
| chr3-141686402-INS-56 | chr3-141686402-INS-56 |
| chr3-141693198-INS-176 | chr3-141693198-INS-177 |
| chr3-141715890-INS-56 | chr3-141715901-INS-73 |
| chr3-141722240-INS-400 | chr3-141722240-INS-400 |
| chr3-141753157-INS-4301 | chr3-141753158-INS-4305 |
| chr3-141775857-INS-77 | chr3-141775857-INS-69 |
| chr3-141794789-INS-54 | chr3-141794789-INS-60 |
| chr3-141824076-INS-54 | chr3-141824076-INS-100 |
| chr3-141834724-INS-124 | chr3-141834724-INS-124 |
| chr3-141878759-INS-84 | chr3-141878777-INS-96 |
| chr3-141887171-INS-312 | chr3-141887171-INS-313 |
| chr3-141894941-INS-5267 | chr3-141894942-INS-5269 |
| chr3-141899177-INS-112 | chr3-141899177-INS-76 |
| chr3-141903553-INS-163 | chr3-141903553-INS-163 |
| chr3-141908902-INS-162 | chr3-141908902-INS-133 |
| chr3-141913498-INS-491 | chr3-141913498-INS-491 |
| chr3-141932899-INS-72 | chr3-141932899-INS-72 |
| chr3-141933171-INS-7557 | chr3-141933172-INS-7560 |
| chr3-141970242-INS-7111 | chr3-141970243-INS-7113 |
| chr3-141992373-INS-108 | chr3-141992404-INS-99 |
| chr3-142002377-INS-92 | chr3-142002377-INS-92 |
| chr3-142034754-INS-70 | chr3-142034754-INS-70 |
| chr3-142041372-INS-64 | chr3-142041372-INS-64 |
| chr3-142067086-INS-4671 | chr3-142067087-INS-4671 |
| chr3-142077935-INS-121 | chr3-142077935-INS-124 |
| chr3-142093016-INS-62 | chr3-142093016-INS-56 |
| chr3-142096753-INS-62 | chr3-142096773-INS-56 |
| chr3-142109908-INS-762 | chr3-142109908-INS-761 |
| chr3-142125047-INS-300 | chr3-142125047-INS-316 |
| chr3-142134369-INS-102 | chr3-142134367-INS-94 |
| chr3-142152167-INS-197 | chr3-142152167-INS-197 |
| chr3-142193128-INS-200 | chr3-142193122-INS-192 |
| chr3-142210254-INS-777 | chr3-142210254-INS-782 |
| chr3-142214714-INS-255 | chr3-142214714-INS-261 |
| chr3-142230541-INS-462 | chr3-142230541-INS-465 |
| chr3-142238861-INS-346 | chr3-142238861-INS-346 |
| chr3-142240856-INS-64 | chr3-142240856-INS-64 |
| chr3-142245102-INS-66 | chr3-142245151-INS-67 |
| chr3-142266031-INS-393 | chr3-142266031-INS-393 |
| chr3-142274327-INS-51 | chr3-142274327-INS-53 |
| chr3-142283458-INS-56 | chr3-142283458-INS-91 |
| chr3-142290886-INS-434 | chr3-142290886-INS-440 |
| chr3-142351840-INS-708 | chr3-142351840-INS-708 |
| chr3-142361551-INS-207 | chr3-142361551-INS-207 |
| chr3-142374240-INS-110 | chr3-142374240-INS-134 |
| chr3-142385168-INS-8986 | chr3-142385169-INS-8986 |
| chr3-142396194-INS-353 | chr3-142396194-INS-353 |
| chr3-142401145-INS-210 | chr3-142401145-INS-210 |
| chr3-142404667-INS-494 | chr3-142404667-INS-494 |
| chr3-142411730-INS-63 | chr3-142411730-INS-65 |
| chr3-142418093-INS-52 | chr3-142418093-INS-60 |
| chr3-142419168-INS-204 | chr3-142419168-INS-203 |
| chr3-142426365-INS-497 | chr3-142426365-INS-496 |
| chr3-142449162-INS-52 | chr3-142449162-INS-54 |
| chr3-142456267-INS-5313 | chr3-142456268-INS-5313 |
| chr3-142474636-INS-216 | chr3-142474636-INS-217 |
| chr3-142475376-INS-801 | chr3-142475376-INS-801 |
| chr3-142478546-INS-969 | chr3-142478546-INS-969 |

| | |
|---|---|
| chr3-142501854-INS-155 | chr3-142501856-INS-159 |
| chr3-142507306-INS-201 | chr3-142507306-INS-201 |
| chr3-142520723-INS-772 | chr3-142520723-INS-772 |
| chr3-142548427-INS-719 | chr3-142548427-INS-718 |
| chr3-142574654-INS-229 | chr3-142574654-INS-229 |
| chr3-142578473-INS-100 | chr3-142578473-INS-84 |
| chr3-142581759-INS-88 | chr3-142581759-INS-92 |
| chr3-142601069-INS-124 | chr3-142601069-INS-123 |
| chr3-142603735-INS-563 | chr3-142603740-INS-559 |
| chr3-142606521-INS-159 | chr3-142606521-INS-159 |
| chr3-142616749-INS-55 | chr3-142616749-INS-55 |
| chr3-142653372-INS-58 | chr3-142653372-INS-58 |
| chr3-142654927-INS-174 | chr3-142654927-INS-174 |
| chr3-142658287-INS-54 | chr3-142658297-INS-52 |
| chr3-142680245-INS-785 | chr3-142680245-INS-779 |
| chr3-142707476-INS-66 | chr3-142707476-INS-88 |
| chr3-142709970-INS-333 | chr3-142709970-INS-333 |
| chr3-142777003-INS-60 | chr3-142777003-INS-50 |
| chr3-142784036-INS-1505 | chr3-142784037-INS-1507 |
| chr3-142792572-INS-651 | chr3-142792572-INS-651 |
| chr3-142828935-INS-202 | chr3-142828935-INS-201 |
| chr3-142829420-INS-1702 | chr3-142829421-INS-1703 |
| chr3-142834133-INS-265 | chr3-142834128-INS-304 |
| chr3-142843191-INS-52 | chr3-142843191-INS-54 |
| chr3-142853867-INS-54 | chr3-142853867-INS-56 |
| chr3-142855209-INS-66 | chr3-142855241-INS-50 |
| chr3-142880269-INS-385 | chr3-142880269-INS-382 |
| chr3-142883872-INS-104 | chr3-142883872-INS-104 |
| chr3-142885642-INS-114 | chr3-142885679-INS-126 |
| chr3-142891294-INS-50 | chr3-142891270-INS-56 |
| chr3-142908756-INS-604 | chr3-142908756-INS-604 |
| chr3-142937392-INS-280 | chr3-142937392-INS-280 |
| chr3-142939863-INS-2975 | chr3-142939864-INS-2975 |
| chr3-142944893-INS-84 | chr3-142944893-INS-84 |
| chr3-143009868-INS-52 | chr3-143009868-INS-50 |
| chr3-143015360-INS-1523 | chr3-143015361-INS-1523 |
| chr3-143016721-INS-231 | chr3-143016721-INS-231 |
| chr3-143055967-INS-84 | chr3-143055967-INS-126 |
| chr3-143058970-INS-237 | chr3-143058970-INS-237 |
| chr3-143073003-INS-93 | chr3-143073003-INS-93 |
| chr3-143075207-INS-110 | chr3-143075207-INS-110 |
| chr3-143093715-INS-82 | chr3-143093724-INS-54 |
| chr3-143108453-INS-64 | chr3-143108453-INS-68 |
| chr3-143118207-INS-1150 | chr3-143118215-INS-1140 |
| chr3-143139625-INS-532 | chr3-143139625-INS-530 |
| chr3-143145556-INS-277 | chr3-143145556-INS-275 |
| chr3-143183455-INS-72 | chr3-143183455-INS-72 |
| chr3-143214487-INS-64 | chr3-143214487-INS-68 |
| chr3-143223004-INS-10417 | chr3-143223004-INS-10415 |
| chr3-143228443-INS-72 | chr3-143228443-INS-72 |
| chr3-143230689-INS-110 | chr3-143230682-INS-76 |
| chr3-143247291-INS-688 | chr3-143247291-INS-688 |
| chr3-143249503-INS-74 | chr3-143249501-INS-76 |
| chr3-143251520-INS-92 | chr3-143251521-INS-96 |
| chr3-143340703-INS-76 | chr3-143340703-INS-76 |
| chr3-143345479-INS-56 | chr3-143345479-INS-56 |
| chr3-143352409-INS-416 | chr3-143352409-INS-416 |
| chr3-143375871-INS-100 | chr3-143375871-INS-96 |
| chr3-143401036-INS-6743 | chr3-143401037-INS-6743 |
| chr3-143478803-INS-104 | chr3-143478816-INS-122 |
| chr3-143489016-INS-80 | chr3-143489016-INS-80 |
| chr3-143500027-INS-194 | chr3-143500027-INS-194 |
| chr3-143511842-INS-64 | chr3-143511842-INS-64 |
| chr3-143525986-INS-463 | chr3-143525986-INS-463 |
| chr3-143539297-INS-54 | chr3-143539297-INS-56 |
| chr3-143542015-INS-108 | chr3-143542015-INS-108 |
| chr3-143588426-INS-716 | chr3-143588426-INS-718 |
| chr3-143604438-INS-60 | chr3-143604438-INS-60 |
| chr3-143611434-INS-56 | chr3-143611434-INS-56 |
| chr3-143622070-INS-75 | chr3-143622070-INS-75 |
| chr3-143628664-INS-50 | chr3-143628629-INS-62 |
| chr3-143634308-INS-110 | chr3-143634308-INS-108 |
| chr3-143636027-INS-251 | chr3-143636027-INS-251 |
| chr3-143641515-INS-58 | chr3-143641515-INS-58 |
| chr3-143659774-INS-2733 | chr3-143659775-INS-2733 |
| chr3-143662891-INS-98 | chr3-143662891-INS-98 |
| chr3-143684429-INS-56 | chr3-143684420-INS-70 |

| | |
|---|---|
| chr3-143692490-INS-130 | chr3-143692490-INS-126 |
| chr3-143697424-INS-55 | chr3-143697427-INS-55 |
| chr3-143711921-INS-77 | chr3-143711921-INS-78 |
| chr3-143720759-INS-62 | chr3-143720769-INS-50 |
| chr3-143758246-INS-100 | chr3-143758246-INS-60 |
| chr3-143777042-INS-335 | chr3-143777042-INS-335 |
| chr3-143781926-INS-54 | chr3-143781926-INS-50 |
| chr3-143796588-INS-1891 | chr3-143796589-INS-1891 |
| chr3-143826800-INS-174 | chr3-143826800-INS-175 |
| chr3-143838540-INS-53 | chr3-143838540-INS-89 |
| chr3-143857627-INS-208 | chr3-143857627-INS-208 |
| chr3-143862993-INS-5885 | chr3-143862994-INS-5887 |
| chr3-143866382-INS-132 | chr3-143866382-INS-132 |
| chr3-143876730-INS-74 | chr3-143876730-INS-56 |
| chr3-143883167-INS-113 | chr3-143883126-INS-148 |
| chr3-143926762-INS-59 | chr3-143926762-INS-60 |
| chr3-143977956-INS-59 | chr3-143977956-INS-59 |
| chr3-143987014-INS-60 | chr3-143986966-INS-72 |
| chr3-144057698-INS-202 | chr3-144057698-INS-202 |
| chr3-144080967-INS-85 | chr3-144080967-INS-85 |
| chr3-144098033-INS-203 | chr3-144098033-INS-203 |
| chr3-144121362-INS-64 | chr3-144121396-INS-50 |
| chr3-144127402-INS-110 | chr3-144127402-INS-110 |
| chr3-144141594-INS-132 | chr3-144141602-INS-141 |
| chr3-144145760-INS-90 | chr3-144145760-INS-90 |
| chr3-144160526-INS-6473 | chr3-144160527-INS-6473 |
| chr3-144163489-INS-58 | chr3-144163489-INS-58 |
| chr3-144179476-INS-7917 | chr3-144179477-INS-7918 |
| chr3-144210516-INS-434 | chr3-144210516-INS-434 |
| chr3-144216065-INS-82 | chr3-144216065-INS-98 |
| chr3-144232789-INS-145 | chr3-144232789-INS-145 |
| chr3-144238050-INS-89 | chr3-144238050-INS-60 |
| chr3-144279122-INS-89 | chr3-144279122-INS-89 |
| chr3-144286604-INS-144 | chr3-144286604-INS-144 |
| chr3-144290268-INS-106 | chr3-144290257-INS-73 |
| chr3-144307958-INS-202 | chr3-144307958-INS-202 |
| chr3-144314510-INS-70 | chr3-144314510-INS-70 |
| chr3-144315503-INS-51 | chr3-144315501-INS-59 |
| chr3-144371507-INS-208 | chr3-144371507-INS-207 |
| chr3-144419284-INS-54 | chr3-144419284-INS-54 |
| chr3-144420906-INS-54 | chr3-144420906-INS-54 |
| chr3-144421470-INS-2017 | chr3-144421471-INS-2017 |
| chr3-144426324-INS-136 | chr3-144426324-INS-136 |
| chr3-144451343-INS-66 | chr3-144451343-INS-66 |
| chr3-144466276-INS-120 | chr3-144466276-INS-120 |
| chr3-144542258-INS-95 | chr3-144542258-INS-105 |
| chr3-144557320-INS-166 | chr3-144557320-INS-165 |
| chr3-144566770-INS-277 | chr3-144566770-INS-277 |
| chr3-144683089-INS-212 | chr3-144683089-INS-212 |
| chr3-144688675-INS-123 | chr3-144688675-INS-91 |
| chr3-144693269-INS-84 | chr3-144693269-INS-84 |
| chr3-144695859-INS-98 | chr3-144695862-INS-86 |
| chr3-144701141-INS-215 | chr3-144701141-INS-215 |
| chr3-144773085-INS-188 | chr3-144773085-INS-191 |
| chr3-144801329-INS-6384 | chr3-144801330-INS-6384 |
| chr3-144857174-INS-53 | chr3-144857174-INS-53 |
| chr3-144902735-INS-76 | chr3-144902735-INS-76 |
| chr3-144903672-INS-6027 | chr3-144903673-INS-6027 |
| chr3-144919689-INS-56 | chr3-144919675-INS-50 |
| chr3-144940320-INS-64 | chr3-144940320-INS-64 |
| chr3-144970356-INS-541 | chr3-144970356-INS-542 |
| chr3-144977119-INS-6619 | chr3-144977120-INS-6623 |
| chr3-145009513-INS-123 | chr3-145009513-INS-109 |
| chr3-145014115-INS-59 | chr3-145014115-INS-54 |
| chr3-145016904-INS-6473 | chr3-145016905-INS-6474 |
| chr3-145070382-INS-195 | chr3-145070382-INS-195 |
| chr3-145079301-INS-54 | chr3-145079301-INS-62 |
| chr3-145093881-INS-57 | chr3-145093881-INS-57 |
| chr3-145124934-INS-779 | chr3-145124934-INS-779 |
| chr3-145148788-INS-1795 | chr3-145148789-INS-1795 |
| chr3-145167524-INS-54 | chr3-145167524-INS-54 |
| chr3-145241873-INS-62 | chr3-145241873-INS-62 |
| chr3-145243756-INS-113 | chr3-145243756-INS-113 |
| chr3-145267221-INS-120 | chr3-145267223-INS-99 |
| chr3-145276492-INS-56 | chr3-145276492-INS-56 |
| chr3-145278829-INS-181 | chr3-145278829-INS-180 |
| chr3-145306374-INS-65 | chr3-145306380-INS-53 |

| | |
|---|---|
| chr3-145361407-INS-78 | chr3-145361407-INS-60 |
| chr3-145379889-INS-7083 | chr3-145379890-INS-7083 |
| chr3-145384060-INS-510 | chr3-145384060-INS-510 |
| chr3-145462911-INS-298 | chr3-145462911-INS-297 |
| chr3-145500567-INS-212 | chr3-145500567-INS-212 |
| chr3-145594615-INS-202 | chr3-145594615-INS-198 |
| chr3-145614370-INS-216 | chr3-145614370-INS-216 |
| chr3-145617409-INS-539 | chr3-145617410-INS-539 |
| chr3-145625961-INS-186 | chr3-145625961-INS-186 |
| chr3-145636640-INS-197 | chr3-145636640-INS-197 |
| chr3-145641669-INS-82 | chr3-145641669-INS-82 |
| chr3-145664995-INS-74 | chr3-145664995-INS-90 |
| chr3-145671767-INS-209 | chr3-145671767-INS-234 |
| chr3-145683326-INS-6014 | chr3-145683327-INS-6014 |
| chr3-145708363-INS-204 | chr3-145708363-INS-204 |
| chr3-145712346-INS-126 | chr3-145712346-INS-126 |
| chr3-145728775-INS-85 | chr3-145728775-INS-85 |
| chr3-145744245-INS-1400 | chr3-145744246-INS-1417 |
| chr3-145778158-INS-53 | chr3-145778158-INS-53 |
| chr3-145785856-INS-216 | chr3-145785851-INS-217 |
| chr3-145796414-INS-56 | chr3-145796418-INS-56 |
| chr3-145812050-INS-68 | chr3-145812000-INS-80 |
| chr3-145817191-INS-66 | chr3-145817191-INS-68 |
| chr3-145832469-INS-91 | chr3-145832471-INS-101 |
| chr3-145858817-INS-57 | chr3-145858817-INS-57 |
| chr3-145864521-INS-58 | chr3-145864522-INS-60 |
| chr3-145906354-INS-132 | chr3-145906356-INS-141 |
| chr3-145931547-INS-92 | chr3-145931547-INS-72 |
| chr3-145960264-INS-68 | chr3-145960264-INS-68 |
| chr3-146006404-INS-62 | chr3-146006404-INS-63 |
| chr3-146011423-INS-57 | chr3-146011423-INS-63 |
| chr3-146029247-INS-66 | chr3-146029247-INS-66 |
| chr3-146031577-INS-153 | chr3-146031577-INS-137 |
| chr3-146072561-INS-78 | chr3-146072561-INS-78 |
| chr3-146086520-INS-60 | chr3-146086520-INS-60 |
| chr3-146093235-INS-136 | chr3-146093235-INS-136 |
| chr3-146096494-INS-328 | chr3-146096494-INS-328 |
| chr3-146123385-INS-52 | chr3-146123385-INS-52 |
| chr3-146131273-INS-891 | chr3-146131273-INS-892 |
| chr3-146148467-INS-1873 | chr3-146148468-INS-1873 |
| chr3-146174406-INS-68 | chr3-146174403-INS-72 |
| chr3-146178954-INS-7314 | chr3-146178955-INS-7314 |
| chr3-146183596-INS-1721 | chr3-146183597-INS-1721 |
| chr3-146184876-INS-205 | chr3-146184876-INS-205 |
| chr3-146197293-INS-74 | chr3-146197293-INS-80 |
| chr3-146231438-INS-192 | chr3-146231438-INS-192 |
| chr3-146232817-INS-70 | chr3-146232817-INS-74 |
| chr3-146243585-INS-4506 | chr3-146243586-INS-4506 |
| chr3-146277452-INS-173 | chr3-146277452-INS-173 |
| chr3-146279083-INS-63 | chr3-146279083-INS-63 |
| chr3-146286719-INS-54 | chr3-146286719-INS-56 |
| chr3-146289892-INS-94 | chr3-146289890-INS-98 |
| chr3-146299022-INS-89 | chr3-146299030-INS-90 |
| chr3-146301638-INS-66 | chr3-146301638-INS-66 |
| chr3-146369209-INS-4202 | chr3-146369210-INS-4202 |
| chr3-146392098-INS-5544 | chr3-146392099-INS-5544 |
| chr3-146416799-INS-62 | chr3-146416799-INS-50 |
| chr3-146424712-INS-126 | chr3-146424712-INS-126 |
| chr3-146431101-INS-949 | chr3-146431101-INS-939 |
| chr3-146448611-INS-225 | chr3-146448611-INS-225 |
| chr3-146543069-INS-75 | chr3-146543067-INS-63 |
| chr3-146545690-INS-52 | chr3-146545690-INS-54 |
| chr3-146561375-INS-187 | chr3-146561376-INS-186 |
| chr3-146566637-INS-7640 | chr3-146566638-INS-7639 |
| chr3-146573432-INS-132 | chr3-146573432-INS-135 |
| chr3-146591050-INS-69 | chr3-146591050-INS-69 |
| chr3-146592395-INS-78 | chr3-146592395-INS-78 |
| chr3-146613554-INS-64 | chr3-146613555-INS-64 |
| chr3-146648099-INS-226 | chr3-146648099-INS-226 |
| chr3-146650110-INS-72 | chr3-146650110-INS-72 |
| chr3-146654047-INS-186 | chr3-146654047-INS-186 |
| chr3-146676968-INS-56 | chr3-146676968-INS-56 |
| chr3-146696340-INS-85 | chr3-146696340-INS-105 |
| chr3-146714988-INS-6646 | chr3-146714989-INS-6648 |
| chr3-146721352-INS-58 | chr3-146721380-INS-62 |
| chr3-146737379-INS-362 | chr3-146737379-INS-365 |
| chr3-146742192-INS-54 | chr3-146742192-INS-56 |

| | |
|---|---|
| chr3-146791850-INS-52 | chr3-146791850-INS-52 |
| chr3-146839828-INS-52 | chr3-146839828-INS-54 |
| chr3-146847025-INS-908 | chr3-146847025-INS-909 |
| chr3-146847052-INS-51 | chr3-146847052-INS-51 |
| chr3-146871565-INS-54 | chr3-146871567-INS-54 |
| chr3-146878825-INS-202 | chr3-146878795-INS-211 |
| chr3-146891617-INS-83 | chr3-146891602-INS-91 |
| chr3-146897572-INS-7157 | chr3-146897573-INS-7157 |
| chr3-146919014-INS-66 | chr3-146919014-INS-69 |
| chr3-146937689-INS-56 | chr3-146937687-INS-52 |
| chr3-146963138-INS-6864 | chr3-146963139-INS-6865 |
| chr3-146998801-INS-58 | chr3-146998801-INS-62 |
| chr3-147033637-INS-116 | chr3-147033637-INS-116 |
| chr3-147039276-INS-50 | chr3-147039277-INS-76 |
| chr3-147057653-INS-82 | chr3-147057653-INS-82 |
| chr3-147068787-INS-72 | chr3-147068787-INS-72 |
| chr3-147080760-INS-212 | chr3-147080760-INS-213 |
| chr3-147121921-INS-52 | chr3-147121921-INS-78 |
| chr3-147123284-INS-64 | chr3-147123328-INS-62 |
| chr3-147149237-INS-122 | chr3-147149239-INS-133 |
| chr3-147159813-INS-6410 | chr3-147159814-INS-6410 |
| chr3-147160134-INS-703 | chr3-147160134-INS-705 |
| chr3-147209613-INS-2783 | chr3-147209614-INS-2784 |
| chr3-147215240-INS-934 | chr3-147215240-INS-941 |
| chr3-147221965-INS-6236 | chr3-147221966-INS-6236 |
| chr3-147250910-INS-4796 | chr3-147250911-INS-4796 |
| chr3-147253931-INS-239 | chr3-147253931-INS-231 |
| chr3-147274297-INS-105 | chr3-147274297-INS-105 |
| chr3-147279943-INS-982 | chr3-147279943-INS-983 |
| chr3-147302647-INS-159 | chr3-147302647-INS-159 |
| chr3-147327113-INS-136 | chr3-147327113-INS-137 |
| chr3-147330640-INS-795 | chr3-147330641-INS-800 |
| chr3-147350241-INS-57 | chr3-147350241-INS-57 |
| chr3-147351935-INS-64 | chr3-147351935-INS-64 |
| chr3-147387218-INS-56 | chr3-147387218-INS-63 |
| chr3-147395997-INS-383 | chr3-147395997-INS-383 |
| chr3-147396553-INS-4135 | chr3-147396554-INS-4135 |
| chr3-147407236-INS-54 | chr3-147407236-INS-52 |
| chr3-147408770-INS-81 | chr3-147408770-INS-84 |
| chr3-147434048-INS-7105 | chr3-147434049-INS-7105 |
| chr3-147439585-INS-126 | chr3-147439585-INS-126 |
| chr3-147444959-INS-1279 | chr3-147444966-INS-1273 |
| chr3-147445993-INS-78 | chr3-147445993-INS-57 |
| chr3-147447156-INS-6407 | chr3-147447157-INS-6407 |
| chr3-147465104-INS-199 | chr3-147465104-INS-201 |
| chr3-147483907-INS-482 | chr3-147483907-INS-482 |
| chr3-147499795-INS-494 | chr3-147499795-INS-494 |
| chr3-147503837-INS-5543 | chr3-147503838-INS-5543 |
| chr3-147533175-INS-99 | chr3-147533175-INS-96 |
| chr3-147534270-INS-99 | chr3-147534271-INS-99 |
| chr3-147542585-INS-80 | chr3-147542585-INS-80 |
| chr3-147563711-INS-1107 | chr3-147563712-INS-1107 |
| chr3-147564808-INS-94 | chr3-147564804-INS-118 |
| chr3-147576106-INS-56 | chr3-147576106-INS-52 |
| chr3-147609016-INS-53 | chr3-147609016-INS-53 |
| chr3-147610884-INS-145 | chr3-147610893-INS-117 |
| chr3-147611791-INS-59 | chr3-147611791-INS-69 |
| chr3-147620433-INS-74 | chr3-147620433-INS-74 |
| chr3-147623713-INS-108 | chr3-147623713-INS-100 |
| chr3-147635799-INS-88 | chr3-147635799-INS-88 |
| chr3-147702352-INS-137 | chr3-147702402-INS-133 |
| chr3-147709377-INS-354 | chr3-147709377-INS-354 |
| chr3-147769331-INS-55 | chr3-147769331-INS-55 |
| chr3-147788726-INS-358 | chr3-147788726-INS-358 |
| chr3-147789541-INS-132 | chr3-147789541-INS-132 |
| chr3-147863158-INS-66 | chr3-147863158-INS-66 |
| chr3-147878595-INS-651 | chr3-147878595-INS-651 |
| chr3-147894656-INS-61 | chr3-147894656-INS-61 |
| chr3-147899441-INS-279 | chr3-147899441-INS-281 |
| chr3-147914732-INS-862 | chr3-147914732-INS-862 |
| chr3-147951571-INS-3156 | chr3-147951572-INS-3156 |
| chr3-147968146-INS-166 | chr3-147968146-INS-118 |
| chr3-148022628-INS-413 | chr3-148022628-INS-418 |
| chr3-148047359-INS-65 | chr3-148047358-INS-50 |
| chr3-148053887-INS-63 | chr3-148053909-INS-71 |
| chr3-148062944-INS-89 | chr3-148062944-INS-87 |
| chr3-148080323-INS-80 | chr3-148080329-INS-78 |

| | |
|---|---|
| chr3-148097098-INS-422 | chr3-148097098-INS-423 |
| chr3-148117765-INS-74 | chr3-148117750-INS-110 |
| chr3-148133401-INS-80 | chr3-148133401-INS-80 |
| chr3-148144705-INS-594 | chr3-148144705-INS-594 |
| chr3-148165619-INS-64 | chr3-148165619-INS-70 |
| chr3-148261920-INS-140 | chr3-148261920-INS-140 |
| chr3-148267787-INS-335 | chr3-148267787-INS-336 |
| chr3-148271083-INS-52 | chr3-148271083-INS-52 |
| chr3-148277283-INS-1701 | chr3-148277284-INS-1698 |
| chr3-148290831-INS-201 | chr3-148290844-INS-220 |
| chr3-148304857-INS-7007 | chr3-148304858-INS-7007 |
| chr3-148331645-INS-152 | chr3-148331645-INS-152 |
| chr3-148358587-INS-56 | chr3-148358587-INS-57 |
| chr3-148363252-INS-57 | chr3-148363252-INS-57 |
| chr3-148370300-INS-70 | chr3-148370300-INS-70 |
| chr3-148374681-INS-1480 | chr3-148374682-INS-1480 |
| chr3-148393683-INS-1634 | chr3-148393684-INS-1634 |
| chr3-148397182-INS-85 | chr3-148397182-INS-85 |
| chr3-148404627-INS-128 | chr3-148404624-INS-110 |
| chr3-148407117-INS-54 | chr3-148407117-INS-54 |
| chr3-148432007-INS-51 | chr3-148432007-INS-51 |
| chr3-148488078-INS-106 | chr3-148488078-INS-110 |
| chr3-148494840-INS-89 | chr3-148494884-INS-55 |
| chr3-148504425-INS-500 | chr3-148504419-INS-491 |
| chr3-148508991-INS-130 | chr3-148508991-INS-138 |
| chr3-148518735-INS-68 | chr3-148518735-INS-68 |
| chr3-148528531-INS-58 | chr3-148528531-INS-58 |
| chr3-148535064-INS-552 | chr3-148535064-INS-548 |
| chr3-148539026-INS-80 | chr3-148539024-INS-78 |
| chr3-148574433-INS-84 | chr3-148574433-INS-84 |
| chr3-148591239-INS-56 | chr3-148591239-INS-58 |
| chr3-148630088-INS-96 | chr3-148630088-INS-72 |
| chr3-148632714-INS-709 | chr3-148632715-INS-701 |
| chr3-148634995-INS-56 | chr3-148634995-INS-56 |
| chr3-148641352-INS-56 | chr3-148641352-INS-56 |
| chr3-148687877-INS-53 | chr3-148687875-INS-54 |
| chr3-148699383-INS-94 | chr3-148699383-INS-94 |
| chr3-148705788-INS-58 | chr3-148705788-INS-58 |
| chr3-148708420-INS-6606 | chr3-148708421-INS-6606 |
| chr3-148743723-INS-59 | chr3-148743723-INS-51 |
| chr3-148766233-INS-138 | chr3-148766233-INS-141 |
| chr3-148850063-INS-61 | chr3-148850084-INS-57 |
| chr3-148855782-INS-237 | chr3-148855782-INS-252 |
| chr3-148863642-INS-475 | chr3-148863642-INS-475 |
| chr3-148877897-INS-64 | chr3-148877897-INS-64 |
| chr3-148893154-INS-66 | chr3-148893154-INS-54 |
| chr3-148906700-INS-6233 | chr3-148906701-INS-6233 |
| chr3-148907457-INS-80 | chr3-148907455-INS-94 |
| chr3-148934510-INS-297 | chr3-148934510-INS-297 |
| chr3-148946277-INS-82 | chr3-148946277-INS-86 |
| chr3-148947949-INS-55 | chr3-148947949-INS-75 |
| chr3-148966127-INS-72 | chr3-148966127-INS-72 |
| chr3-148982075-INS-199 | chr3-148982075-INS-204 |
| chr3-148983235-INS-52 | chr3-148983236-INS-70 |
| chr3-148995392-INS-434 | chr3-148995392-INS-434 |
| chr3-149000627-INS-54 | chr3-149000627-INS-56 |
| chr3-149013133-INS-60 | chr3-149013133-INS-56 |
| chr3-149017819-INS-59 | chr3-149017819-INS-59 |
| chr3-149033210-INS-173 | chr3-149033226-INS-172 |
| chr3-149092589-INS-54 | chr3-149092589-INS-55 |
| chr3-149096683-INS-54 | chr3-149096687-INS-56 |
| chr3-149102056-INS-6793 | chr3-149102057-INS-6806 |
| chr3-149103974-INS-189 | chr3-149103974-INS-189 |
| chr3-149115746-INS-126 | chr3-149115746-INS-108 |
| chr3-149118752-INS-102 | chr3-149118750-INS-82 |
| chr3-149121614-INS-55 | chr3-149121614-INS-61 |
| chr3-149134105-INS-72 | chr3-149134115-INS-84 |
| chr3-149134939-INS-247 | chr3-149134939-INS-248 |
| chr3-149146740-INS-56 | chr3-149146740-INS-56 |
| chr3-149152129-INS-98 | chr3-149152129-INS-98 |
| chr3-149154952-INS-89 | chr3-149154952-INS-93 |
| chr3-149171398-INS-177 | chr3-149171424-INS-159 |
| chr3-149191999-INS-52 | chr3-149191999-INS-52 |
| chr3-149203751-INS-223 | chr3-149203745-INS-251 |
| chr3-149247473-INS-76 | chr3-149247473-INS-80 |
| chr3-149275527-INS-54 | chr3-149275527-INS-54 |
| chr3-149307413-INS-80 | chr3-149307413-INS-80 |

| | |
|---|---|
| chr3-149331604-INS-451 | chr3-149331604-INS-451 |
| chr3-149331887-INS-6423 | chr3-149331888-INS-6423 |
| chr3-149373642-INS-104 | chr3-149373639-INS-116 |
| chr3-149389950-INS-127 | chr3-149389950-INS-132 |
| chr3-149404389-INS-691 | chr3-149404389-INS-691 |
| chr3-149437804-INS-6580 | chr3-149437805-INS-6580 |
| chr3-149453685-INS-72 | chr3-149453685-INS-72 |
| chr3-149459362-INS-305 | chr3-149459362-INS-313 |
| chr3-149488064-INS-596 | chr3-149488064-INS-602 |
| chr3-149535412-INS-1135 | chr3-149535413-INS-1132 |
| chr3-149536623-INS-85 | chr3-149536653-INS-54 |
| chr3-149566706-INS-822 | chr3-149566706-INS-820 |
| chr3-149570140-INS-52 | chr3-149570140-INS-54 |
| chr3-149600425-INS-4838 | chr3-149600426-INS-4838 |
| chr3-149618525-INS-1013 | chr3-149618526-INS-1013 |
| chr3-149664399-INS-457 | chr3-149664393-INS-450 |
| chr3-149672982-INS-298 | chr3-149672982-INS-294 |
| chr3-149702684-INS-297 | chr3-149702684-INS-297 |
| chr3-149711835-INS-223 | chr3-149711835-INS-223 |
| chr3-149743296-INS-58 | chr3-149743254-INS-62 |
| chr3-149750344-INS-200 | chr3-149750344-INS-186 |
| chr3-149800822-INS-89 | chr3-149800823-INS-64 |
| chr3-149802375-INS-331 | chr3-149802375-INS-352 |
| chr3-149825138-INS-58 | chr3-149825127-INS-98 |
| chr3-149859456-INS-62 | chr3-149859456-INS-62 |
| chr3-149874236-INS-6740 | chr3-149874237-INS-6742 |
| chr3-149903120-INS-6466 | chr3-149903121-INS-6467 |
| chr3-149903160-INS-1485 | chr3-149903161-INS-1485 |
| chr3-149945213-INS-52 | chr3-149945213-INS-54 |
| chr3-149958225-INS-223 | chr3-149958225-INS-223 |
| chr3-150004535-INS-1067 | chr3-150004536-INS-1067 |
| chr3-150029581-INS-478 | chr3-150029581-INS-478 |
| chr3-150069212-INS-52 | chr3-150069208-INS-58 |
| chr3-150083500-INS-200 | chr3-150083500-INS-205 |
| chr3-150086754-INS-98 | chr3-150086754-INS-98 |
| chr3-150090768-INS-487 | chr3-150090768-INS-487 |
| chr3-150104304-INS-51 | chr3-150104304-INS-51 |
| chr3-150120777-INS-122 | chr3-150120774-INS-127 |
| chr3-150148421-INS-115 | chr3-150148421-INS-115 |
| chr3-150157342-INS-6507 | chr3-150157343-INS-6507 |
| chr3-150170099-INS-4392 | chr3-150170100-INS-4392 |
| chr3-150174112-INS-54 | chr3-150174113-INS-52 |
| chr3-150202743-INS-1132 | chr3-150202744-INS-1133 |
| chr3-150217718-INS-368 | chr3-150217718-INS-368 |
| chr3-150235464-INS-4851 | chr3-150235465-INS-4851 |
| chr3-150244147-INS-198 | chr3-150244147-INS-198 |
| chr3-150251957-INS-3097 | chr3-150251958-INS-3097 |
| chr3-150270355-INS-95 | chr3-150270355-INS-95 |
| chr3-150282617-INS-98 | chr3-150282617-INS-98 |
| chr3-150285365-INS-6595 | chr3-150285366-INS-6597 |
| chr3-150289145-INS-76 | chr3-150289145-INS-76 |
| chr3-150290816-INS-54 | chr3-150290791-INS-78 |
| chr3-150320990-INS-481 | chr3-150320990-INS-481 |
| chr3-150327818-INS-281 | chr3-150327819-INS-281 |
| chr3-150334316-INS-56 | chr3-150334291-INS-52 |
| chr3-150363973-INS-5251 | chr3-150363974-INS-5251 |
| chr3-150364922-INS-6656 | chr3-150364923-INS-6657 |
| chr3-150410970-INS-1911 | chr3-150410971-INS-1911 |
| chr3-150419871-INS-55 | chr3-150419871-INS-55 |
| chr3-150422298-INS-125 | chr3-150422294-INS-123 |
| chr3-150472162-INS-51 | chr3-150472192-INS-70 |
| chr3-150472866-INS-85 | chr3-150472866-INS-75 |
| chr3-150488527-INS-256 | chr3-150488527-INS-256 |
| chr3-150494900-INS-483 | chr3-150494900-INS-483 |
| chr3-150506692-INS-139 | chr3-150506692-INS-138 |
| chr3-150509109-INS-68 | chr3-150509109-INS-68 |
| chr3-150535104-INS-54 | chr3-150535104-INS-54 |
| chr3-150537316-INS-56 | chr3-150537316-INS-52 |
| chr3-150540351-INS-211 | chr3-150540350-INS-208 |
| chr3-150571640-INS-57 | chr3-150571640-INS-57 |
| chr3-150630951-INS-195 | chr3-150630951-INS-192 |
| chr3-150683012-INS-401 | chr3-150683012-INS-401 |
| chr3-150694337-INS-82 | chr3-150694337-INS-86 |
| chr3-150710447-INS-85 | chr3-150710447-INS-85 |
| chr3-150748512-INS-726 | chr3-150748512-INS-726 |
| chr3-150754983-INS-358 | chr3-150754983-INS-358 |
| chr3-150758434-INS-54 | chr3-150758435-INS-94 |

| | |
|---|---|
| chr3-150769381-INS-395 | chr3-150769381-INS-395 |
| chr3-150790997-INS-5326 | chr3-150790998-INS-5326 |
| chr3-150792583-INS-56 | chr3-150792603-INS-68 |
| chr3-150796042-INS-652 | chr3-150796042-INS-652 |
| chr3-150803127-INS-73 | chr3-150803127-INS-71 |
| chr3-150805168-INS-72 | chr3-150805168-INS-70 |
| chr3-150810825-INS-96 | chr3-150810825-INS-96 |
| chr3-150831588-INS-75 | chr3-150831588-INS-75 |
| chr3-150843027-INS-93 | chr3-150843039-INS-53 |
| chr3-150896897-INS-60 | chr3-150896897-INS-62 |
| chr3-150903121-INS-110 | chr3-150903121-INS-110 |
| chr3-150912131-INS-584 | chr3-150912131-INS-584 |
| chr3-150950033-INS-568 | chr3-150950033-INS-568 |
| chr3-150951686-INS-66 | chr3-150951686-INS-66 |
| chr3-150959562-INS-187 | chr3-150959562-INS-187 |
| chr3-150962382-INS-959 | chr3-150962382-INS-955 |
| chr3-150966502-INS-206 | chr3-150966502-INS-206 |
| chr3-151010902-INS-120 | chr3-151010902-INS-120 |
| chr3-151023540-INS-3048 | chr3-151023541-INS-3048 |
| chr3-151030191-INS-54 | chr3-151030191-INS-50 |
| chr3-151046691-INS-4124 | chr3-151046692-INS-4125 |
| chr3-151049598-INS-130 | chr3-151049598-INS-131 |
| chr3-151051993-INS-51 | chr3-151051993-INS-57 |
| chr3-151052044-INS-617 | chr3-151052045-INS-602 |
| chr3-151068397-INS-7039 | chr3-151068398-INS-7039 |
| chr3-151100223-INS-55 | chr3-151100223-INS-57 |
| chr3-151137840-INS-348 | chr3-151137840-INS-348 |
| chr3-151139182-INS-962 | chr3-151139182-INS-971 |
| chr3-151147708-INS-54 | chr3-151147709-INS-54 |
| chr3-151167360-INS-57 | chr3-151167358-INS-54 |
| chr3-151175148-INS-62 | chr3-151175148-INS-62 |
| chr3-151225394-INS-84 | chr3-151225410-INS-72 |
| chr3-151226660-INS-74 | chr3-151226660-INS-74 |
| chr3-151233570-INS-90 | chr3-151233570-INS-92 |
| chr3-151243945-INS-107 | chr3-151243945-INS-107 |
| chr3-151267290-INS-144 | chr3-151267290-INS-150 |
| chr3-151269001-INS-4837 | chr3-151269002-INS-4837 |
| chr3-151270377-INS-146 | chr3-151270377-INS-154 |
| chr3-151271329-INS-466 | chr3-151271330-INS-467 |
| chr3-151286656-INS-336 | chr3-151286656-INS-336 |
| chr3-151315046-INS-64 | chr3-151315046-INS-70 |
| chr3-151329192-INS-498 | chr3-151329192-INS-498 |
| chr3-151354755-INS-75 | chr3-151354737-INS-60 |
| chr3-151447801-INS-65 | chr3-151447801-INS-65 |
| chr3-151456336-INS-184 | chr3-151456336-INS-184 |
| chr3-151458495-INS-74 | chr3-151458495-INS-86 |
| chr3-151494999-INS-62 | chr3-151494999-INS-64 |
| chr3-151497997-INS-56 | chr3-151497997-INS-56 |
| chr3-151537125-INS-65 | chr3-151537118-INS-68 |
| chr3-151546146-INS-913 | chr3-151546146-INS-913 |
| chr3-151558101-INS-56 | chr3-151558101-INS-60 |
| chr3-151560695-INS-5407 | chr3-151560696-INS-5407 |
| chr3-151564651-INS-461 | chr3-151564651-INS-461 |
| chr3-151573113-INS-117 | chr3-151573089-INS-123 |
| chr3-151604024-INS-147 | chr3-151604024-INS-155 |
| chr3-151618148-INS-56 | chr3-151618148-INS-56 |
| chr3-151629879-INS-454 | chr3-151629879-INS-454 |
| chr3-151687125-INS-954 | chr3-151687125-INS-954 |
| chr3-151691797-INS-2786 | chr3-151691798-INS-2786 |
| chr3-151699069-INS-65 | chr3-151699069-INS-65 |
| chr3-151705240-INS-1069 | chr3-151705241-INS-1069 |
| chr3-151713937-INS-286 | chr3-151713938-INS-287 |
| chr3-151722158-INS-113 | chr3-151722158-INS-113 |
| chr3-151733360-INS-72 | chr3-151733360-INS-72 |
| chr3-151749112-INS-54 | chr3-151749115-INS-67 |
| chr3-151759495-INS-903 | chr3-151759495-INS-903 |
| chr3-151763828-INS-2562 | chr3-151763829-INS-2563 |
| chr3-151764807-INS-343 | chr3-151764807-INS-343 |
| chr3-151780395-INS-1831 | chr3-151780396-INS-1831 |
| chr3-151789035-INS-56 | chr3-151789025-INS-50 |
| chr3-151804562-INS-70 | chr3-151804570-INS-60 |
| chr3-151822931-INS-64 | chr3-151822931-INS-66 |
| chr3-151857899-INS-174 | chr3-151857899-INS-173 |
| chr3-151865632-INS-7174 | chr3-151865633-INS-7173 |
| chr3-151902660-INS-308 | chr3-151902660-INS-308 |
| chr3-151907430-INS-64 | chr3-151907435-INS-71 |
| chr3-151975532-INS-7097 | chr3-151975533-INS-7097 |

| | |
|---|---|
| chr3-151992236-INS-96 | chr3-151992233-INS-96 |
| chr3-151999662-INS-143 | chr3-151999662-INS-143 |
| chr3-152000628-INS-206 | chr3-152000615-INS-206 |
| chr3-152002615-INS-64 | chr3-152002615-INS-64 |
| chr3-152006945-INS-228 | chr3-152006945-INS-228 |
| chr3-152021506-INS-88 | chr3-152021506-INS-88 |
| chr3-152022655-INS-60 | chr3-152022655-INS-60 |
| chr3-152038258-INS-66 | chr3-152038258-INS-66 |
| chr3-152060908-INS-943 | chr3-152060908-INS-953 |
| chr3-152107476-INS-132 | chr3-152107476-INS-132 |
| chr3-152138201-INS-246 | chr3-152138201-INS-246 |
| chr3-152148702-INS-77 | chr3-152148702-INS-77 |
| chr3-152195387-INS-170 | chr3-152195387-INS-166 |
| chr3-152230095-INS-509 | chr3-152230095-INS-509 |
| chr3-152234300-INS-85 | chr3-152234300-INS-85 |
| chr3-152235799-INS-569 | chr3-152235799-INS-569 |
| chr3-152239763-INS-60 | chr3-152239756-INS-101 |
| chr3-152310548-INS-55 | chr3-152310557-INS-56 |
| chr3-152320344-INS-120 | chr3-152320344-INS-120 |
| chr3-152325606-INS-117 | chr3-152325606-INS-117 |
| chr3-152340932-INS-62 | chr3-152340932-INS-62 |
| chr3-152346467-INS-88 | chr3-152346467-INS-88 |
| chr3-152358331-INS-193 | chr3-152358332-INS-194 |
| chr3-152381619-INS-291 | chr3-152381619-INS-291 |
| chr3-152385660-INS-453 | chr3-152385660-INS-453 |
| chr3-152390625-INS-272 | chr3-152390625-INS-269 |
| chr3-152407224-INS-88 | chr3-152407240-INS-52 |
| chr3-152411733-INS-692 | chr3-152411733-INS-692 |
| chr3-152429783-INS-187 | chr3-152429783-INS-189 |
| chr3-152431550-INS-77 | chr3-152431550-INS-103 |
| chr3-152438274-INS-385 | chr3-152438274-INS-385 |
| chr3-152442175-INS-316 | chr3-152442175-INS-316 |
| chr3-152447786-INS-135 | chr3-152447786-INS-135 |
| chr3-152449773-INS-466 | chr3-152449773-INS-466 |
| chr3-152486091-INS-196 | chr3-152486091-INS-196 |
| chr3-152500251-INS-74 | chr3-152500251-INS-74 |
| chr3-152518427-INS-107 | chr3-152518427-INS-107 |
| chr3-152531725-INS-1059 | chr3-152531726-INS-1059 |
| chr3-152560552-INS-60 | chr3-152560552-INS-60 |
| chr3-152569190-INS-370 | chr3-152569190-INS-370 |
| chr3-152596377-INS-54 | chr3-152596407-INS-63 |
| chr3-152598453-INS-96 | chr3-152598453-INS-91 |
| chr3-152612656-INS-3331 | chr3-152612657-INS-3331 |
| chr3-152638582-INS-905 | chr3-152638582-INS-905 |
| chr3-152646639-INS-588 | chr3-152646639-INS-590 |
| chr3-152721746-INS-68 | chr3-152721742-INS-68 |
| chr3-152726533-INS-58 | chr3-152726533-INS-58 |
| chr3-152738962-INS-62 | chr3-152738962-INS-62 |
| chr3-152748211-INS-68 | chr3-152748211-INS-68 |
| chr3-152756877-INS-64 | chr3-152756877-INS-64 |
| chr3-152765852-INS-91 | chr3-152765852-INS-76 |
| chr3-152769019-INS-71 | chr3-152769019-INS-71 |
| chr3-152771033-INS-102 | chr3-152771058-INS-102 |
| chr3-152794792-INS-452 | chr3-152794792-INS-452 |
| chr3-152819753-INS-62 | chr3-152819753-INS-58 |
| chr3-152824851-INS-124 | chr3-152824851-INS-124 |
| chr3-152835274-INS-52 | chr3-152835274-INS-52 |
| chr3-152838744-INS-64 | chr3-152838744-INS-68 |
| chr3-152848246-INS-587 | chr3-152848246-INS-587 |
| chr3-152863153-INS-52 | chr3-152863153-INS-52 |
| chr3-152899851-INS-497 | chr3-152899851-INS-498 |
| chr3-152927009-INS-363 | chr3-152927009-INS-363 |
| chr3-152927952-INS-164 | chr3-152927952-INS-164 |
| chr3-152944772-INS-82 | chr3-152944772-INS-82 |
| chr3-152950975-INS-56 | chr3-152950975-INS-56 |
| chr3-152978415-INS-6544 | chr3-152978416-INS-6544 |
| chr3-152993687-INS-651 | chr3-152993687-INS-651 |
| chr3-153021239-INS-150 | chr3-153021239-INS-150 |
| chr3-153033994-INS-284 | chr3-153033994-INS-269 |
| chr3-153060080-INS-57 | chr3-153060104-INS-57 |
| chr3-153093884-INS-96 | chr3-153093884-INS-96 |
| chr3-153108489-INS-64 | chr3-153108489-INS-64 |
| chr3-153139334-INS-57 | chr3-153139334-INS-59 |
| chr3-153139874-INS-62 | chr3-153139862-INS-62 |
| chr3-153147510-INS-1892 | chr3-153147511-INS-1892 |
| chr3-153154049-INS-150 | chr3-153154049-INS-150 |
| chr3-153155955-INS-96 | chr3-153155955-INS-96 |

| | |
|---|---|
| chr3-153190026-INS-83 | chr3-153190026-INS-87 |
| chr3-153230401-INS-81 | chr3-153230401-INS-81 |
| chr3-153236694-INS-156 | chr3-153236694-INS-158 |
| chr3-153243507-INS-68 | chr3-153243507-INS-68 |
| chr3-153253742-INS-60 | chr3-153253742-INS-60 |
| chr3-153284389-INS-54 | chr3-153284389-INS-52 |
| chr3-153286995-INS-390 | chr3-153286995-INS-400 |
| chr3-153287863-INS-54 | chr3-153287863-INS-69 |
| chr3-153295241-INS-632 | chr3-153295241-INS-632 |
| chr3-153313933-INS-436 | chr3-153313933-INS-450 |
| chr3-153327602-INS-56 | chr3-153327602-INS-56 |
| chr3-153346491-INS-431 | chr3-153346491-INS-432 |
| chr3-153352069-INS-73 | chr3-153352069-INS-73 |
| chr3-153376594-INS-427 | chr3-153376594-INS-427 |
| chr3-153377659-INS-282 | chr3-153377659-INS-283 |
| chr3-153390700-INS-140 | chr3-153390700-INS-140 |
| chr3-153406531-INS-218 | chr3-153406531-INS-218 |
| chr3-153409290-INS-65 | chr3-153409290-INS-92 |
| chr3-153436503-INS-60 | chr3-153436503-INS-60 |
| chr3-153457906-INS-67 | chr3-153457905-INS-64 |
| chr3-153459131-INS-235 | chr3-153459142-INS-213 |
| chr3-153460169-INS-134 | chr3-153460169-INS-134 |
| chr3-153484549-INS-213 | chr3-153484549-INS-214 |
| chr3-153521686-INS-482 | chr3-153521686-INS-482 |
| chr3-153532201-INS-52 | chr3-153532201-INS-54 |
| chr3-153545243-INS-1396 | chr3-153545244-INS-1396 |
| chr3-153551221-INS-254 | chr3-153551224-INS-255 |
| chr3-153562433-INS-197 | chr3-153562424-INS-203 |
| chr3-153581142-INS-464 | chr3-153581142-INS-466 |
| chr3-153583257-INS-61 | chr3-153583257-INS-61 |
| chr3-153589039-INS-820 | chr3-153589039-INS-820 |
| chr3-153590707-INS-202 | chr3-153590707-INS-202 |
| chr3-153596052-INS-7777 | chr3-153596053-INS-7778 |
| chr3-153605974-INS-519 | chr3-153605974-INS-519 |
| chr3-153620918-INS-199 | chr3-153620918-INS-198 |
| chr3-153622920-INS-199 | chr3-153622920-INS-199 |
| chr3-153655712-INS-657 | chr3-153655712-INS-657 |
| chr3-153666204-INS-198 | chr3-153666204-INS-198 |
| chr3-153672245-INS-168 | chr3-153672245-INS-196 |
| chr3-153673687-INS-122 | chr3-153673687-INS-123 |
| chr3-153684879-INS-104 | chr3-153684880-INS-90 |
| chr3-153694835-INS-81 | chr3-153694835-INS-81 |
| chr3-153708332-INS-64 | chr3-153708332-INS-64 |
| chr3-153757430-INS-245 | chr3-153757430-INS-245 |
| chr3-153761438-INS-480 | chr3-153761438-INS-478 |
| chr3-153772052-INS-74 | chr3-153772051-INS-75 |
| chr3-153782183-INS-473 | chr3-153782183-INS-473 |
| chr3-153784715-INS-100 | chr3-153784715-INS-104 |
| chr3-153791318-INS-68 | chr3-153791318-INS-68 |
| chr3-153802053-INS-512 | chr3-153802053-INS-518 |
| chr3-153806042-INS-72 | chr3-153806041-INS-88 |
| chr3-153833122-INS-654 | chr3-153833122-INS-658 |
| chr3-153837896-INS-402 | chr3-153837896-INS-402 |
| chr3-153852551-INS-7289 | chr3-153852552-INS-7289 |
| chr3-153882684-INS-1033 | chr3-153882685-INS-1026 |
| chr3-153892075-INS-107 | chr3-153892075-INS-89 |
| chr3-153900978-INS-60 | chr3-153900978-INS-60 |
| chr3-153945416-INS-72 | chr3-153945416-INS-72 |
| chr3-153959962-INS-120 | chr3-153959962-INS-120 |
| chr3-153982719-INS-68 | chr3-153982719-INS-60 |
| chr3-153987271-INS-100 | chr3-153987271-INS-100 |
| chr3-154097276-INS-124 | chr3-154097276-INS-123 |
| chr3-154105121-INS-520 | chr3-154105121-INS-521 |
| chr3-154117470-INS-74 | chr3-154117470-INS-74 |
| chr3-154123571-INS-132 | chr3-154123571-INS-132 |
| chr3-154159154-INS-144 | chr3-154159154-INS-148 |
| chr3-154184328-INS-194 | chr3-154184328-INS-194 |
| chr3-154195615-INS-3694 | chr3-154195616-INS-3670 |
| chr3-154209844-INS-97 | chr3-154209844-INS-97 |
| chr3-154230709-INS-53 | chr3-154230703-INS-56 |
| chr3-154232187-INS-81 | chr3-154232187-INS-81 |
| chr3-154240994-INS-97 | chr3-154240994-INS-97 |
| chr3-154242870-INS-7160 | chr3-154242871-INS-7160 |
| chr3-154244471-INS-53 | chr3-154244471-INS-53 |
| chr3-154263251-INS-60 | chr3-154263251-INS-60 |
| chr3-154300064-INS-144 | chr3-154300064-INS-144 |
| chr3-154387378-INS-88 | chr3-154387378-INS-88 |

| | |
|---|---|
| chr3-154390372-INS-204 | chr3-154390372-INS-202 |
| chr3-154399420-INS-203 | chr3-154399420-INS-202 |
| chr3-154442671-INS-96 | chr3-154442671-INS-96 |
| chr3-154480914-INS-165 | chr3-154480914-INS-165 |
| chr3-154490362-INS-60 | chr3-154490362-INS-68 |
| chr3-154495079-INS-7122 | chr3-154495080-INS-7122 |
| chr3-154544239-INS-456 | chr3-154544239-INS-453 |
| chr3-154560644-INS-244 | chr3-154560646-INS-264 |
| chr3-154563716-INS-50 | chr3-154563716-INS-50 |
| chr3-154615219-INS-98 | chr3-154615218-INS-95 |
| chr3-154621720-INS-73 | chr3-154621720-INS-75 |
| chr3-154643130-INS-179 | chr3-154643130-INS-179 |
| chr3-154651253-INS-61 | chr3-154651259-INS-89 |
| chr3-154660368-INS-7195 | chr3-154660369-INS-7195 |
| chr3-154685115-INS-54 | chr3-154685113-INS-56 |
| chr3-154696451-INS-72 | chr3-154696414-INS-94 |
| chr3-154715292-INS-54 | chr3-154715292-INS-54 |
| chr3-154716371-INS-76 | chr3-154716361-INS-58 |
| chr3-154722816-INS-52 | chr3-154722816-INS-52 |
| chr3-154727877-INS-86 | chr3-154727880-INS-89 |
| chr3-154798619-INS-146 | chr3-154798619-INS-146 |
| chr3-154825278-INS-57 | chr3-154825278-INS-64 |
| chr3-154835716-INS-73 | chr3-154835716-INS-73 |
| chr3-154844544-INS-7151 | chr3-154844545-INS-7152 |
| chr3-154857581-INS-199 | chr3-154857581-INS-199 |
| chr3-154895487-INS-52 | chr3-154895487-INS-56 |
| chr3-154907468-INS-54 | chr3-154907468-INS-54 |
| chr3-154910787-INS-230 | chr3-154910787-INS-230 |
| chr3-154938958-INS-115 | chr3-154938958-INS-120 |
| chr3-154984894-INS-80 | chr3-154984922-INS-96 |
| chr3-154993357-INS-54 | chr3-154993357-INS-72 |
| chr3-155015568-INS-212 | chr3-155015568-INS-212 |
| chr3-155037307-INS-52 | chr3-155037307-INS-52 |
| chr3-155040134-INS-244 | chr3-155040134-INS-245 |
| chr3-155050165-INS-7170 | chr3-155050166-INS-7170 |
| chr3-155139177-INS-6180 | chr3-155139178-INS-6180 |
| chr3-155148634-INS-819 | chr3-155148634-INS-821 |
| chr3-155149332-INS-76 | chr3-155149332-INS-76 |
| chr3-155157616-INS-115 | chr3-155157616-INS-115 |
| chr3-155171051-INS-68 | chr3-155171038-INS-57 |
| chr3-155172951-INS-67 | chr3-155172951-INS-55 |
| chr3-155175716-INS-54 | chr3-155175716-INS-53 |
| chr3-155176698-INS-585 | chr3-155176698-INS-587 |
| chr3-155186429-INS-384 | chr3-155186429-INS-387 |
| chr3-155203981-INS-61 | chr3-155203981-INS-65 |
| chr3-155219613-INS-132 | chr3-155219618-INS-134 |
| chr3-155286633-INS-497 | chr3-155286633-INS-495 |
| chr3-155294040-INS-84 | chr3-155294042-INS-54 |
| chr3-155341166-INS-171 | chr3-155341166-INS-169 |
| chr3-155352755-INS-76 | chr3-155352755-INS-96 |
| chr3-155382886-INS-593 | chr3-155382886-INS-573 |
| chr3-155384952-INS-68 | chr3-155384952-INS-68 |
| chr3-155386994-INS-136 | chr3-155386994-INS-136 |
| chr3-155406735-INS-236 | chr3-155406735-INS-216 |
| chr3-155419042-INS-230 | chr3-155419042-INS-230 |
| chr3-155419780-INS-6382 | chr3-155419776-INS-6355 |
| chr3-155434806-INS-342 | chr3-155434806-INS-342 |
| chr3-155446246-INS-212 | chr3-155446246-INS-213 |
| chr3-155457703-INS-109 | chr3-155457703-INS-109 |
| chr3-155474196-INS-2866 | chr3-155474197-INS-2866 |
| chr3-155495856-INS-97 | chr3-155495856-INS-104 |
| chr3-155531703-INS-70 | chr3-155531701-INS-86 |
| chr3-155531895-INS-6303 | chr3-155531896-INS-6303 |
| chr3-155537075-INS-8225 | chr3-155537076-INS-8225 |
| chr3-155537076-INS-424 | chr3-155537076-INS-424 |
| chr3-155592854-INS-1423 | chr3-155592855-INS-1421 |
| chr3-155624093-INS-73 | chr3-155624091-INS-65 |
| chr3-155625702-INS-112 | chr3-155625702-INS-117 |
| chr3-155637705-INS-328 | chr3-155637706-INS-328 |
| chr3-155661050-INS-58 | chr3-155661050-INS-58 |
| chr3-155680822-INS-308 | chr3-155680828-INS-306 |
| chr3-155702097-INS-52 | chr3-155702097-INS-50 |
| chr3-155705302-INS-656 | chr3-155705302-INS-656 |
| chr3-155736511-INS-65 | chr3-155736511-INS-65 |
| chr3-155767630-INS-1696 | chr3-155767631-INS-1696 |
| chr3-155769958-INS-64 | chr3-155769958-INS-50 |
| chr3-155781784-INS-540 | chr3-155781784-INS-539 |

| | |
|---|---|
| chr3-155807056-INS-162 | chr3-155807056-INS-164 |
| chr3-155814061-INS-405 | chr3-155814061-INS-405 |
| chr3-155821156-INS-60 | chr3-155821156-INS-60 |
| chr3-155823372-INS-58 | chr3-155823374-INS-56 |
| chr3-155865301-INS-6401 | chr3-155865302-INS-6401 |
| chr3-155904485-INS-828 | chr3-155904485-INS-828 |
| chr3-155917867-INS-74 | chr3-155917867-INS-78 |
| chr3-155919644-INS-85 | chr3-155919644-INS-86 |
| chr3-155934669-INS-114 | chr3-155934669-INS-114 |
| chr3-155935792-INS-6612 | chr3-155935793-INS-6612 |
| chr3-155949636-INS-7103 | chr3-155949637-INS-7103 |
| chr3-155956847-INS-70 | chr3-155956847-INS-114 |
| chr3-155982602-INS-130 | chr3-155982602-INS-130 |
| chr3-155982677-INS-7144 | chr3-155982678-INS-7146 |
| chr3-156057988-INS-81 | chr3-156057988-INS-81 |
| chr3-156075398-INS-455 | chr3-156075398-INS-455 |
| chr3-156082872-INS-88 | chr3-156082872-INS-91 |
| chr3-156086667-INS-52 | chr3-156086667-INS-54 |
| chr3-156088549-INS-4600 | chr3-156088550-INS-4597 |
| chr3-156090967-INS-345 | chr3-156090967-INS-365 |
| chr3-156097662-INS-54 | chr3-156097662-INS-72 |
| chr3-156114603-INS-64 | chr3-156114603-INS-72 |
| chr3-156127373-INS-119 | chr3-156127373-INS-118 |
| chr3-156129257-INS-5191 | chr3-156129258-INS-5190 |
| chr3-156131440-INS-182 | chr3-156131440-INS-182 |
| chr3-156139132-INS-120 | chr3-156139132-INS-120 |
| chr3-156139696-INS-880 | chr3-156139696-INS-880 |
| chr3-156150829-INS-68 | chr3-156150829-INS-110 |
| chr3-156166143-INS-76 | chr3-156166107-INS-90 |
| chr3-156180799-INS-1224 | chr3-156180800-INS-1224 |
| chr3-156188652-INS-58 | chr3-156188652-INS-58 |
| chr3-156220975-INS-6473 | chr3-156220976-INS-6473 |
| chr3-156221221-INS-76 | chr3-156221221-INS-76 |
| chr3-156224086-INS-379 | chr3-156224086-INS-380 |
| chr3-156226529-INS-122 | chr3-156226529-INS-122 |
| chr3-156235768-INS-81 | chr3-156235772-INS-97 |
| chr3-156239163-INS-106 | chr3-156239163-INS-110 |
| chr3-156247455-INS-6178 | chr3-156247456-INS-6180 |
| chr3-156257104-INS-98 | chr3-156257104-INS-98 |
| chr3-156266935-INS-50 | chr3-156266935-INS-55 |
| chr3-156282971-INS-82 | chr3-156282970-INS-75 |
| chr3-156293743-INS-64 | chr3-156293743-INS-64 |
| chr3-156297832-INS-298 | chr3-156297832-INS-298 |
| chr3-156322797-INS-6384 | chr3-156322798-INS-6384 |
| chr3-156345518-INS-130 | chr3-156345511-INS-122 |
| chr3-156369588-INS-6628 | chr3-156369589-INS-6629 |
| chr3-156389765-INS-652 | chr3-156389765-INS-652 |
| chr3-156395616-INS-96 | chr3-156395616-INS-96 |
| chr3-156403056-INS-7171 | chr3-156403057-INS-7172 |
| chr3-156428868-INS-51 | chr3-156428869-INS-51 |
| chr3-156445067-INS-113 | chr3-156445067-INS-113 |
| chr3-156466992-INS-1292 | chr3-156466993-INS-1292 |
| chr3-156474678-INS-52 | chr3-156474678-INS-52 |
| chr3-156501205-INS-142 | chr3-156501205-INS-142 |
| chr3-156549174-INS-98 | chr3-156549178-INS-78 |
| chr3-156581286-INS-52 | chr3-156581286-INS-52 |
| chr3-156634002-INS-75 | chr3-156634002-INS-71 |
| chr3-156688938-INS-227 | chr3-156688938-INS-227 |
| chr3-156720171-INS-223 | chr3-156720171-INS-223 |
| chr3-156730570-INS-7152 | chr3-156730571-INS-7152 |
| chr3-156738812-INS-52 | chr3-156738812-INS-52 |
| chr3-156747499-INS-172 | chr3-156747499-INS-172 |
| chr3-156764164-INS-82 | chr3-156764196-INS-74 |
| chr3-156805112-INS-1032 | chr3-156805113-INS-1032 |
| chr3-156815074-INS-62 | chr3-156815025-INS-66 |
| chr3-156821978-INS-53 | chr3-156821978-INS-53 |
| chr3-156831761-INS-114 | chr3-156831761-INS-115 |
| chr3-156856832-INS-162 | chr3-156856831-INS-164 |
| chr3-156904020-INS-2881 | chr3-156904021-INS-2881 |
| chr3-156912786-INS-270 | chr3-156912786-INS-223 |
| chr3-156921065-INS-248 | chr3-156921065-INS-252 |
| chr3-157015101-INS-66 | chr3-157015101-INS-66 |
| chr3-157083112-INS-441 | chr3-157083112-INS-441 |
| chr3-157104103-INS-85 | chr3-157104115-INS-98 |
| chr3-157105545-INS-71 | chr3-157105542-INS-72 |
| chr3-157147226-INS-1626 | chr3-157147227-INS-1626 |
| chr3-157158443-INS-1506 | chr3-157158444-INS-1508 |

| | |
|---|---|
| chr3-157174837-INS-85 | chr3-157174850-INS-70 |
| chr3-157213635-INS-6896 | chr3-157213636-INS-6896 |
| chr3-157238963-INS-200 | chr3-157238963-INS-200 |
| chr3-157302234-INS-54 | chr3-157302234-INS-54 |
| chr3-157318420-INS-58 | chr3-157318434-INS-78 |
| chr3-157341797-INS-77 | chr3-157341797-INS-77 |
| chr3-157362240-INS-682 | chr3-157362240-INS-686 |
| chr3-157381446-INS-6692 | chr3-157381447-INS-6692 |
| chr3-157389351-INS-110 | chr3-157389351-INS-81 |
| chr3-157392780-INS-3274 | chr3-157392781-INS-3279 |
| chr3-157429595-INS-469 | chr3-157429595-INS-469 |
| chr3-157470532-INS-198 | chr3-157470532-INS-198 |
| chr3-157486160-INS-83 | chr3-157486160-INS-83 |
| chr3-157490735-INS-3642 | chr3-157490736-INS-3642 |
| chr3-157497616-INS-282 | chr3-157497617-INS-282 |
| chr3-157532394-INS-421 | chr3-157532394-INS-421 |
| chr3-157535053-INS-212 | chr3-157535053-INS-212 |
| chr3-157539897-INS-169 | chr3-157539897-INS-169 |
| chr3-157545213-INS-221 | chr3-157545213-INS-223 |
| chr3-157551358-INS-264 | chr3-157551362-INS-265 |
| chr3-157555486-INS-401 | chr3-157555486-INS-401 |
| chr3-157559134-INS-207 | chr3-157559134-INS-207 |
| chr3-157571113-INS-233 | chr3-157571113-INS-233 |
| chr3-157593367-INS-143 | chr3-157593367-INS-143 |
| chr3-157595755-INS-402 | chr3-157595712-INS-426 |
| chr3-157602621-INS-384 | chr3-157602621-INS-384 |
| chr3-157606494-INS-221 | chr3-157606494-INS-221 |
| chr3-157623135-INS-204 | chr3-157623135-INS-204 |
| chr3-157644803-INS-193 | chr3-157644803-INS-193 |
| chr3-157650535-INS-191 | chr3-157650535-INS-191 |
| chr3-157664064-INS-228 | chr3-157664064-INS-228 |
| chr3-157679898-INS-400 | chr3-157679898-INS-400 |
| chr3-157701763-INS-76 | chr3-157701742-INS-52 |
| chr3-157704851-INS-359 | chr3-157704851-INS-359 |
| chr3-157752200-INS-79 | chr3-157752200-INS-79 |
| chr3-157759031-INS-58 | chr3-157759031-INS-62 |
| chr3-157777033-INS-453 | chr3-157777033-INS-453 |
| chr3-157780567-INS-72 | chr3-157780560-INS-52 |
| chr3-157798828-INS-83 | chr3-157798828-INS-83 |
| chr3-157845637-INS-753 | chr3-157845638-INS-754 |
| chr3-157889814-INS-103 | chr3-157889814-INS-108 |
| chr3-157891324-INS-81 | chr3-157891322-INS-102 |
| chr3-157894085-INS-112 | chr3-157894085-INS-121 |
| chr3-157894745-INS-542 | chr3-157894745-INS-542 |
| chr3-157913545-INS-188 | chr3-157913545-INS-188 |
| chr3-157916986-INS-176 | chr3-157916983-INS-176 |
| chr3-157933647-INS-64 | chr3-157933647-INS-64 |
| chr3-157974223-INS-60 | chr3-157974223-INS-60 |
| chr3-157978942-INS-52 | chr3-157978942-INS-54 |
| chr3-158019532-INS-3053 | chr3-158019533-INS-3053 |
| chr3-158059303-INS-219 | chr3-158059303-INS-219 |
| chr3-158070607-INS-6745 | chr3-158070608-INS-6745 |
| chr3-158083461-INS-82 | chr3-158083458-INS-63 |
| chr3-158153793-INS-212 | chr3-158153793-INS-212 |
| chr3-158155935-INS-56 | chr3-158155935-INS-56 |
| chr3-158164747-INS-433 | chr3-158164749-INS-433 |
| chr3-158232237-INS-504 | chr3-158232237-INS-499 |
| chr3-158248626-INS-76 | chr3-158248632-INS-82 |
| chr3-158279320-INS-6475 | chr3-158279321-INS-6475 |
| chr3-158284868-INS-72 | chr3-158284868-INS-72 |
| chr3-158285469-INS-1735 | chr3-158285470-INS-1735 |
| chr3-158297718-INS-494 | chr3-158297718-INS-494 |
| chr3-158298775-INS-441 | chr3-158298775-INS-441 |
| chr3-158314260-INS-126 | chr3-158314260-INS-126 |
| chr3-158361786-INS-6197 | chr3-158361787-INS-6197 |
| chr3-158392217-INS-1058 | chr3-158392218-INS-1058 |
| chr3-158422980-INS-1069 | chr3-158422981-INS-1071 |
| chr3-158462856-INS-1592 | chr3-158462877-INS-1569 |
| chr3-158466284-INS-52 | chr3-158466284-INS-56 |
| chr3-158485532-INS-6601 | chr3-158485533-INS-6601 |
| chr3-158517483-INS-68 | chr3-158517483-INS-72 |
| chr3-158526946-INS-56 | chr3-158526946-INS-56 |
| chr3-158546393-INS-264 | chr3-158546393-INS-264 |
| chr3-158555975-INS-3413 | chr3-158555976-INS-3413 |
| chr3-158556417-INS-998 | chr3-158556417-INS-1004 |
| chr3-158559655-INS-213 | chr3-158559655-INS-213 |
| chr3-158618631-INS-1163 | chr3-158618632-INS-1164 |

| | |
|---|---|
| chr3-158624800-INS-7847 | chr3-158624801-INS-7847 |
| chr3-158636327-INS-1327 | chr3-158636328-INS-1327 |
| chr3-158641521-INS-1398 | chr3-158641517-INS-1377 |
| chr3-158672659-INS-860 | chr3-158672659-INS-861 |
| chr3-158675815-INS-62 | chr3-158675815-INS-64 |
| chr3-158725984-INS-159 | chr3-158725984-INS-159 |
| chr3-158733704-INS-54 | chr3-158733700-INS-56 |
| chr3-158765725-INS-86 | chr3-158765725-INS-86 |
| chr3-158774653-INS-498 | chr3-158774653-INS-498 |
| chr3-158804769-INS-64 | chr3-158804795-INS-71 |
| chr3-158845264-INS-6885 | chr3-158845265-INS-6885 |
| chr3-158904272-INS-186 | chr3-158904272-INS-186 |
| chr3-158914007-INS-138 | chr3-158914007-INS-142 |
| chr3-158944539-INS-208 | chr3-158944539-INS-208 |
| chr3-159001026-INS-134 | chr3-159001026-INS-134 |
| chr3-159002793-INS-486 | chr3-159002793-INS-486 |
| chr3-159038168-INS-967 | chr3-159038168-INS-971 |
| chr3-159049766-INS-475 | chr3-159049766-INS-475 |
| chr3-159057599-INS-187 | chr3-159057599-INS-187 |
| chr3-159076166-INS-977 | chr3-159076166-INS-977 |
| chr3-159112157-INS-545 | chr3-159112157-INS-545 |
| chr3-159124528-INS-1573 | chr3-159124529-INS-1573 |
| chr3-159128489-INS-1718 | chr3-159128490-INS-1718 |
| chr3-159136349-INS-122 | chr3-159136349-INS-122 |
| chr3-159148784-INS-3296 | chr3-159148785-INS-3296 |
| chr3-159159477-INS-5926 | chr3-159159478-INS-5926 |
| chr3-159179920-INS-125 | chr3-159179920-INS-125 |
| chr3-159193952-INS-56 | chr3-159193952-INS-52 |
| chr3-159196534-INS-68 | chr3-159196534-INS-68 |
| chr3-159309167-INS-73 | chr3-159309137-INS-84 |
| chr3-159349204-INS-94 | chr3-159349217-INS-82 |
| chr3-159365016-INS-5509 | chr3-159365017-INS-5509 |
| chr3-159414760-INS-56 | chr3-159414760-INS-68 |
| chr3-159435623-INS-126 | chr3-159435623-INS-121 |
| chr3-159508358-INS-3444 | chr3-159508359-INS-3444 |
| chr3-159509761-INS-222 | chr3-159509761-INS-222 |
| chr3-159520620-INS-190 | chr3-159520620-INS-190 |
| chr3-159543929-INS-220 | chr3-159543929-INS-220 |
| chr3-159554042-INS-3530 | chr3-159554043-INS-3530 |
| chr4-3055957-INS-125 | chr4-3055957-INS-125 |
| chr4-3232406-INS-441 | chr4-3232407-INS-441 |
| chr4-3342298-INS-71 | chr4-3342299-INS-71 |
| chr4-3353112-INS-204 | chr4-3353112-INS-204 |
| chr4-3371215-INS-218 | chr4-3371215-INS-217 |
| chr4-3382788-INS-92 | chr4-3382806-INS-72 |
| chr4-3383730-INS-418 | chr4-3383730-INS-418 |
| chr4-3407686-INS-197 | chr4-3407686-INS-197 |
| chr4-3411336-INS-5202 | chr4-3411337-INS-5202 |
| chr4-3423000-INS-54 | chr4-3423000-INS-54 |
| chr4-3424195-INS-156 | chr4-3424195-INS-156 |
| chr4-3426849-INS-82 | chr4-3426849-INS-82 |
| chr4-3433610-INS-61 | chr4-3433607-INS-58 |
| chr4-3435578-INS-5472 | chr4-3435579-INS-5472 |
| chr4-3451110-INS-56 | chr4-3451110-INS-62 |
| chr4-3456139-INS-95 | chr4-3456139-INS-95 |
| chr4-3489491-INS-65 | chr4-3489491-INS-60 |
| chr4-3490812-INS-377 | chr4-3490812-INS-377 |
| chr4-3499602-INS-242 | chr4-3499602-INS-242 |
| chr4-3521885-INS-6321 | chr4-3521886-INS-6321 |
| chr4-3537275-INS-3960 | chr4-3537276-INS-3960 |
| chr4-3545974-INS-433 | chr4-3545974-INS-433 |
| chr4-3550670-INS-3322 | chr4-3550671-INS-3322 |
| chr4-3550671-INS-230 | chr4-3550671-INS-230 |
| chr4-3553910-INS-68 | chr4-3553910-INS-68 |
| chr4-3574579-INS-752 | chr4-3574533-INS-796 |
| chr4-3595728-INS-57 | chr4-3595728-INS-57 |
| chr4-3640300-INS-497 | chr4-3640300-INS-497 |
| chr4-3672723-INS-177 | chr4-3672723-INS-162 |
| chr4-3684707-INS-1543 | chr4-3684708-INS-1545 |
| chr4-3688223-INS-62 | chr4-3688219-INS-66 |
| chr4-3694124-INS-76 | chr4-3694124-INS-78 |
| chr4-3698686-INS-121 | chr4-3698686-INS-121 |
| chr4-3706161-INS-7068 | chr4-3706162-INS-7068 |
| chr4-3721983-INS-74 | chr4-3721983-INS-82 |
| chr4-3724746-INS-60 | chr4-3724747-INS-82 |
| chr4-3758438-INS-56 | chr4-3758438-INS-64 |
| chr4-3786493-INS-200 | chr4-3786493-INS-200 |

| | |
|---|---|
| chr4-3789260-INS-104 | chr4-3789260-INS-104 |
| chr4-3792176-INS-209 | chr4-3792176-INS-209 |
| chr4-3805869-INS-67 | chr4-3805869-INS-102 |
| chr4-3817896-INS-5543 | chr4-3817897-INS-5543 |
| chr4-3866800-INS-56 | chr4-3866800-INS-56 |
| chr4-3868415-INS-208 | chr4-3868415-INS-208 |
| chr4-3878421-INS-205 | chr4-3878421-INS-205 |
| chr4-3886789-INS-205 | chr4-3886789-INS-205 |
| chr4-3893320-INS-68 | chr4-3893320-INS-68 |
| chr4-3896777-INS-1620 | chr4-3896778-INS-1620 |
| chr4-3908054-INS-125 | chr4-3908089-INS-96 |
| chr4-3914495-INS-60 | chr4-3914487-INS-88 |
| chr4-3915107-INS-703 | chr4-3915107-INS-704 |
| chr4-3960289-INS-78 | chr4-3960328-INS-76 |
| chr4-3963567-INS-194 | chr4-3963567-INS-194 |
| chr4-3997128-INS-179 | chr4-3997128-INS-180 |
| chr4-4001998-INS-2020 | chr4-4001999-INS-2020 |
| chr4-4021543-INS-353 | chr4-4021545-INS-352 |
| chr4-4023939-INS-146 | chr4-4023939-INS-146 |
| chr4-4052904-INS-182 | chr4-4052904-INS-216 |
| chr4-4118206-INS-2817 | chr4-4118207-INS-2817 |
| chr4-4170633-INS-202 | chr4-4170633-INS-202 |
| chr4-4182499-INS-282 | chr4-4182499-INS-301 |
| chr4-4208221-INS-192 | chr4-4208207-INS-198 |
| chr4-4220548-INS-54 | chr4-4220548-INS-54 |
| chr4-4246482-INS-479 | chr4-4246482-INS-479 |
| chr4-4292660-INS-6600 | chr4-4292661-INS-6600 |
| chr4-4328559-INS-54 | chr4-4328559-INS-54 |
| chr4-4331377-INS-196 | chr4-4331377-INS-196 |
| chr4-4362268-INS-818 | chr4-4362268-INS-818 |
| chr4-4368456-INS-813 | chr4-4368456-INS-813 |
| chr4-4374588-INS-929 | chr4-4374588-INS-929 |
| chr4-4410266-INS-8982 | chr4-4410267-INS-8982 |
| chr4-4515411-INS-52 | chr4-4515411-INS-56 |
| chr4-4516597-INS-445 | chr4-4516597-INS-445 |
| chr4-4543051-INS-577 | chr4-4543051-INS-577 |
| chr4-4555176-INS-6677 | chr4-4555177-INS-6680 |
| chr4-4583611-INS-7195 | chr4-4583612-INS-7195 |
| chr4-4625868-INS-213 | chr4-4625868-INS-213 |
| chr4-4628721-INS-63 | chr4-4628721-INS-63 |
| chr4-4631517-INS-6620 | chr4-4631518-INS-6621 |
| chr4-4695271-INS-68 | chr4-4695271-INS-80 |
| chr4-4705761-INS-60 | chr4-4705761-INS-64 |
| chr4-4722754-INS-83 | chr4-4722754-INS-95 |
| chr4-4734147-INS-63 | chr4-4734147-INS-92 |
| chr4-4749511-INS-5026 | chr4-4749512-INS-5026 |
| chr4-4861952-INS-71 | chr4-4861952-INS-71 |
| chr4-4863251-INS-208 | chr4-4863241-INS-208 |
| chr4-4917251-INS-504 | chr4-4917251-INS-504 |
| chr4-4954709-INS-53 | chr4-4954709-INS-53 |
| chr4-4987962-INS-96 | chr4-4987962-INS-96 |
| chr4-4994867-INS-66 | chr4-4994867-INS-66 |
| chr4-4999078-INS-193 | chr4-4999079-INS-193 |
| chr4-5083847-INS-447 | chr4-5083847-INS-447 |
| chr4-5100730-INS-96 | chr4-5100730-INS-107 |
| chr4-5240288-INS-748 | chr4-5240288-INS-755 |
| chr4-5272334-INS-2180 | chr4-5272335-INS-2181 |
| chr4-5281119-INS-6221 | chr4-5281120-INS-6221 |
| chr4-5292867-INS-65 | chr4-5292864-INS-65 |
| chr4-5343156-INS-3118 | chr4-5343157-INS-3118 |
| chr4-5391221-INS-167 | chr4-5391221-INS-167 |
| chr4-5423515-INS-1364 | chr4-5423532-INS-1361 |
| chr4-5471669-INS-6689 | chr4-5471670-INS-6692 |
| chr4-5473587-INS-55 | chr4-5473587-INS-52 |
| chr4-5480455-INS-56 | chr4-5480447-INS-71 |
| chr4-5485015-INS-145 | chr4-5485015-INS-145 |
| chr4-5485167-INS-3294 | chr4-5485168-INS-3294 |
| chr4-5495499-INS-8939 | chr4-5495500-INS-8940 |
| chr4-5535398-INS-69 | chr4-5535398-INS-69 |
| chr4-5552014-INS-70 | chr4-5552014-INS-70 |
| chr4-5594232-INS-320 | chr4-5594232-INS-320 |
| chr4-5661024-INS-288 | chr4-5661024-INS-288 |
| chr4-5679669-INS-82 | chr4-5679677-INS-82 |
| chr4-5704868-INS-90 | chr4-5704868-INS-93 |
| chr4-5714670-INS-212 | chr4-5714670-INS-212 |
| chr4-5783913-INS-442 | chr4-5783882-INS-443 |
| chr4-5900272-INS-100 | chr4-5900272-INS-94 |

| | |
|---|---|
| chr4-5951057-INS-6401 | chr4-5951058-INS-6401 |
| chr4-5981873-INS-56 | chr4-5981873-INS-58 |
| chr4-5989408-INS-5107 | chr4-5989409-INS-5107 |
| chr4-6041054-INS-211 | chr4-6041054-INS-235 |
| chr4-6056875-INS-68 | chr4-6056875-INS-68 |
| chr4-6077946-INS-7395 | chr4-6077947-INS-7395 |
| chr4-6087388-INS-2837 | chr4-6087389-INS-2837 |
| chr4-6091041-INS-2063 | chr4-6091042-INS-2064 |
| chr4-6103345-INS-2220 | chr4-6103346-INS-2220 |
| chr4-6106718-INS-1893 | chr4-6106719-INS-1893 |
| chr4-6112160-INS-74 | chr4-6112160-INS-74 |
| chr4-6159669-INS-63 | chr4-6159669-INS-63 |
| chr4-6162318-INS-422 | chr4-6162318-INS-422 |
| chr4-6208434-INS-152 | chr4-6208434-INS-152 |
| chr4-6230734-INS-168 | chr4-6230734-INS-168 |
| chr4-6242132-INS-164 | chr4-6242132-INS-171 |
| chr4-6249188-INS-740 | chr4-6249188-INS-739 |
| chr4-6270973-INS-177 | chr4-6270973-INS-177 |
| chr4-6277497-INS-270 | chr4-6277497-INS-272 |
| chr4-6283177-INS-601 | chr4-6283177-INS-609 |
| chr4-6329129-INS-376 | chr4-6329129-INS-376 |
| chr4-6343986-INS-82 | chr4-6343986-INS-67 |
| chr4-6351833-INS-196 | chr4-6351833-INS-196 |
| chr4-6381934-INS-219 | chr4-6381934-INS-224 |
| chr4-6383119-INS-51 | chr4-6383119-INS-51 |
| chr4-6391047-INS-223 | chr4-6391047-INS-224 |
| chr4-6415919-INS-75 | chr4-6415919-INS-72 |
| chr4-6422382-INS-80 | chr4-6422382-INS-64 |
| chr4-6463311-INS-1121 | chr4-6463312-INS-1121 |
| chr4-6465815-INS-86 | chr4-6465815-INS-86 |
| chr4-6472476-INS-590 | chr4-6472476-INS-590 |
| chr4-6486192-INS-212 | chr4-6486192-INS-213 |
| chr4-6486739-INS-2861 | chr4-6486740-INS-2861 |
| chr4-6498696-INS-51 | chr4-6498709-INS-83 |
| chr4-6529936-INS-205 | chr4-6529936-INS-205 |
| chr4-6537000-INS-101 | chr4-6536999-INS-99 |
| chr4-6545749-INS-2979 | chr4-6545750-INS-2980 |
| chr4-6575167-INS-70 | chr4-6575167-INS-117 |
| chr4-6581028-INS-53 | chr4-6581028-INS-53 |
| chr4-6602860-INS-182 | chr4-6602860-INS-178 |
| chr4-6620324-INS-7123 | chr4-6620325-INS-7123 |
| chr4-6629314-INS-52 | chr4-6629308-INS-53 |
| chr4-6629359-INS-782 | chr4-6629360-INS-808 |
| chr4-6639558-INS-58 | chr4-6639558-INS-62 |
| chr4-6659754-INS-319 | chr4-6659754-INS-319 |
| chr4-6681130-INS-729 | chr4-6681130-INS-730 |
| chr4-6705714-INS-98 | chr4-6705714-INS-98 |
| chr4-6878427-INS-61 | chr4-6878427-INS-61 |
| chr4-6934087-INS-6693 | chr4-6934088-INS-6693 |
| chr4-6999198-INS-1831 | chr4-6999199-INS-1831 |
| chr4-7016117-INS-1091 | chr4-7016118-INS-1092 |
| chr4-7021468-INS-92 | chr4-7021468-INS-92 |
| chr4-7048354-INS-737 | chr4-7048354-INS-737 |
| chr4-7136260-INS-6627 | chr4-7136261-INS-6627 |
| chr4-7144185-INS-1340 | chr4-7144186-INS-1340 |
| chr4-7160534-INS-68 | chr4-7160535-INS-64 |
| chr4-7163917-INS-51 | chr4-7163917-INS-51 |
| chr4-7174767-INS-199 | chr4-7174767-INS-200 |
| chr4-7186884-INS-1361 | chr4-7186885-INS-1361 |
| chr4-7200524-INS-63 | chr4-7200524-INS-60 |
| chr4-7206273-INS-6802 | chr4-7206274-INS-6802 |
| chr4-7209264-INS-76 | chr4-7209264-INS-76 |
| chr4-7222252-INS-84 | chr4-7222250-INS-87 |
| chr4-7237292-INS-143 | chr4-7237320-INS-142 |
| chr4-7245207-INS-70 | chr4-7245207-INS-68 |
| chr4-7249921-INS-110 | chr4-7249921-INS-110 |
| chr4-7258709-INS-385 | chr4-7258709-INS-385 |
| chr4-7329374-INS-142 | chr4-7329374-INS-142 |
| chr4-7339822-INS-4622 | chr4-7339823-INS-4622 |
| chr4-7392121-INS-78 | chr4-7392121-INS-82 |
| chr4-7538168-INS-6920 | chr4-7538169-INS-6919 |
| chr4-7617789-INS-255 | chr4-7617789-INS-255 |
| chr4-7659461-INS-54 | chr4-7659461-INS-58 |
| chr4-7750508-INS-92 | chr4-7750508-INS-76 |
| chr4-7753074-INS-84 | chr4-7753074-INS-78 |
| chr4-7773554-INS-76 | chr4-7773554-INS-76 |
| chr4-7789421-INS-74 | chr4-7789421-INS-78 |

| | |
|---|---|
| chr4-7856844-INS-65 | chr4-7856844-INS-65 |
| chr4-7868645-INS-62 | chr4-7868645-INS-56 |
| chr4-7911271-INS-98 | chr4-7911272-INS-59 |
| chr4-7911823-INS-51 | chr4-7911823-INS-51 |
| chr4-7927484-INS-476 | chr4-7927484-INS-476 |
| chr4-7930928-INS-6233 | chr4-7930929-INS-6232 |
| chr4-7940457-INS-54 | chr4-7940457-INS-58 |
| chr4-7942888-INS-86 | chr4-7942889-INS-81 |
| chr4-7949538-INS-2190 | chr4-7949539-INS-2191 |
| chr4-7970125-INS-75 | chr4-7970109-INS-85 |
| chr4-7982642-INS-76 | chr4-7982641-INS-81 |
| chr4-7984441-INS-128 | chr4-7984441-INS-128 |
| chr4-8081072-INS-66 | chr4-8081072-INS-66 |
| chr4-8104668-INS-52 | chr4-8104668-INS-58 |
| chr4-8108743-INS-51 | chr4-8108743-INS-51 |
| chr4-8124710-INS-134 | chr4-8124710-INS-136 |
| chr4-8137168-INS-128 | chr4-8137168-INS-128 |
| chr4-8157209-INS-272 | chr4-8157209-INS-272 |
| chr4-8166150-INS-439 | chr4-8166150-INS-439 |
| chr4-8191770-INS-212 | chr4-8191770-INS-212 |
| chr4-8242155-INS-74 | chr4-8242157-INS-73 |
| chr4-8250253-INS-202 | chr4-8250253-INS-202 |
| chr4-8272386-INS-95 | chr4-8272386-INS-95 |
| chr4-8278039-INS-6683 | chr4-8278040-INS-6684 |
| chr4-8338387-INS-52 | chr4-8338387-INS-76 |
| chr4-8342321-INS-140 | chr4-8342321-INS-140 |
| chr4-8348871-INS-120 | chr4-8348871-INS-116 |
| chr4-8359648-INS-52 | chr4-8359650-INS-62 |
| chr4-8365954-INS-2846 | chr4-8365955-INS-2846 |
| chr4-8368530-INS-72 | chr4-8368530-INS-72 |
| chr4-8398117-INS-198 | chr4-8398143-INS-204 |
| chr4-8415282-INS-328 | chr4-8415282-INS-328 |
| chr4-8502739-INS-206 | chr4-8502739-INS-207 |
| chr4-8512226-INS-52 | chr4-8512226-INS-52 |
| chr4-8526933-INS-1043 | chr4-8526934-INS-1043 |
| chr4-8529043-INS-69 | chr4-8529043-INS-69 |
| chr4-8570749-INS-52 | chr4-8570749-INS-52 |
| chr4-8589356-INS-57 | chr4-8589357-INS-67 |
| chr4-8593184-INS-720 | chr4-8593184-INS-720 |
| chr4-8597129-INS-198 | chr4-8597129-INS-198 |
| chr4-8613275-INS-343 | chr4-8613275-INS-343 |
| chr4-8621457-INS-483 | chr4-8621457-INS-483 |
| chr4-8623891-INS-504 | chr4-8623891-INS-504 |
| chr4-8626763-INS-514 | chr4-8626763-INS-514 |
| chr4-8640836-INS-51 | chr4-8640840-INS-50 |
| chr4-8645533-INS-59 | chr4-8645533-INS-59 |
| chr4-8652181-INS-66 | chr4-8652185-INS-63 |
| chr4-8663912-INS-137 | chr4-8663918-INS-133 |
| chr4-8667600-INS-68 | chr4-8667600-INS-72 |
| chr4-8680456-INS-307 | chr4-8680456-INS-307 |
| chr4-8703702-INS-103 | chr4-8703702-INS-103 |
| chr4-8758729-INS-73 | chr4-8758729-INS-73 |
| chr4-8766160-INS-55 | chr4-8766160-INS-55 |
| chr4-8767652-INS-173 | chr4-8767652-INS-168 |
| chr4-8917993-INS-121 | chr4-8917993-INS-114 |
| chr4-8947117-INS-265 | chr4-8947117-INS-267 |
| chr4-8960090-INS-350 | chr4-8960090-INS-342 |
| chr4-8960128-INS-830 | chr4-8960128-INS-830 |
| chr4-8964494-INS-328 | chr4-8964494-INS-328 |
| chr4-8985010-INS-55 | chr4-8985010-INS-55 |
| chr4-9011537-INS-66 | chr4-9011537-INS-69 |
| chr4-9014174-INS-77 | chr4-9014174-INS-60 |
| chr4-9025740-INS-140 | chr4-9025740-INS-140 |
| chr4-9054687-INS-106 | chr4-9054687-INS-106 |
| chr4-9064025-INS-96 | chr4-9064018-INS-110 |
| chr4-9076406-INS-169 | chr4-9076406-INS-169 |
| chr4-9085440-INS-81 | chr4-9085440-INS-81 |
| chr4-9106435-INS-56 | chr4-9106435-INS-60 |
| chr4-9115818-INS-202 | chr4-9115818-INS-202 |
| chr4-9120740-INS-7605 | chr4-9120741-INS-7605 |
| chr4-9123898-INS-54 | chr4-9123898-INS-54 |
| chr4-9125307-INS-7207 | chr4-9125308-INS-7207 |
| chr4-9130463-INS-2605 | chr4-9130464-INS-2605 |
| chr4-9136618-INS-130 | chr4-9136618-INS-132 |
| chr4-9140405-INS-110 | chr4-9140405-INS-110 |
| chr4-9160877-INS-51 | chr4-9160877-INS-51 |
| chr4-9178563-INS-5633 | chr4-9178564-INS-5633 |

| | |
|---|---|
| chr4-9198875-INS-70 | chr4-9198875-INS-70 |
| chr4-9207438-INS-124 | chr4-9207448-INS-117 |
| chr4-9209923-INS-174 | chr4-9209923-INS-174 |
| chr4-9219648-INS-148 | chr4-9219648-INS-146 |
| chr4-9223635-INS-235 | chr4-9223635-INS-235 |
| chr4-9238750-INS-7450 | chr4-9238751-INS-7450 |
| chr4-9245079-INS-67 | chr4-9245079-INS-67 |
| chr4-9249745-INS-146 | chr4-9249745-INS-146 |
| chr4-9264310-INS-80 | chr4-9264310-INS-80 |
| chr4-9268940-INS-52 | chr4-9268940-INS-52 |
| chr4-9287942-INS-196 | chr4-9287942-INS-196 |
| chr4-9321347-INS-136 | chr4-9321347-INS-136 |
| chr4-9324909-INS-153 | chr4-9324909-INS-153 |
| chr4-9328548-INS-939 | chr4-9328548-INS-920 |
| chr4-9342141-INS-503 | chr4-9342141-INS-493 |
| chr4-9344479-INS-337 | chr4-9344479-INS-337 |
| chr4-9371914-INS-440 | chr4-9371914-INS-440 |
| chr4-9383970-INS-79 | chr4-9383970-INS-119 |
| chr4-9388959-INS-63 | chr4-9388959-INS-61 |
| chr4-9401676-INS-98 | chr4-9401676-INS-110 |
| chr4-9432410-INS-60 | chr4-9432410-INS-64 |
| chr4-9454577-INS-130 | chr4-9454607-INS-90 |
| chr4-9459385-INS-99 | chr4-9459408-INS-66 |
| chr4-9469812-INS-199 | chr4-9469812-INS-199 |
| chr4-9477950-INS-54 | chr4-9477950-INS-80 |
| chr4-9492274-INS-106 | chr4-9492274-INS-96 |
| chr4-9502701-INS-134 | chr4-9502701-INS-134 |
| chr4-9514380-INS-62 | chr4-9514380-INS-64 |
| chr4-9522369-INS-245 | chr4-9522369-INS-245 |
| chr4-9523679-INS-5301 | chr4-9523680-INS-5303 |
| chr4-9530661-INS-100 | chr4-9530661-INS-88 |
| chr4-9542318-INS-66 | chr4-9542318-INS-58 |
| chr4-9544332-INS-358 | chr4-9544332-INS-358 |
| chr4-9556052-INS-62 | chr4-9556052-INS-62 |
| chr4-9559360-INS-54 | chr4-9559360-INS-54 |
| chr4-9572019-INS-75 | chr4-9572019-INS-50 |
| chr4-9580066-INS-94 | chr4-9580066-INS-94 |
| chr4-9620218-INS-52 | chr4-9620237-INS-50 |
| chr4-9646296-INS-52 | chr4-9646300-INS-75 |
| chr4-9659320-INS-84 | chr4-9659324-INS-54 |
| chr4-9659858-INS-52 | chr4-9659858-INS-52 |
| chr4-9684575-INS-417 | chr4-9684575-INS-425 |
| chr4-9696658-INS-256 | chr4-9696658-INS-256 |
| chr4-9705235-INS-56 | chr4-9705239-INS-57 |
| chr4-9708007-INS-389 | chr4-9708007-INS-389 |
| chr4-9744168-INS-115 | chr4-9744168-INS-115 |
| chr4-9749035-INS-53 | chr4-9749035-INS-53 |
| chr4-9752131-INS-52 | chr4-9752131-INS-52 |
| chr4-9761381-INS-457 | chr4-9761381-INS-457 |
| chr4-9777698-INS-94 | chr4-9777698-INS-94 |
| chr4-9778725-INS-166 | chr4-9778725-INS-166 |
| chr4-9824475-INS-107 | chr4-9824475-INS-107 |
| chr4-9832796-INS-106 | chr4-9832796-INS-108 |
| chr4-9928047-INS-208 | chr4-9928047-INS-209 |
| chr4-9970962-INS-1888 | chr4-9970963-INS-1888 |
| chr4-9975926-INS-83 | chr4-9975926-INS-83 |
| chr4-9987158-INS-196 | chr4-9987158-INS-197 |
| chr4-10034514-INS-81 | chr4-10034514-INS-81 |
| chr4-10056052-INS-6653 | chr4-10056053-INS-6650 |
| chr4-10061827-INS-3187 | chr4-10061828-INS-3186 |
| chr4-10085128-INS-1140 | chr4-10085129-INS-1140 |
| chr4-10093959-INS-72 | chr4-10093990-INS-62 |
| chr4-10095244-INS-2031 | chr4-10095245-INS-1996 |
| chr4-10131354-INS-474 | chr4-10131354-INS-474 |
| chr4-10149991-INS-3811 | chr4-10149992-INS-3811 |
| chr4-10213299-INS-896 | chr4-10213299-INS-896 |
| chr4-10225580-INS-2642 | chr4-10225581-INS-2642 |
| chr4-10225937-INS-436 | chr4-10225937-INS-436 |
| chr4-10229117-INS-2869 | chr4-10229118-INS-2869 |
| chr4-10229118-INS-459 | chr4-10229118-INS-459 |
| chr4-10238090-INS-87 | chr4-10238090-INS-87 |
| chr4-10246775-INS-4667 | chr4-10246776-INS-4667 |
| chr4-10267120-INS-1166 | chr4-10267121-INS-1166 |
| chr4-10268492-INS-1975 | chr4-10268493-INS-1975 |
| chr4-10277472-INS-104 | chr4-10277472-INS-56 |
| chr4-10286491-INS-88 | chr4-10286491-INS-88 |
| chr4-10344030-INS-198 | chr4-10344030-INS-198 |

| | |
|---|---|
| chr4-10366475-INS-432 | chr4-10366475-INS-432 |
| chr4-10390882-INS-631 | chr4-10390882-INS-631 |
| chr4-10412020-INS-1022 | chr4-10412021-INS-1022 |
| chr4-10425345-INS-4937 | chr4-10425346-INS-4938 |
| chr4-10458606-INS-67 | chr4-10458609-INS-54 |
| chr4-10516829-INS-5601 | chr4-10516830-INS-5601 |
| chr4-10518926-INS-667 | chr4-10518926-INS-667 |
| chr4-10573863-INS-102 | chr4-10573863-INS-105 |
| chr4-10586559-INS-52 | chr4-10586549-INS-96 |
| chr4-10599168-INS-317 | chr4-10599169-INS-310 |
| chr4-10605205-INS-1650 | chr4-10605206-INS-1651 |
| chr4-10621820-INS-69 | chr4-10621820-INS-71 |
| chr4-10633089-INS-7207 | chr4-10633090-INS-7207 |
| chr4-10659338-INS-390 | chr4-10659338-INS-390 |
| chr4-10683976-INS-294 | chr4-10683976-INS-294 |
| chr4-10685138-INS-798 | chr4-10685138-INS-798 |
| chr4-10710984-INS-690 | chr4-10710984-INS-690 |
| chr4-10717138-INS-56 | chr4-10717138-INS-50 |
| chr4-10749580-INS-549 | chr4-10749580-INS-549 |
| chr4-10761400-INS-107 | chr4-10761400-INS-107 |
| chr4-10763482-INS-1403 | chr4-10763483-INS-1403 |
| chr4-10792195-INS-78 | chr4-10792195-INS-78 |
| chr4-10793626-INS-1040 | chr4-10793627-INS-1040 |
| chr4-10821041-INS-56 | chr4-10821041-INS-52 |
| chr4-10889427-INS-238 | chr4-10889427-INS-238 |
| chr4-10892063-INS-70 | chr4-10892063-INS-70 |
| chr4-10911394-INS-148 | chr4-10911394-INS-149 |
| chr4-10917351-INS-396 | chr4-10917352-INS-396 |
| chr4-10928295-INS-722 | chr4-10928295-INS-722 |
| chr4-10931820-INS-70 | chr4-10931820-INS-72 |
| chr4-10948121-INS-4910 | chr4-10948122-INS-4910 |
| chr4-10968855-INS-69 | chr4-10968855-INS-69 |
| chr4-10969560-INS-66 | chr4-10969560-INS-66 |
| chr4-10971830-INS-337 | chr4-10971831-INS-340 |
| chr4-10972627-INS-587 | chr4-10972627-INS-587 |
| chr4-10975346-INS-200 | chr4-10975346-INS-200 |
| chr4-10978670-INS-212 | chr4-10978670-INS-213 |
| chr4-10980077-INS-636 | chr4-10980077-INS-636 |
| chr4-10984286-INS-106 | chr4-10984290-INS-108 |
| chr4-10989522-INS-84 | chr4-10989522-INS-84 |
| chr4-11022212-INS-955 | chr4-11022212-INS-955 |
| chr4-11025640-INS-432 | chr4-11025640-INS-432 |
| chr4-11029640-INS-172 | chr4-11029640-INS-172 |
| chr4-11035601-INS-64 | chr4-11035601-INS-64 |
| chr4-11039437-INS-192 | chr4-11039437-INS-192 |
| chr4-11052015-INS-215 | chr4-11052015-INS-215 |
| chr4-11063264-INS-247 | chr4-11063264-INS-247 |
| chr4-11065402-INS-76 | chr4-11065402-INS-76 |
| chr4-11066441-INS-86 | chr4-11066441-INS-92 |
| chr4-11069383-INS-166 | chr4-11069383-INS-166 |
| chr4-11092971-INS-110 | chr4-11092971-INS-110 |
| chr4-11094585-INS-5305 | chr4-11094586-INS-5305 |
| chr4-11102060-INS-87 | chr4-11102060-INS-87 |
| chr4-11109489-INS-191 | chr4-11109489-INS-207 |
| chr4-11127559-INS-54 | chr4-11127559-INS-52 |
| chr4-11171049-INS-52 | chr4-11171049-INS-52 |
| chr4-11176226-INS-6446 | chr4-11176227-INS-6479 |
| chr4-11221788-INS-214 | chr4-11221788-INS-214 |
| chr4-11226018-INS-74 | chr4-11226018-INS-74 |
| chr4-11237843-INS-226 | chr4-11237843-INS-226 |
| chr4-11269334-INS-1556 | chr4-11269335-INS-1556 |
| chr4-11279073-INS-114 | chr4-11279073-INS-114 |
| chr4-11284401-INS-105 | chr4-11284401-INS-105 |
| chr4-11286724-INS-230 | chr4-11286724-INS-230 |
| chr4-11297735-INS-75 | chr4-11297735-INS-75 |
| chr4-11309693-INS-68 | chr4-11309693-INS-68 |
| chr4-11311972-INS-1890 | chr4-11311973-INS-1890 |
| chr4-11314079-INS-498 | chr4-11314079-INS-498 |
| chr4-11350754-INS-205 | chr4-11350754-INS-205 |
| chr4-11362164-INS-86 | chr4-11362164-INS-86 |
| chr4-11380803-INS-158 | chr4-11380803-INS-158 |
| chr4-11404263-INS-7181 | chr4-11404264-INS-7181 |
| chr4-11472291-INS-60 | chr4-11472291-INS-60 |
| chr4-11477305-INS-131 | chr4-11477305-INS-139 |
| chr4-11496169-INS-60 | chr4-11496169-INS-90 |
| chr4-11519550-INS-108 | chr4-11519550-INS-109 |
| chr4-11535555-INS-380 | chr4-11535556-INS-381 |

| | |
|---|---|
| chr4-11557124-INS-52 | chr4-11557124-INS-56 |
| chr4-11566946-INS-203 | chr4-11566946-INS-203 |
| chr4-11575900-INS-72 | chr4-11575900-INS-72 |
| chr4-11590019-INS-53 | chr4-11590019-INS-53 |
| chr4-11597356-INS-219 | chr4-11597356-INS-218 |
| chr4-11622098-INS-60 | chr4-11622098-INS-60 |
| chr4-11625299-INS-261 | chr4-11625299-INS-272 |
| chr4-11643015-INS-328 | chr4-11643015-INS-323 |
| chr4-11657832-INS-92 | chr4-11657832-INS-89 |
| chr4-11666676-INS-499 | chr4-11666676-INS-499 |
| chr4-11670449-INS-69 | chr4-11670449-INS-69 |
| chr4-11677031-INS-56 | chr4-11677031-INS-60 |
| chr4-11692879-INS-61 | chr4-11692879-INS-61 |
| chr4-11710273-INS-5566 | chr4-11710274-INS-5567 |
| chr4-11740020-INS-56 | chr4-11740020-INS-60 |
| chr4-11742081-INS-6444 | chr4-11742082-INS-6429 |
| chr4-11759640-INS-63 | chr4-11759637-INS-59 |
| chr4-11762727-INS-914 | chr4-11762726-INS-916 |
| chr4-11792246-INS-2987 | chr4-11792247-INS-2988 |
| chr4-11825646-INS-956 | chr4-11825646-INS-956 |
| chr4-11907529-INS-87 | chr4-11907529-INS-87 |
| chr4-11945570-INS-144 | chr4-11945570-INS-144 |
| chr4-11949771-INS-60 | chr4-11949771-INS-60 |
| chr4-12020229-INS-147 | chr4-12020229-INS-149 |
| chr4-12028282-INS-4338 | chr4-12028283-INS-4338 |
| chr4-12028537-INS-202 | chr4-12028537-INS-203 |
| chr4-12034652-INS-1657 | chr4-12034653-INS-1657 |
| chr4-12034790-INS-105 | chr4-12034790-INS-104 |
| chr4-12107360-INS-119 | chr4-12107360-INS-119 |
| chr4-12154993-INS-69 | chr4-12154993-INS-77 |
| chr4-12178951-INS-116 | chr4-12178956-INS-117 |
| chr4-12189512-INS-4082 | chr4-12189513-INS-4082 |
| chr4-12229314-INS-100 | chr4-12229326-INS-102 |
| chr4-12231395-INS-56 | chr4-12231395-INS-56 |
| chr4-12260936-INS-7150 | chr4-12260937-INS-7150 |
| chr4-12281145-INS-234 | chr4-12281145-INS-234 |
| chr4-12312603-INS-95 | chr4-12312603-INS-111 |
| chr4-12330606-INS-56 | chr4-12330606-INS-65 |
| chr4-12338963-INS-84 | chr4-12338963-INS-84 |
| chr4-12341613-INS-215 | chr4-12341615-INS-224 |
| chr4-12345879-INS-391 | chr4-12345879-INS-391 |
| chr4-12364589-INS-7278 | chr4-12364590-INS-7278 |
| chr4-12376003-INS-1469 | chr4-12376004-INS-1461 |
| chr4-12390742-INS-281 | chr4-12390743-INS-280 |
| chr4-12398234-INS-1789 | chr4-12398235-INS-1789 |
| chr4-12439380-INS-285 | chr4-12439380-INS-316 |
| chr4-12459777-INS-201 | chr4-12459777-INS-201 |
| chr4-12545373-INS-1501 | chr4-12545380-INS-1479 |
| chr4-12562991-INS-63 | chr4-12562991-INS-63 |
| chr4-12565402-INS-6583 | chr4-12565403-INS-6583 |
| chr4-12574323-INS-59 | chr4-12574323-INS-59 |
| chr4-12581501-INS-7400 | chr4-12581502-INS-7400 |
| chr4-12610768-INS-86 | chr4-12610768-INS-76 |
| chr4-12627994-INS-309 | chr4-12627994-INS-313 |
| chr4-12661299-INS-6436 | chr4-12661300-INS-6436 |
| chr4-12677142-INS-72 | chr4-12677142-INS-72 |
| chr4-12687062-INS-5019 | chr4-12687063-INS-5019 |
| chr4-12692775-INS-71 | chr4-12692775-INS-71 |
| chr4-12701331-INS-60 | chr4-12701331-INS-60 |
| chr4-12714623-INS-132 | chr4-12714623-INS-132 |
| chr4-12729597-INS-334 | chr4-12729597-INS-334 |
| chr4-12744535-INS-65 | chr4-12744535-INS-65 |
| chr4-12769604-INS-468 | chr4-12769604-INS-468 |
| chr4-12804897-INS-191 | chr4-12804897-INS-191 |
| chr4-12807505-INS-88 | chr4-12807517-INS-97 |
| chr4-12809222-INS-181 | chr4-12809222-INS-183 |
| chr4-12819661-INS-64 | chr4-12819661-INS-64 |
| chr4-12826010-INS-1912 | chr4-12826011-INS-1912 |
| chr4-12843020-INS-202 | chr4-12843020-INS-202 |
| chr4-12857665-INS-281 | chr4-12857665-INS-281 |
| chr4-12873984-INS-69 | chr4-12873984-INS-69 |
| chr4-12913944-INS-82 | chr4-12913944-INS-82 |
| chr4-12919887-INS-93 | chr4-12919887-INS-93 |
| chr4-12929508-INS-6484 | chr4-12929509-INS-6484 |
| chr4-12932303-INS-56 | chr4-12932303-INS-58 |
| chr4-12936574-INS-2998 | chr4-12936575-INS-2998 |
| chr4-12946023-INS-576 | chr4-12946023-INS-576 |

| | |
|---|---|
| chr4-12961426-INS-100 | chr4-12961426-INS-100 |
| chr4-12985120-INS-60 | chr4-12985124-INS-56 |
| chr4-12990592-INS-227 | chr4-12990592-INS-227 |
| chr4-13065123-INS-386 | chr4-13065123-INS-386 |
| chr4-13087055-INS-423 | chr4-13087055-INS-424 |
| chr4-13114902-INS-355 | chr4-13114902-INS-355 |
| chr4-13129155-INS-362 | chr4-13129155-INS-362 |
| chr4-13192015-INS-131 | chr4-13192013-INS-131 |
| chr4-13219346-INS-60 | chr4-13219346-INS-62 |
| chr4-13222463-INS-105 | chr4-13222463-INS-85 |
| chr4-13246263-INS-85 | chr4-13246263-INS-84 |
| chr4-13262403-INS-140 | chr4-13262403-INS-145 |
| chr4-13277685-INS-78 | chr4-13277685-INS-78 |
| chr4-13330774-INS-66 | chr4-13330774-INS-72 |
| chr4-13342410-INS-494 | chr4-13342410-INS-493 |
| chr4-13355197-INS-1488 | chr4-13355198-INS-1488 |
| chr4-13373380-INS-66 | chr4-13373380-INS-78 |
| chr4-13422695-INS-2544 | chr4-13422696-INS-2544 |
| chr4-13434657-INS-507 | chr4-13434657-INS-507 |
| chr4-13484970-INS-781 | chr4-13484970-INS-781 |
| chr4-13514377-INS-6907 | chr4-13514378-INS-6907 |
| chr4-13519475-INS-87 | chr4-13519475-INS-87 |
| chr4-13527754-INS-6521 | chr4-13527755-INS-6521 |
| chr4-13553732-INS-7126 | chr4-13553733-INS-7126 |
| chr4-13555075-INS-58 | chr4-13555075-INS-58 |
| chr4-13590961-INS-689 | chr4-13590967-INS-685 |
| chr4-13602157-INS-378 | chr4-13602157-INS-378 |
| chr4-13604639-INS-100 | chr4-13604639-INS-100 |
| chr4-13612097-INS-60 | chr4-13612097-INS-60 |
| chr4-13622715-INS-120 | chr4-13622713-INS-148 |
| chr4-13626412-INS-92 | chr4-13626412-INS-91 |
| chr4-13655573-INS-87 | chr4-13655573-INS-87 |
| chr4-13686697-INS-128 | chr4-13686662-INS-174 |
| chr4-13709962-INS-2397 | chr4-13709963-INS-2397 |
| chr4-13755155-INS-71 | chr4-13755163-INS-84 |
| chr4-13757638-INS-317 | chr4-13757634-INS-318 |
| chr4-13800004-INS-74 | chr4-13800004-INS-74 |
| chr4-13827407-INS-54 | chr4-13827407-INS-54 |
| chr4-13883071-INS-52 | chr4-13883071-INS-52 |
| chr4-13910198-INS-1691 | chr4-13910199-INS-1692 |
| chr4-13930202-INS-201 | chr4-13930202-INS-201 |
| chr4-13942975-INS-6314 | chr4-13942976-INS-6315 |
| chr4-13949879-INS-376 | chr4-13949880-INS-370 |
| chr4-13962391-INS-62 | chr4-13962392-INS-58 |
| chr4-13969578-INS-96 | chr4-13969578-INS-105 |
| chr4-14004263-INS-1025 | chr4-14004264-INS-1025 |
| chr4-14053717-INS-670 | chr4-14053717-INS-671 |
| chr4-14079845-INS-6623 | chr4-14079846-INS-6623 |
| chr4-14137480-INS-201 | chr4-14137480-INS-201 |
| chr4-14143257-INS-3164 | chr4-14143258-INS-3164 |
| chr4-14150495-INS-53 | chr4-14150517-INS-56 |
| chr4-14165890-INS-6910 | chr4-14165891-INS-6910 |
| chr4-14176989-INS-565 | chr4-14176989-INS-566 |
| chr4-14219411-INS-53 | chr4-14219409-INS-60 |
| chr4-14228807-INS-577 | chr4-14228807-INS-577 |
| chr4-14245580-INS-6643 | chr4-14245581-INS-6643 |
| chr4-14336426-INS-6285 | chr4-14336427-INS-6285 |
| chr4-14348532-INS-6907 | chr4-14348533-INS-6907 |
| chr4-14351978-INS-2158 | chr4-14351979-INS-2160 |
| chr4-14364451-INS-6425 | chr4-14364452-INS-6426 |
| chr4-14378954-INS-52 | chr4-14378955-INS-74 |
| chr4-14394533-INS-67 | chr4-14394537-INS-71 |
| chr4-14396085-INS-56 | chr4-14396098-INS-86 |
| chr4-14419617-INS-6283 | chr4-14419618-INS-6282 |
| chr4-14424070-INS-199 | chr4-14424070-INS-199 |
| chr4-14454443-INS-54 | chr4-14454443-INS-54 |
| chr4-14456929-INS-401 | chr4-14456929-INS-401 |
| chr4-14460234-INS-84 | chr4-14460237-INS-88 |
| chr4-14499396-INS-436 | chr4-14499396-INS-436 |
| chr4-14500396-INS-832 | chr4-14500396-INS-836 |
| chr4-14508645-INS-82 | chr4-14508643-INS-99 |
| chr4-14535223-INS-500 | chr4-14535227-INS-499 |
| chr4-14537692-INS-369 | chr4-14537692-INS-369 |
| chr4-14542099-INS-380 | chr4-14542123-INS-387 |
| chr4-14572067-INS-55 | chr4-14572067-INS-60 |
| chr4-14592851-INS-1869 | chr4-14592852-INS-1869 |
| chr4-14597205-INS-96 | chr4-14597205-INS-96 |

| | |
|---|---|
| chr4-14597256-INS-2008 | chr4-14597257-INS-2009 |
| chr4-14598190-INS-164 | chr4-14598190-INS-164 |
| chr4-14621536-INS-52 | chr4-14621536-INS-54 |
| chr4-14671071-INS-64 | chr4-14671051-INS-56 |
| chr4-14709817-INS-94 | chr4-14709817-INS-94 |
| chr4-14731533-INS-562 | chr4-14731529-INS-569 |
| chr4-14739505-INS-231 | chr4-14739505-INS-231 |
| chr4-14753378-INS-102 | chr4-14753378-INS-102 |
| chr4-14775978-INS-151 | chr4-14775978-INS-151 |
| chr4-14827506-INS-75 | chr4-14827463-INS-77 |
| chr4-14837005-INS-437 | chr4-14837005-INS-437 |
| chr4-14845064-INS-62 | chr4-14845064-INS-62 |
| chr4-14856293-INS-243 | chr4-14856294-INS-243 |
| chr4-14857645-INS-62 | chr4-14857645-INS-62 |
| chr4-14860939-INS-417 | chr4-14860939-INS-458 |
| chr4-14867383-INS-73 | chr4-14867383-INS-73 |
| chr4-14876955-INS-72 | chr4-14876955-INS-72 |
| chr4-14884147-INS-521 | chr4-14884147-INS-521 |
| chr4-14893319-INS-81 | chr4-14893319-INS-77 |
| chr4-14917204-INS-205 | chr4-14917204-INS-205 |
| chr4-14920361-INS-396 | chr4-14920361-INS-429 |
| chr4-14923246-INS-786 | chr4-14923246-INS-786 |
| chr4-14942515-INS-160 | chr4-14942515-INS-162 |
| chr4-14966278-INS-175 | chr4-14966278-INS-175 |
| chr4-14971827-INS-6180 | chr4-14971828-INS-6180 |
| chr4-14973681-INS-200 | chr4-14973681-INS-200 |
| chr4-14978095-INS-52 | chr4-14978095-INS-60 |
| chr4-14994376-INS-6263 | chr4-14994377-INS-6264 |
| chr4-15024013-INS-57 | chr4-15024013-INS-57 |
| chr4-15026446-INS-644 | chr4-15026448-INS-644 |
| chr4-15076245-INS-6984 | chr4-15076246-INS-6984 |
| chr4-15092961-INS-1894 | chr4-15092962-INS-1894 |
| chr4-15127972-INS-95 | chr4-15127972-INS-100 |
| chr4-15152726-INS-206 | chr4-15152726-INS-206 |
| chr4-15160615-INS-6464 | chr4-15160616-INS-6464 |
| chr4-15179136-INS-6291 | chr4-15179137-INS-6291 |
| chr4-15197968-INS-55 | chr4-15197953-INS-69 |
| chr4-15221277-INS-418 | chr4-15221277-INS-418 |
| chr4-15292592-INS-290 | chr4-15292592-INS-284 |
| chr4-15324471-INS-6666 | chr4-15324472-INS-6666 |
| chr4-15334670-INS-54 | chr4-15334670-INS-54 |
| chr4-15364344-INS-61 | chr4-15364344-INS-64 |
| chr4-15369551-INS-58 | chr4-15369551-INS-58 |
| chr4-15371341-INS-66 | chr4-15371341-INS-66 |
| chr4-15387521-INS-50 | chr4-15387521-INS-68 |
| chr4-15389430-INS-6665 | chr4-15389431-INS-6666 |
| chr4-15410045-INS-54 | chr4-15410045-INS-54 |
| chr4-15415629-INS-156 | chr4-15415629-INS-156 |
| chr4-15418410-INS-60 | chr4-15418411-INS-64 |
| chr4-15419951-INS-56 | chr4-15419951-INS-56 |
| chr4-15425203-INS-1069 | chr4-15425204-INS-1069 |
| chr4-15432734-INS-1878 | chr4-15432735-INS-1878 |
| chr4-15451905-INS-134 | chr4-15451905-INS-134 |
| chr4-15477401-INS-963 | chr4-15477401-INS-963 |
| chr4-15496596-INS-198 | chr4-15496596-INS-198 |
| chr4-15512990-INS-432 | chr4-15512990-INS-432 |
| chr4-15516268-INS-186 | chr4-15516268-INS-186 |
| chr4-15520147-INS-138 | chr4-15520149-INS-127 |
| chr4-15529515-INS-68 | chr4-15529515-INS-68 |
| chr4-15536418-INS-2384 | chr4-15536419-INS-2384 |
| chr4-15557704-INS-6908 | chr4-15557705-INS-6908 |
| chr4-15605058-INS-383 | chr4-15605058-INS-383 |
| chr4-15655246-INS-199 | chr4-15655246-INS-199 |
| chr4-15673221-INS-564 | chr4-15673221-INS-564 |
| chr4-15688107-INS-6227 | chr4-15688108-INS-6229 |
| chr4-15702141-INS-255 | chr4-15702141-INS-253 |
| chr4-15724196-INS-6790 | chr4-15724197-INS-6792 |
| chr4-15729716-INS-905 | chr4-15729716-INS-905 |
| chr4-15730894-INS-2554 | chr4-15730896-INS-2555 |
| chr4-15736825-INS-2522 | chr4-15736826-INS-2522 |
| chr4-15746556-INS-1098 | chr4-15746557-INS-1098 |
| chr4-15762733-INS-680 | chr4-15762733-INS-677 |
| chr4-15872074-INS-239 | chr4-15872074-INS-239 |
| chr4-15910018-INS-185 | chr4-15910018-INS-183 |
| chr4-15952249-INS-245 | chr4-15952249-INS-245 |
| chr4-15980160-INS-219 | chr4-15980160-INS-219 |
| chr4-15981104-INS-62 | chr4-15981104-INS-62 |

| | |
|---|---|
| chr4-16010563-INS-146 | chr4-16010563-INS-146 |
| chr4-16066885-INS-61 | chr4-16066885-INS-60 |
| chr4-16087348-INS-334 | chr4-16087348-INS-334 |
| chr4-16103891-INS-93 | chr4-16103894-INS-51 |
| chr4-16127410-INS-61 | chr4-16127405-INS-78 |
| chr4-16175923-INS-1864 | chr4-16175924-INS-1864 |
| chr4-16179112-INS-104 | chr4-16179150-INS-104 |
| chr4-16197954-INS-417 | chr4-16197954-INS-417 |
| chr4-16205369-INS-233 | chr4-16205369-INS-233 |
| chr4-16211942-INS-100 | chr4-16211953-INS-104 |
| chr4-16223026-INS-88 | chr4-16223025-INS-82 |
| chr4-16228551-INS-260 | chr4-16228551-INS-260 |
| chr4-16240602-INS-54 | chr4-16240602-INS-56 |
| chr4-16245662-INS-104 | chr4-16245660-INS-70 |
| chr4-16273486-INS-112 | chr4-16273486-INS-111 |
| chr4-16350298-INS-74 | chr4-16350298-INS-74 |
| chr4-16354907-INS-434 | chr4-16354907-INS-434 |
| chr4-16415773-INS-4726 | chr4-16415774-INS-4726 |
| chr4-16419940-INS-2694 | chr4-16419938-INS-2674 |
| chr4-16459594-INS-5544 | chr4-16459595-INS-5544 |
| chr4-16462443-INS-62 | chr4-16462443-INS-54 |
| chr4-16462812-INS-1620 | chr4-16462813-INS-1621 |
| chr4-16467698-INS-100 | chr4-16467693-INS-85 |
| chr4-16478743-INS-4180 | chr4-16478726-INS-4212 |
| chr4-16480102-INS-401 | chr4-16480102-INS-402 |
| chr4-16510141-INS-60 | chr4-16510141-INS-58 |
| chr4-16561017-INS-140 | chr4-16561017-INS-140 |
| chr4-16563733-INS-57 | chr4-16563733-INS-53 |
| chr4-16568459-INS-78 | chr4-16568459-INS-74 |
| chr4-16569680-INS-110 | chr4-16569680-INS-112 |
| chr4-16599786-INS-741 | chr4-16599786-INS-741 |
| chr4-16600857-INS-5886 | chr4-16600858-INS-5886 |
| chr4-16605595-INS-822 | chr4-16605596-INS-822 |
| chr4-16605797-INS-60 | chr4-16605797-INS-85 |
| chr4-16645396-INS-200 | chr4-16645403-INS-200 |
| chr4-16658961-INS-63 | chr4-16658961-INS-63 |
| chr4-16688148-INS-165 | chr4-16688148-INS-172 |
| chr4-16696511-INS-58 | chr4-16696511-INS-58 |
| chr4-16702045-INS-2704 | chr4-16702046-INS-2704 |
| chr4-16742763-INS-7180 | chr4-16742764-INS-7180 |
| chr4-16753680-INS-245 | chr4-16753680-INS-245 |
| chr4-16757395-INS-4877 | chr4-16757396-INS-4877 |
| chr4-16757688-INS-308 | chr4-16757688-INS-308 |
| chr4-16768021-INS-3182 | chr4-16768022-INS-3182 |
| chr4-16794403-INS-751 | chr4-16794403-INS-751 |
| chr4-16810750-INS-201 | chr4-16810750-INS-201 |
| chr4-16812580-INS-105 | chr4-16812596-INS-78 |
| chr4-16817601-INS-705 | chr4-16817601-INS-709 |
| chr4-16821294-INS-52 | chr4-16821294-INS-54 |
| chr4-16831935-INS-13997 | chr4-16831936-INS-13997 |
| chr4-16844599-INS-4845 | chr4-16844600-INS-4845 |
| chr4-16899948-INS-1601 | chr4-16899949-INS-1601 |
| chr4-16908116-INS-619 | chr4-16908116-INS-621 |
| chr4-16909776-INS-708 | chr4-16909777-INS-709 |
| chr4-16917909-INS-1353 | chr4-16917910-INS-1353 |
| chr4-16957118-INS-53 | chr4-16957118-INS-53 |
| chr4-16995609-INS-431 | chr4-16995609-INS-431 |
| chr4-17002740-INS-96 | chr4-17002735-INS-89 |
| chr4-17005656-INS-3574 | chr4-17005657-INS-3574 |
| chr4-17018068-INS-201 | chr4-17018068-INS-201 |
| chr4-17046131-INS-477 | chr4-17046131-INS-520 |
| chr4-17076785-INS-1864 | chr4-17076786-INS-1864 |
| chr4-17106426-INS-1765 | chr4-17106427-INS-1768 |
| chr4-17107235-INS-76 | chr4-17107235-INS-76 |
| chr4-17115795-INS-645 | chr4-17115795-INS-645 |
| chr4-17132024-INS-164 | chr4-17132024-INS-165 |
| chr4-17137396-INS-65 | chr4-17137396-INS-69 |
| chr4-17167461-INS-4388 | chr4-17167462-INS-4388 |
| chr4-17171802-INS-196 | chr4-17171802-INS-196 |
| chr4-17178628-INS-3794 | chr4-17178629-INS-3794 |
| chr4-17190201-INS-1125 | chr4-17190202-INS-1125 |
| chr4-17234551-INS-81 | chr4-17234530-INS-78 |
| chr4-17263538-INS-2565 | chr4-17263539-INS-2564 |
| chr4-17278888-INS-284 | chr4-17278888-INS-285 |
| chr4-17313105-INS-358 | chr4-17313105-INS-363 |
| chr4-17325099-INS-169 | chr4-17325080-INS-191 |
| chr4-17333931-INS-66 | chr4-17333931-INS-66 |

| | |
|---|---|
| chr4-17339032-INS-64 | chr4-17339040-INS-67 |
| chr4-17343655-INS-236 | chr4-17343655-INS-233 |
| chr4-17348603-INS-417 | chr4-17348603-INS-417 |
| chr4-17350591-INS-93 | chr4-17350598-INS-50 |
| chr4-17360329-INS-523 | chr4-17360329-INS-522 |
| chr4-17362834-INS-341 | chr4-17362834-INS-341 |
| chr4-17436742-INS-170 | chr4-17436742-INS-148 |
| chr4-17439223-INS-101 | chr4-17439223-INS-101 |
| chr4-17476978-INS-76 | chr4-17476978-INS-81 |
| chr4-17492379-INS-60 | chr4-17492379-INS-60 |
| chr4-17506223-INS-109 | chr4-17506214-INS-107 |
| chr4-17515094-INS-52 | chr4-17515094-INS-54 |
| chr4-17554762-INS-8015 | chr4-17554763-INS-8016 |
| chr4-17556944-INS-362 | chr4-17556944-INS-358 |
| chr4-17560483-INS-440 | chr4-17560483-INS-440 |
| chr4-17587085-INS-369 | chr4-17587085-INS-369 |
| chr4-17592892-INS-81 | chr4-17592892-INS-81 |
| chr4-17594028-INS-190 | chr4-17594033-INS-171 |
| chr4-17600491-INS-51 | chr4-17600487-INS-51 |
| chr4-17624800-INS-6599 | chr4-17624801-INS-6601 |
| chr4-17629664-INS-7093 | chr4-17629665-INS-7093 |
| chr4-17634070-INS-1647 | chr4-17634071-INS-1647 |
| chr4-17640741-INS-98 | chr4-17640741-INS-98 |
| chr4-17653576-INS-71 | chr4-17653576-INS-79 |
| chr4-17671098-INS-70 | chr4-17671098-INS-70 |
| chr4-17673650-INS-54 | chr4-17673650-INS-52 |
| chr4-17692406-INS-6455 | chr4-17692407-INS-6455 |
| chr4-17724830-INS-6151 | chr4-17724831-INS-6151 |
| chr4-17734372-INS-98 | chr4-17734372-INS-100 |
| chr4-17767427-INS-7138 | chr4-17767428-INS-7138 |
| chr4-17772313-INS-721 | chr4-17772313-INS-720 |
| chr4-17777346-INS-615 | chr4-17777346-INS-615 |
| chr4-17793550-INS-68 | chr4-17793550-INS-73 |
| chr4-17818232-INS-176 | chr4-17818232-INS-177 |
| chr4-17820412-INS-138 | chr4-17820406-INS-174 |
| chr4-17822383-INS-126 | chr4-17822379-INS-114 |
| chr4-17890671-INS-155 | chr4-17890671-INS-155 |
| chr4-17928260-INS-537 | chr4-17928260-INS-536 |
| chr4-17949858-INS-7416 | chr4-17949859-INS-7416 |
| chr4-17949859-INS-475 | chr4-17949859-INS-475 |
| chr4-17962029-INS-112 | chr4-17962029-INS-110 |
| chr4-17972036-INS-79 | chr4-17972036-INS-79 |
| chr4-18075684-INS-64 | chr4-18075709-INS-88 |
| chr4-18115592-INS-55 | chr4-18115592-INS-50 |
| chr4-18124157-INS-55 | chr4-18124157-INS-55 |
| chr4-18151591-INS-3205 | chr4-18151592-INS-3207 |
| chr4-18168123-INS-589 | chr4-18168123-INS-589 |
| chr4-18193371-INS-54 | chr4-18193371-INS-54 |
| chr4-18198618-INS-6423 | chr4-18198619-INS-6423 |
| chr4-18229957-INS-56 | chr4-18229936-INS-74 |
| chr4-18239588-INS-1374 | chr4-18239589-INS-1374 |
| chr4-18276056-INS-70 | chr4-18276056-INS-70 |
| chr4-18282876-INS-66 | chr4-18282876-INS-66 |
| chr4-18286017-INS-6447 | chr4-18286018-INS-6447 |
| chr4-18310544-INS-692 | chr4-18310544-INS-692 |
| chr4-18331044-INS-90 | chr4-18331044-INS-87 |
| chr4-18332634-INS-107 | chr4-18332634-INS-107 |
| chr4-18349679-INS-2424 | chr4-18349680-INS-2425 |
| chr4-18358602-INS-6251 | chr4-18358603-INS-6252 |
| chr4-18368567-INS-374 | chr4-18368567-INS-374 |
| chr4-18415964-INS-206 | chr4-18415988-INS-160 |
| chr4-18436327-INS-222 | chr4-18436327-INS-222 |
| chr4-18457085-INS-56 | chr4-18457085-INS-56 |
| chr4-18514001-INS-1289 | chr4-18514008-INS-1288 |
| chr4-18519906-INS-79 | chr4-18519906-INS-79 |
| chr4-18550933-INS-165 | chr4-18550933-INS-165 |
| chr4-18569716-INS-440 | chr4-18569716-INS-439 |
| chr4-18583231-INS-158 | chr4-18583231-INS-159 |
| chr4-18629499-INS-2644 | chr4-18629500-INS-2645 |
| chr4-18630107-INS-328 | chr4-18630107-INS-328 |
| chr4-18641250-INS-6242 | chr4-18641251-INS-6242 |
| chr4-18663798-INS-3535 | chr4-18663799-INS-3533 |
| chr4-18670226-INS-655 | chr4-18670209-INS-658 |
| chr4-18671127-INS-1051 | chr4-18671132-INS-1047 |
| chr4-18726149-INS-290 | chr4-18726150-INS-291 |
| chr4-18739937-INS-367 | chr4-18739937-INS-368 |
| chr4-18765232-INS-246 | chr4-18765232-INS-246 |

| | |
|---|---|
| chr4-18805107-INS-106 | chr4-18805107-INS-106 |
| chr4-18840031-INS-101 | chr4-18840031-INS-101 |
| chr4-18898880-INS-64 | chr4-18898880-INS-64 |
| chr4-18901985-INS-108 | chr4-18901985-INS-108 |
| chr4-18925012-INS-87 | chr4-18925012-INS-90 |
| chr4-19005515-INS-128 | chr4-19005515-INS-114 |
| chr4-19015363-INS-67 | chr4-19015363-INS-65 |
| chr4-19033281-INS-704 | chr4-19033281-INS-704 |
| chr4-19071259-INS-85 | chr4-19071230-INS-66 |
| chr4-19084335-INS-256 | chr4-19084336-INS-274 |
| chr4-19094081-INS-112 | chr4-19094081-INS-112 |
| chr4-19107319-INS-541 | chr4-19107319-INS-542 |
| chr4-19134807-INS-55 | chr4-19134803-INS-69 |
| chr4-19164413-INS-369 | chr4-19164413-INS-369 |
| chr4-19165564-INS-521 | chr4-19165564-INS-506 |
| chr4-19177690-INS-248 | chr4-19177690-INS-207 |
| chr4-19202450-INS-129 | chr4-19202450-INS-129 |
| chr4-19274253-INS-216 | chr4-19274250-INS-216 |
| chr4-19282993-INS-53 | chr4-19282993-INS-53 |
| chr4-19307105-INS-163 | chr4-19307094-INS-157 |
| chr4-19347324-INS-1379 | chr4-19347325-INS-1373 |
| chr4-19356921-INS-348 | chr4-19356921-INS-348 |
| chr4-19361039-INS-88 | chr4-19361039-INS-90 |
| chr4-19368590-INS-67 | chr4-19368590-INS-67 |
| chr4-19381656-INS-128 | chr4-19381656-INS-126 |
| chr4-19411203-INS-132 | chr4-19411203-INS-132 |
| chr4-19423850-INS-278 | chr4-19423850-INS-279 |
| chr4-19429565-INS-69 | chr4-19429565-INS-69 |
| chr4-19438602-INS-1869 | chr4-19438603-INS-1870 |
| chr4-19460903-INS-744 | chr4-19460903-INS-744 |
| chr4-19466642-INS-121 | chr4-19466642-INS-121 |
| chr4-19473112-INS-52 | chr4-19473112-INS-53 |
| chr4-19473419-INS-416 | chr4-19473419-INS-416 |
| chr4-19485358-INS-84 | chr4-19485358-INS-78 |
| chr4-19490199-INS-615 | chr4-19490199-INS-615 |
| chr4-19500977-INS-6463 | chr4-19500978-INS-6463 |
| chr4-19501791-INS-212 | chr4-19501791-INS-213 |
| chr4-19514237-INS-359 | chr4-19514202-INS-399 |
| chr4-19524385-INS-129 | chr4-19524385-INS-130 |
| chr4-19527050-INS-392 | chr4-19527050-INS-392 |
| chr4-19542940-INS-73 | chr4-19542940-INS-77 |
| chr4-19552449-INS-3957 | chr4-19552450-INS-3957 |
| chr4-19577649-INS-7149 | chr4-19577650-INS-7167 |
| chr4-19613898-INS-7148 | chr4-19613899-INS-7148 |
| chr4-19621341-INS-173 | chr4-19621341-INS-173 |
| chr4-19636488-INS-121 | chr4-19636488-INS-121 |
| chr4-19694265-INS-135 | chr4-19694265-INS-135 |
| chr4-19729888-INS-104 | chr4-19729888-INS-104 |
| chr4-19777481-INS-124 | chr4-19777481-INS-124 |
| chr4-19801129-INS-3021 | chr4-19801130-INS-3021 |
| chr4-19821707-INS-716 | chr4-19821707-INS-728 |
| chr4-19828836-INS-66 | chr4-19828836-INS-69 |
| chr4-19872592-INS-766 | chr4-19872592-INS-766 |
| chr4-19892278-INS-219 | chr4-19892278-INS-231 |
| chr4-19894553-INS-329 | chr4-19894555-INS-343 |
| chr4-19899305-INS-147 | chr4-19899293-INS-119 |
| chr4-19908340-INS-65 | chr4-19908337-INS-60 |
| chr4-19913079-INS-394 | chr4-19913079-INS-394 |
| chr4-19925324-INS-55 | chr4-19925330-INS-74 |
| chr4-19962064-INS-6448 | chr4-19962065-INS-6448 |
| chr4-19973043-INS-328 | chr4-19973037-INS-331 |
| chr4-19974403-INS-62 | chr4-19974403-INS-62 |
| chr4-19997584-INS-472 | chr4-19997584-INS-472 |
| chr4-19999583-INS-64 | chr4-19999583-INS-64 |
| chr4-20004372-INS-687 | chr4-20004372-INS-688 |
| chr4-20013931-INS-56 | chr4-20013932-INS-50 |
| chr4-20037515-INS-82 | chr4-20037515-INS-82 |
| chr4-20046833-INS-89 | chr4-20046831-INS-92 |
| chr4-20048036-INS-503 | chr4-20048036-INS-508 |
| chr4-20052524-INS-88 | chr4-20052524-INS-88 |
| chr4-20103278-INS-253 | chr4-20103278-INS-253 |
| chr4-20157170-INS-62 | chr4-20157146-INS-97 |
| chr4-20191914-INS-2400 | chr4-20191915-INS-2400 |
| chr4-20203726-INS-620 | chr4-20203726-INS-620 |
| chr4-20220505-INS-112 | chr4-20220505-INS-112 |
| chr4-20256071-INS-8448 | chr4-20256072-INS-8448 |
| chr4-20278246-INS-101 | chr4-20278246-INS-101 |

| | |
|---|---|
| chr4-20306400-INS-97 | chr4-20306400-INS-97 |
| chr4-20420761-INS-424 | chr4-20420761-INS-425 |
| chr4-20462620-INS-206 | chr4-20462663-INS-206 |
| chr4-20488779-INS-481 | chr4-20488779-INS-481 |
| chr4-20509496-INS-67 | chr4-20509496-INS-67 |
| chr4-20530821-INS-110 | chr4-20530823-INS-100 |
| chr4-20541082-INS-222 | chr4-20541082-INS-222 |
| chr4-20542159-INS-6569 | chr4-20542160-INS-6569 |
| chr4-20543196-INS-272 | chr4-20543196-INS-269 |
| chr4-20555791-INS-420 | chr4-20555791-INS-426 |
| chr4-20579848-INS-2986 | chr4-20579849-INS-2986 |
| chr4-20581840-INS-901 | chr4-20581840-INS-901 |
| chr4-20606350-INS-54 | chr4-20606350-INS-54 |
| chr4-20608724-INS-94 | chr4-20608724-INS-94 |
| chr4-20625213-INS-84 | chr4-20625221-INS-75 |
| chr4-20633455-INS-390 | chr4-20633455-INS-391 |
| chr4-20642834-INS-185 | chr4-20642834-INS-185 |
| chr4-20672304-INS-54 | chr4-20672304-INS-56 |
| chr4-20675462-INS-327 | chr4-20675462-INS-327 |
| chr4-20682418-INS-1562 | chr4-20682419-INS-1562 |
| chr4-20688218-INS-78 | chr4-20688255-INS-84 |
| chr4-20688310-INS-6424 | chr4-20688311-INS-6424 |
| chr4-20730080-INS-74 | chr4-20730080-INS-54 |
| chr4-20739731-INS-140 | chr4-20739731-INS-140 |
| chr4-20744578-INS-7133 | chr4-20744579-INS-7134 |
| chr4-20791702-INS-999 | chr4-20791702-INS-999 |
| chr4-20797750-INS-2240 | chr4-20797751-INS-2240 |
| chr4-20816189-INS-69 | chr4-20816189-INS-72 |
| chr4-20836848-INS-73 | chr4-20836846-INS-95 |
| chr4-20874322-INS-130 | chr4-20874322-INS-130 |
| chr4-20895844-INS-108 | chr4-20895844-INS-104 |
| chr4-20898012-INS-221 | chr4-20898012-INS-221 |
| chr4-20907593-INS-80 | chr4-20907594-INS-65 |
| chr4-20923206-INS-6395 | chr4-20923207-INS-6396 |
| chr4-20982518-INS-1865 | chr4-20982526-INS-1855 |
| chr4-21001064-INS-197 | chr4-21001064-INS-197 |
| chr4-21013838-INS-109 | chr4-21013838-INS-109 |
| chr4-21015611-INS-6391 | chr4-21015612-INS-6391 |
| chr4-21025272-INS-56 | chr4-21025272-INS-56 |
| chr4-21043695-INS-54 | chr4-21043695-INS-54 |
| chr4-21087181-INS-58 | chr4-21087181-INS-58 |
| chr4-21098912-INS-941 | chr4-21098912-INS-941 |
| chr4-21116719-INS-124 | chr4-21116715-INS-78 |
| chr4-21130496-INS-5311 | chr4-21130497-INS-5311 |
| chr4-21168430-INS-918 | chr4-21168430-INS-918 |
| chr4-21177870-INS-215 | chr4-21177870-INS-217 |
| chr4-21189732-INS-143 | chr4-21189732-INS-143 |
| chr4-21191719-INS-194 | chr4-21191719-INS-194 |
| chr4-21207332-INS-5266 | chr4-21207333-INS-5266 |
| chr4-21227044-INS-346 | chr4-21227027-INS-359 |
| chr4-21261101-INS-257 | chr4-21261101-INS-257 |
| chr4-21265277-INS-80 | chr4-21265277-INS-80 |
| chr4-21301475-INS-976 | chr4-21301475-INS-979 |
| chr4-21307011-INS-66 | chr4-21307011-INS-66 |
| chr4-21344433-INS-955 | chr4-21344433-INS-976 |
| chr4-21349263-INS-92 | chr4-21349263-INS-87 |
| chr4-21422929-INS-201 | chr4-21422929-INS-201 |
| chr4-21442991-INS-69 | chr4-21442991-INS-69 |
| chr4-21448752-INS-339 | chr4-21448752-INS-344 |
| chr4-21463805-INS-6284 | chr4-21463806-INS-6285 |
| chr4-21521028-INS-54 | chr4-21521028-INS-54 |
| chr4-21534495-INS-7483 | chr4-21534496-INS-7483 |
| chr4-21537034-INS-52 | chr4-21537034-INS-52 |
| chr4-21542827-INS-707 | chr4-21542827-INS-707 |
| chr4-21562149-INS-243 | chr4-21562149-INS-243 |
| chr4-21569140-INS-66 | chr4-21569139-INS-71 |
| chr4-21588019-INS-92 | chr4-21588025-INS-58 |
| chr4-21598455-INS-53 | chr4-21598456-INS-53 |
| chr4-21631759-INS-115 | chr4-21631759-INS-74 |
| chr4-21640135-INS-823 | chr4-21640135-INS-823 |
| chr4-21671675-INS-873 | chr4-21671675-INS-877 |
| chr4-21703051-INS-945 | chr4-21703051-INS-945 |
| chr4-21708468-INS-329 | chr4-21708468-INS-329 |
| chr4-21720218-INS-210 | chr4-21720218-INS-210 |
| chr4-21763678-INS-180 | chr4-21763678-INS-180 |
| chr4-21784538-INS-223 | chr4-21784538-INS-223 |
| chr4-21887099-INS-1721 | chr4-21887100-INS-1725 |

| | |
|---|---|
| chr4-21913961-INS-1545 | chr4-21913962-INS-1546 |
| chr4-21913962-INS-751 | chr4-21913962-INS-751 |
| chr4-21919178-INS-200 | chr4-21919178-INS-200 |
| chr4-21925550-INS-74 | chr4-21925550-INS-78 |
| chr4-21926571-INS-7110 | chr4-21926572-INS-7110 |
| chr4-21934860-INS-68 | chr4-21934834-INS-82 |
| chr4-21936782-INS-509 | chr4-21936782-INS-525 |
| chr4-21986448-INS-66 | chr4-21986448-INS-66 |
| chr4-22047945-INS-71 | chr4-22047945-INS-71 |
| chr4-22053562-INS-54 | chr4-22053562-INS-58 |
| chr4-22066471-INS-4802 | chr4-22066472-INS-4802 |
| chr4-22080307-INS-56 | chr4-22080307-INS-56 |
| chr4-22080492-INS-7112 | chr4-22080493-INS-7112 |
| chr4-22083988-INS-414 | chr4-22083988-INS-412 |
| chr4-22088254-INS-83 | chr4-22088254-INS-87 |
| chr4-22110164-INS-158 | chr4-22110164-INS-158 |
| chr4-22119195-INS-64 | chr4-22119195-INS-64 |
| chr4-22121478-INS-411 | chr4-22121478-INS-404 |
| chr4-22176734-INS-105 | chr4-22176734-INS-105 |
| chr4-22216482-INS-67 | chr4-22216481-INS-75 |
| chr4-22220448-INS-310 | chr4-22220448-INS-310 |
| chr4-22224898-INS-638 | chr4-22224898-INS-638 |
| chr4-22233697-INS-221 | chr4-22233697-INS-221 |
| chr4-22234609-INS-4782 | chr4-22234610-INS-4782 |
| chr4-22242076-INS-177 | chr4-22242076-INS-177 |
| chr4-22243059-INS-120 | chr4-22243057-INS-122 |
| chr4-22249452-INS-62 | chr4-22249484-INS-62 |
| chr4-22257747-INS-195 | chr4-22257747-INS-195 |
| chr4-22277057-INS-72 | chr4-22277057-INS-76 |
| chr4-22278579-INS-1887 | chr4-22278580-INS-1887 |
| chr4-22294667-INS-136 | chr4-22294643-INS-120 |
| chr4-22340709-INS-78 | chr4-22340709-INS-78 |
| chr4-22344107-INS-915 | chr4-22344107-INS-917 |
| chr4-22430550-INS-68 | chr4-22430550-INS-54 |
| chr4-22442633-INS-553 | chr4-22442633-INS-553 |
| chr4-22454927-INS-90 | chr4-22454927-INS-96 |
| chr4-22463242-INS-208 | chr4-22463242-INS-208 |
| chr4-22490919-INS-53 | chr4-22490921-INS-70 |
| chr4-22545594-INS-207 | chr4-22545594-INS-206 |
| chr4-22574638-INS-7072 | chr4-22574639-INS-7072 |
| chr4-22607298-INS-74 | chr4-22607291-INS-54 |
| chr4-22620987-INS-80 | chr4-22620987-INS-80 |
| chr4-22627096-INS-2929 | chr4-22627097-INS-2929 |
| chr4-22647237-INS-70 | chr4-22647237-INS-78 |
| chr4-22684698-INS-2567 | chr4-22684699-INS-2567 |
| chr4-22718178-INS-118 | chr4-22718178-INS-118 |
| chr4-22734499-INS-1560 | chr4-22734500-INS-1562 |
| chr4-22747299-INS-153 | chr4-22747302-INS-154 |
| chr4-22758535-INS-93 | chr4-22758535-INS-93 |
| chr4-22782656-INS-349 | chr4-22782656-INS-356 |
| chr4-22790215-INS-1012 | chr4-22790216-INS-1012 |
| chr4-22794984-INS-7312 | chr4-22794985-INS-7313 |
| chr4-22817053-INS-231 | chr4-22817053-INS-230 |
| chr4-22825314-INS-334 | chr4-22825314-INS-334 |
| chr4-22827609-INS-3624 | chr4-22827610-INS-3646 |
| chr4-22828754-INS-1017 | chr4-22828755-INS-1017 |
| chr4-22830119-INS-62 | chr4-22830119-INS-62 |
| chr4-22847401-INS-90 | chr4-22847401-INS-70 |
| chr4-22872304-INS-309 | chr4-22872304-INS-309 |
| chr4-22898673-INS-477 | chr4-22898673-INS-477 |
| chr4-22899546-INS-60 | chr4-22899546-INS-60 |
| chr4-22945452-INS-454 | chr4-22945452-INS-454 |
| chr4-22952432-INS-57 | chr4-22952432-INS-57 |
| chr4-22957072-INS-52 | chr4-22957072-INS-52 |
| chr4-22979091-INS-5016 | chr4-22979092-INS-5016 |
| chr4-22982829-INS-51 | chr4-22982829-INS-51 |
| chr4-23003623-INS-7446 | chr4-23003624-INS-7447 |
| chr4-23026546-INS-281 | chr4-23026546-INS-281 |
| chr4-23034350-INS-120 | chr4-23034351-INS-120 |
| chr4-23055757-INS-1215 | chr4-23055758-INS-1215 |
| chr4-23095274-INS-826 | chr4-23095274-INS-826 |
| chr4-23155258-INS-3577 | chr4-23155259-INS-3577 |
| chr4-23177679-INS-70 | chr4-23177679-INS-76 |
| chr4-23180854-INS-50 | chr4-23180854-INS-50 |
| chr4-23181075-INS-493 | chr4-23181075-INS-488 |
| chr4-23215736-INS-1393 | chr4-23215737-INS-1395 |
| chr4-23256827-INS-7097 | chr4-23256828-INS-7097 |

| | |
|---|---|
| chr4-23284301-INS-75 | chr4-23284301-INS-75 |
| chr4-23284509-INS-2709 | chr4-23284510-INS-2709 |
| chr4-23286584-INS-58 | chr4-23286584-INS-52 |
| chr4-23303025-INS-340 | chr4-23303025-INS-340 |
| chr4-23322149-INS-200 | chr4-23322149-INS-201 |
| chr4-23331016-INS-258 | chr4-23331010-INS-307 |
| chr4-23345574-INS-6482 | chr4-23345575-INS-6482 |
| chr4-23371713-INS-7641 | chr4-23371714-INS-7641 |
| chr4-23401335-INS-105 | chr4-23401335-INS-108 |
| chr4-23417365-INS-205 | chr4-23417365-INS-205 |
| chr4-23435782-INS-55 | chr4-23435746-INS-60 |
| chr4-23462297-INS-782 | chr4-23462297-INS-784 |
| chr4-23512616-INS-71 | chr4-23512616-INS-71 |
| chr4-23514730-INS-74 | chr4-23514732-INS-74 |
| chr4-23546296-INS-6675 | chr4-23546297-INS-6675 |
| chr4-23579993-INS-80 | chr4-23579993-INS-84 |
| chr4-23597451-INS-3639 | chr4-23597452-INS-3639 |
| chr4-23621002-INS-7442 | chr4-23621003-INS-7442 |
| chr4-23685468-INS-6458 | chr4-23685469-INS-6458 |
| chr4-23694252-INS-52 | chr4-23694252-INS-58 |
| chr4-23741251-INS-1081 | chr4-23741252-INS-1081 |
| chr4-23785150-INS-69 | chr4-23785150-INS-69 |
| chr4-23824002-INS-72 | chr4-23823999-INS-75 |
| chr4-23829002-INS-55 | chr4-23829002-INS-55 |
| chr4-23845942-INS-51 | chr4-23845942-INS-51 |
| chr4-23880771-INS-60 | chr4-23880771-INS-60 |
| chr4-23901671-INS-316 | chr4-23901671-INS-316 |
| chr4-23942439-INS-428 | chr4-23942439-INS-428 |
| chr4-23953722-INS-469 | chr4-23953723-INS-469 |
| chr4-23956085-INS-55 | chr4-23956085-INS-52 |
| chr4-23967949-INS-354 | chr4-23967949-INS-354 |
| chr4-23979115-INS-80 | chr4-23979115-INS-101 |
| chr4-23981094-INS-126 | chr4-23981094-INS-126 |
| chr4-24016138-INS-62 | chr4-24016138-INS-66 |
| chr4-24018031-INS-71 | chr4-24018031-INS-79 |
| chr4-24029951-INS-57 | chr4-24029951-INS-83 |
| chr4-24036337-INS-105 | chr4-24036345-INS-108 |
| chr4-24116286-INS-64 | chr4-24116314-INS-51 |
| chr4-24123839-INS-74 | chr4-24123839-INS-76 |
| chr4-24316527-INS-185 | chr4-24316527-INS-214 |
| chr4-24334753-INS-230 | chr4-24334753-INS-230 |
| chr4-24357350-INS-716 | chr4-24357350-INS-716 |
| chr4-24365795-INS-712 | chr4-24365795-INS-712 |
| chr4-24384930-INS-117 | chr4-24384930-INS-117 |
| chr4-24407472-INS-50 | chr4-24407472-INS-50 |
| chr4-24409387-INS-184 | chr4-24409383-INS-194 |
| chr4-24416316-INS-220 | chr4-24416314-INS-220 |
| chr4-24443637-INS-6674 | chr4-24443638-INS-6674 |
| chr4-24444555-INS-231 | chr4-24444555-INS-231 |
| chr4-24453195-INS-8672 | chr4-24453196-INS-8672 |
| chr4-24481724-INS-6401 | chr4-24481725-INS-6401 |
| chr4-24483347-INS-261 | chr4-24483347-INS-261 |
| chr4-24491162-INS-181 | chr4-24491162-INS-182 |
| chr4-24493091-INS-313 | chr4-24493092-INS-313 |
| chr4-24529685-INS-58 | chr4-24529685-INS-58 |
| chr4-24575565-INS-757 | chr4-24575565-INS-756 |
| chr4-24598415-INS-88 | chr4-24598415-INS-88 |
| chr4-24611148-INS-584 | chr4-24611148-INS-562 |
| chr4-24637946-INS-418 | chr4-24637946-INS-418 |
| chr4-24638508-INS-116 | chr4-24638508-INS-158 |
| chr4-24684801-INS-557 | chr4-24684801-INS-557 |
| chr4-24705945-INS-55 | chr4-24705945-INS-55 |
| chr4-24720360-INS-116 | chr4-24720360-INS-116 |
| chr4-24731531-INS-71 | chr4-24731531-INS-89 |
| chr4-24851605-INS-187 | chr4-24851605-INS-187 |
| chr4-24873111-INS-325 | chr4-24873111-INS-325 |
| chr4-24885800-INS-63 | chr4-24885819-INS-66 |
| chr4-24898771-INS-60 | chr4-24898771-INS-60 |
| chr4-24913585-INS-54 | chr4-24913585-INS-62 |
| chr4-24934989-INS-157 | chr4-24934989-INS-157 |
| chr4-24945711-INS-270 | chr4-24945711-INS-276 |
| chr4-24949530-INS-174 | chr4-24949530-INS-175 |
| chr4-25057997-INS-65 | chr4-25057997-INS-65 |
| chr4-25176200-INS-7695 | chr4-25176201-INS-7696 |
| chr4-25180864-INS-183 | chr4-25180864-INS-183 |
| chr4-25202969-INS-112 | chr4-25202969-INS-129 |
| chr4-25209024-INS-112 | chr4-25209014-INS-86 |

| | |
|---|---|
| chr4-25244127-INS-80 | chr4-25244127-INS-80 |
| chr4-25277091-INS-137 | chr4-25277091-INS-123 |
| chr4-25328880-INS-6404 | chr4-25328881-INS-6404 |
| chr4-25351056-INS-55 | chr4-25351056-INS-55 |
| chr4-25352311-INS-743 | chr4-25352311-INS-743 |
| chr4-25364883-INS-78 | chr4-25364883-INS-78 |
| chr4-25369380-INS-409 | chr4-25369380-INS-415 |
| chr4-25373779-INS-206 | chr4-25373779-INS-206 |
| chr4-25385174-INS-150 | chr4-25385215-INS-125 |
| chr4-25408005-INS-52 | chr4-25408005-INS-52 |
| chr4-25411725-INS-6603 | chr4-25411726-INS-6603 |
| chr4-25419343-INS-1508 | chr4-25419344-INS-1508 |
| chr4-25421994-INS-1715 | chr4-25421990-INS-1691 |
| chr4-25434148-INS-116 | chr4-25434143-INS-126 |
| chr4-25456743-INS-4118 | chr4-25456744-INS-4118 |
| chr4-25457095-INS-210 | chr4-25457095-INS-210 |
| chr4-25477212-INS-6433 | chr4-25477213-INS-6433 |
| chr4-25482549-INS-72 | chr4-25482524-INS-82 |
| chr4-25524272-INS-60 | chr4-25524272-INS-84 |
| chr4-25541092-INS-6678 | chr4-25541093-INS-6678 |
| chr4-25595735-INS-897 | chr4-25595735-INS-897 |
| chr4-25625335-INS-56 | chr4-25625335-INS-56 |
| chr4-25666800-INS-127 | chr4-25666799-INS-96 |
| chr4-25695957-INS-6269 | chr4-25695958-INS-6270 |
| chr4-25758246-INS-76 | chr4-25758246-INS-84 |
| chr4-25765663-INS-93 | chr4-25765663-INS-94 |
| chr4-25779463-INS-54 | chr4-25779463-INS-54 |
| chr4-25813060-INS-491 | chr4-25813060-INS-489 |
| chr4-25821876-INS-158 | chr4-25821887-INS-141 |
| chr4-25868700-INS-572 | chr4-25868700-INS-572 |
| chr4-25882866-INS-54 | chr4-25882863-INS-60 |
| chr4-25894175-INS-61 | chr4-25894139-INS-66 |
| chr4-25911862-INS-520 | chr4-25911862-INS-520 |
| chr4-25920393-INS-92 | chr4-25920393-INS-92 |
| chr4-25926387-INS-6462 | chr4-25926388-INS-6463 |
| chr4-25931310-INS-6467 | chr4-25931311-INS-6467 |
| chr4-25951545-INS-6645 | chr4-25951546-INS-6645 |
| chr4-25964979-INS-80 | chr4-25964979-INS-80 |
| chr4-25988295-INS-480 | chr4-25988295-INS-480 |
| chr4-25990956-INS-1585 | chr4-25990957-INS-1588 |
| chr4-25996642-INS-6179 | chr4-25996643-INS-6179 |
| chr4-26066725-INS-1167 | chr4-26066726-INS-1167 |
| chr4-26082779-INS-5237 | chr4-26082780-INS-5237 |
| chr4-26170493-INS-124 | chr4-26170493-INS-126 |
| chr4-26211661-INS-4678 | chr4-26211662-INS-4679 |
| chr4-26248467-INS-2221 | chr4-26248468-INS-2222 |
| chr4-26291039-INS-5556 | chr4-26291040-INS-5556 |
| chr4-26302343-INS-91 | chr4-26302343-INS-98 |
| chr4-26340095-INS-267 | chr4-26340095-INS-267 |
| chr4-26344766-INS-58 | chr4-26344766-INS-60 |
| chr4-26377560-INS-4690 | chr4-26377561-INS-4690 |
| chr4-26422926-INS-1889 | chr4-26422927-INS-1889 |
| chr4-26423656-INS-587 | chr4-26423656-INS-593 |
| chr4-26424441-INS-65 | chr4-26424441-INS-65 |
| chr4-26430799-INS-191 | chr4-26430799-INS-161 |
| chr4-26433122-INS-210 | chr4-26433122-INS-210 |
| chr4-26440616-INS-117 | chr4-26440616-INS-111 |
| chr4-26446413-INS-58 | chr4-26446413-INS-58 |
| chr4-26480777-INS-842 | chr4-26480777-INS-843 |
| chr4-26485438-INS-208 | chr4-26485438-INS-207 |
| chr4-26486725-INS-96 | chr4-26486725-INS-96 |
| chr4-26492207-INS-172 | chr4-26492207-INS-171 |
| chr4-26510829-INS-128 | chr4-26510829-INS-128 |
| chr4-26605602-INS-96 | chr4-26605602-INS-98 |
| chr4-26615260-INS-188 | chr4-26615260-INS-188 |
| chr4-26622616-INS-137 | chr4-26622616-INS-132 |
| chr4-26672165-INS-205 | chr4-26672165-INS-208 |
| chr4-26691922-INS-62 | chr4-26691908-INS-67 |
| chr4-26701154-INS-656 | chr4-26701154-INS-657 |
| chr4-26724507-INS-83 | chr4-26724499-INS-86 |
| chr4-26783861-INS-1020 | chr4-26783862-INS-1020 |
| chr4-26806331-INS-183 | chr4-26806331-INS-184 |
| chr4-26807199-INS-1645 | chr4-26807200-INS-1645 |
| chr4-26822138-INS-234 | chr4-26822134-INS-258 |
| chr4-26860230-INS-100 | chr4-26860230-INS-104 |
| chr4-26887171-INS-119 | chr4-26887171-INS-128 |
| chr4-26908465-INS-1875 | chr4-26908466-INS-1875 |

| | |
|---|---|
| chr4-26910725-INS-1890 | chr4-26910726-INS-1890 |
| chr4-26929603-INS-62 | chr4-26929603-INS-64 |
| chr4-26932016-INS-67 | chr4-26932016-INS-67 |
| chr4-26933407-INS-129 | chr4-26933407-INS-129 |
| chr4-27033670-INS-83 | chr4-27033670-INS-83 |
| chr4-27066721-INS-6475 | chr4-27066722-INS-6475 |
| chr4-27076824-INS-56 | chr4-27076824-INS-56 |
| chr4-27116154-INS-6338 | chr4-27116155-INS-6338 |
| chr4-27199791-INS-539 | chr4-27199791-INS-539 |
| chr4-27229771-INS-95 | chr4-27229771-INS-102 |
| chr4-27231451-INS-7112 | chr4-27231452-INS-7112 |
| chr4-27265143-INS-383 | chr4-27265143-INS-385 |
| chr4-27266612-INS-727 | chr4-27266612-INS-727 |
| chr4-27271150-INS-4164 | chr4-27271151-INS-4164 |
| chr4-27283828-INS-156 | chr4-27283822-INS-156 |
| chr4-27314182-INS-55 | chr4-27314182-INS-50 |
| chr4-27324690-INS-98 | chr4-27324690-INS-126 |
| chr4-27341082-INS-76 | chr4-27341082-INS-52 |
| chr4-27361878-INS-182 | chr4-27361878-INS-182 |
| chr4-27372439-INS-141 | chr4-27372439-INS-141 |
| chr4-27373673-INS-60 | chr4-27373673-INS-58 |
| chr4-27381003-INS-66 | chr4-27381005-INS-102 |
| chr4-27402681-INS-401 | chr4-27402681-INS-401 |
| chr4-27423178-INS-111 | chr4-27423178-INS-64 |
| chr4-27424824-INS-72 | chr4-27424824-INS-72 |
| chr4-27438947-INS-417 | chr4-27438947-INS-417 |
| chr4-27516709-INS-3234 | chr4-27516710-INS-3235 |
| chr4-27535648-INS-477 | chr4-27535648-INS-477 |
| chr4-27567377-INS-142 | chr4-27567377-INS-142 |
| chr4-27578940-INS-91 | chr4-27578941-INS-90 |
| chr4-27601468-INS-71 | chr4-27601468-INS-70 |
| chr4-27609411-INS-1166 | chr4-27609412-INS-1166 |
| chr4-27616166-INS-6687 | chr4-27616167-INS-6687 |
| chr4-27619288-INS-771 | chr4-27619288-INS-771 |
| chr4-27621214-INS-66 | chr4-27621214-INS-66 |
| chr4-27623273-INS-420 | chr4-27623273-INS-420 |
| chr4-27651651-INS-906 | chr4-27651651-INS-907 |
| chr4-27654593-INS-67 | chr4-27654593-INS-60 |
| chr4-27697257-INS-66 | chr4-27697257-INS-68 |
| chr4-27724241-INS-51 | chr4-27724245-INS-50 |
| chr4-27784643-INS-2953 | chr4-27784644-INS-2954 |
| chr4-27809497-INS-815 | chr4-27809497-INS-815 |
| chr4-27825512-INS-681 | chr4-27825512-INS-681 |
| chr4-27853453-INS-1827 | chr4-27853454-INS-1828 |
| chr4-27878193-INS-2081 | chr4-27878194-INS-2081 |
| chr4-27911897-INS-786 | chr4-27911897-INS-785 |
| chr4-27981987-INS-52 | chr4-27981983-INS-56 |
| chr4-28041185-INS-143 | chr4-28041189-INS-151 |
| chr4-28043469-INS-62 | chr4-28043469-INS-64 |
| chr4-28051555-INS-51 | chr4-28051555-INS-51 |
| chr4-28068964-INS-62 | chr4-28068964-INS-63 |
| chr4-28073131-INS-6452 | chr4-28073132-INS-6452 |
| chr4-28112370-INS-551 | chr4-28112370-INS-558 |
| chr4-28128288-INS-71 | chr4-28128286-INS-50 |
| chr4-28188939-INS-63 | chr4-28188939-INS-54 |
| chr4-28203227-INS-202 | chr4-28203227-INS-217 |
| chr4-28208396-INS-83 | chr4-28208396-INS-84 |
| chr4-28221316-INS-149 | chr4-28221311-INS-148 |
| chr4-28233359-INS-464 | chr4-28233359-INS-464 |
| chr4-28267322-INS-59 | chr4-28267324-INS-60 |
| chr4-28278286-INS-61 | chr4-28278286-INS-61 |
| chr4-28283800-INS-700 | chr4-28283800-INS-700 |
| chr4-28298221-INS-59 | chr4-28298221-INS-59 |
| chr4-28301574-INS-56 | chr4-28301546-INS-89 |
| chr4-28322572-INS-102 | chr4-28322573-INS-102 |
| chr4-28358947-INS-997 | chr4-28358947-INS-999 |
| chr4-28365553-INS-144 | chr4-28365553-INS-144 |
| chr4-28382222-INS-835 | chr4-28382222-INS-836 |
| chr4-28403574-INS-152 | chr4-28403574-INS-150 |
| chr4-28412019-INS-430 | chr4-28412019-INS-438 |
| chr4-28437618-INS-6729 | chr4-28437619-INS-6729 |
| chr4-28438293-INS-2038 | chr4-28438294-INS-2043 |
| chr4-28470586-INS-54 | chr4-28470586-INS-54 |
| chr4-28477697-INS-424 | chr4-28477697-INS-424 |
| chr4-28484827-INS-135 | chr4-28484827-INS-134 |
| chr4-28495468-INS-63 | chr4-28495468-INS-62 |
| chr4-28528909-INS-101 | chr4-28528909-INS-101 |

| | |
|---|---|
| chr4-28532353-INS-407 | chr4-28532353-INS-407 |
| chr4-28559096-INS-73 | chr4-28559096-INS-73 |
| chr4-28569318-INS-325 | chr4-28569318-INS-325 |
| chr4-28572134-INS-415 | chr4-28572134-INS-415 |
| chr4-28604641-INS-56 | chr4-28604641-INS-54 |
| chr4-28629318-INS-222 | chr4-28629318-INS-222 |
| chr4-28651583-INS-220 | chr4-28651583-INS-220 |
| chr4-28698730-INS-80 | chr4-28698734-INS-76 |
| chr4-28707787-INS-72 | chr4-28707787-INS-74 |
| chr4-28718468-INS-54 | chr4-28718468-INS-56 |
| chr4-28725130-INS-1821 | chr4-28725131-INS-1821 |
| chr4-28774148-INS-416 | chr4-28774148-INS-420 |
| chr4-28784267-INS-54 | chr4-28784267-INS-58 |
| chr4-28796926-INS-2004 | chr4-28796927-INS-2021 |
| chr4-28831071-INS-100 | chr4-28831071-INS-104 |
| chr4-28846271-INS-84 | chr4-28846271-INS-84 |
| chr4-28908453-INS-844 | chr4-28908453-INS-842 |
| chr4-28908993-INS-150 | chr4-28908993-INS-150 |
| chr4-28925102-INS-70 | chr4-28925102-INS-70 |
| chr4-28954220-INS-151 | chr4-28954221-INS-166 |
| chr4-28959615-INS-160 | chr4-28959615-INS-160 |
| chr4-29057745-INS-56 | chr4-29057746-INS-56 |
| chr4-29064317-INS-400 | chr4-29064317-INS-396 |
| chr4-29070634-INS-821 | chr4-29070634-INS-821 |
| chr4-29079885-INS-108 | chr4-29079861-INS-132 |
| chr4-29083561-INS-104 | chr4-29083561-INS-108 |
| chr4-29106842-INS-194 | chr4-29106842-INS-194 |
| chr4-29109088-INS-54 | chr4-29109089-INS-58 |
| chr4-29111601-INS-866 | chr4-29111601-INS-867 |
| chr4-29123774-INS-6674 | chr4-29123775-INS-6675 |
| chr4-29133282-INS-51 | chr4-29133282-INS-51 |
| chr4-29158469-INS-90 | chr4-29158469-INS-93 |
| chr4-29173969-INS-64 | chr4-29173969-INS-64 |
| chr4-29207735-INS-6580 | chr4-29207720-INS-6580 |
| chr4-29229403-INS-128 | chr4-29229414-INS-128 |
| chr4-29269114-INS-4896 | chr4-29269115-INS-4897 |
| chr4-29299025-INS-101 | chr4-29299032-INS-107 |
| chr4-29306031-INS-64 | chr4-29306031-INS-64 |
| chr4-29310937-INS-140 | chr4-29310937-INS-140 |
| chr4-29319737-INS-95 | chr4-29319737-INS-90 |
| chr4-29328698-INS-1105 | chr4-29328699-INS-1105 |
| chr4-29331770-INS-194 | chr4-29331751-INS-198 |
| chr4-29337769-INS-60 | chr4-29337769-INS-60 |
| chr4-29359838-INS-112 | chr4-29359838-INS-112 |
| chr4-29401486-INS-84 | chr4-29401486-INS-84 |
| chr4-29404510-INS-341 | chr4-29404510-INS-341 |
| chr4-29412339-INS-76 | chr4-29412339-INS-76 |
| chr4-29427976-INS-54 | chr4-29427976-INS-54 |
| chr4-29431897-INS-149 | chr4-29431897-INS-149 |
| chr4-29434462-INS-114 | chr4-29434463-INS-121 |
| chr4-29445220-INS-203 | chr4-29445220-INS-203 |
| chr4-29448307-INS-119 | chr4-29448307-INS-119 |
| chr4-29449551-INS-62 | chr4-29449551-INS-58 |
| chr4-29452770-INS-995 | chr4-29452770-INS-988 |
| chr4-29475261-INS-231 | chr4-29475261-INS-243 |
| chr4-29485720-INS-52 | chr4-29485720-INS-52 |
| chr4-29492485-INS-6704 | chr4-29492486-INS-6704 |
| chr4-29517840-INS-63 | chr4-29517836-INS-86 |
| chr4-29519982-INS-56 | chr4-29519982-INS-64 |
| chr4-29524653-INS-64 | chr4-29524653-INS-64 |
| chr4-29525426-INS-244 | chr4-29525426-INS-244 |
| chr4-29551437-INS-66 | chr4-29551433-INS-99 |
| chr4-29556946-INS-110 | chr4-29556946-INS-110 |
| chr4-29575409-INS-1392 | chr4-29575410-INS-1392 |
| chr4-29580587-INS-1033 | chr4-29580588-INS-1033 |
| chr4-29585304-INS-2774 | chr4-29585305-INS-2759 |
| chr4-29587124-INS-112 | chr4-29587124-INS-112 |
| chr4-29605309-INS-133 | chr4-29605309-INS-112 |
| chr4-29617106-INS-128 | chr4-29617106-INS-152 |
| chr4-29653780-INS-151 | chr4-29653780-INS-139 |
| chr4-29669396-INS-132 | chr4-29669438-INS-134 |
| chr4-29682192-INS-66 | chr4-29682204-INS-64 |
| chr4-29687890-INS-782 | chr4-29687897-INS-802 |
| chr4-29688015-INS-101 | chr4-29688025-INS-127 |
| chr4-29722475-INS-392 | chr4-29722475-INS-392 |
| chr4-29733003-INS-2883 | chr4-29733004-INS-2883 |
| chr4-29753152-INS-52 | chr4-29753152-INS-54 |

| | |
|---|---|
| chr4-29769953-INS-62 | chr4-29769953-INS-54 |
| chr4-29772769-INS-57 | chr4-29772769-INS-57 |
| chr4-29809937-INS-800 | chr4-29809934-INS-790 |
| chr4-29827333-INS-2040 | chr4-29827334-INS-2040 |
| chr4-29840810-INS-199 | chr4-29840789-INS-199 |
| chr4-29861823-INS-58 | chr4-29861825-INS-66 |
| chr4-29888713-INS-2371 | chr4-29888714-INS-2377 |
| chr4-29893252-INS-1302 | chr4-29893253-INS-1302 |
| chr4-29902750-INS-602 | chr4-29902750-INS-602 |
| chr4-29904732-INS-3115 | chr4-29904733-INS-3114 |
| chr4-29941170-INS-83 | chr4-29941170-INS-81 |
| chr4-29943825-INS-801 | chr4-29943825-INS-800 |
| chr4-29951293-INS-503 | chr4-29951293-INS-508 |
| chr4-29969229-INS-5380 | chr4-29969230-INS-5383 |
| chr4-30079065-INS-184 | chr4-30079065-INS-201 |
| chr4-30119486-INS-70 | chr4-30119486-INS-70 |
| chr4-30137175-INS-51 | chr4-30137175-INS-54 |
| chr4-30151291-INS-75 | chr4-30151296-INS-60 |
| chr4-30164317-INS-56 | chr4-30164317-INS-56 |
| chr4-30170946-INS-212 | chr4-30170946-INS-212 |
| chr4-30181537-INS-78 | chr4-30181537-INS-78 |
| chr4-30197065-INS-68 | chr4-30197096-INS-50 |
| chr4-30304494-INS-82 | chr4-30304494-INS-82 |
| chr4-30315344-INS-86 | chr4-30315361-INS-86 |
| chr4-30318193-INS-54 | chr4-30318193-INS-78 |
| chr4-30319951-INS-58 | chr4-30319951-INS-63 |
| chr4-30339514-INS-107 | chr4-30339513-INS-87 |
| chr4-30355602-INS-564 | chr4-30355602-INS-564 |
| chr4-30367784-INS-88 | chr4-30367784-INS-72 |
| chr4-30377308-INS-1129 | chr4-30377309-INS-1125 |
| chr4-30382748-INS-92 | chr4-30382749-INS-54 |
| chr4-30388996-INS-70 | chr4-30388992-INS-114 |
| chr4-30390548-INS-400 | chr4-30390548-INS-400 |
| chr4-30416825-INS-90 | chr4-30416825-INS-90 |
| chr4-30417960-INS-102 | chr4-30417960-INS-102 |
| chr4-30437075-INS-80 | chr4-30437075-INS-86 |
| chr4-30440703-INS-494 | chr4-30440703-INS-496 |
| chr4-30461619-INS-56 | chr4-30461619-INS-58 |
| chr4-30541333-INS-80 | chr4-30541333-INS-86 |
| chr4-30547746-INS-200 | chr4-30547746-INS-200 |
| chr4-30592311-INS-598 | chr4-30592311-INS-598 |
| chr4-30596736-INS-116 | chr4-30596736-INS-122 |
| chr4-30604020-INS-86 | chr4-30604020-INS-86 |
| chr4-30628812-INS-653 | chr4-30628809-INS-656 |
| chr4-30644055-INS-63 | chr4-30644096-INS-52 |
| chr4-30685326-INS-57 | chr4-30685326-INS-57 |
| chr4-30697028-INS-303 | chr4-30697028-INS-303 |
| chr4-30717524-INS-279 | chr4-30717524-INS-279 |
| chr4-30725187-INS-970 | chr4-30725187-INS-966 |
| chr4-30733967-INS-88 | chr4-30733979-INS-89 |
| chr4-30740038-INS-84 | chr4-30740041-INS-58 |
| chr4-30773924-INS-55 | chr4-30773924-INS-55 |
| chr4-30794376-INS-152 | chr4-30794376-INS-152 |
| chr4-30814110-INS-497 | chr4-30814110-INS-497 |
| chr4-30848395-INS-4039 | chr4-30848396-INS-4039 |
| chr4-30852213-INS-55 | chr4-30852225-INS-90 |
| chr4-30865077-INS-186 | chr4-30865077-INS-186 |
| chr4-30873418-INS-52 | chr4-30873419-INS-58 |
| chr4-30888089-INS-932 | chr4-30888089-INS-928 |
| chr4-30938650-INS-449 | chr4-30938650-INS-449 |
| chr4-30944339-INS-166 | chr4-30944339-INS-166 |
| chr4-30946895-INS-124 | chr4-30946891-INS-110 |
| chr4-30948385-INS-140 | chr4-30948392-INS-95 |
| chr4-30950614-INS-158 | chr4-30950614-INS-158 |
| chr4-30972672-INS-56 | chr4-30972672-INS-56 |
| chr4-31095834-INS-707 | chr4-31095834-INS-707 |
| chr4-31109244-INS-70 | chr4-31109244-INS-99 |
| chr4-31119037-INS-2322 | chr4-31119038-INS-2322 |
| chr4-31141701-INS-51 | chr4-31141701-INS-51 |
| chr4-31151112-INS-54 | chr4-31151112-INS-54 |
| chr4-31180082-INS-143 | chr4-31180082-INS-142 |
| chr4-31192813-INS-87 | chr4-31192813-INS-105 |
| chr4-31193782-INS-500 | chr4-31193782-INS-502 |
| chr4-31217978-INS-94 | chr4-31217978-INS-94 |
| chr4-31222103-INS-55 | chr4-31222103-INS-55 |
| chr4-31252580-INS-132 | chr4-31252613-INS-99 |
| chr4-31260551-INS-799 | chr4-31260551-INS-798 |

| | |
|---|---|
| chr4-31277724-INS-632 | chr4-31277724-INS-643 |
| chr4-31294843-INS-878 | chr4-31294843-INS-877 |
| chr4-31309946-INS-93 | chr4-31309946-INS-93 |
| chr4-31315273-INS-403 | chr4-31315273-INS-396 |
| chr4-31320413-INS-73 | chr4-31320413-INS-68 |
| chr4-31324380-INS-198 | chr4-31324380-INS-198 |
| chr4-31368184-INS-694 | chr4-31368184-INS-694 |
| chr4-31372303-INS-768 | chr4-31372303-INS-767 |
| chr4-31390155-INS-454 | chr4-31390166-INS-471 |
| chr4-31395089-INS-2482 | chr4-31395090-INS-2482 |
| chr4-31427929-INS-71 | chr4-31427929-INS-71 |
| chr4-31430737-INS-94 | chr4-31430737-INS-94 |
| chr4-31465608-INS-99 | chr4-31465608-INS-101 |
| chr4-31478727-INS-707 | chr4-31478727-INS-707 |
| chr4-31488095-INS-5994 | chr4-31488096-INS-5994 |
| chr4-31515803-INS-6485 | chr4-31515804-INS-6488 |
| chr4-31542228-INS-191 | chr4-31542228-INS-191 |
| chr4-31581399-INS-58 | chr4-31581399-INS-58 |
| chr4-31600619-INS-149 | chr4-31600617-INS-164 |
| chr4-31616028-INS-258 | chr4-31616001-INS-267 |
| chr4-31680400-INS-76 | chr4-31680400-INS-84 |
| chr4-31697296-INS-86 | chr4-31697296-INS-90 |
| chr4-31721869-INS-101 | chr4-31721869-INS-100 |
| chr4-31747150-INS-194 | chr4-31747150-INS-199 |
| chr4-31767389-INS-84 | chr4-31767389-INS-84 |
| chr4-31771208-INS-518 | chr4-31771208-INS-518 |
| chr4-31796336-INS-62 | chr4-31796336-INS-62 |
| chr4-31817705-INS-699 | chr4-31817705-INS-695 |
| chr4-31845194-INS-63 | chr4-31845194-INS-63 |
| chr4-31849231-INS-722 | chr4-31849231-INS-722 |
| chr4-31903844-INS-81 | chr4-31903847-INS-66 |
| chr4-31914689-INS-99 | chr4-31914689-INS-75 |
| chr4-31919751-INS-6440 | chr4-31919752-INS-6440 |
| chr4-31928251-INS-99 | chr4-31928251-INS-99 |
| chr4-31943233-INS-286 | chr4-31943233-INS-286 |
| chr4-31960537-INS-56 | chr4-31960537-INS-50 |
| chr4-31979213-INS-88 | chr4-31979215-INS-81 |
| chr4-32092333-INS-54 | chr4-32092333-INS-54 |
| chr4-32094772-INS-234 | chr4-32094772-INS-234 |
| chr4-32113323-INS-170 | chr4-32113302-INS-166 |
| chr4-32136550-INS-52 | chr4-32136550-INS-54 |
| chr4-32144264-INS-64 | chr4-32144264-INS-72 |
| chr4-32145853-INS-54 | chr4-32145863-INS-57 |
| chr4-32156819-INS-645 | chr4-32156819-INS-646 |
| chr4-32165714-INS-497 | chr4-32165714-INS-490 |
| chr4-32169389-INS-239 | chr4-32169389-INS-241 |
| chr4-32184139-INS-212 | chr4-32184139-INS-212 |
| chr4-32196240-INS-62 | chr4-32196240-INS-56 |
| chr4-32249625-INS-198 | chr4-32249625-INS-198 |
| chr4-32260250-INS-61 | chr4-32260250-INS-61 |
| chr4-32295523-INS-79 | chr4-32295530-INS-73 |
| chr4-32320001-INS-109 | chr4-32320001-INS-109 |
| chr4-32345458-INS-61 | chr4-32345458-INS-73 |
| chr4-32356765-INS-1510 | chr4-32356766-INS-1508 |
| chr4-32382936-INS-239 | chr4-32382936-INS-239 |
| chr4-32390426-INS-380 | chr4-32390426-INS-379 |
| chr4-32430583-INS-56 | chr4-32430583-INS-56 |
| chr4-32482126-INS-135 | chr4-32482126-INS-135 |
| chr4-32492224-INS-1823 | chr4-32492225-INS-1823 |
| chr4-32493917-INS-751 | chr4-32493917-INS-751 |
| chr4-32518937-INS-192 | chr4-32518945-INS-208 |
| chr4-32546045-INS-78 | chr4-32546045-INS-78 |
| chr4-32608327-INS-5384 | chr4-32608328-INS-5432 |
| chr4-32610357-INS-316 | chr4-32610382-INS-331 |
| chr4-32632396-INS-52 | chr4-32632396-INS-52 |
| chr4-32697813-INS-220 | chr4-32697813-INS-220 |
| chr4-32798407-INS-78 | chr4-32798407-INS-78 |
| chr4-32811229-INS-250 | chr4-32811229-INS-253 |
| chr4-32825769-INS-55 | chr4-32825769-INS-55 |
| chr4-32844434-INS-58 | chr4-32844434-INS-58 |
| chr4-32862568-INS-75 | chr4-32862568-INS-75 |
| chr4-32869782-INS-126 | chr4-32869782-INS-128 |
| chr4-32873190-INS-274 | chr4-32873190-INS-275 |
| chr4-32881971-INS-440 | chr4-32881971-INS-440 |
| chr4-32900669-INS-184 | chr4-32900669-INS-182 |
| chr4-32917256-INS-193 | chr4-32917256-INS-193 |
| chr4-32946126-INS-853 | chr4-32946126-INS-854 |

| | |
|---|---|
| chr4-32953301-INS-68 | chr4-32953301-INS-82 |
| chr4-32954168-INS-330 | chr4-32954168-INS-331 |
| chr4-32956630-INS-531 | chr4-32956630-INS-534 |
| chr4-32958940-INS-217 | chr4-32958931-INS-217 |
| chr4-32962110-INS-77 | chr4-32962110-INS-75 |
| chr4-32970778-INS-2722 | chr4-32970779-INS-2722 |
| chr4-32972487-INS-422 | chr4-32972487-INS-422 |
| chr4-32976257-INS-258 | chr4-32976257-INS-272 |
| chr4-32978000-INS-505 | chr4-32978000-INS-492 |
| chr4-32986128-INS-262 | chr4-32986128-INS-261 |
| chr4-32999899-INS-252 | chr4-32999899-INS-253 |
| chr4-33005623-INS-108 | chr4-33005587-INS-99 |
| chr4-33023391-INS-60 | chr4-33023391-INS-60 |
| chr4-33038531-INS-154 | chr4-33038528-INS-155 |
| chr4-33045404-INS-83 | chr4-33045404-INS-83 |
| chr4-33064360-INS-110 | chr4-33064391-INS-70 |
| chr4-33073613-INS-76 | chr4-33073613-INS-76 |
| chr4-33083047-INS-835 | chr4-33083047-INS-828 |
| chr4-33098104-INS-91 | chr4-33098104-INS-91 |
| chr4-33108096-INS-287 | chr4-33108096-INS-289 |
| chr4-33116801-INS-63 | chr4-33116801-INS-57 |
| chr4-33125088-INS-197 | chr4-33125088-INS-197 |
| chr4-33148650-INS-171 | chr4-33148650-INS-173 |
| chr4-33150521-INS-96 | chr4-33150521-INS-96 |
| chr4-33154190-INS-64 | chr4-33154191-INS-52 |
| chr4-33162454-INS-672 | chr4-33162454-INS-672 |
| chr4-33196228-INS-3209 | chr4-33196229-INS-3209 |
| chr4-33203404-INS-64 | chr4-33203404-INS-64 |
| chr4-33230199-INS-52 | chr4-33230236-INS-60 |
| chr4-33263993-INS-302 | chr4-33263993-INS-302 |
| chr4-33286173-INS-54 | chr4-33286173-INS-54 |
| chr4-33288571-INS-50 | chr4-33288523-INS-63 |
| chr4-33290446-INS-98 | chr4-33290446-INS-147 |
| chr4-33292206-INS-1468 | chr4-33292207-INS-1462 |
| chr4-33300765-INS-167 | chr4-33300765-INS-165 |
| chr4-33305806-INS-390 | chr4-33305806-INS-390 |
| chr4-33315027-INS-194 | chr4-33315027-INS-194 |
| chr4-33331136-INS-124 | chr4-33331151-INS-104 |
| chr4-33350360-INS-62 | chr4-33350360-INS-62 |
| chr4-33369599-INS-114 | chr4-33369618-INS-78 |
| chr4-33414620-INS-1022 | chr4-33414621-INS-1021 |
| chr4-33466221-INS-276 | chr4-33466221-INS-276 |
| chr4-33469979-INS-63 | chr4-33469971-INS-74 |
| chr4-33479227-INS-206 | chr4-33479227-INS-226 |
| chr4-33490200-INS-1893 | chr4-33490201-INS-1893 |
| chr4-33493379-INS-428 | chr4-33493379-INS-428 |
| chr4-33504039-INS-357 | chr4-33504039-INS-357 |
| chr4-33512221-INS-70 | chr4-33512221-INS-72 |
| chr4-33514306-INS-63 | chr4-33514308-INS-105 |
| chr4-33528848-INS-64 | chr4-33528848-INS-66 |
| chr4-33529740-INS-538 | chr4-33529740-INS-538 |
| chr4-33550882-INS-2966 | chr4-33550883-INS-2966 |
| chr4-33578264-INS-6440 | chr4-33578265-INS-6440 |
| chr4-33603846-INS-4162 | chr4-33603847-INS-4162 |
| chr4-33608180-INS-166 | chr4-33608180-INS-166 |
| chr4-33645972-INS-4203 | chr4-33645973-INS-4203 |
| chr4-33674164-INS-551 | chr4-33674164-INS-543 |
| chr4-33678948-INS-123 | chr4-33678949-INS-123 |
| chr4-33698024-INS-237 | chr4-33698024-INS-237 |
| chr4-33706694-INS-799 | chr4-33706698-INS-799 |
| chr4-33709740-INS-165 | chr4-33709740-INS-164 |
| chr4-33726733-INS-60 | chr4-33726733-INS-51 |
| chr4-33838034-INS-62 | chr4-33838034-INS-62 |
| chr4-33844015-INS-121 | chr4-33844015-INS-121 |
| chr4-33892279-INS-4783 | chr4-33892280-INS-4785 |
| chr4-33920468-INS-329 | chr4-33920468-INS-329 |
| chr4-33950116-INS-3340 | chr4-33950117-INS-3340 |
| chr4-33950117-INS-688 | chr4-33950117-INS-688 |
| chr4-33976137-INS-61 | chr4-33976137-INS-61 |
| chr4-34121478-INS-2804 | chr4-34121479-INS-2798 |
| chr4-34174794-INS-190 | chr4-34174805-INS-208 |
| chr4-34182040-INS-2687 | chr4-34182041-INS-2687 |
| chr4-34186112-INS-54 | chr4-34186112-INS-52 |
| chr4-34205972-INS-7129 | chr4-34205973-INS-7129 |
| chr4-34230563-INS-88 | chr4-34230562-INS-62 |
| chr4-34235954-INS-121 | chr4-34235954-INS-121 |
| chr4-34243513-INS-445 | chr4-34243513-INS-445 |

| | |
|---|---|
| chr4-34265095-INS-133 | chr4-34265095-INS-142 |
| chr4-34271856-INS-198 | chr4-34271856-INS-198 |
| chr4-34276718-INS-107 | chr4-34276710-INS-116 |
| chr4-34278144-INS-96 | chr4-34278144-INS-96 |
| chr4-34310073-INS-51 | chr4-34310073-INS-51 |
| chr4-34311789-INS-300 | chr4-34311804-INS-327 |
| chr4-34316031-INS-114 | chr4-34316031-INS-114 |
| chr4-34317426-INS-7103 | chr4-34317427-INS-7103 |
| chr4-34333500-INS-55 | chr4-34333500-INS-59 |
| chr4-34380345-INS-81 | chr4-34380345-INS-87 |
| chr4-34383076-INS-6054 | chr4-34383077-INS-6054 |
| chr4-34436712-INS-80 | chr4-34436712-INS-80 |
| chr4-34438154-INS-135 | chr4-34438154-INS-135 |
| chr4-34470852-INS-168 | chr4-34470852-INS-168 |
| chr4-34494177-INS-146 | chr4-34494177-INS-146 |
| chr4-34519819-INS-173 | chr4-34519835-INS-168 |
| chr4-34521503-INS-139 | chr4-34521503-INS-145 |
| chr4-34544896-INS-58 | chr4-34544896-INS-62 |
| chr4-34584975-INS-108 | chr4-34584975-INS-108 |
| chr4-34687990-INS-5408 | chr4-34687987-INS-5398 |
| chr4-34696799-INS-56 | chr4-34696799-INS-58 |
| chr4-34728486-INS-187 | chr4-34728486-INS-187 |
| chr4-34731775-INS-532 | chr4-34731775-INS-542 |
| chr4-34743192-INS-259 | chr4-34743192-INS-279 |
| chr4-34748432-INS-200 | chr4-34748444-INS-203 |
| chr4-34751583-INS-75 | chr4-34751601-INS-79 |
| chr4-34782247-INS-197 | chr4-34782248-INS-197 |
| chr4-34798497-INS-203 | chr4-34798498-INS-173 |
| chr4-34812196-INS-228 | chr4-34812196-INS-228 |
| chr4-34820656-INS-51 | chr4-34820656-INS-51 |
| chr4-34831662-INS-82 | chr4-34831662-INS-86 |
| chr4-34837905-INS-57 | chr4-34837905-INS-57 |
| chr4-34845887-INS-212 | chr4-34845887-INS-213 |
| chr4-34862143-INS-200 | chr4-34862143-INS-200 |
| chr4-34873960-INS-1005 | chr4-34873961-INS-1006 |
| chr4-34910893-INS-8986 | chr4-34910894-INS-8986 |
| chr4-34919213-INS-468 | chr4-34919213-INS-468 |
| chr4-34920712-INS-118 | chr4-34920712-INS-157 |
| chr4-34945375-INS-335 | chr4-34945376-INS-335 |
| chr4-34977968-INS-86 | chr4-34977968-INS-86 |
| chr4-34997705-INS-201 | chr4-34997705-INS-201 |
| chr4-35010268-INS-207 | chr4-35010258-INS-228 |
| chr4-35059361-INS-196 | chr4-35059361-INS-199 |
| chr4-35059902-INS-100 | chr4-35059894-INS-111 |
| chr4-35064904-INS-163 | chr4-35064904-INS-164 |
| chr4-35070006-INS-59 | chr4-35069991-INS-60 |
| chr4-35100700-INS-238 | chr4-35100700-INS-192 |
| chr4-35113071-INS-136 | chr4-35113102-INS-90 |
| chr4-35116512-INS-419 | chr4-35116512-INS-419 |
| chr4-35147081-INS-232 | chr4-35147081-INS-229 |
| chr4-35154868-INS-156 | chr4-35154868-INS-160 |
| chr4-35172045-INS-78 | chr4-35172045-INS-58 |
| chr4-35174280-INS-309 | chr4-35174280-INS-309 |
| chr4-35180450-INS-214 | chr4-35180450-INS-214 |
| chr4-35196879-INS-175 | chr4-35196879-INS-180 |
| chr4-35200474-INS-204 | chr4-35200474-INS-207 |
| chr4-35209928-INS-187 | chr4-35209928-INS-187 |
| chr4-35247598-INS-85 | chr4-35247598-INS-98 |
| chr4-35265202-INS-65 | chr4-35265202-INS-65 |
| chr4-35268989-INS-2072 | chr4-35268990-INS-2069 |
| chr4-35283339-INS-413 | chr4-35283341-INS-413 |
| chr4-35310238-INS-68 | chr4-35310238-INS-68 |
| chr4-35317401-INS-229 | chr4-35317401-INS-229 |
| chr4-35319493-INS-3343 | chr4-35319493-INS-3342 |
| chr4-35325704-INS-54 | chr4-35325675-INS-77 |
| chr4-35367878-INS-122 | chr4-35367878-INS-128 |
| chr4-35384858-INS-66 | chr4-35384858-INS-68 |
| chr4-35392482-INS-88 | chr4-35392482-INS-82 |
| chr4-35424108-INS-198 | chr4-35424108-INS-198 |
| chr4-35441354-INS-100 | chr4-35441354-INS-100 |
| chr4-35480167-INS-53 | chr4-35480198-INS-52 |
| chr4-35485665-INS-107 | chr4-35485663-INS-130 |
| chr4-35507435-INS-421 | chr4-35507435-INS-421 |
| chr4-35522746-INS-84 | chr4-35522746-INS-118 |
| chr4-35536277-INS-144 | chr4-35536277-INS-144 |
| chr4-35565524-INS-148 | chr4-35565524-INS-107 |
| chr4-35572497-INS-7131 | chr4-35572498-INS-7132 |

| | |
|---|---|
| chr4-35639093-INS-62 | chr4-35639095-INS-96 |
| chr4-35782116-INS-188 | chr4-35782116-INS-188 |
| chr4-35785295-INS-7010 | chr4-35785296-INS-7010 |
| chr4-35795098-INS-93 | chr4-35795098-INS-93 |
| chr4-35803624-INS-6756 | chr4-35803625-INS-6756 |
| chr4-35835440-INS-64 | chr4-35835440-INS-72 |
| chr4-35867649-INS-832 | chr4-35867649-INS-832 |
| chr4-35926010-INS-1133 | chr4-35926011-INS-1134 |
| chr4-35935457-INS-60 | chr4-35935457-INS-88 |
| chr4-35939069-INS-1131 | chr4-35939070-INS-1130 |
| chr4-35965039-INS-499 | chr4-35965039-INS-499 |
| chr4-36011468-INS-100 | chr4-36011468-INS-100 |
| chr4-36023327-INS-235 | chr4-36023327-INS-235 |
| chr4-36040895-INS-52 | chr4-36040895-INS-54 |
| chr4-36042593-INS-74 | chr4-36042593-INS-76 |
| chr4-36082984-INS-1199 | chr4-36082985-INS-1199 |
| chr4-36093305-INS-6381 | chr4-36093306-INS-6381 |
| chr4-36095799-INS-52 | chr4-36095799-INS-52 |
| chr4-36115881-INS-64 | chr4-36115882-INS-64 |
| chr4-36116386-INS-937 | chr4-36116386-INS-938 |
| chr4-36117721-INS-136 | chr4-36117720-INS-127 |
| chr4-36135172-INS-5902 | chr4-36135172-INS-5902 |
| chr4-36136567-INS-657 | chr4-36136567-INS-658 |
| chr4-36202365-INS-70 | chr4-36202365-INS-95 |
| chr4-36208764-INS-722 | chr4-36208763-INS-733 |
| chr4-36208784-INS-130 | chr4-36208781-INS-139 |
| chr4-36243674-INS-121 | chr4-36243673-INS-138 |
| chr4-36315174-INS-130 | chr4-36315174-INS-95 |
| chr4-36325258-INS-68 | chr4-36325223-INS-56 |
| chr4-36332112-INS-833 | chr4-36332112-INS-833 |
| chr4-36339178-INS-597 | chr4-36339179-INS-595 |
| chr4-36343078-INS-99 | chr4-36343078-INS-99 |
| chr4-36393927-INS-4796 | chr4-36393928-INS-4797 |
| chr4-36398813-INS-898 | chr4-36398813-INS-897 |
| chr4-36400605-INS-84 | chr4-36400605-INS-84 |
| chr4-36459883-INS-1169 | chr4-36459893-INS-1162 |
| chr4-36515864-INS-170 | chr4-36515864-INS-170 |
| chr4-36524723-INS-115 | chr4-36524723-INS-94 |
| chr4-36527422-INS-60 | chr4-36527422-INS-54 |
| chr4-36533503-INS-89 | chr4-36533503-INS-89 |
| chr4-36629507-INS-792 | chr4-36629507-INS-792 |
| chr4-36666502-INS-236 | chr4-36666502-INS-236 |
| chr4-36681930-INS-2989 | chr4-36681931-INS-2989 |
| chr4-36683002-INS-128 | chr4-36683002-INS-83 |
| chr4-36708464-INS-206 | chr4-36708464-INS-206 |
| chr4-36734389-INS-5276 | chr4-36734390-INS-5276 |
| chr4-36823459-INS-6952 | chr4-36823460-INS-6953 |
| chr4-36864366-INS-130 | chr4-36864366-INS-130 |
| chr4-36870894-INS-725 | chr4-36870894-INS-725 |
| chr4-36875046-INS-209 | chr4-36875046-INS-209 |
| chr4-36892979-INS-6404 | chr4-36892980-INS-6398 |
| chr4-36932124-INS-450 | chr4-36932124-INS-449 |
| chr4-36936105-INS-75 | chr4-36936100-INS-50 |
| chr4-36962007-INS-729 | chr4-36962007-INS-729 |
| chr4-36999319-INS-5858 | chr4-36999320-INS-5858 |
| chr4-37008875-INS-6399 | chr4-37008876-INS-6399 |
| chr4-37029569-INS-60 | chr4-37029569-INS-56 |
| chr4-37038121-INS-566 | chr4-37038121-INS-561 |
| chr4-37055939-INS-507 | chr4-37055939-INS-507 |
| chr4-37057422-INS-404 | chr4-37057422-INS-404 |
| chr4-37086511-INS-88 | chr4-37086511-INS-68 |
| chr4-37129439-INS-448 | chr4-37129439-INS-448 |
| chr4-37139865-INS-201 | chr4-37139865-INS-201 |
| chr4-37206789-INS-1077 | chr4-37206790-INS-1077 |
| chr4-37235824-INS-108 | chr4-37235824-INS-108 |
| chr4-37262222-INS-217 | chr4-37262222-INS-217 |
| chr4-37262986-INS-7098 | chr4-37262987-INS-7098 |
| chr4-37284681-INS-114 | chr4-37284681-INS-114 |
| chr4-37293258-INS-76 | chr4-37293258-INS-60 |
| chr4-37341731-INS-1866 | chr4-37341732-INS-1866 |
| chr4-37393371-INS-6549 | chr4-37393372-INS-6550 |
| chr4-37396162-INS-257 | chr4-37396162-INS-257 |
| chr4-37399223-INS-968 | chr4-37399223-INS-978 |
| chr4-37428141-INS-160 | chr4-37428141-INS-160 |
| chr4-37447331-INS-85 | chr4-37447331-INS-85 |
| chr4-37458140-INS-260 | chr4-37458155-INS-228 |
| chr4-37485943-INS-251 | chr4-37485943-INS-252 |

| | |
|---|---|
| chr4-37496013-INS-578 | chr4-37496013-INS-578 |
| chr4-37510946-INS-99 | chr4-37510946-INS-99 |
| chr4-37515278-INS-1303 | chr4-37515279-INS-1303 |
| chr4-37526177-INS-51 | chr4-37526177-INS-51 |
| chr4-37529061-INS-70 | chr4-37529061-INS-70 |
| chr4-37531910-INS-133 | chr4-37531910-INS-133 |
| chr4-37545319-INS-6560 | chr4-37545320-INS-6560 |
| chr4-37561531-INS-102 | chr4-37561531-INS-102 |
| chr4-37563469-INS-74 | chr4-37563469-INS-74 |
| chr4-37654709-INS-4435 | chr4-37654710-INS-4435 |
| chr4-37668737-INS-137 | chr4-37668758-INS-125 |
| chr4-37686727-INS-5369 | chr4-37686728-INS-5369 |
| chr4-37693397-INS-68 | chr4-37693420-INS-65 |
| chr4-37718628-INS-436 | chr4-37718628-INS-439 |
| chr4-37751929-INS-57 | chr4-37751929-INS-53 |
| chr4-37757512-INS-214 | chr4-37757512-INS-214 |
| chr4-37772405-INS-89 | chr4-37772441-INS-80 |
| chr4-37803444-INS-6424 | chr4-37803445-INS-6424 |
| chr4-37839842-INS-81 | chr4-37839842-INS-80 |
| chr4-37851173-INS-368 | chr4-37851173-INS-368 |
| chr4-37853535-INS-77 | chr4-37853535-INS-77 |
| chr4-37916838-INS-1321 | chr4-37916839-INS-1322 |
| chr4-37974613-INS-96 | chr4-37974613-INS-96 |
| chr4-38032354-INS-406 | chr4-38032354-INS-400 |
| chr4-38038179-INS-233 | chr4-38038179-INS-242 |
| chr4-38039023-INS-1206 | chr4-38039024-INS-1207 |
| chr4-38042389-INS-2814 | chr4-38042390-INS-2788 |
| chr4-38057686-INS-381 | chr4-38057688-INS-384 |
| chr4-38091663-INS-96 | chr4-38091663-INS-95 |
| chr4-38097898-INS-2274 | chr4-38097899-INS-2276 |
| chr4-38124802-INS-215 | chr4-38124802-INS-212 |
| chr4-38145768-INS-341 | chr4-38145768-INS-341 |
| chr4-38265354-INS-64 | chr4-38265339-INS-63 |
| chr4-38286782-INS-54 | chr4-38286782-INS-52 |
| chr4-38322141-INS-212 | chr4-38322141-INS-212 |
| chr4-38341369-INS-236 | chr4-38341371-INS-236 |
| chr4-38371913-INS-58 | chr4-38371913-INS-58 |
| chr4-38399000-INS-80 | chr4-38399000-INS-80 |
| chr4-38434870-INS-52 | chr4-38434870-INS-52 |
| chr4-38438313-INS-7217 | chr4-38438314-INS-7217 |
| chr4-38444489-INS-215 | chr4-38444487-INS-225 |
| chr4-38447684-INS-57 | chr4-38447681-INS-60 |
| chr4-38458050-INS-437 | chr4-38458050-INS-437 |
| chr4-38512704-INS-129 | chr4-38512704-INS-129 |
| chr4-38546011-INS-2105 | chr4-38546012-INS-2106 |
| chr4-38549686-INS-214 | chr4-38549686-INS-214 |
| chr4-38578419-INS-884 | chr4-38578419-INS-884 |
| chr4-38586855-INS-761 | chr4-38586855-INS-752 |
| chr4-38599207-INS-3689 | chr4-38599208-INS-3689 |
| chr4-38614983-INS-492 | chr4-38614983-INS-492 |
| chr4-38626497-INS-7346 | chr4-38626498-INS-7347 |
| chr4-38632223-INS-75 | chr4-38632223-INS-75 |
| chr4-38642511-INS-81 | chr4-38642517-INS-62 |
| chr4-38650106-INS-2904 | chr4-38650107-INS-2904 |
| chr4-38666154-INS-6414 | chr4-38666155-INS-6414 |
| chr4-38671656-INS-53 | chr4-38671656-INS-54 |
| chr4-38671948-INS-1542 | chr4-38671947-INS-1540 |
| chr4-38675155-INS-82 | chr4-38675155-INS-88 |
| chr4-38765069-INS-2064 | chr4-38765070-INS-2064 |
| chr4-38767545-INS-323 | chr4-38767545-INS-323 |
| chr4-38783966-INS-55 | chr4-38783966-INS-55 |
| chr4-38805755-INS-8837 | chr4-38805756-INS-8837 |
| chr4-38882249-INS-107 | chr4-38882249-INS-107 |
| chr4-38892858-INS-66 | chr4-38892858-INS-66 |
| chr4-38905331-INS-90 | chr4-38905331-INS-90 |
| chr4-38919322-INS-211 | chr4-38919322-INS-195 |
| chr4-38945399-INS-284 | chr4-38945399-INS-286 |
| chr4-38975102-INS-54 | chr4-38975057-INS-58 |
| chr4-38985632-INS-109 | chr4-38985629-INS-94 |
| chr4-39009206-INS-3562 | chr4-39009207-INS-3562 |
| chr4-39022043-INS-147 | chr4-39022043-INS-147 |
| chr4-39061068-INS-53 | chr4-39061068-INS-53 |
| chr4-39064824-INS-6790 | chr4-39064825-INS-6790 |
| chr4-39126499-INS-6395 | chr4-39126500-INS-6399 |
| chr4-39137599-INS-56 | chr4-39137601-INS-52 |
| chr4-39146598-INS-129 | chr4-39146598-INS-128 |
| chr4-39154552-INS-76 | chr4-39154552-INS-78 |

| | |
|---|---|
| chr4-39174035-INS-60 | chr4-39174035-INS-63 |
| chr4-39193992-INS-85 | chr4-39193992-INS-115 |
| chr4-39288228-INS-58 | chr4-39288228-INS-58 |
| chr4-39291661-INS-6804 | chr4-39291662-INS-6804 |
| chr4-39296397-INS-206 | chr4-39296397-INS-209 |
| chr4-39300445-INS-184 | chr4-39300445-INS-195 |
| chr4-39327330-INS-87 | chr4-39327330-INS-102 |
| chr4-39378787-INS-90 | chr4-39378787-INS-90 |
| chr4-39419618-INS-6472 | chr4-39419619-INS-6475 |
| chr4-39441285-INS-371 | chr4-39441285-INS-405 |
| chr4-39446124-INS-76 | chr4-39446124-INS-79 |
| chr4-39461308-INS-52 | chr4-39461308-INS-54 |
| chr4-39476847-INS-107 | chr4-39476847-INS-107 |
| chr4-39478114-INS-422 | chr4-39478114-INS-422 |
| chr4-39483520-INS-293 | chr4-39483520-INS-293 |
| chr4-39487811-INS-8099 | chr4-39487812-INS-8099 |
| chr4-39504187-INS-63 | chr4-39504187-INS-63 |
| chr4-39525804-INS-486 | chr4-39525804-INS-486 |
| chr4-39557351-INS-391 | chr4-39557351-INS-391 |
| chr4-39570856-INS-110 | chr4-39570856-INS-110 |
| chr4-39579651-INS-680 | chr4-39579651-INS-696 |
| chr4-39591734-INS-130 | chr4-39591734-INS-141 |
| chr4-39660209-INS-52 | chr4-39660209-INS-52 |
| chr4-39687125-INS-180 | chr4-39687126-INS-180 |
| chr4-39712533-INS-497 | chr4-39712533-INS-497 |
| chr4-39720442-INS-72 | chr4-39720449-INS-84 |
| chr4-39727088-INS-131 | chr4-39727088-INS-122 |
| chr4-39747160-INS-70 | chr4-39747160-INS-70 |
| chr4-39763132-INS-120 | chr4-39763125-INS-123 |
| chr4-39813714-INS-65 | chr4-39813714-INS-65 |
| chr4-39917287-INS-175 | chr4-39917287-INS-189 |
| chr4-39929085-INS-60 | chr4-39929085-INS-60 |
| chr4-39944320-INS-102 | chr4-39944320-INS-111 |
| chr4-39994163-INS-1894 | chr4-39994164-INS-1894 |
| chr4-40006426-INS-6664 | chr4-40006427-INS-6661 |
| chr4-40008741-INS-91 | chr4-40008741-INS-104 |
| chr4-40065787-INS-91 | chr4-40065787-INS-91 |
| chr4-40070793-INS-3881 | chr4-40070794-INS-3881 |
| chr4-40109046-INS-151 | chr4-40109046-INS-151 |
| chr4-40186256-INS-106 | chr4-40186256-INS-106 |
| chr4-40214892-INS-217 | chr4-40214892-INS-217 |
| chr4-40226297-INS-709 | chr4-40226297-INS-709 |
| chr4-40247495-INS-67 | chr4-40247495-INS-67 |
| chr4-40248423-INS-207 | chr4-40248423-INS-207 |
| chr4-40290439-INS-88 | chr4-40290439-INS-94 |
| chr4-40295933-INS-197 | chr4-40295933-INS-197 |
| chr4-40308392-INS-66 | chr4-40308392-INS-66 |
| chr4-40309338-INS-6133 | chr4-40309339-INS-6133 |
| chr4-40364976-INS-99 | chr4-40364976-INS-104 |
| chr4-40377116-INS-70 | chr4-40377132-INS-80 |
| chr4-40379503-INS-60 | chr4-40379503-INS-62 |
| chr4-40383192-INS-585 | chr4-40383192-INS-577 |
| chr4-40402689-INS-54 | chr4-40402683-INS-56 |
| chr4-40405783-INS-768 | chr4-40405783-INS-768 |
| chr4-40446263-INS-126 | chr4-40446263-INS-126 |
| chr4-40447852-INS-53 | chr4-40447852-INS-53 |
| chr4-40461525-INS-1383 | chr4-40461526-INS-1383 |
| chr4-40483057-INS-74 | chr4-40483057-INS-74 |
| chr4-40498315-INS-6846 | chr4-40498316-INS-6848 |
| chr4-40525906-INS-528 | chr4-40525906-INS-543 |
| chr4-40537465-INS-496 | chr4-40537465-INS-496 |
| chr4-40584546-INS-168 | chr4-40584546-INS-168 |
| chr4-40622271-INS-215 | chr4-40622271-INS-215 |
| chr4-40629294-INS-213 | chr4-40629294-INS-213 |
| chr4-40633422-INS-60 | chr4-40633422-INS-60 |
| chr4-40640970-INS-52 | chr4-40640970-INS-52 |
| chr4-40644309-INS-227 | chr4-40644309-INS-227 |
| chr4-40646050-INS-152 | chr4-40646050-INS-152 |
| chr4-40648139-INS-53 | chr4-40648139-INS-53 |
| chr4-40660949-INS-57 | chr4-40660949-INS-57 |
| chr4-40672085-INS-204 | chr4-40672085-INS-204 |
| chr4-40673570-INS-240 | chr4-40673570-INS-240 |
| chr4-40680265-INS-73 | chr4-40680265-INS-73 |
| chr4-40688989-INS-189 | chr4-40688989-INS-190 |
| chr4-40690033-INS-153 | chr4-40690037-INS-171 |
| chr4-40696281-INS-106 | chr4-40696301-INS-114 |
| chr4-40705827-INS-56 | chr4-40705827-INS-66 |

| | |
|---|---|
| chr4-40747168-INS-63 | chr4-40747182-INS-51 |
| chr4-40750875-INS-207 | chr4-40750875-INS-207 |
| chr4-40755840-INS-60 | chr4-40755840-INS-60 |
| chr4-40766068-INS-623 | chr4-40766068-INS-626 |
| chr4-40798539-INS-71 | chr4-40798539-INS-54 |
| chr4-40816298-INS-181 | chr4-40816298-INS-181 |
| chr4-40830689-INS-542 | chr4-40830686-INS-545 |
| chr4-40856233-INS-100 | chr4-40856227-INS-85 |
| chr4-40860679-INS-58 | chr4-40860679-INS-58 |
| chr4-40865224-INS-64 | chr4-40865220-INS-68 |
| chr4-40868154-INS-133 | chr4-40868154-INS-133 |
| chr4-40881770-INS-177 | chr4-40881770-INS-177 |
| chr4-40897940-INS-85 | chr4-40897940-INS-61 |
| chr4-41003804-INS-273 | chr4-41003804-INS-273 |
| chr4-41029993-INS-367 | chr4-41029993-INS-367 |
| chr4-41033324-INS-58 | chr4-41033322-INS-66 |
| chr4-41038599-INS-56 | chr4-41038599-INS-56 |
| chr4-41049964-INS-124 | chr4-41049977-INS-124 |
| chr4-41053595-INS-58 | chr4-41053595-INS-78 |
| chr4-41068246-INS-54 | chr4-41068249-INS-93 |
| chr4-41101680-INS-6403 | chr4-41101681-INS-6403 |
| chr4-41102007-INS-111 | chr4-41102010-INS-111 |
| chr4-41113180-INS-202 | chr4-41113180-INS-202 |
| chr4-41126222-INS-205 | chr4-41126222-INS-205 |
| chr4-41127661-INS-1521 | chr4-41127662-INS-1521 |
| chr4-41130584-INS-194 | chr4-41130584-INS-177 |
| chr4-41143739-INS-81 | chr4-41143740-INS-89 |
| chr4-41153575-INS-272 | chr4-41153575-INS-272 |
| chr4-41160896-INS-229 | chr4-41160896-INS-229 |
| chr4-41182791-INS-237 | chr4-41182791-INS-237 |
| chr4-41245975-INS-201 | chr4-41245976-INS-201 |
| chr4-41299102-INS-176 | chr4-41299102-INS-176 |
| chr4-41340556-INS-207 | chr4-41340556-INS-207 |
| chr4-41351179-INS-60 | chr4-41351179-INS-60 |
| chr4-41389867-INS-202 | chr4-41389867-INS-202 |
| chr4-41397265-INS-72 | chr4-41397265-INS-72 |
| chr4-41399255-INS-478 | chr4-41399255-INS-478 |
| chr4-41399865-INS-5149 | chr4-41399866-INS-5149 |
| chr4-41443356-INS-86 | chr4-41443356-INS-90 |
| chr4-41447718-INS-215 | chr4-41447718-INS-215 |
| chr4-41462263-INS-309 | chr4-41462263-INS-309 |
| chr4-41512412-INS-203 | chr4-41512412-INS-203 |
| chr4-41543163-INS-244 | chr4-41543163-INS-244 |
| chr4-41581247-INS-140 | chr4-41581247-INS-140 |
| chr4-41588665-INS-138 | chr4-41588665-INS-144 |
| chr4-41593015-INS-106 | chr4-41593015-INS-106 |
| chr4-41605391-INS-96 | chr4-41605391-INS-96 |
| chr4-41731909-INS-317 | chr4-41731889-INS-331 |
| chr4-41753353-INS-294 | chr4-41753353-INS-294 |
| chr4-41766870-INS-65 | chr4-41766870-INS-51 |
| chr4-41775550-INS-527 | chr4-41775551-INS-526 |
| chr4-41846003-INS-54 | chr4-41845984-INS-55 |
| chr4-42765903-INS-12383 | chr4-42765904-INS-12383 |
| chr4-42778215-INS-7090 | chr4-42778216-INS-7093 |
| chr4-42786231-INS-379 | chr4-42786231-INS-366 |
| chr4-42865136-INS-170 | chr4-42865136-INS-175 |
| chr4-42867586-INS-186 | chr4-42867586-INS-186 |
| chr4-42877063-INS-1974 | chr4-42877064-INS-1974 |
| chr4-42878200-INS-108 | chr4-42878200-INS-106 |
| chr4-42884305-INS-986 | chr4-42884305-INS-986 |
| chr4-42893555-INS-193 | chr4-42893555-INS-193 |
| chr4-42896815-INS-96 | chr4-42896857-INS-90 |
| chr4-42936733-INS-82 | chr4-42936733-INS-108 |
| chr4-42957128-INS-52 | chr4-42957128-INS-54 |
| chr4-42983214-INS-153 | chr4-42983214-INS-152 |
| chr4-43004438-INS-165 | chr4-43004431-INS-143 |
| chr4-43012397-INS-221 | chr4-43012397-INS-221 |
| chr4-43035313-INS-197 | chr4-43035313-INS-197 |
| chr4-43042971-INS-51 | chr4-43042971-INS-63 |
| chr4-43052478-INS-501 | chr4-43052478-INS-501 |
| chr4-43065573-INS-107 | chr4-43065573-INS-100 |
| chr4-43095056-INS-186 | chr4-43095048-INS-192 |
| chr4-43119837-INS-383 | chr4-43119837-INS-383 |
| chr4-43122199-INS-228 | chr4-43122199-INS-228 |
| chr4-43136469-INS-873 | chr4-43136469-INS-873 |
| chr4-43149639-INS-351 | chr4-43149639-INS-351 |
| chr4-43151540-INS-227 | chr4-43151540-INS-227 |

| | |
|---|---|
| chr4-43158502-INS-2852 | chr4-43158503-INS-2852 |
| chr4-43190193-INS-606 | chr4-43190193-INS-605 |
| chr4-43193055-INS-222 | chr4-43193055-INS-222 |
| chr4-43211356-INS-106 | chr4-43211357-INS-108 |
| chr4-43213965-INS-431 | chr4-43213965-INS-431 |
| chr4-43219489-INS-53 | chr4-43219489-INS-53 |
| chr4-43268427-INS-56 | chr4-43268427-INS-53 |
| chr4-43288702-INS-74 | chr4-43288702-INS-74 |
| chr4-43311196-INS-386 | chr4-43311196-INS-386 |
| chr4-43341000-INS-359 | chr4-43341000-INS-359 |
| chr4-43353242-INS-1228 | chr4-43353243-INS-1230 |
| chr4-43362610-INS-54 | chr4-43362610-INS-52 |
| chr4-43365203-INS-72 | chr4-43365203-INS-72 |
| chr4-43374635-INS-66 | chr4-43374635-INS-66 |
| chr4-43392147-INS-215 | chr4-43392147-INS-215 |
| chr4-43414185-INS-64 | chr4-43414168-INS-60 |
| chr4-43420889-INS-385 | chr4-43420889-INS-385 |
| chr4-43438554-INS-214 | chr4-43438554-INS-216 |
| chr4-43450686-INS-169 | chr4-43450686-INS-191 |
| chr4-43456853-INS-52 | chr4-43456853-INS-60 |
| chr4-43469086-INS-85 | chr4-43469102-INS-65 |
| chr4-43479847-INS-75 | chr4-43479844-INS-72 |
| chr4-43502037-INS-1888 | chr4-43502038-INS-1888 |
| chr4-43513460-INS-96 | chr4-43513460-INS-96 |
| chr4-43591076-INS-2846 | chr4-43591077-INS-2846 |
| chr4-43598284-INS-60 | chr4-43598284-INS-60 |
| chr4-43604405-INS-201 | chr4-43604405-INS-201 |
| chr4-43606443-INS-219 | chr4-43606443-INS-214 |
| chr4-43608249-INS-407 | chr4-43608243-INS-413 |
| chr4-43610980-INS-92 | chr4-43610980-INS-110 |
| chr4-43612994-INS-64 | chr4-43612994-INS-64 |
| chr4-43639972-INS-82 | chr4-43639962-INS-100 |
| chr4-43663035-INS-148 | chr4-43663035-INS-148 |
| chr4-43708983-INS-65 | chr4-43708965-INS-52 |
| chr4-43746084-INS-110 | chr4-43746122-INS-60 |
| chr4-43756892-INS-85 | chr4-43756892-INS-95 |
| chr4-43762600-INS-424 | chr4-43762600-INS-441 |
| chr4-43765968-INS-5511 | chr4-43765969-INS-5504 |
| chr4-43765969-INS-328 | chr4-43765969-INS-328 |
| chr4-43769613-INS-360 | chr4-43769613-INS-360 |
| chr4-43797380-INS-106 | chr4-43797380-INS-106 |
| chr4-43801559-INS-207 | chr4-43801559-INS-207 |
| chr4-43804974-INS-65 | chr4-43804979-INS-70 |
| chr4-43806990-INS-203 | chr4-43806990-INS-203 |
| chr4-43808515-INS-633 | chr4-43808515-INS-633 |
| chr4-43832556-INS-972 | chr4-43832556-INS-973 |
| chr4-43842117-INS-130 | chr4-43842118-INS-130 |
| chr4-43856121-INS-52 | chr4-43856121-INS-52 |
| chr4-43862770-INS-400 | chr4-43862770-INS-400 |
| chr4-43894372-INS-68 | chr4-43894372-INS-79 |
| chr4-43905912-INS-88 | chr4-43905912-INS-88 |
| chr4-43941639-INS-218 | chr4-43941639-INS-218 |
| chr4-43953807-INS-232 | chr4-43953807-INS-280 |
| chr4-43962471-INS-58 | chr4-43962451-INS-62 |
| chr4-43967938-INS-141 | chr4-43967938-INS-141 |
| chr4-43974815-INS-68 | chr4-43974795-INS-90 |
| chr4-43976843-INS-96 | chr4-43976843-INS-92 |
| chr4-43989650-INS-244 | chr4-43989650-INS-243 |
| chr4-43995052-INS-685 | chr4-43995052-INS-698 |
| chr4-44000415-INS-244 | chr4-44000423-INS-245 |
| chr4-44002626-INS-148 | chr4-44002660-INS-158 |
| chr4-44005423-INS-52 | chr4-44005423-INS-68 |
| chr4-44036487-INS-219 | chr4-44036487-INS-219 |
| chr4-44046045-INS-100 | chr4-44046045-INS-100 |
| chr4-44061542-INS-92 | chr4-44061542-INS-98 |
| chr4-44073611-INS-61 | chr4-44073611-INS-61 |
| chr4-44077186-INS-66 | chr4-44077171-INS-91 |
| chr4-44086474-INS-190 | chr4-44086474-INS-190 |
| chr4-44096988-INS-118 | chr4-44096989-INS-99 |
| chr4-44105822-INS-62 | chr4-44105822-INS-62 |
| chr4-44112821-INS-1876 | chr4-44112822-INS-1876 |
| chr4-44116609-INS-84 | chr4-44116609-INS-84 |
| chr4-44153803-INS-57 | chr4-44153803-INS-57 |
| chr4-44164924-INS-207 | chr4-44164924-INS-207 |
| chr4-44168867-INS-101 | chr4-44168867-INS-103 |
| chr4-44181234-INS-201 | chr4-44181234-INS-202 |
| chr4-44193779-INS-2156 | chr4-44193780-INS-2151 |

| | |
|---|---|
| chr4-44195344-INS-215 | chr4-44195344-INS-215 |
| chr4-44199271-INS-208 | chr4-44199271-INS-207 |
| chr4-44212214-INS-204 | chr4-44212231-INS-207 |
| chr4-44213916-INS-241 | chr4-44213874-INS-256 |
| chr4-44215677-INS-197 | chr4-44215677-INS-200 |
| chr4-44222708-INS-62 | chr4-44222708-INS-66 |
| chr4-44224571-INS-61 | chr4-44224546-INS-80 |
| chr4-44235637-INS-304 | chr4-44235637-INS-307 |
| chr4-44237397-INS-78 | chr4-44237397-INS-78 |
| chr4-44249531-INS-498 | chr4-44249531-INS-498 |
| chr4-44255657-INS-195 | chr4-44255657-INS-194 |
| chr4-44256801-INS-2355 | chr4-44256802-INS-2355 |
| chr4-44267207-INS-55 | chr4-44267207-INS-59 |
| chr4-44271870-INS-53 | chr4-44271870-INS-50 |
| chr4-44275615-INS-59 | chr4-44275615-INS-68 |
| chr4-44300505-INS-200 | chr4-44300505-INS-200 |
| chr4-44327291-INS-523 | chr4-44327291-INS-523 |
| chr4-44334642-INS-196 | chr4-44334642-INS-194 |
| chr4-44349141-INS-208 | chr4-44349141-INS-204 |
| chr4-44357874-INS-402 | chr4-44357874-INS-402 |
| chr4-44373960-INS-561 | chr4-44373960-INS-562 |
| chr4-44375042-INS-112 | chr4-44375043-INS-112 |
| chr4-44396576-INS-388 | chr4-44396576-INS-388 |
| chr4-44397918-INS-282 | chr4-44397918-INS-289 |
| chr4-44412669-INS-985 | chr4-44412669-INS-985 |
| chr4-44417525-INS-82 | chr4-44417509-INS-92 |
| chr4-44420064-INS-352 | chr4-44420064-INS-352 |
| chr4-44444817-INS-72 | chr4-44444817-INS-72 |
| chr4-44447960-INS-6442 | chr4-44447961-INS-6442 |
| chr4-44453568-INS-1115 | chr4-44453569-INS-1115 |
| chr4-44482336-INS-187 | chr4-44482336-INS-187 |
| chr4-44506530-INS-52 | chr4-44506530-INS-52 |
| chr4-44544705-INS-885 | chr4-44544705-INS-885 |
| chr4-44597428-INS-63 | chr4-44597428-INS-63 |
| chr4-44619988-INS-87 | chr4-44619988-INS-87 |
| chr4-44629160-INS-90 | chr4-44629164-INS-56 |
| chr4-44638234-INS-3063 | chr4-44638235-INS-3063 |
| chr4-44647369-INS-55 | chr4-44647334-INS-67 |
| chr4-44690523-INS-109 | chr4-44690523-INS-89 |
| chr4-44716098-INS-54 | chr4-44716100-INS-62 |
| chr4-44719132-INS-119 | chr4-44719132-INS-119 |
| chr4-44721723-INS-66 | chr4-44721723-INS-78 |
| chr4-44738139-INS-75 | chr4-44738139-INS-73 |
| chr4-44739609-INS-195 | chr4-44739609-INS-197 |
| chr4-44744615-INS-203 | chr4-44744615-INS-203 |
| chr4-44751136-INS-229 | chr4-44751125-INS-233 |
| chr4-44804535-INS-215 | chr4-44804535-INS-215 |
| chr4-44836702-INS-232 | chr4-44836702-INS-232 |
| chr4-44838705-INS-144 | chr4-44838705-INS-144 |
| chr4-44863421-INS-78 | chr4-44863421-INS-78 |
| chr4-44872351-INS-215 | chr4-44872351-INS-215 |
| chr4-44888067-INS-52 | chr4-44888067-INS-50 |
| chr4-44906991-INS-93 | chr4-44906987-INS-92 |
| chr4-44908874-INS-252 | chr4-44908874-INS-252 |
| chr4-44944315-INS-196 | chr4-44944315-INS-196 |
| chr4-44947372-INS-117 | chr4-44947372-INS-117 |
| chr4-44952713-INS-71 | chr4-44952713-INS-71 |
| chr4-44955354-INS-52 | chr4-44955354-INS-52 |
| chr4-45020613-INS-158 | chr4-45020613-INS-158 |
| chr4-45031511-INS-74 | chr4-45031511-INS-75 |
| chr4-45042406-INS-79 | chr4-45042405-INS-67 |
| chr4-45049613-INS-164 | chr4-45049614-INS-164 |
| chr4-45090198-INS-61 | chr4-45090198-INS-61 |
| chr4-45104773-INS-56 | chr4-45104773-INS-56 |
| chr4-45109503-INS-8985 | chr4-45109504-INS-8985 |
| chr4-45117020-INS-210 | chr4-45117020-INS-208 |
| chr4-45120898-INS-59 | chr4-45120913-INS-51 |
| chr4-45129550-INS-59 | chr4-45129550-INS-59 |
| chr4-45137163-INS-128 | chr4-45137163-INS-108 |
| chr4-45145983-INS-87 | chr4-45145973-INS-59 |
| chr4-45165554-INS-762 | chr4-45165554-INS-749 |
| chr4-45175115-INS-61 | chr4-45175115-INS-57 |
| chr4-45209396-INS-1893 | chr4-45209397-INS-1893 |
| chr4-45220823-INS-288 | chr4-45220864-INS-288 |
| chr4-45234584-INS-58 | chr4-45234584-INS-58 |
| chr4-45235372-INS-63 | chr4-45235368-INS-68 |
| chr4-45262484-INS-189 | chr4-45262484-INS-183 |

| | |
|---|---|
| chr4-45318976-INS-193 | chr4-45318963-INS-199 |
| chr4-45346619-INS-205 | chr4-45346619-INS-207 |
| chr4-45369027-INS-193 | chr4-45369027-INS-193 |
| chr4-45388877-INS-624 | chr4-45388877-INS-624 |
| chr4-45397739-INS-116 | chr4-45397739-INS-80 |
| chr4-45400755-INS-497 | chr4-45400755-INS-498 |
| chr4-45409833-INS-150 | chr4-45409833-INS-158 |
| chr4-45417969-INS-69 | chr4-45417969-INS-69 |
| chr4-45442574-INS-498 | chr4-45442574-INS-498 |
| chr4-45455059-INS-555 | chr4-45455059-INS-555 |
| chr4-45463900-INS-663 | chr4-45463900-INS-667 |
| chr4-45464472-INS-117 | chr4-45464472-INS-117 |
| chr4-45469511-INS-120 | chr4-45469531-INS-98 |
| chr4-45493495-INS-68 | chr4-45493495-INS-68 |
| chr4-45496857-INS-252 | chr4-45496857-INS-252 |
| chr4-45566664-INS-358 | chr4-45566664-INS-358 |
| chr4-45570904-INS-462 | chr4-45570904-INS-436 |
| chr4-45574057-INS-71 | chr4-45574057-INS-71 |
| chr4-45582353-INS-4394 | chr4-45582354-INS-4394 |
| chr4-45594779-INS-70 | chr4-45594779-INS-118 |
| chr4-45599695-INS-204 | chr4-45599695-INS-204 |
| chr4-45602387-INS-81 | chr4-45602387-INS-81 |
| chr4-45612694-INS-108 | chr4-45612694-INS-108 |
| chr4-45615861-INS-1876 | chr4-45615862-INS-1876 |
| chr4-45652160-INS-54 | chr4-45652160-INS-54 |
| chr4-45661105-INS-63 | chr4-45661105-INS-63 |
| chr4-45675887-INS-109 | chr4-45675887-INS-107 |
| chr4-45690624-INS-96 | chr4-45690624-INS-96 |
| chr4-45694955-INS-202 | chr4-45694955-INS-202 |
| chr4-45698215-INS-68 | chr4-45698215-INS-74 |
| chr4-45707718-INS-81 | chr4-45707702-INS-88 |
| chr4-45717660-INS-172 | chr4-45717661-INS-174 |
| chr4-45721502-INS-83 | chr4-45721502-INS-83 |
| chr4-45723094-INS-52 | chr4-45723094-INS-52 |
| chr4-45777880-INS-7185 | chr4-45777881-INS-7185 |
| chr4-45793160-INS-194 | chr4-45793160-INS-194 |
| chr4-45818305-INS-68 | chr4-45818305-INS-68 |
| chr4-45850873-INS-66 | chr4-45850873-INS-66 |
| chr4-45853384-INS-214 | chr4-45853384-INS-215 |
| chr4-45858347-INS-106 | chr4-45858347-INS-106 |
| chr4-45858874-INS-202 | chr4-45858874-INS-202 |
| chr4-45888904-INS-88 | chr4-45888904-INS-124 |
| chr4-45980854-INS-1238 | chr4-45980855-INS-1238 |
| chr4-45980855-INS-419 | chr4-45980855-INS-419 |
| chr4-46029530-INS-532 | chr4-46029530-INS-538 |
| chr4-46045756-INS-82 | chr4-46045756-INS-82 |
| chr4-46055957-INS-80 | chr4-46055957-INS-80 |
| chr4-46070636-INS-108 | chr4-46070636-INS-112 |
| chr4-46082251-INS-170 | chr4-46082251-INS-170 |
| chr4-46122254-INS-453 | chr4-46122254-INS-453 |
| chr4-46192334-INS-76 | chr4-46192334-INS-76 |
| chr4-46195452-INS-56 | chr4-46195452-INS-58 |
| chr4-46215374-INS-92 | chr4-46215374-INS-92 |
| chr4-46219707-INS-140 | chr4-46219707-INS-142 |
| chr4-46223477-INS-151 | chr4-46223477-INS-189 |
| chr4-46269564-INS-65 | chr4-46269564-INS-65 |
| chr4-46277986-INS-91 | chr4-46277988-INS-89 |
| chr4-46286424-INS-245 | chr4-46286424-INS-246 |
| chr4-46295670-INS-253 | chr4-46295670-INS-249 |
| chr4-46318325-INS-87 | chr4-46318323-INS-88 |
| chr4-46325841-INS-148 | chr4-46325841-INS-149 |
| chr4-46378927-INS-198 | chr4-46378927-INS-199 |
| chr4-46389227-INS-105 | chr4-46389227-INS-106 |
| chr4-46399597-INS-592 | chr4-46399597-INS-600 |
| chr4-46409992-INS-141 | chr4-46409980-INS-151 |
| chr4-46422335-INS-887 | chr4-46422335-INS-887 |
| chr4-46425728-INS-93 | chr4-46425728-INS-93 |
| chr4-46428545-INS-62 | chr4-46428516-INS-68 |
| chr4-46436519-INS-237 | chr4-46436519-INS-237 |
| chr4-46468863-INS-60 | chr4-46468863-INS-60 |
| chr4-46478432-INS-94 | chr4-46478415-INS-93 |
| chr4-46479287-INS-93 | chr4-46479288-INS-94 |
| chr4-46501620-INS-247 | chr4-46501620-INS-247 |
| chr4-46523284-INS-224 | chr4-46523284-INS-224 |
| chr4-46527804-INS-207 | chr4-46527803-INS-210 |
| chr4-46545934-INS-54 | chr4-46545935-INS-61 |
| chr4-46547097-INS-220 | chr4-46547097-INS-240 |

| | |
|---|---|
| chr4-46553213-INS-134 | chr4-46553213-INS-134 |
| chr4-46557796-INS-90 | chr4-46557796-INS-90 |
| chr4-46559115-INS-64 | chr4-46559115-INS-66 |
| chr4-46570917-INS-65 | chr4-46570918-INS-86 |
| chr4-46572595-INS-64 | chr4-46572595-INS-68 |
| chr4-46612192-INS-255 | chr4-46612192-INS-274 |
| chr4-46615415-INS-116 | chr4-46615415-INS-107 |
| chr4-46619170-INS-223 | chr4-46619170-INS-223 |
| chr4-46633094-INS-209 | chr4-46633093-INS-172 |
| chr4-46645589-INS-74 | chr4-46645589-INS-74 |
| chr4-46661271-INS-352 | chr4-46661271-INS-351 |
| chr4-46676733-INS-218 | chr4-46676733-INS-218 |
| chr4-46682978-INS-111 | chr4-46682978-INS-111 |
| chr4-46693575-INS-78 | chr4-46693573-INS-84 |
| chr4-46705564-INS-117 | chr4-46705564-INS-117 |
| chr4-46743141-INS-63 | chr4-46743141-INS-51 |
| chr4-46750007-INS-68 | chr4-46750008-INS-56 |
| chr4-46766948-INS-95 | chr4-46766948-INS-95 |
| chr4-46777874-INS-1371 | chr4-46777837-INS-1372 |
| chr4-46793756-INS-228 | chr4-46793756-INS-228 |
| chr4-46915770-INS-4208 | chr4-46915771-INS-4208 |
| chr4-46933868-INS-51 | chr4-46933839-INS-60 |
| chr4-46947129-INS-84 | chr4-46947129-INS-80 |
| chr4-46975141-INS-58 | chr4-46975141-INS-58 |
| chr4-46975387-INS-2247 | chr4-46975385-INS-2267 |
| chr4-46985228-INS-57 | chr4-46985228-INS-63 |
| chr4-47042753-INS-150 | chr4-47042755-INS-151 |
| chr4-47058537-INS-1721 | chr4-47058533-INS-1720 |
| chr4-47059253-INS-356 | chr4-47059253-INS-356 |
| chr4-47060697-INS-130 | chr4-47060697-INS-140 |
| chr4-47084150-INS-56 | chr4-47084127-INS-56 |
| chr4-47118921-INS-80 | chr4-47118921-INS-80 |
| chr4-47131830-INS-6917 | chr4-47131831-INS-6917 |
| chr4-47142638-INS-488 | chr4-47142638-INS-489 |
| chr4-47147531-INS-60 | chr4-47147527-INS-58 |
| chr4-47165166-INS-330 | chr4-47165166-INS-327 |
| chr4-47170869-INS-74 | chr4-47170869-INS-74 |
| chr4-47202874-INS-60 | chr4-47202874-INS-68 |
| chr4-47210398-INS-142 | chr4-47210394-INS-138 |
| chr4-47234859-INS-120 | chr4-47234859-INS-120 |
| chr4-47287282-INS-53 | chr4-47287282-INS-70 |
| chr4-47296252-INS-55 | chr4-47296252-INS-55 |
| chr4-47421113-INS-52 | chr4-47421132-INS-56 |
| chr4-47423876-INS-67 | chr4-47423876-INS-67 |
| chr4-47426779-INS-6643 | chr4-47426780-INS-6643 |
| chr4-47437329-INS-54 | chr4-47437329-INS-57 |
| chr4-47452081-INS-353 | chr4-47452081-INS-352 |
| chr4-47455598-INS-624 | chr4-47455598-INS-614 |
| chr4-47458031-INS-218 | chr4-47458031-INS-220 |
| chr4-47485522-INS-222 | chr4-47485522-INS-222 |
| chr4-47516873-INS-194 | chr4-47516873-INS-199 |
| chr4-47518556-INS-78 | chr4-47518557-INS-63 |
| chr4-47526047-INS-176 | chr4-47526047-INS-176 |
| chr4-47533988-INS-186 | chr4-47533989-INS-185 |
| chr4-47563119-INS-5998 | chr4-47563120-INS-5998 |
| chr4-47565012-INS-89 | chr4-47565012-INS-93 |
| chr4-47571490-INS-489 | chr4-47571490-INS-447 |
| chr4-47604872-INS-70 | chr4-47604902-INS-58 |
| chr4-47647150-INS-203 | chr4-47647150-INS-204 |
| chr4-47673079-INS-68 | chr4-47673079-INS-60 |
| chr4-47736431-INS-424 | chr4-47736431-INS-421 |
| chr4-47748072-INS-72 | chr4-47748072-INS-72 |
| chr4-47774509-INS-102 | chr4-47774509-INS-58 |
| chr4-47779870-INS-72 | chr4-47779870-INS-84 |
| chr4-47805097-INS-156 | chr4-47805097-INS-190 |
| chr4-47819918-INS-63 | chr4-47819918-INS-56 |
| chr4-47860531-INS-114 | chr4-47860531-INS-114 |
| chr4-47913303-INS-90 | chr4-47913303-INS-66 |
| chr4-47920996-INS-52 | chr4-47920996-INS-50 |
| chr4-47922473-INS-978 | chr4-47922472-INS-981 |
| chr4-47926795-INS-72 | chr4-47926787-INS-92 |
| chr4-47940676-INS-634 | chr4-47940676-INS-634 |
| chr4-47941496-INS-98 | chr4-47941515-INS-120 |
| chr4-47945043-INS-98 | chr4-47945043-INS-98 |
| chr4-47954015-INS-65 | chr4-47954015-INS-65 |
| chr4-47969871-INS-178 | chr4-47969871-INS-179 |
| chr4-47976369-INS-6440 | chr4-47976370-INS-6440 |

| | |
|---|---|
| chr4-47997896-INS-252 | chr4-47997896-INS-252 |
| chr4-48029455-INS-68 | chr4-48029456-INS-64 |
| chr4-48098543-INS-60 | chr4-48098543-INS-55 |
| chr4-48105770-INS-52 | chr4-48105802-INS-57 |
| chr4-48107089-INS-94 | chr4-48107089-INS-93 |
| chr4-48137607-INS-63 | chr4-48137607-INS-67 |
| chr4-48139568-INS-58 | chr4-48139568-INS-60 |
| chr4-48142413-INS-398 | chr4-48142413-INS-414 |
| chr4-48148931-INS-94 | chr4-48148931-INS-94 |
| chr4-48163612-INS-72 | chr4-48163612-INS-72 |
| chr4-48166313-INS-95 | chr4-48166313-INS-95 |
| chr4-48168715-INS-67 | chr4-48168715-INS-67 |
| chr4-48173585-INS-52 | chr4-48173585-INS-52 |
| chr4-48249726-INS-63 | chr4-48249739-INS-63 |
| chr4-48302604-INS-539 | chr4-48302604-INS-539 |
| chr4-48312233-INS-55 | chr4-48312233-INS-55 |
| chr4-48368837-INS-6874 | chr4-48368838-INS-6873 |
| chr4-48379223-INS-1047 | chr4-48379224-INS-1071 |
| chr4-48390549-INS-99 | chr4-48390549-INS-99 |
| chr4-48391497-INS-55 | chr4-48391497-INS-55 |
| chr4-48400037-INS-56 | chr4-48400037-INS-56 |
| chr4-48442161-INS-78 | chr4-48442161-INS-93 |
| chr4-48486329-INS-169 | chr4-48486329-INS-169 |
| chr4-48509521-INS-59 | chr4-48509521-INS-59 |
| chr4-48510009-INS-5421 | chr4-48510010-INS-5424 |
| chr4-48521652-INS-131 | chr4-48521652-INS-131 |
| chr4-48522837-INS-245 | chr4-48522837-INS-246 |
| chr4-48525978-INS-436 | chr4-48525978-INS-436 |
| chr4-48527958-INS-216 | chr4-48527958-INS-216 |
| chr4-48530840-INS-52 | chr4-48530840-INS-52 |
| chr4-48532550-INS-308 | chr4-48532550-INS-308 |
| chr4-48543537-INS-181 | chr4-48543537-INS-181 |
| chr4-48553376-INS-208 | chr4-48553376-INS-209 |
| chr4-48555567-INS-56 | chr4-48555567-INS-56 |
| chr4-48568398-INS-229 | chr4-48568398-INS-217 |
| chr4-48573701-INS-79 | chr4-48573701-INS-112 |
| chr4-48575781-INS-55 | chr4-48575781-INS-55 |
| chr4-48580216-INS-219 | chr4-48580216-INS-209 |
| chr4-48582296-INS-7182 | chr4-48582297-INS-7182 |
| chr4-48583668-INS-267 | chr4-48583668-INS-266 |
| chr4-48603269-INS-122 | chr4-48603269-INS-122 |
| chr4-48610911-INS-76 | chr4-48610872-INS-88 |
| chr4-48645278-INS-408 | chr4-48645278-INS-408 |
| chr4-48669522-INS-201 | chr4-48669522-INS-202 |
| chr4-48675548-INS-83 | chr4-48675548-INS-79 |
| chr4-48682546-INS-206 | chr4-48682546-INS-206 |
| chr4-48683182-INS-608 | chr4-48683172-INS-603 |
| chr4-48683658-INS-4198 | chr4-48683659-INS-4198 |
| chr4-48689702-INS-146 | chr4-48689702-INS-130 |
| chr4-48711695-INS-66 | chr4-48711700-INS-62 |
| chr4-48712863-INS-270 | chr4-48712863-INS-225 |
| chr4-48753791-INS-475 | chr4-48753791-INS-477 |
| chr4-48757389-INS-120 | chr4-48757389-INS-120 |
| chr4-48763272-INS-1909 | chr4-48763273-INS-1909 |
| chr4-48765543-INS-7476 | chr4-48765544-INS-7503 |
| chr4-48782026-INS-56 | chr4-48782018-INS-92 |
| chr4-48782380-INS-1423 | chr4-48782381-INS-1423 |
| chr4-48783712-INS-533 | chr4-48783712-INS-539 |
| chr4-48784672-INS-115 | chr4-48784672-INS-115 |
| chr4-48789212-INS-63 | chr4-48789212-INS-59 |
| chr4-48809242-INS-82 | chr4-48809242-INS-82 |
| chr4-48810810-INS-498 | chr4-48810810-INS-498 |
| chr4-48836483-INS-56 | chr4-48836519-INS-64 |
| chr4-48868867-INS-201 | chr4-48868867-INS-201 |
| chr4-48887310-INS-1459 | chr4-48887311-INS-1459 |
| chr4-48918511-INS-52 | chr4-48918511-INS-68 |
| chr4-48954995-INS-2105 | chr4-48954996-INS-2118 |
| chr4-48966577-INS-6155 | chr4-48966578-INS-6156 |
| chr4-48977431-INS-52 | chr4-48977431-INS-52 |
| chr4-48983083-INS-102 | chr4-48983082-INS-100 |
| chr4-48986105-INS-93 | chr4-48986125-INS-111 |
| chr4-49008631-INS-53 | chr4-49008631-INS-53 |
| chr4-49040311-INS-115 | chr4-49040357-INS-82 |
| chr4-49046401-INS-1461 | chr4-49046402-INS-1460 |
| chr4-49059422-INS-60 | chr4-49059422-INS-52 |
| chr4-49112690-INS-70 | chr4-49112690-INS-64 |
| chr4-49147611-INS-79 | chr4-49147611-INS-79 |

| | |
|---|---|
| chr4-49153005-INS-230 | chr4-49153005-INS-230 |
| chr4-49155951-INS-3755 | chr4-49155952-INS-3755 |
| chr4-49163952-INS-80 | chr4-49163952-INS-64 |
| chr4-49168583-INS-60 | chr4-49168578-INS-54 |
| chr4-49172984-INS-54 | chr4-49172984-INS-54 |
| chr4-49186742-INS-336 | chr4-49186742-INS-338 |
| chr4-49196436-INS-89 | chr4-49196436-INS-89 |
| chr4-49211163-INS-380 | chr4-49211164-INS-425 |
| chr4-49216903-INS-55 | chr4-49216903-INS-70 |
| chr4-49276857-INS-159 | chr4-49276857-INS-159 |
| chr4-49279307-INS-90 | chr4-49279308-INS-90 |
| chr4-49281672-INS-60 | chr4-49281672-INS-76 |
| chr4-49291239-INS-453 | chr4-49291239-INS-453 |
| chr4-49298178-INS-6614 | chr4-49298179-INS-6614 |
| chr4-49315648-INS-89 | chr4-49315648-INS-107 |
| chr4-49320456-INS-169 | chr4-49320452-INS-192 |
| chr4-49343109-INS-487 | chr4-49343109-INS-487 |
| chr4-49344888-INS-56 | chr4-49344888-INS-52 |
| chr4-49351527-INS-1542 | chr4-49351528-INS-1541 |
| chr4-49398536-INS-57 | chr4-49398536-INS-58 |
| chr4-49406221-INS-88 | chr4-49406228-INS-128 |
| chr4-49415358-INS-72 | chr4-49415358-INS-60 |
| chr4-49426992-INS-401 | chr4-49426992-INS-401 |
| chr4-49443141-INS-56 | chr4-49443141-INS-56 |
| chr4-49485786-INS-344 | chr4-49485786-INS-345 |
| chr4-49602045-INS-72 | chr4-49602045-INS-74 |
| chr4-49610111-INS-60 | chr4-49610096-INS-60 |
| chr4-49617806-INS-85 | chr4-49617806-INS-71 |
| chr4-49633808-INS-72 | chr4-49633791-INS-56 |
| chr4-49675910-INS-146 | chr4-49675910-INS-146 |
| chr4-49677491-INS-149 | chr4-49677491-INS-137 |
| chr4-49685054-INS-284 | chr4-49685075-INS-291 |
| chr4-49687620-INS-253 | chr4-49687610-INS-265 |
| chr4-49728976-INS-557 | chr4-49728976-INS-555 |
| chr4-49740232-INS-144 | chr4-49740234-INS-125 |
| chr4-49759902-INS-62 | chr4-49759902-INS-62 |
| chr4-49764413-INS-94 | chr4-49764413-INS-94 |
| chr4-49767188-INS-509 | chr4-49767188-INS-509 |
| chr4-49776026-INS-6452 | chr4-49776027-INS-6456 |
| chr4-49796046-INS-7028 | chr4-49796047-INS-7028 |
| chr4-49797702-INS-96 | chr4-49797710-INS-97 |
| chr4-49830342-INS-6945 | chr4-49830343-INS-6945 |
| chr4-49895532-INS-609 | chr4-49895532-INS-609 |
| chr4-49900110-INS-62 | chr4-49900110-INS-62 |
| chr4-49904499-INS-56 | chr4-49904499-INS-56 |
| chr4-49910232-INS-6386 | chr4-49910233-INS-6387 |
| chr4-49912281-INS-166 | chr4-49912281-INS-166 |
| chr4-49938943-INS-6705 | chr4-49938944-INS-6704 |
| chr4-49955352-INS-1101 | chr4-49955353-INS-1101 |
| chr4-49969577-INS-58 | chr4-49969577-INS-62 |
| chr4-50012633-INS-6865 | chr4-50012634-INS-6865 |
| chr4-50124235-INS-427 | chr4-50124235-INS-426 |
| chr4-50157841-INS-69 | chr4-50157841-INS-69 |
| chr4-50189269-INS-292 | chr4-50189269-INS-292 |
| chr4-50216127-INS-150 | chr4-50216127-INS-150 |
| chr4-50234909-INS-4816 | chr4-50234910-INS-4816 |
| chr4-50292781-INS-6570 | chr4-50292782-INS-6570 |
| chr4-50312832-INS-1851 | chr4-50312833-INS-1853 |
| chr4-50334523-INS-191 | chr4-50334523-INS-191 |
| chr4-50348037-INS-153 | chr4-50348035-INS-156 |
| chr4-50389412-INS-121 | chr4-50389412-INS-121 |
| chr4-50410368-INS-636 | chr4-50410368-INS-636 |
| chr4-50417032-INS-427 | chr4-50417032-INS-427 |
| chr4-50419861-INS-62 | chr4-50419879-INS-63 |
| chr4-50424284-INS-843 | chr4-50424284-INS-844 |
| chr4-50440784-INS-230 | chr4-50440784-INS-230 |
| chr4-50471381-INS-1493 | chr4-50471382-INS-1493 |
| chr4-50505361-INS-2968 | chr4-50505362-INS-2977 |
| chr4-50505366-INS-6453 | chr4-50505367-INS-6454 |
| chr4-50527490-INS-853 | chr4-50527490-INS-853 |
| chr4-50575342-INS-6288 | chr4-50575343-INS-6288 |
| chr4-50668143-INS-101 | chr4-50668166-INS-127 |
| chr4-50696986-INS-90 | chr4-50696986-INS-90 |
| chr4-50713996-INS-216 | chr4-50713996-INS-216 |
| chr4-50837118-INS-59 | chr4-50837118-INS-59 |
| chr4-50840457-INS-445 | chr4-50840457-INS-445 |
| chr4-50843367-INS-159 | chr4-50843367-INS-164 |

| | |
|---|---|
| chr4-50865737-INS-465 | chr4-50865737-INS-465 |
| chr4-50871521-INS-387 | chr4-50871521-INS-387 |
| chr4-50891231-INS-150 | chr4-50891231-INS-155 |
| chr4-50898843-INS-63 | chr4-50898853-INS-53 |
| chr4-50902701-INS-165 | chr4-50902701-INS-165 |
| chr4-50905880-INS-64 | chr4-50905894-INS-64 |
| chr4-50910504-INS-1607 | chr4-50910504-INS-1621 |
| chr4-50946009-INS-6296 | chr4-50946010-INS-6296 |
| chr4-50986351-INS-63 | chr4-50986351-INS-76 |
| chr4-51003723-INS-390 | chr4-51003723-INS-390 |
| chr4-51020803-INS-6478 | chr4-51020804-INS-6479 |
| chr4-51024209-INS-495 | chr4-51024209-INS-495 |
| chr4-51060231-INS-206 | chr4-51060231-INS-206 |
| chr4-51081318-INS-3241 | chr4-51081319-INS-3241 |
| chr4-51170625-INS-695 | chr4-51170625-INS-695 |
| chr4-51194296-INS-153 | chr4-51194296-INS-153 |
| chr4-51204226-INS-390 | chr4-51204226-INS-390 |
| chr4-51212619-INS-596 | chr4-51212619-INS-596 |
| chr4-51228991-INS-81 | chr4-51228991-INS-81 |
| chr4-51239767-INS-76 | chr4-51239767-INS-77 |
| chr4-51261561-INS-238 | chr4-51261561-INS-238 |
| chr4-51264615-INS-435 | chr4-51264615-INS-435 |
| chr4-51282341-INS-162 | chr4-51282341-INS-163 |
| chr4-51283235-INS-60 | chr4-51283235-INS-60 |
| chr4-51285661-INS-1007 | chr4-51285662-INS-1008 |
| chr4-51293637-INS-268 | chr4-51293637-INS-270 |
| chr4-51345462-INS-170 | chr4-51345462-INS-168 |
| chr4-51346937-INS-88 | chr4-51346937-INS-88 |
| chr4-51355019-INS-538 | chr4-51355019-INS-538 |
| chr4-51356800-INS-769 | chr4-51356800-INS-745 |
| chr4-51373769-INS-501 | chr4-51373769-INS-501 |
| chr4-51396119-INS-1995 | chr4-51396120-INS-2005 |
| chr4-51443595-INS-463 | chr4-51443595-INS-463 |
| chr4-51452239-INS-94 | chr4-51452239-INS-92 |
| chr4-51466112-INS-439 | chr4-51466112-INS-441 |
| chr4-51467441-INS-114 | chr4-51467441-INS-122 |
| chr4-51474892-INS-52 | chr4-51474892-INS-52 |
| chr4-51506273-INS-95 | chr4-51506273-INS-87 |
| chr4-51534668-INS-70 | chr4-51534668-INS-70 |
| chr4-51540592-INS-68 | chr4-51540555-INS-100 |
| chr4-51542552-INS-198 | chr4-51542552-INS-198 |
| chr4-51557432-INS-2610 | chr4-51557433-INS-2608 |
| chr4-51593460-INS-496 | chr4-51593460-INS-496 |
| chr4-51607285-INS-171 | chr4-51607285-INS-167 |
| chr4-51610820-INS-69 | chr4-51610820-INS-69 |
| chr4-51623991-INS-159 | chr4-51623991-INS-157 |
| chr4-51641740-INS-97 | chr4-51641740-INS-97 |
| chr4-51716825-INS-2488 | chr4-51716826-INS-2488 |
| chr4-51727599-INS-903 | chr4-51727599-INS-903 |
| chr4-51755897-INS-2210 | chr4-51755898-INS-2210 |
| chr4-51775219-INS-553 | chr4-51775219-INS-554 |
| chr4-51795732-INS-162 | chr4-51795732-INS-162 |
| chr4-51803735-INS-54 | chr4-51803727-INS-65 |
| chr4-51803951-INS-5994 | chr4-51803952-INS-5993 |
| chr4-51809519-INS-96 | chr4-51809519-INS-96 |
| chr4-51854095-INS-272 | chr4-51854095-INS-272 |
| chr4-51858378-INS-96 | chr4-51858378-INS-132 |
| chr4-51939338-INS-68 | chr4-51939338-INS-54 |
| chr4-51946636-INS-640 | chr4-51946644-INS-642 |
| chr4-51976597-INS-6130 | chr4-51976598-INS-6130 |
| chr4-52010553-INS-712 | chr4-52010553-INS-714 |
| chr4-52013876-INS-6384 | chr4-52013877-INS-6384 |
| chr4-52022561-INS-52 | chr4-52022562-INS-62 |
| chr4-52023769-INS-6475 | chr4-52023770-INS-6475 |
| chr4-52028264-INS-65 | chr4-52028264-INS-70 |
| chr4-52039697-INS-68 | chr4-52039712-INS-96 |
| chr4-52094095-INS-217 | chr4-52094095-INS-217 |
| chr4-52100790-INS-506 | chr4-52100790-INS-506 |
| chr4-52131472-INS-2331 | chr4-52131473-INS-2331 |
| chr4-52147959-INS-66 | chr4-52147959-INS-66 |
| chr4-52149536-INS-65 | chr4-52149536-INS-65 |
| chr4-52159408-INS-52 | chr4-52159408-INS-52 |
| chr4-52203722-INS-91 | chr4-52203722-INS-91 |
| chr4-52207270-INS-67 | chr4-52207270-INS-72 |
| chr4-52220437-INS-1455 | chr4-52220438-INS-1454 |
| chr4-52245409-INS-588 | chr4-52245410-INS-588 |
| chr4-52274075-INS-294 | chr4-52274075-INS-294 |

| | |
|---|---|
| chr4-52280218-INS-6471 | chr4-52280219-INS-6471 |
| chr4-52289713-INS-2706 | chr4-52289714-INS-2708 |
| chr4-52300082-INS-1601 | chr4-52300083-INS-1601 |
| chr4-52335581-INS-102 | chr4-52335581-INS-102 |
| chr4-52389126-INS-660 | chr4-52389126-INS-661 |
| chr4-52390649-INS-53 | chr4-52390642-INS-84 |
| chr4-52431870-INS-636 | chr4-52431870-INS-636 |
| chr4-52437424-INS-424 | chr4-52437424-INS-424 |
| chr4-52459594-INS-160 | chr4-52459594-INS-163 |
| chr4-52504096-INS-324 | chr4-52504096-INS-324 |
| chr4-52506292-INS-1510 | chr4-52506293-INS-1509 |
| chr4-52507317-INS-158 | chr4-52507347-INS-170 |
| chr4-52511701-INS-186 | chr4-52511701-INS-186 |
| chr4-52558910-INS-71 | chr4-52558915-INS-64 |
| chr4-52575120-INS-387 | chr4-52575120-INS-387 |
| chr4-52710459-INS-128 | chr4-52710442-INS-145 |
| chr4-52713107-INS-70 | chr4-52713107-INS-70 |
| chr4-52728901-INS-6788 | chr4-52728902-INS-6788 |
| chr4-52731117-INS-202 | chr4-52731117-INS-202 |
| chr4-52732046-INS-102 | chr4-52732060-INS-70 |
| chr4-52759600-INS-246 | chr4-52759600-INS-248 |
| chr4-52822785-INS-153 | chr4-52822747-INS-152 |
| chr4-52841014-INS-3018 | chr4-52841015-INS-3018 |
| chr4-52847424-INS-618 | chr4-52847424-INS-618 |
| chr4-52851957-INS-549 | chr4-52851957-INS-549 |
| chr4-52853755-INS-65 | chr4-52853755-INS-65 |
| chr4-52860316-INS-128 | chr4-52860290-INS-132 |
| chr4-52876944-INS-329 | chr4-52876944-INS-329 |
| chr4-52881431-INS-54 | chr4-52881432-INS-59 |
| chr4-52917367-INS-531 | chr4-52917367-INS-532 |
| chr4-52919992-INS-65 | chr4-52919992-INS-66 |
| chr4-52928847-INS-60 | chr4-52928871-INS-52 |
| chr4-52940484-INS-67 | chr4-52940484-INS-71 |
| chr4-52949406-INS-89 | chr4-52949406-INS-89 |
| chr4-52952048-INS-404 | chr4-52952048-INS-406 |
| chr4-52985159-INS-65 | chr4-52985159-INS-65 |
| chr4-52988781-INS-53 | chr4-52988781-INS-53 |
| chr4-53013865-INS-58 | chr4-53013881-INS-78 |
| chr4-53014800-INS-310 | chr4-53014800-INS-310 |
| chr4-53037873-INS-55 | chr4-53037887-INS-73 |
| chr4-53040004-INS-115 | chr4-53040008-INS-115 |
| chr4-53043037-INS-64 | chr4-53043037-INS-64 |
| chr4-53062047-INS-60 | chr4-53062037-INS-50 |
| chr4-53115365-INS-60 | chr4-53115365-INS-50 |
| chr4-53115927-INS-67 | chr4-53115927-INS-68 |
| chr4-53119582-INS-54 | chr4-53119582-INS-54 |
| chr4-53132061-INS-56 | chr4-53132056-INS-56 |
| chr4-53149223-INS-64 | chr4-53149223-INS-66 |
| chr4-53245619-INS-78 | chr4-53245619-INS-78 |
| chr4-53284658-INS-119 | chr4-53284656-INS-129 |
| chr4-53306722-INS-98 | chr4-53306686-INS-90 |
| chr4-53310201-INS-206 | chr4-53310199-INS-207 |
| chr4-53320795-INS-56 | chr4-53320778-INS-66 |
| chr4-53336919-INS-318 | chr4-53336919-INS-318 |
| chr4-53338518-INS-233 | chr4-53338518-INS-233 |
| chr4-53362483-INS-137 | chr4-53362483-INS-178 |
| chr4-53371647-INS-76 | chr4-53371647-INS-60 |
| chr4-53378675-INS-60 | chr4-53378664-INS-76 |
| chr4-53383862-INS-56 | chr4-53383862-INS-56 |
| chr4-53399262-INS-266 | chr4-53399278-INS-240 |
| chr4-53406503-INS-200 | chr4-53406503-INS-200 |
| chr4-53416378-INS-58 | chr4-53416378-INS-58 |
| chr4-53422810-INS-5216 | chr4-53422811-INS-5217 |
| chr4-53426546-INS-214 | chr4-53426546-INS-214 |
| chr4-53432381-INS-215 | chr4-53432381-INS-215 |
| chr4-53433749-INS-826 | chr4-53433749-INS-826 |
| chr4-53523892-INS-62 | chr4-53523892-INS-62 |
| chr4-53530619-INS-86 | chr4-53530619-INS-86 |
| chr4-53534281-INS-60 | chr4-53534281-INS-56 |
| chr4-53549296-INS-56 | chr4-53549296-INS-56 |
| chr4-53601261-INS-281 | chr4-53601261-INS-280 |
| chr4-53619381-INS-1166 | chr4-53619382-INS-1166 |
| chr4-53667623-INS-1141 | chr4-53667624-INS-1142 |
| chr4-53680825-INS-78 | chr4-53680825-INS-117 |
| chr4-53703738-INS-172 | chr4-53703738-INS-172 |
| chr4-53711909-INS-102 | chr4-53711909-INS-103 |
| chr4-53718247-INS-374 | chr4-53718247-INS-374 |

| | |
|---|---|
| chr4-53736216-INS-51 | chr4-53736216-INS-51 |
| chr4-53772814-INS-89 | chr4-53772815-INS-89 |
| chr4-53836802-INS-1018 | chr4-53836803-INS-1018 |
| chr4-53840355-INS-77 | chr4-53840332-INS-75 |
| chr4-53881355-INS-52 | chr4-53881355-INS-52 |
| chr4-53912898-INS-123 | chr4-53912899-INS-126 |
| chr4-53925432-INS-199 | chr4-53925432-INS-199 |
| chr4-53933754-INS-252 | chr4-53933754-INS-252 |
| chr4-53947502-INS-6445 | chr4-53947503-INS-6445 |
| chr4-53951416-INS-77 | chr4-53951416-INS-77 |
| chr4-53973590-INS-6440 | chr4-53973591-INS-6440 |
| chr4-53990203-INS-66 | chr4-53990203-INS-68 |
| chr4-54015495-INS-115 | chr4-54015492-INS-118 |
| chr4-54018820-INS-146 | chr4-54018824-INS-168 |
| chr4-54038879-INS-63 | chr4-54038856-INS-62 |
| chr4-54074536-INS-1414 | chr4-54074537-INS-1414 |
| chr4-54086093-INS-195 | chr4-54086093-INS-185 |
| chr4-54088163-INS-221 | chr4-54088163-INS-221 |
| chr4-54090282-INS-52 | chr4-54090282-INS-52 |
| chr4-54095661-INS-152 | chr4-54095661-INS-154 |
| chr4-54104701-INS-498 | chr4-54104701-INS-498 |
| chr4-54128371-INS-1565 | chr4-54128372-INS-1565 |
| chr4-54168834-INS-65 | chr4-54168834-INS-65 |
| chr4-54183326-INS-1056 | chr4-54183327-INS-1056 |
| chr4-54214440-INS-756 | chr4-54214440-INS-756 |
| chr4-54235406-INS-113 | chr4-54235390-INS-135 |
| chr4-54254233-INS-130 | chr4-54254233-INS-132 |
| chr4-54263394-INS-308 | chr4-54263394-INS-308 |
| chr4-54266924-INS-248 | chr4-54266924-INS-248 |
| chr4-54277989-INS-6530 | chr4-54277990-INS-6530 |
| chr4-54281414-INS-51 | chr4-54281414-INS-60 |
| chr4-54284089-INS-2355 | chr4-54284090-INS-2355 |
| chr4-54311434-INS-6539 | chr4-54311435-INS-6534 |
| chr4-54321858-INS-342 | chr4-54321882-INS-348 |
| chr4-54338243-INS-6871 | chr4-54338244-INS-6883 |
| chr4-54338246-INS-175 | chr4-54338249-INS-159 |
| chr4-54376719-INS-57 | chr4-54376715-INS-70 |
| chr4-54403693-INS-2242 | chr4-54403694-INS-2243 |
| chr4-54418698-INS-74 | chr4-54418698-INS-74 |
| chr4-54424421-INS-88 | chr4-54424422-INS-88 |
| chr4-54474841-INS-259 | chr4-54474841-INS-259 |
| chr4-54516563-INS-175 | chr4-54516578-INS-170 |
| chr4-54546300-INS-78 | chr4-54546300-INS-78 |
| chr4-54550916-INS-66 | chr4-54550916-INS-68 |
| chr4-54552962-INS-483 | chr4-54552962-INS-483 |
| chr4-54568757-INS-4455 | chr4-54568758-INS-4455 |
| chr4-54572550-INS-106 | chr4-54572555-INS-85 |
| chr4-54590119-INS-77 | chr4-54590119-INS-77 |
| chr4-54620416-INS-52 | chr4-54620418-INS-52 |
| chr4-54633551-INS-87 | chr4-54633551-INS-87 |
| chr4-54651276-INS-771 | chr4-54651276-INS-771 |
| chr4-54720361-INS-57 | chr4-54720362-INS-93 |
| chr4-54723506-INS-100 | chr4-54723506-INS-102 |
| chr4-54734484-INS-228 | chr4-54734484-INS-227 |
| chr4-54748774-INS-78 | chr4-54748774-INS-70 |
| chr4-54752286-INS-52 | chr4-54752333-INS-50 |
| chr4-54771023-INS-80 | chr4-54771028-INS-56 |
| chr4-54779954-INS-75 | chr4-54779954-INS-79 |
| chr4-54795828-INS-977 | chr4-54795828-INS-977 |
| chr4-54802295-INS-52 | chr4-54802295-INS-52 |
| chr4-54811445-INS-77 | chr4-54811445-INS-77 |
| chr4-54814923-INS-62 | chr4-54814923-INS-66 |
| chr4-54816105-INS-98 | chr4-54816105-INS-102 |
| chr4-54817459-INS-57 | chr4-54817459-INS-58 |
| chr4-54821840-INS-358 | chr4-54821840-INS-358 |
| chr4-54830745-INS-149 | chr4-54830745-INS-149 |
| chr4-54847746-INS-66 | chr4-54847747-INS-54 |
| chr4-54941732-INS-53 | chr4-54941732-INS-53 |
| chr4-54944083-INS-281 | chr4-54944083-INS-284 |
| chr4-54971859-INS-455 | chr4-54971859-INS-455 |
| chr4-54973575-INS-6013 | chr4-54973576-INS-6013 |
| chr4-54998934-INS-54 | chr4-54998934-INS-54 |
| chr4-55045615-INS-57 | chr4-55045617-INS-67 |
| chr4-55050751-INS-54 | chr4-55050751-INS-50 |
| chr4-55092917-INS-56 | chr4-55092917-INS-54 |
| chr4-55105902-INS-7084 | chr4-55105903-INS-7084 |
| chr4-55109570-INS-180 | chr4-55109570-INS-180 |

| | |
|---|---|
| chr4-55188555-INS-593 | chr4-55188555-INS-593 |
| chr4-55191667-INS-123 | chr4-55191667-INS-117 |
| chr4-55204850-INS-63 | chr4-55204850-INS-63 |
| chr4-55255351-INS-129 | chr4-55255373-INS-150 |
| chr4-55285508-INS-312 | chr4-55285508-INS-312 |
| chr4-55297671-INS-236 | chr4-55297672-INS-233 |
| chr4-55301873-INS-198 | chr4-55301873-INS-200 |
| chr4-55319747-INS-126 | chr4-55319755-INS-77 |
| chr4-55320911-INS-498 | chr4-55320911-INS-498 |
| chr4-55337359-INS-196 | chr4-55337359-INS-196 |
| chr4-55340464-INS-180 | chr4-55340464-INS-180 |
| chr4-55362775-INS-164 | chr4-55362775-INS-168 |
| chr4-55388042-INS-209 | chr4-55388043-INS-209 |
| chr4-55395926-INS-214 | chr4-55395926-INS-214 |
| chr4-55402162-INS-53 | chr4-55402162-INS-53 |
| chr4-55405830-INS-222 | chr4-55405830-INS-223 |
| chr4-55412077-INS-205 | chr4-55412077-INS-205 |
| chr4-55416983-INS-77 | chr4-55416985-INS-78 |
| chr4-55429984-INS-412 | chr4-55429984-INS-412 |
| chr4-55430152-INS-1234 | chr4-55430153-INS-1234 |
| chr4-55449084-INS-53 | chr4-55449112-INS-55 |
| chr4-55463036-INS-169 | chr4-55463036-INS-213 |
| chr4-55470205-INS-187 | chr4-55470205-INS-203 |
| chr4-55494246-INS-209 | chr4-55494246-INS-209 |
| chr4-55495973-INS-159 | chr4-55495973-INS-160 |
| chr4-55502778-INS-87 | chr4-55502779-INS-54 |
| chr4-55516676-INS-56 | chr4-55516676-INS-50 |
| chr4-55545948-INS-344 | chr4-55545948-INS-342 |
| chr4-55564628-INS-50 | chr4-55564628-INS-54 |
| chr4-55588510-INS-147 | chr4-55588510-INS-150 |
| chr4-55625748-INS-82 | chr4-55625764-INS-70 |
| chr4-55644893-INS-165 | chr4-55644871-INS-186 |
| chr4-55665257-INS-72 | chr4-55665257-INS-72 |
| chr4-55668087-INS-273 | chr4-55668087-INS-273 |
| chr4-55675982-INS-186 | chr4-55675982-INS-162 |
| chr4-55686498-INS-537 | chr4-55686498-INS-537 |
| chr4-55695770-INS-98 | chr4-55695770-INS-102 |
| chr4-55713853-INS-98 | chr4-55713853-INS-110 |
| chr4-55719968-INS-115 | chr4-55719968-INS-113 |
| chr4-55753931-INS-7660 | chr4-55753932-INS-7661 |
| chr4-55840428-INS-393 | chr4-55840428-INS-393 |
| chr4-55849159-INS-130 | chr4-55849160-INS-92 |
| chr4-55867392-INS-212 | chr4-55867392-INS-212 |
| chr4-55908056-INS-313 | chr4-55908056-INS-313 |
| chr4-55908448-INS-1524 | chr4-55908449-INS-1524 |
| chr4-55924214-INS-1008 | chr4-55924215-INS-1008 |
| chr4-55958296-INS-52 | chr4-55958296-INS-56 |
| chr4-55998312-INS-186 | chr4-55998312-INS-186 |
| chr4-56005320-INS-177 | chr4-56005320-INS-177 |
| chr4-56045793-INS-199 | chr4-56045793-INS-199 |
| chr4-56048221-INS-255 | chr4-56048221-INS-262 |
| chr4-56049860-INS-78 | chr4-56049866-INS-72 |
| chr4-56052352-INS-5655 | chr4-56052353-INS-5655 |
| chr4-56075439-INS-6261 | chr4-56075440-INS-6261 |
| chr4-56082088-INS-206 | chr4-56082088-INS-206 |
| chr4-56117844-INS-190 | chr4-56117844-INS-150 |
| chr4-56158618-INS-50 | chr4-56158618-INS-52 |
| chr4-56166293-INS-80 | chr4-56166293-INS-64 |
| chr4-56201612-INS-114 | chr4-56201612-INS-114 |
| chr4-56234203-INS-596 | chr4-56234203-INS-596 |
| chr4-56251135-INS-198 | chr4-56251135-INS-198 |
| chr4-56259056-INS-78 | chr4-56259050-INS-88 |
| chr4-56279326-INS-58 | chr4-56279326-INS-60 |
| chr4-56284307-INS-6172 | chr4-56284308-INS-6172 |
| chr4-56313786-INS-6452 | chr4-56313787-INS-6451 |
| chr4-56333234-INS-400 | chr4-56333234-INS-400 |
| chr4-56335831-INS-255 | chr4-56335831-INS-247 |
| chr4-56342682-INS-51 | chr4-56342671-INS-60 |
| chr4-56354509-INS-283 | chr4-56354530-INS-284 |
| chr4-56354907-INS-984 | chr4-56354907-INS-984 |
| chr4-56415727-INS-864 | chr4-56415727-INS-864 |
| chr4-56428646-INS-204 | chr4-56428646-INS-200 |
| chr4-56433861-INS-6359 | chr4-56433862-INS-6359 |
| chr4-56440799-INS-505 | chr4-56440799-INS-505 |
| chr4-56447851-INS-338 | chr4-56447852-INS-351 |
| chr4-56481083-INS-71 | chr4-56481083-INS-71 |
| chr4-56505343-INS-282 | chr4-56505343-INS-273 |

| | |
|---|---|
| chr4-56524921-INS-464 | chr4-56524921-INS-464 |
| chr4-56528060-INS-70 | chr4-56528032-INS-74 |
| chr4-56538629-INS-55 | chr4-56538630-INS-51 |
| chr4-56613062-INS-94 | chr4-56613062-INS-78 |
| chr4-56626437-INS-60 | chr4-56626437-INS-76 |
| chr4-56646437-INS-3020 | chr4-56646438-INS-3020 |
| chr4-56646438-INS-480 | chr4-56646438-INS-480 |
| chr4-56649624-INS-56 | chr4-56649624-INS-55 |
| chr4-56752707-INS-68 | chr4-56752671-INS-118 |
| chr4-56754751-INS-108 | chr4-56754751-INS-108 |
| chr4-56757660-INS-595 | chr4-56757660-INS-596 |
| chr4-56763930-INS-217 | chr4-56763930-INS-217 |
| chr4-56814259-INS-402 | chr4-56814259-INS-402 |
| chr4-56822108-INS-671 | chr4-56822108-INS-671 |
| chr4-56836319-INS-74 | chr4-56836319-INS-77 |
| chr4-56839690-INS-104 | chr4-56839716-INS-98 |
| chr4-56841320-INS-211 | chr4-56841320-INS-212 |
| chr4-56862304-INS-202 | chr4-56862304-INS-203 |
| chr4-56877599-INS-156 | chr4-56877599-INS-156 |
| chr4-56898153-INS-62 | chr4-56898151-INS-104 |
| chr4-56902304-INS-217 | chr4-56902304-INS-219 |
| chr4-56918164-INS-55 | chr4-56918164-INS-55 |
| chr4-56919473-INS-94 | chr4-56919473-INS-94 |
| chr4-56925692-INS-666 | chr4-56925692-INS-654 |
| chr4-56931793-INS-86 | chr4-56931754-INS-86 |
| chr4-56942209-INS-73 | chr4-56942209-INS-73 |
| chr4-56954173-INS-52 | chr4-56954173-INS-56 |
| chr4-56965715-INS-376 | chr4-56965715-INS-376 |
| chr4-56975222-INS-256 | chr4-56975222-INS-257 |
| chr4-56981516-INS-349 | chr4-56981516-INS-349 |
| chr4-57006851-INS-150 | chr4-57006851-INS-154 |
| chr4-57036554-INS-607 | chr4-57036554-INS-608 |
| chr4-57046330-INS-1894 | chr4-57046331-INS-1894 |
| chr4-57058229-INS-91 | chr4-57058216-INS-67 |
| chr4-57074275-INS-638 | chr4-57074275-INS-638 |
| chr4-57090744-INS-207 | chr4-57090744-INS-207 |
| chr4-57114572-INS-94 | chr4-57114572-INS-98 |
| chr4-57140234-INS-67 | chr4-57140234-INS-67 |
| chr4-57162570-INS-88 | chr4-57162570-INS-88 |
| chr4-57166111-INS-55 | chr4-57166111-INS-55 |
| chr4-57167752-INS-300 | chr4-57167753-INS-319 |
| chr4-57205521-INS-58 | chr4-57205521-INS-58 |
| chr4-57210949-INS-83 | chr4-57210965-INS-84 |
| chr4-57240041-INS-144 | chr4-57240041-INS-144 |
| chr4-57255352-INS-66 | chr4-57255352-INS-66 |
| chr4-57258274-INS-50 | chr4-57258274-INS-52 |
| chr4-57282663-INS-174 | chr4-57282663-INS-172 |
| chr4-57294604-INS-64 | chr4-57294604-INS-64 |
| chr4-57322139-INS-659 | chr4-57322139-INS-704 |
| chr4-57344848-INS-263 | chr4-57344848-INS-262 |
| chr4-57348098-INS-91 | chr4-57348116-INS-114 |
| chr4-57353273-INS-59 | chr4-57353273-INS-59 |
| chr4-57362227-INS-209 | chr4-57362227-INS-209 |
| chr4-57365328-INS-151 | chr4-57365327-INS-153 |
| chr4-57368174-INS-165 | chr4-57368174-INS-157 |
| chr4-57372890-INS-493 | chr4-57372890-INS-492 |
| chr4-57396131-INS-382 | chr4-57396132-INS-383 |
| chr4-57400559-INS-145 | chr4-57400559-INS-178 |
| chr4-57403613-INS-297 | chr4-57403613-INS-296 |
| chr4-57406810-INS-116 | chr4-57406810-INS-116 |
| chr4-57424757-INS-91 | chr4-57424757-INS-92 |
| chr4-57474335-INS-125 | chr4-57474336-INS-84 |
| chr4-57499649-INS-74 | chr4-57499649-INS-74 |
| chr4-57504242-INS-492 | chr4-57504242-INS-492 |
| chr4-57520710-INS-893 | chr4-57520710-INS-906 |
| chr4-57539004-INS-6371 | chr4-57539005-INS-6380 |
| chr4-57539942-INS-100 | chr4-57539942-INS-100 |
| chr4-57548163-INS-93 | chr4-57548163-INS-93 |
| chr4-57560857-INS-58 | chr4-57560867-INS-72 |
| chr4-57604482-INS-195 | chr4-57604482-INS-196 |
| chr4-57605156-INS-193 | chr4-57605153-INS-195 |
| chr4-57614350-INS-96 | chr4-57614321-INS-96 |
| chr4-57729250-INS-99 | chr4-57729250-INS-99 |
| chr4-57734546-INS-134 | chr4-57734546-INS-134 |
| chr4-57757322-INS-70 | chr4-57757322-INS-71 |
| chr4-57769730-INS-119 | chr4-57769730-INS-119 |
| chr4-57779737-INS-60 | chr4-57779737-INS-80 |

| | |
|---|---|
| chr4-57796761-INS-173 | chr4-57796761-INS-158 |
| chr4-57820349-INS-61 | chr4-57820372-INS-86 |
| chr4-57837699-INS-62 | chr4-57837699-INS-58 |
| chr4-57840019-INS-68 | chr4-57839995-INS-50 |
| chr4-57858994-INS-210 | chr4-57858994-INS-210 |
| chr4-57861665-INS-94 | chr4-57861665-INS-102 |
| chr4-57882732-INS-3835 | chr4-57882733-INS-3835 |
| chr4-57905116-INS-194 | chr4-57905084-INS-232 |
| chr4-57912652-INS-68 | chr4-57912652-INS-68 |
| chr4-57920205-INS-116 | chr4-57920169-INS-114 |
| chr4-57924206-INS-152 | chr4-57924173-INS-174 |
| chr4-57933770-INS-99 | chr4-57933770-INS-99 |
| chr4-57937165-INS-124 | chr4-57937164-INS-166 |
| chr4-57947961-INS-250 | chr4-57947961-INS-250 |
| chr4-58026354-INS-55 | chr4-58026355-INS-50 |
| chr4-58033855-INS-84 | chr4-58033855-INS-84 |
| chr4-58076893-INS-54 | chr4-58076865-INS-60 |
| chr4-58084272-INS-889 | chr4-58084272-INS-889 |
| chr4-58105287-INS-62 | chr4-58105287-INS-70 |
| chr4-58129762-INS-430 | chr4-58129762-INS-430 |
| chr4-58151782-INS-73 | chr4-58151780-INS-63 |
| chr4-58155418-INS-66 | chr4-58155418-INS-60 |
| chr4-58166146-INS-54 | chr4-58166151-INS-56 |
| chr4-58169647-INS-3522 | chr4-58169646-INS-3531 |
| chr4-58169813-INS-54 | chr4-58169813-INS-60 |
| chr4-58170922-INS-7127 | chr4-58170923-INS-7127 |
| chr4-58173837-INS-77 | chr4-58173835-INS-78 |
| chr4-58194467-INS-64 | chr4-58194485-INS-64 |
| chr4-58209065-INS-66 | chr4-58209065-INS-68 |
| chr4-58210336-INS-340 | chr4-58210336-INS-340 |
| chr4-58239777-INS-63 | chr4-58239777-INS-69 |
| chr4-58246844-INS-345 | chr4-58246844-INS-345 |
| chr4-58279360-INS-2621 | chr4-58279361-INS-2620 |
| chr4-58294229-INS-434 | chr4-58294229-INS-442 |
| chr4-58301299-INS-1521 | chr4-58301300-INS-1521 |
| chr4-58304685-INS-208 | chr4-58304685-INS-216 |
| chr4-58309958-INS-52 | chr4-58309958-INS-52 |
| chr4-58311574-INS-89 | chr4-58311607-INS-91 |
| chr4-58314652-INS-105 | chr4-58314635-INS-142 |
| chr4-58319002-INS-55 | chr4-58319002-INS-55 |
| chr4-58348594-INS-170 | chr4-58348594-INS-158 |
| chr4-58376830-INS-72 | chr4-58376837-INS-108 |
| chr4-58379496-INS-114 | chr4-58379497-INS-114 |
| chr4-58380928-INS-85 | chr4-58380889-INS-114 |
| chr4-58382818-INS-58 | chr4-58382818-INS-58 |
| chr4-58402771-INS-125 | chr4-58402781-INS-88 |
| chr4-58413097-INS-1925 | chr4-58413098-INS-1925 |
| chr4-58433101-INS-145 | chr4-58433101-INS-145 |
| chr4-58443165-INS-61 | chr4-58443165-INS-61 |
| chr4-58479009-INS-124 | chr4-58479009-INS-124 |
| chr4-58506910-INS-304 | chr4-58506910-INS-304 |
| chr4-58519322-INS-64 | chr4-58519322-INS-64 |
| chr4-58523463-INS-84 | chr4-58523474-INS-95 |
| chr4-58532868-INS-78 | chr4-58532850-INS-81 |
| chr4-58539319-INS-186 | chr4-58539319-INS-186 |
| chr4-58558067-INS-71 | chr4-58558073-INS-66 |
| chr4-58587169-INS-99 | chr4-58587170-INS-99 |
| chr4-58600377-INS-77 | chr4-58600355-INS-77 |
| chr4-58602585-INS-61 | chr4-58602585-INS-62 |
| chr4-58610371-INS-62 | chr4-58610369-INS-100 |
| chr4-58616055-INS-56 | chr4-58616055-INS-60 |
| chr4-58635351-INS-52 | chr4-58635351-INS-52 |
| chr4-58710902-INS-5668 | chr4-58710903-INS-5668 |
| chr4-58717278-INS-105 | chr4-58717278-INS-110 |
| chr4-58727967-INS-263 | chr4-58727967-INS-263 |
| chr4-58752749-INS-310 | chr4-58752749-INS-310 |
| chr4-58758927-INS-135 | chr4-58758927-INS-132 |
| chr4-58782011-INS-529 | chr4-58782011-INS-529 |
| chr4-58788056-INS-111 | chr4-58788056-INS-140 |
| chr4-58788994-INS-766 | chr4-58788994-INS-766 |
| chr4-58790515-INS-6482 | chr4-58790516-INS-6482 |
| chr4-58821989-INS-98 | chr4-58821989-INS-98 |
| chr4-58866928-INS-384 | chr4-58866928-INS-374 |
| chr4-58899207-INS-78 | chr4-58899221-INS-100 |
| chr4-58899987-INS-272 | chr4-58899987-INS-272 |
| chr4-58925329-INS-64 | chr4-58925329-INS-64 |
| chr4-58926541-INS-1140 | chr4-58926542-INS-1141 |

| | |
|---|---|
| chr4-58940780-INS-221 | chr4-58940780-INS-222 |
| chr4-58951334-INS-129 | chr4-58951334-INS-129 |
| chr4-58987209-INS-64 | chr4-58987209-INS-62 |
| chr4-59000282-INS-88 | chr4-59000313-INS-66 |
| chr4-59007475-INS-105 | chr4-59007475-INS-105 |
| chr4-59037458-INS-162 | chr4-59037458-INS-168 |
| chr4-59072858-INS-698 | chr4-59072858-INS-728 |
| chr4-59102181-INS-74 | chr4-59102179-INS-60 |
| chr4-59109916-INS-590 | chr4-59109913-INS-632 |
| chr4-59121796-INS-99 | chr4-59121796-INS-99 |
| chr4-59125222-INS-180 | chr4-59125222-INS-198 |
| chr4-59140771-INS-61 | chr4-59140771-INS-61 |
| chr4-59144862-INS-210 | chr4-59144862-INS-217 |
| chr4-59147411-INS-226 | chr4-59147411-INS-226 |
| chr4-59149256-INS-87 | chr4-59149256-INS-87 |
| chr4-59154722-INS-195 | chr4-59154722-INS-195 |
| chr4-59156898-INS-95 | chr4-59156898-INS-95 |
| chr4-59169901-INS-125 | chr4-59169901-INS-112 |
| chr4-59178576-INS-173 | chr4-59178576-INS-173 |
| chr4-59188174-INS-120 | chr4-59188174-INS-120 |
| chr4-59246205-INS-5587 | chr4-59246206-INS-5587 |
| chr4-59260153-INS-58 | chr4-59260153-INS-58 |
| chr4-59266147-INS-89 | chr4-59266147-INS-89 |
| chr4-59298096-INS-54 | chr4-59298096-INS-54 |
| chr4-59312151-INS-73 | chr4-59312151-INS-102 |
| chr4-59367707-INS-160 | chr4-59367707-INS-158 |
| chr4-59368860-INS-156 | chr4-59368860-INS-156 |
| chr4-59371269-INS-98 | chr4-59371269-INS-96 |
| chr4-59427121-INS-106 | chr4-59427121-INS-118 |
| chr4-59429778-INS-775 | chr4-59429778-INS-779 |
| chr4-59464066-INS-884 | chr4-59464066-INS-884 |
| chr4-59467235-INS-198 | chr4-59467235-INS-198 |
| chr4-59500993-INS-61 | chr4-59500993-INS-65 |
| chr4-59514973-INS-170 | chr4-59514973-INS-168 |
| chr4-59527905-INS-352 | chr4-59527921-INS-355 |
| chr4-59544884-INS-209 | chr4-59544884-INS-208 |
| chr4-59551322-INS-96 | chr4-59551322-INS-96 |
| chr4-59554637-INS-54 | chr4-59554621-INS-62 |
| chr4-59565049-INS-144 | chr4-59565049-INS-168 |
| chr4-59571731-INS-185 | chr4-59571731-INS-194 |
| chr4-59572666-INS-53 | chr4-59572666-INS-53 |
| chr4-59577987-INS-191 | chr4-59577987-INS-191 |
| chr4-59613312-INS-1491 | chr4-59613313-INS-1540 |
| chr4-59628666-INS-56 | chr4-59628666-INS-52 |
| chr4-59633808-INS-97 | chr4-59633772-INS-127 |
| chr4-59678612-INS-138 | chr4-59678612-INS-138 |
| chr4-59748666-INS-61 | chr4-59748666-INS-61 |
| chr4-59767036-INS-3087 | chr4-59767037-INS-3086 |
| chr4-59772825-INS-419 | chr4-59772814-INS-439 |
| chr4-59786578-INS-88 | chr4-59786578-INS-88 |
| chr4-59787928-INS-84 | chr4-59787936-INS-68 |
| chr4-59801656-INS-55 | chr4-59801656-INS-59 |
| chr4-59803869-INS-131 | chr4-59803881-INS-143 |
| chr4-59850503-INS-123 | chr4-59850504-INS-123 |
| chr4-59858042-INS-80 | chr4-59858042-INS-80 |
| chr4-59861790-INS-6480 | chr4-59861791-INS-6480 |
| chr4-59862267-INS-578 | chr4-59862268-INS-578 |
| chr4-59867365-INS-69 | chr4-59867365-INS-80 |
| chr4-59879157-INS-56 | chr4-59879157-INS-56 |
| chr4-59895507-INS-77 | chr4-59895507-INS-77 |
| chr4-59897725-INS-56 | chr4-59897725-INS-56 |
| chr4-59915731-INS-53 | chr4-59915731-INS-53 |
| chr4-59942619-INS-64 | chr4-59942619-INS-64 |
| chr4-59959128-INS-90 | chr4-59959128-INS-90 |
| chr4-59964491-INS-6475 | chr4-59964492-INS-6475 |
| chr4-59987797-INS-56 | chr4-59987797-INS-56 |
| chr4-60016052-INS-52 | chr4-60016052-INS-52 |
| chr4-60166610-INS-74 | chr4-60166610-INS-52 |
| chr4-60765852-INS-53 | chr4-60765852-INS-53 |
| chr4-61640112-INS-1356 | chr4-61640113-INS-1360 |
| chr4-61711587-INS-66 | chr4-61711587-INS-78 |
| chr4-61757070-INS-60 | chr4-61757070-INS-66 |
| chr4-61770413-INS-488 | chr4-61770413-INS-488 |
| chr4-61771795-INS-70 | chr4-61771795-INS-74 |
| chr4-61774450-INS-102 | chr4-61774450-INS-102 |
| chr4-61805954-INS-1557 | chr4-61805955-INS-1557 |
| chr4-61814642-INS-6603 | chr4-61814643-INS-6603 |

| | |
|---|---|
| chr4-61862542-INS-135 | chr4-61862542-INS-135 |
| chr4-61931191-INS-245 | chr4-61931191-INS-245 |
| chr4-61937885-INS-6543 | chr4-61937886-INS-6544 |
| chr4-62001282-INS-205 | chr4-62001282-INS-205 |
| chr4-62058510-INS-6442 | chr4-62058511-INS-6442 |
| chr4-62077856-INS-2448 | chr4-62077857-INS-2463 |
| chr4-62103894-INS-4421 | chr4-62103895-INS-4421 |
| chr4-62131382-INS-215 | chr4-62131382-INS-215 |
| chr4-62176087-INS-1185 | chr4-62176088-INS-1185 |
| chr4-62199202-INS-525 | chr4-62199202-INS-525 |
| chr4-62232160-INS-61 | chr4-62232160-INS-61 |
| chr4-62285067-INS-56 | chr4-62285068-INS-56 |
| chr4-62324875-INS-146 | chr4-62324875-INS-146 |
| chr4-62336306-INS-219 | chr4-62336306-INS-220 |
| chr4-62351576-INS-113 | chr4-62351576-INS-114 |
| chr4-62393614-INS-1430 | chr4-62393615-INS-1428 |
| chr4-62444388-INS-54 | chr4-62444361-INS-60 |
| chr4-62471419-INS-597 | chr4-62471419-INS-618 |
| chr4-62480082-INS-97 | chr4-62480106-INS-68 |
| chr4-62497874-INS-172 | chr4-62497874-INS-174 |
| chr4-62505759-INS-486 | chr4-62505759-INS-487 |
| chr4-62534454-INS-73 | chr4-62534454-INS-66 |
| chr4-62535691-INS-154 | chr4-62535691-INS-154 |
| chr4-62607859-INS-54 | chr4-62607859-INS-54 |
| chr4-62613006-INS-281 | chr4-62613006-INS-281 |
| chr4-62687716-INS-197 | chr4-62687716-INS-197 |
| chr4-62718268-INS-632 | chr4-62718268-INS-631 |
| chr4-62738198-INS-7466 | chr4-62738199-INS-7466 |
| chr4-62740662-INS-96 | chr4-62740650-INS-134 |
| chr4-62750915-INS-119 | chr4-62750915-INS-119 |
| chr4-62794828-INS-54 | chr4-62794828-INS-54 |
| chr4-62806306-INS-345 | chr4-62806308-INS-371 |
| chr4-62818605-INS-272 | chr4-62818617-INS-283 |
| chr4-62828730-INS-287 | chr4-62828730-INS-294 |
| chr4-62838651-INS-312 | chr4-62838651-INS-312 |
| chr4-62851409-INS-137 | chr4-62851409-INS-137 |
| chr4-62863189-INS-604 | chr4-62863189-INS-604 |
| chr4-62872808-INS-58 | chr4-62872808-INS-58 |
| chr4-62874956-INS-3100 | chr4-62874957-INS-3100 |
| chr4-62879807-INS-7166 | chr4-62879808-INS-7166 |
| chr4-62909247-INS-59 | chr4-62909247-INS-68 |
| chr4-62921454-INS-94 | chr4-62921454-INS-94 |
| chr4-62940050-INS-55 | chr4-62940050-INS-55 |
| chr4-62950062-INS-68 | chr4-62950058-INS-71 |
| chr4-62985825-INS-272 | chr4-62985825-INS-272 |
| chr4-63009478-INS-151 | chr4-63009478-INS-170 |
| chr4-63025446-INS-310 | chr4-63025446-INS-310 |
| chr4-63078238-INS-54 | chr4-63078238-INS-54 |
| chr4-63095090-INS-58 | chr4-63095108-INS-52 |
| chr4-63098284-INS-160 | chr4-63098284-INS-160 |
| chr4-63104106-INS-56 | chr4-63104106-INS-56 |
| chr4-63107557-INS-131 | chr4-63107554-INS-180 |
| chr4-63115252-INS-62 | chr4-63115252-INS-62 |
| chr4-63140507-INS-135 | chr4-63140507-INS-135 |
| chr4-63150628-INS-389 | chr4-63150628-INS-389 |
| chr4-63208973-INS-52 | chr4-63208973-INS-52 |
| chr4-63225901-INS-198 | chr4-63225902-INS-186 |
| chr4-63278549-INS-156 | chr4-63278549-INS-157 |
| chr4-63311572-INS-201 | chr4-63311572-INS-201 |
| chr4-63324579-INS-205 | chr4-63324579-INS-205 |
| chr4-63358846-INS-1083 | chr4-63358847-INS-1083 |
| chr4-63369893-INS-204 | chr4-63369893-INS-204 |
| chr4-63376259-INS-184 | chr4-63376259-INS-183 |
| chr4-63378356-INS-124 | chr4-63378356-INS-149 |
| chr4-63450706-INS-271 | chr4-63450706-INS-271 |
| chr4-63481253-INS-574 | chr4-63481253-INS-525 |
| chr4-63495064-INS-75 | chr4-63495064-INS-75 |
| chr4-63507935-INS-184 | chr4-63507935-INS-188 |
| chr4-63511005-INS-62 | chr4-63511005-INS-62 |
| chr4-63521837-INS-60 | chr4-63521837-INS-60 |
| chr4-63535127-INS-669 | chr4-63535127-INS-669 |
| chr4-63540125-INS-73 | chr4-63540115-INS-53 |
| chr4-63542455-INS-52 | chr4-63542455-INS-56 |
| chr4-63555411-INS-1864 | chr4-63555412-INS-1864 |
| chr4-63591706-INS-52 | chr4-63591706-INS-54 |
| chr4-63602748-INS-72 | chr4-63602748-INS-72 |
| chr4-63619543-INS-421 | chr4-63619543-INS-421 |

| | |
|---|---|
| chr4-63633071-INS-80 | chr4-63633071-INS-56 |
| chr4-63687119-INS-80 | chr4-63687121-INS-86 |
| chr4-63701146-INS-106 | chr4-63701146-INS-106 |
| chr4-63729432-INS-504 | chr4-63729432-INS-504 |
| chr4-63755547-INS-59 | chr4-63755547-INS-51 |
| chr4-63763992-INS-132 | chr4-63763992-INS-132 |
| chr4-63766251-INS-82 | chr4-63766251-INS-76 |
| chr4-63798587-INS-146 | chr4-63798587-INS-126 |
| chr4-63819027-INS-51 | chr4-63819025-INS-68 |
| chr4-63826714-INS-159 | chr4-63826714-INS-159 |
| chr4-63829240-INS-1450 | chr4-63829241-INS-1451 |
| chr4-63832114-INS-3350 | chr4-63832115-INS-3351 |
| chr4-63855455-INS-64 | chr4-63855443-INS-100 |
| chr4-63872831-INS-3096 | chr4-63872832-INS-3096 |
| chr4-63904386-INS-55 | chr4-63904386-INS-51 |
| chr4-63915082-INS-64 | chr4-63915082-INS-68 |
| chr4-63968169-INS-58 | chr4-63968169-INS-57 |
| chr4-63983988-INS-392 | chr4-63983988-INS-390 |
| chr4-63995075-INS-3190 | chr4-63995076-INS-3190 |
| chr4-64006356-INS-54 | chr4-64006359-INS-58 |
| chr4-64010029-INS-230 | chr4-64010029-INS-212 |
| chr4-64025644-INS-6321 | chr4-64025645-INS-6322 |
| chr4-64078688-INS-380 | chr4-64078688-INS-380 |
| chr4-64085652-INS-77 | chr4-64085654-INS-82 |
| chr4-64114915-INS-400 | chr4-64114915-INS-400 |
| chr4-64122229-INS-497 | chr4-64122229-INS-497 |
| chr4-64135499-INS-249 | chr4-64135498-INS-255 |
| chr4-64176040-INS-1420 | chr4-64176041-INS-1420 |
| chr4-64185584-INS-54 | chr4-64185584-INS-54 |
| chr4-64189458-INS-54 | chr4-64189456-INS-56 |
| chr4-64211811-INS-212 | chr4-64211811-INS-212 |
| chr4-64230378-INS-236 | chr4-64230378-INS-236 |
| chr4-64242677-INS-245 | chr4-64242677-INS-245 |
| chr4-64250761-INS-2082 | chr4-64250762-INS-2082 |
| chr4-64287050-INS-52 | chr4-64287050-INS-52 |
| chr4-64291228-INS-77 | chr4-64291228-INS-77 |
| chr4-64296882-INS-429 | chr4-64296882-INS-413 |
| chr4-64297516-INS-57 | chr4-64297516-INS-57 |
| chr4-64301693-INS-7084 | chr4-64301694-INS-7085 |
| chr4-64320218-INS-72 | chr4-64320218-INS-74 |
| chr4-64362512-INS-81 | chr4-64362520-INS-80 |
| chr4-64412580-INS-218 | chr4-64412580-INS-221 |
| chr4-64423534-INS-285 | chr4-64423534-INS-274 |
| chr4-64460313-INS-62 | chr4-64460313-INS-64 |
| chr4-64490910-INS-52 | chr4-64490910-INS-50 |
| chr4-64499388-INS-58 | chr4-64499403-INS-54 |
| chr4-64511655-INS-104 | chr4-64511655-INS-104 |
| chr4-64516503-INS-1309 | chr4-64516504-INS-1309 |
| chr4-64520500-INS-53 | chr4-64520500-INS-55 |
| chr4-64521987-INS-917 | chr4-64521987-INS-917 |
| chr4-64524588-INS-6240 | chr4-64524589-INS-6240 |
| chr4-64544633-INS-5255 | chr4-64544634-INS-5255 |
| chr4-64554850-INS-68 | chr4-64554850-INS-68 |
| chr4-64580641-INS-269 | chr4-64580641-INS-269 |
| chr4-64613943-INS-313 | chr4-64613931-INS-310 |
| chr4-64625372-INS-84 | chr4-64625372-INS-84 |
| chr4-64633385-INS-65 | chr4-64633385-INS-65 |
| chr4-64637209-INS-74 | chr4-64637209-INS-80 |
| chr4-64661109-INS-892 | chr4-64661109-INS-892 |
| chr4-64683521-INS-56 | chr4-64683521-INS-60 |
| chr4-64763772-INS-50 | chr4-64763772-INS-50 |
| chr4-64764597-INS-2411 | chr4-64764598-INS-2415 |
| chr4-64787760-INS-368 | chr4-64787760-INS-368 |
| chr4-64809262-INS-65 | chr4-64809259-INS-64 |
| chr4-64825055-INS-116 | chr4-64825092-INS-125 |
| chr4-64853167-INS-2097 | chr4-64853168-INS-2097 |
| chr4-64870126-INS-1379 | chr4-64870127-INS-1386 |
| chr4-64875398-INS-6299 | chr4-64875399-INS-6299 |
| chr4-64885921-INS-389 | chr4-64885921-INS-389 |
| chr4-64896869-INS-117 | chr4-64896869-INS-117 |
| chr4-64915148-INS-152 | chr4-64915154-INS-138 |
| chr4-64930215-INS-662 | chr4-64930216-INS-687 |
| chr4-64999889-INS-88 | chr4-64999877-INS-100 |
| chr4-65010517-INS-1293 | chr4-65010518-INS-1294 |
| chr4-65012177-INS-6424 | chr4-65012178-INS-6423 |
| chr4-65037706-INS-516 | chr4-65037706-INS-516 |
| chr4-65058774-INS-251 | chr4-65058774-INS-251 |

| | |
|---|---|
| chr4-65079018-INS-54 | chr4-65079018-INS-56 |
| chr4-65164509-INS-58 | chr4-65164509-INS-78 |
| chr4-65186268-INS-108 | chr4-65186268-INS-108 |
| chr4-65229587-INS-132 | chr4-65229587-INS-130 |
| chr4-65259644-INS-52 | chr4-65259644-INS-54 |
| chr4-65261912-INS-60 | chr4-65261912-INS-54 |
| chr4-65274381-INS-76 | chr4-65274377-INS-68 |
| chr4-65329745-INS-748 | chr4-65329745-INS-748 |
| chr4-65330952-INS-85 | chr4-65330952-INS-56 |
| chr4-65398406-INS-77 | chr4-65398406-INS-77 |
| chr4-65405555-INS-171 | chr4-65405555-INS-174 |
| chr4-65436901-INS-92 | chr4-65436901-INS-92 |
| chr4-65469719-INS-756 | chr4-65469719-INS-755 |
| chr4-65491675-INS-198 | chr4-65491640-INS-199 |
| chr4-65505591-INS-432 | chr4-65505598-INS-382 |
| chr4-65507103-INS-628 | chr4-65507103-INS-649 |
| chr4-65558150-INS-818 | chr4-65558150-INS-818 |
| chr4-65561703-INS-105 | chr4-65561703-INS-100 |
| chr4-65580828-INS-224 | chr4-65580828-INS-224 |
| chr4-65600491-INS-56 | chr4-65600492-INS-54 |
| chr4-65621318-INS-62 | chr4-65621318-INS-62 |
| chr4-65643646-INS-60 | chr4-65643646-INS-61 |
| chr4-65663333-INS-72 | chr4-65663333-INS-72 |
| chr4-65702082-INS-106 | chr4-65702082-INS-106 |
| chr4-65728569-INS-409 | chr4-65728569-INS-409 |
| chr4-65739713-INS-249 | chr4-65739713-INS-249 |
| chr4-65765748-INS-164 | chr4-65765748-INS-164 |
| chr4-65802099-INS-590 | chr4-65802092-INS-595 |
| chr4-65825002-INS-66 | chr4-65824969-INS-115 |
| chr4-65843442-INS-3101 | chr4-65843443-INS-3074 |
| chr4-65903325-INS-1329 | chr4-65903326-INS-1329 |
| chr4-65924150-INS-3872 | chr4-65924151-INS-3872 |
| chr4-65951567-INS-101 | chr4-65951567-INS-104 |
| chr4-65989164-INS-212 | chr4-65989164-INS-212 |
| chr4-66008637-INS-87 | chr4-66008594-INS-105 |
| chr4-66039331-INS-785 | chr4-66039331-INS-786 |
| chr4-66055559-INS-6806 | chr4-66055556-INS-6802 |
| chr4-66095546-INS-980 | chr4-66095546-INS-983 |
| chr4-66105932-INS-51 | chr4-66105932-INS-50 |
| chr4-66183410-INS-65 | chr4-66183410-INS-69 |
| chr4-66248637-INS-89 | chr4-66248637-INS-89 |
| chr4-66250816-INS-118 | chr4-66250816-INS-119 |
| chr4-66252669-INS-54 | chr4-66252669-INS-50 |
| chr4-66255799-INS-1245 | chr4-66255800-INS-1245 |
| chr4-66293332-INS-158 | chr4-66293334-INS-128 |
| chr4-66310434-INS-1054 | chr4-66310435-INS-1056 |
| chr4-66315196-INS-60 | chr4-66315209-INS-78 |
| chr4-66333675-INS-182 | chr4-66333659-INS-173 |
| chr4-66343441-INS-6897 | chr4-66343442-INS-6897 |
| chr4-66343503-INS-488 | chr4-66343505-INS-488 |
| chr4-66359312-INS-6380 | chr4-66359309-INS-6381 |
| chr4-66375810-INS-86 | chr4-66375810-INS-94 |
| chr4-66376964-INS-5284 | chr4-66376965-INS-5290 |
| chr4-66378123-INS-298 | chr4-66378123-INS-298 |
| chr4-66503927-INS-351 | chr4-66503927-INS-351 |
| chr4-66508698-INS-52 | chr4-66508698-INS-52 |
| chr4-66530415-INS-1418 | chr4-66530416-INS-1418 |
| chr4-66602991-INS-51 | chr4-66602991-INS-51 |
| chr4-66615190-INS-73 | chr4-66615190-INS-73 |
| chr4-66661535-INS-115 | chr4-66661535-INS-115 |
| chr4-66692562-INS-71 | chr4-66692553-INS-90 |
| chr4-66707426-INS-1186 | chr4-66707427-INS-1186 |
| chr4-66709951-INS-5420 | chr4-66709952-INS-5418 |
| chr4-66733267-INS-329 | chr4-66733267-INS-329 |
| chr4-66769773-INS-131 | chr4-66769773-INS-133 |
| chr4-66787605-INS-55 | chr4-66787606-INS-60 |
| chr4-66859657-INS-58 | chr4-66859657-INS-52 |
| chr4-66861009-INS-1279 | chr4-66861010-INS-1276 |
| chr4-66861716-INS-80 | chr4-66861708-INS-88 |
| chr4-66882519-INS-79 | chr4-66882508-INS-108 |
| chr4-66907674-INS-69 | chr4-66907674-INS-69 |
| chr4-66915992-INS-549 | chr4-66915992-INS-549 |
| chr4-66969166-INS-1450 | chr4-66969167-INS-1450 |
| chr4-66969702-INS-1280 | chr4-66969703-INS-1280 |
| chr4-66987530-INS-401 | chr4-66987530-INS-401 |
| chr4-66993352-INS-6649 | chr4-66993353-INS-6649 |
| chr4-67001487-INS-569 | chr4-67001489-INS-572 |

| | |
|---|---|
| chr4-67005287-INS-392 | chr4-67005287-INS-391 |
| chr4-67021478-INS-52 | chr4-67021478-INS-52 |
| chr4-67029761-INS-6240 | chr4-67029762-INS-6246 |
| chr4-67048755-INS-64 | chr4-67048757-INS-72 |
| chr4-67052861-INS-4347 | chr4-67052862-INS-4346 |
| chr4-67082043-INS-95 | chr4-67082043-INS-95 |
| chr4-67083261-INS-59 | chr4-67083261-INS-61 |
| chr4-67100370-INS-62 | chr4-67100350-INS-82 |
| chr4-67102720-INS-164 | chr4-67102720-INS-144 |
| chr4-67126644-INS-76 | chr4-67126644-INS-55 |
| chr4-67169144-INS-51 | chr4-67169144-INS-51 |
| chr4-67184235-INS-84 | chr4-67184235-INS-60 |
| chr4-67186711-INS-64 | chr4-67186711-INS-64 |
| chr4-67202325-INS-82 | chr4-67202325-INS-84 |
| chr4-67206988-INS-96 | chr4-67206984-INS-95 |
| chr4-67252149-INS-51 | chr4-67252149-INS-51 |
| chr4-67261273-INS-2584 | chr4-67261274-INS-2583 |
| chr4-67264407-INS-7354 | chr4-67264408-INS-7350 |
| chr4-67286995-INS-7303 | chr4-67286996-INS-7303 |
| chr4-67304421-INS-56 | chr4-67304421-INS-56 |
| chr4-67310841-INS-338 | chr4-67310841-INS-338 |
| chr4-67319254-INS-54 | chr4-67319272-INS-84 |
| chr4-67327819-INS-4105 | chr4-67327820-INS-4105 |
| chr4-67340068-INS-55 | chr4-67340068-INS-56 |
| chr4-67341921-INS-113 | chr4-67341919-INS-110 |
| chr4-67350579-INS-68 | chr4-67350579-INS-68 |
| chr4-67353673-INS-6454 | chr4-67353674-INS-6454 |
| chr4-67389909-INS-164 | chr4-67389913-INS-156 |
| chr4-67413622-INS-51 | chr4-67413622-INS-51 |
| chr4-67436027-INS-52 | chr4-67436047-INS-52 |
| chr4-67462326-INS-407 | chr4-67462326-INS-407 |
| chr4-67472934-INS-84 | chr4-67472972-INS-60 |
| chr4-67488231-INS-206 | chr4-67488231-INS-206 |
| chr4-67494617-INS-77 | chr4-67494613-INS-85 |
| chr4-67499099-INS-62 | chr4-67499099-INS-52 |
| chr4-67535154-INS-675 | chr4-67535154-INS-675 |
| chr4-67574377-INS-72 | chr4-67574377-INS-64 |
| chr4-67591502-INS-67 | chr4-67591502-INS-67 |
| chr4-67593571-INS-83 | chr4-67593571-INS-83 |
| chr4-67628091-INS-922 | chr4-67628089-INS-924 |
| chr4-67631264-INS-2036 | chr4-67631269-INS-2035 |
| chr4-67634650-INS-6193 | chr4-67634651-INS-6193 |
| chr4-67670499-INS-1200 | chr4-67670500-INS-1201 |
| chr4-67680875-INS-341 | chr4-67680875-INS-345 |
| chr4-67684322-INS-52 | chr4-67684322-INS-54 |
| chr4-67689170-INS-4282 | chr4-67689162-INS-4277 |
| chr4-67712097-INS-1069 | chr4-67712098-INS-1062 |
| chr4-67721439-INS-589 | chr4-67721441-INS-598 |
| chr4-67723868-INS-506 | chr4-67723857-INS-495 |
| chr4-67724190-INS-1336 | chr4-67724191-INS-1330 |
| chr4-67768266-INS-216 | chr4-67768267-INS-216 |
| chr4-67777078-INS-1811 | chr4-67777079-INS-1811 |
| chr4-67778430-INS-234 | chr4-67778430-INS-234 |
| chr4-67780248-INS-73 | chr4-67780248-INS-73 |
| chr4-67811275-INS-358 | chr4-67811275-INS-358 |
| chr4-67815224-INS-104 | chr4-67815234-INS-80 |
| chr4-67816298-INS-2837 | chr4-67816299-INS-2837 |
| chr4-67819054-INS-391 | chr4-67819054-INS-391 |
| chr4-67824007-INS-55 | chr4-67824007-INS-60 |
| chr4-67846640-INS-78 | chr4-67846640-INS-78 |
| chr4-67865080-INS-179 | chr4-67865080-INS-183 |
| chr4-67927308-INS-52 | chr4-67927308-INS-50 |
| chr4-67931102-INS-673 | chr4-67931102-INS-672 |
| chr4-67936392-INS-54 | chr4-67936392-INS-54 |
| chr4-67954788-INS-178 | chr4-67954788-INS-178 |
| chr4-67979615-INS-1140 | chr4-67979616-INS-1140 |
| chr4-67996207-INS-7171 | chr4-67996208-INS-7172 |
| chr4-68009573-INS-6381 | chr4-68009574-INS-6381 |
| chr4-68020398-INS-52 | chr4-68020410-INS-50 |
| chr4-68030347-INS-83 | chr4-68030347-INS-98 |
| chr4-68076597-INS-696 | chr4-68076597-INS-695 |
| chr4-68101267-INS-51 | chr4-68101267-INS-51 |
| chr4-68138449-INS-52 | chr4-68138449-INS-52 |
| chr4-68139358-INS-583 | chr4-68139358-INS-583 |
| chr4-68141457-INS-59 | chr4-68141457-INS-59 |
| chr4-68169442-INS-64 | chr4-68169442-INS-64 |
| chr4-68172663-INS-99 | chr4-68172663-INS-99 |

| | |
|---|---|
| chr4-68176704-INS-330 | chr4-68176704-INS-330 |
| chr4-68202340-INS-170 | chr4-68202345-INS-132 |
| chr4-68261699-INS-61 | chr4-68261699-INS-57 |
| chr4-68263622-INS-55 | chr4-68263622-INS-57 |
| chr4-68272423-INS-55 | chr4-68272423-INS-51 |
| chr4-68276149-INS-876 | chr4-68276149-INS-876 |
| chr4-68293938-INS-52 | chr4-68293938-INS-52 |
| chr4-68304367-INS-53 | chr4-68304367-INS-63 |
| chr4-68314494-INS-51 | chr4-68314494-INS-55 |
| chr4-68321595-INS-81 | chr4-68321595-INS-56 |
| chr4-68341200-INS-351 | chr4-68341200-INS-351 |
| chr4-68342296-INS-971 | chr4-68342296-INS-971 |
| chr4-68352950-INS-198 | chr4-68352950-INS-198 |
| chr4-68357348-INS-77 | chr4-68357348-INS-76 |
| chr4-68364281-INS-69 | chr4-68364281-INS-71 |
| chr4-68394123-INS-126 | chr4-68394123-INS-126 |
| chr4-68396255-INS-137 | chr4-68396255-INS-137 |
| chr4-68423770-INS-54 | chr4-68423770-INS-52 |
| chr4-68452566-INS-813 | chr4-68452566-INS-812 |
| chr4-68467851-INS-306 | chr4-68467851-INS-309 |
| chr4-68477886-INS-56 | chr4-68477886-INS-58 |
| chr4-68485050-INS-83 | chr4-68485050-INS-108 |
| chr4-68498090-INS-305 | chr4-68498090-INS-305 |
| chr4-68555730-INS-66 | chr4-68555730-INS-66 |
| chr4-68576038-INS-95 | chr4-68576038-INS-96 |
| chr4-68620486-INS-66 | chr4-68620486-INS-66 |
| chr4-68644888-INS-123 | chr4-68644883-INS-126 |
| chr4-68648993-INS-953 | chr4-68648993-INS-954 |
| chr4-68651583-INS-2002 | chr4-68651584-INS-2003 |
| chr4-68653751-INS-6564 | chr4-68653752-INS-6564 |
| chr4-68654979-INS-2267 | chr4-68654980-INS-2267 |
| chr4-68657338-INS-579 | chr4-68657338-INS-579 |
| chr4-68666971-INS-2751 | chr4-68666972-INS-2751 |
| chr4-68679052-INS-58 | chr4-68679052-INS-58 |
| chr4-68682024-INS-52 | chr4-68682024-INS-52 |
| chr4-68682365-INS-1888 | chr4-68682366-INS-1888 |
| chr4-68736658-INS-174 | chr4-68736658-INS-172 |
| chr4-68742652-INS-147 | chr4-68742652-INS-149 |
| chr4-68772933-INS-7728 | chr4-68772934-INS-7728 |
| chr4-68782079-INS-5261 | chr4-68782080-INS-5261 |
| chr4-68794375-INS-51 | chr4-68794375-INS-51 |
| chr4-68810831-INS-6888 | chr4-68810832-INS-6888 |
| chr4-68970401-INS-356 | chr4-68970401-INS-356 |
| chr4-68972789-INS-465 | chr4-68972789-INS-467 |
| chr4-68988902-INS-411 | chr4-68988902-INS-393 |
| chr4-68992137-INS-6685 | chr4-68992138-INS-6685 |
| chr4-68994291-INS-60 | chr4-68994291-INS-92 |
| chr4-69018301-INS-60 | chr4-69018301-INS-50 |
| chr4-69026372-INS-54 | chr4-69026372-INS-58 |
| chr4-69032781-INS-90 | chr4-69032781-INS-93 |
| chr4-69042514-INS-72 | chr4-69042498-INS-75 |
| chr4-69055119-INS-100 | chr4-69055119-INS-91 |
| chr4-69065877-INS-2635 | chr4-69065878-INS-2635 |
| chr4-69071201-INS-219 | chr4-69071201-INS-219 |
| chr4-69095620-INS-214 | chr4-69095620-INS-214 |
| chr4-69122352-INS-179 | chr4-69122352-INS-179 |
| chr4-69155532-INS-447 | chr4-69155532-INS-447 |
| chr4-69193586-INS-68 | chr4-69193588-INS-50 |
| chr4-69231767-INS-1383 | chr4-69231768-INS-1383 |
| chr4-69319880-INS-54 | chr4-69319880-INS-68 |
| chr4-69332315-INS-85 | chr4-69332315-INS-85 |
| chr4-69341091-INS-326 | chr4-69341091-INS-284 |
| chr4-69359958-INS-55 | chr4-69359958-INS-66 |
| chr4-69391086-INS-6449 | chr4-69391087-INS-6449 |
| chr4-69400882-INS-6417 | chr4-69400883-INS-6417 |
| chr4-69444582-INS-63 | chr4-69444582-INS-80 |
| chr4-69450762-INS-529 | chr4-69450762-INS-529 |
| chr4-69452417-INS-65 | chr4-69452458-INS-53 |
| chr4-69469352-INS-6246 | chr4-69469353-INS-6246 |
| chr4-69472578-INS-364 | chr4-69472578-INS-364 |
| chr4-69508948-INS-596 | chr4-69508948-INS-596 |
| chr4-69524746-INS-60 | chr4-69524780-INS-76 |
| chr4-69540531-INS-56 | chr4-69540499-INS-96 |
| chr4-69626067-INS-212 | chr4-69626067-INS-212 |
| chr4-69686795-INS-53 | chr4-69686795-INS-58 |
| chr4-69689180-INS-6538 | chr4-69689181-INS-6538 |
| chr4-69728229-INS-80 | chr4-69728229-INS-80 |

| | |
|---|---|
| chr4-69767282-INS-74 | chr4-69767282-INS-117 |
| chr4-69779592-INS-56 | chr4-69779592-INS-54 |
| chr4-69829957-INS-1172 | chr4-69829958-INS-1172 |
| chr4-69841865-INS-78 | chr4-69841865-INS-78 |
| chr4-69861068-INS-602 | chr4-69861074-INS-611 |
| chr4-69864004-INS-510 | chr4-69864004-INS-510 |
| chr4-69867250-INS-68 | chr4-69867250-INS-70 |
| chr4-69875777-INS-542 | chr4-69875777-INS-542 |
| chr4-69936910-INS-258 | chr4-69936910-INS-210 |
| chr4-69939906-INS-6481 | chr4-69939907-INS-6481 |
| chr4-69975507-INS-5200 | chr4-69975508-INS-5200 |
| chr4-69997694-INS-241 | chr4-69997694-INS-248 |
| chr4-70014430-INS-1072 | chr4-70014431-INS-1072 |
| chr4-70025688-INS-306 | chr4-70025688-INS-306 |
| chr4-70033696-INS-126 | chr4-70033696-INS-114 |
| chr4-70050070-INS-131 | chr4-70050070-INS-131 |
| chr4-70052018-INS-2929 | chr4-70052019-INS-2941 |
| chr4-70052463-INS-66 | chr4-70052463-INS-66 |
| chr4-70063697-INS-55 | chr4-70063697-INS-55 |
| chr4-70075996-INS-7367 | chr4-70075997-INS-7369 |
| chr4-70132557-INS-156 | chr4-70132557-INS-156 |
| chr4-70137888-INS-781 | chr4-70137888-INS-781 |
| chr4-70138495-INS-121 | chr4-70138475-INS-94 |
| chr4-70139317-INS-163 | chr4-70139315-INS-165 |
| chr4-70211947-INS-464 | chr4-70211947-INS-465 |
| chr4-70240714-INS-2848 | chr4-70240715-INS-2845 |
| chr4-70276382-INS-203 | chr4-70276382-INS-203 |
| chr4-70281103-INS-206 | chr4-70281103-INS-217 |
| chr4-70305104-INS-165 | chr4-70305104-INS-165 |
| chr4-70340256-INS-53 | chr4-70340256-INS-53 |
| chr4-70355702-INS-276 | chr4-70355702-INS-276 |
| chr4-70364063-INS-2102 | chr4-70364056-INS-2096 |
| chr4-70369078-INS-2744 | chr4-70369079-INS-2744 |
| chr4-70372141-INS-136 | chr4-70372141-INS-136 |
| chr4-70381922-INS-279 | chr4-70381922-INS-279 |
| chr4-70388505-INS-203 | chr4-70388505-INS-201 |
| chr4-70436788-INS-56 | chr4-70436804-INS-74 |
| chr4-70443283-INS-373 | chr4-70443283-INS-373 |
| chr4-70450784-INS-245 | chr4-70450784-INS-245 |
| chr4-70466827-INS-6421 | chr4-70466828-INS-6421 |
| chr4-70493159-INS-343 | chr4-70493159-INS-343 |
| chr4-70503558-INS-849 | chr4-70503558-INS-848 |
| chr4-70534808-INS-909 | chr4-70534808-INS-895 |
| chr4-70534980-INS-64 | chr4-70534980-INS-52 |
| chr4-70545733-INS-361 | chr4-70545733-INS-361 |
| chr4-70546907-INS-86 | chr4-70546905-INS-88 |
| chr4-70548180-INS-60 | chr4-70548180-INS-72 |
| chr4-70551876-INS-72 | chr4-70551878-INS-73 |
| chr4-70559939-INS-128 | chr4-70559939-INS-128 |
| chr4-70584328-INS-575 | chr4-70584328-INS-575 |
| chr4-70597969-INS-64 | chr4-70597969-INS-68 |
| chr4-70627200-INS-87 | chr4-70627200-INS-87 |
| chr4-70638289-INS-128 | chr4-70638289-INS-145 |
| chr4-70659212-INS-507 | chr4-70659212-INS-507 |
| chr4-70680368-INS-225 | chr4-70680368-INS-225 |
| chr4-70714391-INS-83 | chr4-70714391-INS-83 |
| chr4-70715541-INS-2438 | chr4-70715542-INS-2439 |
| chr4-70723951-INS-6427 | chr4-70723952-INS-6427 |
| chr4-70725984-INS-427 | chr4-70725984-INS-427 |
| chr4-70736237-INS-1494 | chr4-70736238-INS-1494 |
| chr4-70741416-INS-78 | chr4-70741416-INS-82 |
| chr4-70779213-INS-73 | chr4-70779213-INS-73 |
| chr4-70798314-INS-150 | chr4-70798314-INS-150 |
| chr4-70811318-INS-1100 | chr4-70811319-INS-1100 |
| chr4-70820452-INS-610 | chr4-70820452-INS-610 |
| chr4-70827542-INS-69 | chr4-70827542-INS-69 |
| chr4-70903941-INS-450 | chr4-70903941-INS-450 |
| chr4-70906183-INS-1494 | chr4-70906184-INS-1494 |
| chr4-70917172-INS-66 | chr4-70917172-INS-66 |
| chr4-70930213-INS-1039 | chr4-70930222-INS-1028 |
| chr4-70943888-INS-576 | chr4-70943895-INS-577 |
| chr4-70951896-INS-64 | chr4-70951896-INS-60 |
| chr4-70963532-INS-6937 | chr4-70963533-INS-6938 |
| chr4-70984450-INS-292 | chr4-70984448-INS-304 |
| chr4-71030798-INS-111 | chr4-71030770-INS-122 |
| chr4-71033503-INS-131 | chr4-71033503-INS-131 |
| chr4-71044340-INS-56 | chr4-71044340-INS-56 |

| | |
|---|---|
| chr4-71047162-INS-51 | chr4-71047147-INS-64 |
| chr4-71068215-INS-153 | chr4-71068217-INS-154 |
| chr4-71113512-INS-219 | chr4-71113512-INS-218 |
| chr4-71121842-INS-68 | chr4-71121844-INS-50 |
| chr4-71124599-INS-6318 | chr4-71124600-INS-6319 |
| chr4-71139865-INS-78 | chr4-71139865-INS-78 |
| chr4-71149953-INS-64 | chr4-71149953-INS-64 |
| chr4-71153777-INS-204 | chr4-71153792-INS-204 |
| chr4-71163639-INS-88 | chr4-71163639-INS-94 |
| chr4-71189358-INS-51 | chr4-71189352-INS-97 |
| chr4-71197302-INS-172 | chr4-71197298-INS-176 |
| chr4-71211632-INS-159 | chr4-71211632-INS-159 |
| chr4-71230111-INS-224 | chr4-71230110-INS-228 |
| chr4-71281463-INS-66 | chr4-71281463-INS-93 |
| chr4-71293548-INS-61 | chr4-71293548-INS-61 |
| chr4-71309533-INS-432 | chr4-71309534-INS-432 |
| chr4-71336547-INS-64 | chr4-71336547-INS-64 |
| chr4-71352564-INS-428 | chr4-71352564-INS-428 |
| chr4-71360795-INS-71 | chr4-71360795-INS-73 |
| chr4-71382332-INS-72 | chr4-71382332-INS-72 |
| chr4-71390106-INS-90 | chr4-71390134-INS-95 |
| chr4-71395261-INS-114 | chr4-71395261-INS-115 |
| chr4-71400577-INS-56 | chr4-71400536-INS-84 |
| chr4-71453529-INS-52 | chr4-71453561-INS-52 |
| chr4-71465371-INS-2280 | chr4-71465372-INS-2280 |
| chr4-71467045-INS-6019 | chr4-71467046-INS-6019 |
| chr4-71496170-INS-170 | chr4-71496164-INS-168 |
| chr4-71499434-INS-1411 | chr4-71499435-INS-1412 |
| chr4-71512134-INS-467 | chr4-71512134-INS-467 |
| chr4-71527478-INS-2627 | chr4-71527479-INS-2627 |
| chr4-71533415-INS-153 | chr4-71533415-INS-153 |
| chr4-71536609-INS-75 | chr4-71536609-INS-75 |
| chr4-71551754-INS-55 | chr4-71551754-INS-55 |
| chr4-71560749-INS-112 | chr4-71560749-INS-112 |
| chr4-71563151-INS-6757 | chr4-71563152-INS-6757 |
| chr4-71570008-INS-6390 | chr4-71570009-INS-6390 |
| chr4-71581701-INS-6372 | chr4-71581702-INS-6372 |
| chr4-71591965-INS-8176 | chr4-71591966-INS-8176 |
| chr4-71608814-INS-850 | chr4-71608814-INS-850 |
| chr4-71631451-INS-388 | chr4-71631451-INS-393 |
| chr4-71638719-INS-215 | chr4-71638719-INS-215 |
| chr4-71640563-INS-228 | chr4-71640563-INS-228 |
| chr4-71649236-INS-420 | chr4-71649236-INS-420 |
| chr4-71656178-INS-92 | chr4-71656138-INS-103 |
| chr4-71687419-INS-62 | chr4-71687419-INS-62 |
| chr4-71687774-INS-3856 | chr4-71687775-INS-3856 |
| chr4-71697546-INS-71 | chr4-71697546-INS-71 |
| chr4-71741039-INS-127 | chr4-71741039-INS-127 |
| chr4-71774927-INS-889 | chr4-71774927-INS-888 |
| chr4-71778555-INS-400 | chr4-71778555-INS-400 |
| chr4-71786735-INS-60 | chr4-71786735-INS-62 |
| chr4-71796639-INS-263 | chr4-71796639-INS-263 |
| chr4-71857610-INS-54 | chr4-71857592-INS-62 |
| chr4-71873548-INS-114 | chr4-71873548-INS-114 |
| chr4-71952548-INS-55 | chr4-71952548-INS-52 |
| chr4-71969028-INS-176 | chr4-71969028-INS-186 |
| chr4-71972727-INS-183 | chr4-71972727-INS-183 |
| chr4-71989680-INS-293 | chr4-71989680-INS-306 |
| chr4-71998530-INS-212 | chr4-71998530-INS-212 |
| chr4-72002907-INS-496 | chr4-72002907-INS-496 |
| chr4-72023822-INS-76 | chr4-72023822-INS-76 |
| chr4-72037078-INS-58 | chr4-72037078-INS-56 |
| chr4-72044227-INS-1125 | chr4-72044228-INS-1125 |
| chr4-72066096-INS-82 | chr4-72066096-INS-82 |
| chr4-72074443-INS-5231 | chr4-72074444-INS-5232 |
| chr4-72074758-INS-56 | chr4-72074758-INS-64 |
| chr4-72126743-INS-667 | chr4-72126743-INS-667 |
| chr4-72145617-INS-3142 | chr4-72145618-INS-3140 |
| chr4-72152359-INS-53 | chr4-72152359-INS-54 |
| chr4-72156998-INS-134 | chr4-72156998-INS-134 |
| chr4-72159060-INS-3325 | chr4-72159061-INS-3325 |
| chr4-72166860-INS-70 | chr4-72166860-INS-60 |
| chr4-72181575-INS-228 | chr4-72181575-INS-229 |
| chr4-72182918-INS-1894 | chr4-72182919-INS-1894 |
| chr4-72196421-INS-53 | chr4-72196421-INS-53 |
| chr4-72210712-INS-57 | chr4-72210712-INS-57 |
| chr4-72264875-INS-62 | chr4-72264875-INS-52 |

| | |
|---|---|
| chr4-72283272-INS-71 | chr4-72283268-INS-72 |
| chr4-72305554-INS-339 | chr4-72305554-INS-339 |
| chr4-72339803-INS-91 | chr4-72339803-INS-94 |
| chr4-72380046-INS-354 | chr4-72380046-INS-354 |
| chr4-72384025-INS-60 | chr4-72384026-INS-54 |
| chr4-72393913-INS-8341 | chr4-72393914-INS-8342 |
| chr4-72452898-INS-407 | chr4-72452898-INS-407 |
| chr4-72472393-INS-184 | chr4-72472393-INS-184 |
| chr4-72493963-INS-358 | chr4-72493963-INS-358 |
| chr4-72502044-INS-631 | chr4-72502044-INS-632 |
| chr4-72518578-INS-136 | chr4-72518578-INS-136 |
| chr4-72530919-INS-52 | chr4-72530919-INS-52 |
| chr4-72539487-INS-281 | chr4-72539487-INS-281 |
| chr4-72543770-INS-172 | chr4-72543770-INS-172 |
| chr4-72562120-INS-1865 | chr4-72562121-INS-1865 |
| chr4-72569265-INS-51 | chr4-72569265-INS-58 |
| chr4-72574756-INS-1811 | chr4-72574757-INS-1808 |
| chr4-72577460-INS-3976 | chr4-72577461-INS-3977 |
| chr4-72581940-INS-6300 | chr4-72581941-INS-6298 |
| chr4-72659874-INS-62 | chr4-72659874-INS-69 |
| chr4-72663210-INS-800 | chr4-72663210-INS-801 |
| chr4-72670260-INS-106 | chr4-72670260-INS-106 |
| chr4-72679035-INS-2504 | chr4-72679036-INS-2505 |
| chr4-72699124-INS-2680 | chr4-72699125-INS-2680 |
| chr4-72712798-INS-52 | chr4-72712798-INS-52 |
| chr4-72717229-INS-658 | chr4-72717229-INS-658 |
| chr4-72721477-INS-62 | chr4-72721477-INS-62 |
| chr4-72723543-INS-6238 | chr4-72723544-INS-6240 |
| chr4-72731519-INS-6407 | chr4-72731520-INS-6408 |
| chr4-72748194-INS-260 | chr4-72748194-INS-288 |
| chr4-72764595-INS-2385 | chr4-72764596-INS-2392 |
| chr4-72772389-INS-194 | chr4-72772389-INS-186 |
| chr4-72778382-INS-3603 | chr4-72778383-INS-3618 |
| chr4-72794888-INS-77 | chr4-72794888-INS-78 |
| chr4-72800495-INS-211 | chr4-72800495-INS-211 |
| chr4-72809416-INS-72 | chr4-72809416-INS-72 |
| chr4-72818983-INS-57 | chr4-72818983-INS-65 |
| chr4-72822004-INS-408 | chr4-72822004-INS-407 |
| chr4-72831884-INS-371 | chr4-72831884-INS-371 |
| chr4-72839967-INS-158 | chr4-72839999-INS-154 |
| chr4-72842311-INS-98 | chr4-72842312-INS-81 |
| chr4-72842830-INS-129 | chr4-72842827-INS-122 |
| chr4-72856277-INS-56 | chr4-72856277-INS-55 |
| chr4-72857109-INS-504 | chr4-72857109-INS-504 |
| chr4-72908702-INS-4509 | chr4-72908703-INS-4509 |
| chr4-72923559-INS-9149 | chr4-72923559-INS-9149 |
| chr4-72946975-INS-6155 | chr4-72946976-INS-6155 |
| chr4-72949461-INS-55 | chr4-72949455-INS-68 |
| chr4-72954731-INS-54 | chr4-72954731-INS-52 |
| chr4-72964983-INS-2277 | chr4-72964984-INS-2277 |
| chr4-72971790-INS-58 | chr4-72971791-INS-58 |
| chr4-72978057-INS-146 | chr4-72978057-INS-146 |
| chr4-73014963-INS-6292 | chr4-73014964-INS-6292 |
| chr4-73034144-INS-6084 | chr4-73034145-INS-6085 |
| chr4-73054332-INS-101 | chr4-73054332-INS-101 |
| chr4-73060729-INS-402 | chr4-73060729-INS-405 |
| chr4-73111043-INS-3756 | chr4-73111044-INS-3756 |
| chr4-73150525-INS-300 | chr4-73150525-INS-298 |
| chr4-73172369-INS-573 | chr4-73172369-INS-578 |
| chr4-73225906-INS-201 | chr4-73225909-INS-201 |
| chr4-73259719-INS-390 | chr4-73259719-INS-390 |
| chr4-73269556-INS-306 | chr4-73269556-INS-306 |
| chr4-73327494-INS-3611 | chr4-73327495-INS-3611 |
| chr4-73342309-INS-490 | chr4-73342309-INS-490 |
| chr4-73396603-INS-85 | chr4-73396605-INS-61 |
| chr4-73403940-INS-257 | chr4-73403941-INS-256 |
| chr4-73452302-INS-5214 | chr4-73452303-INS-5214 |
| chr4-73475937-INS-102 | chr4-73475952-INS-76 |
| chr4-73487262-INS-56 | chr4-73487262-INS-56 |
| chr4-73673675-INS-275 | chr4-73673675-INS-275 |
| chr4-73682222-INS-409 | chr4-73682222-INS-413 |
| chr4-73694918-INS-84 | chr4-73694918-INS-83 |
| chr4-73713964-INS-4743 | chr4-73713965-INS-4743 |
| chr4-73725650-INS-406 | chr4-73725650-INS-406 |
| chr4-73758326-INS-853 | chr4-73758326-INS-853 |
| chr4-73904303-INS-144 | chr4-73904303-INS-144 |
| chr4-73921670-INS-1627 | chr4-73921671-INS-1627 |

| | |
|---|---|
| chr4-73921948-INS-83 | chr4-73921948-INS-85 |
| chr4-73933226-INS-3537 | chr4-73933227-INS-3536 |
| chr4-73937001-INS-59 | chr4-73936980-INS-55 |
| chr4-73940650-INS-393 | chr4-73940653-INS-394 |
| chr4-73942648-INS-6251 | chr4-73942649-INS-6251 |
| chr4-73945270-INS-506 | chr4-73945270-INS-506 |
| chr4-73957964-INS-54 | chr4-73957964-INS-54 |
| chr4-73965599-INS-792 | chr4-73965599-INS-792 |
| chr4-73971255-INS-69 | chr4-73971255-INS-69 |
| chr4-74002708-INS-88 | chr4-74002708-INS-88 |
| chr4-74005003-INS-383 | chr4-74005003-INS-383 |
| chr4-74006928-INS-1320 | chr4-74006929-INS-1321 |
| chr4-74006929-INS-160 | chr4-74006929-INS-160 |
| chr4-74008842-INS-498 | chr4-74008842-INS-498 |
| chr4-74014785-INS-13597 | chr4-74014786-INS-13596 |
| chr4-74017109-INS-1165 | chr4-74017110-INS-1165 |
| chr4-74025423-INS-60 | chr4-74025429-INS-56 |
| chr4-74034604-INS-438 | chr4-74034604-INS-438 |
| chr4-74037045-INS-74 | chr4-74037045-INS-74 |
| chr4-74048422-INS-512 | chr4-74048422-INS-512 |
| chr4-74061515-INS-225 | chr4-74061515-INS-225 |
| chr4-74062739-INS-549 | chr4-74062739-INS-549 |
| chr4-74063662-INS-265 | chr4-74063662-INS-265 |
| chr4-74063711-INS-834 | chr4-74063711-INS-836 |
| chr4-74073914-INS-234 | chr4-74073914-INS-234 |
| chr4-74075364-INS-717 | chr4-74075364-INS-717 |
| chr4-74084328-INS-142 | chr4-74084328-INS-142 |
| chr4-74088098-INS-204 | chr4-74088098-INS-204 |
| chr4-74094080-INS-512 | chr4-74094080-INS-514 |
| chr4-74109136-INS-57 | chr4-74109144-INS-65 |
| chr4-74114869-INS-71 | chr4-74114842-INS-54 |
| chr4-74118806-INS-71 | chr4-74118807-INS-69 |
| chr4-74121646-INS-231 | chr4-74121646-INS-228 |
| chr4-74124509-INS-201 | chr4-74124509-INS-200 |
| chr4-74209255-INS-187 | chr4-74209255-INS-140 |
| chr4-74221204-INS-255 | chr4-74221204-INS-255 |
| chr4-74262739-INS-212 | chr4-74262739-INS-212 |
| chr4-74300643-INS-69 | chr4-74300643-INS-67 |
| chr4-74324835-INS-60 | chr4-74324835-INS-62 |
| chr4-74363710-INS-191 | chr4-74363709-INS-184 |
| chr4-74384755-INS-732 | chr4-74384755-INS-732 |
| chr4-74413932-INS-1154 | chr4-74413933-INS-1154 |
| chr4-74424899-INS-65 | chr4-74424899-INS-65 |
| chr4-74436900-INS-435 | chr4-74436900-INS-434 |
| chr4-74473821-INS-737 | chr4-74473821-INS-737 |
| chr4-74486347-INS-164 | chr4-74486347-INS-168 |
| chr4-74537094-INS-66 | chr4-74537094-INS-68 |
| chr4-74540410-INS-51 | chr4-74540410-INS-79 |
| chr4-74542356-INS-121 | chr4-74542357-INS-97 |
| chr4-74552436-INS-175 | chr4-74552431-INS-180 |
| chr4-74555101-INS-103 | chr4-74555103-INS-102 |
| chr4-74586585-INS-225 | chr4-74586585-INS-225 |
| chr4-74589443-INS-7784 | chr4-74589444-INS-7785 |
| chr4-74590664-INS-52 | chr4-74590664-INS-54 |
| chr4-74655716-INS-3710 | chr4-74655717-INS-3710 |
| chr4-74660987-INS-54 | chr4-74660987-INS-52 |
| chr4-74661604-INS-2195 | chr4-74661605-INS-2195 |
| chr4-74663216-INS-72 | chr4-74663216-INS-60 |
| chr4-74668369-INS-2016 | chr4-74668370-INS-2016 |
| chr4-74679449-INS-1821 | chr4-74679450-INS-1821 |
| chr4-74724102-INS-512 | chr4-74724102-INS-511 |
| chr4-74726994-INS-523 | chr4-74726994-INS-523 |
| chr4-74742425-INS-60 | chr4-74742425-INS-75 |
| chr4-74764458-INS-1459 | chr4-74764459-INS-1459 |
| chr4-74777169-INS-143 | chr4-74777169-INS-102 |
| chr4-74782814-INS-214 | chr4-74782814-INS-214 |
| chr4-74794613-INS-6857 | chr4-74794614-INS-6858 |
| chr4-74810446-INS-54 | chr4-74810446-INS-54 |
| chr4-74815131-INS-54 | chr4-74815131-INS-56 |
| chr4-74818638-INS-172 | chr4-74818638-INS-180 |
| chr4-74840576-INS-52 | chr4-74840576-INS-52 |
| chr4-74869490-INS-59 | chr4-74869490-INS-59 |
| chr4-74880473-INS-115 | chr4-74880473-INS-115 |
| chr4-74918618-INS-3154 | chr4-74918619-INS-3154 |
| chr4-75071074-INS-6403 | chr4-75071075-INS-6411 |
| chr4-75074508-INS-4851 | chr4-75074509-INS-4851 |
| chr4-75120451-INS-399 | chr4-75120451-INS-399 |

| | |
|---|---|
| chr4-75201096-INS-4196 | chr4-75201097-INS-4195 |
| chr4-75218053-INS-434 | chr4-75218053-INS-434 |
| chr4-75219738-INS-66 | chr4-75219738-INS-60 |
| chr4-75264238-INS-148 | chr4-75264238-INS-148 |
| chr4-75274261-INS-161 | chr4-75274298-INS-133 |
| chr4-75295552-INS-515 | chr4-75295552-INS-515 |
| chr4-75318241-INS-58 | chr4-75318241-INS-58 |
| chr4-75325778-INS-2245 | chr4-75325779-INS-2245 |
| chr4-75329726-INS-84 | chr4-75329726-INS-54 |
| chr4-75340412-INS-7887 | chr4-75340413-INS-7889 |
| chr4-75361566-INS-64 | chr4-75361566-INS-69 |
| chr4-75369213-INS-66 | chr4-75369219-INS-50 |
| chr4-75382587-INS-559 | chr4-75382587-INS-558 |
| chr4-75386874-INS-180 | chr4-75386875-INS-179 |
| chr4-75401305-INS-75 | chr4-75401305-INS-78 |
| chr4-75410672-INS-200 | chr4-75410672-INS-200 |
| chr4-75430868-INS-498 | chr4-75430868-INS-498 |
| chr4-75435026-INS-844 | chr4-75435026-INS-844 |
| chr4-75451172-INS-549 | chr4-75451169-INS-550 |
| chr4-75454534-INS-57 | chr4-75454534-INS-62 |
| chr4-75457765-INS-99 | chr4-75457739-INS-78 |
| chr4-75463091-INS-380 | chr4-75463074-INS-410 |
| chr4-75481909-INS-70 | chr4-75481909-INS-70 |
| chr4-75485306-INS-124 | chr4-75485306-INS-116 |
| chr4-75487257-INS-58 | chr4-75487257-INS-60 |
| chr4-75512846-INS-436 | chr4-75512846-INS-436 |
| chr4-75518303-INS-442 | chr4-75518303-INS-442 |
| chr4-75533378-INS-2446 | chr4-75533379-INS-2440 |
| chr4-75559456-INS-1172 | chr4-75559457-INS-1172 |
| chr4-75562094-INS-146 | chr4-75562094-INS-119 |
| chr4-75576015-INS-823 | chr4-75576015-INS-823 |
| chr4-75586804-INS-2365 | chr4-75586805-INS-2365 |
| chr4-75597126-INS-6650 | chr4-75597127-INS-6654 |
| chr4-75664220-INS-2628 | chr4-75664221-INS-2629 |
| chr4-75679182-INS-60 | chr4-75679182-INS-60 |
| chr4-75694649-INS-6648 | chr4-75694650-INS-6647 |
| chr4-75713898-INS-106 | chr4-75713898-INS-82 |
| chr4-75716997-INS-260 | chr4-75716997-INS-261 |
| chr4-75725549-INS-211 | chr4-75725578-INS-165 |
| chr4-75735402-INS-89 | chr4-75735402-INS-81 |
| chr4-75740151-INS-114 | chr4-75740151-INS-114 |
| chr4-75744502-INS-51 | chr4-75744507-INS-59 |
| chr4-75781142-INS-60 | chr4-75781170-INS-64 |
| chr4-75791645-INS-89 | chr4-75791645-INS-89 |
| chr4-75811314-INS-255 | chr4-75811314-INS-255 |
| chr4-75825774-INS-6705 | chr4-75825775-INS-6706 |
| chr4-75837699-INS-297 | chr4-75837699-INS-297 |
| chr4-75875918-INS-56 | chr4-75875918-INS-56 |
| chr4-75899636-INS-1742 | chr4-75899637-INS-1745 |
| chr4-75915830-INS-64 | chr4-75915830-INS-66 |
| chr4-75917060-INS-246 | chr4-75917060-INS-246 |
| chr4-76024164-INS-66 | chr4-76024164-INS-72 |
| chr4-76030157-INS-206 | chr4-76030157-INS-206 |
| chr4-76032515-INS-7539 | chr4-76032516-INS-7541 |
| chr4-76034945-INS-145 | chr4-76034946-INS-145 |
| chr4-76037489-INS-53 | chr4-76037489-INS-53 |
| chr4-76042015-INS-445 | chr4-76042015-INS-445 |
| chr4-76072258-INS-6502 | chr4-76072259-INS-6504 |
| chr4-76077756-INS-52 | chr4-76077773-INS-60 |
| chr4-76085150-INS-7309 | chr4-76085151-INS-7309 |
| chr4-76091388-INS-183 | chr4-76091354-INS-180 |
| chr4-76118974-INS-83 | chr4-76118974-INS-83 |
| chr4-76125037-INS-54 | chr4-76125037-INS-50 |
| chr4-76143461-INS-178 | chr4-76143461-INS-178 |
| chr4-76154429-INS-84 | chr4-76154429-INS-84 |
| chr4-76163421-INS-54 | chr4-76163421-INS-54 |
| chr4-76173868-INS-361 | chr4-76173868-INS-361 |
| chr4-76211775-INS-199 | chr4-76211775-INS-199 |
| chr4-76223824-INS-59 | chr4-76223824-INS-59 |
| chr4-76272390-INS-1022 | chr4-76272391-INS-1022 |
| chr4-76283341-INS-6758 | chr4-76283342-INS-6758 |
| chr4-76289413-INS-137 | chr4-76289413-INS-151 |
| chr4-76308913-INS-147 | chr4-76308913-INS-147 |
| chr4-76343768-INS-78 | chr4-76343768-INS-78 |
| chr4-76378806-INS-58 | chr4-76378805-INS-67 |
| chr4-76398329-INS-64 | chr4-76398331-INS-52 |
| chr4-76412333-INS-6715 | chr4-76412318-INS-6718 |

| | |
|---|---|
| chr4-76429089-INS-54 | chr4-76429090-INS-54 |
| chr4-76449593-INS-64 | chr4-76449593-INS-64 |
| chr4-76463944-INS-770 | chr4-76463944-INS-771 |
| chr4-76466491-INS-62 | chr4-76466491-INS-56 |
| chr4-76484566-INS-250 | chr4-76484566-INS-250 |
| chr4-76494893-INS-991 | chr4-76494893-INS-991 |
| chr4-76514373-INS-60 | chr4-76514349-INS-58 |
| chr4-76534619-INS-90 | chr4-76534619-INS-100 |
| chr4-76573862-INS-237 | chr4-76573862-INS-241 |
| chr4-76617317-INS-79 | chr4-76617317-INS-72 |
| chr4-76659593-INS-97 | chr4-76659593-INS-97 |
| chr4-76667458-INS-386 | chr4-76667458-INS-386 |
| chr4-76674583-INS-58 | chr4-76674583-INS-58 |
| chr4-76719369-INS-67 | chr4-76719368-INS-74 |
| chr4-76722220-INS-205 | chr4-76722220-INS-205 |
| chr4-76745620-INS-152 | chr4-76745620-INS-142 |
| chr4-76763606-INS-214 | chr4-76763606-INS-214 |
| chr4-76774135-INS-305 | chr4-76774135-INS-299 |
| chr4-76775360-INS-994 | chr4-76775360-INS-994 |
| chr4-76791683-INS-52 | chr4-76791694-INS-80 |
| chr4-76794494-INS-90 | chr4-76794494-INS-124 |
| chr4-76801344-INS-1597 | chr4-76801345-INS-1597 |
| chr4-76805441-INS-103 | chr4-76805441-INS-103 |
| chr4-76840774-INS-721 | chr4-76840774-INS-721 |
| chr4-76842639-INS-52 | chr4-76842639-INS-52 |
| chr4-76844377-INS-344 | chr4-76844377-INS-344 |
| chr4-76863792-INS-55 | chr4-76863792-INS-59 |
| chr4-76875944-INS-403 | chr4-76875944-INS-403 |
| chr4-76898384-INS-110 | chr4-76898384-INS-117 |
| chr4-76905143-INS-405 | chr4-76905143-INS-405 |
| chr4-76942097-INS-154 | chr4-76942097-INS-162 |
| chr4-76943591-INS-60 | chr4-76943594-INS-56 |
| chr4-76953701-INS-89 | chr4-76953701-INS-89 |
| chr4-76959956-INS-6926 | chr4-76959957-INS-6929 |
| chr4-76960131-INS-72 | chr4-76960131-INS-72 |
| chr4-76979139-INS-62 | chr4-76979139-INS-62 |
| chr4-76990196-INS-1195 | chr4-76990197-INS-1195 |
| chr4-77001288-INS-68 | chr4-77001282-INS-78 |
| chr4-77016369-INS-381 | chr4-77016369-INS-381 |
| chr4-77018462-INS-138 | chr4-77018462-INS-138 |
| chr4-77020049-INS-84 | chr4-77020049-INS-91 |
| chr4-77087539-INS-64 | chr4-77087543-INS-78 |
| chr4-77097451-INS-54 | chr4-77097451-INS-72 |
| chr4-77126769-INS-56 | chr4-77126769-INS-56 |
| chr4-77129335-INS-358 | chr4-77129335-INS-334 |
| chr4-77133764-INS-3145 | chr4-77133765-INS-3143 |
| chr4-77138591-INS-61 | chr4-77138591-INS-61 |
| chr4-77153117-INS-432 | chr4-77153117-INS-432 |
| chr4-77162483-INS-542 | chr4-77162483-INS-542 |
| chr4-77190974-INS-180 | chr4-77190952-INS-181 |
| chr4-77201367-INS-328 | chr4-77201367-INS-328 |
| chr4-77212281-INS-51 | chr4-77212281-INS-51 |
| chr4-77213786-INS-6883 | chr4-77213787-INS-6885 |
| chr4-77221928-INS-943 | chr4-77221928-INS-985 |
| chr4-77241219-INS-55 | chr4-77241219-INS-50 |
| chr4-77281236-INS-624 | chr4-77281236-INS-625 |
| chr4-77302902-INS-91 | chr4-77302920-INS-106 |
| chr4-77323040-INS-3094 | chr4-77323041-INS-3086 |
| chr4-77330245-INS-272 | chr4-77330245-INS-272 |
| chr4-77335637-INS-1603 | chr4-77335638-INS-1603 |
| chr4-77346162-INS-984 | chr4-77346162-INS-985 |
| chr4-77348753-INS-498 | chr4-77348753-INS-498 |
| chr4-77387528-INS-188 | chr4-77387528-INS-168 |
| chr4-77402360-INS-53 | chr4-77402360-INS-61 |
| chr4-77411756-INS-584 | chr4-77411756-INS-583 |
| chr4-77456714-INS-1763 | chr4-77456715-INS-1763 |
| chr4-77476470-INS-53 | chr4-77476470-INS-53 |
| chr4-77480735-INS-194 | chr4-77480735-INS-194 |
| chr4-77532312-INS-54 | chr4-77532314-INS-50 |
| chr4-77534330-INS-68 | chr4-77534330-INS-68 |
| chr4-77538209-INS-1583 | chr4-77538210-INS-1583 |
| chr4-77540800-INS-419 | chr4-77540801-INS-418 |
| chr4-77543683-INS-77 | chr4-77543683-INS-77 |
| chr4-77549220-INS-6535 | chr4-77549221-INS-6535 |
| chr4-77575698-INS-396 | chr4-77575698-INS-396 |
| chr4-77603781-INS-6359 | chr4-77603782-INS-6359 |
| chr4-77610603-INS-267 | chr4-77610603-INS-267 |

| | |
|---|---|
| chr4-77628103-INS-54 | chr4-77628103-INS-56 |
| chr4-77644063-INS-59 | chr4-77644063-INS-69 |
| chr4-77695717-INS-833 | chr4-77695717-INS-833 |
| chr4-77703961-INS-4068 | chr4-77703962-INS-4068 |
| chr4-77706687-INS-64 | chr4-77706687-INS-64 |
| chr4-77721455-INS-56 | chr4-77721455-INS-56 |
| chr4-77730162-INS-80 | chr4-77730162-INS-80 |
| chr4-77733421-INS-120 | chr4-77733421-INS-120 |
| chr4-77743863-INS-146 | chr4-77743886-INS-152 |
| chr4-77745036-INS-466 | chr4-77745036-INS-466 |
| chr4-77763814-INS-999 | chr4-77763814-INS-999 |
| chr4-77764533-INS-52 | chr4-77764537-INS-52 |
| chr4-77787054-INS-78 | chr4-77787054-INS-86 |
| chr4-77799608-INS-59 | chr4-77799629-INS-63 |
| chr4-77852580-INS-145 | chr4-77852580-INS-145 |
| chr4-77860232-INS-58 | chr4-77860232-INS-62 |
| chr4-77888400-INS-149 | chr4-77888400-INS-149 |
| chr4-77907993-INS-195 | chr4-77907993-INS-195 |
| chr4-77920834-INS-80 | chr4-77920834-INS-82 |
| chr4-77928336-INS-116 | chr4-77928336-INS-85 |
| chr4-77932528-INS-201 | chr4-77932528-INS-201 |
| chr4-77944280-INS-557 | chr4-77944280-INS-557 |
| chr4-77952927-INS-148 | chr4-77952927-INS-128 |
| chr4-77969921-INS-61 | chr4-77969921-INS-61 |
| chr4-78034123-INS-56 | chr4-78034123-INS-70 |
| chr4-78086213-INS-202 | chr4-78086213-INS-202 |
| chr4-78095786-INS-82 | chr4-78095829-INS-54 |
| chr4-78099867-INS-252 | chr4-78099867-INS-243 |
| chr4-78102061-INS-168 | chr4-78102061-INS-168 |
| chr4-78114186-INS-56 | chr4-78114186-INS-56 |
| chr4-78116087-INS-88 | chr4-78116087-INS-86 |
| chr4-78134235-INS-183 | chr4-78134235-INS-185 |
| chr4-78141179-INS-186 | chr4-78141179-INS-182 |
| chr4-78181137-INS-79 | chr4-78181137-INS-79 |
| chr4-78202116-INS-740 | chr4-78202116-INS-740 |
| chr4-78228165-INS-6955 | chr4-78228166-INS-6955 |
| chr4-78259775-INS-197 | chr4-78259775-INS-198 |
| chr4-78320811-INS-1886 | chr4-78320812-INS-1886 |
| chr4-78345773-INS-327 | chr4-78345773-INS-327 |
| chr4-78347982-INS-68 | chr4-78347954-INS-94 |
| chr4-78354429-INS-561 | chr4-78354462-INS-563 |
| chr4-78362456-INS-52 | chr4-78362456-INS-52 |
| chr4-78378252-INS-118 | chr4-78378252-INS-118 |
| chr4-78394879-INS-64 | chr4-78394879-INS-60 |
| chr4-78444541-INS-154 | chr4-78444541-INS-156 |
| chr4-78480529-INS-123 | chr4-78480531-INS-123 |
| chr4-78480573-INS-3059 | chr4-78480574-INS-3049 |
| chr4-78490971-INS-4378 | chr4-78490972-INS-4379 |
| chr4-78492066-INS-69 | chr4-78492048-INS-101 |
| chr4-78506792-INS-74 | chr4-78506792-INS-98 |
| chr4-78524304-INS-566 | chr4-78524305-INS-566 |
| chr4-78536049-INS-88 | chr4-78536055-INS-76 |
| chr4-78537965-INS-64 | chr4-78537965-INS-56 |
| chr4-78671042-INS-104 | chr4-78671045-INS-105 |
| chr4-78680235-INS-183 | chr4-78680235-INS-192 |
| chr4-78687659-INS-75 | chr4-78687659-INS-75 |
| chr4-78701330-INS-514 | chr4-78701330-INS-514 |
| chr4-78707553-INS-65 | chr4-78707553-INS-65 |
| chr4-78714165-INS-116 | chr4-78714165-INS-116 |
| chr4-78739755-INS-131 | chr4-78739755-INS-131 |
| chr4-78793489-INS-284 | chr4-78793489-INS-284 |
| chr4-78797306-INS-96 | chr4-78797306-INS-96 |
| chr4-78800641-INS-440 | chr4-78800642-INS-439 |
| chr4-78833996-INS-52 | chr4-78833996-INS-52 |
| chr4-78870658-INS-206 | chr4-78870654-INS-210 |
| chr4-78895649-INS-6643 | chr4-78895650-INS-6644 |
| chr4-78953937-INS-140 | chr4-78953937-INS-140 |
| chr4-78996149-INS-434 | chr4-78996149-INS-435 |
| chr4-79046104-INS-12764 | chr4-79046105-INS-12763 |
| chr4-79094874-INS-6584 | chr4-79094875-INS-6584 |
| chr4-79153177-INS-6462 | chr4-79153178-INS-6459 |
| chr4-79153821-INS-665 | chr4-79153819-INS-675 |
| chr4-79197530-INS-350 | chr4-79197530-INS-350 |
| chr4-79206332-INS-601 | chr4-79206332-INS-599 |
| chr4-79245404-INS-52 | chr4-79245404-INS-52 |
| chr4-79279782-INS-385 | chr4-79279782-INS-385 |
| chr4-79281648-INS-124 | chr4-79281648-INS-124 |

| | |
|---|---|
| chr4-79293542-INS-121 | chr4-79293559-INS-146 |
| chr4-79301856-INS-505 | chr4-79301856-INS-505 |
| chr4-79314726-INS-6512 | chr4-79314727-INS-6516 |
| chr4-79330168-INS-6270 | chr4-79330169-INS-6271 |
| chr4-79350605-INS-75 | chr4-79350605-INS-75 |
| chr4-79395631-INS-2144 | chr4-79395632-INS-2145 |
| chr4-79398888-INS-183 | chr4-79398888-INS-183 |
| chr4-79408927-INS-197 | chr4-79408927-INS-197 |
| chr4-79424145-INS-70 | chr4-79424145-INS-74 |
| chr4-79541957-INS-128 | chr4-79541957-INS-120 |
| chr4-79597103-INS-60 | chr4-79597103-INS-57 |
| chr4-79603862-INS-52 | chr4-79603862-INS-54 |
| chr4-79637956-INS-66 | chr4-79637956-INS-66 |
| chr4-79644092-INS-591 | chr4-79644092-INS-592 |
| chr4-79648684-INS-81 | chr4-79648684-INS-81 |
| chr4-79655594-INS-76 | chr4-79655594-INS-76 |
| chr4-79724818-INS-280 | chr4-79724815-INS-285 |
| chr4-79739930-INS-487 | chr4-79739930-INS-501 |
| chr4-79765212-INS-384 | chr4-79765212-INS-384 |
| chr4-79806457-INS-558 | chr4-79806457-INS-558 |
| chr4-79834284-INS-4168 | chr4-79834285-INS-4168 |
| chr4-79842943-INS-5550 | chr4-79842944-INS-5550 |
| chr4-79880043-INS-475 | chr4-79880043-INS-475 |
| chr4-79882420-INS-118 | chr4-79882421-INS-114 |
| chr4-79890012-INS-233 | chr4-79890012-INS-241 |
| chr4-79907566-INS-781 | chr4-79907567-INS-781 |
| chr4-79926785-INS-176 | chr4-79926785-INS-160 |
| chr4-79950053-INS-128 | chr4-79950053-INS-104 |
| chr4-79968369-INS-159 | chr4-79968363-INS-158 |
| chr4-79989561-INS-406 | chr4-79989561-INS-414 |
| chr4-80028601-INS-6277 | chr4-80028602-INS-6278 |
| chr4-80076819-INS-52 | chr4-80076819-INS-52 |
| chr4-80093708-INS-55 | chr4-80093708-INS-70 |
| chr4-80096621-INS-150 | chr4-80096624-INS-148 |
| chr4-80106852-INS-80 | chr4-80106852-INS-82 |
| chr4-80110987-INS-353 | chr4-80110993-INS-366 |
| chr4-80189891-INS-116 | chr4-80189890-INS-113 |
| chr4-80213611-INS-66 | chr4-80213611-INS-66 |
| chr4-80236833-INS-303 | chr4-80236833-INS-303 |
| chr4-80271304-INS-158 | chr4-80271304-INS-158 |
| chr4-80271616-INS-2278 | chr4-80271617-INS-2281 |
| chr4-80272635-INS-2620 | chr4-80272652-INS-2622 |
| chr4-80293138-INS-3938 | chr4-80293139-INS-3933 |
| chr4-80303490-INS-510 | chr4-80303490-INS-517 |
| chr4-80312989-INS-6147 | chr4-80312990-INS-6146 |
| chr4-80325871-INS-401 | chr4-80325871-INS-401 |
| chr4-80333541-INS-71 | chr4-80333541-INS-71 |
| chr4-80356711-INS-64 | chr4-80356712-INS-66 |
| chr4-80363771-INS-677 | chr4-80363771-INS-677 |
| chr4-80366420-INS-831 | chr4-80366420-INS-831 |
| chr4-80371019-INS-401 | chr4-80371019-INS-401 |
| chr4-80421666-INS-1167 | chr4-80421667-INS-1166 |
| chr4-80452687-INS-58 | chr4-80452683-INS-64 |
| chr4-80458074-INS-6581 | chr4-80458075-INS-6581 |
| chr4-80461912-INS-6470 | chr4-80461913-INS-6470 |
| chr4-80472455-INS-1001 | chr4-80472456-INS-999 |
| chr4-80491575-INS-66 | chr4-80491575-INS-54 |
| chr4-80494415-INS-1490 | chr4-80494416-INS-1487 |
| chr4-80527740-INS-1340 | chr4-80527741-INS-1341 |
| chr4-80551226-INS-63 | chr4-80551226-INS-82 |
| chr4-80572643-INS-88 | chr4-80572643-INS-88 |
| chr4-80580890-INS-317 | chr4-80580890-INS-317 |
| chr4-80591814-INS-110 | chr4-80591814-INS-110 |
| chr4-80616096-INS-6640 | chr4-80616097-INS-6643 |
| chr4-80675035-INS-84 | chr4-80675035-INS-84 |
| chr4-80705415-INS-293 | chr4-80705410-INS-293 |
| chr4-80778070-INS-96 | chr4-80778038-INS-105 |
| chr4-80788922-INS-86 | chr4-80788922-INS-86 |
| chr4-80790856-INS-123 | chr4-80790856-INS-123 |
| chr4-80793883-INS-2185 | chr4-80793884-INS-2185 |
| chr4-80797583-INS-81 | chr4-80797583-INS-56 |
| chr4-80839540-INS-72 | chr4-80839545-INS-56 |
| chr4-80853004-INS-119 | chr4-80853014-INS-90 |
| chr4-80861962-INS-218 | chr4-80861962-INS-218 |
| chr4-80869710-INS-78 | chr4-80869710-INS-86 |
| chr4-80878619-INS-7428 | chr4-80878620-INS-7427 |
| chr4-80923287-INS-65 | chr4-80923287-INS-65 |

| | |
|---|---|
| chr4-80942877-INS-54 | chr4-80942877-INS-54 |
| chr4-80946431-INS-653 | chr4-80946431-INS-657 |
| chr4-80949620-INS-637 | chr4-80949620-INS-637 |
| chr4-80950632-INS-394 | chr4-80950632-INS-394 |
| chr4-80954477-INS-309 | chr4-80954480-INS-315 |
| chr4-81029977-INS-6482 | chr4-81029978-INS-6484 |
| chr4-81057024-INS-390 | chr4-81057021-INS-403 |
| chr4-81101176-INS-60 | chr4-81101149-INS-78 |
| chr4-81109506-INS-108 | chr4-81109506-INS-80 |
| chr4-81114831-INS-635 | chr4-81114831-INS-629 |
| chr4-81118452-INS-2653 | chr4-81118453-INS-2653 |
| chr4-81136813-INS-75 | chr4-81136813-INS-78 |
| chr4-81169995-INS-72 | chr4-81169995-INS-76 |
| chr4-81186047-INS-66 | chr4-81186047-INS-51 |
| chr4-81193241-INS-200 | chr4-81193221-INS-186 |
| chr4-81204584-INS-58 | chr4-81204584-INS-58 |
| chr4-81253536-INS-191 | chr4-81253536-INS-195 |
| chr4-81263439-INS-912 | chr4-81263439-INS-913 |
| chr4-81312477-INS-541 | chr4-81312477-INS-541 |
| chr4-81356111-INS-337 | chr4-81356111-INS-337 |
| chr4-81437670-INS-114 | chr4-81437670-INS-126 |
| chr4-81450352-INS-55 | chr4-81450352-INS-55 |
| chr4-81481571-INS-78 | chr4-81481571-INS-80 |
| chr4-81501549-INS-77 | chr4-81501549-INS-70 |
| chr4-81531085-INS-358 | chr4-81531085-INS-358 |
| chr4-81559519-INS-261 | chr4-81559519-INS-261 |
| chr4-81591369-INS-6512 | chr4-81591370-INS-6513 |
| chr4-81604981-INS-118 | chr4-81604981-INS-117 |
| chr4-81628739-INS-157 | chr4-81628739-INS-170 |
| chr4-81661641-INS-99 | chr4-81661628-INS-65 |
| chr4-81707487-INS-62 | chr4-81707487-INS-62 |
| chr4-81731441-INS-6339 | chr4-81731442-INS-6338 |
| chr4-81762688-INS-581 | chr4-81762688-INS-581 |
| chr4-81786016-INS-63 | chr4-81786016-INS-63 |
| chr4-81846631-INS-74 | chr4-81846631-INS-74 |
| chr4-81889592-INS-126 | chr4-81889592-INS-126 |
| chr4-81891640-INS-91 | chr4-81891640-INS-96 |
| chr4-81899460-INS-115 | chr4-81899460-INS-115 |
| chr4-81902120-INS-6673 | chr4-81902121-INS-6673 |
| chr4-81905247-INS-72 | chr4-81905247-INS-72 |
| chr4-81922609-INS-272 | chr4-81922609-INS-272 |
| chr4-81942465-INS-72 | chr4-81942464-INS-75 |
| chr4-81950672-INS-622 | chr4-81950672-INS-619 |
| chr4-81995588-INS-6492 | chr4-81995589-INS-6493 |
| chr4-82003131-INS-3129 | chr4-82003132-INS-3129 |
| chr4-82003134-INS-211 | chr4-82003134-INS-210 |
| chr4-82018056-INS-56 | chr4-82018056-INS-54 |
| chr4-82019573-INS-90 | chr4-82019572-INS-77 |
| chr4-82022302-INS-419 | chr4-82022303-INS-419 |
| chr4-82028062-INS-332 | chr4-82028062-INS-328 |
| chr4-82045788-INS-400 | chr4-82045788-INS-400 |
| chr4-82061580-INS-70 | chr4-82061580-INS-70 |
| chr4-82172467-INS-58 | chr4-82172458-INS-91 |
| chr4-82179817-INS-4189 | chr4-82179818-INS-4189 |
| chr4-82188214-INS-205 | chr4-82188214-INS-206 |
| chr4-82256264-INS-66 | chr4-82256259-INS-76 |
| chr4-82326593-INS-78 | chr4-82326593-INS-83 |
| chr4-82333141-INS-205 | chr4-82333141-INS-205 |
| chr4-82342121-INS-261 | chr4-82342121-INS-261 |
| chr4-82391600-INS-60 | chr4-82391620-INS-60 |
| chr4-82440585-INS-52 | chr4-82440585-INS-54 |
| chr4-82452045-INS-57 | chr4-82452046-INS-60 |
| chr4-82463075-INS-383 | chr4-82463075-INS-383 |
| chr4-82477366-INS-52 | chr4-82477391-INS-68 |
| chr4-82480841-INS-76 | chr4-82480841-INS-80 |
| chr4-82635311-INS-55 | chr4-82635311-INS-55 |
| chr4-82642324-INS-586 | chr4-82642324-INS-585 |
| chr4-82643477-INS-804 | chr4-82643477-INS-793 |
| chr4-82654426-INS-164 | chr4-82654426-INS-160 |
| chr4-82687306-INS-1390 | chr4-82687307-INS-1390 |
| chr4-82733725-INS-489 | chr4-82733725-INS-489 |
| chr4-82749887-INS-189 | chr4-82749887-INS-189 |
| chr4-82755038-INS-75 | chr4-82755038-INS-75 |
| chr4-82803770-INS-88 | chr4-82803768-INS-100 |
| chr4-82806514-INS-59 | chr4-82806529-INS-74 |
| chr4-82846324-INS-3135 | chr4-82846325-INS-3135 |
| chr4-82851728-INS-110 | chr4-82851753-INS-127 |

| | |
|---|---|
| chr4-82874196-INS-56 | chr4-82874196-INS-64 |
| chr4-82895221-INS-76 | chr4-82895222-INS-116 |
| chr4-82937770-INS-68 | chr4-82937770-INS-70 |
| chr4-82938412-INS-7188 | chr4-82938413-INS-7188 |
| chr4-82990644-INS-6442 | chr4-82990645-INS-6442 |
| chr4-83014877-INS-652 | chr4-83014877-INS-652 |
| chr4-83020948-INS-76 | chr4-83020948-INS-78 |
| chr4-83025928-INS-65 | chr4-83025928-INS-60 |
| chr4-83029522-INS-90 | chr4-83029522-INS-90 |
| chr4-83035386-INS-71 | chr4-83035386-INS-71 |
| chr4-83036162-INS-486 | chr4-83036162-INS-510 |
| chr4-83062490-INS-75 | chr4-83062497-INS-72 |
| chr4-83069114-INS-68 | chr4-83069096-INS-72 |
| chr4-83070873-INS-76 | chr4-83070867-INS-88 |
| chr4-83083716-INS-257 | chr4-83083716-INS-257 |
| chr4-83086089-INS-56 | chr4-83086089-INS-56 |
| chr4-83093060-INS-60 | chr4-83093060-INS-62 |
| chr4-83099655-INS-144 | chr4-83099655-INS-144 |
| chr4-83104606-INS-245 | chr4-83104606-INS-246 |
| chr4-83108027-INS-400 | chr4-83108027-INS-400 |
| chr4-83112998-INS-225 | chr4-83112998-INS-225 |
| chr4-83154673-INS-128 | chr4-83154673-INS-128 |
| chr4-83174133-INS-279 | chr4-83174134-INS-269 |
| chr4-83198099-INS-323 | chr4-83198099-INS-321 |
| chr4-83209533-INS-229 | chr4-83209533-INS-232 |
| chr4-83212386-INS-6416 | chr4-83212387-INS-6416 |
| chr4-83230005-INS-208 | chr4-83230005-INS-208 |
| chr4-83237061-INS-327 | chr4-83237073-INS-302 |
| chr4-83239336-INS-206 | chr4-83239336-INS-202 |
| chr4-83247598-INS-196 | chr4-83247598-INS-223 |
| chr4-83266022-INS-56 | chr4-83266022-INS-60 |
| chr4-83276841-INS-206 | chr4-83276841-INS-204 |
| chr4-83296032-INS-217 | chr4-83296032-INS-218 |
| chr4-83300464-INS-221 | chr4-83300464-INS-221 |
| chr4-83302665-INS-1515 | chr4-83302666-INS-1518 |
| chr4-83304107-INS-175 | chr4-83304107-INS-175 |
| chr4-83307984-INS-7091 | chr4-83307985-INS-7091 |
| chr4-83308421-INS-1778 | chr4-83308422-INS-1779 |
| chr4-83310043-INS-191 | chr4-83310043-INS-186 |
| chr4-83310852-INS-64 | chr4-83310852-INS-64 |
| chr4-83316407-INS-230 | chr4-83316407-INS-200 |
| chr4-83317307-INS-127 | chr4-83317307-INS-126 |
| chr4-83322898-INS-78 | chr4-83322908-INS-98 |
| chr4-83375795-INS-220 | chr4-83375795-INS-220 |
| chr4-83385457-INS-121 | chr4-83385457-INS-121 |
| chr4-83408670-INS-57 | chr4-83408671-INS-75 |
| chr4-83427686-INS-562 | chr4-83427686-INS-558 |
| chr4-83441398-INS-51 | chr4-83441398-INS-54 |
| chr4-83449777-INS-188 | chr4-83449777-INS-188 |
| chr4-83467205-INS-201 | chr4-83467205-INS-201 |
| chr4-83483150-INS-463 | chr4-83483150-INS-463 |
| chr4-83486049-INS-58 | chr4-83486049-INS-60 |
| chr4-83492430-INS-71 | chr4-83492425-INS-67 |
| chr4-83497717-INS-90 | chr4-83497717-INS-90 |
| chr4-83538241-INS-1141 | chr4-83538242-INS-1140 |
| chr4-83538743-INS-79 | chr4-83538743-INS-117 |
| chr4-83539890-INS-178 | chr4-83539890-INS-178 |
| chr4-83541492-INS-72 | chr4-83541486-INS-108 |
| chr4-83569517-INS-194 | chr4-83569517-INS-192 |
| chr4-83578908-INS-414 | chr4-83578908-INS-414 |
| chr4-83582725-INS-169 | chr4-83582729-INS-146 |
| chr4-83600978-INS-73 | chr4-83600978-INS-75 |
| chr4-83607745-INS-160 | chr4-83607745-INS-160 |
| chr4-83643320-INS-238 | chr4-83643320-INS-238 |
| chr4-83682620-INS-145 | chr4-83682620-INS-145 |
| chr4-83690977-INS-123 | chr4-83690945-INS-128 |
| chr4-83702496-INS-1239 | chr4-83702497-INS-1238 |
| chr4-83726439-INS-6612 | chr4-83726440-INS-6613 |
| chr4-83727425-INS-132 | chr4-83727425-INS-132 |
| chr4-83742070-INS-102 | chr4-83742070-INS-102 |
| chr4-83756261-INS-227 | chr4-83756261-INS-241 |
| chr4-83758403-INS-80 | chr4-83758403-INS-56 |
| chr4-83760377-INS-240 | chr4-83760377-INS-240 |
| chr4-83768417-INS-56 | chr4-83768417-INS-56 |
| chr4-83779773-INS-323 | chr4-83779773-INS-323 |
| chr4-83791847-INS-445 | chr4-83791847-INS-445 |
| chr4-83818304-INS-162 | chr4-83818294-INS-198 |

| | |
|---|---|
| chr4-83824726-INS-95 | chr4-83824726-INS-93 |
| chr4-83827597-INS-3810 | chr4-83827598-INS-3810 |
| chr4-83830263-INS-6378 | chr4-83830264-INS-6378 |
| chr4-83887721-INS-63 | chr4-83887721-INS-78 |
| chr4-83910401-INS-64 | chr4-83910401-INS-64 |
| chr4-83916898-INS-119 | chr4-83916898-INS-119 |
| chr4-83918443-INS-1642 | chr4-83918444-INS-1642 |
| chr4-83977242-INS-226 | chr4-83977242-INS-221 |
| chr4-83979130-INS-234 | chr4-83979130-INS-236 |
| chr4-83995608-INS-7752 | chr4-83995609-INS-7753 |
| chr4-84007503-INS-64 | chr4-84007504-INS-86 |
| chr4-84012258-INS-60 | chr4-84012298-INS-68 |
| chr4-84029611-INS-93 | chr4-84029611-INS-93 |
| chr4-84060092-INS-178 | chr4-84060092-INS-177 |
| chr4-84099662-INS-54 | chr4-84099661-INS-54 |
| chr4-84154967-INS-237 | chr4-84154967-INS-237 |
| chr4-84160956-INS-196 | chr4-84160956-INS-227 |
| chr4-84165309-INS-6697 | chr4-84165296-INS-6694 |
| chr4-84170802-INS-77 | chr4-84170802-INS-77 |
| chr4-84263244-INS-56 | chr4-84263244-INS-56 |
| chr4-84317562-INS-128 | chr4-84317562-INS-128 |
| chr4-84333748-INS-74 | chr4-84333748-INS-58 |
| chr4-84407855-INS-216 | chr4-84407855-INS-183 |
| chr4-84412037-INS-61 | chr4-84412038-INS-97 |
| chr4-84436227-INS-56 | chr4-84436227-INS-58 |
| chr4-84444831-INS-81 | chr4-84444831-INS-79 |
| chr4-84492901-INS-399 | chr4-84492901-INS-399 |
| chr4-84493902-INS-407 | chr4-84493902-INS-407 |
| chr4-84499056-INS-94 | chr4-84499056-INS-94 |
| chr4-84524812-INS-194 | chr4-84524812-INS-194 |
| chr4-84539000-INS-60 | chr4-84539001-INS-52 |
| chr4-84554649-INS-198 | chr4-84554648-INS-201 |
| chr4-84567167-INS-54 | chr4-84567208-INS-58 |
| chr4-84569211-INS-86 | chr4-84569211-INS-86 |
| chr4-84601929-INS-5552 | chr4-84601930-INS-5552 |
| chr4-84627884-INS-67 | chr4-84627888-INS-68 |
| chr4-84649039-INS-208 | chr4-84649039-INS-207 |
| chr4-84672730-INS-6363 | chr4-84672731-INS-6364 |
| chr4-84730374-INS-79 | chr4-84730374-INS-83 |
| chr4-84746265-INS-299 | chr4-84746265-INS-300 |
| chr4-84747246-INS-115 | chr4-84747243-INS-133 |
| chr4-84776168-INS-56 | chr4-84776168-INS-58 |
| chr4-84784220-INS-56 | chr4-84784220-INS-58 |
| chr4-84786209-INS-1208 | chr4-84786210-INS-1209 |
| chr4-84788252-INS-178 | chr4-84788252-INS-177 |
| chr4-84817362-INS-1608 | chr4-84817363-INS-1608 |
| chr4-84838531-INS-3111 | chr4-84838532-INS-3123 |
| chr4-84847499-INS-341 | chr4-84847499-INS-341 |
| chr4-84857810-INS-1140 | chr4-84857811-INS-1139 |
| chr4-84872464-INS-65 | chr4-84872464-INS-65 |
| chr4-84877802-INS-104 | chr4-84877802-INS-104 |
| chr4-84890387-INS-54 | chr4-84890387-INS-71 |
| chr4-84901595-INS-776 | chr4-84901595-INS-776 |
| chr4-84913335-INS-72 | chr4-84913327-INS-64 |
| chr4-84920794-INS-52 | chr4-84920794-INS-55 |
| chr4-84941747-INS-6336 | chr4-84941748-INS-6337 |
| chr4-84949885-INS-231 | chr4-84949885-INS-231 |
| chr4-84953014-INS-712 | chr4-84953014-INS-713 |
| chr4-84953609-INS-201 | chr4-84953609-INS-201 |
| chr4-84968966-INS-6437 | chr4-84968967-INS-6436 |
| chr4-84971092-INS-127 | chr4-84971092-INS-127 |
| chr4-85000364-INS-213 | chr4-85000364-INS-213 |
| chr4-85051953-INS-328 | chr4-85051953-INS-328 |
| chr4-85060154-INS-126 | chr4-85060154-INS-126 |
| chr4-85111023-INS-1047 | chr4-85111024-INS-1046 |
| chr4-85122195-INS-696 | chr4-85122195-INS-696 |
| chr4-85129638-INS-196 | chr4-85129638-INS-198 |
| chr4-85130827-INS-6707 | chr4-85130828-INS-6709 |
| chr4-85135402-INS-172 | chr4-85135402-INS-172 |
| chr4-85138364-INS-293 | chr4-85138364-INS-297 |
| chr4-85145571-INS-52 | chr4-85145571-INS-52 |
| chr4-85161451-INS-209 | chr4-85161451-INS-204 |
| chr4-85170233-INS-56 | chr4-85170233-INS-102 |
| chr4-85191804-INS-839 | chr4-85191804-INS-839 |
| chr4-85196700-INS-51 | chr4-85196700-INS-51 |
| chr4-85205369-INS-54 | chr4-85205370-INS-68 |
| chr4-85219086-INS-61 | chr4-85219086-INS-61 |

| | |
|---|---|
| chr4-85227455-INS-6275 | chr4-85227456-INS-6276 |
| chr4-85245873-INS-396 | chr4-85245873-INS-396 |
| chr4-85253369-INS-73 | chr4-85253369-INS-73 |
| chr4-85266628-INS-8325 | chr4-85266629-INS-8325 |
| chr4-85267208-INS-71 | chr4-85267208-INS-71 |
| chr4-85286064-INS-57 | chr4-85286021-INS-50 |
| chr4-85288224-INS-583 | chr4-85288225-INS-583 |
| chr4-85303342-INS-66 | chr4-85303353-INS-56 |
| chr4-85314744-INS-243 | chr4-85314744-INS-243 |
| chr4-85316101-INS-265 | chr4-85316099-INS-265 |
| chr4-85336784-INS-193 | chr4-85336784-INS-195 |
| chr4-85344335-INS-64 | chr4-85344343-INS-50 |
| chr4-85360590-INS-6159 | chr4-85360591-INS-6159 |
| chr4-85363532-INS-553 | chr4-85363532-INS-554 |
| chr4-85368661-INS-64 | chr4-85368661-INS-54 |
| chr4-85428224-INS-60 | chr4-85428224-INS-60 |
| chr4-85439947-INS-51 | chr4-85439947-INS-51 |
| chr4-85479000-INS-72 | chr4-85479000-INS-76 |
| chr4-85499726-INS-56 | chr4-85499726-INS-56 |
| chr4-85521012-INS-78 | chr4-85521012-INS-78 |
| chr4-85524192-INS-51 | chr4-85524193-INS-87 |
| chr4-85583799-INS-6466 | chr4-85583800-INS-6466 |
| chr4-85633189-INS-136 | chr4-85633165-INS-125 |
| chr4-85636530-INS-253 | chr4-85636528-INS-259 |
| chr4-85650587-INS-175 | chr4-85650587-INS-175 |
| chr4-85680901-INS-823 | chr4-85680901-INS-823 |
| chr4-85680952-INS-6879 | chr4-85680953-INS-6881 |
| chr4-85686949-INS-52 | chr4-85686949-INS-54 |
| chr4-85701492-INS-52 | chr4-85701492-INS-60 |
| chr4-85706699-INS-2280 | chr4-85706700-INS-2280 |
| chr4-85707479-INS-60 | chr4-85707479-INS-54 |
| chr4-85709179-INS-621 | chr4-85709179-INS-621 |
| chr4-85722942-INS-138 | chr4-85722942-INS-145 |
| chr4-85729216-INS-83 | chr4-85729216-INS-83 |
| chr4-85733077-INS-114 | chr4-85733077-INS-114 |
| chr4-85758444-INS-69 | chr4-85758444-INS-70 |
| chr4-85769594-INS-114 | chr4-85769594-INS-112 |
| chr4-85810096-INS-214 | chr4-85810096-INS-214 |
| chr4-85826830-INS-96 | chr4-85826830-INS-102 |
| chr4-85837168-INS-131 | chr4-85837168-INS-131 |
| chr4-85873554-INS-64 | chr4-85873554-INS-68 |
| chr4-85891282-INS-100 | chr4-85891282-INS-108 |
| chr4-85898887-INS-56 | chr4-85898885-INS-68 |
| chr4-85912450-INS-898 | chr4-85912450-INS-898 |
| chr4-85956299-INS-164 | chr4-85956299-INS-164 |
| chr4-85958414-INS-6657 | chr4-85958415-INS-6656 |
| chr4-86023740-INS-7718 | chr4-86023741-INS-7718 |
| chr4-86051789-INS-159 | chr4-86051789-INS-168 |
| chr4-86055033-INS-4323 | chr4-86055034-INS-4323 |
| chr4-86056345-INS-62 | chr4-86056345-INS-56 |
| chr4-86071660-INS-66 | chr4-86071660-INS-81 |
| chr4-86073582-INS-178 | chr4-86073582-INS-178 |
| chr4-86084025-INS-2150 | chr4-86084026-INS-2150 |
| chr4-86097982-INS-294 | chr4-86097983-INS-294 |
| chr4-86113978-INS-54 | chr4-86113981-INS-55 |
| chr4-86132806-INS-75 | chr4-86132806-INS-75 |
| chr4-86136444-INS-82 | chr4-86136444-INS-82 |
| chr4-86145160-INS-6577 | chr4-86145161-INS-6577 |
| chr4-86147545-INS-114 | chr4-86147545-INS-144 |
| chr4-86180369-INS-357 | chr4-86180369-INS-359 |
| chr4-86187963-INS-110 | chr4-86187989-INS-72 |
| chr4-86235877-INS-554 | chr4-86235877-INS-554 |
| chr4-86270479-INS-495 | chr4-86270479-INS-499 |
| chr4-86295673-INS-66 | chr4-86295673-INS-66 |
| chr4-86299116-INS-78 | chr4-86299116-INS-84 |
| chr4-86318163-INS-61 | chr4-86318206-INS-52 |
| chr4-86354124-INS-642 | chr4-86354124-INS-608 |
| chr4-86396748-INS-281 | chr4-86396748-INS-281 |
| chr4-86398299-INS-57 | chr4-86398299-INS-57 |
| chr4-86413629-INS-6074 | chr4-86413630-INS-6075 |
| chr4-86423675-INS-3640 | chr4-86423676-INS-3638 |
| chr4-86457611-INS-76 | chr4-86457611-INS-80 |
| chr4-86459558-INS-61 | chr4-86459558-INS-61 |
| chr4-86463010-INS-500 | chr4-86463010-INS-501 |
| chr4-86467386-INS-399 | chr4-86467386-INS-399 |
| chr4-86507597-INS-88 | chr4-86507597-INS-89 |
| chr4-86548159-INS-484 | chr4-86548159-INS-484 |

| | |
|---|---|
| chr4-86558518-INS-53 | chr4-86558518-INS-53 |
| chr4-86584884-INS-256 | chr4-86584884-INS-256 |
| chr4-86590521-INS-6457 | chr4-86590522-INS-6457 |
| chr4-86628023-INS-1667 | chr4-86628024-INS-1650 |
| chr4-86629737-INS-6403 | chr4-86629738-INS-6403 |
| chr4-86642154-INS-59 | chr4-86642154-INS-54 |
| chr4-86647634-INS-519 | chr4-86647634-INS-519 |
| chr4-86653936-INS-64 | chr4-86653933-INS-69 |
| chr4-86654879-INS-129 | chr4-86654879-INS-133 |
| chr4-86657247-INS-52 | chr4-86657247-INS-52 |
| chr4-86662265-INS-215 | chr4-86662265-INS-215 |
| chr4-86683136-INS-204 | chr4-86683136-INS-203 |
| chr4-86693609-INS-58 | chr4-86693609-INS-58 |
| chr4-86700432-INS-217 | chr4-86700411-INS-209 |
| chr4-86701640-INS-1498 | chr4-86701641-INS-1499 |
| chr4-86702179-INS-2833 | chr4-86702180-INS-2833 |
| chr4-86713608-INS-211 | chr4-86713608-INS-212 |
| chr4-86724133-INS-216 | chr4-86724133-INS-215 |
| chr4-86748931-INS-85 | chr4-86748931-INS-85 |
| chr4-86749885-INS-228 | chr4-86749885-INS-228 |
| chr4-86750964-INS-334 | chr4-86750964-INS-334 |
| chr4-86761664-INS-587 | chr4-86761664-INS-588 |
| chr4-86762688-INS-202 | chr4-86762688-INS-203 |
| chr4-86771166-INS-58 | chr4-86771166-INS-65 |
| chr4-86801504-INS-606 | chr4-86801504-INS-606 |
| chr4-86809177-INS-186 | chr4-86809177-INS-186 |
| chr4-86810350-INS-65 | chr4-86810350-INS-65 |
| chr4-86826501-INS-200 | chr4-86826501-INS-200 |
| chr4-86828322-INS-206 | chr4-86828322-INS-206 |
| chr4-86831503-INS-453 | chr4-86831503-INS-453 |
| chr4-86832883-INS-74 | chr4-86832883-INS-74 |
| chr4-86839434-INS-138 | chr4-86839434-INS-142 |
| chr4-86853784-INS-79 | chr4-86853784-INS-71 |
| chr4-86856879-INS-774 | chr4-86856852-INS-818 |
| chr4-86858136-INS-161 | chr4-86858136-INS-189 |
| chr4-86860295-INS-509 | chr4-86860295-INS-509 |
| chr4-86864968-INS-530 | chr4-86864968-INS-530 |
| chr4-86880451-INS-985 | chr4-86880451-INS-985 |
| chr4-86910576-INS-84 | chr4-86910584-INS-80 |
| chr4-86932806-INS-203 | chr4-86932806-INS-203 |
| chr4-86976467-INS-65 | chr4-86976467-INS-65 |
| chr4-86982559-INS-105 | chr4-86982544-INS-117 |
| chr4-86989670-INS-80 | chr4-86989670-INS-82 |
| chr4-87008846-INS-79 | chr4-87008846-INS-79 |
| chr4-87017501-INS-6682 | chr4-87017502-INS-6684 |
| chr4-87024451-INS-6722 | chr4-87024452-INS-6722 |
| chr4-87036007-INS-6379 | chr4-87036008-INS-6379 |
| chr4-87059137-INS-7411 | chr4-87059138-INS-7411 |
| chr4-87062711-INS-4862 | chr4-87062712-INS-4862 |
| chr4-87083366-INS-6522 | chr4-87083367-INS-6524 |
| chr4-87087302-INS-766 | chr4-87087302-INS-766 |
| chr4-87087827-INS-109 | chr4-87087808-INS-109 |
| chr4-87090985-INS-56 | chr4-87090985-INS-62 |
| chr4-87135317-INS-3110 | chr4-87135318-INS-3111 |
| chr4-87156102-INS-142 | chr4-87156102-INS-142 |
| chr4-87157426-INS-6675 | chr4-87157427-INS-6675 |
| chr4-87161095-INS-73 | chr4-87161095-INS-73 |
| chr4-87179103-INS-7250 | chr4-87179104-INS-7249 |
| chr4-87180043-INS-1272 | chr4-87180044-INS-1272 |
| chr4-87182824-INS-77 | chr4-87182824-INS-77 |
| chr4-87201495-INS-1283 | chr4-87201496-INS-1285 |
| chr4-87213592-INS-73 | chr4-87213576-INS-78 |
| chr4-87221051-INS-3389 | chr4-87221052-INS-3389 |
| chr4-87252724-INS-6241 | chr4-87252725-INS-6241 |
| chr4-87253304-INS-760 | chr4-87253304-INS-758 |
| chr4-87256177-INS-118 | chr4-87256177-INS-88 |
| chr4-87267785-INS-576 | chr4-87267785-INS-576 |
| chr4-87277551-INS-475 | chr4-87277551-INS-477 |
| chr4-87292108-INS-6238 | chr4-87292109-INS-6239 |
| chr4-87307131-INS-246 | chr4-87307126-INS-264 |
| chr4-87321595-INS-688 | chr4-87321595-INS-688 |
| chr4-87325471-INS-73 | chr4-87325471-INS-89 |
| chr4-87328039-INS-60 | chr4-87328039-INS-56 |
| chr4-87336971-INS-506 | chr4-87336971-INS-506 |
| chr4-87355772-INS-875 | chr4-87355772-INS-876 |
| chr4-87356413-INS-198 | chr4-87356413-INS-198 |
| chr4-87362262-INS-62 | chr4-87362262-INS-62 |

| | |
|---|---|
| chr4-87371715-INS-56 | chr4-87371715-INS-66 |
| chr4-87381354-INS-52 | chr4-87381354-INS-52 |
| chr4-87391365-INS-547 | chr4-87391365-INS-543 |
| chr4-87400025-INS-150 | chr4-87400025-INS-150 |
| chr4-87419848-INS-64 | chr4-87419872-INS-73 |
| chr4-87428137-INS-96 | chr4-87428137-INS-78 |
| chr4-87439143-INS-73 | chr4-87439143-INS-81 |
| chr4-87453140-INS-52 | chr4-87453140-INS-52 |
| chr4-87454472-INS-6310 | chr4-87454473-INS-6310 |
| chr4-87455028-INS-527 | chr4-87455028-INS-528 |
| chr4-87547723-INS-368 | chr4-87547721-INS-356 |
| chr4-87550649-INS-69 | chr4-87550647-INS-70 |
| chr4-87567222-INS-59 | chr4-87567222-INS-62 |
| chr4-87584584-INS-989 | chr4-87584584-INS-989 |
| chr4-87592156-INS-93 | chr4-87592189-INS-87 |
| chr4-87638698-INS-2928 | chr4-87638699-INS-2943 |
| chr4-87674661-INS-51 | chr4-87674662-INS-51 |
| chr4-87682538-INS-51 | chr4-87682538-INS-51 |
| chr4-87714402-INS-6474 | chr4-87714403-INS-6474 |
| chr4-87754070-INS-155 | chr4-87754070-INS-155 |
| chr4-87803942-INS-57 | chr4-87803943-INS-69 |
| chr4-87834670-INS-190 | chr4-87834676-INS-201 |
| chr4-87866417-INS-980 | chr4-87866417-INS-980 |
| chr4-87871420-INS-202 | chr4-87871420-INS-202 |
| chr4-87887112-INS-54 | chr4-87887106-INS-68 |
| chr4-87898815-INS-80 | chr4-87898815-INS-87 |
| chr4-87909957-INS-70 | chr4-87909967-INS-66 |
| chr4-87996584-INS-209 | chr4-87996584-INS-209 |
| chr4-88022237-INS-69 | chr4-88022237-INS-81 |
| chr4-88030771-INS-74 | chr4-88030771-INS-74 |
| chr4-88033936-INS-7115 | chr4-88033937-INS-7115 |
| chr4-88035447-INS-122 | chr4-88035447-INS-122 |
| chr4-88064662-INS-205 | chr4-88064662-INS-205 |
| chr4-88083962-INS-52 | chr4-88083958-INS-56 |
| chr4-88095363-INS-216 | chr4-88095363-INS-217 |
| chr4-88099885-INS-6218 | chr4-88099886-INS-6218 |
| chr4-88122645-INS-202 | chr4-88122645-INS-202 |
| chr4-88149558-INS-2149 | chr4-88149559-INS-2149 |
| chr4-88177705-INS-54 | chr4-88177747-INS-56 |
| chr4-88179855-INS-165 | chr4-88179855-INS-155 |
| chr4-88242817-INS-54 | chr4-88242817-INS-54 |
| chr4-88245667-INS-6632 | chr4-88245668-INS-6634 |
| chr4-88263122-INS-66 | chr4-88263122-INS-66 |
| chr4-88363969-INS-1340 | chr4-88363970-INS-1340 |
| chr4-88366197-INS-80 | chr4-88366197-INS-80 |
| chr4-88382102-INS-130 | chr4-88382100-INS-130 |
| chr4-88392632-INS-59 | chr4-88392649-INS-54 |
| chr4-88451503-INS-127 | chr4-88451503-INS-127 |
| chr4-88465663-INS-1547 | chr4-88465664-INS-1549 |
| chr4-88477287-INS-62 | chr4-88477287-INS-60 |
| chr4-88483196-INS-208 | chr4-88483196-INS-208 |
| chr4-88491019-INS-67 | chr4-88491019-INS-97 |
| chr4-88543137-INS-52 | chr4-88543138-INS-50 |
| chr4-88570996-INS-60 | chr4-88570996-INS-50 |
| chr4-88595836-INS-1043 | chr4-88595831-INS-1053 |
| chr4-88627085-INS-66 | chr4-88627085-INS-54 |
| chr4-88630200-INS-832 | chr4-88630200-INS-832 |
| chr4-88631592-INS-86 | chr4-88631592-INS-86 |
| chr4-88642730-INS-3232 | chr4-88642731-INS-3232 |
| chr4-88654675-INS-93 | chr4-88654675-INS-93 |
| chr4-88736225-INS-3282 | chr4-88736226-INS-3273 |
| chr4-88766557-INS-56 | chr4-88766555-INS-66 |
| chr4-88771700-INS-105 | chr4-88771703-INS-120 |
| chr4-88775868-INS-143 | chr4-88775868-INS-143 |
| chr4-88812851-INS-124 | chr4-88812857-INS-101 |
| chr4-88817359-INS-54 | chr4-88817359-INS-54 |
| chr4-88834328-INS-405 | chr4-88834328-INS-405 |
| chr4-88859691-INS-5269 | chr4-88859691-INS-5269 |
| chr4-88910707-INS-55 | chr4-88910707-INS-55 |
| chr4-88929346-INS-6653 | chr4-88929347-INS-6653 |
| chr4-88939296-INS-2492 | chr4-88939307-INS-2486 |
| chr4-88939476-INS-398 | chr4-88939476-INS-398 |
| chr4-88945400-INS-204 | chr4-88945400-INS-204 |
| chr4-88950473-INS-67 | chr4-88950473-INS-64 |
| chr4-88968766-INS-642 | chr4-88968766-INS-642 |
| chr4-88973361-INS-87 | chr4-88973361-INS-87 |
| chr4-88993074-INS-76 | chr4-88993074-INS-76 |

| | |
|---|---|
| chr4-88994526-INS-55 | chr4-88994526-INS-57 |
| chr4-89000481-INS-66 | chr4-89000481-INS-66 |
| chr4-89001478-INS-59 | chr4-89001478-INS-62 |
| chr4-89003681-INS-75 | chr4-89003681-INS-75 |
| chr4-89016040-INS-1078 | chr4-89016041-INS-1078 |
| chr4-89045366-INS-64 | chr4-89045366-INS-66 |
| chr4-89050006-INS-102 | chr4-89050006-INS-102 |
| chr4-89057623-INS-50 | chr4-89057623-INS-54 |
| chr4-89062150-INS-423 | chr4-89062127-INS-451 |
| chr4-89064502-INS-2677 | chr4-89064491-INS-2697 |
| chr4-89067697-INS-178 | chr4-89067697-INS-178 |
| chr4-89101987-INS-2788 | chr4-89101988-INS-2788 |
| chr4-89113503-INS-327 | chr4-89113503-INS-327 |
| chr4-89136735-INS-111 | chr4-89136735-INS-111 |
| chr4-89152514-INS-768 | chr4-89152514-INS-783 |
| chr4-89156196-INS-152 | chr4-89156196-INS-152 |
| chr4-89171552-INS-4581 | chr4-89171553-INS-4581 |
| chr4-89172964-INS-230 | chr4-89172964-INS-230 |
| chr4-89192054-INS-54 | chr4-89192054-INS-54 |
| chr4-89196871-INS-58 | chr4-89196871-INS-58 |
| chr4-89199375-INS-54 | chr4-89199375-INS-54 |
| chr4-89203313-INS-1353 | chr4-89203314-INS-1353 |
| chr4-89217219-INS-109 | chr4-89217219-INS-109 |
| chr4-89240710-INS-641 | chr4-89240710-INS-641 |
| chr4-89302768-INS-357 | chr4-89302768-INS-358 |
| chr4-89305725-INS-148 | chr4-89305725-INS-148 |
| chr4-89306805-INS-1347 | chr4-89306806-INS-1347 |
| chr4-89312805-INS-238 | chr4-89312805-INS-238 |
| chr4-89325458-INS-118 | chr4-89325458-INS-120 |
| chr4-89340471-INS-7335 | chr4-89340471-INS-7342 |
| chr4-89343915-INS-60 | chr4-89343915-INS-60 |
| chr4-89362775-INS-53 | chr4-89362776-INS-53 |
| chr4-89368079-INS-105 | chr4-89368079-INS-106 |
| chr4-89386498-INS-466 | chr4-89386498-INS-456 |
| chr4-89411131-INS-52 | chr4-89411131-INS-68 |
| chr4-89422585-INS-7059 | chr4-89422586-INS-7059 |
| chr4-89425582-INS-3378 | chr4-89425583-INS-3378 |
| chr4-89434036-INS-59 | chr4-89434032-INS-51 |
| chr4-89444900-INS-58 | chr4-89444900-INS-58 |
| chr4-89449880-INS-59 | chr4-89449880-INS-55 |
| chr4-89463172-INS-78 | chr4-89463172-INS-86 |
| chr4-89470346-INS-105 | chr4-89470346-INS-113 |
| chr4-89484220-INS-549 | chr4-89484220-INS-549 |
| chr4-89484673-INS-132 | chr4-89484673-INS-128 |
| chr4-89520229-INS-6405 | chr4-89520230-INS-6406 |
| chr4-89523418-INS-1435 | chr4-89523419-INS-1435 |
| chr4-89529338-INS-423 | chr4-89529338-INS-423 |
| chr4-89553001-INS-7079 | chr4-89553002-INS-7079 |
| chr4-89596864-INS-201 | chr4-89596864-INS-201 |
| chr4-89600265-INS-435 | chr4-89600265-INS-438 |
| chr4-89623432-INS-4137 | chr4-89623433-INS-4134 |
| chr4-89627759-INS-6287 | chr4-89627760-INS-6287 |
| chr4-89646932-INS-78 | chr4-89646932-INS-52 |
| chr4-89662934-INS-1148 | chr4-89662935-INS-1148 |
| chr4-89665125-INS-6277 | chr4-89665126-INS-6277 |
| chr4-89707782-INS-6193 | chr4-89707783-INS-6191 |
| chr4-89720958-INS-555 | chr4-89720958-INS-555 |
| chr4-89743506-INS-7092 | chr4-89743507-INS-7092 |
| chr4-89790013-INS-73 | chr4-89790060-INS-76 |
| chr4-89790674-INS-61 | chr4-89790674-INS-61 |
| chr4-89795644-INS-58 | chr4-89795644-INS-64 |
| chr4-89798154-INS-853 | chr4-89798154-INS-852 |
| chr4-89820880-INS-6831 | chr4-89820881-INS-6833 |
| chr4-89822437-INS-2336 | chr4-89822438-INS-2335 |
| chr4-89825827-INS-784 | chr4-89825827-INS-784 |
| chr4-89828878-INS-7120 | chr4-89828879-INS-7121 |
| chr4-89846026-INS-188 | chr4-89846026-INS-188 |
| chr4-89882066-INS-66 | chr4-89882066-INS-72 |
| chr4-89911139-INS-6471 | chr4-89911140-INS-6471 |
| chr4-89929275-INS-421 | chr4-89929275-INS-420 |
| chr4-89932792-INS-1088 | chr4-89932793-INS-1089 |
| chr4-89934553-INS-178 | chr4-89934553-INS-179 |
| chr4-89944995-INS-304 | chr4-89944995-INS-304 |
| chr4-89961358-INS-390 | chr4-89961349-INS-386 |
| chr4-89983676-INS-60 | chr4-89983676-INS-62 |
| chr4-89996180-INS-6646 | chr4-89996181-INS-6646 |
| chr4-90047976-INS-1002 | chr4-90048019-INS-1003 |

| | |
|---|---|
| chr4-90054091-INS-210 | chr4-90054091-INS-210 |
| chr4-90105827-INS-7065 | chr4-90105828-INS-7065 |
| chr4-90113937-INS-52 | chr4-90113937-INS-52 |
| chr4-90127706-INS-730 | chr4-90127706-INS-730 |
| chr4-90165417-INS-145 | chr4-90165417-INS-145 |
| chr4-90178572-INS-488 | chr4-90178572-INS-488 |
| chr4-90239875-INS-54 | chr4-90239875-INS-69 |
| chr4-90242599-INS-86 | chr4-90242599-INS-80 |
| chr4-90243729-INS-52 | chr4-90243729-INS-53 |
| chr4-90256068-INS-435 | chr4-90256068-INS-435 |
| chr4-90262503-INS-101 | chr4-90262503-INS-101 |
| chr4-90264446-INS-1036 | chr4-90264447-INS-1036 |
| chr4-90293288-INS-76 | chr4-90293288-INS-74 |
| chr4-90296745-INS-91 | chr4-90296745-INS-86 |
| chr4-90301329-INS-405 | chr4-90301329-INS-405 |
| chr4-90322575-INS-56 | chr4-90322575-INS-56 |
| chr4-90373842-INS-413 | chr4-90373842-INS-415 |
| chr4-90379286-INS-387 | chr4-90379286-INS-388 |
| chr4-90382599-INS-76 | chr4-90382599-INS-64 |
| chr4-90397425-INS-62 | chr4-90397393-INS-60 |
| chr4-90402387-INS-70 | chr4-90402387-INS-70 |
| chr4-90419267-INS-992 | chr4-90419268-INS-992 |
| chr4-90426922-INS-134 | chr4-90426922-INS-134 |
| chr4-90453468-INS-342 | chr4-90453468-INS-342 |
| chr4-90503219-INS-3689 | chr4-90503218-INS-3688 |
| chr4-90514757-INS-453 | chr4-90514757-INS-454 |
| chr4-90529275-INS-102 | chr4-90529275-INS-100 |
| chr4-90555432-INS-223 | chr4-90555432-INS-223 |
| chr4-90571941-INS-421 | chr4-90571941-INS-422 |
| chr4-90579391-INS-646 | chr4-90579397-INS-656 |
| chr4-90580358-INS-498 | chr4-90580358-INS-498 |
| chr4-90598889-INS-78 | chr4-90598889-INS-90 |
| chr4-90607336-INS-453 | chr4-90607338-INS-456 |
| chr4-90644543-INS-56 | chr4-90644543-INS-56 |
| chr4-90666802-INS-6401 | chr4-90666803-INS-6400 |
| chr4-90671548-INS-160 | chr4-90671544-INS-174 |
| chr4-90700245-INS-414 | chr4-90700245-INS-414 |
| chr4-90712522-INS-6365 | chr4-90712523-INS-6366 |
| chr4-90713327-INS-59 | chr4-90713327-INS-59 |
| chr4-90721276-INS-91 | chr4-90721276-INS-91 |
| chr4-90721487-INS-434 | chr4-90721485-INS-438 |
| chr4-90737604-INS-496 | chr4-90737604-INS-496 |
| chr4-90739736-INS-80 | chr4-90739731-INS-76 |
| chr4-90750310-INS-539 | chr4-90750310-INS-539 |
| chr4-90801359-INS-82 | chr4-90801359-INS-76 |
| chr4-90836772-INS-58 | chr4-90836765-INS-73 |
| chr4-90845099-INS-175 | chr4-90845099-INS-175 |
| chr4-90854448-INS-84 | chr4-90854448-INS-53 |
| chr4-90866335-INS-13831 | chr4-90866336-INS-13822 |
| chr4-90896651-INS-1483 | chr4-90896636-INS-1483 |
| chr4-90932963-INS-314 | chr4-90932963-INS-314 |
| chr4-90936719-INS-690 | chr4-90936719-INS-690 |
| chr4-90941272-INS-681 | chr4-90941272-INS-680 |
| chr4-91007903-INS-134 | chr4-91007903-INS-138 |
| chr4-91017409-INS-112 | chr4-91017409-INS-156 |
| chr4-91026994-INS-60 | chr4-91026994-INS-70 |
| chr4-91040915-INS-213 | chr4-91040915-INS-213 |
| chr4-91050893-INS-650 | chr4-91050893-INS-651 |
| chr4-91079218-INS-54 | chr4-91079230-INS-62 |
| chr4-91081789-INS-66 | chr4-91081789-INS-68 |
| chr4-91093816-INS-75 | chr4-91093816-INS-75 |
| chr4-91109742-INS-53 | chr4-91109742-INS-53 |
| chr4-91184556-INS-88 | chr4-91184556-INS-84 |
| chr4-91222470-INS-676 | chr4-91222470-INS-675 |
| chr4-91292581-INS-56 | chr4-91292581-INS-63 |
| chr4-91304066-INS-205 | chr4-91304066-INS-209 |
| chr4-91327870-INS-64 | chr4-91327867-INS-67 |
| chr4-91333520-INS-6442 | chr4-91333521-INS-6442 |
| chr4-91350404-INS-407 | chr4-91350404-INS-407 |
| chr4-91353290-INS-58 | chr4-91353290-INS-60 |
| chr4-91397297-INS-498 | chr4-91397297-INS-498 |
| chr4-91408008-INS-6183 | chr4-91408009-INS-6184 |
| chr4-91435145-INS-1402 | chr4-91435146-INS-1402 |
| chr4-91444858-INS-58 | chr4-91444836-INS-64 |
| chr4-91467414-INS-433 | chr4-91467418-INS-434 |
| chr4-91470215-INS-83 | chr4-91470215-INS-83 |
| chr4-91472525-INS-67 | chr4-91472525-INS-87 |

| | |
|---|---|
| chr4-91480113-INS-114 | chr4-91480100-INS-91 |
| chr4-91493737-INS-6338 | chr4-91493738-INS-6337 |
| chr4-91568461-INS-64 | chr4-91568461-INS-68 |
| chr4-91584031-INS-212 | chr4-91584031-INS-212 |
| chr4-91590150-INS-76 | chr4-91590150-INS-76 |
| chr4-91642475-INS-1890 | chr4-91642476-INS-1890 |
| chr4-91651733-INS-346 | chr4-91651733-INS-345 |
| chr4-91652898-INS-6545 | chr4-91652899-INS-6545 |
| chr4-91670130-INS-127 | chr4-91670130-INS-125 |
| chr4-91680396-INS-2897 | chr4-91680397-INS-2897 |
| chr4-91698403-INS-72 | chr4-91698403-INS-72 |
| chr4-91708573-INS-66 | chr4-91708573-INS-80 |
| chr4-91757536-INS-51 | chr4-91757526-INS-75 |
| chr4-91788664-INS-212 | chr4-91788664-INS-212 |
| chr4-91807814-INS-360 | chr4-91807818-INS-359 |
| chr4-91834274-INS-422 | chr4-91834274-INS-431 |
| chr4-91837998-INS-6164 | chr4-91837999-INS-6164 |
| chr4-91853349-INS-52 | chr4-91853350-INS-56 |
| chr4-91952058-INS-53 | chr4-91952061-INS-54 |
| chr4-91961070-INS-74 | chr4-91961070-INS-74 |
| chr4-92005521-INS-57 | chr4-92005521-INS-57 |
| chr4-92040329-INS-58 | chr4-92040295-INS-78 |
| chr4-92042169-INS-56 | chr4-92042169-INS-56 |
| chr4-92048681-INS-967 | chr4-92048681-INS-967 |
| chr4-92063148-INS-391 | chr4-92063148-INS-391 |
| chr4-92073528-INS-6182 | chr4-92073529-INS-6182 |
| chr4-92110715-INS-462 | chr4-92110737-INS-478 |
| chr4-92115891-INS-73 | chr4-92115891-INS-73 |
| chr4-92133296-INS-226 | chr4-92133301-INS-227 |
| chr4-92138458-INS-6722 | chr4-92138459-INS-6724 |
| chr4-92147202-INS-59 | chr4-92147202-INS-59 |
| chr4-92177363-INS-152 | chr4-92177363-INS-196 |
| chr4-92179109-INS-101 | chr4-92179109-INS-76 |
| chr4-92205412-INS-104 | chr4-92205412-INS-105 |
| chr4-92207608-INS-140 | chr4-92207608-INS-174 |
| chr4-92211345-INS-51 | chr4-92211361-INS-61 |
| chr4-92216936-INS-74 | chr4-92216936-INS-76 |
| chr4-92218283-INS-153 | chr4-92218283-INS-157 |
| chr4-92224883-INS-51 | chr4-92224883-INS-51 |
| chr4-92249415-INS-62 | chr4-92249415-INS-62 |
| chr4-92261113-INS-2478 | chr4-92261114-INS-2492 |
| chr4-92272052-INS-632 | chr4-92272052-INS-632 |
| chr4-92323550-INS-7081 | chr4-92323551-INS-7081 |
| chr4-92385522-INS-60 | chr4-92385522-INS-60 |
| chr4-92394723-INS-91 | chr4-92394723-INS-91 |
| chr4-92412154-INS-4124 | chr4-92412155-INS-4124 |
| chr4-92429735-INS-75 | chr4-92429735-INS-50 |
| chr4-92433180-INS-515 | chr4-92433180-INS-516 |
| chr4-92458187-INS-496 | chr4-92458187-INS-496 |
| chr4-92473055-INS-104 | chr4-92473024-INS-101 |
| chr4-92477644-INS-62 | chr4-92477643-INS-56 |
| chr4-92487067-INS-699 | chr4-92487067-INS-690 |
| chr4-92507038-INS-1066 | chr4-92507039-INS-1054 |
| chr4-92513614-INS-164 | chr4-92513610-INS-186 |
| chr4-92521466-INS-580 | chr4-92521466-INS-580 |
| chr4-92530365-INS-194 | chr4-92530365-INS-194 |
| chr4-92535015-INS-796 | chr4-92535015-INS-796 |
| chr4-92537922-INS-246 | chr4-92537922-INS-246 |
| chr4-92559719-INS-6606 | chr4-92559720-INS-6606 |
| chr4-92560308-INS-518 | chr4-92560308-INS-518 |
| chr4-92589317-INS-52 | chr4-92589318-INS-52 |
| chr4-92589535-INS-6379 | chr4-92589536-INS-6379 |
| chr4-92607015-INS-73 | chr4-92607015-INS-67 |
| chr4-92652242-INS-63 | chr4-92652242-INS-63 |
| chr4-92656134-INS-589 | chr4-92656134-INS-589 |
| chr4-92678189-INS-52 | chr4-92678189-INS-52 |
| chr4-92715887-INS-269 | chr4-92715886-INS-290 |
| chr4-92728233-INS-2968 | chr4-92728234-INS-2968 |
| chr4-92854028-INS-100 | chr4-92854028-INS-100 |
| chr4-92940681-INS-52 | chr4-92940681-INS-52 |
| chr4-92943419-INS-52 | chr4-92943419-INS-58 |
| chr4-92945428-INS-77 | chr4-92945428-INS-89 |
| chr4-92947755-INS-141 | chr4-92947755-INS-147 |
| chr4-92952760-INS-129 | chr4-92952760-INS-129 |
| chr4-92980534-INS-68 | chr4-92980534-INS-50 |
| chr4-93011173-INS-190 | chr4-93011173-INS-190 |
| chr4-93026801-INS-496 | chr4-93026801-INS-496 |

| | |
|---|---|
| chr4-93036653-INS-96 | chr4-93036653-INS-118 |
| chr4-93078198-INS-198 | chr4-93078198-INS-197 |
| chr4-93083501-INS-687 | chr4-93083501-INS-688 |
| chr4-93084094-INS-106 | chr4-93084094-INS-100 |
| chr4-93085550-INS-111 | chr4-93085546-INS-128 |
| chr4-93164312-INS-192 | chr4-93164312-INS-192 |
| chr4-93175592-INS-124 | chr4-93175592-INS-124 |
| chr4-93177626-INS-57 | chr4-93177626-INS-54 |
| chr4-93188107-INS-53 | chr4-93188092-INS-70 |
| chr4-93206441-INS-755 | chr4-93206441-INS-755 |
| chr4-93210410-INS-423 | chr4-93210436-INS-437 |
| chr4-93230913-INS-3687 | chr4-93230914-INS-3687 |
| chr4-93236752-INS-2555 | chr4-93236753-INS-2556 |
| chr4-93266132-INS-1146 | chr4-93266133-INS-1146 |
| chr4-93274259-INS-1561 | chr4-93274297-INS-1541 |
| chr4-93284530-INS-72 | chr4-93284530-INS-72 |
| chr4-93296370-INS-760 | chr4-93296370-INS-761 |
| chr4-93300498-INS-4434 | chr4-93300499-INS-4434 |
| chr4-93330592-INS-141 | chr4-93330592-INS-130 |
| chr4-93334021-INS-7232 | chr4-93334022-INS-7232 |
| chr4-93339004-INS-56 | chr4-93339004-INS-54 |
| chr4-93368348-INS-150 | chr4-93368351-INS-135 |
| chr4-93379970-INS-185 | chr4-93379970-INS-195 |
| chr4-93389585-INS-83 | chr4-93389586-INS-83 |
| chr4-93399343-INS-130 | chr4-93399338-INS-111 |
| chr4-93411390-INS-4380 | chr4-93411391-INS-4380 |
| chr4-93418929-INS-60 | chr4-93418924-INS-77 |
| chr4-93426513-INS-211 | chr4-93426513-INS-226 |
| chr4-93489706-INS-1412 | chr4-93489707-INS-1412 |
| chr4-93506273-INS-102 | chr4-93506270-INS-116 |
| chr4-93536546-INS-3350 | chr4-93536547-INS-3350 |
| chr4-93547547-INS-129 | chr4-93547570-INS-114 |
| chr4-93557088-INS-4240 | chr4-93557089-INS-4236 |
| chr4-93575280-INS-576 | chr4-93575280-INS-599 |
| chr4-93613885-INS-52 | chr4-93613885-INS-52 |
| chr4-93620685-INS-56 | chr4-93620685-INS-50 |
| chr4-93658096-INS-54 | chr4-93658096-INS-58 |
| chr4-93732643-INS-55 | chr4-93732643-INS-55 |
| chr4-93759919-INS-173 | chr4-93759919-INS-151 |
| chr4-93782720-INS-6413 | chr4-93782721-INS-6413 |
| chr4-93803278-INS-3061 | chr4-93803279-INS-3061 |
| chr4-93844667-INS-6521 | chr4-93844668-INS-6521 |
| chr4-93863952-INS-114 | chr4-93863990-INS-112 |
| chr4-93902230-INS-267 | chr4-93902230-INS-267 |
| chr4-93903361-INS-322 | chr4-93903361-INS-321 |
| chr4-93915067-INS-85 | chr4-93915045-INS-114 |
| chr4-93962865-INS-6687 | chr4-93962866-INS-6687 |
| chr4-93966226-INS-156 | chr4-93966226-INS-157 |
| chr4-93984317-INS-56 | chr4-93984316-INS-72 |
| chr4-93995766-INS-261 | chr4-93995766-INS-254 |
| chr4-94129472-INS-62 | chr4-94129450-INS-53 |
| chr4-94155218-INS-509 | chr4-94155218-INS-509 |
| chr4-94240778-INS-70 | chr4-94240778-INS-70 |
| chr4-94247973-INS-54 | chr4-94247973-INS-86 |
| chr4-94253294-INS-74 | chr4-94253294-INS-92 |
| chr4-94275253-INS-123 | chr4-94275290-INS-105 |
| chr4-94281400-INS-712 | chr4-94281400-INS-712 |
| chr4-94324778-INS-7203 | chr4-94324779-INS-7203 |
| chr4-94329544-INS-1941 | chr4-94329545-INS-1941 |
| chr4-94341889-INS-55 | chr4-94341889-INS-55 |
| chr4-94359202-INS-501 | chr4-94359202-INS-501 |
| chr4-94366012-INS-206 | chr4-94366012-INS-206 |
| chr4-94367241-INS-288 | chr4-94367241-INS-252 |
| chr4-94370586-INS-207 | chr4-94370586-INS-207 |
| chr4-94395502-INS-71 | chr4-94395502-INS-66 |
| chr4-94403947-INS-52 | chr4-94403947-INS-52 |
| chr4-94412315-INS-169 | chr4-94412315-INS-169 |
| chr4-94435295-INS-209 | chr4-94435295-INS-211 |
| chr4-94447094-INS-71 | chr4-94447094-INS-71 |
| chr4-94470389-INS-65 | chr4-94470389-INS-65 |
| chr4-94473238-INS-6521 | chr4-94473239-INS-6521 |
| chr4-94492309-INS-498 | chr4-94492309-INS-498 |
| chr4-94493167-INS-195 | chr4-94493167-INS-195 |
| chr4-94508076-INS-592 | chr4-94508076-INS-571 |
| chr4-94571287-INS-100 | chr4-94571287-INS-100 |
| chr4-94574197-INS-182 | chr4-94574211-INS-162 |
| chr4-94584738-INS-409 | chr4-94584738-INS-410 |

| | |
|---|---|
| chr4-94617109-INS-67 | chr4-94617109-INS-67 |
| chr4-94620273-INS-52 | chr4-94620273-INS-52 |
| chr4-94665693-INS-485 | chr4-94665693-INS-485 |
| chr4-94811635-INS-77 | chr4-94811635-INS-95 |
| chr4-94819299-INS-59 | chr4-94819299-INS-71 |
| chr4-94828497-INS-2197 | chr4-94828498-INS-2197 |
| chr4-94828789-INS-212 | chr4-94828789-INS-212 |
| chr4-94829190-INS-530 | chr4-94829190-INS-530 |
| chr4-94863750-INS-54 | chr4-94863750-INS-62 |
| chr4-94865395-INS-718 | chr4-94865395-INS-718 |
| chr4-94874684-INS-520 | chr4-94874685-INS-520 |
| chr4-94884435-INS-53 | chr4-94884438-INS-54 |
| chr4-94908047-INS-130 | chr4-94908015-INS-179 |
| chr4-94909867-INS-76 | chr4-94909861-INS-90 |
| chr4-94965563-INS-60 | chr4-94965563-INS-60 |
| chr4-94968501-INS-78 | chr4-94968502-INS-78 |
| chr4-94972295-INS-96 | chr4-94972295-INS-72 |
| chr4-94997944-INS-67 | chr4-94997944-INS-55 |
| chr4-95014945-INS-108 | chr4-95014945-INS-96 |
| chr4-95023665-INS-5348 | chr4-95023666-INS-5348 |
| chr4-95047757-INS-220 | chr4-95047757-INS-220 |
| chr4-95072129-INS-152 | chr4-95072129-INS-124 |
| chr4-95082217-INS-76 | chr4-95082217-INS-78 |
| chr4-95085929-INS-64 | chr4-95085918-INS-50 |
| chr4-95089516-INS-51 | chr4-95089516-INS-51 |
| chr4-95092253-INS-55 | chr4-95092257-INS-67 |
| chr4-95136962-INS-1845 | chr4-95136963-INS-1845 |
| chr4-95170035-INS-565 | chr4-95170035-INS-565 |
| chr4-95251924-INS-69 | chr4-95251916-INS-101 |
| chr4-95257236-INS-499 | chr4-95257240-INS-504 |
| chr4-95258405-INS-7091 | chr4-95258406-INS-7091 |
| chr4-95298712-INS-52 | chr4-95298712-INS-54 |
| chr4-95331412-INS-192 | chr4-95331416-INS-200 |
| chr4-95354696-INS-61 | chr4-95354696-INS-62 |
| chr4-95371334-INS-60 | chr4-95371334-INS-60 |
| chr4-95394118-INS-62 | chr4-95394102-INS-70 |
| chr4-95394689-INS-1760 | chr4-95394690-INS-1762 |
| chr4-95417090-INS-394 | chr4-95417090-INS-394 |
| chr4-95427118-INS-107 | chr4-95427118-INS-103 |
| chr4-95450140-INS-159 | chr4-95450140-INS-159 |
| chr4-95464556-INS-188 | chr4-95464556-INS-188 |
| chr4-95467866-INS-325 | chr4-95467866-INS-325 |
| chr4-95480559-INS-53 | chr4-95480557-INS-51 |
| chr4-95485701-INS-299 | chr4-95485701-INS-299 |
| chr4-95490607-INS-611 | chr4-95490607-INS-610 |
| chr4-95502064-INS-440 | chr4-95502064-INS-440 |
| chr4-95523132-INS-60 | chr4-95523132-INS-60 |
| chr4-95553277-INS-56 | chr4-95553277-INS-58 |
| chr4-95605761-INS-72 | chr4-95605761-INS-78 |
| chr4-95608682-INS-64 | chr4-95608682-INS-64 |
| chr4-95623049-INS-123 | chr4-95623049-INS-86 |
| chr4-95646447-INS-87 | chr4-95646447-INS-121 |
| chr4-95679453-INS-54 | chr4-95679453-INS-54 |
| chr4-95692801-INS-6925 | chr4-95692802-INS-6934 |
| chr4-95697022-INS-958 | chr4-95697022-INS-960 |
| chr4-95701146-INS-64 | chr4-95701150-INS-64 |
| chr4-95706775-INS-153 | chr4-95706775-INS-153 |
| chr4-95719842-INS-53 | chr4-95719842-INS-53 |
| chr4-95727212-INS-103 | chr4-95727212-INS-153 |
| chr4-95756119-INS-62 | chr4-95756119-INS-64 |
| chr4-95807983-INS-67 | chr4-95807983-INS-67 |
| chr4-95823638-INS-150 | chr4-95823638-INS-150 |
| chr4-95826277-INS-740 | chr4-95826277-INS-738 |
| chr4-95845064-INS-105 | chr4-95845078-INS-107 |
| chr4-95850345-INS-265 | chr4-95850345-INS-265 |
| chr4-95882182-INS-55 | chr4-95882182-INS-51 |
| chr4-95914566-INS-5501 | chr4-95914567-INS-5501 |
| chr4-95921761-INS-477 | chr4-95921761-INS-477 |
| chr4-95927080-INS-514 | chr4-95927080-INS-514 |
| chr4-95950983-INS-816 | chr4-95950983-INS-819 |
| chr4-95954120-INS-100 | chr4-95954120-INS-102 |
| chr4-95964939-INS-55 | chr4-95964891-INS-53 |
| chr4-96010037-INS-68 | chr4-96010004-INS-76 |
| chr4-96021924-INS-499 | chr4-96021924-INS-499 |
| chr4-96031607-INS-197 | chr4-96031607-INS-197 |
| chr4-96050645-INS-332 | chr4-96050645-INS-332 |
| chr4-96059693-INS-875 | chr4-96059693-INS-875 |

| | |
|---|---|
| chr4-96072010-INS-81 | chr4-96072010-INS-81 |
| chr4-96073481-INS-465 | chr4-96073481-INS-465 |
| chr4-96124542-INS-55 | chr4-96124542-INS-62 |
| chr4-96260440-INS-595 | chr4-96260440-INS-599 |
| chr4-96267888-INS-59 | chr4-96267863-INS-54 |
| chr4-96292147-INS-198 | chr4-96292147-INS-198 |
| chr4-96297388-INS-518 | chr4-96297388-INS-518 |
| chr4-96310798-INS-335 | chr4-96310798-INS-347 |
| chr4-96396966-INS-802 | chr4-96396966-INS-801 |
| chr4-96408688-INS-156 | chr4-96408688-INS-172 |
| chr4-96444641-INS-1566 | chr4-96444642-INS-1569 |
| chr4-96450733-INS-6375 | chr4-96450734-INS-6389 |
| chr4-96457896-INS-176 | chr4-96457896-INS-176 |
| chr4-96467951-INS-60 | chr4-96467905-INS-90 |
| chr4-96523793-INS-51 | chr4-96523793-INS-51 |
| chr4-96527386-INS-79 | chr4-96527386-INS-79 |
| chr4-96584926-INS-199 | chr4-96584926-INS-199 |
| chr4-96631056-INS-72 | chr4-96631056-INS-86 |
| chr4-96652891-INS-54 | chr4-96652856-INS-62 |
| chr4-96665112-INS-199 | chr4-96665112-INS-224 |
| chr4-96711190-INS-230 | chr4-96711190-INS-230 |
| chr4-96779786-INS-2819 | chr4-96779787-INS-2819 |
| chr4-96781473-INS-143 | chr4-96781473-INS-145 |
| chr4-96805793-INS-488 | chr4-96805793-INS-488 |
| chr4-96831098-INS-80 | chr4-96831098-INS-80 |
| chr4-96851449-INS-108 | chr4-96851454-INS-122 |
| chr4-96854942-INS-2485 | chr4-96854943-INS-2485 |
| chr4-96860759-INS-58 | chr4-96860759-INS-58 |
| chr4-96903998-INS-6220 | chr4-96903999-INS-6220 |
| chr4-96904674-INS-176 | chr4-96904674-INS-176 |
| chr4-96968380-INS-199 | chr4-96968380-INS-199 |
| chr4-96983175-INS-115 | chr4-96983175-INS-115 |
| chr4-96988988-INS-69 | chr4-96988988-INS-69 |
| chr4-97021789-INS-59 | chr4-97021792-INS-59 |
| chr4-97027760-INS-2585 | chr4-97027761-INS-2585 |
| chr4-97066605-INS-471 | chr4-97066605-INS-423 |
| chr4-97071094-INS-624 | chr4-97071094-INS-626 |
| chr4-97074620-INS-519 | chr4-97074620-INS-512 |
| chr4-97077906-INS-93 | chr4-97077879-INS-124 |
| chr4-97090392-INS-6243 | chr4-97090393-INS-6242 |
| chr4-97090923-INS-799 | chr4-97090923-INS-799 |
| chr4-97098327-INS-123 | chr4-97098328-INS-123 |
| chr4-97125794-INS-1896 | chr4-97125795-INS-1896 |
| chr4-97136574-INS-719 | chr4-97136574-INS-717 |
| chr4-97185343-INS-56 | chr4-97185343-INS-54 |
| chr4-97199733-INS-5405 | chr4-97199734-INS-5404 |
| chr4-97201463-INS-481 | chr4-97201463-INS-481 |
| chr4-97209472-INS-199 | chr4-97209472-INS-199 |
| chr4-97229977-INS-62 | chr4-97229988-INS-68 |
| chr4-97267128-INS-80 | chr4-97267123-INS-56 |
| chr4-97268560-INS-845 | chr4-97268561-INS-845 |
| chr4-97276724-INS-204 | chr4-97276717-INS-204 |
| chr4-97282259-INS-725 | chr4-97282260-INS-741 |
| chr4-97290438-INS-65 | chr4-97290439-INS-54 |
| chr4-97291334-INS-304 | chr4-97291334-INS-304 |
| chr4-97331593-INS-1124 | chr4-97331594-INS-1144 |
| chr4-97339728-INS-98 | chr4-97339728-INS-98 |
| chr4-97343072-INS-390 | chr4-97343072-INS-390 |
| chr4-97374364-INS-114 | chr4-97374362-INS-112 |
| chr4-97389311-INS-918 | chr4-97389311-INS-918 |
| chr4-97406691-INS-377 | chr4-97406691-INS-357 |
| chr4-97446490-INS-76 | chr4-97446488-INS-90 |
| chr4-97454213-INS-6725 | chr4-97454199-INS-6705 |
| chr4-97464685-INS-300 | chr4-97464689-INS-278 |
| chr4-97510775-INS-599 | chr4-97510775-INS-602 |
| chr4-97523270-INS-71 | chr4-97523256-INS-50 |
| chr4-97583108-INS-75 | chr4-97583108-INS-75 |
| chr4-97619646-INS-62 | chr4-97619646-INS-64 |
| chr4-97703634-INS-134 | chr4-97703634-INS-132 |
| chr4-97843063-INS-64 | chr4-97843063-INS-65 |
| chr4-97844494-INS-924 | chr4-97844494-INS-925 |
| chr4-97845723-INS-127 | chr4-97845723-INS-77 |
| chr4-97879088-INS-65 | chr4-97879088-INS-65 |
| chr4-97888225-INS-55 | chr4-97888225-INS-55 |
| chr4-97985161-INS-110 | chr4-97985155-INS-113 |
| chr4-97998620-INS-53 | chr4-97998620-INS-52 |
| chr4-98026291-INS-174 | chr4-98026291-INS-174 |

| | |
|---|---|
| chr4-98054268-INS-202 | chr4-98054268-INS-202 |
| chr4-98059218-INS-51 | chr4-98059218-INS-51 |
| chr4-98081958-INS-192 | chr4-98081959-INS-192 |
| chr4-98092383-INS-54 | chr4-98092383-INS-50 |
| chr4-98093758-INS-429 | chr4-98093758-INS-429 |
| chr4-98096000-INS-56 | chr4-98095998-INS-62 |
| chr4-98122215-INS-100 | chr4-98122215-INS-80 |
| chr4-98130253-INS-63 | chr4-98130253-INS-63 |
| chr4-98135513-INS-1590 | chr4-98135514-INS-1588 |
| chr4-98138804-INS-272 | chr4-98138804-INS-222 |
| chr4-98179444-INS-293 | chr4-98179444-INS-293 |
| chr4-98213113-INS-90 | chr4-98213113-INS-67 |
| chr4-98254926-INS-417 | chr4-98254927-INS-417 |
| chr4-98312921-INS-62 | chr4-98312921-INS-62 |
| chr4-98386202-INS-60 | chr4-98386202-INS-60 |
| chr4-98438902-INS-53 | chr4-98438902-INS-53 |
| chr4-98465711-INS-325 | chr4-98465711-INS-326 |
| chr4-98470774-INS-59 | chr4-98470775-INS-50 |
| chr4-98472046-INS-112 | chr4-98472046-INS-112 |
| chr4-98475401-INS-1177 | chr4-98475402-INS-1177 |
| chr4-98476555-INS-112 | chr4-98476555-INS-112 |
| chr4-98487451-INS-56 | chr4-98487451-INS-58 |
| chr4-98520962-INS-249 | chr4-98520962-INS-249 |
| chr4-98536985-INS-204 | chr4-98536985-INS-204 |
| chr4-98547407-INS-372 | chr4-98547407-INS-373 |
| chr4-98588815-INS-191 | chr4-98588815-INS-195 |
| chr4-98593723-INS-72 | chr4-98593723-INS-88 |
| chr4-98597313-INS-94 | chr4-98597289-INS-101 |
| chr4-98604040-INS-96 | chr4-98604040-INS-80 |
| chr4-98613649-INS-195 | chr4-98613650-INS-195 |
| chr4-98636896-INS-52 | chr4-98636892-INS-84 |
| chr4-98662206-INS-503 | chr4-98662206-INS-503 |
| chr4-98669588-INS-398 | chr4-98669588-INS-398 |
| chr4-98686758-INS-272 | chr4-98686759-INS-270 |
| chr4-98698881-INS-52 | chr4-98698892-INS-80 |
| chr4-98713185-INS-486 | chr4-98713185-INS-486 |
| chr4-98725684-INS-100 | chr4-98725684-INS-100 |
| chr4-98728174-INS-197 | chr4-98728174-INS-197 |
| chr4-98738809-INS-53 | chr4-98738809-INS-53 |
| chr4-98748603-INS-203 | chr4-98748603-INS-203 |
| chr4-98758841-INS-217 | chr4-98758841-INS-213 |
| chr4-98762790-INS-58 | chr4-98762790-INS-54 |
| chr4-98773838-INS-54 | chr4-98773838-INS-54 |
| chr4-98776321-INS-690 | chr4-98776321-INS-690 |
| chr4-98778156-INS-362 | chr4-98778156-INS-362 |
| chr4-98791216-INS-206 | chr4-98791219-INS-203 |
| chr4-98798811-INS-186 | chr4-98798811-INS-186 |
| chr4-98801048-INS-7365 | chr4-98801049-INS-7365 |
| chr4-98815780-INS-210 | chr4-98815780-INS-200 |
| chr4-98827911-INS-68 | chr4-98827911-INS-72 |
| chr4-98831995-INS-201 | chr4-98831996-INS-156 |
| chr4-98833910-INS-54 | chr4-98833910-INS-71 |
| chr4-98858786-INS-109 | chr4-98858786-INS-109 |
| chr4-98875828-INS-165 | chr4-98875828-INS-165 |
| chr4-98878890-INS-640 | chr4-98878890-INS-640 |
| chr4-98898250-INS-71 | chr4-98898250-INS-51 |
| chr4-98898494-INS-516 | chr4-98898493-INS-515 |
| chr4-98937796-INS-152 | chr4-98937796-INS-152 |
| chr4-98961752-INS-1489 | chr4-98961753-INS-1489 |
| chr4-98975013-INS-57 | chr4-98975013-INS-63 |
| chr4-98980487-INS-57 | chr4-98980487-INS-52 |
| chr4-99003034-INS-2890 | chr4-99003035-INS-2890 |
| chr4-99011438-INS-88 | chr4-99011442-INS-56 |
| chr4-99015647-INS-5231 | chr4-99015648-INS-5231 |
| chr4-99024146-INS-959 | chr4-99024156-INS-970 |
| chr4-99025802-INS-911 | chr4-99025802-INS-905 |
| chr4-99026612-INS-63 | chr4-99026612-INS-63 |
| chr4-99028682-INS-77 | chr4-99028682-INS-77 |
| chr4-99040018-INS-54 | chr4-99040018-INS-54 |
| chr4-99053638-INS-52 | chr4-99053638-INS-52 |
| chr4-99054920-INS-660 | chr4-99054924-INS-658 |
| chr4-99055903-INS-471 | chr4-99055903-INS-470 |
| chr4-99063116-INS-52 | chr4-99063113-INS-54 |
| chr4-99064338-INS-205 | chr4-99064338-INS-206 |
| chr4-99071296-INS-189 | chr4-99071296-INS-189 |
| chr4-99075088-INS-52 | chr4-99075088-INS-52 |
| chr4-99080997-INS-232 | chr4-99080997-INS-232 |

| | |
|---|---|
| chr4-99092691-INS-125 | chr4-99092691-INS-125 |
| chr4-99094191-INS-190 | chr4-99094240-INS-190 |
| chr4-99108714-INS-6446 | chr4-99108725-INS-6432 |
| chr4-99114794-INS-332 | chr4-99114794-INS-332 |
| chr4-99130852-INS-56 | chr4-99130852-INS-56 |
| chr4-99140585-INS-58 | chr4-99140585-INS-58 |
| chr4-99142292-INS-55 | chr4-99142255-INS-99 |
| chr4-99147435-INS-94 | chr4-99147424-INS-107 |
| chr4-99147955-INS-218 | chr4-99147955-INS-212 |
| chr4-99169121-INS-53 | chr4-99169080-INS-59 |
| chr4-99181004-INS-60 | chr4-99181036-INS-60 |
| chr4-99184952-INS-52 | chr4-99184953-INS-63 |
| chr4-99197480-INS-80 | chr4-99197480-INS-80 |
| chr4-99203909-INS-238 | chr4-99203909-INS-238 |
| chr4-99223797-INS-217 | chr4-99223797-INS-218 |
| chr4-99235997-INS-462 | chr4-99235997-INS-462 |
| chr4-99245104-INS-54 | chr4-99245110-INS-71 |
| chr4-99368927-INS-94 | chr4-99368927-INS-94 |
| chr4-99416266-INS-954 | chr4-99416266-INS-954 |
| chr4-99455034-INS-220 | chr4-99455034-INS-212 |
| chr4-99475636-INS-253 | chr4-99475636-INS-223 |
| chr4-99509931-INS-68 | chr4-99509931-INS-72 |
| chr4-99512197-INS-53 | chr4-99512202-INS-55 |
| chr4-99515988-INS-166 | chr4-99515988-INS-166 |
| chr4-99520547-INS-163 | chr4-99520547-INS-135 |
| chr4-99527067-INS-127 | chr4-99527067-INS-127 |
| chr4-99530799-INS-54 | chr4-99530799-INS-52 |
| chr4-99561408-INS-54 | chr4-99561408-INS-54 |
| chr4-99570210-INS-290 | chr4-99570210-INS-290 |
| chr4-99583985-INS-80 | chr4-99583985-INS-66 |
| chr4-99598011-INS-60 | chr4-99598020-INS-52 |
| chr4-99602139-INS-215 | chr4-99602139-INS-216 |
| chr4-99602740-INS-251 | chr4-99602740-INS-251 |
| chr4-99636407-INS-125 | chr4-99636376-INS-156 |
| chr4-99680612-INS-781 | chr4-99680612-INS-779 |
| chr4-99756607-INS-56 | chr4-99756607-INS-60 |
| chr4-99813994-INS-103 | chr4-99813994-INS-103 |
| chr4-99837683-INS-86 | chr4-99837683-INS-86 |
| chr4-99842144-INS-6624 | chr4-99842145-INS-6626 |
| chr4-99883250-INS-221 | chr4-99883222-INS-228 |
| chr4-99886591-INS-2656 | chr4-99886592-INS-2648 |
| chr4-99903931-INS-112 | chr4-99903931-INS-112 |
| chr4-99905588-INS-6475 | chr4-99905589-INS-6475 |
| chr4-99910073-INS-56 | chr4-99910050-INS-54 |
| chr4-99929824-INS-129 | chr4-99929832-INS-123 |
| chr4-99932673-INS-308 | chr4-99932673-INS-308 |
| chr4-99966391-INS-85 | chr4-99966391-INS-91 |
| chr4-99968604-INS-84 | chr4-99968604-INS-69 |
| chr4-100016577-INS-64 | chr4-100016577-INS-64 |
| chr4-100025266-INS-54 | chr4-100025266-INS-56 |
| chr4-100026284-INS-54 | chr4-100026284-INS-54 |
| chr4-100038339-INS-105 | chr4-100038295-INS-150 |
| chr4-100041216-INS-67 | chr4-100041202-INS-64 |
| chr4-100097163-INS-76 | chr4-100097174-INS-100 |
| chr4-100114062-INS-2930 | chr4-100114063-INS-2930 |
| chr4-100126809-INS-76 | chr4-100126809-INS-76 |
| chr4-100136118-INS-70 | chr4-100136118-INS-70 |
| chr4-100142375-INS-86 | chr4-100142375-INS-80 |
| chr4-100143904-INS-58 | chr4-100143939-INS-51 |
| chr4-100174396-INS-218 | chr4-100174396-INS-218 |
| chr4-100180196-INS-308 | chr4-100180202-INS-324 |
| chr4-100184126-INS-62 | chr4-100184126-INS-62 |
| chr4-100203289-INS-1262 | chr4-100203290-INS-1264 |
| chr4-100213803-INS-90 | chr4-100213803-INS-90 |
| chr4-100218569-INS-60 | chr4-100218569-INS-54 |
| chr4-100244558-INS-88 | chr4-100244558-INS-88 |
| chr4-100255665-INS-74 | chr4-100255665-INS-74 |
| chr4-100262084-INS-64 | chr4-100262088-INS-59 |
| chr4-100271175-INS-973 | chr4-100271175-INS-972 |
| chr4-100305569-INS-80 | chr4-100305573-INS-68 |
| chr4-100407567-INS-76 | chr4-100407567-INS-76 |
| chr4-100430996-INS-54 | chr4-100430996-INS-54 |
| chr4-100483578-INS-734 | chr4-100483578-INS-731 |
| chr4-100485410-INS-5503 | chr4-100485411-INS-5503 |
| chr4-100490376-INS-1656 | chr4-100490377-INS-1656 |
| chr4-100498027-INS-1132 | chr4-100498028-INS-1156 |
| chr4-100505389-INS-72 | chr4-100505389-INS-62 |

| | |
|---|---|
| chr4-100523716-INS-101 | chr4-100523716-INS-101 |
| chr4-100547426-INS-321 | chr4-100547426-INS-321 |
| chr4-100552007-INS-57 | chr4-100552007-INS-58 |
| chr4-100565877-INS-975 | chr4-100565877-INS-975 |
| chr4-100568142-INS-52 | chr4-100568142-INS-52 |
| chr4-100573932-INS-134 | chr4-100573932-INS-134 |
| chr4-100592026-INS-198 | chr4-100592026-INS-198 |
| chr4-100598115-INS-524 | chr4-100598115-INS-524 |
| chr4-100605926-INS-774 | chr4-100605926-INS-774 |
| chr4-100625424-INS-69 | chr4-100625430-INS-99 |
| chr4-100629335-INS-4252 | chr4-100629336-INS-4252 |
| chr4-100663432-INS-84 | chr4-100663432-INS-84 |
| chr4-100664557-INS-507 | chr4-100664557-INS-507 |
| chr4-100670963-INS-214 | chr4-100670963-INS-214 |
| chr4-100673869-INS-53 | chr4-100673869-INS-75 |
| chr4-100731917-INS-199 | chr4-100731917-INS-198 |
| chr4-100735781-INS-69 | chr4-100735781-INS-69 |
| chr4-100767577-INS-51 | chr4-100767577-INS-50 |
| chr4-100774662-INS-202 | chr4-100774662-INS-218 |
| chr4-100782958-INS-56 | chr4-100782951-INS-104 |
| chr4-100804585-INS-64 | chr4-100804586-INS-64 |
| chr4-100805434-INS-194 | chr4-100805430-INS-230 |
| chr4-100889218-INS-66 | chr4-100889218-INS-51 |
| chr4-100893548-INS-78 | chr4-100893556-INS-72 |
| chr4-100920030-INS-84 | chr4-100920030-INS-84 |
| chr4-100945423-INS-1288 | chr4-100945424-INS-1268 |
| chr4-100947069-INS-59 | chr4-100947069-INS-65 |
| chr4-100971997-INS-242 | chr4-100971997-INS-241 |
| chr4-100973907-INS-109 | chr4-100973907-INS-76 |
| chr4-100975219-INS-267 | chr4-100975219-INS-267 |
| chr4-100995305-INS-159 | chr4-100995305-INS-159 |
| chr4-101039371-INS-353 | chr4-101039371-INS-358 |
| chr4-101045056-INS-95 | chr4-101045056-INS-99 |
| chr4-101067781-INS-1384 | chr4-101067780-INS-1354 |
| chr4-101092274-INS-62 | chr4-101092292-INS-90 |
| chr4-101130100-INS-99 | chr4-101130100-INS-99 |
| chr4-101143119-INS-58 | chr4-101143165-INS-60 |
| chr4-101144675-INS-66 | chr4-101144675-INS-66 |
| chr4-101148191-INS-121 | chr4-101148191-INS-106 |
| chr4-101156737-INS-196 | chr4-101156737-INS-196 |
| chr4-101160941-INS-64 | chr4-101160941-INS-64 |
| chr4-101172365-INS-84 | chr4-101172366-INS-70 |
| chr4-101185016-INS-80 | chr4-101185016-INS-80 |
| chr4-101195140-INS-66 | chr4-101195140-INS-113 |
| chr4-101201844-INS-145 | chr4-101201844-INS-145 |
| chr4-101203692-INS-188 | chr4-101203692-INS-188 |
| chr4-101215846-INS-93 | chr4-101215846-INS-106 |
| chr4-101236787-INS-212 | chr4-101236787-INS-212 |
| chr4-101241239-INS-96 | chr4-101241213-INS-80 |
| chr4-101245353-INS-116 | chr4-101245355-INS-104 |
| chr4-101286444-INS-64 | chr4-101286444-INS-64 |
| chr4-101288645-INS-126 | chr4-101288645-INS-126 |
| chr4-101293509-INS-76 | chr4-101293509-INS-80 |
| chr4-101299566-INS-54 | chr4-101299566-INS-54 |
| chr4-101300638-INS-197 | chr4-101300638-INS-197 |
| chr4-101304638-INS-488 | chr4-101304638-INS-488 |
| chr4-101311774-INS-134 | chr4-101311774-INS-134 |
| chr4-101316964-INS-276 | chr4-101316964-INS-277 |
| chr4-101318806-INS-98 | chr4-101318808-INS-100 |
| chr4-101350523-INS-80 | chr4-101350523-INS-50 |
| chr4-101377632-INS-182 | chr4-101377632-INS-182 |
| chr4-101435259-INS-55 | chr4-101435259-INS-55 |
| chr4-101462749-INS-230 | chr4-101462749-INS-228 |
| chr4-101501008-INS-172 | chr4-101501008-INS-172 |
| chr4-101517917-INS-156 | chr4-101517917-INS-156 |
| chr4-101543447-INS-250 | chr4-101543447-INS-250 |
| chr4-101604223-INS-386 | chr4-101604223-INS-386 |
| chr4-101605276-INS-119 | chr4-101605276-INS-115 |
| chr4-101609623-INS-125 | chr4-101609623-INS-125 |
| chr4-101624044-INS-54 | chr4-101624048-INS-68 |
| chr4-101627783-INS-259 | chr4-101627783-INS-251 |
| chr4-101633325-INS-58 | chr4-101633325-INS-58 |
| chr4-101649608-INS-460 | chr4-101649608-INS-460 |
| chr4-101662559-INS-176 | chr4-101662559-INS-179 |
| chr4-101664373-INS-637 | chr4-101664372-INS-637 |
| chr4-101703656-INS-1419 | chr4-101703657-INS-1419 |
| chr4-101732019-INS-1093 | chr4-101732020-INS-1093 |

| | |
|---|---|
| chr4-101735540-INS-56 | chr4-101735540-INS-58 |
| chr4-101737998-INS-246 | chr4-101737998-INS-246 |
| chr4-101747714-INS-91 | chr4-101747714-INS-91 |
| chr4-101751567-INS-179 | chr4-101751567-INS-179 |
| chr4-101766962-INS-181 | chr4-101766962-INS-186 |
| chr4-101789959-INS-717 | chr4-101789959-INS-717 |
| chr4-101809434-INS-365 | chr4-101809434-INS-364 |
| chr4-101811977-INS-437 | chr4-101811977-INS-436 |
| chr4-101813842-INS-178 | chr4-101813842-INS-178 |
| chr4-101839262-INS-178 | chr4-101839262-INS-178 |
| chr4-101851880-INS-3044 | chr4-101851881-INS-3045 |
| chr4-101867177-INS-7610 | chr4-101867178-INS-7612 |
| chr4-101874578-INS-169 | chr4-101874578-INS-174 |
| chr4-101878652-INS-61 | chr4-101878652-INS-61 |
| chr4-101881644-INS-124 | chr4-101881644-INS-124 |
| chr4-101884123-INS-454 | chr4-101884123-INS-457 |
| chr4-101895357-INS-65 | chr4-101895333-INS-65 |
| chr4-101901686-INS-304 | chr4-101901686-INS-304 |
| chr4-101941107-INS-237 | chr4-101941107-INS-237 |
| chr4-101942791-INS-65 | chr4-101942791-INS-65 |
| chr4-102012665-INS-503 | chr4-102012665-INS-503 |
| chr4-102024447-INS-7084 | chr4-102024448-INS-7084 |
| chr4-102030662-INS-97 | chr4-102030662-INS-97 |
| chr4-102078976-INS-466 | chr4-102078976-INS-466 |
| chr4-102089076-INS-105 | chr4-102089076-INS-90 |
| chr4-102125121-INS-6162 | chr4-102125122-INS-6162 |
| chr4-102135261-INS-6156 | chr4-102135262-INS-6157 |
| chr4-102183537-INS-152 | chr4-102183537-INS-152 |
| chr4-102200652-INS-1719 | chr4-102200653-INS-1721 |
| chr4-102246495-INS-971 | chr4-102246495-INS-971 |
| chr4-102275481-INS-578 | chr4-102275481-INS-585 |
| chr4-102301679-INS-282 | chr4-102301679-INS-283 |
| chr4-102312943-INS-194 | chr4-102312943-INS-231 |
| chr4-102327214-INS-72 | chr4-102327214-INS-72 |
| chr4-102330831-INS-2258 | chr4-102330831-INS-2274 |
| chr4-102347834-INS-6167 | chr4-102347836-INS-6161 |
| chr4-102351691-INS-1582 | chr4-102351702-INS-1592 |
| chr4-102375970-INS-67 | chr4-102375970-INS-67 |
| chr4-102436088-INS-259 | chr4-102436088-INS-259 |
| chr4-102453363-INS-64 | chr4-102453363-INS-64 |
| chr4-102491285-INS-982 | chr4-102491285-INS-988 |
| chr4-102513980-INS-154 | chr4-102513980-INS-154 |
| chr4-102524022-INS-235 | chr4-102524022-INS-235 |
| chr4-102528425-INS-78 | chr4-102528425-INS-68 |
| chr4-102543601-INS-6900 | chr4-102543602-INS-6902 |
| chr4-102567418-INS-63 | chr4-102567418-INS-79 |
| chr4-102571678-INS-84 | chr4-102571678-INS-107 |
| chr4-102601015-INS-908 | chr4-102601015-INS-907 |
| chr4-102612484-INS-57 | chr4-102612484-INS-60 |
| chr4-102634313-INS-52 | chr4-102634315-INS-85 |
| chr4-102641847-INS-379 | chr4-102641847-INS-381 |
| chr4-102660562-INS-63 | chr4-102660562-INS-63 |
| chr4-102662769-INS-55 | chr4-102662769-INS-55 |
| chr4-102684323-INS-74 | chr4-102684355-INS-72 |
| chr4-102687619-INS-419 | chr4-102687619-INS-401 |
| chr4-102692476-INS-52 | chr4-102692476-INS-50 |
| chr4-102712087-INS-177 | chr4-102712088-INS-195 |
| chr4-102730594-INS-90 | chr4-102730594-INS-90 |
| chr4-102845403-INS-59 | chr4-102845403-INS-59 |
| chr4-102860410-INS-177 | chr4-102860410-INS-177 |
| chr4-102861100-INS-747 | chr4-102861100-INS-745 |
| chr4-102892817-INS-58 | chr4-102892817-INS-60 |
| chr4-102895672-INS-82 | chr4-102895672-INS-82 |
| chr4-102905924-INS-68 | chr4-102905924-INS-68 |
| chr4-102909402-INS-56 | chr4-102909402-INS-56 |
| chr4-102936328-INS-413 | chr4-102936328-INS-413 |
| chr4-102955622-INS-528 | chr4-102955622-INS-528 |
| chr4-102960941-INS-3322 | chr4-102960942-INS-3322 |
| chr4-103004914-INS-365 | chr4-103004914-INS-365 |
| chr4-103026096-INS-54 | chr4-103026069-INS-54 |
| chr4-103036582-INS-486 | chr4-103036582-INS-486 |
| chr4-103055202-INS-54 | chr4-103055202-INS-54 |
| chr4-103059885-INS-97 | chr4-103059900-INS-102 |
| chr4-103082338-INS-323 | chr4-103082338-INS-323 |
| chr4-103102462-INS-137 | chr4-103102462-INS-137 |
| chr4-103115994-INS-83 | chr4-103115994-INS-75 |
| chr4-103130520-INS-212 | chr4-103130520-INS-212 |

| | |
|---|---|
| chr4-103146828-INS-165 | chr4-103146828-INS-165 |
| chr4-103153835-INS-57 | chr4-103153835-INS-57 |
| chr4-103167505-INS-620 | chr4-103167505-INS-619 |
| chr4-103175875-INS-138 | chr4-103175910-INS-140 |
| chr4-103181271-INS-52 | chr4-103181271-INS-64 |
| chr4-103217934-INS-73 | chr4-103217935-INS-73 |
| chr4-103237252-INS-292 | chr4-103237252-INS-289 |
| chr4-103244992-INS-6681 | chr4-103244993-INS-6681 |
| chr4-103261219-INS-176 | chr4-103261219-INS-176 |
| chr4-103264629-INS-954 | chr4-103264629-INS-954 |
| chr4-103272418-INS-54 | chr4-103272418-INS-54 |
| chr4-103283225-INS-667 | chr4-103283225-INS-666 |
| chr4-103316699-INS-59 | chr4-103316699-INS-60 |
| chr4-103327682-INS-80 | chr4-103327651-INS-100 |
| chr4-103359138-INS-1803 | chr4-103359139-INS-1803 |
| chr4-103378456-INS-121 | chr4-103378456-INS-121 |
| chr4-103382691-INS-100 | chr4-103382691-INS-102 |
| chr4-103394524-INS-2979 | chr4-103394525-INS-2979 |
| chr4-103407620-INS-74 | chr4-103407620-INS-74 |
| chr4-103451276-INS-1681 | chr4-103451277-INS-1681 |
| chr4-103472282-INS-56 | chr4-103472270-INS-52 |
| chr4-103525490-INS-374 | chr4-103525490-INS-375 |
| chr4-103528354-INS-336 | chr4-103528354-INS-336 |
| chr4-103546398-INS-98 | chr4-103546401-INS-100 |
| chr4-103547906-INS-1582 | chr4-103547907-INS-1582 |
| chr4-103549850-INS-152 | chr4-103549849-INS-120 |
| chr4-103567677-INS-62 | chr4-103567677-INS-50 |
| chr4-103578984-INS-138 | chr4-103578984-INS-142 |
| chr4-103585282-INS-122 | chr4-103585282-INS-165 |
| chr4-103587505-INS-6362 | chr4-103587506-INS-6362 |
| chr4-103597842-INS-4718 | chr4-103597843-INS-4719 |
| chr4-103605391-INS-3275 | chr4-103605392-INS-3275 |
| chr4-103613985-INS-71 | chr4-103613985-INS-90 |
| chr4-103616582-INS-54 | chr4-103616582-INS-52 |
| chr4-103624201-INS-171 | chr4-103624201-INS-181 |
| chr4-103655208-INS-54 | chr4-103655208-INS-60 |
| chr4-103657977-INS-219 | chr4-103657977-INS-225 |
| chr4-103696153-INS-68 | chr4-103696153-INS-68 |
| chr4-103700911-INS-100 | chr4-103700923-INS-50 |
| chr4-103715901-INS-166 | chr4-103715901-INS-165 |
| chr4-103717205-INS-51 | chr4-103717205-INS-86 |
| chr4-103723371-INS-592 | chr4-103723371-INS-590 |
| chr4-103767827-INS-6414 | chr4-103767828-INS-6414 |
| chr4-103808620-INS-3508 | chr4-103808621-INS-3509 |
| chr4-103820216-INS-114 | chr4-103820216-INS-66 |
| chr4-103830417-INS-72 | chr4-103830417-INS-72 |
| chr4-103836697-INS-1314 | chr4-103836698-INS-1314 |
| chr4-103849793-INS-65 | chr4-103849818-INS-55 |
| chr4-103900104-INS-64 | chr4-103900104-INS-64 |
| chr4-103974508-INS-1526 | chr4-103974509-INS-1526 |
| chr4-103994990-INS-388 | chr4-103994990-INS-388 |
| chr4-103996559-INS-78 | chr4-103996559-INS-52 |
| chr4-104004292-INS-77 | chr4-104004292-INS-59 |
| chr4-104014829-INS-4865 | chr4-104014830-INS-4865 |
| chr4-104034467-INS-68 | chr4-104034467-INS-70 |
| chr4-104036629-INS-274 | chr4-104036618-INS-269 |
| chr4-104063942-INS-248 | chr4-104063942-INS-249 |
| chr4-104098996-INS-114 | chr4-104098996-INS-114 |
| chr4-104101928-INS-2248 | chr4-104101929-INS-2248 |
| chr4-104103273-INS-320 | chr4-104103273-INS-320 |
| chr4-104196134-INS-76 | chr4-104196134-INS-76 |
| chr4-104289744-INS-92 | chr4-104289746-INS-82 |
| chr4-104296388-INS-70 | chr4-104296388-INS-71 |
| chr4-104322139-INS-63 | chr4-104322139-INS-57 |
| chr4-104376386-INS-60 | chr4-104376386-INS-60 |
| chr4-104394368-INS-90 | chr4-104394391-INS-66 |
| chr4-104406291-INS-1405 | chr4-104406292-INS-1404 |
| chr4-104420405-INS-6487 | chr4-104420406-INS-6487 |
| chr4-104434863-INS-630 | chr4-104434863-INS-629 |
| chr4-104479816-INS-61 | chr4-104479816-INS-61 |
| chr4-104556141-INS-174 | chr4-104556141-INS-175 |
| chr4-104565055-INS-54 | chr4-104565055-INS-60 |
| chr4-104579231-INS-54 | chr4-104579231-INS-58 |
| chr4-104593129-INS-59 | chr4-104593129-INS-59 |
| chr4-104603398-INS-878 | chr4-104603398-INS-872 |
| chr4-104618391-INS-91 | chr4-104618391-INS-83 |
| chr4-104641437-INS-195 | chr4-104641437-INS-195 |

| | |
|---|---|
| chr4-104724151-INS-78 | chr4-104724151-INS-78 |
| chr4-104727846-INS-114 | chr4-104727846-INS-114 |
| chr4-104731606-INS-98 | chr4-104731606-INS-98 |
| chr4-104744255-INS-460 | chr4-104744255-INS-413 |
| chr4-104746415-INS-138 | chr4-104746415-INS-125 |
| chr4-104750525-INS-140 | chr4-104750493-INS-130 |
| chr4-104766608-INS-70 | chr4-104766608-INS-70 |
| chr4-104779630-INS-51 | chr4-104779630-INS-51 |
| chr4-104797547-INS-98 | chr4-104797547-INS-98 |
| chr4-104826156-INS-573 | chr4-104826156-INS-574 |
| chr4-104831157-INS-545 | chr4-104831157-INS-545 |
| chr4-104879738-INS-122 | chr4-104879738-INS-123 |
| chr4-104895620-INS-51 | chr4-104895622-INS-62 |
| chr4-104920235-INS-60 | chr4-104920235-INS-60 |
| chr4-104931325-INS-132 | chr4-104931320-INS-136 |
| chr4-104939982-INS-174 | chr4-104939982-INS-180 |
| chr4-104945456-INS-60 | chr4-104945456-INS-62 |
| chr4-104994956-INS-60 | chr4-104994956-INS-62 |
| chr4-105004174-INS-130 | chr4-105004174-INS-115 |
| chr4-105034657-INS-99 | chr4-105034657-INS-99 |
| chr4-105072672-INS-61 | chr4-105072672-INS-61 |
| chr4-105093172-INS-52 | chr4-105093172-INS-52 |
| chr4-105102587-INS-6385 | chr4-105102588-INS-6386 |
| chr4-105104166-INS-80 | chr4-105104166-INS-56 |
| chr4-105110619-INS-63 | chr4-105110619-INS-64 |
| chr4-105124987-INS-90 | chr4-105124987-INS-90 |
| chr4-105139929-INS-384 | chr4-105139929-INS-383 |
| chr4-105141982-INS-252 | chr4-105141982-INS-252 |
| chr4-105146012-INS-64 | chr4-105146012-INS-76 |
| chr4-105147913-INS-63 | chr4-105147913-INS-55 |
| chr4-105173591-INS-204 | chr4-105173591-INS-204 |
| chr4-105192881-INS-54 | chr4-105192865-INS-68 |
| chr4-105194696-INS-67 | chr4-105194672-INS-77 |
| chr4-105240232-INS-80 | chr4-105240236-INS-96 |
| chr4-105252667-INS-60 | chr4-105252667-INS-60 |
| chr4-105266054-INS-55 | chr4-105266091-INS-67 |
| chr4-105344777-INS-183 | chr4-105344777-INS-183 |
| chr4-105349155-INS-70 | chr4-105349157-INS-58 |
| chr4-105352172-INS-563 | chr4-105352172-INS-563 |
| chr4-105366436-INS-57 | chr4-105366436-INS-57 |
| chr4-105400531-INS-281 | chr4-105400531-INS-287 |
| chr4-105409868-INS-206 | chr4-105409868-INS-206 |
| chr4-105427937-INS-656 | chr4-105427937-INS-657 |
| chr4-105440513-INS-225 | chr4-105440513-INS-225 |
| chr4-105468978-INS-52 | chr4-105468978-INS-52 |
| chr4-105489195-INS-6738 | chr4-105489196-INS-6741 |
| chr4-105499715-INS-214 | chr4-105499715-INS-211 |
| chr4-105502337-INS-88 | chr4-105502337-INS-88 |
| chr4-105503809-INS-407 | chr4-105503809-INS-399 |
| chr4-105614981-INS-6542 | chr4-105614982-INS-6542 |
| chr4-105629627-INS-72 | chr4-105629627-INS-72 |
| chr4-105645236-INS-234 | chr4-105645236-INS-234 |
| chr4-105682089-INS-4586 | chr4-105682089-INS-4586 |
| chr4-105706333-INS-2646 | chr4-105706345-INS-2649 |
| chr4-105775259-INS-152 | chr4-105775308-INS-118 |
| chr4-105787138-INS-55 | chr4-105787138-INS-55 |
| chr4-105797584-INS-120 | chr4-105797583-INS-147 |
| chr4-105803408-INS-695 | chr4-105803408-INS-695 |
| chr4-105817656-INS-55 | chr4-105817656-INS-55 |
| chr4-105821250-INS-107 | chr4-105821250-INS-107 |
| chr4-105838853-INS-92 | chr4-105838853-INS-92 |
| chr4-105856447-INS-2524 | chr4-105856448-INS-2524 |
| chr4-105866345-INS-59 | chr4-105866345-INS-59 |
| chr4-105882292-INS-6380 | chr4-105882293-INS-6380 |
| chr4-105904113-INS-82 | chr4-105904116-INS-83 |
| chr4-105904689-INS-4097 | chr4-105904690-INS-4097 |
| chr4-105925887-INS-286 | chr4-105925887-INS-286 |
| chr4-105969135-INS-1360 | chr4-105969136-INS-1360 |
| chr4-105976484-INS-6931 | chr4-105976485-INS-6932 |
| chr4-105984328-INS-3461 | chr4-105984329-INS-3452 |
| chr4-106003198-INS-149 | chr4-106003189-INS-164 |
| chr4-106007310-INS-244 | chr4-106007292-INS-236 |
| chr4-106027967-INS-106 | chr4-106027967-INS-111 |
| chr4-106031019-INS-304 | chr4-106031019-INS-303 |
| chr4-106058023-INS-96 | chr4-106058023-INS-96 |
| chr4-106086719-INS-54 | chr4-106086719-INS-54 |
| chr4-106087447-INS-556 | chr4-106087447-INS-557 |

| | |
|---|---|
| chr4-106103763-INS-4199 | chr4-106103764-INS-4200 |
| chr4-106106558-INS-254 | chr4-106106558-INS-255 |
| chr4-106119023-INS-178 | chr4-106119023-INS-167 |
| chr4-106138557-INS-84 | chr4-106138557-INS-84 |
| chr4-106146529-INS-58 | chr4-106146500-INS-60 |
| chr4-106162884-INS-6400 | chr4-106162885-INS-6400 |
| chr4-106167598-INS-58 | chr4-106167598-INS-56 |
| chr4-106169642-INS-54 | chr4-106169642-INS-54 |
| chr4-106170224-INS-182 | chr4-106170224-INS-182 |
| chr4-106171559-INS-67 | chr4-106171559-INS-67 |
| chr4-106259627-INS-70 | chr4-106259664-INS-66 |
| chr4-106322996-INS-353 | chr4-106323026-INS-319 |
| chr4-106326577-INS-138 | chr4-106326568-INS-147 |
| chr4-106344667-INS-54 | chr4-106344656-INS-63 |
| chr4-106348858-INS-72 | chr4-106348858-INS-72 |
| chr4-106351860-INS-2994 | chr4-106351861-INS-2994 |
| chr4-106370244-INS-1895 | chr4-106370245-INS-1895 |
| chr4-106389898-INS-204 | chr4-106389898-INS-204 |
| chr4-106400739-INS-55 | chr4-106400731-INS-62 |
| chr4-106406235-INS-216 | chr4-106406235-INS-216 |
| chr4-106416225-INS-66 | chr4-106416225-INS-66 |
| chr4-106424169-INS-221 | chr4-106424169-INS-221 |
| chr4-106427023-INS-62 | chr4-106427023-INS-62 |
| chr4-106441215-INS-132 | chr4-106441216-INS-132 |
| chr4-106533493-INS-310 | chr4-106533493-INS-310 |
| chr4-106547230-INS-117 | chr4-106547230-INS-125 |
| chr4-106634162-INS-63 | chr4-106634162-INS-63 |
| chr4-106658776-INS-72 | chr4-106658744-INS-56 |
| chr4-106702150-INS-404 | chr4-106702150-INS-404 |
| chr4-106703932-INS-104 | chr4-106703932-INS-99 |
| chr4-106724695-INS-160 | chr4-106724695-INS-180 |
| chr4-106731193-INS-84 | chr4-106731193-INS-84 |
| chr4-106734326-INS-891 | chr4-106734327-INS-911 |
| chr4-106738648-INS-165 | chr4-106738648-INS-165 |
| chr4-106751238-INS-228 | chr4-106751238-INS-236 |
| chr4-106754090-INS-52 | chr4-106754090-INS-52 |
| chr4-106761364-INS-72 | chr4-106761364-INS-74 |
| chr4-106812691-INS-51 | chr4-106812691-INS-51 |
| chr4-106847423-INS-62 | chr4-106847423-INS-58 |
| chr4-106900078-INS-102 | chr4-106900072-INS-144 |
| chr4-106907443-INS-55 | chr4-106907443-INS-55 |
| chr4-106917185-INS-218 | chr4-106917185-INS-243 |
| chr4-106945989-INS-57 | chr4-106945989-INS-50 |
| chr4-106947517-INS-117 | chr4-106947517-INS-118 |
| chr4-106971234-INS-57 | chr4-106971234-INS-57 |
| chr4-107010875-INS-425 | chr4-107010875-INS-425 |
| chr4-107021998-INS-3546 | chr4-107021999-INS-3546 |
| chr4-107023922-INS-73 | chr4-107023922-INS-61 |
| chr4-107030995-INS-7153 | chr4-107030996-INS-7153 |
| chr4-107055723-INS-98 | chr4-107055705-INS-84 |
| chr4-107096611-INS-117 | chr4-107096611-INS-117 |
| chr4-107113020-INS-54 | chr4-107113020-INS-54 |
| chr4-107150571-INS-145 | chr4-107150579-INS-134 |
| chr4-107151875-INS-93 | chr4-107151875-INS-83 |
| chr4-107157681-INS-63 | chr4-107157681-INS-56 |
| chr4-107160814-INS-174 | chr4-107160814-INS-175 |
| chr4-107184416-INS-1037 | chr4-107184417-INS-1037 |
| chr4-107194389-INS-201 | chr4-107194389-INS-205 |
| chr4-107198010-INS-52 | chr4-107198010-INS-52 |
| chr4-107216393-INS-209 | chr4-107216393-INS-208 |
| chr4-107220595-INS-405 | chr4-107220596-INS-405 |
| chr4-107228587-INS-211 | chr4-107228587-INS-210 |
| chr4-107234500-INS-179 | chr4-107234500-INS-179 |
| chr4-107235271-INS-205 | chr4-107235259-INS-200 |
| chr4-107245218-INS-192 | chr4-107245218-INS-190 |
| chr4-107270335-INS-170 | chr4-107270335-INS-170 |
| chr4-107307765-INS-251 | chr4-107307765-INS-253 |
| chr4-107337411-INS-167 | chr4-107337407-INS-214 |
| chr4-107343034-INS-71 | chr4-107343034-INS-71 |
| chr4-107385165-INS-186 | chr4-107385165-INS-187 |
| chr4-107390286-INS-100 | chr4-107390286-INS-100 |
| chr4-107418999-INS-220 | chr4-107418999-INS-220 |
| chr4-107441206-INS-80 | chr4-107441207-INS-52 |
| chr4-107475131-INS-55 | chr4-107475131-INS-53 |
| chr4-107515416-INS-124 | chr4-107515416-INS-160 |
| chr4-107570417-INS-92 | chr4-107570417-INS-92 |
| chr4-107571542-INS-54 | chr4-107571542-INS-55 |

| | |
|---|---|
| chr4-107580016-INS-204 | chr4-107580016-INS-204 |
| chr4-107589756-INS-197 | chr4-107589756-INS-186 |
| chr4-107606234-INS-185 | chr4-107606234-INS-185 |
| chr4-107615460-INS-504 | chr4-107615460-INS-504 |
| chr4-107621119-INS-180 | chr4-107621119-INS-180 |
| chr4-107640606-INS-215 | chr4-107640606-INS-215 |
| chr4-107646819-INS-215 | chr4-107646819-INS-214 |
| chr4-107710003-INS-399 | chr4-107710003-INS-399 |
| chr4-107758728-INS-55 | chr4-107758755-INS-55 |
| chr4-107763105-INS-7116 | chr4-107763106-INS-7116 |
| chr4-107803527-INS-207 | chr4-107803527-INS-207 |
| chr4-107810799-INS-140 | chr4-107810799-INS-140 |
| chr4-107849015-INS-54 | chr4-107849015-INS-54 |
| chr4-107855300-INS-81 | chr4-107855300-INS-87 |
| chr4-107857818-INS-56 | chr4-107857818-INS-53 |
| chr4-107880209-INS-51 | chr4-107880209-INS-51 |
| chr4-107892600-INS-204 | chr4-107892606-INS-198 |
| chr4-107897588-INS-202 | chr4-107897571-INS-204 |
| chr4-107904365-INS-270 | chr4-107904365-INS-283 |
| chr4-107909692-INS-1865 | chr4-107909693-INS-1865 |
| chr4-107922984-INS-200 | chr4-107922984-INS-199 |
| chr4-107932939-INS-166 | chr4-107932941-INS-187 |
| chr4-107940034-INS-95 | chr4-107940034-INS-95 |
| chr4-107951257-INS-219 | chr4-107951257-INS-213 |
| chr4-107953285-INS-443 | chr4-107953285-INS-443 |
| chr4-107960303-INS-124 | chr4-107960288-INS-136 |
| chr4-107962626-INS-112 | chr4-107962626-INS-112 |
| chr4-107964033-INS-109 | chr4-107964033-INS-109 |
| chr4-107977077-INS-204 | chr4-107977069-INS-205 |
| chr4-107986887-INS-68 | chr4-107986887-INS-68 |
| chr4-108017497-INS-170 | chr4-108017492-INS-173 |
| chr4-108019247-INS-183 | chr4-108019245-INS-211 |
| chr4-108027600-INS-140 | chr4-108027600-INS-100 |
| chr4-108030874-INS-68 | chr4-108030874-INS-102 |
| chr4-108033070-INS-52 | chr4-108033070-INS-50 |
| chr4-108034568-INS-182 | chr4-108034568-INS-182 |
| chr4-108038146-INS-62 | chr4-108038146-INS-64 |
| chr4-108063562-INS-60 | chr4-108063562-INS-58 |
| chr4-108076562-INS-208 | chr4-108076562-INS-206 |
| chr4-108081092-INS-427 | chr4-108081092-INS-427 |
| chr4-108083209-INS-63 | chr4-108083209-INS-70 |
| chr4-108086600-INS-199 | chr4-108086600-INS-200 |
| chr4-108149567-INS-124 | chr4-108149567-INS-124 |
| chr4-108172463-INS-61 | chr4-108172463-INS-61 |
| chr4-108224299-INS-74 | chr4-108224254-INS-108 |
| chr4-108228520-INS-205 | chr4-108228520-INS-205 |
| chr4-108229561-INS-66 | chr4-108229561-INS-70 |
| chr4-108232112-INS-236 | chr4-108232112-INS-236 |
| chr4-108238559-INS-6403 | chr4-108238560-INS-6403 |
| chr4-108256697-INS-199 | chr4-108256697-INS-199 |
| chr4-108298089-INS-407 | chr4-108298089-INS-407 |
| chr4-108311216-INS-208 | chr4-108311216-INS-208 |
| chr4-108324602-INS-213 | chr4-108324602-INS-213 |
| chr4-108328958-INS-290 | chr4-108328958-INS-290 |
| chr4-108406049-INS-212 | chr4-108406049-INS-212 |
| chr4-108450893-INS-59 | chr4-108450893-INS-59 |
| chr4-108492756-INS-70 | chr4-108492756-INS-69 |
| chr4-108499679-INS-54 | chr4-108499679-INS-54 |
| chr4-108525179-INS-215 | chr4-108525179-INS-217 |
| chr4-108533237-INS-746 | chr4-108533237-INS-746 |
| chr4-108537444-INS-117 | chr4-108537444-INS-136 |
| chr4-108544714-INS-234 | chr4-108544714-INS-234 |
| chr4-108552620-INS-203 | chr4-108552616-INS-206 |
| chr4-108572726-INS-60 | chr4-108572714-INS-91 |
| chr4-108580067-INS-208 | chr4-108580067-INS-209 |
| chr4-108598000-INS-136 | chr4-108598000-INS-136 |
| chr4-108614320-INS-70 | chr4-108614320-INS-74 |
| chr4-108617857-INS-53 | chr4-108617827-INS-58 |
| chr4-108628134-INS-187 | chr4-108628134-INS-194 |
| chr4-108639235-INS-64 | chr4-108639225-INS-58 |
| chr4-108646080-INS-54 | chr4-108646080-INS-54 |
| chr4-108653193-INS-140 | chr4-108653193-INS-148 |
| chr4-108654424-INS-168 | chr4-108654424-INS-168 |
| chr4-108656939-INS-204 | chr4-108656939-INS-203 |
| chr4-108659058-INS-212 | chr4-108659058-INS-212 |
| chr4-108664237-INS-69 | chr4-108664242-INS-85 |
| chr4-108667596-INS-225 | chr4-108667596-INS-225 |

| | |
|---|---|
| chr4-108668713-INS-201 | chr4-108668713-INS-203 |
| chr4-108677302-INS-896 | chr4-108677312-INS-924 |
| chr4-108722143-INS-114 | chr4-108722143-INS-114 |
| chr4-108738509-INS-153 | chr4-108738509-INS-153 |
| chr4-108746463-INS-100 | chr4-108746463-INS-100 |
| chr4-108754699-INS-776 | chr4-108754699-INS-776 |
| chr4-108758352-INS-175 | chr4-108758352-INS-175 |
| chr4-108778770-INS-5437 | chr4-108778771-INS-5437 |
| chr4-108782268-INS-99 | chr4-108782280-INS-100 |
| chr4-108785420-INS-181 | chr4-108785420-INS-180 |
| chr4-108805276-INS-222 | chr4-108805276-INS-224 |
| chr4-108834973-INS-59 | chr4-108834973-INS-59 |
| chr4-108837411-INS-66 | chr4-108837411-INS-60 |
| chr4-108837794-INS-5586 | chr4-108837795-INS-5586 |
| chr4-108882595-INS-355 | chr4-108882595-INS-355 |
| chr4-108904009-INS-70 | chr4-108904009-INS-70 |
| chr4-108920134-INS-182 | chr4-108920134-INS-183 |
| chr4-108947447-INS-59 | chr4-108947447-INS-59 |
| chr4-108950055-INS-86 | chr4-108950055-INS-86 |
| chr4-108983064-INS-70 | chr4-108983064-INS-70 |
| chr4-109044374-INS-51 | chr4-109044374-INS-51 |
| chr4-109078681-INS-217 | chr4-109078681-INS-218 |
| chr4-109105142-INS-60 | chr4-109105142-INS-58 |
| chr4-109134996-INS-703 | chr4-109134996-INS-704 |
| chr4-109135855-INS-5391 | chr4-109135856-INS-5391 |
| chr4-109148380-INS-242 | chr4-109148381-INS-222 |
| chr4-109165408-INS-94 | chr4-109165408-INS-74 |
| chr4-109189125-INS-54 | chr4-109189125-INS-54 |
| chr4-109221314-INS-155 | chr4-109221319-INS-160 |
| chr4-109234677-INS-127 | chr4-109234677-INS-126 |
| chr4-109246606-INS-3215 | chr4-109246607-INS-3215 |
| chr4-109266192-INS-124 | chr4-109266192-INS-132 |
| chr4-109347253-INS-108 | chr4-109347253-INS-108 |
| chr4-109414232-INS-119 | chr4-109414232-INS-119 |
| chr4-109427680-INS-251 | chr4-109427680-INS-252 |
| chr4-109437321-INS-69 | chr4-109437321-INS-69 |
| chr4-109442497-INS-54 | chr4-109442501-INS-58 |
| chr4-109448249-INS-82 | chr4-109448249-INS-82 |
| chr4-109458959-INS-489 | chr4-109458959-INS-505 |
| chr4-109479456-INS-198 | chr4-109479456-INS-198 |
| chr4-109484416-INS-116 | chr4-109484416-INS-116 |
| chr4-109499351-INS-287 | chr4-109499351-INS-287 |
| chr4-109503777-INS-66 | chr4-109503777-INS-69 |
| chr4-109528474-INS-234 | chr4-109528474-INS-237 |
| chr4-109541828-INS-60 | chr4-109541828-INS-60 |
| chr4-109546318-INS-789 | chr4-109546318-INS-789 |
| chr4-109551763-INS-217 | chr4-109551763-INS-217 |
| chr4-109612299-INS-55 | chr4-109612299-INS-57 |
| chr4-109651070-INS-90 | chr4-109651070-INS-90 |
| chr4-109680178-INS-990 | chr4-109680178-INS-988 |
| chr4-109690208-INS-66 | chr4-109690208-INS-72 |
| chr4-109700553-INS-345 | chr4-109700555-INS-347 |
| chr4-109709036-INS-1855 | chr4-109709037-INS-1854 |
| chr4-109714544-INS-72 | chr4-109714544-INS-54 |
| chr4-109752377-INS-68 | chr4-109752377-INS-76 |
| chr4-109762038-INS-52 | chr4-109762038-INS-52 |
| chr4-109790538-INS-58 | chr4-109790538-INS-58 |
| chr4-109795574-INS-588 | chr4-109795574-INS-588 |
| chr4-109816559-INS-101 | chr4-109816552-INS-74 |
| chr4-109823705-INS-403 | chr4-109823705-INS-403 |
| chr4-109824223-INS-54 | chr4-109824221-INS-53 |
| chr4-109863261-INS-188 | chr4-109863261-INS-188 |
| chr4-109868831-INS-6215 | chr4-109868832-INS-6215 |
| chr4-109886027-INS-174 | chr4-109886028-INS-174 |
| chr4-109900689-INS-89 | chr4-109900689-INS-89 |
| chr4-109907290-INS-6442 | chr4-109907291-INS-6442 |
| chr4-109919642-INS-56 | chr4-109919642-INS-56 |
| chr4-110000835-INS-111 | chr4-110000835-INS-111 |
| chr4-110013031-INS-1181 | chr4-110013032-INS-1181 |
| chr4-110030523-INS-6823 | chr4-110030524-INS-6825 |
| chr4-110057832-INS-199 | chr4-110057832-INS-200 |
| chr4-110059124-INS-56 | chr4-110059124-INS-62 |
| chr4-110080850-INS-57 | chr4-110080815-INS-57 |
| chr4-110106503-INS-140 | chr4-110106503-INS-140 |
| chr4-110145457-INS-68 | chr4-110145445-INS-52 |
| chr4-110216028-INS-96 | chr4-110216028-INS-96 |
| chr4-110229097-INS-471 | chr4-110229097-INS-470 |

| | |
|---|---|
| chr4-110240679-INS-243 | chr4-110240679-INS-243 |
| chr4-110278491-INS-6546 | chr4-110278492-INS-6547 |
| chr4-110300402-INS-1017 | chr4-110300403-INS-1008 |
| chr4-110305347-INS-190 | chr4-110305347-INS-191 |
| chr4-110345412-INS-5530 | chr4-110345413-INS-5530 |
| chr4-110359907-INS-170 | chr4-110359907-INS-170 |
| chr4-110393737-INS-60 | chr4-110393729-INS-64 |
| chr4-110421654-INS-61 | chr4-110421654-INS-61 |
| chr4-110422419-INS-109 | chr4-110422419-INS-109 |
| chr4-110493500-INS-5359 | chr4-110493501-INS-5361 |
| chr4-110499896-INS-2782 | chr4-110499897-INS-2825 |
| chr4-110533705-INS-1885 | chr4-110533706-INS-1885 |
| chr4-110544501-INS-125 | chr4-110544501-INS-125 |
| chr4-110575256-INS-214 | chr4-110575256-INS-214 |
| chr4-110578498-INS-79 | chr4-110578498-INS-79 |
| chr4-110627846-INS-174 | chr4-110627846-INS-181 |
| chr4-110655972-INS-708 | chr4-110655972-INS-708 |
| chr4-110688637-INS-907 | chr4-110688636-INS-912 |
| chr4-110693938-INS-661 | chr4-110693938-INS-661 |
| chr4-110733627-INS-61 | chr4-110733641-INS-80 |
| chr4-110748814-INS-206 | chr4-110748814-INS-230 |
| chr4-110756833-INS-160 | chr4-110756833-INS-119 |
| chr4-110764733-INS-10743 | chr4-110764734-INS-10750 |
| chr4-110779863-INS-83 | chr4-110779863-INS-83 |
| chr4-110790536-INS-82 | chr4-110790536-INS-90 |
| chr4-110806917-INS-884 | chr4-110806917-INS-884 |
| chr4-110863061-INS-231 | chr4-110863061-INS-231 |
| chr4-110919319-INS-152 | chr4-110919319-INS-152 |
| chr4-110920526-INS-240 | chr4-110920526-INS-196 |
| chr4-110954127-INS-6240 | chr4-110954128-INS-6240 |
| chr4-111008715-INS-61 | chr4-111008715-INS-67 |
| chr4-111031520-INS-199 | chr4-111031470-INS-199 |
| chr4-111034046-INS-196 | chr4-111034046-INS-196 |
| chr4-111038247-INS-89 | chr4-111038253-INS-92 |
| chr4-111071201-INS-436 | chr4-111071201-INS-436 |
| chr4-111141500-INS-1078 | chr4-111141501-INS-1078 |
| chr4-111165112-INS-7103 | chr4-111165113-INS-7103 |
| chr4-111175963-INS-302 | chr4-111175963-INS-302 |
| chr4-111198637-INS-60 | chr4-111198660-INS-68 |
| chr4-111245301-INS-201 | chr4-111245301-INS-201 |
| chr4-111262750-INS-1898 | chr4-111262751-INS-1898 |
| chr4-111271986-INS-143 | chr4-111271986-INS-143 |
| chr4-111280597-INS-223 | chr4-111280597-INS-225 |
| chr4-111319081-INS-148 | chr4-111319079-INS-163 |
| chr4-111322675-INS-60 | chr4-111322647-INS-53 |
| chr4-111323905-INS-60 | chr4-111323905-INS-62 |
| chr4-111328222-INS-65 | chr4-111328222-INS-65 |
| chr4-111366865-INS-116 | chr4-111366865-INS-112 |
| chr4-111398276-INS-765 | chr4-111398276-INS-765 |
| chr4-111406146-INS-123 | chr4-111406146-INS-123 |
| chr4-111408055-INS-67 | chr4-111408019-INS-115 |
| chr4-111415972-INS-5432 | chr4-111415973-INS-5432 |
| chr4-111419017-INS-1605 | chr4-111419018-INS-1605 |
| chr4-111449630-INS-60 | chr4-111449630-INS-52 |
| chr4-111461817-INS-169 | chr4-111461817-INS-169 |
| chr4-111471535-INS-54 | chr4-111471535-INS-50 |
| chr4-111478630-INS-60 | chr4-111478630-INS-64 |
| chr4-111505022-INS-67 | chr4-111505022-INS-50 |
| chr4-111529732-INS-60 | chr4-111529732-INS-64 |
| chr4-111553191-INS-73 | chr4-111553221-INS-65 |
| chr4-111561759-INS-55 | chr4-111561759-INS-55 |
| chr4-111567682-INS-56 | chr4-111567682-INS-54 |
| chr4-111571129-INS-56 | chr4-111571129-INS-58 |
| chr4-111572284-INS-3088 | chr4-111572285-INS-3088 |
| chr4-111574929-INS-256 | chr4-111574929-INS-256 |
| chr4-111591610-INS-70 | chr4-111591610-INS-65 |
| chr4-111606530-INS-1395 | chr4-111606531-INS-1396 |
| chr4-111638787-INS-60 | chr4-111638787-INS-60 |
| chr4-111650966-INS-1383 | chr4-111650967-INS-1383 |
| chr4-111660522-INS-582 | chr4-111660522-INS-582 |
| chr4-111704410-INS-176 | chr4-111704410-INS-176 |
| chr4-111745510-INS-74 | chr4-111745510-INS-80 |
| chr4-111820455-INS-322 | chr4-111820455-INS-323 |
| chr4-111833428-INS-331 | chr4-111833428-INS-331 |
| chr4-111867592-INS-175 | chr4-111867566-INS-196 |
| chr4-112142363-INS-1340 | chr4-112142365-INS-1336 |
| chr4-112475236-INS-199 | chr4-112475236-INS-199 |

| | |
|---|---|
| chr4-112507296-INS-57 | chr4-112507296-INS-61 |
| chr4-112515264-INS-246 | chr4-112515264-INS-246 |
| chr4-112577872-INS-6477 | chr4-112577873-INS-6477 |
| chr4-112593264-INS-58 | chr4-112593264-INS-58 |
| chr4-112598924-INS-162 | chr4-112598924-INS-162 |
| chr4-112603339-INS-56 | chr4-112603339-INS-58 |
| chr4-112607177-INS-125 | chr4-112607177-INS-125 |
| chr4-112647347-INS-53 | chr4-112647346-INS-61 |
| chr4-112654524-INS-62 | chr4-112654524-INS-64 |
| chr4-113817208-INS-60 | chr4-113817208-INS-58 |
| chr4-113842871-INS-63 | chr4-113842874-INS-73 |
| chr4-113846668-INS-52 | chr4-113846668-INS-52 |
| chr4-113868871-INS-547 | chr4-113868871-INS-547 |
| chr4-113988108-INS-58 | chr4-113988120-INS-62 |
| chr4-114055147-INS-703 | chr4-114055147-INS-700 |
| chr4-114182614-INS-482 | chr4-114182614-INS-482 |
| chr4-114225797-INS-252 | chr4-114225797-INS-252 |
| chr4-114251455-INS-57 | chr4-114251455-INS-57 |
| chr4-114262034-INS-120 | chr4-114262034-INS-126 |
| chr4-114286121-INS-51 | chr4-114286121-INS-51 |
| chr4-114295176-INS-54 | chr4-114295176-INS-62 |
| chr4-114322727-INS-69 | chr4-114322727-INS-81 |
| chr4-114324620-INS-243 | chr4-114324620-INS-243 |
| chr4-114389846-INS-171 | chr4-114389846-INS-171 |
| chr4-114394522-INS-219 | chr4-114394522-INS-233 |
| chr4-114440247-INS-68 | chr4-114440247-INS-68 |
| chr4-114450033-INS-283 | chr4-114450033-INS-283 |
| chr4-114454605-INS-1035 | chr4-114454606-INS-1039 |
| chr4-114465665-INS-52 | chr4-114465665-INS-56 |
| chr4-114501005-INS-85 | chr4-114501005-INS-85 |
| chr4-114547497-INS-64 | chr4-114547476-INS-55 |
| chr4-114588871-INS-6742 | chr4-114588872-INS-6742 |
| chr4-114594514-INS-1886 | chr4-114594515-INS-1886 |
| chr4-114623563-INS-60 | chr4-114623563-INS-60 |
| chr4-114627294-INS-202 | chr4-114627294-INS-203 |
| chr4-114657894-INS-102 | chr4-114657898-INS-110 |
| chr4-114716806-INS-5281 | chr4-114716807-INS-5281 |
| chr4-114723226-INS-1033 | chr4-114723227-INS-1034 |
| chr4-114743824-INS-74 | chr4-114743810-INS-51 |
| chr4-114776120-INS-1897 | chr4-114776121-INS-1897 |
| chr4-114810955-INS-102 | chr4-114810941-INS-129 |
| chr4-114858954-INS-54 | chr4-114858954-INS-54 |
| chr4-114908931-INS-102 | chr4-114908931-INS-125 |
| chr4-114929334-INS-198 | chr4-114929334-INS-198 |
| chr4-114937730-INS-58 | chr4-114937730-INS-58 |
| chr4-114953529-INS-286 | chr4-114953529-INS-286 |
| chr4-114961340-INS-193 | chr4-114961340-INS-193 |
| chr4-114967684-INS-244 | chr4-114967684-INS-244 |
| chr4-114972341-INS-67 | chr4-114972341-INS-69 |
| chr4-114975366-INS-139 | chr4-114975366-INS-140 |
| chr4-114982584-INS-211 | chr4-114982584-INS-212 |
| chr4-114993556-INS-56 | chr4-114993560-INS-76 |
| chr4-115018868-INS-551 | chr4-115018867-INS-550 |
| chr4-115062048-INS-1983 | chr4-115062049-INS-1984 |
| chr4-115098542-INS-5387 | chr4-115098543-INS-5387 |
| chr4-115194446-INS-212 | chr4-115194446-INS-212 |
| chr4-115216976-INS-214 | chr4-115216976-INS-214 |
| chr4-115228053-INS-85 | chr4-115228053-INS-85 |
| chr4-115243415-INS-2367 | chr4-115243416-INS-2367 |
| chr4-115257929-INS-2876 | chr4-115257930-INS-2878 |
| chr4-115282598-INS-199 | chr4-115282598-INS-199 |
| chr4-115325703-INS-328 | chr4-115325703-INS-328 |
| chr4-115330769-INS-7241 | chr4-115330770-INS-7241 |
| chr4-115336058-INS-53 | chr4-115336065-INS-52 |
| chr4-115353637-INS-223 | chr4-115353637-INS-228 |
| chr4-115363556-INS-7116 | chr4-115363557-INS-7116 |
| chr4-115407386-INS-6465 | chr4-115407387-INS-6477 |
| chr4-115407574-INS-2638 | chr4-115407575-INS-2638 |
| chr4-115464767-INS-183 | chr4-115464767-INS-183 |
| chr4-115478937-INS-82 | chr4-115478937-INS-82 |
| chr4-115484620-INS-1359 | chr4-115484621-INS-1358 |
| chr4-115491626-INS-51 | chr4-115491626-INS-51 |
| chr4-115507486-INS-67 | chr4-115507486-INS-67 |
| chr4-115512970-INS-92 | chr4-115512970-INS-93 |
| chr4-115565342-INS-52 | chr4-115565338-INS-51 |
| chr4-115573755-INS-119 | chr4-115573755-INS-119 |
| chr4-115581375-INS-2490 | chr4-115581376-INS-2490 |

| | |
|---|---|
| chr4-115589730-INS-90 | chr4-115589720-INS-74 |
| chr4-115601358-INS-217 | chr4-115601358-INS-217 |
| chr4-115612153-INS-197 | chr4-115612153-INS-197 |
| chr4-115621414-INS-212 | chr4-115621414-INS-212 |
| chr4-115649539-INS-1890 | chr4-115649540-INS-1890 |
| chr4-115654370-INS-125 | chr4-115654382-INS-85 |
| chr4-115667258-INS-1604 | chr4-115667259-INS-1606 |
| chr4-115669326-INS-234 | chr4-115669326-INS-235 |
| chr4-115760607-INS-71 | chr4-115760607-INS-71 |
| chr4-115792920-INS-56 | chr4-115792920-INS-58 |
| chr4-115816442-INS-57 | chr4-115816442-INS-60 |
| chr4-115822951-INS-271 | chr4-115822951-INS-276 |
| chr4-115835917-INS-84 | chr4-115835916-INS-60 |
| chr4-115838477-INS-72 | chr4-115838477-INS-72 |
| chr4-115849664-INS-295 | chr4-115849664-INS-296 |
| chr4-115863486-INS-55 | chr4-115863487-INS-50 |
| chr4-115864274-INS-202 | chr4-115864274-INS-202 |
| chr4-115866148-INS-6440 | chr4-115866149-INS-6440 |
| chr4-115871489-INS-169 | chr4-115871495-INS-174 |
| chr4-115883916-INS-116 | chr4-115883916-INS-116 |
| chr4-115898999-INS-175 | chr4-115898999-INS-175 |
| chr4-115921551-INS-200 | chr4-115921551-INS-198 |
| chr4-115924900-INS-199 | chr4-115924900-INS-199 |
| chr4-115935323-INS-57 | chr4-115935323-INS-59 |
| chr4-115938999-INS-216 | chr4-115938999-INS-208 |
| chr4-115971249-INS-307 | chr4-115971249-INS-307 |
| chr4-116013570-INS-203 | chr4-116013570-INS-209 |
| chr4-116027234-INS-63 | chr4-116027235-INS-63 |
| chr4-116038321-INS-101 | chr4-116038305-INS-96 |
| chr4-116072274-INS-55 | chr4-116072279-INS-56 |
| chr4-116074885-INS-56 | chr4-116074885-INS-59 |
| chr4-116093963-INS-108 | chr4-116093963-INS-108 |
| chr4-116100200-INS-140 | chr4-116100200-INS-140 |
| chr4-116101223-INS-787 | chr4-116101223-INS-787 |
| chr4-116106008-INS-54 | chr4-116105998-INS-89 |
| chr4-116107321-INS-212 | chr4-116107321-INS-212 |
| chr4-116111605-INS-111 | chr4-116111605-INS-120 |
| chr4-116118857-INS-64 | chr4-116118857-INS-66 |
| chr4-116124850-INS-113 | chr4-116124850-INS-150 |
| chr4-116130827-INS-84 | chr4-116130820-INS-88 |
| chr4-116151959-INS-191 | chr4-116151959-INS-191 |
| chr4-116160980-INS-223 | chr4-116160980-INS-225 |
| chr4-116184191-INS-443 | chr4-116184191-INS-443 |
| chr4-116195251-INS-263 | chr4-116195251-INS-263 |
| chr4-116196508-INS-204 | chr4-116196508-INS-205 |
| chr4-116202352-INS-1146 | chr4-116202353-INS-1191 |
| chr4-116206140-INS-181 | chr4-116206140-INS-182 |
| chr4-116211548-INS-60 | chr4-116211548-INS-66 |
| chr4-116264311-INS-119 | chr4-116264311-INS-119 |
| chr4-116282877-INS-227 | chr4-116282877-INS-206 |
| chr4-116305852-INS-211 | chr4-116305852-INS-211 |
| chr4-116347725-INS-347 | chr4-116347725-INS-347 |
| chr4-116386578-INS-392 | chr4-116386578-INS-392 |
| chr4-116399351-INS-206 | chr4-116399338-INS-222 |
| chr4-116412818-INS-204 | chr4-116412818-INS-200 |
| chr4-116428220-INS-54 | chr4-116428220-INS-54 |
| chr4-116430801-INS-53 | chr4-116430791-INS-51 |
| chr4-116436355-INS-217 | chr4-116436355-INS-217 |
| chr4-116444243-INS-107 | chr4-116444243-INS-111 |
| chr4-116452426-INS-86 | chr4-116452426-INS-86 |
| chr4-116459895-INS-179 | chr4-116459895-INS-178 |
| chr4-116472081-INS-146 | chr4-116472081-INS-145 |
| chr4-116475830-INS-159 | chr4-116475830-INS-159 |
| chr4-116477325-INS-95 | chr4-116477325-INS-95 |
| chr4-116485282-INS-111 | chr4-116485282-INS-111 |
| chr4-116490608-INS-222 | chr4-116490608-INS-222 |
| chr4-116502275-INS-240 | chr4-116502275-INS-240 |
| chr4-116519017-INS-212 | chr4-116519018-INS-212 |
| chr4-116520491-INS-4730 | chr4-116520492-INS-4730 |
| chr4-116520943-INS-191 | chr4-116520943-INS-192 |
| chr4-116526471-INS-82 | chr4-116526499-INS-76 |
| chr4-116528858-INS-123 | chr4-116528858-INS-123 |
| chr4-116529343-INS-204 | chr4-116529349-INS-212 |
| chr4-116535620-INS-233 | chr4-116535620-INS-232 |
| chr4-116541528-INS-200 | chr4-116541528-INS-200 |
| chr4-116543740-INS-218 | chr4-116543740-INS-218 |
| chr4-116545104-INS-325 | chr4-116545105-INS-326 |

| | |
|---|---|
| chr4-116547922-INS-98 | chr4-116547922-INS-98 |
| chr4-116551951-INS-200 | chr4-116551951-INS-200 |
| chr4-116569480-INS-177 | chr4-116569480-INS-177 |
| chr4-116572887-INS-218 | chr4-116572887-INS-218 |
| chr4-116581268-INS-192 | chr4-116581268-INS-193 |
| chr4-116587836-INS-61 | chr4-116587847-INS-50 |
| chr4-116590108-INS-100 | chr4-116590108-INS-100 |
| chr4-116599963-INS-197 | chr4-116599963-INS-197 |
| chr4-116605537-INS-171 | chr4-116605538-INS-155 |
| chr4-116612274-INS-214 | chr4-116612274-INS-214 |
| chr4-116616192-INS-234 | chr4-116616192-INS-234 |
| chr4-116623933-INS-213 | chr4-116623933-INS-216 |
| chr4-116626271-INS-130 | chr4-116626271-INS-130 |
| chr4-116634139-INS-706 | chr4-116634139-INS-706 |
| chr4-116641151-INS-51 | chr4-116641151-INS-51 |
| chr4-116642227-INS-203 | chr4-116642227-INS-211 |
| chr4-116644734-INS-60 | chr4-116644734-INS-68 |
| chr4-116647628-INS-318 | chr4-116647628-INS-318 |
| chr4-116648552-INS-166 | chr4-116648541-INS-166 |
| chr4-116654678-INS-68 | chr4-116654678-INS-68 |
| chr4-116655038-INS-222 | chr4-116655039-INS-208 |
| chr4-116669401-INS-5253 | chr4-116669402-INS-5253 |
| chr4-116691418-INS-180 | chr4-116691418-INS-180 |
| chr4-116694906-INS-230 | chr4-116694906-INS-227 |
| chr4-116698662-INS-218 | chr4-116698662-INS-218 |
| chr4-116702293-INS-319 | chr4-116702293-INS-273 |
| chr4-116740812-INS-123 | chr4-116740812-INS-140 |
| chr4-116748177-INS-198 | chr4-116748177-INS-198 |
| chr4-116750756-INS-520 | chr4-116750756-INS-520 |
| chr4-116754126-INS-55 | chr4-116754126-INS-98 |
| chr4-116772033-INS-1049 | chr4-116772034-INS-1049 |
| chr4-116774330-INS-211 | chr4-116774330-INS-211 |
| chr4-116788096-INS-498 | chr4-116788096-INS-498 |
| chr4-116792412-INS-2935 | chr4-116792413-INS-2935 |
| chr4-116801158-INS-134 | chr4-116801152-INS-148 |
| chr4-116807561-INS-200 | chr4-116807561-INS-201 |
| chr4-116813222-INS-105 | chr4-116813222-INS-67 |
| chr4-116822172-INS-111 | chr4-116822172-INS-111 |
| chr4-116831587-INS-57 | chr4-116831587-INS-57 |
| chr4-116835780-INS-1701 | chr4-116835781-INS-1702 |
| chr4-116867444-INS-204 | chr4-116867432-INS-184 |
| chr4-116874420-INS-210 | chr4-116874420-INS-210 |
| chr4-116877393-INS-196 | chr4-116877393-INS-196 |
| chr4-116886382-INS-211 | chr4-116886382-INS-211 |
| chr4-116895972-INS-170 | chr4-116895972-INS-170 |
| chr4-116899246-INS-247 | chr4-116899292-INS-251 |
| chr4-116902389-INS-204 | chr4-116902389-INS-204 |
| chr4-116908731-INS-220 | chr4-116908731-INS-221 |
| chr4-116922453-INS-203 | chr4-116922453-INS-204 |
| chr4-116924942-INS-67 | chr4-116924942-INS-67 |
| chr4-116926828-INS-216 | chr4-116926829-INS-213 |
| chr4-116931996-INS-203 | chr4-116931996-INS-203 |
| chr4-116943500-INS-320 | chr4-116943500-INS-290 |
| chr4-116950479-INS-98 | chr4-116950470-INS-95 |
| chr4-116960277-INS-250 | chr4-116960277-INS-250 |
| chr4-116964488-INS-68 | chr4-116964488-INS-68 |
| chr4-116969905-INS-211 | chr4-116969901-INS-213 |
| chr4-116972825-INS-202 | chr4-116972827-INS-203 |
| chr4-116994045-INS-193 | chr4-116994045-INS-196 |
| chr4-116998694-INS-212 | chr4-116998694-INS-212 |
| chr4-117000942-INS-55 | chr4-117000942-INS-57 |
| chr4-117007016-INS-209 | chr4-117007016-INS-187 |
| chr4-117012057-INS-110 | chr4-117012057-INS-110 |
| chr4-117014628-INS-205 | chr4-117014628-INS-205 |
| chr4-117018370-INS-212 | chr4-117018370-INS-212 |
| chr4-117061229-INS-204 | chr4-117061229-INS-214 |
| chr4-117063185-INS-149 | chr4-117063185-INS-149 |
| chr4-117064514-INS-204 | chr4-117064515-INS-203 |
| chr4-117069154-INS-68 | chr4-117069153-INS-68 |
| chr4-117080785-INS-209 | chr4-117080785-INS-208 |
| chr4-117094808-INS-58 | chr4-117094808-INS-58 |
| chr4-117097666-INS-215 | chr4-117097666-INS-215 |
| chr4-117103299-INS-219 | chr4-117103299-INS-219 |
| chr4-117113483-INS-200 | chr4-117113483-INS-200 |
| chr4-117116584-INS-88 | chr4-117116584-INS-88 |
| chr4-117125953-INS-67 | chr4-117125917-INS-65 |
| chr4-117165921-INS-208 | chr4-117165921-INS-208 |

| | |
|---|---|
| chr4-117187191-INS-206 | chr4-117187191-INS-206 |
| chr4-117197640-INS-101 | chr4-117197640-INS-101 |
| chr4-117198334-INS-210 | chr4-117198334-INS-211 |
| chr4-117212749-INS-259 | chr4-117212749-INS-241 |
| chr4-117220382-INS-846 | chr4-117220382-INS-849 |
| chr4-117279264-INS-209 | chr4-117279264-INS-209 |
| chr4-117310234-INS-90 | chr4-117310234-INS-94 |
| chr4-117360732-INS-96 | chr4-117360732-INS-102 |
| chr4-117369155-INS-211 | chr4-117369155-INS-212 |
| chr4-117380833-INS-193 | chr4-117380833-INS-193 |
| chr4-117396624-INS-124 | chr4-117396624-INS-124 |
| chr4-117417223-INS-217 | chr4-117417225-INS-220 |
| chr4-117472128-INS-252 | chr4-117472128-INS-252 |
| chr4-117499167-INS-123 | chr4-117499167-INS-154 |
| chr4-117520848-INS-495 | chr4-117520848-INS-511 |
| chr4-117525420-INS-158 | chr4-117525420-INS-158 |
| chr4-117526408-INS-194 | chr4-117526408-INS-195 |
| chr4-117563339-INS-83 | chr4-117563339-INS-79 |
| chr4-117577482-INS-207 | chr4-117577482-INS-208 |
| chr4-117605590-INS-199 | chr4-117605590-INS-199 |
| chr4-117610342-INS-208 | chr4-117610342-INS-208 |
| chr4-117621795-INS-266 | chr4-117621795-INS-266 |
| chr4-117632937-INS-70 | chr4-117632957-INS-74 |
| chr4-117641754-INS-197 | chr4-117641754-INS-197 |
| chr4-117644167-INS-96 | chr4-117644157-INS-62 |
| chr4-117649929-INS-195 | chr4-117649929-INS-195 |
| chr4-117665565-INS-6441 | chr4-117665566-INS-6441 |
| chr4-117675886-INS-187 | chr4-117675886-INS-188 |
| chr4-117684514-INS-69 | chr4-117684515-INS-69 |
| chr4-117728132-INS-64 | chr4-117728132-INS-74 |
| chr4-117746600-INS-57 | chr4-117746600-INS-52 |
| chr4-117767720-INS-56 | chr4-117767720-INS-56 |
| chr4-117778139-INS-446 | chr4-117778139-INS-446 |
| chr4-117795964-INS-94 | chr4-117795964-INS-50 |
| chr4-117803105-INS-175 | chr4-117803105-INS-176 |
| chr4-117803754-INS-95 | chr4-117803754-INS-95 |
| chr4-117826082-INS-426 | chr4-117826082-INS-426 |
| chr4-117844740-INS-117 | chr4-117844740-INS-117 |
| chr4-117866134-INS-161 | chr4-117866134-INS-161 |
| chr4-117886860-INS-69 | chr4-117886861-INS-70 |
| chr4-117888075-INS-54 | chr4-117888075-INS-54 |
| chr4-117888690-INS-54 | chr4-117888690-INS-54 |
| chr4-117897858-INS-513 | chr4-117897858-INS-513 |
| chr4-117927379-INS-201 | chr4-117927379-INS-201 |
| chr4-117930287-INS-203 | chr4-117930287-INS-203 |
| chr4-117941351-INS-4051 | chr4-117941352-INS-4051 |
| chr4-117949368-INS-59 | chr4-117949368-INS-62 |
| chr4-117950818-INS-227 | chr4-117950818-INS-227 |
| chr4-117967442-INS-52 | chr4-117967442-INS-54 |
| chr4-117970446-INS-77 | chr4-117970447-INS-82 |
| chr4-117998188-INS-428 | chr4-117998188-INS-428 |
| chr4-118026112-INS-212 | chr4-118026112-INS-215 |
| chr4-118050302-INS-54 | chr4-118050302-INS-54 |
| chr4-118057144-INS-84 | chr4-118057144-INS-84 |
| chr4-118085346-INS-262 | chr4-118085346-INS-254 |
| chr4-118110775-INS-208 | chr4-118110775-INS-209 |
| chr4-118161991-INS-64 | chr4-118161991-INS-68 |
| chr4-118164055-INS-212 | chr4-118164055-INS-210 |
| chr4-118170681-INS-58 | chr4-118170681-INS-61 |
| chr4-118180472-INS-6452 | chr4-118180473-INS-6452 |
| chr4-118193093-INS-205 | chr4-118193093-INS-200 |
| chr4-118194455-INS-6884 | chr4-118194456-INS-6876 |
| chr4-118194536-INS-203 | chr4-118194536-INS-203 |
| chr4-118206917-INS-139 | chr4-118206917-INS-113 |
| chr4-118212263-INS-60 | chr4-118212263-INS-60 |
| chr4-118214712-INS-221 | chr4-118214712-INS-221 |
| chr4-118232429-INS-207 | chr4-118232429-INS-207 |
| chr4-118234202-INS-217 | chr4-118234202-INS-217 |
| chr4-118277066-INS-169 | chr4-118277066-INS-169 |
| chr4-118299146-INS-187 | chr4-118299146-INS-187 |
| chr4-118371359-INS-113 | chr4-118371359-INS-130 |
| chr4-118373524-INS-339 | chr4-118373528-INS-339 |
| chr4-118382312-INS-56 | chr4-118382312-INS-56 |
| chr4-118389727-INS-216 | chr4-118389727-INS-216 |
| chr4-118391515-INS-211 | chr4-118391515-INS-210 |
| chr4-118428595-INS-52 | chr4-118428587-INS-56 |
| chr4-118435387-INS-172 | chr4-118435384-INS-195 |

| | |
|---|---|
| chr4-118442381-INS-131 | chr4-118442381-INS-131 |
| chr4-118446347-INS-329 | chr4-118446347-INS-301 |
| chr4-118449769-INS-6992 | chr4-118449770-INS-6994 |
| chr4-118450857-INS-100 | chr4-118450857-INS-100 |
| chr4-118469798-INS-235 | chr4-118469798-INS-236 |
| chr4-118476887-INS-994 | chr4-118476887-INS-994 |
| chr4-118480620-INS-57 | chr4-118480620-INS-57 |
| chr4-118486087-INS-78 | chr4-118486087-INS-78 |
| chr4-118490913-INS-51 | chr4-118490909-INS-54 |
| chr4-118513491-INS-202 | chr4-118513491-INS-202 |
| chr4-118527669-INS-236 | chr4-118527669-INS-220 |
| chr4-118563905-INS-84 | chr4-118563905-INS-100 |
| chr4-118574946-INS-51 | chr4-118574922-INS-54 |
| chr4-118579333-INS-221 | chr4-118579333-INS-221 |
| chr4-118608056-INS-148 | chr4-118608056-INS-148 |
| chr4-118612611-INS-1893 | chr4-118612612-INS-1893 |
| chr4-118661911-INS-65 | chr4-118661897-INS-65 |
| chr4-118664701-INS-507 | chr4-118664701-INS-508 |
| chr4-118700946-INS-430 | chr4-118700946-INS-430 |
| chr4-118706156-INS-7081 | chr4-118706157-INS-7081 |
| chr4-118729019-INS-660 | chr4-118729019-INS-660 |
| chr4-118736117-INS-476 | chr4-118736117-INS-477 |
| chr4-118744117-INS-361 | chr4-118744117-INS-361 |
| chr4-118760874-INS-830 | chr4-118760874-INS-831 |
| chr4-118762749-INS-331 | chr4-118762749-INS-331 |
| chr4-118801855-INS-1894 | chr4-118801856-INS-1894 |
| chr4-118802960-INS-6917 | chr4-118802961-INS-6917 |
| chr4-118814021-INS-60 | chr4-118814021-INS-56 |
| chr4-118845581-INS-50 | chr4-118845581-INS-60 |
| chr4-118853517-INS-59 | chr4-118853517-INS-59 |
| chr4-118861020-INS-219 | chr4-118861020-INS-219 |
| chr4-118864577-INS-88 | chr4-118864577-INS-92 |
| chr4-118869805-INS-62 | chr4-118869805-INS-65 |
| chr4-118884590-INS-219 | chr4-118884590-INS-219 |
| chr4-118894677-INS-68 | chr4-118894677-INS-64 |
| chr4-118932488-INS-84 | chr4-118932488-INS-86 |
| chr4-118937048-INS-60 | chr4-118937048-INS-63 |
| chr4-118999336-INS-182 | chr4-118999345-INS-165 |
| chr4-119002389-INS-598 | chr4-119002389-INS-598 |
| chr4-119015684-INS-402 | chr4-119015684-INS-402 |
| chr4-119020282-INS-151 | chr4-119020282-INS-129 |
| chr4-119029326-INS-187 | chr4-119029326-INS-185 |
| chr4-119037988-INS-272 | chr4-119037988-INS-267 |
| chr4-119041031-INS-77 | chr4-119041031-INS-77 |
| chr4-119043717-INS-190 | chr4-119043717-INS-199 |
| chr4-119047406-INS-223 | chr4-119047406-INS-223 |
| chr4-119064549-INS-203 | chr4-119064549-INS-202 |
| chr4-119068866-INS-185 | chr4-119068866-INS-185 |
| chr4-119070476-INS-272 | chr4-119070476-INS-272 |
| chr4-119086889-INS-209 | chr4-119086889-INS-209 |
| chr4-119108065-INS-68 | chr4-119108065-INS-68 |
| chr4-119117520-INS-215 | chr4-119117520-INS-216 |
| chr4-119122299-INS-245 | chr4-119122299-INS-245 |
| chr4-119148861-INS-54 | chr4-119148861-INS-54 |
| chr4-119155421-INS-209 | chr4-119155421-INS-209 |
| chr4-119239074-INS-209 | chr4-119239074-INS-209 |
| chr4-119239862-INS-68 | chr4-119239862-INS-68 |
| chr4-119253221-INS-87 | chr4-119253221-INS-70 |
| chr4-119288644-INS-104 | chr4-119288646-INS-114 |
| chr4-119299977-INS-74 | chr4-119299977-INS-74 |
| chr4-119334594-INS-2833 | chr4-119334595-INS-2833 |
| chr4-119336239-INS-58 | chr4-119336239-INS-62 |
| chr4-119355243-INS-54 | chr4-119355243-INS-56 |
| chr4-119365136-INS-56 | chr4-119365160-INS-68 |
| chr4-119393794-INS-125 | chr4-119393794-INS-125 |
| chr4-119410227-INS-547 | chr4-119410227-INS-547 |
| chr4-119417274-INS-89 | chr4-119417274-INS-89 |
| chr4-119434512-INS-121 | chr4-119434512-INS-121 |
| chr4-119436097-INS-99 | chr4-119436097-INS-99 |
| chr4-119500422-INS-283 | chr4-119500422-INS-270 |
| chr4-119517619-INS-80 | chr4-119517619-INS-80 |
| chr4-119525658-INS-79 | chr4-119525658-INS-64 |
| chr4-119556324-INS-1060 | chr4-119556325-INS-1060 |
| chr4-119562684-INS-52 | chr4-119562684-INS-52 |
| chr4-119565451-INS-58 | chr4-119565451-INS-52 |
| chr4-119570887-INS-52 | chr4-119570887-INS-52 |
| chr4-119572852-INS-69 | chr4-119572852-INS-77 |

| | |
|---|---|
| chr4-119575419-INS-341 | chr4-119575419-INS-341 |
| chr4-119588025-INS-152 | chr4-119588025-INS-152 |
| chr4-119614256-INS-60 | chr4-119614256-INS-50 |
| chr4-119614920-INS-70 | chr4-119614925-INS-89 |
| chr4-119621509-INS-87 | chr4-119621509-INS-87 |
| chr4-119630783-INS-94 | chr4-119630783-INS-98 |
| chr4-119638473-INS-2839 | chr4-119638474-INS-2833 |
| chr4-119641859-INS-1160 | chr4-119641860-INS-1181 |
| chr4-119645554-INS-2994 | chr4-119645555-INS-2994 |
| chr4-119680185-INS-743 | chr4-119680185-INS-743 |
| chr4-119707488-INS-52 | chr4-119707475-INS-73 |
| chr4-119715683-INS-72 | chr4-119715683-INS-108 |
| chr4-119747349-INS-160 | chr4-119747349-INS-161 |
| chr4-119749260-INS-89 | chr4-119749279-INS-104 |
| chr4-119766305-INS-265 | chr4-119766305-INS-265 |
| chr4-119769377-INS-2571 | chr4-119769378-INS-2571 |
| chr4-119781061-INS-229 | chr4-119781061-INS-229 |
| chr4-119804566-INS-400 | chr4-119804566-INS-400 |
| chr4-119874540-INS-1893 | chr4-119874541-INS-1893 |
| chr4-119935231-INS-545 | chr4-119935231-INS-546 |
| chr4-119957705-INS-222 | chr4-119957695-INS-232 |
| chr4-119993915-INS-96 | chr4-119993915-INS-89 |
| chr4-120001097-INS-51 | chr4-120001097-INS-51 |
| chr4-120008619-INS-201 | chr4-120008615-INS-203 |
| chr4-120028348-INS-51 | chr4-120028348-INS-54 |
| chr4-120032951-INS-86 | chr4-120032952-INS-54 |
| chr4-120034975-INS-100 | chr4-120034975-INS-100 |
| chr4-120045112-INS-151 | chr4-120045112-INS-151 |
| chr4-120057642-INS-101 | chr4-120057637-INS-113 |
| chr4-120061216-INS-61 | chr4-120061216-INS-66 |
| chr4-120072769-INS-73 | chr4-120072769-INS-73 |
| chr4-120085491-INS-70 | chr4-120085516-INS-84 |
| chr4-120154906-INS-206 | chr4-120154906-INS-207 |
| chr4-120160179-INS-130 | chr4-120160179-INS-144 |
| chr4-120169062-INS-429 | chr4-120169062-INS-429 |
| chr4-120169733-INS-54 | chr4-120169719-INS-55 |
| chr4-120198544-INS-533 | chr4-120198544-INS-533 |
| chr4-120231595-INS-180 | chr4-120231595-INS-180 |
| chr4-120243438-INS-209 | chr4-120243438-INS-209 |
| chr4-120244600-INS-218 | chr4-120244600-INS-219 |
| chr4-120260283-INS-5152 | chr4-120260284-INS-5152 |
| chr4-120297799-INS-215 | chr4-120297799-INS-215 |
| chr4-120299272-INS-198 | chr4-120299272-INS-198 |
| chr4-120326582-INS-202 | chr4-120326582-INS-202 |
| chr4-120350538-INS-88 | chr4-120350538-INS-88 |
| chr4-120372165-INS-75 | chr4-120372165-INS-75 |
| chr4-120378419-INS-908 | chr4-120378419-INS-908 |
| chr4-120414574-INS-306 | chr4-120414574-INS-306 |
| chr4-120422424-INS-189 | chr4-120422424-INS-190 |
| chr4-120423589-INS-55 | chr4-120423589-INS-55 |
| chr4-120440188-INS-51 | chr4-120440188-INS-51 |
| chr4-120440436-INS-7195 | chr4-120440437-INS-7196 |
| chr4-120443796-INS-193 | chr4-120443796-INS-194 |
| chr4-120444644-INS-1006 | chr4-120444645-INS-966 |
| chr4-120445372-INS-52 | chr4-120445372-INS-80 |
| chr4-120448977-INS-62 | chr4-120448977-INS-62 |
| chr4-120451507-INS-55 | chr4-120451507-INS-58 |
| chr4-120464059-INS-80 | chr4-120464059-INS-80 |
| chr4-120482303-INS-356 | chr4-120482303-INS-356 |
| chr4-120487995-INS-52 | chr4-120487995-INS-52 |
| chr4-120510482-INS-63 | chr4-120510482-INS-70 |
| chr4-120512922-INS-213 | chr4-120512922-INS-205 |
| chr4-120514604-INS-5199 | chr4-120514605-INS-5199 |
| chr4-120534671-INS-84 | chr4-120534671-INS-76 |
| chr4-120537998-INS-106 | chr4-120537994-INS-89 |
| chr4-120546050-INS-112 | chr4-120546050-INS-112 |
| chr4-120684916-INS-7116 | chr4-120684917-INS-7116 |
| chr4-120692718-INS-57 | chr4-120692718-INS-57 |
| chr4-120708208-INS-219 | chr4-120708208-INS-219 |
| chr4-120719694-INS-137 | chr4-120719694-INS-137 |
| chr4-120768871-INS-536 | chr4-120768871-INS-536 |
| chr4-120773010-INS-306 | chr4-120772992-INS-307 |
| chr4-120773463-INS-72 | chr4-120773460-INS-102 |
| chr4-120774547-INS-120 | chr4-120774547-INS-120 |
| chr4-120801215-INS-210 | chr4-120801215-INS-211 |
| chr4-120810874-INS-60 | chr4-120810874-INS-68 |
| chr4-120814735-INS-93 | chr4-120814735-INS-93 |

| | |
|---|---|
| chr4-120816208-INS-203 | chr4-120816208-INS-203 |
| chr4-120818543-INS-183 | chr4-120818543-INS-184 |
| chr4-120822693-INS-71 | chr4-120822693-INS-73 |
| chr4-120823647-INS-185 | chr4-120823647-INS-185 |
| chr4-120829797-INS-244 | chr4-120829797-INS-233 |
| chr4-120843770-INS-98 | chr4-120843770-INS-98 |
| chr4-120864268-INS-153 | chr4-120864268-INS-153 |
| chr4-120878881-INS-51 | chr4-120878891-INS-81 |
| chr4-120886937-INS-338 | chr4-120886937-INS-335 |
| chr4-120889025-INS-164 | chr4-120889025-INS-163 |
| chr4-120891875-INS-208 | chr4-120891875-INS-207 |
| chr4-120895456-INS-208 | chr4-120895456-INS-208 |
| chr4-120918480-INS-176 | chr4-120918480-INS-176 |
| chr4-120921167-INS-382 | chr4-120921167-INS-383 |
| chr4-120930170-INS-197 | chr4-120930170-INS-201 |
| chr4-120972138-INS-9046 | chr4-120972139-INS-9046 |
| chr4-120973756-INS-274 | chr4-120973756-INS-276 |
| chr4-120983983-INS-215 | chr4-120983983-INS-215 |
| chr4-121009535-INS-206 | chr4-121009535-INS-203 |
| chr4-121074436-INS-61 | chr4-121074415-INS-56 |
| chr4-121088877-INS-199 | chr4-121088877-INS-200 |
| chr4-121136259-INS-71 | chr4-121136259-INS-74 |
| chr4-121293569-INS-58 | chr4-121293569-INS-58 |
| chr4-121301467-INS-631 | chr4-121301459-INS-595 |
| chr4-121301516-INS-59 | chr4-121301516-INS-84 |
| chr4-121304598-INS-493 | chr4-121304598-INS-495 |
| chr4-121407087-INS-946 | chr4-121407087-INS-942 |
| chr4-121444804-INS-137 | chr4-121444804-INS-132 |
| chr4-121480509-INS-62 | chr4-121480488-INS-52 |
| chr4-121571838-INS-115 | chr4-121571838-INS-115 |
| chr4-121596765-INS-90 | chr4-121596765-INS-90 |
| chr4-121604305-INS-522 | chr4-121604303-INS-526 |
| chr4-122438218-INS-6459 | chr4-122438219-INS-6460 |
| chr4-122507139-INS-5428 | chr4-122507140-INS-5428 |
| chr4-122513756-INS-6930 | chr4-122513757-INS-6931 |
| chr4-122546423-INS-1890 | chr4-122546424-INS-1890 |
| chr4-122574742-INS-1611 | chr4-122574743-INS-1611 |
| chr4-122579725-INS-268 | chr4-122579725-INS-268 |
| chr4-122607376-INS-424 | chr4-122607377-INS-424 |
| chr4-122610659-INS-327 | chr4-122610659-INS-327 |
| chr4-122611415-INS-1893 | chr4-122611416-INS-1893 |
| chr4-122694442-INS-221 | chr4-122694442-INS-223 |
| chr4-122704682-INS-401 | chr4-122704682-INS-401 |
| chr4-122771077-INS-1863 | chr4-122771078-INS-1863 |
| chr4-122804099-INS-72 | chr4-122804093-INS-52 |
| chr4-122808913-INS-101 | chr4-122808913-INS-113 |
| chr4-122813757-INS-75 | chr4-122813757-INS-75 |
| chr4-122818817-INS-107 | chr4-122818818-INS-84 |
| chr4-122832166-INS-175 | chr4-122832166-INS-168 |
| chr4-122833687-INS-104 | chr4-122833687-INS-152 |
| chr4-122836735-INS-68 | chr4-122836735-INS-68 |
| chr4-122878160-INS-211 | chr4-122878160-INS-208 |
| chr4-122883975-INS-162 | chr4-122883975-INS-161 |
| chr4-122902498-INS-57 | chr4-122902494-INS-85 |
| chr4-122912216-INS-1212 | chr4-122912217-INS-1202 |
| chr4-122917211-INS-53 | chr4-122917211-INS-53 |
| chr4-122964373-INS-277 | chr4-122964373-INS-274 |
| chr4-122967339-INS-87 | chr4-122967339-INS-87 |
| chr4-122970215-INS-64 | chr4-122970215-INS-64 |
| chr4-122983678-INS-60 | chr4-122983678-INS-60 |
| chr4-123012893-INS-64 | chr4-123012893-INS-64 |
| chr4-123019762-INS-67 | chr4-123019762-INS-67 |
| chr4-123024643-INS-172 | chr4-123024643-INS-172 |
| chr4-123036987-INS-596 | chr4-123036987-INS-596 |
| chr4-123060798-INS-120 | chr4-123060799-INS-86 |
| chr4-123062995-INS-343 | chr4-123062995-INS-343 |
| chr4-123071465-INS-55 | chr4-123071465-INS-55 |
| chr4-123083163-INS-316 | chr4-123083163-INS-318 |
| chr4-123091151-INS-605 | chr4-123091151-INS-606 |
| chr4-123116130-INS-173 | chr4-123116154-INS-190 |
| chr4-123145682-INS-58 | chr4-123145682-INS-58 |
| chr4-123148470-INS-159 | chr4-123148470-INS-159 |
| chr4-123155132-INS-169 | chr4-123155132-INS-168 |
| chr4-123165561-INS-193 | chr4-123165561-INS-191 |
| chr4-123168080-INS-72 | chr4-123168080-INS-72 |
| chr4-123196282-INS-181 | chr4-123196282-INS-181 |
| chr4-123202595-INS-283 | chr4-123202595-INS-285 |

| | |
|---|---|
| chr4-123225661-INS-172 | chr4-123225661-INS-175 |
| chr4-123227905-INS-55 | chr4-123227905-INS-54 |
| chr4-123229902-INS-224 | chr4-123229902-INS-213 |
| chr4-123242673-INS-212 | chr4-123242673-INS-212 |
| chr4-123250529-INS-173 | chr4-123250529-INS-174 |
| chr4-123281660-INS-238 | chr4-123281660-INS-238 |
| chr4-123294003-INS-207 | chr4-123294003-INS-204 |
| chr4-123297435-INS-3443 | chr4-123297436-INS-3443 |
| chr4-123298568-INS-213 | chr4-123298568-INS-213 |
| chr4-123300365-INS-212 | chr4-123300365-INS-212 |
| chr4-123333738-INS-205 | chr4-123333738-INS-205 |
| chr4-123358640-INS-376 | chr4-123358640-INS-378 |
| chr4-123406480-INS-288 | chr4-123406480-INS-289 |
| chr4-123440283-INS-59 | chr4-123440283-INS-59 |
| chr4-123464544-INS-220 | chr4-123464544-INS-220 |
| chr4-123474059-INS-64 | chr4-123474038-INS-56 |
| chr4-123479259-INS-116 | chr4-123479259-INS-117 |
| chr4-123499645-INS-221 | chr4-123499645-INS-221 |
| chr4-123506163-INS-216 | chr4-123506163-INS-216 |
| chr4-123518734-INS-72 | chr4-123518734-INS-72 |
| chr4-123519116-INS-427 | chr4-123519116-INS-427 |
| chr4-123522631-INS-52 | chr4-123522635-INS-56 |
| chr4-123541370-INS-196 | chr4-123541370-INS-196 |
| chr4-123544684-INS-223 | chr4-123544684-INS-223 |
| chr4-123559101-INS-56 | chr4-123559101-INS-56 |
| chr4-123589578-INS-179 | chr4-123589579-INS-194 |
| chr4-123594267-INS-64 | chr4-123594221-INS-85 |
| chr4-123594623-INS-441 | chr4-123594623-INS-441 |
| chr4-123603578-INS-58 | chr4-123603578-INS-58 |
| chr4-123606964-INS-190 | chr4-123606964-INS-190 |
| chr4-123616025-INS-178 | chr4-123616025-INS-178 |
| chr4-123625085-INS-197 | chr4-123625085-INS-196 |
| chr4-123684441-INS-5406 | chr4-123684442-INS-5406 |
| chr4-123684450-INS-134 | chr4-123684450-INS-135 |
| chr4-123709319-INS-199 | chr4-123709319-INS-194 |
| chr4-123713834-INS-526 | chr4-123713834-INS-527 |
| chr4-123722670-INS-68 | chr4-123722658-INS-98 |
| chr4-123747316-INS-1890 | chr4-123747317-INS-1890 |
| chr4-123748313-INS-130 | chr4-123748313-INS-130 |
| chr4-123786606-INS-187 | chr4-123786609-INS-188 |
| chr4-123809434-INS-210 | chr4-123809434-INS-210 |
| chr4-123858252-INS-609 | chr4-123858252-INS-610 |
| chr4-123863550-INS-405 | chr4-123863550-INS-405 |
| chr4-123864978-INS-196 | chr4-123864978-INS-196 |
| chr4-123879196-INS-151 | chr4-123879195-INS-178 |
| chr4-123910321-INS-211 | chr4-123910321-INS-211 |
| chr4-123914255-INS-61 | chr4-123914255-INS-61 |
| chr4-123921800-INS-88 | chr4-123921800-INS-88 |
| chr4-123955483-INS-50 | chr4-123955487-INS-53 |
| chr4-123971807-INS-76 | chr4-123971808-INS-76 |
| chr4-123982434-INS-70 | chr4-123982434-INS-70 |
| chr4-123989919-INS-52 | chr4-123989919-INS-52 |
| chr4-124021053-INS-541 | chr4-124021053-INS-541 |
| chr4-124030338-INS-64 | chr4-124030344-INS-76 |
| chr4-124031649-INS-91 | chr4-124031645-INS-75 |
| chr4-124050178-INS-88 | chr4-124050178-INS-88 |
| chr4-124060886-INS-68 | chr4-124060886-INS-68 |
| chr4-124076741-INS-55 | chr4-124076739-INS-54 |
| chr4-124078208-INS-476 | chr4-124078208-INS-476 |
| chr4-124096014-INS-52 | chr4-124096014-INS-55 |
| chr4-124118567-INS-52 | chr4-124118567-INS-50 |
| chr4-124124541-INS-54 | chr4-124124541-INS-54 |
| chr4-124144249-INS-180 | chr4-124144249-INS-177 |
| chr4-124156929-INS-103 | chr4-124156924-INS-115 |
| chr4-124161792-INS-77 | chr4-124161792-INS-77 |
| chr4-124190783-INS-1163 | chr4-124190784-INS-1163 |
| chr4-124191506-INS-55 | chr4-124191506-INS-55 |
| chr4-124194228-INS-55 | chr4-124194228-INS-55 |
| chr4-124252745-INS-68 | chr4-124252741-INS-68 |
| chr4-124256604-INS-100 | chr4-124256604-INS-100 |
| chr4-124259828-INS-66 | chr4-124259828-INS-70 |
| chr4-124273061-INS-156 | chr4-124273061-INS-156 |
| chr4-124276013-INS-74 | chr4-124276013-INS-75 |
| chr4-124293573-INS-255 | chr4-124293573-INS-255 |
| chr4-124306539-INS-231 | chr4-124306539-INS-233 |
| chr4-124309311-INS-224 | chr4-124309311-INS-225 |
| chr4-124318801-INS-124 | chr4-124318801-INS-174 |

| | |
|---|---|
| chr4-124348837-INS-4185 | chr4-124348838-INS-4185 |
| chr4-124351449-INS-126 | chr4-124351447-INS-126 |
| chr4-124362792-INS-56 | chr4-124362792-INS-56 |
| chr4-124384813-INS-184 | chr4-124384813-INS-184 |
| chr4-124416983-INS-98 | chr4-124417004-INS-65 |
| chr4-124453860-INS-331 | chr4-124453860-INS-295 |
| chr4-124457049-INS-219 | chr4-124457049-INS-220 |
| chr4-124466532-INS-85 | chr4-124466532-INS-120 |
| chr4-124473419-INS-84 | chr4-124473419-INS-84 |
| chr4-124523309-INS-204 | chr4-124523309-INS-204 |
| chr4-124526339-INS-68 | chr4-124526339-INS-69 |
| chr4-124544575-INS-206 | chr4-124544575-INS-207 |
| chr4-124545648-INS-108 | chr4-124545648-INS-108 |
| chr4-124562925-INS-115 | chr4-124562925-INS-160 |
| chr4-124572355-INS-58 | chr4-124572355-INS-58 |
| chr4-124582859-INS-247 | chr4-124582859-INS-248 |
| chr4-124595453-INS-74 | chr4-124595460-INS-88 |
| chr4-124603434-INS-74 | chr4-124603433-INS-81 |
| chr4-124610177-INS-6440 | chr4-124610178-INS-6440 |
| chr4-124614349-INS-208 | chr4-124614349-INS-208 |
| chr4-124619427-INS-74 | chr4-124619427-INS-76 |
| chr4-124620280-INS-237 | chr4-124620280-INS-235 |
| chr4-124628728-INS-81 | chr4-124628728-INS-74 |
| chr4-124643918-INS-93 | chr4-124643918-INS-93 |
| chr4-124655838-INS-280 | chr4-124655838-INS-280 |
| chr4-124663810-INS-66 | chr4-124663823-INS-75 |
| chr4-124669060-INS-1241 | chr4-124669061-INS-1241 |
| chr4-124694191-INS-282 | chr4-124694191-INS-282 |
| chr4-124724353-INS-98 | chr4-124724353-INS-98 |
| chr4-124730581-INS-211 | chr4-124730581-INS-212 |
| chr4-124732855-INS-359 | chr4-124732858-INS-376 |
| chr4-124734686-INS-52 | chr4-124734686-INS-52 |
| chr4-124754578-INS-202 | chr4-124754578-INS-204 |
| chr4-124765421-INS-212 | chr4-124765421-INS-212 |
| chr4-124768371-INS-75 | chr4-124768398-INS-95 |
| chr4-124794856-INS-143 | chr4-124794856-INS-145 |
| chr4-124804975-INS-541 | chr4-124804975-INS-562 |
| chr4-124820344-INS-215 | chr4-124820344-INS-215 |
| chr4-124833246-INS-194 | chr4-124833247-INS-185 |
| chr4-124836904-INS-205 | chr4-124836904-INS-205 |
| chr4-124843928-INS-6440 | chr4-124843929-INS-6440 |
| chr4-124860249-INS-182 | chr4-124860251-INS-175 |
| chr4-124872148-INS-68 | chr4-124872148-INS-69 |
| chr4-124898087-INS-574 | chr4-124898087-INS-572 |
| chr4-124916107-INS-71 | chr4-124916110-INS-50 |
| chr4-124925807-INS-504 | chr4-124925807-INS-504 |
| chr4-124955335-INS-232 | chr4-124955335-INS-233 |
| chr4-125006469-INS-57 | chr4-125006469-INS-57 |
| chr4-125035293-INS-57 | chr4-125035293-INS-51 |
| chr4-125060005-INS-3337 | chr4-125060006-INS-3337 |
| chr4-125082791-INS-63 | chr4-125082791-INS-63 |
| chr4-125102682-INS-3414 | chr4-125102683-INS-3413 |
| chr4-125102854-INS-60 | chr4-125102849-INS-63 |
| chr4-125138346-INS-78 | chr4-125138346-INS-78 |
| chr4-125139978-INS-51 | chr4-125139975-INS-54 |
| chr4-125161976-INS-223 | chr4-125161976-INS-223 |
| chr4-125163838-INS-114 | chr4-125163838-INS-114 |
| chr4-125178183-INS-82 | chr4-125178183-INS-86 |
| chr4-125184261-INS-78 | chr4-125184261-INS-74 |
| chr4-125209604-INS-70 | chr4-125209604-INS-84 |
| chr4-125213885-INS-52 | chr4-125213886-INS-52 |
| chr4-125218866-INS-129 | chr4-125218866-INS-141 |
| chr4-125233381-INS-761 | chr4-125233381-INS-761 |
| chr4-125242770-INS-125 | chr4-125242770-INS-75 |
| chr4-125264004-INS-6401 | chr4-125264005-INS-6401 |
| chr4-125267117-INS-71 | chr4-125267117-INS-74 |
| chr4-125281571-INS-103 | chr4-125281572-INS-103 |
| chr4-125309962-INS-96 | chr4-125309962-INS-64 |
| chr4-125312403-INS-643 | chr4-125312403-INS-643 |
| chr4-125380338-INS-96 | chr4-125380338-INS-96 |
| chr4-125393572-INS-52 | chr4-125393572-INS-52 |
| chr4-125415804-INS-60 | chr4-125415804-INS-61 |
| chr4-125433992-INS-165 | chr4-125433992-INS-171 |
| chr4-125465969-INS-302 | chr4-125465999-INS-297 |
| chr4-125466800-INS-71 | chr4-125466801-INS-64 |
| chr4-125529838-INS-206 | chr4-125529838-INS-206 |
| chr4-125582344-INS-52 | chr4-125582344-INS-69 |

| | |
|---|---|
| chr4-125611533-INS-610 | chr4-125611533-INS-610 |
| chr4-125624618-INS-62 | chr4-125624618-INS-62 |
| chr4-125693902-INS-58 | chr4-125693902-INS-58 |
| chr4-125702659-INS-72 | chr4-125702659-INS-96 |
| chr4-125710529-INS-60 | chr4-125710529-INS-60 |
| chr4-125719176-INS-1893 | chr4-125719177-INS-1893 |
| chr4-125719744-INS-70 | chr4-125719744-INS-70 |
| chr4-125721476-INS-641 | chr4-125721476-INS-640 |
| chr4-125758034-INS-250 | chr4-125758035-INS-250 |
| chr4-125780782-INS-205 | chr4-125780782-INS-205 |
| chr4-125849022-INS-158 | chr4-125849014-INS-168 |
| chr4-125855239-INS-87 | chr4-125855253-INS-108 |
| chr4-125859391-INS-162 | chr4-125859391-INS-161 |
| chr4-125860928-INS-91 | chr4-125860928-INS-104 |
| chr4-125862097-INS-220 | chr4-125862097-INS-220 |
| chr4-125869393-INS-172 | chr4-125869394-INS-167 |
| chr4-125872552-INS-400 | chr4-125872552-INS-400 |
| chr4-125876028-INS-57 | chr4-125876028-INS-57 |
| chr4-125889498-INS-204 | chr4-125889498-INS-204 |
| chr4-125904774-INS-64 | chr4-125904774-INS-64 |
| chr4-125907851-INS-208 | chr4-125907847-INS-212 |
| chr4-125922273-INS-140 | chr4-125922274-INS-163 |
| chr4-125930149-INS-213 | chr4-125930149-INS-212 |
| chr4-125934891-INS-177 | chr4-125934891-INS-177 |
| chr4-125950528-INS-253 | chr4-125950528-INS-253 |
| chr4-125983342-INS-54 | chr4-125983342-INS-56 |
| chr4-125985991-INS-75 | chr4-125985991-INS-75 |
| chr4-125989046-INS-978 | chr4-125989046-INS-975 |
| chr4-126004258-INS-215 | chr4-126004258-INS-216 |
| chr4-126025059-INS-133 | chr4-126025059-INS-133 |
| chr4-126035614-INS-194 | chr4-126035614-INS-194 |
| chr4-126067336-INS-203 | chr4-126067336-INS-203 |
| chr4-126075851-INS-203 | chr4-126075851-INS-203 |
| chr4-126077932-INS-207 | chr4-126077932-INS-207 |
| chr4-126089251-INS-226 | chr4-126089251-INS-226 |
| chr4-126103137-INS-204 | chr4-126103135-INS-206 |
| chr4-126105404-INS-178 | chr4-126105404-INS-189 |
| chr4-126108953-INS-56 | chr4-126108954-INS-54 |
| chr4-126109785-INS-81 | chr4-126109785-INS-77 |
| chr4-126110552-INS-3247 | chr4-126110553-INS-3258 |
| chr4-126111898-INS-232 | chr4-126111898-INS-232 |
| chr4-126117153-INS-215 | chr4-126117153-INS-214 |
| chr4-126118561-INS-361 | chr4-126118561-INS-361 |
| chr4-126120242-INS-497 | chr4-126120242-INS-497 |
| chr4-126130586-INS-63 | chr4-126130586-INS-57 |
| chr4-126155504-INS-219 | chr4-126155535-INS-207 |
| chr4-126176490-INS-255 | chr4-126176490-INS-255 |
| chr4-126176920-INS-1048 | chr4-126176921-INS-1048 |
| chr4-126189890-INS-60 | chr4-126189890-INS-60 |
| chr4-126209760-INS-135 | chr4-126209760-INS-135 |
| chr4-126212828-INS-285 | chr4-126212787-INS-316 |
| chr4-126240166-INS-219 | chr4-126240166-INS-220 |
| chr4-126253409-INS-209 | chr4-126253432-INS-225 |
| chr4-126255324-INS-156 | chr4-126255324-INS-156 |
| chr4-126263484-INS-52 | chr4-126263484-INS-68 |
| chr4-126275734-INS-237 | chr4-126275734-INS-238 |
| chr4-126290335-INS-146 | chr4-126290335-INS-146 |
| chr4-126299305-INS-89 | chr4-126299305-INS-94 |
| chr4-126308982-INS-175 | chr4-126308982-INS-175 |
| chr4-126309782-INS-224 | chr4-126309782-INS-224 |
| chr4-126311723-INS-7125 | chr4-126311724-INS-7125 |
| chr4-126312698-INS-59 | chr4-126312698-INS-57 |
| chr4-126315631-INS-208 | chr4-126315631-INS-208 |
| chr4-126334703-INS-53 | chr4-126334704-INS-53 |
| chr4-126343400-INS-214 | chr4-126343400-INS-215 |
| chr4-126363387-INS-336 | chr4-126363387-INS-336 |
| chr4-126365838-INS-207 | chr4-126365838-INS-207 |
| chr4-126369592-INS-118 | chr4-126369592-INS-68 |
| chr4-126396856-INS-120 | chr4-126396856-INS-120 |
| chr4-126407075-INS-51 | chr4-126407074-INS-52 |
| chr4-126415316-INS-824 | chr4-126415317-INS-824 |
| chr4-126423442-INS-327 | chr4-126423442-INS-327 |
| chr4-126424540-INS-252 | chr4-126424540-INS-250 |
| chr4-126429587-INS-473 | chr4-126429587-INS-473 |
| chr4-126440940-INS-185 | chr4-126440944-INS-173 |
| chr4-126442830-INS-213 | chr4-126442831-INS-211 |
| chr4-126443804-INS-283 | chr4-126443805-INS-285 |

| | |
|---|---|
| chr4-126448914-INS-60 | chr4-126448914-INS-76 |
| chr4-126454712-INS-210 | chr4-126454712-INS-210 |
| chr4-126492361-INS-106 | chr4-126492361-INS-90 |
| chr4-126493805-INS-479 | chr4-126493802-INS-475 |
| chr4-126509697-INS-54 | chr4-126509697-INS-54 |
| chr4-126523560-INS-424 | chr4-126523560-INS-426 |
| chr4-126536721-INS-78 | chr4-126536721-INS-78 |
| chr4-126566421-INS-4825 | chr4-126566422-INS-4825 |
| chr4-126589291-INS-61 | chr4-126589291-INS-61 |
| chr4-126592009-INS-315 | chr4-126592009-INS-284 |
| chr4-126602120-INS-286 | chr4-126602120-INS-285 |
| chr4-126607833-INS-102 | chr4-126607833-INS-100 |
| chr4-126608400-INS-210 | chr4-126608400-INS-210 |
| chr4-126619906-INS-123 | chr4-126619906-INS-122 |
| chr4-126669508-INS-1608 | chr4-126669509-INS-1607 |
| chr4-126670071-INS-189 | chr4-126670071-INS-189 |
| chr4-126691804-INS-1594 | chr4-126691805-INS-1581 |
| chr4-126692596-INS-54 | chr4-126692620-INS-52 |
| chr4-126695049-INS-65 | chr4-126695041-INS-79 |
| chr4-126703702-INS-57 | chr4-126703702-INS-57 |
| chr4-126705737-INS-54 | chr4-126705742-INS-63 |
| chr4-126712547-INS-258 | chr4-126712547-INS-259 |
| chr4-126732327-INS-190 | chr4-126732327-INS-192 |
| chr4-126769256-INS-93 | chr4-126769256-INS-63 |
| chr4-126777723-INS-198 | chr4-126777723-INS-197 |
| chr4-126779391-INS-202 | chr4-126779391-INS-201 |
| chr4-126783611-INS-199 | chr4-126783612-INS-200 |
| chr4-126785898-INS-213 | chr4-126785898-INS-215 |
| chr4-126787295-INS-85 | chr4-126787310-INS-61 |
| chr4-126789724-INS-219 | chr4-126789724-INS-220 |
| chr4-126796687-INS-321 | chr4-126796687-INS-325 |
| chr4-126808396-INS-1051 | chr4-126808397-INS-1067 |
| chr4-126812674-INS-1888 | chr4-126812675-INS-1888 |
| chr4-126827512-INS-217 | chr4-126827512-INS-217 |
| chr4-126866520-INS-204 | chr4-126866520-INS-204 |
| chr4-126876819-INS-184 | chr4-126876819-INS-194 |
| chr4-126887140-INS-201 | chr4-126887140-INS-228 |
| chr4-126891707-INS-70 | chr4-126891707-INS-74 |
| chr4-126893746-INS-774 | chr4-126893746-INS-774 |
| chr4-126910092-INS-212 | chr4-126910092-INS-212 |
| chr4-126937347-INS-206 | chr4-126937347-INS-206 |
| chr4-126939156-INS-234 | chr4-126939156-INS-234 |
| chr4-126949186-INS-115 | chr4-126949186-INS-115 |
| chr4-126950592-INS-322 | chr4-126950592-INS-322 |
| chr4-126956212-INS-369 | chr4-126956212-INS-370 |
| chr4-126957306-INS-760 | chr4-126957306-INS-760 |
| chr4-126959743-INS-6442 | chr4-126959744-INS-6442 |
| chr4-126964360-INS-214 | chr4-126964360-INS-214 |
| chr4-126966576-INS-66 | chr4-126966576-INS-66 |
| chr4-126974602-INS-62 | chr4-126974602-INS-62 |
| chr4-126987003-INS-6442 | chr4-126987004-INS-6442 |
| chr4-126995862-INS-76 | chr4-126995866-INS-64 |
| chr4-126998889-INS-1893 | chr4-126998890-INS-1893 |
| chr4-127000780-INS-56 | chr4-127000780-INS-56 |
| chr4-127025787-INS-239 | chr4-127025787-INS-239 |
| chr4-127029899-INS-88 | chr4-127029899-INS-132 |
| chr4-127037049-INS-169 | chr4-127037049-INS-177 |
| chr4-127042740-INS-182 | chr4-127042740-INS-183 |
| chr4-127053212-INS-256 | chr4-127053212-INS-256 |
| chr4-127065826-INS-52 | chr4-127065812-INS-58 |
| chr4-127124554-INS-96 | chr4-127124554-INS-61 |
| chr4-127151514-INS-58 | chr4-127151514-INS-58 |
| chr4-127159675-INS-155 | chr4-127159675-INS-155 |
| chr4-127162941-INS-64 | chr4-127162941-INS-64 |
| chr4-127169368-INS-1268 | chr4-127169369-INS-1226 |
| chr4-127177461-INS-107 | chr4-127177461-INS-103 |
| chr4-127186978-INS-73 | chr4-127187009-INS-79 |
| chr4-127193252-INS-498 | chr4-127193252-INS-498 |
| chr4-127202846-INS-70 | chr4-127202796-INS-74 |
| chr4-127227262-INS-170 | chr4-127227263-INS-169 |
| chr4-127247897-INS-7059 | chr4-127247898-INS-7059 |
| chr4-127268897-INS-53 | chr4-127268901-INS-75 |
| chr4-127271640-INS-64 | chr4-127271626-INS-94 |
| chr4-127304083-INS-123 | chr4-127304083-INS-123 |
| chr4-127305888-INS-208 | chr4-127305888-INS-209 |
| chr4-127312836-INS-106 | chr4-127312836-INS-100 |
| chr4-127326495-INS-104 | chr4-127326451-INS-123 |

| | |
|---|---|
| chr4-127327251-INS-131 | chr4-127327251-INS-136 |
| chr4-127347123-INS-82 | chr4-127347122-INS-68 |
| chr4-127363316-INS-65 | chr4-127363316-INS-65 |
| chr4-127408810-INS-68 | chr4-127408810-INS-70 |
| chr4-127434144-INS-314 | chr4-127434154-INS-297 |
| chr4-127442480-INS-126 | chr4-127442493-INS-150 |
| chr4-127446339-INS-209 | chr4-127446339-INS-215 |
| chr4-127468084-INS-70 | chr4-127468083-INS-86 |
| chr4-127540709-INS-156 | chr4-127540709-INS-156 |
| chr4-127545428-INS-82 | chr4-127545428-INS-97 |
| chr4-127550373-INS-51 | chr4-127550373-INS-51 |
| chr4-127565016-INS-1623 | chr4-127565017-INS-1623 |
| chr4-127578257-INS-52 | chr4-127578257-INS-52 |
| chr4-127615429-INS-939 | chr4-127615431-INS-939 |
| chr4-127631515-INS-883 | chr4-127631515-INS-885 |
| chr4-127657043-INS-104 | chr4-127657043-INS-136 |
| chr4-127660839-INS-7160 | chr4-127660840-INS-7160 |
| chr4-127674610-INS-66 | chr4-127674610-INS-66 |
| chr4-127681274-INS-4664 | chr4-127681275-INS-4641 |
| chr4-127686860-INS-97 | chr4-127686863-INS-92 |
| chr4-127688592-INS-399 | chr4-127688592-INS-391 |
| chr4-127703634-INS-233 | chr4-127703634-INS-236 |
| chr4-127722934-INS-54 | chr4-127722934-INS-74 |
| chr4-127744872-INS-120 | chr4-127744872-INS-93 |
| chr4-127760768-INS-155 | chr4-127760768-INS-155 |
| chr4-127820895-INS-68 | chr4-127820895-INS-68 |
| chr4-127874091-INS-1864 | chr4-127874092-INS-1864 |
| chr4-127904982-INS-72 | chr4-127904982-INS-72 |
| chr4-127921623-INS-55 | chr4-127921623-INS-55 |
| chr4-127967137-INS-57 | chr4-127967137-INS-66 |
| chr4-127974605-INS-51 | chr4-127974638-INS-54 |
| chr4-127976178-INS-132 | chr4-127976178-INS-132 |
| chr4-127983412-INS-141 | chr4-127983420-INS-139 |
| chr4-127985603-INS-5375 | chr4-127985604-INS-5375 |
| chr4-127991175-INS-68 | chr4-127991198-INS-75 |
| chr4-127995398-INS-211 | chr4-127995398-INS-211 |
| chr4-128022189-INS-324 | chr4-128022189-INS-344 |
| chr4-128037420-INS-617 | chr4-128037420-INS-616 |
| chr4-128075117-INS-593 | chr4-128075117-INS-593 |
| chr4-128092988-INS-69 | chr4-128092988-INS-72 |
| chr4-128106204-INS-85 | chr4-128106191-INS-90 |
| chr4-128115333-INS-1305 | chr4-128115334-INS-1260 |
| chr4-128117936-INS-52 | chr4-128117936-INS-58 |
| chr4-128160563-INS-303 | chr4-128160563-INS-303 |
| chr4-128166655-INS-620 | chr4-128166655-INS-620 |
| chr4-128176457-INS-72 | chr4-128176457-INS-58 |
| chr4-128183238-INS-231 | chr4-128183238-INS-231 |
| chr4-128198663-INS-173 | chr4-128198713-INS-173 |
| chr4-128213156-INS-84 | chr4-128213127-INS-89 |
| chr4-128225105-INS-497 | chr4-128225105-INS-497 |
| chr4-128253318-INS-588 | chr4-128253318-INS-588 |
| chr4-128265614-INS-55 | chr4-128265614-INS-60 |
| chr4-128283074-INS-56 | chr4-128283074-INS-58 |
| chr4-128299856-INS-52 | chr4-128299856-INS-56 |
| chr4-128334016-INS-58 | chr4-128334016-INS-60 |
| chr4-128337102-INS-52 | chr4-128337103-INS-62 |
| chr4-128348707-INS-108 | chr4-128348707-INS-60 |
| chr4-128375506-INS-306 | chr4-128375507-INS-306 |
| chr4-128388228-INS-93 | chr4-128388228-INS-93 |
| chr4-128397544-INS-83 | chr4-128397544-INS-83 |
| chr4-128398419-INS-345 | chr4-128398419-INS-391 |
| chr4-128403941-INS-100 | chr4-128403941-INS-100 |
| chr4-128411605-INS-198 | chr4-128411605-INS-198 |
| chr4-128452532-INS-84 | chr4-128452532-INS-84 |
| chr4-128465314-INS-75 | chr4-128465314-INS-75 |
| chr4-128499176-INS-55 | chr4-128499177-INS-51 |
| chr4-128516812-INS-7114 | chr4-128516813-INS-7114 |
| chr4-128524073-INS-120 | chr4-128524114-INS-80 |
| chr4-128524751-INS-65 | chr4-128524751-INS-65 |
| chr4-128558710-INS-218 | chr4-128558710-INS-198 |
| chr4-128560877-INS-55 | chr4-128560877-INS-65 |
| chr4-128571822-INS-50 | chr4-128571822-INS-50 |
| chr4-128575318-INS-59 | chr4-128575340-INS-52 |
| chr4-128591447-INS-109 | chr4-128591447-INS-109 |
| chr4-128596940-INS-70 | chr4-128596940-INS-70 |
| chr4-128603944-INS-135 | chr4-128603944-INS-162 |
| chr4-128613600-INS-183 | chr4-128613600-INS-219 |

| | |
|---|---|
| chr4-128662818-INS-203 | chr4-128662818-INS-204 |
| chr4-128681616-INS-204 | chr4-128681616-INS-205 |
| chr4-128708752-INS-246 | chr4-128708752-INS-246 |
| chr4-128715897-INS-132 | chr4-128715894-INS-122 |
| chr4-128717886-INS-498 | chr4-128717886-INS-498 |
| chr4-128725954-INS-71 | chr4-128725949-INS-61 |
| chr4-128737466-INS-228 | chr4-128737466-INS-228 |
| chr4-128745883-INS-184 | chr4-128745883-INS-167 |
| chr4-128750526-INS-208 | chr4-128750526-INS-208 |
| chr4-128753642-INS-2620 | chr4-128753643-INS-2620 |
| chr4-128759606-INS-93 | chr4-128759606-INS-93 |
| chr4-128761127-INS-190 | chr4-128761127-INS-189 |
| chr4-128769246-INS-64 | chr4-128769246-INS-62 |
| chr4-128770373-INS-120 | chr4-128770373-INS-120 |
| chr4-128776697-INS-239 | chr4-128776697-INS-201 |
| chr4-128796781-INS-62 | chr4-128796781-INS-62 |
| chr4-128822242-INS-633 | chr4-128822242-INS-633 |
| chr4-128826479-INS-61 | chr4-128826479-INS-61 |
| chr4-128830768-INS-52 | chr4-128830755-INS-50 |
| chr4-128848399-INS-65 | chr4-128848378-INS-84 |
| chr4-128862853-INS-508 | chr4-128862853-INS-508 |
| chr4-128872177-INS-66 | chr4-128872186-INS-70 |
| chr4-128873150-INS-126 | chr4-128873150-INS-126 |
| chr4-128877090-INS-71 | chr4-128877090-INS-71 |
| chr4-128885309-INS-176 | chr4-128885309-INS-177 |
| chr4-128888225-INS-97 | chr4-128888225-INS-96 |
| chr4-128892299-INS-208 | chr4-128892299-INS-208 |
| chr4-128903787-INS-65 | chr4-128903787-INS-95 |
| chr4-128919834-INS-204 | chr4-128919834-INS-205 |
| chr4-128944937-INS-169 | chr4-128944937-INS-169 |
| chr4-128946810-INS-4140 | chr4-128946811-INS-4140 |
| chr4-128946811-INS-342 | chr4-128946811-INS-342 |
| chr4-128947703-INS-4834 | chr4-128947704-INS-4834 |
| chr4-128985560-INS-290 | chr4-128985561-INS-284 |
| chr4-128986130-INS-219 | chr4-128986130-INS-219 |
| chr4-128993150-INS-98 | chr4-128993150-INS-86 |
| chr4-128998613-INS-88 | chr4-128998611-INS-58 |
| chr4-129032757-INS-72 | chr4-129032757-INS-73 |
| chr4-129053449-INS-217 | chr4-129053445-INS-239 |
| chr4-129064721-INS-55 | chr4-129064721-INS-55 |
| chr4-129085984-INS-226 | chr4-129085984-INS-227 |
| chr4-129094699-INS-433 | chr4-129094699-INS-433 |
| chr4-129129270-INS-222 | chr4-129129270-INS-223 |
| chr4-129149515-INS-218 | chr4-129149515-INS-224 |
| chr4-129150102-INS-54 | chr4-129150102-INS-54 |
| chr4-129153556-INS-4004 | chr4-129153557-INS-4004 |
| chr4-129155015-INS-160 | chr4-129154986-INS-171 |
| chr4-129160551-INS-202 | chr4-129160551-INS-202 |
| chr4-129161773-INS-415 | chr4-129161773-INS-415 |
| chr4-129164243-INS-227 | chr4-129164243-INS-229 |
| chr4-129172774-INS-182 | chr4-129172774-INS-182 |
| chr4-129184447-INS-165 | chr4-129184448-INS-125 |
| chr4-129193034-INS-176 | chr4-129193034-INS-176 |
| chr4-129207640-INS-374 | chr4-129207640-INS-367 |
| chr4-129225965-INS-80 | chr4-129226001-INS-80 |
| chr4-129245793-INS-181 | chr4-129245793-INS-181 |
| chr4-129247413-INS-202 | chr4-129247413-INS-205 |
| chr4-129251141-INS-159 | chr4-129251147-INS-182 |
| chr4-129258563-INS-215 | chr4-129258563-INS-215 |
| chr4-129259987-INS-218 | chr4-129259987-INS-218 |
| chr4-129262623-INS-193 | chr4-129262623-INS-193 |
| chr4-129264504-INS-410 | chr4-129264504-INS-410 |
| chr4-129265833-INS-206 | chr4-129265833-INS-206 |
| chr4-129269971-INS-174 | chr4-129269971-INS-174 |
| chr4-129279621-INS-78 | chr4-129279621-INS-78 |
| chr4-129285542-INS-151 | chr4-129285542-INS-155 |
| chr4-129287661-INS-64 | chr4-129287652-INS-95 |
| chr4-129289346-INS-199 | chr4-129289331-INS-204 |
| chr4-129295753-INS-198 | chr4-129295753-INS-198 |
| chr4-129308464-INS-2767 | chr4-129308477-INS-2749 |
| chr4-129311316-INS-131 | chr4-129311316-INS-131 |
| chr4-129313793-INS-201 | chr4-129313785-INS-191 |
| chr4-129316419-INS-72 | chr4-129316419-INS-60 |
| chr4-129319133-INS-232 | chr4-129319134-INS-217 |
| chr4-129335058-INS-218 | chr4-129335058-INS-219 |
| chr4-129340666-INS-207 | chr4-129340666-INS-206 |
| chr4-129344826-INS-175 | chr4-129344826-INS-177 |

| | |
|---|---|
| chr4-129357620-INS-68 | chr4-129357620-INS-68 |
| chr4-129387519-INS-175 | chr4-129387519-INS-175 |
| chr4-129421164-INS-169 | chr4-129421168-INS-163 |
| chr4-129438957-INS-197 | chr4-129438957-INS-197 |
| chr4-129445031-INS-103 | chr4-129445032-INS-93 |
| chr4-129453749-INS-202 | chr4-129453749-INS-202 |
| chr4-129456855-INS-191 | chr4-129456856-INS-176 |
| chr4-129458279-INS-213 | chr4-129458279-INS-213 |
| chr4-129462483-INS-206 | chr4-129462483-INS-206 |
| chr4-129476389-INS-175 | chr4-129476389-INS-174 |
| chr4-129482607-INS-71 | chr4-129482607-INS-71 |
| chr4-129495308-INS-207 | chr4-129495308-INS-207 |
| chr4-129503545-INS-52 | chr4-129503545-INS-52 |
| chr4-129556738-INS-1580 | chr4-129556739-INS-1580 |
| chr4-129561783-INS-522 | chr4-129561783-INS-522 |
| chr4-129562743-INS-54 | chr4-129562743-INS-62 |
| chr4-129581460-INS-146 | chr4-129581460-INS-146 |
| chr4-129612546-INS-55 | chr4-129612546-INS-55 |
| chr4-129619018-INS-204 | chr4-129619018-INS-213 |
| chr4-129641881-INS-51 | chr4-129641884-INS-66 |
| chr4-129645214-INS-691 | chr4-129645214-INS-691 |
| chr4-129645916-INS-224 | chr4-129645916-INS-224 |
| chr4-129657070-INS-105 | chr4-129657070-INS-109 |
| chr4-129663235-INS-158 | chr4-129663231-INS-165 |
| chr4-129666457-INS-53 | chr4-129666457-INS-55 |
| chr4-129672922-INS-550 | chr4-129672922-INS-551 |
| chr4-129686094-INS-328 | chr4-129686094-INS-330 |
| chr4-129689669-INS-114 | chr4-129689669-INS-114 |
| chr4-129691430-INS-73 | chr4-129691431-INS-72 |
| chr4-129694988-INS-241 | chr4-129694988-INS-246 |
| chr4-129707887-INS-518 | chr4-129707883-INS-518 |
| chr4-129721931-INS-172 | chr4-129721931-INS-172 |
| chr4-129740197-INS-184 | chr4-129740197-INS-184 |
| chr4-129756394-INS-252 | chr4-129756394-INS-252 |
| chr4-129768264-INS-56 | chr4-129768264-INS-56 |
| chr4-129795396-INS-55 | chr4-129795396-INS-53 |
| chr4-129813661-INS-128 | chr4-129813661-INS-146 |
| chr4-129817334-INS-291 | chr4-129817336-INS-291 |
| chr4-129819812-INS-361 | chr4-129819812-INS-361 |
| chr4-129824517-INS-197 | chr4-129824517-INS-197 |
| chr4-129831350-INS-254 | chr4-129831350-INS-259 |
| chr4-129836473-INS-89 | chr4-129836473-INS-89 |
| chr4-129863511-INS-201 | chr4-129863511-INS-201 |
| chr4-129873765-INS-56 | chr4-129873765-INS-58 |
| chr4-129890055-INS-209 | chr4-129890055-INS-209 |
| chr4-129896578-INS-142 | chr4-129896578-INS-154 |
| chr4-129902695-INS-102 | chr4-129902695-INS-116 |
| chr4-129920233-INS-96 | chr4-129920233-INS-100 |
| chr4-129931573-INS-199 | chr4-129931573-INS-199 |
| chr4-129954207-INS-67 | chr4-129954207-INS-60 |
| chr4-129988506-INS-211 | chr4-129988506-INS-211 |
| chr4-129996871-INS-64 | chr4-129996871-INS-64 |
| chr4-129998403-INS-216 | chr4-129998403-INS-216 |
| chr4-130004602-INS-227 | chr4-130004602-INS-227 |
| chr4-130008025-INS-68 | chr4-130008025-INS-68 |
| chr4-130009593-INS-101 | chr4-130009593-INS-103 |
| chr4-130042089-INS-118 | chr4-130042089-INS-118 |
| chr4-130050592-INS-81 | chr4-130050614-INS-96 |
| chr4-130059031-INS-94 | chr4-130059042-INS-110 |
| chr4-130101047-INS-75 | chr4-130101048-INS-98 |
| chr4-130127621-INS-55 | chr4-130127633-INS-70 |
| chr4-130130996-INS-108 | chr4-130130980-INS-124 |
| chr4-130142153-INS-264 | chr4-130142153-INS-276 |
| chr4-130153129-INS-90 | chr4-130153132-INS-60 |
| chr4-130167541-INS-51 | chr4-130167541-INS-51 |
| chr4-130175454-INS-52 | chr4-130175454-INS-52 |
| chr4-130176388-INS-87 | chr4-130176346-INS-102 |
| chr4-130179692-INS-58 | chr4-130179692-INS-58 |
| chr4-130187128-INS-151 | chr4-130187108-INS-162 |
| chr4-130193036-INS-64 | chr4-130193049-INS-69 |
| chr4-130224395-INS-227 | chr4-130224395-INS-237 |
| chr4-130229940-INS-216 | chr4-130229940-INS-216 |
| chr4-130242555-INS-185 | chr4-130242523-INS-182 |
| chr4-130254918-INS-202 | chr4-130254918-INS-202 |
| chr4-130261028-INS-224 | chr4-130261028-INS-212 |
| chr4-130271645-INS-199 | chr4-130271645-INS-200 |
| chr4-130276953-INS-64 | chr4-130276953-INS-66 |

| | |
|---|---|
| chr4-130280221-INS-211 | chr4-130280221-INS-211 |
| chr4-130290138-INS-62 | chr4-130290140-INS-60 |
| chr4-130322832-INS-163 | chr4-130322831-INS-162 |
| chr4-130326119-INS-62 | chr4-130326151-INS-66 |
| chr4-130335260-INS-177 | chr4-130335260-INS-177 |
| chr4-130346023-INS-193 | chr4-130346023-INS-193 |
| chr4-130363228-INS-183 | chr4-130363228-INS-188 |
| chr4-130378402-INS-205 | chr4-130378402-INS-204 |
| chr4-130393336-INS-103 | chr4-130393336-INS-103 |
| chr4-130416489-INS-203 | chr4-130416489-INS-203 |
| chr4-130452148-INS-57 | chr4-130452148-INS-57 |
| chr4-130470233-INS-80 | chr4-130470233-INS-80 |
| chr4-130486389-INS-111 | chr4-130486389-INS-111 |
| chr4-130571471-INS-53 | chr4-130571471-INS-53 |
| chr4-130574851-INS-384 | chr4-130574869-INS-374 |
| chr4-130582792-INS-139 | chr4-130582769-INS-169 |
| chr4-130603077-INS-269 | chr4-130603077-INS-223 |
| chr4-130616442-INS-72 | chr4-130616442-INS-76 |
| chr4-130622342-INS-58 | chr4-130622342-INS-82 |
| chr4-130624178-INS-110 | chr4-130624136-INS-110 |
| chr4-130627554-INS-312 | chr4-130627554-INS-274 |
| chr4-130631425-INS-500 | chr4-130631425-INS-504 |
| chr4-130660444-INS-130 | chr4-130660444-INS-130 |
| chr4-130681026-INS-52 | chr4-130681026-INS-52 |
| chr4-130689972-INS-100 | chr4-130689972-INS-80 |
| chr4-130691096-INS-300 | chr4-130691096-INS-300 |
| chr4-130715914-INS-80 | chr4-130715914-INS-80 |
| chr4-130722664-INS-160 | chr4-130722673-INS-144 |
| chr4-130735151-INS-66 | chr4-130735151-INS-66 |
| chr4-130744052-INS-192 | chr4-130744052-INS-192 |
| chr4-130786398-INS-125 | chr4-130786398-INS-126 |
| chr4-130787522-INS-110 | chr4-130787522-INS-110 |
| chr4-130790701-INS-385 | chr4-130790701-INS-385 |
| chr4-130797054-INS-58 | chr4-130797054-INS-58 |
| chr4-130799483-INS-493 | chr4-130799483-INS-497 |
| chr4-130807023-INS-117 | chr4-130807023-INS-117 |
| chr4-130815730-INS-119 | chr4-130815726-INS-88 |
| chr4-130823239-INS-75 | chr4-130823239-INS-75 |
| chr4-130858094-INS-305 | chr4-130858094-INS-305 |
| chr4-130882570-INS-160 | chr4-130882570-INS-160 |
| chr4-130899984-INS-55 | chr4-130899994-INS-55 |
| chr4-130928436-INS-66 | chr4-130928436-INS-66 |
| chr4-130989514-INS-81 | chr4-130989514-INS-81 |
| chr4-130998439-INS-100 | chr4-130998439-INS-100 |
| chr4-131010119-INS-66 | chr4-131010119-INS-66 |
| chr4-131019009-INS-96 | chr4-131019009-INS-96 |
| chr4-131027638-INS-72 | chr4-131027630-INS-104 |
| chr4-131030741-INS-92 | chr4-131030726-INS-92 |
| chr4-131035988-INS-72 | chr4-131035988-INS-58 |
| chr4-131040459-INS-96 | chr4-131040459-INS-96 |
| chr4-131067134-INS-55 | chr4-131067134-INS-57 |
| chr4-131069958-INS-66 | chr4-131069941-INS-65 |
| chr4-131081738-INS-54 | chr4-131081735-INS-60 |
| chr4-131089523-INS-72 | chr4-131089523-INS-74 |
| chr4-131111625-INS-50 | chr4-131111625-INS-52 |
| chr4-131129119-INS-57 | chr4-131129097-INS-90 |
| chr4-131140893-INS-80 | chr4-131140893-INS-80 |
| chr4-131142505-INS-73 | chr4-131142507-INS-73 |
| chr4-131165850-INS-55 | chr4-131165850-INS-55 |
| chr4-131169560-INS-104 | chr4-131169560-INS-104 |
| chr4-131177762-INS-4751 | chr4-131177779-INS-4732 |
| chr4-131194766-INS-141 | chr4-131194766-INS-100 |
| chr4-131203036-INS-98 | chr4-131203036-INS-100 |
| chr4-131220265-INS-104 | chr4-131220265-INS-108 |
| chr4-131225048-INS-182 | chr4-131225048-INS-181 |
| chr4-131316386-INS-56 | chr4-131316426-INS-56 |
| chr4-131345239-INS-95 | chr4-131345240-INS-95 |
| chr4-131357468-INS-78 | chr4-131357468-INS-78 |
| chr4-131365207-INS-53 | chr4-131365207-INS-68 |
| chr4-131376438-INS-1888 | chr4-131376439-INS-1888 |
| chr4-131381434-INS-90 | chr4-131381434-INS-85 |
| chr4-131395016-INS-213 | chr4-131395016-INS-213 |
| chr4-131397671-INS-96 | chr4-131397671-INS-96 |
| chr4-131425743-INS-151 | chr4-131425743-INS-152 |
| chr4-131463432-INS-88 | chr4-131463432-INS-53 |
| chr4-131504997-INS-95 | chr4-131504997-INS-59 |
| chr4-131520690-INS-1151 | chr4-131520691-INS-1151 |

| | |
|---|---|
| chr4-131539209-INS-199 | chr4-131539209-INS-199 |
| chr4-131574396-INS-2502 | chr4-131574397-INS-2497 |
| chr4-131646101-INS-58 | chr4-131646103-INS-53 |
| chr4-131662376-INS-534 | chr4-131662376-INS-534 |
| chr4-131701081-INS-84 | chr4-131701081-INS-84 |
| chr4-131707309-INS-257 | chr4-131707309-INS-257 |
| chr4-131708080-INS-216 | chr4-131708080-INS-216 |
| chr4-131712085-INS-105 | chr4-131712085-INS-105 |
| chr4-131720212-INS-228 | chr4-131720212-INS-230 |
| chr4-131722516-INS-213 | chr4-131722516-INS-213 |
| chr4-131735317-INS-168 | chr4-131735317-INS-168 |
| chr4-131746771-INS-214 | chr4-131746771-INS-214 |
| chr4-131747982-INS-106 | chr4-131747982-INS-106 |
| chr4-131753697-INS-919 | chr4-131753722-INS-911 |
| chr4-131785546-INS-215 | chr4-131785546-INS-215 |
| chr4-131808778-INS-54 | chr4-131808775-INS-57 |
| chr4-131818450-INS-168 | chr4-131818450-INS-180 |
| chr4-131842435-INS-236 | chr4-131842435-INS-232 |
| chr4-131872298-INS-106 | chr4-131872298-INS-106 |
| chr4-131892457-INS-718 | chr4-131892457-INS-719 |
| chr4-131908149-INS-204 | chr4-131908149-INS-204 |
| chr4-131962949-INS-204 | chr4-131962949-INS-204 |
| chr4-131967464-INS-241 | chr4-131967464-INS-241 |
| chr4-131972791-INS-121 | chr4-131972791-INS-121 |
| chr4-131991377-INS-175 | chr4-131991377-INS-175 |
| chr4-132000451-INS-215 | chr4-132000451-INS-219 |
| chr4-132003912-INS-205 | chr4-132003912-INS-205 |
| chr4-132007153-INS-338 | chr4-132007153-INS-338 |
| chr4-132019844-INS-247 | chr4-132019844-INS-247 |
| chr4-132026832-INS-77 | chr4-132026832-INS-60 |
| chr4-132027501-INS-206 | chr4-132027501-INS-206 |
| chr4-132030759-INS-205 | chr4-132030759-INS-206 |
| chr4-132050060-INS-221 | chr4-132050060-INS-211 |
| chr4-132074332-INS-185 | chr4-132074332-INS-182 |
| chr4-132079311-INS-209 | chr4-132079311-INS-208 |
| chr4-132107955-INS-359 | chr4-132107955-INS-351 |
| chr4-132112677-INS-197 | chr4-132112677-INS-205 |
| chr4-132116604-INS-199 | chr4-132116604-INS-199 |
| chr4-132120274-INS-202 | chr4-132120274-INS-199 |
| chr4-132120951-INS-113 | chr4-132120951-INS-113 |
| chr4-132128044-INS-212 | chr4-132128044-INS-210 |
| chr4-132128107-INS-7182 | chr4-132128108-INS-7182 |
| chr4-132154545-INS-94 | chr4-132154545-INS-94 |
| chr4-132158185-INS-665 | chr4-132158185-INS-666 |
| chr4-132160923-INS-128 | chr4-132160955-INS-122 |
| chr4-132169193-INS-131 | chr4-132169193-INS-143 |
| chr4-132184899-INS-225 | chr4-132184900-INS-237 |
| chr4-132189917-INS-181 | chr4-132189909-INS-166 |
| chr4-132192808-INS-201 | chr4-132192808-INS-201 |
| chr4-132206469-INS-204 | chr4-132206469-INS-203 |
| chr4-132246781-INS-207 | chr4-132246781-INS-207 |
| chr4-132248831-INS-262 | chr4-132248831-INS-266 |
| chr4-132250067-INS-175 | chr4-132250067-INS-176 |
| chr4-132257264-INS-426 | chr4-132257290-INS-423 |
| chr4-132276340-INS-215 | chr4-132276340-INS-216 |
| chr4-132277343-INS-470 | chr4-132277343-INS-470 |
| chr4-132278541-INS-189 | chr4-132278542-INS-196 |
| chr4-132285810-INS-52 | chr4-132285810-INS-52 |
| chr4-132340170-INS-60 | chr4-132340170-INS-58 |
| chr4-132341665-INS-59 | chr4-132341665-INS-55 |
| chr4-132351351-INS-60 | chr4-132351351-INS-60 |
| chr4-132371618-INS-301 | chr4-132371618-INS-301 |
| chr4-132375976-INS-200 | chr4-132375976-INS-200 |
| chr4-132393705-INS-208 | chr4-132393705-INS-208 |
| chr4-132398007-INS-230 | chr4-132398007-INS-232 |
| chr4-132424130-INS-100 | chr4-132424130-INS-100 |
| chr4-132425803-INS-134 | chr4-132425803-INS-134 |
| chr4-132452249-INS-198 | chr4-132452249-INS-198 |
| chr4-132482737-INS-56 | chr4-132482742-INS-79 |
| chr4-132485788-INS-51 | chr4-132485788-INS-73 |
| chr4-132491697-INS-73 | chr4-132491716-INS-63 |
| chr4-132498669-INS-166 | chr4-132498669-INS-166 |
| chr4-132533223-INS-197 | chr4-132533223-INS-199 |
| chr4-132539820-INS-450 | chr4-132539820-INS-450 |
| chr4-132572195-INS-206 | chr4-132572195-INS-206 |
| chr4-132609778-INS-90 | chr4-132609778-INS-58 |
| chr4-132610696-INS-199 | chr4-132610696-INS-199 |

| | |
|---|---|
| chr4-132636285-INS-226 | chr4-132636285-INS-225 |
| chr4-132642063-INS-4057 | chr4-132642064-INS-4024 |
| chr4-132650861-INS-190 | chr4-132650861-INS-190 |
| chr4-132656734-INS-375 | chr4-132656734-INS-375 |
| chr4-132657323-INS-209 | chr4-132657323-INS-209 |
| chr4-132663363-INS-215 | chr4-132663363-INS-215 |
| chr4-132666488-INS-202 | chr4-132666488-INS-202 |
| chr4-132679374-INS-206 | chr4-132679374-INS-206 |
| chr4-132681311-INS-99 | chr4-132681284-INS-125 |
| chr4-132684871-INS-204 | chr4-132684871-INS-205 |
| chr4-132687433-INS-80 | chr4-132687433-INS-80 |
| chr4-132696598-INS-219 | chr4-132696598-INS-220 |
| chr4-132699175-INS-183 | chr4-132699175-INS-183 |
| chr4-132717475-INS-202 | chr4-132717475-INS-202 |
| chr4-132718209-INS-139 | chr4-132718218-INS-145 |
| chr4-132722451-INS-189 | chr4-132722451-INS-189 |
| chr4-132723942-INS-52 | chr4-132723942-INS-68 |
| chr4-132729192-INS-178 | chr4-132729192-INS-178 |
| chr4-132736453-INS-173 | chr4-132736453-INS-179 |
| chr4-132762394-INS-106 | chr4-132762394-INS-106 |
| chr4-132838320-INS-208 | chr4-132838320-INS-208 |
| chr4-132845764-INS-114 | chr4-132845764-INS-129 |
| chr4-132848724-INS-52 | chr4-132848720-INS-52 |
| chr4-132851378-INS-459 | chr4-132851378-INS-457 |
| chr4-132855699-INS-184 | chr4-132855699-INS-184 |
| chr4-132863492-INS-78 | chr4-132863492-INS-62 |
| chr4-132868154-INS-228 | chr4-132868154-INS-228 |
| chr4-132900098-INS-857 | chr4-132900098-INS-857 |
| chr4-132910727-INS-51 | chr4-132910727-INS-51 |
| chr4-132931621-INS-76 | chr4-132931621-INS-76 |
| chr4-132943543-INS-141 | chr4-132943529-INS-122 |
| chr4-132956154-INS-214 | chr4-132956154-INS-214 |
| chr4-132967036-INS-347 | chr4-132967036-INS-346 |
| chr4-132980354-INS-188 | chr4-132980354-INS-188 |
| chr4-132986993-INS-78 | chr4-132986993-INS-78 |
| chr4-133003339-INS-272 | chr4-133003339-INS-272 |
| chr4-133038292-INS-55 | chr4-133038292-INS-55 |
| chr4-133041972-INS-54 | chr4-133041972-INS-54 |
| chr4-133049287-INS-57 | chr4-133049287-INS-57 |
| chr4-133053307-INS-207 | chr4-133053307-INS-207 |
| chr4-133057141-INS-245 | chr4-133057141-INS-245 |
| chr4-133063341-INS-207 | chr4-133063341-INS-208 |
| chr4-133093924-INS-198 | chr4-133093924-INS-198 |
| chr4-133114046-INS-162 | chr4-133114046-INS-162 |
| chr4-133126156-INS-276 | chr4-133126156-INS-276 |
| chr4-133151957-INS-202 | chr4-133151957-INS-202 |
| chr4-133154126-INS-184 | chr4-133154126-INS-184 |
| chr4-133158181-INS-132 | chr4-133158181-INS-128 |
| chr4-133179380-INS-659 | chr4-133179380-INS-658 |
| chr4-133189265-INS-212 | chr4-133189265-INS-212 |
| chr4-133193521-INS-174 | chr4-133193521-INS-174 |
| chr4-133198718-INS-805 | chr4-133198718-INS-805 |
| chr4-133203133-INS-195 | chr4-133203133-INS-195 |
| chr4-133211215-INS-215 | chr4-133211215-INS-215 |
| chr4-133220538-INS-72 | chr4-133220514-INS-50 |
| chr4-133228754-INS-68 | chr4-133228754-INS-65 |
| chr4-133237402-INS-283 | chr4-133237402-INS-283 |
| chr4-133242606-INS-210 | chr4-133242606-INS-210 |
| chr4-133254049-INS-144 | chr4-133254049-INS-144 |
| chr4-133258729-INS-333 | chr4-133258729-INS-333 |
| chr4-133266473-INS-52 | chr4-133266473-INS-52 |
| chr4-133278267-INS-65 | chr4-133278267-INS-65 |
| chr4-133297837-INS-55 | chr4-133297838-INS-55 |
| chr4-133300960-INS-179 | chr4-133300960-INS-179 |
| chr4-133310249-INS-213 | chr4-133310265-INS-225 |
| chr4-133310853-INS-516 | chr4-133310853-INS-516 |
| chr4-133325565-INS-203 | chr4-133325565-INS-203 |
| chr4-133343119-INS-61 | chr4-133343119-INS-61 |
| chr4-133370318-INS-202 | chr4-133370318-INS-202 |
| chr4-133382303-INS-210 | chr4-133382311-INS-202 |
| chr4-133386702-INS-195 | chr4-133386702-INS-195 |
| chr4-133393248-INS-471 | chr4-133393248-INS-492 |
| chr4-133405181-INS-190 | chr4-133405181-INS-191 |
| chr4-133413231-INS-204 | chr4-133413231-INS-204 |
| chr4-133430095-INS-72 | chr4-133430095-INS-72 |
| chr4-133442278-INS-214 | chr4-133442278-INS-214 |
| chr4-133459001-INS-75 | chr4-133459001-INS-75 |

| | |
|---|---|
| chr4-133494891-INS-408 | chr4-133494891-INS-408 |
| chr4-133498268-INS-187 | chr4-133498268-INS-187 |
| chr4-133499631-INS-175 | chr4-133499631-INS-175 |
| chr4-133507933-INS-394 | chr4-133507934-INS-394 |
| chr4-133513685-INS-179 | chr4-133513685-INS-179 |
| chr4-133516881-INS-59 | chr4-133516881-INS-63 |
| chr4-133536822-INS-64 | chr4-133536810-INS-56 |
| chr4-133550289-INS-582 | chr4-133550289-INS-582 |
| chr4-133554006-INS-343 | chr4-133554006-INS-344 |
| chr4-133562484-INS-206 | chr4-133562485-INS-206 |
| chr4-133582686-INS-96 | chr4-133582683-INS-94 |
| chr4-133611287-INS-199 | chr4-133611287-INS-199 |
| chr4-133616903-INS-144 | chr4-133616903-INS-144 |
| chr4-133621332-INS-206 | chr4-133621332-INS-206 |
| chr4-133623151-INS-199 | chr4-133623151-INS-198 |
| chr4-133624443-INS-163 | chr4-133624443-INS-163 |
| chr4-133631081-INS-1237 | chr4-133631082-INS-1237 |
| chr4-133633684-INS-179 | chr4-133633684-INS-179 |
| chr4-133638989-INS-182 | chr4-133638989-INS-182 |
| chr4-133640582-INS-202 | chr4-133640582-INS-199 |
| chr4-133643808-INS-595 | chr4-133643808-INS-602 |
| chr4-133649369-INS-51 | chr4-133649369-INS-51 |
| chr4-133670017-INS-60 | chr4-133670017-INS-60 |
| chr4-133671400-INS-7081 | chr4-133671401-INS-7081 |
| chr4-133681428-INS-199 | chr4-133681428-INS-199 |
| chr4-133684317-INS-206 | chr4-133684317-INS-205 |
| chr4-133692302-INS-179 | chr4-133692302-INS-179 |
| chr4-133700383-INS-218 | chr4-133700383-INS-218 |
| chr4-133715562-INS-204 | chr4-133715562-INS-204 |
| chr4-133719077-INS-223 | chr4-133719077-INS-223 |
| chr4-133734123-INS-169 | chr4-133734123-INS-183 |
| chr4-133736136-INS-154 | chr4-133736136-INS-154 |
| chr4-133741292-INS-54 | chr4-133741292-INS-54 |
| chr4-133748418-INS-187 | chr4-133748418-INS-187 |
| chr4-133750437-INS-216 | chr4-133750437-INS-216 |
| chr4-133752975-INS-61 | chr4-133752989-INS-74 |
| chr4-133764029-INS-122 | chr4-133764029-INS-116 |
| chr4-133765240-INS-62 | chr4-133765240-INS-62 |
| chr4-133768388-INS-203 | chr4-133768388-INS-203 |
| chr4-133770071-INS-190 | chr4-133770071-INS-190 |
| chr4-133777252-INS-197 | chr4-133777252-INS-207 |
| chr4-133781709-INS-497 | chr4-133781709-INS-497 |
| chr4-133791935-INS-5318 | chr4-133791935-INS-5317 |
| chr4-133812900-INS-199 | chr4-133812901-INS-191 |
| chr4-133831098-INS-53 | chr4-133831098-INS-53 |
| chr4-133832391-INS-554 | chr4-133832391-INS-555 |
| chr4-133838471-INS-65 | chr4-133838471-INS-65 |
| chr4-133871436-INS-722 | chr4-133871436-INS-722 |
| chr4-133904981-INS-92 | chr4-133904981-INS-92 |
| chr4-133908130-INS-399 | chr4-133908130-INS-399 |
| chr4-133926874-INS-63 | chr4-133926874-INS-63 |
| chr4-133928535-INS-205 | chr4-133928535-INS-204 |
| chr4-133933542-INS-90 | chr4-133933542-INS-90 |
| chr4-133941277-INS-184 | chr4-133941277-INS-184 |
| chr4-133944598-INS-129 | chr4-133944598-INS-105 |
| chr4-133968146-INS-400 | chr4-133968146-INS-400 |
| chr4-134019495-INS-164 | chr4-134019495-INS-164 |
| chr4-134024268-INS-62 | chr4-134024268-INS-52 |
| chr4-134031967-INS-80 | chr4-134031967-INS-80 |
| chr4-134033480-INS-74 | chr4-134033480-INS-74 |
| chr4-134051020-INS-149 | chr4-134051012-INS-157 |
| chr4-134065460-INS-416 | chr4-134065460-INS-417 |
| chr4-134085493-INS-59 | chr4-134085493-INS-59 |
| chr4-134103173-INS-123 | chr4-134103173-INS-125 |
| chr4-134134855-INS-201 | chr4-134134855-INS-201 |
| chr4-134136176-INS-480 | chr4-134136176-INS-480 |
| chr4-134150436-INS-61 | chr4-134150436-INS-61 |
| chr4-134169666-INS-6676 | chr4-134169667-INS-6676 |
| chr4-134181344-INS-269 | chr4-134181344-INS-267 |
| chr4-134199219-INS-206 | chr4-134199219-INS-206 |
| chr4-134209349-INS-87 | chr4-134209349-INS-102 |
| chr4-134216172-INS-249 | chr4-134216172-INS-249 |
| chr4-134226332-INS-62 | chr4-134226333-INS-62 |
| chr4-134243582-INS-72 | chr4-134243582-INS-72 |
| chr4-134248178-INS-401 | chr4-134248178-INS-401 |
| chr4-134275174-INS-66 | chr4-134275174-INS-66 |
| chr4-134283301-INS-132 | chr4-134283301-INS-88 |

| | |
|---|---|
| chr4-134322178-INS-124 | chr4-134322178-INS-124 |
| chr4-134378679-INS-52 | chr4-134378679-INS-50 |
| chr4-134383162-INS-100 | chr4-134383163-INS-130 |
| chr4-134386722-INS-548 | chr4-134386722-INS-548 |
| chr4-134391693-INS-132 | chr4-134391693-INS-132 |
| chr4-134398481-INS-374 | chr4-134398481-INS-374 |
| chr4-134407580-INS-67 | chr4-134407580-INS-67 |
| chr4-134413818-INS-215 | chr4-134413818-INS-215 |
| chr4-134414911-INS-390 | chr4-134414911-INS-390 |
| chr4-134433921-INS-56 | chr4-134433921-INS-55 |
| chr4-134438120-INS-73 | chr4-134438120-INS-81 |
| chr4-134459829-INS-55 | chr4-134459829-INS-79 |
| chr4-134485314-INS-403 | chr4-134485314-INS-403 |
| chr4-134500496-INS-3085 | chr4-134500497-INS-3085 |
| chr4-134511822-INS-204 | chr4-134511822-INS-204 |
| chr4-134522650-INS-62 | chr4-134522624-INS-71 |
| chr4-134527862-INS-56 | chr4-134527862-INS-56 |
| chr4-134571977-INS-501 | chr4-134571976-INS-510 |
| chr4-134590917-INS-145 | chr4-134590918-INS-145 |
| chr4-134603372-INS-229 | chr4-134603372-INS-230 |
| chr4-134612958-INS-113 | chr4-134612959-INS-134 |
| chr4-134620020-INS-123 | chr4-134620020-INS-123 |
| chr4-134627035-INS-1908 | chr4-134627036-INS-1908 |
| chr4-134627523-INS-152 | chr4-134627523-INS-152 |
| chr4-134641374-INS-111 | chr4-134641374-INS-112 |
| chr4-134647959-INS-63 | chr4-134647959-INS-63 |
| chr4-134648561-INS-210 | chr4-134648561-INS-210 |
| chr4-134663680-INS-54 | chr4-134663680-INS-54 |
| chr4-134668541-INS-203 | chr4-134668541-INS-203 |
| chr4-134675261-INS-50 | chr4-134675271-INS-62 |
| chr4-134677842-INS-150 | chr4-134677842-INS-152 |
| chr4-134680280-INS-53 | chr4-134680280-INS-53 |
| chr4-134688679-INS-80 | chr4-134688679-INS-80 |
| chr4-134690735-INS-204 | chr4-134690735-INS-204 |
| chr4-134703631-INS-408 | chr4-134703631-INS-412 |
| chr4-134711085-INS-84 | chr4-134711085-INS-83 |
| chr4-134738775-INS-137 | chr4-134738775-INS-139 |
| chr4-134743026-INS-1888 | chr4-134743027-INS-1888 |
| chr4-134761860-INS-120 | chr4-134761860-INS-120 |
| chr4-134767816-INS-54 | chr4-134767816-INS-56 |
| chr4-134779644-INS-135 | chr4-134779642-INS-134 |
| chr4-134796801-INS-52 | chr4-134796801-INS-52 |
| chr4-134817538-INS-138 | chr4-134817538-INS-143 |
| chr4-134824063-INS-58 | chr4-134824063-INS-58 |
| chr4-134835474-INS-79 | chr4-134835479-INS-85 |
| chr4-134837831-INS-104 | chr4-134837829-INS-123 |
| chr4-134877055-INS-238 | chr4-134877055-INS-239 |
| chr4-134881415-INS-70 | chr4-134881415-INS-73 |
| chr4-134885244-INS-213 | chr4-134885244-INS-214 |
| chr4-134908953-INS-216 | chr4-134908953-INS-212 |
| chr4-134911305-INS-151 | chr4-134911305-INS-159 |
| chr4-134935234-INS-87 | chr4-134935234-INS-50 |
| chr4-134948752-INS-56 | chr4-134948752-INS-78 |
| chr4-134957228-INS-222 | chr4-134957228-INS-223 |
| chr4-134968063-INS-199 | chr4-134968063-INS-199 |
| chr4-134972026-INS-296 | chr4-134972026-INS-296 |
| chr4-134993221-INS-223 | chr4-134993221-INS-223 |
| chr4-135009177-INS-70 | chr4-135009175-INS-73 |
| chr4-135012396-INS-406 | chr4-135012396-INS-406 |
| chr4-135018765-INS-70 | chr4-135018765-INS-70 |
| chr4-135020726-INS-82 | chr4-135020739-INS-74 |
| chr4-135031062-INS-167 | chr4-135031062-INS-166 |
| chr4-135031769-INS-176 | chr4-135031769-INS-177 |
| chr4-135068105-INS-209 | chr4-135068105-INS-209 |
| chr4-135094159-INS-72 | chr4-135094159-INS-72 |
| chr4-135119176-INS-63 | chr4-135119176-INS-63 |
| chr4-135126364-INS-115 | chr4-135126364-INS-116 |
| chr4-135138522-INS-221 | chr4-135138522-INS-221 |
| chr4-135151753-INS-78 | chr4-135151753-INS-66 |
| chr4-135166716-INS-89 | chr4-135166716-INS-89 |
| chr4-135167594-INS-177 | chr4-135167616-INS-151 |
| chr4-135168870-INS-52 | chr4-135168870-INS-54 |
| chr4-135169568-INS-551 | chr4-135169568-INS-551 |
| chr4-135196516-INS-56 | chr4-135196516-INS-56 |
| chr4-135198418-INS-226 | chr4-135198418-INS-226 |
| chr4-135204487-INS-216 | chr4-135204487-INS-228 |
| chr4-135207291-INS-174 | chr4-135207294-INS-175 |

| | |
|---|---|
| chr4-135219394-INS-96 | chr4-135219391-INS-103 |
| chr4-135226309-INS-219 | chr4-135226309-INS-220 |
| chr4-135232620-INS-510 | chr4-135232620-INS-510 |
| chr4-135236445-INS-195 | chr4-135236445-INS-195 |
| chr4-135241997-INS-193 | chr4-135241997-INS-193 |
| chr4-135279807-INS-120 | chr4-135279807-INS-122 |
| chr4-135292486-INS-195 | chr4-135292486-INS-195 |
| chr4-135324279-INS-56 | chr4-135324279-INS-60 |
| chr4-135340780-INS-55 | chr4-135340746-INS-76 |
| chr4-135341190-INS-327 | chr4-135341190-INS-327 |
| chr4-135352859-INS-56 | chr4-135352860-INS-60 |
| chr4-135371634-INS-123 | chr4-135371634-INS-123 |
| chr4-135378884-INS-188 | chr4-135378878-INS-194 |
| chr4-135384373-INS-73 | chr4-135384373-INS-73 |
| chr4-135390576-INS-235 | chr4-135390576-INS-247 |
| chr4-135418091-INS-98 | chr4-135418091-INS-98 |
| chr4-135420872-INS-378 | chr4-135420872-INS-378 |
| chr4-135437362-INS-63 | chr4-135437362-INS-62 |
| chr4-135440869-INS-54 | chr4-135440869-INS-56 |
| chr4-135442918-INS-63 | chr4-135442916-INS-56 |
| chr4-135451154-INS-110 | chr4-135451154-INS-115 |
| chr4-135452361-INS-149 | chr4-135452342-INS-143 |
| chr4-135453426-INS-79 | chr4-135453414-INS-84 |
| chr4-135465865-INS-215 | chr4-135465865-INS-215 |
| chr4-135471929-INS-88 | chr4-135471929-INS-82 |
| chr4-135481236-INS-522 | chr4-135481236-INS-538 |
| chr4-135490494-INS-56 | chr4-135490494-INS-68 |
| chr4-135495232-INS-84 | chr4-135495232-INS-84 |
| chr4-135496463-INS-66 | chr4-135496463-INS-62 |
| chr4-135498688-INS-261 | chr4-135498688-INS-260 |
| chr4-135503454-INS-95 | chr4-135503454-INS-95 |
| chr4-135509805-INS-201 | chr4-135509805-INS-201 |
| chr4-135511693-INS-474 | chr4-135511693-INS-480 |
| chr4-135544100-INS-58 | chr4-135544100-INS-58 |
| chr4-135544489-INS-173 | chr4-135544489-INS-173 |
| chr4-135550980-INS-252 | chr4-135550980-INS-252 |
| chr4-135555221-INS-56 | chr4-135555221-INS-56 |
| chr4-135562967-INS-148 | chr4-135562966-INS-147 |
| chr4-135570859-INS-60 | chr4-135570855-INS-64 |
| chr4-135584994-INS-180 | chr4-135584994-INS-180 |
| chr4-135605093-INS-203 | chr4-135605093-INS-203 |
| chr4-135609628-INS-78 | chr4-135609628-INS-78 |
| chr4-135616715-INS-219 | chr4-135616715-INS-219 |
| chr4-135617681-INS-54 | chr4-135617682-INS-60 |
| chr4-135621004-INS-201 | chr4-135621005-INS-201 |
| chr4-135662304-INS-71 | chr4-135662304-INS-71 |
| chr4-135671542-INS-52 | chr4-135671542-INS-52 |
| chr4-135682264-INS-219 | chr4-135682264-INS-219 |
| chr4-135683607-INS-116 | chr4-135683598-INS-100 |
| chr4-135687533-INS-202 | chr4-135687533-INS-202 |
| chr4-135709870-INS-83 | chr4-135709870-INS-51 |
| chr4-135721967-INS-116 | chr4-135721967-INS-116 |
| chr4-135723897-INS-200 | chr4-135723897-INS-200 |
| chr4-135726206-INS-81 | chr4-135726205-INS-82 |
| chr4-135750478-INS-61 | chr4-135750478-INS-61 |
| chr4-135752974-INS-650 | chr4-135752974-INS-650 |
| chr4-135766301-INS-209 | chr4-135766302-INS-209 |
| chr4-135769221-INS-208 | chr4-135769221-INS-208 |
| chr4-135775301-INS-199 | chr4-135775301-INS-199 |
| chr4-135784013-INS-212 | chr4-135784013-INS-212 |
| chr4-135786876-INS-225 | chr4-135786876-INS-225 |
| chr4-135791573-INS-232 | chr4-135791608-INS-225 |
| chr4-135796398-INS-200 | chr4-135796425-INS-175 |
| chr4-135797833-INS-459 | chr4-135797833-INS-466 |
| chr4-135802519-INS-180 | chr4-135802519-INS-180 |
| chr4-135804961-INS-196 | chr4-135804961-INS-200 |
| chr4-135806320-INS-159 | chr4-135806320-INS-159 |
| chr4-135807199-INS-233 | chr4-135807199-INS-234 |
| chr4-135808101-INS-63 | chr4-135808101-INS-63 |
| chr4-135826492-INS-134 | chr4-135826492-INS-135 |
| chr4-135839780-INS-189 | chr4-135839780-INS-187 |
| chr4-135842846-INS-227 | chr4-135842846-INS-227 |
| chr4-135845047-INS-207 | chr4-135845046-INS-188 |
| chr4-135859857-INS-439 | chr4-135859857-INS-439 |
| chr4-135882937-INS-93 | chr4-135882937-INS-62 |
| chr4-135896105-INS-197 | chr4-135896105-INS-197 |
| chr4-135917024-INS-195 | chr4-135917024-INS-195 |

| | |
|---|---|
| chr4-135926460-INS-187 | chr4-135926460-INS-187 |
| chr4-135927758-INS-201 | chr4-135927758-INS-201 |
| chr4-135929157-INS-432 | chr4-135929157-INS-432 |
| chr4-135991461-INS-4370 | chr4-135991462-INS-4378 |
| chr4-135993947-INS-149 | chr4-135993947-INS-159 |
| chr4-135996113-INS-480 | chr4-135996113-INS-480 |
| chr4-135997126-INS-155 | chr4-135997154-INS-162 |
| chr4-135999262-INS-65 | chr4-135999269-INS-82 |
| chr4-136002050-INS-216 | chr4-136002050-INS-217 |
| chr4-136005735-INS-84 | chr4-136005735-INS-88 |
| chr4-136011176-INS-50 | chr4-136011176-INS-54 |
| chr4-136016422-INS-200 | chr4-136016422-INS-200 |
| chr4-136019007-INS-70 | chr4-136019007-INS-70 |
| chr4-136034504-INS-211 | chr4-136034504-INS-211 |
| chr4-136044862-INS-184 | chr4-136044862-INS-184 |
| chr4-136049975-INS-1071 | chr4-136049976-INS-1070 |
| chr4-136053156-INS-203 | chr4-136053156-INS-203 |
| chr4-136057455-INS-221 | chr4-136057455-INS-221 |
| chr4-136059975-INS-107 | chr4-136059975-INS-107 |
| chr4-136081060-INS-59 | chr4-136081060-INS-59 |
| chr4-136086167-INS-473 | chr4-136086167-INS-473 |
| chr4-136093516-INS-52 | chr4-136093516-INS-52 |
| chr4-136116533-INS-876 | chr4-136116533-INS-876 |
| chr4-136120441-INS-200 | chr4-136120441-INS-200 |
| chr4-136123546-INS-212 | chr4-136123546-INS-212 |
| chr4-136125552-INS-67 | chr4-136125543-INS-55 |
| chr4-136149309-INS-100 | chr4-136149266-INS-100 |
| chr4-136164936-INS-56 | chr4-136164932-INS-60 |
| chr4-136167026-INS-485 | chr4-136167027-INS-481 |
| chr4-136168476-INS-166 | chr4-136168476-INS-166 |
| chr4-136174823-INS-64 | chr4-136174823-INS-64 |
| chr4-136178529-INS-228 | chr4-136178529-INS-229 |
| chr4-136201956-INS-221 | chr4-136201956-INS-220 |
| chr4-136207550-INS-210 | chr4-136207550-INS-211 |
| chr4-136216009-INS-117 | chr4-136216009-INS-99 |
| chr4-136217838-INS-208 | chr4-136217838-INS-208 |
| chr4-136221867-INS-1431 | chr4-136221868-INS-1480 |
| chr4-136225749-INS-213 | chr4-136225749-INS-211 |
| chr4-136230560-INS-201 | chr4-136230560-INS-200 |
| chr4-136247092-INS-62 | chr4-136247092-INS-64 |
| chr4-136297182-INS-202 | chr4-136297182-INS-202 |
| chr4-136321215-INS-235 | chr4-136321215-INS-235 |
| chr4-136323077-INS-210 | chr4-136323077-INS-209 |
| chr4-136332840-INS-62 | chr4-136332840-INS-62 |
| chr4-136336481-INS-65 | chr4-136336481-INS-65 |
| chr4-136338980-INS-199 | chr4-136338981-INS-199 |
| chr4-136346303-INS-110 | chr4-136346303-INS-110 |
| chr4-136357823-INS-78 | chr4-136357845-INS-84 |
| chr4-136361115-INS-237 | chr4-136361116-INS-237 |
| chr4-136365877-INS-208 | chr4-136365877-INS-211 |
| chr4-136388577-INS-191 | chr4-136388577-INS-191 |
| chr4-136392828-INS-190 | chr4-136392830-INS-194 |
| chr4-136420262-INS-188 | chr4-136420263-INS-189 |
| chr4-136436991-INS-497 | chr4-136436991-INS-497 |
| chr4-136464918-INS-56 | chr4-136464918-INS-64 |
| chr4-136474927-INS-153 | chr4-136474927-INS-153 |
| chr4-136484832-INS-121 | chr4-136484832-INS-128 |
| chr4-136497745-INS-358 | chr4-136497745-INS-358 |
| chr4-136501615-INS-105 | chr4-136501615-INS-105 |
| chr4-136515583-INS-57 | chr4-136515583-INS-57 |
| chr4-136537898-INS-63 | chr4-136537897-INS-75 |
| chr4-136572836-INS-56 | chr4-136572836-INS-56 |
| chr4-136579837-INS-106 | chr4-136579837-INS-106 |
| chr4-136596685-INS-207 | chr4-136596685-INS-207 |
| chr4-136605985-INS-269 | chr4-136605985-INS-269 |
| chr4-136642600-INS-103 | chr4-136642598-INS-133 |
| chr4-136648317-INS-137 | chr4-136648311-INS-114 |
| chr4-136652082-INS-122 | chr4-136652082-INS-102 |
| chr4-136676045-INS-110 | chr4-136676045-INS-110 |
| chr4-136691371-INS-217 | chr4-136691371-INS-217 |
| chr4-136720489-INS-195 | chr4-136720489-INS-193 |
| chr4-136750601-INS-100 | chr4-136750601-INS-100 |
| chr4-136774472-INS-64 | chr4-136774472-INS-64 |
| chr4-136784101-INS-148 | chr4-136784101-INS-148 |
| chr4-136790631-INS-721 | chr4-136790631-INS-721 |
| chr4-136796676-INS-143 | chr4-136796668-INS-175 |
| chr4-136798497-INS-200 | chr4-136798497-INS-200 |

| | |
|---|---|
| chr4-136802888-INS-198 | chr4-136802888-INS-198 |
| chr4-136810890-INS-215 | chr4-136810890-INS-215 |
| chr4-136830283-INS-216 | chr4-136830283-INS-216 |
| chr4-136868616-INS-60 | chr4-136868616-INS-60 |
| chr4-136895058-INS-60 | chr4-136895058-INS-60 |
| chr4-136929356-INS-63 | chr4-136929356-INS-63 |
| chr4-136931893-INS-105 | chr4-136931846-INS-131 |
| chr4-136958406-INS-164 | chr4-136958406-INS-168 |
| chr4-136971712-INS-61 | chr4-136971697-INS-64 |
| chr4-137016705-INS-81 | chr4-137016705-INS-81 |
| chr4-137019270-INS-52 | chr4-137019270-INS-52 |
| chr4-137038689-INS-166 | chr4-137038689-INS-166 |
| chr4-137051326-INS-75 | chr4-137051326-INS-85 |
| chr4-137057016-INS-182 | chr4-137057016-INS-166 |
| chr4-137064095-INS-8977 | chr4-137064096-INS-8978 |
| chr4-137075331-INS-228 | chr4-137075331-INS-228 |
| chr4-137100175-INS-211 | chr4-137100175-INS-209 |
| chr4-137104747-INS-184 | chr4-137104747-INS-184 |
| chr4-137126093-INS-204 | chr4-137126092-INS-205 |
| chr4-137140282-INS-436 | chr4-137140282-INS-469 |
| chr4-137157287-INS-207 | chr4-137157287-INS-207 |
| chr4-137162416-INS-153 | chr4-137162416-INS-163 |
| chr4-137169534-INS-138 | chr4-137169534-INS-127 |
| chr4-137179943-INS-73 | chr4-137179931-INS-66 |
| chr4-137229570-INS-203 | chr4-137229570-INS-203 |
| chr4-137288290-INS-289 | chr4-137288290-INS-277 |
| chr4-137291954-INS-60 | chr4-137291954-INS-60 |
| chr4-137307949-INS-163 | chr4-137307949-INS-162 |
| chr4-137309119-INS-240 | chr4-137309119-INS-234 |
| chr4-137315358-INS-215 | chr4-137315358-INS-214 |
| chr4-137337691-INS-54 | chr4-137337691-INS-50 |
| chr4-137355040-INS-69 | chr4-137355040-INS-53 |
| chr4-137381098-INS-92 | chr4-137381098-INS-92 |
| chr4-137408315-INS-70 | chr4-137408315-INS-98 |
| chr4-137412005-INS-54 | chr4-137412005-INS-54 |
| chr4-137418178-INS-74 | chr4-137418176-INS-76 |
| chr4-137433204-INS-225 | chr4-137433204-INS-225 |
| chr4-137494297-INS-207 | chr4-137494297-INS-209 |
| chr4-137504025-INS-130 | chr4-137504025-INS-126 |
| chr4-137507445-INS-192 | chr4-137507445-INS-192 |
| chr4-137513704-INS-63 | chr4-137513705-INS-55 |
| chr4-137526665-INS-52 | chr4-137526665-INS-52 |
| chr4-137528178-INS-68 | chr4-137528186-INS-53 |
| chr4-137531446-INS-70 | chr4-137531452-INS-52 |
| chr4-137533560-INS-225 | chr4-137533560-INS-225 |
| chr4-137538071-INS-90 | chr4-137538071-INS-87 |
| chr4-137545484-INS-204 | chr4-137545484-INS-204 |
| chr4-137553995-INS-201 | chr4-137553995-INS-201 |
| chr4-137555376-INS-596 | chr4-137555376-INS-589 |
| chr4-137561805-INS-208 | chr4-137561805-INS-212 |
| chr4-137563365-INS-81 | chr4-137563365-INS-84 |
| chr4-137612796-INS-1880 | chr4-137612797-INS-1880 |
| chr4-137617916-INS-82 | chr4-137617916-INS-84 |
| chr4-137651989-INS-58 | chr4-137651966-INS-87 |
| chr4-137689617-INS-485 | chr4-137689617-INS-486 |
| chr4-137693200-INS-80 | chr4-137693200-INS-80 |
| chr4-137731845-INS-817 | chr4-137731845-INS-816 |
| chr4-137738515-INS-76 | chr4-137738515-INS-67 |
| chr4-137740397-INS-545 | chr4-137740397-INS-544 |
| chr4-137741094-INS-227 | chr4-137741094-INS-225 |
| chr4-137745777-INS-94 | chr4-137745777-INS-85 |
| chr4-137752771-INS-155 | chr4-137752771-INS-155 |
| chr4-137754456-INS-209 | chr4-137754456-INS-210 |
| chr4-137768293-INS-54 | chr4-137768293-INS-54 |
| chr4-137774581-INS-102 | chr4-137774581-INS-102 |
| chr4-137796388-INS-198 | chr4-137796388-INS-199 |
| chr4-137802979-INS-52 | chr4-137802978-INS-52 |
| chr4-137808642-INS-235 | chr4-137808642-INS-235 |
| chr4-137839946-INS-172 | chr4-137839947-INS-172 |
| chr4-137852088-INS-52 | chr4-137852090-INS-53 |
| chr4-137869205-INS-497 | chr4-137869205-INS-497 |
| chr4-137883349-INS-195 | chr4-137883349-INS-195 |
| chr4-137912821-INS-167 | chr4-137912821-INS-167 |
| chr4-137920864-INS-251 | chr4-137920865-INS-257 |
| chr4-137921672-INS-4475 | chr4-137921673-INS-4475 |
| chr4-137945816-INS-212 | chr4-137945816-INS-212 |
| chr4-137952124-INS-210 | chr4-137952124-INS-209 |

| | |
|---|---|
| chr4-137958433-INS-214 | chr4-137958433-INS-215 |
| chr4-137964768-INS-99 | chr4-137964768-INS-105 |
| chr4-137972337-INS-102 | chr4-137972337-INS-114 |
| chr4-137973239-INS-144 | chr4-137973239-INS-150 |
| chr4-137993579-INS-202 | chr4-137993579-INS-203 |
| chr4-138003197-INS-83 | chr4-138003183-INS-65 |
| chr4-138011491-INS-146 | chr4-138011491-INS-146 |
| chr4-138016587-INS-57 | chr4-138016587-INS-57 |
| chr4-138018230-INS-162 | chr4-138018230-INS-179 |
| chr4-138026857-INS-226 | chr4-138026857-INS-226 |
| chr4-138043683-INS-183 | chr4-138043683-INS-184 |
| chr4-138047921-INS-97 | chr4-138047921-INS-97 |
| chr4-138060660-INS-209 | chr4-138060660-INS-205 |
| chr4-138084548-INS-838 | chr4-138084548-INS-838 |
| chr4-138099119-INS-90 | chr4-138099119-INS-90 |
| chr4-138108917-INS-67 | chr4-138108917-INS-67 |
| chr4-138153649-INS-201 | chr4-138153649-INS-205 |
| chr4-138176945-INS-222 | chr4-138176945-INS-221 |
| chr4-138184295-INS-54 | chr4-138184295-INS-54 |
| chr4-138204526-INS-185 | chr4-138204526-INS-188 |
| chr4-138212544-INS-595 | chr4-138212544-INS-593 |
| chr4-138213420-INS-203 | chr4-138213420-INS-203 |
| chr4-138228803-INS-51 | chr4-138228803-INS-53 |
| chr4-138256668-INS-125 | chr4-138256668-INS-138 |
| chr4-138265871-INS-57 | chr4-138265871-INS-57 |
| chr4-138289070-INS-66 | chr4-138289071-INS-74 |
| chr4-138383779-INS-101 | chr4-138383779-INS-101 |
| chr4-138390783-INS-110 | chr4-138390775-INS-117 |
| chr4-138394726-INS-69 | chr4-138394724-INS-69 |
| chr4-138398361-INS-194 | chr4-138398361-INS-195 |
| chr4-138407788-INS-200 | chr4-138407788-INS-200 |
| chr4-138416213-INS-301 | chr4-138416211-INS-301 |
| chr4-138453594-INS-68 | chr4-138453594-INS-68 |
| chr4-138468886-INS-115 | chr4-138468886-INS-116 |
| chr4-138482591-INS-158 | chr4-138482591-INS-158 |
| chr4-138492496-INS-92 | chr4-138492497-INS-92 |
| chr4-138497632-INS-54 | chr4-138497632-INS-54 |
| chr4-138511720-INS-200 | chr4-138511720-INS-201 |
| chr4-138517338-INS-51 | chr4-138517338-INS-53 |
| chr4-138526120-INS-278 | chr4-138526119-INS-294 |
| chr4-138531404-INS-60 | chr4-138531404-INS-60 |
| chr4-138532329-INS-200 | chr4-138532329-INS-200 |
| chr4-138544207-INS-90 | chr4-138544207-INS-90 |
| chr4-138551945-INS-1284 | chr4-138551946-INS-1283 |
| chr4-138558729-INS-93 | chr4-138558729-INS-93 |
| chr4-138565306-INS-790 | chr4-138565306-INS-790 |
| chr4-138611243-INS-60 | chr4-138611243-INS-60 |
| chr4-138614022-INS-267 | chr4-138614022-INS-267 |
| chr4-138627213-INS-98 | chr4-138627213-INS-98 |
| chr4-138635513-INS-119 | chr4-138635513-INS-119 |
| chr4-138644877-INS-182 | chr4-138644877-INS-182 |
| chr4-138648270-INS-65 | chr4-138648270-INS-65 |
| chr4-138667563-INS-60 | chr4-138667563-INS-62 |
| chr4-138673032-INS-879 | chr4-138673032-INS-879 |
| chr4-138730472-INS-109 | chr4-138730480-INS-121 |
| chr4-138730888-INS-5416 | chr4-138730889-INS-5416 |
| chr4-138743664-INS-84 | chr4-138743664-INS-106 |
| chr4-138748567-INS-52 | chr4-138748567-INS-52 |
| chr4-138758420-INS-63 | chr4-138758420-INS-63 |
| chr4-138769720-INS-196 | chr4-138769724-INS-212 |
| chr4-138770713-INS-220 | chr4-138770713-INS-220 |
| chr4-138784293-INS-62 | chr4-138784299-INS-60 |
| chr4-138789616-INS-69 | chr4-138789616-INS-69 |
| chr4-138798980-INS-116 | chr4-138798980-INS-120 |
| chr4-138824619-INS-220 | chr4-138824619-INS-222 |
| chr4-138849712-INS-240 | chr4-138849712-INS-240 |
| chr4-138855551-INS-235 | chr4-138855551-INS-245 |
| chr4-138859959-INS-59 | chr4-138859959-INS-59 |
| chr4-138885516-INS-56 | chr4-138885516-INS-56 |
| chr4-138890615-INS-209 | chr4-138890615-INS-209 |
| chr4-138908911-INS-331 | chr4-138908911-INS-330 |
| chr4-138916358-INS-66 | chr4-138916362-INS-74 |
| chr4-138934194-INS-85 | chr4-138934146-INS-85 |
| chr4-138983909-INS-210 | chr4-138983909-INS-210 |
| chr4-139041036-INS-95 | chr4-139041033-INS-97 |
| chr4-139048256-INS-205 | chr4-139048260-INS-187 |
| chr4-139055703-INS-54 | chr4-139055703-INS-54 |

| | |
|---|---|
| chr4-139072318-INS-206 | chr4-139072318-INS-206 |
| chr4-139121978-INS-54 | chr4-139121978-INS-50 |
| chr4-139144850-INS-308 | chr4-139144850-INS-308 |
| chr4-139193411-INS-171 | chr4-139193411-INS-171 |
| chr4-139205763-INS-108 | chr4-139205763-INS-108 |
| chr4-139228863-INS-51 | chr4-139228863-INS-51 |
| chr4-139251026-INS-2456 | chr4-139251027-INS-2448 |
| chr4-139280618-INS-267 | chr4-139280618-INS-231 |
| chr4-139288791-INS-486 | chr4-139288768-INS-492 |
| chr4-139297045-INS-225 | chr4-139297045-INS-225 |
| chr4-139342946-INS-366 | chr4-139342946-INS-349 |
| chr4-139401705-INS-51 | chr4-139401705-INS-55 |
| chr4-139416158-INS-62 | chr4-139416158-INS-62 |
| chr4-139429431-INS-70 | chr4-139429431-INS-70 |
| chr4-139431459-INS-81 | chr4-139431459-INS-81 |
| chr4-139447787-INS-1958 | chr4-139447788-INS-1970 |
| chr4-139459126-INS-401 | chr4-139459126-INS-401 |
| chr4-139465648-INS-56 | chr4-139465648-INS-56 |
| chr4-139491995-INS-207 | chr4-139491995-INS-207 |
| chr4-139502056-INS-62 | chr4-139502056-INS-68 |
| chr4-139513428-INS-63 | chr4-139513428-INS-59 |
| chr4-139530733-INS-139 | chr4-139530724-INS-127 |
| chr4-139534124-INS-56 | chr4-139534124-INS-50 |
| chr4-139554587-INS-101 | chr4-139554587-INS-101 |
| chr4-139580882-INS-51 | chr4-139580882-INS-53 |
| chr4-139582034-INS-67 | chr4-139582034-INS-67 |
| chr4-139591763-INS-386 | chr4-139591768-INS-395 |
| chr4-139593033-INS-114 | chr4-139593054-INS-114 |
| chr4-139617283-INS-62 | chr4-139617283-INS-69 |
| chr4-139622612-INS-116 | chr4-139622612-INS-116 |
| chr4-139647831-INS-92 | chr4-139647831-INS-97 |
| chr4-139658171-INS-292 | chr4-139658150-INS-279 |
| chr4-139662596-INS-105 | chr4-139662596-INS-105 |
| chr4-139707307-INS-200 | chr4-139707307-INS-200 |
| chr4-139712985-INS-70 | chr4-139712986-INS-70 |
| chr4-139718251-INS-225 | chr4-139718294-INS-189 |
| chr4-139768873-INS-144 | chr4-139768876-INS-140 |
| chr4-139776764-INS-87 | chr4-139776764-INS-87 |
| chr4-139800931-INS-203 | chr4-139800931-INS-204 |
| chr4-139821763-INS-54 | chr4-139821763-INS-54 |
| chr4-139828471-INS-111 | chr4-139828471-INS-111 |
| chr4-139830639-INS-60 | chr4-139830639-INS-50 |
| chr4-139833289-INS-104 | chr4-139833289-INS-104 |
| chr4-139835090-INS-78 | chr4-139835092-INS-76 |
| chr4-139837808-INS-160 | chr4-139837808-INS-158 |
| chr4-139854357-INS-52 | chr4-139854344-INS-50 |
| chr4-139857964-INS-138 | chr4-139857964-INS-137 |
| chr4-139864939-INS-63 | chr4-139864939-INS-63 |
| chr4-139900051-INS-197 | chr4-139900051-INS-197 |
| chr4-139904618-INS-90 | chr4-139904618-INS-90 |
| chr4-139922671-INS-62 | chr4-139922671-INS-62 |
| chr4-139966665-INS-478 | chr4-139966665-INS-480 |
| chr4-139996360-INS-131 | chr4-139996360-INS-131 |
| chr4-140008985-INS-132 | chr4-140008985-INS-132 |
| chr4-140010643-INS-210 | chr4-140010643-INS-210 |
| chr4-140031719-INS-348 | chr4-140031721-INS-352 |
| chr4-140039867-INS-410 | chr4-140039867-INS-410 |
| chr4-140058864-INS-422 | chr4-140058864-INS-422 |
| chr4-140073942-INS-132 | chr4-140073942-INS-110 |
| chr4-140080052-INS-400 | chr4-140080052-INS-400 |
| chr4-140087121-INS-121 | chr4-140087121-INS-121 |
| chr4-140088946-INS-103 | chr4-140088946-INS-102 |
| chr4-140099629-INS-150 | chr4-140099629-INS-149 |
| chr4-140146203-INS-72 | chr4-140146203-INS-72 |
| chr4-140165954-INS-54 | chr4-140165954-INS-54 |
| chr4-140168352-INS-220 | chr4-140168352-INS-220 |
| chr4-140230185-INS-370 | chr4-140230185-INS-370 |
| chr4-140249840-INS-80 | chr4-140249826-INS-76 |
| chr4-140252269-INS-71 | chr4-140252269-INS-53 |
| chr4-140287763-INS-346 | chr4-140287755-INS-356 |
| chr4-140301274-INS-227 | chr4-140301275-INS-228 |
| chr4-140302446-INS-208 | chr4-140302446-INS-208 |
| chr4-140318203-INS-171 | chr4-140318203-INS-171 |
| chr4-140323159-INS-192 | chr4-140323159-INS-178 |
| chr4-140327509-INS-108 | chr4-140327509-INS-108 |
| chr4-140338281-INS-66 | chr4-140338281-INS-66 |
| chr4-140348639-INS-62 | chr4-140348639-INS-62 |

| | |
|---|---|
| chr4-140349243-INS-199 | chr4-140349243-INS-199 |
| chr4-140365480-INS-291 | chr4-140365499-INS-280 |
| chr4-140382121-INS-57 | chr4-140382113-INS-69 |
| chr4-140382985-INS-207 | chr4-140382985-INS-210 |
| chr4-140387805-INS-234 | chr4-140387805-INS-265 |
| chr4-140391279-INS-189 | chr4-140391279-INS-189 |
| chr4-140396754-INS-138 | chr4-140396754-INS-138 |
| chr4-140408207-INS-151 | chr4-140408207-INS-145 |
| chr4-140409546-INS-201 | chr4-140409546-INS-201 |
| chr4-140411087-INS-68 | chr4-140411087-INS-69 |
| chr4-140447771-INS-199 | chr4-140447771-INS-175 |
| chr4-140454264-INS-200 | chr4-140454264-INS-200 |
| chr4-140495129-INS-85 | chr4-140495137-INS-116 |
| chr4-140528362-INS-58 | chr4-140528362-INS-58 |
| chr4-140531446-INS-150 | chr4-140531432-INS-181 |
| chr4-140534183-INS-64 | chr4-140534183-INS-66 |
| chr4-140562975-INS-72 | chr4-140562978-INS-50 |
| chr4-140564652-INS-54 | chr4-140564652-INS-58 |
| chr4-140587574-INS-88 | chr4-140587574-INS-88 |
| chr4-140596616-INS-70 | chr4-140596616-INS-70 |
| chr4-140610410-INS-151 | chr4-140610410-INS-151 |
| chr4-140648338-INS-182 | chr4-140648338-INS-182 |
| chr4-140654138-INS-84 | chr4-140654142-INS-88 |
| chr4-140660008-INS-59 | chr4-140660008-INS-59 |
| chr4-140682924-INS-64 | chr4-140682924-INS-64 |
| chr4-140706059-INS-206 | chr4-140706059-INS-206 |
| chr4-140707485-INS-246 | chr4-140707485-INS-244 |
| chr4-140736753-INS-53 | chr4-140736753-INS-53 |
| chr4-140754647-INS-52 | chr4-140754647-INS-54 |
| chr4-140764897-INS-211 | chr4-140764898-INS-211 |
| chr4-140767418-INS-202 | chr4-140767418-INS-203 |
| chr4-140789175-INS-308 | chr4-140789170-INS-309 |
| chr4-140802096-INS-118 | chr4-140802096-INS-118 |
| chr4-140815066-INS-53 | chr4-140815040-INS-56 |
| chr4-140822073-INS-76 | chr4-140822073-INS-82 |
| chr4-140833120-INS-230 | chr4-140833120-INS-233 |
| chr4-140836874-INS-197 | chr4-140836874-INS-196 |
| chr4-140861159-INS-151 | chr4-140861157-INS-154 |
| chr4-140878685-INS-54 | chr4-140878685-INS-54 |
| chr4-140886456-INS-220 | chr4-140886456-INS-220 |
| chr4-140899746-INS-204 | chr4-140899746-INS-205 |
| chr4-140912369-INS-68 | chr4-140912369-INS-68 |
| chr4-140923054-INS-210 | chr4-140923054-INS-210 |
| chr4-140925187-INS-210 | chr4-140925187-INS-210 |
| chr4-140948490-INS-205 | chr4-140948490-INS-204 |
| chr4-140949728-INS-70 | chr4-140949728-INS-74 |
| chr4-140953591-INS-66 | chr4-140953591-INS-66 |
| chr4-140957789-INS-202 | chr4-140957789-INS-203 |
| chr4-140960936-INS-55 | chr4-140960936-INS-50 |
| chr4-140965022-INS-133 | chr4-140965039-INS-135 |
| chr4-140989015-INS-440 | chr4-140989015-INS-440 |
| chr4-140990611-INS-176 | chr4-140990611-INS-176 |
| chr4-140992284-INS-165 | chr4-140992284-INS-162 |
| chr4-140996029-INS-365 | chr4-140996029-INS-365 |
| chr4-141006407-INS-167 | chr4-141006407-INS-165 |
| chr4-141007597-INS-100 | chr4-141007597-INS-120 |
| chr4-141026070-INS-58 | chr4-141026081-INS-74 |
| chr4-141062486-INS-65 | chr4-141062487-INS-54 |
| chr4-141094976-INS-56 | chr4-141094976-INS-56 |
| chr4-141130353-INS-212 | chr4-141130353-INS-212 |
| chr4-141145507-INS-152 | chr4-141145502-INS-155 |
| chr4-141152825-INS-202 | chr4-141152825-INS-201 |
| chr4-141210549-INS-1891 | chr4-141210553-INS-1881 |
| chr4-141222369-INS-57 | chr4-141222369-INS-57 |
| chr4-141222527-INS-1767 | chr4-141222521-INS-1731 |
| chr4-141233051-INS-199 | chr4-141233051-INS-199 |
| chr4-141238353-INS-120 | chr4-141238353-INS-120 |
| chr4-141246329-INS-213 | chr4-141246329-INS-213 |
| chr4-141251327-INS-240 | chr4-141251327-INS-244 |
| chr4-141255560-INS-118 | chr4-141255560-INS-118 |
| chr4-141257593-INS-75 | chr4-141257593-INS-70 |
| chr4-141279187-INS-209 | chr4-141279187-INS-209 |
| chr4-141291260-INS-206 | chr4-141291260-INS-207 |
| chr4-141292435-INS-169 | chr4-141292436-INS-176 |
| chr4-141294871-INS-184 | chr4-141294871-INS-184 |
| chr4-141306016-INS-205 | chr4-141306016-INS-195 |
| chr4-141306858-INS-57 | chr4-141306858-INS-57 |

| | |
|---|---|
| chr4-141311177-INS-190 | chr4-141311177-INS-190 |
| chr4-141315305-INS-223 | chr4-141315305-INS-224 |
| chr4-141323727-INS-94 | chr4-141323728-INS-52 |
| chr4-141331185-INS-217 | chr4-141331185-INS-218 |
| chr4-141336119-INS-53 | chr4-141336129-INS-61 |
| chr4-141339492-INS-202 | chr4-141339492-INS-202 |
| chr4-141346603-INS-52 | chr4-141346603-INS-51 |
| chr4-141355005-INS-202 | chr4-141355005-INS-202 |
| chr4-141360742-INS-83 | chr4-141360789-INS-53 |
| chr4-141365384-INS-98 | chr4-141365384-INS-98 |
| chr4-141373920-INS-203 | chr4-141373920-INS-203 |
| chr4-141375017-INS-146 | chr4-141375017-INS-145 |
| chr4-141382196-INS-76 | chr4-141382229-INS-52 |
| chr4-141395300-INS-110 | chr4-141395263-INS-110 |
| chr4-141399339-INS-50 | chr4-141399355-INS-60 |
| chr4-141423520-INS-52 | chr4-141423522-INS-53 |
| chr4-141424764-INS-175 | chr4-141424764-INS-175 |
| chr4-141431286-INS-204 | chr4-141431286-INS-211 |
| chr4-141438551-INS-208 | chr4-141438551-INS-208 |
| chr4-141446685-INS-145 | chr4-141446685-INS-144 |
| chr4-141458285-INS-74 | chr4-141458285-INS-74 |
| chr4-141468382-INS-77 | chr4-141468383-INS-66 |
| chr4-141471473-INS-188 | chr4-141471473-INS-187 |
| chr4-141473557-INS-74 | chr4-141473577-INS-71 |
| chr4-141476076-INS-72 | chr4-141476076-INS-72 |
| chr4-141499150-INS-52 | chr4-141499150-INS-52 |
| chr4-141499841-INS-211 | chr4-141499841-INS-211 |
| chr4-141525618-INS-66 | chr4-141525622-INS-54 |
| chr4-141545375-INS-243 | chr4-141545375-INS-244 |
| chr4-141549701-INS-212 | chr4-141549701-INS-212 |
| chr4-141552065-INS-101 | chr4-141552053-INS-145 |
| chr4-141556769-INS-120 | chr4-141556769-INS-120 |
| chr4-141564331-INS-133 | chr4-141564328-INS-144 |
| chr4-141603302-INS-57 | chr4-141603302-INS-57 |
| chr4-141612435-INS-111 | chr4-141612438-INS-120 |
| chr4-141616363-INS-216 | chr4-141616352-INS-217 |
| chr4-141630979-INS-77 | chr4-141630977-INS-68 |
| chr4-141649416-INS-188 | chr4-141649416-INS-188 |
| chr4-141658188-INS-53 | chr4-141658188-INS-53 |
| chr4-141661764-INS-58 | chr4-141661764-INS-58 |
| chr4-141670248-INS-6442 | chr4-141670249-INS-6442 |
| chr4-141674499-INS-101 | chr4-141674499-INS-103 |
| chr4-141692260-INS-84 | chr4-141692260-INS-84 |
| chr4-141703127-INS-250 | chr4-141703127-INS-250 |
| chr4-141708531-INS-112 | chr4-141708531-INS-112 |
| chr4-141713511-INS-94 | chr4-141713511-INS-94 |
| chr4-141725666-INS-247 | chr4-141725666-INS-247 |
| chr4-141739584-INS-64 | chr4-141739584-INS-64 |
| chr4-141760640-INS-210 | chr4-141760640-INS-210 |
| chr4-141801646-INS-178 | chr4-141801646-INS-180 |
| chr4-141805922-INS-66 | chr4-141805922-INS-66 |
| chr4-141816683-INS-60 | chr4-141816680-INS-60 |
| chr4-141817988-INS-68 | chr4-141818017-INS-65 |
| chr4-141821953-INS-7125 | chr4-141821954-INS-7125 |
| chr4-141824608-INS-419 | chr4-141824608-INS-401 |
| chr4-141838438-INS-91 | chr4-141838438-INS-91 |
| chr4-141849439-INS-66 | chr4-141849437-INS-73 |
| chr4-141852844-INS-203 | chr4-141852844-INS-203 |
| chr4-141881324-INS-52 | chr4-141881324-INS-52 |
| chr4-141946363-INS-226 | chr4-141946363-INS-227 |
| chr4-141960822-INS-195 | chr4-141960822-INS-195 |
| chr4-141985857-INS-131 | chr4-141985857-INS-135 |
| chr4-141994501-INS-194 | chr4-141994501-INS-194 |
| chr4-141999034-INS-60 | chr4-141999034-INS-60 |
| chr4-141999887-INS-665 | chr4-141999887-INS-648 |
| chr4-142002848-INS-209 | chr4-142002848-INS-209 |
| chr4-142004637-INS-121 | chr4-142004637-INS-87 |
| chr4-142015717-INS-163 | chr4-142015717-INS-159 |
| chr4-142102372-INS-481 | chr4-142102372-INS-481 |
| chr4-142125527-INS-483 | chr4-142125527-INS-483 |
| chr4-142136453-INS-97 | chr4-142136458-INS-99 |
| chr4-142141666-INS-90 | chr4-142141666-INS-90 |
| chr4-142154727-INS-414 | chr4-142154727-INS-414 |
| chr4-142177397-INS-158 | chr4-142177400-INS-166 |
| chr4-142182898-INS-60 | chr4-142182898-INS-60 |
| chr4-142195890-INS-57 | chr4-142195875-INS-61 |
| chr4-142208494-INS-168 | chr4-142208494-INS-168 |

| | |
|---|---|
| chr4-142209420-INS-385 | chr4-142209420-INS-386 |
| chr4-142212112-INS-54 | chr4-142212112-INS-54 |
| chr4-142221375-INS-56 | chr4-142221375-INS-72 |
| chr4-142229733-INS-358 | chr4-142229733-INS-358 |
| chr4-142236671-INS-66 | chr4-142236671-INS-68 |
| chr4-142240620-INS-66 | chr4-142240620-INS-66 |
| chr4-142251413-INS-74 | chr4-142251413-INS-76 |
| chr4-142255573-INS-62 | chr4-142255567-INS-101 |
| chr4-142276934-INS-64 | chr4-142276934-INS-68 |
| chr4-142280006-INS-64 | chr4-142280006-INS-64 |
| chr4-142314715-INS-52 | chr4-142314715-INS-50 |
| chr4-142316482-INS-60 | chr4-142316482-INS-72 |
| chr4-142318564-INS-4134 | chr4-142318565-INS-4134 |
| chr4-142330025-INS-106 | chr4-142330026-INS-142 |
| chr4-142351230-INS-7163 | chr4-142351231-INS-7163 |
| chr4-142353492-INS-320 | chr4-142353492-INS-320 |
| chr4-142354382-INS-82 | chr4-142354337-INS-127 |
| chr4-142369527-INS-195 | chr4-142369527-INS-195 |
| chr4-142380802-INS-1030 | chr4-142380803-INS-1032 |
| chr4-142393387-INS-72 | chr4-142393388-INS-76 |
| chr4-142405865-INS-60 | chr4-142405865-INS-54 |
| chr4-142419595-INS-311 | chr4-142419595-INS-311 |
| chr4-142420728-INS-923 | chr4-142420728-INS-925 |
| chr4-142493761-INS-90 | chr4-142493761-INS-100 |
| chr4-142495065-INS-52 | chr4-142495065-INS-54 |
| chr4-142502641-INS-214 | chr4-142502641-INS-214 |
| chr4-142515132-INS-603 | chr4-142515132-INS-602 |
| chr4-142547509-INS-211 | chr4-142547509-INS-210 |
| chr4-142555302-INS-122 | chr4-142555302-INS-99 |
| chr4-142563819-INS-127 | chr4-142563835-INS-122 |
| chr4-142577700-INS-132 | chr4-142577700-INS-132 |
| chr4-142598300-INS-142 | chr4-142598263-INS-150 |
| chr4-142605339-INS-289 | chr4-142605339-INS-289 |
| chr4-142615939-INS-161 | chr4-142615939-INS-161 |
| chr4-142622903-INS-101 | chr4-142622903-INS-77 |
| chr4-142686995-INS-122 | chr4-142686995-INS-126 |
| chr4-142694423-INS-60 | chr4-142694423-INS-60 |
| chr4-142712004-INS-187 | chr4-142712009-INS-183 |
| chr4-142713987-INS-163 | chr4-142713987-INS-122 |
| chr4-142719915-INS-56 | chr4-142719915-INS-64 |
| chr4-142736914-INS-128 | chr4-142736914-INS-119 |
| chr4-142743779-INS-139 | chr4-142743779-INS-184 |
| chr4-142746512-INS-61 | chr4-142746512-INS-61 |
| chr4-142790383-INS-80 | chr4-142790383-INS-95 |
| chr4-142806441-INS-52 | chr4-142806441-INS-52 |
| chr4-142807728-INS-54 | chr4-142807728-INS-56 |
| chr4-142810197-INS-85 | chr4-142810197-INS-80 |
| chr4-142817679-INS-1248 | chr4-142817680-INS-1248 |
| chr4-142850958-INS-1026 | chr4-142850959-INS-1028 |
| chr4-142876044-INS-212 | chr4-142876044-INS-211 |
| chr4-142908671-INS-54 | chr4-142908671-INS-52 |
| chr4-142913951-INS-178 | chr4-142913951-INS-188 |
| chr4-142917615-INS-183 | chr4-142917615-INS-183 |
| chr4-142922784-INS-201 | chr4-142922784-INS-202 |
| chr4-142936589-INS-168 | chr4-142936577-INS-168 |
| chr4-142951963-INS-69 | chr4-142951963-INS-85 |
| chr4-142963274-INS-168 | chr4-142963274-INS-168 |
| chr4-142964679-INS-58 | chr4-142964679-INS-58 |
| chr4-142977785-INS-181 | chr4-142977784-INS-163 |
| chr4-142979218-INS-421 | chr4-142979219-INS-421 |
| chr4-142981908-INS-111 | chr4-142981908-INS-110 |
| chr4-142993089-INS-205 | chr4-142993089-INS-205 |
| chr4-142993734-INS-181 | chr4-142993748-INS-181 |
| chr4-142998051-INS-82 | chr4-142998047-INS-88 |
| chr4-143016901-INS-89 | chr4-143016901-INS-89 |
| chr4-143019759-INS-175 | chr4-143019759-INS-172 |
| chr4-143020441-INS-607 | chr4-143020440-INS-601 |
| chr4-143046961-INS-213 | chr4-143046961-INS-213 |
| chr4-143053520-INS-66 | chr4-143053520-INS-66 |
| chr4-143070220-INS-211 | chr4-143070220-INS-212 |
| chr4-143087702-INS-54 | chr4-143087702-INS-62 |
| chr4-143108658-INS-89 | chr4-143108652-INS-65 |
| chr4-143151484-INS-527 | chr4-143151484-INS-527 |
| chr4-143152696-INS-4132 | chr4-143152697-INS-4132 |
| chr4-143187523-INS-466 | chr4-143187523-INS-466 |
| chr4-143193844-INS-1092 | chr4-143193845-INS-1092 |
| chr4-143222491-INS-86 | chr4-143222485-INS-86 |

| | |
|---|---|
| chr4-143239573-INS-80 | chr4-143239581-INS-62 |
| chr4-143240977-INS-106 | chr4-143240977-INS-102 |
| chr4-143250595-INS-204 | chr4-143250595-INS-204 |
| chr4-143266341-INS-204 | chr4-143266341-INS-204 |
| chr4-143352251-INS-519 | chr4-143352251-INS-519 |
| chr4-143359746-INS-199 | chr4-143359746-INS-199 |
| chr4-143363651-INS-51 | chr4-143363651-INS-59 |
| chr4-143422263-INS-767 | chr4-143422263-INS-764 |
| chr4-143433814-INS-53 | chr4-143433814-INS-54 |
| chr4-143434808-INS-388 | chr4-143434808-INS-388 |
| chr4-143450394-INS-57 | chr4-143450394-INS-54 |
| chr4-143460806-INS-885 | chr4-143460806-INS-884 |
| chr4-143466869-INS-185 | chr4-143466889-INS-170 |
| chr4-143483329-INS-66 | chr4-143483334-INS-78 |
| chr4-143522484-INS-210 | chr4-143522452-INS-212 |
| chr4-143560561-INS-571 | chr4-143560561-INS-571 |
| chr4-143644108-INS-70 | chr4-143644108-INS-70 |
| chr4-143688125-INS-264 | chr4-143688125-INS-264 |
| chr4-143740054-INS-214 | chr4-143740054-INS-213 |
| chr4-143765383-INS-55 | chr4-143765383-INS-54 |
| chr4-143770399-INS-669 | chr4-143770399-INS-675 |
| chr4-143894544-INS-53 | chr4-143894526-INS-53 |
| chr4-143930327-INS-6867 | chr4-143930327-INS-6867 |
| chr4-143980729-INS-1894 | chr4-143980730-INS-1893 |
| chr4-143980874-INS-300 | chr4-143980875-INS-300 |
| chr4-143990368-INS-328 | chr4-143990368-INS-330 |
| chr4-143994280-INS-5148 | chr4-143994281-INS-5148 |
| chr4-143994281-INS-471 | chr4-143994281-INS-471 |
| chr4-144037246-INS-180 | chr4-144037246-INS-182 |
| chr4-144047427-INS-220 | chr4-144047427-INS-219 |
| chr4-144048496-INS-79 | chr4-144048496-INS-79 |
| chr4-144073961-INS-204 | chr4-144073961-INS-204 |
| chr4-144097254-INS-215 | chr4-144097254-INS-219 |
| chr4-144116958-INS-52 | chr4-144116958-INS-52 |
| chr4-144162320-INS-60 | chr4-144162320-INS-62 |
| chr4-144169965-INS-52 | chr4-144169965-INS-50 |
| chr4-144187069-INS-4101 | chr4-144187070-INS-4080 |
| chr4-144222755-INS-199 | chr4-144222755-INS-199 |
| chr4-144232072-INS-1425 | chr4-144232073-INS-1425 |
| chr4-144245644-INS-60 | chr4-144245641-INS-77 |
| chr4-144248481-INS-71 | chr4-144248481-INS-71 |
| chr4-144265005-INS-59 | chr4-144265005-INS-54 |
| chr4-144300843-INS-7095 | chr4-144300844-INS-7095 |
| chr4-144318593-INS-217 | chr4-144318593-INS-217 |
| chr4-144355276-INS-329 | chr4-144355276-INS-329 |
| chr4-144400007-INS-7148 | chr4-144400008-INS-7148 |
| chr4-144414193-INS-7416 | chr4-144414194-INS-7417 |
| chr4-144420607-INS-102 | chr4-144420607-INS-98 |
| chr4-144424869-INS-1658 | chr4-144424870-INS-1662 |
| chr4-144441088-INS-218 | chr4-144441088-INS-218 |
| chr4-144456383-INS-203 | chr4-144456383-INS-203 |
| chr4-144458590-INS-65 | chr4-144458590-INS-65 |
| chr4-144461917-INS-823 | chr4-144461917-INS-823 |
| chr4-144483370-INS-248 | chr4-144483370-INS-248 |
| chr4-144488067-INS-1600 | chr4-144488068-INS-1602 |
| chr4-144493122-INS-54 | chr4-144493098-INS-66 |
| chr4-144507845-INS-199 | chr4-144507845-INS-199 |
| chr4-144518365-INS-64 | chr4-144518365-INS-64 |
| chr4-144521265-INS-3150 | chr4-144521266-INS-3150 |
| chr4-144523330-INS-658 | chr4-144523330-INS-658 |
| chr4-144529494-INS-208 | chr4-144529494-INS-208 |
| chr4-144539790-INS-77 | chr4-144539790-INS-106 |
| chr4-144595613-INS-60 | chr4-144595613-INS-60 |
| chr4-144605716-INS-204 | chr4-144605716-INS-202 |
| chr4-144608551-INS-63 | chr4-144608551-INS-63 |
| chr4-144613049-INS-52 | chr4-144613049-INS-54 |
| chr4-144686923-INS-903 | chr4-144686923-INS-906 |
| chr4-144733795-INS-70 | chr4-144733795-INS-115 |
| chr4-144741598-INS-396 | chr4-144741598-INS-396 |
| chr4-144764442-INS-54 | chr4-144764442-INS-54 |
| chr4-144771204-INS-92 | chr4-144771204-INS-96 |
| chr4-144802171-INS-7072 | chr4-144802172-INS-7056 |
| chr4-144803700-INS-201 | chr4-144803700-INS-201 |
| chr4-144817447-INS-93 | chr4-144817447-INS-93 |
| chr4-144855530-INS-181 | chr4-144855530-INS-213 |
| chr4-144878829-INS-179 | chr4-144878829-INS-179 |
| chr4-144892636-INS-62 | chr4-144892636-INS-62 |

| | |
|---|---|
| chr4-144899918-INS-453 | chr4-144899918-INS-453 |
| chr4-144927618-INS-58 | chr4-144927618-INS-52 |
| chr4-144938357-INS-165 | chr4-144938357-INS-195 |
| chr4-144953060-INS-209 | chr4-144953060-INS-208 |
| chr4-145011315-INS-183 | chr4-145011315-INS-183 |
| chr4-145017893-INS-232 | chr4-145017893-INS-233 |
| chr4-145020582-INS-85 | chr4-145020582-INS-85 |
| chr4-145046472-INS-185 | chr4-145046472-INS-190 |
| chr4-145053481-INS-466 | chr4-145053481-INS-466 |
| chr4-145056753-INS-181 | chr4-145056753-INS-181 |
| chr4-145066833-INS-52 | chr4-145066833-INS-52 |
| chr4-145069323-INS-530 | chr4-145069323-INS-523 |
| chr4-145081150-INS-216 | chr4-145081150-INS-215 |
| chr4-145105839-INS-403 | chr4-145105839-INS-405 |
| chr4-145115270-INS-254 | chr4-145115270-INS-254 |
| chr4-145217279-INS-390 | chr4-145217279-INS-390 |
| chr4-145262308-INS-498 | chr4-145262310-INS-500 |
| chr4-145275366-INS-70 | chr4-145275367-INS-71 |
| chr4-145328366-INS-2429 | chr4-145328367-INS-2429 |
| chr4-145350165-INS-84 | chr4-145350166-INS-84 |
| chr4-145453558-INS-367 | chr4-145453558-INS-367 |
| chr4-146061338-INS-84 | chr4-146061341-INS-84 |
| chr4-146291509-INS-4770 | chr4-146291510-INS-4770 |
| chr4-146638154-INS-171 | chr4-146638154-INS-171 |
| chr4-147012523-INS-197 | chr4-147012524-INS-198 |
| chr4-147015690-INS-577 | chr4-147015691-INS-577 |
| chr4-147022329-INS-114 | chr4-147022329-INS-114 |
| chr4-147104459-INS-56 | chr4-147104457-INS-57 |
| chr4-147192523-INS-171 | chr4-147192524-INS-169 |
| chr4-147195645-INS-185 | chr4-147195646-INS-184 |
| chr4-147332410-INS-2922 | chr4-147332403-INS-2913 |
| chr4-147343453-INS-420 | chr4-147343454-INS-420 |
| chr4-147443253-INS-56 | chr4-147443253-INS-56 |
| chr4-147524301-INS-188 | chr4-147524301-INS-188 |
| chr4-147572112-INS-2873 | chr4-147572113-INS-2873 |
| chr4-147698576-INS-310 | chr4-147698576-INS-310 |
| chr4-147729479-INS-56 | chr4-147729479-INS-56 |
| chr4-147770215-INS-291 | chr4-147770215-INS-283 |
| chr4-147906011-INS-231 | chr4-147906011-INS-231 |
| chr4-147926974-INS-199 | chr4-147926974-INS-199 |
| chr4-147931823-INS-255 | chr4-147931823-INS-259 |
| chr4-147944144-INS-224 | chr4-147944144-INS-228 |
| chr4-147945231-INS-394 | chr4-147945231-INS-394 |
| chr4-147958725-INS-64 | chr4-147958725-INS-64 |
| chr4-147963815-INS-55 | chr4-147963815-INS-55 |
| chr4-148035155-INS-57 | chr4-148035155-INS-57 |
| chr4-148038469-INS-211 | chr4-148038469-INS-211 |
| chr4-148045942-INS-78 | chr4-148045942-INS-84 |
| chr4-148049286-INS-177 | chr4-148049286-INS-177 |
| chr4-148063552-INS-271 | chr4-148063552-INS-271 |
| chr4-148075302-INS-210 | chr4-148075302-INS-210 |
| chr4-148106942-INS-86 | chr4-148106942-INS-85 |
| chr4-148118029-INS-208 | chr4-148118029-INS-207 |
| chr4-148121049-INS-173 | chr4-148121049-INS-173 |
| chr4-148148114-INS-206 | chr4-148148114-INS-206 |
| chr4-148157843-INS-187 | chr4-148157843-INS-187 |
| chr4-148163627-INS-53 | chr4-148163625-INS-56 |
| chr4-148164115-INS-225 | chr4-148164115-INS-225 |
| chr4-148168505-INS-1221 | chr4-148168506-INS-1221 |
| chr4-148168506-INS-381 | chr4-148168506-INS-381 |
| chr4-148171289-INS-391 | chr4-148171289-INS-391 |
| chr4-148175840-INS-365 | chr4-148175840-INS-365 |
| chr4-148184523-INS-223 | chr4-148184523-INS-223 |
| chr4-148189033-INS-234 | chr4-148189033-INS-240 |
| chr4-148253432-INS-271 | chr4-148253432-INS-271 |
| chr4-148256537-INS-201 | chr4-148256537-INS-201 |
| chr4-148281425-INS-420 | chr4-148281425-INS-420 |
| chr4-148292426-INS-304 | chr4-148292427-INS-305 |
| chr4-148295068-INS-310 | chr4-148295068-INS-265 |
| chr4-148306659-INS-119 | chr4-148306659-INS-118 |
| chr4-148341510-INS-160 | chr4-148341510-INS-146 |
| chr4-148344630-INS-86 | chr4-148344630-INS-86 |
| chr4-148346006-INS-53 | chr4-148346006-INS-53 |
| chr4-148350609-INS-185 | chr4-148350609-INS-193 |
| chr4-148363910-INS-140 | chr4-148363907-INS-167 |
| chr4-148379791-INS-62 | chr4-148379765-INS-88 |
| chr4-148386435-INS-76 | chr4-148386391-INS-86 |

| | |
|---|---|
| chr4-148434638-INS-51 | chr4-148434650-INS-63 |
| chr4-148440133-INS-136 | chr4-148440133-INS-128 |
| chr4-148453066-INS-104 | chr4-148453066-INS-104 |
| chr4-148475739-INS-60 | chr4-148475739-INS-60 |
| chr4-148477141-INS-62 | chr4-148477141-INS-58 |
| chr4-148480401-INS-68 | chr4-148480401-INS-77 |
| chr4-148486904-INS-84 | chr4-148486904-INS-85 |
| chr4-148494476-INS-105 | chr4-148494521-INS-110 |
| chr4-148496292-INS-504 | chr4-148496292-INS-504 |
| chr4-148503859-INS-58 | chr4-148503859-INS-72 |
| chr4-148523644-INS-5548 | chr4-148523645-INS-5548 |
| chr4-148524150-INS-162 | chr4-148524139-INS-132 |
| chr4-148529778-INS-202 | chr4-148529778-INS-202 |
| chr4-148536514-INS-221 | chr4-148536514-INS-222 |
| chr4-148538880-INS-56 | chr4-148538880-INS-56 |
| chr4-148541575-INS-685 | chr4-148541575-INS-685 |
| chr4-148544923-INS-56 | chr4-148544923-INS-56 |
| chr4-148592811-INS-233 | chr4-148592811-INS-233 |
| chr4-148600638-INS-83 | chr4-148600633-INS-83 |
| chr4-148604971-INS-207 | chr4-148604971-INS-207 |
| chr4-148613269-INS-213 | chr4-148613269-INS-213 |
| chr4-148619898-INS-69 | chr4-148619898-INS-67 |
| chr4-148627212-INS-205 | chr4-148627212-INS-205 |
| chr4-148627836-INS-74 | chr4-148627836-INS-52 |
| chr4-148629113-INS-172 | chr4-148629113-INS-173 |
| chr4-148636760-INS-210 | chr4-148636760-INS-209 |
| chr4-148670996-INS-206 | chr4-148670996-INS-205 |
| chr4-148673229-INS-222 | chr4-148673229-INS-222 |
| chr4-148712852-INS-220 | chr4-148712852-INS-219 |
| chr4-148718587-INS-51 | chr4-148718587-INS-53 |
| chr4-148720126-INS-531 | chr4-148720126-INS-531 |
| chr4-148722484-INS-190 | chr4-148722484-INS-191 |
| chr4-148732659-INS-210 | chr4-148732659-INS-222 |
| chr4-148735789-INS-210 | chr4-148735789-INS-211 |
| chr4-148739754-INS-207 | chr4-148739754-INS-207 |
| chr4-148750043-INS-209 | chr4-148750036-INS-212 |
| chr4-148784693-INS-615 | chr4-148784693-INS-596 |
| chr4-148801471-INS-97 | chr4-148801482-INS-78 |
| chr4-148813616-INS-123 | chr4-148813616-INS-123 |
| chr4-148844346-INS-2415 | chr4-148844347-INS-2419 |
| chr4-148861889-INS-140 | chr4-148861924-INS-148 |
| chr4-148865318-INS-6442 | chr4-148865319-INS-6442 |
| chr4-148909303-INS-54 | chr4-148909303-INS-56 |
| chr4-148928146-INS-126 | chr4-148928146-INS-126 |
| chr4-148996218-INS-80 | chr4-148996218-INS-96 |
| chr4-149065322-INS-169 | chr4-149065322-INS-169 |
| chr4-149087074-INS-82 | chr4-149087074-INS-82 |
| chr4-149087760-INS-222 | chr4-149087760-INS-222 |
| chr4-149094976-INS-168 | chr4-149094976-INS-174 |
| chr4-149116818-INS-3693 | chr4-149116819-INS-3693 |
| chr4-149117157-INS-122 | chr4-149117157-INS-122 |
| chr4-149121867-INS-52 | chr4-149121867-INS-52 |
| chr4-149126573-INS-95 | chr4-149126574-INS-95 |
| chr4-149135745-INS-54 | chr4-149135745-INS-60 |
| chr4-149168518-INS-493 | chr4-149168518-INS-493 |
| chr4-149175842-INS-201 | chr4-149175842-INS-203 |
| chr4-149179975-INS-307 | chr4-149179975-INS-305 |
| chr4-149191228-INS-220 | chr4-149191228-INS-220 |
| chr4-149197685-INS-205 | chr4-149197685-INS-205 |
| chr4-149203374-INS-75 | chr4-149203374-INS-95 |
| chr4-149205265-INS-58 | chr4-149205264-INS-63 |
| chr4-149207550-INS-184 | chr4-149207550-INS-195 |
| chr4-149209632-INS-208 | chr4-149209632-INS-208 |
| chr4-149222878-INS-217 | chr4-149222870-INS-221 |
| chr4-149239233-INS-162 | chr4-149239233-INS-169 |
| chr4-149259780-INS-207 | chr4-149259781-INS-207 |
| chr4-149275950-INS-73 | chr4-149275950-INS-73 |
| chr4-149280362-INS-196 | chr4-149280362-INS-228 |
| chr4-149281119-INS-209 | chr4-149281119-INS-215 |
| chr4-149289255-INS-165 | chr4-149289255-INS-165 |
| chr4-149351391-INS-217 | chr4-149351391-INS-217 |
| chr4-149356364-INS-199 | chr4-149356364-INS-199 |
| chr4-149376555-INS-753 | chr4-149376554-INS-747 |
| chr4-149379367-INS-237 | chr4-149379357-INS-198 |
| chr4-149389944-INS-96 | chr4-149389944-INS-96 |
| chr4-149396824-INS-452 | chr4-149396824-INS-450 |
| chr4-149421463-INS-178 | chr4-149421463-INS-179 |

| | |
|---|---|
| chr4-149428106-INS-487 | chr4-149428106-INS-487 |
| chr4-149437280-INS-174 | chr4-149437280-INS-174 |
| chr4-149445971-INS-51 | chr4-149445989-INS-58 |
| chr4-149461313-INS-159 | chr4-149461313-INS-156 |
| chr4-149468778-INS-204 | chr4-149468778-INS-207 |
| chr4-149501830-INS-107 | chr4-149501830-INS-107 |
| chr4-149505360-INS-52 | chr4-149505360-INS-90 |
| chr4-149511757-INS-184 | chr4-149511757-INS-185 |
| chr4-149533205-INS-130 | chr4-149533206-INS-132 |
| chr4-149568667-INS-57 | chr4-149568667-INS-57 |
| chr4-149571104-INS-150 | chr4-149571104-INS-150 |
| chr4-149577574-INS-291 | chr4-149577574-INS-291 |
| chr4-149612426-INS-203 | chr4-149612426-INS-203 |
| chr4-149613384-INS-216 | chr4-149613384-INS-216 |
| chr4-149615597-INS-380 | chr4-149615598-INS-385 |
| chr4-149618671-INS-52 | chr4-149618671-INS-52 |
| chr4-149686918-INS-59 | chr4-149686916-INS-59 |
| chr4-149703835-INS-62 | chr4-149703835-INS-62 |
| chr4-149722546-INS-178 | chr4-149722546-INS-178 |
| chr4-149747413-INS-54 | chr4-149747414-INS-54 |
| chr4-149762314-INS-58 | chr4-149762314-INS-58 |
| chr4-149789214-INS-184 | chr4-149789214-INS-184 |
| chr4-149790164-INS-345 | chr4-149790164-INS-345 |
| chr4-149795425-INS-608 | chr4-149795423-INS-606 |
| chr4-149805277-INS-4999 | chr4-149805278-INS-4999 |
| chr4-149847070-INS-379 | chr4-149847070-INS-382 |
| chr4-149863962-INS-204 | chr4-149863962-INS-205 |
| chr4-149878391-INS-62 | chr4-149878375-INS-57 |
| chr4-149905741-INS-68 | chr4-149905791-INS-68 |
| chr4-149924148-INS-93 | chr4-149924148-INS-95 |
| chr4-149958489-INS-569 | chr4-149958489-INS-569 |
| chr4-149965818-INS-300 | chr4-149965820-INS-298 |
| chr4-149972453-INS-167 | chr4-149972453-INS-167 |
| chr4-150012764-INS-93 | chr4-150012764-INS-93 |
| chr4-150051217-INS-60 | chr4-150051217-INS-60 |
| chr4-150060686-INS-165 | chr4-150060659-INS-193 |
| chr4-150062810-INS-77 | chr4-150062810-INS-77 |
| chr4-150067438-INS-152 | chr4-150067438-INS-152 |
| chr4-150122770-INS-53 | chr4-150122766-INS-61 |
| chr4-150141571-INS-211 | chr4-150141571-INS-211 |
| chr4-150156835-INS-220 | chr4-150156835-INS-220 |
| chr4-150161095-INS-1009 | chr4-150161096-INS-1009 |
| chr4-150165981-INS-224 | chr4-150165982-INS-201 |
| chr4-150175235-INS-194 | chr4-150175244-INS-191 |
| chr4-150181543-INS-73 | chr4-150181543-INS-73 |
| chr4-150198437-INS-472 | chr4-150198437-INS-472 |
| chr4-150202180-INS-93 | chr4-150202180-INS-95 |
| chr4-150218182-INS-66 | chr4-150218182-INS-70 |
| chr4-150240584-INS-85 | chr4-150240634-INS-64 |
| chr4-150265306-INS-125 | chr4-150265306-INS-125 |
| chr4-150266714-INS-3333 | chr4-150266715-INS-3333 |
| chr4-150268494-INS-100 | chr4-150268494-INS-100 |
| chr4-150273323-INS-916 | chr4-150273313-INS-931 |
| chr4-150277302-INS-61 | chr4-150277293-INS-56 |
| chr4-150286298-INS-68 | chr4-150286282-INS-108 |
| chr4-150289059-INS-244 | chr4-150289059-INS-243 |
| chr4-150293846-INS-71 | chr4-150293846-INS-71 |
| chr4-150296436-INS-61 | chr4-150296436-INS-65 |
| chr4-150312013-INS-204 | chr4-150312013-INS-204 |
| chr4-150313560-INS-55 | chr4-150313546-INS-102 |
| chr4-150319549-INS-201 | chr4-150319549-INS-202 |
| chr4-150348535-INS-205 | chr4-150348535-INS-203 |
| chr4-150359496-INS-248 | chr4-150359496-INS-220 |
| chr4-150368400-INS-52 | chr4-150368400-INS-52 |
| chr4-150378142-INS-216 | chr4-150378142-INS-217 |
| chr4-150386314-INS-167 | chr4-150386313-INS-164 |
| chr4-150398734-INS-64 | chr4-150398734-INS-64 |
| chr4-150402951-INS-86 | chr4-150402951-INS-84 |
| chr4-150417864-INS-207 | chr4-150417864-INS-207 |
| chr4-150419013-INS-229 | chr4-150419013-INS-230 |
| chr4-150421192-INS-214 | chr4-150421193-INS-214 |
| chr4-150430369-INS-215 | chr4-150430369-INS-215 |
| chr4-150443794-INS-56 | chr4-150443794-INS-56 |
| chr4-150444553-INS-75 | chr4-150444553-INS-71 |
| chr4-150449168-INS-62 | chr4-150449154-INS-61 |
| chr4-150457189-INS-106 | chr4-150457149-INS-145 |
| chr4-150459047-INS-194 | chr4-150459047-INS-194 |

| | |
|---|---|
| chr4-150469703-INS-53 | chr4-150469702-INS-51 |
| chr4-150478439-INS-128 | chr4-150478439-INS-131 |
| chr4-150544381-INS-57 | chr4-150544381-INS-72 |
| chr4-150545803-INS-170 | chr4-150545803-INS-170 |
| chr4-150561688-INS-166 | chr4-150561688-INS-166 |
| chr4-150574181-INS-121 | chr4-150574181-INS-121 |
| chr4-150591396-INS-100 | chr4-150591428-INS-84 |
| chr4-150595491-INS-195 | chr4-150595491-INS-197 |
| chr4-150596305-INS-54 | chr4-150596305-INS-52 |
| chr4-150618211-INS-213 | chr4-150618211-INS-213 |
| chr4-150619198-INS-72 | chr4-150619198-INS-56 |
| chr4-150621172-INS-52 | chr4-150621172-INS-52 |
| chr4-150627169-INS-1879 | chr4-150627170-INS-1879 |
| chr4-150631089-INS-465 | chr4-150631089-INS-500 |
| chr4-150631913-INS-67 | chr4-150631913-INS-67 |
| chr4-150633368-INS-120 | chr4-150633355-INS-140 |
| chr4-150634918-INS-438 | chr4-150634918-INS-438 |
| chr4-150657225-INS-193 | chr4-150657225-INS-193 |
| chr4-150663467-INS-200 | chr4-150663467-INS-200 |
| chr4-150665558-INS-144 | chr4-150665559-INS-144 |
| chr4-150665929-INS-504 | chr4-150665929-INS-504 |
| chr4-150671133-INS-72 | chr4-150671133-INS-87 |
| chr4-150673202-INS-228 | chr4-150673202-INS-228 |
| chr4-150685492-INS-53 | chr4-150685492-INS-53 |
| chr4-150716219-INS-61 | chr4-150716221-INS-50 |
| chr4-150730520-INS-54 | chr4-150730520-INS-54 |
| chr4-150738766-INS-76 | chr4-150738766-INS-76 |
| chr4-150743674-INS-80 | chr4-150743706-INS-70 |
| chr4-150745955-INS-173 | chr4-150745955-INS-196 |
| chr4-150749080-INS-108 | chr4-150749083-INS-88 |
| chr4-150756598-INS-141 | chr4-150756598-INS-145 |
| chr4-150762798-INS-89 | chr4-150762827-INS-97 |
| chr4-150768288-INS-267 | chr4-150768288-INS-268 |
| chr4-150769026-INS-212 | chr4-150769026-INS-212 |
| chr4-150786546-INS-968 | chr4-150786546-INS-966 |
| chr4-150795004-INS-161 | chr4-150795004-INS-161 |
| chr4-150803011-INS-64 | chr4-150803011-INS-64 |
| chr4-150811823-INS-88 | chr4-150811851-INS-100 |
| chr4-150814139-INS-100 | chr4-150814139-INS-100 |
| chr4-150820217-INS-133 | chr4-150820217-INS-133 |
| chr4-150834849-INS-57 | chr4-150834849-INS-57 |
| chr4-150849866-INS-391 | chr4-150849866-INS-391 |
| chr4-150850658-INS-70 | chr4-150850658-INS-70 |
| chr4-150877121-INS-111 | chr4-150877124-INS-125 |
| chr4-150885871-INS-204 | chr4-150885871-INS-204 |
| chr4-150895091-INS-98 | chr4-150895078-INS-89 |
| chr4-150903089-INS-213 | chr4-150903089-INS-213 |
| chr4-150926257-INS-60 | chr4-150926238-INS-96 |
| chr4-150928286-INS-135 | chr4-150928286-INS-135 |
| chr4-150959449-INS-155 | chr4-150959449-INS-155 |
| chr4-150980905-INS-209 | chr4-150980905-INS-211 |
| chr4-151001795-INS-69 | chr4-151001795-INS-73 |
| chr4-151005369-INS-60 | chr4-151005369-INS-60 |
| chr4-151025767-INS-56 | chr4-151025767-INS-54 |
| chr4-151060112-INS-92 | chr4-151060112-INS-92 |
| chr4-151087031-INS-200 | chr4-151087031-INS-200 |
| chr4-151096305-INS-96 | chr4-151096305-INS-96 |
| chr4-151099788-INS-133 | chr4-151099778-INS-123 |
| chr4-151110230-INS-228 | chr4-151110230-INS-222 |
| chr4-151141106-INS-201 | chr4-151141106-INS-200 |
| chr4-151182728-INS-211 | chr4-151182728-INS-212 |
| chr4-151219055-INS-74 | chr4-151219042-INS-110 |
| chr4-151251223-INS-372 | chr4-151251223-INS-380 |
| chr4-151260592-INS-360 | chr4-151260592-INS-358 |
| chr4-151264790-INS-51 | chr4-151264790-INS-51 |
| chr4-151285403-INS-54 | chr4-151285403-INS-54 |
| chr4-151291112-INS-6442 | chr4-151291113-INS-6442 |
| chr4-151300765-INS-84 | chr4-151300765-INS-63 |
| chr4-151312157-INS-57 | chr4-151312156-INS-64 |
| chr4-151313401-INS-244 | chr4-151313446-INS-207 |
| chr4-151321259-INS-925 | chr4-151321250-INS-943 |
| chr4-151322929-INS-60 | chr4-151322940-INS-64 |
| chr4-151324664-INS-167 | chr4-151324664-INS-167 |
| chr4-151326158-INS-150 | chr4-151326158-INS-150 |
| chr4-151331174-INS-74 | chr4-151331175-INS-74 |
| chr4-151353685-INS-75 | chr4-151353685-INS-67 |
| chr4-151379388-INS-54 | chr4-151379388-INS-56 |

| | |
|---|---|
| chr4-151414238-INS-120 | chr4-151414252-INS-90 |
| chr4-151418422-INS-1708 | chr4-151418423-INS-1716 |
| chr4-151457475-INS-55 | chr4-151457475-INS-55 |
| chr4-151481649-INS-388 | chr4-151481650-INS-354 |
| chr4-151483824-INS-71 | chr4-151483824-INS-71 |
| chr4-151500928-INS-108 | chr4-151500928-INS-109 |
| chr4-151512512-INS-202 | chr4-151512512-INS-202 |
| chr4-151525541-INS-60 | chr4-151525541-INS-60 |
| chr4-151533339-INS-68 | chr4-151533339-INS-68 |
| chr4-151554018-INS-2702 | chr4-151554019-INS-2698 |
| chr4-151587121-INS-91 | chr4-151587121-INS-91 |
| chr4-151605045-INS-64 | chr4-151605046-INS-64 |
| chr4-151606741-INS-52 | chr4-151606741-INS-52 |
| chr4-151609723-INS-191 | chr4-151609723-INS-191 |
| chr4-151633060-INS-72 | chr4-151633060-INS-78 |
| chr4-151635869-INS-108 | chr4-151635869-INS-108 |
| chr4-151654759-INS-60 | chr4-151654759-INS-60 |
| chr4-151667385-INS-131 | chr4-151667385-INS-120 |
| chr4-151674598-INS-60 | chr4-151674598-INS-60 |
| chr4-151721127-INS-76 | chr4-151721127-INS-76 |
| chr4-151755791-INS-57 | chr4-151755791-INS-61 |
| chr4-151762474-INS-499 | chr4-151762472-INS-499 |
| chr4-151771034-INS-61 | chr4-151771037-INS-63 |
| chr4-151773762-INS-82 | chr4-151773757-INS-104 |
| chr4-151787120-INS-53 | chr4-151787120-INS-53 |
| chr4-151810831-INS-59 | chr4-151810831-INS-100 |
| chr4-151815539-INS-138 | chr4-151815539-INS-138 |
| chr4-151818443-INS-83 | chr4-151818477-INS-52 |
| chr4-151822628-INS-1810 | chr4-151822619-INS-1807 |
| chr4-151843988-INS-258 | chr4-151843988-INS-258 |
| chr4-151885575-INS-743 | chr4-151885575-INS-743 |
| chr4-151890624-INS-200 | chr4-151890624-INS-192 |
| chr4-151924231-INS-125 | chr4-151924231-INS-125 |
| chr4-151955498-INS-140 | chr4-151955498-INS-144 |
| chr4-151958280-INS-176 | chr4-151958280-INS-176 |
| chr4-151990449-INS-322 | chr4-151990449-INS-322 |
| chr4-151999814-INS-66 | chr4-151999800-INS-76 |
| chr4-152025817-INS-533 | chr4-152025817-INS-536 |
| chr4-152030747-INS-174 | chr4-152030748-INS-174 |
| chr4-152032538-INS-202 | chr4-152032538-INS-202 |
| chr4-152038544-INS-51 | chr4-152038544-INS-51 |
| chr4-152060449-INS-208 | chr4-152060449-INS-206 |
| chr4-152083267-INS-55 | chr4-152083267-INS-55 |
| chr4-152087889-INS-177 | chr4-152087889-INS-177 |
| chr4-152089668-INS-226 | chr4-152089668-INS-226 |
| chr4-152089783-INS-7403 | chr4-152089784-INS-7403 |
| chr4-152109033-INS-227 | chr4-152109033-INS-227 |
| chr4-152113870-INS-176 | chr4-152113870-INS-176 |
| chr4-152115363-INS-4396 | chr4-152115364-INS-4396 |
| chr4-152129457-INS-365 | chr4-152129457-INS-365 |
| chr4-152133139-INS-64 | chr4-152133145-INS-54 |
| chr4-152151293-INS-194 | chr4-152151318-INS-196 |
| chr4-152159733-INS-201 | chr4-152159733-INS-201 |
| chr4-152163510-INS-198 | chr4-152163508-INS-185 |
| chr4-152170922-INS-62 | chr4-152170922-INS-62 |
| chr4-152185055-INS-65 | chr4-152185055-INS-65 |
| chr4-152206632-INS-207 | chr4-152206632-INS-208 |
| chr4-152226016-INS-289 | chr4-152226016-INS-292 |
| chr4-152249434-INS-206 | chr4-152249434-INS-211 |
| chr4-152253403-INS-267 | chr4-152253403-INS-268 |
| chr4-152275163-INS-132 | chr4-152275166-INS-135 |
| chr4-152304683-INS-170 | chr4-152304683-INS-170 |
| chr4-152310463-INS-245 | chr4-152310463-INS-245 |
| chr4-152348375-INS-57 | chr4-152348375-INS-57 |
| chr4-152363111-INS-206 | chr4-152363111-INS-206 |
| chr4-152373910-INS-796 | chr4-152373911-INS-796 |
| chr4-152386628-INS-210 | chr4-152386628-INS-211 |
| chr4-152415858-INS-66 | chr4-152415858-INS-66 |
| chr4-152419198-INS-212 | chr4-152419198-INS-215 |
| chr4-152432533-INS-68 | chr4-152432533-INS-72 |
| chr4-152443138-INS-81 | chr4-152443148-INS-75 |
| chr4-152487074-INS-399 | chr4-152487074-INS-399 |
| chr4-152539081-INS-52 | chr4-152539081-INS-54 |
| chr4-152543907-INS-246 | chr4-152543907-INS-246 |
| chr4-152548794-INS-250 | chr4-152548794-INS-250 |
| chr4-152551248-INS-2523 | chr4-152551249-INS-2523 |
| chr4-152563782-INS-311 | chr4-152563782-INS-311 |

| | |
|---|---|
| chr4-152609394-INS-365 | chr4-152609394-INS-365 |
| chr4-152620217-INS-149 | chr4-152620217-INS-149 |
| chr4-152627027-INS-76 | chr4-152627027-INS-76 |
| chr4-152629952-INS-56 | chr4-152629954-INS-56 |
| chr4-152630544-INS-135 | chr4-152630544-INS-135 |
| chr4-152636501-INS-126 | chr4-152636501-INS-130 |
| chr4-152642310-INS-131 | chr4-152642310-INS-131 |
| chr4-152659464-INS-158 | chr4-152659464-INS-157 |
| chr4-152662352-INS-52 | chr4-152662352-INS-52 |
| chr4-152663265-INS-55 | chr4-152663265-INS-53 |
| chr4-152667321-INS-93 | chr4-152667321-INS-94 |
| chr4-152689917-INS-204 | chr4-152689884-INS-249 |
| chr4-152718194-INS-82 | chr4-152718194-INS-82 |
| chr4-152735297-INS-55 | chr4-152735297-INS-55 |
| chr4-152737234-INS-56 | chr4-152737234-INS-63 |
| chr4-152745362-INS-559 | chr4-152745362-INS-553 |
| chr4-152751430-INS-62 | chr4-152751430-INS-70 |
| chr4-152755726-INS-155 | chr4-152755726-INS-155 |
| chr4-152761284-INS-84 | chr4-152761280-INS-120 |
| chr4-152785028-INS-220 | chr4-152785028-INS-220 |
| chr4-152789700-INS-188 | chr4-152789700-INS-193 |
| chr4-152801147-INS-87 | chr4-152801147-INS-87 |
| chr4-152801661-INS-50 | chr4-152801683-INS-56 |
| chr4-152811427-INS-74 | chr4-152811427-INS-58 |
| chr4-152814940-INS-124 | chr4-152814940-INS-124 |
| chr4-152822604-INS-54 | chr4-152822604-INS-56 |
| chr4-152836235-INS-342 | chr4-152836235-INS-342 |
| chr4-152862115-INS-56 | chr4-152862115-INS-58 |
| chr4-152870337-INS-62 | chr4-152870337-INS-66 |
| chr4-152883188-INS-120 | chr4-152883188-INS-104 |
| chr4-152904679-INS-353 | chr4-152904679-INS-353 |
| chr4-152908778-INS-255 | chr4-152908778-INS-259 |
| chr4-152911681-INS-68 | chr4-152911688-INS-54 |
| chr4-152949144-INS-68 | chr4-152949127-INS-85 |
| chr4-152957315-INS-152 | chr4-152957315-INS-139 |
| chr4-152959114-INS-60 | chr4-152959114-INS-60 |
| chr4-152989467-INS-183 | chr4-152989467-INS-182 |
| chr4-152991098-INS-186 | chr4-152991098-INS-186 |
| chr4-153001732-INS-138 | chr4-153001740-INS-152 |
| chr4-153010098-INS-62 | chr4-153010098-INS-62 |
| chr4-153014975-INS-288 | chr4-153014975-INS-288 |
| chr4-153018696-INS-107 | chr4-153018696-INS-107 |
| chr4-153035944-INS-70 | chr4-153035945-INS-70 |
| chr4-153045217-INS-226 | chr4-153045210-INS-186 |
| chr4-153048752-INS-301 | chr4-153048752-INS-297 |
| chr4-153100909-INS-400 | chr4-153100909-INS-400 |
| chr4-153105563-INS-306 | chr4-153105563-INS-306 |
| chr4-153107295-INS-58 | chr4-153107295-INS-58 |
| chr4-153108804-INS-92 | chr4-153108804-INS-124 |
| chr4-153109822-INS-147 | chr4-153109822-INS-147 |
| chr4-153116606-INS-1437 | chr4-153116607-INS-1437 |
| chr4-153123010-INS-90 | chr4-153123010-INS-64 |
| chr4-153133910-INS-92 | chr4-153133910-INS-92 |
| chr4-153165174-INS-108 | chr4-153165174-INS-108 |
| chr4-153180182-INS-58 | chr4-153180183-INS-58 |
| chr4-153217187-INS-66 | chr4-153217186-INS-94 |
| chr4-153224504-INS-80 | chr4-153224504-INS-80 |
| chr4-153230386-INS-74 | chr4-153230386-INS-76 |
| chr4-153238315-INS-470 | chr4-153238315-INS-470 |
| chr4-153245731-INS-52 | chr4-153245731-INS-54 |
| chr4-153271911-INS-76 | chr4-153271903-INS-86 |
| chr4-153274611-INS-58 | chr4-153274611-INS-58 |
| chr4-153277565-INS-675 | chr4-153277565-INS-701 |
| chr4-153296302-INS-147 | chr4-153296301-INS-149 |
| chr4-153308776-INS-205 | chr4-153308776-INS-166 |
| chr4-153339245-INS-112 | chr4-153339245-INS-112 |
| chr4-153353165-INS-466 | chr4-153353165-INS-467 |
| chr4-153387792-INS-54 | chr4-153387792-INS-54 |
| chr4-153397582-INS-54 | chr4-153397594-INS-50 |
| chr4-153404777-INS-411 | chr4-153404777-INS-417 |
| chr4-153408121-INS-104 | chr4-153408121-INS-104 |
| chr4-153446586-INS-54 | chr4-153446586-INS-54 |
| chr4-153472014-INS-6442 | chr4-153472015-INS-6442 |
| chr4-153478814-INS-87 | chr4-153478814-INS-84 |
| chr4-153494407-INS-56 | chr4-153494407-INS-56 |
| chr4-153497477-INS-66 | chr4-153497483-INS-60 |
| chr4-153537686-INS-76 | chr4-153537686-INS-76 |

| | |
|---|---|
| chr4-153548503-INS-122 | chr4-153548546-INS-127 |
| chr4-153577795-INS-61 | chr4-153577795-INS-52 |
| chr4-153580405-INS-56 | chr4-153580405-INS-55 |
| chr4-153582791-INS-57 | chr4-153582796-INS-85 |
| chr4-153584477-INS-66 | chr4-153584466-INS-50 |
| chr4-153605450-INS-7082 | chr4-153605451-INS-7082 |
| chr4-153607277-INS-297 | chr4-153607277-INS-313 |
| chr4-153609793-INS-64 | chr4-153609793-INS-64 |
| chr4-153611905-INS-65 | chr4-153611905-INS-65 |
| chr4-153636083-INS-62 | chr4-153636084-INS-72 |
| chr4-153712724-INS-52 | chr4-153712724-INS-52 |
| chr4-153743067-INS-55 | chr4-153743067-INS-58 |
| chr4-153745535-INS-318 | chr4-153745535-INS-318 |
| chr4-153765855-INS-390 | chr4-153765855-INS-392 |
| chr4-153767730-INS-234 | chr4-153767730-INS-235 |
| chr4-153777810-INS-95 | chr4-153777793-INS-136 |
| chr4-153789753-INS-140 | chr4-153789758-INS-145 |
| chr4-153793488-INS-51 | chr4-153793488-INS-51 |
| chr4-153794544-INS-55 | chr4-153794544-INS-55 |
| chr4-153795861-INS-161 | chr4-153795861-INS-161 |
| chr4-153798066-INS-430 | chr4-153798066-INS-430 |
| chr4-153837165-INS-205 | chr4-153837165-INS-167 |
| chr4-153844513-INS-59 | chr4-153844513-INS-59 |
| chr4-153860138-INS-853 | chr4-153860138-INS-853 |
| chr4-153888942-INS-59 | chr4-153888942-INS-59 |
| chr4-153923043-INS-371 | chr4-153923043-INS-385 |
| chr4-153936797-INS-7294 | chr4-153936798-INS-7294 |
| chr4-153976783-INS-74 | chr4-153976765-INS-64 |
| chr4-153978324-INS-72 | chr4-153978353-INS-60 |
| chr4-153981916-INS-260 | chr4-153981916-INS-260 |
| chr4-153986198-INS-55 | chr4-153986172-INS-54 |
| chr4-154019889-INS-335 | chr4-154019889-INS-334 |
| chr4-154032094-INS-7261 | chr4-154032095-INS-7261 |
| chr4-154044503-INS-62 | chr4-154044503-INS-62 |
| chr4-154055879-INS-2834 | chr4-154055880-INS-2834 |
| chr4-154062949-INS-256 | chr4-154062951-INS-258 |
| chr4-154065269-INS-270 | chr4-154065269-INS-270 |
| chr4-154070536-INS-96 | chr4-154070536-INS-96 |
| chr4-154078994-INS-119 | chr4-154078994-INS-121 |
| chr4-154085689-INS-240 | chr4-154085690-INS-240 |
| chr4-154087326-INS-135 | chr4-154087299-INS-173 |
| chr4-154089675-INS-217 | chr4-154089712-INS-189 |
| chr4-154090910-INS-63 | chr4-154090912-INS-64 |
| chr4-154104501-INS-52 | chr4-154104501-INS-52 |
| chr4-154128543-INS-54 | chr4-154128543-INS-54 |
| chr4-154129247-INS-190 | chr4-154129247-INS-191 |
| chr4-154144105-INS-233 | chr4-154144089-INS-273 |
| chr4-154146061-INS-195 | chr4-154146061-INS-195 |
| chr4-154150163-INS-351 | chr4-154150150-INS-352 |
| chr4-154160230-INS-107 | chr4-154160230-INS-107 |
| chr4-154176348-INS-57 | chr4-154176348-INS-57 |
| chr4-154188960-INS-58 | chr4-154188960-INS-58 |
| chr4-154211763-INS-302 | chr4-154211763-INS-302 |
| chr4-154212471-INS-80 | chr4-154212492-INS-56 |
| chr4-154213990-INS-120 | chr4-154213970-INS-128 |
| chr4-154215298-INS-205 | chr4-154215298-INS-205 |
| chr4-154245940-INS-1585 | chr4-154245941-INS-1585 |
| chr4-154266028-INS-576 | chr4-154266028-INS-584 |
| chr4-154276073-INS-308 | chr4-154276073-INS-308 |
| chr4-154279283-INS-199 | chr4-154279283-INS-191 |
| chr4-154294434-INS-52 | chr4-154294434-INS-52 |
| chr4-154295038-INS-54 | chr4-154295038-INS-80 |
| chr4-154295863-INS-419 | chr4-154295837-INS-399 |
| chr4-154297325-INS-124 | chr4-154297325-INS-124 |
| chr4-154330786-INS-81 | chr4-154330786-INS-81 |
| chr4-154345123-INS-137 | chr4-154345134-INS-140 |
| chr4-154348964-INS-70 | chr4-154348938-INS-69 |
| chr4-154351169-INS-70 | chr4-154351169-INS-105 |
| chr4-154353566-INS-78 | chr4-154353566-INS-78 |
| chr4-154355909-INS-402 | chr4-154355909-INS-402 |
| chr4-154386983-INS-1585 | chr4-154386984-INS-1584 |
| chr4-154390825-INS-513 | chr4-154390825-INS-513 |
| chr4-154427910-INS-120 | chr4-154427910-INS-120 |
| chr4-154450110-INS-1864 | chr4-154450111-INS-1864 |
| chr4-154465102-INS-1820 | chr4-154465103-INS-1821 |
| chr4-154493393-INS-912 | chr4-154493393-INS-912 |
| chr4-154501821-INS-349 | chr4-154501821-INS-350 |

| | |
|---|---|
| chr4-154516036-INS-108 | chr4-154516036-INS-90 |
| chr4-154518480-INS-84 | chr4-154518480-INS-84 |
| chr4-154530639-INS-148 | chr4-154530639-INS-139 |
| chr4-154595381-INS-174 | chr4-154595381-INS-174 |
| chr4-154602326-INS-447 | chr4-154602318-INS-472 |
| chr4-154607393-INS-76 | chr4-154607393-INS-76 |
| chr4-154618544-INS-593 | chr4-154618544-INS-593 |
| chr4-154619471-INS-275 | chr4-154619471-INS-275 |
| chr4-154623344-INS-248 | chr4-154623345-INS-248 |
| chr4-154657022-INS-67 | chr4-154657022-INS-67 |
| chr4-154673065-INS-54 | chr4-154673065-INS-54 |
| chr4-154675314-INS-376 | chr4-154675322-INS-408 |
| chr4-154681394-INS-333 | chr4-154681394-INS-335 |
| chr4-154712881-INS-62 | chr4-154712881-INS-62 |
| chr4-154729483-INS-95 | chr4-154729479-INS-97 |
| chr4-154734238-INS-211 | chr4-154734238-INS-213 |
| chr4-154738031-INS-401 | chr4-154738032-INS-401 |
| chr4-154749210-INS-55 | chr4-154749210-INS-77 |
| chr4-154768347-INS-78 | chr4-154768346-INS-84 |
| chr4-154776198-INS-1148 | chr4-154776199-INS-1152 |
| chr4-154819412-INS-60 | chr4-154819412-INS-60 |
| chr4-154819466-INS-3038 | chr4-154819467-INS-3038 |
| chr4-154822839-INS-574 | chr4-154822839-INS-568 |
| chr4-154843253-INS-78 | chr4-154843253-INS-78 |
| chr4-154847883-INS-376 | chr4-154847879-INS-404 |
| chr4-154861581-INS-78 | chr4-154861581-INS-78 |
| chr4-154883411-INS-451 | chr4-154883411-INS-451 |
| chr4-154937985-INS-208 | chr4-154937985-INS-208 |
| chr4-154943185-INS-104 | chr4-154943185-INS-104 |
| chr4-154950251-INS-344 | chr4-154950251-INS-344 |
| chr4-154962654-INS-97 | chr4-154962654-INS-99 |
| chr4-154966896-INS-189 | chr4-154966896-INS-190 |
| chr4-154969733-INS-193 | chr4-154969733-INS-197 |
| chr4-154973191-INS-161 | chr4-154973191-INS-161 |
| chr4-154995013-INS-96 | chr4-154995057-INS-98 |
| chr4-155026655-INS-537 | chr4-155026642-INS-534 |
| chr4-155061013-INS-63 | chr4-155061013-INS-63 |
| chr4-155074247-INS-5545 | chr4-155074248-INS-5545 |
| chr4-155116549-INS-72 | chr4-155116575-INS-88 |
| chr4-155140003-INS-194 | chr4-155140003-INS-194 |
| chr4-155165038-INS-327 | chr4-155165019-INS-326 |
| chr4-155178414-INS-196 | chr4-155178414-INS-196 |
| chr4-155182722-INS-86 | chr4-155182722-INS-86 |
| chr4-155187916-INS-201 | chr4-155187916-INS-200 |
| chr4-155197840-INS-296 | chr4-155197840-INS-304 |
| chr4-155268107-INS-82 | chr4-155268107-INS-80 |
| chr4-155281707-INS-161 | chr4-155281707-INS-120 |
| chr4-155307261-INS-205 | chr4-155307261-INS-205 |
| chr4-155327708-INS-94 | chr4-155327708-INS-97 |
| chr4-155333404-INS-164 | chr4-155333404-INS-164 |
| chr4-155348882-INS-706 | chr4-155348883-INS-716 |
| chr4-155350651-INS-528 | chr4-155350651-INS-540 |
| chr4-155356619-INS-214 | chr4-155356619-INS-214 |
| chr4-155368731-INS-5630 | chr4-155368732-INS-5630 |
| chr4-155370639-INS-232 | chr4-155370639-INS-233 |
| chr4-155377810-INS-75 | chr4-155377810-INS-75 |
| chr4-155396409-INS-208 | chr4-155396409-INS-208 |
| chr4-155399513-INS-194 | chr4-155399513-INS-196 |
| chr4-155414097-INS-200 | chr4-155414097-INS-198 |
| chr4-155416243-INS-1290 | chr4-155416244-INS-1290 |
| chr4-155421915-INS-5429 | chr4-155421916-INS-5429 |
| chr4-155464693-INS-62 | chr4-155464693-INS-78 |
| chr4-155465258-INS-205 | chr4-155465258-INS-205 |
| chr4-155490255-INS-106 | chr4-155490255-INS-108 |
| chr4-155514793-INS-82 | chr4-155514793-INS-82 |
| chr4-155515506-INS-56 | chr4-155515506-INS-56 |
| chr4-155525400-INS-211 | chr4-155525400-INS-209 |
| chr4-155556466-INS-201 | chr4-155556477-INS-189 |
| chr4-155567050-INS-106 | chr4-155567057-INS-117 |
| chr4-155570908-INS-199 | chr4-155570908-INS-199 |
| chr4-155578401-INS-171 | chr4-155578401-INS-170 |
| chr4-155579248-INS-72 | chr4-155579248-INS-72 |
| chr4-155582911-INS-222 | chr4-155582911-INS-221 |
| chr4-155586319-INS-251 | chr4-155586319-INS-256 |
| chr4-155589729-INS-184 | chr4-155589729-INS-184 |
| chr4-155599463-INS-655 | chr4-155599471-INS-656 |
| chr4-155608096-INS-142 | chr4-155608096-INS-141 |

| | |
|---|---|
| chr4-155610041-INS-249 | chr4-155610041-INS-241 |
| chr4-155615810-INS-78 | chr4-155615807-INS-61 |
| chr4-155620329-INS-257 | chr4-155620329-INS-257 |
| chr4-155626999-INS-200 | chr4-155626999-INS-200 |
| chr4-155631518-INS-68 | chr4-155631518-INS-66 |
| chr4-155640303-INS-202 | chr4-155640303-INS-202 |
| chr4-155654744-INS-354 | chr4-155654744-INS-353 |
| chr4-155677837-INS-204 | chr4-155677837-INS-204 |
| chr4-155683338-INS-177 | chr4-155683338-INS-174 |
| chr4-155685536-INS-191 | chr4-155685536-INS-190 |
| chr4-155691290-INS-57 | chr4-155691290-INS-57 |
| chr4-155693038-INS-215 | chr4-155693038-INS-215 |
| chr4-155703487-INS-63 | chr4-155703487-INS-60 |
| chr4-155707022-INS-210 | chr4-155707022-INS-207 |
| chr4-155713664-INS-177 | chr4-155713664-INS-176 |
| chr4-155722348-INS-198 | chr4-155722348-INS-198 |
| chr4-155727378-INS-74 | chr4-155727378-INS-74 |
| chr4-155730736-INS-2546 | chr4-155730737-INS-2546 |
| chr4-155744830-INS-58 | chr4-155744830-INS-59 |
| chr4-155754692-INS-202 | chr4-155754692-INS-201 |
| chr4-155759111-INS-196 | chr4-155759111-INS-196 |
| chr4-155760423-INS-215 | chr4-155760423-INS-234 |
| chr4-155769004-INS-67 | chr4-155769004-INS-67 |
| chr4-155793069-INS-94 | chr4-155793078-INS-96 |
| chr4-155796406-INS-248 | chr4-155796406-INS-244 |
| chr4-155798989-INS-324 | chr4-155798989-INS-324 |
| chr4-155801075-INS-181 | chr4-155801075-INS-181 |
| chr4-155808822-INS-207 | chr4-155808822-INS-207 |
| chr4-155847863-INS-7184 | chr4-155847864-INS-7184 |
| chr4-155850784-INS-192 | chr4-155850784-INS-192 |
| chr4-155863966-INS-211 | chr4-155863966-INS-211 |
| chr4-155867086-INS-203 | chr4-155867086-INS-203 |
| chr4-155879420-INS-65 | chr4-155879420-INS-55 |
| chr4-155880375-INS-211 | chr4-155880375-INS-211 |
| chr4-155892245-INS-319 | chr4-155892245-INS-319 |
| chr4-155907513-INS-56 | chr4-155907513-INS-50 |
| chr4-155910696-INS-168 | chr4-155910696-INS-168 |
| chr4-155917576-INS-209 | chr4-155917576-INS-209 |
| chr4-155919280-INS-165 | chr4-155919280-INS-165 |
| chr4-155921360-INS-204 | chr4-155921360-INS-204 |
| chr4-155969145-INS-54 | chr4-155969145-INS-50 |
| chr4-155973321-INS-210 | chr4-155973321-INS-210 |
| chr4-155988895-INS-56 | chr4-155988897-INS-54 |
| chr4-156008070-INS-153 | chr4-156008039-INS-159 |
| chr4-156016865-INS-213 | chr4-156016865-INS-213 |
| chr4-156020368-INS-4503 | chr4-156020369-INS-4503 |
| chr4-156037268-INS-214 | chr4-156037268-INS-214 |
| chr4-156045665-INS-58 | chr4-156045665-INS-58 |
| chr4-156052784-INS-96 | chr4-156052784-INS-113 |
| chr4-156067515-INS-208 | chr4-156067515-INS-208 |
| chr4-156069557-INS-200 | chr4-156069557-INS-200 |
| chr4-156070886-INS-115 | chr4-156070888-INS-116 |
| chr4-156082252-INS-156 | chr4-156082273-INS-144 |
| chr4-156087494-INS-200 | chr4-156087494-INS-201 |
| chr4-156103141-INS-197 | chr4-156103141-INS-197 |
| chr4-156119673-INS-54 | chr4-156119673-INS-54 |
| chr4-156123123-INS-123 | chr4-156123123-INS-123 |
| chr4-156123854-INS-231 | chr4-156123854-INS-231 |
| chr4-156152648-INS-60 | chr4-156152627-INS-68 |
| chr4-156155602-INS-67 | chr4-156155602-INS-99 |
| chr4-156172426-INS-245 | chr4-156172426-INS-246 |
| chr4-156186020-INS-211 | chr4-156186020-INS-211 |
| chr4-156192691-INS-64 | chr4-156192691-INS-64 |
| chr4-156196630-INS-222 | chr4-156196630-INS-222 |
| chr4-156199461-INS-199 | chr4-156199472-INS-193 |
| chr4-156231786-INS-143 | chr4-156231786-INS-143 |
| chr4-156234735-INS-75 | chr4-156234735-INS-75 |
| chr4-156291343-INS-199 | chr4-156291343-INS-199 |
| chr4-156297773-INS-55 | chr4-156297773-INS-53 |
| chr4-156332083-INS-187 | chr4-156332083-INS-187 |
| chr4-156381972-INS-384 | chr4-156381974-INS-386 |
| chr4-156386572-INS-112 | chr4-156386572-INS-112 |
| chr4-156389407-INS-582 | chr4-156389407-INS-582 |
| chr4-156399488-INS-101 | chr4-156399488-INS-95 |
| chr4-156456402-INS-56 | chr4-156456402-INS-54 |
| chr4-156467772-INS-199 | chr4-156467772-INS-198 |
| chr4-156476301-INS-601 | chr4-156476302-INS-601 |

| | |
|---|---|
| chr5-3052027-INS-402 | chr5-3052027-INS-402 |
| chr5-3090990-INS-7469 | chr5-3090991-INS-7469 |
| chr5-3097061-INS-2799 | chr5-3097062-INS-2799 |
| chr5-3101393-INS-318 | chr5-3101393-INS-318 |
| chr5-3214032-INS-6475 | chr5-3214033-INS-6475 |
| chr5-3222910-INS-58 | chr5-3222910-INS-58 |
| chr5-3272485-INS-104 | chr5-3272485-INS-104 |
| chr5-3290200-INS-66 | chr5-3290201-INS-70 |
| chr5-3293581-INS-76 | chr5-3293581-INS-76 |
| chr5-3323292-INS-55 | chr5-3323292-INS-55 |
| chr5-3345270-INS-61 | chr5-3345270-INS-61 |
| chr5-3359016-INS-670 | chr5-3359016-INS-670 |
| chr5-3399067-INS-201 | chr5-3399067-INS-201 |
| chr5-3410586-INS-138 | chr5-3410586-INS-138 |
| chr5-3439224-INS-51 | chr5-3439224-INS-51 |
| chr5-3456994-INS-5215 | chr5-3456995-INS-5215 |
| chr5-3464450-INS-120 | chr5-3464450-INS-120 |
| chr5-3520391-INS-199 | chr5-3520391-INS-199 |
| chr5-3526493-INS-137 | chr5-3526492-INS-146 |
| chr5-3540100-INS-195 | chr5-3540100-INS-195 |
| chr5-3545081-INS-363 | chr5-3545081-INS-363 |
| chr5-3559860-INS-65 | chr5-3559860-INS-65 |
| chr5-3562893-INS-53 | chr5-3562891-INS-51 |
| chr5-3576016-INS-64 | chr5-3576004-INS-88 |
| chr5-3577768-INS-70 | chr5-3577768-INS-70 |
| chr5-3581637-INS-102 | chr5-3581637-INS-102 |
| chr5-3588476-INS-331 | chr5-3588476-INS-326 |
| chr5-3622892-INS-2784 | chr5-3622893-INS-2807 |
| chr5-3625025-INS-78 | chr5-3625025-INS-78 |
| chr5-3668011-INS-540 | chr5-3668011-INS-541 |
| chr5-3670789-INS-85 | chr5-3670789-INS-85 |
| chr5-3682826-INS-514 | chr5-3682826-INS-514 |
| chr5-3703856-INS-5249 | chr5-3703857-INS-5249 |
| chr5-3727489-INS-310 | chr5-3727496-INS-284 |
| chr5-3738829-INS-59 | chr5-3738829-INS-59 |
| chr5-3745958-INS-84 | chr5-3745958-INS-84 |
| chr5-3748976-INS-210 | chr5-3748976-INS-210 |
| chr5-3762363-INS-206 | chr5-3762363-INS-206 |
| chr5-3783213-INS-80 | chr5-3783213-INS-88 |
| chr5-3802089-INS-102 | chr5-3802089-INS-102 |
| chr5-3811196-INS-614 | chr5-3811196-INS-614 |
| chr5-3820756-INS-388 | chr5-3820756-INS-387 |
| chr5-3833848-INS-54 | chr5-3833848-INS-54 |
| chr5-3845131-INS-8681 | chr5-3845132-INS-8681 |
| chr5-3850126-INS-78 | chr5-3850121-INS-78 |
| chr5-3870053-INS-346 | chr5-3870053-INS-346 |
| chr5-3886096-INS-203 | chr5-3886096-INS-203 |
| chr5-3896650-INS-6340 | chr5-3896651-INS-6340 |
| chr5-3897493-INS-216 | chr5-3897493-INS-216 |
| chr5-3954577-INS-81 | chr5-3954578-INS-81 |
| chr5-4001184-INS-114 | chr5-4001184-INS-114 |
| chr5-4016321-INS-6405 | chr5-4016322-INS-6405 |
| chr5-4017674-INS-7053 | chr5-4017675-INS-7054 |
| chr5-4022516-INS-247 | chr5-4022516-INS-247 |
| chr5-4052125-INS-216 | chr5-4052115-INS-220 |
| chr5-4061752-INS-150 | chr5-4061752-INS-150 |
| chr5-4101607-INS-52 | chr5-4101607-INS-52 |
| chr5-4160665-INS-2219 | chr5-4160666-INS-2219 |
| chr5-4177246-INS-206 | chr5-4177246-INS-206 |
| chr5-4206052-INS-59 | chr5-4206097-INS-52 |
| chr5-4236048-INS-51 | chr5-4236048-INS-53 |
| chr5-4240569-INS-6423 | chr5-4240570-INS-6424 |
| chr5-4259965-INS-53 | chr5-4259966-INS-59 |
| chr5-4263740-INS-2238 | chr5-4263741-INS-2238 |
| chr5-4273849-INS-67 | chr5-4273850-INS-50 |
| chr5-4279268-INS-1165 | chr5-4279269-INS-1166 |
| chr5-4281027-INS-56 | chr5-4281027-INS-54 |
| chr5-4288709-INS-6607 | chr5-4288710-INS-6607 |
| chr5-4297133-INS-646 | chr5-4297133-INS-646 |
| chr5-4322406-INS-54 | chr5-4322406-INS-54 |
| chr5-4325125-INS-4363 | chr5-4325126-INS-4363 |
| chr5-4337585-INS-6756 | chr5-4337586-INS-6756 |
| chr5-4340910-INS-524 | chr5-4340910-INS-524 |
| chr5-4349156-INS-80 | chr5-4349156-INS-80 |
| chr5-4360967-INS-292 | chr5-4360967-INS-294 |
| chr5-4365793-INS-51 | chr5-4365793-INS-52 |
| chr5-4380044-INS-6120 | chr5-4380045-INS-6120 |

| | |
|---|---|
| chr5-4380566-INS-70 | chr5-4380566-INS-50 |
| chr5-4411234-INS-397 | chr5-4411234-INS-397 |
| chr5-4417914-INS-881 | chr5-4417914-INS-881 |
| chr5-4438772-INS-197 | chr5-4438772-INS-197 |
| chr5-4444064-INS-600 | chr5-4444064-INS-601 |
| chr5-4447227-INS-184 | chr5-4447228-INS-158 |
| chr5-4449566-INS-358 | chr5-4449566-INS-359 |
| chr5-4451876-INS-527 | chr5-4451877-INS-482 |
| chr5-4464924-INS-481 | chr5-4464924-INS-483 |
| chr5-4466119-INS-2817 | chr5-4466120-INS-2817 |
| chr5-4473210-INS-57 | chr5-4473210-INS-61 |
| chr5-4480826-INS-53 | chr5-4480826-INS-53 |
| chr5-4496112-INS-500 | chr5-4496112-INS-500 |
| chr5-4504238-INS-157 | chr5-4504238-INS-157 |
| chr5-4519886-INS-485 | chr5-4519886-INS-489 |
| chr5-4521782-INS-58 | chr5-4521782-INS-58 |
| chr5-4538298-INS-56 | chr5-4538298-INS-64 |
| chr5-4543648-INS-62 | chr5-4543648-INS-62 |
| chr5-4575520-INS-184 | chr5-4575520-INS-184 |
| chr5-4578861-INS-304 | chr5-4578861-INS-310 |
| chr5-4602069-INS-268 | chr5-4602069-INS-268 |
| chr5-4613484-INS-267 | chr5-4613484-INS-255 |
| chr5-4620808-INS-3231 | chr5-4620809-INS-3231 |
| chr5-4651467-INS-212 | chr5-4651467-INS-212 |
| chr5-4661068-INS-104 | chr5-4661068-INS-104 |
| chr5-4664441-INS-72 | chr5-4664445-INS-79 |
| chr5-4770059-INS-2645 | chr5-4770060-INS-2645 |
| chr5-4780615-INS-64 | chr5-4780615-INS-64 |
| chr5-4813343-INS-62 | chr5-4813343-INS-62 |
| chr5-4834707-INS-66 | chr5-4834702-INS-78 |
| chr5-4847260-INS-82 | chr5-4847260-INS-80 |
| chr5-4896850-INS-264 | chr5-4896850-INS-267 |
| chr5-4902242-INS-56 | chr5-4902242-INS-56 |
| chr5-4904239-INS-370 | chr5-4904239-INS-370 |
| chr5-4910519-INS-60 | chr5-4910519-INS-58 |
| chr5-4922263-INS-789 | chr5-4922263-INS-789 |
| chr5-4929566-INS-54 | chr5-4929566-INS-54 |
| chr5-4963539-INS-266 | chr5-4963539-INS-266 |
| chr5-4973316-INS-95 | chr5-4973316-INS-95 |
| chr5-4977709-INS-56 | chr5-4977709-INS-60 |
| chr5-4997108-INS-226 | chr5-4997108-INS-226 |
| chr5-4998759-INS-216 | chr5-4998759-INS-216 |
| chr5-5007762-INS-112 | chr5-5007762-INS-112 |
| chr5-5023791-INS-64 | chr5-5023791-INS-64 |
| chr5-5055162-INS-86 | chr5-5055133-INS-74 |
| chr5-5057402-INS-52 | chr5-5057374-INS-50 |
| chr5-5065819-INS-52 | chr5-5065819-INS-52 |
| chr5-5077381-INS-5108 | chr5-5077382-INS-5107 |
| chr5-5091362-INS-178 | chr5-5091362-INS-179 |
| chr5-5091969-INS-105 | chr5-5091969-INS-108 |
| chr5-5127511-INS-71 | chr5-5127511-INS-71 |
| chr5-5138973-INS-198 | chr5-5138973-INS-198 |
| chr5-5140325-INS-102 | chr5-5140325-INS-102 |
| chr5-5170221-INS-1446 | chr5-5170222-INS-1446 |
| chr5-5175874-INS-1435 | chr5-5175875-INS-1426 |
| chr5-5179438-INS-68 | chr5-5179438-INS-70 |
| chr5-5192826-INS-513 | chr5-5192826-INS-513 |
| chr5-5201638-INS-82 | chr5-5201638-INS-82 |
| chr5-5225907-INS-421 | chr5-5225907-INS-421 |
| chr5-5247396-INS-442 | chr5-5247396-INS-442 |
| chr5-5249184-INS-492 | chr5-5249182-INS-496 |
| chr5-5253000-INS-60 | chr5-5253005-INS-60 |
| chr5-5254004-INS-63 | chr5-5254004-INS-63 |
| chr5-5264930-INS-2862 | chr5-5264931-INS-2862 |
| chr5-5265346-INS-97 | chr5-5265346-INS-97 |
| chr5-5270149-INS-395 | chr5-5270149-INS-395 |
| chr5-5272638-INS-90 | chr5-5272638-INS-90 |
| chr5-5275012-INS-80 | chr5-5275012-INS-80 |
| chr5-5291749-INS-6257 | chr5-5291750-INS-6257 |
| chr5-5300757-INS-55 | chr5-5300757-INS-55 |
| chr5-5324616-INS-218 | chr5-5324616-INS-218 |
| chr5-5349569-INS-96 | chr5-5349569-INS-96 |
| chr5-5374252-INS-701 | chr5-5374252-INS-701 |
| chr5-5399289-INS-730 | chr5-5399289-INS-730 |
| chr5-5407042-INS-545 | chr5-5407042-INS-565 |
| chr5-5410935-INS-51 | chr5-5410927-INS-60 |
| chr5-5413552-INS-441 | chr5-5413552-INS-441 |

| | |
|---|---|
| chr5-5417104-INS-1169 | chr5-5417105-INS-1169 |
| chr5-5451696-INS-498 | chr5-5451696-INS-498 |
| chr5-5465352-INS-666 | chr5-5465352-INS-666 |
| chr5-5470822-INS-52 | chr5-5470813-INS-80 |
| chr5-5491878-INS-106 | chr5-5491878-INS-141 |
| chr5-5503821-INS-773 | chr5-5503821-INS-773 |
| chr5-5509109-INS-791 | chr5-5509109-INS-791 |
| chr5-5540329-INS-63 | chr5-5540329-INS-58 |
| chr5-5570568-INS-1158 | chr5-5570569-INS-1155 |
| chr5-5584767-INS-574 | chr5-5584767-INS-574 |
| chr5-5616456-INS-379 | chr5-5616456-INS-379 |
| chr5-5621727-INS-55 | chr5-5621727-INS-59 |
| chr5-5622358-INS-1364 | chr5-5622359-INS-1364 |
| chr5-5628006-INS-6462 | chr5-5628007-INS-6462 |
| chr5-5661777-INS-1476 | chr5-5661778-INS-1478 |
| chr5-5668637-INS-181 | chr5-5668637-INS-199 |
| chr5-5678609-INS-3905 | chr5-5678610-INS-3905 |
| chr5-5708039-INS-69 | chr5-5708039-INS-69 |
| chr5-5739230-INS-2992 | chr5-5739231-INS-2992 |
| chr5-5789415-INS-219 | chr5-5789415-INS-219 |
| chr5-5791440-INS-94 | chr5-5791440-INS-94 |
| chr5-5801023-INS-108 | chr5-5801023-INS-108 |
| chr5-5813740-INS-192 | chr5-5813740-INS-192 |
| chr5-5825617-INS-112 | chr5-5825617-INS-112 |
| chr5-5840383-INS-584 | chr5-5840383-INS-586 |
| chr5-5856892-INS-128 | chr5-5856892-INS-128 |
| chr5-5858756-INS-504 | chr5-5858756-INS-504 |
| chr5-5889855-INS-2408 | chr5-5889856-INS-2408 |
| chr5-5900912-INS-70 | chr5-5900912-INS-70 |
| chr5-5904102-INS-805 | chr5-5904102-INS-804 |
| chr5-5938662-INS-204 | chr5-5938662-INS-204 |
| chr5-5956087-INS-191 | chr5-5956087-INS-191 |
| chr5-5967575-INS-76 | chr5-5967589-INS-76 |
| chr5-5981587-INS-59 | chr5-5981587-INS-59 |
| chr5-5994874-INS-76 | chr5-5994910-INS-76 |
| chr5-6019796-INS-52 | chr5-6019797-INS-97 |
| chr5-6021175-INS-1063 | chr5-6021176-INS-1063 |
| chr5-6022803-INS-757 | chr5-6022803-INS-757 |
| chr5-6044462-INS-7087 | chr5-6044463-INS-7087 |
| chr5-6056334-INS-208 | chr5-6056334-INS-208 |
| chr5-6057827-INS-97 | chr5-6057827-INS-97 |
| chr5-6065346-INS-592 | chr5-6065346-INS-599 |
| chr5-6065567-INS-6253 | chr5-6065568-INS-6253 |
| chr5-6077796-INS-436 | chr5-6077796-INS-436 |
| chr5-6100188-INS-655 | chr5-6100188-INS-655 |
| chr5-6107470-INS-61 | chr5-6107470-INS-52 |
| chr5-6116550-INS-68 | chr5-6116550-INS-68 |
| chr5-6137312-INS-835 | chr5-6137312-INS-835 |
| chr5-6155193-INS-67 | chr5-6155193-INS-62 |
| chr5-6159136-INS-183 | chr5-6159121-INS-189 |
| chr5-6165481-INS-266 | chr5-6165467-INS-265 |
| chr5-6169543-INS-11191 | chr5-6169544-INS-11193 |
| chr5-6180759-INS-60 | chr5-6180759-INS-64 |
| chr5-6187412-INS-1453 | chr5-6187413-INS-1452 |
| chr5-6192208-INS-208 | chr5-6192208-INS-208 |
| chr5-6194418-INS-669 | chr5-6194423-INS-668 |
| chr5-6204382-INS-1422 | chr5-6204381-INS-1420 |
| chr5-6206036-INS-347 | chr5-6206036-INS-347 |
| chr5-6227718-INS-56 | chr5-6227729-INS-56 |
| chr5-6244185-INS-531 | chr5-6244185-INS-531 |
| chr5-6285637-INS-375 | chr5-6285637-INS-375 |
| chr5-6393155-INS-447 | chr5-6393155-INS-447 |
| chr5-6414895-INS-130 | chr5-6414895-INS-130 |
| chr5-6442477-INS-223 | chr5-6442477-INS-223 |
| chr5-6452501-INS-212 | chr5-6452501-INS-212 |
| chr5-6456336-INS-198 | chr5-6456336-INS-198 |
| chr5-6476989-INS-6228 | chr5-6476990-INS-6228 |
| chr5-6479919-INS-2057 | chr5-6479920-INS-2057 |
| chr5-6489844-INS-6451 | chr5-6489845-INS-6452 |
| chr5-6496163-INS-6262 | chr5-6496164-INS-6262 |
| chr5-6525253-INS-55 | chr5-6525253-INS-55 |
| chr5-6622149-INS-391 | chr5-6622149-INS-391 |
| chr5-6650914-INS-72 | chr5-6650914-INS-74 |
| chr5-6659222-INS-112 | chr5-6659238-INS-107 |
| chr5-6690267-INS-172 | chr5-6690268-INS-170 |
| chr5-6691117-INS-60 | chr5-6691117-INS-60 |
| chr5-6716730-INS-3108 | chr5-6716731-INS-3108 |

| | |
|---|---|
| chr5-6717667-INS-58 | chr5-6717667-INS-58 |
| chr5-6718436-INS-76 | chr5-6718437-INS-79 |
| chr5-6720768-INS-5540 | chr5-6720769-INS-5540 |
| chr5-6721517-INS-53 | chr5-6721517-INS-53 |
| chr5-6724734-INS-71 | chr5-6724725-INS-51 |
| chr5-6749866-INS-52 | chr5-6749866-INS-72 |
| chr5-6767436-INS-1956 | chr5-6767437-INS-1956 |
| chr5-6792600-INS-119 | chr5-6792600-INS-124 |
| chr5-6810883-INS-6331 | chr5-6810884-INS-6331 |
| chr5-6849837-INS-7209 | chr5-6849838-INS-7212 |
| chr5-6869439-INS-51 | chr5-6869439-INS-51 |
| chr5-6893551-INS-406 | chr5-6893551-INS-406 |
| chr5-6895115-INS-222 | chr5-6895115-INS-222 |
| chr5-6915154-INS-81 | chr5-6915154-INS-81 |
| chr5-6941609-INS-73 | chr5-6941609-INS-73 |
| chr5-6951567-INS-272 | chr5-6951567-INS-273 |
| chr5-6966999-INS-128 | chr5-6966999-INS-128 |
| chr5-6967970-INS-88 | chr5-6967970-INS-88 |
| chr5-6977272-INS-53 | chr5-6977272-INS-53 |
| chr5-6986023-INS-130 | chr5-6986023-INS-96 |
| chr5-6996609-INS-1058 | chr5-6996610-INS-1058 |
| chr5-7011632-INS-342 | chr5-7011632-INS-342 |
| chr5-7013772-INS-51 | chr5-7013772-INS-51 |
| chr5-7053550-INS-6552 | chr5-7053551-INS-6551 |
| chr5-7070313-INS-50 | chr5-7070306-INS-52 |
| chr5-7108411-INS-6667 | chr5-7108412-INS-6668 |
| chr5-7125129-INS-142 | chr5-7125129-INS-142 |
| chr5-7149010-INS-268 | chr5-7149010-INS-268 |
| chr5-7170740-INS-54 | chr5-7170733-INS-54 |
| chr5-7175323-INS-64 | chr5-7175323-INS-64 |
| chr5-7182837-INS-194 | chr5-7182838-INS-194 |
| chr5-7190486-INS-69 | chr5-7190486-INS-74 |
| chr5-7224627-INS-250 | chr5-7224627-INS-258 |
| chr5-7235411-INS-220 | chr5-7235411-INS-220 |
| chr5-7279082-INS-223 | chr5-7279082-INS-223 |
| chr5-7286793-INS-52 | chr5-7286793-INS-52 |
| chr5-7288409-INS-107 | chr5-7288409-INS-111 |
| chr5-7292091-INS-3541 | chr5-7292092-INS-3541 |
| chr5-7315533-INS-56 | chr5-7315561-INS-72 |
| chr5-7332156-INS-7357 | chr5-7332157-INS-7364 |
| chr5-7332763-INS-2331 | chr5-7332764-INS-2332 |
| chr5-7368459-INS-63 | chr5-7368459-INS-75 |
| chr5-7380769-INS-113 | chr5-7380765-INS-140 |
| chr5-7408070-INS-211 | chr5-7408070-INS-215 |
| chr5-7427087-INS-56 | chr5-7427043-INS-62 |
| chr5-7443047-INS-198 | chr5-7443047-INS-198 |
| chr5-7472764-INS-943 | chr5-7472764-INS-943 |
| chr5-7496064-INS-54 | chr5-7496064-INS-54 |
| chr5-7510750-INS-56 | chr5-7510750-INS-56 |
| chr5-7523971-INS-4170 | chr5-7523972-INS-4170 |
| chr5-7549394-INS-6675 | chr5-7549395-INS-6676 |
| chr5-7554601-INS-2876 | chr5-7554602-INS-2876 |
| chr5-7574876-INS-5231 | chr5-7574877-INS-5232 |
| chr5-7582879-INS-359 | chr5-7582879-INS-359 |
| chr5-7629570-INS-71 | chr5-7629572-INS-73 |
| chr5-7641114-INS-893 | chr5-7641114-INS-893 |
| chr5-7641572-INS-988 | chr5-7641572-INS-988 |
| chr5-7652425-INS-4144 | chr5-7652426-INS-4103 |
| chr5-7657370-INS-1774 | chr5-7657371-INS-1775 |
| chr5-7697238-INS-316 | chr5-7697238-INS-316 |
| chr5-7706119-INS-68 | chr5-7706119-INS-64 |
| chr5-7714757-INS-84 | chr5-7714757-INS-85 |
| chr5-7723205-INS-682 | chr5-7723205-INS-682 |
| chr5-7730593-INS-58 | chr5-7730593-INS-60 |
| chr5-7750938-INS-354 | chr5-7750938-INS-354 |
| chr5-7778441-INS-184 | chr5-7778441-INS-184 |
| chr5-7790738-INS-642 | chr5-7790738-INS-642 |
| chr5-7791729-INS-92 | chr5-7791729-INS-138 |
| chr5-7804751-INS-156 | chr5-7804767-INS-140 |
| chr5-7811348-INS-85 | chr5-7811348-INS-85 |
| chr5-7816543-INS-397 | chr5-7816543-INS-397 |
| chr5-7841711-INS-6298 | chr5-7841712-INS-6298 |
| chr5-7848382-INS-212 | chr5-7848382-INS-212 |
| chr5-7864699-INS-1968 | chr5-7864700-INS-1968 |
| chr5-7885780-INS-214 | chr5-7885780-INS-214 |
| chr5-7936013-INS-92 | chr5-7936013-INS-92 |
| chr5-7962684-INS-58 | chr5-7962688-INS-104 |

| | |
|---|---|
| chr5-7977551-INS-492 | chr5-7977551-INS-492 |
| chr5-7978789-INS-67 | chr5-7978789-INS-67 |
| chr5-7982117-INS-223 | chr5-7982117-INS-224 |
| chr5-8017475-INS-259 | chr5-8017475-INS-259 |
| chr5-8049657-INS-253 | chr5-8049657-INS-253 |
| chr5-8052508-INS-6905 | chr5-8052509-INS-6906 |
| chr5-8102425-INS-163 | chr5-8102425-INS-163 |
| chr5-8120554-INS-61 | chr5-8120515-INS-93 |
| chr5-8129417-INS-196 | chr5-8129413-INS-206 |
| chr5-8129998-INS-96 | chr5-8129998-INS-101 |
| chr5-8154172-INS-358 | chr5-8154172-INS-358 |
| chr5-8161306-INS-129 | chr5-8161306-INS-139 |
| chr5-8165790-INS-336 | chr5-8165790-INS-336 |
| chr5-8170327-INS-220 | chr5-8170327-INS-220 |
| chr5-8171681-INS-70 | chr5-8171681-INS-104 |
| chr5-8228523-INS-1260 | chr5-8228524-INS-1256 |
| chr5-8230925-INS-297 | chr5-8230926-INS-297 |
| chr5-8277523-INS-6700 | chr5-8277524-INS-6700 |
| chr5-8278687-INS-198 | chr5-8278687-INS-198 |
| chr5-8298313-INS-54 | chr5-8298313-INS-56 |
| chr5-8329996-INS-3050 | chr5-8329997-INS-3050 |
| chr5-8339715-INS-84 | chr5-8339715-INS-70 |
| chr5-8346354-INS-549 | chr5-8346354-INS-549 |
| chr5-8348686-INS-52 | chr5-8348686-INS-52 |
| chr5-8359094-INS-70 | chr5-8359094-INS-80 |
| chr5-8365666-INS-60 | chr5-8365666-INS-60 |
| chr5-8387768-INS-475 | chr5-8387768-INS-475 |
| chr5-8429383-INS-106 | chr5-8429383-INS-106 |
| chr5-8449350-INS-244 | chr5-8449350-INS-236 |
| chr5-8487519-INS-6461 | chr5-8487520-INS-6461 |
| chr5-8492669-INS-65 | chr5-8492669-INS-65 |
| chr5-8508531-INS-1450 | chr5-8508532-INS-1452 |
| chr5-8509186-INS-7080 | chr5-8509187-INS-7080 |
| chr5-8529173-INS-426 | chr5-8529173-INS-426 |
| chr5-8539040-INS-55 | chr5-8539040-INS-55 |
| chr5-8574397-INS-63 | chr5-8574402-INS-66 |
| chr5-8585926-INS-204 | chr5-8585926-INS-204 |
| chr5-8590255-INS-97 | chr5-8590255-INS-96 |
| chr5-8614258-INS-124 | chr5-8614258-INS-124 |
| chr5-8637073-INS-1446 | chr5-8637074-INS-1453 |
| chr5-8639128-INS-1157 | chr5-8639129-INS-1158 |
| chr5-8667128-INS-231 | chr5-8667128-INS-232 |
| chr5-8682518-INS-592 | chr5-8682518-INS-592 |
| chr5-8713779-INS-559 | chr5-8713779-INS-559 |
| chr5-8727287-INS-60 | chr5-8727287-INS-60 |
| chr5-8746318-INS-272 | chr5-8746318-INS-272 |
| chr5-8751924-INS-180 | chr5-8751924-INS-180 |
| chr5-8770715-INS-269 | chr5-8770715-INS-273 |
| chr5-8776719-INS-293 | chr5-8776719-INS-293 |
| chr5-8780912-INS-200 | chr5-8780912-INS-200 |
| chr5-8786026-INS-84 | chr5-8786026-INS-82 |
| chr5-8809512-INS-73 | chr5-8809512-INS-73 |
| chr5-8822811-INS-60 | chr5-8822811-INS-60 |
| chr5-8839016-INS-585 | chr5-8839016-INS-580 |
| chr5-8866782-INS-60 | chr5-8866782-INS-56 |
| chr5-8896445-INS-55 | chr5-8896445-INS-70 |
| chr5-8900711-INS-4502 | chr5-8900712-INS-4502 |
| chr5-8908987-INS-56 | chr5-8908987-INS-76 |
| chr5-8924283-INS-779 | chr5-8924283-INS-779 |
| chr5-8937449-INS-346 | chr5-8937449-INS-346 |
| chr5-8938168-INS-57 | chr5-8938165-INS-61 |
| chr5-8947504-INS-223 | chr5-8947504-INS-224 |
| chr5-8969442-INS-399 | chr5-8969442-INS-399 |
| chr5-8984033-INS-329 | chr5-8984033-INS-329 |
| chr5-9003562-INS-89 | chr5-9003562-INS-89 |
| chr5-9007826-INS-209 | chr5-9007826-INS-209 |
| chr5-9008578-INS-391 | chr5-9008578-INS-391 |
| chr5-9019646-INS-241 | chr5-9019646-INS-241 |
| chr5-9022405-INS-73 | chr5-9022405-INS-73 |
| chr5-9029558-INS-60 | chr5-9029558-INS-65 |
| chr5-9031714-INS-201 | chr5-9031714-INS-201 |
| chr5-9036251-INS-658 | chr5-9036251-INS-663 |
| chr5-9057647-INS-472 | chr5-9057647-INS-472 |
| chr5-9080707-INS-60 | chr5-9080707-INS-64 |
| chr5-9128122-INS-211 | chr5-9128122-INS-211 |
| chr5-9130609-INS-771 | chr5-9130609-INS-771 |
| chr5-9131492-INS-324 | chr5-9131492-INS-324 |

| | |
|---|---|
| chr5-9203890-INS-4927 | chr5-9203891-INS-4927 |
| chr5-9204664-INS-340 | chr5-9204664-INS-340 |
| chr5-9209766-INS-71 | chr5-9209766-INS-71 |
| chr5-9221616-INS-8040 | chr5-9221617-INS-8040 |
| chr5-9236316-INS-379 | chr5-9236316-INS-379 |
| chr5-9244953-INS-1885 | chr5-9244954-INS-1885 |
| chr5-9260260-INS-366 | chr5-9260260-INS-368 |
| chr5-9269372-INS-70 | chr5-9269379-INS-110 |
| chr5-9276491-INS-120 | chr5-9276491-INS-120 |
| chr5-9294664-INS-6231 | chr5-9294665-INS-6232 |
| chr5-9296670-INS-256 | chr5-9296665-INS-266 |
| chr5-9317995-INS-114 | chr5-9317995-INS-128 |
| chr5-9325994-INS-312 | chr5-9325994-INS-312 |
| chr5-9334863-INS-57 | chr5-9334863-INS-57 |
| chr5-9341958-INS-152 | chr5-9341958-INS-152 |
| chr5-9344507-INS-68 | chr5-9344505-INS-58 |
| chr5-9363491-INS-113 | chr5-9363491-INS-113 |
| chr5-9378668-INS-208 | chr5-9378668-INS-208 |
| chr5-9411650-INS-126 | chr5-9411650-INS-126 |
| chr5-9415184-INS-84 | chr5-9415184-INS-84 |
| chr5-9422582-INS-166 | chr5-9422582-INS-166 |
| chr5-9425393-INS-98 | chr5-9425393-INS-90 |
| chr5-9426376-INS-70 | chr5-9426376-INS-72 |
| chr5-9427398-INS-64 | chr5-9427398-INS-64 |
| chr5-9438409-INS-476 | chr5-9438409-INS-476 |
| chr5-9442074-INS-57 | chr5-9442074-INS-62 |
| chr5-9450791-INS-104 | chr5-9450791-INS-60 |
| chr5-9451236-INS-201 | chr5-9451236-INS-201 |
| chr5-9470086-INS-817 | chr5-9470086-INS-817 |
| chr5-9473317-INS-77 | chr5-9473317-INS-77 |
| chr5-9491594-INS-67 | chr5-9491594-INS-67 |
| chr5-9492891-INS-51 | chr5-9492892-INS-53 |
| chr5-9497323-INS-91 | chr5-9497324-INS-52 |
| chr5-9517441-INS-52 | chr5-9517441-INS-52 |
| chr5-9548699-INS-268 | chr5-9548699-INS-268 |
| chr5-9575616-INS-204 | chr5-9575616-INS-204 |
| chr5-9579180-INS-152 | chr5-9579181-INS-152 |
| chr5-9588281-INS-629 | chr5-9588281-INS-629 |
| chr5-9605620-INS-721 | chr5-9605620-INS-721 |
| chr5-9625521-INS-155 | chr5-9625521-INS-155 |
| chr5-9625790-INS-621 | chr5-9625791-INS-613 |
| chr5-9634989-INS-88 | chr5-9634989-INS-86 |
| chr5-9695252-INS-136 | chr5-9695252-INS-137 |
| chr5-9722978-INS-1116 | chr5-9722979-INS-1117 |
| chr5-9754161-INS-265 | chr5-9754175-INS-240 |
| chr5-9758022-INS-505 | chr5-9758012-INS-535 |
| chr5-9779087-INS-164 | chr5-9779087-INS-164 |
| chr5-9785177-INS-460 | chr5-9785182-INS-467 |
| chr5-9789888-INS-94 | chr5-9789889-INS-81 |
| chr5-9796506-INS-62 | chr5-9796506-INS-62 |
| chr5-9797380-INS-54 | chr5-9797380-INS-56 |
| chr5-9805242-INS-132 | chr5-9805244-INS-114 |
| chr5-9815729-INS-62 | chr5-9815729-INS-80 |
| chr5-9848906-INS-180 | chr5-9848906-INS-180 |
| chr5-9851121-INS-1737 | chr5-9851122-INS-1737 |
| chr5-9872910-INS-386 | chr5-9872910-INS-386 |
| chr5-9881480-INS-687 | chr5-9881480-INS-687 |
| chr5-9882157-INS-193 | chr5-9882157-INS-191 |
| chr5-9893030-INS-572 | chr5-9893031-INS-573 |
| chr5-9909758-INS-307 | chr5-9909758-INS-307 |
| chr5-9918810-INS-316 | chr5-9918810-INS-328 |
| chr5-9942841-INS-2631 | chr5-9942842-INS-2631 |
| chr5-9973440-INS-530 | chr5-9973440-INS-532 |
| chr5-10018201-INS-2105 | chr5-10018202-INS-2105 |
| chr5-10049251-INS-939 | chr5-10049251-INS-940 |
| chr5-10062870-INS-6464 | chr5-10062871-INS-6464 |
| chr5-10086302-INS-68 | chr5-10086302-INS-68 |
| chr5-10100639-INS-610 | chr5-10100639-INS-610 |
| chr5-10103977-INS-497 | chr5-10103978-INS-497 |
| chr5-10144803-INS-730 | chr5-10144803-INS-732 |
| chr5-10157324-INS-122 | chr5-10157324-INS-110 |
| chr5-10184176-INS-6401 | chr5-10184177-INS-6401 |
| chr5-10194371-INS-317 | chr5-10194372-INS-317 |
| chr5-10202692-INS-101 | chr5-10202664-INS-144 |
| chr5-10210013-INS-91 | chr5-10210013-INS-91 |
| chr5-10218312-INS-2658 | chr5-10218313-INS-2657 |
| chr5-10224957-INS-204 | chr5-10224957-INS-204 |

| | |
|---|---|
| chr5-10241758-INS-53 | chr5-10241758-INS-53 |
| chr5-10321820-INS-9531 | chr5-10321821-INS-9531 |
| chr5-10363823-INS-166 | chr5-10363808-INS-175 |
| chr5-10363855-INS-658 | chr5-10363858-INS-658 |
| chr5-10390828-INS-189 | chr5-10390828-INS-187 |
| chr5-10395665-INS-90 | chr5-10395665-INS-90 |
| chr5-10414107-INS-75 | chr5-10414107-INS-75 |
| chr5-10436945-INS-2029 | chr5-10436946-INS-2035 |
| chr5-10457117-INS-97 | chr5-10457117-INS-100 |
| chr5-10459220-INS-343 | chr5-10459220-INS-343 |
| chr5-10460153-INS-357 | chr5-10460153-INS-308 |
| chr5-10468447-INS-879 | chr5-10468447-INS-879 |
| chr5-10470385-INS-234 | chr5-10470385-INS-254 |
| chr5-10517470-INS-519 | chr5-10517470-INS-519 |
| chr5-10545133-INS-57 | chr5-10545133-INS-61 |
| chr5-10552448-INS-1095 | chr5-10552449-INS-1095 |
| chr5-10555703-INS-240 | chr5-10555703-INS-240 |
| chr5-10563278-INS-132 | chr5-10563278-INS-132 |
| chr5-10567353-INS-87 | chr5-10567356-INS-88 |
| chr5-10609827-INS-136 | chr5-10609831-INS-132 |
| chr5-10707249-INS-283 | chr5-10707249-INS-281 |
| chr5-10737860-INS-576 | chr5-10737860-INS-560 |
| chr5-10739275-INS-609 | chr5-10739275-INS-612 |
| chr5-10781780-INS-1731 | chr5-10781784-INS-1726 |
| chr5-10792775-INS-1709 | chr5-10792776-INS-1710 |
| chr5-10792925-INS-9318 | chr5-10792925-INS-9317 |
| chr5-10802408-INS-87 | chr5-10802408-INS-130 |
| chr5-10838736-INS-695 | chr5-10838736-INS-695 |
| chr5-10845522-INS-323 | chr5-10845522-INS-329 |
| chr5-10883961-INS-70 | chr5-10883961-INS-92 |
| chr5-10894539-INS-3176 | chr5-10894540-INS-3176 |
| chr5-12037251-INS-443 | chr5-12037251-INS-444 |
| chr5-12044642-INS-90 | chr5-12044642-INS-90 |
| chr5-12047639-INS-143 | chr5-12047639-INS-143 |
| chr5-12051671-INS-795 | chr5-12051671-INS-796 |
| chr5-12074411-INS-452 | chr5-12074412-INS-451 |
| chr5-12155310-INS-52 | chr5-12155307-INS-64 |
| chr5-12202669-INS-146 | chr5-12202661-INS-124 |
| chr5-12203812-INS-68 | chr5-12203804-INS-80 |
| chr5-12244696-INS-154 | chr5-12244696-INS-154 |
| chr5-12258332-INS-96 | chr5-12258332-INS-96 |
| chr5-12273756-INS-133 | chr5-12273756-INS-135 |
| chr5-12288853-INS-110 | chr5-12288853-INS-119 |
| chr5-12293064-INS-56 | chr5-12293064-INS-60 |
| chr5-12339284-INS-54 | chr5-12339334-INS-51 |
| chr5-12349506-INS-61 | chr5-12349505-INS-65 |
| chr5-12364608-INS-92 | chr5-12364608-INS-92 |
| chr5-12375878-INS-103 | chr5-12375878-INS-103 |
| chr5-12409359-INS-6463 | chr5-12409360-INS-6463 |
| chr5-12415854-INS-6947 | chr5-12415855-INS-6948 |
| chr5-12420404-INS-826 | chr5-12420404-INS-832 |
| chr5-12423561-INS-6953 | chr5-12423562-INS-6953 |
| chr5-12459844-INS-143 | chr5-12459844-INS-140 |
| chr5-12468174-INS-222 | chr5-12468174-INS-222 |
| chr5-12470994-INS-187 | chr5-12470994-INS-187 |
| chr5-12474468-INS-4897 | chr5-12474469-INS-4897 |
| chr5-12482206-INS-68 | chr5-12482208-INS-72 |
| chr5-12520496-INS-51 | chr5-12520497-INS-55 |
| chr5-12529095-INS-7010 | chr5-12529096-INS-7010 |
| chr5-12534281-INS-218 | chr5-12534281-INS-228 |
| chr5-12536643-INS-56 | chr5-12536643-INS-56 |
| chr5-12544406-INS-6548 | chr5-12544407-INS-6548 |
| chr5-12546972-INS-373 | chr5-12546972-INS-371 |
| chr5-12558297-INS-197 | chr5-12558297-INS-197 |
| chr5-12565687-INS-172 | chr5-12565698-INS-164 |
| chr5-12580596-INS-56 | chr5-12580596-INS-56 |
| chr5-12621888-INS-52 | chr5-12621888-INS-50 |
| chr5-12648945-INS-6398 | chr5-12648946-INS-6398 |
| chr5-12693085-INS-836 | chr5-12693085-INS-840 |
| chr5-12706854-INS-1084 | chr5-12706839-INS-1073 |
| chr5-12733492-INS-144 | chr5-12733492-INS-141 |
| chr5-12744267-INS-66 | chr5-12744267-INS-66 |
| chr5-12759931-INS-56 | chr5-12759931-INS-56 |
| chr5-12773655-INS-52 | chr5-12773650-INS-57 |
| chr5-12807263-INS-78 | chr5-12807263-INS-78 |
| chr5-12814604-INS-443 | chr5-12814604-INS-439 |
| chr5-12848657-INS-575 | chr5-12848657-INS-575 |

| | |
|---|---|
| chr5-12858581-INS-69 | chr5-12858534-INS-92 |
| chr5-12870814-INS-261 | chr5-12870814-INS-261 |
| chr5-12881143-INS-207 | chr5-12881143-INS-207 |
| chr5-12896818-INS-3020 | chr5-12896819-INS-3016 |
| chr5-12923593-INS-502 | chr5-12923593-INS-503 |
| chr5-12957514-INS-824 | chr5-12957514-INS-824 |
| chr5-13028542-INS-2748 | chr5-13028543-INS-2748 |
| chr5-13038042-INS-64 | chr5-13038043-INS-96 |
| chr5-13046345-INS-58 | chr5-13046345-INS-60 |
| chr5-13072003-INS-206 | chr5-13072003-INS-206 |
| chr5-13076566-INS-180 | chr5-13076566-INS-173 |
| chr5-13092159-INS-60 | chr5-13092159-INS-68 |
| chr5-13140192-INS-68 | chr5-13140192-INS-68 |
| chr5-13147519-INS-265 | chr5-13147519-INS-265 |
| chr5-13156486-INS-6527 | chr5-13156487-INS-6527 |
| chr5-13190598-INS-1027 | chr5-13190599-INS-1027 |
| chr5-13208276-INS-142 | chr5-13208276-INS-142 |
| chr5-13210609-INS-6472 | chr5-13210610-INS-6472 |
| chr5-13213587-INS-222 | chr5-13213587-INS-222 |
| chr5-13226385-INS-92 | chr5-13226404-INS-68 |
| chr5-13240298-INS-156 | chr5-13240298-INS-156 |
| chr5-13242017-INS-1400 | chr5-13242018-INS-1400 |
| chr5-13282353-INS-311 | chr5-13282353-INS-311 |
| chr5-13301030-INS-992 | chr5-13301030-INS-989 |
| chr5-13308823-INS-52 | chr5-13308823-INS-62 |
| chr5-13322002-INS-54 | chr5-13322012-INS-55 |
| chr5-13346915-INS-741 | chr5-13346915-INS-740 |
| chr5-13356633-INS-92 | chr5-13356633-INS-92 |
| chr5-13373986-INS-327 | chr5-13373986-INS-327 |
| chr5-13383510-INS-104 | chr5-13383510-INS-100 |
| chr5-13408441-INS-624 | chr5-13408441-INS-624 |
| chr5-13427336-INS-81 | chr5-13427336-INS-84 |
| chr5-13430105-INS-58 | chr5-13430105-INS-50 |
| chr5-13431693-INS-56 | chr5-13431693-INS-56 |
| chr5-13445851-INS-56 | chr5-13445828-INS-56 |
| chr5-13447166-INS-70 | chr5-13447164-INS-82 |
| chr5-13504404-INS-378 | chr5-13504404-INS-378 |
| chr5-13524685-INS-182 | chr5-13524685-INS-182 |
| chr5-13541150-INS-74 | chr5-13541173-INS-52 |
| chr5-13551817-INS-87 | chr5-13551817-INS-87 |
| chr5-13563928-INS-1533 | chr5-13563929-INS-1544 |
| chr5-13566642-INS-518 | chr5-13566642-INS-519 |
| chr5-13571980-INS-55 | chr5-13571980-INS-55 |
| chr5-13625539-INS-6912 | chr5-13625540-INS-6914 |
| chr5-13626644-INS-526 | chr5-13626644-INS-527 |
| chr5-13628147-INS-56 | chr5-13628145-INS-56 |
| chr5-13632938-INS-103 | chr5-13632952-INS-103 |
| chr5-13640637-INS-177 | chr5-13640637-INS-177 |
| chr5-13745797-INS-108 | chr5-13745844-INS-73 |
| chr5-13769259-INS-4200 | chr5-13769260-INS-4200 |
| chr5-13773514-INS-1197 | chr5-13773515-INS-1197 |
| chr5-13859430-INS-557 | chr5-13859435-INS-542 |
| chr5-13874248-INS-400 | chr5-13874248-INS-400 |
| chr5-13875574-INS-80 | chr5-13875565-INS-60 |
| chr5-13894749-INS-581 | chr5-13894749-INS-602 |
| chr5-13897170-INS-333 | chr5-13897170-INS-332 |
| chr5-13945268-INS-1894 | chr5-13945269-INS-1894 |
| chr5-13948612-INS-91 | chr5-13948616-INS-71 |
| chr5-13954320-INS-930 | chr5-13954320-INS-930 |
| chr5-13978499-INS-1539 | chr5-13978500-INS-1541 |
| chr5-13980947-INS-269 | chr5-13980971-INS-273 |
| chr5-13995928-INS-58 | chr5-13995928-INS-58 |
| chr5-14009768-INS-315 | chr5-14009770-INS-316 |
| chr5-14017539-INS-3355 | chr5-14017540-INS-3355 |
| chr5-14030707-INS-53 | chr5-14030707-INS-53 |
| chr5-14040025-INS-449 | chr5-14040025-INS-450 |
| chr5-14057873-INS-75 | chr5-14057873-INS-75 |
| chr5-14074507-INS-228 | chr5-14074507-INS-228 |
| chr5-14122585-INS-171 | chr5-14122585-INS-172 |
| chr5-14125256-INS-602 | chr5-14125256-INS-602 |
| chr5-14127111-INS-954 | chr5-14127111-INS-954 |
| chr5-14129044-INS-176 | chr5-14129044-INS-176 |
| chr5-14129906-INS-1734 | chr5-14129907-INS-1733 |
| chr5-14134264-INS-74 | chr5-14134264-INS-80 |
| chr5-14161955-INS-6183 | chr5-14161956-INS-6183 |
| chr5-14164635-INS-1011 | chr5-14164643-INS-1006 |
| chr5-14173790-INS-69 | chr5-14173790-INS-69 |

| | |
|---|---|
| chr5-14175022-INS-536 | chr5-14175022-INS-536 |
| chr5-14192352-INS-105 | chr5-14192352-INS-105 |
| chr5-14205422-INS-52 | chr5-14205422-INS-56 |
| chr5-14235254-INS-53 | chr5-14235254-INS-77 |
| chr5-14271156-INS-6580 | chr5-14271157-INS-6580 |
| chr5-14273872-INS-56 | chr5-14273872-INS-56 |
| chr5-14287406-INS-2659 | chr5-14287407-INS-2659 |
| chr5-14297494-INS-343 | chr5-14297494-INS-343 |
| chr5-14301674-INS-6591 | chr5-14301675-INS-6592 |
| chr5-14383028-INS-135 | chr5-14383028-INS-140 |
| chr5-14397816-INS-70 | chr5-14397798-INS-60 |
| chr5-14420033-INS-205 | chr5-14420033-INS-205 |
| chr5-14484757-INS-1454 | chr5-14484758-INS-1454 |
| chr5-14489530-INS-885 | chr5-14489531-INS-895 |
| chr5-14500868-INS-914 | chr5-14500868-INS-914 |
| chr5-14519286-INS-78 | chr5-14519283-INS-72 |
| chr5-14520953-INS-1877 | chr5-14520954-INS-1877 |
| chr5-14528108-INS-6440 | chr5-14528109-INS-6440 |
| chr5-14537427-INS-60 | chr5-14537427-INS-58 |
| chr5-14580826-INS-62 | chr5-14580826-INS-70 |
| chr5-14596267-INS-143 | chr5-14596268-INS-143 |
| chr5-14609054-INS-101 | chr5-14609054-INS-100 |
| chr5-14611538-INS-212 | chr5-14611538-INS-212 |
| chr5-14622794-INS-76 | chr5-14622794-INS-79 |
| chr5-14648139-INS-269 | chr5-14648139-INS-269 |
| chr5-14655668-INS-89 | chr5-14655668-INS-89 |
| chr5-14657648-INS-852 | chr5-14657648-INS-852 |
| chr5-14674235-INS-530 | chr5-14674235-INS-529 |
| chr5-14687760-INS-84 | chr5-14687760-INS-84 |
| chr5-14695173-INS-66 | chr5-14695173-INS-66 |
| chr5-14700437-INS-173 | chr5-14700437-INS-173 |
| chr5-14736334-INS-80 | chr5-14736334-INS-83 |
| chr5-14745437-INS-198 | chr5-14745437-INS-197 |
| chr5-14758673-INS-52 | chr5-14758683-INS-53 |
| chr5-14767024-INS-476 | chr5-14767022-INS-474 |
| chr5-14771780-INS-305 | chr5-14771780-INS-306 |
| chr5-14782726-INS-108 | chr5-14782726-INS-109 |
| chr5-14817252-INS-3279 | chr5-14817270-INS-3306 |
| chr5-14833992-INS-223 | chr5-14833992-INS-222 |
| chr5-14836082-INS-78 | chr5-14836074-INS-88 |
| chr5-14837440-INS-3189 | chr5-14837441-INS-3189 |
| chr5-14837441-INS-394 | chr5-14837441-INS-394 |
| chr5-14875442-INS-99 | chr5-14875442-INS-125 |
| chr5-14882275-INS-73 | chr5-14882275-INS-73 |
| chr5-14915482-INS-204 | chr5-14915482-INS-204 |
| chr5-14919798-INS-130 | chr5-14919798-INS-130 |
| chr5-14936924-INS-63 | chr5-14936924-INS-63 |
| chr5-14941377-INS-978 | chr5-14941377-INS-978 |
| chr5-14947021-INS-212 | chr5-14947021-INS-213 |
| chr5-14953152-INS-778 | chr5-14953152-INS-774 |
| chr5-14972833-INS-101 | chr5-14972833-INS-98 |
| chr5-15067763-INS-200 | chr5-15067756-INS-193 |
| chr5-15070303-INS-123 | chr5-15070305-INS-118 |
| chr5-15841670-INS-392 | chr5-15841670-INS-392 |
| chr5-15927809-INS-108 | chr5-15927809-INS-104 |
| chr5-15934333-INS-89 | chr5-15934336-INS-88 |
| chr5-15939540-INS-82 | chr5-15939534-INS-82 |
| chr5-15954148-INS-486 | chr5-15954148-INS-486 |
| chr5-16004906-INS-55 | chr5-16004906-INS-57 |
| chr5-16007143-INS-96 | chr5-16007182-INS-74 |
| chr5-16063000-INS-6586 | chr5-16063001-INS-6585 |
| chr5-16069875-INS-498 | chr5-16069875-INS-498 |
| chr5-16071732-INS-343 | chr5-16071732-INS-351 |
| chr5-16110259-INS-99 | chr5-16110231-INS-94 |
| chr5-16123408-INS-96 | chr5-16123404-INS-102 |
| chr5-16157354-INS-62 | chr5-16157354-INS-62 |
| chr5-16169217-INS-70 | chr5-16169218-INS-85 |
| chr5-16181674-INS-194 | chr5-16181674-INS-204 |
| chr5-16232243-INS-231 | chr5-16232244-INS-191 |
| chr5-16258398-INS-56 | chr5-16258401-INS-83 |
| chr5-16281740-INS-56 | chr5-16281740-INS-56 |
| chr5-16357048-INS-60 | chr5-16357048-INS-61 |
| chr5-16373342-INS-7816 | chr5-16373343-INS-7816 |
| chr5-16387312-INS-80 | chr5-16387314-INS-72 |
| chr5-16399270-INS-8473 | chr5-16399271-INS-8473 |
| chr5-16412856-INS-53 | chr5-16412856-INS-54 |
| chr5-16414879-INS-580 | chr5-16414879-INS-581 |

| | |
|---|---|
| chr5-16417103-INS-68 | chr5-16417103-INS-68 |
| chr5-16487473-INS-52 | chr5-16487473-INS-56 |
| chr5-16498012-INS-380 | chr5-16498012-INS-389 |
| chr5-16564689-INS-67 | chr5-16564689-INS-67 |
| chr5-16573328-INS-522 | chr5-16573328-INS-522 |
| chr5-16583691-INS-96 | chr5-16583689-INS-57 |
| chr5-16597937-INS-394 | chr5-16597937-INS-394 |
| chr5-16599847-INS-7083 | chr5-16599848-INS-7083 |
| chr5-16599860-INS-201 | chr5-16599860-INS-201 |
| chr5-16603682-INS-60 | chr5-16603694-INS-61 |
| chr5-16657950-INS-103 | chr5-16657950-INS-103 |
| chr5-16718755-INS-88 | chr5-16718755-INS-88 |
| chr5-16731789-INS-632 | chr5-16731789-INS-632 |
| chr5-16755735-INS-5489 | chr5-16755736-INS-5489 |
| chr5-16768179-INS-79 | chr5-16768179-INS-79 |
| chr5-16789985-INS-58 | chr5-16789985-INS-62 |
| chr5-16799255-INS-55 | chr5-16799255-INS-60 |
| chr5-16799907-INS-528 | chr5-16799907-INS-528 |
| chr5-16804346-INS-220 | chr5-16804346-INS-220 |
| chr5-16815704-INS-58 | chr5-16815704-INS-58 |
| chr5-16822564-INS-168 | chr5-16822564-INS-169 |
| chr5-16850707-INS-6281 | chr5-16850708-INS-6281 |
| chr5-16860205-INS-324 | chr5-16860215-INS-342 |
| chr5-16862215-INS-61 | chr5-16862219-INS-53 |
| chr5-16886085-INS-64 | chr5-16886049-INS-82 |
| chr5-16893529-INS-55 | chr5-16893522-INS-54 |
| chr5-16907198-INS-415 | chr5-16907198-INS-415 |
| chr5-16908909-INS-197 | chr5-16908909-INS-197 |
| chr5-16910893-INS-2060 | chr5-16910894-INS-2061 |
| chr5-16916825-INS-61 | chr5-16916831-INS-53 |
| chr5-16974195-INS-96 | chr5-16974207-INS-76 |
| chr5-16984924-INS-74 | chr5-16984936-INS-54 |
| chr5-16991205-INS-2281 | chr5-16991206-INS-2282 |
| chr5-16995398-INS-76 | chr5-16995398-INS-76 |
| chr5-17087935-INS-139 | chr5-17087935-INS-139 |
| chr5-17088257-INS-1469 | chr5-17088258-INS-1470 |
| chr5-17141664-INS-327 | chr5-17141664-INS-327 |
| chr5-17155613-INS-112 | chr5-17155612-INS-101 |
| chr5-17163004-INS-876 | chr5-17163004-INS-876 |
| chr5-17172272-INS-5714 | chr5-17172273-INS-5714 |
| chr5-17174486-INS-92 | chr5-17174486-INS-92 |
| chr5-17257536-INS-112 | chr5-17257536-INS-98 |
| chr5-17266375-INS-100 | chr5-17266375-INS-100 |
| chr5-17270027-INS-552 | chr5-17270006-INS-557 |
| chr5-17278856-INS-89 | chr5-17278856-INS-89 |
| chr5-17281305-INS-343 | chr5-17281311-INS-343 |
| chr5-17287949-INS-1010 | chr5-17287950-INS-1005 |
| chr5-17297337-INS-210 | chr5-17297337-INS-210 |
| chr5-17313041-INS-156 | chr5-17313035-INS-168 |
| chr5-17325626-INS-258 | chr5-17325626-INS-258 |
| chr5-17348642-INS-229 | chr5-17348642-INS-229 |
| chr5-17349448-INS-2515 | chr5-17349449-INS-2515 |
| chr5-17354412-INS-164 | chr5-17354413-INS-164 |
| chr5-17381376-INS-412 | chr5-17381378-INS-411 |
| chr5-17394997-INS-693 | chr5-17394997-INS-693 |
| chr5-17403612-INS-76 | chr5-17403602-INS-88 |
| chr5-17405448-INS-463 | chr5-17405449-INS-463 |
| chr5-17439545-INS-58 | chr5-17439545-INS-58 |
| chr5-17466911-INS-372 | chr5-17466911-INS-372 |
| chr5-17496935-INS-72 | chr5-17496935-INS-72 |
| chr5-17508953-INS-3848 | chr5-17508954-INS-3848 |
| chr5-17509385-INS-69 | chr5-17509385-INS-71 |
| chr5-17523644-INS-7149 | chr5-17523645-INS-7149 |
| chr5-17525351-INS-1527 | chr5-17525352-INS-1529 |
| chr5-17543896-INS-92 | chr5-17543896-INS-80 |
| chr5-17564797-INS-6442 | chr5-17564798-INS-6442 |
| chr5-17574502-INS-765 | chr5-17574502-INS-765 |
| chr5-17606337-INS-1533 | chr5-17606338-INS-1534 |
| chr5-17612594-INS-242 | chr5-17612594-INS-242 |
| chr5-17620782-INS-52 | chr5-17620782-INS-52 |
| chr5-17640881-INS-200 | chr5-17640881-INS-200 |
| chr5-17662781-INS-66 | chr5-17662781-INS-76 |
| chr5-17676456-INS-67 | chr5-17676422-INS-67 |
| chr5-17718885-INS-57 | chr5-17718885-INS-65 |
| chr5-17726150-INS-52 | chr5-17726150-INS-52 |
| chr5-17727333-INS-157 | chr5-17727329-INS-160 |
| chr5-17756190-INS-224 | chr5-17756190-INS-224 |

| | |
|---|---|
| chr5-17757216-INS-68 | chr5-17757216-INS-68 |
| chr5-17767330-INS-413 | chr5-17767330-INS-413 |
| chr5-17768263-INS-52 | chr5-17768290-INS-50 |
| chr5-17770115-INS-266 | chr5-17770115-INS-266 |
| chr5-17836527-INS-57 | chr5-17836527-INS-63 |
| chr5-17847558-INS-346 | chr5-17847558-INS-346 |
| chr5-17861447-INS-232 | chr5-17861447-INS-232 |
| chr5-17865494-INS-56 | chr5-17865494-INS-66 |
| chr5-17899497-INS-899 | chr5-17899497-INS-899 |
| chr5-17908486-INS-177 | chr5-17908486-INS-179 |
| chr5-17920988-INS-717 | chr5-17920988-INS-717 |
| chr5-17975349-INS-144 | chr5-17975346-INS-120 |
| chr5-17999090-INS-200 | chr5-17999090-INS-200 |
| chr5-18014905-INS-474 | chr5-18014905-INS-474 |
| chr5-18017916-INS-126 | chr5-18017916-INS-128 |
| chr5-18050652-INS-95 | chr5-18050651-INS-64 |
| chr5-18070466-INS-67 | chr5-18070478-INS-74 |
| chr5-18073420-INS-290 | chr5-18073420-INS-290 |
| chr5-18086846-INS-78 | chr5-18086846-INS-78 |
| chr5-18091841-INS-6448 | chr5-18091842-INS-6450 |
| chr5-18121914-INS-466 | chr5-18121914-INS-467 |
| chr5-18126128-INS-111 | chr5-18126129-INS-114 |
| chr5-18131112-INS-1685 | chr5-18131113-INS-1685 |
| chr5-18131807-INS-435 | chr5-18131807-INS-435 |
| chr5-18135442-INS-100 | chr5-18135442-INS-115 |
| chr5-18152277-INS-64 | chr5-18152277-INS-52 |
| chr5-18168845-INS-236 | chr5-18168845-INS-236 |
| chr5-18177513-INS-6596 | chr5-18177514-INS-6596 |
| chr5-18214552-INS-90 | chr5-18214552-INS-90 |
| chr5-18290985-INS-171 | chr5-18290985-INS-171 |
| chr5-18312352-INS-1875 | chr5-18312353-INS-1875 |
| chr5-18319884-INS-82 | chr5-18319884-INS-72 |
| chr5-18342100-INS-186 | chr5-18342100-INS-186 |
| chr5-18380676-INS-57 | chr5-18380681-INS-58 |
| chr5-18388669-INS-472 | chr5-18388669-INS-472 |
| chr5-18399578-INS-4901 | chr5-18399579-INS-4901 |
| chr5-18402018-INS-2134 | chr5-18402019-INS-2134 |
| chr5-18405110-INS-6174 | chr5-18405111-INS-6174 |
| chr5-18409557-INS-647 | chr5-18409557-INS-647 |
| chr5-18448353-INS-289 | chr5-18448353-INS-287 |
| chr5-18454920-INS-54 | chr5-18454920-INS-54 |
| chr5-18457147-INS-89 | chr5-18457147-INS-52 |
| chr5-18459248-INS-56 | chr5-18459230-INS-74 |
| chr5-18469514-INS-71 | chr5-18469514-INS-79 |
| chr5-18510754-INS-59 | chr5-18510754-INS-59 |
| chr5-18511983-INS-190 | chr5-18511983-INS-191 |
| chr5-18518946-INS-617 | chr5-18518946-INS-617 |
| chr5-18521918-INS-70 | chr5-18521918-INS-70 |
| chr5-18560532-INS-494 | chr5-18560532-INS-493 |
| chr5-18577585-INS-175 | chr5-18577585-INS-175 |
| chr5-18583713-INS-345 | chr5-18583713-INS-345 |
| chr5-18595898-INS-4403 | chr5-18595899-INS-4403 |
| chr5-18598919-INS-7254 | chr5-18598920-INS-7254 |
| chr5-18612838-INS-91 | chr5-18612838-INS-63 |
| chr5-18617743-INS-251 | chr5-18617743-INS-251 |
| chr5-18629407-INS-6196 | chr5-18629408-INS-6196 |
| chr5-18631000-INS-2376 | chr5-18631001-INS-2376 |
| chr5-18660141-INS-440 | chr5-18660141-INS-440 |
| chr5-18674045-INS-86 | chr5-18674045-INS-108 |
| chr5-18681778-INS-326 | chr5-18681778-INS-326 |
| chr5-18683829-INS-52 | chr5-18683829-INS-56 |
| chr5-18693842-INS-83 | chr5-18693842-INS-71 |
| chr5-18694436-INS-197 | chr5-18694436-INS-197 |
| chr5-18713260-INS-53 | chr5-18713275-INS-54 |
| chr5-18735247-INS-371 | chr5-18735247-INS-371 |
| chr5-18750827-INS-86 | chr5-18750827-INS-86 |
| chr5-18767569-INS-505 | chr5-18767569-INS-505 |
| chr5-18776182-INS-654 | chr5-18776182-INS-654 |
| chr5-18851762-INS-176 | chr5-18851763-INS-176 |
| chr5-18888420-INS-79 | chr5-18888420-INS-58 |
| chr5-18894765-INS-425 | chr5-18894765-INS-426 |
| chr5-18921324-INS-2864 | chr5-18921325-INS-2858 |
| chr5-18925437-INS-337 | chr5-18925437-INS-330 |
| chr5-18951283-INS-577 | chr5-18951283-INS-577 |
| chr5-18972352-INS-390 | chr5-18972352-INS-390 |
| chr5-18993129-INS-187 | chr5-18993129-INS-187 |
| chr5-19003382-INS-435 | chr5-19003382-INS-435 |

| | |
|---|---|
| chr5-19015876-INS-51 | chr5-19015876-INS-51 |
| chr5-19054348-INS-3116 | chr5-19054349-INS-3116 |
| chr5-19054349-INS-726 | chr5-19054349-INS-726 |
| chr5-19058028-INS-214 | chr5-19058028-INS-214 |
| chr5-19071786-INS-633 | chr5-19071794-INS-651 |
| chr5-19089229-INS-609 | chr5-19089229-INS-610 |
| chr5-19090850-INS-2685 | chr5-19090851-INS-2687 |
| chr5-19093719-INS-6458 | chr5-19093720-INS-6459 |
| chr5-19114106-INS-225 | chr5-19114106-INS-224 |
| chr5-19176547-INS-226 | chr5-19176547-INS-222 |
| chr5-19180549-INS-80 | chr5-19180561-INS-56 |
| chr5-19253184-INS-2183 | chr5-19253185-INS-2183 |
| chr5-19275183-INS-75 | chr5-19275183-INS-79 |
| chr5-19280411-INS-68 | chr5-19280411-INS-72 |
| chr5-19296883-INS-1147 | chr5-19296884-INS-1147 |
| chr5-19328821-INS-7153 | chr5-19328822-INS-7156 |
| chr5-19342227-INS-75 | chr5-19342227-INS-68 |
| chr5-19383164-INS-187 | chr5-19383160-INS-186 |
| chr5-19427081-INS-352 | chr5-19427081-INS-361 |
| chr5-19438576-INS-72 | chr5-19438576-INS-72 |
| chr5-19470562-INS-328 | chr5-19470571-INS-356 |
| chr5-19471648-INS-185 | chr5-19471648-INS-186 |
| chr5-19495469-INS-60 | chr5-19495467-INS-76 |
| chr5-19512772-INS-103 | chr5-19512772-INS-72 |
| chr5-19524346-INS-69 | chr5-19524346-INS-73 |
| chr5-19534313-INS-51 | chr5-19534313-INS-51 |
| chr5-19570592-INS-60 | chr5-19570592-INS-60 |
| chr5-19595293-INS-80 | chr5-19595293-INS-56 |
| chr5-19605925-INS-765 | chr5-19605944-INS-776 |
| chr5-19605968-INS-64 | chr5-19605968-INS-66 |
| chr5-19617566-INS-469 | chr5-19617566-INS-469 |
| chr5-19627158-INS-552 | chr5-19627159-INS-544 |
| chr5-19631596-INS-925 | chr5-19631596-INS-924 |
| chr5-19648744-INS-127 | chr5-19648744-INS-127 |
| chr5-19657762-INS-5165 | chr5-19657763-INS-5166 |
| chr5-19658219-INS-54 | chr5-19658219-INS-54 |
| chr5-19660283-INS-200 | chr5-19660282-INS-205 |
| chr5-19663113-INS-68 | chr5-19663113-INS-68 |
| chr5-19663681-INS-3289 | chr5-19663682-INS-3289 |
| chr5-19673026-INS-1158 | chr5-19673027-INS-1159 |
| chr5-19674371-INS-8241 | chr5-19674372-INS-8240 |
| chr5-19691045-INS-651 | chr5-19691045-INS-651 |
| chr5-19692763-INS-72 | chr5-19692763-INS-72 |
| chr5-19738597-INS-203 | chr5-19738597-INS-202 |
| chr5-19786803-INS-6182 | chr5-19786804-INS-6183 |
| chr5-19803871-INS-58 | chr5-19803871-INS-53 |
| chr5-19828074-INS-126 | chr5-19828074-INS-126 |
| chr5-19830900-INS-104 | chr5-19830900-INS-104 |
| chr5-19901434-INS-440 | chr5-19901434-INS-440 |
| chr5-19902292-INS-6517 | chr5-19902293-INS-6519 |
| chr5-19916716-INS-51 | chr5-19916716-INS-53 |
| chr5-19921357-INS-453 | chr5-19921357-INS-453 |
| chr5-19952769-INS-83 | chr5-19952769-INS-83 |
| chr5-19960495-INS-982 | chr5-19960495-INS-983 |
| chr5-19988454-INS-189 | chr5-19988454-INS-162 |
| chr5-19990047-INS-3454 | chr5-19990048-INS-3454 |
| chr5-20025152-INS-4240 | chr5-20025153-INS-4240 |
| chr5-20064939-INS-89 | chr5-20064948-INS-59 |
| chr5-20075628-INS-68 | chr5-20075628-INS-68 |
| chr5-20083409-INS-1418 | chr5-20083410-INS-1418 |
| chr5-20100353-INS-62 | chr5-20100353-INS-62 |
| chr5-20130568-INS-815 | chr5-20130568-INS-814 |
| chr5-20133053-INS-611 | chr5-20133053-INS-610 |
| chr5-20133559-INS-62 | chr5-20133559-INS-86 |
| chr5-20135308-INS-85 | chr5-20135306-INS-56 |
| chr5-20146218-INS-92 | chr5-20146216-INS-94 |
| chr5-20161015-INS-193 | chr5-20161015-INS-194 |
| chr5-20190197-INS-860 | chr5-20190193-INS-863 |
| chr5-20247728-INS-80 | chr5-20247728-INS-81 |
| chr5-20260928-INS-6746 | chr5-20260929-INS-6747 |
| chr5-20301729-INS-189 | chr5-20301729-INS-198 |
| chr5-20333199-INS-56 | chr5-20333199-INS-52 |
| chr5-20351068-INS-3062 | chr5-20351069-INS-3062 |
| chr5-20352081-INS-744 | chr5-20352078-INS-745 |
| chr5-20369449-INS-243 | chr5-20369449-INS-243 |
| chr5-20419447-INS-5501 | chr5-20419448-INS-5501 |
| chr5-20427203-INS-238 | chr5-20427203-INS-238 |

| | |
|---|---|
| chr5-20477112-INS-58 | chr5-20477102-INS-60 |
| chr5-20479684-INS-64 | chr5-20479684-INS-50 |
| chr5-20499632-INS-805 | chr5-20499632-INS-805 |
| chr5-20518736-INS-64 | chr5-20518737-INS-54 |
| chr5-20528513-INS-66 | chr5-20528513-INS-50 |
| chr5-20530600-INS-1542 | chr5-20530601-INS-1541 |
| chr5-20543673-INS-536 | chr5-20543673-INS-536 |
| chr5-20594819-INS-54 | chr5-20594819-INS-54 |
| chr5-20609016-INS-403 | chr5-20609016-INS-403 |
| chr5-20621822-INS-64 | chr5-20621836-INS-76 |
| chr5-20625222-INS-6939 | chr5-20625223-INS-6940 |
| chr5-20626153-INS-329 | chr5-20626153-INS-329 |
| chr5-20636647-INS-92 | chr5-20636647-INS-89 |
| chr5-20649588-INS-65 | chr5-20649589-INS-71 |
| chr5-20678213-INS-463 | chr5-20678213-INS-463 |
| chr5-20682279-INS-52 | chr5-20682279-INS-50 |
| chr5-20720177-INS-438 | chr5-20720178-INS-438 |
| chr5-20720731-INS-425 | chr5-20720731-INS-425 |
| chr5-20722905-INS-229 | chr5-20722905-INS-229 |
| chr5-20727173-INS-102 | chr5-20727173-INS-102 |
| chr5-20757990-INS-203 | chr5-20757990-INS-203 |
| chr5-20779769-INS-353 | chr5-20779769-INS-355 |
| chr5-20781422-INS-182 | chr5-20781422-INS-177 |
| chr5-20783607-INS-54 | chr5-20783607-INS-54 |
| chr5-20818627-INS-56 | chr5-20818627-INS-56 |
| chr5-20821450-INS-2787 | chr5-20821451-INS-2787 |
| chr5-20852626-INS-670 | chr5-20852626-INS-671 |
| chr5-20863537-INS-52 | chr5-20863568-INS-54 |
| chr5-20892705-INS-61 | chr5-20892705-INS-61 |
| chr5-20900315-INS-422 | chr5-20900315-INS-422 |
| chr5-20930151-INS-64 | chr5-20930151-INS-64 |
| chr5-20937899-INS-205 | chr5-20937917-INS-160 |
| chr5-20940470-INS-114 | chr5-20940470-INS-110 |
| chr5-20953062-INS-486 | chr5-20953062-INS-486 |
| chr5-20957439-INS-3247 | chr5-20957440-INS-3247 |
| chr5-20982779-INS-197 | chr5-20982779-INS-197 |
| chr5-20985601-INS-131 | chr5-20985601-INS-131 |
| chr5-20986938-INS-82 | chr5-20986938-INS-82 |
| chr5-21009535-INS-58 | chr5-21009535-INS-58 |
| chr5-21021576-INS-139 | chr5-21021576-INS-141 |
| chr5-21023634-INS-84 | chr5-21023634-INS-84 |
| chr5-21053146-INS-147 | chr5-21053146-INS-147 |
| chr5-21069507-INS-941 | chr5-21069507-INS-946 |
| chr5-21076438-INS-112 | chr5-21076438-INS-111 |
| chr5-21115762-INS-55 | chr5-21115762-INS-55 |
| chr5-21135396-INS-359 | chr5-21135396-INS-359 |
| chr5-21149776-INS-197 | chr5-21149776-INS-197 |
| chr5-21193497-INS-246 | chr5-21193497-INS-198 |
| chr5-21211775-INS-396 | chr5-21211775-INS-397 |
| chr5-21213014-INS-208 | chr5-21213014-INS-209 |
| chr5-21264421-INS-173 | chr5-21264421-INS-173 |
| chr5-21324550-INS-5259 | chr5-21324551-INS-5259 |
| chr5-21357440-INS-64 | chr5-21357440-INS-64 |
| chr5-21365989-INS-449 | chr5-21365989-INS-449 |
| chr5-21372923-INS-497 | chr5-21372923-INS-497 |
| chr5-21381946-INS-203 | chr5-21381946-INS-203 |
| chr5-21386191-INS-82 | chr5-21386191-INS-66 |
| chr5-21448623-INS-107 | chr5-21448623-INS-107 |
| chr5-21452067-INS-218 | chr5-21452067-INS-201 |
| chr5-21471417-INS-999 | chr5-21471418-INS-971 |
| chr5-21486795-INS-140 | chr5-21486795-INS-140 |
| chr5-21501767-INS-211 | chr5-21501767-INS-211 |
| chr5-21509221-INS-5287 | chr5-21509222-INS-5287 |
| chr5-21527195-INS-321 | chr5-21527195-INS-324 |
| chr5-21551161-INS-432 | chr5-21551161-INS-432 |
| chr5-21562452-INS-68 | chr5-21562452-INS-68 |
| chr5-21586315-INS-58 | chr5-21586315-INS-58 |
| chr5-21602291-INS-420 | chr5-21602291-INS-400 |
| chr5-21604334-INS-64 | chr5-21604334-INS-64 |
| chr5-21632553-INS-168 | chr5-21632553-INS-168 |
| chr5-21692479-INS-52 | chr5-21692479-INS-52 |
| chr5-21703728-INS-235 | chr5-21703728-INS-235 |
| chr5-21714158-INS-327 | chr5-21714158-INS-327 |
| chr5-21715140-INS-108 | chr5-21715167-INS-108 |
| chr5-21721393-INS-75 | chr5-21721393-INS-75 |
| chr5-21723944-INS-976 | chr5-21723944-INS-976 |
| chr5-21731281-INS-646 | chr5-21731281-INS-637 |

| | |
|---|---|
| chr5-21737127-INS-587 | chr5-21737127-INS-587 |
| chr5-21760113-INS-102 | chr5-21760113-INS-102 |
| chr5-21773968-INS-60 | chr5-21773968-INS-60 |
| chr5-21775365-INS-116 | chr5-21775365-INS-116 |
| chr5-21791457-INS-250 | chr5-21791457-INS-250 |
| chr5-21797481-INS-205 | chr5-21797481-INS-205 |
| chr5-21800448-INS-66 | chr5-21800448-INS-51 |
| chr5-21822873-INS-142 | chr5-21822873-INS-142 |
| chr5-21832064-INS-60 | chr5-21832064-INS-62 |
| chr5-21841234-INS-4452 | chr5-21841235-INS-4452 |
| chr5-21843401-INS-79 | chr5-21843402-INS-55 |
| chr5-21861741-INS-477 | chr5-21861741-INS-477 |
| chr5-21865663-INS-214 | chr5-21865663-INS-214 |
| chr5-21873514-INS-95 | chr5-21873514-INS-78 |
| chr5-21881563-INS-1077 | chr5-21881564-INS-1081 |
| chr5-21882150-INS-425 | chr5-21882150-INS-425 |
| chr5-21882933-INS-193 | chr5-21882933-INS-194 |
| chr5-21888019-INS-172 | chr5-21888019-INS-185 |
| chr5-21889028-INS-162 | chr5-21889028-INS-144 |
| chr5-21895211-INS-209 | chr5-21895211-INS-209 |
| chr5-21918994-INS-325 | chr5-21918994-INS-325 |
| chr5-21929162-INS-207 | chr5-21929162-INS-207 |
| chr5-21930433-INS-114 | chr5-21930433-INS-114 |
| chr5-21937267-INS-209 | chr5-21937267-INS-209 |
| chr5-21942568-INS-76 | chr5-21942568-INS-84 |
| chr5-21954191-INS-730 | chr5-21954191-INS-730 |
| chr5-21962539-INS-217 | chr5-21962539-INS-217 |
| chr5-21970570-INS-191 | chr5-21970571-INS-205 |
| chr5-21983202-INS-207 | chr5-21983202-INS-207 |
| chr5-21991110-INS-69 | chr5-21991110-INS-69 |
| chr5-22003998-INS-222 | chr5-22003998-INS-224 |
| chr5-22008881-INS-80 | chr5-22008882-INS-52 |
| chr5-22021181-INS-51 | chr5-22021181-INS-51 |
| chr5-22028002-INS-54 | chr5-22028002-INS-54 |
| chr5-22029283-INS-1888 | chr5-22029284-INS-1888 |
| chr5-22033492-INS-55 | chr5-22033492-INS-50 |
| chr5-22048024-INS-96 | chr5-22048024-INS-96 |
| chr5-22049230-INS-98 | chr5-22049232-INS-100 |
| chr5-22051167-INS-245 | chr5-22051167-INS-245 |
| chr5-22062558-INS-101 | chr5-22062543-INS-135 |
| chr5-22078984-INS-322 | chr5-22078984-INS-322 |
| chr5-22080364-INS-433 | chr5-22080364-INS-432 |
| chr5-22111374-INS-164 | chr5-22111334-INS-180 |
| chr5-22112523-INS-78 | chr5-22112523-INS-78 |
| chr5-22113135-INS-251 | chr5-22113135-INS-271 |
| chr5-22117646-INS-73 | chr5-22117646-INS-73 |
| chr5-22130578-INS-56 | chr5-22130578-INS-66 |
| chr5-22133037-INS-680 | chr5-22133037-INS-697 |
| chr5-22133955-INS-64 | chr5-22133955-INS-64 |
| chr5-22134886-INS-7415 | chr5-22134887-INS-7415 |
| chr5-22135863-INS-108 | chr5-22135863-INS-108 |
| chr5-22174696-INS-338 | chr5-22174696-INS-338 |
| chr5-22179059-INS-132 | chr5-22179047-INS-132 |
| chr5-22187350-INS-223 | chr5-22187350-INS-223 |
| chr5-22218680-INS-54 | chr5-22218680-INS-54 |
| chr5-22219622-INS-6539 | chr5-22219623-INS-6539 |
| chr5-22242022-INS-102 | chr5-22242022-INS-102 |
| chr5-22256653-INS-246 | chr5-22256653-INS-245 |
| chr5-22271270-INS-1296 | chr5-22271271-INS-1296 |
| chr5-22303916-INS-895 | chr5-22303916-INS-896 |
| chr5-22305651-INS-350 | chr5-22305651-INS-350 |
| chr5-22306355-INS-81 | chr5-22306355-INS-81 |
| chr5-22316031-INS-395 | chr5-22316031-INS-401 |
| chr5-22325858-INS-174 | chr5-22325858-INS-174 |
| chr5-22329535-INS-136 | chr5-22329535-INS-140 |
| chr5-22350219-INS-53 | chr5-22350219-INS-69 |
| chr5-22375289-INS-345 | chr5-22375289-INS-345 |
| chr5-22381814-INS-264 | chr5-22381814-INS-264 |
| chr5-22384581-INS-73 | chr5-22384574-INS-106 |
| chr5-22426287-INS-640 | chr5-22426287-INS-636 |
| chr5-22435930-INS-65 | chr5-22435930-INS-68 |
| chr5-22467623-INS-68 | chr5-22467623-INS-68 |
| chr5-22491606-INS-63 | chr5-22491606-INS-59 |
| chr5-22495033-INS-938 | chr5-22495033-INS-938 |
| chr5-22497363-INS-56 | chr5-22497363-INS-56 |
| chr5-22522121-INS-82 | chr5-22522121-INS-82 |
| chr5-22572892-INS-1540 | chr5-22572893-INS-1540 |

| | |
|---|---|
| chr5-22589340-INS-94 | chr5-22589376-INS-66 |
| chr5-22608635-INS-129 | chr5-22608635-INS-123 |
| chr5-22630782-INS-174 | chr5-22630782-INS-174 |
| chr5-22659280-INS-70 | chr5-22659280-INS-58 |
| chr5-22731299-INS-51 | chr5-22731299-INS-51 |
| chr5-22757136-INS-220 | chr5-22757136-INS-220 |
| chr5-22758894-INS-54 | chr5-22758894-INS-54 |
| chr5-22770246-INS-577 | chr5-22770246-INS-577 |
| chr5-22788453-INS-336 | chr5-22788458-INS-349 |
| chr5-22795030-INS-356 | chr5-22795030-INS-356 |
| chr5-22819709-INS-7598 | chr5-22819705-INS-7596 |
| chr5-22843694-INS-93 | chr5-22843694-INS-93 |
| chr5-22854141-INS-56 | chr5-22854141-INS-56 |
| chr5-22857061-INS-88 | chr5-22857068-INS-108 |
| chr5-22860081-INS-57 | chr5-22860081-INS-56 |
| chr5-22861296-INS-78 | chr5-22861296-INS-78 |
| chr5-22863139-INS-510 | chr5-22863139-INS-510 |
| chr5-22888094-INS-6478 | chr5-22888095-INS-6478 |
| chr5-22942581-INS-96 | chr5-22942581-INS-92 |
| chr5-22947128-INS-156 | chr5-22947128-INS-156 |
| chr5-22958413-INS-58 | chr5-22958431-INS-60 |
| chr5-22963496-INS-558 | chr5-22963495-INS-556 |
| chr5-23017067-INS-51 | chr5-23017068-INS-51 |
| chr5-23023781-INS-414 | chr5-23023781-INS-418 |
| chr5-23033587-INS-52 | chr5-23033583-INS-56 |
| chr5-23057674-INS-344 | chr5-23057674-INS-344 |
| chr5-23062337-INS-6200 | chr5-23062328-INS-6190 |
| chr5-23066298-INS-66 | chr5-23066298-INS-52 |
| chr5-23094339-INS-498 | chr5-23094339-INS-498 |
| chr5-23125851-INS-74 | chr5-23125851-INS-80 |
| chr5-23146890-INS-56 | chr5-23146890-INS-56 |
| chr5-23154429-INS-5162 | chr5-23154430-INS-5166 |
| chr5-23190140-INS-103 | chr5-23190140-INS-101 |
| chr5-23216472-INS-153 | chr5-23216472-INS-153 |
| chr5-23226857-INS-79 | chr5-23226905-INS-70 |
| chr5-23239996-INS-98 | chr5-23239996-INS-98 |
| chr5-23247167-INS-51 | chr5-23247154-INS-74 |
| chr5-23252250-INS-71 | chr5-23252250-INS-71 |
| chr5-23260117-INS-385 | chr5-23260117-INS-385 |
| chr5-23270124-INS-365 | chr5-23270125-INS-365 |
| chr5-23287467-INS-71 | chr5-23287467-INS-71 |
| chr5-23308718-INS-53 | chr5-23308714-INS-52 |
| chr5-23359537-INS-6485 | chr5-23359538-INS-6486 |
| chr5-23360636-INS-597 | chr5-23360636-INS-597 |
| chr5-23373063-INS-118 | chr5-23373063-INS-118 |
| chr5-23374224-INS-680 | chr5-23374224-INS-680 |
| chr5-23379496-INS-60 | chr5-23379496-INS-62 |
| chr5-23380138-INS-128 | chr5-23380138-INS-128 |
| chr5-23388856-INS-113 | chr5-23388856-INS-113 |
| chr5-23404692-INS-108 | chr5-23404688-INS-76 |
| chr5-23414129-INS-211 | chr5-23414129-INS-212 |
| chr5-23440368-INS-80 | chr5-23440368-INS-80 |
| chr5-23446882-INS-233 | chr5-23446887-INS-237 |
| chr5-23461926-INS-78 | chr5-23461932-INS-72 |
| chr5-23465835-INS-54 | chr5-23465838-INS-70 |
| chr5-23483028-INS-56 | chr5-23483028-INS-56 |
| chr5-23485705-INS-58 | chr5-23485705-INS-58 |
| chr5-23504851-INS-1033 | chr5-23504852-INS-1033 |
| chr5-23514965-INS-94 | chr5-23514965-INS-94 |
| chr5-23533877-INS-908 | chr5-23533877-INS-908 |
| chr5-23549279-INS-182 | chr5-23549279-INS-182 |
| chr5-23567484-INS-738 | chr5-23567485-INS-740 |
| chr5-23580963-INS-53 | chr5-23580963-INS-54 |
| chr5-23581982-INS-3666 | chr5-23581983-INS-3666 |
| chr5-23632056-INS-145 | chr5-23632056-INS-144 |
| chr5-23725248-INS-700 | chr5-23725248-INS-702 |
| chr5-23726750-INS-261 | chr5-23726750-INS-261 |
| chr5-23754906-INS-229 | chr5-23754906-INS-229 |
| chr5-23762632-INS-53 | chr5-23762632-INS-53 |
| chr5-23765876-INS-426 | chr5-23765876-INS-426 |
| chr5-23768781-INS-223 | chr5-23768781-INS-224 |
| chr5-23781301-INS-180 | chr5-23781301-INS-184 |
| chr5-23794705-INS-78 | chr5-23794706-INS-86 |
| chr5-23818702-INS-146 | chr5-23818703-INS-160 |
| chr5-23835835-INS-211 | chr5-23835835-INS-211 |
| chr5-23836863-INS-72 | chr5-23836847-INS-60 |
| chr5-23847978-INS-137 | chr5-23847978-INS-138 |

| | |
|---|---|
| chr5-23848289-INS-365 | chr5-23848289-INS-365 |
| chr5-23853498-INS-211 | chr5-23853498-INS-211 |
| chr5-23866390-INS-230 | chr5-23866391-INS-230 |
| chr5-23868875-INS-51 | chr5-23868875-INS-51 |
| chr5-23869840-INS-510 | chr5-23869840-INS-510 |
| chr5-23870848-INS-218 | chr5-23870848-INS-218 |
| chr5-23871841-INS-118 | chr5-23871841-INS-118 |
| chr5-23888046-INS-195 | chr5-23888046-INS-195 |
| chr5-23895141-INS-83 | chr5-23895141-INS-86 |
| chr5-23899407-INS-5713 | chr5-23899408-INS-5714 |
| chr5-23917020-INS-745 | chr5-23917020-INS-745 |
| chr5-23920446-INS-1524 | chr5-23920456-INS-1513 |
| chr5-23926883-INS-132 | chr5-23926883-INS-133 |
| chr5-23928015-INS-219 | chr5-23928015-INS-219 |
| chr5-23931935-INS-78 | chr5-23931935-INS-78 |
| chr5-23938188-INS-480 | chr5-23938188-INS-480 |
| chr5-23940304-INS-207 | chr5-23940291-INS-177 |
| chr5-23940827-INS-251 | chr5-23940827-INS-246 |
| chr5-23944124-INS-201 | chr5-23944124-INS-201 |
| chr5-23951053-INS-116 | chr5-23951053-INS-116 |
| chr5-23955110-INS-188 | chr5-23955111-INS-156 |
| chr5-23957774-INS-244 | chr5-23957774-INS-244 |
| chr5-23978054-INS-117 | chr5-23978054-INS-117 |
| chr5-23983863-INS-207 | chr5-23983863-INS-207 |
| chr5-23984676-INS-57 | chr5-23984676-INS-57 |
| chr5-23986415-INS-208 | chr5-23986416-INS-176 |
| chr5-24001331-INS-205 | chr5-24001331-INS-203 |
| chr5-24004981-INS-6945 | chr5-24004982-INS-6946 |
| chr5-24017007-INS-2830 | chr5-24017008-INS-2838 |
| chr5-24018171-INS-54 | chr5-24018176-INS-74 |
| chr5-24023757-INS-107 | chr5-24023757-INS-108 |
| chr5-24030809-INS-252 | chr5-24030809-INS-252 |
| chr5-24033029-INS-265 | chr5-24033029-INS-265 |
| chr5-24043677-INS-113 | chr5-24043677-INS-113 |
| chr5-24049760-INS-3166 | chr5-24049761-INS-3177 |
| chr5-24050791-INS-114 | chr5-24050791-INS-117 |
| chr5-24051691-INS-344 | chr5-24051691-INS-341 |
| chr5-24057887-INS-201 | chr5-24057887-INS-201 |
| chr5-24059401-INS-75 | chr5-24059401-INS-75 |
| chr5-24066253-INS-72 | chr5-24066253-INS-76 |
| chr5-24071823-INS-207 | chr5-24071823-INS-207 |
| chr5-24088416-INS-200 | chr5-24088416-INS-202 |
| chr5-24090899-INS-57 | chr5-24090883-INS-57 |
| chr5-24097869-INS-88 | chr5-24097869-INS-88 |
| chr5-24127235-INS-62 | chr5-24127235-INS-60 |
| chr5-24130959-INS-64 | chr5-24130959-INS-64 |
| chr5-24134507-INS-319 | chr5-24134507-INS-319 |
| chr5-24139439-INS-498 | chr5-24139439-INS-498 |
| chr5-24142558-INS-78 | chr5-24142591-INS-53 |
| chr5-24155206-INS-206 | chr5-24155206-INS-205 |
| chr5-24157953-INS-81 | chr5-24157954-INS-51 |
| chr5-24176386-INS-208 | chr5-24176386-INS-208 |
| chr5-24183720-INS-137 | chr5-24183720-INS-137 |
| chr5-24186780-INS-196 | chr5-24186798-INS-206 |
| chr5-24220003-INS-691 | chr5-24220003-INS-691 |
| chr5-24229378-INS-215 | chr5-24229378-INS-215 |
| chr5-24232610-INS-225 | chr5-24232610-INS-204 |
| chr5-24256869-INS-158 | chr5-24256869-INS-158 |
| chr5-24259876-INS-194 | chr5-24259876-INS-194 |
| chr5-24272738-INS-81 | chr5-24272738-INS-81 |
| chr5-24276773-INS-338 | chr5-24276773-INS-338 |
| chr5-24289175-INS-667 | chr5-24289175-INS-667 |
| chr5-24294980-INS-202 | chr5-24294980-INS-202 |
| chr5-24301471-INS-7128 | chr5-24301472-INS-7142 |
| chr5-24301834-INS-58 | chr5-24301834-INS-50 |
| chr5-24314871-INS-384 | chr5-24314871-INS-385 |
| chr5-24316897-INS-214 | chr5-24316897-INS-214 |
| chr5-24325395-INS-218 | chr5-24325395-INS-218 |
| chr5-24351331-INS-5313 | chr5-24351332-INS-5313 |
| chr5-24354935-INS-113 | chr5-24354935-INS-113 |
| chr5-24383666-INS-168 | chr5-24383666-INS-168 |
| chr5-24412740-INS-154 | chr5-24412740-INS-154 |
| chr5-24434642-INS-785 | chr5-24434642-INS-785 |
| chr5-24435570-INS-196 | chr5-24435570-INS-196 |
| chr5-24437083-INS-112 | chr5-24437083-INS-112 |
| chr5-24444421-INS-137 | chr5-24444421-INS-132 |
| chr5-24446758-INS-122 | chr5-24446751-INS-122 |

| | |
|---|---|
| chr5-24447851-INS-509 | chr5-24447851-INS-544 |
| chr5-24451905-INS-52 | chr5-24451905-INS-62 |
| chr5-24473707-INS-201 | chr5-24473707-INS-201 |
| chr5-24484671-INS-400 | chr5-24484671-INS-400 |
| chr5-24487403-INS-57 | chr5-24487403-INS-57 |
| chr5-24502618-INS-115 | chr5-24502618-INS-112 |
| chr5-24510934-INS-72 | chr5-24510934-INS-72 |
| chr5-24583887-INS-86 | chr5-24583887-INS-86 |
| chr5-24587203-INS-58 | chr5-24587180-INS-54 |
| chr5-24659249-INS-52 | chr5-24659249-INS-54 |
| chr5-24664458-INS-172 | chr5-24664458-INS-175 |
| chr5-24725881-INS-55 | chr5-24725881-INS-50 |
| chr5-24728439-INS-65 | chr5-24728439-INS-90 |
| chr5-24752571-INS-116 | chr5-24752571-INS-116 |
| chr5-24769987-INS-224 | chr5-24769987-INS-224 |
| chr5-24779719-INS-205 | chr5-24779719-INS-211 |
| chr5-24790303-INS-6472 | chr5-24790304-INS-6472 |
| chr5-24834323-INS-173 | chr5-24834324-INS-173 |
| chr5-24835748-INS-199 | chr5-24835748-INS-200 |
| chr5-24849186-INS-185 | chr5-24849186-INS-185 |
| chr5-24851612-INS-209 | chr5-24851612-INS-209 |
| chr5-24862140-INS-56 | chr5-24862140-INS-58 |
| chr5-24868746-INS-209 | chr5-24868746-INS-209 |
| chr5-24873543-INS-53 | chr5-24873543-INS-55 |
| chr5-24879320-INS-833 | chr5-24879320-INS-839 |
| chr5-24884079-INS-440 | chr5-24884079-INS-440 |
| chr5-24888530-INS-240 | chr5-24888530-INS-240 |
| chr5-24889431-INS-239 | chr5-24889431-INS-239 |
| chr5-24896681-INS-68 | chr5-24896681-INS-71 |
| chr5-24922528-INS-204 | chr5-24922528-INS-210 |
| chr5-24933570-INS-214 | chr5-24933570-INS-215 |
| chr5-24947807-INS-51 | chr5-24947807-INS-51 |
| chr5-24951859-INS-172 | chr5-24951820-INS-195 |
| chr5-24959400-INS-222 | chr5-24959400-INS-224 |
| chr5-24964881-INS-56 | chr5-24964881-INS-56 |
| chr5-24983498-INS-181 | chr5-24983498-INS-181 |
| chr5-24984683-INS-87 | chr5-24984683-INS-87 |
| chr5-24987102-INS-52 | chr5-24987102-INS-52 |
| chr5-24992776-INS-359 | chr5-24992776-INS-359 |
| chr5-24994634-INS-66 | chr5-24994634-INS-66 |
| chr5-24997254-INS-55 | chr5-24997254-INS-55 |
| chr5-25003356-INS-206 | chr5-25003356-INS-206 |
| chr5-25049420-INS-4145 | chr5-25049421-INS-4145 |
| chr5-25063034-INS-768 | chr5-25063034-INS-768 |
| chr5-25064549-INS-62 | chr5-25064549-INS-62 |
| chr5-25070859-INS-124 | chr5-25070835-INS-100 |
| chr5-25089678-INS-87 | chr5-25089678-INS-87 |
| chr5-25105191-INS-57 | chr5-25105191-INS-62 |
| chr5-25110526-INS-59 | chr5-25110526-INS-59 |
| chr5-25121932-INS-499 | chr5-25121932-INS-498 |
| chr5-25147144-INS-7192 | chr5-25147145-INS-7192 |
| chr5-25171398-INS-1904 | chr5-25171399-INS-1910 |
| chr5-25178091-INS-117 | chr5-25178091-INS-78 |
| chr5-25179972-INS-60 | chr5-25179972-INS-60 |
| chr5-25184087-INS-147 | chr5-25184087-INS-147 |
| chr5-25186286-INS-208 | chr5-25186286-INS-208 |
| chr5-25205036-INS-223 | chr5-25205036-INS-223 |
| chr5-25213400-INS-143 | chr5-25213381-INS-153 |
| chr5-25230575-INS-6504 | chr5-25230576-INS-6506 |
| chr5-25251678-INS-84 | chr5-25251674-INS-88 |
| chr5-25253061-INS-66 | chr5-25253061-INS-69 |
| chr5-25271161-INS-1889 | chr5-25271162-INS-1889 |
| chr5-25278303-INS-563 | chr5-25278303-INS-580 |
| chr5-25281979-INS-96 | chr5-25281980-INS-115 |
| chr5-25292637-INS-609 | chr5-25292637-INS-609 |
| chr5-25293645-INS-69 | chr5-25293645-INS-102 |
| chr5-25311799-INS-84 | chr5-25311799-INS-87 |
| chr5-25336198-INS-208 | chr5-25336198-INS-208 |
| chr5-25345543-INS-56 | chr5-25345544-INS-50 |
| chr5-25367386-INS-209 | chr5-25367386-INS-208 |
| chr5-25380421-INS-197 | chr5-25380428-INS-177 |
| chr5-25384666-INS-111 | chr5-25384678-INS-120 |
| chr5-25398022-INS-212 | chr5-25398022-INS-207 |
| chr5-25414014-INS-242 | chr5-25414014-INS-234 |
| chr5-25447203-INS-225 | chr5-25447203-INS-205 |
| chr5-25447706-INS-64 | chr5-25447706-INS-64 |
| chr5-25484929-INS-181 | chr5-25484929-INS-188 |

| | |
|---|---|
| chr5-25507005-INS-247 | chr5-25507005-INS-250 |
| chr5-25509843-INS-241 | chr5-25509843-INS-241 |
| chr5-25514236-INS-192 | chr5-25514236-INS-192 |
| chr5-25518014-INS-181 | chr5-25518014-INS-181 |
| chr5-25522131-INS-77 | chr5-25522131-INS-77 |
| chr5-25545725-INS-69 | chr5-25545725-INS-79 |
| chr5-25546753-INS-3729 | chr5-25546768-INS-3722 |
| chr5-25598053-INS-55 | chr5-25598058-INS-50 |
| chr5-25605128-INS-120 | chr5-25605128-INS-122 |
| chr5-25613287-INS-601 | chr5-25613287-INS-601 |
| chr5-25623310-INS-55 | chr5-25623310-INS-57 |
| chr5-25626047-INS-5407 | chr5-25626048-INS-5408 |
| chr5-25659207-INS-237 | chr5-25659205-INS-236 |
| chr5-25659813-INS-1050 | chr5-25659814-INS-1097 |
| chr5-25674556-INS-205 | chr5-25674556-INS-205 |
| chr5-25682671-INS-86 | chr5-25682668-INS-86 |
| chr5-25688407-INS-298 | chr5-25688407-INS-298 |
| chr5-25709346-INS-58 | chr5-25709346-INS-58 |
| chr5-25728736-INS-98 | chr5-25728736-INS-98 |
| chr5-25738803-INS-198 | chr5-25738803-INS-198 |
| chr5-25750826-INS-182 | chr5-25750826-INS-182 |
| chr5-25753754-INS-343 | chr5-25753754-INS-325 |
| chr5-25761175-INS-584 | chr5-25761175-INS-584 |
| chr5-25773159-INS-200 | chr5-25773159-INS-200 |
| chr5-25781276-INS-200 | chr5-25781276-INS-200 |
| chr5-25808413-INS-233 | chr5-25808413-INS-245 |
| chr5-25810444-INS-56 | chr5-25810444-INS-56 |
| chr5-25819427-INS-52 | chr5-25819427-INS-52 |
| chr5-25825175-INS-220 | chr5-25825175-INS-221 |
| chr5-25828701-INS-220 | chr5-25828701-INS-221 |
| chr5-25836614-INS-140 | chr5-25836614-INS-136 |
| chr5-25845009-INS-492 | chr5-25845009-INS-494 |
| chr5-25854539-INS-639 | chr5-25854541-INS-645 |
| chr5-25866092-INS-129 | chr5-25866097-INS-95 |
| chr5-25872119-INS-105 | chr5-25872119-INS-115 |
| chr5-25873534-INS-147 | chr5-25873534-INS-147 |
| chr5-25874662-INS-160 | chr5-25874662-INS-160 |
| chr5-25879352-INS-107 | chr5-25879352-INS-89 |
| chr5-25881813-INS-77 | chr5-25881779-INS-77 |
| chr5-25909196-INS-52 | chr5-25909218-INS-62 |
| chr5-25911668-INS-78 | chr5-25911668-INS-84 |
| chr5-25913209-INS-64 | chr5-25913209-INS-70 |
| chr5-25922829-INS-511 | chr5-25922829-INS-524 |
| chr5-25935082-INS-158 | chr5-25935082-INS-131 |
| chr5-25937300-INS-57 | chr5-25937300-INS-57 |
| chr5-25939725-INS-150 | chr5-25939714-INS-150 |
| chr5-25941445-INS-199 | chr5-25941445-INS-199 |
| chr5-25947222-INS-88 | chr5-25947222-INS-92 |
| chr5-25949159-INS-55 | chr5-25949159-INS-55 |
| chr5-25952178-INS-110 | chr5-25952178-INS-110 |
| chr5-25974020-INS-93 | chr5-25974025-INS-133 |
| chr5-25975087-INS-120 | chr5-25975087-INS-120 |
| chr5-25975514-INS-4577 | chr5-25975515-INS-4577 |
| chr5-25976619-INS-72 | chr5-25976619-INS-72 |
| chr5-26005875-INS-58 | chr5-26005875-INS-58 |
| chr5-26008300-INS-163 | chr5-26008300-INS-162 |
| chr5-26036160-INS-74 | chr5-26036161-INS-75 |
| chr5-26038809-INS-109 | chr5-26038809-INS-129 |
| chr5-26048764-INS-54 | chr5-26048740-INS-68 |
| chr5-26067273-INS-60 | chr5-26067273-INS-60 |
| chr5-26070289-INS-107 | chr5-26070281-INS-73 |
| chr5-26080443-INS-391 | chr5-26080441-INS-346 |
| chr5-26106541-INS-72 | chr5-26106541-INS-56 |
| chr5-26113064-INS-90 | chr5-26113064-INS-90 |
| chr5-26349451-INS-1054 | chr5-26349452-INS-1054 |
| chr5-26370836-INS-52 | chr5-26370836-INS-54 |
| chr5-26393398-INS-122 | chr5-26393384-INS-109 |
| chr5-26396732-INS-370 | chr5-26396732-INS-370 |
| chr5-26424233-INS-164 | chr5-26424233-INS-164 |
| chr5-26439077-INS-794 | chr5-26439077-INS-792 |
| chr5-26454494-INS-7111 | chr5-26454495-INS-7111 |
| chr5-26459032-INS-81 | chr5-26459032-INS-81 |
| chr5-26488271-INS-6896 | chr5-26488272-INS-6896 |
| chr5-26492639-INS-153 | chr5-26492639-INS-153 |
| chr5-26495176-INS-151 | chr5-26495176-INS-151 |
| chr5-26518101-INS-4322 | chr5-26518102-INS-4340 |
| chr5-26546531-INS-64 | chr5-26546531-INS-66 |

| | |
|---|---|
| chr5-26563130-INS-164 | chr5-26563130-INS-164 |
| chr5-26580546-INS-52 | chr5-26580546-INS-80 |
| chr5-26595591-INS-145 | chr5-26595592-INS-145 |
| chr5-26603261-INS-164 | chr5-26603261-INS-174 |
| chr5-26609834-INS-290 | chr5-26609834-INS-290 |
| chr5-26631984-INS-60 | chr5-26631984-INS-56 |
| chr5-26645810-INS-80 | chr5-26645810-INS-80 |
| chr5-26647068-INS-66 | chr5-26647068-INS-66 |
| chr5-26655161-INS-88 | chr5-26655163-INS-98 |
| chr5-26668173-INS-1048 | chr5-26668174-INS-1049 |
| chr5-26670906-INS-445 | chr5-26670906-INS-445 |
| chr5-26675100-INS-51 | chr5-26675101-INS-63 |
| chr5-26703676-INS-709 | chr5-26703676-INS-709 |
| chr5-26718860-INS-154 | chr5-26718860-INS-202 |
| chr5-26742218-INS-2092 | chr5-26742219-INS-2093 |
| chr5-26757289-INS-100 | chr5-26757321-INS-115 |
| chr5-26772290-INS-495 | chr5-26772290-INS-492 |
| chr5-26801669-INS-98 | chr5-26801670-INS-56 |
| chr5-26823976-INS-849 | chr5-26823977-INS-856 |
| chr5-26825990-INS-500 | chr5-26825991-INS-500 |
| chr5-26842506-INS-408 | chr5-26842506-INS-363 |
| chr5-26881447-INS-69 | chr5-26881447-INS-73 |
| chr5-26900071-INS-548 | chr5-26900071-INS-548 |
| chr5-26930942-INS-498 | chr5-26930942-INS-498 |
| chr5-26935396-INS-931 | chr5-26935396-INS-931 |
| chr5-26947304-INS-762 | chr5-26947304-INS-762 |
| chr5-26957379-INS-127 | chr5-26957360-INS-127 |
| chr5-26963745-INS-64 | chr5-26963745-INS-64 |
| chr5-26966410-INS-336 | chr5-26966410-INS-336 |
| chr5-26971860-INS-74 | chr5-26971826-INS-68 |
| chr5-26976009-INS-93 | chr5-26976009-INS-99 |
| chr5-26978390-INS-73 | chr5-26978342-INS-72 |
| chr5-26982471-INS-6789 | chr5-26982472-INS-6789 |
| chr5-26983200-INS-176 | chr5-26983200-INS-176 |
| chr5-27000019-INS-6962 | chr5-27000020-INS-6962 |
| chr5-27003890-INS-71 | chr5-27003893-INS-101 |
| chr5-27007869-INS-102 | chr5-27007869-INS-102 |
| chr5-27013807-INS-133 | chr5-27013807-INS-133 |
| chr5-27027129-INS-82 | chr5-27027129-INS-82 |
| chr5-27046381-INS-110 | chr5-27046381-INS-110 |
| chr5-27059001-INS-71 | chr5-27059001-INS-71 |
| chr5-27064883-INS-60 | chr5-27064883-INS-64 |
| chr5-27068827-INS-1840 | chr5-27068828-INS-1852 |
| chr5-27119175-INS-64 | chr5-27119175-INS-64 |
| chr5-27119849-INS-2075 | chr5-27119850-INS-2077 |
| chr5-27127067-INS-76 | chr5-27127110-INS-59 |
| chr5-27161719-INS-1722 | chr5-27161720-INS-1723 |
| chr5-27174220-INS-751 | chr5-27174220-INS-776 |
| chr5-27194361-INS-68 | chr5-27194361-INS-68 |
| chr5-27200264-INS-148 | chr5-27200264-INS-150 |
| chr5-27219450-INS-51 | chr5-27219456-INS-50 |
| chr5-27244140-INS-418 | chr5-27244140-INS-420 |
| chr5-27286839-INS-781 | chr5-27286839-INS-781 |
| chr5-27298996-INS-54 | chr5-27298996-INS-78 |
| chr5-27316780-INS-6360 | chr5-27316781-INS-6360 |
| chr5-27318367-INS-817 | chr5-27318367-INS-817 |
| chr5-27341332-INS-102 | chr5-27341332-INS-102 |
| chr5-27350960-INS-331 | chr5-27350960-INS-331 |
| chr5-27368271-INS-341 | chr5-27368271-INS-341 |
| chr5-27410455-INS-70 | chr5-27410455-INS-70 |
| chr5-27420197-INS-109 | chr5-27420197-INS-109 |
| chr5-27431950-INS-1267 | chr5-27431951-INS-1269 |
| chr5-27451085-INS-197 | chr5-27451110-INS-215 |
| chr5-27526737-INS-56 | chr5-27526737-INS-51 |
| chr5-27585179-INS-64 | chr5-27585179-INS-64 |
| chr5-27611396-INS-947 | chr5-27611396-INS-970 |
| chr5-27615137-INS-939 | chr5-27615137-INS-912 |
| chr5-27670776-INS-64 | chr5-27670776-INS-64 |
| chr5-27680761-INS-202 | chr5-27680761-INS-219 |
| chr5-27683673-INS-116 | chr5-27683673-INS-116 |
| chr5-27694587-INS-266 | chr5-27694587-INS-266 |
| chr5-27704541-INS-104 | chr5-27704541-INS-108 |
| chr5-27720524-INS-59 | chr5-27720524-INS-59 |
| chr5-27735430-INS-63 | chr5-27735430-INS-68 |
| chr5-27757159-INS-95 | chr5-27757159-INS-90 |
| chr5-27762721-INS-88 | chr5-27762721-INS-90 |
| chr5-27782710-INS-2321 | chr5-27782711-INS-2321 |

| | |
|---|---|
| chr5-27789175-INS-60 | chr5-27789175-INS-60 |
| chr5-27790834-INS-54 | chr5-27790834-INS-56 |
| chr5-27795334-INS-2146 | chr5-27795335-INS-2143 |
| chr5-27796016-INS-54 | chr5-27796016-INS-58 |
| chr5-27802622-INS-65 | chr5-27802622-INS-65 |
| chr5-27812298-INS-350 | chr5-27812302-INS-351 |
| chr5-27826400-INS-213 | chr5-27826400-INS-213 |
| chr5-27828557-INS-376 | chr5-27828557-INS-376 |
| chr5-27864783-INS-136 | chr5-27864814-INS-140 |
| chr5-27869938-INS-75 | chr5-27869938-INS-76 |
| chr5-27870990-INS-175 | chr5-27870990-INS-175 |
| chr5-27886559-INS-86 | chr5-27886588-INS-86 |
| chr5-27909400-INS-161 | chr5-27909400-INS-161 |
| chr5-27924498-INS-905 | chr5-27924498-INS-905 |
| chr5-27935451-INS-68 | chr5-27935451-INS-68 |
| chr5-27939336-INS-767 | chr5-27939336-INS-767 |
| chr5-27955827-INS-113 | chr5-27955788-INS-123 |
| chr5-27961374-INS-230 | chr5-27961374-INS-230 |
| chr5-28000370-INS-64 | chr5-28000354-INS-73 |
| chr5-28002993-INS-206 | chr5-28002993-INS-206 |
| chr5-28005654-INS-120 | chr5-28005654-INS-122 |
| chr5-28013034-INS-63 | chr5-28013034-INS-66 |
| chr5-28023175-INS-246 | chr5-28023175-INS-246 |
| chr5-28027707-INS-514 | chr5-28027707-INS-514 |
| chr5-28036185-INS-67 | chr5-28036185-INS-76 |
| chr5-28044034-INS-208 | chr5-28044034-INS-208 |
| chr5-28051074-INS-204 | chr5-28051074-INS-203 |
| chr5-28064408-INS-240 | chr5-28064408-INS-239 |
| chr5-28071498-INS-205 | chr5-28071498-INS-204 |
| chr5-28088721-INS-796 | chr5-28088721-INS-794 |
| chr5-28094061-INS-7224 | chr5-28094062-INS-7224 |
| chr5-28095496-INS-84 | chr5-28095496-INS-84 |
| chr5-28098828-INS-80 | chr5-28098828-INS-80 |
| chr5-28123442-INS-88 | chr5-28123442-INS-129 |
| chr5-28124180-INS-6442 | chr5-28124181-INS-6442 |
| chr5-28130524-INS-565 | chr5-28130524-INS-565 |
| chr5-28132050-INS-6714 | chr5-28132051-INS-6717 |
| chr5-28137041-INS-278 | chr5-28137041-INS-279 |
| chr5-28138510-INS-219 | chr5-28138510-INS-219 |
| chr5-28141814-INS-67 | chr5-28141858-INS-50 |
| chr5-28166373-INS-138 | chr5-28166373-INS-137 |
| chr5-28196085-INS-5021 | chr5-28196086-INS-5021 |
| chr5-28219888-INS-704 | chr5-28219888-INS-704 |
| chr5-28224849-INS-449 | chr5-28224849-INS-449 |
| chr5-28245246-INS-52 | chr5-28245246-INS-52 |
| chr5-28269892-INS-108 | chr5-28269892-INS-60 |
| chr5-28271278-INS-147 | chr5-28271278-INS-147 |
| chr5-28289592-INS-274 | chr5-28289592-INS-274 |
| chr5-28337100-INS-95 | chr5-28337100-INS-95 |
| chr5-28352411-INS-74 | chr5-28352411-INS-74 |
| chr5-28372467-INS-55 | chr5-28372465-INS-52 |
| chr5-28383150-INS-143 | chr5-28383150-INS-129 |
| chr5-28387224-INS-62 | chr5-28387222-INS-64 |
| chr5-28451646-INS-203 | chr5-28451646-INS-202 |
| chr5-28469484-INS-259 | chr5-28469484-INS-253 |
| chr5-28477513-INS-245 | chr5-28477513-INS-245 |
| chr5-28487578-INS-777 | chr5-28487578-INS-777 |
| chr5-28502934-INS-97 | chr5-28502934-INS-76 |
| chr5-28509403-INS-128 | chr5-28509403-INS-128 |
| chr5-28528613-INS-201 | chr5-28528613-INS-201 |
| chr5-28652223-INS-204 | chr5-28652223-INS-204 |
| chr5-28654412-INS-62 | chr5-28654412-INS-64 |
| chr5-28656276-INS-6915 | chr5-28656277-INS-6915 |
| chr5-28678062-INS-201 | chr5-28678062-INS-202 |
| chr5-28686663-INS-4483 | chr5-28686664-INS-4484 |
| chr5-28723338-INS-54 | chr5-28723338-INS-54 |
| chr5-28733899-INS-175 | chr5-28733885-INS-209 |
| chr5-28748852-INS-64 | chr5-28748852-INS-55 |
| chr5-28789262-INS-148 | chr5-28789262-INS-148 |
| chr5-28809293-INS-212 | chr5-28809293-INS-212 |
| chr5-28821792-INS-1682 | chr5-28821793-INS-1682 |
| chr5-28856869-INS-56 | chr5-28856863-INS-56 |
| chr5-28873149-INS-405 | chr5-28873149-INS-405 |
| chr5-28876178-INS-76 | chr5-28876178-INS-76 |
| chr5-28903151-INS-401 | chr5-28903151-INS-401 |
| chr5-28932682-INS-52 | chr5-28932682-INS-54 |
| chr5-28959251-INS-2734 | chr5-28959252-INS-2734 |

| | |
|---|---|
| chr5-29006674-INS-171 | chr5-29006674-INS-171 |
| chr5-29093230-INS-55 | chr5-29093231-INS-56 |
| chr5-29114898-INS-649 | chr5-29114898-INS-649 |
| chr5-29115559-INS-60 | chr5-29115559-INS-60 |
| chr5-29211928-INS-62 | chr5-29211928-INS-62 |
| chr5-29218750-INS-6586 | chr5-29218751-INS-6586 |
| chr5-29223947-INS-55 | chr5-29223947-INS-55 |
| chr5-29226186-INS-383 | chr5-29226186-INS-384 |
| chr5-29254892-INS-101 | chr5-29254892-INS-101 |
| chr5-29258731-INS-70 | chr5-29258732-INS-108 |
| chr5-29268267-INS-7207 | chr5-29268268-INS-7207 |
| chr5-29273858-INS-440 | chr5-29273858-INS-440 |
| chr5-29310987-INS-71 | chr5-29310987-INS-62 |
| chr5-29313836-INS-177 | chr5-29313836-INS-177 |
| chr5-29327340-INS-7098 | chr5-29327341-INS-7098 |
| chr5-29336109-INS-373 | chr5-29336109-INS-373 |
| chr5-29362543-INS-192 | chr5-29362543-INS-192 |
| chr5-29381802-INS-212 | chr5-29381802-INS-212 |
| chr5-29489066-INS-58 | chr5-29489066-INS-64 |
| chr5-29502507-INS-6592 | chr5-29502508-INS-6592 |
| chr5-29516490-INS-60 | chr5-29516490-INS-60 |
| chr5-29539181-INS-260 | chr5-29539181-INS-248 |
| chr5-29542923-INS-54 | chr5-29542915-INS-63 |
| chr5-29553061-INS-166 | chr5-29553061-INS-144 |
| chr5-29554104-INS-196 | chr5-29554104-INS-195 |
| chr5-29587112-INS-234 | chr5-29587112-INS-235 |
| chr5-29618526-INS-361 | chr5-29618526-INS-364 |
| chr5-29622116-INS-197 | chr5-29622116-INS-200 |
| chr5-29625495-INS-118 | chr5-29625495-INS-118 |
| chr5-29700395-INS-52 | chr5-29700352-INS-76 |
| chr5-29739241-INS-188 | chr5-29739241-INS-183 |
| chr5-29741873-INS-131 | chr5-29741847-INS-160 |
| chr5-29762771-INS-215 | chr5-29762771-INS-215 |
| chr5-29764871-INS-616 | chr5-29764871-INS-616 |
| chr5-29769058-INS-7455 | chr5-29769059-INS-7455 |
| chr5-29813566-INS-1026 | chr5-29813567-INS-1008 |
| chr5-29895582-INS-8986 | chr5-29895583-INS-8986 |
| chr5-29896777-INS-178 | chr5-29896777-INS-177 |
| chr5-29910855-INS-99 | chr5-29910853-INS-99 |
| chr5-29947783-INS-134 | chr5-29947784-INS-131 |
| chr5-29997462-INS-210 | chr5-29997462-INS-210 |
| chr5-30001178-INS-288 | chr5-30001178-INS-283 |
| chr5-30008613-INS-60 | chr5-30008613-INS-60 |
| chr5-30017915-INS-96 | chr5-30017915-INS-96 |
| chr5-30059398-INS-340 | chr5-30059398-INS-341 |
| chr5-30092863-INS-400 | chr5-30092863-INS-400 |
| chr5-30096604-INS-178 | chr5-30096604-INS-178 |
| chr5-30099238-INS-85 | chr5-30099238-INS-84 |
| chr5-30130875-INS-70 | chr5-30130875-INS-70 |
| chr5-30131796-INS-184 | chr5-30131796-INS-184 |
| chr5-30138644-INS-412 | chr5-30138644-INS-412 |
| chr5-30147037-INS-200 | chr5-30147037-INS-205 |
| chr5-30167029-INS-237 | chr5-30167029-INS-249 |
| chr5-30168048-INS-206 | chr5-30168048-INS-200 |
| chr5-30192168-INS-77 | chr5-30192168-INS-93 |
| chr5-30196437-INS-99 | chr5-30196416-INS-77 |
| chr5-30225280-INS-184 | chr5-30225280-INS-182 |
| chr5-30245323-INS-53 | chr5-30245323-INS-59 |
| chr5-30252824-INS-202 | chr5-30252824-INS-203 |
| chr5-30253806-INS-180 | chr5-30253806-INS-180 |
| chr5-30264281-INS-7127 | chr5-30264282-INS-7127 |
| chr5-30280007-INS-221 | chr5-30280007-INS-220 |
| chr5-30284035-INS-430 | chr5-30284035-INS-431 |
| chr5-30289751-INS-58 | chr5-30289751-INS-54 |
| chr5-30292234-INS-5056 | chr5-30292235-INS-5056 |
| chr5-30302269-INS-274 | chr5-30302269-INS-274 |
| chr5-30352725-INS-65 | chr5-30352714-INS-69 |
| chr5-30376023-INS-76 | chr5-30376023-INS-56 |
| chr5-30381623-INS-60 | chr5-30381623-INS-69 |
| chr5-30383931-INS-207 | chr5-30383931-INS-200 |
| chr5-30392261-INS-199 | chr5-30392261-INS-199 |
| chr5-30411764-INS-8734 | chr5-30411765-INS-8734 |
| chr5-30414248-INS-207 | chr5-30414248-INS-207 |
| chr5-30434509-INS-118 | chr5-30434509-INS-75 |
| chr5-30502281-INS-63 | chr5-30502281-INS-69 |
| chr5-30507211-INS-201 | chr5-30507197-INS-202 |
| chr5-30512699-INS-54 | chr5-30512670-INS-72 |

| | |
|---|---|
| chr5-30516359-INS-223 | chr5-30516359-INS-216 |
| chr5-30522739-INS-204 | chr5-30522738-INS-194 |
| chr5-30525777-INS-88 | chr5-30525826-INS-77 |
| chr5-30534958-INS-264 | chr5-30534958-INS-264 |
| chr5-30551616-INS-66 | chr5-30551616-INS-66 |
| chr5-30569791-INS-314 | chr5-30569791-INS-314 |
| chr5-30583007-INS-58 | chr5-30583007-INS-58 |
| chr5-30613040-INS-75 | chr5-30613040-INS-75 |
| chr5-30615853-INS-70 | chr5-30615853-INS-70 |
| chr5-30641782-INS-237 | chr5-30641782-INS-246 |
| chr5-30655145-INS-148 | chr5-30655145-INS-147 |
| chr5-30665780-INS-199 | chr5-30665780-INS-207 |
| chr5-30680100-INS-883 | chr5-30680101-INS-886 |
| chr5-30682927-INS-134 | chr5-30682927-INS-134 |
| chr5-30714731-INS-72 | chr5-30714731-INS-72 |
| chr5-30758388-INS-55 | chr5-30758388-INS-52 |
| chr5-30790824-INS-100 | chr5-30790830-INS-56 |
| chr5-30794558-INS-64 | chr5-30794558-INS-85 |
| chr5-30844499-INS-105 | chr5-30844499-INS-89 |
| chr5-30852565-INS-7471 | chr5-30852566-INS-7471 |
| chr5-30865856-INS-55 | chr5-30865856-INS-55 |
| chr5-30886521-INS-56 | chr5-30886521-INS-52 |
| chr5-30889064-INS-68 | chr5-30889064-INS-68 |
| chr5-30893381-INS-84 | chr5-30893381-INS-99 |
| chr5-30901897-INS-413 | chr5-30901898-INS-409 |
| chr5-30914033-INS-137 | chr5-30914033-INS-137 |
| chr5-30915318-INS-52 | chr5-30915318-INS-53 |
| chr5-31041251-INS-230 | chr5-31041261-INS-231 |
| chr5-31095793-INS-197 | chr5-31095793-INS-199 |
| chr5-31108227-INS-421 | chr5-31108227-INS-418 |
| chr5-31114330-INS-179 | chr5-31114330-INS-202 |
| chr5-31119733-INS-208 | chr5-31119733-INS-208 |
| chr5-31133869-INS-430 | chr5-31133869-INS-430 |
| chr5-31142872-INS-61 | chr5-31142872-INS-62 |
| chr5-31150261-INS-142 | chr5-31150261-INS-163 |
| chr5-31158377-INS-227 | chr5-31158377-INS-228 |
| chr5-31158886-INS-755 | chr5-31158886-INS-755 |
| chr5-31173663-INS-161 | chr5-31173663-INS-161 |
| chr5-31186654-INS-202 | chr5-31186654-INS-201 |
| chr5-31199374-INS-123 | chr5-31199374-INS-123 |
| chr5-31207809-INS-176 | chr5-31207809-INS-176 |
| chr5-31220232-INS-110 | chr5-31220226-INS-118 |
| chr5-31221421-INS-217 | chr5-31221421-INS-217 |
| chr5-31223289-INS-247 | chr5-31223289-INS-243 |
| chr5-31238757-INS-180 | chr5-31238757-INS-180 |
| chr5-31240042-INS-59 | chr5-31240042-INS-57 |
| chr5-31241206-INS-54 | chr5-31241206-INS-56 |
| chr5-31251410-INS-74 | chr5-31251410-INS-74 |
| chr5-31262583-INS-64 | chr5-31262583-INS-64 |
| chr5-31277987-INS-570 | chr5-31277987-INS-570 |
| chr5-31301623-INS-216 | chr5-31301623-INS-216 |
| chr5-31367277-INS-189 | chr5-31367277-INS-189 |
| chr5-31381301-INS-203 | chr5-31381301-INS-203 |
| chr5-31387114-INS-217 | chr5-31387114-INS-217 |
| chr5-31394412-INS-165 | chr5-31394412-INS-165 |
| chr5-31432347-INS-60 | chr5-31432347-INS-54 |
| chr5-31446853-INS-110 | chr5-31446853-INS-110 |
| chr5-31457802-INS-177 | chr5-31457802-INS-177 |
| chr5-31468982-INS-121 | chr5-31468985-INS-116 |
| chr5-31511531-INS-197 | chr5-31511531-INS-197 |
| chr5-31522585-INS-6488 | chr5-31522586-INS-6488 |
| chr5-31525616-INS-3908 | chr5-31525617-INS-3908 |
| chr5-31545132-INS-6530 | chr5-31545133-INS-6530 |
| chr5-31560406-INS-213 | chr5-31560453-INS-213 |
| chr5-31561760-INS-274 | chr5-31561760-INS-274 |
| chr5-31647706-INS-203 | chr5-31647706-INS-205 |
| chr5-31660312-INS-175 | chr5-31660327-INS-172 |
| chr5-31662403-INS-216 | chr5-31662403-INS-216 |
| chr5-31668616-INS-51 | chr5-31668616-INS-51 |
| chr5-31670402-INS-307 | chr5-31670402-INS-307 |
| chr5-31679890-INS-483 | chr5-31679890-INS-483 |
| chr5-31680715-INS-213 | chr5-31680715-INS-212 |
| chr5-31681389-INS-66 | chr5-31681389-INS-66 |
| chr5-31693452-INS-52 | chr5-31693452-INS-52 |
| chr5-31703891-INS-53 | chr5-31703891-INS-53 |
| chr5-31713692-INS-263 | chr5-31713692-INS-262 |
| chr5-31729523-INS-67 | chr5-31729523-INS-67 |

| | |
|---|---|
| chr5-31732354-INS-80 | chr5-31732354-INS-68 |
| chr5-31746026-INS-75 | chr5-31746026-INS-75 |
| chr5-31755689-INS-216 | chr5-31755689-INS-213 |
| chr5-31766681-INS-212 | chr5-31766681-INS-214 |
| chr5-31783363-INS-215 | chr5-31783363-INS-215 |
| chr5-31787248-INS-79 | chr5-31787248-INS-52 |
| chr5-31797460-INS-200 | chr5-31797460-INS-200 |
| chr5-31804372-INS-95 | chr5-31804372-INS-89 |
| chr5-31816511-INS-5413 | chr5-31816512-INS-5413 |
| chr5-31825468-INS-129 | chr5-31825468-INS-129 |
| chr5-31839205-INS-211 | chr5-31839205-INS-210 |
| chr5-31868076-INS-198 | chr5-31868076-INS-198 |
| chr5-31881436-INS-4390 | chr5-31881437-INS-4391 |
| chr5-31906215-INS-53 | chr5-31906215-INS-52 |
| chr5-31909881-INS-99 | chr5-31909872-INS-96 |
| chr5-31961676-INS-108 | chr5-31961676-INS-108 |
| chr5-32001055-INS-92 | chr5-32001099-INS-76 |
| chr5-32065328-INS-441 | chr5-32065328-INS-441 |
| chr5-32069940-INS-144 | chr5-32069940-INS-147 |
| chr5-32093419-INS-6436 | chr5-32093420-INS-6436 |
| chr5-32097867-INS-1014 | chr5-32097868-INS-1014 |
| chr5-32146261-INS-83 | chr5-32146261-INS-83 |
| chr5-32185051-INS-84 | chr5-32185051-INS-84 |
| chr5-32328004-INS-2600 | chr5-32328005-INS-2600 |
| chr5-32368976-INS-80 | chr5-32368995-INS-68 |
| chr5-32385540-INS-92 | chr5-32385540-INS-92 |
| chr5-32387333-INS-6444 | chr5-32387334-INS-6444 |
| chr5-32394071-INS-83 | chr5-32394071-INS-90 |
| chr5-32408387-INS-6461 | chr5-32408388-INS-6461 |
| chr5-32408475-INS-98 | chr5-32408475-INS-90 |
| chr5-32411934-INS-198 | chr5-32411893-INS-217 |
| chr5-32441833-INS-117 | chr5-32441833-INS-118 |
| chr5-32445674-INS-1110 | chr5-32445675-INS-1109 |
| chr5-32490439-INS-265 | chr5-32490439-INS-265 |
| chr5-32533641-INS-188 | chr5-32533641-INS-189 |
| chr5-32536991-INS-115 | chr5-32536991-INS-115 |
| chr5-32538202-INS-246 | chr5-32538202-INS-246 |
| chr5-32579793-INS-66 | chr5-32579793-INS-66 |
| chr5-32607489-INS-726 | chr5-32607489-INS-726 |
| chr5-32612957-INS-72 | chr5-32612957-INS-72 |
| chr5-32647009-INS-164 | chr5-32647009-INS-164 |
| chr5-32655577-INS-400 | chr5-32655577-INS-400 |
| chr5-32661307-INS-377 | chr5-32661307-INS-377 |
| chr5-32667137-INS-494 | chr5-32667137-INS-494 |
| chr5-32678870-INS-970 | chr5-32678870-INS-970 |
| chr5-32688140-INS-127 | chr5-32688140-INS-98 |
| chr5-32689964-INS-66 | chr5-32689964-INS-66 |
| chr5-32743876-INS-64 | chr5-32743876-INS-56 |
| chr5-32748444-INS-90 | chr5-32748449-INS-72 |
| chr5-32749912-INS-178 | chr5-32749912-INS-178 |
| chr5-32753734-INS-213 | chr5-32753734-INS-213 |
| chr5-32781808-INS-172 | chr5-32781808-INS-171 |
| chr5-32806566-INS-58 | chr5-32806566-INS-58 |
| chr5-32814123-INS-219 | chr5-32814123-INS-219 |
| chr5-32852609-INS-340 | chr5-32852609-INS-340 |
| chr5-32867141-INS-544 | chr5-32867141-INS-544 |
| chr5-32870947-INS-76 | chr5-32870947-INS-76 |
| chr5-32877127-INS-77 | chr5-32877127-INS-77 |
| chr5-32924808-INS-210 | chr5-32924808-INS-210 |
| chr5-32929720-INS-76 | chr5-32929720-INS-67 |
| chr5-32933852-INS-56 | chr5-32933852-INS-52 |
| chr5-32937467-INS-5499 | chr5-32937468-INS-5499 |
| chr5-32942212-INS-211 | chr5-32942212-INS-211 |
| chr5-32944325-INS-211 | chr5-32944325-INS-205 |
| chr5-32952026-INS-224 | chr5-32952026-INS-224 |
| chr5-32982225-INS-186 | chr5-32982221-INS-236 |
| chr5-32997839-INS-202 | chr5-32997839-INS-202 |
| chr5-33012210-INS-289 | chr5-33012201-INS-254 |
| chr5-33014732-INS-191 | chr5-33014732-INS-191 |
| chr5-33017808-INS-314 | chr5-33017808-INS-319 |
| chr5-33035258-INS-121 | chr5-33035258-INS-121 |
| chr5-33041153-INS-52 | chr5-33041167-INS-77 |
| chr5-33054953-INS-481 | chr5-33054953-INS-490 |
| chr5-33064742-INS-279 | chr5-33064741-INS-312 |
| chr5-33065748-INS-198 | chr5-33065748-INS-198 |
| chr5-33088672-INS-382 | chr5-33088672-INS-382 |
| chr5-33104500-INS-211 | chr5-33104500-INS-213 |

| | |
|---|---|
| chr5-33122803-INS-145 | chr5-33122803-INS-151 |
| chr5-33140393-INS-219 | chr5-33140393-INS-220 |
| chr5-33192446-INS-221 | chr5-33192446-INS-223 |
| chr5-33237542-INS-211 | chr5-33237542-INS-211 |
| chr5-33239461-INS-139 | chr5-33239453-INS-120 |
| chr5-33255463-INS-73 | chr5-33255463-INS-73 |
| chr5-33269319-INS-110 | chr5-33269303-INS-84 |
| chr5-33270292-INS-105 | chr5-33270292-INS-95 |
| chr5-33296299-INS-51 | chr5-33296283-INS-52 |
| chr5-33297912-INS-215 | chr5-33297912-INS-214 |
| chr5-33304792-INS-51 | chr5-33304756-INS-54 |
| chr5-33337965-INS-52 | chr5-33337965-INS-52 |
| chr5-33384929-INS-243 | chr5-33384929-INS-246 |
| chr5-33391535-INS-214 | chr5-33391535-INS-214 |
| chr5-33461133-INS-60 | chr5-33461133-INS-60 |
| chr5-33463866-INS-60 | chr5-33463866-INS-60 |
| chr5-33490931-INS-3269 | chr5-33490932-INS-3269 |
| chr5-33509012-INS-2582 | chr5-33509013-INS-2583 |
| chr5-33534241-INS-498 | chr5-33534241-INS-498 |
| chr5-33548041-INS-60 | chr5-33548041-INS-60 |
| chr5-33574922-INS-120 | chr5-33574922-INS-90 |
| chr5-33636026-INS-889 | chr5-33636026-INS-889 |
| chr5-33651247-INS-59 | chr5-33651286-INS-59 |
| chr5-33709689-INS-333 | chr5-33709690-INS-339 |
| chr5-33734432-INS-81 | chr5-33734438-INS-54 |
| chr5-33736369-INS-743 | chr5-33736369-INS-711 |
| chr5-33744442-INS-220 | chr5-33744442-INS-198 |
| chr5-33748451-INS-3304 | chr5-33748452-INS-3304 |
| chr5-33750094-INS-199 | chr5-33750094-INS-198 |
| chr5-33751889-INS-207 | chr5-33751889-INS-222 |
| chr5-33775393-INS-209 | chr5-33775393-INS-209 |
| chr5-33801682-INS-192 | chr5-33801691-INS-168 |
| chr5-33802679-INS-204 | chr5-33802679-INS-204 |
| chr5-33838096-INS-79 | chr5-33838096-INS-79 |
| chr5-33854336-INS-206 | chr5-33854336-INS-229 |
| chr5-33920309-INS-562 | chr5-33920309-INS-563 |
| chr5-33940841-INS-62 | chr5-33940844-INS-80 |
| chr5-34101612-INS-56 | chr5-34101612-INS-56 |
| chr5-34150438-INS-1892 | chr5-34150439-INS-1892 |
| chr5-34157741-INS-177 | chr5-34157741-INS-177 |
| chr5-34197237-INS-66 | chr5-34197237-INS-66 |
| chr5-34205455-INS-214 | chr5-34205455-INS-214 |
| chr5-34219893-INS-51 | chr5-34219893-INS-52 |
| chr5-34239094-INS-56 | chr5-34239106-INS-55 |
| chr5-34241686-INS-76 | chr5-34241684-INS-82 |
| chr5-34244167-INS-6476 | chr5-34244168-INS-6477 |
| chr5-34246211-INS-63 | chr5-34246211-INS-81 |
| chr5-34253336-INS-179 | chr5-34253336-INS-178 |
| chr5-34266440-INS-355 | chr5-34266440-INS-355 |
| chr5-34287745-INS-137 | chr5-34287745-INS-135 |
| chr5-34296666-INS-148 | chr5-34296666-INS-148 |
| chr5-34329165-INS-7062 | chr5-34329166-INS-7050 |
| chr5-34370631-INS-64 | chr5-34370623-INS-62 |
| chr5-34379496-INS-115 | chr5-34379496-INS-116 |
| chr5-34385112-INS-96 | chr5-34385112-INS-98 |
| chr5-34409116-INS-206 | chr5-34409116-INS-205 |
| chr5-34416977-INS-175 | chr5-34416977-INS-171 |
| chr5-34418285-INS-849 | chr5-34418285-INS-863 |
| chr5-34498026-INS-242 | chr5-34498027-INS-207 |
| chr5-34523900-INS-52 | chr5-34523906-INS-50 |
| chr5-34532272-INS-351 | chr5-34532272-INS-351 |
| chr5-34545747-INS-3626 | chr5-34545748-INS-3626 |
| chr5-34554047-INS-130 | chr5-34554047-INS-130 |
| chr5-34682990-INS-67 | chr5-34682972-INS-76 |
| chr5-34705563-INS-200 | chr5-34705563-INS-201 |
| chr5-34726683-INS-62 | chr5-34726683-INS-66 |
| chr5-34729449-INS-173 | chr5-34729449-INS-173 |
| chr5-34740029-INS-211 | chr5-34740029-INS-211 |
| chr5-34750430-INS-56 | chr5-34750430-INS-56 |
| chr5-34756452-INS-209 | chr5-34756452-INS-210 |
| chr5-34759255-INS-128 | chr5-34759255-INS-128 |
| chr5-34760222-INS-77 | chr5-34760222-INS-77 |
| chr5-34768923-INS-205 | chr5-34768923-INS-200 |
| chr5-34848845-INS-562 | chr5-34848845-INS-562 |
| chr5-34853641-INS-6475 | chr5-34853642-INS-6475 |
| chr5-34874550-INS-75 | chr5-34874547-INS-60 |
| chr5-34914157-INS-213 | chr5-34914157-INS-214 |

| | |
|---|---|
| chr5-35026785-INS-962 | chr5-35026785-INS-962 |
| chr5-35070354-INS-6403 | chr5-35070355-INS-6403 |
| chr5-35095515-INS-220 | chr5-35095515-INS-220 |
| chr5-35098737-INS-53 | chr5-35098737-INS-53 |
| chr5-35109912-INS-52 | chr5-35109912-INS-52 |
| chr5-35114280-INS-70 | chr5-35114280-INS-70 |
| chr5-35129283-INS-202 | chr5-35129283-INS-202 |
| chr5-35163481-INS-544 | chr5-35163481-INS-544 |
| chr5-35195422-INS-64 | chr5-35195422-INS-64 |
| chr5-35273607-INS-199 | chr5-35273607-INS-199 |
| chr5-35276899-INS-409 | chr5-35276899-INS-409 |
| chr5-35278867-INS-184 | chr5-35278867-INS-197 |
| chr5-35300161-INS-52 | chr5-35300161-INS-52 |
| chr5-35326000-INS-83 | chr5-35326000-INS-83 |
| chr5-35327306-INS-109 | chr5-35327306-INS-113 |
| chr5-35344142-INS-77 | chr5-35344123-INS-50 |
| chr5-35379353-INS-89 | chr5-35379353-INS-91 |
| chr5-35399316-INS-414 | chr5-35399316-INS-414 |
| chr5-35416831-INS-76 | chr5-35416831-INS-76 |
| chr5-35445167-INS-142 | chr5-35445167-INS-142 |
| chr5-35478484-INS-201 | chr5-35478485-INS-201 |
| chr5-35479129-INS-131 | chr5-35479130-INS-131 |
| chr5-35491244-INS-167 | chr5-35491244-INS-161 |
| chr5-35514807-INS-176 | chr5-35514807-INS-176 |
| chr5-35581125-INS-1084 | chr5-35581126-INS-1084 |
| chr5-35581981-INS-354 | chr5-35581981-INS-354 |
| chr5-35590583-INS-54 | chr5-35590583-INS-54 |
| chr5-35604333-INS-187 | chr5-35604333-INS-186 |
| chr5-35635624-INS-64 | chr5-35635624-INS-64 |
| chr5-35649790-INS-68 | chr5-35649790-INS-118 |
| chr5-35701668-INS-158 | chr5-35701666-INS-151 |
| chr5-35702774-INS-346 | chr5-35702774-INS-339 |
| chr5-35718098-INS-136 | chr5-35718098-INS-136 |
| chr5-35733300-INS-255 | chr5-35733300-INS-223 |
| chr5-35748551-INS-1257 | chr5-35748552-INS-1257 |
| chr5-35776179-INS-62 | chr5-35776179-INS-82 |
| chr5-35784713-INS-56 | chr5-35784713-INS-56 |
| chr5-35786706-INS-52 | chr5-35786706-INS-54 |
| chr5-35797314-INS-101 | chr5-35797314-INS-110 |
| chr5-35812068-INS-131 | chr5-35812068-INS-132 |
| chr5-35840343-INS-208 | chr5-35840343-INS-208 |
| chr5-35849859-INS-122 | chr5-35849855-INS-128 |
| chr5-35857318-INS-65 | chr5-35857318-INS-65 |
| chr5-35862029-INS-807 | chr5-35862029-INS-807 |
| chr5-35879784-INS-77 | chr5-35879778-INS-111 |
| chr5-35887172-INS-56 | chr5-35887191-INS-64 |
| chr5-35925911-INS-231 | chr5-35925911-INS-231 |
| chr5-35995252-INS-161 | chr5-35995252-INS-162 |
| chr5-36010706-INS-283 | chr5-36010706-INS-283 |
| chr5-36016717-INS-160 | chr5-36016718-INS-157 |
| chr5-36042978-INS-64 | chr5-36042979-INS-72 |
| chr5-36075078-INS-1607 | chr5-36075079-INS-1602 |
| chr5-36087609-INS-335 | chr5-36087609-INS-334 |
| chr5-36096861-INS-52 | chr5-36096845-INS-60 |
| chr5-36129436-INS-53 | chr5-36129429-INS-53 |
| chr5-36135639-INS-52 | chr5-36135639-INS-54 |
| chr5-36148932-INS-144 | chr5-36148932-INS-162 |
| chr5-36200144-INS-64 | chr5-36200144-INS-68 |
| chr5-36200676-INS-76 | chr5-36200676-INS-76 |
| chr5-36205874-INS-440 | chr5-36205874-INS-440 |
| chr5-36223403-INS-140 | chr5-36223403-INS-140 |
| chr5-36271094-INS-97 | chr5-36271094-INS-104 |
| chr5-36280051-INS-7076 | chr5-36280052-INS-7076 |
| chr5-36281372-INS-54 | chr5-36281372-INS-54 |
| chr5-36327489-INS-6440 | chr5-36327490-INS-6440 |
| chr5-36334731-INS-248 | chr5-36334731-INS-248 |
| chr5-36340580-INS-54 | chr5-36340580-INS-54 |
| chr5-36343145-INS-54 | chr5-36343159-INS-57 |
| chr5-36345516-INS-110 | chr5-36345516-INS-110 |
| chr5-36373300-INS-232 | chr5-36373300-INS-234 |
| chr5-36447652-INS-639 | chr5-36447652-INS-639 |
| chr5-36453488-INS-216 | chr5-36453488-INS-216 |
| chr5-36465949-INS-142 | chr5-36465949-INS-142 |
| chr5-36476068-INS-238 | chr5-36476068-INS-252 |
| chr5-36573426-INS-156 | chr5-36573426-INS-156 |
| chr5-36576666-INS-200 | chr5-36576666-INS-196 |
| chr5-36591423-INS-64 | chr5-36591423-INS-64 |

| | |
|---|---|
| chr5-36604814-INS-388 | chr5-36604814-INS-388 |
| chr5-36612515-INS-7070 | chr5-36612516-INS-7022 |
| chr5-36614462-INS-285 | chr5-36614463-INS-285 |
| chr5-36617302-INS-143 | chr5-36617296-INS-150 |
| chr5-36661518-INS-6346 | chr5-36661519-INS-6346 |
| chr5-36679648-INS-276 | chr5-36679648-INS-276 |
| chr5-36686632-INS-6457 | chr5-36686633-INS-6457 |
| chr5-36696808-INS-88 | chr5-36696808-INS-88 |
| chr5-36751431-INS-203 | chr5-36751431-INS-204 |
| chr5-36784953-INS-996 | chr5-36784953-INS-997 |
| chr5-36828057-INS-60 | chr5-36828057-INS-64 |
| chr5-36865793-INS-70 | chr5-36865793-INS-70 |
| chr5-36884813-INS-59 | chr5-36884813-INS-59 |
| chr5-36911215-INS-56 | chr5-36911215-INS-58 |
| chr5-36928559-INS-68 | chr5-36928559-INS-64 |
| chr5-36934848-INS-101 | chr5-36934848-INS-103 |
| chr5-36947677-INS-52 | chr5-36947677-INS-52 |
| chr5-36950729-INS-488 | chr5-36950729-INS-488 |
| chr5-36983302-INS-61 | chr5-36983302-INS-61 |
| chr5-36989380-INS-82 | chr5-36989380-INS-82 |
| chr5-37016213-INS-205 | chr5-37016213-INS-205 |
| chr5-37054639-INS-114 | chr5-37054639-INS-114 |
| chr5-37060487-INS-55 | chr5-37060487-INS-55 |
| chr5-37063679-INS-115 | chr5-37063679-INS-117 |
| chr5-37114242-INS-59 | chr5-37114242-INS-59 |
| chr5-37119278-INS-212 | chr5-37119278-INS-212 |
| chr5-37137588-INS-142 | chr5-37137588-INS-141 |
| chr5-37148724-INS-193 | chr5-37148724-INS-193 |
| chr5-37159372-INS-65 | chr5-37159372-INS-85 |
| chr5-37176885-INS-56 | chr5-37176885-INS-53 |
| chr5-37219018-INS-160 | chr5-37219018-INS-164 |
| chr5-37220111-INS-197 | chr5-37220111-INS-197 |
| chr5-37231242-INS-6855 | chr5-37231243-INS-6857 |
| chr5-37244194-INS-220 | chr5-37244230-INS-216 |
| chr5-37247492-INS-223 | chr5-37247492-INS-223 |
| chr5-37258595-INS-84 | chr5-37258595-INS-86 |
| chr5-37261389-INS-90 | chr5-37261389-INS-90 |
| chr5-37267280-INS-66 | chr5-37267280-INS-66 |
| chr5-37333458-INS-163 | chr5-37333458-INS-163 |
| chr5-37361472-INS-208 | chr5-37361472-INS-208 |
| chr5-37412893-INS-181 | chr5-37412893-INS-182 |
| chr5-37418461-INS-54 | chr5-37418461-INS-54 |
| chr5-37426266-INS-97 | chr5-37426266-INS-97 |
| chr5-37427277-INS-97 | chr5-37427277-INS-94 |
| chr5-37435980-INS-68 | chr5-37435980-INS-68 |
| chr5-37460423-INS-60 | chr5-37460423-INS-106 |
| chr5-37488985-INS-873 | chr5-37488985-INS-873 |
| chr5-37507873-INS-55 | chr5-37507873-INS-60 |
| chr5-37515704-INS-129 | chr5-37515745-INS-130 |
| chr5-37520299-INS-62 | chr5-37520316-INS-50 |
| chr5-37530516-INS-76 | chr5-37530535-INS-91 |
| chr5-37539886-INS-58 | chr5-37539886-INS-58 |
| chr5-37543275-INS-88 | chr5-37543275-INS-88 |
| chr5-37565222-INS-362 | chr5-37565222-INS-370 |
| chr5-37592643-INS-52 | chr5-37592648-INS-56 |
| chr5-37599703-INS-53 | chr5-37599703-INS-53 |
| chr5-37617859-INS-80 | chr5-37617859-INS-80 |
| chr5-37642721-INS-112 | chr5-37642721-INS-107 |
| chr5-37691368-INS-285 | chr5-37691368-INS-285 |
| chr5-37693593-INS-928 | chr5-37693593-INS-931 |
| chr5-37702992-INS-151 | chr5-37702992-INS-141 |
| chr5-37724258-INS-398 | chr5-37724258-INS-398 |
| chr5-37802912-INS-765 | chr5-37802912-INS-754 |
| chr5-37811360-INS-58 | chr5-37811336-INS-64 |
| chr5-37841980-INS-68 | chr5-37841980-INS-68 |
| chr5-37881530-INS-142 | chr5-37881530-INS-142 |
| chr5-37890671-INS-176 | chr5-37890651-INS-182 |
| chr5-37893842-INS-132 | chr5-37893842-INS-132 |
| chr5-37918499-INS-54 | chr5-37918499-INS-55 |
| chr5-37936584-INS-54 | chr5-37936584-INS-54 |
| chr5-37937307-INS-85 | chr5-37937307-INS-85 |
| chr5-37945500-INS-408 | chr5-37945500-INS-408 |
| chr5-37997340-INS-483 | chr5-37997340-INS-479 |
| chr5-38001731-INS-56 | chr5-38001731-INS-68 |
| chr5-38010867-INS-138 | chr5-38010856-INS-171 |
| chr5-38024032-INS-5355 | chr5-38024033-INS-5355 |
| chr5-38030933-INS-87 | chr5-38030933-INS-87 |

| | |
|---|---|
| chr5-38045257-INS-66 | chr5-38045257-INS-50 |
| chr5-38047551-INS-55 | chr5-38047541-INS-83 |
| chr5-38099308-INS-220 | chr5-38099308-INS-249 |
| chr5-38126940-INS-222 | chr5-38126940-INS-222 |
| chr5-38135756-INS-196 | chr5-38135756-INS-196 |
| chr5-38144402-INS-433 | chr5-38144402-INS-433 |
| chr5-38155349-INS-127 | chr5-38155349-INS-127 |
| chr5-38165494-INS-71 | chr5-38165497-INS-55 |
| chr5-38207561-INS-204 | chr5-38207561-INS-203 |
| chr5-38250708-INS-171 | chr5-38250708-INS-188 |
| chr5-38266889-INS-73 | chr5-38266889-INS-73 |
| chr5-38326682-INS-109 | chr5-38326648-INS-107 |
| chr5-38346789-INS-55 | chr5-38346754-INS-60 |
| chr5-38353600-INS-62 | chr5-38353600-INS-64 |
| chr5-38355640-INS-64 | chr5-38355640-INS-64 |
| chr5-38375997-INS-498 | chr5-38375997-INS-498 |
| chr5-38382401-INS-75 | chr5-38382405-INS-75 |
| chr5-38396481-INS-325 | chr5-38396481-INS-332 |
| chr5-38397194-INS-236 | chr5-38397194-INS-237 |
| chr5-38400100-INS-51 | chr5-38400064-INS-58 |
| chr5-38403441-INS-386 | chr5-38403441-INS-386 |
| chr5-38420049-INS-88 | chr5-38420049-INS-88 |
| chr5-38452195-INS-58 | chr5-38452195-INS-58 |
| chr5-38455574-INS-167 | chr5-38455574-INS-168 |
| chr5-38481855-INS-6475 | chr5-38481856-INS-6475 |
| chr5-38517828-INS-94 | chr5-38517839-INS-62 |
| chr5-38528592-INS-54 | chr5-38528592-INS-60 |
| chr5-38529128-INS-68 | chr5-38529128-INS-96 |
| chr5-38530969-INS-92 | chr5-38530994-INS-68 |
| chr5-38531700-INS-66 | chr5-38531700-INS-66 |
| chr5-38532794-INS-63 | chr5-38532794-INS-63 |
| chr5-38540820-INS-180 | chr5-38540820-INS-180 |
| chr5-38593427-INS-258 | chr5-38593427-INS-259 |
| chr5-38620983-INS-206 | chr5-38620983-INS-206 |
| chr5-38640224-INS-57 | chr5-38640224-INS-57 |
| chr5-38693700-INS-101 | chr5-38693700-INS-101 |
| chr5-38720965-INS-4538 | chr5-38720966-INS-4538 |
| chr5-38730820-INS-54 | chr5-38730841-INS-54 |
| chr5-38737614-INS-321 | chr5-38737614-INS-311 |
| chr5-38738069-INS-388 | chr5-38738069-INS-388 |
| chr5-38761602-INS-397 | chr5-38761602-INS-397 |
| chr5-38764802-INS-184 | chr5-38764802-INS-184 |
| chr5-38778659-INS-173 | chr5-38778659-INS-191 |
| chr5-38780507-INS-5476 | chr5-38780508-INS-5476 |
| chr5-38788556-INS-124 | chr5-38788556-INS-127 |
| chr5-38793468-INS-228 | chr5-38793468-INS-227 |
| chr5-38794342-INS-65 | chr5-38794342-INS-83 |
| chr5-38798381-INS-135 | chr5-38798381-INS-107 |
| chr5-38816935-INS-63 | chr5-38816935-INS-63 |
| chr5-38822125-INS-785 | chr5-38822125-INS-785 |
| chr5-38847667-INS-167 | chr5-38847667-INS-160 |
| chr5-38867249-INS-107 | chr5-38867249-INS-112 |
| chr5-38871969-INS-197 | chr5-38871969-INS-197 |
| chr5-38886225-INS-199 | chr5-38886225-INS-192 |
| chr5-38888981-INS-72 | chr5-38888982-INS-75 |
| chr5-38909719-INS-97 | chr5-38909719-INS-97 |
| chr5-38916063-INS-193 | chr5-38916063-INS-193 |
| chr5-38923071-INS-55 | chr5-38923071-INS-71 |
| chr5-38936132-INS-415 | chr5-38936132-INS-415 |
| chr5-38955349-INS-165 | chr5-38955349-INS-169 |
| chr5-39002045-INS-108 | chr5-39002045-INS-63 |
| chr5-39006460-INS-452 | chr5-39006460-INS-452 |
| chr5-39040896-INS-400 | chr5-39040896-INS-400 |
| chr5-39042770-INS-640 | chr5-39042770-INS-654 |
| chr5-39045133-INS-60 | chr5-39045133-INS-64 |
| chr5-39066299-INS-114 | chr5-39066299-INS-121 |
| chr5-39081500-INS-94 | chr5-39081499-INS-94 |
| chr5-39105732-INS-338 | chr5-39105732-INS-337 |
| chr5-39108784-INS-83 | chr5-39108784-INS-83 |
| chr5-39128901-INS-99 | chr5-39128906-INS-99 |
| chr5-39138147-INS-53 | chr5-39138148-INS-63 |
| chr5-39156096-INS-4542 | chr5-39156097-INS-4542 |
| chr5-39162831-INS-88 | chr5-39162787-INS-131 |
| chr5-39166640-INS-54 | chr5-39166640-INS-52 |
| chr5-39216887-INS-376 | chr5-39216897-INS-406 |
| chr5-39245470-INS-80 | chr5-39245471-INS-78 |
| chr5-39251323-INS-401 | chr5-39251323-INS-401 |

| | |
|---|---|
| chr5-39252949-INS-676 | chr5-39252949-INS-680 |
| chr5-39276374-INS-270 | chr5-39276374-INS-268 |
| chr5-39323292-INS-6186 | chr5-39323293-INS-6186 |
| chr5-39338106-INS-146 | chr5-39338106-INS-158 |
| chr5-39343726-INS-272 | chr5-39343726-INS-270 |
| chr5-39359618-INS-967 | chr5-39359618-INS-967 |
| chr5-39378289-INS-72 | chr5-39378289-INS-72 |
| chr5-39407940-INS-645 | chr5-39407940-INS-645 |
| chr5-39418696-INS-52 | chr5-39418696-INS-54 |
| chr5-39479798-INS-86 | chr5-39479798-INS-121 |
| chr5-39502108-INS-1880 | chr5-39502108-INS-1871 |
| chr5-39504936-INS-738 | chr5-39504936-INS-738 |
| chr5-39507527-INS-1341 | chr5-39507528-INS-1341 |
| chr5-39513884-INS-332 | chr5-39513884-INS-332 |
| chr5-39514757-INS-570 | chr5-39514757-INS-573 |
| chr5-39535407-INS-57 | chr5-39535407-INS-57 |
| chr5-39548350-INS-74 | chr5-39548360-INS-98 |
| chr5-39552382-INS-629 | chr5-39552382-INS-629 |
| chr5-39561499-INS-528 | chr5-39561499-INS-528 |
| chr5-39603846-INS-81 | chr5-39603813-INS-71 |
| chr5-39630232-INS-780 | chr5-39630232-INS-780 |
| chr5-39650926-INS-912 | chr5-39650926-INS-910 |
| chr5-39655508-INS-422 | chr5-39655508-INS-423 |
| chr5-39670412-INS-62 | chr5-39670413-INS-66 |
| chr5-39693216-INS-62 | chr5-39693216-INS-62 |
| chr5-39709266-INS-7976 | chr5-39709267-INS-7978 |
| chr5-39713769-INS-123 | chr5-39713769-INS-123 |
| chr5-39717536-INS-2665 | chr5-39717537-INS-2663 |
| chr5-39757547-INS-54 | chr5-39757547-INS-56 |
| chr5-39841913-INS-508 | chr5-39841913-INS-508 |
| chr5-39846162-INS-60 | chr5-39846150-INS-74 |
| chr5-39905446-INS-58 | chr5-39905446-INS-56 |
| chr5-39908767-INS-575 | chr5-39908767-INS-564 |
| chr5-39921010-INS-1292 | chr5-39921011-INS-1293 |
| chr5-39942658-INS-201 | chr5-39942652-INS-202 |
| chr5-39964618-INS-51 | chr5-39964618-INS-51 |
| chr5-39965829-INS-55 | chr5-39965829-INS-55 |
| chr5-39985865-INS-132 | chr5-39985865-INS-132 |
| chr5-40019911-INS-123 | chr5-40019941-INS-124 |
| chr5-40036633-INS-2626 | chr5-40036634-INS-2621 |
| chr5-40039699-INS-174 | chr5-40039702-INS-177 |
| chr5-40102430-INS-56 | chr5-40102430-INS-60 |
| chr5-40109425-INS-499 | chr5-40109425-INS-499 |
| chr5-40111303-INS-58 | chr5-40111303-INS-56 |
| chr5-40131423-INS-54 | chr5-40131423-INS-54 |
| chr5-40173962-INS-54 | chr5-40173962-INS-66 |
| chr5-40180125-INS-6593 | chr5-40180126-INS-6593 |
| chr5-40222591-INS-54 | chr5-40222591-INS-52 |
| chr5-40252625-INS-1696 | chr5-40252626-INS-1695 |
| chr5-40260886-INS-152 | chr5-40260886-INS-187 |
| chr5-40300960-INS-139 | chr5-40300960-INS-139 |
| chr5-40307297-INS-1588 | chr5-40307298-INS-1588 |
| chr5-40310519-INS-4060 | chr5-40310515-INS-4064 |
| chr5-40325802-INS-194 | chr5-40325802-INS-194 |
| chr5-40332109-INS-72 | chr5-40332109-INS-74 |
| chr5-40352013-INS-442 | chr5-40352013-INS-442 |
| chr5-40366226-INS-179 | chr5-40366226-INS-179 |
| chr5-40369005-INS-66 | chr5-40369005-INS-50 |
| chr5-40371430-INS-138 | chr5-40371430-INS-138 |
| chr5-40379951-INS-60 | chr5-40379951-INS-62 |
| chr5-40389238-INS-2051 | chr5-40389239-INS-2052 |
| chr5-40392119-INS-6193 | chr5-40392120-INS-6196 |
| chr5-40397913-INS-163 | chr5-40397913-INS-173 |
| chr5-40403727-INS-687 | chr5-40403745-INS-695 |
| chr5-40412396-INS-6971 | chr5-40412406-INS-6972 |
| chr5-40414972-INS-259 | chr5-40414972-INS-259 |
| chr5-40424232-INS-92 | chr5-40424282-INS-52 |
| chr5-40436586-INS-233 | chr5-40436586-INS-233 |
| chr5-40437486-INS-82 | chr5-40437484-INS-78 |
| chr5-40439573-INS-6792 | chr5-40439574-INS-6792 |
| chr5-40440173-INS-88 | chr5-40440179-INS-87 |
| chr5-40440843-INS-1229 | chr5-40440844-INS-1229 |
| chr5-40453735-INS-4448 | chr5-40453736-INS-4448 |
| chr5-40468077-INS-1943 | chr5-40468089-INS-1936 |
| chr5-40478700-INS-471 | chr5-40478700-INS-471 |
| chr5-40485368-INS-77 | chr5-40485368-INS-77 |
| chr5-40491594-INS-60 | chr5-40491594-INS-62 |

| | |
|---|---|
| chr5-40501380-INS-147 | chr5-40501380-INS-147 |
| chr5-40520362-INS-51 | chr5-40520372-INS-59 |
| chr5-40545673-INS-59 | chr5-40545678-INS-55 |
| chr5-40547877-INS-52 | chr5-40547887-INS-54 |
| chr5-40561246-INS-161 | chr5-40561247-INS-160 |
| chr5-40561300-INS-1152 | chr5-40561308-INS-1147 |
| chr5-40564973-INS-201 | chr5-40564973-INS-201 |
| chr5-40576283-INS-89 | chr5-40576283-INS-81 |
| chr5-40577819-INS-2692 | chr5-40577820-INS-2691 |
| chr5-40621226-INS-92 | chr5-40621226-INS-90 |
| chr5-40622940-INS-52 | chr5-40622940-INS-56 |
| chr5-40678467-INS-493 | chr5-40678467-INS-493 |
| chr5-40685240-INS-5822 | chr5-40685241-INS-5822 |
| chr5-40805915-INS-943 | chr5-40805915-INS-943 |
| chr5-40855991-INS-80 | chr5-40855991-INS-84 |
| chr5-40862971-INS-620 | chr5-40862971-INS-620 |
| chr5-40877881-INS-60 | chr5-40877873-INS-53 |
| chr5-40889193-INS-56 | chr5-40889193-INS-58 |
| chr5-40899903-INS-2042 | chr5-40899904-INS-2042 |
| chr5-40905488-INS-6799 | chr5-40905489-INS-6799 |
| chr5-40912429-INS-566 | chr5-40912429-INS-566 |
| chr5-40946279-INS-86 | chr5-40946279-INS-88 |
| chr5-40985126-INS-519 | chr5-40985126-INS-521 |
| chr5-41002731-INS-92 | chr5-41002751-INS-55 |
| chr5-41015407-INS-66 | chr5-41015407-INS-68 |
| chr5-41031466-INS-2031 | chr5-41031467-INS-2031 |
| chr5-41041462-INS-2214 | chr5-41041468-INS-2210 |
| chr5-41057466-INS-119 | chr5-41057432-INS-160 |
| chr5-41095436-INS-4768 | chr5-41095437-INS-4768 |
| chr5-41104719-INS-7116 | chr5-41104720-INS-7116 |
| chr5-41119973-INS-1896 | chr5-41119974-INS-1896 |
| chr5-41120757-INS-53 | chr5-41120757-INS-53 |
| chr5-41189285-INS-162 | chr5-41189300-INS-191 |
| chr5-41202600-INS-314 | chr5-41202600-INS-314 |
| chr5-41216126-INS-892 | chr5-41216123-INS-901 |
| chr5-41253527-INS-7247 | chr5-41253528-INS-7247 |
| chr5-41263489-INS-64 | chr5-41263489-INS-80 |
| chr5-41272406-INS-203 | chr5-41272406-INS-203 |
| chr5-41314485-INS-889 | chr5-41314485-INS-889 |
| chr5-41321809-INS-7565 | chr5-41321823-INS-7560 |
| chr5-41336283-INS-69 | chr5-41336283-INS-66 |
| chr5-41341159-INS-66 | chr5-41341159-INS-66 |
| chr5-41386788-INS-6263 | chr5-41386789-INS-6260 |
| chr5-41426182-INS-104 | chr5-41426182-INS-104 |
| chr5-41440441-INS-66 | chr5-41440441-INS-66 |
| chr5-41451151-INS-57 | chr5-41451159-INS-71 |
| chr5-41456847-INS-251 | chr5-41456847-INS-251 |
| chr5-41459356-INS-54 | chr5-41459356-INS-54 |
| chr5-41481728-INS-445 | chr5-41481728-INS-445 |
| chr5-41490953-INS-185 | chr5-41490953-INS-185 |
| chr5-41492460-INS-6950 | chr5-41492461-INS-6948 |
| chr5-41493419-INS-276 | chr5-41493419-INS-262 |
| chr5-41502061-INS-2354 | chr5-41502062-INS-2354 |
| chr5-41528111-INS-153 | chr5-41528103-INS-145 |
| chr5-41589045-INS-6231 | chr5-41589046-INS-6230 |
| chr5-41600536-INS-6481 | chr5-41600537-INS-6493 |
| chr5-41622023-INS-98 | chr5-41622023-INS-100 |
| chr5-41660915-INS-52 | chr5-41660924-INS-54 |
| chr5-41667999-INS-96 | chr5-41667999-INS-96 |
| chr5-41680811-INS-96 | chr5-41680811-INS-96 |
| chr5-41693576-INS-3055 | chr5-41693577-INS-3055 |
| chr5-41708428-INS-1561 | chr5-41708429-INS-1563 |
| chr5-41712358-INS-6261 | chr5-41712359-INS-6263 |
| chr5-41738647-INS-363 | chr5-41738647-INS-363 |
| chr5-41749828-INS-944 | chr5-41749828-INS-944 |
| chr5-41769471-INS-53 | chr5-41769484-INS-61 |
| chr5-41778022-INS-6317 | chr5-41778004-INS-6317 |
| chr5-41781361-INS-392 | chr5-41781362-INS-390 |
| chr5-41789338-INS-203 | chr5-41789338-INS-203 |
| chr5-41826063-INS-206 | chr5-41826063-INS-206 |
| chr5-41840353-INS-937 | chr5-41840353-INS-937 |
| chr5-41849948-INS-986 | chr5-41849948-INS-987 |
| chr5-41900006-INS-219 | chr5-41900006-INS-219 |
| chr5-41926331-INS-224 | chr5-41926331-INS-224 |
| chr5-41927452-INS-231 | chr5-41927452-INS-231 |
| chr5-41932168-INS-425 | chr5-41932168-INS-425 |
| chr5-41960676-INS-209 | chr5-41960676-INS-209 |

| | |
|---|---|
| chr5-41986309-INS-1831 | chr5-41986310-INS-1831 |
| chr5-41995134-INS-2316 | chr5-41995107-INS-2317 |
| chr5-42036031-INS-497 | chr5-42036031-INS-497 |
| chr5-42038642-INS-953 | chr5-42038642-INS-953 |
| chr5-42040960-INS-82 | chr5-42040960-INS-82 |
| chr5-42043003-INS-98 | chr5-42043003-INS-99 |
| chr5-42045215-INS-310 | chr5-42045215-INS-310 |
| chr5-42056652-INS-440 | chr5-42056652-INS-440 |
| chr5-42074843-INS-855 | chr5-42074843-INS-855 |
| chr5-42079582-INS-111 | chr5-42079582-INS-111 |
| chr5-42083656-INS-60 | chr5-42083656-INS-93 |
| chr5-42088138-INS-1319 | chr5-42088139-INS-1321 |
| chr5-42091928-INS-6265 | chr5-42091929-INS-6266 |
| chr5-42139207-INS-99 | chr5-42139207-INS-99 |
| chr5-42209964-INS-909 | chr5-42209964-INS-909 |
| chr5-42238237-INS-942 | chr5-42238233-INS-936 |
| chr5-42247561-INS-1669 | chr5-42247562-INS-1669 |
| chr5-42251231-INS-470 | chr5-42251231-INS-470 |
| chr5-42353086-INS-6413 | chr5-42353087-INS-6413 |
| chr5-42366173-INS-2210 | chr5-42366174-INS-2211 |
| chr5-42393099-INS-97 | chr5-42393100-INS-50 |
| chr5-42404760-INS-321 | chr5-42404760-INS-321 |
| chr5-42414212-INS-242 | chr5-42414212-INS-237 |
| chr5-42422612-INS-2367 | chr5-42422613-INS-2367 |
| chr5-42434506-INS-102 | chr5-42434506-INS-102 |
| chr5-42460949-INS-1361 | chr5-42460950-INS-1361 |
| chr5-42472417-INS-96 | chr5-42472417-INS-96 |
| chr5-42472742-INS-6499 | chr5-42472743-INS-6499 |
| chr5-42480811-INS-515 | chr5-42480811-INS-515 |
| chr5-42485276-INS-118 | chr5-42485280-INS-122 |
| chr5-42486755-INS-202 | chr5-42486746-INS-165 |
| chr5-42489966-INS-60 | chr5-42489966-INS-60 |
| chr5-42498925-INS-5846 | chr5-42498926-INS-5846 |
| chr5-42540909-INS-203 | chr5-42540912-INS-203 |
| chr5-42568427-INS-67 | chr5-42568386-INS-77 |
| chr5-42611347-INS-1096 | chr5-42611348-INS-1097 |
| chr5-42617581-INS-70 | chr5-42617613-INS-83 |
| chr5-42619192-INS-152 | chr5-42619162-INS-181 |
| chr5-42628709-INS-196 | chr5-42628726-INS-198 |
| chr5-42690556-INS-60 | chr5-42690553-INS-56 |
| chr5-42692638-INS-55 | chr5-42692638-INS-61 |
| chr5-42732297-INS-73 | chr5-42732297-INS-73 |
| chr5-42767176-INS-541 | chr5-42767176-INS-541 |
| chr5-42781313-INS-72 | chr5-42781340-INS-85 |
| chr5-42798750-INS-322 | chr5-42798750-INS-322 |
| chr5-42800120-INS-54 | chr5-42800120-INS-52 |
| chr5-42853122-INS-6497 | chr5-42853123-INS-6499 |
| chr5-42856495-INS-542 | chr5-42856495-INS-542 |
| chr5-42887236-INS-52 | chr5-42887236-INS-52 |
| chr5-42937706-INS-6496 | chr5-42937707-INS-6496 |
| chr5-42950676-INS-6391 | chr5-42950677-INS-6391 |
| chr5-42995813-INS-2229 | chr5-42995814-INS-2229 |
| chr5-43020826-INS-6320 | chr5-43020827-INS-6320 |
| chr5-43030591-INS-268 | chr5-43030586-INS-297 |
| chr5-43037535-INS-65 | chr5-43037535-INS-65 |
| chr5-43043160-INS-126 | chr5-43043160-INS-126 |
| chr5-43067270-INS-60 | chr5-43067270-INS-51 |
| chr5-43068278-INS-56 | chr5-43068278-INS-60 |
| chr5-43086952-INS-70 | chr5-43086952-INS-65 |
| chr5-43089868-INS-162 | chr5-43089868-INS-162 |
| chr5-43152050-INS-59 | chr5-43152050-INS-57 |
| chr5-43175430-INS-74 | chr5-43175430-INS-104 |
| chr5-43183151-INS-148 | chr5-43183151-INS-148 |
| chr5-43186942-INS-262 | chr5-43186942-INS-262 |
| chr5-43188968-INS-6029 | chr5-43188969-INS-6035 |
| chr5-43193599-INS-503 | chr5-43193599-INS-501 |
| chr5-43197058-INS-59 | chr5-43197060-INS-58 |
| chr5-43233343-INS-194 | chr5-43233343-INS-194 |
| chr5-43246296-INS-54 | chr5-43246296-INS-56 |
| chr5-43251768-INS-131 | chr5-43251768-INS-131 |
| chr5-43310253-INS-110 | chr5-43310251-INS-113 |
| chr5-43315175-INS-4433 | chr5-43315176-INS-4433 |
| chr5-43362955-INS-177 | chr5-43362955-INS-179 |
| chr5-43401206-INS-54 | chr5-43401206-INS-60 |
| chr5-43456710-INS-2029 | chr5-43456711-INS-2020 |
| chr5-43465077-INS-60 | chr5-43465077-INS-60 |
| chr5-43466130-INS-511 | chr5-43466130-INS-511 |

| | |
|---|---|
| chr5-43485578-INS-1117 | chr5-43485579-INS-1117 |
| chr5-43532598-INS-358 | chr5-43532598-INS-358 |
| chr5-43546367-INS-75 | chr5-43546367-INS-87 |
| chr5-43562277-INS-148 | chr5-43562277-INS-148 |
| chr5-43563068-INS-83 | chr5-43563073-INS-66 |
| chr5-43566764-INS-604 | chr5-43566764-INS-604 |
| chr5-43576626-INS-537 | chr5-43576626-INS-511 |
| chr5-43618834-INS-78 | chr5-43618834-INS-82 |
| chr5-43624945-INS-1179 | chr5-43624946-INS-1179 |
| chr5-43633805-INS-136 | chr5-43633803-INS-138 |
| chr5-43679138-INS-51 | chr5-43679138-INS-51 |
| chr5-43680543-INS-75 | chr5-43680543-INS-75 |
| chr5-43683006-INS-1913 | chr5-43683007-INS-1913 |
| chr5-43694812-INS-3878 | chr5-43694813-INS-3878 |
| chr5-43694953-INS-518 | chr5-43694953-INS-519 |
| chr5-43698762-INS-85 | chr5-43698749-INS-65 |
| chr5-43701099-INS-96 | chr5-43701099-INS-111 |
| chr5-43758440-INS-72 | chr5-43758440-INS-72 |
| chr5-43769419-INS-95 | chr5-43769417-INS-58 |
| chr5-43829405-INS-68 | chr5-43829429-INS-88 |
| chr5-43848379-INS-158 | chr5-43848379-INS-158 |
| chr5-43851755-INS-70 | chr5-43851755-INS-70 |
| chr5-43878579-INS-87 | chr5-43878558-INS-81 |
| chr5-43893152-INS-62 | chr5-43893154-INS-50 |
| chr5-43920394-INS-197 | chr5-43920394-INS-197 |
| chr5-43941803-INS-201 | chr5-43941803-INS-201 |
| chr5-43958528-INS-66 | chr5-43958528-INS-76 |
| chr5-43961361-INS-392 | chr5-43961361-INS-392 |
| chr5-43969913-INS-111 | chr5-43969913-INS-111 |
| chr5-43970638-INS-1536 | chr5-43970639-INS-1535 |
| chr5-43971032-INS-299 | chr5-43971032-INS-300 |
| chr5-43972392-INS-428 | chr5-43972392-INS-446 |
| chr5-43973987-INS-198 | chr5-43973987-INS-198 |
| chr5-44005638-INS-1325 | chr5-44005639-INS-1321 |
| chr5-44012969-INS-166 | chr5-44012969-INS-184 |
| chr5-44013529-INS-586 | chr5-44013529-INS-554 |
| chr5-44018933-INS-57 | chr5-44018924-INS-51 |
| chr5-44045146-INS-68 | chr5-44045146-INS-88 |
| chr5-44054749-INS-202 | chr5-44054749-INS-202 |
| chr5-44059774-INS-105 | chr5-44059774-INS-105 |
| chr5-44068957-INS-518 | chr5-44068957-INS-518 |
| chr5-44077314-INS-166 | chr5-44077315-INS-162 |
| chr5-44099819-INS-293 | chr5-44099804-INS-326 |
| chr5-44101158-INS-61 | chr5-44101158-INS-78 |
| chr5-44126509-INS-107 | chr5-44126509-INS-90 |
| chr5-44161100-INS-215 | chr5-44161100-INS-215 |
| chr5-44164847-INS-64 | chr5-44164847-INS-64 |
| chr5-44168143-INS-185 | chr5-44168143-INS-149 |
| chr5-44172818-INS-107 | chr5-44172818-INS-103 |
| chr5-44178317-INS-68 | chr5-44178317-INS-51 |
| chr5-44181741-INS-203 | chr5-44181741-INS-203 |
| chr5-44206037-INS-111 | chr5-44206037-INS-111 |
| chr5-44220427-INS-533 | chr5-44220427-INS-533 |
| chr5-44227879-INS-175 | chr5-44227879-INS-175 |
| chr5-44247417-INS-54 | chr5-44247417-INS-70 |
| chr5-44250548-INS-204 | chr5-44250548-INS-212 |
| chr5-44262918-INS-67 | chr5-44262918-INS-67 |
| chr5-44272284-INS-53 | chr5-44272284-INS-65 |
| chr5-44274432-INS-170 | chr5-44274432-INS-161 |
| chr5-44275992-INS-188 | chr5-44275992-INS-140 |
| chr5-44288632-INS-444 | chr5-44288632-INS-444 |
| chr5-44289872-INS-192 | chr5-44289872-INS-192 |
| chr5-44303502-INS-56 | chr5-44303522-INS-83 |
| chr5-44406639-INS-92 | chr5-44406665-INS-78 |
| chr5-44408540-INS-2701 | chr5-44408541-INS-2701 |
| chr5-44413415-INS-7105 | chr5-44413416-INS-7105 |
| chr5-44416250-INS-74 | chr5-44416250-INS-74 |
| chr5-44422848-INS-226 | chr5-44422848-INS-226 |
| chr5-44424155-INS-492 | chr5-44424155-INS-500 |
| chr5-44429095-INS-63 | chr5-44429095-INS-63 |
| chr5-44431148-INS-195 | chr5-44431148-INS-195 |
| chr5-44436482-INS-196 | chr5-44436482-INS-196 |
| chr5-44439272-INS-213 | chr5-44439272-INS-212 |
| chr5-44447051-INS-223 | chr5-44447051-INS-223 |
| chr5-44465413-INS-53 | chr5-44465413-INS-53 |
| chr5-44486266-INS-6462 | chr5-44486267-INS-6463 |
| chr5-44492545-INS-219 | chr5-44492545-INS-220 |

| | |
|---|---|
| chr5-44499736-INS-200 | chr5-44499736-INS-200 |
| chr5-44503548-INS-129 | chr5-44503548-INS-129 |
| chr5-44526499-INS-95 | chr5-44526503-INS-75 |
| chr5-44542868-INS-375 | chr5-44542868-INS-375 |
| chr5-44556382-INS-414 | chr5-44556382-INS-414 |
| chr5-44564080-INS-64 | chr5-44564080-INS-61 |
| chr5-44585161-INS-56 | chr5-44585161-INS-57 |
| chr5-44587395-INS-115 | chr5-44587399-INS-123 |
| chr5-44597325-INS-5228 | chr5-44597326-INS-5228 |
| chr5-44601215-INS-52 | chr5-44601215-INS-56 |
| chr5-44612914-INS-52 | chr5-44612914-INS-60 |
| chr5-44614102-INS-71 | chr5-44614102-INS-71 |
| chr5-44618308-INS-52 | chr5-44618308-INS-52 |
| chr5-44631920-INS-88 | chr5-44631920-INS-88 |
| chr5-44634071-INS-56 | chr5-44634071-INS-56 |
| chr5-44657539-INS-72 | chr5-44657539-INS-72 |
| chr5-44659960-INS-7388 | chr5-44659961-INS-7389 |
| chr5-44704552-INS-75 | chr5-44704552-INS-75 |
| chr5-44717133-INS-954 | chr5-44717138-INS-957 |
| chr5-44729913-INS-72 | chr5-44729913-INS-50 |
| chr5-44748016-INS-124 | chr5-44748016-INS-124 |
| chr5-44771102-INS-62 | chr5-44771102-INS-62 |
| chr5-44794029-INS-204 | chr5-44794029-INS-202 |
| chr5-44799055-INS-94 | chr5-44799055-INS-94 |
| chr5-44812887-INS-87 | chr5-44812891-INS-72 |
| chr5-44814882-INS-79 | chr5-44814882-INS-79 |
| chr5-44823376-INS-6425 | chr5-44823372-INS-6419 |
| chr5-44837712-INS-4888 | chr5-44837713-INS-4887 |
| chr5-44858958-INS-62 | chr5-44858942-INS-108 |
| chr5-44870187-INS-52 | chr5-44870191-INS-76 |
| chr5-44878448-INS-844 | chr5-44878448-INS-844 |
| chr5-44886851-INS-56 | chr5-44886854-INS-72 |
| chr5-44898737-INS-73 | chr5-44898762-INS-80 |
| chr5-44909269-INS-62 | chr5-44909269-INS-62 |
| chr5-44911741-INS-6403 | chr5-44911742-INS-6403 |
| chr5-44914689-INS-166 | chr5-44914689-INS-174 |
| chr5-44963071-INS-129 | chr5-44963071-INS-129 |
| chr5-44989345-INS-52 | chr5-44989345-INS-52 |
| chr5-45003297-INS-68 | chr5-45003297-INS-68 |
| chr5-45043344-INS-185 | chr5-45043344-INS-186 |
| chr5-45071946-INS-214 | chr5-45071946-INS-213 |
| chr5-45080802-INS-55 | chr5-45080802-INS-55 |
| chr5-45088283-INS-987 | chr5-45088283-INS-987 |
| chr5-45098734-INS-6587 | chr5-45098735-INS-6587 |
| chr5-45113947-INS-380 | chr5-45113947-INS-379 |
| chr5-45124397-INS-174 | chr5-45124397-INS-174 |
| chr5-45145607-INS-6324 | chr5-45145608-INS-6324 |
| chr5-45149449-INS-129 | chr5-45149449-INS-129 |
| chr5-45171809-INS-6727 | chr5-45171810-INS-6727 |
| chr5-45175788-INS-230 | chr5-45175788-INS-230 |
| chr5-45191390-INS-73 | chr5-45191376-INS-84 |
| chr5-45219043-INS-60 | chr5-45219043-INS-83 |
| chr5-45221561-INS-484 | chr5-45221561-INS-482 |
| chr5-45264264-INS-318 | chr5-45264264-INS-323 |
| chr5-45307461-INS-6204 | chr5-45307462-INS-6204 |
| chr5-45329368-INS-68 | chr5-45329368-INS-68 |
| chr5-45435320-INS-60 | chr5-45435320-INS-51 |
| chr5-45454416-INS-70 | chr5-45454416-INS-75 |
| chr5-45516663-INS-144 | chr5-45516663-INS-144 |
| chr5-45517717-INS-84 | chr5-45517713-INS-87 |
| chr5-45525070-INS-565 | chr5-45525070-INS-565 |
| chr5-45549633-INS-732 | chr5-45549633-INS-732 |
| chr5-45587895-INS-55 | chr5-45587895-INS-55 |
| chr5-45593341-INS-209 | chr5-45593341-INS-209 |
| chr5-45603010-INS-178 | chr5-45603010-INS-178 |
| chr5-45620235-INS-91 | chr5-45620217-INS-74 |
| chr5-45631650-INS-86 | chr5-45631650-INS-86 |
| chr5-45648602-INS-214 | chr5-45648602-INS-219 |
| chr5-45662389-INS-128 | chr5-45662389-INS-128 |
| chr5-45676427-INS-591 | chr5-45676425-INS-591 |
| chr5-45684925-INS-61 | chr5-45684925-INS-58 |
| chr5-45705929-INS-89 | chr5-45705937-INS-102 |
| chr5-45744726-INS-89 | chr5-45744726-INS-107 |
| chr5-45758064-INS-52 | chr5-45758064-INS-52 |
| chr5-45772303-INS-309 | chr5-45772303-INS-309 |
| chr5-45773749-INS-217 | chr5-45773726-INS-215 |
| chr5-45778314-INS-137 | chr5-45778314-INS-139 |

| | |
|---|---|
| chr5-45795243-INS-267 | chr5-45795243-INS-267 |
| chr5-45801339-INS-907 | chr5-45801339-INS-907 |
| chr5-45838069-INS-535 | chr5-45838069-INS-566 |
| chr5-45870840-INS-54 | chr5-45870840-INS-54 |
| chr5-45872866-INS-119 | chr5-45872866-INS-118 |
| chr5-45902918-INS-233 | chr5-45902918-INS-232 |
| chr5-45909246-INS-6247 | chr5-45909247-INS-6247 |
| chr5-45914998-INS-581 | chr5-45914998-INS-581 |
| chr5-45936563-INS-222 | chr5-45936563-INS-222 |
| chr5-45972215-INS-128 | chr5-45972215-INS-128 |
| chr5-46022375-INS-4811 | chr5-46022376-INS-4801 |
| chr5-46025523-INS-3166 | chr5-46025524-INS-3166 |
| chr5-46028497-INS-190 | chr5-46028504-INS-187 |
| chr5-46043047-INS-210 | chr5-46043047-INS-210 |
| chr5-46071683-INS-253 | chr5-46071683-INS-253 |
| chr5-46077320-INS-128 | chr5-46077320-INS-128 |
| chr5-46103023-INS-134 | chr5-46103054-INS-151 |
| chr5-46184274-INS-58 | chr5-46184274-INS-72 |
| chr5-46192966-INS-1795 | chr5-46192967-INS-1795 |
| chr5-46196506-INS-6659 | chr5-46196507-INS-6659 |
| chr5-46245794-INS-68 | chr5-46245794-INS-72 |
| chr5-46255883-INS-185 | chr5-46255883-INS-185 |
| chr5-46268682-INS-74 | chr5-46268682-INS-76 |
| chr5-46297182-INS-220 | chr5-46297182-INS-220 |
| chr5-46317069-INS-6400 | chr5-46317070-INS-6400 |
| chr5-46348869-INS-538 | chr5-46348869-INS-539 |
| chr5-46381976-INS-151 | chr5-46381976-INS-152 |
| chr5-46382722-INS-103 | chr5-46382722-INS-105 |
| chr5-46385227-INS-204 | chr5-46385226-INS-204 |
| chr5-46440087-INS-95 | chr5-46440087-INS-95 |
| chr5-46441220-INS-86 | chr5-46441220-INS-86 |
| chr5-46462635-INS-94 | chr5-46462635-INS-94 |
| chr5-46483433-INS-399 | chr5-46483434-INS-399 |
| chr5-46518940-INS-112 | chr5-46518940-INS-100 |
| chr5-46530496-INS-86 | chr5-46530496-INS-86 |
| chr5-46534680-INS-449 | chr5-46534680-INS-449 |
| chr5-46537883-INS-204 | chr5-46537883-INS-204 |
| chr5-46582687-INS-1888 | chr5-46582688-INS-1888 |
| chr5-46593117-INS-89 | chr5-46593111-INS-107 |
| chr5-46601227-INS-2549 | chr5-46601228-INS-2549 |
| chr5-46615819-INS-208 | chr5-46615819-INS-208 |
| chr5-46627060-INS-169 | chr5-46627060-INS-169 |
| chr5-46667494-INS-7238 | chr5-46667495-INS-7238 |
| chr5-46668657-INS-66 | chr5-46668657-INS-66 |
| chr5-46683240-INS-165 | chr5-46683240-INS-152 |
| chr5-46684177-INS-765 | chr5-46684177-INS-775 |
| chr5-46716871-INS-6939 | chr5-46716872-INS-6939 |
| chr5-46774634-INS-67 | chr5-46774634-INS-76 |
| chr5-46788000-INS-736 | chr5-46788000-INS-736 |
| chr5-46790881-INS-211 | chr5-46790881-INS-211 |
| chr5-46804917-INS-784 | chr5-46804917-INS-784 |
| chr5-46810227-INS-76 | chr5-46810227-INS-88 |
| chr5-46810764-INS-549 | chr5-46810764-INS-549 |
| chr5-46833485-INS-577 | chr5-46833485-INS-577 |
| chr5-46850592-INS-70 | chr5-46850592-INS-70 |
| chr5-46871906-INS-2274 | chr5-46871907-INS-2275 |
| chr5-46877915-INS-199 | chr5-46877915-INS-199 |
| chr5-46880243-INS-296 | chr5-46880244-INS-296 |
| chr5-46910719-INS-293 | chr5-46910719-INS-293 |
| chr5-46925860-INS-76 | chr5-46925860-INS-100 |
| chr5-46938230-INS-64 | chr5-46938231-INS-66 |
| chr5-46939416-INS-489 | chr5-46939416-INS-489 |
| chr5-46943498-INS-157 | chr5-46943498-INS-157 |
| chr5-46954272-INS-6480 | chr5-46954273-INS-6480 |
| chr5-46955826-INS-58 | chr5-46955826-INS-69 |
| chr5-46963332-INS-1884 | chr5-46963333-INS-1884 |
| chr5-46977331-INS-968 | chr5-46977331-INS-968 |
| chr5-46985264-INS-505 | chr5-46985264-INS-505 |
| chr5-47001850-INS-7157 | chr5-47001851-INS-7157 |
| chr5-47015540-INS-86 | chr5-47015540-INS-102 |
| chr5-47026659-INS-52 | chr5-47026659-INS-60 |
| chr5-47032958-INS-83 | chr5-47032958-INS-77 |
| chr5-47041802-INS-2266 | chr5-47041803-INS-2266 |
| chr5-47056856-INS-64 | chr5-47056856-INS-64 |
| chr5-47063369-INS-117 | chr5-47063369-INS-117 |
| chr5-47075763-INS-262 | chr5-47075763-INS-266 |
| chr5-47081390-INS-85 | chr5-47081423-INS-100 |

| | |
|---|---|
| chr5-47085371-INS-210 | chr5-47085371-INS-211 |
| chr5-47094494-INS-1229 | chr5-47094493-INS-1226 |
| chr5-47106709-INS-353 | chr5-47106709-INS-353 |
| chr5-47109889-INS-70 | chr5-47109891-INS-57 |
| chr5-47124152-INS-236 | chr5-47124152-INS-235 |
| chr5-47129548-INS-54 | chr5-47129548-INS-54 |
| chr5-47139614-INS-134 | chr5-47139614-INS-128 |
| chr5-47175070-INS-60 | chr5-47175070-INS-60 |
| chr5-47182551-INS-544 | chr5-47182551-INS-543 |
| chr5-47184580-INS-1046 | chr5-47184581-INS-1046 |
| chr5-47187635-INS-6518 | chr5-47187636-INS-6518 |
| chr5-47195544-INS-1737 | chr5-47195545-INS-1736 |
| chr5-47198896-INS-1846 | chr5-47198897-INS-1862 |
| chr5-47204489-INS-6216 | chr5-47204490-INS-6216 |
| chr5-47211605-INS-3134 | chr5-47211606-INS-3134 |
| chr5-47221354-INS-78 | chr5-47221354-INS-78 |
| chr5-47223684-INS-54 | chr5-47223684-INS-56 |
| chr5-47229921-INS-6677 | chr5-47229922-INS-6680 |
| chr5-47244940-INS-1212 | chr5-47244941-INS-1212 |
| chr5-47250361-INS-2892 | chr5-47250362-INS-2892 |
| chr5-47255804-INS-1766 | chr5-47255805-INS-1764 |
| chr5-47272375-INS-72 | chr5-47272373-INS-78 |
| chr5-47302824-INS-52 | chr5-47302824-INS-56 |
| chr5-47331877-INS-62 | chr5-47331881-INS-60 |
| chr5-47378435-INS-112 | chr5-47378435-INS-113 |
| chr5-47389764-INS-672 | chr5-47389764-INS-672 |
| chr5-47398437-INS-2435 | chr5-47398438-INS-2435 |
| chr5-47446518-INS-879 | chr5-47446518-INS-879 |
| chr5-47475142-INS-462 | chr5-47475142-INS-462 |
| chr5-47476267-INS-3348 | chr5-47476268-INS-3348 |
| chr5-47478008-INS-52 | chr5-47478008-INS-52 |
| chr5-47480905-INS-676 | chr5-47480907-INS-667 |
| chr5-47488632-INS-6463 | chr5-47488633-INS-6463 |
| chr5-47498131-INS-96 | chr5-47498131-INS-96 |
| chr5-47515172-INS-72 | chr5-47515172-INS-72 |
| chr5-47518392-INS-56 | chr5-47518392-INS-56 |
| chr5-47532915-INS-195 | chr5-47532905-INS-198 |
| chr5-47559714-INS-62 | chr5-47559730-INS-58 |
| chr5-47568175-INS-53 | chr5-47568175-INS-57 |
| chr5-47576704-INS-6568 | chr5-47576705-INS-6568 |
| chr5-47589670-INS-6607 | chr5-47589671-INS-6607 |
| chr5-47599589-INS-1993 | chr5-47599590-INS-1993 |
| chr5-47621568-INS-358 | chr5-47621568-INS-358 |
| chr5-47646956-INS-1637 | chr5-47646957-INS-1643 |
| chr5-47654286-INS-61 | chr5-47654263-INS-101 |
| chr5-47655864-INS-66 | chr5-47655881-INS-50 |
| chr5-47675141-INS-7082 | chr5-47675142-INS-7082 |
| chr5-47696587-INS-6177 | chr5-47696588-INS-6177 |
| chr5-47700445-INS-84 | chr5-47700443-INS-86 |
| chr5-47711119-INS-1654 | chr5-47711120-INS-1654 |
| chr5-47713977-INS-54 | chr5-47713985-INS-50 |
| chr5-47727282-INS-80 | chr5-47727282-INS-70 |
| chr5-47758378-INS-3864 | chr5-47758379-INS-3864 |
| chr5-47761257-INS-60 | chr5-47761257-INS-50 |
| chr5-47761896-INS-143 | chr5-47761899-INS-148 |
| chr5-47776414-INS-68 | chr5-47776414-INS-72 |
| chr5-47802631-INS-54 | chr5-47802631-INS-54 |
| chr5-47830093-INS-507 | chr5-47830093-INS-507 |
| chr5-47862902-INS-66 | chr5-47862862-INS-58 |
| chr5-47867539-INS-716 | chr5-47867539-INS-716 |
| chr5-47870141-INS-56 | chr5-47870141-INS-82 |
| chr5-47900174-INS-6722 | chr5-47900175-INS-6722 |
| chr5-47916402-INS-219 | chr5-47916402-INS-220 |
| chr5-47935721-INS-1912 | chr5-47935722-INS-1912 |
| chr5-47946088-INS-7313 | chr5-47946089-INS-7314 |
| chr5-47950390-INS-52 | chr5-47950390-INS-50 |
| chr5-47953020-INS-6349 | chr5-47953021-INS-6350 |
| chr5-47953413-INS-545 | chr5-47953413-INS-545 |
| chr5-47956336-INS-3227 | chr5-47956337-INS-3228 |
| chr5-47959896-INS-52 | chr5-47959896-INS-52 |
| chr5-47976562-INS-93 | chr5-47976543-INS-85 |
| chr5-47989828-INS-213 | chr5-47989828-INS-213 |
| chr5-47990930-INS-452 | chr5-47990930-INS-452 |
| chr5-48002587-INS-90 | chr5-48002587-INS-90 |
| chr5-48023389-INS-141 | chr5-48023389-INS-141 |
| chr5-48026244-INS-66 | chr5-48026238-INS-58 |
| chr5-48037678-INS-13569 | chr5-48037679-INS-13570 |

| | |
|---|---|
| chr5-48075301-INS-6017 | chr5-48075302-INS-6017 |
| chr5-48093250-INS-9246 | chr5-48093251-INS-9247 |
| chr5-48103969-INS-52 | chr5-48103969-INS-52 |
| chr5-48116863-INS-246 | chr5-48116863-INS-246 |
| chr5-48118413-INS-64 | chr5-48118411-INS-68 |
| chr5-48122478-INS-1203 | chr5-48122479-INS-1201 |
| chr5-48164891-INS-4420 | chr5-48164892-INS-4419 |
| chr5-48208993-INS-989 | chr5-48208993-INS-992 |
| chr5-48224717-INS-199 | chr5-48224717-INS-199 |
| chr5-48237003-INS-159 | chr5-48237003-INS-159 |
| chr5-48249620-INS-276 | chr5-48249620-INS-276 |
| chr5-48268093-INS-78 | chr5-48268093-INS-70 |
| chr5-48304437-INS-1327 | chr5-48304438-INS-1327 |
| chr5-48310951-INS-77 | chr5-48310951-INS-77 |
| chr5-48315705-INS-57 | chr5-48315705-INS-50 |
| chr5-48330549-INS-480 | chr5-48330549-INS-480 |
| chr5-48347352-INS-57 | chr5-48347349-INS-60 |
| chr5-48357274-INS-772 | chr5-48357274-INS-772 |
| chr5-48360780-INS-52 | chr5-48360780-INS-52 |
| chr5-48397502-INS-56 | chr5-48397502-INS-56 |
| chr5-48404770-INS-4740 | chr5-48404771-INS-4740 |
| chr5-48425395-INS-1133 | chr5-48425396-INS-1135 |
| chr5-48474932-INS-518 | chr5-48474932-INS-518 |
| chr5-48507373-INS-4540 | chr5-48507374-INS-4540 |
| chr5-48517181-INS-82 | chr5-48517181-INS-82 |
| chr5-48517767-INS-1163 | chr5-48517768-INS-1163 |
| chr5-48526982-INS-54 | chr5-48526978-INS-58 |
| chr5-48535892-INS-130 | chr5-48535892-INS-130 |
| chr5-48540963-INS-71 | chr5-48540963-INS-71 |
| chr5-48541796-INS-406 | chr5-48541796-INS-406 |
| chr5-48557568-INS-57 | chr5-48557568-INS-57 |
| chr5-48568344-INS-1301 | chr5-48568345-INS-1302 |
| chr5-48625312-INS-80 | chr5-48625312-INS-80 |
| chr5-48633090-INS-147 | chr5-48633090-INS-147 |
| chr5-48636850-INS-445 | chr5-48636850-INS-445 |
| chr5-48646041-INS-719 | chr5-48646041-INS-720 |
| chr5-48659574-INS-232 | chr5-48659574-INS-232 |
| chr5-48670455-INS-76 | chr5-48670446-INS-81 |
| chr5-48703689-INS-60 | chr5-48703689-INS-54 |
| chr5-48709679-INS-187 | chr5-48709679-INS-187 |
| chr5-48736756-INS-51 | chr5-48736756-INS-52 |
| chr5-48763114-INS-73 | chr5-48763114-INS-88 |
| chr5-48788375-INS-142 | chr5-48788374-INS-144 |
| chr5-48790850-INS-106 | chr5-48790850-INS-144 |
| chr5-48803563-INS-66 | chr5-48803563-INS-66 |
| chr5-48823002-INS-63 | chr5-48823003-INS-61 |
| chr5-48824652-INS-90 | chr5-48824652-INS-102 |
| chr5-48825208-INS-193 | chr5-48825208-INS-196 |
| chr5-48826657-INS-995 | chr5-48826657-INS-995 |
| chr5-48839936-INS-203 | chr5-48839936-INS-203 |
| chr5-48866084-INS-56 | chr5-48866084-INS-56 |
| chr5-48880787-INS-82 | chr5-48880787-INS-82 |
| chr5-48882442-INS-633 | chr5-48882442-INS-633 |
| chr5-48884934-INS-276 | chr5-48884934-INS-275 |
| chr5-48896913-INS-72 | chr5-48896890-INS-88 |
| chr5-48916566-INS-346 | chr5-48916566-INS-346 |
| chr5-48924051-INS-423 | chr5-48924051-INS-423 |
| chr5-48940640-INS-244 | chr5-48940640-INS-244 |
| chr5-48966586-INS-201 | chr5-48966586-INS-201 |
| chr5-48980825-INS-103 | chr5-48980825-INS-103 |
| chr5-48985357-INS-441 | chr5-48985357-INS-441 |
| chr5-49009564-INS-182 | chr5-49009564-INS-182 |
| chr5-49017069-INS-70 | chr5-49017069-INS-54 |
| chr5-49020468-INS-58 | chr5-49020468-INS-58 |
| chr5-49024201-INS-61 | chr5-49024193-INS-60 |
| chr5-49025977-INS-730 | chr5-49025977-INS-730 |
| chr5-49154116-INS-167 | chr5-49154116-INS-176 |
| chr5-49158871-INS-62 | chr5-49158886-INS-59 |
| chr5-49181914-INS-56 | chr5-49181912-INS-50 |
| chr5-49197090-INS-856 | chr5-49197090-INS-856 |
| chr5-49205213-INS-55 | chr5-49205213-INS-55 |
| chr5-49215535-INS-411 | chr5-49215536-INS-411 |
| chr5-49236373-INS-3510 | chr5-49236374-INS-3507 |
| chr5-49240944-INS-52 | chr5-49240938-INS-60 |
| chr5-49248847-INS-460 | chr5-49248847-INS-460 |
| chr5-49253606-INS-70 | chr5-49253606-INS-71 |
| chr5-49277011-INS-55 | chr5-49277011-INS-55 |

| | |
|---|---|
| chr5-49280496-INS-6628 | chr5-49280497-INS-6630 |
| chr5-49290520-INS-537 | chr5-49290520-INS-537 |
| chr5-49301773-INS-66 | chr5-49301791-INS-68 |
| chr5-49358026-INS-60 | chr5-49358018-INS-66 |
| chr5-49403295-INS-154 | chr5-49403295-INS-154 |
| chr5-49453744-INS-52 | chr5-49453744-INS-54 |
| chr5-49459972-INS-372 | chr5-49459972-INS-372 |
| chr5-49480876-INS-1798 | chr5-49480877-INS-1796 |
| chr5-49487466-INS-78 | chr5-49487466-INS-78 |
| chr5-49512561-INS-212 | chr5-49512561-INS-212 |
| chr5-49514203-INS-254 | chr5-49514203-INS-254 |
| chr5-49515247-INS-2335 | chr5-49515248-INS-2335 |
| chr5-49520346-INS-1700 | chr5-49520347-INS-1701 |
| chr5-49521391-INS-180 | chr5-49521383-INS-188 |
| chr5-49535609-INS-202 | chr5-49535609-INS-201 |
| chr5-49559112-INS-296 | chr5-49559112-INS-296 |
| chr5-49568697-INS-359 | chr5-49568697-INS-362 |
| chr5-49596187-INS-3036 | chr5-49596188-INS-3030 |
| chr5-49603090-INS-486 | chr5-49603090-INS-485 |
| chr5-49650265-INS-4361 | chr5-49650266-INS-4361 |
| chr5-49651460-INS-58 | chr5-49651461-INS-58 |
| chr5-49653982-INS-527 | chr5-49653982-INS-540 |
| chr5-49666515-INS-65 | chr5-49666515-INS-93 |
| chr5-49679678-INS-532 | chr5-49679678-INS-532 |
| chr5-49682520-INS-68 | chr5-49682520-INS-60 |
| chr5-49694233-INS-62 | chr5-49694233-INS-62 |
| chr5-49709623-INS-2405 | chr5-49709623-INS-2398 |
| chr5-49713854-INS-184 | chr5-49713856-INS-140 |
| chr5-49739746-INS-107 | chr5-49739746-INS-110 |
| chr5-49743854-INS-59 | chr5-49743855-INS-56 |
| chr5-49755480-INS-1185 | chr5-49755481-INS-1185 |
| chr5-49757437-INS-357 | chr5-49757437-INS-357 |
| chr5-49766910-INS-72 | chr5-49766910-INS-72 |
| chr5-49776484-INS-100 | chr5-49776484-INS-85 |
| chr5-49783338-INS-66 | chr5-49783338-INS-66 |
| chr5-49786410-INS-226 | chr5-49786410-INS-226 |
| chr5-49789347-INS-463 | chr5-49789347-INS-463 |
| chr5-49790447-INS-98 | chr5-49790447-INS-98 |
| chr5-49791853-INS-68 | chr5-49791845-INS-56 |
| chr5-49797055-INS-80 | chr5-49797055-INS-80 |
| chr5-49811947-INS-485 | chr5-49811947-INS-484 |
| chr5-49852828-INS-324 | chr5-49852828-INS-325 |
| chr5-49870008-INS-95 | chr5-49870008-INS-95 |
| chr5-49922364-INS-59 | chr5-49922345-INS-56 |
| chr5-49932630-INS-85 | chr5-49932630-INS-85 |
| chr5-49937703-INS-626 | chr5-49937703-INS-628 |
| chr5-49944943-INS-206 | chr5-49944943-INS-206 |
| chr5-49957495-INS-295 | chr5-49957495-INS-295 |
| chr5-49960743-INS-84 | chr5-49960743-INS-80 |
| chr5-49962228-INS-183 | chr5-49962249-INS-195 |
| chr5-49965882-INS-51 | chr5-49965882-INS-51 |
| chr5-49987516-INS-6401 | chr5-49987517-INS-6401 |
| chr5-50027561-INS-3891 | chr5-50027562-INS-3891 |
| chr5-50029639-INS-255 | chr5-50029639-INS-267 |
| chr5-50052063-INS-66 | chr5-50052063-INS-66 |
| chr5-50055178-INS-597 | chr5-50055178-INS-597 |
| chr5-50056436-INS-87 | chr5-50056436-INS-87 |
| chr5-50073993-INS-64 | chr5-50073971-INS-56 |
| chr5-50077821-INS-108 | chr5-50077821-INS-108 |
| chr5-50098208-INS-387 | chr5-50098208-INS-387 |
| chr5-50145136-INS-206 | chr5-50145136-INS-204 |
| chr5-50155790-INS-262 | chr5-50155790-INS-262 |
| chr5-50191089-INS-68 | chr5-50191089-INS-68 |
| chr5-50237331-INS-611 | chr5-50237331-INS-615 |
| chr5-50250512-INS-1254 | chr5-50250513-INS-1258 |
| chr5-50315802-INS-72 | chr5-50315802-INS-56 |
| chr5-50331867-INS-150 | chr5-50331867-INS-150 |
| chr5-50332830-INS-1553 | chr5-50332831-INS-1553 |
| chr5-50337029-INS-84 | chr5-50337029-INS-84 |
| chr5-50352660-INS-52 | chr5-50352660-INS-50 |
| chr5-50369809-INS-156 | chr5-50369809-INS-157 |
| chr5-50403495-INS-57 | chr5-50403495-INS-60 |
| chr5-50433571-INS-337 | chr5-50433571-INS-337 |
| chr5-50477886-INS-67 | chr5-50477882-INS-56 |
| chr5-50479333-INS-70 | chr5-50479331-INS-72 |
| chr5-50501949-INS-72 | chr5-50501949-INS-76 |
| chr5-50526716-INS-6498 | chr5-50526717-INS-6501 |

| | |
|---|---|
| chr5-50545808-INS-62 | chr5-50545803-INS-62 |
| chr5-50554139-INS-62 | chr5-50554139-INS-62 |
| chr5-50566609-INS-2762 | chr5-50566610-INS-2762 |
| chr5-50587508-INS-597 | chr5-50587508-INS-597 |
| chr5-50619676-INS-179 | chr5-50619676-INS-179 |
| chr5-50640076-INS-425 | chr5-50640076-INS-425 |
| chr5-50642776-INS-1043 | chr5-50642777-INS-1043 |
| chr5-50655469-INS-482 | chr5-50655469-INS-482 |
| chr5-50661728-INS-66 | chr5-50661728-INS-71 |
| chr5-50689808-INS-946 | chr5-50689808-INS-946 |
| chr5-50691336-INS-171 | chr5-50691336-INS-171 |
| chr5-50697984-INS-83 | chr5-50698009-INS-71 |
| chr5-50702364-INS-220 | chr5-50702364-INS-220 |
| chr5-50714078-INS-383 | chr5-50714078-INS-383 |
| chr5-50715874-INS-437 | chr5-50715874-INS-437 |
| chr5-50725171-INS-131 | chr5-50725171-INS-131 |
| chr5-50730262-INS-1398 | chr5-50730263-INS-1398 |
| chr5-50747369-INS-456 | chr5-50747369-INS-458 |
| chr5-50763845-INS-6415 | chr5-50763846-INS-6415 |
| chr5-50821216-INS-77 | chr5-50821216-INS-80 |
| chr5-50837321-INS-1801 | chr5-50837322-INS-1802 |
| chr5-50837977-INS-6462 | chr5-50837978-INS-6461 |
| chr5-50841129-INS-60 | chr5-50841129-INS-52 |
| chr5-50868773-INS-179 | chr5-50868773-INS-185 |
| chr5-50877090-INS-555 | chr5-50877090-INS-555 |
| chr5-50888431-INS-68 | chr5-50888431-INS-76 |
| chr5-50890308-INS-401 | chr5-50890308-INS-401 |
| chr5-50947146-INS-198 | chr5-50947146-INS-198 |
| chr5-50954925-INS-84 | chr5-50954925-INS-84 |
| chr5-50960760-INS-56 | chr5-50960760-INS-56 |
| chr5-50983322-INS-449 | chr5-50983322-INS-449 |
| chr5-51021341-INS-112 | chr5-51021341-INS-118 |
| chr5-51022236-INS-388 | chr5-51022236-INS-388 |
| chr5-51025292-INS-67 | chr5-51025292-INS-67 |
| chr5-51036727-INS-408 | chr5-51036727-INS-408 |
| chr5-51048200-INS-60 | chr5-51048200-INS-51 |
| chr5-51084099-INS-429 | chr5-51084099-INS-436 |
| chr5-51088553-INS-6595 | chr5-51088554-INS-6595 |
| chr5-51098386-INS-54 | chr5-51098385-INS-58 |
| chr5-51106050-INS-7089 | chr5-51106051-INS-7089 |
| chr5-51123261-INS-82 | chr5-51123261-INS-84 |
| chr5-51197985-INS-89 | chr5-51197985-INS-91 |
| chr5-51209286-INS-1100 | chr5-51209287-INS-1100 |
| chr5-51214459-INS-533 | chr5-51214471-INS-545 |
| chr5-51238181-INS-497 | chr5-51238181-INS-497 |
| chr5-51239621-INS-89 | chr5-51239621-INS-90 |
| chr5-51246175-INS-68 | chr5-51246175-INS-70 |
| chr5-51267878-INS-59 | chr5-51267878-INS-59 |
| chr5-51268636-INS-877 | chr5-51268636-INS-877 |
| chr5-51280967-INS-688 | chr5-51280967-INS-678 |
| chr5-51300954-INS-96 | chr5-51300954-INS-98 |
| chr5-51321044-INS-6384 | chr5-51321045-INS-6387 |
| chr5-51332695-INS-60 | chr5-51332695-INS-60 |
| chr5-51334656-INS-212 | chr5-51334656-INS-212 |
| chr5-51335492-INS-801 | chr5-51335492-INS-801 |
| chr5-51360085-INS-193 | chr5-51360084-INS-194 |
| chr5-51384389-INS-96 | chr5-51384389-INS-78 |
| chr5-51400033-INS-66 | chr5-51400033-INS-52 |
| chr5-51413301-INS-370 | chr5-51413301-INS-369 |
| chr5-51421782-INS-357 | chr5-51421782-INS-357 |
| chr5-51439289-INS-504 | chr5-51439279-INS-512 |
| chr5-51441202-INS-314 | chr5-51441202-INS-314 |
| chr5-51458893-INS-119 | chr5-51458893-INS-109 |
| chr5-51470427-INS-6251 | chr5-51470428-INS-6250 |
| chr5-51484489-INS-56 | chr5-51484510-INS-84 |
| chr5-51499317-INS-449 | chr5-51499317-INS-449 |
| chr5-51512739-INS-499 | chr5-51512739-INS-499 |
| chr5-51559367-INS-124 | chr5-51559367-INS-138 |
| chr5-51563662-INS-55 | chr5-51563662-INS-55 |
| chr5-51582931-INS-409 | chr5-51582931-INS-409 |
| chr5-51594045-INS-64 | chr5-51594045-INS-66 |
| chr5-51602927-INS-56 | chr5-51602879-INS-55 |
| chr5-51623202-INS-444 | chr5-51623202-INS-444 |
| chr5-51634148-INS-94 | chr5-51634139-INS-64 |
| chr5-51646475-INS-108 | chr5-51646425-INS-113 |
| chr5-51664229-INS-142 | chr5-51664223-INS-181 |
| chr5-51664825-INS-1891 | chr5-51664826-INS-1891 |

| | |
|---|---|
| chr5-51668148-INS-68 | chr5-51668148-INS-72 |
| chr5-51726516-INS-78 | chr5-51726543-INS-72 |
| chr5-51735791-INS-84 | chr5-51735787-INS-112 |
| chr5-51755338-INS-111 | chr5-51755338-INS-112 |
| chr5-51764374-INS-80 | chr5-51764377-INS-51 |
| chr5-51806543-INS-69 | chr5-51806543-INS-55 |
| chr5-51830456-INS-340 | chr5-51830456-INS-340 |
| chr5-51846575-INS-82 | chr5-51846574-INS-60 |
| chr5-51891654-INS-54 | chr5-51891680-INS-64 |
| chr5-51902207-INS-60 | chr5-51902208-INS-102 |
| chr5-51936268-INS-149 | chr5-51936268-INS-149 |
| chr5-51979427-INS-63 | chr5-51979427-INS-63 |
| chr5-51989172-INS-82 | chr5-51989172-INS-90 |
| chr5-52086441-INS-72 | chr5-52086438-INS-96 |
| chr5-52089991-INS-147 | chr5-52089991-INS-147 |
| chr5-52098520-INS-905 | chr5-52098520-INS-905 |
| chr5-52101824-INS-68 | chr5-52101824-INS-98 |
| chr5-52111660-INS-831 | chr5-52111660-INS-831 |
| chr5-52116668-INS-600 | chr5-52116668-INS-588 |
| chr5-52121799-INS-6684 | chr5-52121800-INS-6685 |
| chr5-52159562-INS-73 | chr5-52159562-INS-59 |
| chr5-52175901-INS-54 | chr5-52175901-INS-50 |
| chr5-52214279-INS-54 | chr5-52214279-INS-56 |
| chr5-52239774-INS-90 | chr5-52239750-INS-64 |
| chr5-52248512-INS-74 | chr5-52248512-INS-74 |
| chr5-52254591-INS-69 | chr5-52254591-INS-69 |
| chr5-52310108-INS-82 | chr5-52310108-INS-82 |
| chr5-52312827-INS-234 | chr5-52312827-INS-234 |
| chr5-52344896-INS-52 | chr5-52344896-INS-56 |
| chr5-52370169-INS-181 | chr5-52370169-INS-174 |
| chr5-52374488-INS-108 | chr5-52374488-INS-108 |
| chr5-52423317-INS-89 | chr5-52423317-INS-87 |
| chr5-52445581-INS-82 | chr5-52445577-INS-80 |
| chr5-52449471-INS-6444 | chr5-52449472-INS-6446 |
| chr5-52519071-INS-493 | chr5-52519071-INS-493 |
| chr5-52529690-INS-78 | chr5-52529702-INS-75 |
| chr5-52546662-INS-65 | chr5-52546662-INS-65 |
| chr5-52553020-INS-53 | chr5-52553020-INS-53 |
| chr5-52554390-INS-442 | chr5-52554417-INS-416 |
| chr5-52564799-INS-62 | chr5-52564799-INS-62 |
| chr5-52588791-INS-542 | chr5-52588791-INS-528 |
| chr5-52592688-INS-792 | chr5-52592688-INS-791 |
| chr5-52613979-INS-59 | chr5-52613979-INS-59 |
| chr5-52623292-INS-76 | chr5-52623292-INS-50 |
| chr5-52643846-INS-78 | chr5-52643846-INS-84 |
| chr5-52654026-INS-1896 | chr5-52654027-INS-1896 |
| chr5-52657715-INS-82 | chr5-52657715-INS-128 |
| chr5-52669070-INS-183 | chr5-52669079-INS-135 |
| chr5-52669755-INS-1893 | chr5-52669756-INS-1893 |
| chr5-52679267-INS-498 | chr5-52679267-INS-501 |
| chr5-52688452-INS-71 | chr5-52688431-INS-86 |
| chr5-52710586-INS-134 | chr5-52710586-INS-133 |
| chr5-52746343-INS-924 | chr5-52746358-INS-923 |
| chr5-52781448-INS-270 | chr5-52781448-INS-270 |
| chr5-52781488-INS-817 | chr5-52781488-INS-818 |
| chr5-52789105-INS-307 | chr5-52789105-INS-299 |
| chr5-52792929-INS-116 | chr5-52792932-INS-95 |
| chr5-52818724-INS-384 | chr5-52818724-INS-384 |
| chr5-52823244-INS-188 | chr5-52823244-INS-188 |
| chr5-52828828-INS-81 | chr5-52828828-INS-81 |
| chr5-52837243-INS-379 | chr5-52837243-INS-379 |
| chr5-52879438-INS-409 | chr5-52879438-INS-409 |
| chr5-52894096-INS-6503 | chr5-52894097-INS-6499 |
| chr5-52903428-INS-69 | chr5-52903428-INS-61 |
| chr5-52909311-INS-477 | chr5-52909311-INS-478 |
| chr5-52925614-INS-209 | chr5-52925614-INS-209 |
| chr5-52929654-INS-64 | chr5-52929654-INS-76 |
| chr5-52967901-INS-141 | chr5-52967901-INS-94 |
| chr5-53019804-INS-58 | chr5-53019804-INS-58 |
| chr5-53035582-INS-119 | chr5-53035582-INS-119 |
| chr5-53042136-INS-405 | chr5-53042148-INS-407 |
| chr5-53044582-INS-971 | chr5-53044582-INS-971 |
| chr5-53061576-INS-184 | chr5-53061576-INS-184 |
| chr5-53065059-INS-139 | chr5-53065012-INS-160 |
| chr5-53111645-INS-64 | chr5-53111645-INS-65 |
| chr5-53117479-INS-83 | chr5-53117479-INS-62 |
| chr5-53121506-INS-160 | chr5-53121506-INS-160 |

| | |
|---|---|
| chr5-53125755-INS-71 | chr5-53125752-INS-50 |
| chr5-53169981-INS-78 | chr5-53169981-INS-89 |
| chr5-53171639-INS-79 | chr5-53171639-INS-79 |
| chr5-53173008-INS-171 | chr5-53173016-INS-173 |
| chr5-53198850-INS-61 | chr5-53198850-INS-61 |
| chr5-53223680-INS-305 | chr5-53223680-INS-300 |
| chr5-53235021-INS-226 | chr5-53235021-INS-211 |
| chr5-53264165-INS-314 | chr5-53264165-INS-314 |
| chr5-53279804-INS-1888 | chr5-53279805-INS-1888 |
| chr5-53347819-INS-60 | chr5-53347819-INS-52 |
| chr5-53353462-INS-696 | chr5-53353462-INS-696 |
| chr5-53355688-INS-539 | chr5-53355688-INS-537 |
| chr5-53392290-INS-6414 | chr5-53392291-INS-6414 |
| chr5-53412759-INS-107 | chr5-53412737-INS-113 |
| chr5-53431281-INS-1594 | chr5-53431282-INS-1594 |
| chr5-53446702-INS-6485 | chr5-53446703-INS-6486 |
| chr5-53462880-INS-348 | chr5-53462880-INS-348 |
| chr5-53471456-INS-364 | chr5-53471456-INS-365 |
| chr5-53474759-INS-116 | chr5-53474759-INS-116 |
| chr5-53493479-INS-59 | chr5-53493479-INS-75 |
| chr5-53507260-INS-509 | chr5-53507260-INS-507 |
| chr5-53516728-INS-112 | chr5-53516728-INS-112 |
| chr5-53528824-INS-111 | chr5-53528831-INS-100 |
| chr5-53569188-INS-68 | chr5-53569188-INS-68 |
| chr5-53608753-INS-204 | chr5-53608754-INS-203 |
| chr5-53636160-INS-104 | chr5-53636162-INS-104 |
| chr5-53654256-INS-70 | chr5-53654255-INS-72 |
| chr5-53669943-INS-5488 | chr5-53669944-INS-5488 |
| chr5-53688350-INS-74 | chr5-53688350-INS-78 |
| chr5-53690550-INS-3279 | chr5-53690551-INS-3279 |
| chr5-53696412-INS-193 | chr5-53696412-INS-193 |
| chr5-53709220-INS-487 | chr5-53709220-INS-487 |
| chr5-53716750-INS-214 | chr5-53716750-INS-214 |
| chr5-53745307-INS-62 | chr5-53745307-INS-62 |
| chr5-53771480-INS-395 | chr5-53771480-INS-395 |
| chr5-53789237-INS-196 | chr5-53789237-INS-196 |
| chr5-53795067-INS-202 | chr5-53795067-INS-201 |
| chr5-53808535-INS-167 | chr5-53808535-INS-168 |
| chr5-53861478-INS-52 | chr5-53861478-INS-52 |
| chr5-53871736-INS-205 | chr5-53871736-INS-203 |
| chr5-53897542-INS-345 | chr5-53897542-INS-345 |
| chr5-53917296-INS-52 | chr5-53917296-INS-52 |
| chr5-53921711-INS-75 | chr5-53921712-INS-52 |
| chr5-53940964-INS-957 | chr5-53940964-INS-957 |
| chr5-53963588-INS-6759 | chr5-53963589-INS-6760 |
| chr5-54000357-INS-3973 | chr5-54000358-INS-3975 |
| chr5-54012609-INS-1407 | chr5-54012610-INS-1407 |
| chr5-54037632-INS-113 | chr5-54037632-INS-112 |
| chr5-54071975-INS-1247 | chr5-54071976-INS-1244 |
| chr5-54073091-INS-66 | chr5-54073091-INS-64 |
| chr5-54174489-INS-628 | chr5-54174489-INS-629 |
| chr5-54196663-INS-281 | chr5-54196663-INS-311 |
| chr5-54224982-INS-204 | chr5-54224982-INS-203 |
| chr5-54244772-INS-60 | chr5-54244772-INS-66 |
| chr5-54253374-INS-273 | chr5-54253374-INS-273 |
| chr5-54258350-INS-333 | chr5-54258350-INS-296 |
| chr5-54288613-INS-52 | chr5-54288616-INS-95 |
| chr5-54310668-INS-54 | chr5-54310658-INS-50 |
| chr5-54332295-INS-486 | chr5-54332295-INS-486 |
| chr5-54346359-INS-67 | chr5-54346359-INS-67 |
| chr5-54401642-INS-6194 | chr5-54401643-INS-6195 |
| chr5-54409551-INS-58 | chr5-54409551-INS-58 |
| chr5-54466369-INS-56 | chr5-54466369-INS-60 |
| chr5-54493318-INS-165 | chr5-54493333-INS-180 |
| chr5-54520874-INS-102 | chr5-54520874-INS-102 |
| chr5-54530908-INS-4693 | chr5-54530909-INS-4693 |
| chr5-54555898-INS-93 | chr5-54555898-INS-93 |
| chr5-54558810-INS-2062 | chr5-54558811-INS-2062 |
| chr5-54584839-INS-747 | chr5-54584839-INS-747 |
| chr5-54594361-INS-213 | chr5-54594361-INS-213 |
| chr5-54604968-INS-74 | chr5-54604968-INS-64 |
| chr5-54651819-INS-65 | chr5-54651844-INS-84 |
| chr5-54669724-INS-64 | chr5-54669720-INS-64 |
| chr5-54691550-INS-6576 | chr5-54691551-INS-6575 |
| chr5-54697446-INS-214 | chr5-54697446-INS-214 |
| chr5-54728757-INS-1680 | chr5-54728758-INS-1680 |
| chr5-54733658-INS-62 | chr5-54733658-INS-62 |

| | |
|---|---|
| chr5-54745583-INS-54 | chr5-54745587-INS-58 |
| chr5-54772325-INS-56 | chr5-54772325-INS-60 |
| chr5-54802572-INS-7586 | chr5-54802573-INS-7587 |
| chr5-54813921-INS-93 | chr5-54813921-INS-97 |
| chr5-54816077-INS-899 | chr5-54816077-INS-899 |
| chr5-54816787-INS-56 | chr5-54816787-INS-56 |
| chr5-54863490-INS-1554 | chr5-54863506-INS-1541 |
| chr5-54893995-INS-76 | chr5-54893995-INS-72 |
| chr5-54894571-INS-6334 | chr5-54894572-INS-6334 |
| chr5-54917531-INS-90 | chr5-54917531-INS-90 |
| chr5-54928914-INS-55 | chr5-54928914-INS-55 |
| chr5-54991516-INS-117 | chr5-54991517-INS-117 |
| chr5-55001785-INS-73 | chr5-55001785-INS-64 |
| chr5-55013576-INS-57 | chr5-55013576-INS-57 |
| chr5-55022352-INS-75 | chr5-55022352-INS-75 |
| chr5-55026219-INS-1844 | chr5-55026220-INS-1845 |
| chr5-55029449-INS-523 | chr5-55029449-INS-523 |
| chr5-55066572-INS-213 | chr5-55066572-INS-213 |
| chr5-55072462-INS-2988 | chr5-55072463-INS-2988 |
| chr5-55090790-INS-6420 | chr5-55090791-INS-6421 |
| chr5-55113608-INS-64 | chr5-55113608-INS-59 |
| chr5-55125437-INS-255 | chr5-55125437-INS-255 |
| chr5-55131989-INS-214 | chr5-55131989-INS-214 |
| chr5-55156723-INS-4851 | chr5-55156724-INS-4852 |
| chr5-55166019-INS-77 | chr5-55166042-INS-70 |
| chr5-55174489-INS-187 | chr5-55174470-INS-193 |
| chr5-55184593-INS-227 | chr5-55184593-INS-227 |
| chr5-55185234-INS-439 | chr5-55185232-INS-434 |
| chr5-55191428-INS-1208 | chr5-55191429-INS-1209 |
| chr5-55200382-INS-6833 | chr5-55200383-INS-6833 |
| chr5-55226568-INS-802 | chr5-55226568-INS-802 |
| chr5-55232936-INS-1958 | chr5-55232937-INS-1958 |
| chr5-55276014-INS-741 | chr5-55276014-INS-741 |
| chr5-55282195-INS-12357 | chr5-55282196-INS-12360 |
| chr5-55298446-INS-6267 | chr5-55298447-INS-6266 |
| chr5-55302022-INS-68 | chr5-55302022-INS-68 |
| chr5-55340669-INS-5470 | chr5-55340670-INS-5470 |
| chr5-55381509-INS-82 | chr5-55381470-INS-76 |
| chr5-55382737-INS-85 | chr5-55382737-INS-85 |
| chr5-55391124-INS-53 | chr5-55391124-INS-53 |
| chr5-55397946-INS-731 | chr5-55397946-INS-731 |
| chr5-55410187-INS-57 | chr5-55410187-INS-57 |
| chr5-55410689-INS-1891 | chr5-55410690-INS-1891 |
| chr5-55410861-INS-548 | chr5-55410861-INS-548 |
| chr5-55437383-INS-956 | chr5-55437384-INS-979 |
| chr5-55440941-INS-75 | chr5-55440941-INS-75 |
| chr5-55478650-INS-5611 | chr5-55478651-INS-5611 |
| chr5-55501540-INS-84 | chr5-55501540-INS-84 |
| chr5-55527266-INS-697 | chr5-55527266-INS-695 |
| chr5-55528409-INS-6660 | chr5-55528410-INS-6660 |
| chr5-55560332-INS-5359 | chr5-55560333-INS-5360 |
| chr5-55561256-INS-1316 | chr5-55561257-INS-1266 |
| chr5-55593607-INS-87 | chr5-55593607-INS-58 |
| chr5-55602609-INS-772 | chr5-55602609-INS-772 |
| chr5-55621172-INS-58 | chr5-55621172-INS-58 |
| chr5-55626781-INS-1887 | chr5-55626782-INS-1887 |
| chr5-55665735-INS-6228 | chr5-55665736-INS-6228 |
| chr5-55665928-INS-60 | chr5-55665928-INS-61 |
| chr5-55781367-INS-74 | chr5-55781368-INS-74 |
| chr5-55789916-INS-6809 | chr5-55789917-INS-6809 |
| chr5-55802736-INS-76 | chr5-55802736-INS-76 |
| chr5-55828276-INS-133 | chr5-55828276-INS-133 |
| chr5-55847775-INS-204 | chr5-55847775-INS-204 |
| chr5-55848358-INS-6785 | chr5-55848359-INS-6784 |
| chr5-55902703-INS-944 | chr5-55902703-INS-940 |
| chr5-55925640-INS-204 | chr5-55925640-INS-204 |
| chr5-55936144-INS-3855 | chr5-55936145-INS-3855 |
| chr5-55945361-INS-724 | chr5-55945361-INS-724 |
| chr5-55945935-INS-88 | chr5-55945935-INS-88 |
| chr5-55961743-INS-171 | chr5-55961743-INS-173 |
| chr5-55963378-INS-112 | chr5-55963378-INS-112 |
| chr5-55977694-INS-103 | chr5-55977694-INS-118 |
| chr5-56007795-INS-59 | chr5-56007795-INS-55 |
| chr5-56017008-INS-104 | chr5-56017008-INS-104 |
| chr5-56021496-INS-136 | chr5-56021500-INS-152 |
| chr5-56022944-INS-79 | chr5-56022957-INS-63 |
| chr5-56040453-INS-756 | chr5-56040453-INS-755 |

| | |
|---|---|
| chr5-56041936-INS-56 | chr5-56041936-INS-56 |
| chr5-56052010-INS-773 | chr5-56052010-INS-773 |
| chr5-56069782-INS-52 | chr5-56069782-INS-51 |
| chr5-56092981-INS-160 | chr5-56092981-INS-160 |
| chr5-56105994-INS-199 | chr5-56105994-INS-199 |
| chr5-56113399-INS-54 | chr5-56113399-INS-56 |
| chr5-56162581-INS-6412 | chr5-56162582-INS-6412 |
| chr5-56178884-INS-2138 | chr5-56178885-INS-2138 |
| chr5-56178885-INS-710 | chr5-56178885-INS-709 |
| chr5-56182337-INS-71 | chr5-56182337-INS-71 |
| chr5-56210968-INS-540 | chr5-56210968-INS-540 |
| chr5-56214745-INS-88 | chr5-56214745-INS-58 |
| chr5-56230514-INS-199 | chr5-56230514-INS-199 |
| chr5-56236115-INS-78 | chr5-56236115-INS-78 |
| chr5-56242204-INS-466 | chr5-56242204-INS-466 |
| chr5-56244132-INS-118 | chr5-56244132-INS-118 |
| chr5-56265089-INS-1995 | chr5-56265090-INS-1995 |
| chr5-56287475-INS-104 | chr5-56287495-INS-98 |
| chr5-56323338-INS-63 | chr5-56323334-INS-71 |
| chr5-56344732-INS-122 | chr5-56344732-INS-122 |
| chr5-56398145-INS-54 | chr5-56398103-INS-70 |
| chr5-56420431-INS-358 | chr5-56420431-INS-358 |
| chr5-56427488-INS-319 | chr5-56427488-INS-322 |
| chr5-56428557-INS-476 | chr5-56428557-INS-427 |
| chr5-56430981-INS-80 | chr5-56430968-INS-74 |
| chr5-56479348-INS-5055 | chr5-56479349-INS-5055 |
| chr5-56488506-INS-77 | chr5-56488506-INS-50 |
| chr5-56492214-INS-154 | chr5-56492214-INS-152 |
| chr5-56520379-INS-511 | chr5-56520379-INS-511 |
| chr5-56535618-INS-502 | chr5-56535618-INS-502 |
| chr5-56538806-INS-457 | chr5-56538806-INS-457 |
| chr5-56551919-INS-167 | chr5-56551919-INS-169 |
| chr5-56554304-INS-504 | chr5-56554304-INS-504 |
| chr5-56568629-INS-2816 | chr5-56568630-INS-2819 |
| chr5-56586238-INS-1821 | chr5-56586239-INS-1821 |
| chr5-56588541-INS-6814 | chr5-56588542-INS-6815 |
| chr5-56596369-INS-56 | chr5-56596369-INS-56 |
| chr5-56601242-INS-408 | chr5-56601242-INS-409 |
| chr5-56629626-INS-212 | chr5-56629626-INS-212 |
| chr5-56637338-INS-173 | chr5-56637339-INS-180 |
| chr5-56641438-INS-54 | chr5-56641438-INS-52 |
| chr5-56649950-INS-108 | chr5-56649945-INS-73 |
| chr5-56650670-INS-424 | chr5-56650670-INS-424 |
| chr5-56667290-INS-803 | chr5-56667292-INS-803 |
| chr5-56690220-INS-58 | chr5-56690220-INS-56 |
| chr5-56699563-INS-209 | chr5-56699563-INS-209 |
| chr5-56720653-INS-390 | chr5-56720653-INS-390 |
| chr5-56724709-INS-58 | chr5-56724709-INS-58 |
| chr5-56726404-INS-160 | chr5-56726404-INS-158 |
| chr5-56755469-INS-59 | chr5-56755421-INS-60 |
| chr5-56762753-INS-408 | chr5-56762753-INS-408 |
| chr5-56763436-INS-57 | chr5-56763436-INS-57 |
| chr5-56772239-INS-356 | chr5-56772239-INS-356 |
| chr5-56791779-INS-692 | chr5-56791779-INS-694 |
| chr5-56802502-INS-1126 | chr5-56802503-INS-1126 |
| chr5-56814159-INS-71 | chr5-56814207-INS-72 |
| chr5-56862388-INS-52 | chr5-56862388-INS-54 |
| chr5-56873042-INS-6525 | chr5-56873043-INS-6526 |
| chr5-56915709-INS-108 | chr5-56915709-INS-108 |
| chr5-56921464-INS-390 | chr5-56921464-INS-390 |
| chr5-56923982-INS-6263 | chr5-56923983-INS-6263 |
| chr5-56947789-INS-1594 | chr5-56947790-INS-1594 |
| chr5-56951180-INS-88 | chr5-56951180-INS-88 |
| chr5-56955749-INS-5749 | chr5-56955750-INS-5749 |
| chr5-56957367-INS-68 | chr5-56957390-INS-82 |
| chr5-56958948-INS-55 | chr5-56958945-INS-51 |
| chr5-56961823-INS-86 | chr5-56961830-INS-81 |
| chr5-56964648-INS-221 | chr5-56964648-INS-222 |
| chr5-56970929-INS-79 | chr5-56970929-INS-81 |
| chr5-56975019-INS-81 | chr5-56975019-INS-81 |
| chr5-56979148-INS-200 | chr5-56979148-INS-200 |
| chr5-56983105-INS-454 | chr5-56983105-INS-454 |
| chr5-57009706-INS-73 | chr5-57009704-INS-73 |
| chr5-57019761-INS-53 | chr5-57019761-INS-53 |
| chr5-57024823-INS-6758 | chr5-57024824-INS-6760 |
| chr5-57043519-INS-60 | chr5-57043519-INS-60 |
| chr5-57105722-INS-68 | chr5-57105722-INS-62 |

| | |
|---|---|
| chr5-57115137-INS-90 | chr5-57115137-INS-87 |
| chr5-57135439-INS-148 | chr5-57135439-INS-108 |
| chr5-57160450-INS-54 | chr5-57160450-INS-54 |
| chr5-57169923-INS-60 | chr5-57169923-INS-76 |
| chr5-57199810-INS-615 | chr5-57199810-INS-615 |
| chr5-57203985-INS-112 | chr5-57204013-INS-86 |
| chr5-57232254-INS-6374 | chr5-57232255-INS-6395 |
| chr5-57233451-INS-2947 | chr5-57233452-INS-2948 |
| chr5-57237649-INS-78 | chr5-57237649-INS-76 |
| chr5-57252271-INS-107 | chr5-57252273-INS-120 |
| chr5-57267271-INS-149 | chr5-57267271-INS-142 |
| chr5-57279315-INS-379 | chr5-57279315-INS-379 |
| chr5-57347083-INS-57 | chr5-57347069-INS-77 |
| chr5-57349549-INS-58 | chr5-57349549-INS-52 |
| chr5-57361232-INS-691 | chr5-57361232-INS-690 |
| chr5-57396670-INS-360 | chr5-57396670-INS-318 |
| chr5-57413055-INS-4493 | chr5-57413056-INS-4493 |
| chr5-57425157-INS-586 | chr5-57425157-INS-583 |
| chr5-57473420-INS-66 | chr5-57473371-INS-113 |
| chr5-57512403-INS-1584 | chr5-57512380-INS-1567 |
| chr5-57520627-INS-755 | chr5-57520627-INS-754 |
| chr5-57548830-INS-56 | chr5-57548830-INS-66 |
| chr5-57648488-INS-55 | chr5-57648488-INS-55 |
| chr5-57693122-INS-6422 | chr5-57693123-INS-6428 |
| chr5-57712080-INS-83 | chr5-57712080-INS-83 |
| chr5-57739238-INS-1820 | chr5-57739239-INS-1820 |
| chr5-57780813-INS-604 | chr5-57780813-INS-603 |
| chr5-57789691-INS-6946 | chr5-57789692-INS-6949 |
| chr5-57819370-INS-1891 | chr5-57819371-INS-1891 |
| chr5-57832508-INS-304 | chr5-57832508-INS-304 |
| chr5-57859906-INS-435 | chr5-57859906-INS-434 |
| chr5-57864327-INS-52 | chr5-57864327-INS-53 |
| chr5-57892736-INS-60 | chr5-57892736-INS-60 |
| chr5-57897951-INS-4086 | chr5-57897952-INS-4086 |
| chr5-57925771-INS-715 | chr5-57925771-INS-717 |
| chr5-57965692-INS-83 | chr5-57965692-INS-82 |
| chr5-57975078-INS-59 | chr5-57975078-INS-59 |
| chr5-58004188-INS-160 | chr5-58004188-INS-160 |
| chr5-58005327-INS-518 | chr5-58005321-INS-528 |
| chr5-58043894-INS-212 | chr5-58043894-INS-209 |
| chr5-58048630-INS-77 | chr5-58048630-INS-77 |
| chr5-58090707-INS-111 | chr5-58090707-INS-133 |
| chr5-58094271-INS-221 | chr5-58094271-INS-222 |
| chr5-58113124-INS-1013 | chr5-58113125-INS-1013 |
| chr5-58122006-INS-68 | chr5-58122006-INS-52 |
| chr5-58128182-INS-160 | chr5-58128182-INS-160 |
| chr5-58177720-INS-54 | chr5-58177720-INS-77 |
| chr5-58191027-INS-549 | chr5-58191027-INS-550 |
| chr5-58221569-INS-2799 | chr5-58221570-INS-2799 |
| chr5-58230361-INS-864 | chr5-58230361-INS-864 |
| chr5-58251045-INS-6532 | chr5-58251046-INS-6532 |
| chr5-58262666-INS-1317 | chr5-58262667-INS-1318 |
| chr5-58264764-INS-624 | chr5-58264764-INS-624 |
| chr5-58280528-INS-6574 | chr5-58280529-INS-6574 |
| chr5-58282454-INS-110 | chr5-58282454-INS-121 |
| chr5-58285371-INS-94 | chr5-58285371-INS-94 |
| chr5-58292728-INS-66 | chr5-58292728-INS-58 |
| chr5-58293519-INS-52 | chr5-58293519-INS-56 |
| chr5-58303327-INS-6335 | chr5-58303328-INS-6335 |
| chr5-58306888-INS-399 | chr5-58306888-INS-399 |
| chr5-58336520-INS-8072 | chr5-58336521-INS-8072 |
| chr5-58355986-INS-82 | chr5-58355986-INS-112 |
| chr5-58359552-INS-696 | chr5-58359547-INS-691 |
| chr5-58367565-INS-71 | chr5-58367566-INS-62 |
| chr5-58372475-INS-74 | chr5-58372475-INS-74 |
| chr5-58377013-INS-68 | chr5-58377013-INS-68 |
| chr5-58390290-INS-3384 | chr5-58390291-INS-3384 |
| chr5-58391738-INS-1544 | chr5-58391739-INS-1544 |
| chr5-58417366-INS-237 | chr5-58417366-INS-205 |
| chr5-58468280-INS-190 | chr5-58468280-INS-190 |
| chr5-58486204-INS-1012 | chr5-58486205-INS-1012 |
| chr5-58497407-INS-91 | chr5-58497407-INS-91 |
| chr5-58506773-INS-55 | chr5-58506773-INS-55 |
| chr5-58549271-INS-54 | chr5-58549271-INS-60 |
| chr5-58606757-INS-65 | chr5-58606757-INS-65 |
| chr5-58653216-INS-51 | chr5-58653216-INS-51 |
| chr5-58667142-INS-6156 | chr5-58667143-INS-6157 |

| | |
|---|---|
| chr5-58697823-INS-243 | chr5-58697823-INS-244 |
| chr5-58709115-INS-64 | chr5-58709115-INS-71 |
| chr5-58722279-INS-94 | chr5-58722295-INS-55 |
| chr5-58755772-INS-472 | chr5-58755772-INS-473 |
| chr5-58767719-INS-57 | chr5-58767719-INS-69 |
| chr5-58790306-INS-569 | chr5-58790306-INS-569 |
| chr5-58805489-INS-406 | chr5-58805499-INS-409 |
| chr5-58809456-INS-105 | chr5-58809456-INS-105 |
| chr5-58840400-INS-761 | chr5-58840400-INS-763 |
| chr5-58904248-INS-180 | chr5-58904248-INS-180 |
| chr5-58915184-INS-1893 | chr5-58915185-INS-1893 |
| chr5-58926504-INS-56 | chr5-58926504-INS-58 |
| chr5-58946468-INS-121 | chr5-58946468-INS-121 |
| chr5-58955076-INS-112 | chr5-58955067-INS-141 |
| chr5-58967183-INS-83 | chr5-58967183-INS-83 |
| chr5-58968101-INS-328 | chr5-58968101-INS-328 |
| chr5-59080500-INS-1859 | chr5-59080501-INS-1859 |
| chr5-59105251-INS-1891 | chr5-59105252-INS-1891 |
| chr5-59109321-INS-188 | chr5-59109321-INS-188 |
| chr5-59173638-INS-327 | chr5-59173638-INS-327 |
| chr5-59189188-INS-1645 | chr5-59189185-INS-1656 |
| chr5-59189284-INS-87 | chr5-59189264-INS-114 |
| chr5-59191886-INS-72 | chr5-59191886-INS-72 |
| chr5-59219602-INS-6389 | chr5-59219603-INS-6390 |
| chr5-59232805-INS-6385 | chr5-59232806-INS-6385 |
| chr5-59236587-INS-205 | chr5-59236587-INS-242 |
| chr5-59249284-INS-212 | chr5-59249284-INS-212 |
| chr5-59259093-INS-791 | chr5-59259093-INS-791 |
| chr5-59290393-INS-242 | chr5-59290393-INS-239 |
| chr5-59305389-INS-135 | chr5-59305389-INS-135 |
| chr5-59310538-INS-6217 | chr5-59310559-INS-6207 |
| chr5-59310617-INS-60 | chr5-59310617-INS-62 |
| chr5-59316147-INS-137 | chr5-59316147-INS-149 |
| chr5-59349695-INS-2824 | chr5-59349696-INS-2824 |
| chr5-59356971-INS-7671 | chr5-59356974-INS-7672 |
| chr5-59370242-INS-6894 | chr5-59370243-INS-6894 |
| chr5-59384961-INS-398 | chr5-59384961-INS-398 |
| chr5-59393207-INS-72 | chr5-59393207-INS-54 |
| chr5-59425461-INS-265 | chr5-59425461-INS-265 |
| chr5-59453699-INS-53 | chr5-59453699-INS-77 |
| chr5-59465010-INS-72 | chr5-59465010-INS-72 |
| chr5-59494685-INS-52 | chr5-59494685-INS-52 |
| chr5-59500249-INS-71 | chr5-59500246-INS-51 |
| chr5-59520555-INS-60 | chr5-59520555-INS-70 |
| chr5-59525683-INS-993 | chr5-59525683-INS-994 |
| chr5-59543459-INS-255 | chr5-59543459-INS-241 |
| chr5-59604380-INS-177 | chr5-59604380-INS-177 |
| chr5-59630583-INS-94 | chr5-59630583-INS-85 |
| chr5-59640998-INS-55 | chr5-59640998-INS-55 |
| chr5-59655348-INS-4503 | chr5-59655349-INS-4503 |
| chr5-59670851-INS-130 | chr5-59670851-INS-139 |
| chr5-59686974-INS-156 | chr5-59686967-INS-129 |
| chr5-59697046-INS-122 | chr5-59697059-INS-126 |
| chr5-59699232-INS-63 | chr5-59699232-INS-66 |
| chr5-59730783-INS-7134 | chr5-59730784-INS-7142 |
| chr5-59748103-INS-55 | chr5-59748103-INS-55 |
| chr5-59772260-INS-84 | chr5-59772260-INS-68 |
| chr5-59791489-INS-52 | chr5-59791489-INS-54 |
| chr5-59836377-INS-59 | chr5-59836377-INS-59 |
| chr5-59858226-INS-68 | chr5-59858226-INS-60 |
| chr5-59935109-INS-272 | chr5-59935109-INS-273 |
| chr5-59942138-INS-58 | chr5-59942138-INS-60 |
| chr5-59968309-INS-6676 | chr5-59968310-INS-6676 |
| chr5-59970219-INS-2955 | chr5-59970220-INS-2955 |
| chr5-59978862-INS-65 | chr5-59978839-INS-90 |
| chr5-59992078-INS-2944 | chr5-59992079-INS-2944 |
| chr5-60019472-INS-122 | chr5-60019473-INS-78 |
| chr5-60023477-INS-4172 | chr5-60023478-INS-4172 |
| chr5-60026060-INS-6472 | chr5-60026061-INS-6475 |
| chr5-60034748-INS-228 | chr5-60034748-INS-233 |
| chr5-60058215-INS-6572 | chr5-60058216-INS-6573 |
| chr5-60076188-INS-87 | chr5-60076216-INS-72 |
| chr5-60088548-INS-1080 | chr5-60088549-INS-1064 |
| chr5-60127449-INS-197 | chr5-60127449-INS-197 |
| chr5-60169219-INS-6885 | chr5-60169220-INS-6887 |
| chr5-60202638-INS-5263 | chr5-60202639-INS-5263 |
| chr5-60210320-INS-422 | chr5-60210326-INS-423 |

| | |
|---|---|
| chr5-60280925-INS-707 | chr5-60280925-INS-706 |
| chr5-60285430-INS-1326 | chr5-60285431-INS-1303 |
| chr5-60286552-INS-2973 | chr5-60286553-INS-2963 |
| chr5-60301580-INS-457 | chr5-60301580-INS-456 |
| chr5-60313238-INS-232 | chr5-60313238-INS-232 |
| chr5-60326592-INS-88 | chr5-60326619-INS-86 |
| chr5-60357595-INS-56 | chr5-60357595-INS-60 |
| chr5-60369400-INS-6453 | chr5-60369401-INS-6453 |
| chr5-60398470-INS-268 | chr5-60398455-INS-301 |
| chr5-60404291-INS-97 | chr5-60404291-INS-97 |
| chr5-60411570-INS-70 | chr5-60411570-INS-68 |
| chr5-60419919-INS-2980 | chr5-60419920-INS-2980 |
| chr5-60430655-INS-118 | chr5-60430655-INS-118 |
| chr5-60439541-INS-7075 | chr5-60439542-INS-7077 |
| chr5-60442361-INS-132 | chr5-60442361-INS-132 |
| chr5-60450239-INS-173 | chr5-60450239-INS-173 |
| chr5-60450469-INS-4301 | chr5-60450470-INS-4301 |
| chr5-60468727-INS-52 | chr5-60468727-INS-54 |
| chr5-60472986-INS-259 | chr5-60472984-INS-264 |
| chr5-60513073-INS-400 | chr5-60513073-INS-400 |
| chr5-60517607-INS-219 | chr5-60517607-INS-225 |
| chr5-60523076-INS-69 | chr5-60523116-INS-60 |
| chr5-60547825-INS-1844 | chr5-60547826-INS-1844 |
| chr5-60569180-INS-686 | chr5-60569180-INS-686 |
| chr5-60585378-INS-223 | chr5-60585378-INS-223 |
| chr5-60603819-INS-64 | chr5-60603819-INS-64 |
| chr5-60609868-INS-179 | chr5-60609898-INS-178 |
| chr5-60615160-INS-54 | chr5-60615155-INS-57 |
| chr5-60645049-INS-556 | chr5-60645049-INS-557 |
| chr5-60652041-INS-59 | chr5-60652041-INS-62 |
| chr5-60663789-INS-5286 | chr5-60663790-INS-5288 |
| chr5-60668668-INS-53 | chr5-60668668-INS-53 |
| chr5-60694105-INS-60 | chr5-60694105-INS-65 |
| chr5-60704355-INS-63 | chr5-60704355-INS-63 |
| chr5-60753919-INS-492 | chr5-60753919-INS-492 |
| chr5-60756410-INS-67 | chr5-60756410-INS-67 |
| chr5-60835296-INS-148 | chr5-60835301-INS-147 |
| chr5-60840806-INS-174 | chr5-60840806-INS-174 |
| chr5-60857733-INS-638 | chr5-60857733-INS-638 |
| chr5-60866467-INS-90 | chr5-60866467-INS-90 |
| chr5-60867728-INS-888 | chr5-60867730-INS-884 |
| chr5-60886467-INS-208 | chr5-60886468-INS-208 |
| chr5-60951454-INS-95 | chr5-60951454-INS-100 |
| chr5-60973509-INS-56 | chr5-60973509-INS-54 |
| chr5-60973978-INS-1809 | chr5-60973978-INS-1806 |
| chr5-60976164-INS-980 | chr5-60976164-INS-992 |
| chr5-60984571-INS-160 | chr5-60984571-INS-142 |
| chr5-61007862-INS-101 | chr5-61007862-INS-68 |
| chr5-61031944-INS-220 | chr5-61031944-INS-220 |
| chr5-61041383-INS-434 | chr5-61041383-INS-434 |
| chr5-61058567-INS-55 | chr5-61058567-INS-55 |
| chr5-61081765-INS-6801 | chr5-61081766-INS-6801 |
| chr5-61176406-INS-58 | chr5-61176406-INS-58 |
| chr5-61183921-INS-426 | chr5-61183921-INS-426 |
| chr5-61198377-INS-1149 | chr5-61198378-INS-1144 |
| chr5-61238729-INS-68 | chr5-61238729-INS-80 |
| chr5-61247211-INS-582 | chr5-61247212-INS-583 |
| chr5-61250130-INS-89 | chr5-61250156-INS-76 |
| chr5-61327152-INS-7432 | chr5-61327153-INS-7432 |
| chr5-61368249-INS-2891 | chr5-61368250-INS-2905 |
| chr5-61370305-INS-212 | chr5-61370305-INS-212 |
| chr5-61378844-INS-771 | chr5-61378844-INS-771 |
| chr5-61381904-INS-296 | chr5-61381904-INS-296 |
| chr5-61400836-INS-3672 | chr5-61400837-INS-3672 |
| chr5-61423979-INS-934 | chr5-61423979-INS-932 |
| chr5-61476377-INS-429 | chr5-61476377-INS-429 |
| chr5-61521209-INS-229 | chr5-61521209-INS-230 |
| chr5-61522666-INS-65 | chr5-61522670-INS-77 |
| chr5-61541297-INS-7086 | chr5-61541298-INS-7087 |
| chr5-61575753-INS-609 | chr5-61575753-INS-610 |
| chr5-61620332-INS-54 | chr5-61620332-INS-54 |
| chr5-61624321-INS-846 | chr5-61624321-INS-846 |
| chr5-61648923-INS-6418 | chr5-61648924-INS-6419 |
| chr5-61660340-INS-94 | chr5-61660340-INS-94 |
| chr5-61671507-INS-1988 | chr5-61671508-INS-1988 |
| chr5-61673146-INS-594 | chr5-61673146-INS-594 |
| chr5-61677612-INS-160 | chr5-61677612-INS-160 |

| | |
|---|---|
| chr5-61679381-INS-88 | chr5-61679381-INS-88 |
| chr5-61680789-INS-74 | chr5-61680789-INS-50 |
| chr5-61685989-INS-395 | chr5-61685989-INS-398 |
| chr5-61695647-INS-326 | chr5-61695648-INS-326 |
| chr5-61720668-INS-107 | chr5-61720662-INS-111 |
| chr5-61722751-INS-370 | chr5-61722751-INS-366 |
| chr5-61743682-INS-61 | chr5-61743682-INS-61 |
| chr5-61771537-INS-4546 | chr5-61771538-INS-4546 |
| chr5-61777229-INS-6198 | chr5-61777230-INS-6226 |
| chr5-61785487-INS-498 | chr5-61785487-INS-498 |
| chr5-61806926-INS-814 | chr5-61806926-INS-814 |
| chr5-61812392-INS-6656 | chr5-61812393-INS-6656 |
| chr5-61815798-INS-72 | chr5-61815798-INS-72 |
| chr5-61855073-INS-206 | chr5-61855073-INS-206 |
| chr5-61858754-INS-335 | chr5-61858754-INS-336 |
| chr5-61871474-INS-6513 | chr5-61871475-INS-6513 |
| chr5-61876080-INS-55 | chr5-61876080-INS-55 |
| chr5-61876913-INS-307 | chr5-61876913-INS-308 |
| chr5-61997884-INS-53 | chr5-61997877-INS-68 |
| chr5-62071346-INS-208 | chr5-62071346-INS-208 |
| chr5-62082924-INS-7295 | chr5-62082925-INS-7295 |
| chr5-62124899-INS-80 | chr5-62124899-INS-80 |
| chr5-62180315-INS-83 | chr5-62180303-INS-88 |
| chr5-62239336-INS-665 | chr5-62239342-INS-663 |
| chr5-62266034-INS-410 | chr5-62266026-INS-408 |
| chr5-62310302-INS-117 | chr5-62310302-INS-96 |
| chr5-62317651-INS-6444 | chr5-62317652-INS-6444 |
| chr5-62336850-INS-394 | chr5-62336849-INS-394 |
| chr5-62380861-INS-2635 | chr5-62380862-INS-2635 |
| chr5-62410070-INS-3037 | chr5-62410071-INS-3037 |
| chr5-62410472-INS-503 | chr5-62410472-INS-503 |
| chr5-62413013-INS-1366 | chr5-62413014-INS-1366 |
| chr5-62439752-INS-79 | chr5-62439753-INS-76 |
| chr5-62441856-INS-2286 | chr5-62441862-INS-2301 |
| chr5-62457026-INS-203 | chr5-62457026-INS-200 |
| chr5-62492918-INS-59 | chr5-62492918-INS-59 |
| chr5-62520526-INS-1546 | chr5-62520527-INS-1546 |
| chr5-62549734-INS-1087 | chr5-62549735-INS-1087 |
| chr5-62574106-INS-129 | chr5-62574106-INS-129 |
| chr5-62583704-INS-3653 | chr5-62583705-INS-3649 |
| chr5-62630083-INS-201 | chr5-62630083-INS-201 |
| chr5-62636975-INS-51 | chr5-62636975-INS-52 |
| chr5-62692440-INS-95 | chr5-62692440-INS-54 |
| chr5-62705156-INS-507 | chr5-62705156-INS-508 |
| chr5-62730744-INS-6860 | chr5-62730745-INS-6860 |
| chr5-62774257-INS-91 | chr5-62774257-INS-124 |
| chr5-62781945-INS-108 | chr5-62781945-INS-108 |
| chr5-62805584-INS-102 | chr5-62805584-INS-84 |
| chr5-62817006-INS-4408 | chr5-62817007-INS-4410 |
| chr5-62850684-INS-64 | chr5-62850684-INS-64 |
| chr5-62865026-INS-114 | chr5-62865026-INS-117 |
| chr5-62964708-INS-1729 | chr5-62964709-INS-1728 |
| chr5-62979723-INS-6859 | chr5-62979724-INS-6859 |
| chr5-62981313-INS-55 | chr5-62981313-INS-58 |
| chr5-62986759-INS-6134 | chr5-62986760-INS-6135 |
| chr5-63005202-INS-558 | chr5-63005201-INS-556 |
| chr5-63034146-INS-943 | chr5-63034146-INS-943 |
| chr5-63044687-INS-769 | chr5-63044675-INS-778 |
| chr5-63053904-INS-1127 | chr5-63053905-INS-1127 |
| chr5-63065698-INS-6260 | chr5-63065699-INS-6260 |
| chr5-63069567-INS-64 | chr5-63069567-INS-51 |
| chr5-63081737-INS-594 | chr5-63081737-INS-595 |
| chr5-63122838-INS-115 | chr5-63122838-INS-115 |
| chr5-63158781-INS-60 | chr5-63158781-INS-60 |
| chr5-63162416-INS-63 | chr5-63162416-INS-63 |
| chr5-63175287-INS-68 | chr5-63175287-INS-68 |
| chr5-63205418-INS-736 | chr5-63205418-INS-736 |
| chr5-63209513-INS-2598 | chr5-63209514-INS-2600 |
| chr5-63213718-INS-698 | chr5-63213718-INS-698 |
| chr5-63215275-INS-591 | chr5-63215275-INS-591 |
| chr5-63229448-INS-710 | chr5-63229448-INS-711 |
| chr5-63249543-INS-6447 | chr5-63249544-INS-6447 |
| chr5-63281647-INS-6440 | chr5-63281648-INS-6440 |
| chr5-63288566-INS-58 | chr5-63288566-INS-60 |
| chr5-63295158-INS-6170 | chr5-63295159-INS-6170 |
| chr5-63365852-INS-234 | chr5-63365852-INS-237 |
| chr5-63410863-INS-51 | chr5-63410863-INS-51 |

| | |
|---|---|
| chr5-63416991-INS-78 | chr5-63416991-INS-72 |
| chr5-63421819-INS-464 | chr5-63421819-INS-465 |
| chr5-63425877-INS-82 | chr5-63425885-INS-105 |
| chr5-63435379-INS-75 | chr5-63435379-INS-103 |
| chr5-63444447-INS-60 | chr5-63444447-INS-60 |
| chr5-63448927-INS-482 | chr5-63448931-INS-517 |
| chr5-63449993-INS-6526 | chr5-63449994-INS-6527 |
| chr5-63458887-INS-1873 | chr5-63458888-INS-1873 |
| chr5-63465948-INS-64 | chr5-63465948-INS-76 |
| chr5-63466671-INS-675 | chr5-63466671-INS-674 |
| chr5-63474435-INS-1389 | chr5-63474436-INS-1391 |
| chr5-63502817-INS-85 | chr5-63502817-INS-85 |
| chr5-63510370-INS-1860 | chr5-63510371-INS-1860 |
| chr5-63525987-INS-214 | chr5-63525959-INS-214 |
| chr5-63553323-INS-183 | chr5-63553323-INS-184 |
| chr5-63567906-INS-58 | chr5-63567899-INS-50 |
| chr5-63582988-INS-4649 | chr5-63582989-INS-4649 |
| chr5-63594559-INS-7088 | chr5-63594560-INS-7088 |
| chr5-63630047-INS-1842 | chr5-63630048-INS-1842 |
| chr5-63646704-INS-3712 | chr5-63646705-INS-3721 |
| chr5-63683257-INS-6138 | chr5-63683258-INS-6132 |
| chr5-63754381-INS-205 | chr5-63754381-INS-205 |
| chr5-63791454-INS-84 | chr5-63791454-INS-68 |
| chr5-63793108-INS-1632 | chr5-63793109-INS-1638 |
| chr5-63797713-INS-343 | chr5-63797713-INS-343 |
| chr5-63800278-INS-54 | chr5-63800294-INS-86 |
| chr5-63814892-INS-212 | chr5-63814892-INS-212 |
| chr5-63849941-INS-223 | chr5-63849941-INS-222 |
| chr5-63891359-INS-55 | chr5-63891364-INS-65 |
| chr5-63894615-INS-2765 | chr5-63894616-INS-2764 |
| chr5-63922575-INS-1890 | chr5-63922576-INS-1890 |
| chr5-63924334-INS-52 | chr5-63924334-INS-52 |
| chr5-63926933-INS-751 | chr5-63926933-INS-751 |
| chr5-63942633-INS-92 | chr5-63942633-INS-92 |
| chr5-63997050-INS-410 | chr5-63997050-INS-406 |
| chr5-64030958-INS-79 | chr5-64030958-INS-79 |
| chr5-64075388-INS-94 | chr5-64075388-INS-94 |
| chr5-64088956-INS-614 | chr5-64088956-INS-607 |
| chr5-64092935-INS-53 | chr5-64092935-INS-53 |
| chr5-64103321-INS-5105 | chr5-64103322-INS-5105 |
| chr5-64112045-INS-71 | chr5-64112070-INS-60 |
| chr5-64115649-INS-57 | chr5-64115649-INS-57 |
| chr5-64120866-INS-277 | chr5-64120866-INS-277 |
| chr5-64123055-INS-219 | chr5-64123055-INS-219 |
| chr5-64123982-INS-78 | chr5-64123982-INS-78 |
| chr5-64139023-INS-7108 | chr5-64139024-INS-7108 |
| chr5-64146556-INS-88 | chr5-64146556-INS-76 |
| chr5-64154411-INS-243 | chr5-64154411-INS-243 |
| chr5-64166120-INS-65 | chr5-64166120-INS-65 |
| chr5-64169489-INS-325 | chr5-64169490-INS-326 |
| chr5-64184530-INS-397 | chr5-64184530-INS-397 |
| chr5-64202999-INS-64 | chr5-64202999-INS-70 |
| chr5-64223659-INS-246 | chr5-64223672-INS-223 |
| chr5-64234361-INS-54 | chr5-64234361-INS-54 |
| chr5-64240444-INS-7105 | chr5-64240445-INS-7105 |
| chr5-64242622-INS-196 | chr5-64242622-INS-196 |
| chr5-64245698-INS-76 | chr5-64245708-INS-73 |
| chr5-64258913-INS-98 | chr5-64258913-INS-98 |
| chr5-64297193-INS-5086 | chr5-64297194-INS-5086 |
| chr5-64302501-INS-2921 | chr5-64302502-INS-2921 |
| chr5-64319029-INS-85 | chr5-64319029-INS-85 |
| chr5-64322306-INS-1890 | chr5-64322307-INS-1890 |
| chr5-64342207-INS-518 | chr5-64342207-INS-523 |
| chr5-64349275-INS-58 | chr5-64349275-INS-58 |
| chr5-64356418-INS-1888 | chr5-64356419-INS-1888 |
| chr5-64427006-INS-7150 | chr5-64427007-INS-7150 |
| chr5-64487405-INS-285 | chr5-64487405-INS-285 |
| chr5-64488394-INS-80 | chr5-64488394-INS-82 |
| chr5-64493848-INS-207 | chr5-64493848-INS-207 |
| chr5-64497637-INS-74 | chr5-64497637-INS-74 |
| chr5-64498852-INS-2321 | chr5-64498853-INS-2321 |
| chr5-64545383-INS-77 | chr5-64545375-INS-68 |
| chr5-64573457-INS-93 | chr5-64573457-INS-93 |
| chr5-64601044-INS-100 | chr5-64601050-INS-94 |
| chr5-64617004-INS-403 | chr5-64617004-INS-403 |
| chr5-64624527-INS-105 | chr5-64624527-INS-111 |
| chr5-64672897-INS-212 | chr5-64672897-INS-211 |

| | |
|---|---|
| chr5-64677388-INS-549 | chr5-64677374-INS-563 |
| chr5-64682410-INS-96 | chr5-64682405-INS-106 |
| chr5-64683450-INS-219 | chr5-64683450-INS-219 |
| chr5-64698631-INS-90 | chr5-64698631-INS-90 |
| chr5-64728121-INS-205 | chr5-64728121-INS-205 |
| chr5-64788645-INS-663 | chr5-64788645-INS-663 |
| chr5-64796473-INS-4754 | chr5-64796474-INS-4754 |
| chr5-64802568-INS-401 | chr5-64802568-INS-401 |
| chr5-64827805-INS-134 | chr5-64827795-INS-147 |
| chr5-64837181-INS-4708 | chr5-64837182-INS-4708 |
| chr5-64843099-INS-72 | chr5-64843099-INS-72 |
| chr5-64850822-INS-93 | chr5-64850823-INS-93 |
| chr5-64866334-INS-53 | chr5-64866337-INS-61 |
| chr5-64870798-INS-100 | chr5-64870798-INS-71 |
| chr5-64878096-INS-1493 | chr5-64878097-INS-1493 |
| chr5-64881233-INS-7140 | chr5-64881245-INS-7126 |
| chr5-64899105-INS-8403 | chr5-64899106-INS-8404 |
| chr5-64905618-INS-292 | chr5-64905618-INS-275 |
| chr5-64921273-INS-214 | chr5-64921298-INS-207 |
| chr5-64930444-INS-66 | chr5-64930432-INS-50 |
| chr5-64933827-INS-138 | chr5-64933827-INS-139 |
| chr5-64953602-INS-54 | chr5-64953553-INS-54 |
| chr5-64967734-INS-640 | chr5-64967734-INS-640 |
| chr5-64999687-INS-399 | chr5-64999687-INS-399 |
| chr5-65004439-INS-6556 | chr5-65004440-INS-6558 |
| chr5-65009361-INS-69 | chr5-65009361-INS-68 |
| chr5-65067895-INS-352 | chr5-65067895-INS-352 |
| chr5-65075434-INS-170 | chr5-65075434-INS-172 |
| chr5-65080338-INS-243 | chr5-65080338-INS-243 |
| chr5-65081373-INS-1705 | chr5-65081374-INS-1706 |
| chr5-65086723-INS-92 | chr5-65086723-INS-100 |
| chr5-65088016-INS-56 | chr5-65088016-INS-58 |
| chr5-65092148-INS-54 | chr5-65092148-INS-51 |
| chr5-65118921-INS-79 | chr5-65118921-INS-80 |
| chr5-65123045-INS-66 | chr5-65123045-INS-66 |
| chr5-65125478-INS-143 | chr5-65125453-INS-157 |
| chr5-65140397-INS-76 | chr5-65140409-INS-92 |
| chr5-65158396-INS-195 | chr5-65158396-INS-195 |
| chr5-65170896-INS-74 | chr5-65170896-INS-62 |
| chr5-65181946-INS-65 | chr5-65181946-INS-65 |
| chr5-65183323-INS-502 | chr5-65183323-INS-502 |
| chr5-65184814-INS-306 | chr5-65184814-INS-295 |
| chr5-65188864-INS-87 | chr5-65188853-INS-72 |
| chr5-65195327-INS-209 | chr5-65195327-INS-209 |
| chr5-65199722-INS-441 | chr5-65199722-INS-441 |
| chr5-65281490-INS-146 | chr5-65281490-INS-146 |
| chr5-65312606-INS-65 | chr5-65312595-INS-52 |
| chr5-65328178-INS-128 | chr5-65328178-INS-134 |
| chr5-65369266-INS-152 | chr5-65369266-INS-152 |
| chr5-65391765-INS-56 | chr5-65391765-INS-56 |
| chr5-65395111-INS-230 | chr5-65395111-INS-247 |
| chr5-65491100-INS-193 | chr5-65491100-INS-193 |
| chr5-65499400-INS-192 | chr5-65499400-INS-192 |
| chr5-65512777-INS-151 | chr5-65512777-INS-151 |
| chr5-65519537-INS-55 | chr5-65519537-INS-55 |
| chr5-65525675-INS-234 | chr5-65525675-INS-234 |
| chr5-65552560-INS-214 | chr5-65552560-INS-214 |
| chr5-65564735-INS-88 | chr5-65564735-INS-88 |
| chr5-65575111-INS-58 | chr5-65575114-INS-64 |
| chr5-65596533-INS-67 | chr5-65596533-INS-62 |
| chr5-65631362-INS-218 | chr5-65631362-INS-218 |
| chr5-65634919-INS-208 | chr5-65634919-INS-208 |
| chr5-65658913-INS-562 | chr5-65658913-INS-567 |
| chr5-65669148-INS-221 | chr5-65669148-INS-221 |
| chr5-65670780-INS-55 | chr5-65670780-INS-55 |
| chr5-65675261-INS-3856 | chr5-65675264-INS-3868 |
| chr5-65740638-INS-96 | chr5-65740638-INS-64 |
| chr5-65803426-INS-214 | chr5-65803426-INS-214 |
| chr5-65808101-INS-212 | chr5-65808101-INS-225 |
| chr5-65862296-INS-211 | chr5-65862296-INS-211 |
| chr5-65864936-INS-283 | chr5-65864936-INS-283 |
| chr5-65869311-INS-216 | chr5-65869311-INS-216 |
| chr5-65884687-INS-83 | chr5-65884677-INS-78 |
| chr5-65892026-INS-2135 | chr5-65892027-INS-2136 |
| chr5-65898660-INS-424 | chr5-65898660-INS-424 |
| chr5-65903551-INS-60 | chr5-65903551-INS-64 |
| chr5-65910977-INS-218 | chr5-65910977-INS-217 |

| | |
|---|---|
| chr5-65935392-INS-54 | chr5-65935411-INS-64 |
| chr5-65948661-INS-248 | chr5-65948661-INS-250 |
| chr5-65951634-INS-295 | chr5-65951634-INS-301 |
| chr5-65957221-INS-70 | chr5-65957221-INS-70 |
| chr5-65996886-INS-87 | chr5-65996875-INS-60 |
| chr5-66035427-INS-69 | chr5-66035427-INS-69 |
| chr5-66043204-INS-198 | chr5-66043204-INS-198 |
| chr5-66071270-INS-122 | chr5-66071253-INS-120 |
| chr5-66107665-INS-80 | chr5-66107666-INS-98 |
| chr5-66119023-INS-201 | chr5-66119023-INS-184 |
| chr5-66125368-INS-324 | chr5-66125368-INS-324 |
| chr5-66146544-INS-177 | chr5-66146544-INS-177 |
| chr5-66154403-INS-60 | chr5-66154403-INS-50 |
| chr5-66181141-INS-117 | chr5-66181148-INS-127 |
| chr5-66192001-INS-209 | chr5-66192001-INS-209 |
| chr5-66209379-INS-100 | chr5-66209379-INS-80 |
| chr5-66210771-INS-57 | chr5-66210771-INS-57 |
| chr5-66221335-INS-104 | chr5-66221335-INS-100 |
| chr5-66239457-INS-69 | chr5-66239457-INS-69 |
| chr5-66253154-INS-6456 | chr5-66253155-INS-6456 |
| chr5-66275568-INS-105 | chr5-66275568-INS-108 |
| chr5-66281996-INS-53 | chr5-66281996-INS-54 |
| chr5-66286788-INS-360 | chr5-66286788-INS-372 |
| chr5-66294203-INS-709 | chr5-66294204-INS-697 |
| chr5-66295836-INS-206 | chr5-66295836-INS-206 |
| chr5-66314699-INS-252 | chr5-66314699-INS-252 |
| chr5-66346762-INS-108 | chr5-66346763-INS-112 |
| chr5-66351400-INS-218 | chr5-66351400-INS-209 |
| chr5-66366213-INS-75 | chr5-66366213-INS-75 |
| chr5-66381035-INS-64 | chr5-66381035-INS-64 |
| chr5-66382370-INS-418 | chr5-66382370-INS-418 |
| chr5-66385989-INS-51 | chr5-66385989-INS-53 |
| chr5-66418079-INS-65 | chr5-66418067-INS-111 |
| chr5-66424962-INS-1890 | chr5-66424963-INS-1890 |
| chr5-66428945-INS-735 | chr5-66428945-INS-734 |
| chr5-66449600-INS-55 | chr5-66449600-INS-55 |
| chr5-66476814-INS-234 | chr5-66476814-INS-234 |
| chr5-66513061-INS-226 | chr5-66513061-INS-226 |
| chr5-66553683-INS-152 | chr5-66553683-INS-152 |
| chr5-66564463-INS-50 | chr5-66564457-INS-67 |
| chr5-66585320-INS-52 | chr5-66585320-INS-54 |
| chr5-66619357-INS-218 | chr5-66619357-INS-218 |
| chr5-66635683-INS-120 | chr5-66635683-INS-136 |
| chr5-66683969-INS-51 | chr5-66683965-INS-59 |
| chr5-66718964-INS-88 | chr5-66718964-INS-93 |
| chr5-66723970-INS-54 | chr5-66723992-INS-64 |
| chr5-66724505-INS-66 | chr5-66724505-INS-66 |
| chr5-66776899-INS-278 | chr5-66776899-INS-278 |
| chr5-66780389-INS-7093 | chr5-66780390-INS-7093 |
| chr5-66791528-INS-117 | chr5-66791525-INS-107 |
| chr5-66796490-INS-54 | chr5-66796490-INS-101 |
| chr5-66797655-INS-210 | chr5-66797655-INS-210 |
| chr5-66812139-INS-126 | chr5-66812139-INS-126 |
| chr5-66838063-INS-160 | chr5-66838063-INS-160 |
| chr5-66844746-INS-222 | chr5-66844746-INS-222 |
| chr5-66855586-INS-314 | chr5-66855586-INS-310 |
| chr5-66872631-INS-84 | chr5-66872648-INS-87 |
| chr5-66899729-INS-703 | chr5-66899729-INS-703 |
| chr5-66968476-INS-663 | chr5-66968476-INS-640 |
| chr5-66985696-INS-67 | chr5-66985696-INS-63 |
| chr5-66999754-INS-56 | chr5-66999754-INS-56 |
| chr5-67001491-INS-83 | chr5-67001491-INS-82 |
| chr5-67001989-INS-5246 | chr5-67001990-INS-5246 |
| chr5-67021010-INS-94 | chr5-67021010-INS-87 |
| chr5-67023463-INS-3089 | chr5-67023464-INS-3089 |
| chr5-67029730-INS-64 | chr5-67029730-INS-64 |
| chr5-67052139-INS-386 | chr5-67052139-INS-386 |
| chr5-67112417-INS-121 | chr5-67112417-INS-120 |
| chr5-67137273-INS-77 | chr5-67137273-INS-77 |
| chr5-67144028-INS-151 | chr5-67144028-INS-151 |
| chr5-67156822-INS-63 | chr5-67156822-INS-62 |
| chr5-67223075-INS-238 | chr5-67223075-INS-238 |
| chr5-67233335-INS-56 | chr5-67233335-INS-52 |
| chr5-67246140-INS-52 | chr5-67246140-INS-56 |
| chr5-67299369-INS-113 | chr5-67299369-INS-116 |
| chr5-67314691-INS-870 | chr5-67314691-INS-870 |
| chr5-67350645-INS-3235 | chr5-67350646-INS-3235 |

| | |
|---|---|
| chr5-67402717-INS-185 | chr5-67402717-INS-178 |
| chr5-67407731-INS-86 | chr5-67407735-INS-58 |
| chr5-67411977-INS-89 | chr5-67411975-INS-91 |
| chr5-67477582-INS-280 | chr5-67477582-INS-280 |
| chr5-67536024-INS-54 | chr5-67536024-INS-64 |
| chr5-67537134-INS-246 | chr5-67537134-INS-259 |
| chr5-67552082-INS-58 | chr5-67552082-INS-58 |
| chr5-67579254-INS-144 | chr5-67579254-INS-144 |
| chr5-67597597-INS-188 | chr5-67597597-INS-189 |
| chr5-67599728-INS-90 | chr5-67599728-INS-91 |
| chr5-67601974-INS-248 | chr5-67601974-INS-251 |
| chr5-67602858-INS-90 | chr5-67602878-INS-115 |
| chr5-67606100-INS-5401 | chr5-67606101-INS-5399 |
| chr5-67642011-INS-60 | chr5-67642011-INS-64 |
| chr5-67653048-INS-259 | chr5-67653048-INS-259 |
| chr5-67661287-INS-52 | chr5-67661287-INS-58 |
| chr5-67666672-INS-302 | chr5-67666672-INS-302 |
| chr5-67672449-INS-58 | chr5-67672449-INS-58 |
| chr5-67680306-INS-65 | chr5-67680306-INS-65 |
| chr5-67739973-INS-157 | chr5-67739967-INS-172 |
| chr5-67788168-INS-60 | chr5-67788168-INS-60 |
| chr5-67814740-INS-81 | chr5-67814742-INS-82 |
| chr5-67827493-INS-212 | chr5-67827493-INS-212 |
| chr5-67854984-INS-62 | chr5-67854980-INS-63 |
| chr5-67930795-INS-60 | chr5-67930795-INS-64 |
| chr5-67956157-INS-64 | chr5-67956157-INS-64 |
| chr5-67968078-INS-124 | chr5-67968078-INS-128 |
| chr5-67995570-INS-714 | chr5-67995570-INS-714 |
| chr5-68012049-INS-128 | chr5-68012049-INS-128 |
| chr5-68014623-INS-54 | chr5-68014634-INS-57 |
| chr5-68015949-INS-55 | chr5-68015949-INS-55 |
| chr5-68028755-INS-939 | chr5-68028755-INS-939 |
| chr5-68035594-INS-67 | chr5-68035594-INS-67 |
| chr5-68054075-INS-671 | chr5-68054080-INS-715 |
| chr5-68108045-INS-7079 | chr5-68108046-INS-7080 |
| chr5-68115938-INS-7136 | chr5-68115939-INS-7136 |
| chr5-68124626-INS-150 | chr5-68124626-INS-150 |
| chr5-68160369-INS-131 | chr5-68160366-INS-144 |
| chr5-68167427-INS-71 | chr5-68167427-INS-71 |
| chr5-68177262-INS-111 | chr5-68177262-INS-153 |
| chr5-68183301-INS-92 | chr5-68183303-INS-90 |
| chr5-68210134-INS-99 | chr5-68210134-INS-109 |
| chr5-68219323-INS-176 | chr5-68219323-INS-176 |
| chr5-68230290-INS-76 | chr5-68230290-INS-76 |
| chr5-68250493-INS-419 | chr5-68250493-INS-427 |
| chr5-68263686-INS-58 | chr5-68263686-INS-62 |
| chr5-68281546-INS-82 | chr5-68281542-INS-84 |
| chr5-68309217-INS-106 | chr5-68309217-INS-106 |
| chr5-68364176-INS-440 | chr5-68364176-INS-440 |
| chr5-68390919-INS-850 | chr5-68390904-INS-857 |
| chr5-68420463-INS-90 | chr5-68420463-INS-90 |
| chr5-68435406-INS-6886 | chr5-68435407-INS-6886 |
| chr5-68437053-INS-52 | chr5-68437053-INS-54 |
| chr5-68440769-INS-167 | chr5-68440769-INS-194 |
| chr5-68444065-INS-486 | chr5-68444065-INS-486 |
| chr5-68511685-INS-56 | chr5-68511685-INS-56 |
| chr5-68513518-INS-63 | chr5-68513519-INS-63 |
| chr5-68523288-INS-54 | chr5-68523288-INS-54 |
| chr5-68559061-INS-753 | chr5-68559061-INS-753 |
| chr5-68578046-INS-232 | chr5-68578050-INS-237 |
| chr5-68581220-INS-473 | chr5-68581220-INS-473 |
| chr5-68582041-INS-291 | chr5-68582041-INS-282 |
| chr5-68612045-INS-201 | chr5-68612045-INS-201 |
| chr5-68641320-INS-76 | chr5-68641320-INS-76 |
| chr5-68643443-INS-72 | chr5-68643443-INS-72 |
| chr5-68757049-INS-561 | chr5-68757049-INS-561 |
| chr5-68770255-INS-88 | chr5-68770255-INS-88 |
| chr5-68772466-INS-529 | chr5-68772467-INS-529 |
| chr5-68781746-INS-418 | chr5-68781746-INS-418 |
| chr5-68799163-INS-62 | chr5-68799165-INS-70 |
| chr5-68813816-INS-105 | chr5-68813815-INS-112 |
| chr5-68815704-INS-104 | chr5-68815680-INS-80 |
| chr5-68823814-INS-70 | chr5-68823814-INS-65 |
| chr5-68857129-INS-74 | chr5-68857129-INS-75 |
| chr5-68892243-INS-6401 | chr5-68892244-INS-6401 |
| chr5-68915981-INS-772 | chr5-68915981-INS-770 |
| chr5-68975999-INS-6265 | chr5-68976000-INS-6265 |

| | |
|---|---|
| chr5-68996479-INS-100 | chr5-68996479-INS-100 |
| chr5-69003136-INS-772 | chr5-69003136-INS-772 |
| chr5-69039187-INS-1091 | chr5-69039188-INS-1091 |
| chr5-69043348-INS-81 | chr5-69043348-INS-84 |
| chr5-69088323-INS-74 | chr5-69088323-INS-74 |
| chr5-69132164-INS-517 | chr5-69132164-INS-517 |
| chr5-69156911-INS-238 | chr5-69156911-INS-238 |
| chr5-69162824-INS-555 | chr5-69162824-INS-574 |
| chr5-69200970-INS-74 | chr5-69200991-INS-66 |
| chr5-69210669-INS-322 | chr5-69210669-INS-322 |
| chr5-69248482-INS-86 | chr5-69248482-INS-86 |
| chr5-69260597-INS-587 | chr5-69260597-INS-587 |
| chr5-69293454-INS-6495 | chr5-69293455-INS-6495 |
| chr5-69322647-INS-70 | chr5-69322648-INS-61 |
| chr5-69359329-INS-431 | chr5-69359329-INS-431 |
| chr5-69407284-INS-324 | chr5-69407288-INS-331 |
| chr5-69424861-INS-106 | chr5-69424861-INS-106 |
| chr5-69429057-INS-847 | chr5-69429057-INS-853 |
| chr5-69456798-INS-734 | chr5-69456798-INS-734 |
| chr5-69459741-INS-826 | chr5-69459739-INS-819 |
| chr5-69479529-INS-66 | chr5-69479529-INS-66 |
| chr5-69510240-INS-6226 | chr5-69510241-INS-6226 |
| chr5-69515030-INS-55 | chr5-69515030-INS-55 |
| chr5-69543242-INS-288 | chr5-69543242-INS-288 |
| chr5-69561842-INS-162 | chr5-69561833-INS-165 |
| chr5-69571805-INS-75 | chr5-69571805-INS-90 |
| chr5-69621245-INS-220 | chr5-69621245-INS-220 |
| chr5-69711706-INS-356 | chr5-69711706-INS-356 |
| chr5-69735614-INS-1146 | chr5-69735615-INS-1146 |
| chr5-69736552-INS-444 | chr5-69736553-INS-450 |
| chr5-69744901-INS-317 | chr5-69744901-INS-317 |
| chr5-69817990-INS-3023 | chr5-69817991-INS-3023 |
| chr5-69918972-INS-490 | chr5-69918972-INS-490 |
| chr5-69966442-INS-683 | chr5-69966442-INS-686 |
| chr5-69968137-INS-242 | chr5-69968137-INS-242 |
| chr5-69983375-INS-6641 | chr5-69983376-INS-6641 |
| chr5-69994282-INS-51 | chr5-69994287-INS-50 |
| chr5-70012466-INS-265 | chr5-70012467-INS-300 |
| chr5-70020880-INS-54 | chr5-70020879-INS-88 |
| chr5-70085464-INS-53 | chr5-70085464-INS-53 |
| chr5-70086810-INS-215 | chr5-70086771-INS-204 |
| chr5-70089811-INS-725 | chr5-70089811-INS-724 |
| chr5-70095218-INS-200 | chr5-70095219-INS-200 |
| chr5-70101770-INS-264 | chr5-70101770-INS-264 |
| chr5-70110285-INS-58 | chr5-70110285-INS-58 |
| chr5-70114281-INS-4005 | chr5-70114282-INS-4005 |
| chr5-70121818-INS-62 | chr5-70121818-INS-62 |
| chr5-70167427-INS-379 | chr5-70167427-INS-379 |
| chr5-70223119-INS-485 | chr5-70223119-INS-485 |
| chr5-70267183-INS-80 | chr5-70267182-INS-88 |
| chr5-70298321-INS-76 | chr5-70298321-INS-90 |
| chr5-70349426-INS-56 | chr5-70349426-INS-56 |
| chr5-70371389-INS-100 | chr5-70371379-INS-94 |
| chr5-70381479-INS-50 | chr5-70381479-INS-54 |
| chr5-70458197-INS-1003 | chr5-70458198-INS-1003 |
| chr5-70516004-INS-3723 | chr5-70516005-INS-3723 |
| chr5-70517267-INS-56 | chr5-70517267-INS-52 |
| chr5-70520571-INS-153 | chr5-70520571-INS-153 |
| chr5-70524083-INS-66 | chr5-70524083-INS-62 |
| chr5-70562827-INS-505 | chr5-70562827-INS-505 |
| chr5-70569802-INS-73 | chr5-70569802-INS-73 |
| chr5-70603913-INS-68 | chr5-70603913-INS-68 |
| chr5-70627916-INS-75 | chr5-70627916-INS-79 |
| chr5-70632910-INS-112 | chr5-70632910-INS-112 |
| chr5-70730154-INS-486 | chr5-70730154-INS-486 |
| chr5-70745498-INS-6684 | chr5-70745499-INS-6688 |
| chr5-70746014-INS-478 | chr5-70746014-INS-478 |
| chr5-70757521-INS-338 | chr5-70757521-INS-340 |
| chr5-70796721-INS-57 | chr5-70796721-INS-57 |
| chr5-70849255-INS-54 | chr5-70849257-INS-76 |
| chr5-70859115-INS-96 | chr5-70859142-INS-75 |
| chr5-70862784-INS-205 | chr5-70862786-INS-237 |
| chr5-70864612-INS-6483 | chr5-70864613-INS-6483 |
| chr5-70890465-INS-64 | chr5-70890465-INS-79 |
| chr5-70895301-INS-192 | chr5-70895301-INS-192 |
| chr5-70962740-INS-407 | chr5-70962741-INS-407 |
| chr5-71002945-INS-118 | chr5-71002945-INS-125 |

| | |
|---|---|
| chr5-71065458-INS-60 | chr5-71065458-INS-60 |
| chr5-71070422-INS-114 | chr5-71070422-INS-114 |
| chr5-71082220-INS-2498 | chr5-71082221-INS-2498 |
| chr5-71091006-INS-54 | chr5-71091006-INS-52 |
| chr5-71096766-INS-52 | chr5-71096766-INS-54 |
| chr5-71101244-INS-737 | chr5-71101244-INS-737 |
| chr5-71140976-INS-62 | chr5-71140976-INS-86 |
| chr5-71195396-INS-151 | chr5-71195396-INS-151 |
| chr5-71219394-INS-118 | chr5-71219398-INS-120 |
| chr5-71242513-INS-220 | chr5-71242513-INS-220 |
| chr5-71283344-INS-126 | chr5-71283342-INS-130 |
| chr5-71303342-INS-976 | chr5-71303343-INS-976 |
| chr5-71308863-INS-74 | chr5-71308863-INS-74 |
| chr5-71333153-INS-110 | chr5-71333153-INS-108 |
| chr5-71338742-INS-398 | chr5-71338742-INS-397 |
| chr5-71348944-INS-443 | chr5-71348944-INS-443 |
| chr5-71361729-INS-316 | chr5-71361729-INS-302 |
| chr5-71586751-INS-5893 | chr5-71586752-INS-5893 |
| chr5-71587066-INS-75 | chr5-71587066-INS-78 |
| chr5-71608884-INS-6135 | chr5-71608885-INS-6135 |
| chr5-71612770-INS-62 | chr5-71612770-INS-58 |
| chr5-71637887-INS-431 | chr5-71637887-INS-432 |
| chr5-71703756-INS-595 | chr5-71703756-INS-596 |
| chr5-71743515-INS-102 | chr5-71743515-INS-118 |
| chr5-71751617-INS-330 | chr5-71751617-INS-330 |
| chr5-71830115-INS-858 | chr5-71830115-INS-857 |
| chr5-71849316-INS-395 | chr5-71849316-INS-395 |
| chr5-71930821-INS-132 | chr5-71930821-INS-132 |
| chr5-71936931-INS-6376 | chr5-71936932-INS-6376 |
| chr5-71978491-INS-139 | chr5-71978491-INS-139 |
| chr5-71982955-INS-274 | chr5-71982955-INS-274 |
| chr5-72020458-INS-70 | chr5-72020458-INS-70 |
| chr5-72041042-INS-6264 | chr5-72041043-INS-6265 |
| chr5-72064460-INS-2376 | chr5-72064461-INS-2376 |
| chr5-72075892-INS-479 | chr5-72075892-INS-479 |
| chr5-72087151-INS-56 | chr5-72087172-INS-65 |
| chr5-72102418-INS-6712 | chr5-72102419-INS-6712 |
| chr5-72131358-INS-53 | chr5-72131358-INS-53 |
| chr5-72161473-INS-741 | chr5-72161473-INS-741 |
| chr5-72167961-INS-77 | chr5-72167961-INS-77 |
| chr5-72172418-INS-64 | chr5-72172418-INS-76 |
| chr5-72177829-INS-77 | chr5-72177829-INS-77 |
| chr5-72191288-INS-54 | chr5-72191288-INS-54 |
| chr5-72202196-INS-90 | chr5-72202196-INS-90 |
| chr5-72261094-INS-4510 | chr5-72261093-INS-4506 |
| chr5-72287348-INS-130 | chr5-72287348-INS-130 |
| chr5-72323920-INS-208 | chr5-72323920-INS-205 |
| chr5-72326502-INS-144 | chr5-72326502-INS-156 |
| chr5-72327520-INS-89 | chr5-72327520-INS-89 |
| chr5-72330676-INS-257 | chr5-72330676-INS-257 |
| chr5-72338269-INS-52 | chr5-72338269-INS-56 |
| chr5-72341011-INS-84 | chr5-72341011-INS-84 |
| chr5-72366216-INS-204 | chr5-72366216-INS-204 |
| chr5-72381508-INS-63 | chr5-72381508-INS-73 |
| chr5-72409552-INS-54 | chr5-72409552-INS-56 |
| chr5-72411798-INS-189 | chr5-72411785-INS-179 |
| chr5-72446940-INS-6877 | chr5-72446941-INS-6877 |
| chr5-72483106-INS-677 | chr5-72483106-INS-677 |
| chr5-72522552-INS-173 | chr5-72522552-INS-167 |
| chr5-72544797-INS-344 | chr5-72544797-INS-345 |
| chr5-72577266-INS-187 | chr5-72577266-INS-187 |
| chr5-72579885-INS-76 | chr5-72579867-INS-93 |
| chr5-72588970-INS-286 | chr5-72588970-INS-286 |
| chr5-72599690-INS-191 | chr5-72599690-INS-191 |
| chr5-72607902-INS-96 | chr5-72607902-INS-96 |
| chr5-72615781-INS-277 | chr5-72615781-INS-285 |
| chr5-72637542-INS-53 | chr5-72637542-INS-57 |
| chr5-72644792-INS-6597 | chr5-72644793-INS-6595 |
| chr5-72649305-INS-59 | chr5-72649305-INS-59 |
| chr5-72711497-INS-165 | chr5-72711497-INS-147 |
| chr5-72724019-INS-490 | chr5-72724019-INS-490 |
| chr5-72754307-INS-62 | chr5-72754307-INS-62 |
| chr5-72789350-INS-138 | chr5-72789350-INS-138 |
| chr5-72800494-INS-195 | chr5-72800497-INS-192 |
| chr5-72806171-INS-180 | chr5-72806171-INS-174 |
| chr5-72818996-INS-76 | chr5-72818996-INS-76 |
| chr5-72836847-INS-505 | chr5-72836847-INS-505 |

| | |
|---|---|
| chr5-72839582-INS-200 | chr5-72839582-INS-200 |
| chr5-72868757-INS-150 | chr5-72868757-INS-150 |
| chr5-72876501-INS-560 | chr5-72876501-INS-560 |
| chr5-72880534-INS-200 | chr5-72880535-INS-200 |
| chr5-72892032-INS-379 | chr5-72892032-INS-380 |
| chr5-72922128-INS-915 | chr5-72922128-INS-915 |
| chr5-72937795-INS-116 | chr5-72937795-INS-117 |
| chr5-72974181-INS-242 | chr5-72974181-INS-252 |
| chr5-72975888-INS-69 | chr5-72975888-INS-69 |
| chr5-72984564-INS-721 | chr5-72984564-INS-727 |
| chr5-73036411-INS-93 | chr5-73036415-INS-87 |
| chr5-73047365-INS-70 | chr5-73047341-INS-72 |
| chr5-73055181-INS-55 | chr5-73055186-INS-56 |
| chr5-73058572-INS-7095 | chr5-73058573-INS-7095 |
| chr5-73090077-INS-214 | chr5-73090077-INS-215 |
| chr5-73096454-INS-195 | chr5-73096464-INS-207 |
| chr5-73116049-INS-58 | chr5-73116049-INS-58 |
| chr5-73132261-INS-572 | chr5-73132261-INS-572 |
| chr5-73137613-INS-238 | chr5-73137613-INS-245 |
| chr5-73138254-INS-7106 | chr5-73138255-INS-7106 |
| chr5-73140165-INS-210 | chr5-73140165-INS-211 |
| chr5-73218830-INS-885 | chr5-73218830-INS-885 |
| chr5-73225871-INS-640 | chr5-73225871-INS-643 |
| chr5-73265163-INS-148 | chr5-73265163-INS-154 |
| chr5-73271175-INS-58 | chr5-73271175-INS-58 |
| chr5-73290406-INS-96 | chr5-73290406-INS-96 |
| chr5-73324404-INS-223 | chr5-73324404-INS-225 |
| chr5-73346319-INS-2079 | chr5-73346320-INS-2080 |
| chr5-73347915-INS-52 | chr5-73347916-INS-84 |
| chr5-73370999-INS-104 | chr5-73370999-INS-104 |
| chr5-73373413-INS-658 | chr5-73373413-INS-665 |
| chr5-73395965-INS-886 | chr5-73395965-INS-885 |
| chr5-73429178-INS-4729 | chr5-73429179-INS-4729 |
| chr5-73440756-INS-187 | chr5-73440756-INS-187 |
| chr5-73469597-INS-250 | chr5-73469597-INS-250 |
| chr5-73497953-INS-204 | chr5-73497953-INS-204 |
| chr5-73507770-INS-1897 | chr5-73507771-INS-1897 |
| chr5-73509000-INS-131 | chr5-73509000-INS-131 |
| chr5-73522072-INS-201 | chr5-73522072-INS-202 |
| chr5-73561325-INS-151 | chr5-73561325-INS-151 |
| chr5-73565271-INS-144 | chr5-73565271-INS-144 |
| chr5-73572954-INS-54 | chr5-73572954-INS-60 |
| chr5-73581401-INS-56 | chr5-73581401-INS-56 |
| chr5-73600285-INS-900 | chr5-73600285-INS-918 |
| chr5-73605707-INS-138 | chr5-73605712-INS-157 |
| chr5-73632315-INS-52 | chr5-73632315-INS-56 |
| chr5-73689304-INS-1796 | chr5-73689305-INS-1797 |
| chr5-73692263-INS-5388 | chr5-73692264-INS-5388 |
| chr5-73709760-INS-120 | chr5-73709760-INS-120 |
| chr5-73730180-INS-206 | chr5-73730180-INS-206 |
| chr5-73756119-INS-56 | chr5-73756119-INS-62 |
| chr5-73770016-INS-65 | chr5-73770027-INS-61 |
| chr5-73785762-INS-93 | chr5-73785762-INS-93 |
| chr5-73800965-INS-88 | chr5-73800955-INS-100 |
| chr5-73812303-INS-51 | chr5-73812304-INS-51 |
| chr5-73823309-INS-67 | chr5-73823309-INS-67 |
| chr5-73845920-INS-473 | chr5-73845920-INS-472 |
| chr5-73852181-INS-3854 | chr5-73852182-INS-3856 |
| chr5-73856391-INS-591 | chr5-73856392-INS-593 |
| chr5-73886378-INS-106 | chr5-73886378-INS-106 |
| chr5-73888581-INS-74 | chr5-73888581-INS-74 |
| chr5-73917776-INS-104 | chr5-73917730-INS-114 |
| chr5-73921230-INS-191 | chr5-73921230-INS-191 |
| chr5-73931443-INS-151 | chr5-73931443-INS-151 |
| chr5-73959364-INS-102 | chr5-73959364-INS-102 |
| chr5-73961314-INS-60 | chr5-73961314-INS-60 |
| chr5-73995944-INS-205 | chr5-73995944-INS-206 |
| chr5-74011528-INS-213 | chr5-74011528-INS-214 |
| chr5-74046414-INS-569 | chr5-74046416-INS-566 |
| chr5-74071537-INS-110 | chr5-74071537-INS-110 |
| chr5-74084345-INS-6596 | chr5-74084346-INS-6596 |
| chr5-74161669-INS-52 | chr5-74161669-INS-76 |
| chr5-74167058-INS-53 | chr5-74167058-INS-59 |
| chr5-74192782-INS-208 | chr5-74192782-INS-209 |
| chr5-74195953-INS-54 | chr5-74195954-INS-54 |
| chr5-74206651-INS-198 | chr5-74206651-INS-200 |
| chr5-74209842-INS-7001 | chr5-74209843-INS-7001 |

| | |
|---|---|
| chr5-74261666-INS-88 | chr5-74261666-INS-88 |
| chr5-74285539-INS-174 | chr5-74285539-INS-190 |
| chr5-74289214-INS-105 | chr5-74289214-INS-105 |
| chr5-74291238-INS-291 | chr5-74291238-INS-280 |
| chr5-74294744-INS-525 | chr5-74294744-INS-525 |
| chr5-74296166-INS-76 | chr5-74296166-INS-85 |
| chr5-74314575-INS-56 | chr5-74314575-INS-56 |
| chr5-74317454-INS-188 | chr5-74317454-INS-188 |
| chr5-74324376-INS-96 | chr5-74324367-INS-94 |
| chr5-74332847-INS-74 | chr5-74332847-INS-72 |
| chr5-74345063-INS-56 | chr5-74345063-INS-52 |
| chr5-74345718-INS-51 | chr5-74345718-INS-51 |
| chr5-74353992-INS-150 | chr5-74353997-INS-106 |
| chr5-74389620-INS-95 | chr5-74389620-INS-123 |
| chr5-74402870-INS-205 | chr5-74402870-INS-203 |
| chr5-74407836-INS-201 | chr5-74407836-INS-201 |
| chr5-74418078-INS-1131 | chr5-74418079-INS-1092 |
| chr5-74426051-INS-170 | chr5-74426051-INS-170 |
| chr5-74449216-INS-122 | chr5-74449216-INS-122 |
| chr5-74459676-INS-175 | chr5-74459676-INS-175 |
| chr5-74469191-INS-144 | chr5-74469191-INS-144 |
| chr5-74528974-INS-835 | chr5-74528974-INS-832 |
| chr5-74537164-INS-265 | chr5-74537164-INS-265 |
| chr5-74561557-INS-203 | chr5-74561557-INS-203 |
| chr5-74577461-INS-146 | chr5-74577511-INS-144 |
| chr5-74582751-INS-445 | chr5-74582751-INS-445 |
| chr5-74606010-INS-275 | chr5-74606010-INS-275 |
| chr5-74611070-INS-64 | chr5-74611070-INS-64 |
| chr5-74621912-INS-53 | chr5-74621912-INS-53 |
| chr5-74652887-INS-131 | chr5-74652887-INS-117 |
| chr5-74654719-INS-121 | chr5-74654719-INS-132 |
| chr5-74656396-INS-229 | chr5-74656396-INS-229 |
| chr5-74660776-INS-198 | chr5-74660776-INS-198 |
| chr5-74675905-INS-771 | chr5-74675905-INS-774 |
| chr5-74683422-INS-7119 | chr5-74683423-INS-7119 |
| chr5-74710938-INS-96 | chr5-74710939-INS-96 |
| chr5-74785668-INS-610 | chr5-74785668-INS-614 |
| chr5-74787585-INS-50 | chr5-74787544-INS-100 |
| chr5-74810377-INS-58 | chr5-74810377-INS-58 |
| chr5-74813895-INS-4325 | chr5-74813896-INS-4325 |
| chr5-74855826-INS-192 | chr5-74855826-INS-192 |
| chr5-74880968-INS-124 | chr5-74880968-INS-104 |
| chr5-74882997-INS-85 | chr5-74882997-INS-88 |
| chr5-74898490-INS-171 | chr5-74898480-INS-169 |
| chr5-74927989-INS-82 | chr5-74927989-INS-82 |
| chr5-74929196-INS-60 | chr5-74929196-INS-56 |
| chr5-74933215-INS-93 | chr5-74933215-INS-93 |
| chr5-74965250-INS-77 | chr5-74965250-INS-71 |
| chr5-74981153-INS-129 | chr5-74981153-INS-129 |
| chr5-74988291-INS-58 | chr5-74988309-INS-85 |
| chr5-75028119-INS-1288 | chr5-75028120-INS-1288 |
| chr5-75045104-INS-113 | chr5-75045104-INS-63 |
| chr5-75063861-INS-72 | chr5-75063861-INS-80 |
| chr5-75076426-INS-72 | chr5-75076426-INS-116 |
| chr5-75084798-INS-7106 | chr5-75084799-INS-7105 |
| chr5-75089235-INS-60 | chr5-75089235-INS-60 |
| chr5-75107639-INS-151 | chr5-75107639-INS-159 |
| chr5-75111141-INS-57 | chr5-75111142-INS-57 |
| chr5-75153361-INS-203 | chr5-75153361-INS-203 |
| chr5-75201694-INS-81 | chr5-75201706-INS-86 |
| chr5-75222676-INS-464 | chr5-75222676-INS-464 |
| chr5-75247653-INS-1890 | chr5-75247654-INS-1890 |
| chr5-75253496-INS-144 | chr5-75253493-INS-117 |
| chr5-75262562-INS-79 | chr5-75262562-INS-90 |
| chr5-75265598-INS-189 | chr5-75265598-INS-189 |
| chr5-75275694-INS-312 | chr5-75275693-INS-308 |
| chr5-75279109-INS-198 | chr5-75279109-INS-198 |
| chr5-75281504-INS-104 | chr5-75281504-INS-104 |
| chr5-75286717-INS-400 | chr5-75286717-INS-400 |
| chr5-75290699-INS-51 | chr5-75290699-INS-51 |
| chr5-75295756-INS-58 | chr5-75295756-INS-58 |
| chr5-75300910-INS-134 | chr5-75300911-INS-127 |
| chr5-75306348-INS-209 | chr5-75306348-INS-209 |
| chr5-75318154-INS-84 | chr5-75318169-INS-63 |
| chr5-75332961-INS-400 | chr5-75332961-INS-400 |
| chr5-75336280-INS-75 | chr5-75336300-INS-69 |
| chr5-75337910-INS-62 | chr5-75337898-INS-78 |

| | |
|---|---|
| chr5-75373229-INS-52 | chr5-75373229-INS-58 |
| chr5-75378314-INS-52 | chr5-75378314-INS-52 |
| chr5-75380323-INS-827 | chr5-75380323-INS-827 |
| chr5-75387745-INS-114 | chr5-75387745-INS-120 |
| chr5-75425611-INS-199 | chr5-75425611-INS-198 |
| chr5-75504339-INS-222 | chr5-75504339-INS-222 |
| chr5-75527371-INS-210 | chr5-75527371-INS-210 |
| chr5-75540741-INS-72 | chr5-75540741-INS-72 |
| chr5-75564559-INS-64 | chr5-75564559-INS-64 |
| chr5-75613228-INS-4539 | chr5-75613229-INS-4539 |
| chr5-75640167-INS-55 | chr5-75640167-INS-55 |
| chr5-75669159-INS-66 | chr5-75669165-INS-75 |
| chr5-75680237-INS-56 | chr5-75680251-INS-72 |
| chr5-75698070-INS-6442 | chr5-75698071-INS-6443 |
| chr5-75718888-INS-54 | chr5-75718888-INS-54 |
| chr5-75731470-INS-937 | chr5-75731470-INS-937 |
| chr5-75788422-INS-215 | chr5-75788422-INS-215 |
| chr5-75804487-INS-55 | chr5-75804487-INS-55 |
| chr5-75820990-INS-64 | chr5-75820991-INS-56 |
| chr5-75846252-INS-125 | chr5-75846252-INS-102 |
| chr5-75849465-INS-7124 | chr5-75849466-INS-7124 |
| chr5-75851989-INS-56 | chr5-75851989-INS-56 |
| chr5-75860642-INS-88 | chr5-75860642-INS-88 |
| chr5-75869422-INS-440 | chr5-75869422-INS-440 |
| chr5-75880266-INS-127 | chr5-75880265-INS-85 |
| chr5-75889636-INS-64 | chr5-75889636-INS-62 |
| chr5-75890174-INS-76 | chr5-75890180-INS-60 |
| chr5-75967413-INS-7134 | chr5-75967414-INS-7134 |
| chr5-75983171-INS-6579 | chr5-75983172-INS-6579 |
| chr5-76007781-INS-227 | chr5-76007781-INS-228 |
| chr5-76018355-INS-58 | chr5-76018355-INS-77 |
| chr5-76027964-INS-98 | chr5-76027964-INS-83 |
| chr5-76046687-INS-569 | chr5-76046687-INS-569 |
| chr5-76049012-INS-58 | chr5-76049013-INS-50 |
| chr5-76074258-INS-6614 | chr5-76074259-INS-6614 |
| chr5-76089029-INS-438 | chr5-76089033-INS-416 |
| chr5-76092457-INS-70 | chr5-76092433-INS-62 |
| chr5-76092553-INS-338 | chr5-76092553-INS-338 |
| chr5-76171100-INS-772 | chr5-76171100-INS-771 |
| chr5-76196237-INS-103 | chr5-76196213-INS-115 |
| chr5-76243050-INS-54 | chr5-76243050-INS-54 |
| chr5-76279820-INS-65 | chr5-76279820-INS-65 |
| chr5-76349086-INS-1247 | chr5-76349087-INS-1247 |
| chr5-76405595-INS-201 | chr5-76405595-INS-200 |
| chr5-76415760-INS-7214 | chr5-76415761-INS-7214 |
| chr5-76476838-INS-62 | chr5-76476838-INS-62 |
| chr5-76495457-INS-115 | chr5-76495457-INS-115 |
| chr5-76498738-INS-54 | chr5-76498738-INS-54 |
| chr5-76501980-INS-201 | chr5-76501980-INS-201 |
| chr5-76502999-INS-144 | chr5-76502999-INS-147 |
| chr5-76510177-INS-118 | chr5-76510175-INS-134 |
| chr5-76529984-INS-7124 | chr5-76529984-INS-7123 |
| chr5-76577189-INS-51 | chr5-76577189-INS-81 |
| chr5-76583872-INS-85 | chr5-76583872-INS-87 |
| chr5-76587856-INS-1492 | chr5-76587857-INS-1492 |
| chr5-76594393-INS-61 | chr5-76594393-INS-61 |
| chr5-76595991-INS-547 | chr5-76595991-INS-547 |
| chr5-76614356-INS-54 | chr5-76614356-INS-54 |
| chr5-76615614-INS-328 | chr5-76615616-INS-366 |
| chr5-76632096-INS-86 | chr5-76632096-INS-89 |
| chr5-76635073-INS-58 | chr5-76635073-INS-60 |
| chr5-76651263-INS-78 | chr5-76651264-INS-57 |
| chr5-76658319-INS-391 | chr5-76658319-INS-391 |
| chr5-76661494-INS-732 | chr5-76661494-INS-732 |
| chr5-76689225-INS-249 | chr5-76689225-INS-249 |
| chr5-76724190-INS-65 | chr5-76724190-INS-66 |
| chr5-76728751-INS-52 | chr5-76728751-INS-52 |
| chr5-76729451-INS-224 | chr5-76729451-INS-224 |
| chr5-76731686-INS-156 | chr5-76731686-INS-156 |
| chr5-76742125-INS-68 | chr5-76742125-INS-68 |
| chr5-76766401-INS-220 | chr5-76766401-INS-221 |
| chr5-76801429-INS-59 | chr5-76801429-INS-75 |
| chr5-76821957-INS-96 | chr5-76821957-INS-96 |
| chr5-76829770-INS-75 | chr5-76829770-INS-75 |
| chr5-76847962-INS-113 | chr5-76847962-INS-113 |
| chr5-76854349-INS-177 | chr5-76854349-INS-178 |
| chr5-76878368-INS-558 | chr5-76878368-INS-558 |

| | |
|---|---|
| chr5-76882170-INS-153 | chr5-76882170-INS-161 |
| chr5-76888029-INS-67 | chr5-76888029-INS-68 |
| chr5-76889680-INS-63 | chr5-76889680-INS-63 |
| chr5-76897663-INS-232 | chr5-76897663-INS-232 |
| chr5-76939912-INS-56 | chr5-76939892-INS-56 |
| chr5-76953062-INS-500 | chr5-76953062-INS-499 |
| chr5-76961296-INS-59 | chr5-76961296-INS-59 |
| chr5-76985829-INS-1360 | chr5-76985830-INS-1361 |
| chr5-76989343-INS-70 | chr5-76989343-INS-70 |
| chr5-77008628-INS-146 | chr5-77008628-INS-147 |
| chr5-77023019-INS-53 | chr5-77023019-INS-53 |
| chr5-77028175-INS-123 | chr5-77028175-INS-122 |
| chr5-77031605-INS-72 | chr5-77031637-INS-72 |
| chr5-77038707-INS-76 | chr5-77038707-INS-76 |
| chr5-77043136-INS-1156 | chr5-77043137-INS-1157 |
| chr5-77047739-INS-184 | chr5-77047739-INS-185 |
| chr5-77057448-INS-51 | chr5-77057448-INS-51 |
| chr5-77060499-INS-61 | chr5-77060499-INS-61 |
| chr5-77108660-INS-90 | chr5-77108660-INS-90 |
| chr5-77145736-INS-204 | chr5-77145736-INS-204 |
| chr5-77150338-INS-57 | chr5-77150338-INS-57 |
| chr5-77175364-INS-55 | chr5-77175364-INS-55 |
| chr5-77179983-INS-129 | chr5-77179980-INS-127 |
| chr5-77180489-INS-74 | chr5-77180478-INS-111 |
| chr5-77209956-INS-90 | chr5-77209942-INS-68 |
| chr5-77213091-INS-623 | chr5-77213089-INS-624 |
| chr5-77231722-INS-287 | chr5-77231722-INS-287 |
| chr5-77254941-INS-52 | chr5-77254941-INS-52 |
| chr5-77258050-INS-125 | chr5-77258050-INS-125 |
| chr5-77260591-INS-58 | chr5-77260591-INS-58 |
| chr5-77264788-INS-72 | chr5-77264788-INS-68 |
| chr5-77278340-INS-56 | chr5-77278340-INS-56 |
| chr5-77304631-INS-440 | chr5-77304643-INS-447 |
| chr5-77309099-INS-179 | chr5-77309099-INS-179 |
| chr5-77310159-INS-1602 | chr5-77310160-INS-1602 |
| chr5-77320242-INS-68 | chr5-77320242-INS-68 |
| chr5-77350793-INS-64 | chr5-77350793-INS-96 |
| chr5-77355815-INS-272 | chr5-77355815-INS-272 |
| chr5-77364553-INS-178 | chr5-77364553-INS-174 |
| chr5-77365621-INS-56 | chr5-77365621-INS-56 |
| chr5-77370427-INS-1146 | chr5-77370419-INS-1152 |
| chr5-77373953-INS-189 | chr5-77373953-INS-181 |
| chr5-77374953-INS-350 | chr5-77374949-INS-375 |
| chr5-77376031-INS-207 | chr5-77376031-INS-204 |
| chr5-77380606-INS-92 | chr5-77380610-INS-126 |
| chr5-77461738-INS-69 | chr5-77461738-INS-69 |
| chr5-77466297-INS-96 | chr5-77466315-INS-95 |
| chr5-77469169-INS-51 | chr5-77469162-INS-95 |
| chr5-77479949-INS-205 | chr5-77479949-INS-205 |
| chr5-77496367-INS-180 | chr5-77496367-INS-180 |
| chr5-77503880-INS-69 | chr5-77503880-INS-69 |
| chr5-77516840-INS-162 | chr5-77516840-INS-162 |
| chr5-77530955-INS-191 | chr5-77530955-INS-191 |
| chr5-77543959-INS-592 | chr5-77543959-INS-587 |
| chr5-77582830-INS-6440 | chr5-77582831-INS-6440 |
| chr5-77599750-INS-7117 | chr5-77599751-INS-7117 |
| chr5-77602855-INS-1004 | chr5-77602856-INS-1004 |
| chr5-77611308-INS-4460 | chr5-77611309-INS-4460 |
| chr5-77618803-INS-128 | chr5-77618803-INS-128 |
| chr5-77654399-INS-60 | chr5-77654402-INS-86 |
| chr5-77674512-INS-89 | chr5-77674512-INS-95 |
| chr5-77693575-INS-98 | chr5-77693575-INS-98 |
| chr5-77698784-INS-65 | chr5-77698774-INS-60 |
| chr5-77719363-INS-4131 | chr5-77719364-INS-4131 |
| chr5-77742346-INS-86 | chr5-77742344-INS-103 |
| chr5-77755763-INS-220 | chr5-77755763-INS-220 |
| chr5-77762738-INS-179 | chr5-77762738-INS-174 |
| chr5-77771573-INS-424 | chr5-77771573-INS-424 |
| chr5-77779673-INS-1561 | chr5-77779674-INS-1561 |
| chr5-77788031-INS-406 | chr5-77788031-INS-406 |
| chr5-77798004-INS-119 | chr5-77797999-INS-153 |
| chr5-77815407-INS-1451 | chr5-77815412-INS-1446 |
| chr5-77815762-INS-429 | chr5-77815762-INS-429 |
| chr5-77841095-INS-140 | chr5-77841095-INS-105 |
| chr5-77851811-INS-68 | chr5-77851811-INS-68 |
| chr5-77857437-INS-63 | chr5-77857437-INS-63 |
| chr5-77864461-INS-531 | chr5-77864462-INS-539 |

| | |
|---|---|
| chr5-77879242-INS-331 | chr5-77879242-INS-331 |
| chr5-77884430-INS-3894 | chr5-77884431-INS-3894 |
| chr5-77954055-INS-52 | chr5-77954055-INS-50 |
| chr5-77987715-INS-4497 | chr5-77987716-INS-4497 |
| chr5-77997312-INS-214 | chr5-77997312-INS-214 |
| chr5-78005667-INS-313 | chr5-78005672-INS-308 |
| chr5-78024118-INS-1779 | chr5-78024119-INS-1779 |
| chr5-78063848-INS-66 | chr5-78063848-INS-66 |
| chr5-78071224-INS-313 | chr5-78071224-INS-313 |
| chr5-78098937-INS-58 | chr5-78098935-INS-64 |
| chr5-78127073-INS-3992 | chr5-78127074-INS-3992 |
| chr5-78131269-INS-52 | chr5-78131270-INS-100 |
| chr5-78161669-INS-106 | chr5-78161670-INS-107 |
| chr5-78198857-INS-64 | chr5-78198857-INS-66 |
| chr5-78199501-INS-6328 | chr5-78199502-INS-6328 |
| chr5-78223616-INS-577 | chr5-78223613-INS-582 |
| chr5-78229256-INS-444 | chr5-78229256-INS-444 |
| chr5-78231188-INS-576 | chr5-78231188-INS-578 |
| chr5-78245123-INS-197 | chr5-78245142-INS-197 |
| chr5-78270008-INS-84 | chr5-78270008-INS-84 |
| chr5-78315428-INS-99 | chr5-78315428-INS-99 |
| chr5-78348904-INS-2451 | chr5-78348905-INS-2440 |
| chr5-78384327-INS-60 | chr5-78384327-INS-60 |
| chr5-78413927-INS-5197 | chr5-78413928-INS-5197 |
| chr5-78424413-INS-52 | chr5-78424413-INS-52 |
| chr5-78452598-INS-102 | chr5-78452598-INS-102 |
| chr5-78459564-INS-350 | chr5-78459564-INS-350 |
| chr5-78467228-INS-65 | chr5-78467227-INS-87 |
| chr5-78510456-INS-341 | chr5-78510426-INS-335 |
| chr5-78518373-INS-1463 | chr5-78518374-INS-1463 |
| chr5-78540903-INS-110 | chr5-78540903-INS-110 |
| chr5-78543238-INS-389 | chr5-78543238-INS-389 |
| chr5-78563808-INS-817 | chr5-78563808-INS-820 |
| chr5-78595516-INS-224 | chr5-78595516-INS-224 |
| chr5-78608910-INS-75 | chr5-78608913-INS-72 |
| chr5-78650079-INS-61 | chr5-78650079-INS-57 |
| chr5-78661557-INS-78 | chr5-78661557-INS-54 |
| chr5-78685958-INS-57 | chr5-78685958-INS-57 |
| chr5-78688151-INS-503 | chr5-78688152-INS-496 |
| chr5-78694213-INS-93 | chr5-78694220-INS-51 |
| chr5-78702389-INS-971 | chr5-78702389-INS-972 |
| chr5-78727535-INS-1703 | chr5-78727534-INS-1703 |
| chr5-78753242-INS-408 | chr5-78753242-INS-408 |
| chr5-78768445-INS-653 | chr5-78768445-INS-653 |
| chr5-78776775-INS-3871 | chr5-78776776-INS-3871 |
| chr5-78857803-INS-200 | chr5-78857803-INS-200 |
| chr5-78885717-INS-104 | chr5-78885717-INS-104 |
| chr5-78924101-INS-392 | chr5-78924101-INS-392 |
| chr5-78942494-INS-82 | chr5-78942488-INS-122 |
| chr5-78969238-INS-489 | chr5-78969238-INS-489 |
| chr5-78971309-INS-172 | chr5-78971309-INS-172 |
| chr5-78979772-INS-7096 | chr5-78979773-INS-7098 |
| chr5-79041734-INS-62 | chr5-79041734-INS-68 |
| chr5-79050077-INS-150 | chr5-79050077-INS-150 |
| chr5-79058719-INS-6695 | chr5-79058720-INS-6694 |
| chr5-79076623-INS-64 | chr5-79076623-INS-64 |
| chr5-79080850-INS-172 | chr5-79080852-INS-173 |
| chr5-79118852-INS-54 | chr5-79118852-INS-52 |
| chr5-79122314-INS-6386 | chr5-79122315-INS-6387 |
| chr5-79130453-INS-381 | chr5-79130453-INS-381 |
| chr5-79174863-INS-94 | chr5-79174861-INS-94 |
| chr5-79176058-INS-1958 | chr5-79176059-INS-1959 |
| chr5-79224231-INS-106 | chr5-79224238-INS-133 |
| chr5-79235588-INS-86 | chr5-79235588-INS-86 |
| chr5-79243334-INS-1902 | chr5-79243335-INS-1903 |
| chr5-79269725-INS-2307 | chr5-79269726-INS-2307 |
| chr5-79280856-INS-59 | chr5-79280856-INS-59 |
| chr5-79282099-INS-114 | chr5-79282099-INS-68 |
| chr5-79330175-INS-535 | chr5-79330175-INS-536 |
| chr5-79373467-INS-203 | chr5-79373467-INS-204 |
| chr5-79395274-INS-206 | chr5-79395274-INS-206 |
| chr5-79408275-INS-67 | chr5-79408275-INS-73 |
| chr5-79416369-INS-543 | chr5-79416369-INS-551 |
| chr5-79454702-INS-159 | chr5-79454702-INS-159 |
| chr5-79474380-INS-54 | chr5-79474380-INS-52 |
| chr5-79492231-INS-1145 | chr5-79492232-INS-1154 |
| chr5-79495828-INS-98 | chr5-79495828-INS-98 |

| | |
|---|---|
| chr5-79499803-INS-61 | chr5-79499821-INS-56 |
| chr5-79509667-INS-76 | chr5-79509632-INS-112 |
| chr5-79527862-INS-122 | chr5-79527862-INS-132 |
| chr5-79578895-INS-180 | chr5-79578927-INS-138 |
| chr5-79585208-INS-6175 | chr5-79585209-INS-6175 |
| chr5-79606214-INS-1222 | chr5-79606215-INS-1222 |
| chr5-79697105-INS-60 | chr5-79697106-INS-65 |
| chr5-79705182-INS-338 | chr5-79705182-INS-326 |
| chr5-79712835-INS-72 | chr5-79712866-INS-64 |
| chr5-79740724-INS-80 | chr5-79740745-INS-72 |
| chr5-79742143-INS-217 | chr5-79742143-INS-217 |
| chr5-79756581-INS-6817 | chr5-79756582-INS-6818 |
| chr5-79824746-INS-512 | chr5-79824746-INS-512 |
| chr5-79833390-INS-3064 | chr5-79833391-INS-3064 |
| chr5-79835775-INS-63 | chr5-79835775-INS-63 |
| chr5-79856774-INS-82 | chr5-79856774-INS-50 |
| chr5-79871378-INS-68 | chr5-79871378-INS-68 |
| chr5-79872444-INS-68 | chr5-79872444-INS-70 |
| chr5-79890760-INS-208 | chr5-79890760-INS-208 |
| chr5-79902104-INS-6476 | chr5-79902105-INS-6478 |
| chr5-79906974-INS-4640 | chr5-79906975-INS-4640 |
| chr5-79915536-INS-444 | chr5-79915536-INS-444 |
| chr5-79934091-INS-1290 | chr5-79934092-INS-1290 |
| chr5-79938750-INS-6453 | chr5-79938751-INS-6453 |
| chr5-79957932-INS-69 | chr5-79957932-INS-69 |
| chr5-79992434-INS-66 | chr5-79992434-INS-64 |
| chr5-79995422-INS-208 | chr5-79995422-INS-210 |
| chr5-80014332-INS-1700 | chr5-80014333-INS-1700 |
| chr5-80067983-INS-172 | chr5-80067983-INS-172 |
| chr5-80074729-INS-68 | chr5-80074729-INS-70 |
| chr5-80099012-INS-501 | chr5-80099012-INS-502 |
| chr5-80175601-INS-611 | chr5-80175601-INS-611 |
| chr5-80248232-INS-158 | chr5-80248232-INS-158 |
| chr5-80303197-INS-68 | chr5-80303197-INS-75 |
| chr5-80329023-INS-52 | chr5-80329023-INS-60 |
| chr5-80331381-INS-56 | chr5-80331371-INS-60 |
| chr5-80365846-INS-234 | chr5-80365846-INS-234 |
| chr5-80376588-INS-199 | chr5-80376613-INS-199 |
| chr5-80426617-INS-58 | chr5-80426617-INS-50 |
| chr5-80441696-INS-523 | chr5-80441696-INS-532 |
| chr5-80444384-INS-186 | chr5-80444384-INS-186 |
| chr5-80460598-INS-770 | chr5-80460598-INS-770 |
| chr5-80493092-INS-4876 | chr5-80493093-INS-4876 |
| chr5-80513972-INS-332 | chr5-80513972-INS-332 |
| chr5-80540150-INS-84 | chr5-80540150-INS-84 |
| chr5-80555696-INS-52 | chr5-80555687-INS-68 |
| chr5-80573047-INS-6460 | chr5-80573048-INS-6462 |
| chr5-80581630-INS-168 | chr5-80581636-INS-173 |
| chr5-80596110-INS-504 | chr5-80596110-INS-504 |
| chr5-80629098-INS-799 | chr5-80629098-INS-799 |
| chr5-80635081-INS-215 | chr5-80635081-INS-219 |
| chr5-80691665-INS-92 | chr5-80691665-INS-92 |
| chr5-80696902-INS-63 | chr5-80696902-INS-56 |
| chr5-80712304-INS-63 | chr5-80712304-INS-63 |
| chr5-80723876-INS-2063 | chr5-80723877-INS-2063 |
| chr5-80726212-INS-200 | chr5-80726212-INS-200 |
| chr5-80742093-INS-209 | chr5-80742093-INS-209 |
| chr5-80744079-INS-70 | chr5-80744079-INS-70 |
| chr5-80755506-INS-81 | chr5-80755506-INS-81 |
| chr5-80771704-INS-3590 | chr5-80771705-INS-3590 |
| chr5-80772519-INS-78 | chr5-80772519-INS-78 |
| chr5-80777570-INS-514 | chr5-80777570-INS-514 |
| chr5-80779730-INS-72 | chr5-80779730-INS-72 |
| chr5-80800246-INS-56 | chr5-80800246-INS-52 |
| chr5-80828295-INS-495 | chr5-80828295-INS-495 |
| chr5-80832782-INS-119 | chr5-80832782-INS-119 |
| chr5-80865262-INS-2856 | chr5-80865263-INS-2856 |
| chr5-80867774-INS-52 | chr5-80867774-INS-53 |
| chr5-80874585-INS-105 | chr5-80874585-INS-105 |
| chr5-80889155-INS-83 | chr5-80889142-INS-67 |
| chr5-80949682-INS-94 | chr5-80949682-INS-96 |
| chr5-80956640-INS-56 | chr5-80956640-INS-56 |
| chr5-80975622-INS-157 | chr5-80975622-INS-157 |
| chr5-81036835-INS-1269 | chr5-81036836-INS-1270 |
| chr5-81049651-INS-75 | chr5-81049651-INS-60 |
| chr5-81065631-INS-793 | chr5-81065629-INS-790 |
| chr5-81072537-INS-146 | chr5-81072534-INS-146 |

| | |
|---|---|
| chr5-81082199-INS-64 | chr5-81082198-INS-62 |
| chr5-81094086-INS-228 | chr5-81094086-INS-229 |
| chr5-81101747-INS-7186 | chr5-81101748-INS-7185 |
| chr5-81108616-INS-100 | chr5-81108616-INS-100 |
| chr5-81118102-INS-98 | chr5-81118103-INS-98 |
| chr5-81158482-INS-223 | chr5-81158482-INS-223 |
| chr5-81159514-INS-92 | chr5-81159506-INS-52 |
| chr5-81163182-INS-63 | chr5-81163182-INS-68 |
| chr5-81270091-INS-71 | chr5-81270095-INS-92 |
| chr5-81284206-INS-1469 | chr5-81284207-INS-1469 |
| chr5-81289678-INS-7335 | chr5-81289679-INS-7336 |
| chr5-81290047-INS-58 | chr5-81290047-INS-50 |
| chr5-81299218-INS-597 | chr5-81299218-INS-598 |
| chr5-81365602-INS-471 | chr5-81365602-INS-470 |
| chr5-81368152-INS-288 | chr5-81368152-INS-300 |
| chr5-81417860-INS-606 | chr5-81417855-INS-609 |
| chr5-81420931-INS-378 | chr5-81420931-INS-377 |
| chr5-81423934-INS-88 | chr5-81423934-INS-96 |
| chr5-81497875-INS-58 | chr5-81497875-INS-58 |
| chr5-81499638-INS-177 | chr5-81499638-INS-177 |
| chr5-81505843-INS-2215 | chr5-81505844-INS-2213 |
| chr5-81582520-INS-62 | chr5-81582520-INS-64 |
| chr5-81592380-INS-645 | chr5-81592380-INS-653 |
| chr5-81602927-INS-256 | chr5-81602927-INS-256 |
| chr5-81606302-INS-56 | chr5-81606302-INS-56 |
| chr5-81637362-INS-55 | chr5-81637362-INS-55 |
| chr5-81640984-INS-1592 | chr5-81640985-INS-1596 |
| chr5-81650566-INS-61 | chr5-81650566-INS-61 |
| chr5-81689100-INS-579 | chr5-81689100-INS-579 |
| chr5-81699731-INS-58 | chr5-81699731-INS-60 |
| chr5-81747592-INS-60 | chr5-81747592-INS-60 |
| chr5-81765770-INS-90 | chr5-81765770-INS-92 |
| chr5-81798357-INS-127 | chr5-81798357-INS-123 |
| chr5-81814877-INS-74 | chr5-81814877-INS-74 |
| chr5-81845836-INS-52 | chr5-81845836-INS-50 |
| chr5-81902066-INS-58 | chr5-81902066-INS-58 |
| chr5-81909055-INS-90 | chr5-81909055-INS-91 |
| chr5-81956601-INS-511 | chr5-81956601-INS-511 |
| chr5-82004268-INS-148 | chr5-82004268-INS-148 |
| chr5-82037452-INS-64 | chr5-82037452-INS-66 |
| chr5-82155422-INS-204 | chr5-82155422-INS-204 |
| chr5-82197174-INS-815 | chr5-82197174-INS-815 |
| chr5-82199489-INS-497 | chr5-82199489-INS-497 |
| chr5-82203776-INS-4010 | chr5-82203777-INS-4010 |
| chr5-82213442-INS-6229 | chr5-82213443-INS-6229 |
| chr5-82218418-INS-7287 | chr5-82218419-INS-7287 |
| chr5-82297284-INS-2680 | chr5-82297285-INS-2680 |
| chr5-82316363-INS-1892 | chr5-82316364-INS-1892 |
| chr5-82322942-INS-203 | chr5-82322942-INS-203 |
| chr5-82324864-INS-6461 | chr5-82324865-INS-6461 |
| chr5-82352880-INS-575 | chr5-82352880-INS-574 |
| chr5-82353644-INS-74 | chr5-82353644-INS-92 |
| chr5-82396028-INS-67 | chr5-82396028-INS-68 |
| chr5-82402278-INS-62 | chr5-82402282-INS-62 |
| chr5-82457045-INS-565 | chr5-82457045-INS-564 |
| chr5-82480949-INS-6596 | chr5-82480950-INS-6596 |
| chr5-82510280-INS-6602 | chr5-82510281-INS-6597 |
| chr5-82548216-INS-304 | chr5-82548216-INS-303 |
| chr5-82553182-INS-580 | chr5-82553182-INS-580 |
| chr5-82584661-INS-771 | chr5-82584661-INS-771 |
| chr5-82598685-INS-199 | chr5-82598685-INS-199 |
| chr5-82602649-INS-66 | chr5-82602649-INS-58 |
| chr5-82609196-INS-53 | chr5-82609196-INS-53 |
| chr5-82616411-INS-120 | chr5-82616411-INS-120 |
| chr5-82620246-INS-431 | chr5-82620246-INS-432 |
| chr5-82624190-INS-52 | chr5-82624190-INS-50 |
| chr5-82630465-INS-56 | chr5-82630465-INS-54 |
| chr5-82633744-INS-55 | chr5-82633744-INS-70 |
| chr5-82672087-INS-668 | chr5-82672087-INS-664 |
| chr5-82678454-INS-6532 | chr5-82678455-INS-6532 |
| chr5-82723144-INS-6229 | chr5-82723145-INS-6235 |
| chr5-82751892-INS-60 | chr5-82751892-INS-58 |
| chr5-82761627-INS-208 | chr5-82761627-INS-208 |
| chr5-82797795-INS-416 | chr5-82797795-INS-416 |
| chr5-82797860-INS-1040 | chr5-82797861-INS-1040 |
| chr5-82802495-INS-54 | chr5-82802495-INS-54 |
| chr5-82844724-INS-7082 | chr5-82844725-INS-7082 |

| | |
|---|---|
| chr5-82846019-INS-76 | chr5-82846019-INS-50 |
| chr5-82868781-INS-70 | chr5-82868780-INS-83 |
| chr5-82899771-INS-7070 | chr5-82899772-INS-7070 |
| chr5-82929352-INS-731 | chr5-82929352-INS-731 |
| chr5-82953231-INS-58 | chr5-82953231-INS-58 |
| chr5-82970228-INS-553 | chr5-82970228-INS-554 |
| chr5-83035832-INS-96 | chr5-83035830-INS-98 |
| chr5-83038716-INS-835 | chr5-83038716-INS-835 |
| chr5-83049719-INS-1513 | chr5-83049720-INS-1512 |
| chr5-83066917-INS-95 | chr5-83066917-INS-95 |
| chr5-83079582-INS-559 | chr5-83079582-INS-558 |
| chr5-83119563-INS-75 | chr5-83119563-INS-85 |
| chr5-83177441-INS-195 | chr5-83177441-INS-195 |
| chr5-83205130-INS-64 | chr5-83205130-INS-75 |
| chr5-83207236-INS-5668 | chr5-83207237-INS-5668 |
| chr5-83213391-INS-54 | chr5-83213391-INS-57 |
| chr5-83237653-INS-70 | chr5-83237653-INS-70 |
| chr5-83292880-INS-53 | chr5-83292881-INS-51 |
| chr5-83299610-INS-164 | chr5-83299610-INS-165 |
| chr5-83314000-INS-117 | chr5-83314000-INS-117 |
| chr5-83343929-INS-6168 | chr5-83343930-INS-6168 |
| chr5-83347931-INS-96 | chr5-83347935-INS-92 |
| chr5-83361334-INS-5071 | chr5-83361335-INS-5071 |
| chr5-83375466-INS-842 | chr5-83375466-INS-844 |
| chr5-83414540-INS-7323 | chr5-83414541-INS-7323 |
| chr5-83426966-INS-1537 | chr5-83426967-INS-1543 |
| chr5-83438180-INS-56 | chr5-83438182-INS-52 |
| chr5-83465740-INS-864 | chr5-83465740-INS-864 |
| chr5-83487131-INS-635 | chr5-83487131-INS-622 |
| chr5-83497746-INS-154 | chr5-83497746-INS-154 |
| chr5-83500722-INS-871 | chr5-83500722-INS-871 |
| chr5-83661323-INS-78 | chr5-83661323-INS-92 |
| chr5-83664120-INS-71 | chr5-83664099-INS-64 |
| chr5-83726380-INS-90 | chr5-83726380-INS-90 |
| chr5-83767833-INS-1673 | chr5-83767834-INS-1674 |
| chr5-83771856-INS-1781 | chr5-83771857-INS-1782 |
| chr5-83824406-INS-253 | chr5-83824406-INS-253 |
| chr5-83866074-INS-469 | chr5-83866074-INS-469 |
| chr5-83882555-INS-175 | chr5-83882555-INS-181 |
| chr5-83894862-INS-66 | chr5-83894862-INS-84 |
| chr5-83920661-INS-449 | chr5-83920661-INS-449 |
| chr5-83927181-INS-99 | chr5-83927181-INS-99 |
| chr5-83956787-INS-151 | chr5-83956830-INS-147 |
| chr5-84036710-INS-450 | chr5-84036710-INS-450 |
| chr5-84077629-INS-223 | chr5-84077629-INS-223 |
| chr5-84214579-INS-68 | chr5-84214579-INS-77 |
| chr5-84233435-INS-940 | chr5-84233437-INS-947 |
| chr5-84244807-INS-104 | chr5-84244807-INS-99 |
| chr5-84276588-INS-785 | chr5-84276588-INS-785 |
| chr5-84375763-INS-805 | chr5-84375763-INS-805 |
| chr5-84378720-INS-834 | chr5-84378720-INS-832 |
| chr5-84448731-INS-1744 | chr5-84448732-INS-1741 |
| chr5-84567315-INS-208 | chr5-84567315-INS-208 |
| chr5-84577189-INS-59 | chr5-84577189-INS-84 |
| chr5-84761375-INS-6847 | chr5-84761376-INS-6847 |
| chr5-84779547-INS-52 | chr5-84779551-INS-52 |
| chr5-84783229-INS-231 | chr5-84783229-INS-231 |
| chr5-84795680-INS-55 | chr5-84795680-INS-55 |
| chr5-84821450-INS-67 | chr5-84821450-INS-73 |
| chr5-84833307-INS-630 | chr5-84833307-INS-631 |
| chr5-84890138-INS-368 | chr5-84890138-INS-375 |
| chr5-84943675-INS-71 | chr5-84943675-INS-69 |
| chr5-85035182-INS-624 | chr5-85035182-INS-623 |
| chr5-85060721-INS-56 | chr5-85060743-INS-56 |
| chr5-85084755-INS-1071 | chr5-85084756-INS-1072 |
| chr5-85131913-INS-76 | chr5-85131913-INS-76 |
| chr5-85142505-INS-150 | chr5-85142505-INS-150 |
| chr5-85144889-INS-68 | chr5-85144889-INS-56 |
| chr5-85253415-INS-6452 | chr5-85253428-INS-6446 |
| chr5-85285889-INS-57 | chr5-85285889-INS-57 |
| chr5-85335990-INS-189 | chr5-85335990-INS-189 |
| chr5-85389837-INS-222 | chr5-85389839-INS-222 |
| chr5-85464100-INS-52 | chr5-85464098-INS-50 |
| chr5-85471586-INS-84 | chr5-85471586-INS-57 |
| chr5-85495283-INS-56 | chr5-85495280-INS-65 |
| chr5-85504231-INS-194 | chr5-85504231-INS-193 |
| chr5-85532818-INS-400 | chr5-85532818-INS-400 |

| | |
|---|---|
| chr5-85582369-INS-58 | chr5-85582369-INS-88 |
| chr5-85584431-INS-234 | chr5-85584432-INS-233 |
| chr5-85842067-INS-485 | chr5-85842067-INS-485 |
| chr5-85950073-INS-53 | chr5-85950073-INS-53 |
| chr5-85955776-INS-72 | chr5-85955766-INS-111 |
| chr5-85996775-INS-424 | chr5-85996775-INS-424 |
| chr5-86028207-INS-615 | chr5-86028207-INS-615 |
| chr5-86046394-INS-1617 | chr5-86046395-INS-1617 |
| chr5-86069503-INS-56 | chr5-86069503-INS-80 |
| chr5-86073147-INS-354 | chr5-86073147-INS-353 |
| chr5-86076454-INS-489 | chr5-86076454-INS-492 |
| chr5-86093511-INS-64 | chr5-86093523-INS-58 |
| chr5-86096341-INS-7919 | chr5-86096342-INS-7920 |
| chr5-86101738-INS-117 | chr5-86101738-INS-123 |
| chr5-86114676-INS-8852 | chr5-86114660-INS-8825 |
| chr5-86117607-INS-639 | chr5-86117610-INS-643 |
| chr5-86169634-INS-1864 | chr5-86169635-INS-1864 |
| chr5-86172913-INS-504 | chr5-86172913-INS-504 |
| chr5-86192973-INS-52 | chr5-86192973-INS-52 |
| chr5-86213635-INS-56 | chr5-86213635-INS-56 |
| chr5-86223862-INS-72 | chr5-86223862-INS-72 |
| chr5-86225453-INS-334 | chr5-86225453-INS-337 |
| chr5-86233918-INS-136 | chr5-86233918-INS-141 |
| chr5-86324722-INS-448 | chr5-86324722-INS-448 |
| chr5-86330486-INS-108 | chr5-86330486-INS-108 |
| chr5-86338142-INS-401 | chr5-86338142-INS-401 |
| chr5-86347775-INS-1780 | chr5-86347776-INS-1780 |
| chr5-86375344-INS-3214 | chr5-86375345-INS-3214 |
| chr5-86391974-INS-109 | chr5-86391974-INS-109 |
| chr5-86416132-INS-1108 | chr5-86416133-INS-1109 |
| chr5-86418834-INS-895 | chr5-86418834-INS-897 |
| chr5-86420172-INS-58 | chr5-86420172-INS-58 |
| chr5-86437023-INS-90 | chr5-86437023-INS-90 |
| chr5-86441031-INS-52 | chr5-86441031-INS-54 |
| chr5-86449631-INS-52 | chr5-86449631-INS-61 |
| chr5-86453150-INS-855 | chr5-86453148-INS-840 |
| chr5-86504435-INS-134 | chr5-86504435-INS-134 |
| chr5-86505194-INS-424 | chr5-86505194-INS-424 |
| chr5-86529557-INS-82 | chr5-86529557-INS-82 |
| chr5-86546039-INS-408 | chr5-86546039-INS-408 |
| chr5-86548197-INS-210 | chr5-86548197-INS-210 |
| chr5-86598332-INS-214 | chr5-86598332-INS-214 |
| chr5-86608959-INS-1724 | chr5-86608960-INS-1727 |
| chr5-86609765-INS-56 | chr5-86609773-INS-79 |
| chr5-86617268-INS-56 | chr5-86617268-INS-52 |
| chr5-86657568-INS-201 | chr5-86657568-INS-201 |
| chr5-86678157-INS-227 | chr5-86678154-INS-230 |
| chr5-86686082-INS-144 | chr5-86686082-INS-144 |
| chr5-86696026-INS-204 | chr5-86696026-INS-204 |
| chr5-86714358-INS-142 | chr5-86714358-INS-145 |
| chr5-86778052-INS-125 | chr5-86778052-INS-125 |
| chr5-86794489-INS-6648 | chr5-86794489-INS-6647 |
| chr5-86847355-INS-185 | chr5-86847355-INS-185 |
| chr5-86849114-INS-161 | chr5-86849114-INS-143 |
| chr5-86857426-INS-68 | chr5-86857419-INS-110 |
| chr5-86870423-INS-60 | chr5-86870423-INS-96 |
| chr5-86872914-INS-752 | chr5-86872914-INS-752 |
| chr5-86879847-INS-7184 | chr5-86879848-INS-7184 |
| chr5-86890583-INS-119 | chr5-86890582-INS-120 |
| chr5-86913618-INS-183 | chr5-86913618-INS-183 |
| chr5-86933081-INS-241 | chr5-86933081-INS-241 |
| chr5-86937439-INS-50 | chr5-86937429-INS-61 |
| chr5-86950124-INS-1119 | chr5-86950125-INS-1123 |
| chr5-86950429-INS-344 | chr5-86950429-INS-344 |
| chr5-87039782-INS-495 | chr5-87039782-INS-499 |
| chr5-87043118-INS-695 | chr5-87043119-INS-734 |
| chr5-87159512-INS-62 | chr5-87159512-INS-62 |
| chr5-87200383-INS-56 | chr5-87200383-INS-64 |
| chr5-87227517-INS-426 | chr5-87227517-INS-426 |
| chr5-87246422-INS-460 | chr5-87246422-INS-460 |
| chr5-87255788-INS-399 | chr5-87255788-INS-399 |
| chr5-87270247-INS-139 | chr5-87270247-INS-139 |
| chr5-87289748-INS-90 | chr5-87289776-INS-98 |
| chr5-87323902-INS-76 | chr5-87323902-INS-76 |
| chr5-87334058-INS-639 | chr5-87334046-INS-642 |
| chr5-87335462-INS-76 | chr5-87335462-INS-76 |
| chr5-87375859-INS-547 | chr5-87375859-INS-546 |

| | |
|---|---|
| chr5-87426861-INS-459 | chr5-87426862-INS-457 |
| chr5-87445930-INS-278 | chr5-87445930-INS-278 |
| chr5-87475719-INS-96 | chr5-87475719-INS-96 |
| chr5-87494551-INS-170 | chr5-87494551-INS-170 |
| chr5-87499329-INS-1812 | chr5-87499330-INS-1814 |
| chr5-87513848-INS-3635 | chr5-87513849-INS-3635 |
| chr5-87584743-INS-1695 | chr5-87584744-INS-1695 |
| chr5-87599835-INS-329 | chr5-87599835-INS-329 |
| chr5-87609485-INS-717 | chr5-87609485-INS-718 |
| chr5-87635944-INS-76 | chr5-87635945-INS-76 |
| chr5-87653497-INS-205 | chr5-87653497-INS-205 |
| chr5-87662205-INS-452 | chr5-87662188-INS-482 |
| chr5-87671987-INS-62 | chr5-87671987-INS-62 |
| chr5-87746923-INS-54 | chr5-87746923-INS-76 |
| chr5-87750178-INS-52 | chr5-87750178-INS-56 |
| chr5-87804739-INS-58 | chr5-87804731-INS-59 |
| chr5-87807577-INS-179 | chr5-87807545-INS-167 |
| chr5-87817077-INS-51 | chr5-87817077-INS-51 |
| chr5-87826076-INS-56 | chr5-87826076-INS-56 |
| chr5-87828250-INS-121 | chr5-87828250-INS-121 |
| chr5-87906750-INS-571 | chr5-87906750-INS-571 |
| chr5-87925046-INS-52 | chr5-87925055-INS-58 |
| chr5-87925975-INS-1889 | chr5-87925976-INS-1889 |
| chr5-87949570-INS-66 | chr5-87949570-INS-66 |
| chr5-87969561-INS-57 | chr5-87969561-INS-103 |
| chr5-87994446-INS-4005 | chr5-87994447-INS-4006 |
| chr5-88020530-INS-51 | chr5-88020530-INS-53 |
| chr5-88023698-INS-209 | chr5-88023698-INS-209 |
| chr5-88043302-INS-6390 | chr5-88043303-INS-6390 |
| chr5-88044577-INS-80 | chr5-88044577-INS-80 |
| chr5-88070007-INS-371 | chr5-88070007-INS-371 |
| chr5-88134524-INS-86 | chr5-88134523-INS-87 |
| chr5-88173563-INS-412 | chr5-88173563-INS-413 |
| chr5-88189759-INS-896 | chr5-88189759-INS-896 |
| chr5-88222916-INS-57 | chr5-88222902-INS-60 |
| chr5-88231886-INS-447 | chr5-88231886-INS-447 |
| chr5-88239904-INS-73 | chr5-88239904-INS-70 |
| chr5-88243632-INS-132 | chr5-88243629-INS-122 |
| chr5-88248231-INS-13560 | chr5-88248232-INS-13562 |
| chr5-88253452-INS-179 | chr5-88253452-INS-181 |
| chr5-88258418-INS-265 | chr5-88258418-INS-265 |
| chr5-88266431-INS-7311 | chr5-88266432-INS-7311 |
| chr5-88290123-INS-54 | chr5-88290123-INS-54 |
| chr5-88325718-INS-1040 | chr5-88325719-INS-1041 |
| chr5-88348892-INS-131 | chr5-88348887-INS-131 |
| chr5-88407302-INS-214 | chr5-88407302-INS-214 |
| chr5-88427037-INS-6493 | chr5-88427038-INS-6491 |
| chr5-88463100-INS-63 | chr5-88463100-INS-78 |
| chr5-88464368-INS-244 | chr5-88464368-INS-244 |
| chr5-88468688-INS-121 | chr5-88468689-INS-77 |
| chr5-88469420-INS-165 | chr5-88469420-INS-165 |
| chr5-88474012-INS-213 | chr5-88474012-INS-213 |
| chr5-88483590-INS-72 | chr5-88483590-INS-78 |
| chr5-88512457-INS-57 | chr5-88512445-INS-69 |
| chr5-88590007-INS-185 | chr5-88590007-INS-184 |
| chr5-88591169-INS-54 | chr5-88591169-INS-50 |
| chr5-88593592-INS-808 | chr5-88593592-INS-808 |
| chr5-88623442-INS-377 | chr5-88623442-INS-382 |
| chr5-88649969-INS-99 | chr5-88649969-INS-99 |
| chr5-88662139-INS-281 | chr5-88662139-INS-283 |
| chr5-88663929-INS-537 | chr5-88663929-INS-529 |
| chr5-88668150-INS-85 | chr5-88668150-INS-85 |
| chr5-88669568-INS-738 | chr5-88669568-INS-738 |
| chr5-88691978-INS-52 | chr5-88691978-INS-52 |
| chr5-88730957-INS-85 | chr5-88730957-INS-93 |
| chr5-88767828-INS-189 | chr5-88767827-INS-197 |
| chr5-88784691-INS-68 | chr5-88784691-INS-68 |
| chr5-88797467-INS-217 | chr5-88797467-INS-217 |
| chr5-88802825-INS-263 | chr5-88802825-INS-278 |
| chr5-88846064-INS-477 | chr5-88846064-INS-469 |
| chr5-88893177-INS-68 | chr5-88893149-INS-76 |
| chr5-88894520-INS-2793 | chr5-88894521-INS-2793 |
| chr5-88896326-INS-279 | chr5-88896326-INS-279 |
| chr5-88918387-INS-205 | chr5-88918387-INS-204 |
| chr5-88929556-INS-82 | chr5-88929556-INS-82 |
| chr5-88932333-INS-332 | chr5-88932333-INS-332 |
| chr5-88966151-INS-127 | chr5-88966151-INS-127 |

| | |
|---|---|
| chr5-88967495-INS-52 | chr5-88967495-INS-54 |
| chr5-88991929-INS-266 | chr5-88991930-INS-280 |
| chr5-88993342-INS-815 | chr5-88993342-INS-815 |
| chr5-89025223-INS-504 | chr5-89025223-INS-504 |
| chr5-89035306-INS-180 | chr5-89035306-INS-190 |
| chr5-89054886-INS-70 | chr5-89054886-INS-70 |
| chr5-89067123-INS-95 | chr5-89067117-INS-76 |
| chr5-89127331-INS-972 | chr5-89127331-INS-973 |
| chr5-89168712-INS-2109 | chr5-89168732-INS-2090 |
| chr5-89213411-INS-100 | chr5-89213411-INS-100 |
| chr5-89242803-INS-68 | chr5-89242803-INS-68 |
| chr5-89286911-INS-399 | chr5-89286911-INS-399 |
| chr5-89292519-INS-6420 | chr5-89292520-INS-6420 |
| chr5-89321995-INS-138 | chr5-89321995-INS-138 |
| chr5-89366953-INS-68 | chr5-89366953-INS-68 |
| chr5-89398816-INS-6340 | chr5-89398817-INS-6339 |
| chr5-89409648-INS-6322 | chr5-89409649-INS-6320 |
| chr5-89412313-INS-2941 | chr5-89412314-INS-2941 |
| chr5-89427703-INS-51 | chr5-89427703-INS-51 |
| chr5-89436875-INS-58 | chr5-89436875-INS-62 |
| chr5-89491734-INS-1856 | chr5-89491735-INS-1853 |
| chr5-89508771-INS-90 | chr5-89508802-INS-71 |
| chr5-89512562-INS-770 | chr5-89512562-INS-770 |
| chr5-89550467-INS-104 | chr5-89550467-INS-89 |
| chr5-89557604-INS-123 | chr5-89557604-INS-123 |
| chr5-89585368-INS-80 | chr5-89585380-INS-68 |
| chr5-89637973-INS-81 | chr5-89637973-INS-81 |
| chr5-89639615-INS-169 | chr5-89639615-INS-169 |
| chr5-89691204-INS-776 | chr5-89691204-INS-779 |
| chr5-89693526-INS-6881 | chr5-89693527-INS-6881 |
| chr5-89698490-INS-1931 | chr5-89698491-INS-1931 |
| chr5-89699798-INS-66 | chr5-89699798-INS-66 |
| chr5-89703559-INS-240 | chr5-89703559-INS-240 |
| chr5-89708673-INS-67 | chr5-89708673-INS-69 |
| chr5-89757641-INS-401 | chr5-89757641-INS-401 |
| chr5-89798513-INS-105 | chr5-89798513-INS-105 |
| chr5-89802168-INS-1176 | chr5-89802169-INS-1176 |
| chr5-89806977-INS-51 | chr5-89806977-INS-53 |
| chr5-89835057-INS-70 | chr5-89835057-INS-70 |
| chr5-89867476-INS-111 | chr5-89867476-INS-67 |
| chr5-89874623-INS-56 | chr5-89874623-INS-56 |
| chr5-89914556-INS-69 | chr5-89914556-INS-69 |
| chr5-89949123-INS-790 | chr5-89949124-INS-790 |
| chr5-90001128-INS-62 | chr5-90001128-INS-65 |
| chr5-90005879-INS-107 | chr5-90005879-INS-99 |
| chr5-90051525-INS-75 | chr5-90051525-INS-75 |
| chr5-90060169-INS-108 | chr5-90060169-INS-108 |
| chr5-90067056-INS-63 | chr5-90067092-INS-70 |
| chr5-90075322-INS-295 | chr5-90075322-INS-297 |
| chr5-90077434-INS-55 | chr5-90077434-INS-55 |
| chr5-90093973-INS-487 | chr5-90093973-INS-487 |
| chr5-90171095-INS-61 | chr5-90171095-INS-61 |
| chr5-90193202-INS-59 | chr5-90193198-INS-63 |
| chr5-90210381-INS-121 | chr5-90210378-INS-128 |
| chr5-90222698-INS-375 | chr5-90222698-INS-367 |
| chr5-90255910-INS-470 | chr5-90255910-INS-470 |
| chr5-90290442-INS-52 | chr5-90290442-INS-72 |
| chr5-90296535-INS-54 | chr5-90296520-INS-61 |
| chr5-90333147-INS-1891 | chr5-90333148-INS-1891 |
| chr5-90333475-INS-51 | chr5-90333470-INS-54 |
| chr5-90362537-INS-100 | chr5-90362541-INS-56 |
| chr5-90397904-INS-68 | chr5-90397904-INS-70 |
| chr5-90406921-INS-481 | chr5-90406921-INS-483 |
| chr5-90417591-INS-215 | chr5-90417591-INS-215 |
| chr5-90429326-INS-211 | chr5-90429326-INS-212 |
| chr5-90443830-INS-57 | chr5-90443830-INS-57 |
| chr5-90471077-INS-211 | chr5-90471077-INS-211 |
| chr5-90528977-INS-53 | chr5-90528977-INS-60 |
| chr5-90530577-INS-77 | chr5-90530577-INS-79 |
| chr5-90542223-INS-58 | chr5-90542223-INS-58 |
| chr5-90557710-INS-150 | chr5-90557710-INS-150 |
| chr5-90574593-INS-205 | chr5-90574593-INS-205 |
| chr5-90611373-INS-188 | chr5-90611373-INS-188 |
| chr5-90618920-INS-211 | chr5-90618920-INS-214 |
| chr5-90661703-INS-277 | chr5-90661703-INS-277 |
| chr5-90703625-INS-76 | chr5-90703625-INS-76 |
| chr5-90715965-INS-384 | chr5-90715966-INS-412 |

| | |
|---|---|
| chr5-90734221-INS-306 | chr5-90734221-INS-304 |
| chr5-90737835-INS-67 | chr5-90737835-INS-67 |
| chr5-90746020-INS-6769 | chr5-90746021-INS-6768 |
| chr5-90801966-INS-94 | chr5-90801966-INS-94 |
| chr5-90822755-INS-6613 | chr5-90822756-INS-6612 |
| chr5-90867768-INS-68 | chr5-90867768-INS-68 |
| chr5-90888535-INS-159 | chr5-90888535-INS-159 |
| chr5-90908769-INS-274 | chr5-90908769-INS-280 |
| chr5-90954615-INS-112 | chr5-90954615-INS-112 |
| chr5-91003653-INS-6171 | chr5-91003654-INS-6169 |
| chr5-91021723-INS-59 | chr5-91021723-INS-63 |
| chr5-91082944-INS-72 | chr5-91082945-INS-96 |
| chr5-91128516-INS-683 | chr5-91128516-INS-686 |
| chr5-91145943-INS-491 | chr5-91145944-INS-491 |
| chr5-91160412-INS-375 | chr5-91160412-INS-375 |
| chr5-91167590-INS-112 | chr5-91167590-INS-112 |
| chr5-91217424-INS-78 | chr5-91217424-INS-108 |
| chr5-91218095-INS-847 | chr5-91218095-INS-847 |
| chr5-91224282-INS-60 | chr5-91224282-INS-68 |
| chr5-91229136-INS-60 | chr5-91229119-INS-93 |
| chr5-91269871-INS-819 | chr5-91269871-INS-819 |
| chr5-91278990-INS-96 | chr5-91278990-INS-72 |
| chr5-91280102-INS-105 | chr5-91280095-INS-112 |
| chr5-91286332-INS-90 | chr5-91286332-INS-125 |
| chr5-91351451-INS-90 | chr5-91351454-INS-102 |
| chr5-91363334-INS-87 | chr5-91363334-INS-104 |
| chr5-91369887-INS-4471 | chr5-91369888-INS-4471 |
| chr5-91395156-INS-858 | chr5-91395156-INS-858 |
| chr5-91438994-INS-7109 | chr5-91438995-INS-7109 |
| chr5-91448498-INS-464 | chr5-91448498-INS-464 |
| chr5-91489156-INS-80 | chr5-91489156-INS-80 |
| chr5-91527790-INS-74 | chr5-91527782-INS-60 |
| chr5-91557802-INS-99 | chr5-91557802-INS-50 |
| chr5-91581470-INS-798 | chr5-91581470-INS-748 |
| chr5-91584327-INS-78 | chr5-91584295-INS-60 |
| chr5-91598254-INS-162 | chr5-91598254-INS-162 |
| chr5-91630209-INS-52 | chr5-91630210-INS-56 |
| chr5-91640917-INS-308 | chr5-91640917-INS-308 |
| chr5-91649624-INS-118 | chr5-91649622-INS-78 |
| chr5-91651236-INS-52 | chr5-91651236-INS-52 |
| chr5-91682314-INS-269 | chr5-91682306-INS-271 |
| chr5-91697062-INS-62 | chr5-91697062-INS-72 |
| chr5-91729932-INS-6336 | chr5-91729933-INS-6336 |
| chr5-91738134-INS-1151 | chr5-91738133-INS-1136 |
| chr5-91739530-INS-153 | chr5-91739530-INS-133 |
| chr5-91805884-INS-6222 | chr5-91805885-INS-6223 |
| chr5-91818960-INS-2412 | chr5-91818961-INS-2414 |
| chr5-91846982-INS-72 | chr5-91846982-INS-72 |
| chr5-91869401-INS-78 | chr5-91869401-INS-78 |
| chr5-91880630-INS-61 | chr5-91880630-INS-61 |
| chr5-91907445-INS-80 | chr5-91907445-INS-80 |
| chr5-91917862-INS-51 | chr5-91917862-INS-52 |
| chr5-91922693-INS-54 | chr5-91922694-INS-94 |
| chr5-91928463-INS-945 | chr5-91928463-INS-952 |
| chr5-91953102-INS-1284 | chr5-91953103-INS-1284 |
| chr5-91973693-INS-304 | chr5-91973693-INS-304 |
| chr5-91986661-INS-169 | chr5-91986661-INS-169 |
| chr5-92036374-INS-6799 | chr5-92036375-INS-6799 |
| chr5-92047254-INS-217 | chr5-92047254-INS-216 |
| chr5-92047783-INS-207 | chr5-92047783-INS-207 |
| chr5-92056069-INS-51 | chr5-92056069-INS-50 |
| chr5-92080498-INS-257 | chr5-92080498-INS-249 |
| chr5-92081062-INS-58 | chr5-92081062-INS-58 |
| chr5-92087487-INS-143 | chr5-92087456-INS-135 |
| chr5-92096514-INS-343 | chr5-92096514-INS-344 |
| chr5-92098617-INS-365 | chr5-92098617-INS-380 |
| chr5-92114955-INS-183 | chr5-92114955-INS-183 |
| chr5-92116054-INS-297 | chr5-92116054-INS-297 |
| chr5-92120797-INS-1487 | chr5-92120798-INS-1487 |
| chr5-92122442-INS-4891 | chr5-92122443-INS-4891 |
| chr5-92132471-INS-59 | chr5-92132467-INS-58 |
| chr5-92153513-INS-93 | chr5-92153513-INS-93 |
| chr5-92169112-INS-1864 | chr5-92169113-INS-1864 |
| chr5-92172917-INS-211 | chr5-92172917-INS-211 |
| chr5-92178005-INS-65 | chr5-92178005-INS-65 |
| chr5-92218145-INS-227 | chr5-92218145-INS-227 |
| chr5-92220002-INS-135 | chr5-92220002-INS-135 |

| | |
|---|---|
| chr5-92228065-INS-63 | chr5-92228019-INS-67 |
| chr5-92239547-INS-213 | chr5-92239547-INS-213 |
| chr5-92263317-INS-483 | chr5-92263317-INS-485 |
| chr5-92294653-INS-218 | chr5-92294653-INS-218 |
| chr5-92297735-INS-275 | chr5-92297735-INS-275 |
| chr5-92299969-INS-210 | chr5-92299969-INS-210 |
| chr5-92343890-INS-152 | chr5-92343890-INS-120 |
| chr5-92353081-INS-197 | chr5-92353081-INS-197 |
| chr5-92366124-INS-6903 | chr5-92366125-INS-6903 |
| chr5-92372495-INS-97 | chr5-92372495-INS-99 |
| chr5-92378918-INS-51 | chr5-92378929-INS-55 |
| chr5-92382606-INS-92 | chr5-92382596-INS-82 |
| chr5-92389693-INS-58 | chr5-92389693-INS-60 |
| chr5-92409664-INS-81 | chr5-92409664-INS-83 |
| chr5-92419525-INS-121 | chr5-92419525-INS-121 |
| chr5-92435528-INS-1322 | chr5-92435529-INS-1319 |
| chr5-92445241-INS-225 | chr5-92445241-INS-224 |
| chr5-92447149-INS-129 | chr5-92447149-INS-129 |
| chr5-92458831-INS-396 | chr5-92458827-INS-400 |
| chr5-92469026-INS-72 | chr5-92469016-INS-56 |
| chr5-92485113-INS-80 | chr5-92485113-INS-80 |
| chr5-92488040-INS-308 | chr5-92488040-INS-308 |
| chr5-92491866-INS-152 | chr5-92491866-INS-156 |
| chr5-92503820-INS-57 | chr5-92503831-INS-57 |
| chr5-92511884-INS-64 | chr5-92511884-INS-58 |
| chr5-92542387-INS-88 | chr5-92542377-INS-71 |
| chr5-92544654-INS-5537 | chr5-92544655-INS-5537 |
| chr5-92548554-INS-850 | chr5-92548545-INS-860 |
| chr5-92552849-INS-56 | chr5-92552849-INS-60 |
| chr5-92558532-INS-62 | chr5-92558532-INS-62 |
| chr5-92561042-INS-54 | chr5-92561060-INS-70 |
| chr5-92566532-INS-217 | chr5-92566532-INS-217 |
| chr5-92571474-INS-219 | chr5-92571474-INS-220 |
| chr5-92574419-INS-170 | chr5-92574419-INS-170 |
| chr5-92603776-INS-216 | chr5-92603776-INS-215 |
| chr5-92607107-INS-90 | chr5-92607107-INS-100 |
| chr5-92635749-INS-86 | chr5-92635749-INS-86 |
| chr5-92637945-INS-199 | chr5-92637945-INS-199 |
| chr5-92647749-INS-86 | chr5-92647729-INS-108 |
| chr5-92648983-INS-84 | chr5-92648958-INS-60 |
| chr5-92654902-INS-217 | chr5-92654900-INS-243 |
| chr5-92668907-INS-184 | chr5-92668907-INS-184 |
| chr5-92718687-INS-51 | chr5-92718687-INS-89 |
| chr5-92726022-INS-722 | chr5-92726022-INS-722 |
| chr5-92777764-INS-172 | chr5-92777808-INS-174 |
| chr5-92784974-INS-208 | chr5-92784974-INS-209 |
| chr5-92788009-INS-538 | chr5-92788009-INS-538 |
| chr5-92797592-INS-369 | chr5-92797592-INS-366 |
| chr5-92811928-INS-108 | chr5-92811928-INS-108 |
| chr5-92825289-INS-54 | chr5-92825288-INS-55 |
| chr5-92841602-INS-196 | chr5-92841602-INS-196 |
| chr5-92905736-INS-221 | chr5-92905736-INS-222 |
| chr5-92906272-INS-205 | chr5-92906272-INS-205 |
| chr5-92907393-INS-1719 | chr5-92907394-INS-1719 |
| chr5-92907394-INS-357 | chr5-92907394-INS-357 |
| chr5-92913143-INS-207 | chr5-92913143-INS-193 |
| chr5-92930221-INS-194 | chr5-92930221-INS-153 |
| chr5-92948353-INS-54 | chr5-92948369-INS-62 |
| chr5-92953497-INS-1894 | chr5-92953498-INS-1894 |
| chr5-92953895-INS-509 | chr5-92953895-INS-512 |
| chr5-92956693-INS-87 | chr5-92956693-INS-83 |
| chr5-92963651-INS-200 | chr5-92963651-INS-200 |
| chr5-92974361-INS-174 | chr5-92974361-INS-174 |
| chr5-92977882-INS-216 | chr5-92977889-INS-200 |
| chr5-92981404-INS-204 | chr5-92981404-INS-204 |
| chr5-93015602-INS-213 | chr5-93015602-INS-213 |
| chr5-93019480-INS-487 | chr5-93019480-INS-474 |
| chr5-93025507-INS-55 | chr5-93025507-INS-55 |
| chr5-93027153-INS-197 | chr5-93027149-INS-202 |
| chr5-93033588-INS-68 | chr5-93033588-INS-68 |
| chr5-93039949-INS-214 | chr5-93039949-INS-214 |
| chr5-93052724-INS-52 | chr5-93052724-INS-52 |
| chr5-93058445-INS-310 | chr5-93058445-INS-310 |
| chr5-93080313-INS-295 | chr5-93080313-INS-293 |
| chr5-93082944-INS-63 | chr5-93082944-INS-63 |
| chr5-93141296-INS-202 | chr5-93141296-INS-202 |
| chr5-93144918-INS-206 | chr5-93144918-INS-206 |

| | |
|---|---|
| chr5-93155637-INS-74 | chr5-93155637-INS-74 |
| chr5-93169255-INS-193 | chr5-93169255-INS-200 |
| chr5-93178323-INS-701 | chr5-93178323-INS-701 |
| chr5-93183426-INS-77 | chr5-93183424-INS-81 |
| chr5-93189123-INS-203 | chr5-93189123-INS-203 |
| chr5-93255941-INS-154 | chr5-93255941-INS-160 |
| chr5-93290949-INS-53 | chr5-93290949-INS-53 |
| chr5-93296711-INS-79 | chr5-93296711-INS-79 |
| chr5-93315004-INS-279 | chr5-93315025-INS-280 |
| chr5-93356285-INS-198 | chr5-93356285-INS-198 |
| chr5-93364904-INS-498 | chr5-93364904-INS-498 |
| chr5-93368646-INS-62 | chr5-93368646-INS-62 |
| chr5-93374011-INS-61 | chr5-93374012-INS-54 |
| chr5-93378414-INS-2837 | chr5-93378415-INS-2837 |
| chr5-93427486-INS-246 | chr5-93427486-INS-241 |
| chr5-93482144-INS-210 | chr5-93482144-INS-210 |
| chr5-93505192-INS-641 | chr5-93505192-INS-641 |
| chr5-93514526-INS-738 | chr5-93514526-INS-739 |
| chr5-93520566-INS-1667 | chr5-93520567-INS-1676 |
| chr5-93553461-INS-641 | chr5-93553461-INS-641 |
| chr5-93605340-INS-209 | chr5-93605340-INS-210 |
| chr5-93611077-INS-199 | chr5-93611077-INS-199 |
| chr5-93638584-INS-195 | chr5-93638584-INS-196 |
| chr5-93697347-INS-170 | chr5-93697347-INS-170 |
| chr5-93744530-INS-194 | chr5-93744530-INS-194 |
| chr5-93900496-INS-1538 | chr5-93900497-INS-1538 |
| chr5-96017166-INS-633 | chr5-96017166-INS-638 |
| chr5-96048818-INS-84 | chr5-96048818-INS-68 |
| chr5-96056228-INS-6358 | chr5-96056229-INS-6360 |
| chr5-96097001-INS-389 | chr5-96097001-INS-390 |
| chr5-96101073-INS-643 | chr5-96101073-INS-644 |
| chr5-96257199-INS-60 | chr5-96257196-INS-66 |
| chr5-96300984-INS-661 | chr5-96300984-INS-661 |
| chr5-96324523-INS-52 | chr5-96324523-INS-54 |
| chr5-96332235-INS-747 | chr5-96332235-INS-747 |
| chr5-96332252-INS-1866 | chr5-96332253-INS-1866 |
| chr5-96363512-INS-64 | chr5-96363512-INS-64 |
| chr5-96366086-INS-51 | chr5-96366086-INS-51 |
| chr5-96400139-INS-200 | chr5-96400139-INS-200 |
| chr5-96459015-INS-191 | chr5-96459015-INS-191 |
| chr5-96502881-INS-88 | chr5-96502881-INS-86 |
| chr5-96514946-INS-68 | chr5-96514946-INS-68 |
| chr5-96519282-INS-52 | chr5-96519251-INS-92 |
| chr5-96577137-INS-348 | chr5-96577137-INS-351 |
| chr5-96610088-INS-110 | chr5-96610088-INS-110 |
| chr5-96619459-INS-52 | chr5-96619460-INS-82 |
| chr5-96641077-INS-118 | chr5-96641077-INS-119 |
| chr5-96662272-INS-70 | chr5-96662272-INS-75 |
| chr5-96740044-INS-55 | chr5-96740044-INS-55 |
| chr5-96763558-INS-54 | chr5-96763550-INS-56 |
| chr5-96779916-INS-53 | chr5-96779916-INS-53 |
| chr5-96782852-INS-1498 | chr5-96782853-INS-1519 |
| chr5-96804842-INS-124 | chr5-96804842-INS-124 |
| chr5-96859615-INS-77 | chr5-96859615-INS-77 |
| chr5-96871659-INS-56 | chr5-96871659-INS-54 |
| chr5-96876946-INS-51 | chr5-96876947-INS-71 |
| chr5-96882579-INS-590 | chr5-96882579-INS-592 |
| chr5-96885063-INS-171 | chr5-96885063-INS-171 |
| chr5-96892344-INS-62 | chr5-96892352-INS-66 |
| chr5-96903824-INS-57 | chr5-96903824-INS-57 |
| chr5-96922724-INS-73 | chr5-96922724-INS-73 |
| chr5-96944252-INS-59 | chr5-96944252-INS-63 |
| chr5-96959936-INS-201 | chr5-96959936-INS-201 |
| chr5-96985242-INS-72 | chr5-96985242-INS-76 |
| chr5-96989818-INS-117 | chr5-96989777-INS-118 |
| chr5-97009051-INS-83 | chr5-97009054-INS-103 |
| chr5-97012928-INS-7082 | chr5-97012929-INS-7082 |
| chr5-97026378-INS-96 | chr5-97026378-INS-100 |
| chr5-97063366-INS-789 | chr5-97063366-INS-789 |
| chr5-97073263-INS-64 | chr5-97073247-INS-76 |
| chr5-97112222-INS-113 | chr5-97112222-INS-114 |
| chr5-97114016-INS-53 | chr5-97114016-INS-97 |
| chr5-97115306-INS-209 | chr5-97115306-INS-210 |
| chr5-97121690-INS-222 | chr5-97121690-INS-222 |
| chr5-97124442-INS-163 | chr5-97124442-INS-163 |
| chr5-97136124-INS-151 | chr5-97136124-INS-151 |
| chr5-97161462-INS-145 | chr5-97161462-INS-145 |

| | |
|---|---|
| chr5-97171620-INS-348 | chr5-97171620-INS-347 |
| chr5-97209949-INS-74 | chr5-97209949-INS-74 |
| chr5-97224680-INS-476 | chr5-97224680-INS-476 |
| chr5-97230339-INS-99 | chr5-97230339-INS-99 |
| chr5-97248438-INS-55 | chr5-97248438-INS-55 |
| chr5-97288701-INS-865 | chr5-97288701-INS-866 |
| chr5-97308466-INS-1780 | chr5-97308467-INS-1779 |
| chr5-97322420-INS-111 | chr5-97322420-INS-111 |
| chr5-97332959-INS-200 | chr5-97332959-INS-200 |
| chr5-97342471-INS-208 | chr5-97342471-INS-208 |
| chr5-97353029-INS-486 | chr5-97353029-INS-487 |
| chr5-97368018-INS-75 | chr5-97368018-INS-75 |
| chr5-97376544-INS-73 | chr5-97376544-INS-73 |
| chr5-97377268-INS-939 | chr5-97377268-INS-939 |
| chr5-97384227-INS-616 | chr5-97384227-INS-616 |
| chr5-97388510-INS-92 | chr5-97388510-INS-109 |
| chr5-97392579-INS-830 | chr5-97392579-INS-829 |
| chr5-97430983-INS-246 | chr5-97430983-INS-244 |
| chr5-97434934-INS-172 | chr5-97434956-INS-184 |
| chr5-97440390-INS-60 | chr5-97440350-INS-52 |
| chr5-97443981-INS-90 | chr5-97443981-INS-90 |
| chr5-97526855-INS-6879 | chr5-97526856-INS-6880 |
| chr5-97528127-INS-52 | chr5-97528123-INS-60 |
| chr5-97537333-INS-129 | chr5-97537333-INS-129 |
| chr5-97548823-INS-209 | chr5-97548823-INS-209 |
| chr5-97573290-INS-53 | chr5-97573290-INS-53 |
| chr5-97581047-INS-424 | chr5-97581047-INS-424 |
| chr5-97582079-INS-6270 | chr5-97582080-INS-6270 |
| chr5-97609561-INS-120 | chr5-97609561-INS-124 |
| chr5-97625496-INS-169 | chr5-97625494-INS-169 |
| chr5-97629184-INS-4764 | chr5-97629185-INS-4764 |
| chr5-97691521-INS-66 | chr5-97691496-INS-66 |
| chr5-97708920-INS-147 | chr5-97708917-INS-138 |
| chr5-97710544-INS-70 | chr5-97710544-INS-70 |
| chr5-97714444-INS-56 | chr5-97714444-INS-56 |
| chr5-97750154-INS-1893 | chr5-97750155-INS-1893 |
| chr5-97770587-INS-56 | chr5-97770587-INS-83 |
| chr5-97779086-INS-63 | chr5-97779086-INS-67 |
| chr5-97795001-INS-2271 | chr5-97795002-INS-2224 |
| chr5-97803840-INS-52 | chr5-97803840-INS-52 |
| chr5-97810756-INS-66 | chr5-97810756-INS-66 |
| chr5-97822067-INS-187 | chr5-97822067-INS-187 |
| chr5-97823940-INS-4997 | chr5-97823941-INS-4997 |
| chr5-97842379-INS-6354 | chr5-97842380-INS-6354 |
| chr5-97863986-INS-134 | chr5-97863986-INS-134 |
| chr5-97876327-INS-6161 | chr5-97876328-INS-6161 |
| chr5-97892242-INS-64 | chr5-97892248-INS-54 |
| chr5-97930415-INS-282 | chr5-97930415-INS-286 |
| chr5-97931579-INS-217 | chr5-97931583-INS-213 |
| chr5-97937806-INS-62 | chr5-97937807-INS-60 |
| chr5-97940304-INS-2682 | chr5-97940305-INS-2682 |
| chr5-97943836-INS-1297 | chr5-97943837-INS-1317 |
| chr5-97981245-INS-1272 | chr5-97981246-INS-1272 |
| chr5-97986003-INS-60 | chr5-97986003-INS-60 |
| chr5-97991269-INS-1609 | chr5-97991270-INS-1609 |
| chr5-98003429-INS-209 | chr5-98003429-INS-209 |
| chr5-98028172-INS-494 | chr5-98028172-INS-494 |
| chr5-98072223-INS-6215 | chr5-98072224-INS-6215 |
| chr5-98114833-INS-126 | chr5-98114833-INS-131 |
| chr5-98125119-INS-52 | chr5-98125119-INS-56 |
| chr5-98132658-INS-90 | chr5-98132659-INS-53 |
| chr5-98185722-INS-311 | chr5-98185722-INS-310 |
| chr5-98215062-INS-54 | chr5-98215062-INS-50 |
| chr5-98221048-INS-84 | chr5-98221048-INS-84 |
| chr5-98251356-INS-368 | chr5-98251356-INS-359 |
| chr5-98258736-INS-4089 | chr5-98258737-INS-4089 |
| chr5-98338393-INS-74 | chr5-98338393-INS-62 |
| chr5-98350553-INS-116 | chr5-98350553-INS-116 |
| chr5-98373038-INS-239 | chr5-98373038-INS-241 |
| chr5-98389637-INS-66 | chr5-98389637-INS-66 |
| chr5-98413217-INS-194 | chr5-98413217-INS-194 |
| chr5-98427226-INS-112 | chr5-98427226-INS-112 |
| chr5-98502870-INS-72 | chr5-98502886-INS-76 |
| chr5-98516387-INS-120 | chr5-98516403-INS-109 |
| chr5-98541202-INS-55 | chr5-98541202-INS-50 |
| chr5-98543742-INS-146 | chr5-98543742-INS-145 |
| chr5-98577040-INS-72 | chr5-98577040-INS-72 |

| | |
|---|---|
| chr5-98595853-INS-87 | chr5-98595853-INS-89 |
| chr5-98615898-INS-58 | chr5-98615898-INS-58 |
| chr5-98632774-INS-53 | chr5-98632774-INS-56 |
| chr5-98644112-INS-218 | chr5-98644112-INS-218 |
| chr5-98722187-INS-455 | chr5-98722187-INS-455 |
| chr5-98728654-INS-59 | chr5-98728654-INS-59 |
| chr5-98734540-INS-278 | chr5-98734531-INS-292 |
| chr5-98744696-INS-188 | chr5-98744696-INS-187 |
| chr5-98804128-INS-58 | chr5-98804129-INS-84 |
| chr5-98831801-INS-6587 | chr5-98831802-INS-6587 |
| chr5-98842300-INS-63 | chr5-98842300-INS-63 |
| chr5-98854137-INS-6344 | chr5-98854138-INS-6344 |
| chr5-98865287-INS-986 | chr5-98865314-INS-988 |
| chr5-98921894-INS-55 | chr5-98921894-INS-55 |
| chr5-98933179-INS-242 | chr5-98933179-INS-218 |
| chr5-98945483-INS-79 | chr5-98945478-INS-76 |
| chr5-98957382-INS-7143 | chr5-98957383-INS-7143 |
| chr5-98965259-INS-52 | chr5-98965240-INS-62 |
| chr5-98970866-INS-74 | chr5-98970866-INS-74 |
| chr5-98973924-INS-70 | chr5-98973949-INS-91 |
| chr5-98982416-INS-157 | chr5-98982412-INS-154 |
| chr5-99036569-INS-64 | chr5-99036569-INS-64 |
| chr5-99038414-INS-127 | chr5-99038414-INS-78 |
| chr5-99127126-INS-269 | chr5-99127126-INS-269 |
| chr5-99201220-INS-1147 | chr5-99201221-INS-1150 |
| chr5-99206144-INS-93 | chr5-99206144-INS-92 |
| chr5-99224399-INS-340 | chr5-99224399-INS-340 |
| chr5-99242853-INS-88 | chr5-99242853-INS-88 |
| chr5-99244803-INS-478 | chr5-99244803-INS-478 |
| chr5-99272206-INS-481 | chr5-99272172-INS-481 |
| chr5-99273075-INS-57 | chr5-99273075-INS-57 |
| chr5-99289635-INS-543 | chr5-99289635-INS-543 |
| chr5-99296120-INS-61 | chr5-99296120-INS-61 |
| chr5-99298555-INS-52 | chr5-99298555-INS-58 |
| chr5-99306162-INS-58 | chr5-99306162-INS-54 |
| chr5-99390870-INS-259 | chr5-99390870-INS-259 |
| chr5-99396171-INS-60 | chr5-99396197-INS-60 |
| chr5-99418070-INS-115 | chr5-99418070-INS-115 |
| chr5-99438490-INS-126 | chr5-99438490-INS-126 |
| chr5-99445711-INS-58 | chr5-99445711-INS-60 |
| chr5-99509756-INS-67 | chr5-99509756-INS-69 |
| chr5-99532390-INS-198 | chr5-99532390-INS-198 |
| chr5-99607411-INS-66 | chr5-99607394-INS-105 |
| chr5-99615853-INS-68 | chr5-99615839-INS-65 |
| chr5-99620091-INS-1889 | chr5-99620092-INS-1889 |
| chr5-99682111-INS-64 | chr5-99682111-INS-62 |
| chr5-99686383-INS-67 | chr5-99686383-INS-67 |
| chr5-99709661-INS-5308 | chr5-99709662-INS-5308 |
| chr5-99734580-INS-80 | chr5-99734630-INS-62 |
| chr5-99740817-INS-66 | chr5-99740817-INS-66 |
| chr5-99766095-INS-91 | chr5-99766122-INS-51 |
| chr5-99826930-INS-54 | chr5-99826930-INS-55 |
| chr5-99841708-INS-78 | chr5-99841727-INS-85 |
| chr5-99860557-INS-52 | chr5-99860557-INS-52 |
| chr5-99918452-INS-56 | chr5-99918439-INS-60 |
| chr5-99932584-INS-6437 | chr5-99932585-INS-6439 |
| chr5-99969824-INS-83 | chr5-99969794-INS-100 |
| chr5-99970626-INS-110 | chr5-99970627-INS-66 |
| chr5-99979647-INS-135 | chr5-99979647-INS-135 |
| chr5-99995535-INS-75 | chr5-99995535-INS-75 |
| chr5-100003002-INS-116 | chr5-100003002-INS-95 |
| chr5-100003488-INS-178 | chr5-100003488-INS-178 |
| chr5-100011081-INS-76 | chr5-100011081-INS-76 |
| chr5-100012599-INS-136 | chr5-100012599-INS-136 |
| chr5-100014442-INS-312 | chr5-100014442-INS-312 |
| chr5-100022800-INS-6273 | chr5-100022801-INS-6273 |
| chr5-100045714-INS-442 | chr5-100045714-INS-442 |
| chr5-100066597-INS-639 | chr5-100066597-INS-656 |
| chr5-100083194-INS-438 | chr5-100083183-INS-415 |
| chr5-100093184-INS-53 | chr5-100093184-INS-53 |
| chr5-100152649-INS-204 | chr5-100152649-INS-204 |
| chr5-100168816-INS-408 | chr5-100168816-INS-409 |
| chr5-100216368-INS-71 | chr5-100216368-INS-71 |
| chr5-100225521-INS-60 | chr5-100225521-INS-60 |
| chr5-100285432-INS-644 | chr5-100285432-INS-645 |
| chr5-100285567-INS-1508 | chr5-100285568-INS-1508 |
| chr5-100289152-INS-56 | chr5-100289164-INS-63 |

| | |
|---|---|
| chr5-100291586-INS-70 | chr5-100291586-INS-70 |
| chr5-100295121-INS-679 | chr5-100295121-INS-681 |
| chr5-100312967-INS-64 | chr5-100312931-INS-64 |
| chr5-100315801-INS-101 | chr5-100315801-INS-101 |
| chr5-100326338-INS-7539 | chr5-100326339-INS-7554 |
| chr5-100342616-INS-85 | chr5-100342616-INS-85 |
| chr5-100350854-INS-52 | chr5-100350854-INS-56 |
| chr5-100360190-INS-73 | chr5-100360190-INS-80 |
| chr5-100361157-INS-5548 | chr5-100361158-INS-5548 |
| chr5-100371573-INS-1115 | chr5-100371574-INS-1133 |
| chr5-100392135-INS-227 | chr5-100392123-INS-209 |
| chr5-100401112-INS-2730 | chr5-100401113-INS-2730 |
| chr5-100410087-INS-1038 | chr5-100410088-INS-1038 |
| chr5-100415135-INS-64 | chr5-100415135-INS-64 |
| chr5-100425018-INS-56 | chr5-100425018-INS-56 |
| chr5-100428452-INS-72 | chr5-100428465-INS-84 |
| chr5-100448046-INS-6403 | chr5-100448047-INS-6403 |
| chr5-100471312-INS-237 | chr5-100471312-INS-238 |
| chr5-100489753-INS-193 | chr5-100489749-INS-187 |
| chr5-100500563-INS-7579 | chr5-100500564-INS-7579 |
| chr5-100526270-INS-123 | chr5-100526270-INS-123 |
| chr5-100643721-INS-58 | chr5-100643721-INS-58 |
| chr5-100651919-INS-65 | chr5-100651915-INS-65 |
| chr5-100665308-INS-402 | chr5-100665308-INS-402 |
| chr5-100673917-INS-5445 | chr5-100673918-INS-5445 |
| chr5-100688163-INS-76 | chr5-100688163-INS-76 |
| chr5-100704199-INS-202 | chr5-100704199-INS-201 |
| chr5-100705394-INS-210 | chr5-100705394-INS-214 |
| chr5-100717671-INS-343 | chr5-100717671-INS-342 |
| chr5-100751612-INS-111 | chr5-100751604-INS-104 |
| chr5-100762609-INS-145 | chr5-100762609-INS-157 |
| chr5-100786835-INS-54 | chr5-100786835-INS-63 |
| chr5-100791436-INS-210 | chr5-100791436-INS-210 |
| chr5-100800071-INS-104 | chr5-100800071-INS-104 |
| chr5-100817147-INS-206 | chr5-100817147-INS-206 |
| chr5-100838244-INS-423 | chr5-100838244-INS-423 |
| chr5-100850018-INS-55 | chr5-100850005-INS-55 |
| chr5-100851170-INS-580 | chr5-100851170-INS-620 |
| chr5-100852197-INS-180 | chr5-100852197-INS-181 |
| chr5-100864474-INS-116 | chr5-100864440-INS-110 |
| chr5-100883264-INS-98 | chr5-100883265-INS-94 |
| chr5-100886949-INS-68 | chr5-100886949-INS-68 |
| chr5-100925288-INS-52 | chr5-100925288-INS-52 |
| chr5-100948252-INS-648 | chr5-100948253-INS-648 |
| chr5-100956515-INS-103 | chr5-100956538-INS-96 |
| chr5-100982940-INS-74 | chr5-100982940-INS-77 |
| chr5-100986136-INS-201 | chr5-100986136-INS-201 |
| chr5-100998023-INS-165 | chr5-100998023-INS-167 |
| chr5-101002289-INS-66 | chr5-101002289-INS-66 |
| chr5-101005908-INS-61 | chr5-101005906-INS-78 |
| chr5-101007050-INS-92 | chr5-101007050-INS-84 |
| chr5-101015584-INS-126 | chr5-101015631-INS-84 |
| chr5-101025301-INS-205 | chr5-101025301-INS-207 |
| chr5-101039750-INS-102 | chr5-101039750-INS-102 |
| chr5-101043068-INS-217 | chr5-101043068-INS-217 |
| chr5-101052600-INS-70 | chr5-101052584-INS-63 |
| chr5-101072807-INS-301 | chr5-101072807-INS-320 |
| chr5-101134839-INS-63 | chr5-101134839-INS-63 |
| chr5-101138099-INS-426 | chr5-101138099-INS-426 |
| chr5-101146489-INS-52 | chr5-101146489-INS-54 |
| chr5-101173444-INS-201 | chr5-101173444-INS-201 |
| chr5-101175183-INS-144 | chr5-101175183-INS-144 |
| chr5-101194964-INS-827 | chr5-101194964-INS-827 |
| chr5-101221716-INS-361 | chr5-101221716-INS-361 |
| chr5-101262642-INS-61 | chr5-101262643-INS-61 |
| chr5-101284794-INS-159 | chr5-101284794-INS-159 |
| chr5-101292129-INS-7080 | chr5-101292130-INS-7081 |
| chr5-101302540-INS-52 | chr5-101302540-INS-70 |
| chr5-101331170-INS-96 | chr5-101331196-INS-100 |
| chr5-101336943-INS-99 | chr5-101336916-INS-105 |
| chr5-101340318-INS-222 | chr5-101340318-INS-222 |
| chr5-101344132-INS-69 | chr5-101344132-INS-69 |
| chr5-101391617-INS-200 | chr5-101391617-INS-200 |
| chr5-101446233-INS-179 | chr5-101446233-INS-180 |
| chr5-101467146-INS-76 | chr5-101467146-INS-76 |
| chr5-101484854-INS-61 | chr5-101484854-INS-61 |
| chr5-101507607-INS-60 | chr5-101507607-INS-58 |

| | |
|---|---|
| chr5-101539468-INS-156 | chr5-101539468-INS-168 |
| chr5-101563918-INS-125 | chr5-101563918-INS-125 |
| chr5-101565695-INS-401 | chr5-101565653-INS-401 |
| chr5-101590214-INS-72 | chr5-101590214-INS-60 |
| chr5-101601570-INS-61 | chr5-101601570-INS-61 |
| chr5-101605885-INS-57 | chr5-101605885-INS-57 |
| chr5-101614465-INS-84 | chr5-101614465-INS-84 |
| chr5-101704093-INS-124 | chr5-101704095-INS-76 |
| chr5-101712249-INS-66 | chr5-101712249-INS-66 |
| chr5-101720511-INS-493 | chr5-101720511-INS-492 |
| chr5-101722198-INS-60 | chr5-101722198-INS-60 |
| chr5-101724875-INS-1998 | chr5-101724876-INS-1998 |
| chr5-101728190-INS-51 | chr5-101728181-INS-57 |
| chr5-101759419-INS-184 | chr5-101759419-INS-178 |
| chr5-101783706-INS-108 | chr5-101783706-INS-108 |
| chr5-101785539-INS-79 | chr5-101785539-INS-75 |
| chr5-101789920-INS-52 | chr5-101789920-INS-52 |
| chr5-101790913-INS-193 | chr5-101790913-INS-193 |
| chr5-101803160-INS-53 | chr5-101803160-INS-56 |
| chr5-101850244-INS-116 | chr5-101850244-INS-118 |
| chr5-101851370-INS-198 | chr5-101851370-INS-199 |
| chr5-101852741-INS-213 | chr5-101852740-INS-220 |
| chr5-101854530-INS-78 | chr5-101854530-INS-78 |
| chr5-101857376-INS-201 | chr5-101857376-INS-219 |
| chr5-101858869-INS-51 | chr5-101858877-INS-86 |
| chr5-101864079-INS-76 | chr5-101864079-INS-76 |
| chr5-101895750-INS-76 | chr5-101895725-INS-90 |
| chr5-101927749-INS-63 | chr5-101927749-INS-63 |
| chr5-101929639-INS-1895 | chr5-101929668-INS-1857 |
| chr5-101954473-INS-272 | chr5-101954473-INS-272 |
| chr5-101972621-INS-106 | chr5-101972622-INS-94 |
| chr5-101995802-INS-112 | chr5-101995802-INS-112 |
| chr5-102043380-INS-85 | chr5-102043380-INS-87 |
| chr5-102093335-INS-86 | chr5-102093335-INS-86 |
| chr5-102097359-INS-85 | chr5-102097359-INS-85 |
| chr5-102117210-INS-1890 | chr5-102117211-INS-1890 |
| chr5-102154865-INS-116 | chr5-102154865-INS-116 |
| chr5-102168520-INS-165 | chr5-102168520-INS-165 |
| chr5-102180675-INS-57 | chr5-102180675-INS-57 |
| chr5-102182775-INS-431 | chr5-102182775-INS-431 |
| chr5-102190839-INS-201 | chr5-102190839-INS-201 |
| chr5-102205612-INS-52 | chr5-102205612-INS-52 |
| chr5-102283575-INS-114 | chr5-102283590-INS-124 |
| chr5-102305873-INS-4762 | chr5-102305874-INS-4762 |
| chr5-102321628-INS-144 | chr5-102321637-INS-141 |
| chr5-102334568-INS-56 | chr5-102334555-INS-50 |
| chr5-102349473-INS-63 | chr5-102349473-INS-66 |
| chr5-102358269-INS-493 | chr5-102358269-INS-495 |
| chr5-102376272-INS-98 | chr5-102376272-INS-98 |
| chr5-102377067-INS-68 | chr5-102377067-INS-85 |
| chr5-102379874-INS-71 | chr5-102379874-INS-71 |
| chr5-102392460-INS-291 | chr5-102392460-INS-291 |
| chr5-102402636-INS-65 | chr5-102402636-INS-55 |
| chr5-102407183-INS-6622 | chr5-102407184-INS-6622 |
| chr5-102428316-INS-174 | chr5-102428310-INS-182 |
| chr5-102446056-INS-81 | chr5-102446061-INS-87 |
| chr5-102459347-INS-56 | chr5-102459347-INS-60 |
| chr5-102465364-INS-173 | chr5-102465364-INS-173 |
| chr5-102467458-INS-177 | chr5-102467458-INS-179 |
| chr5-102469488-INS-752 | chr5-102469488-INS-749 |
| chr5-102487686-INS-53 | chr5-102487641-INS-99 |
| chr5-102499652-INS-77 | chr5-102499652-INS-77 |
| chr5-102505345-INS-623 | chr5-102505345-INS-623 |
| chr5-102513782-INS-70 | chr5-102513782-INS-78 |
| chr5-102529210-INS-74 | chr5-102529210-INS-74 |
| chr5-102624485-INS-340 | chr5-102624486-INS-340 |
| chr5-102640667-INS-80 | chr5-102640667-INS-52 |
| chr5-102656179-INS-67 | chr5-102656188-INS-68 |
| chr5-102705383-INS-65 | chr5-102705383-INS-65 |
| chr5-102708237-INS-147 | chr5-102708237-INS-147 |
| chr5-102709775-INS-60 | chr5-102709775-INS-56 |
| chr5-102711863-INS-67 | chr5-102711863-INS-67 |
| chr5-102747071-INS-115 | chr5-102747071-INS-114 |
| chr5-102753548-INS-214 | chr5-102753548-INS-214 |
| chr5-102787122-INS-110 | chr5-102787122-INS-80 |
| chr5-102806915-INS-61 | chr5-102806915-INS-53 |
| chr5-102815914-INS-59 | chr5-102815914-INS-59 |

| | |
|---|---|
| chr5-102824320-INS-90 | chr5-102824365-INS-90 |
| chr5-102843478-INS-264 | chr5-102843478-INS-264 |
| chr5-102854825-INS-170 | chr5-102854825-INS-171 |
| chr5-102880797-INS-299 | chr5-102880797-INS-299 |
| chr5-102896716-INS-94 | chr5-102896716-INS-51 |
| chr5-102935980-INS-130 | chr5-102935980-INS-166 |
| chr5-102949330-INS-130 | chr5-102949330-INS-156 |
| chr5-103042920-INS-759 | chr5-103042920-INS-759 |
| chr5-103049087-INS-60 | chr5-103049104-INS-52 |
| chr5-103074676-INS-62 | chr5-103074676-INS-62 |
| chr5-103115860-INS-78 | chr5-103115835-INS-104 |
| chr5-103122746-INS-62 | chr5-103122746-INS-60 |
| chr5-103130441-INS-74 | chr5-103130464-INS-80 |
| chr5-103168489-INS-124 | chr5-103168489-INS-124 |
| chr5-103170532-INS-74 | chr5-103170532-INS-74 |
| chr5-103206542-INS-71 | chr5-103206542-INS-71 |
| chr5-103246961-INS-67 | chr5-103246962-INS-51 |
| chr5-103261876-INS-218 | chr5-103261876-INS-218 |
| chr5-103271880-INS-1991 | chr5-103271880-INS-1991 |
| chr5-103276283-INS-283 | chr5-103276283-INS-284 |
| chr5-103361171-INS-98 | chr5-103361171-INS-98 |
| chr5-103388236-INS-204 | chr5-103388236-INS-204 |
| chr5-103399227-INS-67 | chr5-103399227-INS-67 |
| chr5-103411154-INS-228 | chr5-103411154-INS-228 |
| chr5-103415555-INS-499 | chr5-103415555-INS-499 |
| chr5-103428782-INS-64 | chr5-103428782-INS-64 |
| chr5-103443544-INS-181 | chr5-103443544-INS-180 |
| chr5-103546759-INS-211 | chr5-103546759-INS-211 |
| chr5-103591448-INS-72 | chr5-103591448-INS-70 |
| chr5-103614169-INS-65 | chr5-103614169-INS-67 |
| chr5-103615401-INS-157 | chr5-103615401-INS-157 |
| chr5-103641083-INS-52 | chr5-103641083-INS-52 |
| chr5-103642174-INS-499 | chr5-103642174-INS-505 |
| chr5-103691916-INS-97 | chr5-103691916-INS-97 |
| chr5-103692538-INS-66 | chr5-103692538-INS-66 |
| chr5-103701756-INS-1749 | chr5-103701757-INS-1751 |
| chr5-103705076-INS-396 | chr5-103705076-INS-396 |
| chr5-103722799-INS-104 | chr5-103722799-INS-104 |
| chr5-103727742-INS-53 | chr5-103727742-INS-53 |
| chr5-103755547-INS-107 | chr5-103755547-INS-107 |
| chr5-103801006-INS-234 | chr5-103801006-INS-234 |
| chr5-103807181-INS-103 | chr5-103807181-INS-103 |
| chr5-103810614-INS-193 | chr5-103810614-INS-193 |
| chr5-103814350-INS-240 | chr5-103814350-INS-240 |
| chr5-103819618-INS-175 | chr5-103819618-INS-175 |
| chr5-103843303-INS-280 | chr5-103843303-INS-280 |
| chr5-103858857-INS-55 | chr5-103858857-INS-50 |
| chr5-103865137-INS-214 | chr5-103865137-INS-214 |
| chr5-103873886-INS-215 | chr5-103873886-INS-215 |
| chr5-103875317-INS-919 | chr5-103875311-INS-915 |
| chr5-103876734-INS-218 | chr5-103876734-INS-219 |
| chr5-103879005-INS-115 | chr5-103879005-INS-115 |
| chr5-103913764-INS-204 | chr5-103913764-INS-204 |
| chr5-103925870-INS-211 | chr5-103925870-INS-214 |
| chr5-103940272-INS-215 | chr5-103940272-INS-215 |
| chr5-103941193-INS-211 | chr5-103941193-INS-212 |
| chr5-103942439-INS-193 | chr5-103942440-INS-193 |
| chr5-103961460-INS-52 | chr5-103961460-INS-52 |
| chr5-103980286-INS-59 | chr5-103980286-INS-59 |
| chr5-103992021-INS-588 | chr5-103992021-INS-582 |
| chr5-103993784-INS-217 | chr5-103993784-INS-216 |
| chr5-104030616-INS-332 | chr5-104030650-INS-330 |
| chr5-104044549-INS-228 | chr5-104044549-INS-197 |
| chr5-104046730-INS-220 | chr5-104046730-INS-220 |
| chr5-104049368-INS-81 | chr5-104049368-INS-81 |
| chr5-104077537-INS-52 | chr5-104077537-INS-56 |
| chr5-104079730-INS-51 | chr5-104079730-INS-51 |
| chr5-104089948-INS-56 | chr5-104089948-INS-56 |
| chr5-104113288-INS-111 | chr5-104113288-INS-82 |
| chr5-104119333-INS-117 | chr5-104119333-INS-117 |
| chr5-104119858-INS-320 | chr5-104119858-INS-333 |
| chr5-104152272-INS-763 | chr5-104152272-INS-763 |
| chr5-104167149-INS-60 | chr5-104167106-INS-53 |
| chr5-104191009-INS-74 | chr5-104191010-INS-74 |
| chr5-104197207-INS-60 | chr5-104197192-INS-65 |
| chr5-104236861-INS-88 | chr5-104236861-INS-88 |
| chr5-104249035-INS-152 | chr5-104249036-INS-152 |

| | |
|---|---|
| chr5-104254275-INS-73 | chr5-104254275-INS-73 |
| chr5-104269980-INS-130 | chr5-104269980-INS-130 |
| chr5-104273356-INS-109 | chr5-104273356-INS-109 |
| chr5-104285658-INS-122 | chr5-104285658-INS-124 |
| chr5-104288159-INS-5544 | chr5-104288160-INS-5544 |
| chr5-104289669-INS-1285 | chr5-104289670-INS-1287 |
| chr5-104290411-INS-211 | chr5-104290411-INS-211 |
| chr5-104299447-INS-211 | chr5-104299447-INS-211 |
| chr5-104308765-INS-266 | chr5-104308777-INS-277 |
| chr5-104331310-INS-982 | chr5-104331310-INS-982 |
| chr5-104355900-INS-71 | chr5-104355900-INS-76 |
| chr5-104357487-INS-68 | chr5-104357487-INS-68 |
| chr5-104359244-INS-69 | chr5-104359244-INS-70 |
| chr5-104364009-INS-141 | chr5-104364010-INS-141 |
| chr5-104368679-INS-56 | chr5-104368708-INS-64 |
| chr5-104369907-INS-172 | chr5-104369907-INS-172 |
| chr5-104375636-INS-74 | chr5-104375638-INS-77 |
| chr5-104387448-INS-95 | chr5-104387448-INS-95 |
| chr5-104397125-INS-89 | chr5-104397125-INS-89 |
| chr5-104411179-INS-87 | chr5-104411179-INS-87 |
| chr5-104441568-INS-102 | chr5-104441568-INS-102 |
| chr5-104457417-INS-56 | chr5-104457417-INS-62 |
| chr5-104460075-INS-80 | chr5-104460075-INS-80 |
| chr5-104461819-INS-423 | chr5-104461819-INS-458 |
| chr5-104490250-INS-391 | chr5-104490250-INS-391 |
| chr5-104492190-INS-176 | chr5-104492190-INS-177 |
| chr5-104543484-INS-204 | chr5-104543484-INS-205 |
| chr5-104554836-INS-71 | chr5-104554836-INS-71 |
| chr5-104561624-INS-74 | chr5-104561624-INS-74 |
| chr5-104565297-INS-6777 | chr5-104565298-INS-6779 |
| chr5-104579385-INS-114 | chr5-104579385-INS-114 |
| chr5-104583314-INS-52 | chr5-104583337-INS-59 |
| chr5-104586681-INS-202 | chr5-104586681-INS-202 |
| chr5-104598322-INS-164 | chr5-104598322-INS-160 |
| chr5-104608429-INS-182 | chr5-104608429-INS-184 |
| chr5-104609671-INS-94 | chr5-104609671-INS-92 |
| chr5-104644958-INS-7087 | chr5-104644959-INS-7087 |
| chr5-104660128-INS-199 | chr5-104660128-INS-199 |
| chr5-104673296-INS-2326 | chr5-104673297-INS-2326 |
| chr5-104688538-INS-180 | chr5-104688538-INS-180 |
| chr5-104696429-INS-218 | chr5-104696429-INS-218 |
| chr5-104699955-INS-213 | chr5-104699955-INS-213 |
| chr5-104707351-INS-206 | chr5-104707351-INS-206 |
| chr5-104721539-INS-178 | chr5-104721540-INS-177 |
| chr5-104730384-INS-252 | chr5-104730384-INS-252 |
| chr5-104736049-INS-13214 | chr5-104736050-INS-13257 |
| chr5-104739042-INS-5249 | chr5-104739043-INS-5249 |
| chr5-104766717-INS-504 | chr5-104766717-INS-506 |
| chr5-104819643-INS-541 | chr5-104819663-INS-534 |
| chr5-104845708-INS-80 | chr5-104845708-INS-80 |
| chr5-104850403-INS-170 | chr5-104850393-INS-171 |
| chr5-104872606-INS-12919 | chr5-104872606-INS-12920 |
| chr5-104945013-INS-201 | chr5-104945013-INS-201 |
| chr5-104980295-INS-130 | chr5-104980293-INS-84 |
| chr5-104981306-INS-185 | chr5-104981306-INS-185 |
| chr5-104992615-INS-519 | chr5-104992615-INS-525 |
| chr5-104996284-INS-160 | chr5-104996284-INS-160 |
| chr5-104999974-INS-211 | chr5-104999974-INS-211 |
| chr5-105111956-INS-10178 | chr5-105111957-INS-10178 |
| chr5-105113470-INS-399 | chr5-105113470-INS-399 |
| chr5-105123411-INS-208 | chr5-105123411-INS-208 |
| chr5-105146981-INS-78 | chr5-105146985-INS-58 |
| chr5-105151634-INS-397 | chr5-105151634-INS-397 |
| chr5-105153254-INS-6633 | chr5-105153255-INS-6634 |
| chr5-105186414-INS-66 | chr5-105186411-INS-63 |
| chr5-105221448-INS-215 | chr5-105221448-INS-215 |
| chr5-105250146-INS-331 | chr5-105250146-INS-331 |
| chr5-105255018-INS-1888 | chr5-105255019-INS-1888 |
| chr5-105298873-INS-7501 | chr5-105298874-INS-7500 |
| chr5-105351823-INS-1893 | chr5-105351824-INS-1893 |
| chr5-105411888-INS-297 | chr5-105411888-INS-297 |
| chr5-105413842-INS-5964 | chr5-105413843-INS-5965 |
| chr5-105451040-INS-6205 | chr5-105451041-INS-6205 |
| chr5-105454569-INS-54 | chr5-105454569-INS-54 |
| chr5-105466242-INS-108 | chr5-105466250-INS-69 |
| chr5-105485381-INS-7075 | chr5-105485382-INS-7077 |
| chr5-105503137-INS-131 | chr5-105503137-INS-131 |

| | |
|---|---|
| chr5-105560312-INS-63 | chr5-105560312-INS-63 |
| chr5-105580591-INS-138 | chr5-105580587-INS-146 |
| chr5-105583133-INS-143 | chr5-105583133-INS-143 |
| chr5-105601129-INS-72 | chr5-105601126-INS-72 |
| chr5-105631061-INS-240 | chr5-105631061-INS-240 |
| chr5-105642905-INS-64 | chr5-105642905-INS-51 |
| chr5-105646078-INS-134 | chr5-105646078-INS-139 |
| chr5-105659442-INS-56 | chr5-105659442-INS-58 |
| chr5-105687739-INS-604 | chr5-105687739-INS-604 |
| chr5-105698899-INS-54 | chr5-105698899-INS-56 |
| chr5-105706583-INS-200 | chr5-105706583-INS-200 |
| chr5-105748137-INS-342 | chr5-105748137-INS-342 |
| chr5-105751268-INS-54 | chr5-105751268-INS-54 |
| chr5-105784093-INS-53 | chr5-105784092-INS-53 |
| chr5-105786483-INS-72 | chr5-105786483-INS-72 |
| chr5-105792309-INS-523 | chr5-105792309-INS-524 |
| chr5-105798993-INS-90 | chr5-105798993-INS-104 |
| chr5-105807121-INS-119 | chr5-105807121-INS-119 |
| chr5-105807985-INS-268 | chr5-105807985-INS-268 |
| chr5-105810248-INS-197 | chr5-105810248-INS-197 |
| chr5-105811990-INS-193 | chr5-105811990-INS-197 |
| chr5-105812891-INS-115 | chr5-105812889-INS-97 |
| chr5-105814117-INS-157 | chr5-105814117-INS-157 |
| chr5-105819373-INS-52 | chr5-105819373-INS-52 |
| chr5-105826704-INS-168 | chr5-105826705-INS-131 |
| chr5-105829612-INS-185 | chr5-105829612-INS-185 |
| chr5-105841332-INS-189 | chr5-105841332-INS-189 |
| chr5-105869313-INS-909 | chr5-105869313-INS-909 |
| chr5-105886558-INS-80 | chr5-105886563-INS-55 |
| chr5-105893546-INS-84 | chr5-105893522-INS-125 |
| chr5-105916687-INS-70 | chr5-105916687-INS-70 |
| chr5-105932232-INS-80 | chr5-105932232-INS-80 |
| chr5-105943889-INS-57 | chr5-105943889-INS-57 |
| chr5-106000780-INS-164 | chr5-106000780-INS-183 |
| chr5-106006270-INS-156 | chr5-106006270-INS-156 |
| chr5-106022506-INS-72 | chr5-106022506-INS-72 |
| chr5-106066805-INS-3244 | chr5-106066806-INS-3244 |
| chr5-106093690-INS-307 | chr5-106093691-INS-319 |
| chr5-106100188-INS-246 | chr5-106100188-INS-235 |
| chr5-106105697-INS-214 | chr5-106105697-INS-214 |
| chr5-106106205-INS-56 | chr5-106106205-INS-56 |
| chr5-106122152-INS-126 | chr5-106122152-INS-126 |
| chr5-106156151-INS-643 | chr5-106156151-INS-645 |
| chr5-106163666-INS-170 | chr5-106163666-INS-169 |
| chr5-106229291-INS-171 | chr5-106229283-INS-192 |
| chr5-106255655-INS-51 | chr5-106255663-INS-61 |
| chr5-106264439-INS-186 | chr5-106264439-INS-186 |
| chr5-106280251-INS-70 | chr5-106280251-INS-75 |
| chr5-106280600-INS-968 | chr5-106280600-INS-968 |
| chr5-106283138-INS-67 | chr5-106283135-INS-58 |
| chr5-106314475-INS-557 | chr5-106314475-INS-562 |
| chr5-106358715-INS-56 | chr5-106358715-INS-56 |
| chr5-106379786-INS-7160 | chr5-106379787-INS-7160 |
| chr5-106393605-INS-52 | chr5-106393605-INS-52 |
| chr5-106395345-INS-2836 | chr5-106395346-INS-2836 |
| chr5-106397051-INS-1290 | chr5-106397047-INS-1287 |
| chr5-106407661-INS-75 | chr5-106407661-INS-64 |
| chr5-106411615-INS-479 | chr5-106411615-INS-479 |
| chr5-106413109-INS-60 | chr5-106413109-INS-62 |
| chr5-106435342-INS-96 | chr5-106435342-INS-96 |
| chr5-106441490-INS-7096 | chr5-106441491-INS-7096 |
| chr5-106461001-INS-67 | chr5-106461002-INS-67 |
| chr5-106501919-INS-98 | chr5-106501919-INS-102 |
| chr5-106513861-INS-233 | chr5-106513861-INS-233 |
| chr5-106517968-INS-115 | chr5-106517968-INS-132 |
| chr5-106530907-INS-219 | chr5-106530907-INS-220 |
| chr5-106547288-INS-504 | chr5-106547288-INS-504 |
| chr5-106551862-INS-56 | chr5-106551862-INS-56 |
| chr5-106573296-INS-75 | chr5-106573293-INS-68 |
| chr5-106574231-INS-173 | chr5-106574231-INS-168 |
| chr5-106575605-INS-373 | chr5-106575605-INS-373 |
| chr5-106578044-INS-81 | chr5-106578044-INS-81 |
| chr5-106582943-INS-54 | chr5-106582943-INS-60 |
| chr5-106628069-INS-570 | chr5-106628069-INS-569 |
| chr5-106634391-INS-211 | chr5-106634391-INS-211 |
| chr5-106663827-INS-7165 | chr5-106663828-INS-7165 |
| chr5-106666111-INS-222 | chr5-106666111-INS-222 |

| | |
|---|---|
| chr5-106667491-INS-211 | chr5-106667495-INS-207 |
| chr5-106670535-INS-106 | chr5-106670540-INS-105 |
| chr5-106687474-INS-260 | chr5-106687474-INS-257 |
| chr5-106742891-INS-60 | chr5-106742913-INS-60 |
| chr5-106811169-INS-557 | chr5-106811169-INS-557 |
| chr5-106822366-INS-667 | chr5-106822366-INS-667 |
| chr5-106849021-INS-69 | chr5-106849021-INS-69 |
| chr5-106850325-INS-239 | chr5-106850325-INS-238 |
| chr5-106860518-INS-79 | chr5-106860518-INS-79 |
| chr5-106866927-INS-90 | chr5-106866927-INS-90 |
| chr5-106872235-INS-7101 | chr5-106872236-INS-7101 |
| chr5-106877924-INS-1014 | chr5-106877925-INS-1010 |
| chr5-106884376-INS-93 | chr5-106884376-INS-93 |
| chr5-106886631-INS-428 | chr5-106886631-INS-396 |
| chr5-106887429-INS-106 | chr5-106887420-INS-117 |
| chr5-106910027-INS-7106 | chr5-106910028-INS-7106 |
| chr5-106910560-INS-89 | chr5-106910574-INS-54 |
| chr5-106924541-INS-214 | chr5-106924541-INS-214 |
| chr5-106930241-INS-438 | chr5-106930241-INS-445 |
| chr5-106972112-INS-70 | chr5-106972112-INS-72 |
| chr5-106974428-INS-202 | chr5-106974428-INS-206 |
| chr5-107051882-INS-100 | chr5-107051882-INS-50 |
| chr5-107093215-INS-62 | chr5-107093215-INS-50 |
| chr5-107098292-INS-55 | chr5-107098275-INS-52 |
| chr5-107151716-INS-210 | chr5-107151716-INS-210 |
| chr5-107157435-INS-350 | chr5-107157435-INS-350 |
| chr5-107169572-INS-959 | chr5-107169572-INS-960 |
| chr5-107176452-INS-53 | chr5-107176452-INS-53 |
| chr5-107180359-INS-78 | chr5-107180359-INS-78 |
| chr5-107188668-INS-670 | chr5-107188668-INS-670 |
| chr5-107189498-INS-70 | chr5-107189498-INS-70 |
| chr5-107209896-INS-65 | chr5-107209896-INS-65 |
| chr5-107215271-INS-236 | chr5-107215271-INS-236 |
| chr5-107242081-INS-62 | chr5-107242061-INS-78 |
| chr5-107281613-INS-2874 | chr5-107281614-INS-2874 |
| chr5-107322827-INS-62 | chr5-107322827-INS-62 |
| chr5-107337987-INS-1112 | chr5-107337988-INS-1112 |
| chr5-107387982-INS-109 | chr5-107387982-INS-109 |
| chr5-107411175-INS-68 | chr5-107411176-INS-56 |
| chr5-107423253-INS-105 | chr5-107423253-INS-105 |
| chr5-107446583-INS-1383 | chr5-107446584-INS-1365 |
| chr5-107454296-INS-107 | chr5-107454296-INS-107 |
| chr5-107480650-INS-60 | chr5-107480650-INS-60 |
| chr5-107519674-INS-60 | chr5-107519675-INS-52 |
| chr5-107529203-INS-1864 | chr5-107529204-INS-1864 |
| chr5-107534561-INS-81 | chr5-107534561-INS-81 |
| chr5-107553256-INS-159 | chr5-107553256-INS-159 |
| chr5-107577110-INS-79 | chr5-107577115-INS-79 |
| chr5-107628138-INS-51 | chr5-107628138-INS-51 |
| chr5-107631378-INS-7109 | chr5-107631379-INS-7108 |
| chr5-107644835-INS-199 | chr5-107644835-INS-200 |
| chr5-107647680-INS-214 | chr5-107647680-INS-214 |
| chr5-107700305-INS-113 | chr5-107700305-INS-113 |
| chr5-107717014-INS-6489 | chr5-107717015-INS-6489 |
| chr5-107801137-INS-4453 | chr5-107801138-INS-4453 |
| chr5-107803307-INS-95 | chr5-107803307-INS-95 |
| chr5-107819929-INS-202 | chr5-107819929-INS-203 |
| chr5-107840303-INS-115 | chr5-107840303-INS-126 |
| chr5-107881327-INS-4663 | chr5-107881328-INS-4663 |
| chr5-107923401-INS-184 | chr5-107923401-INS-184 |
| chr5-107935033-INS-6652 | chr5-107935034-INS-6652 |
| chr5-107967130-INS-1862 | chr5-107967131-INS-1862 |
| chr5-107982397-INS-208 | chr5-107982397-INS-208 |
| chr5-107992506-INS-56 | chr5-107992506-INS-56 |
| chr5-108034619-INS-106 | chr5-108034620-INS-80 |
| chr5-108048018-INS-75 | chr5-108048018-INS-75 |
| chr5-108065971-INS-204 | chr5-108065971-INS-204 |
| chr5-108069582-INS-217 | chr5-108069582-INS-217 |
| chr5-108072112-INS-215 | chr5-108072112-INS-216 |
| chr5-108075528-INS-105 | chr5-108075528-INS-105 |
| chr5-108085012-INS-282 | chr5-108085012-INS-282 |
| chr5-108105538-INS-210 | chr5-108105538-INS-211 |
| chr5-108124253-INS-162 | chr5-108124253-INS-162 |
| chr5-108166511-INS-222 | chr5-108166511-INS-214 |
| chr5-108172341-INS-205 | chr5-108172341-INS-205 |
| chr5-108181286-INS-71 | chr5-108181286-INS-71 |
| chr5-108189685-INS-653 | chr5-108189685-INS-653 |

| | |
|---|---|
| chr5-108196127-INS-1266 | chr5-108196128-INS-1265 |
| chr5-108197624-INS-57 | chr5-108197626-INS-81 |
| chr5-108199822-INS-231 | chr5-108199822-INS-234 |
| chr5-108202750-INS-188 | chr5-108202750-INS-188 |
| chr5-108205452-INS-198 | chr5-108205452-INS-198 |
| chr5-108209570-INS-205 | chr5-108209570-INS-205 |
| chr5-108219129-INS-206 | chr5-108219129-INS-206 |
| chr5-108219708-INS-72 | chr5-108219708-INS-72 |
| chr5-108235076-INS-211 | chr5-108235076-INS-211 |
| chr5-108243038-INS-211 | chr5-108243038-INS-211 |
| chr5-108248501-INS-212 | chr5-108248501-INS-212 |
| chr5-108295139-INS-195 | chr5-108295139-INS-191 |
| chr5-108296991-INS-54 | chr5-108296991-INS-54 |
| chr5-108312015-INS-128 | chr5-108312015-INS-126 |
| chr5-108339311-INS-4506 | chr5-108339312-INS-4506 |
| chr5-108346025-INS-58 | chr5-108346025-INS-61 |
| chr5-108347446-INS-221 | chr5-108347446-INS-221 |
| chr5-108351941-INS-160 | chr5-108351941-INS-163 |
| chr5-108356753-INS-69 | chr5-108356753-INS-69 |
| chr5-108364102-INS-216 | chr5-108364102-INS-216 |
| chr5-108376354-INS-196 | chr5-108376354-INS-196 |
| chr5-108399686-INS-639 | chr5-108399686-INS-639 |
| chr5-108402601-INS-102 | chr5-108402599-INS-104 |
| chr5-108431816-INS-216 | chr5-108431816-INS-233 |
| chr5-108455493-INS-278 | chr5-108455493-INS-278 |
| chr5-108468002-INS-203 | chr5-108468002-INS-204 |
| chr5-108470613-INS-295 | chr5-108470613-INS-295 |
| chr5-108477575-INS-193 | chr5-108477575-INS-193 |
| chr5-108538277-INS-8645 | chr5-108538278-INS-8645 |
| chr5-108542504-INS-60 | chr5-108542504-INS-60 |
| chr5-108559021-INS-57 | chr5-108559021-INS-60 |
| chr5-108563941-INS-116 | chr5-108563941-INS-124 |
| chr5-108566969-INS-170 | chr5-108566969-INS-170 |
| chr5-108573042-INS-612 | chr5-108573042-INS-611 |
| chr5-108574751-INS-69 | chr5-108574751-INS-69 |
| chr5-108588426-INS-491 | chr5-108588427-INS-491 |
| chr5-108599630-INS-1152 | chr5-108599631-INS-1153 |
| chr5-108608176-INS-68 | chr5-108608176-INS-68 |
| chr5-108664102-INS-78 | chr5-108664102-INS-77 |
| chr5-108691005-INS-222 | chr5-108691005-INS-222 |
| chr5-108725369-INS-228 | chr5-108725369-INS-228 |
| chr5-108753067-INS-384 | chr5-108753067-INS-383 |
| chr5-108763396-INS-60 | chr5-108763396-INS-56 |
| chr5-108774448-INS-247 | chr5-108774448-INS-248 |
| chr5-108781052-INS-220 | chr5-108781052-INS-221 |
| chr5-108810608-INS-216 | chr5-108810608-INS-217 |
| chr5-108849841-INS-351 | chr5-108849841-INS-351 |
| chr5-108863209-INS-3034 | chr5-108863210-INS-3034 |
| chr5-108868216-INS-52 | chr5-108868216-INS-56 |
| chr5-108868855-INS-51 | chr5-108868855-INS-54 |
| chr5-108890822-INS-301 | chr5-108890822-INS-302 |
| chr5-108906708-INS-6402 | chr5-108906709-INS-6402 |
| chr5-108919604-INS-68 | chr5-108919604-INS-68 |
| chr5-108936575-INS-76 | chr5-108936575-INS-78 |
| chr5-108958311-INS-4257 | chr5-108958312-INS-4249 |
| chr5-108967600-INS-3182 | chr5-108967601-INS-3182 |
| chr5-109007795-INS-1132 | chr5-109007796-INS-1132 |
| chr5-109026900-INS-4335 | chr5-109026901-INS-4335 |
| chr5-109026901-INS-345 | chr5-109026901-INS-345 |
| chr5-109047555-INS-117 | chr5-109047555-INS-117 |
| chr5-109107943-INS-7115 | chr5-109107944-INS-7115 |
| chr5-109173637-INS-428 | chr5-109173637-INS-428 |
| chr5-109222085-INS-1100 | chr5-109222086-INS-1094 |
| chr5-109247583-INS-7210 | chr5-109247584-INS-7210 |
| chr5-109344891-INS-522 | chr5-109344891-INS-522 |
| chr5-109372256-INS-56 | chr5-109372256-INS-64 |
| chr5-109374514-INS-8346 | chr5-109374515-INS-8346 |
| chr5-109407869-INS-469 | chr5-109407869-INS-469 |
| chr5-109462249-INS-7122 | chr5-109462250-INS-7122 |
| chr5-109475267-INS-960 | chr5-109475267-INS-960 |
| chr5-109484995-INS-2952 | chr5-109484996-INS-2954 |
| chr5-109490272-INS-6452 | chr5-109490273-INS-6455 |
| chr5-109499858-INS-543 | chr5-109499858-INS-543 |
| chr5-109521139-INS-11588 | chr5-109521140-INS-11588 |
| chr5-109528110-INS-112 | chr5-109528110-INS-112 |
| chr5-109563098-INS-6403 | chr5-109563099-INS-6403 |
| chr5-109610379-INS-384 | chr5-109610379-INS-384 |

| | |
|---|---|
| chr5-109650458-INS-103 | chr5-109650458-INS-103 |
| chr5-109660863-INS-58 | chr5-109660864-INS-58 |
| chr5-109697313-INS-53 | chr5-109697313-INS-53 |
| chr5-109706978-INS-454 | chr5-109706982-INS-443 |
| chr5-109713202-INS-208 | chr5-109713202-INS-208 |
| chr5-109725349-INS-6212 | chr5-109725350-INS-6212 |
| chr5-109731952-INS-6403 | chr5-109731953-INS-6403 |
| chr5-109750064-INS-204 | chr5-109750064-INS-204 |
| chr5-109751748-INS-60 | chr5-109751748-INS-60 |
| chr5-109753862-INS-63 | chr5-109753861-INS-100 |
| chr5-109756189-INS-86 | chr5-109756193-INS-90 |
| chr5-109760621-INS-74 | chr5-109760610-INS-68 |
| chr5-109805136-INS-52 | chr5-109805132-INS-52 |
| chr5-109809214-INS-655 | chr5-109809214-INS-657 |
| chr5-109832674-INS-7732 | chr5-109832675-INS-7734 |
| chr5-109832833-INS-57 | chr5-109832833-INS-57 |
| chr5-109841141-INS-6446 | chr5-109841142-INS-6447 |
| chr5-109841538-INS-1308 | chr5-109841545-INS-1304 |
| chr5-109886154-INS-299 | chr5-109886154-INS-299 |
| chr5-109901677-INS-53 | chr5-109901670-INS-53 |
| chr5-109902709-INS-410 | chr5-109902710-INS-413 |
| chr5-109904412-INS-203 | chr5-109904412-INS-202 |
| chr5-109921729-INS-7160 | chr5-109921730-INS-7160 |
| chr5-109953658-INS-183 | chr5-109953658-INS-183 |
| chr5-109963436-INS-268 | chr5-109963436-INS-268 |
| chr5-109968114-INS-434 | chr5-109968114-INS-434 |
| chr5-109970306-INS-215 | chr5-109970306-INS-216 |
| chr5-109980920-INS-85 | chr5-109980920-INS-85 |
| chr5-109986723-INS-76 | chr5-109986723-INS-76 |
| chr5-110002512-INS-193 | chr5-110002512-INS-193 |
| chr5-110016420-INS-58 | chr5-110016420-INS-58 |
| chr5-110025794-INS-100 | chr5-110025794-INS-100 |
| chr5-110034890-INS-51 | chr5-110034890-INS-51 |
| chr5-110047171-INS-93 | chr5-110047171-INS-93 |
| chr5-110057909-INS-84 | chr5-110057909-INS-50 |
| chr5-110078732-INS-172 | chr5-110078732-INS-179 |
| chr5-110119682-INS-78 | chr5-110119682-INS-78 |
| chr5-110133532-INS-559 | chr5-110133532-INS-559 |
| chr5-110139639-INS-6664 | chr5-110139640-INS-6664 |
| chr5-110175841-INS-63 | chr5-110175841-INS-67 |
| chr5-110177202-INS-62 | chr5-110177202-INS-63 |
| chr5-110179052-INS-3588 | chr5-110179053-INS-3590 |
| chr5-110191584-INS-705 | chr5-110191584-INS-705 |
| chr5-110195927-INS-173 | chr5-110195927-INS-173 |
| chr5-110197373-INS-213 | chr5-110197373-INS-213 |
| chr5-110224606-INS-210 | chr5-110224606-INS-210 |
| chr5-110260060-INS-171 | chr5-110260060-INS-171 |
| chr5-110261353-INS-52 | chr5-110261353-INS-52 |
| chr5-110265875-INS-210 | chr5-110265875-INS-210 |
| chr5-110272152-INS-205 | chr5-110272152-INS-205 |
| chr5-110275243-INS-337 | chr5-110275243-INS-336 |
| chr5-110279591-INS-101 | chr5-110279591-INS-101 |
| chr5-110287507-INS-128 | chr5-110287507-INS-128 |
| chr5-110292019-INS-181 | chr5-110292019-INS-182 |
| chr5-110296901-INS-209 | chr5-110296901-INS-209 |
| chr5-110299276-INS-194 | chr5-110299276-INS-194 |
| chr5-110301963-INS-198 | chr5-110301963-INS-198 |
| chr5-110311994-INS-191 | chr5-110311995-INS-190 |
| chr5-110328469-INS-190 | chr5-110328469-INS-183 |
| chr5-110364095-INS-245 | chr5-110364095-INS-245 |
| chr5-110376672-INS-85 | chr5-110376672-INS-81 |
| chr5-110382190-INS-227 | chr5-110382190-INS-228 |
| chr5-110393829-INS-8557 | chr5-110393830-INS-8556 |
| chr5-110396745-INS-88 | chr5-110396745-INS-89 |
| chr5-110399866-INS-76 | chr5-110399866-INS-76 |
| chr5-110410997-INS-224 | chr5-110410997-INS-225 |
| chr5-110412814-INS-170 | chr5-110412814-INS-170 |
| chr5-110421579-INS-217 | chr5-110421579-INS-217 |
| chr5-110428797-INS-199 | chr5-110428797-INS-199 |
| chr5-110438571-INS-215 | chr5-110438571-INS-216 |
| chr5-110445983-INS-211 | chr5-110445983-INS-211 |
| chr5-110498177-INS-208 | chr5-110498177-INS-208 |
| chr5-110504802-INS-157 | chr5-110504802-INS-179 |
| chr5-110542538-INS-57 | chr5-110542522-INS-86 |
| chr5-110550292-INS-1747 | chr5-110550293-INS-1739 |
| chr5-110553647-INS-1430 | chr5-110553648-INS-1430 |
| chr5-110558272-INS-181 | chr5-110558272-INS-182 |

| | |
|---|---|
| chr5-110560613-INS-217 | chr5-110560613-INS-217 |
| chr5-110612070-INS-188 | chr5-110612070-INS-188 |
| chr5-110626702-INS-227 | chr5-110626702-INS-227 |
| chr5-110652755-INS-150 | chr5-110652759-INS-158 |
| chr5-110657400-INS-501 | chr5-110657399-INS-502 |
| chr5-110664931-INS-68 | chr5-110664931-INS-68 |
| chr5-110672205-INS-190 | chr5-110672205-INS-190 |
| chr5-110758822-INS-87 | chr5-110758820-INS-93 |
| chr5-110760317-INS-70 | chr5-110760299-INS-66 |
| chr5-110772138-INS-120 | chr5-110772138-INS-120 |
| chr5-110781459-INS-212 | chr5-110781459-INS-212 |
| chr5-110789946-INS-222 | chr5-110789946-INS-222 |
| chr5-110830059-INS-75 | chr5-110830059-INS-75 |
| chr5-110857644-INS-217 | chr5-110857644-INS-217 |
| chr5-110860644-INS-86 | chr5-110860644-INS-50 |
| chr5-110915146-INS-224 | chr5-110915146-INS-224 |
| chr5-110945230-INS-170 | chr5-110945258-INS-166 |
| chr5-110951807-INS-145 | chr5-110951793-INS-111 |
| chr5-110958650-INS-55 | chr5-110958650-INS-55 |
| chr5-110964055-INS-128 | chr5-110964051-INS-151 |
| chr5-110967652-INS-189 | chr5-110967646-INS-176 |
| chr5-110968962-INS-90 | chr5-110968963-INS-56 |
| chr5-110979408-INS-205 | chr5-110979408-INS-205 |
| chr5-110982249-INS-232 | chr5-110982249-INS-232 |
| chr5-110992005-INS-606 | chr5-110992005-INS-606 |
| chr5-110999909-INS-211 | chr5-110999909-INS-211 |
| chr5-111033157-INS-64 | chr5-111033129-INS-54 |
| chr5-111049151-INS-74 | chr5-111049151-INS-74 |
| chr5-111061818-INS-198 | chr5-111061818-INS-199 |
| chr5-111077227-INS-262 | chr5-111077225-INS-264 |
| chr5-111083269-INS-199 | chr5-111083269-INS-199 |
| chr5-111096163-INS-218 | chr5-111096163-INS-219 |
| chr5-111136528-INS-67 | chr5-111136529-INS-54 |
| chr5-111164276-INS-132 | chr5-111164276-INS-132 |
| chr5-111220845-INS-460 | chr5-111220845-INS-460 |
| chr5-111243913-INS-102 | chr5-111243913-INS-104 |
| chr5-111266731-INS-840 | chr5-111266731-INS-840 |
| chr5-111271812-INS-621 | chr5-111271812-INS-614 |
| chr5-111282732-INS-217 | chr5-111282732-INS-217 |
| chr5-111304609-INS-136 | chr5-111304609-INS-136 |
| chr5-111359213-INS-112 | chr5-111359210-INS-112 |
| chr5-111376931-INS-624 | chr5-111376931-INS-626 |
| chr5-111394944-INS-147 | chr5-111394944-INS-147 |
| chr5-111399732-INS-60 | chr5-111399732-INS-60 |
| chr5-111420419-INS-95 | chr5-111420419-INS-95 |
| chr5-111423246-INS-98 | chr5-111423246-INS-98 |
| chr5-111437201-INS-56 | chr5-111437201-INS-56 |
| chr5-111438883-INS-5301 | chr5-111438884-INS-5301 |
| chr5-111439965-INS-62 | chr5-111439965-INS-70 |
| chr5-111444026-INS-234 | chr5-111444026-INS-235 |
| chr5-111445096-INS-1700 | chr5-111445097-INS-1700 |
| chr5-111445635-INS-68 | chr5-111445635-INS-69 |
| chr5-111448294-INS-72 | chr5-111448294-INS-72 |
| chr5-111463430-INS-939 | chr5-111463430-INS-941 |
| chr5-111468202-INS-205 | chr5-111468202-INS-205 |
| chr5-111497984-INS-161 | chr5-111497984-INS-156 |
| chr5-111541668-INS-57 | chr5-111541668-INS-57 |
| chr5-111589037-INS-82 | chr5-111589037-INS-82 |
| chr5-111636996-INS-272 | chr5-111636996-INS-274 |
| chr5-111638299-INS-76 | chr5-111638343-INS-52 |
| chr5-111665787-INS-85 | chr5-111665787-INS-85 |
| chr5-111671527-INS-120 | chr5-111671527-INS-120 |
| chr5-111676603-INS-51 | chr5-111676603-INS-51 |
| chr5-111722844-INS-106 | chr5-111722844-INS-106 |
| chr5-111736136-INS-60 | chr5-111736136-INS-60 |
| chr5-111737921-INS-100 | chr5-111737921-INS-100 |
| chr5-111748232-INS-1891 | chr5-111748233-INS-1891 |
| chr5-111777012-INS-189 | chr5-111777012-INS-189 |
| chr5-111795076-INS-124 | chr5-111795076-INS-124 |
| chr5-111886305-INS-148 | chr5-111886305-INS-148 |
| chr5-111896929-INS-84 | chr5-111896929-INS-84 |
| chr5-111904701-INS-56 | chr5-111904701-INS-62 |
| chr5-111905718-INS-151 | chr5-111905718-INS-151 |
| chr5-111917352-INS-549 | chr5-111917352-INS-549 |
| chr5-111918707-INS-129 | chr5-111918708-INS-145 |
| chr5-111933520-INS-98 | chr5-111933519-INS-120 |
| chr5-111948999-INS-142 | chr5-111948999-INS-142 |

| | |
|---|---|
| chr5-111985577-INS-2007 | chr5-111985578-INS-2009 |
| chr5-112052945-INS-242 | chr5-112052945-INS-242 |
| chr5-112075509-INS-92 | chr5-112075509-INS-92 |
| chr5-112106632-INS-52 | chr5-112106632-INS-56 |
| chr5-112137543-INS-80 | chr5-112137540-INS-68 |
| chr5-112154525-INS-749 | chr5-112154525-INS-768 |
| chr5-112171499-INS-200 | chr5-112171499-INS-200 |
| chr5-112181528-INS-54 | chr5-112181528-INS-58 |
| chr5-112214312-INS-72 | chr5-112214300-INS-54 |
| chr5-112219830-INS-380 | chr5-112219830-INS-380 |
| chr5-112220435-INS-57 | chr5-112220435-INS-77 |
| chr5-112235372-INS-118 | chr5-112235372-INS-118 |
| chr5-112245475-INS-63 | chr5-112245475-INS-63 |
| chr5-112275540-INS-149 | chr5-112275540-INS-149 |
| chr5-112303359-INS-136 | chr5-112303359-INS-137 |
| chr5-112331842-INS-60 | chr5-112331842-INS-60 |
| chr5-112342305-INS-55 | chr5-112342310-INS-59 |
| chr5-112349017-INS-196 | chr5-112349017-INS-196 |
| chr5-112350250-INS-54 | chr5-112350250-INS-54 |
| chr5-112350739-INS-100 | chr5-112350739-INS-52 |
| chr5-112360392-INS-52 | chr5-112360392-INS-50 |
| chr5-112401533-INS-62 | chr5-112401531-INS-90 |
| chr5-112412913-INS-277 | chr5-112412913-INS-277 |
| chr5-112437175-INS-150 | chr5-112437175-INS-150 |
| chr5-112443768-INS-93 | chr5-112443768-INS-93 |
| chr5-112451397-INS-150 | chr5-112451397-INS-149 |
| chr5-112465631-INS-69 | chr5-112465639-INS-82 |
| chr5-112467427-INS-157 | chr5-112467427-INS-157 |
| chr5-112469928-INS-60 | chr5-112469928-INS-60 |
| chr5-112530230-INS-57 | chr5-112530230-INS-67 |
| chr5-112563553-INS-52 | chr5-112563553-INS-54 |
| chr5-112578322-INS-614 | chr5-112578322-INS-615 |
| chr5-112606679-INS-64 | chr5-112606679-INS-68 |
| chr5-112617794-INS-68 | chr5-112617794-INS-68 |
| chr5-112655276-INS-55 | chr5-112655265-INS-90 |
| chr5-112694901-INS-118 | chr5-112694901-INS-118 |
| chr5-112710538-INS-234 | chr5-112710538-INS-234 |
| chr5-112711966-INS-54 | chr5-112711966-INS-54 |
| chr5-112714422-INS-52 | chr5-112714423-INS-52 |
| chr5-112741850-INS-56 | chr5-112741850-INS-56 |
| chr5-112754736-INS-81 | chr5-112754736-INS-84 |
| chr5-112768289-INS-60 | chr5-112768289-INS-60 |
| chr5-112769335-INS-68 | chr5-112769335-INS-70 |
| chr5-112780418-INS-2169 | chr5-112780419-INS-2167 |
| chr5-112805827-INS-132 | chr5-112805827-INS-132 |
| chr5-112808479-INS-1761 | chr5-112808480-INS-1756 |
| chr5-112820051-INS-159 | chr5-112820049-INS-159 |
| chr5-112854960-INS-99 | chr5-112854961-INS-97 |
| chr5-112869082-INS-7087 | chr5-112869083-INS-7087 |
| chr5-112886166-INS-60 | chr5-112886189-INS-64 |
| chr5-112894413-INS-57 | chr5-112894413-INS-57 |
| chr5-112896606-INS-633 | chr5-112896606-INS-647 |
| chr5-112900256-INS-88 | chr5-112900256-INS-84 |
| chr5-112911436-INS-94 | chr5-112911434-INS-104 |
| chr5-112923410-INS-94 | chr5-112923407-INS-94 |
| chr5-112977956-INS-70 | chr5-112977956-INS-70 |
| chr5-112983549-INS-55 | chr5-112983549-INS-55 |
| chr5-112990397-INS-97 | chr5-112990367-INS-97 |
| chr5-113001487-INS-108 | chr5-113001487-INS-107 |
| chr5-113028943-INS-68 | chr5-113028943-INS-74 |
| chr5-113046599-INS-398 | chr5-113046599-INS-398 |
| chr5-113055975-INS-150 | chr5-113055928-INS-162 |
| chr5-113068473-INS-52 | chr5-113068492-INS-55 |
| chr5-113072484-INS-65 | chr5-113072532-INS-55 |
| chr5-113099254-INS-60 | chr5-113099254-INS-60 |
| chr5-113105502-INS-58 | chr5-113105502-INS-58 |
| chr5-113117897-INS-104 | chr5-113117897-INS-104 |
| chr5-113128184-INS-204 | chr5-113128184-INS-204 |
| chr5-113165572-INS-6442 | chr5-113165573-INS-6442 |
| chr5-113175885-INS-338 | chr5-113175885-INS-338 |
| chr5-113241507-INS-60 | chr5-113241509-INS-65 |
| chr5-113247228-INS-141 | chr5-113247228-INS-141 |
| chr5-113309928-INS-77 | chr5-113309931-INS-75 |
| chr5-113312641-INS-57 | chr5-113312641-INS-56 |
| chr5-113337754-INS-64 | chr5-113337754-INS-64 |
| chr5-113365786-INS-87 | chr5-113365786-INS-87 |
| chr5-113393937-INS-69 | chr5-113393937-INS-69 |

| | |
|---|---|
| chr5-113395272-INS-66 | chr5-113395272-INS-66 |
| chr5-113416247-INS-465 | chr5-113416247-INS-465 |
| chr5-113467474-INS-183 | chr5-113467474-INS-171 |
| chr5-113484300-INS-223 | chr5-113484300-INS-232 |
| chr5-113550384-INS-71 | chr5-113550384-INS-71 |
| chr5-113565831-INS-52 | chr5-113565812-INS-64 |
| chr5-113569042-INS-66 | chr5-113569042-INS-54 |
| chr5-113574125-INS-235 | chr5-113574125-INS-235 |
| chr5-113588741-INS-144 | chr5-113588742-INS-144 |
| chr5-113594993-INS-146 | chr5-113594993-INS-145 |
| chr5-113611354-INS-1889 | chr5-113611355-INS-1889 |
| chr5-113628738-INS-59 | chr5-113628738-INS-64 |
| chr5-113650443-INS-78 | chr5-113650455-INS-88 |
| chr5-113655574-INS-55 | chr5-113655563-INS-104 |
| chr5-113675775-INS-55 | chr5-113675775-INS-55 |
| chr5-113699800-INS-60 | chr5-113699800-INS-60 |
| chr5-113703124-INS-223 | chr5-113703124-INS-225 |
| chr5-113714708-INS-80 | chr5-113714708-INS-84 |
| chr5-113736160-INS-54 | chr5-113736160-INS-54 |
| chr5-113740093-INS-60 | chr5-113740093-INS-70 |
| chr5-113749098-INS-66 | chr5-113749090-INS-71 |
| chr5-113765870-INS-213 | chr5-113765870-INS-213 |
| chr5-113786229-INS-76 | chr5-113786229-INS-81 |
| chr5-113817338-INS-116 | chr5-113817338-INS-116 |
| chr5-113825051-INS-440 | chr5-113825051-INS-440 |
| chr5-113826389-INS-91 | chr5-113826398-INS-66 |
| chr5-113828052-INS-56 | chr5-113828052-INS-52 |
| chr5-113830549-INS-74 | chr5-113830549-INS-74 |
| chr5-113832438-INS-66 | chr5-113832438-INS-61 |
| chr5-113838538-INS-54 | chr5-113838538-INS-54 |
| chr5-113844758-INS-66 | chr5-113844758-INS-72 |
| chr5-113861463-INS-65 | chr5-113861461-INS-89 |
| chr5-113867713-INS-65 | chr5-113867716-INS-62 |
| chr5-113871825-INS-56 | chr5-113871780-INS-83 |
| chr5-113872707-INS-281 | chr5-113872707-INS-281 |
| chr5-113879321-INS-61 | chr5-113879321-INS-61 |
| chr5-113888168-INS-3771 | chr5-113888169-INS-3777 |
| chr5-113892718-INS-53 | chr5-113892718-INS-53 |
| chr5-113895146-INS-354 | chr5-113895146-INS-354 |
| chr5-113912911-INS-56 | chr5-113912909-INS-76 |
| chr5-113929238-INS-122 | chr5-113929272-INS-80 |
| chr5-113939985-INS-572 | chr5-113939984-INS-569 |
| chr5-113949645-INS-60 | chr5-113949630-INS-69 |
| chr5-113966923-INS-60 | chr5-113966923-INS-60 |
| chr5-114005771-INS-104 | chr5-114005771-INS-108 |
| chr5-114010449-INS-1004 | chr5-114010450-INS-1004 |
| chr5-114013411-INS-248 | chr5-114013411-INS-248 |
| chr5-114016076-INS-163 | chr5-114016076-INS-163 |
| chr5-114030003-INS-54 | chr5-114030003-INS-54 |
| chr5-114058524-INS-55 | chr5-114058520-INS-51 |
| chr5-114071166-INS-199 | chr5-114071166-INS-199 |
| chr5-114149892-INS-60 | chr5-114149919-INS-60 |
| chr5-114156184-INS-226 | chr5-114156184-INS-226 |
| chr5-114164753-INS-373 | chr5-114164738-INS-375 |
| chr5-114186964-INS-54 | chr5-114186964-INS-54 |
| chr5-114199226-INS-159 | chr5-114199247-INS-159 |
| chr5-114200589-INS-58 | chr5-114200613-INS-58 |
| chr5-114233014-INS-168 | chr5-114233014-INS-168 |
| chr5-114234231-INS-880 | chr5-114234182-INS-881 |
| chr5-114263851-INS-232 | chr5-114263851-INS-232 |
| chr5-114278794-INS-91 | chr5-114278787-INS-103 |
| chr5-114279085-INS-7206 | chr5-114279086-INS-7206 |
| chr5-114282430-INS-118 | chr5-114282430-INS-119 |
| chr5-114307732-INS-225 | chr5-114307732-INS-225 |
| chr5-114320670-INS-173 | chr5-114320670-INS-173 |
| chr5-114336638-INS-112 | chr5-114336638-INS-114 |
| chr5-114340743-INS-197 | chr5-114340768-INS-211 |
| chr5-114346178-INS-64 | chr5-114346178-INS-64 |
| chr5-114346599-INS-164 | chr5-114346599-INS-164 |
| chr5-114355293-INS-56 | chr5-114355297-INS-53 |
| chr5-114359841-INS-201 | chr5-114359841-INS-201 |
| chr5-114362725-INS-223 | chr5-114362725-INS-224 |
| chr5-114384725-INS-65 | chr5-114384690-INS-86 |
| chr5-114402913-INS-621 | chr5-114402913-INS-621 |
| chr5-114507177-INS-89 | chr5-114507177-INS-89 |
| chr5-114546368-INS-1145 | chr5-114546369-INS-1145 |
| chr5-114554824-INS-216 | chr5-114554824-INS-216 |

| | |
|---|---|
| chr5-114587454-INS-177 | chr5-114587454-INS-139 |
| chr5-114604852-INS-71 | chr5-114604820-INS-63 |
| chr5-114614243-INS-155 | chr5-114614243-INS-155 |
| chr5-114626708-INS-170 | chr5-114626704-INS-174 |
| chr5-114627250-INS-56 | chr5-114627248-INS-51 |
| chr5-114635883-INS-6401 | chr5-114635884-INS-6401 |
| chr5-114656226-INS-60 | chr5-114656226-INS-60 |
| chr5-114660624-INS-209 | chr5-114660624-INS-209 |
| chr5-114664654-INS-850 | chr5-114664654-INS-871 |
| chr5-114668008-INS-218 | chr5-114668008-INS-218 |
| chr5-114679017-INS-433 | chr5-114679017-INS-433 |
| chr5-114700215-INS-215 | chr5-114700215-INS-215 |
| chr5-114718657-INS-646 | chr5-114718657-INS-646 |
| chr5-114732878-INS-218 | chr5-114732878-INS-219 |
| chr5-114738237-INS-214 | chr5-114738237-INS-214 |
| chr5-114743230-INS-106 | chr5-114743230-INS-112 |
| chr5-114775054-INS-147 | chr5-114775054-INS-147 |
| chr5-114778636-INS-279 | chr5-114778648-INS-271 |
| chr5-114810803-INS-274 | chr5-114810780-INS-270 |
| chr5-114824522-INS-51 | chr5-114824522-INS-50 |
| chr5-114880573-INS-70 | chr5-114880573-INS-70 |
| chr5-114926615-INS-228 | chr5-114926615-INS-230 |
| chr5-114946515-INS-53 | chr5-114946515-INS-56 |
| chr5-114948315-INS-211 | chr5-114948315-INS-208 |
| chr5-114970601-INS-340 | chr5-114970601-INS-341 |
| chr5-114972182-INS-212 | chr5-114972182-INS-212 |
| chr5-114979539-INS-207 | chr5-114979539-INS-207 |
| chr5-114993675-INS-181 | chr5-114993676-INS-181 |
| chr5-115005254-INS-202 | chr5-115005254-INS-202 |
| chr5-115037628-INS-176 | chr5-115037628-INS-175 |
| chr5-115039937-INS-202 | chr5-115039937-INS-202 |
| chr5-115057429-INS-636 | chr5-115057429-INS-636 |
| chr5-115072386-INS-239 | chr5-115072386-INS-240 |
| chr5-115098509-INS-84 | chr5-115098509-INS-84 |
| chr5-115113284-INS-117 | chr5-115113285-INS-68 |
| chr5-115122182-INS-161 | chr5-115122182-INS-161 |
| chr5-115127587-INS-414 | chr5-115127587-INS-414 |
| chr5-115135199-INS-106 | chr5-115135199-INS-106 |
| chr5-115143087-INS-192 | chr5-115143087-INS-185 |
| chr5-115170108-INS-193 | chr5-115170108-INS-193 |
| chr5-115193459-INS-210 | chr5-115193459-INS-207 |
| chr5-115222037-INS-217 | chr5-115222037-INS-217 |
| chr5-115230251-INS-174 | chr5-115230251-INS-174 |
| chr5-115259947-INS-100 | chr5-115259947-INS-50 |
| chr5-115300647-INS-84 | chr5-115300649-INS-55 |
| chr5-115317017-INS-211 | chr5-115317017-INS-212 |
| chr5-115337932-INS-76 | chr5-115337932-INS-104 |
| chr5-115339126-INS-399 | chr5-115339126-INS-399 |
| chr5-115351974-INS-120 | chr5-115351974-INS-120 |
| chr5-115357650-INS-51 | chr5-115357642-INS-54 |
| chr5-115378168-INS-225 | chr5-115378168-INS-213 |
| chr5-115402455-INS-217 | chr5-115402455-INS-217 |
| chr5-115407188-INS-227 | chr5-115407188-INS-228 |
| chr5-115411126-INS-209 | chr5-115411126-INS-209 |
| chr5-115412881-INS-217 | chr5-115412881-INS-217 |
| chr5-115416542-INS-399 | chr5-115416542-INS-399 |
| chr5-115418994-INS-209 | chr5-115418994-INS-209 |
| chr5-115423295-INS-214 | chr5-115423295-INS-215 |
| chr5-115426999-INS-231 | chr5-115426999-INS-231 |
| chr5-115444334-INS-5537 | chr5-115444335-INS-5537 |
| chr5-115449021-INS-204 | chr5-115449021-INS-204 |
| chr5-115451393-INS-307 | chr5-115451393-INS-307 |
| chr5-115459040-INS-1152 | chr5-115459041-INS-1153 |
| chr5-115460150-INS-207 | chr5-115460150-INS-200 |
| chr5-115476680-INS-178 | chr5-115476680-INS-179 |
| chr5-115490808-INS-7167 | chr5-115490809-INS-7167 |
| chr5-115494915-INS-61 | chr5-115494918-INS-100 |
| chr5-115500608-INS-55 | chr5-115500608-INS-55 |
| chr5-115504518-INS-251 | chr5-115504518-INS-254 |
| chr5-115511438-INS-213 | chr5-115511438-INS-211 |
| chr5-115516436-INS-139 | chr5-115516436-INS-139 |
| chr5-115556157-INS-79 | chr5-115556157-INS-84 |
| chr5-115584758-INS-215 | chr5-115584758-INS-216 |
| chr5-115611043-INS-203 | chr5-115611043-INS-203 |
| chr5-115616639-INS-94 | chr5-115616639-INS-95 |
| chr5-115620166-INS-217 | chr5-115620166-INS-217 |
| chr5-115626136-INS-429 | chr5-115626136-INS-427 |

| | |
|---|---|
| chr5-115631156-INS-109 | chr5-115631156-INS-109 |
| chr5-115631681-INS-670 | chr5-115631681-INS-672 |
| chr5-115633351-INS-198 | chr5-115633351-INS-198 |
| chr5-115636076-INS-194 | chr5-115636077-INS-194 |
| chr5-115638925-INS-198 | chr5-115638925-INS-198 |
| chr5-115640607-INS-192 | chr5-115640607-INS-191 |
| chr5-115647735-INS-88 | chr5-115647735-INS-78 |
| chr5-115657835-INS-185 | chr5-115657835-INS-185 |
| chr5-115665785-INS-328 | chr5-115665785-INS-328 |
| chr5-115672725-INS-220 | chr5-115672725-INS-221 |
| chr5-115708584-INS-204 | chr5-115708584-INS-204 |
| chr5-115712122-INS-180 | chr5-115712122-INS-187 |
| chr5-115713395-INS-61 | chr5-115713395-INS-61 |
| chr5-115724819-INS-139 | chr5-115724808-INS-119 |
| chr5-115725636-INS-51 | chr5-115725636-INS-53 |
| chr5-115743858-INS-1158 | chr5-115743859-INS-1163 |
| chr5-115766176-INS-235 | chr5-115766176-INS-235 |
| chr5-115793877-INS-138 | chr5-115793877-INS-140 |
| chr5-115799793-INS-237 | chr5-115799793-INS-236 |
| chr5-115827449-INS-52 | chr5-115827449-INS-52 |
| chr5-115838358-INS-386 | chr5-115838358-INS-386 |
| chr5-115847942-INS-234 | chr5-115847942-INS-234 |
| chr5-115855664-INS-259 | chr5-115855664-INS-259 |
| chr5-115877852-INS-222 | chr5-115877852-INS-222 |
| chr5-115885334-INS-82 | chr5-115885334-INS-82 |
| chr5-115890532-INS-385 | chr5-115890532-INS-385 |
| chr5-115897087-INS-449 | chr5-115897082-INS-485 |
| chr5-115914827-INS-220 | chr5-115914827-INS-220 |
| chr5-115924029-INS-568 | chr5-115924030-INS-563 |
| chr5-115943698-INS-216 | chr5-115943698-INS-215 |
| chr5-115957274-INS-360 | chr5-115957274-INS-360 |
| chr5-115963109-INS-204 | chr5-115963109-INS-205 |
| chr5-115968395-INS-515 | chr5-115968395-INS-516 |
| chr5-115969861-INS-204 | chr5-115969857-INS-186 |
| chr5-115977100-INS-67 | chr5-115977100-INS-67 |
| chr5-116019673-INS-71 | chr5-116019673-INS-71 |
| chr5-116027752-INS-189 | chr5-116027752-INS-189 |
| chr5-116037732-INS-54 | chr5-116037713-INS-60 |
| chr5-116097397-INS-104 | chr5-116097397-INS-104 |
| chr5-116132989-INS-433 | chr5-116132989-INS-433 |
| chr5-116149455-INS-119 | chr5-116149455-INS-90 |
| chr5-116154201-INS-156 | chr5-116154201-INS-156 |
| chr5-116177512-INS-171 | chr5-116177512-INS-171 |
| chr5-116179304-INS-317 | chr5-116179297-INS-313 |
| chr5-116193906-INS-105 | chr5-116193906-INS-102 |
| chr5-116211367-INS-179 | chr5-116211367-INS-179 |
| chr5-116222024-INS-2145 | chr5-116222038-INS-2129 |
| chr5-116227190-INS-196 | chr5-116227190-INS-196 |
| chr5-116233490-INS-55 | chr5-116233490-INS-52 |
| chr5-116256388-INS-91 | chr5-116256397-INS-86 |
| chr5-116277483-INS-343 | chr5-116277483-INS-345 |
| chr5-116301219-INS-276 | chr5-116301219-INS-276 |
| chr5-116351537-INS-352 | chr5-116351537-INS-351 |
| chr5-116354334-INS-88 | chr5-116354295-INS-77 |
| chr5-116356847-INS-55 | chr5-116356847-INS-65 |
| chr5-116359347-INS-100 | chr5-116359347-INS-106 |
| chr5-116368517-INS-343 | chr5-116368517-INS-339 |
| chr5-116435009-INS-710 | chr5-116435009-INS-720 |
| chr5-116437863-INS-64 | chr5-116437863-INS-66 |
| chr5-116439391-INS-58 | chr5-116439391-INS-58 |
| chr5-116449519-INS-86 | chr5-116449523-INS-88 |
| chr5-116478928-INS-440 | chr5-116478913-INS-438 |
| chr5-116488210-INS-4306 | chr5-116488211-INS-4306 |
| chr5-116495757-INS-52 | chr5-116495757-INS-52 |
| chr5-116512047-INS-74 | chr5-116512032-INS-91 |
| chr5-116516112-INS-76 | chr5-116516120-INS-72 |
| chr5-116521250-INS-211 | chr5-116521250-INS-215 |
| chr5-116541310-INS-4584 | chr5-116541311-INS-4584 |
| chr5-116542616-INS-60 | chr5-116542616-INS-64 |
| chr5-116660969-INS-52 | chr5-116660969-INS-52 |
| chr5-116682937-INS-94 | chr5-116682937-INS-94 |
| chr5-116688589-INS-108 | chr5-116688589-INS-114 |
| chr5-116721946-INS-55 | chr5-116721946-INS-58 |
| chr5-116749971-INS-108 | chr5-116749970-INS-114 |
| chr5-116815760-INS-66 | chr5-116815760-INS-66 |
| chr5-116819807-INS-52 | chr5-116819807-INS-52 |
| chr5-116820937-INS-124 | chr5-116820937-INS-126 |

| | |
|---|---|
| chr5-116824358-INS-76 | chr5-116824358-INS-76 |
| chr5-116848183-INS-5541 | chr5-116848184-INS-5541 |
| chr5-116854214-INS-72 | chr5-116854214-INS-72 |
| chr5-116869339-INS-109 | chr5-116869339-INS-108 |
| chr5-116877885-INS-96 | chr5-116877885-INS-96 |
| chr5-116880512-INS-108 | chr5-116880512-INS-108 |
| chr5-116886734-INS-117 | chr5-116886734-INS-117 |
| chr5-116904056-INS-155 | chr5-116904056-INS-169 |
| chr5-116925326-INS-68 | chr5-116925326-INS-72 |
| chr5-116942641-INS-187 | chr5-116942661-INS-198 |
| chr5-116967534-INS-711 | chr5-116967534-INS-711 |
| chr5-116969220-INS-62 | chr5-116969220-INS-60 |
| chr5-116992485-INS-54 | chr5-116992485-INS-54 |
| chr5-117019183-INS-77 | chr5-117019183-INS-77 |
| chr5-117041616-INS-115 | chr5-117041616-INS-110 |
| chr5-117063292-INS-51 | chr5-117063292-INS-51 |
| chr5-117076011-INS-64 | chr5-117076001-INS-64 |
| chr5-117086409-INS-495 | chr5-117086409-INS-495 |
| chr5-117097582-INS-55 | chr5-117097605-INS-65 |
| chr5-117144984-INS-1893 | chr5-117144985-INS-1893 |
| chr5-117191817-INS-124 | chr5-117191817-INS-124 |
| chr5-117214150-INS-58 | chr5-117214150-INS-58 |
| chr5-117222543-INS-328 | chr5-117222543-INS-325 |
| chr5-117236914-INS-72 | chr5-117236914-INS-72 |
| chr5-117249200-INS-115 | chr5-117249200-INS-115 |
| chr5-117288725-INS-69 | chr5-117288727-INS-69 |
| chr5-117291399-INS-173 | chr5-117291399-INS-174 |
| chr5-117295211-INS-755 | chr5-117295212-INS-750 |
| chr5-117305457-INS-204 | chr5-117305458-INS-204 |
| chr5-117332472-INS-56 | chr5-117332472-INS-56 |
| chr5-117343463-INS-312 | chr5-117343461-INS-309 |
| chr5-117364333-INS-54 | chr5-117364334-INS-54 |
| chr5-117395794-INS-195 | chr5-117395795-INS-175 |
| chr5-117411512-INS-607 | chr5-117411512-INS-596 |
| chr5-117414771-INS-99 | chr5-117414771-INS-92 |
| chr5-117426908-INS-226 | chr5-117426908-INS-226 |
| chr5-117450991-INS-178 | chr5-117450991-INS-178 |
| chr5-117469685-INS-192 | chr5-117469685-INS-192 |
| chr5-117472376-INS-92 | chr5-117472344-INS-90 |
| chr5-117488666-INS-163 | chr5-117488666-INS-163 |
| chr5-117492869-INS-181 | chr5-117492869-INS-181 |
| chr5-117511363-INS-390 | chr5-117511363-INS-390 |
| chr5-117532688-INS-203 | chr5-117532688-INS-203 |
| chr5-117533097-INS-519 | chr5-117533097-INS-519 |
| chr5-117568641-INS-506 | chr5-117568641-INS-522 |
| chr5-117569911-INS-108 | chr5-117569911-INS-108 |
| chr5-117592234-INS-212 | chr5-117592234-INS-212 |
| chr5-117612786-INS-100 | chr5-117612786-INS-100 |
| chr5-117639495-INS-93 | chr5-117639515-INS-112 |
| chr5-117641765-INS-253 | chr5-117641765-INS-255 |
| chr5-117645419-INS-89 | chr5-117645419-INS-102 |
| chr5-117685570-INS-73 | chr5-117685570-INS-95 |
| chr5-117692796-INS-67 | chr5-117692796-INS-67 |
| chr5-117696554-INS-100 | chr5-117696554-INS-100 |
| chr5-117703008-INS-51 | chr5-117703001-INS-76 |
| chr5-117725329-INS-144 | chr5-117725329-INS-144 |
| chr5-117727244-INS-52 | chr5-117727231-INS-76 |
| chr5-117735495-INS-68 | chr5-117735476-INS-66 |
| chr5-117737563-INS-72 | chr5-117737563-INS-72 |
| chr5-117739905-INS-143 | chr5-117739905-INS-143 |
| chr5-117760998-INS-92 | chr5-117761000-INS-66 |
| chr5-117789931-INS-54 | chr5-117789931-INS-81 |
| chr5-117843813-INS-68 | chr5-117843801-INS-62 |
| chr5-117854863-INS-515 | chr5-117854863-INS-515 |
| chr5-117857340-INS-7002 | chr5-117857341-INS-7002 |
| chr5-117860175-INS-194 | chr5-117860175-INS-194 |
| chr5-117867098-INS-84 | chr5-117867098-INS-70 |
| chr5-117872827-INS-147 | chr5-117872827-INS-147 |
| chr5-117885575-INS-69 | chr5-117885575-INS-69 |
| chr5-117912382-INS-56 | chr5-117912382-INS-56 |
| chr5-117951283-INS-198 | chr5-117951283-INS-198 |
| chr5-118002713-INS-151 | chr5-118002713-INS-162 |
| chr5-118007736-INS-58 | chr5-118007747-INS-88 |
| chr5-118065448-INS-70 | chr5-118065448-INS-56 |
| chr5-118089317-INS-93 | chr5-118089317-INS-93 |
| chr5-118104869-INS-70 | chr5-118104869-INS-55 |
| chr5-118122255-INS-77 | chr5-118122255-INS-77 |

| | |
|---|---|
| chr5-118159945-INS-54 | chr5-118159945-INS-54 |
| chr5-118180423-INS-63 | chr5-118180403-INS-110 |
| chr5-118187010-INS-101 | chr5-118187010-INS-101 |
| chr5-118240869-INS-159 | chr5-118240869-INS-165 |
| chr5-118242034-INS-466 | chr5-118242034-INS-466 |
| chr5-118248039-INS-7155 | chr5-118248040-INS-7155 |
| chr5-118272546-INS-55 | chr5-118272546-INS-55 |
| chr5-118277641-INS-413 | chr5-118277641-INS-447 |
| chr5-118282912-INS-218 | chr5-118282912-INS-217 |
| chr5-118330475-INS-60 | chr5-118330475-INS-60 |
| chr5-118389875-INS-137 | chr5-118389875-INS-137 |
| chr5-118396107-INS-86 | chr5-118396107-INS-86 |
| chr5-118403031-INS-91 | chr5-118403038-INS-70 |
| chr5-118415762-INS-209 | chr5-118415762-INS-212 |
| chr5-118424597-INS-186 | chr5-118424597-INS-186 |
| chr5-118431628-INS-56 | chr5-118431628-INS-76 |
| chr5-118446580-INS-519 | chr5-118446580-INS-519 |
| chr5-118451444-INS-68 | chr5-118451444-INS-60 |
| chr5-118456446-INS-58 | chr5-118456446-INS-52 |
| chr5-118539662-INS-204 | chr5-118539662-INS-204 |
| chr5-118558638-INS-97 | chr5-118558639-INS-112 |
| chr5-118583104-INS-191 | chr5-118583104-INS-191 |
| chr5-118601443-INS-101 | chr5-118601443-INS-53 |
| chr5-118637356-INS-6216 | chr5-118637357-INS-6216 |
| chr5-118661034-INS-63 | chr5-118661019-INS-58 |
| chr5-118685779-INS-221 | chr5-118685779-INS-222 |
| chr5-118686666-INS-52 | chr5-118686666-INS-54 |
| chr5-118705900-INS-58 | chr5-118705900-INS-60 |
| chr5-118720420-INS-2987 | chr5-118720421-INS-2999 |
| chr5-118747518-INS-93 | chr5-118747518-INS-94 |
| chr5-118767240-INS-84 | chr5-118767240-INS-92 |
| chr5-118864486-INS-68 | chr5-118864486-INS-68 |
| chr5-118929663-INS-674 | chr5-118929663-INS-674 |
| chr5-118960243-INS-51 | chr5-118960243-INS-51 |
| chr5-118975621-INS-400 | chr5-118975621-INS-400 |
| chr5-118996778-INS-83 | chr5-118996778-INS-56 |
| chr5-119003320-INS-2303 | chr5-119003321-INS-2303 |
| chr5-119051686-INS-257 | chr5-119051686-INS-257 |
| chr5-119092761-INS-57 | chr5-119092720-INS-72 |
| chr5-119106602-INS-54 | chr5-119106602-INS-78 |
| chr5-119120974-INS-61 | chr5-119120974-INS-55 |
| chr5-119131935-INS-1062 | chr5-119131936-INS-1062 |
| chr5-119132131-INS-105 | chr5-119132131-INS-105 |
| chr5-119151966-INS-79 | chr5-119151966-INS-61 |
| chr5-119163728-INS-76 | chr5-119163728-INS-50 |
| chr5-119165742-INS-90 | chr5-119165742-INS-90 |
| chr5-119183889-INS-2214 | chr5-119183879-INS-2214 |
| chr5-119186705-INS-62 | chr5-119186705-INS-62 |
| chr5-119212966-INS-202 | chr5-119212966-INS-200 |
| chr5-119216994-INS-202 | chr5-119216994-INS-212 |
| chr5-119219282-INS-116 | chr5-119219282-INS-132 |
| chr5-119261789-INS-549 | chr5-119261789-INS-549 |
| chr5-119263513-INS-94 | chr5-119263513-INS-93 |
| chr5-119284224-INS-74 | chr5-119284224-INS-78 |
| chr5-119288476-INS-64 | chr5-119288476-INS-68 |
| chr5-119319900-INS-235 | chr5-119319878-INS-223 |
| chr5-119338932-INS-178 | chr5-119338932-INS-178 |
| chr5-119360156-INS-308 | chr5-119360156-INS-275 |
| chr5-119418208-INS-60 | chr5-119418208-INS-50 |
| chr5-119427372-INS-81 | chr5-119427371-INS-76 |
| chr5-119435527-INS-7101 | chr5-119435525-INS-7098 |
| chr5-119438617-INS-53 | chr5-119438617-INS-50 |
| chr5-119471712-INS-90 | chr5-119471712-INS-91 |
| chr5-119472988-INS-159 | chr5-119472988-INS-159 |
| chr5-119479840-INS-75 | chr5-119479840-INS-83 |
| chr5-119488128-INS-64 | chr5-119488168-INS-56 |
| chr5-119489804-INS-58 | chr5-119489826-INS-52 |
| chr5-119490842-INS-170 | chr5-119490842-INS-170 |
| chr5-119508031-INS-57 | chr5-119508031-INS-52 |
| chr5-119510057-INS-86 | chr5-119510057-INS-85 |
| chr5-119511011-INS-66 | chr5-119511011-INS-66 |
| chr5-119514077-INS-74 | chr5-119514077-INS-74 |
| chr5-119555770-INS-334 | chr5-119555767-INS-314 |
| chr5-119592065-INS-77 | chr5-119592065-INS-77 |
| chr5-119606041-INS-1231 | chr5-119606042-INS-1231 |
| chr5-119633797-INS-118 | chr5-119633790-INS-92 |
| chr5-119635495-INS-1545 | chr5-119635496-INS-1545 |

| | |
|---|---|
| chr5-119644460-INS-78 | chr5-119644501-INS-78 |
| chr5-119654361-INS-775 | chr5-119654361-INS-776 |
| chr5-119658021-INS-63 | chr5-119658021-INS-63 |
| chr5-119671574-INS-73 | chr5-119671574-INS-71 |
| chr5-119691348-INS-74 | chr5-119691348-INS-74 |
| chr5-119696981-INS-123 | chr5-119696981-INS-123 |
| chr5-119704949-INS-59 | chr5-119704928-INS-51 |
| chr5-119755792-INS-572 | chr5-119755792-INS-555 |
| chr5-119756965-INS-99 | chr5-119756942-INS-90 |
| chr5-119786414-INS-177 | chr5-119786414-INS-178 |
| chr5-119788690-INS-90 | chr5-119788690-INS-90 |
| chr5-119873952-INS-68 | chr5-119873952-INS-68 |
| chr5-119879508-INS-148 | chr5-119879508-INS-198 |
| chr5-119908209-INS-83 | chr5-119908208-INS-92 |
| chr5-119913870-INS-169 | chr5-119913870-INS-175 |
| chr5-119961604-INS-59 | chr5-119961604-INS-54 |
| chr5-119988972-INS-64 | chr5-119988972-INS-64 |
| chr5-120005765-INS-52 | chr5-120005765-INS-62 |
| chr5-120028266-INS-110 | chr5-120028266-INS-116 |
| chr5-120050916-INS-68 | chr5-120050916-INS-113 |
| chr5-120054407-INS-183 | chr5-120054407-INS-162 |
| chr5-120100337-INS-115 | chr5-120100337-INS-131 |
| chr5-120103638-INS-101 | chr5-120103638-INS-81 |
| chr5-120110490-INS-107 | chr5-120110490-INS-107 |
| chr5-120138915-INS-1885 | chr5-120138916-INS-1885 |
| chr5-120151100-INS-203 | chr5-120151100-INS-203 |
| chr5-120158845-INS-105 | chr5-120158845-INS-105 |
| chr5-120163023-INS-63 | chr5-120163044-INS-63 |
| chr5-120190186-INS-2808 | chr5-120190187-INS-2808 |
| chr5-120274204-INS-66 | chr5-120274204-INS-66 |
| chr5-120299827-INS-53 | chr5-120299827-INS-53 |
| chr5-120374610-INS-54 | chr5-120374610-INS-54 |
| chr5-120396768-INS-224 | chr5-120396768-INS-224 |
| chr5-120400598-INS-62 | chr5-120400598-INS-74 |
| chr5-120425805-INS-214 | chr5-120425806-INS-216 |
| chr5-120437411-INS-157 | chr5-120437411-INS-132 |
| chr5-120441968-INS-184 | chr5-120441968-INS-184 |
| chr5-120448837-INS-160 | chr5-120448837-INS-140 |
| chr5-120462251-INS-63 | chr5-120462251-INS-58 |
| chr5-120466837-INS-312 | chr5-120466849-INS-311 |
| chr5-120487215-INS-735 | chr5-120487215-INS-735 |
| chr5-120499532-INS-52 | chr5-120499548-INS-64 |
| chr5-120509226-INS-155 | chr5-120509225-INS-139 |
| chr5-120529362-INS-208 | chr5-120529362-INS-208 |
| chr5-120568539-INS-56 | chr5-120568515-INS-68 |
| chr5-120578757-INS-69 | chr5-120578772-INS-94 |
| chr5-120595197-INS-79 | chr5-120595221-INS-60 |
| chr5-120608834-INS-155 | chr5-120608834-INS-155 |
| chr5-120643141-INS-193 | chr5-120643141-INS-194 |
| chr5-120669671-INS-70 | chr5-120669647-INS-76 |
| chr5-120677761-INS-92 | chr5-120677757-INS-60 |
| chr5-120682970-INS-135 | chr5-120682964-INS-127 |
| chr5-120687946-INS-115 | chr5-120687946-INS-116 |
| chr5-120697840-INS-205 | chr5-120697840-INS-197 |
| chr5-120710196-INS-60 | chr5-120710171-INS-68 |
| chr5-120719450-INS-168 | chr5-120719450-INS-168 |
| chr5-120728942-INS-54 | chr5-120728942-INS-56 |
| chr5-120730316-INS-222 | chr5-120730316-INS-222 |
| chr5-120753007-INS-223 | chr5-120753007-INS-225 |
| chr5-120784405-INS-67 | chr5-120784405-INS-67 |
| chr5-120799464-INS-56 | chr5-120799464-INS-80 |
| chr5-120812557-INS-64 | chr5-120812557-INS-65 |
| chr5-120858947-INS-112 | chr5-120858941-INS-152 |
| chr5-120866668-INS-76 | chr5-120866668-INS-76 |
| chr5-120878330-INS-95 | chr5-120878336-INS-93 |
| chr5-120886306-INS-95 | chr5-120886306-INS-138 |
| chr5-120903816-INS-196 | chr5-120903822-INS-212 |
| chr5-120913105-INS-56 | chr5-120913105-INS-56 |
| chr5-120916850-INS-55 | chr5-120916852-INS-56 |
| chr5-120923008-INS-112 | chr5-120923008-INS-112 |
| chr5-120968836-INS-194 | chr5-120968836-INS-196 |
| chr5-120998491-INS-102 | chr5-120998491-INS-98 |
| chr5-121056022-INS-60 | chr5-121056022-INS-60 |
| chr5-121097620-INS-60 | chr5-121097614-INS-68 |
| chr5-121111265-INS-162 | chr5-121111288-INS-120 |
| chr5-121132492-INS-238 | chr5-121132492-INS-238 |
| chr5-121171134-INS-141 | chr5-121171134-INS-141 |

| | |
|---|---|
| chr5-121185117-INS-95 | chr5-121185117-INS-95 |
| chr5-121193566-INS-52 | chr5-121193567-INS-52 |
| chr5-121199247-INS-240 | chr5-121199247-INS-240 |
| chr5-121201793-INS-86 | chr5-121201793-INS-86 |
| chr5-121208562-INS-62 | chr5-121208562-INS-62 |
| chr5-121222858-INS-149 | chr5-121222858-INS-149 |
| chr5-121231174-INS-218 | chr5-121231174-INS-218 |
| chr5-121233576-INS-84 | chr5-121233570-INS-84 |
| chr5-121235180-INS-54 | chr5-121235180-INS-54 |
| chr5-121242580-INS-52 | chr5-121242580-INS-50 |
| chr5-121244669-INS-1395 | chr5-121244670-INS-1395 |
| chr5-121251112-INS-77 | chr5-121251112-INS-77 |
| chr5-121282172-INS-221 | chr5-121282185-INS-228 |
| chr5-121288068-INS-171 | chr5-121288069-INS-200 |
| chr5-121291018-INS-209 | chr5-121291018-INS-209 |
| chr5-121298853-INS-226 | chr5-121298853-INS-214 |
| chr5-121302380-INS-67 | chr5-121302380-INS-71 |
| chr5-121313156-INS-460 | chr5-121313156-INS-460 |
| chr5-121313933-INS-169 | chr5-121313933-INS-169 |
| chr5-121350185-INS-206 | chr5-121350184-INS-206 |
| chr5-121385635-INS-206 | chr5-121385635-INS-206 |
| chr5-121388313-INS-217 | chr5-121388313-INS-214 |
| chr5-121389049-INS-181 | chr5-121389049-INS-184 |
| chr5-121391738-INS-584 | chr5-121391738-INS-586 |
| chr5-121422612-INS-218 | chr5-121422612-INS-218 |
| chr5-121425599-INS-111 | chr5-121425599-INS-123 |
| chr5-121450022-INS-74 | chr5-121450022-INS-74 |
| chr5-121480176-INS-534 | chr5-121480176-INS-534 |
| chr5-121489249-INS-357 | chr5-121489249-INS-357 |
| chr5-121506668-INS-108 | chr5-121506668-INS-108 |
| chr5-121514348-INS-91 | chr5-121514348-INS-92 |
| chr5-121520284-INS-92 | chr5-121520284-INS-92 |
| chr5-121555202-INS-74 | chr5-121555202-INS-74 |
| chr5-121562037-INS-7135 | chr5-121562038-INS-7135 |
| chr5-121597677-INS-210 | chr5-121597677-INS-208 |
| chr5-121603344-INS-170 | chr5-121603344-INS-172 |
| chr5-121642329-INS-1890 | chr5-121642330-INS-1890 |
| chr5-121672914-INS-287 | chr5-121672914-INS-279 |
| chr5-121690594-INS-415 | chr5-121690586-INS-411 |
| chr5-121692830-INS-65 | chr5-121692830-INS-65 |
| chr5-121695170-INS-186 | chr5-121695170-INS-186 |
| chr5-121696812-INS-57 | chr5-121696812-INS-60 |
| chr5-121700114-INS-66 | chr5-121700114-INS-66 |
| chr5-121710932-INS-56 | chr5-121710935-INS-57 |
| chr5-121713615-INS-67 | chr5-121713615-INS-67 |
| chr5-121726019-INS-194 | chr5-121726019-INS-193 |
| chr5-121735584-INS-61 | chr5-121735584-INS-61 |
| chr5-121751984-INS-61 | chr5-121751984-INS-61 |
| chr5-121762910-INS-226 | chr5-121762910-INS-226 |
| chr5-121792425-INS-183 | chr5-121792425-INS-183 |
| chr5-121807987-INS-65 | chr5-121807987-INS-55 |
| chr5-121812480-INS-529 | chr5-121812480-INS-529 |
| chr5-121817844-INS-207 | chr5-121817844-INS-207 |
| chr5-121820699-INS-203 | chr5-121820699-INS-201 |
| chr5-121827868-INS-209 | chr5-121827868-INS-209 |
| chr5-121880837-INS-175 | chr5-121880838-INS-179 |
| chr5-121894460-INS-224 | chr5-121894460-INS-224 |
| chr5-121902205-INS-51 | chr5-121902205-INS-50 |
| chr5-121907562-INS-81 | chr5-121907562-INS-80 |
| chr5-121963967-INS-249 | chr5-121963967-INS-251 |
| chr5-122059516-INS-84 | chr5-122059534-INS-94 |
| chr5-122134464-INS-66 | chr5-122134464-INS-66 |
| chr5-122195479-INS-209 | chr5-122195479-INS-209 |
| chr5-122216645-INS-666 | chr5-122216645-INS-667 |
| chr5-122220287-INS-214 | chr5-122220287-INS-214 |
| chr5-122228621-INS-206 | chr5-122228621-INS-206 |
| chr5-122262424-INS-68 | chr5-122262414-INS-68 |
| chr5-122264245-INS-75 | chr5-122264245-INS-63 |
| chr5-122266092-INS-54 | chr5-122266092-INS-54 |
| chr5-122278539-INS-130 | chr5-122278550-INS-140 |
| chr5-122316439-INS-6464 | chr5-122316440-INS-6464 |
| chr5-122319356-INS-223 | chr5-122319356-INS-223 |
| chr5-122367255-INS-158 | chr5-122367211-INS-190 |
| chr5-122401820-INS-205 | chr5-122401820-INS-205 |
| chr5-122403731-INS-191 | chr5-122403731-INS-194 |
| chr5-122421393-INS-96 | chr5-122421391-INS-74 |
| chr5-122437002-INS-122 | chr5-122437002-INS-126 |

| | |
|---|---|
| chr5-122453751-INS-217 | chr5-122453751-INS-217 |
| chr5-122467321-INS-206 | chr5-122467321-INS-209 |
| chr5-122473252-INS-88 | chr5-122473252-INS-88 |
| chr5-122506813-INS-56 | chr5-122506813-INS-55 |
| chr5-122508144-INS-204 | chr5-122508144-INS-204 |
| chr5-122535622-INS-198 | chr5-122535622-INS-198 |
| chr5-122554397-INS-327 | chr5-122554397-INS-327 |
| chr5-122556773-INS-4145 | chr5-122556774-INS-4135 |
| chr5-122562908-INS-207 | chr5-122562908-INS-208 |
| chr5-122679192-INS-70 | chr5-122679192-INS-70 |
| chr5-122682484-INS-144 | chr5-122682484-INS-144 |
| chr5-122689842-INS-81 | chr5-122689842-INS-91 |
| chr5-122699064-INS-68 | chr5-122699064-INS-68 |
| chr5-122714439-INS-66 | chr5-122714470-INS-56 |
| chr5-122717125-INS-84 | chr5-122717125-INS-108 |
| chr5-122736912-INS-201 | chr5-122736912-INS-201 |
| chr5-122746910-INS-4725 | chr5-122746911-INS-4725 |
| chr5-122754849-INS-105 | chr5-122754849-INS-105 |
| chr5-122762288-INS-222 | chr5-122762288-INS-223 |
| chr5-122772234-INS-64 | chr5-122772234-INS-64 |
| chr5-122776621-INS-93 | chr5-122776621-INS-93 |
| chr5-122787019-INS-311 | chr5-122787020-INS-311 |
| chr5-122809531-INS-121 | chr5-122809531-INS-156 |
| chr5-122841307-INS-131 | chr5-122841341-INS-105 |
| chr5-122887173-INS-182 | chr5-122887173-INS-168 |
| chr5-122889561-INS-828 | chr5-122889561-INS-828 |
| chr5-122890112-INS-164 | chr5-122890099-INS-179 |
| chr5-122893918-INS-233 | chr5-122893918-INS-235 |
| chr5-122895945-INS-201 | chr5-122895945-INS-202 |
| chr5-122923401-INS-559 | chr5-122923401-INS-596 |
| chr5-122947836-INS-214 | chr5-122947836-INS-214 |
| chr5-122966356-INS-199 | chr5-122966356-INS-199 |
| chr5-122969648-INS-203 | chr5-122969644-INS-191 |
| chr5-122978296-INS-186 | chr5-122978296-INS-187 |
| chr5-122981819-INS-155 | chr5-122981819-INS-155 |
| chr5-122990130-INS-213 | chr5-122990129-INS-228 |
| chr5-122991793-INS-209 | chr5-122991793-INS-208 |
| chr5-123000548-INS-209 | chr5-123000548-INS-209 |
| chr5-123004780-INS-204 | chr5-123004780-INS-205 |
| chr5-123008495-INS-185 | chr5-123008495-INS-185 |
| chr5-123028712-INS-61 | chr5-123028712-INS-61 |
| chr5-123034975-INS-93 | chr5-123034975-INS-90 |
| chr5-123048195-INS-355 | chr5-123048195-INS-355 |
| chr5-123050403-INS-139 | chr5-123050403-INS-140 |
| chr5-123050714-INS-5918 | chr5-123050715-INS-5918 |
| chr5-123051521-INS-157 | chr5-123051521-INS-157 |
| chr5-123055317-INS-208 | chr5-123055317-INS-208 |
| chr5-123056122-INS-178 | chr5-123056122-INS-178 |
| chr5-123066465-INS-54 | chr5-123066465-INS-51 |
| chr5-123067266-INS-74 | chr5-123067267-INS-74 |
| chr5-123081138-INS-210 | chr5-123081138-INS-210 |
| chr5-123082754-INS-160 | chr5-123082754-INS-128 |
| chr5-123084952-INS-56 | chr5-123084952-INS-52 |
| chr5-123101249-INS-98 | chr5-123101227-INS-102 |
| chr5-123111166-INS-590 | chr5-123111166-INS-591 |
| chr5-123116602-INS-52 | chr5-123116602-INS-52 |
| chr5-123135134-INS-219 | chr5-123135134-INS-219 |
| chr5-123137937-INS-730 | chr5-123137937-INS-730 |
| chr5-123163684-INS-210 | chr5-123163684-INS-225 |
| chr5-123187002-INS-80 | chr5-123187002-INS-80 |
| chr5-123195154-INS-52 | chr5-123195117-INS-68 |
| chr5-123206042-INS-56 | chr5-123206043-INS-64 |
| chr5-123209605-INS-207 | chr5-123209605-INS-207 |
| chr5-123211428-INS-275 | chr5-123211428-INS-289 |
| chr5-123216298-INS-120 | chr5-123216318-INS-125 |
| chr5-123258304-INS-150 | chr5-123258319-INS-150 |
| chr5-123263837-INS-56 | chr5-123263837-INS-52 |
| chr5-123362198-INS-152 | chr5-123362198-INS-144 |
| chr5-123365899-INS-622 | chr5-123365899-INS-622 |
| chr5-123369345-INS-498 | chr5-123369345-INS-498 |
| chr5-123382033-INS-574 | chr5-123382027-INS-575 |
| chr5-123384959-INS-164 | chr5-123384960-INS-164 |
| chr5-123394533-INS-64 | chr5-123394529-INS-64 |
| chr5-123452820-INS-57 | chr5-123452818-INS-94 |
| chr5-123467931-INS-208 | chr5-123467931-INS-208 |
| chr5-123547317-INS-62 | chr5-123547317-INS-62 |
| chr5-123557628-INS-67 | chr5-123557628-INS-54 |

| | |
|---|---|
| chr5-123563076-INS-62 | chr5-123563074-INS-63 |
| chr5-123564024-INS-124 | chr5-123564024-INS-90 |
| chr5-123566690-INS-171 | chr5-123566690-INS-172 |
| chr5-123567407-INS-89 | chr5-123567440-INS-52 |
| chr5-123584316-INS-232 | chr5-123584316-INS-231 |
| chr5-123602600-INS-198 | chr5-123602600-INS-198 |
| chr5-123603806-INS-132 | chr5-123603806-INS-132 |
| chr5-123604517-INS-754 | chr5-123604517-INS-744 |
| chr5-123610189-INS-683 | chr5-123610189-INS-675 |
| chr5-123621907-INS-2339 | chr5-123621907-INS-2323 |
| chr5-123642659-INS-73 | chr5-123642659-INS-73 |
| chr5-123674298-INS-62 | chr5-123674298-INS-62 |
| chr5-123696107-INS-56 | chr5-123696107-INS-54 |
| chr5-123699096-INS-166 | chr5-123699096-INS-166 |
| chr5-123701172-INS-110 | chr5-123701172-INS-110 |
| chr5-123707092-INS-56 | chr5-123707087-INS-84 |
| chr5-123715069-INS-213 | chr5-123715069-INS-214 |
| chr5-123735530-INS-81 | chr5-123735530-INS-81 |
| chr5-123754079-INS-73 | chr5-123754079-INS-73 |
| chr5-123775708-INS-83 | chr5-123775708-INS-131 |
| chr5-123790489-INS-220 | chr5-123790489-INS-219 |
| chr5-123795249-INS-189 | chr5-123795249-INS-189 |
| chr5-123801235-INS-222 | chr5-123801235-INS-236 |
| chr5-123810615-INS-213 | chr5-123810615-INS-213 |
| chr5-123817576-INS-224 | chr5-123817576-INS-250 |
| chr5-123841628-INS-213 | chr5-123841645-INS-198 |
| chr5-123852503-INS-202 | chr5-123852470-INS-205 |
| chr5-123856428-INS-65 | chr5-123856428-INS-69 |
| chr5-123871330-INS-209 | chr5-123871330-INS-209 |
| chr5-123884780-INS-213 | chr5-123884780-INS-214 |
| chr5-123921700-INS-53 | chr5-123921700-INS-53 |
| chr5-123943563-INS-88 | chr5-123943563-INS-88 |
| chr5-123966239-INS-68 | chr5-123966239-INS-64 |
| chr5-123967207-INS-207 | chr5-123967207-INS-209 |
| chr5-123971295-INS-7139 | chr5-123971296-INS-7139 |
| chr5-123977226-INS-92 | chr5-123977226-INS-63 |
| chr5-123985059-INS-200 | chr5-123985059-INS-199 |
| chr5-123987371-INS-250 | chr5-123987371-INS-298 |
| chr5-123990575-INS-75 | chr5-123990575-INS-74 |
| chr5-124020184-INS-158 | chr5-124020184-INS-158 |
| chr5-124028525-INS-344 | chr5-124028523-INS-377 |
| chr5-124041239-INS-159 | chr5-124041239-INS-162 |
| chr5-124058500-INS-63 | chr5-124058500-INS-63 |
| chr5-124064282-INS-217 | chr5-124064282-INS-237 |
| chr5-124065228-INS-188 | chr5-124065228-INS-189 |
| chr5-124068384-INS-192 | chr5-124068384-INS-192 |
| chr5-124095350-INS-673 | chr5-124095350-INS-660 |
| chr5-124102034-INS-202 | chr5-124102034-INS-202 |
| chr5-124155070-INS-270 | chr5-124155070-INS-274 |
| chr5-124166038-INS-217 | chr5-124166038-INS-217 |
| chr5-124172384-INS-186 | chr5-124172384-INS-187 |
| chr5-124184246-INS-75 | chr5-124184246-INS-75 |
| chr5-124186119-INS-221 | chr5-124186119-INS-221 |
| chr5-124189021-INS-202 | chr5-124189021-INS-206 |
| chr5-124201937-INS-184 | chr5-124201937-INS-184 |
| chr5-124210362-INS-203 | chr5-124210362-INS-202 |
| chr5-124233004-INS-52 | chr5-124233004-INS-54 |
| chr5-124238604-INS-957 | chr5-124238604-INS-958 |
| chr5-124290826-INS-52 | chr5-124290826-INS-52 |
| chr5-124307755-INS-686 | chr5-124307755-INS-686 |
| chr5-124363209-INS-64 | chr5-124363209-INS-66 |
| chr5-124416414-INS-210 | chr5-124416414-INS-209 |
| chr5-124426853-INS-207 | chr5-124426853-INS-207 |
| chr5-124448873-INS-168 | chr5-124448873-INS-168 |
| chr5-124455614-INS-363 | chr5-124455614-INS-363 |
| chr5-124483261-INS-390 | chr5-124483265-INS-379 |
| chr5-124485633-INS-212 | chr5-124485633-INS-215 |
| chr5-124495180-INS-206 | chr5-124495180-INS-206 |
| chr5-124609979-INS-198 | chr5-124609974-INS-197 |
| chr5-124614594-INS-346 | chr5-124614594-INS-348 |
| chr5-124673669-INS-97 | chr5-124673649-INS-115 |
| chr5-124685477-INS-102 | chr5-124685477-INS-102 |
| chr5-124710041-INS-202 | chr5-124710041-INS-203 |
| chr5-124729309-INS-104 | chr5-124729309-INS-104 |
| chr5-124734957-INS-187 | chr5-124734957-INS-187 |
| chr5-124754982-INS-65 | chr5-124754982-INS-51 |
| chr5-124781265-INS-205 | chr5-124781265-INS-207 |

| | |
|---|---|
| chr5-124782653-INS-168 | chr5-124782660-INS-184 |
| chr5-124786778-INS-828 | chr5-124786779-INS-828 |
| chr5-124808062-INS-144 | chr5-124808062-INS-144 |
| chr5-124827718-INS-211 | chr5-124827718-INS-212 |
| chr5-124844866-INS-6440 | chr5-124844867-INS-6440 |
| chr5-124855401-INS-54 | chr5-124855401-INS-54 |
| chr5-124860163-INS-113 | chr5-124860163-INS-114 |
| chr5-124869258-INS-2920 | chr5-124869259-INS-2920 |
| chr5-124893959-INS-66 | chr5-124893959-INS-56 |
| chr5-124948727-INS-210 | chr5-124948727-INS-210 |
| chr5-124991228-INS-69 | chr5-124991228-INS-54 |
| chr5-124993568-INS-58 | chr5-124993568-INS-58 |
| chr5-125006402-INS-962 | chr5-125006404-INS-963 |
| chr5-125015682-INS-70 | chr5-125015694-INS-70 |
| chr5-125019358-INS-364 | chr5-125019358-INS-364 |
| chr5-125034299-INS-92 | chr5-125034295-INS-58 |
| chr5-125061432-INS-213 | chr5-125061432-INS-207 |
| chr5-125072229-INS-55 | chr5-125072229-INS-55 |
| chr5-125075801-INS-54 | chr5-125075801-INS-50 |
| chr5-125089976-INS-66 | chr5-125089976-INS-66 |
| chr5-125105287-INS-60 | chr5-125105299-INS-80 |
| chr5-125189595-INS-56 | chr5-125189595-INS-68 |
| chr5-125203488-INS-207 | chr5-125203488-INS-207 |
| chr5-125212736-INS-133 | chr5-125212743-INS-139 |
| chr5-125220092-INS-214 | chr5-125220092-INS-214 |
| chr5-125241237-INS-735 | chr5-125241237-INS-749 |
| chr5-125266174-INS-197 | chr5-125266174-INS-198 |
| chr5-125270295-INS-215 | chr5-125270295-INS-214 |
| chr5-125285452-INS-54 | chr5-125285452-INS-54 |
| chr5-125311022-INS-51 | chr5-125311022-INS-51 |
| chr5-125318842-INS-63 | chr5-125318840-INS-53 |
| chr5-125332651-INS-209 | chr5-125332651-INS-209 |
| chr5-125333558-INS-324 | chr5-125333558-INS-324 |
| chr5-125348636-INS-198 | chr5-125348636-INS-197 |
| chr5-125353534-INS-160 | chr5-125353534-INS-160 |
| chr5-125380802-INS-508 | chr5-125380802-INS-510 |
| chr5-125400528-INS-215 | chr5-125400528-INS-215 |
| chr5-125407493-INS-4518 | chr5-125407494-INS-4518 |
| chr5-125442635-INS-51 | chr5-125442636-INS-57 |
| chr5-125459774-INS-728 | chr5-125459774-INS-728 |
| chr5-125484904-INS-64 | chr5-125484934-INS-52 |
| chr5-125489515-INS-84 | chr5-125489514-INS-100 |
| chr5-125544138-INS-76 | chr5-125544138-INS-76 |
| chr5-125545055-INS-114 | chr5-125545055-INS-114 |
| chr5-125546476-INS-100 | chr5-125546476-INS-95 |
| chr5-125549844-INS-51 | chr5-125549845-INS-77 |
| chr5-125563439-INS-68 | chr5-125563439-INS-56 |
| chr5-125576257-INS-51 | chr5-125576257-INS-59 |
| chr5-125615280-INS-52 | chr5-125615280-INS-54 |
| chr5-125618607-INS-228 | chr5-125618608-INS-228 |
| chr5-125668588-INS-58 | chr5-125668588-INS-58 |
| chr5-125675801-INS-67 | chr5-125675802-INS-63 |
| chr5-125688557-INS-76 | chr5-125688557-INS-76 |
| chr5-125690191-INS-56 | chr5-125690191-INS-52 |
| chr5-125747421-INS-99 | chr5-125747421-INS-99 |
| chr5-125788916-INS-53 | chr5-125788916-INS-53 |
| chr5-125790143-INS-155 | chr5-125790143-INS-117 |
| chr5-125800954-INS-130 | chr5-125800954-INS-130 |
| chr5-125818945-INS-78 | chr5-125818900-INS-128 |
| chr5-125830267-INS-57 | chr5-125830267-INS-69 |
| chr5-125851845-INS-314 | chr5-125851845-INS-314 |
| chr5-125854715-INS-493 | chr5-125854715-INS-493 |
| chr5-125868893-INS-56 | chr5-125868893-INS-76 |
| chr5-125873374-INS-129 | chr5-125873373-INS-113 |
| chr5-125885561-INS-70 | chr5-125885561-INS-81 |
| chr5-125897802-INS-191 | chr5-125897802-INS-189 |
| chr5-125909022-INS-97 | chr5-125909022-INS-87 |
| chr5-125912933-INS-65 | chr5-125912933-INS-61 |
| chr5-125919797-INS-392 | chr5-125919798-INS-392 |
| chr5-125921997-INS-118 | chr5-125921997-INS-124 |
| chr5-125932372-INS-180 | chr5-125932372-INS-181 |
| chr5-125937018-INS-179 | chr5-125937018-INS-179 |
| chr5-125939950-INS-6935 | chr5-125939951-INS-6935 |
| chr5-125965250-INS-365 | chr5-125965250-INS-365 |
| chr5-125975115-INS-399 | chr5-125975125-INS-419 |
| chr5-125980227-INS-185 | chr5-125980227-INS-185 |
| chr5-125988102-INS-500 | chr5-125988102-INS-500 |

| | |
|---|---|
| chr5-125991637-INS-658 | chr5-125991637-INS-657 |
| chr5-125992392-INS-78 | chr5-125992392-INS-78 |
| chr5-126002259-INS-108 | chr5-126002247-INS-124 |
| chr5-126026510-INS-68 | chr5-126026510-INS-68 |
| chr5-126037361-INS-73 | chr5-126037361-INS-75 |
| chr5-126059653-INS-246 | chr5-126059653-INS-246 |
| chr5-126070359-INS-889 | chr5-126070357-INS-888 |
| chr5-126082657-INS-224 | chr5-126082657-INS-224 |
| chr5-126083999-INS-357 | chr5-126083999-INS-358 |
| chr5-126160126-INS-84 | chr5-126160126-INS-84 |
| chr5-126169415-INS-144 | chr5-126169415-INS-162 |
| chr5-126172194-INS-6327 | chr5-126172195-INS-6327 |
| chr5-126174585-INS-180 | chr5-126174585-INS-184 |
| chr5-126192286-INS-510 | chr5-126192286-INS-510 |
| chr5-126194384-INS-212 | chr5-126194388-INS-222 |
| chr5-126211891-INS-72 | chr5-126211889-INS-72 |
| chr5-126215942-INS-134 | chr5-126215942-INS-134 |
| chr5-126230002-INS-179 | chr5-126230002-INS-179 |
| chr5-126251704-INS-931 | chr5-126251704-INS-927 |
| chr5-126269528-INS-112 | chr5-126269528-INS-112 |
| chr5-126274992-INS-100 | chr5-126274992-INS-112 |
| chr5-126286805-INS-78 | chr5-126286805-INS-78 |
| chr5-126324232-INS-60 | chr5-126324232-INS-60 |
| chr5-126333444-INS-212 | chr5-126333444-INS-192 |
| chr5-126337144-INS-59 | chr5-126337144-INS-62 |
| chr5-126354633-INS-432 | chr5-126354633-INS-433 |
| chr5-126365022-INS-8184 | chr5-126365023-INS-8191 |
| chr5-126372104-INS-1563 | chr5-126372105-INS-1561 |
| chr5-126373642-INS-129 | chr5-126373640-INS-133 |
| chr5-126378106-INS-10795 | chr5-126378107-INS-10795 |
| chr5-126386532-INS-149 | chr5-126386532-INS-149 |
| chr5-126388744-INS-1062 | chr5-126388745-INS-1062 |
| chr5-126391414-INS-129 | chr5-126391434-INS-136 |
| chr5-126395393-INS-51 | chr5-126395393-INS-51 |
| chr5-126404067-INS-65 | chr5-126404067-INS-65 |
| chr5-126417818-INS-189 | chr5-126417818-INS-189 |
| chr5-126428439-INS-60 | chr5-126428439-INS-98 |
| chr5-126481937-INS-3115 | chr5-126481920-INS-3116 |
| chr5-126483614-INS-193 | chr5-126483614-INS-199 |
| chr5-126484892-INS-58 | chr5-126484892-INS-54 |
| chr5-126486069-INS-265 | chr5-126486069-INS-265 |
| chr5-126504791-INS-2613 | chr5-126504792-INS-2613 |
| chr5-126510997-INS-1153 | chr5-126510998-INS-1155 |
| chr5-126533267-INS-788 | chr5-126533267-INS-788 |
| chr5-126577786-INS-77 | chr5-126577786-INS-88 |
| chr5-126580690-INS-498 | chr5-126580690-INS-498 |
| chr5-126585472-INS-7392 | chr5-126585473-INS-7392 |
| chr5-126602507-INS-60 | chr5-126602507-INS-60 |
| chr5-126603905-INS-141 | chr5-126603905-INS-141 |
| chr5-126626694-INS-106 | chr5-126626694-INS-102 |
| chr5-126652741-INS-7158 | chr5-126652742-INS-7158 |
| chr5-126677198-INS-61 | chr5-126677200-INS-63 |
| chr5-126713862-INS-6395 | chr5-126713863-INS-6395 |
| chr5-126736321-INS-183 | chr5-126736327-INS-187 |
| chr5-126745803-INS-199 | chr5-126745803-INS-199 |
| chr5-126754634-INS-2046 | chr5-126754635-INS-2047 |
| chr5-126770660-INS-82 | chr5-126770660-INS-84 |
| chr5-126800154-INS-98 | chr5-126800178-INS-52 |
| chr5-126827161-INS-154 | chr5-126827161-INS-154 |
| chr5-126834317-INS-102 | chr5-126834297-INS-130 |
| chr5-126841906-INS-133 | chr5-126841895-INS-127 |
| chr5-126893432-INS-56 | chr5-126893432-INS-60 |
| chr5-126895295-INS-56 | chr5-126895295-INS-58 |
| chr5-126905833-INS-255 | chr5-126905833-INS-255 |
| chr5-126928154-INS-151 | chr5-126928154-INS-150 |
| chr5-126940629-INS-128 | chr5-126940629-INS-130 |
| chr5-126950514-INS-6453 | chr5-126950515-INS-6454 |
| chr5-126955516-INS-63 | chr5-126955516-INS-59 |
| chr5-126960100-INS-505 | chr5-126960100-INS-505 |
| chr5-126969644-INS-64 | chr5-126969644-INS-62 |
| chr5-126975128-INS-111 | chr5-126975128-INS-117 |
| chr5-126976273-INS-58 | chr5-126976272-INS-66 |
| chr5-126997589-INS-936 | chr5-126997604-INS-940 |
| chr5-127016207-INS-86 | chr5-127016207-INS-86 |
| chr5-127029879-INS-67 | chr5-127029879-INS-67 |
| chr5-127030646-INS-100 | chr5-127030646-INS-102 |
| chr5-127048386-INS-78 | chr5-127048386-INS-84 |

| | |
|---|---|
| chr5-127063704-INS-91 | chr5-127063704-INS-91 |
| chr5-127095755-INS-60 | chr5-127095755-INS-60 |
| chr5-127109223-INS-123 | chr5-127109223-INS-123 |
| chr5-127113201-INS-78 | chr5-127113201-INS-78 |
| chr5-127118360-INS-71 | chr5-127118360-INS-57 |
| chr5-127150097-INS-291 | chr5-127150097-INS-293 |
| chr5-127176809-INS-61 | chr5-127176809-INS-51 |
| chr5-127193951-INS-90 | chr5-127193951-INS-90 |
| chr5-127207247-INS-7095 | chr5-127207248-INS-7095 |
| chr5-127209143-INS-101 | chr5-127209143-INS-100 |
| chr5-127210197-INS-196 | chr5-127210197-INS-196 |
| chr5-127225854-INS-800 | chr5-127225854-INS-800 |
| chr5-127243251-INS-59 | chr5-127243251-INS-59 |
| chr5-127254152-INS-61 | chr5-127254152-INS-66 |
| chr5-127288329-INS-6195 | chr5-127288330-INS-6195 |
| chr5-127303268-INS-445 | chr5-127303268-INS-443 |
| chr5-127307900-INS-479 | chr5-127307900-INS-478 |
| chr5-127311022-INS-66 | chr5-127311022-INS-68 |
| chr5-127323839-INS-127 | chr5-127323839-INS-95 |
| chr5-127359008-INS-62 | chr5-127359008-INS-62 |
| chr5-127382767-INS-84 | chr5-127382767-INS-84 |
| chr5-127396840-INS-82 | chr5-127396840-INS-82 |
| chr5-127399128-INS-80 | chr5-127399128-INS-82 |
| chr5-127410758-INS-81 | chr5-127410738-INS-55 |
| chr5-127468682-INS-198 | chr5-127468682-INS-199 |
| chr5-127489511-INS-4072 | chr5-127489512-INS-4072 |
| chr5-127501844-INS-226 | chr5-127501844-INS-192 |
| chr5-127505955-INS-60 | chr5-127505955-INS-60 |
| chr5-127507281-INS-70 | chr5-127507244-INS-80 |
| chr5-127532810-INS-86 | chr5-127532811-INS-62 |
| chr5-127534644-INS-150 | chr5-127534632-INS-156 |
| chr5-127560051-INS-202 | chr5-127560051-INS-201 |
| chr5-127599704-INS-70 | chr5-127599704-INS-70 |
| chr5-127600656-INS-447 | chr5-127600656-INS-447 |
| chr5-127610872-INS-90 | chr5-127610872-INS-84 |
| chr5-127625851-INS-55 | chr5-127625889-INS-53 |
| chr5-127658501-INS-169 | chr5-127658501-INS-143 |
| chr5-127667509-INS-182 | chr5-127667509-INS-182 |
| chr5-127670020-INS-200 | chr5-127670020-INS-200 |
| chr5-127686257-INS-71 | chr5-127686257-INS-71 |
| chr5-127689948-INS-194 | chr5-127689953-INS-189 |
| chr5-127691397-INS-69 | chr5-127691397-INS-68 |
| chr5-127696245-INS-204 | chr5-127696245-INS-201 |
| chr5-127699705-INS-206 | chr5-127699705-INS-206 |
| chr5-127706428-INS-226 | chr5-127706428-INS-226 |
| chr5-127712846-INS-85 | chr5-127712846-INS-85 |
| chr5-127745863-INS-419 | chr5-127745863-INS-419 |
| chr5-127777814-INS-60 | chr5-127777814-INS-64 |
| chr5-127793842-INS-84 | chr5-127793844-INS-98 |
| chr5-127798610-INS-58 | chr5-127798610-INS-56 |
| chr5-127812294-INS-1892 | chr5-127812295-INS-1892 |
| chr5-127863755-INS-229 | chr5-127863755-INS-260 |
| chr5-127894786-INS-1652 | chr5-127894787-INS-1651 |
| chr5-127904657-INS-150 | chr5-127904679-INS-100 |
| chr5-127912184-INS-146 | chr5-127912177-INS-139 |
| chr5-127919620-INS-494 | chr5-127919618-INS-492 |
| chr5-127927003-INS-632 | chr5-127927003-INS-628 |
| chr5-127938508-INS-65 | chr5-127938508-INS-70 |
| chr5-127976650-INS-137 | chr5-127976658-INS-102 |
| chr5-127981694-INS-484 | chr5-127981694-INS-483 |
| chr5-128005589-INS-688 | chr5-128005589-INS-688 |
| chr5-128019917-INS-72 | chr5-128019917-INS-72 |
| chr5-128057322-INS-105 | chr5-128057322-INS-105 |
| chr5-128076351-INS-80 | chr5-128076351-INS-79 |
| chr5-128114583-INS-1784 | chr5-128114584-INS-1786 |
| chr5-128117980-INS-515 | chr5-128117980-INS-539 |
| chr5-128137325-INS-167 | chr5-128137300-INS-179 |
| chr5-128138020-INS-217 | chr5-128138020-INS-217 |
| chr5-128146673-INS-1194 | chr5-128146674-INS-1200 |
| chr5-128154907-INS-1135 | chr5-128154908-INS-1136 |
| chr5-128157658-INS-1602 | chr5-128157659-INS-1596 |
| chr5-128163203-INS-328 | chr5-128163203-INS-328 |
| chr5-128192371-INS-391 | chr5-128192371-INS-391 |
| chr5-128197783-INS-165 | chr5-128197783-INS-126 |
| chr5-128203966-INS-852 | chr5-128203934-INS-865 |
| chr5-128211577-INS-99 | chr5-128211577-INS-97 |
| chr5-128242490-INS-56 | chr5-128242488-INS-57 |

| | |
|---|---|
| chr5-128245493-INS-128 | chr5-128245493-INS-128 |
| chr5-128257755-INS-54 | chr5-128257755-INS-54 |
| chr5-128336253-INS-497 | chr5-128336253-INS-497 |
| chr5-128361828-INS-70 | chr5-128361824-INS-68 |
| chr5-128371799-INS-70 | chr5-128371797-INS-70 |
| chr5-128380674-INS-176 | chr5-128380674-INS-176 |
| chr5-128384501-INS-302 | chr5-128384501-INS-302 |
| chr5-128389669-INS-1892 | chr5-128389670-INS-1892 |
| chr5-128391895-INS-82 | chr5-128391895-INS-74 |
| chr5-128407944-INS-419 | chr5-128407944-INS-420 |
| chr5-128421960-INS-171 | chr5-128421960-INS-171 |
| chr5-128426159-INS-540 | chr5-128426159-INS-540 |
| chr5-128427134-INS-625 | chr5-128427134-INS-625 |
| chr5-128434924-INS-77 | chr5-128434924-INS-77 |
| chr5-128447921-INS-338 | chr5-128447921-INS-338 |
| chr5-128450406-INS-81 | chr5-128450406-INS-84 |
| chr5-128456436-INS-77 | chr5-128456425-INS-63 |
| chr5-128462893-INS-54 | chr5-128462893-INS-50 |
| chr5-128500695-INS-381 | chr5-128500695-INS-381 |
| chr5-128518003-INS-75 | chr5-128518003-INS-75 |
| chr5-128529937-INS-135 | chr5-128529938-INS-118 |
| chr5-128530710-INS-172 | chr5-128530711-INS-160 |
| chr5-128543098-INS-2953 | chr5-128543099-INS-2953 |
| chr5-128547615-INS-54 | chr5-128547615-INS-62 |
| chr5-128564666-INS-80 | chr5-128564666-INS-80 |
| chr5-128567581-INS-249 | chr5-128567581-INS-249 |
| chr5-128576380-INS-84 | chr5-128576380-INS-87 |
| chr5-128580075-INS-102 | chr5-128580073-INS-61 |
| chr5-128582895-INS-56 | chr5-128582895-INS-56 |
| chr5-128584088-INS-89 | chr5-128584088-INS-84 |
| chr5-128587912-INS-56 | chr5-128587912-INS-60 |
| chr5-128604148-INS-189 | chr5-128604148-INS-173 |
| chr5-128608600-INS-316 | chr5-128608600-INS-316 |
| chr5-128612118-INS-206 | chr5-128612118-INS-208 |
| chr5-128621796-INS-217 | chr5-128621796-INS-216 |
| chr5-128627081-INS-51 | chr5-128627081-INS-51 |
| chr5-128660921-INS-81 | chr5-128660921-INS-81 |
| chr5-128682580-INS-52 | chr5-128682580-INS-52 |
| chr5-128699460-INS-139 | chr5-128699458-INS-139 |
| chr5-128714760-INS-360 | chr5-128714760-INS-360 |
| chr5-128717218-INS-401 | chr5-128717218-INS-400 |
| chr5-128728455-INS-785 | chr5-128728455-INS-785 |
| chr5-128728995-INS-97 | chr5-128728995-INS-97 |
| chr5-128740768-INS-89 | chr5-128740768-INS-91 |
| chr5-128756862-INS-52 | chr5-128756862-INS-54 |
| chr5-128757452-INS-210 | chr5-128757452-INS-210 |
| chr5-128771720-INS-967 | chr5-128771720-INS-967 |
| chr5-128793847-INS-342 | chr5-128793847-INS-342 |
| chr5-128798732-INS-51 | chr5-128798732-INS-51 |
| chr5-128803865-INS-254 | chr5-128803865-INS-254 |
| chr5-128815621-INS-134 | chr5-128815621-INS-136 |
| chr5-128843031-INS-67 | chr5-128843031-INS-67 |
| chr5-128867646-INS-477 | chr5-128867646-INS-480 |
| chr5-128867996-INS-726 | chr5-128867973-INS-729 |
| chr5-128871721-INS-78 | chr5-128871745-INS-68 |
| chr5-128875601-INS-64 | chr5-128875601-INS-64 |
| chr5-128899791-INS-51 | chr5-128899791-INS-55 |
| chr5-128906617-INS-83 | chr5-128906617-INS-53 |
| chr5-128910901-INS-313 | chr5-128910901-INS-313 |
| chr5-128933397-INS-54 | chr5-128933397-INS-56 |
| chr5-128940851-INS-199 | chr5-128940851-INS-197 |
| chr5-128958021-INS-159 | chr5-128958018-INS-167 |
| chr5-128965944-INS-767 | chr5-128965944-INS-768 |
| chr5-128968265-INS-58 | chr5-128968284-INS-61 |
| chr5-128969288-INS-127 | chr5-128969288-INS-122 |
| chr5-128974241-INS-264 | chr5-128974241-INS-276 |
| chr5-128989037-INS-63 | chr5-128989037-INS-73 |
| chr5-129025441-INS-52 | chr5-129025441-INS-76 |
| chr5-129062083-INS-126 | chr5-129062083-INS-105 |
| chr5-129078651-INS-52 | chr5-129078651-INS-52 |
| chr5-129093037-INS-88 | chr5-129093037-INS-88 |
| chr5-129113035-INS-107 | chr5-129113035-INS-104 |
| chr5-129127278-INS-263 | chr5-129127278-INS-264 |
| chr5-129128902-INS-53 | chr5-129128871-INS-98 |
| chr5-129131661-INS-86 | chr5-129131661-INS-110 |
| chr5-129137401-INS-207 | chr5-129137401-INS-207 |
| chr5-129146481-INS-104 | chr5-129146481-INS-104 |

| | |
|---|---|
| chr5-129157371-INS-402 | chr5-129157371-INS-390 |
| chr5-129166638-INS-132 | chr5-129166638-INS-136 |
| chr5-129169344-INS-112 | chr5-129169344-INS-112 |
| chr5-129172021-INS-52 | chr5-129172021-INS-56 |
| chr5-129187401-INS-73 | chr5-129187401-INS-54 |
| chr5-129196679-INS-64 | chr5-129196680-INS-64 |
| chr5-129234247-INS-10859 | chr5-129234248-INS-10850 |
| chr5-129250388-INS-69 | chr5-129250388-INS-69 |
| chr5-129276633-INS-172 | chr5-129276623-INS-155 |
| chr5-129305135-INS-139 | chr5-129305135-INS-139 |
| chr5-129307059-INS-82 | chr5-129307059-INS-82 |
| chr5-129318109-INS-84 | chr5-129318109-INS-84 |
| chr5-129361787-INS-106 | chr5-129361763-INS-156 |
| chr5-129363636-INS-255 | chr5-129363636-INS-255 |
| chr5-129378351-INS-60 | chr5-129378351-INS-80 |
| chr5-129385512-INS-94 | chr5-129385512-INS-94 |
| chr5-129392747-INS-84 | chr5-129392747-INS-84 |
| chr5-129399751-INS-114 | chr5-129399754-INS-75 |
| chr5-129403047-INS-207 | chr5-129403047-INS-208 |
| chr5-129406424-INS-72 | chr5-129406424-INS-68 |
| chr5-129490880-INS-91 | chr5-129490880-INS-91 |
| chr5-129498369-INS-55 | chr5-129498369-INS-55 |
| chr5-129499756-INS-72 | chr5-129499761-INS-80 |
| chr5-129508912-INS-52 | chr5-129508912-INS-52 |
| chr5-129543264-INS-68 | chr5-129543301-INS-58 |
| chr5-129561054-INS-1863 | chr5-129561055-INS-1863 |
| chr5-129568775-INS-477 | chr5-129568775-INS-477 |
| chr5-129583404-INS-205 | chr5-129583404-INS-205 |
| chr5-129592224-INS-116 | chr5-129592238-INS-116 |
| chr5-129608028-INS-75 | chr5-129608028-INS-73 |
| chr5-129643168-INS-67 | chr5-129643168-INS-67 |
| chr5-129645556-INS-61 | chr5-129645556-INS-61 |
| chr5-129654440-INS-222 | chr5-129654441-INS-222 |
| chr5-129656417-INS-93 | chr5-129656417-INS-93 |
| chr5-129665549-INS-88 | chr5-129665549-INS-88 |
| chr5-129677199-INS-84 | chr5-129677199-INS-84 |
| chr5-129696811-INS-359 | chr5-129696811-INS-387 |
| chr5-129709392-INS-170 | chr5-129709391-INS-155 |
| chr5-129713410-INS-643 | chr5-129713410-INS-643 |
| chr5-129751690-INS-964 | chr5-129751690-INS-964 |
| chr5-129755703-INS-205 | chr5-129755703-INS-205 |
| chr5-129778412-INS-158 | chr5-129778443-INS-109 |
| chr5-129798448-INS-58 | chr5-129798449-INS-58 |
| chr5-129801310-INS-186 | chr5-129801310-INS-186 |
| chr5-129835578-INS-91 | chr5-129835578-INS-91 |
| chr5-129838231-INS-52 | chr5-129838264-INS-67 |
| chr5-129842975-INS-89 | chr5-129842975-INS-111 |
| chr5-129845063-INS-56 | chr5-129845063-INS-56 |
| chr5-129852181-INS-130 | chr5-129852181-INS-132 |
| chr5-129865873-INS-212 | chr5-129865872-INS-214 |
| chr5-129870540-INS-5715 | chr5-129870541-INS-5715 |
| chr5-129877383-INS-100 | chr5-129877383-INS-100 |
| chr5-129879821-INS-216 | chr5-129879821-INS-216 |
| chr5-129888274-INS-354 | chr5-129888275-INS-354 |
| chr5-129892794-INS-219 | chr5-129892795-INS-219 |
| chr5-129894649-INS-183 | chr5-129894649-INS-183 |
| chr5-129896898-INS-54 | chr5-129896896-INS-54 |
| chr5-129906544-INS-198 | chr5-129906555-INS-196 |
| chr5-129908114-INS-215 | chr5-129908114-INS-215 |
| chr5-129921656-INS-186 | chr5-129921656-INS-186 |
| chr5-129926884-INS-170 | chr5-129926886-INS-184 |
| chr5-129941223-INS-204 | chr5-129941223-INS-200 |
| chr5-129948842-INS-85 | chr5-129948842-INS-85 |
| chr5-129952234-INS-199 | chr5-129952234-INS-200 |
| chr5-129959308-INS-58 | chr5-129959308-INS-58 |
| chr5-130026036-INS-209 | chr5-130026036-INS-211 |
| chr5-130028471-INS-206 | chr5-130028471-INS-206 |
| chr5-130035464-INS-1045 | chr5-130035465-INS-1045 |
| chr5-130046015-INS-111 | chr5-130046019-INS-149 |
| chr5-130059719-INS-126 | chr5-130059719-INS-126 |
| chr5-130062566-INS-944 | chr5-130062591-INS-940 |
| chr5-130063319-INS-2934 | chr5-130063320-INS-2930 |
| chr5-130110984-INS-218 | chr5-130110984-INS-218 |
| chr5-130125743-INS-61 | chr5-130125743-INS-61 |
| chr5-130147957-INS-5797 | chr5-130147958-INS-5797 |
| chr5-130154682-INS-114 | chr5-130154668-INS-130 |
| chr5-130164873-INS-70 | chr5-130164860-INS-76 |

| | |
|---|---|
| chr5-130167472-INS-175 | chr5-130167473-INS-162 |
| chr5-130170427-INS-99 | chr5-130170427-INS-99 |
| chr5-130177272-INS-85 | chr5-130177272-INS-87 |
| chr5-130206183-INS-7110 | chr5-130206184-INS-7110 |
| chr5-130217990-INS-65 | chr5-130217988-INS-67 |
| chr5-130218497-INS-360 | chr5-130218497-INS-360 |
| chr5-130228261-INS-60 | chr5-130228248-INS-78 |
| chr5-130230711-INS-193 | chr5-130230711-INS-193 |
| chr5-130337510-INS-157 | chr5-130337510-INS-157 |
| chr5-130341648-INS-95 | chr5-130341654-INS-111 |
| chr5-130368036-INS-9045 | chr5-130368037-INS-9045 |
| chr5-130392159-INS-55 | chr5-130392159-INS-97 |
| chr5-130397872-INS-58 | chr5-130397833-INS-52 |
| chr5-130399476-INS-59 | chr5-130399476-INS-59 |
| chr5-130409208-INS-436 | chr5-130409203-INS-420 |
| chr5-130418093-INS-208 | chr5-130418093-INS-247 |
| chr5-130423680-INS-85 | chr5-130423680-INS-85 |
| chr5-130454520-INS-311 | chr5-130454520-INS-317 |
| chr5-130492906-INS-53 | chr5-130492906-INS-53 |
| chr5-130509862-INS-53 | chr5-130509862-INS-53 |
| chr5-130517106-INS-715 | chr5-130517103-INS-714 |
| chr5-130521155-INS-91 | chr5-130521155-INS-88 |
| chr5-130522699-INS-100 | chr5-130522699-INS-116 |
| chr5-130527488-INS-226 | chr5-130527488-INS-226 |
| chr5-130538571-INS-68 | chr5-130538548-INS-50 |
| chr5-130562553-INS-110 | chr5-130562553-INS-110 |
| chr5-130566867-INS-56 | chr5-130566867-INS-56 |
| chr5-130583985-INS-1258 | chr5-130583986-INS-1258 |
| chr5-130586243-INS-63 | chr5-130586243-INS-73 |
| chr5-130599397-INS-114 | chr5-130599397-INS-156 |
| chr5-130604015-INS-52 | chr5-130604015-INS-50 |
| chr5-130618584-INS-207 | chr5-130618584-INS-207 |
| chr5-130645700-INS-78 | chr5-130645709-INS-68 |
| chr5-130655583-INS-135 | chr5-130655608-INS-134 |
| chr5-130662415-INS-67 | chr5-130662428-INS-66 |
| chr5-130665005-INS-357 | chr5-130665005-INS-357 |
| chr5-130683154-INS-53 | chr5-130683154-INS-53 |
| chr5-130698085-INS-53 | chr5-130698115-INS-53 |
| chr5-130704146-INS-77 | chr5-130704156-INS-111 |
| chr5-130708825-INS-196 | chr5-130708825-INS-196 |
| chr5-130719796-INS-215 | chr5-130719796-INS-211 |
| chr5-130725574-INS-53 | chr5-130725574-INS-54 |
| chr5-130726649-INS-115 | chr5-130726649-INS-115 |
| chr5-130729768-INS-562 | chr5-130729768-INS-562 |
| chr5-130730590-INS-53 | chr5-130730596-INS-81 |
| chr5-130738586-INS-76 | chr5-130738594-INS-70 |
| chr5-130750108-INS-842 | chr5-130750108-INS-842 |
| chr5-130755870-INS-7723 | chr5-130755871-INS-7726 |
| chr5-130777895-INS-105 | chr5-130777847-INS-139 |
| chr5-130780049-INS-182 | chr5-130780049-INS-194 |
| chr5-130813053-INS-1188 | chr5-130813054-INS-1188 |
| chr5-130818803-INS-142 | chr5-130818803-INS-142 |
| chr5-130830938-INS-109 | chr5-130830944-INS-119 |
| chr5-130841351-INS-64 | chr5-130841351-INS-52 |
| chr5-130843583-INS-108 | chr5-130843583-INS-111 |
| chr5-130856023-INS-462 | chr5-130856023-INS-462 |
| chr5-130884464-INS-65 | chr5-130884445-INS-50 |
| chr5-130893004-INS-181 | chr5-130893004-INS-181 |
| chr5-130899654-INS-52 | chr5-130899654-INS-52 |
| chr5-130903567-INS-74 | chr5-130903567-INS-72 |
| chr5-130914040-INS-58 | chr5-130914040-INS-60 |
| chr5-130921417-INS-54 | chr5-130921417-INS-54 |
| chr5-130927390-INS-99 | chr5-130927380-INS-63 |
| chr5-130960391-INS-124 | chr5-130960391-INS-104 |
| chr5-130964451-INS-6223 | chr5-130964452-INS-6223 |
| chr5-130970169-INS-53 | chr5-130970169-INS-53 |
| chr5-130985021-INS-90 | chr5-130985021-INS-89 |
| chr5-130999818-INS-52 | chr5-130999818-INS-52 |
| chr5-131044870-INS-58 | chr5-131044870-INS-64 |
| chr5-131049358-INS-60 | chr5-131049354-INS-66 |
| chr5-131092509-INS-60 | chr5-131092509-INS-60 |
| chr5-131132992-INS-228 | chr5-131132992-INS-228 |
| chr5-131134086-INS-53 | chr5-131134086-INS-53 |
| chr5-131186190-INS-81 | chr5-131186190-INS-81 |
| chr5-131204109-INS-201 | chr5-131204109-INS-180 |
| chr5-131212978-INS-125 | chr5-131212979-INS-105 |
| chr5-131214710-INS-52 | chr5-131214674-INS-56 |

| | |
|---|---|
| chr5-131221560-INS-68 | chr5-131221560-INS-68 |
| chr5-131224846-INS-64 | chr5-131224846-INS-64 |
| chr5-131242569-INS-54 | chr5-131242569-INS-56 |
| chr5-131259184-INS-221 | chr5-131259184-INS-221 |
| chr5-131262098-INS-295 | chr5-131262098-INS-295 |
| chr5-131264048-INS-64 | chr5-131264048-INS-64 |
| chr5-131296641-INS-58 | chr5-131296639-INS-68 |
| chr5-131300763-INS-80 | chr5-131300763-INS-93 |
| chr5-131340888-INS-236 | chr5-131340888-INS-236 |
| chr5-131352225-INS-66 | chr5-131352225-INS-62 |
| chr5-131358289-INS-58 | chr5-131358289-INS-50 |
| chr5-131364120-INS-100 | chr5-131364160-INS-92 |
| chr5-131387578-INS-112 | chr5-131387563-INS-86 |
| chr5-131397990-INS-93 | chr5-131397991-INS-93 |
| chr5-131440098-INS-203 | chr5-131440098-INS-203 |
| chr5-131444696-INS-68 | chr5-131444696-INS-68 |
| chr5-131451598-INS-81 | chr5-131451598-INS-83 |
| chr5-131471463-INS-66 | chr5-131471463-INS-56 |
| chr5-131477209-INS-70 | chr5-131477221-INS-101 |
| chr5-131480636-INS-981 | chr5-131480636-INS-983 |
| chr5-131482965-INS-52 | chr5-131482965-INS-56 |
| chr5-131578568-INS-60 | chr5-131578564-INS-88 |
| chr5-131593944-INS-65 | chr5-131593944-INS-65 |
| chr5-131596910-INS-206 | chr5-131596910-INS-206 |
| chr5-131612714-INS-879 | chr5-131612714-INS-879 |
| chr5-131622835-INS-213 | chr5-131622835-INS-213 |
| chr5-131650796-INS-344 | chr5-131650798-INS-350 |
| chr5-131662497-INS-141 | chr5-131662498-INS-141 |
| chr5-131677378-INS-53 | chr5-131677378-INS-53 |
| chr5-131684806-INS-64 | chr5-131684811-INS-64 |
| chr5-131690469-INS-511 | chr5-131690470-INS-466 |
| chr5-131697637-INS-57 | chr5-131697637-INS-75 |
| chr5-131714919-INS-405 | chr5-131714916-INS-377 |
| chr5-131766906-INS-99 | chr5-131766906-INS-99 |
| chr5-131827967-INS-1219 | chr5-131827968-INS-1203 |
| chr5-131828509-INS-1425 | chr5-131828510-INS-1430 |
| chr5-131850752-INS-103 | chr5-131850752-INS-103 |
| chr5-131882226-INS-53 | chr5-131882226-INS-53 |
| chr5-131883587-INS-53 | chr5-131883587-INS-53 |
| chr5-131884884-INS-52 | chr5-131884884-INS-52 |
| chr5-131897802-INS-62 | chr5-131897802-INS-62 |
| chr5-131914142-INS-76 | chr5-131914142-INS-52 |
| chr5-131947740-INS-130 | chr5-131947740-INS-130 |
| chr5-131974550-INS-112 | chr5-131974550-INS-112 |
| chr5-131989584-INS-6401 | chr5-131989585-INS-6401 |
| chr5-132009332-INS-172 | chr5-132009332-INS-172 |
| chr5-132038222-INS-336 | chr5-132038222-INS-336 |
| chr5-132041443-INS-176 | chr5-132041430-INS-181 |
| chr5-132064570-INS-106 | chr5-132064577-INS-101 |
| chr5-132116457-INS-66 | chr5-132116457-INS-66 |
| chr5-132146980-INS-94 | chr5-132146980-INS-68 |
| chr5-132155072-INS-75 | chr5-132155072-INS-77 |
| chr5-132272787-INS-496 | chr5-132272787-INS-496 |
| chr5-132300693-INS-75 | chr5-132300693-INS-71 |
| chr5-132305958-INS-202 | chr5-132305958-INS-206 |
| chr5-132423315-INS-59 | chr5-132423315-INS-52 |
| chr5-132452328-INS-88 | chr5-132452328-INS-90 |
| chr5-132459556-INS-496 | chr5-132459556-INS-496 |
| chr5-132463852-INS-117 | chr5-132463852-INS-117 |
| chr5-132566847-INS-72 | chr5-132566847-INS-60 |
| chr5-132593114-INS-198 | chr5-132593114-INS-196 |
| chr5-132616011-INS-81 | chr5-132616011-INS-57 |
| chr5-132647399-INS-358 | chr5-132647399-INS-358 |
| chr5-132650027-INS-53 | chr5-132650027-INS-53 |
| chr5-132680881-INS-420 | chr5-132680881-INS-420 |
| chr5-132741915-INS-82 | chr5-132741925-INS-65 |
| chr5-132749986-INS-51 | chr5-132749986-INS-68 |
| chr5-132755717-INS-92 | chr5-132755669-INS-98 |
| chr5-132780134-INS-53 | chr5-132780134-INS-53 |
| chr5-132787013-INS-57 | chr5-132787013-INS-57 |
| chr5-132790516-INS-59 | chr5-132790516-INS-91 |
| chr5-132832144-INS-64 | chr5-132832144-INS-68 |
| chr5-132836827-INS-61 | chr5-132836840-INS-54 |
| chr5-132845908-INS-88 | chr5-132845908-INS-88 |
| chr5-132855872-INS-300 | chr5-132855850-INS-312 |
| chr5-132862448-INS-865 | chr5-132862448-INS-865 |
| chr5-132870986-INS-72 | chr5-132870986-INS-72 |

| | |
|---|---|
| chr5-132873108-INS-80 | chr5-132873102-INS-94 |
| chr5-132889319-INS-54 | chr5-132889319-INS-54 |
| chr5-132891136-INS-51 | chr5-132891150-INS-54 |
| chr5-132924149-INS-72 | chr5-132924150-INS-58 |
| chr5-132929025-INS-106 | chr5-132929025-INS-106 |
| chr5-132932500-INS-53 | chr5-132932500-INS-53 |
| chr5-132960831-INS-499 | chr5-132960831-INS-499 |
| chr5-132965290-INS-56 | chr5-132965290-INS-76 |
| chr5-132975979-INS-58 | chr5-132975979-INS-58 |
| chr5-132991899-INS-110 | chr5-132991900-INS-94 |
| chr5-133009656-INS-68 | chr5-133009648-INS-56 |
| chr5-133010149-INS-6310 | chr5-133010150-INS-6310 |
| chr5-133014814-INS-97 | chr5-133014784-INS-120 |
| chr5-133023823-INS-114 | chr5-133023823-INS-108 |
| chr5-133041446-INS-97 | chr5-133041446-INS-139 |
| chr5-133045323-INS-90 | chr5-133045317-INS-74 |
| chr5-133071995-INS-198 | chr5-133071995-INS-198 |
| chr5-133082379-INS-850 | chr5-133082378-INS-864 |
| chr5-133105160-INS-135 | chr5-133105156-INS-95 |
| chr5-133111974-INS-90 | chr5-133111974-INS-92 |
| chr5-133139975-INS-68 | chr5-133139993-INS-67 |
| chr5-133172604-INS-737 | chr5-133172586-INS-739 |
| chr5-133177248-INS-179 | chr5-133177250-INS-164 |
| chr5-133178333-INS-95 | chr5-133178333-INS-100 |
| chr5-133180657-INS-132 | chr5-133180657-INS-132 |
| chr5-133195204-INS-360 | chr5-133195204-INS-360 |
| chr5-133211286-INS-72 | chr5-133211286-INS-106 |
| chr5-133213489-INS-123 | chr5-133213489-INS-123 |
| chr5-133225260-INS-74 | chr5-133225260-INS-74 |
| chr5-133226725-INS-119 | chr5-133226725-INS-91 |
| chr5-133248308-INS-63 | chr5-133248308-INS-63 |
| chr5-133260151-INS-56 | chr5-133260151-INS-56 |
| chr5-133266676-INS-6362 | chr5-133266677-INS-6362 |
| chr5-133300958-INS-91 | chr5-133300960-INS-112 |
| chr5-133311114-INS-310 | chr5-133311117-INS-316 |
| chr5-133312578-INS-78 | chr5-133312578-INS-64 |
| chr5-133319424-INS-91 | chr5-133319424-INS-91 |
| chr5-133334246-INS-278 | chr5-133334246-INS-277 |
| chr5-133338929-INS-78 | chr5-133338929-INS-106 |
| chr5-133349796-INS-51 | chr5-133349796-INS-51 |
| chr5-133369898-INS-443 | chr5-133369898-INS-443 |
| chr5-133378381-INS-71 | chr5-133378381-INS-70 |
| chr5-133442139-INS-940 | chr5-133442138-INS-941 |
| chr5-133537869-INS-54 | chr5-133537869-INS-54 |
| chr5-133606096-INS-53 | chr5-133606096-INS-53 |
| chr5-133618376-INS-105 | chr5-133618376-INS-105 |
| chr5-133624374-INS-73 | chr5-133624374-INS-76 |
| chr5-133628533-INS-165 | chr5-133628517-INS-133 |
| chr5-133634872-INS-100 | chr5-133634872-INS-131 |
| chr5-133639559-INS-230 | chr5-133639559-INS-226 |
| chr5-133640934-INS-140 | chr5-133640934-INS-140 |
| chr5-133659386-INS-325 | chr5-133659375-INS-354 |
| chr5-133671344-INS-295 | chr5-133671344-INS-259 |
| chr5-133685485-INS-228 | chr5-133685486-INS-189 |
| chr5-133697490-INS-135 | chr5-133697490-INS-135 |
| chr5-133698330-INS-327 | chr5-133698330-INS-327 |
| chr5-133701103-INS-106 | chr5-133701103-INS-106 |
| chr5-133711569-INS-58 | chr5-133711569-INS-60 |
| chr5-133712166-INS-232 | chr5-133712166-INS-229 |
| chr5-133736184-INS-54 | chr5-133736184-INS-53 |
| chr5-133758947-INS-106 | chr5-133758947-INS-108 |
| chr5-133765252-INS-123 | chr5-133765252-INS-123 |
| chr5-133767638-INS-74 | chr5-133767612-INS-74 |
| chr5-133782367-INS-66 | chr5-133782367-INS-62 |
| chr5-133785068-INS-361 | chr5-133785068-INS-361 |
| chr5-133791127-INS-56 | chr5-133791127-INS-56 |
| chr5-133798133-INS-111 | chr5-133798138-INS-119 |
| chr5-133806248-INS-64 | chr5-133806253-INS-58 |
| chr5-133822188-INS-557 | chr5-133822188-INS-557 |
| chr5-133842089-INS-53 | chr5-133842089-INS-53 |
| chr5-133859080-INS-54 | chr5-133859074-INS-64 |
| chr5-133872125-INS-56 | chr5-133872125-INS-58 |
| chr5-133905798-INS-63 | chr5-133905802-INS-62 |
| chr5-133915887-INS-2029 | chr5-133915888-INS-2024 |
| chr5-133923268-INS-112 | chr5-133923268-INS-87 |
| chr5-133938538-INS-2271 | chr5-133938544-INS-2270 |
| chr5-133949192-INS-124 | chr5-133949192-INS-124 |

| | |
|---|---|
| chr5-133950643-INS-53 | chr5-133950643-INS-53 |
| chr5-133988695-INS-114 | chr5-133988695-INS-126 |
| chr5-133993200-INS-66 | chr5-133993200-INS-66 |
| chr5-134003997-INS-172 | chr5-134003997-INS-164 |
| chr5-134006108-INS-538 | chr5-134006108-INS-539 |
| chr5-134009912-INS-68 | chr5-134009912-INS-69 |
| chr5-134021632-INS-394 | chr5-134021632-INS-394 |
| chr5-134030436-INS-316 | chr5-134030436-INS-317 |
| chr5-134050293-INS-53 | chr5-134050294-INS-53 |
| chr5-134069877-INS-56 | chr5-134069878-INS-53 |
| chr5-134070461-INS-59 | chr5-134070462-INS-62 |
| chr5-134077615-INS-57 | chr5-134077615-INS-57 |
| chr5-134086665-INS-155 | chr5-134086665-INS-180 |
| chr5-134091946-INS-76 | chr5-134091946-INS-59 |
| chr5-134099325-INS-454 | chr5-134099325-INS-453 |
| chr5-134102046-INS-184 | chr5-134102046-INS-183 |
| chr5-134105305-INS-54 | chr5-134105305-INS-54 |
| chr5-134108269-INS-53 | chr5-134108269-INS-53 |
| chr5-134114373-INS-224 | chr5-134114373-INS-224 |
| chr5-134121667-INS-77 | chr5-134121667-INS-54 |
| chr5-134124336-INS-52 | chr5-134124329-INS-52 |
| chr5-134136770-INS-599 | chr5-134136770-INS-599 |
| chr5-134138775-INS-174 | chr5-134138775-INS-175 |
| chr5-134156369-INS-196 | chr5-134156369-INS-196 |
| chr5-134157266-INS-214 | chr5-134157266-INS-214 |
| chr5-134158711-INS-53 | chr5-134158711-INS-53 |
| chr5-134163178-INS-196 | chr5-134163178-INS-196 |
| chr5-134179268-INS-65 | chr5-134179268-INS-65 |
| chr5-134183646-INS-106 | chr5-134183646-INS-106 |
| chr5-134198861-INS-78 | chr5-134198896-INS-81 |
| chr5-134233837-INS-219 | chr5-134233837-INS-219 |
| chr5-134259357-INS-172 | chr5-134259357-INS-172 |
| chr5-134297365-INS-204 | chr5-134297365-INS-204 |
| chr5-134346570-INS-126 | chr5-134346570-INS-127 |
| chr5-134348685-INS-440 | chr5-134348685-INS-440 |
| chr5-134353608-INS-212 | chr5-134353608-INS-213 |
| chr5-134366877-INS-205 | chr5-134366877-INS-205 |
| chr5-134368771-INS-197 | chr5-134368771-INS-199 |
| chr5-134369845-INS-102 | chr5-134369845-INS-102 |
| chr5-134380308-INS-251 | chr5-134380308-INS-253 |
| chr5-134385472-INS-194 | chr5-134385465-INS-193 |
| chr5-134412042-INS-1114 | chr5-134412043-INS-1116 |
| chr5-134419309-INS-53 | chr5-134419309-INS-53 |
| chr5-134428900-INS-218 | chr5-134428900-INS-217 |
| chr5-134430362-INS-108 | chr5-134430362-INS-107 |
| chr5-134458148-INS-54 | chr5-134458148-INS-50 |
| chr5-134482890-INS-64 | chr5-134482895-INS-50 |
| chr5-134486736-INS-64 | chr5-134486736-INS-64 |
| chr5-134489675-INS-192 | chr5-134489675-INS-189 |
| chr5-134501200-INS-104 | chr5-134501200-INS-104 |
| chr5-134508333-INS-204 | chr5-134508333-INS-204 |
| chr5-134539404-INS-53 | chr5-134539404-INS-53 |
| chr5-134544178-INS-195 | chr5-134544178-INS-197 |
| chr5-134565267-INS-53 | chr5-134565267-INS-53 |
| chr5-134567940-INS-199 | chr5-134567940-INS-201 |
| chr5-134572596-INS-199 | chr5-134572596-INS-199 |
| chr5-134574801-INS-60 | chr5-134574830-INS-60 |
| chr5-134577553-INS-207 | chr5-134577553-INS-207 |
| chr5-134586918-INS-217 | chr5-134586918-INS-217 |
| chr5-134606149-INS-144 | chr5-134606149-INS-104 |
| chr5-134608571-INS-348 | chr5-134608571-INS-347 |
| chr5-134617651-INS-150 | chr5-134617651-INS-150 |
| chr5-134633245-INS-116 | chr5-134633252-INS-152 |
| chr5-134678917-INS-53 | chr5-134678920-INS-57 |
| chr5-134684354-INS-241 | chr5-134684354-INS-241 |
| chr5-134712622-INS-57 | chr5-134712622-INS-58 |
| chr5-134730831-INS-158 | chr5-134730831-INS-158 |
| chr5-134754816-INS-52 | chr5-134754787-INS-53 |
| chr5-134768641-INS-54 | chr5-134768641-INS-54 |
| chr5-134777922-INS-99 | chr5-134777922-INS-111 |
| chr5-134780243-INS-125 | chr5-134780243-INS-125 |
| chr5-134782461-INS-104 | chr5-134782461-INS-104 |
| chr5-134785738-INS-68 | chr5-134785738-INS-52 |
| chr5-134798859-INS-55 | chr5-134798859-INS-55 |
| chr5-134812751-INS-157 | chr5-134812751-INS-157 |
| chr5-134827434-INS-78 | chr5-134827434-INS-56 |
| chr5-134834608-INS-120 | chr5-134834608-INS-120 |

| | |
|---|---|
| chr5-134871418-INS-211 | chr5-134871418-INS-211 |
| chr5-134884117-INS-106 | chr5-134884117-INS-106 |
| chr5-134905122-INS-357 | chr5-134905122-INS-358 |
| chr5-134910450-INS-92 | chr5-134910493-INS-53 |
| chr5-134914804-INS-228 | chr5-134914804-INS-233 |
| chr5-134936669-INS-192 | chr5-134936669-INS-192 |
| chr5-134939835-INS-70 | chr5-134939833-INS-76 |
| chr5-134941429-INS-103 | chr5-134941405-INS-107 |
| chr5-134941541-INS-848 | chr5-134941542-INS-848 |
| chr5-134962624-INS-53 | chr5-134962624-INS-53 |
| chr5-134965534-INS-58 | chr5-134965534-INS-58 |
| chr5-134966514-INS-197 | chr5-134966514-INS-197 |
| chr5-134987382-INS-64 | chr5-134987382-INS-52 |
| chr5-135002946-INS-53 | chr5-135002946-INS-53 |
| chr5-135026352-INS-73 | chr5-135026352-INS-77 |
| chr5-135035691-INS-54 | chr5-135035691-INS-58 |
| chr5-135055713-INS-208 | chr5-135055713-INS-207 |
| chr5-135057125-INS-94 | chr5-135057165-INS-64 |
| chr5-135084315-INS-60 | chr5-135084315-INS-60 |
| chr5-135087524-INS-209 | chr5-135087524-INS-209 |
| chr5-135114431-INS-155 | chr5-135114431-INS-132 |
| chr5-135131839-INS-673 | chr5-135131839-INS-710 |
| chr5-135148538-INS-203 | chr5-135148539-INS-203 |
| chr5-135153192-INS-159 | chr5-135153192-INS-159 |
| chr5-135158112-INS-209 | chr5-135158112-INS-209 |
| chr5-135159741-INS-200 | chr5-135159741-INS-200 |
| chr5-135171215-INS-159 | chr5-135171208-INS-159 |
| chr5-135175103-INS-213 | chr5-135175103-INS-213 |
| chr5-135180533-INS-52 | chr5-135180533-INS-52 |
| chr5-135226914-INS-88 | chr5-135226942-INS-81 |
| chr5-135230775-INS-205 | chr5-135230775-INS-205 |
| chr5-135232374-INS-401 | chr5-135232374-INS-402 |
| chr5-135233123-INS-131 | chr5-135233123-INS-131 |
| chr5-135253503-INS-664 | chr5-135253503-INS-664 |
| chr5-135259958-INS-54 | chr5-135259958-INS-52 |
| chr5-135284822-INS-75 | chr5-135284812-INS-87 |
| chr5-135296097-INS-1088 | chr5-135296098-INS-1088 |
| chr5-135306318-INS-116 | chr5-135306277-INS-114 |
| chr5-135320231-INS-52 | chr5-135320231-INS-54 |
| chr5-135329410-INS-69 | chr5-135329417-INS-73 |
| chr5-135331522-INS-4945 | chr5-135331523-INS-4945 |
| chr5-135332049-INS-1284 | chr5-135332050-INS-1284 |
| chr5-135333675-INS-210 | chr5-135333675-INS-210 |
| chr5-135347877-INS-92 | chr5-135347877-INS-96 |
| chr5-135351066-INS-268 | chr5-135351066-INS-268 |
| chr5-135365891-INS-118 | chr5-135365889-INS-120 |
| chr5-135370626-INS-233 | chr5-135370626-INS-197 |
| chr5-135386228-INS-80 | chr5-135386228-INS-80 |
| chr5-135388266-INS-53 | chr5-135388266-INS-53 |
| chr5-135423682-INS-127 | chr5-135423682-INS-139 |
| chr5-135424609-INS-201 | chr5-135424609-INS-201 |
| chr5-135434359-INS-56 | chr5-135434359-INS-56 |
| chr5-135449707-INS-62 | chr5-135449670-INS-66 |
| chr5-135455398-INS-218 | chr5-135455398-INS-217 |
| chr5-135466750-INS-202 | chr5-135466751-INS-202 |
| chr5-135469799-INS-181 | chr5-135469799-INS-182 |
| chr5-135481419-INS-69 | chr5-135481419-INS-53 |
| chr5-135485037-INS-235 | chr5-135485037-INS-215 |
| chr5-135499314-INS-192 | chr5-135499314-INS-192 |
| chr5-135500315-INS-158 | chr5-135500315-INS-165 |
| chr5-135505636-INS-62 | chr5-135505636-INS-62 |
| chr5-135531419-INS-64 | chr5-135531421-INS-75 |
| chr5-135538612-INS-52 | chr5-135538612-INS-52 |
| chr5-135539652-INS-131 | chr5-135539652-INS-131 |
| chr5-135548744-INS-68 | chr5-135548744-INS-68 |
| chr5-135550219-INS-206 | chr5-135550219-INS-205 |
| chr5-135552033-INS-156 | chr5-135552033-INS-155 |
| chr5-135558798-INS-294 | chr5-135558798-INS-294 |
| chr5-135566053-INS-138 | chr5-135566053-INS-132 |
| chr5-135567930-INS-70 | chr5-135567896-INS-68 |
| chr5-135577149-INS-174 | chr5-135577149-INS-174 |
| chr5-135578325-INS-196 | chr5-135578325-INS-196 |
| chr5-135583086-INS-51 | chr5-135583095-INS-68 |
| chr5-135587354-INS-84 | chr5-135587354-INS-56 |
| chr5-135602637-INS-60 | chr5-135602637-INS-60 |
| chr5-135609742-INS-215 | chr5-135609742-INS-197 |
| chr5-135613074-INS-106 | chr5-135613074-INS-106 |

| | |
|---|---|
| chr5-135616550-INS-106 | chr5-135616550-INS-105 |
| chr5-135621271-INS-950 | chr5-135621271-INS-955 |
| chr5-135624290-INS-488 | chr5-135624290-INS-488 |
| chr5-135639792-INS-56 | chr5-135639792-INS-56 |
| chr5-135658071-INS-102 | chr5-135658072-INS-98 |
| chr5-135660714-INS-172 | chr5-135660714-INS-168 |
| chr5-135666138-INS-1501 | chr5-135666139-INS-1502 |
| chr5-135668515-INS-159 | chr5-135668515-INS-159 |
| chr5-135699774-INS-107 | chr5-135699774-INS-107 |
| chr5-135708683-INS-197 | chr5-135708683-INS-197 |
| chr5-135709462-INS-185 | chr5-135709462-INS-186 |
| chr5-135711122-INS-200 | chr5-135711122-INS-200 |
| chr5-135723863-INS-52 | chr5-135723863-INS-52 |
| chr5-135726894-INS-143 | chr5-135726894-INS-143 |
| chr5-135728348-INS-56 | chr5-135728348-INS-56 |
| chr5-135769242-INS-203 | chr5-135769242-INS-205 |
| chr5-135777130-INS-177 | chr5-135777115-INS-170 |
| chr5-135781186-INS-157 | chr5-135781186-INS-157 |
| chr5-135799063-INS-152 | chr5-135799063-INS-152 |
| chr5-135813453-INS-58 | chr5-135813453-INS-58 |
| chr5-135832282-INS-51 | chr5-135832286-INS-52 |
| chr5-135834492-INS-96 | chr5-135834492-INS-79 |
| chr5-135843478-INS-54 | chr5-135843478-INS-54 |
| chr5-135848238-INS-196 | chr5-135848238-INS-196 |
| chr5-135851930-INS-57 | chr5-135851930-INS-53 |
| chr5-135862798-INS-210 | chr5-135862798-INS-210 |
| chr5-135901119-INS-206 | chr5-135901119-INS-205 |
| chr5-135905330-INS-590 | chr5-135905330-INS-586 |
| chr5-135916396-INS-208 | chr5-135916396-INS-208 |
| chr5-135924201-INS-73 | chr5-135924201-INS-68 |
| chr5-135931495-INS-95 | chr5-135931495-INS-98 |
| chr5-135957553-INS-72 | chr5-135957553-INS-72 |
| chr5-135964463-INS-202 | chr5-135964463-INS-202 |
| chr5-135970093-INS-208 | chr5-135970093-INS-224 |
| chr5-135976277-INS-5565 | chr5-135976278-INS-5565 |
| chr5-135978929-INS-53 | chr5-135978929-INS-53 |
| chr5-135993667-INS-548 | chr5-135993664-INS-553 |
| chr5-136008088-INS-55 | chr5-136008088-INS-70 |
| chr5-136014135-INS-247 | chr5-136014135-INS-247 |
| chr5-136017521-INS-96 | chr5-136017521-INS-96 |
| chr5-136019191-INS-223 | chr5-136019191-INS-223 |
| chr5-136019852-INS-204 | chr5-136019852-INS-207 |
| chr5-136029567-INS-208 | chr5-136029567-INS-208 |
| chr5-136047045-INS-210 | chr5-136047045-INS-210 |
| chr5-136052218-INS-119 | chr5-136052192-INS-146 |
| chr5-136070963-INS-87 | chr5-136070963-INS-87 |
| chr5-136096381-INS-938 | chr5-136096381-INS-942 |
| chr5-136098202-INS-62 | chr5-136098200-INS-54 |
| chr5-136103066-INS-160 | chr5-136103066-INS-160 |
| chr5-136119046-INS-73 | chr5-136119034-INS-91 |
| chr5-136125691-INS-66 | chr5-136125659-INS-57 |
| chr5-136126738-INS-174 | chr5-136126738-INS-134 |
| chr5-136133937-INS-53 | chr5-136133937-INS-53 |
| chr5-136152880-INS-187 | chr5-136152880-INS-187 |
| chr5-136180164-INS-53 | chr5-136180164-INS-53 |
| chr5-136195727-INS-55 | chr5-136195748-INS-58 |
| chr5-136197898-INS-128 | chr5-136197898-INS-160 |
| chr5-136207651-INS-169 | chr5-136207651-INS-174 |
| chr5-136245724-INS-472 | chr5-136245724-INS-472 |
| chr5-136256710-INS-53 | chr5-136256710-INS-53 |
| chr5-136258434-INS-56 | chr5-136258434-INS-60 |
| chr5-136276688-INS-53 | chr5-136276693-INS-53 |
| chr5-136352708-INS-51 | chr5-136352712-INS-53 |
| chr5-136364591-INS-55 | chr5-136364591-INS-55 |
| chr5-136378297-INS-57 | chr5-136378297-INS-57 |
| chr5-136380116-INS-80 | chr5-136380116-INS-80 |
| chr5-136426440-INS-56 | chr5-136426440-INS-56 |
| chr5-136428491-INS-676 | chr5-136428491-INS-662 |
| chr5-136461568-INS-97 | chr5-136461568-INS-92 |
| chr5-136472383-INS-97 | chr5-136472383-INS-102 |
| chr5-136530591-INS-51 | chr5-136530591-INS-51 |
| chr5-136538225-INS-192 | chr5-136538225-INS-192 |
| chr5-136546910-INS-6403 | chr5-136546911-INS-6403 |
| chr5-136551216-INS-169 | chr5-136551221-INS-130 |
| chr5-136565263-INS-84 | chr5-136565267-INS-54 |
| chr5-136576345-INS-99 | chr5-136576341-INS-85 |
| chr5-136606608-INS-54 | chr5-136606614-INS-61 |

| | |
|---|---|
| chr5-136614357-INS-105 | chr5-136614357-INS-105 |
| chr5-136626447-INS-116 | chr5-136626447-INS-76 |
| chr5-136648094-INS-56 | chr5-136648109-INS-53 |
| chr5-136668102-INS-279 | chr5-136668102-INS-279 |
| chr5-136670352-INS-53 | chr5-136670352-INS-53 |
| chr5-136683502-INS-91 | chr5-136683502-INS-104 |
| chr5-136687609-INS-444 | chr5-136687609-INS-445 |
| chr5-136689711-INS-103 | chr5-136689744-INS-106 |
| chr5-136709507-INS-207 | chr5-136709507-INS-207 |
| chr5-136719907-INS-60 | chr5-136719907-INS-60 |
| chr5-136722892-INS-58 | chr5-136722892-INS-58 |
| chr5-136726611-INS-53 | chr5-136726611-INS-53 |
| chr5-136729019-INS-53 | chr5-136729057-INS-53 |
| chr5-136756161-INS-110 | chr5-136756161-INS-110 |
| chr5-136757454-INS-197 | chr5-136757454-INS-197 |
| chr5-136779546-INS-53 | chr5-136779546-INS-53 |
| chr5-136784639-INS-193 | chr5-136784639-INS-193 |
| chr5-136786200-INS-201 | chr5-136786200-INS-201 |
| chr5-136789516-INS-112 | chr5-136789521-INS-152 |
| chr5-136792305-INS-57 | chr5-136792305-INS-57 |
| chr5-136806247-INS-237 | chr5-136806247-INS-239 |
| chr5-136808031-INS-158 | chr5-136808032-INS-158 |
| chr5-136810041-INS-204 | chr5-136810041-INS-204 |
| chr5-136812759-INS-79 | chr5-136812759-INS-79 |
| chr5-136818713-INS-522 | chr5-136818713-INS-522 |
| chr5-136825530-INS-56 | chr5-136825488-INS-105 |
| chr5-136833563-INS-112 | chr5-136833563-INS-112 |
| chr5-136852967-INS-150 | chr5-136852970-INS-140 |
| chr5-136854270-INS-152 | chr5-136854271-INS-104 |
| chr5-136858591-INS-105 | chr5-136858591-INS-105 |
| chr5-136862225-INS-206 | chr5-136862225-INS-208 |
| chr5-136865995-INS-107 | chr5-136865995-INS-107 |
| chr5-136867088-INS-145 | chr5-136867088-INS-162 |
| chr5-136895197-INS-381 | chr5-136895198-INS-381 |
| chr5-136899334-INS-105 | chr5-136899352-INS-106 |
| chr5-136913633-INS-52 | chr5-136913633-INS-52 |
| chr5-136922564-INS-52 | chr5-136922564-INS-52 |
| chr5-136939139-INS-58 | chr5-136939174-INS-70 |
| chr5-136940311-INS-53 | chr5-136940311-INS-53 |
| chr5-136941022-INS-53 | chr5-136941059-INS-53 |
| chr5-136945677-INS-58 | chr5-136945685-INS-76 |
| chr5-136946878-INS-570 | chr5-136946878-INS-568 |
| chr5-136949415-INS-176 | chr5-136949415-INS-176 |
| chr5-136952178-INS-55 | chr5-136952153-INS-53 |
| chr5-136966967-INS-107 | chr5-136966967-INS-107 |
| chr5-137005697-INS-213 | chr5-137005697-INS-204 |
| chr5-137032200-INS-78 | chr5-137032200-INS-72 |
| chr5-137033540-INS-102 | chr5-137033540-INS-136 |
| chr5-137036868-INS-205 | chr5-137036868-INS-205 |
| chr5-137092663-INS-106 | chr5-137092663-INS-106 |
| chr5-137102440-INS-146 | chr5-137102450-INS-116 |
| chr5-137112867-INS-692 | chr5-137112867-INS-694 |
| chr5-137115810-INS-101 | chr5-137115810-INS-101 |
| chr5-137120232-INS-203 | chr5-137120232-INS-204 |
| chr5-137146012-INS-426 | chr5-137146012-INS-426 |
| chr5-137160173-INS-53 | chr5-137160173-INS-53 |
| chr5-137178802-INS-90 | chr5-137178802-INS-90 |
| chr5-137191776-INS-52 | chr5-137191776-INS-52 |
| chr5-137196359-INS-115 | chr5-137196359-INS-115 |
| chr5-137201768-INS-97 | chr5-137201775-INS-69 |
| chr5-137210609-INS-52 | chr5-137210609-INS-52 |
| chr5-137212146-INS-6440 | chr5-137212147-INS-6440 |
| chr5-137249159-INS-53 | chr5-137249159-INS-53 |
| chr5-137252763-INS-54 | chr5-137252763-INS-54 |
| chr5-137263568-INS-56 | chr5-137263568-INS-54 |
| chr5-137265030-INS-80 | chr5-137265030-INS-80 |
| chr5-137278527-INS-90 | chr5-137278527-INS-80 |
| chr5-137282779-INS-212 | chr5-137282780-INS-212 |
| chr5-137284176-INS-238 | chr5-137284176-INS-238 |
| chr5-137291237-INS-54 | chr5-137291237-INS-58 |
| chr5-137317845-INS-265 | chr5-137317845-INS-265 |
| chr5-137318666-INS-54 | chr5-137318666-INS-54 |
| chr5-137320231-INS-53 | chr5-137320231-INS-53 |
| chr5-137325322-INS-128 | chr5-137325322-INS-128 |
| chr5-137328778-INS-201 | chr5-137328778-INS-201 |
| chr5-137332661-INS-203 | chr5-137332661-INS-202 |
| chr5-137334344-INS-76 | chr5-137334344-INS-76 |

| | |
|---|---|
| chr5-137358871-INS-177 | chr5-137358871-INS-178 |
| chr5-137364702-INS-68 | chr5-137364702-INS-70 |
| chr5-137374495-INS-91 | chr5-137374495-INS-91 |
| chr5-137380113-INS-206 | chr5-137380113-INS-205 |
| chr5-137385615-INS-62 | chr5-137385615-INS-62 |
| chr5-137388443-INS-222 | chr5-137388430-INS-223 |
| chr5-137390868-INS-190 | chr5-137390868-INS-190 |
| chr5-137395315-INS-56 | chr5-137395315-INS-56 |
| chr5-137409193-INS-53 | chr5-137409193-INS-55 |
| chr5-137425661-INS-221 | chr5-137425661-INS-222 |
| chr5-137449418-INS-215 | chr5-137449418-INS-214 |
| chr5-137456471-INS-57 | chr5-137456471-INS-57 |
| chr5-137462190-INS-99 | chr5-137462190-INS-99 |
| chr5-137467754-INS-125 | chr5-137467760-INS-76 |
| chr5-137473555-INS-58 | chr5-137473583-INS-58 |
| chr5-137475989-INS-86 | chr5-137476032-INS-58 |
| chr5-137490468-INS-56 | chr5-137490468-INS-56 |
| chr5-137493177-INS-86 | chr5-137493180-INS-90 |
| chr5-137534168-INS-215 | chr5-137534168-INS-215 |
| chr5-137542682-INS-176 | chr5-137542686-INS-144 |
| chr5-137548993-INS-103 | chr5-137549043-INS-80 |
| chr5-137560696-INS-54 | chr5-137560696-INS-58 |
| chr5-137574327-INS-66 | chr5-137574328-INS-93 |
| chr5-137582947-INS-53 | chr5-137582932-INS-53 |
| chr5-137590492-INS-52 | chr5-137590492-INS-52 |
| chr5-137597013-INS-448 | chr5-137597013-INS-459 |
| chr5-137604767-INS-72 | chr5-137604767-INS-76 |
| chr5-137605961-INS-64 | chr5-137605961-INS-66 |
| chr5-137651179-INS-171 | chr5-137651140-INS-165 |
| chr5-137653165-INS-170 | chr5-137653158-INS-179 |
| chr5-137659199-INS-213 | chr5-137659199-INS-213 |
| chr5-137677059-INS-574 | chr5-137677059-INS-585 |
| chr5-137699807-INS-475 | chr5-137699807-INS-475 |
| chr5-137726484-INS-104 | chr5-137726484-INS-104 |
| chr5-137797568-INS-55 | chr5-137797568-INS-55 |
| chr5-137804377-INS-72 | chr5-137804377-INS-71 |
| chr5-137830705-INS-68 | chr5-137830705-INS-70 |
| chr5-137843339-INS-199 | chr5-137843325-INS-203 |
| chr5-137867804-INS-58 | chr5-137867804-INS-56 |
| chr5-137884787-INS-54 | chr5-137884787-INS-56 |
| chr5-137927600-INS-421 | chr5-137927600-INS-421 |
| chr5-137962046-INS-256 | chr5-137962084-INS-216 |
| chr5-138012441-INS-204 | chr5-138012441-INS-204 |
| chr5-138015643-INS-58 | chr5-138015598-INS-66 |
| chr5-138022531-INS-129 | chr5-138022531-INS-130 |
| chr5-138053513-INS-1292 | chr5-138053514-INS-1292 |
| chr5-138079436-INS-598 | chr5-138079436-INS-597 |
| chr5-138085953-INS-199 | chr5-138085953-INS-199 |
| chr5-138087786-INS-203 | chr5-138087786-INS-202 |
| chr5-138089473-INS-292 | chr5-138089474-INS-292 |
| chr5-138090870-INS-207 | chr5-138090870-INS-207 |
| chr5-138106231-INS-206 | chr5-138106231-INS-206 |
| chr5-138108623-INS-59 | chr5-138108642-INS-59 |
| chr5-138112222-INS-64 | chr5-138112222-INS-66 |
| chr5-138118281-INS-401 | chr5-138118276-INS-398 |
| chr5-138125469-INS-56 | chr5-138125469-INS-56 |
| chr5-138147930-INS-521 | chr5-138147931-INS-521 |
| chr5-138152552-INS-76 | chr5-138152552-INS-76 |
| chr5-138155129-INS-198 | chr5-138155129-INS-198 |
| chr5-138189017-INS-197 | chr5-138189017-INS-197 |
| chr5-138192168-INS-201 | chr5-138192168-INS-201 |
| chr5-138195766-INS-203 | chr5-138195766-INS-203 |
| chr5-138230914-INS-55 | chr5-138230914-INS-55 |
| chr5-138250128-INS-59 | chr5-138250128-INS-59 |
| chr5-138312441-INS-172 | chr5-138312441-INS-172 |
| chr5-138314385-INS-197 | chr5-138314385-INS-198 |
| chr5-138348842-INS-234 | chr5-138348842-INS-235 |
| chr5-138362310-INS-477 | chr5-138362310-INS-477 |
| chr5-138371761-INS-66 | chr5-138371761-INS-66 |
| chr5-138383227-INS-80 | chr5-138383227-INS-82 |
| chr5-138386459-INS-5507 | chr5-138386460-INS-5507 |
| chr5-138405181-INS-144 | chr5-138405181-INS-116 |
| chr5-138408128-INS-165 | chr5-138408128-INS-165 |
| chr5-138431658-INS-3327 | chr5-138431659-INS-3327 |
| chr5-138434627-INS-126 | chr5-138434627-INS-78 |
| chr5-138435883-INS-54 | chr5-138435883-INS-54 |
| chr5-138465864-INS-394 | chr5-138465864-INS-394 |

| | |
|---|---|
| chr5-138470962-INS-671 | chr5-138470962-INS-671 |
| chr5-138475624-INS-449 | chr5-138475624-INS-449 |
| chr5-138515909-INS-1394 | chr5-138515910-INS-1394 |
| chr5-138515966-INS-1891 | chr5-138515967-INS-1891 |
| chr5-138533801-INS-3246 | chr5-138533802-INS-3247 |
| chr5-138556851-INS-280 | chr5-138556851-INS-280 |
| chr5-138558186-INS-74 | chr5-138558186-INS-76 |
| chr5-138575241-INS-689 | chr5-138575241-INS-689 |
| chr5-138600447-INS-81 | chr5-138600447-INS-81 |
| chr5-138601485-INS-4713 | chr5-138601486-INS-4713 |
| chr5-138606433-INS-83 | chr5-138606433-INS-83 |
| chr5-138608303-INS-195 | chr5-138608303-INS-195 |
| chr5-138632532-INS-61 | chr5-138632532-INS-61 |
| chr5-138640217-INS-153 | chr5-138640217-INS-153 |
| chr5-138646714-INS-481 | chr5-138646714-INS-481 |
| chr5-138650987-INS-89 | chr5-138651001-INS-54 |
| chr5-138658660-INS-160 | chr5-138658660-INS-160 |
| chr5-138664057-INS-166 | chr5-138664057-INS-166 |
| chr5-138668277-INS-54 | chr5-138668277-INS-58 |
| chr5-138702038-INS-53 | chr5-138702038-INS-53 |
| chr5-138704380-INS-70 | chr5-138704362-INS-70 |
| chr5-138713374-INS-162 | chr5-138713374-INS-162 |
| chr5-138720507-INS-61 | chr5-138720507-INS-61 |
| chr5-138765168-INS-143 | chr5-138765180-INS-137 |
| chr5-138823159-INS-200 | chr5-138823159-INS-192 |
| chr5-138828471-INS-159 | chr5-138828471-INS-159 |
| chr5-138841879-INS-62 | chr5-138841879-INS-62 |
| chr5-138862552-INS-124 | chr5-138862552-INS-124 |
| chr5-138891045-INS-95 | chr5-138891045-INS-95 |
| chr5-138894317-INS-158 | chr5-138894317-INS-158 |
| chr5-138898281-INS-88 | chr5-138898281-INS-54 |
| chr5-138907585-INS-68 | chr5-138907580-INS-104 |
| chr5-138910186-INS-60 | chr5-138910187-INS-62 |
| chr5-138912169-INS-53 | chr5-138912169-INS-53 |
| chr5-138912431-INS-528 | chr5-138912431-INS-528 |
| chr5-138940935-INS-425 | chr5-138940936-INS-425 |
| chr5-138969248-INS-265 | chr5-138969248-INS-264 |
| chr5-139042609-INS-86 | chr5-139042609-INS-86 |
| chr5-139065721-INS-7319 | chr5-139065722-INS-7319 |
| chr5-139072087-INS-106 | chr5-139072092-INS-110 |
| chr5-139144275-INS-777 | chr5-139144275-INS-778 |
| chr5-139160528-INS-84 | chr5-139160527-INS-80 |
| chr5-139162566-INS-70 | chr5-139162566-INS-70 |
| chr5-139169544-INS-64 | chr5-139169544-INS-64 |
| chr5-139171554-INS-61 | chr5-139171554-INS-61 |
| chr5-139174572-INS-243 | chr5-139174572-INS-243 |
| chr5-139175582-INS-61 | chr5-139175582-INS-64 |
| chr5-139177124-INS-56 | chr5-139177124-INS-56 |
| chr5-139179611-INS-204 | chr5-139179611-INS-205 |
| chr5-139180786-INS-66 | chr5-139180786-INS-66 |
| chr5-139185737-INS-204 | chr5-139185737-INS-205 |
| chr5-139194708-INS-232 | chr5-139194691-INS-240 |
| chr5-139217094-INS-173 | chr5-139217098-INS-211 |
| chr5-139233036-INS-52 | chr5-139233036-INS-52 |
| chr5-139262960-INS-54 | chr5-139262960-INS-54 |
| chr5-139266600-INS-54 | chr5-139266600-INS-56 |
| chr5-139267532-INS-52 | chr5-139267532-INS-52 |
| chr5-139273386-INS-54 | chr5-139273386-INS-54 |
| chr5-139279432-INS-191 | chr5-139279432-INS-191 |
| chr5-139289015-INS-52 | chr5-139289015-INS-52 |
| chr5-139299218-INS-196 | chr5-139299212-INS-219 |
| chr5-139329880-INS-211 | chr5-139329880-INS-212 |
| chr5-139338857-INS-96 | chr5-139338857-INS-108 |
| chr5-139353521-INS-181 | chr5-139353521-INS-177 |
| chr5-139358291-INS-202 | chr5-139358336-INS-188 |
| chr5-139359419-INS-56 | chr5-139359419-INS-84 |
| chr5-139378141-INS-73 | chr5-139378141-INS-73 |
| chr5-139381580-INS-204 | chr5-139381580-INS-204 |
| chr5-139423987-INS-105 | chr5-139423987-INS-105 |
| chr5-139441729-INS-211 | chr5-139441729-INS-210 |
| chr5-139444216-INS-231 | chr5-139444216-INS-216 |
| chr5-139450349-INS-234 | chr5-139450349-INS-235 |
| chr5-139467868-INS-57 | chr5-139467868-INS-57 |
| chr5-139476629-INS-104 | chr5-139476629-INS-104 |
| chr5-139478091-INS-52 | chr5-139478091-INS-68 |
| chr5-139489180-INS-166 | chr5-139489180-INS-168 |
| chr5-139493543-INS-230 | chr5-139493555-INS-215 |

| | |
|---|---|
| chr5-139497973-INS-126 | chr5-139497973-INS-117 |
| chr5-139503887-INS-1895 | chr5-139503888-INS-1895 |
| chr5-139507102-INS-96 | chr5-139507100-INS-76 |
| chr5-139508557-INS-52 | chr5-139508557-INS-52 |
| chr5-139513866-INS-53 | chr5-139513866-INS-53 |
| chr5-139516051-INS-62 | chr5-139516057-INS-54 |
| chr5-139560474-INS-52 | chr5-139560474-INS-52 |
| chr5-139566588-INS-88 | chr5-139566588-INS-92 |
| chr5-139574228-INS-105 | chr5-139574228-INS-105 |
| chr5-139595078-INS-67 | chr5-139595078-INS-67 |
| chr5-139616738-INS-62 | chr5-139616734-INS-50 |
| chr5-139620105-INS-211 | chr5-139620105-INS-211 |
| chr5-139636148-INS-69 | chr5-139636122-INS-60 |
| chr5-139675525-INS-206 | chr5-139675534-INS-208 |
| chr5-139705925-INS-159 | chr5-139705925-INS-159 |
| chr5-139714479-INS-90 | chr5-139714479-INS-50 |
| chr5-139739113-INS-52 | chr5-139739113-INS-54 |
| chr5-139741577-INS-84 | chr5-139741577-INS-86 |
| chr5-139749452-INS-67 | chr5-139749452-INS-67 |
| chr5-139765699-INS-233 | chr5-139765699-INS-204 |
| chr5-139806272-INS-286 | chr5-139806274-INS-299 |
| chr5-139818254-INS-58 | chr5-139818254-INS-58 |
| chr5-139819209-INS-122 | chr5-139819240-INS-87 |
| chr5-139823308-INS-62 | chr5-139823308-INS-61 |
| chr5-139837082-INS-158 | chr5-139837082-INS-158 |
| chr5-139840496-INS-98 | chr5-139840490-INS-100 |
| chr5-139851296-INS-194 | chr5-139851296-INS-194 |
| chr5-139883835-INS-52 | chr5-139883835-INS-53 |
| chr5-139890489-INS-53 | chr5-139890489-INS-53 |
| chr5-139907335-INS-205 | chr5-139907335-INS-205 |
| chr5-139914682-INS-131 | chr5-139914660-INS-171 |
| chr5-139922868-INS-211 | chr5-139922868-INS-211 |
| chr5-139972010-INS-62 | chr5-139972028-INS-80 |
| chr5-139972665-INS-59 | chr5-139972665-INS-65 |
| chr5-139975114-INS-205 | chr5-139975114-INS-205 |
| chr5-139979427-INS-54 | chr5-139979427-INS-54 |
| chr5-139980278-INS-60 | chr5-139980278-INS-60 |
| chr5-139983181-INS-154 | chr5-139983181-INS-166 |
| chr5-139992435-INS-54 | chr5-139992435-INS-54 |
| chr5-140025849-INS-215 | chr5-140025849-INS-215 |
| chr5-140033059-INS-96 | chr5-140033059-INS-96 |
| chr5-140052359-INS-62 | chr5-140052359-INS-64 |
| chr5-140094589-INS-195 | chr5-140094589-INS-195 |
| chr5-140112961-INS-520 | chr5-140112961-INS-520 |
| chr5-140139908-INS-52 | chr5-140139908-INS-52 |
| chr5-140142236-INS-56 | chr5-140142236-INS-50 |
| chr5-140158757-INS-62 | chr5-140158757-INS-62 |
| chr5-140188319-INS-674 | chr5-140188331-INS-684 |
| chr5-140189937-INS-390 | chr5-140189937-INS-370 |
| chr5-140191844-INS-68 | chr5-140191844-INS-60 |
| chr5-140222986-INS-206 | chr5-140222986-INS-207 |
| chr5-140233404-INS-518 | chr5-140233404-INS-518 |
| chr5-140252101-INS-129 | chr5-140252101-INS-129 |
| chr5-140295204-INS-74 | chr5-140295204-INS-50 |
| chr5-140299720-INS-80 | chr5-140299753-INS-56 |
| chr5-140308044-INS-204 | chr5-140308044-INS-204 |
| chr5-140310527-INS-211 | chr5-140310527-INS-212 |
| chr5-140333972-INS-165 | chr5-140333972-INS-116 |
| chr5-140344523-INS-209 | chr5-140344523-INS-216 |
| chr5-140352204-INS-208 | chr5-140352205-INS-191 |
| chr5-140363321-INS-74 | chr5-140363321-INS-74 |
| chr5-140370310-INS-212 | chr5-140370311-INS-209 |
| chr5-140386644-INS-64 | chr5-140386644-INS-64 |
| chr5-140390079-INS-202 | chr5-140390079-INS-203 |
| chr5-140390853-INS-55 | chr5-140390853-INS-66 |
| chr5-140398787-INS-4818 | chr5-140398788-INS-4818 |
| chr5-140405782-INS-66 | chr5-140405782-INS-67 |
| chr5-140410634-INS-106 | chr5-140410634-INS-106 |
| chr5-140420364-INS-93 | chr5-140420364-INS-56 |
| chr5-140423555-INS-63 | chr5-140423563-INS-71 |
| chr5-140424608-INS-198 | chr5-140424608-INS-198 |
| chr5-140433402-INS-456 | chr5-140433402-INS-456 |
| chr5-140446172-INS-252 | chr5-140446172-INS-251 |
| chr5-140448607-INS-64 | chr5-140448607-INS-70 |
| chr5-140458042-INS-147 | chr5-140458042-INS-147 |
| chr5-140463805-INS-60 | chr5-140463805-INS-60 |
| chr5-140465471-INS-87 | chr5-140465471-INS-97 |

| | |
|---|---|
| chr5-140469900-INS-435 | chr5-140469900-INS-435 |
| chr5-140478522-INS-236 | chr5-140478522-INS-243 |
| chr5-140513324-INS-52 | chr5-140513282-INS-52 |
| chr5-140521221-INS-59 | chr5-140521221-INS-71 |
| chr5-140523583-INS-177 | chr5-140523583-INS-177 |
| chr5-140532596-INS-200 | chr5-140532596-INS-200 |
| chr5-140540420-INS-72 | chr5-140540420-INS-72 |
| chr5-140567967-INS-51 | chr5-140567967-INS-51 |
| chr5-140576649-INS-52 | chr5-140576649-INS-52 |
| chr5-140606021-INS-241 | chr5-140606021-INS-247 |
| chr5-140625172-INS-198 | chr5-140625172-INS-197 |
| chr5-140641169-INS-222 | chr5-140641169-INS-222 |
| chr5-140665759-INS-209 | chr5-140665759-INS-207 |
| chr5-140667394-INS-51 | chr5-140667394-INS-51 |
| chr5-140667445-INS-580 | chr5-140667445-INS-580 |
| chr5-140684564-INS-104 | chr5-140684564-INS-104 |
| chr5-140701414-INS-220 | chr5-140701414-INS-220 |
| chr5-140725096-INS-53 | chr5-140725096-INS-53 |
| chr5-140728432-INS-98 | chr5-140728432-INS-84 |
| chr5-140729387-INS-207 | chr5-140729387-INS-207 |
| chr5-140731325-INS-53 | chr5-140731325-INS-53 |
| chr5-140731743-INS-189 | chr5-140731743-INS-191 |
| chr5-140773279-INS-217 | chr5-140773279-INS-217 |
| chr5-140773332-INS-6452 | chr5-140773333-INS-6452 |
| chr5-140779166-INS-180 | chr5-140779166-INS-177 |
| chr5-140833333-INS-56 | chr5-140833333-INS-58 |
| chr5-140872019-INS-83 | chr5-140872004-INS-57 |
| chr5-140874555-INS-104 | chr5-140874555-INS-104 |
| chr5-140884403-INS-53 | chr5-140884403-INS-53 |
| chr5-140892149-INS-82 | chr5-140892153-INS-116 |
| chr5-140897439-INS-53 | chr5-140897439-INS-53 |
| chr5-140903979-INS-209 | chr5-140903979-INS-228 |
| chr5-140931661-INS-59 | chr5-140931661-INS-50 |
| chr5-140936793-INS-56 | chr5-140936794-INS-60 |
| chr5-140941687-INS-70 | chr5-140941687-INS-70 |
| chr5-140943574-INS-60 | chr5-140943570-INS-60 |
| chr5-140951987-INS-53 | chr5-140951987-INS-53 |
| chr5-140969446-INS-79 | chr5-140969468-INS-106 |
| chr5-140983010-INS-106 | chr5-140983010-INS-106 |
| chr5-140997055-INS-52 | chr5-140997055-INS-50 |
| chr5-141014674-INS-124 | chr5-141014675-INS-96 |
| chr5-141065980-INS-99 | chr5-141065972-INS-100 |
| chr5-141066985-INS-127 | chr5-141066984-INS-89 |
| chr5-141079034-INS-53 | chr5-141079034-INS-53 |
| chr5-141084211-INS-55 | chr5-141084208-INS-53 |
| chr5-141088759-INS-264 | chr5-141088759-INS-264 |
| chr5-141090001-INS-4877 | chr5-141090002-INS-4881 |
| chr5-141112151-INS-582 | chr5-141112161-INS-577 |
| chr5-141124142-INS-53 | chr5-141124142-INS-53 |
| chr5-141128214-INS-121 | chr5-141128223-INS-93 |
| chr5-141164101-INS-162 | chr5-141164095-INS-175 |
| chr5-141172196-INS-119 | chr5-141172196-INS-127 |
| chr5-141177975-INS-137 | chr5-141177974-INS-114 |
| chr5-141185770-INS-60 | chr5-141185770-INS-60 |
| chr5-141241684-INS-1060 | chr5-141241685-INS-1060 |
| chr5-141246171-INS-6365 | chr5-141246172-INS-6365 |
| chr5-141272304-INS-1659 | chr5-141272305-INS-1659 |
| chr5-141291037-INS-84 | chr5-141291063-INS-76 |
| chr5-141306020-INS-73 | chr5-141306020-INS-73 |
| chr5-141380714-INS-78 | chr5-141380714-INS-62 |
| chr5-141412239-INS-311 | chr5-141412239-INS-311 |
| chr5-141413325-INS-58 | chr5-141413325-INS-68 |
| chr5-141431464-INS-228 | chr5-141431464-INS-231 |
| chr5-141441186-INS-138 | chr5-141441184-INS-140 |
| chr5-141463527-INS-73 | chr5-141463527-INS-71 |
| chr5-141493945-INS-191 | chr5-141493945-INS-191 |
| chr5-141537330-INS-229 | chr5-141537330-INS-234 |
| chr5-141543302-INS-68 | chr5-141543293-INS-64 |
| chr5-141567034-INS-208 | chr5-141567033-INS-182 |
| chr5-141594959-INS-54 | chr5-141594959-INS-54 |
| chr5-141608000-INS-168 | chr5-141608000-INS-181 |
| chr5-141619778-INS-498 | chr5-141619778-INS-505 |
| chr5-141622361-INS-142 | chr5-141622366-INS-108 |
| chr5-141624505-INS-60 | chr5-141624505-INS-65 |
| chr5-141686318-INS-52 | chr5-141686318-INS-52 |
| chr5-141693832-INS-52 | chr5-141693834-INS-50 |
| chr5-141699931-INS-56 | chr5-141699931-INS-66 |

| | |
|---|---|
| chr5-141723289-INS-135 | chr5-141723289-INS-135 |
| chr5-141750234-INS-164 | chr5-141750234-INS-164 |
| chr5-141754072-INS-350 | chr5-141754071-INS-301 |
| chr5-141783088-INS-64 | chr5-141783088-INS-60 |
| chr5-141801412-INS-378 | chr5-141801412-INS-389 |
| chr5-141802292-INS-54 | chr5-141802305-INS-54 |
| chr5-141808086-INS-58 | chr5-141808084-INS-62 |
| chr5-141828616-INS-58 | chr5-141828621-INS-50 |
| chr5-141859328-INS-173 | chr5-141859328-INS-173 |
| chr5-141886275-INS-210 | chr5-141886321-INS-210 |
| chr5-141915556-INS-201 | chr5-141915556-INS-201 |
| chr5-141920788-INS-7108 | chr5-141920789-INS-7087 |
| chr5-141956674-INS-78 | chr5-141956674-INS-60 |
| chr5-141972675-INS-52 | chr5-141972632-INS-102 |
| chr5-141973833-INS-124 | chr5-141973833-INS-128 |
| chr5-141983239-INS-54 | chr5-141983239-INS-54 |
| chr5-142027939-INS-52 | chr5-142027939-INS-54 |
| chr5-142056018-INS-416 | chr5-142056018-INS-416 |
| chr5-142069087-INS-53 | chr5-142069087-INS-53 |
| chr5-142094875-INS-107 | chr5-142094875-INS-107 |
| chr5-142126807-INS-66 | chr5-142126807-INS-54 |
| chr5-142133287-INS-73 | chr5-142133287-INS-73 |
| chr5-142143479-INS-52 | chr5-142143479-INS-52 |
| chr5-142144873-INS-63 | chr5-142144873-INS-61 |
| chr5-142145156-INS-179 | chr5-142145156-INS-179 |
| chr5-142154530-INS-52 | chr5-142154530-INS-52 |
| chr5-142163085-INS-323 | chr5-142163085-INS-322 |
| chr5-142182979-INS-200 | chr5-142182979-INS-200 |
| chr5-142186973-INS-69 | chr5-142186973-INS-69 |
| chr5-142190357-INS-53 | chr5-142190366-INS-65 |
| chr5-142197150-INS-499 | chr5-142197150-INS-499 |
| chr5-142211793-INS-498 | chr5-142211793-INS-498 |
| chr5-142226066-INS-407 | chr5-142226066-INS-408 |
| chr5-142237839-INS-77 | chr5-142237839-INS-77 |
| chr5-142244997-INS-53 | chr5-142244997-INS-53 |
| chr5-142245946-INS-165 | chr5-142245947-INS-132 |
| chr5-142257675-INS-522 | chr5-142257675-INS-522 |
| chr5-142272847-INS-136 | chr5-142272847-INS-136 |
| chr5-142279173-INS-106 | chr5-142279173-INS-106 |
| chr5-142312921-INS-52 | chr5-142312921-INS-56 |
| chr5-142332990-INS-53 | chr5-142332990-INS-51 |
| chr5-142338143-INS-51 | chr5-142338143-INS-51 |
| chr5-142340741-INS-823 | chr5-142340741-INS-826 |
| chr5-142365223-INS-58 | chr5-142365223-INS-58 |
| chr5-142390661-INS-59 | chr5-142390652-INS-59 |
| chr5-142391826-INS-53 | chr5-142391826-INS-53 |
| chr5-142396887-INS-5549 | chr5-142396888-INS-5549 |
| chr5-142409969-INS-213 | chr5-142409969-INS-214 |
| chr5-142412559-INS-58 | chr5-142412559-INS-54 |
| chr5-142417056-INS-108 | chr5-142417056-INS-108 |
| chr5-142430853-INS-373 | chr5-142430854-INS-375 |
| chr5-142473588-INS-136 | chr5-142473588-INS-128 |
| chr5-142484518-INS-105 | chr5-142484518-INS-105 |
| chr5-142503262-INS-53 | chr5-142503262-INS-53 |
| chr5-142507707-INS-56 | chr5-142507707-INS-56 |
| chr5-142520769-INS-498 | chr5-142520769-INS-514 |
| chr5-142523105-INS-243 | chr5-142523093-INS-250 |
| chr5-142546754-INS-107 | chr5-142546754-INS-107 |
| chr5-142569135-INS-54 | chr5-142569135-INS-54 |
| chr5-142612156-INS-5549 | chr5-142612157-INS-5549 |
| chr5-142632087-INS-324 | chr5-142632087-INS-324 |
| chr5-142644412-INS-70 | chr5-142644364-INS-72 |
| chr5-142657182-INS-177 | chr5-142657182-INS-185 |
| chr5-142663011-INS-57 | chr5-142663011-INS-50 |
| chr5-142674466-INS-106 | chr5-142674466-INS-106 |
| chr5-142677493-INS-53 | chr5-142677493-INS-53 |
| chr5-142705219-INS-51 | chr5-142705219-INS-51 |
| chr5-142718974-INS-475 | chr5-142718974-INS-475 |
| chr5-142727300-INS-211 | chr5-142727300-INS-211 |
| chr5-142728508-INS-800 | chr5-142728508-INS-800 |
| chr5-142728527-INS-208 | chr5-142728526-INS-202 |
| chr5-142752231-INS-221 | chr5-142752231-INS-219 |
| chr5-142754177-INS-181 | chr5-142754177-INS-182 |
| chr5-142771566-INS-54 | chr5-142771566-INS-54 |
| chr5-142789403-INS-190 | chr5-142789403-INS-190 |
| chr5-142792452-INS-196 | chr5-142792452-INS-196 |
| chr5-142806592-INS-56 | chr5-142806592-INS-56 |

| | |
|---|---|
| chr5-142816156-INS-58 | chr5-142816156-INS-77 |
| chr5-142821650-INS-70 | chr5-142821637-INS-88 |
| chr5-142822477-INS-194 | chr5-142822477-INS-194 |
| chr5-142825689-INS-108 | chr5-142825689-INS-108 |
| chr5-142832340-INS-203 | chr5-142832340-INS-203 |
| chr5-142834544-INS-66 | chr5-142834544-INS-66 |
| chr5-142839446-INS-70 | chr5-142839446-INS-63 |
| chr5-142844749-INS-52 | chr5-142844749-INS-52 |
| chr5-142846712-INS-55 | chr5-142846712-INS-63 |
| chr5-142851842-INS-211 | chr5-142851842-INS-211 |
| chr5-142883651-INS-220 | chr5-142883651-INS-221 |
| chr5-142915507-INS-51 | chr5-142915507-INS-51 |
| chr5-142917707-INS-157 | chr5-142917707-INS-157 |
| chr5-142921114-INS-198 | chr5-142921114-INS-198 |
| chr5-142924706-INS-54 | chr5-142924706-INS-50 |
| chr5-142929524-INS-235 | chr5-142929525-INS-198 |
| chr5-142932665-INS-207 | chr5-142932665-INS-207 |
| chr5-142942283-INS-176 | chr5-142942283-INS-177 |
| chr5-142951170-INS-52 | chr5-142951170-INS-52 |
| chr5-142957159-INS-53 | chr5-142957159-INS-53 |
| chr5-142961951-INS-180 | chr5-142961951-INS-182 |
| chr5-142969255-INS-131 | chr5-142969255-INS-142 |
| chr5-142984735-INS-230 | chr5-142984735-INS-228 |
| chr5-142997590-INS-234 | chr5-142997589-INS-221 |
| chr5-143005087-INS-209 | chr5-143005087-INS-209 |
| chr5-143025112-INS-67 | chr5-143025112-INS-67 |
| chr5-143042921-INS-57 | chr5-143042921-INS-57 |
| chr5-143049811-INS-210 | chr5-143049811-INS-210 |
| chr5-143112287-INS-63 | chr5-143112292-INS-67 |
| chr5-143112949-INS-64 | chr5-143112949-INS-64 |
| chr5-143135974-INS-154 | chr5-143135974-INS-156 |
| chr5-143141540-INS-559 | chr5-143141540-INS-559 |
| chr5-143145018-INS-113 | chr5-143145018-INS-113 |
| chr5-143161250-INS-54 | chr5-143161250-INS-54 |
| chr5-143180259-INS-60 | chr5-143180259-INS-62 |
| chr5-143182975-INS-453 | chr5-143182975-INS-435 |
| chr5-143186546-INS-430 | chr5-143186546-INS-430 |
| chr5-143193129-INS-3454 | chr5-143193130-INS-3454 |
| chr5-143200062-INS-203 | chr5-143200062-INS-203 |
| chr5-143218602-INS-59 | chr5-143218602-INS-59 |
| chr5-143251815-INS-201 | chr5-143251815-INS-201 |
| chr5-143280038-INS-255 | chr5-143280038-INS-256 |
| chr5-143317746-INS-204 | chr5-143317746-INS-203 |
| chr5-143319745-INS-60 | chr5-143319745-INS-60 |
| chr5-143335552-INS-61 | chr5-143335552-INS-61 |
| chr5-143368880-INS-72 | chr5-143368880-INS-72 |
| chr5-143378827-INS-52 | chr5-143378827-INS-58 |
| chr5-143432616-INS-224 | chr5-143432616-INS-224 |
| chr5-143455807-INS-97 | chr5-143455807-INS-97 |
| chr5-143459368-INS-336 | chr5-143459368-INS-336 |
| chr5-143462533-INS-117 | chr5-143462527-INS-139 |
| chr5-143463222-INS-264 | chr5-143463222-INS-264 |
| chr5-143484228-INS-71 | chr5-143484228-INS-71 |
| chr5-143555429-INS-392 | chr5-143555429-INS-392 |
| chr5-143562912-INS-232 | chr5-143562912-INS-244 |
| chr5-143567284-INS-52 | chr5-143567284-INS-52 |
| chr5-143568432-INS-52 | chr5-143568420-INS-53 |
| chr5-143585890-INS-226 | chr5-143585890-INS-228 |
| chr5-143588196-INS-70 | chr5-143588196-INS-64 |
| chr5-143596963-INS-238 | chr5-143596963-INS-238 |
| chr5-143605113-INS-101 | chr5-143605113-INS-133 |
| chr5-143632727-INS-177 | chr5-143632727-INS-180 |
| chr5-143644068-INS-234 | chr5-143644068-INS-238 |
| chr5-143652761-INS-220 | chr5-143652761-INS-220 |
| chr5-143654446-INS-1018 | chr5-143654447-INS-1008 |
| chr5-143681535-INS-6568 | chr5-143681536-INS-6568 |
| chr5-143694717-INS-52 | chr5-143694717-INS-50 |
| chr5-143719248-INS-200 | chr5-143719248-INS-200 |
| chr5-143733493-INS-80 | chr5-143733493-INS-65 |
| chr5-143741215-INS-225 | chr5-143741216-INS-185 |
| chr5-143747127-INS-90 | chr5-143747127-INS-90 |
| chr5-143751509-INS-57 | chr5-143751509-INS-57 |
| chr5-143758304-INS-54 | chr5-143758320-INS-64 |
| chr5-143807417-INS-50 | chr5-143807417-INS-50 |
| chr5-143815328-INS-204 | chr5-143815328-INS-202 |
| chr5-143830239-INS-162 | chr5-143830239-INS-156 |
| chr5-143845341-INS-218 | chr5-143845341-INS-218 |

| | |
|---|---|
| chr5-143862922-INS-124 | chr5-143862922-INS-124 |
| chr5-143879613-INS-168 | chr5-143879613-INS-124 |
| chr5-143889140-INS-200 | chr5-143889140-INS-200 |
| chr5-143906914-INS-217 | chr5-143906914-INS-214 |
| chr5-143931255-INS-59 | chr5-143931255-INS-59 |
| chr5-143944451-INS-186 | chr5-143944451-INS-175 |
| chr5-143949613-INS-216 | chr5-143949613-INS-214 |
| chr5-143952059-INS-170 | chr5-143952059-INS-169 |
| chr5-143967754-INS-180 | chr5-143967754-INS-180 |
| chr5-143971250-INS-104 | chr5-143971250-INS-104 |
| chr5-143982460-INS-193 | chr5-143982460-INS-195 |
| chr5-143986727-INS-167 | chr5-143986727-INS-167 |
| chr5-143987845-INS-99 | chr5-143987845-INS-119 |
| chr5-144006301-INS-217 | chr5-144006301-INS-217 |
| chr5-144008631-INS-78 | chr5-144008631-INS-56 |
| chr5-144020953-INS-187 | chr5-144020953-INS-197 |
| chr5-144033547-INS-547 | chr5-144033547-INS-548 |
| chr5-144057836-INS-207 | chr5-144057836-INS-222 |
| chr5-144070393-INS-124 | chr5-144070393-INS-115 |
| chr5-144073084-INS-52 | chr5-144073084-INS-52 |
| chr5-144074759-INS-123 | chr5-144074759-INS-123 |
| chr5-144079742-INS-81 | chr5-144079742-INS-81 |
| chr5-144094963-INS-400 | chr5-144094963-INS-393 |
| chr5-144098698-INS-178 | chr5-144098698-INS-178 |
| chr5-144168096-INS-408 | chr5-144168096-INS-409 |
| chr5-144170179-INS-65 | chr5-144170179-INS-65 |
| chr5-144172953-INS-363 | chr5-144172953-INS-363 |
| chr5-144184694-INS-213 | chr5-144184694-INS-213 |
| chr5-144189000-INS-86 | chr5-144189000-INS-86 |
| chr5-144204242-INS-148 | chr5-144204242-INS-99 |
| chr5-144207928-INS-127 | chr5-144207928-INS-131 |
| chr5-144226542-INS-375 | chr5-144226542-INS-375 |
| chr5-144244609-INS-7134 | chr5-144244610-INS-7134 |
| chr5-144259953-INS-576 | chr5-144259953-INS-577 |
| chr5-144275595-INS-156 | chr5-144275595-INS-156 |
| chr5-144287901-INS-66 | chr5-144287901-INS-66 |
| chr5-144303382-INS-110 | chr5-144303382-INS-110 |
| chr5-144311187-INS-210 | chr5-144311187-INS-210 |
| chr5-144314028-INS-60 | chr5-144314045-INS-60 |
| chr5-144325012-INS-79 | chr5-144325010-INS-83 |
| chr5-144340103-INS-221 | chr5-144340103-INS-221 |
| chr5-144361515-INS-160 | chr5-144361515-INS-160 |
| chr5-144368870-INS-96 | chr5-144368870-INS-80 |
| chr5-144370200-INS-68 | chr5-144370180-INS-96 |
| chr5-144387217-INS-53 | chr5-144387217-INS-53 |
| chr5-144407841-INS-54 | chr5-144407841-INS-88 |
| chr5-144412306-INS-63 | chr5-144412306-INS-63 |
| chr5-144426133-INS-78 | chr5-144426127-INS-84 |
| chr5-144447442-INS-51 | chr5-144447442-INS-50 |
| chr5-144457869-INS-92 | chr5-144457889-INS-80 |
| chr5-144461193-INS-72 | chr5-144461189-INS-68 |
| chr5-144470326-INS-635 | chr5-144470342-INS-638 |
| chr5-144482111-INS-89 | chr5-144482109-INS-84 |
| chr5-144486739-INS-5173 | chr5-144486740-INS-5173 |
| chr5-144491811-INS-178 | chr5-144491811-INS-182 |
| chr5-144508163-INS-204 | chr5-144508163-INS-204 |
| chr5-144516396-INS-217 | chr5-144516396-INS-217 |
| chr5-144528876-INS-148 | chr5-144528876-INS-148 |
| chr5-144529010-INS-518 | chr5-144529010-INS-518 |
| chr5-144530159-INS-354 | chr5-144530173-INS-361 |
| chr5-144546258-INS-53 | chr5-144546247-INS-62 |
| chr5-144553283-INS-264 | chr5-144553283-INS-264 |
| chr5-144556330-INS-189 | chr5-144556349-INS-182 |
| chr5-144585671-INS-138 | chr5-144585671-INS-143 |
| chr5-144585807-INS-504 | chr5-144585806-INS-517 |
| chr5-144592740-INS-57 | chr5-144592731-INS-57 |
| chr5-144595974-INS-65 | chr5-144595974-INS-70 |
| chr5-144630344-INS-72 | chr5-144630344-INS-73 |
| chr5-144634353-INS-90 | chr5-144634353-INS-78 |
| chr5-144639935-INS-52 | chr5-144639923-INS-53 |
| chr5-144678607-INS-195 | chr5-144678607-INS-199 |
| chr5-144688718-INS-266 | chr5-144688718-INS-266 |
| chr5-144701923-INS-204 | chr5-144701923-INS-204 |
| chr5-144720961-INS-225 | chr5-144720961-INS-225 |
| chr5-144724912-INS-117 | chr5-144724912-INS-117 |
| chr5-144732637-INS-64 | chr5-144732637-INS-64 |
| chr5-144784171-INS-213 | chr5-144784171-INS-213 |

| | |
|---|---|
| chr5-144805778-INS-203 | chr5-144805778-INS-203 |
| chr5-144812264-INS-5452 | chr5-144812265-INS-5452 |
| chr5-144859799-INS-67 | chr5-144859799-INS-67 |
| chr5-144863657-INS-57 | chr5-144863657-INS-57 |
| chr5-144864680-INS-79 | chr5-144864630-INS-89 |
| chr5-144868132-INS-64 | chr5-144868132-INS-54 |
| chr5-144878760-INS-71 | chr5-144878760-INS-71 |
| chr5-144892079-INS-53 | chr5-144892079-INS-53 |
| chr5-144896628-INS-60 | chr5-144896628-INS-60 |
| chr5-144925936-INS-88 | chr5-144925926-INS-70 |
| chr5-144927254-INS-207 | chr5-144927283-INS-204 |
| chr5-144933462-INS-228 | chr5-144933462-INS-238 |
| chr5-144935822-INS-71 | chr5-144935822-INS-68 |
| chr5-144938974-INS-130 | chr5-144938973-INS-128 |
| chr5-145003794-INS-65 | chr5-145003793-INS-57 |
| chr5-145004664-INS-3210 | chr5-145004665-INS-3210 |
| chr5-145009496-INS-74 | chr5-145009505-INS-50 |
| chr5-145034498-INS-141 | chr5-145034498-INS-141 |
| chr5-145044503-INS-54 | chr5-145044506-INS-50 |
| chr5-145046840-INS-461 | chr5-145046840-INS-461 |
| chr5-145054323-INS-105 | chr5-145054323-INS-105 |
| chr5-145062981-INS-603 | chr5-145062981-INS-605 |
| chr5-145090806-INS-357 | chr5-145090806-INS-355 |
| chr5-145092656-INS-205 | chr5-145092656-INS-205 |
| chr5-145116896-INS-82 | chr5-145116896-INS-82 |
| chr5-145126736-INS-229 | chr5-145126736-INS-229 |
| chr5-145148487-INS-190 | chr5-145148487-INS-190 |
| chr5-145149742-INS-207 | chr5-145149742-INS-207 |
| chr5-145156556-INS-217 | chr5-145156556-INS-210 |
| chr5-145161773-INS-65 | chr5-145161728-INS-113 |
| chr5-145165568-INS-358 | chr5-145165568-INS-358 |
| chr5-145177521-INS-212 | chr5-145177521-INS-212 |
| chr5-145184802-INS-53 | chr5-145184802-INS-53 |
| chr5-145202131-INS-941 | chr5-145202131-INS-941 |
| chr5-145221017-INS-110 | chr5-145221027-INS-78 |
| chr5-145235283-INS-5258 | chr5-145235284-INS-5258 |
| chr5-145238069-INS-63 | chr5-145238066-INS-60 |
| chr5-145243470-INS-7160 | chr5-145243471-INS-7160 |
| chr5-145246826-INS-167 | chr5-145246826-INS-167 |
| chr5-145288502-INS-74 | chr5-145288502-INS-74 |
| chr5-145289624-INS-339 | chr5-145289624-INS-339 |
| chr5-145297676-INS-175 | chr5-145297676-INS-175 |
| chr5-145302794-INS-765 | chr5-145302794-INS-765 |
| chr5-145315015-INS-56 | chr5-145315015-INS-56 |
| chr5-145330971-INS-1367 | chr5-145330972-INS-1366 |
| chr5-145361092-INS-547 | chr5-145361092-INS-547 |
| chr5-145529675-INS-58 | chr5-145529675-INS-58 |
| chr5-145670417-INS-234 | chr5-145670420-INS-234 |
| chr5-145689980-INS-2983 | chr5-145689981-INS-2983 |
| chr5-145724389-INS-379 | chr5-145724389-INS-379 |
| chr5-145818969-INS-1145 | chr5-145818970-INS-1145 |
| chr5-145861851-INS-209 | chr5-145861851-INS-209 |
| chr5-145884264-INS-505 | chr5-145884264-INS-505 |
| chr5-145891806-INS-80 | chr5-145891806-INS-82 |
| chr5-145897204-INS-1886 | chr5-145897205-INS-1886 |
| chr5-145909906-INS-71 | chr5-145909905-INS-76 |
| chr5-145921092-INS-180 | chr5-145921092-INS-182 |
| chr5-145930987-INS-587 | chr5-145930987-INS-587 |
| chr5-145945729-INS-58 | chr5-145945729-INS-50 |
| chr5-145948679-INS-52 | chr5-145948679-INS-52 |
| chr5-146010524-INS-166 | chr5-146010526-INS-153 |
| chr5-146041277-INS-54 | chr5-146041278-INS-54 |
| chr5-146042489-INS-85 | chr5-146042491-INS-122 |
| chr5-146044158-INS-5775 | chr5-146044158-INS-5775 |
| chr5-146044600-INS-56 | chr5-146044606-INS-82 |
| chr5-146072656-INS-6235 | chr5-146072657-INS-6235 |
| chr5-146076512-INS-58 | chr5-146076512-INS-60 |
| chr5-146081105-INS-62 | chr5-146081105-INS-62 |
| chr5-146089734-INS-132 | chr5-146089734-INS-132 |
| chr5-146128765-INS-2686 | chr5-146128766-INS-2665 |
| chr5-146137201-INS-176 | chr5-146137201-INS-176 |
| chr5-146225810-INS-56 | chr5-146225800-INS-58 |
| chr5-146230651-INS-113 | chr5-146230647-INS-68 |
| chr5-146264806-INS-116 | chr5-146264801-INS-106 |
| chr5-146282152-INS-135 | chr5-146282152-INS-135 |
| chr5-146284431-INS-54 | chr5-146284431-INS-54 |
| chr5-146292903-INS-67 | chr5-146292906-INS-86 |

| | |
|---|---|
| chr5-146296739-INS-6951 | chr5-146296740-INS-6951 |
| chr5-146300018-INS-60 | chr5-146300018-INS-60 |
| chr5-146303380-INS-60 | chr5-146303380-INS-75 |
| chr5-146314087-INS-54 | chr5-146314087-INS-54 |
| chr5-146324900-INS-57 | chr5-146324901-INS-60 |
| chr5-146336467-INS-228 | chr5-146336464-INS-219 |
| chr5-146374107-INS-379 | chr5-146374107-INS-379 |
| chr5-146396653-INS-92 | chr5-146396653-INS-92 |
| chr5-146409596-INS-61 | chr5-146409604-INS-61 |
| chr5-146492810-INS-97 | chr5-146492810-INS-97 |
| chr5-146498676-INS-64 | chr5-146498676-INS-64 |
| chr5-146538052-INS-72 | chr5-146538052-INS-72 |
| chr5-146542450-INS-152 | chr5-146542450-INS-164 |
| chr5-146552858-INS-90 | chr5-146552862-INS-114 |
| chr5-146556482-INS-1886 | chr5-146556483-INS-1886 |
| chr5-146556944-INS-53 | chr5-146556944-INS-53 |
| chr5-146559466-INS-135 | chr5-146559481-INS-136 |
| chr5-146565328-INS-80 | chr5-146565328-INS-80 |
| chr5-146572035-INS-5158 | chr5-146572036-INS-5158 |
| chr5-146582321-INS-408 | chr5-146582321-INS-412 |
| chr5-146599979-INS-113 | chr5-146599979-INS-113 |
| chr5-146605438-INS-310 | chr5-146605438-INS-310 |
| chr5-146606850-INS-349 | chr5-146606827-INS-332 |
| chr5-146607409-INS-162 | chr5-146607409-INS-162 |
| chr5-146611443-INS-177 | chr5-146611443-INS-146 |
| chr5-146621040-INS-66 | chr5-146621040-INS-90 |
| chr5-146641613-INS-51 | chr5-146641634-INS-50 |
| chr5-146649039-INS-206 | chr5-146649039-INS-207 |
| chr5-146655439-INS-70 | chr5-146655439-INS-64 |
| chr5-146658358-INS-100 | chr5-146658352-INS-140 |
| chr5-146685668-INS-57 | chr5-146685668-INS-50 |
| chr5-146691791-INS-345 | chr5-146691791-INS-345 |
| chr5-146695026-INS-119 | chr5-146695027-INS-160 |
| chr5-146696942-INS-358 | chr5-146696942-INS-358 |
| chr5-146708281-INS-223 | chr5-146708281-INS-223 |
| chr5-146717878-INS-199 | chr5-146717878-INS-195 |
| chr5-146718941-INS-171 | chr5-146718941-INS-171 |
| chr5-146733573-INS-61 | chr5-146733573-INS-61 |
| chr5-146738122-INS-507 | chr5-146738122-INS-507 |
| chr5-146739668-INS-90 | chr5-146739668-INS-90 |
| chr5-146747073-INS-216 | chr5-146747073-INS-215 |
| chr5-146748471-INS-189 | chr5-146748475-INS-162 |
| chr5-146750871-INS-52 | chr5-146750871-INS-52 |
| chr5-146752664-INS-98 | chr5-146752664-INS-146 |
| chr5-146778893-INS-100 | chr5-146778893-INS-100 |
| chr5-146785245-INS-101 | chr5-146785245-INS-101 |
| chr5-146786604-INS-99 | chr5-146786604-INS-108 |
| chr5-146791659-INS-212 | chr5-146791659-INS-212 |
| chr5-146796802-INS-57 | chr5-146796772-INS-57 |
| chr5-146808965-INS-210 | chr5-146808965-INS-210 |
| chr5-146826380-INS-216 | chr5-146826380-INS-209 |
| chr5-146827935-INS-72 | chr5-146827935-INS-99 |
| chr5-146835177-INS-397 | chr5-146835177-INS-397 |
| chr5-146835960-INS-84 | chr5-146835960-INS-66 |
| chr5-146842749-INS-76 | chr5-146842749-INS-59 |
| chr5-146871922-INS-228 | chr5-146871922-INS-229 |
| chr5-146872578-INS-122 | chr5-146872578-INS-122 |
| chr5-146878326-INS-56 | chr5-146878326-INS-60 |
| chr5-146909346-INS-217 | chr5-146909341-INS-220 |
| chr5-146918721-INS-79 | chr5-146918721-INS-79 |
| chr5-146921766-INS-52 | chr5-146921766-INS-52 |
| chr5-146947016-INS-3586 | chr5-146947017-INS-3586 |
| chr5-146969504-INS-718 | chr5-146969504-INS-718 |
| chr5-147027376-INS-55 | chr5-147027376-INS-57 |
| chr5-147034624-INS-53 | chr5-147034624-INS-53 |
| chr5-147056225-INS-60 | chr5-147056225-INS-75 |
| chr5-147064342-INS-150 | chr5-147064332-INS-140 |
| chr5-147077187-INS-64 | chr5-147077187-INS-58 |
| chr5-147082169-INS-228 | chr5-147082169-INS-229 |
| chr5-147084997-INS-115 | chr5-147084997-INS-85 |
| chr5-147099766-INS-165 | chr5-147099766-INS-165 |
| chr5-147101066-INS-64 | chr5-147101066-INS-64 |
| chr5-147102998-INS-138 | chr5-147102998-INS-114 |
| chr5-147144787-INS-68 | chr5-147144787-INS-68 |
| chr5-147147807-INS-148 | chr5-147147807-INS-152 |
| chr5-147149214-INS-90 | chr5-147149215-INS-76 |
| chr5-147151057-INS-442 | chr5-147151066-INS-440 |

| | |
|---|---|
| chr5-147151929-INS-106 | chr5-147151929-INS-130 |
| chr5-147161700-INS-54 | chr5-147161747-INS-55 |
| chr5-147173505-INS-88 | chr5-147173515-INS-114 |
| chr5-147175983-INS-7103 | chr5-147175984-INS-7103 |
| chr5-147186054-INS-217 | chr5-147186054-INS-217 |
| chr5-147210780-INS-52 | chr5-147210780-INS-56 |
| chr5-147218141-INS-52 | chr5-147218141-INS-52 |
| chr5-147222249-INS-63 | chr5-147222242-INS-85 |
| chr5-147226993-INS-177 | chr5-147226993-INS-178 |
| chr5-147270070-INS-173 | chr5-147270079-INS-176 |
| chr5-147285458-INS-56 | chr5-147285458-INS-52 |
| chr5-147301656-INS-52 | chr5-147301656-INS-52 |
| chr5-147318583-INS-1358 | chr5-147318584-INS-1360 |
| chr5-147319861-INS-219 | chr5-147319861-INS-219 |
| chr5-147333257-INS-218 | chr5-147333257-INS-210 |
| chr5-147351053-INS-74 | chr5-147351053-INS-74 |
| chr5-147363771-INS-185 | chr5-147363771-INS-183 |
| chr5-147374859-INS-107 | chr5-147374859-INS-106 |
| chr5-147383331-INS-492 | chr5-147383331-INS-492 |
| chr5-147387999-INS-113 | chr5-147388022-INS-68 |
| chr5-147405812-INS-102 | chr5-147405787-INS-79 |
| chr5-147407335-INS-79 | chr5-147407334-INS-68 |
| chr5-147423912-INS-201 | chr5-147423912-INS-199 |
| chr5-147433657-INS-256 | chr5-147433657-INS-214 |
| chr5-147437312-INS-55 | chr5-147437311-INS-56 |
| chr5-147489964-INS-54 | chr5-147489964-INS-54 |
| chr5-147498981-INS-148 | chr5-147498981-INS-123 |
| chr5-147502540-INS-70 | chr5-147502540-INS-70 |
| chr5-147516065-INS-214 | chr5-147516065-INS-214 |
| chr5-147528516-INS-76 | chr5-147528531-INS-76 |
| chr5-147542703-INS-314 | chr5-147542703-INS-320 |
| chr5-147556891-INS-64 | chr5-147556891-INS-64 |
| chr5-147558491-INS-103 | chr5-147558492-INS-84 |
| chr5-147563299-INS-71 | chr5-147563299-INS-69 |
| chr5-147565572-INS-52 | chr5-147565573-INS-52 |
| chr5-147586567-INS-67 | chr5-147586567-INS-67 |
| chr5-147596566-INS-100 | chr5-147596566-INS-100 |
| chr5-147615563-INS-51 | chr5-147615563-INS-55 |
| chr5-147633409-INS-201 | chr5-147633409-INS-200 |
| chr5-147634544-INS-138 | chr5-147634544-INS-143 |
| chr5-147637057-INS-103 | chr5-147637057-INS-103 |
| chr5-147675785-INS-53 | chr5-147675785-INS-52 |
| chr5-147697585-INS-51 | chr5-147697585-INS-51 |
| chr5-147702633-INS-155 | chr5-147702601-INS-163 |
| chr5-147714530-INS-76 | chr5-147714547-INS-70 |
| chr5-147721823-INS-71 | chr5-147721823-INS-70 |
| chr5-147727197-INS-78 | chr5-147727197-INS-78 |
| chr5-147748387-INS-130 | chr5-147748390-INS-99 |
| chr5-147768156-INS-80 | chr5-147768156-INS-80 |
| chr5-147771244-INS-63 | chr5-147771244-INS-60 |
| chr5-147811507-INS-200 | chr5-147811507-INS-201 |
| chr5-147826555-INS-72 | chr5-147826555-INS-72 |
| chr5-147834127-INS-204 | chr5-147834109-INS-223 |
| chr5-147835371-INS-56 | chr5-147835371-INS-60 |
| chr5-147840898-INS-91 | chr5-147840904-INS-85 |
| chr5-147848419-INS-65 | chr5-147848419-INS-65 |
| chr5-147855679-INS-212 | chr5-147855679-INS-212 |
| chr5-147856180-INS-197 | chr5-147856180-INS-199 |
| chr5-147865105-INS-386 | chr5-147865105-INS-400 |
| chr5-147872478-INS-56 | chr5-147872478-INS-56 |
| chr5-147910090-INS-211 | chr5-147910090-INS-210 |
| chr5-147929003-INS-83 | chr5-147929000-INS-82 |
| chr5-147935280-INS-96 | chr5-147935280-INS-98 |
| chr5-147956812-INS-89 | chr5-147956821-INS-56 |
| chr5-147961731-INS-64 | chr5-147961731-INS-64 |
| chr5-147971148-INS-71 | chr5-147971148-INS-71 |
| chr5-147986969-INS-75 | chr5-147986969-INS-75 |
| chr5-148007705-INS-105 | chr5-148007705-INS-105 |
| chr5-148020342-INS-125 | chr5-148020342-INS-125 |
| chr5-148033423-INS-116 | chr5-148033465-INS-98 |
| chr5-148040074-INS-56 | chr5-148040049-INS-78 |
| chr5-148046069-INS-120 | chr5-148046069-INS-131 |
| chr5-148074322-INS-173 | chr5-148074327-INS-178 |
| chr5-148102900-INS-70 | chr5-148102900-INS-64 |
| chr5-148109763-INS-295 | chr5-148109763-INS-295 |
| chr5-148115831-INS-74 | chr5-148115831-INS-66 |
| chr5-148122469-INS-325 | chr5-148122469-INS-325 |

| | |
|---|---|
| chr5-148131937-INS-54 | chr5-148131937-INS-50 |
| chr5-148143043-INS-57 | chr5-148143043-INS-56 |
| chr5-148144798-INS-400 | chr5-148144798-INS-400 |
| chr5-148159412-INS-117 | chr5-148159412-INS-117 |
| chr5-148175083-INS-55 | chr5-148175085-INS-56 |
| chr5-148189960-INS-164 | chr5-148189960-INS-164 |
| chr5-148218017-INS-70 | chr5-148218020-INS-56 |
| chr5-148243167-INS-52 | chr5-148243167-INS-52 |
| chr5-148252153-INS-58 | chr5-148252153-INS-58 |
| chr5-148255293-INS-351 | chr5-148255293-INS-350 |
| chr5-148283452-INS-64 | chr5-148283440-INS-68 |
| chr5-148285360-INS-62 | chr5-148285360-INS-64 |
| chr5-148309206-INS-209 | chr5-148309206-INS-209 |
| chr5-148337661-INS-52 | chr5-148337661-INS-53 |
| chr5-148339926-INS-58 | chr5-148339926-INS-78 |
| chr5-148379372-INS-106 | chr5-148379372-INS-111 |
| chr5-148404529-INS-171 | chr5-148404529-INS-171 |
| chr5-148462036-INS-212 | chr5-148462036-INS-212 |
| chr5-148526765-INS-133 | chr5-148526753-INS-151 |
| chr5-148531310-INS-52 | chr5-148531310-INS-52 |
| chr5-148537766-INS-54 | chr5-148537766-INS-64 |
| chr5-148542347-INS-62 | chr5-148542347-INS-70 |
| chr5-148544744-INS-364 | chr5-148544744-INS-364 |
| chr5-148554944-INS-217 | chr5-148554936-INS-224 |
| chr5-148557685-INS-65 | chr5-148557685-INS-65 |
| chr5-148561179-INS-68 | chr5-148561179-INS-68 |
| chr5-148562887-INS-440 | chr5-148562887-INS-440 |
| chr5-148579534-INS-1474 | chr5-148579535-INS-1474 |
| chr5-148607279-INS-65 | chr5-148607264-INS-81 |
| chr5-148614809-INS-51 | chr5-148614809-INS-56 |
| chr5-148617082-INS-4546 | chr5-148617083-INS-4546 |
| chr5-148635651-INS-7160 | chr5-148635652-INS-7160 |
| chr5-148650485-INS-52 | chr5-148650485-INS-64 |
| chr5-148653310-INS-118 | chr5-148653310-INS-120 |
| chr5-148656811-INS-91 | chr5-148656811-INS-76 |
| chr5-148662413-INS-92 | chr5-148662415-INS-94 |
| chr5-148674375-INS-84 | chr5-148674375-INS-85 |
| chr5-148698617-INS-327 | chr5-148698617-INS-327 |
| chr5-148706194-INS-84 | chr5-148706194-INS-86 |
| chr5-148707072-INS-136 | chr5-148707072-INS-116 |
| chr5-148730958-INS-375 | chr5-148730958-INS-375 |
| chr5-148736712-INS-117 | chr5-148736712-INS-114 |
| chr5-148739834-INS-813 | chr5-148739833-INS-811 |
| chr5-148740951-INS-55 | chr5-148740951-INS-55 |
| chr5-148742357-INS-649 | chr5-148742357-INS-649 |
| chr5-148747568-INS-761 | chr5-148747568-INS-761 |
| chr5-148769713-INS-86 | chr5-148769713-INS-68 |
| chr5-148777674-INS-445 | chr5-148777674-INS-400 |
| chr5-148779469-INS-756 | chr5-148779469-INS-755 |
| chr5-148792596-INS-192 | chr5-148792596-INS-190 |
| chr5-148794640-INS-175 | chr5-148794640-INS-175 |
| chr5-148806306-INS-391 | chr5-148806306-INS-391 |
| chr5-148814664-INS-96 | chr5-148814664-INS-96 |
| chr5-148842654-INS-72 | chr5-148842656-INS-68 |
| chr5-148844040-INS-714 | chr5-148844041-INS-698 |
| chr5-148857171-INS-1660 | chr5-148857172-INS-1660 |
| chr5-148894531-INS-180 | chr5-148894531-INS-180 |
| chr5-148911165-INS-60 | chr5-148911160-INS-80 |
| chr5-148917170-INS-384 | chr5-148917170-INS-382 |
| chr5-148937482-INS-58 | chr5-148937482-INS-58 |
| chr5-148974274-INS-254 | chr5-148974274-INS-253 |
| chr5-148982171-INS-144 | chr5-148982171-INS-149 |
| chr5-148992227-INS-60 | chr5-148992236-INS-92 |
| chr5-149002730-INS-58 | chr5-149002734-INS-62 |
| chr5-149004093-INS-59 | chr5-149004093-INS-69 |
| chr5-149009134-INS-56 | chr5-149009134-INS-71 |
| chr5-149024375-INS-90 | chr5-149024375-INS-90 |
| chr5-149063181-INS-506 | chr5-149063181-INS-506 |
| chr5-149063839-INS-54 | chr5-149063839-INS-54 |
| chr5-149075500-INS-354 | chr5-149075500-INS-354 |
| chr5-149076663-INS-58 | chr5-149076663-INS-77 |
| chr5-149088888-INS-124 | chr5-149088863-INS-156 |
| chr5-149097638-INS-139 | chr5-149097638-INS-139 |
| chr5-149098894-INS-92 | chr5-149098894-INS-92 |
| chr5-149117169-INS-98 | chr5-149117169-INS-84 |
| chr5-149156559-INS-5529 | chr5-149156560-INS-5527 |
| chr5-149173298-INS-72 | chr5-149173301-INS-68 |

| | |
|---|---|
| chr5-149177648-INS-75 | chr5-149177648-INS-108 |
| chr5-149188037-INS-155 | chr5-149188039-INS-150 |
| chr5-149193605-INS-87 | chr5-149193605-INS-57 |
| chr5-149223773-INS-55 | chr5-149223773-INS-55 |
| chr5-149235875-INS-202 | chr5-149235875-INS-202 |
| chr5-149242924-INS-646 | chr5-149242924-INS-661 |
| chr5-149247615-INS-58 | chr5-149247599-INS-63 |
| chr5-149248736-INS-119 | chr5-149248734-INS-109 |
| chr5-149253960-INS-244 | chr5-149253986-INS-219 |
| chr5-149261912-INS-157 | chr5-149261912-INS-158 |
| chr5-149264395-INS-110 | chr5-149264440-INS-85 |
| chr5-149280135-INS-608 | chr5-149280135-INS-608 |
| chr5-149296383-INS-7120 | chr5-149296384-INS-7120 |
| chr5-149324974-INS-252 | chr5-149324974-INS-244 |
| chr5-149327791-INS-138 | chr5-149327744-INS-182 |
| chr5-149330775-INS-85 | chr5-149330756-INS-104 |
| chr5-149332517-INS-102 | chr5-149332517-INS-102 |
| chr5-149339679-INS-74 | chr5-149339679-INS-76 |
| chr5-149341259-INS-409 | chr5-149341259-INS-409 |
| chr5-149358369-INS-216 | chr5-149358369-INS-217 |
| chr5-149361603-INS-70 | chr5-149361603-INS-74 |
| chr5-149372798-INS-64 | chr5-149372798-INS-64 |
| chr5-149376142-INS-211 | chr5-149376142-INS-210 |
| chr5-149404666-INS-234 | chr5-149404666-INS-234 |
| chr5-149445622-INS-110 | chr5-149445622-INS-111 |
| chr5-149446403-INS-2295 | chr5-149446404-INS-2307 |
| chr5-149453484-INS-52 | chr5-149453484-INS-52 |
| chr5-149470170-INS-67 | chr5-149470170-INS-67 |
| chr5-149473797-INS-168 | chr5-149473797-INS-167 |
| chr5-149480263-INS-213 | chr5-149480263-INS-213 |
| chr5-149486439-INS-173 | chr5-149486434-INS-176 |
| chr5-149504857-INS-207 | chr5-149504876-INS-213 |
| chr5-149509962-INS-215 | chr5-149509962-INS-215 |
| chr5-149518240-INS-200 | chr5-149518240-INS-200 |
| chr5-149588320-INS-196 | chr5-149588320-INS-201 |
| chr5-149620078-INS-73 | chr5-149620084-INS-58 |
| chr5-149629799-INS-162 | chr5-149629799-INS-162 |
| chr5-149641886-INS-68 | chr5-149641886-INS-50 |
| chr5-149662810-INS-140 | chr5-149662810-INS-140 |
| chr5-149669077-INS-116 | chr5-149669077-INS-104 |
| chr5-149669737-INS-349 | chr5-149669737-INS-351 |
| chr5-149689617-INS-54 | chr5-149689617-INS-54 |
| chr5-149761399-INS-54 | chr5-149761403-INS-56 |
| chr5-149798075-INS-90 | chr5-149798079-INS-122 |
| chr5-149804247-INS-98 | chr5-149804247-INS-100 |
| chr5-149807023-INS-56 | chr5-149807023-INS-60 |
| chr5-149809248-INS-60 | chr5-149809237-INS-55 |
| chr5-149833392-INS-204 | chr5-149833392-INS-204 |
| chr5-149850297-INS-53 | chr5-149850297-INS-53 |
| chr5-149854193-INS-82 | chr5-149854193-INS-82 |
| chr5-149855240-INS-828 | chr5-149855240-INS-828 |
| chr5-149856798-INS-381 | chr5-149856798-INS-379 |
| chr5-149858489-INS-64 | chr5-149858489-INS-56 |
| chr5-149866864-INS-81 | chr5-149866864-INS-56 |
| chr5-149879715-INS-97 | chr5-149879713-INS-74 |
| chr5-149906966-INS-97 | chr5-149906960-INS-103 |
| chr5-149937822-INS-100 | chr5-149937818-INS-86 |
| chr5-149957885-INS-86 | chr5-149957885-INS-86 |
| chr5-149975105-INS-112 | chr5-149975105-INS-80 |
| chr5-150008746-INS-6440 | chr5-150008747-INS-6440 |
| chr5-150014917-INS-895 | chr5-150014917-INS-895 |
| chr5-150029290-INS-60 | chr5-150029289-INS-67 |
| chr5-150060929-INS-62 | chr5-150060929-INS-60 |
| chr5-150094667-INS-64 | chr5-150094667-INS-64 |
| chr5-150149512-INS-61 | chr5-150149512-INS-65 |
| chr5-150151261-INS-7079 | chr5-150151262-INS-7079 |
| chr5-150154883-INS-128 | chr5-150154883-INS-128 |
| chr5-150167761-INS-125 | chr5-150167761-INS-134 |
| chr5-150183923-INS-64 | chr5-150183923-INS-68 |
| chr5-150192426-INS-203 | chr5-150192426-INS-203 |
| chr5-150201887-INS-431 | chr5-150201887-INS-431 |
| chr5-150213749-INS-62 | chr5-150213749-INS-66 |
| chr5-150232003-INS-2356 | chr5-150232004-INS-2356 |
| chr5-150248195-INS-179 | chr5-150248196-INS-191 |
| chr5-150264035-INS-88 | chr5-150264069-INS-77 |
| chr5-150270700-INS-374 | chr5-150270700-INS-370 |
| chr5-150285026-INS-416 | chr5-150285026-INS-416 |

| | |
|---|---|
| chr5-150302085-INS-185 | chr5-150302085-INS-185 |
| chr5-150313029-INS-257 | chr5-150313029-INS-257 |
| chr5-150314927-INS-96 | chr5-150314927-INS-96 |
| chr5-150344040-INS-138 | chr5-150344040-INS-138 |
| chr5-150348495-INS-572 | chr5-150348495-INS-572 |
| chr5-150361495-INS-121 | chr5-150361495-INS-121 |
| chr5-150378227-INS-54 | chr5-150378227-INS-56 |
| chr5-150380185-INS-88 | chr5-150380185-INS-88 |
| chr5-150404147-INS-271 | chr5-150404147-INS-271 |
| chr5-150414269-INS-56 | chr5-150414269-INS-54 |
| chr5-150436195-INS-54 | chr5-150436195-INS-50 |
| chr5-150454946-INS-205 | chr5-150454946-INS-204 |
| chr5-150458718-INS-202 | chr5-150458718-INS-202 |
| chr5-150488451-INS-54 | chr5-150488467-INS-66 |
| chr5-150547524-INS-76 | chr5-150547524-INS-63 |
| chr5-150552556-INS-218 | chr5-150552556-INS-218 |
| chr5-150558487-INS-210 | chr5-150558487-INS-211 |
| chr5-150606960-INS-60 | chr5-150606960-INS-60 |
| chr5-150608182-INS-3153 | chr5-150608183-INS-3153 |
| chr5-150612603-INS-83 | chr5-150612603-INS-78 |
| chr5-150632263-INS-219 | chr5-150632263-INS-220 |
| chr5-150634063-INS-155 | chr5-150634063-INS-155 |
| chr5-150639595-INS-80 | chr5-150639595-INS-60 |
| chr5-150645295-INS-1877 | chr5-150645296-INS-1877 |
| chr5-150665498-INS-114 | chr5-150665498-INS-114 |
| chr5-150668881-INS-410 | chr5-150668881-INS-411 |
| chr5-150681176-INS-56 | chr5-150681176-INS-56 |
| chr5-150683941-INS-58 | chr5-150683941-INS-62 |
| chr5-150685743-INS-112 | chr5-150685743-INS-112 |
| chr5-150717289-INS-200 | chr5-150717289-INS-200 |
| chr5-150743208-INS-700 | chr5-150743208-INS-699 |
| chr5-150745162-INS-155 | chr5-150745162-INS-164 |
| chr5-150758737-INS-58 | chr5-150758737-INS-59 |
| chr5-150775051-INS-181 | chr5-150775052-INS-181 |
| chr5-150786881-INS-450 | chr5-150786881-INS-450 |
| chr5-150794859-INS-191 | chr5-150794859-INS-191 |
| chr5-150795954-INS-116 | chr5-150795954-INS-124 |
| chr5-150818603-INS-53 | chr5-150818603-INS-53 |
| chr5-150824279-INS-85 | chr5-150824279-INS-85 |
| chr5-150831642-INS-60 | chr5-150831642-INS-64 |
| chr5-150834475-INS-67 | chr5-150834475-INS-62 |
| chr5-150841705-INS-217 | chr5-150841705-INS-215 |
| chr5-150846550-INS-236 | chr5-150846551-INS-236 |
| chr5-150848884-INS-138 | chr5-150848884-INS-153 |
| chr5-150858748-INS-312 | chr5-150858765-INS-294 |
| chr5-150868450-INS-100 | chr5-150868450-INS-129 |
| chr5-150881988-INS-226 | chr5-150881988-INS-226 |
| chr5-150888887-INS-122 | chr5-150888887-INS-122 |
| chr5-150909316-INS-58 | chr5-150909318-INS-51 |
| chr5-150921603-INS-58 | chr5-150921603-INS-58 |
| chr5-150955926-INS-191 | chr5-150955926-INS-191 |
| chr5-150957152-INS-73 | chr5-150957152-INS-73 |
| chr5-150962420-INS-139 | chr5-150962420-INS-139 |
| chr5-150994220-INS-3412 | chr5-150994221-INS-3412 |
| chr5-151044199-INS-54 | chr5-151044199-INS-54 |
| chr5-151051881-INS-1902 | chr5-151051882-INS-1901 |
| chr5-151075665-INS-135 | chr5-151075665-INS-135 |
| chr5-151117328-INS-7096 | chr5-151117329-INS-7096 |
| chr5-151141107-INS-84 | chr5-151141107-INS-80 |
| chr5-151141270-INS-7067 | chr5-151141271-INS-7067 |
| chr5-151159310-INS-72 | chr5-151159310-INS-75 |
| chr5-151189368-INS-58 | chr5-151189367-INS-50 |
| chr5-151226823-INS-8136 | chr5-151226824-INS-8136 |
| chr5-151271604-INS-64 | chr5-151271604-INS-64 |
| chr5-151327457-INS-61 | chr5-151327457-INS-61 |
| chr5-151342952-INS-408 | chr5-151342952-INS-408 |
| chr5-151353734-INS-60 | chr5-151353734-INS-60 |
| chr5-151427053-INS-218 | chr5-151427053-INS-218 |
| chr5-151432361-INS-129 | chr5-151432361-INS-129 |
| chr5-151491402-INS-6418 | chr5-151491403-INS-6421 |
| chr5-151538213-INS-99 | chr5-151538213-INS-99 |
| chr5-151547356-INS-119 | chr5-151547332-INS-111 |
| chr5-151560560-INS-506 | chr5-151560560-INS-506 |
| chr5-151575139-INS-100 | chr5-151575139-INS-100 |
| chr5-151631726-INS-6825 | chr5-151631726-INS-6825 |
| chr5-151651646-INS-1890 | chr5-151651647-INS-1890 |
| chr6-3069709-INS-52 | chr6-3069709-INS-52 |

| | |
|---|---|
| chr6-3139732-INS-66 | chr6-3139732-INS-66 |
| chr6-3168707-INS-353 | chr6-3168707-INS-353 |
| chr6-3208856-INS-4786 | chr6-3208857-INS-4786 |
| chr6-3231711-INS-58 | chr6-3231715-INS-54 |
| chr6-3260082-INS-55 | chr6-3260082-INS-55 |
| chr6-3273937-INS-81 | chr6-3273912-INS-81 |
| chr6-3276895-INS-211 | chr6-3276895-INS-211 |
| chr6-3286204-INS-1118 | chr6-3286205-INS-1118 |
| chr6-3298204-INS-4937 | chr6-3298205-INS-4938 |
| chr6-3319609-INS-270 | chr6-3319618-INS-243 |
| chr6-3322691-INS-272 | chr6-3322691-INS-272 |
| chr6-3341603-INS-1458 | chr6-3341604-INS-1458 |
| chr6-3350359-INS-7477 | chr6-3350360-INS-7477 |
| chr6-3358567-INS-7089 | chr6-3358568-INS-7089 |
| chr6-3390965-INS-268 | chr6-3390965-INS-267 |
| chr6-3394141-INS-72 | chr6-3394157-INS-54 |
| chr6-3425873-INS-549 | chr6-3425873-INS-549 |
| chr6-3429998-INS-109 | chr6-3429998-INS-109 |
| chr6-3434420-INS-60 | chr6-3434420-INS-60 |
| chr6-3437146-INS-53 | chr6-3437146-INS-53 |
| chr6-3446393-INS-200 | chr6-3446393-INS-200 |
| chr6-3448529-INS-657 | chr6-3448529-INS-657 |
| chr6-3477593-INS-143 | chr6-3477593-INS-143 |
| chr6-3489024-INS-156 | chr6-3489024-INS-160 |
| chr6-3521085-INS-327 | chr6-3521085-INS-324 |
| chr6-3560076-INS-78 | chr6-3560076-INS-78 |
| chr6-3566901-INS-215 | chr6-3566901-INS-211 |
| chr6-3574019-INS-247 | chr6-3574019-INS-270 |
| chr6-3577433-INS-52 | chr6-3577437-INS-60 |
| chr6-3595963-INS-84 | chr6-3595963-INS-82 |
| chr6-3638241-INS-972 | chr6-3638241-INS-973 |
| chr6-3673209-INS-519 | chr6-3673209-INS-519 |
| chr6-3678094-INS-182 | chr6-3678094-INS-182 |
| chr6-3703769-INS-128 | chr6-3703769-INS-110 |
| chr6-3725031-INS-191 | chr6-3725031-INS-190 |
| chr6-3780038-INS-63 | chr6-3780038-INS-63 |
| chr6-3820696-INS-212 | chr6-3820696-INS-212 |
| chr6-3824594-INS-116 | chr6-3824594-INS-117 |
| chr6-3826709-INS-163 | chr6-3826702-INS-168 |
| chr6-3886342-INS-120 | chr6-3886340-INS-114 |
| chr6-3908223-INS-64 | chr6-3908223-INS-72 |
| chr6-3918120-INS-3739 | chr6-3918121-INS-3739 |
| chr6-3919475-INS-319 | chr6-3919475-INS-319 |
| chr6-3921324-INS-815 | chr6-3921324-INS-816 |
| chr6-3929629-INS-58 | chr6-3929632-INS-50 |
| chr6-3940781-INS-62 | chr6-3940781-INS-62 |
| chr6-3964647-INS-378 | chr6-3964647-INS-410 |
| chr6-3970061-INS-6682 | chr6-3970062-INS-6682 |
| chr6-3981240-INS-372 | chr6-3981240-INS-372 |
| chr6-4059895-INS-449 | chr6-4059895-INS-451 |
| chr6-4078632-INS-106 | chr6-4078620-INS-83 |
| chr6-4078847-INS-6930 | chr6-4078848-INS-6931 |
| chr6-4119655-INS-72 | chr6-4119655-INS-72 |
| chr6-4138052-INS-4677 | chr6-4138053-INS-4677 |
| chr6-4140485-INS-152 | chr6-4140485-INS-152 |
| chr6-4146329-INS-6592 | chr6-4146330-INS-6592 |
| chr6-4196367-INS-75 | chr6-4196367-INS-55 |
| chr6-4203258-INS-134 | chr6-4203258-INS-134 |
| chr6-4215051-INS-5277 | chr6-4215052-INS-5277 |
| chr6-4218899-INS-3672 | chr6-4218900-INS-3672 |
| chr6-4245072-INS-1324 | chr6-4245073-INS-1323 |
| chr6-4246823-INS-550 | chr6-4246823-INS-550 |
| chr6-4282422-INS-212 | chr6-4282422-INS-212 |
| chr6-4297243-INS-748 | chr6-4297243-INS-749 |
| chr6-4297804-INS-52 | chr6-4297804-INS-82 |
| chr6-4346990-INS-58 | chr6-4346990-INS-60 |
| chr6-4360746-INS-219 | chr6-4360746-INS-220 |
| chr6-4378104-INS-106 | chr6-4378104-INS-106 |
| chr6-4385426-INS-203 | chr6-4385426-INS-203 |
| chr6-4389812-INS-6428 | chr6-4389813-INS-6428 |
| chr6-4406085-INS-261 | chr6-4406107-INS-212 |
| chr6-4416333-INS-568 | chr6-4416333-INS-568 |
| chr6-4448178-INS-112 | chr6-4448178-INS-112 |
| chr6-4450163-INS-498 | chr6-4450163-INS-498 |
| chr6-4545519-INS-59 | chr6-4545499-INS-62 |
| chr6-4547546-INS-54 | chr6-4547546-INS-54 |
| chr6-4573855-INS-56 | chr6-4573861-INS-62 |

| | |
|---|---|
| chr6-4577415-INS-52 | chr6-4577415-INS-92 |
| chr6-4596213-INS-52 | chr6-4596213-INS-52 |
| chr6-4603948-INS-62 | chr6-4603948-INS-66 |
| chr6-4646161-INS-108 | chr6-4646161-INS-108 |
| chr6-4688543-INS-181 | chr6-4688535-INS-171 |
| chr6-4716854-INS-102 | chr6-4716854-INS-102 |
| chr6-4735973-INS-182 | chr6-4735987-INS-188 |
| chr6-4756453-INS-81 | chr6-4756453-INS-81 |
| chr6-4789356-INS-90 | chr6-4789360-INS-84 |
| chr6-4797590-INS-64 | chr6-4797590-INS-66 |
| chr6-4839072-INS-66 | chr6-4839072-INS-60 |
| chr6-4839371-INS-260 | chr6-4839371-INS-260 |
| chr6-4853085-INS-200 | chr6-4853085-INS-200 |
| chr6-4865271-INS-170 | chr6-4865271-INS-168 |
| chr6-4867197-INS-198 | chr6-4867198-INS-199 |
| chr6-4879635-INS-146 | chr6-4879635-INS-150 |
| chr6-4881658-INS-54 | chr6-4881658-INS-54 |
| chr6-4887031-INS-268 | chr6-4887031-INS-268 |
| chr6-4916433-INS-68 | chr6-4916431-INS-72 |
| chr6-4917502-INS-52 | chr6-4917502-INS-52 |
| chr6-4919789-INS-220 | chr6-4919789-INS-220 |
| chr6-4925841-INS-718 | chr6-4925838-INS-736 |
| chr6-4930404-INS-70 | chr6-4930404-INS-70 |
| chr6-4930404-INS-898 | chr6-4930404-INS-922 |
| chr6-4931520-INS-200 | chr6-4931520-INS-200 |
| chr6-4935475-INS-327 | chr6-4935499-INS-331 |
| chr6-4958746-INS-210 | chr6-4958746-INS-209 |
| chr6-4967654-INS-51 | chr6-4967610-INS-54 |
| chr6-5012050-INS-5548 | chr6-5012051-INS-5548 |
| chr6-5034939-INS-211 | chr6-5034939-INS-211 |
| chr6-5055279-INS-129 | chr6-5055279-INS-129 |
| chr6-5056871-INS-84 | chr6-5056871-INS-84 |
| chr6-5081904-INS-147 | chr6-5081894-INS-147 |
| chr6-5090622-INS-84 | chr6-5090622-INS-66 |
| chr6-5106413-INS-182 | chr6-5106413-INS-182 |
| chr6-5113887-INS-61 | chr6-5113887-INS-61 |
| chr6-5142119-INS-126 | chr6-5142119-INS-144 |
| chr6-5146689-INS-77 | chr6-5146690-INS-85 |
| chr6-5195739-INS-153 | chr6-5195739-INS-153 |
| chr6-5200935-INS-89 | chr6-5200935-INS-89 |
| chr6-5208105-INS-7111 | chr6-5208106-INS-7111 |
| chr6-5248492-INS-215 | chr6-5248492-INS-215 |
| chr6-5253320-INS-210 | chr6-5253320-INS-210 |
| chr6-5271289-INS-54 | chr6-5271289-INS-54 |
| chr6-5275611-INS-2513 | chr6-5275612-INS-2513 |
| chr6-5295762-INS-96 | chr6-5295762-INS-125 |
| chr6-5296997-INS-75 | chr6-5297026-INS-75 |
| chr6-5311761-INS-5271 | chr6-5311762-INS-5271 |
| chr6-5346521-INS-58 | chr6-5346521-INS-58 |
| chr6-5356792-INS-111 | chr6-5356790-INS-112 |
| chr6-5362067-INS-210 | chr6-5362067-INS-210 |
| chr6-5364807-INS-445 | chr6-5364807-INS-445 |
| chr6-5383011-INS-1873 | chr6-5383012-INS-1873 |
| chr6-5418184-INS-63 | chr6-5418185-INS-58 |
| chr6-5426954-INS-178 | chr6-5426958-INS-182 |
| chr6-5429658-INS-470 | chr6-5429658-INS-470 |
| chr6-5437319-INS-2508 | chr6-5437320-INS-2508 |
| chr6-5439144-INS-463 | chr6-5439144-INS-463 |
| chr6-5447000-INS-60 | chr6-5447000-INS-80 |
| chr6-5476970-INS-2897 | chr6-5476971-INS-2891 |
| chr6-5532572-INS-108 | chr6-5532572-INS-108 |
| chr6-5584533-INS-505 | chr6-5584533-INS-505 |
| chr6-5591858-INS-62 | chr6-5591858-INS-54 |
| chr6-5602013-INS-62 | chr6-5602013-INS-62 |
| chr6-5604063-INS-508 | chr6-5604063-INS-508 |
| chr6-5617538-INS-95 | chr6-5617538-INS-104 |
| chr6-5632224-INS-342 | chr6-5632224-INS-360 |
| chr6-5638648-INS-104 | chr6-5638632-INS-114 |
| chr6-5663147-INS-179 | chr6-5663147-INS-223 |
| chr6-5664964-INS-81 | chr6-5664964-INS-126 |
| chr6-5672541-INS-487 | chr6-5672541-INS-487 |
| chr6-5684503-INS-62 | chr6-5684503-INS-64 |
| chr6-5691608-INS-192 | chr6-5691608-INS-179 |
| chr6-5700265-INS-206 | chr6-5700265-INS-204 |
| chr6-5705611-INS-1122 | chr6-5705612-INS-1122 |
| chr6-5709485-INS-150 | chr6-5709485-INS-200 |
| chr6-5712137-INS-208 | chr6-5712137-INS-208 |

| | |
|---|---|
| chr6-5716445-INS-54 | chr6-5716445-INS-68 |
| chr6-5717146-INS-83 | chr6-5717146-INS-83 |
| chr6-5732016-INS-77 | chr6-5732016-INS-77 |
| chr6-5737643-INS-3210 | chr6-5737644-INS-3210 |
| chr6-5785407-INS-183 | chr6-5785407-INS-185 |
| chr6-5788753-INS-69 | chr6-5788753-INS-69 |
| chr6-5793300-INS-509 | chr6-5793297-INS-514 |
| chr6-5807699-INS-176 | chr6-5807697-INS-213 |
| chr6-5819372-INS-198 | chr6-5819372-INS-198 |
| chr6-5835516-INS-6402 | chr6-5835517-INS-6405 |
| chr6-5873608-INS-279 | chr6-5873608-INS-279 |
| chr6-5891337-INS-56 | chr6-5891337-INS-56 |
| chr6-5955694-INS-68 | chr6-5955694-INS-68 |
| chr6-5986734-INS-54 | chr6-5986734-INS-56 |
| chr6-6050576-INS-70 | chr6-6050576-INS-70 |
| chr6-6063510-INS-51 | chr6-6063510-INS-51 |
| chr6-6118810-INS-311 | chr6-6118810-INS-310 |
| chr6-6126546-INS-140 | chr6-6126546-INS-152 |
| chr6-6154327-INS-72 | chr6-6154327-INS-72 |
| chr6-6188808-INS-210 | chr6-6188808-INS-210 |
| chr6-6192430-INS-62 | chr6-6192430-INS-62 |
| chr6-6199723-INS-504 | chr6-6199723-INS-504 |
| chr6-6213117-INS-52 | chr6-6213117-INS-52 |
| chr6-6222637-INS-116 | chr6-6222637-INS-90 |
| chr6-6239412-INS-202 | chr6-6239412-INS-202 |
| chr6-6265654-INS-60 | chr6-6265654-INS-60 |
| chr6-6267050-INS-65 | chr6-6267050-INS-65 |
| chr6-6268162-INS-223 | chr6-6268162-INS-225 |
| chr6-6293092-INS-68 | chr6-6293092-INS-68 |
| chr6-6295042-INS-71 | chr6-6295042-INS-76 |
| chr6-6307137-INS-60 | chr6-6307137-INS-60 |
| chr6-6353312-INS-112 | chr6-6353312-INS-113 |
| chr6-6379444-INS-68 | chr6-6379444-INS-68 |
| chr6-6388073-INS-57 | chr6-6388073-INS-57 |
| chr6-6415984-INS-249 | chr6-6415984-INS-243 |
| chr6-6438527-INS-119 | chr6-6438527-INS-119 |
| chr6-6447256-INS-56 | chr6-6447256-INS-56 |
| chr6-6543545-INS-70 | chr6-6543545-INS-70 |
| chr6-6563351-INS-65 | chr6-6563351-INS-65 |
| chr6-6571691-INS-1219 | chr6-6571692-INS-1219 |
| chr6-6572966-INS-79 | chr6-6572966-INS-79 |
| chr6-6595185-INS-5878 | chr6-6595186-INS-5878 |
| chr6-6598211-INS-151 | chr6-6598167-INS-189 |
| chr6-6604990-INS-87 | chr6-6604990-INS-73 |
| chr6-6606707-INS-227 | chr6-6606707-INS-227 |
| chr6-6612121-INS-218 | chr6-6612121-INS-218 |
| chr6-6619759-INS-60 | chr6-6619759-INS-70 |
| chr6-6645758-INS-98 | chr6-6645746-INS-75 |
| chr6-6657148-INS-100 | chr6-6657147-INS-67 |
| chr6-6661908-INS-6233 | chr6-6661909-INS-6233 |
| chr6-6674913-INS-136 | chr6-6674913-INS-160 |
| chr6-6697123-INS-87 | chr6-6697101-INS-65 |
| chr6-6710223-INS-103 | chr6-6710225-INS-103 |
| chr6-6748013-INS-88 | chr6-6748003-INS-92 |
| chr6-6750333-INS-4070 | chr6-6750334-INS-4070 |
| chr6-6787329-INS-2343 | chr6-6787330-INS-2342 |
| chr6-6882350-INS-56 | chr6-6882350-INS-70 |
| chr6-6896562-INS-7084 | chr6-6896563-INS-7084 |
| chr6-6899022-INS-93 | chr6-6899022-INS-93 |
| chr6-6924063-INS-116 | chr6-6924063-INS-119 |
| chr6-6934635-INS-425 | chr6-6934635-INS-425 |
| chr6-6938156-INS-476 | chr6-6938156-INS-476 |
| chr6-6958194-INS-459 | chr6-6958192-INS-463 |
| chr6-6960285-INS-82 | chr6-6960285-INS-52 |
| chr6-6966188-INS-138 | chr6-6966188-INS-138 |
| chr6-6968559-INS-399 | chr6-6968559-INS-399 |
| chr6-6983294-INS-338 | chr6-6983271-INS-347 |
| chr6-7012049-INS-66 | chr6-7012049-INS-66 |
| chr6-7047834-INS-68 | chr6-7047834-INS-68 |
| chr6-7052505-INS-226 | chr6-7052505-INS-231 |
| chr6-7065933-INS-65 | chr6-7065933-INS-65 |
| chr6-7090267-INS-334 | chr6-7090267-INS-334 |
| chr6-7115071-INS-506 | chr6-7115071-INS-495 |
| chr6-7193975-INS-702 | chr6-7193975-INS-695 |
| chr6-7201089-INS-56 | chr6-7201089-INS-58 |
| chr6-7210282-INS-68 | chr6-7210242-INS-66 |
| chr6-7232796-INS-58 | chr6-7232768-INS-75 |

| | |
|---|---|
| chr6-7237931-INS-328 | chr6-7237931-INS-328 |
| chr6-7264750-INS-316 | chr6-7264750-INS-316 |
| chr6-7269517-INS-80 | chr6-7269517-INS-100 |
| chr6-7306789-INS-51 | chr6-7306791-INS-83 |
| chr6-7315251-INS-3018 | chr6-7315252-INS-3018 |
| chr6-7346373-INS-56 | chr6-7346412-INS-52 |
| chr6-7355243-INS-2777 | chr6-7355244-INS-2781 |
| chr6-7372713-INS-66 | chr6-7372713-INS-79 |
| chr6-7402631-INS-54 | chr6-7402631-INS-60 |
| chr6-7416715-INS-811 | chr6-7416718-INS-814 |
| chr6-7419413-INS-888 | chr6-7419413-INS-888 |
| chr6-7450219-INS-1318 | chr6-7450220-INS-1320 |
| chr6-7459706-INS-56 | chr6-7459706-INS-56 |
| chr6-7468672-INS-141 | chr6-7468676-INS-147 |
| chr6-7491640-INS-1777 | chr6-7491641-INS-1788 |
| chr6-7531474-INS-484 | chr6-7531474-INS-484 |
| chr6-7554257-INS-92 | chr6-7554258-INS-92 |
| chr6-7570274-INS-6262 | chr6-7570275-INS-6262 |
| chr6-7572367-INS-107 | chr6-7572367-INS-107 |
| chr6-7602828-INS-58 | chr6-7602828-INS-62 |
| chr6-7607523-INS-223 | chr6-7607523-INS-223 |
| chr6-7614175-INS-193 | chr6-7614175-INS-193 |
| chr6-7629098-INS-84 | chr6-7629095-INS-95 |
| chr6-7660642-INS-52 | chr6-7660614-INS-50 |
| chr6-7681572-INS-98 | chr6-7681588-INS-100 |
| chr6-7690569-INS-100 | chr6-7690569-INS-100 |
| chr6-7693417-INS-72 | chr6-7693417-INS-72 |
| chr6-7696201-INS-59 | chr6-7696232-INS-70 |
| chr6-7718735-INS-237 | chr6-7718735-INS-237 |
| chr6-7737945-INS-413 | chr6-7737945-INS-413 |
| chr6-7741390-INS-54 | chr6-7741390-INS-52 |
| chr6-7755543-INS-167 | chr6-7755543-INS-191 |
| chr6-7775208-INS-108 | chr6-7775208-INS-108 |
| chr6-7776716-INS-148 | chr6-7776716-INS-148 |
| chr6-7778459-INS-84 | chr6-7778459-INS-84 |
| chr6-7790556-INS-1859 | chr6-7790557-INS-1859 |
| chr6-7793583-INS-142 | chr6-7793583-INS-140 |
| chr6-7796395-INS-867 | chr6-7796395-INS-868 |
| chr6-7805395-INS-125 | chr6-7805395-INS-150 |
| chr6-7806574-INS-84 | chr6-7806574-INS-84 |
| chr6-7808318-INS-223 | chr6-7808318-INS-222 |
| chr6-7810927-INS-441 | chr6-7810927-INS-441 |
| chr6-7827258-INS-112 | chr6-7827220-INS-150 |
| chr6-7837731-INS-60 | chr6-7837731-INS-60 |
| chr6-7856260-INS-1976 | chr6-7856261-INS-1973 |
| chr6-7860368-INS-52 | chr6-7860378-INS-53 |
| chr6-7865380-INS-98 | chr6-7865380-INS-98 |
| chr6-7883245-INS-52 | chr6-7883233-INS-64 |
| chr6-7893480-INS-1307 | chr6-7893481-INS-1307 |
| chr6-7906775-INS-2886 | chr6-7906776-INS-2886 |
| chr6-7911640-INS-1354 | chr6-7911641-INS-1335 |
| chr6-7916756-INS-78 | chr6-7916756-INS-78 |
| chr6-7953583-INS-327 | chr6-7953583-INS-327 |
| chr6-7963008-INS-217 | chr6-7963008-INS-217 |
| chr6-7985451-INS-69 | chr6-7985451-INS-69 |
| chr6-7993059-INS-76 | chr6-7993059-INS-76 |
| chr6-7994232-INS-97 | chr6-7994232-INS-81 |
| chr6-8004482-INS-86 | chr6-8004482-INS-86 |
| chr6-8010977-INS-82 | chr6-8010977-INS-98 |
| chr6-8027651-INS-482 | chr6-8027651-INS-483 |
| chr6-8040515-INS-183 | chr6-8040515-INS-183 |
| chr6-8048331-INS-278 | chr6-8048332-INS-277 |
| chr6-8048579-INS-1713 | chr6-8048580-INS-1747 |
| chr6-8050984-INS-101 | chr6-8050973-INS-101 |
| chr6-8064117-INS-100 | chr6-8064117-INS-100 |
| chr6-8067038-INS-432 | chr6-8067038-INS-432 |
| chr6-8084396-INS-68 | chr6-8084396-INS-66 |
| chr6-8097946-INS-51 | chr6-8097946-INS-51 |
| chr6-8131971-INS-81 | chr6-8131976-INS-72 |
| chr6-8136168-INS-1370 | chr6-8136169-INS-1370 |
| chr6-8138845-INS-254 | chr6-8138845-INS-250 |
| chr6-8150190-INS-72 | chr6-8150190-INS-72 |
| chr6-8152256-INS-88 | chr6-8152256-INS-70 |
| chr6-8161260-INS-139 | chr6-8161260-INS-139 |
| chr6-8181759-INS-61 | chr6-8181794-INS-52 |
| chr6-8188699-INS-94 | chr6-8188700-INS-64 |
| chr6-8192217-INS-280 | chr6-8192218-INS-235 |

| | |
|---|---|
| chr6-8193393-INS-52 | chr6-8193393-INS-54 |
| chr6-8198567-INS-207 | chr6-8198567-INS-207 |
| chr6-8199510-INS-6868 | chr6-8199511-INS-6868 |
| chr6-8216598-INS-190 | chr6-8216598-INS-190 |
| chr6-8220002-INS-52 | chr6-8220002-INS-52 |
| chr6-8230529-INS-485 | chr6-8230529-INS-485 |
| chr6-8238824-INS-207 | chr6-8238824-INS-196 |
| chr6-8261388-INS-96 | chr6-8261388-INS-96 |
| chr6-8266753-INS-98 | chr6-8266753-INS-98 |
| chr6-8270901-INS-204 | chr6-8270901-INS-209 |
| chr6-8271888-INS-644 | chr6-8271888-INS-645 |
| chr6-8307138-INS-6612 | chr6-8307139-INS-6612 |
| chr6-8344632-INS-206 | chr6-8344632-INS-205 |
| chr6-8355649-INS-6944 | chr6-8355650-INS-6947 |
| chr6-8369371-INS-58 | chr6-8369371-INS-58 |
| chr6-8375002-INS-314 | chr6-8374953-INS-314 |
| chr6-8376673-INS-896 | chr6-8376673-INS-896 |
| chr6-8385032-INS-6687 | chr6-8385033-INS-6690 |
| chr6-8415003-INS-179 | chr6-8415003-INS-180 |
| chr6-8452080-INS-186 | chr6-8452080-INS-192 |
| chr6-8454677-INS-781 | chr6-8454677-INS-779 |
| chr6-8477311-INS-69 | chr6-8477311-INS-69 |
| chr6-8478938-INS-567 | chr6-8478939-INS-567 |
| chr6-8490642-INS-323 | chr6-8490652-INS-326 |
| chr6-8560570-INS-62 | chr6-8560570-INS-62 |
| chr6-8641961-INS-74 | chr6-8641961-INS-74 |
| chr6-8646693-INS-96 | chr6-8646693-INS-88 |
| chr6-8709609-INS-170 | chr6-8709609-INS-172 |
| chr6-8728743-INS-132 | chr6-8728767-INS-103 |
| chr6-8747656-INS-7165 | chr6-8747657-INS-7165 |
| chr6-8753565-INS-90 | chr6-8753565-INS-90 |
| chr6-8760316-INS-193 | chr6-8760316-INS-191 |
| chr6-8762800-INS-2696 | chr6-8762801-INS-2697 |
| chr6-8776886-INS-96 | chr6-8776886-INS-104 |
| chr6-8790446-INS-66 | chr6-8790446-INS-68 |
| chr6-8800318-INS-53 | chr6-8800318-INS-53 |
| chr6-8816056-INS-324 | chr6-8816057-INS-325 |
| chr6-8818714-INS-6377 | chr6-8818715-INS-6377 |
| chr6-8819194-INS-59 | chr6-8819194-INS-59 |
| chr6-8832290-INS-85 | chr6-8832290-INS-85 |
| chr6-8866132-INS-62 | chr6-8866132-INS-64 |
| chr6-8872899-INS-400 | chr6-8872899-INS-400 |
| chr6-8886527-INS-54 | chr6-8886527-INS-54 |
| chr6-8927786-INS-64 | chr6-8927786-INS-64 |
| chr6-8928727-INS-7131 | chr6-8928728-INS-7132 |
| chr6-8963978-INS-57 | chr6-8963978-INS-76 |
| chr6-8991740-INS-90 | chr6-8991740-INS-90 |
| chr6-8993299-INS-386 | chr6-8993313-INS-385 |
| chr6-8997901-INS-52 | chr6-8997901-INS-52 |
| chr6-9003329-INS-617 | chr6-9003329-INS-617 |
| chr6-9016304-INS-55 | chr6-9016304-INS-55 |
| chr6-9026333-INS-866 | chr6-9026333-INS-866 |
| chr6-9031567-INS-579 | chr6-9031567-INS-579 |
| chr6-9039109-INS-587 | chr6-9039109-INS-587 |
| chr6-9089245-INS-6479 | chr6-9089246-INS-6479 |
| chr6-9110186-INS-745 | chr6-9110186-INS-744 |
| chr6-9222314-INS-65 | chr6-9222353-INS-60 |
| chr6-9245667-INS-97 | chr6-9245667-INS-97 |
| chr6-9291510-INS-2296 | chr6-9291511-INS-2296 |
| chr6-9317322-INS-52 | chr6-9317321-INS-56 |
| chr6-9320784-INS-346 | chr6-9320784-INS-309 |
| chr6-9338168-INS-7509 | chr6-9338169-INS-7510 |
| chr6-9369110-INS-331 | chr6-9369110-INS-329 |
| chr6-9426799-INS-1787 | chr6-9426800-INS-1787 |
| chr6-9475628-INS-52 | chr6-9475628-INS-56 |
| chr6-9488002-INS-497 | chr6-9488002-INS-497 |
| chr6-9499709-INS-120 | chr6-9499709-INS-120 |
| chr6-9513276-INS-59 | chr6-9513276-INS-61 |
| chr6-9524893-INS-747 | chr6-9524893-INS-747 |
| chr6-9540487-INS-371 | chr6-9540487-INS-371 |
| chr6-9543806-INS-676 | chr6-9543806-INS-676 |
| chr6-9576286-INS-72 | chr6-9576319-INS-54 |
| chr6-9596560-INS-101 | chr6-9596561-INS-76 |
| chr6-9601903-INS-78 | chr6-9601903-INS-78 |
| chr6-9654079-INS-659 | chr6-9654079-INS-659 |
| chr6-9661134-INS-116 | chr6-9661135-INS-116 |
| chr6-9673743-INS-208 | chr6-9673743-INS-208 |

| | |
|---|---|
| chr6-9676438-INS-72 | chr6-9676438-INS-64 |
| chr6-9796735-INS-65 | chr6-9796735-INS-71 |
| chr6-9817151-INS-637 | chr6-9817151-INS-636 |
| chr6-9818553-INS-402 | chr6-9818553-INS-402 |
| chr6-9833117-INS-3419 | chr6-9833122-INS-3412 |
| chr6-9907180-INS-1443 | chr6-9907181-INS-1443 |
| chr6-9929153-INS-105 | chr6-9929153-INS-100 |
| chr6-9934916-INS-58 | chr6-9934878-INS-57 |
| chr6-9973489-INS-1098 | chr6-9973490-INS-1098 |
| chr6-9986602-INS-64 | chr6-9986602-INS-62 |
| chr6-9988666-INS-258 | chr6-9988666-INS-258 |
| chr6-9989870-INS-64 | chr6-9989870-INS-64 |
| chr6-10000430-INS-433 | chr6-10000430-INS-441 |
| chr6-10083607-INS-68 | chr6-10083607-INS-68 |
| chr6-10094107-INS-83 | chr6-10094066-INS-96 |
| chr6-10167831-INS-563 | chr6-10167831-INS-564 |
| chr6-10204646-INS-473 | chr6-10204646-INS-473 |
| chr6-10204997-INS-1179 | chr6-10204998-INS-1179 |
| chr6-10206613-INS-475 | chr6-10206613-INS-475 |
| chr6-10213816-INS-1889 | chr6-10213821-INS-1878 |
| chr6-10260276-INS-216 | chr6-10260276-INS-216 |
| chr6-10303023-INS-251 | chr6-10303023-INS-252 |
| chr6-10320653-INS-1334 | chr6-10320654-INS-1334 |
| chr6-10321628-INS-348 | chr6-10321626-INS-351 |
| chr6-10341089-INS-281 | chr6-10341089-INS-281 |
| chr6-10351282-INS-6403 | chr6-10351283-INS-6403 |
| chr6-10359694-INS-75 | chr6-10359694-INS-75 |
| chr6-10361986-INS-86 | chr6-10361986-INS-86 |
| chr6-10369596-INS-1495 | chr6-10369597-INS-1495 |
| chr6-10385960-INS-87 | chr6-10385960-INS-87 |
| chr6-10445453-INS-67 | chr6-10445453-INS-67 |
| chr6-10456565-INS-1946 | chr6-10456566-INS-1945 |
| chr6-10485650-INS-1612 | chr6-10485651-INS-1612 |
| chr6-10543266-INS-54 | chr6-10543266-INS-60 |
| chr6-10592451-INS-286 | chr6-10592441-INS-294 |
| chr6-10664667-INS-67 | chr6-10664667-INS-64 |
| chr6-10678289-INS-803 | chr6-10678289-INS-803 |
| chr6-10685736-INS-161 | chr6-10685736-INS-161 |
| chr6-10725700-INS-95 | chr6-10725700-INS-92 |
| chr6-10744778-INS-81 | chr6-10744817-INS-73 |
| chr6-10762527-INS-236 | chr6-10762529-INS-280 |
| chr6-10764812-INS-55 | chr6-10764808-INS-59 |
| chr6-10765367-INS-5039 | chr6-10765368-INS-5039 |
| chr6-10784246-INS-358 | chr6-10784246-INS-358 |
| chr6-10841083-INS-67 | chr6-10841083-INS-69 |
| chr6-10889442-INS-105 | chr6-10889442-INS-85 |
| chr6-10900087-INS-100 | chr6-10900093-INS-104 |
| chr6-10930201-INS-201 | chr6-10930201-INS-201 |
| chr6-10999641-INS-6472 | chr6-10999642-INS-6472 |
| chr6-11020081-INS-289 | chr6-11020081-INS-289 |
| chr6-11048829-INS-166 | chr6-11048829-INS-166 |
| chr6-11056646-INS-2953 | chr6-11056647-INS-2953 |
| chr6-11095909-INS-67 | chr6-11095909-INS-60 |
| chr6-11170123-INS-402 | chr6-11170124-INS-402 |
| chr6-11170761-INS-58 | chr6-11170761-INS-58 |
| chr6-11184684-INS-483 | chr6-11184684-INS-483 |
| chr6-11190607-INS-249 | chr6-11190607-INS-249 |
| chr6-11203936-INS-599 | chr6-11203936-INS-599 |
| chr6-11250937-INS-201 | chr6-11250937-INS-201 |
| chr6-11266672-INS-404 | chr6-11266672-INS-403 |
| chr6-11278016-INS-198 | chr6-11278016-INS-198 |
| chr6-11287105-INS-52 | chr6-11287105-INS-64 |
| chr6-11291322-INS-4740 | chr6-11291323-INS-4740 |
| chr6-11296660-INS-1305 | chr6-11296661-INS-1305 |
| chr6-11320142-INS-6488 | chr6-11320143-INS-6487 |
| chr6-11335846-INS-55 | chr6-11335846-INS-63 |
| chr6-11350339-INS-216 | chr6-11350338-INS-215 |
| chr6-11366523-INS-124 | chr6-11366523-INS-124 |
| chr6-11389949-INS-99 | chr6-11389949-INS-91 |
| chr6-11470799-INS-2818 | chr6-11470800-INS-2818 |
| chr6-11476121-INS-58 | chr6-11476121-INS-60 |
| chr6-11488237-INS-64 | chr6-11488241-INS-60 |
| chr6-11527919-INS-64 | chr6-11527919-INS-66 |
| chr6-11538011-INS-814 | chr6-11538015-INS-829 |
| chr6-11553627-INS-1120 | chr6-11553628-INS-1120 |
| chr6-11581420-INS-93 | chr6-11581420-INS-84 |
| chr6-11585694-INS-327 | chr6-11585694-INS-327 |

| | |
|---|---|
| chr6-11621036-INS-92 | chr6-11621036-INS-105 |
| chr6-11653035-INS-55 | chr6-11653035-INS-65 |
| chr6-11673888-INS-122 | chr6-11673888-INS-122 |
| chr6-11708228-INS-2351 | chr6-11708229-INS-2351 |
| chr6-11713011-INS-77 | chr6-11713011-INS-82 |
| chr6-11718957-INS-90 | chr6-11718947-INS-90 |
| chr6-11742261-INS-78 | chr6-11742261-INS-74 |
| chr6-11748080-INS-1894 | chr6-11748081-INS-1894 |
| chr6-11748656-INS-424 | chr6-11748656-INS-424 |
| chr6-11808249-INS-1266 | chr6-11808250-INS-1266 |
| chr6-11818390-INS-51 | chr6-11818396-INS-69 |
| chr6-11871829-INS-213 | chr6-11871829-INS-213 |
| chr6-11914224-INS-315 | chr6-11914224-INS-315 |
| chr6-11914965-INS-83 | chr6-11914953-INS-54 |
| chr6-11949141-INS-7099 | chr6-11949142-INS-7099 |
| chr6-12005399-INS-87 | chr6-12005399-INS-87 |
| chr6-12018829-INS-377 | chr6-12018829-INS-377 |
| chr6-12023816-INS-226 | chr6-12023816-INS-226 |
| chr6-12030679-INS-225 | chr6-12030679-INS-225 |
| chr6-12048193-INS-1128 | chr6-12048194-INS-1128 |
| chr6-12050139-INS-57 | chr6-12050139-INS-62 |
| chr6-12076020-INS-6461 | chr6-12076021-INS-6464 |
| chr6-12099510-INS-76 | chr6-12099510-INS-72 |
| chr6-12103264-INS-54 | chr6-12103262-INS-80 |
| chr6-12109790-INS-86 | chr6-12109790-INS-88 |
| chr6-12130055-INS-539 | chr6-12130055-INS-538 |
| chr6-12145924-INS-64 | chr6-12145924-INS-68 |
| chr6-12162915-INS-2020 | chr6-12162916-INS-2020 |
| chr6-12163907-INS-358 | chr6-12163907-INS-358 |
| chr6-12174677-INS-76 | chr6-12174677-INS-92 |
| chr6-12185553-INS-109 | chr6-12185558-INS-117 |
| chr6-12200624-INS-53 | chr6-12200624-INS-53 |
| chr6-12217405-INS-6330 | chr6-12217406-INS-6333 |
| chr6-12260680-INS-117 | chr6-12260688-INS-115 |
| chr6-12263462-INS-103 | chr6-12263462-INS-143 |
| chr6-12269482-INS-76 | chr6-12269484-INS-72 |
| chr6-12274299-INS-76 | chr6-12274327-INS-52 |
| chr6-12308823-INS-129 | chr6-12308823-INS-129 |
| chr6-12319370-INS-292 | chr6-12319363-INS-306 |
| chr6-12368605-INS-66 | chr6-12368605-INS-66 |
| chr6-12385988-INS-6483 | chr6-12385989-INS-6482 |
| chr6-12393726-INS-110 | chr6-12393726-INS-92 |
| chr6-12427056-INS-107 | chr6-12427056-INS-107 |
| chr6-12472762-INS-360 | chr6-12472762-INS-360 |
| chr6-12477632-INS-162 | chr6-12477632-INS-156 |
| chr6-12486388-INS-118 | chr6-12486388-INS-118 |
| chr6-12500361-INS-78 | chr6-12500361-INS-78 |
| chr6-12556588-INS-80 | chr6-12556588-INS-76 |
| chr6-12581129-INS-58 | chr6-12581129-INS-60 |
| chr6-12589269-INS-72 | chr6-12589269-INS-58 |
| chr6-12618300-INS-98 | chr6-12618308-INS-60 |
| chr6-12648861-INS-72 | chr6-12648861-INS-72 |
| chr6-12689679-INS-487 | chr6-12689679-INS-487 |
| chr6-12696291-INS-52 | chr6-12696291-INS-59 |
| chr6-12711581-INS-72 | chr6-12711581-INS-60 |
| chr6-12727436-INS-51 | chr6-12727436-INS-52 |
| chr6-12734879-INS-3291 | chr6-12734880-INS-3291 |
| chr6-12738172-INS-89 | chr6-12738170-INS-76 |
| chr6-12746773-INS-61 | chr6-12746774-INS-67 |
| chr6-12779629-INS-2090 | chr6-12779630-INS-2128 |
| chr6-12796096-INS-67 | chr6-12796096-INS-67 |
| chr6-12825003-INS-52 | chr6-12825010-INS-59 |
| chr6-12838490-INS-323 | chr6-12838491-INS-323 |
| chr6-12844992-INS-965 | chr6-12844992-INS-964 |
| chr6-12851445-INS-391 | chr6-12851445-INS-392 |
| chr6-12853236-INS-115 | chr6-12853236-INS-115 |
| chr6-12855114-INS-66 | chr6-12855114-INS-66 |
| chr6-12879134-INS-419 | chr6-12879134-INS-419 |
| chr6-12897507-INS-368 | chr6-12897507-INS-369 |
| chr6-12905696-INS-401 | chr6-12905696-INS-401 |
| chr6-12905708-INS-3153 | chr6-12905709-INS-3153 |
| chr6-12983673-INS-56 | chr6-12983673-INS-56 |
| chr6-13003495-INS-202 | chr6-13003495-INS-202 |
| chr6-13004291-INS-494 | chr6-13004291-INS-494 |
| chr6-13015692-INS-6373 | chr6-13015693-INS-6373 |
| chr6-13032769-INS-466 | chr6-13032769-INS-466 |
| chr6-13046190-INS-89 | chr6-13046190-INS-89 |

| | |
|---|---|
| chr6-13050940-INS-187 | chr6-13050940-INS-187 |
| chr6-13051935-INS-6403 | chr6-13051936-INS-6403 |
| chr6-13061108-INS-184 | chr6-13061114-INS-136 |
| chr6-13102370-INS-375 | chr6-13102370-INS-375 |
| chr6-13107805-INS-62 | chr6-13107805-INS-51 |
| chr6-13115046-INS-66 | chr6-13115045-INS-72 |
| chr6-13117648-INS-81 | chr6-13117678-INS-81 |
| chr6-13133218-INS-72 | chr6-13133231-INS-56 |
| chr6-13137617-INS-1893 | chr6-13137618-INS-1893 |
| chr6-13165011-INS-70 | chr6-13165015-INS-94 |
| chr6-13178658-INS-1464 | chr6-13178659-INS-1465 |
| chr6-13179860-INS-89 | chr6-13179865-INS-89 |
| chr6-13188842-INS-80 | chr6-13188842-INS-54 |
| chr6-13191239-INS-443 | chr6-13191239-INS-443 |
| chr6-13206114-INS-60 | chr6-13206133-INS-72 |
| chr6-13274239-INS-4105 | chr6-13274240-INS-4105 |
| chr6-13280165-INS-8230 | chr6-13280166-INS-8228 |
| chr6-13282370-INS-54 | chr6-13282370-INS-60 |
| chr6-13306908-INS-76 | chr6-13306908-INS-54 |
| chr6-13309895-INS-81 | chr6-13309895-INS-79 |
| chr6-13338172-INS-129 | chr6-13338172-INS-129 |
| chr6-13401412-INS-2956 | chr6-13401413-INS-2956 |
| chr6-13421707-INS-211 | chr6-13421676-INS-212 |
| chr6-13465661-INS-2456 | chr6-13465662-INS-2456 |
| chr6-13518851-INS-1091 | chr6-13518852-INS-1091 |
| chr6-13521254-INS-192 | chr6-13521269-INS-183 |
| chr6-13548540-INS-2433 | chr6-13548541-INS-2433 |
| chr6-13556221-INS-148 | chr6-13556214-INS-168 |
| chr6-13563156-INS-179 | chr6-13563156-INS-180 |
| chr6-13584271-INS-193 | chr6-13584271-INS-193 |
| chr6-13600089-INS-110 | chr6-13600089-INS-89 |
| chr6-13648794-INS-4557 | chr6-13648795-INS-4557 |
| chr6-13706991-INS-79 | chr6-13706991-INS-79 |
| chr6-13731623-INS-805 | chr6-13731623-INS-803 |
| chr6-13753507-INS-52 | chr6-13753501-INS-56 |
| chr6-13757682-INS-54 | chr6-13757682-INS-54 |
| chr6-13779172-INS-322 | chr6-13779172-INS-322 |
| chr6-13856780-INS-69 | chr6-13856780-INS-56 |
| chr6-13867645-INS-6413 | chr6-13867646-INS-6413 |
| chr6-13904338-INS-6413 | chr6-13904339-INS-6413 |
| chr6-13912099-INS-859 | chr6-13912099-INS-861 |
| chr6-13915844-INS-542 | chr6-13915844-INS-542 |
| chr6-13929044-INS-68 | chr6-13929044-INS-64 |
| chr6-13931987-INS-54 | chr6-13931987-INS-62 |
| chr6-13963973-INS-80 | chr6-13963972-INS-60 |
| chr6-13986133-INS-86 | chr6-13986133-INS-86 |
| chr6-13988993-INS-60 | chr6-13988993-INS-60 |
| chr6-14000596-INS-132 | chr6-14000612-INS-108 |
| chr6-14007998-INS-184 | chr6-14007998-INS-184 |
| chr6-14019845-INS-1802 | chr6-14019846-INS-1802 |
| chr6-14028885-INS-241 | chr6-14028885-INS-241 |
| chr6-14033416-INS-1886 | chr6-14033417-INS-1886 |
| chr6-14033581-INS-60 | chr6-14033581-INS-54 |
| chr6-14041954-INS-1204 | chr6-14041955-INS-1204 |
| chr6-14043432-INS-62 | chr6-14043432-INS-62 |
| chr6-14101540-INS-595 | chr6-14101540-INS-596 |
| chr6-14103785-INS-52 | chr6-14103785-INS-52 |
| chr6-14123932-INS-1183 | chr6-14123933-INS-1184 |
| chr6-14124194-INS-2061 | chr6-14124195-INS-2061 |
| chr6-14129343-INS-75 | chr6-14129343-INS-75 |
| chr6-14145544-INS-102 | chr6-14145542-INS-76 |
| chr6-14151108-INS-83 | chr6-14151108-INS-89 |
| chr6-14155591-INS-441 | chr6-14155591-INS-441 |
| chr6-14161748-INS-142 | chr6-14161749-INS-143 |
| chr6-14162322-INS-4671 | chr6-14162323-INS-4668 |
| chr6-14167313-INS-101 | chr6-14167313-INS-119 |
| chr6-14171810-INS-6595 | chr6-14171811-INS-6595 |
| chr6-14193525-INS-534 | chr6-14193525-INS-534 |
| chr6-14199597-INS-7634 | chr6-14199598-INS-7637 |
| chr6-14202573-INS-629 | chr6-14202573-INS-629 |
| chr6-14205276-INS-51 | chr6-14205276-INS-51 |
| chr6-14215143-INS-7283 | chr6-14215144-INS-7283 |
| chr6-14223034-INS-240 | chr6-14223035-INS-228 |
| chr6-14238505-INS-58 | chr6-14238505-INS-54 |
| chr6-14260751-INS-383 | chr6-14260751-INS-383 |
| chr6-14262006-INS-557 | chr6-14262007-INS-557 |
| chr6-14302255-INS-6438 | chr6-14302256-INS-6438 |

| | |
|---|---|
| chr6-14330216-INS-67 | chr6-14330216-INS-65 |
| chr6-14340042-INS-89 | chr6-14340042-INS-89 |
| chr6-14353211-INS-66 | chr6-14353211-INS-66 |
| chr6-14362407-INS-99 | chr6-14362423-INS-118 |
| chr6-14366521-INS-146 | chr6-14366521-INS-147 |
| chr6-14371474-INS-56 | chr6-14371474-INS-52 |
| chr6-14386208-INS-58 | chr6-14386208-INS-58 |
| chr6-14397489-INS-210 | chr6-14397489-INS-210 |
| chr6-14404045-INS-54 | chr6-14404045-INS-52 |
| chr6-14410557-INS-75 | chr6-14410557-INS-75 |
| chr6-14416848-INS-391 | chr6-14416848-INS-391 |
| chr6-14426785-INS-3236 | chr6-14426786-INS-3236 |
| chr6-14429493-INS-3121 | chr6-14429494-INS-3121 |
| chr6-14469089-INS-329 | chr6-14469089-INS-328 |
| chr6-14480321-INS-50 | chr6-14480329-INS-76 |
| chr6-14481606-INS-2584 | chr6-14481607-INS-2577 |
| chr6-14487304-INS-116 | chr6-14487304-INS-132 |
| chr6-14498483-INS-244 | chr6-14498497-INS-267 |
| chr6-14501363-INS-55 | chr6-14501363-INS-55 |
| chr6-14507246-INS-272 | chr6-14507246-INS-272 |
| chr6-14520641-INS-53 | chr6-14520603-INS-51 |
| chr6-14542080-INS-118 | chr6-14542080-INS-118 |
| chr6-14594033-INS-71 | chr6-14594033-INS-71 |
| chr6-14616232-INS-70 | chr6-14616237-INS-115 |
| chr6-14625200-INS-6367 | chr6-14625201-INS-6367 |
| chr6-14629557-INS-303 | chr6-14629601-INS-304 |
| chr6-14654461-INS-208 | chr6-14654461-INS-208 |
| chr6-14685267-INS-51 | chr6-14685267-INS-51 |
| chr6-14689526-INS-4797 | chr6-14689527-INS-4797 |
| chr6-14697225-INS-211 | chr6-14697225-INS-211 |
| chr6-14699013-INS-1189 | chr6-14699014-INS-1189 |
| chr6-14710118-INS-79 | chr6-14710118-INS-79 |
| chr6-14810132-INS-418 | chr6-14810132-INS-417 |
| chr6-14816993-INS-93 | chr6-14816993-INS-93 |
| chr6-14867554-INS-327 | chr6-14867554-INS-327 |
| chr6-14872068-INS-64 | chr6-14872065-INS-60 |
| chr6-14880512-INS-62 | chr6-14880522-INS-66 |
| chr6-14886060-INS-371 | chr6-14886060-INS-371 |
| chr6-14943263-INS-55 | chr6-14943264-INS-64 |
| chr6-14945940-INS-364 | chr6-14945940-INS-368 |
| chr6-14966748-INS-214 | chr6-14966748-INS-214 |
| chr6-14978281-INS-54 | chr6-14978299-INS-62 |
| chr6-14982378-INS-55 | chr6-14982378-INS-61 |
| chr6-15013215-INS-70 | chr6-15013215-INS-78 |
| chr6-15019643-INS-7342 | chr6-15019644-INS-7343 |
| chr6-15028377-INS-52 | chr6-15028377-INS-54 |
| chr6-15033814-INS-1889 | chr6-15033815-INS-1889 |
| chr6-15140090-INS-71 | chr6-15140095-INS-71 |
| chr6-15147109-INS-226 | chr6-15147109-INS-226 |
| chr6-15164237-INS-72 | chr6-15164237-INS-72 |
| chr6-15168293-INS-59 | chr6-15168293-INS-59 |
| chr6-15169910-INS-135 | chr6-15169866-INS-139 |
| chr6-15170745-INS-738 | chr6-15170745-INS-735 |
| chr6-15174612-INS-69 | chr6-15174612-INS-51 |
| chr6-15179741-INS-615 | chr6-15179741-INS-617 |
| chr6-15227601-INS-165 | chr6-15227601-INS-165 |
| chr6-15232449-INS-54 | chr6-15232449-INS-50 |
| chr6-15257186-INS-6902 | chr6-15257187-INS-6905 |
| chr6-15295997-INS-76 | chr6-15296010-INS-77 |
| chr6-15355382-INS-62 | chr6-15355382-INS-62 |
| chr6-15394016-INS-52 | chr6-15394016-INS-55 |
| chr6-15426195-INS-569 | chr6-15426195-INS-570 |
| chr6-15463666-INS-213 | chr6-15463666-INS-213 |
| chr6-15480174-INS-158 | chr6-15480174-INS-158 |
| chr6-15495607-INS-56 | chr6-15495607-INS-56 |
| chr6-15508577-INS-1656 | chr6-15508578-INS-1656 |
| chr6-15541708-INS-213 | chr6-15541708-INS-220 |
| chr6-15607374-INS-214 | chr6-15607386-INS-214 |
| chr6-15621711-INS-96 | chr6-15621711-INS-96 |
| chr6-15629361-INS-95 | chr6-15629361-INS-95 |
| chr6-15645736-INS-576 | chr6-15645736-INS-576 |
| chr6-15646706-INS-53 | chr6-15646706-INS-56 |
| chr6-15662795-INS-60 | chr6-15662795-INS-60 |
| chr6-15706429-INS-192 | chr6-15706429-INS-192 |
| chr6-15735760-INS-59 | chr6-15735760-INS-60 |
| chr6-15755956-INS-96 | chr6-15755913-INS-119 |
| chr6-15770578-INS-51 | chr6-15770578-INS-50 |

| | |
|---|---|
| chr6-15778847-INS-111 | chr6-15778847-INS-111 |
| chr6-15794554-INS-94 | chr6-15794558-INS-103 |
| chr6-15826673-INS-63 | chr6-15826679-INS-65 |
| chr6-15844194-INS-186 | chr6-15844163-INS-176 |
| chr6-15845255-INS-826 | chr6-15845255-INS-826 |
| chr6-15886294-INS-101 | chr6-15886294-INS-101 |
| chr6-15917536-INS-314 | chr6-15917536-INS-316 |
| chr6-15927735-INS-169 | chr6-15927735-INS-169 |
| chr6-15940383-INS-6542 | chr6-15940384-INS-6540 |
| chr6-15941181-INS-549 | chr6-15941181-INS-552 |
| chr6-15960651-INS-343 | chr6-15960651-INS-343 |
| chr6-15972282-INS-667 | chr6-15972282-INS-664 |
| chr6-15991413-INS-117 | chr6-15991413-INS-117 |
| chr6-16024552-INS-280 | chr6-16024552-INS-280 |
| chr6-16132917-INS-1087 | chr6-16132918-INS-1087 |
| chr6-16174680-INS-188 | chr6-16174680-INS-186 |
| chr6-16219208-INS-120 | chr6-16219208-INS-120 |
| chr6-16223476-INS-66 | chr6-16223493-INS-69 |
| chr6-16226200-INS-94 | chr6-16226200-INS-94 |
| chr6-16243320-INS-2227 | chr6-16243321-INS-2235 |
| chr6-16319789-INS-181 | chr6-16319791-INS-173 |
| chr6-16351606-INS-122 | chr6-16351586-INS-108 |
| chr6-16387499-INS-1107 | chr6-16387500-INS-1107 |
| chr6-16391070-INS-212 | chr6-16391070-INS-212 |
| chr6-16431672-INS-102 | chr6-16431672-INS-117 |
| chr6-16434156-INS-66 | chr6-16434128-INS-56 |
| chr6-16450569-INS-68 | chr6-16450569-INS-102 |
| chr6-16459007-INS-2248 | chr6-16459008-INS-2246 |
| chr6-16474893-INS-461 | chr6-16474893-INS-462 |
| chr6-16520669-INS-517 | chr6-16520669-INS-517 |
| chr6-16534280-INS-87 | chr6-16534261-INS-135 |
| chr6-16543013-INS-116 | chr6-16543031-INS-74 |
| chr6-16567348-INS-166 | chr6-16567348-INS-211 |
| chr6-16573848-INS-70 | chr6-16573839-INS-80 |
| chr6-16615603-INS-66 | chr6-16615603-INS-66 |
| chr6-16620297-INS-62 | chr6-16620297-INS-64 |
| chr6-16694408-INS-57 | chr6-16694408-INS-57 |
| chr6-16707770-INS-4496 | chr6-16707771-INS-4496 |
| chr6-16716169-INS-6512 | chr6-16716170-INS-6513 |
| chr6-16771848-INS-72 | chr6-16771848-INS-72 |
| chr6-16785657-INS-80 | chr6-16785657-INS-64 |
| chr6-16810166-INS-249 | chr6-16810166-INS-249 |
| chr6-16814458-INS-84 | chr6-16814458-INS-85 |
| chr6-16852296-INS-603 | chr6-16852296-INS-603 |
| chr6-16858953-INS-105 | chr6-16858953-INS-99 |
| chr6-16902418-INS-738 | chr6-16902418-INS-738 |
| chr6-16950109-INS-252 | chr6-16950110-INS-253 |
| chr6-16979145-INS-64 | chr6-16979145-INS-64 |
| chr6-16997595-INS-7119 | chr6-16997596-INS-7120 |
| chr6-17002363-INS-203 | chr6-17002363-INS-203 |
| chr6-17004240-INS-204 | chr6-17004240-INS-204 |
| chr6-17100115-INS-6617 | chr6-17100116-INS-6617 |
| chr6-17103169-INS-640 | chr6-17103169-INS-639 |
| chr6-17112008-INS-213 | chr6-17112008-INS-224 |
| chr6-17117340-INS-78 | chr6-17117364-INS-78 |
| chr6-17123225-INS-51 | chr6-17123222-INS-90 |
| chr6-17124189-INS-6377 | chr6-17124190-INS-6377 |
| chr6-17140461-INS-132 | chr6-17140461-INS-132 |
| chr6-17147073-INS-57 | chr6-17147073-INS-60 |
| chr6-17150095-INS-77 | chr6-17150095-INS-80 |
| chr6-17152588-INS-96 | chr6-17152588-INS-100 |
| chr6-17163863-INS-93 | chr6-17163863-INS-93 |
| chr6-17165849-INS-1688 | chr6-17165850-INS-1690 |
| chr6-17166672-INS-217 | chr6-17166672-INS-217 |
| chr6-17171853-INS-94 | chr6-17171853-INS-94 |
| chr6-17188851-INS-66 | chr6-17188851-INS-66 |
| chr6-17195569-INS-1345 | chr6-17195570-INS-1345 |
| chr6-17200727-INS-52 | chr6-17200727-INS-52 |
| chr6-17211799-INS-4403 | chr6-17211800-INS-4403 |
| chr6-17221103-INS-52 | chr6-17221103-INS-58 |
| chr6-17225763-INS-115 | chr6-17225794-INS-102 |
| chr6-17229516-INS-94 | chr6-17229521-INS-97 |
| chr6-17233994-INS-6420 | chr6-17233995-INS-6420 |
| chr6-17243827-INS-356 | chr6-17243827-INS-356 |
| chr6-17335983-INS-509 | chr6-17335984-INS-507 |
| chr6-17365656-INS-55 | chr6-17365656-INS-55 |
| chr6-17399980-INS-401 | chr6-17399980-INS-401 |

| | |
|---|---|
| chr6-17413350-INS-67 | chr6-17413350-INS-67 |
| chr6-17439757-INS-2657 | chr6-17439758-INS-2657 |
| chr6-17448667-INS-58 | chr6-17448667-INS-62 |
| chr6-17457612-INS-115 | chr6-17457597-INS-136 |
| chr6-17501290-INS-611 | chr6-17501293-INS-612 |
| chr6-17565726-INS-58 | chr6-17565727-INS-62 |
| chr6-17570292-INS-460 | chr6-17570298-INS-461 |
| chr6-17599062-INS-2737 | chr6-17599063-INS-2737 |
| chr6-17600756-INS-66 | chr6-17600756-INS-64 |
| chr6-17602844-INS-202 | chr6-17602844-INS-201 |
| chr6-17615330-INS-383 | chr6-17615330-INS-383 |
| chr6-17624689-INS-420 | chr6-17624689-INS-420 |
| chr6-17627134-INS-210 | chr6-17627134-INS-210 |
| chr6-17640477-INS-205 | chr6-17640477-INS-204 |
| chr6-17661312-INS-53 | chr6-17661312-INS-53 |
| chr6-17669519-INS-206 | chr6-17669519-INS-206 |
| chr6-17673175-INS-204 | chr6-17673175-INS-205 |
| chr6-17675809-INS-194 | chr6-17675809-INS-194 |
| chr6-17772578-INS-790 | chr6-17772578-INS-790 |
| chr6-17810265-INS-381 | chr6-17810265-INS-381 |
| chr6-17817786-INS-118 | chr6-17817786-INS-145 |
| chr6-17823152-INS-85 | chr6-17823147-INS-90 |
| chr6-17869290-INS-238 | chr6-17869290-INS-240 |
| chr6-17883279-INS-434 | chr6-17883279-INS-434 |
| chr6-17886236-INS-400 | chr6-17886236-INS-400 |
| chr6-17898951-INS-65 | chr6-17898951-INS-65 |
| chr6-17924832-INS-73 | chr6-17924832-INS-77 |
| chr6-17988932-INS-281 | chr6-17988932-INS-286 |
| chr6-18035275-INS-192 | chr6-18035275-INS-192 |
| chr6-18038828-INS-65 | chr6-18038828-INS-65 |
| chr6-18070037-INS-800 | chr6-18070037-INS-800 |
| chr6-18101460-INS-7008 | chr6-18101461-INS-7007 |
| chr6-18105178-INS-1712 | chr6-18105179-INS-1712 |
| chr6-18118862-INS-390 | chr6-18118856-INS-354 |
| chr6-18126373-INS-2018 | chr6-18126374-INS-2018 |
| chr6-18131274-INS-80 | chr6-18131274-INS-80 |
| chr6-18133105-INS-60 | chr6-18133105-INS-100 |
| chr6-18156718-INS-1189 | chr6-18156719-INS-1192 |
| chr6-18188934-INS-182 | chr6-18188934-INS-181 |
| chr6-18191830-INS-307 | chr6-18191830-INS-307 |
| chr6-18201538-INS-76 | chr6-18201538-INS-76 |
| chr6-18251896-INS-159 | chr6-18251896-INS-159 |
| chr6-18254820-INS-86 | chr6-18254820-INS-86 |
| chr6-18327761-INS-746 | chr6-18327761-INS-747 |
| chr6-18337985-INS-839 | chr6-18337985-INS-844 |
| chr6-18345295-INS-446 | chr6-18345295-INS-442 |
| chr6-18373174-INS-2189 | chr6-18373175-INS-2189 |
| chr6-18386520-INS-56 | chr6-18386520-INS-56 |
| chr6-18390221-INS-6593 | chr6-18390222-INS-6594 |
| chr6-18401999-INS-57 | chr6-18401999-INS-57 |
| chr6-18443454-INS-120 | chr6-18443412-INS-112 |
| chr6-18445778-INS-434 | chr6-18445778-INS-434 |
| chr6-18449633-INS-88 | chr6-18449633-INS-88 |
| chr6-18453234-INS-57 | chr6-18453234-INS-57 |
| chr6-18475305-INS-127 | chr6-18475305-INS-135 |
| chr6-18480780-INS-56 | chr6-18480780-INS-58 |
| chr6-18489756-INS-7105 | chr6-18489757-INS-7105 |
| chr6-18512792-INS-61 | chr6-18512792-INS-59 |
| chr6-18527168-INS-531 | chr6-18527168-INS-531 |
| chr6-18538179-INS-66 | chr6-18538179-INS-66 |
| chr6-18538191-INS-869 | chr6-18538191-INS-869 |
| chr6-18552296-INS-56 | chr6-18552296-INS-54 |
| chr6-18557249-INS-68 | chr6-18557249-INS-70 |
| chr6-18566923-INS-69 | chr6-18566925-INS-70 |
| chr6-18600069-INS-710 | chr6-18600069-INS-710 |
| chr6-18615099-INS-113 | chr6-18615099-INS-113 |
| chr6-18616654-INS-58 | chr6-18616654-INS-58 |
| chr6-18616936-INS-282 | chr6-18616936-INS-282 |
| chr6-18627667-INS-791 | chr6-18627668-INS-783 |
| chr6-18628209-INS-1667 | chr6-18628210-INS-1667 |
| chr6-18628321-INS-83 | chr6-18628325-INS-75 |
| chr6-18635177-INS-106 | chr6-18635177-INS-106 |
| chr6-18636429-INS-199 | chr6-18636429-INS-199 |
| chr6-18640668-INS-64 | chr6-18640668-INS-64 |
| chr6-18643759-INS-76 | chr6-18643759-INS-60 |
| chr6-18656105-INS-73 | chr6-18656105-INS-75 |
| chr6-18675423-INS-195 | chr6-18675377-INS-230 |

| | |
|---|---|
| chr6-18687552-INS-50 | chr6-18687552-INS-58 |
| chr6-18717379-INS-90 | chr6-18717379-INS-90 |
| chr6-18763516-INS-111 | chr6-18763516-INS-111 |
| chr6-18765487-INS-620 | chr6-18765487-INS-619 |
| chr6-18813830-INS-300 | chr6-18813831-INS-290 |
| chr6-18835955-INS-1143 | chr6-18835956-INS-1143 |
| chr6-18836645-INS-2152 | chr6-18836648-INS-2149 |
| chr6-18859794-INS-59 | chr6-18859794-INS-59 |
| chr6-18867736-INS-265 | chr6-18867736-INS-234 |
| chr6-18873080-INS-99 | chr6-18873066-INS-127 |
| chr6-18888748-INS-493 | chr6-18888748-INS-493 |
| chr6-18891138-INS-2724 | chr6-18891139-INS-2724 |
| chr6-18900208-INS-214 | chr6-18900208-INS-214 |
| chr6-18909170-INS-3045 | chr6-18909171-INS-3044 |
| chr6-18944688-INS-5302 | chr6-18944689-INS-5302 |
| chr6-18952042-INS-78 | chr6-18952042-INS-78 |
| chr6-18958862-INS-1557 | chr6-18958864-INS-1555 |
| chr6-19006336-INS-94 | chr6-19006336-INS-58 |
| chr6-19034950-INS-54 | chr6-19034950-INS-54 |
| chr6-19051857-INS-71 | chr6-19051857-INS-71 |
| chr6-19068150-INS-9907 | chr6-19068151-INS-9907 |
| chr6-19089445-INS-6187 | chr6-19089446-INS-6188 |
| chr6-19090985-INS-76 | chr6-19090980-INS-80 |
| chr6-19095016-INS-400 | chr6-19095016-INS-400 |
| chr6-19102875-INS-3112 | chr6-19102876-INS-3112 |
| chr6-19136516-INS-514 | chr6-19136516-INS-514 |
| chr6-19137891-INS-2584 | chr6-19137892-INS-2583 |
| chr6-19161040-INS-90 | chr6-19161040-INS-90 |
| chr6-19178752-INS-426 | chr6-19178752-INS-419 |
| chr6-19182486-INS-473 | chr6-19182486-INS-473 |
| chr6-19203963-INS-79 | chr6-19203963-INS-79 |
| chr6-19205790-INS-426 | chr6-19205790-INS-426 |
| chr6-19207832-INS-56 | chr6-19207832-INS-56 |
| chr6-19216647-INS-2872 | chr6-19216648-INS-2872 |
| chr6-19223265-INS-130 | chr6-19223265-INS-110 |
| chr6-19228246-INS-1944 | chr6-19228247-INS-1945 |
| chr6-19233496-INS-79 | chr6-19233510-INS-76 |
| chr6-19239479-INS-2302 | chr6-19239480-INS-2302 |
| chr6-19256952-INS-4016 | chr6-19256953-INS-4016 |
| chr6-19258430-INS-4179 | chr6-19258431-INS-4178 |
| chr6-19259666-INS-2417 | chr6-19259667-INS-2429 |
| chr6-19274211-INS-141 | chr6-19274211-INS-141 |
| chr6-19276489-INS-6323 | chr6-19276490-INS-6323 |
| chr6-19282707-INS-102 | chr6-19282707-INS-84 |
| chr6-19286431-INS-72 | chr6-19286431-INS-72 |
| chr6-19296854-INS-113 | chr6-19296854-INS-113 |
| chr6-19297129-INS-1524 | chr6-19297130-INS-1523 |
| chr6-19302608-INS-69 | chr6-19302607-INS-117 |
| chr6-19341586-INS-6316 | chr6-19341587-INS-6316 |
| chr6-19342271-INS-65 | chr6-19342271-INS-68 |
| chr6-19346448-INS-51 | chr6-19346448-INS-51 |
| chr6-19356391-INS-240 | chr6-19356391-INS-241 |
| chr6-19388288-INS-5250 | chr6-19388289-INS-5250 |
| chr6-19395894-INS-1953 | chr6-19395895-INS-1953 |
| chr6-19405076-INS-466 | chr6-19405076-INS-466 |
| chr6-19411678-INS-68 | chr6-19411678-INS-68 |
| chr6-19415274-INS-393 | chr6-19415274-INS-393 |
| chr6-19451647-INS-170 | chr6-19451647-INS-170 |
| chr6-19471210-INS-334 | chr6-19471210-INS-334 |
| chr6-19475030-INS-655 | chr6-19475030-INS-655 |
| chr6-19515834-INS-60 | chr6-19515834-INS-68 |
| chr6-19570996-INS-3641 | chr6-19570997-INS-3641 |
| chr6-19579158-INS-78 | chr6-19579160-INS-76 |
| chr6-19583099-INS-158 | chr6-19583105-INS-154 |
| chr6-19589334-INS-197 | chr6-19589347-INS-204 |
| chr6-19598942-INS-326 | chr6-19598942-INS-326 |
| chr6-19700151-INS-464 | chr6-19700151-INS-464 |
| chr6-19718768-INS-2364 | chr6-19718769-INS-2363 |
| chr6-19731267-INS-56 | chr6-19731265-INS-50 |
| chr6-19736995-INS-1939 | chr6-19736996-INS-1939 |
| chr6-19746677-INS-120 | chr6-19746677-INS-126 |
| chr6-19764233-INS-332 | chr6-19764233-INS-332 |
| chr6-19772235-INS-134 | chr6-19772235-INS-134 |
| chr6-19807870-INS-320 | chr6-19807870-INS-320 |
| chr6-19817739-INS-272 | chr6-19817739-INS-272 |
| chr6-19820586-INS-56 | chr6-19820587-INS-78 |
| chr6-19836516-INS-56 | chr6-19836516-INS-52 |

| | |
|---|---|
| chr6-19849602-INS-431 | chr6-19849602-INS-431 |
| chr6-19851556-INS-692 | chr6-19851556-INS-692 |
| chr6-19859777-INS-6374 | chr6-19859778-INS-6374 |
| chr6-19861080-INS-488 | chr6-19861080-INS-488 |
| chr6-19874272-INS-7111 | chr6-19874273-INS-7111 |
| chr6-19875813-INS-112 | chr6-19875813-INS-96 |
| chr6-19879164-INS-84 | chr6-19879164-INS-84 |
| chr6-19895032-INS-60 | chr6-19895032-INS-60 |
| chr6-19896756-INS-62 | chr6-19896756-INS-62 |
| chr6-19909918-INS-59 | chr6-19909916-INS-63 |
| chr6-19913250-INS-82 | chr6-19913250-INS-100 |
| chr6-19933519-INS-84 | chr6-19933519-INS-64 |
| chr6-19939353-INS-1018 | chr6-19939354-INS-1018 |
| chr6-20002108-INS-4742 | chr6-20002109-INS-4742 |
| chr6-20034386-INS-217 | chr6-20034386-INS-214 |
| chr6-20045266-INS-151 | chr6-20045266-INS-151 |
| chr6-20051996-INS-160 | chr6-20051996-INS-190 |
| chr6-20067139-INS-62 | chr6-20067140-INS-71 |
| chr6-20071265-INS-74 | chr6-20071265-INS-72 |
| chr6-20078465-INS-56 | chr6-20078461-INS-58 |
| chr6-20091009-INS-191 | chr6-20091009-INS-191 |
| chr6-20111197-INS-743 | chr6-20111197-INS-743 |
| chr6-20122499-INS-4226 | chr6-20122500-INS-4228 |
| chr6-20159606-INS-2321 | chr6-20159607-INS-2321 |
| chr6-20179960-INS-56 | chr6-20179960-INS-56 |
| chr6-20207040-INS-4795 | chr6-20207041-INS-4795 |
| chr6-20214819-INS-6802 | chr6-20214820-INS-6802 |
| chr6-20218934-INS-78 | chr6-20218934-INS-78 |
| chr6-20237606-INS-485 | chr6-20237606-INS-485 |
| chr6-20247037-INS-464 | chr6-20247037-INS-464 |
| chr6-20247210-INS-4169 | chr6-20247211-INS-4169 |
| chr6-20254553-INS-334 | chr6-20254553-INS-334 |
| chr6-20258750-INS-127 | chr6-20258747-INS-129 |
| chr6-20270801-INS-4177 | chr6-20270802-INS-4177 |
| chr6-20306049-INS-1165 | chr6-20306050-INS-1165 |
| chr6-20334097-INS-408 | chr6-20334097-INS-408 |
| chr6-20337892-INS-6347 | chr6-20337893-INS-6347 |
| chr6-20342115-INS-2961 | chr6-20342116-INS-2961 |
| chr6-20342116-INS-576 | chr6-20342116-INS-576 |
| chr6-20347988-INS-51 | chr6-20347988-INS-51 |
| chr6-20351706-INS-52 | chr6-20351706-INS-52 |
| chr6-20361573-INS-1302 | chr6-20361574-INS-1302 |
| chr6-20368673-INS-584 | chr6-20368673-INS-585 |
| chr6-20387240-INS-66 | chr6-20387226-INS-96 |
| chr6-20398180-INS-435 | chr6-20398180-INS-435 |
| chr6-20414825-INS-114 | chr6-20414825-INS-94 |
| chr6-20420943-INS-51 | chr6-20420943-INS-51 |
| chr6-20424687-INS-202 | chr6-20424687-INS-203 |
| chr6-20459440-INS-4491 | chr6-20459441-INS-4491 |
| chr6-20551937-INS-105 | chr6-20551937-INS-93 |
| chr6-20560187-INS-7469 | chr6-20560188-INS-7469 |
| chr6-20595427-INS-266 | chr6-20595427-INS-268 |
| chr6-20622728-INS-381 | chr6-20622728-INS-381 |
| chr6-20648337-INS-65 | chr6-20648337-INS-65 |
| chr6-20651813-INS-586 | chr6-20651813-INS-586 |
| chr6-20679882-INS-440 | chr6-20679882-INS-440 |
| chr6-20679971-INS-6847 | chr6-20679972-INS-6847 |
| chr6-20680466-INS-355 | chr6-20680466-INS-355 |
| chr6-20703762-INS-156 | chr6-20703762-INS-162 |
| chr6-20721829-INS-3230 | chr6-20721830-INS-3230 |
| chr6-20743898-INS-55 | chr6-20743898-INS-55 |
| chr6-20758879-INS-441 | chr6-20758879-INS-441 |
| chr6-20760454-INS-76 | chr6-20760454-INS-76 |
| chr6-20761825-INS-595 | chr6-20761820-INS-555 |
| chr6-20764862-INS-2212 | chr6-20764863-INS-2212 |
| chr6-20771687-INS-139 | chr6-20771687-INS-129 |
| chr6-20781366-INS-583 | chr6-20781366-INS-583 |
| chr6-20808966-INS-106 | chr6-20808969-INS-84 |
| chr6-20818262-INS-74 | chr6-20818236-INS-72 |
| chr6-20835771-INS-525 | chr6-20835771-INS-524 |
| chr6-20851859-INS-806 | chr6-20851859-INS-806 |
| chr6-20852534-INS-437 | chr6-20852534-INS-438 |
| chr6-20866605-INS-61 | chr6-20866605-INS-63 |
| chr6-20878398-INS-92 | chr6-20878398-INS-92 |
| chr6-20937697-INS-52 | chr6-20937697-INS-61 |
| chr6-20955968-INS-498 | chr6-20955968-INS-498 |
| chr6-20961495-INS-6784 | chr6-20961496-INS-6784 |

| | |
|---|---|
| chr6-20970093-INS-390 | chr6-20970093-INS-398 |
| chr6-20978381-INS-64 | chr6-20978381-INS-64 |
| chr6-20982020-INS-526 | chr6-20982020-INS-520 |
| chr6-20995001-INS-500 | chr6-20995001-INS-499 |
| chr6-21006519-INS-430 | chr6-21006519-INS-430 |
| chr6-21042877-INS-221 | chr6-21042877-INS-217 |
| chr6-21060006-INS-54 | chr6-21060006-INS-56 |
| chr6-21114773-INS-183 | chr6-21114768-INS-195 |
| chr6-21119951-INS-463 | chr6-21119933-INS-448 |
| chr6-21125499-INS-59 | chr6-21125499-INS-59 |
| chr6-21155202-INS-72 | chr6-21155202-INS-72 |
| chr6-21164080-INS-54 | chr6-21164080-INS-56 |
| chr6-21181733-INS-1086 | chr6-21181734-INS-1082 |
| chr6-21221969-INS-51 | chr6-21221968-INS-70 |
| chr6-21228419-INS-852 | chr6-21228408-INS-849 |
| chr6-21252598-INS-96 | chr6-21252598-INS-96 |
| chr6-21285102-INS-724 | chr6-21285102-INS-724 |
| chr6-21289046-INS-608 | chr6-21289046-INS-608 |
| chr6-21309625-INS-231 | chr6-21309625-INS-231 |
| chr6-21312436-INS-158 | chr6-21312436-INS-156 |
| chr6-21334340-INS-190 | chr6-21334340-INS-190 |
| chr6-21340925-INS-1938 | chr6-21340926-INS-1939 |
| chr6-21367916-INS-184 | chr6-21367916-INS-184 |
| chr6-21368451-INS-10701 | chr6-21368452-INS-10702 |
| chr6-21383799-INS-52 | chr6-21383788-INS-57 |
| chr6-21418915-INS-249 | chr6-21418915-INS-250 |
| chr6-21443301-INS-120 | chr6-21443299-INS-72 |
| chr6-21444553-INS-797 | chr6-21444553-INS-798 |
| chr6-21448277-INS-51 | chr6-21448255-INS-64 |
| chr6-21459167-INS-338 | chr6-21459167-INS-338 |
| chr6-21463521-INS-827 | chr6-21463521-INS-828 |
| chr6-21465995-INS-796 | chr6-21465995-INS-796 |
| chr6-21479538-INS-91 | chr6-21479538-INS-91 |
| chr6-21491664-INS-303 | chr6-21491664-INS-303 |
| chr6-21499465-INS-479 | chr6-21499465-INS-479 |
| chr6-21502478-INS-122 | chr6-21502478-INS-122 |
| chr6-21555275-INS-6407 | chr6-21555276-INS-6408 |
| chr6-21647050-INS-2417 | chr6-21647051-INS-2418 |
| chr6-21649467-INS-355 | chr6-21649467-INS-355 |
| chr6-21651064-INS-201 | chr6-21651064-INS-201 |
| chr6-21671520-INS-64 | chr6-21671520-INS-64 |
| chr6-21675171-INS-59 | chr6-21675134-INS-91 |
| chr6-21735597-INS-90 | chr6-21735597-INS-90 |
| chr6-21738064-INS-150 | chr6-21738064-INS-150 |
| chr6-21761293-INS-56 | chr6-21761293-INS-70 |
| chr6-21794122-INS-98 | chr6-21794122-INS-98 |
| chr6-21808981-INS-419 | chr6-21808981-INS-419 |
| chr6-21848068-INS-88 | chr6-21848068-INS-50 |
| chr6-21859852-INS-433 | chr6-21859852-INS-432 |
| chr6-21871558-INS-117 | chr6-21871557-INS-69 |
| chr6-21905824-INS-72 | chr6-21905824-INS-88 |
| chr6-21907917-INS-148 | chr6-21907917-INS-151 |
| chr6-21991814-INS-130 | chr6-21991814-INS-130 |
| chr6-22010917-INS-76 | chr6-22010917-INS-76 |
| chr6-22031804-INS-61 | chr6-22031804-INS-61 |
| chr6-22057600-INS-520 | chr6-22057600-INS-513 |
| chr6-22068617-INS-723 | chr6-22068617-INS-724 |
| chr6-22073126-INS-10103 | chr6-22073127-INS-10104 |
| chr6-22075119-INS-119 | chr6-22075119-INS-119 |
| chr6-22077928-INS-378 | chr6-22077928-INS-378 |
| chr6-22138137-INS-254 | chr6-22138137-INS-260 |
| chr6-22144957-INS-388 | chr6-22144957-INS-387 |
| chr6-22166909-INS-206 | chr6-22166910-INS-209 |
| chr6-22168216-INS-151 | chr6-22168220-INS-145 |
| chr6-22182496-INS-228 | chr6-22182496-INS-233 |
| chr6-22195954-INS-66 | chr6-22195954-INS-66 |
| chr6-22216287-INS-68 | chr6-22216263-INS-76 |
| chr6-22253862-INS-66 | chr6-22253862-INS-66 |
| chr6-22263265-INS-76 | chr6-22263265-INS-80 |
| chr6-22269761-INS-54 | chr6-22269761-INS-54 |
| chr6-22312230-INS-148 | chr6-22312230-INS-148 |
| chr6-22319824-INS-72 | chr6-22319824-INS-104 |
| chr6-22334432-INS-62 | chr6-22334432-INS-62 |
| chr6-22335217-INS-83 | chr6-22335217-INS-83 |
| chr6-22340419-INS-122 | chr6-22340419-INS-122 |
| chr6-22357833-INS-906 | chr6-22357833-INS-906 |
| chr6-22364167-INS-209 | chr6-22364167-INS-210 |

| | |
|---|---|
| chr6-22371109-INS-51 | chr6-22371109-INS-53 |
| chr6-22376545-INS-209 | chr6-22376545-INS-209 |
| chr6-22383509-INS-195 | chr6-22383509-INS-195 |
| chr6-22386061-INS-207 | chr6-22386070-INS-200 |
| chr6-22389627-INS-172 | chr6-22389627-INS-173 |
| chr6-22395005-INS-430 | chr6-22395006-INS-430 |
| chr6-22396867-INS-184 | chr6-22396867-INS-182 |
| chr6-22398490-INS-220 | chr6-22398490-INS-220 |
| chr6-22408820-INS-251 | chr6-22408820-INS-238 |
| chr6-22419207-INS-100 | chr6-22419171-INS-124 |
| chr6-22425576-INS-66 | chr6-22425576-INS-66 |
| chr6-22431794-INS-995 | chr6-22431799-INS-996 |
| chr6-22433625-INS-101 | chr6-22433627-INS-64 |
| chr6-22445662-INS-84 | chr6-22445662-INS-68 |
| chr6-22474418-INS-188 | chr6-22474413-INS-197 |
| chr6-22482358-INS-248 | chr6-22482373-INS-260 |
| chr6-22487371-INS-622 | chr6-22487371-INS-622 |
| chr6-22515273-INS-6687 | chr6-22515274-INS-6689 |
| chr6-22528247-INS-198 | chr6-22528247-INS-198 |
| chr6-22557175-INS-683 | chr6-22557175-INS-683 |
| chr6-22559403-INS-82 | chr6-22559404-INS-54 |
| chr6-22566375-INS-427 | chr6-22566381-INS-426 |
| chr6-22568290-INS-236 | chr6-22568290-INS-236 |
| chr6-22609372-INS-4822 | chr6-22609373-INS-4822 |
| chr6-22614276-INS-199 | chr6-22614276-INS-199 |
| chr6-22618246-INS-6598 | chr6-22618247-INS-6598 |
| chr6-22619683-INS-214 | chr6-22619683-INS-213 |
| chr6-22640132-INS-417 | chr6-22640132-INS-417 |
| chr6-22663309-INS-517 | chr6-22663309-INS-517 |
| chr6-22666956-INS-674 | chr6-22666956-INS-675 |
| chr6-22667565-INS-70 | chr6-22667566-INS-72 |
| chr6-22670086-INS-6209 | chr6-22670087-INS-6209 |
| chr6-22676413-INS-4160 | chr6-22676414-INS-4160 |
| chr6-22697876-INS-1593 | chr6-22697877-INS-1593 |
| chr6-22719397-INS-64 | chr6-22719397-INS-64 |
| chr6-22723947-INS-2860 | chr6-22723948-INS-2860 |
| chr6-22735015-INS-1081 | chr6-22735016-INS-1084 |
| chr6-22757715-INS-325 | chr6-22757715-INS-325 |
| chr6-22766295-INS-116 | chr6-22766295-INS-78 |
| chr6-22768332-INS-607 | chr6-22768332-INS-607 |
| chr6-22785784-INS-203 | chr6-22785784-INS-203 |
| chr6-22803188-INS-217 | chr6-22803188-INS-221 |
| chr6-22804384-INS-213 | chr6-22804384-INS-214 |
| chr6-22806580-INS-589 | chr6-22806580-INS-589 |
| chr6-22825817-INS-5141 | chr6-22825818-INS-5142 |
| chr6-22829402-INS-79 | chr6-22829402-INS-79 |
| chr6-22843120-INS-2012 | chr6-22843121-INS-2011 |
| chr6-22849250-INS-2134 | chr6-22849251-INS-2137 |
| chr6-22850674-INS-6290 | chr6-22850675-INS-6294 |
| chr6-22861455-INS-4105 | chr6-22861456-INS-4106 |
| chr6-22864907-INS-6470 | chr6-22864908-INS-6470 |
| chr6-22925442-INS-143 | chr6-22925442-INS-144 |
| chr6-22927818-INS-136 | chr6-22927798-INS-111 |
| chr6-22953063-INS-85 | chr6-22953063-INS-85 |
| chr6-23025887-INS-80 | chr6-23025882-INS-75 |
| chr6-23028224-INS-1888 | chr6-23028225-INS-1888 |
| chr6-23060066-INS-62 | chr6-23060070-INS-56 |
| chr6-23068768-INS-125 | chr6-23068768-INS-135 |
| chr6-23073548-INS-247 | chr6-23073548-INS-247 |
| chr6-23078213-INS-80 | chr6-23078213-INS-51 |
| chr6-23099154-INS-171 | chr6-23099154-INS-171 |
| chr6-23111786-INS-80 | chr6-23111773-INS-116 |
| chr6-23145631-INS-172 | chr6-23145631-INS-172 |
| chr6-23192571-INS-198 | chr6-23192571-INS-199 |
| chr6-23197607-INS-62 | chr6-23197607-INS-62 |
| chr6-23218792-INS-416 | chr6-23218792-INS-424 |
| chr6-23220264-INS-91 | chr6-23220264-INS-91 |
| chr6-23220733-INS-93 | chr6-23220733-INS-93 |
| chr6-23229918-INS-607 | chr6-23229955-INS-608 |
| chr6-23243557-INS-66 | chr6-23243557-INS-66 |
| chr6-23274453-INS-58 | chr6-23274453-INS-58 |
| chr6-23296488-INS-132 | chr6-23296488-INS-170 |
| chr6-23302141-INS-148 | chr6-23302137-INS-128 |
| chr6-23313831-INS-55 | chr6-23313831-INS-55 |
| chr6-23328237-INS-52 | chr6-23328237-INS-52 |
| chr6-23346453-INS-56 | chr6-23346453-INS-72 |
| chr6-23352069-INS-52 | chr6-23352069-INS-52 |

| | |
|---|---|
| chr6-23362995-INS-172 | chr6-23362995-INS-172 |
| chr6-23374138-INS-270 | chr6-23374138-INS-270 |
| chr6-23415851-INS-72 | chr6-23415851-INS-72 |
| chr6-23421426-INS-729 | chr6-23421426-INS-729 |
| chr6-23488291-INS-590 | chr6-23488292-INS-592 |
| chr6-23535924-INS-58 | chr6-23535924-INS-58 |
| chr6-23571079-INS-584 | chr6-23571098-INS-585 |
| chr6-23593344-INS-990 | chr6-23593344-INS-990 |
| chr6-23609592-INS-4510 | chr6-23609593-INS-4515 |
| chr6-23622734-INS-71 | chr6-23622734-INS-104 |
| chr6-23634391-INS-60 | chr6-23634391-INS-72 |
| chr6-23645178-INS-65 | chr6-23645176-INS-74 |
| chr6-23667328-INS-525 | chr6-23667328-INS-525 |
| chr6-23680636-INS-160 | chr6-23680636-INS-154 |
| chr6-23686570-INS-96 | chr6-23686570-INS-96 |
| chr6-23714959-INS-51 | chr6-23714959-INS-51 |
| chr6-23730442-INS-60 | chr6-23730442-INS-62 |
| chr6-23764433-INS-64 | chr6-23764433-INS-66 |
| chr6-23767010-INS-51 | chr6-23767010-INS-69 |
| chr6-23778117-INS-390 | chr6-23778120-INS-386 |
| chr6-23780627-INS-114 | chr6-23780627-INS-114 |
| chr6-23785563-INS-918 | chr6-23785563-INS-918 |
| chr6-23786088-INS-105 | chr6-23786088-INS-105 |
| chr6-23796677-INS-54 | chr6-23796677-INS-54 |
| chr6-23846114-INS-51 | chr6-23846114-INS-52 |
| chr6-23897934-INS-6585 | chr6-23897935-INS-6586 |
| chr6-23945949-INS-77 | chr6-23945949-INS-77 |
| chr6-23947689-INS-4104 | chr6-23947690-INS-4104 |
| chr6-23955930-INS-93 | chr6-23955930-INS-67 |
| chr6-23956981-INS-66 | chr6-23956981-INS-66 |
| chr6-24023482-INS-66 | chr6-24023482-INS-75 |
| chr6-24024804-INS-590 | chr6-24024804-INS-610 |
| chr6-24077587-INS-57 | chr6-24077587-INS-51 |
| chr6-24082130-INS-504 | chr6-24082130-INS-505 |
| chr6-24105117-INS-62 | chr6-24105117-INS-52 |
| chr6-24113304-INS-318 | chr6-24113304-INS-318 |
| chr6-24125537-INS-62 | chr6-24125526-INS-74 |
| chr6-24130171-INS-103 | chr6-24130172-INS-107 |
| chr6-24131897-INS-109 | chr6-24131897-INS-112 |
| chr6-24140825-INS-108 | chr6-24140825-INS-102 |
| chr6-24181364-INS-165 | chr6-24181364-INS-166 |
| chr6-24199272-INS-377 | chr6-24199272-INS-377 |
| chr6-24235784-INS-52 | chr6-24235784-INS-50 |
| chr6-24263553-INS-58 | chr6-24263553-INS-58 |
| chr6-24277974-INS-93 | chr6-24277974-INS-93 |
| chr6-24301577-INS-55 | chr6-24301602-INS-74 |
| chr6-24329329-INS-75 | chr6-24329336-INS-102 |
| chr6-24352473-INS-400 | chr6-24352473-INS-400 |
| chr6-24354320-INS-996 | chr6-24354313-INS-968 |
| chr6-24355904-INS-1570 | chr6-24355905-INS-1571 |
| chr6-24357682-INS-2350 | chr6-24357681-INS-2336 |
| chr6-24366112-INS-538 | chr6-24366112-INS-538 |
| chr6-24373172-INS-90 | chr6-24373157-INS-110 |
| chr6-24375170-INS-6283 | chr6-24375171-INS-6278 |
| chr6-24386879-INS-1068 | chr6-24386880-INS-1068 |
| chr6-24422817-INS-6826 | chr6-24422818-INS-6826 |
| chr6-24474199-INS-91 | chr6-24474199-INS-132 |
| chr6-24491900-INS-212 | chr6-24491900-INS-212 |
| chr6-24511152-INS-66 | chr6-24511157-INS-71 |
| chr6-24523690-INS-94 | chr6-24523690-INS-94 |
| chr6-24610106-INS-55 | chr6-24610106-INS-55 |
| chr6-24644472-INS-158 | chr6-24644472-INS-158 |
| chr6-24683896-INS-398 | chr6-24683896-INS-398 |
| chr6-24696720-INS-249 | chr6-24696720-INS-241 |
| chr6-24707333-INS-948 | chr6-24707333-INS-948 |
| chr6-24724469-INS-76 | chr6-24724469-INS-76 |
| chr6-24737324-INS-71 | chr6-24737324-INS-75 |
| chr6-24758659-INS-85 | chr6-24758659-INS-75 |
| chr6-24771082-INS-169 | chr6-24771078-INS-173 |
| chr6-24772129-INS-62 | chr6-24772129-INS-70 |
| chr6-24805073-INS-599 | chr6-24805073-INS-599 |
| chr6-24824242-INS-388 | chr6-24824242-INS-381 |
| chr6-24826303-INS-302 | chr6-24826303-INS-302 |
| chr6-24839267-INS-182 | chr6-24839268-INS-162 |
| chr6-24880213-INS-202 | chr6-24880213-INS-202 |
| chr6-24921750-INS-487 | chr6-24921750-INS-487 |
| chr6-24970097-INS-92 | chr6-24970097-INS-92 |

| | |
|---|---|
| chr6-24989263-INS-203 | chr6-24989263-INS-204 |
| chr6-24999910-INS-647 | chr6-24999910-INS-649 |
| chr6-25024597-INS-244 | chr6-25024597-INS-244 |
| chr6-25065453-INS-173 | chr6-25065453-INS-176 |
| chr6-25130435-INS-250 | chr6-25130435-INS-250 |
| chr6-25138581-INS-74 | chr6-25138581-INS-76 |
| chr6-25143856-INS-3440 | chr6-25143857-INS-3440 |
| chr6-25150959-INS-400 | chr6-25150959-INS-400 |
| chr6-25166085-INS-134 | chr6-25166085-INS-130 |
| chr6-25174917-INS-504 | chr6-25174917-INS-504 |
| chr6-25181264-INS-76 | chr6-25181264-INS-80 |
| chr6-25195638-INS-53 | chr6-25195638-INS-53 |
| chr6-25197435-INS-184 | chr6-25197435-INS-184 |
| chr6-25220509-INS-5410 | chr6-25220510-INS-5410 |
| chr6-25244439-INS-209 | chr6-25244439-INS-209 |
| chr6-25254320-INS-3312 | chr6-25254321-INS-3312 |
| chr6-25275238-INS-6527 | chr6-25275239-INS-6527 |
| chr6-25301508-INS-83 | chr6-25301508-INS-83 |
| chr6-25318752-INS-487 | chr6-25318752-INS-487 |
| chr6-25335996-INS-1154 | chr6-25335997-INS-1149 |
| chr6-25373923-INS-96 | chr6-25373923-INS-56 |
| chr6-25409686-INS-119 | chr6-25409686-INS-119 |
| chr6-25411241-INS-220 | chr6-25411241-INS-220 |
| chr6-25457443-INS-374 | chr6-25457443-INS-373 |
| chr6-25500770-INS-150 | chr6-25500768-INS-154 |
| chr6-25587143-INS-1095 | chr6-25587144-INS-1095 |
| chr6-25602553-INS-59 | chr6-25602553-INS-60 |
| chr6-25604501-INS-70 | chr6-25604501-INS-66 |
| chr6-25607412-INS-54 | chr6-25607406-INS-56 |
| chr6-25608354-INS-74 | chr6-25608354-INS-74 |
| chr6-25615947-INS-90 | chr6-25615947-INS-90 |
| chr6-25627602-INS-801 | chr6-25627602-INS-803 |
| chr6-25635475-INS-1153 | chr6-25635476-INS-1153 |
| chr6-25652887-INS-179 | chr6-25652887-INS-183 |
| chr6-25666442-INS-95 | chr6-25666442-INS-100 |
| chr6-25675685-INS-6722 | chr6-25675686-INS-6724 |
| chr6-25693526-INS-643 | chr6-25693526-INS-638 |
| chr6-25696650-INS-1270 | chr6-25696651-INS-1273 |
| chr6-25713181-INS-580 | chr6-25713181-INS-580 |
| chr6-25742102-INS-60 | chr6-25742102-INS-60 |
| chr6-25755227-INS-53 | chr6-25755209-INS-57 |
| chr6-25776462-INS-3282 | chr6-25776463-INS-3282 |
| chr6-25776515-INS-66 | chr6-25776515-INS-66 |
| chr6-25793879-INS-6760 | chr6-25793880-INS-6760 |
| chr6-25811977-INS-70 | chr6-25811977-INS-70 |
| chr6-25816941-INS-1888 | chr6-25816942-INS-1888 |
| chr6-25843390-INS-98 | chr6-25843390-INS-98 |
| chr6-25856939-INS-6996 | chr6-25856940-INS-6998 |
| chr6-25860170-INS-53 | chr6-25860170-INS-51 |
| chr6-25870172-INS-7191 | chr6-25870173-INS-7191 |
| chr6-25895450-INS-74 | chr6-25895450-INS-123 |
| chr6-25902699-INS-259 | chr6-25902699-INS-271 |
| chr6-25923123-INS-130 | chr6-25923126-INS-90 |
| chr6-25940869-INS-88 | chr6-25940869-INS-73 |
| chr6-25975971-INS-449 | chr6-25975971-INS-449 |
| chr6-25999440-INS-1031 | chr6-25999441-INS-1031 |
| chr6-26001890-INS-654 | chr6-26001890-INS-654 |
| chr6-26054504-INS-61 | chr6-26054498-INS-77 |
| chr6-26082417-INS-56 | chr6-26082417-INS-56 |
| chr6-26154034-INS-150 | chr6-26154035-INS-156 |
| chr6-26171975-INS-233 | chr6-26171975-INS-240 |
| chr6-26179055-INS-71 | chr6-26179051-INS-110 |
| chr6-26210240-INS-117 | chr6-26210240-INS-117 |
| chr6-26211423-INS-178 | chr6-26211423-INS-158 |
| chr6-26222649-INS-1108 | chr6-26222650-INS-1108 |
| chr6-26234688-INS-205 | chr6-26234688-INS-205 |
| chr6-26280307-INS-201 | chr6-26280307-INS-201 |
| chr6-26290940-INS-608 | chr6-26290940-INS-608 |
| chr6-26306638-INS-4756 | chr6-26306639-INS-4756 |
| chr6-26312416-INS-72 | chr6-26312416-INS-75 |
| chr6-26325747-INS-109 | chr6-26325747-INS-109 |
| chr6-26328315-INS-6522 | chr6-26328316-INS-6522 |
| chr6-26360017-INS-6567 | chr6-26360027-INS-6566 |
| chr6-26366464-INS-720 | chr6-26366464-INS-720 |
| chr6-26381963-INS-1912 | chr6-26381964-INS-1912 |
| chr6-26385749-INS-56 | chr6-26385753-INS-56 |
| chr6-26391033-INS-2865 | chr6-26391034-INS-2865 |

| | |
|---|---|
| chr6-26430079-INS-93 | chr6-26430079-INS-103 |
| chr6-26431433-INS-71 | chr6-26431422-INS-87 |
| chr6-26442717-INS-734 | chr6-26442717-INS-734 |
| chr6-26447983-INS-193 | chr6-26447983-INS-193 |
| chr6-26450844-INS-54 | chr6-26450844-INS-54 |
| chr6-26460771-INS-242 | chr6-26460771-INS-242 |
| chr6-26464760-INS-192 | chr6-26464760-INS-193 |
| chr6-26494403-INS-64 | chr6-26494403-INS-68 |
| chr6-26539794-INS-984 | chr6-26539794-INS-983 |
| chr6-26553385-INS-4830 | chr6-26553386-INS-4830 |
| chr6-26558030-INS-69 | chr6-26558030-INS-69 |
| chr6-26585759-INS-53 | chr6-26585759-INS-53 |
| chr6-26613930-INS-197 | chr6-26613930-INS-198 |
| chr6-26632664-INS-968 | chr6-26632664-INS-968 |
| chr6-26658418-INS-53 | chr6-26658418-INS-53 |
| chr6-26668202-INS-88 | chr6-26668202-INS-60 |
| chr6-26725472-INS-707 | chr6-26725474-INS-711 |
| chr6-26802718-INS-822 | chr6-26802718-INS-811 |
| chr6-26819808-INS-325 | chr6-26819808-INS-325 |
| chr6-26823261-INS-586 | chr6-26823261-INS-588 |
| chr6-26897540-INS-212 | chr6-26897540-INS-200 |
| chr6-26902424-INS-81 | chr6-26902416-INS-67 |
| chr6-26934360-INS-60 | chr6-26934370-INS-50 |
| chr6-26955125-INS-86 | chr6-26955129-INS-87 |
| chr6-26974039-INS-889 | chr6-26974039-INS-892 |
| chr6-26975697-INS-150 | chr6-26975697-INS-168 |
| chr6-26981985-INS-68 | chr6-26981980-INS-58 |
| chr6-27002008-INS-103 | chr6-27002008-INS-108 |
| chr6-27006784-INS-72 | chr6-27006754-INS-78 |
| chr6-27019037-INS-6483 | chr6-27019038-INS-6483 |
| chr6-27049446-INS-721 | chr6-27049446-INS-720 |
| chr6-27053408-INS-97 | chr6-27053408-INS-97 |
| chr6-27059594-INS-341 | chr6-27059594-INS-313 |
| chr6-27060201-INS-82 | chr6-27060201-INS-84 |
| chr6-27185204-INS-329 | chr6-27185204-INS-329 |
| chr6-27257556-INS-6163 | chr6-27257543-INS-6162 |
| chr6-27281515-INS-172 | chr6-27281515-INS-172 |
| chr6-27282696-INS-243 | chr6-27282696-INS-244 |
| chr6-27287906-INS-221 | chr6-27287908-INS-224 |
| chr6-27299506-INS-65 | chr6-27299506-INS-62 |
| chr6-27303103-INS-137 | chr6-27303103-INS-137 |
| chr6-27357200-INS-330 | chr6-27357200-INS-329 |
| chr6-27365498-INS-52 | chr6-27365498-INS-52 |
| chr6-27374583-INS-559 | chr6-27374583-INS-558 |
| chr6-27375335-INS-1769 | chr6-27375336-INS-1769 |
| chr6-27395510-INS-6908 | chr6-27395511-INS-6908 |
| chr6-27425794-INS-152 | chr6-27425794-INS-153 |
| chr6-27427105-INS-8943 | chr6-27427106-INS-8943 |
| chr6-27439675-INS-6446 | chr6-27439676-INS-6446 |
| chr6-27449816-INS-358 | chr6-27449816-INS-358 |
| chr6-27469710-INS-225 | chr6-27469710-INS-225 |
| chr6-27470079-INS-607 | chr6-27470079-INS-607 |
| chr6-27485059-INS-94 | chr6-27485059-INS-140 |
| chr6-27493889-INS-102 | chr6-27493931-INS-82 |
| chr6-27564701-INS-244 | chr6-27564701-INS-244 |
| chr6-27576470-INS-390 | chr6-27576470-INS-390 |
| chr6-27608134-INS-102 | chr6-27608134-INS-102 |
| chr6-27619754-INS-62 | chr6-27619754-INS-58 |
| chr6-27622467-INS-708 | chr6-27622467-INS-706 |
| chr6-27632050-INS-145 | chr6-27632050-INS-147 |
| chr6-27657430-INS-6730 | chr6-27657431-INS-6731 |
| chr6-27695907-INS-75 | chr6-27695907-INS-73 |
| chr6-27696935-INS-386 | chr6-27696924-INS-394 |
| chr6-27711525-INS-434 | chr6-27711527-INS-467 |
| chr6-27720379-INS-192 | chr6-27720379-INS-192 |
| chr6-27768342-INS-2596 | chr6-27768343-INS-2596 |
| chr6-27874450-INS-212 | chr6-27874450-INS-213 |
| chr6-27882434-INS-129 | chr6-27882434-INS-129 |
| chr6-27895973-INS-60 | chr6-27895973-INS-60 |
| chr6-27917777-INS-74 | chr6-27917777-INS-74 |
| chr6-27935499-INS-7937 | chr6-27935500-INS-7933 |
| chr6-27938333-INS-187 | chr6-27938333-INS-187 |
| chr6-27958767-INS-66 | chr6-27958767-INS-66 |
| chr6-27963769-INS-62 | chr6-27963769-INS-62 |
| chr6-27977960-INS-197 | chr6-27977963-INS-214 |
| chr6-27989509-INS-1764 | chr6-27989510-INS-1765 |
| chr6-27989718-INS-65 | chr6-27989690-INS-55 |

| | |
|---|---|
| chr6-28010690-INS-73 | chr6-28010676-INS-50 |
| chr6-28030711-INS-1361 | chr6-28030712-INS-1362 |
| chr6-28038636-INS-352 | chr6-28038636-INS-348 |
| chr6-28070740-INS-602 | chr6-28070740-INS-597 |
| chr6-28104706-INS-486 | chr6-28104706-INS-486 |
| chr6-28123112-INS-1070 | chr6-28123113-INS-1071 |
| chr6-28135443-INS-58 | chr6-28135443-INS-57 |
| chr6-28143795-INS-668 | chr6-28143795-INS-671 |
| chr6-28160149-INS-461 | chr6-28160149-INS-461 |
| chr6-28169074-INS-70 | chr6-28169074-INS-72 |
| chr6-28170358-INS-54 | chr6-28170358-INS-52 |
| chr6-28185632-INS-53 | chr6-28185637-INS-56 |
| chr6-28189847-INS-1109 | chr6-28189848-INS-1109 |
| chr6-28194499-INS-96 | chr6-28194499-INS-56 |
| chr6-28237319-INS-150 | chr6-28237319-INS-150 |
| chr6-28267389-INS-282 | chr6-28267389-INS-282 |
| chr6-28281845-INS-811 | chr6-28281845-INS-812 |
| chr6-28301473-INS-211 | chr6-28301473-INS-212 |
| chr6-28365324-INS-7221 | chr6-28365325-INS-7201 |
| chr6-28414241-INS-75 | chr6-28414241-INS-75 |
| chr6-28430852-INS-51 | chr6-28430852-INS-51 |
| chr6-28439192-INS-56 | chr6-28439192-INS-56 |
| chr6-28443406-INS-82 | chr6-28443407-INS-55 |
| chr6-28444268-INS-150 | chr6-28444268-INS-150 |
| chr6-28445650-INS-465 | chr6-28445650-INS-465 |
| chr6-28453070-INS-96 | chr6-28453070-INS-71 |
| chr6-28454041-INS-2771 | chr6-28454042-INS-2771 |
| chr6-28459544-INS-6159 | chr6-28459545-INS-6159 |
| chr6-28530939-INS-94 | chr6-28530939-INS-94 |
| chr6-28579523-INS-97 | chr6-28579523-INS-97 |
| chr6-28683349-INS-66 | chr6-28683325-INS-52 |
| chr6-28685086-INS-90 | chr6-28685086-INS-85 |
| chr6-28690782-INS-71 | chr6-28690782-INS-71 |
| chr6-28718627-INS-212 | chr6-28718627-INS-208 |
| chr6-28769916-INS-84 | chr6-28769916-INS-79 |
| chr6-28810672-INS-79 | chr6-28810672-INS-79 |
| chr6-28870080-INS-143 | chr6-28870080-INS-143 |
| chr6-28912122-INS-328 | chr6-28912122-INS-328 |
| chr6-28942481-INS-218 | chr6-28942481-INS-218 |
| chr6-28944959-INS-54 | chr6-28944959-INS-54 |
| chr6-28949828-INS-68 | chr6-28949828-INS-68 |
| chr6-28957374-INS-64 | chr6-28957372-INS-62 |
| chr6-28962581-INS-6672 | chr6-28962582-INS-6672 |
| chr6-29003151-INS-199 | chr6-29003151-INS-199 |
| chr6-29014071-INS-76 | chr6-29014071-INS-76 |
| chr6-29034638-INS-82 | chr6-29034671-INS-56 |
| chr6-29037274-INS-64 | chr6-29037274-INS-64 |
| chr6-29063791-INS-90 | chr6-29063791-INS-90 |
| chr6-29089345-INS-65 | chr6-29089345-INS-65 |
| chr6-29090169-INS-159 | chr6-29090169-INS-159 |
| chr6-29118047-INS-62 | chr6-29118047-INS-76 |
| chr6-29142748-INS-210 | chr6-29142748-INS-210 |
| chr6-29180548-INS-60 | chr6-29180548-INS-60 |
| chr6-29196773-INS-193 | chr6-29196773-INS-191 |
| chr6-29211000-INS-194 | chr6-29211000-INS-195 |
| chr6-29220684-INS-296 | chr6-29220684-INS-297 |
| chr6-29224510-INS-406 | chr6-29224510-INS-406 |
| chr6-29228899-INS-68 | chr6-29228899-INS-58 |
| chr6-29233649-INS-72 | chr6-29233649-INS-72 |
| chr6-29238883-INS-203 | chr6-29238883-INS-203 |
| chr6-29259057-INS-206 | chr6-29259057-INS-204 |
| chr6-29268617-INS-226 | chr6-29268617-INS-226 |
| chr6-29284699-INS-63 | chr6-29284699-INS-63 |
| chr6-29324921-INS-54 | chr6-29324917-INS-80 |
| chr6-29364918-INS-295 | chr6-29364908-INS-293 |
| chr6-29369639-INS-176 | chr6-29369639-INS-186 |
| chr6-29468559-INS-79 | chr6-29468559-INS-79 |
| chr6-29480413-INS-70 | chr6-29480425-INS-58 |
| chr6-29497130-INS-54 | chr6-29497130-INS-54 |
| chr6-29503254-INS-66 | chr6-29503254-INS-54 |
| chr6-29513915-INS-71 | chr6-29513915-INS-72 |
| chr6-29568200-INS-63 | chr6-29568200-INS-51 |
| chr6-29626214-INS-96 | chr6-29626214-INS-72 |
| chr6-29627792-INS-3006 | chr6-29627793-INS-3006 |
| chr6-29631663-INS-139 | chr6-29631657-INS-106 |
| chr6-29632336-INS-645 | chr6-29632336-INS-645 |
| chr6-29637479-INS-200 | chr6-29637479-INS-199 |

| | |
|---|---|
| chr6-29641217-INS-486 | chr6-29641217-INS-486 |
| chr6-29646039-INS-96 | chr6-29646039-INS-95 |
| chr6-29647529-INS-356 | chr6-29647529-INS-356 |
| chr6-29654730-INS-78 | chr6-29654723-INS-117 |
| chr6-29692457-INS-124 | chr6-29692457-INS-107 |
| chr6-29712281-INS-762 | chr6-29712281-INS-762 |
| chr6-29727693-INS-195 | chr6-29727693-INS-199 |
| chr6-29738585-INS-58 | chr6-29738570-INS-70 |
| chr6-29747210-INS-226 | chr6-29747211-INS-224 |
| chr6-29750407-INS-206 | chr6-29750407-INS-206 |
| chr6-29775247-INS-229 | chr6-29775247-INS-229 |
| chr6-29832273-INS-190 | chr6-29832273-INS-190 |
| chr6-29842667-INS-222 | chr6-29842667-INS-222 |
| chr6-29844227-INS-3161 | chr6-29844228-INS-3139 |
| chr6-29844228-INS-525 | chr6-29844228-INS-525 |
| chr6-29873071-INS-125 | chr6-29873072-INS-125 |
| chr6-29959675-INS-214 | chr6-29959675-INS-214 |
| chr6-29965683-INS-790 | chr6-29965683-INS-774 |
| chr6-29988710-INS-220 | chr6-29988710-INS-220 |
| chr6-29996278-INS-205 | chr6-29996278-INS-205 |
| chr6-30003154-INS-201 | chr6-30003154-INS-192 |
| chr6-30007701-INS-186 | chr6-30007701-INS-186 |
| chr6-30010292-INS-80 | chr6-30010292-INS-76 |
| chr6-30011413-INS-225 | chr6-30011413-INS-225 |
| chr6-30015901-INS-402 | chr6-30015901-INS-401 |
| chr6-30034682-INS-66 | chr6-30034681-INS-75 |
| chr6-30071878-INS-109 | chr6-30071883-INS-112 |
| chr6-30090504-INS-184 | chr6-30090504-INS-184 |
| chr6-30100126-INS-207 | chr6-30100126-INS-207 |
| chr6-30107464-INS-285 | chr6-30107464-INS-285 |
| chr6-30131645-INS-57 | chr6-30131645-INS-71 |
| chr6-30163170-INS-209 | chr6-30163170-INS-208 |
| chr6-30196968-INS-179 | chr6-30196976-INS-209 |
| chr6-30202446-INS-211 | chr6-30202446-INS-210 |
| chr6-30216160-INS-198 | chr6-30216160-INS-199 |
| chr6-30219374-INS-237 | chr6-30219374-INS-237 |
| chr6-30254091-INS-65 | chr6-30254091-INS-66 |
| chr6-30262958-INS-1414 | chr6-30262959-INS-1415 |
| chr6-30294158-INS-80 | chr6-30294158-INS-80 |
| chr6-30319752-INS-125 | chr6-30319762-INS-85 |
| chr6-30333423-INS-200 | chr6-30333424-INS-168 |
| chr6-30361195-INS-122 | chr6-30361195-INS-122 |
| chr6-30363933-INS-189 | chr6-30363933-INS-196 |
| chr6-30366197-INS-142 | chr6-30366187-INS-183 |
| chr6-30387961-INS-113 | chr6-30387961-INS-113 |
| chr6-30412181-INS-255 | chr6-30412181-INS-255 |
| chr6-30416439-INS-217 | chr6-30416439-INS-217 |
| chr6-30490010-INS-213 | chr6-30490010-INS-213 |
| chr6-30565795-INS-63 | chr6-30565795-INS-52 |
| chr6-30577808-INS-138 | chr6-30577789-INS-138 |
| chr6-30728284-INS-270 | chr6-30728284-INS-270 |
| chr6-30809862-INS-759 | chr6-30809862-INS-760 |
| chr6-30816412-INS-343 | chr6-30816412-INS-343 |
| chr6-30820574-INS-99 | chr6-30820574-INS-99 |
| chr6-31007246-INS-357 | chr6-31007242-INS-360 |
| chr6-31024836-INS-510 | chr6-31024836-INS-514 |
| chr6-31154794-INS-432 | chr6-31154794-INS-432 |
| chr6-31200690-INS-68 | chr6-31200690-INS-84 |
| chr6-31207211-INS-100 | chr6-31207206-INS-98 |
| chr6-31223958-INS-100 | chr6-31223958-INS-96 |
| chr6-31246518-INS-148 | chr6-31246518-INS-153 |
| chr6-31251603-INS-1845 | chr6-31251604-INS-1854 |
| chr6-31252355-INS-54 | chr6-31252355-INS-54 |
| chr6-31281100-INS-120 | chr6-31281100-INS-122 |
| chr6-31288312-INS-58 | chr6-31288284-INS-72 |
| chr6-31309102-INS-117 | chr6-31309128-INS-120 |
| chr6-31315763-INS-62 | chr6-31315763-INS-62 |
| chr6-31369779-INS-74 | chr6-31369779-INS-74 |
| chr6-31409405-INS-128 | chr6-31409405-INS-84 |
| chr6-31414052-INS-486 | chr6-31414052-INS-486 |
| chr6-31426304-INS-90 | chr6-31426304-INS-70 |
| chr6-31429602-INS-1185 | chr6-31429603-INS-1191 |
| chr6-31430033-INS-498 | chr6-31430033-INS-498 |
| chr6-31433409-INS-219 | chr6-31433409-INS-220 |
| chr6-31459608-INS-156 | chr6-31459608-INS-156 |
| chr6-31512455-INS-126 | chr6-31512455-INS-152 |
| chr6-31541028-INS-52 | chr6-31541032-INS-60 |

| | |
|---|---|
| chr6-31558334-INS-102 | chr6-31558334-INS-102 |
| chr6-31612795-INS-92 | chr6-31612793-INS-114 |
| chr6-31616842-INS-845 | chr6-31616842-INS-846 |
| chr6-31627823-INS-55 | chr6-31627823-INS-55 |
| chr6-31633121-INS-7114 | chr6-31633122-INS-7114 |
| chr6-31665530-INS-56 | chr6-31665515-INS-53 |
| chr6-31666764-INS-198 | chr6-31666764-INS-198 |
| chr6-31679833-INS-52 | chr6-31679833-INS-50 |
| chr6-31683033-INS-173 | chr6-31683033-INS-171 |
| chr6-31701974-INS-54 | chr6-31701988-INS-50 |
| chr6-31730329-INS-225 | chr6-31730329-INS-220 |
| chr6-31760171-INS-59 | chr6-31760171-INS-60 |
| chr6-31760928-INS-72 | chr6-31760898-INS-54 |
| chr6-31776270-INS-66 | chr6-31776270-INS-67 |
| chr6-31785595-INS-148 | chr6-31785595-INS-148 |
| chr6-31858925-INS-234 | chr6-31858925-INS-241 |
| chr6-31876586-INS-53 | chr6-31876573-INS-51 |
| chr6-31882192-INS-58 | chr6-31882192-INS-52 |
| chr6-31906966-INS-78 | chr6-31906974-INS-70 |
| chr6-31914951-INS-423 | chr6-31914951-INS-423 |
| chr6-31943681-INS-104 | chr6-31943681-INS-76 |
| chr6-31952337-INS-66 | chr6-31952337-INS-66 |
| chr6-31954708-INS-65 | chr6-31954709-INS-65 |
| chr6-31980849-INS-6912 | chr6-31980850-INS-6913 |
| chr6-32070528-INS-424 | chr6-32070528-INS-424 |
| chr6-32109713-INS-144 | chr6-32109715-INS-124 |
| chr6-32184199-INS-92 | chr6-32184207-INS-80 |
| chr6-32261946-INS-2694 | chr6-32261947-INS-2694 |
| chr6-32265313-INS-60 | chr6-32265313-INS-58 |
| chr6-32300756-INS-597 | chr6-32300756-INS-598 |
| chr6-32368268-INS-157 | chr6-32368268-INS-133 |
| chr6-32404041-INS-58 | chr6-32404034-INS-54 |
| chr6-32413862-INS-391 | chr6-32413862-INS-391 |
| chr6-32434718-INS-310 | chr6-32434718-INS-310 |
| chr6-32436955-INS-66 | chr6-32436951-INS-56 |
| chr6-32459199-INS-70 | chr6-32459199-INS-74 |
| chr6-32487529-INS-54 | chr6-32487515-INS-50 |
| chr6-32492641-INS-52 | chr6-32492641-INS-52 |
| chr6-32553369-INS-51 | chr6-32553369-INS-53 |
| chr6-32588528-INS-75 | chr6-32588528-INS-75 |
| chr6-32629992-INS-183 | chr6-32629992-INS-182 |
| chr6-32644673-INS-76 | chr6-32644673-INS-76 |
| chr6-32659681-INS-334 | chr6-32659681-INS-334 |
| chr6-32703278-INS-54 | chr6-32703278-INS-54 |
| chr6-32730880-INS-90 | chr6-32730880-INS-90 |
| chr6-32740270-INS-346 | chr6-32740270-INS-347 |
| chr6-32776718-INS-53 | chr6-32776719-INS-51 |
| chr6-32789995-INS-106 | chr6-32789996-INS-108 |
| chr6-32807592-INS-1557 | chr6-32807593-INS-1553 |
| chr6-32852932-INS-240 | chr6-32852932-INS-240 |
| chr6-32867513-INS-2362 | chr6-32867514-INS-2364 |
| chr6-32868402-INS-139 | chr6-32868402-INS-139 |
| chr6-32879811-INS-60 | chr6-32879786-INS-60 |
| chr6-32883201-INS-84 | chr6-32883201-INS-84 |
| chr6-32909970-INS-61 | chr6-32909970-INS-61 |
| chr6-32926016-INS-119 | chr6-32926008-INS-127 |
| chr6-32994404-INS-1725 | chr6-32994405-INS-1725 |
| chr6-32994474-INS-210 | chr6-32994474-INS-210 |
| chr6-33032392-INS-166 | chr6-33032392-INS-166 |
| chr6-33056533-INS-83 | chr6-33056533-INS-79 |
| chr6-33060981-INS-120 | chr6-33060981-INS-120 |
| chr6-33066357-INS-76 | chr6-33066357-INS-72 |
| chr6-33076877-INS-65 | chr6-33076877-INS-79 |
| chr6-33087059-INS-146 | chr6-33087059-INS-146 |
| chr6-33093230-INS-1506 | chr6-33093231-INS-1509 |
| chr6-33187971-INS-483 | chr6-33187971-INS-471 |
| chr6-33225465-INS-64 | chr6-33225465-INS-64 |
| chr6-33229257-INS-52 | chr6-33229257-INS-52 |
| chr6-33233103-INS-177 | chr6-33233103-INS-178 |
| chr6-33288202-INS-76 | chr6-33288202-INS-104 |
| chr6-33294541-INS-6872 | chr6-33294542-INS-6872 |
| chr6-33302577-INS-52 | chr6-33302577-INS-54 |
| chr6-33439375-INS-219 | chr6-33439372-INS-223 |
| chr6-33441045-INS-69 | chr6-33441045-INS-69 |
| chr6-33504807-INS-102 | chr6-33504807-INS-60 |
| chr6-33555053-INS-71 | chr6-33555053-INS-71 |
| chr6-33561627-INS-52 | chr6-33561627-INS-52 |

| | |
|---|---|
| chr6-33574773-INS-96 | chr6-33574787-INS-50 |
| chr6-33578945-INS-1404 | chr6-33578946-INS-1402 |
| chr6-33589279-INS-150 | chr6-33589310-INS-159 |
| chr6-33595665-INS-333 | chr6-33595665-INS-333 |
| chr6-33627154-INS-88 | chr6-33627154-INS-88 |
| chr6-33661406-INS-59 | chr6-33661406-INS-59 |
| chr6-33666851-INS-208 | chr6-33666851-INS-209 |
| chr6-33702047-INS-403 | chr6-33702047-INS-403 |
| chr6-33734305-INS-4597 | chr6-33734305-INS-4579 |
| chr6-33754136-INS-56 | chr6-33754136-INS-56 |
| chr6-33768620-INS-1979 | chr6-33768621-INS-1979 |
| chr6-33776689-INS-93 | chr6-33776689-INS-93 |
| chr6-33779125-INS-176 | chr6-33779125-INS-176 |
| chr6-33787166-INS-3034 | chr6-33787167-INS-3034 |
| chr6-33794639-INS-237 | chr6-33794639-INS-235 |
| chr6-33807450-INS-6386 | chr6-33807451-INS-6386 |
| chr6-33811750-INS-60 | chr6-33811750-INS-86 |
| chr6-33817006-INS-139 | chr6-33817012-INS-94 |
| chr6-33840974-INS-52 | chr6-33840974-INS-52 |
| chr6-33858560-INS-88 | chr6-33858564-INS-92 |
| chr6-33866553-INS-87 | chr6-33866553-INS-87 |
| chr6-33877101-INS-2911 | chr6-33877102-INS-2926 |
| chr6-33878501-INS-6687 | chr6-33878502-INS-6687 |
| chr6-33922981-INS-212 | chr6-33922981-INS-212 |
| chr6-33924543-INS-111 | chr6-33924543-INS-111 |
| chr6-33944769-INS-98 | chr6-33944770-INS-98 |
| chr6-33960304-INS-186 | chr6-33960304-INS-188 |
| chr6-33971454-INS-132 | chr6-33971454-INS-132 |
| chr6-33974409-INS-234 | chr6-33974409-INS-234 |
| chr6-33979063-INS-60 | chr6-33979063-INS-60 |
| chr6-33983814-INS-88 | chr6-33983814-INS-70 |
| chr6-33989203-INS-124 | chr6-33989204-INS-93 |
| chr6-34034604-INS-56 | chr6-34034604-INS-56 |
| chr6-34083063-INS-73 | chr6-34083063-INS-74 |
| chr6-34089982-INS-53 | chr6-34089982-INS-73 |
| chr6-34094625-INS-177 | chr6-34094625-INS-179 |
| chr6-34106435-INS-73 | chr6-34106435-INS-65 |
| chr6-34113944-INS-6448 | chr6-34113945-INS-6448 |
| chr6-34159842-INS-118 | chr6-34159853-INS-106 |
| chr6-34183265-INS-64 | chr6-34183265-INS-66 |
| chr6-34188237-INS-59 | chr6-34188237-INS-59 |
| chr6-34190888-INS-754 | chr6-34190888-INS-754 |
| chr6-34228110-INS-144 | chr6-34228130-INS-136 |
| chr6-34231548-INS-102 | chr6-34231548-INS-102 |
| chr6-34242646-INS-1889 | chr6-34242647-INS-1888 |
| chr6-34278409-INS-7111 | chr6-34278410-INS-7111 |
| chr6-34296391-INS-171 | chr6-34296391-INS-136 |
| chr6-34311527-INS-5322 | chr6-34311528-INS-5322 |
| chr6-34313435-INS-55 | chr6-34313435-INS-50 |
| chr6-34328243-INS-53 | chr6-34328243-INS-61 |
| chr6-34330785-INS-60 | chr6-34330785-INS-60 |
| chr6-34339371-INS-148 | chr6-34339371-INS-140 |
| chr6-34398911-INS-3979 | chr6-34398912-INS-3979 |
| chr6-34402812-INS-6402 | chr6-34402813-INS-6402 |
| chr6-34418777-INS-538 | chr6-34418777-INS-538 |
| chr6-34428390-INS-626 | chr6-34428390-INS-626 |
| chr6-34429121-INS-85 | chr6-34429121-INS-85 |
| chr6-34436451-INS-60 | chr6-34436473-INS-54 |
| chr6-34449749-INS-66 | chr6-34449749-INS-92 |
| chr6-34471422-INS-376 | chr6-34471422-INS-376 |
| chr6-34497296-INS-5354 | chr6-34497297-INS-5354 |
| chr6-34511322-INS-64 | chr6-34511322-INS-65 |
| chr6-34518807-INS-187 | chr6-34518807-INS-187 |
| chr6-34526384-INS-64 | chr6-34526384-INS-64 |
| chr6-34541248-INS-74 | chr6-34541248-INS-74 |
| chr6-34566791-INS-52 | chr6-34566791-INS-50 |
| chr6-34584640-INS-193 | chr6-34584640-INS-193 |
| chr6-34624791-INS-77 | chr6-34624791-INS-77 |
| chr6-34659983-INS-122 | chr6-34659983-INS-122 |
| chr6-34663790-INS-337 | chr6-34663790-INS-333 |
| chr6-34735774-INS-1508 | chr6-34735775-INS-1515 |
| chr6-34754112-INS-56 | chr6-34754112-INS-82 |
| chr6-34783911-INS-252 | chr6-34783911-INS-252 |
| chr6-34785938-INS-2133 | chr6-34785934-INS-2128 |
| chr6-34898445-INS-155 | chr6-34898445-INS-124 |
| chr6-34899485-INS-60 | chr6-34899485-INS-60 |
| chr6-34909227-INS-80 | chr6-34909227-INS-80 |

| | |
|---|---|
| chr6-34911784-INS-168 | chr6-34911796-INS-180 |
| chr6-34932666-INS-70 | chr6-34932652-INS-56 |
| chr6-34981982-INS-182 | chr6-34981961-INS-187 |
| chr6-35011105-INS-232 | chr6-35011105-INS-232 |
| chr6-35031761-INS-665 | chr6-35031761-INS-618 |
| chr6-35038530-INS-96 | chr6-35038527-INS-96 |
| chr6-35039620-INS-1143 | chr6-35039621-INS-1141 |
| chr6-35053612-INS-198 | chr6-35053612-INS-199 |
| chr6-35067493-INS-57 | chr6-35067494-INS-105 |
| chr6-35088216-INS-206 | chr6-35088216-INS-206 |
| chr6-35119365-INS-182 | chr6-35119365-INS-182 |
| chr6-35122814-INS-210 | chr6-35122814-INS-210 |
| chr6-35124741-INS-181 | chr6-35124741-INS-181 |
| chr6-35135836-INS-1914 | chr6-35135837-INS-1915 |
| chr6-35192683-INS-226 | chr6-35192683-INS-228 |
| chr6-35270759-INS-272 | chr6-35270759-INS-251 |
| chr6-35277403-INS-462 | chr6-35277403-INS-462 |
| chr6-35280456-INS-66 | chr6-35280505-INS-66 |
| chr6-35294683-INS-104 | chr6-35294683-INS-104 |
| chr6-35325790-INS-438 | chr6-35325792-INS-435 |
| chr6-35363227-INS-141 | chr6-35363227-INS-138 |
| chr6-35410450-INS-208 | chr6-35410450-INS-208 |
| chr6-35460029-INS-56 | chr6-35460029-INS-56 |
| chr6-35473540-INS-52 | chr6-35473540-INS-52 |
| chr6-35522567-INS-104 | chr6-35522567-INS-100 |
| chr6-35557163-INS-52 | chr6-35557163-INS-64 |
| chr6-35578253-INS-344 | chr6-35578253-INS-344 |
| chr6-35613402-INS-82 | chr6-35613380-INS-84 |
| chr6-35684705-INS-53 | chr6-35684705-INS-56 |
| chr6-35687008-INS-523 | chr6-35687009-INS-525 |
| chr6-35709368-INS-798 | chr6-35709368-INS-800 |
| chr6-35741522-INS-68 | chr6-35741522-INS-64 |
| chr6-35764709-INS-58 | chr6-35764709-INS-76 |
| chr6-35775086-INS-58 | chr6-35775078-INS-52 |
| chr6-35812426-INS-1854 | chr6-35812427-INS-1853 |
| chr6-35858151-INS-6119 | chr6-35858152-INS-6119 |
| chr6-35878799-INS-292 | chr6-35878799-INS-292 |
| chr6-35881016-INS-217 | chr6-35881016-INS-218 |
| chr6-35916199-INS-429 | chr6-35916199-INS-429 |
| chr6-35969453-INS-6440 | chr6-35969454-INS-6440 |
| chr6-35977501-INS-717 | chr6-35977501-INS-717 |
| chr6-35996104-INS-462 | chr6-35996104-INS-462 |
| chr6-36002116-INS-105 | chr6-36002116-INS-102 |
| chr6-36029561-INS-7086 | chr6-36029562-INS-7084 |
| chr6-36031492-INS-58 | chr6-36031502-INS-63 |
| chr6-36043891-INS-1762 | chr6-36043892-INS-1762 |
| chr6-36051939-INS-89 | chr6-36051939-INS-91 |
| chr6-36069662-INS-56 | chr6-36069662-INS-56 |
| chr6-36082049-INS-96 | chr6-36082049-INS-57 |
| chr6-36104778-INS-96 | chr6-36104778-INS-96 |
| chr6-36120959-INS-6662 | chr6-36120960-INS-6662 |
| chr6-36192151-INS-747 | chr6-36192151-INS-747 |
| chr6-36224723-INS-138 | chr6-36224723-INS-138 |
| chr6-36226396-INS-2303 | chr6-36226397-INS-2303 |
| chr6-36237609-INS-86 | chr6-36237588-INS-65 |
| chr6-36241526-INS-60 | chr6-36241526-INS-64 |
| chr6-36242486-INS-896 | chr6-36242486-INS-897 |
| chr6-36250561-INS-316 | chr6-36250561-INS-312 |
| chr6-36252574-INS-6324 | chr6-36252575-INS-6329 |
| chr6-36281202-INS-56 | chr6-36281202-INS-56 |
| chr6-36309214-INS-117 | chr6-36309214-INS-117 |
| chr6-36354961-INS-327 | chr6-36354961-INS-327 |
| chr6-36454230-INS-52 | chr6-36454230-INS-52 |
| chr6-36459322-INS-1057 | chr6-36459323-INS-1060 |
| chr6-36516443-INS-400 | chr6-36516454-INS-380 |
| chr6-36519317-INS-270 | chr6-36519316-INS-271 |
| chr6-36530379-INS-82 | chr6-36530379-INS-90 |
| chr6-36582274-INS-85 | chr6-36582274-INS-78 |
| chr6-36590985-INS-60 | chr6-36590985-INS-60 |
| chr6-36634909-INS-64 | chr6-36634909-INS-64 |
| chr6-36641380-INS-1042 | chr6-36641381-INS-1042 |
| chr6-36642745-INS-6880 | chr6-36642746-INS-6880 |
| chr6-36661584-INS-52 | chr6-36661584-INS-52 |
| chr6-36681586-INS-962 | chr6-36681586-INS-962 |
| chr6-36688368-INS-201 | chr6-36688368-INS-201 |
| chr6-36738431-INS-58 | chr6-36738431-INS-62 |
| chr6-36747421-INS-79 | chr6-36747431-INS-50 |

| | |
|---|---|
| chr6-36798681-INS-85 | chr6-36798705-INS-63 |
| chr6-36806553-INS-6657 | chr6-36806554-INS-6658 |
| chr6-36808211-INS-57 | chr6-36808211-INS-57 |
| chr6-36832120-INS-595 | chr6-36832120-INS-594 |
| chr6-36907917-INS-6442 | chr6-36907918-INS-6442 |
| chr6-37058370-INS-53 | chr6-37058375-INS-50 |
| chr6-37072003-INS-146 | chr6-37072016-INS-173 |
| chr6-37098171-INS-84 | chr6-37098171-INS-72 |
| chr6-37110264-INS-173 | chr6-37110264-INS-173 |
| chr6-37132005-INS-5541 | chr6-37132006-INS-5541 |
| chr6-37155902-INS-183 | chr6-37155903-INS-176 |
| chr6-37161931-INS-67 | chr6-37161944-INS-60 |
| chr6-37198842-INS-56 | chr6-37198833-INS-68 |
| chr6-37203982-INS-213 | chr6-37203983-INS-213 |
| chr6-37253700-INS-75 | chr6-37253700-INS-75 |
| chr6-37261736-INS-113 | chr6-37261736-INS-113 |
| chr6-37262475-INS-1156 | chr6-37262476-INS-1156 |
| chr6-37302826-INS-441 | chr6-37302826-INS-441 |
| chr6-37316167-INS-440 | chr6-37316167-INS-451 |
| chr6-37324014-INS-77 | chr6-37324014-INS-77 |
| chr6-37349355-INS-74 | chr6-37349355-INS-74 |
| chr6-37381836-INS-58 | chr6-37381822-INS-68 |
| chr6-37414956-INS-1213 | chr6-37414957-INS-1223 |
| chr6-37434372-INS-92 | chr6-37434372-INS-63 |
| chr6-37437775-INS-325 | chr6-37437775-INS-325 |
| chr6-37457331-INS-2878 | chr6-37457332-INS-2876 |
| chr6-37462735-INS-197 | chr6-37462735-INS-197 |
| chr6-37504701-INS-149 | chr6-37504701-INS-145 |
| chr6-37506651-INS-103 | chr6-37506651-INS-103 |
| chr6-37511059-INS-92 | chr6-37511059-INS-82 |
| chr6-37536632-INS-1889 | chr6-37536633-INS-1889 |
| chr6-37538027-INS-192 | chr6-37538028-INS-192 |
| chr6-37580920-INS-79 | chr6-37580920-INS-55 |
| chr6-37593935-INS-136 | chr6-37593935-INS-138 |
| chr6-37601577-INS-428 | chr6-37601577-INS-428 |
| chr6-37606722-INS-263 | chr6-37606722-INS-259 |
| chr6-37614704-INS-6938 | chr6-37614705-INS-6937 |
| chr6-37615562-INS-1427 | chr6-37615563-INS-1427 |
| chr6-37617302-INS-172 | chr6-37617302-INS-175 |
| chr6-37638958-INS-3581 | chr6-37638959-INS-3581 |
| chr6-37638959-INS-920 | chr6-37638959-INS-920 |
| chr6-37643987-INS-62 | chr6-37643983-INS-70 |
| chr6-37645050-INS-99 | chr6-37645050-INS-99 |
| chr6-37647475-INS-52 | chr6-37647475-INS-62 |
| chr6-37653938-INS-119 | chr6-37653938-INS-107 |
| chr6-37656294-INS-58 | chr6-37656294-INS-64 |
| chr6-37666728-INS-191 | chr6-37666728-INS-193 |
| chr6-37672543-INS-1888 | chr6-37672544-INS-1888 |
| chr6-37693277-INS-216 | chr6-37693277-INS-216 |
| chr6-37694253-INS-325 | chr6-37694253-INS-328 |
| chr6-37710818-INS-120 | chr6-37710818-INS-126 |
| chr6-37718296-INS-94 | chr6-37718296-INS-80 |
| chr6-37730250-INS-209 | chr6-37730250-INS-209 |
| chr6-37736478-INS-222 | chr6-37736478-INS-218 |
| chr6-37770074-INS-908 | chr6-37770074-INS-908 |
| chr6-37779580-INS-192 | chr6-37779580-INS-185 |
| chr6-37789327-INS-129 | chr6-37789328-INS-100 |
| chr6-37826919-INS-7080 | chr6-37826920-INS-7080 |
| chr6-37829597-INS-1871 | chr6-37829598-INS-1871 |
| chr6-37867815-INS-10242 | chr6-37867816-INS-10242 |
| chr6-37881462-INS-205 | chr6-37881462-INS-211 |
| chr6-37887614-INS-54 | chr6-37887614-INS-54 |
| chr6-37915695-INS-204 | chr6-37915695-INS-204 |
| chr6-37919699-INS-99 | chr6-37919699-INS-99 |
| chr6-37965714-INS-209 | chr6-37965714-INS-209 |
| chr6-37978046-INS-628 | chr6-37978046-INS-628 |
| chr6-37982672-INS-77 | chr6-37982672-INS-77 |
| chr6-37994769-INS-460 | chr6-37994769-INS-415 |
| chr6-37996211-INS-167 | chr6-37996211-INS-173 |
| chr6-38037329-INS-93 | chr6-38037329-INS-70 |
| chr6-38055528-INS-739 | chr6-38055528-INS-757 |
| chr6-38079861-INS-88 | chr6-38079834-INS-66 |
| chr6-38114162-INS-52 | chr6-38114162-INS-52 |
| chr6-38115233-INS-73 | chr6-38115233-INS-73 |
| chr6-38116655-INS-60 | chr6-38116655-INS-92 |
| chr6-38121390-INS-176 | chr6-38121390-INS-176 |
| chr6-38123479-INS-54 | chr6-38123465-INS-50 |

| | |
|---|---|
| chr6-38134403-INS-207 | chr6-38134403-INS-219 |
| chr6-38135264-INS-184 | chr6-38135264-INS-184 |
| chr6-38152728-INS-142 | chr6-38152728-INS-142 |
| chr6-38167191-INS-88 | chr6-38167191-INS-88 |
| chr6-38188229-INS-173 | chr6-38188229-INS-210 |
| chr6-38202727-INS-78 | chr6-38202727-INS-78 |
| chr6-38203377-INS-73 | chr6-38203377-INS-73 |
| chr6-38204117-INS-54 | chr6-38204117-INS-54 |
| chr6-38219295-INS-60 | chr6-38219295-INS-66 |
| chr6-38227611-INS-51 | chr6-38227605-INS-54 |
| chr6-38239097-INS-224 | chr6-38239097-INS-225 |
| chr6-38268735-INS-5319 | chr6-38268736-INS-5319 |
| chr6-38286855-INS-60 | chr6-38286855-INS-60 |
| chr6-38289729-INS-197 | chr6-38289729-INS-202 |
| chr6-38290820-INS-171 | chr6-38290820-INS-171 |
| chr6-38303707-INS-127 | chr6-38303707-INS-127 |
| chr6-38313034-INS-171 | chr6-38313034-INS-174 |
| chr6-38318348-INS-203 | chr6-38318346-INS-214 |
| chr6-38326533-INS-188 | chr6-38326533-INS-190 |
| chr6-38327589-INS-171 | chr6-38327589-INS-171 |
| chr6-38335324-INS-226 | chr6-38335332-INS-177 |
| chr6-38352593-INS-204 | chr6-38352593-INS-204 |
| chr6-38355788-INS-182 | chr6-38355788-INS-182 |
| chr6-38361244-INS-209 | chr6-38361244-INS-209 |
| chr6-38372760-INS-240 | chr6-38372760-INS-242 |
| chr6-38436093-INS-120 | chr6-38436093-INS-120 |
| chr6-38502081-INS-52 | chr6-38502069-INS-50 |
| chr6-38506478-INS-207 | chr6-38506478-INS-207 |
| chr6-38512369-INS-54 | chr6-38512369-INS-54 |
| chr6-38519951-INS-190 | chr6-38519951-INS-190 |
| chr6-38530649-INS-404 | chr6-38530649-INS-405 |
| chr6-38533094-INS-186 | chr6-38533094-INS-186 |
| chr6-38589258-INS-201 | chr6-38589258-INS-201 |
| chr6-38601791-INS-341 | chr6-38601791-INS-341 |
| chr6-38608244-INS-238 | chr6-38608244-INS-224 |
| chr6-38609240-INS-215 | chr6-38609240-INS-212 |
| chr6-38620238-INS-60 | chr6-38620238-INS-60 |
| chr6-38626491-INS-77 | chr6-38626491-INS-77 |
| chr6-38638437-INS-330 | chr6-38638437-INS-330 |
| chr6-38655756-INS-84 | chr6-38655764-INS-88 |
| chr6-38683982-INS-165 | chr6-38683982-INS-165 |
| chr6-38691804-INS-101 | chr6-38691804-INS-101 |
| chr6-38706116-INS-64 | chr6-38706130-INS-80 |
| chr6-38717694-INS-209 | chr6-38717694-INS-209 |
| chr6-38752391-INS-51 | chr6-38752391-INS-51 |
| chr6-38754458-INS-88 | chr6-38754458-INS-70 |
| chr6-38786547-INS-139 | chr6-38786547-INS-139 |
| chr6-38875358-INS-314 | chr6-38875358-INS-332 |
| chr6-38897820-INS-60 | chr6-38897840-INS-62 |
| chr6-38903019-INS-54 | chr6-38903039-INS-56 |
| chr6-38911078-INS-158 | chr6-38911078-INS-156 |
| chr6-38930761-INS-81 | chr6-38930761-INS-112 |
| chr6-38935918-INS-80 | chr6-38935919-INS-61 |
| chr6-38942959-INS-1892 | chr6-38942960-INS-1892 |
| chr6-38973063-INS-131 | chr6-38973063-INS-131 |
| chr6-38998776-INS-150 | chr6-38998776-INS-135 |
| chr6-39069434-INS-917 | chr6-39069450-INS-925 |
| chr6-39071282-INS-2382 | chr6-39071283-INS-2382 |
| chr6-39106122-INS-516 | chr6-39106122-INS-516 |
| chr6-39160439-INS-202 | chr6-39160439-INS-201 |
| chr6-39204242-INS-82 | chr6-39204288-INS-70 |
| chr6-39242576-INS-117 | chr6-39242576-INS-143 |
| chr6-39258809-INS-172 | chr6-39258809-INS-180 |
| chr6-39278171-INS-909 | chr6-39278170-INS-922 |
| chr6-39281167-INS-176 | chr6-39281167-INS-176 |
| chr6-39283465-INS-162 | chr6-39283465-INS-163 |
| chr6-39285792-INS-65 | chr6-39285787-INS-65 |
| chr6-39296001-INS-60 | chr6-39296001-INS-62 |
| chr6-39403369-INS-163 | chr6-39403362-INS-164 |
| chr6-39410187-INS-441 | chr6-39410187-INS-445 |
| chr6-39413808-INS-231 | chr6-39413808-INS-215 |
| chr6-39415256-INS-56 | chr6-39415256-INS-85 |
| chr6-39418546-INS-203 | chr6-39418544-INS-194 |
| chr6-39421648-INS-208 | chr6-39421664-INS-216 |
| chr6-39425034-INS-117 | chr6-39425030-INS-119 |
| chr6-39428700-INS-356 | chr6-39428700-INS-356 |
| chr6-39448960-INS-212 | chr6-39448960-INS-213 |

| | |
|---|---|
| chr6-39456641-INS-215 | chr6-39456641-INS-199 |
| chr6-39462864-INS-78 | chr6-39462864-INS-92 |
| chr6-39463968-INS-142 | chr6-39463969-INS-131 |
| chr6-39464439-INS-136 | chr6-39464427-INS-146 |
| chr6-39467480-INS-72 | chr6-39467512-INS-66 |
| chr6-39469384-INS-196 | chr6-39469384-INS-197 |
| chr6-39511306-INS-373 | chr6-39511306-INS-370 |
| chr6-39538403-INS-157 | chr6-39538403-INS-169 |
| chr6-39542792-INS-191 | chr6-39542792-INS-193 |
| chr6-39546966-INS-198 | chr6-39546966-INS-195 |
| chr6-39593478-INS-65 | chr6-39593478-INS-65 |
| chr6-39594902-INS-223 | chr6-39594902-INS-223 |
| chr6-39604267-INS-168 | chr6-39604267-INS-169 |
| chr6-39611808-INS-209 | chr6-39611808-INS-209 |
| chr6-39615892-INS-329 | chr6-39615892-INS-327 |
| chr6-39631169-INS-341 | chr6-39631169-INS-341 |
| chr6-39637080-INS-76 | chr6-39637080-INS-82 |
| chr6-39655306-INS-671 | chr6-39655303-INS-676 |
| chr6-39656473-INS-93 | chr6-39656473-INS-93 |
| chr6-39662042-INS-198 | chr6-39662042-INS-198 |
| chr6-39663407-INS-198 | chr6-39663407-INS-196 |
| chr6-39667424-INS-199 | chr6-39667424-INS-203 |
| chr6-39687315-INS-218 | chr6-39687315-INS-218 |
| chr6-39725562-INS-79 | chr6-39725536-INS-69 |
| chr6-39737040-INS-198 | chr6-39737040-INS-198 |
| chr6-39755778-INS-54 | chr6-39755778-INS-54 |
| chr6-39757299-INS-76 | chr6-39757295-INS-84 |
| chr6-39814710-INS-68 | chr6-39814710-INS-68 |
| chr6-39869896-INS-56 | chr6-39869896-INS-83 |
| chr6-39925480-INS-200 | chr6-39925480-INS-201 |
| chr6-39938537-INS-76 | chr6-39938537-INS-76 |
| chr6-39985039-INS-53 | chr6-39985039-INS-53 |
| chr6-40012586-INS-63 | chr6-40012586-INS-65 |
| chr6-40047540-INS-53 | chr6-40047531-INS-57 |
| chr6-40050939-INS-53 | chr6-40050974-INS-60 |
| chr6-40101844-INS-210 | chr6-40101844-INS-210 |
| chr6-40136371-INS-200 | chr6-40136371-INS-200 |
| chr6-40206571-INS-227 | chr6-40206571-INS-229 |
| chr6-40212077-INS-210 | chr6-40212077-INS-207 |
| chr6-40275338-INS-54 | chr6-40275338-INS-68 |
| chr6-40293013-INS-91 | chr6-40293013-INS-91 |
| chr6-40294972-INS-159 | chr6-40294972-INS-159 |
| chr6-40337281-INS-181 | chr6-40337281-INS-181 |
| chr6-40354676-INS-60 | chr6-40354676-INS-90 |
| chr6-40390049-INS-331 | chr6-40390049-INS-329 |
| chr6-40409013-INS-88 | chr6-40409013-INS-88 |
| chr6-40414476-INS-84 | chr6-40414483-INS-60 |
| chr6-40450572-INS-61 | chr6-40450572-INS-61 |
| chr6-40456487-INS-331 | chr6-40456487-INS-331 |
| chr6-40488673-INS-1090 | chr6-40488674-INS-1091 |
| chr6-40488807-INS-207 | chr6-40488807-INS-207 |
| chr6-40491404-INS-194 | chr6-40491405-INS-199 |
| chr6-40545171-INS-58 | chr6-40545154-INS-64 |
| chr6-40548384-INS-7102 | chr6-40548385-INS-7102 |
| chr6-40549878-INS-64 | chr6-40549878-INS-64 |
| chr6-40570891-INS-160 | chr6-40570891-INS-160 |
| chr6-40623735-INS-998 | chr6-40623735-INS-997 |
| chr6-40654305-INS-6320 | chr6-40654306-INS-6315 |
| chr6-40742083-INS-385 | chr6-40742083-INS-384 |
| chr6-40760931-INS-56 | chr6-40760931-INS-56 |
| chr6-40785680-INS-6478 | chr6-40785681-INS-6477 |
| chr6-40793329-INS-68 | chr6-40793310-INS-63 |
| chr6-40883933-INS-409 | chr6-40883933-INS-409 |
| chr6-40925082-INS-441 | chr6-40925082-INS-441 |
| chr6-40972165-INS-2830 | chr6-40972166-INS-2830 |
| chr6-40996715-INS-121 | chr6-40996715-INS-121 |
| chr6-41025143-INS-599 | chr6-41025143-INS-602 |
| chr6-41051303-INS-86 | chr6-41051303-INS-86 |
| chr6-41056785-INS-329 | chr6-41056785-INS-333 |
| chr6-41095956-INS-52 | chr6-41095956-INS-52 |
| chr6-41115993-INS-74 | chr6-41116029-INS-54 |
| chr6-41141910-INS-437 | chr6-41141901-INS-440 |
| chr6-41145657-INS-68 | chr6-41145657-INS-68 |
| chr6-41146705-INS-76 | chr6-41146705-INS-68 |
| chr6-41180227-INS-88 | chr6-41180239-INS-50 |
| chr6-41213657-INS-430 | chr6-41213657-INS-432 |
| chr6-41225612-INS-51 | chr6-41225609-INS-52 |

| | |
|---|---|
| chr6-41235441-INS-380 | chr6-41235441-INS-380 |
| chr6-41240894-INS-87 | chr6-41240894-INS-88 |
| chr6-41293420-INS-78 | chr6-41293420-INS-78 |
| chr6-41298940-INS-116 | chr6-41298940-INS-116 |
| chr6-41333434-INS-267 | chr6-41333434-INS-267 |
| chr6-41423778-INS-657 | chr6-41423778-INS-659 |
| chr6-41439765-INS-97 | chr6-41439771-INS-99 |
| chr6-41458972-INS-370 | chr6-41458972-INS-366 |
| chr6-41470194-INS-91 | chr6-41470194-INS-91 |
| chr6-41510484-INS-53 | chr6-41510484-INS-53 |
| chr6-41514643-INS-656 | chr6-41514644-INS-656 |
| chr6-41542800-INS-203 | chr6-41542800-INS-202 |
| chr6-41544195-INS-95 | chr6-41544205-INS-107 |
| chr6-41548802-INS-70 | chr6-41548802-INS-110 |
| chr6-41554184-INS-103 | chr6-41554178-INS-91 |
| chr6-41559843-INS-4005 | chr6-41559844-INS-4005 |
| chr6-41670448-INS-71 | chr6-41670448-INS-71 |
| chr6-41715907-INS-64 | chr6-41715907-INS-71 |
| chr6-41747084-INS-96 | chr6-41747084-INS-96 |
| chr6-41777046-INS-206 | chr6-41777046-INS-207 |
| chr6-41800192-INS-177 | chr6-41800192-INS-177 |
| chr6-41834565-INS-399 | chr6-41834565-INS-401 |
| chr6-41842610-INS-204 | chr6-41842610-INS-204 |
| chr6-41866703-INS-244 | chr6-41866706-INS-247 |
| chr6-41891341-INS-64 | chr6-41891327-INS-53 |
| chr6-41916242-INS-78 | chr6-41916242-INS-70 |
| chr6-41929958-INS-7102 | chr6-41929959-INS-7102 |
| chr6-42006800-INS-547 | chr6-42006800-INS-547 |
| chr6-42018809-INS-946 | chr6-42018809-INS-944 |
| chr6-42061404-INS-52 | chr6-42061438-INS-52 |
| chr6-42080492-INS-202 | chr6-42080541-INS-203 |
| chr6-42112281-INS-505 | chr6-42112281-INS-520 |
| chr6-42155998-INS-80 | chr6-42155998-INS-104 |
| chr6-42191629-INS-58 | chr6-42191630-INS-64 |
| chr6-42199847-INS-172 | chr6-42199800-INS-215 |
| chr6-42204257-INS-500 | chr6-42204257-INS-501 |
| chr6-42239752-INS-121 | chr6-42239752-INS-122 |
| chr6-42265113-INS-60 | chr6-42265113-INS-99 |
| chr6-42283299-INS-394 | chr6-42283299-INS-396 |
| chr6-42305661-INS-222 | chr6-42305661-INS-182 |
| chr6-42363276-INS-64 | chr6-42363276-INS-72 |
| chr6-42400604-INS-116 | chr6-42400604-INS-122 |
| chr6-42409355-INS-54 | chr6-42409361-INS-50 |
| chr6-42418917-INS-61 | chr6-42418917-INS-85 |
| chr6-42421616-INS-131 | chr6-42421616-INS-132 |
| chr6-42427687-INS-214 | chr6-42427687-INS-214 |
| chr6-42443734-INS-713 | chr6-42443734-INS-713 |
| chr6-42529658-INS-60 | chr6-42529658-INS-60 |
| chr6-42532155-INS-1210 | chr6-42532156-INS-1210 |
| chr6-42537122-INS-1223 | chr6-42537123-INS-1224 |
| chr6-42540232-INS-6329 | chr6-42540233-INS-6330 |
| chr6-42583737-INS-173 | chr6-42583737-INS-173 |
| chr6-42616000-INS-224 | chr6-42616002-INS-256 |
| chr6-42621214-INS-895 | chr6-42621214-INS-895 |
| chr6-42660662-INS-62 | chr6-42660660-INS-66 |
| chr6-42747264-INS-83 | chr6-42747264-INS-83 |
| chr6-42756526-INS-70 | chr6-42756526-INS-74 |
| chr6-42770418-INS-463 | chr6-42770418-INS-463 |
| chr6-42820191-INS-174 | chr6-42820193-INS-176 |
| chr6-42845081-INS-113 | chr6-42845081-INS-113 |
| chr6-42867644-INS-60 | chr6-42867644-INS-60 |
| chr6-42868423-INS-1130 | chr6-42868424-INS-1130 |
| chr6-42910756-INS-1210 | chr6-42910757-INS-1210 |
| chr6-42931066-INS-76 | chr6-42931066-INS-77 |
| chr6-42936244-INS-1727 | chr6-42936245-INS-1729 |
| chr6-42936861-INS-1536 | chr6-42936862-INS-1536 |
| chr6-42992672-INS-232 | chr6-42992688-INS-216 |
| chr6-43030268-INS-112 | chr6-43030268-INS-112 |
| chr6-43037600-INS-121 | chr6-43037594-INS-123 |
| chr6-43114285-INS-400 | chr6-43114285-INS-400 |
| chr6-43126221-INS-103 | chr6-43126221-INS-105 |
| chr6-43159566-INS-298 | chr6-43159566-INS-286 |
| chr6-43205659-INS-2893 | chr6-43205660-INS-2893 |
| chr6-43227547-INS-203 | chr6-43227547-INS-201 |
| chr6-43279269-INS-1678 | chr6-43279270-INS-1679 |
| chr6-43315273-INS-645 | chr6-43315273-INS-646 |
| chr6-43337583-INS-3589 | chr6-43337584-INS-3589 |

| | |
|---|---|
| chr6-43341101-INS-312 | chr6-43341101-INS-311 |
| chr6-43342111-INS-597 | chr6-43342111-INS-597 |
| chr6-43403953-INS-613 | chr6-43403953-INS-614 |
| chr6-43406731-INS-347 | chr6-43406731-INS-347 |
| chr6-43472193-INS-103 | chr6-43472193-INS-103 |
| chr6-43478903-INS-439 | chr6-43478903-INS-436 |
| chr6-43501653-INS-77 | chr6-43501657-INS-77 |
| chr6-43509261-INS-154 | chr6-43509261-INS-156 |
| chr6-43516051-INS-1346 | chr6-43516052-INS-1346 |
| chr6-43517640-INS-60 | chr6-43517622-INS-84 |
| chr6-43548549-INS-367 | chr6-43548549-INS-367 |
| chr6-43581902-INS-60 | chr6-43581902-INS-71 |
| chr6-43620285-INS-110 | chr6-43620285-INS-110 |
| chr6-43702864-INS-385 | chr6-43702864-INS-385 |
| chr6-43730337-INS-778 | chr6-43730337-INS-780 |
| chr6-43795644-INS-62 | chr6-43795644-INS-70 |
| chr6-43814932-INS-677 | chr6-43814932-INS-677 |
| chr6-43814953-INS-53 | chr6-43814953-INS-53 |
| chr6-43827645-INS-56 | chr6-43827645-INS-62 |
| chr6-43829959-INS-359 | chr6-43829959-INS-359 |
| chr6-43841520-INS-186 | chr6-43841520-INS-186 |
| chr6-43869593-INS-2019 | chr6-43869594-INS-2019 |
| chr6-43887781-INS-6401 | chr6-43887782-INS-6401 |
| chr6-43902245-INS-376 | chr6-43902276-INS-329 |
| chr6-43905368-INS-81 | chr6-43905368-INS-86 |
| chr6-43948468-INS-68 | chr6-43948468-INS-68 |
| chr6-43992773-INS-82 | chr6-43992773-INS-82 |
| chr6-44020465-INS-68 | chr6-44020465-INS-72 |
| chr6-44065220-INS-79 | chr6-44065221-INS-64 |
| chr6-44074205-INS-57 | chr6-44074205-INS-66 |
| chr6-44102241-INS-2297 | chr6-44102242-INS-2298 |
| chr6-44110171-INS-498 | chr6-44110171-INS-492 |
| chr6-44120506-INS-6419 | chr6-44120507-INS-6419 |
| chr6-44154271-INS-215 | chr6-44154271-INS-215 |
| chr6-44155662-INS-222 | chr6-44155626-INS-210 |
| chr6-44160445-INS-116 | chr6-44160445-INS-120 |
| chr6-44163911-INS-96 | chr6-44163916-INS-88 |
| chr6-44164529-INS-1441 | chr6-44164530-INS-1442 |
| chr6-44172144-INS-52 | chr6-44172144-INS-50 |
| chr6-44185176-INS-76 | chr6-44185176-INS-76 |
| chr6-44197044-INS-66 | chr6-44197044-INS-66 |
| chr6-44203213-INS-80 | chr6-44203213-INS-80 |
| chr6-44205984-INS-550 | chr6-44205984-INS-550 |
| chr6-44238985-INS-110 | chr6-44238985-INS-68 |
| chr6-44270547-INS-63 | chr6-44270547-INS-67 |
| chr6-44282064-INS-70 | chr6-44282064-INS-70 |
| chr6-44326160-INS-7122 | chr6-44326161-INS-7122 |
| chr6-44409534-INS-215 | chr6-44409534-INS-215 |
| chr6-44415108-INS-2425 | chr6-44415109-INS-2425 |
| chr6-44438292-INS-54 | chr6-44438292-INS-54 |
| chr6-44446435-INS-62 | chr6-44446435-INS-54 |
| chr6-44486067-INS-69 | chr6-44486076-INS-87 |
| chr6-44524770-INS-52 | chr6-44524743-INS-50 |
| chr6-44543755-INS-56 | chr6-44543755-INS-52 |
| chr6-44577443-INS-4537 | chr6-44577444-INS-4537 |
| chr6-44582535-INS-390 | chr6-44582535-INS-390 |
| chr6-44585952-INS-54 | chr6-44585948-INS-52 |
| chr6-44586674-INS-2932 | chr6-44586672-INS-2937 |
| chr6-44602470-INS-185 | chr6-44602470-INS-190 |
| chr6-44656762-INS-836 | chr6-44656762-INS-836 |
| chr6-44681584-INS-425 | chr6-44681584-INS-424 |
| chr6-44693595-INS-3384 | chr6-44693596-INS-3384 |
| chr6-44728729-INS-6217 | chr6-44728730-INS-6217 |
| chr6-44729288-INS-2404 | chr6-44729289-INS-2405 |
| chr6-44742582-INS-1446 | chr6-44742583-INS-1446 |
| chr6-44753092-INS-358 | chr6-44753092-INS-358 |
| chr6-44785403-INS-6701 | chr6-44785404-INS-6701 |
| chr6-44815113-INS-96 | chr6-44815113-INS-96 |
| chr6-44850934-INS-112 | chr6-44850934-INS-112 |
| chr6-44872009-INS-307 | chr6-44872009-INS-312 |
| chr6-44885789-INS-773 | chr6-44885793-INS-780 |
| chr6-44938849-INS-266 | chr6-44938849-INS-234 |
| chr6-44958962-INS-131 | chr6-44958967-INS-132 |
| chr6-44972228-INS-170 | chr6-44972228-INS-174 |
| chr6-45026269-INS-53 | chr6-45026269-INS-54 |
| chr6-45074990-INS-137 | chr6-45075010-INS-115 |
| chr6-45077449-INS-128 | chr6-45077449-INS-128 |

| | |
|---|---|
| chr6-45078175-INS-5444 | chr6-45078176-INS-5444 |
| chr6-45097167-INS-1671 | chr6-45097168-INS-1672 |
| chr6-45116816-INS-90 | chr6-45116838-INS-89 |
| chr6-45172999-INS-85 | chr6-45172999-INS-85 |
| chr6-45176472-INS-73 | chr6-45176471-INS-84 |
| chr6-45224062-INS-463 | chr6-45224062-INS-463 |
| chr6-45282564-INS-79 | chr6-45282566-INS-85 |
| chr6-45292299-INS-1771 | chr6-45292300-INS-1771 |
| chr6-45300475-INS-6723 | chr6-45300476-INS-6723 |
| chr6-45310348-INS-52 | chr6-45310348-INS-56 |
| chr6-45331440-INS-6444 | chr6-45331441-INS-6445 |
| chr6-45332447-INS-203 | chr6-45332447-INS-197 |
| chr6-45337719-INS-4307 | chr6-45337720-INS-4308 |
| chr6-45339308-INS-53 | chr6-45339308-INS-57 |
| chr6-45345247-INS-72 | chr6-45345247-INS-72 |
| chr6-45361660-INS-6202 | chr6-45361661-INS-6203 |
| chr6-45366933-INS-1874 | chr6-45366934-INS-1875 |
| chr6-45383265-INS-147 | chr6-45383265-INS-148 |
| chr6-45413706-INS-826 | chr6-45413706-INS-826 |
| chr6-45425008-INS-65 | chr6-45425003-INS-71 |
| chr6-45428616-INS-54 | chr6-45428615-INS-52 |
| chr6-45435593-INS-232 | chr6-45435593-INS-232 |
| chr6-45443866-INS-189 | chr6-45443866-INS-189 |
| chr6-45451905-INS-234 | chr6-45451905-INS-234 |
| chr6-45473455-INS-179 | chr6-45473455-INS-181 |
| chr6-45480544-INS-2813 | chr6-45480545-INS-2813 |
| chr6-45483020-INS-89 | chr6-45483020-INS-99 |
| chr6-45490582-INS-63 | chr6-45490582-INS-63 |
| chr6-45497692-INS-181 | chr6-45497673-INS-167 |
| chr6-45503888-INS-147 | chr6-45503888-INS-147 |
| chr6-45508488-INS-224 | chr6-45508488-INS-224 |
| chr6-45550578-INS-78 | chr6-45550578-INS-68 |
| chr6-45558770-INS-639 | chr6-45558770-INS-639 |
| chr6-45559625-INS-81 | chr6-45559625-INS-71 |
| chr6-45560690-INS-199 | chr6-45560690-INS-194 |
| chr6-45587712-INS-517 | chr6-45587712-INS-517 |
| chr6-45590975-INS-635 | chr6-45590975-INS-635 |
| chr6-45612164-INS-203 | chr6-45612164-INS-203 |
| chr6-45675644-INS-707 | chr6-45675644-INS-708 |
| chr6-45688892-INS-70 | chr6-45688893-INS-64 |
| chr6-45703683-INS-934 | chr6-45703683-INS-934 |
| chr6-45713581-INS-2524 | chr6-45713582-INS-2526 |
| chr6-45737033-INS-74 | chr6-45737033-INS-74 |
| chr6-45776269-INS-1589 | chr6-45776270-INS-1590 |
| chr6-45782165-INS-6696 | chr6-45782166-INS-6697 |
| chr6-45784633-INS-172 | chr6-45784633-INS-184 |
| chr6-45825874-INS-2848 | chr6-45825875-INS-2848 |
| chr6-45929434-INS-59 | chr6-45929434-INS-59 |
| chr6-45941877-INS-54 | chr6-45941915-INS-66 |
| chr6-45988629-INS-118 | chr6-45988636-INS-126 |
| chr6-45991020-INS-52 | chr6-45991024-INS-53 |
| chr6-45997804-INS-52 | chr6-45997804-INS-68 |
| chr6-46037515-INS-1272 | chr6-46037516-INS-1268 |
| chr6-46086112-INS-52 | chr6-46086124-INS-52 |
| chr6-46088650-INS-96 | chr6-46088654-INS-98 |
| chr6-46090508-INS-123 | chr6-46090508-INS-123 |
| chr6-46103370-INS-71 | chr6-46103370-INS-72 |
| chr6-46147388-INS-339 | chr6-46147388-INS-337 |
| chr6-46161944-INS-1556 | chr6-46161945-INS-1557 |
| chr6-46173339-INS-64 | chr6-46173339-INS-66 |
| chr6-46181775-INS-1395 | chr6-46181776-INS-1395 |
| chr6-46209544-INS-323 | chr6-46209544-INS-323 |
| chr6-46231795-INS-1395 | chr6-46231796-INS-1395 |
| chr6-46242422-INS-746 | chr6-46242422-INS-746 |
| chr6-46256718-INS-686 | chr6-46256720-INS-687 |
| chr6-46257768-INS-54 | chr6-46257769-INS-71 |
| chr6-46289367-INS-201 | chr6-46289367-INS-201 |
| chr6-46306002-INS-161 | chr6-46306002-INS-160 |
| chr6-46306718-INS-60 | chr6-46306718-INS-64 |
| chr6-46313619-INS-65 | chr6-46313619-INS-73 |
| chr6-46313837-INS-6575 | chr6-46313838-INS-6575 |
| chr6-46315573-INS-704 | chr6-46315573-INS-706 |
| chr6-46359254-INS-660 | chr6-46359254-INS-662 |
| chr6-46408165-INS-402 | chr6-46408165-INS-402 |
| chr6-46434240-INS-6524 | chr6-46434241-INS-6524 |
| chr6-46436719-INS-53 | chr6-46436713-INS-56 |
| chr6-46446826-INS-76 | chr6-46446826-INS-80 |

| | |
|---|---|
| chr6-46451043-INS-333 | chr6-46451043-INS-341 |
| chr6-46454136-INS-219 | chr6-46454136-INS-219 |
| chr6-46460168-INS-84 | chr6-46460168-INS-84 |
| chr6-46467589-INS-51 | chr6-46467589-INS-55 |
| chr6-46517486-INS-267 | chr6-46517486-INS-268 |
| chr6-46525720-INS-3952 | chr6-46525721-INS-3953 |
| chr6-46597630-INS-54 | chr6-46597630-INS-56 |
| chr6-46605526-INS-75 | chr6-46605526-INS-78 |
| chr6-46634747-INS-60 | chr6-46634754-INS-98 |
| chr6-46636450-INS-54 | chr6-46636450-INS-52 |
| chr6-46664210-INS-60 | chr6-46664210-INS-50 |
| chr6-46736453-INS-764 | chr6-46736453-INS-764 |
| chr6-46812129-INS-450 | chr6-46812129-INS-450 |
| chr6-46851210-INS-70 | chr6-46851210-INS-76 |
| chr6-46863820-INS-1494 | chr6-46863821-INS-1494 |
| chr6-46870372-INS-54 | chr6-46870372-INS-56 |
| chr6-46875645-INS-191 | chr6-46875645-INS-191 |
| chr6-46888387-INS-186 | chr6-46888387-INS-186 |
| chr6-46909093-INS-486 | chr6-46909093-INS-486 |
| chr6-46929311-INS-198 | chr6-46929311-INS-234 |
| chr6-46969887-INS-460 | chr6-46969887-INS-465 |
| chr6-46992501-INS-55 | chr6-46992501-INS-55 |
| chr6-46996972-INS-1904 | chr6-46996973-INS-1904 |
| chr6-46999710-INS-7088 | chr6-46999711-INS-7088 |
| chr6-47016493-INS-1451 | chr6-47016494-INS-1451 |
| chr6-47039099-INS-430 | chr6-47039099-INS-432 |
| chr6-47050338-INS-649 | chr6-47050338-INS-650 |
| chr6-47052922-INS-6382 | chr6-47052923-INS-6382 |
| chr6-47063211-INS-358 | chr6-47063211-INS-358 |
| chr6-47090651-INS-62 | chr6-47090610-INS-84 |
| chr6-47095653-INS-190 | chr6-47095653-INS-190 |
| chr6-47101938-INS-76 | chr6-47101938-INS-76 |
| chr6-47135656-INS-58 | chr6-47135657-INS-58 |
| chr6-47143606-INS-586 | chr6-47143606-INS-587 |
| chr6-47158090-INS-488 | chr6-47158090-INS-488 |
| chr6-47178872-INS-138 | chr6-47178877-INS-126 |
| chr6-47180446-INS-75 | chr6-47180446-INS-67 |
| chr6-47189080-INS-142 | chr6-47189080-INS-142 |
| chr6-47194378-INS-285 | chr6-47194378-INS-280 |
| chr6-47224005-INS-81 | chr6-47224005-INS-81 |
| chr6-47238662-INS-54 | chr6-47238662-INS-60 |
| chr6-47273998-INS-717 | chr6-47273998-INS-717 |
| chr6-47290354-INS-66 | chr6-47290354-INS-66 |
| chr6-47305721-INS-132 | chr6-47305721-INS-133 |
| chr6-47313266-INS-3879 | chr6-47313267-INS-3880 |
| chr6-47316482-INS-200 | chr6-47316482-INS-200 |
| chr6-47321919-INS-209 | chr6-47321927-INS-244 |
| chr6-47323696-INS-74 | chr6-47323696-INS-82 |
| chr6-47339226-INS-97 | chr6-47339233-INS-98 |
| chr6-47342582-INS-80 | chr6-47342576-INS-52 |
| chr6-47354146-INS-207 | chr6-47354146-INS-209 |
| chr6-47363250-INS-72 | chr6-47363292-INS-71 |
| chr6-47379387-INS-528 | chr6-47379387-INS-528 |
| chr6-47396244-INS-83 | chr6-47396244-INS-83 |
| chr6-47413062-INS-201 | chr6-47413062-INS-201 |
| chr6-47421824-INS-614 | chr6-47421824-INS-614 |
| chr6-47444182-INS-223 | chr6-47444182-INS-225 |
| chr6-47541104-INS-218 | chr6-47541104-INS-218 |
| chr6-47545318-INS-247 | chr6-47545318-INS-247 |
| chr6-47556488-INS-439 | chr6-47556488-INS-439 |
| chr6-47565582-INS-55 | chr6-47565582-INS-64 |
| chr6-47595059-INS-1084 | chr6-47595060-INS-1084 |
| chr6-47758223-INS-230 | chr6-47758223-INS-231 |
| chr6-47775064-INS-294 | chr6-47775064-INS-294 |
| chr6-47775889-INS-100 | chr6-47775889-INS-100 |
| chr6-47801781-INS-54 | chr6-47801781-INS-62 |
| chr6-47805678-INS-177 | chr6-47805678-INS-178 |
| chr6-47809183-INS-3842 | chr6-47809184-INS-3842 |
| chr6-47850800-INS-55 | chr6-47850800-INS-55 |
| chr6-47886765-INS-84 | chr6-47886797-INS-80 |
| chr6-47893661-INS-207 | chr6-47893661-INS-207 |
| chr6-47904676-INS-158 | chr6-47904676-INS-158 |
| chr6-47910271-INS-2459 | chr6-47910272-INS-2459 |
| chr6-47920792-INS-646 | chr6-47920792-INS-646 |
| chr6-47941607-INS-58 | chr6-47941607-INS-58 |
| chr6-47953739-INS-242 | chr6-47953739-INS-243 |
| chr6-47961191-INS-84 | chr6-47961191-INS-84 |

| | |
|---|---|
| chr6-47962837-INS-213 | chr6-47962837-INS-212 |
| chr6-48009524-INS-360 | chr6-48009524-INS-360 |
| chr6-48033599-INS-58 | chr6-48033599-INS-58 |
| chr6-48037388-INS-517 | chr6-48037388-INS-517 |
| chr6-48101714-INS-374 | chr6-48101714-INS-375 |
| chr6-48168353-INS-90 | chr6-48168353-INS-93 |
| chr6-48232888-INS-58 | chr6-48232888-INS-58 |
| chr6-48241064-INS-689 | chr6-48241064-INS-689 |
| chr6-48311311-INS-389 | chr6-48311311-INS-392 |
| chr6-48336721-INS-219 | chr6-48336721-INS-219 |
| chr6-48361445-INS-3355 | chr6-48361446-INS-3355 |
| chr6-48367934-INS-701 | chr6-48367933-INS-699 |
| chr6-48391296-INS-218 | chr6-48391296-INS-218 |
| chr6-48436210-INS-127 | chr6-48436210-INS-119 |
| chr6-48454214-INS-168 | chr6-48454216-INS-163 |
| chr6-48466523-INS-56 | chr6-48466523-INS-56 |
| chr6-48502327-INS-195 | chr6-48502327-INS-195 |
| chr6-48502929-INS-108 | chr6-48502919-INS-72 |
| chr6-48534212-INS-165 | chr6-48534207-INS-158 |
| chr6-48555486-INS-379 | chr6-48555486-INS-377 |
| chr6-48580469-INS-81 | chr6-48580469-INS-105 |
| chr6-48619534-INS-1890 | chr6-48619535-INS-1890 |
| chr6-48621913-INS-66 | chr6-48621913-INS-69 |
| chr6-48669721-INS-1880 | chr6-48669722-INS-1880 |
| chr6-48688796-INS-64 | chr6-48688819-INS-65 |
| chr6-48693607-INS-566 | chr6-48693607-INS-566 |
| chr6-48699628-INS-69 | chr6-48699628-INS-69 |
| chr6-48725682-INS-75 | chr6-48725693-INS-55 |
| chr6-48739487-INS-179 | chr6-48739487-INS-179 |
| chr6-48746885-INS-55 | chr6-48746885-INS-55 |
| chr6-48836027-INS-455 | chr6-48836027-INS-455 |
| chr6-48840754-INS-88 | chr6-48840752-INS-81 |
| chr6-48851882-INS-453 | chr6-48851883-INS-453 |
| chr6-48857424-INS-58 | chr6-48857420-INS-58 |
| chr6-48877721-INS-125 | chr6-48877717-INS-93 |
| chr6-48928016-INS-404 | chr6-48928016-INS-404 |
| chr6-48933511-INS-614 | chr6-48933511-INS-614 |
| chr6-48936539-INS-59 | chr6-48936539-INS-59 |
| chr6-48946618-INS-93 | chr6-48946618-INS-93 |
| chr6-48964956-INS-54 | chr6-48964956-INS-54 |
| chr6-48983325-INS-52 | chr6-48983325-INS-52 |
| chr6-48997504-INS-100 | chr6-48997504-INS-100 |
| chr6-49019356-INS-72 | chr6-49019356-INS-72 |
| chr6-49037508-INS-68 | chr6-49037508-INS-70 |
| chr6-49038926-INS-199 | chr6-49038926-INS-199 |
| chr6-49045418-INS-118 | chr6-49045418-INS-117 |
| chr6-49082142-INS-97 | chr6-49082142-INS-97 |
| chr6-49087231-INS-128 | chr6-49087231-INS-156 |
| chr6-49093499-INS-80 | chr6-49093499-INS-68 |
| chr6-49095450-INS-56 | chr6-49095450-INS-56 |
| chr6-49111053-INS-54 | chr6-49111053-INS-58 |
| chr6-49174670-INS-93 | chr6-49174670-INS-111 |
| chr6-49180946-INS-216 | chr6-49180946-INS-216 |
| chr6-49202258-INS-70 | chr6-49202258-INS-70 |
| chr6-49211948-INS-76 | chr6-49211932-INS-56 |
| chr6-49220439-INS-64 | chr6-49220439-INS-64 |
| chr6-49229147-INS-196 | chr6-49229145-INS-197 |
| chr6-49233552-INS-219 | chr6-49233552-INS-224 |
| chr6-49234407-INS-200 | chr6-49234407-INS-199 |
| chr6-49235605-INS-205 | chr6-49235605-INS-203 |
| chr6-49246937-INS-208 | chr6-49246937-INS-207 |
| chr6-49256200-INS-53 | chr6-49256200-INS-53 |
| chr6-49258728-INS-56 | chr6-49258728-INS-84 |
| chr6-49275845-INS-109 | chr6-49275845-INS-109 |
| chr6-49278308-INS-320 | chr6-49278308-INS-319 |
| chr6-49279610-INS-601 | chr6-49279610-INS-587 |
| chr6-49287807-INS-205 | chr6-49287807-INS-207 |
| chr6-49293510-INS-1476 | chr6-49293511-INS-1476 |
| chr6-49297643-INS-80 | chr6-49297643-INS-80 |
| chr6-49302200-INS-242 | chr6-49302200-INS-248 |
| chr6-49323995-INS-189 | chr6-49323995-INS-189 |
| chr6-49336050-INS-231 | chr6-49336050-INS-231 |
| chr6-49339694-INS-218 | chr6-49339694-INS-222 |
| chr6-49356672-INS-309 | chr6-49356672-INS-309 |
| chr6-49357664-INS-217 | chr6-49357664-INS-218 |
| chr6-49362842-INS-6443 | chr6-49362843-INS-6443 |
| chr6-49377580-INS-5544 | chr6-49377581-INS-5544 |

| | |
|---|---|
| chr6-49383333-INS-316 | chr6-49383336-INS-321 |
| chr6-49440591-INS-2587 | chr6-49440592-INS-2587 |
| chr6-49461149-INS-59 | chr6-49461136-INS-79 |
| chr6-49486174-INS-66 | chr6-49486184-INS-67 |
| chr6-49491855-INS-463 | chr6-49491855-INS-463 |
| chr6-49516354-INS-260 | chr6-49516354-INS-265 |
| chr6-49524013-INS-80 | chr6-49524013-INS-81 |
| chr6-49549555-INS-210 | chr6-49549555-INS-210 |
| chr6-49558002-INS-515 | chr6-49558002-INS-515 |
| chr6-49575227-INS-522 | chr6-49575227-INS-522 |
| chr6-49579682-INS-399 | chr6-49579682-INS-399 |
| chr6-49591864-INS-85 | chr6-49591864-INS-85 |
| chr6-49593719-INS-206 | chr6-49593719-INS-206 |
| chr6-49603817-INS-78 | chr6-49603817-INS-78 |
| chr6-49612441-INS-5560 | chr6-49612442-INS-5560 |
| chr6-49613264-INS-2865 | chr6-49613265-INS-2865 |
| chr6-49618817-INS-213 | chr6-49618812-INS-216 |
| chr6-49652178-INS-299 | chr6-49652171-INS-282 |
| chr6-49657665-INS-1285 | chr6-49657669-INS-1263 |
| chr6-49660445-INS-69 | chr6-49660445-INS-54 |
| chr6-49664930-INS-6764 | chr6-49664931-INS-6764 |
| chr6-49670968-INS-1956 | chr6-49670969-INS-1956 |
| chr6-49675722-INS-503 | chr6-49675722-INS-503 |
| chr6-49678189-INS-128 | chr6-49678189-INS-130 |
| chr6-49697374-INS-193 | chr6-49697374-INS-194 |
| chr6-49701641-INS-86 | chr6-49701641-INS-88 |
| chr6-49710909-INS-62 | chr6-49710909-INS-62 |
| chr6-49723890-INS-108 | chr6-49723890-INS-108 |
| chr6-49763613-INS-654 | chr6-49763613-INS-654 |
| chr6-49767777-INS-54 | chr6-49767777-INS-50 |
| chr6-49778930-INS-737 | chr6-49778930-INS-737 |
| chr6-49796768-INS-498 | chr6-49796768-INS-498 |
| chr6-49818206-INS-395 | chr6-49818206-INS-395 |
| chr6-49820581-INS-497 | chr6-49820580-INS-543 |
| chr6-49826517-INS-1864 | chr6-49826518-INS-1864 |
| chr6-49847007-INS-58 | chr6-49847022-INS-52 |
| chr6-49847583-INS-164 | chr6-49847583-INS-164 |
| chr6-49867042-INS-59 | chr6-49867042-INS-59 |
| chr6-49877138-INS-68 | chr6-49877138-INS-68 |
| chr6-49880994-INS-804 | chr6-49880994-INS-804 |
| chr6-49883175-INS-59 | chr6-49883175-INS-59 |
| chr6-49888981-INS-6440 | chr6-49888982-INS-6440 |
| chr6-49892959-INS-52 | chr6-49892939-INS-50 |
| chr6-49895008-INS-61 | chr6-49895012-INS-100 |
| chr6-49905670-INS-78 | chr6-49905676-INS-54 |
| chr6-49922132-INS-4213 | chr6-49922133-INS-4213 |
| chr6-49941180-INS-60 | chr6-49941180-INS-62 |
| chr6-49964572-INS-68 | chr6-49964572-INS-72 |
| chr6-50001093-INS-66 | chr6-50001093-INS-66 |
| chr6-50007961-INS-402 | chr6-50007961-INS-402 |
| chr6-50083169-INS-51 | chr6-50083169-INS-51 |
| chr6-50114611-INS-54 | chr6-50114611-INS-58 |
| chr6-50147145-INS-1236 | chr6-50147160-INS-1220 |
| chr6-50172768-INS-60 | chr6-50172768-INS-60 |
| chr6-50183751-INS-7158 | chr6-50183752-INS-7158 |
| chr6-50185971-INS-171 | chr6-50185971-INS-171 |
| chr6-50195053-INS-160 | chr6-50195036-INS-200 |
| chr6-50208840-INS-227 | chr6-50208840-INS-227 |
| chr6-50234984-INS-68 | chr6-50234984-INS-68 |
| chr6-50243236-INS-148 | chr6-50243236-INS-148 |
| chr6-50249935-INS-409 | chr6-50249935-INS-409 |
| chr6-50261234-INS-222 | chr6-50261234-INS-223 |
| chr6-50269165-INS-61 | chr6-50269165-INS-61 |
| chr6-50293744-INS-183 | chr6-50293744-INS-183 |
| chr6-50306852-INS-116 | chr6-50306852-INS-116 |
| chr6-50327106-INS-2990 | chr6-50327107-INS-2990 |
| chr6-50335750-INS-73 | chr6-50335744-INS-55 |
| chr6-50341983-INS-68 | chr6-50341983-INS-68 |
| chr6-50408345-INS-76 | chr6-50408345-INS-76 |
| chr6-50428430-INS-338 | chr6-50428420-INS-316 |
| chr6-50431355-INS-294 | chr6-50431355-INS-304 |
| chr6-50434029-INS-78 | chr6-50434031-INS-96 |
| chr6-50446767-INS-992 | chr6-50446767-INS-992 |
| chr6-50467337-INS-98 | chr6-50467337-INS-79 |
| chr6-50468506-INS-117 | chr6-50468507-INS-98 |
| chr6-50478397-INS-123 | chr6-50478397-INS-123 |
| chr6-50503972-INS-86 | chr6-50503972-INS-68 |

| | |
|---|---|
| chr6-50509369-INS-662 | chr6-50509365-INS-673 |
| chr6-50515097-INS-64 | chr6-50515097-INS-64 |
| chr6-50527479-INS-83 | chr6-50527459-INS-113 |
| chr6-50528868-INS-205 | chr6-50528868-INS-200 |
| chr6-50530365-INS-6401 | chr6-50530366-INS-6401 |
| chr6-50544888-INS-61 | chr6-50544888-INS-61 |
| chr6-50651536-INS-102 | chr6-50651536-INS-102 |
| chr6-50651957-INS-332 | chr6-50651955-INS-335 |
| chr6-50673986-INS-668 | chr6-50673986-INS-668 |
| chr6-50701092-INS-427 | chr6-50701092-INS-427 |
| chr6-50722608-INS-58 | chr6-50722608-INS-58 |
| chr6-50731745-INS-176 | chr6-50731745-INS-176 |
| chr6-50748769-INS-86 | chr6-50748769-INS-86 |
| chr6-50755665-INS-74 | chr6-50755665-INS-74 |
| chr6-50765136-INS-96 | chr6-50765136-INS-96 |
| chr6-50793408-INS-87 | chr6-50793408-INS-58 |
| chr6-50801685-INS-140 | chr6-50801685-INS-140 |
| chr6-50815502-INS-60 | chr6-50815500-INS-62 |
| chr6-50820327-INS-203 | chr6-50820327-INS-203 |
| chr6-50830920-INS-83 | chr6-50830902-INS-61 |
| chr6-50832202-INS-55 | chr6-50832209-INS-50 |
| chr6-50843290-INS-376 | chr6-50843290-INS-419 |
| chr6-50844579-INS-4408 | chr6-50844580-INS-4408 |
| chr6-50847884-INS-2478 | chr6-50847885-INS-2478 |
| chr6-50851522-INS-103 | chr6-50851513-INS-64 |
| chr6-50859101-INS-111 | chr6-50859100-INS-111 |
| chr6-50899973-INS-246 | chr6-50899967-INS-241 |
| chr6-50904239-INS-53 | chr6-50904239-INS-54 |
| chr6-50909286-INS-7182 | chr6-50909287-INS-7182 |
| chr6-50933200-INS-56 | chr6-50933200-INS-56 |
| chr6-50955493-INS-75 | chr6-50955477-INS-80 |
| chr6-50957766-INS-153 | chr6-50957766-INS-153 |
| chr6-50962334-INS-256 | chr6-50962334-INS-256 |
| chr6-50968545-INS-5522 | chr6-50968546-INS-5522 |
| chr6-50984835-INS-457 | chr6-50984835-INS-458 |
| chr6-51006823-INS-453 | chr6-51006823-INS-453 |
| chr6-51012372-INS-74 | chr6-51012372-INS-75 |
| chr6-51032223-INS-273 | chr6-51032223-INS-279 |
| chr6-51062148-INS-63 | chr6-51062168-INS-83 |
| chr6-51086708-INS-149 | chr6-51086735-INS-114 |
| chr6-51090279-INS-56 | chr6-51090279-INS-64 |
| chr6-51094811-INS-75 | chr6-51094783-INS-118 |
| chr6-51099271-INS-72 | chr6-51099271-INS-80 |
| chr6-51116184-INS-52 | chr6-51116184-INS-76 |
| chr6-51118082-INS-106 | chr6-51118082-INS-111 |
| chr6-51182338-INS-6431 | chr6-51182339-INS-6431 |
| chr6-51191689-INS-865 | chr6-51191689-INS-864 |
| chr6-51198314-INS-125 | chr6-51198337-INS-141 |
| chr6-51198971-INS-62 | chr6-51198992-INS-62 |
| chr6-51204757-INS-315 | chr6-51204757-INS-315 |
| chr6-51230692-INS-2847 | chr6-51230693-INS-2846 |
| chr6-51239121-INS-68 | chr6-51239122-INS-86 |
| chr6-51241785-INS-80 | chr6-51241785-INS-80 |
| chr6-51261348-INS-128 | chr6-51261348-INS-128 |
| chr6-51271301-INS-63 | chr6-51271301-INS-63 |
| chr6-51291334-INS-5314 | chr6-51291335-INS-5314 |
| chr6-51327597-INS-6401 | chr6-51327598-INS-6401 |
| chr6-51329288-INS-64 | chr6-51329288-INS-56 |
| chr6-51380552-INS-251 | chr6-51380553-INS-248 |
| chr6-51386619-INS-140 | chr6-51386619-INS-140 |
| chr6-51403277-INS-2885 | chr6-51403278-INS-2875 |
| chr6-51420044-INS-95 | chr6-51420044-INS-83 |
| chr6-51425728-INS-55 | chr6-51425728-INS-55 |
| chr6-51442916-INS-199 | chr6-51442916-INS-199 |
| chr6-51480896-INS-97 | chr6-51480896-INS-101 |
| chr6-51481393-INS-210 | chr6-51481393-INS-210 |
| chr6-51489585-INS-479 | chr6-51489585-INS-478 |
| chr6-51492280-INS-207 | chr6-51492280-INS-209 |
| chr6-51493790-INS-184 | chr6-51493790-INS-191 |
| chr6-51499406-INS-232 | chr6-51499406-INS-232 |
| chr6-51503801-INS-96 | chr6-51503778-INS-72 |
| chr6-51521165-INS-58 | chr6-51521165-INS-56 |
| chr6-51525552-INS-212 | chr6-51525553-INS-199 |
| chr6-51538467-INS-5035 | chr6-51538468-INS-5035 |
| chr6-51551206-INS-5225 | chr6-51551207-INS-5225 |
| chr6-51575251-INS-52 | chr6-51575251-INS-56 |
| chr6-51590099-INS-55 | chr6-51590095-INS-78 |

| | |
|---|---|
| chr6-51597997-INS-117 | chr6-51597997-INS-88 |
| chr6-51605967-INS-3102 | chr6-51605968-INS-3102 |
| chr6-51614527-INS-52 | chr6-51614527-INS-52 |
| chr6-51619835-INS-77 | chr6-51619835-INS-77 |
| chr6-51623616-INS-4540 | chr6-51623617-INS-4540 |
| chr6-51632306-INS-257 | chr6-51632306-INS-280 |
| chr6-51640623-INS-105 | chr6-51640623-INS-105 |
| chr6-51648954-INS-68 | chr6-51648949-INS-81 |
| chr6-51681728-INS-87 | chr6-51681728-INS-87 |
| chr6-51714708-INS-72 | chr6-51714708-INS-100 |
| chr6-51724878-INS-57 | chr6-51724878-INS-57 |
| chr6-51728341-INS-328 | chr6-51728341-INS-328 |
| chr6-51731748-INS-167 | chr6-51731748-INS-145 |
| chr6-51758363-INS-596 | chr6-51758363-INS-596 |
| chr6-51770638-INS-60 | chr6-51770638-INS-68 |
| chr6-51782327-INS-66 | chr6-51782359-INS-66 |
| chr6-51786764-INS-1836 | chr6-51786765-INS-1860 |
| chr6-51793822-INS-1935 | chr6-51793823-INS-1930 |
| chr6-51798354-INS-53 | chr6-51798333-INS-80 |
| chr6-51803504-INS-480 | chr6-51803504-INS-480 |
| chr6-51829017-INS-575 | chr6-51829017-INS-575 |
| chr6-51831718-INS-91 | chr6-51831718-INS-91 |
| chr6-51902784-INS-116 | chr6-51902785-INS-114 |
| chr6-51903523-INS-119 | chr6-51903524-INS-119 |
| chr6-51918203-INS-151 | chr6-51918203-INS-151 |
| chr6-51922822-INS-71 | chr6-51922792-INS-85 |
| chr6-51926872-INS-76 | chr6-51926872-INS-76 |
| chr6-51974082-INS-226 | chr6-51974082-INS-226 |
| chr6-52033632-INS-141 | chr6-52033632-INS-102 |
| chr6-52035935-INS-84 | chr6-52035935-INS-71 |
| chr6-52039937-INS-2906 | chr6-52039938-INS-2906 |
| chr6-52048108-INS-80 | chr6-52048153-INS-77 |
| chr6-52067470-INS-52 | chr6-52067470-INS-52 |
| chr6-52070286-INS-1735 | chr6-52070287-INS-1718 |
| chr6-52074868-INS-108 | chr6-52074851-INS-93 |
| chr6-52118010-INS-122 | chr6-52118019-INS-74 |
| chr6-52122956-INS-63 | chr6-52122957-INS-63 |
| chr6-52123869-INS-672 | chr6-52123869-INS-673 |
| chr6-52247175-INS-70 | chr6-52247175-INS-70 |
| chr6-52265104-INS-218 | chr6-52265104-INS-218 |
| chr6-52318130-INS-59 | chr6-52318133-INS-63 |
| chr6-52342715-INS-108 | chr6-52342715-INS-108 |
| chr6-52351565-INS-277 | chr6-52351565-INS-277 |
| chr6-52428803-INS-140 | chr6-52428803-INS-96 |
| chr6-52476916-INS-64 | chr6-52476916-INS-64 |
| chr6-52487194-INS-7132 | chr6-52487195-INS-7132 |
| chr6-52511073-INS-417 | chr6-52511073-INS-417 |
| chr6-52548130-INS-1661 | chr6-52548131-INS-1661 |
| chr6-52557707-INS-2890 | chr6-52557708-INS-2890 |
| chr6-52567283-INS-183 | chr6-52567283-INS-184 |
| chr6-52568263-INS-87 | chr6-52568263-INS-87 |
| chr6-52573976-INS-51 | chr6-52573976-INS-51 |
| chr6-52586502-INS-60 | chr6-52586502-INS-60 |
| chr6-52602632-INS-61 | chr6-52602632-INS-61 |
| chr6-52630207-INS-532 | chr6-52630207-INS-531 |
| chr6-52643772-INS-63 | chr6-52643762-INS-64 |
| chr6-52647184-INS-714 | chr6-52647184-INS-714 |
| chr6-52722932-INS-124 | chr6-52722932-INS-124 |
| chr6-52728915-INS-1576 | chr6-52728916-INS-1593 |
| chr6-52729787-INS-1400 | chr6-52729788-INS-1400 |
| chr6-52739186-INS-721 | chr6-52739186-INS-735 |
| chr6-52784159-INS-2098 | chr6-52784165-INS-2107 |
| chr6-52792335-INS-63 | chr6-52792335-INS-63 |
| chr6-52858606-INS-305 | chr6-52858578-INS-283 |
| chr6-52877112-INS-87 | chr6-52877112-INS-99 |
| chr6-52897783-INS-92 | chr6-52897822-INS-92 |
| chr6-52904697-INS-92 | chr6-52904697-INS-92 |
| chr6-52940054-INS-978 | chr6-52940054-INS-980 |
| chr6-52947943-INS-94 | chr6-52947943-INS-94 |
| chr6-52999662-INS-145 | chr6-52999662-INS-145 |
| chr6-53061513-INS-442 | chr6-53061520-INS-477 |
| chr6-53108767-INS-347 | chr6-53108768-INS-334 |
| chr6-53110349-INS-1156 | chr6-53110350-INS-1155 |
| chr6-53128778-INS-270 | chr6-53128815-INS-270 |
| chr6-53130446-INS-97 | chr6-53130449-INS-97 |
| chr6-53133654-INS-249 | chr6-53133655-INS-264 |
| chr6-53139262-INS-71 | chr6-53139279-INS-75 |

| | |
|---|---|
| chr6-53156608-INS-466 | chr6-53156608-INS-466 |
| chr6-53187725-INS-212 | chr6-53187725-INS-212 |
| chr6-53196786-INS-156 | chr6-53196786-INS-174 |
| chr6-53214807-INS-996 | chr6-53214807-INS-995 |
| chr6-53217806-INS-125 | chr6-53217806-INS-123 |
| chr6-53231018-INS-62 | chr6-53231018-INS-62 |
| chr6-53232910-INS-76 | chr6-53232910-INS-77 |
| chr6-53234785-INS-54 | chr6-53234785-INS-53 |
| chr6-53235962-INS-226 | chr6-53235962-INS-226 |
| chr6-53249794-INS-86 | chr6-53249794-INS-86 |
| chr6-53258987-INS-107 | chr6-53258987-INS-109 |
| chr6-53268673-INS-76 | chr6-53268673-INS-77 |
| chr6-53273358-INS-59 | chr6-53273362-INS-55 |
| chr6-53298437-INS-169 | chr6-53298437-INS-169 |
| chr6-53336747-INS-652 | chr6-53336747-INS-652 |
| chr6-53371661-INS-885 | chr6-53371661-INS-885 |
| chr6-53403881-INS-631 | chr6-53403881-INS-631 |
| chr6-53471173-INS-93 | chr6-53471177-INS-78 |
| chr6-53482124-INS-94 | chr6-53482124-INS-75 |
| chr6-53485753-INS-63 | chr6-53485753-INS-63 |
| chr6-53519427-INS-84 | chr6-53519427-INS-112 |
| chr6-53536248-INS-58 | chr6-53536260-INS-58 |
| chr6-53543217-INS-63 | chr6-53543217-INS-57 |
| chr6-53590215-INS-691 | chr6-53590215-INS-690 |
| chr6-53625027-INS-123 | chr6-53625027-INS-123 |
| chr6-53625878-INS-507 | chr6-53625878-INS-507 |
| chr6-53656854-INS-60 | chr6-53656854-INS-52 |
| chr6-53680334-INS-497 | chr6-53680334-INS-497 |
| chr6-53688978-INS-68 | chr6-53688978-INS-68 |
| chr6-53691010-INS-56 | chr6-53691010-INS-56 |
| chr6-53759529-INS-327 | chr6-53759529-INS-340 |
| chr6-53762408-INS-78 | chr6-53762408-INS-88 |
| chr6-53776908-INS-86 | chr6-53776908-INS-86 |
| chr6-53779893-INS-51 | chr6-53779893-INS-51 |
| chr6-53781585-INS-100 | chr6-53781585-INS-100 |
| chr6-53837551-INS-55 | chr6-53837521-INS-53 |
| chr6-53863246-INS-407 | chr6-53863246-INS-407 |
| chr6-53912481-INS-409 | chr6-53912481-INS-409 |
| chr6-53917657-INS-1332 | chr6-53917658-INS-1332 |
| chr6-53921541-INS-5273 | chr6-53921542-INS-5273 |
| chr6-53922225-INS-73 | chr6-53922225-INS-73 |
| chr6-53936804-INS-56 | chr6-53936804-INS-52 |
| chr6-53967973-INS-168 | chr6-53968009-INS-164 |
| chr6-53970605-INS-328 | chr6-53970605-INS-328 |
| chr6-53972022-INS-63 | chr6-53972022-INS-67 |
| chr6-53975983-INS-400 | chr6-53975983-INS-400 |
| chr6-54003562-INS-86 | chr6-54003562-INS-88 |
| chr6-54013819-INS-161 | chr6-54013819-INS-161 |
| chr6-54094856-INS-60 | chr6-54094856-INS-60 |
| chr6-54120134-INS-64 | chr6-54120134-INS-68 |
| chr6-54136178-INS-735 | chr6-54136178-INS-735 |
| chr6-54162052-INS-67 | chr6-54162058-INS-61 |
| chr6-54185206-INS-234 | chr6-54185206-INS-234 |
| chr6-54270198-INS-68 | chr6-54270198-INS-68 |
| chr6-54286262-INS-235 | chr6-54286262-INS-250 |
| chr6-54311726-INS-72 | chr6-54311726-INS-72 |
| chr6-54337255-INS-171 | chr6-54337274-INS-152 |
| chr6-54345002-INS-241 | chr6-54345002-INS-243 |
| chr6-54350923-INS-260 | chr6-54350906-INS-268 |
| chr6-54359599-INS-2877 | chr6-54359600-INS-2877 |
| chr6-54361275-INS-82 | chr6-54361275-INS-96 |
| chr6-54372968-INS-109 | chr6-54372968-INS-110 |
| chr6-54378418-INS-64 | chr6-54378418-INS-50 |
| chr6-54392455-INS-80 | chr6-54392455-INS-80 |
| chr6-54395362-INS-806 | chr6-54395362-INS-806 |
| chr6-54401689-INS-202 | chr6-54401689-INS-203 |
| chr6-54423347-INS-52 | chr6-54423347-INS-56 |
| chr6-54453862-INS-71 | chr6-54453862-INS-71 |
| chr6-54490828-INS-124 | chr6-54490828-INS-136 |
| chr6-54498682-INS-891 | chr6-54498682-INS-891 |
| chr6-54510491-INS-116 | chr6-54510491-INS-116 |
| chr6-54513736-INS-1618 | chr6-54513737-INS-1618 |
| chr6-54537701-INS-201 | chr6-54537701-INS-201 |
| chr6-54544118-INS-62 | chr6-54544120-INS-71 |
| chr6-54550227-INS-62 | chr6-54550225-INS-73 |
| chr6-54554861-INS-185 | chr6-54554861-INS-185 |
| chr6-54582863-INS-5513 | chr6-54582864-INS-5513 |

| | |
|---|---|
| chr6-54589888-INS-2643 | chr6-54589889-INS-2643 |
| chr6-54596707-INS-53 | chr6-54596707-INS-53 |
| chr6-54600269-INS-54 | chr6-54600269-INS-54 |
| chr6-54608462-INS-232 | chr6-54608462-INS-232 |
| chr6-54630658-INS-151 | chr6-54630648-INS-129 |
| chr6-54637000-INS-345 | chr6-54637000-INS-345 |
| chr6-54723101-INS-179 | chr6-54723101-INS-180 |
| chr6-54724305-INS-75 | chr6-54724305-INS-100 |
| chr6-54738358-INS-64 | chr6-54738358-INS-56 |
| chr6-54745874-INS-213 | chr6-54745874-INS-213 |
| chr6-54751958-INS-420 | chr6-54751958-INS-420 |
| chr6-54753165-INS-56 | chr6-54753165-INS-56 |
| chr6-54771840-INS-1265 | chr6-54771841-INS-1265 |
| chr6-54814722-INS-207 | chr6-54814722-INS-207 |
| chr6-54816774-INS-492 | chr6-54816774-INS-492 |
| chr6-54826605-INS-56 | chr6-54826605-INS-58 |
| chr6-54827603-INS-203 | chr6-54827603-INS-205 |
| chr6-54858989-INS-52 | chr6-54858989-INS-52 |
| chr6-54861989-INS-193 | chr6-54861989-INS-194 |
| chr6-54877219-INS-216 | chr6-54877220-INS-209 |
| chr6-54878080-INS-2615 | chr6-54878081-INS-2616 |
| chr6-54878443-INS-377 | chr6-54878444-INS-377 |
| chr6-54894868-INS-200 | chr6-54894868-INS-200 |
| chr6-54899256-INS-56 | chr6-54899256-INS-56 |
| chr6-54908848-INS-213 | chr6-54908848-INS-213 |
| chr6-54932014-INS-7185 | chr6-54932015-INS-7185 |
| chr6-54932700-INS-2087 | chr6-54932701-INS-2087 |
| chr6-54939875-INS-64 | chr6-54939875-INS-60 |
| chr6-54954870-INS-338 | chr6-54954870-INS-338 |
| chr6-54970938-INS-478 | chr6-54970938-INS-478 |
| chr6-54998811-INS-67 | chr6-54998811-INS-67 |
| chr6-55009797-INS-109 | chr6-55009797-INS-109 |
| chr6-55013719-INS-61 | chr6-55013719-INS-61 |
| chr6-55020468-INS-54 | chr6-55020458-INS-62 |
| chr6-55102925-INS-229 | chr6-55102925-INS-216 |
| chr6-55108210-INS-116 | chr6-55108210-INS-98 |
| chr6-55125879-INS-111 | chr6-55125850-INS-122 |
| chr6-55149079-INS-60 | chr6-55149079-INS-62 |
| chr6-55158045-INS-55 | chr6-55158045-INS-70 |
| chr6-55187118-INS-100 | chr6-55187132-INS-52 |
| chr6-55223283-INS-108 | chr6-55223278-INS-101 |
| chr6-55237251-INS-203 | chr6-55237251-INS-203 |
| chr6-55299607-INS-734 | chr6-55299607-INS-734 |
| chr6-55300873-INS-54 | chr6-55300873-INS-54 |
| chr6-55317461-INS-101 | chr6-55317461-INS-101 |
| chr6-55327341-INS-90 | chr6-55327341-INS-90 |
| chr6-55406184-INS-6088 | chr6-55406185-INS-6088 |
| chr6-55422218-INS-56 | chr6-55422218-INS-56 |
| chr6-55484445-INS-2709 | chr6-55484446-INS-2709 |
| chr6-55487792-INS-3387 | chr6-55487793-INS-3387 |
| chr6-55516836-INS-103 | chr6-55516827-INS-106 |
| chr6-55525427-INS-96 | chr6-55525427-INS-72 |
| chr6-55534284-INS-70 | chr6-55534272-INS-82 |
| chr6-55595685-INS-61 | chr6-55595685-INS-68 |
| chr6-55597809-INS-98 | chr6-55597809-INS-98 |
| chr6-55649009-INS-66 | chr6-55649009-INS-66 |
| chr6-55667870-INS-112 | chr6-55667865-INS-70 |
| chr6-55669335-INS-64 | chr6-55669335-INS-64 |
| chr6-55671614-INS-62 | chr6-55671614-INS-56 |
| chr6-55683496-INS-66 | chr6-55683496-INS-51 |
| chr6-55686663-INS-2425 | chr6-55686664-INS-2426 |
| chr6-55698170-INS-1433 | chr6-55698171-INS-1434 |
| chr6-55712437-INS-958 | chr6-55712437-INS-915 |
| chr6-55720597-INS-2882 | chr6-55720598-INS-2882 |
| chr6-55721781-INS-53 | chr6-55721781-INS-53 |
| chr6-55732138-INS-4439 | chr6-55732139-INS-4439 |
| chr6-55749217-INS-74 | chr6-55749217-INS-74 |
| chr6-55766571-INS-118 | chr6-55766589-INS-78 |
| chr6-55806634-INS-921 | chr6-55806634-INS-921 |
| chr6-55808928-INS-622 | chr6-55808928-INS-622 |
| chr6-55821717-INS-111 | chr6-55821676-INS-138 |
| chr6-55844177-INS-184 | chr6-55844177-INS-183 |
| chr6-55855638-INS-168 | chr6-55855635-INS-150 |
| chr6-55883412-INS-56 | chr6-55883406-INS-70 |
| chr6-55904085-INS-77 | chr6-55904123-INS-56 |
| chr6-55943064-INS-78 | chr6-55943064-INS-78 |
| chr6-55944423-INS-201 | chr6-55944423-INS-201 |

| | |
|---|---|
| chr6-55969189-INS-161 | chr6-55969189-INS-161 |
| chr6-55972649-INS-56 | chr6-55972650-INS-56 |
| chr6-55985511-INS-55 | chr6-55985511-INS-51 |
| chr6-55991687-INS-85 | chr6-55991687-INS-97 |
| chr6-56009754-INS-76 | chr6-56009756-INS-110 |
| chr6-56027724-INS-107 | chr6-56027724-INS-117 |
| chr6-56036998-INS-462 | chr6-56036998-INS-468 |
| chr6-56049959-INS-275 | chr6-56049959-INS-275 |
| chr6-56074742-INS-256 | chr6-56074742-INS-256 |
| chr6-56105758-INS-1979 | chr6-56105759-INS-1979 |
| chr6-56194236-INS-526 | chr6-56194236-INS-526 |
| chr6-56208085-INS-2650 | chr6-56208086-INS-2650 |
| chr6-56210001-INS-100 | chr6-56210001-INS-100 |
| chr6-56240885-INS-1558 | chr6-56240886-INS-1558 |
| chr6-56241628-INS-3569 | chr6-56241629-INS-3569 |
| chr6-56253931-INS-241 | chr6-56253931-INS-245 |
| chr6-56257057-INS-291 | chr6-56257057-INS-295 |
| chr6-56262554-INS-964 | chr6-56262554-INS-965 |
| chr6-56339659-INS-77 | chr6-56339659-INS-127 |
| chr6-56347429-INS-77 | chr6-56347429-INS-60 |
| chr6-56364233-INS-1103 | chr6-56364234-INS-1103 |
| chr6-56372616-INS-2086 | chr6-56372617-INS-2079 |
| chr6-56388819-INS-52 | chr6-56388819-INS-60 |
| chr6-56400601-INS-2731 | chr6-56400602-INS-2731 |
| chr6-56415875-INS-231 | chr6-56415875-INS-235 |
| chr6-56448273-INS-367 | chr6-56448273-INS-367 |
| chr6-56454115-INS-398 | chr6-56454115-INS-398 |
| chr6-56462581-INS-374 | chr6-56462581-INS-374 |
| chr6-56481215-INS-220 | chr6-56481214-INS-227 |
| chr6-56495630-INS-59 | chr6-56495627-INS-56 |
| chr6-56496121-INS-716 | chr6-56496121-INS-717 |
| chr6-56514866-INS-76 | chr6-56514866-INS-63 |
| chr6-56516328-INS-75 | chr6-56516328-INS-65 |
| chr6-56523681-INS-664 | chr6-56523681-INS-664 |
| chr6-56541673-INS-294 | chr6-56541673-INS-292 |
| chr6-56546747-INS-248 | chr6-56546747-INS-248 |
| chr6-56552295-INS-72 | chr6-56552295-INS-74 |
| chr6-56557273-INS-6627 | chr6-56557274-INS-6628 |
| chr6-56561593-INS-347 | chr6-56561589-INS-347 |
| chr6-56567887-INS-191 | chr6-56567887-INS-191 |
| chr6-56575993-INS-948 | chr6-56575993-INS-948 |
| chr6-56577208-INS-691 | chr6-56577208-INS-691 |
| chr6-56580171-INS-54 | chr6-56580171-INS-54 |
| chr6-56584472-INS-347 | chr6-56584472-INS-347 |
| chr6-56594586-INS-1394 | chr6-56594587-INS-1393 |
| chr6-56606126-INS-205 | chr6-56606126-INS-205 |
| chr6-56626782-INS-52 | chr6-56626756-INS-61 |
| chr6-56641508-INS-283 | chr6-56641508-INS-284 |
| chr6-56652514-INS-260 | chr6-56652514-INS-260 |
| chr6-56664991-INS-53 | chr6-56664992-INS-53 |
| chr6-56666188-INS-748 | chr6-56666188-INS-748 |
| chr6-56670611-INS-52 | chr6-56670611-INS-52 |
| chr6-56677228-INS-103 | chr6-56677228-INS-103 |
| chr6-56699454-INS-177 | chr6-56699454-INS-177 |
| chr6-56731034-INS-57 | chr6-56731034-INS-57 |
| chr6-56744034-INS-129 | chr6-56744034-INS-129 |
| chr6-56749998-INS-89 | chr6-56749998-INS-87 |
| chr6-56780087-INS-55 | chr6-56780087-INS-55 |
| chr6-56782222-INS-201 | chr6-56782222-INS-201 |
| chr6-56786527-INS-118 | chr6-56786527-INS-128 |
| chr6-56788215-INS-3300 | chr6-56788216-INS-3300 |
| chr6-56791798-INS-203 | chr6-56791798-INS-203 |
| chr6-56794170-INS-321 | chr6-56794170-INS-321 |
| chr6-56821233-INS-206 | chr6-56821233-INS-247 |
| chr6-56831521-INS-201 | chr6-56831521-INS-201 |
| chr6-56839214-INS-762 | chr6-56839214-INS-796 |
| chr6-56840158-INS-126 | chr6-56840164-INS-139 |
| chr6-56874258-INS-347 | chr6-56874258-INS-347 |
| chr6-56885930-INS-65 | chr6-56885930-INS-60 |
| chr6-56950318-INS-149 | chr6-56950318-INS-149 |
| chr6-56950739-INS-457 | chr6-56950739-INS-457 |
| chr6-56956910-INS-1273 | chr6-56956911-INS-1274 |
| chr6-56973601-INS-204 | chr6-56973601-INS-201 |
| chr6-56983797-INS-79 | chr6-56983797-INS-79 |
| chr6-57012057-INS-1105 | chr6-57012058-INS-1105 |
| chr6-57039027-INS-85 | chr6-57039027-INS-85 |
| chr6-57046064-INS-56 | chr6-57046064-INS-55 |

| | |
|---|---|
| chr6-57051489-INS-1887 | chr6-57051490-INS-1887 |
| chr6-57082043-INS-83 | chr6-57082043-INS-83 |
| chr6-57095824-INS-163 | chr6-57095844-INS-171 |
| chr6-57117226-INS-63 | chr6-57117226-INS-63 |
| chr6-57141949-INS-6326 | chr6-57141950-INS-6326 |
| chr6-57182929-INS-280 | chr6-57182930-INS-280 |
| chr6-57222202-INS-984 | chr6-57222202-INS-984 |
| chr6-57230180-INS-457 | chr6-57230180-INS-457 |
| chr6-57232761-INS-53 | chr6-57232808-INS-55 |
| chr6-57234376-INS-245 | chr6-57234376-INS-254 |
| chr6-57242711-INS-328 | chr6-57242711-INS-322 |
| chr6-57247355-INS-61 | chr6-57247355-INS-61 |
| chr6-57254876-INS-98 | chr6-57254876-INS-98 |
| chr6-57287624-INS-275 | chr6-57287624-INS-275 |
| chr6-57304459-INS-101 | chr6-57304459-INS-83 |
| chr6-57320369-INS-206 | chr6-57320369-INS-206 |
| chr6-57336134-INS-98 | chr6-57336134-INS-98 |
| chr6-57364776-INS-200 | chr6-57364776-INS-200 |
| chr6-57375488-INS-6194 | chr6-57375489-INS-6194 |
| chr6-57382791-INS-452 | chr6-57382791-INS-452 |
| chr6-57384472-INS-70 | chr6-57384472-INS-92 |
| chr6-57390589-INS-2138 | chr6-57390590-INS-2138 |
| chr6-57423468-INS-99 | chr6-57423468-INS-108 |
| chr6-57428217-INS-56 | chr6-57428217-INS-100 |
| chr6-57459072-INS-7045 | chr6-57459073-INS-7045 |
| chr6-57463821-INS-75 | chr6-57463821-INS-75 |
| chr6-57471356-INS-312 | chr6-57471362-INS-275 |
| chr6-57490007-INS-124 | chr6-57490007-INS-124 |
| chr6-57500891-INS-54 | chr6-57500891-INS-56 |
| chr6-57528011-INS-54 | chr6-57528011-INS-54 |
| chr6-57536009-INS-55 | chr6-57536009-INS-63 |
| chr6-57540396-INS-140 | chr6-57540396-INS-96 |
| chr6-57549351-INS-6703 | chr6-57549352-INS-6704 |
| chr6-57561494-INS-213 | chr6-57561494-INS-213 |
| chr6-57572068-INS-55 | chr6-57572068-INS-55 |
| chr6-57580436-INS-66 | chr6-57580436-INS-50 |
| chr6-57589295-INS-90 | chr6-57589295-INS-92 |
| chr6-57602085-INS-159 | chr6-57602085-INS-159 |
| chr6-57613885-INS-330 | chr6-57613885-INS-344 |
| chr6-57620008-INS-6745 | chr6-57620009-INS-6745 |
| chr6-57650609-INS-212 | chr6-57650609-INS-212 |
| chr6-57659878-INS-51 | chr6-57659878-INS-51 |
| chr6-57673523-INS-137 | chr6-57673523-INS-137 |
| chr6-57676589-INS-76 | chr6-57676589-INS-76 |
| chr6-57682027-INS-331 | chr6-57682027-INS-331 |
| chr6-57708248-INS-1677 | chr6-57708249-INS-1677 |
| chr6-57721759-INS-57 | chr6-57721759-INS-57 |
| chr6-57739757-INS-6031 | chr6-57739758-INS-6032 |
| chr6-57749476-INS-56 | chr6-57749476-INS-58 |
| chr6-57757907-INS-55 | chr6-57757907-INS-55 |
| chr6-57794449-INS-6495 | chr6-57794450-INS-6492 |
| chr6-57831277-INS-109 | chr6-57831277-INS-113 |
| chr6-57837586-INS-65 | chr6-57837590-INS-95 |
| chr6-57850689-INS-895 | chr6-57850689-INS-895 |
| chr6-57851598-INS-255 | chr6-57851598-INS-255 |
| chr6-57896335-INS-6947 | chr6-57896336-INS-6948 |
| chr6-57920592-INS-643 | chr6-57920592-INS-643 |
| chr6-57928261-INS-460 | chr6-57928261-INS-460 |
| chr6-57950640-INS-4392 | chr6-57950641-INS-4389 |
| chr6-57956654-INS-342 | chr6-57956654-INS-343 |
| chr6-57960920-INS-497 | chr6-57960920-INS-505 |
| chr6-57988362-INS-74 | chr6-57988362-INS-54 |
| chr6-58000846-INS-6865 | chr6-58000846-INS-6863 |
| chr6-58065125-INS-4745 | chr6-58065126-INS-4745 |
| chr6-58080046-INS-90 | chr6-58080046-INS-90 |
| chr6-58084793-INS-242 | chr6-58084793-INS-242 |
| chr6-58118020-INS-553 | chr6-58118020-INS-589 |
| chr6-58141254-INS-81 | chr6-58141254-INS-81 |
| chr6-58158977-INS-203 | chr6-58158977-INS-203 |
| chr6-58168254-INS-296 | chr6-58168254-INS-298 |
| chr6-58175789-INS-568 | chr6-58175789-INS-568 |
| chr6-58179464-INS-72 | chr6-58179464-INS-80 |
| chr6-58189618-INS-59 | chr6-58189618-INS-56 |
| chr6-58244644-INS-8474 | chr6-58244631-INS-8470 |
| chr6-58275474-INS-117 | chr6-58275474-INS-117 |
| chr6-58293271-INS-921 | chr6-58293271-INS-921 |
| chr6-58320250-INS-64 | chr6-58320250-INS-52 |

| | |
|---|---|
| chr6-58339675-INS-112 | chr6-58339675-INS-112 |
| chr6-58437817-INS-56 | chr6-58437817-INS-56 |
| chr6-58456381-INS-112 | chr6-58456381-INS-112 |
| chr6-58503513-INS-317 | chr6-58503513-INS-317 |
| chr6-58555975-INS-54 | chr6-58555975-INS-66 |
| chr6-58575856-INS-666 | chr6-58575856-INS-666 |
| chr6-58593914-INS-405 | chr6-58593915-INS-405 |
| chr6-58605750-INS-2011 | chr6-58605745-INS-2022 |
| chr6-58611405-INS-1083 | chr6-58611406-INS-1115 |
| chr6-58614175-INS-1604 | chr6-58614176-INS-1604 |
| chr6-58637289-INS-52 | chr6-58637286-INS-52 |
| chr6-58648090-INS-150 | chr6-58648090-INS-150 |
| chr6-58676393-INS-1455 | chr6-58676394-INS-1455 |
| chr6-58708453-INS-62 | chr6-58708464-INS-94 |
| chr6-58711955-INS-173 | chr6-58711989-INS-126 |
| chr6-58727137-INS-1554 | chr6-58727138-INS-1558 |
| chr6-58749758-INS-7271 | chr6-58749759-INS-7271 |
| chr6-58752121-INS-56 | chr6-58752121-INS-56 |
| chr6-58786737-INS-679 | chr6-58786737-INS-678 |
| chr6-58837581-INS-96 | chr6-58837576-INS-95 |
| chr6-58890911-INS-201 | chr6-58890911-INS-243 |
| chr6-58892235-INS-326 | chr6-58892235-INS-326 |
| chr6-58948428-INS-332 | chr6-58948426-INS-330 |
| chr6-59016600-INS-332 | chr6-59016600-INS-332 |
| chr6-59028399-INS-6214 | chr6-59028400-INS-6214 |
| chr6-59039104-INS-6744 | chr6-59039105-INS-6744 |
| chr6-59054590-INS-300 | chr6-59054590-INS-300 |
| chr6-59058055-INS-1783 | chr6-59058056-INS-1784 |
| chr6-59080720-INS-718 | chr6-59080720-INS-732 |
| chr6-59099046-INS-119 | chr6-59099046-INS-147 |
| chr6-59111072-INS-1124 | chr6-59111073-INS-1124 |
| chr6-59130311-INS-821 | chr6-59130311-INS-821 |
| chr6-59141416-INS-781 | chr6-59141416-INS-781 |
| chr6-59149300-INS-56 | chr6-59149300-INS-64 |
| chr6-59167859-INS-258 | chr6-59167859-INS-258 |
| chr6-59177081-INS-212 | chr6-59177081-INS-212 |
| chr6-59199188-INS-412 | chr6-59199188-INS-412 |
| chr6-59243121-INS-1461 | chr6-59243122-INS-1461 |
| chr6-59246882-INS-217 | chr6-59246882-INS-217 |
| chr6-59380516-INS-51 | chr6-59380516-INS-54 |
| chr6-59407037-INS-6622 | chr6-59407038-INS-6622 |
| chr6-59408565-INS-267 | chr6-59408559-INS-280 |
| chr6-59409785-INS-95 | chr6-59409791-INS-98 |
| chr6-59427039-INS-56 | chr6-59427039-INS-56 |
| chr6-59471107-INS-6171 | chr6-59471108-INS-6173 |
| chr6-59503112-INS-121 | chr6-59503112-INS-124 |
| chr6-59508313-INS-614 | chr6-59508313-INS-614 |
| chr6-59534700-INS-512 | chr6-59534700-INS-513 |
| chr6-59540807-INS-1768 | chr6-59540808-INS-1769 |
| chr6-59542316-INS-70 | chr6-59542316-INS-72 |
| chr6-59545964-INS-1848 | chr6-59545965-INS-1848 |
| chr6-59582079-INS-830 | chr6-59582079-INS-875 |
| chr6-59595420-INS-254 | chr6-59595420-INS-254 |
| chr6-59604925-INS-498 | chr6-59604925-INS-498 |
| chr6-59619255-INS-2870 | chr6-59619256-INS-2870 |
| chr6-59681039-INS-302 | chr6-59681039-INS-302 |
| chr6-59727076-INS-567 | chr6-59727076-INS-552 |
| chr6-59735357-INS-614 | chr6-59735357-INS-613 |
| chr6-59883733-INS-697 | chr6-59883733-INS-695 |
| chr6-60070332-INS-53 | chr6-60070332-INS-56 |
| chr6-60314108-INS-202 | chr6-60314108-INS-203 |
| chr6-60358962-INS-954 | chr6-60358962-INS-955 |
| chr6-60364169-INS-4837 | chr6-60364170-INS-4837 |
| chr6-60368896-INS-114 | chr6-60368896-INS-112 |
| chr6-60378973-INS-323 | chr6-60378973-INS-323 |
| chr6-60382386-INS-55 | chr6-60382397-INS-51 |
| chr6-60466375-INS-377 | chr6-60466393-INS-330 |
| chr6-60469415-INS-232 | chr6-60469397-INS-212 |
| chr6-60525507-INS-284 | chr6-60525507-INS-284 |
| chr6-60528051-INS-923 | chr6-60528051-INS-924 |
| chr6-60538304-INS-94 | chr6-60538296-INS-72 |
| chr6-60553794-INS-190 | chr6-60553794-INS-232 |
| chr6-60567610-INS-82 | chr6-60567610-INS-68 |
| chr6-60575961-INS-199 | chr6-60575961-INS-199 |
| chr6-60577264-INS-73 | chr6-60577264-INS-73 |
| chr6-60581590-INS-203 | chr6-60581590-INS-203 |
| chr6-60590828-INS-204 | chr6-60590828-INS-201 |

| | |
|---|---|
| chr6-60611721-INS-91 | chr6-60611720-INS-113 |
| chr6-60622948-INS-52 | chr6-60622948-INS-54 |
| chr6-60623843-INS-334 | chr6-60623843-INS-334 |
| chr6-60698184-INS-272 | chr6-60698184-INS-272 |
| chr6-60708208-INS-62 | chr6-60708210-INS-58 |
| chr6-60714747-INS-922 | chr6-60714747-INS-921 |
| chr6-60729031-INS-89 | chr6-60729031-INS-88 |
| chr6-60730725-INS-93 | chr6-60730727-INS-57 |
| chr6-60738154-INS-238 | chr6-60738154-INS-238 |
| chr6-60756541-INS-7013 | chr6-60756542-INS-7015 |
| chr6-60759694-INS-1321 | chr6-60759695-INS-1321 |
| chr6-60765456-INS-54 | chr6-60765456-INS-64 |
| chr6-60798882-INS-280 | chr6-60798882-INS-280 |
| chr6-60819523-INS-339 | chr6-60819523-INS-338 |
| chr6-60847239-INS-68 | chr6-60847239-INS-68 |
| chr6-60851976-INS-980 | chr6-60851976-INS-980 |
| chr6-60879198-INS-63 | chr6-60879198-INS-63 |
| chr6-60906870-INS-175 | chr6-60906870-INS-175 |
| chr6-60909887-INS-336 | chr6-60909898-INS-333 |
| chr6-60913163-INS-69 | chr6-60913163-INS-69 |
| chr6-60932277-INS-991 | chr6-60932277-INS-989 |
| chr6-60963914-INS-167 | chr6-60963914-INS-139 |
| chr6-60970023-INS-63 | chr6-60970023-INS-63 |
| chr6-60976954-INS-348 | chr6-60976954-INS-375 |
| chr6-60988387-INS-2036 | chr6-60988388-INS-2033 |
| chr6-61012361-INS-476 | chr6-61012361-INS-475 |
| chr6-61015615-INS-492 | chr6-61015615-INS-492 |
| chr6-61016537-INS-73 | chr6-61016537-INS-82 |
| chr6-61033727-INS-68 | chr6-61033727-INS-70 |
| chr6-61057299-INS-174 | chr6-61057299-INS-174 |
| chr6-61065465-INS-214 | chr6-61065433-INS-214 |
| chr6-61078423-INS-60 | chr6-61078423-INS-58 |
| chr6-61098595-INS-402 | chr6-61098595-INS-396 |
| chr6-61099078-INS-3695 | chr6-61099079-INS-3695 |
| chr6-61144437-INS-64 | chr6-61144437-INS-64 |
| chr6-61150259-INS-409 | chr6-61150259-INS-409 |
| chr6-61231722-INS-65 | chr6-61231722-INS-65 |
| chr6-61241365-INS-7356 | chr6-61241366-INS-7351 |
| chr6-61242088-INS-84 | chr6-61242088-INS-88 |
| chr6-61290143-INS-187 | chr6-61290143-INS-188 |
| chr6-61291953-INS-488 | chr6-61291953-INS-488 |
| chr6-61293703-INS-431 | chr6-61293703-INS-431 |
| chr6-61301079-INS-94 | chr6-61301079-INS-94 |
| chr6-61348882-INS-600 | chr6-61348882-INS-600 |
| chr6-61350008-INS-1791 | chr6-61350006-INS-1764 |
| chr6-61385404-INS-1538 | chr6-61385405-INS-1538 |
| chr6-61385511-INS-344 | chr6-61385511-INS-344 |
| chr6-61407168-INS-3737 | chr6-61407169-INS-3737 |
| chr6-61476363-INS-1779 | chr6-61476364-INS-1778 |
| chr6-61497323-INS-86 | chr6-61497323-INS-86 |
| chr6-61504678-INS-411 | chr6-61504678-INS-411 |
| chr6-61524822-INS-52 | chr6-61524822-INS-96 |
| chr6-61531465-INS-6628 | chr6-61531466-INS-6629 |
| chr6-61543743-INS-6410 | chr6-61543744-INS-6410 |
| chr6-61545806-INS-128 | chr6-61545806-INS-130 |
| chr6-61667442-INS-6593 | chr6-61667443-INS-6593 |
| chr6-61669581-INS-58 | chr6-61669581-INS-60 |
| chr6-61675738-INS-2686 | chr6-61675739-INS-2686 |
| chr6-61681166-INS-6800 | chr6-61681167-INS-6800 |
| chr6-61731172-INS-6209 | chr6-61731173-INS-6209 |
| chr6-61815344-INS-104 | chr6-61815355-INS-109 |
| chr6-61837926-INS-294 | chr6-61837928-INS-295 |
| chr6-61849425-INS-91 | chr6-61849425-INS-91 |
| chr6-61882344-INS-490 | chr6-61882344-INS-490 |
| chr6-61891877-INS-64 | chr6-61891877-INS-68 |
| chr6-61902196-INS-438 | chr6-61902196-INS-440 |
| chr6-61903627-INS-81 | chr6-61903623-INS-98 |
| chr6-61905945-INS-102 | chr6-61905945-INS-105 |
| chr6-61924445-INS-1894 | chr6-61924446-INS-1894 |
| chr6-61925215-INS-51 | chr6-61925215-INS-53 |
| chr6-61949500-INS-68 | chr6-61949500-INS-68 |
| chr6-61963028-INS-318 | chr6-61963028-INS-318 |
| chr6-61985824-INS-172 | chr6-61985824-INS-173 |
| chr6-61994580-INS-85 | chr6-61994580-INS-85 |
| chr6-62001984-INS-417 | chr6-62001984-INS-417 |
| chr6-62008644-INS-200 | chr6-62008644-INS-200 |
| chr6-62029061-INS-74 | chr6-62029061-INS-74 |

| | |
|---|---|
| chr6-62103365-INS-476 | chr6-62103365-INS-476 |
| chr6-62147305-INS-105 | chr6-62147305-INS-111 |
| chr6-62173463-INS-202 | chr6-62173463-INS-202 |
| chr6-62206946-INS-6694 | chr6-62206947-INS-6695 |
| chr6-62209733-INS-705 | chr6-62209733-INS-686 |
| chr6-62243814-INS-68 | chr6-62243814-INS-70 |
| chr6-62244407-INS-836 | chr6-62244407-INS-853 |
| chr6-62345340-INS-51 | chr6-62345340-INS-51 |
| chr6-62368124-INS-212 | chr6-62368124-INS-212 |
| chr6-62381234-INS-912 | chr6-62381234-INS-912 |
| chr6-62449659-INS-109 | chr6-62449659-INS-109 |
| chr6-62461602-INS-104 | chr6-62461602-INS-106 |
| chr6-62463264-INS-387 | chr6-62463268-INS-389 |
| chr6-62493572-INS-80 | chr6-62493572-INS-82 |
| chr6-62510435-INS-652 | chr6-62510435-INS-646 |
| chr6-62511372-INS-238 | chr6-62511372-INS-238 |
| chr6-62518157-INS-445 | chr6-62518157-INS-445 |
| chr6-62542537-INS-201 | chr6-62542496-INS-201 |
| chr6-62549515-INS-1887 | chr6-62549516-INS-1887 |
| chr6-62561985-INS-1483 | chr6-62561986-INS-1483 |
| chr6-62578413-INS-156 | chr6-62578413-INS-156 |
| chr6-62591396-INS-57 | chr6-62591396-INS-57 |
| chr6-62632353-INS-134 | chr6-62632353-INS-134 |
| chr6-62694877-INS-1940 | chr6-62694878-INS-1940 |
| chr6-62706794-INS-142 | chr6-62706796-INS-133 |
| chr6-62710939-INS-58 | chr6-62710939-INS-58 |
| chr6-62738086-INS-132 | chr6-62738086-INS-121 |
| chr6-62748141-INS-504 | chr6-62748140-INS-510 |
| chr6-62749551-INS-714 | chr6-62749551-INS-714 |
| chr6-62752068-INS-958 | chr6-62752068-INS-958 |
| chr6-62787890-INS-64 | chr6-62787890-INS-68 |
| chr6-62935264-INS-92 | chr6-62935260-INS-96 |
| chr6-62991149-INS-260 | chr6-62991149-INS-261 |
| chr6-63013010-INS-76 | chr6-63013010-INS-77 |
| chr6-63019334-INS-88 | chr6-63019332-INS-92 |
| chr6-63034125-INS-1958 | chr6-63034126-INS-1959 |
| chr6-63040572-INS-2485 | chr6-63040573-INS-2485 |
| chr6-63049287-INS-120 | chr6-63049287-INS-120 |
| chr6-63057872-INS-174 | chr6-63057872-INS-174 |
| chr6-63077362-INS-158 | chr6-63077362-INS-162 |
| chr6-63125643-INS-7086 | chr6-63125644-INS-7086 |
| chr6-63136775-INS-146 | chr6-63136775-INS-134 |
| chr6-63163679-INS-55 | chr6-63163679-INS-55 |
| chr6-63268092-INS-97 | chr6-63268080-INS-79 |
| chr6-63269900-INS-446 | chr6-63269900-INS-446 |
| chr6-63288536-INS-64 | chr6-63288536-INS-64 |
| chr6-63306783-INS-4132 | chr6-63306784-INS-4134 |
| chr6-63331994-INS-198 | chr6-63331994-INS-198 |
| chr6-63338278-INS-60 | chr6-63338278-INS-66 |
| chr6-63371048-INS-184 | chr6-63371048-INS-184 |
| chr6-63383362-INS-113 | chr6-63383362-INS-113 |
| chr6-63500879-INS-185 | chr6-63500879-INS-185 |
| chr6-63501784-INS-6857 | chr6-63501785-INS-6858 |
| chr6-63504315-INS-757 | chr6-63504315-INS-757 |
| chr6-63534392-INS-110 | chr6-63534392-INS-111 |
| chr6-63575019-INS-5127 | chr6-63575020-INS-5095 |
| chr6-63607770-INS-64 | chr6-63607770-INS-64 |
| chr6-63619639-INS-1253 | chr6-63619640-INS-1253 |
| chr6-63678095-INS-199 | chr6-63678095-INS-199 |
| chr6-63727716-INS-488 | chr6-63727716-INS-488 |
| chr6-63735309-INS-365 | chr6-63735309-INS-365 |
| chr6-63749507-INS-3372 | chr6-63749508-INS-3372 |
| chr6-63752384-INS-2999 | chr6-63752385-INS-2999 |
| chr6-63760402-INS-204 | chr6-63760402-INS-204 |
| chr6-63823174-INS-381 | chr6-63823175-INS-381 |
| chr6-63831188-INS-128 | chr6-63831188-INS-133 |
| chr6-63862559-INS-112 | chr6-63862561-INS-113 |
| chr6-63932652-INS-456 | chr6-63932652-INS-456 |
| chr6-63978714-INS-52 | chr6-63978714-INS-56 |
| chr6-64041710-INS-272 | chr6-64041710-INS-254 |
| chr6-64047744-INS-1000 | chr6-64047744-INS-1001 |
| chr6-64059101-INS-1208 | chr6-64059102-INS-1208 |
| chr6-64091988-INS-100 | chr6-64091988-INS-106 |
| chr6-64094590-INS-102 | chr6-64094590-INS-72 |
| chr6-64159551-INS-54 | chr6-64159551-INS-56 |
| chr6-64172045-INS-66 | chr6-64172035-INS-62 |
| chr6-64215953-INS-64 | chr6-64215928-INS-88 |

| | |
|---|---|
| chr6-64235755-INS-682 | chr6-64235755-INS-680 |
| chr6-64236770-INS-85 | chr6-64236770-INS-62 |
| chr6-64243503-INS-69 | chr6-64243503-INS-69 |
| chr6-64256662-INS-66 | chr6-64256662-INS-68 |
| chr6-64325532-INS-52 | chr6-64325532-INS-54 |
| chr6-64332569-INS-279 | chr6-64332569-INS-279 |
| chr6-64382837-INS-64 | chr6-64382826-INS-52 |
| chr6-64386239-INS-740 | chr6-64386239-INS-746 |
| chr6-64392947-INS-112 | chr6-64392947-INS-114 |
| chr6-64409036-INS-62 | chr6-64409036-INS-62 |
| chr6-64459572-INS-1193 | chr6-64459573-INS-1194 |
| chr6-64462710-INS-156 | chr6-64462710-INS-156 |
| chr6-64493986-INS-6277 | chr6-64493987-INS-6277 |
| chr6-64498745-INS-6277 | chr6-64498746-INS-6276 |
| chr6-64501855-INS-1342 | chr6-64501856-INS-1341 |
| chr6-64507758-INS-564 | chr6-64507758-INS-564 |
| chr6-64512834-INS-1018 | chr6-64512835-INS-1016 |
| chr6-64530791-INS-92 | chr6-64530791-INS-92 |
| chr6-64538336-INS-6587 | chr6-64538337-INS-6587 |
| chr6-64557543-INS-50 | chr6-64557543-INS-67 |
| chr6-64577506-INS-6992 | chr6-64577507-INS-6992 |
| chr6-64599506-INS-103 | chr6-64599504-INS-58 |
| chr6-64617755-INS-1909 | chr6-64617756-INS-1932 |
| chr6-64655694-INS-60 | chr6-64655694-INS-60 |
| chr6-64672177-INS-50 | chr6-64672177-INS-58 |
| chr6-64723832-INS-61 | chr6-64723865-INS-78 |
| chr6-64752962-INS-65 | chr6-64752962-INS-69 |
| chr6-64766249-INS-870 | chr6-64766249-INS-870 |
| chr6-64772009-INS-66 | chr6-64772009-INS-89 |
| chr6-64795859-INS-71 | chr6-64795862-INS-70 |
| chr6-64807869-INS-75 | chr6-64807869-INS-79 |
| chr6-64808549-INS-55 | chr6-64808549-INS-55 |
| chr6-64813790-INS-207 | chr6-64813790-INS-209 |
| chr6-64815873-INS-54 | chr6-64815831-INS-60 |
| chr6-64828405-INS-2561 | chr6-64828406-INS-2562 |
| chr6-64830641-INS-141 | chr6-64830674-INS-138 |
| chr6-64832252-INS-5396 | chr6-64832253-INS-5395 |
| chr6-64836613-INS-166 | chr6-64836613-INS-166 |
| chr6-64853887-INS-898 | chr6-64853887-INS-898 |
| chr6-64858866-INS-208 | chr6-64858866-INS-208 |
| chr6-64866179-INS-183 | chr6-64866179-INS-178 |
| chr6-64885779-INS-51 | chr6-64885780-INS-51 |
| chr6-64891185-INS-101 | chr6-64891185-INS-97 |
| chr6-64898243-INS-209 | chr6-64898236-INS-215 |
| chr6-64913791-INS-1888 | chr6-64913792-INS-1888 |
| chr6-64916677-INS-739 | chr6-64916676-INS-741 |
| chr6-64958693-INS-89 | chr6-64958694-INS-89 |
| chr6-64962712-INS-126 | chr6-64962712-INS-126 |
| chr6-64970279-INS-105 | chr6-64970279-INS-113 |
| chr6-64978039-INS-244 | chr6-64978039-INS-250 |
| chr6-64983425-INS-203 | chr6-64983425-INS-203 |
| chr6-64997524-INS-195 | chr6-64997524-INS-195 |
| chr6-64998389-INS-101 | chr6-64998389-INS-99 |
| chr6-65000122-INS-99 | chr6-65000122-INS-99 |
| chr6-65006134-INS-179 | chr6-65006135-INS-181 |
| chr6-65011004-INS-70 | chr6-65011004-INS-55 |
| chr6-65016070-INS-98 | chr6-65016070-INS-98 |
| chr6-65045240-INS-291 | chr6-65045240-INS-291 |
| chr6-65083336-INS-215 | chr6-65083336-INS-215 |
| chr6-65102598-INS-87 | chr6-65102598-INS-87 |
| chr6-65118773-INS-210 | chr6-65118772-INS-169 |
| chr6-65125722-INS-539 | chr6-65125722-INS-541 |
| chr6-65139315-INS-82 | chr6-65139306-INS-107 |
| chr6-65143043-INS-66 | chr6-65143043-INS-66 |
| chr6-65217813-INS-232 | chr6-65217813-INS-232 |
| chr6-65237934-INS-208 | chr6-65237934-INS-208 |
| chr6-65240354-INS-492 | chr6-65240354-INS-495 |
| chr6-65245282-INS-95 | chr6-65245303-INS-72 |
| chr6-65257626-INS-3008 | chr6-65257611-INS-2991 |
| chr6-65268791-INS-471 | chr6-65268791-INS-471 |
| chr6-65288579-INS-95 | chr6-65288579-INS-85 |
| chr6-65323422-INS-64 | chr6-65323422-INS-72 |
| chr6-65331233-INS-204 | chr6-65331223-INS-203 |
| chr6-65336778-INS-554 | chr6-65336778-INS-554 |
| chr6-65389040-INS-90 | chr6-65389040-INS-65 |
| chr6-65403083-INS-6565 | chr6-65403084-INS-6566 |
| chr6-65422454-INS-123 | chr6-65422451-INS-132 |

| | |
|---|---|
| chr6-65429155-INS-169 | chr6-65429155-INS-170 |
| chr6-65433857-INS-146 | chr6-65433857-INS-146 |
| chr6-65460388-INS-197 | chr6-65460388-INS-197 |
| chr6-65500819-INS-72 | chr6-65500819-INS-72 |
| chr6-65563249-INS-552 | chr6-65563249-INS-552 |
| chr6-65584076-INS-990 | chr6-65584076-INS-990 |
| chr6-65605044-INS-485 | chr6-65605044-INS-486 |
| chr6-65607944-INS-162 | chr6-65607944-INS-162 |
| chr6-65613570-INS-6448 | chr6-65613571-INS-6454 |
| chr6-65628018-INS-261 | chr6-65628023-INS-295 |
| chr6-65647675-INS-56 | chr6-65647675-INS-68 |
| chr6-65693220-INS-8680 | chr6-65693221-INS-8680 |
| chr6-65696529-INS-60 | chr6-65696529-INS-60 |
| chr6-65706479-INS-59 | chr6-65706479-INS-70 |
| chr6-65725239-INS-169 | chr6-65725239-INS-169 |
| chr6-65730858-INS-52 | chr6-65730811-INS-69 |
| chr6-65737935-INS-52 | chr6-65737935-INS-52 |
| chr6-65752669-INS-946 | chr6-65752669-INS-946 |
| chr6-65765459-INS-63 | chr6-65765459-INS-63 |
| chr6-65790373-INS-75 | chr6-65790373-INS-74 |
| chr6-65807542-INS-342 | chr6-65807542-INS-336 |
| chr6-65810507-INS-201 | chr6-65810507-INS-200 |
| chr6-65815017-INS-6323 | chr6-65815018-INS-6319 |
| chr6-65821785-INS-72 | chr6-65821785-INS-76 |
| chr6-65824566-INS-1719 | chr6-65824567-INS-1721 |
| chr6-65872643-INS-649 | chr6-65872643-INS-649 |
| chr6-65960592-INS-74 | chr6-65960590-INS-79 |
| chr6-65975552-INS-2079 | chr6-65975553-INS-2079 |
| chr6-65994192-INS-6571 | chr6-65994193-INS-6572 |
| chr6-66002374-INS-55 | chr6-66002374-INS-55 |
| chr6-66122152-INS-988 | chr6-66122152-INS-993 |
| chr6-66131976-INS-78 | chr6-66131976-INS-78 |
| chr6-66162050-INS-78 | chr6-66162001-INS-82 |
| chr6-66175551-INS-77 | chr6-66175551-INS-77 |
| chr6-66181565-INS-4012 | chr6-66181566-INS-4013 |
| chr6-66247528-INS-466 | chr6-66247528-INS-472 |
| chr6-66311874-INS-63 | chr6-66311866-INS-60 |
| chr6-66315916-INS-6897 | chr6-66315917-INS-6899 |
| chr6-66320880-INS-7528 | chr6-66320881-INS-7529 |
| chr6-66361335-INS-6070 | chr6-66361336-INS-6079 |
| chr6-66386560-INS-803 | chr6-66386560-INS-801 |
| chr6-66410443-INS-64 | chr6-66410443-INS-64 |
| chr6-66421666-INS-2652 | chr6-66421667-INS-2652 |
| chr6-66476276-INS-172 | chr6-66476276-INS-172 |
| chr6-66498503-INS-251 | chr6-66498503-INS-242 |
| chr6-66508689-INS-128 | chr6-66508689-INS-128 |
| chr6-66542216-INS-95 | chr6-66542223-INS-80 |
| chr6-66563950-INS-1372 | chr6-66563951-INS-1360 |
| chr6-66598440-INS-5518 | chr6-66598441-INS-5545 |
| chr6-66599099-INS-2438 | chr6-66599100-INS-2432 |
| chr6-66599100-INS-299 | chr6-66599100-INS-299 |
| chr6-66622969-INS-2453 | chr6-66622970-INS-2453 |
| chr6-66638189-INS-52 | chr6-66638189-INS-61 |
| chr6-66665500-INS-53 | chr6-66665500-INS-53 |
| chr6-66672250-INS-534 | chr6-66672250-INS-534 |
| chr6-66678379-INS-4867 | chr6-66678380-INS-4846 |
| chr6-66683658-INS-2534 | chr6-66683659-INS-2534 |
| chr6-66750180-INS-55 | chr6-66750180-INS-55 |
| chr6-66762050-INS-208 | chr6-66762051-INS-213 |
| chr6-66784877-INS-205 | chr6-66784877-INS-203 |
| chr6-66838794-INS-82 | chr6-66838794-INS-82 |
| chr6-66861988-INS-3071 | chr6-66861989-INS-3071 |
| chr6-66910272-INS-83 | chr6-66910272-INS-78 |
| chr6-66917803-INS-200 | chr6-66917803-INS-200 |
| chr6-66932203-INS-66 | chr6-66932203-INS-66 |
| chr6-66947305-INS-384 | chr6-66947306-INS-384 |
| chr6-66993987-INS-76 | chr6-66993987-INS-76 |
| chr6-67011175-INS-68 | chr6-67011175-INS-68 |
| chr6-67030600-INS-156 | chr6-67030600-INS-160 |
| chr6-67058816-INS-93 | chr6-67058816-INS-93 |
| chr6-67060340-INS-444 | chr6-67060340-INS-445 |
| chr6-67074348-INS-87 | chr6-67074351-INS-66 |
| chr6-67079169-INS-260 | chr6-67079166-INS-294 |
| chr6-67083489-INS-60 | chr6-67083445-INS-58 |
| chr6-67144475-INS-52 | chr6-67144475-INS-52 |
| chr6-67146556-INS-100 | chr6-67146556-INS-100 |
| chr6-67152517-INS-54 | chr6-67152517-INS-54 |

| | |
|---|---|
| chr6-67184550-INS-305 | chr6-67184550-INS-338 |
| chr6-67186781-INS-245 | chr6-67186781-INS-245 |
| chr6-67191428-INS-54 | chr6-67191421-INS-58 |
| chr6-67204754-INS-192 | chr6-67204754-INS-192 |
| chr6-67208277-INS-149 | chr6-67208278-INS-145 |
| chr6-67221888-INS-100 | chr6-67221888-INS-100 |
| chr6-67239861-INS-159 | chr6-67239861-INS-161 |
| chr6-67242615-INS-1585 | chr6-67242616-INS-1586 |
| chr6-67251308-INS-182 | chr6-67251308-INS-183 |
| chr6-67256322-INS-198 | chr6-67256322-INS-198 |
| chr6-67291203-INS-81 | chr6-67291209-INS-105 |
| chr6-67292756-INS-219 | chr6-67292756-INS-219 |
| chr6-67299976-INS-191 | chr6-67299976-INS-191 |
| chr6-67330851-INS-456 | chr6-67330851-INS-456 |
| chr6-67341662-INS-201 | chr6-67341662-INS-201 |
| chr6-67348574-INS-60 | chr6-67348574-INS-96 |
| chr6-67349950-INS-1502 | chr6-67349951-INS-1503 |
| chr6-67361844-INS-761 | chr6-67361844-INS-761 |
| chr6-67371714-INS-155 | chr6-67371714-INS-155 |
| chr6-67418195-INS-85 | chr6-67418195-INS-85 |
| chr6-67434597-INS-209 | chr6-67434597-INS-209 |
| chr6-67435755-INS-79 | chr6-67435755-INS-79 |
| chr6-67441237-INS-63 | chr6-67441237-INS-63 |
| chr6-67446248-INS-337 | chr6-67446248-INS-333 |
| chr6-67447259-INS-51 | chr6-67447259-INS-51 |
| chr6-67458426-INS-64 | chr6-67458426-INS-64 |
| chr6-67496449-INS-436 | chr6-67496449-INS-436 |
| chr6-67502938-INS-79 | chr6-67502938-INS-79 |
| chr6-67504797-INS-67 | chr6-67504797-INS-67 |
| chr6-67520391-INS-143 | chr6-67520391-INS-143 |
| chr6-67527737-INS-81 | chr6-67527737-INS-81 |
| chr6-67540135-INS-999 | chr6-67540135-INS-999 |
| chr6-67540789-INS-84 | chr6-67540790-INS-84 |
| chr6-67659008-INS-63 | chr6-67659008-INS-65 |
| chr6-67700857-INS-73 | chr6-67700857-INS-73 |
| chr6-67716783-INS-58 | chr6-67716783-INS-58 |
| chr6-67726383-INS-2767 | chr6-67726384-INS-2767 |
| chr6-67727950-INS-91 | chr6-67727950-INS-91 |
| chr6-67757504-INS-56 | chr6-67757504-INS-56 |
| chr6-67776972-INS-146 | chr6-67776972-INS-146 |
| chr6-67807975-INS-168 | chr6-67807976-INS-156 |
| chr6-67840588-INS-205 | chr6-67840588-INS-206 |
| chr6-67865239-INS-58 | chr6-67865239-INS-58 |
| chr6-67885928-INS-6383 | chr6-67885929-INS-6382 |
| chr6-67895134-INS-57 | chr6-67895134-INS-57 |
| chr6-67896956-INS-232 | chr6-67896956-INS-231 |
| chr6-67898905-INS-12053 | chr6-67898906-INS-12040 |
| chr6-67899431-INS-560 | chr6-67899431-INS-560 |
| chr6-67947945-INS-53 | chr6-67947945-INS-53 |
| chr6-67955524-INS-87 | chr6-67955524-INS-87 |
| chr6-67968088-INS-399 | chr6-67968088-INS-399 |
| chr6-67976685-INS-63 | chr6-67976685-INS-68 |
| chr6-67993330-INS-488 | chr6-67993330-INS-488 |
| chr6-68023374-INS-80 | chr6-68023374-INS-80 |
| chr6-68058248-INS-329 | chr6-68058248-INS-329 |
| chr6-68158692-INS-1206 | chr6-68158693-INS-1206 |
| chr6-68191166-INS-318 | chr6-68191162-INS-319 |
| chr6-68232580-INS-928 | chr6-68232580-INS-928 |
| chr6-68235180-INS-109 | chr6-68235180-INS-109 |
| chr6-68289109-INS-84 | chr6-68289109-INS-84 |
| chr6-68363724-INS-1895 | chr6-68363725-INS-1895 |
| chr6-68400706-INS-849 | chr6-68400706-INS-846 |
| chr6-68419029-INS-54 | chr6-68419027-INS-62 |
| chr6-68421816-INS-1314 | chr6-68421817-INS-1304 |
| chr6-68435889-INS-74 | chr6-68435889-INS-74 |
| chr6-68449091-INS-54 | chr6-68449091-INS-54 |
| chr6-68454695-INS-51 | chr6-68454697-INS-58 |
| chr6-68486992-INS-956 | chr6-68486992-INS-956 |
| chr6-68502712-INS-72 | chr6-68502712-INS-54 |
| chr6-68507866-INS-60 | chr6-68507889-INS-50 |
| chr6-68511859-INS-120 | chr6-68511859-INS-120 |
| chr6-68516189-INS-79 | chr6-68516189-INS-79 |
| chr6-68522265-INS-95 | chr6-68522253-INS-140 |
| chr6-68533902-INS-416 | chr6-68533902-INS-416 |
| chr6-68544882-INS-52 | chr6-68544882-INS-52 |
| chr6-68547067-INS-422 | chr6-68547067-INS-422 |
| chr6-68556964-INS-342 | chr6-68556964-INS-350 |

| | |
|---|---|
| chr6-68612484-INS-504 | chr6-68612484-INS-504 |
| chr6-68613528-INS-119 | chr6-68613528-INS-115 |
| chr6-68632463-INS-927 | chr6-68632463-INS-919 |
| chr6-68678922-INS-61 | chr6-68678922-INS-61 |
| chr6-68695945-INS-68 | chr6-68695945-INS-68 |
| chr6-68747446-INS-464 | chr6-68747446-INS-488 |
| chr6-68802667-INS-52 | chr6-68802667-INS-52 |
| chr6-68807342-INS-84 | chr6-68807342-INS-86 |
| chr6-68843424-INS-73 | chr6-68843424-INS-73 |
| chr6-68867409-INS-95 | chr6-68867409-INS-102 |
| chr6-68889755-INS-102 | chr6-68889755-INS-103 |
| chr6-68916101-INS-484 | chr6-68916101-INS-485 |
| chr6-68937204-INS-60 | chr6-68937208-INS-76 |
| chr6-68963809-INS-199 | chr6-68963809-INS-199 |
| chr6-69028301-INS-531 | chr6-69028301-INS-531 |
| chr6-69036017-INS-56 | chr6-69036017-INS-84 |
| chr6-69068646-INS-1994 | chr6-69068647-INS-1994 |
| chr6-69113046-INS-54 | chr6-69113046-INS-54 |
| chr6-69116490-INS-528 | chr6-69116490-INS-528 |
| chr6-69278752-INS-685 | chr6-69278752-INS-685 |
| chr6-69295989-INS-91 | chr6-69295978-INS-87 |
| chr6-69301949-INS-117 | chr6-69301949-INS-117 |
| chr6-69315412-INS-129 | chr6-69315412-INS-129 |
| chr6-69315607-INS-5550 | chr6-69315608-INS-5550 |
| chr6-69323715-INS-330 | chr6-69323715-INS-316 |
| chr6-69327962-INS-90 | chr6-69327962-INS-90 |
| chr6-69329476-INS-68 | chr6-69329476-INS-68 |
| chr6-69350011-INS-209 | chr6-69350011-INS-209 |
| chr6-69352962-INS-198 | chr6-69352962-INS-198 |
| chr6-69370188-INS-199 | chr6-69370188-INS-199 |
| chr6-69388519-INS-7187 | chr6-69388520-INS-7187 |
| chr6-69632290-INS-57 | chr6-69632290-INS-50 |
| chr6-69645675-INS-841 | chr6-69645675-INS-841 |
| chr6-69680359-INS-712 | chr6-69680359-INS-712 |
| chr6-69697079-INS-488 | chr6-69697078-INS-494 |
| chr6-69700774-INS-484 | chr6-69700774-INS-484 |
| chr6-69707185-INS-322 | chr6-69707224-INS-308 |
| chr6-69730781-INS-1246 | chr6-69730782-INS-1247 |
| chr6-69731568-INS-318 | chr6-69731568-INS-322 |
| chr6-69735155-INS-253 | chr6-69735155-INS-253 |
| chr6-69735868-INS-8282 | chr6-69735869-INS-8282 |
| chr6-69735872-INS-3878 | chr6-69735873-INS-3878 |
| chr6-69751987-INS-158 | chr6-69751987-INS-156 |
| chr6-69756691-INS-704 | chr6-69756691-INS-703 |
| chr6-69831295-INS-240 | chr6-69831295-INS-240 |
| chr6-69834516-INS-64 | chr6-69834516-INS-69 |
| chr6-69846174-INS-319 | chr6-69846179-INS-322 |
| chr6-69851615-INS-458 | chr6-69851615-INS-458 |
| chr6-69855515-INS-56 | chr6-69855516-INS-56 |
| chr6-69862438-INS-2141 | chr6-69862439-INS-2129 |
| chr6-69863942-INS-892 | chr6-69863943-INS-875 |
| chr6-69887654-INS-54 | chr6-69887654-INS-54 |
| chr6-69890374-INS-105 | chr6-69890366-INS-111 |
| chr6-69897192-INS-237 | chr6-69897192-INS-236 |
| chr6-69917928-INS-158 | chr6-69917928-INS-158 |
| chr6-69921620-INS-72 | chr6-69921620-INS-56 |
| chr6-69944463-INS-153 | chr6-69944463-INS-151 |
| chr6-69945667-INS-5548 | chr6-69945667-INS-5548 |
| chr6-69960342-INS-60 | chr6-69960342-INS-60 |
| chr6-69998746-INS-498 | chr6-69998746-INS-498 |
| chr6-70002316-INS-136 | chr6-70002316-INS-136 |
| chr6-70084400-INS-892 | chr6-70084400-INS-892 |
| chr6-70094855-INS-233 | chr6-70094855-INS-233 |
| chr6-70123447-INS-460 | chr6-70123447-INS-464 |
| chr6-70171412-INS-189 | chr6-70171412-INS-189 |
| chr6-70193375-INS-136 | chr6-70193375-INS-136 |
| chr6-70195438-INS-94 | chr6-70195438-INS-94 |
| chr6-70291079-INS-77 | chr6-70291079-INS-77 |
| chr6-70398721-INS-107 | chr6-70398722-INS-107 |
| chr6-70443445-INS-82 | chr6-70443445-INS-68 |
| chr6-70497104-INS-63 | chr6-70497104-INS-65 |
| chr6-70554172-INS-51 | chr6-70554172-INS-51 |
| chr6-70558547-INS-486 | chr6-70558547-INS-486 |
| chr6-70591665-INS-476 | chr6-70591665-INS-476 |
| chr6-70683656-INS-965 | chr6-70683656-INS-965 |
| chr6-70773399-INS-112 | chr6-70773399-INS-136 |
| chr6-70780236-INS-212 | chr6-70780252-INS-218 |

| | |
|---|---|
| chr6-70797480-INS-82 | chr6-70797480-INS-82 |
| chr6-70813725-INS-91 | chr6-70813725-INS-95 |
| chr6-70815376-INS-54 | chr6-70815376-INS-54 |
| chr6-70817963-INS-7113 | chr6-70817964-INS-7113 |
| chr6-70830262-INS-55 | chr6-70830261-INS-71 |
| chr6-70894440-INS-220 | chr6-70894440-INS-220 |
| chr6-70899509-INS-67 | chr6-70899509-INS-67 |
| chr6-70912683-INS-160 | chr6-70912683-INS-172 |
| chr6-70913429-INS-973 | chr6-70913429-INS-973 |
| chr6-70935479-INS-56 | chr6-70935479-INS-59 |
| chr6-70949412-INS-54 | chr6-70949413-INS-51 |
| chr6-70962650-INS-56 | chr6-70962607-INS-96 |
| chr6-70972163-INS-966 | chr6-70972163-INS-966 |
| chr6-70972696-INS-130 | chr6-70972698-INS-128 |
| chr6-70988097-INS-207 | chr6-70988097-INS-207 |
| chr6-71005103-INS-183 | chr6-71005103-INS-183 |
| chr6-71008606-INS-201 | chr6-71008608-INS-201 |
| chr6-71017906-INS-64 | chr6-71017906-INS-105 |
| chr6-71022138-INS-200 | chr6-71022138-INS-200 |
| chr6-71031734-INS-552 | chr6-71031734-INS-553 |
| chr6-71032781-INS-441 | chr6-71032781-INS-441 |
| chr6-71037770-INS-183 | chr6-71037770-INS-177 |
| chr6-71044870-INS-152 | chr6-71044870-INS-152 |
| chr6-71067316-INS-249 | chr6-71067316-INS-249 |
| chr6-71076636-INS-60 | chr6-71076636-INS-56 |
| chr6-71087612-INS-78 | chr6-71087612-INS-78 |
| chr6-71089844-INS-634 | chr6-71089844-INS-643 |
| chr6-71097396-INS-375 | chr6-71097396-INS-375 |
| chr6-71107188-INS-100 | chr6-71107188-INS-100 |
| chr6-71114946-INS-67 | chr6-71114939-INS-60 |
| chr6-71123598-INS-159 | chr6-71123598-INS-160 |
| chr6-71141325-INS-52 | chr6-71141325-INS-54 |
| chr6-71142477-INS-73 | chr6-71142478-INS-66 |
| chr6-71144400-INS-58 | chr6-71144401-INS-58 |
| chr6-71155348-INS-411 | chr6-71155346-INS-411 |
| chr6-71199541-INS-987 | chr6-71199541-INS-1000 |
| chr6-71200284-INS-201 | chr6-71200284-INS-200 |
| chr6-71207677-INS-100 | chr6-71207688-INS-50 |
| chr6-71209452-INS-112 | chr6-71209452-INS-114 |
| chr6-71228901-INS-110 | chr6-71228901-INS-82 |
| chr6-71230798-INS-52 | chr6-71230798-INS-64 |
| chr6-71231248-INS-701 | chr6-71231248-INS-701 |
| chr6-71234781-INS-207 | chr6-71234782-INS-214 |
| chr6-71266753-INS-193 | chr6-71266751-INS-187 |
| chr6-71275663-INS-95 | chr6-71275659-INS-96 |
| chr6-71292717-INS-135 | chr6-71292718-INS-135 |
| chr6-71295191-INS-223 | chr6-71295191-INS-209 |
| chr6-71295457-INS-4174 | chr6-71295458-INS-4174 |
| chr6-71322873-INS-54 | chr6-71322873-INS-54 |
| chr6-71336000-INS-290 | chr6-71336001-INS-300 |
| chr6-71358101-INS-6485 | chr6-71358102-INS-6485 |
| chr6-71408332-INS-3404 | chr6-71408334-INS-3396 |
| chr6-71410032-INS-215 | chr6-71410032-INS-216 |
| chr6-71414843-INS-350 | chr6-71414843-INS-353 |
| chr6-71424835-INS-196 | chr6-71424835-INS-198 |
| chr6-71430633-INS-72 | chr6-71430633-INS-72 |
| chr6-71463853-INS-4984 | chr6-71463854-INS-4984 |
| chr6-71483365-INS-911 | chr6-71483394-INS-928 |
| chr6-71488381-INS-100 | chr6-71488381-INS-100 |
| chr6-71498652-INS-200 | chr6-71498652-INS-200 |
| chr6-71559233-INS-54 | chr6-71559233-INS-62 |
| chr6-71572792-INS-204 | chr6-71572792-INS-203 |
| chr6-71597606-INS-62 | chr6-71597606-INS-62 |
| chr6-71619575-INS-68 | chr6-71619575-INS-56 |
| chr6-71650642-INS-123 | chr6-71650653-INS-75 |
| chr6-71658195-INS-108 | chr6-71658191-INS-156 |
| chr6-71665136-INS-169 | chr6-71665136-INS-169 |
| chr6-71676508-INS-254 | chr6-71676508-INS-213 |
| chr6-71694510-INS-55 | chr6-71694510-INS-55 |
| chr6-71698880-INS-66 | chr6-71698880-INS-66 |
| chr6-71702811-INS-809 | chr6-71702811-INS-809 |
| chr6-71718409-INS-208 | chr6-71718409-INS-208 |
| chr6-71727911-INS-253 | chr6-71727911-INS-254 |
| chr6-71749029-INS-54 | chr6-71749029-INS-56 |
| chr6-71759831-INS-53 | chr6-71759831-INS-53 |
| chr6-71766341-INS-85 | chr6-71766341-INS-65 |
| chr6-71769272-INS-264 | chr6-71769272-INS-261 |

| | |
|---|---|
| chr6-71773404-INS-354 | chr6-71773404-INS-354 |
| chr6-71774442-INS-250 | chr6-71774436-INS-247 |
| chr6-71778665-INS-55 | chr6-71778665-INS-55 |
| chr6-71803630-INS-74 | chr6-71803600-INS-82 |
| chr6-71810520-INS-112 | chr6-71810520-INS-108 |
| chr6-71813801-INS-6954 | chr6-71813802-INS-6954 |
| chr6-71815002-INS-1006 | chr6-71815003-INS-1006 |
| chr6-71820156-INS-237 | chr6-71820156-INS-231 |
| chr6-71824715-INS-60 | chr6-71824715-INS-60 |
| chr6-71845925-INS-56 | chr6-71845925-INS-52 |
| chr6-71859870-INS-86 | chr6-71859870-INS-50 |
| chr6-71869131-INS-246 | chr6-71869131-INS-246 |
| chr6-71877769-INS-51 | chr6-71877769-INS-51 |
| chr6-71899548-INS-101 | chr6-71899548-INS-101 |
| chr6-71904880-INS-262 | chr6-71904880-INS-263 |
| chr6-71906933-INS-399 | chr6-71906933-INS-399 |
| chr6-71939785-INS-208 | chr6-71939785-INS-208 |
| chr6-71966036-INS-57 | chr6-71966036-INS-55 |
| chr6-71969988-INS-189 | chr6-71969993-INS-190 |
| chr6-71972521-INS-194 | chr6-71972521-INS-194 |
| chr6-71980557-INS-72 | chr6-71980557-INS-72 |
| chr6-71983651-INS-985 | chr6-71983651-INS-985 |
| chr6-71984049-INS-64 | chr6-71984020-INS-69 |
| chr6-71993646-INS-127 | chr6-71993680-INS-111 |
| chr6-71998375-INS-891 | chr6-71998375-INS-891 |
| chr6-72018110-INS-72 | chr6-72018110-INS-72 |
| chr6-72023321-INS-54 | chr6-72023325-INS-56 |
| chr6-72024345-INS-63 | chr6-72024346-INS-53 |
| chr6-72026603-INS-354 | chr6-72026603-INS-354 |
| chr6-72032078-INS-495 | chr6-72032079-INS-495 |
| chr6-72035549-INS-519 | chr6-72035549-INS-499 |
| chr6-72044875-INS-2785 | chr6-72044876-INS-2785 |
| chr6-72044876-INS-410 | chr6-72044876-INS-410 |
| chr6-72054293-INS-710 | chr6-72054293-INS-711 |
| chr6-72061319-INS-764 | chr6-72061320-INS-754 |
| chr6-72070040-INS-110 | chr6-72070040-INS-110 |
| chr6-72111022-INS-72 | chr6-72111022-INS-72 |
| chr6-72126993-INS-111 | chr6-72126993-INS-128 |
| chr6-72127989-INS-56 | chr6-72127989-INS-54 |
| chr6-72139186-INS-175 | chr6-72139186-INS-175 |
| chr6-72151733-INS-2932 | chr6-72151734-INS-2932 |
| chr6-72155984-INS-115 | chr6-72155981-INS-111 |
| chr6-72172367-INS-51 | chr6-72172368-INS-75 |
| chr6-72186167-INS-975 | chr6-72186167-INS-965 |
| chr6-72204629-INS-150 | chr6-72204629-INS-150 |
| chr6-72214328-INS-1286 | chr6-72214329-INS-1287 |
| chr6-72228862-INS-98 | chr6-72228904-INS-52 |
| chr6-72241374-INS-533 | chr6-72241374-INS-530 |
| chr6-72247928-INS-200 | chr6-72247928-INS-200 |
| chr6-72255099-INS-172 | chr6-72255099-INS-174 |
| chr6-72256210-INS-737 | chr6-72256210-INS-740 |
| chr6-72257060-INS-6348 | chr6-72257061-INS-6348 |
| chr6-72259101-INS-66 | chr6-72259101-INS-66 |
| chr6-72269153-INS-57 | chr6-72269153-INS-57 |
| chr6-72273873-INS-58 | chr6-72273873-INS-60 |
| chr6-72285180-INS-368 | chr6-72285180-INS-369 |
| chr6-72297425-INS-362 | chr6-72297425-INS-362 |
| chr6-72305733-INS-61 | chr6-72305733-INS-61 |
| chr6-72306380-INS-103 | chr6-72306380-INS-103 |
| chr6-72344034-INS-190 | chr6-72344034-INS-190 |
| chr6-72349788-INS-104 | chr6-72349793-INS-121 |
| chr6-72351732-INS-63 | chr6-72351732-INS-63 |
| chr6-72361362-INS-224 | chr6-72361362-INS-237 |
| chr6-72363483-INS-185 | chr6-72363483-INS-186 |
| chr6-72383381-INS-68 | chr6-72383381-INS-68 |
| chr6-72386546-INS-70 | chr6-72386546-INS-70 |
| chr6-72407385-INS-59 | chr6-72407385-INS-62 |
| chr6-72409683-INS-427 | chr6-72409683-INS-427 |
| chr6-72422744-INS-54 | chr6-72422744-INS-58 |
| chr6-72426680-INS-69 | chr6-72426707-INS-81 |
| chr6-72435073-INS-181 | chr6-72435073-INS-180 |
| chr6-72436938-INS-54 | chr6-72436947-INS-66 |
| chr6-72438641-INS-182 | chr6-72438641-INS-191 |
| chr6-72443825-INS-53 | chr6-72443845-INS-52 |
| chr6-72461531-INS-175 | chr6-72461531-INS-205 |
| chr6-72465049-INS-174 | chr6-72465049-INS-170 |
| chr6-72469361-INS-384 | chr6-72469361-INS-384 |

| | |
|---|---|
| chr6-72477643-INS-68 | chr6-72477643-INS-68 |
| chr6-72484413-INS-205 | chr6-72484413-INS-199 |
| chr6-72488308-INS-205 | chr6-72488308-INS-205 |
| chr6-72494948-INS-196 | chr6-72494948-INS-199 |
| chr6-72509099-INS-54 | chr6-72509099-INS-56 |
| chr6-72510982-INS-59 | chr6-72510982-INS-59 |
| chr6-72514280-INS-311 | chr6-72514281-INS-311 |
| chr6-72548660-INS-108 | chr6-72548676-INS-60 |
| chr6-72551201-INS-69 | chr6-72551201-INS-69 |
| chr6-72565530-INS-73 | chr6-72565530-INS-73 |
| chr6-72572468-INS-56 | chr6-72572468-INS-56 |
| chr6-72581737-INS-476 | chr6-72581737-INS-476 |
| chr6-72584962-INS-227 | chr6-72584962-INS-228 |
| chr6-72621923-INS-55 | chr6-72621923-INS-55 |
| chr6-72636847-INS-206 | chr6-72636847-INS-206 |
| chr6-72639174-INS-201 | chr6-72639174-INS-201 |
| chr6-72673573-INS-75 | chr6-72673530-INS-80 |
| chr6-72693752-INS-193 | chr6-72693752-INS-193 |
| chr6-72749880-INS-207 | chr6-72749880-INS-207 |
| chr6-72771433-INS-52 | chr6-72771433-INS-52 |
| chr6-72776235-INS-64 | chr6-72776235-INS-62 |
| chr6-72777102-INS-178 | chr6-72777102-INS-179 |
| chr6-72778964-INS-206 | chr6-72778964-INS-206 |
| chr6-72785681-INS-100 | chr6-72785726-INS-104 |
| chr6-72788639-INS-61 | chr6-72788639-INS-62 |
| chr6-72796482-INS-200 | chr6-72796482-INS-200 |
| chr6-72806646-INS-414 | chr6-72806646-INS-414 |
| chr6-72807825-INS-218 | chr6-72807825-INS-218 |
| chr6-72809695-INS-208 | chr6-72809695-INS-208 |
| chr6-72849141-INS-214 | chr6-72849141-INS-211 |
| chr6-72862453-INS-63 | chr6-72862453-INS-60 |
| chr6-72878922-INS-315 | chr6-72878925-INS-315 |
| chr6-72883784-INS-263 | chr6-72883784-INS-219 |
| chr6-72898391-INS-60 | chr6-72898391-INS-60 |
| chr6-72909061-INS-88 | chr6-72909046-INS-121 |
| chr6-72927891-INS-53 | chr6-72927891-INS-53 |
| chr6-72928780-INS-94 | chr6-72928780-INS-94 |
| chr6-72943109-INS-216 | chr6-72943109-INS-216 |
| chr6-72951373-INS-204 | chr6-72951373-INS-204 |
| chr6-72964309-INS-76 | chr6-72964309-INS-76 |
| chr6-72965187-INS-214 | chr6-72965187-INS-215 |
| chr6-72979595-INS-77 | chr6-72979595-INS-77 |
| chr6-73000132-INS-54 | chr6-73000131-INS-96 |
| chr6-73006937-INS-68 | chr6-73006937-INS-74 |
| chr6-73009416-INS-542 | chr6-73009416-INS-543 |
| chr6-73020868-INS-78 | chr6-73020868-INS-84 |
| chr6-73035803-INS-99 | chr6-73035803-INS-99 |
| chr6-73054502-INS-901 | chr6-73054502-INS-901 |
| chr6-73092592-INS-2330 | chr6-73092593-INS-2334 |
| chr6-73098133-INS-3132 | chr6-73098134-INS-3132 |
| chr6-73102029-INS-208 | chr6-73102029-INS-208 |
| chr6-73105368-INS-56 | chr6-73105368-INS-56 |
| chr6-73110960-INS-6584 | chr6-73110961-INS-6584 |
| chr6-73122068-INS-88 | chr6-73122068-INS-88 |
| chr6-73139319-INS-269 | chr6-73139319-INS-270 |
| chr6-73143690-INS-198 | chr6-73143690-INS-198 |
| chr6-73152766-INS-139 | chr6-73152766-INS-154 |
| chr6-73159156-INS-51 | chr6-73159151-INS-53 |
| chr6-73164525-INS-89 | chr6-73164525-INS-89 |
| chr6-73181060-INS-66 | chr6-73181060-INS-66 |
| chr6-73187726-INS-52 | chr6-73187726-INS-52 |
| chr6-73203774-INS-60 | chr6-73203774-INS-60 |
| chr6-73205309-INS-84 | chr6-73205309-INS-122 |
| chr6-73234635-INS-520 | chr6-73234635-INS-520 |
| chr6-73249304-INS-262 | chr6-73249304-INS-263 |
| chr6-73251448-INS-6441 | chr6-73251449-INS-6442 |
| chr6-73259585-INS-677 | chr6-73259585-INS-679 |
| chr6-73290726-INS-199 | chr6-73290726-INS-199 |
| chr6-73291107-INS-1895 | chr6-73291108-INS-1895 |
| chr6-73307184-INS-65 | chr6-73307184-INS-55 |
| chr6-73316432-INS-120 | chr6-73316456-INS-76 |
| chr6-73316478-INS-6355 | chr6-73316472-INS-6367 |
| chr6-73340502-INS-129 | chr6-73340502-INS-129 |
| chr6-73340576-INS-508 | chr6-73340561-INS-514 |
| chr6-73344364-INS-76 | chr6-73344364-INS-65 |
| chr6-73347016-INS-65 | chr6-73347016-INS-111 |
| chr6-73365667-INS-128 | chr6-73365667-INS-104 |

| | |
|---|---|
| chr6-73366815-INS-68 | chr6-73366815-INS-68 |
| chr6-73372576-INS-66 | chr6-73372576-INS-96 |
| chr6-73374054-INS-56 | chr6-73374054-INS-56 |
| chr6-73382419-INS-168 | chr6-73382419-INS-168 |
| chr6-73399606-INS-217 | chr6-73399606-INS-217 |
| chr6-73404220-INS-3235 | chr6-73404221-INS-3235 |
| chr6-73406804-INS-122 | chr6-73406804-INS-122 |
| chr6-73410342-INS-66 | chr6-73410342-INS-66 |
| chr6-73447366-INS-60 | chr6-73447366-INS-80 |
| chr6-73449059-INS-90 | chr6-73449059-INS-90 |
| chr6-73449654-INS-1822 | chr6-73449655-INS-1822 |
| chr6-73497782-INS-155 | chr6-73497782-INS-159 |
| chr6-73506273-INS-176 | chr6-73506273-INS-176 |
| chr6-73509216-INS-803 | chr6-73509216-INS-799 |
| chr6-73520991-INS-446 | chr6-73520991-INS-448 |
| chr6-73540334-INS-52 | chr6-73540334-INS-68 |
| chr6-73541857-INS-112 | chr6-73541856-INS-100 |
| chr6-73544355-INS-136 | chr6-73544355-INS-136 |
| chr6-73546355-INS-60 | chr6-73546356-INS-90 |
| chr6-73550768-INS-76 | chr6-73550774-INS-78 |
| chr6-73563051-INS-72 | chr6-73563051-INS-72 |
| chr6-73570171-INS-2582 | chr6-73570172-INS-2582 |
| chr6-73587791-INS-55 | chr6-73587791-INS-50 |
| chr6-73601122-INS-6595 | chr6-73601123-INS-6595 |
| chr6-73614053-INS-56 | chr6-73614053-INS-56 |
| chr6-73621709-INS-70 | chr6-73621709-INS-70 |
| chr6-73644035-INS-96 | chr6-73644040-INS-106 |
| chr6-73653193-INS-166 | chr6-73653193-INS-150 |
| chr6-73662122-INS-362 | chr6-73662122-INS-362 |
| chr6-73665643-INS-511 | chr6-73665644-INS-511 |
| chr6-73690692-INS-2323 | chr6-73690693-INS-2324 |
| chr6-73738222-INS-131 | chr6-73738222-INS-131 |
| chr6-73743390-INS-374 | chr6-73743390-INS-374 |
| chr6-73751376-INS-6662 | chr6-73751377-INS-6662 |
| chr6-73784610-INS-835 | chr6-73784610-INS-835 |
| chr6-73791147-INS-1985 | chr6-73791148-INS-1985 |
| chr6-73800451-INS-103 | chr6-73800451-INS-103 |
| chr6-73826107-INS-6674 | chr6-73826104-INS-6706 |
| chr6-73829571-INS-64 | chr6-73829571-INS-100 |
| chr6-73835184-INS-2423 | chr6-73835185-INS-2423 |
| chr6-73851628-INS-167 | chr6-73851628-INS-167 |
| chr6-73863033-INS-57 | chr6-73863033-INS-59 |
| chr6-73876150-INS-6470 | chr6-73876151-INS-6470 |
| chr6-73904743-INS-5456 | chr6-73904744-INS-5457 |
| chr6-73909875-INS-387 | chr6-73909875-INS-387 |
| chr6-73922733-INS-501 | chr6-73922733-INS-501 |
| chr6-73924462-INS-145 | chr6-73924462-INS-144 |
| chr6-73954617-INS-93 | chr6-73954617-INS-93 |
| chr6-73955934-INS-829 | chr6-73955934-INS-829 |
| chr6-73962322-INS-209 | chr6-73962320-INS-248 |
| chr6-73979569-INS-762 | chr6-73979569-INS-763 |
| chr6-73980236-INS-53 | chr6-73980236-INS-59 |
| chr6-73984586-INS-84 | chr6-73984586-INS-84 |
| chr6-73992311-INS-56 | chr6-73992302-INS-72 |
| chr6-74001845-INS-1430 | chr6-74001846-INS-1430 |
| chr6-74049119-INS-77 | chr6-74049093-INS-60 |
| chr6-74057273-INS-125 | chr6-74057273-INS-97 |
| chr6-74062973-INS-121 | chr6-74062973-INS-122 |
| chr6-74081101-INS-142 | chr6-74081101-INS-143 |
| chr6-74083379-INS-337 | chr6-74083379-INS-338 |
| chr6-74092942-INS-499 | chr6-74092944-INS-503 |
| chr6-74110512-INS-52 | chr6-74110512-INS-52 |
| chr6-74140752-INS-6687 | chr6-74140753-INS-6687 |
| chr6-74147632-INS-56 | chr6-74147632-INS-56 |
| chr6-74169146-INS-688 | chr6-74169146-INS-688 |
| chr6-74186626-INS-6442 | chr6-74186627-INS-6442 |
| chr6-74193399-INS-64 | chr6-74193395-INS-82 |
| chr6-74199620-INS-65 | chr6-74199585-INS-58 |
| chr6-74208101-INS-283 | chr6-74208101-INS-283 |
| chr6-74229638-INS-92 | chr6-74229638-INS-88 |
| chr6-74237136-INS-120 | chr6-74237136-INS-122 |
| chr6-74248218-INS-52 | chr6-74248218-INS-56 |
| chr6-74262154-INS-4845 | chr6-74262155-INS-4845 |
| chr6-74263126-INS-85 | chr6-74263123-INS-95 |
| chr6-74268911-INS-6436 | chr6-74268912-INS-6436 |
| chr6-74281071-INS-483 | chr6-74281072-INS-483 |
| chr6-74282743-INS-209 | chr6-74282743-INS-209 |

| | |
|---|---|
| chr6-74290335-INS-62 | chr6-74290335-INS-63 |
| chr6-74307175-INS-53 | chr6-74307175-INS-53 |
| chr6-74322138-INS-2196 | chr6-74322139-INS-2196 |
| chr6-74322905-INS-536 | chr6-74322905-INS-536 |
| chr6-74326761-INS-130 | chr6-74326761-INS-130 |
| chr6-74351204-INS-146 | chr6-74351174-INS-181 |
| chr6-74353837-INS-64 | chr6-74353837-INS-64 |
| chr6-74365547-INS-6712 | chr6-74365548-INS-6713 |
| chr6-74368888-INS-52 | chr6-74368891-INS-59 |
| chr6-74378540-INS-79 | chr6-74378577-INS-78 |
| chr6-74395674-INS-179 | chr6-74395689-INS-150 |
| chr6-74411592-INS-434 | chr6-74411592-INS-430 |
| chr6-74435833-INS-787 | chr6-74435833-INS-788 |
| chr6-74453804-INS-104 | chr6-74453804-INS-112 |
| chr6-74466241-INS-6902 | chr6-74466242-INS-6903 |
| chr6-74473554-INS-132 | chr6-74473554-INS-130 |
| chr6-74475269-INS-80 | chr6-74475269-INS-92 |
| chr6-74481801-INS-735 | chr6-74481801-INS-735 |
| chr6-74492102-INS-261 | chr6-74492102-INS-261 |
| chr6-74494748-INS-482 | chr6-74494748-INS-482 |
| chr6-74497256-INS-7202 | chr6-74497257-INS-7202 |
| chr6-74504656-INS-1594 | chr6-74504657-INS-1594 |
| chr6-74509152-INS-69 | chr6-74509152-INS-73 |
| chr6-74516095-INS-320 | chr6-74516095-INS-320 |
| chr6-74521715-INS-102 | chr6-74521715-INS-102 |
| chr6-74530756-INS-2823 | chr6-74530757-INS-2823 |
| chr6-74532575-INS-102 | chr6-74532575-INS-102 |
| chr6-74570872-INS-95 | chr6-74570872-INS-107 |
| chr6-74598807-INS-862 | chr6-74598807-INS-862 |
| chr6-74615572-INS-52 | chr6-74615586-INS-80 |
| chr6-74616347-INS-294 | chr6-74616347-INS-309 |
| chr6-74636665-INS-76 | chr6-74636665-INS-74 |
| chr6-74650099-INS-390 | chr6-74650099-INS-390 |
| chr6-74683112-INS-424 | chr6-74683112-INS-424 |
| chr6-74686242-INS-65 | chr6-74686245-INS-61 |
| chr6-74692368-INS-410 | chr6-74692368-INS-410 |
| chr6-74723260-INS-6779 | chr6-74723261-INS-6777 |
| chr6-74748421-INS-56 | chr6-74748421-INS-56 |
| chr6-74776663-INS-500 | chr6-74776663-INS-500 |
| chr6-74793970-INS-1450 | chr6-74793971-INS-1450 |
| chr6-74801836-INS-95 | chr6-74801836-INS-99 |
| chr6-74814017-INS-211 | chr6-74814017-INS-211 |
| chr6-74820907-INS-66 | chr6-74820907-INS-66 |
| chr6-74823264-INS-471 | chr6-74823264-INS-471 |
| chr6-74869041-INS-1888 | chr6-74869042-INS-1888 |
| chr6-74885270-INS-364 | chr6-74885270-INS-364 |
| chr6-74889884-INS-62 | chr6-74889885-INS-62 |
| chr6-74890644-INS-2897 | chr6-74890645-INS-2897 |
| chr6-74895138-INS-176 | chr6-74895138-INS-176 |
| chr6-74916805-INS-940 | chr6-74916805-INS-941 |
| chr6-74917856-INS-71 | chr6-74917856-INS-71 |
| chr6-74948410-INS-54 | chr6-74948410-INS-92 |
| chr6-74953661-INS-3018 | chr6-74953662-INS-3018 |
| chr6-74956712-INS-984 | chr6-74956712-INS-986 |
| chr6-74964315-INS-60 | chr6-74964315-INS-64 |
| chr6-75004469-INS-784 | chr6-75004469-INS-779 |
| chr6-75010897-INS-233 | chr6-75010897-INS-233 |
| chr6-75014686-INS-401 | chr6-75014686-INS-401 |
| chr6-75030290-INS-187 | chr6-75030290-INS-187 |
| chr6-75077478-INS-74 | chr6-75077478-INS-74 |
| chr6-75095647-INS-216 | chr6-75095647-INS-216 |
| chr6-75098801-INS-4907 | chr6-75098802-INS-4907 |
| chr6-75099588-INS-218 | chr6-75099588-INS-218 |
| chr6-75169899-INS-72 | chr6-75169899-INS-72 |
| chr6-75190463-INS-260 | chr6-75190464-INS-259 |
| chr6-75194098-INS-210 | chr6-75194098-INS-210 |
| chr6-75220047-INS-1658 | chr6-75220048-INS-1658 |
| chr6-75221267-INS-5548 | chr6-75221268-INS-5548 |
| chr6-75241137-INS-65 | chr6-75241143-INS-50 |
| chr6-75245674-INS-444 | chr6-75245674-INS-444 |
| chr6-75258340-INS-56 | chr6-75258340-INS-58 |
| chr6-75262145-INS-6462 | chr6-75262146-INS-6462 |
| chr6-75281127-INS-52 | chr6-75281127-INS-52 |
| chr6-75318748-INS-431 | chr6-75318748-INS-431 |
| chr6-75339781-INS-84 | chr6-75339781-INS-96 |
| chr6-75341346-INS-56 | chr6-75341346-INS-57 |
| chr6-75343443-INS-52 | chr6-75343443-INS-52 |

| | |
|---|---|
| chr6-75346261-INS-61 | chr6-75346262-INS-75 |
| chr6-75358221-INS-3370 | chr6-75358222-INS-3370 |
| chr6-75367432-INS-7143 | chr6-75367433-INS-7143 |
| chr6-75368613-INS-52 | chr6-75368613-INS-60 |
| chr6-75369560-INS-114 | chr6-75369560-INS-114 |
| chr6-75376796-INS-236 | chr6-75376796-INS-237 |
| chr6-75380158-INS-441 | chr6-75380158-INS-441 |
| chr6-75388552-INS-57 | chr6-75388553-INS-52 |
| chr6-75390709-INS-134 | chr6-75390709-INS-134 |
| chr6-75393392-INS-483 | chr6-75393387-INS-486 |
| chr6-75395549-INS-290 | chr6-75395549-INS-292 |
| chr6-75402389-INS-6444 | chr6-75402390-INS-6444 |
| chr6-75406846-INS-98 | chr6-75406846-INS-98 |
| chr6-75431027-INS-60 | chr6-75431027-INS-60 |
| chr6-75455858-INS-3246 | chr6-75455859-INS-3246 |
| chr6-75460787-INS-139 | chr6-75460791-INS-182 |
| chr6-75501268-INS-176 | chr6-75501268-INS-176 |
| chr6-75510411-INS-56 | chr6-75510411-INS-60 |
| chr6-75512550-INS-6172 | chr6-75512551-INS-6172 |
| chr6-75523763-INS-114 | chr6-75523763-INS-114 |
| chr6-75531199-INS-763 | chr6-75531199-INS-763 |
| chr6-75547756-INS-2141 | chr6-75547757-INS-2141 |
| chr6-75551458-INS-2066 | chr6-75551459-INS-2106 |
| chr6-75575745-INS-53 | chr6-75575745-INS-53 |
| chr6-75588282-INS-325 | chr6-75588282-INS-325 |
| chr6-75595657-INS-849 | chr6-75595657-INS-849 |
| chr6-75598271-INS-6963 | chr6-75598272-INS-6963 |
| chr6-75605394-INS-374 | chr6-75605394-INS-374 |
| chr6-75620875-INS-1891 | chr6-75620876-INS-1891 |
| chr6-75627082-INS-52 | chr6-75627083-INS-52 |
| chr6-75646145-INS-61 | chr6-75646145-INS-59 |
| chr6-75647878-INS-6692 | chr6-75647879-INS-6692 |
| chr6-75670615-INS-54 | chr6-75670597-INS-60 |
| chr6-75674571-INS-62 | chr6-75674571-INS-66 |
| chr6-75679913-INS-64 | chr6-75679913-INS-90 |
| chr6-75704510-INS-217 | chr6-75704515-INS-217 |
| chr6-75719488-INS-1331 | chr6-75719489-INS-1334 |
| chr6-75725175-INS-122 | chr6-75725172-INS-126 |
| chr6-75727956-INS-65 | chr6-75727956-INS-65 |
| chr6-75728135-INS-1163 | chr6-75728136-INS-1150 |
| chr6-75730410-INS-357 | chr6-75730410-INS-357 |
| chr6-75738847-INS-98 | chr6-75738847-INS-102 |
| chr6-75748343-INS-65 | chr6-75748343-INS-55 |
| chr6-75751983-INS-464 | chr6-75751983-INS-464 |
| chr6-75760181-INS-251 | chr6-75760181-INS-259 |
| chr6-75766810-INS-3039 | chr6-75766811-INS-3039 |
| chr6-75779364-INS-369 | chr6-75779364-INS-369 |
| chr6-75807027-INS-363 | chr6-75807027-INS-363 |
| chr6-75807345-INS-671 | chr6-75807345-INS-671 |
| chr6-75819988-INS-232 | chr6-75819988-INS-233 |
| chr6-75828558-INS-66 | chr6-75828558-INS-66 |
| chr6-75849286-INS-691 | chr6-75849286-INS-691 |
| chr6-75857314-INS-59 | chr6-75857314-INS-59 |
| chr6-75860580-INS-1413 | chr6-75860590-INS-1379 |
| chr6-75868436-INS-1379 | chr6-75868437-INS-1379 |
| chr6-75870421-INS-169 | chr6-75870421-INS-169 |
| chr6-75876623-INS-65 | chr6-75876576-INS-66 |
| chr6-75903603-INS-57 | chr6-75903594-INS-53 |
| chr6-75916839-INS-64 | chr6-75916839-INS-82 |
| chr6-75919709-INS-651 | chr6-75919711-INS-658 |
| chr6-75934404-INS-204 | chr6-75934404-INS-204 |
| chr6-75940662-INS-75 | chr6-75940662-INS-110 |
| chr6-75955014-INS-198 | chr6-75955049-INS-198 |
| chr6-75958239-INS-51 | chr6-75958239-INS-51 |
| chr6-75964657-INS-5830 | chr6-75964658-INS-5830 |
| chr6-75968079-INS-1703 | chr6-75968080-INS-1703 |
| chr6-75973070-INS-900 | chr6-75973070-INS-900 |
| chr6-75973681-INS-54 | chr6-75973687-INS-62 |
| chr6-75974940-INS-1124 | chr6-75974941-INS-1124 |
| chr6-75979027-INS-76 | chr6-75979027-INS-76 |
| chr6-75986669-INS-2624 | chr6-75986670-INS-2624 |
| chr6-76022811-INS-117 | chr6-76022811-INS-117 |
| chr6-76028869-INS-222 | chr6-76028869-INS-222 |
| chr6-76030490-INS-53 | chr6-76030491-INS-54 |
| chr6-76032340-INS-599 | chr6-76032340-INS-599 |
| chr6-76047622-INS-245 | chr6-76047622-INS-246 |
| chr6-76069793-INS-77 | chr6-76069793-INS-78 |

| | |
|---|---|
| chr6-76072229-INS-466 | chr6-76072229-INS-494 |
| chr6-76083608-INS-479 | chr6-76083608-INS-478 |
| chr6-76092192-INS-80 | chr6-76092192-INS-56 |
| chr6-76102727-INS-114 | chr6-76102727-INS-127 |
| chr6-76141688-INS-6381 | chr6-76141689-INS-6381 |
| chr6-76150382-INS-877 | chr6-76150382-INS-877 |
| chr6-76158004-INS-7939 | chr6-76158005-INS-7939 |
| chr6-76171806-INS-62 | chr6-76171806-INS-62 |
| chr6-76179297-INS-106 | chr6-76179297-INS-108 |
| chr6-76182870-INS-753 | chr6-76182870-INS-753 |
| chr6-76186724-INS-176 | chr6-76186724-INS-176 |
| chr6-76189355-INS-59 | chr6-76189355-INS-59 |
| chr6-76202161-INS-134 | chr6-76202137-INS-115 |
| chr6-76204648-INS-183 | chr6-76204648-INS-183 |
| chr6-76220542-INS-54 | chr6-76220542-INS-56 |
| chr6-76223230-INS-72 | chr6-76223230-INS-76 |
| chr6-76233977-INS-6384 | chr6-76233978-INS-6385 |
| chr6-76237296-INS-962 | chr6-76237296-INS-962 |
| chr6-76244676-INS-428 | chr6-76244676-INS-428 |
| chr6-76256043-INS-4759 | chr6-76256044-INS-4766 |
| chr6-76276349-INS-57 | chr6-76276350-INS-52 |
| chr6-76282959-INS-80 | chr6-76282959-INS-80 |
| chr6-76299313-INS-68 | chr6-76299310-INS-55 |
| chr6-76311476-INS-659 | chr6-76311476-INS-659 |
| chr6-76322499-INS-133 | chr6-76322499-INS-163 |
| chr6-76325146-INS-359 | chr6-76325146-INS-359 |
| chr6-76332392-INS-160 | chr6-76332392-INS-163 |
| chr6-76342754-INS-138 | chr6-76342754-INS-138 |
| chr6-76348295-INS-397 | chr6-76348295-INS-404 |
| chr6-76363445-INS-96 | chr6-76363445-INS-96 |
| chr6-76374593-INS-529 | chr6-76374593-INS-529 |
| chr6-76384203-INS-1153 | chr6-76384204-INS-1154 |
| chr6-76399422-INS-235 | chr6-76399422-INS-235 |
| chr6-76420494-INS-50 | chr6-76420494-INS-74 |
| chr6-76428317-INS-86 | chr6-76428317-INS-58 |
| chr6-76435692-INS-52 | chr6-76435692-INS-95 |
| chr6-76457497-INS-2909 | chr6-76457498-INS-2910 |
| chr6-76479948-INS-136 | chr6-76479972-INS-140 |
| chr6-76519233-INS-505 | chr6-76519233-INS-505 |
| chr6-76526108-INS-283 | chr6-76526108-INS-283 |
| chr6-76527946-INS-6944 | chr6-76527947-INS-6944 |
| chr6-76533511-INS-57 | chr6-76533511-INS-57 |
| chr6-76575612-INS-2821 | chr6-76575613-INS-2821 |
| chr6-76589914-INS-814 | chr6-76589914-INS-814 |
| chr6-76595337-INS-204 | chr6-76595345-INS-203 |
| chr6-76722754-INS-120 | chr6-76722754-INS-120 |
| chr6-76729063-INS-57 | chr6-76729063-INS-51 |
| chr6-76747527-INS-663 | chr6-76747527-INS-671 |
| chr6-76751107-INS-174 | chr6-76751107-INS-174 |
| chr6-76752467-INS-77 | chr6-76752467-INS-77 |
| chr6-76759628-INS-180 | chr6-76759638-INS-216 |
| chr6-76777835-INS-133 | chr6-76777835-INS-133 |
| chr6-76786615-INS-3491 | chr6-76786616-INS-3491 |
| chr6-76804003-INS-146 | chr6-76804003-INS-146 |
| chr6-76832249-INS-115 | chr6-76832249-INS-101 |
| chr6-76838116-INS-3286 | chr6-76838117-INS-3286 |
| chr6-76874746-INS-66 | chr6-76874746-INS-66 |
| chr6-76886584-INS-116 | chr6-76886559-INS-106 |
| chr6-76889130-INS-81 | chr6-76889130-INS-81 |
| chr6-76905134-INS-176 | chr6-76905170-INS-137 |
| chr6-76918872-INS-60 | chr6-76918872-INS-60 |
| chr6-76963453-INS-63 | chr6-76963453-INS-57 |
| chr6-76986532-INS-74 | chr6-76986532-INS-76 |
| chr6-76988629-INS-110 | chr6-76988629-INS-91 |
| chr6-77013164-INS-56 | chr6-77013164-INS-54 |
| chr6-77028269-INS-55 | chr6-77028270-INS-56 |
| chr6-77033134-INS-424 | chr6-77033134-INS-424 |
| chr6-77069928-INS-592 | chr6-77069928-INS-592 |
| chr6-77081738-INS-54 | chr6-77081738-INS-52 |
| chr6-77120197-INS-53 | chr6-77120219-INS-53 |
| chr6-77126177-INS-1332 | chr6-77126178-INS-1332 |
| chr6-77131043-INS-70 | chr6-77131043-INS-70 |
| chr6-77139724-INS-52 | chr6-77139724-INS-54 |
| chr6-77157951-INS-60 | chr6-77157951-INS-60 |
| chr6-77168198-INS-54 | chr6-77168198-INS-54 |
| chr6-77184143-INS-413 | chr6-77184142-INS-419 |
| chr6-77201852-INS-56 | chr6-77201884-INS-52 |

| | |
|---|---|
| chr6-77217317-INS-60 | chr6-77217317-INS-52 |
| chr6-77235482-INS-252 | chr6-77235482-INS-265 |
| chr6-77275068-INS-51 | chr6-77275072-INS-94 |
| chr6-77278745-INS-66 | chr6-77278761-INS-66 |
| chr6-77279436-INS-486 | chr6-77279436-INS-483 |
| chr6-77346653-INS-101 | chr6-77346653-INS-101 |
| chr6-77370779-INS-266 | chr6-77370779-INS-273 |
| chr6-77373620-INS-512 | chr6-77373620-INS-512 |
| chr6-77393449-INS-94 | chr6-77393445-INS-76 |
| chr6-77399995-INS-302 | chr6-77399995-INS-303 |
| chr6-77405609-INS-92 | chr6-77405609-INS-92 |
| chr6-77417010-INS-76 | chr6-77417010-INS-60 |
| chr6-77480965-INS-821 | chr6-77480965-INS-818 |
| chr6-77492580-INS-149 | chr6-77492580-INS-149 |
| chr6-77500987-INS-256 | chr6-77500988-INS-259 |
| chr6-77512534-INS-74 | chr6-77512534-INS-74 |
| chr6-77524438-INS-106 | chr6-77524434-INS-118 |
| chr6-77531425-INS-1795 | chr6-77531426-INS-1792 |
| chr6-77552795-INS-60 | chr6-77552795-INS-60 |
| chr6-77572775-INS-3035 | chr6-77572776-INS-3035 |
| chr6-77584486-INS-454 | chr6-77584486-INS-454 |
| chr6-77589819-INS-546 | chr6-77589819-INS-546 |
| chr6-77593054-INS-160 | chr6-77593031-INS-151 |
| chr6-77625096-INS-138 | chr6-77625096-INS-138 |
| chr6-77632817-INS-80 | chr6-77632817-INS-56 |
| chr6-77650431-INS-68 | chr6-77650431-INS-68 |
| chr6-77654007-INS-52 | chr6-77654020-INS-58 |
| chr6-77655595-INS-322 | chr6-77655595-INS-322 |
| chr6-77664553-INS-59 | chr6-77664553-INS-54 |
| chr6-77668304-INS-785 | chr6-77668304-INS-785 |
| chr6-77692275-INS-498 | chr6-77692275-INS-498 |
| chr6-77721136-INS-58 | chr6-77721136-INS-53 |
| chr6-77723841-INS-411 | chr6-77723844-INS-412 |
| chr6-77731211-INS-109 | chr6-77731211-INS-109 |
| chr6-77733181-INS-165 | chr6-77733181-INS-165 |
| chr6-77734578-INS-62 | chr6-77734578-INS-62 |
| chr6-77791752-INS-61 | chr6-77791768-INS-76 |
| chr6-77797499-INS-62 | chr6-77797499-INS-68 |
| chr6-77800119-INS-65 | chr6-77800119-INS-80 |
| chr6-77802142-INS-266 | chr6-77802142-INS-266 |
| chr6-77804181-INS-91 | chr6-77804181-INS-91 |
| chr6-77843818-INS-107 | chr6-77843818-INS-107 |
| chr6-77851095-INS-56 | chr6-77851095-INS-52 |
| chr6-77870177-INS-642 | chr6-77870177-INS-642 |
| chr6-77872722-INS-319 | chr6-77872722-INS-318 |
| chr6-77873988-INS-220 | chr6-77873982-INS-228 |
| chr6-77875413-INS-358 | chr6-77875413-INS-358 |
| chr6-77883587-INS-66 | chr6-77883587-INS-66 |
| chr6-77944340-INS-56 | chr6-77944340-INS-56 |
| chr6-77967331-INS-1978 | chr6-77967332-INS-1984 |
| chr6-77971770-INS-1179 | chr6-77971771-INS-1178 |
| chr6-77985757-INS-260 | chr6-77985757-INS-260 |
| chr6-77989949-INS-128 | chr6-77989949-INS-128 |
| chr6-77998681-INS-181 | chr6-77998681-INS-180 |
| chr6-78041693-INS-5858 | chr6-78041693-INS-5860 |
| chr6-78069216-INS-64 | chr6-78069216-INS-64 |
| chr6-78080057-INS-3152 | chr6-78080058-INS-3151 |
| chr6-78087264-INS-7004 | chr6-78087265-INS-7004 |
| chr6-78114144-INS-219 | chr6-78114144-INS-219 |
| chr6-78126910-INS-3439 | chr6-78126911-INS-3439 |
| chr6-78132062-INS-212 | chr6-78132062-INS-212 |
| chr6-78142267-INS-52 | chr6-78142267-INS-52 |
| chr6-78149784-INS-463 | chr6-78149784-INS-463 |
| chr6-78158155-INS-784 | chr6-78158155-INS-785 |
| chr6-78166555-INS-372 | chr6-78166554-INS-372 |
| chr6-78168827-INS-67 | chr6-78168827-INS-65 |
| chr6-78177951-INS-50 | chr6-78177951-INS-50 |
| chr6-78203548-INS-66 | chr6-78203564-INS-82 |
| chr6-78225532-INS-4725 | chr6-78225533-INS-4725 |
| chr6-78226346-INS-538 | chr6-78226346-INS-538 |
| chr6-78247619-INS-52 | chr6-78247619-INS-52 |
| chr6-78251166-INS-75 | chr6-78251166-INS-75 |
| chr6-78266143-INS-256 | chr6-78266145-INS-258 |
| chr6-78312663-INS-168 | chr6-78312663-INS-168 |
| chr6-78314203-INS-52 | chr6-78314206-INS-58 |
| chr6-78359830-INS-525 | chr6-78359830-INS-525 |
| chr6-78399894-INS-518 | chr6-78399894-INS-518 |

| | |
|---|---|
| chr6-78406833-INS-328 | chr6-78406833-INS-328 |
| chr6-78427133-INS-380 | chr6-78427133-INS-380 |
| chr6-78477769-INS-144 | chr6-78477769-INS-142 |
| chr6-78488791-INS-504 | chr6-78488792-INS-504 |
| chr6-78538485-INS-62 | chr6-78538485-INS-62 |
| chr6-78579450-INS-7314 | chr6-78579451-INS-7314 |
| chr6-78601732-INS-858 | chr6-78601732-INS-858 |
| chr6-78606753-INS-187 | chr6-78606753-INS-187 |
| chr6-78632484-INS-557 | chr6-78632484-INS-557 |
| chr6-78654698-INS-76 | chr6-78654698-INS-76 |
| chr6-78675371-INS-118 | chr6-78675372-INS-128 |
| chr6-78731247-INS-61 | chr6-78731247-INS-63 |
| chr6-78769649-INS-394 | chr6-78769649-INS-394 |
| chr6-78770577-INS-667 | chr6-78770563-INS-665 |
| chr6-78801557-INS-62 | chr6-78801557-INS-64 |
| chr6-78811268-INS-82 | chr6-78811268-INS-107 |
| chr6-78928302-INS-2128 | chr6-78928303-INS-2128 |
| chr6-78948897-INS-70 | chr6-78948897-INS-65 |
| chr6-78975109-INS-6718 | chr6-78975110-INS-6718 |
| chr6-78984311-INS-51 | chr6-78984311-INS-51 |
| chr6-78992697-INS-1253 | chr6-78992691-INS-1250 |
| chr6-79053630-INS-2071 | chr6-79053631-INS-2071 |
| chr6-79054130-INS-8026 | chr6-79054131-INS-8027 |
| chr6-79096924-INS-872 | chr6-79096924-INS-872 |
| chr6-79123524-INS-54 | chr6-79123524-INS-54 |
| chr6-79162477-INS-1899 | chr6-79162478-INS-1900 |
| chr6-79165862-INS-150 | chr6-79165862-INS-150 |
| chr6-79175062-INS-240 | chr6-79175093-INS-250 |
| chr6-79201521-INS-356 | chr6-79201508-INS-357 |
| chr6-79206789-INS-58 | chr6-79206789-INS-56 |
| chr6-79210417-INS-606 | chr6-79210418-INS-587 |
| chr6-79216918-INS-6317 | chr6-79216919-INS-6317 |
| chr6-79219834-INS-79 | chr6-79219834-INS-95 |
| chr6-79219991-INS-3149 | chr6-79219992-INS-3149 |
| chr6-79285069-INS-66 | chr6-79285069-INS-66 |
| chr6-79303011-INS-102 | chr6-79303012-INS-102 |
| chr6-79304934-INS-510 | chr6-79304940-INS-513 |
| chr6-79329477-INS-1357 | chr6-79329478-INS-1359 |
| chr6-79357229-INS-231 | chr6-79357229-INS-231 |
| chr6-79361493-INS-4786 | chr6-79361494-INS-4786 |
| chr6-79363531-INS-548 | chr6-79363531-INS-548 |
| chr6-79455118-INS-303 | chr6-79455118-INS-304 |
| chr6-79465136-INS-1403 | chr6-79465137-INS-1403 |
| chr6-79473321-INS-52 | chr6-79473321-INS-52 |
| chr6-79488601-INS-1690 | chr6-79488602-INS-1689 |
| chr6-79507769-INS-4039 | chr6-79507770-INS-4041 |
| chr6-79513810-INS-424 | chr6-79513810-INS-424 |
| chr6-79525420-INS-86 | chr6-79525420-INS-89 |
| chr6-79533776-INS-6372 | chr6-79533777-INS-6372 |
| chr6-79543531-INS-79 | chr6-79543531-INS-79 |
| chr6-79546975-INS-82 | chr6-79546975-INS-82 |
| chr6-79568067-INS-125 | chr6-79568067-INS-126 |
| chr6-79571539-INS-2651 | chr6-79571540-INS-2651 |
| chr6-79583343-INS-60 | chr6-79583355-INS-57 |
| chr6-79583367-INS-240 | chr6-79583334-INS-252 |
| chr6-79653066-INS-55 | chr6-79653066-INS-57 |
| chr6-79680885-INS-74 | chr6-79680885-INS-74 |
| chr6-79684714-INS-70 | chr6-79684714-INS-70 |
| chr6-79713893-INS-67 | chr6-79713893-INS-78 |
| chr6-79751863-INS-96 | chr6-79751863-INS-92 |
| chr6-79788778-INS-5764 | chr6-79788779-INS-5763 |
| chr6-79795866-INS-60 | chr6-79795835-INS-68 |
| chr6-79798323-INS-641 | chr6-79798323-INS-642 |
| chr6-79814795-INS-143 | chr6-79814795-INS-143 |
| chr6-79820736-INS-2362 | chr6-79820737-INS-2364 |
| chr6-79834980-INS-6378 | chr6-79834981-INS-6379 |
| chr6-79837589-INS-2801 | chr6-79837590-INS-2801 |
| chr6-79843265-INS-321 | chr6-79843265-INS-322 |
| chr6-79844849-INS-83 | chr6-79844849-INS-83 |
| chr6-79914887-INS-500 | chr6-79914887-INS-502 |
| chr6-79945588-INS-64 | chr6-79945588-INS-64 |
| chr6-79960636-INS-2501 | chr6-79960637-INS-2501 |
| chr6-79965113-INS-54 | chr6-79965113-INS-52 |
| chr6-79973599-INS-170 | chr6-79973599-INS-170 |
| chr6-80014536-INS-54 | chr6-80014536-INS-54 |
| chr6-80022159-INS-2450 | chr6-80022160-INS-2450 |
| chr6-80077144-INS-996 | chr6-80077144-INS-996 |

| | |
|---|---|
| chr6-80078495-INS-147 | chr6-80078489-INS-159 |
| chr6-80089807-INS-56 | chr6-80089807-INS-56 |
| chr6-80092788-INS-58 | chr6-80092788-INS-64 |
| chr6-80118718-INS-6461 | chr6-80118719-INS-6493 |
| chr6-80132006-INS-254 | chr6-80132006-INS-254 |
| chr6-80142494-INS-292 | chr6-80142494-INS-290 |
| chr6-80179115-INS-1774 | chr6-80179116-INS-1777 |
| chr6-80179558-INS-125 | chr6-80179558-INS-125 |
| chr6-80194769-INS-63 | chr6-80194769-INS-63 |
| chr6-80231492-INS-94 | chr6-80231499-INS-58 |
| chr6-80330082-INS-6982 | chr6-80330083-INS-6983 |
| chr6-80332858-INS-68 | chr6-80332858-INS-68 |
| chr6-80335976-INS-94 | chr6-80335976-INS-98 |
| chr6-80342180-INS-176 | chr6-80342180-INS-176 |
| chr6-80354560-INS-744 | chr6-80354560-INS-747 |
| chr6-80378863-INS-60 | chr6-80378863-INS-95 |
| chr6-80389206-INS-56 | chr6-80389206-INS-55 |
| chr6-80394443-INS-52 | chr6-80394444-INS-52 |
| chr6-80420520-INS-6403 | chr6-80420521-INS-6403 |
| chr6-80426008-INS-189 | chr6-80426008-INS-189 |
| chr6-80435668-INS-6506 | chr6-80435669-INS-6507 |
| chr6-80468807-INS-108 | chr6-80468807-INS-108 |
| chr6-80475569-INS-204 | chr6-80475569-INS-205 |
| chr6-80476952-INS-191 | chr6-80476952-INS-188 |
| chr6-80491327-INS-199 | chr6-80491327-INS-201 |
| chr6-80494461-INS-67 | chr6-80494462-INS-71 |
| chr6-80505457-INS-730 | chr6-80505457-INS-731 |
| chr6-80542856-INS-59 | chr6-80542849-INS-68 |
| chr6-80554698-INS-54 | chr6-80554696-INS-98 |
| chr6-80558815-INS-496 | chr6-80558815-INS-496 |
| chr6-80565626-INS-2199 | chr6-80565627-INS-2194 |
| chr6-80568650-INS-412 | chr6-80568650-INS-412 |
| chr6-80574210-INS-89 | chr6-80574210-INS-89 |
| chr6-80584775-INS-64 | chr6-80584775-INS-61 |
| chr6-80591212-INS-271 | chr6-80591212-INS-271 |
| chr6-80651990-INS-88 | chr6-80651990-INS-90 |
| chr6-80692107-INS-82 | chr6-80692107-INS-98 |
| chr6-80709518-INS-273 | chr6-80709518-INS-273 |
| chr6-80725764-INS-111 | chr6-80725768-INS-104 |
| chr6-80737182-INS-3924 | chr6-80737183-INS-3923 |
| chr6-80739585-INS-304 | chr6-80739585-INS-304 |
| chr6-80741183-INS-983 | chr6-80741183-INS-984 |
| chr6-80756125-INS-79 | chr6-80756125-INS-79 |
| chr6-80758260-INS-294 | chr6-80758260-INS-294 |
| chr6-80783508-INS-464 | chr6-80783500-INS-482 |
| chr6-80796058-INS-199 | chr6-80796058-INS-199 |
| chr6-80819149-INS-727 | chr6-80819149-INS-727 |
| chr6-80828525-INS-728 | chr6-80828525-INS-729 |
| chr6-80844411-INS-607 | chr6-80844411-INS-607 |
| chr6-80845930-INS-62 | chr6-80845930-INS-62 |
| chr6-80847854-INS-1599 | chr6-80847867-INS-1586 |
| chr6-80864102-INS-54 | chr6-80864102-INS-66 |
| chr6-80888314-INS-1040 | chr6-80888315-INS-1040 |
| chr6-80938524-INS-6418 | chr6-80938525-INS-6418 |
| chr6-80946487-INS-74 | chr6-80946487-INS-73 |
| chr6-80948138-INS-7304 | chr6-80948139-INS-7305 |
| chr6-80956569-INS-181 | chr6-80956569-INS-176 |
| chr6-80979058-INS-355 | chr6-80979058-INS-355 |
| chr6-80990338-INS-326 | chr6-80990338-INS-326 |
| chr6-80999702-INS-120 | chr6-80999702-INS-120 |
| chr6-81005162-INS-64 | chr6-81005162-INS-64 |
| chr6-81007562-INS-66 | chr6-81007562-INS-66 |
| chr6-81026275-INS-432 | chr6-81026275-INS-432 |
| chr6-81063654-INS-599 | chr6-81063654-INS-604 |
| chr6-81071960-INS-7080 | chr6-81071961-INS-7081 |
| chr6-81076799-INS-6512 | chr6-81076800-INS-6510 |
| chr6-81080372-INS-66 | chr6-81080372-INS-84 |
| chr6-81081359-INS-2416 | chr6-81081360-INS-2416 |
| chr6-81122749-INS-86 | chr6-81122749-INS-84 |
| chr6-81124817-INS-293 | chr6-81124817-INS-291 |
| chr6-81134629-INS-53 | chr6-81134629-INS-53 |
| chr6-81139826-INS-103 | chr6-81139826-INS-101 |
| chr6-81195233-INS-111 | chr6-81195233-INS-111 |
| chr6-81199783-INS-953 | chr6-81199783-INS-956 |
| chr6-81216061-INS-56 | chr6-81216061-INS-56 |
| chr6-81217375-INS-53 | chr6-81217375-INS-53 |
| chr6-81220607-INS-441 | chr6-81220607-INS-440 |

| | |
|---|---|
| chr6-81223286-INS-58 | chr6-81223266-INS-58 |
| chr6-81247337-INS-1223 | chr6-81247338-INS-1225 |
| chr6-81261552-INS-61 | chr6-81261555-INS-62 |
| chr6-81264932-INS-56 | chr6-81264932-INS-88 |
| chr6-81268056-INS-73 | chr6-81268027-INS-78 |
| chr6-81301958-INS-56 | chr6-81301958-INS-66 |
| chr6-81323694-INS-131 | chr6-81323694-INS-131 |
| chr6-81367519-INS-64 | chr6-81367519-INS-64 |
| chr6-81380761-INS-469 | chr6-81380761-INS-468 |
| chr6-81399863-INS-54 | chr6-81399863-INS-54 |
| chr6-81403725-INS-90 | chr6-81403729-INS-89 |
| chr6-81431085-INS-58 | chr6-81431081-INS-66 |
| chr6-81453880-INS-92 | chr6-81453880-INS-92 |
| chr6-81466420-INS-54 | chr6-81466420-INS-54 |
| chr6-81472477-INS-58 | chr6-81472481-INS-66 |
| chr6-81473612-INS-400 | chr6-81473612-INS-400 |
| chr6-81537530-INS-108 | chr6-81537530-INS-106 |
| chr6-81539664-INS-85 | chr6-81539664-INS-85 |
| chr6-81547725-INS-1867 | chr6-81547726-INS-1862 |
| chr6-81572712-INS-66 | chr6-81572712-INS-50 |
| chr6-81582466-INS-70 | chr6-81582466-INS-66 |
| chr6-81598903-INS-69 | chr6-81598895-INS-74 |
| chr6-81603177-INS-124 | chr6-81603177-INS-132 |
| chr6-81649768-INS-66 | chr6-81649768-INS-86 |
| chr6-81664633-INS-6586 | chr6-81664634-INS-6586 |
| chr6-81668911-INS-52 | chr6-81668911-INS-52 |
| chr6-81671820-INS-7198 | chr6-81671821-INS-7199 |
| chr6-81696879-INS-98 | chr6-81696879-INS-122 |
| chr6-81709466-INS-4704 | chr6-81709467-INS-4704 |
| chr6-81723550-INS-64 | chr6-81723550-INS-60 |
| chr6-81734762-INS-59 | chr6-81734762-INS-59 |
| chr6-81783136-INS-70 | chr6-81783136-INS-70 |
| chr6-81798463-INS-173 | chr6-81798463-INS-173 |
| chr6-81821854-INS-195 | chr6-81821854-INS-196 |
| chr6-81893425-INS-66 | chr6-81893425-INS-66 |
| chr6-81896905-INS-1156 | chr6-81896906-INS-1156 |
| chr6-81909000-INS-192 | chr6-81909004-INS-193 |
| chr6-81931017-INS-120 | chr6-81931017-INS-120 |
| chr6-81950806-INS-72 | chr6-81950811-INS-58 |
| chr6-81954228-INS-400 | chr6-81954229-INS-400 |
| chr6-81966555-INS-52 | chr6-81966555-INS-50 |
| chr6-81989255-INS-62 | chr6-81989255-INS-64 |
| chr6-81994814-INS-2288 | chr6-81994815-INS-2288 |
| chr6-81997478-INS-58 | chr6-81997478-INS-72 |
| chr6-82000043-INS-66 | chr6-82000050-INS-104 |
| chr6-82025992-INS-97 | chr6-82025992-INS-97 |
| chr6-82034920-INS-63 | chr6-82034920-INS-63 |
| chr6-82048577-INS-90 | chr6-82048577-INS-52 |
| chr6-82058513-INS-205 | chr6-82058513-INS-207 |
| chr6-82064975-INS-134 | chr6-82064971-INS-136 |
| chr6-82066513-INS-774 | chr6-82066513-INS-779 |
| chr6-82083321-INS-51 | chr6-82083321-INS-72 |
| chr6-82100832-INS-80 | chr6-82100832-INS-84 |
| chr6-82114977-INS-58 | chr6-82114966-INS-84 |
| chr6-82122552-INS-96 | chr6-82122552-INS-96 |
| chr6-82141719-INS-93 | chr6-82141719-INS-93 |
| chr6-82148248-INS-64 | chr6-82148248-INS-64 |
| chr6-82170590-INS-471 | chr6-82170590-INS-471 |
| chr6-82204376-INS-108 | chr6-82204376-INS-72 |
| chr6-82212083-INS-202 | chr6-82212083-INS-202 |
| chr6-82218427-INS-68 | chr6-82218427-INS-68 |
| chr6-82220830-INS-54 | chr6-82220829-INS-87 |
| chr6-82242284-INS-68 | chr6-82242284-INS-68 |
| chr6-82249955-INS-136 | chr6-82249955-INS-107 |
| chr6-82257483-INS-52 | chr6-82257483-INS-52 |
| chr6-82261978-INS-58 | chr6-82261978-INS-62 |
| chr6-82265604-INS-5549 | chr6-82265605-INS-5549 |
| chr6-82286319-INS-90 | chr6-82286319-INS-86 |
| chr6-82304621-INS-51 | chr6-82304608-INS-57 |
| chr6-82309803-INS-68 | chr6-82309803-INS-68 |
| chr6-82313953-INS-399 | chr6-82313953-INS-400 |
| chr6-82315334-INS-529 | chr6-82315334-INS-528 |
| chr6-82323846-INS-212 | chr6-82323846-INS-212 |
| chr6-82342809-INS-60 | chr6-82342809-INS-52 |
| chr6-82343373-INS-124 | chr6-82343373-INS-124 |
| chr6-82346479-INS-6324 | chr6-82346484-INS-6347 |
| chr6-82346905-INS-123 | chr6-82346892-INS-110 |

| | |
|---|---|
| chr6-82351898-INS-5234 | chr6-82351899-INS-5235 |
| chr6-82355471-INS-1303 | chr6-82355472-INS-1303 |
| chr6-82371432-INS-1373 | chr6-82371433-INS-1373 |
| chr6-82394281-INS-530 | chr6-82394281-INS-530 |
| chr6-82396256-INS-60 | chr6-82396256-INS-96 |
| chr6-82416793-INS-139 | chr6-82416793-INS-139 |
| chr6-82417590-INS-6211 | chr6-82417591-INS-6211 |
| chr6-82479496-INS-114 | chr6-82479496-INS-118 |
| chr6-82481791-INS-60 | chr6-82481791-INS-88 |
| chr6-82483876-INS-56 | chr6-82483876-INS-56 |
| chr6-82509976-INS-328 | chr6-82509976-INS-328 |
| chr6-82513648-INS-5236 | chr6-82513649-INS-5236 |
| chr6-82531750-INS-178 | chr6-82531750-INS-178 |
| chr6-82533259-INS-73 | chr6-82533260-INS-76 |
| chr6-82535164-INS-62 | chr6-82535170-INS-54 |
| chr6-82540068-INS-73 | chr6-82540068-INS-73 |
| chr6-82557409-INS-51 | chr6-82557427-INS-63 |
| chr6-82558204-INS-887 | chr6-82558204-INS-887 |
| chr6-82571708-INS-1527 | chr6-82571709-INS-1527 |
| chr6-82581082-INS-133 | chr6-82581085-INS-89 |
| chr6-82610432-INS-142 | chr6-82610432-INS-146 |
| chr6-82612057-INS-211 | chr6-82612057-INS-211 |
| chr6-82613903-INS-56 | chr6-82613903-INS-55 |
| chr6-82631647-INS-602 | chr6-82631647-INS-597 |
| chr6-82636485-INS-10710 | chr6-82636486-INS-10711 |
| chr6-82646280-INS-68 | chr6-82646280-INS-68 |
| chr6-82666235-INS-64 | chr6-82666235-INS-64 |
| chr6-82690757-INS-82 | chr6-82690757-INS-72 |
| chr6-82698497-INS-81 | chr6-82698497-INS-81 |
| chr6-82699633-INS-443 | chr6-82699633-INS-449 |
| chr6-82717299-INS-245 | chr6-82717299-INS-245 |
| chr6-82724682-INS-203 | chr6-82724682-INS-203 |
| chr6-82753547-INS-64 | chr6-82753558-INS-65 |
| chr6-82758452-INS-120 | chr6-82758452-INS-120 |
| chr6-82772129-INS-292 | chr6-82772125-INS-323 |
| chr6-82773924-INS-174 | chr6-82773924-INS-128 |
| chr6-82778820-INS-82 | chr6-82778836-INS-71 |
| chr6-82782543-INS-106 | chr6-82782543-INS-112 |
| chr6-82788126-INS-406 | chr6-82788126-INS-406 |
| chr6-82789522-INS-56 | chr6-82789522-INS-52 |
| chr6-82794759-INS-66 | chr6-82794759-INS-66 |
| chr6-82809057-INS-54 | chr6-82809057-INS-60 |
| chr6-82813773-INS-576 | chr6-82813773-INS-576 |
| chr6-82825757-INS-52 | chr6-82825757-INS-50 |
| chr6-82830793-INS-282 | chr6-82830793-INS-282 |
| chr6-82833335-INS-194 | chr6-82833335-INS-194 |
| chr6-82870673-INS-85 | chr6-82870673-INS-71 |
| chr6-82883508-INS-206 | chr6-82883508-INS-209 |
| chr6-82931764-INS-68 | chr6-82931764-INS-68 |
| chr6-82959663-INS-346 | chr6-82959693-INS-297 |
| chr6-82963753-INS-185 | chr6-82963753-INS-185 |
| chr6-82972632-INS-69 | chr6-82972632-INS-66 |
| chr6-82978194-INS-132 | chr6-82978194-INS-182 |
| chr6-82988710-INS-72 | chr6-82988710-INS-76 |
| chr6-82990431-INS-184 | chr6-82990431-INS-184 |
| chr6-82996058-INS-87 | chr6-82996050-INS-119 |
| chr6-83018651-INS-193 | chr6-83018651-INS-191 |
| chr6-83023485-INS-150 | chr6-83023485-INS-156 |
| chr6-83051887-INS-78 | chr6-83051887-INS-78 |
| chr6-83066579-INS-54 | chr6-83066573-INS-50 |
| chr6-83081572-INS-84 | chr6-83081572-INS-85 |
| chr6-83088599-INS-146 | chr6-83088599-INS-146 |
| chr6-83103335-INS-205 | chr6-83103335-INS-205 |
| chr6-83110094-INS-91 | chr6-83110094-INS-91 |
| chr6-83127309-INS-124 | chr6-83127309-INS-124 |
| chr6-83144709-INS-239 | chr6-83144709-INS-238 |
| chr6-83147544-INS-70 | chr6-83147555-INS-70 |
| chr6-83177750-INS-66 | chr6-83177750-INS-66 |
| chr6-83204183-INS-68 | chr6-83204183-INS-68 |
| chr6-83212067-INS-102 | chr6-83212067-INS-102 |
| chr6-83216792-INS-210 | chr6-83216792-INS-210 |
| chr6-83218767-INS-64 | chr6-83218765-INS-60 |
| chr6-83224932-INS-86 | chr6-83224932-INS-86 |
| chr6-83253311-INS-190 | chr6-83253311-INS-190 |
| chr6-83255910-INS-107 | chr6-83255910-INS-107 |
| chr6-83260750-INS-70 | chr6-83260752-INS-84 |
| chr6-83264383-INS-91 | chr6-83264383-INS-94 |

| | |
|---|---|
| chr6-83270436-INS-124 | chr6-83270436-INS-124 |
| chr6-83302439-INS-207 | chr6-83302439-INS-207 |
| chr6-83310296-INS-80 | chr6-83310326-INS-64 |
| chr6-83355013-INS-243 | chr6-83355013-INS-244 |
| chr6-83365945-INS-95 | chr6-83365989-INS-95 |
| chr6-83394154-INS-80 | chr6-83394154-INS-80 |
| chr6-83396279-INS-72 | chr6-83396279-INS-72 |
| chr6-83430466-INS-61 | chr6-83430466-INS-61 |
| chr6-83438812-INS-287 | chr6-83438812-INS-287 |
| chr6-83450263-INS-68 | chr6-83450263-INS-70 |
| chr6-83455795-INS-178 | chr6-83455785-INS-201 |
| chr6-83472085-INS-112 | chr6-83472085-INS-112 |
| chr6-83480432-INS-82 | chr6-83480432-INS-83 |
| chr6-83494877-INS-134 | chr6-83494877-INS-146 |
| chr6-83512315-INS-70 | chr6-83512282-INS-60 |
| chr6-83521951-INS-68 | chr6-83521951-INS-70 |
| chr6-83523826-INS-78 | chr6-83523826-INS-78 |
| chr6-83544113-INS-56 | chr6-83544113-INS-56 |
| chr6-83556030-INS-129 | chr6-83556030-INS-129 |
| chr6-83561280-INS-73 | chr6-83561280-INS-51 |
| chr6-83579528-INS-52 | chr6-83579528-INS-52 |
| chr6-83600371-INS-3713 | chr6-83600372-INS-3730 |
| chr6-83613743-INS-60 | chr6-83613716-INS-87 |
| chr6-83627993-INS-148 | chr6-83627994-INS-98 |
| chr6-83635811-INS-138 | chr6-83635811-INS-138 |
| chr6-83638328-INS-203 | chr6-83638328-INS-203 |
| chr6-83655997-INS-56 | chr6-83655998-INS-60 |
| chr6-83658507-INS-6777 | chr6-83658508-INS-6777 |
| chr6-83669156-INS-61 | chr6-83669156-INS-61 |
| chr6-83676704-INS-185 | chr6-83676704-INS-186 |
| chr6-83685838-INS-212 | chr6-83685838-INS-213 |
| chr6-83700804-INS-111 | chr6-83700804-INS-111 |
| chr6-83702326-INS-215 | chr6-83702326-INS-215 |
| chr6-83708898-INS-113 | chr6-83708898-INS-116 |
| chr6-83730973-INS-1888 | chr6-83730974-INS-1888 |
| chr6-83793713-INS-74 | chr6-83793713-INS-74 |
| chr6-83816712-INS-119 | chr6-83816712-INS-121 |
| chr6-83821215-INS-56 | chr6-83821215-INS-68 |
| chr6-83849562-INS-216 | chr6-83849562-INS-217 |
| chr6-83865437-INS-65 | chr6-83865440-INS-53 |
| chr6-83896288-INS-176 | chr6-83896288-INS-176 |
| chr6-83916507-INS-235 | chr6-83916507-INS-244 |
| chr6-83979084-INS-310 | chr6-83979084-INS-310 |
| chr6-83981829-INS-80 | chr6-83981829-INS-83 |
| chr6-83986441-INS-57 | chr6-83986441-INS-60 |
| chr6-83997963-INS-73 | chr6-83997963-INS-117 |
| chr6-84001680-INS-156 | chr6-84001680-INS-122 |
| chr6-84005094-INS-54 | chr6-84005091-INS-52 |
| chr6-84013658-INS-173 | chr6-84013658-INS-185 |
| chr6-84017124-INS-58 | chr6-84017124-INS-54 |
| chr6-84018258-INS-82 | chr6-84018258-INS-87 |
| chr6-84035994-INS-5549 | chr6-84035995-INS-5549 |
| chr6-84037232-INS-518 | chr6-84037232-INS-539 |
| chr6-84055197-INS-80 | chr6-84055197-INS-80 |
| chr6-84056495-INS-62 | chr6-84056495-INS-58 |
| chr6-84067373-INS-53 | chr6-84067388-INS-77 |
| chr6-84069170-INS-182 | chr6-84069170-INS-182 |
| chr6-84081653-INS-222 | chr6-84081661-INS-199 |
| chr6-84091180-INS-199 | chr6-84091180-INS-199 |
| chr6-84104351-INS-275 | chr6-84104349-INS-259 |
| chr6-84117430-INS-77 | chr6-84117430-INS-77 |
| chr6-84120580-INS-157 | chr6-84120579-INS-157 |
| chr6-84149833-INS-68 | chr6-84149833-INS-68 |
| chr6-84154951-INS-78 | chr6-84154951-INS-82 |
| chr6-84170113-INS-397 | chr6-84170113-INS-396 |
| chr6-84170835-INS-60 | chr6-84170835-INS-66 |
| chr6-84189108-INS-52 | chr6-84189108-INS-52 |
| chr6-84219493-INS-787 | chr6-84219493-INS-787 |
| chr6-84234132-INS-57 | chr6-84234132-INS-57 |
| chr6-84240567-INS-141 | chr6-84240567-INS-136 |
| chr6-84248437-INS-305 | chr6-84248437-INS-305 |
| chr6-84250977-INS-159 | chr6-84250978-INS-159 |
| chr6-84256058-INS-2399 | chr6-84256059-INS-2399 |
| chr6-84283203-INS-70 | chr6-84283203-INS-72 |
| chr6-84289306-INS-88 | chr6-84289306-INS-87 |
| chr6-84304061-INS-94 | chr6-84304061-INS-100 |
| chr6-84316986-INS-216 | chr6-84316986-INS-215 |

| | |
|---|---|
| chr6-84341865-INS-88 | chr6-84341865-INS-88 |
| chr6-84346509-INS-1293 | chr6-84346510-INS-1293 |
| chr6-84369718-INS-422 | chr6-84369718-INS-422 |
| chr6-84374842-INS-812 | chr6-84374842-INS-811 |
| chr6-84392031-INS-235 | chr6-84392031-INS-235 |
| chr6-84399116-INS-116 | chr6-84399116-INS-116 |
| chr6-84437290-INS-83 | chr6-84437290-INS-83 |
| chr6-84450448-INS-52 | chr6-84450448-INS-52 |
| chr6-84499288-INS-104 | chr6-84499288-INS-104 |
| chr6-84501497-INS-138 | chr6-84501498-INS-146 |
| chr6-84513165-INS-196 | chr6-84513165-INS-196 |
| chr6-84521529-INS-225 | chr6-84521529-INS-225 |
| chr6-84535887-INS-98 | chr6-84535887-INS-105 |
| chr6-84537787-INS-83 | chr6-84537788-INS-75 |
| chr6-84579303-INS-82 | chr6-84579313-INS-52 |
| chr6-84584536-INS-382 | chr6-84584536-INS-382 |
| chr6-84640492-INS-5457 | chr6-84640493-INS-5457 |
| chr6-84683505-INS-2529 | chr6-84683506-INS-2528 |
| chr6-84687350-INS-60 | chr6-84687304-INS-75 |
| chr6-84846191-INS-7323 | chr6-84846192-INS-7324 |
| chr6-84907578-INS-289 | chr6-84907578-INS-290 |
| chr6-84909948-INS-63 | chr6-84909948-INS-51 |
| chr6-84922713-INS-517 | chr6-84922713-INS-521 |
| chr6-84924825-INS-6965 | chr6-84924826-INS-6971 |
| chr6-85047645-INS-52 | chr6-85047652-INS-54 |
| chr6-85066668-INS-2076 | chr6-85066669-INS-2034 |
| chr6-85141315-INS-131 | chr6-85141312-INS-125 |
| chr6-85162516-INS-82 | chr6-85162516-INS-82 |
| chr6-85251520-INS-69 | chr6-85251520-INS-69 |
| chr6-85296904-INS-173 | chr6-85296904-INS-169 |
| chr6-85359631-INS-209 | chr6-85359631-INS-209 |
| chr6-85370057-INS-104 | chr6-85370057-INS-104 |
| chr6-85390700-INS-214 | chr6-85390700-INS-232 |
| chr6-85397473-INS-183 | chr6-85397473-INS-185 |
| chr6-85407397-INS-117 | chr6-85407397-INS-117 |
| chr6-85484522-INS-158 | chr6-85484522-INS-160 |
| chr6-85492242-INS-58 | chr6-85492242-INS-58 |
| chr6-85498332-INS-100 | chr6-85498332-INS-99 |
| chr6-85505851-INS-64 | chr6-85505851-INS-66 |
| chr6-85517271-INS-117 | chr6-85517271-INS-117 |
| chr6-85536689-INS-6440 | chr6-85536690-INS-6440 |
| chr6-85546198-INS-55 | chr6-85546198-INS-55 |
| chr6-85555579-INS-105 | chr6-85555577-INS-109 |
| chr6-85576038-INS-55 | chr6-85576038-INS-57 |
| chr6-85580319-INS-713 | chr6-85580319-INS-712 |
| chr6-85610142-INS-60 | chr6-85610156-INS-56 |
| chr6-85612859-INS-226 | chr6-85612872-INS-263 |
| chr6-85725078-INS-54 | chr6-85725078-INS-84 |
| chr6-85772510-INS-168 | chr6-85772510-INS-168 |
| chr6-85777523-INS-56 | chr6-85777508-INS-56 |
| chr6-85816687-INS-72 | chr6-85816687-INS-56 |
| chr6-85820281-INS-60 | chr6-85820308-INS-52 |
| chr6-85839426-INS-72 | chr6-85839426-INS-72 |
| chr6-85865259-INS-195 | chr6-85865259-INS-187 |
| chr6-85865793-INS-355 | chr6-85865793-INS-355 |
| chr6-85908097-INS-54 | chr6-85908097-INS-52 |
| chr6-85972779-INS-86 | chr6-85972779-INS-86 |
| chr6-85994108-INS-62 | chr6-85994108-INS-62 |
| chr6-86012301-INS-106 | chr6-86012302-INS-66 |
| chr6-86023223-INS-1314 | chr6-86023224-INS-1314 |
| chr6-86025597-INS-55 | chr6-86025597-INS-55 |
| chr6-86037564-INS-237 | chr6-86037564-INS-237 |
| chr6-86044562-INS-230 | chr6-86044562-INS-230 |
| chr6-86052975-INS-65 | chr6-86052975-INS-58 |
| chr6-86058554-INS-56 | chr6-86058554-INS-60 |
| chr6-86067931-INS-75 | chr6-86067931-INS-75 |
| chr6-86072356-INS-63 | chr6-86072356-INS-67 |
| chr6-86081869-INS-508 | chr6-86081869-INS-508 |
| chr6-86103934-INS-65 | chr6-86103934-INS-65 |
| chr6-86126884-INS-106 | chr6-86126884-INS-82 |
| chr6-86165772-INS-325 | chr6-86165772-INS-372 |
| chr6-86177561-INS-62 | chr6-86177561-INS-63 |
| chr6-86187258-INS-6352 | chr6-86187259-INS-6353 |
| chr6-86193921-INS-158 | chr6-86193918-INS-154 |
| chr6-86233522-INS-211 | chr6-86233522-INS-213 |
| chr6-86288630-INS-3249 | chr6-86288631-INS-3251 |
| chr6-86301516-INS-9046 | chr6-86301517-INS-9046 |

| | |
|---|---|
| chr6-86305075-INS-283 | chr6-86305075-INS-298 |
| chr6-86314852-INS-56 | chr6-86314852-INS-56 |
| chr6-86331568-INS-128 | chr6-86331568-INS-120 |
| chr6-86337064-INS-144 | chr6-86337084-INS-138 |
| chr6-86350684-INS-206 | chr6-86350684-INS-206 |
| chr6-86377565-INS-102 | chr6-86377565-INS-108 |
| chr6-86389219-INS-58 | chr6-86389219-INS-60 |
| chr6-86399015-INS-219 | chr6-86399015-INS-220 |
| chr6-86424009-INS-70 | chr6-86424009-INS-70 |
| chr6-86446652-INS-201 | chr6-86446652-INS-200 |
| chr6-86457978-INS-172 | chr6-86457978-INS-172 |
| chr6-86461367-INS-209 | chr6-86461367-INS-209 |
| chr6-86471497-INS-164 | chr6-86471497-INS-164 |
| chr6-86474590-INS-88 | chr6-86474590-INS-88 |
| chr6-86480798-INS-171 | chr6-86480798-INS-169 |
| chr6-86495316-INS-56 | chr6-86495316-INS-88 |
| chr6-86530429-INS-92 | chr6-86530429-INS-90 |
| chr6-86542031-INS-75 | chr6-86542031-INS-75 |
| chr6-86549278-INS-98 | chr6-86549228-INS-116 |
| chr6-86557113-INS-81 | chr6-86557113-INS-81 |
| chr6-86559458-INS-51 | chr6-86559458-INS-55 |
| chr6-86567193-INS-167 | chr6-86567193-INS-166 |
| chr6-86571051-INS-100 | chr6-86571068-INS-114 |
| chr6-86579975-INS-66 | chr6-86579975-INS-74 |
| chr6-86611839-INS-82 | chr6-86611839-INS-82 |
| chr6-86617522-INS-102 | chr6-86617522-INS-102 |
| chr6-86618412-INS-177 | chr6-86618412-INS-175 |
| chr6-86658510-INS-218 | chr6-86658510-INS-217 |
| chr6-86665599-INS-445 | chr6-86665599-INS-445 |
| chr6-86680777-INS-206 | chr6-86680777-INS-213 |
| chr6-86690290-INS-70 | chr6-86690290-INS-70 |
| chr6-86691585-INS-201 | chr6-86691585-INS-201 |
| chr6-86715115-INS-76 | chr6-86715112-INS-52 |
| chr6-86719248-INS-52 | chr6-86719248-INS-52 |
| chr6-86726208-INS-86 | chr6-86726208-INS-86 |
| chr6-86739726-INS-63 | chr6-86739726-INS-63 |
| chr6-86783854-INS-163 | chr6-86783854-INS-205 |
| chr6-86784043-INS-507 | chr6-86784044-INS-507 |
| chr6-86793998-INS-60 | chr6-86794014-INS-63 |
| chr6-86801084-INS-273 | chr6-86801084-INS-273 |
| chr6-86857690-INS-196 | chr6-86857690-INS-196 |
| chr6-86874340-INS-222 | chr6-86874340-INS-222 |
| chr6-86893945-INS-67 | chr6-86893945-INS-67 |
| chr6-86910227-INS-90 | chr6-86910189-INS-90 |
| chr6-86926017-INS-209 | chr6-86926017-INS-209 |
| chr6-86941493-INS-168 | chr6-86941493-INS-168 |
| chr6-86953723-INS-233 | chr6-86953723-INS-233 |
| chr6-86959988-INS-203 | chr6-86959988-INS-203 |
| chr6-86986044-INS-774 | chr6-86986044-INS-775 |
| chr6-87008173-INS-74 | chr6-87008173-INS-74 |
| chr6-87009394-INS-215 | chr6-87009394-INS-215 |
| chr6-87012601-INS-209 | chr6-87012601-INS-209 |
| chr6-87020726-INS-184 | chr6-87020726-INS-184 |
| chr6-87024192-INS-54 | chr6-87024188-INS-66 |
| chr6-87026064-INS-445 | chr6-87026064-INS-445 |
| chr6-87027809-INS-170 | chr6-87027809-INS-171 |
| chr6-87037114-INS-753 | chr6-87037114-INS-753 |
| chr6-87054491-INS-66 | chr6-87054491-INS-66 |
| chr6-87088195-INS-102 | chr6-87088195-INS-54 |
| chr6-87098843-INS-440 | chr6-87098843-INS-440 |
| chr6-87099887-INS-108 | chr6-87099887-INS-140 |
| chr6-87137488-INS-63 | chr6-87137488-INS-63 |
| chr6-87141972-INS-72 | chr6-87141972-INS-72 |
| chr6-87146973-INS-282 | chr6-87146973-INS-283 |
| chr6-87151244-INS-57 | chr6-87151244-INS-66 |
| chr6-87160045-INS-221 | chr6-87160045-INS-224 |
| chr6-87188513-INS-134 | chr6-87188513-INS-136 |
| chr6-87193858-INS-85 | chr6-87193858-INS-85 |
| chr6-87199093-INS-71 | chr6-87199093-INS-71 |
| chr6-87202021-INS-659 | chr6-87202021-INS-660 |
| chr6-87205472-INS-636 | chr6-87205472-INS-636 |
| chr6-87252063-INS-63 | chr6-87252063-INS-63 |
| chr6-87279367-INS-84 | chr6-87279357-INS-91 |
| chr6-87297437-INS-563 | chr6-87297437-INS-563 |
| chr6-87300045-INS-61 | chr6-87300045-INS-53 |
| chr6-87302176-INS-170 | chr6-87302176-INS-171 |
| chr6-87307090-INS-60 | chr6-87307090-INS-60 |

| | |
|---|---|
| chr6-87307974-INS-68 | chr6-87307974-INS-68 |
| chr6-87309029-INS-325 | chr6-87309029-INS-326 |
| chr6-87335230-INS-54 | chr6-87335232-INS-54 |
| chr6-87337010-INS-860 | chr6-87337008-INS-863 |
| chr6-87346856-INS-110 | chr6-87346856-INS-110 |
| chr6-87359921-INS-114 | chr6-87359921-INS-76 |
| chr6-87374707-INS-57 | chr6-87374691-INS-90 |
| chr6-87413046-INS-84 | chr6-87413046-INS-50 |
| chr6-87434365-INS-1578 | chr6-87434366-INS-1578 |
| chr6-87434888-INS-2887 | chr6-87434907-INS-2869 |
| chr6-87434899-INS-662 | chr6-87434914-INS-663 |
| chr6-87456280-INS-234 | chr6-87456280-INS-234 |
| chr6-87459686-INS-2362 | chr6-87459687-INS-2380 |
| chr6-87498931-INS-70 | chr6-87498931-INS-70 |
| chr6-87524316-INS-526 | chr6-87524317-INS-526 |
| chr6-87550988-INS-53 | chr6-87550988-INS-53 |
| chr6-87560872-INS-64 | chr6-87560872-INS-64 |
| chr6-87573205-INS-135 | chr6-87573205-INS-135 |
| chr6-87576360-INS-92 | chr6-87576359-INS-51 |
| chr6-87593158-INS-118 | chr6-87593158-INS-126 |
| chr6-87607283-INS-58 | chr6-87607283-INS-58 |
| chr6-87623364-INS-454 | chr6-87623364-INS-454 |
| chr6-87629744-INS-57 | chr6-87629744-INS-57 |
| chr6-87660689-INS-285 | chr6-87660683-INS-329 |
| chr6-87667385-INS-70 | chr6-87667385-INS-70 |
| chr6-87672790-INS-202 | chr6-87672790-INS-202 |
| chr6-87686662-INS-208 | chr6-87686662-INS-208 |
| chr6-87702533-INS-1963 | chr6-87702534-INS-1963 |
| chr6-87704246-INS-124 | chr6-87704246-INS-124 |
| chr6-87711845-INS-110 | chr6-87711845-INS-154 |
| chr6-87844133-INS-68 | chr6-87844133-INS-68 |
| chr6-87858980-INS-136 | chr6-87858980-INS-142 |
| chr6-87862720-INS-60 | chr6-87862720-INS-60 |
| chr6-87864501-INS-204 | chr6-87864500-INS-197 |
| chr6-87865564-INS-142 | chr6-87865564-INS-135 |
| chr6-87873207-INS-67 | chr6-87873207-INS-67 |
| chr6-87878218-INS-187 | chr6-87878218-INS-205 |
| chr6-87923072-INS-115 | chr6-87923072-INS-115 |
| chr6-87932332-INS-58 | chr6-87932331-INS-51 |
| chr6-87936738-INS-209 | chr6-87936738-INS-209 |
| chr6-87940543-INS-4697 | chr6-87940544-INS-4697 |
| chr6-87974438-INS-101 | chr6-87974438-INS-106 |
| chr6-87985467-INS-169 | chr6-87985467-INS-169 |
| chr6-87989614-INS-62 | chr6-87989614-INS-58 |
| chr6-87993409-INS-210 | chr6-87993409-INS-210 |
| chr6-88012145-INS-63 | chr6-88012145-INS-63 |
| chr6-88015188-INS-186 | chr6-88015188-INS-182 |
| chr6-88034409-INS-52 | chr6-88034409-INS-52 |
| chr6-88035666-INS-200 | chr6-88035666-INS-200 |
| chr6-88038366-INS-856 | chr6-88038366-INS-856 |
| chr6-88040389-INS-127 | chr6-88040389-INS-127 |
| chr6-88042970-INS-200 | chr6-88042970-INS-200 |
| chr6-88044571-INS-76 | chr6-88044571-INS-88 |
| chr6-88046290-INS-92 | chr6-88046324-INS-67 |
| chr6-88055408-INS-117 | chr6-88055439-INS-114 |
| chr6-88115917-INS-72 | chr6-88115917-INS-72 |
| chr6-88134606-INS-7004 | chr6-88134607-INS-7004 |
| chr6-88160791-INS-2533 | chr6-88160792-INS-2529 |
| chr6-88161758-INS-109 | chr6-88161758-INS-109 |
| chr6-88164579-INS-284 | chr6-88164580-INS-306 |
| chr6-88220433-INS-93 | chr6-88220433-INS-62 |
| chr6-88267798-INS-1240 | chr6-88267799-INS-1240 |
| chr6-88325292-INS-65 | chr6-88325292-INS-65 |
| chr6-88357430-INS-217 | chr6-88357430-INS-219 |
| chr6-88364535-INS-58 | chr6-88364535-INS-58 |
| chr6-88407775-INS-1270 | chr6-88407776-INS-1270 |
| chr6-88429508-INS-5338 | chr6-88429509-INS-5338 |
| chr6-88432026-INS-175 | chr6-88432026-INS-175 |
| chr6-88435078-INS-74 | chr6-88435078-INS-50 |
| chr6-88444068-INS-194 | chr6-88444068-INS-195 |
| chr6-88446510-INS-206 | chr6-88446510-INS-206 |
| chr6-88452883-INS-93 | chr6-88452883-INS-93 |
| chr6-88493516-INS-66 | chr6-88493516-INS-96 |
| chr6-88503626-INS-202 | chr6-88503626-INS-202 |
| chr6-88514745-INS-54 | chr6-88514772-INS-71 |
| chr6-88537659-INS-50 | chr6-88537659-INS-52 |
| chr6-88560313-INS-356 | chr6-88560313-INS-356 |

| | |
|---|---|
| chr6-88590336-INS-76 | chr6-88590336-INS-76 |
| chr6-88612505-INS-6366 | chr6-88612506-INS-6366 |
| chr6-88621312-INS-633 | chr6-88621305-INS-635 |
| chr6-88638866-INS-57 | chr6-88638866-INS-57 |
| chr6-88644330-INS-189 | chr6-88644330-INS-189 |
| chr6-88668885-INS-72 | chr6-88668885-INS-73 |
| chr6-88673970-INS-83 | chr6-88673970-INS-83 |
| chr6-88693317-INS-2627 | chr6-88693318-INS-2632 |
| chr6-88741943-INS-63 | chr6-88741943-INS-51 |
| chr6-88756694-INS-197 | chr6-88756694-INS-205 |
| chr6-88761663-INS-6413 | chr6-88761664-INS-6413 |
| chr6-88782927-INS-201 | chr6-88782927-INS-201 |
| chr6-88793548-INS-58 | chr6-88793548-INS-58 |
| chr6-88823653-INS-260 | chr6-88823653-INS-258 |
| chr6-88832659-INS-90 | chr6-88832659-INS-94 |
| chr6-88849701-INS-128 | chr6-88849691-INS-121 |
| chr6-88865022-INS-142 | chr6-88865022-INS-142 |
| chr6-88885349-INS-145 | chr6-88885349-INS-145 |
| chr6-88954912-INS-80 | chr6-88954912-INS-75 |
| chr6-88959010-INS-100 | chr6-88959010-INS-80 |
| chr6-88967943-INS-250 | chr6-88967973-INS-232 |
| chr6-89001906-INS-434 | chr6-89001906-INS-434 |
| chr6-89011732-INS-2805 | chr6-89011733-INS-2805 |
| chr6-89038391-INS-70 | chr6-89038391-INS-64 |
| chr6-89082230-INS-65 | chr6-89082232-INS-57 |
| chr6-89183810-INS-681 | chr6-89183810-INS-681 |
| chr6-89207004-INS-414 | chr6-89207004-INS-415 |
| chr6-89230331-INS-81 | chr6-89230331-INS-96 |
| chr6-89254002-INS-222 | chr6-89254003-INS-227 |
| chr6-89255808-INS-51 | chr6-89255808-INS-55 |
| chr6-89263131-INS-234 | chr6-89263131-INS-232 |
| chr6-89264267-INS-70 | chr6-89264267-INS-62 |
| chr6-89267298-INS-205 | chr6-89267298-INS-205 |
| chr6-89283702-INS-1889 | chr6-89283703-INS-1889 |
| chr6-89297352-INS-167 | chr6-89297352-INS-167 |
| chr6-89372900-INS-65 | chr6-89372900-INS-65 |
| chr6-89388853-INS-51 | chr6-89388853-INS-51 |
| chr6-89420169-INS-52 | chr6-89420169-INS-52 |
| chr6-89481662-INS-87 | chr6-89481668-INS-63 |
| chr6-89579604-INS-173 | chr6-89579604-INS-177 |
| chr6-89657878-INS-96 | chr6-89657878-INS-96 |
| chr6-89691868-INS-415 | chr6-89691868-INS-417 |
| chr6-89740161-INS-6564 | chr6-89740162-INS-6564 |
| chr6-89774829-INS-70 | chr6-89774833-INS-71 |
| chr6-89786714-INS-847 | chr6-89786714-INS-847 |
| chr6-89797076-INS-669 | chr6-89797076-INS-669 |
| chr6-89852771-INS-324 | chr6-89852771-INS-324 |
| chr6-89902568-INS-4398 | chr6-89902569-INS-4398 |
| chr6-89919546-INS-4620 | chr6-89919547-INS-4620 |
| chr6-89923608-INS-5255 | chr6-89923609-INS-5255 |
| chr6-89933224-INS-7782 | chr6-89933225-INS-7784 |
| chr6-89956581-INS-1495 | chr6-89956582-INS-1495 |
| chr6-89981034-INS-203 | chr6-89981038-INS-203 |
| chr6-90032714-INS-80 | chr6-90032714-INS-80 |
| chr6-90037084-INS-917 | chr6-90037084-INS-924 |
| chr6-90083609-INS-67 | chr6-90083610-INS-50 |
| chr6-90089000-INS-198 | chr6-90089000-INS-201 |
| chr6-90128379-INS-65 | chr6-90128379-INS-65 |
| chr6-90162563-INS-7440 | chr6-90162564-INS-7440 |
| chr6-90247433-INS-66 | chr6-90247433-INS-58 |
| chr6-90264780-INS-107 | chr6-90264780-INS-108 |
| chr6-90277507-INS-1974 | chr6-90277508-INS-1973 |
| chr6-90285570-INS-87 | chr6-90285570-INS-87 |
| chr6-90294778-INS-102 | chr6-90294777-INS-58 |
| chr6-90330316-INS-55 | chr6-90330315-INS-56 |
| chr6-90465957-INS-56 | chr6-90465990-INS-61 |
| chr6-90493982-INS-105 | chr6-90493982-INS-111 |
| chr6-90530237-INS-56 | chr6-90530237-INS-54 |
| chr6-90541615-INS-127 | chr6-90541615-INS-127 |
| chr6-90543870-INS-222 | chr6-90543870-INS-222 |
| chr6-90569272-INS-126 | chr6-90569272-INS-126 |
| chr6-90609789-INS-115 | chr6-90609818-INS-83 |
| chr6-90650980-INS-64 | chr6-90650980-INS-60 |
| chr6-90660975-INS-6625 | chr6-90660976-INS-6625 |
| chr6-90727466-INS-202 | chr6-90727466-INS-202 |
| chr6-90767519-INS-60 | chr6-90767509-INS-60 |
| chr6-90787640-INS-564 | chr6-90787640-INS-566 |

| | |
|---|---|
| chr6-90809443-INS-292 | chr6-90809443-INS-292 |
| chr6-90812171-INS-52 | chr6-90812171-INS-54 |
| chr6-90991843-INS-406 | chr6-90991843-INS-406 |
| chr6-91004429-INS-207 | chr6-91004429-INS-207 |
| chr6-91026069-INS-127 | chr6-91026074-INS-124 |
| chr6-91038487-INS-222 | chr6-91038487-INS-219 |
| chr6-91066793-INS-217 | chr6-91066793-INS-217 |
| chr6-91094694-INS-815 | chr6-91094694-INS-815 |
| chr6-91142061-INS-98 | chr6-91142061-INS-110 |
| chr6-91152164-INS-78 | chr6-91152152-INS-70 |
| chr6-91192052-INS-172 | chr6-91192052-INS-172 |
| chr6-91250327-INS-54 | chr6-91250328-INS-54 |
| chr6-91275371-INS-1377 | chr6-91275372-INS-1382 |
| chr6-91373569-INS-60 | chr6-91373569-INS-60 |
| chr6-91398644-INS-247 | chr6-91398644-INS-247 |
| chr6-91419946-INS-411 | chr6-91419946-INS-407 |
| chr6-91420976-INS-252 | chr6-91420976-INS-252 |
| chr6-91429833-INS-459 | chr6-91429833-INS-454 |
| chr6-91437729-INS-203 | chr6-91437729-INS-203 |
| chr6-91441264-INS-215 | chr6-91441264-INS-216 |
| chr6-91511276-INS-54 | chr6-91511278-INS-50 |
| chr6-91551953-INS-66 | chr6-91551953-INS-58 |
| chr6-91564253-INS-134 | chr6-91564244-INS-104 |
| chr6-91596053-INS-446 | chr6-91596066-INS-433 |
| chr6-91605242-INS-342 | chr6-91605242-INS-342 |
| chr6-91614541-INS-71 | chr6-91614541-INS-69 |
| chr6-91626011-INS-126 | chr6-91626011-INS-90 |
| chr6-91690579-INS-7959 | chr6-91690580-INS-7959 |
| chr6-91795899-INS-97 | chr6-91795899-INS-105 |
| chr6-91796767-INS-102 | chr6-91796767-INS-90 |
| chr6-91802048-INS-335 | chr6-91802048-INS-343 |
| chr6-91810667-INS-69 | chr6-91810643-INS-107 |
| chr6-91831824-INS-1210 | chr6-91831825-INS-1209 |
| chr6-91846194-INS-62 | chr6-91846194-INS-76 |
| chr6-91847859-INS-326 | chr6-91847859-INS-342 |
| chr6-91853626-INS-108 | chr6-91853626-INS-110 |
| chr6-91888509-INS-88 | chr6-91888509-INS-75 |
| chr6-91914657-INS-517 | chr6-91914660-INS-518 |
| chr6-91970281-INS-57 | chr6-91970281-INS-63 |
| chr6-91977755-INS-68 | chr6-91977755-INS-68 |
| chr6-92002536-INS-109 | chr6-92002536-INS-110 |
| chr6-92002553-INS-1494 | chr6-92002554-INS-1494 |
| chr6-92025152-INS-97 | chr6-92025152-INS-81 |
| chr6-92073763-INS-1888 | chr6-92073764-INS-1888 |
| chr6-92077734-INS-213 | chr6-92077734-INS-208 |
| chr6-92116610-INS-136 | chr6-92116610-INS-171 |
| chr6-92119128-INS-200 | chr6-92119135-INS-204 |
| chr6-92159599-INS-106 | chr6-92159615-INS-80 |
| chr6-92162567-INS-1067 | chr6-92162577-INS-1049 |
| chr6-92182208-INS-7135 | chr6-92182209-INS-7135 |
| chr6-92191049-INS-150 | chr6-92191049-INS-120 |
| chr6-92196367-INS-82 | chr6-92196367-INS-83 |
| chr6-92270297-INS-60 | chr6-92270298-INS-76 |
| chr6-92272430-INS-57 | chr6-92272437-INS-56 |
| chr6-92278419-INS-641 | chr6-92278419-INS-634 |
| chr6-92304787-INS-208 | chr6-92304787-INS-208 |
| chr6-92309156-INS-844 | chr6-92309156-INS-844 |
| chr6-92321310-INS-391 | chr6-92321310-INS-391 |
| chr6-92324505-INS-210 | chr6-92324505-INS-211 |
| chr6-92331979-INS-1104 | chr6-92331980-INS-1104 |
| chr6-92365359-INS-1867 | chr6-92365360-INS-1867 |
| chr6-92366299-INS-51 | chr6-92366298-INS-51 |
| chr6-92383113-INS-52 | chr6-92383113-INS-52 |
| chr6-92419609-INS-54 | chr6-92419609-INS-58 |
| chr6-92452107-INS-96 | chr6-92452107-INS-94 |
| chr6-92484136-INS-191 | chr6-92484136-INS-191 |
| chr6-92494671-INS-219 | chr6-92494671-INS-219 |
| chr6-92516565-INS-52 | chr6-92516575-INS-78 |
| chr6-92542987-INS-1318 | chr6-92542988-INS-1318 |
| chr6-92551903-INS-6279 | chr6-92551904-INS-6278 |
| chr6-92566392-INS-63 | chr6-92566400-INS-52 |
| chr6-92576436-INS-64 | chr6-92576440-INS-60 |
| chr6-92593864-INS-60 | chr6-92593864-INS-64 |
| chr6-92613790-INS-4817 | chr6-92613791-INS-4817 |
| chr6-92630346-INS-199 | chr6-92630346-INS-199 |
| chr6-92650948-INS-186 | chr6-92650963-INS-215 |
| chr6-92720332-INS-116 | chr6-92720332-INS-116 |

| | |
|---|---|
| chr6-92724788-INS-121 | chr6-92724788-INS-121 |
| chr6-92749607-INS-52 | chr6-92749607-INS-52 |
| chr6-92764871-INS-96 | chr6-92764871-INS-120 |
| chr6-92767426-INS-51 | chr6-92767434-INS-83 |
| chr6-92770705-INS-63 | chr6-92770696-INS-67 |
| chr6-92844267-INS-85 | chr6-92844267-INS-90 |
| chr6-92856318-INS-138 | chr6-92856318-INS-138 |
| chr6-92869968-INS-112 | chr6-92869959-INS-118 |
| chr6-92875583-INS-134 | chr6-92875583-INS-134 |
| chr6-92885971-INS-53 | chr6-92885971-INS-53 |
| chr6-92890253-INS-81 | chr6-92890253-INS-85 |
| chr6-92895223-INS-104 | chr6-92895223-INS-102 |
| chr6-92916686-INS-54 | chr6-92916703-INS-57 |
| chr6-92951891-INS-208 | chr6-92951886-INS-171 |
| chr6-92984010-INS-402 | chr6-92984010-INS-402 |
| chr6-92995479-INS-52 | chr6-92995483-INS-57 |
| chr6-92999563-INS-51 | chr6-92999563-INS-51 |
| chr6-93004535-INS-87 | chr6-93004535-INS-87 |
| chr6-93018592-INS-280 | chr6-93018592-INS-280 |
| chr6-93031557-INS-352 | chr6-93031557-INS-352 |
| chr6-93056248-INS-59 | chr6-93056248-INS-60 |
| chr6-93069987-INS-66 | chr6-93069975-INS-68 |
| chr6-93089808-INS-266 | chr6-93089808-INS-252 |
| chr6-93093172-INS-63 | chr6-93093172-INS-63 |
| chr6-93120971-INS-71 | chr6-93120971-INS-74 |
| chr6-93124461-INS-159 | chr6-93124455-INS-160 |
| chr6-93128979-INS-103 | chr6-93129010-INS-80 |
| chr6-93129908-INS-324 | chr6-93129908-INS-324 |
| chr6-93137106-INS-76 | chr6-93137106-INS-76 |
| chr6-93152199-INS-203 | chr6-93152199-INS-203 |
| chr6-93180524-INS-6383 | chr6-93180525-INS-6383 |
| chr6-93184135-INS-52 | chr6-93184135-INS-58 |
| chr6-93192128-INS-110 | chr6-93192115-INS-90 |
| chr6-93208038-INS-140 | chr6-93208038-INS-140 |
| chr6-93211136-INS-366 | chr6-93211136-INS-390 |
| chr6-93230136-INS-52 | chr6-93230136-INS-68 |
| chr6-93232740-INS-101 | chr6-93232740-INS-107 |
| chr6-93255870-INS-100 | chr6-93255858-INS-136 |
| chr6-93269020-INS-167 | chr6-93269020-INS-165 |
| chr6-93273829-INS-6677 | chr6-93273830-INS-6674 |
| chr6-93299794-INS-143 | chr6-93299794-INS-123 |
| chr6-93301638-INS-84 | chr6-93301638-INS-86 |
| chr6-93316709-INS-54 | chr6-93316709-INS-54 |
| chr6-93324632-INS-1268 | chr6-93324633-INS-1268 |
| chr6-93336853-INS-1115 | chr6-93336854-INS-1115 |
| chr6-93348008-INS-66 | chr6-93348008-INS-62 |
| chr6-93377930-INS-473 | chr6-93377930-INS-473 |
| chr6-93395935-INS-62 | chr6-93395935-INS-58 |
| chr6-93407897-INS-55 | chr6-93407898-INS-54 |
| chr6-93412649-INS-2573 | chr6-93412650-INS-2573 |
| chr6-93424146-INS-70 | chr6-93424146-INS-77 |
| chr6-93444147-INS-58 | chr6-93444147-INS-58 |
| chr6-93471274-INS-56 | chr6-93471246-INS-56 |
| chr6-93493294-INS-382 | chr6-93493294-INS-382 |
| chr6-93497943-INS-270 | chr6-93497943-INS-270 |
| chr6-93507184-INS-95 | chr6-93507182-INS-92 |
| chr6-93509669-INS-6601 | chr6-93509670-INS-6601 |
| chr6-93511466-INS-57 | chr6-93511466-INS-57 |
| chr6-93516546-INS-197 | chr6-93516546-INS-197 |
| chr6-93522423-INS-68 | chr6-93522423-INS-64 |
| chr6-93523655-INS-59 | chr6-93523631-INS-80 |
| chr6-93527703-INS-197 | chr6-93527702-INS-190 |
| chr6-93548072-INS-106 | chr6-93548072-INS-112 |
| chr6-93561996-INS-6081 | chr6-93561997-INS-6081 |
| chr6-93569077-INS-253 | chr6-93569077-INS-249 |
| chr6-93573773-INS-86 | chr6-93573773-INS-84 |
| chr6-93589042-INS-6662 | chr6-93589043-INS-6662 |
| chr6-93608286-INS-70 | chr6-93608286-INS-66 |
| chr6-93635494-INS-125 | chr6-93635494-INS-120 |
| chr6-93641128-INS-533 | chr6-93641128-INS-533 |
| chr6-93653633-INS-167 | chr6-93653633-INS-167 |
| chr6-93657029-INS-187 | chr6-93657029-INS-188 |
| chr6-93696771-INS-69 | chr6-93696771-INS-69 |
| chr6-93711097-INS-217 | chr6-93711097-INS-218 |
| chr6-93730050-INS-96 | chr6-93730050-INS-96 |
| chr6-93741800-INS-52 | chr6-93741800-INS-52 |
| chr6-93744116-INS-52 | chr6-93744116-INS-52 |

| | |
|---|---|
| chr6-93753801-INS-954 | chr6-93753801-INS-954 |
| chr6-93760714-INS-70 | chr6-93760714-INS-70 |
| chr6-93767571-INS-7086 | chr6-93767572-INS-7086 |
| chr6-93791358-INS-153 | chr6-93791358-INS-153 |
| chr6-93800913-INS-58 | chr6-93800888-INS-59 |
| chr6-93825727-INS-329 | chr6-93825727-INS-329 |
| chr6-93826160-INS-5180 | chr6-93826161-INS-5180 |
| chr6-93832087-INS-344 | chr6-93832087-INS-344 |
| chr6-93834948-INS-3056 | chr6-93834949-INS-3056 |
| chr6-93843451-INS-60 | chr6-93843451-INS-60 |
| chr6-93846014-INS-407 | chr6-93846015-INS-407 |
| chr6-93864876-INS-1890 | chr6-93864877-INS-1890 |
| chr6-93893168-INS-54 | chr6-93893168-INS-54 |
| chr6-93894917-INS-51 | chr6-93894917-INS-51 |
| chr6-93900894-INS-61 | chr6-93900894-INS-61 |
| chr6-93908207-INS-55 | chr6-93908207-INS-90 |
| chr6-93921800-INS-58 | chr6-93921800-INS-58 |
| chr6-93958940-INS-52 | chr6-93958940-INS-56 |
| chr6-93965744-INS-88 | chr6-93965747-INS-64 |
| chr6-93977759-INS-180 | chr6-93977759-INS-180 |
| chr6-93982906-INS-154 | chr6-93982906-INS-139 |
| chr6-94004245-INS-229 | chr6-94004245-INS-229 |
| chr6-94006004-INS-54 | chr6-94006004-INS-54 |
| chr6-94015621-INS-262 | chr6-94015621-INS-262 |
| chr6-94023746-INS-52 | chr6-94023746-INS-54 |
| chr6-94027450-INS-60 | chr6-94027450-INS-60 |
| chr6-94073934-INS-139 | chr6-94073934-INS-139 |
| chr6-94075209-INS-1033 | chr6-94075210-INS-1033 |
| chr6-94075316-INS-58 | chr6-94075316-INS-58 |
| chr6-94077378-INS-195 | chr6-94077378-INS-195 |
| chr6-94118064-INS-60 | chr6-94118070-INS-66 |
| chr6-94197207-INS-207 | chr6-94197207-INS-204 |
| chr6-94205646-INS-98 | chr6-94205647-INS-98 |
| chr6-94206630-INS-205 | chr6-94206630-INS-206 |
| chr6-94210794-INS-953 | chr6-94210794-INS-954 |
| chr6-94226351-INS-209 | chr6-94226351-INS-208 |
| chr6-94257063-INS-81 | chr6-94257063-INS-81 |
| chr6-94262531-INS-574 | chr6-94262531-INS-574 |
| chr6-94284876-INS-104 | chr6-94284876-INS-104 |
| chr6-94289189-INS-90 | chr6-94289190-INS-94 |
| chr6-94291342-INS-66 | chr6-94291342-INS-66 |
| chr6-94306892-INS-234 | chr6-94306892-INS-232 |
| chr6-94315096-INS-61 | chr6-94315096-INS-63 |
| chr6-94323049-INS-91 | chr6-94323094-INS-86 |
| chr6-94329465-INS-215 | chr6-94329465-INS-216 |
| chr6-94331755-INS-109 | chr6-94331756-INS-108 |
| chr6-94336559-INS-56 | chr6-94336559-INS-60 |
| chr6-94348538-INS-74 | chr6-94348512-INS-62 |
| chr6-94354912-INS-62 | chr6-94354912-INS-70 |
| chr6-94359336-INS-180 | chr6-94359336-INS-180 |
| chr6-94368894-INS-75 | chr6-94368867-INS-81 |
| chr6-94379228-INS-63 | chr6-94379228-INS-63 |
| chr6-94393040-INS-185 | chr6-94393040-INS-185 |
| chr6-94402207-INS-234 | chr6-94402207-INS-235 |
| chr6-94409617-INS-211 | chr6-94409617-INS-210 |
| chr6-94412936-INS-132 | chr6-94412936-INS-138 |
| chr6-94424769-INS-55 | chr6-94424769-INS-56 |
| chr6-94437748-INS-88 | chr6-94437769-INS-54 |
| chr6-94446172-INS-170 | chr6-94446172-INS-170 |
| chr6-94460904-INS-60 | chr6-94460904-INS-60 |
| chr6-94469385-INS-358 | chr6-94469385-INS-358 |
| chr6-94481484-INS-201 | chr6-94481484-INS-200 |
| chr6-94490111-INS-217 | chr6-94490101-INS-191 |
| chr6-94492165-INS-129 | chr6-94492165-INS-129 |
| chr6-94495281-INS-60 | chr6-94495281-INS-60 |
| chr6-94522426-INS-135 | chr6-94522426-INS-135 |
| chr6-94555971-INS-194 | chr6-94555971-INS-194 |
| chr6-94640547-INS-208 | chr6-94640547-INS-204 |
| chr6-94642805-INS-51 | chr6-94642805-INS-53 |
| chr6-94697115-INS-430 | chr6-94697115-INS-430 |
| chr6-94706112-INS-61 | chr6-94706151-INS-54 |
| chr6-94735211-INS-56 | chr6-94735211-INS-56 |
| chr6-94748336-INS-204 | chr6-94748336-INS-203 |
| chr6-94781196-INS-123 | chr6-94781196-INS-127 |
| chr6-94826954-INS-135 | chr6-94826955-INS-115 |
| chr6-94832936-INS-579 | chr6-94832930-INS-576 |
| chr6-94841758-INS-51 | chr6-94841758-INS-51 |

| | |
|---|---|
| chr6-94852801-INS-75 | chr6-94852801-INS-78 |
| chr6-94874195-INS-220 | chr6-94874195-INS-220 |
| chr6-94894684-INS-50 | chr6-94894684-INS-52 |
| chr6-94897068-INS-86 | chr6-94897060-INS-96 |
| chr6-94918345-INS-91 | chr6-94918345-INS-109 |
| chr6-94930025-INS-52 | chr6-94930011-INS-58 |
| chr6-94934000-INS-6266 | chr6-94934001-INS-6268 |
| chr6-94942212-INS-626 | chr6-94942212-INS-626 |
| chr6-94950131-INS-561 | chr6-94950131-INS-562 |
| chr6-94956937-INS-3297 | chr6-94956938-INS-3297 |
| chr6-94994785-INS-63 | chr6-94994785-INS-62 |
| chr6-94997452-INS-152 | chr6-94997452-INS-152 |
| chr6-95029407-INS-226 | chr6-95029407-INS-224 |
| chr6-95037298-INS-219 | chr6-95037298-INS-220 |
| chr6-95048524-INS-1545 | chr6-95048525-INS-1545 |
| chr6-95074061-INS-60 | chr6-95074061-INS-63 |
| chr6-95093584-INS-54 | chr6-95093584-INS-68 |
| chr6-95117662-INS-95 | chr6-95117662-INS-103 |
| chr6-95125390-INS-83 | chr6-95125390-INS-83 |
| chr6-95126706-INS-656 | chr6-95126706-INS-656 |
| chr6-95128458-INS-269 | chr6-95128458-INS-269 |
| chr6-95131312-INS-7166 | chr6-95131313-INS-7166 |
| chr6-95134666-INS-57 | chr6-95134666-INS-61 |
| chr6-95147158-INS-65 | chr6-95147158-INS-65 |
| chr6-95152822-INS-120 | chr6-95152822-INS-120 |
| chr6-95168254-INS-978 | chr6-95168257-INS-982 |
| chr6-95235457-INS-188 | chr6-95235432-INS-192 |
| chr6-95251961-INS-114 | chr6-95251961-INS-130 |
| chr6-95260972-INS-228 | chr6-95260972-INS-229 |
| chr6-95261672-INS-174 | chr6-95261672-INS-174 |
| chr6-95264728-INS-235 | chr6-95264728-INS-209 |
| chr6-95271566-INS-828 | chr6-95271566-INS-829 |
| chr6-95308874-INS-7779 | chr6-95308875-INS-7779 |
| chr6-95323940-INS-88 | chr6-95323940-INS-88 |
| chr6-95367473-INS-64 | chr6-95367473-INS-69 |
| chr6-95370962-INS-6181 | chr6-95370963-INS-6181 |
| chr6-95394714-INS-102 | chr6-95394711-INS-64 |
| chr6-95421524-INS-73 | chr6-95421524-INS-73 |
| chr6-95430831-INS-6241 | chr6-95430835-INS-6233 |
| chr6-95474991-INS-453 | chr6-95474991-INS-453 |
| chr6-95477446-INS-4825 | chr6-95477447-INS-4825 |
| chr6-95482605-INS-598 | chr6-95482605-INS-605 |
| chr6-95487913-INS-66 | chr6-95487913-INS-66 |
| chr6-95540424-INS-3771 | chr6-95540425-INS-3765 |
| chr6-95542152-INS-396 | chr6-95542152-INS-396 |
| chr6-95553804-INS-398 | chr6-95553804-INS-398 |
| chr6-95579202-INS-108 | chr6-95579202-INS-108 |
| chr6-95595777-INS-827 | chr6-95595774-INS-831 |
| chr6-95608028-INS-56 | chr6-95608028-INS-56 |
| chr6-95616318-INS-146 | chr6-95616318-INS-146 |
| chr6-95630776-INS-55 | chr6-95630776-INS-55 |
| chr6-95655364-INS-188 | chr6-95655364-INS-188 |
| chr6-95655847-INS-5546 | chr6-95655848-INS-5546 |
| chr6-95659072-INS-83 | chr6-95659103-INS-51 |
| chr6-95667039-INS-79 | chr6-95667039-INS-79 |
| chr6-95712573-INS-80 | chr6-95712585-INS-108 |
| chr6-95715116-INS-57 | chr6-95715121-INS-57 |
| chr6-95723550-INS-894 | chr6-95723550-INS-898 |
| chr6-95734512-INS-113 | chr6-95734512-INS-64 |
| chr6-95740648-INS-167 | chr6-95740652-INS-168 |
| chr6-95746865-INS-225 | chr6-95746865-INS-209 |
| chr6-95770007-INS-661 | chr6-95770009-INS-672 |
| chr6-95798424-INS-6867 | chr6-95798425-INS-6867 |
| chr6-95838014-INS-216 | chr6-95838014-INS-216 |
| chr6-95838396-INS-7148 | chr6-95838397-INS-7148 |
| chr6-95870401-INS-204 | chr6-95870401-INS-204 |
| chr6-95915393-INS-196 | chr6-95915393-INS-184 |
| chr6-95925488-INS-87 | chr6-95925488-INS-87 |
| chr6-95979131-INS-172 | chr6-95979131-INS-172 |
| chr6-96006700-INS-104 | chr6-96006678-INS-126 |
| chr6-96019716-INS-61 | chr6-96019690-INS-69 |
| chr6-96043296-INS-54 | chr6-96043296-INS-52 |
| chr6-96127287-INS-192 | chr6-96127287-INS-192 |
| chr6-96146806-INS-6352 | chr6-96146807-INS-6352 |
| chr6-96146807-INS-215 | chr6-96146807-INS-213 |
| chr6-96153190-INS-71 | chr6-96153190-INS-58 |
| chr6-96158624-INS-86 | chr6-96158602-INS-90 |

| | |
|---|---|
| chr6-96164052-INS-55 | chr6-96164052-INS-60 |
| chr6-96197980-INS-553 | chr6-96197980-INS-551 |
| chr6-96208255-INS-106 | chr6-96208255-INS-106 |
| chr6-96212727-INS-83 | chr6-96212727-INS-83 |
| chr6-96221966-INS-342 | chr6-96221966-INS-338 |
| chr6-96252087-INS-68 | chr6-96252087-INS-68 |
| chr6-96290834-INS-715 | chr6-96290834-INS-714 |
| chr6-96300235-INS-524 | chr6-96300235-INS-525 |
| chr6-96332153-INS-133 | chr6-96332140-INS-111 |
| chr6-96366703-INS-66 | chr6-96366703-INS-66 |
| chr6-96375632-INS-64 | chr6-96375632-INS-60 |
| chr6-96394560-INS-9030 | chr6-96394561-INS-9031 |
| chr6-96435952-INS-52 | chr6-96435936-INS-50 |
| chr6-96437601-INS-1358 | chr6-96437602-INS-1356 |
| chr6-96445155-INS-697 | chr6-96445155-INS-697 |
| chr6-96477787-INS-73 | chr6-96477787-INS-73 |
| chr6-96485223-INS-101 | chr6-96485223-INS-101 |
| chr6-96486216-INS-1188 | chr6-96486217-INS-1188 |
| chr6-96499053-INS-281 | chr6-96499053-INS-281 |
| chr6-96517171-INS-214 | chr6-96517171-INS-215 |
| chr6-96519522-INS-77 | chr6-96519522-INS-77 |
| chr6-96520107-INS-4362 | chr6-96520108-INS-4362 |
| chr6-96542391-INS-645 | chr6-96542391-INS-645 |
| chr6-96546536-INS-60 | chr6-96546545-INS-61 |
| chr6-96615131-INS-133 | chr6-96615129-INS-131 |
| chr6-96616383-INS-1214 | chr6-96616384-INS-1215 |
| chr6-96638217-INS-6361 | chr6-96638218-INS-6362 |
| chr6-96646673-INS-78 | chr6-96646669-INS-84 |
| chr6-96664423-INS-100 | chr6-96664422-INS-65 |
| chr6-96685655-INS-412 | chr6-96685655-INS-412 |
| chr6-96692276-INS-167 | chr6-96692287-INS-135 |
| chr6-96708193-INS-52 | chr6-96708193-INS-52 |
| chr6-96712382-INS-80 | chr6-96712382-INS-80 |
| chr6-96728421-INS-59 | chr6-96728432-INS-53 |
| chr6-96763947-INS-64 | chr6-96763947-INS-64 |
| chr6-96823838-INS-56 | chr6-96823838-INS-56 |
| chr6-96852431-INS-69 | chr6-96852431-INS-69 |
| chr6-96895351-INS-52 | chr6-96895351-INS-52 |
| chr6-96905880-INS-179 | chr6-96905877-INS-177 |
| chr6-96911098-INS-604 | chr6-96911098-INS-604 |
| chr6-96978118-INS-479 | chr6-96978118-INS-479 |
| chr6-96981445-INS-283 | chr6-96981428-INS-308 |
| chr6-97026113-INS-63 | chr6-97026090-INS-106 |
| chr6-97026875-INS-72 | chr6-97026876-INS-66 |
| chr6-97032124-INS-1229 | chr6-97032125-INS-1231 |
| chr6-97033343-INS-133 | chr6-97033344-INS-111 |
| chr6-97055031-INS-400 | chr6-97055031-INS-400 |
| chr6-97092539-INS-56 | chr6-97092539-INS-50 |
| chr6-97104076-INS-104 | chr6-97104076-INS-107 |
| chr6-97105778-INS-68 | chr6-97105778-INS-68 |
| chr6-97113199-INS-292 | chr6-97113199-INS-288 |
| chr6-97118672-INS-208 | chr6-97118672-INS-208 |
| chr6-97156599-INS-200 | chr6-97156599-INS-203 |
| chr6-97164551-INS-197 | chr6-97164551-INS-197 |
| chr6-97193445-INS-166 | chr6-97193445-INS-166 |
| chr6-97195292-INS-57 | chr6-97195277-INS-59 |
| chr6-97202800-INS-85 | chr6-97202800-INS-115 |
| chr6-97213936-INS-62 | chr6-97213953-INS-88 |
| chr6-97221447-INS-285 | chr6-97221447-INS-285 |
| chr6-97235366-INS-110 | chr6-97235361-INS-123 |
| chr6-97240627-INS-68 | chr6-97240627-INS-68 |
| chr6-97260832-INS-369 | chr6-97260832-INS-369 |
| chr6-97282412-INS-210 | chr6-97282412-INS-214 |
| chr6-97291913-INS-58 | chr6-97291913-INS-73 |
| chr6-97312857-INS-78 | chr6-97312830-INS-82 |
| chr6-97339230-INS-54 | chr6-97339230-INS-58 |
| chr6-97347390-INS-54 | chr6-97347390-INS-58 |
| chr6-97379469-INS-67 | chr6-97379469-INS-55 |
| chr6-97422690-INS-53 | chr6-97422685-INS-52 |
| chr6-97433776-INS-77 | chr6-97433776-INS-77 |
| chr6-97452073-INS-6473 | chr6-97452074-INS-6473 |
| chr6-97452830-INS-174 | chr6-97452831-INS-134 |
| chr6-97458596-INS-213 | chr6-97458596-INS-206 |
| chr6-97472817-INS-79 | chr6-97472817-INS-79 |
| chr6-97516340-INS-58 | chr6-97516340-INS-58 |
| chr6-97550401-INS-6403 | chr6-97550402-INS-6403 |
| chr6-97576857-INS-91 | chr6-97576857-INS-99 |

| | |
|---|---|
| chr6-97581642-INS-102 | chr6-97581642-INS-102 |
| chr6-97663148-INS-140 | chr6-97663148-INS-144 |
| chr6-97664525-INS-507 | chr6-97664525-INS-507 |
| chr6-97688192-INS-64 | chr6-97688200-INS-60 |
| chr6-97726366-INS-114 | chr6-97726377-INS-114 |
| chr6-97735693-INS-64 | chr6-97735693-INS-62 |
| chr6-97747195-INS-146 | chr6-97747195-INS-120 |
| chr6-97763289-INS-1326 | chr6-97763290-INS-1326 |
| chr6-97861720-INS-89 | chr6-97861720-INS-89 |
| chr6-97892294-INS-56 | chr6-97892294-INS-56 |
| chr6-97899402-INS-1878 | chr6-97899403-INS-1878 |
| chr6-97902152-INS-225 | chr6-97902152-INS-225 |
| chr6-97908227-INS-528 | chr6-97908227-INS-529 |
| chr6-97913777-INS-54 | chr6-97913777-INS-58 |
| chr6-97957294-INS-86 | chr6-97957294-INS-86 |
| chr6-97986291-INS-88 | chr6-97986318-INS-85 |
| chr6-98011440-INS-87 | chr6-98011440-INS-89 |
| chr6-98013447-INS-931 | chr6-98013447-INS-931 |
| chr6-98035179-INS-181 | chr6-98035179-INS-185 |
| chr6-98039805-INS-95 | chr6-98039805-INS-95 |
| chr6-98052961-INS-101 | chr6-98052961-INS-110 |
| chr6-98063413-INS-4089 | chr6-98063414-INS-4089 |
| chr6-98069162-INS-53 | chr6-98069162-INS-53 |
| chr6-98097028-INS-78 | chr6-98097028-INS-60 |
| chr6-98118139-INS-72 | chr6-98118128-INS-56 |
| chr6-98176803-INS-71 | chr6-98176803-INS-51 |
| chr6-98191372-INS-294 | chr6-98191372-INS-294 |
| chr6-98196957-INS-190 | chr6-98196997-INS-193 |
| chr6-98226152-INS-367 | chr6-98226153-INS-380 |
| chr6-98285282-INS-52 | chr6-98285282-INS-50 |
| chr6-98297676-INS-57 | chr6-98297676-INS-54 |
| chr6-98308329-INS-6541 | chr6-98308330-INS-6541 |
| chr6-98309596-INS-53 | chr6-98309598-INS-51 |
| chr6-98334301-INS-50 | chr6-98334315-INS-80 |
| chr6-98350490-INS-150 | chr6-98350490-INS-150 |
| chr6-98357324-INS-126 | chr6-98357324-INS-126 |
| chr6-98380851-INS-179 | chr6-98380851-INS-179 |
| chr6-98382692-INS-742 | chr6-98382692-INS-742 |
| chr6-98412220-INS-228 | chr6-98412220-INS-228 |
| chr6-98419505-INS-349 | chr6-98419505-INS-349 |
| chr6-98429841-INS-52 | chr6-98429841-INS-52 |
| chr6-98445706-INS-828 | chr6-98445706-INS-828 |
| chr6-98464269-INS-54 | chr6-98464269-INS-54 |
| chr6-98520698-INS-180 | chr6-98520698-INS-181 |
| chr6-98571959-INS-108 | chr6-98571942-INS-84 |
| chr6-98606522-INS-63 | chr6-98606522-INS-62 |
| chr6-98625236-INS-288 | chr6-98625236-INS-299 |
| chr6-98629760-INS-52 | chr6-98629760-INS-52 |
| chr6-98658588-INS-72 | chr6-98658588-INS-68 |
| chr6-98671626-INS-72 | chr6-98671625-INS-76 |
| chr6-98743293-INS-70 | chr6-98743321-INS-75 |
| chr6-98750663-INS-52 | chr6-98750663-INS-50 |
| chr6-98788548-INS-52 | chr6-98788549-INS-52 |
| chr6-98813613-INS-53 | chr6-98813613-INS-53 |
| chr6-98822451-INS-203 | chr6-98822451-INS-203 |
| chr6-98839521-INS-2050 | chr6-98839522-INS-2047 |
| chr6-98864628-INS-498 | chr6-98864628-INS-498 |
| chr6-98866446-INS-144 | chr6-98866446-INS-108 |
| chr6-98867473-INS-52 | chr6-98867466-INS-80 |
| chr6-98961881-INS-67 | chr6-98961881-INS-67 |
| chr6-98967670-INS-55 | chr6-98967670-INS-55 |
| chr6-99036969-INS-187 | chr6-99036969-INS-167 |
| chr6-99043941-INS-188 | chr6-99043941-INS-191 |
| chr6-99083320-INS-3243 | chr6-99083321-INS-3244 |
| chr6-99124477-INS-67 | chr6-99124477-INS-66 |
| chr6-99216234-INS-242 | chr6-99216232-INS-244 |
| chr6-99240443-INS-66 | chr6-99240455-INS-52 |
| chr6-99264167-INS-60 | chr6-99264167-INS-60 |
| chr6-99277889-INS-1231 | chr6-99277890-INS-1245 |
| chr6-99315192-INS-126 | chr6-99315192-INS-126 |
| chr6-99324239-INS-159 | chr6-99324239-INS-159 |
| chr6-99366010-INS-54 | chr6-99366029-INS-55 |
| chr6-99442283-INS-346 | chr6-99442283-INS-366 |
| chr6-99453819-INS-100 | chr6-99453819-INS-100 |
| chr6-99511238-INS-69 | chr6-99511238-INS-69 |
| chr6-99593978-INS-99 | chr6-99593978-INS-51 |
| chr6-99628795-INS-100 | chr6-99628795-INS-50 |

| | |
|---|---|
| chr6-99695494-INS-590 | chr6-99695494-INS-590 |
| chr6-99729674-INS-78 | chr6-99729674-INS-90 |
| chr6-99732181-INS-236 | chr6-99732181-INS-241 |
| chr6-99735518-INS-106 | chr6-99735518-INS-108 |
| chr6-99762769-INS-974 | chr6-99762769-INS-976 |
| chr6-99766994-INS-120 | chr6-99766994-INS-115 |
| chr6-99785936-INS-82 | chr6-99785936-INS-88 |
| chr6-99803146-INS-58 | chr6-99803146-INS-58 |
| chr6-99808064-INS-59 | chr6-99808064-INS-59 |
| chr6-99815331-INS-52 | chr6-99815331-INS-65 |
| chr6-99838263-INS-358 | chr6-99838263-INS-358 |
| chr6-99845615-INS-153 | chr6-99845615-INS-153 |
| chr6-99863735-INS-217 | chr6-99863736-INS-218 |
| chr6-99881371-INS-232 | chr6-99881371-INS-241 |
| chr6-99883683-INS-110 | chr6-99883683-INS-113 |
| chr6-99903003-INS-82 | chr6-99903003-INS-83 |
| chr6-99914934-INS-184 | chr6-99914934-INS-184 |
| chr6-99918381-INS-503 | chr6-99918381-INS-503 |
| chr6-99925303-INS-132 | chr6-99925303-INS-132 |
| chr6-99938530-INS-99 | chr6-99938510-INS-106 |
| chr6-99951933-INS-112 | chr6-99951933-INS-114 |
| chr6-99961359-INS-116 | chr6-99961359-INS-116 |
| chr6-99993208-INS-80 | chr6-99993208-INS-64 |
| chr6-100004472-INS-118 | chr6-100004472-INS-121 |
| chr6-100017177-INS-94 | chr6-100017175-INS-78 |
| chr6-100177978-INS-520 | chr6-100177978-INS-520 |
| chr6-100187992-INS-222 | chr6-100187992-INS-222 |
| chr6-100191020-INS-50 | chr6-100191020-INS-54 |
| chr6-100202588-INS-67 | chr6-100202578-INS-52 |
| chr6-100222505-INS-56 | chr6-100222505-INS-54 |
| chr6-100224400-INS-1009 | chr6-100224396-INS-1002 |
| chr6-100229525-INS-90 | chr6-100229525-INS-90 |
| chr6-100238154-INS-62 | chr6-100238154-INS-62 |
| chr6-100259421-INS-378 | chr6-100259421-INS-396 |
| chr6-100267496-INS-200 | chr6-100267496-INS-200 |
| chr6-100271312-INS-58 | chr6-100271312-INS-58 |
| chr6-100285289-INS-207 | chr6-100285289-INS-207 |
| chr6-100287835-INS-96 | chr6-100287835-INS-102 |
| chr6-100302893-INS-320 | chr6-100302893-INS-321 |
| chr6-100306262-INS-198 | chr6-100306262-INS-200 |
| chr6-100312054-INS-199 | chr6-100312054-INS-199 |
| chr6-100327194-INS-60 | chr6-100327194-INS-60 |
| chr6-100331831-INS-177 | chr6-100331831-INS-177 |
| chr6-100338478-INS-387 | chr6-100338478-INS-389 |
| chr6-100339595-INS-55 | chr6-100339592-INS-62 |
| chr6-100346975-INS-56 | chr6-100346975-INS-58 |
| chr6-100348253-INS-68 | chr6-100348253-INS-68 |
| chr6-100370476-INS-212 | chr6-100370476-INS-212 |
| chr6-100374541-INS-206 | chr6-100374541-INS-206 |
| chr6-100376786-INS-54 | chr6-100376787-INS-54 |
| chr6-100379000-INS-80 | chr6-100379000-INS-80 |
| chr6-100389772-INS-84 | chr6-100389772-INS-84 |
| chr6-100419044-INS-244 | chr6-100419044-INS-244 |
| chr6-100424907-INS-54 | chr6-100424907-INS-54 |
| chr6-100425175-INS-3086 | chr6-100425176-INS-3111 |
| chr6-100430779-INS-375 | chr6-100430779-INS-375 |
| chr6-100432525-INS-68 | chr6-100432524-INS-112 |
| chr6-100440026-INS-64 | chr6-100440027-INS-64 |
| chr6-100442483-INS-64 | chr6-100442483-INS-80 |
| chr6-100443645-INS-171 | chr6-100443645-INS-171 |
| chr6-100465312-INS-752 | chr6-100465312-INS-749 |
| chr6-100474145-INS-98 | chr6-100474145-INS-98 |
| chr6-100477198-INS-52 | chr6-100477198-INS-75 |
| chr6-100479701-INS-52 | chr6-100479701-INS-52 |
| chr6-100483737-INS-87 | chr6-100483737-INS-87 |
| chr6-100491736-INS-194 | chr6-100491736-INS-194 |
| chr6-100506447-INS-105 | chr6-100506447-INS-134 |
| chr6-100526693-INS-182 | chr6-100526739-INS-184 |
| chr6-100532900-INS-60 | chr6-100532890-INS-84 |
| chr6-100566883-INS-129 | chr6-100566883-INS-130 |
| chr6-100570676-INS-104 | chr6-100570676-INS-104 |
| chr6-100574857-INS-54 | chr6-100574857-INS-54 |
| chr6-100602258-INS-429 | chr6-100602258-INS-429 |
| chr6-100611438-INS-180 | chr6-100611438-INS-180 |
| chr6-100635047-INS-62 | chr6-100635047-INS-62 |
| chr6-100643474-INS-80 | chr6-100643474-INS-80 |
| chr6-100690873-INS-150 | chr6-100690873-INS-150 |

| | |
|---|---|
| chr6-100697634-INS-100 | chr6-100697634-INS-100 |
| chr6-100705734-INS-200 | chr6-100705734-INS-199 |
| chr6-100736684-INS-640 | chr6-100736684-INS-638 |
| chr6-100784144-INS-58 | chr6-100784157-INS-65 |
| chr6-100832565-INS-125 | chr6-100832565-INS-127 |
| chr6-100834744-INS-954 | chr6-100834744-INS-954 |
| chr6-100853138-INS-88 | chr6-100853152-INS-89 |
| chr6-100853519-INS-436 | chr6-100853533-INS-424 |
| chr6-100858861-INS-622 | chr6-100858861-INS-622 |
| chr6-100878564-INS-59 | chr6-100878564-INS-59 |
| chr6-100905406-INS-57 | chr6-100905406-INS-69 |
| chr6-100922133-INS-52 | chr6-100922133-INS-68 |
| chr6-100923328-INS-165 | chr6-100923328-INS-171 |
| chr6-100935055-INS-54 | chr6-100935055-INS-60 |
| chr6-100944040-INS-92 | chr6-100944044-INS-56 |
| chr6-100955762-INS-77 | chr6-100955762-INS-69 |
| chr6-100959497-INS-288 | chr6-100959514-INS-297 |
| chr6-100969811-INS-122 | chr6-100969811-INS-122 |
| chr6-100992718-INS-183 | chr6-100992718-INS-183 |
| chr6-101003092-INS-342 | chr6-101003092-INS-342 |
| chr6-101022946-INS-84 | chr6-101022938-INS-80 |
| chr6-101032858-INS-57 | chr6-101032858-INS-51 |
| chr6-101034812-INS-295 | chr6-101034812-INS-281 |
| chr6-101050271-INS-306 | chr6-101050272-INS-302 |
| chr6-101064023-INS-106 | chr6-101064023-INS-106 |
| chr6-101096416-INS-80 | chr6-101096416-INS-80 |
| chr6-101097971-INS-225 | chr6-101097971-INS-223 |
| chr6-101109377-INS-249 | chr6-101109377-INS-228 |
| chr6-101217842-INS-888 | chr6-101217842-INS-888 |
| chr6-101249673-INS-79 | chr6-101249673-INS-79 |
| chr6-101251289-INS-54 | chr6-101251290-INS-54 |
| chr6-101280922-INS-404 | chr6-101280922-INS-404 |
| chr6-101283724-INS-64 | chr6-101283724-INS-64 |
| chr6-101301901-INS-67 | chr6-101301901-INS-67 |
| chr6-101353000-INS-88 | chr6-101353000-INS-88 |
| chr6-101374429-INS-771 | chr6-101374429-INS-771 |
| chr6-101378195-INS-81 | chr6-101378195-INS-81 |
| chr6-101410558-INS-59 | chr6-101410558-INS-61 |
| chr6-101413914-INS-1156 | chr6-101413915-INS-1156 |
| chr6-101438631-INS-394 | chr6-101438631-INS-395 |
| chr6-101440277-INS-175 | chr6-101440277-INS-175 |
| chr6-101448127-INS-102 | chr6-101448128-INS-132 |
| chr6-101467378-INS-52 | chr6-101467378-INS-50 |
| chr6-101480439-INS-166 | chr6-101480439-INS-165 |
| chr6-101488891-INS-53 | chr6-101488891-INS-57 |
| chr6-101491779-INS-527 | chr6-101491779-INS-527 |
| chr6-101513673-INS-278 | chr6-101513673-INS-278 |
| chr6-101524540-INS-51 | chr6-101524540-INS-51 |
| chr6-101543402-INS-69 | chr6-101543402-INS-69 |
| chr6-101554028-INS-204 | chr6-101554028-INS-204 |
| chr6-101563252-INS-207 | chr6-101563252-INS-207 |
| chr6-101585884-INS-51 | chr6-101585884-INS-51 |
| chr6-101634233-INS-53 | chr6-101634233-INS-58 |
| chr6-101645417-INS-109 | chr6-101645390-INS-129 |
| chr6-101675357-INS-781 | chr6-101675357-INS-783 |
| chr6-101680332-INS-170 | chr6-101680332-INS-164 |
| chr6-101744186-INS-1228 | chr6-101744187-INS-1228 |
| chr6-101744381-INS-175 | chr6-101744381-INS-219 |
| chr6-101756689-INS-52 | chr6-101756689-INS-60 |
| chr6-101777951-INS-1318 | chr6-101777952-INS-1319 |
| chr6-101788904-INS-56 | chr6-101788904-INS-56 |
| chr6-101793192-INS-3345 | chr6-101793193-INS-3346 |
| chr6-101793471-INS-195 | chr6-101793471-INS-195 |
| chr6-101812252-INS-61 | chr6-101812252-INS-66 |
| chr6-101816973-INS-529 | chr6-101816973-INS-529 |
| chr6-101827510-INS-60 | chr6-101827510-INS-72 |
| chr6-101842960-INS-85 | chr6-101842960-INS-50 |
| chr6-101847708-INS-406 | chr6-101847708-INS-406 |
| chr6-101880433-INS-81 | chr6-101880433-INS-81 |
| chr6-101893778-INS-531 | chr6-101893778-INS-531 |
| chr6-101905106-INS-142 | chr6-101905089-INS-167 |
| chr6-101907127-INS-671 | chr6-101907127-INS-676 |
| chr6-101908678-INS-1303 | chr6-101908679-INS-1304 |
| chr6-101911483-INS-60 | chr6-101911483-INS-60 |
| chr6-101949036-INS-692 | chr6-101949036-INS-702 |
| chr6-101977668-INS-111 | chr6-101977682-INS-72 |
| chr6-101985945-INS-133 | chr6-101985945-INS-133 |

| | |
|---|---|
| chr6-101993703-INS-60 | chr6-101993703-INS-60 |
| chr6-102030426-INS-98 | chr6-102030401-INS-87 |
| chr6-102037722-INS-3983 | chr6-102037723-INS-3983 |
| chr6-102045524-INS-132 | chr6-102045524-INS-132 |
| chr6-102056392-INS-6904 | chr6-102056393-INS-6905 |
| chr6-102068854-INS-2891 | chr6-102068855-INS-2891 |
| chr6-102075541-INS-6620 | chr6-102075542-INS-6620 |
| chr6-102085517-INS-212 | chr6-102085517-INS-212 |
| chr6-102091133-INS-67 | chr6-102091137-INS-59 |
| chr6-102099955-INS-309 | chr6-102099955-INS-309 |
| chr6-102116356-INS-123 | chr6-102116356-INS-123 |
| chr6-102120532-INS-199 | chr6-102120532-INS-199 |
| chr6-102166069-INS-87 | chr6-102166069-INS-87 |
| chr6-102171374-INS-75 | chr6-102171374-INS-75 |
| chr6-102173316-INS-86 | chr6-102173317-INS-86 |
| chr6-102242555-INS-90 | chr6-102242524-INS-91 |
| chr6-102284338-INS-74 | chr6-102284339-INS-53 |
| chr6-102339427-INS-54 | chr6-102339413-INS-58 |
| chr6-102350716-INS-472 | chr6-102350710-INS-493 |
| chr6-102354597-INS-5347 | chr6-102354598-INS-5347 |
| chr6-102389009-INS-203 | chr6-102389009-INS-203 |
| chr6-102498717-INS-498 | chr6-102498717-INS-498 |
| chr6-102509783-INS-366 | chr6-102509784-INS-366 |
| chr6-102511986-INS-81 | chr6-102511981-INS-97 |
| chr6-102558267-INS-352 | chr6-102558267-INS-353 |
| chr6-102583318-INS-132 | chr6-102583318-INS-132 |
| chr6-102598830-INS-280 | chr6-102598830-INS-281 |
| chr6-102628421-INS-605 | chr6-102628421-INS-605 |
| chr6-102632905-INS-62 | chr6-102632905-INS-64 |
| chr6-102644189-INS-212 | chr6-102644189-INS-212 |
| chr6-102667808-INS-233 | chr6-102667828-INS-247 |
| chr6-102669771-INS-370 | chr6-102669771-INS-367 |
| chr6-102680654-INS-4488 | chr6-102680655-INS-4488 |
| chr6-102726263-INS-84 | chr6-102726263-INS-86 |
| chr6-102727872-INS-101 | chr6-102727877-INS-60 |
| chr6-102738565-INS-852 | chr6-102738566-INS-852 |
| chr6-102759045-INS-199 | chr6-102759045-INS-199 |
| chr6-102783704-INS-2350 | chr6-102783705-INS-2349 |
| chr6-102786346-INS-6362 | chr6-102786347-INS-6363 |
| chr6-102813172-INS-427 | chr6-102813172-INS-434 |
| chr6-102820197-INS-86 | chr6-102820197-INS-100 |
| chr6-102823479-INS-6394 | chr6-102823480-INS-6396 |
| chr6-102836807-INS-279 | chr6-102836807-INS-280 |
| chr6-102842771-INS-1239 | chr6-102842772-INS-1209 |
| chr6-102857491-INS-2730 | chr6-102857492-INS-2730 |
| chr6-102874156-INS-888 | chr6-102874156-INS-889 |
| chr6-102878780-INS-727 | chr6-102878781-INS-727 |
| chr6-102879653-INS-1241 | chr6-102879654-INS-1241 |
| chr6-102882018-INS-315 | chr6-102882018-INS-315 |
| chr6-102912358-INS-440 | chr6-102912358-INS-440 |
| chr6-102921397-INS-649 | chr6-102921397-INS-649 |
| chr6-102943644-INS-53 | chr6-102943644-INS-53 |
| chr6-102954352-INS-7140 | chr6-102954353-INS-7140 |
| chr6-102956466-INS-574 | chr6-102956466-INS-574 |
| chr6-102971124-INS-781 | chr6-102971124-INS-781 |
| chr6-102975913-INS-6568 | chr6-102975914-INS-6568 |
| chr6-102978463-INS-52 | chr6-102978463-INS-58 |
| chr6-102981187-INS-11325 | chr6-102981188-INS-11325 |
| chr6-102986595-INS-201 | chr6-102986595-INS-201 |
| chr6-102986681-INS-647 | chr6-102986681-INS-647 |
| chr6-102987259-INS-547 | chr6-102987259-INS-548 |
| chr6-102996398-INS-1183 | chr6-102996399-INS-1184 |
| chr6-102996624-INS-94 | chr6-102996623-INS-52 |
| chr6-103016915-INS-127 | chr6-103016915-INS-127 |
| chr6-103035496-INS-419 | chr6-103035496-INS-419 |
| chr6-103062784-INS-65 | chr6-103062778-INS-52 |
| chr6-103066399-INS-51 | chr6-103066399-INS-51 |
| chr6-103083278-INS-211 | chr6-103083278-INS-211 |
| chr6-103086152-INS-60 | chr6-103086152-INS-60 |
| chr6-103089336-INS-52 | chr6-103089322-INS-57 |
| chr6-103125706-INS-64 | chr6-103125706-INS-56 |
| chr6-103138302-INS-96 | chr6-103138302-INS-102 |
| chr6-103145119-INS-70 | chr6-103145119-INS-60 |
| chr6-103176715-INS-92 | chr6-103176715-INS-94 |
| chr6-103184990-INS-6390 | chr6-103184991-INS-6390 |
| chr6-103231732-INS-308 | chr6-103231732-INS-309 |
| chr6-103242682-INS-98 | chr6-103242682-INS-52 |

| | |
|---|---|
| chr6-103260759-INS-3059 | chr6-103260760-INS-3059 |
| chr6-103264047-INS-499 | chr6-103264047-INS-499 |
| chr6-103278236-INS-83 | chr6-103278236-INS-81 |
| chr6-103301755-INS-118 | chr6-103301755-INS-133 |
| chr6-103305769-INS-60 | chr6-103305769-INS-60 |
| chr6-103320258-INS-85 | chr6-103320259-INS-95 |
| chr6-103345123-INS-338 | chr6-103345123-INS-332 |
| chr6-103350615-INS-100 | chr6-103350613-INS-102 |
| chr6-103356329-INS-62 | chr6-103356329-INS-62 |
| chr6-103366469-INS-566 | chr6-103366469-INS-564 |
| chr6-103378228-INS-528 | chr6-103378228-INS-528 |
| chr6-103415616-INS-897 | chr6-103415616-INS-897 |
| chr6-103432603-INS-54 | chr6-103432603-INS-54 |
| chr6-103433841-INS-67 | chr6-103433841-INS-67 |
| chr6-103434478-INS-94 | chr6-103434478-INS-96 |
| chr6-103435613-INS-395 | chr6-103435613-INS-395 |
| chr6-103443219-INS-588 | chr6-103443220-INS-591 |
| chr6-103462034-INS-80 | chr6-103462020-INS-68 |
| chr6-103481452-INS-57 | chr6-103481452-INS-57 |
| chr6-103516389-INS-126 | chr6-103516389-INS-126 |
| chr6-103523858-INS-95 | chr6-103523858-INS-50 |
| chr6-103531911-INS-215 | chr6-103531911-INS-217 |
| chr6-103535082-INS-60 | chr6-103535088-INS-63 |
| chr6-103568666-INS-83 | chr6-103568666-INS-83 |
| chr6-103585020-INS-72 | chr6-103585020-INS-66 |
| chr6-103633322-INS-67 | chr6-103633322-INS-67 |
| chr6-103653425-INS-6265 | chr6-103653426-INS-6266 |
| chr6-103663868-INS-88 | chr6-103663868-INS-88 |
| chr6-103664972-INS-488 | chr6-103664972-INS-488 |
| chr6-103680061-INS-58 | chr6-103680061-INS-58 |
| chr6-103694983-INS-667 | chr6-103694983-INS-667 |
| chr6-103707086-INS-52 | chr6-103707086-INS-60 |
| chr6-103721205-INS-53 | chr6-103721245-INS-50 |
| chr6-103754678-INS-522 | chr6-103754678-INS-523 |
| chr6-103777344-INS-439 | chr6-103777345-INS-439 |
| chr6-103786052-INS-76 | chr6-103786056-INS-56 |
| chr6-103826693-INS-400 | chr6-103826693-INS-400 |
| chr6-103834020-INS-237 | chr6-103834020-INS-237 |
| chr6-103866079-INS-70 | chr6-103866054-INS-50 |
| chr6-103876736-INS-91 | chr6-103876737-INS-56 |
| chr6-103888300-INS-2231 | chr6-103888307-INS-2254 |
| chr6-104020024-INS-70 | chr6-104020024-INS-75 |
| chr6-104020855-INS-318 | chr6-104020855-INS-318 |
| chr6-104027822-INS-98 | chr6-104027822-INS-105 |
| chr6-104066863-INS-82 | chr6-104066863-INS-66 |
| chr6-104073861-INS-5283 | chr6-104073862-INS-5283 |
| chr6-104082647-INS-126 | chr6-104082647-INS-162 |
| chr6-104088093-INS-56 | chr6-104088093-INS-56 |
| chr6-104095816-INS-72 | chr6-104095805-INS-54 |
| chr6-104108057-INS-347 | chr6-104108057-INS-346 |
| chr6-104131904-INS-3593 | chr6-104131905-INS-3593 |
| chr6-104143010-INS-6137 | chr6-104143011-INS-6137 |
| chr6-104172913-INS-1888 | chr6-104172914-INS-1888 |
| chr6-104177966-INS-911 | chr6-104177966-INS-911 |
| chr6-104193882-INS-52 | chr6-104193882-INS-52 |
| chr6-104204609-INS-60 | chr6-104204609-INS-92 |
| chr6-104208771-INS-121 | chr6-104208764-INS-84 |
| chr6-104220503-INS-78 | chr6-104220503-INS-78 |
| chr6-104224376-INS-238 | chr6-104224376-INS-239 |
| chr6-104247594-INS-359 | chr6-104247594-INS-360 |
| chr6-104263419-INS-56 | chr6-104263420-INS-56 |
| chr6-104267381-INS-55 | chr6-104267381-INS-55 |
| chr6-104289561-INS-70 | chr6-104289561-INS-62 |
| chr6-104311558-INS-598 | chr6-104311558-INS-599 |
| chr6-104446498-INS-101 | chr6-104446498-INS-101 |
| chr6-104448327-INS-313 | chr6-104448327-INS-313 |
| chr6-104452178-INS-498 | chr6-104452178-INS-498 |
| chr6-104499218-INS-878 | chr6-104499218-INS-878 |
| chr6-104533754-INS-136 | chr6-104533754-INS-136 |
| chr6-104567909-INS-108 | chr6-104567909-INS-113 |
| chr6-104596095-INS-545 | chr6-104596095-INS-545 |
| chr6-104601456-INS-56 | chr6-104601456-INS-76 |
| chr6-104645118-INS-1227 | chr6-104645119-INS-1226 |
| chr6-104665394-INS-127 | chr6-104665394-INS-127 |
| chr6-104683155-INS-6878 | chr6-104683156-INS-6877 |
| chr6-104706982-INS-1890 | chr6-104706983-INS-1891 |
| chr6-104717415-INS-84 | chr6-104717415-INS-88 |

| | |
|---|---|
| chr6-104803961-INS-67 | chr6-104803961-INS-67 |
| chr6-104827196-INS-54 | chr6-104827197-INS-54 |
| chr6-104830269-INS-712 | chr6-104830262-INS-726 |
| chr6-104831378-INS-54 | chr6-104831378-INS-54 |
| chr6-104848332-INS-57 | chr6-104848332-INS-59 |
| chr6-104884258-INS-80 | chr6-104884258-INS-82 |
| chr6-104885689-INS-336 | chr6-104885689-INS-336 |
| chr6-104901832-INS-1353 | chr6-104901833-INS-1353 |
| chr6-104916418-INS-401 | chr6-104916418-INS-400 |
| chr6-104917889-INS-3226 | chr6-104917890-INS-3226 |
| chr6-104921366-INS-6366 | chr6-104921367-INS-6366 |
| chr6-104925608-INS-98 | chr6-104925608-INS-93 |
| chr6-104953721-INS-494 | chr6-104953721-INS-494 |
| chr6-104956922-INS-6441 | chr6-104956923-INS-6442 |
| chr6-104958245-INS-91 | chr6-104958245-INS-105 |
| chr6-104978849-INS-5029 | chr6-104978850-INS-5029 |
| chr6-105016557-INS-7087 | chr6-105016558-INS-7087 |
| chr6-105029867-INS-162 | chr6-105029867-INS-162 |
| chr6-105031125-INS-94 | chr6-105031125-INS-96 |
| chr6-105039995-INS-398 | chr6-105039995-INS-398 |
| chr6-105069224-INS-61 | chr6-105069224-INS-61 |
| chr6-105079502-INS-6592 | chr6-105079503-INS-6592 |
| chr6-105095588-INS-199 | chr6-105095581-INS-199 |
| chr6-105096059-INS-4221 | chr6-105096060-INS-4221 |
| chr6-105177485-INS-201 | chr6-105177485-INS-201 |
| chr6-105199832-INS-56 | chr6-105199834-INS-52 |
| chr6-105239669-INS-1044 | chr6-105239670-INS-1036 |
| chr6-105285045-INS-124 | chr6-105285041-INS-128 |
| chr6-105301153-INS-56 | chr6-105301157-INS-57 |
| chr6-105310628-INS-66 | chr6-105310600-INS-53 |
| chr6-105329098-INS-198 | chr6-105329098-INS-199 |
| chr6-105331308-INS-62 | chr6-105331308-INS-60 |
| chr6-105339266-INS-321 | chr6-105339266-INS-321 |
| chr6-105351300-INS-1448 | chr6-105351301-INS-1448 |
| chr6-105361544-INS-102 | chr6-105361538-INS-68 |
| chr6-105376910-INS-160 | chr6-105376906-INS-120 |
| chr6-105405951-INS-174 | chr6-105405951-INS-174 |
| chr6-105502022-INS-52 | chr6-105502022-INS-52 |
| chr6-105503827-INS-80 | chr6-105503827-INS-112 |
| chr6-105618337-INS-8311 | chr6-105618338-INS-8312 |
| chr6-105633525-INS-135 | chr6-105633525-INS-132 |
| chr6-105640405-INS-206 | chr6-105640406-INS-206 |
| chr6-105658919-INS-56 | chr6-105658911-INS-94 |
| chr6-105697677-INS-7779 | chr6-105697678-INS-7779 |
| chr6-105740841-INS-66 | chr6-105740841-INS-50 |
| chr6-105758422-INS-76 | chr6-105758415-INS-60 |
| chr6-105800294-INS-205 | chr6-105800294-INS-204 |
| chr6-105830003-INS-76 | chr6-105830003-INS-74 |
| chr6-105839552-INS-470 | chr6-105839557-INS-466 |
| chr6-105853758-INS-661 | chr6-105853758-INS-660 |
| chr6-105854192-INS-6286 | chr6-105854193-INS-6286 |
| chr6-105854866-INS-803 | chr6-105854866-INS-803 |
| chr6-105908575-INS-95 | chr6-105908575-INS-95 |
| chr6-105924144-INS-226 | chr6-105924144-INS-226 |
| chr6-105937033-INS-208 | chr6-105937033-INS-208 |
| chr6-105972879-INS-607 | chr6-105972879-INS-607 |
| chr6-105979947-INS-52 | chr6-105979947-INS-52 |
| chr6-106029300-INS-60 | chr6-106029300-INS-60 |
| chr6-106044848-INS-1325 | chr6-106044849-INS-1325 |
| chr6-106058756-INS-98 | chr6-106058756-INS-102 |
| chr6-106119078-INS-80 | chr6-106119123-INS-50 |
| chr6-106126016-INS-90 | chr6-106125996-INS-75 |
| chr6-106128843-INS-54 | chr6-106128843-INS-54 |
| chr6-106136561-INS-251 | chr6-106136562-INS-251 |
| chr6-106147482-INS-63 | chr6-106147482-INS-63 |
| chr6-106148620-INS-6678 | chr6-106148621-INS-6679 |
| chr6-106169023-INS-52 | chr6-106169023-INS-52 |
| chr6-106201538-INS-63 | chr6-106201538-INS-105 |
| chr6-106202306-INS-167 | chr6-106202306-INS-167 |
| chr6-106221901-INS-100 | chr6-106221928-INS-69 |
| chr6-106237560-INS-189 | chr6-106237560-INS-189 |
| chr6-106249740-INS-54 | chr6-106249740-INS-52 |
| chr6-106269587-INS-173 | chr6-106269587-INS-173 |
| chr6-106280241-INS-67 | chr6-106280241-INS-52 |
| chr6-106287099-INS-1319 | chr6-106287100-INS-1319 |
| chr6-106292272-INS-204 | chr6-106292272-INS-204 |
| chr6-106347520-INS-112 | chr6-106347520-INS-112 |

| | |
|---|---|
| chr6-106354624-INS-311 | chr6-106354624-INS-309 |
| chr6-106382477-INS-90 | chr6-106382480-INS-129 |
| chr6-106385657-INS-72 | chr6-106385657-INS-78 |
| chr6-106403992-INS-613 | chr6-106403992-INS-613 |
| chr6-106404065-INS-6602 | chr6-106404066-INS-6602 |
| chr6-106407060-INS-549 | chr6-106407060-INS-549 |
| chr6-106410058-INS-69 | chr6-106410058-INS-60 |
| chr6-106419476-INS-7808 | chr6-106419477-INS-7809 |
| chr6-106425076-INS-382 | chr6-106425076-INS-386 |
| chr6-106431620-INS-66 | chr6-106431620-INS-66 |
| chr6-106433514-INS-3389 | chr6-106433515-INS-3389 |
| chr6-106434541-INS-379 | chr6-106434541-INS-379 |
| chr6-106437117-INS-58 | chr6-106437117-INS-58 |
| chr6-106437214-INS-7705 | chr6-106437215-INS-7706 |
| chr6-106508912-INS-2690 | chr6-106508913-INS-2690 |
| chr6-106529720-INS-457 | chr6-106529720-INS-472 |
| chr6-106544992-INS-452 | chr6-106544988-INS-468 |
| chr6-106546783-INS-85 | chr6-106546783-INS-70 |
| chr6-106556408-INS-54 | chr6-106556408-INS-54 |
| chr6-106582197-INS-259 | chr6-106582202-INS-260 |
| chr6-106609146-INS-52 | chr6-106609146-INS-52 |
| chr6-106612509-INS-3614 | chr6-106612510-INS-3615 |
| chr6-106621778-INS-672 | chr6-106621778-INS-665 |
| chr6-106624984-INS-198 | chr6-106624983-INS-156 |
| chr6-106633213-INS-64 | chr6-106633213-INS-64 |
| chr6-106681139-INS-51 | chr6-106681139-INS-52 |
| chr6-106687367-INS-161 | chr6-106687367-INS-167 |
| chr6-106745502-INS-96 | chr6-106745499-INS-62 |
| chr6-106769120-INS-60 | chr6-106769120-INS-60 |
| chr6-106770177-INS-126 | chr6-106770176-INS-147 |
| chr6-106779630-INS-72 | chr6-106779630-INS-72 |
| chr6-106781010-INS-52 | chr6-106780992-INS-54 |
| chr6-106820991-INS-428 | chr6-106820991-INS-428 |
| chr6-106824338-INS-1276 | chr6-106824339-INS-1276 |
| chr6-106831642-INS-84 | chr6-106831646-INS-96 |
| chr6-106835308-INS-601 | chr6-106835308-INS-601 |
| chr6-106837700-INS-153 | chr6-106837700-INS-153 |
| chr6-106845201-INS-182 | chr6-106845201-INS-182 |
| chr6-106903584-INS-73 | chr6-106903584-INS-77 |
| chr6-106912919-INS-192 | chr6-106912919-INS-192 |
| chr6-106930258-INS-736 | chr6-106930258-INS-736 |
| chr6-106948270-INS-247 | chr6-106948270-INS-250 |
| chr6-106962054-INS-1892 | chr6-106962055-INS-1892 |
| chr6-106972859-INS-60 | chr6-106972859-INS-50 |
| chr6-106984463-INS-427 | chr6-106984463-INS-427 |
| chr6-107017695-INS-232 | chr6-107017695-INS-232 |
| chr6-107022581-INS-356 | chr6-107022581-INS-356 |
| chr6-107034006-INS-396 | chr6-107034010-INS-363 |
| chr6-107036151-INS-65 | chr6-107036133-INS-70 |
| chr6-107047122-INS-52 | chr6-107047122-INS-52 |
| chr6-107048372-INS-8834 | chr6-107048373-INS-8834 |
| chr6-107059104-INS-83 | chr6-107059104-INS-83 |
| chr6-107061208-INS-178 | chr6-107061208-INS-178 |
| chr6-107082549-INS-58 | chr6-107082549-INS-62 |
| chr6-107083190-INS-62 | chr6-107083190-INS-50 |
| chr6-107091379-INS-70 | chr6-107091379-INS-64 |
| chr6-107103374-INS-3143 | chr6-107103375-INS-3143 |
| chr6-107130988-INS-58 | chr6-107130988-INS-62 |
| chr6-107136963-INS-69 | chr6-107136943-INS-72 |
| chr6-107190996-INS-4545 | chr6-107190997-INS-4549 |
| chr6-107208266-INS-54 | chr6-107208266-INS-50 |
| chr6-107218446-INS-182 | chr6-107218446-INS-183 |
| chr6-107288969-INS-73 | chr6-107288969-INS-75 |
| chr6-107306761-INS-208 | chr6-107306761-INS-208 |
| chr6-107336253-INS-65 | chr6-107336253-INS-67 |
| chr6-107364755-INS-88 | chr6-107364755-INS-94 |
| chr6-107366052-INS-71 | chr6-107366053-INS-98 |
| chr6-107368098-INS-6228 | chr6-107368099-INS-6229 |
| chr6-107381343-INS-2170 | chr6-107381344-INS-2170 |
| chr6-107387065-INS-8732 | chr6-107387066-INS-8732 |
| chr6-107411565-INS-6445 | chr6-107411566-INS-6446 |
| chr6-107413359-INS-735 | chr6-107413360-INS-735 |
| chr6-107417651-INS-6982 | chr6-107417638-INS-6984 |
| chr6-107420199-INS-54 | chr6-107420169-INS-94 |
| chr6-107435146-INS-8484 | chr6-107435147-INS-8485 |
| chr6-107439833-INS-150 | chr6-107439836-INS-114 |
| chr6-107454472-INS-111 | chr6-107454473-INS-126 |

| | |
|---|---|
| chr6-107474712-INS-399 | chr6-107474712-INS-399 |
| chr6-107505866-INS-68 | chr6-107505866-INS-62 |
| chr6-107519750-INS-871 | chr6-107519750-INS-871 |
| chr6-107526728-INS-84 | chr6-107526728-INS-50 |
| chr6-107529887-INS-319 | chr6-107529887-INS-320 |
| chr6-107550509-INS-6424 | chr6-107550510-INS-6424 |
| chr6-107554749-INS-368 | chr6-107554741-INS-335 |
| chr6-107570278-INS-56 | chr6-107570278-INS-60 |
| chr6-107617463-INS-1141 | chr6-107617464-INS-1144 |
| chr6-107624549-INS-54 | chr6-107624549-INS-54 |
| chr6-107625865-INS-4454 | chr6-107625866-INS-4454 |
| chr6-107682694-INS-6887 | chr6-107682695-INS-6888 |
| chr6-107708607-INS-68 | chr6-107708597-INS-50 |
| chr6-107714233-INS-6440 | chr6-107714234-INS-6440 |
| chr6-107727395-INS-6777 | chr6-107727396-INS-6777 |
| chr6-107728246-INS-180 | chr6-107728243-INS-183 |
| chr6-107745410-INS-204 | chr6-107745410-INS-204 |
| chr6-107751912-INS-1005 | chr6-107751913-INS-1005 |
| chr6-107779626-INS-7664 | chr6-107779627-INS-7665 |
| chr6-107803521-INS-175 | chr6-107803521-INS-175 |
| chr6-107811351-INS-347 | chr6-107811351-INS-347 |
| chr6-107823245-INS-148 | chr6-107823245-INS-140 |
| chr6-107824245-INS-192 | chr6-107824245-INS-192 |
| chr6-107839568-INS-2454 | chr6-107839569-INS-2454 |
| chr6-107866153-INS-319 | chr6-107866153-INS-323 |
| chr6-107868293-INS-596 | chr6-107868302-INS-595 |
| chr6-107884776-INS-1047 | chr6-107884777-INS-1045 |
| chr6-107906836-INS-567 | chr6-107906836-INS-567 |
| chr6-107912430-INS-295 | chr6-107912430-INS-297 |
| chr6-107958151-INS-704 | chr6-107958151-INS-700 |
| chr6-107998297-INS-52 | chr6-107998297-INS-55 |
| chr6-108007466-INS-105 | chr6-108007466-INS-130 |
| chr6-108016854-INS-785 | chr6-108016854-INS-785 |
| chr6-108024117-INS-6436 | chr6-108024118-INS-6436 |
| chr6-108041258-INS-974 | chr6-108041258-INS-972 |
| chr6-108055936-INS-214 | chr6-108055936-INS-220 |
| chr6-108065789-INS-56 | chr6-108065789-INS-54 |
| chr6-108068396-INS-249 | chr6-108068396-INS-249 |
| chr6-108076109-INS-243 | chr6-108076110-INS-198 |
| chr6-108078664-INS-281 | chr6-108078664-INS-283 |
| chr6-108108917-INS-275 | chr6-108108917-INS-269 |
| chr6-108110149-INS-160 | chr6-108110149-INS-160 |
| chr6-108151646-INS-81 | chr6-108151646-INS-81 |
| chr6-108161259-INS-202 | chr6-108161259-INS-201 |
| chr6-108167249-INS-477 | chr6-108167249-INS-477 |
| chr6-108253274-INS-197 | chr6-108253274-INS-197 |
| chr6-108256263-INS-62 | chr6-108256263-INS-56 |
| chr6-108262519-INS-54 | chr6-108262520-INS-51 |
| chr6-108322623-INS-134 | chr6-108322623-INS-113 |
| chr6-108327603-INS-58 | chr6-108327603-INS-58 |
| chr6-108331061-INS-54 | chr6-108331061-INS-54 |
| chr6-108354235-INS-95 | chr6-108354235-INS-95 |
| chr6-108373550-INS-65 | chr6-108373547-INS-108 |
| chr6-108378958-INS-63 | chr6-108378958-INS-63 |
| chr6-108382513-INS-52 | chr6-108382513-INS-52 |
| chr6-108395981-INS-65 | chr6-108395981-INS-61 |
| chr6-108472764-INS-2943 | chr6-108472765-INS-2943 |
| chr6-108485727-INS-226 | chr6-108485727-INS-199 |
| chr6-108497382-INS-56 | chr6-108497345-INS-60 |
| chr6-108536590-INS-120 | chr6-108536591-INS-120 |
| chr6-108547193-INS-60 | chr6-108547202-INS-69 |
| chr6-108553871-INS-263 | chr6-108553871-INS-264 |
| chr6-108609787-INS-7096 | chr6-108609788-INS-7096 |
| chr6-108615098-INS-7397 | chr6-108615099-INS-7397 |
| chr6-108620429-INS-54 | chr6-108620445-INS-64 |
| chr6-108622157-INS-57 | chr6-108622157-INS-57 |
| chr6-108627888-INS-206 | chr6-108627888-INS-208 |
| chr6-108647091-INS-51 | chr6-108647091-INS-51 |
| chr6-108667155-INS-168 | chr6-108667155-INS-178 |
| chr6-108671838-INS-53 | chr6-108671838-INS-53 |
| chr6-108674003-INS-134 | chr6-108674003-INS-134 |
| chr6-108683444-INS-56 | chr6-108683444-INS-64 |
| chr6-108689836-INS-63 | chr6-108689836-INS-63 |
| chr6-108693324-INS-98 | chr6-108693324-INS-54 |
| chr6-108695338-INS-6441 | chr6-108695339-INS-6439 |
| chr6-108715058-INS-73 | chr6-108715058-INS-73 |
| chr6-108727740-INS-181 | chr6-108727740-INS-182 |

| | |
|---|---|
| chr6-108744387-INS-54 | chr6-108744385-INS-92 |
| chr6-108747773-INS-199 | chr6-108747773-INS-199 |
| chr6-108749155-INS-73 | chr6-108749155-INS-122 |
| chr6-108756929-INS-351 | chr6-108756929-INS-351 |
| chr6-108778668-INS-4957 | chr6-108778669-INS-4957 |
| chr6-108785490-INS-762 | chr6-108785490-INS-731 |
| chr6-108821630-INS-163 | chr6-108821630-INS-164 |
| chr6-108830941-INS-52 | chr6-108830966-INS-56 |
| chr6-108859981-INS-330 | chr6-108859981-INS-332 |
| chr6-108861903-INS-2221 | chr6-108861904-INS-2211 |
| chr6-108865575-INS-87 | chr6-108865575-INS-100 |
| chr6-108866029-INS-2432 | chr6-108866030-INS-2432 |
| chr6-108876256-INS-333 | chr6-108876256-INS-333 |
| chr6-108877441-INS-6923 | chr6-108877442-INS-6925 |
| chr6-108895874-INS-70 | chr6-108895876-INS-80 |
| chr6-108901395-INS-60 | chr6-108901395-INS-69 |
| chr6-108905040-INS-468 | chr6-108905078-INS-478 |
| chr6-108912861-INS-830 | chr6-108912861-INS-830 |
| chr6-108916208-INS-207 | chr6-108916208-INS-207 |
| chr6-108930315-INS-121 | chr6-108930315-INS-111 |
| chr6-108931172-INS-7050 | chr6-108931173-INS-7049 |
| chr6-108939584-INS-60 | chr6-108939584-INS-64 |
| chr6-108965202-INS-200 | chr6-108965202-INS-201 |
| chr6-108972312-INS-70 | chr6-108972312-INS-70 |
| chr6-109002548-INS-199 | chr6-109002548-INS-199 |
| chr6-109003252-INS-220 | chr6-109003252-INS-220 |
| chr6-109005874-INS-393 | chr6-109005874-INS-393 |
| chr6-109040494-INS-54 | chr6-109040493-INS-55 |
| chr6-109103420-INS-201 | chr6-109103420-INS-200 |
| chr6-109133340-INS-64 | chr6-109133340-INS-64 |
| chr6-109135075-INS-1169 | chr6-109135076-INS-1164 |
| chr6-109138077-INS-121 | chr6-109138077-INS-121 |
| chr6-109148598-INS-81 | chr6-109148595-INS-85 |
| chr6-109171523-INS-663 | chr6-109171523-INS-663 |
| chr6-109207193-INS-66 | chr6-109207193-INS-66 |
| chr6-109240128-INS-4013 | chr6-109240129-INS-4013 |
| chr6-109251931-INS-1889 | chr6-109251932-INS-1893 |
| chr6-109254809-INS-190 | chr6-109254809-INS-190 |
| chr6-109287084-INS-1256 | chr6-109287085-INS-1260 |
| chr6-109306176-INS-515 | chr6-109306176-INS-515 |
| chr6-109309806-INS-1886 | chr6-109309807-INS-1886 |
| chr6-109318877-INS-402 | chr6-109318877-INS-402 |
| chr6-109352907-INS-64 | chr6-109352907-INS-66 |
| chr6-109358876-INS-6456 | chr6-109358877-INS-6457 |
| chr6-109360073-INS-1933 | chr6-109360074-INS-1932 |
| chr6-109360937-INS-2810 | chr6-109360938-INS-2810 |
| chr6-109370229-INS-198 | chr6-109370229-INS-197 |
| chr6-109374039-INS-2820 | chr6-109374040-INS-2833 |
| chr6-109385940-INS-83 | chr6-109385940-INS-79 |
| chr6-109392535-INS-4702 | chr6-109392536-INS-4702 |
| chr6-109406719-INS-60 | chr6-109406719-INS-62 |
| chr6-109411299-INS-332 | chr6-109411299-INS-333 |
| chr6-109414401-INS-6187 | chr6-109414402-INS-6187 |
| chr6-109416280-INS-54 | chr6-109416280-INS-54 |
| chr6-109447386-INS-137 | chr6-109447386-INS-137 |
| chr6-109452333-INS-65 | chr6-109452329-INS-77 |
| chr6-109460447-INS-57 | chr6-109460447-INS-57 |
| chr6-109474942-INS-70 | chr6-109474942-INS-69 |
| chr6-109481742-INS-228 | chr6-109481742-INS-228 |
| chr6-109486743-INS-54 | chr6-109486743-INS-56 |
| chr6-109538194-INS-56 | chr6-109538194-INS-56 |
| chr6-109609374-INS-1148 | chr6-109609375-INS-1148 |
| chr6-109617941-INS-102 | chr6-109617941-INS-106 |
| chr6-109618874-INS-208 | chr6-109618874-INS-208 |
| chr6-109631018-INS-1518 | chr6-109631019-INS-1519 |
| chr6-109637732-INS-102 | chr6-109637734-INS-52 |
| chr6-109643602-INS-6412 | chr6-109643603-INS-6412 |
| chr6-109661272-INS-205 | chr6-109661272-INS-205 |
| chr6-109663572-INS-58 | chr6-109663572-INS-58 |
| chr6-109671639-INS-582 | chr6-109671639-INS-591 |
| chr6-109673260-INS-592 | chr6-109673260-INS-592 |
| chr6-109701523-INS-3586 | chr6-109701524-INS-3586 |
| chr6-109741521-INS-6177 | chr6-109741522-INS-6177 |
| chr6-109758645-INS-6167 | chr6-109758646-INS-6167 |
| chr6-109765976-INS-57 | chr6-109765976-INS-65 |
| chr6-109767309-INS-72 | chr6-109767309-INS-72 |
| chr6-109769863-INS-70 | chr6-109769863-INS-76 |

| | |
|---|---|
| chr6-109777575-INS-106 | chr6-109777549-INS-156 |
| chr6-109820177-INS-80 | chr6-109820177-INS-80 |
| chr6-109831641-INS-308 | chr6-109831641-INS-308 |
| chr6-109834181-INS-93 | chr6-109834181-INS-93 |
| chr6-109838011-INS-196 | chr6-109838011-INS-193 |
| chr6-109842558-INS-134 | chr6-109842557-INS-150 |
| chr6-109849117-INS-54 | chr6-109849117-INS-56 |
| chr6-109858424-INS-70 | chr6-109858424-INS-67 |
| chr6-109876116-INS-87 | chr6-109876116-INS-87 |
| chr6-109900082-INS-784 | chr6-109900082-INS-784 |
| chr6-109924220-INS-75 | chr6-109924220-INS-63 |
| chr6-109973244-INS-198 | chr6-109973244-INS-198 |
| chr6-110032061-INS-82 | chr6-110032061-INS-82 |
| chr6-110044021-INS-64 | chr6-110044014-INS-70 |
| chr6-110087039-INS-52 | chr6-110087039-INS-52 |
| chr6-110091410-INS-52 | chr6-110091410-INS-54 |
| chr6-110094026-INS-385 | chr6-110094026-INS-385 |
| chr6-110127248-INS-67 | chr6-110127255-INS-70 |
| chr6-110201538-INS-378 | chr6-110201554-INS-389 |
| chr6-110206366-INS-78 | chr6-110206372-INS-72 |
| chr6-110207124-INS-53 | chr6-110207124-INS-53 |
| chr6-110210827-INS-66 | chr6-110210827-INS-78 |
| chr6-110215690-INS-231 | chr6-110215690-INS-231 |
| chr6-110218431-INS-60 | chr6-110218431-INS-60 |
| chr6-110233741-INS-187 | chr6-110233741-INS-187 |
| chr6-110257826-INS-54 | chr6-110257828-INS-56 |
| chr6-110262334-INS-340 | chr6-110262333-INS-378 |
| chr6-110275391-INS-68 | chr6-110275392-INS-52 |
| chr6-110278617-INS-6845 | chr6-110278618-INS-6844 |
| chr6-110319323-INS-98 | chr6-110319323-INS-98 |
| chr6-110325448-INS-6360 | chr6-110325449-INS-6360 |
| chr6-110338939-INS-271 | chr6-110338945-INS-231 |
| chr6-110339977-INS-56 | chr6-110340007-INS-50 |
| chr6-110345914-INS-460 | chr6-110345914-INS-460 |
| chr6-110375345-INS-65 | chr6-110375345-INS-65 |
| chr6-110376382-INS-642 | chr6-110376382-INS-642 |
| chr6-110377691-INS-84 | chr6-110377655-INS-84 |
| chr6-110389494-INS-186 | chr6-110389494-INS-186 |
| chr6-110391086-INS-700 | chr6-110391086-INS-699 |
| chr6-110412417-INS-6308 | chr6-110412418-INS-6308 |
| chr6-110429020-INS-6783 | chr6-110429021-INS-6783 |
| chr6-110445826-INS-428 | chr6-110445826-INS-428 |
| chr6-110472258-INS-52 | chr6-110472258-INS-52 |
| chr6-110517948-INS-166 | chr6-110517948-INS-175 |
| chr6-110521365-INS-464 | chr6-110521365-INS-464 |
| chr6-110559298-INS-219 | chr6-110559298-INS-219 |
| chr6-110563956-INS-681 | chr6-110563956-INS-681 |
| chr6-110578787-INS-51 | chr6-110578787-INS-51 |
| chr6-110598799-INS-64 | chr6-110598799-INS-64 |
| chr6-110613384-INS-125 | chr6-110613384-INS-125 |
| chr6-110653460-INS-832 | chr6-110653465-INS-842 |
| chr6-110664766-INS-90 | chr6-110664766-INS-94 |
| chr6-110666645-INS-86 | chr6-110666645-INS-78 |
| chr6-110701787-INS-1465 | chr6-110701788-INS-1457 |
| chr6-110713199-INS-258 | chr6-110713199-INS-216 |
| chr6-110723527-INS-6487 | chr6-110723528-INS-6486 |
| chr6-110738001-INS-118 | chr6-110738001-INS-119 |
| chr6-110738617-INS-66 | chr6-110738617-INS-59 |
| chr6-110746598-INS-172 | chr6-110746598-INS-168 |
| chr6-110757721-INS-165 | chr6-110757721-INS-165 |
| chr6-110781181-INS-72 | chr6-110781181-INS-75 |
| chr6-110785909-INS-62 | chr6-110785917-INS-92 |
| chr6-110791014-INS-876 | chr6-110791014-INS-875 |
| chr6-110812070-INS-114 | chr6-110812070-INS-117 |
| chr6-110817002-INS-197 | chr6-110817002-INS-197 |
| chr6-110835239-INS-61 | chr6-110835239-INS-52 |
| chr6-110847625-INS-56 | chr6-110847626-INS-94 |
| chr6-110849800-INS-459 | chr6-110849800-INS-464 |
| chr6-110874963-INS-237 | chr6-110874971-INS-229 |
| chr6-110881495-INS-51 | chr6-110881495-INS-51 |
| chr6-110907800-INS-642 | chr6-110907800-INS-644 |
| chr6-110910905-INS-1450 | chr6-110910906-INS-1450 |
| chr6-110957120-INS-52 | chr6-110957120-INS-51 |
| chr6-111002756-INS-232 | chr6-111002756-INS-236 |
| chr6-111011662-INS-96 | chr6-111011662-INS-64 |
| chr6-111043623-INS-60 | chr6-111043623-INS-60 |
| chr6-111055495-INS-509 | chr6-111055495-INS-509 |

| | |
|---|---|
| chr6-111059715-INS-781 | chr6-111059715-INS-785 |
| chr6-111072728-INS-52 | chr6-111072728-INS-84 |
| chr6-111111625-INS-56 | chr6-111111625-INS-60 |
| chr6-111117334-INS-90 | chr6-111117334-INS-90 |
| chr6-111149431-INS-60 | chr6-111149435-INS-52 |
| chr6-111155321-INS-207 | chr6-111155321-INS-208 |
| chr6-111167891-INS-455 | chr6-111167891-INS-450 |
| chr6-111184578-INS-134 | chr6-111184582-INS-86 |
| chr6-111207006-INS-445 | chr6-111207006-INS-445 |
| chr6-111209153-INS-425 | chr6-111209153-INS-425 |
| chr6-111253463-INS-357 | chr6-111253463-INS-356 |
| chr6-111263565-INS-554 | chr6-111263565-INS-556 |
| chr6-111284039-INS-196 | chr6-111284039-INS-195 |
| chr6-111296282-INS-267 | chr6-111296282-INS-265 |
| chr6-111298413-INS-2356 | chr6-111298414-INS-2367 |
| chr6-111313814-INS-141 | chr6-111313814-INS-141 |
| chr6-111334811-INS-56 | chr6-111334811-INS-60 |
| chr6-111338083-INS-7148 | chr6-111338084-INS-7124 |
| chr6-111346684-INS-119 | chr6-111346684-INS-154 |
| chr6-111366499-INS-84 | chr6-111366499-INS-84 |
| chr6-111461313-INS-137 | chr6-111461313-INS-90 |
| chr6-111463656-INS-54 | chr6-111463656-INS-72 |
| chr6-111474659-INS-60 | chr6-111474659-INS-60 |
| chr6-111501569-INS-7148 | chr6-111501570-INS-7148 |
| chr6-111503399-INS-351 | chr6-111503401-INS-352 |
| chr6-111553868-INS-502 | chr6-111553868-INS-502 |
| chr6-111578779-INS-397 | chr6-111578779-INS-397 |
| chr6-111595454-INS-6816 | chr6-111595455-INS-6816 |
| chr6-111601804-INS-63 | chr6-111601804-INS-63 |
| chr6-111603164-INS-687 | chr6-111603173-INS-677 |
| chr6-111620581-INS-56 | chr6-111620549-INS-66 |
| chr6-111635374-INS-278 | chr6-111635374-INS-280 |
| chr6-111638281-INS-2442 | chr6-111638282-INS-2442 |
| chr6-111680689-INS-51 | chr6-111680689-INS-52 |
| chr6-111689599-INS-667 | chr6-111689599-INS-671 |
| chr6-111717504-INS-2890 | chr6-111717505-INS-2901 |
| chr6-111730566-INS-54 | chr6-111730566-INS-54 |
| chr6-111736829-INS-85 | chr6-111736829-INS-55 |
| chr6-111786694-INS-461 | chr6-111786694-INS-462 |
| chr6-111795111-INS-73 | chr6-111795111-INS-103 |
| chr6-111810091-INS-68 | chr6-111810091-INS-52 |
| chr6-111816181-INS-188 | chr6-111816181-INS-188 |
| chr6-111820070-INS-242 | chr6-111820070-INS-237 |
| chr6-111826285-INS-60 | chr6-111826285-INS-52 |
| chr6-111882771-INS-60 | chr6-111882771-INS-64 |
| chr6-111884117-INS-68 | chr6-111884119-INS-51 |
| chr6-111889486-INS-110 | chr6-111889478-INS-101 |
| chr6-111909271-INS-134 | chr6-111909271-INS-95 |
| chr6-111981672-INS-70 | chr6-111981672-INS-60 |
| chr6-111999203-INS-68 | chr6-111999203-INS-68 |
| chr6-112011304-INS-93 | chr6-112011304-INS-103 |
| chr6-112032425-INS-52 | chr6-112032431-INS-66 |
| chr6-112065438-INS-52 | chr6-112065438-INS-56 |
| chr6-112066728-INS-92 | chr6-112066728-INS-94 |
| chr6-112089327-INS-2414 | chr6-112089328-INS-2415 |
| chr6-112097652-INS-515 | chr6-112097652-INS-515 |
| chr6-112128397-INS-6485 | chr6-112128398-INS-6485 |
| chr6-112136456-INS-75 | chr6-112136456-INS-111 |
| chr6-112137486-INS-1475 | chr6-112137487-INS-1471 |
| chr6-112174183-INS-538 | chr6-112174183-INS-542 |
| chr6-112190413-INS-78 | chr6-112190413-INS-72 |
| chr6-112193839-INS-204 | chr6-112193839-INS-204 |
| chr6-112211854-INS-71 | chr6-112211854-INS-99 |
| chr6-112228676-INS-426 | chr6-112228676-INS-426 |
| chr6-112278858-INS-61 | chr6-112278858-INS-61 |
| chr6-112323633-INS-252 | chr6-112323633-INS-249 |
| chr6-112340850-INS-64 | chr6-112340850-INS-61 |
| chr6-112393816-INS-443 | chr6-112393816-INS-443 |
| chr6-112426390-INS-55 | chr6-112426353-INS-53 |
| chr6-112432967-INS-82 | chr6-112432956-INS-87 |
| chr6-112438023-INS-68 | chr6-112438014-INS-57 |
| chr6-112476247-INS-185 | chr6-112476211-INS-192 |
| chr6-112484255-INS-401 | chr6-112484255-INS-371 |
| chr6-112489159-INS-862 | chr6-112489159-INS-873 |
| chr6-112537828-INS-75 | chr6-112537828-INS-75 |
| chr6-112545339-INS-551 | chr6-112545339-INS-552 |
| chr6-112546978-INS-484 | chr6-112546978-INS-511 |

| | |
|---|---|
| chr6-112554973-INS-115 | chr6-112554973-INS-109 |
| chr6-112562493-INS-972 | chr6-112562493-INS-973 |
| chr6-112594496-INS-150 | chr6-112594504-INS-153 |
| chr6-112605596-INS-105 | chr6-112605596-INS-105 |
| chr6-112617093-INS-3063 | chr6-112617094-INS-3063 |
| chr6-112625778-INS-243 | chr6-112625778-INS-243 |
| chr6-112649492-INS-67 | chr6-112649492-INS-84 |
| chr6-112657628-INS-853 | chr6-112657628-INS-853 |
| chr6-112690027-INS-500 | chr6-112690027-INS-500 |
| chr6-112735217-INS-91 | chr6-112735206-INS-104 |
| chr6-112744625-INS-62 | chr6-112744616-INS-62 |
| chr6-112784395-INS-60 | chr6-112784395-INS-60 |
| chr6-112808607-INS-54 | chr6-112808613-INS-51 |
| chr6-112815272-INS-199 | chr6-112815272-INS-199 |
| chr6-112846295-INS-103 | chr6-112846295-INS-103 |
| chr6-112855764-INS-60 | chr6-112855764-INS-55 |
| chr6-112863088-INS-182 | chr6-112863088-INS-199 |
| chr6-112864871-INS-73 | chr6-112864870-INS-74 |
| chr6-112869965-INS-210 | chr6-112869990-INS-168 |
| chr6-112923325-INS-84 | chr6-112923325-INS-84 |
| chr6-112927279-INS-212 | chr6-112927279-INS-212 |
| chr6-112936855-INS-171 | chr6-112936844-INS-171 |
| chr6-112973690-INS-1896 | chr6-112973691-INS-1896 |
| chr6-112978983-INS-943 | chr6-112978983-INS-943 |
| chr6-112982133-INS-190 | chr6-112982133-INS-194 |
| chr6-112984649-INS-56 | chr6-112984649-INS-62 |
| chr6-112990754-INS-671 | chr6-112990754-INS-670 |
| chr6-113000737-INS-398 | chr6-113000737-INS-398 |
| chr6-113015861-INS-497 | chr6-113015861-INS-497 |
| chr6-113016580-INS-1032 | chr6-113016581-INS-1032 |
| chr6-113058832-INS-64 | chr6-113058832-INS-68 |
| chr6-113075038-INS-76 | chr6-113075038-INS-80 |
| chr6-113110216-INS-71 | chr6-113110216-INS-77 |
| chr6-113133597-INS-216 | chr6-113133597-INS-216 |
| chr6-113147543-INS-54 | chr6-113147543-INS-56 |
| chr6-113179971-INS-90 | chr6-113179971-INS-90 |
| chr6-113183082-INS-125 | chr6-113183082-INS-125 |
| chr6-113220702-INS-156 | chr6-113220702-INS-156 |
| chr6-113326560-INS-204 | chr6-113326560-INS-205 |
| chr6-113331817-INS-53 | chr6-113331818-INS-95 |
| chr6-113333517-INS-63 | chr6-113333517-INS-63 |
| chr6-113338007-INS-202 | chr6-113338007-INS-202 |
| chr6-113347498-INS-74 | chr6-113347498-INS-74 |
| chr6-113349012-INS-96 | chr6-113349014-INS-92 |
| chr6-113350306-INS-214 | chr6-113350306-INS-215 |
| chr6-113359249-INS-217 | chr6-113359249-INS-218 |
| chr6-113360269-INS-2815 | chr6-113360270-INS-2815 |
| chr6-113408349-INS-78 | chr6-113408339-INS-96 |
| chr6-113430488-INS-235 | chr6-113430488-INS-235 |
| chr6-113431308-INS-206 | chr6-113431308-INS-206 |
| chr6-113482134-INS-5568 | chr6-113482135-INS-5568 |
| chr6-113483919-INS-390 | chr6-113483919-INS-390 |
| chr6-113526059-INS-78 | chr6-113526059-INS-82 |
| chr6-113555970-INS-117 | chr6-113556000-INS-86 |
| chr6-113558288-INS-202 | chr6-113558288-INS-202 |
| chr6-113591763-INS-61 | chr6-113591763-INS-61 |
| chr6-113603335-INS-182 | chr6-113603335-INS-181 |
| chr6-113624823-INS-147 | chr6-113624823-INS-147 |
| chr6-113630398-INS-129 | chr6-113630399-INS-141 |
| chr6-113640999-INS-86 | chr6-113640999-INS-86 |
| chr6-113651645-INS-5230 | chr6-113651646-INS-5230 |
| chr6-113657775-INS-203 | chr6-113657775-INS-212 |
| chr6-113708707-INS-182 | chr6-113708707-INS-193 |
| chr6-113746538-INS-123 | chr6-113746538-INS-123 |
| chr6-113748704-INS-135 | chr6-113748704-INS-147 |
| chr6-113774707-INS-62 | chr6-113774707-INS-64 |
| chr6-113777312-INS-104 | chr6-113777308-INS-67 |
| chr6-113793261-INS-735 | chr6-113793261-INS-736 |
| chr6-113809637-INS-116 | chr6-113809643-INS-116 |
| chr6-113906822-INS-170 | chr6-113906850-INS-145 |
| chr6-113926074-INS-82 | chr6-113926072-INS-68 |
| chr6-113968139-INS-1151 | chr6-113968140-INS-1151 |
| chr6-113991458-INS-60 | chr6-113991458-INS-62 |
| chr6-113993981-INS-244 | chr6-113993981-INS-244 |
| chr6-114033730-INS-179 | chr6-114033730-INS-179 |
| chr6-114041570-INS-88 | chr6-114041570-INS-88 |
| chr6-114089313-INS-65 | chr6-114089313-INS-62 |

| | |
|---|---|
| chr6-114090682-INS-148 | chr6-114090696-INS-104 |
| chr6-114107166-INS-226 | chr6-114107166-INS-226 |
| chr6-114114295-INS-64 | chr6-114114295-INS-56 |
| chr6-114147425-INS-400 | chr6-114147425-INS-400 |
| chr6-114187802-INS-1893 | chr6-114187803-INS-1893 |
| chr6-114198223-INS-1030 | chr6-114198228-INS-1031 |
| chr6-114223567-INS-400 | chr6-114223567-INS-400 |
| chr6-114237808-INS-116 | chr6-114237808-INS-121 |
| chr6-114245016-INS-65 | chr6-114245016-INS-64 |
| chr6-114251701-INS-54 | chr6-114251701-INS-55 |
| chr6-114296612-INS-103 | chr6-114296612-INS-123 |
| chr6-114325849-INS-92 | chr6-114325849-INS-92 |
| chr6-114338026-INS-591 | chr6-114338026-INS-589 |
| chr6-114344251-INS-80 | chr6-114344271-INS-72 |
| chr6-114356071-INS-174 | chr6-114356089-INS-190 |
| chr6-114367667-INS-150 | chr6-114367667-INS-150 |
| chr6-114370800-INS-54 | chr6-114370800-INS-72 |
| chr6-114390895-INS-137 | chr6-114390871-INS-184 |
| chr6-114423316-INS-72 | chr6-114423316-INS-72 |
| chr6-114440936-INS-479 | chr6-114440936-INS-479 |
| chr6-114470854-INS-54 | chr6-114470854-INS-54 |
| chr6-114473355-INS-85 | chr6-114473355-INS-88 |
| chr6-114509380-INS-76 | chr6-114509381-INS-116 |
| chr6-114677357-INS-102 | chr6-114677357-INS-102 |
| chr6-114706426-INS-77 | chr6-114706426-INS-77 |
| chr6-114709930-INS-1033 | chr6-114709931-INS-1033 |
| chr6-114764503-INS-86 | chr6-114764500-INS-93 |
| chr6-114784494-INS-265 | chr6-114784494-INS-265 |
| chr6-114791934-INS-52 | chr6-114791940-INS-52 |
| chr6-114812939-INS-60 | chr6-114812939-INS-60 |
| chr6-114819530-INS-80 | chr6-114819530-INS-88 |
| chr6-114822425-INS-58 | chr6-114822420-INS-58 |
| chr6-114881070-INS-741 | chr6-114881070-INS-709 |
| chr6-114923277-INS-78 | chr6-114923277-INS-78 |
| chr6-114925349-INS-200 | chr6-114925349-INS-199 |
| chr6-114927466-INS-110 | chr6-114927467-INS-107 |
| chr6-114929930-INS-53 | chr6-114929930-INS-54 |
| chr6-114950631-INS-91 | chr6-114950644-INS-50 |
| chr6-114975296-INS-318 | chr6-114975296-INS-324 |
| chr6-115010339-INS-337 | chr6-115010339-INS-337 |
| chr6-115045464-INS-142 | chr6-115045464-INS-152 |
| chr6-115073356-INS-922 | chr6-115073356-INS-922 |
| chr6-115075099-INS-126 | chr6-115075099-INS-90 |
| chr6-115077567-INS-523 | chr6-115077567-INS-523 |
| chr6-115080666-INS-200 | chr6-115080666-INS-200 |
| chr6-115109281-INS-2657 | chr6-115109282-INS-2657 |
| chr6-115139450-INS-2682 | chr6-115139451-INS-2682 |
| chr6-115147835-INS-52 | chr6-115147835-INS-52 |
| chr6-115163119-INS-1506 | chr6-115163120-INS-1506 |
| chr6-115165527-INS-333 | chr6-115165527-INS-333 |
| chr6-115171570-INS-110 | chr6-115171570-INS-110 |
| chr6-115185083-INS-1108 | chr6-115185084-INS-1067 |
| chr6-115212676-INS-54 | chr6-115212676-INS-54 |
| chr6-115217546-INS-1721 | chr6-115217519-INS-1722 |
| chr6-115242316-INS-399 | chr6-115242316-INS-399 |
| chr6-115270696-INS-619 | chr6-115270696-INS-619 |
| chr6-115282671-INS-157 | chr6-115282671-INS-157 |
| chr6-115328229-INS-242 | chr6-115328229-INS-238 |
| chr6-115334665-INS-104 | chr6-115334665-INS-92 |
| chr6-115421301-INS-55 | chr6-115421341-INS-56 |
| chr6-115458418-INS-84 | chr6-115458418-INS-108 |
| chr6-115459705-INS-54 | chr6-115459705-INS-58 |
| chr6-115468678-INS-194 | chr6-115468676-INS-197 |
| chr6-115475512-INS-52 | chr6-115475512-INS-52 |
| chr6-115483463-INS-56 | chr6-115483463-INS-56 |
| chr6-115509795-INS-120 | chr6-115509795-INS-112 |
| chr6-115520552-INS-6442 | chr6-115520553-INS-6442 |
| chr6-115526670-INS-56 | chr6-115526670-INS-56 |
| chr6-115529070-INS-232 | chr6-115529070-INS-233 |
| chr6-115535804-INS-58 | chr6-115535804-INS-58 |
| chr6-115550503-INS-168 | chr6-115550503-INS-168 |
| chr6-115559411-INS-1886 | chr6-115559412-INS-1886 |
| chr6-115568436-INS-72 | chr6-115568436-INS-72 |
| chr6-115570240-INS-184 | chr6-115570240-INS-184 |
| chr6-115599045-INS-690 | chr6-115599045-INS-690 |
| chr6-115602447-INS-177 | chr6-115602447-INS-177 |
| chr6-115610279-INS-88 | chr6-115610279-INS-110 |

| | |
|---|---|
| chr6-115616877-INS-92 | chr6-115616877-INS-67 |
| chr6-115619504-INS-65 | chr6-115619484-INS-74 |
| chr6-115622571-INS-56 | chr6-115622571-INS-56 |
| chr6-115625419-INS-82 | chr6-115625420-INS-128 |
| chr6-115629457-INS-208 | chr6-115629457-INS-208 |
| chr6-115635203-INS-205 | chr6-115635203-INS-206 |
| chr6-115666064-INS-243 | chr6-115666064-INS-249 |
| chr6-115675511-INS-231 | chr6-115675511-INS-231 |
| chr6-115680701-INS-200 | chr6-115680701-INS-200 |
| chr6-115688874-INS-217 | chr6-115688874-INS-218 |
| chr6-115690417-INS-331 | chr6-115690417-INS-331 |
| chr6-115703141-INS-215 | chr6-115703141-INS-215 |
| chr6-115719771-INS-168 | chr6-115719771-INS-168 |
| chr6-115725625-INS-94 | chr6-115725633-INS-99 |
| chr6-115730194-INS-282 | chr6-115730190-INS-280 |
| chr6-115731682-INS-210 | chr6-115731682-INS-211 |
| chr6-115733197-INS-52 | chr6-115733197-INS-52 |
| chr6-115735437-INS-105 | chr6-115735437-INS-105 |
| chr6-115739928-INS-818 | chr6-115739928-INS-850 |
| chr6-115744388-INS-210 | chr6-115744388-INS-210 |
| chr6-115771421-INS-59 | chr6-115771461-INS-60 |
| chr6-115811317-INS-214 | chr6-115811317-INS-214 |
| chr6-115816082-INS-136 | chr6-115816074-INS-146 |
| chr6-115886561-INS-59 | chr6-115886561-INS-55 |
| chr6-115905258-INS-73 | chr6-115905258-INS-73 |
| chr6-115926396-INS-268 | chr6-115926396-INS-258 |
| chr6-115980170-INS-56 | chr6-115980170-INS-52 |
| chr6-115993812-INS-52 | chr6-115993818-INS-80 |
| chr6-116000037-INS-113 | chr6-116000033-INS-118 |
| chr6-116011862-INS-208 | chr6-116011862-INS-208 |
| chr6-116048703-INS-201 | chr6-116048703-INS-201 |
| chr6-116116268-INS-3072 | chr6-116116269-INS-3072 |
| chr6-116126477-INS-59 | chr6-116126477-INS-59 |
| chr6-116129341-INS-216 | chr6-116129341-INS-214 |
| chr6-116166033-INS-112 | chr6-116166033-INS-112 |
| chr6-116205233-INS-59 | chr6-116205233-INS-59 |
| chr6-116214911-INS-125 | chr6-116214911-INS-144 |
| chr6-116217368-INS-207 | chr6-116217368-INS-207 |
| chr6-116219136-INS-289 | chr6-116219148-INS-282 |
| chr6-116226237-INS-208 | chr6-116226237-INS-211 |
| chr6-116228701-INS-868 | chr6-116228702-INS-868 |
| chr6-116236798-INS-54 | chr6-116236757-INS-50 |
| chr6-116261269-INS-209 | chr6-116261269-INS-209 |
| chr6-116275356-INS-62 | chr6-116275356-INS-62 |
| chr6-116279240-INS-331 | chr6-116279240-INS-331 |
| chr6-116288866-INS-198 | chr6-116288866-INS-202 |
| chr6-116294992-INS-424 | chr6-116294992-INS-436 |
| chr6-116310545-INS-51 | chr6-116310545-INS-51 |
| chr6-116312889-INS-1253 | chr6-116312890-INS-1262 |
| chr6-116319392-INS-676 | chr6-116319392-INS-676 |
| chr6-116322218-INS-380 | chr6-116322218-INS-392 |
| chr6-116352686-INS-53 | chr6-116352686-INS-53 |
| chr6-116357406-INS-98 | chr6-116357406-INS-98 |
| chr6-116377730-INS-60 | chr6-116377730-INS-60 |
| chr6-116393676-INS-73 | chr6-116393676-INS-108 |
| chr6-116396442-INS-223 | chr6-116396442-INS-223 |
| chr6-116397918-INS-70 | chr6-116397922-INS-64 |
| chr6-116408626-INS-66 | chr6-116408626-INS-66 |
| chr6-116419643-INS-63 | chr6-116419643-INS-53 |
| chr6-116429606-INS-75 | chr6-116429609-INS-70 |
| chr6-116460478-INS-197 | chr6-116460478-INS-198 |
| chr6-116496503-INS-122 | chr6-116496505-INS-122 |
| chr6-116524647-INS-210 | chr6-116524647-INS-210 |
| chr6-116537959-INS-328 | chr6-116537959-INS-329 |
| chr6-116539582-INS-484 | chr6-116539583-INS-477 |
| chr6-116562974-INS-573 | chr6-116562974-INS-573 |
| chr6-116597534-INS-436 | chr6-116597535-INS-436 |
| chr6-116626506-INS-6711 | chr6-116626507-INS-6711 |
| chr6-116633468-INS-60 | chr6-116633468-INS-60 |
| chr6-116634454-INS-124 | chr6-116634454-INS-122 |
| chr6-116635133-INS-238 | chr6-116635133-INS-238 |
| chr6-116637892-INS-175 | chr6-116637892-INS-178 |
| chr6-116641265-INS-571 | chr6-116641265-INS-571 |
| chr6-116659873-INS-51 | chr6-116659873-INS-51 |
| chr6-116668790-INS-88 | chr6-116668790-INS-66 |
| chr6-116679668-INS-296 | chr6-116679668-INS-296 |
| chr6-116682012-INS-96 | chr6-116682012-INS-96 |

| | |
|---|---|
| chr6-116683501-INS-310 | chr6-116683501-INS-310 |
| chr6-116686546-INS-7460 | chr6-116686547-INS-7460 |
| chr6-116688605-INS-54 | chr6-116688604-INS-61 |
| chr6-116729942-INS-50 | chr6-116729942-INS-52 |
| chr6-116742177-INS-433 | chr6-116742177-INS-433 |
| chr6-116746728-INS-7085 | chr6-116746729-INS-7085 |
| chr6-116751307-INS-68 | chr6-116751305-INS-72 |
| chr6-116762982-INS-116 | chr6-116762986-INS-84 |
| chr6-116785713-INS-1907 | chr6-116785714-INS-1908 |
| chr6-116796672-INS-100 | chr6-116796676-INS-100 |
| chr6-116808266-INS-80 | chr6-116808266-INS-86 |
| chr6-116865088-INS-393 | chr6-116865088-INS-393 |
| chr6-116868241-INS-63 | chr6-116868241-INS-71 |
| chr6-116875967-INS-4374 | chr6-116875968-INS-4374 |
| chr6-116889882-INS-290 | chr6-116889882-INS-290 |
| chr6-116927006-INS-205 | chr6-116927006-INS-205 |
| chr6-116935476-INS-300 | chr6-116935476-INS-298 |
| chr6-116941164-INS-555 | chr6-116941164-INS-560 |
| chr6-116948534-INS-66 | chr6-116948534-INS-66 |
| chr6-116960137-INS-317 | chr6-116960133-INS-318 |
| chr6-116974324-INS-456 | chr6-116974324-INS-461 |
| chr6-117013402-INS-467 | chr6-117013402-INS-466 |
| chr6-117032489-INS-773 | chr6-117032511-INS-774 |
| chr6-117045730-INS-51 | chr6-117045730-INS-51 |
| chr6-117046443-INS-1894 | chr6-117046444-INS-1894 |
| chr6-117053005-INS-528 | chr6-117053006-INS-566 |
| chr6-117054896-INS-61 | chr6-117054896-INS-65 |
| chr6-117075399-INS-182 | chr6-117075399-INS-174 |
| chr6-117128083-INS-57 | chr6-117128109-INS-81 |
| chr6-117177166-INS-65 | chr6-117177162-INS-58 |
| chr6-117194313-INS-65 | chr6-117194328-INS-50 |
| chr6-117205651-INS-53 | chr6-117205652-INS-61 |
| chr6-117207302-INS-4828 | chr6-117207303-INS-4828 |
| chr6-117252592-INS-64 | chr6-117252592-INS-64 |
| chr6-117281393-INS-201 | chr6-117281393-INS-201 |
| chr6-117295261-INS-328 | chr6-117295261-INS-328 |
| chr6-117306814-INS-91 | chr6-117306813-INS-88 |
| chr6-117332849-INS-146 | chr6-117332849-INS-146 |
| chr6-117335866-INS-419 | chr6-117335866-INS-419 |
| chr6-117348141-INS-62 | chr6-117348141-INS-62 |
| chr6-117359185-INS-81 | chr6-117359184-INS-101 |
| chr6-117369703-INS-67 | chr6-117369703-INS-67 |
| chr6-117377622-INS-70 | chr6-117377622-INS-66 |
| chr6-117388571-INS-213 | chr6-117388571-INS-214 |
| chr6-117393392-INS-74 | chr6-117393392-INS-81 |
| chr6-117452611-INS-51 | chr6-117452611-INS-58 |
| chr6-117464598-INS-63 | chr6-117464598-INS-76 |
| chr6-117488003-INS-60 | chr6-117488003-INS-62 |
| chr6-117493452-INS-171 | chr6-117493452-INS-171 |
| chr6-117517774-INS-229 | chr6-117517774-INS-221 |
| chr6-117519023-INS-58 | chr6-117519023-INS-82 |
| chr6-117521397-INS-378 | chr6-117521397-INS-378 |
| chr6-117526850-INS-639 | chr6-117526850-INS-637 |
| chr6-117548561-INS-60 | chr6-117548561-INS-60 |
| chr6-117572152-INS-672 | chr6-117572152-INS-672 |
| chr6-117581832-INS-65 | chr6-117581832-INS-63 |
| chr6-117585256-INS-53 | chr6-117585284-INS-55 |
| chr6-117588095-INS-6568 | chr6-117588096-INS-6568 |
| chr6-117593287-INS-72 | chr6-117593287-INS-52 |
| chr6-117604984-INS-504 | chr6-117604984-INS-504 |
| chr6-117629542-INS-72 | chr6-117629542-INS-76 |
| chr6-117645150-INS-104 | chr6-117645152-INS-105 |
| chr6-117650031-INS-92 | chr6-117650009-INS-92 |
| chr6-117651833-INS-397 | chr6-117651833-INS-397 |
| chr6-117697309-INS-114 | chr6-117697286-INS-161 |
| chr6-117697823-INS-167 | chr6-117697823-INS-167 |
| chr6-117715655-INS-185 | chr6-117715642-INS-173 |
| chr6-117716808-INS-58 | chr6-117716808-INS-58 |
| chr6-117741962-INS-127 | chr6-117741962-INS-127 |
| chr6-117742666-INS-678 | chr6-117742666-INS-689 |
| chr6-117751367-INS-56 | chr6-117751367-INS-56 |
| chr6-117760523-INS-204 | chr6-117760523-INS-207 |
| chr6-117787409-INS-240 | chr6-117787409-INS-243 |
| chr6-117807682-INS-1092 | chr6-117807690-INS-1085 |
| chr6-117827768-INS-458 | chr6-117827768-INS-460 |
| chr6-117834094-INS-526 | chr6-117834094-INS-533 |
| chr6-117839118-INS-221 | chr6-117839119-INS-194 |

| | |
|---|---|
| chr6-117840477-INS-207 | chr6-117840478-INS-207 |
| chr6-117846479-INS-58 | chr6-117846479-INS-52 |
| chr6-117851218-INS-191 | chr6-117851218-INS-192 |
| chr6-117863279-INS-209 | chr6-117863279-INS-210 |
| chr6-117867686-INS-447 | chr6-117867682-INS-451 |
| chr6-117872444-INS-210 | chr6-117872444-INS-210 |
| chr6-117876556-INS-60 | chr6-117876556-INS-60 |
| chr6-117879105-INS-110 | chr6-117879105-INS-114 |
| chr6-117881916-INS-232 | chr6-117881912-INS-228 |
| chr6-117892940-INS-226 | chr6-117892940-INS-226 |
| chr6-117908903-INS-214 | chr6-117908903-INS-214 |
| chr6-117911791-INS-187 | chr6-117911792-INS-184 |
| chr6-117918962-INS-203 | chr6-117918962-INS-202 |
| chr6-117921351-INS-58 | chr6-117921351-INS-90 |
| chr6-117936914-INS-77 | chr6-117936914-INS-77 |
| chr6-117938073-INS-604 | chr6-117938084-INS-616 |
| chr6-117939047-INS-109 | chr6-117939047-INS-109 |
| chr6-117943759-INS-209 | chr6-117943759-INS-208 |
| chr6-117945825-INS-214 | chr6-117945825-INS-206 |
| chr6-117946895-INS-92 | chr6-117946895-INS-142 |
| chr6-117951516-INS-400 | chr6-117951516-INS-400 |
| chr6-117961565-INS-87 | chr6-117961574-INS-69 |
| chr6-117970089-INS-293 | chr6-117970089-INS-294 |
| chr6-117978074-INS-72 | chr6-117978074-INS-72 |
| chr6-117992653-INS-327 | chr6-117992653-INS-329 |
| chr6-117994417-INS-6487 | chr6-117994418-INS-6487 |
| chr6-118030374-INS-4642 | chr6-118030375-INS-4642 |
| chr6-118036123-INS-173 | chr6-118036120-INS-165 |
| chr6-118056800-INS-60 | chr6-118056800-INS-69 |
| chr6-118057759-INS-80 | chr6-118057759-INS-80 |
| chr6-118075247-INS-2879 | chr6-118075248-INS-2879 |
| chr6-118075961-INS-548 | chr6-118075961-INS-536 |
| chr6-118102042-INS-198 | chr6-118102042-INS-198 |
| chr6-118124746-INS-146 | chr6-118124746-INS-123 |
| chr6-118125896-INS-126 | chr6-118125896-INS-95 |
| chr6-118193340-INS-133 | chr6-118193341-INS-123 |
| chr6-118216278-INS-110 | chr6-118216290-INS-62 |
| chr6-118237172-INS-5349 | chr6-118237173-INS-5349 |
| chr6-118240021-INS-6930 | chr6-118240022-INS-6931 |
| chr6-118250460-INS-207 | chr6-118250460-INS-207 |
| chr6-118261772-INS-276 | chr6-118261772-INS-276 |
| chr6-118271429-INS-114 | chr6-118271422-INS-117 |
| chr6-118328012-INS-168 | chr6-118328012-INS-168 |
| chr6-118350932-INS-175 | chr6-118350932-INS-170 |
| chr6-118375837-INS-61 | chr6-118375837-INS-61 |
| chr6-118400973-INS-1912 | chr6-118400974-INS-1912 |
| chr6-118402317-INS-68 | chr6-118402327-INS-52 |
| chr6-118411362-INS-51 | chr6-118411362-INS-52 |
| chr6-118445274-INS-120 | chr6-118445274-INS-120 |
| chr6-118447933-INS-177 | chr6-118447933-INS-176 |
| chr6-118460812-INS-255 | chr6-118460812-INS-251 |
| chr6-118486509-INS-2765 | chr6-118486510-INS-2770 |
| chr6-118496070-INS-442 | chr6-118496070-INS-398 |
| chr6-118507553-INS-169 | chr6-118507547-INS-199 |
| chr6-118511827-INS-559 | chr6-118511827-INS-559 |
| chr6-118516632-INS-60 | chr6-118516632-INS-60 |
| chr6-118524511-INS-60 | chr6-118524511-INS-60 |
| chr6-118527531-INS-207 | chr6-118527531-INS-207 |
| chr6-118533806-INS-1140 | chr6-118533807-INS-1140 |
| chr6-118537591-INS-70 | chr6-118537591-INS-70 |
| chr6-118552598-INS-466 | chr6-118552598-INS-466 |
| chr6-118556309-INS-61 | chr6-118556309-INS-62 |
| chr6-118559272-INS-123 | chr6-118559272-INS-94 |
| chr6-118559453-INS-6419 | chr6-118559454-INS-6419 |
| chr6-118574462-INS-54 | chr6-118574462-INS-54 |
| chr6-118585738-INS-607 | chr6-118585738-INS-607 |
| chr6-118627974-INS-56 | chr6-118627974-INS-56 |
| chr6-118643868-INS-68 | chr6-118643868-INS-98 |
| chr6-118694995-INS-370 | chr6-118694995-INS-372 |
| chr6-118703970-INS-61 | chr6-118703970-INS-91 |
| chr6-118745405-INS-105 | chr6-118745405-INS-111 |
| chr6-118781164-INS-111 | chr6-118781164-INS-111 |
| chr6-118791967-INS-105 | chr6-118791965-INS-97 |
| chr6-118794301-INS-181 | chr6-118794301-INS-181 |
| chr6-118822659-INS-100 | chr6-118822659-INS-70 |
| chr6-118878354-INS-51 | chr6-118878354-INS-51 |
| chr6-118908819-INS-218 | chr6-118908819-INS-224 |

| | |
|---|---|
| chr6-118933677-INS-277 | chr6-118933677-INS-277 |
| chr6-118947621-INS-401 | chr6-118947621-INS-401 |
| chr6-118961933-INS-140 | chr6-118961933-INS-140 |
| chr6-118972206-INS-174 | chr6-118972206-INS-174 |
| chr6-118976114-INS-56 | chr6-118976074-INS-68 |
| chr6-118998669-INS-70 | chr6-118998669-INS-64 |
| chr6-119043547-INS-298 | chr6-119043547-INS-298 |
| chr6-119064178-INS-288 | chr6-119064179-INS-288 |
| chr6-119078591-INS-74 | chr6-119078593-INS-72 |
| chr6-119079034-INS-130 | chr6-119079052-INS-96 |
| chr6-119079475-INS-198 | chr6-119079475-INS-186 |
| chr6-119081952-INS-56 | chr6-119081952-INS-56 |
| chr6-119088581-INS-268 | chr6-119088581-INS-270 |
| chr6-119097559-INS-115 | chr6-119097559-INS-118 |
| chr6-119163425-INS-53 | chr6-119163425-INS-53 |
| chr6-119212703-INS-67 | chr6-119212703-INS-67 |
| chr6-119224132-INS-99 | chr6-119224132-INS-99 |
| chr6-119225643-INS-432 | chr6-119225643-INS-432 |
| chr6-119227875-INS-274 | chr6-119227875-INS-273 |
| chr6-119238211-INS-189 | chr6-119238212-INS-231 |
| chr6-119289025-INS-138 | chr6-119289025-INS-138 |
| chr6-119319964-INS-253 | chr6-119319965-INS-253 |
| chr6-119354327-INS-198 | chr6-119354327-INS-198 |
| chr6-119377456-INS-536 | chr6-119377436-INS-536 |
| chr6-119379694-INS-199 | chr6-119379694-INS-199 |
| chr6-119389909-INS-86 | chr6-119389909-INS-90 |
| chr6-119406696-INS-113 | chr6-119406711-INS-83 |
| chr6-119424042-INS-86 | chr6-119424042-INS-86 |
| chr6-119430359-INS-444 | chr6-119430358-INS-445 |
| chr6-119434872-INS-514 | chr6-119434872-INS-514 |
| chr6-119443924-INS-398 | chr6-119443924-INS-398 |
| chr6-119447199-INS-140 | chr6-119447199-INS-140 |
| chr6-119461144-INS-80 | chr6-119461144-INS-77 |
| chr6-119483704-INS-62 | chr6-119483704-INS-51 |
| chr6-119535481-INS-70 | chr6-119535471-INS-106 |
| chr6-119540185-INS-54 | chr6-119540185-INS-54 |
| chr6-119561713-INS-205 | chr6-119561713-INS-204 |
| chr6-119602809-INS-212 | chr6-119602809-INS-211 |
| chr6-119617201-INS-2565 | chr6-119617202-INS-2565 |
| chr6-119620208-INS-80 | chr6-119620208-INS-80 |
| chr6-119621206-INS-1285 | chr6-119621207-INS-1286 |
| chr6-119652599-INS-79 | chr6-119652599-INS-79 |
| chr6-119670776-INS-211 | chr6-119670776-INS-211 |
| chr6-119693035-INS-120 | chr6-119693025-INS-114 |
| chr6-119732128-INS-749 | chr6-119732128-INS-750 |
| chr6-119744937-INS-291 | chr6-119744937-INS-252 |
| chr6-119749794-INS-126 | chr6-119749794-INS-126 |
| chr6-119808193-INS-256 | chr6-119808193-INS-256 |
| chr6-119817194-INS-53 | chr6-119817194-INS-50 |
| chr6-119823208-INS-220 | chr6-119823208-INS-220 |
| chr6-119861372-INS-88 | chr6-119861376-INS-56 |
| chr6-119863101-INS-208 | chr6-119863101-INS-209 |
| chr6-119871293-INS-225 | chr6-119871293-INS-226 |
| chr6-119876546-INS-72 | chr6-119876546-INS-68 |
| chr6-119893843-INS-56 | chr6-119893842-INS-88 |
| chr6-119948819-INS-214 | chr6-119948819-INS-216 |
| chr6-119972721-INS-6237 | chr6-119972722-INS-6237 |
| chr6-120026337-INS-438 | chr6-120026338-INS-438 |
| chr6-120037232-INS-60 | chr6-120037232-INS-50 |
| chr6-120044991-INS-64 | chr6-120044991-INS-64 |
| chr6-120067999-INS-77 | chr6-120067999-INS-77 |
| chr6-120075699-INS-229 | chr6-120075699-INS-229 |
| chr6-120088283-INS-206 | chr6-120088283-INS-205 |
| chr6-120090029-INS-64 | chr6-120090033-INS-59 |
| chr6-120100451-INS-175 | chr6-120100451-INS-176 |
| chr6-120102470-INS-498 | chr6-120102470-INS-498 |
| chr6-120117249-INS-112 | chr6-120117249-INS-95 |
| chr6-120121557-INS-60 | chr6-120121533-INS-52 |
| chr6-120153155-INS-85 | chr6-120153155-INS-89 |
| chr6-120166997-INS-371 | chr6-120166997-INS-371 |
| chr6-120222539-INS-72 | chr6-120222544-INS-75 |
| chr6-120226988-INS-77 | chr6-120226988-INS-77 |
| chr6-120228139-INS-399 | chr6-120228139-INS-399 |
| chr6-120246734-INS-1380 | chr6-120246735-INS-1380 |
| chr6-120254645-INS-75 | chr6-120254645-INS-84 |
| chr6-120259768-INS-189 | chr6-120259768-INS-188 |
| chr6-120299799-INS-2243 | chr6-120299800-INS-2246 |

| | |
|---|---|
| chr6-120349374-INS-197 | chr6-120349374-INS-196 |
| chr6-120357608-INS-206 | chr6-120357607-INS-200 |
| chr6-120381509-INS-656 | chr6-120381509-INS-666 |
| chr6-120383131-INS-217 | chr6-120383132-INS-208 |
| chr6-120387837-INS-211 | chr6-120387837-INS-210 |
| chr6-120408448-INS-625 | chr6-120408448-INS-625 |
| chr6-120423728-INS-209 | chr6-120423728-INS-209 |
| chr6-120430764-INS-445 | chr6-120430766-INS-484 |
| chr6-120432050-INS-103 | chr6-120432050-INS-103 |
| chr6-120435237-INS-202 | chr6-120435237-INS-201 |
| chr6-120445505-INS-208 | chr6-120445505-INS-209 |
| chr6-120476722-INS-339 | chr6-120476722-INS-339 |
| chr6-120480896-INS-92 | chr6-120480896-INS-133 |
| chr6-120504469-INS-51 | chr6-120504469-INS-51 |
| chr6-120521268-INS-209 | chr6-120521268-INS-209 |
| chr6-120526806-INS-208 | chr6-120526806-INS-208 |
| chr6-120531009-INS-188 | chr6-120531009-INS-188 |
| chr6-120536903-INS-211 | chr6-120536903-INS-211 |
| chr6-120542220-INS-190 | chr6-120542220-INS-195 |
| chr6-120554272-INS-102 | chr6-120554272-INS-108 |
| chr6-120560778-INS-55 | chr6-120560778-INS-55 |
| chr6-120571319-INS-135 | chr6-120571323-INS-138 |
| chr6-120579080-INS-135 | chr6-120579081-INS-135 |
| chr6-120586124-INS-285 | chr6-120586124-INS-285 |
| chr6-120590745-INS-51 | chr6-120590745-INS-51 |
| chr6-120626813-INS-215 | chr6-120626813-INS-215 |
| chr6-120635286-INS-213 | chr6-120635286-INS-213 |
| chr6-120658220-INS-65 | chr6-120658220-INS-65 |
| chr6-120665124-INS-167 | chr6-120665124-INS-143 |
| chr6-120716362-INS-218 | chr6-120716362-INS-218 |
| chr6-120730597-INS-207 | chr6-120730597-INS-207 |
| chr6-120740411-INS-551 | chr6-120740416-INS-552 |
| chr6-120818651-INS-223 | chr6-120818651-INS-270 |
| chr6-120830696-INS-197 | chr6-120830696-INS-199 |
| chr6-120841713-INS-214 | chr6-120841698-INS-216 |
| chr6-120862061-INS-207 | chr6-120862061-INS-206 |
| chr6-120883289-INS-68 | chr6-120883293-INS-64 |
| chr6-120897494-INS-167 | chr6-120897494-INS-159 |
| chr6-120956516-INS-92 | chr6-120956516-INS-96 |
| chr6-120971379-INS-60 | chr6-120971379-INS-60 |
| chr6-121026330-INS-69 | chr6-121026330-INS-69 |
| chr6-121056167-INS-202 | chr6-121056167-INS-202 |
| chr6-121069536-INS-205 | chr6-121069536-INS-205 |
| chr6-121075063-INS-67 | chr6-121075063-INS-67 |
| chr6-121089992-INS-98 | chr6-121089988-INS-101 |
| chr6-121092278-INS-84 | chr6-121092278-INS-86 |
| chr6-121100928-INS-53 | chr6-121100946-INS-73 |
| chr6-121102389-INS-64 | chr6-121102388-INS-96 |
| chr6-121104421-INS-56 | chr6-121104421-INS-60 |
| chr6-121165637-INS-199 | chr6-121165637-INS-199 |
| chr6-121180884-INS-80 | chr6-121180884-INS-80 |
| chr6-121183969-INS-84 | chr6-121183969-INS-72 |
| chr6-121195744-INS-102 | chr6-121195744-INS-102 |
| chr6-121201677-INS-192 | chr6-121201677-INS-224 |
| chr6-121228526-INS-70 | chr6-121228538-INS-54 |
| chr6-121259433-INS-6887 | chr6-121259434-INS-6887 |
| chr6-121271396-INS-196 | chr6-121271396-INS-196 |
| chr6-121284807-INS-53 | chr6-121284807-INS-51 |
| chr6-121286477-INS-53 | chr6-121286455-INS-51 |
| chr6-121287436-INS-2044 | chr6-121287437-INS-2044 |
| chr6-121287437-INS-368 | chr6-121287437-INS-370 |
| chr6-121316119-INS-4847 | chr6-121316120-INS-4847 |
| chr6-121354447-INS-130 | chr6-121354447-INS-130 |
| chr6-121358535-INS-94 | chr6-121358535-INS-94 |
| chr6-121372582-INS-537 | chr6-121372582-INS-537 |
| chr6-121381735-INS-65 | chr6-121381715-INS-113 |
| chr6-121424897-INS-51 | chr6-121424897-INS-54 |
| chr6-121465383-INS-54 | chr6-121465384-INS-54 |
| chr6-121473463-INS-972 | chr6-121473463-INS-973 |
| chr6-121489935-INS-232 | chr6-121489935-INS-232 |
| chr6-121508608-INS-70 | chr6-121508608-INS-71 |
| chr6-121515955-INS-332 | chr6-121515955-INS-331 |
| chr6-121523318-INS-1519 | chr6-121523319-INS-1519 |
| chr6-121523936-INS-211 | chr6-121523936-INS-211 |
| chr6-121538014-INS-558 | chr6-121538014-INS-558 |
| chr6-121544758-INS-450 | chr6-121544758-INS-450 |
| chr6-121553476-INS-7168 | chr6-121553477-INS-7169 |

| | |
|---|---|
| chr6-121582246-INS-114 | chr6-121582246-INS-114 |
| chr6-121604537-INS-77 | chr6-121604537-INS-77 |
| chr6-121613865-INS-104 | chr6-121613865-INS-104 |
| chr6-121617166-INS-80 | chr6-121617164-INS-88 |
| chr6-121626005-INS-64 | chr6-121626005-INS-56 |
| chr6-121693149-INS-72 | chr6-121693149-INS-72 |
| chr6-121694775-INS-339 | chr6-121694775-INS-339 |
| chr6-121699123-INS-189 | chr6-121699123-INS-189 |
| chr6-121702756-INS-94 | chr6-121702754-INS-97 |
| chr6-121711309-INS-506 | chr6-121711309-INS-506 |
| chr6-121719028-INS-84 | chr6-121719028-INS-84 |
| chr6-121761782-INS-71 | chr6-121761782-INS-71 |
| chr6-121823745-INS-141 | chr6-121823745-INS-123 |
| chr6-121852309-INS-52 | chr6-121852309-INS-52 |
| chr6-121867636-INS-509 | chr6-121867636-INS-508 |
| chr6-121904813-INS-60 | chr6-121904813-INS-60 |
| chr6-121905604-INS-564 | chr6-121905604-INS-564 |
| chr6-121941733-INS-662 | chr6-121941733-INS-659 |
| chr6-122018487-INS-5445 | chr6-122018488-INS-5445 |
| chr6-122083208-INS-62 | chr6-122083208-INS-62 |
| chr6-122099978-INS-202 | chr6-122099978-INS-203 |
| chr6-122115750-INS-120 | chr6-122115750-INS-108 |
| chr6-122125329-INS-88 | chr6-122125329-INS-88 |
| chr6-122181128-INS-670 | chr6-122181128-INS-670 |
| chr6-122213519-INS-1078 | chr6-122213520-INS-1078 |
| chr6-122234485-INS-84 | chr6-122234485-INS-81 |
| chr6-122274449-INS-57 | chr6-122274449-INS-58 |
| chr6-122311010-INS-206 | chr6-122311010-INS-208 |
| chr6-122323307-INS-205 | chr6-122323307-INS-203 |
| chr6-122328186-INS-420 | chr6-122328186-INS-414 |
| chr6-122337084-INS-215 | chr6-122337077-INS-210 |
| chr6-122349627-INS-4335 | chr6-122349628-INS-4298 |
| chr6-122363264-INS-101 | chr6-122363267-INS-111 |
| chr6-122367099-INS-848 | chr6-122367085-INS-876 |
| chr6-122384393-INS-344 | chr6-122384393-INS-344 |
| chr6-122387063-INS-137 | chr6-122387063-INS-137 |
| chr6-122401415-INS-64 | chr6-122401396-INS-64 |
| chr6-122449927-INS-196 | chr6-122449927-INS-196 |
| chr6-122456486-INS-54 | chr6-122456486-INS-54 |
| chr6-122488756-INS-218 | chr6-122488756-INS-211 |
| chr6-122496171-INS-300 | chr6-122496171-INS-300 |
| chr6-122523179-INS-57 | chr6-122523179-INS-57 |
| chr6-122529809-INS-468 | chr6-122529788-INS-471 |
| chr6-122545299-INS-256 | chr6-122545299-INS-244 |
| chr6-122547478-INS-142 | chr6-122547478-INS-130 |
| chr6-122556344-INS-52 | chr6-122556336-INS-54 |
| chr6-122561078-INS-595 | chr6-122561078-INS-596 |
| chr6-122582462-INS-286 | chr6-122582463-INS-286 |
| chr6-122588967-INS-56 | chr6-122588967-INS-56 |
| chr6-122590756-INS-62 | chr6-122590756-INS-50 |
| chr6-122596579-INS-192 | chr6-122596565-INS-158 |
| chr6-122601114-INS-56 | chr6-122601136-INS-82 |
| chr6-122609179-INS-207 | chr6-122609179-INS-206 |
| chr6-122612031-INS-56 | chr6-122612014-INS-65 |
| chr6-122647825-INS-209 | chr6-122647825-INS-209 |
| chr6-122649037-INS-106 | chr6-122649037-INS-107 |
| chr6-122652420-INS-218 | chr6-122652420-INS-218 |
| chr6-122658119-INS-223 | chr6-122658119-INS-223 |
| chr6-122661810-INS-59 | chr6-122661810-INS-59 |
| chr6-122667454-INS-223 | chr6-122667454-INS-204 |
| chr6-122670502-INS-74 | chr6-122670507-INS-69 |
| chr6-122679198-INS-196 | chr6-122679198-INS-196 |
| chr6-122680727-INS-117 | chr6-122680727-INS-103 |
| chr6-122687993-INS-78 | chr6-122687993-INS-72 |
| chr6-122728728-INS-54 | chr6-122728728-INS-52 |
| chr6-122748022-INS-64 | chr6-122748022-INS-64 |
| chr6-122750435-INS-149 | chr6-122750431-INS-155 |
| chr6-122763388-INS-206 | chr6-122763388-INS-207 |
| chr6-122770296-INS-55 | chr6-122770296-INS-55 |
| chr6-122774410-INS-75 | chr6-122774387-INS-76 |
| chr6-122775299-INS-218 | chr6-122775299-INS-218 |
| chr6-122786859-INS-203 | chr6-122786859-INS-200 |
| chr6-122792900-INS-202 | chr6-122792900-INS-202 |
| chr6-122793652-INS-5519 | chr6-122793653-INS-5519 |
| chr6-122794919-INS-198 | chr6-122794919-INS-198 |
| chr6-122813586-INS-202 | chr6-122813586-INS-202 |
| chr6-122814509-INS-90 | chr6-122814509-INS-91 |

| | |
|---|---|
| chr6-122823871-INS-6401 | chr6-122823872-INS-6401 |
| chr6-122873582-INS-105 | chr6-122873576-INS-147 |
| chr6-122880124-INS-6538 | chr6-122880125-INS-6539 |
| chr6-122907400-INS-406 | chr6-122907413-INS-415 |
| chr6-122908342-INS-165 | chr6-122908342-INS-166 |
| chr6-122915789-INS-99 | chr6-122915789-INS-66 |
| chr6-122944749-INS-61 | chr6-122944749-INS-91 |
| chr6-122946761-INS-52 | chr6-122946761-INS-54 |
| chr6-122952033-INS-212 | chr6-122952033-INS-212 |
| chr6-122960420-INS-1061 | chr6-122960421-INS-1062 |
| chr6-122990637-INS-622 | chr6-122990637-INS-635 |
| chr6-123000197-INS-215 | chr6-123000196-INS-215 |
| chr6-123001484-INS-95 | chr6-123001484-INS-90 |
| chr6-123004062-INS-1160 | chr6-123004063-INS-1160 |
| chr6-123005098-INS-414 | chr6-123005098-INS-443 |
| chr6-123008140-INS-198 | chr6-123008140-INS-198 |
| chr6-123028970-INS-107 | chr6-123028970-INS-92 |
| chr6-123031630-INS-123 | chr6-123031611-INS-95 |
| chr6-123039966-INS-708 | chr6-123039966-INS-704 |
| chr6-123078649-INS-478 | chr6-123078649-INS-478 |
| chr6-123087492-INS-60 | chr6-123087492-INS-60 |
| chr6-123104540-INS-130 | chr6-123104540-INS-130 |
| chr6-123115294-INS-196 | chr6-123115294-INS-196 |
| chr6-123119769-INS-54 | chr6-123119768-INS-50 |
| chr6-123154016-INS-344 | chr6-123154016-INS-354 |
| chr6-123157169-INS-780 | chr6-123157169-INS-780 |
| chr6-123163023-INS-137 | chr6-123163023-INS-147 |
| chr6-123169909-INS-649 | chr6-123169909-INS-648 |
| chr6-123193574-INS-325 | chr6-123193574-INS-325 |
| chr6-123212638-INS-283 | chr6-123212638-INS-286 |
| chr6-123214706-INS-1140 | chr6-123214707-INS-1136 |
| chr6-123226877-INS-87 | chr6-123226877-INS-76 |
| chr6-123232734-INS-56 | chr6-123232734-INS-58 |
| chr6-123238824-INS-302 | chr6-123238824-INS-303 |
| chr6-123239384-INS-56 | chr6-123239376-INS-52 |
| chr6-123241196-INS-710 | chr6-123241196-INS-710 |
| chr6-123284208-INS-52 | chr6-123284208-INS-56 |
| chr6-123287806-INS-271 | chr6-123287803-INS-271 |
| chr6-123311660-INS-338 | chr6-123311660-INS-338 |
| chr6-123312434-INS-92 | chr6-123312434-INS-92 |
| chr6-123321423-INS-952 | chr6-123321423-INS-953 |
| chr6-123361261-INS-485 | chr6-123361261-INS-485 |
| chr6-123589154-INS-106 | chr6-123589154-INS-107 |
| chr6-123602269-INS-1775 | chr6-123602270-INS-1763 |
| chr6-123628558-INS-369 | chr6-123628558-INS-346 |
| chr6-123684065-INS-352 | chr6-123684065-INS-351 |
| chr6-123704283-INS-153 | chr6-123704283-INS-153 |
| chr6-123709737-INS-62 | chr6-123709736-INS-68 |
| chr6-123725269-INS-809 | chr6-123725270-INS-808 |
| chr6-123728603-INS-545 | chr6-123728603-INS-544 |
| chr6-123761389-INS-175 | chr6-123761389-INS-175 |
| chr6-123917918-INS-204 | chr6-123917918-INS-204 |
| chr6-123953617-INS-2386 | chr6-123953618-INS-2386 |
| chr6-123993728-INS-2804 | chr6-123993689-INS-2804 |
| chr6-123999862-INS-1483 | chr6-123999863-INS-1483 |
| chr6-124005052-INS-6390 | chr6-124005053-INS-6390 |
| chr6-124019531-INS-57 | chr6-124019545-INS-53 |
| chr6-124039205-INS-4257 | chr6-124039206-INS-4256 |
| chr6-124065559-INS-183 | chr6-124065559-INS-183 |
| chr6-124076845-INS-110 | chr6-124076847-INS-106 |
| chr6-124095052-INS-3659 | chr6-124095054-INS-3655 |
| chr6-124097759-INS-185 | chr6-124097752-INS-185 |
| chr6-124133650-INS-140 | chr6-124133650-INS-140 |
| chr6-124147766-INS-439 | chr6-124147767-INS-419 |
| chr6-124216559-INS-1904 | chr6-124216560-INS-1904 |
| chr6-124249680-INS-6442 | chr6-124249681-INS-6442 |
| chr6-124293597-INS-147 | chr6-124293595-INS-110 |
| chr6-124331525-INS-76 | chr6-124331525-INS-72 |
| chr6-124342802-INS-1891 | chr6-124342803-INS-1891 |
| chr6-124348259-INS-69 | chr6-124348259-INS-70 |
| chr6-124352159-INS-422 | chr6-124352159-INS-422 |
| chr6-124353724-INS-53 | chr6-124353724-INS-61 |
| chr6-124366691-INS-216 | chr6-124366692-INS-208 |
| chr6-124369261-INS-1701 | chr6-124369262-INS-1701 |
| chr6-124370374-INS-60 | chr6-124370374-INS-60 |
| chr6-124404159-INS-199 | chr6-124404159-INS-199 |
| chr6-124429758-INS-172 | chr6-124429785-INS-172 |

| | |
|---|---|
| chr6-124431712-INS-225 | chr6-124431712-INS-214 |
| chr6-124447705-INS-68 | chr6-124447705-INS-68 |
| chr6-124452888-INS-200 | chr6-124452888-INS-203 |
| chr6-124455337-INS-186 | chr6-124455337-INS-187 |
| chr6-124469504-INS-2884 | chr6-124469505-INS-2884 |
| chr6-124475126-INS-199 | chr6-124475126-INS-198 |
| chr6-124512470-INS-82 | chr6-124512472-INS-70 |
| chr6-124538053-INS-1054 | chr6-124538054-INS-1054 |
| chr6-124577222-INS-4960 | chr6-124577223-INS-4960 |
| chr6-124589352-INS-5382 | chr6-124589353-INS-5382 |
| chr6-124607573-INS-56 | chr6-124607551-INS-60 |
| chr6-124637536-INS-4010 | chr6-124637537-INS-4010 |
| chr6-124687993-INS-52 | chr6-124687993-INS-52 |
| chr6-124690465-INS-99 | chr6-124690465-INS-99 |
| chr6-124699840-INS-70 | chr6-124699844-INS-72 |
| chr6-124757339-INS-52 | chr6-124757339-INS-56 |
| chr6-124759005-INS-53 | chr6-124759005-INS-53 |
| chr6-124768340-INS-206 | chr6-124768340-INS-210 |
| chr6-124783940-INS-268 | chr6-124783940-INS-230 |
| chr6-124818943-INS-138 | chr6-124818943-INS-138 |
| chr6-124821634-INS-207 | chr6-124821634-INS-204 |
| chr6-124824316-INS-93 | chr6-124824318-INS-93 |
| chr6-124904979-INS-208 | chr6-124904979-INS-208 |
| chr6-124922576-INS-220 | chr6-124922576-INS-222 |
| chr6-124928109-INS-325 | chr6-124928109-INS-323 |
| chr6-124971671-INS-222 | chr6-124971671-INS-222 |
| chr6-124975709-INS-6442 | chr6-124975710-INS-6442 |
| chr6-124988658-INS-213 | chr6-124988658-INS-214 |
| chr6-124996941-INS-104 | chr6-124996941-INS-104 |
| chr6-125003636-INS-252 | chr6-125003636-INS-252 |
| chr6-125029896-INS-196 | chr6-125029895-INS-200 |
| chr6-125081537-INS-70 | chr6-125081523-INS-74 |
| chr6-125082633-INS-78 | chr6-125082633-INS-78 |
| chr6-125118762-INS-103 | chr6-125118749-INS-140 |
| chr6-125127545-INS-88 | chr6-125127545-INS-88 |
| chr6-125134644-INS-177 | chr6-125134644-INS-176 |
| chr6-125164175-INS-72 | chr6-125164175-INS-72 |
| chr6-125169870-INS-84 | chr6-125169880-INS-63 |
| chr6-125176165-INS-7155 | chr6-125176166-INS-7155 |
| chr6-125187701-INS-181 | chr6-125187701-INS-189 |
| chr6-125189408-INS-207 | chr6-125189408-INS-212 |
| chr6-125232511-INS-249 | chr6-125232511-INS-249 |
| chr6-125236286-INS-105 | chr6-125236286-INS-105 |
| chr6-125237493-INS-353 | chr6-125237489-INS-378 |
| chr6-125251904-INS-130 | chr6-125251904-INS-138 |
| chr6-125253192-INS-822 | chr6-125253193-INS-799 |
| chr6-125253960-INS-106 | chr6-125253960-INS-78 |
| chr6-125267556-INS-52 | chr6-125267556-INS-52 |
| chr6-125317768-INS-56 | chr6-125317768-INS-56 |
| chr6-125325137-INS-196 | chr6-125325144-INS-168 |
| chr6-125342559-INS-68 | chr6-125342566-INS-80 |
| chr6-125351222-INS-60 | chr6-125351222-INS-60 |
| chr6-125353188-INS-104 | chr6-125353188-INS-104 |
| chr6-125384143-INS-67 | chr6-125384143-INS-67 |
| chr6-125413580-INS-167 | chr6-125413597-INS-199 |
| chr6-125418921-INS-221 | chr6-125418921-INS-221 |
| chr6-125464309-INS-965 | chr6-125464309-INS-965 |
| chr6-125476517-INS-51 | chr6-125476517-INS-51 |
| chr6-125478453-INS-212 | chr6-125478453-INS-211 |
| chr6-125484782-INS-252 | chr6-125484782-INS-252 |
| chr6-125509765-INS-69 | chr6-125509815-INS-50 |
| chr6-125540047-INS-125 | chr6-125540047-INS-125 |
| chr6-125594961-INS-64 | chr6-125594963-INS-55 |
| chr6-125610019-INS-54 | chr6-125610046-INS-74 |
| chr6-125639226-INS-52 | chr6-125639228-INS-69 |
| chr6-125701729-INS-1892 | chr6-125701730-INS-1892 |
| chr6-125727876-INS-325 | chr6-125727869-INS-325 |
| chr6-125742003-INS-69 | chr6-125742003-INS-63 |
| chr6-125748325-INS-66 | chr6-125748325-INS-71 |
| chr6-125762459-INS-129 | chr6-125762459-INS-119 |
| chr6-125807718-INS-229 | chr6-125807718-INS-229 |
| chr6-125814406-INS-379 | chr6-125814406-INS-379 |
| chr6-125854587-INS-552 | chr6-125854587-INS-551 |
| chr6-125914303-INS-65 | chr6-125914278-INS-75 |
| chr6-125923893-INS-152 | chr6-125923893-INS-152 |
| chr6-125942972-INS-1811 | chr6-125942973-INS-1810 |
| chr6-125962338-INS-72 | chr6-125962338-INS-72 |

| | |
|---|---|
| chr6-125964224-INS-6181 | chr6-125964225-INS-6182 |
| chr6-125989934-INS-171 | chr6-125989932-INS-136 |
| chr6-126017699-INS-404 | chr6-126017699-INS-404 |
| chr6-126018912-INS-392 | chr6-126018912-INS-393 |
| chr6-126028658-INS-755 | chr6-126028658-INS-756 |
| chr6-126043954-INS-436 | chr6-126043954-INS-436 |
| chr6-126086790-INS-72 | chr6-126086790-INS-72 |
| chr6-126118361-INS-63 | chr6-126118361-INS-93 |
| chr6-126122936-INS-84 | chr6-126122893-INS-104 |
| chr6-126138294-INS-52 | chr6-126138294-INS-52 |
| chr6-126141405-INS-51 | chr6-126141405-INS-51 |
| chr6-126166277-INS-58 | chr6-126166277-INS-74 |
| chr6-126179282-INS-245 | chr6-126179282-INS-205 |
| chr6-126199938-INS-291 | chr6-126199938-INS-292 |
| chr6-126206843-INS-114 | chr6-126206843-INS-114 |
| chr6-126210514-INS-68 | chr6-126210514-INS-68 |
| chr6-126216699-INS-58 | chr6-126216711-INS-64 |
| chr6-126222802-INS-78 | chr6-126222802-INS-78 |
| chr6-126226834-INS-72 | chr6-126226834-INS-72 |
| chr6-126240242-INS-201 | chr6-126240242-INS-201 |
| chr6-126269900-INS-122 | chr6-126269900-INS-122 |
| chr6-126270923-INS-83 | chr6-126270957-INS-54 |
| chr6-126287843-INS-1375 | chr6-126287844-INS-1375 |
| chr6-126325165-INS-104 | chr6-126325158-INS-104 |
| chr6-126326471-INS-290 | chr6-126326471-INS-294 |
| chr6-126327488-INS-133 | chr6-126327488-INS-133 |
| chr6-126330767-INS-149 | chr6-126330768-INS-128 |
| chr6-126360916-INS-159 | chr6-126360916-INS-138 |
| chr6-126370838-INS-114 | chr6-126370838-INS-112 |
| chr6-126377240-INS-71 | chr6-126377240-INS-72 |
| chr6-126396172-INS-55 | chr6-126396172-INS-50 |
| chr6-126400431-INS-579 | chr6-126400431-INS-581 |
| chr6-126443757-INS-401 | chr6-126443757-INS-401 |
| chr6-126447118-INS-210 | chr6-126447115-INS-242 |
| chr6-126452262-INS-86 | chr6-126452261-INS-53 |
| chr6-126454864-INS-181 | chr6-126454864-INS-181 |
| chr6-126491166-INS-627 | chr6-126491206-INS-624 |
| chr6-126494902-INS-102 | chr6-126494902-INS-111 |
| chr6-126500292-INS-57 | chr6-126500292-INS-57 |
| chr6-126509828-INS-55 | chr6-126509828-INS-55 |
| chr6-126529475-INS-524 | chr6-126529475-INS-525 |
| chr6-126540228-INS-58 | chr6-126540228-INS-66 |
| chr6-126546628-INS-634 | chr6-126546628-INS-635 |
| chr6-126591511-INS-236 | chr6-126591511-INS-262 |
| chr6-126639434-INS-461 | chr6-126639434-INS-461 |
| chr6-126664746-INS-256 | chr6-126664746-INS-263 |
| chr6-126710563-INS-103 | chr6-126710588-INS-76 |
| chr6-126728070-INS-108 | chr6-126728070-INS-107 |
| chr6-126757147-INS-65 | chr6-126757147-INS-65 |
| chr6-126776281-INS-388 | chr6-126776281-INS-388 |
| chr6-126813003-INS-68 | chr6-126813003-INS-68 |
| chr6-126833286-INS-217 | chr6-126833286-INS-219 |
| chr6-126883515-INS-440 | chr6-126883516-INS-440 |
| chr6-126898327-INS-103 | chr6-126898327-INS-103 |
| chr6-126910486-INS-4740 | chr6-126910487-INS-4740 |
| chr6-127044027-INS-52 | chr6-127044029-INS-50 |
| chr6-127046218-INS-182 | chr6-127046219-INS-182 |
| chr6-127047047-INS-80 | chr6-127047048-INS-75 |
| chr6-127048122-INS-525 | chr6-127048122-INS-525 |
| chr6-127061509-INS-185 | chr6-127061509-INS-185 |
| chr6-127068252-INS-52 | chr6-127068252-INS-52 |
| chr6-127070959-INS-58 | chr6-127070959-INS-58 |
| chr6-127100973-INS-81 | chr6-127100979-INS-65 |
| chr6-127195900-INS-57 | chr6-127195899-INS-60 |
| chr6-127206894-INS-157 | chr6-127206894-INS-157 |
| chr6-127247775-INS-106 | chr6-127247775-INS-82 |
| chr6-127259533-INS-1738 | chr6-127259534-INS-1737 |
| chr6-127260042-INS-178 | chr6-127260042-INS-177 |
| chr6-127274583-INS-63 | chr6-127274577-INS-78 |
| chr6-127286988-INS-146 | chr6-127286988-INS-146 |
| chr6-127292672-INS-282 | chr6-127292672-INS-294 |
| chr6-127299184-INS-55 | chr6-127299184-INS-76 |
| chr6-127312167-INS-52 | chr6-127312167-INS-52 |
| chr6-127316018-INS-2733 | chr6-127316019-INS-2733 |
| chr6-127377668-INS-193 | chr6-127377668-INS-193 |
| chr6-127389215-INS-89 | chr6-127389215-INS-89 |
| chr6-127394000-INS-81 | chr6-127394000-INS-81 |

| | |
|---|---|
| chr6-127414901-INS-6481 | chr6-127414902-INS-6481 |
| chr6-127437892-INS-5433 | chr6-127437893-INS-5433 |
| chr6-127445869-INS-94 | chr6-127445869-INS-82 |
| chr6-127450711-INS-111 | chr6-127450715-INS-63 |
| chr6-127461353-INS-233 | chr6-127461353-INS-229 |
| chr6-127465271-INS-54 | chr6-127465256-INS-53 |
| chr6-127486127-INS-68 | chr6-127486127-INS-72 |
| chr6-127501843-INS-74 | chr6-127501842-INS-78 |
| chr6-127522319-INS-86 | chr6-127522319-INS-86 |
| chr6-127548975-INS-73 | chr6-127548975-INS-73 |
| chr6-127551581-INS-56 | chr6-127551570-INS-74 |
| chr6-127618038-INS-399 | chr6-127618038-INS-399 |
| chr6-127685015-INS-77 | chr6-127685015-INS-59 |
| chr6-127702338-INS-144 | chr6-127702338-INS-144 |
| chr6-127703651-INS-178 | chr6-127703651-INS-184 |
| chr6-127821787-INS-56 | chr6-127821787-INS-52 |
| chr6-127841383-INS-415 | chr6-127841383-INS-422 |
| chr6-127859515-INS-64 | chr6-127859515-INS-56 |
| chr6-127877491-INS-62 | chr6-127877491-INS-62 |
| chr6-127898374-INS-90 | chr6-127898374-INS-70 |
| chr6-127926058-INS-282 | chr6-127926058-INS-282 |
| chr6-127951605-INS-251 | chr6-127951605-INS-264 |
| chr6-128114673-INS-205 | chr6-128114691-INS-210 |
| chr6-128117638-INS-227 | chr6-128117638-INS-220 |
| chr6-128175859-INS-476 | chr6-128175859-INS-475 |
| chr6-128183859-INS-70 | chr6-128183859-INS-70 |
| chr6-128212880-INS-193 | chr6-128212880-INS-193 |
| chr6-128218854-INS-211 | chr6-128218854-INS-211 |
| chr6-128254274-INS-60 | chr6-128254274-INS-60 |
| chr6-128257136-INS-112 | chr6-128257136-INS-111 |
| chr6-128281714-INS-58 | chr6-128281694-INS-70 |
| chr6-128285587-INS-88 | chr6-128285587-INS-84 |
| chr6-128296101-INS-735 | chr6-128296101-INS-735 |
| chr6-128357684-INS-78 | chr6-128357684-INS-78 |
| chr6-128383538-INS-165 | chr6-128383538-INS-164 |
| chr6-128390734-INS-112 | chr6-128390734-INS-91 |
| chr6-128391849-INS-202 | chr6-128391849-INS-204 |
| chr6-128396142-INS-7087 | chr6-128396143-INS-7087 |
| chr6-128405425-INS-385 | chr6-128405425-INS-385 |
| chr6-128420685-INS-129 | chr6-128420685-INS-129 |
| chr6-128428831-INS-389 | chr6-128428831-INS-390 |
| chr6-128443960-INS-205 | chr6-128443960-INS-204 |
| chr6-128448254-INS-60 | chr6-128448254-INS-60 |
| chr6-128451488-INS-79 | chr6-128451488-INS-82 |
| chr6-128456722-INS-97 | chr6-128456722-INS-103 |
| chr6-128490203-INS-69 | chr6-128490200-INS-84 |
| chr6-128496768-INS-60 | chr6-128496768-INS-64 |
| chr6-128541400-INS-143 | chr6-128541400-INS-143 |
| chr6-128548419-INS-486 | chr6-128548419-INS-486 |
| chr6-128562199-INS-98 | chr6-128562199-INS-98 |
| chr6-128571023-INS-829 | chr6-128571023-INS-829 |
| chr6-128602468-INS-112 | chr6-128602471-INS-112 |
| chr6-128620850-INS-228 | chr6-128620850-INS-227 |
| chr6-128624955-INS-5440 | chr6-128624956-INS-5440 |
| chr6-128685259-INS-341 | chr6-128685260-INS-341 |
| chr6-128713116-INS-635 | chr6-128713116-INS-635 |
| chr6-128743244-INS-202 | chr6-128743244-INS-202 |
| chr6-128744331-INS-754 | chr6-128744331-INS-790 |
| chr6-128767723-INS-448 | chr6-128767723-INS-448 |
| chr6-128770051-INS-64 | chr6-128770051-INS-64 |
| chr6-128778093-INS-107 | chr6-128778093-INS-107 |
| chr6-128788184-INS-860 | chr6-128788184-INS-860 |
| chr6-128791418-INS-205 | chr6-128791418-INS-204 |
| chr6-128796563-INS-91 | chr6-128796563-INS-91 |
| chr6-128798361-INS-288 | chr6-128798361-INS-322 |
| chr6-128829562-INS-74 | chr6-128829562-INS-74 |
| chr6-128846022-INS-94 | chr6-128846022-INS-94 |
| chr6-128862976-INS-235 | chr6-128862974-INS-236 |
| chr6-128927657-INS-55 | chr6-128927657-INS-50 |
| chr6-128940528-INS-51 | chr6-128940528-INS-51 |
| chr6-128983434-INS-64 | chr6-128983434-INS-64 |
| chr6-129011452-INS-458 | chr6-129011452-INS-457 |
| chr6-129013083-INS-462 | chr6-129013083-INS-462 |
| chr6-129020021-INS-232 | chr6-129020021-INS-232 |
| chr6-129044982-INS-160 | chr6-129044982-INS-160 |
| chr6-129058282-INS-108 | chr6-129058282-INS-108 |
| chr6-129061265-INS-3127 | chr6-129061266-INS-3126 |

| | |
|---|---|
| chr6-129061266-INS-530 | chr6-129061266-INS-530 |
| chr6-129063036-INS-5719 | chr6-129063037-INS-5717 |
| chr6-129072440-INS-747 | chr6-129072440-INS-747 |
| chr6-129090703-INS-76 | chr6-129090703-INS-78 |
| chr6-129105675-INS-64 | chr6-129105675-INS-68 |
| chr6-129110373-INS-331 | chr6-129110374-INS-331 |
| chr6-129116778-INS-2897 | chr6-129116779-INS-2897 |
| chr6-129128361-INS-5102 | chr6-129128362-INS-5108 |
| chr6-129128362-INS-529 | chr6-129128362-INS-529 |
| chr6-129130901-INS-965 | chr6-129130901-INS-965 |
| chr6-129134566-INS-3729 | chr6-129134567-INS-3729 |
| chr6-129151285-INS-185 | chr6-129151285-INS-185 |
| chr6-129153586-INS-6440 | chr6-129153587-INS-6440 |
| chr6-129173715-INS-7093 | chr6-129173716-INS-7093 |
| chr6-129201641-INS-131 | chr6-129201641-INS-131 |
| chr6-129205650-INS-96 | chr6-129205650-INS-66 |
| chr6-129220637-INS-2362 | chr6-129220638-INS-2362 |
| chr6-129240208-INS-193 | chr6-129240208-INS-175 |
| chr6-129246440-INS-69 | chr6-129246448-INS-72 |
| chr6-129258563-INS-209 | chr6-129258563-INS-209 |
| chr6-129259169-INS-55 | chr6-129259169-INS-56 |
| chr6-129264201-INS-486 | chr6-129264201-INS-488 |
| chr6-129268683-INS-73 | chr6-129268679-INS-53 |
| chr6-129272134-INS-68 | chr6-129272132-INS-51 |
| chr6-129310844-INS-1843 | chr6-129310845-INS-1843 |
| chr6-129328871-INS-68 | chr6-129328871-INS-116 |
| chr6-129333486-INS-160 | chr6-129333486-INS-160 |
| chr6-129338241-INS-55 | chr6-129338242-INS-50 |
| chr6-129340045-INS-104 | chr6-129340045-INS-104 |
| chr6-129347413-INS-103 | chr6-129347413-INS-103 |
| chr6-129355301-INS-676 | chr6-129355301-INS-676 |
| chr6-129359925-INS-220 | chr6-129359925-INS-220 |
| chr6-129381533-INS-80 | chr6-129381535-INS-127 |
| chr6-129391591-INS-60 | chr6-129391591-INS-64 |
| chr6-129393817-INS-60 | chr6-129393816-INS-90 |
| chr6-129410417-INS-105 | chr6-129410417-INS-95 |
| chr6-129421921-INS-60 | chr6-129421921-INS-56 |
| chr6-129463734-INS-178 | chr6-129463734-INS-172 |
| chr6-129483276-INS-57 | chr6-129483276-INS-61 |
| chr6-129489229-INS-78 | chr6-129489261-INS-64 |
| chr6-129544895-INS-127 | chr6-129544895-INS-127 |
| chr6-129560581-INS-96 | chr6-129560581-INS-92 |
| chr6-129562195-INS-105 | chr6-129562199-INS-97 |
| chr6-129582788-INS-433 | chr6-129582788-INS-434 |
| chr6-129608105-INS-78 | chr6-129608145-INS-60 |
| chr6-129609429-INS-722 | chr6-129609429-INS-722 |
| chr6-129621006-INS-684 | chr6-129620992-INS-685 |
| chr6-129625027-INS-270 | chr6-129625027-INS-282 |
| chr6-129661672-INS-2799 | chr6-129661673-INS-2799 |
| chr6-129666578-INS-717 | chr6-129666578-INS-717 |
| chr6-129678976-INS-81 | chr6-129678976-INS-81 |
| chr6-129681300-INS-668 | chr6-129681300-INS-667 |
| chr6-129694694-INS-73 | chr6-129694698-INS-110 |
| chr6-129712672-INS-248 | chr6-129712672-INS-276 |
| chr6-129720735-INS-90 | chr6-129720727-INS-80 |
| chr6-129732276-INS-339 | chr6-129732276-INS-345 |
| chr6-129732920-INS-1073 | chr6-129732921-INS-1073 |
| chr6-129744568-INS-54 | chr6-129744568-INS-54 |
| chr6-129750735-INS-800 | chr6-129750735-INS-811 |
| chr6-129752051-INS-72 | chr6-129752051-INS-72 |
| chr6-129766354-INS-84 | chr6-129766354-INS-84 |
| chr6-129770186-INS-204 | chr6-129770186-INS-204 |
| chr6-129772431-INS-288 | chr6-129772431-INS-289 |
| chr6-129797934-INS-61 | chr6-129797934-INS-58 |
| chr6-129803091-INS-7087 | chr6-129803092-INS-7087 |
| chr6-129804488-INS-390 | chr6-129804488-INS-390 |
| chr6-129806497-INS-54 | chr6-129806457-INS-74 |
| chr6-129810410-INS-6378 | chr6-129810411-INS-6379 |
| chr6-129926283-INS-602 | chr6-129926283-INS-603 |
| chr6-129968894-INS-214 | chr6-129968894-INS-212 |
| chr6-129978047-INS-322 | chr6-129978047-INS-322 |
| chr6-129978836-INS-6263 | chr6-129978837-INS-6263 |
| chr6-129982698-INS-55 | chr6-129982698-INS-55 |
| chr6-130066420-INS-227 | chr6-130066422-INS-232 |
| chr6-130112574-INS-54 | chr6-130112555-INS-82 |
| chr6-130116133-INS-59 | chr6-130116133-INS-60 |
| chr6-130117225-INS-573 | chr6-130117225-INS-573 |

| | |
|---|---|
| chr6-130123357-INS-64 | chr6-130123357-INS-64 |
| chr6-130153198-INS-76 | chr6-130153225-INS-62 |
| chr6-130174243-INS-282 | chr6-130174237-INS-288 |
| chr6-130192771-INS-5645 | chr6-130192774-INS-5645 |
| chr6-130194856-INS-84 | chr6-130194856-INS-61 |
| chr6-130195395-INS-52 | chr6-130195395-INS-52 |
| chr6-130197529-INS-2290 | chr6-130197530-INS-2293 |
| chr6-130197848-INS-102 | chr6-130197848-INS-94 |
| chr6-130214466-INS-60 | chr6-130214450-INS-50 |
| chr6-130234799-INS-499 | chr6-130234800-INS-498 |
| chr6-130235320-INS-207 | chr6-130235320-INS-207 |
| chr6-130236356-INS-355 | chr6-130236356-INS-355 |
| chr6-130238701-INS-67 | chr6-130238701-INS-76 |
| chr6-130262736-INS-52 | chr6-130262730-INS-68 |
| chr6-130267263-INS-84 | chr6-130267263-INS-96 |
| chr6-130276422-INS-53 | chr6-130276422-INS-52 |
| chr6-130297447-INS-88 | chr6-130297497-INS-56 |
| chr6-130340307-INS-114 | chr6-130340351-INS-80 |
| chr6-130354140-INS-52 | chr6-130354140-INS-56 |
| chr6-130407937-INS-1260 | chr6-130407938-INS-1260 |
| chr6-130408109-INS-54 | chr6-130408110-INS-72 |
| chr6-130423736-INS-7261 | chr6-130423736-INS-7261 |
| chr6-130455650-INS-64 | chr6-130455650-INS-64 |
| chr6-130463233-INS-58 | chr6-130463233-INS-60 |
| chr6-130492292-INS-52 | chr6-130492290-INS-50 |
| chr6-130500842-INS-6388 | chr6-130500843-INS-6388 |
| chr6-130519575-INS-4527 | chr6-130519576-INS-4527 |
| chr6-130520677-INS-227 | chr6-130520677-INS-227 |
| chr6-130542065-INS-2812 | chr6-130542066-INS-2812 |
| chr6-130558421-INS-194 | chr6-130558421-INS-217 |
| chr6-130579770-INS-201 | chr6-130579818-INS-201 |
| chr6-130582298-INS-660 | chr6-130582298-INS-661 |
| chr6-130595582-INS-55 | chr6-130595582-INS-55 |
| chr6-130600376-INS-1787 | chr6-130600377-INS-1788 |
| chr6-130601142-INS-108 | chr6-130601142-INS-105 |
| chr6-130652910-INS-252 | chr6-130652910-INS-252 |
| chr6-130659190-INS-51 | chr6-130659150-INS-54 |
| chr6-130707937-INS-138 | chr6-130707938-INS-138 |
| chr6-130732807-INS-214 | chr6-130732807-INS-214 |
| chr6-130752629-INS-220 | chr6-130752629-INS-219 |
| chr6-130755294-INS-131 | chr6-130755304-INS-134 |
| chr6-130764624-INS-84 | chr6-130764624-INS-84 |
| chr6-130787848-INS-204 | chr6-130787848-INS-204 |
| chr6-130803954-INS-110 | chr6-130803949-INS-120 |
| chr6-130809469-INS-89 | chr6-130809475-INS-59 |
| chr6-130819386-INS-1673 | chr6-130819387-INS-1672 |
| chr6-130866595-INS-4334 | chr6-130866596-INS-4334 |
| chr6-130867679-INS-101 | chr6-130867679-INS-100 |
| chr6-130884819-INS-60 | chr6-130884819-INS-56 |
| chr6-130973941-INS-7172 | chr6-130973942-INS-7172 |
| chr6-131019555-INS-55 | chr6-131019555-INS-55 |
| chr6-131043606-INS-442 | chr6-131043606-INS-442 |
| chr6-131044783-INS-68 | chr6-131044783-INS-68 |
| chr6-131051341-INS-517 | chr6-131051341-INS-517 |
| chr6-131052247-INS-499 | chr6-131052247-INS-499 |
| chr6-131092140-INS-428 | chr6-131092140-INS-428 |
| chr6-131106205-INS-473 | chr6-131106205-INS-473 |
| chr6-131110685-INS-85 | chr6-131110685-INS-95 |
| chr6-131121722-INS-222 | chr6-131121722-INS-222 |
| chr6-131165350-INS-66 | chr6-131165350-INS-66 |
| chr6-131234876-INS-2959 | chr6-131234877-INS-2959 |
| chr6-131238299-INS-365 | chr6-131238299-INS-371 |
| chr6-131241453-INS-420 | chr6-131241453-INS-408 |
| chr6-131250933-INS-114 | chr6-131250933-INS-114 |
| chr6-131261388-INS-201 | chr6-131261388-INS-201 |
| chr6-131266225-INS-139 | chr6-131266225-INS-139 |
| chr6-131267708-INS-66 | chr6-131267708-INS-70 |
| chr6-131275419-INS-560 | chr6-131275419-INS-560 |
| chr6-131280355-INS-184 | chr6-131280355-INS-184 |
| chr6-131287221-INS-178 | chr6-131287221-INS-178 |
| chr6-131303376-INS-73 | chr6-131303376-INS-73 |
| chr6-131311387-INS-772 | chr6-131311387-INS-772 |
| chr6-131322216-INS-222 | chr6-131322216-INS-222 |
| chr6-131331174-INS-204 | chr6-131331174-INS-204 |
| chr6-131339226-INS-59 | chr6-131339226-INS-80 |
| chr6-131352481-INS-90 | chr6-131352481-INS-91 |
| chr6-131354045-INS-202 | chr6-131354045-INS-200 |

| | |
|---|---|
| chr6-131371655-INS-102 | chr6-131371655-INS-102 |
| chr6-131404310-INS-530 | chr6-131404310-INS-530 |
| chr6-131412595-INS-207 | chr6-131412595-INS-215 |
| chr6-131433702-INS-94 | chr6-131433702-INS-94 |
| chr6-131435691-INS-210 | chr6-131435691-INS-210 |
| chr6-131444438-INS-102 | chr6-131444438-INS-96 |
| chr6-131446535-INS-230 | chr6-131446535-INS-230 |
| chr6-131450694-INS-364 | chr6-131450694-INS-364 |
| chr6-131466829-INS-215 | chr6-131466829-INS-215 |
| chr6-131504773-INS-251 | chr6-131504773-INS-251 |
| chr6-131506690-INS-119 | chr6-131506690-INS-119 |
| chr6-131551930-INS-206 | chr6-131551930-INS-206 |
| chr6-131600977-INS-186 | chr6-131600977-INS-186 |
| chr6-131629317-INS-67 | chr6-131629317-INS-71 |
| chr6-131750719-INS-207 | chr6-131750719-INS-207 |
| chr6-131758920-INS-517 | chr6-131758920-INS-517 |
| chr6-131764728-INS-111 | chr6-131764728-INS-111 |
| chr6-131774501-INS-416 | chr6-131774501-INS-416 |
| chr6-131801493-INS-423 | chr6-131801494-INS-423 |
| chr6-131802444-INS-59 | chr6-131802467-INS-61 |
| chr6-131880634-INS-82 | chr6-131880634-INS-82 |
| chr6-131883733-INS-6214 | chr6-131883734-INS-6214 |
| chr6-131930516-INS-76 | chr6-131930488-INS-98 |
| chr6-131964873-INS-66 | chr6-131964873-INS-66 |
| chr6-131985471-INS-56 | chr6-131985471-INS-58 |
| chr6-131997191-INS-351 | chr6-131997191-INS-351 |
| chr6-132006528-INS-65 | chr6-132006528-INS-65 |
| chr6-132008562-INS-52 | chr6-132008562-INS-53 |
| chr6-132048115-INS-72 | chr6-132048115-INS-70 |
| chr6-132062060-INS-104 | chr6-132062073-INS-60 |
| chr6-132066566-INS-1749 | chr6-132066567-INS-1749 |
| chr6-132071107-INS-1887 | chr6-132071111-INS-1888 |
| chr6-132081609-INS-709 | chr6-132081609-INS-709 |
| chr6-132091656-INS-54 | chr6-132091657-INS-54 |
| chr6-132110586-INS-441 | chr6-132110586-INS-441 |
| chr6-132118088-INS-573 | chr6-132118088-INS-573 |
| chr6-132139770-INS-476 | chr6-132139770-INS-476 |
| chr6-132160040-INS-95 | chr6-132160040-INS-95 |
| chr6-132317094-INS-68 | chr6-132317095-INS-63 |
| chr6-132329003-INS-375 | chr6-132329003-INS-375 |
| chr6-132338666-INS-356 | chr6-132338666-INS-327 |
| chr6-132344327-INS-581 | chr6-132344323-INS-584 |
| chr6-132384843-INS-197 | chr6-132384843-INS-197 |
| chr6-132392435-INS-6486 | chr6-132392435-INS-6486 |
| chr6-132407994-INS-3303 | chr6-132407995-INS-3303 |
| chr6-132453071-INS-96 | chr6-132453071-INS-116 |
| chr6-132456848-INS-76 | chr6-132456820-INS-108 |
| chr6-132469173-INS-426 | chr6-132469173-INS-426 |
| chr6-132487915-INS-70 | chr6-132487915-INS-70 |
| chr6-132488151-INS-4275 | chr6-132488152-INS-4276 |
| chr6-132520079-INS-575 | chr6-132520079-INS-575 |
| chr6-132530843-INS-118 | chr6-132530843-INS-118 |
| chr6-132549261-INS-123 | chr6-132549261-INS-123 |
| chr6-132579934-INS-173 | chr6-132579934-INS-165 |
| chr6-132638300-INS-945 | chr6-132638301-INS-944 |
| chr6-132642061-INS-84 | chr6-132642061-INS-93 |
| chr6-132696817-INS-56 | chr6-132696817-INS-62 |
| chr6-132708563-INS-7786 | chr6-132708564-INS-7804 |
| chr6-132725486-INS-6247 | chr6-132725487-INS-6247 |
| chr6-132727275-INS-63 | chr6-132727275-INS-63 |
| chr6-132736223-INS-106 | chr6-132736223-INS-106 |
| chr6-132750697-INS-61 | chr6-132750697-INS-61 |
| chr6-132755520-INS-306 | chr6-132755521-INS-307 |
| chr6-132759523-INS-79 | chr6-132759523-INS-85 |
| chr6-132771607-INS-166 | chr6-132771607-INS-174 |
| chr6-132775029-INS-56 | chr6-132775029-INS-80 |
| chr6-132844553-INS-56 | chr6-132844553-INS-56 |
| chr6-132852824-INS-208 | chr6-132852824-INS-208 |
| chr6-132861157-INS-92 | chr6-132861157-INS-92 |
| chr6-132873607-INS-560 | chr6-132873607-INS-560 |
| chr6-132875937-INS-161 | chr6-132875937-INS-175 |
| chr6-132912118-INS-55 | chr6-132912118-INS-55 |
| chr6-132931693-INS-50 | chr6-132931693-INS-67 |
| chr6-132932824-INS-498 | chr6-132932824-INS-498 |
| chr6-132936203-INS-310 | chr6-132936203-INS-310 |
| chr6-132979728-INS-1612 | chr6-132979729-INS-1612 |
| chr6-132991824-INS-434 | chr6-132991824-INS-434 |

| | |
|---|---|
| chr6-133012383-INS-197 | chr6-133012383-INS-197 |
| chr6-133015554-INS-87 | chr6-133015555-INS-103 |
| chr6-133016023-INS-717 | chr6-133016023-INS-719 |
| chr6-133022289-INS-242 | chr6-133022289-INS-242 |
| chr6-133025688-INS-4530 | chr6-133025689-INS-4530 |
| chr6-133027492-INS-384 | chr6-133027492-INS-384 |
| chr6-133054049-INS-64 | chr6-133054047-INS-84 |
| chr6-133075598-INS-2676 | chr6-133075599-INS-2677 |
| chr6-133081816-INS-54 | chr6-133081816-INS-54 |
| chr6-133083395-INS-103 | chr6-133083395-INS-103 |
| chr6-133101719-INS-95 | chr6-133101719-INS-83 |
| chr6-133153025-INS-67 | chr6-133153024-INS-55 |
| chr6-133179148-INS-447 | chr6-133179148-INS-447 |
| chr6-133180024-INS-1403 | chr6-133180025-INS-1403 |
| chr6-133219564-INS-54 | chr6-133219569-INS-51 |
| chr6-133228512-INS-366 | chr6-133228512-INS-357 |
| chr6-133241088-INS-2861 | chr6-133241089-INS-2861 |
| chr6-133268231-INS-66 | chr6-133268231-INS-60 |
| chr6-133297080-INS-98 | chr6-133297080-INS-99 |
| chr6-133313243-INS-2664 | chr6-133313244-INS-2664 |
| chr6-133336712-INS-90 | chr6-133336679-INS-93 |
| chr6-133353668-INS-90 | chr6-133353668-INS-90 |
| chr6-133376232-INS-388 | chr6-133376232-INS-388 |
| chr6-133378089-INS-65 | chr6-133378089-INS-65 |
| chr6-133398643-INS-60 | chr6-133398643-INS-56 |
| chr6-133416418-INS-97 | chr6-133416418-INS-98 |
| chr6-133460285-INS-53 | chr6-133460286-INS-58 |
| chr6-133469323-INS-186 | chr6-133469345-INS-202 |
| chr6-133491528-INS-872 | chr6-133491528-INS-872 |
| chr6-133569021-INS-141 | chr6-133569015-INS-148 |
| chr6-133570658-INS-52 | chr6-133570659-INS-52 |
| chr6-133576748-INS-52 | chr6-133576748-INS-52 |
| chr6-133578175-INS-53 | chr6-133578175-INS-53 |
| chr6-133579533-INS-72 | chr6-133579533-INS-74 |
| chr6-133586682-INS-122 | chr6-133586682-INS-122 |
| chr6-133595152-INS-491 | chr6-133595152-INS-491 |
| chr6-133599869-INS-72 | chr6-133599869-INS-76 |
| chr6-133600462-INS-702 | chr6-133600462-INS-700 |
| chr6-133621841-INS-4525 | chr6-133621842-INS-4525 |
| chr6-133641608-INS-6629 | chr6-133641609-INS-6630 |
| chr6-133645672-INS-77 | chr6-133645672-INS-77 |
| chr6-133657306-INS-54 | chr6-133657306-INS-72 |
| chr6-133664875-INS-6387 | chr6-133664876-INS-6387 |
| chr6-133673244-INS-58 | chr6-133673244-INS-56 |
| chr6-133703639-INS-79 | chr6-133703639-INS-79 |
| chr6-133705863-INS-567 | chr6-133705864-INS-567 |
| chr6-133721255-INS-3132 | chr6-133721256-INS-3132 |
| chr6-133737665-INS-122 | chr6-133737665-INS-122 |
| chr6-133746173-INS-109 | chr6-133746173-INS-106 |
| chr6-133757423-INS-8323 | chr6-133757424-INS-8323 |
| chr6-133775475-INS-79 | chr6-133775475-INS-85 |
| chr6-133789928-INS-122 | chr6-133789928-INS-122 |
| chr6-133831235-INS-1178 | chr6-133831236-INS-1178 |
| chr6-133837358-INS-6305 | chr6-133837359-INS-6305 |
| chr6-133846255-INS-388 | chr6-133846255-INS-388 |
| chr6-133875987-INS-127 | chr6-133875987-INS-127 |
| chr6-133881391-INS-75 | chr6-133881391-INS-75 |
| chr6-133909464-INS-447 | chr6-133909464-INS-447 |
| chr6-133914763-INS-54 | chr6-133914763-INS-56 |
| chr6-133915045-INS-343 | chr6-133915045-INS-343 |
| chr6-133918035-INS-487 | chr6-133918035-INS-487 |
| chr6-133936299-INS-54 | chr6-133936299-INS-62 |
| chr6-133948062-INS-52 | chr6-133948062-INS-56 |
| chr6-133963679-INS-78 | chr6-133963679-INS-75 |
| chr6-133976683-INS-1795 | chr6-133976684-INS-1796 |
| chr6-133978343-INS-201 | chr6-133978343-INS-201 |
| chr6-133984718-INS-289 | chr6-133984718-INS-293 |
| chr6-134002453-INS-264 | chr6-134002453-INS-264 |
| chr6-134005116-INS-179 | chr6-134005116-INS-180 |
| chr6-134031189-INS-72 | chr6-134031189-INS-72 |
| chr6-134038869-INS-52 | chr6-134038867-INS-52 |
| chr6-134048238-INS-165 | chr6-134048238-INS-166 |
| chr6-134052869-INS-209 | chr6-134052869-INS-210 |
| chr6-134056934-INS-66 | chr6-134056934-INS-78 |
| chr6-134066251-INS-70 | chr6-134066251-INS-70 |
| chr6-134093673-INS-58 | chr6-134093671-INS-58 |
| chr6-134096547-INS-98 | chr6-134096547-INS-64 |

| | |
|---|---|
| chr6-134108933-INS-86 | chr6-134108933-INS-86 |
| chr6-134165337-INS-64 | chr6-134165337-INS-55 |
| chr6-134214385-INS-54 | chr6-134214385-INS-52 |
| chr6-134258601-INS-67 | chr6-134258601-INS-66 |
| chr6-134260133-INS-684 | chr6-134260133-INS-684 |
| chr6-134311803-INS-90 | chr6-134311803-INS-90 |
| chr6-134317717-INS-1961 | chr6-134317718-INS-1962 |
| chr6-134324158-INS-93 | chr6-134324158-INS-93 |
| chr6-134344352-INS-114 | chr6-134344352-INS-114 |
| chr6-134347351-INS-193 | chr6-134347351-INS-187 |
| chr6-134356870-INS-55 | chr6-134356870-INS-63 |
| chr6-134361476-INS-384 | chr6-134361448-INS-376 |
| chr6-134385545-INS-97 | chr6-134385545-INS-97 |
| chr6-134401630-INS-64 | chr6-134401629-INS-76 |
| chr6-134412240-INS-52 | chr6-134412240-INS-54 |
| chr6-134413752-INS-168 | chr6-134413752-INS-168 |
| chr6-134425852-INS-244 | chr6-134425852-INS-242 |
| chr6-134428988-INS-217 | chr6-134428988-INS-217 |
| chr6-134438791-INS-298 | chr6-134438791-INS-298 |
| chr6-134441750-INS-782 | chr6-134441750-INS-782 |
| chr6-134459060-INS-51 | chr6-134459060-INS-51 |
| chr6-134461871-INS-145 | chr6-134461871-INS-145 |
| chr6-134485794-INS-72 | chr6-134485795-INS-64 |
| chr6-134496168-INS-212 | chr6-134496168-INS-213 |
| chr6-134497601-INS-218 | chr6-134497601-INS-218 |
| chr6-134524087-INS-147 | chr6-134524087-INS-147 |
| chr6-134531346-INS-67 | chr6-134531346-INS-67 |
| chr6-134548357-INS-192 | chr6-134548357-INS-192 |
| chr6-134601289-INS-56 | chr6-134601289-INS-56 |
| chr6-134610886-INS-52 | chr6-134610886-INS-52 |
| chr6-134622262-INS-102 | chr6-134622262-INS-102 |
| chr6-134639571-INS-198 | chr6-134639571-INS-198 |
| chr6-134648504-INS-84 | chr6-134648504-INS-90 |
| chr6-134651127-INS-78 | chr6-134651127-INS-78 |
| chr6-134656908-INS-215 | chr6-134656908-INS-216 |
| chr6-134661151-INS-200 | chr6-134661151-INS-200 |
| chr6-134681038-INS-240 | chr6-134681038-INS-240 |
| chr6-134727002-INS-210 | chr6-134727002-INS-210 |
| chr6-134736507-INS-100 | chr6-134736507-INS-104 |
| chr6-134747536-INS-194 | chr6-134747536-INS-194 |
| chr6-134765829-INS-193 | chr6-134765829-INS-193 |
| chr6-134769868-INS-64 | chr6-134769868-INS-64 |
| chr6-134788969-INS-3740 | chr6-134788970-INS-3738 |
| chr6-134831084-INS-57 | chr6-134831084-INS-57 |
| chr6-134851618-INS-216 | chr6-134851618-INS-216 |
| chr6-134854528-INS-125 | chr6-134854528-INS-92 |
| chr6-134855134-INS-118 | chr6-134855134-INS-114 |
| chr6-134866857-INS-200 | chr6-134866857-INS-208 |
| chr6-134886086-INS-213 | chr6-134886086-INS-212 |
| chr6-134942136-INS-171 | chr6-134942138-INS-175 |
| chr6-134947077-INS-83 | chr6-134947081-INS-79 |
| chr6-134950886-INS-79 | chr6-134950886-INS-79 |
| chr6-134959797-INS-51 | chr6-134959797-INS-51 |
| chr6-134969137-INS-101 | chr6-134969143-INS-99 |
| chr6-134982921-INS-52 | chr6-134982921-INS-52 |
| chr6-134997488-INS-201 | chr6-134997488-INS-201 |
| chr6-135016696-INS-66 | chr6-135016696-INS-68 |
| chr6-135019776-INS-96 | chr6-135019776-INS-96 |
| chr6-135039134-INS-303 | chr6-135039116-INS-321 |
| chr6-135045242-INS-104 | chr6-135045242-INS-106 |
| chr6-135065922-INS-51 | chr6-135065923-INS-51 |
| chr6-135067630-INS-75 | chr6-135067609-INS-84 |
| chr6-135068352-INS-97 | chr6-135068348-INS-53 |
| chr6-135081085-INS-92 | chr6-135081109-INS-69 |
| chr6-135109285-INS-52 | chr6-135109285-INS-52 |
| chr6-135127648-INS-137 | chr6-135127648-INS-140 |
| chr6-135129276-INS-56 | chr6-135129276-INS-64 |
| chr6-135136888-INS-202 | chr6-135136888-INS-207 |
| chr6-135137807-INS-106 | chr6-135137807-INS-106 |
| chr6-135162057-INS-532 | chr6-135162057-INS-534 |
| chr6-135166960-INS-97 | chr6-135166960-INS-97 |
| chr6-135189183-INS-60 | chr6-135189183-INS-60 |
| chr6-135208214-INS-77 | chr6-135208214-INS-77 |
| chr6-135219378-INS-52 | chr6-135219378-INS-52 |
| chr6-135233101-INS-81 | chr6-135233101-INS-81 |
| chr6-135239067-INS-84 | chr6-135239067-INS-84 |
| chr6-135245012-INS-72 | chr6-135245016-INS-86 |

| | |
|---|---|
| chr6-135253363-INS-60 | chr6-135253360-INS-65 |
| chr6-135257414-INS-51 | chr6-135257414-INS-51 |
| chr6-135276821-INS-62 | chr6-135276821-INS-62 |
| chr6-135293561-INS-79 | chr6-135293561-INS-103 |
| chr6-135294843-INS-224 | chr6-135294843-INS-224 |
| chr6-135316010-INS-63 | chr6-135316010-INS-84 |
| chr6-135325159-INS-69 | chr6-135325159-INS-69 |
| chr6-135333402-INS-52 | chr6-135333403-INS-66 |
| chr6-135344820-INS-54 | chr6-135344820-INS-66 |
| chr6-135347382-INS-51 | chr6-135347382-INS-60 |
| chr6-135389768-INS-52 | chr6-135389768-INS-52 |
| chr6-135430568-INS-241 | chr6-135430568-INS-241 |
| chr6-135473329-INS-6829 | chr6-135473330-INS-6828 |
| chr6-135485007-INS-154 | chr6-135485007-INS-154 |
| chr6-135520079-INS-70 | chr6-135520079-INS-70 |
| chr6-135527760-INS-55 | chr6-135527760-INS-50 |
| chr6-135532668-INS-457 | chr6-135532668-INS-474 |
| chr6-135541291-INS-268 | chr6-135541291-INS-231 |
| chr6-135545663-INS-589 | chr6-135545663-INS-589 |
| chr6-135563852-INS-231 | chr6-135563850-INS-231 |
| chr6-135572832-INS-52 | chr6-135572833-INS-53 |
| chr6-135586298-INS-117 | chr6-135586316-INS-92 |
| chr6-135622884-INS-355 | chr6-135622884-INS-356 |
| chr6-135625086-INS-66 | chr6-135625086-INS-66 |
| chr6-135625575-INS-80 | chr6-135625575-INS-104 |
| chr6-135656328-INS-72 | chr6-135656328-INS-78 |
| chr6-135666661-INS-54 | chr6-135666679-INS-52 |
| chr6-135668609-INS-167 | chr6-135668609-INS-167 |
| chr6-135698056-INS-190 | chr6-135698056-INS-190 |
| chr6-135705656-INS-61 | chr6-135705662-INS-54 |
| chr6-135724484-INS-123 | chr6-135724484-INS-123 |
| chr6-135730879-INS-107 | chr6-135730879-INS-107 |
| chr6-135741373-INS-128 | chr6-135741373-INS-128 |
| chr6-135776466-INS-72 | chr6-135776463-INS-78 |
| chr6-135841730-INS-126 | chr6-135841730-INS-126 |
| chr6-135862984-INS-108 | chr6-135862984-INS-108 |
| chr6-135890504-INS-130 | chr6-135890495-INS-90 |
| chr6-135893823-INS-114 | chr6-135893823-INS-107 |
| chr6-135897698-INS-60 | chr6-135897702-INS-60 |
| chr6-135906392-INS-183 | chr6-135906392-INS-183 |
| chr6-135914928-INS-58 | chr6-135914928-INS-60 |
| chr6-135927812-INS-55 | chr6-135927812-INS-55 |
| chr6-135933108-INS-52 | chr6-135933108-INS-54 |
| chr6-135959849-INS-129 | chr6-135959849-INS-123 |
| chr6-135962873-INS-56 | chr6-135962873-INS-56 |
| chr6-135964611-INS-74 | chr6-135964611-INS-74 |
| chr6-135973410-INS-85 | chr6-135973410-INS-85 |
| chr6-135977069-INS-252 | chr6-135977069-INS-250 |
| chr6-135978285-INS-6472 | chr6-135978286-INS-6472 |
| chr6-136026567-INS-128 | chr6-136026567-INS-128 |
| chr6-136029142-INS-80 | chr6-136029142-INS-80 |
| chr6-136032354-INS-120 | chr6-136032310-INS-160 |
| chr6-136032564-INS-1893 | chr6-136032565-INS-1893 |
| chr6-136050513-INS-6707 | chr6-136050514-INS-6693 |
| chr6-136052861-INS-56 | chr6-136052861-INS-58 |
| chr6-136075934-INS-651 | chr6-136075934-INS-651 |
| chr6-136100179-INS-57 | chr6-136100180-INS-63 |
| chr6-136101322-INS-212 | chr6-136101322-INS-229 |
| chr6-136111645-INS-55 | chr6-136111645-INS-55 |
| chr6-136137238-INS-95 | chr6-136137238-INS-95 |
| chr6-136145590-INS-60 | chr6-136145590-INS-50 |
| chr6-136176644-INS-140 | chr6-136176644-INS-152 |
| chr6-136202844-INS-252 | chr6-136202844-INS-263 |
| chr6-136247577-INS-923 | chr6-136247577-INS-966 |
| chr6-136251128-INS-97 | chr6-136251128-INS-97 |
| chr6-136255740-INS-2555 | chr6-136255741-INS-2554 |
| chr6-136273951-INS-6523 | chr6-136273952-INS-6524 |
| chr6-136274142-INS-147 | chr6-136274142-INS-147 |
| chr6-136275811-INS-104 | chr6-136275808-INS-154 |
| chr6-136290698-INS-185 | chr6-136290698-INS-190 |
| chr6-136308468-INS-56 | chr6-136308469-INS-56 |
| chr6-136310657-INS-64 | chr6-136310657-INS-64 |
| chr6-136314584-INS-168 | chr6-136314537-INS-189 |
| chr6-136337218-INS-159 | chr6-136337218-INS-159 |
| chr6-136350019-INS-108 | chr6-136350019-INS-108 |
| chr6-136356489-INS-617 | chr6-136356489-INS-617 |
| chr6-136358016-INS-206 | chr6-136358016-INS-206 |

| | |
|---|---|
| chr6-136358609-INS-201 | chr6-136358609-INS-200 |
| chr6-136368863-INS-196 | chr6-136368872-INS-188 |
| chr6-136371955-INS-90 | chr6-136371955-INS-92 |
| chr6-136375341-INS-287 | chr6-136375341-INS-287 |
| chr6-136385749-INS-76 | chr6-136385749-INS-88 |
| chr6-136400794-INS-213 | chr6-136400794-INS-213 |
| chr6-136401526-INS-60 | chr6-136401526-INS-60 |
| chr6-136426095-INS-207 | chr6-136426095-INS-214 |
| chr6-136456001-INS-197 | chr6-136456001-INS-197 |
| chr6-136467474-INS-54 | chr6-136467474-INS-54 |
| chr6-136474748-INS-203 | chr6-136474748-INS-203 |
| chr6-136503087-INS-200 | chr6-136503087-INS-200 |
| chr6-136507833-INS-81 | chr6-136507833-INS-81 |
| chr6-136554682-INS-209 | chr6-136554682-INS-209 |
| chr6-136563991-INS-251 | chr6-136563991-INS-251 |
| chr6-136575269-INS-200 | chr6-136575269-INS-199 |
| chr6-136577439-INS-176 | chr6-136577439-INS-176 |
| chr6-136609002-INS-62 | chr6-136609002-INS-62 |
| chr6-136616840-INS-283 | chr6-136616840-INS-283 |
| chr6-136620150-INS-404 | chr6-136620150-INS-406 |
| chr6-136641051-INS-62 | chr6-136641051-INS-62 |
| chr6-136644405-INS-208 | chr6-136644405-INS-194 |
| chr6-136646884-INS-58 | chr6-136646884-INS-66 |
| chr6-136664799-INS-418 | chr6-136664799-INS-416 |
| chr6-136708168-INS-312 | chr6-136708168-INS-308 |
| chr6-136721899-INS-99 | chr6-136721899-INS-99 |
| chr6-136727447-INS-62 | chr6-136727457-INS-59 |
| chr6-136729828-INS-85 | chr6-136729828-INS-85 |
| chr6-136740257-INS-82 | chr6-136740261-INS-73 |
| chr6-136763865-INS-184 | chr6-136763865-INS-216 |
| chr6-136764412-INS-2905 | chr6-136764413-INS-2878 |
| chr6-136765409-INS-167 | chr6-136765409-INS-144 |
| chr6-136767633-INS-204 | chr6-136767633-INS-204 |
| chr6-136771487-INS-152 | chr6-136771487-INS-152 |
| chr6-136774842-INS-52 | chr6-136774842-INS-52 |
| chr6-136784519-INS-60 | chr6-136784498-INS-72 |
| chr6-136796578-INS-98 | chr6-136796566-INS-102 |
| chr6-136799303-INS-52 | chr6-136799303-INS-52 |
| chr6-136823746-INS-101 | chr6-136823727-INS-105 |
| chr6-136850667-INS-72 | chr6-136850667-INS-72 |
| chr6-136863198-INS-1390 | chr6-136863199-INS-1385 |
| chr6-136863285-INS-300 | chr6-136863286-INS-300 |
| chr6-136875287-INS-99 | chr6-136875287-INS-99 |
| chr6-136878921-INS-152 | chr6-136878921-INS-152 |
| chr6-136884223-INS-168 | chr6-136884223-INS-174 |
| chr6-136890263-INS-332 | chr6-136890244-INS-315 |
| chr6-136901450-INS-55 | chr6-136901450-INS-56 |
| chr6-136903537-INS-72 | chr6-136903537-INS-72 |
| chr6-136906836-INS-68 | chr6-136906803-INS-68 |
| chr6-136911014-INS-115 | chr6-136911042-INS-113 |
| chr6-136916349-INS-94 | chr6-136916349-INS-89 |
| chr6-136927561-INS-2078 | chr6-136927562-INS-2081 |
| chr6-136943892-INS-52 | chr6-136943892-INS-52 |
| chr6-136956037-INS-120 | chr6-136956037-INS-122 |
| chr6-136984911-INS-390 | chr6-136984911-INS-390 |
| chr6-136986912-INS-60 | chr6-136986916-INS-80 |
| chr6-136992808-INS-106 | chr6-136992809-INS-106 |
| chr6-137003089-INS-57 | chr6-137003089-INS-63 |
| chr6-137003397-INS-998 | chr6-137003389-INS-999 |
| chr6-137020829-INS-67 | chr6-137020830-INS-67 |
| chr6-137030169-INS-57 | chr6-137030169-INS-57 |
| chr6-137057062-INS-107 | chr6-137057062-INS-108 |
| chr6-137064859-INS-82 | chr6-137064877-INS-106 |
| chr6-137076324-INS-145 | chr6-137076324-INS-146 |
| chr6-137093010-INS-1729 | chr6-137093011-INS-1729 |
| chr6-137112129-INS-254 | chr6-137112129-INS-227 |
| chr6-137126209-INS-84 | chr6-137126206-INS-83 |
| chr6-137155047-INS-196 | chr6-137155047-INS-196 |
| chr6-137161594-INS-100 | chr6-137161594-INS-100 |
| chr6-137163394-INS-88 | chr6-137163394-INS-90 |
| chr6-137166771-INS-269 | chr6-137166771-INS-269 |
| chr6-137167919-INS-454 | chr6-137167919-INS-454 |
| chr6-137185827-INS-52 | chr6-137185827-INS-52 |
| chr6-137192788-INS-314 | chr6-137192788-INS-325 |
| chr6-137195961-INS-2190 | chr6-137195962-INS-2163 |
| chr6-137225957-INS-66 | chr6-137225957-INS-66 |
| chr6-137227940-INS-54 | chr6-137227940-INS-62 |

| | |
|---|---|
| chr6-137252462-INS-53 | chr6-137252462-INS-53 |
| chr6-137286419-INS-208 | chr6-137286419-INS-192 |
| chr6-137291306-INS-52 | chr6-137291306-INS-52 |
| chr6-137296754-INS-69 | chr6-137296755-INS-69 |
| chr6-137303285-INS-90 | chr6-137303285-INS-57 |
| chr6-137306661-INS-275 | chr6-137306661-INS-275 |
| chr6-137327339-INS-934 | chr6-137327339-INS-935 |
| chr6-137350539-INS-103 | chr6-137350539-INS-103 |
| chr6-137364296-INS-119 | chr6-137364296-INS-117 |
| chr6-137387573-INS-60 | chr6-137387564-INS-51 |
| chr6-137423218-INS-106 | chr6-137423218-INS-106 |
| chr6-137474373-INS-89 | chr6-137474373-INS-89 |
| chr6-137497796-INS-168 | chr6-137497796-INS-168 |
| chr6-137503897-INS-204 | chr6-137503897-INS-204 |
| chr6-137505976-INS-207 | chr6-137505976-INS-245 |
| chr6-137532830-INS-438 | chr6-137532830-INS-438 |
| chr6-137535794-INS-198 | chr6-137535794-INS-198 |
| chr6-137550028-INS-217 | chr6-137550028-INS-218 |
| chr6-137556092-INS-199 | chr6-137556092-INS-199 |
| chr6-137570686-INS-58 | chr6-137570687-INS-58 |
| chr6-137576962-INS-552 | chr6-137576962-INS-552 |
| chr6-137583831-INS-109 | chr6-137583831-INS-109 |
| chr6-137629132-INS-172 | chr6-137629132-INS-177 |
| chr6-137636041-INS-199 | chr6-137636042-INS-199 |
| chr6-137658890-INS-164 | chr6-137658890-INS-164 |
| chr6-137669518-INS-174 | chr6-137669518-INS-195 |
| chr6-137697796-INS-52 | chr6-137697796-INS-52 |
| chr6-137704517-INS-476 | chr6-137704517-INS-476 |
| chr6-137706940-INS-170 | chr6-137706962-INS-122 |
| chr6-137721929-INS-54 | chr6-137721929-INS-54 |
| chr6-137766123-INS-64 | chr6-137766123-INS-64 |
| chr6-137790989-INS-447 | chr6-137790989-INS-447 |
| chr6-137795969-INS-58 | chr6-137795969-INS-56 |
| chr6-137797132-INS-53 | chr6-137797132-INS-53 |
| chr6-137801969-INS-75 | chr6-137801969-INS-75 |
| chr6-137813222-INS-87 | chr6-137813222-INS-87 |
| chr6-137815202-INS-176 | chr6-137815202-INS-176 |
| chr6-137820578-INS-525 | chr6-137820578-INS-525 |
| chr6-137834963-INS-60 | chr6-137834963-INS-64 |
| chr6-137843013-INS-172 | chr6-137843013-INS-172 |
| chr6-137844662-INS-104 | chr6-137844662-INS-100 |
| chr6-137851953-INS-923 | chr6-137851958-INS-924 |
| chr6-137857721-INS-501 | chr6-137857726-INS-534 |
| chr6-137872700-INS-54 | chr6-137872700-INS-54 |
| chr6-137899109-INS-72 | chr6-137899107-INS-70 |
| chr6-137911626-INS-165 | chr6-137911626-INS-201 |
| chr6-137921616-INS-62 | chr6-137921616-INS-56 |
| chr6-137928581-INS-202 | chr6-137928581-INS-201 |
| chr6-137935512-INS-641 | chr6-137935512-INS-642 |
| chr6-137944537-INS-463 | chr6-137944537-INS-463 |
| chr6-137971086-INS-105 | chr6-137971086-INS-107 |
| chr6-137988627-INS-51 | chr6-137988627-INS-51 |
| chr6-138017682-INS-200 | chr6-138017682-INS-200 |
| chr6-138025713-INS-74 | chr6-138025713-INS-74 |
| chr6-138037494-INS-813 | chr6-138037494-INS-813 |
| chr6-138089856-INS-144 | chr6-138089857-INS-138 |
| chr6-138097744-INS-75 | chr6-138097744-INS-61 |
| chr6-138102046-INS-51 | chr6-138102046-INS-53 |
| chr6-138106858-INS-54 | chr6-138106858-INS-54 |
| chr6-138108585-INS-70 | chr6-138108585-INS-70 |
| chr6-138114288-INS-222 | chr6-138114288-INS-222 |
| chr6-138120854-INS-57 | chr6-138120854-INS-68 |
| chr6-138142490-INS-98 | chr6-138142480-INS-62 |
| chr6-138146628-INS-295 | chr6-138146628-INS-295 |
| chr6-138161874-INS-93 | chr6-138161873-INS-88 |
| chr6-138180784-INS-557 | chr6-138180784-INS-558 |
| chr6-138203341-INS-67 | chr6-138203341-INS-67 |
| chr6-138213208-INS-53 | chr6-138213208-INS-55 |
| chr6-138250086-INS-56 | chr6-138250086-INS-60 |
| chr6-138267835-INS-105 | chr6-138267835-INS-105 |
| chr6-138279310-INS-206 | chr6-138279310-INS-195 |
| chr6-138297969-INS-56 | chr6-138297969-INS-56 |
| chr6-138302294-INS-522 | chr6-138302294-INS-522 |
| chr6-138327490-INS-151 | chr6-138327523-INS-139 |
| chr6-138331606-INS-66 | chr6-138331606-INS-66 |
| chr6-138367991-INS-131 | chr6-138367991-INS-131 |
| chr6-138389333-INS-60 | chr6-138389333-INS-62 |

| | |
|---|---|
| chr6-138400060-INS-81 | chr6-138400060-INS-114 |
| chr6-138407226-INS-392 | chr6-138407226-INS-402 |
| chr6-138475025-INS-130 | chr6-138475025-INS-138 |
| chr6-138477899-INS-76 | chr6-138477900-INS-76 |
| chr6-138488323-INS-431 | chr6-138488323-INS-432 |
| chr6-138493044-INS-6660 | chr6-138493045-INS-6663 |
| chr6-138505616-INS-58 | chr6-138505616-INS-50 |
| chr6-138518246-INS-174 | chr6-138518246-INS-175 |
| chr6-138533275-INS-84 | chr6-138533275-INS-84 |
| chr6-138562780-INS-681 | chr6-138562780-INS-681 |
| chr6-138575386-INS-498 | chr6-138575386-INS-498 |
| chr6-138585610-INS-247 | chr6-138585610-INS-250 |
| chr6-138596626-INS-807 | chr6-138596626-INS-807 |
| chr6-138620291-INS-56 | chr6-138620291-INS-56 |
| chr6-138622850-INS-267 | chr6-138622851-INS-267 |
| chr6-138646909-INS-54 | chr6-138646909-INS-60 |
| chr6-138648091-INS-52 | chr6-138648091-INS-52 |
| chr6-138672209-INS-54 | chr6-138672170-INS-50 |
| chr6-138695877-INS-199 | chr6-138695877-INS-195 |
| chr6-138715135-INS-63 | chr6-138715135-INS-67 |
| chr6-138745488-INS-149 | chr6-138745488-INS-149 |
| chr6-138752733-INS-64 | chr6-138752726-INS-64 |
| chr6-138758209-INS-119 | chr6-138758209-INS-119 |
| chr6-138773008-INS-368 | chr6-138773008-INS-368 |
| chr6-138788494-INS-508 | chr6-138788494-INS-508 |
| chr6-138809549-INS-58 | chr6-138809549-INS-58 |
| chr6-138814736-INS-61 | chr6-138814736-INS-61 |
| chr6-138861614-INS-601 | chr6-138861614-INS-601 |
| chr6-138881452-INS-60 | chr6-138881417-INS-63 |
| chr6-138901037-INS-65 | chr6-138901037-INS-65 |
| chr6-138908609-INS-58 | chr6-138908609-INS-60 |
| chr6-138923335-INS-59 | chr6-138923300-INS-68 |
| chr6-138964048-INS-67 | chr6-138964065-INS-62 |
| chr6-138989693-INS-52 | chr6-138989693-INS-54 |
| chr6-138993056-INS-390 | chr6-138993056-INS-390 |
| chr6-139037676-INS-52 | chr6-139037677-INS-52 |
| chr6-139042788-INS-5458 | chr6-139042789-INS-5458 |
| chr6-139081328-INS-65 | chr6-139081328-INS-65 |
| chr6-139092897-INS-478 | chr6-139092897-INS-478 |
| chr6-139107898-INS-96 | chr6-139107898-INS-96 |
| chr6-139185701-INS-94 | chr6-139185701-INS-94 |
| chr6-139189150-INS-1219 | chr6-139189151-INS-1223 |
| chr6-139200798-INS-100 | chr6-139200798-INS-100 |
| chr6-139228503-INS-93 | chr6-139228503-INS-93 |
| chr6-139249918-INS-179 | chr6-139249918-INS-179 |
| chr6-139250496-INS-344 | chr6-139250446-INS-354 |
| chr6-139276290-INS-54 | chr6-139276290-INS-54 |
| chr6-139280524-INS-181 | chr6-139280524-INS-181 |
| chr6-139285890-INS-52 | chr6-139285890-INS-56 |
| chr6-139302865-INS-7111 | chr6-139302866-INS-7111 |
| chr6-139305852-INS-64 | chr6-139305852-INS-92 |
| chr6-139325804-INS-806 | chr6-139325804-INS-806 |
| chr6-139361467-INS-56 | chr6-139361467-INS-62 |
| chr6-139369587-INS-391 | chr6-139369587-INS-391 |
| chr6-139391295-INS-697 | chr6-139391295-INS-697 |
| chr6-139396545-INS-134 | chr6-139396545-INS-93 |
| chr6-139404620-INS-52 | chr6-139404620-INS-52 |
| chr6-139409303-INS-431 | chr6-139409303-INS-431 |
| chr6-139431045-INS-51 | chr6-139431045-INS-51 |
| chr6-139434276-INS-191 | chr6-139434290-INS-205 |
| chr6-139435050-INS-54 | chr6-139435050-INS-54 |
| chr6-139439050-INS-116 | chr6-139439044-INS-162 |
| chr6-139484072-INS-52 | chr6-139484072-INS-50 |
| chr6-139486627-INS-52 | chr6-139486627-INS-54 |
| chr6-139497482-INS-6664 | chr6-139497483-INS-6664 |
| chr6-139528112-INS-1146 | chr6-139528113-INS-1146 |
| chr6-139534977-INS-64 | chr6-139534977-INS-50 |
| chr6-139539814-INS-314 | chr6-139539814-INS-314 |
| chr6-139540865-INS-78 | chr6-139540865-INS-78 |
| chr6-139541811-INS-321 | chr6-139541811-INS-321 |
| chr6-139574387-INS-86 | chr6-139574387-INS-98 |
| chr6-139664850-INS-144 | chr6-139664859-INS-164 |
| chr6-139671701-INS-62 | chr6-139671701-INS-62 |
| chr6-139706297-INS-898 | chr6-139706297-INS-898 |
| chr6-139721902-INS-56 | chr6-139721902-INS-52 |
| chr6-139726113-INS-100 | chr6-139726113-INS-100 |
| chr6-139748785-INS-178 | chr6-139748785-INS-178 |

| | |
|---|---|
| chr6-139794512-INS-74 | chr6-139794512-INS-84 |
| chr6-139799457-INS-57 | chr6-139799457-INS-57 |
| chr6-139804357-INS-59 | chr6-139804357-INS-59 |
| chr6-139809292-INS-118 | chr6-139809292-INS-118 |
| chr6-139822857-INS-55 | chr6-139822857-INS-60 |
| chr6-139825770-INS-205 | chr6-139825770-INS-205 |
| chr6-139829285-INS-842 | chr6-139829285-INS-842 |
| chr6-139835398-INS-68 | chr6-139835398-INS-66 |
| chr6-139838709-INS-250 | chr6-139838709-INS-250 |
| chr6-139846307-INS-280 | chr6-139846307-INS-280 |
| chr6-139864469-INS-179 | chr6-139864469-INS-180 |
| chr6-139881232-INS-61 | chr6-139881232-INS-61 |
| chr6-139888718-INS-73 | chr6-139888718-INS-73 |
| chr6-139894631-INS-548 | chr6-139894631-INS-548 |
| chr6-139907079-INS-69 | chr6-139907080-INS-69 |
| chr6-139910145-INS-56 | chr6-139910145-INS-52 |
| chr6-139943095-INS-352 | chr6-139943095-INS-350 |
| chr6-139949843-INS-6750 | chr6-139949844-INS-6750 |
| chr6-139951076-INS-135 | chr6-139951076-INS-135 |
| chr6-139965768-INS-111 | chr6-139965768-INS-111 |
| chr6-139969598-INS-6202 | chr6-139969599-INS-6202 |
| chr6-139981093-INS-138 | chr6-139981123-INS-143 |
| chr6-139991110-INS-120 | chr6-139991110-INS-84 |
| chr6-139998765-INS-56 | chr6-139998741-INS-56 |
| chr6-140000805-INS-465 | chr6-140000805-INS-466 |
| chr6-140001589-INS-931 | chr6-140001589-INS-931 |
| chr6-140007600-INS-2778 | chr6-140007601-INS-2778 |
| chr6-140008200-INS-56 | chr6-140008200-INS-56 |
| chr6-140019051-INS-90 | chr6-140019051-INS-90 |
| chr6-140020938-INS-183 | chr6-140020938-INS-189 |
| chr6-140039014-INS-71 | chr6-140039014-INS-71 |
| chr6-140049514-INS-117 | chr6-140049514-INS-117 |
| chr6-140058910-INS-55 | chr6-140058910-INS-55 |
| chr6-140077360-INS-520 | chr6-140077363-INS-521 |
| chr6-140087893-INS-91 | chr6-140087893-INS-91 |
| chr6-140090845-INS-487 | chr6-140090845-INS-487 |
| chr6-140109907-INS-159 | chr6-140109907-INS-159 |
| chr6-140116830-INS-89 | chr6-140116830-INS-67 |
| chr6-140117641-INS-1698 | chr6-140117642-INS-1698 |
| chr6-140120951-INS-87 | chr6-140120951-INS-87 |
| chr6-140128708-INS-7786 | chr6-140128709-INS-7787 |
| chr6-140151649-INS-91 | chr6-140151649-INS-103 |
| chr6-140159347-INS-90 | chr6-140159347-INS-122 |
| chr6-140189984-INS-68 | chr6-140189984-INS-76 |
| chr6-140204622-INS-90 | chr6-140204622-INS-94 |
| chr6-140210370-INS-246 | chr6-140210370-INS-252 |
| chr6-140211219-INS-556 | chr6-140211219-INS-572 |
| chr6-140223613-INS-234 | chr6-140223613-INS-218 |
| chr6-140234222-INS-408 | chr6-140234222-INS-407 |
| chr6-140236252-INS-62 | chr6-140236252-INS-62 |
| chr6-140242431-INS-108 | chr6-140242431-INS-60 |
| chr6-140247005-INS-111 | chr6-140247005-INS-97 |
| chr6-140275536-INS-68 | chr6-140275536-INS-51 |
| chr6-140278405-INS-238 | chr6-140278406-INS-232 |
| chr6-140294402-INS-87 | chr6-140294399-INS-128 |
| chr6-140316974-INS-58 | chr6-140316974-INS-58 |
| chr6-140325477-INS-290 | chr6-140325477-INS-290 |
| chr6-140325477-INS-640 | chr6-140325477-INS-640 |
| chr6-140333823-INS-94 | chr6-140333823-INS-94 |
| chr6-140340785-INS-51 | chr6-140340761-INS-99 |
| chr6-140359371-INS-70 | chr6-140359371-INS-74 |
| chr6-140373916-INS-257 | chr6-140373916-INS-257 |
| chr6-140391760-INS-1464 | chr6-140391761-INS-1422 |
| chr6-140395143-INS-204 | chr6-140395143-INS-202 |
| chr6-140467410-INS-98 | chr6-140467410-INS-98 |
| chr6-140493276-INS-215 | chr6-140493276-INS-215 |
| chr6-140517588-INS-2438 | chr6-140517589-INS-2452 |
| chr6-140535484-INS-217 | chr6-140535484-INS-217 |
| chr6-140537001-INS-1386 | chr6-140537002-INS-1384 |
| chr6-140553029-INS-232 | chr6-140553029-INS-229 |
| chr6-140558115-INS-508 | chr6-140558115-INS-510 |
| chr6-140558664-INS-1194 | chr6-140558665-INS-1194 |
| chr6-140561806-INS-296 | chr6-140561807-INS-297 |
| chr6-140562540-INS-234 | chr6-140562540-INS-236 |
| chr6-140583286-INS-207 | chr6-140583286-INS-208 |
| chr6-140588715-INS-724 | chr6-140588715-INS-724 |
| chr6-140590913-INS-69 | chr6-140590913-INS-69 |

| | |
|---|---|
| chr6-140612078-INS-210 | chr6-140612078-INS-210 |
| chr6-140627516-INS-162 | chr6-140627516-INS-174 |
| chr6-140643761-INS-204 | chr6-140643761-INS-204 |
| chr6-140703854-INS-330 | chr6-140703855-INS-330 |
| chr6-140707610-INS-249 | chr6-140707610-INS-251 |
| chr6-140713280-INS-185 | chr6-140713280-INS-207 |
| chr6-140721503-INS-201 | chr6-140721503-INS-201 |
| chr6-140723742-INS-99 | chr6-140723749-INS-50 |
| chr6-140731674-INS-58 | chr6-140731674-INS-50 |
| chr6-140736242-INS-302 | chr6-140736242-INS-304 |
| chr6-140761070-INS-138 | chr6-140761070-INS-138 |
| chr6-140773494-INS-347 | chr6-140773494-INS-348 |
| chr6-140778604-INS-938 | chr6-140778604-INS-938 |
| chr6-140795175-INS-239 | chr6-140795175-INS-244 |
| chr6-140796340-INS-486 | chr6-140796340-INS-486 |
| chr6-140857632-INS-114 | chr6-140857632-INS-116 |
| chr6-140874381-INS-200 | chr6-140874375-INS-202 |
| chr6-140927266-INS-52 | chr6-140927238-INS-60 |
| chr6-140949898-INS-168 | chr6-140949898-INS-180 |
| chr6-140998663-INS-224 | chr6-140998663-INS-224 |
| chr6-141020159-INS-52 | chr6-141020159-INS-52 |
| chr6-141029644-INS-136 | chr6-141029644-INS-136 |
| chr6-141037815-INS-95 | chr6-141037815-INS-95 |
| chr6-141053531-INS-78 | chr6-141053531-INS-78 |
| chr6-141059627-INS-129 | chr6-141059627-INS-129 |
| chr6-141061646-INS-57 | chr6-141061646-INS-57 |
| chr6-141067986-INS-796 | chr6-141067986-INS-796 |
| chr6-141073894-INS-79 | chr6-141073894-INS-79 |
| chr6-141079467-INS-72 | chr6-141079467-INS-72 |
| chr6-141105622-INS-102 | chr6-141105622-INS-96 |
| chr6-141108329-INS-201 | chr6-141108329-INS-201 |
| chr6-141115272-INS-261 | chr6-141115269-INS-268 |
| chr6-141118505-INS-455 | chr6-141118505-INS-455 |
| chr6-141123864-INS-475 | chr6-141123864-INS-473 |
| chr6-141153523-INS-61 | chr6-141153523-INS-61 |
| chr6-141174340-INS-107 | chr6-141174340-INS-107 |
| chr6-141185156-INS-163 | chr6-141185156-INS-163 |
| chr6-141238600-INS-190 | chr6-141238600-INS-190 |
| chr6-141270921-INS-52 | chr6-141270921-INS-52 |
| chr6-141292665-INS-725 | chr6-141292665-INS-725 |
| chr6-141294272-INS-191 | chr6-141294272-INS-191 |
| chr6-141295206-INS-198 | chr6-141295199-INS-162 |
| chr6-141300340-INS-893 | chr6-141300340-INS-893 |
| chr6-141312224-INS-92 | chr6-141312244-INS-97 |
| chr6-141333357-INS-133 | chr6-141333357-INS-146 |
| chr6-141364297-INS-51 | chr6-141364297-INS-51 |
| chr6-141383042-INS-60 | chr6-141383042-INS-60 |
| chr6-141386773-INS-209 | chr6-141386773-INS-206 |
| chr6-141400117-INS-236 | chr6-141400111-INS-280 |
| chr6-141406197-INS-226 | chr6-141406196-INS-226 |
| chr6-141411786-INS-60 | chr6-141411786-INS-60 |
| chr6-141421144-INS-68 | chr6-141421151-INS-70 |
| chr6-141423055-INS-78 | chr6-141423055-INS-78 |
| chr6-141425781-INS-181 | chr6-141425781-INS-180 |
| chr6-141428751-INS-74 | chr6-141428729-INS-52 |
| chr6-141477421-INS-83 | chr6-141477421-INS-83 |
| chr6-141481291-INS-52 | chr6-141481291-INS-52 |
| chr6-141491768-INS-136 | chr6-141491768-INS-136 |
| chr6-141507321-INS-638 | chr6-141507321-INS-638 |
| chr6-141521165-INS-66 | chr6-141521165-INS-66 |
| chr6-141532109-INS-58 | chr6-141532109-INS-74 |
| chr6-141581957-INS-275 | chr6-141581957-INS-276 |
| chr6-141615837-INS-298 | chr6-141615838-INS-319 |
| chr6-141637992-INS-127 | chr6-141637989-INS-147 |
| chr6-141641930-INS-1191 | chr6-141641932-INS-1190 |
| chr6-141648058-INS-2159 | chr6-141648059-INS-2159 |
| chr6-141648911-INS-77 | chr6-141648911-INS-77 |
| chr6-141656837-INS-1340 | chr6-141656838-INS-1326 |
| chr6-141673980-INS-150 | chr6-141674001-INS-103 |
| chr6-141683382-INS-63 | chr6-141683382-INS-66 |
| chr6-141688635-INS-66 | chr6-141688635-INS-66 |
| chr6-141709681-INS-448 | chr6-141709681-INS-448 |
| chr6-141721383-INS-451 | chr6-141721384-INS-451 |
| chr6-141723565-INS-52 | chr6-141723565-INS-52 |
| chr6-141725980-INS-56 | chr6-141725980-INS-56 |
| chr6-141760563-INS-385 | chr6-141760563-INS-393 |
| chr6-141798200-INS-196 | chr6-141798200-INS-196 |

| | |
|---|---|
| chr6-141820866-INS-4125 | chr6-141820867-INS-4125 |
| chr6-141823829-INS-583 | chr6-141823853-INS-596 |
| chr6-141827010-INS-52 | chr6-141827010-INS-54 |
| chr6-141827513-INS-57 | chr6-141827513-INS-57 |
| chr6-141836927-INS-165 | chr6-141836927-INS-165 |
| chr6-141845880-INS-55 | chr6-141845865-INS-60 |
| chr6-141861592-INS-126 | chr6-141861587-INS-132 |
| chr6-141876855-INS-81 | chr6-141876859-INS-54 |
| chr6-141900974-INS-297 | chr6-141900972-INS-300 |
| chr6-141910556-INS-345 | chr6-141910556-INS-345 |
| chr6-141916383-INS-365 | chr6-141916383-INS-365 |
| chr6-141928250-INS-60 | chr6-141928250-INS-60 |
| chr6-141931145-INS-220 | chr6-141931145-INS-220 |
| chr6-141953235-INS-2651 | chr6-141953236-INS-2653 |
| chr6-141973544-INS-1594 | chr6-141973545-INS-1594 |
| chr6-141975097-INS-60 | chr6-141975097-INS-60 |
| chr6-141994925-INS-280 | chr6-141994922-INS-283 |
| chr6-142018585-INS-128 | chr6-142018585-INS-128 |
| chr6-142039383-INS-147 | chr6-142039383-INS-152 |
| chr6-142045570-INS-221 | chr6-142045562-INS-269 |
| chr6-142123165-INS-87 | chr6-142123165-INS-91 |
| chr6-142134869-INS-56 | chr6-142134869-INS-52 |
| chr6-142153833-INS-1936 | chr6-142153834-INS-1937 |
| chr6-142157944-INS-54 | chr6-142157944-INS-54 |
| chr6-142159193-INS-332 | chr6-142159193-INS-332 |
| chr6-142176265-INS-91 | chr6-142176234-INS-114 |
| chr6-142182206-INS-85 | chr6-142182202-INS-82 |
| chr6-142197845-INS-458 | chr6-142197846-INS-459 |
| chr6-142233127-INS-230 | chr6-142233127-INS-230 |
| chr6-142238939-INS-539 | chr6-142238939-INS-539 |
| chr6-142240818-INS-58 | chr6-142240843-INS-70 |
| chr6-142255516-INS-432 | chr6-142255516-INS-432 |
| chr6-142286634-INS-148 | chr6-142286634-INS-149 |
| chr6-142328777-INS-2903 | chr6-142328778-INS-2903 |
| chr6-142350581-INS-120 | chr6-142350581-INS-120 |
| chr6-142358456-INS-71 | chr6-142358454-INS-77 |
| chr6-142405833-INS-64 | chr6-142405833-INS-64 |
| chr6-142415878-INS-73 | chr6-142415878-INS-84 |
| chr6-142425146-INS-222 | chr6-142425146-INS-222 |
| chr6-142432971-INS-206 | chr6-142432955-INS-213 |
| chr6-142437310-INS-124 | chr6-142437310-INS-124 |
| chr6-142465681-INS-324 | chr6-142465681-INS-324 |
| chr6-142467792-INS-92 | chr6-142467793-INS-92 |
| chr6-142469900-INS-82 | chr6-142469923-INS-66 |
| chr6-142475479-INS-612 | chr6-142475505-INS-607 |
| chr6-142482251-INS-78 | chr6-142482249-INS-77 |
| chr6-142527731-INS-227 | chr6-142527731-INS-227 |
| chr6-142552488-INS-73 | chr6-142552488-INS-64 |
| chr6-142570152-INS-114 | chr6-142570152-INS-114 |
| chr6-142588267-INS-60 | chr6-142588251-INS-57 |
| chr6-142608628-INS-60 | chr6-142608628-INS-60 |
| chr6-142624565-INS-206 | chr6-142624565-INS-206 |
| chr6-142644355-INS-55 | chr6-142644353-INS-51 |
| chr6-142650276-INS-64 | chr6-142650276-INS-64 |
| chr6-142663095-INS-103 | chr6-142663095-INS-103 |
| chr6-142670714-INS-3807 | chr6-142670715-INS-3807 |
| chr6-142684778-INS-320 | chr6-142684778-INS-320 |
| chr6-142715037-INS-57 | chr6-142715037-INS-57 |
| chr6-142746178-INS-61 | chr6-142746178-INS-67 |
| chr6-142784891-INS-53 | chr6-142784891-INS-53 |
| chr6-142819433-INS-51 | chr6-142819433-INS-51 |
| chr6-142833148-INS-59 | chr6-142833146-INS-65 |
| chr6-142879777-INS-96 | chr6-142879777-INS-96 |
| chr6-142882683-INS-64 | chr6-142882683-INS-64 |
| chr6-142884938-INS-88 | chr6-142884938-INS-88 |
| chr6-142886235-INS-150 | chr6-142886236-INS-150 |
| chr6-142888445-INS-64 | chr6-142888451-INS-72 |
| chr6-142913600-INS-112 | chr6-142913555-INS-123 |
| chr6-142929696-INS-166 | chr6-142929716-INS-190 |
| chr6-142932445-INS-198 | chr6-142932445-INS-200 |
| chr6-142959452-INS-80 | chr6-142959452-INS-120 |
| chr6-142963302-INS-221 | chr6-142963302-INS-221 |
| chr6-142967162-INS-202 | chr6-142967162-INS-202 |
| chr6-142979926-INS-919 | chr6-142979926-INS-920 |
| chr6-142993810-INS-71 | chr6-142993796-INS-86 |
| chr6-143026352-INS-80 | chr6-143026352-INS-84 |
| chr6-143028167-INS-210 | chr6-143028167-INS-209 |

| | |
|---|---|
| chr6-143040304-INS-74 | chr6-143040304-INS-74 |
| chr6-143049601-INS-207 | chr6-143049601-INS-207 |
| chr6-143064458-INS-56 | chr6-143064458-INS-88 |
| chr6-143081862-INS-172 | chr6-143081862-INS-174 |
| chr6-143084504-INS-86 | chr6-143084504-INS-82 |
| chr6-143106383-INS-191 | chr6-143106383-INS-189 |
| chr6-143108203-INS-52 | chr6-143108203-INS-53 |
| chr6-143109677-INS-66 | chr6-143109677-INS-66 |
| chr6-143113069-INS-116 | chr6-143113069-INS-116 |
| chr6-143169361-INS-204 | chr6-143169361-INS-204 |
| chr6-143174703-INS-58 | chr6-143174703-INS-58 |
| chr6-143223771-INS-54 | chr6-143223771-INS-54 |
| chr6-143273526-INS-56 | chr6-143273526-INS-58 |
| chr6-143275487-INS-52 | chr6-143275487-INS-50 |
| chr6-143277682-INS-117 | chr6-143277682-INS-117 |
| chr6-143284699-INS-52 | chr6-143284699-INS-52 |
| chr6-143306752-INS-65 | chr6-143306752-INS-90 |
| chr6-143351572-INS-52 | chr6-143351572-INS-91 |
| chr6-143373615-INS-86 | chr6-143373612-INS-113 |
| chr6-143396735-INS-86 | chr6-143396735-INS-86 |
| chr6-143419521-INS-92 | chr6-143419500-INS-127 |
| chr6-143430476-INS-397 | chr6-143430476-INS-397 |
| chr6-143442233-INS-62 | chr6-143442233-INS-60 |
| chr6-143470134-INS-63 | chr6-143470139-INS-66 |
| chr6-143472256-INS-115 | chr6-143472256-INS-147 |
| chr6-143556140-INS-129 | chr6-143556141-INS-101 |
| chr6-143571885-INS-86 | chr6-143571885-INS-86 |
| chr6-143596015-INS-52 | chr6-143596015-INS-54 |
| chr6-143607171-INS-339 | chr6-143607171-INS-339 |
| chr6-143614430-INS-54 | chr6-143614441-INS-56 |
| chr6-143624589-INS-77 | chr6-143624589-INS-91 |
| chr6-143677250-INS-340 | chr6-143677250-INS-341 |
| chr6-143689602-INS-491 | chr6-143689602-INS-491 |
| chr6-143704843-INS-157 | chr6-143704843-INS-157 |
| chr6-143728594-INS-164 | chr6-143728594-INS-170 |
| chr6-143740350-INS-6587 | chr6-143740351-INS-6587 |
| chr6-143768403-INS-108 | chr6-143768403-INS-102 |
| chr6-143803693-INS-54 | chr6-143803693-INS-54 |
| chr6-143804932-INS-72 | chr6-143804932-INS-72 |
| chr6-143834020-INS-69 | chr6-143834020-INS-69 |
| chr6-143937399-INS-434 | chr6-143937399-INS-438 |
| chr6-143984697-INS-54 | chr6-143984697-INS-51 |
| chr6-143993888-INS-61 | chr6-143993888-INS-62 |
| chr6-144008905-INS-98 | chr6-144008905-INS-98 |
| chr6-144011378-INS-56 | chr6-144011378-INS-56 |
| chr6-144075298-INS-118 | chr6-144075295-INS-118 |
| chr6-144155263-INS-58 | chr6-144155263-INS-56 |
| chr6-144165048-INS-60 | chr6-144165058-INS-60 |
| chr6-144169291-INS-103 | chr6-144169291-INS-54 |
| chr6-144181293-INS-147 | chr6-144181293-INS-147 |
| chr6-144209820-INS-455 | chr6-144209820-INS-455 |
| chr6-144237928-INS-145 | chr6-144237928-INS-144 |
| chr6-144239165-INS-59 | chr6-144239165-INS-61 |
| chr6-144267140-INS-150 | chr6-144267140-INS-150 |
| chr6-144273453-INS-52 | chr6-144273453-INS-50 |
| chr6-144275161-INS-124 | chr6-144275161-INS-124 |
| chr6-144278951-INS-413 | chr6-144278960-INS-423 |
| chr6-144306065-INS-4831 | chr6-144306066-INS-4831 |
| chr6-144307893-INS-193 | chr6-144307893-INS-191 |
| chr6-144316590-INS-74 | chr6-144316590-INS-74 |
| chr6-144334226-INS-54 | chr6-144334237-INS-58 |
| chr6-144338717-INS-145 | chr6-144338717-INS-145 |
| chr6-144345329-INS-54 | chr6-144345329-INS-56 |
| chr6-144347776-INS-56 | chr6-144347776-INS-52 |
| chr6-144362483-INS-52 | chr6-144362472-INS-64 |
| chr6-144368763-INS-145 | chr6-144368763-INS-122 |
| chr6-144377416-INS-244 | chr6-144377416-INS-244 |
| chr6-144381330-INS-473 | chr6-144381330-INS-466 |
| chr6-144397137-INS-337 | chr6-144397137-INS-337 |
| chr6-144404301-INS-486 | chr6-144404336-INS-486 |
| chr6-144414436-INS-466 | chr6-144414436-INS-467 |
| chr6-144431239-INS-147 | chr6-144431239-INS-148 |
| chr6-144443224-INS-6557 | chr6-144443225-INS-6557 |
| chr6-144484279-INS-135 | chr6-144484279-INS-135 |
| chr6-144488672-INS-89 | chr6-144488669-INS-96 |
| chr6-144489975-INS-52 | chr6-144489983-INS-52 |
| chr6-144497257-INS-304 | chr6-144497258-INS-304 |

| | |
|---|---|
| chr6-144498315-INS-466 | chr6-144498315-INS-466 |
| chr6-144502649-INS-82 | chr6-144502649-INS-82 |
| chr6-144521895-INS-68 | chr6-144521895-INS-70 |
| chr6-144529423-INS-169 | chr6-144529414-INS-192 |
| chr6-144539593-INS-316 | chr6-144539593-INS-316 |
| chr6-144567644-INS-52 | chr6-144567644-INS-56 |
| chr6-144574920-INS-76 | chr6-144574920-INS-52 |
| chr6-144581229-INS-54 | chr6-144581229-INS-54 |
| chr6-144592562-INS-168 | chr6-144592562-INS-168 |
| chr6-144593401-INS-196 | chr6-144593427-INS-168 |
| chr6-144635987-INS-126 | chr6-144635987-INS-114 |
| chr6-144665684-INS-61 | chr6-144665713-INS-73 |
| chr6-144698315-INS-71 | chr6-144698315-INS-71 |
| chr6-144706790-INS-52 | chr6-144706793-INS-52 |
| chr6-144708880-INS-111 | chr6-144708880-INS-100 |
| chr6-144713053-INS-107 | chr6-144713053-INS-83 |
| chr6-144739967-INS-549 | chr6-144739967-INS-547 |
| chr6-144742053-INS-58 | chr6-144742053-INS-58 |
| chr6-144746766-INS-70 | chr6-144746766-INS-70 |
| chr6-144767903-INS-76 | chr6-144767903-INS-76 |
| chr6-144771355-INS-97 | chr6-144771355-INS-97 |
| chr6-144786015-INS-82 | chr6-144786019-INS-82 |
| chr6-144800606-INS-107 | chr6-144800606-INS-107 |
| chr6-144806858-INS-212 | chr6-144806858-INS-212 |
| chr6-144815858-INS-53 | chr6-144815858-INS-59 |
| chr6-144833066-INS-126 | chr6-144833066-INS-127 |
| chr6-144837026-INS-51 | chr6-144837026-INS-51 |
| chr6-144844256-INS-1430 | chr6-144844257-INS-1430 |
| chr6-144845473-INS-54 | chr6-144845473-INS-54 |
| chr6-144852858-INS-110 | chr6-144852858-INS-130 |
| chr6-144866950-INS-101 | chr6-144866938-INS-107 |
| chr6-144872733-INS-95 | chr6-144872733-INS-75 |
| chr6-144874706-INS-137 | chr6-144874706-INS-137 |
| chr6-144886923-INS-57 | chr6-144886926-INS-81 |
| chr6-144915652-INS-116 | chr6-144915652-INS-116 |
| chr6-144917819-INS-447 | chr6-144917819-INS-447 |
| chr6-144967533-INS-627 | chr6-144967534-INS-631 |
| chr6-144980979-INS-51 | chr6-144980980-INS-53 |
| chr6-145003090-INS-179 | chr6-145003090-INS-179 |
| chr6-145012895-INS-634 | chr6-145012896-INS-630 |
| chr6-145018530-INS-746 | chr6-145018530-INS-747 |
| chr6-145026076-INS-78 | chr6-145026076-INS-79 |
| chr6-145029424-INS-213 | chr6-145029424-INS-214 |
| chr6-145037472-INS-110 | chr6-145037472-INS-110 |
| chr6-145040888-INS-61 | chr6-145040888-INS-61 |
| chr6-145049438-INS-199 | chr6-145049437-INS-205 |
| chr6-145054216-INS-203 | chr6-145054216-INS-203 |
| chr6-145078734-INS-51 | chr6-145078734-INS-51 |
| chr6-145084478-INS-521 | chr6-145084478-INS-521 |
| chr6-145106859-INS-389 | chr6-145106846-INS-387 |
| chr6-145111489-INS-222 | chr6-145111489-INS-222 |
| chr6-145137605-INS-72 | chr6-145137605-INS-96 |
| chr6-145141462-INS-202 | chr6-145141462-INS-202 |
| chr6-145152934-INS-121 | chr6-145152934-INS-121 |
| chr6-145155475-INS-467 | chr6-145155475-INS-467 |
| chr6-145203999-INS-96 | chr6-145203999-INS-96 |
| chr6-145208827-INS-120 | chr6-145208827-INS-120 |
| chr6-145219843-INS-100 | chr6-145219843-INS-101 |
| chr6-145245482-INS-87 | chr6-145245482-INS-92 |
| chr6-145264689-INS-400 | chr6-145264689-INS-400 |
| chr6-145269662-INS-56 | chr6-145269662-INS-56 |
| chr6-145276793-INS-224 | chr6-145276793-INS-230 |
| chr6-145295892-INS-218 | chr6-145295892-INS-218 |
| chr6-145300805-INS-60 | chr6-145300805-INS-60 |
| chr6-145317024-INS-62 | chr6-145317024-INS-62 |
| chr6-145320960-INS-58 | chr6-145320960-INS-58 |
| chr6-145351285-INS-201 | chr6-145351285-INS-201 |
| chr6-145364100-INS-64 | chr6-145364100-INS-64 |
| chr6-145385074-INS-261 | chr6-145385077-INS-262 |
| chr6-145388539-INS-512 | chr6-145388539-INS-512 |
| chr6-145390452-INS-112 | chr6-145390448-INS-112 |
| chr6-145395116-INS-72 | chr6-145395116-INS-114 |
| chr6-145430722-INS-96 | chr6-145430722-INS-96 |
| chr6-145448522-INS-90 | chr6-145448522-INS-135 |
| chr6-145474299-INS-60 | chr6-145474299-INS-60 |
| chr6-145483247-INS-114 | chr6-145483247-INS-108 |
| chr6-145492310-INS-648 | chr6-145492310-INS-648 |

| | |
|---|---|
| chr6-145496342-INS-92 | chr6-145496346-INS-86 |
| chr6-145501744-INS-530 | chr6-145501744-INS-527 |
| chr6-145512156-INS-154 | chr6-145512156-INS-157 |
| chr6-145520296-INS-401 | chr6-145520297-INS-401 |
| chr6-145526839-INS-422 | chr6-145526839-INS-422 |
| chr6-145529128-INS-71 | chr6-145529125-INS-73 |
| chr6-145533382-INS-55 | chr6-145533382-INS-55 |
| chr6-145562173-INS-64 | chr6-145562173-INS-64 |
| chr6-145575482-INS-62 | chr6-145575482-INS-64 |
| chr6-145585881-INS-7135 | chr6-145585882-INS-7135 |
| chr6-145599166-INS-132 | chr6-145599166-INS-132 |
| chr6-145600298-INS-54 | chr6-145600280-INS-86 |
| chr6-145607315-INS-214 | chr6-145607315-INS-214 |
| chr6-145618728-INS-7235 | chr6-145618729-INS-7235 |
| chr6-145641600-INS-54 | chr6-145641600-INS-52 |
| chr6-145653005-INS-62 | chr6-145653005-INS-62 |
| chr6-145669648-INS-1892 | chr6-145669649-INS-1892 |
| chr6-145670204-INS-1892 | chr6-145670205-INS-1892 |
| chr6-145670862-INS-520 | chr6-145670862-INS-520 |
| chr6-145719107-INS-90 | chr6-145719107-INS-90 |
| chr6-145741359-INS-52 | chr6-145741359-INS-52 |
| chr6-145749648-INS-72 | chr6-145749648-INS-72 |
| chr6-145753872-INS-58 | chr6-145753872-INS-58 |
| chr6-145757459-INS-64 | chr6-145757459-INS-70 |
| chr6-145769862-INS-506 | chr6-145769862-INS-506 |
| chr6-145775739-INS-245 | chr6-145775739-INS-231 |
| chr6-145777990-INS-79 | chr6-145777991-INS-83 |
| chr6-145787667-INS-62 | chr6-145787667-INS-62 |
| chr6-145827066-INS-92 | chr6-145827066-INS-92 |
| chr6-145832822-INS-86 | chr6-145832822-INS-87 |
| chr6-145834046-INS-4800 | chr6-145834047-INS-4762 |
| chr6-145864342-INS-92 | chr6-145864342-INS-92 |
| chr6-145872019-INS-136 | chr6-145872019-INS-138 |
| chr6-145916666-INS-72 | chr6-145916666-INS-72 |
| chr6-145920118-INS-282 | chr6-145920119-INS-281 |
| chr6-145924201-INS-60 | chr6-145924201-INS-60 |
| chr6-145930458-INS-198 | chr6-145930458-INS-199 |
| chr6-145946875-INS-78 | chr6-145946875-INS-78 |
| chr6-145947781-INS-504 | chr6-145947781-INS-504 |
| chr6-146007205-INS-735 | chr6-146007205-INS-729 |
| chr6-146021224-INS-3191 | chr6-146021225-INS-3191 |
| chr6-146029882-INS-4777 | chr6-146029883-INS-4779 |
| chr6-146036639-INS-1369 | chr6-146036640-INS-1377 |
| chr6-146044536-INS-332 | chr6-146044537-INS-333 |
| chr6-146065394-INS-68 | chr6-146065394-INS-68 |
| chr6-146084738-INS-106 | chr6-146084738-INS-68 |
| chr6-146086053-INS-7082 | chr6-146086054-INS-7082 |
| chr6-146119948-INS-67 | chr6-146119948-INS-67 |
| chr6-146131534-INS-62 | chr6-146131553-INS-66 |
| chr6-146153544-INS-122 | chr6-146153544-INS-122 |
| chr6-146163340-INS-575 | chr6-146163340-INS-578 |
| chr6-146193521-INS-79 | chr6-146193521-INS-79 |
| chr6-146208640-INS-145 | chr6-146208640-INS-145 |
| chr6-146210163-INS-189 | chr6-146210163-INS-189 |
| chr6-146222817-INS-96 | chr6-146222817-INS-96 |
| chr6-146227072-INS-58 | chr6-146227072-INS-58 |
| chr6-146248480-INS-132 | chr6-146248480-INS-132 |
| chr6-146253973-INS-6401 | chr6-146253974-INS-6401 |
| chr6-146261839-INS-69 | chr6-146261839-INS-69 |
| chr6-146265579-INS-65 | chr6-146265578-INS-69 |
| chr6-146268745-INS-60 | chr6-146268745-INS-80 |
| chr6-146271653-INS-497 | chr6-146271653-INS-497 |
| chr6-146272171-INS-1511 | chr6-146272172-INS-1511 |
| chr6-146293177-INS-88 | chr6-146293177-INS-90 |
| chr6-146295016-INS-95 | chr6-146295016-INS-98 |
| chr6-146315435-INS-757 | chr6-146315435-INS-757 |
| chr6-146364534-INS-74 | chr6-146364537-INS-80 |
| chr6-146377302-INS-300 | chr6-146377302-INS-300 |
| chr6-146379700-INS-161 | chr6-146379700-INS-170 |
| chr6-146383860-INS-60 | chr6-146383860-INS-60 |
| chr6-146408278-INS-93 | chr6-146408278-INS-93 |
| chr6-146419036-INS-54 | chr6-146419036-INS-54 |
| chr6-146433568-INS-61 | chr6-146433578-INS-54 |
| chr6-146439400-INS-182 | chr6-146439400-INS-182 |
| chr6-146441550-INS-60 | chr6-146441550-INS-78 |
| chr6-146481919-INS-203 | chr6-146481919-INS-203 |
| chr6-146488632-INS-205 | chr6-146488632-INS-205 |

| | |
|---|---|
| chr6-146516115-INS-262 | chr6-146516145-INS-264 |
| chr6-146517704-INS-58 | chr6-146517704-INS-58 |
| chr6-146539074-INS-206 | chr6-146539074-INS-205 |
| chr6-146560094-INS-224 | chr6-146560094-INS-224 |
| chr6-146565457-INS-194 | chr6-146565457-INS-194 |
| chr6-146573683-INS-214 | chr6-146573683-INS-216 |
| chr6-146579337-INS-57 | chr6-146579336-INS-66 |
| chr6-146588119-INS-203 | chr6-146588110-INS-165 |
| chr6-146604593-INS-179 | chr6-146604593-INS-179 |
| chr6-146630864-INS-210 | chr6-146630864-INS-210 |
| chr6-146642654-INS-368 | chr6-146642654-INS-368 |
| chr6-146665648-INS-205 | chr6-146665648-INS-205 |
| chr6-146676048-INS-210 | chr6-146676048-INS-210 |
| chr6-146684021-INS-60 | chr6-146684021-INS-60 |
| chr6-146730544-INS-70 | chr6-146730546-INS-66 |
| chr6-146732568-INS-242 | chr6-146732568-INS-244 |
| chr6-146734471-INS-129 | chr6-146734507-INS-127 |
| chr6-146735526-INS-196 | chr6-146735526-INS-196 |
| chr6-146741684-INS-136 | chr6-146741684-INS-136 |
| chr6-146753425-INS-387 | chr6-146753425-INS-385 |
| chr6-146762933-INS-127 | chr6-146762937-INS-113 |
| chr6-146765713-INS-201 | chr6-146765713-INS-202 |
| chr6-146770755-INS-158 | chr6-146770756-INS-113 |
| chr6-146810347-INS-789 | chr6-146810347-INS-789 |
| chr6-146811300-INS-540 | chr6-146811300-INS-541 |
| chr6-146813838-INS-82 | chr6-146813837-INS-73 |
| chr6-146827823-INS-437 | chr6-146827823-INS-437 |
| chr6-146832589-INS-53 | chr6-146832589-INS-54 |
| chr6-146869217-INS-220 | chr6-146869217-INS-217 |
| chr6-146872337-INS-3155 | chr6-146872338-INS-3155 |
| chr6-146877622-INS-2166 | chr6-146877623-INS-2166 |
| chr6-146877623-INS-285 | chr6-146877623-INS-285 |
| chr6-146883848-INS-101 | chr6-146883848-INS-101 |
| chr6-146884226-INS-692 | chr6-146884226-INS-709 |
| chr6-146889377-INS-6676 | chr6-146889378-INS-6676 |
| chr6-146899134-INS-56 | chr6-146899134-INS-56 |
| chr6-146906863-INS-72 | chr6-146906863-INS-72 |
| chr6-146937772-INS-272 | chr6-146937772-INS-272 |
| chr6-146955120-INS-60 | chr6-146955120-INS-60 |
| chr6-146955190-INS-562 | chr6-146955190-INS-562 |
| chr6-147005975-INS-57 | chr6-147005975-INS-57 |
| chr6-147011951-INS-196 | chr6-147011952-INS-196 |
| chr6-147025808-INS-90 | chr6-147025809-INS-92 |
| chr6-147066677-INS-56 | chr6-147066677-INS-56 |
| chr6-147070251-INS-731 | chr6-147070251-INS-729 |
| chr6-147093915-INS-420 | chr6-147093915-INS-419 |
| chr6-147099167-INS-68 | chr6-147099167-INS-70 |
| chr6-147099535-INS-225 | chr6-147099535-INS-206 |
| chr6-147120235-INS-680 | chr6-147120235-INS-685 |
| chr6-147123725-INS-207 | chr6-147123725-INS-209 |
| chr6-147129758-INS-265 | chr6-147129758-INS-265 |
| chr6-147141756-INS-76 | chr6-147141747-INS-62 |
| chr6-147243907-INS-102 | chr6-147243889-INS-84 |
| chr6-147253428-INS-225 | chr6-147253428-INS-180 |
| chr6-147287182-INS-72 | chr6-147287182-INS-50 |
| chr6-147298656-INS-384 | chr6-147298657-INS-384 |
| chr6-147303768-INS-56 | chr6-147303793-INS-54 |
| chr6-147324752-INS-194 | chr6-147324752-INS-194 |
| chr6-147342811-INS-1908 | chr6-147342812-INS-1908 |
| chr6-147347229-INS-88 | chr6-147347242-INS-68 |
| chr6-147349076-INS-346 | chr6-147349076-INS-346 |
| chr6-147370189-INS-1473 | chr6-147370190-INS-1474 |
| chr6-147381571-INS-225 | chr6-147381571-INS-226 |
| chr6-147389904-INS-1472 | chr6-147389905-INS-1474 |
| chr6-147423790-INS-58 | chr6-147423791-INS-58 |
| chr6-147427407-INS-220 | chr6-147427407-INS-220 |
| chr6-147432642-INS-2385 | chr6-147432643-INS-2385 |
| chr6-147442398-INS-6451 | chr6-147442399-INS-6451 |
| chr6-147501060-INS-1876 | chr6-147501061-INS-1876 |
| chr6-147518406-INS-301 | chr6-147518406-INS-301 |
| chr6-147523078-INS-64 | chr6-147523078-INS-64 |
| chr6-147527971-INS-82 | chr6-147527971-INS-82 |
| chr6-147551469-INS-506 | chr6-147551469-INS-506 |
| chr6-147563647-INS-495 | chr6-147563647-INS-495 |
| chr6-147568446-INS-103 | chr6-147568446-INS-103 |
| chr6-147590190-INS-2193 | chr6-147590191-INS-2193 |
| chr6-147626632-INS-179 | chr6-147626593-INS-200 |

| | |
|---|---|
| chr6-147631617-INS-1152 | chr6-147631618-INS-1152 |
| chr6-147635739-INS-82 | chr6-147635739-INS-79 |
| chr6-147664508-INS-197 | chr6-147664508-INS-196 |
| chr6-147688074-INS-83 | chr6-147688074-INS-83 |
| chr6-147689628-INS-160 | chr6-147689628-INS-160 |
| chr6-147694941-INS-75 | chr6-147694941-INS-75 |
| chr6-147725103-INS-6426 | chr6-147725104-INS-6426 |
| chr6-147729259-INS-57 | chr6-147729259-INS-57 |
| chr6-147732817-INS-70 | chr6-147732829-INS-60 |
| chr6-147737009-INS-346 | chr6-147737009-INS-346 |
| chr6-147753276-INS-82 | chr6-147753276-INS-98 |
| chr6-147756833-INS-71 | chr6-147756833-INS-70 |
| chr6-147774502-INS-58 | chr6-147774502-INS-60 |
| chr6-147778957-INS-2404 | chr6-147778958-INS-2404 |
| chr6-147781513-INS-1875 | chr6-147781514-INS-1875 |
| chr6-147792086-INS-64 | chr6-147792086-INS-64 |
| chr6-147794026-INS-52 | chr6-147794027-INS-53 |
| chr6-147817456-INS-72 | chr6-147817451-INS-52 |
| chr6-147845252-INS-128 | chr6-147845252-INS-128 |
| chr6-147847286-INS-1509 | chr6-147847287-INS-1528 |
| chr6-147867471-INS-246 | chr6-147867471-INS-246 |
| chr6-147871649-INS-997 | chr6-147871649-INS-998 |
| chr6-147871914-INS-1771 | chr6-147871915-INS-1769 |
| chr6-147873470-INS-6453 | chr6-147873471-INS-6453 |
| chr6-147891159-INS-1456 | chr6-147891160-INS-1457 |
| chr6-147891749-INS-289 | chr6-147891748-INS-293 |
| chr6-147905761-INS-100 | chr6-147905761-INS-112 |
| chr6-147915610-INS-96 | chr6-147915610-INS-96 |
| chr6-147921013-INS-5497 | chr6-147921014-INS-5497 |
| chr6-147944406-INS-352 | chr6-147944406-INS-352 |
| chr6-148037312-INS-77 | chr6-148037312-INS-104 |
| chr6-148062681-INS-675 | chr6-148062680-INS-657 |
| chr6-148129628-INS-704 | chr6-148129626-INS-715 |
| chr6-148135941-INS-376 | chr6-148135941-INS-376 |
| chr6-148190986-INS-711 | chr6-148190986-INS-711 |
| chr6-148200497-INS-243 | chr6-148200497-INS-243 |
| chr6-148205410-INS-89 | chr6-148205410-INS-79 |
| chr6-148267130-INS-636 | chr6-148267130-INS-636 |
| chr6-148314832-INS-1886 | chr6-148314833-INS-1886 |
| chr6-148320949-INS-68 | chr6-148320949-INS-80 |
| chr6-148333639-INS-56 | chr6-148333680-INS-58 |
| chr6-148338675-INS-52 | chr6-148338675-INS-52 |
| chr6-148369821-INS-5247 | chr6-148369822-INS-5247 |
| chr6-148393355-INS-588 | chr6-148393355-INS-588 |
| chr6-148433773-INS-195 | chr6-148433773-INS-196 |
| chr6-148449910-INS-327 | chr6-148449910-INS-327 |
| chr6-148454606-INS-52 | chr6-148454594-INS-70 |
| chr6-148469979-INS-6248 | chr6-148469980-INS-6248 |
| chr6-148482841-INS-62 | chr6-148482841-INS-62 |
| chr6-148501506-INS-53 | chr6-148501506-INS-52 |
| chr6-148506635-INS-448 | chr6-148506635-INS-452 |
| chr6-148514857-INS-54 | chr6-148514857-INS-61 |
| chr6-148524687-INS-6376 | chr6-148524688-INS-6377 |
| chr6-148561023-INS-421 | chr6-148561023-INS-421 |
| chr6-148573380-INS-207 | chr6-148573380-INS-205 |
| chr6-148573669-INS-441 | chr6-148573669-INS-441 |
| chr6-148586833-INS-1892 | chr6-148586834-INS-1892 |
| chr6-148590661-INS-485 | chr6-148590661-INS-485 |
| chr6-148594263-INS-7308 | chr6-148594264-INS-7309 |
| chr6-148597616-INS-123 | chr6-148597616-INS-123 |
| chr6-148627007-INS-504 | chr6-148627007-INS-504 |
| chr6-148637547-INS-54 | chr6-148637547-INS-60 |
| chr6-148649744-INS-102 | chr6-148649744-INS-102 |
| chr6-148657994-INS-54 | chr6-148657982-INS-54 |
| chr6-148659606-INS-128 | chr6-148659606-INS-133 |
| chr6-148660591-INS-56 | chr6-148660591-INS-52 |
| chr6-148669778-INS-64 | chr6-148669777-INS-54 |
| chr6-148701508-INS-125 | chr6-148701508-INS-125 |
| chr6-148704113-INS-1005 | chr6-148704114-INS-992 |
| chr6-148707295-INS-209 | chr6-148707295-INS-208 |
| chr6-148715071-INS-62 | chr6-148715071-INS-61 |
| chr6-148747786-INS-181 | chr6-148747786-INS-171 |
| chr6-148756320-INS-60 | chr6-148756320-INS-68 |
| chr6-148773813-INS-73 | chr6-148773813-INS-57 |
| chr6-148795290-INS-69 | chr6-148795290-INS-67 |
| chr6-148798472-INS-333 | chr6-148798472-INS-333 |
| chr6-148804249-INS-414 | chr6-148804249-INS-414 |

| | |
|---|---|
| chr6-148825375-INS-218 | chr6-148825375-INS-218 |
| chr6-148826840-INS-217 | chr6-148826840-INS-201 |
| chr6-148830208-INS-83 | chr6-148830208-INS-83 |
| chr6-148837155-INS-190 | chr6-148837155-INS-190 |
| chr6-148861319-INS-211 | chr6-148861319-INS-211 |
| chr6-148868625-INS-272 | chr6-148868625-INS-272 |
| chr6-148878285-INS-56 | chr6-148878285-INS-52 |
| chr6-148886507-INS-148 | chr6-148886508-INS-163 |
| chr6-148887475-INS-7328 | chr6-148887476-INS-7328 |
| chr6-148904891-INS-202 | chr6-148904891-INS-203 |
| chr6-148907753-INS-240 | chr6-148907767-INS-248 |
| chr6-148908877-INS-210 | chr6-148908877-INS-208 |
| chr6-148916163-INS-103 | chr6-148916155-INS-92 |
| chr6-148917789-INS-52 | chr6-148917789-INS-52 |
| chr6-148919489-INS-210 | chr6-148919489-INS-211 |
| chr6-148922733-INS-122 | chr6-148922733-INS-123 |
| chr6-148944272-INS-253 | chr6-148944283-INS-252 |
| chr6-149014714-INS-209 | chr6-149014714-INS-209 |
| chr6-149027000-INS-212 | chr6-149027001-INS-196 |
| chr6-149034627-INS-86 | chr6-149034616-INS-71 |
| chr6-149040839-INS-3175 | chr6-149040840-INS-3172 |
| chr6-149041507-INS-52 | chr6-149041507-INS-68 |
| chr6-149046056-INS-213 | chr6-149046056-INS-213 |
| chr6-149055257-INS-203 | chr6-149055257-INS-204 |
| chr6-149056770-INS-788 | chr6-149056770-INS-789 |
| chr6-149060635-INS-173 | chr6-149060635-INS-175 |
| chr6-149062288-INS-60 | chr6-149062271-INS-54 |
| chr6-149070605-INS-801 | chr6-149070605-INS-801 |
| chr6-149082348-INS-195 | chr6-149082348-INS-196 |
| chr6-149099620-INS-312 | chr6-149099620-INS-293 |
| chr6-149112448-INS-60 | chr6-149112448-INS-51 |
| chr6-149168630-INS-3281 | chr6-149168631-INS-3281 |
| chr6-149182949-INS-535 | chr6-149182947-INS-540 |
| chr6-149191525-INS-323 | chr6-149191525-INS-323 |
| chr6-149194994-INS-5360 | chr6-149194995-INS-5360 |
| chr6-149224730-INS-562 | chr6-149224730-INS-562 |
| chr6-149252246-INS-95 | chr6-149252246-INS-102 |
| chr6-149276882-INS-213 | chr6-149276882-INS-213 |
| chr6-149281361-INS-331 | chr6-149281361-INS-331 |
| chr6-149304048-INS-63 | chr6-149304048-INS-63 |
| chr6-149335346-INS-7129 | chr6-149335347-INS-7129 |
| chr6-149358424-INS-7792 | chr6-149358425-INS-7792 |
| chr6-149366293-INS-63 | chr6-149366293-INS-63 |
| chr6-149370115-INS-205 | chr6-149370115-INS-212 |
| chr6-149420325-INS-209 | chr6-149420325-INS-209 |
| chr6-149447521-INS-204 | chr6-149447521-INS-204 |
| chr6-149474103-INS-195 | chr6-149474103-INS-195 |
| chr6-149476060-INS-219 | chr6-149476060-INS-221 |
| chr7-3058947-INS-182 | chr7-3058947-INS-174 |
| chr7-3089632-INS-270 | chr7-3089632-INS-258 |
| chr7-3113887-INS-58 | chr7-3113887-INS-60 |
| chr7-3135478-INS-498 | chr7-3135478-INS-498 |
| chr7-3154900-INS-54 | chr7-3154901-INS-62 |
| chr7-3170789-INS-6856 | chr7-3170790-INS-6857 |
| chr7-3193648-INS-196 | chr7-3193648-INS-197 |
| chr7-3210876-INS-133 | chr7-3210876-INS-144 |
| chr7-3213452-INS-52 | chr7-3213452-INS-52 |
| chr7-3223824-INS-51 | chr7-3223824-INS-51 |
| chr7-3233689-INS-203 | chr7-3233689-INS-203 |
| chr7-3239999-INS-199 | chr7-3239999-INS-224 |
| chr7-3243495-INS-219 | chr7-3243495-INS-220 |
| chr7-3244564-INS-416 | chr7-3244565-INS-416 |
| chr7-3251751-INS-181 | chr7-3251747-INS-185 |
| chr7-3276523-INS-184 | chr7-3276485-INS-184 |
| chr7-3278696-INS-175 | chr7-3278696-INS-176 |
| chr7-3279420-INS-230 | chr7-3279406-INS-215 |
| chr7-3291591-INS-99 | chr7-3291543-INS-99 |
| chr7-3309002-INS-100 | chr7-3309002-INS-100 |
| chr7-3312173-INS-180 | chr7-3312173-INS-180 |
| chr7-3313293-INS-146 | chr7-3313293-INS-138 |
| chr7-3315841-INS-203 | chr7-3315841-INS-204 |
| chr7-3317169-INS-171 | chr7-3317162-INS-182 |
| chr7-3317817-INS-86 | chr7-3317789-INS-76 |
| chr7-3319842-INS-56 | chr7-3319842-INS-56 |
| chr7-3320645-INS-206 | chr7-3320645-INS-206 |
| chr7-3327054-INS-181 | chr7-3327054-INS-181 |
| chr7-3332474-INS-147 | chr7-3332474-INS-147 |

| | |
|---|---|
| chr7-3337184-INS-63 | chr7-3337179-INS-51 |
| chr7-3344572-INS-188 | chr7-3344572-INS-188 |
| chr7-3365112-INS-64 | chr7-3365068-INS-60 |
| chr7-3372302-INS-227 | chr7-3372302-INS-227 |
| chr7-3373505-INS-102 | chr7-3373505-INS-53 |
| chr7-3375615-INS-190 | chr7-3375615-INS-186 |
| chr7-3387668-INS-80 | chr7-3387668-INS-80 |
| chr7-3392889-INS-63 | chr7-3392889-INS-63 |
| chr7-3428429-INS-210 | chr7-3428429-INS-219 |
| chr7-3429579-INS-356 | chr7-3429579-INS-355 |
| chr7-3440414-INS-51 | chr7-3440414-INS-56 |
| chr7-3451107-INS-72 | chr7-3451102-INS-67 |
| chr7-3452917-INS-60 | chr7-3452917-INS-64 |
| chr7-3457416-INS-216 | chr7-3457408-INS-235 |
| chr7-3468895-INS-192 | chr7-3468895-INS-192 |
| chr7-3478539-INS-91 | chr7-3478539-INS-94 |
| chr7-3493996-INS-223 | chr7-3493996-INS-222 |
| chr7-3498570-INS-124 | chr7-3498547-INS-119 |
| chr7-3499993-INS-511 | chr7-3499993-INS-511 |
| chr7-3507593-INS-207 | chr7-3507593-INS-207 |
| chr7-3520831-INS-205 | chr7-3520831-INS-204 |
| chr7-3522062-INS-196 | chr7-3522062-INS-196 |
| chr7-3537506-INS-693 | chr7-3537506-INS-693 |
| chr7-3553037-INS-212 | chr7-3553037-INS-212 |
| chr7-3557117-INS-96 | chr7-3557117-INS-102 |
| chr7-3561124-INS-202 | chr7-3561124-INS-203 |
| chr7-3562883-INS-204 | chr7-3562883-INS-204 |
| chr7-3587241-INS-479 | chr7-3587254-INS-472 |
| chr7-3608777-INS-155 | chr7-3608777-INS-107 |
| chr7-3659765-INS-212 | chr7-3659765-INS-213 |
| chr7-3670685-INS-216 | chr7-3670685-INS-216 |
| chr7-3693352-INS-233 | chr7-3693352-INS-233 |
| chr7-3702267-INS-101 | chr7-3702267-INS-101 |
| chr7-3741114-INS-167 | chr7-3741112-INS-167 |
| chr7-3792773-INS-214 | chr7-3792773-INS-214 |
| chr7-3865514-INS-360 | chr7-3865514-INS-360 |
| chr7-3988617-INS-115 | chr7-3988617-INS-115 |
| chr7-4106704-INS-78 | chr7-4106704-INS-78 |
| chr7-4117262-INS-95 | chr7-4117262-INS-98 |
| chr7-4128941-INS-115 | chr7-4128941-INS-115 |
| chr7-4130096-INS-195 | chr7-4130096-INS-199 |
| chr7-4161176-INS-212 | chr7-4161176-INS-212 |
| chr7-4186009-INS-681 | chr7-4186009-INS-681 |
| chr7-4207386-INS-97 | chr7-4207386-INS-97 |
| chr7-4213940-INS-415 | chr7-4213939-INS-408 |
| chr7-4255323-INS-190 | chr7-4255323-INS-190 |
| chr7-4287426-INS-1906 | chr7-4287427-INS-1905 |
| chr7-4292271-INS-5586 | chr7-4292272-INS-5595 |
| chr7-4303826-INS-200 | chr7-4303826-INS-200 |
| chr7-4328448-INS-96 | chr7-4328448-INS-96 |
| chr7-4336447-INS-248 | chr7-4336447-INS-249 |
| chr7-4343309-INS-73 | chr7-4343302-INS-54 |
| chr7-4360458-INS-56 | chr7-4360458-INS-56 |
| chr7-4371488-INS-280 | chr7-4371488-INS-280 |
| chr7-4385384-INS-70 | chr7-4385384-INS-70 |
| chr7-4399957-INS-63 | chr7-4399957-INS-51 |
| chr7-4401506-INS-54 | chr7-4401506-INS-55 |
| chr7-4410969-INS-230 | chr7-4410950-INS-219 |
| chr7-4416224-INS-203 | chr7-4416224-INS-205 |
| chr7-4417412-INS-7077 | chr7-4417413-INS-7077 |
| chr7-4419002-INS-1373 | chr7-4419003-INS-1375 |
| chr7-4433548-INS-197 | chr7-4433548-INS-177 |
| chr7-4441948-INS-187 | chr7-4441949-INS-186 |
| chr7-4450863-INS-216 | chr7-4450863-INS-215 |
| chr7-4496420-INS-123 | chr7-4496420-INS-123 |
| chr7-4527467-INS-80 | chr7-4527467-INS-54 |
| chr7-4598478-INS-475 | chr7-4598478-INS-475 |
| chr7-4613869-INS-57 | chr7-4613869-INS-57 |
| chr7-4672613-INS-276 | chr7-4672613-INS-276 |
| chr7-4679499-INS-57 | chr7-4679477-INS-76 |
| chr7-4681716-INS-68 | chr7-4681716-INS-68 |
| chr7-4688343-INS-102 | chr7-4688342-INS-126 |
| chr7-4695285-INS-115 | chr7-4695285-INS-115 |
| chr7-4699268-INS-252 | chr7-4699269-INS-294 |
| chr7-4721509-INS-244 | chr7-4721509-INS-245 |
| chr7-4745898-INS-98 | chr7-4745898-INS-98 |
| chr7-4750936-INS-54 | chr7-4750936-INS-54 |

| | |
|---|---|
| chr7-4766453-INS-71 | chr7-4766453-INS-77 |
| chr7-4781847-INS-116 | chr7-4781847-INS-115 |
| chr7-4806059-INS-202 | chr7-4806059-INS-202 |
| chr7-4812502-INS-205 | chr7-4812502-INS-205 |
| chr7-4822013-INS-56 | chr7-4822013-INS-55 |
| chr7-4824064-INS-204 | chr7-4824064-INS-204 |
| chr7-4838532-INS-227 | chr7-4838532-INS-214 |
| chr7-4840693-INS-202 | chr7-4840693-INS-202 |
| chr7-4842770-INS-56 | chr7-4842770-INS-56 |
| chr7-4843700-INS-176 | chr7-4843700-INS-176 |
| chr7-4850367-INS-248 | chr7-4850367-INS-248 |
| chr7-4858785-INS-75 | chr7-4858785-INS-54 |
| chr7-4860532-INS-64 | chr7-4860532-INS-64 |
| chr7-4863581-INS-590 | chr7-4863581-INS-590 |
| chr7-4870464-INS-225 | chr7-4870464-INS-224 |
| chr7-4872439-INS-205 | chr7-4872432-INS-199 |
| chr7-4906529-INS-120 | chr7-4906529-INS-124 |
| chr7-4942852-INS-88 | chr7-4942852-INS-88 |
| chr7-4957886-INS-122 | chr7-4957887-INS-122 |
| chr7-4968267-INS-174 | chr7-4968267-INS-174 |
| chr7-4984615-INS-141 | chr7-4984609-INS-142 |
| chr7-5012444-INS-361 | chr7-5012444-INS-359 |
| chr7-5013503-INS-98 | chr7-5013503-INS-98 |
| chr7-5021767-INS-212 | chr7-5021767-INS-212 |
| chr7-5030770-INS-58 | chr7-5030770-INS-58 |
| chr7-5047866-INS-222 | chr7-5047866-INS-222 |
| chr7-5050693-INS-156 | chr7-5050693-INS-155 |
| chr7-5053128-INS-197 | chr7-5053128-INS-195 |
| chr7-5103016-INS-86 | chr7-5103016-INS-86 |
| chr7-5162192-INS-848 | chr7-5162192-INS-848 |
| chr7-5179824-INS-1446 | chr7-5179825-INS-1446 |
| chr7-5185040-INS-5407 | chr7-5185041-INS-5407 |
| chr7-5190172-INS-52 | chr7-5190193-INS-58 |
| chr7-5288053-INS-110 | chr7-5288070-INS-117 |
| chr7-5336479-INS-3161 | chr7-5336480-INS-3161 |
| chr7-5336907-INS-260 | chr7-5336907-INS-260 |
| chr7-5356628-INS-83 | chr7-5356628-INS-113 |
| chr7-5364927-INS-397 | chr7-5364927-INS-397 |
| chr7-5390533-INS-4781 | chr7-5390534-INS-4779 |
| chr7-5404412-INS-58 | chr7-5404412-INS-58 |
| chr7-5435537-INS-89 | chr7-5435537-INS-89 |
| chr7-5480399-INS-180 | chr7-5480399-INS-180 |
| chr7-5481819-INS-6187 | chr7-5481820-INS-6187 |
| chr7-5489765-INS-62 | chr7-5489765-INS-64 |
| chr7-5494980-INS-155 | chr7-5495025-INS-159 |
| chr7-5495626-INS-62 | chr7-5495626-INS-62 |
| chr7-5582908-INS-7215 | chr7-5582909-INS-7215 |
| chr7-5852618-INS-2250 | chr7-5852619-INS-2250 |
| chr7-5873538-INS-117 | chr7-5873549-INS-117 |
| chr7-5885505-INS-1747 | chr7-5885506-INS-1766 |
| chr7-5930013-INS-200 | chr7-5930013-INS-205 |
| chr7-5931124-INS-355 | chr7-5931124-INS-355 |
| chr7-5939973-INS-141 | chr7-5939973-INS-141 |
| chr7-6023867-INS-65 | chr7-6023867-INS-66 |
| chr7-6030941-INS-6954 | chr7-6030942-INS-6957 |
| chr7-6032224-INS-82 | chr7-6032195-INS-91 |
| chr7-6051137-INS-56 | chr7-6051137-INS-58 |
| chr7-6078014-INS-66 | chr7-6077998-INS-110 |
| chr7-6137410-INS-52 | chr7-6137410-INS-52 |
| chr7-6144190-INS-68 | chr7-6144190-INS-88 |
| chr7-6165004-INS-440 | chr7-6165004-INS-440 |
| chr7-6168695-INS-95 | chr7-6168688-INS-93 |
| chr7-6188036-INS-720 | chr7-6188068-INS-715 |
| chr7-6205530-INS-55 | chr7-6205530-INS-55 |
| chr7-6214864-INS-94 | chr7-6214864-INS-96 |
| chr7-6225830-INS-178 | chr7-6225830-INS-165 |
| chr7-6239109-INS-149 | chr7-6239109-INS-149 |
| chr7-6240856-INS-100 | chr7-6240854-INS-102 |
| chr7-6243351-INS-177 | chr7-6243347-INS-149 |
| chr7-6257410-INS-156 | chr7-6257410-INS-156 |
| chr7-6261308-INS-196 | chr7-6261308-INS-196 |
| chr7-6265181-INS-76 | chr7-6265181-INS-76 |
| chr7-6266150-INS-75 | chr7-6266139-INS-75 |
| chr7-6267449-INS-52 | chr7-6267449-INS-52 |
| chr7-6273521-INS-207 | chr7-6273521-INS-203 |
| chr7-6292625-INS-63 | chr7-6292625-INS-63 |
| chr7-6318286-INS-586 | chr7-6318286-INS-589 |

| | |
|---|---|
| chr7-6318921-INS-52 | chr7-6318932-INS-87 |
| chr7-6324826-INS-158 | chr7-6324826-INS-158 |
| chr7-6335473-INS-108 | chr7-6335473-INS-108 |
| chr7-6339999-INS-351 | chr7-6339999-INS-351 |
| chr7-6345535-INS-399 | chr7-6345535-INS-399 |
| chr7-6353405-INS-90 | chr7-6353405-INS-90 |
| chr7-6358672-INS-58 | chr7-6358668-INS-50 |
| chr7-6399923-INS-108 | chr7-6399923-INS-108 |
| chr7-6402373-INS-419 | chr7-6402373-INS-419 |
| chr7-6450133-INS-52 | chr7-6450133-INS-54 |
| chr7-6493621-INS-240 | chr7-6493621-INS-260 |
| chr7-6498642-INS-124 | chr7-6498642-INS-124 |
| chr7-6508078-INS-53 | chr7-6508078-INS-53 |
| chr7-6511510-INS-56 | chr7-6511510-INS-56 |
| chr7-6606142-INS-66 | chr7-6606142-INS-66 |
| chr7-6617111-INS-261 | chr7-6617111-INS-261 |
| chr7-6628297-INS-53 | chr7-6628311-INS-52 |
| chr7-6645688-INS-108 | chr7-6645688-INS-108 |
| chr7-6647500-INS-120 | chr7-6647500-INS-120 |
| chr7-6674784-INS-64 | chr7-6674784-INS-64 |
| chr7-6735897-INS-75 | chr7-6735897-INS-75 |
| chr7-6768174-INS-60 | chr7-6768175-INS-88 |
| chr7-6784255-INS-60 | chr7-6784255-INS-62 |
| chr7-6792469-INS-64 | chr7-6792469-INS-58 |
| chr7-6841765-INS-77 | chr7-6841765-INS-87 |
| chr7-6857357-INS-57 | chr7-6857337-INS-58 |
| chr7-6858251-INS-287 | chr7-6858251-INS-247 |
| chr7-6866497-INS-782 | chr7-6866497-INS-784 |
| chr7-6870022-INS-54 | chr7-6870021-INS-63 |
| chr7-6878512-INS-96 | chr7-6878512-INS-90 |
| chr7-6886488-INS-55 | chr7-6886488-INS-55 |
| chr7-6891124-INS-83 | chr7-6891124-INS-73 |
| chr7-6956108-INS-229 | chr7-6956108-INS-229 |
| chr7-6962557-INS-82 | chr7-6962517-INS-82 |
| chr7-7005002-INS-153 | chr7-7005002-INS-153 |
| chr7-7006673-INS-192 | chr7-7006673-INS-192 |
| chr7-7019318-INS-209 | chr7-7019318-INS-209 |
| chr7-7034081-INS-52 | chr7-7034052-INS-50 |
| chr7-7036389-INS-201 | chr7-7036389-INS-201 |
| chr7-7043513-INS-60 | chr7-7043513-INS-64 |
| chr7-7048743-INS-211 | chr7-7048743-INS-210 |
| chr7-7099910-INS-231 | chr7-7099910-INS-232 |
| chr7-7106297-INS-160 | chr7-7106297-INS-164 |
| chr7-7112769-INS-195 | chr7-7112769-INS-208 |
| chr7-7141432-INS-101 | chr7-7141432-INS-88 |
| chr7-7190098-INS-5288 | chr7-7190099-INS-5288 |
| chr7-7225688-INS-214 | chr7-7225688-INS-214 |
| chr7-8553408-INS-84 | chr7-8553424-INS-84 |
| chr7-9069715-INS-59 | chr7-9069715-INS-61 |
| chr7-9739077-INS-881 | chr7-9739077-INS-877 |
| chr7-9860268-INS-6583 | chr7-9860269-INS-6583 |
| chr7-9894494-INS-61 | chr7-9894494-INS-61 |
| chr7-9897734-INS-3117 | chr7-9897735-INS-3115 |
| chr7-9958441-INS-84 | chr7-9958441-INS-80 |
| chr7-10007619-INS-54 | chr7-10007619-INS-51 |
| chr7-10018932-INS-54 | chr7-10018932-INS-60 |
| chr7-10071849-INS-54 | chr7-10071849-INS-54 |
| chr7-10074056-INS-634 | chr7-10074056-INS-634 |
| chr7-10166584-INS-87 | chr7-10166584-INS-95 |
| chr7-10213834-INS-121 | chr7-10213832-INS-123 |
| chr7-10237117-INS-605 | chr7-10237117-INS-605 |
| chr7-10254849-INS-7009 | chr7-10254850-INS-7009 |
| chr7-10280769-INS-55 | chr7-10280769-INS-54 |
| chr7-10292680-INS-52 | chr7-10292680-INS-54 |
| chr7-10320566-INS-95 | chr7-10320566-INS-95 |
| chr7-10327083-INS-974 | chr7-10327085-INS-977 |
| chr7-10343801-INS-584 | chr7-10343801-INS-583 |
| chr7-10406877-INS-285 | chr7-10406877-INS-285 |
| chr7-10412221-INS-3369 | chr7-10412222-INS-3369 |
| chr7-10419217-INS-7081 | chr7-10419218-INS-7082 |
| chr7-10471316-INS-54 | chr7-10471345-INS-62 |
| chr7-10473971-INS-50 | chr7-10473971-INS-50 |
| chr7-10477751-INS-63 | chr7-10477751-INS-66 |
| chr7-10486790-INS-776 | chr7-10486790-INS-775 |
| chr7-10488357-INS-85 | chr7-10488357-INS-85 |
| chr7-10865230-INS-238 | chr7-10865230-INS-238 |
| chr7-11026211-INS-185 | chr7-11026211-INS-186 |

| | |
|---|---|
| chr7-11340075-INS-77 | chr7-11340075-INS-77 |
| chr7-11341334-INS-72 | chr7-11341342-INS-87 |
| chr7-11367073-INS-2234 | chr7-11367074-INS-2234 |
| chr7-11368160-INS-852 | chr7-11368160-INS-852 |
| chr7-11422613-INS-401 | chr7-11422613-INS-401 |
| chr7-11431269-INS-3082 | chr7-11431270-INS-3082 |
| chr7-11485530-INS-93 | chr7-11485530-INS-93 |
| chr7-11538981-INS-111 | chr7-11538981-INS-111 |
| chr7-11551881-INS-386 | chr7-11551881-INS-402 |
| chr7-11583302-INS-90 | chr7-11583302-INS-90 |
| chr7-11587182-INS-1538 | chr7-11587183-INS-1536 |
| chr7-11596773-INS-588 | chr7-11596773-INS-588 |
| chr7-11622197-INS-1139 | chr7-11622198-INS-1139 |
| chr7-11626578-INS-56 | chr7-11626578-INS-52 |
| chr7-11637214-INS-7094 | chr7-11637215-INS-7093 |
| chr7-11641253-INS-82 | chr7-11641266-INS-104 |
| chr7-11680784-INS-334 | chr7-11680784-INS-339 |
| chr7-11681004-INS-1582 | chr7-11681005-INS-1584 |
| chr7-11695126-INS-50 | chr7-11695122-INS-58 |
| chr7-11782418-INS-744 | chr7-11782418-INS-749 |
| chr7-11809874-INS-199 | chr7-11809874-INS-199 |
| chr7-11824440-INS-128 | chr7-11824440-INS-126 |
| chr7-11838722-INS-95 | chr7-11838722-INS-96 |
| chr7-11841081-INS-65 | chr7-11841081-INS-65 |
| chr7-11871691-INS-166 | chr7-11871691-INS-166 |
| chr7-11879476-INS-314 | chr7-11879476-INS-314 |
| chr7-11883441-INS-249 | chr7-11883477-INS-250 |
| chr7-11884509-INS-112 | chr7-11884509-INS-132 |
| chr7-11887617-INS-62 | chr7-11887617-INS-52 |
| chr7-11889432-INS-51 | chr7-11889410-INS-51 |
| chr7-11894547-INS-6302 | chr7-11894548-INS-6302 |
| chr7-11985708-INS-509 | chr7-11985708-INS-492 |
| chr7-11992270-INS-172 | chr7-11992270-INS-172 |
| chr7-12000781-INS-671 | chr7-12000780-INS-671 |
| chr7-12009447-INS-57 | chr7-12009447-INS-60 |
| chr7-12035904-INS-56 | chr7-12035904-INS-50 |
| chr7-12075486-INS-341 | chr7-12075486-INS-341 |
| chr7-12079039-INS-57 | chr7-12079055-INS-55 |
| chr7-12134845-INS-57 | chr7-12134845-INS-57 |
| chr7-12140653-INS-55 | chr7-12140653-INS-55 |
| chr7-12157145-INS-56 | chr7-12157145-INS-58 |
| chr7-12164588-INS-369 | chr7-12164588-INS-369 |
| chr7-12192211-INS-208 | chr7-12192211-INS-208 |
| chr7-12212914-INS-93 | chr7-12212882-INS-105 |
| chr7-12224532-INS-353 | chr7-12224532-INS-352 |
| chr7-12225141-INS-825 | chr7-12225141-INS-834 |
| chr7-12253272-INS-341 | chr7-12253272-INS-341 |
| chr7-12283308-INS-830 | chr7-12283308-INS-830 |
| chr7-12294401-INS-233 | chr7-12294401-INS-233 |
| chr7-12313921-INS-4844 | chr7-12313922-INS-4844 |
| chr7-12343122-INS-425 | chr7-12343122-INS-425 |
| chr7-12364928-INS-94 | chr7-12364938-INS-63 |
| chr7-12372401-INS-64 | chr7-12372405-INS-64 |
| chr7-12389632-INS-7316 | chr7-12389633-INS-7313 |
| chr7-12433210-INS-564 | chr7-12433210-INS-564 |
| chr7-12471480-INS-135 | chr7-12471480-INS-139 |
| chr7-12487015-INS-204 | chr7-12487015-INS-204 |
| chr7-12508438-INS-72 | chr7-12508424-INS-91 |
| chr7-12515289-INS-266 | chr7-12515289-INS-266 |
| chr7-12524779-INS-6597 | chr7-12524780-INS-6599 |
| chr7-12525339-INS-213 | chr7-12525339-INS-213 |
| chr7-12534291-INS-196 | chr7-12534291-INS-196 |
| chr7-12585030-INS-104 | chr7-12585030-INS-104 |
| chr7-12591442-INS-195 | chr7-12591442-INS-192 |
| chr7-12602358-INS-237 | chr7-12602358-INS-237 |
| chr7-12602851-INS-5132 | chr7-12602852-INS-5132 |
| chr7-12622997-INS-7234 | chr7-12622998-INS-7240 |
| chr7-12635095-INS-309 | chr7-12635095-INS-309 |
| chr7-12666655-INS-202 | chr7-12666655-INS-202 |
| chr7-12682461-INS-207 | chr7-12682461-INS-209 |
| chr7-12684277-INS-9083 | chr7-12684278-INS-9083 |
| chr7-12686449-INS-242 | chr7-12686449-INS-243 |
| chr7-12692358-INS-206 | chr7-12692359-INS-178 |
| chr7-12700450-INS-110 | chr7-12700449-INS-114 |
| chr7-12704114-INS-122 | chr7-12704099-INS-147 |
| chr7-12709006-INS-234 | chr7-12709006-INS-234 |
| chr7-12721252-INS-56 | chr7-12721234-INS-56 |

| | |
|---|---|
| chr7-12724764-INS-60 | chr7-12724764-INS-78 |
| chr7-12728221-INS-410 | chr7-12728229-INS-403 |
| chr7-12736154-INS-200 | chr7-12736154-INS-199 |
| chr7-12744092-INS-67 | chr7-12744092-INS-52 |
| chr7-12748769-INS-172 | chr7-12748769-INS-173 |
| chr7-12775096-INS-132 | chr7-12775096-INS-132 |
| chr7-12780907-INS-51 | chr7-12780908-INS-53 |
| chr7-12800599-INS-71 | chr7-12800599-INS-68 |
| chr7-12838918-INS-552 | chr7-12838918-INS-552 |
| chr7-12854200-INS-107 | chr7-12854200-INS-107 |
| chr7-12869164-INS-54 | chr7-12869164-INS-54 |
| chr7-12885699-INS-1925 | chr7-12885700-INS-1926 |
| chr7-12891050-INS-651 | chr7-12891044-INS-623 |
| chr7-12893958-INS-116 | chr7-12893958-INS-116 |
| chr7-12911808-INS-554 | chr7-12911808-INS-554 |
| chr7-12928307-INS-96 | chr7-12928307-INS-101 |
| chr7-12930409-INS-464 | chr7-12930409-INS-464 |
| chr7-12935332-INS-7125 | chr7-12935333-INS-7125 |
| chr7-12951497-INS-191 | chr7-12951497-INS-190 |
| chr7-12976346-INS-53 | chr7-12976346-INS-53 |
| chr7-13021481-INS-217 | chr7-13021481-INS-217 |
| chr7-13047459-INS-5211 | chr7-13047460-INS-5211 |
| chr7-13086760-INS-558 | chr7-13086760-INS-552 |
| chr7-13093681-INS-172 | chr7-13093681-INS-172 |
| chr7-13115265-INS-52 | chr7-13115265-INS-52 |
| chr7-13124994-INS-160 | chr7-13124994-INS-160 |
| chr7-13177102-INS-80 | chr7-13177102-INS-80 |
| chr7-13190457-INS-144 | chr7-13190457-INS-142 |
| chr7-13192215-INS-299 | chr7-13192215-INS-299 |
| chr7-13220904-INS-118 | chr7-13220904-INS-118 |
| chr7-13229803-INS-452 | chr7-13229803-INS-452 |
| chr7-13242208-INS-640 | chr7-13242208-INS-641 |
| chr7-13271232-INS-6269 | chr7-13271233-INS-6269 |
| chr7-13276822-INS-53 | chr7-13276834-INS-63 |
| chr7-13305366-INS-666 | chr7-13305366-INS-666 |
| chr7-13331309-INS-416 | chr7-13331310-INS-416 |
| chr7-13393997-INS-3197 | chr7-13393998-INS-3197 |
| chr7-13409032-INS-2907 | chr7-13409033-INS-2907 |
| chr7-13433352-INS-118 | chr7-13433352-INS-70 |
| chr7-13579639-INS-64 | chr7-13579635-INS-66 |
| chr7-13587640-INS-907 | chr7-13587640-INS-907 |
| chr7-13601148-INS-360 | chr7-13601134-INS-394 |
| chr7-13622698-INS-1752 | chr7-13622699-INS-1752 |
| chr7-13644769-INS-54 | chr7-13644769-INS-54 |
| chr7-13656537-INS-62 | chr7-13656499-INS-64 |
| chr7-13723471-INS-4756 | chr7-13723472-INS-4756 |
| chr7-13804935-INS-7067 | chr7-13804936-INS-7068 |
| chr7-13917507-INS-204 | chr7-13917507-INS-204 |
| chr7-13937902-INS-798 | chr7-13937886-INS-797 |
| chr7-13972824-INS-6828 | chr7-13972824-INS-6829 |
| chr7-13974907-INS-164 | chr7-13974907-INS-147 |
| chr7-13995705-INS-1429 | chr7-13995706-INS-1429 |
| chr7-13996928-INS-97 | chr7-13996928-INS-99 |
| chr7-14006967-INS-444 | chr7-14006967-INS-444 |
| chr7-14011901-INS-60 | chr7-14011901-INS-60 |
| chr7-14048733-INS-52 | chr7-14048733-INS-52 |
| chr7-14058721-INS-195 | chr7-14058721-INS-196 |
| chr7-14071882-INS-65 | chr7-14071882-INS-67 |
| chr7-14104201-INS-201 | chr7-14104201-INS-201 |
| chr7-14162960-INS-4459 | chr7-14162961-INS-4460 |
| chr7-14175685-INS-56 | chr7-14175685-INS-56 |
| chr7-14201519-INS-976 | chr7-14201519-INS-978 |
| chr7-14207186-INS-59 | chr7-14207186-INS-59 |
| chr7-14248654-INS-54 | chr7-14248654-INS-58 |
| chr7-14253193-INS-65 | chr7-14253193-INS-65 |
| chr7-14277234-INS-451 | chr7-14277234-INS-458 |
| chr7-14284763-INS-399 | chr7-14284763-INS-399 |
| chr7-14301463-INS-61 | chr7-14301463-INS-50 |
| chr7-14301552-INS-681 | chr7-14301552-INS-679 |
| chr7-14317846-INS-6845 | chr7-14317847-INS-6845 |
| chr7-14345661-INS-154 | chr7-14345661-INS-154 |
| chr7-14348093-INS-4820 | chr7-14348094-INS-4820 |
| chr7-14388939-INS-86 | chr7-14388899-INS-80 |
| chr7-14393069-INS-577 | chr7-14393069-INS-577 |
| chr7-14440781-INS-158 | chr7-14440781-INS-157 |
| chr7-14553482-INS-148 | chr7-14553482-INS-148 |
| chr7-14611127-INS-214 | chr7-14611127-INS-214 |

| | |
|---|---|
| chr7-14643008-INS-1892 | chr7-14643009-INS-1892 |
| chr7-14691422-INS-204 | chr7-14691422-INS-204 |
| chr7-14750608-INS-201 | chr7-14750608-INS-201 |
| chr7-14795116-INS-56 | chr7-14795080-INS-52 |
| chr7-15013758-INS-81 | chr7-15013758-INS-80 |
| chr7-15018776-INS-368 | chr7-15018776-INS-368 |
| chr7-15291684-INS-344 | chr7-15291684-INS-345 |
| chr7-15388126-INS-91 | chr7-15388126-INS-91 |
| chr7-15424272-INS-69 | chr7-15424272-INS-82 |
| chr7-15479469-INS-75 | chr7-15479469-INS-80 |
| chr7-15511248-INS-852 | chr7-15511248-INS-853 |
| chr7-15516796-INS-85 | chr7-15516800-INS-130 |
| chr7-15552819-INS-60 | chr7-15552827-INS-54 |
| chr7-15575767-INS-197 | chr7-15575767-INS-197 |
| chr7-15590218-INS-5336 | chr7-15590219-INS-5336 |
| chr7-15604418-INS-73 | chr7-15604430-INS-52 |
| chr7-15625390-INS-5363 | chr7-15625391-INS-5363 |
| chr7-15635002-INS-72 | chr7-15635003-INS-58 |
| chr7-15642239-INS-80 | chr7-15642239-INS-80 |
| chr7-15652082-INS-53 | chr7-15652082-INS-53 |
| chr7-15662111-INS-97 | chr7-15662104-INS-95 |
| chr7-15669741-INS-378 | chr7-15669732-INS-384 |
| chr7-15671590-INS-159 | chr7-15671603-INS-158 |
| chr7-15695914-INS-53 | chr7-15695914-INS-53 |
| chr7-15735575-INS-522 | chr7-15735575-INS-522 |
| chr7-15745275-INS-159 | chr7-15745275-INS-159 |
| chr7-15755284-INS-83 | chr7-15755284-INS-90 |
| chr7-15759689-INS-191 | chr7-15759689-INS-191 |
| chr7-15764862-INS-207 | chr7-15764862-INS-213 |
| chr7-15783608-INS-67 | chr7-15783608-INS-67 |
| chr7-15787945-INS-51 | chr7-15787945-INS-51 |
| chr7-15871925-INS-212 | chr7-15871925-INS-211 |
| chr7-15884802-INS-192 | chr7-15884803-INS-193 |
| chr7-15888946-INS-151 | chr7-15888925-INS-155 |
| chr7-15897271-INS-51 | chr7-15897271-INS-53 |
| chr7-15898091-INS-265 | chr7-15898091-INS-265 |
| chr7-15904495-INS-86 | chr7-15904493-INS-100 |
| chr7-15915163-INS-54 | chr7-15915163-INS-54 |
| chr7-15919260-INS-216 | chr7-15919260-INS-216 |
| chr7-15928480-INS-54 | chr7-15928480-INS-54 |
| chr7-15929909-INS-116 | chr7-15929909-INS-116 |
| chr7-15942258-INS-114 | chr7-15942248-INS-130 |
| chr7-15959899-INS-172 | chr7-15959899-INS-172 |
| chr7-15966369-INS-202 | chr7-15966369-INS-203 |
| chr7-15968971-INS-60 | chr7-15968971-INS-72 |
| chr7-15970294-INS-82 | chr7-15970294-INS-82 |
| chr7-15975438-INS-1539 | chr7-15975439-INS-1542 |
| chr7-15977021-INS-204 | chr7-15977022-INS-204 |
| chr7-15985214-INS-208 | chr7-15985214-INS-209 |
| chr7-15991092-INS-53 | chr7-15991092-INS-53 |
| chr7-15998108-INS-205 | chr7-15998101-INS-206 |
| chr7-16019143-INS-167 | chr7-16019143-INS-168 |
| chr7-16023256-INS-714 | chr7-16023256-INS-714 |
| chr7-16025798-INS-81 | chr7-16025798-INS-81 |
| chr7-16038491-INS-197 | chr7-16038491-INS-197 |
| chr7-16062105-INS-70 | chr7-16062105-INS-70 |
| chr7-16063808-INS-509 | chr7-16063808-INS-509 |
| chr7-16104641-INS-207 | chr7-16104641-INS-207 |
| chr7-16116689-INS-231 | chr7-16116689-INS-227 |
| chr7-16125420-INS-204 | chr7-16125420-INS-204 |
| chr7-16127161-INS-185 | chr7-16127161-INS-185 |
| chr7-16131805-INS-56 | chr7-16131805-INS-56 |
| chr7-16144029-INS-121 | chr7-16144029-INS-109 |
| chr7-16147829-INS-186 | chr7-16147829-INS-186 |
| chr7-16176973-INS-359 | chr7-16176973-INS-359 |
| chr7-16177602-INS-52 | chr7-16177602-INS-52 |
| chr7-16188619-INS-106 | chr7-16188619-INS-106 |
| chr7-16190385-INS-52 | chr7-16190385-INS-52 |
| chr7-16195197-INS-564 | chr7-16195197-INS-564 |
| chr7-16199917-INS-53 | chr7-16199917-INS-53 |
| chr7-16205471-INS-155 | chr7-16205460-INS-168 |
| chr7-16221207-INS-140 | chr7-16221207-INS-140 |
| chr7-16240060-INS-221 | chr7-16240060-INS-222 |
| chr7-16245040-INS-198 | chr7-16245053-INS-192 |
| chr7-16250062-INS-58 | chr7-16250062-INS-52 |
| chr7-16268786-INS-152 | chr7-16268794-INS-132 |
| chr7-16333553-INS-52 | chr7-16333554-INS-52 |

| | |
|---|---|
| chr7-16370969-INS-50 | chr7-16371000-INS-62 |
| chr7-16373882-INS-339 | chr7-16373882-INS-339 |
| chr7-16377181-INS-86 | chr7-16377181-INS-86 |
| chr7-16377705-INS-197 | chr7-16377705-INS-196 |
| chr7-16379227-INS-54 | chr7-16379231-INS-55 |
| chr7-16388012-INS-2627 | chr7-16388013-INS-2627 |
| chr7-16391437-INS-219 | chr7-16391437-INS-219 |
| chr7-16421058-INS-134 | chr7-16421058-INS-134 |
| chr7-16436942-INS-501 | chr7-16436942-INS-501 |
| chr7-16444377-INS-56 | chr7-16444377-INS-56 |
| chr7-16449008-INS-194 | chr7-16449008-INS-194 |
| chr7-16450043-INS-145 | chr7-16450043-INS-145 |
| chr7-16451115-INS-477 | chr7-16451115-INS-477 |
| chr7-16466435-INS-59 | chr7-16466435-INS-59 |
| chr7-16474913-INS-215 | chr7-16474913-INS-216 |
| chr7-16486578-INS-162 | chr7-16486578-INS-162 |
| chr7-16492289-INS-180 | chr7-16492289-INS-181 |
| chr7-16511112-INS-74 | chr7-16511112-INS-101 |
| chr7-16518283-INS-53 | chr7-16518283-INS-53 |
| chr7-16524772-INS-705 | chr7-16524772-INS-705 |
| chr7-16524773-INS-92 | chr7-16524773-INS-92 |
| chr7-16553364-INS-72 | chr7-16553364-INS-72 |
| chr7-16568965-INS-207 | chr7-16568965-INS-208 |
| chr7-16570006-INS-76 | chr7-16570006-INS-56 |
| chr7-16585119-INS-122 | chr7-16585106-INS-165 |
| chr7-16591417-INS-213 | chr7-16591417-INS-213 |
| chr7-16592777-INS-64 | chr7-16592777-INS-60 |
| chr7-16594178-INS-192 | chr7-16594178-INS-191 |
| chr7-16598421-INS-52 | chr7-16598421-INS-52 |
| chr7-16612599-INS-264 | chr7-16612599-INS-264 |
| chr7-16639236-INS-208 | chr7-16639236-INS-208 |
| chr7-16666947-INS-54 | chr7-16666985-INS-62 |
| chr7-16692230-INS-635 | chr7-16692230-INS-635 |
| chr7-16706531-INS-218 | chr7-16706531-INS-218 |
| chr7-16709137-INS-208 | chr7-16709137-INS-209 |
| chr7-16713724-INS-53 | chr7-16713724-INS-53 |
| chr7-16725057-INS-175 | chr7-16725057-INS-175 |
| chr7-16735318-INS-53 | chr7-16735318-INS-53 |
| chr7-16759496-INS-63 | chr7-16759496-INS-63 |
| chr7-16791637-INS-234 | chr7-16791637-INS-237 |
| chr7-16800723-INS-58 | chr7-16800723-INS-56 |
| chr7-16814669-INS-222 | chr7-16814669-INS-222 |
| chr7-16823826-INS-408 | chr7-16823826-INS-409 |
| chr7-16871354-INS-139 | chr7-16871354-INS-144 |
| chr7-16874836-INS-669 | chr7-16874836-INS-669 |
| chr7-16903580-INS-67 | chr7-16903616-INS-51 |
| chr7-16910625-INS-241 | chr7-16910625-INS-253 |
| chr7-16943760-INS-99 | chr7-16943760-INS-97 |
| chr7-16959074-INS-56 | chr7-16959076-INS-85 |
| chr7-16963519-INS-866 | chr7-16963519-INS-866 |
| chr7-16969519-INS-103 | chr7-16969519-INS-103 |
| chr7-16973253-INS-75 | chr7-16973253-INS-75 |
| chr7-16985594-INS-328 | chr7-16985594-INS-327 |
| chr7-17008399-INS-201 | chr7-17008399-INS-201 |
| chr7-17057569-INS-542 | chr7-17057569-INS-542 |
| chr7-17296698-INS-136 | chr7-17296695-INS-138 |
| chr7-17337666-INS-51 | chr7-17337666-INS-70 |
| chr7-17339920-INS-210 | chr7-17339920-INS-210 |
| chr7-17355251-INS-6505 | chr7-17355252-INS-6505 |
| chr7-17389759-INS-180 | chr7-17389759-INS-184 |
| chr7-17422551-INS-6674 | chr7-17422552-INS-6674 |
| chr7-17432401-INS-2023 | chr7-17432402-INS-2023 |
| chr7-17458850-INS-1819 | chr7-17458851-INS-1819 |
| chr7-17468590-INS-6397 | chr7-17468591-INS-6396 |
| chr7-17471666-INS-55 | chr7-17471667-INS-50 |
| chr7-17505183-INS-54 | chr7-17505183-INS-60 |
| chr7-17505947-INS-6253 | chr7-17505948-INS-6253 |
| chr7-17515261-INS-1596 | chr7-17515262-INS-1596 |
| chr7-17534272-INS-208 | chr7-17534272-INS-208 |
| chr7-17616160-INS-6868 | chr7-17616161-INS-6868 |
| chr7-17642101-INS-68 | chr7-17642101-INS-68 |
| chr7-17644113-INS-62 | chr7-17644113-INS-100 |
| chr7-17656075-INS-144 | chr7-17656075-INS-144 |
| chr7-17674617-INS-201 | chr7-17674617-INS-201 |
| chr7-17723113-INS-86 | chr7-17723113-INS-75 |
| chr7-17734403-INS-76 | chr7-17734403-INS-82 |
| chr7-17769015-INS-196 | chr7-17769015-INS-196 |

| | |
|---|---|
| chr7-17786407-INS-5404 | chr7-17786408-INS-5404 |
| chr7-17790351-INS-104 | chr7-17790351-INS-106 |
| chr7-17828285-INS-218 | chr7-17828285-INS-218 |
| chr7-17858412-INS-61 | chr7-17858412-INS-61 |
| chr7-17879530-INS-72 | chr7-17879531-INS-96 |
| chr7-17894827-INS-59 | chr7-17894833-INS-60 |
| chr7-17909618-INS-376 | chr7-17909615-INS-375 |
| chr7-17915573-INS-77 | chr7-17915573-INS-79 |
| chr7-17923250-INS-5049 | chr7-17923251-INS-5049 |
| chr7-17942297-INS-5145 | chr7-17942298-INS-5145 |
| chr7-17945343-INS-51 | chr7-17945367-INS-59 |
| chr7-17957839-INS-101 | chr7-17957839-INS-102 |
| chr7-17970789-INS-8421 | chr7-17970790-INS-8421 |
| chr7-18018388-INS-7115 | chr7-18018389-INS-7117 |
| chr7-18052389-INS-428 | chr7-18052405-INS-429 |
| chr7-18080307-INS-458 | chr7-18080307-INS-458 |
| chr7-18101184-INS-84 | chr7-18101184-INS-92 |
| chr7-18158852-INS-1890 | chr7-18158853-INS-1890 |
| chr7-18197844-INS-3296 | chr7-18197845-INS-3297 |
| chr7-18207894-INS-4161 | chr7-18207895-INS-4161 |
| chr7-18239632-INS-268 | chr7-18239632-INS-268 |
| chr7-18243918-INS-233 | chr7-18243918-INS-233 |
| chr7-18246941-INS-76 | chr7-18246941-INS-92 |
| chr7-18252144-INS-3083 | chr7-18252145-INS-3082 |
| chr7-18268804-INS-54 | chr7-18268804-INS-54 |
| chr7-18278165-INS-52 | chr7-18278165-INS-52 |
| chr7-18281996-INS-80 | chr7-18281996-INS-80 |
| chr7-18333849-INS-82 | chr7-18333849-INS-102 |
| chr7-18430398-INS-431 | chr7-18430398-INS-431 |
| chr7-18440601-INS-146 | chr7-18440601-INS-152 |
| chr7-18482945-INS-82 | chr7-18482945-INS-84 |
| chr7-18485441-INS-53 | chr7-18485441-INS-53 |
| chr7-18493815-INS-542 | chr7-18493815-INS-542 |
| chr7-18494746-INS-187 | chr7-18494746-INS-173 |
| chr7-18514062-INS-52 | chr7-18514062-INS-54 |
| chr7-18515333-INS-101 | chr7-18515333-INS-107 |
| chr7-18515997-INS-96 | chr7-18515997-INS-96 |
| chr7-18521559-INS-112 | chr7-18521559-INS-112 |
| chr7-18531610-INS-52 | chr7-18531610-INS-52 |
| chr7-18543964-INS-95 | chr7-18543949-INS-96 |
| chr7-18546177-INS-8844 | chr7-18546178-INS-8844 |
| chr7-18557561-INS-435 | chr7-18557561-INS-435 |
| chr7-18559217-INS-218 | chr7-18559217-INS-218 |
| chr7-18560028-INS-479 | chr7-18560028-INS-479 |
| chr7-18562054-INS-5509 | chr7-18562055-INS-5509 |
| chr7-18588611-INS-53 | chr7-18588611-INS-57 |
| chr7-18596531-INS-224 | chr7-18596531-INS-225 |
| chr7-18606556-INS-180 | chr7-18606522-INS-182 |
| chr7-18609145-INS-107 | chr7-18609109-INS-106 |
| chr7-18610919-INS-400 | chr7-18610919-INS-400 |
| chr7-18616499-INS-58 | chr7-18616490-INS-84 |
| chr7-18622097-INS-514 | chr7-18622097-INS-514 |
| chr7-18645779-INS-105 | chr7-18645779-INS-105 |
| chr7-18648923-INS-64 | chr7-18648897-INS-58 |
| chr7-18654064-INS-62 | chr7-18654064-INS-64 |
| chr7-18655825-INS-199 | chr7-18655825-INS-199 |
| chr7-18663330-INS-52 | chr7-18663330-INS-52 |
| chr7-18664211-INS-199 | chr7-18664211-INS-199 |
| chr7-18678355-INS-3113 | chr7-18678356-INS-3113 |
| chr7-18700673-INS-105 | chr7-18700673-INS-105 |
| chr7-18705963-INS-52 | chr7-18705964-INS-60 |
| chr7-18707050-INS-225 | chr7-18707050-INS-225 |
| chr7-18708360-INS-64 | chr7-18708360-INS-56 |
| chr7-18710884-INS-105 | chr7-18710884-INS-105 |
| chr7-18762331-INS-189 | chr7-18762331-INS-189 |
| chr7-18798283-INS-60 | chr7-18798283-INS-60 |
| chr7-18801602-INS-217 | chr7-18801602-INS-217 |
| chr7-18805507-INS-53 | chr7-18805507-INS-53 |
| chr7-18806483-INS-105 | chr7-18806483-INS-105 |
| chr7-18823355-INS-53 | chr7-18823356-INS-56 |
| chr7-18848015-INS-208 | chr7-18848015-INS-208 |
| chr7-18878324-INS-187 | chr7-18878324-INS-187 |
| chr7-18901196-INS-165 | chr7-18901197-INS-166 |
| chr7-18907145-INS-6463 | chr7-18907145-INS-6463 |
| chr7-18917782-INS-95 | chr7-18917782-INS-95 |
| chr7-18924345-INS-76 | chr7-18924345-INS-76 |
| chr7-18933249-INS-53 | chr7-18933249-INS-53 |

| | |
|---|---|
| chr7-18934795-INS-170 | chr7-18934795-INS-171 |
| chr7-18945069-INS-52 | chr7-18945070-INS-52 |
| chr7-18949950-INS-190 | chr7-18949950-INS-190 |
| chr7-18952840-INS-55 | chr7-18952840-INS-55 |
| chr7-18958097-INS-214 | chr7-18958097-INS-215 |
| chr7-18974267-INS-201 | chr7-18974260-INS-179 |
| chr7-18984617-INS-215 | chr7-18984617-INS-215 |
| chr7-19000982-INS-60 | chr7-19000992-INS-54 |
| chr7-19011473-INS-173 | chr7-19011473-INS-173 |
| chr7-19014701-INS-168 | chr7-19014701-INS-168 |
| chr7-19023684-INS-197 | chr7-19023684-INS-197 |
| chr7-19030569-INS-61 | chr7-19030569-INS-61 |
| chr7-19032626-INS-209 | chr7-19032626-INS-209 |
| chr7-19046398-INS-228 | chr7-19046398-INS-228 |
| chr7-19047111-INS-122 | chr7-19047111-INS-122 |
| chr7-19054238-INS-249 | chr7-19054238-INS-246 |
| chr7-19056266-INS-53 | chr7-19056266-INS-53 |
| chr7-19058161-INS-202 | chr7-19058161-INS-200 |
| chr7-19059017-INS-53 | chr7-19059017-INS-53 |
| chr7-19067976-INS-54 | chr7-19067976-INS-54 |
| chr7-19078091-INS-184 | chr7-19078092-INS-184 |
| chr7-19078720-INS-64 | chr7-19078720-INS-96 |
| chr7-19082578-INS-100 | chr7-19082579-INS-58 |
| chr7-19102077-INS-53 | chr7-19102077-INS-53 |
| chr7-19113557-INS-194 | chr7-19113557-INS-194 |
| chr7-19148579-INS-53 | chr7-19148579-INS-53 |
| chr7-19164290-INS-196 | chr7-19164290-INS-196 |
| chr7-19169175-INS-469 | chr7-19169175-INS-469 |
| chr7-19175329-INS-210 | chr7-19175329-INS-210 |
| chr7-19184915-INS-52 | chr7-19184915-INS-50 |
| chr7-19188429-INS-105 | chr7-19188429-INS-105 |
| chr7-19204287-INS-64 | chr7-19204287-INS-50 |
| chr7-19224570-INS-208 | chr7-19224570-INS-205 |
| chr7-19264437-INS-53 | chr7-19264437-INS-53 |
| chr7-19267626-INS-215 | chr7-19267626-INS-215 |
| chr7-19272914-INS-159 | chr7-19272914-INS-159 |
| chr7-19273609-INS-3770 | chr7-19273610-INS-3770 |
| chr7-19276645-INS-4718 | chr7-19276646-INS-4718 |
| chr7-19302129-INS-66 | chr7-19302129-INS-70 |
| chr7-19304755-INS-51 | chr7-19304755-INS-51 |
| chr7-19308751-INS-86 | chr7-19308751-INS-86 |
| chr7-19331054-INS-55 | chr7-19331054-INS-56 |
| chr7-19347393-INS-53 | chr7-19347393-INS-53 |
| chr7-19355267-INS-159 | chr7-19355267-INS-159 |
| chr7-19400508-INS-216 | chr7-19400508-INS-215 |
| chr7-19418598-INS-162 | chr7-19418599-INS-160 |
| chr7-19422926-INS-88 | chr7-19422926-INS-88 |
| chr7-19435870-INS-185 | chr7-19435870-INS-185 |
| chr7-19453471-INS-65 | chr7-19453471-INS-65 |
| chr7-19459306-INS-159 | chr7-19459306-INS-159 |
| chr7-19491469-INS-105 | chr7-19491469-INS-105 |
| chr7-19501200-INS-106 | chr7-19501200-INS-105 |
| chr7-19509139-INS-169 | chr7-19509139-INS-169 |
| chr7-19514359-INS-122 | chr7-19514347-INS-141 |
| chr7-19523371-INS-52 | chr7-19523371-INS-52 |
| chr7-19527426-INS-53 | chr7-19527426-INS-53 |
| chr7-19540824-INS-108 | chr7-19540824-INS-108 |
| chr7-19546833-INS-105 | chr7-19546833-INS-105 |
| chr7-19571458-INS-53 | chr7-19571458-INS-53 |
| chr7-19579753-INS-52 | chr7-19579753-INS-52 |
| chr7-19583340-INS-53 | chr7-19583352-INS-54 |
| chr7-19584434-INS-64 | chr7-19584434-INS-64 |
| chr7-19603922-INS-64 | chr7-19603922-INS-80 |
| chr7-19623401-INS-62 | chr7-19623401-INS-62 |
| chr7-19647697-INS-106 | chr7-19647697-INS-106 |
| chr7-19654830-INS-56 | chr7-19654830-INS-57 |
| chr7-19658507-INS-200 | chr7-19658507-INS-201 |
| chr7-19663997-INS-137 | chr7-19663997-INS-137 |
| chr7-19676199-INS-498 | chr7-19676199-INS-498 |
| chr7-19683496-INS-1061 | chr7-19683497-INS-1059 |
| chr7-19685927-INS-70 | chr7-19685927-INS-70 |
| chr7-19690950-INS-413 | chr7-19690950-INS-413 |
| chr7-19697194-INS-85 | chr7-19697194-INS-88 |
| chr7-19704755-INS-146 | chr7-19704755-INS-129 |
| chr7-19708706-INS-210 | chr7-19708706-INS-211 |
| chr7-19721023-INS-52 | chr7-19721023-INS-58 |
| chr7-19722693-INS-100 | chr7-19722693-INS-100 |

| | |
|---|---|
| chr7-19724354-INS-300 | chr7-19724354-INS-319 |
| chr7-19727135-INS-142 | chr7-19727135-INS-141 |
| chr7-19746018-INS-127 | chr7-19746018-INS-127 |
| chr7-19760816-INS-57 | chr7-19760801-INS-63 |
| chr7-19801058-INS-90 | chr7-19801058-INS-75 |
| chr7-20694949-INS-195 | chr7-20694950-INS-198 |
| chr7-21162123-INS-498 | chr7-21162123-INS-498 |
| chr7-22995746-INS-158 | chr7-22995746-INS-161 |
| chr7-23058854-INS-52 | chr7-23058854-INS-52 |
| chr7-23076922-INS-61 | chr7-23076922-INS-108 |
| chr7-23082133-INS-5340 | chr7-23082134-INS-5340 |
| chr7-23101120-INS-59 | chr7-23101120-INS-59 |
| chr7-23136568-INS-69 | chr7-23136568-INS-72 |
| chr7-23154072-INS-498 | chr7-23154080-INS-524 |
| chr7-23170609-INS-8849 | chr7-23170610-INS-8842 |
| chr7-23188397-INS-5149 | chr7-23188398-INS-5149 |
| chr7-23191569-INS-53 | chr7-23191569-INS-52 |
| chr7-23203747-INS-94 | chr7-23203747-INS-94 |
| chr7-23210504-INS-60 | chr7-23210504-INS-65 |
| chr7-23219051-INS-10268 | chr7-23219052-INS-10268 |
| chr7-23226845-INS-3035 | chr7-23226846-INS-3035 |
| chr7-23277823-INS-54 | chr7-23277823-INS-54 |
| chr7-23320839-INS-90 | chr7-23320839-INS-90 |
| chr7-23335659-INS-186 | chr7-23335658-INS-205 |
| chr7-23342005-INS-156 | chr7-23342005-INS-162 |
| chr7-23461891-INS-7795 | chr7-23461891-INS-7797 |
| chr7-23485658-INS-6454 | chr7-23485659-INS-6454 |
| chr7-23498680-INS-1236 | chr7-23498681-INS-1236 |
| chr7-23504982-INS-471 | chr7-23504982-INS-471 |
| chr7-23523942-INS-89 | chr7-23523942-INS-87 |
| chr7-23536610-INS-569 | chr7-23536610-INS-569 |
| chr7-23545285-INS-148 | chr7-23545285-INS-148 |
| chr7-23553970-INS-72 | chr7-23553970-INS-72 |
| chr7-23557311-INS-5442 | chr7-23557307-INS-5451 |
| chr7-23575193-INS-197 | chr7-23575193-INS-198 |
| chr7-23580517-INS-1016 | chr7-23580518-INS-1016 |
| chr7-23580798-INS-3660 | chr7-23580799-INS-3660 |
| chr7-23585687-INS-7098 | chr7-23585688-INS-7098 |
| chr7-23587890-INS-108 | chr7-23587890-INS-112 |
| chr7-23618879-INS-160 | chr7-23618879-INS-132 |
| chr7-23632899-INS-62 | chr7-23632904-INS-64 |
| chr7-23640085-INS-6452 | chr7-23640086-INS-6452 |
| chr7-23644322-INS-397 | chr7-23644322-INS-397 |
| chr7-23654121-INS-1222 | chr7-23654122-INS-1223 |
| chr7-23677346-INS-169 | chr7-23677346-INS-168 |
| chr7-23706229-INS-340 | chr7-23706229-INS-340 |
| chr7-23758811-INS-54 | chr7-23758811-INS-52 |
| chr7-23763068-INS-60 | chr7-23763068-INS-55 |
| chr7-23776384-INS-53 | chr7-23776384-INS-53 |
| chr7-23778006-INS-87 | chr7-23778019-INS-72 |
| chr7-23786629-INS-111 | chr7-23786629-INS-111 |
| chr7-23789183-INS-214 | chr7-23789227-INS-204 |
| chr7-23798693-INS-159 | chr7-23798693-INS-159 |
| chr7-23823087-INS-68 | chr7-23823060-INS-88 |
| chr7-23828326-INS-58 | chr7-23828314-INS-70 |
| chr7-23829316-INS-189 | chr7-23829316-INS-210 |
| chr7-23846897-INS-360 | chr7-23846897-INS-360 |
| chr7-23849570-INS-130 | chr7-23849570-INS-110 |
| chr7-23851495-INS-117 | chr7-23851484-INS-90 |
| chr7-23880563-INS-75 | chr7-23880563-INS-75 |
| chr7-23916758-INS-88 | chr7-23916758-INS-117 |
| chr7-23938120-INS-53 | chr7-23938120-INS-53 |
| chr7-23943703-INS-52 | chr7-23943703-INS-50 |
| chr7-23967357-INS-99 | chr7-23967357-INS-99 |
| chr7-23984611-INS-6551 | chr7-23984612-INS-6551 |
| chr7-23989891-INS-79 | chr7-23989869-INS-55 |
| chr7-23996099-INS-308 | chr7-23996099-INS-308 |
| chr7-23999171-INS-100 | chr7-23999171-INS-96 |
| chr7-24003810-INS-98 | chr7-24003810-INS-97 |
| chr7-24009432-INS-66 | chr7-24009432-INS-66 |
| chr7-24009932-INS-959 | chr7-24009932-INS-954 |
| chr7-24037728-INS-55 | chr7-24037733-INS-93 |
| chr7-24039576-INS-59 | chr7-24039576-INS-59 |
| chr7-24044455-INS-383 | chr7-24044455-INS-383 |
| chr7-24046590-INS-94 | chr7-24046590-INS-84 |
| chr7-24050668-INS-332 | chr7-24050668-INS-332 |
| chr7-24062384-INS-557 | chr7-24062412-INS-539 |

| | |
|---|---|
| chr7-24066447-INS-107 | chr7-24066447-INS-107 |
| chr7-24074993-INS-477 | chr7-24074994-INS-477 |
| chr7-24077369-INS-62 | chr7-24077376-INS-56 |
| chr7-24085270-INS-52 | chr7-24085270-INS-56 |
| chr7-24091813-INS-109 | chr7-24091813-INS-109 |
| chr7-24127090-INS-7087 | chr7-24127091-INS-7087 |
| chr7-24140978-INS-7122 | chr7-24140979-INS-7122 |
| chr7-24149530-INS-8675 | chr7-24149531-INS-8675 |
| chr7-24154072-INS-114 | chr7-24154072-INS-100 |
| chr7-24164522-INS-193 | chr7-24164522-INS-192 |
| chr7-24170446-INS-53 | chr7-24170446-INS-53 |
| chr7-24189816-INS-71 | chr7-24189816-INS-71 |
| chr7-24195205-INS-107 | chr7-24195205-INS-107 |
| chr7-24227640-INS-52 | chr7-24227640-INS-52 |
| chr7-24228685-INS-198 | chr7-24228685-INS-199 |
| chr7-24238438-INS-53 | chr7-24238438-INS-53 |
| chr7-24248190-INS-411 | chr7-24248190-INS-411 |
| chr7-24254900-INS-194 | chr7-24254903-INS-187 |
| chr7-24262087-INS-51 | chr7-24262087-INS-51 |
| chr7-24289446-INS-54 | chr7-24289448-INS-56 |
| chr7-24346395-INS-54 | chr7-24346395-INS-54 |
| chr7-24357633-INS-297 | chr7-24357633-INS-297 |
| chr7-24374102-INS-60 | chr7-24374102-INS-60 |
| chr7-24390646-INS-122 | chr7-24390646-INS-122 |
| chr7-24407759-INS-55 | chr7-24407725-INS-85 |
| chr7-24408893-INS-214 | chr7-24408893-INS-214 |
| chr7-24409953-INS-90 | chr7-24409953-INS-90 |
| chr7-24415491-INS-65 | chr7-24415491-INS-65 |
| chr7-24420650-INS-120 | chr7-24420650-INS-120 |
| chr7-24455121-INS-105 | chr7-24455121-INS-105 |
| chr7-24457569-INS-62 | chr7-24457569-INS-50 |
| chr7-24467935-INS-84 | chr7-24467935-INS-84 |
| chr7-24470965-INS-7140 | chr7-24470966-INS-7140 |
| chr7-24492897-INS-70 | chr7-24492897-INS-70 |
| chr7-24494014-INS-55 | chr7-24494014-INS-55 |
| chr7-24533500-INS-56 | chr7-24533533-INS-64 |
| chr7-24539979-INS-67 | chr7-24539979-INS-67 |
| chr7-24552833-INS-61 | chr7-24552833-INS-63 |
| chr7-24555302-INS-430 | chr7-24555302-INS-413 |
| chr7-24567250-INS-162 | chr7-24567250-INS-162 |
| chr7-24581221-INS-193 | chr7-24581221-INS-193 |
| chr7-24615582-INS-68 | chr7-24615578-INS-102 |
| chr7-24645228-INS-111 | chr7-24645228-INS-111 |
| chr7-24669041-INS-67 | chr7-24669041-INS-67 |
| chr7-24676392-INS-53 | chr7-24676381-INS-53 |
| chr7-24730962-INS-61 | chr7-24730962-INS-63 |
| chr7-24764073-INS-123 | chr7-24764073-INS-122 |
| chr7-24801397-INS-101 | chr7-24801397-INS-104 |
| chr7-24815314-INS-108 | chr7-24815314-INS-112 |
| chr7-24816543-INS-56 | chr7-24816543-INS-56 |
| chr7-24823545-INS-413 | chr7-24823545-INS-412 |
| chr7-24842378-INS-160 | chr7-24842378-INS-160 |
| chr7-24885780-INS-72 | chr7-24885780-INS-72 |
| chr7-24891193-INS-358 | chr7-24891193-INS-358 |
| chr7-24898251-INS-119 | chr7-24898255-INS-126 |
| chr7-24916491-INS-208 | chr7-24916491-INS-208 |
| chr7-24941519-INS-260 | chr7-24941519-INS-259 |
| chr7-24952823-INS-192 | chr7-24952823-INS-192 |
| chr7-24957341-INS-84 | chr7-24957341-INS-51 |
| chr7-24959553-INS-60 | chr7-24959553-INS-60 |
| chr7-24965163-INS-122 | chr7-24965157-INS-132 |
| chr7-24965631-INS-205 | chr7-24965631-INS-207 |
| chr7-24996137-INS-165 | chr7-24996116-INS-193 |
| chr7-25000462-INS-158 | chr7-25000462-INS-164 |
| chr7-25032134-INS-102 | chr7-25032134-INS-102 |
| chr7-25039786-INS-128 | chr7-25039786-INS-128 |
| chr7-25043253-INS-182 | chr7-25043253-INS-185 |
| chr7-25050641-INS-159 | chr7-25050679-INS-159 |
| chr7-25082359-INS-76 | chr7-25082359-INS-81 |
| chr7-25112944-INS-58 | chr7-25112944-INS-73 |
| chr7-25119629-INS-106 | chr7-25119629-INS-111 |
| chr7-25144548-INS-194 | chr7-25144548-INS-194 |
| chr7-25182376-INS-292 | chr7-25182376-INS-292 |
| chr7-25203476-INS-200 | chr7-25203476-INS-199 |
| chr7-25225012-INS-178 | chr7-25225012-INS-178 |
| chr7-25232677-INS-1081 | chr7-25232678-INS-1081 |
| chr7-25235993-INS-174 | chr7-25235993-INS-175 |

| | |
|---|---|
| chr7-25258277-INS-4896 | chr7-25258278-INS-4896 |
| chr7-25309502-INS-66 | chr7-25309502-INS-66 |
| chr7-25312766-INS-168 | chr7-25312766-INS-210 |
| chr7-25318323-INS-54 | chr7-25318323-INS-64 |
| chr7-25333291-INS-206 | chr7-25333291-INS-206 |
| chr7-25343402-INS-54 | chr7-25343402-INS-84 |
| chr7-25371564-INS-139 | chr7-25371540-INS-182 |
| chr7-25382442-INS-179 | chr7-25382442-INS-163 |
| chr7-25408796-INS-223 | chr7-25408796-INS-210 |
| chr7-25467106-INS-106 | chr7-25467106-INS-106 |
| chr7-25474196-INS-58 | chr7-25474196-INS-56 |
| chr7-25494039-INS-171 | chr7-25494039-INS-182 |
| chr7-25523736-INS-56 | chr7-25523736-INS-56 |
| chr7-25534372-INS-189 | chr7-25534372-INS-189 |
| chr7-25540917-INS-103 | chr7-25540917-INS-103 |
| chr7-25565863-INS-54 | chr7-25565852-INS-59 |
| chr7-25571397-INS-431 | chr7-25571397-INS-407 |
| chr7-25575200-INS-125 | chr7-25575217-INS-81 |
| chr7-25591886-INS-90 | chr7-25591886-INS-105 |
| chr7-25593795-INS-304 | chr7-25593796-INS-305 |
| chr7-25600734-INS-1910 | chr7-25600735-INS-1911 |
| chr7-25610222-INS-193 | chr7-25610244-INS-203 |
| chr7-25617979-INS-58 | chr7-25617979-INS-58 |
| chr7-25662000-INS-52 | chr7-25662000-INS-52 |
| chr7-25671148-INS-484 | chr7-25671148-INS-484 |
| chr7-25713091-INS-100 | chr7-25713089-INS-102 |
| chr7-25751804-INS-177 | chr7-25751804-INS-177 |
| chr7-25783833-INS-6585 | chr7-25783834-INS-6585 |
| chr7-25816068-INS-354 | chr7-25816068-INS-354 |
| chr7-25887430-INS-156 | chr7-25887430-INS-156 |
| chr7-25902197-INS-68 | chr7-25902197-INS-68 |
| chr7-25924944-INS-7175 | chr7-25924945-INS-7175 |
| chr7-25953538-INS-1709 | chr7-25953539-INS-1709 |
| chr7-25967615-INS-159 | chr7-25967615-INS-159 |
| chr7-25972121-INS-1886 | chr7-25972122-INS-1886 |
| chr7-25977166-INS-51 | chr7-25977166-INS-51 |
| chr7-25987831-INS-6832 | chr7-25987832-INS-6833 |
| chr7-25999254-INS-52 | chr7-25999254-INS-50 |
| chr7-26000262-INS-594 | chr7-26000262-INS-600 |
| chr7-26039281-INS-3023 | chr7-26039282-INS-3022 |
| chr7-26100231-INS-1682 | chr7-26100232-INS-1685 |
| chr7-26115784-INS-137 | chr7-26115834-INS-131 |
| chr7-26126474-INS-55 | chr7-26126474-INS-55 |
| chr7-26132259-INS-6415 | chr7-26132260-INS-6415 |
| chr7-26140500-INS-187 | chr7-26140500-INS-193 |
| chr7-26150505-INS-61 | chr7-26150505-INS-65 |
| chr7-26206153-INS-150 | chr7-26206153-INS-150 |
| chr7-26213348-INS-73 | chr7-26213348-INS-73 |
| chr7-26217948-INS-6764 | chr7-26217949-INS-6764 |
| chr7-26243721-INS-62 | chr7-26243721-INS-62 |
| chr7-26244246-INS-164 | chr7-26244246-INS-164 |
| chr7-26269559-INS-95 | chr7-26269559-INS-77 |
| chr7-26271092-INS-6758 | chr7-26271093-INS-6757 |
| chr7-26274488-INS-291 | chr7-26274488-INS-330 |
| chr7-26347822-INS-1889 | chr7-26347823-INS-1889 |
| chr7-26377539-INS-328 | chr7-26377539-INS-328 |
| chr7-26380647-INS-65 | chr7-26380645-INS-64 |
| chr7-26383925-INS-343 | chr7-26383925-INS-343 |
| chr7-26384983-INS-6231 | chr7-26384984-INS-6231 |
| chr7-26417879-INS-779 | chr7-26417879-INS-795 |
| chr7-26425754-INS-5487 | chr7-26425755-INS-5487 |
| chr7-26434538-INS-60 | chr7-26434492-INS-110 |
| chr7-26443055-INS-102 | chr7-26443055-INS-96 |
| chr7-26451917-INS-446 | chr7-26451917-INS-446 |
| chr7-26463201-INS-501 | chr7-26463201-INS-501 |
| chr7-26467572-INS-106 | chr7-26467572-INS-106 |
| chr7-26469346-INS-72 | chr7-26469348-INS-71 |
| chr7-26479833-INS-90 | chr7-26479833-INS-72 |
| chr7-26527219-INS-76 | chr7-26527219-INS-82 |
| chr7-26599112-INS-58 | chr7-26599112-INS-58 |
| chr7-26600143-INS-401 | chr7-26600143-INS-401 |
| chr7-26640225-INS-52 | chr7-26640225-INS-54 |
| chr7-26694826-INS-4959 | chr7-26694827-INS-4959 |
| chr7-26697086-INS-567 | chr7-26697086-INS-567 |
| chr7-26703061-INS-6512 | chr7-26703062-INS-6514 |
| chr7-26735526-INS-55 | chr7-26735526-INS-55 |
| chr7-26760551-INS-52 | chr7-26760551-INS-50 |

| | |
|---|---|
| chr7-26771349-INS-53 | chr7-26771349-INS-53 |
| chr7-26774714-INS-69 | chr7-26774714-INS-63 |
| chr7-26793200-INS-159 | chr7-26793200-INS-159 |
| chr7-26824182-INS-210 | chr7-26824182-INS-210 |
| chr7-26834145-INS-436 | chr7-26834145-INS-436 |
| chr7-26839830-INS-5586 | chr7-26839831-INS-5586 |
| chr7-26840029-INS-1838 | chr7-26840030-INS-1838 |
| chr7-26898759-INS-99 | chr7-26898759-INS-82 |
| chr7-26900995-INS-156 | chr7-26901005-INS-161 |
| chr7-26915660-INS-280 | chr7-26915660-INS-284 |
| chr7-26921930-INS-106 | chr7-26921930-INS-106 |
| chr7-26923851-INS-557 | chr7-26923851-INS-557 |
| chr7-26929118-INS-76 | chr7-26929118-INS-60 |
| chr7-26942468-INS-99 | chr7-26942468-INS-94 |
| chr7-26947088-INS-195 | chr7-26947088-INS-195 |
| chr7-26954798-INS-54 | chr7-26954798-INS-54 |
| chr7-26975568-INS-117 | chr7-26975568-INS-117 |
| chr7-26989679-INS-179 | chr7-26989679-INS-180 |
| chr7-26999886-INS-173 | chr7-26999887-INS-173 |
| chr7-27000104-INS-1500 | chr7-27000105-INS-1488 |
| chr7-27000104-INS-84 | chr7-27000105-INS-120 |
| chr7-27000918-INS-7103 | chr7-27000919-INS-7103 |
| chr7-27043134-INS-98 | chr7-27043116-INS-77 |
| chr7-27045989-INS-216 | chr7-27045989-INS-216 |
| chr7-27070483-INS-70 | chr7-27070483-INS-70 |
| chr7-27074623-INS-197 | chr7-27074623-INS-197 |
| chr7-27089748-INS-54 | chr7-27089748-INS-54 |
| chr7-27091242-INS-440 | chr7-27091249-INS-441 |
| chr7-27113313-INS-53 | chr7-27113313-INS-53 |
| chr7-27131944-INS-191 | chr7-27131901-INS-217 |
| chr7-27136792-INS-80 | chr7-27136792-INS-56 |
| chr7-27143152-INS-391 | chr7-27143152-INS-391 |
| chr7-27149829-INS-226 | chr7-27149829-INS-226 |
| chr7-27152984-INS-217 | chr7-27152985-INS-194 |
| chr7-27159617-INS-53 | chr7-27159617-INS-53 |
| chr7-27163842-INS-186 | chr7-27163842-INS-187 |
| chr7-27164588-INS-208 | chr7-27164588-INS-208 |
| chr7-27174881-INS-56 | chr7-27174881-INS-56 |
| chr7-27179084-INS-201 | chr7-27179084-INS-201 |
| chr7-27181345-INS-62 | chr7-27181345-INS-68 |
| chr7-27194224-INS-53 | chr7-27194224-INS-53 |
| chr7-27206924-INS-210 | chr7-27206924-INS-209 |
| chr7-27221563-INS-88 | chr7-27221563-INS-89 |
| chr7-27238074-INS-219 | chr7-27238074-INS-220 |
| chr7-27263092-INS-82 | chr7-27263092-INS-82 |
| chr7-27270074-INS-81 | chr7-27270096-INS-82 |
| chr7-27271375-INS-83 | chr7-27271375-INS-83 |
| chr7-27278235-INS-51 | chr7-27278233-INS-57 |
| chr7-27284026-INS-58 | chr7-27284027-INS-58 |
| chr7-27285473-INS-190 | chr7-27285474-INS-190 |
| chr7-27312803-INS-238 | chr7-27312803-INS-239 |
| chr7-27317110-INS-217 | chr7-27317110-INS-217 |
| chr7-27323644-INS-177 | chr7-27323644-INS-178 |
| chr7-27334720-INS-270 | chr7-27334720-INS-270 |
| chr7-27344705-INS-82 | chr7-27344708-INS-68 |
| chr7-27349424-INS-207 | chr7-27349424-INS-207 |
| chr7-27360196-INS-212 | chr7-27360198-INS-203 |
| chr7-27363094-INS-232 | chr7-27363094-INS-238 |
| chr7-27363094-INS-75 | chr7-27363094-INS-85 |
| chr7-27367136-INS-53 | chr7-27367136-INS-53 |
| chr7-27367754-INS-57 | chr7-27367754-INS-57 |
| chr7-27376886-INS-138 | chr7-27376906-INS-106 |
| chr7-27382504-INS-176 | chr7-27382504-INS-176 |
| chr7-27383966-INS-70 | chr7-27383969-INS-52 |
| chr7-27385661-INS-488 | chr7-27385661-INS-488 |
| chr7-27386341-INS-8143 | chr7-27386342-INS-8143 |
| chr7-27387081-INS-113 | chr7-27387081-INS-113 |
| chr7-27388592-INS-157 | chr7-27388590-INS-163 |
| chr7-27399479-INS-63 | chr7-27399479-INS-63 |
| chr7-27406613-INS-364 | chr7-27406613-INS-364 |
| chr7-27423820-INS-148 | chr7-27423828-INS-120 |
| chr7-27424551-INS-410 | chr7-27424551-INS-410 |
| chr7-27426643-INS-5544 | chr7-27426644-INS-5544 |
| chr7-27444285-INS-230 | chr7-27444285-INS-230 |
| chr7-27471797-INS-223 | chr7-27471797-INS-222 |
| chr7-27476839-INS-195 | chr7-27476839-INS-196 |
| chr7-27477613-INS-243 | chr7-27477613-INS-244 |

| | |
|---|---|
| chr7-27479716-INS-179 | chr7-27479716-INS-174 |
| chr7-27483417-INS-51 | chr7-27483417-INS-51 |
| chr7-27487052-INS-65 | chr7-27487062-INS-70 |
| chr7-27487756-INS-506 | chr7-27487756-INS-506 |
| chr7-27492931-INS-176 | chr7-27492931-INS-172 |
| chr7-27498114-INS-71 | chr7-27498114-INS-75 |
| chr7-27499478-INS-508 | chr7-27499478-INS-508 |
| chr7-27501004-INS-183 | chr7-27501004-INS-187 |
| chr7-27502042-INS-7772 | chr7-27502043-INS-7772 |
| chr7-27503442-INS-1181 | chr7-27503443-INS-1181 |
| chr7-27514857-INS-238 | chr7-27514857-INS-238 |
| chr7-27534606-INS-892 | chr7-27534601-INS-883 |
| chr7-27536376-INS-52 | chr7-27536376-INS-52 |
| chr7-27556977-INS-2533 | chr7-27556978-INS-2533 |
| chr7-27556978-INS-349 | chr7-27556978-INS-349 |
| chr7-27565380-INS-194 | chr7-27565380-INS-194 |
| chr7-27586758-INS-147 | chr7-27586728-INS-153 |
| chr7-27588785-INS-545 | chr7-27588785-INS-544 |
| chr7-27598662-INS-607 | chr7-27598672-INS-571 |
| chr7-27615470-INS-210 | chr7-27615470-INS-210 |
| chr7-27618226-INS-54 | chr7-27618226-INS-54 |
| chr7-27632574-INS-533 | chr7-27632574-INS-536 |
| chr7-27642331-INS-59 | chr7-27642331-INS-59 |
| chr7-27651691-INS-193 | chr7-27651691-INS-193 |
| chr7-27653801-INS-205 | chr7-27653801-INS-205 |
| chr7-27672397-INS-3274 | chr7-27672398-INS-3274 |
| chr7-27695960-INS-211 | chr7-27695960-INS-211 |
| chr7-27717576-INS-987 | chr7-27717576-INS-984 |
| chr7-27728945-INS-852 | chr7-27728945-INS-852 |
| chr7-27756643-INS-1258 | chr7-27756644-INS-1258 |
| chr7-27764423-INS-74 | chr7-27764463-INS-80 |
| chr7-27781466-INS-282 | chr7-27781466-INS-282 |
| chr7-27822422-INS-69 | chr7-27822422-INS-50 |
| chr7-27851280-INS-63 | chr7-27851280-INS-63 |
| chr7-27927903-INS-74 | chr7-27927915-INS-78 |
| chr7-27937177-INS-1921 | chr7-27937178-INS-1922 |
| chr7-27941038-INS-216 | chr7-27941038-INS-216 |
| chr7-27948390-INS-72 | chr7-27948390-INS-72 |
| chr7-27970006-INS-54 | chr7-27970006-INS-54 |
| chr7-27984303-INS-65 | chr7-27984309-INS-54 |
| chr7-27989634-INS-64 | chr7-27989637-INS-75 |
| chr7-28036565-INS-60 | chr7-28036565-INS-108 |
| chr7-28047228-INS-218 | chr7-28047228-INS-220 |
| chr7-28082191-INS-8413 | chr7-28082192-INS-8413 |
| chr7-28138582-INS-193 | chr7-28138582-INS-193 |
| chr7-28139255-INS-187 | chr7-28139255-INS-186 |
| chr7-28175384-INS-218 | chr7-28175384-INS-218 |
| chr7-28178889-INS-138 | chr7-28178889-INS-138 |
| chr7-28181667-INS-60 | chr7-28181667-INS-76 |
| chr7-28204585-INS-60 | chr7-28204585-INS-61 |
| chr7-28206327-INS-112 | chr7-28206327-INS-112 |
| chr7-28223983-INS-60 | chr7-28223983-INS-60 |
| chr7-28232895-INS-2387 | chr7-28232896-INS-2362 |
| chr7-28245181-INS-63 | chr7-28245181-INS-62 |
| chr7-28248898-INS-53 | chr7-28248865-INS-53 |
| chr7-28279632-INS-207 | chr7-28279632-INS-207 |
| chr7-28334082-INS-246 | chr7-28334082-INS-252 |
| chr7-28336759-INS-94 | chr7-28336759-INS-94 |
| chr7-28338077-INS-106 | chr7-28338077-INS-106 |
| chr7-28340023-INS-402 | chr7-28340023-INS-402 |
| chr7-28353824-INS-173 | chr7-28353824-INS-173 |
| chr7-28360862-INS-210 | chr7-28360862-INS-210 |
| chr7-28362234-INS-96 | chr7-28362234-INS-96 |
| chr7-28362318-INS-5262 | chr7-28362319-INS-5262 |
| chr7-28364414-INS-260 | chr7-28364414-INS-259 |
| chr7-28366056-INS-60 | chr7-28366050-INS-54 |
| chr7-28405538-INS-156 | chr7-28405538-INS-157 |
| chr7-28409272-INS-262 | chr7-28409272-INS-262 |
| chr7-28415491-INS-210 | chr7-28415491-INS-208 |
| chr7-28418318-INS-446 | chr7-28418319-INS-446 |
| chr7-28419944-INS-72 | chr7-28419940-INS-80 |
| chr7-28421743-INS-54 | chr7-28421743-INS-54 |
| chr7-28425255-INS-125 | chr7-28425255-INS-127 |
| chr7-28426654-INS-85 | chr7-28426654-INS-85 |
| chr7-28429698-INS-310 | chr7-28429698-INS-318 |
| chr7-28431657-INS-206 | chr7-28431657-INS-206 |
| chr7-28437723-INS-56 | chr7-28437723-INS-56 |

| | |
|---|---|
| chr7-28440065-INS-87 | chr7-28440065-INS-87 |
| chr7-28442976-INS-170 | chr7-28442990-INS-165 |
| chr7-28457662-INS-204 | chr7-28457662-INS-204 |
| chr7-28473156-INS-106 | chr7-28473152-INS-90 |
| chr7-28477071-INS-87 | chr7-28477071-INS-51 |
| chr7-28493697-INS-542 | chr7-28493697-INS-542 |
| chr7-28503383-INS-354 | chr7-28503383-INS-356 |
| chr7-28505112-INS-661 | chr7-28505112-INS-662 |
| chr7-28519267-INS-246 | chr7-28519267-INS-246 |
| chr7-28537691-INS-227 | chr7-28537691-INS-228 |
| chr7-28546964-INS-211 | chr7-28546964-INS-210 |
| chr7-28550163-INS-211 | chr7-28550163-INS-211 |
| chr7-28553919-INS-183 | chr7-28553919-INS-183 |
| chr7-28555531-INS-206 | chr7-28555536-INS-210 |
| chr7-28569279-INS-203 | chr7-28569279-INS-203 |
| chr7-28586630-INS-220 | chr7-28586630-INS-220 |
| chr7-28599642-INS-170 | chr7-28599642-INS-173 |
| chr7-28622800-INS-72 | chr7-28622776-INS-50 |
| chr7-28641937-INS-56 | chr7-28641937-INS-60 |
| chr7-28653764-INS-173 | chr7-28653765-INS-174 |
| chr7-28656191-INS-216 | chr7-28656191-INS-211 |
| chr7-28666942-INS-174 | chr7-28666942-INS-191 |
| chr7-28678230-INS-216 | chr7-28678230-INS-216 |
| chr7-28691772-INS-255 | chr7-28691772-INS-255 |
| chr7-28693258-INS-52 | chr7-28693258-INS-52 |
| chr7-28703622-INS-160 | chr7-28703622-INS-160 |
| chr7-28724475-INS-190 | chr7-28724475-INS-192 |
| chr7-28730816-INS-96 | chr7-28730816-INS-96 |
| chr7-28739153-INS-199 | chr7-28739153-INS-199 |
| chr7-28740601-INS-77 | chr7-28740630-INS-74 |
| chr7-28742127-INS-99 | chr7-28742128-INS-54 |
| chr7-28763289-INS-53 | chr7-28763289-INS-53 |
| chr7-28764592-INS-187 | chr7-28764592-INS-187 |
| chr7-28765447-INS-53 | chr7-28765447-INS-53 |
| chr7-28774778-INS-57 | chr7-28774778-INS-57 |
| chr7-28800445-INS-52 | chr7-28800445-INS-52 |
| chr7-28807400-INS-174 | chr7-28807400-INS-174 |
| chr7-28807767-INS-212 | chr7-28807772-INS-195 |
| chr7-28826880-INS-97 | chr7-28826868-INS-134 |
| chr7-28828904-INS-156 | chr7-28828904-INS-156 |
| chr7-28840176-INS-223 | chr7-28840176-INS-225 |
| chr7-28842860-INS-213 | chr7-28842863-INS-210 |
| chr7-28844994-INS-159 | chr7-28844994-INS-159 |
| chr7-28880377-INS-945 | chr7-28880377-INS-948 |
| chr7-28890628-INS-204 | chr7-28890628-INS-206 |
| chr7-28915593-INS-1297 | chr7-28915594-INS-1297 |
| chr7-28920492-INS-144 | chr7-28920489-INS-164 |
| chr7-28922343-INS-133 | chr7-28922343-INS-132 |
| chr7-28932969-INS-241 | chr7-28932969-INS-233 |
| chr7-28936677-INS-198 | chr7-28936677-INS-198 |
| chr7-28958503-INS-206 | chr7-28958503-INS-205 |
| chr7-28976741-INS-206 | chr7-28976741-INS-206 |
| chr7-28983055-INS-212 | chr7-28983055-INS-212 |
| chr7-28986388-INS-73 | chr7-28986385-INS-96 |
| chr7-29001260-INS-1888 | chr7-29001261-INS-1888 |
| chr7-29011619-INS-153 | chr7-29011619-INS-153 |
| chr7-29032843-INS-228 | chr7-29032843-INS-229 |
| chr7-29035324-INS-90 | chr7-29035324-INS-65 |
| chr7-29037648-INS-54 | chr7-29037630-INS-53 |
| chr7-29055942-INS-201 | chr7-29055942-INS-201 |
| chr7-29064566-INS-214 | chr7-29064566-INS-214 |
| chr7-29065899-INS-62 | chr7-29065899-INS-58 |
| chr7-29113808-INS-152 | chr7-29113808-INS-152 |
| chr7-29122279-INS-151 | chr7-29122279-INS-155 |
| chr7-29136755-INS-59 | chr7-29136755-INS-59 |
| chr7-29145026-INS-53 | chr7-29145026-INS-53 |
| chr7-29159897-INS-53 | chr7-29159897-INS-53 |
| chr7-29178593-INS-120 | chr7-29178613-INS-136 |
| chr7-29182877-INS-72 | chr7-29182877-INS-72 |
| chr7-29202066-INS-105 | chr7-29202066-INS-105 |
| chr7-29209829-INS-194 | chr7-29209829-INS-194 |
| chr7-29211429-INS-72 | chr7-29211423-INS-50 |
| chr7-29242003-INS-212 | chr7-29242003-INS-210 |
| chr7-29253665-INS-171 | chr7-29253671-INS-168 |
| chr7-29272498-INS-174 | chr7-29272498-INS-174 |
| chr7-29290678-INS-6401 | chr7-29290679-INS-6401 |
| chr7-29302692-INS-146 | chr7-29302693-INS-140 |

| | |
|---|---|
| chr7-29425109-INS-96 | chr7-29425112-INS-96 |
| chr7-29441453-INS-147 | chr7-29441453-INS-147 |
| chr7-29452564-INS-200 | chr7-29452564-INS-200 |
| chr7-29454360-INS-264 | chr7-29454360-INS-264 |
| chr7-29484201-INS-125 | chr7-29484201-INS-120 |
| chr7-29484719-INS-231 | chr7-29484719-INS-231 |
| chr7-29514064-INS-212 | chr7-29514064-INS-209 |
| chr7-29519986-INS-234 | chr7-29519986-INS-234 |
| chr7-29577573-INS-54 | chr7-29577573-INS-54 |
| chr7-29581778-INS-203 | chr7-29581778-INS-203 |
| chr7-29583142-INS-66 | chr7-29583142-INS-66 |
| chr7-29603878-INS-209 | chr7-29603878-INS-209 |
| chr7-29617060-INS-88 | chr7-29617060-INS-88 |
| chr7-29618048-INS-115 | chr7-29618048-INS-115 |
| chr7-29618849-INS-225 | chr7-29618849-INS-228 |
| chr7-29622340-INS-420 | chr7-29622340-INS-420 |
| chr7-29637951-INS-118 | chr7-29637952-INS-86 |
| chr7-29640001-INS-68 | chr7-29640001-INS-68 |
| chr7-29652256-INS-84 | chr7-29652256-INS-86 |
| chr7-29654195-INS-199 | chr7-29654195-INS-199 |
| chr7-29661072-INS-164 | chr7-29661072-INS-164 |
| chr7-29663741-INS-215 | chr7-29663741-INS-214 |
| chr7-29665452-INS-210 | chr7-29665452-INS-210 |
| chr7-29668492-INS-780 | chr7-29668492-INS-780 |
| chr7-29675955-INS-202 | chr7-29675955-INS-201 |
| chr7-29682947-INS-203 | chr7-29682947-INS-203 |
| chr7-29696147-INS-205 | chr7-29696147-INS-205 |
| chr7-29698540-INS-206 | chr7-29698541-INS-206 |
| chr7-29708542-INS-172 | chr7-29708542-INS-172 |
| chr7-29711403-INS-205 | chr7-29711403-INS-205 |
| chr7-29720919-INS-61 | chr7-29720919-INS-61 |
| chr7-29727805-INS-296 | chr7-29727805-INS-296 |
| chr7-29731629-INS-399 | chr7-29731629-INS-399 |
| chr7-29735557-INS-74 | chr7-29735557-INS-94 |
| chr7-29739781-INS-99 | chr7-29739781-INS-94 |
| chr7-29749551-INS-881 | chr7-29749551-INS-881 |
| chr7-29759236-INS-1826 | chr7-29759237-INS-1826 |
| chr7-29762778-INS-196 | chr7-29762779-INS-196 |
| chr7-29769992-INS-202 | chr7-29769992-INS-202 |
| chr7-29774250-INS-172 | chr7-29774250-INS-174 |
| chr7-29775443-INS-372 | chr7-29775443-INS-402 |
| chr7-29781015-INS-74 | chr7-29781015-INS-74 |
| chr7-29783506-INS-432 | chr7-29783506-INS-432 |
| chr7-29793057-INS-68 | chr7-29793057-INS-52 |
| chr7-29812287-INS-401 | chr7-29812287-INS-401 |
| chr7-29815322-INS-2635 | chr7-29815323-INS-2637 |
| chr7-29816395-INS-208 | chr7-29816395-INS-214 |
| chr7-29819598-INS-52 | chr7-29819598-INS-56 |
| chr7-29836201-INS-194 | chr7-29836200-INS-194 |
| chr7-29846455-INS-207 | chr7-29846455-INS-207 |
| chr7-29847455-INS-138 | chr7-29847455-INS-139 |
| chr7-29863511-INS-61 | chr7-29863511-INS-61 |
| chr7-29871245-INS-204 | chr7-29871245-INS-204 |
| chr7-29874545-INS-203 | chr7-29874545-INS-203 |
| chr7-29875552-INS-636 | chr7-29875553-INS-636 |
| chr7-29878435-INS-212 | chr7-29878435-INS-212 |
| chr7-29880975-INS-212 | chr7-29880975-INS-212 |
| chr7-29898745-INS-203 | chr7-29898745-INS-202 |
| chr7-29902124-INS-210 | chr7-29902124-INS-210 |
| chr7-29903235-INS-397 | chr7-29903235-INS-397 |
| chr7-29905453-INS-80 | chr7-29905453-INS-80 |
| chr7-29909683-INS-180 | chr7-29909651-INS-182 |
| chr7-29913800-INS-221 | chr7-29913800-INS-223 |
| chr7-29924994-INS-51 | chr7-29924994-INS-51 |
| chr7-29957913-INS-81 | chr7-29957913-INS-81 |
| chr7-29959187-INS-183 | chr7-29959168-INS-204 |
| chr7-29960156-INS-80 | chr7-29960156-INS-80 |
| chr7-29967934-INS-219 | chr7-29967930-INS-203 |
| chr7-29974173-INS-209 | chr7-29974173-INS-192 |
| chr7-29984696-INS-406 | chr7-29984696-INS-406 |
| chr7-29987576-INS-231 | chr7-29987576-INS-231 |
| chr7-30030451-INS-199 | chr7-30030451-INS-199 |
| chr7-30037634-INS-198 | chr7-30037634-INS-198 |
| chr7-30065503-INS-343 | chr7-30065503-INS-310 |
| chr7-30111355-INS-108 | chr7-30111355-INS-108 |
| chr7-30116310-INS-209 | chr7-30116310-INS-209 |
| chr7-30142372-INS-53 | chr7-30142372-INS-53 |

| | |
|---|---|
| chr7-30163949-INS-215 | chr7-30163949-INS-215 |
| chr7-30172981-INS-62 | chr7-30172981-INS-70 |
| chr7-30177476-INS-83 | chr7-30177476-INS-83 |
| chr7-30182282-INS-202 | chr7-30182282-INS-201 |
| chr7-30188674-INS-64 | chr7-30188674-INS-64 |
| chr7-30201658-INS-91 | chr7-30201658-INS-91 |
| chr7-30211560-INS-89 | chr7-30211560-INS-89 |
| chr7-30230672-INS-112 | chr7-30230672-INS-106 |
| chr7-30245401-INS-134 | chr7-30245401-INS-134 |
| chr7-30259584-INS-228 | chr7-30259584-INS-228 |
| chr7-30294306-INS-55 | chr7-30294306-INS-57 |
| chr7-30305555-INS-87 | chr7-30305555-INS-87 |
| chr7-30308315-INS-68 | chr7-30308326-INS-97 |
| chr7-30315188-INS-200 | chr7-30315188-INS-204 |
| chr7-30315967-INS-256 | chr7-30315967-INS-256 |
| chr7-30328275-INS-604 | chr7-30328275-INS-604 |
| chr7-30332457-INS-202 | chr7-30332457-INS-202 |
| chr7-30342628-INS-188 | chr7-30342628-INS-180 |
| chr7-30361700-INS-175 | chr7-30361700-INS-127 |
| chr7-30380373-INS-4920 | chr7-30380371-INS-4921 |
| chr7-30384405-INS-195 | chr7-30384405-INS-195 |
| chr7-30407162-INS-52 | chr7-30407162-INS-52 |
| chr7-30410416-INS-171 | chr7-30410416-INS-171 |
| chr7-30423211-INS-54 | chr7-30423211-INS-58 |
| chr7-30442999-INS-4277 | chr7-30443000-INS-4278 |
| chr7-30467259-INS-52 | chr7-30467259-INS-52 |
| chr7-30494353-INS-213 | chr7-30494353-INS-216 |
| chr7-30503445-INS-59 | chr7-30503445-INS-60 |
| chr7-30506723-INS-88 | chr7-30506723-INS-88 |
| chr7-30508799-INS-213 | chr7-30508799-INS-213 |
| chr7-30529825-INS-66 | chr7-30529825-INS-66 |
| chr7-30535403-INS-52 | chr7-30535403-INS-51 |
| chr7-30540825-INS-255 | chr7-30540825-INS-255 |
| chr7-30546667-INS-58 | chr7-30546687-INS-82 |
| chr7-30564310-INS-69 | chr7-30564310-INS-68 |
| chr7-30570258-INS-64 | chr7-30570258-INS-66 |
| chr7-30577837-INS-794 | chr7-30577837-INS-794 |
| chr7-30579959-INS-164 | chr7-30579959-INS-166 |
| chr7-30596151-INS-212 | chr7-30596160-INS-213 |
| chr7-30598125-INS-54 | chr7-30598125-INS-60 |
| chr7-30601538-INS-63 | chr7-30601538-INS-63 |
| chr7-30618697-INS-178 | chr7-30618649-INS-213 |
| chr7-30619725-INS-211 | chr7-30619725-INS-211 |
| chr7-30622785-INS-80 | chr7-30622785-INS-85 |
| chr7-30638426-INS-60 | chr7-30638426-INS-60 |
| chr7-30678763-INS-185 | chr7-30678763-INS-185 |
| chr7-30680280-INS-203 | chr7-30680280-INS-205 |
| chr7-30705452-INS-53 | chr7-30705452-INS-52 |
| chr7-30711835-INS-211 | chr7-30711835-INS-211 |
| chr7-30737170-INS-80 | chr7-30737178-INS-101 |
| chr7-30774625-INS-96 | chr7-30774631-INS-97 |
| chr7-30780642-INS-389 | chr7-30780644-INS-410 |
| chr7-30787544-INS-52 | chr7-30787544-INS-52 |
| chr7-30805317-INS-53 | chr7-30805317-INS-53 |
| chr7-30830990-INS-53 | chr7-30830993-INS-53 |
| chr7-30863469-INS-85 | chr7-30863457-INS-101 |
| chr7-30897285-INS-58 | chr7-30897285-INS-59 |
| chr7-30923983-INS-214 | chr7-30923983-INS-214 |
| chr7-31030477-INS-82 | chr7-31030490-INS-77 |
| chr7-31037028-INS-58 | chr7-31037028-INS-58 |
| chr7-31039339-INS-373 | chr7-31039339-INS-373 |
| chr7-31111248-INS-54 | chr7-31111248-INS-52 |
| chr7-31121486-INS-578 | chr7-31121486-INS-578 |
| chr7-31193991-INS-6507 | chr7-31193992-INS-6507 |
| chr7-31219689-INS-56 | chr7-31219689-INS-52 |
| chr7-31293858-INS-935 | chr7-31293858-INS-935 |
| chr7-31296864-INS-201 | chr7-31296886-INS-199 |
| chr7-31449030-INS-52 | chr7-31449030-INS-52 |
| chr7-31562395-INS-200 | chr7-31562395-INS-200 |
| chr7-31635181-INS-131 | chr7-31635181-INS-131 |
| chr7-31657231-INS-53 | chr7-31657231-INS-53 |
| chr7-31726192-INS-241 | chr7-31726192-INS-239 |
| chr7-31786796-INS-80 | chr7-31786796-INS-80 |
| chr7-32869821-INS-2388 | chr7-32869822-INS-2391 |
| chr7-32901015-INS-3703 | chr7-32901016-INS-3703 |
| chr7-33040061-INS-1776 | chr7-33040062-INS-1790 |
| chr7-33110953-INS-68 | chr7-33110969-INS-80 |

| | |
|---|---|
| chr7-33127649-INS-5265 | chr7-33127650-INS-5265 |
| chr7-33158523-INS-56 | chr7-33158523-INS-56 |
| chr7-33185985-INS-728 | chr7-33185985-INS-728 |
| chr7-33223594-INS-165 | chr7-33223594-INS-164 |
| chr7-33246002-INS-131 | chr7-33246002-INS-127 |
| chr7-33262733-INS-200 | chr7-33262733-INS-201 |
| chr7-33270085-INS-105 | chr7-33270085-INS-109 |
| chr7-33329343-INS-78 | chr7-33329343-INS-78 |
| chr7-33341297-INS-62 | chr7-33341301-INS-63 |
| chr7-33343860-INS-1493 | chr7-33343861-INS-1493 |
| chr7-33350271-INS-58 | chr7-33350271-INS-60 |
| chr7-33356152-INS-55 | chr7-33356152-INS-55 |
| chr7-33357139-INS-441 | chr7-33357139-INS-441 |
| chr7-33360338-INS-1486 | chr7-33360339-INS-1486 |
| chr7-33415942-INS-128 | chr7-33415942-INS-131 |
| chr7-33427375-INS-663 | chr7-33427375-INS-663 |
| chr7-33471577-INS-572 | chr7-33471577-INS-572 |
| chr7-33482908-INS-6673 | chr7-33482909-INS-6673 |
| chr7-33484136-INS-56 | chr7-33484135-INS-66 |
| chr7-33497777-INS-70 | chr7-33497777-INS-76 |
| chr7-33520688-INS-934 | chr7-33520688-INS-934 |
| chr7-33593048-INS-6410 | chr7-33593049-INS-6410 |
| chr7-33608135-INS-1846 | chr7-33608136-INS-1846 |
| chr7-33696539-INS-114 | chr7-33696539-INS-114 |
| chr7-33719562-INS-559 | chr7-33719562-INS-559 |
| chr7-33726333-INS-256 | chr7-33726333-INS-256 |
| chr7-33803488-INS-6597 | chr7-33803489-INS-6598 |
| chr7-33806573-INS-76 | chr7-33806572-INS-76 |
| chr7-33825762-INS-765 | chr7-33825766-INS-766 |
| chr7-33828177-INS-98 | chr7-33828177-INS-98 |
| chr7-33828435-INS-7127 | chr7-33828436-INS-7128 |
| chr7-33848450-INS-64 | chr7-33848450-INS-66 |
| chr7-33850770-INS-215 | chr7-33850770-INS-215 |
| chr7-33854658-INS-190 | chr7-33854658-INS-190 |
| chr7-33858699-INS-207 | chr7-33858699-INS-208 |
| chr7-33861602-INS-56 | chr7-33861602-INS-56 |
| chr7-33865710-INS-197 | chr7-33865710-INS-197 |
| chr7-33894539-INS-201 | chr7-33894539-INS-201 |
| chr7-33907548-INS-102 | chr7-33907548-INS-102 |
| chr7-33910715-INS-80 | chr7-33910715-INS-80 |
| chr7-33912033-INS-69 | chr7-33912033-INS-65 |
| chr7-33913379-INS-216 | chr7-33913379-INS-216 |
| chr7-33936117-INS-205 | chr7-33936117-INS-205 |
| chr7-33947127-INS-913 | chr7-33947127-INS-912 |
| chr7-33976227-INS-209 | chr7-33976227-INS-209 |
| chr7-33979081-INS-203 | chr7-33979081-INS-203 |
| chr7-33986100-INS-93 | chr7-33986100-INS-81 |
| chr7-33994317-INS-68 | chr7-33994317-INS-66 |
| chr7-34015208-INS-178 | chr7-34015208-INS-178 |
| chr7-34018994-INS-57 | chr7-34018996-INS-51 |
| chr7-34027262-INS-417 | chr7-34027262-INS-417 |
| chr7-34028216-INS-160 | chr7-34028252-INS-112 |
| chr7-34046720-INS-1195 | chr7-34046721-INS-1195 |
| chr7-34046721-INS-169 | chr7-34046722-INS-170 |
| chr7-34061731-INS-62 | chr7-34061731-INS-62 |
| chr7-34072082-INS-244 | chr7-34072082-INS-245 |
| chr7-34074316-INS-64 | chr7-34074321-INS-74 |
| chr7-34080354-INS-178 | chr7-34080354-INS-178 |
| chr7-34091034-INS-207 | chr7-34091034-INS-207 |
| chr7-34097014-INS-4727 | chr7-34097015-INS-4727 |
| chr7-34101899-INS-58 | chr7-34101899-INS-58 |
| chr7-34103564-INS-181 | chr7-34103564-INS-181 |
| chr7-34110691-INS-549 | chr7-34110691-INS-549 |
| chr7-34121578-INS-75 | chr7-34121578-INS-75 |
| chr7-34135524-INS-53 | chr7-34135500-INS-60 |
| chr7-34139734-INS-203 | chr7-34139734-INS-203 |
| chr7-34143699-INS-58 | chr7-34143699-INS-53 |
| chr7-34155514-INS-207 | chr7-34155514-INS-207 |
| chr7-34157172-INS-400 | chr7-34157173-INS-414 |
| chr7-34163730-INS-71 | chr7-34163730-INS-71 |
| chr7-34178627-INS-195 | chr7-34178630-INS-195 |
| chr7-34201338-INS-215 | chr7-34201338-INS-214 |
| chr7-34205159-INS-72 | chr7-34205159-INS-72 |
| chr7-34232244-INS-173 | chr7-34232244-INS-173 |
| chr7-34233738-INS-208 | chr7-34233738-INS-208 |
| chr7-34262329-INS-53 | chr7-34262329-INS-55 |
| chr7-34265987-INS-209 | chr7-34265987-INS-209 |

| | |
|---|---|
| chr7-34267592-INS-90 | chr7-34267595-INS-98 |
| chr7-34275627-INS-65 | chr7-34275627-INS-65 |
| chr7-34290531-INS-292 | chr7-34290522-INS-300 |
| chr7-34298475-INS-94 | chr7-34298455-INS-76 |
| chr7-34319563-INS-53 | chr7-34319563-INS-53 |
| chr7-34322745-INS-70 | chr7-34322746-INS-70 |
| chr7-34326791-INS-77 | chr7-34326791-INS-57 |
| chr7-34335309-INS-82 | chr7-34335304-INS-80 |
| chr7-34341990-INS-94 | chr7-34341990-INS-56 |
| chr7-34350471-INS-56 | chr7-34350471-INS-60 |
| chr7-34382644-INS-86 | chr7-34382644-INS-72 |
| chr7-34399782-INS-109 | chr7-34399782-INS-109 |
| chr7-34400390-INS-91 | chr7-34400390-INS-95 |
| chr7-34405324-INS-493 | chr7-34405324-INS-493 |
| chr7-34411615-INS-1816 | chr7-34411616-INS-1813 |
| chr7-34428510-INS-106 | chr7-34428510-INS-106 |
| chr7-34444196-INS-68 | chr7-34444196-INS-68 |
| chr7-34446277-INS-418 | chr7-34446277-INS-419 |
| chr7-34469715-INS-70 | chr7-34469715-INS-50 |
| chr7-34483289-INS-6434 | chr7-34483290-INS-6434 |
| chr7-34491000-INS-1893 | chr7-34491001-INS-1899 |
| chr7-34499079-INS-72 | chr7-34499079-INS-72 |
| chr7-34533240-INS-97 | chr7-34533240-INS-97 |
| chr7-34543990-INS-131 | chr7-34543990-INS-178 |
| chr7-34546380-INS-171 | chr7-34546380-INS-173 |
| chr7-34550554-INS-518 | chr7-34550554-INS-518 |
| chr7-34551762-INS-139 | chr7-34551762-INS-120 |
| chr7-34558590-INS-51 | chr7-34558590-INS-51 |
| chr7-34572237-INS-125 | chr7-34572237-INS-125 |
| chr7-34587595-INS-371 | chr7-34587595-INS-371 |
| chr7-34639883-INS-6452 | chr7-34639884-INS-6452 |
| chr7-34640400-INS-115 | chr7-34640400-INS-115 |
| chr7-34655451-INS-76 | chr7-34655451-INS-76 |
| chr7-34720727-INS-86 | chr7-34720747-INS-76 |
| chr7-34726137-INS-503 | chr7-34726137-INS-502 |
| chr7-34787462-INS-253 | chr7-34787462-INS-253 |
| chr7-34789286-INS-269 | chr7-34789286-INS-269 |
| chr7-34804692-INS-56 | chr7-34804692-INS-56 |
| chr7-34809297-INS-268 | chr7-34809297-INS-268 |
| chr7-34854625-INS-64 | chr7-34854625-INS-66 |
| chr7-34858467-INS-204 | chr7-34858467-INS-204 |
| chr7-34863604-INS-87 | chr7-34863604-INS-61 |
| chr7-34870387-INS-120 | chr7-34870387-INS-120 |
| chr7-34878654-INS-424 | chr7-34878654-INS-424 |
| chr7-34906956-INS-105 | chr7-34907001-INS-80 |
| chr7-34922626-INS-1035 | chr7-34922634-INS-1032 |
| chr7-34952032-INS-217 | chr7-34952032-INS-217 |
| chr7-34954006-INS-125 | chr7-34954006-INS-125 |
| chr7-34956489-INS-106 | chr7-34956461-INS-106 |
| chr7-34958923-INS-69 | chr7-34958920-INS-76 |
| chr7-34978149-INS-918 | chr7-34978149-INS-918 |
| chr7-34980617-INS-7258 | chr7-34980618-INS-7258 |
| chr7-34996039-INS-61 | chr7-34996039-INS-81 |
| chr7-35001767-INS-56 | chr7-35001767-INS-60 |
| chr7-35006841-INS-52 | chr7-35006852-INS-62 |
| chr7-35013262-INS-74 | chr7-35013262-INS-74 |
| chr7-35015383-INS-68 | chr7-35015387-INS-84 |
| chr7-35018964-INS-76 | chr7-35018964-INS-75 |
| chr7-35028566-INS-179 | chr7-35028566-INS-179 |
| chr7-35030370-INS-210 | chr7-35030370-INS-210 |
| chr7-35056208-INS-88 | chr7-35056158-INS-95 |
| chr7-35059450-INS-64 | chr7-35059450-INS-66 |
| chr7-35067442-INS-167 | chr7-35067442-INS-167 |
| chr7-35072044-INS-54 | chr7-35072045-INS-54 |
| chr7-35086400-INS-62 | chr7-35086400-INS-62 |
| chr7-35097303-INS-76 | chr7-35097303-INS-78 |
| chr7-35100793-INS-280 | chr7-35100802-INS-264 |
| chr7-35104684-INS-214 | chr7-35104684-INS-211 |
| chr7-35105283-INS-101 | chr7-35105283-INS-101 |
| chr7-35110490-INS-197 | chr7-35110490-INS-197 |
| chr7-35117487-INS-84 | chr7-35117487-INS-84 |
| chr7-35119246-INS-60 | chr7-35119246-INS-55 |
| chr7-35122646-INS-66 | chr7-35122646-INS-66 |
| chr7-35126486-INS-409 | chr7-35126486-INS-409 |
| chr7-35139769-INS-264 | chr7-35139769-INS-215 |
| chr7-35146599-INS-54 | chr7-35146599-INS-87 |
| chr7-35181602-INS-778 | chr7-35181602-INS-779 |

| | |
|---|---|
| chr7-35184121-INS-216 | chr7-35184121-INS-218 |
| chr7-35192826-INS-102 | chr7-35192852-INS-98 |
| chr7-35197363-INS-240 | chr7-35197363-INS-240 |
| chr7-35216921-INS-56 | chr7-35216921-INS-56 |
| chr7-35221851-INS-206 | chr7-35221852-INS-206 |
| chr7-35222405-INS-192 | chr7-35222405-INS-193 |
| chr7-35229192-INS-177 | chr7-35229192-INS-178 |
| chr7-35233831-INS-52 | chr7-35233832-INS-68 |
| chr7-35246858-INS-175 | chr7-35246858-INS-175 |
| chr7-35261639-INS-199 | chr7-35261639-INS-200 |
| chr7-35268192-INS-199 | chr7-35268192-INS-202 |
| chr7-35273445-INS-66 | chr7-35273453-INS-51 |
| chr7-35293974-INS-53 | chr7-35293974-INS-53 |
| chr7-35296369-INS-268 | chr7-35296369-INS-268 |
| chr7-35298317-INS-56 | chr7-35298317-INS-50 |
| chr7-35299577-INS-56 | chr7-35299577-INS-50 |
| chr7-35313212-INS-399 | chr7-35313213-INS-399 |
| chr7-35318485-INS-212 | chr7-35318485-INS-212 |
| chr7-35323177-INS-73 | chr7-35323177-INS-73 |
| chr7-35327031-INS-6442 | chr7-35327032-INS-6442 |
| chr7-35331397-INS-51 | chr7-35331397-INS-51 |
| chr7-35335357-INS-54 | chr7-35335353-INS-58 |
| chr7-35339916-INS-216 | chr7-35339916-INS-216 |
| chr7-35344637-INS-205 | chr7-35344637-INS-205 |
| chr7-35350911-INS-391 | chr7-35350911-INS-391 |
| chr7-35364171-INS-202 | chr7-35364171-INS-202 |
| chr7-35366230-INS-196 | chr7-35366230-INS-172 |
| chr7-35368176-INS-185 | chr7-35368176-INS-185 |
| chr7-35373073-INS-205 | chr7-35373087-INS-204 |
| chr7-35382848-INS-55 | chr7-35382848-INS-57 |
| chr7-35410927-INS-112 | chr7-35410926-INS-114 |
| chr7-35419573-INS-6453 | chr7-35419574-INS-6453 |
| chr7-35420128-INS-181 | chr7-35420128-INS-181 |
| chr7-35448452-INS-276 | chr7-35448452-INS-276 |
| chr7-35461399-INS-53 | chr7-35461399-INS-53 |
| chr7-35465696-INS-56 | chr7-35465697-INS-57 |
| chr7-35469681-INS-204 | chr7-35469681-INS-204 |
| chr7-35517756-INS-196 | chr7-35517756-INS-202 |
| chr7-35543602-INS-551 | chr7-35543602-INS-551 |
| chr7-35549664-INS-53 | chr7-35549664-INS-53 |
| chr7-35562061-INS-54 | chr7-35562062-INS-78 |
| chr7-35562566-INS-60 | chr7-35562566-INS-60 |
| chr7-35571749-INS-52 | chr7-35571749-INS-52 |
| chr7-35587524-INS-58 | chr7-35587524-INS-50 |
| chr7-35631744-INS-213 | chr7-35631744-INS-214 |
| chr7-35698959-INS-84 | chr7-35698959-INS-84 |
| chr7-35701385-INS-53 | chr7-35701385-INS-53 |
| chr7-35709567-INS-73 | chr7-35709567-INS-73 |
| chr7-35741502-INS-60 | chr7-35741502-INS-64 |
| chr7-35757254-INS-336 | chr7-35757254-INS-336 |
| chr7-35782049-INS-84 | chr7-35782050-INS-70 |
| chr7-35782977-INS-84 | chr7-35782978-INS-59 |
| chr7-35793685-INS-613 | chr7-35793685-INS-613 |
| chr7-35842032-INS-197 | chr7-35842032-INS-197 |
| chr7-35845578-INS-121 | chr7-35845595-INS-126 |
| chr7-35857467-INS-78 | chr7-35857467-INS-120 |
| chr7-35865952-INS-98 | chr7-35865952-INS-100 |
| chr7-35870871-INS-226 | chr7-35870871-INS-220 |
| chr7-35873628-INS-53 | chr7-35873628-INS-53 |
| chr7-35882360-INS-62 | chr7-35882360-INS-77 |
| chr7-35894849-INS-153 | chr7-35894849-INS-104 |
| chr7-35913803-INS-50 | chr7-35913803-INS-56 |
| chr7-35930037-INS-58 | chr7-35930037-INS-50 |
| chr7-35944788-INS-91 | chr7-35944788-INS-91 |
| chr7-35948301-INS-62 | chr7-35948299-INS-80 |
| chr7-35959798-INS-91 | chr7-35959802-INS-87 |
| chr7-35962107-INS-115 | chr7-35962107-INS-120 |
| chr7-35970154-INS-74 | chr7-35970154-INS-74 |
| chr7-35984188-INS-825 | chr7-35984188-INS-825 |
| chr7-35986080-INS-115 | chr7-35986081-INS-97 |
| chr7-35994448-INS-53 | chr7-35994448-INS-53 |
| chr7-35999704-INS-51 | chr7-35999705-INS-51 |
| chr7-36018543-INS-52 | chr7-36018543-INS-60 |
| chr7-36038444-INS-202 | chr7-36038444-INS-202 |
| chr7-36041923-INS-54 | chr7-36041923-INS-56 |
| chr7-36060670-INS-131 | chr7-36060668-INS-127 |
| chr7-36084384-INS-76 | chr7-36084393-INS-68 |

| | |
|---|---|
| chr7-36094114-INS-1395 | chr7-36094115-INS-1395 |
| chr7-36111607-INS-151 | chr7-36111605-INS-145 |
| chr7-36127260-INS-56 | chr7-36127274-INS-74 |
| chr7-36166582-INS-6906 | chr7-36166583-INS-6907 |
| chr7-36192748-INS-165 | chr7-36192748-INS-165 |
| chr7-36198359-INS-123 | chr7-36198361-INS-112 |
| chr7-36205386-INS-57 | chr7-36205386-INS-65 |
| chr7-36241817-INS-286 | chr7-36241817-INS-286 |
| chr7-36244921-INS-52 | chr7-36244921-INS-52 |
| chr7-36255956-INS-60 | chr7-36255956-INS-62 |
| chr7-36276061-INS-645 | chr7-36276054-INS-619 |
| chr7-36276061-INS-96 | chr7-36276061-INS-96 |
| chr7-36278622-INS-154 | chr7-36278622-INS-154 |
| chr7-36279511-INS-400 | chr7-36279511-INS-400 |
| chr7-36295408-INS-66 | chr7-36295408-INS-54 |
| chr7-36298146-INS-524 | chr7-36298146-INS-524 |
| chr7-36314331-INS-130 | chr7-36314329-INS-132 |
| chr7-36367716-INS-72 | chr7-36367710-INS-57 |
| chr7-36369101-INS-53 | chr7-36369101-INS-53 |
| chr7-36374446-INS-122 | chr7-36374476-INS-129 |
| chr7-36452735-INS-102 | chr7-36452731-INS-103 |
| chr7-36458767-INS-106 | chr7-36458767-INS-106 |
| chr7-36459329-INS-98 | chr7-36459329-INS-107 |
| chr7-36490441-INS-57 | chr7-36490441-INS-57 |
| chr7-36494087-INS-177 | chr7-36494087-INS-177 |
| chr7-36501614-INS-309 | chr7-36501614-INS-309 |
| chr7-36505096-INS-58 | chr7-36505096-INS-87 |
| chr7-36535164-INS-72 | chr7-36535164-INS-70 |
| chr7-36538365-INS-68 | chr7-36538348-INS-87 |
| chr7-36546208-INS-52 | chr7-36546208-INS-52 |
| chr7-36571668-INS-128 | chr7-36571668-INS-128 |
| chr7-36581465-INS-160 | chr7-36581465-INS-160 |
| chr7-36585880-INS-172 | chr7-36585880-INS-174 |
| chr7-36600410-INS-92 | chr7-36600410-INS-92 |
| chr7-36704012-INS-116 | chr7-36704012-INS-137 |
| chr7-36705457-INS-220 | chr7-36705474-INS-240 |
| chr7-36719065-INS-106 | chr7-36719065-INS-106 |
| chr7-36725516-INS-88 | chr7-36725516-INS-88 |
| chr7-36750038-INS-1146 | chr7-36750039-INS-1146 |
| chr7-36753562-INS-62 | chr7-36753556-INS-66 |
| chr7-36758617-INS-72 | chr7-36758617-INS-66 |
| chr7-36760419-INS-100 | chr7-36760419-INS-102 |
| chr7-36765235-INS-54 | chr7-36765235-INS-60 |
| chr7-36768041-INS-61 | chr7-36768041-INS-71 |
| chr7-36788093-INS-99 | chr7-36788093-INS-99 |
| chr7-36828596-INS-868 | chr7-36828596-INS-868 |
| chr7-36835435-INS-86 | chr7-36835439-INS-64 |
| chr7-36855802-INS-94 | chr7-36855802-INS-94 |
| chr7-36868146-INS-61 | chr7-36868146-INS-61 |
| chr7-36886246-INS-66 | chr7-36886246-INS-66 |
| chr7-36888735-INS-64 | chr7-36888735-INS-66 |
| chr7-36889707-INS-224 | chr7-36889707-INS-224 |
| chr7-36895467-INS-66 | chr7-36895467-INS-66 |
| chr7-36911406-INS-57 | chr7-36911406-INS-60 |
| chr7-36916845-INS-52 | chr7-36916845-INS-52 |
| chr7-36961659-INS-60 | chr7-36961675-INS-60 |
| chr7-36968089-INS-84 | chr7-36968089-INS-78 |
| chr7-36968119-INS-566 | chr7-36968119-INS-548 |
| chr7-36973225-INS-7294 | chr7-36973226-INS-7294 |
| chr7-36986773-INS-81 | chr7-36986776-INS-97 |
| chr7-36987919-INS-71 | chr7-36987919-INS-71 |
| chr7-37020957-INS-409 | chr7-37020957-INS-413 |
| chr7-37034552-INS-508 | chr7-37034552-INS-508 |
| chr7-37076772-INS-62 | chr7-37076772-INS-62 |
| chr7-37082187-INS-88 | chr7-37082223-INS-95 |
| chr7-37110616-INS-53 | chr7-37110616-INS-53 |
| chr7-37115202-INS-87 | chr7-37115202-INS-116 |
| chr7-37116356-INS-200 | chr7-37116356-INS-200 |
| chr7-37117731-INS-58 | chr7-37117697-INS-58 |
| chr7-37137927-INS-106 | chr7-37137924-INS-118 |
| chr7-37147743-INS-52 | chr7-37147743-INS-52 |
| chr7-37161673-INS-130 | chr7-37161673-INS-130 |
| chr7-37174532-INS-224 | chr7-37174532-INS-223 |
| chr7-37192922-INS-52 | chr7-37192922-INS-52 |
| chr7-37252197-INS-630 | chr7-37252227-INS-639 |
| chr7-37253003-INS-68 | chr7-37253003-INS-64 |
| chr7-37279629-INS-84 | chr7-37279629-INS-84 |

| | |
|---|---|
| chr7-37291088-INS-223 | chr7-37291088-INS-223 |
| chr7-37292013-INS-87 | chr7-37292004-INS-81 |
| chr7-37298209-INS-201 | chr7-37298181-INS-235 |
| chr7-37325981-INS-54 | chr7-37325981-INS-54 |
| chr7-37346913-INS-330 | chr7-37346913-INS-330 |
| chr7-37381627-INS-120 | chr7-37381627-INS-120 |
| chr7-37397088-INS-53 | chr7-37397088-INS-53 |
| chr7-37407161-INS-61 | chr7-37407164-INS-58 |
| chr7-37408778-INS-88 | chr7-37408778-INS-66 |
| chr7-37438386-INS-92 | chr7-37438386-INS-92 |
| chr7-37498170-INS-57 | chr7-37498170-INS-57 |
| chr7-37503913-INS-110 | chr7-37503913-INS-111 |
| chr7-37521835-INS-108 | chr7-37521800-INS-117 |
| chr7-37553087-INS-104 | chr7-37553087-INS-105 |
| chr7-37554406-INS-51 | chr7-37554405-INS-50 |
| chr7-37566633-INS-66 | chr7-37566633-INS-60 |
| chr7-37582519-INS-120 | chr7-37582519-INS-120 |
| chr7-37645383-INS-233 | chr7-37645383-INS-233 |
| chr7-37650825-INS-242 | chr7-37650825-INS-241 |
| chr7-37706348-INS-167 | chr7-37706348-INS-175 |
| chr7-37710492-INS-127 | chr7-37710492-INS-127 |
| chr7-37715415-INS-135 | chr7-37715415-INS-139 |
| chr7-37733742-INS-82 | chr7-37733742-INS-82 |
| chr7-37739517-INS-179 | chr7-37739517-INS-179 |
| chr7-37754773-INS-434 | chr7-37754773-INS-434 |
| chr7-37765225-INS-196 | chr7-37765225-INS-197 |
| chr7-37771553-INS-210 | chr7-37771553-INS-211 |
| chr7-37788766-INS-72 | chr7-37788811-INS-72 |
| chr7-37804627-INS-208 | chr7-37804627-INS-208 |
| chr7-37813741-INS-237 | chr7-37813742-INS-237 |
| chr7-37815456-INS-67 | chr7-37815464-INS-71 |
| chr7-37831453-INS-132 | chr7-37831453-INS-132 |
| chr7-37839445-INS-225 | chr7-37839445-INS-188 |
| chr7-37852157-INS-106 | chr7-37852157-INS-106 |
| chr7-37857141-INS-64 | chr7-37857141-INS-50 |
| chr7-37859920-INS-56 | chr7-37859918-INS-50 |
| chr7-37866478-INS-168 | chr7-37866478-INS-168 |
| chr7-37939031-INS-200 | chr7-37939031-INS-200 |
| chr7-37955058-INS-388 | chr7-37955058-INS-388 |
| chr7-38098178-INS-228 | chr7-38098141-INS-229 |
| chr7-38176859-INS-6239 | chr7-38176860-INS-6239 |
| chr7-38239497-INS-71 | chr7-38239497-INS-71 |
| chr7-38504882-INS-199 | chr7-38504883-INS-199 |
| chr7-38917087-INS-5709 | chr7-38917088-INS-5707 |
| chr7-38997405-INS-62 | chr7-38997405-INS-63 |
| chr7-39083486-INS-182 | chr7-39083486-INS-183 |
| chr7-39096241-INS-166 | chr7-39096241-INS-166 |
| chr7-39111274-INS-54 | chr7-39111274-INS-57 |
| chr7-39115932-INS-254 | chr7-39115932-INS-254 |
| chr7-39127316-INS-52 | chr7-39127309-INS-50 |
| chr7-39135325-INS-59 | chr7-39135325-INS-59 |
| chr7-39138579-INS-104 | chr7-39138579-INS-104 |
| chr7-39143941-INS-427 | chr7-39143941-INS-427 |
| chr7-39171315-INS-181 | chr7-39171315-INS-181 |
| chr7-39172254-INS-150 | chr7-39172254-INS-150 |
| chr7-39179003-INS-61 | chr7-39179003-INS-51 |
| chr7-39223299-INS-881 | chr7-39223299-INS-881 |
| chr7-39224992-INS-200 | chr7-39224992-INS-204 |
| chr7-39243847-INS-389 | chr7-39243847-INS-389 |
| chr7-39250345-INS-88 | chr7-39250345-INS-88 |
| chr7-39317583-INS-116 | chr7-39317587-INS-108 |
| chr7-39329231-INS-68 | chr7-39329231-INS-68 |
| chr7-39403652-INS-60 | chr7-39403652-INS-60 |
| chr7-39405675-INS-116 | chr7-39405675-INS-116 |
| chr7-39456032-INS-1724 | chr7-39456033-INS-1724 |
| chr7-39457394-INS-97 | chr7-39457394-INS-97 |
| chr7-39465067-INS-1826 | chr7-39465068-INS-1826 |
| chr7-39465161-INS-671 | chr7-39465161-INS-672 |
| chr7-39466965-INS-183 | chr7-39466965-INS-184 |
| chr7-39483252-INS-77 | chr7-39483287-INS-81 |
| chr7-39507362-INS-6453 | chr7-39507363-INS-6453 |
| chr7-39554501-INS-180 | chr7-39554501-INS-175 |
| chr7-39624764-INS-616 | chr7-39624764-INS-616 |
| chr7-39627927-INS-315 | chr7-39627961-INS-317 |
| chr7-39628812-INS-59 | chr7-39628783-INS-56 |
| chr7-39668294-INS-59 | chr7-39668297-INS-91 |
| chr7-39673596-INS-915 | chr7-39673596-INS-915 |

| | |
|---|---|
| chr7-39676744-INS-77 | chr7-39676744-INS-77 |
| chr7-39677748-INS-1827 | chr7-39677749-INS-1869 |
| chr7-39685035-INS-57 | chr7-39685035-INS-57 |
| chr7-39702788-INS-87 | chr7-39702788-INS-89 |
| chr7-39706827-INS-639 | chr7-39706827-INS-639 |
| chr7-39721207-INS-989 | chr7-39721207-INS-988 |
| chr7-39751413-INS-76 | chr7-39751413-INS-76 |
| chr7-39789271-INS-100 | chr7-39789271-INS-100 |
| chr7-39812923-INS-65 | chr7-39812923-INS-69 |
| chr7-39830376-INS-82 | chr7-39830376-INS-82 |
| chr7-39850418-INS-50 | chr7-39850418-INS-50 |
| chr7-39854043-INS-64 | chr7-39854044-INS-63 |
| chr7-39854261-INS-399 | chr7-39854261-INS-399 |
| chr7-39856139-INS-497 | chr7-39856139-INS-497 |
| chr7-39864800-INS-140 | chr7-39864801-INS-179 |
| chr7-39873353-INS-55 | chr7-39873353-INS-63 |
| chr7-39892220-INS-575 | chr7-39892220-INS-579 |
| chr7-39901698-INS-52 | chr7-39901699-INS-56 |
| chr7-39927245-INS-175 | chr7-39927245-INS-139 |
| chr7-39946333-INS-56 | chr7-39946333-INS-56 |
| chr7-39978303-INS-568 | chr7-39978303-INS-568 |
| chr7-39981227-INS-126 | chr7-39981227-INS-132 |
| chr7-40003950-INS-626 | chr7-40003950-INS-621 |
| chr7-40028628-INS-115 | chr7-40028617-INS-103 |
| chr7-40031221-INS-97 | chr7-40031221-INS-97 |
| chr7-40035715-INS-106 | chr7-40035751-INS-113 |
| chr7-40078833-INS-425 | chr7-40078833-INS-425 |
| chr7-40092556-INS-69 | chr7-40092551-INS-64 |
| chr7-40092924-INS-9785 | chr7-40092925-INS-9792 |
| chr7-40095491-INS-52 | chr7-40095491-INS-54 |
| chr7-40147656-INS-145 | chr7-40147657-INS-104 |
| chr7-40158469-INS-52 | chr7-40158469-INS-52 |
| chr7-40164830-INS-98 | chr7-40164831-INS-78 |
| chr7-40167933-INS-468 | chr7-40167933-INS-468 |
| chr7-40169451-INS-58 | chr7-40169452-INS-70 |
| chr7-40201772-INS-956 | chr7-40201772-INS-956 |
| chr7-40206104-INS-86 | chr7-40206057-INS-104 |
| chr7-40206901-INS-92 | chr7-40206917-INS-68 |
| chr7-40233327-INS-2451 | chr7-40233328-INS-2451 |
| chr7-40233679-INS-61 | chr7-40233679-INS-61 |
| chr7-40251042-INS-65 | chr7-40251042-INS-65 |
| chr7-40264768-INS-86 | chr7-40264772-INS-60 |
| chr7-40268014-INS-84 | chr7-40268017-INS-64 |
| chr7-40278319-INS-6176 | chr7-40278320-INS-6176 |
| chr7-40296426-INS-5032 | chr7-40296427-INS-5032 |
| chr7-40296561-INS-78 | chr7-40296561-INS-78 |
| chr7-40328774-INS-100 | chr7-40328774-INS-100 |
| chr7-40333512-INS-57 | chr7-40333512-INS-57 |
| chr7-40339903-INS-83 | chr7-40339891-INS-129 |
| chr7-40397889-INS-284 | chr7-40397889-INS-284 |
| chr7-40442866-INS-74 | chr7-40442866-INS-79 |
| chr7-40470254-INS-1113 | chr7-40470255-INS-1115 |
| chr7-40475857-INS-65 | chr7-40475857-INS-65 |
| chr7-40484149-INS-7078 | chr7-40484150-INS-7078 |
| chr7-40490881-INS-3004 | chr7-40490882-INS-3004 |
| chr7-40595513-INS-69 | chr7-40595508-INS-82 |
| chr7-40603482-INS-52 | chr7-40603480-INS-52 |
| chr7-40617045-INS-668 | chr7-40617045-INS-668 |
| chr7-40620717-INS-321 | chr7-40620719-INS-324 |
| chr7-40652111-INS-130 | chr7-40652111-INS-130 |
| chr7-40663572-INS-231 | chr7-40663572-INS-231 |
| chr7-40744355-INS-58 | chr7-40744310-INS-79 |
| chr7-40901275-INS-571 | chr7-40901275-INS-570 |
| chr7-40929392-INS-174 | chr7-40929392-INS-170 |
| chr7-40931222-INS-125 | chr7-40931211-INS-95 |
| chr7-40940514-INS-121 | chr7-40940514-INS-125 |
| chr7-40970493-INS-1086 | chr7-40970494-INS-1086 |
| chr7-40989071-INS-70 | chr7-40989071-INS-62 |
| chr7-40998562-INS-57 | chr7-40998562-INS-84 |
| chr7-41017709-INS-3416 | chr7-41017710-INS-3415 |
| chr7-41077861-INS-60 | chr7-41077861-INS-60 |
| chr7-41091650-INS-109 | chr7-41091650-INS-109 |
| chr7-41125984-INS-209 | chr7-41125984-INS-210 |
| chr7-41153202-INS-99 | chr7-41153202-INS-99 |
| chr7-41161387-INS-546 | chr7-41161387-INS-542 |
| chr7-41166207-INS-52 | chr7-41166207-INS-56 |
| chr7-41194699-INS-241 | chr7-41194699-INS-242 |

| | |
|---|---|
| chr7-41200321-INS-203 | chr7-41200321-INS-201 |
| chr7-41211485-INS-103 | chr7-41211485-INS-103 |
| chr7-41223421-INS-111 | chr7-41223412-INS-120 |
| chr7-41229168-INS-200 | chr7-41229168-INS-208 |
| chr7-41231713-INS-209 | chr7-41231713-INS-210 |
| chr7-41235437-INS-203 | chr7-41235437-INS-203 |
| chr7-41237320-INS-112 | chr7-41237320-INS-112 |
| chr7-41238371-INS-491 | chr7-41238371-INS-491 |
| chr7-41238916-INS-105 | chr7-41238916-INS-105 |
| chr7-41246349-INS-56 | chr7-41246362-INS-80 |
| chr7-41247357-INS-1745 | chr7-41247358-INS-1725 |
| chr7-41261393-INS-234 | chr7-41261393-INS-234 |
| chr7-41269278-INS-207 | chr7-41269278-INS-206 |
| chr7-41273836-INS-462 | chr7-41273836-INS-473 |
| chr7-41288717-INS-204 | chr7-41288714-INS-188 |
| chr7-41294288-INS-199 | chr7-41294288-INS-200 |
| chr7-41300737-INS-178 | chr7-41300736-INS-176 |
| chr7-41316413-INS-4960 | chr7-41316414-INS-4960 |
| chr7-41324227-INS-1532 | chr7-41324228-INS-1533 |
| chr7-41334215-INS-54 | chr7-41334215-INS-54 |
| chr7-41339950-INS-419 | chr7-41339950-INS-419 |
| chr7-41350849-INS-6363 | chr7-41350851-INS-6364 |
| chr7-41388314-INS-1604 | chr7-41388315-INS-1606 |
| chr7-41488608-INS-116 | chr7-41488608-INS-121 |
| chr7-41524227-INS-68 | chr7-41524227-INS-56 |
| chr7-41539024-INS-1788 | chr7-41539025-INS-1788 |
| chr7-41554997-INS-342 | chr7-41554997-INS-343 |
| chr7-41573600-INS-401 | chr7-41573600-INS-401 |
| chr7-41604916-INS-187 | chr7-41604920-INS-207 |
| chr7-41606759-INS-6582 | chr7-41606760-INS-6582 |
| chr7-41608850-INS-238 | chr7-41608850-INS-238 |
| chr7-41616601-INS-613 | chr7-41616602-INS-619 |
| chr7-41674131-INS-6419 | chr7-41674132-INS-6420 |
| chr7-41727211-INS-100 | chr7-41727178-INS-108 |
| chr7-41730661-INS-3588 | chr7-41730662-INS-3616 |
| chr7-41734163-INS-235 | chr7-41734163-INS-220 |
| chr7-41741686-INS-6363 | chr7-41741687-INS-6363 |
| chr7-41747222-INS-1889 | chr7-41747223-INS-1889 |
| chr7-41751098-INS-2826 | chr7-41751099-INS-2822 |
| chr7-41752026-INS-589 | chr7-41752026-INS-589 |
| chr7-41809650-INS-608 | chr7-41809650-INS-609 |
| chr7-41833296-INS-200 | chr7-41833296-INS-201 |
| chr7-41834876-INS-106 | chr7-41834876-INS-72 |
| chr7-41905494-INS-98 | chr7-41905494-INS-98 |
| chr7-41908959-INS-367 | chr7-41908959-INS-367 |
| chr7-41914820-INS-590 | chr7-41914820-INS-590 |
| chr7-41935797-INS-1571 | chr7-41935798-INS-1575 |
| chr7-41975839-INS-2891 | chr7-41975840-INS-2891 |
| chr7-41986052-INS-210 | chr7-41986052-INS-210 |
| chr7-41990779-INS-174 | chr7-41990777-INS-133 |
| chr7-41992015-INS-111 | chr7-41992041-INS-63 |
| chr7-42080876-INS-717 | chr7-42080876-INS-717 |
| chr7-42104636-INS-327 | chr7-42104636-INS-327 |
| chr7-42106163-INS-194 | chr7-42106163-INS-195 |
| chr7-42114211-INS-58 | chr7-42114211-INS-58 |
| chr7-42123693-INS-516 | chr7-42123693-INS-516 |
| chr7-42129562-INS-99 | chr7-42129562-INS-95 |
| chr7-42150717-INS-237 | chr7-42150717-INS-237 |
| chr7-42189400-INS-52 | chr7-42189400-INS-50 |
| chr7-42192409-INS-368 | chr7-42192409-INS-368 |
| chr7-42214754-INS-209 | chr7-42214754-INS-209 |
| chr7-42276604-INS-2732 | chr7-42276605-INS-2744 |
| chr7-42276605-INS-455 | chr7-42276605-INS-457 |
| chr7-42299132-INS-202 | chr7-42299132-INS-203 |
| chr7-42319063-INS-108 | chr7-42319063-INS-108 |
| chr7-42331374-INS-5114 | chr7-42331375-INS-5114 |
| chr7-42337164-INS-211 | chr7-42337164-INS-210 |
| chr7-42347949-INS-208 | chr7-42347949-INS-208 |
| chr7-42380709-INS-54 | chr7-42380709-INS-54 |
| chr7-42396717-INS-1604 | chr7-42396718-INS-1599 |
| chr7-42469023-INS-168 | chr7-42469024-INS-168 |
| chr7-42526223-INS-107 | chr7-42526223-INS-102 |
| chr7-42529647-INS-66 | chr7-42529647-INS-76 |
| chr7-42763768-INS-389 | chr7-42763768-INS-388 |
| chr7-42794713-INS-8756 | chr7-42794714-INS-8757 |
| chr7-42881749-INS-52 | chr7-42881749-INS-52 |
| chr7-42893401-INS-104 | chr7-42893401-INS-114 |

| | |
|---|---|
| chr7-42909105-INS-71 | chr7-42909105-INS-71 |
| chr7-42923668-INS-52 | chr7-42923668-INS-52 |
| chr7-42925723-INS-91 | chr7-42925723-INS-91 |
| chr7-42930373-INS-116 | chr7-42930330-INS-146 |
| chr7-42937016-INS-120 | chr7-42937016-INS-120 |
| chr7-42937671-INS-3548 | chr7-42937672-INS-3538 |
| chr7-42938961-INS-78 | chr7-42938961-INS-86 |
| chr7-42964564-INS-108 | chr7-42964564-INS-112 |
| chr7-42979689-INS-4186 | chr7-42979690-INS-4187 |
| chr7-42983029-INS-63 | chr7-42983029-INS-60 |
| chr7-42987036-INS-3105 | chr7-42987037-INS-3105 |
| chr7-42995644-INS-327 | chr7-42995644-INS-327 |
| chr7-43004276-INS-62 | chr7-43004276-INS-64 |
| chr7-43007047-INS-107 | chr7-43007047-INS-108 |
| chr7-43026986-INS-2950 | chr7-43026987-INS-2950 |
| chr7-43027865-INS-381 | chr7-43027866-INS-378 |
| chr7-43032843-INS-6285 | chr7-43032844-INS-6285 |
| chr7-43051061-INS-535 | chr7-43051061-INS-535 |
| chr7-43073996-INS-63 | chr7-43073980-INS-106 |
| chr7-43081608-INS-254 | chr7-43081608-INS-251 |
| chr7-43116396-INS-53 | chr7-43116396-INS-58 |
| chr7-43121762-INS-107 | chr7-43121762-INS-107 |
| chr7-43137272-INS-76 | chr7-43137272-INS-88 |
| chr7-43159383-INS-94 | chr7-43159376-INS-110 |
| chr7-43162497-INS-206 | chr7-43162497-INS-207 |
| chr7-43168528-INS-127 | chr7-43168528-INS-130 |
| chr7-43173121-INS-204 | chr7-43173121-INS-203 |
| chr7-43191438-INS-52 | chr7-43191435-INS-62 |
| chr7-43207271-INS-198 | chr7-43207271-INS-197 |
| chr7-43216616-INS-130 | chr7-43216616-INS-130 |
| chr7-43221106-INS-296 | chr7-43221106-INS-284 |
| chr7-43225635-INS-201 | chr7-43225635-INS-200 |
| chr7-43227619-INS-91 | chr7-43227619-INS-91 |
| chr7-43252537-INS-195 | chr7-43252537-INS-195 |
| chr7-43258033-INS-127 | chr7-43258033-INS-127 |
| chr7-43279854-INS-1466 | chr7-43279855-INS-1466 |
| chr7-43284568-INS-72 | chr7-43284576-INS-58 |
| chr7-43290052-INS-458 | chr7-43290052-INS-458 |
| chr7-43312199-INS-206 | chr7-43312199-INS-207 |
| chr7-43332296-INS-107 | chr7-43332296-INS-113 |
| chr7-43334156-INS-211 | chr7-43334158-INS-212 |
| chr7-43339320-INS-56 | chr7-43339320-INS-56 |
| chr7-43344596-INS-124 | chr7-43344590-INS-133 |
| chr7-43345187-INS-197 | chr7-43345187-INS-197 |
| chr7-43356786-INS-94 | chr7-43356786-INS-74 |
| chr7-43365789-INS-51 | chr7-43365789-INS-51 |
| chr7-43380403-INS-60 | chr7-43380403-INS-64 |
| chr7-43387497-INS-609 | chr7-43387497-INS-609 |
| chr7-43415574-INS-59 | chr7-43415574-INS-68 |
| chr7-43440950-INS-58 | chr7-43440950-INS-56 |
| chr7-43444791-INS-54 | chr7-43444757-INS-80 |
| chr7-43452218-INS-52 | chr7-43452218-INS-52 |
| chr7-43455407-INS-68 | chr7-43455407-INS-66 |
| chr7-43469444-INS-56 | chr7-43469444-INS-67 |
| chr7-43504866-INS-59 | chr7-43504866-INS-61 |
| chr7-43544924-INS-146 | chr7-43544924-INS-146 |
| chr7-43570164-INS-218 | chr7-43570164-INS-218 |
| chr7-43601866-INS-72 | chr7-43601866-INS-72 |
| chr7-43644295-INS-281 | chr7-43644295-INS-282 |
| chr7-43670419-INS-340 | chr7-43670419-INS-340 |
| chr7-43683914-INS-182 | chr7-43683914-INS-182 |
| chr7-43704771-INS-82 | chr7-43704771-INS-78 |
| chr7-43762067-INS-5313 | chr7-43762068-INS-5313 |
| chr7-43768769-INS-61 | chr7-43768769-INS-62 |
| chr7-43815423-INS-58 | chr7-43815423-INS-58 |
| chr7-43845554-INS-146 | chr7-43845557-INS-146 |
| chr7-43868905-INS-79 | chr7-43868905-INS-54 |
| chr7-43898363-INS-113 | chr7-43898363-INS-113 |
| chr7-43919762-INS-69 | chr7-43919762-INS-87 |
| chr7-43952237-INS-71 | chr7-43952237-INS-71 |
| chr7-43959688-INS-333 | chr7-43959688-INS-333 |
| chr7-43990855-INS-209 | chr7-43990855-INS-209 |
| chr7-43999883-INS-86 | chr7-43999879-INS-50 |
| chr7-44013710-INS-180 | chr7-44013710-INS-181 |
| chr7-44014500-INS-54 | chr7-44014500-INS-56 |
| chr7-44029936-INS-130 | chr7-44029936-INS-130 |
| chr7-44030774-INS-76 | chr7-44030774-INS-82 |

| | |
|---|---|
| chr7-44048713-INS-60 | chr7-44048713-INS-60 |
| chr7-44054936-INS-145 | chr7-44054936-INS-145 |
| chr7-44061041-INS-63 | chr7-44061041-INS-69 |
| chr7-44068960-INS-344 | chr7-44068960-INS-344 |
| chr7-44069634-INS-55 | chr7-44069634-INS-55 |
| chr7-44076420-INS-242 | chr7-44076420-INS-206 |
| chr7-44083961-INS-5549 | chr7-44083962-INS-5549 |
| chr7-44086321-INS-166 | chr7-44086321-INS-121 |
| chr7-44126876-INS-224 | chr7-44126876-INS-224 |
| chr7-44132942-INS-70 | chr7-44132942-INS-70 |
| chr7-44148098-INS-196 | chr7-44148098-INS-194 |
| chr7-44159796-INS-431 | chr7-44159796-INS-431 |
| chr7-44161104-INS-684 | chr7-44161101-INS-700 |
| chr7-44169982-INS-106 | chr7-44169995-INS-98 |
| chr7-44171651-INS-66 | chr7-44171651-INS-66 |
| chr7-44210904-INS-56 | chr7-44210904-INS-56 |
| chr7-44213139-INS-196 | chr7-44213138-INS-187 |
| chr7-44216233-INS-66 | chr7-44216233-INS-66 |
| chr7-44222563-INS-118 | chr7-44222563-INS-118 |
| chr7-44225417-INS-92 | chr7-44225419-INS-62 |
| chr7-44254849-INS-77 | chr7-44254849-INS-77 |
| chr7-44264610-INS-88 | chr7-44264610-INS-88 |
| chr7-44270654-INS-84 | chr7-44270654-INS-112 |
| chr7-44293401-INS-203 | chr7-44293401-INS-203 |
| chr7-44315754-INS-643 | chr7-44315720-INS-663 |
| chr7-44331628-INS-112 | chr7-44331607-INS-85 |
| chr7-44333883-INS-52 | chr7-44333883-INS-52 |
| chr7-44338850-INS-55 | chr7-44338850-INS-55 |
| chr7-44362063-INS-53 | chr7-44362063-INS-53 |
| chr7-44362867-INS-470 | chr7-44362867-INS-470 |
| chr7-44370243-INS-208 | chr7-44370243-INS-208 |
| chr7-44379204-INS-107 | chr7-44379203-INS-110 |
| chr7-44388440-INS-1836 | chr7-44388418-INS-1834 |
| chr7-44390439-INS-162 | chr7-44390439-INS-172 |
| chr7-44401391-INS-209 | chr7-44401391-INS-209 |
| chr7-44402067-INS-204 | chr7-44402067-INS-204 |
| chr7-44411667-INS-173 | chr7-44411667-INS-173 |
| chr7-44412162-INS-234 | chr7-44412162-INS-235 |
| chr7-44412994-INS-87 | chr7-44412994-INS-87 |
| chr7-44424364-INS-52 | chr7-44424364-INS-52 |
| chr7-44438082-INS-124 | chr7-44438082-INS-124 |
| chr7-44443304-INS-6413 | chr7-44443305-INS-6413 |
| chr7-44449591-INS-206 | chr7-44449591-INS-204 |
| chr7-44582472-INS-203 | chr7-44582472-INS-203 |
| chr7-44587521-INS-179 | chr7-44587521-INS-180 |
| chr7-44598326-INS-1307 | chr7-44598327-INS-1307 |
| chr7-44640231-INS-437 | chr7-44640231-INS-396 |
| chr7-44641580-INS-51 | chr7-44641568-INS-55 |
| chr7-44649020-INS-228 | chr7-44649020-INS-228 |
| chr7-44672017-INS-82 | chr7-44672017-INS-82 |
| chr7-44691835-INS-211 | chr7-44691835-INS-211 |
| chr7-44727578-INS-209 | chr7-44727578-INS-209 |
| chr7-44730104-INS-209 | chr7-44730104-INS-209 |
| chr7-44742246-INS-72 | chr7-44742255-INS-55 |
| chr7-44745361-INS-169 | chr7-44745361-INS-174 |
| chr7-44748287-INS-53 | chr7-44748287-INS-53 |
| chr7-44752833-INS-58 | chr7-44752833-INS-58 |
| chr7-44758035-INS-68 | chr7-44758035-INS-68 |
| chr7-44759963-INS-173 | chr7-44759963-INS-173 |
| chr7-44764560-INS-203 | chr7-44764560-INS-203 |
| chr7-44787659-INS-204 | chr7-44787659-INS-205 |
| chr7-44790122-INS-2881 | chr7-44790123-INS-2866 |
| chr7-44799673-INS-56 | chr7-44799673-INS-58 |
| chr7-44803308-INS-78 | chr7-44803308-INS-84 |
| chr7-44817044-INS-70 | chr7-44817042-INS-74 |
| chr7-44852369-INS-54 | chr7-44852369-INS-54 |
| chr7-44863137-INS-6462 | chr7-44863138-INS-6462 |
| chr7-44875762-INS-143 | chr7-44875762-INS-143 |
| chr7-44877240-INS-68 | chr7-44877260-INS-72 |
| chr7-44893163-INS-200 | chr7-44893163-INS-199 |
| chr7-44898211-INS-202 | chr7-44898211-INS-209 |
| chr7-44924549-INS-200 | chr7-44924549-INS-200 |
| chr7-44977724-INS-102 | chr7-44977699-INS-96 |
| chr7-45000845-INS-76 | chr7-45000861-INS-60 |
| chr7-45042062-INS-222 | chr7-45042062-INS-222 |
| chr7-45045749-INS-215 | chr7-45045749-INS-213 |
| chr7-45051589-INS-127 | chr7-45051589-INS-127 |

| | |
|---|---|
| chr7-45053900-INS-56 | chr7-45053900-INS-56 |
| chr7-45112711-INS-102 | chr7-45112711-INS-103 |
| chr7-45122165-INS-217 | chr7-45122165-INS-218 |
| chr7-45135071-INS-197 | chr7-45135072-INS-196 |
| chr7-45150966-INS-65 | chr7-45150966-INS-76 |
| chr7-45168582-INS-63 | chr7-45168582-INS-63 |
| chr7-45180856-INS-183 | chr7-45180856-INS-184 |
| chr7-45182767-INS-75 | chr7-45182767-INS-74 |
| chr7-45189047-INS-56 | chr7-45189047-INS-50 |
| chr7-45191872-INS-71 | chr7-45191872-INS-73 |
| chr7-45193728-INS-214 | chr7-45193728-INS-214 |
| chr7-45207127-INS-208 | chr7-45207127-INS-208 |
| chr7-45212946-INS-153 | chr7-45212946-INS-137 |
| chr7-45215273-INS-55 | chr7-45215273-INS-55 |
| chr7-45228714-INS-163 | chr7-45228714-INS-163 |
| chr7-45234240-INS-1866 | chr7-45234241-INS-1866 |
| chr7-45235976-INS-72 | chr7-45235976-INS-80 |
| chr7-45263099-INS-236 | chr7-45263083-INS-221 |
| chr7-45312160-INS-948 | chr7-45312160-INS-949 |
| chr7-45318111-INS-88 | chr7-45318111-INS-88 |
| chr7-45326634-INS-56 | chr7-45326644-INS-56 |
| chr7-45335159-INS-82 | chr7-45335159-INS-82 |
| chr7-45351522-INS-221 | chr7-45351522-INS-221 |
| chr7-45379320-INS-98 | chr7-45379300-INS-108 |
| chr7-45400488-INS-75 | chr7-45400488-INS-75 |
| chr7-45414714-INS-204 | chr7-45414714-INS-204 |
| chr7-45496296-INS-140 | chr7-45496296-INS-148 |
| chr7-45498514-INS-72 | chr7-45498514-INS-64 |
| chr7-45498983-INS-160 | chr7-45498983-INS-160 |
| chr7-45584759-INS-59 | chr7-45584759-INS-59 |
| chr7-45601524-INS-63 | chr7-45601520-INS-79 |
| chr7-45602772-INS-100 | chr7-45602772-INS-105 |
| chr7-45604205-INS-325 | chr7-45604205-INS-325 |
| chr7-45606296-INS-341 | chr7-45606296-INS-341 |
| chr7-45607209-INS-989 | chr7-45607209-INS-989 |
| chr7-45618891-INS-222 | chr7-45618891-INS-222 |
| chr7-45640901-INS-64 | chr7-45640901-INS-64 |
| chr7-45669872-INS-144 | chr7-45669872-INS-144 |
| chr7-45689556-INS-306 | chr7-45689556-INS-306 |
| chr7-45691222-INS-365 | chr7-45691222-INS-365 |
| chr7-45701869-INS-651 | chr7-45701869-INS-651 |
| chr7-45712292-INS-54 | chr7-45712292-INS-56 |
| chr7-45714522-INS-326 | chr7-45714508-INS-336 |
| chr7-45720617-INS-148 | chr7-45720617-INS-148 |
| chr7-45731615-INS-54 | chr7-45731615-INS-52 |
| chr7-45738958-INS-63 | chr7-45738971-INS-65 |
| chr7-45756540-INS-5340 | chr7-45756541-INS-5340 |
| chr7-45783955-INS-68 | chr7-45783958-INS-74 |
| chr7-45790485-INS-73 | chr7-45790485-INS-73 |
| chr7-45810913-INS-153 | chr7-45810913-INS-153 |
| chr7-45828275-INS-132 | chr7-45828275-INS-104 |
| chr7-45835642-INS-1895 | chr7-45835643-INS-1895 |
| chr7-45842507-INS-70 | chr7-45842507-INS-100 |
| chr7-45850973-INS-90 | chr7-45850971-INS-92 |
| chr7-45854289-INS-115 | chr7-45854289-INS-115 |
| chr7-45873279-INS-120 | chr7-45873279-INS-155 |
| chr7-45891926-INS-52 | chr7-45891926-INS-52 |
| chr7-45933728-INS-734 | chr7-45933728-INS-734 |
| chr7-45950412-INS-90 | chr7-45950412-INS-90 |
| chr7-45965756-INS-67 | chr7-45965780-INS-79 |
| chr7-45975132-INS-477 | chr7-45975132-INS-477 |
| chr7-45978157-INS-154 | chr7-45978157-INS-154 |
| chr7-45978748-INS-826 | chr7-45978739-INS-828 |
| chr7-46021268-INS-60 | chr7-46021268-INS-68 |
| chr7-46105120-INS-539 | chr7-46105120-INS-540 |
| chr7-46116632-INS-386 | chr7-46116632-INS-386 |
| chr7-46166423-INS-60 | chr7-46166415-INS-66 |
| chr7-46205557-INS-97 | chr7-46205558-INS-105 |
| chr7-46210291-INS-363 | chr7-46210291-INS-363 |
| chr7-46221213-INS-176 | chr7-46221213-INS-176 |
| chr7-46270266-INS-850 | chr7-46270266-INS-850 |
| chr7-46367598-INS-520 | chr7-46367598-INS-520 |
| chr7-46397593-INS-80 | chr7-46397593-INS-80 |
| chr7-46407326-INS-441 | chr7-46407327-INS-441 |
| chr7-46419347-INS-312 | chr7-46419345-INS-326 |
| chr7-46426563-INS-392 | chr7-46426563-INS-400 |
| chr7-46431781-INS-84 | chr7-46431781-INS-84 |

| | |
|---|---|
| chr7-46442659-INS-82 | chr7-46442659-INS-65 |
| chr7-46447262-INS-263 | chr7-46447262-INS-267 |
| chr7-46449603-INS-207 | chr7-46449603-INS-207 |
| chr7-46451933-INS-207 | chr7-46451933-INS-207 |
| chr7-46455647-INS-220 | chr7-46455647-INS-220 |
| chr7-46464174-INS-204 | chr7-46464174-INS-204 |
| chr7-46486200-INS-228 | chr7-46486200-INS-227 |
| chr7-46489629-INS-62 | chr7-46489629-INS-64 |
| chr7-46492219-INS-60 | chr7-46492219-INS-54 |
| chr7-46497586-INS-189 | chr7-46497586-INS-189 |
| chr7-46514015-INS-233 | chr7-46513996-INS-202 |
| chr7-46518165-INS-195 | chr7-46518165-INS-213 |
| chr7-46541266-INS-204 | chr7-46541266-INS-204 |
| chr7-46551989-INS-215 | chr7-46551989-INS-217 |
| chr7-46552831-INS-66 | chr7-46552817-INS-52 |
| chr7-46572709-INS-188 | chr7-46572708-INS-192 |
| chr7-46574276-INS-426 | chr7-46574276-INS-427 |
| chr7-46579615-INS-193 | chr7-46579615-INS-191 |
| chr7-46643527-INS-93 | chr7-46643527-INS-93 |
| chr7-46670431-INS-209 | chr7-46670431-INS-209 |
| chr7-46742889-INS-80 | chr7-46742889-INS-60 |
| chr7-46770195-INS-378 | chr7-46770195-INS-378 |
| chr7-46770514-INS-643 | chr7-46770514-INS-644 |
| chr7-46793535-INS-84 | chr7-46793535-INS-67 |
| chr7-46794760-INS-111 | chr7-46794760-INS-117 |
| chr7-46804027-INS-192 | chr7-46804027-INS-192 |
| chr7-46806337-INS-208 | chr7-46806335-INS-204 |
| chr7-46817651-INS-118 | chr7-46817651-INS-118 |
| chr7-46822848-INS-66 | chr7-46822848-INS-68 |
| chr7-46830772-INS-204 | chr7-46830757-INS-204 |
| chr7-46957157-INS-180 | chr7-46957154-INS-183 |
| chr7-46967374-INS-185 | chr7-46967374-INS-186 |
| chr7-46971410-INS-90 | chr7-46971410-INS-90 |
| chr7-47002344-INS-94 | chr7-47002344-INS-94 |
| chr7-47008369-INS-101 | chr7-47008368-INS-101 |
| chr7-47043361-INS-63 | chr7-47043361-INS-80 |
| chr7-47153048-INS-312 | chr7-47153048-INS-315 |
| chr7-47186995-INS-56 | chr7-47186991-INS-56 |
| chr7-47190590-INS-326 | chr7-47190590-INS-326 |
| chr7-47200829-INS-4500 | chr7-47200830-INS-4500 |
| chr7-47330710-INS-74 | chr7-47330710-INS-74 |
| chr7-47398001-INS-170 | chr7-47398002-INS-120 |
| chr7-47410882-INS-120 | chr7-47410880-INS-131 |
| chr7-47481690-INS-55 | chr7-47481690-INS-53 |
| chr7-47507113-INS-507 | chr7-47507113-INS-507 |
| chr7-47512733-INS-2296 | chr7-47512734-INS-2297 |
| chr7-47525134-INS-2929 | chr7-47525135-INS-2929 |
| chr7-47593323-INS-86 | chr7-47593327-INS-56 |
| chr7-47632617-INS-894 | chr7-47632617-INS-894 |
| chr7-47647048-INS-199 | chr7-47647048-INS-199 |
| chr7-47656071-INS-68 | chr7-47656071-INS-68 |
| chr7-47673334-INS-54 | chr7-47673327-INS-74 |
| chr7-47678984-INS-963 | chr7-47678984-INS-963 |
| chr7-47685816-INS-73 | chr7-47685816-INS-118 |
| chr7-47691705-INS-1872 | chr7-47691706-INS-1876 |
| chr7-47702845-INS-6193 | chr7-47702846-INS-6192 |
| chr7-47710305-INS-64 | chr7-47710339-INS-65 |
| chr7-47722646-INS-2749 | chr7-47722647-INS-2748 |
| chr7-47760585-INS-2617 | chr7-47760586-INS-2628 |
| chr7-47815152-INS-101 | chr7-47815152-INS-101 |
| chr7-47820606-INS-679 | chr7-47820606-INS-678 |
| chr7-47830697-INS-216 | chr7-47830697-INS-216 |
| chr7-47855026-INS-382 | chr7-47855026-INS-383 |
| chr7-47860403-INS-610 | chr7-47860403-INS-610 |
| chr7-47884328-INS-52 | chr7-47884328-INS-57 |
| chr7-47891425-INS-62 | chr7-47891425-INS-62 |
| chr7-47913814-INS-68 | chr7-47913817-INS-62 |
| chr7-47973013-INS-954 | chr7-47973013-INS-954 |
| chr7-47989657-INS-54 | chr7-47989657-INS-54 |
| chr7-48001281-INS-65 | chr7-48001281-INS-75 |
| chr7-48030348-INS-51 | chr7-48030348-INS-51 |
| chr7-48074641-INS-66 | chr7-48074641-INS-66 |
| chr7-48081857-INS-2505 | chr7-48081858-INS-2505 |
| chr7-48089412-INS-78 | chr7-48089412-INS-80 |
| chr7-48101461-INS-203 | chr7-48101461-INS-203 |
| chr7-48156509-INS-395 | chr7-48156509-INS-395 |
| chr7-48187494-INS-1964 | chr7-48187495-INS-1964 |

| | |
|---|---|
| chr7-48225759-INS-2086 | chr7-48225760-INS-2084 |
| chr7-48248339-INS-95 | chr7-48248339-INS-95 |
| chr7-48367480-INS-70 | chr7-48367480-INS-68 |
| chr7-48371630-INS-158 | chr7-48371634-INS-131 |
| chr7-48471966-INS-101 | chr7-48471966-INS-101 |
| chr7-48479658-INS-207 | chr7-48479658-INS-207 |
| chr7-48481377-INS-58 | chr7-48481361-INS-60 |
| chr7-48536527-INS-504 | chr7-48536527-INS-504 |
| chr7-48551390-INS-144 | chr7-48551390-INS-144 |
| chr7-48552924-INS-935 | chr7-48552924-INS-918 |
| chr7-48565389-INS-65 | chr7-48565389-INS-53 |
| chr7-48578661-INS-220 | chr7-48578661-INS-220 |
| chr7-48581674-INS-304 | chr7-48581674-INS-302 |
| chr7-48583370-INS-4531 | chr7-48583371-INS-4531 |
| chr7-48587868-INS-528 | chr7-48587868-INS-528 |
| chr7-48642313-INS-1183 | chr7-48642314-INS-1211 |
| chr7-48655422-INS-66 | chr7-48655422-INS-56 |
| chr7-48662019-INS-6478 | chr7-48662020-INS-6480 |
| chr7-48673277-INS-51 | chr7-48673275-INS-50 |
| chr7-48699931-INS-84 | chr7-48699901-INS-104 |
| chr7-48765349-INS-97 | chr7-48765349-INS-101 |
| chr7-48780448-INS-571 | chr7-48780449-INS-571 |
| chr7-48785434-INS-60 | chr7-48785434-INS-50 |
| chr7-48804340-INS-137 | chr7-48804338-INS-159 |
| chr7-48814119-INS-83 | chr7-48814119-INS-77 |
| chr7-48818923-INS-52 | chr7-48818923-INS-60 |
| chr7-48832230-INS-54 | chr7-48832230-INS-54 |
| chr7-48849881-INS-116 | chr7-48849881-INS-116 |
| chr7-48855078-INS-57 | chr7-48855119-INS-54 |
| chr7-48856523-INS-86 | chr7-48856523-INS-88 |
| chr7-48858017-INS-179 | chr7-48858017-INS-179 |
| chr7-48879275-INS-78 | chr7-48879275-INS-63 |
| chr7-48883478-INS-282 | chr7-48883478-INS-294 |
| chr7-48887381-INS-4030 | chr7-48887382-INS-4032 |
| chr7-48889423-INS-104 | chr7-48889425-INS-103 |
| chr7-48895207-INS-66 | chr7-48895207-INS-66 |
| chr7-48917132-INS-76 | chr7-48917127-INS-79 |
| chr7-48924503-INS-58 | chr7-48924503-INS-70 |
| chr7-48926597-INS-173 | chr7-48926610-INS-173 |
| chr7-48953515-INS-59 | chr7-48953517-INS-64 |
| chr7-48992593-INS-69 | chr7-48992593-INS-69 |
| chr7-49010857-INS-155 | chr7-49010857-INS-155 |
| chr7-49116247-INS-64 | chr7-49116247-INS-64 |
| chr7-49119900-INS-181 | chr7-49119900-INS-182 |
| chr7-49121477-INS-305 | chr7-49121477-INS-305 |
| chr7-49136731-INS-210 | chr7-49136731-INS-222 |
| chr7-49154005-INS-51 | chr7-49154005-INS-54 |
| chr7-49158539-INS-236 | chr7-49158539-INS-240 |
| chr7-49159102-INS-211 | chr7-49159102-INS-211 |
| chr7-49169955-INS-166 | chr7-49169955-INS-166 |
| chr7-49182776-INS-83 | chr7-49182763-INS-64 |
| chr7-49196262-INS-73 | chr7-49196261-INS-50 |
| chr7-49199517-INS-60 | chr7-49199512-INS-63 |
| chr7-49218160-INS-52 | chr7-49218160-INS-56 |
| chr7-49218803-INS-1892 | chr7-49218804-INS-1892 |
| chr7-49246181-INS-62 | chr7-49246181-INS-62 |
| chr7-49258159-INS-55 | chr7-49258159-INS-53 |
| chr7-49260702-INS-626 | chr7-49260702-INS-626 |
| chr7-49265643-INS-81 | chr7-49265643-INS-81 |
| chr7-49296497-INS-132 | chr7-49296468-INS-162 |
| chr7-49321844-INS-60 | chr7-49321844-INS-60 |
| chr7-49322963-INS-340 | chr7-49322963-INS-342 |
| chr7-49340852-INS-323 | chr7-49340852-INS-323 |
| chr7-49360079-INS-70 | chr7-49360049-INS-50 |
| chr7-49383743-INS-90 | chr7-49383743-INS-70 |
| chr7-49388268-INS-62 | chr7-49388268-INS-66 |
| chr7-49403872-INS-406 | chr7-49403872-INS-406 |
| chr7-49423922-INS-217 | chr7-49423922-INS-217 |
| chr7-49430726-INS-217 | chr7-49430726-INS-218 |
| chr7-49474351-INS-209 | chr7-49474351-INS-209 |
| chr7-49484455-INS-608 | chr7-49484455-INS-608 |
| chr7-49525603-INS-210 | chr7-49525603-INS-210 |
| chr7-49535293-INS-804 | chr7-49535293-INS-804 |
| chr7-49536438-INS-205 | chr7-49536438-INS-206 |
| chr7-49550495-INS-401 | chr7-49550495-INS-401 |
| chr7-49565277-INS-57 | chr7-49565277-INS-57 |
| chr7-49605600-INS-156 | chr7-49605608-INS-134 |

| | |
|---|---|
| chr7-49614410-INS-68 | chr7-49614410-INS-68 |
| chr7-49622240-INS-52 | chr7-49622238-INS-50 |
| chr7-49636001-INS-56 | chr7-49635967-INS-104 |
| chr7-49647295-INS-521 | chr7-49647300-INS-516 |
| chr7-49647900-INS-85 | chr7-49647900-INS-85 |
| chr7-49669501-INS-3163 | chr7-49669502-INS-3162 |
| chr7-49671680-INS-64 | chr7-49671680-INS-60 |
| chr7-49680936-INS-338 | chr7-49680936-INS-338 |
| chr7-49682694-INS-226 | chr7-49682700-INS-225 |
| chr7-49693161-INS-374 | chr7-49693161-INS-382 |
| chr7-49715141-INS-62 | chr7-49715141-INS-62 |
| chr7-49733300-INS-524 | chr7-49733300-INS-524 |
| chr7-49734575-INS-102 | chr7-49734575-INS-92 |
| chr7-49736321-INS-52 | chr7-49736321-INS-52 |
| chr7-49737152-INS-501 | chr7-49737139-INS-508 |
| chr7-49737158-INS-1030 | chr7-49737159-INS-1030 |
| chr7-49772504-INS-434 | chr7-49772504-INS-434 |
| chr7-49789301-INS-121 | chr7-49789316-INS-86 |
| chr7-49827374-INS-6524 | chr7-49827375-INS-6524 |
| chr7-49838859-INS-156 | chr7-49838811-INS-166 |
| chr7-49905586-INS-97 | chr7-49905542-INS-95 |
| chr7-49906083-INS-291 | chr7-49906083-INS-293 |
| chr7-49954764-INS-212 | chr7-49954764-INS-212 |
| chr7-49957141-INS-1853 | chr7-49957142-INS-1853 |
| chr7-49978116-INS-67 | chr7-49978113-INS-65 |
| chr7-49997881-INS-684 | chr7-49997881-INS-684 |
| chr7-50061288-INS-230 | chr7-50061288-INS-231 |
| chr7-50062466-INS-539 | chr7-50062466-INS-536 |
| chr7-50063360-INS-119 | chr7-50063360-INS-119 |
| chr7-50085852-INS-55 | chr7-50085859-INS-94 |
| chr7-50135036-INS-508 | chr7-50135036-INS-507 |
| chr7-50135922-INS-146 | chr7-50135921-INS-138 |
| chr7-50138015-INS-100 | chr7-50138016-INS-101 |
| chr7-50160586-INS-6272 | chr7-50160587-INS-6272 |
| chr7-50189800-INS-65 | chr7-50189812-INS-72 |
| chr7-50226138-INS-709 | chr7-50226138-INS-710 |
| chr7-50253953-INS-61 | chr7-50253953-INS-64 |
| chr7-50256235-INS-96 | chr7-50256247-INS-116 |
| chr7-50263576-INS-67 | chr7-50263576-INS-67 |
| chr7-50292344-INS-196 | chr7-50292344-INS-196 |
| chr7-50318704-INS-78 | chr7-50318704-INS-78 |
| chr7-50359071-INS-113 | chr7-50359079-INS-96 |
| chr7-50370243-INS-889 | chr7-50370243-INS-889 |
| chr7-50371901-INS-198 | chr7-50371902-INS-198 |
| chr7-50443967-INS-1891 | chr7-50443968-INS-1891 |
| chr7-50496858-INS-8575 | chr7-50496859-INS-8575 |
| chr7-50502984-INS-2075 | chr7-50502985-INS-2075 |
| chr7-50508694-INS-187 | chr7-50508694-INS-187 |
| chr7-50511509-INS-509 | chr7-50511509-INS-548 |
| chr7-50550170-INS-291 | chr7-50550170-INS-291 |
| chr7-50570450-INS-128 | chr7-50570450-INS-128 |
| chr7-50589672-INS-562 | chr7-50589672-INS-562 |
| chr7-50595066-INS-150 | chr7-50595066-INS-150 |
| chr7-50603817-INS-150 | chr7-50603818-INS-150 |
| chr7-50605034-INS-85 | chr7-50605035-INS-114 |
| chr7-50612563-INS-150 | chr7-50612564-INS-150 |
| chr7-50621317-INS-150 | chr7-50621317-INS-150 |
| chr7-50630070-INS-150 | chr7-50630070-INS-150 |
| chr7-50657660-INS-98 | chr7-50657661-INS-112 |
| chr7-50717927-INS-55 | chr7-50717927-INS-55 |
| chr7-50743432-INS-306 | chr7-50743432-INS-306 |
| chr7-50764581-INS-145 | chr7-50764581-INS-149 |
| chr7-50823778-INS-65 | chr7-50823778-INS-74 |
| chr7-50827507-INS-70 | chr7-50827507-INS-82 |
| chr7-50870998-INS-195 | chr7-50870998-INS-195 |
| chr7-50894496-INS-7380 | chr7-50894497-INS-7384 |
| chr7-50907356-INS-52 | chr7-50907328-INS-76 |
| chr7-50918424-INS-240 | chr7-50918424-INS-240 |
| chr7-50945500-INS-222 | chr7-50945500-INS-222 |
| chr7-50956342-INS-152 | chr7-50956342-INS-152 |
| chr7-50992749-INS-6503 | chr7-50992750-INS-6504 |
| chr7-50994206-INS-522 | chr7-50994206-INS-527 |
| chr7-51002052-INS-682 | chr7-51002052-INS-682 |
| chr7-51004823-INS-6281 | chr7-51004824-INS-6282 |
| chr7-51050030-INS-120 | chr7-51050030-INS-128 |
| chr7-51056114-INS-88 | chr7-51056114-INS-88 |
| chr7-51087812-INS-75 | chr7-51087810-INS-95 |

| | |
|---|---|
| chr7-51136917-INS-61 | chr7-51136917-INS-61 |
| chr7-51144722-INS-1894 | chr7-51144723-INS-1894 |
| chr7-51171315-INS-1198 | chr7-51171316-INS-1199 |
| chr7-51189475-INS-657 | chr7-51189475-INS-657 |
| chr7-51208707-INS-52 | chr7-51208707-INS-50 |
| chr7-51227462-INS-642 | chr7-51227462-INS-644 |
| chr7-51231633-INS-76 | chr7-51231629-INS-52 |
| chr7-51234078-INS-82 | chr7-51234078-INS-100 |
| chr7-51237774-INS-1888 | chr7-51237775-INS-1888 |
| chr7-51252975-INS-56 | chr7-51252975-INS-56 |
| chr7-51268820-INS-1889 | chr7-51268821-INS-1889 |
| chr7-51285637-INS-81 | chr7-51285628-INS-63 |
| chr7-51298859-INS-120 | chr7-51298859-INS-96 |
| chr7-51327715-INS-105 | chr7-51327693-INS-93 |
| chr7-51361623-INS-1551 | chr7-51361624-INS-1551 |
| chr7-51377649-INS-663 | chr7-51377649-INS-663 |
| chr7-51381967-INS-1570 | chr7-51381968-INS-1573 |
| chr7-51457843-INS-840 | chr7-51457843-INS-840 |
| chr7-51493828-INS-374 | chr7-51493828-INS-374 |
| chr7-51515821-INS-70 | chr7-51515821-INS-75 |
| chr7-51525411-INS-7089 | chr7-51525412-INS-7091 |
| chr7-51542724-INS-51 | chr7-51542724-INS-51 |
| chr7-51554403-INS-162 | chr7-51554403-INS-162 |
| chr7-51619990-INS-62 | chr7-51619990-INS-62 |
| chr7-51631767-INS-78 | chr7-51631767-INS-78 |
| chr7-51664422-INS-58 | chr7-51664422-INS-58 |
| chr7-51669177-INS-66 | chr7-51669163-INS-62 |
| chr7-51698767-INS-56 | chr7-51698767-INS-56 |
| chr7-51701140-INS-59 | chr7-51701126-INS-50 |
| chr7-51711736-INS-615 | chr7-51711736-INS-615 |
| chr7-51760758-INS-53 | chr7-51760756-INS-51 |
| chr7-51762357-INS-70 | chr7-51762357-INS-86 |
| chr7-51843515-INS-2249 | chr7-51843516-INS-2249 |
| chr7-51848980-INS-73 | chr7-51848980-INS-73 |
| chr7-51853770-INS-120 | chr7-51853774-INS-152 |
| chr7-51869058-INS-778 | chr7-51869058-INS-779 |
| chr7-51871123-INS-190 | chr7-51871123-INS-190 |
| chr7-51895010-INS-78 | chr7-51895014-INS-74 |
| chr7-51901208-INS-767 | chr7-51901208-INS-768 |
| chr7-51908811-INS-56 | chr7-51908811-INS-70 |
| chr7-51910144-INS-90 | chr7-51910144-INS-90 |
| chr7-51921659-INS-271 | chr7-51921659-INS-271 |
| chr7-51949104-INS-60 | chr7-51949104-INS-64 |
| chr7-51960298-INS-453 | chr7-51960298-INS-453 |
| chr7-52001388-INS-1189 | chr7-52001389-INS-1189 |
| chr7-52028128-INS-98 | chr7-52028128-INS-98 |
| chr7-52045570-INS-154 | chr7-52045570-INS-154 |
| chr7-52068106-INS-104 | chr7-52068106-INS-76 |
| chr7-52080052-INS-87 | chr7-52080049-INS-59 |
| chr7-52092895-INS-785 | chr7-52092895-INS-783 |
| chr7-52095563-INS-401 | chr7-52095563-INS-402 |
| chr7-52101788-INS-350 | chr7-52101788-INS-350 |
| chr7-52110642-INS-119 | chr7-52110642-INS-119 |
| chr7-52120095-INS-337 | chr7-52120095-INS-337 |
| chr7-52129531-INS-71 | chr7-52129531-INS-72 |
| chr7-52131400-INS-121 | chr7-52131401-INS-120 |
| chr7-52139251-INS-52 | chr7-52139251-INS-58 |
| chr7-52177661-INS-54 | chr7-52177661-INS-54 |
| chr7-52204017-INS-7145 | chr7-52204018-INS-7145 |
| chr7-52221000-INS-801 | chr7-52221000-INS-801 |
| chr7-52222087-INS-2063 | chr7-52222088-INS-2063 |
| chr7-52255323-INS-883 | chr7-52255323-INS-883 |
| chr7-52256686-INS-59 | chr7-52256698-INS-64 |
| chr7-52319015-INS-124 | chr7-52319015-INS-140 |
| chr7-52332664-INS-254 | chr7-52332664-INS-254 |
| chr7-52334459-INS-70 | chr7-52334497-INS-60 |
| chr7-52335654-INS-58 | chr7-52335654-INS-58 |
| chr7-52346554-INS-114 | chr7-52346554-INS-114 |
| chr7-52371480-INS-72 | chr7-52371480-INS-72 |
| chr7-52374517-INS-6212 | chr7-52374517-INS-6211 |
| chr7-52374940-INS-199 | chr7-52374940-INS-199 |
| chr7-52405423-INS-1034 | chr7-52405424-INS-1035 |
| chr7-52411340-INS-853 | chr7-52411352-INS-852 |
| chr7-52411352-INS-287 | chr7-52411353-INS-256 |
| chr7-52431384-INS-6452 | chr7-52431385-INS-6452 |
| chr7-52433581-INS-1524 | chr7-52433582-INS-1524 |
| chr7-52508151-INS-849 | chr7-52508151-INS-847 |

| | |
|---|---|
| chr7-52548977-INS-7091 | chr7-52548978-INS-7090 |
| chr7-52575169-INS-56 | chr7-52575169-INS-58 |
| chr7-52577135-INS-2042 | chr7-52577136-INS-2041 |
| chr7-52596443-INS-57 | chr7-52596474-INS-52 |
| chr7-52596777-INS-954 | chr7-52596781-INS-952 |
| chr7-52608612-INS-758 | chr7-52608612-INS-758 |
| chr7-52612117-INS-57 | chr7-52612117-INS-57 |
| chr7-52621908-INS-6379 | chr7-52621909-INS-6379 |
| chr7-52622610-INS-188 | chr7-52622610-INS-188 |
| chr7-52627898-INS-115 | chr7-52627898-INS-115 |
| chr7-52630675-INS-84 | chr7-52630675-INS-92 |
| chr7-52640051-INS-461 | chr7-52640051-INS-461 |
| chr7-52645371-INS-6577 | chr7-52645372-INS-6577 |
| chr7-52652228-INS-854 | chr7-52652228-INS-854 |
| chr7-52656092-INS-396 | chr7-52656092-INS-396 |
| chr7-52662201-INS-106 | chr7-52662201-INS-105 |
| chr7-52675840-INS-55 | chr7-52675840-INS-55 |
| chr7-52696897-INS-87 | chr7-52696897-INS-127 |
| chr7-52703314-INS-710 | chr7-52703314-INS-710 |
| chr7-52710684-INS-115 | chr7-52710684-INS-115 |
| chr7-52717051-INS-115 | chr7-52717070-INS-116 |
| chr7-52726057-INS-171 | chr7-52726057-INS-179 |
| chr7-52732342-INS-89 | chr7-52732342-INS-89 |
| chr7-52735978-INS-1128 | chr7-52735979-INS-1118 |
| chr7-52748311-INS-52 | chr7-52748311-INS-54 |
| chr7-52750784-INS-69 | chr7-52750815-INS-84 |
| chr7-52784632-INS-70 | chr7-52784613-INS-52 |
| chr7-52801809-INS-719 | chr7-52801812-INS-708 |
| chr7-52823655-INS-64 | chr7-52823655-INS-80 |
| chr7-52844125-INS-101 | chr7-52844125-INS-105 |
| chr7-52852789-INS-58 | chr7-52852787-INS-56 |
| chr7-52899808-INS-989 | chr7-52899808-INS-988 |
| chr7-52908508-INS-68 | chr7-52908508-INS-56 |
| chr7-52918101-INS-92 | chr7-52918092-INS-108 |
| chr7-52922255-INS-81 | chr7-52922255-INS-81 |
| chr7-52961870-INS-81 | chr7-52961870-INS-81 |
| chr7-52971933-INS-380 | chr7-52971933-INS-380 |
| chr7-52975272-INS-447 | chr7-52975272-INS-447 |
| chr7-52985316-INS-91 | chr7-52985316-INS-90 |
| chr7-53020524-INS-88 | chr7-53020524-INS-88 |
| chr7-53037415-INS-177 | chr7-53037417-INS-174 |
| chr7-53066068-INS-63 | chr7-53066064-INS-74 |
| chr7-53085445-INS-303 | chr7-53085445-INS-303 |
| chr7-53088656-INS-60 | chr7-53088656-INS-60 |
| chr7-53102833-INS-63 | chr7-53102833-INS-51 |
| chr7-53105422-INS-1176 | chr7-53105423-INS-1176 |
| chr7-53108947-INS-637 | chr7-53108931-INS-650 |
| chr7-53117804-INS-80 | chr7-53117805-INS-70 |
| chr7-53129288-INS-52 | chr7-53129288-INS-52 |
| chr7-53131407-INS-421 | chr7-53131408-INS-416 |
| chr7-53138337-INS-590 | chr7-53138333-INS-583 |
| chr7-53148298-INS-69 | chr7-53148293-INS-60 |
| chr7-53189254-INS-74 | chr7-53189254-INS-74 |
| chr7-53195258-INS-62 | chr7-53195259-INS-62 |
| chr7-53196247-INS-852 | chr7-53196247-INS-852 |
| chr7-53200577-INS-504 | chr7-53200577-INS-518 |
| chr7-53205102-INS-315 | chr7-53205102-INS-315 |
| chr7-53232785-INS-57 | chr7-53232785-INS-69 |
| chr7-53265847-INS-82 | chr7-53265838-INS-68 |
| chr7-53301253-INS-88 | chr7-53301278-INS-104 |
| chr7-53302048-INS-575 | chr7-53302048-INS-591 |
| chr7-53306255-INS-928 | chr7-53306255-INS-932 |
| chr7-53325738-INS-186 | chr7-53325741-INS-182 |
| chr7-53341991-INS-341 | chr7-53341991-INS-379 |
| chr7-53366922-INS-627 | chr7-53366922-INS-628 |
| chr7-53370020-INS-119 | chr7-53370020-INS-84 |
| chr7-53380785-INS-76 | chr7-53380785-INS-76 |
| chr7-53382642-INS-60 | chr7-53382642-INS-54 |
| chr7-53397266-INS-112 | chr7-53397266-INS-116 |
| chr7-53398352-INS-54 | chr7-53398352-INS-54 |
| chr7-53404420-INS-449 | chr7-53404420-INS-449 |
| chr7-53428895-INS-73 | chr7-53428895-INS-73 |
| chr7-53457866-INS-61 | chr7-53457851-INS-53 |
| chr7-53459558-INS-3516 | chr7-53459559-INS-3513 |
| chr7-53460899-INS-68 | chr7-53460899-INS-68 |
| chr7-53482765-INS-54 | chr7-53482765-INS-58 |
| chr7-53502098-INS-61 | chr7-53502098-INS-55 |

| | |
|---|---|
| chr7-53532232-INS-3520 | chr7-53532233-INS-3520 |
| chr7-53540544-INS-75 | chr7-53540529-INS-60 |
| chr7-53548565-INS-279 | chr7-53548571-INS-279 |
| chr7-53568471-INS-103 | chr7-53568471-INS-95 |
| chr7-53621254-INS-156 | chr7-53621254-INS-158 |
| chr7-53626170-INS-72 | chr7-53626170-INS-72 |
| chr7-53634509-INS-164 | chr7-53634509-INS-166 |
| chr7-53648891-INS-663 | chr7-53648891-INS-663 |
| chr7-53653495-INS-54 | chr7-53653495-INS-54 |
| chr7-53662487-INS-1241 | chr7-53662488-INS-1242 |
| chr7-53685355-INS-134 | chr7-53685355-INS-134 |
| chr7-53694155-INS-451 | chr7-53694147-INS-462 |
| chr7-53743593-INS-100 | chr7-53743593-INS-100 |
| chr7-53758684-INS-201 | chr7-53758684-INS-201 |
| chr7-53771024-INS-6575 | chr7-53771025-INS-6575 |
| chr7-53776022-INS-6453 | chr7-53776023-INS-6451 |
| chr7-53826503-INS-2610 | chr7-53826504-INS-2610 |
| chr7-53837781-INS-6778 | chr7-53837782-INS-6778 |
| chr7-53846874-INS-103 | chr7-53846874-INS-99 |
| chr7-53851458-INS-1127 | chr7-53851459-INS-1115 |
| chr7-53877128-INS-2257 | chr7-53877129-INS-2257 |
| chr7-53878498-INS-197 | chr7-53878498-INS-197 |
| chr7-53883761-INS-102 | chr7-53883761-INS-102 |
| chr7-53889499-INS-80 | chr7-53889499-INS-68 |
| chr7-53894298-INS-427 | chr7-53894299-INS-427 |
| chr7-53897182-INS-60 | chr7-53897182-INS-60 |
| chr7-53900754-INS-1583 | chr7-53900755-INS-1587 |
| chr7-53934977-INS-217 | chr7-53934977-INS-219 |
| chr7-53949779-INS-194 | chr7-53949779-INS-194 |
| chr7-53974713-INS-96 | chr7-53974713-INS-96 |
| chr7-53978715-INS-60 | chr7-53978715-INS-60 |
| chr7-53979900-INS-301 | chr7-53979900-INS-301 |
| chr7-53981073-INS-95 | chr7-53981069-INS-85 |
| chr7-54016513-INS-84 | chr7-54016513-INS-94 |
| chr7-54046699-INS-321 | chr7-54046699-INS-321 |
| chr7-54055966-INS-252 | chr7-54055967-INS-252 |
| chr7-54060247-INS-131 | chr7-54060248-INS-137 |
| chr7-54085602-INS-1624 | chr7-54085603-INS-1635 |
| chr7-54090663-INS-75 | chr7-54090663-INS-75 |
| chr7-54094717-INS-141 | chr7-54094717-INS-141 |
| chr7-54112036-INS-4801 | chr7-54112037-INS-4801 |
| chr7-54125356-INS-714 | chr7-54125356-INS-714 |
| chr7-54141790-INS-211 | chr7-54141781-INS-183 |
| chr7-54147625-INS-62 | chr7-54147625-INS-62 |
| chr7-54155469-INS-485 | chr7-54155469-INS-485 |
| chr7-54206704-INS-830 | chr7-54206704-INS-830 |
| chr7-54215078-INS-60 | chr7-54215078-INS-64 |
| chr7-54223192-INS-751 | chr7-54223192-INS-753 |
| chr7-54297708-INS-72 | chr7-54297705-INS-68 |
| chr7-54314350-INS-252 | chr7-54314350-INS-252 |
| chr7-54357690-INS-122 | chr7-54357691-INS-116 |
| chr7-54383254-INS-485 | chr7-54383254-INS-485 |
| chr7-54386287-INS-1923 | chr7-54386288-INS-1923 |
| chr7-54388038-INS-206 | chr7-54388038-INS-207 |
| chr7-54392384-INS-440 | chr7-54392384-INS-440 |
| chr7-54407131-INS-54 | chr7-54407159-INS-60 |
| chr7-54429398-INS-1244 | chr7-54429399-INS-1244 |
| chr7-54464248-INS-216 | chr7-54464248-INS-217 |
| chr7-54475735-INS-151 | chr7-54475735-INS-111 |
| chr7-54500210-INS-773 | chr7-54500210-INS-776 |
| chr7-54515745-INS-77 | chr7-54515745-INS-77 |
| chr7-54538316-INS-53 | chr7-54538316-INS-53 |
| chr7-54541608-INS-994 | chr7-54541608-INS-986 |
| chr7-54545675-INS-52 | chr7-54545675-INS-50 |
| chr7-54631342-INS-2368 | chr7-54631343-INS-2368 |
| chr7-54643177-INS-1736 | chr7-54643178-INS-1736 |
| chr7-54646396-INS-98 | chr7-54646396-INS-98 |
| chr7-54720645-INS-183 | chr7-54720665-INS-191 |
| chr7-54721285-INS-81 | chr7-54721285-INS-56 |
| chr7-54736706-INS-62 | chr7-54736706-INS-62 |
| chr7-54738139-INS-64 | chr7-54738139-INS-66 |
| chr7-54813060-INS-304 | chr7-54813060-INS-328 |
| chr7-54813814-INS-108 | chr7-54813815-INS-121 |
| chr7-54831577-INS-75 | chr7-54831577-INS-76 |
| chr7-54853644-INS-402 | chr7-54853644-INS-402 |
| chr7-54862797-INS-52 | chr7-54862798-INS-68 |
| chr7-54883856-INS-370 | chr7-54883856-INS-370 |

| | |
|---|---|
| chr7-54903374-INS-810 | chr7-54903374-INS-810 |
| chr7-54920361-INS-72 | chr7-54920361-INS-60 |
| chr7-54928300-INS-75 | chr7-54928268-INS-117 |
| chr7-54954698-INS-431 | chr7-54954698-INS-431 |
| chr7-54992133-INS-238 | chr7-54992133-INS-238 |
| chr7-55018300-INS-2542 | chr7-55018313-INS-2540 |
| chr7-55022996-INS-6288 | chr7-55022997-INS-6294 |
| chr7-55034966-INS-6667 | chr7-55034967-INS-6668 |
| chr7-55041504-INS-1803 | chr7-55041505-INS-1800 |
| chr7-55064665-INS-60 | chr7-55064665-INS-60 |
| chr7-55065268-INS-189 | chr7-55065268-INS-189 |
| chr7-55089416-INS-89 | chr7-55089416-INS-89 |
| chr7-55130107-INS-462 | chr7-55130107-INS-463 |
| chr7-55143309-INS-551 | chr7-55143309-INS-552 |
| chr7-55156628-INS-3643 | chr7-55156629-INS-3642 |
| chr7-55158091-INS-6324 | chr7-55158092-INS-6327 |
| chr7-55160144-INS-483 | chr7-55160146-INS-459 |
| chr7-55163763-INS-1213 | chr7-55163764-INS-1213 |
| chr7-55168140-INS-57 | chr7-55168140-INS-57 |
| chr7-55180342-INS-52 | chr7-55180348-INS-82 |
| chr7-55180865-INS-6590 | chr7-55180866-INS-6591 |
| chr7-55184180-INS-130 | chr7-55184180-INS-127 |
| chr7-55319099-INS-95 | chr7-55319099-INS-95 |
| chr7-55361692-INS-644 | chr7-55361692-INS-644 |
| chr7-55373662-INS-148 | chr7-55373662-INS-132 |
| chr7-55391994-INS-66 | chr7-55391995-INS-72 |
| chr7-55412809-INS-52 | chr7-55412811-INS-54 |
| chr7-55428213-INS-162 | chr7-55428213-INS-162 |
| chr7-55431502-INS-538 | chr7-55431502-INS-538 |
| chr7-55434823-INS-75 | chr7-55434823-INS-75 |
| chr7-55462560-INS-123 | chr7-55462550-INS-148 |
| chr7-55505889-INS-68 | chr7-55505889-INS-86 |
| chr7-55508974-INS-218 | chr7-55508974-INS-218 |
| chr7-55518748-INS-255 | chr7-55518748-INS-235 |
| chr7-55519588-INS-104 | chr7-55519588-INS-104 |
| chr7-55534776-INS-125 | chr7-55534776-INS-125 |
| chr7-55554864-INS-102 | chr7-55554864-INS-80 |
| chr7-55592070-INS-214 | chr7-55592070-INS-214 |
| chr7-55615560-INS-255 | chr7-55615564-INS-256 |
| chr7-55619954-INS-55 | chr7-55619971-INS-84 |
| chr7-55620967-INS-67 | chr7-55620968-INS-71 |
| chr7-55622083-INS-512 | chr7-55622083-INS-525 |
| chr7-55631134-INS-62 | chr7-55631154-INS-62 |
| chr7-55634886-INS-497 | chr7-55634886-INS-502 |
| chr7-55671472-INS-488 | chr7-55671468-INS-490 |
| chr7-55674372-INS-1048 | chr7-55674373-INS-1049 |
| chr7-55676677-INS-428 | chr7-55676677-INS-428 |
| chr7-55677999-INS-206 | chr7-55677999-INS-206 |
| chr7-55707462-INS-207 | chr7-55707462-INS-212 |
| chr7-55748377-INS-99 | chr7-55748377-INS-100 |
| chr7-55755561-INS-78 | chr7-55755561-INS-78 |
| chr7-55761031-INS-338 | chr7-55761029-INS-317 |
| chr7-55774972-INS-6639 | chr7-55774973-INS-6639 |
| chr7-55777768-INS-60 | chr7-55777768-INS-60 |
| chr7-55808915-INS-191 | chr7-55808938-INS-195 |
| chr7-55815182-INS-4739 | chr7-55815183-INS-4739 |
| chr7-55826147-INS-69 | chr7-55826168-INS-60 |
| chr7-55830516-INS-53 | chr7-55830516-INS-53 |
| chr7-55831021-INS-479 | chr7-55831021-INS-482 |
| chr7-55850739-INS-130 | chr7-55850739-INS-130 |
| chr7-55897317-INS-6485 | chr7-55897318-INS-6484 |
| chr7-55948548-INS-1494 | chr7-55948549-INS-1481 |
| chr7-55952347-INS-237 | chr7-55952347-INS-237 |
| chr7-55954994-INS-59 | chr7-55954958-INS-82 |
| chr7-55970720-INS-6453 | chr7-55970721-INS-6453 |
| chr7-56010779-INS-139 | chr7-56010779-INS-139 |
| chr7-56010886-INS-1371 | chr7-56010878-INS-1370 |
| chr7-56030833-INS-54 | chr7-56030833-INS-53 |
| chr7-56033509-INS-752 | chr7-56033509-INS-751 |
| chr7-56036780-INS-62 | chr7-56036780-INS-72 |
| chr7-56084614-INS-621 | chr7-56084611-INS-620 |
| chr7-56092517-INS-60 | chr7-56092517-INS-50 |
| chr7-56095586-INS-52 | chr7-56095587-INS-60 |
| chr7-56096562-INS-136 | chr7-56096562-INS-132 |
| chr7-56119042-INS-2304 | chr7-56119043-INS-2304 |
| chr7-56148878-INS-6836 | chr7-56148879-INS-6836 |
| chr7-56159393-INS-191 | chr7-56159393-INS-190 |

| | |
|---|---|
| chr7-56173120-INS-1895 | chr7-56173121-INS-1894 |
| chr7-56185816-INS-60 | chr7-56185816-INS-60 |
| chr7-56191329-INS-313 | chr7-56191329-INS-313 |
| chr7-56210464-INS-60 | chr7-56210436-INS-54 |
| chr7-56230446-INS-252 | chr7-56230446-INS-252 |
| chr7-56257956-INS-62 | chr7-56257956-INS-62 |
| chr7-56259710-INS-68 | chr7-56259710-INS-68 |
| chr7-56280721-INS-7407 | chr7-56280722-INS-7407 |
| chr7-56281394-INS-186 | chr7-56281394-INS-186 |
| chr7-56293191-INS-51 | chr7-56293191-INS-51 |
| chr7-56316847-INS-6414 | chr7-56316848-INS-6414 |
| chr7-56322786-INS-59 | chr7-56322786-INS-52 |
| chr7-56325216-INS-1490 | chr7-56325217-INS-1491 |
| chr7-56345943-INS-165 | chr7-56345943-INS-165 |
| chr7-56351751-INS-621 | chr7-56351751-INS-621 |
| chr7-56356769-INS-110 | chr7-56356769-INS-78 |
| chr7-56358186-INS-741 | chr7-56358186-INS-741 |
| chr7-56381566-INS-7368 | chr7-56381567-INS-7374 |
| chr7-56390141-INS-2863 | chr7-56390142-INS-2863 |
| chr7-56392984-INS-452 | chr7-56392984-INS-455 |
| chr7-56449021-INS-429 | chr7-56449021-INS-428 |
| chr7-56464885-INS-1044 | chr7-56464886-INS-1045 |
| chr7-56480678-INS-135 | chr7-56480678-INS-135 |
| chr7-56496051-INS-84 | chr7-56496051-INS-132 |
| chr7-56498836-INS-52 | chr7-56498837-INS-52 |
| chr7-56499701-INS-56 | chr7-56499702-INS-54 |
| chr7-56522774-INS-239 | chr7-56522774-INS-249 |
| chr7-56523957-INS-312 | chr7-56523957-INS-320 |
| chr7-56528074-INS-200 | chr7-56528074-INS-200 |
| chr7-56541195-INS-60 | chr7-56541195-INS-72 |
| chr7-56546610-INS-62 | chr7-56546603-INS-108 |
| chr7-56583364-INS-3308 | chr7-56583365-INS-3308 |
| chr7-56585522-INS-130 | chr7-56585522-INS-130 |
| chr7-56592987-INS-820 | chr7-56592987-INS-820 |
| chr7-56613205-INS-688 | chr7-56613205-INS-688 |
| chr7-56623362-INS-399 | chr7-56623362-INS-386 |
| chr7-56633601-INS-93 | chr7-56633601-INS-93 |
| chr7-56651837-INS-2776 | chr7-56651838-INS-2776 |
| chr7-56660178-INS-56 | chr7-56660178-INS-58 |
| chr7-56663687-INS-379 | chr7-56663685-INS-383 |
| chr7-56680841-INS-595 | chr7-56680841-INS-596 |
| chr7-56696596-INS-64 | chr7-56696604-INS-52 |
| chr7-56719702-INS-1436 | chr7-56719703-INS-1434 |
| chr7-56742851-INS-2967 | chr7-56742852-INS-2967 |
| chr7-56746350-INS-164 | chr7-56746350-INS-164 |
| chr7-56776961-INS-97 | chr7-56776960-INS-56 |
| chr7-56781430-INS-1892 | chr7-56781431-INS-1892 |
| chr7-56800919-INS-452 | chr7-56800919-INS-453 |
| chr7-56807176-INS-91 | chr7-56807176-INS-93 |
| chr7-56808618-INS-374 | chr7-56808618-INS-374 |
| chr7-56813674-INS-3237 | chr7-56813675-INS-3237 |
| chr7-56815160-INS-7483 | chr7-56815161-INS-7485 |
| chr7-56900164-INS-52 | chr7-56900152-INS-54 |
| chr7-56901646-INS-64 | chr7-56901646-INS-64 |
| chr7-56918967-INS-82 | chr7-56918967-INS-82 |
| chr7-56922928-INS-176 | chr7-56922887-INS-178 |
| chr7-56927293-INS-420 | chr7-56927293-INS-420 |
| chr7-56935280-INS-54 | chr7-56935286-INS-58 |
| chr7-56953896-INS-100 | chr7-56953897-INS-81 |
| chr7-56966564-INS-242 | chr7-56966564-INS-242 |
| chr7-56981404-INS-550 | chr7-56981403-INS-584 |
| chr7-56999070-INS-137 | chr7-56999070-INS-147 |
| chr7-56999378-INS-832 | chr7-56999378-INS-832 |
| chr7-57009940-INS-52 | chr7-57009940-INS-74 |
| chr7-57012065-INS-58 | chr7-57012065-INS-76 |
| chr7-57025034-INS-150 | chr7-57025028-INS-109 |
| chr7-57029766-INS-55 | chr7-57029765-INS-59 |
| chr7-57048031-INS-91 | chr7-57048031-INS-91 |
| chr7-57048936-INS-212 | chr7-57048936-INS-212 |
| chr7-57054296-INS-68 | chr7-57054327-INS-72 |
| chr7-57059887-INS-55 | chr7-57059890-INS-56 |
| chr7-57089312-INS-206 | chr7-57089289-INS-254 |
| chr7-57105747-INS-269 | chr7-57105747-INS-269 |
| chr7-57114419-INS-55 | chr7-57114419-INS-57 |
| chr7-57124806-INS-809 | chr7-57124806-INS-809 |
| chr7-57142180-INS-120 | chr7-57142180-INS-120 |
| chr7-57170188-INS-1413 | chr7-57170189-INS-1412 |

| | |
|---|---|
| chr7-57176052-INS-100 | chr7-57176052-INS-64 |
| chr7-57186430-INS-124 | chr7-57186430-INS-108 |
| chr7-57189549-INS-299 | chr7-57189549-INS-299 |
| chr7-57192546-INS-61 | chr7-57192563-INS-80 |
| chr7-57213468-INS-387 | chr7-57213468-INS-391 |
| chr7-57226332-INS-52 | chr7-57226332-INS-54 |
| chr7-57285895-INS-64 | chr7-57285893-INS-100 |
| chr7-57300806-INS-4715 | chr7-57300807-INS-4715 |
| chr7-57318591-INS-63 | chr7-57318591-INS-63 |
| chr7-57320773-INS-829 | chr7-57320773-INS-829 |
| chr7-57323413-INS-58 | chr7-57323413-INS-50 |
| chr7-57332340-INS-57 | chr7-57332340-INS-62 |
| chr7-57340503-INS-54 | chr7-57340503-INS-58 |
| chr7-57343428-INS-7077 | chr7-57343429-INS-7078 |
| chr7-57346298-INS-189 | chr7-57346298-INS-189 |
| chr7-57354272-INS-663 | chr7-57354276-INS-694 |
| chr7-57375345-INS-238 | chr7-57375345-INS-237 |
| chr7-57377495-INS-412 | chr7-57377495-INS-414 |
| chr7-57386017-INS-386 | chr7-57386017-INS-386 |
| chr7-57406997-INS-171 | chr7-57406997-INS-171 |
| chr7-57412037-INS-403 | chr7-57412037-INS-403 |
| chr7-57421576-INS-56 | chr7-57421576-INS-60 |
| chr7-57436969-INS-2613 | chr7-57436970-INS-2613 |
| chr7-57448059-INS-6404 | chr7-57448060-INS-6404 |
| chr7-57450663-INS-136 | chr7-57450634-INS-157 |
| chr7-57533307-INS-357 | chr7-57533307-INS-358 |
| chr7-57551736-INS-550 | chr7-57551736-INS-550 |
| chr7-57572534-INS-80 | chr7-57572534-INS-82 |
| chr7-57594407-INS-59 | chr7-57594407-INS-59 |
| chr7-57596427-INS-106 | chr7-57596428-INS-99 |
| chr7-57606150-INS-56 | chr7-57606150-INS-58 |
| chr7-57606689-INS-538 | chr7-57606689-INS-538 |
| chr7-57668000-INS-437 | chr7-57668000-INS-436 |
| chr7-57669973-INS-62 | chr7-57669973-INS-62 |
| chr7-57683351-INS-177 | chr7-57683351-INS-176 |
| chr7-57775981-INS-72 | chr7-57775981-INS-72 |
| chr7-57781418-INS-52 | chr7-57781406-INS-56 |
| chr7-57784963-INS-150 | chr7-57784963-INS-149 |
| chr7-57790740-INS-58 | chr7-57790734-INS-82 |
| chr7-57831758-INS-1894 | chr7-57831759-INS-1896 |
| chr7-57862413-INS-410 | chr7-57862413-INS-409 |
| chr7-57891573-INS-412 | chr7-57891573-INS-412 |
| chr7-57902397-INS-6440 | chr7-57902398-INS-6440 |
| chr7-58034474-INS-6677 | chr7-58034475-INS-6677 |
| chr7-58046195-INS-53 | chr7-58046178-INS-84 |
| chr7-58056610-INS-112 | chr7-58056612-INS-112 |
| chr7-58062315-INS-195 | chr7-58062315-INS-195 |
| chr7-58081490-INS-55 | chr7-58081497-INS-52 |
| chr7-58094436-INS-1255 | chr7-58094437-INS-1258 |
| chr7-58096743-INS-56 | chr7-58096703-INS-72 |
| chr7-58097656-INS-58 | chr7-58097668-INS-58 |
| chr7-58100309-INS-204 | chr7-58100309-INS-204 |
| chr7-58116296-INS-758 | chr7-58116296-INS-752 |
| chr7-58138444-INS-460 | chr7-58138444-INS-460 |
| chr7-58160257-INS-1593 | chr7-58160258-INS-1593 |
| chr7-58168276-INS-148 | chr7-58168276-INS-148 |
| chr7-58221792-INS-427 | chr7-58221792-INS-430 |
| chr7-58281727-INS-56 | chr7-58281727-INS-60 |
| chr7-58283846-INS-882 | chr7-58283846-INS-882 |
| chr7-58290489-INS-411 | chr7-58290489-INS-411 |
| chr7-58297689-INS-72 | chr7-58297689-INS-84 |
| chr7-58299858-INS-6334 | chr7-58299859-INS-6333 |
| chr7-58306615-INS-58 | chr7-58306614-INS-84 |
| chr7-58313019-INS-56 | chr7-58313019-INS-68 |
| chr7-58325382-INS-174 | chr7-58325382-INS-202 |
| chr7-58378034-INS-97 | chr7-58378034-INS-97 |
| chr7-58384899-INS-108 | chr7-58384902-INS-138 |
| chr7-58406631-INS-149 | chr7-58406631-INS-149 |
| chr7-58407799-INS-203 | chr7-58407799-INS-201 |
| chr7-58408940-INS-375 | chr7-58408940-INS-376 |
| chr7-58413971-INS-213 | chr7-58413971-INS-213 |
| chr7-58414797-INS-7111 | chr7-58414798-INS-7111 |
| chr7-58418803-INS-78 | chr7-58418803-INS-105 |
| chr7-58440061-INS-92 | chr7-58440061-INS-92 |
| chr7-58458306-INS-66 | chr7-58458306-INS-70 |
| chr7-58550366-INS-214 | chr7-58550366-INS-214 |
| chr7-58553436-INS-54 | chr7-58553432-INS-58 |

| | |
|---|---|
| chr7-58555069-INS-344 | chr7-58555069-INS-344 |
| chr7-58583239-INS-1299 | chr7-58583240-INS-1299 |
| chr7-58603251-INS-55 | chr7-58603251-INS-59 |
| chr7-58612419-INS-6796 | chr7-58612420-INS-6796 |
| chr7-58623627-INS-58 | chr7-58623623-INS-62 |
| chr7-58633890-INS-6930 | chr7-58633891-INS-6931 |
| chr7-58639680-INS-62 | chr7-58639680-INS-52 |
| chr7-58645598-INS-7222 | chr7-58645599-INS-7222 |
| chr7-58652375-INS-7222 | chr7-58652376-INS-7222 |
| chr7-58682290-INS-6648 | chr7-58682291-INS-6648 |
| chr7-58707075-INS-758 | chr7-58707075-INS-758 |
| chr7-58771662-INS-50 | chr7-58771662-INS-51 |
| chr7-58792768-INS-115 | chr7-58792813-INS-80 |
| chr7-58817501-INS-60 | chr7-58817493-INS-64 |
| chr7-58822094-INS-175 | chr7-58822094-INS-191 |
| chr7-58927001-INS-215 | chr7-58927001-INS-216 |
| chr7-58932635-INS-945 | chr7-58932635-INS-945 |
| chr7-58973985-INS-52 | chr7-58974010-INS-56 |
| chr7-59043918-INS-215 | chr7-59043939-INS-215 |
| chr7-59277617-INS-4792 | chr7-59277618-INS-4794 |
| chr7-59280099-INS-52 | chr7-59280099-INS-56 |
| chr7-59285377-INS-75 | chr7-59285377-INS-66 |
| chr7-59290273-INS-297 | chr7-59290273-INS-297 |
| chr7-59304489-INS-6596 | chr7-59304490-INS-6596 |
| chr7-59308731-INS-198 | chr7-59308731-INS-198 |
| chr7-59326892-INS-66 | chr7-59326939-INS-54 |
| chr7-59580402-INS-68 | chr7-59580361-INS-60 |
| chr7-59585156-INS-208 | chr7-59585156-INS-208 |
| chr7-59610706-INS-114 | chr7-59610678-INS-144 |
| chr7-59636798-INS-627 | chr7-59636798-INS-627 |
| chr7-59663785-INS-2694 | chr7-59663786-INS-2688 |
| chr7-59667496-INS-5317 | chr7-59667497-INS-5318 |
| chr7-59682423-INS-486 | chr7-59682423-INS-480 |
| chr7-59688677-INS-1270 | chr7-59688682-INS-1263 |
| chr7-59690381-INS-488 | chr7-59690381-INS-488 |
| chr7-59697894-INS-157 | chr7-59697894-INS-158 |
| chr7-59722303-INS-710 | chr7-59722303-INS-710 |
| chr7-59726935-INS-2110 | chr7-59726936-INS-2110 |
| chr7-59728685-INS-205 | chr7-59728685-INS-205 |
| chr7-59767532-INS-206 | chr7-59767532-INS-206 |
| chr7-59787563-INS-499 | chr7-59787563-INS-493 |
| chr7-59795974-INS-52 | chr7-59795974-INS-52 |
| chr7-59817113-INS-241 | chr7-59817113-INS-241 |
| chr7-59837805-INS-296 | chr7-59837805-INS-296 |
| chr7-59843242-INS-4711 | chr7-59843243-INS-4711 |
| chr7-59937610-INS-627 | chr7-59937610-INS-627 |
| chr7-59967906-INS-2443 | chr7-59967907-INS-2443 |
| chr7-60001091-INS-214 | chr7-60001091-INS-214 |
| chr7-60045868-INS-6652 | chr7-60045869-INS-6652 |
| chr7-60047742-INS-54 | chr7-60047742-INS-54 |
| chr7-60060003-INS-377 | chr7-60060003-INS-377 |
| chr7-60066877-INS-8751 | chr7-60066878-INS-8751 |
| chr7-60080990-INS-119 | chr7-60080990-INS-110 |
| chr7-60137266-INS-1060 | chr7-60137267-INS-1063 |
| chr7-60157470-INS-6456 | chr7-60157471-INS-6458 |
| chr7-60168867-INS-871 | chr7-60168877-INS-869 |
| chr7-60210137-INS-108 | chr7-60210127-INS-80 |
| chr7-60246055-INS-88 | chr7-60246055-INS-86 |
| chr7-60266241-INS-56 | chr7-60266241-INS-57 |
| chr7-60274223-INS-3213 | chr7-60274224-INS-3213 |
| chr7-60274674-INS-76 | chr7-60274674-INS-80 |
| chr7-60280346-INS-1797 | chr7-60280347-INS-1797 |
| chr7-60290784-INS-54 | chr7-60290784-INS-58 |
| chr7-60369148-INS-924 | chr7-60369148-INS-924 |
| chr7-60462552-INS-6592 | chr7-60462553-INS-6590 |
| chr7-60471138-INS-52 | chr7-60471138-INS-54 |
| chr7-60483249-INS-75 | chr7-60483249-INS-75 |
| chr7-60486199-INS-399 | chr7-60486199-INS-399 |
| chr7-60486532-INS-4670 | chr7-60486533-INS-4670 |
| chr7-60491921-INS-1328 | chr7-60491922-INS-1328 |
| chr7-60497905-INS-2428 | chr7-60497906-INS-2428 |
| chr7-60502581-INS-1553 | chr7-60502582-INS-1553 |
| chr7-60504736-INS-159 | chr7-60504736-INS-159 |
| chr7-60536165-INS-3207 | chr7-60536166-INS-3207 |
| chr7-60548744-INS-3934 | chr7-60548745-INS-3915 |
| chr7-60549881-INS-55 | chr7-60549881-INS-51 |
| chr7-60563445-INS-114 | chr7-60563445-INS-134 |

| | |
|---|---|
| chr7-60569396-INS-1247 | chr7-60569397-INS-1248 |
| chr7-60579860-INS-7494 | chr7-60579861-INS-7490 |
| chr7-60582060-INS-510 | chr7-60582060-INS-510 |
| chr7-60643665-INS-938 | chr7-60643665-INS-938 |
| chr7-60647804-INS-67 | chr7-60647804-INS-68 |
| chr7-60662329-INS-1513 | chr7-60662330-INS-1513 |
| chr7-60668870-INS-6600 | chr7-60668871-INS-6600 |
| chr7-60674786-INS-620 | chr7-60674786-INS-621 |
| chr7-60706634-INS-6395 | chr7-60706635-INS-6395 |
| chr7-60713474-INS-308 | chr7-60713474-INS-307 |
| chr7-60725432-INS-145 | chr7-60725432-INS-154 |
| chr7-60729310-INS-291 | chr7-60729310-INS-261 |
| chr7-60762999-INS-102 | chr7-60762975-INS-135 |
| chr7-60766384-INS-618 | chr7-60766384-INS-619 |
| chr7-60768748-INS-6447 | chr7-60768749-INS-6447 |
| chr7-60780554-INS-491 | chr7-60780554-INS-491 |
| chr7-60783434-INS-52 | chr7-60783434-INS-54 |
| chr7-60853277-INS-61 | chr7-60853277-INS-61 |
| chr7-60874037-INS-70 | chr7-60874037-INS-75 |
| chr7-60876923-INS-56 | chr7-60876923-INS-58 |
| chr7-60888161-INS-3526 | chr7-60888162-INS-3526 |
| chr7-60907783-INS-5124 | chr7-60907784-INS-5124 |
| chr7-60953363-INS-225 | chr7-60953345-INS-273 |
| chr7-60984923-INS-70 | chr7-60984916-INS-88 |
| chr7-61011789-INS-51 | chr7-61011790-INS-53 |
| chr7-61054881-INS-6369 | chr7-61054882-INS-6369 |
| chr7-61063651-INS-56 | chr7-61063651-INS-56 |
| chr7-61079892-INS-3244 | chr7-61079893-INS-3243 |
| chr7-61090457-INS-2272 | chr7-61090458-INS-2272 |
| chr7-61107917-INS-426 | chr7-61107917-INS-426 |
| chr7-61142311-INS-2677 | chr7-61142312-INS-2691 |
| chr7-61150786-INS-187 | chr7-61150786-INS-187 |
| chr7-61269262-INS-907 | chr7-61269262-INS-907 |
| chr7-61359869-INS-1890 | chr7-61359870-INS-1890 |
| chr7-61369069-INS-58 | chr7-61369093-INS-56 |
| chr7-61408994-INS-322 | chr7-61408994-INS-322 |
| chr7-61428416-INS-1938 | chr7-61428417-INS-1938 |
| chr7-61434901-INS-101 | chr7-61434902-INS-100 |
| chr7-61452976-INS-91 | chr7-61452976-INS-58 |
| chr7-61469506-INS-109 | chr7-61469512-INS-64 |
| chr7-61470806-INS-121 | chr7-61470806-INS-84 |
| chr7-61476614-INS-124 | chr7-61476588-INS-112 |
| chr7-61518547-INS-57 | chr7-61518547-INS-73 |
| chr7-61540511-INS-59 | chr7-61540511-INS-59 |
| chr7-61541259-INS-68 | chr7-61541259-INS-59 |
| chr7-61550305-INS-156 | chr7-61550305-INS-176 |
| chr7-61558165-INS-504 | chr7-61558165-INS-504 |
| chr7-61565696-INS-58 | chr7-61565696-INS-58 |
| chr7-61571418-INS-52 | chr7-61571418-INS-58 |
| chr7-61578692-INS-56 | chr7-61578692-INS-54 |
| chr7-61579452-INS-65 | chr7-61579452-INS-65 |
| chr7-61597069-INS-56 | chr7-61597069-INS-60 |
| chr7-61603282-INS-192 | chr7-61603282-INS-192 |
| chr7-61621901-INS-248 | chr7-61621901-INS-250 |
| chr7-61624631-INS-53 | chr7-61624631-INS-53 |
| chr7-61639494-INS-62 | chr7-61639494-INS-64 |
| chr7-61672981-INS-586 | chr7-61672981-INS-586 |
| chr7-61683705-INS-68 | chr7-61683705-INS-68 |
| chr7-61692490-INS-536 | chr7-61692490-INS-541 |
| chr7-61701475-INS-1408 | chr7-61701476-INS-1408 |
| chr7-61704441-INS-184 | chr7-61704441-INS-184 |
| chr7-61734415-INS-81 | chr7-61734411-INS-52 |
| chr7-61738527-INS-64 | chr7-61738527-INS-64 |
| chr7-61741161-INS-86 | chr7-61741150-INS-68 |
| chr7-61760439-INS-4367 | chr7-61760440-INS-4366 |
| chr7-61769634-INS-200 | chr7-61769635-INS-200 |
| chr7-61827322-INS-476 | chr7-61827322-INS-476 |
| chr7-61840211-INS-3664 | chr7-61840212-INS-3656 |
| chr7-61852040-INS-72 | chr7-61852040-INS-72 |
| chr7-61864875-INS-543 | chr7-61864875-INS-543 |
| chr7-61871030-INS-65 | chr7-61871036-INS-53 |
| chr7-61875458-INS-65 | chr7-61875458-INS-65 |
| chr7-61889304-INS-6473 | chr7-61889305-INS-6474 |
| chr7-61921782-INS-54 | chr7-61921795-INS-60 |
| chr7-61943480-INS-6451 | chr7-61943481-INS-6450 |
| chr7-61955846-INS-1894 | chr7-61955847-INS-1894 |
| chr7-61968065-INS-75 | chr7-61968070-INS-90 |

| | |
|---|---|
| chr7-61984995-INS-212 | chr7-61984995-INS-212 |
| chr7-62001582-INS-56 | chr7-62001582-INS-56 |
| chr7-62020358-INS-6405 | chr7-62020359-INS-6408 |
| chr7-62024272-INS-6400 | chr7-62024273-INS-6400 |
| chr7-62051595-INS-78 | chr7-62051608-INS-75 |
| chr7-62163619-INS-7582 | chr7-62163620-INS-7582 |
| chr7-62168967-INS-938 | chr7-62168943-INS-938 |
| chr7-62201644-INS-201 | chr7-62201644-INS-201 |
| chr7-62216711-INS-7581 | chr7-62216712-INS-7581 |
| chr7-62218653-INS-1896 | chr7-62218654-INS-1896 |
| chr7-62224307-INS-54 | chr7-62224307-INS-57 |
| chr7-62232947-INS-73 | chr7-62232947-INS-75 |
| chr7-62249851-INS-827 | chr7-62249851-INS-827 |
| chr7-62261358-INS-215 | chr7-62261356-INS-227 |
| chr7-62268872-INS-82 | chr7-62268872-INS-82 |
| chr7-62321119-INS-418 | chr7-62321119-INS-418 |
| chr7-62387287-INS-283 | chr7-62387287-INS-283 |
| chr7-62493308-INS-1895 | chr7-62493309-INS-1895 |
| chr7-62521233-INS-109 | chr7-62521234-INS-109 |
| chr7-62526666-INS-491 | chr7-62526663-INS-490 |
| chr7-62559065-INS-60 | chr7-62559065-INS-60 |
| chr7-62566479-INS-326 | chr7-62566479-INS-327 |
| chr7-62571890-INS-71 | chr7-62571890-INS-62 |
| chr7-62611267-INS-407 | chr7-62611267-INS-404 |
| chr7-62633466-INS-343 | chr7-62633468-INS-343 |
| chr7-62645828-INS-498 | chr7-62645828-INS-498 |
| chr7-62686180-INS-52 | chr7-62686180-INS-87 |
| chr7-62723928-INS-59 | chr7-62723942-INS-61 |
| chr7-62733800-INS-6638 | chr7-62733801-INS-6639 |
| chr7-62735335-INS-740 | chr7-62735336-INS-741 |
| chr7-62764383-INS-212 | chr7-62764383-INS-212 |
| chr7-62854240-INS-6444 | chr7-62854241-INS-6446 |
| chr7-62958644-INS-509 | chr7-62958644-INS-509 |
| chr7-62979088-INS-82 | chr7-62979072-INS-94 |
| chr7-62991966-INS-102 | chr7-62991966-INS-80 |
| chr7-63017717-INS-2534 | chr7-63017711-INS-2534 |
| chr7-63029700-INS-3087 | chr7-63029701-INS-3087 |
| chr7-63039528-INS-7111 | chr7-63039529-INS-7111 |
| chr7-63073647-INS-179 | chr7-63073647-INS-179 |
| chr7-63078027-INS-155 | chr7-63078052-INS-112 |
| chr7-63089174-INS-68 | chr7-63089174-INS-65 |
| chr7-63119100-INS-60 | chr7-63119100-INS-60 |
| chr7-63128518-INS-218 | chr7-63128518-INS-218 |
| chr7-63135740-INS-245 | chr7-63135740-INS-245 |
| chr7-63143417-INS-66 | chr7-63143417-INS-66 |
| chr7-63149465-INS-136 | chr7-63149465-INS-138 |
| chr7-63160165-INS-6657 | chr7-63160166-INS-6659 |
| chr7-63178117-INS-155 | chr7-63178117-INS-155 |
| chr7-63196435-INS-6871 | chr7-63196436-INS-6872 |
| chr7-63200588-INS-1344 | chr7-63200589-INS-1344 |
| chr7-63208585-INS-533 | chr7-63208585-INS-534 |
| chr7-63218235-INS-168 | chr7-63218235-INS-168 |
| chr7-63237820-INS-480 | chr7-63237820-INS-481 |
| chr7-63241802-INS-7315 | chr7-63241803-INS-7315 |
| chr7-63247108-INS-765 | chr7-63247108-INS-764 |
| chr7-63262473-INS-60 | chr7-63262453-INS-57 |
| chr7-63270316-INS-593 | chr7-63270316-INS-593 |
| chr7-63273672-INS-66 | chr7-63273672-INS-66 |
| chr7-63277386-INS-4646 | chr7-63277387-INS-4646 |
| chr7-63310400-INS-178 | chr7-63310400-INS-178 |
| chr7-63389686-INS-244 | chr7-63389686-INS-246 |
| chr7-63395015-INS-67 | chr7-63395015-INS-67 |
| chr7-63421298-INS-75 | chr7-63421298-INS-75 |
| chr7-63435943-INS-180 | chr7-63435943-INS-188 |
| chr7-63437513-INS-75 | chr7-63437513-INS-75 |
| chr7-63469801-INS-96 | chr7-63469801-INS-92 |
| chr7-63498829-INS-205 | chr7-63498829-INS-205 |
| chr7-63523443-INS-72 | chr7-63523443-INS-54 |
| chr7-63583991-INS-304 | chr7-63584008-INS-306 |
| chr7-63596911-INS-52 | chr7-63596911-INS-50 |
| chr7-63613409-INS-208 | chr7-63613409-INS-208 |
| chr7-63616606-INS-124 | chr7-63616601-INS-135 |
| chr7-63617071-INS-261 | chr7-63617072-INS-261 |
| chr7-63629817-INS-76 | chr7-63629817-INS-57 |
| chr7-63630760-INS-160 | chr7-63630795-INS-131 |
| chr7-63631964-INS-59 | chr7-63631928-INS-84 |
| chr7-63653133-INS-51 | chr7-63653133-INS-54 |

| | |
|---|---|
| chr7-63654734-INS-180 | chr7-63654734-INS-181 |
| chr7-63659466-INS-519 | chr7-63659466-INS-519 |
| chr7-63668060-INS-189 | chr7-63668060-INS-188 |
| chr7-63677342-INS-51 | chr7-63677342-INS-51 |
| chr7-63681244-INS-144 | chr7-63681243-INS-105 |
| chr7-63686144-INS-120 | chr7-63686144-INS-120 |
| chr7-63703954-INS-119 | chr7-63703954-INS-119 |
| chr7-63707917-INS-413 | chr7-63707917-INS-413 |
| chr7-63708710-INS-130 | chr7-63708710-INS-130 |
| chr7-63718807-INS-89 | chr7-63718807-INS-91 |
| chr7-63720543-INS-56 | chr7-63720543-INS-56 |
| chr7-63789073-INS-118 | chr7-63789073-INS-118 |
| chr7-63797106-INS-55 | chr7-63797106-INS-55 |
| chr7-63799865-INS-53 | chr7-63799912-INS-50 |
| chr7-63806814-INS-206 | chr7-63806814-INS-206 |
| chr7-63814332-INS-182 | chr7-63814332-INS-182 |
| chr7-63819013-INS-230 | chr7-63819013-INS-242 |
| chr7-63853966-INS-70 | chr7-63853966-INS-61 |
| chr7-63855187-INS-558 | chr7-63855187-INS-558 |
| chr7-63855854-INS-84 | chr7-63855836-INS-56 |
| chr7-63892952-INS-72 | chr7-63892952-INS-67 |
| chr7-63899944-INS-354 | chr7-63899944-INS-354 |
| chr7-63937066-INS-242 | chr7-63937066-INS-238 |
| chr7-63941175-INS-67 | chr7-63941175-INS-59 |
| chr7-63949908-INS-72 | chr7-63949908-INS-72 |
| chr7-63969389-INS-78 | chr7-63969389-INS-82 |
| chr7-63984582-INS-338 | chr7-63984582-INS-338 |
| chr7-63991636-INS-63 | chr7-63991637-INS-54 |
| chr7-63993419-INS-175 | chr7-63993419-INS-175 |
| chr7-63995029-INS-77 | chr7-63995029-INS-77 |
| chr7-63997664-INS-157 | chr7-63997670-INS-157 |
| chr7-64001643-INS-236 | chr7-64001643-INS-228 |
| chr7-64014002-INS-51 | chr7-64014002-INS-51 |
| chr7-64016216-INS-176 | chr7-64016216-INS-176 |
| chr7-64021741-INS-92 | chr7-64021741-INS-98 |
| chr7-64028935-INS-224 | chr7-64028935-INS-225 |
| chr7-64052889-INS-114 | chr7-64052889-INS-115 |
| chr7-64059630-INS-673 | chr7-64059630-INS-673 |
| chr7-64065432-INS-52 | chr7-64065432-INS-52 |
| chr7-64092893-INS-645 | chr7-64092893-INS-644 |
| chr7-64097207-INS-7138 | chr7-64097208-INS-7138 |
| chr7-64153958-INS-51 | chr7-64153951-INS-72 |
| chr7-64158489-INS-78 | chr7-64158505-INS-86 |
| chr7-64169686-INS-167 | chr7-64169651-INS-162 |
| chr7-64184586-INS-123 | chr7-64184586-INS-124 |
| chr7-64184991-INS-943 | chr7-64184991-INS-943 |
| chr7-64187412-INS-102 | chr7-64187412-INS-60 |
| chr7-64193833-INS-112 | chr7-64193833-INS-112 |
| chr7-64239904-INS-62 | chr7-64239904-INS-62 |
| chr7-64246026-INS-194 | chr7-64246026-INS-198 |
| chr7-64279320-INS-99 | chr7-64279320-INS-99 |
| chr7-64307090-INS-6701 | chr7-64307091-INS-6700 |
| chr7-64331920-INS-401 | chr7-64331920-INS-401 |
| chr7-64354874-INS-69 | chr7-64354874-INS-65 |
| chr7-64361247-INS-54 | chr7-64361247-INS-52 |
| chr7-64390052-INS-78 | chr7-64390053-INS-78 |
| chr7-64492847-INS-504 | chr7-64492847-INS-490 |
| chr7-64509623-INS-542 | chr7-64509623-INS-550 |
| chr7-64525939-INS-211 | chr7-64525939-INS-193 |
| chr7-64553256-INS-67 | chr7-64553256-INS-79 |
| chr7-64563348-INS-101 | chr7-64563348-INS-105 |
| chr7-64637665-INS-107 | chr7-64637665-INS-107 |
| chr7-64642113-INS-903 | chr7-64642113-INS-903 |
| chr7-64649013-INS-400 | chr7-64649013-INS-399 |
| chr7-64666763-INS-54 | chr7-64666763-INS-54 |
| chr7-64728024-INS-84 | chr7-64728024-INS-84 |
| chr7-64776732-INS-53 | chr7-64776732-INS-57 |
| chr7-64778936-INS-73 | chr7-64778936-INS-73 |
| chr7-64789907-INS-277 | chr7-64789907-INS-279 |
| chr7-64795845-INS-56 | chr7-64795845-INS-76 |
| chr7-64809660-INS-56 | chr7-64809660-INS-64 |
| chr7-64810930-INS-66 | chr7-64810930-INS-72 |
| chr7-64819793-INS-82 | chr7-64819793-INS-52 |
| chr7-64824660-INS-400 | chr7-64824668-INS-399 |
| chr7-64829596-INS-68 | chr7-64829592-INS-72 |
| chr7-64837082-INS-204 | chr7-64837082-INS-204 |
| chr7-64874525-INS-358 | chr7-64874526-INS-358 |

| | |
|---|---|
| chr7-64895101-INS-788 | chr7-64895101-INS-788 |
| chr7-64914565-INS-6403 | chr7-64914566-INS-6403 |
| chr7-64919107-INS-51 | chr7-64919107-INS-50 |
| chr7-64926361-INS-207 | chr7-64926361-INS-207 |
| chr7-64934849-INS-953 | chr7-64934848-INS-960 |
| chr7-64948902-INS-206 | chr7-64948902-INS-206 |
| chr7-64982858-INS-68 | chr7-64982860-INS-65 |
| chr7-65030481-INS-197 | chr7-65030481-INS-199 |
| chr7-65050212-INS-221 | chr7-65050212-INS-222 |
| chr7-65054331-INS-180 | chr7-65054331-INS-180 |
| chr7-65059914-INS-162 | chr7-65059914-INS-160 |
| chr7-65078919-INS-207 | chr7-65078919-INS-207 |
| chr7-65082636-INS-175 | chr7-65082636-INS-176 |
| chr7-65101719-INS-217 | chr7-65101719-INS-217 |
| chr7-65135143-INS-211 | chr7-65135144-INS-212 |
| chr7-65144740-INS-84 | chr7-65144740-INS-84 |
| chr7-65164401-INS-57 | chr7-65164401-INS-69 |
| chr7-65186399-INS-57 | chr7-65186399-INS-54 |
| chr7-65195184-INS-103 | chr7-65195184-INS-56 |
| chr7-65223192-INS-94 | chr7-65223192-INS-65 |
| chr7-65242895-INS-51 | chr7-65242895-INS-55 |
| chr7-65246613-INS-62 | chr7-65246609-INS-59 |
| chr7-65248619-INS-51 | chr7-65248619-INS-51 |
| chr7-65256752-INS-63 | chr7-65256752-INS-63 |
| chr7-65279243-INS-77 | chr7-65279270-INS-93 |
| chr7-65286710-INS-68 | chr7-65286710-INS-68 |
| chr7-65298323-INS-364 | chr7-65298323-INS-364 |
| chr7-65299181-INS-145 | chr7-65299181-INS-145 |
| chr7-65302903-INS-95 | chr7-65302903-INS-95 |
| chr7-65313030-INS-88 | chr7-65313073-INS-88 |
| chr7-65345477-INS-751 | chr7-65345477-INS-776 |
| chr7-65349302-INS-56 | chr7-65349302-INS-56 |
| chr7-65354575-INS-177 | chr7-65354575-INS-177 |
| chr7-65358362-INS-957 | chr7-65358362-INS-969 |
| chr7-65365203-INS-52 | chr7-65365203-INS-52 |
| chr7-65365692-INS-734 | chr7-65365692-INS-734 |
| chr7-65378952-INS-113 | chr7-65378952-INS-91 |
| chr7-65391457-INS-88 | chr7-65391460-INS-64 |
| chr7-65407195-INS-112 | chr7-65407199-INS-119 |
| chr7-65414941-INS-92 | chr7-65414941-INS-100 |
| chr7-65417653-INS-90 | chr7-65417653-INS-90 |
| chr7-65428247-INS-202 | chr7-65428247-INS-202 |
| chr7-65437262-INS-52 | chr7-65437262-INS-50 |
| chr7-65438540-INS-210 | chr7-65438540-INS-211 |
| chr7-65451819-INS-227 | chr7-65451819-INS-227 |
| chr7-65478968-INS-195 | chr7-65478955-INS-204 |
| chr7-65482162-INS-61 | chr7-65482162-INS-52 |
| chr7-65491102-INS-100 | chr7-65491102-INS-106 |
| chr7-65492226-INS-968 | chr7-65492226-INS-968 |
| chr7-65497949-INS-91 | chr7-65497949-INS-97 |
| chr7-65508697-INS-116 | chr7-65508697-INS-116 |
| chr7-65516665-INS-7502 | chr7-65516666-INS-7502 |
| chr7-65527817-INS-69 | chr7-65527817-INS-51 |
| chr7-65540721-INS-211 | chr7-65540721-INS-213 |
| chr7-65543128-INS-5225 | chr7-65543129-INS-5224 |
| chr7-65547270-INS-203 | chr7-65547270-INS-203 |
| chr7-65561352-INS-168 | chr7-65561352-INS-156 |
| chr7-65583773-INS-222 | chr7-65583773-INS-222 |
| chr7-65589141-INS-87 | chr7-65589141-INS-87 |
| chr7-65600989-INS-58 | chr7-65600989-INS-58 |
| chr7-65606446-INS-62 | chr7-65606447-INS-62 |
| chr7-65610312-INS-58 | chr7-65610312-INS-53 |
| chr7-65617209-INS-117 | chr7-65617209-INS-133 |
| chr7-65618911-INS-195 | chr7-65618911-INS-195 |
| chr7-65627407-INS-86 | chr7-65627407-INS-86 |
| chr7-65636092-INS-54 | chr7-65636094-INS-54 |
| chr7-65664323-INS-77 | chr7-65664323-INS-77 |
| chr7-65665363-INS-213 | chr7-65665363-INS-213 |
| chr7-65678270-INS-51 | chr7-65678223-INS-56 |
| chr7-65691149-INS-550 | chr7-65691149-INS-550 |
| chr7-65692398-INS-892 | chr7-65692398-INS-892 |
| chr7-65704699-INS-195 | chr7-65704699-INS-195 |
| chr7-65713745-INS-254 | chr7-65713745-INS-254 |
| chr7-65741819-INS-58 | chr7-65741819-INS-58 |
| chr7-65756014-INS-93 | chr7-65756014-INS-95 |
| chr7-65784999-INS-199 | chr7-65784999-INS-201 |
| chr7-65794325-INS-94 | chr7-65794325-INS-94 |

| | |
|---|---|
| chr7-65795621-INS-739 | chr7-65795622-INS-739 |
| chr7-65805145-INS-56 | chr7-65805145-INS-56 |
| chr7-65815796-INS-187 | chr7-65815800-INS-176 |
| chr7-65817734-INS-152 | chr7-65817730-INS-156 |
| chr7-65819511-INS-179 | chr7-65819511-INS-177 |
| chr7-65840993-INS-107 | chr7-65840993-INS-107 |
| chr7-65842252-INS-137 | chr7-65842252-INS-137 |
| chr7-65878977-INS-128 | chr7-65878971-INS-130 |
| chr7-65881264-INS-62 | chr7-65881264-INS-62 |
| chr7-65890043-INS-170 | chr7-65890043-INS-171 |
| chr7-65895421-INS-800 | chr7-65895421-INS-836 |
| chr7-65897946-INS-259 | chr7-65897946-INS-259 |
| chr7-65914745-INS-130 | chr7-65914745-INS-130 |
| chr7-65926396-INS-261 | chr7-65926396-INS-261 |
| chr7-65941862-INS-74 | chr7-65941862-INS-74 |
| chr7-65950932-INS-57 | chr7-65950932-INS-57 |
| chr7-65957308-INS-73 | chr7-65957305-INS-77 |
| chr7-65959927-INS-571 | chr7-65959927-INS-571 |
| chr7-65963808-INS-603 | chr7-65963808-INS-604 |
| chr7-65969141-INS-116 | chr7-65969174-INS-124 |
| chr7-65971704-INS-263 | chr7-65971704-INS-253 |
| chr7-65983151-INS-448 | chr7-65983151-INS-446 |
| chr7-65989855-INS-64 | chr7-65989855-INS-64 |
| chr7-66010045-INS-227 | chr7-66010045-INS-231 |
| chr7-66024070-INS-80 | chr7-66024070-INS-102 |
| chr7-66058853-INS-60 | chr7-66058853-INS-60 |
| chr7-66065765-INS-7165 | chr7-66065766-INS-7165 |
| chr7-66089418-INS-135 | chr7-66089418-INS-135 |
| chr7-66133346-INS-90 | chr7-66133346-INS-58 |
| chr7-66135818-INS-50 | chr7-66135818-INS-52 |
| chr7-66162145-INS-1713 | chr7-66162146-INS-1713 |
| chr7-66163530-INS-56 | chr7-66163530-INS-56 |
| chr7-66184715-INS-54 | chr7-66184715-INS-54 |
| chr7-66219004-INS-75 | chr7-66219004-INS-76 |
| chr7-66247001-INS-899 | chr7-66247001-INS-899 |
| chr7-66255034-INS-314 | chr7-66255034-INS-314 |
| chr7-66269552-INS-68 | chr7-66269552-INS-66 |
| chr7-66274890-INS-886 | chr7-66274890-INS-886 |
| chr7-66281520-INS-88 | chr7-66281520-INS-88 |
| chr7-66282429-INS-2605 | chr7-66282430-INS-2605 |
| chr7-66283721-INS-78 | chr7-66283747-INS-78 |
| chr7-66356170-INS-56 | chr7-66356132-INS-87 |
| chr7-66356805-INS-1759 | chr7-66356806-INS-1769 |
| chr7-66419932-INS-833 | chr7-66419932-INS-831 |
| chr7-66483167-INS-1941 | chr7-66483168-INS-1940 |
| chr7-66484096-INS-111 | chr7-66484096-INS-111 |
| chr7-66506835-INS-62 | chr7-66506835-INS-62 |
| chr7-66527712-INS-487 | chr7-66527712-INS-488 |
| chr7-66535532-INS-80 | chr7-66535532-INS-81 |
| chr7-66542376-INS-60 | chr7-66542376-INS-80 |
| chr7-66546030-INS-65 | chr7-66546030-INS-74 |
| chr7-66548171-INS-60 | chr7-66548171-INS-60 |
| chr7-66554812-INS-6650 | chr7-66554813-INS-6649 |
| chr7-66563557-INS-96 | chr7-66563557-INS-96 |
| chr7-66600689-INS-6409 | chr7-66600690-INS-6415 |
| chr7-66620354-INS-80 | chr7-66620337-INS-108 |
| chr7-66653376-INS-60 | chr7-66653376-INS-56 |
| chr7-66682863-INS-155 | chr7-66682864-INS-152 |
| chr7-66691845-INS-2136 | chr7-66691846-INS-2118 |
| chr7-66695625-INS-280 | chr7-66695625-INS-326 |
| chr7-66709357-INS-72 | chr7-66709357-INS-84 |
| chr7-66715912-INS-106 | chr7-66715912-INS-114 |
| chr7-66734687-INS-80 | chr7-66734687-INS-54 |
| chr7-66744512-INS-267 | chr7-66744551-INS-267 |
| chr7-66756340-INS-80 | chr7-66756340-INS-76 |
| chr7-66834497-INS-132 | chr7-66834497-INS-132 |
| chr7-66845313-INS-342 | chr7-66845313-INS-353 |
| chr7-66849171-INS-53 | chr7-66849171-INS-59 |
| chr7-66853035-INS-401 | chr7-66853035-INS-401 |
| chr7-66854677-INS-346 | chr7-66854677-INS-351 |
| chr7-66909059-INS-523 | chr7-66909059-INS-523 |
| chr7-66913937-INS-82 | chr7-66913937-INS-82 |
| chr7-66923995-INS-212 | chr7-66923995-INS-212 |
| chr7-66970690-INS-52 | chr7-66970690-INS-52 |
| chr7-66985594-INS-6389 | chr7-66985595-INS-6389 |
| chr7-66989198-INS-3013 | chr7-66989199-INS-3014 |
| chr7-67075855-INS-66 | chr7-67075855-INS-54 |

| | |
|---|---|
| chr7-67079770-INS-60 | chr7-67079771-INS-88 |
| chr7-67084024-INS-77 | chr7-67084019-INS-52 |
| chr7-67135045-INS-491 | chr7-67135045-INS-447 |
| chr7-67144996-INS-114 | chr7-67144996-INS-120 |
| chr7-67162323-INS-249 | chr7-67162323-INS-248 |
| chr7-67180779-INS-51 | chr7-67180779-INS-51 |
| chr7-67184128-INS-289 | chr7-67184128-INS-283 |
| chr7-67281954-INS-209 | chr7-67281954-INS-207 |
| chr7-67289443-INS-213 | chr7-67289443-INS-195 |
| chr7-67291220-INS-122 | chr7-67291220-INS-108 |
| chr7-67316690-INS-207 | chr7-67316690-INS-207 |
| chr7-67327452-INS-324 | chr7-67327461-INS-346 |
| chr7-67408162-INS-100 | chr7-67408162-INS-50 |
| chr7-67417508-INS-327 | chr7-67417508-INS-327 |
| chr7-67426032-INS-72 | chr7-67425994-INS-87 |
| chr7-67429414-INS-83 | chr7-67429414-INS-83 |
| chr7-67463955-INS-62 | chr7-67463957-INS-76 |
| chr7-67597752-INS-101 | chr7-67597752-INS-101 |
| chr7-67656647-INS-4812 | chr7-67656648-INS-4812 |
| chr7-67717603-INS-350 | chr7-67717618-INS-351 |
| chr7-67726624-INS-276 | chr7-67726603-INS-253 |
| chr7-67737026-INS-84 | chr7-67737026-INS-86 |
| chr7-67756321-INS-122 | chr7-67756321-INS-122 |
| chr7-67834423-INS-788 | chr7-67834423-INS-788 |
| chr7-67890618-INS-5559 | chr7-67890617-INS-5561 |
| chr7-67898529-INS-256 | chr7-67898529-INS-257 |
| chr7-67902166-INS-5665 | chr7-67902167-INS-5665 |
| chr7-67905745-INS-54 | chr7-67905762-INS-60 |
| chr7-67909407-INS-7137 | chr7-67909408-INS-7137 |
| chr7-67915145-INS-54 | chr7-67915145-INS-54 |
| chr7-67921382-INS-56 | chr7-67921382-INS-56 |
| chr7-67923403-INS-183 | chr7-67923403-INS-186 |
| chr7-67934688-INS-329 | chr7-67934688-INS-329 |
| chr7-67937079-INS-68 | chr7-67937075-INS-76 |
| chr7-67937785-INS-7181 | chr7-67937786-INS-7181 |
| chr7-67958148-INS-383 | chr7-67958148-INS-383 |
| chr7-67983818-INS-1412 | chr7-67983819-INS-1412 |
| chr7-68004649-INS-178 | chr7-68004649-INS-175 |
| chr7-68008857-INS-66 | chr7-68008846-INS-68 |
| chr7-68013755-INS-124 | chr7-68013749-INS-145 |
| chr7-68022616-INS-297 | chr7-68022616-INS-297 |
| chr7-68031970-INS-55 | chr7-68031970-INS-56 |
| chr7-68033006-INS-260 | chr7-68033006-INS-268 |
| chr7-68044685-INS-56 | chr7-68044685-INS-54 |
| chr7-68067870-INS-266 | chr7-68067870-INS-266 |
| chr7-68076747-INS-229 | chr7-68076747-INS-229 |
| chr7-68077720-INS-94 | chr7-68077720-INS-130 |
| chr7-68083646-INS-410 | chr7-68083646-INS-410 |
| chr7-68087018-INS-58 | chr7-68087018-INS-104 |
| chr7-68110661-INS-358 | chr7-68110661-INS-358 |
| chr7-68111927-INS-554 | chr7-68111927-INS-554 |
| chr7-68145493-INS-2854 | chr7-68145494-INS-2847 |
| chr7-68155058-INS-860 | chr7-68155058-INS-861 |
| chr7-68156999-INS-1241 | chr7-68157000-INS-1241 |
| chr7-68169005-INS-54 | chr7-68169005-INS-73 |
| chr7-68177824-INS-51 | chr7-68177824-INS-51 |
| chr7-68184145-INS-54 | chr7-68184145-INS-54 |
| chr7-68187537-INS-64 | chr7-68187537-INS-64 |
| chr7-68190894-INS-379 | chr7-68190894-INS-383 |
| chr7-68199729-INS-89 | chr7-68199729-INS-89 |
| chr7-68204353-INS-86 | chr7-68204353-INS-92 |
| chr7-68206773-INS-116 | chr7-68206803-INS-80 |
| chr7-68216263-INS-198 | chr7-68216263-INS-198 |
| chr7-68218290-INS-6438 | chr7-68218291-INS-6442 |
| chr7-68231064-INS-91 | chr7-68231064-INS-91 |
| chr7-68235539-INS-216 | chr7-68235539-INS-239 |
| chr7-68250409-INS-6611 | chr7-68250410-INS-6611 |
| chr7-68254545-INS-80 | chr7-68254545-INS-82 |
| chr7-68307561-INS-192 | chr7-68307561-INS-193 |
| chr7-68310995-INS-115 | chr7-68310990-INS-123 |
| chr7-68401333-INS-1893 | chr7-68401334-INS-1893 |
| chr7-68414559-INS-490 | chr7-68414559-INS-490 |
| chr7-68420509-INS-76 | chr7-68420509-INS-76 |
| chr7-68438084-INS-53 | chr7-68438084-INS-89 |
| chr7-68445931-INS-78 | chr7-68445931-INS-72 |
| chr7-68450508-INS-54 | chr7-68450508-INS-50 |
| chr7-68458586-INS-85 | chr7-68458586-INS-85 |

| | |
|---|---|
| chr7-68473734-INS-70 | chr7-68473734-INS-70 |
| chr7-68485189-INS-68 | chr7-68485189-INS-70 |
| chr7-68498423-INS-74 | chr7-68498423-INS-60 |
| chr7-68507849-INS-193 | chr7-68507849-INS-193 |
| chr7-68535576-INS-212 | chr7-68535574-INS-182 |
| chr7-68542892-INS-669 | chr7-68542892-INS-670 |
| chr7-68548326-INS-595 | chr7-68548326-INS-595 |
| chr7-68554655-INS-310 | chr7-68554655-INS-310 |
| chr7-68574828-INS-320 | chr7-68574828-INS-323 |
| chr7-68616416-INS-7052 | chr7-68616417-INS-7052 |
| chr7-68617171-INS-92 | chr7-68617171-INS-52 |
| chr7-68671328-INS-80 | chr7-68671281-INS-84 |
| chr7-68682523-INS-806 | chr7-68682523-INS-806 |
| chr7-68702130-INS-6631 | chr7-68702131-INS-6632 |
| chr7-68714536-INS-198 | chr7-68714533-INS-198 |
| chr7-68730977-INS-1002 | chr7-68730978-INS-999 |
| chr7-68745068-INS-166 | chr7-68745068-INS-165 |
| chr7-68747161-INS-60 | chr7-68747161-INS-69 |
| chr7-68796909-INS-4523 | chr7-68796910-INS-4523 |
| chr7-68808320-INS-54 | chr7-68808321-INS-56 |
| chr7-68831768-INS-88 | chr7-68831768-INS-88 |
| chr7-68834107-INS-76 | chr7-68834107-INS-76 |
| chr7-68858083-INS-6403 | chr7-68858084-INS-6403 |
| chr7-68871646-INS-1919 | chr7-68871647-INS-1919 |
| chr7-68874448-INS-400 | chr7-68874448-INS-400 |
| chr7-68880094-INS-53 | chr7-68880094-INS-53 |
| chr7-68889177-INS-52 | chr7-68889177-INS-52 |
| chr7-68891600-INS-85 | chr7-68891593-INS-80 |
| chr7-68896806-INS-504 | chr7-68896806-INS-504 |
| chr7-68902956-INS-852 | chr7-68902956-INS-844 |
| chr7-68946922-INS-586 | chr7-68946922-INS-586 |
| chr7-68980400-INS-66 | chr7-68980400-INS-69 |
| chr7-68993618-INS-79 | chr7-68993631-INS-111 |
| chr7-68994834-INS-80 | chr7-68994834-INS-92 |
| chr7-69065187-INS-742 | chr7-69065187-INS-742 |
| chr7-69071307-INS-147 | chr7-69071307-INS-147 |
| chr7-69090575-INS-57 | chr7-69090575-INS-65 |
| chr7-69097899-INS-69 | chr7-69097899-INS-79 |
| chr7-69121211-INS-56 | chr7-69121211-INS-58 |
| chr7-69174350-INS-516 | chr7-69174350-INS-516 |
| chr7-69240574-INS-634 | chr7-69240574-INS-633 |
| chr7-69260935-INS-96 | chr7-69260935-INS-98 |
| chr7-69290462-INS-52 | chr7-69290462-INS-52 |
| chr7-69290965-INS-775 | chr7-69290965-INS-775 |
| chr7-69321954-INS-58 | chr7-69321954-INS-58 |
| chr7-69332204-INS-81 | chr7-69332204-INS-77 |
| chr7-69333129-INS-3443 | chr7-69333130-INS-3442 |
| chr7-69340404-INS-101 | chr7-69340407-INS-90 |
| chr7-69348792-INS-335 | chr7-69348792-INS-335 |
| chr7-69366826-INS-67 | chr7-69366826-INS-67 |
| chr7-69369898-INS-130 | chr7-69369894-INS-106 |
| chr7-69372729-INS-502 | chr7-69372729-INS-511 |
| chr7-69439241-INS-270 | chr7-69439241-INS-270 |
| chr7-69453437-INS-865 | chr7-69453437-INS-866 |
| chr7-69455178-INS-118 | chr7-69455179-INS-68 |
| chr7-69461983-INS-63 | chr7-69461983-INS-63 |
| chr7-69467410-INS-65 | chr7-69467410-INS-68 |
| chr7-69474690-INS-153 | chr7-69474690-INS-155 |
| chr7-69486407-INS-514 | chr7-69486407-INS-514 |
| chr7-69525018-INS-319 | chr7-69525018-INS-319 |
| chr7-69552489-INS-63 | chr7-69552489-INS-66 |
| chr7-69623304-INS-6697 | chr7-69623305-INS-6684 |
| chr7-69646828-INS-6421 | chr7-69646829-INS-6422 |
| chr7-69650465-INS-195 | chr7-69650477-INS-187 |
| chr7-69663334-INS-58 | chr7-69663334-INS-50 |
| chr7-69677966-INS-975 | chr7-69677966-INS-975 |
| chr7-69714442-INS-60 | chr7-69714442-INS-60 |
| chr7-69732447-INS-327 | chr7-69732447-INS-327 |
| chr7-69746849-INS-172 | chr7-69746849-INS-172 |
| chr7-69747875-INS-1741 | chr7-69747876-INS-1741 |
| chr7-69748679-INS-73 | chr7-69748696-INS-50 |
| chr7-69763206-INS-54 | chr7-69763206-INS-56 |
| chr7-69784963-INS-52 | chr7-69784963-INS-52 |
| chr7-69855537-INS-83 | chr7-69855537-INS-68 |
| chr7-69860186-INS-213 | chr7-69860186-INS-213 |
| chr7-69868445-INS-66 | chr7-69868445-INS-66 |
| chr7-69880311-INS-130 | chr7-69880348-INS-130 |

| | |
|---|---|
| chr7-69883924-INS-396 | chr7-69883924-INS-383 |
| chr7-69911046-INS-143 | chr7-69911046-INS-143 |
| chr7-69916517-INS-164 | chr7-69916517-INS-165 |
| chr7-69949597-INS-102 | chr7-69949598-INS-102 |
| chr7-69988794-INS-63 | chr7-69988822-INS-60 |
| chr7-70029219-INS-89 | chr7-70029173-INS-94 |
| chr7-70096803-INS-630 | chr7-70096803-INS-609 |
| chr7-70112519-INS-52 | chr7-70112517-INS-78 |
| chr7-70114171-INS-52 | chr7-70114171-INS-56 |
| chr7-70135075-INS-880 | chr7-70135075-INS-879 |
| chr7-70152137-INS-55 | chr7-70152137-INS-80 |
| chr7-70167321-INS-139 | chr7-70167321-INS-123 |
| chr7-70173708-INS-78 | chr7-70173710-INS-55 |
| chr7-70182292-INS-86 | chr7-70182304-INS-52 |
| chr7-70209297-INS-2787 | chr7-70209298-INS-2787 |
| chr7-70212481-INS-400 | chr7-70212481-INS-400 |
| chr7-70247702-INS-51 | chr7-70247702-INS-51 |
| chr7-70262755-INS-178 | chr7-70262755-INS-170 |
| chr7-70284568-INS-4091 | chr7-70284569-INS-4091 |
| chr7-70292080-INS-55 | chr7-70292081-INS-50 |
| chr7-70344691-INS-237 | chr7-70344691-INS-237 |
| chr7-70353772-INS-87 | chr7-70353772-INS-93 |
| chr7-70358913-INS-88 | chr7-70358913-INS-91 |
| chr7-70384048-INS-54 | chr7-70384048-INS-56 |
| chr7-70385850-INS-58 | chr7-70385850-INS-58 |
| chr7-70395518-INS-66 | chr7-70395517-INS-74 |
| chr7-70466781-INS-52 | chr7-70466781-INS-50 |
| chr7-70475577-INS-64 | chr7-70475577-INS-54 |
| chr7-70487666-INS-106 | chr7-70487666-INS-100 |
| chr7-70492560-INS-85 | chr7-70492560-INS-85 |
| chr7-70498918-INS-390 | chr7-70498918-INS-381 |
| chr7-70543226-INS-215 | chr7-70543226-INS-215 |
| chr7-70553784-INS-52 | chr7-70553752-INS-76 |
| chr7-70555552-INS-6789 | chr7-70555553-INS-6790 |
| chr7-70566666-INS-2229 | chr7-70566667-INS-2218 |
| chr7-70573393-INS-6359 | chr7-70573394-INS-6359 |
| chr7-70585867-INS-2946 | chr7-70585868-INS-2946 |
| chr7-70597569-INS-51 | chr7-70597569-INS-51 |
| chr7-70601815-INS-442 | chr7-70601815-INS-442 |
| chr7-70606714-INS-56 | chr7-70606715-INS-96 |
| chr7-70611675-INS-1888 | chr7-70611676-INS-1888 |
| chr7-70622708-INS-105 | chr7-70622708-INS-105 |
| chr7-70643388-INS-440 | chr7-70643388-INS-440 |
| chr7-70650888-INS-264 | chr7-70650888-INS-258 |
| chr7-70669617-INS-948 | chr7-70669617-INS-948 |
| chr7-70730393-INS-63 | chr7-70730393-INS-63 |
| chr7-70760925-INS-61 | chr7-70760925-INS-61 |
| chr7-70789393-INS-334 | chr7-70789393-INS-334 |
| chr7-70802835-INS-52 | chr7-70802835-INS-52 |
| chr7-70805932-INS-6463 | chr7-70805933-INS-6463 |
| chr7-70808230-INS-64 | chr7-70808230-INS-69 |
| chr7-70817726-INS-52 | chr7-70817726-INS-52 |
| chr7-70818922-INS-830 | chr7-70818922-INS-836 |
| chr7-70895466-INS-56 | chr7-70895466-INS-60 |
| chr7-70909141-INS-57 | chr7-70909141-INS-57 |
| chr7-70926018-INS-57 | chr7-70926018-INS-57 |
| chr7-70930681-INS-59 | chr7-70930693-INS-70 |
| chr7-70941295-INS-84 | chr7-70941295-INS-84 |
| chr7-70944877-INS-51 | chr7-70944874-INS-51 |
| chr7-70954556-INS-176 | chr7-70954556-INS-176 |
| chr7-71002023-INS-241 | chr7-71002023-INS-244 |
| chr7-71024479-INS-149 | chr7-71024479-INS-152 |
| chr7-71027410-INS-87 | chr7-71027407-INS-81 |
| chr7-71046912-INS-68 | chr7-71046913-INS-52 |
| chr7-71053758-INS-98 | chr7-71053758-INS-98 |
| chr7-71069923-INS-1054 | chr7-71069924-INS-1074 |
| chr7-71094860-INS-53 | chr7-71094860-INS-53 |
| chr7-71159350-INS-168 | chr7-71159350-INS-170 |
| chr7-71170813-INS-306 | chr7-71170813-INS-306 |
| chr7-71197680-INS-6605 | chr7-71197681-INS-6605 |
| chr7-71248581-INS-84 | chr7-71248581-INS-84 |
| chr7-71257220-INS-58 | chr7-71257220-INS-60 |
| chr7-71277413-INS-66 | chr7-71277413-INS-64 |
| chr7-71279425-INS-4331 | chr7-71279426-INS-4331 |
| chr7-71280949-INS-629 | chr7-71280949-INS-629 |
| chr7-71302001-INS-3928 | chr7-71302002-INS-3928 |
| chr7-71318047-INS-56 | chr7-71318047-INS-55 |

| | |
|---|---|
| chr7-71352220-INS-186 | chr7-71352220-INS-189 |
| chr7-71360317-INS-1203 | chr7-71360318-INS-1203 |
| chr7-71361373-INS-65 | chr7-71361377-INS-65 |
| chr7-71371541-INS-658 | chr7-71371541-INS-659 |
| chr7-71374048-INS-54 | chr7-71374048-INS-72 |
| chr7-71374973-INS-568 | chr7-71374973-INS-568 |
| chr7-71392281-INS-194 | chr7-71392281-INS-195 |
| chr7-71405938-INS-6381 | chr7-71405939-INS-6382 |
| chr7-71427669-INS-52 | chr7-71427669-INS-52 |
| chr7-71506436-INS-129 | chr7-71506388-INS-169 |
| chr7-71507793-INS-60 | chr7-71507793-INS-57 |
| chr7-71509810-INS-117 | chr7-71509810-INS-117 |
| chr7-71525551-INS-232 | chr7-71525551-INS-232 |
| chr7-71528163-INS-88 | chr7-71528163-INS-52 |
| chr7-71570991-INS-62 | chr7-71570991-INS-50 |
| chr7-71601007-INS-7237 | chr7-71601008-INS-7237 |
| chr7-71612034-INS-132 | chr7-71612034-INS-136 |
| chr7-71625530-INS-52 | chr7-71625530-INS-88 |
| chr7-71634036-INS-208 | chr7-71634036-INS-208 |
| chr7-71634609-INS-116 | chr7-71634609-INS-124 |
| chr7-71647493-INS-269 | chr7-71647481-INS-240 |
| chr7-71649297-INS-1358 | chr7-71649298-INS-1360 |
| chr7-71652189-INS-193 | chr7-71652188-INS-192 |
| chr7-71664488-INS-436 | chr7-71664488-INS-436 |
| chr7-71678028-INS-127 | chr7-71678028-INS-127 |
| chr7-71682975-INS-796 | chr7-71682975-INS-796 |
| chr7-71695817-INS-161 | chr7-71695817-INS-161 |
| chr7-71699115-INS-62 | chr7-71699115-INS-62 |
| chr7-71723025-INS-191 | chr7-71723025-INS-192 |
| chr7-71738182-INS-130 | chr7-71738182-INS-99 |
| chr7-71744077-INS-74 | chr7-71744095-INS-78 |
| chr7-71751761-INS-265 | chr7-71751761-INS-242 |
| chr7-71756762-INS-7075 | chr7-71756763-INS-7075 |
| chr7-71805385-INS-68 | chr7-71805385-INS-68 |
| chr7-71813807-INS-60 | chr7-71813807-INS-72 |
| chr7-71872906-INS-54 | chr7-71872906-INS-54 |
| chr7-71886149-INS-58 | chr7-71886168-INS-67 |
| chr7-71888933-INS-206 | chr7-71888933-INS-205 |
| chr7-71902784-INS-358 | chr7-71902784-INS-360 |
| chr7-71907791-INS-6270 | chr7-71907792-INS-6270 |
| chr7-71936197-INS-65 | chr7-71936197-INS-65 |
| chr7-71937644-INS-125 | chr7-71937644-INS-126 |
| chr7-71961466-INS-72 | chr7-71961466-INS-72 |
| chr7-71966018-INS-106 | chr7-71966018-INS-108 |
| chr7-71971352-INS-1785 | chr7-71971353-INS-1785 |
| chr7-71980375-INS-153 | chr7-71980384-INS-117 |
| chr7-71999314-INS-55 | chr7-71999290-INS-52 |
| chr7-72000602-INS-822 | chr7-72000603-INS-822 |
| chr7-72003961-INS-951 | chr7-72003961-INS-951 |
| chr7-72016921-INS-50 | chr7-72016904-INS-74 |
| chr7-72039938-INS-1797 | chr7-72039939-INS-1801 |
| chr7-72052956-INS-101 | chr7-72052957-INS-101 |
| chr7-72062293-INS-239 | chr7-72062293-INS-238 |
| chr7-72080883-INS-79 | chr7-72080881-INS-76 |
| chr7-72100958-INS-1003 | chr7-72100959-INS-1003 |
| chr7-72118021-INS-669 | chr7-72118021-INS-669 |
| chr7-72123858-INS-2842 | chr7-72123859-INS-2842 |
| chr7-72181887-INS-477 | chr7-72181878-INS-496 |
| chr7-72190525-INS-84 | chr7-72190525-INS-90 |
| chr7-72205044-INS-4875 | chr7-72205042-INS-4877 |
| chr7-72215682-INS-819 | chr7-72215682-INS-810 |
| chr7-72222987-INS-54 | chr7-72222987-INS-70 |
| chr7-72273137-INS-735 | chr7-72273137-INS-736 |
| chr7-72283580-INS-758 | chr7-72283580-INS-760 |
| chr7-72319651-INS-51 | chr7-72319651-INS-51 |
| chr7-72321305-INS-268 | chr7-72321305-INS-284 |
| chr7-72325286-INS-386 | chr7-72325286-INS-389 |
| chr7-72330637-INS-984 | chr7-72330637-INS-984 |
| chr7-72339358-INS-110 | chr7-72339358-INS-66 |
| chr7-72344466-INS-2483 | chr7-72344467-INS-2483 |
| chr7-72380907-INS-1622 | chr7-72380908-INS-1621 |
| chr7-72382568-INS-318 | chr7-72382568-INS-342 |
| chr7-72399130-INS-7117 | chr7-72399130-INS-7118 |
| chr7-72400166-INS-163 | chr7-72400164-INS-165 |
| chr7-72404889-INS-92 | chr7-72404889-INS-92 |
| chr7-72430176-INS-122 | chr7-72430176-INS-122 |
| chr7-72437826-INS-74 | chr7-72437826-INS-72 |

| | |
|---|---|
| chr7-72442414-INS-59 | chr7-72442414-INS-59 |
| chr7-72443928-INS-259 | chr7-72443928-INS-258 |
| chr7-72460797-INS-399 | chr7-72460797-INS-399 |
| chr7-72502129-INS-433 | chr7-72502129-INS-433 |
| chr7-72514746-INS-56 | chr7-72514746-INS-56 |
| chr7-72544571-INS-132 | chr7-72544534-INS-150 |
| chr7-72556215-INS-54 | chr7-72556214-INS-103 |
| chr7-72564792-INS-97 | chr7-72564792-INS-89 |
| chr7-72589630-INS-441 | chr7-72589630-INS-448 |
| chr7-72596997-INS-82 | chr7-72596997-INS-104 |
| chr7-72626097-INS-1073 | chr7-72626098-INS-1071 |
| chr7-72678712-INS-60 | chr7-72678712-INS-62 |
| chr7-72721893-INS-60 | chr7-72721893-INS-60 |
| chr7-72740551-INS-90 | chr7-72740551-INS-90 |
| chr7-72785329-INS-107 | chr7-72785329-INS-80 |
| chr7-72793520-INS-54 | chr7-72793520-INS-54 |
| chr7-72799838-INS-74 | chr7-72799838-INS-74 |
| chr7-72805070-INS-204 | chr7-72805070-INS-220 |
| chr7-72859097-INS-73 | chr7-72859097-INS-73 |
| chr7-72861886-INS-628 | chr7-72861886-INS-628 |
| chr7-72872710-INS-106 | chr7-72872684-INS-120 |
| chr7-72879406-INS-205 | chr7-72879406-INS-220 |
| chr7-72886041-INS-5217 | chr7-72886042-INS-5216 |
| chr7-72888842-INS-174 | chr7-72888842-INS-177 |
| chr7-72888842-INS-742 | chr7-72888842-INS-704 |
| chr7-72898285-INS-96 | chr7-72898285-INS-96 |
| chr7-72908255-INS-801 | chr7-72908255-INS-801 |
| chr7-72919030-INS-71 | chr7-72919030-INS-71 |
| chr7-72945694-INS-219 | chr7-72945694-INS-220 |
| chr7-73002556-INS-60 | chr7-73002525-INS-86 |
| chr7-73123374-INS-247 | chr7-73123374-INS-253 |
| chr7-73162006-INS-208 | chr7-73162006-INS-209 |
| chr7-73173310-INS-52 | chr7-73173310-INS-60 |
| chr7-73177308-INS-207 | chr7-73177308-INS-210 |
| chr7-73181347-INS-54 | chr7-73181347-INS-54 |
| chr7-73186265-INS-193 | chr7-73186265-INS-194 |
| chr7-73212066-INS-52 | chr7-73212086-INS-52 |
| chr7-73213980-INS-208 | chr7-73213980-INS-209 |
| chr7-73226748-INS-514 | chr7-73226748-INS-508 |
| chr7-73228052-INS-72 | chr7-73228052-INS-72 |
| chr7-73229245-INS-201 | chr7-73229245-INS-201 |
| chr7-73254651-INS-52 | chr7-73254651-INS-54 |
| chr7-73260591-INS-193 | chr7-73260604-INS-200 |
| chr7-73261978-INS-356 | chr7-73261978-INS-359 |
| chr7-73272798-INS-54 | chr7-73272778-INS-54 |
| chr7-73273620-INS-180 | chr7-73273620-INS-177 |
| chr7-73275842-INS-157 | chr7-73275842-INS-127 |
| chr7-73284152-INS-226 | chr7-73284152-INS-226 |
| chr7-73288248-INS-66 | chr7-73288248-INS-70 |
| chr7-73291482-INS-230 | chr7-73291482-INS-235 |
| chr7-73301580-INS-400 | chr7-73301580-INS-400 |
| chr7-73303802-INS-6492 | chr7-73303803-INS-6490 |
| chr7-73311527-INS-325 | chr7-73311527-INS-325 |
| chr7-73313335-INS-679 | chr7-73313335-INS-679 |
| chr7-73348145-INS-214 | chr7-73348145-INS-214 |
| chr7-73395592-INS-51 | chr7-73395592-INS-51 |
| chr7-73409363-INS-517 | chr7-73409363-INS-517 |
| chr7-73415689-INS-52 | chr7-73415697-INS-82 |
| chr7-73428285-INS-311 | chr7-73428285-INS-311 |
| chr7-73490897-INS-163 | chr7-73490901-INS-193 |
| chr7-73509878-INS-649 | chr7-73509878-INS-650 |
| chr7-73522669-INS-84 | chr7-73522669-INS-100 |
| chr7-73523829-INS-64 | chr7-73523829-INS-62 |
| chr7-73525108-INS-1968 | chr7-73525109-INS-1968 |
| chr7-73539904-INS-3703 | chr7-73539905-INS-3703 |
| chr7-73564918-INS-6455 | chr7-73564919-INS-6458 |
| chr7-73603223-INS-52 | chr7-73603223-INS-52 |
| chr7-73607046-INS-140 | chr7-73607046-INS-140 |
| chr7-73646159-INS-110 | chr7-73646159-INS-110 |
| chr7-73650286-INS-6667 | chr7-73650287-INS-6667 |
| chr7-73660397-INS-127 | chr7-73660391-INS-117 |
| chr7-73721974-INS-89 | chr7-73721974-INS-84 |
| chr7-73725359-INS-186 | chr7-73725359-INS-186 |
| chr7-73735921-INS-297 | chr7-73735921-INS-297 |
| chr7-73743452-INS-52 | chr7-73743452-INS-52 |
| chr7-73746177-INS-6506 | chr7-73746178-INS-6506 |
| chr7-73746696-INS-96 | chr7-73746680-INS-120 |

| | |
|---|---|
| chr7-73759721-INS-125 | chr7-73759710-INS-120 |
| chr7-73777445-INS-6590 | chr7-73777446-INS-6590 |
| chr7-73814258-INS-6668 | chr7-73814259-INS-6651 |
| chr7-73826845-INS-54 | chr7-73826845-INS-54 |
| chr7-73846933-INS-147 | chr7-73846933-INS-147 |
| chr7-73874297-INS-391 | chr7-73874297-INS-388 |
| chr7-73894856-INS-720 | chr7-73894856-INS-720 |
| chr7-73921614-INS-52 | chr7-73921600-INS-80 |
| chr7-73983358-INS-6702 | chr7-73983359-INS-6702 |
| chr7-73984140-INS-59 | chr7-73984140-INS-59 |
| chr7-73995849-INS-226 | chr7-73995893-INS-224 |
| chr7-74004929-INS-58 | chr7-74004929-INS-58 |
| chr7-74052105-INS-376 | chr7-74052105-INS-391 |
| chr7-74071093-INS-122 | chr7-74071093-INS-122 |
| chr7-74072907-INS-88 | chr7-74072907-INS-88 |
| chr7-74082221-INS-6454 | chr7-74082222-INS-6454 |
| chr7-74092033-INS-144 | chr7-74092033-INS-144 |
| chr7-74095632-INS-928 | chr7-74095632-INS-922 |
| chr7-74097931-INS-70 | chr7-74097920-INS-76 |
| chr7-74110479-INS-1666 | chr7-74110480-INS-1666 |
| chr7-74112743-INS-85 | chr7-74112714-INS-80 |
| chr7-74129079-INS-60 | chr7-74129079-INS-60 |
| chr7-74132236-INS-96 | chr7-74132236-INS-86 |
| chr7-74136207-INS-52 | chr7-74136207-INS-54 |
| chr7-74152578-INS-186 | chr7-74152578-INS-186 |
| chr7-74177152-INS-65 | chr7-74177152-INS-65 |
| chr7-74178846-INS-5685 | chr7-74178847-INS-5679 |
| chr7-74198208-INS-118 | chr7-74198220-INS-96 |
| chr7-74234923-INS-59 | chr7-74234922-INS-66 |
| chr7-74246938-INS-443 | chr7-74246939-INS-445 |
| chr7-74261964-INS-738 | chr7-74261964-INS-738 |
| chr7-74291877-INS-84 | chr7-74291845-INS-96 |
| chr7-74297595-INS-78 | chr7-74297595-INS-110 |
| chr7-74308974-INS-56 | chr7-74308974-INS-58 |
| chr7-74319001-INS-7570 | chr7-74319002-INS-7563 |
| chr7-74319613-INS-539 | chr7-74319613-INS-539 |
| chr7-74330293-INS-865 | chr7-74330293-INS-863 |
| chr7-74336633-INS-89 | chr7-74336633-INS-90 |
| chr7-74342293-INS-528 | chr7-74342293-INS-528 |
| chr7-74348309-INS-54 | chr7-74348309-INS-54 |
| chr7-74354033-INS-70 | chr7-74354033-INS-70 |
| chr7-74419945-INS-6529 | chr7-74419946-INS-6529 |
| chr7-74423295-INS-128 | chr7-74423299-INS-129 |
| chr7-74475171-INS-198 | chr7-74475175-INS-202 |
| chr7-74477160-INS-152 | chr7-74477160-INS-152 |
| chr7-74502602-INS-72 | chr7-74502582-INS-70 |
| chr7-74520368-INS-6434 | chr7-74520369-INS-6434 |
| chr7-74528260-INS-173 | chr7-74528260-INS-173 |
| chr7-74537432-INS-7144 | chr7-74537433-INS-7144 |
| chr7-74550565-INS-194 | chr7-74550541-INS-218 |
| chr7-74598009-INS-60 | chr7-74598009-INS-56 |
| chr7-74614963-INS-103 | chr7-74614963-INS-103 |
| chr7-74644207-INS-292 | chr7-74644208-INS-288 |
| chr7-74644530-INS-822 | chr7-74644532-INS-842 |
| chr7-74673020-INS-90 | chr7-74673020-INS-88 |
| chr7-74685707-INS-2253 | chr7-74685708-INS-2253 |
| chr7-74692576-INS-54 | chr7-74692576-INS-52 |
| chr7-74700257-INS-538 | chr7-74700255-INS-542 |
| chr7-74740242-INS-84 | chr7-74740240-INS-84 |
| chr7-74772527-INS-176 | chr7-74772517-INS-182 |
| chr7-74797222-INS-141 | chr7-74797214-INS-132 |
| chr7-74807985-INS-790 | chr7-74807985-INS-790 |
| chr7-74815043-INS-75 | chr7-74815043-INS-75 |
| chr7-74821777-INS-58 | chr7-74821771-INS-58 |
| chr7-74824485-INS-60 | chr7-74824485-INS-64 |
| chr7-74859501-INS-273 | chr7-74859501-INS-273 |
| chr7-74879577-INS-3529 | chr7-74879578-INS-3531 |
| chr7-74882016-INS-72 | chr7-74882002-INS-50 |
| chr7-74884194-INS-103 | chr7-74884195-INS-95 |
| chr7-74889422-INS-57 | chr7-74889422-INS-57 |
| chr7-74907823-INS-72 | chr7-74907825-INS-90 |
| chr7-74914283-INS-321 | chr7-74914283-INS-320 |
| chr7-74927483-INS-50 | chr7-74927483-INS-94 |
| chr7-74931257-INS-92 | chr7-74931257-INS-120 |
| chr7-74947984-INS-882 | chr7-74947984-INS-881 |
| chr7-74949484-INS-296 | chr7-74949484-INS-298 |
| chr7-74959786-INS-6471 | chr7-74959787-INS-6475 |

| | |
|---|---|
| chr7-74960318-INS-701 | chr7-74960318-INS-710 |
| chr7-74999728-INS-76 | chr7-74999716-INS-107 |
| chr7-75000954-INS-56 | chr7-75000954-INS-58 |
| chr7-75004108-INS-80 | chr7-75004108-INS-50 |
| chr7-75006221-INS-70 | chr7-75006221-INS-68 |
| chr7-75007289-INS-61 | chr7-75007289-INS-56 |
| chr7-75019645-INS-618 | chr7-75019645-INS-620 |
| chr7-75025517-INS-52 | chr7-75025517-INS-54 |
| chr7-75026954-INS-66 | chr7-75026954-INS-66 |
| chr7-75036666-INS-62 | chr7-75036671-INS-70 |
| chr7-75048516-INS-211 | chr7-75048516-INS-228 |
| chr7-75067056-INS-51 | chr7-75067056-INS-66 |
| chr7-75074773-INS-54 | chr7-75074773-INS-54 |
| chr7-75080190-INS-52 | chr7-75080190-INS-64 |
| chr7-75116835-INS-705 | chr7-75116835-INS-706 |
| chr7-75122690-INS-65 | chr7-75122690-INS-66 |
| chr7-75156086-INS-154 | chr7-75156086-INS-193 |
| chr7-75164056-INS-93 | chr7-75164056-INS-71 |
| chr7-75168065-INS-55 | chr7-75168065-INS-57 |
| chr7-75228281-INS-150 | chr7-75228309-INS-100 |
| chr7-75243789-INS-125 | chr7-75243829-INS-118 |
| chr7-75280904-INS-69 | chr7-75280865-INS-114 |
| chr7-75301291-INS-6886 | chr7-75301292-INS-6887 |
| chr7-75309011-INS-247 | chr7-75309021-INS-252 |
| chr7-75312298-INS-182 | chr7-75312298-INS-182 |
| chr7-75405337-INS-210 | chr7-75405338-INS-212 |
| chr7-75429883-INS-96 | chr7-75429883-INS-96 |
| chr7-75431097-INS-121 | chr7-75431097-INS-130 |
| chr7-75437516-INS-80 | chr7-75437516-INS-80 |
| chr7-75484930-INS-178 | chr7-75484930-INS-167 |
| chr7-75490166-INS-677 | chr7-75490166-INS-670 |
| chr7-75509970-INS-2572 | chr7-75509971-INS-2572 |
| chr7-75517332-INS-229 | chr7-75517332-INS-229 |
| chr7-75529053-INS-6012 | chr7-75529054-INS-6012 |
| chr7-75546472-INS-268 | chr7-75546472-INS-268 |
| chr7-75559999-INS-117 | chr7-75559999-INS-102 |
| chr7-75568962-INS-157 | chr7-75568962-INS-157 |
| chr7-75589925-INS-60 | chr7-75589925-INS-60 |
| chr7-75594541-INS-261 | chr7-75594541-INS-260 |
| chr7-75604707-INS-58 | chr7-75604723-INS-76 |
| chr7-75631366-INS-282 | chr7-75631366-INS-282 |
| chr7-75647502-INS-80 | chr7-75647502-INS-80 |
| chr7-75651293-INS-215 | chr7-75651293-INS-215 |
| chr7-75683146-INS-202 | chr7-75683146-INS-202 |
| chr7-75695707-INS-6462 | chr7-75695708-INS-6462 |
| chr7-75697086-INS-1246 | chr7-75697087-INS-1246 |
| chr7-75729988-INS-338 | chr7-75729958-INS-377 |
| chr7-75736152-INS-102 | chr7-75736152-INS-104 |
| chr7-75737154-INS-200 | chr7-75737154-INS-200 |
| chr7-75740898-INS-6508 | chr7-75740899-INS-6510 |
| chr7-75745456-INS-111 | chr7-75745456-INS-111 |
| chr7-75752374-INS-213 | chr7-75752374-INS-213 |
| chr7-75783571-INS-62 | chr7-75783571-INS-66 |
| chr7-75799245-INS-92 | chr7-75799260-INS-92 |
| chr7-75811142-INS-102 | chr7-75811167-INS-53 |
| chr7-75840189-INS-72 | chr7-75840189-INS-104 |
| chr7-75860273-INS-65 | chr7-75860273-INS-59 |
| chr7-75860777-INS-78 | chr7-75860777-INS-81 |
| chr7-75884032-INS-68 | chr7-75884032-INS-68 |
| chr7-75924316-INS-181 | chr7-75924316-INS-187 |
| chr7-75945516-INS-97 | chr7-75945517-INS-132 |
| chr7-76030871-INS-88 | chr7-76030871-INS-88 |
| chr7-76050620-INS-59 | chr7-76050620-INS-59 |
| chr7-76065315-INS-403 | chr7-76065315-INS-404 |
| chr7-76066573-INS-482 | chr7-76066573-INS-482 |
| chr7-76086672-INS-7172 | chr7-76086673-INS-7172 |
| chr7-76089635-INS-129 | chr7-76089635-INS-135 |
| chr7-76105055-INS-993 | chr7-76105055-INS-993 |
| chr7-76116079-INS-120 | chr7-76116073-INS-126 |
| chr7-76128349-INS-109 | chr7-76128370-INS-118 |
| chr7-76135734-INS-63 | chr7-76135734-INS-55 |
| chr7-76173835-INS-68 | chr7-76173835-INS-58 |
| chr7-76177448-INS-418 | chr7-76177449-INS-403 |
| chr7-76251587-INS-252 | chr7-76251587-INS-252 |
| chr7-76275563-INS-400 | chr7-76275563-INS-400 |
| chr7-76320598-INS-102 | chr7-76320598-INS-103 |
| chr7-76335389-INS-85 | chr7-76335389-INS-85 |

| | |
|---|---|
| chr7-76336724-INS-57 | chr7-76336724-INS-62 |
| chr7-76346097-INS-574 | chr7-76346097-INS-574 |
| chr7-76390080-INS-396 | chr7-76390080-INS-396 |
| chr7-76409156-INS-68 | chr7-76409156-INS-68 |
| chr7-76446897-INS-105 | chr7-76446897-INS-103 |
| chr7-76453371-INS-2301 | chr7-76453372-INS-2301 |
| chr7-76491377-INS-55 | chr7-76491377-INS-55 |
| chr7-76493985-INS-86 | chr7-76493985-INS-86 |
| chr7-76500356-INS-51 | chr7-76500351-INS-63 |
| chr7-76513304-INS-5551 | chr7-76513305-INS-5551 |
| chr7-76535442-INS-168 | chr7-76535442-INS-168 |
| chr7-76545447-INS-1233 | chr7-76545448-INS-1234 |
| chr7-76551548-INS-60 | chr7-76551548-INS-60 |
| chr7-76554102-INS-7559 | chr7-76554103-INS-7560 |
| chr7-76561845-INS-257 | chr7-76561845-INS-257 |
| chr7-76563751-INS-109 | chr7-76563751-INS-109 |
| chr7-76567076-INS-75 | chr7-76567076-INS-75 |
| chr7-76575838-INS-6869 | chr7-76575839-INS-6869 |
| chr7-76587484-INS-993 | chr7-76587484-INS-993 |
| chr7-76607404-INS-7572 | chr7-76607405-INS-7574 |
| chr7-76613013-INS-1939 | chr7-76613014-INS-1939 |
| chr7-76678627-INS-80 | chr7-76678635-INS-70 |
| chr7-76731095-INS-5339 | chr7-76731096-INS-5339 |
| chr7-76732696-INS-100 | chr7-76732696-INS-100 |
| chr7-76738740-INS-3273 | chr7-76738741-INS-3274 |
| chr7-76757836-INS-1720 | chr7-76757837-INS-1720 |
| chr7-76765433-INS-52 | chr7-76765433-INS-52 |
| chr7-76784981-INS-733 | chr7-76784981-INS-733 |
| chr7-76870360-INS-278 | chr7-76870360-INS-278 |
| chr7-76878645-INS-60 | chr7-76878645-INS-56 |
| chr7-76896265-INS-66 | chr7-76896265-INS-66 |
| chr7-76914195-INS-78 | chr7-76914164-INS-60 |
| chr7-76962922-INS-1833 | chr7-76962923-INS-1833 |
| chr7-76967261-INS-54 | chr7-76967261-INS-54 |
| chr7-77007687-INS-6501 | chr7-77007688-INS-6501 |
| chr7-77017819-INS-126 | chr7-77017832-INS-111 |
| chr7-77038283-INS-64 | chr7-77038298-INS-63 |
| chr7-77044218-INS-155 | chr7-77044218-INS-158 |
| chr7-77048046-INS-124 | chr7-77048050-INS-161 |
| chr7-77057341-INS-458 | chr7-77057341-INS-458 |
| chr7-77064829-INS-6749 | chr7-77064830-INS-6750 |
| chr7-77096793-INS-460 | chr7-77096793-INS-455 |
| chr7-77133871-INS-575 | chr7-77133871-INS-575 |
| chr7-77137882-INS-533 | chr7-77137882-INS-540 |
| chr7-77142891-INS-203 | chr7-77142891-INS-203 |
| chr7-77189692-INS-124 | chr7-77189687-INS-116 |
| chr7-77190927-INS-81 | chr7-77190925-INS-66 |
| chr7-77207410-INS-122 | chr7-77207423-INS-84 |
| chr7-77224869-INS-681 | chr7-77224869-INS-681 |
| chr7-77232415-INS-56 | chr7-77232377-INS-76 |
| chr7-77241331-INS-279 | chr7-77241331-INS-279 |
| chr7-77252744-INS-137 | chr7-77252744-INS-180 |
| chr7-77304821-INS-636 | chr7-77304822-INS-636 |
| chr7-77377056-INS-56 | chr7-77377020-INS-104 |
| chr7-77381019-INS-6198 | chr7-77381020-INS-6232 |
| chr7-77390137-INS-135 | chr7-77390137-INS-135 |
| chr7-77392639-INS-7487 | chr7-77392640-INS-7487 |
| chr7-77398504-INS-136 | chr7-77398508-INS-152 |
| chr7-77408696-INS-218 | chr7-77408696-INS-217 |
| chr7-77432045-INS-105 | chr7-77432045-INS-105 |
| chr7-77440640-INS-235 | chr7-77440640-INS-263 |
| chr7-77467528-INS-110 | chr7-77467528-INS-118 |
| chr7-77477741-INS-169 | chr7-77477741-INS-169 |
| chr7-77487419-INS-1899 | chr7-77487420-INS-1899 |
| chr7-77490009-INS-6315 | chr7-77490010-INS-6315 |
| chr7-77502088-INS-94 | chr7-77502086-INS-86 |
| chr7-77516651-INS-91 | chr7-77516669-INS-51 |
| chr7-77517904-INS-230 | chr7-77517904-INS-230 |
| chr7-77550571-INS-127 | chr7-77550581-INS-133 |
| chr7-77552808-INS-484 | chr7-77552808-INS-484 |
| chr7-77571828-INS-344 | chr7-77571828-INS-346 |
| chr7-77586481-INS-65 | chr7-77586481-INS-75 |
| chr7-77593205-INS-920 | chr7-77593205-INS-920 |
| chr7-77610599-INS-1193 | chr7-77610600-INS-1199 |
| chr7-77627729-INS-52 | chr7-77627697-INS-80 |
| chr7-77629998-INS-231 | chr7-77629998-INS-231 |
| chr7-77630531-INS-55 | chr7-77630531-INS-55 |

| | |
|---|---|
| chr7-77644004-INS-361 | chr7-77644004-INS-361 |
| chr7-77658039-INS-101 | chr7-77658039-INS-89 |
| chr7-77667129-INS-6213 | chr7-77667130-INS-6213 |
| chr7-77678834-INS-5311 | chr7-77678835-INS-5311 |
| chr7-77682312-INS-989 | chr7-77682312-INS-989 |
| chr7-77734418-INS-465 | chr7-77734418-INS-447 |
| chr7-77741092-INS-1269 | chr7-77741093-INS-1269 |
| chr7-77760488-INS-4037 | chr7-77760489-INS-4036 |
| chr7-77797021-INS-307 | chr7-77797021-INS-307 |
| chr7-77818664-INS-1915 | chr7-77818665-INS-1913 |
| chr7-77827556-INS-1651 | chr7-77827557-INS-1653 |
| chr7-77844733-INS-427 | chr7-77844733-INS-443 |
| chr7-77846878-INS-54 | chr7-77846847-INS-64 |
| chr7-77861861-INS-224 | chr7-77861861-INS-225 |
| chr7-77882263-INS-66 | chr7-77882263-INS-66 |
| chr7-77898820-INS-195 | chr7-77898820-INS-195 |
| chr7-77915000-INS-96 | chr7-77915000-INS-138 |
| chr7-77982791-INS-55 | chr7-77982791-INS-55 |
| chr7-77983093-INS-6895 | chr7-77983094-INS-6895 |
| chr7-78027026-INS-56 | chr7-78027026-INS-82 |
| chr7-78036345-INS-1866 | chr7-78036346-INS-1866 |
| chr7-78046546-INS-403 | chr7-78046546-INS-401 |
| chr7-78048335-INS-469 | chr7-78048335-INS-469 |
| chr7-78094181-INS-54 | chr7-78094181-INS-54 |
| chr7-78120869-INS-239 | chr7-78120869-INS-233 |
| chr7-78122462-INS-7315 | chr7-78122463-INS-7316 |
| chr7-78129178-INS-103 | chr7-78129178-INS-83 |
| chr7-78134029-INS-60 | chr7-78134043-INS-68 |
| chr7-78147725-INS-76 | chr7-78147725-INS-76 |
| chr7-78200592-INS-66 | chr7-78200592-INS-66 |
| chr7-78215130-INS-162 | chr7-78215130-INS-162 |
| chr7-78260726-INS-114 | chr7-78260729-INS-81 |
| chr7-78263564-INS-102 | chr7-78263564-INS-102 |
| chr7-78299148-INS-294 | chr7-78299148-INS-295 |
| chr7-78312765-INS-123 | chr7-78312745-INS-140 |
| chr7-78332025-INS-84 | chr7-78332025-INS-84 |
| chr7-78341645-INS-880 | chr7-78341645-INS-880 |
| chr7-78351929-INS-60 | chr7-78351929-INS-62 |
| chr7-78362560-INS-2406 | chr7-78362561-INS-2404 |
| chr7-78420916-INS-118 | chr7-78420916-INS-118 |
| chr7-78502032-INS-217 | chr7-78502032-INS-217 |
| chr7-78512148-INS-64 | chr7-78512159-INS-100 |
| chr7-78524111-INS-188 | chr7-78524111-INS-144 |
| chr7-78540658-INS-205 | chr7-78540658-INS-205 |
| chr7-78584198-INS-409 | chr7-78584198-INS-411 |
| chr7-78593306-INS-57 | chr7-78593306-INS-57 |
| chr7-78600621-INS-4352 | chr7-78600622-INS-4353 |
| chr7-78611027-INS-70 | chr7-78611027-INS-72 |
| chr7-78634048-INS-644 | chr7-78634048-INS-644 |
| chr7-78634050-INS-52 | chr7-78634050-INS-52 |
| chr7-78647676-INS-57 | chr7-78647677-INS-53 |
| chr7-78648300-INS-70 | chr7-78648300-INS-70 |
| chr7-78655164-INS-131 | chr7-78655164-INS-131 |
| chr7-78660048-INS-131 | chr7-78660055-INS-111 |
| chr7-78674477-INS-203 | chr7-78674477-INS-203 |
| chr7-78717101-INS-387 | chr7-78717101-INS-382 |
| chr7-78717908-INS-99 | chr7-78717907-INS-60 |
| chr7-78724038-INS-199 | chr7-78724041-INS-200 |
| chr7-78752779-INS-60 | chr7-78752779-INS-56 |
| chr7-78775871-INS-68 | chr7-78775871-INS-68 |
| chr7-78797995-INS-57 | chr7-78797995-INS-78 |
| chr7-78811158-INS-161 | chr7-78811159-INS-161 |
| chr7-78812687-INS-198 | chr7-78812687-INS-199 |
| chr7-78815211-INS-498 | chr7-78815211-INS-498 |
| chr7-78819056-INS-54 | chr7-78819056-INS-54 |
| chr7-78824456-INS-109 | chr7-78824456-INS-97 |
| chr7-78830916-INS-55 | chr7-78830916-INS-80 |
| chr7-78835239-INS-197 | chr7-78835240-INS-197 |
| chr7-78837550-INS-227 | chr7-78837550-INS-227 |
| chr7-78850232-INS-52 | chr7-78850232-INS-52 |
| chr7-78851510-INS-56 | chr7-78851510-INS-72 |
| chr7-78859841-INS-220 | chr7-78859841-INS-220 |
| chr7-78860580-INS-209 | chr7-78860580-INS-209 |
| chr7-78869987-INS-73 | chr7-78870028-INS-56 |
| chr7-78873492-INS-413 | chr7-78873492-INS-413 |
| chr7-78892539-INS-195 | chr7-78892539-INS-175 |
| chr7-78893500-INS-54 | chr7-78893500-INS-56 |

| | |
|---|---|
| chr7-78896779-INS-130 | chr7-78896765-INS-164 |
| chr7-78907593-INS-71 | chr7-78907593-INS-60 |
| chr7-78917410-INS-72 | chr7-78917411-INS-58 |
| chr7-78955614-INS-387 | chr7-78955614-INS-387 |
| chr7-79007401-INS-100 | chr7-79007397-INS-104 |
| chr7-79023817-INS-214 | chr7-79023818-INS-208 |
| chr7-79026088-INS-195 | chr7-79026088-INS-192 |
| chr7-79027602-INS-55 | chr7-79027602-INS-55 |
| chr7-79033075-INS-217 | chr7-79033075-INS-217 |
| chr7-79037614-INS-210 | chr7-79037614-INS-207 |
| chr7-79069473-INS-111 | chr7-79069473-INS-112 |
| chr7-79072130-INS-210 | chr7-79072130-INS-210 |
| chr7-79087629-INS-206 | chr7-79087629-INS-206 |
| chr7-79122282-INS-436 | chr7-79122282-INS-436 |
| chr7-79125128-INS-64 | chr7-79125128-INS-52 |
| chr7-79133037-INS-93 | chr7-79133037-INS-97 |
| chr7-79141702-INS-3023 | chr7-79141703-INS-3067 |
| chr7-79143429-INS-52 | chr7-79143429-INS-52 |
| chr7-79170658-INS-54 | chr7-79170658-INS-54 |
| chr7-79177176-INS-209 | chr7-79177176-INS-209 |
| chr7-79213889-INS-54 | chr7-79213891-INS-82 |
| chr7-79231866-INS-254 | chr7-79231866-INS-253 |
| chr7-79246700-INS-56 | chr7-79246700-INS-56 |
| chr7-79287214-INS-91 | chr7-79287214-INS-91 |
| chr7-79295899-INS-124 | chr7-79295875-INS-135 |
| chr7-79297819-INS-204 | chr7-79297819-INS-204 |
| chr7-79302518-INS-172 | chr7-79302483-INS-176 |
| chr7-79305763-INS-51 | chr7-79305763-INS-50 |
| chr7-79314922-INS-273 | chr7-79314922-INS-273 |
| chr7-79373889-INS-102 | chr7-79373889-INS-69 |
| chr7-79409722-INS-189 | chr7-79409722-INS-189 |
| chr7-79539008-INS-189 | chr7-79539010-INS-189 |
| chr7-79572864-INS-207 | chr7-79572864-INS-208 |
| chr7-79602364-INS-234 | chr7-79602365-INS-226 |
| chr7-79605722-INS-111 | chr7-79605722-INS-111 |
| chr7-79610859-INS-127 | chr7-79610861-INS-160 |
| chr7-79629884-INS-200 | chr7-79629884-INS-197 |
| chr7-79642448-INS-194 | chr7-79642445-INS-177 |
| chr7-79649762-INS-208 | chr7-79649762-INS-209 |
| chr7-79650667-INS-86 | chr7-79650667-INS-90 |
| chr7-79651219-INS-155 | chr7-79651219-INS-154 |
| chr7-79656740-INS-161 | chr7-79656741-INS-165 |
| chr7-79662094-INS-72 | chr7-79662094-INS-72 |
| chr7-79704538-INS-87 | chr7-79704538-INS-79 |
| chr7-79756627-INS-203 | chr7-79756627-INS-203 |
| chr7-79758832-INS-200 | chr7-79758832-INS-200 |
| chr7-79760903-INS-226 | chr7-79760903-INS-221 |
| chr7-79783992-INS-207 | chr7-79783992-INS-208 |
| chr7-79794161-INS-1397 | chr7-79794162-INS-1404 |
| chr7-79811110-INS-66 | chr7-79811110-INS-54 |
| chr7-79818736-INS-209 | chr7-79818736-INS-209 |
| chr7-79824842-INS-138 | chr7-79824842-INS-138 |
| chr7-79830292-INS-199 | chr7-79830292-INS-199 |
| chr7-79884834-INS-71 | chr7-79884834-INS-71 |
| chr7-79887685-INS-404 | chr7-79887685-INS-403 |
| chr7-79899723-INS-458 | chr7-79899723-INS-458 |
| chr7-79916912-INS-58 | chr7-79916912-INS-58 |
| chr7-79919203-INS-71 | chr7-79919203-INS-86 |
| chr7-79921450-INS-61 | chr7-79921450-INS-57 |
| chr7-79973424-INS-213 | chr7-79973424-INS-212 |
| chr7-79998404-INS-55 | chr7-79998404-INS-60 |
| chr7-80009790-INS-63 | chr7-80009790-INS-63 |
| chr7-80024696-INS-968 | chr7-80024692-INS-971 |
| chr7-80086217-INS-90 | chr7-80086217-INS-91 |
| chr7-80087530-INS-229 | chr7-80087530-INS-229 |
| chr7-80091819-INS-214 | chr7-80091819-INS-214 |
| chr7-80097888-INS-135 | chr7-80097888-INS-135 |
| chr7-80104626-INS-235 | chr7-80104626-INS-235 |
| chr7-80111720-INS-102 | chr7-80111720-INS-102 |
| chr7-80115902-INS-488 | chr7-80115902-INS-488 |
| chr7-80127365-INS-214 | chr7-80127365-INS-217 |
| chr7-80129892-INS-128 | chr7-80129892-INS-128 |
| chr7-80144497-INS-198 | chr7-80144497-INS-198 |
| chr7-80177999-INS-96 | chr7-80177977-INS-96 |
| chr7-80179522-INS-2164 | chr7-80179523-INS-2163 |
| chr7-80190814-INS-64 | chr7-80190814-INS-66 |
| chr7-80193904-INS-4207 | chr7-80193905-INS-4207 |

| | |
|---|---|
| chr7-80196215-INS-395 | chr7-80196215-INS-395 |
| chr7-80199023-INS-86 | chr7-80199023-INS-86 |
| chr7-80206409-INS-794 | chr7-80206409-INS-788 |
| chr7-80212215-INS-60 | chr7-80212215-INS-60 |
| chr7-80216797-INS-111 | chr7-80216797-INS-114 |
| chr7-80220996-INS-165 | chr7-80220996-INS-165 |
| chr7-80224403-INS-278 | chr7-80224404-INS-274 |
| chr7-80243402-INS-81 | chr7-80243402-INS-81 |
| chr7-80252702-INS-56 | chr7-80252702-INS-64 |
| chr7-80295250-INS-120 | chr7-80295250-INS-120 |
| chr7-80299159-INS-1891 | chr7-80299160-INS-1891 |
| chr7-80304451-INS-54 | chr7-80304451-INS-54 |
| chr7-80324863-INS-1397 | chr7-80324864-INS-1401 |
| chr7-80325983-INS-215 | chr7-80325983-INS-214 |
| chr7-80333187-INS-128 | chr7-80333187-INS-130 |
| chr7-80350580-INS-54 | chr7-80350580-INS-52 |
| chr7-80367092-INS-1890 | chr7-80367093-INS-1890 |
| chr7-80385621-INS-78 | chr7-80385621-INS-90 |
| chr7-80442818-INS-219 | chr7-80442818-INS-222 |
| chr7-80443901-INS-110 | chr7-80443926-INS-129 |
| chr7-80455363-INS-237 | chr7-80455363-INS-237 |
| chr7-80465728-INS-214 | chr7-80465728-INS-214 |
| chr7-80470857-INS-220 | chr7-80470857-INS-219 |
| chr7-80491704-INS-202 | chr7-80491704-INS-202 |
| chr7-80497322-INS-69 | chr7-80497322-INS-75 |
| chr7-80499985-INS-203 | chr7-80499985-INS-203 |
| chr7-80501859-INS-222 | chr7-80501862-INS-232 |
| chr7-80507211-INS-228 | chr7-80507211-INS-228 |
| chr7-80521617-INS-240 | chr7-80521617-INS-240 |
| chr7-80524487-INS-190 | chr7-80524487-INS-165 |
| chr7-80532276-INS-180 | chr7-80532289-INS-180 |
| chr7-80533315-INS-922 | chr7-80533315-INS-922 |
| chr7-80534986-INS-130 | chr7-80534986-INS-127 |
| chr7-80535758-INS-54 | chr7-80535781-INS-51 |
| chr7-80540114-INS-618 | chr7-80540114-INS-584 |
| chr7-80558899-INS-219 | chr7-80558899-INS-219 |
| chr7-80562854-INS-51 | chr7-80562854-INS-51 |
| chr7-80570656-INS-208 | chr7-80570656-INS-208 |
| chr7-80578741-INS-209 | chr7-80578741-INS-209 |
| chr7-80597750-INS-174 | chr7-80597750-INS-174 |
| chr7-80624486-INS-183 | chr7-80624486-INS-183 |
| chr7-80627978-INS-114 | chr7-80627978-INS-114 |
| chr7-80629921-INS-171 | chr7-80629921-INS-170 |
| chr7-80638495-INS-164 | chr7-80638495-INS-164 |
| chr7-80656117-INS-55 | chr7-80656117-INS-60 |
| chr7-80669479-INS-59 | chr7-80669479-INS-56 |
| chr7-80701833-INS-65 | chr7-80701833-INS-65 |
| chr7-80708498-INS-79 | chr7-80708498-INS-79 |
| chr7-80711398-INS-63 | chr7-80711398-INS-75 |
| chr7-80723740-INS-76 | chr7-80723740-INS-76 |
| chr7-80731775-INS-76 | chr7-80731775-INS-98 |
| chr7-80737594-INS-52 | chr7-80737593-INS-52 |
| chr7-80741443-INS-52 | chr7-80741443-INS-52 |
| chr7-80769262-INS-259 | chr7-80769262-INS-259 |
| chr7-80817481-INS-239 | chr7-80817485-INS-237 |
| chr7-80822397-INS-198 | chr7-80822397-INS-198 |
| chr7-80844140-INS-70 | chr7-80844138-INS-64 |
| chr7-80853270-INS-283 | chr7-80853270-INS-237 |
| chr7-80854700-INS-4400 | chr7-80854701-INS-4399 |
| chr7-80867871-INS-628 | chr7-80867872-INS-626 |
| chr7-80871711-INS-220 | chr7-80871711-INS-212 |
| chr7-80888619-INS-90 | chr7-80888619-INS-90 |
| chr7-80890992-INS-57 | chr7-80890992-INS-57 |
| chr7-80896451-INS-474 | chr7-80896451-INS-474 |
| chr7-80906276-INS-196 | chr7-80906276-INS-197 |
| chr7-80947911-INS-160 | chr7-80947911-INS-165 |
| chr7-80964657-INS-228 | chr7-80964657-INS-225 |
| chr7-80965633-INS-327 | chr7-80965633-INS-327 |
| chr7-80984858-INS-186 | chr7-80984858-INS-187 |
| chr7-81040534-INS-201 | chr7-81040534-INS-200 |
| chr7-81067666-INS-76 | chr7-81067666-INS-76 |
| chr7-81075164-INS-94 | chr7-81075164-INS-94 |
| chr7-81090700-INS-204 | chr7-81090700-INS-204 |
| chr7-81093753-INS-208 | chr7-81093753-INS-208 |
| chr7-81132588-INS-499 | chr7-81132586-INS-497 |
| chr7-81164603-INS-87 | chr7-81164603-INS-87 |
| chr7-81171896-INS-102 | chr7-81171896-INS-102 |

| | |
|---|---|
| chr7-81191477-INS-76 | chr7-81191477-INS-76 |
| chr7-81203897-INS-231 | chr7-81203897-INS-232 |
| chr7-81219393-INS-203 | chr7-81219394-INS-173 |
| chr7-81220141-INS-497 | chr7-81220141-INS-492 |
| chr7-81269190-INS-129 | chr7-81269190-INS-129 |
| chr7-81271040-INS-120 | chr7-81271040-INS-112 |
| chr7-81282827-INS-2468 | chr7-81282828-INS-2468 |
| chr7-81287121-INS-52 | chr7-81287121-INS-52 |
| chr7-81297610-INS-162 | chr7-81297618-INS-175 |
| chr7-81310606-INS-210 | chr7-81310558-INS-222 |
| chr7-81311994-INS-319 | chr7-81311994-INS-319 |
| chr7-81364533-INS-6403 | chr7-81364534-INS-6403 |
| chr7-81373175-INS-440 | chr7-81373176-INS-440 |
| chr7-81424665-INS-96 | chr7-81424665-INS-96 |
| chr7-81439566-INS-66 | chr7-81439566-INS-67 |
| chr7-81486070-INS-59 | chr7-81486070-INS-59 |
| chr7-81560019-INS-53 | chr7-81560019-INS-53 |
| chr7-81562188-INS-190 | chr7-81562188-INS-185 |
| chr7-81573819-INS-210 | chr7-81573819-INS-211 |
| chr7-81657755-INS-476 | chr7-81657751-INS-477 |
| chr7-81664684-INS-191 | chr7-81664684-INS-193 |
| chr7-81664684-INS-705 | chr7-81664684-INS-711 |
| chr7-81685557-INS-145 | chr7-81685557-INS-145 |
| chr7-81725394-INS-86 | chr7-81725394-INS-86 |
| chr7-81730408-INS-325 | chr7-81730408-INS-325 |
| chr7-81759144-INS-132 | chr7-81759144-INS-132 |
| chr7-81760325-INS-211 | chr7-81760325-INS-211 |
| chr7-81771268-INS-70 | chr7-81771268-INS-70 |
| chr7-81782824-INS-52 | chr7-81782824-INS-52 |
| chr7-81787323-INS-146 | chr7-81787321-INS-146 |
| chr7-81790608-INS-79 | chr7-81790608-INS-115 |
| chr7-81794275-INS-52 | chr7-81794275-INS-52 |
| chr7-81799517-INS-92 | chr7-81799517-INS-92 |
| chr7-81843871-INS-96 | chr7-81843871-INS-96 |
| chr7-81891297-INS-201 | chr7-81891297-INS-201 |
| chr7-81892411-INS-168 | chr7-81892413-INS-174 |
| chr7-81899913-INS-94 | chr7-81899913-INS-94 |
| chr7-81920694-INS-60 | chr7-81920694-INS-60 |
| chr7-81925067-INS-138 | chr7-81925067-INS-138 |
| chr7-81926754-INS-110 | chr7-81926754-INS-110 |
| chr7-81930470-INS-102 | chr7-81930473-INS-117 |
| chr7-81939415-INS-211 | chr7-81939415-INS-212 |
| chr7-81943825-INS-54 | chr7-81943825-INS-54 |
| chr7-81954909-INS-93 | chr7-81954909-INS-82 |
| chr7-81957288-INS-78 | chr7-81957288-INS-78 |
| chr7-81958558-INS-898 | chr7-81958558-INS-897 |
| chr7-82007242-INS-51 | chr7-82007258-INS-76 |
| chr7-82011276-INS-205 | chr7-82011276-INS-204 |
| chr7-82026250-INS-6271 | chr7-82026251-INS-6273 |
| chr7-82069904-INS-56 | chr7-82069904-INS-56 |
| chr7-82087212-INS-677 | chr7-82087212-INS-676 |
| chr7-82095234-INS-99 | chr7-82095229-INS-100 |
| chr7-82129601-INS-265 | chr7-82129601-INS-265 |
| chr7-82154110-INS-60 | chr7-82154110-INS-54 |
| chr7-82159355-INS-6777 | chr7-82159356-INS-6778 |
| chr7-82174226-INS-632 | chr7-82174220-INS-623 |
| chr7-82213199-INS-196 | chr7-82213199-INS-196 |
| chr7-82251952-INS-57 | chr7-82251986-INS-56 |
| chr7-82260132-INS-103 | chr7-82260132-INS-103 |
| chr7-82273443-INS-99 | chr7-82273443-INS-88 |
| chr7-82274468-INS-103 | chr7-82274468-INS-105 |
| chr7-82287037-INS-6213 | chr7-82287038-INS-6213 |
| chr7-82300607-INS-212 | chr7-82300607-INS-211 |
| chr7-82363166-INS-222 | chr7-82363166-INS-223 |
| chr7-82417167-INS-1518 | chr7-82417168-INS-1554 |
| chr7-82427508-INS-204 | chr7-82427508-INS-203 |
| chr7-82432916-INS-207 | chr7-82432916-INS-207 |
| chr7-82438584-INS-215 | chr7-82438584-INS-217 |
| chr7-82442306-INS-51 | chr7-82442306-INS-67 |
| chr7-82451946-INS-58 | chr7-82451946-INS-76 |
| chr7-82453635-INS-277 | chr7-82453635-INS-277 |
| chr7-82453676-INS-6577 | chr7-82453677-INS-6577 |
| chr7-82478428-INS-181 | chr7-82478428-INS-181 |
| chr7-82485629-INS-79 | chr7-82485637-INS-87 |
| chr7-82502804-INS-99 | chr7-82502801-INS-107 |
| chr7-82510602-INS-5354 | chr7-82510603-INS-5354 |
| chr7-82521761-INS-163 | chr7-82521795-INS-173 |

| | |
|---|---|
| chr7-82527660-INS-213 | chr7-82527660-INS-212 |
| chr7-82550714-INS-72 | chr7-82550724-INS-106 |
| chr7-82555892-INS-56 | chr7-82555893-INS-51 |
| chr7-82576575-INS-52 | chr7-82576575-INS-52 |
| chr7-82599631-INS-234 | chr7-82599631-INS-234 |
| chr7-82607241-INS-52 | chr7-82607241-INS-52 |
| chr7-82618950-INS-318 | chr7-82618950-INS-318 |
| chr7-82670956-INS-69 | chr7-82670956-INS-69 |
| chr7-82701349-INS-83 | chr7-82701349-INS-84 |
| chr7-82739543-INS-6502 | chr7-82739544-INS-6503 |
| chr7-82758796-INS-54 | chr7-82758823-INS-66 |
| chr7-82763887-INS-57 | chr7-82763888-INS-54 |
| chr7-82774336-INS-59 | chr7-82774336-INS-63 |
| chr7-82788411-INS-6400 | chr7-82788414-INS-6399 |
| chr7-82801036-INS-324 | chr7-82801036-INS-324 |
| chr7-82808310-INS-85 | chr7-82808310-INS-85 |
| chr7-82815316-INS-190 | chr7-82815322-INS-191 |
| chr7-82905609-INS-143 | chr7-82905609-INS-143 |
| chr7-82923338-INS-62 | chr7-82923348-INS-60 |
| chr7-82941639-INS-592 | chr7-82941639-INS-592 |
| chr7-82951764-INS-54 | chr7-82951764-INS-56 |
| chr7-83012354-INS-210 | chr7-83012341-INS-226 |
| chr7-83017622-INS-70 | chr7-83017622-INS-74 |
| chr7-83025775-INS-370 | chr7-83025775-INS-370 |
| chr7-83030683-INS-6347 | chr7-83030684-INS-6352 |
| chr7-83044954-INS-128 | chr7-83044949-INS-94 |
| chr7-83104243-INS-69 | chr7-83104257-INS-56 |
| chr7-83108392-INS-695 | chr7-83108392-INS-715 |
| chr7-83112589-INS-119 | chr7-83112583-INS-90 |
| chr7-83116537-INS-203 | chr7-83116537-INS-203 |
| chr7-83126066-INS-72 | chr7-83126066-INS-100 |
| chr7-83136504-INS-192 | chr7-83136504-INS-192 |
| chr7-83147401-INS-198 | chr7-83147401-INS-198 |
| chr7-83159610-INS-6376 | chr7-83159611-INS-6376 |
| chr7-83165389-INS-89 | chr7-83165389-INS-99 |
| chr7-83175626-INS-223 | chr7-83175626-INS-222 |
| chr7-83180504-INS-79 | chr7-83180504-INS-71 |
| chr7-83225176-INS-482 | chr7-83225176-INS-482 |
| chr7-83232517-INS-5182 | chr7-83232518-INS-5182 |
| chr7-83237088-INS-774 | chr7-83237088-INS-816 |
| chr7-83239677-INS-69 | chr7-83239677-INS-69 |
| chr7-83245913-INS-6626 | chr7-83245914-INS-6626 |
| chr7-83260073-INS-57 | chr7-83260066-INS-53 |
| chr7-83260188-INS-1165 | chr7-83260189-INS-1165 |
| chr7-83270024-INS-59 | chr7-83270027-INS-51 |
| chr7-83276643-INS-198 | chr7-83276643-INS-202 |
| chr7-83283959-INS-174 | chr7-83283959-INS-174 |
| chr7-83346250-INS-133 | chr7-83346250-INS-133 |
| chr7-83397181-INS-6616 | chr7-83397182-INS-6615 |
| chr7-83428829-INS-217 | chr7-83428829-INS-218 |
| chr7-83454836-INS-193 | chr7-83454836-INS-221 |
| chr7-83474658-INS-303 | chr7-83474658-INS-310 |
| chr7-83481983-INS-56 | chr7-83481987-INS-60 |
| chr7-83489016-INS-93 | chr7-83489022-INS-94 |
| chr7-83522123-INS-280 | chr7-83522123-INS-280 |
| chr7-83525518-INS-218 | chr7-83525518-INS-224 |
| chr7-83535434-INS-226 | chr7-83535434-INS-226 |
| chr7-83547698-INS-168 | chr7-83547698-INS-168 |
| chr7-83563783-INS-225 | chr7-83563783-INS-267 |
| chr7-83587092-INS-101 | chr7-83587093-INS-80 |
| chr7-83622365-INS-425 | chr7-83622365-INS-425 |
| chr7-83641528-INS-81 | chr7-83641528-INS-72 |
| chr7-83724296-INS-245 | chr7-83724296-INS-245 |
| chr7-83727304-INS-519 | chr7-83727304-INS-520 |
| chr7-83729350-INS-62 | chr7-83729350-INS-56 |
| chr7-83753319-INS-6229 | chr7-83753320-INS-6242 |
| chr7-83776541-INS-205 | chr7-83776541-INS-205 |
| chr7-83802946-INS-86 | chr7-83802946-INS-86 |
| chr7-83819488-INS-58 | chr7-83819488-INS-58 |
| chr7-83839152-INS-78 | chr7-83839152-INS-82 |
| chr7-83852781-INS-113 | chr7-83852753-INS-125 |
| chr7-83861280-INS-6487 | chr7-83861281-INS-6487 |
| chr7-83879292-INS-108 | chr7-83879292-INS-108 |
| chr7-83881721-INS-1761 | chr7-83881722-INS-1759 |
| chr7-83888586-INS-1043 | chr7-83888587-INS-1043 |
| chr7-83890575-INS-158 | chr7-83890571-INS-160 |
| chr7-83967453-INS-611 | chr7-83967453-INS-611 |

| | |
|---|---|
| chr7-83996479-INS-92 | chr7-83996479-INS-92 |
| chr7-84027064-INS-52 | chr7-84027064-INS-64 |
| chr7-84036622-INS-157 | chr7-84036622-INS-159 |
| chr7-84047988-INS-52 | chr7-84047988-INS-58 |
| chr7-84074189-INS-422 | chr7-84074189-INS-422 |
| chr7-84088746-INS-285 | chr7-84088746-INS-289 |
| chr7-84113624-INS-687 | chr7-84113624-INS-687 |
| chr7-84149005-INS-58 | chr7-84149005-INS-58 |
| chr7-84159969-INS-477 | chr7-84159955-INS-475 |
| chr7-84168057-INS-710 | chr7-84168057-INS-708 |
| chr7-84173431-INS-54 | chr7-84173431-INS-56 |
| chr7-84173940-INS-54 | chr7-84173915-INS-79 |
| chr7-84211157-INS-617 | chr7-84211161-INS-619 |
| chr7-84240864-INS-4208 | chr7-84240865-INS-4208 |
| chr7-84243821-INS-137 | chr7-84243821-INS-137 |
| chr7-84251719-INS-96 | chr7-84251727-INS-59 |
| chr7-84258104-INS-68 | chr7-84258104-INS-55 |
| chr7-84282504-INS-68 | chr7-84282504-INS-68 |
| chr7-84288645-INS-95 | chr7-84288645-INS-95 |
| chr7-84305432-INS-1325 | chr7-84305433-INS-1326 |
| chr7-84311138-INS-6511 | chr7-84311139-INS-6511 |
| chr7-84317263-INS-294 | chr7-84317263-INS-294 |
| chr7-84342656-INS-6420 | chr7-84342657-INS-6420 |
| chr7-84355418-INS-60 | chr7-84355418-INS-60 |
| chr7-84449737-INS-110 | chr7-84449737-INS-110 |
| chr7-84459588-INS-102 | chr7-84459588-INS-114 |
| chr7-84469763-INS-3863 | chr7-84469764-INS-3864 |
| chr7-84473751-INS-951 | chr7-84473751-INS-950 |
| chr7-84475878-INS-6470 | chr7-84475879-INS-6470 |
| chr7-84480663-INS-6072 | chr7-84480664-INS-6072 |
| chr7-84520530-INS-773 | chr7-84520530-INS-773 |
| chr7-84540174-INS-60 | chr7-84540174-INS-55 |
| chr7-84577653-INS-4379 | chr7-84577654-INS-4379 |
| chr7-84600516-INS-60 | chr7-84600516-INS-80 |
| chr7-84608274-INS-156 | chr7-84608274-INS-156 |
| chr7-84634030-INS-78 | chr7-84634030-INS-65 |
| chr7-84648780-INS-827 | chr7-84648780-INS-831 |
| chr7-84658072-INS-65 | chr7-84658072-INS-57 |
| chr7-84658745-INS-239 | chr7-84658745-INS-236 |
| chr7-84692728-INS-102 | chr7-84692728-INS-102 |
| chr7-84717541-INS-100 | chr7-84717541-INS-96 |
| chr7-84727227-INS-2007 | chr7-84727228-INS-2007 |
| chr7-84740800-INS-1588 | chr7-84740801-INS-1587 |
| chr7-84811179-INS-3008 | chr7-84811180-INS-3008 |
| chr7-84909085-INS-78 | chr7-84909097-INS-86 |
| chr7-84914961-INS-458 | chr7-84914961-INS-458 |
| chr7-84923883-INS-82 | chr7-84923883-INS-82 |
| chr7-84943143-INS-112 | chr7-84943143-INS-112 |
| chr7-84948890-INS-494 | chr7-84948896-INS-493 |
| chr7-84996464-INS-825 | chr7-84996464-INS-825 |
| chr7-85010004-INS-92 | chr7-85010004-INS-92 |
| chr7-85036965-INS-424 | chr7-85036965-INS-424 |
| chr7-85070639-INS-452 | chr7-85070639-INS-452 |
| chr7-85092488-INS-201 | chr7-85092488-INS-201 |
| chr7-85096207-INS-825 | chr7-85096207-INS-825 |
| chr7-85105638-INS-825 | chr7-85105638-INS-825 |
| chr7-85232623-INS-115 | chr7-85232623-INS-115 |
| chr7-85248399-INS-64 | chr7-85248399-INS-109 |
| chr7-85262265-INS-235 | chr7-85262265-INS-237 |
| chr7-85266857-INS-82 | chr7-85266857-INS-82 |
| chr7-85268213-INS-129 | chr7-85268213-INS-129 |
| chr7-85271301-INS-63 | chr7-85271301-INS-63 |
| chr7-85298088-INS-6228 | chr7-85298089-INS-6228 |
| chr7-85310741-INS-475 | chr7-85310741-INS-475 |
| chr7-85313270-INS-3848 | chr7-85313271-INS-3843 |
| chr7-85389893-INS-55 | chr7-85389893-INS-70 |
| chr7-85468056-INS-528 | chr7-85468056-INS-528 |
| chr7-85481225-INS-7082 | chr7-85481226-INS-7082 |
| chr7-85486491-INS-63 | chr7-85486479-INS-53 |
| chr7-85488311-INS-4799 | chr7-85488312-INS-4799 |
| chr7-85496750-INS-481 | chr7-85496750-INS-481 |
| chr7-85554287-INS-84 | chr7-85554287-INS-54 |
| chr7-85592406-INS-236 | chr7-85592406-INS-237 |
| chr7-85597826-INS-929 | chr7-85597830-INS-933 |
| chr7-85599024-INS-64 | chr7-85599024-INS-64 |
| chr7-85603035-INS-88 | chr7-85603031-INS-96 |
| chr7-85633080-INS-61 | chr7-85633069-INS-81 |

| | |
|---|---|
| chr7-85655282-INS-101 | chr7-85655282-INS-101 |
| chr7-85698837-INS-101 | chr7-85698837-INS-101 |
| chr7-85707019-INS-707 | chr7-85707019-INS-707 |
| chr7-85710405-INS-56 | chr7-85710405-INS-60 |
| chr7-85767084-INS-1898 | chr7-85767085-INS-1898 |
| chr7-85826838-INS-10462 | chr7-85826839-INS-10462 |
| chr7-85837281-INS-172 | chr7-85837281-INS-207 |
| chr7-85844631-INS-6456 | chr7-85844632-INS-6457 |
| chr7-85851863-INS-76 | chr7-85851851-INS-100 |
| chr7-85900823-INS-113 | chr7-85900825-INS-111 |
| chr7-85914432-INS-300 | chr7-85914432-INS-288 |
| chr7-85966823-INS-1265 | chr7-85966824-INS-1269 |
| chr7-85976741-INS-1864 | chr7-85976742-INS-1864 |
| chr7-85983563-INS-55 | chr7-85983563-INS-57 |
| chr7-86040864-INS-58 | chr7-86040912-INS-60 |
| chr7-86049629-INS-75 | chr7-86049614-INS-55 |
| chr7-86080838-INS-384 | chr7-86080838-INS-385 |
| chr7-86083590-INS-135 | chr7-86083590-INS-135 |
| chr7-86089046-INS-8985 | chr7-86089047-INS-8985 |
| chr7-86114569-INS-64 | chr7-86114569-INS-62 |
| chr7-86177103-INS-1688 | chr7-86177104-INS-1688 |
| chr7-86241587-INS-194 | chr7-86241587-INS-194 |
| chr7-86243864-INS-67 | chr7-86243863-INS-68 |
| chr7-86285960-INS-87 | chr7-86285960-INS-94 |
| chr7-86295192-INS-5518 | chr7-86295193-INS-5518 |
| chr7-86303732-INS-5348 | chr7-86303733-INS-5348 |
| chr7-86312122-INS-232 | chr7-86312122-INS-232 |
| chr7-86327032-INS-1667 | chr7-86327033-INS-1667 |
| chr7-86335924-INS-931 | chr7-86335924-INS-931 |
| chr7-86384035-INS-170 | chr7-86384035-INS-170 |
| chr7-86402694-INS-78 | chr7-86402694-INS-78 |
| chr7-86422101-INS-105 | chr7-86422101-INS-105 |
| chr7-86453547-INS-56 | chr7-86453547-INS-70 |
| chr7-86494141-INS-56 | chr7-86494141-INS-56 |
| chr7-86495122-INS-198 | chr7-86495122-INS-198 |
| chr7-86512343-INS-482 | chr7-86512343-INS-482 |
| chr7-86517271-INS-699 | chr7-86517271-INS-666 |
| chr7-86561317-INS-328 | chr7-86561317-INS-328 |
| chr7-86632214-INS-452 | chr7-86632214-INS-453 |
| chr7-86643688-INS-83 | chr7-86643688-INS-68 |
| chr7-86702315-INS-174 | chr7-86702315-INS-175 |
| chr7-86712089-INS-2544 | chr7-86712090-INS-2544 |
| chr7-86715188-INS-3323 | chr7-86715189-INS-3323 |
| chr7-86736592-INS-295 | chr7-86736592-INS-295 |
| chr7-86785577-INS-807 | chr7-86785577-INS-807 |
| chr7-86835508-INS-54 | chr7-86835512-INS-65 |
| chr7-86883748-INS-200 | chr7-86883749-INS-169 |
| chr7-86886272-INS-6217 | chr7-86886273-INS-6216 |
| chr7-86900930-INS-64 | chr7-86900930-INS-64 |
| chr7-86943136-INS-231 | chr7-86943136-INS-231 |
| chr7-86961478-INS-249 | chr7-86961478-INS-249 |
| chr7-86974012-INS-231 | chr7-86974012-INS-231 |
| chr7-86979437-INS-184 | chr7-86979433-INS-198 |
| chr7-86985995-INS-702 | chr7-86985995-INS-702 |
| chr7-87000948-INS-56 | chr7-87000948-INS-62 |
| chr7-87042612-INS-576 | chr7-87042612-INS-576 |
| chr7-87052933-INS-165 | chr7-87052933-INS-165 |
| chr7-87058141-INS-56 | chr7-87058141-INS-58 |
| chr7-87058852-INS-8419 | chr7-87058853-INS-8419 |
| chr7-87090550-INS-2314 | chr7-87090551-INS-2314 |
| chr7-87097043-INS-60 | chr7-87097043-INS-64 |
| chr7-87100201-INS-1101 | chr7-87100202-INS-1101 |
| chr7-87126761-INS-6680 | chr7-87126762-INS-6680 |
| chr7-87171561-INS-60 | chr7-87171561-INS-60 |
| chr7-87177874-INS-155 | chr7-87177874-INS-114 |
| chr7-87190487-INS-1464 | chr7-87190488-INS-1465 |
| chr7-87205485-INS-68 | chr7-87205476-INS-50 |
| chr7-87244340-INS-504 | chr7-87244340-INS-504 |
| chr7-87262810-INS-86 | chr7-87262810-INS-68 |
| chr7-87290417-INS-84 | chr7-87290417-INS-84 |
| chr7-87298701-INS-423 | chr7-87298701-INS-422 |
| chr7-87314420-INS-472 | chr7-87314420-INS-473 |
| chr7-87325158-INS-516 | chr7-87325158-INS-516 |
| chr7-87338209-INS-699 | chr7-87338209-INS-701 |
| chr7-87343361-INS-63 | chr7-87343379-INS-51 |
| chr7-87375355-INS-68 | chr7-87375355-INS-53 |
| chr7-87427411-INS-116 | chr7-87427411-INS-126 |

| | |
|---|---|
| chr7-87469755-INS-135 | chr7-87469755-INS-99 |
| chr7-87490703-INS-84 | chr7-87490703-INS-86 |
| chr7-87505528-INS-124 | chr7-87505528-INS-78 |
| chr7-87506445-INS-208 | chr7-87506445-INS-208 |
| chr7-87517974-INS-1569 | chr7-87517975-INS-1569 |
| chr7-87520560-INS-58 | chr7-87520560-INS-74 |
| chr7-87540582-INS-52 | chr7-87540580-INS-72 |
| chr7-87589673-INS-4839 | chr7-87589674-INS-4839 |
| chr7-87609420-INS-56 | chr7-87609424-INS-52 |
| chr7-87703765-INS-6229 | chr7-87703766-INS-6229 |
| chr7-87732944-INS-349 | chr7-87732944-INS-346 |
| chr7-87862008-INS-186 | chr7-87862008-INS-185 |
| chr7-87878575-INS-582 | chr7-87878575-INS-582 |
| chr7-87948380-INS-674 | chr7-87948380-INS-674 |
| chr7-87953905-INS-418 | chr7-87953905-INS-418 |
| chr7-87993904-INS-809 | chr7-87993904-INS-809 |
| chr7-88017073-INS-104 | chr7-88017077-INS-74 |
| chr7-88027778-INS-426 | chr7-88027778-INS-444 |
| chr7-88029675-INS-424 | chr7-88029675-INS-417 |
| chr7-88058587-INS-54 | chr7-88058587-INS-56 |
| chr7-88068630-INS-111 | chr7-88068630-INS-111 |
| chr7-88096261-INS-440 | chr7-88096261-INS-440 |
| chr7-88119427-INS-238 | chr7-88119427-INS-240 |
| chr7-88189215-INS-82 | chr7-88189215-INS-83 |
| chr7-88250367-INS-519 | chr7-88250367-INS-519 |
| chr7-88251836-INS-160 | chr7-88251814-INS-171 |
| chr7-88268167-INS-197 | chr7-88268167-INS-197 |
| chr7-88296156-INS-188 | chr7-88296156-INS-188 |
| chr7-88316186-INS-71 | chr7-88316186-INS-82 |
| chr7-88384560-INS-469 | chr7-88384560-INS-466 |
| chr7-88389153-INS-494 | chr7-88389153-INS-493 |
| chr7-88420955-INS-2248 | chr7-88420956-INS-2252 |
| chr7-88436056-INS-533 | chr7-88436056-INS-533 |
| chr7-88450130-INS-145 | chr7-88450130-INS-175 |
| chr7-88499448-INS-3025 | chr7-88499449-INS-3025 |
| chr7-88534841-INS-546 | chr7-88534841-INS-552 |
| chr7-88568952-INS-60 | chr7-88568952-INS-68 |
| chr7-88576542-INS-367 | chr7-88576542-INS-359 |
| chr7-88621244-INS-55 | chr7-88621244-INS-55 |
| chr7-88626663-INS-235 | chr7-88626663-INS-235 |
| chr7-88679931-INS-68 | chr7-88679931-INS-68 |
| chr7-88690532-INS-2246 | chr7-88690533-INS-2246 |
| chr7-88721273-INS-147 | chr7-88721317-INS-148 |
| chr7-88725402-INS-91 | chr7-88725402-INS-91 |
| chr7-88729292-INS-6861 | chr7-88729293-INS-6861 |
| chr7-88744201-INS-51 | chr7-88744201-INS-54 |
| chr7-88812816-INS-405 | chr7-88812816-INS-405 |
| chr7-88816238-INS-7169 | chr7-88816239-INS-7169 |
| chr7-88840002-INS-2296 | chr7-88840003-INS-2296 |
| chr7-88865826-INS-67 | chr7-88865826-INS-67 |
| chr7-88887921-INS-1012 | chr7-88887922-INS-1012 |
| chr7-88915713-INS-141 | chr7-88915713-INS-141 |
| chr7-88953365-INS-74 | chr7-88953365-INS-78 |
| chr7-89074273-INS-64 | chr7-89074273-INS-54 |
| chr7-89089452-INS-54 | chr7-89089452-INS-54 |
| chr7-89131334-INS-142 | chr7-89131334-INS-122 |
| chr7-89134170-INS-629 | chr7-89134170-INS-631 |
| chr7-89192375-INS-80 | chr7-89192375-INS-52 |
| chr7-89193584-INS-177 | chr7-89193585-INS-217 |
| chr7-89194640-INS-128 | chr7-89194687-INS-127 |
| chr7-89199752-INS-70 | chr7-89199752-INS-50 |
| chr7-89249842-INS-344 | chr7-89249842-INS-344 |
| chr7-89250993-INS-308 | chr7-89250994-INS-271 |
| chr7-89269422-INS-444 | chr7-89269422-INS-444 |
| chr7-89279372-INS-77 | chr7-89279372-INS-77 |
| chr7-89295540-INS-57 | chr7-89295540-INS-57 |
| chr7-89318416-INS-123 | chr7-89318422-INS-123 |
| chr7-89319702-INS-57 | chr7-89319702-INS-57 |
| chr7-89370702-INS-77 | chr7-89370701-INS-69 |
| chr7-89371796-INS-75 | chr7-89371794-INS-92 |
| chr7-89384636-INS-116 | chr7-89384636-INS-116 |
| chr7-89390915-INS-506 | chr7-89390916-INS-506 |
| chr7-89393837-INS-718 | chr7-89393837-INS-719 |
| chr7-89395787-INS-64 | chr7-89395787-INS-62 |
| chr7-89438585-INS-1781 | chr7-89438600-INS-1779 |
| chr7-89457942-INS-629 | chr7-89457942-INS-630 |
| chr7-89477580-INS-257 | chr7-89477580-INS-257 |

| | |
|---|---|
| chr7-89498361-INS-205 | chr7-89498361-INS-205 |
| chr7-89524288-INS-55 | chr7-89524288-INS-54 |
| chr7-89546206-INS-199 | chr7-89546206-INS-199 |
| chr7-89555647-INS-99 | chr7-89555647-INS-99 |
| chr7-89556871-INS-51 | chr7-89556871-INS-54 |
| chr7-89560037-INS-205 | chr7-89560037-INS-205 |
| chr7-89569550-INS-353 | chr7-89569550-INS-353 |
| chr7-89576754-INS-215 | chr7-89576755-INS-200 |
| chr7-89578802-INS-74 | chr7-89578802-INS-74 |
| chr7-89584487-INS-51 | chr7-89584445-INS-50 |
| chr7-89608093-INS-51 | chr7-89608093-INS-51 |
| chr7-89616665-INS-179 | chr7-89616665-INS-203 |
| chr7-89625851-INS-207 | chr7-89625851-INS-207 |
| chr7-89632392-INS-204 | chr7-89632392-INS-204 |
| chr7-89634170-INS-5193 | chr7-89634171-INS-5193 |
| chr7-89641659-INS-356 | chr7-89641659-INS-356 |
| chr7-89651158-INS-212 | chr7-89651158-INS-219 |
| chr7-89668008-INS-57 | chr7-89668050-INS-52 |
| chr7-89685885-INS-222 | chr7-89685885-INS-222 |
| chr7-89702204-INS-2110 | chr7-89702205-INS-2110 |
| chr7-89715722-INS-202 | chr7-89715722-INS-202 |
| chr7-89725005-INS-1888 | chr7-89725006-INS-1888 |
| chr7-89747089-INS-205 | chr7-89747089-INS-205 |
| chr7-89757797-INS-1894 | chr7-89757798-INS-1894 |
| chr7-89760430-INS-1703 | chr7-89760431-INS-1705 |
| chr7-89770107-INS-91 | chr7-89770119-INS-102 |
| chr7-89797108-INS-606 | chr7-89797108-INS-606 |
| chr7-89815457-INS-55 | chr7-89815457-INS-55 |
| chr7-89844958-INS-202 | chr7-89844958-INS-204 |
| chr7-89871228-INS-2509 | chr7-89871229-INS-2509 |
| chr7-89876696-INS-60 | chr7-89876696-INS-60 |
| chr7-89878432-INS-53 | chr7-89878432-INS-53 |
| chr7-89922102-INS-188 | chr7-89922102-INS-188 |
| chr7-89923525-INS-75 | chr7-89923525-INS-75 |
| chr7-89938265-INS-53 | chr7-89938265-INS-64 |
| chr7-89940640-INS-381 | chr7-89940640-INS-384 |
| chr7-89949845-INS-6442 | chr7-89949846-INS-6442 |
| chr7-89951234-INS-72 | chr7-89951234-INS-80 |
| chr7-89964319-INS-58 | chr7-89964319-INS-58 |
| chr7-89976830-INS-115 | chr7-89976830-INS-118 |
| chr7-89992401-INS-6542 | chr7-89992402-INS-6542 |
| chr7-90046890-INS-206 | chr7-90046890-INS-210 |
| chr7-90066581-INS-147 | chr7-90066581-INS-145 |
| chr7-90074740-INS-69 | chr7-90074743-INS-72 |
| chr7-90081235-INS-145 | chr7-90081216-INS-181 |
| chr7-90084309-INS-108 | chr7-90084309-INS-108 |
| chr7-90097432-INS-229 | chr7-90097432-INS-247 |
| chr7-90109102-INS-90 | chr7-90109090-INS-93 |
| chr7-90118696-INS-58 | chr7-90118696-INS-62 |
| chr7-90136744-INS-110 | chr7-90136745-INS-107 |
| chr7-90143770-INS-1320 | chr7-90143765-INS-1356 |
| chr7-90148176-INS-590 | chr7-90148176-INS-590 |
| chr7-90148986-INS-56 | chr7-90148941-INS-71 |
| chr7-90151146-INS-58 | chr7-90151146-INS-58 |
| chr7-90195210-INS-216 | chr7-90195210-INS-216 |
| chr7-90210924-INS-447 | chr7-90210925-INS-446 |
| chr7-90216685-INS-338 | chr7-90216685-INS-340 |
| chr7-90225447-INS-56 | chr7-90225447-INS-56 |
| chr7-90242724-INS-1730 | chr7-90242725-INS-1729 |
| chr7-90246299-INS-315 | chr7-90246299-INS-315 |
| chr7-90248317-INS-60 | chr7-90248312-INS-65 |
| chr7-90287204-INS-6267 | chr7-90287205-INS-6267 |
| chr7-90293881-INS-232 | chr7-90293881-INS-231 |
| chr7-90310333-INS-54 | chr7-90310333-INS-54 |
| chr7-90332185-INS-75 | chr7-90332161-INS-83 |
| chr7-90333069-INS-7085 | chr7-90333070-INS-7085 |
| chr7-90355646-INS-52 | chr7-90355646-INS-52 |
| chr7-90383071-INS-101 | chr7-90383071-INS-58 |
| chr7-90394687-INS-123 | chr7-90394685-INS-122 |
| chr7-90422435-INS-6292 | chr7-90422436-INS-6295 |
| chr7-90422732-INS-752 | chr7-90422732-INS-754 |
| chr7-90432500-INS-431 | chr7-90432500-INS-431 |
| chr7-90438239-INS-6588 | chr7-90438240-INS-6588 |
| chr7-90451141-INS-84 | chr7-90451141-INS-84 |
| chr7-90503058-INS-56 | chr7-90503057-INS-52 |
| chr7-90553494-INS-255 | chr7-90553494-INS-255 |
| chr7-90558331-INS-60 | chr7-90558312-INS-89 |

| | |
|---|---|
| chr7-90567126-INS-82 | chr7-90567126-INS-82 |
| chr7-90572630-INS-5735 | chr7-90572631-INS-5737 |
| chr7-90604138-INS-55 | chr7-90604138-INS-50 |
| chr7-90615685-INS-60 | chr7-90615685-INS-60 |
| chr7-90635576-INS-288 | chr7-90635576-INS-288 |
| chr7-90649205-INS-56 | chr7-90649205-INS-56 |
| chr7-90656380-INS-576 | chr7-90656380-INS-576 |
| chr7-90662787-INS-73 | chr7-90662787-INS-73 |
| chr7-90669482-INS-294 | chr7-90669482-INS-294 |
| chr7-90686212-INS-534 | chr7-90686251-INS-487 |
| chr7-90696226-INS-372 | chr7-90696226-INS-370 |
| chr7-90739278-INS-58 | chr7-90739278-INS-62 |
| chr7-90754281-INS-874 | chr7-90754281-INS-873 |
| chr7-90756200-INS-59 | chr7-90756200-INS-59 |
| chr7-90758129-INS-118 | chr7-90758126-INS-124 |
| chr7-90774392-INS-178 | chr7-90774392-INS-178 |
| chr7-90823485-INS-1545 | chr7-90823486-INS-1545 |
| chr7-90836343-INS-76 | chr7-90836325-INS-54 |
| chr7-90840091-INS-660 | chr7-90840092-INS-660 |
| chr7-90871579-INS-201 | chr7-90871580-INS-201 |
| chr7-90872360-INS-102 | chr7-90872396-INS-108 |
| chr7-90876271-INS-58 | chr7-90876271-INS-58 |
| chr7-90885299-INS-68 | chr7-90885299-INS-81 |
| chr7-90887792-INS-52 | chr7-90887792-INS-52 |
| chr7-91013938-INS-445 | chr7-91013938-INS-448 |
| chr7-91022013-INS-6477 | chr7-91022014-INS-6475 |
| chr7-91030668-INS-326 | chr7-91030668-INS-326 |
| chr7-91043641-INS-4507 | chr7-91043641-INS-4500 |
| chr7-91045294-INS-797 | chr7-91045294-INS-794 |
| chr7-91051278-INS-1441 | chr7-91051279-INS-1441 |
| chr7-91062216-INS-970 | chr7-91062216-INS-970 |
| chr7-91066823-INS-202 | chr7-91066823-INS-202 |
| chr7-91069039-INS-236 | chr7-91069039-INS-236 |
| chr7-91073547-INS-92 | chr7-91073557-INS-86 |
| chr7-91129699-INS-1398 | chr7-91129700-INS-1398 |
| chr7-91140878-INS-1546 | chr7-91140879-INS-1545 |
| chr7-91217664-INS-95 | chr7-91217644-INS-129 |
| chr7-91222480-INS-762 | chr7-91222480-INS-762 |
| chr7-91223889-INS-272 | chr7-91223889-INS-272 |
| chr7-91230187-INS-107 | chr7-91230175-INS-114 |
| chr7-91248784-INS-62 | chr7-91248771-INS-104 |
| chr7-91267376-INS-1446 | chr7-91267377-INS-1446 |
| chr7-91280378-INS-64 | chr7-91280372-INS-64 |
| chr7-91290612-INS-162 | chr7-91290612-INS-190 |
| chr7-91299435-INS-72 | chr7-91299435-INS-80 |
| chr7-91306669-INS-386 | chr7-91306669-INS-389 |
| chr7-91310854-INS-137 | chr7-91310854-INS-110 |
| chr7-91311807-INS-81 | chr7-91311807-INS-81 |
| chr7-91331666-INS-56 | chr7-91331666-INS-56 |
| chr7-91344534-INS-555 | chr7-91344534-INS-555 |
| chr7-91398083-INS-61 | chr7-91398083-INS-57 |
| chr7-91443635-INS-1785 | chr7-91443636-INS-1786 |
| chr7-91506045-INS-54 | chr7-91506023-INS-52 |
| chr7-91511927-INS-109 | chr7-91511927-INS-117 |
| chr7-91514459-INS-490 | chr7-91514459-INS-490 |
| chr7-91533070-INS-72 | chr7-91533070-INS-54 |
| chr7-91534931-INS-66 | chr7-91534928-INS-51 |
| chr7-91545200-INS-105 | chr7-91545200-INS-108 |
| chr7-91548642-INS-756 | chr7-91548642-INS-741 |
| chr7-91555731-INS-68 | chr7-91555710-INS-87 |
| chr7-91604880-INS-64 | chr7-91604880-INS-64 |
| chr7-91632565-INS-1511 | chr7-91632566-INS-1514 |
| chr7-91634178-INS-570 | chr7-91634179-INS-570 |
| chr7-91640131-INS-104 | chr7-91640131-INS-104 |
| chr7-91650554-INS-619 | chr7-91650554-INS-620 |
| chr7-91671569-INS-2234 | chr7-91671570-INS-2234 |
| chr7-91687318-INS-636 | chr7-91687318-INS-661 |
| chr7-91702102-INS-643 | chr7-91702102-INS-644 |
| chr7-91743223-INS-671 | chr7-91743234-INS-685 |
| chr7-91795710-INS-582 | chr7-91795708-INS-580 |
| chr7-91801780-INS-528 | chr7-91801780-INS-528 |
| chr7-91856488-INS-176 | chr7-91856488-INS-179 |
| chr7-91869692-INS-527 | chr7-91869692-INS-528 |
| chr7-91882035-INS-82 | chr7-91882035-INS-82 |
| chr7-91932696-INS-163 | chr7-91932696-INS-163 |
| chr7-91933146-INS-6443 | chr7-91933147-INS-6443 |
| chr7-91946612-INS-153 | chr7-91946612-INS-155 |

| | |
|---|---|
| chr7-91949689-INS-156 | chr7-91949689-INS-157 |
| chr7-91952673-INS-90 | chr7-91952676-INS-55 |
| chr7-91954291-INS-97 | chr7-91954295-INS-101 |
| chr7-91959024-INS-76 | chr7-91959026-INS-78 |
| chr7-91973761-INS-53 | chr7-91973761-INS-53 |
| chr7-91977806-INS-760 | chr7-91977806-INS-756 |
| chr7-91979551-INS-58 | chr7-91979551-INS-58 |
| chr7-91981820-INS-203 | chr7-91981820-INS-197 |
| chr7-91983342-INS-348 | chr7-91983342-INS-348 |
| chr7-91992793-INS-541 | chr7-91992793-INS-541 |
| chr7-92015619-INS-98 | chr7-92015610-INS-105 |
| chr7-92051451-INS-109 | chr7-92051440-INS-105 |
| chr7-92053720-INS-68 | chr7-92053720-INS-52 |
| chr7-92057557-INS-4464 | chr7-92057558-INS-4464 |
| chr7-92069002-INS-100 | chr7-92069002-INS-100 |
| chr7-92160362-INS-88 | chr7-92160362-INS-88 |
| chr7-92176850-INS-53 | chr7-92176850-INS-53 |
| chr7-92224329-INS-207 | chr7-92224329-INS-207 |
| chr7-92224923-INS-788 | chr7-92224923-INS-788 |
| chr7-92277166-INS-53 | chr7-92277166-INS-53 |
| chr7-92312265-INS-52 | chr7-92312255-INS-70 |
| chr7-92332217-INS-60 | chr7-92332217-INS-62 |
| chr7-92342141-INS-150 | chr7-92342145-INS-178 |
| chr7-92352758-INS-1888 | chr7-92352759-INS-1888 |
| chr7-92362452-INS-6859 | chr7-92362453-INS-6860 |
| chr7-92372154-INS-189 | chr7-92372154-INS-189 |
| chr7-92375700-INS-328 | chr7-92375700-INS-329 |
| chr7-92385681-INS-243 | chr7-92385681-INS-234 |
| chr7-92406737-INS-1891 | chr7-92406738-INS-1891 |
| chr7-92411240-INS-454 | chr7-92411240-INS-454 |
| chr7-92421713-INS-54 | chr7-92421713-INS-58 |
| chr7-92433500-INS-532 | chr7-92433500-INS-523 |
| chr7-92454436-INS-58 | chr7-92454436-INS-102 |
| chr7-92457740-INS-52 | chr7-92457741-INS-56 |
| chr7-92514228-INS-346 | chr7-92514228-INS-347 |
| chr7-92518711-INS-536 | chr7-92518711-INS-536 |
| chr7-92520224-INS-691 | chr7-92520224-INS-692 |
| chr7-92539365-INS-3197 | chr7-92539366-INS-3199 |
| chr7-92546087-INS-173 | chr7-92546087-INS-171 |
| chr7-92556249-INS-52 | chr7-92556249-INS-52 |
| chr7-92583086-INS-5272 | chr7-92583087-INS-5272 |
| chr7-92587805-INS-6404 | chr7-92587806-INS-6404 |
| chr7-92589401-INS-1609 | chr7-92589402-INS-1613 |
| chr7-92648994-INS-1056 | chr7-92648995-INS-1056 |
| chr7-92653251-INS-320 | chr7-92653251-INS-320 |
| chr7-92700226-INS-477 | chr7-92700226-INS-477 |
| chr7-92702539-INS-54 | chr7-92702539-INS-56 |
| chr7-92714005-INS-162 | chr7-92714005-INS-172 |
| chr7-92734458-INS-535 | chr7-92734458-INS-536 |
| chr7-92735631-INS-338 | chr7-92735628-INS-333 |
| chr7-92748327-INS-5083 | chr7-92748328-INS-5083 |
| chr7-92761390-INS-75 | chr7-92761390-INS-83 |
| chr7-92768222-INS-380 | chr7-92768222-INS-377 |
| chr7-92777462-INS-63 | chr7-92777454-INS-55 |
| chr7-92828812-INS-124 | chr7-92828812-INS-124 |
| chr7-92868775-INS-90 | chr7-92868821-INS-58 |
| chr7-92879817-INS-400 | chr7-92879817-INS-400 |
| chr7-92880720-INS-52 | chr7-92880720-INS-52 |
| chr7-92892630-INS-55 | chr7-92892630-INS-51 |
| chr7-92908306-INS-4600 | chr7-92908307-INS-4600 |
| chr7-92933641-INS-343 | chr7-92933641-INS-343 |
| chr7-92948508-INS-5194 | chr7-92948509-INS-5194 |
| chr7-92951610-INS-1234 | chr7-92951611-INS-1238 |
| chr7-92970675-INS-6731 | chr7-92970676-INS-6733 |
| chr7-92994340-INS-6388 | chr7-92994340-INS-6385 |
| chr7-93003816-INS-2480 | chr7-93003817-INS-2480 |
| chr7-93025013-INS-58 | chr7-93025013-INS-50 |
| chr7-93027317-INS-6582 | chr7-93027318-INS-6582 |
| chr7-93041392-INS-275 | chr7-93041420-INS-293 |
| chr7-93086955-INS-390 | chr7-93086955-INS-390 |
| chr7-93091281-INS-1060 | chr7-93091282-INS-1077 |
| chr7-93106529-INS-76 | chr7-93106529-INS-85 |
| chr7-93109767-INS-464 | chr7-93109767-INS-464 |
| chr7-93150258-INS-1316 | chr7-93150259-INS-1319 |
| chr7-93165599-INS-1108 | chr7-93165600-INS-1108 |
| chr7-93170773-INS-201 | chr7-93170773-INS-201 |
| chr7-93214955-INS-431 | chr7-93214955-INS-432 |

| | |
|---|---|
| chr7-93216359-INS-71 | chr7-93216359-INS-89 |
| chr7-93241528-INS-99 | chr7-93241528-INS-100 |
| chr7-93242507-INS-60 | chr7-93242507-INS-60 |
| chr7-93272802-INS-155 | chr7-93272802-INS-155 |
| chr7-93279744-INS-78 | chr7-93279748-INS-64 |
| chr7-93345931-INS-202 | chr7-93345931-INS-202 |
| chr7-93355392-INS-6716 | chr7-93355393-INS-6717 |
| chr7-93356635-INS-210 | chr7-93356635-INS-210 |
| chr7-93368888-INS-418 | chr7-93368888-INS-418 |
| chr7-93383176-INS-467 | chr7-93383176-INS-464 |
| chr7-93408473-INS-92 | chr7-93408516-INS-52 |
| chr7-93422525-INS-194 | chr7-93422526-INS-194 |
| chr7-93425946-INS-68 | chr7-93425945-INS-63 |
| chr7-93440738-INS-423 | chr7-93440738-INS-375 |
| chr7-93481472-INS-56 | chr7-93481480-INS-52 |
| chr7-93488472-INS-225 | chr7-93488472-INS-225 |
| chr7-93492198-INS-50 | chr7-93492198-INS-58 |
| chr7-93498968-INS-394 | chr7-93498945-INS-384 |
| chr7-93525684-INS-7367 | chr7-93525685-INS-7368 |
| chr7-93569836-INS-7070 | chr7-93569837-INS-7070 |
| chr7-93581141-INS-7153 | chr7-93581142-INS-7153 |
| chr7-93593447-INS-485 | chr7-93593447-INS-484 |
| chr7-93611004-INS-62 | chr7-93611004-INS-56 |
| chr7-93627347-INS-3103 | chr7-93627348-INS-3103 |
| chr7-93633477-INS-62 | chr7-93633477-INS-68 |
| chr7-93661996-INS-56 | chr7-93661996-INS-56 |
| chr7-93670870-INS-169 | chr7-93670870-INS-169 |
| chr7-93675562-INS-59 | chr7-93675558-INS-66 |
| chr7-93691333-INS-121 | chr7-93691333-INS-119 |
| chr7-93717445-INS-67 | chr7-93717445-INS-66 |
| chr7-93717707-INS-1180 | chr7-93717708-INS-1181 |
| chr7-93720621-INS-7102 | chr7-93720622-INS-7102 |
| chr7-93720833-INS-326 | chr7-93720833-INS-326 |
| chr7-93725119-INS-76 | chr7-93725119-INS-82 |
| chr7-93745368-INS-2598 | chr7-93745369-INS-2598 |
| chr7-93747245-INS-503 | chr7-93747245-INS-503 |
| chr7-93754866-INS-1460 | chr7-93754867-INS-1460 |
| chr7-93769210-INS-147 | chr7-93769210-INS-147 |
| chr7-93773843-INS-52 | chr7-93773841-INS-56 |
| chr7-93794930-INS-161 | chr7-93794930-INS-162 |
| chr7-93806517-INS-54 | chr7-93806518-INS-55 |
| chr7-93821973-INS-76 | chr7-93821973-INS-60 |
| chr7-93864785-INS-51 | chr7-93864785-INS-51 |
| chr7-93872941-INS-1136 | chr7-93872940-INS-1109 |
| chr7-93887678-INS-52 | chr7-93887678-INS-56 |
| chr7-93913324-INS-7509 | chr7-93913325-INS-7510 |
| chr7-93922166-INS-68 | chr7-93922166-INS-58 |
| chr7-93940107-INS-128 | chr7-93940107-INS-128 |
| chr7-93949811-INS-138 | chr7-93949810-INS-160 |
| chr7-93974772-INS-988 | chr7-93974772-INS-975 |
| chr7-93975336-INS-2161 | chr7-93975337-INS-2161 |
| chr7-93975489-INS-78 | chr7-93975489-INS-80 |
| chr7-93991044-INS-66 | chr7-93991017-INS-66 |
| chr7-93991207-INS-3868 | chr7-93991208-INS-3865 |
| chr7-93997336-INS-156 | chr7-93997336-INS-164 |
| chr7-94002284-INS-3027 | chr7-94002285-INS-3023 |
| chr7-94008354-INS-72 | chr7-94008354-INS-70 |
| chr7-94057376-INS-85 | chr7-94057375-INS-119 |
| chr7-94058315-INS-1553 | chr7-94058321-INS-1544 |
| chr7-94070512-INS-98 | chr7-94070512-INS-68 |
| chr7-94079431-INS-108 | chr7-94079431-INS-108 |
| chr7-94093592-INS-117 | chr7-94093592-INS-117 |
| chr7-94121609-INS-159 | chr7-94121609-INS-161 |
| chr7-94122624-INS-438 | chr7-94122624-INS-438 |
| chr7-94126716-INS-3436 | chr7-94126717-INS-3436 |
| chr7-94131789-INS-443 | chr7-94131789-INS-442 |
| chr7-94133050-INS-5548 | chr7-94133051-INS-5548 |
| chr7-94145599-INS-1892 | chr7-94145600-INS-1892 |
| chr7-94148833-INS-6275 | chr7-94148834-INS-6278 |
| chr7-94154343-INS-68 | chr7-94154343-INS-68 |
| chr7-94163330-INS-63 | chr7-94163331-INS-98 |
| chr7-94186136-INS-60 | chr7-94186136-INS-60 |
| chr7-94194053-INS-491 | chr7-94194053-INS-491 |
| chr7-94202710-INS-655 | chr7-94202711-INS-655 |
| chr7-94241467-INS-60 | chr7-94241467-INS-66 |
| chr7-94283674-INS-367 | chr7-94283674-INS-367 |
| chr7-94291586-INS-6172 | chr7-94291587-INS-6172 |

| | |
|---|---|
| chr7-94304090-INS-58 | chr7-94304090-INS-72 |
| chr7-94308589-INS-1126 | chr7-94308589-INS-1142 |
| chr7-94332384-INS-3081 | chr7-94332385-INS-3081 |
| chr7-94384710-INS-7205 | chr7-94384711-INS-7202 |
| chr7-94395295-INS-230 | chr7-94395295-INS-228 |
| chr7-94404773-INS-52 | chr7-94404773-INS-60 |
| chr7-94409830-INS-569 | chr7-94409830-INS-570 |
| chr7-94410496-INS-56 | chr7-94410497-INS-50 |
| chr7-94417690-INS-202 | chr7-94417690-INS-202 |
| chr7-94424054-INS-55 | chr7-94424054-INS-55 |
| chr7-94444943-INS-334 | chr7-94444943-INS-331 |
| chr7-94448187-INS-190 | chr7-94448187-INS-190 |
| chr7-94457878-INS-72 | chr7-94457878-INS-72 |
| chr7-94462589-INS-407 | chr7-94462589-INS-407 |
| chr7-94468190-INS-56 | chr7-94468190-INS-68 |
| chr7-94511755-INS-1503 | chr7-94511756-INS-1503 |
| chr7-94518513-INS-951 | chr7-94518516-INS-952 |
| chr7-94519296-INS-6539 | chr7-94519297-INS-6539 |
| chr7-94527671-INS-271 | chr7-94527671-INS-271 |
| chr7-94533093-INS-58 | chr7-94533093-INS-54 |
| chr7-94543764-INS-89 | chr7-94543764-INS-89 |
| chr7-94548597-INS-1863 | chr7-94548598-INS-1863 |
| chr7-94548598-INS-394 | chr7-94548598-INS-394 |
| chr7-94576660-INS-76 | chr7-94576660-INS-76 |
| chr7-94598769-INS-1487 | chr7-94598770-INS-1450 |
| chr7-94599118-INS-6809 | chr7-94599119-INS-6809 |
| chr7-94611692-INS-104 | chr7-94611692-INS-104 |
| chr7-94674341-INS-81 | chr7-94674341-INS-72 |
| chr7-94676006-INS-5393 | chr7-94676007-INS-5393 |
| chr7-94677908-INS-68 | chr7-94677908-INS-66 |
| chr7-94680930-INS-553 | chr7-94680930-INS-554 |
| chr7-94690188-INS-4042 | chr7-94690189-INS-4042 |
| chr7-94696326-INS-128 | chr7-94696326-INS-129 |
| chr7-94727335-INS-832 | chr7-94727336-INS-832 |
| chr7-94729299-INS-1216 | chr7-94729300-INS-1216 |
| chr7-94730632-INS-51 | chr7-94730632-INS-51 |
| chr7-94742370-INS-71 | chr7-94742370-INS-69 |
| chr7-94778206-INS-2001 | chr7-94778207-INS-2001 |
| chr7-94781881-INS-58 | chr7-94781849-INS-60 |
| chr7-94819548-INS-533 | chr7-94819548-INS-533 |
| chr7-94849410-INS-150 | chr7-94849410-INS-160 |
| chr7-94864320-INS-88 | chr7-94864320-INS-95 |
| chr7-94869639-INS-59 | chr7-94869639-INS-59 |
| chr7-94914892-INS-82 | chr7-94914892-INS-82 |
| chr7-94928970-INS-61 | chr7-94928970-INS-59 |
| chr7-94943942-INS-59 | chr7-94943942-INS-59 |
| chr7-94952869-INS-4770 | chr7-94952870-INS-4770 |
| chr7-94957207-INS-319 | chr7-94957207-INS-319 |
| chr7-94976086-INS-4980 | chr7-94976087-INS-4967 |
| chr7-94992751-INS-291 | chr7-94992751-INS-291 |
| chr7-95010252-INS-75 | chr7-95010252-INS-75 |
| chr7-95033567-INS-3883 | chr7-95033568-INS-3882 |
| chr7-95061844-INS-60 | chr7-95061844-INS-64 |
| chr7-95066016-INS-4658 | chr7-95066017-INS-4651 |
| chr7-95071869-INS-75 | chr7-95071869-INS-75 |
| chr7-95084416-INS-56 | chr7-95084416-INS-56 |
| chr7-95092985-INS-120 | chr7-95092985-INS-120 |
| chr7-95103087-INS-114 | chr7-95103087-INS-148 |
| chr7-95103978-INS-54 | chr7-95103962-INS-54 |
| chr7-95113213-INS-51 | chr7-95113213-INS-53 |
| chr7-95125710-INS-6719 | chr7-95125711-INS-6720 |
| chr7-95155633-INS-66 | chr7-95155633-INS-66 |
| chr7-95157644-INS-693 | chr7-95157644-INS-693 |
| chr7-95161179-INS-176 | chr7-95161179-INS-164 |
| chr7-95174318-INS-241 | chr7-95174318-INS-242 |
| chr7-95186408-INS-68 | chr7-95186433-INS-61 |
| chr7-95206127-INS-157 | chr7-95206127-INS-157 |
| chr7-95230157-INS-7091 | chr7-95230158-INS-7091 |
| chr7-95233627-INS-60 | chr7-95233627-INS-64 |
| chr7-95244550-INS-6594 | chr7-95244551-INS-6595 |
| chr7-95249880-INS-89 | chr7-95249880-INS-89 |
| chr7-95251787-INS-1664 | chr7-95251788-INS-1665 |
| chr7-95266887-INS-840 | chr7-95266887-INS-838 |
| chr7-95272003-INS-7145 | chr7-95272004-INS-7145 |
| chr7-95292541-INS-99 | chr7-95292541-INS-117 |
| chr7-95293550-INS-1442 | chr7-95293551-INS-1444 |
| chr7-95293978-INS-192 | chr7-95293978-INS-198 |

| | |
|---|---|
| chr7-95297957-INS-100 | chr7-95297957-INS-100 |
| chr7-95320114-INS-54 | chr7-95320114-INS-52 |
| chr7-95331621-INS-144 | chr7-95331621-INS-144 |
| chr7-95332735-INS-264 | chr7-95332735-INS-266 |
| chr7-95345925-INS-1970 | chr7-95345926-INS-1970 |
| chr7-95355704-INS-122 | chr7-95355704-INS-123 |
| chr7-95359947-INS-52 | chr7-95359947-INS-58 |
| chr7-95375351-INS-178 | chr7-95375351-INS-178 |
| chr7-95382570-INS-1260 | chr7-95382571-INS-1261 |
| chr7-95383332-INS-431 | chr7-95383332-INS-432 |
| chr7-95408194-INS-79 | chr7-95408210-INS-55 |
| chr7-95409116-INS-181 | chr7-95409116-INS-181 |
| chr7-95424916-INS-85 | chr7-95424916-INS-52 |
| chr7-95432716-INS-292 | chr7-95432716-INS-260 |
| chr7-95437893-INS-6421 | chr7-95437894-INS-6422 |
| chr7-95441410-INS-53 | chr7-95441410-INS-58 |
| chr7-95444108-INS-452 | chr7-95444108-INS-452 |
| chr7-95461836-INS-734 | chr7-95461836-INS-733 |
| chr7-95506780-INS-245 | chr7-95506780-INS-245 |
| chr7-95518582-INS-64 | chr7-95518560-INS-80 |
| chr7-95533172-INS-6370 | chr7-95533173-INS-6370 |
| chr7-95547075-INS-52 | chr7-95547075-INS-51 |
| chr7-95554496-INS-6613 | chr7-95554497-INS-6613 |
| chr7-95561952-INS-185 | chr7-95561952-INS-185 |
| chr7-95621752-INS-80 | chr7-95621752-INS-80 |
| chr7-95637375-INS-76 | chr7-95637375-INS-88 |
| chr7-95643310-INS-167 | chr7-95643310-INS-167 |
| chr7-95670727-INS-86 | chr7-95670720-INS-79 |
| chr7-95707313-INS-1045 | chr7-95707314-INS-1045 |
| chr7-95708192-INS-1089 | chr7-95708193-INS-1089 |
| chr7-95725540-INS-1224 | chr7-95725541-INS-1224 |
| chr7-95773297-INS-389 | chr7-95773297-INS-389 |
| chr7-95780046-INS-70 | chr7-95780046-INS-72 |
| chr7-95810057-INS-782 | chr7-95810057-INS-785 |
| chr7-95810646-INS-64 | chr7-95810663-INS-80 |
| chr7-95835084-INS-233 | chr7-95835084-INS-233 |
| chr7-95848613-INS-101 | chr7-95848613-INS-101 |
| chr7-95857586-INS-52 | chr7-95857586-INS-52 |
| chr7-95871540-INS-64 | chr7-95871541-INS-52 |
| chr7-95882081-INS-52 | chr7-95882096-INS-50 |
| chr7-95915820-INS-56 | chr7-95915820-INS-56 |
| chr7-95942604-INS-105 | chr7-95942604-INS-105 |
| chr7-95947003-INS-222 | chr7-95947003-INS-234 |
| chr7-95951193-INS-56 | chr7-95951193-INS-56 |
| chr7-95977796-INS-133 | chr7-95977796-INS-137 |
| chr7-96006510-INS-304 | chr7-96006510-INS-302 |
| chr7-96007325-INS-52 | chr7-96007313-INS-54 |
| chr7-96024246-INS-6472 | chr7-96024247-INS-6472 |
| chr7-96027354-INS-92 | chr7-96027354-INS-94 |
| chr7-96051934-INS-349 | chr7-96051934-INS-349 |
| chr7-96062994-INS-91 | chr7-96062990-INS-97 |
| chr7-96080187-INS-52 | chr7-96080187-INS-52 |
| chr7-96084682-INS-118 | chr7-96084682-INS-126 |
| chr7-96099552-INS-68 | chr7-96099552-INS-68 |
| chr7-96162683-INS-222 | chr7-96162683-INS-222 |
| chr7-96164024-INS-179 | chr7-96164024-INS-136 |
| chr7-96186333-INS-82 | chr7-96186380-INS-82 |
| chr7-96252737-INS-67 | chr7-96252737-INS-67 |
| chr7-96256175-INS-58 | chr7-96256175-INS-58 |
| chr7-96270848-INS-202 | chr7-96270848-INS-202 |
| chr7-96271995-INS-93 | chr7-96271995-INS-94 |
| chr7-96273656-INS-397 | chr7-96273656-INS-397 |
| chr7-96289297-INS-54 | chr7-96289297-INS-52 |
| chr7-96307382-INS-54 | chr7-96307382-INS-54 |
| chr7-96312205-INS-78 | chr7-96312207-INS-60 |
| chr7-96318903-INS-71 | chr7-96318903-INS-67 |
| chr7-96333826-INS-67 | chr7-96333826-INS-69 |
| chr7-96344707-INS-2153 | chr7-96344708-INS-2135 |
| chr7-96363394-INS-52 | chr7-96363394-INS-52 |
| chr7-96413477-INS-256 | chr7-96413477-INS-263 |
| chr7-96421049-INS-70 | chr7-96421049-INS-120 |
| chr7-96424938-INS-6644 | chr7-96424939-INS-6645 |
| chr7-96486025-INS-130 | chr7-96486045-INS-94 |
| chr7-96502587-INS-52 | chr7-96502599-INS-52 |
| chr7-96506646-INS-176 | chr7-96506646-INS-176 |
| chr7-96520645-INS-113 | chr7-96520626-INS-126 |
| chr7-96530161-INS-200 | chr7-96530161-INS-199 |

| | |
|---|---|
| chr7-96543032-INS-144 | chr7-96543067-INS-108 |
| chr7-96550619-INS-98 | chr7-96550619-INS-98 |
| chr7-96568379-INS-1365 | chr7-96568380-INS-1403 |
| chr7-96575212-INS-6221 | chr7-96575213-INS-6222 |
| chr7-96588766-INS-61 | chr7-96588766-INS-61 |
| chr7-96612650-INS-410 | chr7-96612650-INS-410 |
| chr7-96648386-INS-195 | chr7-96648386-INS-195 |
| chr7-96656201-INS-62 | chr7-96656201-INS-62 |
| chr7-96656729-INS-98 | chr7-96656729-INS-97 |
| chr7-96657871-INS-144 | chr7-96657871-INS-144 |
| chr7-96663040-INS-105 | chr7-96663040-INS-105 |
| chr7-96666989-INS-119 | chr7-96666989-INS-119 |
| chr7-96687300-INS-394 | chr7-96687300-INS-394 |
| chr7-96692311-INS-57 | chr7-96692311-INS-57 |
| chr7-96727716-INS-78 | chr7-96727716-INS-78 |
| chr7-96732393-INS-199 | chr7-96732393-INS-199 |
| chr7-96740129-INS-76 | chr7-96740129-INS-68 |
| chr7-96772533-INS-898 | chr7-96772533-INS-898 |
| chr7-96803809-INS-60 | chr7-96803809-INS-62 |
| chr7-96832324-INS-883 | chr7-96832324-INS-885 |
| chr7-96834338-INS-6482 | chr7-96834339-INS-6482 |
| chr7-96849126-INS-66 | chr7-96849123-INS-66 |
| chr7-96871388-INS-242 | chr7-96871388-INS-242 |
| chr7-96910146-INS-202 | chr7-96910146-INS-204 |
| chr7-96914425-INS-206 | chr7-96914425-INS-169 |
| chr7-96943546-INS-269 | chr7-96943546-INS-269 |
| chr7-96972212-INS-110 | chr7-96972212-INS-110 |
| chr7-96989346-INS-159 | chr7-96989346-INS-163 |
| chr7-97007383-INS-368 | chr7-97007334-INS-410 |
| chr7-97011682-INS-1199 | chr7-97011683-INS-1199 |
| chr7-97014772-INS-506 | chr7-97014773-INS-528 |
| chr7-97019675-INS-120 | chr7-97019675-INS-145 |
| chr7-97032657-INS-60 | chr7-97032656-INS-56 |
| chr7-97044609-INS-1864 | chr7-97044610-INS-1864 |
| chr7-97052389-INS-108 | chr7-97052393-INS-112 |
| chr7-97061653-INS-54 | chr7-97061653-INS-54 |
| chr7-97064097-INS-52 | chr7-97064097-INS-60 |
| chr7-97080622-INS-57 | chr7-97080620-INS-50 |
| chr7-97088042-INS-94 | chr7-97088042-INS-94 |
| chr7-97091257-INS-80 | chr7-97091233-INS-75 |
| chr7-97092144-INS-246 | chr7-97092144-INS-246 |
| chr7-97100578-INS-179 | chr7-97100578-INS-179 |
| chr7-97104673-INS-171 | chr7-97104673-INS-140 |
| chr7-97108129-INS-183 | chr7-97108129-INS-183 |
| chr7-97109106-INS-52 | chr7-97109106-INS-50 |
| chr7-97112423-INS-249 | chr7-97112423-INS-252 |
| chr7-97114134-INS-327 | chr7-97114134-INS-327 |
| chr7-97132550-INS-73 | chr7-97132551-INS-95 |
| chr7-97150511-INS-70 | chr7-97150511-INS-70 |
| chr7-97159227-INS-121 | chr7-97159227-INS-121 |
| chr7-97172152-INS-192 | chr7-97172152-INS-192 |
| chr7-97180890-INS-83 | chr7-97180858-INS-84 |
| chr7-97183816-INS-7724 | chr7-97183817-INS-7724 |
| chr7-97191130-INS-130 | chr7-97191130-INS-130 |
| chr7-97196757-INS-74 | chr7-97196757-INS-74 |
| chr7-97207935-INS-868 | chr7-97207935-INS-868 |
| chr7-97217935-INS-130 | chr7-97217935-INS-131 |
| chr7-97220266-INS-206 | chr7-97220266-INS-206 |
| chr7-97222381-INS-212 | chr7-97222381-INS-218 |
| chr7-97236557-INS-53 | chr7-97236573-INS-86 |
| chr7-97237390-INS-223 | chr7-97237390-INS-222 |
| chr7-97295058-INS-206 | chr7-97295057-INS-204 |
| chr7-97306917-INS-54 | chr7-97306917-INS-50 |
| chr7-97321484-INS-407 | chr7-97321484-INS-407 |
| chr7-97354368-INS-56 | chr7-97354368-INS-58 |
| chr7-97358967-INS-165 | chr7-97358967-INS-165 |
| chr7-97370905-INS-230 | chr7-97370907-INS-235 |
| chr7-97377402-INS-5431 | chr7-97377403-INS-5431 |
| chr7-97393292-INS-205 | chr7-97393292-INS-205 |
| chr7-97398786-INS-218 | chr7-97398786-INS-218 |
| chr7-97400839-INS-196 | chr7-97400839-INS-204 |
| chr7-97402487-INS-191 | chr7-97402487-INS-191 |
| chr7-97405259-INS-7085 | chr7-97405260-INS-7085 |
| chr7-97406123-INS-325 | chr7-97406123-INS-326 |
| chr7-97409022-INS-200 | chr7-97409022-INS-200 |
| chr7-97423864-INS-95 | chr7-97423864-INS-95 |
| chr7-97445594-INS-1491 | chr7-97445595-INS-1491 |

| | |
|---|---|
| chr7-97471767-INS-114 | chr7-97471767-INS-114 |
| chr7-97485219-INS-222 | chr7-97485219-INS-222 |
| chr7-97494065-INS-205 | chr7-97494065-INS-209 |
| chr7-97496344-INS-202 | chr7-97496344-INS-202 |
| chr7-97497480-INS-206 | chr7-97497480-INS-241 |
| chr7-97525891-INS-119 | chr7-97525891-INS-119 |
| chr7-97546283-INS-76 | chr7-97546283-INS-76 |
| chr7-97555179-INS-220 | chr7-97555179-INS-220 |
| chr7-97607705-INS-60 | chr7-97607705-INS-56 |
| chr7-97631830-INS-70 | chr7-97631830-INS-74 |
| chr7-97632901-INS-7444 | chr7-97632902-INS-7457 |
| chr7-97640725-INS-498 | chr7-97640725-INS-498 |
| chr7-97641507-INS-967 | chr7-97641507-INS-967 |
| chr7-97669103-INS-100 | chr7-97669118-INS-111 |
| chr7-97680304-INS-241 | chr7-97680304-INS-235 |
| chr7-97730931-INS-78 | chr7-97730931-INS-78 |
| chr7-97742452-INS-89 | chr7-97742452-INS-93 |
| chr7-97757000-INS-200 | chr7-97757000-INS-204 |
| chr7-97763494-INS-87 | chr7-97763494-INS-102 |
| chr7-97773609-INS-138 | chr7-97773609-INS-89 |
| chr7-97780612-INS-78 | chr7-97780612-INS-78 |
| chr7-97798647-INS-199 | chr7-97798647-INS-199 |
| chr7-97810345-INS-552 | chr7-97810345-INS-552 |
| chr7-97819924-INS-94 | chr7-97819924-INS-94 |
| chr7-97848830-INS-929 | chr7-97848830-INS-928 |
| chr7-97856119-INS-5077 | chr7-97856120-INS-5076 |
| chr7-97871508-INS-2096 | chr7-97871509-INS-2082 |
| chr7-97893323-INS-124 | chr7-97893323-INS-96 |
| chr7-97899194-INS-62 | chr7-97899194-INS-62 |
| chr7-97917945-INS-52 | chr7-97917945-INS-54 |
| chr7-97919546-INS-175 | chr7-97919547-INS-175 |
| chr7-97925569-INS-100 | chr7-97925562-INS-88 |
| chr7-97932028-INS-525 | chr7-97932028-INS-525 |
| chr7-97941449-INS-288 | chr7-97941449-INS-289 |
| chr7-97964003-INS-384 | chr7-97964004-INS-384 |
| chr7-97970236-INS-52 | chr7-97970236-INS-52 |
| chr7-97978927-INS-179 | chr7-97978927-INS-180 |
| chr7-97981811-INS-78 | chr7-97981811-INS-70 |
| chr7-98014978-INS-310 | chr7-98014978-INS-310 |
| chr7-98016278-INS-50 | chr7-98016283-INS-94 |
| chr7-98020125-INS-270 | chr7-98020125-INS-239 |
| chr7-98021045-INS-55 | chr7-98021045-INS-58 |
| chr7-98039784-INS-181 | chr7-98039784-INS-181 |
| chr7-98062325-INS-449 | chr7-98062342-INS-454 |
| chr7-98069682-INS-185 | chr7-98069682-INS-185 |
| chr7-98075131-INS-508 | chr7-98075131-INS-507 |
| chr7-98088707-INS-172 | chr7-98088707-INS-162 |
| chr7-98127105-INS-146 | chr7-98127105-INS-156 |
| chr7-98153863-INS-60 | chr7-98153851-INS-76 |
| chr7-98163585-INS-193 | chr7-98163585-INS-193 |
| chr7-98216424-INS-200 | chr7-98216424-INS-205 |
| chr7-98227260-INS-468 | chr7-98227260-INS-474 |
| chr7-98303261-INS-1883 | chr7-98303262-INS-1883 |
| chr7-98332607-INS-196 | chr7-98332607-INS-196 |
| chr7-98343712-INS-376 | chr7-98343712-INS-376 |
| chr7-98344630-INS-239 | chr7-98344630-INS-231 |
| chr7-98351064-INS-2392 | chr7-98351065-INS-2392 |
| chr7-98387641-INS-120 | chr7-98387641-INS-120 |
| chr7-98403159-INS-96 | chr7-98403159-INS-96 |
| chr7-98407565-INS-67 | chr7-98407565-INS-67 |
| chr7-98411199-INS-64 | chr7-98411203-INS-68 |
| chr7-98415837-INS-58 | chr7-98415837-INS-58 |
| chr7-98419680-INS-296 | chr7-98419680-INS-296 |
| chr7-98421701-INS-109 | chr7-98421701-INS-109 |
| chr7-98471817-INS-102 | chr7-98471815-INS-108 |
| chr7-98504902-INS-70 | chr7-98504902-INS-70 |
| chr7-98553037-INS-56 | chr7-98553038-INS-74 |
| chr7-98583390-INS-80 | chr7-98583390-INS-68 |
| chr7-98617836-INS-252 | chr7-98617836-INS-252 |
| chr7-98636836-INS-132 | chr7-98636836-INS-132 |
| chr7-98644760-INS-51 | chr7-98644760-INS-51 |
| chr7-98646345-INS-1211 | chr7-98646346-INS-1211 |
| chr7-98673756-INS-73 | chr7-98673756-INS-78 |
| chr7-98708170-INS-56 | chr7-98708134-INS-76 |
| chr7-98708754-INS-54 | chr7-98708754-INS-56 |
| chr7-98724065-INS-68 | chr7-98724065-INS-68 |
| chr7-98739397-INS-60 | chr7-98739397-INS-60 |

| | |
|---|---|
| chr7-98745168-INS-54 | chr7-98745168-INS-54 |
| chr7-98753258-INS-52 | chr7-98753258-INS-56 |
| chr7-98760325-INS-7139 | chr7-98760326-INS-7139 |
| chr7-98771687-INS-52 | chr7-98771687-INS-56 |
| chr7-98775038-INS-501 | chr7-98775038-INS-501 |
| chr7-98779058-INS-133 | chr7-98779082-INS-107 |
| chr7-98782544-INS-245 | chr7-98782544-INS-241 |
| chr7-98784913-INS-70 | chr7-98784913-INS-70 |
| chr7-98832499-INS-59 | chr7-98832499-INS-50 |
| chr7-98859823-INS-233 | chr7-98859823-INS-213 |
| chr7-98877372-INS-90 | chr7-98877372-INS-89 |
| chr7-98879490-INS-163 | chr7-98879490-INS-198 |
| chr7-98889403-INS-63 | chr7-98889363-INS-72 |
| chr7-98893452-INS-193 | chr7-98893452-INS-192 |
| chr7-98896370-INS-430 | chr7-98896370-INS-430 |
| chr7-98898290-INS-51 | chr7-98898290-INS-51 |
| chr7-98910221-INS-70 | chr7-98910221-INS-58 |
| chr7-98937593-INS-64 | chr7-98937593-INS-64 |
| chr7-98951126-INS-6685 | chr7-98951127-INS-6685 |
| chr7-98953713-INS-136 | chr7-98953708-INS-140 |
| chr7-98955111-INS-233 | chr7-98955111-INS-223 |
| chr7-98963727-INS-52 | chr7-98963727-INS-52 |
| chr7-98969814-INS-68 | chr7-98969814-INS-68 |
| chr7-98980750-INS-130 | chr7-98980750-INS-158 |
| chr7-98983020-INS-109 | chr7-98983020-INS-137 |
| chr7-98996727-INS-214 | chr7-98996727-INS-214 |
| chr7-99009790-INS-74 | chr7-99009790-INS-70 |
| chr7-99012363-INS-52 | chr7-99012363-INS-56 |
| chr7-99015613-INS-55 | chr7-99015613-INS-55 |
| chr7-99016786-INS-210 | chr7-99016799-INS-181 |
| chr7-99020587-INS-403 | chr7-99020587-INS-403 |
| chr7-99028622-INS-211 | chr7-99028622-INS-211 |
| chr7-99049599-INS-247 | chr7-99049599-INS-247 |
| chr7-99066896-INS-70 | chr7-99066897-INS-82 |
| chr7-99086870-INS-70 | chr7-99086870-INS-70 |
| chr7-99111963-INS-480 | chr7-99111963-INS-480 |
| chr7-99121891-INS-76 | chr7-99121891-INS-76 |
| chr7-99130233-INS-94 | chr7-99130233-INS-94 |
| chr7-99136672-INS-66 | chr7-99136675-INS-56 |
| chr7-99151401-INS-58 | chr7-99151401-INS-50 |
| chr7-99166698-INS-210 | chr7-99166700-INS-205 |
| chr7-99167139-INS-400 | chr7-99167139-INS-398 |
| chr7-99180586-INS-153 | chr7-99180586-INS-153 |
| chr7-99193338-INS-84 | chr7-99193339-INS-106 |
| chr7-99200849-INS-205 | chr7-99200849-INS-205 |
| chr7-99213708-INS-52 | chr7-99213708-INS-52 |
| chr7-99235121-INS-238 | chr7-99235121-INS-238 |
| chr7-99258994-INS-68 | chr7-99258994-INS-68 |
| chr7-99284153-INS-74 | chr7-99284153-INS-74 |
| chr7-99290275-INS-53 | chr7-99290276-INS-56 |
| chr7-99292383-INS-53 | chr7-99292383-INS-60 |
| chr7-99301859-INS-79 | chr7-99301859-INS-105 |
| chr7-99302883-INS-127 | chr7-99302884-INS-81 |
| chr7-99306027-INS-401 | chr7-99306027-INS-401 |
| chr7-99326702-INS-110 | chr7-99326702-INS-110 |
| chr7-99369255-INS-82 | chr7-99369255-INS-82 |
| chr7-99373908-INS-66 | chr7-99373908-INS-68 |
| chr7-99375779-INS-4114 | chr7-99375780-INS-4114 |
| chr7-99377533-INS-391 | chr7-99377533-INS-391 |
| chr7-99391072-INS-81 | chr7-99391072-INS-81 |
| chr7-99405274-INS-129 | chr7-99405301-INS-133 |
| chr7-99430928-INS-54 | chr7-99430928-INS-54 |
| chr7-99489573-INS-198 | chr7-99489573-INS-198 |
| chr7-99514183-INS-60 | chr7-99514147-INS-62 |
| chr7-99517098-INS-1169 | chr7-99517099-INS-1169 |
| chr7-99539156-INS-190 | chr7-99539156-INS-179 |
| chr7-99552400-INS-490 | chr7-99552400-INS-492 |
| chr7-99586116-INS-57 | chr7-99586116-INS-57 |
| chr7-99614453-INS-253 | chr7-99614453-INS-253 |
| chr7-99615635-INS-4620 | chr7-99615636-INS-4648 |
| chr7-99619936-INS-54 | chr7-99619936-INS-54 |
| chr7-99624843-INS-5363 | chr7-99624844-INS-5363 |
| chr7-99631046-INS-55 | chr7-99631046-INS-52 |
| chr7-99634435-INS-88 | chr7-99634435-INS-80 |
| chr7-99638635-INS-143 | chr7-99638635-INS-143 |
| chr7-99643444-INS-53 | chr7-99643444-INS-59 |
| chr7-99651708-INS-445 | chr7-99651708-INS-445 |

| | |
|---|---|
| chr7-99661231-INS-122 | chr7-99661231-INS-122 |
| chr7-99662235-INS-657 | chr7-99662245-INS-659 |
| chr7-99715625-INS-58 | chr7-99715625-INS-58 |
| chr7-99761412-INS-190 | chr7-99761412-INS-189 |
| chr7-99769492-INS-135 | chr7-99769492-INS-135 |
| chr7-99775656-INS-57 | chr7-99775657-INS-54 |
| chr7-99782652-INS-1450 | chr7-99782653-INS-1450 |
| chr7-99802179-INS-196 | chr7-99802179-INS-196 |
| chr7-99812574-INS-57 | chr7-99812574-INS-57 |
| chr7-99814815-INS-1889 | chr7-99814816-INS-1889 |
| chr7-99817349-INS-489 | chr7-99817342-INS-484 |
| chr7-99842993-INS-78 | chr7-99842987-INS-52 |
| chr7-99844911-INS-68 | chr7-99844911-INS-64 |
| chr7-99846324-INS-247 | chr7-99846324-INS-247 |
| chr7-99859499-INS-78 | chr7-99859499-INS-78 |
| chr7-99875437-INS-194 | chr7-99875437-INS-194 |
| chr7-99894834-INS-82 | chr7-99894834-INS-82 |
| chr7-99900118-INS-7202 | chr7-99900119-INS-7202 |
| chr7-99902902-INS-76 | chr7-99902902-INS-84 |
| chr7-99918453-INS-63 | chr7-99918453-INS-57 |
| chr7-99926175-INS-52 | chr7-99926176-INS-52 |
| chr7-99954050-INS-986 | chr7-99954050-INS-986 |
| chr7-99961829-INS-79 | chr7-99961848-INS-80 |
| chr7-99969499-INS-202 | chr7-99969499-INS-202 |
| chr7-99983621-INS-78 | chr7-99983621-INS-78 |
| chr7-100022730-INS-1079 | chr7-100022731-INS-1079 |
| chr7-100038330-INS-5616 | chr7-100038331-INS-5617 |
| chr7-100039843-INS-52 | chr7-100039843-INS-52 |
| chr7-100070200-INS-90 | chr7-100070224-INS-91 |
| chr7-100075361-INS-703 | chr7-100075361-INS-712 |
| chr7-100098823-INS-62 | chr7-100098819-INS-78 |
| chr7-100112283-INS-383 | chr7-100112283-INS-384 |
| chr7-100126673-INS-56 | chr7-100126673-INS-56 |
| chr7-100138002-INS-430 | chr7-100138002-INS-431 |
| chr7-100158395-INS-59 | chr7-100158395-INS-55 |
| chr7-100164534-INS-123 | chr7-100164534-INS-124 |
| chr7-100174342-INS-53 | chr7-100174342-INS-53 |
| chr7-100200459-INS-189 | chr7-100200459-INS-189 |
| chr7-100205647-INS-4004 | chr7-100205648-INS-4036 |
| chr7-100214152-INS-498 | chr7-100214152-INS-498 |
| chr7-100218445-INS-196 | chr7-100218445-INS-196 |
| chr7-100219995-INS-215 | chr7-100219995-INS-215 |
| chr7-100230900-INS-498 | chr7-100230900-INS-498 |
| chr7-100240896-INS-60 | chr7-100240896-INS-54 |
| chr7-100253243-INS-149 | chr7-100253259-INS-137 |
| chr7-100274058-INS-235 | chr7-100274058-INS-236 |
| chr7-100276997-INS-64 | chr7-100277020-INS-69 |
| chr7-100312879-INS-226 | chr7-100312879-INS-226 |
| chr7-100315176-INS-248 | chr7-100315176-INS-248 |
| chr7-100317536-INS-68 | chr7-100317536-INS-56 |
| chr7-100321119-INS-58 | chr7-100321119-INS-58 |
| chr7-100324295-INS-412 | chr7-100324295-INS-412 |
| chr7-100327015-INS-192 | chr7-100327015-INS-199 |
| chr7-100332954-INS-227 | chr7-100332954-INS-227 |
| chr7-100336031-INS-667 | chr7-100336031-INS-669 |
| chr7-100344029-INS-147 | chr7-100344029-INS-159 |
| chr7-100347291-INS-197 | chr7-100347291-INS-181 |
| chr7-100359969-INS-196 | chr7-100359969-INS-196 |
| chr7-100369654-INS-360 | chr7-100369654-INS-405 |
| chr7-100375761-INS-219 | chr7-100375761-INS-219 |
| chr7-100381563-INS-210 | chr7-100381563-INS-210 |
| chr7-100383101-INS-56 | chr7-100383101-INS-56 |
| chr7-100405990-INS-68 | chr7-100405990-INS-68 |
| chr7-100412814-INS-85 | chr7-100412800-INS-112 |
| chr7-100424985-INS-202 | chr7-100424985-INS-206 |
| chr7-100438915-INS-62 | chr7-100438915-INS-62 |
| chr7-100443487-INS-196 | chr7-100443487-INS-196 |
| chr7-100451379-INS-150 | chr7-100451379-INS-150 |
| chr7-100452325-INS-231 | chr7-100452325-INS-232 |
| chr7-100455470-INS-220 | chr7-100455470-INS-220 |
| chr7-100456668-INS-67 | chr7-100456668-INS-68 |
| chr7-100495015-INS-211 | chr7-100495015-INS-211 |
| chr7-100524452-INS-261 | chr7-100524452-INS-246 |
| chr7-100571695-INS-56 | chr7-100571695-INS-56 |
| chr7-100584787-INS-98 | chr7-100584787-INS-98 |
| chr7-100597881-INS-74 | chr7-100597881-INS-74 |
| chr7-100610032-INS-61 | chr7-100610032-INS-61 |

| | |
|---|---|
| chr7-100620776-INS-3425 | chr7-100620777-INS-3425 |
| chr7-100626045-INS-66 | chr7-100626045-INS-68 |
| chr7-100656873-INS-239 | chr7-100656873-INS-239 |
| chr7-100674304-INS-114 | chr7-100674304-INS-114 |
| chr7-100696449-INS-116 | chr7-100696449-INS-116 |
| chr7-100710947-INS-58 | chr7-100710947-INS-56 |
| chr7-100719658-INS-106 | chr7-100719658-INS-113 |
| chr7-100735836-INS-57 | chr7-100735837-INS-51 |
| chr7-100745493-INS-79 | chr7-100745493-INS-63 |
| chr7-100751870-INS-88 | chr7-100751866-INS-54 |
| chr7-100825048-INS-113 | chr7-100825052-INS-131 |
| chr7-100845644-INS-197 | chr7-100845644-INS-197 |
| chr7-100865527-INS-58 | chr7-100865527-INS-62 |
| chr7-100918113-INS-207 | chr7-100918113-INS-207 |
| chr7-100930140-INS-1875 | chr7-100930141-INS-1861 |
| chr7-100940436-INS-164 | chr7-100940439-INS-165 |
| chr7-100944499-INS-89 | chr7-100944535-INS-52 |
| chr7-100959202-INS-198 | chr7-100959202-INS-198 |
| chr7-100961253-INS-210 | chr7-100961253-INS-210 |
| chr7-101067535-INS-118 | chr7-101067495-INS-119 |
| chr7-101078227-INS-306 | chr7-101078227-INS-276 |
| chr7-101081102-INS-64 | chr7-101081102-INS-64 |
| chr7-101096826-INS-54 | chr7-101096826-INS-54 |
| chr7-101116400-INS-287 | chr7-101116400-INS-287 |
| chr7-101127044-INS-64 | chr7-101127045-INS-68 |
| chr7-101142914-INS-64 | chr7-101142914-INS-56 |
| chr7-101193234-INS-56 | chr7-101193226-INS-64 |
| chr7-101194518-INS-63 | chr7-101194519-INS-51 |
| chr7-101206117-INS-144 | chr7-101206117-INS-144 |
| chr7-101213385-INS-347 | chr7-101213385-INS-347 |
| chr7-101231453-INS-134 | chr7-101231453-INS-108 |
| chr7-101234459-INS-210 | chr7-101234459-INS-212 |
| chr7-101255986-INS-215 | chr7-101255986-INS-217 |
| chr7-101260225-INS-91 | chr7-101260225-INS-91 |
| chr7-101264582-INS-218 | chr7-101264582-INS-219 |
| chr7-101276379-INS-61 | chr7-101276379-INS-61 |
| chr7-101277506-INS-135 | chr7-101277506-INS-135 |
| chr7-101291341-INS-79 | chr7-101291355-INS-90 |
| chr7-101300070-INS-195 | chr7-101300070-INS-195 |
| chr7-101303982-INS-60 | chr7-101303982-INS-60 |
| chr7-101308002-INS-7093 | chr7-101308003-INS-7093 |
| chr7-101310100-INS-193 | chr7-101310100-INS-195 |
| chr7-101321944-INS-89 | chr7-101321944-INS-89 |
| chr7-101328040-INS-147 | chr7-101328040-INS-147 |
| chr7-101397995-INS-148 | chr7-101397995-INS-148 |
| chr7-101402650-INS-346 | chr7-101402650-INS-346 |
| chr7-101409648-INS-69 | chr7-101409648-INS-69 |
| chr7-101435668-INS-67 | chr7-101435668-INS-67 |
| chr7-101436369-INS-65 | chr7-101436369-INS-65 |
| chr7-101439977-INS-55 | chr7-101439977-INS-55 |
| chr7-101449875-INS-97 | chr7-101449875-INS-97 |
| chr7-101455188-INS-68 | chr7-101455188-INS-68 |
| chr7-101456265-INS-53 | chr7-101456265-INS-53 |
| chr7-101459244-INS-92 | chr7-101459234-INS-88 |
| chr7-101463924-INS-198 | chr7-101463923-INS-196 |
| chr7-101466441-INS-68 | chr7-101466434-INS-84 |
| chr7-101471993-INS-87 | chr7-101471995-INS-98 |
| chr7-101498881-INS-70 | chr7-101498857-INS-71 |
| chr7-101502762-INS-65 | chr7-101502762-INS-75 |
| chr7-101507334-INS-52 | chr7-101507334-INS-52 |
| chr7-101521218-INS-104 | chr7-101521222-INS-80 |
| chr7-101526259-INS-58 | chr7-101526259-INS-70 |
| chr7-101528559-INS-134 | chr7-101528559-INS-164 |
| chr7-101529636-INS-190 | chr7-101529636-INS-189 |
| chr7-101530539-INS-363 | chr7-101530539-INS-335 |
| chr7-101536521-INS-52 | chr7-101536521-INS-52 |
| chr7-101545174-INS-212 | chr7-101545174-INS-212 |
| chr7-101596566-INS-177 | chr7-101596566-INS-178 |
| chr7-101611173-INS-210 | chr7-101611173-INS-212 |
| chr7-101612415-INS-195 | chr7-101612415-INS-195 |
| chr7-101668195-INS-99 | chr7-101668195-INS-67 |
| chr7-101674697-INS-687 | chr7-101674697-INS-687 |
| chr7-101684473-INS-193 | chr7-101684473-INS-193 |
| chr7-101690414-INS-170 | chr7-101690414-INS-170 |
| chr7-101697268-INS-85 | chr7-101697268-INS-50 |
| chr7-101703710-INS-206 | chr7-101703710-INS-205 |
| chr7-101710730-INS-208 | chr7-101710730-INS-208 |

| | |
|---|---|
| chr7-101717920-INS-200 | chr7-101717920-INS-207 |
| chr7-101720283-INS-440 | chr7-101720283-INS-440 |
| chr7-101723246-INS-60 | chr7-101723256-INS-65 |
| chr7-101738588-INS-99 | chr7-101738588-INS-54 |
| chr7-101744740-INS-408 | chr7-101744740-INS-398 |
| chr7-101753930-INS-130 | chr7-101753930-INS-130 |
| chr7-101771942-INS-1892 | chr7-101771943-INS-1892 |
| chr7-101777250-INS-236 | chr7-101777250-INS-236 |
| chr7-101778764-INS-218 | chr7-101778764-INS-218 |
| chr7-101799731-INS-64 | chr7-101799731-INS-64 |
| chr7-101813770-INS-321 | chr7-101813770-INS-324 |
| chr7-101816176-INS-200 | chr7-101816169-INS-200 |
| chr7-101858486-INS-248 | chr7-101858486-INS-250 |
| chr7-101861509-INS-211 | chr7-101861509-INS-209 |
| chr7-101865854-INS-204 | chr7-101865854-INS-205 |
| chr7-101894863-INS-496 | chr7-101894813-INS-537 |
| chr7-101912430-INS-54 | chr7-101912430-INS-54 |
| chr7-101915250-INS-309 | chr7-101915248-INS-312 |
| chr7-101930468-INS-832 | chr7-101930469-INS-832 |
| chr7-101935929-INS-187 | chr7-101935929-INS-190 |
| chr7-101943208-INS-98 | chr7-101943208-INS-98 |
| chr7-101948881-INS-56 | chr7-101948880-INS-50 |
| chr7-101970800-INS-205 | chr7-101970800-INS-205 |
| chr7-101972820-INS-504 | chr7-101972820-INS-507 |
| chr7-101973606-INS-75 | chr7-101973607-INS-75 |
| chr7-101987582-INS-84 | chr7-101987582-INS-90 |
| chr7-101990595-INS-966 | chr7-101990595-INS-966 |
| chr7-101993161-INS-1008 | chr7-101993162-INS-1008 |
| chr7-102009018-INS-310 | chr7-102009018-INS-311 |
| chr7-102023456-INS-52 | chr7-102023456-INS-52 |
| chr7-102031676-INS-231 | chr7-102031676-INS-208 |
| chr7-102036564-INS-212 | chr7-102036564-INS-212 |
| chr7-102042276-INS-71 | chr7-102042261-INS-119 |
| chr7-102083019-INS-197 | chr7-102083019-INS-197 |
| chr7-102090577-INS-174 | chr7-102090578-INS-174 |
| chr7-102102437-INS-220 | chr7-102102437-INS-220 |
| chr7-102107073-INS-208 | chr7-102107073-INS-208 |
| chr7-102137838-INS-178 | chr7-102137838-INS-178 |
| chr7-102141979-INS-180 | chr7-102141979-INS-180 |
| chr7-102150270-INS-144 | chr7-102150270-INS-144 |
| chr7-102165351-INS-51 | chr7-102165351-INS-51 |
| chr7-102175807-INS-942 | chr7-102175808-INS-942 |
| chr7-102176956-INS-62 | chr7-102176956-INS-62 |
| chr7-102181260-INS-68 | chr7-102181260-INS-70 |
| chr7-102182763-INS-414 | chr7-102182764-INS-417 |
| chr7-102204962-INS-204 | chr7-102204962-INS-204 |
| chr7-102223446-INS-60 | chr7-102223480-INS-52 |
| chr7-102248607-INS-113 | chr7-102248579-INS-127 |
| chr7-102255617-INS-283 | chr7-102255617-INS-279 |
| chr7-102266893-INS-69 | chr7-102266893-INS-74 |
| chr7-102286191-INS-108 | chr7-102286194-INS-104 |
| chr7-102309403-INS-57 | chr7-102309403-INS-57 |
| chr7-102339311-INS-6454 | chr7-102339312-INS-6454 |
| chr7-102342185-INS-54 | chr7-102342185-INS-54 |
| chr7-102352340-INS-104 | chr7-102352338-INS-108 |
| chr7-102369932-INS-6309 | chr7-102369933-INS-6309 |
| chr7-102375042-INS-132 | chr7-102375042-INS-132 |
| chr7-102388572-INS-1148 | chr7-102388573-INS-1147 |
| chr7-102389211-INS-142 | chr7-102389209-INS-150 |
| chr7-102399311-INS-952 | chr7-102399311-INS-952 |
| chr7-102400318-INS-409 | chr7-102400318-INS-409 |
| chr7-102404618-INS-59 | chr7-102404618-INS-59 |
| chr7-102411317-INS-103 | chr7-102411300-INS-92 |
| chr7-102420707-INS-55 | chr7-102420707-INS-55 |
| chr7-102426377-INS-53 | chr7-102426377-INS-53 |
| chr7-102427367-INS-100 | chr7-102427367-INS-100 |
| chr7-102430052-INS-55 | chr7-102430048-INS-59 |
| chr7-102465590-INS-51 | chr7-102465590-INS-51 |
| chr7-102469424-INS-198 | chr7-102469401-INS-197 |
| chr7-102517223-INS-6665 | chr7-102517224-INS-6670 |
| chr7-102518568-INS-64 | chr7-102518568-INS-64 |
| chr7-102531273-INS-55 | chr7-102531273-INS-59 |
| chr7-102538531-INS-117 | chr7-102538531-INS-117 |
| chr7-102567058-INS-185 | chr7-102567058-INS-185 |
| chr7-102571777-INS-60 | chr7-102571762-INS-92 |
| chr7-102572346-INS-356 | chr7-102572346-INS-366 |
| chr7-102581424-INS-54 | chr7-102581430-INS-77 |

| | |
|---|---|
| chr7-102582441-INS-1994 | chr7-102582442-INS-1994 |
| chr7-102609539-INS-67 | chr7-102609539-INS-67 |
| chr7-102635404-INS-95 | chr7-102635380-INS-76 |
| chr7-102638374-INS-106 | chr7-102638374-INS-106 |
| chr7-102642115-INS-386 | chr7-102642115-INS-386 |
| chr7-102645137-INS-224 | chr7-102645137-INS-224 |
| chr7-102664268-INS-76 | chr7-102664268-INS-80 |
| chr7-102697762-INS-7734 | chr7-102697763-INS-7736 |
| chr7-102704564-INS-527 | chr7-102704564-INS-527 |
| chr7-102705747-INS-99 | chr7-102705735-INS-102 |
| chr7-102719065-INS-131 | chr7-102719065-INS-115 |
| chr7-102743205-INS-265 | chr7-102743205-INS-264 |
| chr7-102747915-INS-442 | chr7-102747915-INS-440 |
| chr7-102749945-INS-7164 | chr7-102749946-INS-7164 |
| chr7-102766959-INS-65 | chr7-102766959-INS-65 |
| chr7-102788441-INS-1890 | chr7-102788442-INS-1890 |
| chr7-102815948-INS-6341 | chr7-102815949-INS-6341 |
| chr7-102843836-INS-51 | chr7-102843838-INS-86 |
| chr7-102889413-INS-401 | chr7-102889413-INS-401 |
| chr7-102902220-INS-67 | chr7-102902220-INS-55 |
| chr7-102919167-INS-370 | chr7-102919167-INS-370 |
| chr7-102931871-INS-208 | chr7-102931871-INS-208 |
| chr7-102955782-INS-196 | chr7-102955782-INS-196 |
| chr7-103018030-INS-717 | chr7-103018030-INS-723 |
| chr7-103026021-INS-181 | chr7-103026021-INS-181 |
| chr7-103034938-INS-960 | chr7-103034938-INS-960 |
| chr7-103045335-INS-63 | chr7-103045335-INS-67 |
| chr7-103048671-INS-977 | chr7-103048671-INS-977 |
| chr7-103061870-INS-2375 | chr7-103061871-INS-2367 |
| chr7-103063892-INS-70 | chr7-103063888-INS-50 |
| chr7-103081602-INS-715 | chr7-103081602-INS-715 |
| chr7-103101635-INS-539 | chr7-103101635-INS-539 |
| chr7-103112636-INS-120 | chr7-103112636-INS-120 |
| chr7-103114325-INS-52 | chr7-103114325-INS-54 |
| chr7-103146807-INS-59 | chr7-103146799-INS-51 |
| chr7-103150354-INS-65 | chr7-103150354-INS-65 |
| chr7-103164366-INS-3386 | chr7-103164367-INS-3386 |
| chr7-103177888-INS-204 | chr7-103177888-INS-204 |
| chr7-103186506-INS-52 | chr7-103186506-INS-52 |
| chr7-103192977-INS-128 | chr7-103192977-INS-128 |
| chr7-103262535-INS-96 | chr7-103262531-INS-96 |
| chr7-103269115-INS-8854 | chr7-103269116-INS-8854 |
| chr7-103281838-INS-74 | chr7-103281838-INS-70 |
| chr7-103284551-INS-339 | chr7-103284551-INS-339 |
| chr7-103290159-INS-161 | chr7-103290159-INS-161 |
| chr7-103305728-INS-54 | chr7-103305728-INS-64 |
| chr7-103326725-INS-565 | chr7-103326725-INS-565 |
| chr7-103327655-INS-64 | chr7-103327655-INS-64 |
| chr7-103328077-INS-828 | chr7-103328077-INS-828 |
| chr7-103371585-INS-97 | chr7-103371585-INS-97 |
| chr7-103377299-INS-571 | chr7-103377300-INS-571 |
| chr7-103377440-INS-5568 | chr7-103377441-INS-5568 |
| chr7-103393187-INS-52 | chr7-103393187-INS-52 |
| chr7-103398541-INS-101 | chr7-103398541-INS-101 |
| chr7-103401559-INS-61 | chr7-103401559-INS-53 |
| chr7-103407451-INS-342 | chr7-103407451-INS-342 |
| chr7-103429813-INS-133 | chr7-103429812-INS-88 |
| chr7-103430911-INS-964 | chr7-103430911-INS-964 |
| chr7-103438512-INS-53 | chr7-103438511-INS-55 |
| chr7-103443160-INS-63 | chr7-103443159-INS-95 |
| chr7-103451670-INS-938 | chr7-103451670-INS-938 |
| chr7-103479946-INS-92 | chr7-103479953-INS-95 |
| chr7-103505343-INS-52 | chr7-103505343-INS-56 |
| chr7-103520256-INS-70 | chr7-103520249-INS-52 |
| chr7-103527924-INS-141 | chr7-103527924-INS-141 |
| chr7-103549936-INS-90 | chr7-103549936-INS-90 |
| chr7-103560060-INS-150 | chr7-103560020-INS-161 |
| chr7-103563781-INS-518 | chr7-103563781-INS-518 |
| chr7-103567847-INS-78 | chr7-103567847-INS-79 |
| chr7-103580850-INS-544 | chr7-103580850-INS-544 |
| chr7-103594042-INS-61 | chr7-103594039-INS-57 |
| chr7-103595519-INS-88 | chr7-103595519-INS-112 |
| chr7-103613525-INS-5495 | chr7-103613526-INS-5495 |
| chr7-103662439-INS-997 | chr7-103662439-INS-1008 |
| chr7-103696576-INS-122 | chr7-103696576-INS-116 |
| chr7-103710435-INS-65 | chr7-103710435-INS-52 |
| chr7-103723072-INS-84 | chr7-103723072-INS-76 |

| | |
|---|---|
| chr7-103724121-INS-56 | chr7-103724122-INS-56 |
| chr7-103732677-INS-433 | chr7-103732677-INS-432 |
| chr7-103742238-INS-155 | chr7-103742238-INS-155 |
| chr7-103765423-INS-121 | chr7-103765423-INS-121 |
| chr7-103787726-INS-57 | chr7-103787727-INS-55 |
| chr7-103804137-INS-62 | chr7-103804137-INS-62 |
| chr7-103814896-INS-372 | chr7-103814896-INS-372 |
| chr7-103842558-INS-80 | chr7-103842558-INS-80 |
| chr7-103848657-INS-82 | chr7-103848657-INS-66 |
| chr7-103853610-INS-87 | chr7-103853610-INS-82 |
| chr7-103863375-INS-54 | chr7-103863343-INS-52 |
| chr7-103872952-INS-220 | chr7-103872954-INS-195 |
| chr7-103874370-INS-176 | chr7-103874370-INS-176 |
| chr7-103878539-INS-103 | chr7-103878539-INS-103 |
| chr7-103882736-INS-87 | chr7-103882699-INS-90 |
| chr7-103888549-INS-78 | chr7-103888549-INS-121 |
| chr7-103890982-INS-82 | chr7-103890982-INS-82 |
| chr7-103907123-INS-66 | chr7-103907123-INS-66 |
| chr7-103913718-INS-78 | chr7-103913768-INS-52 |
| chr7-103963207-INS-359 | chr7-103963207-INS-352 |
| chr7-103980248-INS-176 | chr7-103980248-INS-177 |
| chr7-103997139-INS-7065 | chr7-103997140-INS-7065 |
| chr7-103999249-INS-1444 | chr7-103999250-INS-1444 |
| chr7-104012324-INS-381 | chr7-104012324-INS-381 |
| chr7-104051866-INS-371 | chr7-104051866-INS-371 |
| chr7-104093558-INS-194 | chr7-104093558-INS-194 |
| chr7-104152652-INS-7115 | chr7-104152653-INS-7115 |
| chr7-104156210-INS-251 | chr7-104156213-INS-242 |
| chr7-104207064-INS-727 | chr7-104207064-INS-727 |
| chr7-104233857-INS-140 | chr7-104233857-INS-140 |
| chr7-104239446-INS-1311 | chr7-104239447-INS-1311 |
| chr7-104249244-INS-233 | chr7-104249244-INS-233 |
| chr7-104255484-INS-4217 | chr7-104255485-INS-4218 |
| chr7-104261833-INS-328 | chr7-104261833-INS-328 |
| chr7-104263832-INS-365 | chr7-104263862-INS-379 |
| chr7-104275095-INS-231 | chr7-104275095-INS-234 |
| chr7-104295063-INS-90 | chr7-104295063-INS-90 |
| chr7-104307902-INS-694 | chr7-104307902-INS-694 |
| chr7-104312501-INS-192 | chr7-104312501-INS-192 |
| chr7-104318017-INS-1850 | chr7-104318018-INS-1847 |
| chr7-104344478-INS-7400 | chr7-104344479-INS-7400 |
| chr7-104370462-INS-427 | chr7-104370462-INS-427 |
| chr7-104375597-INS-368 | chr7-104375597-INS-368 |
| chr7-104460379-INS-80 | chr7-104460379-INS-77 |
| chr7-104465608-INS-56 | chr7-104465598-INS-66 |
| chr7-104474163-INS-213 | chr7-104474163-INS-213 |
| chr7-104476391-INS-70 | chr7-104476389-INS-68 |
| chr7-104483278-INS-6440 | chr7-104483279-INS-6436 |
| chr7-104495797-INS-5132 | chr7-104495798-INS-5132 |
| chr7-104516282-INS-5449 | chr7-104516283-INS-5449 |
| chr7-104522070-INS-6369 | chr7-104522071-INS-6369 |
| chr7-104537913-INS-163 | chr7-104537913-INS-163 |
| chr7-104582699-INS-96 | chr7-104582711-INS-68 |
| chr7-104610795-INS-75 | chr7-104610795-INS-70 |
| chr7-104669774-INS-414 | chr7-104669774-INS-406 |
| chr7-104699423-INS-66 | chr7-104699423-INS-63 |
| chr7-104701106-INS-7110 | chr7-104701107-INS-7110 |
| chr7-104702948-INS-60 | chr7-104702948-INS-60 |
| chr7-104708830-INS-56 | chr7-104708830-INS-68 |
| chr7-104711351-INS-111 | chr7-104711351-INS-117 |
| chr7-104740662-INS-172 | chr7-104740662-INS-171 |
| chr7-104759154-INS-185 | chr7-104759152-INS-194 |
| chr7-104809106-INS-6827 | chr7-104809107-INS-6830 |
| chr7-104826082-INS-56 | chr7-104826086-INS-52 |
| chr7-104846780-INS-1502 | chr7-104846781-INS-1502 |
| chr7-104857503-INS-467 | chr7-104857503-INS-467 |
| chr7-104882801-INS-6356 | chr7-104882802-INS-6356 |
| chr7-104926441-INS-51 | chr7-104926441-INS-60 |
| chr7-104927634-INS-56 | chr7-104927634-INS-54 |
| chr7-104947412-INS-3182 | chr7-104947413-INS-3185 |
| chr7-104978342-INS-62 | chr7-104978349-INS-74 |
| chr7-105052545-INS-2027 | chr7-105052546-INS-2011 |
| chr7-105080555-INS-97 | chr7-105080555-INS-93 |
| chr7-105090822-INS-58 | chr7-105090822-INS-58 |
| chr7-105094236-INS-525 | chr7-105094236-INS-527 |
| chr7-105095393-INS-6433 | chr7-105095394-INS-6433 |
| chr7-105097368-INS-740 | chr7-105097368-INS-782 |

| | |
|---|---|
| chr7-105099457-INS-269 | chr7-105099457-INS-269 |
| chr7-105100711-INS-334 | chr7-105100711-INS-334 |
| chr7-105106613-INS-881 | chr7-105106613-INS-881 |
| chr7-105114337-INS-53 | chr7-105114347-INS-70 |
| chr7-105157553-INS-483 | chr7-105157553-INS-483 |
| chr7-105160257-INS-243 | chr7-105160263-INS-250 |
| chr7-105161602-INS-84 | chr7-105161602-INS-84 |
| chr7-105210088-INS-94 | chr7-105210092-INS-118 |
| chr7-105235600-INS-78 | chr7-105235588-INS-99 |
| chr7-105237903-INS-123 | chr7-105237903-INS-123 |
| chr7-105259731-INS-219 | chr7-105259731-INS-212 |
| chr7-105292476-INS-60 | chr7-105292481-INS-50 |
| chr7-105299538-INS-78 | chr7-105299538-INS-78 |
| chr7-105419177-INS-7074 | chr7-105419178-INS-7074 |
| chr7-105462147-INS-1407 | chr7-105462144-INS-1398 |
| chr7-105467388-INS-172 | chr7-105467388-INS-173 |
| chr7-105485110-INS-572 | chr7-105485110-INS-573 |
| chr7-105488799-INS-57 | chr7-105488799-INS-54 |
| chr7-105588459-INS-105 | chr7-105588445-INS-105 |
| chr7-105833331-INS-4777 | chr7-105833332-INS-4777 |
| chr7-105983914-INS-404 | chr7-105983914-INS-408 |
| chr7-105995444-INS-326 | chr7-105995444-INS-325 |
| chr7-106007556-INS-2777 | chr7-106007557-INS-2777 |
| chr7-106015651-INS-2104 | chr7-106015652-INS-2104 |
| chr7-106016665-INS-920 | chr7-106016665-INS-919 |
| chr7-106027640-INS-931 | chr7-106027640-INS-931 |
| chr7-106028276-INS-82 | chr7-106028276-INS-82 |
| chr7-106072149-INS-81 | chr7-106072149-INS-81 |
| chr7-106073263-INS-2075 | chr7-106073264-INS-2075 |
| chr7-106084866-INS-234 | chr7-106084866-INS-230 |
| chr7-106099527-INS-56 | chr7-106099527-INS-56 |
| chr7-106102374-INS-62 | chr7-106102374-INS-62 |
| chr7-106123497-INS-14336 | chr7-106123498-INS-14340 |
| chr7-106139627-INS-1492 | chr7-106139628-INS-1493 |
| chr7-106144002-INS-140 | chr7-106144002-INS-144 |
| chr7-106161024-INS-6607 | chr7-106161025-INS-6606 |
| chr7-106176869-INS-324 | chr7-106176869-INS-324 |
| chr7-106180338-INS-5696 | chr7-106180339-INS-5696 |
| chr7-106183964-INS-231 | chr7-106183964-INS-231 |
| chr7-106192373-INS-124 | chr7-106192373-INS-124 |
| chr7-106194381-INS-964 | chr7-106194381-INS-964 |
| chr7-106200106-INS-206 | chr7-106200106-INS-206 |
| chr7-106221221-INS-6207 | chr7-106221222-INS-6208 |
| chr7-106222797-INS-64 | chr7-106222807-INS-68 |
| chr7-106242959-INS-7386 | chr7-106242960-INS-7386 |
| chr7-106272804-INS-963 | chr7-106272804-INS-963 |
| chr7-106296971-INS-3098 | chr7-106296972-INS-3098 |
| chr7-106303045-INS-1234 | chr7-106303046-INS-1233 |
| chr7-106399009-INS-6399 | chr7-106399010-INS-6399 |
| chr7-106412452-INS-475 | chr7-106412452-INS-475 |
| chr7-106421373-INS-6641 | chr7-106421374-INS-6641 |
| chr7-106454439-INS-6652 | chr7-106454440-INS-6652 |
| chr7-106463211-INS-263 | chr7-106463211-INS-262 |
| chr7-106477455-INS-7078 | chr7-106477456-INS-7079 |
| chr7-106500860-INS-374 | chr7-106500860-INS-355 |
| chr7-106514740-INS-6485 | chr7-106514741-INS-6485 |
| chr7-106527091-INS-68 | chr7-106527111-INS-78 |
| chr7-106533133-INS-500 | chr7-106533133-INS-503 |
| chr7-106550677-INS-143 | chr7-106550677-INS-143 |
| chr7-106558913-INS-67 | chr7-106558918-INS-72 |
| chr7-106558922-INS-9114 | chr7-106558923-INS-9116 |
| chr7-106567037-INS-52 | chr7-106567037-INS-52 |
| chr7-106572863-INS-337 | chr7-106572863-INS-337 |
| chr7-106605797-INS-64 | chr7-106605782-INS-70 |
| chr7-106616528-INS-1309 | chr7-106616529-INS-1310 |
| chr7-106630362-INS-430 | chr7-106630362-INS-430 |
| chr7-106636676-INS-71 | chr7-106636676-INS-71 |
| chr7-106651379-INS-330 | chr7-106651379-INS-330 |
| chr7-106659189-INS-2613 | chr7-106659190-INS-2613 |
| chr7-106664291-INS-80 | chr7-106664291-INS-80 |
| chr7-106711577-INS-76 | chr7-106711577-INS-79 |
| chr7-106732155-INS-54 | chr7-106732155-INS-59 |
| chr7-106749941-INS-155 | chr7-106749941-INS-156 |
| chr7-106750248-INS-7517 | chr7-106750249-INS-7517 |
| chr7-106761608-INS-744 | chr7-106761604-INS-744 |
| chr7-106778137-INS-188 | chr7-106778138-INS-191 |
| chr7-106794198-INS-668 | chr7-106794198-INS-668 |

| | |
|---|---|
| chr7-106800033-INS-331 | chr7-106800033-INS-331 |
| chr7-106801453-INS-1146 | chr7-106801454-INS-1146 |
| chr7-106851944-INS-223 | chr7-106851944-INS-223 |
| chr7-106909654-INS-176 | chr7-106909654-INS-176 |
| chr7-106911395-INS-202 | chr7-106911395-INS-202 |
| chr7-106917061-INS-231 | chr7-106917061-INS-233 |
| chr7-106928621-INS-1894 | chr7-106928622-INS-1894 |
| chr7-106929932-INS-56 | chr7-106929932-INS-52 |
| chr7-106945905-INS-64 | chr7-106945905-INS-55 |
| chr7-106948715-INS-181 | chr7-106948715-INS-182 |
| chr7-106966687-INS-58 | chr7-106966687-INS-60 |
| chr7-106975473-INS-68 | chr7-106975479-INS-71 |
| chr7-106994051-INS-178 | chr7-106994051-INS-179 |
| chr7-107000322-INS-190 | chr7-107000322-INS-188 |
| chr7-107019100-INS-515 | chr7-107019100-INS-515 |
| chr7-107020446-INS-815 | chr7-107020446-INS-814 |
| chr7-107021837-INS-79 | chr7-107021837-INS-83 |
| chr7-107023694-INS-156 | chr7-107023694-INS-154 |
| chr7-107043398-INS-52 | chr7-107043398-INS-52 |
| chr7-107057398-INS-6405 | chr7-107057399-INS-6404 |
| chr7-107072384-INS-2918 | chr7-107072384-INS-2918 |
| chr7-107082244-INS-126 | chr7-107082244-INS-126 |
| chr7-107095398-INS-75 | chr7-107095398-INS-50 |
| chr7-107102422-INS-65 | chr7-107102422-INS-65 |
| chr7-107115789-INS-52 | chr7-107115792-INS-78 |
| chr7-107124149-INS-193 | chr7-107124149-INS-183 |
| chr7-107171385-INS-607 | chr7-107171385-INS-607 |
| chr7-107172612-INS-87 | chr7-107172612-INS-89 |
| chr7-107179903-INS-107 | chr7-107179915-INS-116 |
| chr7-107209658-INS-112 | chr7-107209658-INS-111 |
| chr7-107221665-INS-170 | chr7-107221665-INS-170 |
| chr7-107231148-INS-104 | chr7-107231148-INS-104 |
| chr7-107256727-INS-195 | chr7-107256728-INS-195 |
| chr7-107271120-INS-84 | chr7-107271120-INS-84 |
| chr7-107275613-INS-97 | chr7-107275613-INS-97 |
| chr7-107291453-INS-89 | chr7-107291453-INS-89 |
| chr7-107300761-INS-422 | chr7-107300761-INS-422 |
| chr7-107336414-INS-181 | chr7-107336414-INS-181 |
| chr7-107353649-INS-58 | chr7-107353649-INS-58 |
| chr7-107356762-INS-58 | chr7-107356761-INS-58 |
| chr7-107365130-INS-562 | chr7-107365125-INS-562 |
| chr7-107366469-INS-377 | chr7-107366469-INS-377 |
| chr7-107366760-INS-6593 | chr7-107366761-INS-6593 |
| chr7-107371665-INS-178 | chr7-107371665-INS-179 |
| chr7-107374697-INS-68 | chr7-107374697-INS-68 |
| chr7-107379264-INS-50 | chr7-107379264-INS-50 |
| chr7-107381650-INS-63 | chr7-107381650-INS-66 |
| chr7-107381650-INS-846 | chr7-107381650-INS-844 |
| chr7-107387740-INS-72 | chr7-107387740-INS-73 |
| chr7-107394704-INS-187 | chr7-107394704-INS-171 |
| chr7-107407981-INS-137 | chr7-107407981-INS-154 |
| chr7-107430566-INS-195 | chr7-107430569-INS-195 |
| chr7-107436251-INS-637 | chr7-107436252-INS-637 |
| chr7-107485776-INS-285 | chr7-107485776-INS-285 |
| chr7-107495898-INS-918 | chr7-107495898-INS-918 |
| chr7-107550118-INS-120 | chr7-107550118-INS-128 |
| chr7-107577567-INS-162 | chr7-107577567-INS-132 |
| chr7-107581612-INS-53 | chr7-107581612-INS-53 |
| chr7-107600090-INS-187 | chr7-107600090-INS-188 |
| chr7-107601693-INS-122 | chr7-107601710-INS-142 |
| chr7-107604534-INS-809 | chr7-107604534-INS-810 |
| chr7-107613888-INS-641 | chr7-107613888-INS-644 |
| chr7-107618724-INS-280 | chr7-107618725-INS-280 |
| chr7-107628184-INS-503 | chr7-107628184-INS-503 |
| chr7-107630450-INS-80 | chr7-107630450-INS-81 |
| chr7-107678533-INS-105 | chr7-107678533-INS-105 |
| chr7-107679407-INS-579 | chr7-107679407-INS-580 |
| chr7-107681963-INS-100 | chr7-107681963-INS-100 |
| chr7-107685925-INS-402 | chr7-107685925-INS-415 |
| chr7-107695653-INS-63 | chr7-107695653-INS-57 |
| chr7-107696973-INS-1877 | chr7-107696974-INS-1877 |
| chr7-107712666-INS-63 | chr7-107712666-INS-63 |
| chr7-107721160-INS-787 | chr7-107721160-INS-787 |
| chr7-107723055-INS-350 | chr7-107723054-INS-350 |
| chr7-107802940-INS-296 | chr7-107802940-INS-296 |
| chr7-107851327-INS-195 | chr7-107851327-INS-197 |
| chr7-107869658-INS-106 | chr7-107869659-INS-114 |

| | |
|---|---|
| chr7-107880520-INS-84 | chr7-107880520-INS-60 |
| chr7-107892651-INS-86 | chr7-107892651-INS-85 |
| chr7-107894227-INS-497 | chr7-107894227-INS-497 |
| chr7-107898933-INS-4710 | chr7-107898934-INS-4710 |
| chr7-107902732-INS-201 | chr7-107902730-INS-201 |
| chr7-107918020-INS-88 | chr7-107918020-INS-82 |
| chr7-107921922-INS-62 | chr7-107921922-INS-66 |
| chr7-107933064-INS-2337 | chr7-107933065-INS-2336 |
| chr7-107935593-INS-2970 | chr7-107935594-INS-2954 |
| chr7-107940447-INS-111 | chr7-107940447-INS-83 |
| chr7-107955159-INS-853 | chr7-107955159-INS-853 |
| chr7-107958765-INS-2967 | chr7-107958766-INS-2968 |
| chr7-107963034-INS-498 | chr7-107963034-INS-498 |
| chr7-107979771-INS-548 | chr7-107979771-INS-548 |
| chr7-108010719-INS-1851 | chr7-108010720-INS-1851 |
| chr7-108020819-INS-217 | chr7-108020819-INS-217 |
| chr7-108048273-INS-52 | chr7-108048273-INS-54 |
| chr7-108056852-INS-86 | chr7-108056852-INS-86 |
| chr7-108137528-INS-451 | chr7-108137528-INS-451 |
| chr7-108138713-INS-51 | chr7-108138713-INS-51 |
| chr7-108142801-INS-52 | chr7-108142801-INS-50 |
| chr7-108157695-INS-385 | chr7-108157694-INS-385 |
| chr7-108158712-INS-199 | chr7-108158712-INS-199 |
| chr7-108166555-INS-1154 | chr7-108166556-INS-1154 |
| chr7-108183999-INS-175 | chr7-108183999-INS-175 |
| chr7-108190717-INS-134 | chr7-108190717-INS-136 |
| chr7-108205121-INS-335 | chr7-108205121-INS-335 |
| chr7-108252504-INS-6475 | chr7-108252505-INS-6475 |
| chr7-108271873-INS-161 | chr7-108271873-INS-161 |
| chr7-108294904-INS-6019 | chr7-108294905-INS-6036 |
| chr7-108300776-INS-709 | chr7-108300776-INS-709 |
| chr7-108329603-INS-1747 | chr7-108329604-INS-1747 |
| chr7-108358202-INS-117 | chr7-108358202-INS-129 |
| chr7-108360602-INS-95 | chr7-108360602-INS-99 |
| chr7-108363940-INS-95 | chr7-108363960-INS-74 |
| chr7-108390443-INS-60 | chr7-108390443-INS-60 |
| chr7-108411469-INS-5046 | chr7-108411469-INS-5042 |
| chr7-108418941-INS-51 | chr7-108418941-INS-64 |
| chr7-108430966-INS-171 | chr7-108430966-INS-183 |
| chr7-108435615-INS-4671 | chr7-108435616-INS-4671 |
| chr7-108446350-INS-833 | chr7-108446350-INS-828 |
| chr7-108452322-INS-209 | chr7-108452322-INS-212 |
| chr7-108461979-INS-486 | chr7-108462005-INS-494 |
| chr7-108482831-INS-502 | chr7-108482836-INS-509 |
| chr7-108495632-INS-443 | chr7-108495632-INS-451 |
| chr7-108501175-INS-66 | chr7-108501182-INS-70 |
| chr7-108525729-INS-90 | chr7-108525729-INS-90 |
| chr7-108552841-INS-221 | chr7-108552841-INS-208 |
| chr7-108576958-INS-68 | chr7-108576958-INS-68 |
| chr7-108588622-INS-80 | chr7-108588622-INS-119 |
| chr7-108594678-INS-224 | chr7-108594678-INS-224 |
| chr7-108669286-INS-70 | chr7-108669286-INS-70 |
| chr7-108730669-INS-52 | chr7-108730672-INS-64 |
| chr7-108747295-INS-65 | chr7-108747295-INS-65 |
| chr7-108748121-INS-421 | chr7-108748121-INS-423 |
| chr7-108756473-INS-1451 | chr7-108756474-INS-1451 |
| chr7-108764640-INS-79 | chr7-108764639-INS-79 |
| chr7-108788608-INS-86 | chr7-108788608-INS-86 |
| chr7-108811181-INS-69 | chr7-108811181-INS-72 |
| chr7-108823326-INS-52 | chr7-108823326-INS-52 |
| chr7-108830395-INS-457 | chr7-108830365-INS-438 |
| chr7-108838819-INS-54 | chr7-108838819-INS-56 |
| chr7-108902495-INS-505 | chr7-108902495-INS-505 |
| chr7-108915773-INS-62 | chr7-108915773-INS-62 |
| chr7-108915946-INS-3745 | chr7-108915947-INS-3746 |
| chr7-108947321-INS-80 | chr7-108947298-INS-56 |
| chr7-109026639-INS-177 | chr7-109026639-INS-177 |
| chr7-109029687-INS-52 | chr7-109029687-INS-58 |
| chr7-109036693-INS-52 | chr7-109036719-INS-52 |
| chr7-109044908-INS-434 | chr7-109044908-INS-434 |
| chr7-109061714-INS-60 | chr7-109061714-INS-60 |
| chr7-109070749-INS-811 | chr7-109070749-INS-828 |
| chr7-109081050-INS-60 | chr7-109081050-INS-60 |
| chr7-109095196-INS-69 | chr7-109095196-INS-76 |
| chr7-109146006-INS-563 | chr7-109146006-INS-563 |
| chr7-109200323-INS-70 | chr7-109200323-INS-70 |
| chr7-109241044-INS-201 | chr7-109241044-INS-201 |

| | |
|---|---|
| chr7-109249278-INS-214 | chr7-109249278-INS-214 |
| chr7-109275802-INS-135 | chr7-109275802-INS-135 |
| chr7-109280293-INS-146 | chr7-109280293-INS-146 |
| chr7-109288187-INS-173 | chr7-109288190-INS-176 |
| chr7-109334015-INS-196 | chr7-109334015-INS-191 |
| chr7-109352556-INS-94 | chr7-109352556-INS-94 |
| chr7-109381511-INS-80 | chr7-109381511-INS-50 |
| chr7-109386794-INS-86 | chr7-109386794-INS-92 |
| chr7-109410164-INS-1738 | chr7-109410165-INS-1738 |
| chr7-109415738-INS-82 | chr7-109415738-INS-82 |
| chr7-109455342-INS-147 | chr7-109455342-INS-147 |
| chr7-109461725-INS-225 | chr7-109461709-INS-226 |
| chr7-109472011-INS-67 | chr7-109471993-INS-61 |
| chr7-109476898-INS-54 | chr7-109476898-INS-54 |
| chr7-109484631-INS-202 | chr7-109484638-INS-199 |
| chr7-109492469-INS-53 | chr7-109492469-INS-53 |
| chr7-109494962-INS-111 | chr7-109494962-INS-111 |
| chr7-109494962-INS-761 | chr7-109494962-INS-762 |
| chr7-109505868-INS-135 | chr7-109505868-INS-140 |
| chr7-109510452-INS-182 | chr7-109510452-INS-173 |
| chr7-109520774-INS-208 | chr7-109520774-INS-208 |
| chr7-109526431-INS-217 | chr7-109526431-INS-217 |
| chr7-109549632-INS-194 | chr7-109549633-INS-187 |
| chr7-109556165-INS-205 | chr7-109556165-INS-206 |
| chr7-109563518-INS-199 | chr7-109563523-INS-208 |
| chr7-109564044-INS-222 | chr7-109564044-INS-222 |
| chr7-109579531-INS-195 | chr7-109579544-INS-201 |
| chr7-109601789-INS-1018 | chr7-109601790-INS-1002 |
| chr7-109605069-INS-62 | chr7-109605069-INS-62 |
| chr7-109615995-INS-213 | chr7-109615995-INS-213 |
| chr7-109627900-INS-221 | chr7-109627900-INS-221 |
| chr7-109631225-INS-380 | chr7-109631225-INS-381 |
| chr7-109642864-INS-216 | chr7-109642864-INS-217 |
| chr7-109644921-INS-183 | chr7-109644921-INS-184 |
| chr7-109646931-INS-161 | chr7-109646931-INS-161 |
| chr7-109663120-INS-141 | chr7-109663120-INS-141 |
| chr7-109687699-INS-182 | chr7-109687699-INS-182 |
| chr7-109702490-INS-62 | chr7-109702476-INS-50 |
| chr7-109708598-INS-51 | chr7-109708599-INS-51 |
| chr7-109715444-INS-108 | chr7-109715444-INS-59 |
| chr7-109732953-INS-206 | chr7-109732953-INS-206 |
| chr7-109739432-INS-202 | chr7-109739432-INS-202 |
| chr7-109743231-INS-339 | chr7-109743231-INS-340 |
| chr7-109746499-INS-213 | chr7-109746499-INS-213 |
| chr7-109757658-INS-108 | chr7-109757658-INS-108 |
| chr7-109761836-INS-146 | chr7-109761836-INS-132 |
| chr7-109769103-INS-126 | chr7-109769109-INS-122 |
| chr7-109775388-INS-204 | chr7-109775388-INS-203 |
| chr7-109779883-INS-56 | chr7-109779883-INS-68 |
| chr7-109781129-INS-176 | chr7-109781129-INS-193 |
| chr7-109789707-INS-66 | chr7-109789707-INS-66 |
| chr7-109790211-INS-52 | chr7-109790211-INS-56 |
| chr7-109791735-INS-52 | chr7-109791735-INS-68 |
| chr7-109794896-INS-212 | chr7-109794896-INS-212 |
| chr7-109822631-INS-100 | chr7-109822631-INS-68 |
| chr7-109837029-INS-54 | chr7-109837029-INS-56 |
| chr7-109838439-INS-201 | chr7-109838439-INS-201 |
| chr7-109845064-INS-213 | chr7-109845064-INS-213 |
| chr7-109850709-INS-52 | chr7-109850709-INS-52 |
| chr7-109852346-INS-169 | chr7-109852346-INS-168 |
| chr7-109861456-INS-167 | chr7-109861456-INS-186 |
| chr7-109865486-INS-203 | chr7-109865486-INS-200 |
| chr7-109896127-INS-201 | chr7-109896127-INS-201 |
| chr7-109902787-INS-508 | chr7-109902787-INS-507 |
| chr7-109907897-INS-76 | chr7-109907899-INS-77 |
| chr7-109908781-INS-62 | chr7-109908781-INS-62 |
| chr7-109919101-INS-185 | chr7-109919101-INS-180 |
| chr7-109935404-INS-820 | chr7-109935404-INS-820 |
| chr7-109937695-INS-381 | chr7-109937695-INS-381 |
| chr7-109962321-INS-1005 | chr7-109962322-INS-1006 |
| chr7-109964455-INS-569 | chr7-109964455-INS-569 |
| chr7-109973843-INS-168 | chr7-109973870-INS-168 |
| chr7-109982899-INS-6443 | chr7-109982900-INS-6444 |
| chr7-109996498-INS-192 | chr7-109996498-INS-192 |
| chr7-110022815-INS-1645 | chr7-110022816-INS-1645 |
| chr7-110023432-INS-450 | chr7-110023432-INS-450 |
| chr7-110045099-INS-3269 | chr7-110045100-INS-3269 |

| | |
|---|---|
| chr7-110046112-INS-90 | chr7-110046112-INS-80 |
| chr7-110074164-INS-561 | chr7-110074164-INS-562 |
| chr7-110134879-INS-6658 | chr7-110134880-INS-6658 |
| chr7-110190491-INS-54 | chr7-110190491-INS-54 |
| chr7-110207602-INS-105 | chr7-110207602-INS-105 |
| chr7-110209366-INS-114 | chr7-110209366-INS-122 |
| chr7-110239471-INS-57 | chr7-110239471-INS-87 |
| chr7-110252336-INS-3476 | chr7-110252347-INS-3472 |
| chr7-110278606-INS-463 | chr7-110278601-INS-463 |
| chr7-110313548-INS-56 | chr7-110313548-INS-56 |
| chr7-110343071-INS-169 | chr7-110343071-INS-165 |
| chr7-110344549-INS-1335 | chr7-110344550-INS-1335 |
| chr7-110350218-INS-51 | chr7-110350218-INS-51 |
| chr7-110382059-INS-188 | chr7-110382059-INS-188 |
| chr7-110384079-INS-333 | chr7-110384079-INS-333 |
| chr7-110516691-INS-58 | chr7-110516691-INS-50 |
| chr7-110526715-INS-140 | chr7-110526715-INS-140 |
| chr7-110558691-INS-52 | chr7-110558691-INS-56 |
| chr7-110565375-INS-257 | chr7-110565375-INS-255 |
| chr7-110575386-INS-56 | chr7-110575386-INS-52 |
| chr7-110602679-INS-177 | chr7-110602679-INS-177 |
| chr7-110609233-INS-88 | chr7-110609233-INS-88 |
| chr7-110612248-INS-204 | chr7-110612248-INS-209 |
| chr7-110655290-INS-55 | chr7-110655290-INS-90 |
| chr7-110666932-INS-1731 | chr7-110666933-INS-1732 |
| chr7-110707727-INS-95 | chr7-110707737-INS-80 |
| chr7-110714744-INS-72 | chr7-110714744-INS-88 |
| chr7-110731171-INS-218 | chr7-110731171-INS-219 |
| chr7-110796285-INS-571 | chr7-110796285-INS-577 |
| chr7-110797107-INS-51 | chr7-110797107-INS-57 |
| chr7-110808806-INS-56 | chr7-110808806-INS-58 |
| chr7-110839307-INS-335 | chr7-110839307-INS-334 |
| chr7-110843778-INS-119 | chr7-110843762-INS-120 |
| chr7-110849612-INS-299 | chr7-110849612-INS-299 |
| chr7-110862477-INS-178 | chr7-110862477-INS-178 |
| chr7-110895519-INS-342 | chr7-110895519-INS-342 |
| chr7-110911718-INS-114 | chr7-110911718-INS-104 |
| chr7-110913353-INS-91 | chr7-110913353-INS-91 |
| chr7-110916565-INS-68 | chr7-110916572-INS-52 |
| chr7-110933029-INS-172 | chr7-110933029-INS-171 |
| chr7-110938088-INS-171 | chr7-110938094-INS-184 |
| chr7-110941079-INS-227 | chr7-110941079-INS-227 |
| chr7-110945281-INS-447 | chr7-110945281-INS-447 |
| chr7-110957183-INS-215 | chr7-110957182-INS-222 |
| chr7-110971116-INS-807 | chr7-110971116-INS-807 |
| chr7-110973916-INS-96 | chr7-110973916-INS-96 |
| chr7-110985440-INS-1147 | chr7-110985441-INS-1135 |
| chr7-111034603-INS-440 | chr7-111034603-INS-440 |
| chr7-111036647-INS-186 | chr7-111036647-INS-187 |
| chr7-111060887-INS-68 | chr7-111060887-INS-72 |
| chr7-111086287-INS-83 | chr7-111086287-INS-84 |
| chr7-111111908-INS-179 | chr7-111111908-INS-179 |
| chr7-111121546-INS-449 | chr7-111121547-INS-450 |
| chr7-111136507-INS-173 | chr7-111136510-INS-181 |
| chr7-111143882-INS-61 | chr7-111143882-INS-61 |
| chr7-111175090-INS-53 | chr7-111175086-INS-58 |
| chr7-111180054-INS-60 | chr7-111180054-INS-72 |
| chr7-111190179-INS-66 | chr7-111190214-INS-60 |
| chr7-111191481-INS-75 | chr7-111191481-INS-75 |
| chr7-111201296-INS-54 | chr7-111201296-INS-60 |
| chr7-111216794-INS-160 | chr7-111216794-INS-160 |
| chr7-111252587-INS-6604 | chr7-111252588-INS-6604 |
| chr7-111253107-INS-131 | chr7-111253107-INS-131 |
| chr7-111261060-INS-213 | chr7-111261060-INS-210 |
| chr7-111313903-INS-125 | chr7-111313903-INS-135 |
| chr7-111317750-INS-328 | chr7-111317750-INS-329 |
| chr7-111318556-INS-74 | chr7-111318556-INS-74 |
| chr7-111325749-INS-97 | chr7-111325749-INS-89 |
| chr7-111334658-INS-76 | chr7-111334658-INS-76 |
| chr7-111361467-INS-97 | chr7-111361497-INS-87 |
| chr7-111408617-INS-368 | chr7-111408617-INS-336 |
| chr7-111414178-INS-253 | chr7-111414178-INS-253 |
| chr7-111473410-INS-53 | chr7-111473367-INS-67 |
| chr7-111490619-INS-2137 | chr7-111490620-INS-2137 |
| chr7-111494026-INS-6752 | chr7-111494027-INS-6752 |
| chr7-111502199-INS-60 | chr7-111502199-INS-56 |
| chr7-111534432-INS-216 | chr7-111534432-INS-216 |

| | |
|---|---|
| chr7-111567059-INS-451 | chr7-111567059-INS-451 |
| chr7-111578236-INS-178 | chr7-111578236-INS-174 |
| chr7-111583564-INS-60 | chr7-111583564-INS-60 |
| chr7-111590172-INS-83 | chr7-111590172-INS-84 |
| chr7-111596260-INS-194 | chr7-111596260-INS-194 |
| chr7-111600535-INS-151 | chr7-111600534-INS-158 |
| chr7-111609492-INS-52 | chr7-111609493-INS-72 |
| chr7-111634893-INS-62 | chr7-111634893-INS-61 |
| chr7-111685821-INS-166 | chr7-111685821-INS-166 |
| chr7-111759284-INS-971 | chr7-111759284-INS-962 |
| chr7-111782869-INS-88 | chr7-111782868-INS-96 |
| chr7-111790415-INS-91 | chr7-111790415-INS-93 |
| chr7-111808624-INS-64 | chr7-111808624-INS-64 |
| chr7-111817199-INS-877 | chr7-111817199-INS-844 |
| chr7-111949094-INS-3200 | chr7-111949095-INS-3201 |
| chr7-111986934-INS-58 | chr7-111986934-INS-50 |
| chr7-111988540-INS-65 | chr7-111988563-INS-66 |
| chr7-112023098-INS-2898 | chr7-112023099-INS-2898 |
| chr7-112026566-INS-103 | chr7-112026566-INS-103 |
| chr7-112066534-INS-96 | chr7-112066534-INS-102 |
| chr7-112075499-INS-1890 | chr7-112075500-INS-1890 |
| chr7-112076921-INS-94 | chr7-112076941-INS-86 |
| chr7-112083008-INS-156 | chr7-112083009-INS-162 |
| chr7-112101901-INS-52 | chr7-112101901-INS-56 |
| chr7-112147063-INS-230 | chr7-112147063-INS-232 |
| chr7-112183646-INS-181 | chr7-112183646-INS-181 |
| chr7-112193803-INS-105 | chr7-112193803-INS-142 |
| chr7-112196271-INS-58 | chr7-112196271-INS-58 |
| chr7-112199297-INS-58 | chr7-112199345-INS-60 |
| chr7-112218205-INS-182 | chr7-112218205-INS-183 |
| chr7-112220761-INS-64 | chr7-112220759-INS-82 |
| chr7-112221488-INS-70 | chr7-112221454-INS-88 |
| chr7-112233870-INS-443 | chr7-112233870-INS-448 |
| chr7-112235698-INS-508 | chr7-112235699-INS-508 |
| chr7-112251851-INS-208 | chr7-112251851-INS-208 |
| chr7-112256934-INS-60 | chr7-112256934-INS-68 |
| chr7-112263459-INS-70 | chr7-112263459-INS-68 |
| chr7-112268637-INS-213 | chr7-112268637-INS-213 |
| chr7-112302483-INS-54 | chr7-112302483-INS-54 |
| chr7-112310781-INS-219 | chr7-112310781-INS-218 |
| chr7-112340169-INS-80 | chr7-112340169-INS-80 |
| chr7-112352958-INS-62 | chr7-112352958-INS-62 |
| chr7-112363228-INS-1869 | chr7-112363229-INS-1869 |
| chr7-112371746-INS-110 | chr7-112371746-INS-110 |
| chr7-112380597-INS-91 | chr7-112380597-INS-91 |
| chr7-112383769-INS-207 | chr7-112383769-INS-207 |
| chr7-112403621-INS-92 | chr7-112403621-INS-92 |
| chr7-112432377-INS-52 | chr7-112432377-INS-50 |
| chr7-112437142-INS-53 | chr7-112437142-INS-53 |
| chr7-112469717-INS-213 | chr7-112469717-INS-213 |
| chr7-112486994-INS-203 | chr7-112486994-INS-203 |
| chr7-112489804-INS-214 | chr7-112489804-INS-217 |
| chr7-112525424-INS-157 | chr7-112525428-INS-167 |
| chr7-112532371-INS-64 | chr7-112532371-INS-64 |
| chr7-112538736-INS-60 | chr7-112538736-INS-60 |
| chr7-112546579-INS-123 | chr7-112546579-INS-123 |
| chr7-112558968-INS-90 | chr7-112558968-INS-90 |
| chr7-112584025-INS-66 | chr7-112584025-INS-58 |
| chr7-112593557-INS-269 | chr7-112593557-INS-269 |
| chr7-112597973-INS-51 | chr7-112597973-INS-51 |
| chr7-112604545-INS-71 | chr7-112604545-INS-69 |
| chr7-112620276-INS-70 | chr7-112620276-INS-73 |
| chr7-112637860-INS-174 | chr7-112637860-INS-175 |
| chr7-112660758-INS-60 | chr7-112660758-INS-60 |
| chr7-112676179-INS-64 | chr7-112676177-INS-62 |
| chr7-112699228-INS-112 | chr7-112699228-INS-112 |
| chr7-112713313-INS-465 | chr7-112713313-INS-465 |
| chr7-112715748-INS-72 | chr7-112715748-INS-108 |
| chr7-112722284-INS-66 | chr7-112722284-INS-111 |
| chr7-112735663-INS-88 | chr7-112735656-INS-64 |
| chr7-112736295-INS-56 | chr7-112736300-INS-50 |
| chr7-112771085-INS-92 | chr7-112771090-INS-75 |
| chr7-112775486-INS-2570 | chr7-112775487-INS-2565 |
| chr7-112797817-INS-418 | chr7-112797818-INS-418 |
| chr7-112798602-INS-170 | chr7-112798597-INS-188 |
| chr7-112800674-INS-132 | chr7-112800674-INS-120 |
| chr7-112860619-INS-110 | chr7-112860619-INS-110 |

| | |
|---|---|
| chr7-112862919-INS-76 | chr7-112862878-INS-68 |
| chr7-112865091-INS-199 | chr7-112865091-INS-200 |
| chr7-112871360-INS-6816 | chr7-112871361-INS-6817 |
| chr7-112877716-INS-61 | chr7-112877716-INS-61 |
| chr7-112894426-INS-89 | chr7-112894426-INS-89 |
| chr7-112920888-INS-110 | chr7-112920888-INS-110 |
| chr7-112970274-INS-188 | chr7-112970274-INS-190 |
| chr7-112974629-INS-58 | chr7-112974629-INS-58 |
| chr7-112979677-INS-70 | chr7-112979654-INS-70 |
| chr7-112984425-INS-90 | chr7-112984425-INS-90 |
| chr7-112989982-INS-154 | chr7-112989963-INS-150 |
| chr7-113002154-INS-804 | chr7-113002154-INS-799 |
| chr7-113009384-INS-297 | chr7-113009384-INS-297 |
| chr7-113011421-INS-150 | chr7-113011417-INS-153 |
| chr7-113014861-INS-58 | chr7-113014861-INS-58 |
| chr7-113019914-INS-58 | chr7-113019914-INS-52 |
| chr7-113020991-INS-460 | chr7-113020992-INS-460 |
| chr7-113031885-INS-612 | chr7-113031885-INS-612 |
| chr7-113044279-INS-54 | chr7-113044279-INS-52 |
| chr7-113060392-INS-60 | chr7-113060392-INS-60 |
| chr7-113072603-INS-62 | chr7-113072603-INS-56 |
| chr7-113080425-INS-867 | chr7-113080425-INS-867 |
| chr7-113118935-INS-6451 | chr7-113118936-INS-6451 |
| chr7-113157177-INS-1410 | chr7-113157178-INS-1409 |
| chr7-113177952-INS-88 | chr7-113177952-INS-86 |
| chr7-113232418-INS-5504 | chr7-113232419-INS-5504 |
| chr7-113249927-INS-4533 | chr7-113249928-INS-4533 |
| chr7-113273724-INS-65 | chr7-113273724-INS-65 |
| chr7-113278020-INS-4679 | chr7-113278021-INS-4678 |
| chr7-113292318-INS-87 | chr7-113292318-INS-63 |
| chr7-113297509-INS-287 | chr7-113297509-INS-293 |
| chr7-113298841-INS-486 | chr7-113298841-INS-504 |
| chr7-113330000-INS-2492 | chr7-113330001-INS-2488 |
| chr7-113352490-INS-317 | chr7-113352490-INS-307 |
| chr7-113361018-INS-207 | chr7-113361034-INS-198 |
| chr7-113366157-INS-52 | chr7-113366157-INS-54 |
| chr7-113390314-INS-116 | chr7-113390318-INS-88 |
| chr7-113405202-INS-92 | chr7-113405202-INS-72 |
| chr7-113412295-INS-444 | chr7-113412295-INS-444 |
| chr7-113442883-INS-122 | chr7-113442859-INS-131 |
| chr7-113506842-INS-57 | chr7-113506840-INS-66 |
| chr7-113511324-INS-2213 | chr7-113511325-INS-2214 |
| chr7-113537407-INS-531 | chr7-113537407-INS-531 |
| chr7-113583926-INS-938 | chr7-113583926-INS-938 |
| chr7-113616518-INS-56 | chr7-113616564-INS-50 |
| chr7-113627837-INS-212 | chr7-113627837-INS-212 |
| chr7-113631226-INS-115 | chr7-113631226-INS-115 |
| chr7-113637959-INS-75 | chr7-113637959-INS-80 |
| chr7-113656221-INS-647 | chr7-113656221-INS-648 |
| chr7-113657372-INS-84 | chr7-113657372-INS-85 |
| chr7-113662548-INS-315 | chr7-113662548-INS-315 |
| chr7-113683937-INS-69 | chr7-113683937-INS-69 |
| chr7-113732512-INS-61 | chr7-113732512-INS-61 |
| chr7-113743424-INS-5218 | chr7-113743425-INS-5218 |
| chr7-113748302-INS-80 | chr7-113748313-INS-50 |
| chr7-113764799-INS-60 | chr7-113764800-INS-108 |
| chr7-113767222-INS-4533 | chr7-113767223-INS-4532 |
| chr7-113769799-INS-214 | chr7-113769799-INS-215 |
| chr7-113774457-INS-200 | chr7-113774457-INS-201 |
| chr7-113789516-INS-639 | chr7-113789516-INS-640 |
| chr7-113794407-INS-109 | chr7-113794371-INS-144 |
| chr7-113799434-INS-51 | chr7-113799434-INS-71 |
| chr7-113806093-INS-58 | chr7-113806089-INS-82 |
| chr7-113821414-INS-51 | chr7-113821414-INS-51 |
| chr7-113847976-INS-135 | chr7-113847976-INS-135 |
| chr7-113851201-INS-410 | chr7-113851202-INS-410 |
| chr7-113855211-INS-213 | chr7-113855211-INS-213 |
| chr7-113881826-INS-65 | chr7-113881826-INS-64 |
| chr7-113909314-INS-132 | chr7-113909314-INS-132 |
| chr7-113947128-INS-876 | chr7-113947128-INS-875 |
| chr7-113950737-INS-88 | chr7-113950748-INS-62 |
| chr7-113954582-INS-2718 | chr7-113954583-INS-2718 |
| chr7-113963537-INS-67 | chr7-113963554-INS-55 |
| chr7-114000483-INS-52 | chr7-114000479-INS-52 |
| chr7-114026619-INS-655 | chr7-114026619-INS-656 |
| chr7-114034315-INS-76 | chr7-114034315-INS-76 |
| chr7-114045876-INS-347 | chr7-114045876-INS-347 |

| | |
|---|---|
| chr7-114059726-INS-140 | chr7-114059726-INS-145 |
| chr7-114076758-INS-77 | chr7-114076758-INS-81 |
| chr7-114095552-INS-297 | chr7-114095552-INS-300 |
| chr7-114101605-INS-60 | chr7-114101605-INS-60 |
| chr7-114141826-INS-2887 | chr7-114141827-INS-2885 |
| chr7-114147912-INS-310 | chr7-114147912-INS-311 |
| chr7-114162591-INS-209 | chr7-114162591-INS-209 |
| chr7-114178090-INS-493 | chr7-114178090-INS-501 |
| chr7-114193536-INS-67 | chr7-114193536-INS-73 |
| chr7-114204922-INS-91 | chr7-114204922-INS-91 |
| chr7-114213399-INS-414 | chr7-114213399-INS-413 |
| chr7-114220084-INS-6367 | chr7-114220085-INS-6370 |
| chr7-114221210-INS-91 | chr7-114221210-INS-93 |
| chr7-114255675-INS-309 | chr7-114255675-INS-304 |
| chr7-114276029-INS-116 | chr7-114276025-INS-105 |
| chr7-114276512-INS-7075 | chr7-114276513-INS-7075 |
| chr7-114293522-INS-55 | chr7-114293522-INS-54 |
| chr7-114315699-INS-52 | chr7-114315700-INS-54 |
| chr7-114335078-INS-125 | chr7-114335076-INS-119 |
| chr7-114417592-INS-128 | chr7-114417634-INS-122 |
| chr7-114447295-INS-65 | chr7-114447295-INS-60 |
| chr7-114519780-INS-198 | chr7-114519780-INS-198 |
| chr7-114546741-INS-156 | chr7-114546741-INS-161 |
| chr7-114624858-INS-54 | chr7-114624828-INS-56 |
| chr7-114628396-INS-91 | chr7-114628396-INS-91 |
| chr7-114640374-INS-69 | chr7-114640374-INS-70 |
| chr7-114744603-INS-90 | chr7-114744604-INS-60 |
| chr7-114745496-INS-71 | chr7-114745497-INS-71 |
| chr7-114765470-INS-52 | chr7-114765470-INS-54 |
| chr7-114782806-INS-139 | chr7-114782793-INS-104 |
| chr7-114789552-INS-171 | chr7-114789552-INS-171 |
| chr7-114791706-INS-72 | chr7-114791706-INS-72 |
| chr7-114793512-INS-52 | chr7-114793512-INS-56 |
| chr7-114796920-INS-436 | chr7-114796920-INS-436 |
| chr7-114801909-INS-88 | chr7-114801909-INS-90 |
| chr7-114813968-INS-54 | chr7-114813968-INS-54 |
| chr7-114816966-INS-121 | chr7-114816957-INS-130 |
| chr7-114835863-INS-58 | chr7-114835862-INS-82 |
| chr7-114840817-INS-63 | chr7-114840800-INS-62 |
| chr7-114850504-INS-55 | chr7-114850504-INS-59 |
| chr7-114856596-INS-58 | chr7-114856596-INS-58 |
| chr7-114895490-INS-100 | chr7-114895490-INS-91 |
| chr7-114913711-INS-164 | chr7-114913712-INS-169 |
| chr7-114936991-INS-217 | chr7-114936991-INS-248 |
| chr7-114938838-INS-68 | chr7-114938838-INS-70 |
| chr7-114954632-INS-55 | chr7-114954617-INS-65 |
| chr7-114964492-INS-319 | chr7-114964492-INS-319 |
| chr7-114968537-INS-52 | chr7-114968512-INS-62 |
| chr7-114985134-INS-308 | chr7-114985134-INS-326 |
| chr7-115005706-INS-585 | chr7-115005706-INS-586 |
| chr7-115015400-INS-72 | chr7-115015400-INS-72 |
| chr7-115023522-INS-164 | chr7-115023541-INS-168 |
| chr7-115035209-INS-52 | chr7-115035209-INS-54 |
| chr7-115040801-INS-517 | chr7-115040801-INS-517 |
| chr7-115049138-INS-65 | chr7-115049138-INS-85 |
| chr7-115059339-INS-1210 | chr7-115059340-INS-1210 |
| chr7-115079485-INS-64 | chr7-115079485-INS-68 |
| chr7-115086220-INS-60 | chr7-115086211-INS-68 |
| chr7-115098695-INS-76 | chr7-115098695-INS-76 |
| chr7-115121703-INS-1552 | chr7-115121704-INS-1552 |
| chr7-115130874-INS-63 | chr7-115130874-INS-63 |
| chr7-115134037-INS-259 | chr7-115134037-INS-259 |
| chr7-115136507-INS-57 | chr7-115136507-INS-57 |
| chr7-115156850-INS-130 | chr7-115156861-INS-136 |
| chr7-115162949-INS-52 | chr7-115162949-INS-54 |
| chr7-115173386-INS-54 | chr7-115173386-INS-54 |
| chr7-115218746-INS-54 | chr7-115218746-INS-54 |
| chr7-115238856-INS-252 | chr7-115238856-INS-252 |
| chr7-115239818-INS-55 | chr7-115239818-INS-60 |
| chr7-115247886-INS-1907 | chr7-115247887-INS-1907 |
| chr7-115281364-INS-6860 | chr7-115281365-INS-6861 |
| chr7-115296584-INS-68 | chr7-115296584-INS-64 |
| chr7-115298903-INS-66 | chr7-115298903-INS-66 |
| chr7-115373928-INS-214 | chr7-115373928-INS-214 |
| chr7-115402566-INS-70 | chr7-115402566-INS-70 |
| chr7-115428840-INS-56 | chr7-115428840-INS-64 |
| chr7-115437814-INS-77 | chr7-115437790-INS-102 |

| | |
|---|---|
| chr7-115443004-INS-63 | chr7-115443004-INS-62 |
| chr7-115467629-INS-93 | chr7-115467622-INS-94 |
| chr7-115486406-INS-60 | chr7-115486406-INS-60 |
| chr7-115490977-INS-88 | chr7-115490977-INS-88 |
| chr7-115508475-INS-64 | chr7-115508475-INS-52 |
| chr7-115515107-INS-162 | chr7-115515107-INS-153 |
| chr7-115523534-INS-54 | chr7-115523534-INS-56 |
| chr7-115527255-INS-92 | chr7-115527261-INS-84 |
| chr7-115576912-INS-50 | chr7-115576912-INS-50 |
| chr7-115579092-INS-54 | chr7-115579092-INS-54 |
| chr7-115588227-INS-844 | chr7-115588227-INS-850 |
| chr7-115603405-INS-6498 | chr7-115603406-INS-6499 |
| chr7-115616263-INS-52 | chr7-115616275-INS-55 |
| chr7-115621173-INS-66 | chr7-115621173-INS-66 |
| chr7-115632095-INS-51 | chr7-115632095-INS-57 |
| chr7-115637040-INS-51 | chr7-115637041-INS-54 |
| chr7-115664560-INS-211 | chr7-115664560-INS-211 |
| chr7-115665249-INS-8457 | chr7-115665250-INS-8457 |
| chr7-115683496-INS-5539 | chr7-115683497-INS-5539 |
| chr7-115700150-INS-211 | chr7-115700150-INS-211 |
| chr7-115722057-INS-477 | chr7-115722057-INS-477 |
| chr7-115730236-INS-587 | chr7-115730236-INS-587 |
| chr7-115745581-INS-60 | chr7-115745581-INS-50 |
| chr7-115766532-INS-98 | chr7-115766532-INS-102 |
| chr7-115828719-INS-52 | chr7-115828719-INS-54 |
| chr7-115829009-INS-891 | chr7-115828990-INS-885 |
| chr7-115833035-INS-210 | chr7-115833035-INS-210 |
| chr7-115840087-INS-199 | chr7-115840087-INS-197 |
| chr7-115842906-INS-112 | chr7-115842906-INS-112 |
| chr7-115857388-INS-176 | chr7-115857388-INS-176 |
| chr7-115872705-INS-224 | chr7-115872705-INS-224 |
| chr7-115881010-INS-110 | chr7-115881010-INS-110 |
| chr7-115892109-INS-111 | chr7-115892110-INS-111 |
| chr7-115909436-INS-215 | chr7-115909436-INS-214 |
| chr7-115918375-INS-202 | chr7-115918375-INS-202 |
| chr7-115922127-INS-199 | chr7-115922127-INS-199 |
| chr7-115985919-INS-400 | chr7-115985919-INS-400 |
| chr7-116000706-INS-222 | chr7-116000706-INS-232 |
| chr7-116024139-INS-216 | chr7-116024139-INS-216 |
| chr7-116055534-INS-213 | chr7-116055534-INS-211 |
| chr7-116064099-INS-92 | chr7-116064099-INS-92 |
| chr7-116088177-INS-54 | chr7-116088177-INS-56 |
| chr7-116092708-INS-59 | chr7-116092708-INS-59 |
| chr7-116105517-INS-256 | chr7-116105517-INS-261 |
| chr7-116145877-INS-130 | chr7-116145877-INS-180 |
| chr7-116150707-INS-194 | chr7-116150715-INS-195 |
| chr7-116187298-INS-72 | chr7-116187298-INS-60 |
| chr7-116193297-INS-200 | chr7-116193297-INS-200 |
| chr7-116204244-INS-156 | chr7-116204244-INS-150 |
| chr7-116204968-INS-6403 | chr7-116204969-INS-6403 |
| chr7-116218568-INS-79 | chr7-116218568-INS-86 |
| chr7-116221509-INS-144 | chr7-116221509-INS-144 |
| chr7-116224153-INS-2065 | chr7-116224154-INS-2065 |
| chr7-116227937-INS-667 | chr7-116227937-INS-667 |
| chr7-116267326-INS-6452 | chr7-116267327-INS-6452 |
| chr7-116270925-INS-1138 | chr7-116270926-INS-1138 |
| chr7-116288314-INS-325 | chr7-116288314-INS-325 |
| chr7-116291862-INS-6594 | chr7-116291863-INS-6598 |
| chr7-116364143-INS-51 | chr7-116364143-INS-51 |
| chr7-116365417-INS-290 | chr7-116365417-INS-290 |
| chr7-116366303-INS-728 | chr7-116366303-INS-728 |
| chr7-116396710-INS-149 | chr7-116396710-INS-149 |
| chr7-116397733-INS-242 | chr7-116397733-INS-250 |
| chr7-116401591-INS-75 | chr7-116401591-INS-75 |
| chr7-116416179-INS-230 | chr7-116416179-INS-232 |
| chr7-116429926-INS-410 | chr7-116429926-INS-423 |
| chr7-116444471-INS-6671 | chr7-116444472-INS-6671 |
| chr7-116487033-INS-52 | chr7-116487033-INS-52 |
| chr7-116495824-INS-1943 | chr7-116495825-INS-1956 |
| chr7-116498113-INS-54 | chr7-116498113-INS-57 |
| chr7-116516697-INS-313 | chr7-116516698-INS-313 |
| chr7-116526494-INS-71 | chr7-116526494-INS-71 |
| chr7-116568686-INS-76 | chr7-116568686-INS-76 |
| chr7-116575001-INS-120 | chr7-116575001-INS-123 |
| chr7-116584597-INS-64 | chr7-116584597-INS-64 |
| chr7-116585979-INS-475 | chr7-116585977-INS-471 |
| chr7-116596614-INS-100 | chr7-116596615-INS-100 |

| | |
|---|---|
| chr7-116608559-INS-739 | chr7-116608559-INS-739 |
| chr7-116686322-INS-1894 | chr7-116686323-INS-1894 |
| chr7-116692025-INS-110 | chr7-116692025-INS-110 |
| chr7-116713401-INS-486 | chr7-116713401-INS-486 |
| chr7-116724199-INS-256 | chr7-116724200-INS-256 |
| chr7-116737693-INS-70 | chr7-116737693-INS-74 |
| chr7-116773782-INS-59 | chr7-116773746-INS-58 |
| chr7-116792358-INS-86 | chr7-116792358-INS-90 |
| chr7-116797016-INS-63 | chr7-116797016-INS-60 |
| chr7-116828666-INS-147 | chr7-116828666-INS-147 |
| chr7-116837318-INS-124 | chr7-116837318-INS-124 |
| chr7-116838268-INS-3429 | chr7-116838269-INS-3431 |
| chr7-116855032-INS-76 | chr7-116855032-INS-60 |
| chr7-116858319-INS-155 | chr7-116858336-INS-157 |
| chr7-116868448-INS-74 | chr7-116868448-INS-79 |
| chr7-116900355-INS-75 | chr7-116900355-INS-75 |
| chr7-116903413-INS-220 | chr7-116903413-INS-220 |
| chr7-116905950-INS-72 | chr7-116905950-INS-72 |
| chr7-116931665-INS-671 | chr7-116931665-INS-671 |
| chr7-116948845-INS-80 | chr7-116948845-INS-62 |
| chr7-116951305-INS-54 | chr7-116951305-INS-58 |
| chr7-116961346-INS-100 | chr7-116961346-INS-100 |
| chr7-116962915-INS-156 | chr7-116962915-INS-156 |
| chr7-116965472-INS-60 | chr7-116965472-INS-64 |
| chr7-116974052-INS-204 | chr7-116974052-INS-205 |
| chr7-117019566-INS-57 | chr7-117019568-INS-53 |
| chr7-117024698-INS-56 | chr7-117024699-INS-56 |
| chr7-117027212-INS-62 | chr7-117027212-INS-62 |
| chr7-117035839-INS-191 | chr7-117035839-INS-191 |
| chr7-117044454-INS-85 | chr7-117044454-INS-85 |
| chr7-117056632-INS-64 | chr7-117056632-INS-64 |
| chr7-117058892-INS-711 | chr7-117058892-INS-711 |
| chr7-117066568-INS-51 | chr7-117066568-INS-51 |
| chr7-117075372-INS-241 | chr7-117075372-INS-241 |
| chr7-117163648-INS-52 | chr7-117163648-INS-82 |
| chr7-117166441-INS-62 | chr7-117166441-INS-60 |
| chr7-117183164-INS-163 | chr7-117183164-INS-163 |
| chr7-117191452-INS-74 | chr7-117191452-INS-76 |
| chr7-117223699-INS-1990 | chr7-117223700-INS-1990 |
| chr7-117234296-INS-159 | chr7-117234296-INS-137 |
| chr7-117243172-INS-101 | chr7-117243172-INS-104 |
| chr7-117249227-INS-153 | chr7-117249227-INS-154 |
| chr7-117253879-INS-1890 | chr7-117253880-INS-1890 |
| chr7-117274719-INS-76 | chr7-117274719-INS-74 |
| chr7-117336995-INS-275 | chr7-117336995-INS-275 |
| chr7-117353410-INS-132 | chr7-117353445-INS-132 |
| chr7-117366096-INS-200 | chr7-117366096-INS-200 |
| chr7-117369980-INS-64 | chr7-117369980-INS-64 |
| chr7-117384645-INS-197 | chr7-117384645-INS-197 |
| chr7-117388632-INS-89 | chr7-117388627-INS-94 |
| chr7-117392217-INS-84 | chr7-117392241-INS-77 |
| chr7-117411512-INS-52 | chr7-117411512-INS-62 |
| chr7-117453393-INS-6599 | chr7-117453394-INS-6599 |
| chr7-117482490-INS-82 | chr7-117482490-INS-82 |
| chr7-117484740-INS-554 | chr7-117484752-INS-565 |
| chr7-117487481-INS-186 | chr7-117487478-INS-195 |
| chr7-117494873-INS-51 | chr7-117494868-INS-53 |
| chr7-117530298-INS-212 | chr7-117530298-INS-208 |
| chr7-117545407-INS-525 | chr7-117545407-INS-525 |
| chr7-117565698-INS-57 | chr7-117565698-INS-57 |
| chr7-117573831-INS-84 | chr7-117573833-INS-94 |
| chr7-117580542-INS-5362 | chr7-117580543-INS-5362 |
| chr7-117587226-INS-88 | chr7-117587214-INS-116 |
| chr7-117589164-INS-104 | chr7-117589164-INS-104 |
| chr7-117592411-INS-54 | chr7-117592411-INS-56 |
| chr7-117606778-INS-60 | chr7-117606778-INS-60 |
| chr7-117628028-INS-52 | chr7-117628028-INS-52 |
| chr7-117636490-INS-100 | chr7-117636490-INS-100 |
| chr7-117648453-INS-218 | chr7-117648453-INS-218 |
| chr7-117661589-INS-53 | chr7-117661589-INS-57 |
| chr7-117668454-INS-209 | chr7-117668454-INS-209 |
| chr7-117677087-INS-138 | chr7-117677087-INS-138 |
| chr7-117681318-INS-108 | chr7-117681318-INS-108 |
| chr7-117685469-INS-189 | chr7-117685469-INS-189 |
| chr7-117693961-INS-56 | chr7-117693964-INS-69 |
| chr7-117700810-INS-59 | chr7-117700808-INS-59 |
| chr7-117716196-INS-222 | chr7-117716195-INS-219 |

| | |
|---|---|
| chr7-117721123-INS-56 | chr7-117721123-INS-52 |
| chr7-117757990-INS-184 | chr7-117757990-INS-184 |
| chr7-117763915-INS-57 | chr7-117763915-INS-72 |
| chr7-117777254-INS-240 | chr7-117777254-INS-246 |
| chr7-117798715-INS-69 | chr7-117798708-INS-60 |
| chr7-117850893-INS-217 | chr7-117850894-INS-217 |
| chr7-117865192-INS-896 | chr7-117865192-INS-896 |
| chr7-117882068-INS-55 | chr7-117882060-INS-56 |
| chr7-117902229-INS-175 | chr7-117902225-INS-141 |
| chr7-117907402-INS-169 | chr7-117907403-INS-168 |
| chr7-117910499-INS-78 | chr7-117910499-INS-78 |
| chr7-117924708-INS-84 | chr7-117924708-INS-84 |
| chr7-117937807-INS-5914 | chr7-117937808-INS-5914 |
| chr7-117942246-INS-81 | chr7-117942246-INS-81 |
| chr7-117947980-INS-101 | chr7-117947980-INS-93 |
| chr7-117950645-INS-126 | chr7-117950645-INS-132 |
| chr7-117956294-INS-65 | chr7-117956294-INS-50 |
| chr7-117971484-INS-1890 | chr7-117971485-INS-1890 |
| chr7-117984211-INS-108 | chr7-117984211-INS-108 |
| chr7-117993592-INS-338 | chr7-117993592-INS-338 |
| chr7-117995973-INS-55 | chr7-117995973-INS-55 |
| chr7-118010771-INS-239 | chr7-118010757-INS-240 |
| chr7-118031282-INS-68 | chr7-118031282-INS-68 |
| chr7-118042417-INS-92 | chr7-118042409-INS-90 |
| chr7-118072572-INS-98 | chr7-118072572-INS-96 |
| chr7-118080927-INS-206 | chr7-118080927-INS-206 |
| chr7-118096963-INS-581 | chr7-118096963-INS-581 |
| chr7-118101486-INS-50 | chr7-118101507-INS-50 |
| chr7-118104696-INS-273 | chr7-118104696-INS-236 |
| chr7-118115769-INS-423 | chr7-118115769-INS-423 |
| chr7-118117708-INS-267 | chr7-118117708-INS-267 |
| chr7-118127110-INS-202 | chr7-118127110-INS-206 |
| chr7-118156570-INS-463 | chr7-118156570-INS-463 |
| chr7-118162157-INS-106 | chr7-118162157-INS-106 |
| chr7-118192544-INS-72 | chr7-118192544-INS-72 |
| chr7-118217930-INS-58 | chr7-118217930-INS-60 |
| chr7-118226695-INS-55 | chr7-118226691-INS-73 |
| chr7-118236697-INS-77 | chr7-118236691-INS-72 |
| chr7-118238920-INS-205 | chr7-118238920-INS-205 |
| chr7-118243754-INS-197 | chr7-118243754-INS-197 |
| chr7-118255837-INS-68 | chr7-118255837-INS-64 |
| chr7-118259201-INS-77 | chr7-118259201-INS-77 |
| chr7-118279435-INS-200 | chr7-118279435-INS-200 |
| chr7-118279643-INS-512 | chr7-118279643-INS-512 |
| chr7-118282348-INS-82 | chr7-118282348-INS-82 |
| chr7-118285273-INS-294 | chr7-118285273-INS-294 |
| chr7-118291709-INS-55 | chr7-118291709-INS-55 |
| chr7-118293506-INS-5543 | chr7-118293507-INS-5544 |
| chr7-118317039-INS-209 | chr7-118317039-INS-210 |
| chr7-118342716-INS-764 | chr7-118342716-INS-764 |
| chr7-118344351-INS-177 | chr7-118344352-INS-140 |
| chr7-118358904-INS-433 | chr7-118358904-INS-433 |
| chr7-118365413-INS-52 | chr7-118365414-INS-50 |
| chr7-118371286-INS-542 | chr7-118371286-INS-542 |
| chr7-118376635-INS-188 | chr7-118376635-INS-187 |
| chr7-118386314-INS-132 | chr7-118386314-INS-132 |
| chr7-118389087-INS-89 | chr7-118389087-INS-89 |
| chr7-118392355-INS-202 | chr7-118392355-INS-202 |
| chr7-118410090-INS-90 | chr7-118410090-INS-92 |
| chr7-118412498-INS-152 | chr7-118412494-INS-192 |
| chr7-118426769-INS-186 | chr7-118426769-INS-186 |
| chr7-118429317-INS-50 | chr7-118429317-INS-62 |
| chr7-118430452-INS-568 | chr7-118430451-INS-570 |
| chr7-118451481-INS-76 | chr7-118451481-INS-76 |
| chr7-118453818-INS-273 | chr7-118453818-INS-304 |
| chr7-118460981-INS-187 | chr7-118460981-INS-188 |
| chr7-118470710-INS-186 | chr7-118470694-INS-182 |
| chr7-118473522-INS-186 | chr7-118473522-INS-186 |
| chr7-118511969-INS-64 | chr7-118511969-INS-64 |
| chr7-118512518-INS-67 | chr7-118512518-INS-67 |
| chr7-118524824-INS-90 | chr7-118524824-INS-90 |
| chr7-118526215-INS-125 | chr7-118526215-INS-125 |
| chr7-118535604-INS-114 | chr7-118535604-INS-114 |
| chr7-118551468-INS-402 | chr7-118551468-INS-402 |
| chr7-118607389-INS-217 | chr7-118607389-INS-217 |
| chr7-118614370-INS-72 | chr7-118614370-INS-72 |
| chr7-118615249-INS-579 | chr7-118615235-INS-587 |

| | |
|---|---|
| chr7-118619627-INS-84 | chr7-118619627-INS-86 |
| chr7-118624625-INS-810 | chr7-118624625-INS-810 |
| chr7-118625945-INS-54 | chr7-118625945-INS-56 |
| chr7-118627174-INS-280 | chr7-118627174-INS-284 |
| chr7-118643138-INS-53 | chr7-118643138-INS-53 |
| chr7-118663110-INS-99 | chr7-118663110-INS-95 |
| chr7-118670895-INS-4998 | chr7-118670896-INS-4998 |
| chr7-118729827-INS-159 | chr7-118729811-INS-145 |
| chr7-118734803-INS-108 | chr7-118734803-INS-108 |
| chr7-118752907-INS-75 | chr7-118752901-INS-71 |
| chr7-118788064-INS-58 | chr7-118788075-INS-54 |
| chr7-118794320-INS-115 | chr7-118794312-INS-111 |
| chr7-118804900-INS-348 | chr7-118804901-INS-371 |
| chr7-118840815-INS-80 | chr7-118840827-INS-70 |
| chr7-118883658-INS-62 | chr7-118883658-INS-68 |
| chr7-118900004-INS-344 | chr7-118900004-INS-344 |
| chr7-118901100-INS-206 | chr7-118901100-INS-206 |
| chr7-118907916-INS-62 | chr7-118907916-INS-60 |
| chr7-118912736-INS-6175 | chr7-118912737-INS-6175 |
| chr7-118933701-INS-51 | chr7-118933701-INS-51 |
| chr7-118945515-INS-54 | chr7-118945515-INS-54 |
| chr7-118948195-INS-213 | chr7-118948195-INS-215 |
| chr7-118951809-INS-66 | chr7-118951809-INS-66 |
| chr7-118960835-INS-110 | chr7-118960836-INS-106 |
| chr7-118968181-INS-59 | chr7-118968181-INS-57 |
| chr7-118970193-INS-408 | chr7-118970193-INS-408 |
| chr7-118973569-INS-90 | chr7-118973569-INS-88 |
| chr7-118989549-INS-60 | chr7-118989549-INS-58 |
| chr7-119021477-INS-68 | chr7-119021517-INS-70 |
| chr7-119071881-INS-211 | chr7-119071881-INS-209 |
| chr7-119079992-INS-480 | chr7-119079992-INS-481 |
| chr7-119090420-INS-98 | chr7-119090420-INS-98 |
| chr7-119100490-INS-561 | chr7-119100490-INS-563 |
| chr7-119126228-INS-240 | chr7-119126228-INS-240 |
| chr7-119140798-INS-83 | chr7-119140798-INS-63 |
| chr7-119146188-INS-206 | chr7-119146188-INS-206 |
| chr7-119149004-INS-226 | chr7-119149004-INS-226 |
| chr7-119154364-INS-64 | chr7-119154364-INS-108 |
| chr7-119168533-INS-661 | chr7-119168533-INS-661 |
| chr7-119222142-INS-416 | chr7-119222141-INS-415 |
| chr7-119224493-INS-670 | chr7-119224493-INS-669 |
| chr7-119225684-INS-64 | chr7-119225684-INS-64 |
| chr7-119230274-INS-60 | chr7-119230274-INS-60 |
| chr7-119238027-INS-52 | chr7-119238026-INS-56 |
| chr7-119251080-INS-881 | chr7-119251080-INS-881 |
| chr7-119264563-INS-531 | chr7-119264563-INS-531 |
| chr7-119279220-INS-81 | chr7-119279220-INS-81 |
| chr7-119315944-INS-54 | chr7-119315944-INS-54 |
| chr7-119325140-INS-156 | chr7-119325140-INS-156 |
| chr7-119352241-INS-70 | chr7-119352241-INS-70 |
| chr7-119354458-INS-192 | chr7-119354458-INS-192 |
| chr7-119394854-INS-56 | chr7-119394854-INS-64 |
| chr7-119404990-INS-212 | chr7-119404990-INS-212 |
| chr7-119408249-INS-73 | chr7-119408249-INS-74 |
| chr7-119413543-INS-209 | chr7-119413543-INS-209 |
| chr7-119458508-INS-560 | chr7-119458508-INS-560 |
| chr7-119522734-INS-53 | chr7-119522734-INS-53 |
| chr7-119529085-INS-225 | chr7-119529085-INS-225 |
| chr7-119591046-INS-255 | chr7-119591046-INS-256 |
| chr7-119593921-INS-52 | chr7-119593921-INS-56 |
| chr7-119631083-INS-69 | chr7-119631083-INS-69 |
| chr7-119661584-INS-54 | chr7-119661584-INS-50 |
| chr7-119674399-INS-586 | chr7-119674400-INS-586 |
| chr7-119684982-INS-72 | chr7-119684982-INS-73 |
| chr7-119716564-INS-173 | chr7-119716564-INS-173 |
| chr7-119718288-INS-141 | chr7-119718288-INS-145 |
| chr7-119724058-INS-80 | chr7-119724048-INS-110 |
| chr7-119736907-INS-122 | chr7-119736907-INS-122 |
| chr7-119776105-INS-209 | chr7-119776105-INS-209 |
| chr7-119783295-INS-51 | chr7-119783295-INS-57 |
| chr7-119796465-INS-60 | chr7-119796465-INS-56 |
| chr7-119803064-INS-556 | chr7-119803064-INS-556 |
| chr7-119810051-INS-78 | chr7-119810051-INS-66 |
| chr7-119811768-INS-160 | chr7-119811769-INS-163 |
| chr7-119837226-INS-112 | chr7-119837226-INS-112 |
| chr7-119843174-INS-76 | chr7-119843174-INS-77 |
| chr7-119859544-INS-5423 | chr7-119859544-INS-5423 |

| | |
|---|---|
| chr7-119878086-INS-74 | chr7-119878086-INS-74 |
| chr7-119892052-INS-650 | chr7-119892052-INS-650 |
| chr7-119908058-INS-135 | chr7-119908061-INS-143 |
| chr7-119914023-INS-159 | chr7-119914023-INS-158 |
| chr7-119945622-INS-77 | chr7-119945648-INS-50 |
| chr7-120048038-INS-652 | chr7-120048038-INS-652 |
| chr7-120071703-INS-187 | chr7-120071703-INS-187 |
| chr7-120127260-INS-60 | chr7-120127260-INS-63 |
| chr7-120160091-INS-69 | chr7-120160091-INS-69 |
| chr7-120172374-INS-816 | chr7-120172374-INS-817 |
| chr7-120181341-INS-440 | chr7-120181341-INS-440 |
| chr7-120189666-INS-337 | chr7-120189666-INS-336 |
| chr7-120190931-INS-62 | chr7-120190931-INS-62 |
| chr7-120209120-INS-88 | chr7-120209120-INS-88 |
| chr7-120213438-INS-63 | chr7-120213434-INS-72 |
| chr7-120214765-INS-199 | chr7-120214765-INS-199 |
| chr7-120230336-INS-283 | chr7-120230357-INS-270 |
| chr7-120242554-INS-221 | chr7-120242554-INS-222 |
| chr7-120294851-INS-220 | chr7-120294851-INS-220 |
| chr7-120361295-INS-108 | chr7-120361296-INS-81 |
| chr7-120370915-INS-73 | chr7-120370915-INS-73 |
| chr7-120388521-INS-6440 | chr7-120388522-INS-6440 |
| chr7-120393142-INS-51 | chr7-120393140-INS-71 |
| chr7-120396842-INS-110 | chr7-120396842-INS-111 |
| chr7-120405104-INS-96 | chr7-120405104-INS-96 |
| chr7-120408802-INS-54 | chr7-120408802-INS-102 |
| chr7-120413074-INS-76 | chr7-120413074-INS-70 |
| chr7-120423631-INS-209 | chr7-120423631-INS-209 |
| chr7-120424754-INS-115 | chr7-120424763-INS-115 |
| chr7-120440484-INS-344 | chr7-120440484-INS-344 |
| chr7-120458689-INS-873 | chr7-120458690-INS-874 |
| chr7-120467450-INS-197 | chr7-120467450-INS-197 |
| chr7-120569209-INS-1942 | chr7-120569210-INS-1942 |
| chr7-120610513-INS-193 | chr7-120610513-INS-190 |
| chr7-120633087-INS-266 | chr7-120633087-INS-267 |
| chr7-120660424-INS-66 | chr7-120660450-INS-55 |
| chr7-120688730-INS-91 | chr7-120688730-INS-91 |
| chr7-120699295-INS-101 | chr7-120699312-INS-80 |
| chr7-120729856-INS-100 | chr7-120729846-INS-74 |
| chr7-120745255-INS-2376 | chr7-120745256-INS-2381 |
| chr7-120766938-INS-538 | chr7-120766942-INS-550 |
| chr7-120770034-INS-2757 | chr7-120770035-INS-2757 |
| chr7-120779946-INS-58 | chr7-120779946-INS-62 |
| chr7-120790598-INS-93 | chr7-120790598-INS-53 |
| chr7-120796207-INS-52 | chr7-120796207-INS-54 |
| chr7-120821386-INS-60 | chr7-120821384-INS-60 |
| chr7-120827938-INS-60 | chr7-120827938-INS-64 |
| chr7-120833931-INS-1883 | chr7-120833932-INS-1882 |
| chr7-120841992-INS-205 | chr7-120841988-INS-190 |
| chr7-120854519-INS-63 | chr7-120854520-INS-50 |
| chr7-120864939-INS-120 | chr7-120864939-INS-120 |
| chr7-120882440-INS-215 | chr7-120882440-INS-214 |
| chr7-120942943-INS-298 | chr7-120942943-INS-306 |
| chr7-120947196-INS-100 | chr7-120947202-INS-66 |
| chr7-120971674-INS-134 | chr7-120971674-INS-134 |
| chr7-120984427-INS-130 | chr7-120984436-INS-129 |
| chr7-121020691-INS-384 | chr7-121020691-INS-342 |
| chr7-121046995-INS-268 | chr7-121046995-INS-256 |
| chr7-121066538-INS-143 | chr7-121066538-INS-143 |
| chr7-121090130-INS-55 | chr7-121090130-INS-55 |
| chr7-121090550-INS-730 | chr7-121090550-INS-736 |
| chr7-121095192-INS-68 | chr7-121095192-INS-56 |
| chr7-121104644-INS-124 | chr7-121104644-INS-136 |
| chr7-121122616-INS-399 | chr7-121122616-INS-399 |
| chr7-121138139-INS-520 | chr7-121138139-INS-521 |
| chr7-121150081-INS-111 | chr7-121150088-INS-91 |
| chr7-121212402-INS-399 | chr7-121212402-INS-399 |
| chr7-121273102-INS-151 | chr7-121273102-INS-151 |
| chr7-121279019-INS-144 | chr7-121279019-INS-143 |
| chr7-121282707-INS-6421 | chr7-121282708-INS-6422 |
| chr7-121288716-INS-107 | chr7-121288700-INS-117 |
| chr7-121335113-INS-207 | chr7-121335105-INS-220 |
| chr7-121340603-INS-1620 | chr7-121340604-INS-1621 |
| chr7-121428471-INS-61 | chr7-121428427-INS-61 |
| chr7-121436912-INS-91 | chr7-121436912-INS-84 |
| chr7-121445965-INS-56 | chr7-121445965-INS-60 |
| chr7-121447482-INS-56 | chr7-121447482-INS-58 |

| | |
|---|---|
| chr7-121467267-INS-72 | chr7-121467261-INS-52 |
| chr7-121493895-INS-5395 | chr7-121493896-INS-5395 |
| chr7-121503206-INS-55 | chr7-121503212-INS-78 |
| chr7-121507384-INS-56 | chr7-121507384-INS-56 |
| chr7-121509211-INS-199 | chr7-121509222-INS-197 |
| chr7-121514713-INS-234 | chr7-121514713-INS-234 |
| chr7-121517787-INS-133 | chr7-121517787-INS-133 |
| chr7-121544598-INS-170 | chr7-121544598-INS-170 |
| chr7-121558107-INS-206 | chr7-121558107-INS-207 |
| chr7-121573461-INS-68 | chr7-121573461-INS-68 |
| chr7-121578083-INS-52 | chr7-121578083-INS-52 |
| chr7-121593385-INS-241 | chr7-121593385-INS-241 |
| chr7-121603970-INS-153 | chr7-121603970-INS-153 |
| chr7-121617326-INS-99 | chr7-121617326-INS-76 |
| chr7-121626377-INS-207 | chr7-121626377-INS-207 |
| chr7-121627139-INS-183 | chr7-121627140-INS-183 |
| chr7-121654676-INS-5425 | chr7-121654677-INS-5425 |
| chr7-121660556-INS-213 | chr7-121660556-INS-211 |
| chr7-121700717-INS-216 | chr7-121700717-INS-216 |
| chr7-121706328-INS-54 | chr7-121706328-INS-59 |
| chr7-121707397-INS-64 | chr7-121707397-INS-66 |
| chr7-121711926-INS-220 | chr7-121711926-INS-242 |
| chr7-121724551-INS-204 | chr7-121724551-INS-204 |
| chr7-121735590-INS-80 | chr7-121735590-INS-80 |
| chr7-121738060-INS-203 | chr7-121738060-INS-203 |
| chr7-121742156-INS-128 | chr7-121742156-INS-128 |
| chr7-121746058-INS-58 | chr7-121746058-INS-58 |
| chr7-121749852-INS-86 | chr7-121749852-INS-70 |
| chr7-121764179-INS-228 | chr7-121764179-INS-228 |
| chr7-121781491-INS-84 | chr7-121781491-INS-62 |
| chr7-121797626-INS-99 | chr7-121797626-INS-99 |
| chr7-121817194-INS-54 | chr7-121817167-INS-71 |
| chr7-121824520-INS-91 | chr7-121824514-INS-99 |
| chr7-121833474-INS-201 | chr7-121833474-INS-183 |
| chr7-121842705-INS-75 | chr7-121842705-INS-87 |
| chr7-121857051-INS-72 | chr7-121857051-INS-72 |
| chr7-121942421-INS-183 | chr7-121942421-INS-175 |
| chr7-122023457-INS-51 | chr7-122023452-INS-50 |
| chr7-122065450-INS-57 | chr7-122065450-INS-51 |
| chr7-122068823-INS-140 | chr7-122068823-INS-152 |
| chr7-122073010-INS-176 | chr7-122073010-INS-177 |
| chr7-122083095-INS-60 | chr7-122083095-INS-58 |
| chr7-122085601-INS-58 | chr7-122085601-INS-82 |
| chr7-122130785-INS-78 | chr7-122130785-INS-78 |
| chr7-122133516-INS-135 | chr7-122133516-INS-135 |
| chr7-122167787-INS-140 | chr7-122167823-INS-90 |
| chr7-122188501-INS-52 | chr7-122188522-INS-53 |
| chr7-122189966-INS-109 | chr7-122189966-INS-108 |
| chr7-122198620-INS-60 | chr7-122198620-INS-64 |
| chr7-122211376-INS-87 | chr7-122211376-INS-60 |
| chr7-122232463-INS-83 | chr7-122232513-INS-83 |
| chr7-122242278-INS-111 | chr7-122242278-INS-111 |
| chr7-122251031-INS-99 | chr7-122251031-INS-99 |
| chr7-122292237-INS-108 | chr7-122292250-INS-144 |
| chr7-122297764-INS-58 | chr7-122297793-INS-74 |
| chr7-122300216-INS-125 | chr7-122300216-INS-126 |
| chr7-122305683-INS-55 | chr7-122305683-INS-55 |
| chr7-122313762-INS-95 | chr7-122313762-INS-95 |
| chr7-122319699-INS-1891 | chr7-122319700-INS-1891 |
| chr7-122329463-INS-96 | chr7-122329463-INS-96 |
| chr7-122351364-INS-66 | chr7-122351364-INS-70 |
| chr7-122358855-INS-96 | chr7-122358858-INS-111 |
| chr7-122381178-INS-185 | chr7-122381179-INS-185 |
| chr7-122381785-INS-115 | chr7-122381797-INS-65 |
| chr7-122391177-INS-71 | chr7-122391175-INS-75 |
| chr7-122438329-INS-6442 | chr7-122438330-INS-6442 |
| chr7-122448175-INS-104 | chr7-122448175-INS-104 |
| chr7-122452956-INS-5239 | chr7-122452957-INS-5239 |
| chr7-122453760-INS-326 | chr7-122453760-INS-326 |
| chr7-122455810-INS-93 | chr7-122455810-INS-71 |
| chr7-122469745-INS-88 | chr7-122469761-INS-76 |
| chr7-122478714-INS-54 | chr7-122478714-INS-56 |
| chr7-122483103-INS-54 | chr7-122483102-INS-92 |
| chr7-122503930-INS-431 | chr7-122503930-INS-435 |
| chr7-122539653-INS-121 | chr7-122539653-INS-120 |
| chr7-122542715-INS-54 | chr7-122542716-INS-54 |
| chr7-122544887-INS-105 | chr7-122544887-INS-105 |

| | |
|---|---|
| chr7-122546398-INS-57 | chr7-122546398-INS-57 |
| chr7-122554068-INS-66 | chr7-122554112-INS-71 |
| chr7-122564545-INS-206 | chr7-122564545-INS-206 |
| chr7-122583034-INS-183 | chr7-122583034-INS-176 |
| chr7-122590996-INS-244 | chr7-122590996-INS-244 |
| chr7-122593185-INS-161 | chr7-122593185-INS-161 |
| chr7-122606375-INS-165 | chr7-122606375-INS-165 |
| chr7-122615417-INS-63 | chr7-122615417-INS-67 |
| chr7-122616734-INS-212 | chr7-122616734-INS-212 |
| chr7-122617792-INS-56 | chr7-122617792-INS-56 |
| chr7-122636811-INS-162 | chr7-122636814-INS-170 |
| chr7-122644212-INS-204 | chr7-122644212-INS-200 |
| chr7-122656279-INS-113 | chr7-122656279-INS-113 |
| chr7-122674518-INS-70 | chr7-122674518-INS-70 |
| chr7-122681810-INS-55 | chr7-122681810-INS-54 |
| chr7-122685060-INS-94 | chr7-122685045-INS-74 |
| chr7-122686844-INS-656 | chr7-122686844-INS-657 |
| chr7-122700108-INS-60 | chr7-122700108-INS-55 |
| chr7-122705034-INS-61 | chr7-122705034-INS-65 |
| chr7-122743828-INS-62 | chr7-122743828-INS-62 |
| chr7-122761277-INS-202 | chr7-122761277-INS-201 |
| chr7-122773129-INS-217 | chr7-122773129-INS-219 |
| chr7-122808353-INS-203 | chr7-122808353-INS-204 |
| chr7-122812259-INS-126 | chr7-122812290-INS-135 |
| chr7-122812806-INS-102 | chr7-122812806-INS-66 |
| chr7-122826267-INS-100 | chr7-122826267-INS-100 |
| chr7-122836218-INS-53 | chr7-122836218-INS-53 |
| chr7-122840383-INS-209 | chr7-122840383-INS-209 |
| chr7-122842134-INS-86 | chr7-122842134-INS-66 |
| chr7-122862224-INS-177 | chr7-122862224-INS-177 |
| chr7-122899587-INS-118 | chr7-122899587-INS-118 |
| chr7-122931194-INS-363 | chr7-122931194-INS-363 |
| chr7-122938766-INS-491 | chr7-122938766-INS-492 |
| chr7-122974405-INS-216 | chr7-122974405-INS-216 |
| chr7-122981803-INS-71 | chr7-122981803-INS-74 |
| chr7-122983298-INS-230 | chr7-122983298-INS-230 |
| chr7-122992235-INS-66 | chr7-122992235-INS-66 |
| chr7-123008740-INS-160 | chr7-123008740-INS-192 |
| chr7-123015293-INS-288 | chr7-123015293-INS-288 |
| chr7-123033289-INS-512 | chr7-123033289-INS-515 |
| chr7-123042670-INS-7114 | chr7-123042671-INS-7114 |
| chr7-123043633-INS-222 | chr7-123043633-INS-223 |
| chr7-123046778-INS-200 | chr7-123046778-INS-200 |
| chr7-123047652-INS-66 | chr7-123047652-INS-60 |
| chr7-123063837-INS-111 | chr7-123063837-INS-110 |
| chr7-123072522-INS-5338 | chr7-123072523-INS-5338 |
| chr7-123102589-INS-66 | chr7-123102589-INS-66 |
| chr7-123105006-INS-505 | chr7-123105006-INS-505 |
| chr7-123115563-INS-204 | chr7-123115563-INS-204 |
| chr7-123144752-INS-94 | chr7-123144752-INS-133 |
| chr7-123154454-INS-5406 | chr7-123154455-INS-5406 |
| chr7-123162456-INS-98 | chr7-123162456-INS-76 |
| chr7-123169481-INS-58 | chr7-123169482-INS-53 |
| chr7-123197492-INS-695 | chr7-123197492-INS-696 |
| chr7-123229810-INS-8128 | chr7-123229811-INS-8129 |
| chr7-123247756-INS-234 | chr7-123247756-INS-234 |
| chr7-123275484-INS-96 | chr7-123275484-INS-121 |
| chr7-123358914-INS-103 | chr7-123358914-INS-103 |
| chr7-123360167-INS-80 | chr7-123360167-INS-78 |
| chr7-123389602-INS-75 | chr7-123389602-INS-76 |
| chr7-123390652-INS-902 | chr7-123390652-INS-901 |
| chr7-123391757-INS-497 | chr7-123391757-INS-468 |
| chr7-123400842-INS-439 | chr7-123400842-INS-456 |
| chr7-123403385-INS-64 | chr7-123403385-INS-64 |
| chr7-123483239-INS-86 | chr7-123483239-INS-94 |
| chr7-123486924-INS-659 | chr7-123486924-INS-659 |
| chr7-123491843-INS-69 | chr7-123491843-INS-69 |
| chr7-123512415-INS-554 | chr7-123512415-INS-554 |
| chr7-123519672-INS-364 | chr7-123519672-INS-364 |
| chr7-123551760-INS-2469 | chr7-123551761-INS-2471 |
| chr7-123578264-INS-676 | chr7-123578264-INS-676 |
| chr7-123599231-INS-235 | chr7-123599229-INS-236 |
| chr7-123600701-INS-53 | chr7-123600700-INS-80 |
| chr7-123601329-INS-54 | chr7-123601329-INS-58 |
| chr7-123605403-INS-69 | chr7-123605403-INS-69 |
| chr7-123656426-INS-1698 | chr7-123656443-INS-1697 |
| chr7-123669479-INS-6429 | chr7-123669480-INS-6429 |

| | |
|---|---|
| chr7-123682927-INS-62 | chr7-123682907-INS-78 |
| chr7-123699401-INS-54 | chr7-123699401-INS-54 |
| chr7-123710669-INS-75 | chr7-123710669-INS-75 |
| chr7-123724243-INS-6252 | chr7-123724244-INS-6252 |
| chr7-123761193-INS-381 | chr7-123761194-INS-383 |
| chr7-123767440-INS-920 | chr7-123767439-INS-926 |
| chr7-123771627-INS-6264 | chr7-123771628-INS-6265 |
| chr7-123772988-INS-162 | chr7-123772988-INS-162 |
| chr7-123783858-INS-81 | chr7-123783858-INS-51 |
| chr7-123823729-INS-59 | chr7-123823730-INS-74 |
| chr7-123898398-INS-257 | chr7-123898398-INS-239 |
| chr7-123972710-INS-364 | chr7-123972710-INS-364 |
| chr7-124000531-INS-57 | chr7-124000531-INS-57 |
| chr7-124009426-INS-194 | chr7-124009426-INS-194 |
| chr7-124011686-INS-454 | chr7-124011686-INS-454 |
| chr7-124024363-INS-59 | chr7-124024363-INS-59 |
| chr7-124028852-INS-58 | chr7-124028852-INS-58 |
| chr7-124056962-INS-84 | chr7-124056962-INS-102 |
| chr7-124070701-INS-185 | chr7-124070686-INS-215 |
| chr7-124083963-INS-404 | chr7-124083963-INS-404 |
| chr7-124139183-INS-6857 | chr7-124139184-INS-6859 |
| chr7-124145042-INS-101 | chr7-124145042-INS-101 |
| chr7-124157107-INS-5384 | chr7-124157108-INS-5384 |
| chr7-124158812-INS-329 | chr7-124158812-INS-329 |
| chr7-124162535-INS-64 | chr7-124162536-INS-67 |
| chr7-124167785-INS-131 | chr7-124167785-INS-147 |
| chr7-124173344-INS-6746 | chr7-124173345-INS-6747 |
| chr7-124184256-INS-506 | chr7-124184256-INS-484 |
| chr7-124205520-INS-55 | chr7-124205520-INS-55 |
| chr7-124210779-INS-197 | chr7-124210779-INS-197 |
| chr7-124214205-INS-72 | chr7-124214203-INS-62 |
| chr7-124256929-INS-86 | chr7-124256929-INS-71 |
| chr7-124268814-INS-6947 | chr7-124268815-INS-6951 |
| chr7-124270741-INS-51 | chr7-124270741-INS-54 |
| chr7-124272004-INS-82 | chr7-124272004-INS-66 |
| chr7-124286525-INS-51 | chr7-124286525-INS-51 |
| chr7-124322145-INS-110 | chr7-124322147-INS-96 |
| chr7-124332302-INS-56 | chr7-124332302-INS-56 |
| chr7-124372403-INS-397 | chr7-124372403-INS-397 |
| chr7-124373646-INS-516 | chr7-124373646-INS-517 |
| chr7-124376184-INS-204 | chr7-124376230-INS-204 |
| chr7-124398853-INS-144 | chr7-124398853-INS-148 |
| chr7-124425719-INS-58 | chr7-124425719-INS-62 |
| chr7-124463159-INS-102 | chr7-124463159-INS-104 |
| chr7-124467941-INS-6669 | chr7-124467942-INS-6670 |
| chr7-124480684-INS-146 | chr7-124480684-INS-158 |
| chr7-124485667-INS-50 | chr7-124485667-INS-64 |
| chr7-124516621-INS-104 | chr7-124516613-INS-116 |
| chr7-124521182-INS-5547 | chr7-124521183-INS-5547 |
| chr7-124541400-INS-518 | chr7-124541400-INS-518 |
| chr7-124549771-INS-169 | chr7-124549771-INS-169 |
| chr7-124566528-INS-62 | chr7-124566528-INS-64 |
| chr7-124569854-INS-92 | chr7-124569854-INS-56 |
| chr7-124587963-INS-116 | chr7-124587963-INS-116 |
| chr7-124589670-INS-104 | chr7-124589707-INS-56 |
| chr7-124651350-INS-177 | chr7-124651356-INS-215 |
| chr7-124661097-INS-3359 | chr7-124661098-INS-3359 |
| chr7-124725699-INS-215 | chr7-124725699-INS-214 |
| chr7-124741652-INS-895 | chr7-124741652-INS-895 |
| chr7-124759434-INS-1019 | chr7-124759435-INS-1022 |
| chr7-124762349-INS-51 | chr7-124762349-INS-51 |
| chr7-124782357-INS-61 | chr7-124782357-INS-61 |
| chr7-124796711-INS-51 | chr7-124796711-INS-53 |
| chr7-124806073-INS-7144 | chr7-124806074-INS-7144 |
| chr7-124809512-INS-58 | chr7-124809512-INS-60 |
| chr7-124835833-INS-152 | chr7-124835833-INS-152 |
| chr7-124853914-INS-106 | chr7-124853914-INS-110 |
| chr7-124873473-INS-59 | chr7-124873480-INS-83 |
| chr7-124882678-INS-60 | chr7-124882678-INS-60 |
| chr7-124888054-INS-87 | chr7-124888054-INS-87 |
| chr7-124938859-INS-6260 | chr7-124938860-INS-6260 |
| chr7-124961131-INS-101 | chr7-124961131-INS-105 |
| chr7-124964347-INS-1016 | chr7-124964345-INS-1012 |
| chr7-124978156-INS-99 | chr7-124978156-INS-52 |
| chr7-124997634-INS-74 | chr7-124997634-INS-72 |
| chr7-124998831-INS-67 | chr7-124998831-INS-70 |
| chr7-125009947-INS-89 | chr7-125009944-INS-89 |

| | |
|---|---|
| chr7-125015137-INS-88 | chr7-125015137-INS-88 |
| chr7-125020381-INS-170 | chr7-125020379-INS-145 |
| chr7-125038081-INS-169 | chr7-125038081-INS-169 |
| chr7-125051362-INS-68 | chr7-125051358-INS-63 |
| chr7-125056425-INS-567 | chr7-125056425-INS-567 |
| chr7-125056964-INS-51 | chr7-125056965-INS-51 |
| chr7-125066604-INS-101 | chr7-125066604-INS-101 |
| chr7-125074781-INS-54 | chr7-125074781-INS-54 |
| chr7-125085085-INS-86 | chr7-125085085-INS-87 |
| chr7-125086995-INS-215 | chr7-125086995-INS-215 |
| chr7-125098929-INS-6348 | chr7-125098930-INS-6349 |
| chr7-125121021-INS-67 | chr7-125121003-INS-56 |
| chr7-125127892-INS-66 | chr7-125127886-INS-52 |
| chr7-125135255-INS-97 | chr7-125135255-INS-97 |
| chr7-125163305-INS-84 | chr7-125163305-INS-96 |
| chr7-125165580-INS-57 | chr7-125165580-INS-70 |
| chr7-125179684-INS-141 | chr7-125179684-INS-141 |
| chr7-125181030-INS-211 | chr7-125181030-INS-210 |
| chr7-125192668-INS-147 | chr7-125192668-INS-148 |
| chr7-125193698-INS-80 | chr7-125193698-INS-50 |
| chr7-125233910-INS-201 | chr7-125233910-INS-207 |
| chr7-125242902-INS-59 | chr7-125242902-INS-59 |
| chr7-125249967-INS-895 | chr7-125249973-INS-905 |
| chr7-125270068-INS-60 | chr7-125270068-INS-60 |
| chr7-125287563-INS-440 | chr7-125287563-INS-440 |
| chr7-125297759-INS-269 | chr7-125297759-INS-268 |
| chr7-125328221-INS-7359 | chr7-125328222-INS-7359 |
| chr7-125342612-INS-420 | chr7-125342612-INS-420 |
| chr7-125351572-INS-163 | chr7-125351572-INS-173 |
| chr7-125381099-INS-54 | chr7-125381099-INS-56 |
| chr7-125388124-INS-116 | chr7-125388124-INS-116 |
| chr7-125406680-INS-1989 | chr7-125406681-INS-1989 |
| chr7-125420505-INS-55 | chr7-125420505-INS-53 |
| chr7-125447035-INS-140 | chr7-125447035-INS-140 |
| chr7-125467295-INS-110 | chr7-125467291-INS-85 |
| chr7-125639699-INS-138 | chr7-125639699-INS-138 |
| chr7-125652333-INS-3537 | chr7-125652334-INS-3544 |
| chr7-125654198-INS-177 | chr7-125654198-INS-177 |
| chr7-125727118-INS-1538 | chr7-125727119-INS-1538 |
| chr7-125731701-INS-52 | chr7-125731701-INS-52 |
| chr7-125782607-INS-197 | chr7-125782607-INS-197 |
| chr7-125785119-INS-239 | chr7-125785119-INS-240 |
| chr7-125800361-INS-269 | chr7-125800361-INS-270 |
| chr7-125804245-INS-175 | chr7-125804245-INS-177 |
| chr7-125805236-INS-283 | chr7-125805236-INS-270 |
| chr7-125810504-INS-183 | chr7-125810505-INS-188 |
| chr7-125814843-INS-7914 | chr7-125814844-INS-7914 |
| chr7-125840594-INS-187 | chr7-125840594-INS-187 |
| chr7-125842893-INS-204 | chr7-125842893-INS-204 |
| chr7-125850317-INS-118 | chr7-125850317-INS-118 |
| chr7-125856697-INS-1888 | chr7-125856698-INS-1888 |
| chr7-125864530-INS-71 | chr7-125864530-INS-71 |
| chr7-125869871-INS-213 | chr7-125869871-INS-213 |
| chr7-125874736-INS-111 | chr7-125874736-INS-111 |
| chr7-125898004-INS-56 | chr7-125898004-INS-52 |
| chr7-125903840-INS-205 | chr7-125903840-INS-206 |
| chr7-125910044-INS-105 | chr7-125910044-INS-121 |
| chr7-125915797-INS-69 | chr7-125915797-INS-69 |
| chr7-125917516-INS-103 | chr7-125917507-INS-108 |
| chr7-125926570-INS-193 | chr7-125926570-INS-193 |
| chr7-125929866-INS-65 | chr7-125929828-INS-70 |
| chr7-125932446-INS-223 | chr7-125932446-INS-223 |
| chr7-125933551-INS-239 | chr7-125933551-INS-240 |
| chr7-125936493-INS-184 | chr7-125936493-INS-186 |
| chr7-125950495-INS-94 | chr7-125950495-INS-94 |
| chr7-125952023-INS-201 | chr7-125952023-INS-202 |
| chr7-125953139-INS-414 | chr7-125953139-INS-413 |
| chr7-125977451-INS-179 | chr7-125977451-INS-179 |
| chr7-125979107-INS-211 | chr7-125979107-INS-212 |
| chr7-125980443-INS-177 | chr7-125980443-INS-177 |
| chr7-125987119-INS-59 | chr7-125987099-INS-62 |
| chr7-126006887-INS-3101 | chr7-126006888-INS-3093 |
| chr7-126019271-INS-57 | chr7-126019272-INS-67 |
| chr7-126026564-INS-203 | chr7-126026564-INS-203 |
| chr7-126041989-INS-210 | chr7-126041989-INS-211 |
| chr7-126076087-INS-176 | chr7-126076087-INS-176 |
| chr7-126104649-INS-189 | chr7-126104649-INS-189 |

| | |
|---|---|
| chr7-126111374-INS-199 | chr7-126111386-INS-197 |
| chr7-126113787-INS-971 | chr7-126113787-INS-978 |
| chr7-126120830-INS-200 | chr7-126120830-INS-200 |
| chr7-126122789-INS-58 | chr7-126122789-INS-58 |
| chr7-126134745-INS-189 | chr7-126134745-INS-199 |
| chr7-126137408-INS-206 | chr7-126137407-INS-210 |
| chr7-126143034-INS-173 | chr7-126143034-INS-173 |
| chr7-126147592-INS-170 | chr7-126147592-INS-195 |
| chr7-126148711-INS-216 | chr7-126148722-INS-212 |
| chr7-126151104-INS-191 | chr7-126151117-INS-187 |
| chr7-126153598-INS-256 | chr7-126153598-INS-243 |
| chr7-126160230-INS-465 | chr7-126160230-INS-475 |
| chr7-126163369-INS-57 | chr7-126163369-INS-59 |
| chr7-126167690-INS-1890 | chr7-126167691-INS-1890 |
| chr7-126181520-INS-67 | chr7-126181520-INS-67 |
| chr7-126186855-INS-66 | chr7-126186855-INS-66 |
| chr7-126195547-INS-162 | chr7-126195545-INS-204 |
| chr7-126221439-INS-212 | chr7-126221439-INS-205 |
| chr7-126228051-INS-484 | chr7-126228051-INS-484 |
| chr7-126241317-INS-211 | chr7-126241317-INS-211 |
| chr7-126242303-INS-780 | chr7-126242304-INS-787 |
| chr7-126245716-INS-214 | chr7-126245716-INS-182 |
| chr7-126268988-INS-212 | chr7-126268988-INS-210 |
| chr7-126285079-INS-60 | chr7-126285068-INS-52 |
| chr7-126310031-INS-242 | chr7-126310031-INS-242 |
| chr7-126333658-INS-398 | chr7-126333658-INS-398 |
| chr7-126338112-INS-188 | chr7-126338112-INS-188 |
| chr7-126340752-INS-230 | chr7-126340752-INS-230 |
| chr7-126366016-INS-7253 | chr7-126366017-INS-7253 |
| chr7-126369476-INS-97 | chr7-126369476-INS-97 |
| chr7-126372890-INS-5283 | chr7-126372891-INS-5283 |
| chr7-126405300-INS-64 | chr7-126405300-INS-58 |
| chr7-126410315-INS-70 | chr7-126410315-INS-70 |
| chr7-126439193-INS-545 | chr7-126439193-INS-545 |
| chr7-126443306-INS-54 | chr7-126443306-INS-54 |
| chr7-126472458-INS-215 | chr7-126472458-INS-216 |
| chr7-126489932-INS-200 | chr7-126489932-INS-208 |
| chr7-126508246-INS-51 | chr7-126508256-INS-51 |
| chr7-126524925-INS-70 | chr7-126524925-INS-70 |
| chr7-126557288-INS-217 | chr7-126557288-INS-217 |
| chr7-126599433-INS-60 | chr7-126599409-INS-88 |
| chr7-126656943-INS-212 | chr7-126656943-INS-212 |
| chr7-126659553-INS-57 | chr7-126659553-INS-60 |
| chr7-126660480-INS-227 | chr7-126660480-INS-215 |
| chr7-126666419-INS-203 | chr7-126666419-INS-203 |
| chr7-126719141-INS-116 | chr7-126719141-INS-122 |
| chr7-126751814-INS-69 | chr7-126751814-INS-69 |
| chr7-126758069-INS-177 | chr7-126758069-INS-179 |
| chr7-126761500-INS-177 | chr7-126761482-INS-204 |
| chr7-126764083-INS-212 | chr7-126764083-INS-212 |
| chr7-126766581-INS-59 | chr7-126766581-INS-59 |
| chr7-126774270-INS-81 | chr7-126774270-INS-73 |
| chr7-126775589-INS-177 | chr7-126775589-INS-177 |
| chr7-126787382-INS-179 | chr7-126787389-INS-187 |
| chr7-126790339-INS-596 | chr7-126790339-INS-595 |
| chr7-126824498-INS-200 | chr7-126824498-INS-200 |
| chr7-126863172-INS-1110 | chr7-126863173-INS-1110 |
| chr7-126863247-INS-60 | chr7-126863240-INS-50 |
| chr7-126867481-INS-434 | chr7-126867481-INS-444 |
| chr7-126868534-INS-1042 | chr7-126868535-INS-1042 |
| chr7-126868805-INS-180 | chr7-126868805-INS-180 |
| chr7-126874662-INS-451 | chr7-126874662-INS-453 |
| chr7-126884557-INS-539 | chr7-126884557-INS-539 |
| chr7-126886424-INS-176 | chr7-126886424-INS-180 |
| chr7-126887956-INS-170 | chr7-126887961-INS-169 |
| chr7-126890788-INS-199 | chr7-126890788-INS-199 |
| chr7-126902791-INS-75 | chr7-126902792-INS-102 |
| chr7-126903177-INS-295 | chr7-126903177-INS-297 |
| chr7-126930600-INS-196 | chr7-126930600-INS-196 |
| chr7-126938685-INS-212 | chr7-126938685-INS-211 |
| chr7-126941472-INS-65 | chr7-126941472-INS-79 |
| chr7-126947946-INS-172 | chr7-126947946-INS-156 |
| chr7-126950701-INS-253 | chr7-126950701-INS-254 |
| chr7-126965916-INS-70 | chr7-126965916-INS-106 |
| chr7-126994952-INS-208 | chr7-126994952-INS-209 |
| chr7-127011002-INS-69 | chr7-127011002-INS-69 |
| chr7-127016974-INS-215 | chr7-127016974-INS-217 |

| | |
|---|---|
| chr7-127030626-INS-213 | chr7-127030626-INS-213 |
| chr7-127033287-INS-168 | chr7-127033287-INS-171 |
| chr7-127050210-INS-203 | chr7-127050210-INS-203 |
| chr7-127051143-INS-103 | chr7-127051143-INS-103 |
| chr7-127054735-INS-191 | chr7-127054735-INS-191 |
| chr7-127064045-INS-214 | chr7-127064045-INS-206 |
| chr7-127199619-INS-138 | chr7-127199619-INS-92 |
| chr7-127233257-INS-55 | chr7-127233257-INS-63 |
| chr7-127239477-INS-189 | chr7-127239475-INS-186 |
| chr7-127243204-INS-188 | chr7-127243204-INS-184 |
| chr7-127252478-INS-7280 | chr7-127252479-INS-7280 |
| chr7-127280791-INS-212 | chr7-127280791-INS-212 |
| chr7-127305452-INS-208 | chr7-127305453-INS-208 |
| chr7-127319452-INS-88 | chr7-127319451-INS-94 |
| chr7-127320906-INS-207 | chr7-127320907-INS-214 |
| chr7-127321804-INS-203 | chr7-127321804-INS-203 |
| chr7-127323446-INS-91 | chr7-127323447-INS-92 |
| chr7-127411166-INS-207 | chr7-127411166-INS-216 |
| chr7-127433387-INS-181 | chr7-127433374-INS-181 |
| chr7-127493297-INS-68 | chr7-127493297-INS-68 |
| chr7-127496491-INS-63 | chr7-127496487-INS-64 |
| chr7-127521888-INS-7166 | chr7-127521889-INS-7166 |
| chr7-127540795-INS-131 | chr7-127540795-INS-89 |
| chr7-127541181-INS-211 | chr7-127541183-INS-217 |
| chr7-127559906-INS-681 | chr7-127559906-INS-681 |
| chr7-127563083-INS-60 | chr7-127563083-INS-60 |
| chr7-127601888-INS-52 | chr7-127601888-INS-52 |
| chr7-127605349-INS-269 | chr7-127605349-INS-269 |
| chr7-127610881-INS-145 | chr7-127610881-INS-145 |
| chr7-127614320-INS-801 | chr7-127614320-INS-794 |
| chr7-127642518-INS-168 | chr7-127642519-INS-156 |
| chr7-127650710-INS-186 | chr7-127650710-INS-186 |
| chr7-127656629-INS-52 | chr7-127656629-INS-54 |
| chr7-127660737-INS-94 | chr7-127660737-INS-97 |
| chr7-127665225-INS-212 | chr7-127665225-INS-212 |
| chr7-127677066-INS-103 | chr7-127677066-INS-91 |
| chr7-127703585-INS-316 | chr7-127703585-INS-316 |
| chr7-127711063-INS-71 | chr7-127711064-INS-68 |
| chr7-127726433-INS-214 | chr7-127726433-INS-214 |
| chr7-127743060-INS-52 | chr7-127743060-INS-52 |
| chr7-127755011-INS-731 | chr7-127755013-INS-737 |
| chr7-127759185-INS-75 | chr7-127759185-INS-75 |
| chr7-127766809-INS-54 | chr7-127766809-INS-56 |
| chr7-127771553-INS-67 | chr7-127771553-INS-73 |
| chr7-127778968-INS-121 | chr7-127778968-INS-120 |
| chr7-127791131-INS-506 | chr7-127791131-INS-506 |
| chr7-127800020-INS-118 | chr7-127800020-INS-116 |
| chr7-127824704-INS-201 | chr7-127824704-INS-201 |
| chr7-127829286-INS-128 | chr7-127829286-INS-128 |
| chr7-127849958-INS-72 | chr7-127849958-INS-54 |
| chr7-127856095-INS-408 | chr7-127856095-INS-408 |
| chr7-127866967-INS-75 | chr7-127866967-INS-75 |
| chr7-127868967-INS-137 | chr7-127868967-INS-186 |
| chr7-127886862-INS-63 | chr7-127886864-INS-65 |
| chr7-127907799-INS-375 | chr7-127907799-INS-376 |
| chr7-127917425-INS-53 | chr7-127917425-INS-53 |
| chr7-127948206-INS-6272 | chr7-127948207-INS-6272 |
| chr7-127953364-INS-148 | chr7-127953369-INS-148 |
| chr7-127984845-INS-74 | chr7-127984831-INS-52 |
| chr7-128036542-INS-97 | chr7-128036540-INS-112 |
| chr7-128069517-INS-144 | chr7-128069517-INS-144 |
| chr7-128079025-INS-106 | chr7-128079039-INS-117 |
| chr7-128106463-INS-1515 | chr7-128106463-INS-1515 |
| chr7-128113262-INS-242 | chr7-128113262-INS-242 |
| chr7-128114165-INS-112 | chr7-128114165-INS-108 |
| chr7-128128016-INS-236 | chr7-128128016-INS-236 |
| chr7-128136691-INS-220 | chr7-128136691-INS-205 |
| chr7-128146111-INS-201 | chr7-128146111-INS-201 |
| chr7-128157552-INS-140 | chr7-128157552-INS-140 |
| chr7-128174348-INS-200 | chr7-128174348-INS-200 |
| chr7-128185495-INS-108 | chr7-128185495-INS-108 |
| chr7-128196695-INS-203 | chr7-128196691-INS-196 |
| chr7-128223259-INS-207 | chr7-128223259-INS-207 |
| chr7-128226862-INS-164 | chr7-128226862-INS-164 |
| chr7-128245140-INS-100 | chr7-128245140-INS-100 |
| chr7-128257038-INS-181 | chr7-128257038-INS-182 |
| chr7-128263749-INS-94 | chr7-128263749-INS-96 |

| | |
|---|---|
| chr7-128267491-INS-73 | chr7-128267491-INS-73 |
| chr7-128276705-INS-56 | chr7-128276705-INS-56 |
| chr7-128278088-INS-70 | chr7-128278088-INS-70 |
| chr7-128284637-INS-222 | chr7-128284637-INS-222 |
| chr7-128286509-INS-85 | chr7-128286509-INS-85 |
| chr7-128291745-INS-56 | chr7-128291729-INS-60 |
| chr7-128301637-INS-207 | chr7-128301637-INS-207 |
| chr7-128306564-INS-201 | chr7-128306564-INS-203 |
| chr7-128312516-INS-77 | chr7-128312505-INS-82 |
| chr7-128340367-INS-71 | chr7-128340367-INS-71 |
| chr7-128347301-INS-147 | chr7-128347301-INS-147 |
| chr7-128375426-INS-120 | chr7-128375426-INS-120 |
| chr7-128387878-INS-231 | chr7-128387878-INS-231 |
| chr7-128391159-INS-6423 | chr7-128391160-INS-6423 |
| chr7-128402528-INS-71 | chr7-128402528-INS-75 |
| chr7-128406982-INS-212 | chr7-128406982-INS-212 |
| chr7-128409919-INS-233 | chr7-128409919-INS-233 |
| chr7-128413997-INS-172 | chr7-128413997-INS-173 |
| chr7-128425167-INS-514 | chr7-128425167-INS-514 |
| chr7-128450017-INS-120 | chr7-128450017-INS-112 |
| chr7-128463557-INS-117 | chr7-128463557-INS-82 |
| chr7-128475313-INS-90 | chr7-128475313-INS-90 |
| chr7-128481380-INS-112 | chr7-128481380-INS-116 |
| chr7-128524599-INS-102 | chr7-128524599-INS-106 |
| chr7-128526387-INS-3348 | chr7-128526388-INS-3348 |
| chr7-128532295-INS-87 | chr7-128532295-INS-87 |
| chr7-128558402-INS-62 | chr7-128558399-INS-69 |
| chr7-128561141-INS-80 | chr7-128561142-INS-80 |
| chr7-128562206-INS-72 | chr7-128562170-INS-73 |
| chr7-128563478-INS-74 | chr7-128563429-INS-102 |
| chr7-128572297-INS-136 | chr7-128572290-INS-156 |
| chr7-128573348-INS-257 | chr7-128573348-INS-257 |
| chr7-128608645-INS-802 | chr7-128608645-INS-802 |
| chr7-128620172-INS-89 | chr7-128620172-INS-89 |
| chr7-128625656-INS-226 | chr7-128625656-INS-226 |
| chr7-128654697-INS-75 | chr7-128654697-INS-75 |
| chr7-128690654-INS-66 | chr7-128690654-INS-66 |
| chr7-128696803-INS-1446 | chr7-128696804-INS-1449 |
| chr7-128699283-INS-60 | chr7-128699281-INS-62 |
| chr7-128715127-INS-1372 | chr7-128715128-INS-1370 |
| chr7-128725558-INS-148 | chr7-128725556-INS-152 |
| chr7-128738037-INS-620 | chr7-128738037-INS-620 |
| chr7-128764467-INS-6402 | chr7-128764468-INS-6402 |
| chr7-128774508-INS-2970 | chr7-128774509-INS-2970 |
| chr7-128787735-INS-212 | chr7-128787735-INS-212 |
| chr7-128795964-INS-149 | chr7-128795964-INS-149 |
| chr7-128802296-INS-478 | chr7-128802296-INS-478 |
| chr7-128810007-INS-83 | chr7-128810007-INS-83 |
| chr7-128824252-INS-3004 | chr7-128824253-INS-3004 |
| chr7-128831064-INS-372 | chr7-128831064-INS-372 |
| chr7-128832086-INS-449 | chr7-128832086-INS-449 |
| chr7-128841416-INS-62 | chr7-128841416-INS-78 |
| chr7-128842635-INS-132 | chr7-128842635-INS-132 |
| chr7-128852020-INS-142 | chr7-128852020-INS-142 |
| chr7-128854532-INS-50 | chr7-128854532-INS-54 |
| chr7-128873003-INS-498 | chr7-128873003-INS-498 |
| chr7-128897882-INS-94 | chr7-128897882-INS-94 |
| chr7-128945202-INS-63 | chr7-128945182-INS-98 |
| chr7-128965923-INS-77 | chr7-128965923-INS-79 |
| chr7-128967371-INS-72 | chr7-128967372-INS-71 |
| chr7-128993789-INS-7055 | chr7-128993790-INS-7055 |
| chr7-129003494-INS-250 | chr7-129003494-INS-250 |
| chr7-129007415-INS-97 | chr7-129007415-INS-97 |
| chr7-129008208-INS-6385 | chr7-129008209-INS-6385 |
| chr7-129013712-INS-6492 | chr7-129013713-INS-6492 |
| chr7-129018954-INS-6708 | chr7-129018955-INS-6708 |
| chr7-129040005-INS-78 | chr7-129040005-INS-78 |
| chr7-129046242-INS-72 | chr7-129046242-INS-70 |
| chr7-129072507-INS-89 | chr7-129072507-INS-113 |
| chr7-129114412-INS-57 | chr7-129114412-INS-57 |
| chr7-129117500-INS-425 | chr7-129117500-INS-427 |
| chr7-129134683-INS-1865 | chr7-129134684-INS-1865 |
| chr7-129140904-INS-58 | chr7-129140904-INS-58 |
| chr7-129142562-INS-59 | chr7-129142562-INS-59 |
| chr7-129188366-INS-323 | chr7-129188366-INS-330 |
| chr7-129200153-INS-852 | chr7-129200153-INS-853 |
| chr7-129201525-INS-81 | chr7-129201526-INS-63 |

| | |
|---|---|
| chr7-129242498-INS-402 | chr7-129242503-INS-388 |
| chr7-129244762-INS-60 | chr7-129244762-INS-60 |
| chr7-129246452-INS-110 | chr7-129246490-INS-118 |
| chr7-129259484-INS-124 | chr7-129259476-INS-128 |
| chr7-129265146-INS-2948 | chr7-129265147-INS-2948 |
| chr7-129329757-INS-479 | chr7-129329757-INS-479 |
| chr7-129355860-INS-90 | chr7-129355860-INS-90 |
| chr7-129382009-INS-83 | chr7-129382009-INS-77 |
| chr7-129394452-INS-162 | chr7-129394452-INS-162 |
| chr7-129406742-INS-85 | chr7-129406742-INS-80 |
| chr7-129408055-INS-229 | chr7-129408055-INS-229 |
| chr7-129420336-INS-194 | chr7-129420336-INS-194 |
| chr7-129435887-INS-51 | chr7-129435887-INS-51 |
| chr7-129489499-INS-1705 | chr7-129489500-INS-1707 |
| chr7-129493433-INS-343 | chr7-129493433-INS-343 |
| chr7-129499032-INS-61 | chr7-129499032-INS-61 |
| chr7-129527407-INS-62 | chr7-129527407-INS-66 |
| chr7-129534257-INS-498 | chr7-129534257-INS-498 |
| chr7-129534818-INS-6206 | chr7-129534819-INS-6207 |
| chr7-129538365-INS-60 | chr7-129538365-INS-60 |
| chr7-129545254-INS-442 | chr7-129545254-INS-442 |
| chr7-129557843-INS-58 | chr7-129557843-INS-58 |
| chr7-129567303-INS-72 | chr7-129567303-INS-74 |
| chr7-129587343-INS-81 | chr7-129587343-INS-81 |
| chr7-129607814-INS-169 | chr7-129607814-INS-169 |
| chr7-129609350-INS-84 | chr7-129609350-INS-84 |
| chr7-129641952-INS-98 | chr7-129641960-INS-102 |
| chr7-129642491-INS-77 | chr7-129642491-INS-77 |
| chr7-129673576-INS-140 | chr7-129673576-INS-140 |
| chr7-129692582-INS-578 | chr7-129692583-INS-578 |
| chr7-129733130-INS-209 | chr7-129733131-INS-209 |
| chr7-129755773-INS-187 | chr7-129755773-INS-162 |
| chr7-129769767-INS-51 | chr7-129769767-INS-51 |
| chr7-129789675-INS-98 | chr7-129789675-INS-104 |
| chr7-129816340-INS-57 | chr7-129816340-INS-54 |
| chr7-129820383-INS-181 | chr7-129820383-INS-181 |
| chr7-129820383-INS-781 | chr7-129820383-INS-781 |
| chr7-129883363-INS-52 | chr7-129883363-INS-54 |
| chr7-129897046-INS-60 | chr7-129897046-INS-60 |
| chr7-129930945-INS-73 | chr7-129930962-INS-86 |
| chr7-129936075-INS-184 | chr7-129936075-INS-184 |
| chr7-129958574-INS-125 | chr7-129958574-INS-84 |
| chr7-129959584-INS-62 | chr7-129959584-INS-63 |
| chr7-129965446-INS-857 | chr7-129965446-INS-857 |
| chr7-129970427-INS-76 | chr7-129970427-INS-76 |
| chr7-129973633-INS-178 | chr7-129973634-INS-187 |
| chr7-129985572-INS-59 | chr7-129985572-INS-59 |
| chr7-129992786-INS-206 | chr7-129992786-INS-206 |
| chr7-130000002-INS-163 | chr7-130000002-INS-164 |
| chr7-130007426-INS-516 | chr7-130007426-INS-515 |
| chr7-130012356-INS-1455 | chr7-130012357-INS-1455 |
| chr7-130018448-INS-191 | chr7-130018448-INS-192 |
| chr7-130036883-INS-59 | chr7-130036883-INS-59 |
| chr7-130064075-INS-81 | chr7-130064075-INS-99 |
| chr7-130091942-INS-154 | chr7-130091942-INS-155 |
| chr7-130101170-INS-209 | chr7-130101170-INS-209 |
| chr7-130101897-INS-144 | chr7-130101897-INS-120 |
| chr7-130108820-INS-6439 | chr7-130108821-INS-6439 |
| chr7-130110591-INS-110 | chr7-130110591-INS-65 |
| chr7-130125167-INS-218 | chr7-130125167-INS-214 |
| chr7-130127855-INS-168 | chr7-130127855-INS-168 |
| chr7-130128764-INS-178 | chr7-130128764-INS-179 |
| chr7-130142742-INS-295 | chr7-130142742-INS-297 |
| chr7-130196389-INS-84 | chr7-130196415-INS-108 |
| chr7-130205896-INS-190 | chr7-130205896-INS-228 |
| chr7-130207558-INS-145 | chr7-130207558-INS-140 |
| chr7-130210669-INS-81 | chr7-130210670-INS-78 |
| chr7-130231148-INS-192 | chr7-130231148-INS-192 |
| chr7-130241027-INS-6441 | chr7-130241028-INS-6441 |
| chr7-130253416-INS-76 | chr7-130253416-INS-78 |
| chr7-130262674-INS-197 | chr7-130262674-INS-196 |
| chr7-130264434-INS-299 | chr7-130264434-INS-299 |
| chr7-130265321-INS-206 | chr7-130265321-INS-204 |
| chr7-130270303-INS-195 | chr7-130270304-INS-196 |
| chr7-130286896-INS-188 | chr7-130286896-INS-188 |
| chr7-130302101-INS-211 | chr7-130302101-INS-208 |
| chr7-130302793-INS-408 | chr7-130302793-INS-407 |

| | |
|---|---|
| chr7-130309651-INS-120 | chr7-130309651-INS-120 |
| chr7-130327688-INS-498 | chr7-130327688-INS-498 |
| chr7-130337962-INS-56 | chr7-130337962-INS-69 |
| chr7-130351615-INS-195 | chr7-130351615-INS-195 |
| chr7-130354536-INS-357 | chr7-130354536-INS-362 |
| chr7-130372301-INS-201 | chr7-130372289-INS-201 |
| chr7-130409042-INS-399 | chr7-130409042-INS-399 |
| chr7-130423479-INS-216 | chr7-130423478-INS-213 |
| chr7-130435296-INS-175 | chr7-130435296-INS-175 |
| chr7-130451816-INS-83 | chr7-130451816-INS-83 |
| chr7-130460418-INS-169 | chr7-130460418-INS-184 |
| chr7-130475691-INS-70 | chr7-130475691-INS-70 |
| chr7-130477262-INS-81 | chr7-130477263-INS-81 |
| chr7-130507220-INS-293 | chr7-130507220-INS-293 |
| chr7-130528880-INS-213 | chr7-130528880-INS-213 |
| chr7-130535959-INS-203 | chr7-130535959-INS-206 |
| chr7-130558496-INS-168 | chr7-130558497-INS-168 |
| chr7-130561439-INS-81 | chr7-130561439-INS-81 |
| chr7-130599596-INS-62 | chr7-130599596-INS-62 |
| chr7-130621904-INS-214 | chr7-130621904-INS-213 |
| chr7-130636158-INS-399 | chr7-130636158-INS-399 |
| chr7-130640693-INS-239 | chr7-130640693-INS-241 |
| chr7-130657231-INS-52 | chr7-130657211-INS-99 |
| chr7-130746531-INS-82 | chr7-130746537-INS-55 |
| chr7-130764814-INS-116 | chr7-130764810-INS-119 |
| chr7-130782853-INS-68 | chr7-130782853-INS-68 |
| chr7-130784694-INS-55 | chr7-130784694-INS-55 |
| chr7-130806256-INS-56 | chr7-130806256-INS-50 |
| chr7-130813055-INS-74 | chr7-130813056-INS-74 |
| chr7-130829641-INS-853 | chr7-130829641-INS-854 |
| chr7-130834889-INS-65 | chr7-130834904-INS-60 |
| chr7-130841589-INS-51 | chr7-130841589-INS-51 |
| chr7-130858381-INS-102 | chr7-130858381-INS-102 |
| chr7-130861828-INS-230 | chr7-130861828-INS-230 |
| chr7-130878969-INS-755 | chr7-130878969-INS-755 |
| chr7-130920156-INS-89 | chr7-130920156-INS-89 |
| chr7-130940306-INS-71 | chr7-130940306-INS-71 |
| chr7-130954460-INS-112 | chr7-130954460-INS-112 |
| chr7-130956912-INS-64 | chr7-130956912-INS-80 |
| chr7-130971587-INS-146 | chr7-130971585-INS-150 |
| chr7-131016191-INS-3154 | chr7-131016192-INS-3154 |
| chr7-131059052-INS-81 | chr7-131059052-INS-97 |
| chr7-131117933-INS-971 | chr7-131117933-INS-981 |
| chr7-131150841-INS-82 | chr7-131150841-INS-83 |
| chr7-131178282-INS-108 | chr7-131178282-INS-108 |
| chr7-131183186-INS-97 | chr7-131183186-INS-104 |
| chr7-131191047-INS-80 | chr7-131191047-INS-80 |
| chr7-131197836-INS-470 | chr7-131197836-INS-469 |
| chr7-131213512-INS-81 | chr7-131213512-INS-81 |
| chr7-131215313-INS-100 | chr7-131215313-INS-100 |
| chr7-131217748-INS-133 | chr7-131217752-INS-129 |
| chr7-131250654-INS-76 | chr7-131250654-INS-70 |
| chr7-131275857-INS-216 | chr7-131275865-INS-247 |
| chr7-131315984-INS-87 | chr7-131315985-INS-60 |
| chr7-131366173-INS-72 | chr7-131366173-INS-72 |
| chr7-131370163-INS-53 | chr7-131370163-INS-53 |
| chr7-131379223-INS-77 | chr7-131379223-INS-77 |
| chr7-131381694-INS-60 | chr7-131381694-INS-60 |
| chr7-131408155-INS-60 | chr7-131408159-INS-62 |
| chr7-131443965-INS-1054 | chr7-131443966-INS-1054 |
| chr7-131475037-INS-82 | chr7-131475039-INS-123 |
| chr7-131503514-INS-63 | chr7-131503514-INS-63 |
| chr7-131506233-INS-116 | chr7-131506233-INS-152 |
| chr7-131527997-INS-127 | chr7-131527997-INS-129 |
| chr7-131637194-INS-56 | chr7-131637188-INS-56 |
| chr7-131651623-INS-220 | chr7-131651623-INS-220 |
| chr7-131679031-INS-529 | chr7-131679029-INS-526 |
| chr7-131722144-INS-94 | chr7-131722144-INS-94 |
| chr7-131763737-INS-101 | chr7-131763737-INS-104 |
| chr7-131765869-INS-1704 | chr7-131765870-INS-1704 |
| chr7-131804665-INS-434 | chr7-131804665-INS-434 |
| chr7-131805613-INS-96 | chr7-131805613-INS-56 |
| chr7-131814840-INS-52 | chr7-131814840-INS-52 |
| chr7-131825181-INS-53 | chr7-131825169-INS-54 |
| chr7-131836813-INS-190 | chr7-131836814-INS-230 |
| chr7-131841699-INS-114 | chr7-131841656-INS-108 |
| chr7-131856000-INS-140 | chr7-131856000-INS-140 |

| | |
|---|---|
| chr7-131930939-INS-95 | chr7-131930939-INS-95 |
| chr7-131940513-INS-58 | chr7-131940513-INS-62 |
| chr7-131962096-INS-56 | chr7-131962098-INS-80 |
| chr7-132007453-INS-155 | chr7-132007453-INS-145 |
| chr7-132008816-INS-158 | chr7-132008816-INS-158 |
| chr7-132029562-INS-82 | chr7-132029562-INS-113 |
| chr7-132054779-INS-62 | chr7-132054779-INS-62 |
| chr7-132095032-INS-94 | chr7-132095032-INS-59 |
| chr7-132144622-INS-970 | chr7-132144622-INS-970 |
| chr7-132208554-INS-132 | chr7-132208554-INS-132 |
| chr7-132231042-INS-180 | chr7-132231042-INS-194 |
| chr7-132233789-INS-72 | chr7-132233783-INS-56 |
| chr7-132236983-INS-418 | chr7-132236983-INS-410 |
| chr7-132241543-INS-52 | chr7-132241543-INS-52 |
| chr7-132251494-INS-58 | chr7-132251494-INS-74 |
| chr7-132262995-INS-537 | chr7-132262995-INS-537 |
| chr7-132282635-INS-190 | chr7-132282635-INS-163 |
| chr7-132287274-INS-7509 | chr7-132287276-INS-7505 |
| chr7-132299825-INS-249 | chr7-132299825-INS-249 |
| chr7-132326055-INS-61 | chr7-132326055-INS-61 |
| chr7-132419669-INS-186 | chr7-132419669-INS-183 |
| chr7-132430162-INS-203 | chr7-132430162-INS-205 |
| chr7-132473846-INS-252 | chr7-132473846-INS-259 |
| chr7-132487053-INS-64 | chr7-132487053-INS-64 |
| chr7-132509179-INS-186 | chr7-132509179-INS-178 |
| chr7-132512504-INS-203 | chr7-132512504-INS-211 |
| chr7-132520310-INS-220 | chr7-132520311-INS-220 |
| chr7-132526898-INS-162 | chr7-132526898-INS-149 |
| chr7-132534738-INS-214 | chr7-132534738-INS-214 |
| chr7-132609818-INS-77 | chr7-132609818-INS-77 |
| chr7-132641136-INS-60 | chr7-132641136-INS-60 |
| chr7-132645446-INS-7227 | chr7-132645447-INS-7227 |
| chr7-132669436-INS-60 | chr7-132669459-INS-54 |
| chr7-132737974-INS-66 | chr7-132737974-INS-50 |
| chr7-132743295-INS-395 | chr7-132743295-INS-395 |
| chr7-132750701-INS-204 | chr7-132750673-INS-212 |
| chr7-132772151-INS-709 | chr7-132772151-INS-709 |
| chr7-132829607-INS-5706 | chr7-132829608-INS-5706 |
| chr7-132887607-INS-108 | chr7-132887599-INS-112 |
| chr7-132896786-INS-52 | chr7-132896786-INS-52 |
| chr7-132902555-INS-60 | chr7-132902555-INS-60 |
| chr7-132915734-INS-115 | chr7-132915734-INS-121 |
| chr7-132918648-INS-87 | chr7-132918649-INS-81 |
| chr7-132942398-INS-201 | chr7-132942392-INS-163 |
| chr7-132944044-INS-58 | chr7-132944044-INS-58 |
| chr7-132969027-INS-66 | chr7-132969027-INS-68 |
| chr7-132972405-INS-530 | chr7-132972405-INS-532 |
| chr7-132982562-INS-72 | chr7-132982562-INS-72 |
| chr7-133026747-INS-1562 | chr7-133026748-INS-1563 |
| chr7-133034023-INS-240 | chr7-133034023-INS-243 |
| chr7-133039115-INS-56 | chr7-133039115-INS-54 |
| chr7-133041352-INS-83 | chr7-133041352-INS-83 |
| chr7-133043562-INS-437 | chr7-133043562-INS-438 |
| chr7-133055126-INS-55 | chr7-133055126-INS-50 |
| chr7-133063441-INS-140 | chr7-133063453-INS-118 |
| chr7-133074005-INS-218 | chr7-133074005-INS-218 |
| chr7-133077907-INS-71 | chr7-133077907-INS-71 |
| chr7-133092654-INS-195 | chr7-133092656-INS-178 |
| chr7-133098917-INS-107 | chr7-133098911-INS-113 |
| chr7-133111978-INS-69 | chr7-133111967-INS-98 |
| chr7-133140309-INS-523 | chr7-133140309-INS-523 |
| chr7-133149056-INS-61 | chr7-133149056-INS-65 |
| chr7-133168725-INS-416 | chr7-133168725-INS-417 |
| chr7-133183426-INS-191 | chr7-133183426-INS-191 |
| chr7-133199932-INS-1895 | chr7-133199933-INS-1895 |
| chr7-133223719-INS-66 | chr7-133223752-INS-55 |
| chr7-133230409-INS-85 | chr7-133230393-INS-54 |
| chr7-133238315-INS-201 | chr7-133238315-INS-201 |
| chr7-133281586-INS-121 | chr7-133281578-INS-136 |
| chr7-133291977-INS-65 | chr7-133291977-INS-65 |
| chr7-133314921-INS-54 | chr7-133314921-INS-56 |
| chr7-133329975-INS-192 | chr7-133329975-INS-192 |
| chr7-133384222-INS-346 | chr7-133384227-INS-356 |
| chr7-133392375-INS-431 | chr7-133392375-INS-431 |
| chr7-133404863-INS-210 | chr7-133404863-INS-211 |
| chr7-133429940-INS-72 | chr7-133429940-INS-67 |
| chr7-133436279-INS-329 | chr7-133436279-INS-329 |

| | |
|---|---|
| chr7-133446047-INS-7169 | chr7-133446048-INS-7169 |
| chr7-133447439-INS-3880 | chr7-133447440-INS-3858 |
| chr7-133464606-INS-424 | chr7-133464606-INS-424 |
| chr7-133466340-INS-1900 | chr7-133466341-INS-1901 |
| chr7-133481283-INS-740 | chr7-133481283-INS-738 |
| chr7-133493085-INS-97 | chr7-133493085-INS-97 |
| chr7-133494814-INS-455 | chr7-133494814-INS-455 |
| chr7-133578942-INS-62 | chr7-133578942-INS-62 |
| chr7-133593558-INS-133 | chr7-133593568-INS-136 |
| chr7-133625854-INS-84 | chr7-133625854-INS-84 |
| chr7-133630763-INS-171 | chr7-133630716-INS-176 |
| chr7-133655083-INS-55 | chr7-133655083-INS-55 |
| chr7-133677483-INS-137 | chr7-133677483-INS-124 |
| chr7-133682860-INS-60 | chr7-133682860-INS-50 |
| chr7-133696695-INS-70 | chr7-133696695-INS-66 |
| chr7-133759332-INS-88 | chr7-133759329-INS-106 |
| chr7-133765862-INS-90 | chr7-133765862-INS-90 |
| chr7-133797029-INS-498 | chr7-133797029-INS-498 |
| chr7-133854244-INS-75 | chr7-133854244-INS-73 |
| chr7-133885524-INS-52 | chr7-133885524-INS-52 |
| chr7-133905964-INS-698 | chr7-133905959-INS-666 |
| chr7-133913294-INS-128 | chr7-133913294-INS-147 |
| chr7-133925486-INS-7168 | chr7-133925492-INS-7166 |
| chr7-133927896-INS-78 | chr7-133927896-INS-66 |
| chr7-133951646-INS-211 | chr7-133951646-INS-212 |
| chr7-133998033-INS-2871 | chr7-133998034-INS-2871 |
| chr7-133998034-INS-392 | chr7-133998034-INS-392 |
| chr7-134052418-INS-1607 | chr7-134052419-INS-1607 |
| chr7-134056291-INS-5170 | chr7-134056292-INS-5171 |
| chr7-134093271-INS-88 | chr7-134093271-INS-76 |
| chr7-134095794-INS-59 | chr7-134095793-INS-107 |
| chr7-134116813-INS-698 | chr7-134116813-INS-698 |
| chr7-134140276-INS-60 | chr7-134140276-INS-58 |
| chr7-134156619-INS-404 | chr7-134156619-INS-406 |
| chr7-134160357-INS-181 | chr7-134160357-INS-181 |
| chr7-134171046-INS-474 | chr7-134171047-INS-482 |
| chr7-134177195-INS-64 | chr7-134177195-INS-58 |
| chr7-134181273-INS-663 | chr7-134181273-INS-663 |
| chr7-134191502-INS-161 | chr7-134191497-INS-132 |
| chr7-134193137-INS-156 | chr7-134193137-INS-120 |
| chr7-134195180-INS-6334 | chr7-134195181-INS-6333 |
| chr7-134206835-INS-337 | chr7-134206836-INS-338 |
| chr7-134262829-INS-94 | chr7-134262829-INS-94 |
| chr7-134283054-INS-391 | chr7-134283055-INS-392 |
| chr7-134328156-INS-77 | chr7-134328144-INS-84 |
| chr7-134388340-INS-90 | chr7-134388340-INS-90 |
| chr7-134396495-INS-62 | chr7-134396482-INS-52 |
| chr7-134429786-INS-53 | chr7-134429798-INS-54 |
| chr7-134434725-INS-149 | chr7-134434737-INS-128 |
| chr7-134440835-INS-52 | chr7-134440835-INS-54 |
| chr7-134490512-INS-66 | chr7-134490512-INS-66 |
| chr7-134541282-INS-186 | chr7-134541282-INS-186 |
| chr7-134554068-INS-58 | chr7-134554068-INS-56 |
| chr7-134555911-INS-224 | chr7-134555900-INS-256 |
| chr7-134573005-INS-83 | chr7-134572993-INS-119 |
| chr7-134609888-INS-165 | chr7-134609889-INS-167 |
| chr7-134615616-INS-344 | chr7-134615616-INS-344 |
| chr7-134632406-INS-121 | chr7-134632406-INS-121 |
| chr7-134637409-INS-80 | chr7-134637409-INS-74 |
| chr7-134643542-INS-170 | chr7-134643546-INS-170 |
| chr7-134658943-INS-1517 | chr7-134658944-INS-1516 |
| chr7-134693489-INS-1065 | chr7-134693490-INS-1066 |
| chr7-134771947-INS-456 | chr7-134771947-INS-455 |
| chr7-134787236-INS-66 | chr7-134787212-INS-72 |
| chr7-134790843-INS-859 | chr7-134790840-INS-863 |
| chr7-134801960-INS-78 | chr7-134801960-INS-78 |
| chr7-134807757-INS-104 | chr7-134807757-INS-74 |
| chr7-134809174-INS-6655 | chr7-134809175-INS-6655 |
| chr7-134850769-INS-63 | chr7-134850769-INS-57 |
| chr7-134898931-INS-446 | chr7-134898931-INS-448 |
| chr7-134906762-INS-187 | chr7-134906762-INS-187 |
| chr7-134911084-INS-103 | chr7-134911084-INS-103 |
| chr7-134949104-INS-2397 | chr7-134949105-INS-2396 |
| chr7-134949637-INS-65 | chr7-134949637-INS-65 |
| chr7-134953527-INS-484 | chr7-134953527-INS-484 |
| chr7-134961215-INS-62 | chr7-134961214-INS-108 |
| chr7-134976327-INS-131 | chr7-134976368-INS-110 |

| | |
|---|---|
| chr7-134987122-INS-521 | chr7-134987122-INS-529 |
| chr7-135027840-INS-2807 | chr7-135027841-INS-2807 |
| chr7-135038136-INS-133 | chr7-135038136-INS-133 |
| chr7-135082496-INS-87 | chr7-135082496-INS-94 |
| chr7-135085115-INS-94 | chr7-135085115-INS-77 |
| chr7-135118010-INS-310 | chr7-135118010-INS-306 |
| chr7-135157752-INS-607 | chr7-135157752-INS-607 |
| chr7-135160675-INS-186 | chr7-135160675-INS-192 |
| chr7-135197299-INS-264 | chr7-135197299-INS-265 |
| chr7-135217334-INS-84 | chr7-135217370-INS-63 |
| chr7-135222705-INS-58 | chr7-135222674-INS-57 |
| chr7-135240164-INS-57 | chr7-135240164-INS-57 |
| chr7-135241527-INS-208 | chr7-135241527-INS-208 |
| chr7-135281166-INS-58 | chr7-135281166-INS-50 |
| chr7-135286041-INS-118 | chr7-135286041-INS-108 |
| chr7-135325658-INS-440 | chr7-135325658-INS-440 |
| chr7-135326626-INS-439 | chr7-135326626-INS-452 |
| chr7-135356014-INS-55 | chr7-135356014-INS-79 |
| chr7-135362419-INS-57 | chr7-135362419-INS-57 |
| chr7-135374442-INS-332 | chr7-135374442-INS-334 |
| chr7-135425251-INS-59 | chr7-135425251-INS-58 |
| chr7-135430178-INS-1517 | chr7-135430179-INS-1517 |
| chr7-135474991-INS-125 | chr7-135474976-INS-114 |
| chr7-135517938-INS-553 | chr7-135517938-INS-553 |
| chr7-135576072-INS-58 | chr7-135576072-INS-51 |
| chr7-135577746-INS-80 | chr7-135577754-INS-69 |
| chr7-135586252-INS-92 | chr7-135586253-INS-67 |
| chr7-135614055-INS-2233 | chr7-135614056-INS-2234 |
| chr7-135649976-INS-125 | chr7-135649976-INS-126 |
| chr7-135681602-INS-441 | chr7-135681602-INS-441 |
| chr7-135693065-INS-81 | chr7-135693065-INS-84 |
| chr7-135716354-INS-203 | chr7-135716354-INS-206 |
| chr7-135767339-INS-2647 | chr7-135767340-INS-2647 |
| chr7-135774333-INS-135 | chr7-135774333-INS-170 |
| chr7-135792512-INS-84 | chr7-135792512-INS-84 |
| chr7-135803201-INS-111 | chr7-135803201-INS-95 |
| chr7-135860006-INS-92 | chr7-135860006-INS-76 |
| chr7-135863023-INS-402 | chr7-135863023-INS-402 |
| chr7-135897623-INS-103 | chr7-135897623-INS-103 |
| chr7-135919455-INS-6487 | chr7-135919456-INS-6487 |
| chr7-135926630-INS-2070 | chr7-135926631-INS-2067 |
| chr7-135966204-INS-72 | chr7-135966206-INS-76 |
| chr7-135984014-INS-78 | chr7-135984013-INS-86 |
| chr7-135985777-INS-169 | chr7-135985777-INS-170 |
| chr7-136001831-INS-855 | chr7-136001831-INS-863 |
| chr7-136037457-INS-60 | chr7-136037457-INS-60 |
| chr7-136049036-INS-56 | chr7-136049036-INS-56 |
| chr7-136143230-INS-593 | chr7-136143230-INS-593 |
| chr7-136152500-INS-381 | chr7-136152500-INS-382 |
| chr7-136177927-INS-72 | chr7-136177927-INS-72 |
| chr7-136206620-INS-80 | chr7-136206620-INS-80 |
| chr7-136225501-INS-388 | chr7-136225501-INS-394 |
| chr7-136262405-INS-509 | chr7-136262405-INS-509 |
| chr7-136266215-INS-54 | chr7-136266215-INS-54 |
| chr7-136325322-INS-193 | chr7-136325322-INS-193 |
| chr7-136345233-INS-815 | chr7-136345233-INS-834 |
| chr7-136368398-INS-175 | chr7-136368398-INS-171 |
| chr7-136398327-INS-59 | chr7-136398327-INS-59 |
| chr7-136401387-INS-94 | chr7-136401387-INS-94 |
| chr7-136415106-INS-50 | chr7-136415106-INS-50 |
| chr7-136421201-INS-901 | chr7-136421203-INS-899 |
| chr7-136440548-INS-60 | chr7-136440538-INS-58 |
| chr7-136489301-INS-56 | chr7-136489301-INS-56 |
| chr7-136517898-INS-344 | chr7-136517898-INS-346 |
| chr7-136529499-INS-73 | chr7-136529499-INS-71 |
| chr7-136564457-INS-289 | chr7-136564457-INS-289 |
| chr7-136616942-INS-121 | chr7-136616942-INS-121 |
| chr7-136619223-INS-76 | chr7-136619223-INS-57 |
| chr7-136643692-INS-65 | chr7-136643692-INS-65 |
| chr7-136660281-INS-171 | chr7-136660281-INS-170 |
| chr7-136661098-INS-68 | chr7-136661073-INS-88 |
| chr7-136669942-INS-455 | chr7-136669942-INS-455 |
| chr7-136720309-INS-2885 | chr7-136720310-INS-2885 |
| chr7-136743651-INS-85 | chr7-136743649-INS-77 |
| chr7-136757401-INS-132 | chr7-136757401-INS-136 |
| chr7-136815019-INS-347 | chr7-136815019-INS-348 |
| chr7-136825155-INS-350 | chr7-136825155-INS-350 |

| | |
|---|---|
| chr7-136830212-INS-97 | chr7-136830212-INS-97 |
| chr7-136869768-INS-69 | chr7-136869768-INS-69 |
| chr7-136882563-INS-395 | chr7-136882563-INS-395 |
| chr7-136927487-INS-120 | chr7-136927480-INS-80 |
| chr7-136939004-INS-96 | chr7-136938980-INS-80 |
| chr7-136940676-INS-88 | chr7-136940676-INS-68 |
| chr7-136947344-INS-188 | chr7-136947344-INS-194 |
| chr7-136949096-INS-198 | chr7-136949096-INS-197 |
| chr7-136952576-INS-62 | chr7-136952576-INS-66 |
| chr7-136964359-INS-244 | chr7-136964359-INS-240 |
| chr7-136965496-INS-59 | chr7-136965496-INS-59 |
| chr7-136973758-INS-100 | chr7-136973758-INS-100 |
| chr7-136988930-INS-166 | chr7-136988930-INS-166 |
| chr7-136992332-INS-202 | chr7-136992332-INS-198 |
| chr7-137000440-INS-53 | chr7-137000440-INS-103 |
| chr7-137002381-INS-299 | chr7-137002381-INS-299 |
| chr7-137007825-INS-201 | chr7-137007825-INS-200 |
| chr7-137028451-INS-72 | chr7-137028436-INS-81 |
| chr7-137046868-INS-216 | chr7-137046868-INS-216 |
| chr7-137065013-INS-222 | chr7-137065013-INS-222 |
| chr7-137071596-INS-201 | chr7-137071555-INS-201 |
| chr7-137073575-INS-176 | chr7-137073575-INS-176 |
| chr7-137076983-INS-89 | chr7-137076984-INS-88 |
| chr7-137082414-INS-51 | chr7-137082404-INS-52 |
| chr7-137084379-INS-185 | chr7-137084379-INS-185 |
| chr7-137096149-INS-71 | chr7-137096153-INS-60 |
| chr7-137102494-INS-571 | chr7-137102494-INS-571 |
| chr7-137107701-INS-515 | chr7-137107701-INS-515 |
| chr7-137110377-INS-96 | chr7-137110377-INS-96 |
| chr7-137112713-INS-544 | chr7-137112713-INS-548 |
| chr7-137117714-INS-64 | chr7-137117714-INS-64 |
| chr7-137118441-INS-68 | chr7-137118395-INS-68 |
| chr7-137121862-INS-53 | chr7-137121861-INS-64 |
| chr7-137129054-INS-63 | chr7-137129054-INS-63 |
| chr7-137142519-INS-611 | chr7-137142519-INS-612 |
| chr7-137155189-INS-62 | chr7-137155189-INS-66 |
| chr7-137157175-INS-137 | chr7-137157175-INS-139 |
| chr7-137162963-INS-312 | chr7-137162974-INS-313 |
| chr7-137174969-INS-58 | chr7-137174969-INS-52 |
| chr7-137181433-INS-70 | chr7-137181433-INS-70 |
| chr7-137182901-INS-54 | chr7-137182901-INS-54 |
| chr7-137240330-INS-95 | chr7-137240330-INS-95 |
| chr7-137267579-INS-204 | chr7-137267579-INS-204 |
| chr7-137293036-INS-136 | chr7-137293045-INS-172 |
| chr7-137309975-INS-107 | chr7-137309975-INS-107 |
| chr7-137312494-INS-123 | chr7-137312494-INS-136 |
| chr7-137338559-INS-56 | chr7-137338559-INS-56 |
| chr7-137345674-INS-1892 | chr7-137345675-INS-1892 |
| chr7-137376550-INS-55 | chr7-137376550-INS-55 |
| chr7-137382072-INS-88 | chr7-137382072-INS-71 |
| chr7-137393002-INS-95 | chr7-137393002-INS-95 |
| chr7-137401050-INS-86 | chr7-137401061-INS-81 |
| chr7-137455444-INS-760 | chr7-137455444-INS-761 |
| chr7-137506226-INS-62 | chr7-137506226-INS-64 |
| chr7-137520904-INS-2453 | chr7-137520905-INS-2439 |
| chr7-137535486-INS-62 | chr7-137535486-INS-62 |
| chr7-137573732-INS-88 | chr7-137573732-INS-88 |
| chr7-137574493-INS-205 | chr7-137574493-INS-205 |
| chr7-137582615-INS-78 | chr7-137582615-INS-78 |
| chr7-137586165-INS-74 | chr7-137586158-INS-58 |
| chr7-137592560-INS-72 | chr7-137592560-INS-76 |
| chr7-137625971-INS-80 | chr7-137625971-INS-80 |
| chr7-137626831-INS-54 | chr7-137626831-INS-56 |
| chr7-137643474-INS-54 | chr7-137643474-INS-55 |
| chr7-137659421-INS-7793 | chr7-137659422-INS-7794 |
| chr7-137764355-INS-433 | chr7-137764357-INS-448 |
| chr7-137792620-INS-114 | chr7-137792620-INS-112 |
| chr7-137798364-INS-112 | chr7-137798364-INS-112 |
| chr7-137840572-INS-348 | chr7-137840572-INS-348 |
| chr7-137877580-INS-304 | chr7-137877580-INS-304 |
| chr7-137884344-INS-52 | chr7-137884344-INS-52 |
| chr7-137896891-INS-74 | chr7-137896896-INS-94 |
| chr7-137900878-INS-52 | chr7-137900877-INS-52 |
| chr7-137905081-INS-151 | chr7-137905089-INS-136 |
| chr7-137931243-INS-86 | chr7-137931209-INS-94 |
| chr7-137935612-INS-67 | chr7-137935612-INS-67 |
| chr7-137955751-INS-108 | chr7-137955751-INS-112 |

| | |
|---|---|
| chr7-137969387-INS-1035 | chr7-137969388-INS-1035 |
| chr7-138040313-INS-72 | chr7-138040313-INS-72 |
| chr7-138044826-INS-52 | chr7-138044826-INS-54 |
| chr7-138048768-INS-86 | chr7-138048787-INS-90 |
| chr7-138079241-INS-216 | chr7-138079241-INS-217 |
| chr7-138083103-INS-56 | chr7-138083089-INS-57 |
| chr7-138109240-INS-262 | chr7-138109240-INS-265 |
| chr7-138112318-INS-1705 | chr7-138112319-INS-1706 |
| chr7-138125000-INS-205 | chr7-138125000-INS-201 |
| chr7-138165898-INS-154 | chr7-138165899-INS-154 |
| chr7-138194937-INS-93 | chr7-138194937-INS-93 |
| chr7-138197335-INS-76 | chr7-138197335-INS-76 |
| chr7-138224257-INS-3333 | chr7-138224258-INS-3333 |
| chr7-138231835-INS-247 | chr7-138231835-INS-247 |
| chr7-138238281-INS-151 | chr7-138238281-INS-151 |
| chr7-138246183-INS-330 | chr7-138246183-INS-331 |
| chr7-138253681-INS-201 | chr7-138253681-INS-201 |
| chr7-138272606-INS-51 | chr7-138272588-INS-55 |
| chr7-138280285-INS-68 | chr7-138280286-INS-66 |
| chr7-138301738-INS-92 | chr7-138301693-INS-88 |
| chr7-138307734-INS-150 | chr7-138307734-INS-108 |
| chr7-138314339-INS-147 | chr7-138314339-INS-147 |
| chr7-138340260-INS-54 | chr7-138340236-INS-68 |
| chr7-138345142-INS-103 | chr7-138345142-INS-103 |
| chr7-138379787-INS-76 | chr7-138379790-INS-78 |
| chr7-138384554-INS-309 | chr7-138384554-INS-309 |
| chr7-138396261-INS-74 | chr7-138396261-INS-80 |
| chr7-138421204-INS-60 | chr7-138421204-INS-90 |
| chr7-138445402-INS-51 | chr7-138445402-INS-51 |
| chr7-138449462-INS-64 | chr7-138449465-INS-65 |
| chr7-138458087-INS-85 | chr7-138458074-INS-85 |
| chr7-138463502-INS-66 | chr7-138463490-INS-52 |
| chr7-138465075-INS-83 | chr7-138465075-INS-83 |
| chr7-138471656-INS-130 | chr7-138471656-INS-130 |
| chr7-138495888-INS-53 | chr7-138495888-INS-53 |
| chr7-138522440-INS-60 | chr7-138522436-INS-92 |
| chr7-138524112-INS-90 | chr7-138524112-INS-105 |
| chr7-138532029-INS-54 | chr7-138532037-INS-62 |
| chr7-138549261-INS-93 | chr7-138549261-INS-93 |
| chr7-138562212-INS-207 | chr7-138562212-INS-207 |
| chr7-138584609-INS-60 | chr7-138584609-INS-60 |
| chr7-138613231-INS-360 | chr7-138613231-INS-359 |
| chr7-138622717-INS-95 | chr7-138622717-INS-95 |
| chr7-138640356-INS-144 | chr7-138640353-INS-192 |
| chr7-138646534-INS-88 | chr7-138646529-INS-65 |
| chr7-138677151-INS-75 | chr7-138677141-INS-67 |
| chr7-138679348-INS-390 | chr7-138679348-INS-390 |
| chr7-138719510-INS-399 | chr7-138719510-INS-399 |
| chr7-138738213-INS-146 | chr7-138738213-INS-144 |
| chr7-138742592-INS-767 | chr7-138742604-INS-766 |
| chr7-138747892-INS-119 | chr7-138747892-INS-111 |
| chr7-138760600-INS-143 | chr7-138760600-INS-143 |
| chr7-138775301-INS-108 | chr7-138775301-INS-108 |
| chr7-138777658-INS-288 | chr7-138777658-INS-288 |
| chr7-138800861-INS-213 | chr7-138800861-INS-213 |
| chr7-138811610-INS-55 | chr7-138811610-INS-81 |
| chr7-138814120-INS-134 | chr7-138814122-INS-140 |
| chr7-138815331-INS-80 | chr7-138815331-INS-84 |
| chr7-138821518-INS-282 | chr7-138821514-INS-288 |
| chr7-138858562-INS-56 | chr7-138858566-INS-52 |
| chr7-138870523-INS-102 | chr7-138870523-INS-102 |
| chr7-138876527-INS-71 | chr7-138876527-INS-70 |
| chr7-138886498-INS-92 | chr7-138886498-INS-104 |
| chr7-138905835-INS-68 | chr7-138905835-INS-68 |
| chr7-138922371-INS-58 | chr7-138922371-INS-58 |
| chr7-138951043-INS-84 | chr7-138951043-INS-84 |
| chr7-138961866-INS-86 | chr7-138961866-INS-86 |
| chr7-138976916-INS-68 | chr7-138976896-INS-68 |
| chr7-138979373-INS-309 | chr7-138979373-INS-309 |
| chr7-138988471-INS-61 | chr7-138988469-INS-85 |
| chr7-139009345-INS-67 | chr7-139009345-INS-107 |
| chr7-139011430-INS-2760 | chr7-139011431-INS-2761 |
| chr7-139015659-INS-189 | chr7-139015659-INS-189 |
| chr7-139019408-INS-56 | chr7-139019410-INS-70 |
| chr7-139049521-INS-268 | chr7-139049521-INS-268 |
| chr7-139082660-INS-58 | chr7-139082660-INS-58 |
| chr7-139090866-INS-126 | chr7-139090860-INS-126 |

| | |
|---|---|
| chr7-139098164-INS-54 | chr7-139098164-INS-53 |
| chr7-139103466-INS-6405 | chr7-139103467-INS-6405 |
| chr7-139109491-INS-74 | chr7-139109491-INS-50 |
| chr7-139119370-INS-68 | chr7-139119370-INS-68 |
| chr7-139150341-INS-242 | chr7-139150342-INS-230 |
| chr7-139165715-INS-816 | chr7-139165715-INS-816 |
| chr7-139167307-INS-170 | chr7-139167307-INS-170 |
| chr7-139182802-INS-170 | chr7-139182804-INS-174 |
| chr7-139204245-INS-497 | chr7-139204245-INS-497 |
| chr7-139247214-INS-220 | chr7-139247214-INS-221 |
| chr7-139251133-INS-210 | chr7-139251133-INS-210 |
| chr7-139252920-INS-272 | chr7-139252903-INS-302 |
| chr7-139256503-INS-52 | chr7-139256503-INS-52 |
| chr7-139280529-INS-506 | chr7-139280529-INS-506 |
| chr7-139293967-INS-273 | chr7-139293967-INS-273 |
| chr7-139295318-INS-58 | chr7-139295318-INS-58 |
| chr7-139319041-INS-53 | chr7-139319041-INS-53 |
| chr7-139321858-INS-137 | chr7-139321858-INS-139 |
| chr7-139366372-INS-95 | chr7-139366372-INS-95 |
| chr7-139379781-INS-5548 | chr7-139379782-INS-5548 |
| chr7-139400608-INS-54 | chr7-139400590-INS-60 |
| chr7-139423376-INS-57 | chr7-139423376-INS-57 |
| chr7-139431722-INS-52 | chr7-139431722-INS-52 |
| chr7-139438511-INS-60 | chr7-139438511-INS-60 |
| chr7-139448020-INS-6178 | chr7-139448021-INS-6144 |
| chr7-139471707-INS-126 | chr7-139471707-INS-126 |
| chr7-139523080-INS-198 | chr7-139523080-INS-198 |
| chr7-139580870-INS-190 | chr7-139580870-INS-190 |
| chr7-139612976-INS-193 | chr7-139612976-INS-197 |
| chr7-139621673-INS-410 | chr7-139621673-INS-410 |
| chr7-139678428-INS-221 | chr7-139678428-INS-220 |
| chr7-139681890-INS-106 | chr7-139681890-INS-66 |
| chr7-139710503-INS-58 | chr7-139710503-INS-59 |
| chr7-139718199-INS-7364 | chr7-139718200-INS-7364 |
| chr7-139739207-INS-346 | chr7-139739207-INS-346 |
| chr7-139746220-INS-97 | chr7-139746221-INS-82 |
| chr7-139746931-INS-980 | chr7-139746933-INS-986 |
| chr7-139755226-INS-114 | chr7-139755226-INS-114 |
| chr7-139773562-INS-1221 | chr7-139773563-INS-1221 |
| chr7-139775084-INS-197 | chr7-139775084-INS-197 |
| chr7-139778018-INS-216 | chr7-139778017-INS-200 |
| chr7-139787705-INS-492 | chr7-139787705-INS-492 |
| chr7-139789470-INS-352 | chr7-139789470-INS-352 |
| chr7-139794636-INS-74 | chr7-139794636-INS-74 |
| chr7-139797537-INS-511 | chr7-139797537-INS-512 |
| chr7-139834526-INS-1153 | chr7-139834527-INS-1153 |
| chr7-139835394-INS-844 | chr7-139835394-INS-844 |
| chr7-139842806-INS-1792 | chr7-139842807-INS-1792 |
| chr7-139843922-INS-90 | chr7-139843934-INS-118 |
| chr7-139862417-INS-137 | chr7-139862417-INS-177 |
| chr7-139887235-INS-201 | chr7-139887235-INS-201 |
| chr7-139891451-INS-66 | chr7-139891462-INS-56 |
| chr7-139910610-INS-724 | chr7-139910610-INS-724 |
| chr7-139938854-INS-165 | chr7-139938854-INS-166 |
| chr7-139946877-INS-179 | chr7-139946877-INS-212 |
| chr7-139952248-INS-195 | chr7-139952248-INS-196 |
| chr7-139968172-INS-6474 | chr7-139968173-INS-6478 |
| chr7-139989579-INS-4723 | chr7-139989580-INS-4723 |
| chr7-140022913-INS-1240 | chr7-140022914-INS-1242 |
| chr7-140022914-INS-507 | chr7-140022914-INS-507 |
| chr7-140039993-INS-401 | chr7-140039993-INS-401 |
| chr7-140041168-INS-80 | chr7-140041168-INS-76 |
| chr7-140053024-INS-160 | chr7-140053024-INS-160 |
| chr7-140057468-INS-72 | chr7-140057468-INS-72 |
| chr7-140103230-INS-6452 | chr7-140103231-INS-6452 |
| chr7-140133689-INS-204 | chr7-140133689-INS-204 |
| chr7-140148723-INS-1275 | chr7-140148724-INS-1275 |
| chr7-140153526-INS-2241 | chr7-140153527-INS-2250 |
| chr7-140157018-INS-848 | chr7-140157020-INS-846 |
| chr7-140178923-INS-6121 | chr7-140178924-INS-6121 |
| chr7-140205962-INS-7075 | chr7-140205963-INS-7075 |
| chr7-140213389-INS-368 | chr7-140213389-INS-369 |
| chr7-140234061-INS-204 | chr7-140234061-INS-204 |
| chr7-140270159-INS-279 | chr7-140270159-INS-279 |
| chr7-140291953-INS-873 | chr7-140291953-INS-873 |
| chr7-140312775-INS-344 | chr7-140312775-INS-344 |
| chr7-140315271-INS-115 | chr7-140315271-INS-116 |

| | |
|---|---|
| chr7-140326102-INS-93 | chr7-140326102-INS-93 |
| chr7-140329395-INS-65 | chr7-140329395-INS-65 |
| chr7-140335794-INS-270 | chr7-140335797-INS-269 |
| chr7-140340433-INS-95 | chr7-140340433-INS-50 |
| chr7-140346320-INS-152 | chr7-140346335-INS-160 |
| chr7-140368907-INS-130 | chr7-140368907-INS-130 |
| chr7-140406016-INS-60 | chr7-140406016-INS-62 |
| chr7-140406887-INS-2152 | chr7-140406888-INS-2153 |
| chr7-140450255-INS-221 | chr7-140450255-INS-220 |
| chr7-140483391-INS-54 | chr7-140483391-INS-54 |
| chr7-140486845-INS-196 | chr7-140486845-INS-215 |
| chr7-140507818-INS-56 | chr7-140507818-INS-56 |
| chr7-140510902-INS-223 | chr7-140510902-INS-223 |
| chr7-140513527-INS-226 | chr7-140513527-INS-226 |
| chr7-140516738-INS-266 | chr7-140516738-INS-266 |
| chr7-140561380-INS-77 | chr7-140561363-INS-96 |
| chr7-140585023-INS-232 | chr7-140585023-INS-232 |
| chr7-140599586-INS-334 | chr7-140599586-INS-334 |
| chr7-140604911-INS-219 | chr7-140604911-INS-220 |
| chr7-140607982-INS-187 | chr7-140607983-INS-186 |
| chr7-140621259-INS-362 | chr7-140621253-INS-367 |
| chr7-140649134-INS-117 | chr7-140649134-INS-117 |
| chr7-140681232-INS-124 | chr7-140681232-INS-124 |
| chr7-140692880-INS-174 | chr7-140692880-INS-174 |
| chr7-140694095-INS-183 | chr7-140694095-INS-184 |
| chr7-140696062-INS-145 | chr7-140696062-INS-145 |
| chr7-140708129-INS-94 | chr7-140708129-INS-100 |
| chr7-140714511-INS-74 | chr7-140714511-INS-62 |
| chr7-140741467-INS-54 | chr7-140741467-INS-54 |
| chr7-140756097-INS-69 | chr7-140756085-INS-72 |
| chr7-140759138-INS-204 | chr7-140759139-INS-210 |
| chr7-140765084-INS-397 | chr7-140765084-INS-397 |
| chr7-140766245-INS-259 | chr7-140766251-INS-300 |
| chr7-140816606-INS-109 | chr7-140816579-INS-115 |
| chr7-140824775-INS-630 | chr7-140824775-INS-630 |
| chr7-140846856-INS-215 | chr7-140846856-INS-211 |
| chr7-140854254-INS-63 | chr7-140854254-INS-63 |
| chr7-140865043-INS-100 | chr7-140865043-INS-100 |
| chr7-140867638-INS-219 | chr7-140867638-INS-220 |
| chr7-140884853-INS-124 | chr7-140884853-INS-124 |
| chr7-140892180-INS-70 | chr7-140892180-INS-58 |
| chr7-140914838-INS-70 | chr7-140914838-INS-70 |
| chr7-140948699-INS-63 | chr7-140948738-INS-63 |
| chr7-140952697-INS-220 | chr7-140952697-INS-220 |
| chr7-140955698-INS-220 | chr7-140955698-INS-220 |
| chr7-140966743-INS-181 | chr7-140966743-INS-181 |
| chr7-140976638-INS-205 | chr7-140976638-INS-205 |
| chr7-140987306-INS-238 | chr7-140987306-INS-238 |
| chr7-140991094-INS-246 | chr7-140991094-INS-247 |
| chr7-140992847-INS-214 | chr7-140992847-INS-214 |
| chr7-140997540-INS-192 | chr7-140997528-INS-196 |
| chr7-141033958-INS-168 | chr7-141033958-INS-168 |
| chr7-141079437-INS-57 | chr7-141079437-INS-52 |
| chr7-141083805-INS-200 | chr7-141083805-INS-200 |
| chr7-141086120-INS-414 | chr7-141086121-INS-398 |
| chr7-141088215-INS-79 | chr7-141088216-INS-59 |
| chr7-141093927-INS-193 | chr7-141093927-INS-193 |
| chr7-141105136-INS-227 | chr7-141105136-INS-227 |
| chr7-141121778-INS-206 | chr7-141121778-INS-206 |
| chr7-141123673-INS-174 | chr7-141123673-INS-174 |
| chr7-141124220-INS-73 | chr7-141124220-INS-73 |
| chr7-141127553-INS-64 | chr7-141127554-INS-60 |
| chr7-141171992-INS-140 | chr7-141171992-INS-140 |
| chr7-141181464-INS-62 | chr7-141181464-INS-62 |
| chr7-141194478-INS-207 | chr7-141194478-INS-207 |
| chr7-141196021-INS-64 | chr7-141196021-INS-64 |
| chr7-141197554-INS-51 | chr7-141197554-INS-51 |
| chr7-141206707-INS-64 | chr7-141206707-INS-68 |
| chr7-141249838-INS-193 | chr7-141249848-INS-197 |
| chr7-141255327-INS-486 | chr7-141255327-INS-486 |
| chr7-141264296-INS-60 | chr7-141264316-INS-56 |
| chr7-141301986-INS-54 | chr7-141301986-INS-55 |
| chr7-141306792-INS-573 | chr7-141306792-INS-573 |
| chr7-141313174-INS-119 | chr7-141313205-INS-118 |
| chr7-141314876-INS-54 | chr7-141314877-INS-54 |
| chr7-141319136-INS-68 | chr7-141319136-INS-68 |
| chr7-141326573-INS-307 | chr7-141326573-INS-308 |

| | |
|---|---|
| chr7-141363453-INS-192 | chr7-141363453-INS-192 |
| chr7-141440708-INS-701 | chr7-141440708-INS-701 |
| chr7-141524523-INS-51 | chr7-141524523-INS-56 |
| chr7-141607139-INS-55 | chr7-141607139-INS-55 |
| chr7-141631295-INS-74 | chr7-141631281-INS-93 |
| chr7-141681757-INS-60 | chr7-141681757-INS-71 |
| chr7-141684420-INS-127 | chr7-141684420-INS-105 |
| chr7-141691454-INS-363 | chr7-141691454-INS-362 |
| chr7-141691465-INS-906 | chr7-141691465-INS-906 |
| chr7-141697858-INS-54 | chr7-141697858-INS-54 |
| chr7-141712069-INS-396 | chr7-141712069-INS-402 |
| chr7-141752373-INS-174 | chr7-141752373-INS-174 |
| chr7-141776838-INS-53 | chr7-141776838-INS-53 |
| chr7-141785633-INS-8457 | chr7-141785634-INS-8457 |
| chr7-141789711-INS-192 | chr7-141789711-INS-192 |
| chr7-141808107-INS-54 | chr7-141808107-INS-54 |
| chr7-141811427-INS-60 | chr7-141811427-INS-60 |
| chr7-141827637-INS-114 | chr7-141827637-INS-84 |
| chr7-141831852-INS-460 | chr7-141831852-INS-460 |
| chr7-141834108-INS-180 | chr7-141834108-INS-180 |
| chr7-141847714-INS-124 | chr7-141847714-INS-130 |
| chr7-141858704-INS-499 | chr7-141858704-INS-499 |
| chr7-141860182-INS-399 | chr7-141860182-INS-399 |
| chr7-141863733-INS-765 | chr7-141863733-INS-763 |
| chr7-141869382-INS-169 | chr7-141869382-INS-161 |
| chr7-141878743-INS-226 | chr7-141878743-INS-226 |
| chr7-141893585-INS-507 | chr7-141893585-INS-507 |
| chr7-141899290-INS-76 | chr7-141899261-INS-58 |
| chr7-141915490-INS-60 | chr7-141915490-INS-80 |
| chr7-141918810-INS-53 | chr7-141918810-INS-56 |
| chr7-141958907-INS-209 | chr7-141958907-INS-209 |
| chr7-141961720-INS-208 | chr7-141961720-INS-208 |
| chr7-141964641-INS-108 | chr7-141964641-INS-108 |
| chr7-142008271-INS-63 | chr7-142008271-INS-63 |
| chr7-142066847-INS-3676 | chr7-142066848-INS-3676 |
| chr7-142137840-INS-61 | chr7-142137840-INS-61 |
| chr7-142240573-INS-400 | chr7-142240573-INS-400 |
| chr7-142248361-INS-96 | chr7-142248361-INS-97 |
| chr7-142252588-INS-56 | chr7-142252588-INS-56 |
| chr7-142255599-INS-492 | chr7-142255599-INS-492 |
| chr7-142265723-INS-98 | chr7-142265723-INS-98 |
| chr7-142269673-INS-71 | chr7-142269673-INS-71 |
| chr7-142271937-INS-81 | chr7-142271937-INS-82 |
| chr7-142273440-INS-234 | chr7-142273440-INS-220 |
| chr7-142294394-INS-940 | chr7-142294394-INS-940 |
| chr7-142295012-INS-208 | chr7-142295012-INS-208 |
| chr7-142301920-INS-1146 | chr7-142301921-INS-1152 |
| chr7-142310817-INS-66 | chr7-142310817-INS-62 |
| chr7-142313212-INS-266 | chr7-142313212-INS-266 |
| chr7-142353608-INS-54 | chr7-142353608-INS-54 |
| chr7-142357875-INS-66 | chr7-142357876-INS-86 |
| chr7-142367602-INS-179 | chr7-142367602-INS-179 |
| chr7-142398222-INS-78 | chr7-142398188-INS-78 |
| chr7-142415871-INS-70 | chr7-142415840-INS-55 |
| chr7-142420858-INS-1861 | chr7-142420859-INS-1870 |
| chr7-142439837-INS-57 | chr7-142439850-INS-86 |
| chr7-142491435-INS-67 | chr7-142491435-INS-85 |
| chr7-142534939-INS-162 | chr7-142534934-INS-179 |
| chr7-142548786-INS-110 | chr7-142548786-INS-110 |
| chr7-142611284-INS-136 | chr7-142611293-INS-136 |
| chr7-142614133-INS-439 | chr7-142614142-INS-433 |
| chr7-142714687-INS-60 | chr7-142714687-INS-60 |
| chr7-142725853-INS-51 | chr7-142725847-INS-54 |
| chr7-142728245-INS-227 | chr7-142728242-INS-230 |
| chr7-142729927-INS-62 | chr7-142729927-INS-62 |
| chr7-142779938-INS-63 | chr7-142779938-INS-61 |
| chr7-142795128-INS-6440 | chr7-142795129-INS-6440 |
| chr7-142811178-INS-155 | chr7-142811178-INS-158 |
| chr7-142830780-INS-51 | chr7-142830790-INS-74 |
| chr7-142920857-INS-178 | chr7-142920857-INS-228 |
| chr7-143002201-INS-73 | chr7-143002201-INS-73 |
| chr7-143004500-INS-81 | chr7-143004500-INS-89 |
| chr7-143041682-INS-51 | chr7-143041682-INS-51 |
| chr7-143042871-INS-54 | chr7-143042871-INS-54 |
| chr7-143062959-INS-229 | chr7-143062968-INS-237 |
| chr7-143154576-INS-58 | chr7-143154576-INS-62 |
| chr7-143170215-INS-81 | chr7-143170216-INS-54 |

| | |
|---|---|
| chr7-143202704-INS-103 | chr7-143202704-INS-103 |
| chr7-143253940-INS-60 | chr7-143253968-INS-72 |
| chr7-143271104-INS-52 | chr7-143271144-INS-52 |
| chr7-143324641-INS-121 | chr7-143324641-INS-113 |
| chr7-143358298-INS-52 | chr7-143358298-INS-52 |
| chr7-143455901-INS-6211 | chr7-143455902-INS-6212 |
| chr7-143462270-INS-98 | chr7-143462270-INS-98 |
| chr7-143474464-INS-53 | chr7-143474464-INS-56 |
| chr7-143477672-INS-341 | chr7-143477672-INS-341 |
| chr7-143500583-INS-150 | chr7-143500583-INS-138 |
| chr7-143506344-INS-50 | chr7-143506359-INS-53 |
| chr7-143509622-INS-60 | chr7-143509622-INS-63 |
| chr7-143554201-INS-510 | chr7-143554201-INS-479 |
| chr7-143558204-INS-75 | chr7-143558204-INS-76 |
| chr7-143576172-INS-76 | chr7-143576172-INS-76 |
| chr7-143588856-INS-66 | chr7-143588849-INS-62 |
| chr7-143633642-INS-212 | chr7-143633642-INS-220 |
| chr7-143635353-INS-135 | chr7-143635353-INS-135 |
| chr7-143644697-INS-68 | chr7-143644683-INS-51 |
| chr7-143647012-INS-223 | chr7-143647012-INS-223 |
| chr7-143719908-INS-598 | chr7-143719908-INS-596 |
| chr7-143765289-INS-196 | chr7-143765289-INS-196 |
| chr7-143766336-INS-52 | chr7-143766336-INS-56 |
| chr7-143770918-INS-367 | chr7-143770918-INS-367 |
| chr7-143834401-INS-98 | chr7-143834401-INS-98 |
| chr7-143854668-INS-69 | chr7-143854668-INS-69 |
| chr7-143918057-INS-198 | chr7-143918057-INS-201 |
| chr7-143919492-INS-132 | chr7-143919492-INS-132 |
| chr7-143921510-INS-399 | chr7-143921510-INS-399 |
| chr7-143925826-INS-740 | chr7-143925826-INS-740 |
| chr7-143953266-INS-1328 | chr7-143953267-INS-1328 |
| chr7-143968491-INS-69 | chr7-143968502-INS-107 |
| chr7-144000613-INS-53 | chr7-144000613-INS-63 |
| chr7-144018232-INS-110 | chr7-144018232-INS-110 |
| chr7-144077996-INS-119 | chr7-144078036-INS-115 |
| chr7-144121920-INS-54 | chr7-144121920-INS-56 |
| chr7-144146048-INS-64 | chr7-144146054-INS-54 |
| chr7-144148696-INS-59 | chr7-144148696-INS-61 |
| chr7-144152689-INS-64 | chr7-144152689-INS-76 |
| chr7-144170655-INS-63 | chr7-144170681-INS-65 |
| chr7-144178000-INS-230 | chr7-144178000-INS-236 |
| chr7-144246306-INS-74 | chr7-144246317-INS-61 |
| chr7-144315655-INS-132 | chr7-144315655-INS-116 |
| chr7-144326483-INS-209 | chr7-144326483-INS-209 |
| chr7-144330854-INS-67 | chr7-144330854-INS-72 |
| chr7-144340957-INS-84 | chr7-144340957-INS-83 |
| chr7-144342203-INS-82 | chr7-144342203-INS-76 |
| chr7-144352476-INS-109 | chr7-144352504-INS-73 |
| chr7-144354326-INS-205 | chr7-144354326-INS-201 |
| chr7-144371834-INS-64 | chr7-144371834-INS-80 |
| chr7-144412954-INS-52 | chr7-144412954-INS-57 |
| chr7-144506178-INS-214 | chr7-144506178-INS-215 |
| chr7-144536068-INS-74 | chr7-144536061-INS-52 |
| chr7-144554722-INS-1635 | chr7-144554727-INS-1623 |
| chr7-144557015-INS-1777 | chr7-144557007-INS-1773 |
| chr7-144558922-INS-143 | chr7-144558922-INS-141 |
| chr7-144575651-INS-52 | chr7-144575651-INS-52 |
| chr7-144576850-INS-1893 | chr7-144576851-INS-1893 |
| chr7-144625195-INS-144 | chr7-144625195-INS-144 |
| chr7-144632431-INS-212 | chr7-144632431-INS-212 |
| chr7-144638955-INS-448 | chr7-144638955-INS-448 |
| chr7-144670346-INS-122 | chr7-144670346-INS-122 |
| chr7-144694702-INS-236 | chr7-144694702-INS-237 |
| chr7-144696435-INS-466 | chr7-144696435-INS-466 |
| chr7-144747043-INS-51 | chr7-144747043-INS-56 |
| chr7-144766686-INS-54 | chr7-144766686-INS-51 |
| chr7-144776930-INS-4862 | chr7-144776931-INS-4870 |
| chr7-144827016-INS-147 | chr7-144827016-INS-147 |
| chr7-144843723-INS-96 | chr7-144843723-INS-94 |
| chr7-144851545-INS-200 | chr7-144851545-INS-202 |
| chr7-144860827-INS-51 | chr7-144860825-INS-57 |
| chr7-144863972-INS-470 | chr7-144863972-INS-470 |
| chr7-144870212-INS-327 | chr7-144870212-INS-327 |
| chr8-3188417-INS-6165 | chr8-3188418-INS-6165 |
| chr8-3190790-INS-3197 | chr8-3190791-INS-3197 |
| chr8-3226765-INS-120 | chr8-3226775-INS-120 |
| chr8-3232362-INS-54 | chr8-3232362-INS-56 |

| | |
|---|---|
| chr8-3269330-INS-51 | chr8-3269330-INS-59 |
| chr8-3272597-INS-70 | chr8-3272611-INS-70 |
| chr8-3278517-INS-54 | chr8-3278517-INS-54 |
| chr8-3288252-INS-1040 | chr8-3288253-INS-1039 |
| chr8-3302335-INS-765 | chr8-3302335-INS-765 |
| chr8-3313876-INS-51 | chr8-3313876-INS-51 |
| chr8-3317230-INS-212 | chr8-3317231-INS-212 |
| chr8-3346096-INS-220 | chr8-3346096-INS-220 |
| chr8-3356097-INS-55 | chr8-3356095-INS-84 |
| chr8-3358293-INS-4734 | chr8-3358294-INS-4734 |
| chr8-3379008-INS-206 | chr8-3379008-INS-206 |
| chr8-3380480-INS-455 | chr8-3380480-INS-455 |
| chr8-3383506-INS-157 | chr8-3383506-INS-157 |
| chr8-3408549-INS-57 | chr8-3408549-INS-57 |
| chr8-3424740-INS-76 | chr8-3424740-INS-76 |
| chr8-3433072-INS-60 | chr8-3433067-INS-90 |
| chr8-3463614-INS-208 | chr8-3463614-INS-208 |
| chr8-3474417-INS-217 | chr8-3474417-INS-217 |
| chr8-3474812-INS-6399 | chr8-3474813-INS-6399 |
| chr8-3495230-INS-120 | chr8-3495192-INS-128 |
| chr8-3497644-INS-115 | chr8-3497644-INS-115 |
| chr8-3514285-INS-60 | chr8-3514285-INS-60 |
| chr8-3548263-INS-96 | chr8-3548263-INS-96 |
| chr8-3553422-INS-257 | chr8-3553422-INS-257 |
| chr8-3554343-INS-62 | chr8-3554343-INS-66 |
| chr8-3600769-INS-175 | chr8-3600769-INS-175 |
| chr8-3604213-INS-59 | chr8-3604213-INS-59 |
| chr8-3608685-INS-197 | chr8-3608685-INS-197 |
| chr8-3612301-INS-232 | chr8-3612302-INS-232 |
| chr8-3613624-INS-60 | chr8-3613624-INS-68 |
| chr8-3618335-INS-108 | chr8-3618335-INS-108 |
| chr8-3619158-INS-90 | chr8-3619158-INS-90 |
| chr8-3627162-INS-210 | chr8-3627162-INS-210 |
| chr8-3776279-INS-120 | chr8-3776279-INS-108 |
| chr8-3791181-INS-3578 | chr8-3791182-INS-3577 |
| chr8-3804349-INS-140 | chr8-3804349-INS-144 |
| chr8-3888918-INS-447 | chr8-3888918-INS-452 |
| chr8-3903124-INS-115 | chr8-3903124-INS-115 |
| chr8-3905696-INS-56 | chr8-3905696-INS-58 |
| chr8-3929188-INS-1862 | chr8-3929189-INS-1862 |
| chr8-3932497-INS-344 | chr8-3932497-INS-344 |
| chr8-3939352-INS-245 | chr8-3939352-INS-245 |
| chr8-3941696-INS-96 | chr8-3941676-INS-97 |
| chr8-3942816-INS-68 | chr8-3942827-INS-72 |
| chr8-3982283-INS-6785 | chr8-3982284-INS-6785 |
| chr8-3990092-INS-82 | chr8-3990092-INS-106 |
| chr8-4019852-INS-569 | chr8-4019852-INS-569 |
| chr8-4024913-INS-6409 | chr8-4024914-INS-6409 |
| chr8-4033339-INS-322 | chr8-4033339-INS-322 |
| chr8-4038360-INS-6583 | chr8-4038361-INS-6584 |
| chr8-4068044-INS-52 | chr8-4068044-INS-50 |
| chr8-4083317-INS-520 | chr8-4083317-INS-520 |
| chr8-4101470-INS-54 | chr8-4101456-INS-70 |
| chr8-4117595-INS-70 | chr8-4117595-INS-78 |
| chr8-4117985-INS-442 | chr8-4117985-INS-442 |
| chr8-4173375-INS-8818 | chr8-4173376-INS-8818 |
| chr8-4190063-INS-57 | chr8-4190063-INS-50 |
| chr8-4240035-INS-165 | chr8-4240035-INS-165 |
| chr8-4250501-INS-301 | chr8-4250501-INS-301 |
| chr8-4262032-INS-58 | chr8-4262014-INS-86 |
| chr8-4275918-INS-207 | chr8-4275918-INS-207 |
| chr8-4280486-INS-228 | chr8-4280486-INS-228 |
| chr8-4314739-INS-58 | chr8-4314739-INS-58 |
| chr8-4322074-INS-212 | chr8-4322074-INS-212 |
| chr8-4323330-INS-223 | chr8-4323330-INS-223 |
| chr8-4384373-INS-166 | chr8-4384373-INS-166 |
| chr8-4395420-INS-63 | chr8-4395420-INS-59 |
| chr8-4408453-INS-134 | chr8-4408453-INS-134 |
| chr8-4442582-INS-215 | chr8-4442582-INS-215 |
| chr8-4457983-INS-231 | chr8-4457983-INS-231 |
| chr8-4463137-INS-59 | chr8-4463137-INS-59 |
| chr8-4465275-INS-56 | chr8-4465275-INS-56 |
| chr8-4527368-INS-640 | chr8-4527368-INS-640 |
| chr8-4540327-INS-171 | chr8-4540327-INS-181 |
| chr8-4551625-INS-65 | chr8-4551625-INS-65 |
| chr8-4555918-INS-81 | chr8-4555878-INS-95 |
| chr8-4596718-INS-614 | chr8-4596718-INS-614 |

| | |
|---|---|
| chr8-4694411-INS-671 | chr8-4694411-INS-670 |
| chr8-4708986-INS-7099 | chr8-4708987-INS-7099 |
| chr8-4717956-INS-79 | chr8-4717956-INS-82 |
| chr8-4729147-INS-102 | chr8-4729147-INS-102 |
| chr8-4730565-INS-64 | chr8-4730565-INS-64 |
| chr8-4766228-INS-174 | chr8-4766228-INS-174 |
| chr8-4778827-INS-118 | chr8-4778827-INS-118 |
| chr8-4789969-INS-60 | chr8-4789969-INS-60 |
| chr8-4791688-INS-197 | chr8-4791688-INS-197 |
| chr8-4850303-INS-78 | chr8-4850303-INS-84 |
| chr8-4892966-INS-7208 | chr8-4892967-INS-7208 |
| chr8-4900577-INS-184 | chr8-4900577-INS-184 |
| chr8-4915798-INS-96 | chr8-4915798-INS-104 |
| chr8-4919757-INS-213 | chr8-4919758-INS-214 |
| chr8-4961837-INS-995 | chr8-4961837-INS-995 |
| chr8-5010259-INS-1171 | chr8-5010278-INS-1150 |
| chr8-5022127-INS-470 | chr8-5022126-INS-475 |
| chr8-5062183-INS-6222 | chr8-5062184-INS-6223 |
| chr8-5073203-INS-175 | chr8-5073203-INS-175 |
| chr8-5079099-INS-75 | chr8-5079101-INS-67 |
| chr8-5096150-INS-390 | chr8-5096150-INS-390 |
| chr8-5103964-INS-832 | chr8-5103964-INS-832 |
| chr8-5119801-INS-58 | chr8-5119802-INS-80 |
| chr8-5122223-INS-208 | chr8-5122223-INS-208 |
| chr8-5126594-INS-241 | chr8-5126594-INS-241 |
| chr8-5128467-INS-6589 | chr8-5128468-INS-6589 |
| chr8-5152371-INS-92 | chr8-5152371-INS-92 |
| chr8-5168169-INS-51 | chr8-5168169-INS-51 |
| chr8-5186042-INS-1818 | chr8-5186043-INS-1818 |
| chr8-5186927-INS-1402 | chr8-5186928-INS-1398 |
| chr8-5218720-INS-89 | chr8-5218720-INS-89 |
| chr8-5292264-INS-214 | chr8-5292264-INS-214 |
| chr8-5312730-INS-52 | chr8-5312730-INS-52 |
| chr8-5327601-INS-1087 | chr8-5327602-INS-1088 |
| chr8-5338926-INS-1740 | chr8-5338923-INS-1724 |
| chr8-5377953-INS-53 | chr8-5377959-INS-50 |
| chr8-5382683-INS-93 | chr8-5382683-INS-97 |
| chr8-5425748-INS-504 | chr8-5425748-INS-504 |
| chr8-5435156-INS-538 | chr8-5435156-INS-538 |
| chr8-5448050-INS-54 | chr8-5448050-INS-54 |
| chr8-5485914-INS-190 | chr8-5485914-INS-190 |
| chr8-5508763-INS-2455 | chr8-5508764-INS-2455 |
| chr8-5513377-INS-168 | chr8-5513377-INS-189 |
| chr8-5522204-INS-199 | chr8-5522204-INS-199 |
| chr8-5527240-INS-4867 | chr8-5527241-INS-4866 |
| chr8-5537125-INS-2267 | chr8-5537126-INS-2267 |
| chr8-5539336-INS-105 | chr8-5539307-INS-125 |
| chr8-5548227-INS-132 | chr8-5548227-INS-89 |
| chr8-5549664-INS-109 | chr8-5549632-INS-120 |
| chr8-5560868-INS-426 | chr8-5560899-INS-425 |
| chr8-5585277-INS-262 | chr8-5585277-INS-262 |
| chr8-5654856-INS-217 | chr8-5654857-INS-213 |
| chr8-5678419-INS-507 | chr8-5678419-INS-507 |
| chr8-5698435-INS-66 | chr8-5698435-INS-100 |
| chr8-5704313-INS-270 | chr8-5704313-INS-270 |
| chr8-5733923-INS-56 | chr8-5733923-INS-56 |
| chr8-5736234-INS-478 | chr8-5736234-INS-478 |
| chr8-5759719-INS-752 | chr8-5759719-INS-752 |
| chr8-5844194-INS-730 | chr8-5844194-INS-730 |
| chr8-5847288-INS-81 | chr8-5847288-INS-81 |
| chr8-5855848-INS-80 | chr8-5855848-INS-82 |
| chr8-5863408-INS-74 | chr8-5863366-INS-68 |
| chr8-5881757-INS-64 | chr8-5881757-INS-64 |
| chr8-5912491-INS-90 | chr8-5912497-INS-68 |
| chr8-5936788-INS-885 | chr8-5936788-INS-881 |
| chr8-5965825-INS-68 | chr8-5965825-INS-66 |
| chr8-5989772-INS-174 | chr8-5989772-INS-174 |
| chr8-5994725-INS-948 | chr8-5994725-INS-948 |
| chr8-5999019-INS-6377 | chr8-5999020-INS-6378 |
| chr8-6005902-INS-2563 | chr8-6005903-INS-2563 |
| chr8-6024995-INS-3536 | chr8-6024996-INS-3536 |
| chr8-6042734-INS-317 | chr8-6042734-INS-317 |
| chr8-6057702-INS-53 | chr8-6057703-INS-63 |
| chr8-6064559-INS-408 | chr8-6064559-INS-408 |
| chr8-6078242-INS-61 | chr8-6078242-INS-65 |
| chr8-6082816-INS-4440 | chr8-6082817-INS-4440 |
| chr8-6084470-INS-595 | chr8-6084470-INS-595 |

| | |
|---|---|
| chr8-6093090-INS-146 | chr8-6093097-INS-147 |
| chr8-6126646-INS-67 | chr8-6126646-INS-71 |
| chr8-6131876-INS-254 | chr8-6131876-INS-254 |
| chr8-6139315-INS-64 | chr8-6139289-INS-54 |
| chr8-6162562-INS-63 | chr8-6162562-INS-62 |
| chr8-6184626-INS-702 | chr8-6184626-INS-702 |
| chr8-6187437-INS-80 | chr8-6187437-INS-88 |
| chr8-6195213-INS-144 | chr8-6195213-INS-144 |
| chr8-6196931-INS-558 | chr8-6196931-INS-558 |
| chr8-6201383-INS-325 | chr8-6201383-INS-325 |
| chr8-6212646-INS-90 | chr8-6212646-INS-92 |
| chr8-6213830-INS-2687 | chr8-6213831-INS-2687 |
| chr8-6224723-INS-176 | chr8-6224723-INS-176 |
| chr8-6229411-INS-63 | chr8-6229411-INS-80 |
| chr8-6234457-INS-4977 | chr8-6234458-INS-4977 |
| chr8-6239142-INS-833 | chr8-6239142-INS-833 |
| chr8-6239971-INS-3436 | chr8-6239972-INS-3436 |
| chr8-6240583-INS-6373 | chr8-6240584-INS-6374 |
| chr8-6240650-INS-52 | chr8-6240650-INS-80 |
| chr8-6246721-INS-187 | chr8-6246721-INS-187 |
| chr8-6258868-INS-97 | chr8-6258868-INS-97 |
| chr8-6261691-INS-6897 | chr8-6261692-INS-6896 |
| chr8-6265274-INS-51 | chr8-6265274-INS-51 |
| chr8-6267220-INS-234 | chr8-6267220-INS-234 |
| chr8-6304687-INS-6324 | chr8-6304688-INS-6326 |
| chr8-6312549-INS-7149 | chr8-6312550-INS-7149 |
| chr8-6318246-INS-505 | chr8-6318246-INS-508 |
| chr8-6324333-INS-2606 | chr8-6324334-INS-2606 |
| chr8-6325143-INS-1797 | chr8-6325144-INS-1797 |
| chr8-6348515-INS-193 | chr8-6348515-INS-193 |
| chr8-6373195-INS-77 | chr8-6373195-INS-77 |
| chr8-6382990-INS-2405 | chr8-6382991-INS-2405 |
| chr8-6433462-INS-6389 | chr8-6433463-INS-6389 |
| chr8-6437091-INS-265 | chr8-6437091-INS-265 |
| chr8-6453351-INS-75 | chr8-6453351-INS-74 |
| chr8-6456222-INS-128 | chr8-6456228-INS-120 |
| chr8-6459834-INS-93 | chr8-6459834-INS-96 |
| chr8-6486020-INS-86 | chr8-6486020-INS-86 |
| chr8-6486772-INS-6547 | chr8-6486773-INS-6547 |
| chr8-6491753-INS-3798 | chr8-6491754-INS-3786 |
| chr8-6514489-INS-3092 | chr8-6514490-INS-3092 |
| chr8-6517512-INS-81 | chr8-6517512-INS-81 |
| chr8-6537063-INS-328 | chr8-6537063-INS-328 |
| chr8-6560100-INS-198 | chr8-6560100-INS-198 |
| chr8-6562203-INS-89 | chr8-6562203-INS-87 |
| chr8-6594076-INS-208 | chr8-6594076-INS-184 |
| chr8-6594646-INS-2862 | chr8-6594647-INS-2862 |
| chr8-6598621-INS-990 | chr8-6598621-INS-996 |
| chr8-6600236-INS-6614 | chr8-6600237-INS-6614 |
| chr8-6605010-INS-118 | chr8-6605010-INS-118 |
| chr8-6624732-INS-59 | chr8-6624732-INS-59 |
| chr8-6638466-INS-240 | chr8-6638466-INS-240 |
| chr8-6651114-INS-2886 | chr8-6651115-INS-2886 |
| chr8-6652954-INS-2098 | chr8-6652955-INS-2091 |
| chr8-6670732-INS-433 | chr8-6670733-INS-435 |
| chr8-6700280-INS-59 | chr8-6700280-INS-59 |
| chr8-6714886-INS-55 | chr8-6714886-INS-56 |
| chr8-6717932-INS-6753 | chr8-6717933-INS-6755 |
| chr8-6717967-INS-417 | chr8-6717967-INS-417 |
| chr8-6722799-INS-63 | chr8-6722799-INS-63 |
| chr8-6726243-INS-4105 | chr8-6726244-INS-4105 |
| chr8-6756716-INS-243 | chr8-6756716-INS-243 |
| chr8-6766314-INS-514 | chr8-6766314-INS-514 |
| chr8-6788705-INS-99 | chr8-6788717-INS-88 |
| chr8-6800338-INS-78 | chr8-6800342-INS-60 |
| chr8-6801665-INS-201 | chr8-6801665-INS-202 |
| chr8-6822246-INS-1895 | chr8-6822247-INS-1895 |
| chr8-6840387-INS-6279 | chr8-6840388-INS-6279 |
| chr8-6848401-INS-7459 | chr8-6848402-INS-7459 |
| chr8-6880697-INS-80 | chr8-6880701-INS-124 |
| chr8-6895374-INS-901 | chr8-6895374-INS-901 |
| chr8-6899222-INS-60 | chr8-6899222-INS-60 |
| chr8-6916212-INS-1903 | chr8-6916213-INS-1903 |
| chr8-6927426-INS-101 | chr8-6927426-INS-101 |
| chr8-6958066-INS-61 | chr8-6958066-INS-73 |
| chr8-6968625-INS-2820 | chr8-6968626-INS-2820 |
| chr8-6994465-INS-486 | chr8-6994465-INS-486 |

| | |
|---|---|
| chr8-6995249-INS-68 | chr8-6995249-INS-58 |
| chr8-7025880-INS-82 | chr8-7025880-INS-82 |
| chr8-7045961-INS-56 | chr8-7045961-INS-50 |
| chr8-7074405-INS-791 | chr8-7074405-INS-793 |
| chr8-7096856-INS-228 | chr8-7096856-INS-228 |
| chr8-7147985-INS-60 | chr8-7147985-INS-65 |
| chr8-7153991-INS-68 | chr8-7153975-INS-68 |
| chr8-7171436-INS-6355 | chr8-7171437-INS-6355 |
| chr8-7208958-INS-68 | chr8-7208958-INS-68 |
| chr8-7221593-INS-63 | chr8-7221593-INS-63 |
| chr8-7227171-INS-405 | chr8-7227171-INS-405 |
| chr8-7243566-INS-60 | chr8-7243549-INS-60 |
| chr8-7321206-INS-72 | chr8-7321206-INS-73 |
| chr8-7340414-INS-6370 | chr8-7340415-INS-6370 |
| chr8-7347608-INS-88 | chr8-7347608-INS-88 |
| chr8-7368066-INS-170 | chr8-7368066-INS-171 |
| chr8-7371578-INS-53 | chr8-7371578-INS-53 |
| chr8-7380464-INS-208 | chr8-7380464-INS-208 |
| chr8-7386365-INS-2413 | chr8-7386366-INS-2413 |
| chr8-7396344-INS-127 | chr8-7396344-INS-127 |
| chr8-7425053-INS-195 | chr8-7425053-INS-189 |
| chr8-7426108-INS-68 | chr8-7426108-INS-68 |
| chr8-7431768-INS-73 | chr8-7431768-INS-75 |
| chr8-7457340-INS-56 | chr8-7457340-INS-62 |
| chr8-7467851-INS-7180 | chr8-7467852-INS-7180 |
| chr8-7470118-INS-302 | chr8-7470118-INS-303 |
| chr8-7472989-INS-1613 | chr8-7472990-INS-1614 |
| chr8-7478983-INS-6574 | chr8-7478984-INS-6574 |
| chr8-7486974-INS-7034 | chr8-7486975-INS-7034 |
| chr8-7514005-INS-261 | chr8-7514005-INS-261 |
| chr8-7529521-INS-60 | chr8-7529521-INS-64 |
| chr8-7545307-INS-54 | chr8-7545307-INS-54 |
| chr8-7551642-INS-309 | chr8-7551642-INS-307 |
| chr8-7560917-INS-58 | chr8-7560917-INS-58 |
| chr8-7561688-INS-497 | chr8-7561688-INS-497 |
| chr8-7569886-INS-62 | chr8-7569859-INS-87 |
| chr8-7570494-INS-9907 | chr8-7570495-INS-9909 |
| chr8-7581858-INS-4477 | chr8-7581859-INS-4477 |
| chr8-7638805-INS-583 | chr8-7638805-INS-583 |
| chr8-7650651-INS-117 | chr8-7650651-INS-117 |
| chr8-7664822-INS-233 | chr8-7664829-INS-232 |
| chr8-7705953-INS-1121 | chr8-7705954-INS-1121 |
| chr8-7708276-INS-1064 | chr8-7708277-INS-1064 |
| chr8-7718516-INS-60 | chr8-7718516-INS-60 |
| chr8-7729403-INS-60 | chr8-7729403-INS-62 |
| chr8-7741095-INS-252 | chr8-7741095-INS-252 |
| chr8-7744112-INS-141 | chr8-7744112-INS-141 |
| chr8-7765726-INS-195 | chr8-7765726-INS-195 |
| chr8-7789161-INS-162 | chr8-7789161-INS-166 |
| chr8-7798291-INS-77 | chr8-7798251-INS-111 |
| chr8-7879766-INS-7081 | chr8-7879767-INS-7081 |
| chr8-7896789-INS-593 | chr8-7896789-INS-593 |
| chr8-7899042-INS-546 | chr8-7899042-INS-546 |
| chr8-7901990-INS-68 | chr8-7901990-INS-68 |
| chr8-7927085-INS-69 | chr8-7927085-INS-69 |
| chr8-7935084-INS-58 | chr8-7935084-INS-58 |
| chr8-7951361-INS-203 | chr8-7951361-INS-203 |
| chr8-7967514-INS-188 | chr8-7967514-INS-192 |
| chr8-7985733-INS-171 | chr8-7985733-INS-170 |
| chr8-7998343-INS-84 | chr8-7998337-INS-80 |
| chr8-8005107-INS-1882 | chr8-8005108-INS-1871 |
| chr8-8021278-INS-7257 | chr8-8021279-INS-7256 |
| chr8-8021279-INS-462 | chr8-8021279-INS-462 |
| chr8-8031736-INS-482 | chr8-8031736-INS-482 |
| chr8-8049431-INS-54 | chr8-8049427-INS-58 |
| chr8-8140186-INS-185 | chr8-8140186-INS-185 |
| chr8-8162592-INS-56 | chr8-8162583-INS-75 |
| chr8-8172303-INS-2671 | chr8-8172304-INS-2671 |
| chr8-8181333-INS-64 | chr8-8181327-INS-64 |
| chr8-8186455-INS-57 | chr8-8186456-INS-66 |
| chr8-8245900-INS-411 | chr8-8245918-INS-422 |
| chr8-8253037-INS-645 | chr8-8253030-INS-649 |
| chr8-8264487-INS-55 | chr8-8264487-INS-55 |
| chr8-8267145-INS-4938 | chr8-8267146-INS-4938 |
| chr8-8269099-INS-116 | chr8-8269099-INS-116 |
| chr8-8271290-INS-307 | chr8-8271290-INS-307 |
| chr8-8313227-INS-76 | chr8-8313215-INS-76 |

| | |
|---|---|
| chr8-8342562-INS-94 | chr8-8342562-INS-79 |
| chr8-8352837-INS-56 | chr8-8352837-INS-56 |
| chr8-8356801-INS-51 | chr8-8356801-INS-51 |
| chr8-8360027-INS-111 | chr8-8360027-INS-114 |
| chr8-8361393-INS-215 | chr8-8361393-INS-215 |
| chr8-8373020-INS-255 | chr8-8373020-INS-255 |
| chr8-8387729-INS-56 | chr8-8387729-INS-56 |
| chr8-8402856-INS-6636 | chr8-8402857-INS-6637 |
| chr8-8403831-INS-83 | chr8-8403831-INS-83 |
| chr8-8418012-INS-645 | chr8-8418012-INS-646 |
| chr8-8422943-INS-54 | chr8-8422943-INS-54 |
| chr8-8432567-INS-92 | chr8-8432567-INS-92 |
| chr8-8436815-INS-546 | chr8-8436815-INS-546 |
| chr8-8445852-INS-844 | chr8-8445852-INS-845 |
| chr8-8492406-INS-7208 | chr8-8492407-INS-7208 |
| chr8-8577480-INS-61 | chr8-8577457-INS-60 |
| chr8-8585365-INS-110 | chr8-8585373-INS-120 |
| chr8-8586369-INS-76 | chr8-8586370-INS-54 |
| chr8-8596507-INS-207 | chr8-8596512-INS-209 |
| chr8-8596911-INS-61 | chr8-8596911-INS-61 |
| chr8-8615510-INS-70 | chr8-8615510-INS-100 |
| chr8-8634739-INS-106 | chr8-8634739-INS-67 |
| chr8-8690566-INS-53 | chr8-8690566-INS-53 |
| chr8-8692134-INS-56 | chr8-8692134-INS-58 |
| chr8-8694643-INS-88 | chr8-8694643-INS-88 |
| chr8-8712745-INS-179 | chr8-8712745-INS-187 |
| chr8-8719074-INS-181 | chr8-8719074-INS-181 |
| chr8-8724648-INS-208 | chr8-8724648-INS-208 |
| chr8-8759796-INS-54 | chr8-8759797-INS-56 |
| chr8-8763681-INS-509 | chr8-8763681-INS-509 |
| chr8-8768759-INS-235 | chr8-8768759-INS-235 |
| chr8-8799520-INS-85 | chr8-8799520-INS-85 |
| chr8-8806424-INS-88 | chr8-8806424-INS-70 |
| chr8-8809133-INS-58 | chr8-8809133-INS-58 |
| chr8-8827384-INS-213 | chr8-8827384-INS-213 |
| chr8-8835851-INS-70 | chr8-8835851-INS-60 |
| chr8-8874480-INS-78 | chr8-8874480-INS-74 |
| chr8-8923955-INS-215 | chr8-8923955-INS-211 |
| chr8-8938517-INS-3691 | chr8-8938518-INS-3691 |
| chr8-8941570-INS-1892 | chr8-8941571-INS-1892 |
| chr8-8951838-INS-3343 | chr8-8951839-INS-3343 |
| chr8-8968260-INS-76 | chr8-8968260-INS-77 |
| chr8-8975490-INS-78 | chr8-8975484-INS-72 |
| chr8-8978114-INS-76 | chr8-8978114-INS-92 |
| chr8-8992142-INS-334 | chr8-8992142-INS-334 |
| chr8-9073954-INS-673 | chr8-9073954-INS-673 |
| chr8-9077331-INS-53 | chr8-9077331-INS-53 |
| chr8-9130426-INS-81 | chr8-9130426-INS-81 |
| chr8-9139139-INS-99 | chr8-9139139-INS-105 |
| chr8-9164245-INS-497 | chr8-9164245-INS-497 |
| chr8-9169297-INS-130 | chr8-9169297-INS-130 |
| chr8-9192025-INS-187 | chr8-9192025-INS-187 |
| chr8-9217795-INS-340 | chr8-9217795-INS-341 |
| chr8-9229368-INS-288 | chr8-9229361-INS-288 |
| chr8-9267299-INS-60 | chr8-9267299-INS-60 |
| chr8-9274341-INS-5319 | chr8-9274342-INS-5320 |
| chr8-9278917-INS-61 | chr8-9278915-INS-51 |
| chr8-9282294-INS-56 | chr8-9282294-INS-56 |
| chr8-9284996-INS-306 | chr8-9284996-INS-306 |
| chr8-9312307-INS-88 | chr8-9312307-INS-76 |
| chr8-9313390-INS-88 | chr8-9313411-INS-109 |
| chr8-9328440-INS-165 | chr8-9328436-INS-165 |
| chr8-9346875-INS-243 | chr8-9346876-INS-270 |
| chr8-9350132-INS-169 | chr8-9350132-INS-169 |
| chr8-9388831-INS-84 | chr8-9388832-INS-52 |
| chr8-9408131-INS-214 | chr8-9408131-INS-214 |
| chr8-9435980-INS-383 | chr8-9435980-INS-386 |
| chr8-9439219-INS-69 | chr8-9439219-INS-74 |
| chr8-9493797-INS-111 | chr8-9493797-INS-111 |
| chr8-9528963-INS-87 | chr8-9528963-INS-87 |
| chr8-9539982-INS-60 | chr8-9539974-INS-52 |
| chr8-9566873-INS-52 | chr8-9566854-INS-72 |
| chr8-9569035-INS-54 | chr8-9569035-INS-54 |
| chr8-9587767-INS-2299 | chr8-9587768-INS-2299 |
| chr8-9594339-INS-114 | chr8-9594339-INS-114 |
| chr8-9609121-INS-372 | chr8-9609121-INS-372 |
| chr8-9662758-INS-70 | chr8-9662758-INS-71 |

| | |
|---|---|
| chr8-9697814-INS-145 | chr8-9697809-INS-155 |
| chr8-9758737-INS-270 | chr8-9758737-INS-275 |
| chr8-9766057-INS-63 | chr8-9766052-INS-63 |
| chr8-9768520-INS-162 | chr8-9768520-INS-162 |
| chr8-9776970-INS-75 | chr8-9776970-INS-75 |
| chr8-9781201-INS-65 | chr8-9781201-INS-65 |
| chr8-9787010-INS-68 | chr8-9787010-INS-76 |
| chr8-9815965-INS-76 | chr8-9815965-INS-76 |
| chr8-9825198-INS-69 | chr8-9825198-INS-72 |
| chr8-9843682-INS-99 | chr8-9843682-INS-99 |
| chr8-9856786-INS-52 | chr8-9856794-INS-60 |
| chr8-9881716-INS-310 | chr8-9881715-INS-290 |
| chr8-9884356-INS-2812 | chr8-9884357-INS-2812 |
| chr8-9902078-INS-246 | chr8-9902078-INS-253 |
| chr8-9903198-INS-71 | chr8-9903198-INS-106 |
| chr8-9916344-INS-254 | chr8-9916344-INS-254 |
| chr8-9921101-INS-56 | chr8-9921101-INS-66 |
| chr8-9941423-INS-127 | chr8-9941465-INS-78 |
| chr8-9944082-INS-137 | chr8-9944082-INS-137 |
| chr8-9945423-INS-106 | chr8-9945429-INS-58 |
| chr8-9964751-INS-68 | chr8-9964751-INS-68 |
| chr8-9966070-INS-54 | chr8-9966070-INS-54 |
| chr8-9981418-INS-71 | chr8-9981418-INS-79 |
| chr8-9984735-INS-54 | chr8-9984735-INS-54 |
| chr8-9992836-INS-416 | chr8-9992836-INS-416 |
| chr8-10037239-INS-1889 | chr8-10037240-INS-1889 |
| chr8-10067737-INS-172 | chr8-10067737-INS-183 |
| chr8-10078122-INS-57 | chr8-10078122-INS-57 |
| chr8-10116779-INS-227 | chr8-10116783-INS-230 |
| chr8-10151465-INS-144 | chr8-10151465-INS-148 |
| chr8-10154272-INS-325 | chr8-10154272-INS-325 |
| chr8-10190310-INS-177 | chr8-10190310-INS-193 |
| chr8-10205495-INS-59 | chr8-10205495-INS-59 |
| chr8-10217664-INS-7838 | chr8-10217665-INS-7840 |
| chr8-10235561-INS-1089 | chr8-10235562-INS-1089 |
| chr8-10240061-INS-94 | chr8-10240061-INS-107 |
| chr8-10258841-INS-6568 | chr8-10258838-INS-6566 |
| chr8-10262229-INS-779 | chr8-10262230-INS-780 |
| chr8-10285429-INS-162 | chr8-10285430-INS-164 |
| chr8-10303319-INS-96 | chr8-10303319-INS-96 |
| chr8-10307107-INS-2095 | chr8-10307108-INS-2095 |
| chr8-10312478-INS-1440 | chr8-10312479-INS-1440 |
| chr8-10316673-INS-236 | chr8-10316673-INS-236 |
| chr8-10322370-INS-52 | chr8-10322381-INS-52 |
| chr8-10327481-INS-293 | chr8-10327481-INS-293 |
| chr8-10341885-INS-118 | chr8-10341885-INS-118 |
| chr8-10346799-INS-64 | chr8-10346798-INS-71 |
| chr8-10350191-INS-81 | chr8-10350191-INS-81 |
| chr8-10351358-INS-54 | chr8-10351358-INS-54 |
| chr8-10355543-INS-91 | chr8-10355543-INS-89 |
| chr8-10362862-INS-295 | chr8-10362862-INS-295 |
| chr8-10370818-INS-126 | chr8-10370818-INS-126 |
| chr8-10381965-INS-481 | chr8-10381965-INS-503 |
| chr8-10427441-INS-60 | chr8-10427442-INS-56 |
| chr8-10440627-INS-115 | chr8-10440627-INS-115 |
| chr8-10441605-INS-619 | chr8-10441605-INS-612 |
| chr8-10476417-INS-63 | chr8-10476417-INS-63 |
| chr8-10495703-INS-74 | chr8-10495703-INS-69 |
| chr8-10506562-INS-119 | chr8-10506562-INS-142 |
| chr8-10515340-INS-58 | chr8-10515347-INS-66 |
| chr8-10545534-INS-64 | chr8-10545534-INS-52 |
| chr8-10554051-INS-67 | chr8-10554051-INS-69 |
| chr8-10557323-INS-1321 | chr8-10557324-INS-1298 |
| chr8-10570665-INS-56 | chr8-10570667-INS-54 |
| chr8-10586540-INS-129 | chr8-10586548-INS-84 |
| chr8-10593633-INS-114 | chr8-10593633-INS-152 |
| chr8-10601725-INS-112 | chr8-10601725-INS-112 |
| chr8-10609905-INS-72 | chr8-10609905-INS-72 |
| chr8-10620272-INS-63 | chr8-10620272-INS-63 |
| chr8-10622655-INS-67 | chr8-10622662-INS-53 |
| chr8-10630792-INS-91 | chr8-10630792-INS-91 |
| chr8-10658556-INS-579 | chr8-10658556-INS-579 |
| chr8-10661026-INS-58 | chr8-10661026-INS-54 |
| chr8-10689372-INS-671 | chr8-10689372-INS-665 |
| chr8-10697332-INS-60 | chr8-10697347-INS-81 |
| chr8-10710587-INS-55 | chr8-10710587-INS-55 |
| chr8-10748986-INS-62 | chr8-10748991-INS-80 |

| | |
|---|---|
| chr8-10756255-INS-906 | chr8-10756255-INS-905 |
| chr8-10763296-INS-326 | chr8-10763295-INS-329 |
| chr8-10804930-INS-96 | chr8-10804930-INS-96 |
| chr8-10808181-INS-54 | chr8-10808181-INS-54 |
| chr8-10838459-INS-74 | chr8-10838459-INS-75 |
| chr8-10850622-INS-634 | chr8-10850627-INS-634 |
| chr8-10852626-INS-54 | chr8-10852626-INS-54 |
| chr8-10864544-INS-336 | chr8-10864544-INS-336 |
| chr8-10868990-INS-225 | chr8-10868990-INS-257 |
| chr8-10881131-INS-260 | chr8-10881127-INS-226 |
| chr8-10895293-INS-556 | chr8-10895293-INS-555 |
| chr8-10896649-INS-125 | chr8-10896649-INS-125 |
| chr8-10908831-INS-647 | chr8-10908831-INS-647 |
| chr8-10911341-INS-731 | chr8-10911341-INS-725 |
| chr8-10915206-INS-66 | chr8-10915206-INS-52 |
| chr8-10927353-INS-58 | chr8-10927353-INS-64 |
| chr8-10955190-INS-109 | chr8-10955191-INS-82 |
| chr8-10972515-INS-71 | chr8-10972515-INS-75 |
| chr8-11005734-INS-683 | chr8-11005734-INS-683 |
| chr8-11066828-INS-60 | chr8-11066828-INS-60 |
| chr8-11069371-INS-204 | chr8-11069371-INS-204 |
| chr8-11084950-INS-54 | chr8-11084950-INS-54 |
| chr8-11085011-INS-7098 | chr8-11085012-INS-7098 |
| chr8-11087918-INS-60 | chr8-11087918-INS-52 |
| chr8-11140475-INS-224 | chr8-11140475-INS-225 |
| chr8-11167138-INS-636 | chr8-11167138-INS-635 |
| chr8-11196616-INS-84 | chr8-11196628-INS-50 |
| chr8-11204446-INS-85 | chr8-11204446-INS-85 |
| chr8-11236564-INS-85 | chr8-11236564-INS-86 |
| chr8-11273231-INS-91 | chr8-11273231-INS-91 |
| chr8-11274699-INS-98 | chr8-11274699-INS-98 |
| chr8-11280703-INS-85 | chr8-11280703-INS-90 |
| chr8-11292633-INS-206 | chr8-11292647-INS-203 |
| chr8-11330447-INS-201 | chr8-11330447-INS-201 |
| chr8-11334728-INS-399 | chr8-11334728-INS-399 |
| chr8-11348315-INS-54 | chr8-11348313-INS-58 |
| chr8-11353968-INS-70 | chr8-11353968-INS-70 |
| chr8-11378779-INS-108 | chr8-11378779-INS-108 |
| chr8-11402217-INS-70 | chr8-11402215-INS-82 |
| chr8-11438699-INS-231 | chr8-11438699-INS-231 |
| chr8-11446452-INS-56 | chr8-11446458-INS-50 |
| chr8-11449181-INS-116 | chr8-11449181-INS-116 |
| chr8-11453015-INS-120 | chr8-11453015-INS-120 |
| chr8-11468694-INS-55 | chr8-11468694-INS-55 |
| chr8-11471877-INS-52 | chr8-11471877-INS-52 |
| chr8-11482297-INS-194 | chr8-11482297-INS-193 |
| chr8-11499501-INS-192 | chr8-11499501-INS-192 |
| chr8-11517947-INS-207 | chr8-11517947-INS-206 |
| chr8-11518600-INS-220 | chr8-11518600-INS-222 |
| chr8-11552349-INS-134 | chr8-11552353-INS-130 |
| chr8-11563469-INS-78 | chr8-11563469-INS-78 |
| chr8-11569693-INS-133 | chr8-11569693-INS-133 |
| chr8-11574799-INS-649 | chr8-11574799-INS-649 |
| chr8-11577057-INS-7136 | chr8-11577058-INS-7136 |
| chr8-11577473-INS-56 | chr8-11577473-INS-51 |
| chr8-11588128-INS-66 | chr8-11588128-INS-66 |
| chr8-11589270-INS-333 | chr8-11589270-INS-332 |
| chr8-11597233-INS-756 | chr8-11597233-INS-751 |
| chr8-11622824-INS-86 | chr8-11622824-INS-86 |
| chr8-11627027-INS-396 | chr8-11627027-INS-396 |
| chr8-11659487-INS-191 | chr8-11659487-INS-192 |
| chr8-11662311-INS-70 | chr8-11662311-INS-70 |
| chr8-11679553-INS-116 | chr8-11679553-INS-116 |
| chr8-11688886-INS-882 | chr8-11688886-INS-882 |
| chr8-11700965-INS-215 | chr8-11700965-INS-217 |
| chr8-11709021-INS-52 | chr8-11709021-INS-50 |
| chr8-11720630-INS-104 | chr8-11720630-INS-104 |
| chr8-11726264-INS-51 | chr8-11726265-INS-52 |
| chr8-11729909-INS-188 | chr8-11729909-INS-188 |
| chr8-11731000-INS-58 | chr8-11731000-INS-58 |
| chr8-11733992-INS-215 | chr8-11733992-INS-216 |
| chr8-11736585-INS-198 | chr8-11736585-INS-198 |
| chr8-11741651-INS-51 | chr8-11741617-INS-98 |
| chr8-11768750-INS-125 | chr8-11768753-INS-102 |
| chr8-11786508-INS-54 | chr8-11786508-INS-80 |
| chr8-11787094-INS-52 | chr8-11787094-INS-52 |
| chr8-11885634-INS-74 | chr8-11885634-INS-76 |

| | |
|---|---|
| chr8-11885960-INS-766 | chr8-11885960-INS-765 |
| chr8-11908658-INS-116 | chr8-11908683-INS-87 |
| chr8-11942723-INS-8827 | chr8-11942724-INS-8827 |
| chr8-11955865-INS-69 | chr8-11955865-INS-68 |
| chr8-11958544-INS-330 | chr8-11958544-INS-332 |
| chr8-11967062-INS-54 | chr8-11967094-INS-67 |
| chr8-11969609-INS-65 | chr8-11969609-INS-65 |
| chr8-11977162-INS-167 | chr8-11977157-INS-167 |
| chr8-11979455-INS-211 | chr8-11979455-INS-212 |
| chr8-11988083-INS-64 | chr8-11988083-INS-59 |
| chr8-11998176-INS-206 | chr8-11998176-INS-206 |
| chr8-11998659-INS-615 | chr8-11998659-INS-615 |
| chr8-12000711-INS-195 | chr8-12000712-INS-194 |
| chr8-12007009-INS-248 | chr8-12007050-INS-203 |
| chr8-12009523-INS-205 | chr8-12009523-INS-205 |
| chr8-12048414-INS-56 | chr8-12048414-INS-60 |
| chr8-12062129-INS-74 | chr8-12062129-INS-75 |
| chr8-12063691-INS-154 | chr8-12063690-INS-177 |
| chr8-12095278-INS-56 | chr8-12095272-INS-90 |
| chr8-12100219-INS-66 | chr8-12100235-INS-66 |
| chr8-12102035-INS-107 | chr8-12102035-INS-107 |
| chr8-12121549-INS-64 | chr8-12121549-INS-64 |
| chr8-12183256-INS-822 | chr8-12183257-INS-822 |
| chr8-12195755-INS-94 | chr8-12195747-INS-68 |
| chr8-12201516-INS-228 | chr8-12201516-INS-204 |
| chr8-12205595-INS-169 | chr8-12205592-INS-172 |
| chr8-12215129-INS-56 | chr8-12215129-INS-54 |
| chr8-12257827-INS-64 | chr8-12257827-INS-64 |
| chr8-12270365-INS-60 | chr8-12270371-INS-56 |
| chr8-12298746-INS-666 | chr8-12298746-INS-666 |
| chr8-12299418-INS-101 | chr8-12299418-INS-101 |
| chr8-12304392-INS-216 | chr8-12304392-INS-222 |
| chr8-12339050-INS-62 | chr8-12339010-INS-86 |
| chr8-12349719-INS-457 | chr8-12349719-INS-457 |
| chr8-12391475-INS-80 | chr8-12391475-INS-80 |
| chr8-12396704-INS-255 | chr8-12396704-INS-256 |
| chr8-12406465-INS-588 | chr8-12406465-INS-627 |
| chr8-12414682-INS-2309 | chr8-12414683-INS-2309 |
| chr8-12421172-INS-93 | chr8-12421172-INS-100 |
| chr8-12422443-INS-66 | chr8-12422443-INS-73 |
| chr8-12470073-INS-214 | chr8-12470073-INS-213 |
| chr8-12490687-INS-1237 | chr8-12490692-INS-1229 |
| chr8-12506752-INS-78 | chr8-12506752-INS-78 |
| chr8-12524725-INS-81 | chr8-12524725-INS-81 |
| chr8-12550240-INS-150 | chr8-12550240-INS-150 |
| chr8-12566772-INS-711 | chr8-12566772-INS-711 |
| chr8-12575048-INS-51 | chr8-12575048-INS-51 |
| chr8-12582418-INS-104 | chr8-12582418-INS-100 |
| chr8-12596597-INS-62 | chr8-12596597-INS-62 |
| chr8-12602480-INS-80 | chr8-12602480-INS-82 |
| chr8-12627998-INS-68 | chr8-12627986-INS-112 |
| chr8-12633396-INS-124 | chr8-12633396-INS-155 |
| chr8-12638192-INS-55 | chr8-12638192-INS-50 |
| chr8-12640633-INS-144 | chr8-12640633-INS-144 |
| chr8-12642199-INS-432 | chr8-12642199-INS-432 |
| chr8-12643062-INS-118 | chr8-12643062-INS-118 |
| chr8-12652657-INS-52 | chr8-12652681-INS-59 |
| chr8-12657166-INS-68 | chr8-12657166-INS-56 |
| chr8-12701760-INS-69 | chr8-12701760-INS-69 |
| chr8-12702785-INS-64 | chr8-12702785-INS-64 |
| chr8-12714431-INS-66 | chr8-12714433-INS-56 |
| chr8-12723444-INS-155 | chr8-12723432-INS-168 |
| chr8-12726631-INS-757 | chr8-12726631-INS-753 |
| chr8-12769622-INS-98 | chr8-12769622-INS-98 |
| chr8-12776961-INS-81 | chr8-12776961-INS-81 |
| chr8-12787755-INS-196 | chr8-12787755-INS-196 |
| chr8-12816476-INS-108 | chr8-12816476-INS-144 |
| chr8-12829886-INS-79 | chr8-12829886-INS-79 |
| chr8-12830552-INS-55 | chr8-12830552-INS-57 |
| chr8-12837345-INS-71 | chr8-12837345-INS-71 |
| chr8-12917049-INS-91 | chr8-12917049-INS-91 |
| chr8-12921367-INS-67 | chr8-12921367-INS-72 |
| chr8-12944314-INS-64 | chr8-12944318-INS-96 |
| chr8-12950024-INS-6013 | chr8-12950025-INS-6013 |
| chr8-12957832-INS-248 | chr8-12957832-INS-243 |
| chr8-12974457-INS-69 | chr8-12974457-INS-75 |
| chr8-12986899-INS-115 | chr8-12986892-INS-165 |

| | |
|---|---|
| chr8-13010618-INS-52 | chr8-13010612-INS-62 |
| chr8-13027067-INS-257 | chr8-13027067-INS-260 |
| chr8-13063361-INS-82 | chr8-13063361-INS-89 |
| chr8-13068432-INS-98 | chr8-13068432-INS-98 |
| chr8-13096764-INS-349 | chr8-13096764-INS-349 |
| chr8-13103589-INS-70 | chr8-13103589-INS-70 |
| chr8-13106216-INS-60 | chr8-13106216-INS-64 |
| chr8-13112095-INS-87 | chr8-13112095-INS-88 |
| chr8-13117405-INS-183 | chr8-13117405-INS-183 |
| chr8-13172748-INS-54 | chr8-13172766-INS-54 |
| chr8-13180124-INS-200 | chr8-13180124-INS-200 |
| chr8-13181801-INS-54 | chr8-13181801-INS-54 |
| chr8-13183414-INS-59 | chr8-13183414-INS-59 |
| chr8-13189934-INS-108 | chr8-13189934-INS-108 |
| chr8-13194452-INS-62 | chr8-13194452-INS-62 |
| chr8-13200441-INS-180 | chr8-13200441-INS-181 |
| chr8-13244069-INS-95 | chr8-13244063-INS-132 |
| chr8-13258715-INS-52 | chr8-13258715-INS-50 |
| chr8-13261747-INS-462 | chr8-13261747-INS-462 |
| chr8-13264437-INS-161 | chr8-13264437-INS-167 |
| chr8-13267748-INS-100 | chr8-13267749-INS-51 |
| chr8-13269422-INS-189 | chr8-13269422-INS-199 |
| chr8-13283336-INS-98 | chr8-13283336-INS-102 |
| chr8-13323727-INS-314 | chr8-13323727-INS-314 |
| chr8-13325710-INS-347 | chr8-13325710-INS-347 |
| chr8-13340017-INS-84 | chr8-13340051-INS-60 |
| chr8-13344590-INS-204 | chr8-13344590-INS-205 |
| chr8-13350138-INS-306 | chr8-13350136-INS-286 |
| chr8-13354300-INS-118 | chr8-13354300-INS-82 |
| chr8-13356182-INS-57 | chr8-13356168-INS-53 |
| chr8-13413736-INS-442 | chr8-13413736-INS-442 |
| chr8-13416591-INS-73 | chr8-13416591-INS-73 |
| chr8-13424331-INS-561 | chr8-13424331-INS-563 |
| chr8-13434376-INS-77 | chr8-13434379-INS-52 |
| chr8-13437331-INS-60 | chr8-13437331-INS-55 |
| chr8-13447366-INS-58 | chr8-13447366-INS-50 |
| chr8-13449343-INS-112 | chr8-13449343-INS-112 |
| chr8-13451034-INS-201 | chr8-13451034-INS-202 |
| chr8-13452073-INS-93 | chr8-13452073-INS-78 |
| chr8-13453244-INS-208 | chr8-13453244-INS-210 |
| chr8-13459578-INS-66 | chr8-13459578-INS-66 |
| chr8-13461999-INS-159 | chr8-13462002-INS-160 |
| chr8-13472052-INS-87 | chr8-13472052-INS-87 |
| chr8-13563369-INS-220 | chr8-13563392-INS-225 |
| chr8-13568531-INS-139 | chr8-13568531-INS-135 |
| chr8-13570335-INS-65 | chr8-13570337-INS-102 |
| chr8-13607437-INS-56 | chr8-13607439-INS-54 |
| chr8-13639504-INS-57 | chr8-13639504-INS-57 |
| chr8-13641680-INS-54 | chr8-13641680-INS-54 |
| chr8-13681954-INS-66 | chr8-13681940-INS-84 |
| chr8-13688312-INS-400 | chr8-13688312-INS-400 |
| chr8-13693233-INS-144 | chr8-13693233-INS-144 |
| chr8-13694577-INS-133 | chr8-13694582-INS-176 |
| chr8-13699639-INS-85 | chr8-13699592-INS-83 |
| chr8-13740430-INS-51 | chr8-13740442-INS-68 |
| chr8-13740958-INS-84 | chr8-13740958-INS-84 |
| chr8-13743117-INS-124 | chr8-13743117-INS-124 |
| chr8-13775545-INS-79 | chr8-13775544-INS-87 |
| chr8-13778227-INS-76 | chr8-13778227-INS-107 |
| chr8-13840360-INS-217 | chr8-13840360-INS-217 |
| chr8-13841478-INS-192 | chr8-13841478-INS-192 |
| chr8-13866313-INS-63 | chr8-13866311-INS-67 |
| chr8-13878427-INS-498 | chr8-13878427-INS-498 |
| chr8-13895531-INS-6442 | chr8-13895532-INS-6442 |
| chr8-13899018-INS-128 | chr8-13899018-INS-128 |
| chr8-13902639-INS-55 | chr8-13902639-INS-72 |
| chr8-13904929-INS-1882 | chr8-13904930-INS-1854 |
| chr8-13944208-INS-74 | chr8-13944210-INS-74 |
| chr8-13949992-INS-174 | chr8-13949992-INS-179 |
| chr8-13951813-INS-64 | chr8-13951813-INS-64 |
| chr8-14000795-INS-372 | chr8-14000795-INS-370 |
| chr8-14017509-INS-1280 | chr8-14017510-INS-1280 |
| chr8-14017871-INS-372 | chr8-14017871-INS-372 |
| chr8-14020458-INS-205 | chr8-14020458-INS-208 |
| chr8-14049934-INS-166 | chr8-14049934-INS-166 |
| chr8-14055027-INS-3447 | chr8-14055034-INS-3434 |
| chr8-14059400-INS-202 | chr8-14059400-INS-202 |

| | |
|---|---|
| chr8-14068681-INS-59 | chr8-14068681-INS-61 |
| chr8-14134321-INS-92 | chr8-14134321-INS-72 |
| chr8-14139081-INS-1243 | chr8-14139082-INS-1258 |
| chr8-14156258-INS-53 | chr8-14156258-INS-50 |
| chr8-14159975-INS-707 | chr8-14159975-INS-707 |
| chr8-14165080-INS-64 | chr8-14165080-INS-56 |
| chr8-14168837-INS-92 | chr8-14168837-INS-92 |
| chr8-14231923-INS-549 | chr8-14231923-INS-549 |
| chr8-14296509-INS-7483 | chr8-14296510-INS-7483 |
| chr8-14309742-INS-55 | chr8-14309742-INS-55 |
| chr8-14311821-INS-53 | chr8-14311821-INS-53 |
| chr8-14318947-INS-3012 | chr8-14318948-INS-3012 |
| chr8-14322445-INS-94 | chr8-14322445-INS-94 |
| chr8-14325668-INS-88 | chr8-14325668-INS-88 |
| chr8-14329240-INS-119 | chr8-14329240-INS-126 |
| chr8-14346057-INS-5549 | chr8-14346058-INS-5549 |
| chr8-14357843-INS-113 | chr8-14357843-INS-117 |
| chr8-14367848-INS-2706 | chr8-14367849-INS-2716 |
| chr8-14408856-INS-489 | chr8-14408856-INS-489 |
| chr8-14414723-INS-176 | chr8-14414723-INS-178 |
| chr8-14417143-INS-69 | chr8-14417143-INS-69 |
| chr8-14465672-INS-6443 | chr8-14465673-INS-6443 |
| chr8-14482556-INS-105 | chr8-14482555-INS-105 |
| chr8-14487239-INS-60 | chr8-14487279-INS-60 |
| chr8-14503106-INS-92 | chr8-14503106-INS-90 |
| chr8-14516762-INS-300 | chr8-14516762-INS-300 |
| chr8-14542453-INS-2006 | chr8-14542454-INS-2006 |
| chr8-14543921-INS-125 | chr8-14543921-INS-125 |
| chr8-14555709-INS-78 | chr8-14555709-INS-117 |
| chr8-14617479-INS-195 | chr8-14617468-INS-196 |
| chr8-14641350-INS-64 | chr8-14641350-INS-64 |
| chr8-14650498-INS-342 | chr8-14650498-INS-342 |
| chr8-14659521-INS-66 | chr8-14659526-INS-66 |
| chr8-14683102-INS-228 | chr8-14683102-INS-228 |
| chr8-14693461-INS-361 | chr8-14693461-INS-361 |
| chr8-14706584-INS-168 | chr8-14706584-INS-168 |
| chr8-14783945-INS-64 | chr8-14783943-INS-54 |
| chr8-14808349-INS-55 | chr8-14808349-INS-55 |
| chr8-14811352-INS-53 | chr8-14811352-INS-53 |
| chr8-14850180-INS-58 | chr8-14850180-INS-58 |
| chr8-14869402-INS-199 | chr8-14869402-INS-225 |
| chr8-14870961-INS-154 | chr8-14870961-INS-154 |
| chr8-14887688-INS-56 | chr8-14887688-INS-56 |
| chr8-14908949-INS-406 | chr8-14908949-INS-406 |
| chr8-14925883-INS-358 | chr8-14925883-INS-358 |
| chr8-14934477-INS-6440 | chr8-14934478-INS-6440 |
| chr8-14959459-INS-231 | chr8-14959459-INS-231 |
| chr8-14967162-INS-8945 | chr8-14967163-INS-8945 |
| chr8-14977691-INS-225 | chr8-14977691-INS-219 |
| chr8-15001779-INS-257 | chr8-15001779-INS-260 |
| chr8-15003219-INS-52 | chr8-15003177-INS-52 |
| chr8-15014518-INS-123 | chr8-15014518-INS-123 |
| chr8-15025771-INS-120 | chr8-15025785-INS-96 |
| chr8-15031589-INS-691 | chr8-15031589-INS-685 |
| chr8-15036425-INS-51 | chr8-15036425-INS-51 |
| chr8-15066315-INS-100 | chr8-15066315-INS-105 |
| chr8-15085729-INS-3436 | chr8-15085730-INS-3436 |
| chr8-15091988-INS-84 | chr8-15091988-INS-84 |
| chr8-15108028-INS-139 | chr8-15108014-INS-144 |
| chr8-15110851-INS-56 | chr8-15110851-INS-56 |
| chr8-15114940-INS-89 | chr8-15114940-INS-91 |
| chr8-15118164-INS-68 | chr8-15118164-INS-56 |
| chr8-15121527-INS-93 | chr8-15121527-INS-93 |
| chr8-15148094-INS-66 | chr8-15148094-INS-66 |
| chr8-15156138-INS-247 | chr8-15156138-INS-247 |
| chr8-15169860-INS-59 | chr8-15169860-INS-60 |
| chr8-15176518-INS-65 | chr8-15176518-INS-65 |
| chr8-15196414-INS-325 | chr8-15196414-INS-325 |
| chr8-15197818-INS-403 | chr8-15197818-INS-403 |
| chr8-15222100-INS-242 | chr8-15222100-INS-242 |
| chr8-15235666-INS-843 | chr8-15235666-INS-843 |
| chr8-15250906-INS-256 | chr8-15250906-INS-256 |
| chr8-15262445-INS-1099 | chr8-15262446-INS-1099 |
| chr8-15325563-INS-64 | chr8-15325563-INS-64 |
| chr8-15334225-INS-637 | chr8-15334225-INS-637 |
| chr8-15334359-INS-6471 | chr8-15334360-INS-6472 |
| chr8-15339115-INS-66 | chr8-15339115-INS-85 |

| | |
|---|---|
| chr8-15381139-INS-523 | chr8-15381139-INS-523 |
| chr8-15399061-INS-72 | chr8-15399061-INS-66 |
| chr8-15428089-INS-104 | chr8-15428089-INS-92 |
| chr8-15433493-INS-70 | chr8-15433473-INS-76 |
| chr8-15436830-INS-79 | chr8-15436830-INS-77 |
| chr8-15456412-INS-2144 | chr8-15456413-INS-2144 |
| chr8-15488682-INS-184 | chr8-15488682-INS-184 |
| chr8-15492718-INS-103 | chr8-15492718-INS-103 |
| chr8-15540279-INS-68 | chr8-15540253-INS-57 |
| chr8-15555713-INS-62 | chr8-15555671-INS-60 |
| chr8-15581158-INS-600 | chr8-15581158-INS-600 |
| chr8-15583042-INS-91 | chr8-15583061-INS-71 |
| chr8-15585844-INS-270 | chr8-15585844-INS-278 |
| chr8-15589027-INS-112 | chr8-15589027-INS-108 |
| chr8-15613776-INS-106 | chr8-15613776-INS-106 |
| chr8-15662426-INS-990 | chr8-15662426-INS-990 |
| chr8-15682339-INS-64 | chr8-15682339-INS-64 |
| chr8-15691676-INS-224 | chr8-15691676-INS-224 |
| chr8-15710747-INS-201 | chr8-15710747-INS-201 |
| chr8-15711658-INS-433 | chr8-15711658-INS-433 |
| chr8-15714326-INS-859 | chr8-15714326-INS-859 |
| chr8-15731541-INS-434 | chr8-15731541-INS-434 |
| chr8-15737039-INS-391 | chr8-15737039-INS-391 |
| chr8-15783755-INS-104 | chr8-15783755-INS-104 |
| chr8-15800817-INS-170 | chr8-15800817-INS-170 |
| chr8-15842900-INS-355 | chr8-15842900-INS-358 |
| chr8-15846836-INS-56 | chr8-15846836-INS-56 |
| chr8-15853118-INS-204 | chr8-15853118-INS-204 |
| chr8-15867886-INS-6663 | chr8-15867887-INS-6663 |
| chr8-15878665-INS-58 | chr8-15878666-INS-82 |
| chr8-15885452-INS-133 | chr8-15885452-INS-133 |
| chr8-15895697-INS-155 | chr8-15895697-INS-155 |
| chr8-15896616-INS-434 | chr8-15896616-INS-434 |
| chr8-15901828-INS-216 | chr8-15901832-INS-212 |
| chr8-15928204-INS-224 | chr8-15928204-INS-224 |
| chr8-15930567-INS-6230 | chr8-15930568-INS-6230 |
| chr8-15939749-INS-167 | chr8-15939796-INS-125 |
| chr8-15948290-INS-60 | chr8-15948290-INS-62 |
| chr8-15950921-INS-687 | chr8-15950921-INS-687 |
| chr8-15951457-INS-2715 | chr8-15951458-INS-2715 |
| chr8-15952265-INS-138 | chr8-15952276-INS-137 |
| chr8-15983979-INS-514 | chr8-15983979-INS-514 |
| chr8-16012581-INS-58 | chr8-16012615-INS-50 |
| chr8-16028384-INS-1894 | chr8-16028385-INS-1894 |
| chr8-16028385-INS-401 | chr8-16028385-INS-401 |
| chr8-16034354-INS-667 | chr8-16034354-INS-667 |
| chr8-16039789-INS-62 | chr8-16039789-INS-62 |
| chr8-16047882-INS-80 | chr8-16047882-INS-80 |
| chr8-16054049-INS-82 | chr8-16054049-INS-79 |
| chr8-16055774-INS-2433 | chr8-16055775-INS-2433 |
| chr8-16057442-INS-6418 | chr8-16057443-INS-6418 |
| chr8-16069543-INS-168 | chr8-16069543-INS-168 |
| chr8-16072213-INS-93 | chr8-16072213-INS-93 |
| chr8-16087910-INS-867 | chr8-16087910-INS-867 |
| chr8-16102700-INS-58 | chr8-16102713-INS-58 |
| chr8-16114914-INS-113 | chr8-16114914-INS-116 |
| chr8-16126421-INS-55 | chr8-16126421-INS-55 |
| chr8-16137786-INS-84 | chr8-16137786-INS-72 |
| chr8-16146295-INS-52 | chr8-16146308-INS-59 |
| chr8-16159442-INS-464 | chr8-16159442-INS-464 |
| chr8-16171890-INS-97 | chr8-16171890-INS-97 |
| chr8-16179610-INS-53 | chr8-16179606-INS-53 |
| chr8-16227660-INS-377 | chr8-16227660-INS-377 |
| chr8-16231858-INS-68 | chr8-16231858-INS-76 |
| chr8-16263938-INS-90 | chr8-16263938-INS-80 |
| chr8-16279896-INS-706 | chr8-16279896-INS-706 |
| chr8-16290967-INS-102 | chr8-16290968-INS-96 |
| chr8-16324242-INS-66 | chr8-16324242-INS-66 |
| chr8-16334319-INS-147 | chr8-16334330-INS-157 |
| chr8-16337156-INS-58 | chr8-16337156-INS-58 |
| chr8-16345424-INS-63 | chr8-16345432-INS-59 |
| chr8-16353063-INS-52 | chr8-16353076-INS-72 |
| chr8-16362502-INS-82 | chr8-16362473-INS-125 |
| chr8-16375091-INS-461 | chr8-16375091-INS-461 |
| chr8-16404866-INS-166 | chr8-16404866-INS-166 |
| chr8-16406228-INS-1674 | chr8-16406229-INS-1665 |
| chr8-16432683-INS-236 | chr8-16432683-INS-238 |

| | |
|---|---|
| chr8-16442999-INS-52 | chr8-16442999-INS-54 |
| chr8-16468310-INS-95 | chr8-16468310-INS-95 |
| chr8-16480236-INS-68 | chr8-16480236-INS-68 |
| chr8-16487070-INS-1771 | chr8-16487071-INS-1772 |
| chr8-16497305-INS-1899 | chr8-16497306-INS-1899 |
| chr8-16522749-INS-1074 | chr8-16522762-INS-1075 |
| chr8-16561045-INS-180 | chr8-16561045-INS-180 |
| chr8-16563479-INS-115 | chr8-16563479-INS-115 |
| chr8-16594455-INS-1673 | chr8-16594456-INS-1673 |
| chr8-16637249-INS-186 | chr8-16637249-INS-186 |
| chr8-16653532-INS-1229 | chr8-16653533-INS-1229 |
| chr8-16667323-INS-297 | chr8-16667323-INS-297 |
| chr8-16673078-INS-647 | chr8-16673078-INS-647 |
| chr8-16674303-INS-601 | chr8-16674303-INS-616 |
| chr8-16688052-INS-109 | chr8-16688052-INS-109 |
| chr8-16697726-INS-143 | chr8-16697724-INS-153 |
| chr8-16706133-INS-52 | chr8-16706133-INS-54 |
| chr8-16735164-INS-58 | chr8-16735164-INS-58 |
| chr8-16736388-INS-96 | chr8-16736388-INS-96 |
| chr8-16740638-INS-115 | chr8-16740638-INS-115 |
| chr8-16746481-INS-2041 | chr8-16746482-INS-2041 |
| chr8-16790103-INS-53 | chr8-16790103-INS-53 |
| chr8-16805357-INS-287 | chr8-16805357-INS-287 |
| chr8-16825543-INS-66 | chr8-16825543-INS-66 |
| chr8-16847677-INS-220 | chr8-16847677-INS-220 |
| chr8-16869637-INS-376 | chr8-16869637-INS-334 |
| chr8-16876787-INS-64 | chr8-16876779-INS-76 |
| chr8-16889074-INS-6384 | chr8-16889075-INS-6384 |
| chr8-16923594-INS-56 | chr8-16923594-INS-56 |
| chr8-16941093-INS-96 | chr8-16941093-INS-104 |
| chr8-16981737-INS-138 | chr8-16981737-INS-137 |
| chr8-16994714-INS-89 | chr8-16994742-INS-96 |
| chr8-17000563-INS-64 | chr8-17000563-INS-64 |
| chr8-17019732-INS-249 | chr8-17019732-INS-249 |
| chr8-17024769-INS-4126 | chr8-17024770-INS-4126 |
| chr8-17032989-INS-136 | chr8-17032989-INS-140 |
| chr8-17038890-INS-384 | chr8-17038890-INS-384 |
| chr8-17047539-INS-696 | chr8-17047539-INS-696 |
| chr8-17054923-INS-54 | chr8-17054920-INS-52 |
| chr8-17061514-INS-52 | chr8-17061514-INS-52 |
| chr8-17113764-INS-54 | chr8-17113764-INS-54 |
| chr8-17127433-INS-911 | chr8-17127433-INS-911 |
| chr8-17140473-INS-678 | chr8-17140473-INS-678 |
| chr8-17147724-INS-89 | chr8-17147724-INS-108 |
| chr8-17151406-INS-67 | chr8-17151411-INS-60 |
| chr8-17160181-INS-95 | chr8-17160181-INS-98 |
| chr8-17175697-INS-92 | chr8-17175697-INS-92 |
| chr8-17176713-INS-134 | chr8-17176713-INS-138 |
| chr8-17186708-INS-84 | chr8-17186692-INS-84 |
| chr8-17188774-INS-239 | chr8-17188774-INS-244 |
| chr8-17209062-INS-60 | chr8-17209064-INS-62 |
| chr8-17238195-INS-416 | chr8-17238195-INS-416 |
| chr8-17247460-INS-62 | chr8-17247460-INS-62 |
| chr8-17271301-INS-219 | chr8-17271267-INS-213 |
| chr8-17280522-INS-107 | chr8-17280522-INS-138 |
| chr8-17287008-INS-662 | chr8-17287008-INS-662 |
| chr8-17307217-INS-60 | chr8-17307217-INS-58 |
| chr8-17344719-INS-52 | chr8-17344719-INS-52 |
| chr8-17350714-INS-178 | chr8-17350714-INS-178 |
| chr8-17358433-INS-55 | chr8-17358433-INS-55 |
| chr8-17373195-INS-6452 | chr8-17373196-INS-6453 |
| chr8-17384949-INS-296 | chr8-17384949-INS-296 |
| chr8-17387370-INS-120 | chr8-17387369-INS-147 |
| chr8-17396016-INS-69 | chr8-17396016-INS-69 |
| chr8-17396610-INS-70 | chr8-17396583-INS-58 |
| chr8-17413996-INS-62 | chr8-17413996-INS-65 |
| chr8-17436040-INS-60 | chr8-17436040-INS-65 |
| chr8-17458144-INS-231 | chr8-17458144-INS-231 |
| chr8-17462374-INS-166 | chr8-17462374-INS-166 |
| chr8-17476621-INS-71 | chr8-17476621-INS-94 |
| chr8-17488627-INS-64 | chr8-17488627-INS-56 |
| chr8-17520946-INS-6683 | chr8-17520947-INS-6683 |
| chr8-17527322-INS-156 | chr8-17527322-INS-120 |
| chr8-17530335-INS-199 | chr8-17530335-INS-199 |
| chr8-17570607-INS-53 | chr8-17570607-INS-54 |
| chr8-17576710-INS-144 | chr8-17576710-INS-138 |
| chr8-17614557-INS-320 | chr8-17614557-INS-320 |

| | |
|---|---|
| chr8-17681583-INS-76 | chr8-17681583-INS-76 |
| chr8-17729026-INS-574 | chr8-17729026-INS-574 |
| chr8-17751706-INS-5395 | chr8-17751707-INS-5401 |
| chr8-17787193-INS-429 | chr8-17787193-INS-429 |
| chr8-17791972-INS-71 | chr8-17791972-INS-71 |
| chr8-17809334-INS-55 | chr8-17809342-INS-52 |
| chr8-17821554-INS-60 | chr8-17821554-INS-68 |
| chr8-17850087-INS-175 | chr8-17850087-INS-174 |
| chr8-17857653-INS-89 | chr8-17857653-INS-89 |
| chr8-17858389-INS-313 | chr8-17858389-INS-313 |
| chr8-17858906-INS-4476 | chr8-17858901-INS-4482 |
| chr8-17884093-INS-401 | chr8-17884093-INS-401 |
| chr8-17890225-INS-56 | chr8-17890225-INS-56 |
| chr8-17913889-INS-1050 | chr8-17913890-INS-1050 |
| chr8-17925711-INS-52 | chr8-17925711-INS-52 |
| chr8-17927048-INS-56 | chr8-17927048-INS-56 |
| chr8-17951911-INS-165 | chr8-17951911-INS-160 |
| chr8-17974973-INS-1862 | chr8-17974974-INS-1862 |
| chr8-17989705-INS-135 | chr8-17989705-INS-135 |
| chr8-17995555-INS-571 | chr8-17995555-INS-572 |
| chr8-18002859-INS-4572 | chr8-18002860-INS-4572 |
| chr8-18006341-INS-240 | chr8-18006341-INS-240 |
| chr8-18035132-INS-694 | chr8-18035132-INS-659 |
| chr8-18041749-INS-520 | chr8-18041749-INS-520 |
| chr8-18045083-INS-842 | chr8-18045083-INS-843 |
| chr8-18069266-INS-1768 | chr8-18069267-INS-1768 |
| chr8-18084451-INS-428 | chr8-18084451-INS-428 |
| chr8-18093304-INS-98 | chr8-18093304-INS-106 |
| chr8-18131247-INS-322 | chr8-18131247-INS-323 |
| chr8-18137587-INS-178 | chr8-18137587-INS-178 |
| chr8-18167642-INS-78 | chr8-18167613-INS-117 |
| chr8-18174944-INS-124 | chr8-18174944-INS-124 |
| chr8-18178539-INS-6027 | chr8-18178540-INS-6027 |
| chr8-18186439-INS-79 | chr8-18186440-INS-72 |
| chr8-18194791-INS-52 | chr8-18194801-INS-66 |
| chr8-18215322-INS-52 | chr8-18215322-INS-52 |
| chr8-18216208-INS-142 | chr8-18216208-INS-152 |
| chr8-18216223-INS-987 | chr8-18216223-INS-988 |
| chr8-18220077-INS-64 | chr8-18220063-INS-52 |
| chr8-18241877-INS-181 | chr8-18241877-INS-185 |
| chr8-18244821-INS-1274 | chr8-18244822-INS-1275 |
| chr8-18250709-INS-76 | chr8-18250709-INS-76 |
| chr8-18259760-INS-216 | chr8-18259755-INS-218 |
| chr8-18261072-INS-6772 | chr8-18261073-INS-6772 |
| chr8-18269479-INS-7207 | chr8-18269480-INS-7207 |
| chr8-18276823-INS-314 | chr8-18276823-INS-314 |
| chr8-18293154-INS-199 | chr8-18293154-INS-202 |
| chr8-18354054-INS-70 | chr8-18354054-INS-70 |
| chr8-18367565-INS-52 | chr8-18367541-INS-60 |
| chr8-18380658-INS-84 | chr8-18380658-INS-100 |
| chr8-18386185-INS-96 | chr8-18386185-INS-96 |
| chr8-18395845-INS-174 | chr8-18395845-INS-174 |
| chr8-18421857-INS-69 | chr8-18421857-INS-86 |
| chr8-18428017-INS-444 | chr8-18428017-INS-445 |
| chr8-18440906-INS-134 | chr8-18440900-INS-163 |
| chr8-18443602-INS-256 | chr8-18443602-INS-260 |
| chr8-18444362-INS-416 | chr8-18444362-INS-416 |
| chr8-18460552-INS-263 | chr8-18460552-INS-297 |
| chr8-18465581-INS-6186 | chr8-18465582-INS-6186 |
| chr8-18466210-INS-80 | chr8-18466210-INS-76 |
| chr8-18501838-INS-288 | chr8-18501837-INS-287 |
| chr8-18515684-INS-62 | chr8-18515684-INS-68 |
| chr8-18520788-INS-276 | chr8-18520788-INS-276 |
| chr8-18523054-INS-96 | chr8-18523068-INS-110 |
| chr8-18526881-INS-103 | chr8-18526888-INS-107 |
| chr8-18537783-INS-465 | chr8-18537783-INS-465 |
| chr8-18557447-INS-285 | chr8-18557447-INS-285 |
| chr8-18579221-INS-1442 | chr8-18579222-INS-1442 |
| chr8-18586950-INS-64 | chr8-18586950-INS-64 |
| chr8-18590599-INS-842 | chr8-18590599-INS-832 |
| chr8-18637799-INS-66 | chr8-18637799-INS-66 |
| chr8-18862582-INS-209 | chr8-18862568-INS-205 |
| chr8-18868998-INS-131 | chr8-18868998-INS-133 |
| chr8-18888130-INS-199 | chr8-18888130-INS-199 |
| chr8-18898796-INS-142 | chr8-18898796-INS-142 |
| chr8-18917008-INS-222 | chr8-18917008-INS-223 |
| chr8-18930048-INS-401 | chr8-18930048-INS-401 |

| | |
|---|---|
| chr8-18945328-INS-71 | chr8-18945324-INS-64 |
| chr8-19003171-INS-566 | chr8-19003219-INS-565 |
| chr8-19004822-INS-60 | chr8-19004822-INS-60 |
| chr8-19013651-INS-87 | chr8-19013651-INS-87 |
| chr8-19047857-INS-52 | chr8-19047857-INS-50 |
| chr8-19089586-INS-469 | chr8-19089586-INS-469 |
| chr8-19097068-INS-57 | chr8-19097051-INS-54 |
| chr8-19110124-INS-201 | chr8-19110124-INS-201 |
| chr8-19116531-INS-67 | chr8-19116538-INS-52 |
| chr8-19155473-INS-54 | chr8-19155478-INS-54 |
| chr8-19156136-INS-465 | chr8-19156136-INS-464 |
| chr8-19163209-INS-360 | chr8-19163209-INS-360 |
| chr8-19187574-INS-220 | chr8-19187574-INS-220 |
| chr8-19210036-INS-202 | chr8-19210036-INS-204 |
| chr8-19237843-INS-114 | chr8-19237841-INS-115 |
| chr8-19243108-INS-1388 | chr8-19243109-INS-1388 |
| chr8-19248404-INS-359 | chr8-19248404-INS-359 |
| chr8-19318734-INS-129 | chr8-19318734-INS-129 |
| chr8-19323842-INS-333 | chr8-19323842-INS-330 |
| chr8-19351419-INS-463 | chr8-19351419-INS-463 |
| chr8-19375779-INS-88 | chr8-19375779-INS-90 |
| chr8-19461561-INS-169 | chr8-19461561-INS-168 |
| chr8-19477961-INS-979 | chr8-19477961-INS-988 |
| chr8-19509647-INS-334 | chr8-19509647-INS-334 |
| chr8-19558886-INS-7105 | chr8-19558887-INS-7106 |
| chr8-19568915-INS-108 | chr8-19568915-INS-105 |
| chr8-19594317-INS-127 | chr8-19594317-INS-132 |
| chr8-19628219-INS-69 | chr8-19628178-INS-85 |
| chr8-19652760-INS-187 | chr8-19652768-INS-192 |
| chr8-19671644-INS-169 | chr8-19671629-INS-176 |
| chr8-19711346-INS-60 | chr8-19711353-INS-56 |
| chr8-19720557-INS-66 | chr8-19720557-INS-66 |
| chr8-19725980-INS-72 | chr8-19725977-INS-93 |
| chr8-20522056-INS-2174 | chr8-20522057-INS-2174 |
| chr8-21336626-INS-4766 | chr8-21336626-INS-4759 |
| chr8-21389913-INS-55 | chr8-21389908-INS-90 |
| chr8-21402988-INS-54 | chr8-21402988-INS-52 |
| chr8-21469458-INS-5251 | chr8-21469459-INS-5251 |
| chr8-21501352-INS-1503 | chr8-21501353-INS-1489 |
| chr8-21568613-INS-82 | chr8-21568613-INS-82 |
| chr8-21832077-INS-68 | chr8-21832077-INS-68 |
| chr8-21962743-INS-68 | chr8-21962743-INS-66 |
| chr8-22197872-INS-344 | chr8-22197872-INS-344 |
| chr8-22239171-INS-7482 | chr8-22239172-INS-7482 |
| chr8-22247754-INS-319 | chr8-22247754-INS-319 |
| chr8-22257657-INS-52 | chr8-22257657-INS-52 |
| chr8-22354258-INS-6701 | chr8-22354259-INS-6698 |
| chr8-22389666-INS-150 | chr8-22389666-INS-148 |
| chr8-22394653-INS-54 | chr8-22394653-INS-52 |
| chr8-22405765-INS-478 | chr8-22405765-INS-478 |
| chr8-22432913-INS-3722 | chr8-22432914-INS-3722 |
| chr8-22443494-INS-210 | chr8-22443494-INS-210 |
| chr8-22466804-INS-780 | chr8-22466804-INS-772 |
| chr8-22489908-INS-242 | chr8-22489908-INS-242 |
| chr8-22492294-INS-599 | chr8-22492294-INS-599 |
| chr8-22504114-INS-709 | chr8-22504114-INS-709 |
| chr8-22543186-INS-60 | chr8-22543156-INS-60 |
| chr8-22565986-INS-54 | chr8-22565986-INS-54 |
| chr8-22567531-INS-491 | chr8-22567532-INS-540 |
| chr8-22582342-INS-113 | chr8-22582342-INS-77 |
| chr8-22609126-INS-68 | chr8-22609084-INS-68 |
| chr8-22611415-INS-87 | chr8-22611391-INS-78 |
| chr8-22614496-INS-207 | chr8-22614496-INS-207 |
| chr8-22624639-INS-341 | chr8-22624639-INS-341 |
| chr8-22640868-INS-156 | chr8-22640870-INS-156 |
| chr8-22653711-INS-111 | chr8-22653711-INS-111 |
| chr8-22687151-INS-181 | chr8-22687151-INS-181 |
| chr8-22746852-INS-688 | chr8-22746854-INS-655 |
| chr8-22784856-INS-498 | chr8-22784856-INS-498 |
| chr8-22792901-INS-462 | chr8-22792901-INS-462 |
| chr8-22793869-INS-67 | chr8-22793869-INS-65 |
| chr8-22805270-INS-60 | chr8-22805234-INS-56 |
| chr8-22820994-INS-116 | chr8-22820994-INS-116 |
| chr8-22849169-INS-55 | chr8-22849169-INS-55 |
| chr8-22850788-INS-159 | chr8-22850788-INS-159 |
| chr8-22882330-INS-52 | chr8-22882330-INS-52 |
| chr8-22887688-INS-81 | chr8-22887688-INS-81 |

| | |
|---|---|
| chr8-22891291-INS-200 | chr8-22891291-INS-200 |
| chr8-22929937-INS-1481 | chr8-22929938-INS-1481 |
| chr8-22930507-INS-228 | chr8-22930507-INS-228 |
| chr8-22959948-INS-175 | chr8-22959948-INS-175 |
| chr8-22987060-INS-4293 | chr8-22987061-INS-4293 |
| chr8-22988080-INS-60 | chr8-22988082-INS-62 |
| chr8-23006442-INS-200 | chr8-23006442-INS-201 |
| chr8-23011402-INS-85 | chr8-23011402-INS-55 |
| chr8-23017477-INS-107 | chr8-23017477-INS-107 |
| chr8-23020410-INS-243 | chr8-23020410-INS-246 |
| chr8-23113055-INS-88 | chr8-23113055-INS-91 |
| chr8-23127959-INS-71 | chr8-23127931-INS-72 |
| chr8-23136674-INS-188 | chr8-23136674-INS-188 |
| chr8-23146944-INS-53 | chr8-23146952-INS-54 |
| chr8-23147380-INS-171 | chr8-23147380-INS-170 |
| chr8-23195697-INS-90 | chr8-23195698-INS-117 |
| chr8-23208458-INS-1110 | chr8-23208459-INS-1107 |
| chr8-23225544-INS-4403 | chr8-23225545-INS-4403 |
| chr8-23242729-INS-208 | chr8-23242729-INS-208 |
| chr8-23254366-INS-83 | chr8-23254366-INS-83 |
| chr8-23275257-INS-223 | chr8-23275258-INS-252 |
| chr8-23280308-INS-56 | chr8-23280308-INS-58 |
| chr8-23304118-INS-56 | chr8-23304118-INS-56 |
| chr8-23305170-INS-67 | chr8-23305162-INS-105 |
| chr8-23315286-INS-77 | chr8-23315302-INS-91 |
| chr8-23333644-INS-60 | chr8-23333645-INS-54 |
| chr8-23454226-INS-71 | chr8-23454226-INS-71 |
| chr8-23457713-INS-62 | chr8-23457713-INS-64 |
| chr8-23476694-INS-64 | chr8-23476694-INS-64 |
| chr8-23530263-INS-69 | chr8-23530263-INS-60 |
| chr8-23562458-INS-139 | chr8-23562499-INS-134 |
| chr8-23566620-INS-169 | chr8-23566620-INS-178 |
| chr8-23574154-INS-94 | chr8-23574157-INS-100 |
| chr8-23575979-INS-111 | chr8-23575979-INS-111 |
| chr8-23595074-INS-2752 | chr8-23595075-INS-2752 |
| chr8-23620987-INS-58 | chr8-23620987-INS-70 |
| chr8-23642910-INS-58 | chr8-23642910-INS-52 |
| chr8-23667770-INS-225 | chr8-23667770-INS-225 |
| chr8-23702668-INS-75 | chr8-23702668-INS-75 |
| chr8-23710854-INS-206 | chr8-23710854-INS-210 |
| chr8-23717592-INS-53 | chr8-23717592-INS-55 |
| chr8-23720981-INS-103 | chr8-23720981-INS-103 |
| chr8-23723985-INS-504 | chr8-23723985-INS-504 |
| chr8-23725261-INS-55 | chr8-23725261-INS-55 |
| chr8-23734849-INS-53 | chr8-23734849-INS-53 |
| chr8-23749584-INS-54 | chr8-23749584-INS-54 |
| chr8-23752418-INS-86 | chr8-23752409-INS-68 |
| chr8-23760130-INS-121 | chr8-23760132-INS-122 |
| chr8-23790069-INS-97 | chr8-23790069-INS-96 |
| chr8-23865754-INS-57 | chr8-23865754-INS-59 |
| chr8-23866694-INS-962 | chr8-23866694-INS-961 |
| chr8-23869377-INS-77 | chr8-23869373-INS-69 |
| chr8-23871037-INS-82 | chr8-23871085-INS-50 |
| chr8-23875643-INS-113 | chr8-23875616-INS-108 |
| chr8-23880915-INS-157 | chr8-23880917-INS-185 |
| chr8-23887180-INS-66 | chr8-23887180-INS-57 |
| chr8-23910472-INS-60 | chr8-23910472-INS-60 |
| chr8-23916533-INS-58 | chr8-23916533-INS-58 |
| chr8-23924741-INS-54 | chr8-23924741-INS-51 |
| chr8-23944699-INS-94 | chr8-23944699-INS-90 |
| chr8-23945969-INS-122 | chr8-23945969-INS-125 |
| chr8-23962976-INS-1864 | chr8-23962977-INS-1864 |
| chr8-23998094-INS-109 | chr8-23998094-INS-109 |
| chr8-24010289-INS-1223 | chr8-24010290-INS-1210 |
| chr8-24016267-INS-166 | chr8-24016268-INS-167 |
| chr8-24019098-INS-55 | chr8-24019098-INS-55 |
| chr8-24059773-INS-182 | chr8-24059773-INS-183 |
| chr8-24069639-INS-73 | chr8-24069639-INS-75 |
| chr8-24079706-INS-154 | chr8-24079706-INS-154 |
| chr8-24082915-INS-179 | chr8-24082948-INS-194 |
| chr8-24091761-INS-68 | chr8-24091761-INS-56 |
| chr8-24102046-INS-54 | chr8-24102046-INS-58 |
| chr8-24130290-INS-107 | chr8-24130290-INS-107 |
| chr8-24139971-INS-221 | chr8-24139971-INS-219 |
| chr8-24156155-INS-51 | chr8-24156155-INS-75 |
| chr8-24206106-INS-2915 | chr8-24206107-INS-2915 |
| chr8-24235445-INS-74 | chr8-24235444-INS-73 |

| | |
|---|---|
| chr8-24241388-INS-70 | chr8-24241387-INS-59 |
| chr8-24247016-INS-92 | chr8-24247016-INS-92 |
| chr8-24266211-INS-62 | chr8-24266211-INS-62 |
| chr8-24292461-INS-75 | chr8-24292461-INS-75 |
| chr8-24310654-INS-1044 | chr8-24310655-INS-1043 |
| chr8-24318287-INS-122 | chr8-24318306-INS-107 |
| chr8-24335492-INS-215 | chr8-24335492-INS-215 |
| chr8-24385759-INS-85 | chr8-24385759-INS-69 |
| chr8-24456010-INS-86 | chr8-24456010-INS-86 |
| chr8-24472980-INS-111 | chr8-24472980-INS-83 |
| chr8-24498589-INS-65 | chr8-24498589-INS-55 |
| chr8-24503572-INS-63 | chr8-24503572-INS-63 |
| chr8-24506946-INS-8787 | chr8-24506947-INS-8788 |
| chr8-24539381-INS-118 | chr8-24539357-INS-134 |
| chr8-24568691-INS-128 | chr8-24568691-INS-123 |
| chr8-24646031-INS-413 | chr8-24646031-INS-413 |
| chr8-24694904-INS-52 | chr8-24694904-INS-52 |
| chr8-24702260-INS-1877 | chr8-24702261-INS-1877 |
| chr8-24758641-INS-221 | chr8-24758641-INS-221 |
| chr8-24768176-INS-147 | chr8-24768176-INS-145 |
| chr8-24778599-INS-433 | chr8-24778599-INS-433 |
| chr8-24790165-INS-7117 | chr8-24790166-INS-7117 |
| chr8-24792315-INS-56 | chr8-24792315-INS-56 |
| chr8-24793270-INS-618 | chr8-24793270-INS-598 |
| chr8-24817282-INS-143 | chr8-24817283-INS-143 |
| chr8-24833828-INS-60 | chr8-24833828-INS-53 |
| chr8-24848075-INS-503 | chr8-24848075-INS-504 |
| chr8-24868858-INS-514 | chr8-24868858-INS-514 |
| chr8-24884142-INS-159 | chr8-24884142-INS-157 |
| chr8-24893072-INS-267 | chr8-24893072-INS-267 |
| chr8-24946623-INS-119 | chr8-24946624-INS-83 |
| chr8-24970328-INS-161 | chr8-24970331-INS-120 |
| chr8-24979601-INS-110 | chr8-24979601-INS-110 |
| chr8-24980361-INS-112 | chr8-24980361-INS-112 |
| chr8-24998562-INS-201 | chr8-24998562-INS-203 |
| chr8-25002917-INS-61 | chr8-25002917-INS-62 |
| chr8-25012243-INS-1893 | chr8-25012244-INS-1893 |
| chr8-25013659-INS-105 | chr8-25013659-INS-100 |
| chr8-25033499-INS-117 | chr8-25033498-INS-131 |
| chr8-25052887-INS-56 | chr8-25052883-INS-56 |
| chr8-25058134-INS-346 | chr8-25058134-INS-351 |
| chr8-25061719-INS-55 | chr8-25061689-INS-80 |
| chr8-25076201-INS-54 | chr8-25076196-INS-66 |
| chr8-25084290-INS-102 | chr8-25084291-INS-102 |
| chr8-25091357-INS-431 | chr8-25091357-INS-473 |
| chr8-25094129-INS-97 | chr8-25094129-INS-97 |
| chr8-25128933-INS-200 | chr8-25128933-INS-200 |
| chr8-25142637-INS-55 | chr8-25142673-INS-56 |
| chr8-25144176-INS-60 | chr8-25144176-INS-78 |
| chr8-25160416-INS-115 | chr8-25160416-INS-120 |
| chr8-25169167-INS-57 | chr8-25169167-INS-57 |
| chr8-25176727-INS-883 | chr8-25176734-INS-885 |
| chr8-25181654-INS-70 | chr8-25181668-INS-52 |
| chr8-25204490-INS-315 | chr8-25204491-INS-314 |
| chr8-25218468-INS-680 | chr8-25218468-INS-680 |
| chr8-25221445-INS-55 | chr8-25221406-INS-68 |
| chr8-25315192-INS-51 | chr8-25315147-INS-83 |
| chr8-25329423-INS-86 | chr8-25329423-INS-86 |
| chr8-25364056-INS-505 | chr8-25364056-INS-505 |
| chr8-25366388-INS-58 | chr8-25366388-INS-58 |
| chr8-25386625-INS-180 | chr8-25386625-INS-180 |
| chr8-25388381-INS-195 | chr8-25388381-INS-223 |
| chr8-25402873-INS-56 | chr8-25402873-INS-56 |
| chr8-25408721-INS-60 | chr8-25408721-INS-60 |
| chr8-25414839-INS-128 | chr8-25414833-INS-129 |
| chr8-25417054-INS-185 | chr8-25417054-INS-182 |
| chr8-25424356-INS-54 | chr8-25424344-INS-52 |
| chr8-25432814-INS-213 | chr8-25432814-INS-209 |
| chr8-25433687-INS-74 | chr8-25433659-INS-70 |
| chr8-25448983-INS-203 | chr8-25448983-INS-203 |
| chr8-25467154-INS-66 | chr8-25467154-INS-66 |
| chr8-25478683-INS-565 | chr8-25478683-INS-565 |
| chr8-25534459-INS-284 | chr8-25534459-INS-252 |
| chr8-25554329-INS-97 | chr8-25554329-INS-97 |
| chr8-25578864-INS-171 | chr8-25578864-INS-173 |
| chr8-25612060-INS-96 | chr8-25612060-INS-55 |
| chr8-25636628-INS-89 | chr8-25636628-INS-89 |

| | |
|---|---|
| chr8-25693709-INS-57 | chr8-25693709-INS-57 |
| chr8-25707656-INS-60 | chr8-25707651-INS-81 |
| chr8-25715843-INS-104 | chr8-25715843-INS-104 |
| chr8-25717781-INS-64 | chr8-25717780-INS-71 |
| chr8-25725671-INS-136 | chr8-25725671-INS-136 |
| chr8-25770228-INS-183 | chr8-25770228-INS-156 |
| chr8-25785792-INS-110 | chr8-25785798-INS-108 |
| chr8-25795908-INS-96 | chr8-25795871-INS-112 |
| chr8-25799355-INS-202 | chr8-25799355-INS-202 |
| chr8-25838923-INS-91 | chr8-25838923-INS-96 |
| chr8-25842270-INS-445 | chr8-25842270-INS-441 |
| chr8-25873741-INS-174 | chr8-25873741-INS-203 |
| chr8-25891234-INS-350 | chr8-25891234-INS-350 |
| chr8-25892031-INS-222 | chr8-25892031-INS-222 |
| chr8-25907242-INS-58 | chr8-25907242-INS-68 |
| chr8-25907765-INS-708 | chr8-25907765-INS-709 |
| chr8-25909235-INS-217 | chr8-25909235-INS-219 |
| chr8-25924447-INS-99 | chr8-25924447-INS-99 |
| chr8-25936820-INS-335 | chr8-25936820-INS-338 |
| chr8-25970170-INS-232 | chr8-25970172-INS-206 |
| chr8-26012387-INS-88 | chr8-26012387-INS-60 |
| chr8-26016803-INS-148 | chr8-26016803-INS-148 |
| chr8-26036719-INS-164 | chr8-26036719-INS-164 |
| chr8-26039412-INS-51 | chr8-26039412-INS-51 |
| chr8-26048780-INS-83 | chr8-26048789-INS-87 |
| chr8-26061234-INS-93 | chr8-26061234-INS-114 |
| chr8-26105073-INS-93 | chr8-26105073-INS-93 |
| chr8-26105169-INS-832 | chr8-26105153-INS-858 |
| chr8-26109355-INS-216 | chr8-26109355-INS-217 |
| chr8-26128205-INS-391 | chr8-26128205-INS-391 |
| chr8-26130505-INS-197 | chr8-26130518-INS-228 |
| chr8-26134471-INS-58 | chr8-26134462-INS-78 |
| chr8-26150888-INS-221 | chr8-26150892-INS-198 |
| chr8-26152202-INS-66 | chr8-26152202-INS-60 |
| chr8-26193935-INS-109 | chr8-26193935-INS-107 |
| chr8-26240279-INS-221 | chr8-26240279-INS-221 |
| chr8-26312953-INS-1864 | chr8-26312954-INS-1864 |
| chr8-26334694-INS-93 | chr8-26334694-INS-93 |
| chr8-26338371-INS-66 | chr8-26338340-INS-84 |
| chr8-26358081-INS-102 | chr8-26358081-INS-102 |
| chr8-26403119-INS-652 | chr8-26403119-INS-652 |
| chr8-26404537-INS-65 | chr8-26404537-INS-65 |
| chr8-26440615-INS-86 | chr8-26440612-INS-69 |
| chr8-26450218-INS-199 | chr8-26450218-INS-198 |
| chr8-26469221-INS-168 | chr8-26469221-INS-169 |
| chr8-26501256-INS-273 | chr8-26501256-INS-273 |
| chr8-26503720-INS-250 | chr8-26503720-INS-250 |
| chr8-26504727-INS-68 | chr8-26504727-INS-58 |
| chr8-26506803-INS-83 | chr8-26506853-INS-76 |
| chr8-26510235-INS-204 | chr8-26510235-INS-204 |
| chr8-26516864-INS-54 | chr8-26516864-INS-54 |
| chr8-26518839-INS-169 | chr8-26518839-INS-169 |
| chr8-26523550-INS-74 | chr8-26523550-INS-74 |
| chr8-26540194-INS-129 | chr8-26540195-INS-129 |
| chr8-26548197-INS-70 | chr8-26548197-INS-70 |
| chr8-26549526-INS-195 | chr8-26549526-INS-198 |
| chr8-26551494-INS-85 | chr8-26551494-INS-85 |
| chr8-26557150-INS-226 | chr8-26557150-INS-226 |
| chr8-26580261-INS-1367 | chr8-26580262-INS-1370 |
| chr8-26663023-INS-73 | chr8-26662991-INS-77 |
| chr8-26675240-INS-238 | chr8-26675240-INS-238 |
| chr8-26684658-INS-187 | chr8-26684658-INS-187 |
| chr8-26690878-INS-127 | chr8-26690878-INS-148 |
| chr8-26695118-INS-1480 | chr8-26695119-INS-1480 |
| chr8-26699138-INS-54 | chr8-26699128-INS-61 |
| chr8-26725040-INS-199 | chr8-26725040-INS-223 |
| chr8-26736676-INS-1055 | chr8-26736677-INS-1055 |
| chr8-26737278-INS-6474 | chr8-26737279-INS-6474 |
| chr8-26738296-INS-54 | chr8-26738298-INS-64 |
| chr8-26742775-INS-1371 | chr8-26742776-INS-1372 |
| chr8-26751917-INS-401 | chr8-26751917-INS-401 |
| chr8-26765869-INS-113 | chr8-26765885-INS-104 |
| chr8-26781260-INS-612 | chr8-26781260-INS-612 |
| chr8-26790698-INS-149 | chr8-26790698-INS-149 |
| chr8-26799534-INS-613 | chr8-26799534-INS-613 |
| chr8-26800541-INS-62 | chr8-26800541-INS-62 |
| chr8-26834955-INS-60 | chr8-26834955-INS-60 |

| | |
|---|---|
| chr8-26850574-INS-140 | chr8-26850574-INS-146 |
| chr8-26862093-INS-592 | chr8-26862093-INS-592 |
| chr8-26866386-INS-54 | chr8-26866386-INS-52 |
| chr8-26888670-INS-7259 | chr8-26888671-INS-7259 |
| chr8-26889103-INS-192 | chr8-26889103-INS-192 |
| chr8-26893296-INS-1254 | chr8-26893297-INS-1254 |
| chr8-26898329-INS-54 | chr8-26898329-INS-54 |
| chr8-26908549-INS-6852 | chr8-26908550-INS-6852 |
| chr8-26911534-INS-647 | chr8-26911534-INS-645 |
| chr8-26912486-INS-74 | chr8-26912486-INS-56 |
| chr8-26947223-INS-60 | chr8-26947223-INS-60 |
| chr8-26950480-INS-1265 | chr8-26950481-INS-1265 |
| chr8-26950965-INS-56 | chr8-26950965-INS-56 |
| chr8-26963159-INS-51 | chr8-26963159-INS-51 |
| chr8-26968981-INS-4187 | chr8-26968982-INS-4187 |
| chr8-27000294-INS-160 | chr8-27000279-INS-155 |
| chr8-27011545-INS-129 | chr8-27011528-INS-140 |
| chr8-27045297-INS-1892 | chr8-27045298-INS-1892 |
| chr8-27077586-INS-424 | chr8-27077586-INS-424 |
| chr8-27145537-INS-75 | chr8-27145537-INS-75 |
| chr8-27161488-INS-52 | chr8-27161494-INS-52 |
| chr8-27173215-INS-56 | chr8-27173215-INS-58 |
| chr8-27191736-INS-503 | chr8-27191734-INS-495 |
| chr8-27208228-INS-470 | chr8-27208228-INS-470 |
| chr8-27210382-INS-6804 | chr8-27210383-INS-6804 |
| chr8-27229365-INS-62 | chr8-27229365-INS-62 |
| chr8-27238410-INS-73 | chr8-27238399-INS-90 |
| chr8-27282732-INS-56 | chr8-27282732-INS-101 |
| chr8-27337884-INS-90 | chr8-27337884-INS-90 |
| chr8-27352429-INS-60 | chr8-27352391-INS-68 |
| chr8-27353935-INS-209 | chr8-27353935-INS-209 |
| chr8-27355292-INS-474 | chr8-27355292-INS-475 |
| chr8-27373318-INS-62 | chr8-27373318-INS-64 |
| chr8-27384182-INS-55 | chr8-27384182-INS-53 |
| chr8-27388700-INS-6563 | chr8-27388701-INS-6563 |
| chr8-27390816-INS-56 | chr8-27390816-INS-68 |
| chr8-27393184-INS-133 | chr8-27393184-INS-133 |
| chr8-27421812-INS-118 | chr8-27421802-INS-134 |
| chr8-27437920-INS-138 | chr8-27437920-INS-138 |
| chr8-27440470-INS-93 | chr8-27440470-INS-93 |
| chr8-27476474-INS-110 | chr8-27476474-INS-110 |
| chr8-27491139-INS-70 | chr8-27491165-INS-74 |
| chr8-27538594-INS-74 | chr8-27538595-INS-100 |
| chr8-27563963-INS-694 | chr8-27563964-INS-694 |
| chr8-27582566-INS-98 | chr8-27582566-INS-98 |
| chr8-27602209-INS-211 | chr8-27602209-INS-212 |
| chr8-27610216-INS-121 | chr8-27610230-INS-110 |
| chr8-27620436-INS-52 | chr8-27620436-INS-52 |
| chr8-27635683-INS-114 | chr8-27635683-INS-114 |
| chr8-27645263-INS-171 | chr8-27645214-INS-205 |
| chr8-27654682-INS-78 | chr8-27654642-INS-104 |
| chr8-27659362-INS-217 | chr8-27659362-INS-219 |
| chr8-27666305-INS-58 | chr8-27666305-INS-66 |
| chr8-27677470-INS-318 | chr8-27677470-INS-343 |
| chr8-27682516-INS-88 | chr8-27682516-INS-88 |
| chr8-27684240-INS-208 | chr8-27684240-INS-208 |
| chr8-27687273-INS-218 | chr8-27687273-INS-219 |
| chr8-27742541-INS-63 | chr8-27742541-INS-63 |
| chr8-27758209-INS-178 | chr8-27758209-INS-136 |
| chr8-27761952-INS-139 | chr8-27761952-INS-150 |
| chr8-27784483-INS-61 | chr8-27784475-INS-97 |
| chr8-27804159-INS-53 | chr8-27804159-INS-53 |
| chr8-27818863-INS-51 | chr8-27818901-INS-55 |
| chr8-27820868-INS-215 | chr8-27820868-INS-211 |
| chr8-27863227-INS-89 | chr8-27863227-INS-81 |
| chr8-27866544-INS-70 | chr8-27866544-INS-100 |
| chr8-27906507-INS-61 | chr8-27906507-INS-69 |
| chr8-27910654-INS-69 | chr8-27910654-INS-69 |
| chr8-27914661-INS-460 | chr8-27914661-INS-460 |
| chr8-27924361-INS-7117 | chr8-27924362-INS-7117 |
| chr8-27937193-INS-6881 | chr8-27937194-INS-6894 |
| chr8-27976528-INS-61 | chr8-27976528-INS-62 |
| chr8-28013635-INS-386 | chr8-28013635-INS-400 |
| chr8-28042865-INS-4201 | chr8-28042866-INS-4202 |
| chr8-28086229-INS-126 | chr8-28086229-INS-108 |
| chr8-28116742-INS-199 | chr8-28116770-INS-199 |
| chr8-28124088-INS-180 | chr8-28124111-INS-157 |

| | |
|---|---|
| chr8-28127462-INS-58 | chr8-28127461-INS-58 |
| chr8-28143655-INS-943 | chr8-28143655-INS-942 |
| chr8-28153341-INS-198 | chr8-28153341-INS-198 |
| chr8-28161151-INS-54 | chr8-28161151-INS-57 |
| chr8-28170131-INS-65 | chr8-28170131-INS-61 |
| chr8-28181085-INS-519 | chr8-28181085-INS-520 |
| chr8-28183594-INS-62 | chr8-28183594-INS-58 |
| chr8-28203636-INS-134 | chr8-28203656-INS-151 |
| chr8-28205062-INS-999 | chr8-28205062-INS-999 |
| chr8-28210976-INS-720 | chr8-28210984-INS-728 |
| chr8-28235988-INS-300 | chr8-28235988-INS-301 |
| chr8-28280440-INS-102 | chr8-28280440-INS-102 |
| chr8-28341141-INS-199 | chr8-28341141-INS-201 |
| chr8-28347294-INS-400 | chr8-28347294-INS-399 |
| chr8-28353239-INS-173 | chr8-28353240-INS-173 |
| chr8-28373658-INS-61 | chr8-28373658-INS-61 |
| chr8-28381926-INS-122 | chr8-28381926-INS-129 |
| chr8-28386269-INS-501 | chr8-28386269-INS-502 |
| chr8-28406233-INS-80 | chr8-28406233-INS-81 |
| chr8-28408144-INS-74 | chr8-28408144-INS-76 |
| chr8-28440007-INS-96 | chr8-28439989-INS-82 |
| chr8-28473554-INS-55 | chr8-28473560-INS-52 |
| chr8-28478602-INS-1625 | chr8-28478601-INS-1634 |
| chr8-28482396-INS-155 | chr8-28482396-INS-155 |
| chr8-28494425-INS-459 | chr8-28494425-INS-459 |
| chr8-28533503-INS-218 | chr8-28533504-INS-223 |
| chr8-28536377-INS-233 | chr8-28536377-INS-233 |
| chr8-28573181-INS-101 | chr8-28573210-INS-78 |
| chr8-28584504-INS-52 | chr8-28584504-INS-54 |
| chr8-28599852-INS-209 | chr8-28599852-INS-221 |
| chr8-28643956-INS-246 | chr8-28643956-INS-237 |
| chr8-28667238-INS-98 | chr8-28667238-INS-98 |
| chr8-28677495-INS-52 | chr8-28677495-INS-53 |
| chr8-28706324-INS-242 | chr8-28706324-INS-242 |
| chr8-28718752-INS-53 | chr8-28718753-INS-57 |
| chr8-28778639-INS-83 | chr8-28778639-INS-83 |
| chr8-28782106-INS-937 | chr8-28782106-INS-937 |
| chr8-28783921-INS-68 | chr8-28783921-INS-70 |
| chr8-28788327-INS-115 | chr8-28788327-INS-115 |
| chr8-28803853-INS-116 | chr8-28803853-INS-116 |
| chr8-28811456-INS-6748 | chr8-28811457-INS-6748 |
| chr8-28814381-INS-57 | chr8-28814381-INS-55 |
| chr8-28834530-INS-54 | chr8-28834530-INS-60 |
| chr8-28847333-INS-145 | chr8-28847333-INS-145 |
| chr8-28850345-INS-62 | chr8-28850345-INS-68 |
| chr8-28851904-INS-155 | chr8-28851904-INS-157 |
| chr8-28874078-INS-559 | chr8-28874078-INS-562 |
| chr8-28886667-INS-324 | chr8-28886667-INS-362 |
| chr8-28897825-INS-145 | chr8-28897825-INS-174 |
| chr8-28901925-INS-1327 | chr8-28901926-INS-1329 |
| chr8-28930304-INS-214 | chr8-28930304-INS-214 |
| chr8-28936246-INS-146 | chr8-28936245-INS-160 |
| chr8-28957520-INS-52 | chr8-28957520-INS-54 |
| chr8-29014209-INS-109 | chr8-29014209-INS-109 |
| chr8-29044440-INS-5959 | chr8-29044441-INS-5942 |
| chr8-29090297-INS-85 | chr8-29090308-INS-71 |
| chr8-29095571-INS-66 | chr8-29095571-INS-66 |
| chr8-29097613-INS-164 | chr8-29097613-INS-164 |
| chr8-29118272-INS-400 | chr8-29118267-INS-402 |
| chr8-29134409-INS-647 | chr8-29134409-INS-647 |
| chr8-29148694-INS-147 | chr8-29148694-INS-169 |
| chr8-29153303-INS-76 | chr8-29153303-INS-76 |
| chr8-29187391-INS-69 | chr8-29187391-INS-71 |
| chr8-29205357-INS-531 | chr8-29205357-INS-526 |
| chr8-29255975-INS-60 | chr8-29255975-INS-72 |
| chr8-29270016-INS-68 | chr8-29270016-INS-72 |
| chr8-29279192-INS-313 | chr8-29279192-INS-313 |
| chr8-29323436-INS-68 | chr8-29323445-INS-70 |
| chr8-29340306-INS-70 | chr8-29340306-INS-68 |
| chr8-29340448-INS-514 | chr8-29340448-INS-514 |
| chr8-29346350-INS-498 | chr8-29346350-INS-498 |
| chr8-29362549-INS-507 | chr8-29362549-INS-507 |
| chr8-29363963-INS-412 | chr8-29363963-INS-412 |
| chr8-29422473-INS-178 | chr8-29422473-INS-178 |
| chr8-29480774-INS-198 | chr8-29480774-INS-197 |
| chr8-29503295-INS-283 | chr8-29503295-INS-280 |
| chr8-29506386-INS-70 | chr8-29506384-INS-90 |

| | |
|---|---|
| chr8-29545703-INS-57 | chr8-29545703-INS-58 |
| chr8-29572031-INS-87 | chr8-29572031-INS-87 |
| chr8-29579082-INS-253 | chr8-29579082-INS-253 |
| chr8-29618159-INS-87 | chr8-29618159-INS-52 |
| chr8-29622101-INS-66 | chr8-29622101-INS-86 |
| chr8-29628604-INS-76 | chr8-29628604-INS-76 |
| chr8-29642083-INS-80 | chr8-29642083-INS-80 |
| chr8-29643941-INS-185 | chr8-29643941-INS-185 |
| chr8-29646194-INS-54 | chr8-29646194-INS-54 |
| chr8-29672410-INS-74 | chr8-29672410-INS-74 |
| chr8-29717050-INS-554 | chr8-29717050-INS-554 |
| chr8-29718235-INS-6447 | chr8-29718236-INS-6449 |
| chr8-29720155-INS-202 | chr8-29720155-INS-202 |
| chr8-29721770-INS-59 | chr8-29721770-INS-59 |
| chr8-29725350-INS-152 | chr8-29725351-INS-136 |
| chr8-29728657-INS-808 | chr8-29728657-INS-796 |
| chr8-29734853-INS-54 | chr8-29734853-INS-54 |
| chr8-29742179-INS-57 | chr8-29742186-INS-72 |
| chr8-29761504-INS-131 | chr8-29761504-INS-133 |
| chr8-29773018-INS-57 | chr8-29773018-INS-106 |
| chr8-29788002-INS-317 | chr8-29788002-INS-317 |
| chr8-29793026-INS-72 | chr8-29793026-INS-72 |
| chr8-29804121-INS-272 | chr8-29804121-INS-290 |
| chr8-29824503-INS-60 | chr8-29824507-INS-64 |
| chr8-29869290-INS-236 | chr8-29869290-INS-236 |
| chr8-29878218-INS-1515 | chr8-29878219-INS-1515 |
| chr8-29891116-INS-87 | chr8-29891116-INS-81 |
| chr8-29914296-INS-226 | chr8-29914296-INS-202 |
| chr8-29919997-INS-64 | chr8-29919997-INS-58 |
| chr8-29935684-INS-92 | chr8-29935684-INS-50 |
| chr8-29954355-INS-60 | chr8-29954355-INS-55 |
| chr8-29958955-INS-61 | chr8-29958955-INS-61 |
| chr8-29962317-INS-8596 | chr8-29962318-INS-8596 |
| chr8-29972581-INS-94 | chr8-29972581-INS-55 |
| chr8-29974300-INS-204 | chr8-29974300-INS-204 |
| chr8-30003778-INS-262 | chr8-30003778-INS-262 |
| chr8-30016668-INS-500 | chr8-30016668-INS-536 |
| chr8-30035397-INS-54 | chr8-30035397-INS-58 |
| chr8-30037776-INS-133 | chr8-30037776-INS-133 |
| chr8-30046274-INS-342 | chr8-30046275-INS-342 |
| chr8-30048990-INS-111 | chr8-30048984-INS-109 |
| chr8-30085254-INS-66 | chr8-30085254-INS-70 |
| chr8-30091927-INS-136 | chr8-30091927-INS-136 |
| chr8-30169307-INS-605 | chr8-30169307-INS-605 |
| chr8-30174497-INS-287 | chr8-30174497-INS-287 |
| chr8-30181094-INS-65 | chr8-30181094-INS-63 |
| chr8-30184717-INS-199 | chr8-30184717-INS-199 |
| chr8-30185507-INS-136 | chr8-30185482-INS-138 |
| chr8-30191320-INS-55 | chr8-30191320-INS-55 |
| chr8-30213976-INS-596 | chr8-30213977-INS-596 |
| chr8-30245979-INS-134 | chr8-30245977-INS-135 |
| chr8-30250626-INS-52 | chr8-30250626-INS-56 |
| chr8-30265299-INS-68 | chr8-30265299-INS-68 |
| chr8-30294637-INS-107 | chr8-30294639-INS-151 |
| chr8-30299464-INS-64 | chr8-30299464-INS-84 |
| chr8-30300707-INS-53 | chr8-30300707-INS-53 |
| chr8-30312114-INS-58 | chr8-30312115-INS-60 |
| chr8-30313470-INS-58 | chr8-30313469-INS-58 |
| chr8-30359007-INS-52 | chr8-30359007-INS-88 |
| chr8-30377468-INS-322 | chr8-30377468-INS-322 |
| chr8-30386815-INS-70 | chr8-30386815-INS-70 |
| chr8-30391050-INS-503 | chr8-30391050-INS-503 |
| chr8-30398322-INS-113 | chr8-30398322-INS-113 |
| chr8-30400399-INS-484 | chr8-30400399-INS-484 |
| chr8-30403972-INS-72 | chr8-30403972-INS-76 |
| chr8-30415902-INS-325 | chr8-30415902-INS-325 |
| chr8-30464110-INS-114 | chr8-30464110-INS-114 |
| chr8-30481795-INS-75 | chr8-30481798-INS-75 |
| chr8-30483713-INS-89 | chr8-30483713-INS-89 |
| chr8-30488574-INS-60 | chr8-30488574-INS-50 |
| chr8-30493274-INS-100 | chr8-30493274-INS-100 |
| chr8-30494011-INS-76 | chr8-30494011-INS-81 |
| chr8-30501551-INS-74 | chr8-30501551-INS-74 |
| chr8-30503861-INS-192 | chr8-30503861-INS-201 |
| chr8-30546339-INS-181 | chr8-30546339-INS-182 |
| chr8-30562518-INS-1358 | chr8-30562519-INS-1358 |
| chr8-30595744-INS-110 | chr8-30595744-INS-110 |

| | |
|---|---|
| chr8-30607879-INS-93 | chr8-30607879-INS-93 |
| chr8-30622629-INS-1931 | chr8-30622630-INS-1930 |
| chr8-30665587-INS-794 | chr8-30665587-INS-795 |
| chr8-30666772-INS-92 | chr8-30666771-INS-93 |
| chr8-30669873-INS-654 | chr8-30669873-INS-654 |
| chr8-30670673-INS-956 | chr8-30670673-INS-956 |
| chr8-30673856-INS-146 | chr8-30673856-INS-144 |
| chr8-30707295-INS-129 | chr8-30707295-INS-140 |
| chr8-30709950-INS-7157 | chr8-30709951-INS-7157 |
| chr8-30714307-INS-94 | chr8-30714307-INS-94 |
| chr8-30725732-INS-150 | chr8-30725732-INS-150 |
| chr8-30733869-INS-96 | chr8-30733867-INS-103 |
| chr8-30767663-INS-68 | chr8-30767663-INS-68 |
| chr8-30779124-INS-52 | chr8-30779124-INS-52 |
| chr8-30786499-INS-4600 | chr8-30786500-INS-4600 |
| chr8-30794682-INS-225 | chr8-30794682-INS-225 |
| chr8-30807875-INS-94 | chr8-30807893-INS-86 |
| chr8-30815623-INS-66 | chr8-30815623-INS-66 |
| chr8-30818682-INS-198 | chr8-30818682-INS-198 |
| chr8-30863103-INS-52 | chr8-30863103-INS-54 |
| chr8-30911558-INS-474 | chr8-30911558-INS-474 |
| chr8-30914795-INS-439 | chr8-30914796-INS-440 |
| chr8-30925614-INS-51 | chr8-30925614-INS-55 |
| chr8-30950014-INS-4190 | chr8-30950015-INS-4192 |
| chr8-30958069-INS-706 | chr8-30958069-INS-706 |
| chr8-30979152-INS-54 | chr8-30979152-INS-57 |
| chr8-31013254-INS-125 | chr8-31013254-INS-125 |
| chr8-31014196-INS-6179 | chr8-31014197-INS-6179 |
| chr8-31031832-INS-484 | chr8-31031832-INS-484 |
| chr8-31033576-INS-210 | chr8-31033576-INS-209 |
| chr8-31054676-INS-54 | chr8-31054676-INS-52 |
| chr8-31057682-INS-140 | chr8-31057682-INS-140 |
| chr8-31111422-INS-64 | chr8-31111422-INS-64 |
| chr8-31118401-INS-333 | chr8-31118401-INS-333 |
| chr8-31136325-INS-79 | chr8-31136325-INS-88 |
| chr8-31157646-INS-62 | chr8-31157639-INS-52 |
| chr8-31180081-INS-93 | chr8-31180081-INS-85 |
| chr8-31205254-INS-166 | chr8-31205255-INS-162 |
| chr8-31226858-INS-872 | chr8-31226853-INS-878 |
| chr8-31246543-INS-64 | chr8-31246543-INS-64 |
| chr8-31249869-INS-62 | chr8-31249869-INS-60 |
| chr8-31264182-INS-8753 | chr8-31264183-INS-8753 |
| chr8-31290095-INS-52 | chr8-31290095-INS-52 |
| chr8-31294929-INS-148 | chr8-31294929-INS-148 |
| chr8-31304297-INS-880 | chr8-31304297-INS-880 |
| chr8-31336049-INS-58 | chr8-31336049-INS-58 |
| chr8-31341287-INS-56 | chr8-31341287-INS-58 |
| chr8-31345448-INS-940 | chr8-31345448-INS-940 |
| chr8-31349893-INS-313 | chr8-31349890-INS-316 |
| chr8-31367895-INS-268 | chr8-31367895-INS-268 |
| chr8-31375282-INS-151 | chr8-31375282-INS-151 |
| chr8-31405208-INS-69 | chr8-31405208-INS-75 |
| chr8-31423425-INS-7973 | chr8-31423426-INS-7974 |
| chr8-31447420-INS-8633 | chr8-31447421-INS-8634 |
| chr8-31455303-INS-74 | chr8-31455303-INS-74 |
| chr8-31456921-INS-54 | chr8-31456922-INS-69 |
| chr8-31459205-INS-306 | chr8-31459205-INS-306 |
| chr8-31469277-INS-301 | chr8-31469282-INS-303 |
| chr8-31477355-INS-72 | chr8-31477355-INS-106 |
| chr8-31479889-INS-163 | chr8-31479889-INS-168 |
| chr8-31501702-INS-400 | chr8-31501702-INS-400 |
| chr8-31510453-INS-118 | chr8-31510453-INS-149 |
| chr8-31525295-INS-102 | chr8-31525295-INS-102 |
| chr8-31537083-INS-683 | chr8-31537083-INS-683 |
| chr8-31540960-INS-921 | chr8-31540960-INS-921 |
| chr8-31545576-INS-87 | chr8-31545576-INS-87 |
| chr8-31557531-INS-313 | chr8-31557524-INS-319 |
| chr8-31567895-INS-210 | chr8-31567895-INS-210 |
| chr8-31607048-INS-382 | chr8-31607048-INS-382 |
| chr8-31610024-INS-80 | chr8-31610024-INS-59 |
| chr8-31615604-INS-134 | chr8-31615604-INS-135 |
| chr8-31617503-INS-393 | chr8-31617503-INS-393 |
| chr8-31634501-INS-185 | chr8-31634502-INS-185 |
| chr8-31681343-INS-350 | chr8-31681343-INS-350 |
| chr8-31723718-INS-163 | chr8-31723718-INS-164 |
| chr8-31738380-INS-262 | chr8-31738380-INS-227 |
| chr8-31741066-INS-54 | chr8-31741040-INS-60 |

| | |
|---|---|
| chr8-31743609-INS-378 | chr8-31743609-INS-378 |
| chr8-31750969-INS-84 | chr8-31750969-INS-100 |
| chr8-31752849-INS-265 | chr8-31752849-INS-265 |
| chr8-31755301-INS-288 | chr8-31755301-INS-288 |
| chr8-31771024-INS-59 | chr8-31771025-INS-62 |
| chr8-31787273-INS-54 | chr8-31787253-INS-70 |
| chr8-31794749-INS-71 | chr8-31794749-INS-91 |
| chr8-31800690-INS-191 | chr8-31800690-INS-190 |
| chr8-31810129-INS-7225 | chr8-31810130-INS-7226 |
| chr8-31814438-INS-213 | chr8-31814438-INS-213 |
| chr8-31823556-INS-212 | chr8-31823556-INS-212 |
| chr8-31827150-INS-136 | chr8-31827150-INS-136 |
| chr8-31837222-INS-74 | chr8-31837222-INS-74 |
| chr8-31852547-INS-602 | chr8-31852547-INS-602 |
| chr8-31907484-INS-52 | chr8-31907484-INS-52 |
| chr8-31918205-INS-534 | chr8-31918205-INS-532 |
| chr8-31937271-INS-242 | chr8-31937278-INS-224 |
| chr8-31953825-INS-56 | chr8-31953825-INS-60 |
| chr8-31958744-INS-52 | chr8-31958744-INS-54 |
| chr8-31962045-INS-124 | chr8-31962045-INS-124 |
| chr8-31966110-INS-152 | chr8-31966110-INS-152 |
| chr8-31971258-INS-51 | chr8-31971259-INS-78 |
| chr8-31978666-INS-168 | chr8-31978666-INS-168 |
| chr8-31988787-INS-542 | chr8-31988787-INS-543 |
| chr8-32007811-INS-57 | chr8-32007811-INS-57 |
| chr8-32049945-INS-1133 | chr8-32049946-INS-1133 |
| chr8-32074927-INS-561 | chr8-32074927-INS-563 |
| chr8-32085197-INS-174 | chr8-32085197-INS-185 |
| chr8-32123476-INS-201 | chr8-32123476-INS-201 |
| chr8-32157777-INS-122 | chr8-32157777-INS-122 |
| chr8-32182619-INS-67 | chr8-32182619-INS-72 |
| chr8-32199395-INS-52 | chr8-32199395-INS-50 |
| chr8-32209580-INS-69 | chr8-32209581-INS-70 |
| chr8-32225488-INS-89 | chr8-32225488-INS-77 |
| chr8-32262127-INS-648 | chr8-32262127-INS-653 |
| chr8-32269598-INS-472 | chr8-32269598-INS-476 |
| chr8-32299246-INS-1611 | chr8-32299247-INS-1612 |
| chr8-32329843-INS-119 | chr8-32329843-INS-119 |
| chr8-32375286-INS-415 | chr8-32375286-INS-415 |
| chr8-32389377-INS-56 | chr8-32389377-INS-50 |
| chr8-32395742-INS-6504 | chr8-32395743-INS-6506 |
| chr8-32416433-INS-119 | chr8-32416434-INS-125 |
| chr8-32440394-INS-69 | chr8-32440394-INS-73 |
| chr8-32455939-INS-63 | chr8-32455940-INS-59 |
| chr8-32475506-INS-548 | chr8-32475506-INS-549 |
| chr8-32488268-INS-56 | chr8-32488268-INS-56 |
| chr8-32526257-INS-113 | chr8-32526240-INS-98 |
| chr8-32541828-INS-922 | chr8-32541829-INS-922 |
| chr8-32552225-INS-60 | chr8-32552225-INS-60 |
| chr8-32560252-INS-108 | chr8-32560249-INS-111 |
| chr8-32566291-INS-249 | chr8-32566291-INS-229 |
| chr8-32569196-INS-243 | chr8-32569196-INS-243 |
| chr8-32578154-INS-52 | chr8-32578154-INS-52 |
| chr8-32607299-INS-130 | chr8-32607299-INS-130 |
| chr8-32633946-INS-59 | chr8-32633946-INS-59 |
| chr8-32638208-INS-74 | chr8-32638214-INS-70 |
| chr8-32651334-INS-118 | chr8-32651334-INS-118 |
| chr8-32656980-INS-61 | chr8-32656980-INS-61 |
| chr8-32663481-INS-565 | chr8-32663481-INS-565 |
| chr8-32664042-INS-112 | chr8-32664042-INS-112 |
| chr8-32672127-INS-242 | chr8-32672127-INS-242 |
| chr8-32696028-INS-188 | chr8-32696028-INS-195 |
| chr8-32705267-INS-58 | chr8-32705230-INS-57 |
| chr8-32730237-INS-555 | chr8-32730237-INS-550 |
| chr8-32737508-INS-114 | chr8-32737508-INS-114 |
| chr8-32758330-INS-413 | chr8-32758330-INS-413 |
| chr8-32759900-INS-60 | chr8-32759900-INS-61 |
| chr8-32773031-INS-59 | chr8-32773031-INS-59 |
| chr8-32785065-INS-72 | chr8-32785065-INS-72 |
| chr8-32791719-INS-458 | chr8-32791720-INS-457 |
| chr8-32803713-INS-52 | chr8-32803713-INS-50 |
| chr8-32828015-INS-59 | chr8-32828017-INS-76 |
| chr8-32838767-INS-175 | chr8-32838767-INS-127 |
| chr8-32843211-INS-2785 | chr8-32843212-INS-2785 |
| chr8-32855666-INS-499 | chr8-32855666-INS-499 |
| chr8-32870304-INS-247 | chr8-32870305-INS-243 |
| chr8-32883021-INS-176 | chr8-32883061-INS-135 |

| | |
|---|---|
| chr8-32909969-INS-205 | chr8-32909969-INS-205 |
| chr8-32923124-INS-1474 | chr8-32923125-INS-1474 |
| chr8-32926147-INS-2084 | chr8-32926156-INS-2071 |
| chr8-32940703-INS-756 | chr8-32940703-INS-756 |
| chr8-32950308-INS-76 | chr8-32950308-INS-76 |
| chr8-32951297-INS-390 | chr8-32951297-INS-390 |
| chr8-32970316-INS-545 | chr8-32970316-INS-546 |
| chr8-32985088-INS-89 | chr8-32985088-INS-50 |
| chr8-32987598-INS-76 | chr8-32987598-INS-80 |
| chr8-33074334-INS-67 | chr8-33074334-INS-73 |
| chr8-33082629-INS-605 | chr8-33082629-INS-605 |
| chr8-33102827-INS-1197 | chr8-33102828-INS-1194 |
| chr8-33106396-INS-74 | chr8-33106396-INS-74 |
| chr8-33139132-INS-709 | chr8-33139132-INS-710 |
| chr8-33142579-INS-205 | chr8-33142579-INS-205 |
| chr8-33198633-INS-56 | chr8-33198633-INS-58 |
| chr8-33215132-INS-67 | chr8-33215132-INS-67 |
| chr8-33220735-INS-248 | chr8-33220735-INS-248 |
| chr8-33245576-INS-302 | chr8-33245596-INS-329 |
| chr8-33249578-INS-53 | chr8-33249578-INS-53 |
| chr8-33269388-INS-68 | chr8-33269388-INS-68 |
| chr8-33273212-INS-184 | chr8-33273212-INS-185 |
| chr8-33282281-INS-58 | chr8-33282281-INS-58 |
| chr8-33284166-INS-52 | chr8-33284166-INS-52 |
| chr8-33291449-INS-711 | chr8-33291450-INS-711 |
| chr8-33382364-INS-53 | chr8-33382364-INS-53 |
| chr8-33391067-INS-184 | chr8-33391067-INS-178 |
| chr8-33394075-INS-85 | chr8-33394075-INS-65 |
| chr8-33414650-INS-619 | chr8-33414650-INS-637 |
| chr8-33447769-INS-58 | chr8-33447769-INS-60 |
| chr8-33456306-INS-866 | chr8-33456306-INS-866 |
| chr8-33466660-INS-6193 | chr8-33466661-INS-6193 |
| chr8-33475536-INS-780 | chr8-33475536-INS-780 |
| chr8-33504955-INS-132 | chr8-33504955-INS-135 |
| chr8-33505639-INS-441 | chr8-33505639-INS-441 |
| chr8-33509845-INS-6323 | chr8-33509846-INS-6323 |
| chr8-33510194-INS-462 | chr8-33510194-INS-462 |
| chr8-33554713-INS-113 | chr8-33554714-INS-107 |
| chr8-33569572-INS-120 | chr8-33569572-INS-116 |
| chr8-33582640-INS-129 | chr8-33582640-INS-129 |
| chr8-33637210-INS-82 | chr8-33637210-INS-50 |
| chr8-33675874-INS-345 | chr8-33675874-INS-345 |
| chr8-33684196-INS-262 | chr8-33684203-INS-272 |
| chr8-33687589-INS-206 | chr8-33687589-INS-206 |
| chr8-33701295-INS-351 | chr8-33701295-INS-351 |
| chr8-33730112-INS-205 | chr8-33730112-INS-205 |
| chr8-33751882-INS-518 | chr8-33751882-INS-518 |
| chr8-33779318-INS-201 | chr8-33779318-INS-201 |
| chr8-33787981-INS-2841 | chr8-33787972-INS-2833 |
| chr8-33798109-INS-56 | chr8-33798109-INS-61 |
| chr8-33812039-INS-991 | chr8-33812039-INS-993 |
| chr8-33832907-INS-6389 | chr8-33832908-INS-6387 |
| chr8-33835670-INS-54 | chr8-33835670-INS-54 |
| chr8-33839039-INS-400 | chr8-33839039-INS-400 |
| chr8-33841302-INS-128 | chr8-33841302-INS-128 |
| chr8-33888796-INS-206 | chr8-33888796-INS-204 |
| chr8-33910433-INS-432 | chr8-33910433-INS-432 |
| chr8-33934597-INS-145 | chr8-33934597-INS-173 |
| chr8-34008356-INS-161 | chr8-34008356-INS-163 |
| chr8-34021808-INS-114 | chr8-34021808-INS-116 |
| chr8-34068514-INS-60 | chr8-34068496-INS-50 |
| chr8-34077973-INS-3230 | chr8-34077974-INS-3230 |
| chr8-34086781-INS-198 | chr8-34086781-INS-198 |
| chr8-34089066-INS-110 | chr8-34089060-INS-118 |
| chr8-34110054-INS-81 | chr8-34110054-INS-81 |
| chr8-34128904-INS-212 | chr8-34128904-INS-212 |
| chr8-34130570-INS-68 | chr8-34130560-INS-55 |
| chr8-34136620-INS-210 | chr8-34136620-INS-210 |
| chr8-34137751-INS-187 | chr8-34137742-INS-192 |
| chr8-34141782-INS-110 | chr8-34141782-INS-110 |
| chr8-34182409-INS-210 | chr8-34182409-INS-210 |
| chr8-34190503-INS-88 | chr8-34190507-INS-72 |
| chr8-34194430-INS-138 | chr8-34194434-INS-146 |
| chr8-34211126-INS-212 | chr8-34211126-INS-212 |
| chr8-34218335-INS-214 | chr8-34218335-INS-214 |
| chr8-34223926-INS-245 | chr8-34223926-INS-245 |
| chr8-34248053-INS-57 | chr8-34248053-INS-57 |

| | |
|---|---|
| chr8-34253092-INS-127 | chr8-34253092-INS-125 |
| chr8-34267959-INS-210 | chr8-34267959-INS-210 |
| chr8-34271539-INS-176 | chr8-34271539-INS-176 |
| chr8-34299243-INS-225 | chr8-34299243-INS-225 |
| chr8-34304455-INS-57 | chr8-34304467-INS-54 |
| chr8-34310878-INS-105 | chr8-34310878-INS-99 |
| chr8-34336945-INS-104 | chr8-34336945-INS-110 |
| chr8-34339114-INS-203 | chr8-34339114-INS-203 |
| chr8-34352557-INS-1144 | chr8-34352558-INS-1148 |
| chr8-34373870-INS-203 | chr8-34373870-INS-203 |
| chr8-34383754-INS-2125 | chr8-34383755-INS-2125 |
| chr8-34402965-INS-189 | chr8-34402961-INS-177 |
| chr8-34411274-INS-178 | chr8-34411274-INS-178 |
| chr8-34424500-INS-229 | chr8-34424500-INS-230 |
| chr8-34427117-INS-230 | chr8-34427117-INS-230 |
| chr8-34430591-INS-229 | chr8-34430591-INS-229 |
| chr8-34439367-INS-72 | chr8-34439389-INS-84 |
| chr8-34443695-INS-211 | chr8-34443695-INS-211 |
| chr8-34448270-INS-52 | chr8-34448269-INS-50 |
| chr8-34472319-INS-75 | chr8-34472319-INS-75 |
| chr8-34490436-INS-192 | chr8-34490436-INS-192 |
| chr8-34492085-INS-247 | chr8-34492086-INS-247 |
| chr8-34507913-INS-150 | chr8-34507913-INS-150 |
| chr8-34512408-INS-252 | chr8-34512408-INS-253 |
| chr8-34517787-INS-200 | chr8-34517787-INS-201 |
| chr8-34520266-INS-65 | chr8-34520266-INS-65 |
| chr8-34524435-INS-58 | chr8-34524410-INS-84 |
| chr8-34527304-INS-103 | chr8-34527304-INS-104 |
| chr8-34537982-INS-202 | chr8-34537983-INS-202 |
| chr8-34548771-INS-203 | chr8-34548757-INS-215 |
| chr8-34558046-INS-54 | chr8-34558047-INS-60 |
| chr8-34577799-INS-111 | chr8-34577799-INS-110 |
| chr8-34579451-INS-202 | chr8-34579451-INS-201 |
| chr8-34581184-INS-451 | chr8-34581184-INS-451 |
| chr8-34582951-INS-212 | chr8-34582951-INS-212 |
| chr8-34584281-INS-56 | chr8-34584281-INS-56 |
| chr8-34585330-INS-240 | chr8-34585330-INS-240 |
| chr8-34590366-INS-65 | chr8-34590366-INS-53 |
| chr8-34593060-INS-325 | chr8-34593060-INS-325 |
| chr8-34593656-INS-235 | chr8-34593656-INS-236 |
| chr8-34598664-INS-88 | chr8-34598664-INS-90 |
| chr8-34606855-INS-65 | chr8-34606855-INS-65 |
| chr8-34618665-INS-56 | chr8-34618666-INS-54 |
| chr8-34620417-INS-902 | chr8-34620417-INS-902 |
| chr8-34630260-INS-7116 | chr8-34630261-INS-7116 |
| chr8-34637775-INS-128 | chr8-34637775-INS-128 |
| chr8-34638497-INS-73 | chr8-34638497-INS-77 |
| chr8-34643335-INS-197 | chr8-34643335-INS-197 |
| chr8-34653731-INS-207 | chr8-34653731-INS-208 |
| chr8-34655446-INS-57 | chr8-34655446-INS-55 |
| chr8-34657882-INS-320 | chr8-34657882-INS-320 |
| chr8-34664782-INS-962 | chr8-34664782-INS-962 |
| chr8-34671212-INS-178 | chr8-34671212-INS-178 |
| chr8-34722029-INS-52 | chr8-34722019-INS-66 |
| chr8-34726676-INS-72 | chr8-34726676-INS-76 |
| chr8-34729631-INS-108 | chr8-34729631-INS-108 |
| chr8-34773953-INS-67 | chr8-34773953-INS-65 |
| chr8-34786372-INS-54 | chr8-34786372-INS-51 |
| chr8-34789403-INS-348 | chr8-34789403-INS-356 |
| chr8-34795484-INS-51 | chr8-34795484-INS-51 |
| chr8-34816341-INS-383 | chr8-34816341-INS-383 |
| chr8-34840945-INS-95 | chr8-34840945-INS-95 |
| chr8-34846519-INS-138 | chr8-34846519-INS-138 |
| chr8-34894382-INS-152 | chr8-34894382-INS-152 |
| chr8-34908772-INS-153 | chr8-34908809-INS-120 |
| chr8-34940198-INS-100 | chr8-34940198-INS-102 |
| chr8-34951050-INS-376 | chr8-34951050-INS-376 |
| chr8-34958236-INS-81 | chr8-34958236-INS-81 |
| chr8-34959358-INS-52 | chr8-34959358-INS-54 |
| chr8-34960587-INS-71 | chr8-34960587-INS-71 |
| chr8-34981177-INS-107 | chr8-34981181-INS-111 |
| chr8-34985485-INS-51 | chr8-34985486-INS-51 |
| chr8-34996542-INS-63 | chr8-34996542-INS-57 |
| chr8-34997890-INS-60 | chr8-34997890-INS-60 |
| chr8-35026624-INS-54 | chr8-35026624-INS-58 |
| chr8-35030899-INS-375 | chr8-35030899-INS-375 |
| chr8-35068533-INS-56 | chr8-35068537-INS-52 |

| | |
|---|---|
| chr8-35097922-INS-172 | chr8-35097922-INS-172 |
| chr8-35108948-INS-57 | chr8-35108952-INS-69 |
| chr8-35115573-INS-100 | chr8-35115573-INS-100 |
| chr8-35146674-INS-55 | chr8-35146644-INS-70 |
| chr8-35163978-INS-76 | chr8-35163978-INS-76 |
| chr8-35188391-INS-307 | chr8-35188391-INS-307 |
| chr8-35203954-INS-921 | chr8-35203954-INS-913 |
| chr8-35209884-INS-112 | chr8-35209884-INS-112 |
| chr8-35235223-INS-311 | chr8-35235223-INS-311 |
| chr8-35269017-INS-255 | chr8-35269017-INS-255 |
| chr8-35290980-INS-66 | chr8-35290980-INS-66 |
| chr8-35310922-INS-57 | chr8-35310922-INS-62 |
| chr8-35315384-INS-69 | chr8-35315385-INS-69 |
| chr8-35375371-INS-1072 | chr8-35375372-INS-1072 |
| chr8-35393798-INS-592 | chr8-35393798-INS-592 |
| chr8-35405538-INS-197 | chr8-35405538-INS-196 |
| chr8-35409674-INS-59 | chr8-35409674-INS-59 |
| chr8-35434266-INS-1979 | chr8-35434267-INS-1983 |
| chr8-35449813-INS-3609 | chr8-35449814-INS-3609 |
| chr8-35457895-INS-431 | chr8-35457895-INS-431 |
| chr8-35512983-INS-191 | chr8-35512983-INS-198 |
| chr8-35518452-INS-359 | chr8-35518452-INS-363 |
| chr8-35519154-INS-58 | chr8-35519155-INS-62 |
| chr8-35525862-INS-104 | chr8-35525862-INS-104 |
| chr8-35539467-INS-1503 | chr8-35539468-INS-1512 |
| chr8-35556193-INS-6960 | chr8-35556194-INS-6960 |
| chr8-35581951-INS-63 | chr8-35581951-INS-63 |
| chr8-35587868-INS-207 | chr8-35587868-INS-207 |
| chr8-35596819-INS-533 | chr8-35596819-INS-533 |
| chr8-35611571-INS-976 | chr8-35611571-INS-976 |
| chr8-35618513-INS-64 | chr8-35618513-INS-64 |
| chr8-35630825-INS-52 | chr8-35630825-INS-52 |
| chr8-35635738-INS-68 | chr8-35635738-INS-63 |
| chr8-35646039-INS-52 | chr8-35646039-INS-52 |
| chr8-35653166-INS-62 | chr8-35653166-INS-62 |
| chr8-35725404-INS-162 | chr8-35725404-INS-164 |
| chr8-35731934-INS-925 | chr8-35731934-INS-925 |
| chr8-35745253-INS-344 | chr8-35745253-INS-344 |
| chr8-35752945-INS-2760 | chr8-35752946-INS-2760 |
| chr8-35753634-INS-469 | chr8-35753634-INS-469 |
| chr8-35754941-INS-246 | chr8-35754945-INS-249 |
| chr8-35762388-INS-839 | chr8-35762388-INS-861 |
| chr8-35769706-INS-52 | chr8-35769706-INS-53 |
| chr8-35774170-INS-435 | chr8-35774170-INS-435 |
| chr8-35777715-INS-116 | chr8-35777718-INS-122 |
| chr8-35783969-INS-569 | chr8-35783969-INS-569 |
| chr8-35793462-INS-75 | chr8-35793462-INS-75 |
| chr8-35803278-INS-108 | chr8-35803278-INS-99 |
| chr8-35807321-INS-92 | chr8-35807321-INS-94 |
| chr8-35813539-INS-100 | chr8-35813539-INS-52 |
| chr8-35822992-INS-54 | chr8-35822992-INS-56 |
| chr8-35846722-INS-145 | chr8-35846748-INS-141 |
| chr8-35850672-INS-180 | chr8-35850672-INS-195 |
| chr8-35857102-INS-268 | chr8-35857102-INS-268 |
| chr8-35888092-INS-150 | chr8-35888092-INS-150 |
| chr8-35894607-INS-3599 | chr8-35894608-INS-3599 |
| chr8-35897298-INS-244 | chr8-35897288-INS-252 |
| chr8-35910105-INS-84 | chr8-35910105-INS-84 |
| chr8-35916703-INS-267 | chr8-35916703-INS-267 |
| chr8-35920958-INS-172 | chr8-35920958-INS-172 |
| chr8-35926052-INS-54 | chr8-35926052-INS-54 |
| chr8-35927039-INS-4581 | chr8-35927040-INS-4581 |
| chr8-35977423-INS-54 | chr8-35977423-INS-54 |
| chr8-36005344-INS-1038 | chr8-36005345-INS-1038 |
| chr8-36008067-INS-7160 | chr8-36008068-INS-7160 |
| chr8-36018018-INS-51 | chr8-36018018-INS-51 |
| chr8-36027531-INS-50 | chr8-36027531-INS-56 |
| chr8-36034065-INS-84 | chr8-36034066-INS-106 |
| chr8-36052343-INS-56 | chr8-36052343-INS-56 |
| chr8-36071372-INS-58 | chr8-36071372-INS-58 |
| chr8-36078680-INS-216 | chr8-36078680-INS-215 |
| chr8-36086089-INS-57 | chr8-36086089-INS-57 |
| chr8-36124055-INS-66 | chr8-36124055-INS-66 |
| chr8-36132843-INS-52 | chr8-36132843-INS-58 |
| chr8-36145482-INS-60 | chr8-36145482-INS-62 |
| chr8-36147530-INS-246 | chr8-36147530-INS-246 |
| chr8-36176816-INS-96 | chr8-36176791-INS-78 |

| | |
|---|---|
| chr8-36178544-INS-407 | chr8-36178544-INS-407 |
| chr8-36189080-INS-185 | chr8-36189080-INS-185 |
| chr8-36197117-INS-198 | chr8-36197117-INS-198 |
| chr8-36197651-INS-125 | chr8-36197651-INS-125 |
| chr8-36216181-INS-137 | chr8-36216181-INS-127 |
| chr8-36218070-INS-66 | chr8-36218070-INS-66 |
| chr8-36234905-INS-59 | chr8-36234905-INS-59 |
| chr8-36284182-INS-58 | chr8-36284145-INS-56 |
| chr8-36288841-INS-85 | chr8-36288841-INS-85 |
| chr8-36295524-INS-153 | chr8-36295560-INS-156 |
| chr8-36308074-INS-78 | chr8-36308074-INS-78 |
| chr8-36313394-INS-144 | chr8-36313394-INS-144 |
| chr8-36320164-INS-1183 | chr8-36320165-INS-1183 |
| chr8-36334395-INS-319 | chr8-36334395-INS-319 |
| chr8-36348060-INS-82 | chr8-36348060-INS-82 |
| chr8-36363840-INS-71 | chr8-36363840-INS-81 |
| chr8-36371949-INS-74 | chr8-36371957-INS-66 |
| chr8-36403054-INS-89 | chr8-36403054-INS-109 |
| chr8-36408500-INS-557 | chr8-36408498-INS-554 |
| chr8-36425382-INS-84 | chr8-36425382-INS-84 |
| chr8-36432974-INS-871 | chr8-36432974-INS-871 |
| chr8-36433168-INS-77 | chr8-36433214-INS-78 |
| chr8-36436516-INS-54 | chr8-36436516-INS-60 |
| chr8-36437152-INS-59 | chr8-36437152-INS-64 |
| chr8-36442773-INS-3347 | chr8-36442774-INS-3347 |
| chr8-36450017-INS-6379 | chr8-36450018-INS-6379 |
| chr8-36474132-INS-112 | chr8-36474132-INS-112 |
| chr8-36495131-INS-54 | chr8-36495128-INS-57 |
| chr8-36527946-INS-91 | chr8-36527946-INS-98 |
| chr8-36601162-INS-2795 | chr8-36601163-INS-2795 |
| chr8-36635030-INS-70 | chr8-36635030-INS-72 |
| chr8-36656881-INS-90 | chr8-36656897-INS-50 |
| chr8-36671838-INS-119 | chr8-36671855-INS-132 |
| chr8-36689040-INS-56 | chr8-36689040-INS-58 |
| chr8-36690190-INS-189 | chr8-36690190-INS-189 |
| chr8-36720741-INS-82 | chr8-36720741-INS-84 |
| chr8-36721874-INS-57 | chr8-36721874-INS-58 |
| chr8-36835653-INS-272 | chr8-36835653-INS-272 |
| chr8-36841384-INS-687 | chr8-36841384-INS-688 |
| chr8-36846962-INS-81 | chr8-36846962-INS-81 |
| chr8-36863922-INS-66 | chr8-36863943-INS-88 |
| chr8-36867876-INS-55 | chr8-36867876-INS-55 |
| chr8-36871304-INS-457 | chr8-36871304-INS-456 |
| chr8-36890301-INS-234 | chr8-36890301-INS-234 |
| chr8-36891814-INS-844 | chr8-36891814-INS-844 |
| chr8-36893677-INS-160 | chr8-36893677-INS-160 |
| chr8-36902750-INS-57 | chr8-36902756-INS-70 |
| chr8-36921216-INS-51 | chr8-36921216-INS-51 |
| chr8-36926626-INS-116 | chr8-36926626-INS-116 |
| chr8-36937982-INS-211 | chr8-36937982-INS-211 |
| chr8-36953876-INS-193 | chr8-36953876-INS-193 |
| chr8-36960641-INS-71 | chr8-36960641-INS-72 |
| chr8-36988150-INS-87 | chr8-36988150-INS-87 |
| chr8-36993992-INS-72 | chr8-36993992-INS-76 |
| chr8-36997550-INS-965 | chr8-36997550-INS-954 |
| chr8-36999590-INS-52 | chr8-36999590-INS-56 |
| chr8-37023152-INS-360 | chr8-37023152-INS-360 |
| chr8-37031658-INS-74 | chr8-37031621-INS-78 |
| chr8-37065314-INS-281 | chr8-37065314-INS-284 |
| chr8-37068298-INS-75 | chr8-37068297-INS-75 |
| chr8-37103263-INS-67 | chr8-37103263-INS-55 |
| chr8-37106214-INS-174 | chr8-37106214-INS-174 |
| chr8-37157261-INS-98 | chr8-37157259-INS-102 |
| chr8-37167086-INS-668 | chr8-37167086-INS-668 |
| chr8-37167701-INS-100 | chr8-37167701-INS-126 |
| chr8-37176743-INS-182 | chr8-37176752-INS-191 |
| chr8-37191720-INS-112 | chr8-37191722-INS-118 |
| chr8-37222056-INS-60 | chr8-37222042-INS-56 |
| chr8-37275153-INS-1859 | chr8-37275154-INS-1859 |
| chr8-37290338-INS-53 | chr8-37290338-INS-53 |
| chr8-37307782-INS-116 | chr8-37307782-INS-83 |
| chr8-37310393-INS-52 | chr8-37310393-INS-50 |
| chr8-37326648-INS-79 | chr8-37326648-INS-55 |
| chr8-37334385-INS-85 | chr8-37334385-INS-85 |
| chr8-37347974-INS-116 | chr8-37347974-INS-128 |
| chr8-37372481-INS-498 | chr8-37372481-INS-497 |
| chr8-37402030-INS-92 | chr8-37402030-INS-90 |

| | |
|---|---|
| chr8-37487392-INS-64 | chr8-37487392-INS-112 |
| chr8-37518694-INS-970 | chr8-37518694-INS-970 |
| chr8-37521215-INS-184 | chr8-37521215-INS-184 |
| chr8-37526552-INS-106 | chr8-37526552-INS-112 |
| chr8-37531902-INS-97 | chr8-37531902-INS-54 |
| chr8-37560229-INS-58 | chr8-37560240-INS-64 |
| chr8-37573648-INS-400 | chr8-37573649-INS-400 |
| chr8-37576277-INS-70 | chr8-37576277-INS-70 |
| chr8-37592671-INS-64 | chr8-37592671-INS-57 |
| chr8-37599929-INS-7095 | chr8-37599930-INS-7095 |
| chr8-37604439-INS-123 | chr8-37604439-INS-126 |
| chr8-37648599-INS-239 | chr8-37648599-INS-239 |
| chr8-37653284-INS-70 | chr8-37653291-INS-69 |
| chr8-37677273-INS-946 | chr8-37677273-INS-946 |
| chr8-37704968-INS-66 | chr8-37704968-INS-68 |
| chr8-37723864-INS-1890 | chr8-37723865-INS-1890 |
| chr8-37737378-INS-59 | chr8-37737377-INS-61 |
| chr8-37751037-INS-69 | chr8-37751026-INS-56 |
| chr8-37752114-INS-7162 | chr8-37752115-INS-7162 |
| chr8-37769479-INS-138 | chr8-37769479-INS-138 |
| chr8-37798197-INS-448 | chr8-37798197-INS-448 |
| chr8-37832540-INS-54 | chr8-37832540-INS-60 |
| chr8-37842880-INS-95 | chr8-37842876-INS-96 |
| chr8-37846908-INS-165 | chr8-37846908-INS-165 |
| chr8-37857702-INS-64 | chr8-37857699-INS-52 |
| chr8-37886260-INS-292 | chr8-37886260-INS-292 |
| chr8-37939796-INS-354 | chr8-37939796-INS-354 |
| chr8-37957361-INS-5051 | chr8-37957362-INS-5051 |
| chr8-37964352-INS-1073 | chr8-37964353-INS-1073 |
| chr8-37972252-INS-93 | chr8-37972252-INS-85 |
| chr8-38000702-INS-109 | chr8-38000702-INS-109 |
| chr8-38011474-INS-891 | chr8-38011472-INS-892 |
| chr8-38034433-INS-210 | chr8-38034433-INS-210 |
| chr8-38051178-INS-1288 | chr8-38051179-INS-1288 |
| chr8-38081757-INS-6206 | chr8-38081761-INS-6190 |
| chr8-38127235-INS-67 | chr8-38127235-INS-68 |
| chr8-38187932-INS-817 | chr8-38187932-INS-816 |
| chr8-38196105-INS-388 | chr8-38196105-INS-387 |
| chr8-38226775-INS-185 | chr8-38226775-INS-185 |
| chr8-38234719-INS-98 | chr8-38234719-INS-96 |
| chr8-38263067-INS-123 | chr8-38263067-INS-123 |
| chr8-38285085-INS-283 | chr8-38285085-INS-283 |
| chr8-38298874-INS-52 | chr8-38298876-INS-60 |
| chr8-38332984-INS-823 | chr8-38332984-INS-820 |
| chr8-38346820-INS-1942 | chr8-38346821-INS-1941 |
| chr8-38357739-INS-156 | chr8-38357739-INS-156 |
| chr8-38367091-INS-371 | chr8-38367091-INS-371 |
| chr8-38395205-INS-51 | chr8-38395205-INS-51 |
| chr8-38400213-INS-63 | chr8-38400213-INS-63 |
| chr8-38408113-INS-370 | chr8-38408111-INS-371 |
| chr8-38416023-INS-490 | chr8-38416023-INS-490 |
| chr8-38437871-INS-115 | chr8-38437871-INS-115 |
| chr8-38472025-INS-61 | chr8-38472026-INS-65 |
| chr8-38503091-INS-975 | chr8-38503091-INS-975 |
| chr8-38504477-INS-84 | chr8-38504477-INS-84 |
| chr8-38508225-INS-70 | chr8-38508225-INS-70 |
| chr8-38510889-INS-525 | chr8-38510889-INS-525 |
| chr8-38514017-INS-174 | chr8-38514017-INS-174 |
| chr8-38525951-INS-449 | chr8-38525951-INS-449 |
| chr8-38531739-INS-723 | chr8-38531739-INS-723 |
| chr8-38559830-INS-400 | chr8-38559830-INS-400 |
| chr8-38563530-INS-498 | chr8-38563530-INS-498 |
| chr8-38565877-INS-7791 | chr8-38565878-INS-7791 |
| chr8-38570393-INS-153 | chr8-38570395-INS-158 |
| chr8-38587353-INS-81 | chr8-38587353-INS-81 |
| chr8-38595183-INS-6161 | chr8-38595184-INS-6161 |
| chr8-38599268-INS-56 | chr8-38599268-INS-56 |
| chr8-38613774-INS-60 | chr8-38613774-INS-60 |
| chr8-38617104-INS-52 | chr8-38617104-INS-52 |
| chr8-38623314-INS-550 | chr8-38623314-INS-557 |
| chr8-38624638-INS-164 | chr8-38624638-INS-164 |
| chr8-38634997-INS-60 | chr8-38634997-INS-60 |
| chr8-38646328-INS-58 | chr8-38646318-INS-51 |
| chr8-38669463-INS-52 | chr8-38669463-INS-52 |
| chr8-38701801-INS-7200 | chr8-38701802-INS-7200 |
| chr8-38711625-INS-150 | chr8-38711576-INS-197 |
| chr8-38713748-INS-394 | chr8-38713748-INS-394 |

| | |
|---|---|
| chr8-38745817-INS-54 | chr8-38745817-INS-63 |
| chr8-38750099-INS-1971 | chr8-38750100-INS-1971 |
| chr8-38772449-INS-51 | chr8-38772450-INS-53 |
| chr8-38779727-INS-2305 | chr8-38779727-INS-2293 |
| chr8-38783008-INS-111 | chr8-38783007-INS-131 |
| chr8-38821048-INS-5381 | chr8-38821049-INS-5381 |
| chr8-38821141-INS-208 | chr8-38821141-INS-192 |
| chr8-38824445-INS-945 | chr8-38824445-INS-945 |
| chr8-38826873-INS-210 | chr8-38826873-INS-211 |
| chr8-38855113-INS-53 | chr8-38855114-INS-70 |
| chr8-38915563-INS-52 | chr8-38915563-INS-58 |
| chr8-38945002-INS-284 | chr8-38945002-INS-284 |
| chr8-38947126-INS-821 | chr8-38947126-INS-821 |
| chr8-38961922-INS-151 | chr8-38961922-INS-151 |
| chr8-38963367-INS-6805 | chr8-38963368-INS-6805 |
| chr8-38998717-INS-6482 | chr8-38998718-INS-6483 |
| chr8-39045458-INS-66 | chr8-39045458-INS-66 |
| chr8-39119178-INS-60 | chr8-39119178-INS-60 |
| chr8-39122886-INS-58 | chr8-39122886-INS-62 |
| chr8-39129812-INS-64 | chr8-39129812-INS-64 |
| chr8-39135857-INS-54 | chr8-39135857-INS-55 |
| chr8-39137890-INS-171 | chr8-39137890-INS-171 |
| chr8-39161724-INS-152 | chr8-39161724-INS-152 |
| chr8-39198026-INS-2634 | chr8-39198027-INS-2634 |
| chr8-39212825-INS-3049 | chr8-39212826-INS-3049 |
| chr8-39225089-INS-717 | chr8-39225089-INS-717 |
| chr8-39231417-INS-54 | chr8-39231417-INS-52 |
| chr8-39253651-INS-64 | chr8-39253651-INS-60 |
| chr8-39254713-INS-52 | chr8-39254721-INS-51 |
| chr8-39262122-INS-136 | chr8-39262122-INS-136 |
| chr8-39324651-INS-503 | chr8-39324651-INS-502 |
| chr8-39350793-INS-54 | chr8-39350793-INS-70 |
| chr8-39363278-INS-1171 | chr8-39363279-INS-1171 |
| chr8-39368283-INS-109 | chr8-39368283-INS-109 |
| chr8-39369501-INS-6244 | chr8-39369502-INS-6244 |
| chr8-39374256-INS-66 | chr8-39374256-INS-66 |
| chr8-39392045-INS-6290 | chr8-39392046-INS-6290 |
| chr8-39411099-INS-193 | chr8-39411099-INS-194 |
| chr8-39411243-INS-583 | chr8-39411243-INS-585 |
| chr8-39442524-INS-6396 | chr8-39442525-INS-6396 |
| chr8-39456811-INS-1488 | chr8-39456812-INS-1487 |
| chr8-39471142-INS-204 | chr8-39471142-INS-204 |
| chr8-39472780-INS-218 | chr8-39472780-INS-218 |
| chr8-39496367-INS-63 | chr8-39496367-INS-63 |
| chr8-39552103-INS-2293 | chr8-39552104-INS-2293 |
| chr8-39554269-INS-60 | chr8-39554269-INS-60 |
| chr8-39567829-INS-293 | chr8-39567829-INS-293 |
| chr8-39576476-INS-76 | chr8-39576476-INS-76 |
| chr8-39594383-INS-368 | chr8-39594383-INS-368 |
| chr8-39670163-INS-1276 | chr8-39670164-INS-1276 |
| chr8-39672482-INS-6664 | chr8-39672483-INS-6664 |
| chr8-39676461-INS-56 | chr8-39676461-INS-56 |
| chr8-39709750-INS-490 | chr8-39709750-INS-490 |
| chr8-39714373-INS-77 | chr8-39714373-INS-77 |
| chr8-39723003-INS-225 | chr8-39723003-INS-225 |
| chr8-39728287-INS-6474 | chr8-39728288-INS-6474 |
| chr8-39743032-INS-94 | chr8-39743050-INS-85 |
| chr8-39760299-INS-69 | chr8-39760299-INS-67 |
| chr8-39762324-INS-70 | chr8-39762324-INS-70 |
| chr8-39765070-INS-52 | chr8-39765070-INS-56 |
| chr8-39779319-INS-417 | chr8-39779319-INS-416 |
| chr8-39813608-INS-197 | chr8-39813608-INS-197 |
| chr8-39826086-INS-52 | chr8-39826086-INS-55 |
| chr8-39853087-INS-68 | chr8-39853075-INS-84 |
| chr8-39897042-INS-142 | chr8-39897042-INS-142 |
| chr8-39901255-INS-78 | chr8-39901255-INS-74 |
| chr8-39931648-INS-142 | chr8-39931648-INS-120 |
| chr8-39945471-INS-540 | chr8-39945471-INS-540 |
| chr8-40014375-INS-120 | chr8-40014375-INS-118 |
| chr8-40017502-INS-66 | chr8-40017502-INS-66 |
| chr8-40023279-INS-195 | chr8-40023279-INS-195 |
| chr8-40027924-INS-540 | chr8-40027924-INS-540 |
| chr8-40044419-INS-835 | chr8-40044415-INS-838 |
| chr8-40050649-INS-940 | chr8-40050649-INS-940 |
| chr8-40088715-INS-68 | chr8-40088707-INS-99 |
| chr8-40089948-INS-57 | chr8-40089948-INS-60 |
| chr8-40137105-INS-174 | chr8-40137105-INS-173 |

| | |
|---|---|
| chr8-40140911-INS-125 | chr8-40140911-INS-133 |
| chr8-40148415-INS-210 | chr8-40148415-INS-211 |
| chr8-40236507-INS-81 | chr8-40236507-INS-81 |
| chr8-40237833-INS-203 | chr8-40237833-INS-203 |
| chr8-40244222-INS-7125 | chr8-40244223-INS-7125 |
| chr8-40253217-INS-124 | chr8-40253217-INS-124 |
| chr8-40275703-INS-329 | chr8-40275703-INS-329 |
| chr8-40276492-INS-58 | chr8-40276492-INS-58 |
| chr8-40314854-INS-184 | chr8-40314854-INS-184 |
| chr8-40333046-INS-53 | chr8-40333036-INS-53 |
| chr8-40336001-INS-120 | chr8-40336005-INS-110 |
| chr8-40378717-INS-6467 | chr8-40378718-INS-6468 |
| chr8-40392394-INS-304 | chr8-40392394-INS-304 |
| chr8-40433290-INS-51 | chr8-40433283-INS-83 |
| chr8-40436948-INS-214 | chr8-40436948-INS-214 |
| chr8-40440632-INS-352 | chr8-40440632-INS-352 |
| chr8-40453290-INS-952 | chr8-40453290-INS-952 |
| chr8-40461736-INS-93 | chr8-40461736-INS-93 |
| chr8-40464187-INS-385 | chr8-40464187-INS-385 |
| chr8-40467986-INS-107 | chr8-40467986-INS-109 |
| chr8-40471021-INS-714 | chr8-40471022-INS-702 |
| chr8-40498447-INS-82 | chr8-40498447-INS-78 |
| chr8-40505714-INS-100 | chr8-40505714-INS-100 |
| chr8-40518640-INS-91 | chr8-40518640-INS-91 |
| chr8-40542198-INS-62 | chr8-40542198-INS-64 |
| chr8-40545217-INS-107 | chr8-40545217-INS-107 |
| chr8-40555233-INS-58 | chr8-40555229-INS-66 |
| chr8-40556256-INS-57 | chr8-40556255-INS-59 |
| chr8-40599475-INS-981 | chr8-40599475-INS-982 |
| chr8-40662969-INS-207 | chr8-40662969-INS-207 |
| chr8-40669150-INS-68 | chr8-40669150-INS-68 |
| chr8-40670392-INS-400 | chr8-40670392-INS-400 |
| chr8-40671188-INS-479 | chr8-40671188-INS-479 |
| chr8-40683641-INS-52 | chr8-40683641-INS-52 |
| chr8-40694606-INS-968 | chr8-40694606-INS-968 |
| chr8-40718607-INS-82 | chr8-40718607-INS-88 |
| chr8-40719449-INS-493 | chr8-40719449-INS-493 |
| chr8-40722193-INS-319 | chr8-40722193-INS-319 |
| chr8-40731425-INS-105 | chr8-40731425-INS-90 |
| chr8-40754948-INS-1789 | chr8-40754949-INS-1790 |
| chr8-40758106-INS-5495 | chr8-40758107-INS-5493 |
| chr8-40764540-INS-165 | chr8-40764540-INS-165 |
| chr8-40780245-INS-6664 | chr8-40780246-INS-6664 |
| chr8-40783106-INS-430 | chr8-40783106-INS-430 |
| chr8-40794085-INS-1354 | chr8-40794086-INS-1353 |
| chr8-40851558-INS-289 | chr8-40851558-INS-290 |
| chr8-40856902-INS-182 | chr8-40856902-INS-190 |
| chr8-40883147-INS-147 | chr8-40883147-INS-147 |
| chr8-40885314-INS-61 | chr8-40885314-INS-61 |
| chr8-40936211-INS-55 | chr8-40936211-INS-60 |
| chr8-40942686-INS-52 | chr8-40942686-INS-52 |
| chr8-41017534-INS-207 | chr8-41017534-INS-207 |
| chr8-41042704-INS-236 | chr8-41042704-INS-236 |
| chr8-41055888-INS-92 | chr8-41055888-INS-92 |
| chr8-41076290-INS-98 | chr8-41076290-INS-97 |
| chr8-41096175-INS-59 | chr8-41096159-INS-62 |
| chr8-41100280-INS-336 | chr8-41100280-INS-336 |
| chr8-41120233-INS-545 | chr8-41120233-INS-545 |
| chr8-41150108-INS-58 | chr8-41150108-INS-58 |
| chr8-41184913-INS-3039 | chr8-41184914-INS-3040 |
| chr8-41192106-INS-203 | chr8-41192106-INS-203 |
| chr8-41193709-INS-1165 | chr8-41193710-INS-1166 |
| chr8-41201997-INS-7046 | chr8-41201998-INS-7047 |
| chr8-41236841-INS-63 | chr8-41236841-INS-55 |
| chr8-41256386-INS-121 | chr8-41256382-INS-120 |
| chr8-41307666-INS-2972 | chr8-41307667-INS-2972 |
| chr8-41345582-INS-99 | chr8-41345582-INS-99 |
| chr8-41349541-INS-108 | chr8-41349541-INS-108 |
| chr8-41357168-INS-70 | chr8-41357192-INS-68 |
| chr8-41362672-INS-139 | chr8-41362672-INS-139 |
| chr8-41366754-INS-206 | chr8-41366754-INS-207 |
| chr8-41370766-INS-62 | chr8-41370766-INS-64 |
| chr8-41378998-INS-395 | chr8-41378998-INS-395 |
| chr8-41415430-INS-438 | chr8-41415430-INS-438 |
| chr8-41418265-INS-52 | chr8-41418285-INS-72 |
| chr8-41422123-INS-56 | chr8-41422123-INS-56 |
| chr8-41449975-INS-399 | chr8-41449975-INS-399 |

| | |
|---|---|
| chr8-41461320-INS-66 | chr8-41461320-INS-66 |
| chr8-41478911-INS-69 | chr8-41478911-INS-69 |
| chr8-41480578-INS-57 | chr8-41480579-INS-64 |
| chr8-41495505-INS-55 | chr8-41495505-INS-55 |
| chr8-41509135-INS-226 | chr8-41509135-INS-227 |
| chr8-41520461-INS-69 | chr8-41520461-INS-69 |
| chr8-41544650-INS-374 | chr8-41544650-INS-374 |
| chr8-41545593-INS-60 | chr8-41545593-INS-60 |
| chr8-41581091-INS-193 | chr8-41581091-INS-194 |
| chr8-41595618-INS-70 | chr8-41595622-INS-74 |
| chr8-41598266-INS-896 | chr8-41598266-INS-896 |
| chr8-41624588-INS-127 | chr8-41624588-INS-127 |
| chr8-41659095-INS-327 | chr8-41659095-INS-327 |
| chr8-41672417-INS-493 | chr8-41672417-INS-494 |
| chr8-41678478-INS-207 | chr8-41678478-INS-207 |
| chr8-41679000-INS-52 | chr8-41679000-INS-56 |
| chr8-41688652-INS-692 | chr8-41688652-INS-692 |
| chr8-41696011-INS-236 | chr8-41696011-INS-236 |
| chr8-41724496-INS-134 | chr8-41724496-INS-134 |
| chr8-41754552-INS-837 | chr8-41754552-INS-837 |
| chr8-41851625-INS-156 | chr8-41851625-INS-156 |
| chr8-41860650-INS-392 | chr8-41860650-INS-392 |
| chr8-41868487-INS-61 | chr8-41868487-INS-61 |
| chr8-41894224-INS-715 | chr8-41894224-INS-715 |
| chr8-41906329-INS-655 | chr8-41906329-INS-654 |
| chr8-41907247-INS-401 | chr8-41907247-INS-401 |
| chr8-41914145-INS-62 | chr8-41914137-INS-64 |
| chr8-41918129-INS-204 | chr8-41918129-INS-204 |
| chr8-41934263-INS-129 | chr8-41934263-INS-129 |
| chr8-41968879-INS-326 | chr8-41968879-INS-326 |
| chr8-41981632-INS-574 | chr8-41981632-INS-574 |
| chr8-42022049-INS-87 | chr8-42022049-INS-87 |
| chr8-42041540-INS-68 | chr8-42041540-INS-68 |
| chr8-42044642-INS-116 | chr8-42044642-INS-116 |
| chr8-42070274-INS-302 | chr8-42070274-INS-302 |
| chr8-42095714-INS-2685 | chr8-42095715-INS-2685 |
| chr8-42103170-INS-485 | chr8-42103170-INS-479 |
| chr8-42104511-INS-752 | chr8-42104511-INS-756 |
| chr8-42124291-INS-640 | chr8-42124291-INS-640 |
| chr8-42125066-INS-92 | chr8-42125066-INS-92 |
| chr8-42126131-INS-6239 | chr8-42126132-INS-6239 |
| chr8-42133461-INS-4229 | chr8-42133462-INS-4229 |
| chr8-42146570-INS-303 | chr8-42146570-INS-303 |
| chr8-42153974-INS-4104 | chr8-42153975-INS-4104 |
| chr8-42188036-INS-709 | chr8-42188036-INS-709 |
| chr8-42190195-INS-432 | chr8-42190195-INS-432 |
| chr8-42198248-INS-1740 | chr8-42198256-INS-1749 |
| chr8-42214070-INS-56 | chr8-42214070-INS-60 |
| chr8-42220401-INS-590 | chr8-42220401-INS-590 |
| chr8-42223232-INS-7856 | chr8-42223233-INS-7857 |
| chr8-42237647-INS-58 | chr8-42237647-INS-58 |
| chr8-42239181-INS-177 | chr8-42239181-INS-177 |
| chr8-42243083-INS-60 | chr8-42243083-INS-60 |
| chr8-42254747-INS-110 | chr8-42254747-INS-116 |
| chr8-42261647-INS-81 | chr8-42261647-INS-81 |
| chr8-42272824-INS-82 | chr8-42272824-INS-82 |
| chr8-42307295-INS-7132 | chr8-42307296-INS-7132 |
| chr8-42317892-INS-6590 | chr8-42317893-INS-6590 |
| chr8-42362388-INS-201 | chr8-42362388-INS-201 |
| chr8-42372368-INS-75 | chr8-42372368-INS-75 |
| chr8-42406322-INS-908 | chr8-42406322-INS-908 |
| chr8-42409384-INS-450 | chr8-42409384-INS-450 |
| chr8-42436746-INS-56 | chr8-42436746-INS-62 |
| chr8-42438056-INS-517 | chr8-42438056-INS-517 |
| chr8-42452540-INS-78 | chr8-42452540-INS-78 |
| chr8-42455449-INS-154 | chr8-42455449-INS-154 |
| chr8-42456997-INS-781 | chr8-42456997-INS-782 |
| chr8-42468578-INS-1490 | chr8-42468579-INS-1490 |
| chr8-42479342-INS-359 | chr8-42479342-INS-359 |
| chr8-42509004-INS-54 | chr8-42509004-INS-54 |
| chr8-42515369-INS-920 | chr8-42515369-INS-920 |
| chr8-42536414-INS-84 | chr8-42536414-INS-84 |
| chr8-42545026-INS-7089 | chr8-42545027-INS-7090 |
| chr8-42549934-INS-322 | chr8-42549934-INS-322 |
| chr8-42557945-INS-82 | chr8-42557945-INS-82 |
| chr8-42559138-INS-2013 | chr8-42559139-INS-2013 |
| chr8-42562647-INS-540 | chr8-42562647-INS-550 |

| | |
|---|---|
| chr8-42571111-INS-526 | chr8-42571111-INS-526 |
| chr8-42572615-INS-155 | chr8-42572615-INS-155 |
| chr8-42576528-INS-138 | chr8-42576528-INS-152 |
| chr8-42579774-INS-239 | chr8-42579774-INS-239 |
| chr8-42606579-INS-182 | chr8-42606579-INS-184 |
| chr8-42613596-INS-534 | chr8-42613596-INS-534 |
| chr8-42618506-INS-73 | chr8-42618506-INS-73 |
| chr8-42632471-INS-1610 | chr8-42632472-INS-1611 |
| chr8-42664580-INS-6472 | chr8-42664581-INS-6472 |
| chr8-42675604-INS-265 | chr8-42675604-INS-266 |
| chr8-42678304-INS-271 | chr8-42678304-INS-271 |
| chr8-42682925-INS-6169 | chr8-42682926-INS-6169 |
| chr8-42683437-INS-55 | chr8-42683437-INS-57 |
| chr8-42688745-INS-226 | chr8-42688745-INS-227 |
| chr8-42690107-INS-395 | chr8-42690107-INS-395 |
| chr8-42706108-INS-58 | chr8-42706108-INS-50 |
| chr8-42718083-INS-70 | chr8-42718061-INS-68 |
| chr8-42732639-INS-85 | chr8-42732639-INS-73 |
| chr8-42766890-INS-862 | chr8-42766890-INS-862 |
| chr8-42783424-INS-6371 | chr8-42783425-INS-6372 |
| chr8-42788915-INS-217 | chr8-42788915-INS-205 |
| chr8-42799142-INS-6236 | chr8-42799143-INS-6236 |
| chr8-42800597-INS-136 | chr8-42800597-INS-140 |
| chr8-42876360-INS-875 | chr8-42876360-INS-875 |
| chr8-42886916-INS-66 | chr8-42886916-INS-66 |
| chr8-42893200-INS-178 | chr8-42893200-INS-178 |
| chr8-42954578-INS-580 | chr8-42954578-INS-580 |
| chr8-42961485-INS-398 | chr8-42961487-INS-398 |
| chr8-42993669-INS-7947 | chr8-42993670-INS-7948 |
| chr8-43001404-INS-1901 | chr8-43001406-INS-1906 |
| chr8-43003675-INS-117 | chr8-43003675-INS-85 |
| chr8-43007786-INS-83 | chr8-43007817-INS-87 |
| chr8-43056314-INS-313 | chr8-43056314-INS-313 |
| chr8-43106987-INS-219 | chr8-43106987-INS-240 |
| chr8-43114328-INS-341 | chr8-43114328-INS-341 |
| chr8-43186538-INS-2743 | chr8-43186539-INS-2743 |
| chr8-43208354-INS-6450 | chr8-43208355-INS-6450 |
| chr8-43216165-INS-776 | chr8-43216165-INS-774 |
| chr8-43228768-INS-626 | chr8-43228768-INS-626 |
| chr8-43283977-INS-164 | chr8-43283977-INS-163 |
| chr8-43341839-INS-115 | chr8-43341839-INS-115 |
| chr8-43361695-INS-6559 | chr8-43361696-INS-6559 |
| chr8-43388452-INS-56 | chr8-43388452-INS-50 |
| chr8-43438628-INS-1810 | chr8-43438635-INS-1801 |
| chr8-43457811-INS-327 | chr8-43457811-INS-327 |
| chr8-43481273-INS-380 | chr8-43481273-INS-380 |
| chr8-43500906-INS-206 | chr8-43500906-INS-206 |
| chr8-43505170-INS-62 | chr8-43505170-INS-72 |
| chr8-43505338-INS-6652 | chr8-43505339-INS-6652 |
| chr8-43506587-INS-60 | chr8-43506587-INS-64 |
| chr8-43507891-INS-206 | chr8-43507893-INS-207 |
| chr8-43508926-INS-655 | chr8-43508926-INS-652 |
| chr8-43518791-INS-87 | chr8-43518791-INS-87 |
| chr8-43537890-INS-359 | chr8-43537890-INS-359 |
| chr8-43539156-INS-72 | chr8-43539156-INS-50 |
| chr8-43553960-INS-69 | chr8-43553960-INS-69 |
| chr8-43563773-INS-685 | chr8-43563773-INS-685 |
| chr8-43591873-INS-58 | chr8-43591873-INS-58 |
| chr8-43626406-INS-294 | chr8-43626406-INS-295 |
| chr8-43653010-INS-6452 | chr8-43653011-INS-6452 |
| chr8-43664887-INS-685 | chr8-43664872-INS-678 |
| chr8-43671093-INS-863 | chr8-43671093-INS-863 |
| chr8-43677069-INS-56 | chr8-43677069-INS-60 |
| chr8-43684956-INS-74 | chr8-43684956-INS-74 |
| chr8-43702935-INS-955 | chr8-43702935-INS-955 |
| chr8-43716208-INS-204 | chr8-43716208-INS-204 |
| chr8-43726907-INS-212 | chr8-43726907-INS-213 |
| chr8-43736412-INS-219 | chr8-43736412-INS-218 |
| chr8-43741495-INS-215 | chr8-43741495-INS-215 |
| chr8-43748052-INS-213 | chr8-43748040-INS-210 |
| chr8-43757945-INS-213 | chr8-43757945-INS-213 |
| chr8-43782593-INS-223 | chr8-43782593-INS-224 |
| chr8-43787664-INS-199 | chr8-43787664-INS-199 |
| chr8-43796826-INS-192 | chr8-43796826-INS-192 |
| chr8-43800934-INS-211 | chr8-43800934-INS-211 |
| chr8-43804453-INS-232 | chr8-43804453-INS-233 |
| chr8-43812848-INS-199 | chr8-43812848-INS-200 |

| | |
|---|---|
| chr8-43829174-INS-72 | chr8-43829174-INS-50 |
| chr8-43832241-INS-458 | chr8-43832241-INS-458 |
| chr8-43836485-INS-220 | chr8-43836485-INS-219 |
| chr8-43838062-INS-53 | chr8-43838063-INS-50 |
| chr8-43849822-INS-358 | chr8-43849822-INS-358 |
| chr8-43886391-INS-135 | chr8-43886391-INS-135 |
| chr8-43894893-INS-275 | chr8-43894893-INS-275 |
| chr8-43900517-INS-57 | chr8-43900517-INS-75 |
| chr8-43914259-INS-868 | chr8-43914259-INS-870 |
| chr8-43937369-INS-54 | chr8-43937371-INS-57 |
| chr8-43940465-INS-2165 | chr8-43940466-INS-2169 |
| chr8-43942099-INS-158 | chr8-43942096-INS-192 |
| chr8-43976450-INS-510 | chr8-43976450-INS-510 |
| chr8-44048008-INS-208 | chr8-44048008-INS-208 |
| chr8-44059703-INS-4663 | chr8-44059704-INS-4663 |
| chr8-44071029-INS-90 | chr8-44071029-INS-90 |
| chr8-44080772-INS-133 | chr8-44080772-INS-133 |
| chr8-44081606-INS-7028 | chr8-44081607-INS-7029 |
| chr8-44099438-INS-137 | chr8-44099438-INS-137 |
| chr8-44114204-INS-6458 | chr8-44114205-INS-6459 |
| chr8-44115777-INS-203 | chr8-44115777-INS-203 |
| chr8-44143854-INS-201 | chr8-44143854-INS-201 |
| chr8-44148783-INS-54 | chr8-44148783-INS-54 |
| chr8-44181253-INS-268 | chr8-44181253-INS-269 |
| chr8-44198946-INS-229 | chr8-44198946-INS-229 |
| chr8-44223967-INS-982 | chr8-44223967-INS-983 |
| chr8-44236145-INS-268 | chr8-44236145-INS-268 |
| chr8-44244740-INS-214 | chr8-44244740-INS-214 |
| chr8-44272006-INS-969 | chr8-44272006-INS-969 |
| chr8-44287163-INS-87 | chr8-44287163-INS-86 |
| chr8-44384871-INS-337 | chr8-44384871-INS-337 |
| chr8-44501662-INS-262 | chr8-44501662-INS-262 |
| chr8-44526715-INS-543 | chr8-44526715-INS-545 |
| chr8-44534370-INS-88 | chr8-44534370-INS-88 |
| chr8-44567265-INS-265 | chr8-44567265-INS-265 |
| chr8-44572439-INS-636 | chr8-44572439-INS-635 |
| chr8-44579082-INS-1388 | chr8-44579068-INS-1395 |
| chr8-44584152-INS-60 | chr8-44584152-INS-62 |
| chr8-44586995-INS-72 | chr8-44586995-INS-60 |
| chr8-44598466-INS-563 | chr8-44598466-INS-563 |
| chr8-44606061-INS-609 | chr8-44606061-INS-608 |
| chr8-44623290-INS-2458 | chr8-44623291-INS-2458 |
| chr8-44652054-INS-6209 | chr8-44652054-INS-6213 |
| chr8-44734329-INS-63 | chr8-44734318-INS-54 |
| chr8-44741310-INS-70 | chr8-44741310-INS-70 |
| chr8-44761623-INS-6404 | chr8-44761624-INS-6403 |
| chr8-44764286-INS-81 | chr8-44764286-INS-81 |
| chr8-44773057-INS-2947 | chr8-44773058-INS-2947 |
| chr8-44777516-INS-70 | chr8-44777516-INS-60 |
| chr8-44782081-INS-82 | chr8-44782081-INS-66 |
| chr8-44812656-INS-4334 | chr8-44812657-INS-4334 |
| chr8-44815158-INS-54 | chr8-44815156-INS-98 |
| chr8-44840882-INS-6409 | chr8-44840883-INS-6409 |
| chr8-44872372-INS-84 | chr8-44872372-INS-86 |
| chr8-44884503-INS-60 | chr8-44884503-INS-62 |
| chr8-44890252-INS-231 | chr8-44890252-INS-232 |
| chr8-44895140-INS-2088 | chr8-44895141-INS-2088 |
| chr8-44903273-INS-58 | chr8-44903273-INS-58 |
| chr8-44918562-INS-619 | chr8-44918562-INS-620 |
| chr8-44920848-INS-188 | chr8-44920848-INS-188 |
| chr8-44939939-INS-90 | chr8-44939939-INS-90 |
| chr8-44942804-INS-78 | chr8-44942804-INS-78 |
| chr8-44957283-INS-52 | chr8-44957237-INS-57 |
| chr8-44963450-INS-56 | chr8-44963450-INS-60 |
| chr8-44970767-INS-967 | chr8-44970769-INS-970 |
| chr8-44974862-INS-122 | chr8-44974862-INS-130 |
| chr8-44998190-INS-774 | chr8-44998190-INS-774 |
| chr8-45000559-INS-70 | chr8-45000563-INS-62 |
| chr8-45020353-INS-281 | chr8-45020353-INS-325 |
| chr8-45042791-INS-165 | chr8-45042791-INS-165 |
| chr8-45046676-INS-345 | chr8-45046676-INS-345 |
| chr8-45056421-INS-400 | chr8-45056421-INS-400 |
| chr8-45057849-INS-56 | chr8-45057849-INS-60 |
| chr8-45079747-INS-83 | chr8-45079747-INS-118 |
| chr8-45097840-INS-60 | chr8-45097840-INS-66 |
| chr8-45106564-INS-97 | chr8-45106564-INS-97 |
| chr8-45108474-INS-224 | chr8-45108474-INS-222 |

| | |
|---|---|
| chr8-45155839-INS-381 | chr8-45155839-INS-379 |
| chr8-45166665-INS-598 | chr8-45166665-INS-599 |
| chr8-45174838-INS-102 | chr8-45174845-INS-118 |
| chr8-45212838-INS-63 | chr8-45212840-INS-62 |
| chr8-45247162-INS-89 | chr8-45247162-INS-89 |
| chr8-45270500-INS-399 | chr8-45270500-INS-399 |
| chr8-45281544-INS-71 | chr8-45281544-INS-65 |
| chr8-45300223-INS-178 | chr8-45300251-INS-196 |
| chr8-45309463-INS-199 | chr8-45309463-INS-199 |
| chr8-45316344-INS-162 | chr8-45316394-INS-116 |
| chr8-45327847-INS-77 | chr8-45327847-INS-76 |
| chr8-45331720-INS-90 | chr8-45331720-INS-94 |
| chr8-45350947-INS-55 | chr8-45350947-INS-55 |
| chr8-45355636-INS-55 | chr8-45355636-INS-55 |
| chr8-45418048-INS-55 | chr8-45418048-INS-55 |
| chr8-45418082-INS-5219 | chr8-45418083-INS-5219 |
| chr8-45440861-INS-902 | chr8-45440861-INS-881 |
| chr8-45511387-INS-6550 | chr8-45511388-INS-6550 |
| chr8-45511993-INS-2776 | chr8-45511994-INS-2776 |
| chr8-45515434-INS-2331 | chr8-45515435-INS-2331 |
| chr8-45527890-INS-434 | chr8-45527890-INS-434 |
| chr8-45533933-INS-148 | chr8-45533933-INS-149 |
| chr8-45549348-INS-185 | chr8-45549348-INS-185 |
| chr8-45550792-INS-80 | chr8-45550783-INS-72 |
| chr8-45555266-INS-432 | chr8-45555266-INS-432 |
| chr8-45569741-INS-68 | chr8-45569741-INS-68 |
| chr8-45577420-INS-54 | chr8-45577396-INS-90 |
| chr8-45578557-INS-59 | chr8-45578557-INS-59 |
| chr8-45601847-INS-108 | chr8-45601852-INS-60 |
| chr8-45640525-INS-72 | chr8-45640525-INS-74 |
| chr8-45701309-INS-74 | chr8-45701309-INS-78 |
| chr8-45712547-INS-947 | chr8-45712547-INS-948 |
| chr8-45757987-INS-351 | chr8-45757987-INS-351 |
| chr8-45794161-INS-361 | chr8-45794161-INS-361 |
| chr8-45809971-INS-269 | chr8-45809971-INS-269 |
| chr8-45815159-INS-70 | chr8-45815145-INS-76 |
| chr8-45824375-INS-204 | chr8-45824375-INS-204 |
| chr8-45826289-INS-622 | chr8-45826284-INS-662 |
| chr8-45832135-INS-100 | chr8-45832139-INS-76 |
| chr8-45847846-INS-56 | chr8-45847846-INS-56 |
| chr8-45867840-INS-74 | chr8-45867840-INS-74 |
| chr8-45926454-INS-56 | chr8-45926454-INS-56 |
| chr8-45927967-INS-165 | chr8-45927967-INS-165 |
| chr8-45935019-INS-482 | chr8-45935019-INS-482 |
| chr8-45936835-INS-62 | chr8-45936834-INS-94 |
| chr8-45949426-INS-66 | chr8-45949426-INS-66 |
| chr8-45954657-INS-401 | chr8-45954657-INS-397 |
| chr8-45966205-INS-121 | chr8-45966214-INS-94 |
| chr8-45983654-INS-60 | chr8-45983654-INS-58 |
| chr8-46013178-INS-66 | chr8-46013178-INS-70 |
| chr8-46024618-INS-168 | chr8-46024622-INS-156 |
| chr8-46135527-INS-140 | chr8-46135527-INS-138 |
| chr8-46143993-INS-564 | chr8-46143993-INS-564 |
| chr8-46179665-INS-65 | chr8-46179660-INS-60 |
| chr8-46202587-INS-118 | chr8-46202586-INS-71 |
| chr8-46270895-INS-130 | chr8-46270895-INS-129 |
| chr8-46291078-INS-54 | chr8-46291078-INS-54 |
| chr8-46305335-INS-160 | chr8-46305335-INS-160 |
| chr8-46309276-INS-165 | chr8-46309277-INS-165 |
| chr8-46318123-INS-196 | chr8-46318123-INS-196 |
| chr8-46323918-INS-94 | chr8-46323903-INS-130 |
| chr8-46343588-INS-81 | chr8-46343588-INS-81 |
| chr8-46353163-INS-87 | chr8-46353163-INS-87 |
| chr8-46407334-INS-62 | chr8-46407334-INS-62 |
| chr8-46415762-INS-118 | chr8-46415762-INS-95 |
| chr8-46439687-INS-119 | chr8-46439687-INS-118 |
| chr8-46472923-INS-140 | chr8-46472923-INS-140 |
| chr8-46547591-INS-209 | chr8-46547591-INS-209 |
| chr8-46556458-INS-67 | chr8-46556474-INS-76 |
| chr8-46572938-INS-839 | chr8-46572938-INS-839 |
| chr8-46581439-INS-462 | chr8-46581438-INS-462 |
| chr8-46589697-INS-1773 | chr8-46589698-INS-1773 |
| chr8-46643275-INS-1128 | chr8-46643276-INS-1133 |
| chr8-46671971-INS-62 | chr8-46671971-INS-62 |
| chr8-46672371-INS-505 | chr8-46672371-INS-505 |
| chr8-46677895-INS-201 | chr8-46677895-INS-200 |
| chr8-46688774-INS-109 | chr8-46688770-INS-116 |

| | |
|---|---|
| chr8-46693587-INS-73 | chr8-46693587-INS-72 |
| chr8-46700566-INS-90 | chr8-46700566-INS-60 |
| chr8-46725018-INS-60 | chr8-46725018-INS-60 |
| chr8-46726109-INS-109 | chr8-46726109-INS-109 |
| chr8-46783436-INS-111 | chr8-46783436-INS-112 |
| chr8-46787932-INS-71 | chr8-46787932-INS-78 |
| chr8-46796426-INS-80 | chr8-46796426-INS-80 |
| chr8-46838552-INS-96 | chr8-46838552-INS-96 |
| chr8-46876942-INS-64 | chr8-46876942-INS-62 |
| chr8-46879633-INS-136 | chr8-46879633-INS-92 |
| chr8-46898105-INS-88 | chr8-46898104-INS-88 |
| chr8-46918677-INS-1268 | chr8-46918678-INS-1271 |
| chr8-46934521-INS-206 | chr8-46934521-INS-206 |
| chr8-46935993-INS-171 | chr8-46935993-INS-171 |
| chr8-46963144-INS-88 | chr8-46963144-INS-72 |
| chr8-47004379-INS-72 | chr8-47004381-INS-62 |
| chr8-47009024-INS-163 | chr8-47009019-INS-164 |
| chr8-47019706-INS-62 | chr8-47019706-INS-76 |
| chr8-47025158-INS-1890 | chr8-47025159-INS-1890 |
| chr8-47044219-INS-94 | chr8-47044219-INS-94 |
| chr8-47067172-INS-72 | chr8-47067172-INS-72 |
| chr8-47087984-INS-225 | chr8-47087984-INS-225 |
| chr8-47096230-INS-74 | chr8-47096230-INS-74 |
| chr8-47099442-INS-504 | chr8-47099442-INS-504 |
| chr8-47139314-INS-153 | chr8-47139314-INS-182 |
| chr8-47149048-INS-60 | chr8-47149048-INS-56 |
| chr8-47161584-INS-190 | chr8-47161582-INS-190 |
| chr8-47199365-INS-441 | chr8-47199365-INS-441 |
| chr8-47217765-INS-55 | chr8-47217765-INS-55 |
| chr8-47287338-INS-56 | chr8-47287338-INS-56 |
| chr8-47290067-INS-461 | chr8-47290067-INS-454 |
| chr8-47329672-INS-94 | chr8-47329672-INS-94 |
| chr8-47338269-INS-52 | chr8-47338269-INS-52 |
| chr8-47340403-INS-2185 | chr8-47340404-INS-2185 |
| chr8-47342313-INS-7067 | chr8-47342314-INS-7067 |
| chr8-47342585-INS-62 | chr8-47342603-INS-60 |
| chr8-47348954-INS-51 | chr8-47348955-INS-57 |
| chr8-47390742-INS-69 | chr8-47390742-INS-84 |
| chr8-47400115-INS-77 | chr8-47400115-INS-77 |
| chr8-47406627-INS-54 | chr8-47406628-INS-55 |
| chr8-47420382-INS-59 | chr8-47420387-INS-84 |
| chr8-47434126-INS-337 | chr8-47434126-INS-337 |
| chr8-47447349-INS-50 | chr8-47447344-INS-62 |
| chr8-47455528-INS-450 | chr8-47455528-INS-442 |
| chr8-47465918-INS-164 | chr8-47465918-INS-164 |
| chr8-47495744-INS-51 | chr8-47495744-INS-51 |
| chr8-47508501-INS-90 | chr8-47508501-INS-90 |
| chr8-47510117-INS-56 | chr8-47510109-INS-50 |
| chr8-47511386-INS-55 | chr8-47511402-INS-65 |
| chr8-47544310-INS-97 | chr8-47544325-INS-94 |
| chr8-47559282-INS-70 | chr8-47559282-INS-70 |
| chr8-47563065-INS-72 | chr8-47563065-INS-72 |
| chr8-47567064-INS-642 | chr8-47567064-INS-642 |
| chr8-47581798-INS-191 | chr8-47581798-INS-191 |
| chr8-47587771-INS-54 | chr8-47587789-INS-60 |
| chr8-47592703-INS-469 | chr8-47592703-INS-469 |
| chr8-47608849-INS-138 | chr8-47608849-INS-138 |
| chr8-47619610-INS-128 | chr8-47619610-INS-128 |
| chr8-47635353-INS-56 | chr8-47635353-INS-56 |
| chr8-47652006-INS-72 | chr8-47652006-INS-76 |
| chr8-47656108-INS-53 | chr8-47656108-INS-53 |
| chr8-47657615-INS-171 | chr8-47657615-INS-186 |
| chr8-47659028-INS-115 | chr8-47659028-INS-117 |
| chr8-47659824-INS-54 | chr8-47659824-INS-54 |
| chr8-47773802-INS-197 | chr8-47773802-INS-198 |
| chr8-47810565-INS-59 | chr8-47810579-INS-53 |
| chr8-47811970-INS-53 | chr8-47811970-INS-53 |
| chr8-47813528-INS-175 | chr8-47813528-INS-195 |
| chr8-47821470-INS-120 | chr8-47821470-INS-120 |
| chr8-47825440-INS-591 | chr8-47825440-INS-591 |
| chr8-47847189-INS-166 | chr8-47847189-INS-166 |
| chr8-47869576-INS-423 | chr8-47869576-INS-423 |
| chr8-47879328-INS-87 | chr8-47879328-INS-87 |
| chr8-47896024-INS-60 | chr8-47896024-INS-52 |
| chr8-47912691-INS-71 | chr8-47912699-INS-59 |
| chr8-47913744-INS-372 | chr8-47913742-INS-377 |
| chr8-47936287-INS-67 | chr8-47936278-INS-68 |

| | |
|---|---|
| chr8-47942573-INS-718 | chr8-47942573-INS-713 |
| chr8-47964107-INS-240 | chr8-47964107-INS-240 |
| chr8-47975908-INS-60 | chr8-47975918-INS-74 |
| chr8-47993553-INS-63 | chr8-47993552-INS-65 |
| chr8-48000052-INS-56 | chr8-48000052-INS-56 |
| chr8-48038235-INS-79 | chr8-48038235-INS-75 |
| chr8-48085090-INS-68 | chr8-48085090-INS-68 |
| chr8-48095042-INS-116 | chr8-48095042-INS-116 |
| chr8-48114518-INS-253 | chr8-48114518-INS-253 |
| chr8-48139440-INS-163 | chr8-48139440-INS-164 |
| chr8-48141096-INS-57 | chr8-48141086-INS-94 |
| chr8-48144211-INS-551 | chr8-48144211-INS-552 |
| chr8-48146014-INS-79 | chr8-48145991-INS-102 |
| chr8-48148451-INS-80 | chr8-48148451-INS-60 |
| chr8-48149777-INS-119 | chr8-48149777-INS-143 |
| chr8-48160183-INS-203 | chr8-48160183-INS-199 |
| chr8-48178227-INS-207 | chr8-48178227-INS-208 |
| chr8-48204760-INS-71 | chr8-48204760-INS-102 |
| chr8-48226450-INS-70 | chr8-48226450-INS-80 |
| chr8-48227320-INS-88 | chr8-48227320-INS-88 |
| chr8-48229315-INS-60 | chr8-48229315-INS-89 |
| chr8-48229878-INS-1196 | chr8-48229879-INS-1201 |
| chr8-48242086-INS-106 | chr8-48242086-INS-119 |
| chr8-48247428-INS-68 | chr8-48247428-INS-68 |
| chr8-48271300-INS-92 | chr8-48271276-INS-92 |
| chr8-48290974-INS-52 | chr8-48290974-INS-52 |
| chr8-48296736-INS-141 | chr8-48296736-INS-141 |
| chr8-48302789-INS-218 | chr8-48302789-INS-218 |
| chr8-48308125-INS-200 | chr8-48308125-INS-200 |
| chr8-48318376-INS-168 | chr8-48318376-INS-168 |
| chr8-48372865-INS-141 | chr8-48372865-INS-96 |
| chr8-48380134-INS-755 | chr8-48380134-INS-755 |
| chr8-48390095-INS-205 | chr8-48390095-INS-203 |
| chr8-48401866-INS-201 | chr8-48401866-INS-198 |
| chr8-48467613-INS-100 | chr8-48467613-INS-100 |
| chr8-48473734-INS-51 | chr8-48473734-INS-51 |
| chr8-48479361-INS-3036 | chr8-48479362-INS-3036 |
| chr8-48482104-INS-64 | chr8-48482104-INS-66 |
| chr8-48483154-INS-868 | chr8-48483154-INS-871 |
| chr8-48487540-INS-66 | chr8-48487540-INS-54 |
| chr8-48494453-INS-177 | chr8-48494453-INS-177 |
| chr8-48494453-INS-888 | chr8-48494453-INS-885 |
| chr8-48500432-INS-52 | chr8-48500432-INS-52 |
| chr8-48600114-INS-103 | chr8-48600114-INS-115 |
| chr8-48623212-INS-52 | chr8-48623213-INS-89 |
| chr8-48638414-INS-473 | chr8-48638414-INS-473 |
| chr8-48641756-INS-67 | chr8-48641755-INS-73 |
| chr8-48673522-INS-217 | chr8-48673522-INS-219 |
| chr8-48695423-INS-100 | chr8-48695423-INS-100 |
| chr8-48706400-INS-58 | chr8-48706400-INS-58 |
| chr8-48727439-INS-135 | chr8-48727439-INS-135 |
| chr8-48775686-INS-435 | chr8-48775686-INS-435 |
| chr8-48779586-INS-578 | chr8-48779586-INS-617 |
| chr8-48808978-INS-159 | chr8-48808978-INS-150 |
| chr8-48810192-INS-69 | chr8-48810192-INS-69 |
| chr8-48821803-INS-110 | chr8-48821803-INS-110 |
| chr8-48846073-INS-600 | chr8-48846073-INS-599 |
| chr8-48850485-INS-81 | chr8-48850485-INS-81 |
| chr8-48851659-INS-75 | chr8-48851659-INS-55 |
| chr8-48913934-INS-62 | chr8-48913934-INS-64 |
| chr8-48922210-INS-269 | chr8-48922210-INS-269 |
| chr8-48924292-INS-123 | chr8-48924285-INS-122 |
| chr8-48937669-INS-62 | chr8-48937669-INS-62 |
| chr8-49016960-INS-172 | chr8-49016960-INS-167 |
| chr8-49020690-INS-52 | chr8-49020690-INS-54 |
| chr8-49069144-INS-160 | chr8-49069144-INS-160 |
| chr8-49085735-INS-6782 | chr8-49085736-INS-6782 |
| chr8-49089413-INS-94 | chr8-49089413-INS-100 |
| chr8-49092811-INS-90 | chr8-49092811-INS-92 |
| chr8-49150504-INS-52 | chr8-49150465-INS-52 |
| chr8-49164322-INS-69 | chr8-49164322-INS-69 |
| chr8-49173491-INS-80 | chr8-49173491-INS-78 |
| chr8-49178167-INS-54 | chr8-49178167-INS-78 |
| chr8-49185736-INS-221 | chr8-49185736-INS-221 |
| chr8-49200523-INS-562 | chr8-49200525-INS-562 |
| chr8-49250735-INS-118 | chr8-49250735-INS-110 |
| chr8-49265779-INS-62 | chr8-49265779-INS-62 |

| | |
|---|---|
| chr8-49285725-INS-54 | chr8-49285725-INS-54 |
| chr8-49306327-INS-56 | chr8-49306327-INS-56 |
| chr8-49319095-INS-82 | chr8-49319094-INS-57 |
| chr8-49335118-INS-64 | chr8-49335099-INS-52 |
| chr8-49345557-INS-112 | chr8-49345571-INS-88 |
| chr8-49389841-INS-53 | chr8-49389841-INS-72 |
| chr8-49404997-INS-67 | chr8-49405017-INS-64 |
| chr8-49406931-INS-201 | chr8-49406931-INS-223 |
| chr8-49418624-INS-60 | chr8-49418648-INS-52 |
| chr8-49418843-INS-468 | chr8-49418843-INS-468 |
| chr8-49429504-INS-57 | chr8-49429504-INS-57 |
| chr8-49431238-INS-84 | chr8-49431260-INS-109 |
| chr8-49467777-INS-81 | chr8-49467814-INS-55 |
| chr8-49478277-INS-55 | chr8-49478277-INS-50 |
| chr8-49480643-INS-6488 | chr8-49480644-INS-6488 |
| chr8-49496217-INS-114 | chr8-49496217-INS-114 |
| chr8-49503657-INS-52 | chr8-49503657-INS-52 |
| chr8-49513260-INS-145 | chr8-49513260-INS-147 |
| chr8-49521198-INS-58 | chr8-49521192-INS-76 |
| chr8-49537013-INS-53 | chr8-49537013-INS-51 |
| chr8-49598127-INS-109 | chr8-49598111-INS-141 |
| chr8-49600873-INS-431 | chr8-49600873-INS-432 |
| chr8-49609188-INS-7840 | chr8-49609189-INS-7841 |
| chr8-49625412-INS-339 | chr8-49625412-INS-340 |
| chr8-49632764-INS-60 | chr8-49632764-INS-50 |
| chr8-49657476-INS-97 | chr8-49657470-INS-96 |
| chr8-49666277-INS-56 | chr8-49666278-INS-87 |
| chr8-49679557-INS-423 | chr8-49679557-INS-380 |
| chr8-49691417-INS-301 | chr8-49691415-INS-331 |
| chr8-49699843-INS-121 | chr8-49699843-INS-89 |
| chr8-49730887-INS-92 | chr8-49730882-INS-94 |
| chr8-49780607-INS-130 | chr8-49780634-INS-151 |
| chr8-49795177-INS-96 | chr8-49795177-INS-99 |
| chr8-49806308-INS-56 | chr8-49806308-INS-56 |
| chr8-49863392-INS-88 | chr8-49863392-INS-88 |
| chr8-49898659-INS-60 | chr8-49898659-INS-60 |
| chr8-49964342-INS-103 | chr8-49964349-INS-69 |
| chr8-49965281-INS-52 | chr8-49965281-INS-56 |
| chr8-49968347-INS-6473 | chr8-49968348-INS-6473 |
| chr8-49970017-INS-51 | chr8-49970017-INS-51 |
| chr8-50002103-INS-67 | chr8-50002103-INS-84 |
| chr8-50038491-INS-71 | chr8-50038491-INS-71 |
| chr8-50088044-INS-64 | chr8-50088044-INS-66 |
| chr8-50090464-INS-307 | chr8-50090464-INS-307 |
| chr8-50143683-INS-723 | chr8-50143683-INS-723 |
| chr8-50149530-INS-59 | chr8-50149530-INS-62 |
| chr8-50151004-INS-4935 | chr8-50151005-INS-4935 |
| chr8-50154213-INS-414 | chr8-50154210-INS-417 |
| chr8-50157903-INS-75 | chr8-50157869-INS-59 |
| chr8-50160690-INS-66 | chr8-50160690-INS-64 |
| chr8-50162019-INS-1547 | chr8-50162020-INS-1548 |
| chr8-50163877-INS-98 | chr8-50163877-INS-101 |
| chr8-50170894-INS-6464 | chr8-50170904-INS-6447 |
| chr8-50173074-INS-72 | chr8-50173074-INS-105 |
| chr8-50204068-INS-98 | chr8-50204068-INS-96 |
| chr8-50213663-INS-118 | chr8-50213663-INS-117 |
| chr8-50220219-INS-132 | chr8-50220222-INS-170 |
| chr8-50266855-INS-1592 | chr8-50266856-INS-1596 |
| chr8-50301772-INS-121 | chr8-50301772-INS-122 |
| chr8-50306729-INS-199 | chr8-50306729-INS-199 |
| chr8-50315944-INS-60 | chr8-50315944-INS-60 |
| chr8-50332222-INS-6081 | chr8-50332223-INS-6081 |
| chr8-50353755-INS-56 | chr8-50353755-INS-59 |
| chr8-50360993-INS-74 | chr8-50360993-INS-78 |
| chr8-50363116-INS-461 | chr8-50363116-INS-461 |
| chr8-50380956-INS-150 | chr8-50380956-INS-151 |
| chr8-50415817-INS-3808 | chr8-50415818-INS-3806 |
| chr8-50417603-INS-6946 | chr8-50417604-INS-6947 |
| chr8-50462537-INS-713 | chr8-50462537-INS-712 |
| chr8-50474524-INS-77 | chr8-50474524-INS-52 |
| chr8-50475932-INS-51 | chr8-50475933-INS-61 |
| chr8-50506625-INS-260 | chr8-50506625-INS-260 |
| chr8-50566027-INS-174 | chr8-50566027-INS-175 |
| chr8-50576854-INS-300 | chr8-50576860-INS-310 |
| chr8-50580796-INS-56 | chr8-50580796-INS-58 |
| chr8-50600211-INS-190 | chr8-50600211-INS-190 |
| chr8-50619734-INS-686 | chr8-50619734-INS-686 |

| | |
|---|---|
| chr8-50635785-INS-76 | chr8-50635797-INS-76 |
| chr8-50645194-INS-56 | chr8-50645198-INS-52 |
| chr8-50665704-INS-2055 | chr8-50665705-INS-2055 |
| chr8-50695157-INS-241 | chr8-50695157-INS-241 |
| chr8-50704785-INS-204 | chr8-50704785-INS-203 |
| chr8-50724372-INS-68 | chr8-50724383-INS-73 |
| chr8-50725686-INS-213 | chr8-50725686-INS-213 |
| chr8-50752275-INS-52 | chr8-50752275-INS-52 |
| chr8-50768915-INS-52 | chr8-50768915-INS-52 |
| chr8-50809616-INS-853 | chr8-50809616-INS-863 |
| chr8-50840924-INS-52 | chr8-50840924-INS-50 |
| chr8-50882873-INS-88 | chr8-50882873-INS-88 |
| chr8-50886246-INS-78 | chr8-50886214-INS-100 |
| chr8-50886909-INS-805 | chr8-50886909-INS-812 |
| chr8-50888016-INS-600 | chr8-50888016-INS-598 |
| chr8-50890962-INS-805 | chr8-50890962-INS-806 |
| chr8-50928814-INS-384 | chr8-50928814-INS-384 |
| chr8-50948541-INS-6587 | chr8-50948542-INS-6589 |
| chr8-50952776-INS-1311 | chr8-50952774-INS-1308 |
| chr8-50964631-INS-75 | chr8-50964631-INS-75 |
| chr8-50965550-INS-1320 | chr8-50965551-INS-1320 |
| chr8-51015074-INS-2581 | chr8-51015075-INS-2581 |
| chr8-51059022-INS-368 | chr8-51059022-INS-368 |
| chr8-51062259-INS-1707 | chr8-51062260-INS-1707 |
| chr8-51087185-INS-60 | chr8-51087185-INS-50 |
| chr8-51105507-INS-1611 | chr8-51105508-INS-1610 |
| chr8-51157852-INS-100 | chr8-51157880-INS-52 |
| chr8-51167047-INS-677 | chr8-51167047-INS-677 |
| chr8-51192933-INS-76 | chr8-51192933-INS-56 |
| chr8-51196093-INS-64 | chr8-51196094-INS-66 |
| chr8-51214868-INS-88 | chr8-51214868-INS-64 |
| chr8-51230534-INS-217 | chr8-51230534-INS-212 |
| chr8-51247152-INS-72 | chr8-51247152-INS-56 |
| chr8-51263726-INS-70 | chr8-51263726-INS-70 |
| chr8-51267250-INS-829 | chr8-51267250-INS-829 |
| chr8-51274493-INS-78 | chr8-51274457-INS-99 |
| chr8-51308797-INS-58 | chr8-51308797-INS-58 |
| chr8-51327086-INS-82 | chr8-51327086-INS-82 |
| chr8-51327348-INS-830 | chr8-51327348-INS-830 |
| chr8-51344051-INS-2342 | chr8-51344052-INS-2337 |
| chr8-51345694-INS-87 | chr8-51345699-INS-102 |
| chr8-51351713-INS-1435 | chr8-51351714-INS-1437 |
| chr8-51358409-INS-220 | chr8-51358409-INS-220 |
| chr8-51371913-INS-54 | chr8-51371913-INS-56 |
| chr8-51382069-INS-52 | chr8-51382069-INS-52 |
| chr8-51393986-INS-2799 | chr8-51393987-INS-2798 |
| chr8-51432887-INS-572 | chr8-51432887-INS-572 |
| chr8-51434610-INS-4449 | chr8-51434611-INS-4449 |
| chr8-51435341-INS-84 | chr8-51435310-INS-104 |
| chr8-51472196-INS-552 | chr8-51472196-INS-552 |
| chr8-51511198-INS-4376 | chr8-51511199-INS-4376 |
| chr8-51511298-INS-58 | chr8-51511304-INS-60 |
| chr8-51520027-INS-56 | chr8-51520027-INS-72 |
| chr8-51549784-INS-356 | chr8-51549785-INS-400 |
| chr8-51566271-INS-90 | chr8-51566263-INS-84 |
| chr8-51567679-INS-60 | chr8-51567701-INS-50 |
| chr8-51570384-INS-6620 | chr8-51570385-INS-6621 |
| chr8-51591608-INS-673 | chr8-51591608-INS-673 |
| chr8-51600261-INS-627 | chr8-51600261-INS-627 |
| chr8-51613938-INS-120 | chr8-51613938-INS-152 |
| chr8-51645069-INS-54 | chr8-51645069-INS-50 |
| chr8-51646463-INS-176 | chr8-51646463-INS-176 |
| chr8-51649989-INS-2085 | chr8-51649990-INS-2085 |
| chr8-51651003-INS-935 | chr8-51651003-INS-936 |
| chr8-51652428-INS-138 | chr8-51652428-INS-122 |
| chr8-51661045-INS-143 | chr8-51661045-INS-142 |
| chr8-51693395-INS-373 | chr8-51693395-INS-373 |
| chr8-51706160-INS-100 | chr8-51706160-INS-100 |
| chr8-51745706-INS-268 | chr8-51745706-INS-268 |
| chr8-51750275-INS-1457 | chr8-51750276-INS-1457 |
| chr8-51759057-INS-59 | chr8-51759057-INS-59 |
| chr8-51781709-INS-168 | chr8-51781709-INS-168 |
| chr8-51799088-INS-282 | chr8-51799088-INS-282 |
| chr8-51856602-INS-325 | chr8-51856601-INS-326 |
| chr8-51875709-INS-2929 | chr8-51875710-INS-2930 |
| chr8-51896232-INS-302 | chr8-51896232-INS-304 |
| chr8-51897324-INS-61 | chr8-51897324-INS-72 |

| | |
|---|---|
| chr8-51898686-INS-313 | chr8-51898675-INS-319 |
| chr8-51948764-INS-57 | chr8-51948759-INS-63 |
| chr8-51963134-INS-87 | chr8-51963134-INS-62 |
| chr8-51972371-INS-95 | chr8-51972371-INS-87 |
| chr8-52054610-INS-226 | chr8-52054610-INS-226 |
| chr8-52106362-INS-13851 | chr8-52106363-INS-13855 |
| chr8-52114412-INS-70 | chr8-52114413-INS-78 |
| chr8-52131527-INS-84 | chr8-52131540-INS-56 |
| chr8-52133955-INS-1530 | chr8-52133956-INS-1529 |
| chr8-52150817-INS-6153 | chr8-52150818-INS-6152 |
| chr8-52161727-INS-4550 | chr8-52161728-INS-4550 |
| chr8-52181637-INS-52 | chr8-52181637-INS-64 |
| chr8-52220710-INS-376 | chr8-52220710-INS-376 |
| chr8-52241255-INS-92 | chr8-52241234-INS-63 |
| chr8-52251554-INS-58 | chr8-52251554-INS-62 |
| chr8-52253254-INS-73 | chr8-52253254-INS-79 |
| chr8-52258770-INS-58 | chr8-52258770-INS-58 |
| chr8-52270571-INS-652 | chr8-52270571-INS-652 |
| chr8-52271331-INS-91 | chr8-52271331-INS-91 |
| chr8-52276694-INS-1344 | chr8-52276695-INS-1344 |
| chr8-52294069-INS-302 | chr8-52294069-INS-301 |
| chr8-52307059-INS-80 | chr8-52307059-INS-80 |
| chr8-52312747-INS-69 | chr8-52312747-INS-63 |
| chr8-52314933-INS-608 | chr8-52314933-INS-608 |
| chr8-52334798-INS-68 | chr8-52334790-INS-80 |
| chr8-52375483-INS-341 | chr8-52375483-INS-341 |
| chr8-52400154-INS-434 | chr8-52400154-INS-434 |
| chr8-52404499-INS-88 | chr8-52404499-INS-88 |
| chr8-52447287-INS-6401 | chr8-52447288-INS-6401 |
| chr8-52471524-INS-52 | chr8-52471524-INS-52 |
| chr8-52472597-INS-544 | chr8-52472597-INS-544 |
| chr8-52493240-INS-724 | chr8-52493240-INS-723 |
| chr8-52503551-INS-53 | chr8-52503561-INS-50 |
| chr8-52540969-INS-6566 | chr8-52540970-INS-6566 |
| chr8-52541296-INS-1787 | chr8-52541297-INS-1788 |
| chr8-52555966-INS-172 | chr8-52555966-INS-172 |
| chr8-52556890-INS-2361 | chr8-52556891-INS-2359 |
| chr8-52566852-INS-52 | chr8-52566852-INS-52 |
| chr8-52572169-INS-75 | chr8-52572169-INS-59 |
| chr8-52583639-INS-53 | chr8-52583612-INS-55 |
| chr8-52588264-INS-90 | chr8-52588266-INS-90 |
| chr8-52609021-INS-3084 | chr8-52609022-INS-3084 |
| chr8-52619427-INS-121 | chr8-52619427-INS-121 |
| chr8-52634189-INS-6013 | chr8-52634190-INS-6013 |
| chr8-52636281-INS-102 | chr8-52636281-INS-94 |
| chr8-52664324-INS-58 | chr8-52664324-INS-62 |
| chr8-52679740-INS-142 | chr8-52679740-INS-144 |
| chr8-52694529-INS-497 | chr8-52694529-INS-497 |
| chr8-52712894-INS-172 | chr8-52712894-INS-168 |
| chr8-52716469-INS-54 | chr8-52716467-INS-86 |
| chr8-52736338-INS-3433 | chr8-52736339-INS-3432 |
| chr8-52747464-INS-261 | chr8-52747464-INS-261 |
| chr8-52749659-INS-1253 | chr8-52749662-INS-1234 |
| chr8-52751851-INS-74 | chr8-52751851-INS-78 |
| chr8-52792886-INS-52 | chr8-52792886-INS-54 |
| chr8-52821553-INS-691 | chr8-52821551-INS-692 |
| chr8-52840145-INS-718 | chr8-52840145-INS-717 |
| chr8-52895648-INS-124 | chr8-52895648-INS-124 |
| chr8-52912419-INS-445 | chr8-52912419-INS-445 |
| chr8-52920801-INS-88 | chr8-52920801-INS-88 |
| chr8-52929908-INS-120 | chr8-52929895-INS-126 |
| chr8-53000379-INS-1004 | chr8-53000380-INS-1000 |
| chr8-53015096-INS-144 | chr8-53015111-INS-103 |
| chr8-53023546-INS-463 | chr8-53023546-INS-464 |
| chr8-53032829-INS-618 | chr8-53032832-INS-622 |
| chr8-53063913-INS-86 | chr8-53063913-INS-62 |
| chr8-53101701-INS-1242 | chr8-53101702-INS-1222 |
| chr8-53117096-INS-390 | chr8-53117096-INS-390 |
| chr8-53135445-INS-72 | chr8-53135441-INS-56 |
| chr8-53153775-INS-59 | chr8-53153775-INS-56 |
| chr8-53186379-INS-472 | chr8-53186379-INS-472 |
| chr8-53201788-INS-269 | chr8-53201788-INS-267 |
| chr8-53233965-INS-1968 | chr8-53233966-INS-1968 |
| chr8-53244705-INS-80 | chr8-53244713-INS-69 |
| chr8-53261579-INS-1088 | chr8-53261586-INS-1069 |
| chr8-53265716-INS-113 | chr8-53265716-INS-113 |
| chr8-53286357-INS-829 | chr8-53286357-INS-828 |

| | |
|---|---|
| chr8-53373957-INS-59 | chr8-53373964-INS-56 |
| chr8-53375732-INS-63 | chr8-53375733-INS-54 |
| chr8-53382931-INS-74 | chr8-53382939-INS-94 |
| chr8-53391784-INS-183 | chr8-53391784-INS-184 |
| chr8-53394481-INS-200 | chr8-53394481-INS-201 |
| chr8-53404551-INS-60 | chr8-53404551-INS-62 |
| chr8-53471889-INS-670 | chr8-53471889-INS-670 |
| chr8-53497658-INS-423 | chr8-53497658-INS-423 |
| chr8-53514465-INS-1861 | chr8-53514466-INS-1859 |
| chr8-53530239-INS-54 | chr8-53530239-INS-54 |
| chr8-53562363-INS-136 | chr8-53562367-INS-128 |
| chr8-53580879-INS-65 | chr8-53580891-INS-92 |
| chr8-53622786-INS-6857 | chr8-53622787-INS-6857 |
| chr8-53628522-INS-6539 | chr8-53628523-INS-6540 |
| chr8-53637739-INS-3175 | chr8-53637740-INS-3126 |
| chr8-53644488-INS-959 | chr8-53644488-INS-959 |
| chr8-53645118-INS-56 | chr8-53645118-INS-56 |
| chr8-53668877-INS-448 | chr8-53668877-INS-448 |
| chr8-53670034-INS-187 | chr8-53670034-INS-187 |
| chr8-53706289-INS-56 | chr8-53706289-INS-56 |
| chr8-53714250-INS-177 | chr8-53714250-INS-186 |
| chr8-53733273-INS-810 | chr8-53733278-INS-791 |
| chr8-53748718-INS-108 | chr8-53748718-INS-108 |
| chr8-53756081-INS-410 | chr8-53756081-INS-410 |
| chr8-53757439-INS-965 | chr8-53757439-INS-965 |
| chr8-53769438-INS-79 | chr8-53769438-INS-80 |
| chr8-53784240-INS-6437 | chr8-53784241-INS-6437 |
| chr8-53804870-INS-52 | chr8-53804870-INS-100 |
| chr8-53849164-INS-70 | chr8-53849164-INS-72 |
| chr8-53855854-INS-102 | chr8-53855854-INS-100 |
| chr8-53870675-INS-52 | chr8-53870678-INS-69 |
| chr8-53888196-INS-280 | chr8-53888199-INS-287 |
| chr8-53900927-INS-59 | chr8-53900927-INS-51 |
| chr8-53922162-INS-91 | chr8-53922162-INS-91 |
| chr8-53927720-INS-56 | chr8-53927720-INS-60 |
| chr8-53936183-INS-489 | chr8-53936183-INS-489 |
| chr8-53941734-INS-78 | chr8-53941734-INS-50 |
| chr8-53976104-INS-122 | chr8-53976106-INS-131 |
| chr8-53988594-INS-74 | chr8-53988594-INS-74 |
| chr8-54011703-INS-98 | chr8-54011703-INS-98 |
| chr8-54020900-INS-73 | chr8-54020898-INS-58 |
| chr8-54072759-INS-75 | chr8-54072759-INS-75 |
| chr8-54105242-INS-66 | chr8-54105213-INS-60 |
| chr8-54107758-INS-654 | chr8-54107758-INS-654 |
| chr8-54115584-INS-56 | chr8-54115584-INS-56 |
| chr8-54118095-INS-81 | chr8-54118095-INS-105 |
| chr8-54143802-INS-6462 | chr8-54143803-INS-6458 |
| chr8-54153200-INS-54 | chr8-54153200-INS-54 |
| chr8-54196673-INS-201 | chr8-54196673-INS-202 |
| chr8-54202857-INS-1199 | chr8-54202858-INS-1198 |
| chr8-54218382-INS-52 | chr8-54218373-INS-60 |
| chr8-54237685-INS-64 | chr8-54237685-INS-64 |
| chr8-54266723-INS-186 | chr8-54266723-INS-216 |
| chr8-54282726-INS-76 | chr8-54282726-INS-76 |
| chr8-54307097-INS-80 | chr8-54307109-INS-82 |
| chr8-54309379-INS-729 | chr8-54309379-INS-729 |
| chr8-54321949-INS-74 | chr8-54321949-INS-74 |
| chr8-54408377-INS-105 | chr8-54408372-INS-80 |
| chr8-54419382-INS-78 | chr8-54419382-INS-78 |
| chr8-54438387-INS-54 | chr8-54438387-INS-78 |
| chr8-54439970-INS-6373 | chr8-54439971-INS-6370 |
| chr8-54449704-INS-369 | chr8-54449704-INS-371 |
| chr8-54472737-INS-190 | chr8-54472737-INS-190 |
| chr8-54475562-INS-667 | chr8-54475562-INS-666 |
| chr8-54549927-INS-52 | chr8-54549927-INS-52 |
| chr8-54568641-INS-68 | chr8-54568641-INS-68 |
| chr8-54578596-INS-236 | chr8-54578596-INS-236 |
| chr8-54648554-INS-82 | chr8-54648554-INS-82 |
| chr8-54667555-INS-467 | chr8-54667555-INS-469 |
| chr8-54677460-INS-54 | chr8-54677478-INS-60 |
| chr8-54681047-INS-58 | chr8-54681047-INS-62 |
| chr8-54689541-INS-4423 | chr8-54689542-INS-4423 |
| chr8-54693413-INS-167 | chr8-54693413-INS-167 |
| chr8-54697350-INS-2884 | chr8-54697351-INS-2884 |
| chr8-54701952-INS-62 | chr8-54701952-INS-62 |
| chr8-54706882-INS-78 | chr8-54706882-INS-74 |
| chr8-54722181-INS-65 | chr8-54722197-INS-72 |

| | |
|---|---|
| chr8-54793340-INS-171 | chr8-54793340-INS-171 |
| chr8-54800711-INS-390 | chr8-54800711-INS-390 |
| chr8-54812199-INS-462 | chr8-54812199-INS-462 |
| chr8-54830426-INS-778 | chr8-54830426-INS-779 |
| chr8-54832995-INS-144 | chr8-54832995-INS-166 |
| chr8-54836919-INS-599 | chr8-54836919-INS-598 |
| chr8-54850810-INS-91 | chr8-54850774-INS-120 |
| chr8-54856938-INS-709 | chr8-54856938-INS-709 |
| chr8-54872094-INS-366 | chr8-54872094-INS-368 |
| chr8-54877997-INS-60 | chr8-54877997-INS-50 |
| chr8-54946178-INS-671 | chr8-54946178-INS-671 |
| chr8-54947544-INS-140 | chr8-54947544-INS-104 |
| chr8-54961590-INS-82 | chr8-54961590-INS-56 |
| chr8-54971207-INS-1189 | chr8-54971208-INS-1189 |
| chr8-54972759-INS-1321 | chr8-54972760-INS-1321 |
| chr8-54994705-INS-1761 | chr8-54994706-INS-1762 |
| chr8-55022275-INS-3258 | chr8-55022276-INS-3258 |
| chr8-55022990-INS-577 | chr8-55022990-INS-577 |
| chr8-55037293-INS-120 | chr8-55037308-INS-88 |
| chr8-55045952-INS-204 | chr8-55045952-INS-199 |
| chr8-55061273-INS-1902 | chr8-55061274-INS-1902 |
| chr8-55120173-INS-92 | chr8-55120193-INS-102 |
| chr8-55129150-INS-56 | chr8-55129150-INS-56 |
| chr8-55144264-INS-55 | chr8-55144264-INS-60 |
| chr8-55195539-INS-155 | chr8-55195539-INS-155 |
| chr8-55318306-INS-76 | chr8-55318306-INS-76 |
| chr8-55423457-INS-354 | chr8-55423457-INS-354 |
| chr8-55461806-INS-75 | chr8-55461806-INS-75 |
| chr8-55470740-INS-462 | chr8-55470741-INS-462 |
| chr8-55496909-INS-7186 | chr8-55496910-INS-7186 |
| chr8-55557030-INS-92 | chr8-55557031-INS-92 |
| chr8-55563511-INS-54 | chr8-55563523-INS-57 |
| chr8-55564761-INS-56 | chr8-55564757-INS-52 |
| chr8-56060503-INS-68 | chr8-56060531-INS-64 |
| chr8-56102684-INS-52 | chr8-56102685-INS-74 |
| chr8-56123687-INS-356 | chr8-56123687-INS-356 |
| chr8-56171037-INS-116 | chr8-56171037-INS-96 |
| chr8-56171186-INS-1549 | chr8-56171187-INS-1549 |
| chr8-56217840-INS-163 | chr8-56217840-INS-163 |
| chr8-56235338-INS-506 | chr8-56235338-INS-506 |
| chr8-56258781-INS-145 | chr8-56258782-INS-145 |
| chr8-56275299-INS-1112 | chr8-56275300-INS-1144 |
| chr8-56311037-INS-2747 | chr8-56311038-INS-2747 |
| chr8-56340928-INS-111 | chr8-56340925-INS-119 |
| chr8-56343737-INS-194 | chr8-56343738-INS-191 |
| chr8-56372305-INS-416 | chr8-56372305-INS-417 |
| chr8-56380454-INS-544 | chr8-56380454-INS-544 |
| chr8-56419240-INS-68 | chr8-56419241-INS-76 |
| chr8-56496445-INS-52 | chr8-56496445-INS-56 |
| chr8-56525727-INS-100 | chr8-56525727-INS-102 |
| chr8-56612032-INS-6456 | chr8-56612032-INS-6458 |
| chr8-56640001-INS-53 | chr8-56640001-INS-53 |
| chr8-56642018-INS-68 | chr8-56642018-INS-68 |
| chr8-56673264-INS-129 | chr8-56673266-INS-81 |
| chr8-56686167-INS-90 | chr8-56686165-INS-80 |
| chr8-56743211-INS-61 | chr8-56743211-INS-61 |
| chr8-56763456-INS-92 | chr8-56763477-INS-68 |
| chr8-56787960-INS-59 | chr8-56787948-INS-60 |
| chr8-56802130-INS-51 | chr8-56802134-INS-56 |
| chr8-56839473-INS-52 | chr8-56839473-INS-50 |
| chr8-56870643-INS-86 | chr8-56870643-INS-86 |
| chr8-56879789-INS-52 | chr8-56879789-INS-52 |
| chr8-56900270-INS-62 | chr8-56900270-INS-60 |
| chr8-56904426-INS-175 | chr8-56904426-INS-175 |
| chr8-56909024-INS-6193 | chr8-56909029-INS-6199 |
| chr8-56912274-INS-6644 | chr8-56912275-INS-6644 |
| chr8-56917638-INS-62 | chr8-56917615-INS-66 |
| chr8-56937444-INS-3183 | chr8-56937445-INS-3183 |
| chr8-56942130-INS-70 | chr8-56942131-INS-105 |
| chr8-56995455-INS-690 | chr8-56995455-INS-691 |
| chr8-57000358-INS-121 | chr8-57000358-INS-85 |
| chr8-57001555-INS-154 | chr8-57001555-INS-114 |
| chr8-57006966-INS-1453 | chr8-57006967-INS-1451 |
| chr8-57009989-INS-75 | chr8-57009989-INS-63 |
| chr8-57034104-INS-3321 | chr8-57034105-INS-3321 |
| chr8-57056761-INS-102 | chr8-57056761-INS-104 |
| chr8-57068536-INS-6327 | chr8-57068537-INS-6336 |

| | |
|---|---|
| chr8-57078296-INS-116 | chr8-57078293-INS-165 |
| chr8-57090207-INS-62 | chr8-57090207-INS-62 |
| chr8-57096649-INS-52 | chr8-57096649-INS-52 |
| chr8-57100762-INS-54 | chr8-57100762-INS-58 |
| chr8-57110039-INS-55 | chr8-57110068-INS-56 |
| chr8-57131623-INS-160 | chr8-57131623-INS-168 |
| chr8-57134645-INS-140 | chr8-57134645-INS-140 |
| chr8-57171886-INS-101 | chr8-57171886-INS-100 |
| chr8-57177688-INS-1857 | chr8-57177689-INS-1857 |
| chr8-57204074-INS-68 | chr8-57204074-INS-68 |
| chr8-57213134-INS-94 | chr8-57213119-INS-79 |
| chr8-57219224-INS-155 | chr8-57219224-INS-155 |
| chr8-57228238-INS-140 | chr8-57228238-INS-92 |
| chr8-57244190-INS-984 | chr8-57244190-INS-954 |
| chr8-57273042-INS-349 | chr8-57273042-INS-349 |
| chr8-57305384-INS-555 | chr8-57305384-INS-589 |
| chr8-57315942-INS-104 | chr8-57315952-INS-56 |
| chr8-57344560-INS-146 | chr8-57344560-INS-143 |
| chr8-57423295-INS-97 | chr8-57423297-INS-102 |
| chr8-57429518-INS-61 | chr8-57429518-INS-61 |
| chr8-57457686-INS-634 | chr8-57457687-INS-635 |
| chr8-57465275-INS-7179 | chr8-57465276-INS-7180 |
| chr8-57514486-INS-51 | chr8-57514486-INS-53 |
| chr8-57514631-INS-446 | chr8-57514631-INS-448 |
| chr8-57514902-INS-2854 | chr8-57514903-INS-2854 |
| chr8-57516791-INS-164 | chr8-57516791-INS-164 |
| chr8-57572538-INS-54 | chr8-57572538-INS-56 |
| chr8-57586951-INS-60 | chr8-57586951-INS-86 |
| chr8-57622436-INS-1316 | chr8-57622437-INS-1324 |
| chr8-57636086-INS-212 | chr8-57636086-INS-212 |
| chr8-57650925-INS-248 | chr8-57650925-INS-200 |
| chr8-57662199-INS-6736 | chr8-57662200-INS-6736 |
| chr8-57681528-INS-2620 | chr8-57681529-INS-2620 |
| chr8-57681755-INS-171 | chr8-57681754-INS-171 |
| chr8-57683515-INS-64 | chr8-57683515-INS-64 |
| chr8-57685799-INS-76 | chr8-57685799-INS-76 |
| chr8-57686681-INS-582 | chr8-57686681-INS-578 |
| chr8-57736228-INS-146 | chr8-57736237-INS-112 |
| chr8-57758187-INS-54 | chr8-57758187-INS-54 |
| chr8-57762005-INS-55 | chr8-57762005-INS-57 |
| chr8-57780040-INS-54 | chr8-57780008-INS-50 |
| chr8-57804307-INS-130 | chr8-57804310-INS-126 |
| chr8-57826389-INS-199 | chr8-57826390-INS-199 |
| chr8-57828204-INS-85 | chr8-57828193-INS-65 |
| chr8-57836355-INS-196 | chr8-57836355-INS-196 |
| chr8-57838713-INS-76 | chr8-57838713-INS-73 |
| chr8-57852258-INS-75 | chr8-57852258-INS-75 |
| chr8-57861918-INS-134 | chr8-57861918-INS-166 |
| chr8-57864879-INS-74 | chr8-57864879-INS-74 |
| chr8-57895360-INS-250 | chr8-57895361-INS-200 |
| chr8-57908540-INS-60 | chr8-57908540-INS-64 |
| chr8-57941896-INS-191 | chr8-57941898-INS-188 |
| chr8-57971816-INS-90 | chr8-57971816-INS-96 |
| chr8-57972486-INS-66 | chr8-57972486-INS-66 |
| chr8-57992089-INS-187 | chr8-57992089-INS-187 |
| chr8-57997819-INS-263 | chr8-57997819-INS-263 |
| chr8-58006329-INS-56 | chr8-58006329-INS-56 |
| chr8-58054618-INS-58 | chr8-58054618-INS-64 |
| chr8-58066978-INS-54 | chr8-58066978-INS-58 |
| chr8-58068029-INS-54 | chr8-58068029-INS-54 |
| chr8-58099365-INS-76 | chr8-58099365-INS-57 |
| chr8-58103809-INS-175 | chr8-58103809-INS-176 |
| chr8-58122695-INS-51 | chr8-58122695-INS-51 |
| chr8-58170385-INS-459 | chr8-58170386-INS-459 |
| chr8-58184580-INS-990 | chr8-58184580-INS-991 |
| chr8-58202541-INS-6298 | chr8-58202542-INS-6298 |
| chr8-58215594-INS-169 | chr8-58215594-INS-169 |
| chr8-58247423-INS-52 | chr8-58247423-INS-56 |
| chr8-58265944-INS-60 | chr8-58265944-INS-60 |
| chr8-58269756-INS-53 | chr8-58269756-INS-84 |
| chr8-58272230-INS-214 | chr8-58272230-INS-214 |
| chr8-58276545-INS-121 | chr8-58276545-INS-122 |
| chr8-58285710-INS-83 | chr8-58285710-INS-92 |
| chr8-58299507-INS-52 | chr8-58299507-INS-52 |
| chr8-58307825-INS-100 | chr8-58307799-INS-120 |
| chr8-58341625-INS-211 | chr8-58341625-INS-211 |
| chr8-58342181-INS-53 | chr8-58342182-INS-50 |

| | |
|---|---|
| chr8-58343500-INS-100 | chr8-58343500-INS-102 |
| chr8-58347824-INS-2794 | chr8-58347825-INS-2794 |
| chr8-58355355-INS-51 | chr8-58355353-INS-56 |
| chr8-58372123-INS-82 | chr8-58372131-INS-55 |
| chr8-58390217-INS-65 | chr8-58390217-INS-65 |
| chr8-58395241-INS-222 | chr8-58395241-INS-222 |
| chr8-58395698-INS-4094 | chr8-58395699-INS-4094 |
| chr8-58399862-INS-55 | chr8-58399862-INS-55 |
| chr8-58406818-INS-7235 | chr8-58406819-INS-7237 |
| chr8-58407375-INS-65 | chr8-58407328-INS-105 |
| chr8-58425234-INS-425 | chr8-58425234-INS-425 |
| chr8-58433529-INS-7387 | chr8-58433530-INS-7389 |
| chr8-58438970-INS-208 | chr8-58438970-INS-208 |
| chr8-58439530-INS-395 | chr8-58439531-INS-389 |
| chr8-58442143-INS-54 | chr8-58442143-INS-54 |
| chr8-58452746-INS-51 | chr8-58452746-INS-54 |
| chr8-58456954-INS-1340 | chr8-58456956-INS-1339 |
| chr8-58475253-INS-400 | chr8-58475253-INS-400 |
| chr8-58484838-INS-67 | chr8-58484838-INS-84 |
| chr8-58496726-INS-53 | chr8-58496726-INS-50 |
| chr8-58500034-INS-1401 | chr8-58500035-INS-1401 |
| chr8-58502609-INS-661 | chr8-58502609-INS-661 |
| chr8-58507538-INS-865 | chr8-58507538-INS-865 |
| chr8-58510400-INS-51 | chr8-58510401-INS-55 |
| chr8-58520215-INS-53 | chr8-58520238-INS-62 |
| chr8-58541149-INS-120 | chr8-58541149-INS-120 |
| chr8-58560253-INS-60 | chr8-58560253-INS-62 |
| chr8-58565014-INS-98 | chr8-58565014-INS-100 |
| chr8-58605066-INS-60 | chr8-58605068-INS-50 |
| chr8-58606135-INS-58 | chr8-58606135-INS-53 |
| chr8-58633034-INS-74 | chr8-58633034-INS-70 |
| chr8-58658958-INS-292 | chr8-58658943-INS-291 |
| chr8-58686776-INS-66 | chr8-58686776-INS-62 |
| chr8-58710295-INS-660 | chr8-58710296-INS-663 |
| chr8-58717597-INS-58 | chr8-58717597-INS-58 |
| chr8-58734433-INS-198 | chr8-58734433-INS-198 |
| chr8-58736187-INS-103 | chr8-58736187-INS-101 |
| chr8-58740790-INS-1640 | chr8-58740791-INS-1645 |
| chr8-58751234-INS-208 | chr8-58751235-INS-208 |
| chr8-58769840-INS-1536 | chr8-58769841-INS-1536 |
| chr8-58783435-INS-84 | chr8-58783435-INS-76 |
| chr8-58794807-INS-70 | chr8-58794807-INS-72 |
| chr8-58808086-INS-56 | chr8-58808086-INS-56 |
| chr8-58810194-INS-110 | chr8-58810194-INS-110 |
| chr8-58812825-INS-68 | chr8-58812825-INS-68 |
| chr8-58838555-INS-88 | chr8-58838542-INS-96 |
| chr8-58839254-INS-136 | chr8-58839224-INS-140 |
| chr8-58888948-INS-191 | chr8-58888957-INS-191 |
| chr8-58903874-INS-604 | chr8-58903874-INS-602 |
| chr8-58910945-INS-412 | chr8-58910945-INS-412 |
| chr8-58914526-INS-83 | chr8-58914526-INS-92 |
| chr8-58927993-INS-4354 | chr8-58927994-INS-4354 |
| chr8-58945058-INS-1844 | chr8-58945059-INS-1844 |
| chr8-58966646-INS-6586 | chr8-58966647-INS-6586 |
| chr8-58974849-INS-241 | chr8-58974849-INS-241 |
| chr8-58976084-INS-72 | chr8-58976084-INS-72 |
| chr8-58986637-INS-826 | chr8-58986630-INS-837 |
| chr8-59006796-INS-66 | chr8-59006796-INS-64 |
| chr8-59013692-INS-5020 | chr8-59013693-INS-5020 |
| chr8-59017836-INS-6461 | chr8-59017837-INS-6461 |
| chr8-59022362-INS-1647 | chr8-59022363-INS-1647 |
| chr8-59052869-INS-108 | chr8-59052869-INS-108 |
| chr8-59087278-INS-786 | chr8-59087278-INS-762 |
| chr8-59106621-INS-88 | chr8-59106621-INS-84 |
| chr8-59111682-INS-496 | chr8-59111682-INS-496 |
| chr8-59126129-INS-991 | chr8-59126129-INS-1008 |
| chr8-59135502-INS-114 | chr8-59135517-INS-118 |
| chr8-59170916-INS-52 | chr8-59170916-INS-52 |
| chr8-59176036-INS-469 | chr8-59176036-INS-469 |
| chr8-59197704-INS-7311 | chr8-59197705-INS-7311 |
| chr8-59200222-INS-502 | chr8-59200222-INS-502 |
| chr8-59215031-INS-609 | chr8-59215031-INS-616 |
| chr8-59224103-INS-94 | chr8-59224103-INS-92 |
| chr8-59236044-INS-398 | chr8-59236044-INS-418 |
| chr8-59251963-INS-111 | chr8-59251963-INS-111 |
| chr8-59258299-INS-6658 | chr8-59258300-INS-6659 |
| chr8-59261255-INS-102 | chr8-59261255-INS-102 |

| | |
|---|---|
| chr8-59273563-INS-6576 | chr8-59273564-INS-6576 |
| chr8-59289306-INS-1702 | chr8-59289307-INS-1702 |
| chr8-59307766-INS-52 | chr8-59307766-INS-52 |
| chr8-59383279-INS-677 | chr8-59383279-INS-678 |
| chr8-59405697-INS-267 | chr8-59405697-INS-267 |
| chr8-59415413-INS-166 | chr8-59415413-INS-166 |
| chr8-59436914-INS-193 | chr8-59436909-INS-205 |
| chr8-59467823-INS-67 | chr8-59467823-INS-67 |
| chr8-59472803-INS-6609 | chr8-59472804-INS-6609 |
| chr8-59486687-INS-1640 | chr8-59486688-INS-1640 |
| chr8-59496126-INS-423 | chr8-59496127-INS-424 |
| chr8-59519889-INS-90 | chr8-59519889-INS-90 |
| chr8-59521779-INS-69 | chr8-59521779-INS-69 |
| chr8-59564412-INS-68 | chr8-59564406-INS-71 |
| chr8-59601712-INS-228 | chr8-59601712-INS-228 |
| chr8-59611023-INS-85 | chr8-59611023-INS-85 |
| chr8-59660056-INS-1520 | chr8-59660057-INS-1520 |
| chr8-59662375-INS-60 | chr8-59662375-INS-60 |
| chr8-59699446-INS-62 | chr8-59699446-INS-64 |
| chr8-59702246-INS-6594 | chr8-59702247-INS-6596 |
| chr8-59713194-INS-67 | chr8-59713194-INS-67 |
| chr8-59721541-INS-769 | chr8-59721541-INS-772 |
| chr8-59733637-INS-6381 | chr8-59733638-INS-6383 |
| chr8-59742734-INS-193 | chr8-59742734-INS-194 |
| chr8-59765093-INS-75 | chr8-59765093-INS-75 |
| chr8-59798390-INS-69 | chr8-59798387-INS-69 |
| chr8-59801348-INS-476 | chr8-59801354-INS-491 |
| chr8-59810215-INS-210 | chr8-59810215-INS-210 |
| chr8-59814154-INS-52 | chr8-59814154-INS-52 |
| chr8-59822723-INS-522 | chr8-59822723-INS-522 |
| chr8-59844948-INS-69 | chr8-59844947-INS-77 |
| chr8-59846820-INS-60 | chr8-59846820-INS-76 |
| chr8-59858943-INS-140 | chr8-59858943-INS-140 |
| chr8-59890217-INS-65 | chr8-59890215-INS-66 |
| chr8-59898371-INS-91 | chr8-59898371-INS-91 |
| chr8-59906265-INS-1285 | chr8-59906266-INS-1286 |
| chr8-59913852-INS-677 | chr8-59913866-INS-655 |
| chr8-59916180-INS-56 | chr8-59916180-INS-86 |
| chr8-59933575-INS-62 | chr8-59933575-INS-58 |
| chr8-59963715-INS-272 | chr8-59963713-INS-298 |
| chr8-59979070-INS-54 | chr8-59979070-INS-58 |
| chr8-59980171-INS-1453 | chr8-59980172-INS-1453 |
| chr8-59981602-INS-632 | chr8-59981602-INS-632 |
| chr8-60038469-INS-80 | chr8-60038469-INS-98 |
| chr8-60039738-INS-92 | chr8-60039738-INS-98 |
| chr8-60065182-INS-274 | chr8-60065182-INS-273 |
| chr8-60079417-INS-56 | chr8-60079417-INS-56 |
| chr8-60088449-INS-824 | chr8-60088449-INS-824 |
| chr8-60119361-INS-84 | chr8-60119374-INS-62 |
| chr8-60131595-INS-214 | chr8-60131595-INS-214 |
| chr8-60148602-INS-118 | chr8-60148618-INS-92 |
| chr8-60158797-INS-463 | chr8-60158797-INS-509 |
| chr8-60160562-INS-105 | chr8-60160562-INS-112 |
| chr8-60164047-INS-262 | chr8-60164047-INS-262 |
| chr8-60169202-INS-74 | chr8-60169198-INS-70 |
| chr8-60171726-INS-94 | chr8-60171726-INS-94 |
| chr8-60175768-INS-173 | chr8-60175768-INS-201 |
| chr8-60196766-INS-166 | chr8-60196766-INS-166 |
| chr8-60350810-INS-71 | chr8-60350821-INS-58 |
| chr8-60351484-INS-1863 | chr8-60351485-INS-1863 |
| chr8-60365783-INS-167 | chr8-60365786-INS-155 |
| chr8-60368603-INS-1889 | chr8-60368604-INS-1889 |
| chr8-60430678-INS-77 | chr8-60430678-INS-72 |
| chr8-60452684-INS-213 | chr8-60452684-INS-213 |
| chr8-60457209-INS-54 | chr8-60457212-INS-66 |
| chr8-60469330-INS-1683 | chr8-60469331-INS-1683 |
| chr8-60478824-INS-217 | chr8-60478824-INS-217 |
| chr8-60483355-INS-218 | chr8-60483355-INS-218 |
| chr8-60485726-INS-1092 | chr8-60485727-INS-1095 |
| chr8-60486280-INS-61 | chr8-60486275-INS-55 |
| chr8-60496126-INS-54 | chr8-60496126-INS-56 |
| chr8-60504523-INS-1526 | chr8-60504524-INS-1524 |
| chr8-60509415-INS-212 | chr8-60509415-INS-212 |
| chr8-60532337-INS-68 | chr8-60532337-INS-55 |
| chr8-60546007-INS-329 | chr8-60546007-INS-329 |
| chr8-60548616-INS-415 | chr8-60548616-INS-369 |
| chr8-60565149-INS-56 | chr8-60565144-INS-100 |

| | |
|---|---|
| chr8-60571590-INS-81 | chr8-60571593-INS-81 |
| chr8-60616168-INS-202 | chr8-60616168-INS-202 |
| chr8-60636014-INS-604 | chr8-60636014-INS-605 |
| chr8-60642845-INS-70 | chr8-60642825-INS-74 |
| chr8-60657131-INS-82 | chr8-60657131-INS-82 |
| chr8-60661373-INS-68 | chr8-60661373-INS-75 |
| chr8-60666217-INS-3110 | chr8-60666218-INS-3111 |
| chr8-60672261-INS-6903 | chr8-60672262-INS-6904 |
| chr8-60678702-INS-1658 | chr8-60678703-INS-1658 |
| chr8-60698849-INS-104 | chr8-60698850-INS-54 |
| chr8-60709746-INS-2117 | chr8-60709747-INS-2100 |
| chr8-60719678-INS-104 | chr8-60719678-INS-104 |
| chr8-60753507-INS-351 | chr8-60753507-INS-351 |
| chr8-60758655-INS-5518 | chr8-60758656-INS-5518 |
| chr8-60758656-INS-664 | chr8-60758656-INS-664 |
| chr8-60760364-INS-6675 | chr8-60760365-INS-6675 |
| chr8-60766482-INS-88 | chr8-60766482-INS-88 |
| chr8-60791490-INS-276 | chr8-60791490-INS-276 |
| chr8-60797530-INS-172 | chr8-60797518-INS-182 |
| chr8-60812329-INS-6442 | chr8-60812330-INS-6442 |
| chr8-60857308-INS-7350 | chr8-60857309-INS-7355 |
| chr8-60857887-INS-60 | chr8-60857887-INS-60 |
| chr8-60910694-INS-93 | chr8-60910694-INS-93 |
| chr8-60911204-INS-170 | chr8-60911204-INS-170 |
| chr8-60920862-INS-99 | chr8-60920862-INS-99 |
| chr8-60930196-INS-496 | chr8-60930196-INS-496 |
| chr8-60932609-INS-84 | chr8-60932610-INS-80 |
| chr8-60941407-INS-492 | chr8-60941407-INS-470 |
| chr8-60964879-INS-7086 | chr8-60964880-INS-7086 |
| chr8-60990981-INS-630 | chr8-60990981-INS-630 |
| chr8-61023039-INS-58 | chr8-61023036-INS-60 |
| chr8-61063715-INS-84 | chr8-61063715-INS-84 |
| chr8-61077575-INS-63 | chr8-61077575-INS-72 |
| chr8-61078001-INS-440 | chr8-61078001-INS-440 |
| chr8-61082049-INS-128 | chr8-61082019-INS-128 |
| chr8-61105436-INS-229 | chr8-61105436-INS-230 |
| chr8-61115714-INS-6442 | chr8-61115715-INS-6442 |
| chr8-61155203-INS-125 | chr8-61155204-INS-125 |
| chr8-61156234-INS-151 | chr8-61156234-INS-151 |
| chr8-61159647-INS-267 | chr8-61159647-INS-280 |
| chr8-61163595-INS-196 | chr8-61163595-INS-196 |
| chr8-61169162-INS-64 | chr8-61169117-INS-65 |
| chr8-61190675-INS-129 | chr8-61190675-INS-105 |
| chr8-61190891-INS-508 | chr8-61190891-INS-507 |
| chr8-61200451-INS-768 | chr8-61200452-INS-768 |
| chr8-61208006-INS-440 | chr8-61208006-INS-440 |
| chr8-61209578-INS-411 | chr8-61209578-INS-411 |
| chr8-61237415-INS-305 | chr8-61237415-INS-305 |
| chr8-61241221-INS-138 | chr8-61241237-INS-146 |
| chr8-61251386-INS-103 | chr8-61251391-INS-111 |
| chr8-61268903-INS-170 | chr8-61268903-INS-170 |
| chr8-61288007-INS-218 | chr8-61288007-INS-218 |
| chr8-61290029-INS-224 | chr8-61290029-INS-225 |
| chr8-61292954-INS-213 | chr8-61292954-INS-213 |
| chr8-61296638-INS-84 | chr8-61296638-INS-66 |
| chr8-61333088-INS-76 | chr8-61333088-INS-76 |
| chr8-61383167-INS-992 | chr8-61383167-INS-972 |
| chr8-61383177-INS-3994 | chr8-61383178-INS-3998 |
| chr8-61397524-INS-208 | chr8-61397524-INS-208 |
| chr8-61402481-INS-436 | chr8-61402481-INS-410 |
| chr8-61465498-INS-1221 | chr8-61465499-INS-1235 |
| chr8-61469487-INS-106 | chr8-61469487-INS-106 |
| chr8-61475785-INS-146 | chr8-61475785-INS-146 |
| chr8-61489961-INS-57 | chr8-61489961-INS-50 |
| chr8-61524480-INS-52 | chr8-61524480-INS-52 |
| chr8-61542193-INS-1892 | chr8-61542194-INS-1892 |
| chr8-61565508-INS-218 | chr8-61565508-INS-218 |
| chr8-61574592-INS-68 | chr8-61574593-INS-68 |
| chr8-61587415-INS-1861 | chr8-61587416-INS-1861 |
| chr8-61587877-INS-76 | chr8-61587877-INS-76 |
| chr8-61603988-INS-86 | chr8-61603988-INS-79 |
| chr8-61622196-INS-240 | chr8-61622196-INS-240 |
| chr8-61642985-INS-282 | chr8-61642985-INS-282 |
| chr8-61666758-INS-96 | chr8-61666758-INS-56 |
| chr8-61673380-INS-352 | chr8-61673380-INS-358 |
| chr8-61709633-INS-143 | chr8-61709633-INS-143 |
| chr8-61723022-INS-54 | chr8-61723022-INS-54 |

| | |
|---|---|
| chr8-61744189-INS-60 | chr8-61744194-INS-65 |
| chr8-61752984-INS-671 | chr8-61752984-INS-671 |
| chr8-61787713-INS-67 | chr8-61787713-INS-67 |
| chr8-61799942-INS-249 | chr8-61799942-INS-265 |
| chr8-61824894-INS-67 | chr8-61824875-INS-67 |
| chr8-61861940-INS-54 | chr8-61861942-INS-66 |
| chr8-61871040-INS-82 | chr8-61871041-INS-84 |
| chr8-61883401-INS-189 | chr8-61883401-INS-189 |
| chr8-61885406-INS-54 | chr8-61885406-INS-54 |
| chr8-61887269-INS-585 | chr8-61887271-INS-592 |
| chr8-61895065-INS-72 | chr8-61895065-INS-74 |
| chr8-61903333-INS-6612 | chr8-61903334-INS-6612 |
| chr8-61910317-INS-220 | chr8-61910317-INS-220 |
| chr8-61923955-INS-776 | chr8-61923955-INS-776 |
| chr8-61931439-INS-72 | chr8-61931439-INS-72 |
| chr8-61933987-INS-372 | chr8-61933987-INS-342 |
| chr8-61939203-INS-221 | chr8-61939203-INS-221 |
| chr8-61946184-INS-158 | chr8-61946184-INS-158 |
| chr8-61954033-INS-208 | chr8-61954033-INS-209 |
| chr8-61961493-INS-588 | chr8-61961493-INS-588 |
| chr8-62014333-INS-66 | chr8-62014333-INS-60 |
| chr8-62033981-INS-190 | chr8-62033981-INS-190 |
| chr8-62069782-INS-77 | chr8-62069782-INS-68 |
| chr8-62076337-INS-372 | chr8-62076337-INS-373 |
| chr8-62091046-INS-74 | chr8-62091076-INS-80 |
| chr8-62106576-INS-440 | chr8-62106576-INS-440 |
| chr8-62112666-INS-946 | chr8-62112663-INS-921 |
| chr8-62168946-INS-57 | chr8-62168946-INS-57 |
| chr8-62172991-INS-210 | chr8-62172991-INS-210 |
| chr8-62186579-INS-51 | chr8-62186579-INS-85 |
| chr8-62208888-INS-246 | chr8-62208888-INS-246 |
| chr8-62215173-INS-56 | chr8-62215141-INS-68 |
| chr8-62219477-INS-626 | chr8-62219477-INS-626 |
| chr8-62303344-INS-76 | chr8-62303344-INS-76 |
| chr8-62345163-INS-53 | chr8-62345156-INS-59 |
| chr8-62350356-INS-1353 | chr8-62350362-INS-1349 |
| chr8-62365421-INS-93 | chr8-62365373-INS-111 |
| chr8-62366724-INS-80 | chr8-62366685-INS-120 |
| chr8-62400865-INS-60 | chr8-62400835-INS-105 |
| chr8-62411446-INS-71 | chr8-62411446-INS-70 |
| chr8-62420483-INS-53 | chr8-62420485-INS-96 |
| chr8-62449248-INS-6778 | chr8-62449249-INS-6778 |
| chr8-62512781-INS-1553 | chr8-62512782-INS-1553 |
| chr8-62513247-INS-743 | chr8-62513247-INS-744 |
| chr8-62523150-INS-52 | chr8-62523150-INS-52 |
| chr8-62581210-INS-6150 | chr8-62581211-INS-6176 |
| chr8-62581435-INS-81 | chr8-62581438-INS-73 |
| chr8-62598947-INS-131 | chr8-62598948-INS-82 |
| chr8-62629502-INS-7858 | chr8-62629503-INS-7859 |
| chr8-62630206-INS-204 | chr8-62630187-INS-203 |
| chr8-62647444-INS-4229 | chr8-62647445-INS-4229 |
| chr8-62648315-INS-458 | chr8-62648315-INS-458 |
| chr8-62662972-INS-103 | chr8-62662955-INS-71 |
| chr8-62720982-INS-70 | chr8-62720987-INS-66 |
| chr8-62743398-INS-71 | chr8-62743380-INS-71 |
| chr8-62775861-INS-76 | chr8-62775893-INS-80 |
| chr8-62786081-INS-56 | chr8-62786081-INS-56 |
| chr8-62804672-INS-66 | chr8-62804672-INS-50 |
| chr8-62815460-INS-65 | chr8-62815465-INS-57 |
| chr8-62827925-INS-9242 | chr8-62827926-INS-9244 |
| chr8-62830368-INS-76 | chr8-62830372-INS-76 |
| chr8-62847352-INS-56 | chr8-62847353-INS-57 |
| chr8-62854830-INS-556 | chr8-62854830-INS-556 |
| chr8-62874871-INS-502 | chr8-62874871-INS-502 |
| chr8-62875985-INS-7235 | chr8-62875986-INS-7235 |
| chr8-62876962-INS-71 | chr8-62876962-INS-72 |
| chr8-62904079-INS-98 | chr8-62904079-INS-104 |
| chr8-62906938-INS-682 | chr8-62906938-INS-666 |
| chr8-62907030-INS-56 | chr8-62907030-INS-56 |
| chr8-62914161-INS-113 | chr8-62914161-INS-110 |
| chr8-62936171-INS-426 | chr8-62936171-INS-426 |
| chr8-62962742-INS-707 | chr8-62962742-INS-706 |
| chr8-62971563-INS-184 | chr8-62971563-INS-184 |
| chr8-62975899-INS-134 | chr8-62975899-INS-136 |
| chr8-62981262-INS-93 | chr8-62981262-INS-93 |
| chr8-62989066-INS-427 | chr8-62989066-INS-426 |
| chr8-63066055-INS-1101 | chr8-63066056-INS-1102 |

| | |
|---|---|
| chr8-63067121-INS-341 | chr8-63067115-INS-348 |
| chr8-63070531-INS-138 | chr8-63070531-INS-138 |
| chr8-63089878-INS-82 | chr8-63089878-INS-86 |
| chr8-63109309-INS-416 | chr8-63109309-INS-416 |
| chr8-63112373-INS-164 | chr8-63112373-INS-158 |
| chr8-63146205-INS-72 | chr8-63146184-INS-66 |
| chr8-63147682-INS-120 | chr8-63147694-INS-123 |
| chr8-63160471-INS-1180 | chr8-63160472-INS-1180 |
| chr8-63170273-INS-56 | chr8-63170273-INS-52 |
| chr8-63183495-INS-228 | chr8-63183495-INS-228 |
| chr8-63193001-INS-125 | chr8-63193000-INS-133 |
| chr8-63232730-INS-166 | chr8-63232730-INS-166 |
| chr8-63240803-INS-1556 | chr8-63240804-INS-1556 |
| chr8-63249161-INS-155 | chr8-63249161-INS-154 |
| chr8-63273754-INS-111 | chr8-63273766-INS-111 |
| chr8-63299669-INS-52 | chr8-63299669-INS-52 |
| chr8-63335664-INS-61 | chr8-63335664-INS-72 |
| chr8-63395984-INS-858 | chr8-63395985-INS-858 |
| chr8-63426737-INS-7177 | chr8-63426738-INS-7184 |
| chr8-63430365-INS-73 | chr8-63430354-INS-59 |
| chr8-63430860-INS-1506 | chr8-63430861-INS-1509 |
| chr8-63445761-INS-280 | chr8-63445761-INS-281 |
| chr8-63464656-INS-210 | chr8-63464656-INS-210 |
| chr8-63479520-INS-337 | chr8-63479520-INS-337 |
| chr8-63498784-INS-217 | chr8-63498784-INS-215 |
| chr8-63509648-INS-54 | chr8-63509648-INS-54 |
| chr8-63535412-INS-66 | chr8-63535410-INS-68 |
| chr8-63548448-INS-6235 | chr8-63548449-INS-6235 |
| chr8-63596832-INS-248 | chr8-63596832-INS-252 |
| chr8-63601137-INS-2653 | chr8-63601138-INS-2653 |
| chr8-63633162-INS-130 | chr8-63633162-INS-130 |
| chr8-63703350-INS-51 | chr8-63703351-INS-51 |
| chr8-63714229-INS-551 | chr8-63714229-INS-552 |
| chr8-63719164-INS-58 | chr8-63719164-INS-74 |
| chr8-63720860-INS-130 | chr8-63720860-INS-151 |
| chr8-63723567-INS-54 | chr8-63723567-INS-54 |
| chr8-63727106-INS-211 | chr8-63727106-INS-220 |
| chr8-63728932-INS-58 | chr8-63728903-INS-52 |
| chr8-63780469-INS-400 | chr8-63780469-INS-400 |
| chr8-63795295-INS-54 | chr8-63795295-INS-52 |
| chr8-63839764-INS-138 | chr8-63839764-INS-138 |
| chr8-63860811-INS-279 | chr8-63860811-INS-280 |
| chr8-63870762-INS-1346 | chr8-63870763-INS-1346 |
| chr8-63884409-INS-405 | chr8-63884409-INS-406 |
| chr8-63891132-INS-56 | chr8-63891132-INS-58 |
| chr8-63900336-INS-6074 | chr8-63900337-INS-6072 |
| chr8-63984157-INS-62 | chr8-63984157-INS-51 |
| chr8-64012608-INS-399 | chr8-64012608-INS-399 |
| chr8-64025171-INS-74 | chr8-64025171-INS-78 |
| chr8-64043138-INS-182 | chr8-64043138-INS-185 |
| chr8-64045014-INS-56 | chr8-64045014-INS-70 |
| chr8-64051950-INS-252 | chr8-64051950-INS-252 |
| chr8-64066129-INS-7139 | chr8-64066130-INS-7139 |
| chr8-64079075-INS-385 | chr8-64079075-INS-385 |
| chr8-64087077-INS-60 | chr8-64087095-INS-52 |
| chr8-64092363-INS-215 | chr8-64092339-INS-255 |
| chr8-64093269-INS-54 | chr8-64093269-INS-54 |
| chr8-64093584-INS-318 | chr8-64093584-INS-312 |
| chr8-64100325-INS-355 | chr8-64100325-INS-355 |
| chr8-64103429-INS-467 | chr8-64103429-INS-467 |
| chr8-64110547-INS-6681 | chr8-64110548-INS-6681 |
| chr8-64116366-INS-179 | chr8-64116366-INS-193 |
| chr8-64190311-INS-68 | chr8-64190311-INS-68 |
| chr8-64202659-INS-539 | chr8-64202659-INS-538 |
| chr8-64205334-INS-53 | chr8-64205334-INS-53 |
| chr8-64226252-INS-150 | chr8-64226252-INS-178 |
| chr8-64248583-INS-2249 | chr8-64248584-INS-2250 |
| chr8-64258211-INS-6404 | chr8-64258212-INS-6404 |
| chr8-64258972-INS-375 | chr8-64258972-INS-360 |
| chr8-64296573-INS-1042 | chr8-64296574-INS-1044 |
| chr8-64299964-INS-6736 | chr8-64299965-INS-6737 |
| chr8-64333272-INS-431 | chr8-64333272-INS-431 |
| chr8-64333961-INS-113 | chr8-64333953-INS-112 |
| chr8-64347936-INS-116 | chr8-64347936-INS-116 |
| chr8-64348899-INS-211 | chr8-64348899-INS-211 |
| chr8-64350492-INS-187 | chr8-64350492-INS-190 |
| chr8-64359813-INS-69 | chr8-64359813-INS-69 |

| | |
|---|---|
| chr8-64375166-INS-136 | chr8-64375166-INS-136 |
| chr8-64376944-INS-52 | chr8-64376944-INS-52 |
| chr8-64385349-INS-76 | chr8-64385349-INS-78 |
| chr8-64389376-INS-741 | chr8-64389377-INS-776 |
| chr8-64395184-INS-122 | chr8-64395170-INS-116 |
| chr8-64417988-INS-734 | chr8-64417988-INS-734 |
| chr8-64421664-INS-182 | chr8-64421664-INS-163 |
| chr8-64433609-INS-1602 | chr8-64433610-INS-1602 |
| chr8-64443388-INS-138 | chr8-64443396-INS-121 |
| chr8-64464978-INS-6801 | chr8-64464979-INS-6801 |
| chr8-64493730-INS-63 | chr8-64493730-INS-54 |
| chr8-64506815-INS-57 | chr8-64506815-INS-97 |
| chr8-64511308-INS-294 | chr8-64511308-INS-296 |
| chr8-64512385-INS-74 | chr8-64512399-INS-80 |
| chr8-64513993-INS-472 | chr8-64514001-INS-474 |
| chr8-64521138-INS-239 | chr8-64521138-INS-239 |
| chr8-64538196-INS-60 | chr8-64538186-INS-54 |
| chr8-64603658-INS-1848 | chr8-64603663-INS-1843 |
| chr8-64605099-INS-72 | chr8-64605110-INS-50 |
| chr8-64612052-INS-405 | chr8-64612052-INS-405 |
| chr8-64614760-INS-50 | chr8-64614760-INS-52 |
| chr8-64653823-INS-91 | chr8-64653823-INS-84 |
| chr8-64656639-INS-60 | chr8-64656639-INS-60 |
| chr8-64661380-INS-304 | chr8-64661380-INS-304 |
| chr8-64667556-INS-429 | chr8-64667556-INS-432 |
| chr8-64668986-INS-90 | chr8-64668986-INS-58 |
| chr8-64678588-INS-6353 | chr8-64678589-INS-6353 |
| chr8-64720169-INS-78 | chr8-64720169-INS-108 |
| chr8-64726935-INS-52 | chr8-64726935-INS-60 |
| chr8-64768240-INS-78 | chr8-64768234-INS-76 |
| chr8-64776559-INS-6943 | chr8-64776567-INS-6939 |
| chr8-64818592-INS-159 | chr8-64818592-INS-177 |
| chr8-64822776-INS-53 | chr8-64822776-INS-59 |
| chr8-64824014-INS-1890 | chr8-64824015-INS-1890 |
| chr8-64824966-INS-52 | chr8-64824966-INS-52 |
| chr8-64829904-INS-4993 | chr8-64829905-INS-4993 |
| chr8-64842966-INS-80 | chr8-64842966-INS-80 |
| chr8-64856404-INS-2688 | chr8-64856411-INS-2683 |
| chr8-64859603-INS-584 | chr8-64859603-INS-584 |
| chr8-64870879-INS-596 | chr8-64870872-INS-604 |
| chr8-64884956-INS-58 | chr8-64884957-INS-58 |
| chr8-64940727-INS-7122 | chr8-64940728-INS-7124 |
| chr8-64953039-INS-360 | chr8-64953039-INS-360 |
| chr8-64961381-INS-315 | chr8-64961382-INS-315 |
| chr8-64962125-INS-993 | chr8-64962125-INS-993 |
| chr8-64977530-INS-1138 | chr8-64977531-INS-1138 |
| chr8-64988739-INS-6199 | chr8-64988740-INS-6199 |
| chr8-65003029-INS-4607 | chr8-65003030-INS-4607 |
| chr8-65034489-INS-122 | chr8-65034489-INS-126 |
| chr8-65045483-INS-66 | chr8-65045471-INS-70 |
| chr8-65053064-INS-1079 | chr8-65053065-INS-1081 |
| chr8-65062524-INS-60 | chr8-65062524-INS-60 |
| chr8-65064750-INS-78 | chr8-65064750-INS-82 |
| chr8-65068564-INS-239 | chr8-65068564-INS-233 |
| chr8-65147086-INS-110 | chr8-65147086-INS-100 |
| chr8-65175248-INS-61 | chr8-65175252-INS-65 |
| chr8-65181304-INS-56 | chr8-65181304-INS-58 |
| chr8-65194683-INS-53 | chr8-65194683-INS-53 |
| chr8-65210837-INS-54 | chr8-65210837-INS-54 |
| chr8-65233267-INS-55 | chr8-65233267-INS-55 |
| chr8-65247082-INS-54 | chr8-65247075-INS-54 |
| chr8-65276143-INS-228 | chr8-65276143-INS-227 |
| chr8-65295440-INS-88 | chr8-65295440-INS-60 |
| chr8-65304754-INS-1885 | chr8-65304755-INS-1885 |
| chr8-65334078-INS-57 | chr8-65334078-INS-59 |
| chr8-65339182-INS-163 | chr8-65339182-INS-163 |
| chr8-65350403-INS-103 | chr8-65350403-INS-103 |
| chr8-65352847-INS-93 | chr8-65352836-INS-77 |
| chr8-65359032-INS-419 | chr8-65359032-INS-419 |
| chr8-65374309-INS-64 | chr8-65374309-INS-64 |
| chr8-65375192-INS-52 | chr8-65375192-INS-56 |
| chr8-65402438-INS-76 | chr8-65402438-INS-76 |
| chr8-65422691-INS-187 | chr8-65422691-INS-187 |
| chr8-65424347-INS-439 | chr8-65424347-INS-436 |
| chr8-65437260-INS-71 | chr8-65437274-INS-79 |
| chr8-65440130-INS-135 | chr8-65440130-INS-134 |
| chr8-65461803-INS-154 | chr8-65461803-INS-157 |

| | |
|---|---|
| chr8-65465819-INS-64 | chr8-65465819-INS-64 |
| chr8-65483185-INS-52 | chr8-65483185-INS-54 |
| chr8-65499328-INS-107 | chr8-65499316-INS-119 |
| chr8-65502312-INS-126 | chr8-65502312-INS-129 |
| chr8-65508126-INS-204 | chr8-65508126-INS-207 |
| chr8-65525274-INS-62 | chr8-65525254-INS-52 |
| chr8-65534709-INS-55 | chr8-65534728-INS-57 |
| chr8-65537998-INS-69 | chr8-65537998-INS-69 |
| chr8-65548260-INS-132 | chr8-65548260-INS-132 |
| chr8-65551725-INS-84 | chr8-65551725-INS-90 |
| chr8-65555469-INS-837 | chr8-65555469-INS-837 |
| chr8-65557277-INS-195 | chr8-65557277-INS-196 |
| chr8-65562039-INS-479 | chr8-65562039-INS-479 |
| chr8-65564520-INS-71 | chr8-65564521-INS-62 |
| chr8-65574097-INS-161 | chr8-65574115-INS-170 |
| chr8-65575358-INS-74 | chr8-65575377-INS-70 |
| chr8-65581248-INS-493 | chr8-65581248-INS-495 |
| chr8-65592457-INS-60 | chr8-65592457-INS-60 |
| chr8-65625825-INS-74 | chr8-65625825-INS-58 |
| chr8-65633801-INS-56 | chr8-65633801-INS-54 |
| chr8-65645661-INS-179 | chr8-65645661-INS-179 |
| chr8-65646865-INS-173 | chr8-65646865-INS-176 |
| chr8-65652798-INS-463 | chr8-65652798-INS-463 |
| chr8-65674779-INS-53 | chr8-65674779-INS-53 |
| chr8-65692939-INS-163 | chr8-65692939-INS-163 |
| chr8-65724562-INS-155 | chr8-65724562-INS-157 |
| chr8-65734469-INS-62 | chr8-65734470-INS-65 |
| chr8-65739971-INS-7088 | chr8-65739972-INS-7088 |
| chr8-65740504-INS-54 | chr8-65740505-INS-64 |
| chr8-65749788-INS-1616 | chr8-65749789-INS-1616 |
| chr8-65755001-INS-2649 | chr8-65755002-INS-2652 |
| chr8-65801255-INS-526 | chr8-65801255-INS-526 |
| chr8-65816268-INS-1666 | chr8-65816267-INS-1659 |
| chr8-65818852-INS-79 | chr8-65818852-INS-89 |
| chr8-65834131-INS-252 | chr8-65834131-INS-252 |
| chr8-65877770-INS-1033 | chr8-65877771-INS-1033 |
| chr8-65878336-INS-498 | chr8-65878337-INS-498 |
| chr8-65932888-INS-70 | chr8-65932888-INS-70 |
| chr8-65942245-INS-6385 | chr8-65942246-INS-6385 |
| chr8-65946210-INS-7111 | chr8-65946211-INS-7110 |
| chr8-65946781-INS-249 | chr8-65946781-INS-249 |
| chr8-65958746-INS-199 | chr8-65958746-INS-199 |
| chr8-65969384-INS-441 | chr8-65969384-INS-441 |
| chr8-65976331-INS-6436 | chr8-65976332-INS-6437 |
| chr8-66002805-INS-117 | chr8-66002784-INS-137 |
| chr8-66010999-INS-108 | chr8-66010999-INS-108 |
| chr8-66016382-INS-59 | chr8-66016382-INS-67 |
| chr8-66027908-INS-53 | chr8-66027908-INS-55 |
| chr8-66029172-INS-400 | chr8-66029172-INS-400 |
| chr8-66044098-INS-424 | chr8-66044098-INS-424 |
| chr8-66064613-INS-115 | chr8-66064613-INS-116 |
| chr8-66077538-INS-969 | chr8-66077538-INS-971 |
| chr8-66078181-INS-55 | chr8-66078181-INS-55 |
| chr8-66086629-INS-656 | chr8-66086629-INS-656 |
| chr8-66087601-INS-959 | chr8-66087609-INS-980 |
| chr8-66104488-INS-60 | chr8-66104488-INS-60 |
| chr8-66118249-INS-314 | chr8-66118249-INS-312 |
| chr8-66134790-INS-72 | chr8-66134790-INS-64 |
| chr8-66144281-INS-414 | chr8-66144281-INS-416 |
| chr8-66146096-INS-863 | chr8-66146096-INS-863 |
| chr8-66153628-INS-63 | chr8-66153628-INS-54 |
| chr8-66155464-INS-100 | chr8-66155476-INS-95 |
| chr8-66156492-INS-75 | chr8-66156492-INS-75 |
| chr8-66176845-INS-4874 | chr8-66176846-INS-4874 |
| chr8-66193584-INS-58 | chr8-66193584-INS-50 |
| chr8-66202463-INS-100 | chr8-66202463-INS-86 |
| chr8-66214425-INS-6274 | chr8-66214426-INS-6276 |
| chr8-66220972-INS-81 | chr8-66220972-INS-85 |
| chr8-66227798-INS-81 | chr8-66227802-INS-61 |
| chr8-66241884-INS-572 | chr8-66241884-INS-574 |
| chr8-66263104-INS-151 | chr8-66263114-INS-122 |
| chr8-66295382-INS-194 | chr8-66295382-INS-193 |
| chr8-66302886-INS-60 | chr8-66302886-INS-55 |
| chr8-66315148-INS-678 | chr8-66315148-INS-678 |
| chr8-66341083-INS-7070 | chr8-66341084-INS-7070 |
| chr8-66361394-INS-84 | chr8-66361394-INS-84 |
| chr8-66391281-INS-100 | chr8-66391281-INS-114 |

| | |
|---|---|
| chr8-66392439-INS-1196 | chr8-66392440-INS-1196 |
| chr8-66397932-INS-52 | chr8-66397948-INS-56 |
| chr8-66401128-INS-1858 | chr8-66401129-INS-1858 |
| chr8-66408522-INS-84 | chr8-66408526-INS-88 |
| chr8-66434248-INS-125 | chr8-66434248-INS-125 |
| chr8-66437587-INS-218 | chr8-66437587-INS-218 |
| chr8-66472380-INS-94 | chr8-66472380-INS-95 |
| chr8-66473407-INS-693 | chr8-66473407-INS-693 |
| chr8-66478286-INS-68 | chr8-66478286-INS-68 |
| chr8-66481607-INS-58 | chr8-66481607-INS-58 |
| chr8-66491467-INS-4034 | chr8-66491468-INS-4050 |
| chr8-66506634-INS-64 | chr8-66506634-INS-56 |
| chr8-66533859-INS-72 | chr8-66533859-INS-58 |
| chr8-66543082-INS-927 | chr8-66543082-INS-923 |
| chr8-66549783-INS-6811 | chr8-66549784-INS-6811 |
| chr8-66565651-INS-4264 | chr8-66565652-INS-4270 |
| chr8-66613158-INS-235 | chr8-66613158-INS-235 |
| chr8-66622553-INS-6895 | chr8-66622554-INS-6893 |
| chr8-66626975-INS-1867 | chr8-66626976-INS-1867 |
| chr8-66639086-INS-606 | chr8-66639088-INS-607 |
| chr8-66655078-INS-126 | chr8-66655078-INS-126 |
| chr8-66662712-INS-803 | chr8-66662712-INS-807 |
| chr8-66672508-INS-90 | chr8-66672503-INS-80 |
| chr8-66681348-INS-57 | chr8-66681348-INS-61 |
| chr8-66694016-INS-723 | chr8-66694016-INS-725 |
| chr8-66701943-INS-8944 | chr8-66701944-INS-8945 |
| chr8-66706902-INS-336 | chr8-66706902-INS-377 |
| chr8-66727568-INS-69 | chr8-66727568-INS-54 |
| chr8-66741538-INS-4146 | chr8-66741539-INS-4146 |
| chr8-66743448-INS-202 | chr8-66743448-INS-202 |
| chr8-66759810-INS-69 | chr8-66759810-INS-66 |
| chr8-66765250-INS-99 | chr8-66765255-INS-75 |
| chr8-66766375-INS-96 | chr8-66766375-INS-80 |
| chr8-66777873-INS-97 | chr8-66777873-INS-97 |
| chr8-66781789-INS-426 | chr8-66781789-INS-426 |
| chr8-66789405-INS-461 | chr8-66789405-INS-461 |
| chr8-66807734-INS-3006 | chr8-66807735-INS-3006 |
| chr8-66851516-INS-92 | chr8-66851528-INS-84 |
| chr8-66859504-INS-3217 | chr8-66859505-INS-3217 |
| chr8-66865143-INS-64 | chr8-66865143-INS-66 |
| chr8-66894476-INS-76 | chr8-66894476-INS-68 |
| chr8-66913021-INS-381 | chr8-66913021-INS-380 |
| chr8-66934055-INS-1468 | chr8-66934056-INS-1469 |
| chr8-66983746-INS-6491 | chr8-66983747-INS-6491 |
| chr8-66987694-INS-335 | chr8-66987694-INS-335 |
| chr8-67011042-INS-75 | chr8-67011042-INS-62 |
| chr8-67015425-INS-104 | chr8-67015470-INS-60 |
| chr8-67024189-INS-64 | chr8-67024189-INS-64 |
| chr8-67035363-INS-56 | chr8-67035363-INS-68 |
| chr8-67037687-INS-6229 | chr8-67037688-INS-6230 |
| chr8-67068798-INS-1268 | chr8-67068792-INS-1264 |
| chr8-67086132-INS-1630 | chr8-67086133-INS-1629 |
| chr8-67088458-INS-342 | chr8-67088458-INS-342 |
| chr8-67089718-INS-52 | chr8-67089718-INS-56 |
| chr8-67101043-INS-64 | chr8-67101043-INS-64 |
| chr8-67111238-INS-268 | chr8-67111238-INS-268 |
| chr8-67139821-INS-83 | chr8-67139813-INS-126 |
| chr8-67177580-INS-96 | chr8-67177576-INS-100 |
| chr8-67182724-INS-6189 | chr8-67182725-INS-6190 |
| chr8-67197012-INS-116 | chr8-67197012-INS-116 |
| chr8-67204159-INS-933 | chr8-67204159-INS-934 |
| chr8-67206275-INS-54 | chr8-67206275-INS-50 |
| chr8-67209407-INS-1802 | chr8-67209408-INS-1802 |
| chr8-67216403-INS-67 | chr8-67216440-INS-64 |
| chr8-67231151-INS-119 | chr8-67231151-INS-76 |
| chr8-67242073-INS-1343 | chr8-67242074-INS-1343 |
| chr8-67247946-INS-123 | chr8-67247946-INS-123 |
| chr8-67257266-INS-5714 | chr8-67257267-INS-5714 |
| chr8-67268501-INS-263 | chr8-67268501-INS-265 |
| chr8-67308465-INS-76 | chr8-67308465-INS-76 |
| chr8-67349318-INS-53 | chr8-67349313-INS-70 |
| chr8-67369605-INS-55 | chr8-67369605-INS-55 |
| chr8-67385896-INS-194 | chr8-67385896-INS-194 |
| chr8-67388172-INS-70 | chr8-67388172-INS-70 |
| chr8-67422758-INS-210 | chr8-67422758-INS-210 |
| chr8-67468634-INS-92 | chr8-67468666-INS-68 |
| chr8-67483582-INS-1890 | chr8-67483583-INS-1890 |

| | |
|---|---|
| chr8-67487838-INS-7244 | chr8-67487839-INS-7245 |
| chr8-67508799-INS-133 | chr8-67508799-INS-133 |
| chr8-67513569-INS-201 | chr8-67513569-INS-200 |
| chr8-67522645-INS-6159 | chr8-67522646-INS-6159 |
| chr8-67570745-INS-186 | chr8-67570740-INS-197 |
| chr8-67582816-INS-214 | chr8-67582816-INS-214 |
| chr8-67590429-INS-232 | chr8-67590429-INS-232 |
| chr8-67601304-INS-331 | chr8-67601304-INS-331 |
| chr8-67602023-INS-91 | chr8-67602069-INS-58 |
| chr8-67645688-INS-116 | chr8-67645688-INS-118 |
| chr8-67660596-INS-52 | chr8-67660596-INS-52 |
| chr8-67662980-INS-3302 | chr8-67662981-INS-3302 |
| chr8-67679200-INS-56 | chr8-67679222-INS-52 |
| chr8-67681772-INS-168 | chr8-67681769-INS-132 |
| chr8-67712569-INS-852 | chr8-67712569-INS-854 |
| chr8-67714137-INS-1892 | chr8-67714138-INS-1892 |
| chr8-67749062-INS-7079 | chr8-67749063-INS-7079 |
| chr8-67750149-INS-203 | chr8-67750149-INS-203 |
| chr8-67765454-INS-125 | chr8-67765454-INS-130 |
| chr8-67771305-INS-468 | chr8-67771305-INS-469 |
| chr8-67794305-INS-203 | chr8-67794305-INS-203 |
| chr8-67814146-INS-187 | chr8-67814146-INS-187 |
| chr8-67819878-INS-203 | chr8-67819878-INS-203 |
| chr8-67824500-INS-324 | chr8-67824500-INS-324 |
| chr8-67853734-INS-134 | chr8-67853734-INS-103 |
| chr8-67909784-INS-56 | chr8-67909804-INS-58 |
| chr8-67924389-INS-129 | chr8-67924395-INS-164 |
| chr8-67964393-INS-169 | chr8-67964393-INS-169 |
| chr8-67986477-INS-79 | chr8-67986476-INS-60 |
| chr8-68022891-INS-7597 | chr8-68022892-INS-7597 |
| chr8-68037692-INS-53 | chr8-68037692-INS-53 |
| chr8-68042136-INS-7454 | chr8-68042137-INS-7454 |
| chr8-68044033-INS-206 | chr8-68044033-INS-206 |
| chr8-68048380-INS-58 | chr8-68048380-INS-60 |
| chr8-68056231-INS-51 | chr8-68056227-INS-60 |
| chr8-68062630-INS-134 | chr8-68062630-INS-134 |
| chr8-68076573-INS-60 | chr8-68076573-INS-60 |
| chr8-68084010-INS-185 | chr8-68084010-INS-186 |
| chr8-68097959-INS-63 | chr8-68097959-INS-63 |
| chr8-68099780-INS-102 | chr8-68099780-INS-102 |
| chr8-68100882-INS-63 | chr8-68100882-INS-63 |
| chr8-68112867-INS-424 | chr8-68112867-INS-424 |
| chr8-68113945-INS-372 | chr8-68113945-INS-370 |
| chr8-68138692-INS-73 | chr8-68138692-INS-70 |
| chr8-68189212-INS-379 | chr8-68189212-INS-378 |
| chr8-68278462-INS-93 | chr8-68278462-INS-78 |
| chr8-68282089-INS-70 | chr8-68282089-INS-70 |
| chr8-68289450-INS-52 | chr8-68289450-INS-92 |
| chr8-68296449-INS-139 | chr8-68296449-INS-139 |
| chr8-68305057-INS-98 | chr8-68305054-INS-52 |
| chr8-68308011-INS-79 | chr8-68308011-INS-79 |
| chr8-68312595-INS-1953 | chr8-68312596-INS-1953 |
| chr8-68345951-INS-112 | chr8-68345947-INS-116 |
| chr8-68384447-INS-65 | chr8-68384447-INS-62 |
| chr8-68396371-INS-52 | chr8-68396371-INS-52 |
| chr8-68414757-INS-112 | chr8-68414757-INS-112 |
| chr8-68434595-INS-79 | chr8-68434595-INS-79 |
| chr8-68440444-INS-304 | chr8-68440444-INS-304 |
| chr8-68447305-INS-1249 | chr8-68447306-INS-1252 |
| chr8-68450586-INS-2779 | chr8-68450587-INS-2779 |
| chr8-68460125-INS-432 | chr8-68460125-INS-408 |
| chr8-68469278-INS-142 | chr8-68469278-INS-145 |
| chr8-68476516-INS-173 | chr8-68476516-INS-173 |
| chr8-68480212-INS-138 | chr8-68480200-INS-102 |
| chr8-68494359-INS-68 | chr8-68494332-INS-76 |
| chr8-68502962-INS-4835 | chr8-68502963-INS-4835 |
| chr8-68506084-INS-62 | chr8-68506049-INS-108 |
| chr8-68510678-INS-309 | chr8-68510678-INS-311 |
| chr8-68541611-INS-839 | chr8-68541611-INS-840 |
| chr8-68553285-INS-6413 | chr8-68553286-INS-6413 |
| chr8-68561205-INS-73 | chr8-68561205-INS-73 |
| chr8-68571683-INS-6037 | chr8-68571684-INS-6037 |
| chr8-68584876-INS-291 | chr8-68584878-INS-291 |
| chr8-68593709-INS-176 | chr8-68593709-INS-178 |
| chr8-68610933-INS-74 | chr8-68610933-INS-58 |
| chr8-68613969-INS-2070 | chr8-68613970-INS-2070 |
| chr8-68628493-INS-416 | chr8-68628493-INS-416 |

| | |
|---|---|
| chr8-68643377-INS-104 | chr8-68643371-INS-108 |
| chr8-68681267-INS-142 | chr8-68681267-INS-142 |
| chr8-68682286-INS-202 | chr8-68682286-INS-202 |
| chr8-68688830-INS-122 | chr8-68688830-INS-123 |
| chr8-68692585-INS-96 | chr8-68692585-INS-96 |
| chr8-68698070-INS-6247 | chr8-68698071-INS-6247 |
| chr8-68705364-INS-63 | chr8-68705364-INS-63 |
| chr8-68715191-INS-463 | chr8-68715191-INS-465 |
| chr8-68743111-INS-6253 | chr8-68743099-INS-6247 |
| chr8-68758363-INS-198 | chr8-68758363-INS-198 |
| chr8-68777510-INS-420 | chr8-68777510-INS-417 |
| chr8-68780201-INS-136 | chr8-68780201-INS-170 |
| chr8-68803145-INS-79 | chr8-68803145-INS-79 |
| chr8-68832371-INS-135 | chr8-68832371-INS-135 |
| chr8-68833562-INS-87 | chr8-68833562-INS-53 |
| chr8-68906014-INS-101 | chr8-68906014-INS-101 |
| chr8-68920862-INS-178 | chr8-68920863-INS-176 |
| chr8-68942431-INS-242 | chr8-68942431-INS-240 |
| chr8-68943231-INS-64 | chr8-68943231-INS-64 |
| chr8-68944715-INS-3112 | chr8-68944716-INS-3112 |
| chr8-68965717-INS-408 | chr8-68965717-INS-408 |
| chr8-68974353-INS-6680 | chr8-68974354-INS-6680 |
| chr8-68996795-INS-70 | chr8-68996795-INS-54 |
| chr8-69021532-INS-107 | chr8-69021532-INS-119 |
| chr8-69030941-INS-52 | chr8-69030941-INS-50 |
| chr8-69032317-INS-63 | chr8-69032317-INS-65 |
| chr8-69034681-INS-951 | chr8-69034681-INS-951 |
| chr8-69081025-INS-400 | chr8-69081025-INS-400 |
| chr8-69096601-INS-63 | chr8-69096563-INS-75 |
| chr8-69102400-INS-59 | chr8-69102400-INS-59 |
| chr8-69127766-INS-360 | chr8-69127766-INS-360 |
| chr8-69135616-INS-416 | chr8-69135616-INS-416 |
| chr8-69138742-INS-52 | chr8-69138743-INS-58 |
| chr8-69146555-INS-55 | chr8-69146555-INS-55 |
| chr8-69158374-INS-54 | chr8-69158374-INS-57 |
| chr8-69199367-INS-7218 | chr8-69199368-INS-7218 |
| chr8-69215089-INS-528 | chr8-69215089-INS-528 |
| chr8-69236082-INS-58 | chr8-69236082-INS-62 |
| chr8-69267789-INS-229 | chr8-69267789-INS-229 |
| chr8-69292875-INS-52 | chr8-69292875-INS-52 |
| chr8-69300701-INS-80 | chr8-69300701-INS-80 |
| chr8-69337356-INS-62 | chr8-69337356-INS-60 |
| chr8-69373035-INS-77 | chr8-69373035-INS-77 |
| chr8-69375646-INS-1083 | chr8-69375647-INS-1085 |
| chr8-69386602-INS-716 | chr8-69386602-INS-717 |
| chr8-69388328-INS-1658 | chr8-69388324-INS-1658 |
| chr8-69389491-INS-307 | chr8-69389492-INS-313 |
| chr8-69401652-INS-52 | chr8-69401652-INS-91 |
| chr8-69403870-INS-3252 | chr8-69403871-INS-3252 |
| chr8-69412134-INS-823 | chr8-69412134-INS-824 |
| chr8-69412668-INS-53 | chr8-69412668-INS-53 |
| chr8-69481130-INS-178 | chr8-69481130-INS-178 |
| chr8-69488929-INS-546 | chr8-69488929-INS-546 |
| chr8-69501873-INS-56 | chr8-69501873-INS-56 |
| chr8-69504896-INS-202 | chr8-69504896-INS-202 |
| chr8-69515849-INS-75 | chr8-69515850-INS-75 |
| chr8-69531056-INS-221 | chr8-69531043-INS-227 |
| chr8-69543498-INS-220 | chr8-69543498-INS-221 |
| chr8-69546103-INS-60 | chr8-69546103-INS-62 |
| chr8-69549904-INS-59 | chr8-69549904-INS-59 |
| chr8-69557253-INS-121 | chr8-69557253-INS-140 |
| chr8-69571978-INS-65 | chr8-69571976-INS-65 |
| chr8-69580700-INS-219 | chr8-69580700-INS-219 |
| chr8-69581575-INS-58 | chr8-69581555-INS-50 |
| chr8-69594742-INS-208 | chr8-69594742-INS-208 |
| chr8-69607658-INS-4345 | chr8-69607659-INS-4344 |
| chr8-69607758-INS-55 | chr8-69607758-INS-55 |
| chr8-69657821-INS-210 | chr8-69657821-INS-210 |
| chr8-69659043-INS-210 | chr8-69659043-INS-209 |
| chr8-69664817-INS-68 | chr8-69664821-INS-56 |
| chr8-69665984-INS-215 | chr8-69665984-INS-215 |
| chr8-69677458-INS-52 | chr8-69677459-INS-68 |
| chr8-69681772-INS-210 | chr8-69681772-INS-209 |
| chr8-69693997-INS-220 | chr8-69693997-INS-216 |
| chr8-69697388-INS-341 | chr8-69697388-INS-341 |
| chr8-69701708-INS-53 | chr8-69701672-INS-74 |
| chr8-69705360-INS-169 | chr8-69705360-INS-169 |

| | |
|---|---|
| chr8-69707642-INS-717 | chr8-69707642-INS-726 |
| chr8-69713731-INS-134 | chr8-69713731-INS-137 |
| chr8-69718265-INS-207 | chr8-69718265-INS-207 |
| chr8-69721900-INS-210 | chr8-69721900-INS-210 |
| chr8-69733874-INS-171 | chr8-69733874-INS-171 |
| chr8-69737532-INS-196 | chr8-69737532-INS-196 |
| chr8-69741699-INS-120 | chr8-69741699-INS-115 |
| chr8-69744545-INS-3236 | chr8-69744546-INS-3236 |
| chr8-69760818-INS-205 | chr8-69760818-INS-205 |
| chr8-69774235-INS-341 | chr8-69774235-INS-341 |
| chr8-69783799-INS-52 | chr8-69783799-INS-52 |
| chr8-69789658-INS-495 | chr8-69789658-INS-495 |
| chr8-69810167-INS-75 | chr8-69810167-INS-74 |
| chr8-69820870-INS-425 | chr8-69820870-INS-425 |
| chr8-69823455-INS-62 | chr8-69823455-INS-62 |
| chr8-69823978-INS-653 | chr8-69823978-INS-653 |
| chr8-69827532-INS-210 | chr8-69827532-INS-210 |
| chr8-69843430-INS-6428 | chr8-69843431-INS-6428 |
| chr8-69846424-INS-257 | chr8-69846424-INS-257 |
| chr8-69849638-INS-6252 | chr8-69849639-INS-6252 |
| chr8-69863085-INS-498 | chr8-69863085-INS-498 |
| chr8-69877903-INS-143 | chr8-69877889-INS-161 |
| chr8-69884576-INS-359 | chr8-69884576-INS-359 |
| chr8-69997949-INS-475 | chr8-69997949-INS-461 |
| chr8-70007996-INS-201 | chr8-70007997-INS-201 |
| chr8-70021142-INS-211 | chr8-70021142-INS-211 |
| chr8-70041323-INS-210 | chr8-70041323-INS-210 |
| chr8-70056119-INS-3801 | chr8-70056120-INS-3801 |
| chr8-70057713-INS-737 | chr8-70057713-INS-739 |
| chr8-70072966-INS-216 | chr8-70072966-INS-217 |
| chr8-70081123-INS-226 | chr8-70081123-INS-226 |
| chr8-70139565-INS-400 | chr8-70139565-INS-400 |
| chr8-70140426-INS-194 | chr8-70140426-INS-195 |
| chr8-70142525-INS-111 | chr8-70142541-INS-88 |
| chr8-70153510-INS-387 | chr8-70153510-INS-387 |
| chr8-70164504-INS-431 | chr8-70164504-INS-430 |
| chr8-70188243-INS-161 | chr8-70188243-INS-161 |
| chr8-70200154-INS-401 | chr8-70200154-INS-401 |
| chr8-70209684-INS-368 | chr8-70209684-INS-369 |
| chr8-70229888-INS-95 | chr8-70229888-INS-95 |
| chr8-70234801-INS-212 | chr8-70234801-INS-212 |
| chr8-70237428-INS-923 | chr8-70237428-INS-923 |
| chr8-70240929-INS-782 | chr8-70240929-INS-782 |
| chr8-70260581-INS-83 | chr8-70260581-INS-83 |
| chr8-70280950-INS-202 | chr8-70280950-INS-202 |
| chr8-70287295-INS-232 | chr8-70287295-INS-232 |
| chr8-70294192-INS-54 | chr8-70294170-INS-56 |
| chr8-70295575-INS-108 | chr8-70295580-INS-111 |
| chr8-70307210-INS-195 | chr8-70307210-INS-195 |
| chr8-70318734-INS-92 | chr8-70318740-INS-96 |
| chr8-70358891-INS-202 | chr8-70358891-INS-203 |
| chr8-70360078-INS-109 | chr8-70360078-INS-109 |
| chr8-70433502-INS-219 | chr8-70433502-INS-219 |
| chr8-70450140-INS-219 | chr8-70450140-INS-182 |
| chr8-70451875-INS-204 | chr8-70451875-INS-204 |
| chr8-70458816-INS-206 | chr8-70458816-INS-206 |
| chr8-70459281-INS-1415 | chr8-70459282-INS-1372 |
| chr8-70462407-INS-269 | chr8-70462407-INS-270 |
| chr8-70488477-INS-180 | chr8-70488477-INS-183 |
| chr8-70499516-INS-223 | chr8-70499516-INS-222 |
| chr8-70513983-INS-211 | chr8-70513983-INS-211 |
| chr8-70521507-INS-221 | chr8-70521507-INS-220 |
| chr8-70580901-INS-86 | chr8-70580901-INS-82 |
| chr8-70590294-INS-52 | chr8-70590294-INS-50 |
| chr8-70595824-INS-87 | chr8-70595824-INS-87 |
| chr8-70602678-INS-116 | chr8-70602682-INS-108 |
| chr8-70607824-INS-218 | chr8-70607824-INS-218 |
| chr8-70619105-INS-65 | chr8-70619105-INS-84 |
| chr8-70623205-INS-251 | chr8-70623205-INS-251 |
| chr8-70631085-INS-217 | chr8-70631086-INS-217 |
| chr8-70648869-INS-94 | chr8-70648869-INS-100 |
| chr8-70698388-INS-180 | chr8-70698388-INS-217 |
| chr8-70723348-INS-204 | chr8-70723348-INS-204 |
| chr8-70727745-INS-203 | chr8-70727745-INS-203 |
| chr8-70761863-INS-106 | chr8-70761863-INS-112 |
| chr8-70780243-INS-202 | chr8-70780243-INS-202 |
| chr8-70796457-INS-214 | chr8-70796457-INS-214 |

| | |
|---|---|
| chr8-70821378-INS-205 | chr8-70821378-INS-205 |
| chr8-70826743-INS-67 | chr8-70826743-INS-67 |
| chr8-70852783-INS-82 | chr8-70852783-INS-82 |
| chr8-70858552-INS-165 | chr8-70858552-INS-165 |
| chr8-70875479-INS-70 | chr8-70875479-INS-70 |
| chr8-70926670-INS-59 | chr8-70926666-INS-59 |
| chr8-70959987-INS-186 | chr8-70959987-INS-196 |
| chr8-70964861-INS-211 | chr8-70964861-INS-215 |
| chr8-71008532-INS-190 | chr8-71008532-INS-190 |
| chr8-71021033-INS-164 | chr8-71021033-INS-164 |
| chr8-71078109-INS-87 | chr8-71078106-INS-90 |
| chr8-71085384-INS-243 | chr8-71085384-INS-243 |
| chr8-71096648-INS-213 | chr8-71096648-INS-213 |
| chr8-71171619-INS-164 | chr8-71171619-INS-160 |
| chr8-71204387-INS-195 | chr8-71204387-INS-195 |
| chr8-71290823-INS-217 | chr8-71290823-INS-217 |
| chr8-71314210-INS-51 | chr8-71314210-INS-51 |
| chr8-71322809-INS-214 | chr8-71322809-INS-214 |
| chr8-71328009-INS-213 | chr8-71328009-INS-213 |
| chr8-71347378-INS-58 | chr8-71347378-INS-58 |
| chr8-71382291-INS-56 | chr8-71382287-INS-56 |
| chr8-71386540-INS-3849 | chr8-71386541-INS-3849 |
| chr8-71388444-INS-6606 | chr8-71388445-INS-6606 |
| chr8-71400354-INS-193 | chr8-71400354-INS-193 |
| chr8-71435488-INS-76 | chr8-71435488-INS-76 |
| chr8-71437740-INS-189 | chr8-71437740-INS-189 |
| chr8-71446691-INS-114 | chr8-71446729-INS-76 |
| chr8-71451313-INS-77 | chr8-71451313-INS-77 |
| chr8-71455604-INS-75 | chr8-71455604-INS-75 |
| chr8-71508934-INS-1000 | chr8-71508934-INS-999 |
| chr8-71545749-INS-6404 | chr8-71545750-INS-6404 |
| chr8-71658802-INS-51 | chr8-71658802-INS-51 |
| chr8-71687493-INS-56 | chr8-71687493-INS-56 |
| chr8-71701537-INS-213 | chr8-71701537-INS-217 |
| chr8-71711017-INS-491 | chr8-71711017-INS-498 |
| chr8-71721604-INS-78 | chr8-71721604-INS-80 |
| chr8-71765081-INS-7436 | chr8-71765082-INS-7434 |
| chr8-71823017-INS-201 | chr8-71823017-INS-200 |
| chr8-71845415-INS-208 | chr8-71845415-INS-207 |
| chr8-71865983-INS-195 | chr8-71865983-INS-195 |
| chr8-71868471-INS-125 | chr8-71868471-INS-127 |
| chr8-71871067-INS-5348 | chr8-71871068-INS-5348 |
| chr8-71871589-INS-202 | chr8-71871589-INS-202 |
| chr8-71872951-INS-451 | chr8-71872948-INS-456 |
| chr8-71881955-INS-970 | chr8-71881956-INS-970 |
| chr8-71887699-INS-207 | chr8-71887699-INS-208 |
| chr8-71891131-INS-52 | chr8-71891131-INS-52 |
| chr8-71919197-INS-161 | chr8-71919197-INS-161 |
| chr8-71952408-INS-504 | chr8-71952408-INS-505 |
| chr8-71956268-INS-82 | chr8-71956268-INS-82 |
| chr8-71980360-INS-250 | chr8-71980360-INS-225 |
| chr8-71994525-INS-664 | chr8-71994525-INS-663 |
| chr8-72050200-INS-249 | chr8-72050200-INS-249 |
| chr8-72070474-INS-6005 | chr8-72070475-INS-6005 |
| chr8-72116455-INS-63 | chr8-72116455-INS-63 |
| chr8-72187439-INS-207 | chr8-72187439-INS-208 |
| chr8-72198743-INS-5546 | chr8-72198749-INS-5544 |
| chr8-72202369-INS-1109 | chr8-72202370-INS-1109 |
| chr8-72218254-INS-3878 | chr8-72218255-INS-3880 |
| chr8-72218255-INS-452 | chr8-72218255-INS-451 |
| chr8-72600469-INS-1752 | chr8-72600470-INS-1747 |
| chr8-72651303-INS-4837 | chr8-72651304-INS-4837 |
| chr8-72708478-INS-210 | chr8-72708478-INS-210 |
| chr8-72713032-INS-212 | chr8-72713032-INS-212 |
| chr8-72737259-INS-109 | chr8-72737259-INS-109 |
| chr8-72757936-INS-201 | chr8-72757936-INS-201 |
| chr8-72766191-INS-201 | chr8-72766191-INS-202 |
| chr8-72806119-INS-200 | chr8-72806119-INS-200 |
| chr8-72815933-INS-207 | chr8-72815933-INS-207 |
| chr8-72860407-INS-186 | chr8-72860407-INS-187 |
| chr8-72862214-INS-118 | chr8-72862214-INS-114 |
| chr8-72863104-INS-971 | chr8-72863104-INS-967 |
| chr8-72866431-INS-6464 | chr8-72866432-INS-6464 |
| chr8-72876886-INS-433 | chr8-72876886-INS-433 |
| chr8-72884162-INS-60 | chr8-72884162-INS-60 |
| chr8-72909799-INS-66 | chr8-72909799-INS-66 |
| chr8-72917818-INS-170 | chr8-72917818-INS-170 |

| | |
|---|---|
| chr8-72946409-INS-198 | chr8-72946409-INS-199 |
| chr8-72955236-INS-59 | chr8-72955237-INS-52 |
| chr8-72963136-INS-52 | chr8-72963136-INS-52 |
| chr8-72971923-INS-300 | chr8-72971923-INS-301 |
| chr8-73005431-INS-180 | chr8-73005431-INS-180 |
| chr8-73021249-INS-217 | chr8-73021249-INS-217 |
| chr8-73035920-INS-62 | chr8-73035920-INS-62 |
| chr8-73043980-INS-221 | chr8-73043980-INS-221 |
| chr8-73066292-INS-61 | chr8-73066292-INS-62 |
| chr8-73154357-INS-2090 | chr8-73154358-INS-2090 |
| chr8-73171293-INS-219 | chr8-73171293-INS-215 |
| chr8-73176542-INS-178 | chr8-73176542-INS-178 |
| chr8-73179130-INS-223 | chr8-73179127-INS-221 |
| chr8-73186076-INS-55 | chr8-73186077-INS-100 |
| chr8-73207739-INS-948 | chr8-73207739-INS-951 |
| chr8-73211626-INS-1505 | chr8-73211622-INS-1522 |
| chr8-73213263-INS-240 | chr8-73213263-INS-240 |
| chr8-73225698-INS-219 | chr8-73225698-INS-219 |
| chr8-73323794-INS-213 | chr8-73323794-INS-212 |
| chr8-73330987-INS-236 | chr8-73330987-INS-236 |
| chr8-73332185-INS-223 | chr8-73332185-INS-223 |
| chr8-73379138-INS-53 | chr8-73379138-INS-53 |
| chr8-73389374-INS-91 | chr8-73389362-INS-125 |
| chr8-73418352-INS-68 | chr8-73418352-INS-68 |
| chr8-73503984-INS-59 | chr8-73503984-INS-59 |
| chr8-73580589-INS-4032 | chr8-73580590-INS-4028 |
| chr8-73601885-INS-83 | chr8-73601887-INS-89 |
| chr8-73633694-INS-368 | chr8-73633694-INS-361 |
| chr8-73641109-INS-110 | chr8-73641109-INS-109 |
| chr8-73645389-INS-56 | chr8-73645389-INS-58 |
| chr8-73661759-INS-195 | chr8-73661763-INS-186 |
| chr8-73671601-INS-114 | chr8-73671601-INS-114 |
| chr8-73709997-INS-171 | chr8-73709997-INS-171 |
| chr8-73761668-INS-68 | chr8-73761668-INS-76 |
| chr8-73771654-INS-59 | chr8-73771658-INS-59 |
| chr8-73774021-INS-70 | chr8-73774021-INS-58 |
| chr8-73779993-INS-357 | chr8-73779993-INS-357 |
| chr8-73788555-INS-62 | chr8-73788553-INS-60 |
| chr8-73789762-INS-68 | chr8-73789762-INS-68 |
| chr8-73795613-INS-84 | chr8-73795613-INS-88 |
| chr8-73828769-INS-102 | chr8-73828769-INS-102 |
| chr8-73834422-INS-100 | chr8-73834422-INS-101 |
| chr8-73856736-INS-907 | chr8-73856736-INS-907 |
| chr8-73885937-INS-462 | chr8-73885937-INS-462 |
| chr8-73897008-INS-553 | chr8-73897008-INS-554 |
| chr8-73898821-INS-252 | chr8-73898821-INS-252 |
| chr8-73911460-INS-434 | chr8-73911460-INS-434 |
| chr8-73912127-INS-823 | chr8-73912127-INS-822 |
| chr8-73916384-INS-142 | chr8-73916384-INS-182 |
| chr8-73935192-INS-184 | chr8-73935192-INS-191 |
| chr8-74006741-INS-109 | chr8-74006754-INS-100 |
| chr8-74008219-INS-61 | chr8-74008219-INS-53 |
| chr8-74019348-INS-56 | chr8-74019377-INS-63 |
| chr8-74085966-INS-237 | chr8-74085966-INS-237 |
| chr8-74113786-INS-9438 | chr8-74113787-INS-9438 |
| chr8-74124558-INS-434 | chr8-74124558-INS-434 |
| chr8-74131308-INS-60 | chr8-74131308-INS-50 |
| chr8-74133660-INS-3918 | chr8-74133661-INS-3918 |
| chr8-74170295-INS-1305 | chr8-74170296-INS-1306 |
| chr8-74217132-INS-83 | chr8-74217132-INS-83 |
| chr8-74270479-INS-398 | chr8-74270479-INS-399 |
| chr8-74291254-INS-229 | chr8-74291254-INS-232 |
| chr8-74292889-INS-219 | chr8-74292889-INS-209 |
| chr8-74324859-INS-54 | chr8-74324859-INS-54 |
| chr8-74331712-INS-276 | chr8-74331712-INS-276 |
| chr8-74345693-INS-62 | chr8-74345693-INS-62 |
| chr8-74359264-INS-137 | chr8-74359265-INS-90 |
| chr8-74372072-INS-57 | chr8-74372072-INS-63 |
| chr8-74383856-INS-58 | chr8-74383856-INS-54 |
| chr8-74391065-INS-4842 | chr8-74391066-INS-4830 |
| chr8-74395207-INS-72 | chr8-74395207-INS-72 |
| chr8-74399119-INS-61 | chr8-74399119-INS-61 |
| chr8-74440734-INS-62 | chr8-74440734-INS-64 |
| chr8-74496349-INS-1890 | chr8-74496350-INS-1890 |
| chr8-74550884-INS-113 | chr8-74550884-INS-113 |
| chr8-74552806-INS-6990 | chr8-74552807-INS-6990 |
| chr8-74565766-INS-556 | chr8-74565766-INS-556 |

| | |
|---|---|
| chr8-74587887-INS-111 | chr8-74587887-INS-111 |
| chr8-74593768-INS-170 | chr8-74593768-INS-170 |
| chr8-74603078-INS-518 | chr8-74603078-INS-519 |
| chr8-74629753-INS-56 | chr8-74629753-INS-60 |
| chr8-74647006-INS-451 | chr8-74647006-INS-451 |
| chr8-74651569-INS-84 | chr8-74651573-INS-80 |
| chr8-74652305-INS-6776 | chr8-74652306-INS-6774 |
| chr8-74670621-INS-7033 | chr8-74670622-INS-7033 |
| chr8-74681643-INS-2951 | chr8-74681644-INS-2951 |
| chr8-74691122-INS-6251 | chr8-74691123-INS-6257 |
| chr8-74706665-INS-122 | chr8-74706665-INS-79 |
| chr8-74747954-INS-1432 | chr8-74747955-INS-1432 |
| chr8-74759608-INS-2477 | chr8-74759609-INS-2477 |
| chr8-74797550-INS-106 | chr8-74797550-INS-106 |
| chr8-74848845-INS-399 | chr8-74848845-INS-398 |
| chr8-74906064-INS-399 | chr8-74906064-INS-399 |
| chr8-74933000-INS-73 | chr8-74933000-INS-73 |
| chr8-74962939-INS-52 | chr8-74962939-INS-56 |
| chr8-74963809-INS-66 | chr8-74963809-INS-66 |
| chr8-74986842-INS-152 | chr8-74986842-INS-145 |
| chr8-74993419-INS-70 | chr8-74993390-INS-72 |
| chr8-75000908-INS-179 | chr8-75000908-INS-200 |
| chr8-75048570-INS-1069 | chr8-75048571-INS-1069 |
| chr8-75049799-INS-501 | chr8-75049799-INS-501 |
| chr8-75053039-INS-89 | chr8-75053039-INS-89 |
| chr8-75064899-INS-167 | chr8-75064899-INS-167 |
| chr8-75071798-INS-1407 | chr8-75071799-INS-1409 |
| chr8-75089285-INS-1756 | chr8-75089286-INS-1758 |
| chr8-75091763-INS-51 | chr8-75091763-INS-55 |
| chr8-75128380-INS-189 | chr8-75128380-INS-189 |
| chr8-75141505-INS-107 | chr8-75141505-INS-76 |
| chr8-75179814-INS-6213 | chr8-75179815-INS-6213 |
| chr8-75203366-INS-70 | chr8-75203366-INS-72 |
| chr8-75206513-INS-645 | chr8-75206513-INS-645 |
| chr8-75211514-INS-642 | chr8-75211514-INS-642 |
| chr8-75253906-INS-64 | chr8-75253906-INS-64 |
| chr8-75268933-INS-1593 | chr8-75268934-INS-1593 |
| chr8-75274000-INS-6212 | chr8-75274001-INS-6212 |
| chr8-75279862-INS-82 | chr8-75279904-INS-64 |
| chr8-75308002-INS-94 | chr8-75308006-INS-70 |
| chr8-75335262-INS-2582 | chr8-75335274-INS-2569 |
| chr8-75355924-INS-88 | chr8-75355922-INS-117 |
| chr8-75374836-INS-180 | chr8-75374836-INS-181 |
| chr8-75388634-INS-143 | chr8-75388634-INS-143 |
| chr8-75393977-INS-105 | chr8-75393977-INS-105 |
| chr8-75400200-INS-1415 | chr8-75400201-INS-1415 |
| chr8-75400201-INS-567 | chr8-75400201-INS-569 |
| chr8-75408094-INS-5374 | chr8-75408095-INS-5374 |
| chr8-75420614-INS-80 | chr8-75420614-INS-84 |
| chr8-75422094-INS-56 | chr8-75422094-INS-56 |
| chr8-75430792-INS-98 | chr8-75430792-INS-92 |
| chr8-75462221-INS-665 | chr8-75462221-INS-665 |
| chr8-75537514-INS-50 | chr8-75537477-INS-58 |
| chr8-75553198-INS-4856 | chr8-75553199-INS-4856 |
| chr8-75579053-INS-591 | chr8-75579053-INS-590 |
| chr8-75582920-INS-76 | chr8-75582918-INS-59 |
| chr8-75584374-INS-118 | chr8-75584374-INS-118 |
| chr8-75608660-INS-807 | chr8-75608660-INS-822 |
| chr8-75662961-INS-196 | chr8-75662961-INS-196 |
| chr8-75663672-INS-406 | chr8-75663672-INS-406 |
| chr8-75670919-INS-193 | chr8-75670908-INS-209 |
| chr8-75694458-INS-103 | chr8-75694458-INS-103 |
| chr8-75710663-INS-400 | chr8-75710663-INS-400 |
| chr8-75786877-INS-210 | chr8-75786877-INS-210 |
| chr8-75794513-INS-214 | chr8-75794513-INS-210 |
| chr8-75804283-INS-66 | chr8-75804283-INS-66 |
| chr8-75841630-INS-241 | chr8-75841630-INS-243 |
| chr8-75856802-INS-4368 | chr8-75856803-INS-4369 |
| chr8-75866105-INS-163 | chr8-75866105-INS-169 |
| chr8-75873715-INS-113 | chr8-75873722-INS-72 |
| chr8-75880483-INS-205 | chr8-75880483-INS-206 |
| chr8-75916110-INS-90 | chr8-75916120-INS-99 |
| chr8-75922556-INS-289 | chr8-75922540-INS-294 |
| chr8-75937308-INS-105 | chr8-75937308-INS-105 |
| chr8-75997943-INS-58 | chr8-75997929-INS-66 |
| chr8-76054656-INS-62 | chr8-76054656-INS-62 |
| chr8-76066190-INS-213 | chr8-76066190-INS-213 |

| | |
|---|---|
| chr8-76081623-INS-216 | chr8-76081623-INS-210 |
| chr8-76092381-INS-6683 | chr8-76092382-INS-6683 |
| chr8-76134479-INS-1079 | chr8-76134480-INS-1086 |
| chr8-76178220-INS-62 | chr8-76178220-INS-66 |
| chr8-76212684-INS-6473 | chr8-76212685-INS-6473 |
| chr8-76214506-INS-55 | chr8-76214506-INS-54 |
| chr8-76219969-INS-61 | chr8-76219969-INS-50 |
| chr8-76227666-INS-942 | chr8-76227666-INS-942 |
| chr8-76231540-INS-6214 | chr8-76231541-INS-6214 |
| chr8-76239623-INS-730 | chr8-76239624-INS-736 |
| chr8-76255796-INS-202 | chr8-76255796-INS-202 |
| chr8-76288273-INS-96 | chr8-76288273-INS-96 |
| chr8-76334669-INS-84 | chr8-76334669-INS-84 |
| chr8-76347132-INS-6347 | chr8-76347133-INS-6346 |
| chr8-76351822-INS-353 | chr8-76351824-INS-357 |
| chr8-76359720-INS-485 | chr8-76359720-INS-485 |
| chr8-76387046-INS-161 | chr8-76387046-INS-152 |
| chr8-76475802-INS-359 | chr8-76475802-INS-359 |
| chr8-76477507-INS-329 | chr8-76477507-INS-329 |
| chr8-76494690-INS-128 | chr8-76494690-INS-128 |
| chr8-76542730-INS-107 | chr8-76542730-INS-95 |
| chr8-76545391-INS-182 | chr8-76545391-INS-173 |
| chr8-76590083-INS-94 | chr8-76590083-INS-93 |
| chr8-76761935-INS-6841 | chr8-76761936-INS-6840 |
| chr8-76778095-INS-67 | chr8-76778091-INS-56 |
| chr8-76799218-INS-83 | chr8-76799265-INS-59 |
| chr8-76824026-INS-56 | chr8-76824026-INS-58 |
| chr8-76833667-INS-165 | chr8-76833667-INS-165 |
| chr8-76834958-INS-53 | chr8-76834958-INS-54 |
| chr8-76960764-INS-7168 | chr8-76960765-INS-7168 |
| chr8-76981058-INS-730 | chr8-76981064-INS-729 |
| chr8-76997533-INS-473 | chr8-76997533-INS-473 |
| chr8-77034259-INS-158 | chr8-77034259-INS-158 |
| chr8-77056159-INS-541 | chr8-77056160-INS-494 |
| chr8-77070078-INS-52 | chr8-77070078-INS-76 |
| chr8-77075318-INS-59 | chr8-77075318-INS-60 |
| chr8-77088154-INS-66 | chr8-77088154-INS-60 |
| chr8-77089004-INS-306 | chr8-77089004-INS-306 |
| chr8-77108688-INS-184 | chr8-77108688-INS-184 |
| chr8-77121676-INS-86 | chr8-77121676-INS-86 |
| chr8-77155594-INS-198 | chr8-77155594-INS-198 |
| chr8-77183085-INS-520 | chr8-77183085-INS-520 |
| chr8-77220629-INS-52 | chr8-77220629-INS-52 |
| chr8-77223389-INS-1417 | chr8-77223390-INS-1412 |
| chr8-77234382-INS-148 | chr8-77234382-INS-148 |
| chr8-77239404-INS-77 | chr8-77239404-INS-77 |
| chr8-77278682-INS-57 | chr8-77278657-INS-84 |
| chr8-77308242-INS-51 | chr8-77308242-INS-54 |
| chr8-77325684-INS-56 | chr8-77325684-INS-58 |
| chr8-77326004-INS-1026 | chr8-77326005-INS-1027 |
| chr8-77346559-INS-78 | chr8-77346559-INS-78 |
| chr8-77381116-INS-132 | chr8-77381116-INS-136 |
| chr8-77393717-INS-187 | chr8-77393717-INS-187 |
| chr8-77399434-INS-393 | chr8-77399434-INS-393 |
| chr8-77421671-INS-196 | chr8-77421662-INS-212 |
| chr8-77431748-INS-56 | chr8-77431755-INS-52 |
| chr8-77433008-INS-66 | chr8-77433008-INS-66 |
| chr8-77433592-INS-54 | chr8-77433592-INS-54 |
| chr8-77435313-INS-258 | chr8-77435313-INS-260 |
| chr8-77450131-INS-738 | chr8-77450131-INS-738 |
| chr8-77452241-INS-1502 | chr8-77452242-INS-1502 |
| chr8-77460981-INS-467 | chr8-77460981-INS-466 |
| chr8-77462933-INS-469 | chr8-77462933-INS-469 |
| chr8-77467965-INS-70 | chr8-77467965-INS-70 |
| chr8-77481982-INS-915 | chr8-77481982-INS-915 |
| chr8-77485391-INS-6865 | chr8-77485392-INS-6866 |
| chr8-77490467-INS-84 | chr8-77490483-INS-68 |
| chr8-77508412-INS-312 | chr8-77508415-INS-353 |
| chr8-77525619-INS-53 | chr8-77525606-INS-66 |
| chr8-77526364-INS-477 | chr8-77526364-INS-477 |
| chr8-77589739-INS-7098 | chr8-77589740-INS-7098 |
| chr8-77595154-INS-95 | chr8-77595153-INS-128 |
| chr8-77608228-INS-59 | chr8-77608228-INS-59 |
| chr8-77610346-INS-71 | chr8-77610361-INS-52 |
| chr8-77612118-INS-135 | chr8-77612118-INS-134 |
| chr8-77647281-INS-104 | chr8-77647281-INS-104 |
| chr8-77683739-INS-54 | chr8-77683739-INS-54 |

| | |
|---|---|
| chr8-77690739-INS-70 | chr8-77690739-INS-54 |
| chr8-77706187-INS-2014 | chr8-77706186-INS-2027 |
| chr8-77722278-INS-54 | chr8-77722278-INS-54 |
| chr8-77728306-INS-832 | chr8-77728308-INS-831 |
| chr8-77814856-INS-165 | chr8-77814856-INS-165 |
| chr8-77829933-INS-54 | chr8-77829933-INS-54 |
| chr8-77843800-INS-274 | chr8-77843800-INS-274 |
| chr8-77844662-INS-1211 | chr8-77844663-INS-1211 |
| chr8-77854507-INS-2267 | chr8-77854508-INS-2267 |
| chr8-77862027-INS-454 | chr8-77862027-INS-454 |
| chr8-77874016-INS-63 | chr8-77874016-INS-83 |
| chr8-77876520-INS-776 | chr8-77876520-INS-775 |
| chr8-77878440-INS-61 | chr8-77878440-INS-53 |
| chr8-77900184-INS-175 | chr8-77900185-INS-161 |
| chr8-77928192-INS-1509 | chr8-77928193-INS-1509 |
| chr8-77944328-INS-86 | chr8-77944328-INS-86 |
| chr8-77948885-INS-354 | chr8-77948845-INS-364 |
| chr8-77955726-INS-2586 | chr8-77955727-INS-2587 |
| chr8-77979530-INS-1212 | chr8-77979531-INS-1213 |
| chr8-77979948-INS-165 | chr8-77979948-INS-165 |
| chr8-77981142-INS-452 | chr8-77981142-INS-446 |
| chr8-77993344-INS-203 | chr8-77993344-INS-203 |
| chr8-78009824-INS-252 | chr8-78009824-INS-243 |
| chr8-78014566-INS-70 | chr8-78014546-INS-83 |
| chr8-78065179-INS-260 | chr8-78065152-INS-261 |
| chr8-78067735-INS-5402 | chr8-78067736-INS-5402 |
| chr8-78089934-INS-478 | chr8-78089923-INS-508 |
| chr8-78096239-INS-658 | chr8-78096239-INS-658 |
| chr8-78100059-INS-207 | chr8-78100059-INS-207 |
| chr8-78101344-INS-164 | chr8-78101344-INS-164 |
| chr8-78108795-INS-194 | chr8-78108797-INS-195 |
| chr8-78129615-INS-129 | chr8-78129615-INS-172 |
| chr8-78137621-INS-312 | chr8-78137621-INS-313 |
| chr8-78149456-INS-1290 | chr8-78149457-INS-1290 |
| chr8-78158678-INS-117 | chr8-78158679-INS-117 |
| chr8-78164158-INS-186 | chr8-78164158-INS-162 |
| chr8-78169799-INS-67 | chr8-78169799-INS-67 |
| chr8-78170495-INS-54 | chr8-78170512-INS-52 |
| chr8-78181675-INS-1053 | chr8-78181678-INS-1047 |
| chr8-78192776-INS-176 | chr8-78192776-INS-176 |
| chr8-78196588-INS-215 | chr8-78196588-INS-215 |
| chr8-78198790-INS-60 | chr8-78198787-INS-53 |
| chr8-78204510-INS-52 | chr8-78204519-INS-52 |
| chr8-78230232-INS-67 | chr8-78230232-INS-67 |
| chr8-78231266-INS-68 | chr8-78231266-INS-68 |
| chr8-78235629-INS-74 | chr8-78235629-INS-76 |
| chr8-78254437-INS-82 | chr8-78254437-INS-82 |
| chr8-78292129-INS-397 | chr8-78292129-INS-399 |
| chr8-78311115-INS-55 | chr8-78311115-INS-55 |
| chr8-78313250-INS-6821 | chr8-78313251-INS-6823 |
| chr8-78318352-INS-260 | chr8-78318352-INS-260 |
| chr8-78344152-INS-406 | chr8-78344152-INS-406 |
| chr8-78346360-INS-51 | chr8-78346359-INS-87 |
| chr8-78356545-INS-134 | chr8-78356546-INS-134 |
| chr8-78362243-INS-64 | chr8-78362243-INS-64 |
| chr8-78369644-INS-195 | chr8-78369644-INS-195 |
| chr8-78393921-INS-55 | chr8-78393921-INS-55 |
| chr8-78410951-INS-1550 | chr8-78410952-INS-1550 |
| chr8-78414903-INS-100 | chr8-78414904-INS-101 |
| chr8-78417552-INS-528 | chr8-78417552-INS-528 |
| chr8-78435830-INS-70 | chr8-78435830-INS-75 |
| chr8-78437781-INS-107 | chr8-78437781-INS-107 |
| chr8-78441509-INS-91 | chr8-78441509-INS-58 |
| chr8-78455519-INS-60 | chr8-78455519-INS-60 |
| chr8-78457700-INS-58 | chr8-78457669-INS-86 |
| chr8-78462693-INS-52 | chr8-78462693-INS-57 |
| chr8-78469398-INS-145 | chr8-78469398-INS-145 |
| chr8-78472209-INS-80 | chr8-78472209-INS-80 |
| chr8-78475277-INS-240 | chr8-78475277-INS-240 |
| chr8-78487763-INS-58 | chr8-78487761-INS-62 |
| chr8-78508349-INS-216 | chr8-78508349-INS-216 |
| chr8-78510589-INS-5442 | chr8-78510590-INS-5442 |
| chr8-78510900-INS-67 | chr8-78510900-INS-73 |
| chr8-78514826-INS-92 | chr8-78514826-INS-100 |
| chr8-78515124-INS-6651 | chr8-78515125-INS-6651 |
| chr8-78529524-INS-126 | chr8-78529524-INS-126 |
| chr8-78530738-INS-95 | chr8-78530738-INS-95 |

| | |
|---|---|
| chr8-78550813-INS-82 | chr8-78550813-INS-82 |
| chr8-78568317-INS-432 | chr8-78568318-INS-414 |
| chr8-78584707-INS-94 | chr8-78584707-INS-94 |
| chr8-78601465-INS-222 | chr8-78601465-INS-223 |
| chr8-78608235-INS-377 | chr8-78608230-INS-360 |
| chr8-78609015-INS-141 | chr8-78608991-INS-126 |
| chr8-78661939-INS-52 | chr8-78661939-INS-52 |
| chr8-78671781-INS-63 | chr8-78671781-INS-63 |
| chr8-78672551-INS-6265 | chr8-78672552-INS-6265 |
| chr8-78681222-INS-54 | chr8-78681222-INS-54 |
| chr8-78685570-INS-56 | chr8-78685570-INS-56 |
| chr8-78699096-INS-60 | chr8-78699097-INS-66 |
| chr8-78704263-INS-74 | chr8-78704263-INS-76 |
| chr8-78714288-INS-198 | chr8-78714288-INS-198 |
| chr8-78726929-INS-72 | chr8-78726932-INS-70 |
| chr8-78733233-INS-71 | chr8-78733233-INS-71 |
| chr8-78750042-INS-62 | chr8-78750040-INS-64 |
| chr8-78753224-INS-107 | chr8-78753224-INS-107 |
| chr8-78781163-INS-738 | chr8-78781163-INS-736 |
| chr8-78781191-INS-95 | chr8-78781191-INS-95 |
| chr8-78783454-INS-53 | chr8-78783449-INS-72 |
| chr8-78793910-INS-95 | chr8-78793910-INS-95 |
| chr8-78821816-INS-6452 | chr8-78821817-INS-6452 |
| chr8-78825722-INS-196 | chr8-78825722-INS-196 |
| chr8-78834974-INS-68 | chr8-78834974-INS-72 |
| chr8-78847639-INS-93 | chr8-78847639-INS-93 |
| chr8-78849760-INS-89 | chr8-78849760-INS-91 |
| chr8-78854841-INS-213 | chr8-78854841-INS-213 |
| chr8-78861105-INS-109 | chr8-78861105-INS-109 |
| chr8-78884034-INS-319 | chr8-78884034-INS-319 |
| chr8-78968935-INS-62 | chr8-78968935-INS-62 |
| chr8-78972565-INS-853 | chr8-78972565-INS-853 |
| chr8-78974990-INS-236 | chr8-78974978-INS-236 |
| chr8-78979537-INS-117 | chr8-78979537-INS-117 |
| chr8-79008785-INS-78 | chr8-79008785-INS-80 |
| chr8-79013100-INS-54 | chr8-79013103-INS-54 |
| chr8-79057161-INS-54 | chr8-79057153-INS-56 |
| chr8-79070534-INS-186 | chr8-79070534-INS-186 |
| chr8-79076211-INS-54 | chr8-79076211-INS-60 |
| chr8-79081587-INS-69 | chr8-79081587-INS-69 |
| chr8-79090894-INS-221 | chr8-79090894-INS-221 |
| chr8-79113633-INS-54 | chr8-79113633-INS-54 |
| chr8-79193324-INS-195 | chr8-79193324-INS-195 |
| chr8-79202127-INS-222 | chr8-79202127-INS-222 |
| chr8-79222096-INS-233 | chr8-79222096-INS-232 |
| chr8-79223703-INS-78 | chr8-79223727-INS-88 |
| chr8-79246582-INS-82 | chr8-79246582-INS-82 |
| chr8-79269426-INS-198 | chr8-79269426-INS-198 |
| chr8-79278316-INS-60 | chr8-79278316-INS-70 |
| chr8-79283142-INS-62 | chr8-79283142-INS-62 |
| chr8-79300588-INS-64 | chr8-79300605-INS-65 |
| chr8-79302523-INS-51 | chr8-79302523-INS-55 |
| chr8-79309066-INS-214 | chr8-79309066-INS-214 |
| chr8-79309993-INS-56 | chr8-79309993-INS-56 |
| chr8-79319062-INS-114 | chr8-79319062-INS-114 |
| chr8-79341695-INS-344 | chr8-79341695-INS-344 |
| chr8-79350275-INS-791 | chr8-79350275-INS-791 |
| chr8-79362995-INS-215 | chr8-79362995-INS-215 |
| chr8-79378596-INS-75 | chr8-79378596-INS-75 |
| chr8-79403330-INS-60 | chr8-79403330-INS-62 |
| chr8-79418402-INS-182 | chr8-79418402-INS-179 |
| chr8-79437981-INS-58 | chr8-79437962-INS-85 |
| chr8-79449215-INS-224 | chr8-79449215-INS-230 |
| chr8-79455713-INS-90 | chr8-79455713-INS-90 |
| chr8-79478637-INS-67 | chr8-79478637-INS-67 |
| chr8-79497798-INS-141 | chr8-79497798-INS-141 |
| chr8-79520746-INS-447 | chr8-79520746-INS-447 |
| chr8-79559365-INS-52 | chr8-79559365-INS-62 |
| chr8-79580213-INS-55 | chr8-79580213-INS-55 |
| chr8-79618918-INS-91 | chr8-79618918-INS-91 |
| chr8-79629001-INS-498 | chr8-79629001-INS-498 |
| chr8-79630119-INS-51 | chr8-79630119-INS-51 |
| chr8-79677465-INS-204 | chr8-79677468-INS-211 |
| chr8-79690252-INS-56 | chr8-79690252-INS-52 |
| chr8-79720770-INS-7120 | chr8-79720771-INS-7120 |
| chr8-79746563-INS-58 | chr8-79746564-INS-64 |
| chr8-79760196-INS-69 | chr8-79760205-INS-50 |

| | |
|---|---|
| chr8-79773779-INS-95 | chr8-79773779-INS-95 |
| chr8-79814401-INS-96 | chr8-79814422-INS-64 |
| chr8-79822307-INS-129 | chr8-79822356-INS-130 |
| chr8-79857012-INS-103 | chr8-79857012-INS-146 |
| chr8-79857295-INS-6396 | chr8-79857296-INS-6396 |
| chr8-79866853-INS-243 | chr8-79866853-INS-243 |
| chr8-79981113-INS-52 | chr8-79981113-INS-52 |
| chr8-80030483-INS-82 | chr8-80030484-INS-54 |
| chr8-80036688-INS-52 | chr8-80036688-INS-52 |
| chr8-80066739-INS-88 | chr8-80066739-INS-88 |
| chr8-80068725-INS-159 | chr8-80068725-INS-159 |
| chr8-80081694-INS-102 | chr8-80081694-INS-68 |
| chr8-80100845-INS-75 | chr8-80100845-INS-75 |
| chr8-80103347-INS-60 | chr8-80103347-INS-72 |
| chr8-80104511-INS-66 | chr8-80104508-INS-74 |
| chr8-80147815-INS-66 | chr8-80147770-INS-96 |
| chr8-80164172-INS-167 | chr8-80164172-INS-162 |
| chr8-80170646-INS-54 | chr8-80170646-INS-81 |
| chr8-80185331-INS-121 | chr8-80185327-INS-131 |
| chr8-80190205-INS-5288 | chr8-80190206-INS-5288 |
| chr8-80191047-INS-6966 | chr8-80191048-INS-6968 |
| chr8-80207759-INS-65 | chr8-80207800-INS-50 |
| chr8-80213849-INS-138 | chr8-80213849-INS-138 |
| chr8-80251426-INS-552 | chr8-80251426-INS-549 |
| chr8-80260071-INS-117 | chr8-80260071-INS-117 |
| chr8-80261553-INS-72 | chr8-80261529-INS-80 |
| chr8-80278242-INS-52 | chr8-80278247-INS-68 |
| chr8-80284737-INS-155 | chr8-80284738-INS-114 |
| chr8-80297842-INS-207 | chr8-80297842-INS-210 |
| chr8-80319262-INS-58 | chr8-80319263-INS-52 |
| chr8-80329206-INS-414 | chr8-80329206-INS-415 |
| chr8-80334288-INS-168 | chr8-80334288-INS-168 |
| chr8-80349497-INS-58 | chr8-80349497-INS-69 |
| chr8-80436770-INS-66 | chr8-80436770-INS-66 |
| chr8-80467911-INS-207 | chr8-80467911-INS-204 |
| chr8-80518890-INS-92 | chr8-80518920-INS-55 |
| chr8-80521863-INS-175 | chr8-80521863-INS-175 |
| chr8-80543252-INS-58 | chr8-80543248-INS-66 |
| chr8-80544813-INS-379 | chr8-80544813-INS-378 |
| chr8-80551147-INS-6441 | chr8-80551148-INS-6441 |
| chr8-80554971-INS-641 | chr8-80554971-INS-649 |
| chr8-80570218-INS-220 | chr8-80570218-INS-219 |
| chr8-80583807-INS-5022 | chr8-80583808-INS-5022 |
| chr8-80591011-INS-105 | chr8-80591011-INS-105 |
| chr8-80594649-INS-507 | chr8-80594649-INS-507 |
| chr8-80619011-INS-6703 | chr8-80619012-INS-6704 |
| chr8-80625406-INS-2871 | chr8-80625407-INS-2871 |
| chr8-80629429-INS-3376 | chr8-80629430-INS-3376 |
| chr8-80643162-INS-78 | chr8-80643139-INS-90 |
| chr8-80647339-INS-360 | chr8-80647339-INS-360 |
| chr8-80649518-INS-698 | chr8-80649518-INS-698 |
| chr8-80656562-INS-6954 | chr8-80656563-INS-6954 |
| chr8-80709677-INS-52 | chr8-80709677-INS-65 |
| chr8-80718256-INS-63 | chr8-80718246-INS-53 |
| chr8-80729696-INS-384 | chr8-80729696-INS-384 |
| chr8-80750982-INS-203 | chr8-80750982-INS-203 |
| chr8-80761045-INS-132 | chr8-80761001-INS-144 |
| chr8-80774064-INS-90 | chr8-80774064-INS-90 |
| chr8-80808426-INS-208 | chr8-80808426-INS-208 |
| chr8-80873251-INS-52 | chr8-80873251-INS-52 |
| chr8-80900654-INS-1608 | chr8-80900655-INS-1608 |
| chr8-80915320-INS-159 | chr8-80915361-INS-159 |
| chr8-80920356-INS-56 | chr8-80920356-INS-56 |
| chr8-80944160-INS-379 | chr8-80944160-INS-372 |
| chr8-80954022-INS-6431 | chr8-80954023-INS-6431 |
| chr8-80967547-INS-268 | chr8-80967547-INS-268 |
| chr8-81002520-INS-76 | chr8-81002520-INS-76 |
| chr8-81004881-INS-110 | chr8-81004881-INS-110 |
| chr8-81013928-INS-141 | chr8-81013928-INS-141 |
| chr8-81048843-INS-62 | chr8-81048843-INS-62 |
| chr8-81097489-INS-70 | chr8-81097526-INS-60 |
| chr8-81125878-INS-336 | chr8-81125878-INS-336 |
| chr8-81130382-INS-62 | chr8-81130378-INS-68 |
| chr8-81131335-INS-175 | chr8-81131335-INS-175 |
| chr8-81148357-INS-2379 | chr8-81148358-INS-2371 |
| chr8-81234451-INS-60 | chr8-81234451-INS-71 |
| chr8-81237045-INS-213 | chr8-81237045-INS-213 |

| | |
|---|---|
| chr8-81246122-INS-232 | chr8-81246122-INS-232 |
| chr8-81259236-INS-199 | chr8-81259236-INS-199 |
| chr8-81278683-INS-3192 | chr8-81278684-INS-3192 |
| chr8-81280572-INS-90 | chr8-81280572-INS-105 |
| chr8-81285316-INS-209 | chr8-81285316-INS-214 |
| chr8-81298708-INS-2002 | chr8-81298709-INS-1992 |
| chr8-81311544-INS-365 | chr8-81311544-INS-365 |
| chr8-81312416-INS-77 | chr8-81312419-INS-81 |
| chr8-81315431-INS-197 | chr8-81315431-INS-197 |
| chr8-81339751-INS-105 | chr8-81339764-INS-68 |
| chr8-81362009-INS-71 | chr8-81362009-INS-71 |
| chr8-81376607-INS-472 | chr8-81376607-INS-471 |
| chr8-81379855-INS-518 | chr8-81379855-INS-511 |
| chr8-81393773-INS-142 | chr8-81393773-INS-142 |
| chr8-81396869-INS-225 | chr8-81396869-INS-237 |
| chr8-81402838-INS-115 | chr8-81402838-INS-120 |
| chr8-81434598-INS-102 | chr8-81434598-INS-116 |
| chr8-81443414-INS-92 | chr8-81443436-INS-71 |
| chr8-81462877-INS-244 | chr8-81462877-INS-244 |
| chr8-81470829-INS-54 | chr8-81470840-INS-54 |
| chr8-81493788-INS-67 | chr8-81493788-INS-67 |
| chr8-81504691-INS-142 | chr8-81504683-INS-166 |
| chr8-81508613-INS-80 | chr8-81508613-INS-92 |
| chr8-81581983-INS-54 | chr8-81581983-INS-54 |
| chr8-81588954-INS-71 | chr8-81588969-INS-68 |
| chr8-81620612-INS-441 | chr8-81620612-INS-441 |
| chr8-81628162-INS-390 | chr8-81628162-INS-390 |
| chr8-81636371-INS-59 | chr8-81636385-INS-76 |
| chr8-81644153-INS-64 | chr8-81644147-INS-52 |
| chr8-81648542-INS-236 | chr8-81648542-INS-237 |
| chr8-81660576-INS-4703 | chr8-81660577-INS-4703 |
| chr8-81661157-INS-82 | chr8-81661157-INS-82 |
| chr8-81666225-INS-52 | chr8-81666228-INS-50 |
| chr8-81678203-INS-197 | chr8-81678203-INS-197 |
| chr8-81680953-INS-225 | chr8-81680953-INS-225 |
| chr8-81692217-INS-223 | chr8-81692217-INS-224 |
| chr8-81702148-INS-793 | chr8-81702148-INS-793 |
| chr8-81713032-INS-80 | chr8-81713004-INS-92 |
| chr8-81715555-INS-1158 | chr8-81715556-INS-1159 |
| chr8-81747093-INS-56 | chr8-81747093-INS-56 |
| chr8-81751837-INS-2827 | chr8-81751838-INS-2827 |
| chr8-81753738-INS-8119 | chr8-81753739-INS-8119 |
| chr8-81756414-INS-4016 | chr8-81756415-INS-4016 |
| chr8-81772341-INS-549 | chr8-81772341-INS-561 |
| chr8-81778953-INS-60 | chr8-81778953-INS-60 |
| chr8-81787979-INS-64 | chr8-81787979-INS-84 |
| chr8-81792291-INS-58 | chr8-81792291-INS-56 |
| chr8-81825154-INS-75 | chr8-81825154-INS-75 |
| chr8-81830417-INS-589 | chr8-81830417-INS-589 |
| chr8-81832191-INS-60 | chr8-81832191-INS-60 |
| chr8-81844589-INS-91 | chr8-81844596-INS-109 |
| chr8-81855155-INS-595 | chr8-81855155-INS-599 |
| chr8-81861004-INS-766 | chr8-81861004-INS-766 |
| chr8-81872146-INS-670 | chr8-81872146-INS-670 |
| chr8-81885357-INS-65 | chr8-81885357-INS-65 |
| chr8-81896485-INS-597 | chr8-81896485-INS-597 |
| chr8-81916409-INS-66 | chr8-81916409-INS-66 |
| chr8-81932749-INS-106 | chr8-81932749-INS-106 |
| chr8-81948297-INS-173 | chr8-81948297-INS-173 |
| chr8-81959074-INS-52 | chr8-81959074-INS-52 |
| chr8-81961964-INS-68 | chr8-81961964-INS-68 |
| chr8-81994159-INS-910 | chr8-81994159-INS-913 |
| chr8-82001759-INS-494 | chr8-82001759-INS-494 |
| chr8-82008489-INS-54 | chr8-82008489-INS-66 |
| chr8-82043185-INS-2364 | chr8-82043186-INS-2364 |
| chr8-82065556-INS-7155 | chr8-82065557-INS-7159 |
| chr8-82074458-INS-437 | chr8-82074458-INS-437 |
| chr8-82086039-INS-60 | chr8-82086049-INS-60 |
| chr8-82095936-INS-266 | chr8-82095917-INS-242 |
| chr8-82104235-INS-939 | chr8-82104235-INS-939 |
| chr8-82110451-INS-66 | chr8-82110447-INS-64 |
| chr8-82124802-INS-52 | chr8-82124802-INS-54 |
| chr8-82149216-INS-475 | chr8-82149216-INS-476 |
| chr8-82178077-INS-560 | chr8-82178077-INS-560 |
| chr8-82179539-INS-1642 | chr8-82179540-INS-1636 |
| chr8-82192005-INS-51 | chr8-82192005-INS-51 |
| chr8-82192877-INS-6677 | chr8-82192883-INS-6679 |

| | |
|---|---|
| chr8-82238502-INS-259 | chr8-82238502-INS-259 |
| chr8-82244020-INS-70 | chr8-82244016-INS-78 |
| chr8-82262951-INS-277 | chr8-82262951-INS-277 |
| chr8-82281858-INS-135 | chr8-82281858-INS-135 |
| chr8-82284903-INS-79 | chr8-82284903-INS-79 |
| chr8-82286102-INS-205 | chr8-82286102-INS-205 |
| chr8-82290692-INS-60 | chr8-82290692-INS-61 |
| chr8-82304575-INS-62 | chr8-82304575-INS-62 |
| chr8-82322256-INS-52 | chr8-82322256-INS-52 |
| chr8-82325433-INS-55 | chr8-82325433-INS-69 |
| chr8-82344230-INS-64 | chr8-82344230-INS-64 |
| chr8-82348041-INS-788 | chr8-82348041-INS-788 |
| chr8-82364694-INS-75 | chr8-82364694-INS-77 |
| chr8-82372450-INS-1474 | chr8-82372451-INS-1441 |
| chr8-82390582-INS-83 | chr8-82390566-INS-88 |
| chr8-82401130-INS-1205 | chr8-82401131-INS-1205 |
| chr8-82427821-INS-73 | chr8-82427821-INS-73 |
| chr8-82446989-INS-135 | chr8-82446989-INS-135 |
| chr8-82488027-INS-6230 | chr8-82488028-INS-6231 |
| chr8-82494962-INS-72 | chr8-82494962-INS-68 |
| chr8-82497627-INS-820 | chr8-82497627-INS-823 |
| chr8-82502636-INS-102 | chr8-82502636-INS-102 |
| chr8-82510730-INS-62 | chr8-82510730-INS-51 |
| chr8-82514777-INS-55 | chr8-82514777-INS-54 |
| chr8-82541860-INS-90 | chr8-82541860-INS-91 |
| chr8-82543493-INS-202 | chr8-82543493-INS-202 |
| chr8-82549612-INS-167 | chr8-82549612-INS-167 |
| chr8-82551541-INS-77 | chr8-82551496-INS-111 |
| chr8-82560807-INS-181 | chr8-82560807-INS-181 |
| chr8-82566510-INS-55 | chr8-82566510-INS-55 |
| chr8-82577929-INS-801 | chr8-82577947-INS-810 |
| chr8-82584351-INS-84 | chr8-82584351-INS-86 |
| chr8-82650190-INS-2336 | chr8-82650191-INS-2336 |
| chr8-82665296-INS-52 | chr8-82665296-INS-56 |
| chr8-82673772-INS-103 | chr8-82673772-INS-103 |
| chr8-82691221-INS-519 | chr8-82691221-INS-517 |
| chr8-82692695-INS-175 | chr8-82692695-INS-173 |
| chr8-82697713-INS-287 | chr8-82697707-INS-296 |
| chr8-82705876-INS-62 | chr8-82705880-INS-67 |
| chr8-82716680-INS-78 | chr8-82716680-INS-78 |
| chr8-82779444-INS-301 | chr8-82779444-INS-301 |
| chr8-82780911-INS-4088 | chr8-82780912-INS-4088 |
| chr8-82783632-INS-489 | chr8-82783632-INS-495 |
| chr8-82797146-INS-6461 | chr8-82797147-INS-6462 |
| chr8-82859215-INS-88 | chr8-82859215-INS-93 |
| chr8-82866461-INS-153 | chr8-82866461-INS-153 |
| chr8-82875270-INS-747 | chr8-82875270-INS-747 |
| chr8-82900406-INS-310 | chr8-82900406-INS-310 |
| chr8-82907043-INS-782 | chr8-82907043-INS-782 |
| chr8-82911830-INS-67 | chr8-82911830-INS-67 |
| chr8-82920542-INS-1469 | chr8-82920542-INS-1437 |
| chr8-82939786-INS-56 | chr8-82939780-INS-64 |
| chr8-82942793-INS-156 | chr8-82942793-INS-158 |
| chr8-82977797-INS-86 | chr8-82977797-INS-86 |
| chr8-83007371-INS-100 | chr8-83007371-INS-100 |
| chr8-83008959-INS-1681 | chr8-83008960-INS-1681 |
| chr8-83036747-INS-934 | chr8-83036747-INS-934 |
| chr8-83041420-INS-6824 | chr8-83041421-INS-6825 |
| chr8-83041438-INS-246 | chr8-83041438-INS-246 |
| chr8-83064988-INS-60 | chr8-83064988-INS-60 |
| chr8-83074958-INS-6342 | chr8-83074959-INS-6342 |
| chr8-83135068-INS-54 | chr8-83135068-INS-51 |
| chr8-83169274-INS-1134 | chr8-83169275-INS-1134 |
| chr8-83183241-INS-87 | chr8-83183241-INS-87 |
| chr8-83199542-INS-50 | chr8-83199532-INS-98 |
| chr8-83251108-INS-62 | chr8-83251108-INS-62 |
| chr8-83272697-INS-236 | chr8-83272697-INS-235 |
| chr8-83276798-INS-58 | chr8-83276798-INS-58 |
| chr8-83288447-INS-309 | chr8-83288447-INS-309 |
| chr8-83290146-INS-1019 | chr8-83290147-INS-1019 |
| chr8-83295966-INS-7191 | chr8-83295967-INS-7191 |
| chr8-83295967-INS-423 | chr8-83295967-INS-423 |
| chr8-83301041-INS-1046 | chr8-83301048-INS-1041 |
| chr8-83315209-INS-857 | chr8-83315209-INS-865 |
| chr8-83354445-INS-4073 | chr8-83354446-INS-4088 |
| chr8-83398826-INS-112 | chr8-83398826-INS-124 |
| chr8-83400668-INS-122 | chr8-83400668-INS-128 |

| | |
|---|---|
| chr8-83412164-INS-81 | chr8-83412163-INS-52 |
| chr8-83414424-INS-793 | chr8-83414424-INS-793 |
| chr8-83464307-INS-2615 | chr8-83464308-INS-2614 |
| chr8-83466215-INS-60 | chr8-83466171-INS-60 |
| chr8-83476805-INS-98 | chr8-83476805-INS-98 |
| chr8-83526518-INS-562 | chr8-83526518-INS-562 |
| chr8-83579905-INS-1893 | chr8-83579906-INS-1893 |
| chr8-83596612-INS-82 | chr8-83596612-INS-82 |
| chr8-83613102-INS-6439 | chr8-83613103-INS-6439 |
| chr8-83618641-INS-62 | chr8-83618641-INS-62 |
| chr8-83657117-INS-91 | chr8-83657117-INS-91 |
| chr8-83681785-INS-69 | chr8-83681785-INS-69 |
| chr8-83721114-INS-1276 | chr8-83721115-INS-1276 |
| chr8-83731351-INS-73 | chr8-83731316-INS-71 |
| chr8-83755456-INS-89 | chr8-83755456-INS-91 |
| chr8-83788516-INS-96 | chr8-83788512-INS-88 |
| chr8-83798063-INS-821 | chr8-83798063-INS-824 |
| chr8-83802189-INS-224 | chr8-83802190-INS-224 |
| chr8-83802754-INS-244 | chr8-83802754-INS-244 |
| chr8-83818290-INS-56 | chr8-83818290-INS-54 |
| chr8-83829426-INS-1351 | chr8-83829427-INS-1359 |
| chr8-83835586-INS-209 | chr8-83835586-INS-209 |
| chr8-83880302-INS-102 | chr8-83880302-INS-108 |
| chr8-83887329-INS-331 | chr8-83887329-INS-331 |
| chr8-83899855-INS-422 | chr8-83899855-INS-422 |
| chr8-83947846-INS-692 | chr8-83947844-INS-696 |
| chr8-83958438-INS-54 | chr8-83958438-INS-54 |
| chr8-83960028-INS-213 | chr8-83960028-INS-231 |
| chr8-83991321-INS-86 | chr8-83991321-INS-90 |
| chr8-84026330-INS-100 | chr8-84026340-INS-110 |
| chr8-84029106-INS-680 | chr8-84029106-INS-679 |
| chr8-84061869-INS-74 | chr8-84061869-INS-74 |
| chr8-84065277-INS-56 | chr8-84065277-INS-56 |
| chr8-84075648-INS-59 | chr8-84075648-INS-60 |
| chr8-84076878-INS-120 | chr8-84076903-INS-82 |
| chr8-84080736-INS-240 | chr8-84080736-INS-240 |
| chr8-84081918-INS-6331 | chr8-84081919-INS-6331 |
| chr8-84090185-INS-273 | chr8-84090185-INS-277 |
| chr8-84105103-INS-64 | chr8-84105103-INS-64 |
| chr8-84106394-INS-211 | chr8-84106394-INS-214 |
| chr8-84107697-INS-50 | chr8-84107697-INS-50 |
| chr8-84119639-INS-160 | chr8-84119654-INS-166 |
| chr8-84166901-INS-58 | chr8-84166901-INS-58 |
| chr8-84172060-INS-76 | chr8-84172060-INS-79 |
| chr8-84181826-INS-204 | chr8-84181826-INS-204 |
| chr8-84188502-INS-68 | chr8-84188498-INS-84 |
| chr8-84190236-INS-963 | chr8-84190236-INS-963 |
| chr8-84213490-INS-120 | chr8-84213490-INS-111 |
| chr8-84223460-INS-1930 | chr8-84223461-INS-1932 |
| chr8-84246796-INS-399 | chr8-84246796-INS-399 |
| chr8-84270472-INS-52 | chr8-84270472-INS-52 |
| chr8-84292021-INS-99 | chr8-84292021-INS-127 |
| chr8-84297671-INS-120 | chr8-84297671-INS-120 |
| chr8-84313254-INS-179 | chr8-84313254-INS-179 |
| chr8-84322225-INS-1141 | chr8-84322226-INS-1145 |
| chr8-84326551-INS-216 | chr8-84326551-INS-216 |
| chr8-84344355-INS-92 | chr8-84344355-INS-91 |
| chr8-84359284-INS-229 | chr8-84359284-INS-230 |
| chr8-84359683-INS-4813 | chr8-84359684-INS-4813 |
| chr8-84376298-INS-200 | chr8-84376298-INS-200 |
| chr8-84405325-INS-70 | chr8-84405325-INS-70 |
| chr8-84440655-INS-661 | chr8-84440653-INS-661 |
| chr8-84481196-INS-94 | chr8-84481196-INS-128 |
| chr8-84484142-INS-54 | chr8-84484142-INS-52 |
| chr8-84487848-INS-200 | chr8-84487851-INS-198 |
| chr8-84495052-INS-64 | chr8-84495052-INS-64 |
| chr8-84501770-INS-233 | chr8-84501770-INS-233 |
| chr8-84522254-INS-151 | chr8-84522254-INS-149 |
| chr8-84526646-INS-214 | chr8-84526646-INS-214 |
| chr8-84534562-INS-232 | chr8-84534562-INS-232 |
| chr8-84538466-INS-87 | chr8-84538466-INS-87 |
| chr8-84565922-INS-200 | chr8-84565922-INS-200 |
| chr8-84568501-INS-5552 | chr8-84568502-INS-5552 |
| chr8-84582008-INS-210 | chr8-84582008-INS-210 |
| chr8-84598699-INS-125 | chr8-84598699-INS-125 |
| chr8-84604483-INS-98 | chr8-84604483-INS-101 |
| chr8-84606413-INS-203 | chr8-84606413-INS-205 |

| | |
|---|---|
| chr8-84608350-INS-134 | chr8-84608350-INS-134 |
| chr8-84616981-INS-207 | chr8-84616981-INS-207 |
| chr8-84676758-INS-242 | chr8-84676758-INS-256 |
| chr8-84712531-INS-63 | chr8-84712531-INS-60 |
| chr8-84732104-INS-209 | chr8-84732104-INS-209 |
| chr8-84741245-INS-120 | chr8-84741240-INS-138 |
| chr8-84761286-INS-183 | chr8-84761286-INS-183 |
| chr8-84761931-INS-139 | chr8-84761932-INS-137 |
| chr8-84773598-INS-69 | chr8-84773598-INS-61 |
| chr8-84774674-INS-199 | chr8-84774674-INS-221 |
| chr8-84784540-INS-390 | chr8-84784540-INS-390 |
| chr8-84807260-INS-206 | chr8-84807260-INS-206 |
| chr8-84809688-INS-67 | chr8-84809688-INS-67 |
| chr8-84849468-INS-164 | chr8-84849468-INS-164 |
| chr8-84868971-INS-116 | chr8-84868971-INS-118 |
| chr8-84889433-INS-85 | chr8-84889433-INS-107 |
| chr8-84901041-INS-214 | chr8-84901041-INS-227 |
| chr8-84914663-INS-245 | chr8-84914663-INS-245 |
| chr8-84936582-INS-72 | chr8-84936582-INS-72 |
| chr8-84940874-INS-201 | chr8-84940874-INS-201 |
| chr8-84945094-INS-153 | chr8-84945094-INS-153 |
| chr8-84946144-INS-5358 | chr8-84946145-INS-5358 |
| chr8-84946668-INS-207 | chr8-84946668-INS-207 |
| chr8-84954638-INS-215 | chr8-84954638-INS-216 |
| chr8-84959339-INS-65 | chr8-84959339-INS-65 |
| chr8-84971405-INS-278 | chr8-84971405-INS-278 |
| chr8-85001141-INS-227 | chr8-85001141-INS-227 |
| chr8-85003063-INS-60 | chr8-85003063-INS-56 |
| chr8-85010413-INS-205 | chr8-85010413-INS-205 |
| chr8-85012917-INS-382 | chr8-85012911-INS-381 |
| chr8-85016597-INS-94 | chr8-85016598-INS-94 |
| chr8-85025953-INS-200 | chr8-85025953-INS-200 |
| chr8-85045868-INS-63 | chr8-85045868-INS-63 |
| chr8-85048263-INS-240 | chr8-85048263-INS-212 |
| chr8-85069765-INS-200 | chr8-85069765-INS-200 |
| chr8-85071727-INS-148 | chr8-85071727-INS-148 |
| chr8-85078827-INS-183 | chr8-85078827-INS-183 |
| chr8-85082564-INS-62 | chr8-85082564-INS-74 |
| chr8-85100871-INS-78 | chr8-85100871-INS-78 |
| chr8-85103234-INS-66 | chr8-85103234-INS-66 |
| chr8-85123089-INS-60 | chr8-85123089-INS-56 |
| chr8-85152611-INS-147 | chr8-85152611-INS-147 |
| chr8-85198631-INS-379 | chr8-85198649-INS-394 |
| chr8-85200760-INS-187 | chr8-85200760-INS-187 |
| chr8-85211944-INS-72 | chr8-85211946-INS-52 |
| chr8-85219442-INS-78 | chr8-85219443-INS-102 |
| chr8-85222179-INS-126 | chr8-85222179-INS-126 |
| chr8-85226943-INS-54 | chr8-85226943-INS-54 |
| chr8-85289733-INS-53 | chr8-85289733-INS-53 |
| chr8-85333127-INS-94 | chr8-85333145-INS-98 |
| chr8-85355618-INS-56 | chr8-85355619-INS-54 |
| chr8-85357764-INS-201 | chr8-85357766-INS-201 |
| chr8-85364897-INS-170 | chr8-85364897-INS-160 |
| chr8-85378364-INS-173 | chr8-85378364-INS-173 |
| chr8-85418752-INS-183 | chr8-85418752-INS-184 |
| chr8-85420234-INS-317 | chr8-85420234-INS-317 |
| chr8-85470491-INS-172 | chr8-85470491-INS-172 |
| chr8-85488795-INS-2401 | chr8-85488796-INS-2401 |
| chr8-85530691-INS-111 | chr8-85530674-INS-111 |
| chr8-85550660-INS-206 | chr8-85550661-INS-206 |
| chr8-85559574-INS-209 | chr8-85559574-INS-209 |
| chr8-85606876-INS-197 | chr8-85606876-INS-197 |
| chr8-85612064-INS-194 | chr8-85612064-INS-194 |
| chr8-85625330-INS-59 | chr8-85625330-INS-59 |
| chr8-85632522-INS-58 | chr8-85632522-INS-60 |
| chr8-85665886-INS-55 | chr8-85665844-INS-93 |
| chr8-85669328-INS-197 | chr8-85669328-INS-197 |
| chr8-85726638-INS-198 | chr8-85726638-INS-198 |
| chr8-85736984-INS-549 | chr8-85736984-INS-534 |
| chr8-85741384-INS-85 | chr8-85741399-INS-50 |
| chr8-85826785-INS-972 | chr8-85826785-INS-972 |
| chr8-85828616-INS-218 | chr8-85828616-INS-218 |
| chr8-85838008-INS-537 | chr8-85838008-INS-537 |
| chr8-85843159-INS-211 | chr8-85843159-INS-211 |
| chr8-85848026-INS-475 | chr8-85848026-INS-475 |
| chr8-85870590-INS-506 | chr8-85870590-INS-506 |
| chr8-85873048-INS-144 | chr8-85873048-INS-145 |

| | |
|---|---|
| chr8-85875806-INS-55 | chr8-85875806-INS-55 |
| chr8-85881938-INS-54 | chr8-85881938-INS-54 |
| chr8-85898990-INS-239 | chr8-85898990-INS-235 |
| chr8-85908330-INS-488 | chr8-85908330-INS-484 |
| chr8-85944298-INS-201 | chr8-85944298-INS-201 |
| chr8-86003944-INS-396 | chr8-86003944-INS-397 |
| chr8-86035966-INS-7117 | chr8-86035967-INS-7117 |
| chr8-86037699-INS-155 | chr8-86037699-INS-155 |
| chr8-86047017-INS-199 | chr8-86047017-INS-199 |
| chr8-86063444-INS-219 | chr8-86063456-INS-214 |
| chr8-86066070-INS-597 | chr8-86066070-INS-597 |
| chr8-86089871-INS-59 | chr8-86089871-INS-59 |
| chr8-86090551-INS-936 | chr8-86090551-INS-936 |
| chr8-86101578-INS-567 | chr8-86101578-INS-574 |
| chr8-86114469-INS-825 | chr8-86114469-INS-825 |
| chr8-86120721-INS-74 | chr8-86120733-INS-83 |
| chr8-86122068-INS-195 | chr8-86122068-INS-195 |
| chr8-86126304-INS-231 | chr8-86126301-INS-234 |
| chr8-86129006-INS-534 | chr8-86129006-INS-534 |
| chr8-86132384-INS-52 | chr8-86132384-INS-52 |
| chr8-86151559-INS-101 | chr8-86151559-INS-101 |
| chr8-86159976-INS-331 | chr8-86159976-INS-331 |
| chr8-86162250-INS-208 | chr8-86162288-INS-162 |
| chr8-86165074-INS-195 | chr8-86165074-INS-194 |
| chr8-86194159-INS-88 | chr8-86194159-INS-88 |
| chr8-86195528-INS-64 | chr8-86195528-INS-64 |
| chr8-86206319-INS-773 | chr8-86206319-INS-762 |
| chr8-86214329-INS-132 | chr8-86214329-INS-131 |
| chr8-86257835-INS-779 | chr8-86257835-INS-779 |
| chr8-86262496-INS-164 | chr8-86262506-INS-173 |
| chr8-86275455-INS-112 | chr8-86275455-INS-112 |
| chr8-86313733-INS-7120 | chr8-86313734-INS-7135 |
| chr8-86314028-INS-56 | chr8-86314028-INS-56 |
| chr8-86329429-INS-441 | chr8-86329429-INS-441 |
| chr8-86332408-INS-63 | chr8-86332408-INS-64 |
| chr8-86338129-INS-298 | chr8-86338129-INS-298 |
| chr8-86340845-INS-64 | chr8-86340845-INS-68 |
| chr8-86342241-INS-556 | chr8-86342247-INS-524 |
| chr8-86380646-INS-240 | chr8-86380646-INS-240 |
| chr8-86387986-INS-142 | chr8-86387986-INS-142 |
| chr8-86398755-INS-162 | chr8-86398755-INS-162 |
| chr8-86415418-INS-175 | chr8-86415418-INS-176 |
| chr8-86432177-INS-146 | chr8-86432177-INS-146 |
| chr8-86456395-INS-96 | chr8-86456395-INS-96 |
| chr8-86473949-INS-989 | chr8-86473949-INS-990 |
| chr8-86493706-INS-66 | chr8-86493709-INS-66 |
| chr8-86523612-INS-454 | chr8-86523612-INS-454 |
| chr8-86531646-INS-53 | chr8-86531646-INS-53 |
| chr8-86543710-INS-7295 | chr8-86543711-INS-7295 |
| chr8-86548689-INS-52 | chr8-86548689-INS-52 |
| chr8-86573797-INS-190 | chr8-86573797-INS-190 |
| chr8-86613550-INS-74 | chr8-86613550-INS-62 |
| chr8-86638703-INS-125 | chr8-86638703-INS-128 |
| chr8-86664537-INS-162 | chr8-86664584-INS-126 |
| chr8-86691221-INS-1389 | chr8-86691222-INS-1389 |
| chr8-86709959-INS-821 | chr8-86709959-INS-822 |
| chr8-86711585-INS-170 | chr8-86711576-INS-191 |
| chr8-86723902-INS-272 | chr8-86723902-INS-304 |
| chr8-86731689-INS-6472 | chr8-86731690-INS-6474 |
| chr8-86739030-INS-59 | chr8-86739030-INS-59 |
| chr8-86739057-INS-647 | chr8-86739057-INS-647 |
| chr8-86749006-INS-2918 | chr8-86749007-INS-2918 |
| chr8-86754351-INS-199 | chr8-86754351-INS-199 |
| chr8-86766399-INS-4716 | chr8-86766400-INS-4716 |
| chr8-86773954-INS-136 | chr8-86773931-INS-125 |
| chr8-86806381-INS-241 | chr8-86806381-INS-277 |
| chr8-86872116-INS-213 | chr8-86872116-INS-213 |
| chr8-86884204-INS-7161 | chr8-86884205-INS-7161 |
| chr8-86886590-INS-68 | chr8-86886553-INS-73 |
| chr8-86898870-INS-64 | chr8-86898855-INS-114 |
| chr8-86915426-INS-81 | chr8-86915426-INS-81 |
| chr8-86925977-INS-135 | chr8-86925977-INS-125 |
| chr8-86927745-INS-53 | chr8-86927745-INS-53 |
| chr8-86928838-INS-474 | chr8-86928838-INS-474 |
| chr8-86931804-INS-61 | chr8-86931804-INS-53 |
| chr8-86933083-INS-1493 | chr8-86933084-INS-1494 |
| chr8-86939150-INS-2848 | chr8-86939151-INS-2849 |

| | |
|---|---|
| chr8-86942803-INS-212 | chr8-86942803-INS-212 |
| chr8-86958733-INS-282 | chr8-86958733-INS-282 |
| chr8-86985236-INS-126 | chr8-86985240-INS-138 |
| chr8-86989535-INS-68 | chr8-86989538-INS-82 |
| chr8-86993193-INS-69 | chr8-86993193-INS-84 |
| chr8-86999259-INS-242 | chr8-86999260-INS-199 |
| chr8-87009486-INS-66 | chr8-87009486-INS-68 |
| chr8-87029171-INS-199 | chr8-87029171-INS-199 |
| chr8-87032559-INS-100 | chr8-87032583-INS-68 |
| chr8-87047916-INS-94 | chr8-87047955-INS-52 |
| chr8-87107881-INS-53 | chr8-87107840-INS-62 |
| chr8-87126427-INS-5891 | chr8-87126428-INS-5891 |
| chr8-87133973-INS-67 | chr8-87133953-INS-67 |
| chr8-87152188-INS-99 | chr8-87152188-INS-99 |
| chr8-87170161-INS-58 | chr8-87170161-INS-50 |
| chr8-87201666-INS-80 | chr8-87201689-INS-55 |
| chr8-87245787-INS-54 | chr8-87245787-INS-54 |
| chr8-87269840-INS-498 | chr8-87269840-INS-498 |
| chr8-87274779-INS-59 | chr8-87274799-INS-58 |
| chr8-87290651-INS-71 | chr8-87290651-INS-71 |
| chr8-87303054-INS-71 | chr8-87303054-INS-59 |
| chr8-87344796-INS-66 | chr8-87344796-INS-66 |
| chr8-87347808-INS-57 | chr8-87347808-INS-57 |
| chr8-87367765-INS-857 | chr8-87367765-INS-857 |
| chr8-87428529-INS-399 | chr8-87428529-INS-399 |
| chr8-87435970-INS-76 | chr8-87435977-INS-76 |
| chr8-87477846-INS-83 | chr8-87477850-INS-61 |
| chr8-87490077-INS-1076 | chr8-87490078-INS-1072 |
| chr8-87501234-INS-88 | chr8-87501233-INS-71 |
| chr8-87508728-INS-86 | chr8-87508731-INS-102 |
| chr8-87521692-INS-72 | chr8-87521692-INS-64 |
| chr8-87522859-INS-84 | chr8-87522859-INS-84 |
| chr8-87525091-INS-5302 | chr8-87525092-INS-5302 |
| chr8-87533219-INS-78 | chr8-87533219-INS-78 |
| chr8-87541349-INS-335 | chr8-87541397-INS-333 |
| chr8-87551757-INS-75 | chr8-87551757-INS-117 |
| chr8-87561423-INS-208 | chr8-87561429-INS-179 |
| chr8-87564409-INS-80 | chr8-87564391-INS-66 |
| chr8-87589841-INS-221 | chr8-87589841-INS-221 |
| chr8-87591349-INS-68 | chr8-87591349-INS-60 |
| chr8-87595343-INS-211 | chr8-87595343-INS-211 |
| chr8-87615590-INS-205 | chr8-87615590-INS-205 |
| chr8-87630923-INS-69 | chr8-87630923-INS-74 |
| chr8-87641488-INS-392 | chr8-87641488-INS-392 |
| chr8-87642831-INS-6402 | chr8-87642832-INS-6402 |
| chr8-87649497-INS-87 | chr8-87649497-INS-87 |
| chr8-87654761-INS-87 | chr8-87654761-INS-87 |
| chr8-87673346-INS-244 | chr8-87673344-INS-215 |
| chr8-87686980-INS-402 | chr8-87686980-INS-402 |
| chr8-87691903-INS-84 | chr8-87691933-INS-84 |
| chr8-87697293-INS-59 | chr8-87697293-INS-59 |
| chr8-87730519-INS-573 | chr8-87730519-INS-573 |
| chr8-87731324-INS-1893 | chr8-87731325-INS-1894 |
| chr8-87750351-INS-5243 | chr8-87750352-INS-5243 |
| chr8-87754420-INS-125 | chr8-87754420-INS-125 |
| chr8-87784919-INS-55 | chr8-87784919-INS-55 |
| chr8-87892532-INS-176 | chr8-87892532-INS-176 |
| chr8-87894233-INS-136 | chr8-87894239-INS-152 |
| chr8-87928673-INS-859 | chr8-87928673-INS-859 |
| chr8-87949716-INS-128 | chr8-87949716-INS-128 |
| chr8-88005269-INS-72 | chr8-88005269-INS-52 |
| chr8-88006666-INS-214 | chr8-88006666-INS-209 |
| chr8-88057998-INS-231 | chr8-88057998-INS-231 |
| chr8-88063376-INS-126 | chr8-88063376-INS-126 |
| chr8-88073913-INS-477 | chr8-88073913-INS-476 |
| chr8-88145344-INS-88 | chr8-88145346-INS-76 |
| chr8-88168579-INS-711 | chr8-88168579-INS-715 |
| chr8-88193887-INS-74 | chr8-88193888-INS-70 |
| chr8-88209707-INS-60 | chr8-88209707-INS-74 |
| chr8-88275257-INS-445 | chr8-88275257-INS-445 |
| chr8-88287553-INS-217 | chr8-88287553-INS-218 |
| chr8-88301423-INS-184 | chr8-88301423-INS-184 |
| chr8-88309673-INS-63 | chr8-88309673-INS-77 |
| chr8-88320945-INS-53 | chr8-88320945-INS-58 |
| chr8-88323280-INS-56 | chr8-88323280-INS-58 |
| chr8-88336046-INS-486 | chr8-88336046-INS-486 |
| chr8-88341326-INS-5278 | chr8-88341327-INS-5278 |

| | |
|---|---|
| chr8-88358966-INS-6375 | chr8-88358965-INS-6364 |
| chr8-88394281-INS-58 | chr8-88394281-INS-58 |
| chr8-88480703-INS-163 | chr8-88480694-INS-183 |
| chr8-88487134-INS-61 | chr8-88487139-INS-67 |
| chr8-88506479-INS-69 | chr8-88506479-INS-69 |
| chr8-88515963-INS-205 | chr8-88515963-INS-205 |
| chr8-88526539-INS-51 | chr8-88526499-INS-73 |
| chr8-88537192-INS-66 | chr8-88537144-INS-70 |
| chr8-88540353-INS-132 | chr8-88540353-INS-136 |
| chr8-88582252-INS-65 | chr8-88582252-INS-66 |
| chr8-88628995-INS-136 | chr8-88628995-INS-118 |
| chr8-88632865-INS-278 | chr8-88632865-INS-278 |
| chr8-88636791-INS-680 | chr8-88636791-INS-676 |
| chr8-88650513-INS-81 | chr8-88650513-INS-81 |
| chr8-88662264-INS-92 | chr8-88662264-INS-80 |
| chr8-88692909-INS-68 | chr8-88692909-INS-68 |
| chr8-88697360-INS-143 | chr8-88697360-INS-143 |
| chr8-88712228-INS-52 | chr8-88712228-INS-52 |
| chr8-88722403-INS-207 | chr8-88722403-INS-207 |
| chr8-88726448-INS-615 | chr8-88726448-INS-615 |
| chr8-88768820-INS-134 | chr8-88768820-INS-134 |
| chr8-88776990-INS-68 | chr8-88776990-INS-68 |
| chr8-88779639-INS-62 | chr8-88779639-INS-62 |
| chr8-88779825-INS-762 | chr8-88779825-INS-762 |
| chr8-88790245-INS-194 | chr8-88790245-INS-193 |
| chr8-88813418-INS-245 | chr8-88813418-INS-245 |
| chr8-88835676-INS-58 | chr8-88835676-INS-57 |
| chr8-88848547-INS-197 | chr8-88848547-INS-199 |
| chr8-88851997-INS-205 | chr8-88851997-INS-205 |
| chr8-88859798-INS-209 | chr8-88859798-INS-210 |
| chr8-88889751-INS-88 | chr8-88889751-INS-92 |
| chr8-88892648-INS-220 | chr8-88892648-INS-221 |
| chr8-88896767-INS-51 | chr8-88896767-INS-64 |
| chr8-88898585-INS-67 | chr8-88898585-INS-50 |
| chr8-88920331-INS-182 | chr8-88920331-INS-183 |
| chr8-88925231-INS-174 | chr8-88925231-INS-175 |
| chr8-88929075-INS-95 | chr8-88929070-INS-85 |
| chr8-88936871-INS-202 | chr8-88936871-INS-205 |
| chr8-88940264-INS-112 | chr8-88940264-INS-147 |
| chr8-88942708-INS-213 | chr8-88942708-INS-217 |
| chr8-88948086-INS-195 | chr8-88948081-INS-196 |
| chr8-88959745-INS-188 | chr8-88959745-INS-202 |
| chr8-88977552-INS-226 | chr8-88977551-INS-219 |
| chr8-88997541-INS-223 | chr8-88997541-INS-224 |
| chr8-89029001-INS-6413 | chr8-89029002-INS-6413 |
| chr8-89032125-INS-72 | chr8-89032090-INS-107 |
| chr8-89038296-INS-872 | chr8-89038296-INS-873 |
| chr8-89039448-INS-389 | chr8-89039446-INS-397 |
| chr8-89047615-INS-102 | chr8-89047599-INS-92 |
| chr8-89069969-INS-57 | chr8-89069969-INS-57 |
| chr8-89099302-INS-58 | chr8-89099302-INS-61 |
| chr8-89108138-INS-82 | chr8-89108138-INS-82 |
| chr8-89113610-INS-297 | chr8-89113610-INS-297 |
| chr8-89127490-INS-160 | chr8-89127490-INS-202 |
| chr8-89147242-INS-55 | chr8-89147242-INS-55 |
| chr8-89159929-INS-60 | chr8-89159921-INS-68 |
| chr8-89186509-INS-211 | chr8-89186509-INS-211 |
| chr8-89195756-INS-80 | chr8-89195735-INS-53 |
| chr8-89205156-INS-57 | chr8-89205156-INS-60 |
| chr8-89244340-INS-52 | chr8-89244340-INS-52 |
| chr8-89255454-INS-1536 | chr8-89255455-INS-1537 |
| chr8-89280797-INS-236 | chr8-89280797-INS-236 |
| chr8-89291863-INS-55 | chr8-89291863-INS-55 |
| chr8-89340742-INS-80 | chr8-89340742-INS-80 |
| chr8-89378525-INS-54 | chr8-89378525-INS-55 |
| chr8-89384624-INS-69 | chr8-89384624-INS-69 |
| chr8-89425599-INS-62 | chr8-89425599-INS-62 |
| chr8-89442401-INS-161 | chr8-89442401-INS-161 |
| chr8-89483472-INS-58 | chr8-89483471-INS-62 |
| chr8-89529797-INS-71 | chr8-89529797-INS-71 |
| chr8-89540341-INS-52 | chr8-89540341-INS-56 |
| chr8-89550448-INS-51 | chr8-89550449-INS-52 |
| chr8-89563691-INS-59 | chr8-89563691-INS-63 |
| chr8-89566312-INS-95 | chr8-89566312-INS-87 |
| chr8-89594885-INS-67 | chr8-89594885-INS-57 |
| chr8-89598732-INS-61 | chr8-89598732-INS-61 |
| chr8-89613078-INS-110 | chr8-89613078-INS-110 |

| | |
|---|---|
| chr8-89636482-INS-258 | chr8-89636482-INS-258 |
| chr8-89643069-INS-56 | chr8-89643069-INS-56 |
| chr8-89647238-INS-549 | chr8-89647238-INS-549 |
| chr8-89653037-INS-73 | chr8-89653035-INS-95 |
| chr8-89662173-INS-56 | chr8-89662173-INS-50 |
| chr8-89673588-INS-86 | chr8-89673588-INS-86 |
| chr8-89677249-INS-434 | chr8-89677249-INS-434 |
| chr8-89683371-INS-182 | chr8-89683371-INS-181 |
| chr8-89688728-INS-310 | chr8-89688728-INS-318 |
| chr8-89693656-INS-204 | chr8-89693656-INS-204 |
| chr8-89695704-INS-339 | chr8-89695704-INS-339 |
| chr8-89721941-INS-448 | chr8-89721941-INS-448 |
| chr8-89727725-INS-165 | chr8-89727725-INS-165 |
| chr8-89765776-INS-52 | chr8-89765776-INS-52 |
| chr8-89796212-INS-66 | chr8-89796212-INS-66 |
| chr8-89831905-INS-613 | chr8-89831929-INS-604 |
| chr8-89831932-INS-75 | chr8-89831905-INS-57 |
| chr8-89841837-INS-203 | chr8-89841837-INS-203 |
| chr8-89864891-INS-711 | chr8-89864891-INS-711 |
| chr8-89891238-INS-400 | chr8-89891238-INS-400 |
| chr8-89897882-INS-68 | chr8-89897882-INS-72 |
| chr8-89902352-INS-126 | chr8-89902351-INS-87 |
| chr8-89905034-INS-413 | chr8-89905034-INS-413 |
| chr8-89914805-INS-144 | chr8-89914806-INS-146 |
| chr8-89917259-INS-141 | chr8-89917259-INS-141 |
| chr8-89949975-INS-445 | chr8-89949975-INS-445 |
| chr8-89952759-INS-102 | chr8-89952773-INS-56 |
| chr8-89953002-INS-1381 | chr8-89953003-INS-1378 |
| chr8-89953544-INS-81 | chr8-89953544-INS-80 |
| chr8-89955287-INS-397 | chr8-89955287-INS-436 |
| chr8-89973702-INS-147 | chr8-89973702-INS-148 |
| chr8-90015064-INS-104 | chr8-90015064-INS-80 |
| chr8-90049530-INS-84 | chr8-90049530-INS-84 |
| chr8-90122539-INS-160 | chr8-90122522-INS-142 |
| chr8-90130201-INS-60 | chr8-90130191-INS-54 |
| chr8-90161476-INS-64 | chr8-90161476-INS-54 |
| chr8-90200804-INS-60 | chr8-90200806-INS-90 |
| chr8-90216825-INS-509 | chr8-90216825-INS-509 |
| chr8-90236375-INS-54 | chr8-90236381-INS-55 |
| chr8-90237432-INS-200 | chr8-90237432-INS-200 |
| chr8-90313603-INS-229 | chr8-90313603-INS-229 |
| chr8-90366591-INS-142 | chr8-90366591-INS-152 |
| chr8-90388518-INS-74 | chr8-90388546-INS-90 |
| chr8-90418522-INS-228 | chr8-90418523-INS-224 |
| chr8-90429666-INS-54 | chr8-90429666-INS-54 |
| chr8-90434993-INS-61 | chr8-90434993-INS-61 |
| chr8-90454879-INS-56 | chr8-90454868-INS-57 |
| chr8-90456701-INS-6835 | chr8-90456702-INS-6836 |
| chr8-90467791-INS-51 | chr8-90467791-INS-51 |
| chr8-90482356-INS-75 | chr8-90482356-INS-75 |
| chr8-90499065-INS-86 | chr8-90499065-INS-86 |
| chr8-90515107-INS-266 | chr8-90515107-INS-266 |
| chr8-90519168-INS-6418 | chr8-90519169-INS-6418 |
| chr8-90531881-INS-3469 | chr8-90531882-INS-3469 |
| chr8-90532886-INS-1160 | chr8-90532887-INS-1160 |
| chr8-90553974-INS-54 | chr8-90553974-INS-60 |
| chr8-90573194-INS-113 | chr8-90573194-INS-113 |
| chr8-90586799-INS-554 | chr8-90586799-INS-554 |
| chr8-90618713-INS-116 | chr8-90618761-INS-72 |
| chr8-90691608-INS-54 | chr8-90691608-INS-86 |
| chr8-90694832-INS-54 | chr8-90694832-INS-78 |
| chr8-90696564-INS-934 | chr8-90696564-INS-934 |
| chr8-90697808-INS-6859 | chr8-90697809-INS-6859 |
| chr8-90707513-INS-81 | chr8-90707515-INS-63 |
| chr8-90711132-INS-100 | chr8-90711132-INS-100 |
| chr8-90745721-INS-99 | chr8-90745715-INS-90 |
| chr8-90750295-INS-104 | chr8-90750295-INS-104 |
| chr8-90770234-INS-86 | chr8-90770234-INS-86 |
| chr8-90777363-INS-56 | chr8-90777366-INS-59 |
| chr8-90782264-INS-59 | chr8-90782264-INS-59 |
| chr8-90788501-INS-54 | chr8-90788501-INS-60 |
| chr8-90805437-INS-87 | chr8-90805437-INS-87 |
| chr8-90873762-INS-132 | chr8-90873762-INS-132 |
| chr8-90880908-INS-399 | chr8-90880908-INS-399 |
| chr8-90904793-INS-104 | chr8-90904793-INS-104 |
| chr8-90911678-INS-152 | chr8-90911678-INS-152 |
| chr8-90928453-INS-705 | chr8-90928453-INS-705 |

| | |
|---|---|
| chr8-91011539-INS-229 | chr8-91011539-INS-235 |
| chr8-91037033-INS-98 | chr8-91037033-INS-94 |
| chr8-91101745-INS-168 | chr8-91101745-INS-168 |
| chr8-91106841-INS-499 | chr8-91106841-INS-499 |
| chr8-91122119-INS-54 | chr8-91122120-INS-54 |
| chr8-91159693-INS-139 | chr8-91159685-INS-149 |
| chr8-91165486-INS-56 | chr8-91165486-INS-60 |
| chr8-91174971-INS-64 | chr8-91174971-INS-64 |
| chr8-91182364-INS-213 | chr8-91182364-INS-213 |
| chr8-91186582-INS-784 | chr8-91186582-INS-784 |
| chr8-91217705-INS-51 | chr8-91217705-INS-60 |
| chr8-91236540-INS-56 | chr8-91236540-INS-56 |
| chr8-91258281-INS-129 | chr8-91258329-INS-117 |
| chr8-91302231-INS-1274 | chr8-91302232-INS-1305 |
| chr8-91305425-INS-166 | chr8-91305425-INS-166 |
| chr8-91309847-INS-52 | chr8-91309847-INS-52 |
| chr8-91325301-INS-237 | chr8-91325301-INS-259 |
| chr8-91331713-INS-84 | chr8-91331713-INS-84 |
| chr8-91333864-INS-739 | chr8-91333864-INS-740 |
| chr8-91335234-INS-195 | chr8-91335234-INS-195 |
| chr8-91360229-INS-64 | chr8-91360229-INS-70 |
| chr8-91362477-INS-68 | chr8-91362477-INS-56 |
| chr8-91367010-INS-52 | chr8-91367010-INS-52 |
| chr8-91379599-INS-61 | chr8-91379603-INS-61 |
| chr8-91385154-INS-60 | chr8-91385154-INS-60 |
| chr8-91406950-INS-166 | chr8-91406946-INS-165 |
| chr8-91409541-INS-69 | chr8-91409541-INS-69 |
| chr8-91410671-INS-436 | chr8-91410671-INS-436 |
| chr8-91414819-INS-69 | chr8-91414819-INS-69 |
| chr8-91424591-INS-171 | chr8-91424591-INS-173 |
| chr8-91435517-INS-128 | chr8-91435517-INS-128 |
| chr8-91440102-INS-109 | chr8-91440102-INS-137 |
| chr8-91452505-INS-72 | chr8-91452505-INS-72 |
| chr8-91459262-INS-190 | chr8-91459263-INS-190 |
| chr8-91460539-INS-210 | chr8-91460540-INS-188 |
| chr8-91475617-INS-3238 | chr8-91475618-INS-3238 |
| chr8-91494992-INS-84 | chr8-91494992-INS-72 |
| chr8-91496989-INS-2016 | chr8-91496990-INS-2016 |
| chr8-91507503-INS-102 | chr8-91507503-INS-54 |
| chr8-91530229-INS-415 | chr8-91530229-INS-416 |
| chr8-91539203-INS-6464 | chr8-91539204-INS-6478 |
| chr8-91539204-INS-498 | chr8-91539204-INS-498 |
| chr8-91541722-INS-209 | chr8-91541722-INS-210 |
| chr8-91550383-INS-212 | chr8-91550383-INS-212 |
| chr8-91557622-INS-207 | chr8-91557622-INS-206 |
| chr8-91560549-INS-209 | chr8-91560549-INS-210 |
| chr8-91573270-INS-62 | chr8-91573270-INS-63 |
| chr8-91583403-INS-236 | chr8-91583403-INS-237 |
| chr8-91590190-INS-350 | chr8-91590190-INS-350 |
| chr8-91592366-INS-100 | chr8-91592360-INS-78 |
| chr8-91599260-INS-196 | chr8-91599260-INS-198 |
| chr8-91608509-INS-212 | chr8-91608509-INS-212 |
| chr8-91612220-INS-122 | chr8-91612224-INS-90 |
| chr8-91646781-INS-3737 | chr8-91646782-INS-3737 |
| chr8-91651640-INS-201 | chr8-91651641-INS-201 |
| chr8-91652713-INS-58 | chr8-91652714-INS-58 |
| chr8-91667843-INS-56 | chr8-91667865-INS-73 |
| chr8-91677335-INS-191 | chr8-91677335-INS-191 |
| chr8-91690344-INS-89 | chr8-91690344-INS-87 |
| chr8-91695032-INS-104 | chr8-91695032-INS-104 |
| chr8-91707910-INS-7427 | chr8-91707911-INS-7428 |
| chr8-91715123-INS-56 | chr8-91715139-INS-58 |
| chr8-91741033-INS-53 | chr8-91741033-INS-56 |
| chr8-91765017-INS-52 | chr8-91765017-INS-50 |
| chr8-91773671-INS-228 | chr8-91773671-INS-224 |
| chr8-91785312-INS-201 | chr8-91785312-INS-212 |
| chr8-91807553-INS-194 | chr8-91807553-INS-194 |
| chr8-91809577-INS-198 | chr8-91809577-INS-198 |
| chr8-91843354-INS-62 | chr8-91843354-INS-62 |
| chr8-91848589-INS-346 | chr8-91848589-INS-332 |
| chr8-91872294-INS-272 | chr8-91872294-INS-268 |
| chr8-91896258-INS-63 | chr8-91896257-INS-61 |
| chr8-91903848-INS-52 | chr8-91903848-INS-52 |
| chr8-91907851-INS-54 | chr8-91907851-INS-54 |
| chr8-91935650-INS-53 | chr8-91935650-INS-53 |
| chr8-91938480-INS-78 | chr8-91938480-INS-78 |
| chr8-91963614-INS-58 | chr8-91963614-INS-56 |

| | |
|---|---|
| chr8-91970643-INS-55 | chr8-91970648-INS-96 |
| chr8-91982213-INS-506 | chr8-91982213-INS-507 |
| chr8-92063573-INS-86 | chr8-92063573-INS-86 |
| chr8-92071283-INS-55 | chr8-92071283-INS-55 |
| chr8-92085730-INS-185 | chr8-92085730-INS-186 |
| chr8-92096527-INS-88 | chr8-92096527-INS-52 |
| chr8-92111952-INS-77 | chr8-92111952-INS-77 |
| chr8-92113139-INS-52 | chr8-92113139-INS-57 |
| chr8-92127055-INS-77 | chr8-92127055-INS-77 |
| chr8-92141004-INS-99 | chr8-92141004-INS-99 |
| chr8-92142249-INS-59 | chr8-92142249-INS-59 |
| chr8-92153742-INS-97 | chr8-92153742-INS-99 |
| chr8-92166282-INS-54 | chr8-92166282-INS-54 |
| chr8-92167246-INS-6413 | chr8-92167247-INS-6413 |
| chr8-92171248-INS-199 | chr8-92171248-INS-199 |
| chr8-92194325-INS-200 | chr8-92194325-INS-200 |
| chr8-92195543-INS-56 | chr8-92195543-INS-76 |
| chr8-92206288-INS-104 | chr8-92206334-INS-104 |
| chr8-92211269-INS-74 | chr8-92211269-INS-74 |
| chr8-92217004-INS-210 | chr8-92217005-INS-217 |
| chr8-92233196-INS-67 | chr8-92233196-INS-73 |
| chr8-92238077-INS-213 | chr8-92238077-INS-212 |
| chr8-92253515-INS-53 | chr8-92253515-INS-53 |
| chr8-92291510-INS-7419 | chr8-92291511-INS-7419 |
| chr8-92294304-INS-217 | chr8-92294304-INS-217 |
| chr8-92295021-INS-221 | chr8-92295021-INS-221 |
| chr8-92297429-INS-6442 | chr8-92297430-INS-6442 |
| chr8-92301270-INS-223 | chr8-92301270-INS-225 |
| chr8-92328230-INS-216 | chr8-92328230-INS-216 |
| chr8-92390429-INS-57 | chr8-92390429-INS-57 |
| chr8-92400542-INS-287 | chr8-92400542-INS-288 |
| chr8-92401789-INS-199 | chr8-92401789-INS-201 |
| chr8-92451623-INS-140 | chr8-92451623-INS-140 |
| chr8-92496253-INS-86 | chr8-92496253-INS-88 |
| chr8-92499796-INS-84 | chr8-92499803-INS-54 |
| chr8-92500828-INS-54 | chr8-92500828-INS-60 |
| chr8-92502036-INS-98 | chr8-92502036-INS-68 |
| chr8-92510199-INS-53 | chr8-92510199-INS-51 |
| chr8-92513247-INS-367 | chr8-92513247-INS-339 |
| chr8-92520446-INS-61 | chr8-92520446-INS-66 |
| chr8-92538923-INS-54 | chr8-92538920-INS-56 |
| chr8-92542745-INS-81 | chr8-92542745-INS-81 |
| chr8-92559635-INS-57 | chr8-92559634-INS-62 |
| chr8-92570497-INS-68 | chr8-92570497-INS-68 |
| chr8-92594455-INS-74 | chr8-92594463-INS-62 |
| chr8-92597880-INS-84 | chr8-92597880-INS-84 |
| chr8-92621173-INS-60 | chr8-92621173-INS-60 |
| chr8-92671908-INS-100 | chr8-92671913-INS-55 |
| chr8-92673338-INS-54 | chr8-92673357-INS-53 |
| chr8-92692297-INS-185 | chr8-92692297-INS-187 |
| chr8-92736765-INS-60 | chr8-92736765-INS-60 |
| chr8-92742578-INS-60 | chr8-92742578-INS-60 |
| chr8-92751385-INS-168 | chr8-92751387-INS-166 |
| chr8-92788552-INS-116 | chr8-92788552-INS-118 |
| chr8-92800948-INS-125 | chr8-92800948-INS-126 |
| chr8-92828924-INS-54 | chr8-92828922-INS-58 |
| chr8-92843092-INS-76 | chr8-92843092-INS-76 |
| chr8-92846115-INS-56 | chr8-92846077-INS-52 |
| chr8-92852595-INS-130 | chr8-92852589-INS-98 |
| chr8-92866705-INS-174 | chr8-92866705-INS-174 |
| chr8-92867202-INS-234 | chr8-92867202-INS-234 |
| chr8-92876926-INS-183 | chr8-92876963-INS-196 |
| chr8-92877963-INS-54 | chr8-92877967-INS-72 |
| chr8-92892563-INS-110 | chr8-92892563-INS-106 |
| chr8-92895233-INS-380 | chr8-92895230-INS-394 |
| chr8-92934366-INS-66 | chr8-92934377-INS-62 |
| chr8-92968461-INS-710 | chr8-92968463-INS-708 |
| chr8-92974763-INS-119 | chr8-92974763-INS-119 |
| chr8-92979233-INS-60 | chr8-92979233-INS-60 |
| chr8-92980854-INS-141 | chr8-92980854-INS-141 |
| chr8-92990683-INS-68 | chr8-92990683-INS-68 |
| chr8-93065389-INS-336 | chr8-93065389-INS-336 |
| chr8-93122160-INS-72 | chr8-93122162-INS-70 |
| chr8-93130553-INS-52 | chr8-93130529-INS-50 |
| chr8-93155050-INS-56 | chr8-93155051-INS-64 |
| chr8-93157013-INS-113 | chr8-93157013-INS-113 |
| chr8-93162411-INS-187 | chr8-93162411-INS-186 |

| | |
|---|---|
| chr8-93189631-INS-2687 | chr8-93189632-INS-2683 |
| chr8-93234124-INS-54 | chr8-93234128-INS-50 |
| chr8-93241999-INS-344 | chr8-93241999-INS-344 |
| chr8-93260149-INS-82 | chr8-93260149-INS-70 |
| chr8-93267478-INS-319 | chr8-93267478-INS-319 |
| chr8-93310705-INS-558 | chr8-93310705-INS-561 |
| chr8-93342020-INS-456 | chr8-93342020-INS-457 |
| chr8-93383418-INS-76 | chr8-93383414-INS-110 |
| chr8-93420824-INS-493 | chr8-93420824-INS-492 |
| chr8-93431110-INS-102 | chr8-93431152-INS-74 |
| chr8-93438311-INS-61 | chr8-93438306-INS-57 |
| chr8-93518217-INS-400 | chr8-93518217-INS-400 |
| chr8-93536201-INS-104 | chr8-93536201-INS-100 |
| chr8-93545265-INS-135 | chr8-93545265-INS-154 |
| chr8-93571338-INS-66 | chr8-93571342-INS-106 |
| chr8-93586835-INS-69 | chr8-93586835-INS-69 |
| chr8-93591731-INS-156 | chr8-93591731-INS-156 |
| chr8-93638540-INS-56 | chr8-93638540-INS-54 |
| chr8-93643057-INS-299 | chr8-93643057-INS-299 |
| chr8-93649578-INS-51 | chr8-93649578-INS-51 |
| chr8-93700137-INS-273 | chr8-93700137-INS-273 |
| chr8-93708425-INS-55 | chr8-93708425-INS-55 |
| chr8-93713072-INS-499 | chr8-93713072-INS-499 |
| chr8-93730833-INS-149 | chr8-93730845-INS-162 |
| chr8-93743102-INS-560 | chr8-93743102-INS-560 |
| chr8-93760118-INS-60 | chr8-93760118-INS-60 |
| chr8-93787236-INS-486 | chr8-93787236-INS-486 |
| chr8-93788331-INS-56 | chr8-93788331-INS-56 |
| chr8-93794907-INS-77 | chr8-93794895-INS-80 |
| chr8-93808702-INS-56 | chr8-93808702-INS-66 |
| chr8-93814311-INS-54 | chr8-93814311-INS-54 |
| chr8-93818933-INS-67 | chr8-93818933-INS-63 |
| chr8-93828501-INS-58 | chr8-93828501-INS-58 |
| chr8-93866174-INS-3545 | chr8-93866175-INS-3547 |
| chr8-93878737-INS-182 | chr8-93878737-INS-182 |
| chr8-93881576-INS-106 | chr8-93881576-INS-70 |
| chr8-93898036-INS-218 | chr8-93898036-INS-218 |
| chr8-93956987-INS-56 | chr8-93956987-INS-51 |
| chr8-93978893-INS-204 | chr8-93978893-INS-203 |
| chr8-93985585-INS-80 | chr8-93985583-INS-86 |
| chr8-94029858-INS-107 | chr8-94029845-INS-121 |
| chr8-94063431-INS-83 | chr8-94063431-INS-71 |
| chr8-94109500-INS-2122 | chr8-94109501-INS-2124 |
| chr8-94125785-INS-65 | chr8-94125750-INS-66 |
| chr8-94133265-INS-217 | chr8-94133265-INS-218 |
| chr8-94166546-INS-519 | chr8-94166546-INS-519 |
| chr8-94169535-INS-238 | chr8-94169535-INS-195 |
| chr8-94195614-INS-6933 | chr8-94195615-INS-6933 |
| chr8-94196820-INS-219 | chr8-94196820-INS-223 |
| chr8-94199643-INS-81 | chr8-94199647-INS-82 |
| chr8-94212096-INS-214 | chr8-94212096-INS-214 |
| chr8-94229671-INS-120 | chr8-94229671-INS-122 |
| chr8-94280463-INS-203 | chr8-94280463-INS-203 |
| chr8-94288659-INS-52 | chr8-94288659-INS-68 |
| chr8-94289969-INS-6539 | chr8-94289970-INS-6538 |
| chr8-94302742-INS-78 | chr8-94302785-INS-76 |
| chr8-94318499-INS-113 | chr8-94318499-INS-120 |
| chr8-94331616-INS-87 | chr8-94331616-INS-88 |
| chr8-94334963-INS-729 | chr8-94334963-INS-730 |
| chr8-94379691-INS-63 | chr8-94379691-INS-63 |
| chr8-94380077-INS-496 | chr8-94380077-INS-505 |
| chr8-94396157-INS-78 | chr8-94396157-INS-72 |
| chr8-94406285-INS-80 | chr8-94406285-INS-104 |
| chr8-94423644-INS-318 | chr8-94423644-INS-323 |
| chr8-94450167-INS-57 | chr8-94450197-INS-51 |
| chr8-94453386-INS-7110 | chr8-94453387-INS-7110 |
| chr8-94460661-INS-97 | chr8-94460661-INS-89 |
| chr8-94468328-INS-4712 | chr8-94468329-INS-4712 |
| chr8-94488820-INS-4817 | chr8-94488821-INS-4819 |
| chr8-94519689-INS-6421 | chr8-94519690-INS-6434 |
| chr8-94530274-INS-55 | chr8-94530281-INS-80 |
| chr8-94532417-INS-601 | chr8-94532417-INS-601 |
| chr8-94555216-INS-58 | chr8-94555216-INS-58 |
| chr8-94561513-INS-1777 | chr8-94561514-INS-1776 |
| chr8-94596436-INS-75 | chr8-94596436-INS-77 |
| chr8-94639866-INS-75 | chr8-94639881-INS-61 |
| chr8-94640890-INS-225 | chr8-94640902-INS-213 |

| | |
|---|---|
| chr8-94657363-INS-844 | chr8-94657363-INS-870 |
| chr8-94673933-INS-82 | chr8-94673933-INS-82 |
| chr8-94676190-INS-75 | chr8-94676190-INS-60 |
| chr8-94751665-INS-844 | chr8-94751665-INS-844 |
| chr8-94770225-INS-1086 | chr8-94770226-INS-1083 |
| chr8-94770401-INS-173 | chr8-94770401-INS-173 |
| chr8-94799179-INS-214 | chr8-94799179-INS-214 |
| chr8-94801698-INS-387 | chr8-94801696-INS-386 |
| chr8-94804223-INS-315 | chr8-94804223-INS-316 |
| chr8-94822357-INS-199 | chr8-94822357-INS-199 |
| chr8-94823307-INS-3305 | chr8-94823308-INS-3301 |
| chr8-94827713-INS-212 | chr8-94827713-INS-212 |
| chr8-94833872-INS-287 | chr8-94833872-INS-289 |
| chr8-94853429-INS-244 | chr8-94853429-INS-245 |
| chr8-94856822-INS-197 | chr8-94856822-INS-197 |
| chr8-94859885-INS-2028 | chr8-94859886-INS-2028 |
| chr8-94878994-INS-54 | chr8-94878994-INS-57 |
| chr8-94895196-INS-66 | chr8-94895196-INS-66 |
| chr8-94901984-INS-283 | chr8-94901984-INS-283 |
| chr8-94910772-INS-194 | chr8-94910772-INS-214 |
| chr8-94921532-INS-178 | chr8-94921533-INS-192 |
| chr8-94933845-INS-432 | chr8-94933845-INS-420 |
| chr8-94943786-INS-539 | chr8-94943790-INS-529 |
| chr8-94947955-INS-106 | chr8-94947955-INS-98 |
| chr8-94952206-INS-1315 | chr8-94952207-INS-1321 |
| chr8-94962948-INS-188 | chr8-94962948-INS-187 |
| chr8-94971717-INS-216 | chr8-94971717-INS-217 |
| chr8-94979712-INS-202 | chr8-94979712-INS-203 |
| chr8-95004551-INS-151 | chr8-95004551-INS-151 |
| chr8-95058933-INS-184 | chr8-95058933-INS-186 |
| chr8-95095184-INS-5375 | chr8-95095185-INS-5375 |
| chr8-95100741-INS-205 | chr8-95100741-INS-205 |
| chr8-95104055-INS-65 | chr8-95104055-INS-65 |
| chr8-95120149-INS-71 | chr8-95120147-INS-78 |
| chr8-95122363-INS-5193 | chr8-95122364-INS-5193 |
| chr8-95124871-INS-327 | chr8-95124871-INS-327 |
| chr8-95126404-INS-703 | chr8-95126404-INS-703 |
| chr8-95126913-INS-50 | chr8-95126894-INS-57 |
| chr8-95138890-INS-52 | chr8-95138890-INS-52 |
| chr8-95143239-INS-141 | chr8-95143239-INS-141 |
| chr8-95150222-INS-236 | chr8-95150222-INS-236 |
| chr8-95153898-INS-4350 | chr8-95153899-INS-4350 |
| chr8-95158628-INS-214 | chr8-95158628-INS-215 |
| chr8-95179286-INS-6401 | chr8-95179287-INS-6401 |
| chr8-95184687-INS-1227 | chr8-95184688-INS-1244 |
| chr8-95190121-INS-7729 | chr8-95190122-INS-7729 |
| chr8-95369664-INS-1568 | chr8-95369665-INS-1568 |
| chr8-95374809-INS-82 | chr8-95374809-INS-82 |
| chr8-95396081-INS-232 | chr8-95396081-INS-231 |
| chr8-95430165-INS-80 | chr8-95430165-INS-54 |
| chr8-95436929-INS-67 | chr8-95436929-INS-69 |
| chr8-95440395-INS-167 | chr8-95440395-INS-168 |
| chr8-95442125-INS-212 | chr8-95442125-INS-212 |
| chr8-95443528-INS-54 | chr8-95443533-INS-55 |
| chr8-95464796-INS-76 | chr8-95464796-INS-76 |
| chr8-95470472-INS-77 | chr8-95470472-INS-79 |
| chr8-95492327-INS-54 | chr8-95492327-INS-54 |
| chr8-95533678-INS-60 | chr8-95533679-INS-50 |
| chr8-95534645-INS-118 | chr8-95534645-INS-118 |
| chr8-95542663-INS-67 | chr8-95542663-INS-67 |
| chr8-95561839-INS-198 | chr8-95561839-INS-198 |
| chr8-95568098-INS-56 | chr8-95568098-INS-56 |
| chr8-95577605-INS-136 | chr8-95577605-INS-136 |
| chr8-95581880-INS-57 | chr8-95581882-INS-58 |
| chr8-95583443-INS-208 | chr8-95583443-INS-208 |
| chr8-95587947-INS-318 | chr8-95587947-INS-318 |
| chr8-95603244-INS-100 | chr8-95603244-INS-100 |
| chr8-95610213-INS-241 | chr8-95610213-INS-241 |
| chr8-95651560-INS-476 | chr8-95651560-INS-476 |
| chr8-95672469-INS-68 | chr8-95672469-INS-51 |
| chr8-95743710-INS-62 | chr8-95743710-INS-62 |
| chr8-95750699-INS-60 | chr8-95750699-INS-60 |
| chr8-95763873-INS-60 | chr8-95763873-INS-60 |
| chr8-95770317-INS-105 | chr8-95770326-INS-66 |
| chr8-95883568-INS-58 | chr8-95883568-INS-58 |
| chr8-95917613-INS-379 | chr8-95917622-INS-392 |
| chr8-95925174-INS-89 | chr8-95925174-INS-80 |

| | |
|---|---|
| chr8-95943255-INS-1058 | chr8-95943256-INS-1058 |
| chr8-95947835-INS-179 | chr8-95947835-INS-169 |
| chr8-95956121-INS-5264 | chr8-95956122-INS-5264 |
| chr8-95972353-INS-6442 | chr8-95972354-INS-6442 |
| chr8-95973274-INS-217 | chr8-95973274-INS-217 |
| chr8-95989933-INS-104 | chr8-95989933-INS-104 |
| chr8-96010049-INS-55 | chr8-96010049-INS-51 |
| chr8-96027735-INS-1020 | chr8-96027736-INS-1020 |
| chr8-96028832-INS-50 | chr8-96028832-INS-50 |
| chr8-96111519-INS-52 | chr8-96111519-INS-50 |
| chr8-96158347-INS-160 | chr8-96158347-INS-208 |
| chr8-96168776-INS-5252 | chr8-96168777-INS-5252 |
| chr8-96180535-INS-200 | chr8-96180535-INS-201 |
| chr8-96276373-INS-63 | chr8-96276393-INS-51 |
| chr8-96289670-INS-112 | chr8-96289671-INS-63 |
| chr8-96320678-INS-64 | chr8-96320678-INS-64 |
| chr8-96326799-INS-80 | chr8-96326799-INS-68 |
| chr8-96342091-INS-114 | chr8-96342091-INS-114 |
| chr8-96346908-INS-1893 | chr8-96346909-INS-1893 |
| chr8-96347138-INS-53 | chr8-96347138-INS-53 |
| chr8-96357728-INS-165 | chr8-96357728-INS-165 |
| chr8-96362247-INS-171 | chr8-96362247-INS-171 |
| chr8-96394317-INS-2880 | chr8-96394318-INS-2880 |
| chr8-96444333-INS-220 | chr8-96444333-INS-220 |
| chr8-96529430-INS-218 | chr8-96529430-INS-217 |
| chr8-96543914-INS-253 | chr8-96543914-INS-212 |
| chr8-96557196-INS-171 | chr8-96557196-INS-156 |
| chr8-96576715-INS-206 | chr8-96576719-INS-226 |
| chr8-96577562-INS-182 | chr8-96577562-INS-182 |
| chr8-96585060-INS-662 | chr8-96585054-INS-660 |
| chr8-96606860-INS-213 | chr8-96606860-INS-213 |
| chr8-96616624-INS-73 | chr8-96616624-INS-82 |
| chr8-96617869-INS-227 | chr8-96617869-INS-227 |
| chr8-96654268-INS-142 | chr8-96654267-INS-169 |
| chr8-96667755-INS-456 | chr8-96667755-INS-456 |
| chr8-96672854-INS-74 | chr8-96672824-INS-58 |
| chr8-96677782-INS-855 | chr8-96677782-INS-855 |
| chr8-96683998-INS-167 | chr8-96683998-INS-132 |
| chr8-96697237-INS-90 | chr8-96697226-INS-87 |
| chr8-96712699-INS-128 | chr8-96712732-INS-128 |
| chr8-96761851-INS-52 | chr8-96761851-INS-52 |
| chr8-96773884-INS-55 | chr8-96773884-INS-95 |
| chr8-96805665-INS-74 | chr8-96805667-INS-52 |
| chr8-96825050-INS-941 | chr8-96825043-INS-948 |
| chr8-96836849-INS-57 | chr8-96836853-INS-54 |
| chr8-96842343-INS-79 | chr8-96842343-INS-79 |
| chr8-96848002-INS-60 | chr8-96848002-INS-60 |
| chr8-96897606-INS-6688 | chr8-96897607-INS-6688 |
| chr8-96927201-INS-58 | chr8-96927201-INS-58 |
| chr8-96963599-INS-6485 | chr8-96963600-INS-6485 |
| chr8-96972986-INS-65 | chr8-96972986-INS-65 |
| chr8-96979571-INS-576 | chr8-96979571-INS-576 |
| chr8-96999319-INS-639 | chr8-96999319-INS-642 |
| chr8-97014155-INS-7152 | chr8-97014156-INS-7152 |
| chr8-97026976-INS-190 | chr8-97026976-INS-189 |
| chr8-97081809-INS-400 | chr8-97081809-INS-401 |
| chr8-97083773-INS-107 | chr8-97083775-INS-58 |
| chr8-97094112-INS-52 | chr8-97094112-INS-54 |
| chr8-97096876-INS-59 | chr8-97096876-INS-51 |
| chr8-97098379-INS-643 | chr8-97098379-INS-643 |
| chr8-97165088-INS-60 | chr8-97165088-INS-60 |
| chr8-97179463-INS-785 | chr8-97179463-INS-783 |
| chr8-97186919-INS-394 | chr8-97186919-INS-394 |
| chr8-97189036-INS-2058 | chr8-97189037-INS-2061 |
| chr8-97194231-INS-64 | chr8-97194231-INS-76 |
| chr8-97210212-INS-458 | chr8-97210212-INS-458 |
| chr8-97222686-INS-1566 | chr8-97222687-INS-1565 |
| chr8-97266793-INS-3045 | chr8-97266794-INS-3045 |
| chr8-97286861-INS-796 | chr8-97286861-INS-796 |
| chr8-97296331-INS-508 | chr8-97296331-INS-508 |
| chr8-97306365-INS-7009 | chr8-97306366-INS-7009 |
| chr8-97330715-INS-56 | chr8-97330748-INS-73 |
| chr8-97345071-INS-103 | chr8-97345072-INS-80 |
| chr8-97351406-INS-79 | chr8-97351406-INS-62 |
| chr8-97351512-INS-1056 | chr8-97351514-INS-1058 |
| chr8-97355659-INS-498 | chr8-97355659-INS-498 |
| chr8-97364460-INS-433 | chr8-97364460-INS-433 |

| | |
|---|---|
| chr8-97375214-INS-52 | chr8-97375215-INS-50 |
| chr8-97396528-INS-14480 | chr8-97396528-INS-14480 |
| chr8-97401217-INS-62 | chr8-97401217-INS-63 |
| chr8-97416652-INS-51 | chr8-97416653-INS-57 |
| chr8-97467353-INS-93 | chr8-97467401-INS-79 |
| chr8-97469706-INS-190 | chr8-97469695-INS-191 |
| chr8-97478908-INS-9320 | chr8-97478909-INS-9321 |
| chr8-97484668-INS-78 | chr8-97484669-INS-78 |
| chr8-97502356-INS-245 | chr8-97502356-INS-245 |
| chr8-97511324-INS-1420 | chr8-97511325-INS-1421 |
| chr8-97524797-INS-125 | chr8-97524797-INS-142 |
| chr8-97531631-INS-131 | chr8-97531632-INS-131 |
| chr8-97538929-INS-70 | chr8-97538929-INS-86 |
| chr8-97543297-INS-4561 | chr8-97543298-INS-4561 |
| chr8-97556607-INS-327 | chr8-97556607-INS-327 |
| chr8-97562395-INS-462 | chr8-97562395-INS-463 |
| chr8-97566978-INS-1973 | chr8-97566979-INS-1972 |
| chr8-97588666-INS-199 | chr8-97588666-INS-199 |
| chr8-97595192-INS-56 | chr8-97595192-INS-58 |
| chr8-97596248-INS-6651 | chr8-97596249-INS-6652 |
| chr8-97598743-INS-55 | chr8-97598743-INS-55 |
| chr8-97602108-INS-6086 | chr8-97602109-INS-6088 |
| chr8-97614179-INS-54 | chr8-97614179-INS-58 |
| chr8-97655655-INS-236 | chr8-97655653-INS-244 |
| chr8-97692214-INS-63 | chr8-97692263-INS-55 |
| chr8-97701993-INS-533 | chr8-97701993-INS-534 |
| chr8-97755026-INS-52 | chr8-97755026-INS-52 |
| chr8-97781640-INS-128 | chr8-97781640-INS-128 |
| chr8-97818799-INS-67 | chr8-97818800-INS-91 |
| chr8-97820269-INS-1478 | chr8-97820270-INS-1476 |
| chr8-97832273-INS-51 | chr8-97832273-INS-53 |
| chr8-97854969-INS-235 | chr8-97854969-INS-235 |
| chr8-97865667-INS-392 | chr8-97865667-INS-392 |
| chr8-97870429-INS-113 | chr8-97870429-INS-113 |
| chr8-97887198-INS-52 | chr8-97887198-INS-50 |
| chr8-97923939-INS-376 | chr8-97923939-INS-376 |
| chr8-97937023-INS-54 | chr8-97937023-INS-54 |
| chr8-97954908-INS-60 | chr8-97954908-INS-66 |
| chr8-97999182-INS-75 | chr8-97999182-INS-67 |
| chr8-98002287-INS-343 | chr8-98002287-INS-344 |
| chr8-98009054-INS-1730 | chr8-98009055-INS-1730 |
| chr8-98016709-INS-53 | chr8-98016709-INS-53 |
| chr8-98036601-INS-2089 | chr8-98036602-INS-2088 |
| chr8-98048914-INS-208 | chr8-98048914-INS-213 |
| chr8-98086869-INS-86 | chr8-98086869-INS-86 |
| chr8-98118499-INS-51 | chr8-98118499-INS-81 |
| chr8-98122092-INS-371 | chr8-98122092-INS-371 |
| chr8-98146490-INS-7406 | chr8-98146491-INS-7406 |
| chr8-98150443-INS-315 | chr8-98150443-INS-315 |
| chr8-98152311-INS-1775 | chr8-98152312-INS-1775 |
| chr8-98170827-INS-56 | chr8-98170787-INS-86 |
| chr8-98173390-INS-2013 | chr8-98173391-INS-2012 |
| chr8-98177783-INS-62 | chr8-98177779-INS-68 |
| chr8-98183513-INS-1225 | chr8-98183514-INS-1225 |
| chr8-98227373-INS-1613 | chr8-98227374-INS-1613 |
| chr8-98241470-INS-7168 | chr8-98241471-INS-7168 |
| chr8-98330034-INS-189 | chr8-98330034-INS-189 |
| chr8-98333276-INS-3935 | chr8-98333277-INS-3935 |
| chr8-98333280-INS-458 | chr8-98333280-INS-458 |
| chr8-98336794-INS-776 | chr8-98336794-INS-785 |
| chr8-98374936-INS-212 | chr8-98374936-INS-216 |
| chr8-98380200-INS-674 | chr8-98380200-INS-673 |
| chr8-98391039-INS-88 | chr8-98391039-INS-90 |
| chr8-98392528-INS-119 | chr8-98392536-INS-91 |
| chr8-98395727-INS-80 | chr8-98395727-INS-110 |
| chr8-98407455-INS-64 | chr8-98407433-INS-64 |
| chr8-98417898-INS-60 | chr8-98417923-INS-52 |
| chr8-98433272-INS-101 | chr8-98433272-INS-101 |
| chr8-98436685-INS-281 | chr8-98436686-INS-279 |
| chr8-98487896-INS-913 | chr8-98487896-INS-913 |
| chr8-98510151-INS-63 | chr8-98510151-INS-69 |
| chr8-98533611-INS-283 | chr8-98533611-INS-284 |
| chr8-98547799-INS-56 | chr8-98547800-INS-68 |
| chr8-98565122-INS-6392 | chr8-98565123-INS-6392 |
| chr8-98655961-INS-94 | chr8-98655967-INS-112 |
| chr8-98660226-INS-139 | chr8-98660207-INS-187 |
| chr8-98661309-INS-76 | chr8-98661309-INS-78 |

| | |
|---|---|
| chr8-98666201-INS-78 | chr8-98666201-INS-78 |
| chr8-98680189-INS-119 | chr8-98680189-INS-83 |
| chr8-98706854-INS-96 | chr8-98706853-INS-58 |
| chr8-98737042-INS-443 | chr8-98737042-INS-444 |
| chr8-98739258-INS-108 | chr8-98739256-INS-110 |
| chr8-98799423-INS-226 | chr8-98799423-INS-231 |
| chr8-98809951-INS-60 | chr8-98809951-INS-60 |
| chr8-98828557-INS-96 | chr8-98828580-INS-85 |
| chr8-98831524-INS-72 | chr8-98831524-INS-52 |
| chr8-98861918-INS-86 | chr8-98861918-INS-98 |
| chr8-98862944-INS-6279 | chr8-98862945-INS-6279 |
| chr8-98865995-INS-100 | chr8-98865955-INS-108 |
| chr8-98895491-INS-180 | chr8-98895491-INS-180 |
| chr8-98900411-INS-68 | chr8-98900407-INS-56 |
| chr8-98918654-INS-69 | chr8-98918654-INS-63 |
| chr8-98949752-INS-339 | chr8-98949752-INS-339 |
| chr8-98953681-INS-620 | chr8-98953681-INS-620 |
| chr8-98959090-INS-52 | chr8-98959090-INS-72 |
| chr8-98968151-INS-60 | chr8-98968182-INS-68 |
| chr8-98985089-INS-74 | chr8-98985089-INS-76 |
| chr8-99079400-INS-52 | chr8-99079400-INS-52 |
| chr8-99086195-INS-90 | chr8-99086195-INS-90 |
| chr8-99086712-INS-199 | chr8-99086713-INS-199 |
| chr8-99088615-INS-62 | chr8-99088615-INS-62 |
| chr8-99089900-INS-358 | chr8-99089900-INS-358 |
| chr8-99127363-INS-58 | chr8-99127363-INS-50 |
| chr8-99129163-INS-162 | chr8-99129163-INS-162 |
| chr8-99145446-INS-2578 | chr8-99145447-INS-2578 |
| chr8-99162924-INS-54 | chr8-99162877-INS-53 |
| chr8-99164369-INS-73 | chr8-99164369-INS-75 |
| chr8-99169865-INS-2230 | chr8-99169871-INS-2216 |
| chr8-99175720-INS-84 | chr8-99175720-INS-84 |
| chr8-99190534-INS-66 | chr8-99190574-INS-70 |
| chr8-99195171-INS-59 | chr8-99195171-INS-65 |
| chr8-99207482-INS-5252 | chr8-99207483-INS-5253 |
| chr8-99208625-INS-88 | chr8-99208625-INS-80 |
| chr8-99225287-INS-65 | chr8-99225287-INS-65 |
| chr8-99239968-INS-421 | chr8-99239968-INS-420 |
| chr8-99264046-INS-132 | chr8-99264012-INS-124 |
| chr8-99286371-INS-2822 | chr8-99286372-INS-2822 |
| chr8-99291470-INS-213 | chr8-99291470-INS-216 |
| chr8-99322771-INS-108 | chr8-99322771-INS-100 |
| chr8-99332655-INS-51 | chr8-99332653-INS-54 |
| chr8-99334617-INS-66 | chr8-99334583-INS-62 |
| chr8-99338106-INS-97 | chr8-99338106-INS-102 |
| chr8-99341031-INS-72 | chr8-99341041-INS-105 |
| chr8-99408011-INS-118 | chr8-99408013-INS-119 |
| chr8-99425941-INS-212 | chr8-99425941-INS-212 |
| chr8-99468399-INS-99 | chr8-99468399-INS-99 |
| chr8-99475059-INS-78 | chr8-99475059-INS-78 |
| chr8-99494142-INS-76 | chr8-99494144-INS-64 |
| chr8-99497624-INS-318 | chr8-99497624-INS-318 |
| chr8-99527384-INS-129 | chr8-99527384-INS-129 |
| chr8-99530975-INS-64 | chr8-99530975-INS-58 |
| chr8-99550779-INS-62 | chr8-99550779-INS-65 |
| chr8-99590893-INS-5345 | chr8-99590894-INS-5345 |
| chr8-99608084-INS-59 | chr8-99608085-INS-50 |
| chr8-99611871-INS-962 | chr8-99611871-INS-964 |
| chr8-99651224-INS-1011 | chr8-99651225-INS-1011 |
| chr8-99658319-INS-719 | chr8-99658319-INS-719 |
| chr8-99668198-INS-72 | chr8-99668198-INS-72 |
| chr8-99671605-INS-54 | chr8-99671605-INS-54 |
| chr8-99676320-INS-361 | chr8-99676320-INS-361 |
| chr8-99693577-INS-53 | chr8-99693577-INS-53 |
| chr8-99694561-INS-54 | chr8-99694561-INS-54 |
| chr8-99701071-INS-93 | chr8-99701071-INS-93 |
| chr8-99705736-INS-82 | chr8-99705736-INS-72 |
| chr8-99729821-INS-575 | chr8-99729821-INS-585 |
| chr8-99732062-INS-509 | chr8-99732062-INS-509 |
| chr8-99787942-INS-135 | chr8-99787942-INS-135 |
| chr8-99794085-INS-76 | chr8-99794085-INS-84 |
| chr8-99808861-INS-173 | chr8-99808861-INS-177 |
| chr8-99815619-INS-177 | chr8-99815613-INS-192 |
| chr8-99831829-INS-68 | chr8-99831829-INS-72 |
| chr8-99854572-INS-356 | chr8-99854572-INS-357 |
| chr8-99883683-INS-57 | chr8-99883699-INS-58 |
| chr8-99894668-INS-52 | chr8-99894666-INS-72 |

| | |
|---|---|
| chr8-99913351-INS-5038 | chr8-99913352-INS-5041 |
| chr8-99935704-INS-52 | chr8-99935704-INS-52 |
| chr8-99941097-INS-63 | chr8-99941097-INS-63 |
| chr8-99954931-INS-56 | chr8-99954931-INS-58 |
| chr8-99960340-INS-556 | chr8-99960340-INS-556 |
| chr8-99962586-INS-66 | chr8-99962631-INS-66 |
| chr8-99978738-INS-620 | chr8-99978738-INS-620 |
| chr8-100007034-INS-330 | chr8-100007034-INS-330 |
| chr8-100015572-INS-56 | chr8-100015593-INS-53 |
| chr8-100020454-INS-108 | chr8-100020454-INS-113 |
| chr8-100079819-INS-164 | chr8-100079837-INS-156 |
| chr8-100138802-INS-400 | chr8-100138802-INS-402 |
| chr8-100191657-INS-633 | chr8-100191656-INS-637 |
| chr8-100263010-INS-93 | chr8-100263011-INS-111 |
| chr8-100293400-INS-103 | chr8-100293404-INS-107 |
| chr8-100362440-INS-101 | chr8-100362440-INS-101 |
| chr8-100367029-INS-390 | chr8-100367029-INS-390 |
| chr8-100383206-INS-76 | chr8-100383206-INS-80 |
| chr8-100419181-INS-3382 | chr8-100419182-INS-3388 |
| chr8-100423143-INS-77 | chr8-100423143-INS-77 |
| chr8-100441304-INS-6189 | chr8-100441305-INS-6174 |
| chr8-100442697-INS-1025 | chr8-100442699-INS-1069 |
| chr8-100468796-INS-286 | chr8-100468796-INS-287 |
| chr8-100476417-INS-623 | chr8-100476380-INS-639 |
| chr8-100486827-INS-52 | chr8-100486827-INS-52 |
| chr8-100533799-INS-2403 | chr8-100533800-INS-2403 |
| chr8-100551942-INS-90 | chr8-100551922-INS-115 |
| chr8-100575710-INS-6596 | chr8-100575711-INS-6596 |
| chr8-100642731-INS-70 | chr8-100642731-INS-70 |
| chr8-100653064-INS-921 | chr8-100653064-INS-908 |
| chr8-100663876-INS-615 | chr8-100663876-INS-615 |
| chr8-100690141-INS-64 | chr8-100690141-INS-66 |
| chr8-100760362-INS-100 | chr8-100760358-INS-64 |
| chr8-100765203-INS-68 | chr8-100765203-INS-76 |
| chr8-100771309-INS-663 | chr8-100771309-INS-663 |
| chr8-100814118-INS-506 | chr8-100814118-INS-506 |
| chr8-100832135-INS-436 | chr8-100832135-INS-436 |
| chr8-100884573-INS-988 | chr8-100884573-INS-990 |
| chr8-100903139-INS-63 | chr8-100903141-INS-74 |
| chr8-100918563-INS-51 | chr8-100918551-INS-59 |
| chr8-100925197-INS-174 | chr8-100925197-INS-175 |
| chr8-100949870-INS-116 | chr8-100949870-INS-109 |
| chr8-100999824-INS-214 | chr8-100999824-INS-215 |
| chr8-101005804-INS-5244 | chr8-101005805-INS-5244 |
| chr8-101050674-INS-71 | chr8-101050674-INS-71 |
| chr8-101057892-INS-72 | chr8-101057892-INS-62 |
| chr8-101068852-INS-56 | chr8-101068852-INS-56 |
| chr8-101071108-INS-1812 | chr8-101071109-INS-1811 |
| chr8-101077853-INS-436 | chr8-101077853-INS-439 |
| chr8-101087395-INS-188 | chr8-101087395-INS-188 |
| chr8-101108694-INS-496 | chr8-101108690-INS-465 |
| chr8-101141965-INS-75 | chr8-101141965-INS-75 |
| chr8-101146089-INS-1544 | chr8-101146090-INS-1544 |
| chr8-101148952-INS-64 | chr8-101148952-INS-65 |
| chr8-101150129-INS-6914 | chr8-101150130-INS-6925 |
| chr8-101152170-INS-206 | chr8-101152170-INS-160 |
| chr8-101169425-INS-6547 | chr8-101169426-INS-6547 |
| chr8-101169900-INS-80 | chr8-101169900-INS-84 |
| chr8-101170176-INS-679 | chr8-101170176-INS-679 |
| chr8-101181210-INS-647 | chr8-101181210-INS-647 |
| chr8-101205536-INS-156 | chr8-101205536-INS-156 |
| chr8-101257247-INS-653 | chr8-101257247-INS-681 |
| chr8-101259486-INS-6368 | chr8-101259487-INS-6370 |
| chr8-101268768-INS-61 | chr8-101268768-INS-97 |
| chr8-101271078-INS-3918 | chr8-101271079-INS-3918 |
| chr8-101276021-INS-83 | chr8-101276017-INS-129 |
| chr8-101300719-INS-96 | chr8-101300720-INS-80 |
| chr8-101303618-INS-844 | chr8-101303618-INS-812 |
| chr8-101369209-INS-62 | chr8-101369209-INS-50 |
| chr8-101375873-INS-193 | chr8-101375873-INS-193 |
| chr8-101461435-INS-74 | chr8-101461435-INS-65 |
| chr8-101466328-INS-835 | chr8-101466328-INS-835 |
| chr8-101468245-INS-7122 | chr8-101468246-INS-7122 |
| chr8-101531798-INS-52 | chr8-101531798-INS-52 |
| chr8-101534676-INS-442 | chr8-101534676-INS-442 |
| chr8-101535489-INS-193 | chr8-101535489-INS-193 |
| chr8-101539947-INS-55 | chr8-101539947-INS-59 |

| | |
|---|---|
| chr8-101544417-INS-55 | chr8-101544417-INS-55 |
| chr8-101546387-INS-12302 | chr8-101546388-INS-12304 |
| chr8-101569627-INS-168 | chr8-101569649-INS-168 |
| chr8-101584043-INS-81 | chr8-101584043-INS-83 |
| chr8-101614965-INS-222 | chr8-101614965-INS-222 |
| chr8-101621788-INS-100 | chr8-101621788-INS-100 |
| chr8-101623804-INS-55 | chr8-101623804-INS-55 |
| chr8-101642350-INS-6875 | chr8-101642351-INS-6876 |
| chr8-101676836-INS-64 | chr8-101676810-INS-50 |
| chr8-101692250-INS-7147 | chr8-101692251-INS-7147 |
| chr8-101739209-INS-824 | chr8-101739209-INS-824 |
| chr8-101768427-INS-160 | chr8-101768427-INS-160 |
| chr8-101774607-INS-397 | chr8-101774607-INS-397 |
| chr8-101823281-INS-80 | chr8-101823281-INS-91 |
| chr8-101833925-INS-194 | chr8-101833925-INS-194 |
| chr8-101877257-INS-89 | chr8-101877257-INS-91 |
| chr8-101880221-INS-1138 | chr8-101880222-INS-1138 |
| chr8-101884316-INS-92 | chr8-101884316-INS-68 |
| chr8-101887806-INS-56 | chr8-101887761-INS-56 |
| chr8-101893579-INS-90 | chr8-101893579-INS-90 |
| chr8-101896602-INS-72 | chr8-101896602-INS-72 |
| chr8-101904287-INS-4752 | chr8-101904288-INS-4752 |
| chr8-101929922-INS-202 | chr8-101929922-INS-198 |
| chr8-101959999-INS-1515 | chr8-101960000-INS-1520 |
| chr8-101990510-INS-1536 | chr8-101990511-INS-1536 |
| chr8-102015916-INS-66 | chr8-102015916-INS-64 |
| chr8-102016975-INS-54 | chr8-102016975-INS-56 |
| chr8-102067136-INS-62 | chr8-102067136-INS-62 |
| chr8-102100073-INS-83 | chr8-102100073-INS-83 |
| chr8-102119754-INS-174 | chr8-102119755-INS-174 |
| chr8-102131144-INS-2805 | chr8-102131145-INS-2802 |
| chr8-102143585-INS-64 | chr8-102143594-INS-65 |
| chr8-102164681-INS-882 | chr8-102164681-INS-882 |
| chr8-102210746-INS-55 | chr8-102210750-INS-56 |
| chr8-102250413-INS-77 | chr8-102250458-INS-58 |
| chr8-102262812-INS-52 | chr8-102262812-INS-64 |
| chr8-102283386-INS-72 | chr8-102283387-INS-70 |
| chr8-102304855-INS-369 | chr8-102304855-INS-414 |
| chr8-102308329-INS-512 | chr8-102308329-INS-512 |
| chr8-102358102-INS-110 | chr8-102358102-INS-114 |
| chr8-102365892-INS-5337 | chr8-102365893-INS-5337 |
| chr8-102382104-INS-85 | chr8-102382064-INS-85 |
| chr8-102409163-INS-52 | chr8-102409163-INS-52 |
| chr8-102432880-INS-257 | chr8-102432880-INS-257 |
| chr8-102437587-INS-64 | chr8-102437587-INS-64 |
| chr8-102463670-INS-100 | chr8-102463670-INS-85 |
| chr8-102480237-INS-63 | chr8-102480269-INS-70 |
| chr8-102487139-INS-183 | chr8-102487139-INS-155 |
| chr8-102496720-INS-392 | chr8-102496720-INS-392 |
| chr8-102501084-INS-365 | chr8-102501084-INS-364 |
| chr8-102507880-INS-7077 | chr8-102507881-INS-7077 |
| chr8-102525515-INS-60 | chr8-102525510-INS-62 |
| chr8-102528182-INS-114 | chr8-102528138-INS-131 |
| chr8-102547598-INS-79 | chr8-102547599-INS-60 |
| chr8-102552997-INS-70 | chr8-102552997-INS-86 |
| chr8-102587567-INS-171 | chr8-102587567-INS-171 |
| chr8-102605402-INS-6458 | chr8-102605403-INS-6458 |
| chr8-102605651-INS-108 | chr8-102605651-INS-108 |
| chr8-102657805-INS-106 | chr8-102657805-INS-152 |
| chr8-102679478-INS-74 | chr8-102679520-INS-63 |
| chr8-102773113-INS-199 | chr8-102773113-INS-199 |
| chr8-102778490-INS-685 | chr8-102778490-INS-685 |
| chr8-102798071-INS-595 | chr8-102798071-INS-597 |
| chr8-102809968-INS-101 | chr8-102809968-INS-98 |
| chr8-102826433-INS-64 | chr8-102826433-INS-54 |
| chr8-102848131-INS-1064 | chr8-102848132-INS-1064 |
| chr8-102849312-INS-277 | chr8-102849313-INS-249 |
| chr8-102880733-INS-2167 | chr8-102880734-INS-2168 |
| chr8-102890367-INS-579 | chr8-102890367-INS-579 |
| chr8-102897602-INS-384 | chr8-102897602-INS-384 |
| chr8-102899045-INS-2699 | chr8-102899046-INS-2699 |
| chr8-102947224-INS-444 | chr8-102947224-INS-444 |
| chr8-102970198-INS-282 | chr8-102970198-INS-282 |
| chr8-102991285-INS-64 | chr8-102991285-INS-98 |
| chr8-102999291-INS-52 | chr8-102999291-INS-60 |
| chr8-103005452-INS-1866 | chr8-103005453-INS-1866 |
| chr8-103009685-INS-6439 | chr8-103009686-INS-6439 |

| | |
|---|---|
| chr8-103030258-INS-323 | chr8-103030258-INS-352 |
| chr8-103085463-INS-55 | chr8-103085463-INS-52 |
| chr8-103104666-INS-268 | chr8-103104666-INS-268 |
| chr8-103108807-INS-204 | chr8-103108808-INS-204 |
| chr8-103110194-INS-52 | chr8-103110194-INS-52 |
| chr8-103111403-INS-5439 | chr8-103111404-INS-5439 |
| chr8-103115282-INS-1530 | chr8-103115283-INS-1530 |
| chr8-103127319-INS-58 | chr8-103127319-INS-58 |
| chr8-103159002-INS-2257 | chr8-103159003-INS-2261 |
| chr8-103165074-INS-677 | chr8-103165074-INS-677 |
| chr8-103210549-INS-78 | chr8-103210549-INS-80 |
| chr8-103221185-INS-52 | chr8-103221185-INS-52 |
| chr8-103231548-INS-52 | chr8-103231548-INS-57 |
| chr8-103248528-INS-128 | chr8-103248528-INS-128 |
| chr8-103254870-INS-601 | chr8-103254874-INS-597 |
| chr8-103281565-INS-183 | chr8-103281565-INS-183 |
| chr8-103287227-INS-942 | chr8-103287227-INS-942 |
| chr8-103289601-INS-52 | chr8-103289585-INS-54 |
| chr8-103302995-INS-197 | chr8-103302995-INS-187 |
| chr8-103309819-INS-100 | chr8-103309819-INS-104 |
| chr8-103314919-INS-3221 | chr8-103314920-INS-3223 |
| chr8-103342773-INS-6191 | chr8-103342774-INS-6191 |
| chr8-103352381-INS-723 | chr8-103352381-INS-724 |
| chr8-103380290-INS-938 | chr8-103380291-INS-940 |
| chr8-103388663-INS-68 | chr8-103388663-INS-68 |
| chr8-103392890-INS-65 | chr8-103392890-INS-65 |
| chr8-103410873-INS-119 | chr8-103410873-INS-106 |
| chr8-103444731-INS-52 | chr8-103444693-INS-58 |
| chr8-103464006-INS-54 | chr8-103464006-INS-56 |
| chr8-103491165-INS-67 | chr8-103491185-INS-65 |
| chr8-103518676-INS-68 | chr8-103518696-INS-52 |
| chr8-103524224-INS-6166 | chr8-103524225-INS-6185 |
| chr8-103559440-INS-299 | chr8-103559440-INS-299 |
| chr8-103578175-INS-469 | chr8-103578175-INS-473 |
| chr8-103580144-INS-50 | chr8-103580144-INS-51 |
| chr8-103580643-INS-5223 | chr8-103580644-INS-5223 |
| chr8-103581513-INS-56 | chr8-103581513-INS-56 |
| chr8-103590372-INS-474 | chr8-103590369-INS-477 |
| chr8-103598652-INS-68 | chr8-103598652-INS-50 |
| chr8-103601164-INS-167 | chr8-103601164-INS-167 |
| chr8-103626598-INS-56 | chr8-103626598-INS-56 |
| chr8-103652082-INS-129 | chr8-103652082-INS-145 |
| chr8-103671936-INS-80 | chr8-103671954-INS-55 |
| chr8-103688825-INS-921 | chr8-103688825-INS-921 |
| chr8-103704185-INS-88 | chr8-103704185-INS-88 |
| chr8-103743271-INS-58 | chr8-103743271-INS-60 |
| chr8-103748376-INS-65 | chr8-103748367-INS-88 |
| chr8-103758923-INS-121 | chr8-103758932-INS-89 |
| chr8-103762179-INS-345 | chr8-103762179-INS-346 |
| chr8-103773258-INS-3073 | chr8-103773259-INS-3073 |
| chr8-103811968-INS-168 | chr8-103811968-INS-168 |
| chr8-103835105-INS-1535 | chr8-103835106-INS-1559 |
| chr8-103853784-INS-266 | chr8-103853784-INS-266 |
| chr8-103889195-INS-66 | chr8-103889195-INS-76 |
| chr8-103892260-INS-66 | chr8-103892287-INS-65 |
| chr8-103899650-INS-150 | chr8-103899650-INS-150 |
| chr8-103942219-INS-462 | chr8-103942218-INS-464 |
| chr8-103986670-INS-324 | chr8-103986670-INS-324 |
| chr8-103992027-INS-162 | chr8-103992027-INS-174 |
| chr8-104025275-INS-322 | chr8-104025275-INS-322 |
| chr8-104050656-INS-65 | chr8-104050656-INS-71 |
| chr8-104061221-INS-1410 | chr8-104061222-INS-1411 |
| chr8-104135671-INS-52 | chr8-104135671-INS-62 |
| chr8-104139940-INS-55 | chr8-104139936-INS-59 |
| chr8-104148920-INS-233 | chr8-104148920-INS-227 |
| chr8-104225671-INS-53 | chr8-104225671-INS-53 |
| chr8-104226433-INS-417 | chr8-104226433-INS-417 |
| chr8-104258084-INS-57 | chr8-104258085-INS-80 |
| chr8-104294266-INS-6777 | chr8-104294267-INS-6766 |
| chr8-104322977-INS-60 | chr8-104322977-INS-60 |
| chr8-104342716-INS-5297 | chr8-104342717-INS-5297 |
| chr8-104367247-INS-61 | chr8-104367246-INS-99 |
| chr8-104386634-INS-6241 | chr8-104386635-INS-6242 |
| chr8-104393785-INS-244 | chr8-104393785-INS-216 |
| chr8-104421436-INS-102 | chr8-104421437-INS-83 |
| chr8-104429643-INS-56 | chr8-104429643-INS-56 |
| chr8-104452857-INS-566 | chr8-104452857-INS-566 |

| | |
|---|---|
| chr8-104512034-INS-124 | chr8-104512034-INS-124 |
| chr8-104553052-INS-129 | chr8-104553054-INS-125 |
| chr8-104684433-INS-1199 | chr8-104684434-INS-1198 |
| chr8-104689964-INS-1157 | chr8-104689965-INS-1156 |
| chr8-104721234-INS-66 | chr8-104721231-INS-53 |
| chr8-104732370-INS-65 | chr8-104732370-INS-65 |
| chr8-104740716-INS-52 | chr8-104740716-INS-52 |
| chr8-104756836-INS-60 | chr8-104756845-INS-67 |
| chr8-104767543-INS-157 | chr8-104767543-INS-157 |
| chr8-104791222-INS-439 | chr8-104791213-INS-463 |
| chr8-104798979-INS-7057 | chr8-104798980-INS-7057 |
| chr8-104805744-INS-462 | chr8-104805744-INS-462 |
| chr8-104825338-INS-80 | chr8-104825338-INS-80 |
| chr8-104834747-INS-62 | chr8-104834747-INS-54 |
| chr8-104849611-INS-53 | chr8-104849646-INS-65 |
| chr8-104860318-INS-107 | chr8-104860327-INS-72 |
| chr8-104866456-INS-54 | chr8-104866476-INS-82 |
| chr8-104890321-INS-77 | chr8-104890320-INS-122 |
| chr8-104919182-INS-51 | chr8-104919182-INS-54 |
| chr8-104938679-INS-79 | chr8-104938639-INS-79 |
| chr8-104956046-INS-400 | chr8-104956046-INS-398 |
| chr8-104967914-INS-6475 | chr8-104967915-INS-6475 |
| chr8-104970831-INS-1148 | chr8-104970832-INS-1172 |
| chr8-104984701-INS-284 | chr8-104984701-INS-287 |
| chr8-104986442-INS-60 | chr8-104986442-INS-60 |
| chr8-105005029-INS-326 | chr8-105005029-INS-326 |
| chr8-105023515-INS-85 | chr8-105023511-INS-61 |
| chr8-105076889-INS-69 | chr8-105076889-INS-69 |
| chr8-105093954-INS-111 | chr8-105093952-INS-135 |
| chr8-105098419-INS-58 | chr8-105098419-INS-58 |
| chr8-105106521-INS-471 | chr8-105106520-INS-458 |
| chr8-105122815-INS-462 | chr8-105122815-INS-428 |
| chr8-105141194-INS-113 | chr8-105141194-INS-113 |
| chr8-105170240-INS-54 | chr8-105170240-INS-54 |
| chr8-105200857-INS-401 | chr8-105200857-INS-400 |
| chr8-105202922-INS-1943 | chr8-105202923-INS-1942 |
| chr8-105204928-INS-179 | chr8-105204928-INS-179 |
| chr8-105208105-INS-116 | chr8-105208105-INS-116 |
| chr8-105262662-INS-87 | chr8-105262662-INS-87 |
| chr8-105263461-INS-84 | chr8-105263461-INS-82 |
| chr8-105278385-INS-63 | chr8-105278393-INS-54 |
| chr8-105286781-INS-95 | chr8-105286741-INS-129 |
| chr8-105289356-INS-202 | chr8-105289356-INS-202 |
| chr8-105295491-INS-209 | chr8-105295491-INS-209 |
| chr8-105296575-INS-53 | chr8-105296573-INS-85 |
| chr8-105299171-INS-85 | chr8-105299171-INS-79 |
| chr8-105301382-INS-174 | chr8-105301382-INS-177 |
| chr8-105307187-INS-206 | chr8-105307187-INS-206 |
| chr8-105310075-INS-54 | chr8-105310075-INS-54 |
| chr8-105331236-INS-670 | chr8-105331236-INS-678 |
| chr8-105338892-INS-52 | chr8-105338892-INS-54 |
| chr8-105340229-INS-205 | chr8-105340229-INS-205 |
| chr8-105399732-INS-210 | chr8-105399732-INS-210 |
| chr8-105409750-INS-138 | chr8-105409750-INS-136 |
| chr8-105516317-INS-208 | chr8-105516317-INS-208 |
| chr8-105526162-INS-124 | chr8-105526162-INS-124 |
| chr8-105554622-INS-54 | chr8-105554622-INS-62 |
| chr8-105571990-INS-206 | chr8-105571990-INS-206 |
| chr8-105640965-INS-6744 | chr8-105640966-INS-6744 |
| chr8-105698812-INS-143 | chr8-105698812-INS-142 |
| chr8-105700669-INS-54 | chr8-105700669-INS-84 |
| chr8-105768976-INS-510 | chr8-105768976-INS-510 |
| chr8-105791641-INS-6455 | chr8-105791642-INS-6455 |
| chr8-105799439-INS-183 | chr8-105799439-INS-182 |
| chr8-105826398-INS-60 | chr8-105826398-INS-60 |
| chr8-105841787-INS-132 | chr8-105841767-INS-127 |
| chr8-105850123-INS-221 | chr8-105850123-INS-221 |
| chr8-105865625-INS-210 | chr8-105865625-INS-203 |
| chr8-105877093-INS-51 | chr8-105877097-INS-52 |
| chr8-105883486-INS-231 | chr8-105883486-INS-234 |
| chr8-105892336-INS-112 | chr8-105892336-INS-112 |
| chr8-105900341-INS-92 | chr8-105900341-INS-92 |
| chr8-105931497-INS-55 | chr8-105931497-INS-55 |
| chr8-105941267-INS-161 | chr8-105941266-INS-163 |
| chr8-105963387-INS-199 | chr8-105963401-INS-187 |
| chr8-105989312-INS-73 | chr8-105989314-INS-87 |
| chr8-106070826-INS-136 | chr8-106070826-INS-152 |

| | |
|---|---|
| chr8-106079423-INS-60 | chr8-106079423-INS-64 |
| chr8-106087141-INS-292 | chr8-106087141-INS-292 |
| chr8-106119769-INS-72 | chr8-106119769-INS-72 |
| chr8-106128918-INS-64 | chr8-106128916-INS-68 |
| chr8-106204156-INS-400 | chr8-106204156-INS-400 |
| chr8-106208194-INS-212 | chr8-106208194-INS-212 |
| chr8-106225788-INS-64 | chr8-106225772-INS-84 |
| chr8-106236471-INS-66 | chr8-106236481-INS-76 |
| chr8-106239864-INS-180 | chr8-106239865-INS-179 |
| chr8-106283296-INS-100 | chr8-106283316-INS-52 |
| chr8-106284678-INS-205 | chr8-106284678-INS-205 |
| chr8-106285310-INS-87 | chr8-106285310-INS-87 |
| chr8-106286736-INS-66 | chr8-106286736-INS-66 |
| chr8-106289286-INS-211 | chr8-106289286-INS-208 |
| chr8-106315440-INS-61 | chr8-106315440-INS-90 |
| chr8-106319993-INS-54 | chr8-106319993-INS-54 |
| chr8-106369728-INS-177 | chr8-106369728-INS-177 |
| chr8-106413941-INS-123 | chr8-106413941-INS-100 |
| chr8-106437110-INS-217 | chr8-106437110-INS-218 |
| chr8-106448458-INS-89 | chr8-106448504-INS-87 |
| chr8-106449840-INS-1018 | chr8-106449841-INS-1018 |
| chr8-106458252-INS-210 | chr8-106458252-INS-211 |
| chr8-106465653-INS-175 | chr8-106465653-INS-175 |
| chr8-106472729-INS-106 | chr8-106472729-INS-84 |
| chr8-106476543-INS-188 | chr8-106476543-INS-188 |
| chr8-106488986-INS-440 | chr8-106488988-INS-436 |
| chr8-106512041-INS-859 | chr8-106512041-INS-859 |
| chr8-106515254-INS-208 | chr8-106515254-INS-206 |
| chr8-106516350-INS-60 | chr8-106516350-INS-60 |
| chr8-106521394-INS-76 | chr8-106521394-INS-108 |
| chr8-106524804-INS-238 | chr8-106524804-INS-237 |
| chr8-106555105-INS-168 | chr8-106555105-INS-168 |
| chr8-106567070-INS-207 | chr8-106567070-INS-208 |
| chr8-106574156-INS-215 | chr8-106574156-INS-212 |
| chr8-106578859-INS-164 | chr8-106578843-INS-163 |
| chr8-106597871-INS-61 | chr8-106597871-INS-81 |
| chr8-106625277-INS-197 | chr8-106625276-INS-198 |
| chr8-106632534-INS-205 | chr8-106632534-INS-205 |
| chr8-106644633-INS-205 | chr8-106644633-INS-197 |
| chr8-106647654-INS-208 | chr8-106647650-INS-232 |
| chr8-106684962-INS-197 | chr8-106684962-INS-211 |
| chr8-106708068-INS-217 | chr8-106708068-INS-221 |
| chr8-106729771-INS-192 | chr8-106729771-INS-193 |
| chr8-106778196-INS-294 | chr8-106778196-INS-334 |
| chr8-106782341-INS-151 | chr8-106782341-INS-151 |
| chr8-106827410-INS-621 | chr8-106827410-INS-617 |
| chr8-106874257-INS-114 | chr8-106874283-INS-113 |
| chr8-106886656-INS-197 | chr8-106886656-INS-198 |
| chr8-106901820-INS-226 | chr8-106901820-INS-224 |
| chr8-106908272-INS-72 | chr8-106908272-INS-72 |
| chr8-106910710-INS-52 | chr8-106910710-INS-64 |
| chr8-106930885-INS-191 | chr8-106930885-INS-191 |
| chr8-106940976-INS-231 | chr8-106940976-INS-230 |
| chr8-106947427-INS-74 | chr8-106947434-INS-50 |
| chr8-106949523-INS-199 | chr8-106949523-INS-199 |
| chr8-106956878-INS-51 | chr8-106956880-INS-51 |
| chr8-107016156-INS-76 | chr8-107016156-INS-66 |
| chr8-107052905-INS-90 | chr8-107052905-INS-70 |
| chr8-107055297-INS-92 | chr8-107055297-INS-92 |
| chr8-107071900-INS-68 | chr8-107071902-INS-50 |
| chr8-107095618-INS-3129 | chr8-107095619-INS-3129 |
| chr8-107112216-INS-95 | chr8-107112216-INS-94 |
| chr8-107113633-INS-382 | chr8-107113633-INS-382 |
| chr8-107121869-INS-104 | chr8-107121870-INS-56 |
| chr8-107129354-INS-63 | chr8-107129354-INS-63 |
| chr8-107164669-INS-389 | chr8-107164669-INS-389 |
| chr8-107178486-INS-65 | chr8-107178486-INS-65 |
| chr8-107182058-INS-183 | chr8-107182058-INS-173 |
| chr8-107203359-INS-199 | chr8-107203353-INS-208 |
| chr8-107221882-INS-561 | chr8-107221882-INS-562 |
| chr8-107250482-INS-60 | chr8-107250474-INS-50 |
| chr8-107262660-INS-204 | chr8-107262660-INS-204 |
| chr8-107269147-INS-329 | chr8-107269148-INS-329 |
| chr8-107278292-INS-226 | chr8-107278292-INS-226 |
| chr8-107282983-INS-77 | chr8-107282987-INS-83 |
| chr8-107285169-INS-193 | chr8-107285169-INS-191 |
| chr8-107310366-INS-201 | chr8-107310366-INS-201 |

| | |
|---|---|
| chr8-107316211-INS-217 | chr8-107316211-INS-217 |
| chr8-107322818-INS-178 | chr8-107322818-INS-178 |
| chr8-107324973-INS-8418 | chr8-107324974-INS-8418 |
| chr8-107349284-INS-294 | chr8-107349284-INS-306 |
| chr8-107391837-INS-199 | chr8-107391837-INS-199 |
| chr8-107393806-INS-150 | chr8-107393806-INS-150 |
| chr8-107413148-INS-180 | chr8-107413148-INS-180 |
| chr8-107418064-INS-189 | chr8-107418064-INS-189 |
| chr8-107429468-INS-412 | chr8-107429453-INS-405 |
| chr8-107430455-INS-270 | chr8-107430455-INS-270 |
| chr8-107439923-INS-279 | chr8-107439923-INS-279 |
| chr8-107446276-INS-67 | chr8-107446276-INS-67 |
| chr8-107452191-INS-107 | chr8-107452191-INS-108 |
| chr8-107458183-INS-171 | chr8-107458193-INS-170 |
| chr8-107464034-INS-162 | chr8-107464032-INS-158 |
| chr8-107465813-INS-65 | chr8-107465813-INS-85 |
| chr8-107477528-INS-56 | chr8-107477494-INS-52 |
| chr8-107479583-INS-219 | chr8-107479589-INS-225 |
| chr8-107498054-INS-53 | chr8-107498054-INS-53 |
| chr8-107510000-INS-231 | chr8-107510000-INS-231 |
| chr8-107521906-INS-161 | chr8-107521906-INS-161 |
| chr8-107531638-INS-193 | chr8-107531638-INS-163 |
| chr8-107534942-INS-237 | chr8-107534942-INS-237 |
| chr8-107548401-INS-284 | chr8-107548427-INS-284 |
| chr8-107549820-INS-51 | chr8-107549820-INS-51 |
| chr8-107554764-INS-189 | chr8-107554764-INS-176 |
| chr8-107637248-INS-80 | chr8-107637248-INS-80 |
| chr8-107655093-INS-5012 | chr8-107655094-INS-5012 |
| chr8-107669153-INS-214 | chr8-107669153-INS-214 |
| chr8-107701433-INS-83 | chr8-107701433-INS-83 |
| chr8-107703063-INS-230 | chr8-107703063-INS-221 |
| chr8-107710713-INS-6868 | chr8-107710714-INS-6868 |
| chr8-107715407-INS-214 | chr8-107715407-INS-214 |
| chr8-107720138-INS-209 | chr8-107720139-INS-170 |
| chr8-107737624-INS-204 | chr8-107737624-INS-201 |
| chr8-107746904-INS-122 | chr8-107746904-INS-118 |
| chr8-107784353-INS-209 | chr8-107784353-INS-209 |
| chr8-107817558-INS-178 | chr8-107817550-INS-169 |
| chr8-107832640-INS-208 | chr8-107832640-INS-208 |
| chr8-107844028-INS-73 | chr8-107844024-INS-83 |
| chr8-107847346-INS-162 | chr8-107847350-INS-168 |
| chr8-107858118-INS-55 | chr8-107858118-INS-55 |
| chr8-107866681-INS-62 | chr8-107866681-INS-50 |
| chr8-107871616-INS-209 | chr8-107871616-INS-209 |
| chr8-107874144-INS-206 | chr8-107874144-INS-214 |
| chr8-107874776-INS-235 | chr8-107874787-INS-211 |
| chr8-107878866-INS-215 | chr8-107878866-INS-217 |
| chr8-107883276-INS-60 | chr8-107883279-INS-60 |
| chr8-107892672-INS-58 | chr8-107892672-INS-58 |
| chr8-107919140-INS-406 | chr8-107919140-INS-406 |
| chr8-107941328-INS-202 | chr8-107941312-INS-207 |
| chr8-107974239-INS-53 | chr8-107974234-INS-54 |
| chr8-107984525-INS-56 | chr8-107984525-INS-60 |
| chr8-108002550-INS-79 | chr8-108002550-INS-65 |
| chr8-108012144-INS-117 | chr8-108012144-INS-130 |
| chr8-108028343-INS-140 | chr8-108028343-INS-141 |
| chr8-108049850-INS-241 | chr8-108049850-INS-237 |
| chr8-108086373-INS-104 | chr8-108086373-INS-104 |
| chr8-108111098-INS-846 | chr8-108111098-INS-847 |
| chr8-108112531-INS-194 | chr8-108112531-INS-194 |
| chr8-108127676-INS-160 | chr8-108127718-INS-118 |
| chr8-108130489-INS-2682 | chr8-108130490-INS-2682 |
| chr8-108133194-INS-240 | chr8-108133194-INS-240 |
| chr8-108134601-INS-165 | chr8-108134601-INS-168 |
| chr8-108149483-INS-213 | chr8-108149483-INS-213 |
| chr8-108208066-INS-210 | chr8-108208066-INS-210 |
| chr8-108225847-INS-140 | chr8-108225847-INS-137 |
| chr8-108232343-INS-117 | chr8-108232343-INS-156 |
| chr8-108243211-INS-212 | chr8-108243211-INS-212 |
| chr8-108300434-INS-187 | chr8-108300434-INS-186 |
| chr8-108313962-INS-326 | chr8-108313962-INS-325 |
| chr8-108369964-INS-470 | chr8-108369964-INS-470 |
| chr8-108370837-INS-64 | chr8-108370837-INS-68 |
| chr8-108386028-INS-60 | chr8-108386026-INS-62 |
| chr8-108453585-INS-389 | chr8-108453585-INS-389 |
| chr8-108474284-INS-352 | chr8-108474282-INS-352 |
| chr8-108485133-INS-62 | chr8-108485131-INS-96 |

| | |
|---|---|
| chr8-108489592-INS-122 | chr8-108489592-INS-73 |
| chr8-108491025-INS-73 | chr8-108491025-INS-71 |
| chr8-108493417-INS-65 | chr8-108493417-INS-76 |
| chr8-108494828-INS-59 | chr8-108494829-INS-55 |
| chr8-108503103-INS-55 | chr8-108503102-INS-94 |
| chr8-108506324-INS-56 | chr8-108506324-INS-58 |
| chr8-108539264-INS-86 | chr8-108539264-INS-108 |
| chr8-108547503-INS-80 | chr8-108547503-INS-64 |
| chr8-108549807-INS-457 | chr8-108549807-INS-457 |
| chr8-108561194-INS-54 | chr8-108561194-INS-54 |
| chr8-108564646-INS-208 | chr8-108564646-INS-208 |
| chr8-108616143-INS-135 | chr8-108616143-INS-145 |
| chr8-108621966-INS-52 | chr8-108621966-INS-58 |
| chr8-108654228-INS-103 | chr8-108654228-INS-108 |
| chr8-108696566-INS-236 | chr8-108696566-INS-236 |
| chr8-108716714-INS-198 | chr8-108716714-INS-198 |
| chr8-108722276-INS-106 | chr8-108722272-INS-114 |
| chr8-108732716-INS-54 | chr8-108732716-INS-54 |
| chr8-108747975-INS-111 | chr8-108747975-INS-66 |
| chr8-108751444-INS-61 | chr8-108751452-INS-50 |
| chr8-108760819-INS-56 | chr8-108760819-INS-56 |
| chr8-108830852-INS-101 | chr8-108830838-INS-91 |
| chr8-108855205-INS-387 | chr8-108855226-INS-386 |
| chr8-108868974-INS-63 | chr8-108868974-INS-63 |
| chr8-108955403-INS-62 | chr8-108955403-INS-62 |
| chr8-108985145-INS-72 | chr8-108985173-INS-70 |
| chr8-108996965-INS-149 | chr8-108996965-INS-162 |
| chr8-109003204-INS-56 | chr8-109003208-INS-63 |
| chr8-109042066-INS-70 | chr8-109042066-INS-74 |
| chr8-109062506-INS-114 | chr8-109062518-INS-118 |
| chr8-109089589-INS-70 | chr8-109089589-INS-68 |
| chr8-109107425-INS-80 | chr8-109107425-INS-84 |
| chr8-109109648-INS-54 | chr8-109109648-INS-54 |
| chr8-109112394-INS-60 | chr8-109112394-INS-60 |
| chr8-109148370-INS-4772 | chr8-109148371-INS-4772 |
| chr8-109165254-INS-72 | chr8-109165254-INS-72 |
| chr8-109206440-INS-52 | chr8-109206419-INS-98 |
| chr8-109225392-INS-109 | chr8-109225392-INS-109 |
| chr8-109251153-INS-575 | chr8-109251154-INS-583 |
| chr8-109278804-INS-70 | chr8-109278804-INS-70 |
| chr8-109289990-INS-608 | chr8-109289990-INS-609 |
| chr8-109315725-INS-88 | chr8-109315725-INS-72 |
| chr8-109349723-INS-227 | chr8-109349723-INS-227 |
| chr8-109352093-INS-71 | chr8-109352083-INS-87 |
| chr8-109377223-INS-72 | chr8-109377223-INS-72 |
| chr8-109391915-INS-7075 | chr8-109391916-INS-7075 |
| chr8-109418288-INS-1889 | chr8-109418289-INS-1889 |
| chr8-109473389-INS-220 | chr8-109473389-INS-220 |
| chr8-109544207-INS-175 | chr8-109544202-INS-152 |
| chr8-109550436-INS-96 | chr8-109550436-INS-96 |
| chr8-109567432-INS-192 | chr8-109567432-INS-192 |
| chr8-109619622-INS-102 | chr8-109619647-INS-88 |
| chr8-109636557-INS-56 | chr8-109636557-INS-56 |
| chr8-109641710-INS-121 | chr8-109641756-INS-72 |
| chr8-109646268-INS-232 | chr8-109646268-INS-230 |
| chr8-109667725-INS-54 | chr8-109667725-INS-54 |
| chr8-109735244-INS-722 | chr8-109735244-INS-722 |
| chr8-109757716-INS-515 | chr8-109757716-INS-515 |
| chr8-109760400-INS-57 | chr8-109760395-INS-70 |
| chr8-109799279-INS-724 | chr8-109799279-INS-724 |
| chr8-109800280-INS-58 | chr8-109800280-INS-92 |
| chr8-109803560-INS-7199 | chr8-109803561-INS-7198 |
| chr8-109848767-INS-52 | chr8-109848767-INS-52 |
| chr8-109849860-INS-92 | chr8-109849893-INS-50 |
| chr8-109860290-INS-52 | chr8-109860251-INS-58 |
| chr8-109881295-INS-74 | chr8-109881295-INS-74 |
| chr8-109884795-INS-207 | chr8-109884795-INS-207 |
| chr8-109891587-INS-417 | chr8-109891587-INS-441 |
| chr8-109894614-INS-199 | chr8-109894615-INS-178 |
| chr8-109904604-INS-1297 | chr8-109904605-INS-1296 |
| chr8-109908361-INS-173 | chr8-109908361-INS-171 |
| chr8-109953555-INS-68 | chr8-109953555-INS-64 |
| chr8-109967763-INS-65 | chr8-109967763-INS-55 |
| chr8-109993531-INS-70 | chr8-109993531-INS-70 |
| chr8-110012160-INS-57 | chr8-110012160-INS-57 |
| chr8-110013422-INS-52 | chr8-110013427-INS-54 |
| chr8-110043672-INS-156 | chr8-110043672-INS-167 |

| | |
|---|---|
| chr8-110066019-INS-424 | chr8-110066019-INS-438 |
| chr8-110077274-INS-81 | chr8-110077298-INS-68 |
| chr8-110099471-INS-156 | chr8-110099471-INS-163 |
| chr8-110140519-INS-240 | chr8-110140519-INS-235 |
| chr8-110167924-INS-183 | chr8-110167924-INS-200 |
| chr8-110173559-INS-72 | chr8-110173559-INS-72 |
| chr8-110192941-INS-51 | chr8-110192952-INS-54 |
| chr8-110207234-INS-54 | chr8-110207234-INS-66 |
| chr8-110270487-INS-368 | chr8-110270488-INS-368 |
| chr8-110309636-INS-162 | chr8-110309636-INS-177 |
| chr8-110312602-INS-178 | chr8-110312602-INS-180 |
| chr8-110317491-INS-6475 | chr8-110317492-INS-6475 |
| chr8-110325560-INS-103 | chr8-110325560-INS-107 |
| chr8-110365231-INS-353 | chr8-110365231-INS-350 |
| chr8-110369682-INS-89 | chr8-110369643-INS-128 |
| chr8-110376280-INS-207 | chr8-110376280-INS-207 |
| chr8-110391636-INS-72 | chr8-110391636-INS-78 |
| chr8-110394721-INS-64 | chr8-110394721-INS-70 |
| chr8-110414536-INS-213 | chr8-110414536-INS-210 |
| chr8-110447020-INS-238 | chr8-110447020-INS-239 |
| chr8-110493902-INS-56 | chr8-110493902-INS-60 |
| chr8-110509545-INS-228 | chr8-110509546-INS-218 |
| chr8-110524389-INS-239 | chr8-110524389-INS-239 |
| chr8-110533507-INS-97 | chr8-110533535-INS-53 |
| chr8-110560921-INS-61 | chr8-110560921-INS-61 |
| chr8-110562046-INS-68 | chr8-110562046-INS-68 |
| chr8-110575950-INS-287 | chr8-110575950-INS-287 |
| chr8-110613873-INS-203 | chr8-110613873-INS-206 |
| chr8-110622664-INS-951 | chr8-110622664-INS-964 |
| chr8-110636260-INS-396 | chr8-110636260-INS-396 |
| chr8-110660903-INS-112 | chr8-110660903-INS-110 |
| chr8-110686150-INS-61 | chr8-110686142-INS-94 |
| chr8-110693190-INS-198 | chr8-110693190-INS-198 |
| chr8-110694511-INS-99 | chr8-110694533-INS-92 |
| chr8-110701045-INS-68 | chr8-110701053-INS-56 |
| chr8-110726445-INS-794 | chr8-110726445-INS-800 |
| chr8-110736809-INS-108 | chr8-110736799-INS-107 |
| chr8-110737795-INS-579 | chr8-110737795-INS-578 |
| chr8-110751896-INS-169 | chr8-110751923-INS-162 |
| chr8-110758854-INS-195 | chr8-110758850-INS-203 |
| chr8-110771248-INS-52 | chr8-110771248-INS-74 |
| chr8-110800592-INS-247 | chr8-110800592-INS-250 |
| chr8-110814561-INS-145 | chr8-110814558-INS-174 |
| chr8-110833225-INS-204 | chr8-110833223-INS-206 |
| chr8-110853965-INS-6282 | chr8-110853966-INS-6285 |
| chr8-110863370-INS-154 | chr8-110863370-INS-154 |
| chr8-110884593-INS-169 | chr8-110884593-INS-168 |
| chr8-110898238-INS-383 | chr8-110898253-INS-384 |
| chr8-110953978-INS-6402 | chr8-110953979-INS-6401 |
| chr8-110990014-INS-116 | chr8-110990014-INS-116 |
| chr8-111000642-INS-599 | chr8-111000670-INS-604 |
| chr8-111020360-INS-160 | chr8-111020360-INS-160 |
| chr8-111026899-INS-92 | chr8-111026890-INS-90 |
| chr8-111106386-INS-78 | chr8-111106386-INS-78 |
| chr8-111126316-INS-108 | chr8-111126322-INS-105 |
| chr8-111170161-INS-5381 | chr8-111170162-INS-5381 |
| chr8-111182814-INS-58 | chr8-111182814-INS-58 |
| chr8-111227148-INS-63 | chr8-111227148-INS-63 |
| chr8-111297725-INS-108 | chr8-111297721-INS-116 |
| chr8-111298115-INS-435 | chr8-111298135-INS-435 |
| chr8-111502953-INS-65 | chr8-111502953-INS-64 |
| chr8-111510347-INS-206 | chr8-111510347-INS-207 |
| chr8-111523607-INS-352 | chr8-111523607-INS-352 |
| chr8-111560146-INS-130 | chr8-111560144-INS-146 |
| chr8-111575399-INS-207 | chr8-111575399-INS-207 |
| chr8-111576796-INS-349 | chr8-111576796-INS-337 |
| chr8-111580047-INS-216 | chr8-111580047-INS-216 |
| chr8-111590621-INS-332 | chr8-111590621-INS-330 |
| chr8-111602034-INS-207 | chr8-111602034-INS-213 |
| chr8-111626307-INS-230 | chr8-111626307-INS-234 |
| chr8-111634531-INS-65 | chr8-111634531-INS-65 |
| chr8-111636153-INS-221 | chr8-111636153-INS-221 |
| chr8-111638596-INS-238 | chr8-111638596-INS-239 |
| chr8-111657698-INS-204 | chr8-111657698-INS-205 |
| chr8-111659683-INS-206 | chr8-111659685-INS-213 |
| chr8-111664907-INS-170 | chr8-111664907-INS-170 |
| chr8-111674569-INS-174 | chr8-111674569-INS-195 |

| | |
|---|---|
| chr8-111698804-INS-181 | chr8-111698804-INS-182 |
| chr8-111722034-INS-95 | chr8-111722034-INS-95 |
| chr8-111729259-INS-194 | chr8-111729259-INS-196 |
| chr8-111732359-INS-180 | chr8-111732359-INS-182 |
| chr8-111750980-INS-218 | chr8-111750980-INS-219 |
| chr8-111770877-INS-214 | chr8-111770877-INS-213 |
| chr8-111781110-INS-197 | chr8-111781110-INS-197 |
| chr8-111804847-INS-234 | chr8-111804847-INS-234 |
| chr8-111822785-INS-180 | chr8-111822786-INS-174 |
| chr8-111831028-INS-409 | chr8-111831028-INS-409 |
| chr8-111851587-INS-63 | chr8-111851587-INS-63 |
| chr8-111866983-INS-60 | chr8-111866983-INS-60 |
| chr8-111929482-INS-919 | chr8-111929483-INS-922 |
| chr8-111931869-INS-100 | chr8-111931869-INS-142 |
| chr8-111938727-INS-203 | chr8-111938727-INS-206 |
| chr8-111940295-INS-225 | chr8-111940295-INS-222 |
| chr8-111945143-INS-397 | chr8-111945143-INS-400 |
| chr8-111965386-INS-54 | chr8-111965386-INS-56 |
| chr8-111971127-INS-191 | chr8-111971127-INS-191 |
| chr8-111981009-INS-209 | chr8-111981009-INS-208 |
| chr8-111991598-INS-60 | chr8-111991598-INS-62 |
| chr8-112001983-INS-132 | chr8-112001976-INS-136 |
| chr8-112002755-INS-3028 | chr8-112002756-INS-3028 |
| chr8-112013656-INS-767 | chr8-112013656-INS-770 |
| chr8-112048053-INS-94 | chr8-112048054-INS-94 |
| chr8-112052780-INS-203 | chr8-112052780-INS-204 |
| chr8-112071277-INS-352 | chr8-112071277-INS-352 |
| chr8-112103595-INS-72 | chr8-112103595-INS-72 |
| chr8-112117808-INS-471 | chr8-112117808-INS-490 |
| chr8-112135192-INS-2962 | chr8-112135193-INS-2962 |
| chr8-112148862-INS-80 | chr8-112148862-INS-62 |
| chr8-112152923-INS-197 | chr8-112152915-INS-197 |
| chr8-112160319-INS-6140 | chr8-112160320-INS-6143 |
| chr8-112182716-INS-52 | chr8-112182716-INS-56 |
| chr8-112213421-INS-120 | chr8-112213421-INS-80 |
| chr8-112261320-INS-185 | chr8-112261324-INS-188 |
| chr8-112265284-INS-136 | chr8-112265284-INS-136 |
| chr8-112271958-INS-171 | chr8-112271958-INS-172 |
| chr8-112281329-INS-114 | chr8-112281329-INS-78 |
| chr8-112286891-INS-192 | chr8-112286891-INS-192 |
| chr8-112334425-INS-204 | chr8-112334425-INS-202 |
| chr8-112365726-INS-415 | chr8-112365726-INS-415 |
| chr8-112379010-INS-232 | chr8-112379010-INS-221 |
| chr8-112384542-INS-239 | chr8-112384542-INS-237 |
| chr8-112395404-INS-59 | chr8-112395410-INS-67 |
| chr8-112400164-INS-127 | chr8-112400164-INS-127 |
| chr8-112401502-INS-153 | chr8-112401502-INS-155 |
| chr8-112408505-INS-84 | chr8-112408505-INS-84 |
| chr8-112416429-INS-745 | chr8-112416429-INS-746 |
| chr8-112432928-INS-249 | chr8-112432928-INS-250 |
| chr8-112434281-INS-88 | chr8-112434281-INS-88 |
| chr8-112445136-INS-203 | chr8-112445136-INS-203 |
| chr8-112447091-INS-76 | chr8-112447091-INS-76 |
| chr8-112449311-INS-551 | chr8-112449311-INS-553 |
| chr8-112484776-INS-82 | chr8-112484776-INS-81 |
| chr8-112487638-INS-60 | chr8-112487638-INS-60 |
| chr8-112504704-INS-186 | chr8-112504704-INS-187 |
| chr8-112513841-INS-110 | chr8-112513840-INS-110 |
| chr8-112515499-INS-91 | chr8-112515499-INS-88 |
| chr8-112535338-INS-125 | chr8-112535338-INS-125 |
| chr8-112544959-INS-109 | chr8-112544959-INS-109 |
| chr8-112561608-INS-229 | chr8-112561608-INS-229 |
| chr8-112562995-INS-70 | chr8-112562995-INS-108 |
| chr8-112578214-INS-59 | chr8-112578214-INS-59 |
| chr8-112583674-INS-196 | chr8-112583674-INS-196 |
| chr8-112585432-INS-220 | chr8-112585433-INS-221 |
| chr8-112587153-INS-70 | chr8-112587153-INS-81 |
| chr8-112596849-INS-202 | chr8-112596849-INS-202 |
| chr8-112605701-INS-187 | chr8-112605701-INS-186 |
| chr8-112634450-INS-153 | chr8-112634450-INS-177 |
| chr8-112641871-INS-258 | chr8-112641871-INS-258 |
| chr8-112644673-INS-64 | chr8-112644641-INS-71 |
| chr8-112646985-INS-217 | chr8-112646986-INS-203 |
| chr8-112655046-INS-52 | chr8-112655046-INS-70 |
| chr8-112657206-INS-96 | chr8-112657206-INS-96 |
| chr8-112669435-INS-55 | chr8-112669435-INS-55 |
| chr8-112681318-INS-202 | chr8-112681318-INS-202 |

| | |
|---|---|
| chr8-112682860-INS-185 | chr8-112682861-INS-185 |
| chr8-112687110-INS-126 | chr8-112687110-INS-126 |
| chr8-112701047-INS-249 | chr8-112701047-INS-241 |
| chr8-112727675-INS-178 | chr8-112727675-INS-179 |
| chr8-112728682-INS-195 | chr8-112728694-INS-196 |
| chr8-112742367-INS-711 | chr8-112742367-INS-752 |
| chr8-112751179-INS-209 | chr8-112751179-INS-209 |
| chr8-112756436-INS-1742 | chr8-112756437-INS-1742 |
| chr8-112758239-INS-144 | chr8-112758239-INS-146 |
| chr8-112763250-INS-111 | chr8-112763253-INS-105 |
| chr8-112773265-INS-51 | chr8-112773253-INS-57 |
| chr8-112783540-INS-224 | chr8-112783540-INS-224 |
| chr8-112787045-INS-66 | chr8-112787045-INS-66 |
| chr8-112789934-INS-390 | chr8-112789934-INS-390 |
| chr8-112837695-INS-206 | chr8-112837695-INS-204 |
| chr8-112840672-INS-479 | chr8-112840672-INS-479 |
| chr8-112846071-INS-213 | chr8-112846072-INS-213 |
| chr8-112846837-INS-208 | chr8-112846837-INS-204 |
| chr8-112856549-INS-152 | chr8-112856557-INS-122 |
| chr8-112875981-INS-90 | chr8-112875981-INS-93 |
| chr8-112879605-INS-66 | chr8-112879605-INS-60 |
| chr8-112884792-INS-68 | chr8-112884792-INS-68 |
| chr8-112898599-INS-97 | chr8-112898599-INS-98 |
| chr8-112909698-INS-70 | chr8-112909698-INS-70 |
| chr8-112927827-INS-56 | chr8-112927827-INS-62 |
| chr8-112929359-INS-66 | chr8-112929359-INS-68 |
| chr8-112933801-INS-6412 | chr8-112933802-INS-6412 |
| chr8-112936475-INS-53 | chr8-112936475-INS-71 |
| chr8-112942717-INS-72 | chr8-112942717-INS-72 |
| chr8-112943699-INS-6960 | chr8-112943700-INS-6960 |
| chr8-112951622-INS-63 | chr8-112951622-INS-63 |
| chr8-112959802-INS-6646 | chr8-112959803-INS-6646 |
| chr8-112964555-INS-64 | chr8-112964555-INS-56 |
| chr8-112972429-INS-109 | chr8-112972429-INS-109 |
| chr8-112977214-INS-136 | chr8-112977214-INS-110 |
| chr8-113017580-INS-51 | chr8-113017533-INS-89 |
| chr8-113031816-INS-63 | chr8-113031811-INS-60 |
| chr8-113033390-INS-52 | chr8-113033390-INS-52 |
| chr8-113040914-INS-60 | chr8-113040914-INS-60 |
| chr8-113045891-INS-122 | chr8-113045891-INS-122 |
| chr8-113052692-INS-91 | chr8-113052708-INS-57 |
| chr8-113086166-INS-484 | chr8-113086166-INS-487 |
| chr8-113092474-INS-311 | chr8-113092474-INS-311 |
| chr8-113117354-INS-58 | chr8-113117354-INS-58 |
| chr8-113122980-INS-71 | chr8-113122978-INS-113 |
| chr8-113127372-INS-66 | chr8-113127372-INS-66 |
| chr8-113127376-INS-1408 | chr8-113127373-INS-1403 |
| chr8-113174903-INS-3329 | chr8-113174904-INS-3341 |
| chr8-113200417-INS-67 | chr8-113200417-INS-54 |
| chr8-113221556-INS-126 | chr8-113221556-INS-126 |
| chr8-113230091-INS-51 | chr8-113230081-INS-67 |
| chr8-113251980-INS-65 | chr8-113252002-INS-83 |
| chr8-113255357-INS-217 | chr8-113255356-INS-222 |
| chr8-113256781-INS-498 | chr8-113256781-INS-498 |
| chr8-113260682-INS-718 | chr8-113260682-INS-718 |
| chr8-113268611-INS-102 | chr8-113268611-INS-102 |
| chr8-113274031-INS-142 | chr8-113274031-INS-142 |
| chr8-113305321-INS-151 | chr8-113305321-INS-149 |
| chr8-113316061-INS-104 | chr8-113316061-INS-104 |
| chr8-113333125-INS-1180 | chr8-113333126-INS-1180 |
| chr8-113335042-INS-102 | chr8-113335064-INS-76 |
| chr8-113335801-INS-54 | chr8-113335781-INS-58 |
| chr8-113364550-INS-56 | chr8-113364550-INS-52 |
| chr8-113378463-INS-208 | chr8-113378463-INS-208 |
| chr8-113405565-INS-98 | chr8-113405565-INS-97 |
| chr8-113417049-INS-94 | chr8-113417049-INS-94 |
| chr8-113462981-INS-329 | chr8-113462981-INS-329 |
| chr8-113505639-INS-198 | chr8-113505639-INS-198 |
| chr8-113509572-INS-76 | chr8-113509572-INS-84 |
| chr8-113622397-INS-7075 | chr8-113622398-INS-7075 |
| chr8-113654835-INS-60 | chr8-113654834-INS-56 |
| chr8-113657719-INS-7550 | chr8-113657720-INS-7550 |
| chr8-113714064-INS-50 | chr8-113714064-INS-55 |
| chr8-113731265-INS-382 | chr8-113731265-INS-382 |
| chr8-113737618-INS-70 | chr8-113737618-INS-70 |
| chr8-113759990-INS-157 | chr8-113759990-INS-157 |
| chr8-113805484-INS-694 | chr8-113805484-INS-700 |

| | |
|---|---|
| chr8-113806149-INS-71 | chr8-113806149-INS-71 |
| chr8-113807852-INS-68 | chr8-113807852-INS-68 |
| chr8-113821806-INS-68 | chr8-113821804-INS-54 |
| chr8-113824955-INS-6760 | chr8-113824956-INS-6761 |
| chr8-113836088-INS-108 | chr8-113836088-INS-108 |
| chr8-113837772-INS-114 | chr8-113837796-INS-96 |
| chr8-113848155-INS-6305 | chr8-113848156-INS-6302 |
| chr8-113856843-INS-86 | chr8-113856843-INS-86 |
| chr8-113859184-INS-85 | chr8-113859184-INS-81 |
| chr8-113867559-INS-56 | chr8-113867559-INS-56 |
| chr8-113873881-INS-4575 | chr8-113873886-INS-4566 |
| chr8-113876337-INS-6493 | chr8-113876338-INS-6494 |
| chr8-113887210-INS-475 | chr8-113887210-INS-475 |
| chr8-113895565-INS-5981 | chr8-113895566-INS-5979 |
| chr8-113911322-INS-343 | chr8-113911322-INS-343 |
| chr8-113934857-INS-88 | chr8-113934841-INS-70 |
| chr8-113969800-INS-6607 | chr8-113969801-INS-6609 |
| chr8-113978474-INS-735 | chr8-113978474-INS-735 |
| chr8-114005918-INS-51 | chr8-114005918-INS-55 |
| chr8-114006630-INS-482 | chr8-114006630-INS-482 |
| chr8-114078196-INS-77 | chr8-114078196-INS-77 |
| chr8-114089909-INS-164 | chr8-114089909-INS-166 |
| chr8-114091808-INS-67 | chr8-114091833-INS-80 |
| chr8-114106012-INS-206 | chr8-114106012-INS-206 |
| chr8-114135332-INS-936 | chr8-114135332-INS-935 |
| chr8-114140276-INS-833 | chr8-114140277-INS-828 |
| chr8-114158290-INS-72 | chr8-114158290-INS-72 |
| chr8-114162214-INS-74 | chr8-114162214-INS-74 |
| chr8-114217250-INS-2621 | chr8-114217251-INS-2621 |
| chr8-114224603-INS-538 | chr8-114224603-INS-538 |
| chr8-114226162-INS-93 | chr8-114226162-INS-105 |
| chr8-114227634-INS-74 | chr8-114227634-INS-72 |
| chr8-114235448-INS-575 | chr8-114235448-INS-575 |
| chr8-114253722-INS-97 | chr8-114253722-INS-101 |
| chr8-114256279-INS-92 | chr8-114256279-INS-64 |
| chr8-114260324-INS-71 | chr8-114260324-INS-71 |
| chr8-114327950-INS-134 | chr8-114327937-INS-135 |
| chr8-114341397-INS-205 | chr8-114341405-INS-204 |
| chr8-114376371-INS-64 | chr8-114376371-INS-52 |
| chr8-114377345-INS-116 | chr8-114377345-INS-116 |
| chr8-114380805-INS-91 | chr8-114380805-INS-91 |
| chr8-114406724-INS-192 | chr8-114406733-INS-168 |
| chr8-114438552-INS-60 | chr8-114438552-INS-60 |
| chr8-114444236-INS-52 | chr8-114444230-INS-50 |
| chr8-114449679-INS-67 | chr8-114449678-INS-59 |
| chr8-114452785-INS-208 | chr8-114452786-INS-208 |
| chr8-114456866-INS-51 | chr8-114456866-INS-51 |
| chr8-114487074-INS-93 | chr8-114487074-INS-93 |
| chr8-114510048-INS-67 | chr8-114510038-INS-62 |
| chr8-114511717-INS-54 | chr8-114511717-INS-54 |
| chr8-114525927-INS-754 | chr8-114525927-INS-754 |
| chr8-114533391-INS-213 | chr8-114533391-INS-213 |
| chr8-114546012-INS-179 | chr8-114546012-INS-181 |
| chr8-114578909-INS-222 | chr8-114578909-INS-222 |
| chr8-114586170-INS-68 | chr8-114586171-INS-68 |
| chr8-114608005-INS-729 | chr8-114608005-INS-729 |
| chr8-114610708-INS-257 | chr8-114610708-INS-257 |
| chr8-114624284-INS-56 | chr8-114624268-INS-64 |
| chr8-114625301-INS-56 | chr8-114625301-INS-60 |
| chr8-114647844-INS-80 | chr8-114647844-INS-52 |
| chr8-114648384-INS-81 | chr8-114648384-INS-126 |
| chr8-114665114-INS-7068 | chr8-114665115-INS-7070 |
| chr8-114708842-INS-89 | chr8-114708842-INS-99 |
| chr8-114735238-INS-102 | chr8-114735238-INS-122 |
| chr8-114748794-INS-545 | chr8-114748794-INS-545 |
| chr8-114767224-INS-61 | chr8-114767224-INS-59 |
| chr8-114768499-INS-80 | chr8-114768500-INS-90 |
| chr8-114771459-INS-52 | chr8-114771459-INS-52 |
| chr8-114783939-INS-51 | chr8-114783939-INS-51 |
| chr8-114810798-INS-54 | chr8-114810798-INS-58 |
| chr8-114822266-INS-110 | chr8-114822266-INS-114 |
| chr8-114824196-INS-88 | chr8-114824196-INS-88 |
| chr8-114842164-INS-87 | chr8-114842187-INS-58 |
| chr8-114845483-INS-376 | chr8-114845483-INS-376 |
| chr8-114881786-INS-297 | chr8-114881786-INS-299 |
| chr8-114886008-INS-58 | chr8-114886039-INS-72 |
| chr8-114895897-INS-728 | chr8-114895897-INS-714 |

| | |
|---|---|
| chr8-114911186-INS-148 | chr8-114911186-INS-148 |
| chr8-114931693-INS-137 | chr8-114931693-INS-88 |
| chr8-114945613-INS-90 | chr8-114945613-INS-91 |
| chr8-114958665-INS-81 | chr8-114958665-INS-81 |
| chr8-115001979-INS-91 | chr8-115001979-INS-97 |
| chr8-115015682-INS-73 | chr8-115015682-INS-73 |
| chr8-115025963-INS-178 | chr8-115025963-INS-178 |
| chr8-115039005-INS-340 | chr8-115039017-INS-345 |
| chr8-115045693-INS-56 | chr8-115045693-INS-60 |
| chr8-115058553-INS-61 | chr8-115058593-INS-51 |
| chr8-115079645-INS-56 | chr8-115079645-INS-58 |
| chr8-115109873-INS-109 | chr8-115109873-INS-109 |
| chr8-115131653-INS-54 | chr8-115131653-INS-54 |
| chr8-115138665-INS-174 | chr8-115138665-INS-170 |
| chr8-115156019-INS-248 | chr8-115156053-INS-256 |
| chr8-115160281-INS-105 | chr8-115160277-INS-117 |
| chr8-115173331-INS-387 | chr8-115173331-INS-387 |
| chr8-115199305-INS-59 | chr8-115199305-INS-95 |
| chr8-115203084-INS-168 | chr8-115203084-INS-168 |
| chr8-115205865-INS-176 | chr8-115205865-INS-179 |
| chr8-115214394-INS-108 | chr8-115214394-INS-108 |
| chr8-115216527-INS-52 | chr8-115216527-INS-52 |
| chr8-115249355-INS-58 | chr8-115249355-INS-58 |
| chr8-115311208-INS-287 | chr8-115311208-INS-311 |
| chr8-115316546-INS-244 | chr8-115316555-INS-282 |
| chr8-115345552-INS-60 | chr8-115345552-INS-60 |
| chr8-115347008-INS-88 | chr8-115347006-INS-122 |
| chr8-115348221-INS-80 | chr8-115348221-INS-80 |
| chr8-115351563-INS-1587 | chr8-115351564-INS-1587 |
| chr8-115354384-INS-162 | chr8-115354384-INS-162 |
| chr8-115358270-INS-386 | chr8-115358298-INS-386 |
| chr8-115386434-INS-134 | chr8-115386434-INS-112 |
| chr8-115396646-INS-78 | chr8-115396646-INS-78 |
| chr8-115399770-INS-58 | chr8-115399770-INS-54 |
| chr8-115417066-INS-2154 | chr8-115417067-INS-2155 |
| chr8-115419430-INS-219 | chr8-115419429-INS-222 |
| chr8-115430051-INS-458 | chr8-115430051-INS-450 |
| chr8-115441240-INS-846 | chr8-115441240-INS-850 |
| chr8-115444219-INS-125 | chr8-115444195-INS-106 |
| chr8-115451320-INS-191 | chr8-115451320-INS-225 |
| chr8-115474001-INS-72 | chr8-115474001-INS-72 |
| chr8-115488840-INS-70 | chr8-115488840-INS-70 |
| chr8-115527258-INS-87 | chr8-115527258-INS-85 |
| chr8-115547602-INS-52 | chr8-115547602-INS-50 |
| chr8-115578524-INS-127 | chr8-115578525-INS-88 |
| chr8-115596850-INS-65 | chr8-115596892-INS-57 |
| chr8-115602616-INS-73 | chr8-115602616-INS-74 |
| chr8-115615496-INS-60 | chr8-115615492-INS-68 |
| chr8-115673299-INS-168 | chr8-115673299-INS-168 |
| chr8-115689810-INS-5859 | chr8-115689811-INS-5859 |
| chr8-115710286-INS-54 | chr8-115710286-INS-56 |
| chr8-115741170-INS-119 | chr8-115741170-INS-119 |
| chr8-115784404-INS-954 | chr8-115784404-INS-954 |
| chr8-115796476-INS-56 | chr8-115796476-INS-56 |
| chr8-115803119-INS-105 | chr8-115803119-INS-63 |
| chr8-115814723-INS-410 | chr8-115814723-INS-410 |
| chr8-115817975-INS-155 | chr8-115817956-INS-168 |
| chr8-115838828-INS-223 | chr8-115838828-INS-229 |
| chr8-115864560-INS-112 | chr8-115864560-INS-116 |
| chr8-115894266-INS-457 | chr8-115894266-INS-457 |
| chr8-115906967-INS-87 | chr8-115906998-INS-52 |
| chr8-115942969-INS-75 | chr8-115942924-INS-94 |
| chr8-116043605-INS-70 | chr8-116043592-INS-108 |
| chr8-116046979-INS-94 | chr8-116046979-INS-94 |
| chr8-116106437-INS-593 | chr8-116106437-INS-597 |
| chr8-116121531-INS-150 | chr8-116121531-INS-120 |
| chr8-116154135-INS-123 | chr8-116154135-INS-113 |
| chr8-116168613-INS-80 | chr8-116168613-INS-84 |
| chr8-116184523-INS-260 | chr8-116184523-INS-254 |
| chr8-116187722-INS-206 | chr8-116187717-INS-205 |
| chr8-116226588-INS-64 | chr8-116226588-INS-64 |
| chr8-116227870-INS-187 | chr8-116227886-INS-207 |
| chr8-116230933-INS-64 | chr8-116230933-INS-64 |
| chr8-116259744-INS-157 | chr8-116259744-INS-157 |
| chr8-116263252-INS-367 | chr8-116263252-INS-367 |
| chr8-116319918-INS-62 | chr8-116319905-INS-74 |
| chr8-116347643-INS-6440 | chr8-116347644-INS-6440 |

| | |
|---|---|
| chr8-116407028-INS-418 | chr8-116407028-INS-418 |
| chr8-116435536-INS-51 | chr8-116435536-INS-55 |
| chr8-116442532-INS-86 | chr8-116442533-INS-85 |
| chr8-116449489-INS-55 | chr8-116449489-INS-59 |
| chr8-116457708-INS-965 | chr8-116457708-INS-965 |
| chr8-116480394-INS-423 | chr8-116480394-INS-423 |
| chr8-116482159-INS-69 | chr8-116482159-INS-69 |
| chr8-116485118-INS-81 | chr8-116485102-INS-87 |
| chr8-116504458-INS-393 | chr8-116504456-INS-391 |
| chr8-116532649-INS-68 | chr8-116532649-INS-68 |
| chr8-116536036-INS-406 | chr8-116536036-INS-406 |
| chr8-116550285-INS-134 | chr8-116550285-INS-134 |
| chr8-116552572-INS-57 | chr8-116552572-INS-55 |
| chr8-116563319-INS-7158 | chr8-116563320-INS-7158 |
| chr8-116566702-INS-58 | chr8-116566702-INS-60 |
| chr8-116569295-INS-58 | chr8-116569299-INS-60 |
| chr8-116575843-INS-65 | chr8-116575843-INS-65 |
| chr8-116585508-INS-132 | chr8-116585533-INS-133 |
| chr8-116600665-INS-96 | chr8-116600665-INS-133 |
| chr8-116619420-INS-341 | chr8-116619420-INS-342 |
| chr8-116620850-INS-84 | chr8-116620850-INS-84 |
| chr8-116641033-INS-805 | chr8-116641037-INS-807 |
| chr8-116642282-INS-54 | chr8-116642278-INS-53 |
| chr8-116657295-INS-204 | chr8-116657303-INS-169 |
| chr8-116705179-INS-77 | chr8-116705179-INS-69 |
| chr8-116711975-INS-65 | chr8-116711964-INS-66 |
| chr8-116716351-INS-54 | chr8-116716351-INS-54 |
| chr8-116733305-INS-65 | chr8-116733305-INS-66 |
| chr8-116735217-INS-135 | chr8-116735217-INS-130 |
| chr8-116765633-INS-87 | chr8-116765588-INS-126 |
| chr8-116770443-INS-75 | chr8-116770443-INS-67 |
| chr8-116781258-INS-103 | chr8-116781258-INS-111 |
| chr8-116794931-INS-58 | chr8-116794942-INS-58 |
| chr8-116800273-INS-142 | chr8-116800273-INS-142 |
| chr8-116808393-INS-69 | chr8-116808393-INS-69 |
| chr8-116817526-INS-122 | chr8-116817526-INS-122 |
| chr8-116829615-INS-8156 | chr8-116829616-INS-8157 |
| chr8-116843199-INS-77 | chr8-116843199-INS-78 |
| chr8-116861431-INS-89 | chr8-116861416-INS-110 |
| chr8-116883652-INS-56 | chr8-116883652-INS-56 |
| chr8-116900485-INS-222 | chr8-116900485-INS-186 |
| chr8-116906775-INS-451 | chr8-116906775-INS-498 |
| chr8-116910061-INS-54 | chr8-116910061-INS-90 |
| chr8-116948050-INS-66 | chr8-116948050-INS-69 |
| chr8-116970123-INS-110 | chr8-116970123-INS-110 |
| chr8-116976987-INS-60 | chr8-116976984-INS-52 |
| chr8-117028283-INS-131 | chr8-117028283-INS-127 |
| chr8-117087407-INS-52 | chr8-117087407-INS-54 |
| chr8-117104647-INS-77 | chr8-117104647-INS-107 |
| chr8-117118100-INS-156 | chr8-117118100-INS-156 |
| chr8-117124535-INS-57 | chr8-117124529-INS-60 |
| chr8-117161525-INS-100 | chr8-117161525-INS-100 |
| chr8-117166607-INS-58 | chr8-117166607-INS-58 |
| chr8-117167463-INS-307 | chr8-117167463-INS-307 |
| chr8-117176716-INS-1517 | chr8-117176717-INS-1517 |
| chr8-117205664-INS-6475 | chr8-117205665-INS-6475 |
| chr8-117229868-INS-6467 | chr8-117229869-INS-6468 |
| chr8-117241053-INS-70 | chr8-117241053-INS-70 |
| chr8-117251548-INS-52 | chr8-117251548-INS-52 |
| chr8-117275745-INS-83 | chr8-117275745-INS-103 |
| chr8-117278638-INS-100 | chr8-117278626-INS-148 |
| chr8-117317413-INS-210 | chr8-117317413-INS-232 |
| chr8-117324637-INS-51 | chr8-117324638-INS-55 |
| chr8-117340743-INS-381 | chr8-117340743-INS-381 |
| chr8-117378438-INS-80 | chr8-117378438-INS-81 |
| chr8-117394806-INS-219 | chr8-117394806-INS-220 |
| chr8-117429490-INS-474 | chr8-117429495-INS-467 |
| chr8-117433874-INS-170 | chr8-117433874-INS-167 |
| chr8-117439545-INS-129 | chr8-117439545-INS-137 |
| chr8-117449227-INS-106 | chr8-117449213-INS-112 |
| chr8-117509432-INS-590 | chr8-117509432-INS-590 |
| chr8-117516020-INS-433 | chr8-117516020-INS-433 |
| chr8-117523194-INS-2225 | chr8-117523208-INS-2226 |
| chr8-117523921-INS-207 | chr8-117523921-INS-207 |
| chr8-117534908-INS-63 | chr8-117534908-INS-63 |
| chr8-117537928-INS-52 | chr8-117537928-INS-50 |
| chr8-117549589-INS-6705 | chr8-117549590-INS-6698 |

| | |
|---|---|
| chr8-117554139-INS-394 | chr8-117554142-INS-396 |
| chr8-117561542-INS-408 | chr8-117561542-INS-408 |
| chr8-117573730-INS-52 | chr8-117573730-INS-52 |
| chr8-117576649-INS-56 | chr8-117576649-INS-56 |
| chr8-117582442-INS-505 | chr8-117582442-INS-504 |
| chr8-117590738-INS-53 | chr8-117590736-INS-55 |
| chr8-117595130-INS-74 | chr8-117595131-INS-99 |
| chr8-117595621-INS-414 | chr8-117595621-INS-414 |
| chr8-117612992-INS-100 | chr8-117612992-INS-100 |
| chr8-117621645-INS-176 | chr8-117621647-INS-224 |
| chr8-117630593-INS-104 | chr8-117630593-INS-128 |
| chr8-117639625-INS-174 | chr8-117639625-INS-174 |
| chr8-117653467-INS-147 | chr8-117653467-INS-143 |
| chr8-117655591-INS-245 | chr8-117655591-INS-245 |
| chr8-117704275-INS-186 | chr8-117704275-INS-186 |
| chr8-117708133-INS-310 | chr8-117708133-INS-310 |
| chr8-117710178-INS-345 | chr8-117710178-INS-345 |
| chr8-117720809-INS-168 | chr8-117720809-INS-170 |
| chr8-117735723-INS-71 | chr8-117735723-INS-69 |
| chr8-117751765-INS-73 | chr8-117751761-INS-66 |
| chr8-117773218-INS-213 | chr8-117773218-INS-213 |
| chr8-117804962-INS-71 | chr8-117804962-INS-71 |
| chr8-117813074-INS-214 | chr8-117813074-INS-214 |
| chr8-117857581-INS-6442 | chr8-117857582-INS-6442 |
| chr8-117858034-INS-109 | chr8-117858034-INS-109 |
| chr8-117860182-INS-56 | chr8-117860231-INS-56 |
| chr8-117862065-INS-99 | chr8-117862065-INS-87 |
| chr8-117863087-INS-67 | chr8-117863072-INS-65 |
| chr8-117865666-INS-468 | chr8-117865666-INS-468 |
| chr8-117873505-INS-115 | chr8-117873505-INS-114 |
| chr8-117921111-INS-64 | chr8-117921111-INS-64 |
| chr8-117935940-INS-221 | chr8-117935940-INS-221 |
| chr8-117954889-INS-978 | chr8-117954889-INS-978 |
| chr8-117956699-INS-63 | chr8-117956719-INS-64 |
| chr8-117957853-INS-54 | chr8-117957853-INS-54 |
| chr8-118018571-INS-92 | chr8-118018571-INS-91 |
| chr8-118029782-INS-96 | chr8-118029778-INS-59 |
| chr8-118064888-INS-443 | chr8-118064888-INS-410 |
| chr8-118084458-INS-91 | chr8-118084459-INS-85 |
| chr8-118135767-INS-129 | chr8-118135787-INS-123 |
| chr8-118165499-INS-201 | chr8-118165499-INS-201 |
| chr8-118171126-INS-60 | chr8-118171126-INS-62 |
| chr8-118196143-INS-95 | chr8-118196143-INS-124 |
| chr8-118210877-INS-197 | chr8-118210877-INS-197 |
| chr8-118234148-INS-211 | chr8-118234148-INS-212 |
| chr8-118263141-INS-158 | chr8-118263141-INS-158 |
| chr8-118284463-INS-260 | chr8-118284463-INS-260 |
| chr8-118294772-INS-139 | chr8-118294772-INS-172 |
| chr8-118296782-INS-55 | chr8-118296773-INS-62 |
| chr8-118299094-INS-222 | chr8-118299094-INS-208 |
| chr8-118315416-INS-105 | chr8-118315416-INS-105 |
| chr8-118336701-INS-54 | chr8-118336678-INS-61 |
| chr8-118344879-INS-54 | chr8-118344879-INS-56 |
| chr8-118356693-INS-830 | chr8-118356693-INS-822 |
| chr8-118362487-INS-161 | chr8-118362487-INS-153 |
| chr8-118370173-INS-366 | chr8-118370173-INS-363 |
| chr8-118414182-INS-66 | chr8-118414171-INS-52 |
| chr8-118451726-INS-69 | chr8-118451775-INS-52 |
| chr8-118464007-INS-141 | chr8-118464007-INS-143 |
| chr8-118526581-INS-1424 | chr8-118526582-INS-1424 |
| chr8-118578590-INS-55 | chr8-118578590-INS-55 |
| chr8-118587181-INS-77 | chr8-118587181-INS-77 |
| chr8-118627178-INS-167 | chr8-118627189-INS-143 |
| chr8-118650379-INS-444 | chr8-118650379-INS-445 |
| chr8-118665915-INS-82 | chr8-118665915-INS-82 |
| chr8-118679193-INS-226 | chr8-118679193-INS-221 |
| chr8-118685635-INS-161 | chr8-118685635-INS-177 |
| chr8-118725121-INS-94 | chr8-118725121-INS-100 |
| chr8-118746137-INS-91 | chr8-118746137-INS-91 |
| chr8-118782381-INS-85 | chr8-118782359-INS-57 |
| chr8-118784684-INS-138 | chr8-118784684-INS-134 |
| chr8-118785951-INS-165 | chr8-118785940-INS-203 |
| chr8-118797111-INS-64 | chr8-118797111-INS-64 |
| chr8-118823677-INS-132 | chr8-118823720-INS-133 |
| chr8-118826828-INS-66 | chr8-118826828-INS-54 |
| chr8-118858826-INS-72 | chr8-118858837-INS-60 |
| chr8-118870787-INS-66 | chr8-118870776-INS-114 |

| | |
|---|---|
| chr8-118919316-INS-118 | chr8-118919313-INS-111 |
| chr8-118923416-INS-54 | chr8-118923416-INS-57 |
| chr8-118925236-INS-63 | chr8-118925236-INS-63 |
| chr8-118935436-INS-96 | chr8-118935436-INS-96 |
| chr8-118956465-INS-60 | chr8-118956465-INS-64 |
| chr8-118967264-INS-52 | chr8-118967264-INS-52 |
| chr8-118983856-INS-359 | chr8-118983864-INS-349 |
| chr8-118990721-INS-52 | chr8-118990721-INS-60 |
| chr8-118998546-INS-68 | chr8-118998546-INS-82 |
| chr8-119027493-INS-96 | chr8-119027493-INS-96 |
| chr8-119061600-INS-66 | chr8-119061600-INS-54 |
| chr8-119076169-INS-120 | chr8-119076183-INS-110 |
| chr8-119077038-INS-56 | chr8-119077038-INS-56 |
| chr8-119089073-INS-53 | chr8-119089073-INS-81 |
| chr8-119146289-INS-99 | chr8-119146289-INS-99 |
| chr8-119196730-INS-1029 | chr8-119196722-INS-1025 |
| chr8-119199084-INS-72 | chr8-119199084-INS-72 |
| chr8-119256659-INS-77 | chr8-119256660-INS-50 |
| chr8-119257697-INS-96 | chr8-119257705-INS-131 |
| chr8-119276229-INS-120 | chr8-119276229-INS-120 |
| chr8-119298296-INS-4063 | chr8-119298297-INS-4108 |
| chr8-119318988-INS-67 | chr8-119318988-INS-75 |
| chr8-119346402-INS-59 | chr8-119346418-INS-50 |
| chr8-119350477-INS-876 | chr8-119350477-INS-877 |
| chr8-119395298-INS-66 | chr8-119395298-INS-66 |
| chr8-119418243-INS-65 | chr8-119418243-INS-65 |
| chr8-119432220-INS-64 | chr8-119432220-INS-64 |
| chr8-119463165-INS-176 | chr8-119463165-INS-176 |
| chr8-119495971-INS-261 | chr8-119495971-INS-261 |
| chr8-119511652-INS-68 | chr8-119511652-INS-114 |
| chr8-119553890-INS-78 | chr8-119553885-INS-84 |
| chr8-119565578-INS-90 | chr8-119565578-INS-90 |
| chr8-119582354-INS-440 | chr8-119582354-INS-440 |
| chr8-119599835-INS-104 | chr8-119599835-INS-104 |
| chr8-119623747-INS-52 | chr8-119623749-INS-97 |
| chr8-119635915-INS-106 | chr8-119635915-INS-106 |
| chr8-119652036-INS-114 | chr8-119652036-INS-114 |
| chr8-119664359-INS-580 | chr8-119664359-INS-580 |
| chr8-119667731-INS-83 | chr8-119667731-INS-83 |
| chr8-119690895-INS-79 | chr8-119690883-INS-74 |
| chr8-119710267-INS-76 | chr8-119710267-INS-56 |
| chr8-119753021-INS-288 | chr8-119753021-INS-288 |
| chr8-119763548-INS-90 | chr8-119763540-INS-98 |
| chr8-119789831-INS-377 | chr8-119789831-INS-327 |
| chr8-119795283-INS-67 | chr8-119795283-INS-67 |
| chr8-119840213-INS-51 | chr8-119840213-INS-57 |
| chr8-119854140-INS-132 | chr8-119854142-INS-102 |
| chr8-119859687-INS-953 | chr8-119859687-INS-953 |
| chr8-119891254-INS-488 | chr8-119891254-INS-513 |
| chr8-119900528-INS-64 | chr8-119900561-INS-64 |
| chr8-119906192-INS-837 | chr8-119906191-INS-818 |
| chr8-119919577-INS-136 | chr8-119919577-INS-136 |
| chr8-119921211-INS-389 | chr8-119921211-INS-389 |
| chr8-119921793-INS-116 | chr8-119921790-INS-117 |
| chr8-119923158-INS-63 | chr8-119923158-INS-63 |
| chr8-119927669-INS-55 | chr8-119927669-INS-55 |
| chr8-119991189-INS-73 | chr8-119991175-INS-70 |
| chr8-120026316-INS-82 | chr8-120026316-INS-82 |
| chr8-120042065-INS-253 | chr8-120042053-INS-269 |
| chr8-120117875-INS-192 | chr8-120117875-INS-191 |
| chr8-120119470-INS-64 | chr8-120119498-INS-60 |
| chr8-120129100-INS-5313 | chr8-120129101-INS-5313 |
| chr8-120129484-INS-125 | chr8-120129484-INS-118 |
| chr8-120131536-INS-51 | chr8-120131536-INS-51 |
| chr8-120199774-INS-117 | chr8-120199774-INS-156 |
| chr8-120203068-INS-118 | chr8-120203065-INS-105 |
| chr8-120208470-INS-212 | chr8-120208474-INS-208 |
| chr8-120217082-INS-173 | chr8-120217057-INS-209 |
| chr8-120249994-INS-204 | chr8-120249995-INS-204 |
| chr8-120254134-INS-195 | chr8-120254134-INS-195 |
| chr8-120283025-INS-213 | chr8-120283025-INS-213 |
| chr8-120301190-INS-211 | chr8-120301190-INS-211 |
| chr8-120336796-INS-298 | chr8-120336796-INS-298 |
| chr8-120341115-INS-60 | chr8-120341115-INS-60 |
| chr8-120358745-INS-56 | chr8-120358745-INS-56 |
| chr8-120379410-INS-73 | chr8-120379438-INS-91 |
| chr8-120381696-INS-81 | chr8-120381696-INS-81 |

| | |
|---|---|
| chr8-120388626-INS-6409 | chr8-120388627-INS-6409 |
| chr8-120398375-INS-211 | chr8-120398375-INS-211 |
| chr8-120410215-INS-265 | chr8-120410243-INS-225 |
| chr8-120414764-INS-81 | chr8-120414764-INS-71 |
| chr8-120430586-INS-83 | chr8-120430589-INS-57 |
| chr8-120436047-INS-60 | chr8-120436047-INS-62 |
| chr8-120449601-INS-52 | chr8-120449601-INS-52 |
| chr8-120453881-INS-138 | chr8-120453881-INS-101 |
| chr8-120491199-INS-78 | chr8-120491198-INS-72 |
| chr8-120511874-INS-80 | chr8-120511870-INS-73 |
| chr8-120535836-INS-202 | chr8-120535836-INS-202 |
| chr8-120542691-INS-55 | chr8-120542691-INS-69 |
| chr8-120550166-INS-275 | chr8-120550166-INS-274 |
| chr8-120553510-INS-376 | chr8-120553510-INS-376 |
| chr8-120586780-INS-63 | chr8-120586814-INS-66 |
| chr8-120595691-INS-192 | chr8-120595691-INS-192 |
| chr8-120608515-INS-986 | chr8-120608529-INS-986 |
| chr8-120611662-INS-66 | chr8-120611666-INS-65 |
| chr8-120626249-INS-203 | chr8-120626249-INS-205 |
| chr8-120644305-INS-72 | chr8-120644306-INS-72 |
| chr8-120652189-INS-140 | chr8-120652189-INS-140 |
| chr8-120663052-INS-63 | chr8-120663052-INS-63 |
| chr8-120666061-INS-137 | chr8-120666061-INS-137 |
| chr8-120676911-INS-904 | chr8-120676911-INS-901 |
| chr8-120710578-INS-206 | chr8-120710578-INS-207 |
| chr8-120714085-INS-88 | chr8-120714085-INS-76 |
| chr8-120756192-INS-104 | chr8-120756192-INS-104 |
| chr8-120782004-INS-60 | chr8-120782004-INS-60 |
| chr8-120835206-INS-430 | chr8-120835206-INS-431 |
| chr8-120866831-INS-58 | chr8-120866831-INS-58 |
| chr8-120878832-INS-135 | chr8-120878832-INS-135 |
| chr8-120893979-INS-88 | chr8-120893979-INS-88 |
| chr8-120900456-INS-425 | chr8-120900456-INS-427 |
| chr8-120939203-INS-197 | chr8-120939203-INS-197 |
| chr8-120940600-INS-112 | chr8-120940600-INS-112 |
| chr8-120950559-INS-79 | chr8-120950559-INS-79 |
| chr8-120972980-INS-912 | chr8-120972980-INS-912 |
| chr8-120996323-INS-190 | chr8-120996323-INS-189 |
| chr8-120997936-INS-214 | chr8-120997936-INS-211 |
| chr8-120999007-INS-70 | chr8-120999007-INS-58 |
| chr8-121023210-INS-146 | chr8-121023210-INS-146 |
| chr8-121069068-INS-400 | chr8-121069068-INS-400 |
| chr8-121091691-INS-387 | chr8-121091691-INS-386 |
| chr8-121116171-INS-555 | chr8-121116171-INS-555 |
| chr8-121121818-INS-216 | chr8-121121818-INS-216 |
| chr8-121127376-INS-195 | chr8-121127376-INS-197 |
| chr8-121170233-INS-96 | chr8-121170233-INS-96 |
| chr8-121184140-INS-109 | chr8-121184143-INS-80 |
| chr8-121194385-INS-85 | chr8-121194385-INS-85 |
| chr8-121195972-INS-136 | chr8-121195934-INS-130 |
| chr8-121250270-INS-216 | chr8-121250270-INS-214 |
| chr8-121314319-INS-191 | chr8-121314319-INS-192 |
| chr8-121320616-INS-206 | chr8-121320616-INS-185 |
| chr8-121350075-INS-188 | chr8-121350075-INS-188 |
| chr8-121352785-INS-202 | chr8-121352785-INS-202 |
| chr8-121365184-INS-208 | chr8-121365184-INS-205 |
| chr8-121376272-INS-141 | chr8-121376272-INS-141 |
| chr8-121416198-INS-458 | chr8-121416198-INS-458 |
| chr8-121427984-INS-410 | chr8-121427984-INS-410 |
| chr8-121449037-INS-396 | chr8-121449035-INS-406 |
| chr8-121490696-INS-152 | chr8-121490662-INS-164 |
| chr8-121496517-INS-57 | chr8-121496517-INS-50 |
| chr8-121505505-INS-199 | chr8-121505505-INS-200 |
| chr8-121543364-INS-539 | chr8-121543364-INS-539 |
| chr8-121562228-INS-57 | chr8-121562228-INS-57 |
| chr8-121564727-INS-140 | chr8-121564716-INS-136 |
| chr8-121571343-INS-59 | chr8-121571340-INS-67 |
| chr8-121589836-INS-170 | chr8-121589836-INS-170 |
| chr8-121601576-INS-76 | chr8-121601576-INS-76 |
| chr8-121608073-INS-268 | chr8-121608073-INS-272 |
| chr8-121643332-INS-75 | chr8-121643352-INS-87 |
| chr8-121650503-INS-160 | chr8-121650511-INS-140 |
| chr8-121657280-INS-212 | chr8-121657280-INS-212 |
| chr8-121659628-INS-68 | chr8-121659628-INS-68 |
| chr8-121661475-INS-168 | chr8-121661475-INS-168 |
| chr8-121697735-INS-511 | chr8-121697735-INS-510 |
| chr8-121700670-INS-149 | chr8-121700628-INS-149 |

| | |
|---|---|
| chr8-121707170-INS-156 | chr8-121707194-INS-106 |
| chr8-121716373-INS-255 | chr8-121716373-INS-255 |
| chr8-121741102-INS-62 | chr8-121741102-INS-50 |
| chr8-121771515-INS-610 | chr8-121771515-INS-600 |
| chr8-121824908-INS-70 | chr8-121824908-INS-70 |
| chr8-121835709-INS-74 | chr8-121835709-INS-72 |
| chr8-121865963-INS-99 | chr8-121865966-INS-57 |
| chr8-121878539-INS-152 | chr8-121878540-INS-106 |
| chr8-121902656-INS-328 | chr8-121902657-INS-336 |
| chr8-121904308-INS-65 | chr8-121904308-INS-65 |
| chr8-121931530-INS-2516 | chr8-121931531-INS-2516 |
| chr8-121941078-INS-4390 | chr8-121941079-INS-4390 |
| chr8-121942798-INS-110 | chr8-121942798-INS-110 |
| chr8-121948634-INS-5096 | chr8-121948635-INS-5096 |
| chr8-121958083-INS-439 | chr8-121958083-INS-439 |
| chr8-121971827-INS-468 | chr8-121971827-INS-476 |
| chr8-121976405-INS-117 | chr8-121976405-INS-117 |
| chr8-121977092-INS-81 | chr8-121977077-INS-113 |
| chr8-121994593-INS-143 | chr8-121994567-INS-180 |
| chr8-121999728-INS-120 | chr8-121999728-INS-146 |
| chr8-122045792-INS-99 | chr8-122045792-INS-113 |
| chr8-122064308-INS-220 | chr8-122064308-INS-220 |
| chr8-122090038-INS-117 | chr8-122090038-INS-117 |
| chr8-122092666-INS-69 | chr8-122092666-INS-69 |
| chr8-122127918-INS-396 | chr8-122127918-INS-396 |
| chr8-122147027-INS-69 | chr8-122147027-INS-69 |
| chr8-122149124-INS-81 | chr8-122149105-INS-82 |
| chr8-122169594-INS-98 | chr8-122169594-INS-98 |
| chr8-122193000-INS-73 | chr8-122193000-INS-73 |
| chr8-122200403-INS-103 | chr8-122200403-INS-79 |
| chr8-122205515-INS-56 | chr8-122205515-INS-56 |
| chr8-122206320-INS-78 | chr8-122206276-INS-126 |
| chr8-122212006-INS-469 | chr8-122212006-INS-472 |
| chr8-122235120-INS-69 | chr8-122235120-INS-72 |
| chr8-122247534-INS-421 | chr8-122247535-INS-421 |
| chr8-122256831-INS-173 | chr8-122256827-INS-188 |
| chr8-122295033-INS-57 | chr8-122295033-INS-57 |
| chr8-122300722-INS-98 | chr8-122300702-INS-114 |
| chr8-122318669-INS-332 | chr8-122318669-INS-332 |
| chr8-122339397-INS-204 | chr8-122339397-INS-204 |
| chr8-122386631-INS-592 | chr8-122386631-INS-594 |
| chr8-122413585-INS-160 | chr8-122413585-INS-154 |
| chr8-122417761-INS-89 | chr8-122417761-INS-89 |
| chr8-122437646-INS-208 | chr8-122437646-INS-208 |
| chr8-122441719-INS-144 | chr8-122441715-INS-113 |
| chr8-122447394-INS-165 | chr8-122447394-INS-181 |
| chr8-122461523-INS-172 | chr8-122461523-INS-172 |
| chr8-122495171-INS-117 | chr8-122495171-INS-117 |
| chr8-122533206-INS-372 | chr8-122533206-INS-372 |
| chr8-122537955-INS-214 | chr8-122537955-INS-214 |
| chr8-122547398-INS-139 | chr8-122547398-INS-139 |
| chr8-122585730-INS-205 | chr8-122585730-INS-205 |
| chr8-122594026-INS-192 | chr8-122594026-INS-187 |
| chr8-122596179-INS-94 | chr8-122596179-INS-94 |
| chr8-122605323-INS-150 | chr8-122605323-INS-150 |
| chr8-122624157-INS-84 | chr8-122624157-INS-84 |
| chr8-122629046-INS-170 | chr8-122629046-INS-136 |
| chr8-122630516-INS-96 | chr8-122630516-INS-96 |
| chr8-122647494-INS-60 | chr8-122647495-INS-60 |
| chr8-122648539-INS-498 | chr8-122648539-INS-498 |
| chr8-122651409-INS-464 | chr8-122651409-INS-464 |
| chr8-122655619-INS-108 | chr8-122655619-INS-108 |
| chr8-122656518-INS-179 | chr8-122656518-INS-179 |
| chr8-122713900-INS-328 | chr8-122713900-INS-328 |
| chr8-122731549-INS-320 | chr8-122731549-INS-320 |
| chr8-122732451-INS-88 | chr8-122732456-INS-101 |
| chr8-122783808-INS-7156 | chr8-122783809-INS-7156 |
| chr8-122786690-INS-77 | chr8-122786690-INS-77 |
| chr8-122819943-INS-271 | chr8-122819943-INS-261 |
| chr8-122854955-INS-70 | chr8-122854955-INS-70 |
| chr8-122857493-INS-154 | chr8-122857493-INS-154 |
| chr8-122865799-INS-284 | chr8-122865829-INS-304 |
| chr8-122889440-INS-1876 | chr8-122889441-INS-1876 |
| chr8-122914671-INS-206 | chr8-122914655-INS-178 |
| chr8-122936683-INS-60 | chr8-122936683-INS-60 |
| chr8-122977804-INS-54 | chr8-122977788-INS-72 |
| chr8-123038781-INS-59 | chr8-123038781-INS-59 |

| | |
|---|---|
| chr8-123044155-INS-64 | chr8-123044156-INS-53 |
| chr8-123045800-INS-223 | chr8-123045800-INS-223 |
| chr8-123052428-INS-58 | chr8-123052428-INS-90 |
| chr8-123074114-INS-335 | chr8-123074114-INS-335 |
| chr8-123084862-INS-98 | chr8-123084882-INS-98 |
| chr8-123097901-INS-52 | chr8-123097901-INS-54 |
| chr8-123109830-INS-208 | chr8-123109830-INS-208 |
| chr8-123123664-INS-136 | chr8-123123664-INS-136 |
| chr8-123173309-INS-265 | chr8-123173309-INS-266 |
| chr8-123198398-INS-99 | chr8-123198398-INS-99 |
| chr8-123209338-INS-64 | chr8-123209338-INS-64 |
| chr8-123233929-INS-199 | chr8-123233929-INS-199 |
| chr8-123235185-INS-60 | chr8-123235185-INS-72 |
| chr8-123255679-INS-293 | chr8-123255679-INS-292 |
| chr8-123281206-INS-232 | chr8-123281206-INS-232 |
| chr8-123289535-INS-174 | chr8-123289535-INS-174 |
| chr8-123290314-INS-194 | chr8-123290317-INS-204 |
| chr8-123300177-INS-185 | chr8-123300177-INS-173 |
| chr8-123320991-INS-352 | chr8-123320991-INS-352 |
| chr8-123354402-INS-68 | chr8-123354402-INS-68 |
| chr8-123454987-INS-126 | chr8-123454987-INS-125 |
| chr8-123457071-INS-104 | chr8-123457071-INS-104 |
| chr8-123458000-INS-80 | chr8-123458000-INS-82 |
| chr8-123463545-INS-140 | chr8-123463537-INS-150 |
| chr8-123484039-INS-74 | chr8-123484031-INS-111 |
| chr8-123499888-INS-110 | chr8-123499888-INS-110 |
| chr8-123508860-INS-62 | chr8-123508837-INS-81 |
| chr8-123526757-INS-7285 | chr8-123526758-INS-7285 |
| chr8-123556999-INS-56 | chr8-123557000-INS-50 |
| chr8-123563526-INS-72 | chr8-123563526-INS-72 |
| chr8-123586705-INS-160 | chr8-123586705-INS-159 |
| chr8-123589410-INS-130 | chr8-123589410-INS-130 |
| chr8-123601570-INS-59 | chr8-123601570-INS-59 |
| chr8-123605460-INS-52 | chr8-123605460-INS-52 |
| chr8-123611336-INS-52 | chr8-123611336-INS-54 |
| chr8-123639482-INS-252 | chr8-123639482-INS-254 |
| chr8-123657448-INS-185 | chr8-123657448-INS-185 |
| chr8-123662802-INS-251 | chr8-123662802-INS-252 |
| chr8-123668385-INS-54 | chr8-123668385-INS-54 |
| chr8-123685035-INS-52 | chr8-123685035-INS-52 |
| chr8-123690468-INS-60 | chr8-123690468-INS-62 |
| chr8-123727822-INS-136 | chr8-123727843-INS-94 |
| chr8-123739434-INS-54 | chr8-123739434-INS-54 |
| chr8-123744336-INS-471 | chr8-123744336-INS-472 |
| chr8-123751824-INS-87 | chr8-123751842-INS-84 |
| chr8-123756322-INS-65 | chr8-123756322-INS-65 |
| chr8-123760049-INS-59 | chr8-123760049-INS-50 |
| chr8-123765690-INS-175 | chr8-123765690-INS-175 |
| chr8-123771128-INS-204 | chr8-123771128-INS-202 |
| chr8-123772906-INS-220 | chr8-123772906-INS-220 |
| chr8-123773449-INS-253 | chr8-123773449-INS-255 |
| chr8-123779222-INS-73 | chr8-123779223-INS-73 |
| chr8-123785345-INS-180 | chr8-123785345-INS-179 |
| chr8-123786773-INS-211 | chr8-123786773-INS-200 |
| chr8-123788525-INS-144 | chr8-123788525-INS-144 |
| chr8-123791023-INS-83 | chr8-123791023-INS-83 |
| chr8-123797096-INS-260 | chr8-123797096-INS-260 |
| chr8-123820242-INS-353 | chr8-123820242-INS-352 |
| chr8-123827035-INS-149 | chr8-123827035-INS-149 |
| chr8-123833337-INS-236 | chr8-123833337-INS-236 |
| chr8-123842297-INS-61 | chr8-123842330-INS-59 |
| chr8-123859144-INS-212 | chr8-123859144-INS-212 |
| chr8-123866983-INS-4827 | chr8-123866984-INS-4827 |
| chr8-123872746-INS-206 | chr8-123872746-INS-206 |
| chr8-123874372-INS-172 | chr8-123874372-INS-174 |
| chr8-123877057-INS-181 | chr8-123877057-INS-181 |
| chr8-123879075-INS-121 | chr8-123879075-INS-111 |
| chr8-123882265-INS-1865 | chr8-123882266-INS-1865 |
| chr8-123887396-INS-60 | chr8-123887396-INS-65 |
| chr8-123903022-INS-203 | chr8-123903023-INS-203 |
| chr8-123922875-INS-189 | chr8-123922875-INS-189 |
| chr8-123945489-INS-209 | chr8-123945489-INS-209 |
| chr8-123980400-INS-68 | chr8-123980400-INS-58 |
| chr8-123988424-INS-125 | chr8-123988424-INS-125 |
| chr8-124002924-INS-134 | chr8-124002924-INS-134 |
| chr8-124006362-INS-210 | chr8-124006362-INS-209 |
| chr8-124008609-INS-216 | chr8-124008609-INS-216 |

| | |
|---|---|
| chr8-124010486-INS-221 | chr8-124010486-INS-222 |
| chr8-124015979-INS-144 | chr8-124015979-INS-144 |
| chr8-124017976-INS-218 | chr8-124017976-INS-219 |
| chr8-124019425-INS-155 | chr8-124019425-INS-157 |
| chr8-124022866-INS-6442 | chr8-124022867-INS-6442 |
| chr8-124033296-INS-258 | chr8-124033296-INS-215 |
| chr8-124041116-INS-209 | chr8-124041116-INS-209 |
| chr8-124043437-INS-92 | chr8-124043437-INS-114 |
| chr8-124046935-INS-92 | chr8-124046935-INS-92 |
| chr8-124052934-INS-635 | chr8-124052934-INS-656 |
| chr8-124067685-INS-172 | chr8-124067685-INS-172 |
| chr8-124076858-INS-205 | chr8-124076858-INS-205 |
| chr8-124083958-INS-73 | chr8-124083958-INS-73 |
| chr8-124086759-INS-66 | chr8-124086758-INS-66 |
| chr8-124087976-INS-53 | chr8-124087976-INS-53 |
| chr8-124095233-INS-54 | chr8-124095233-INS-78 |
| chr8-124098112-INS-190 | chr8-124098112-INS-190 |
| chr8-124105458-INS-954 | chr8-124105458-INS-954 |
| chr8-124113166-INS-248 | chr8-124113166-INS-249 |
| chr8-124119080-INS-215 | chr8-124119080-INS-215 |
| chr8-124142243-INS-203 | chr8-124142243-INS-203 |
| chr8-124146412-INS-203 | chr8-124146412-INS-205 |
| chr8-124164739-INS-224 | chr8-124164739-INS-224 |
| chr8-124165817-INS-96 | chr8-124165814-INS-52 |
| chr8-124178010-INS-208 | chr8-124178010-INS-208 |
| chr8-124237519-INS-127 | chr8-124237519-INS-127 |
| chr8-124245068-INS-109 | chr8-124245058-INS-77 |
| chr8-124253047-INS-145 | chr8-124253057-INS-185 |
| chr8-124255700-INS-209 | chr8-124255700-INS-216 |
| chr8-124367201-INS-988 | chr8-124367201-INS-988 |
| chr8-124408187-INS-646 | chr8-124408187-INS-643 |
| chr8-124408984-INS-81 | chr8-124408989-INS-62 |
| chr8-124413208-INS-66 | chr8-124413208-INS-66 |
| chr8-124422901-INS-88 | chr8-124422901-INS-88 |
| chr8-124424918-INS-80 | chr8-124424918-INS-80 |
| chr8-124429091-INS-599 | chr8-124429091-INS-599 |
| chr8-124432037-INS-52 | chr8-124432037-INS-52 |
| chr8-124483782-INS-68 | chr8-124483782-INS-70 |
| chr8-124497078-INS-98 | chr8-124497078-INS-98 |
| chr8-124501428-INS-66 | chr8-124501405-INS-69 |
| chr8-124504930-INS-136 | chr8-124504930-INS-136 |
| chr8-124510052-INS-111 | chr8-124510052-INS-113 |
| chr8-124518956-INS-110 | chr8-124518956-INS-110 |
| chr8-124519947-INS-95 | chr8-124519947-INS-95 |
| chr8-124522238-INS-212 | chr8-124522238-INS-212 |
| chr8-124526629-INS-188 | chr8-124526601-INS-210 |
| chr8-124543342-INS-64 | chr8-124543340-INS-66 |
| chr8-124548250-INS-117 | chr8-124548250-INS-117 |
| chr8-124593462-INS-405 | chr8-124593451-INS-371 |
| chr8-124598220-INS-192 | chr8-124598220-INS-192 |
| chr8-124600575-INS-80 | chr8-124600606-INS-67 |
| chr8-124627835-INS-217 | chr8-124627835-INS-217 |
| chr8-124628788-INS-52 | chr8-124628834-INS-52 |
| chr8-124631546-INS-195 | chr8-124631546-INS-195 |
| chr8-124632446-INS-300 | chr8-124632446-INS-300 |
| chr8-124641598-INS-95 | chr8-124641598-INS-95 |
| chr8-124646196-INS-225 | chr8-124646196-INS-225 |
| chr8-124651916-INS-212 | chr8-124651916-INS-212 |
| chr8-124689326-INS-72 | chr8-124689326-INS-72 |
| chr8-124690773-INS-209 | chr8-124690773-INS-208 |
| chr8-124693887-INS-72 | chr8-124693887-INS-68 |
| chr8-124716958-INS-245 | chr8-124716959-INS-200 |
| chr8-124719540-INS-52 | chr8-124719540-INS-52 |
| chr8-124722585-INS-90 | chr8-124722585-INS-84 |
| chr8-124731569-INS-112 | chr8-124731569-INS-112 |
| chr8-124740389-INS-326 | chr8-124740385-INS-325 |
| chr8-124748609-INS-54 | chr8-124748609-INS-51 |
| chr8-124751672-INS-201 | chr8-124751672-INS-201 |
| chr8-124771436-INS-62 | chr8-124771436-INS-62 |
| chr8-124791837-INS-58 | chr8-124791847-INS-61 |
| chr8-124794964-INS-196 | chr8-124794964-INS-196 |
| chr8-124796128-INS-63 | chr8-124796128-INS-63 |
| chr8-124812687-INS-64 | chr8-124812687-INS-68 |
| chr8-124880038-INS-325 | chr8-124880038-INS-325 |
| chr8-124881162-INS-57 | chr8-124881162-INS-60 |
| chr8-124893010-INS-80 | chr8-124893010-INS-80 |
| chr8-124926557-INS-126 | chr8-124926557-INS-126 |

| | |
|---|---|
| chr8-124952177-INS-104 | chr8-124952177-INS-104 |
| chr8-124957198-INS-85 | chr8-124957198-INS-85 |
| chr8-124967112-INS-87 | chr8-124967112-INS-87 |
| chr8-124986623-INS-80 | chr8-124986626-INS-68 |
| chr8-124989225-INS-525 | chr8-124989225-INS-525 |
| chr8-124993467-INS-55 | chr8-124993467-INS-91 |
| chr8-125041333-INS-238 | chr8-125041333-INS-238 |
| chr8-125062845-INS-76 | chr8-125062845-INS-76 |
| chr8-125063886-INS-186 | chr8-125063886-INS-186 |
| chr8-125088825-INS-64 | chr8-125088810-INS-64 |
| chr8-125098495-INS-55 | chr8-125098479-INS-71 |
| chr8-125105488-INS-400 | chr8-125105488-INS-400 |
| chr8-125112134-INS-5292 | chr8-125112135-INS-5292 |
| chr8-125122138-INS-209 | chr8-125122138-INS-209 |
| chr8-125123398-INS-62 | chr8-125123398-INS-62 |
| chr8-125127203-INS-210 | chr8-125127203-INS-210 |
| chr8-125133215-INS-100 | chr8-125133215-INS-122 |
| chr8-125141945-INS-82 | chr8-125141945-INS-82 |
| chr8-125153410-INS-329 | chr8-125153410-INS-329 |
| chr8-125173558-INS-96 | chr8-125173558-INS-96 |
| chr8-125177385-INS-114 | chr8-125177385-INS-76 |
| chr8-125183220-INS-90 | chr8-125183220-INS-90 |
| chr8-125184223-INS-150 | chr8-125184226-INS-150 |
| chr8-125186038-INS-169 | chr8-125186038-INS-169 |
| chr8-125199293-INS-218 | chr8-125199293-INS-218 |
| chr8-125202592-INS-488 | chr8-125202592-INS-489 |
| chr8-125204884-INS-124 | chr8-125204884-INS-124 |
| chr8-125210614-INS-96 | chr8-125210614-INS-98 |
| chr8-125225216-INS-55 | chr8-125225216-INS-55 |
| chr8-125238544-INS-216 | chr8-125238544-INS-216 |
| chr8-125258248-INS-260 | chr8-125258248-INS-260 |
| chr8-125292305-INS-7198 | chr8-125292306-INS-7198 |
| chr8-125313730-INS-55 | chr8-125313730-INS-55 |
| chr8-125316939-INS-55 | chr8-125316939-INS-50 |
| chr8-125336435-INS-400 | chr8-125336435-INS-400 |
| chr8-125341088-INS-71 | chr8-125341088-INS-75 |
| chr8-125381951-INS-550 | chr8-125381951-INS-552 |
| chr8-125390822-INS-58 | chr8-125390822-INS-58 |
| chr8-125393439-INS-67 | chr8-125393427-INS-70 |
| chr8-125414647-INS-76 | chr8-125414647-INS-76 |
| chr8-125417772-INS-299 | chr8-125417772-INS-299 |
| chr8-125421411-INS-56 | chr8-125421411-INS-56 |
| chr8-125422116-INS-207 | chr8-125422116-INS-208 |
| chr8-125437146-INS-56 | chr8-125437146-INS-56 |
| chr8-125446935-INS-81 | chr8-125446935-INS-81 |
| chr8-125449761-INS-180 | chr8-125449761-INS-195 |
| chr8-125456524-INS-76 | chr8-125456524-INS-76 |
| chr8-125484351-INS-102 | chr8-125484351-INS-101 |
| chr8-125496294-INS-158 | chr8-125496277-INS-158 |
| chr8-125498081-INS-152 | chr8-125498081-INS-152 |
| chr8-125502369-INS-83 | chr8-125502369-INS-89 |
| chr8-125503251-INS-211 | chr8-125503251-INS-211 |
| chr8-125516343-INS-236 | chr8-125516343-INS-241 |
| chr8-125524602-INS-92 | chr8-125524625-INS-100 |
| chr8-125527069-INS-206 | chr8-125527069-INS-206 |
| chr8-125527602-INS-6403 | chr8-125527603-INS-6403 |
| chr8-125537669-INS-58 | chr8-125537669-INS-60 |
| chr8-125549541-INS-71 | chr8-125549541-INS-73 |
| chr8-125551698-INS-218 | chr8-125551698-INS-219 |
| chr8-125555589-INS-147 | chr8-125555589-INS-147 |
| chr8-125608507-INS-56 | chr8-125608498-INS-62 |
| chr8-125631096-INS-56 | chr8-125631096-INS-52 |
| chr8-125633913-INS-150 | chr8-125633913-INS-150 |
| chr8-125662285-INS-215 | chr8-125662285-INS-196 |
| chr8-125673082-INS-78 | chr8-125673082-INS-78 |
| chr8-125682620-INS-95 | chr8-125682620-INS-95 |
| chr8-125688344-INS-196 | chr8-125688344-INS-196 |
| chr8-125691767-INS-442 | chr8-125691767-INS-442 |
| chr8-125701379-INS-52 | chr8-125701379-INS-52 |
| chr8-125712728-INS-614 | chr8-125712766-INS-617 |
| chr8-125722166-INS-51 | chr8-125722166-INS-60 |
| chr8-125743984-INS-232 | chr8-125743984-INS-232 |
| chr8-125751710-INS-264 | chr8-125751710-INS-264 |
| chr8-125773100-INS-330 | chr8-125773100-INS-320 |
| chr8-125775571-INS-213 | chr8-125775571-INS-213 |
| chr8-125789494-INS-60 | chr8-125789494-INS-60 |
| chr8-125805466-INS-267 | chr8-125805466-INS-267 |

| | |
|---|---|
| chr8-125810971-INS-840 | chr8-125810969-INS-842 |
| chr8-125819266-INS-639 | chr8-125819266-INS-639 |
| chr8-125836668-INS-275 | chr8-125836668-INS-275 |
| chr8-125849536-INS-78 | chr8-125849536-INS-78 |
| chr8-125852537-INS-107 | chr8-125852537-INS-113 |
| chr8-125862786-INS-84 | chr8-125862786-INS-84 |
| chr8-125871286-INS-72 | chr8-125871286-INS-74 |
| chr8-125875021-INS-96 | chr8-125875021-INS-96 |
| chr8-125880179-INS-60 | chr8-125880179-INS-60 |
| chr8-125882207-INS-89 | chr8-125882207-INS-89 |
| chr8-125898052-INS-64 | chr8-125898053-INS-64 |
| chr8-125920636-INS-77 | chr8-125920636-INS-66 |
| chr8-125948194-INS-64 | chr8-125948197-INS-77 |
| chr8-125968023-INS-102 | chr8-125968023-INS-102 |
| chr8-125972157-INS-5393 | chr8-125972158-INS-5393 |
| chr8-125978242-INS-139 | chr8-125978242-INS-139 |
| chr8-125996810-INS-52 | chr8-125996810-INS-52 |
| chr8-126001576-INS-53 | chr8-126001575-INS-64 |
| chr8-126003142-INS-340 | chr8-126003142-INS-340 |
| chr8-126013875-INS-108 | chr8-126013875-INS-99 |
| chr8-126020827-INS-56 | chr8-126020827-INS-64 |
| chr8-126035727-INS-981 | chr8-126035727-INS-981 |
| chr8-126074572-INS-475 | chr8-126074572-INS-475 |
| chr8-126084818-INS-220 | chr8-126084818-INS-206 |
| chr8-126086543-INS-201 | chr8-126086543-INS-201 |
| chr8-126092404-INS-199 | chr8-126092404-INS-199 |
| chr8-126100929-INS-128 | chr8-126100929-INS-126 |
| chr8-126109338-INS-189 | chr8-126109348-INS-200 |
| chr8-126120548-INS-512 | chr8-126120548-INS-512 |
| chr8-126122639-INS-7116 | chr8-126122640-INS-7116 |
| chr8-126124902-INS-59 | chr8-126124902-INS-61 |
| chr8-126141051-INS-117 | chr8-126141051-INS-117 |
| chr8-126156880-INS-312 | chr8-126156880-INS-312 |
| chr8-126170261-INS-58 | chr8-126170261-INS-58 |
| chr8-126205836-INS-171 | chr8-126205830-INS-205 |
| chr8-126210774-INS-56 | chr8-126210774-INS-58 |
| chr8-126213282-INS-56 | chr8-126213282-INS-56 |
| chr8-126215356-INS-61 | chr8-126215356-INS-61 |
| chr8-126220898-INS-485 | chr8-126220898-INS-485 |
| chr8-126256292-INS-120 | chr8-126256293-INS-120 |
| chr8-126273701-INS-209 | chr8-126273701-INS-209 |
| chr8-126299012-INS-66 | chr8-126299012-INS-66 |
| chr8-126310843-INS-54 | chr8-126310843-INS-54 |
| chr8-126316540-INS-54 | chr8-126316541-INS-66 |
| chr8-126330949-INS-52 | chr8-126330949-INS-50 |
| chr8-126333009-INS-51 | chr8-126333027-INS-60 |
| chr8-126367870-INS-359 | chr8-126367870-INS-359 |
| chr8-126394110-INS-82 | chr8-126394110-INS-84 |
| chr8-126401083-INS-62 | chr8-126401053-INS-90 |
| chr8-126419585-INS-641 | chr8-126419585-INS-642 |
| chr8-126423798-INS-73 | chr8-126423798-INS-73 |
| chr8-126428771-INS-72 | chr8-126428771-INS-72 |
| chr8-126438061-INS-193 | chr8-126438061-INS-193 |
| chr8-126441583-INS-2552 | chr8-126441584-INS-2552 |
| chr8-126507879-INS-90 | chr8-126507879-INS-90 |
| chr8-126520587-INS-427 | chr8-126520587-INS-427 |
| chr8-126523046-INS-62 | chr8-126523046-INS-62 |
| chr8-126528170-INS-145 | chr8-126528170-INS-143 |
| chr8-126541028-INS-111 | chr8-126541028-INS-110 |
| chr8-126547074-INS-5313 | chr8-126547075-INS-5313 |
| chr8-126548557-INS-121 | chr8-126548539-INS-147 |
| chr8-126558092-INS-121 | chr8-126558092-INS-121 |
| chr8-126560072-INS-76 | chr8-126560072-INS-96 |
| chr8-126569908-INS-218 | chr8-126569908-INS-218 |
| chr8-126577925-INS-52 | chr8-126577925-INS-52 |
| chr8-126595145-INS-57 | chr8-126595109-INS-64 |
| chr8-126600403-INS-60 | chr8-126600403-INS-60 |
| chr8-126620677-INS-514 | chr8-126620677-INS-513 |
| chr8-126632844-INS-65 | chr8-126632844-INS-65 |
| chr8-126633654-INS-711 | chr8-126633654-INS-711 |
| chr8-126635429-INS-66 | chr8-126635429-INS-64 |
| chr8-126664288-INS-63 | chr8-126664288-INS-63 |
| chr8-126670821-INS-62 | chr8-126670821-INS-52 |
| chr8-126685778-INS-99 | chr8-126685789-INS-63 |
| chr8-126715803-INS-87 | chr8-126715799-INS-95 |
| chr8-126788019-INS-68 | chr8-126788019-INS-76 |
| chr8-126790026-INS-64 | chr8-126790026-INS-56 |

| | |
|---|---|
| chr8-126810254-INS-54 | chr8-126810268-INS-55 |
| chr8-126819268-INS-224 | chr8-126819268-INS-224 |
| chr8-126822708-INS-244 | chr8-126822708-INS-244 |
| chr8-126878445-INS-128 | chr8-126878451-INS-160 |
| chr8-126882432-INS-88 | chr8-126882383-INS-106 |
| chr8-126905197-INS-93 | chr8-126905197-INS-56 |
| chr8-126906339-INS-104 | chr8-126906339-INS-104 |
| chr8-126909055-INS-89 | chr8-126909055-INS-59 |
| chr8-126943631-INS-160 | chr8-126943631-INS-191 |
| chr8-126957419-INS-58 | chr8-126957419-INS-60 |
| chr8-126984875-INS-753 | chr8-126984871-INS-728 |
| chr8-126997347-INS-200 | chr8-126997347-INS-236 |
| chr8-127002362-INS-113 | chr8-127002355-INS-119 |
| chr8-127005299-INS-426 | chr8-127005299-INS-426 |
| chr8-127018055-INS-62 | chr8-127018055-INS-60 |
| chr8-127032693-INS-269 | chr8-127032694-INS-282 |
| chr8-127058003-INS-50 | chr8-127057973-INS-54 |
| chr8-127088825-INS-108 | chr8-127088825-INS-60 |
| chr8-127103804-INS-112 | chr8-127103804-INS-71 |
| chr8-127108784-INS-78 | chr8-127108782-INS-69 |
| chr8-127121152-INS-55 | chr8-127121152-INS-55 |
| chr8-127167853-INS-106 | chr8-127167853-INS-106 |
| chr8-127181029-INS-154 | chr8-127181029-INS-154 |
| chr8-127184779-INS-449 | chr8-127184779-INS-449 |
| chr8-127186228-INS-193 | chr8-127186245-INS-212 |
| chr8-127189650-INS-422 | chr8-127189640-INS-400 |
| chr8-127229289-INS-66 | chr8-127229289-INS-64 |
| chr8-127235611-INS-217 | chr8-127235611-INS-217 |
| chr8-127253460-INS-85 | chr8-127253460-INS-85 |
| chr8-127302500-INS-57 | chr8-127302499-INS-59 |
| chr8-127331052-INS-66 | chr8-127331035-INS-64 |
| chr8-127334745-INS-58 | chr8-127334745-INS-62 |
| chr8-127336234-INS-11520 | chr8-127336231-INS-11563 |
| chr8-127340308-INS-164 | chr8-127340308-INS-164 |
| chr8-127355325-INS-238 | chr8-127355325-INS-234 |
| chr8-127360481-INS-120 | chr8-127360478-INS-132 |
| chr8-127372694-INS-57 | chr8-127372694-INS-57 |
| chr8-127385700-INS-106 | chr8-127385700-INS-104 |
| chr8-127416576-INS-445 | chr8-127416576-INS-445 |
| chr8-127420683-INS-68 | chr8-127420683-INS-68 |
| chr8-127427366-INS-84 | chr8-127427370-INS-80 |
| chr8-127427977-INS-80 | chr8-127427977-INS-80 |
| chr8-127443560-INS-90 | chr8-127443560-INS-90 |
| chr8-127458666-INS-189 | chr8-127458655-INS-201 |
| chr8-127466295-INS-1890 | chr8-127466296-INS-1890 |
| chr8-127482247-INS-4034 | chr8-127482248-INS-4034 |
| chr8-127490117-INS-719 | chr8-127490117-INS-719 |
| chr8-127495576-INS-126 | chr8-127495571-INS-108 |
| chr8-127537109-INS-96 | chr8-127537109-INS-96 |
| chr8-127543517-INS-157 | chr8-127543517-INS-154 |
| chr8-127644996-INS-132 | chr8-127644994-INS-146 |
| chr8-127722824-INS-66 | chr8-127722824-INS-93 |
| chr8-127726638-INS-230 | chr8-127726638-INS-231 |
| chr8-127729471-INS-192 | chr8-127729471-INS-192 |
| chr8-127733163-INS-51 | chr8-127733163-INS-51 |
| chr8-127747545-INS-82 | chr8-127747545-INS-82 |
| chr8-127748604-INS-196 | chr8-127748604-INS-199 |
| chr8-127754841-INS-209 | chr8-127754841-INS-210 |
| chr8-127759540-INS-207 | chr8-127759540-INS-210 |
| chr8-127760965-INS-64 | chr8-127760965-INS-52 |
| chr8-127777732-INS-202 | chr8-127777732-INS-202 |
| chr8-127781038-INS-7119 | chr8-127781039-INS-7119 |
| chr8-127796122-INS-53 | chr8-127796122-INS-54 |
| chr8-127838390-INS-125 | chr8-127838390-INS-125 |
| chr8-127841704-INS-60 | chr8-127841704-INS-60 |
| chr8-127844683-INS-181 | chr8-127844683-INS-181 |
| chr8-127859600-INS-199 | chr8-127859558-INS-210 |
| chr8-127863587-INS-108 | chr8-127863587-INS-108 |
| chr8-127913281-INS-55 | chr8-127913281-INS-50 |
| chr8-127961287-INS-154 | chr8-127961287-INS-154 |
| chr8-127971505-INS-75 | chr8-127971505-INS-60 |
| chr8-127972957-INS-188 | chr8-127972957-INS-187 |
| chr8-127981826-INS-684 | chr8-127981827-INS-685 |
| chr8-128001342-INS-63 | chr8-128001342-INS-63 |
| chr8-128024572-INS-170 | chr8-128024572-INS-169 |
| chr8-128051645-INS-64 | chr8-128051645-INS-65 |
| chr8-128072866-INS-7478 | chr8-128072867-INS-7478 |

| | |
|---|---|
| chr8-128091791-INS-51 | chr8-128091772-INS-54 |
| chr8-128098850-INS-70 | chr8-128098845-INS-60 |
| chr8-128120924-INS-125 | chr8-128120924-INS-125 |
| chr8-128155827-INS-58 | chr8-128155826-INS-64 |
| chr8-128186231-INS-1265 | chr8-128186232-INS-1265 |
| chr8-128246054-INS-100 | chr8-128246054-INS-100 |
| chr8-128246582-INS-74 | chr8-128246582-INS-76 |
| chr8-128254948-INS-148 | chr8-128254990-INS-146 |
| chr8-128259042-INS-576 | chr8-128259042-INS-576 |
| chr8-128260119-INS-60 | chr8-128260119-INS-54 |
| chr8-128289559-INS-372 | chr8-128289559-INS-372 |
| chr8-128307604-INS-198 | chr8-128307604-INS-198 |
| chr8-128312512-INS-58 | chr8-128312512-INS-60 |
| chr8-128340600-INS-65 | chr8-128340600-INS-65 |
| chr8-128352934-INS-1074 | chr8-128352935-INS-1070 |
| chr8-128356711-INS-107 | chr8-128356711-INS-105 |
| chr8-128359934-INS-52 | chr8-128359934-INS-52 |
| chr8-128373466-INS-208 | chr8-128373466-INS-204 |
| chr8-128384223-INS-65 | chr8-128384223-INS-69 |
| chr8-128437354-INS-60 | chr8-128437356-INS-54 |
| chr8-128445809-INS-173 | chr8-128445809-INS-172 |
| chr8-128476019-INS-175 | chr8-128476019-INS-175 |
| chr8-128489375-INS-1890 | chr8-128489376-INS-1890 |
| chr8-128520560-INS-1756 | chr8-128520561-INS-1757 |
| chr8-128555257-INS-384 | chr8-128555279-INS-402 |
| chr8-128561293-INS-56 | chr8-128561293-INS-71 |
| chr8-128585282-INS-111 | chr8-128585281-INS-86 |
| chr8-128590179-INS-201 | chr8-128590179-INS-201 |
| chr8-128616039-INS-92 | chr8-128616039-INS-94 |
| chr8-128639663-INS-785 | chr8-128639663-INS-788 |
| chr8-128655596-INS-128 | chr8-128655596-INS-132 |
| chr8-128669146-INS-116 | chr8-128669146-INS-98 |
| chr8-128671119-INS-354 | chr8-128671119-INS-392 |
| chr8-128678078-INS-6655 | chr8-128678079-INS-6655 |
| chr8-128696848-INS-59 | chr8-128696848-INS-59 |
| chr8-128700625-INS-94 | chr8-128700626-INS-123 |
| chr8-128705122-INS-1609 | chr8-128705123-INS-1610 |
| chr8-128710511-INS-304 | chr8-128710511-INS-305 |
| chr8-128718247-INS-783 | chr8-128718247-INS-783 |
| chr8-128727195-INS-64 | chr8-128727195-INS-78 |
| chr8-128751647-INS-54 | chr8-128751647-INS-58 |
| chr8-128752811-INS-60 | chr8-128752811-INS-68 |
| chr8-128766968-INS-202 | chr8-128766968-INS-203 |
| chr8-128791487-INS-60 | chr8-128791488-INS-54 |
| chr8-128799849-INS-63 | chr8-128799849-INS-63 |
| chr8-128830252-INS-1284 | chr8-128830253-INS-1284 |
| chr8-128875476-INS-92 | chr8-128875476-INS-71 |
| chr8-128879454-INS-7084 | chr8-128879455-INS-7084 |
| chr8-128881716-INS-278 | chr8-128881716-INS-280 |
| chr8-128887916-INS-108 | chr8-128887937-INS-132 |
| chr8-128892319-INS-6618 | chr8-128892320-INS-6619 |
| chr8-128906459-INS-96 | chr8-128906453-INS-104 |
| chr8-128949130-INS-97 | chr8-128949168-INS-60 |
| chr8-128975104-INS-71 | chr8-128975104-INS-71 |
| chr8-129025269-INS-885 | chr8-129025269-INS-903 |
| chr8-129028347-INS-79 | chr8-129028347-INS-81 |
| chr8-129031361-INS-139 | chr8-129031355-INS-130 |
| chr8-129046877-INS-66 | chr8-129046864-INS-70 |
| chr8-129058224-INS-6431 | chr8-129058225-INS-6431 |
| chr8-129157812-INS-52 | chr8-129157812-INS-54 |
| chr8-129212257-INS-58 | chr8-129212257-INS-62 |
| chr8-129241716-INS-53 | chr8-129241716-INS-53 |
| chr8-129245090-INS-178 | chr8-129245090-INS-178 |
| chr8-129277622-INS-69 | chr8-129277624-INS-57 |
| chr8-129287962-INS-108 | chr8-129287962-INS-118 |
| chr8-129309197-INS-949 | chr8-129309197-INS-948 |
| chr8-129337809-INS-7566 | chr8-129337810-INS-7566 |
| chr8-129343106-INS-60 | chr8-129343106-INS-60 |
| chr8-129369345-INS-66 | chr8-129369345-INS-66 |
| chr8-129371098-INS-1894 | chr8-129371099-INS-1894 |
| chr8-129378082-INS-53 | chr8-129378082-INS-53 |
| chr8-129387973-INS-61 | chr8-129387973-INS-66 |
| chr8-129460689-INS-189 | chr8-129460689-INS-191 |
| chr8-129489473-INS-3234 | chr8-129489474-INS-3234 |
| chr8-129526916-INS-333 | chr8-129526916-INS-333 |
| chr8-129557139-INS-57 | chr8-129557139-INS-69 |
| chr8-129575538-INS-3287 | chr8-129575539-INS-3289 |

| | |
|---|---|
| chr8-129579320-INS-62 | chr8-129579320-INS-54 |
| chr8-129579892-INS-6669 | chr8-129579893-INS-6669 |
| chr8-129589236-INS-700 | chr8-129589236-INS-701 |
| chr8-129683035-INS-3178 | chr8-129683036-INS-3178 |
| chr8-129717984-INS-67 | chr8-129717984-INS-67 |
| chr8-129749566-INS-364 | chr8-129749566-INS-364 |
| chr8-129790989-INS-220 | chr8-129790989-INS-220 |
| chr8-129831960-INS-4927 | chr8-129831961-INS-4927 |
| chr8-129850032-INS-66 | chr8-129850032-INS-66 |
| chr8-129861038-INS-208 | chr8-129861038-INS-208 |
| chr8-129874989-INS-2244 | chr8-129874990-INS-2247 |
| chr8-129880791-INS-7430 | chr8-129880792-INS-7430 |
| chr8-129910351-INS-6594 | chr8-129910352-INS-6594 |
| chr8-129911207-INS-209 | chr8-129911207-INS-209 |
| chr8-129938777-INS-66 | chr8-129938777-INS-66 |
| chr8-129947925-INS-493 | chr8-129947925-INS-493 |
| chr8-129972063-INS-54 | chr8-129972063-INS-57 |
| chr8-129987073-INS-6413 | chr8-129987074-INS-6413 |
| chr9-3054054-INS-372 | chr9-3054054-INS-403 |
| chr9-3057410-INS-58 | chr9-3057410-INS-58 |
| chr9-3237367-INS-198 | chr9-3237367-INS-198 |
| chr9-3255070-INS-64 | chr9-3255071-INS-76 |
| chr9-3327796-INS-543 | chr9-3327796-INS-543 |
| chr9-3330727-INS-78 | chr9-3330727-INS-78 |
| chr9-3339990-INS-392 | chr9-3339990-INS-393 |
| chr9-3378776-INS-1887 | chr9-3378777-INS-1886 |
| chr9-3407942-INS-401 | chr9-3407942-INS-401 |
| chr9-3429551-INS-87 | chr9-3429547-INS-88 |
| chr9-3489888-INS-2878 | chr9-3489889-INS-2880 |
| chr9-3493537-INS-83 | chr9-3493539-INS-84 |
| chr9-3505305-INS-4733 | chr9-3505306-INS-4733 |
| chr9-3533765-INS-312 | chr9-3533788-INS-320 |
| chr9-3573057-INS-70 | chr9-3573057-INS-102 |
| chr9-3591756-INS-334 | chr9-3591756-INS-334 |
| chr9-3596782-INS-155 | chr9-3596782-INS-155 |
| chr9-3692153-INS-798 | chr9-3692153-INS-798 |
| chr9-3713852-INS-1410 | chr9-3713853-INS-1410 |
| chr9-3714165-INS-2489 | chr9-3714166-INS-2495 |
| chr9-3860771-INS-7038 | chr9-3860772-INS-7038 |
| chr9-3878048-INS-1505 | chr9-3878049-INS-1506 |
| chr9-3900314-INS-261 | chr9-3900314-INS-261 |
| chr9-3904382-INS-52 | chr9-3904382-INS-52 |
| chr9-3972834-INS-64 | chr9-3972788-INS-72 |
| chr9-3984180-INS-64 | chr9-3984180-INS-80 |
| chr9-3985762-INS-50 | chr9-3985794-INS-51 |
| chr9-4028566-INS-416 | chr9-4028566-INS-416 |
| chr9-4032146-INS-906 | chr9-4032146-INS-909 |
| chr9-4034704-INS-448 | chr9-4034704-INS-448 |
| chr9-4047462-INS-340 | chr9-4047462-INS-337 |
| chr9-4055677-INS-393 | chr9-4055677-INS-393 |
| chr9-4068471-INS-2123 | chr9-4068472-INS-2123 |
| chr9-4069505-INS-1638 | chr9-4069506-INS-1638 |
| chr9-4131340-INS-73 | chr9-4131321-INS-72 |
| chr9-4142731-INS-2324 | chr9-4142732-INS-2324 |
| chr9-4256365-INS-70 | chr9-4256371-INS-78 |
| chr9-4339045-INS-62 | chr9-4339045-INS-58 |
| chr9-4341135-INS-5332 | chr9-4341136-INS-5332 |
| chr9-4387823-INS-483 | chr9-4387823-INS-483 |
| chr9-4401207-INS-234 | chr9-4401207-INS-234 |
| chr9-4406329-INS-555 | chr9-4406329-INS-555 |
| chr9-4423508-INS-5407 | chr9-4423509-INS-5407 |
| chr9-4466662-INS-52 | chr9-4466662-INS-60 |
| chr9-4472936-INS-127 | chr9-4472936-INS-130 |
| chr9-4476300-INS-55 | chr9-4476280-INS-52 |
| chr9-4524423-INS-948 | chr9-4524423-INS-946 |
| chr9-4526010-INS-52 | chr9-4526010-INS-52 |
| chr9-4571718-INS-1637 | chr9-4571719-INS-1665 |
| chr9-4572292-INS-74 | chr9-4572292-INS-55 |
| chr9-4665912-INS-132 | chr9-4665882-INS-140 |
| chr9-4722846-INS-99 | chr9-4722888-INS-78 |
| chr9-4740957-INS-1857 | chr9-4740958-INS-1857 |
| chr9-4776003-INS-3234 | chr9-4776004-INS-3234 |
| chr9-4786875-INS-609 | chr9-4786875-INS-604 |
| chr9-4825306-INS-392 | chr9-4825306-INS-392 |
| chr9-4848722-INS-1717 | chr9-4848723-INS-1718 |
| chr9-4869130-INS-126 | chr9-4869130-INS-147 |
| chr9-4876501-INS-7485 | chr9-4876502-INS-7485 |

| | |
|---|---|
| chr9-4881562-INS-107 | chr9-4881588-INS-118 |
| chr9-4884341-INS-4470 | chr9-4884342-INS-4471 |
| chr9-4944042-INS-2833 | chr9-4944043-INS-2833 |
| chr9-4949225-INS-1894 | chr9-4949226-INS-1894 |
| chr9-4977694-INS-364 | chr9-4977694-INS-355 |
| chr9-5002848-INS-1134 | chr9-5002846-INS-1088 |
| chr9-5056687-INS-2943 | chr9-5056688-INS-2943 |
| chr9-5071556-INS-505 | chr9-5071556-INS-505 |
| chr9-5101198-INS-503 | chr9-5101198-INS-503 |
| chr9-5116960-INS-62 | chr9-5116960-INS-64 |
| chr9-5126721-INS-6868 | chr9-5126722-INS-6868 |
| chr9-5142027-INS-99 | chr9-5142019-INS-75 |
| chr9-5143034-INS-545 | chr9-5143034-INS-545 |
| chr9-5146044-INS-84 | chr9-5146021-INS-68 |
| chr9-5152994-INS-60 | chr9-5152998-INS-52 |
| chr9-5175702-INS-84 | chr9-5175702-INS-56 |
| chr9-5185709-INS-319 | chr9-5185709-INS-321 |
| chr9-5215383-INS-4766 | chr9-5215384-INS-4756 |
| chr9-5222645-INS-390 | chr9-5222645-INS-390 |
| chr9-5233139-INS-92 | chr9-5233133-INS-96 |
| chr9-5371354-INS-61 | chr9-5371357-INS-71 |
| chr9-5373265-INS-1246 | chr9-5373266-INS-1249 |
| chr9-5415125-INS-536 | chr9-5415125-INS-536 |
| chr9-5433599-INS-2848 | chr9-5433600-INS-2848 |
| chr9-5468529-INS-513 | chr9-5468530-INS-513 |
| chr9-5485383-INS-599 | chr9-5485383-INS-599 |
| chr9-5506416-INS-118 | chr9-5506416-INS-118 |
| chr9-5537928-INS-98 | chr9-5537928-INS-98 |
| chr9-5587116-INS-6238 | chr9-5587117-INS-6238 |
| chr9-5615944-INS-54 | chr9-5615944-INS-54 |
| chr9-5672483-INS-4702 | chr9-5672484-INS-4702 |
| chr9-5711050-INS-6548 | chr9-5711051-INS-6548 |
| chr9-5717790-INS-90 | chr9-5717790-INS-80 |
| chr9-5729316-INS-3959 | chr9-5729317-INS-3955 |
| chr9-5730734-INS-218 | chr9-5730729-INS-253 |
| chr9-5738004-INS-5466 | chr9-5738005-INS-5466 |
| chr9-5768342-INS-268 | chr9-5768342-INS-268 |
| chr9-5769372-INS-348 | chr9-5769345-INS-330 |
| chr9-5805119-INS-7491 | chr9-5805120-INS-7485 |
| chr9-5805321-INS-358 | chr9-5805321-INS-358 |
| chr9-5809036-INS-231 | chr9-5809036-INS-240 |
| chr9-5822196-INS-50 | chr9-5822196-INS-60 |
| chr9-5829920-INS-457 | chr9-5829918-INS-461 |
| chr9-5831874-INS-1102 | chr9-5831875-INS-1102 |
| chr9-5841538-INS-730 | chr9-5841538-INS-730 |
| chr9-5843747-INS-266 | chr9-5843747-INS-266 |
| chr9-5848620-INS-67 | chr9-5848636-INS-73 |
| chr9-5848656-INS-6320 | chr9-5848657-INS-6315 |
| chr9-5852018-INS-101 | chr9-5852018-INS-102 |
| chr9-5865497-INS-401 | chr9-5865497-INS-401 |
| chr9-5900264-INS-1891 | chr9-5900265-INS-1891 |
| chr9-5903619-INS-430 | chr9-5903619-INS-431 |
| chr9-5928546-INS-242 | chr9-5928546-INS-242 |
| chr9-5932803-INS-416 | chr9-5932803-INS-416 |
| chr9-6016921-INS-518 | chr9-6016921-INS-518 |
| chr9-6022509-INS-285 | chr9-6022509-INS-285 |
| chr9-6024772-INS-54 | chr9-6024772-INS-54 |
| chr9-6037381-INS-493 | chr9-6037381-INS-493 |
| chr9-6077112-INS-170 | chr9-6077112-INS-170 |
| chr9-6097179-INS-6439 | chr9-6097180-INS-6436 |
| chr9-6103686-INS-1589 | chr9-6103687-INS-1589 |
| chr9-6113167-INS-3799 | chr9-6113168-INS-3799 |
| chr9-6129508-INS-586 | chr9-6129503-INS-587 |
| chr9-6168086-INS-152 | chr9-6168086-INS-160 |
| chr9-6171564-INS-100 | chr9-6171540-INS-108 |
| chr9-6213954-INS-721 | chr9-6213954-INS-768 |
| chr9-6246337-INS-85 | chr9-6246337-INS-85 |
| chr9-6270361-INS-76 | chr9-6270340-INS-53 |
| chr9-6282883-INS-76 | chr9-6282883-INS-56 |
| chr9-6312027-INS-98 | chr9-6312027-INS-97 |
| chr9-6312728-INS-113 | chr9-6312728-INS-109 |
| chr9-6330695-INS-66 | chr9-6330725-INS-50 |
| chr9-6359028-INS-268 | chr9-6359028-INS-268 |
| chr9-6366610-INS-77 | chr9-6366610-INS-88 |
| chr9-6392264-INS-3630 | chr9-6392265-INS-3630 |
| chr9-6435579-INS-7112 | chr9-6435580-INS-7112 |
| chr9-6496604-INS-111 | chr9-6496604-INS-111 |

| | |
|---|---|
| chr9-6506449-INS-4898 | chr9-6506450-INS-4898 |
| chr9-6517888-INS-128 | chr9-6517888-INS-128 |
| chr9-6521057-INS-432 | chr9-6521057-INS-432 |
| chr9-6537047-INS-58 | chr9-6537070-INS-78 |
| chr9-6540952-INS-85 | chr9-6540952-INS-90 |
| chr9-6558541-INS-54 | chr9-6558535-INS-62 |
| chr9-6579638-INS-78 | chr9-6579638-INS-78 |
| chr9-6602929-INS-84 | chr9-6602933-INS-56 |
| chr9-6627042-INS-332 | chr9-6627042-INS-332 |
| chr9-6656633-INS-486 | chr9-6656633-INS-487 |
| chr9-6678104-INS-70 | chr9-6678104-INS-74 |
| chr9-6715788-INS-489 | chr9-6715788-INS-489 |
| chr9-6778271-INS-461 | chr9-6778271-INS-461 |
| chr9-6792133-INS-85 | chr9-6792133-INS-85 |
| chr9-6814470-INS-801 | chr9-6814470-INS-801 |
| chr9-6844299-INS-69 | chr9-6844299-INS-69 |
| chr9-6848280-INS-66 | chr9-6848280-INS-72 |
| chr9-6868462-INS-68 | chr9-6868462-INS-68 |
| chr9-6893900-INS-6403 | chr9-6893901-INS-6403 |
| chr9-6924058-INS-210 | chr9-6924058-INS-211 |
| chr9-6925106-INS-202 | chr9-6925106-INS-202 |
| chr9-6926149-INS-1607 | chr9-6926150-INS-1598 |
| chr9-6961967-INS-231 | chr9-6961967-INS-231 |
| chr9-6986207-INS-593 | chr9-6986207-INS-593 |
| chr9-6993742-INS-185 | chr9-6993742-INS-185 |
| chr9-7000695-INS-1250 | chr9-7000696-INS-1250 |
| chr9-7061134-INS-199 | chr9-7061134-INS-199 |
| chr9-7143641-INS-76 | chr9-7143641-INS-76 |
| chr9-7259365-INS-502 | chr9-7259366-INS-502 |
| chr9-7260558-INS-59 | chr9-7260559-INS-80 |
| chr9-7271539-INS-595 | chr9-7271539-INS-595 |
| chr9-7280628-INS-76 | chr9-7280628-INS-72 |
| chr9-7315045-INS-61 | chr9-7315045-INS-61 |
| chr9-7354488-INS-108 | chr9-7354488-INS-108 |
| chr9-7356607-INS-1303 | chr9-7356608-INS-1303 |
| chr9-7384439-INS-164 | chr9-7384439-INS-165 |
| chr9-7440676-INS-56 | chr9-7440676-INS-60 |
| chr9-7443588-INS-252 | chr9-7443588-INS-252 |
| chr9-7471411-INS-86 | chr9-7471403-INS-90 |
| chr9-7484663-INS-65 | chr9-7484663-INS-65 |
| chr9-7539494-INS-697 | chr9-7539494-INS-698 |
| chr9-7554645-INS-645 | chr9-7554645-INS-645 |
| chr9-7556806-INS-64 | chr9-7556806-INS-66 |
| chr9-7593916-INS-60 | chr9-7593916-INS-60 |
| chr9-7617077-INS-94 | chr9-7617077-INS-91 |
| chr9-7624995-INS-528 | chr9-7624992-INS-531 |
| chr9-7629455-INS-60 | chr9-7629455-INS-60 |
| chr9-7646054-INS-65 | chr9-7646054-INS-81 |
| chr9-7657807-INS-160 | chr9-7657837-INS-120 |
| chr9-7672403-INS-84 | chr9-7672403-INS-84 |
| chr9-7672552-INS-664 | chr9-7672552-INS-664 |
| chr9-7706063-INS-195 | chr9-7706063-INS-195 |
| chr9-7797351-INS-156 | chr9-7797347-INS-168 |
| chr9-7798632-INS-238 | chr9-7798632-INS-239 |
| chr9-7816138-INS-156 | chr9-7816131-INS-123 |
| chr9-7857174-INS-56 | chr9-7857175-INS-57 |
| chr9-7867305-INS-186 | chr9-7867305-INS-186 |
| chr9-7868137-INS-160 | chr9-7868137-INS-128 |
| chr9-7873884-INS-212 | chr9-7873884-INS-208 |
| chr9-7875215-INS-249 | chr9-7875215-INS-249 |
| chr9-7887026-INS-372 | chr9-7887026-INS-372 |
| chr9-7946409-INS-69 | chr9-7946403-INS-55 |
| chr9-7972330-INS-119 | chr9-7972330-INS-119 |
| chr9-7990656-INS-201 | chr9-7990656-INS-200 |
| chr9-8010567-INS-165 | chr9-8010567-INS-165 |
| chr9-8035378-INS-77 | chr9-8035378-INS-69 |
| chr9-8057150-INS-101 | chr9-8057150-INS-101 |
| chr9-8058899-INS-64 | chr9-8058888-INS-92 |
| chr9-8069436-INS-107 | chr9-8069436-INS-108 |
| chr9-8076104-INS-92 | chr9-8076104-INS-88 |
| chr9-8084766-INS-203 | chr9-8084766-INS-203 |
| chr9-8085570-INS-220 | chr9-8085570-INS-221 |
| chr9-8094430-INS-214 | chr9-8094430-INS-214 |
| chr9-8120867-INS-3720 | chr9-8120868-INS-3720 |
| chr9-8121971-INS-1818 | chr9-8121962-INS-1817 |
| chr9-8125828-INS-54 | chr9-8125828-INS-54 |
| chr9-8182817-INS-127 | chr9-8182817-INS-127 |

| | |
|---|---|
| chr9-8183938-INS-148 | chr9-8183938-INS-148 |
| chr9-8187191-INS-178 | chr9-8187191-INS-178 |
| chr9-8192639-INS-2657 | chr9-8192645-INS-2657 |
| chr9-8215277-INS-481 | chr9-8215277-INS-481 |
| chr9-8228224-INS-90 | chr9-8228224-INS-90 |
| chr9-8231132-INS-3954 | chr9-8231133-INS-3955 |
| chr9-8237446-INS-201 | chr9-8237446-INS-201 |
| chr9-8248618-INS-846 | chr9-8248618-INS-846 |
| chr9-8260348-INS-54 | chr9-8260348-INS-66 |
| chr9-8267237-INS-86 | chr9-8267237-INS-126 |
| chr9-8274408-INS-74 | chr9-8274408-INS-82 |
| chr9-8324425-INS-602 | chr9-8324425-INS-606 |
| chr9-8385675-INS-352 | chr9-8385675-INS-352 |
| chr9-8412509-INS-64 | chr9-8412505-INS-104 |
| chr9-8442471-INS-282 | chr9-8442471-INS-282 |
| chr9-8446635-INS-839 | chr9-8446635-INS-838 |
| chr9-8458330-INS-66 | chr9-8458330-INS-66 |
| chr9-8461314-INS-356 | chr9-8461314-INS-356 |
| chr9-8492858-INS-523 | chr9-8492859-INS-523 |
| chr9-8546637-INS-200 | chr9-8546637-INS-200 |
| chr9-8548250-INS-54 | chr9-8548250-INS-54 |
| chr9-8591619-INS-100 | chr9-8591619-INS-56 |
| chr9-8611558-INS-213 | chr9-8611558-INS-215 |
| chr9-8619157-INS-110 | chr9-8619145-INS-131 |
| chr9-8641328-INS-100 | chr9-8641281-INS-100 |
| chr9-8650512-INS-58 | chr9-8650512-INS-78 |
| chr9-8663672-INS-200 | chr9-8663672-INS-200 |
| chr9-8717604-INS-6507 | chr9-8717605-INS-6507 |
| chr9-8719512-INS-6496 | chr9-8719513-INS-6496 |
| chr9-8814723-INS-218 | chr9-8814723-INS-219 |
| chr9-8833303-INS-71 | chr9-8833306-INS-74 |
| chr9-8835820-INS-1042 | chr9-8835821-INS-1042 |
| chr9-8839332-INS-103 | chr9-8839332-INS-103 |
| chr9-8847193-INS-701 | chr9-8847193-INS-701 |
| chr9-8857002-INS-550 | chr9-8857002-INS-550 |
| chr9-8872932-INS-182 | chr9-8872938-INS-187 |
| chr9-8918591-INS-74 | chr9-8918587-INS-52 |
| chr9-8926571-INS-896 | chr9-8926573-INS-894 |
| chr9-8932907-INS-273 | chr9-8932907-INS-255 |
| chr9-8938022-INS-57 | chr9-8938022-INS-57 |
| chr9-8938575-INS-57 | chr9-8938542-INS-63 |
| chr9-8950208-INS-277 | chr9-8950208-INS-278 |
| chr9-8961920-INS-2103 | chr9-8961921-INS-2103 |
| chr9-8962429-INS-108 | chr9-8962429-INS-109 |
| chr9-9033048-INS-58 | chr9-9033045-INS-75 |
| chr9-9034823-INS-77 | chr9-9034823-INS-73 |
| chr9-9085495-INS-354 | chr9-9085495-INS-354 |
| chr9-9090953-INS-103 | chr9-9090953-INS-59 |
| chr9-9098341-INS-388 | chr9-9098341-INS-391 |
| chr9-9113324-INS-55 | chr9-9113324-INS-55 |
| chr9-9156847-INS-722 | chr9-9156888-INS-721 |
| chr9-9157591-INS-92 | chr9-9157548-INS-86 |
| chr9-9183343-INS-358 | chr9-9183343-INS-358 |
| chr9-9201261-INS-68 | chr9-9201266-INS-68 |
| chr9-9222122-INS-56 | chr9-9222117-INS-78 |
| chr9-9227466-INS-204 | chr9-9227466-INS-204 |
| chr9-9236528-INS-92 | chr9-9236528-INS-76 |
| chr9-9246658-INS-120 | chr9-9246658-INS-120 |
| chr9-9260228-INS-68 | chr9-9260228-INS-80 |
| chr9-9266106-INS-75 | chr9-9266106-INS-75 |
| chr9-9271377-INS-53 | chr9-9271362-INS-100 |
| chr9-9318423-INS-82 | chr9-9318423-INS-82 |
| chr9-9339950-INS-97 | chr9-9339950-INS-97 |
| chr9-9355257-INS-264 | chr9-9355257-INS-264 |
| chr9-9356066-INS-1496 | chr9-9356067-INS-1493 |
| chr9-9361408-INS-1482 | chr9-9361409-INS-1482 |
| chr9-9370505-INS-3011 | chr9-9370506-INS-3012 |
| chr9-9390889-INS-6379 | chr9-9390890-INS-6379 |
| chr9-9492540-INS-56 | chr9-9492540-INS-56 |
| chr9-9553155-INS-91 | chr9-9553155-INS-91 |
| chr9-9572625-INS-6274 | chr9-9572626-INS-6274 |
| chr9-9573460-INS-68 | chr9-9573453-INS-52 |
| chr9-9575012-INS-491 | chr9-9575013-INS-488 |
| chr9-9599370-INS-75 | chr9-9599370-INS-75 |
| chr9-9606554-INS-6400 | chr9-9606555-INS-6401 |
| chr9-9629551-INS-78 | chr9-9629529-INS-80 |
| chr9-9633760-INS-400 | chr9-9633760-INS-400 |

| | |
|---|---|
| chr9-9643750-INS-433 | chr9-9643750-INS-433 |
| chr9-9651645-INS-64 | chr9-9651645-INS-71 |
| chr9-9665757-INS-54 | chr9-9665757-INS-54 |
| chr9-9670947-INS-68 | chr9-9670985-INS-66 |
| chr9-9678033-INS-58 | chr9-9678033-INS-62 |
| chr9-9681460-INS-509 | chr9-9681460-INS-487 |
| chr9-9685747-INS-71 | chr9-9685747-INS-71 |
| chr9-9687917-INS-226 | chr9-9687917-INS-238 |
| chr9-9710206-INS-2904 | chr9-9710207-INS-2904 |
| chr9-9718287-INS-205 | chr9-9718287-INS-207 |
| chr9-9730743-INS-51 | chr9-9730745-INS-52 |
| chr9-9754676-INS-65 | chr9-9754676-INS-65 |
| chr9-9775630-INS-3321 | chr9-9775631-INS-3326 |
| chr9-9814738-INS-72 | chr9-9814738-INS-50 |
| chr9-9856030-INS-1950 | chr9-9856031-INS-1952 |
| chr9-9930912-INS-177 | chr9-9930912-INS-184 |
| chr9-9942790-INS-195 | chr9-9942790-INS-195 |
| chr9-9982832-INS-156 | chr9-9982832-INS-140 |
| chr9-10046563-INS-146 | chr9-10046563-INS-146 |
| chr9-10048140-INS-201 | chr9-10048140-INS-200 |
| chr9-10105426-INS-199 | chr9-10105426-INS-200 |
| chr9-10122805-INS-410 | chr9-10122805-INS-410 |
| chr9-10144399-INS-90 | chr9-10144399-INS-89 |
| chr9-10158773-INS-199 | chr9-10158773-INS-199 |
| chr9-10160213-INS-858 | chr9-10160213-INS-858 |
| chr9-10234252-INS-4147 | chr9-10234253-INS-4147 |
| chr9-10261874-INS-58 | chr9-10261874-INS-58 |
| chr9-10266544-INS-360 | chr9-10266544-INS-360 |
| chr9-10297837-INS-71 | chr9-10297837-INS-71 |
| chr9-10303132-INS-136 | chr9-10303132-INS-114 |
| chr9-10314603-INS-54 | chr9-10314562-INS-58 |
| chr9-10349882-INS-81 | chr9-10349882-INS-95 |
| chr9-10362549-INS-56 | chr9-10362549-INS-59 |
| chr9-10372782-INS-736 | chr9-10372782-INS-726 |
| chr9-10383747-INS-68 | chr9-10383736-INS-60 |
| chr9-10420538-INS-60 | chr9-10420539-INS-69 |
| chr9-10423122-INS-458 | chr9-10423122-INS-462 |
| chr9-10442810-INS-52 | chr9-10442810-INS-62 |
| chr9-10451051-INS-52 | chr9-10451047-INS-60 |
| chr9-10523000-INS-353 | chr9-10523000-INS-352 |
| chr9-10571842-INS-6828 | chr9-10571843-INS-6829 |
| chr9-10586157-INS-64 | chr9-10586157-INS-88 |
| chr9-10669104-INS-890 | chr9-10669104-INS-892 |
| chr9-10732783-INS-160 | chr9-10732783-INS-160 |
| chr9-10737172-INS-60 | chr9-10737172-INS-60 |
| chr9-10747707-INS-266 | chr9-10747707-INS-266 |
| chr9-10755443-INS-53 | chr9-10755443-INS-53 |
| chr9-10768602-INS-2852 | chr9-10768603-INS-2852 |
| chr9-10812232-INS-176 | chr9-10812232-INS-177 |
| chr9-10878493-INS-104 | chr9-10878493-INS-104 |
| chr9-10913622-INS-232 | chr9-10913622-INS-233 |
| chr9-10936489-INS-7041 | chr9-10936490-INS-7041 |
| chr9-10942364-INS-72 | chr9-10942364-INS-88 |
| chr9-10952165-INS-1348 | chr9-10952166-INS-1348 |
| chr9-11013796-INS-103 | chr9-11013796-INS-103 |
| chr9-11076320-INS-332 | chr9-11076320-INS-332 |
| chr9-11091165-INS-89 | chr9-11091172-INS-91 |
| chr9-11098846-INS-315 | chr9-11098860-INS-314 |
| chr9-11126461-INS-91 | chr9-11126461-INS-91 |
| chr9-11140495-INS-214 | chr9-11140495-INS-214 |
| chr9-11141728-INS-653 | chr9-11141728-INS-651 |
| chr9-11241553-INS-6534 | chr9-11241554-INS-6534 |
| chr9-11264747-INS-7071 | chr9-11264748-INS-7071 |
| chr9-11265833-INS-769 | chr9-11265833-INS-771 |
| chr9-11269987-INS-93 | chr9-11269987-INS-93 |
| chr9-11286189-INS-72 | chr9-11286189-INS-72 |
| chr9-11299256-INS-172 | chr9-11299256-INS-172 |
| chr9-11324804-INS-9176 | chr9-11324805-INS-9195 |
| chr9-11328444-INS-3037 | chr9-11328444-INS-3036 |
| chr9-11342119-INS-214 | chr9-11342119-INS-214 |
| chr9-11348966-INS-87 | chr9-11348966-INS-87 |
| chr9-11383916-INS-409 | chr9-11383916-INS-423 |
| chr9-11471705-INS-2797 | chr9-11471706-INS-2780 |
| chr9-11577387-INS-6583 | chr9-11577388-INS-6583 |
| chr9-11588903-INS-753 | chr9-11588903-INS-753 |
| chr9-11604189-INS-68 | chr9-11604190-INS-72 |
| chr9-11663355-INS-4103 | chr9-11663356-INS-4103 |

| | |
|---|---|
| chr9-11685759-INS-2851 | chr9-11685760-INS-2851 |
| chr9-11701377-INS-198 | chr9-11701377-INS-194 |
| chr9-11722590-INS-213 | chr9-11722590-INS-213 |
| chr9-11925251-INS-110 | chr9-11925251-INS-148 |
| chr9-11979749-INS-115 | chr9-11979773-INS-76 |
| chr9-11990653-INS-6024 | chr9-11990654-INS-6024 |
| chr9-12071234-INS-90 | chr9-12071234-INS-90 |
| chr9-12088171-INS-191 | chr9-12088171-INS-212 |
| chr9-12107019-INS-256 | chr9-12107019-INS-252 |
| chr9-12158083-INS-74 | chr9-12158083-INS-58 |
| chr9-12158916-INS-132 | chr9-12158924-INS-104 |
| chr9-12249168-INS-739 | chr9-12249168-INS-737 |
| chr9-12278944-INS-2028 | chr9-12278944-INS-2023 |
| chr9-12286150-INS-309 | chr9-12286150-INS-310 |
| chr9-12345455-INS-271 | chr9-12345455-INS-271 |
| chr9-12390727-INS-6401 | chr9-12390728-INS-6401 |
| chr9-12396800-INS-201 | chr9-12396800-INS-199 |
| chr9-12500705-INS-1000 | chr9-12500705-INS-1001 |
| chr9-12558059-INS-827 | chr9-12558060-INS-829 |
| chr9-12614162-INS-652 | chr9-12614162-INS-658 |
| chr9-12618659-INS-521 | chr9-12618659-INS-509 |
| chr9-12646935-INS-2349 | chr9-12646936-INS-2349 |
| chr9-12650816-INS-54 | chr9-12650816-INS-54 |
| chr9-12714400-INS-944 | chr9-12714400-INS-943 |
| chr9-12736887-INS-79 | chr9-12736885-INS-57 |
| chr9-12747400-INS-97 | chr9-12747400-INS-97 |
| chr9-12831278-INS-6607 | chr9-12831279-INS-6607 |
| chr9-12942918-INS-345 | chr9-12942905-INS-362 |
| chr9-13018168-INS-1852 | chr9-13018169-INS-1851 |
| chr9-13026888-INS-516 | chr9-13026888-INS-517 |
| chr9-13048236-INS-486 | chr9-13048236-INS-486 |
| chr9-13064093-INS-141 | chr9-13064059-INS-149 |
| chr9-13118897-INS-7385 | chr9-13118898-INS-7373 |
| chr9-13141829-INS-280 | chr9-13141829-INS-280 |
| chr9-13161310-INS-459 | chr9-13161310-INS-459 |
| chr9-13167577-INS-399 | chr9-13167577-INS-399 |
| chr9-13172688-INS-1871 | chr9-13172689-INS-1871 |
| chr9-13184604-INS-404 | chr9-13184604-INS-404 |
| chr9-13214737-INS-1492 | chr9-13214738-INS-1492 |
| chr9-13215135-INS-198 | chr9-13215135-INS-192 |
| chr9-13237797-INS-6647 | chr9-13237798-INS-6647 |
| chr9-13263421-INS-64 | chr9-13263422-INS-62 |
| chr9-13325619-INS-51 | chr9-13325619-INS-51 |
| chr9-13339045-INS-477 | chr9-13339045-INS-466 |
| chr9-13389557-INS-116 | chr9-13389557-INS-116 |
| chr9-13410598-INS-6271 | chr9-13410599-INS-6278 |
| chr9-13445970-INS-72 | chr9-13445970-INS-72 |
| chr9-13446078-INS-1620 | chr9-13446079-INS-1620 |
| chr9-13458074-INS-56 | chr9-13458074-INS-56 |
| chr9-13472565-INS-97 | chr9-13472565-INS-97 |
| chr9-13482271-INS-125 | chr9-13482271-INS-125 |
| chr9-13490761-INS-233 | chr9-13490761-INS-238 |
| chr9-13505334-INS-87 | chr9-13505334-INS-87 |
| chr9-13538951-INS-98 | chr9-13538951-INS-98 |
| chr9-13545280-INS-120 | chr9-13545306-INS-132 |
| chr9-13557360-INS-73 | chr9-13557360-INS-73 |
| chr9-13566510-INS-7090 | chr9-13566511-INS-7090 |
| chr9-13567828-INS-7096 | chr9-13567829-INS-7096 |
| chr9-13707610-INS-434 | chr9-13707610-INS-434 |
| chr9-13719637-INS-225 | chr9-13719637-INS-226 |
| chr9-13728916-INS-341 | chr9-13728917-INS-338 |
| chr9-13749118-INS-53 | chr9-13749118-INS-53 |
| chr9-13758456-INS-448 | chr9-13758456-INS-463 |
| chr9-13771098-INS-388 | chr9-13771098-INS-387 |
| chr9-13829407-INS-6438 | chr9-13829408-INS-6448 |
| chr9-13837475-INS-197 | chr9-13837475-INS-197 |
| chr9-13877138-INS-150 | chr9-13877138-INS-150 |
| chr9-13891584-INS-79 | chr9-13891590-INS-89 |
| chr9-13902160-INS-68 | chr9-13902147-INS-72 |
| chr9-13962845-INS-3192 | chr9-13962846-INS-3192 |
| chr9-13989944-INS-66 | chr9-13989944-INS-66 |
| chr9-14038598-INS-3336 | chr9-14038599-INS-3337 |
| chr9-14051113-INS-632 | chr9-14051113-INS-632 |
| chr9-14053912-INS-120 | chr9-14053912-INS-120 |
| chr9-14071363-INS-3906 | chr9-14071364-INS-3906 |
| chr9-14101387-INS-51 | chr9-14101387-INS-51 |
| chr9-14293544-INS-53 | chr9-14293544-INS-53 |

| | |
|---|---|
| chr9-14294556-INS-277 | chr9-14294556-INS-277 |
| chr9-14310306-INS-211 | chr9-14310306-INS-212 |
| chr9-14323039-INS-57 | chr9-14323039-INS-58 |
| chr9-14391907-INS-1888 | chr9-14391908-INS-1888 |
| chr9-14422383-INS-198 | chr9-14422383-INS-199 |
| chr9-14432857-INS-262 | chr9-14432857-INS-263 |
| chr9-14434730-INS-85 | chr9-14434730-INS-82 |
| chr9-14452575-INS-100 | chr9-14452575-INS-100 |
| chr9-14508670-INS-2876 | chr9-14508671-INS-2876 |
| chr9-14534739-INS-5219 | chr9-14534740-INS-5219 |
| chr9-14563262-INS-138 | chr9-14563262-INS-138 |
| chr9-14580562-INS-565 | chr9-14580562-INS-567 |
| chr9-14582114-INS-124 | chr9-14582114-INS-124 |
| chr9-14594172-INS-53 | chr9-14594172-INS-51 |
| chr9-14599590-INS-72 | chr9-14599588-INS-60 |
| chr9-14610842-INS-248 | chr9-14610841-INS-291 |
| chr9-14616136-INS-599 | chr9-14616136-INS-599 |
| chr9-14647834-INS-504 | chr9-14647834-INS-504 |
| chr9-14649115-INS-1068 | chr9-14649116-INS-1068 |
| chr9-14657368-INS-64 | chr9-14657368-INS-64 |
| chr9-14661930-INS-177 | chr9-14661930-INS-177 |
| chr9-14709134-INS-128 | chr9-14709132-INS-128 |
| chr9-14713057-INS-213 | chr9-14713057-INS-214 |
| chr9-14719718-INS-113 | chr9-14719715-INS-115 |
| chr9-14757751-INS-1840 | chr9-14757752-INS-1840 |
| chr9-14780837-INS-100 | chr9-14780837-INS-110 |
| chr9-14785393-INS-207 | chr9-14785393-INS-207 |
| chr9-14794237-INS-95 | chr9-14794237-INS-95 |
| chr9-14799784-INS-116 | chr9-14799784-INS-116 |
| chr9-14836125-INS-72 | chr9-14836125-INS-72 |
| chr9-14877454-INS-211 | chr9-14877454-INS-211 |
| chr9-14883725-INS-136 | chr9-14883725-INS-148 |
| chr9-14893954-INS-361 | chr9-14893954-INS-376 |
| chr9-14899740-INS-64 | chr9-14899740-INS-64 |
| chr9-14916712-INS-62 | chr9-14916712-INS-62 |
| chr9-14931096-INS-209 | chr9-14931096-INS-186 |
| chr9-14932038-INS-650 | chr9-14932038-INS-650 |
| chr9-14933176-INS-54 | chr9-14933176-INS-54 |
| chr9-14956860-INS-116 | chr9-14956860-INS-78 |
| chr9-14975934-INS-206 | chr9-14975934-INS-205 |
| chr9-15005553-INS-631 | chr9-15005553-INS-637 |
| chr9-15007860-INS-117 | chr9-15007860-INS-96 |
| chr9-15018834-INS-68 | chr9-15018834-INS-68 |
| chr9-15039639-INS-215 | chr9-15039626-INS-230 |
| chr9-15099479-INS-255 | chr9-15099479-INS-255 |
| chr9-15101594-INS-87 | chr9-15101594-INS-81 |
| chr9-15115330-INS-61 | chr9-15115353-INS-68 |
| chr9-15126112-INS-52 | chr9-15126111-INS-58 |
| chr9-15129258-INS-96 | chr9-15129258-INS-96 |
| chr9-15130917-INS-83 | chr9-15130917-INS-87 |
| chr9-15138101-INS-983 | chr9-15138101-INS-983 |
| chr9-15152909-INS-180 | chr9-15152909-INS-181 |
| chr9-15167229-INS-214 | chr9-15167229-INS-214 |
| chr9-15182791-INS-222 | chr9-15182791-INS-222 |
| chr9-15192071-INS-203 | chr9-15192071-INS-203 |
| chr9-15205088-INS-216 | chr9-15205088-INS-216 |
| chr9-15212064-INS-686 | chr9-15212064-INS-687 |
| chr9-15231887-INS-200 | chr9-15231887-INS-200 |
| chr9-15236795-INS-197 | chr9-15236795-INS-197 |
| chr9-15269991-INS-229 | chr9-15269991-INS-230 |
| chr9-15278458-INS-188 | chr9-15278458-INS-188 |
| chr9-15300528-INS-62 | chr9-15300529-INS-62 |
| chr9-15313570-INS-577 | chr9-15313570-INS-577 |
| chr9-15321655-INS-86 | chr9-15321657-INS-56 |
| chr9-15340193-INS-2019 | chr9-15340194-INS-2019 |
| chr9-15346680-INS-515 | chr9-15346680-INS-515 |
| chr9-15350690-INS-283 | chr9-15350690-INS-297 |
| chr9-15353923-INS-361 | chr9-15353923-INS-361 |
| chr9-15361012-INS-259 | chr9-15361012-INS-260 |
| chr9-15376995-INS-56 | chr9-15376995-INS-56 |
| chr9-15383223-INS-337 | chr9-15383223-INS-338 |
| chr9-15398692-INS-158 | chr9-15398692-INS-158 |
| chr9-15430242-INS-87 | chr9-15430242-INS-58 |
| chr9-15438566-INS-66 | chr9-15438566-INS-68 |
| chr9-15451699-INS-187 | chr9-15451699-INS-188 |
| chr9-15457500-INS-79 | chr9-15457502-INS-60 |
| chr9-15479939-INS-5373 | chr9-15479940-INS-5373 |

| | |
|---|---|
| chr9-15484511-INS-112 | chr9-15484511-INS-112 |
| chr9-15496127-INS-76 | chr9-15496123-INS-116 |
| chr9-15523981-INS-450 | chr9-15523981-INS-450 |
| chr9-15537294-INS-129 | chr9-15537285-INS-110 |
| chr9-15541802-INS-218 | chr9-15541802-INS-216 |
| chr9-15550095-INS-254 | chr9-15550095-INS-250 |
| chr9-15567300-INS-210 | chr9-15567300-INS-210 |
| chr9-15584359-INS-139 | chr9-15584352-INS-145 |
| chr9-15657243-INS-1901 | chr9-15657244-INS-1901 |
| chr9-15661036-INS-666 | chr9-15661036-INS-668 |
| chr9-15689329-INS-216 | chr9-15689329-INS-216 |
| chr9-15701978-INS-66 | chr9-15701978-INS-50 |
| chr9-15705232-INS-145 | chr9-15705232-INS-150 |
| chr9-15713180-INS-82 | chr9-15713180-INS-72 |
| chr9-15715280-INS-3959 | chr9-15715281-INS-3950 |
| chr9-15718665-INS-6277 | chr9-15718666-INS-6278 |
| chr9-15733415-INS-56 | chr9-15733415-INS-57 |
| chr9-15733880-INS-166 | chr9-15733876-INS-181 |
| chr9-15744115-INS-64 | chr9-15744162-INS-50 |
| chr9-15758580-INS-418 | chr9-15758580-INS-418 |
| chr9-15797153-INS-6364 | chr9-15797154-INS-6364 |
| chr9-15804662-INS-535 | chr9-15804662-INS-535 |
| chr9-15810147-INS-180 | chr9-15810147-INS-180 |
| chr9-15813715-INS-201 | chr9-15813715-INS-201 |
| chr9-15821085-INS-7120 | chr9-15821086-INS-7122 |
| chr9-15821086-INS-60 | chr9-15821086-INS-56 |
| chr9-15835781-INS-62 | chr9-15835789-INS-56 |
| chr9-15840272-INS-2784 | chr9-15840273-INS-2788 |
| chr9-15913364-INS-76 | chr9-15913364-INS-80 |
| chr9-15915518-INS-126 | chr9-15915518-INS-125 |
| chr9-15961936-INS-218 | chr9-15961936-INS-218 |
| chr9-15975911-INS-498 | chr9-15975911-INS-498 |
| chr9-15989165-INS-184 | chr9-15989165-INS-184 |
| chr9-16002465-INS-57 | chr9-16002465-INS-55 |
| chr9-16018272-INS-914 | chr9-16018272-INS-919 |
| chr9-16037456-INS-438 | chr9-16037456-INS-438 |
| chr9-16064006-INS-6452 | chr9-16064007-INS-6452 |
| chr9-16095949-INS-220 | chr9-16095949-INS-220 |
| chr9-16138990-INS-67 | chr9-16138970-INS-56 |
| chr9-16154110-INS-86 | chr9-16154116-INS-52 |
| chr9-16173716-INS-1350 | chr9-16173717-INS-1350 |
| chr9-16180234-INS-83 | chr9-16180231-INS-84 |
| chr9-16188099-INS-5394 | chr9-16188100-INS-5394 |
| chr9-16225973-INS-874 | chr9-16225973-INS-873 |
| chr9-16232562-INS-200 | chr9-16232556-INS-201 |
| chr9-16236430-INS-52 | chr9-16236430-INS-82 |
| chr9-16243135-INS-168 | chr9-16243131-INS-198 |
| chr9-16245635-INS-91 | chr9-16245684-INS-66 |
| chr9-16273587-INS-2185 | chr9-16273588-INS-2202 |
| chr9-16328101-INS-104 | chr9-16328101-INS-80 |
| chr9-16355533-INS-120 | chr9-16355533-INS-114 |
| chr9-16395792-INS-138 | chr9-16395792-INS-138 |
| chr9-16413834-INS-80 | chr9-16413834-INS-75 |
| chr9-16447084-INS-63 | chr9-16447107-INS-68 |
| chr9-16479260-INS-147 | chr9-16479260-INS-147 |
| chr9-16499263-INS-6412 | chr9-16499264-INS-6412 |
| chr9-16546728-INS-90 | chr9-16546728-INS-90 |
| chr9-16565331-INS-195 | chr9-16565295-INS-196 |
| chr9-16623748-INS-64 | chr9-16623748-INS-64 |
| chr9-16638311-INS-51 | chr9-16638332-INS-52 |
| chr9-16658743-INS-452 | chr9-16658743-INS-452 |
| chr9-16660111-INS-99 | chr9-16660111-INS-90 |
| chr9-16698876-INS-192 | chr9-16698874-INS-196 |
| chr9-16705825-INS-878 | chr9-16705823-INS-871 |
| chr9-16724116-INS-65 | chr9-16724116-INS-65 |
| chr9-16778058-INS-91 | chr9-16778058-INS-91 |
| chr9-16788444-INS-212 | chr9-16788444-INS-213 |
| chr9-16790932-INS-254 | chr9-16790931-INS-258 |
| chr9-16792359-INS-61 | chr9-16792359-INS-61 |
| chr9-16821001-INS-148 | chr9-16820993-INS-152 |
| chr9-16827704-INS-60 | chr9-16827704-INS-60 |
| chr9-16869449-INS-78 | chr9-16869448-INS-70 |
| chr9-16873838-INS-471 | chr9-16873838-INS-471 |
| chr9-16909735-INS-780 | chr9-16909735-INS-780 |
| chr9-16918586-INS-3747 | chr9-16918587-INS-3748 |
| chr9-16922289-INS-548 | chr9-16922289-INS-548 |
| chr9-16935122-INS-147 | chr9-16935122-INS-147 |

| | |
|---|---|
| chr9-16964949-INS-52 | chr9-16964949-INS-52 |
| chr9-16984187-INS-2439 | chr9-16984188-INS-2444 |
| chr9-16984960-INS-66 | chr9-16984960-INS-66 |
| chr9-16987002-INS-68 | chr9-16987002-INS-68 |
| chr9-17024760-INS-272 | chr9-17024760-INS-273 |
| chr9-17030326-INS-51 | chr9-17030321-INS-89 |
| chr9-17030621-INS-786 | chr9-17030621-INS-795 |
| chr9-17044874-INS-146 | chr9-17044874-INS-188 |
| chr9-17046335-INS-873 | chr9-17046335-INS-867 |
| chr9-17058764-INS-1492 | chr9-17058753-INS-1484 |
| chr9-17061312-INS-71 | chr9-17061312-INS-61 |
| chr9-17076123-INS-130 | chr9-17076123-INS-138 |
| chr9-17076783-INS-1597 | chr9-17076784-INS-1596 |
| chr9-17083043-INS-4336 | chr9-17083044-INS-4336 |
| chr9-17120943-INS-84 | chr9-17120943-INS-88 |
| chr9-17149807-INS-56 | chr9-17149805-INS-68 |
| chr9-17184385-INS-139 | chr9-17184374-INS-155 |
| chr9-17197363-INS-820 | chr9-17197363-INS-821 |
| chr9-17198186-INS-90 | chr9-17198186-INS-90 |
| chr9-17205721-INS-60 | chr9-17205721-INS-62 |
| chr9-17231634-INS-117 | chr9-17231634-INS-117 |
| chr9-17266468-INS-2313 | chr9-17266469-INS-2313 |
| chr9-17305773-INS-193 | chr9-17305773-INS-193 |
| chr9-17337440-INS-1478 | chr9-17337441-INS-1478 |
| chr9-17345835-INS-52 | chr9-17345835-INS-54 |
| chr9-17390482-INS-1894 | chr9-17390483-INS-1894 |
| chr9-17411052-INS-80 | chr9-17411052-INS-80 |
| chr9-17415344-INS-324 | chr9-17415344-INS-327 |
| chr9-17420206-INS-541 | chr9-17420206-INS-512 |
| chr9-17428144-INS-754 | chr9-17428144-INS-756 |
| chr9-17452663-INS-375 | chr9-17452663-INS-375 |
| chr9-17468512-INS-118 | chr9-17468512-INS-120 |
| chr9-17484597-INS-98 | chr9-17484597-INS-97 |
| chr9-17509853-INS-109 | chr9-17509853-INS-94 |
| chr9-17515734-INS-3383 | chr9-17515735-INS-3383 |
| chr9-17518157-INS-366 | chr9-17518157-INS-389 |
| chr9-17520030-INS-672 | chr9-17520030-INS-672 |
| chr9-17523077-INS-503 | chr9-17523077-INS-503 |
| chr9-17524464-INS-73 | chr9-17524472-INS-78 |
| chr9-17538531-INS-521 | chr9-17538529-INS-506 |
| chr9-17567248-INS-1406 | chr9-17567249-INS-1405 |
| chr9-17570131-INS-559 | chr9-17570131-INS-559 |
| chr9-17584490-INS-64 | chr9-17584490-INS-64 |
| chr9-17585787-INS-506 | chr9-17585787-INS-506 |
| chr9-17589307-INS-2931 | chr9-17589308-INS-2931 |
| chr9-17609808-INS-258 | chr9-17609808-INS-260 |
| chr9-17611310-INS-593 | chr9-17611310-INS-592 |
| chr9-17620012-INS-54 | chr9-17620012-INS-55 |
| chr9-17633396-INS-240 | chr9-17633396-INS-240 |
| chr9-17639534-INS-379 | chr9-17639534-INS-375 |
| chr9-17674242-INS-574 | chr9-17674242-INS-574 |
| chr9-17677548-INS-68 | chr9-17677548-INS-68 |
| chr9-17678045-INS-964 | chr9-17678045-INS-962 |
| chr9-17682760-INS-429 | chr9-17682760-INS-429 |
| chr9-17691034-INS-347 | chr9-17691034-INS-347 |
| chr9-17694427-INS-507 | chr9-17694429-INS-515 |
| chr9-17695196-INS-74 | chr9-17695196-INS-56 |
| chr9-17697915-INS-68 | chr9-17697920-INS-52 |
| chr9-17702894-INS-64 | chr9-17702894-INS-60 |
| chr9-17751404-INS-1163 | chr9-17751405-INS-1161 |
| chr9-17757519-INS-1312 | chr9-17757520-INS-1312 |
| chr9-17773519-INS-683 | chr9-17773519-INS-698 |
| chr9-17780799-INS-57 | chr9-17780786-INS-52 |
| chr9-17783161-INS-432 | chr9-17783161-INS-432 |
| chr9-17906985-INS-54 | chr9-17906985-INS-55 |
| chr9-17910546-INS-532 | chr9-17910546-INS-533 |
| chr9-17939938-INS-111 | chr9-17939938-INS-63 |
| chr9-17946256-INS-56 | chr9-17946256-INS-52 |
| chr9-17960269-INS-368 | chr9-17960269-INS-368 |
| chr9-17968200-INS-68 | chr9-17968208-INS-96 |
| chr9-18016150-INS-493 | chr9-18016144-INS-497 |
| chr9-18094326-INS-92 | chr9-18094326-INS-94 |
| chr9-18257641-INS-206 | chr9-18257641-INS-206 |
| chr9-18276684-INS-467 | chr9-18276684-INS-467 |
| chr9-18283246-INS-409 | chr9-18283246-INS-409 |
| chr9-18290194-INS-191 | chr9-18290194-INS-201 |
| chr9-18305322-INS-52 | chr9-18305322-INS-54 |

| | |
|---|---|
| chr9-18331629-INS-210 | chr9-18331629-INS-177 |
| chr9-18344885-INS-208 | chr9-18344876-INS-192 |
| chr9-18347639-INS-241 | chr9-18347639-INS-227 |
| chr9-18370762-INS-328 | chr9-18370762-INS-328 |
| chr9-18374090-INS-468 | chr9-18374090-INS-468 |
| chr9-18483629-INS-1003 | chr9-18483630-INS-1003 |
| chr9-18583411-INS-152 | chr9-18583411-INS-152 |
| chr9-18591588-INS-90 | chr9-18591598-INS-114 |
| chr9-18595223-INS-86 | chr9-18595223-INS-50 |
| chr9-18611226-INS-181 | chr9-18611226-INS-180 |
| chr9-18620715-INS-207 | chr9-18620713-INS-203 |
| chr9-18623620-INS-5307 | chr9-18623621-INS-5307 |
| chr9-18635390-INS-440 | chr9-18635390-INS-454 |
| chr9-18641056-INS-52 | chr9-18641056-INS-54 |
| chr9-18648916-INS-211 | chr9-18648916-INS-209 |
| chr9-18667171-INS-6884 | chr9-18667172-INS-6891 |
| chr9-18762947-INS-98 | chr9-18762947-INS-100 |
| chr9-18803578-INS-114 | chr9-18803578-INS-114 |
| chr9-18821131-INS-119 | chr9-18821131-INS-155 |
| chr9-18831171-INS-2129 | chr9-18831172-INS-2132 |
| chr9-18834575-INS-8985 | chr9-18834576-INS-8985 |
| chr9-18834867-INS-199 | chr9-18834867-INS-199 |
| chr9-18836910-INS-109 | chr9-18836910-INS-109 |
| chr9-18845276-INS-699 | chr9-18845276-INS-699 |
| chr9-18892712-INS-1484 | chr9-18892713-INS-1484 |
| chr9-18938803-INS-1272 | chr9-18938804-INS-1272 |
| chr9-18965218-INS-6139 | chr9-18965219-INS-6139 |
| chr9-18978525-INS-60 | chr9-18978525-INS-60 |
| chr9-19006553-INS-5587 | chr9-19006554-INS-5566 |
| chr9-19018169-INS-55 | chr9-19018169-INS-55 |
| chr9-19031899-INS-57 | chr9-19031899-INS-61 |
| chr9-19033550-INS-217 | chr9-19033550-INS-216 |
| chr9-19102598-INS-500 | chr9-19102598-INS-499 |
| chr9-19112199-INS-362 | chr9-19112219-INS-362 |
| chr9-19152299-INS-67 | chr9-19152299-INS-64 |
| chr9-19159635-INS-52 | chr9-19159635-INS-56 |
| chr9-19183585-INS-136 | chr9-19183585-INS-115 |
| chr9-19184255-INS-425 | chr9-19184255-INS-425 |
| chr9-19185011-INS-178 | chr9-19185011-INS-177 |
| chr9-19209190-INS-94 | chr9-19209190-INS-94 |
| chr9-19240864-INS-3806 | chr9-19240865-INS-3807 |
| chr9-19267191-INS-58 | chr9-19267191-INS-58 |
| chr9-19272757-INS-56 | chr9-19272757-INS-54 |
| chr9-19323643-INS-6401 | chr9-19323644-INS-6401 |
| chr9-19336565-INS-195 | chr9-19336565-INS-195 |
| chr9-19355883-INS-390 | chr9-19355883-INS-390 |
| chr9-19385804-INS-118 | chr9-19385804-INS-117 |
| chr9-19395711-INS-204 | chr9-19395711-INS-204 |
| chr9-19401249-INS-926 | chr9-19401249-INS-935 |
| chr9-19406067-INS-214 | chr9-19406067-INS-214 |
| chr9-19439321-INS-72 | chr9-19439321-INS-72 |
| chr9-19447563-INS-396 | chr9-19447563-INS-396 |
| chr9-19454355-INS-52 | chr9-19454355-INS-50 |
| chr9-19472394-INS-1877 | chr9-19472395-INS-1862 |
| chr9-19482056-INS-145 | chr9-19482056-INS-105 |
| chr9-19483357-INS-513 | chr9-19483357-INS-513 |
| chr9-19534752-INS-145 | chr9-19534752-INS-138 |
| chr9-19540785-INS-433 | chr9-19540785-INS-435 |
| chr9-19575172-INS-87 | chr9-19575180-INS-81 |
| chr9-19575209-INS-6433 | chr9-19575210-INS-6433 |
| chr9-19578100-INS-211 | chr9-19578101-INS-211 |
| chr9-19580689-INS-2246 | chr9-19580690-INS-2247 |
| chr9-19583854-INS-89 | chr9-19583854-INS-89 |
| chr9-19610003-INS-460 | chr9-19610003-INS-460 |
| chr9-19629052-INS-108 | chr9-19629052-INS-108 |
| chr9-19645971-INS-70 | chr9-19645971-INS-71 |
| chr9-19646886-INS-1367 | chr9-19646887-INS-1367 |
| chr9-19655598-INS-397 | chr9-19655598-INS-397 |
| chr9-19662212-INS-6379 | chr9-19662213-INS-6379 |
| chr9-19665895-INS-57 | chr9-19665895-INS-59 |
| chr9-19694280-INS-338 | chr9-19694280-INS-326 |
| chr9-19718383-INS-332 | chr9-19718383-INS-332 |
| chr9-19774024-INS-616 | chr9-19774024-INS-616 |
| chr9-19776555-INS-58 | chr9-19776555-INS-58 |
| chr9-19796066-INS-100 | chr9-19796066-INS-100 |
| chr9-19806207-INS-971 | chr9-19806207-INS-971 |
| chr9-19806794-INS-61 | chr9-19806794-INS-61 |

| | |
|---|---|
| chr9-19854731-INS-62 | chr9-19854731-INS-64 |
| chr9-19858580-INS-52 | chr9-19858580-INS-52 |
| chr9-19932689-INS-343 | chr9-19932689-INS-344 |
| chr9-19955454-INS-67 | chr9-19955454-INS-67 |
| chr9-19984795-INS-71 | chr9-19984813-INS-68 |
| chr9-20016989-INS-72 | chr9-20016989-INS-56 |
| chr9-20026800-INS-5721 | chr9-20026800-INS-5721 |
| chr9-20031556-INS-5970 | chr9-20031557-INS-5972 |
| chr9-20070953-INS-60 | chr9-20070953-INS-62 |
| chr9-20106580-INS-469 | chr9-20106580-INS-469 |
| chr9-20113254-INS-75 | chr9-20113270-INS-78 |
| chr9-20131351-INS-308 | chr9-20131351-INS-309 |
| chr9-20146376-INS-68 | chr9-20146370-INS-69 |
| chr9-20152212-INS-80 | chr9-20152212-INS-88 |
| chr9-20183619-INS-4376 | chr9-20183620-INS-4376 |
| chr9-20187123-INS-103 | chr9-20187123-INS-102 |
| chr9-20196832-INS-319 | chr9-20196832-INS-319 |
| chr9-20252530-INS-352 | chr9-20252530-INS-352 |
| chr9-20264295-INS-68 | chr9-20264295-INS-70 |
| chr9-20337766-INS-4119 | chr9-20337767-INS-4119 |
| chr9-20343416-INS-344 | chr9-20343416-INS-353 |
| chr9-20358935-INS-59 | chr9-20358935-INS-59 |
| chr9-20362366-INS-203 | chr9-20362366-INS-203 |
| chr9-20422762-INS-661 | chr9-20422762-INS-661 |
| chr9-20429355-INS-220 | chr9-20429355-INS-220 |
| chr9-20440566-INS-72 | chr9-20440566-INS-72 |
| chr9-20450052-INS-208 | chr9-20450052-INS-208 |
| chr9-20464304-INS-201 | chr9-20464304-INS-201 |
| chr9-20465916-INS-60 | chr9-20465916-INS-60 |
| chr9-20466328-INS-7218 | chr9-20466329-INS-7218 |
| chr9-20470704-INS-111 | chr9-20470704-INS-111 |
| chr9-20516804-INS-7040 | chr9-20516805-INS-7040 |
| chr9-20541924-INS-200 | chr9-20541924-INS-200 |
| chr9-20546399-INS-62 | chr9-20546389-INS-72 |
| chr9-20564838-INS-70 | chr9-20564838-INS-114 |
| chr9-20588866-INS-88 | chr9-20588866-INS-88 |
| chr9-20593462-INS-159 | chr9-20593462-INS-159 |
| chr9-20597611-INS-625 | chr9-20597611-INS-624 |
| chr9-20625927-INS-56 | chr9-20625927-INS-72 |
| chr9-20639570-INS-82 | chr9-20639570-INS-82 |
| chr9-20667821-INS-67 | chr9-20667821-INS-67 |
| chr9-20680267-INS-66 | chr9-20680290-INS-54 |
| chr9-20688620-INS-119 | chr9-20688652-INS-94 |
| chr9-20695717-INS-91 | chr9-20695717-INS-91 |
| chr9-20698371-INS-85 | chr9-20698375-INS-67 |
| chr9-20719179-INS-54 | chr9-20719179-INS-58 |
| chr9-20750047-INS-225 | chr9-20750047-INS-224 |
| chr9-20752485-INS-60 | chr9-20752485-INS-66 |
| chr9-20759237-INS-57 | chr9-20759237-INS-57 |
| chr9-20774790-INS-105 | chr9-20774790-INS-75 |
| chr9-20792484-INS-114 | chr9-20792484-INS-114 |
| chr9-20841640-INS-6404 | chr9-20841641-INS-6404 |
| chr9-20853807-INS-77 | chr9-20853818-INS-78 |
| chr9-20889143-INS-52 | chr9-20889143-INS-58 |
| chr9-20897510-INS-104 | chr9-20897510-INS-117 |
| chr9-20907039-INS-253 | chr9-20907039-INS-252 |
| chr9-20917105-INS-62 | chr9-20917105-INS-50 |
| chr9-20921107-INS-493 | chr9-20921107-INS-498 |
| chr9-20922123-INS-199 | chr9-20922123-INS-199 |
| chr9-20924195-INS-207 | chr9-20924195-INS-206 |
| chr9-21045769-INS-108 | chr9-21045769-INS-104 |
| chr9-21058307-INS-75 | chr9-21058307-INS-75 |
| chr9-21074011-INS-105 | chr9-21074011-INS-105 |
| chr9-21089012-INS-211 | chr9-21089012-INS-210 |
| chr9-21094101-INS-90 | chr9-21094101-INS-52 |
| chr9-21099445-INS-66 | chr9-21099445-INS-66 |
| chr9-21110449-INS-179 | chr9-21110449-INS-170 |
| chr9-21113198-INS-424 | chr9-21113198-INS-424 |
| chr9-21120635-INS-194 | chr9-21120635-INS-156 |
| chr9-21128440-INS-194 | chr9-21128440-INS-192 |
| chr9-21130088-INS-215 | chr9-21130088-INS-215 |
| chr9-21148177-INS-183 | chr9-21148177-INS-184 |
| chr9-21152900-INS-107 | chr9-21152900-INS-122 |
| chr9-21169361-INS-182 | chr9-21169361-INS-182 |
| chr9-21172609-INS-202 | chr9-21172609-INS-202 |
| chr9-21173704-INS-6751 | chr9-21173705-INS-6751 |
| chr9-21185625-INS-237 | chr9-21185625-INS-212 |

| | |
|---|---|
| chr9-21211425-INS-166 | chr9-21211457-INS-184 |
| chr9-21212312-INS-665 | chr9-21212313-INS-669 |
| chr9-21229068-INS-255 | chr9-21229068-INS-255 |
| chr9-21235220-INS-193 | chr9-21235220-INS-193 |
| chr9-21247289-INS-216 | chr9-21247289-INS-215 |
| chr9-21267465-INS-77 | chr9-21267465-INS-77 |
| chr9-21269866-INS-206 | chr9-21269866-INS-197 |
| chr9-21329517-INS-177 | chr9-21329517-INS-181 |
| chr9-21369436-INS-144 | chr9-21369436-INS-144 |
| chr9-21383659-INS-1947 | chr9-21383660-INS-1937 |
| chr9-21384095-INS-231 | chr9-21384095-INS-231 |
| chr9-21423864-INS-1010 | chr9-21423865-INS-1011 |
| chr9-21434190-INS-8782 | chr9-21434191-INS-8782 |
| chr9-21438059-INS-199 | chr9-21438059-INS-200 |
| chr9-21454293-INS-185 | chr9-21454307-INS-187 |
| chr9-21471491-INS-121 | chr9-21471493-INS-120 |
| chr9-21495416-INS-1678 | chr9-21495417-INS-1678 |
| chr9-21514507-INS-228 | chr9-21514507-INS-229 |
| chr9-21543307-INS-177 | chr9-21543342-INS-172 |
| chr9-21609301-INS-213 | chr9-21609301-INS-213 |
| chr9-21618870-INS-938 | chr9-21618870-INS-961 |
| chr9-21620391-INS-234 | chr9-21620391-INS-241 |
| chr9-21628669-INS-59 | chr9-21628669-INS-79 |
| chr9-21636780-INS-202 | chr9-21636780-INS-202 |
| chr9-21638707-INS-217 | chr9-21638707-INS-217 |
| chr9-21642838-INS-79 | chr9-21642838-INS-79 |
| chr9-21647781-INS-184 | chr9-21647780-INS-184 |
| chr9-21663365-INS-880 | chr9-21663365-INS-880 |
| chr9-21670485-INS-880 | chr9-21670485-INS-880 |
| chr9-21681730-INS-84 | chr9-21681730-INS-84 |
| chr9-21683184-INS-216 | chr9-21683184-INS-239 |
| chr9-21693773-INS-84 | chr9-21693773-INS-84 |
| chr9-21694802-INS-205 | chr9-21694802-INS-205 |
| chr9-21697483-INS-52 | chr9-21697483-INS-52 |
| chr9-21698231-INS-104 | chr9-21698231-INS-104 |
| chr9-21699227-INS-216 | chr9-21699227-INS-216 |
| chr9-21713186-INS-70 | chr9-21713186-INS-78 |
| chr9-21721557-INS-226 | chr9-21721557-INS-226 |
| chr9-21727124-INS-238 | chr9-21727124-INS-238 |
| chr9-21728012-INS-90 | chr9-21728012-INS-90 |
| chr9-21733903-INS-92 | chr9-21733903-INS-92 |
| chr9-21757539-INS-98 | chr9-21757539-INS-98 |
| chr9-21770739-INS-253 | chr9-21770739-INS-256 |
| chr9-21790048-INS-60 | chr9-21790048-INS-60 |
| chr9-21813197-INS-6965 | chr9-21813198-INS-6965 |
| chr9-21823481-INS-372 | chr9-21823481-INS-364 |
| chr9-21833116-INS-148 | chr9-21833116-INS-148 |
| chr9-21835863-INS-72 | chr9-21835863-INS-72 |
| chr9-21850391-INS-97 | chr9-21850391-INS-53 |
| chr9-21933160-INS-221 | chr9-21933142-INS-252 |
| chr9-21938574-INS-86 | chr9-21938582-INS-59 |
| chr9-21950186-INS-68 | chr9-21950186-INS-74 |
| chr9-21958290-INS-84 | chr9-21958290-INS-84 |
| chr9-21980561-INS-54 | chr9-21980564-INS-50 |
| chr9-21990487-INS-197 | chr9-21990487-INS-197 |
| chr9-21999142-INS-214 | chr9-21999142-INS-214 |
| chr9-21999742-INS-239 | chr9-21999742-INS-239 |
| chr9-22002445-INS-62 | chr9-22002445-INS-62 |
| chr9-22004840-INS-183 | chr9-22004840-INS-164 |
| chr9-22026719-INS-430 | chr9-22026719-INS-431 |
| chr9-22027876-INS-88 | chr9-22027876-INS-130 |
| chr9-22040193-INS-72 | chr9-22040193-INS-72 |
| chr9-22044512-INS-150 | chr9-22044512-INS-150 |
| chr9-22049460-INS-133 | chr9-22049462-INS-150 |
| chr9-22050672-INS-120 | chr9-22050671-INS-120 |
| chr9-22056359-INS-137 | chr9-22056359-INS-137 |
| chr9-22058106-INS-189 | chr9-22058106-INS-190 |
| chr9-22062822-INS-216 | chr9-22062822-INS-216 |
| chr9-22067606-INS-198 | chr9-22067606-INS-198 |
| chr9-22069663-INS-245 | chr9-22069663-INS-246 |
| chr9-22081810-INS-234 | chr9-22081810-INS-235 |
| chr9-22084496-INS-915 | chr9-22084497-INS-939 |
| chr9-22085329-INS-221 | chr9-22085329-INS-221 |
| chr9-22101614-INS-442 | chr9-22101615-INS-442 |
| chr9-22177830-INS-208 | chr9-22177830-INS-207 |
| chr9-22179862-INS-378 | chr9-22179862-INS-380 |
| chr9-22181393-INS-76 | chr9-22181393-INS-76 |

| | |
|---|---|
| chr9-22199734-INS-80 | chr9-22199734-INS-100 |
| chr9-22203326-INS-66 | chr9-22203326-INS-66 |
| chr9-22205749-INS-931 | chr9-22205749-INS-932 |
| chr9-22211672-INS-64 | chr9-22211672-INS-64 |
| chr9-22217990-INS-167 | chr9-22217990-INS-168 |
| chr9-22255327-INS-206 | chr9-22255327-INS-206 |
| chr9-22263985-INS-198 | chr9-22263985-INS-200 |
| chr9-22276934-INS-176 | chr9-22276934-INS-176 |
| chr9-22309687-INS-5548 | chr9-22309688-INS-5548 |
| chr9-22328370-INS-400 | chr9-22328370-INS-400 |
| chr9-22352034-INS-55 | chr9-22352034-INS-82 |
| chr9-22352482-INS-715 | chr9-22352482-INS-715 |
| chr9-22353615-INS-258 | chr9-22353615-INS-258 |
| chr9-22389923-INS-56 | chr9-22389921-INS-100 |
| chr9-22394671-INS-51 | chr9-22394671-INS-51 |
| chr9-22397856-INS-654 | chr9-22397856-INS-654 |
| chr9-22400912-INS-156 | chr9-22400912-INS-160 |
| chr9-22413624-INS-4744 | chr9-22413625-INS-4744 |
| chr9-22414869-INS-447 | chr9-22414869-INS-443 |
| chr9-22424990-INS-90 | chr9-22424990-INS-90 |
| chr9-22429638-INS-64 | chr9-22429638-INS-74 |
| chr9-22440481-INS-118 | chr9-22440493-INS-82 |
| chr9-22445685-INS-55 | chr9-22445685-INS-76 |
| chr9-22467697-INS-6401 | chr9-22467698-INS-6401 |
| chr9-22507391-INS-76 | chr9-22507391-INS-76 |
| chr9-22516224-INS-66 | chr9-22516224-INS-66 |
| chr9-22522757-INS-58 | chr9-22522757-INS-52 |
| chr9-22547349-INS-88 | chr9-22547349-INS-88 |
| chr9-22550144-INS-54 | chr9-22550144-INS-54 |
| chr9-22622140-INS-61 | chr9-22622140-INS-61 |
| chr9-22623043-INS-565 | chr9-22623043-INS-565 |
| chr9-22642675-INS-212 | chr9-22642675-INS-212 |
| chr9-22650536-INS-146 | chr9-22650536-INS-147 |
| chr9-22661383-INS-76 | chr9-22661383-INS-76 |
| chr9-22669793-INS-211 | chr9-22669793-INS-211 |
| chr9-22671495-INS-102 | chr9-22671495-INS-102 |
| chr9-22684223-INS-71 | chr9-22684223-INS-83 |
| chr9-22698886-INS-135 | chr9-22698887-INS-136 |
| chr9-22707525-INS-100 | chr9-22707525-INS-100 |
| chr9-22708324-INS-200 | chr9-22708324-INS-200 |
| chr9-22709650-INS-52 | chr9-22709650-INS-64 |
| chr9-22778579-INS-279 | chr9-22778574-INS-252 |
| chr9-22796552-INS-216 | chr9-22796552-INS-216 |
| chr9-22862755-INS-111 | chr9-22862792-INS-74 |
| chr9-22869014-INS-56 | chr9-22869014-INS-72 |
| chr9-22873730-INS-51 | chr9-22873730-INS-51 |
| chr9-22880072-INS-3692 | chr9-22880073-INS-3693 |
| chr9-22935803-INS-74 | chr9-22935803-INS-74 |
| chr9-22991207-INS-99 | chr9-22991207-INS-75 |
| chr9-23001273-INS-61 | chr9-23001273-INS-61 |
| chr9-23029931-INS-450 | chr9-23029931-INS-450 |
| chr9-23069761-INS-51 | chr9-23069761-INS-51 |
| chr9-23121335-INS-64 | chr9-23121335-INS-64 |
| chr9-23155912-INS-78 | chr9-23155912-INS-74 |
| chr9-23179753-INS-455 | chr9-23179753-INS-455 |
| chr9-23195548-INS-212 | chr9-23195548-INS-212 |
| chr9-23204316-INS-214 | chr9-23204316-INS-214 |
| chr9-23228255-INS-78 | chr9-23228261-INS-82 |
| chr9-23240422-INS-292 | chr9-23240422-INS-292 |
| chr9-23248975-INS-368 | chr9-23248975-INS-372 |
| chr9-23264883-INS-74 | chr9-23264883-INS-74 |
| chr9-23282986-INS-64 | chr9-23282986-INS-64 |
| chr9-23389185-INS-533 | chr9-23389185-INS-533 |
| chr9-23391829-INS-83 | chr9-23391829-INS-83 |
| chr9-23400613-INS-802 | chr9-23400613-INS-799 |
| chr9-23407983-INS-112 | chr9-23407983-INS-112 |
| chr9-23419309-INS-72 | chr9-23419309-INS-52 |
| chr9-23427752-INS-53 | chr9-23427752-INS-53 |
| chr9-23463568-INS-52 | chr9-23463568-INS-54 |
| chr9-23463658-INS-6413 | chr9-23463659-INS-6411 |
| chr9-23467043-INS-108 | chr9-23467043-INS-108 |
| chr9-23480800-INS-201 | chr9-23480800-INS-201 |
| chr9-23482056-INS-58 | chr9-23482056-INS-58 |
| chr9-23514552-INS-120 | chr9-23514552-INS-122 |
| chr9-23534240-INS-231 | chr9-23534240-INS-231 |
| chr9-23542037-INS-61 | chr9-23542037-INS-61 |
| chr9-23549107-INS-2722 | chr9-23549108-INS-2723 |

| | |
|---|---|
| chr9-23611062-INS-251 | chr9-23611061-INS-241 |
| chr9-23611874-INS-228 | chr9-23611874-INS-228 |
| chr9-23622667-INS-194 | chr9-23622667-INS-194 |
| chr9-23634441-INS-104 | chr9-23634441-INS-72 |
| chr9-23644394-INS-2933 | chr9-23644395-INS-2933 |
| chr9-23647842-INS-2908 | chr9-23647843-INS-2908 |
| chr9-23661600-INS-97 | chr9-23661600-INS-136 |
| chr9-23667892-INS-317 | chr9-23667892-INS-317 |
| chr9-23707669-INS-403 | chr9-23707669-INS-403 |
| chr9-23719141-INS-7137 | chr9-23719142-INS-7137 |
| chr9-23745075-INS-6768 | chr9-23745076-INS-6768 |
| chr9-23751831-INS-204 | chr9-23751831-INS-204 |
| chr9-23776154-INS-86 | chr9-23776154-INS-86 |
| chr9-23780888-INS-134 | chr9-23780888-INS-132 |
| chr9-23783797-INS-53 | chr9-23783797-INS-54 |
| chr9-23832400-INS-214 | chr9-23832400-INS-214 |
| chr9-23835479-INS-80 | chr9-23835479-INS-82 |
| chr9-23837021-INS-72 | chr9-23837021-INS-76 |
| chr9-23875907-INS-342 | chr9-23875907-INS-342 |
| chr9-23880175-INS-52 | chr9-23880175-INS-70 |
| chr9-23919717-INS-180 | chr9-23919717-INS-180 |
| chr9-23978212-INS-65 | chr9-23978212-INS-97 |
| chr9-23990555-INS-218 | chr9-23990555-INS-218 |
| chr9-23993482-INS-82 | chr9-23993481-INS-86 |
| chr9-23993933-INS-3196 | chr9-23993934-INS-3193 |
| chr9-24022009-INS-167 | chr9-24021998-INS-158 |
| chr9-24040788-INS-52 | chr9-24040788-INS-52 |
| chr9-24042202-INS-82 | chr9-24042202-INS-82 |
| chr9-24046882-INS-98 | chr9-24046882-INS-108 |
| chr9-24078722-INS-480 | chr9-24078722-INS-480 |
| chr9-24081625-INS-56 | chr9-24081625-INS-56 |
| chr9-24083977-INS-51 | chr9-24083977-INS-52 |
| chr9-24084889-INS-3509 | chr9-24084890-INS-3510 |
| chr9-24105181-INS-7524 | chr9-24105182-INS-7527 |
| chr9-24115020-INS-6402 | chr9-24115021-INS-6402 |
| chr9-24120086-INS-196 | chr9-24120086-INS-197 |
| chr9-24123713-INS-100 | chr9-24123713-INS-112 |
| chr9-24128534-INS-315 | chr9-24128534-INS-315 |
| chr9-24142696-INS-401 | chr9-24142696-INS-401 |
| chr9-24143820-INS-73 | chr9-24143820-INS-73 |
| chr9-24145678-INS-75 | chr9-24145671-INS-84 |
| chr9-24169863-INS-90 | chr9-24169863-INS-90 |
| chr9-24211368-INS-81 | chr9-24211336-INS-108 |
| chr9-24216043-INS-80 | chr9-24216052-INS-70 |
| chr9-24240342-INS-348 | chr9-24240342-INS-348 |
| chr9-24242534-INS-76 | chr9-24242534-INS-96 |
| chr9-24246735-INS-682 | chr9-24246735-INS-682 |
| chr9-24251898-INS-55 | chr9-24251898-INS-55 |
| chr9-24257796-INS-360 | chr9-24257796-INS-360 |
| chr9-24258936-INS-1994 | chr9-24258937-INS-1971 |
| chr9-24271693-INS-144 | chr9-24271693-INS-144 |
| chr9-24282983-INS-585 | chr9-24282982-INS-567 |
| chr9-24314509-INS-60 | chr9-24314509-INS-55 |
| chr9-24318171-INS-56 | chr9-24318171-INS-56 |
| chr9-24331415-INS-1893 | chr9-24331416-INS-1893 |
| chr9-24349140-INS-2275 | chr9-24349141-INS-2258 |
| chr9-24382691-INS-65 | chr9-24382691-INS-65 |
| chr9-24385992-INS-122 | chr9-24385992-INS-122 |
| chr9-24388467-INS-163 | chr9-24388429-INS-157 |
| chr9-24429248-INS-438 | chr9-24429248-INS-438 |
| chr9-24437027-INS-214 | chr9-24437027-INS-214 |
| chr9-24479267-INS-81 | chr9-24479267-INS-81 |
| chr9-24488818-INS-436 | chr9-24488818-INS-437 |
| chr9-24500411-INS-526 | chr9-24500411-INS-526 |
| chr9-24501700-INS-6402 | chr9-24501701-INS-6402 |
| chr9-24504997-INS-4046 | chr9-24504998-INS-4047 |
| chr9-24521059-INS-1884 | chr9-24521055-INS-1882 |
| chr9-24532355-INS-2750 | chr9-24532356-INS-2752 |
| chr9-24538713-INS-682 | chr9-24538713-INS-682 |
| chr9-24557180-INS-71 | chr9-24557180-INS-71 |
| chr9-24639694-INS-3246 | chr9-24639695-INS-3246 |
| chr9-24640456-INS-99 | chr9-24640456-INS-99 |
| chr9-24647789-INS-90 | chr9-24647789-INS-94 |
| chr9-24659010-INS-58 | chr9-24659010-INS-58 |
| chr9-24698081-INS-52 | chr9-24698080-INS-74 |
| chr9-24705097-INS-55 | chr9-24705097-INS-65 |
| chr9-24721117-INS-1894 | chr9-24721118-INS-1894 |

| | |
|---|---|
| chr9-24723564-INS-6401 | chr9-24723565-INS-6401 |
| chr9-24733222-INS-210 | chr9-24733222-INS-202 |
| chr9-24736787-INS-61 | chr9-24736787-INS-61 |
| chr9-24751899-INS-209 | chr9-24751899-INS-209 |
| chr9-24755919-INS-214 | chr9-24755919-INS-214 |
| chr9-24770641-INS-80 | chr9-24770641-INS-80 |
| chr9-24784187-INS-1048 | chr9-24784188-INS-1051 |
| chr9-24788268-INS-284 | chr9-24788268-INS-282 |
| chr9-24792080-INS-7314 | chr9-24792081-INS-7314 |
| chr9-24793468-INS-1107 | chr9-24793469-INS-1105 |
| chr9-24798322-INS-3217 | chr9-24798323-INS-3217 |
| chr9-24798997-INS-163 | chr9-24798997-INS-163 |
| chr9-24843456-INS-65 | chr9-24843456-INS-65 |
| chr9-24848279-INS-6473 | chr9-24848280-INS-6473 |
| chr9-24853601-INS-64 | chr9-24853601-INS-61 |
| chr9-24856276-INS-72 | chr9-24856276-INS-72 |
| chr9-24860055-INS-53 | chr9-24860055-INS-79 |
| chr9-24887914-INS-74 | chr9-24887914-INS-74 |
| chr9-24891993-INS-216 | chr9-24891993-INS-216 |
| chr9-24903756-INS-63 | chr9-24903756-INS-63 |
| chr9-24914152-INS-6219 | chr9-24914153-INS-6219 |
| chr9-24924062-INS-69 | chr9-24924062-INS-69 |
| chr9-24939419-INS-119 | chr9-24939419-INS-119 |
| chr9-24956585-INS-58 | chr9-24956581-INS-62 |
| chr9-24969944-INS-2734 | chr9-24969945-INS-2734 |
| chr9-24991066-INS-267 | chr9-24991066-INS-267 |
| chr9-24999541-INS-552 | chr9-24999541-INS-552 |
| chr9-25000726-INS-76 | chr9-25000729-INS-64 |
| chr9-25010297-INS-358 | chr9-25010297-INS-358 |
| chr9-25067805-INS-235 | chr9-25067805-INS-233 |
| chr9-25068085-INS-6279 | chr9-25068086-INS-6279 |
| chr9-25087261-INS-129 | chr9-25087261-INS-129 |
| chr9-25095165-INS-80 | chr9-25095165-INS-80 |
| chr9-25131252-INS-722 | chr9-25131252-INS-723 |
| chr9-25141096-INS-458 | chr9-25141096-INS-458 |
| chr9-25170521-INS-1124 | chr9-25170522-INS-1119 |
| chr9-25172569-INS-1067 | chr9-25172570-INS-1067 |
| chr9-25188910-INS-120 | chr9-25188910-INS-120 |
| chr9-25202806-INS-91 | chr9-25202806-INS-91 |
| chr9-25216145-INS-188 | chr9-25216145-INS-190 |
| chr9-25231381-INS-71 | chr9-25231380-INS-109 |
| chr9-25236334-INS-3627 | chr9-25236335-INS-3627 |
| chr9-25252573-INS-61 | chr9-25252574-INS-56 |
| chr9-25257733-INS-382 | chr9-25257733-INS-381 |
| chr9-25270463-INS-286 | chr9-25270475-INS-307 |
| chr9-25275764-INS-75 | chr9-25275813-INS-54 |
| chr9-25280987-INS-214 | chr9-25280986-INS-242 |
| chr9-25300227-INS-61 | chr9-25300222-INS-95 |
| chr9-25307645-INS-1968 | chr9-25307646-INS-1967 |
| chr9-25322317-INS-198 | chr9-25322317-INS-201 |
| chr9-25325542-INS-65 | chr9-25325542-INS-65 |
| chr9-25357788-INS-7080 | chr9-25357789-INS-7080 |
| chr9-25363652-INS-60 | chr9-25363652-INS-65 |
| chr9-25364347-INS-546 | chr9-25364347-INS-546 |
| chr9-25365912-INS-3208 | chr9-25365913-INS-3208 |
| chr9-25414196-INS-109 | chr9-25414196-INS-108 |
| chr9-25434178-INS-3165 | chr9-25434179-INS-3165 |
| chr9-25458157-INS-132 | chr9-25458157-INS-111 |
| chr9-25458611-INS-6651 | chr9-25458612-INS-6651 |
| chr9-25481102-INS-937 | chr9-25481102-INS-938 |
| chr9-25490859-INS-75 | chr9-25490859-INS-75 |
| chr9-25530449-INS-1384 | chr9-25530450-INS-1384 |
| chr9-25532772-INS-6426 | chr9-25532773-INS-6426 |
| chr9-25535755-INS-182 | chr9-25535755-INS-182 |
| chr9-25540887-INS-4428 | chr9-25540888-INS-4428 |
| chr9-25585018-INS-89 | chr9-25585018-INS-89 |
| chr9-25588108-INS-2830 | chr9-25588109-INS-2830 |
| chr9-25592400-INS-55 | chr9-25592400-INS-55 |
| chr9-25608619-INS-360 | chr9-25608619-INS-360 |
| chr9-25614150-INS-1851 | chr9-25614151-INS-1851 |
| chr9-25620347-INS-214 | chr9-25620347-INS-214 |
| chr9-25624623-INS-67 | chr9-25624623-INS-53 |
| chr9-25644771-INS-2163 | chr9-25644772-INS-2163 |
| chr9-25679827-INS-145 | chr9-25679827-INS-145 |
| chr9-25715062-INS-150 | chr9-25715062-INS-150 |
| chr9-25717162-INS-435 | chr9-25717158-INS-435 |
| chr9-25720151-INS-178 | chr9-25720151-INS-130 |

| | |
|---|---|
| chr9-25738050-INS-102 | chr9-25738053-INS-103 |
| chr9-25787819-INS-226 | chr9-25787819-INS-226 |
| chr9-25811251-INS-105 | chr9-25811251-INS-105 |
| chr9-25830040-INS-54 | chr9-25830040-INS-58 |
| chr9-25857980-INS-102 | chr9-25857980-INS-102 |
| chr9-25870685-INS-52 | chr9-25870685-INS-54 |
| chr9-25895301-INS-72 | chr9-25895301-INS-70 |
| chr9-25971639-INS-3737 | chr9-25971640-INS-3738 |
| chr9-25991361-INS-52 | chr9-25991355-INS-50 |
| chr9-26019081-INS-62 | chr9-26019065-INS-68 |
| chr9-26024592-INS-94 | chr9-26024592-INS-52 |
| chr9-26024877-INS-847 | chr9-26024877-INS-847 |
| chr9-26026343-INS-193 | chr9-26026343-INS-193 |
| chr9-26026653-INS-1426 | chr9-26026654-INS-1426 |
| chr9-26037339-INS-116 | chr9-26037339-INS-116 |
| chr9-26040406-INS-258 | chr9-26040398-INS-264 |
| chr9-26056120-INS-56 | chr9-26056120-INS-58 |
| chr9-26058777-INS-4703 | chr9-26058778-INS-4703 |
| chr9-26079676-INS-73 | chr9-26079680-INS-104 |
| chr9-26090805-INS-579 | chr9-26090805-INS-579 |
| chr9-26106362-INS-6162 | chr9-26106363-INS-6160 |
| chr9-26193231-INS-60 | chr9-26193231-INS-50 |
| chr9-26259601-INS-382 | chr9-26259601-INS-382 |
| chr9-26265698-INS-216 | chr9-26265698-INS-215 |
| chr9-26343682-INS-185 | chr9-26343682-INS-171 |
| chr9-26347449-INS-59 | chr9-26347449-INS-59 |
| chr9-26395428-INS-156 | chr9-26395383-INS-155 |
| chr9-26443818-INS-129 | chr9-26443818-INS-129 |
| chr9-26444826-INS-214 | chr9-26444828-INS-182 |
| chr9-26454005-INS-511 | chr9-26454005-INS-510 |
| chr9-26534843-INS-93 | chr9-26534843-INS-105 |
| chr9-26549970-INS-60 | chr9-26549962-INS-50 |
| chr9-26594344-INS-475 | chr9-26594344-INS-480 |
| chr9-26617954-INS-4076 | chr9-26617955-INS-4078 |
| chr9-26649812-INS-60 | chr9-26649812-INS-90 |
| chr9-26707492-INS-98 | chr9-26707528-INS-108 |
| chr9-26710354-INS-52 | chr9-26710354-INS-52 |
| chr9-26795298-INS-101 | chr9-26795298-INS-106 |
| chr9-26838746-INS-191 | chr9-26838746-INS-203 |
| chr9-26842183-INS-737 | chr9-26842183-INS-740 |
| chr9-26862569-INS-66 | chr9-26862569-INS-66 |
| chr9-26870099-INS-7214 | chr9-26870100-INS-7214 |
| chr9-26874386-INS-1869 | chr9-26874387-INS-1869 |
| chr9-26893754-INS-214 | chr9-26893754-INS-214 |
| chr9-26912247-INS-328 | chr9-26912247-INS-328 |
| chr9-26913313-INS-1336 | chr9-26913314-INS-1336 |
| chr9-26913476-INS-205 | chr9-26913476-INS-205 |
| chr9-26921578-INS-72 | chr9-26921578-INS-72 |
| chr9-26977414-INS-711 | chr9-26977414-INS-701 |
| chr9-27030768-INS-3303 | chr9-27030769-INS-3304 |
| chr9-27045072-INS-84 | chr9-27045072-INS-84 |
| chr9-27088030-INS-66 | chr9-27088030-INS-69 |
| chr9-27101791-INS-4920 | chr9-27101792-INS-4920 |
| chr9-27109172-INS-59 | chr9-27109172-INS-57 |
| chr9-27123030-INS-58 | chr9-27123030-INS-60 |
| chr9-27175165-INS-352 | chr9-27175165-INS-352 |
| chr9-27177920-INS-6895 | chr9-27177921-INS-6896 |
| chr9-27190230-INS-6638 | chr9-27190231-INS-6638 |
| chr9-27289241-INS-6440 | chr9-27289242-INS-6440 |
| chr9-27332100-INS-476 | chr9-27332100-INS-476 |
| chr9-27342670-INS-96 | chr9-27342670-INS-104 |
| chr9-27355578-INS-69 | chr9-27355578-INS-69 |
| chr9-27405807-INS-54 | chr9-27405807-INS-54 |
| chr9-27460091-INS-97 | chr9-27460088-INS-54 |
| chr9-27465105-INS-70 | chr9-27465109-INS-59 |
| chr9-27474483-INS-355 | chr9-27474483-INS-355 |
| chr9-27529251-INS-64 | chr9-27529251-INS-64 |
| chr9-27553436-INS-390 | chr9-27553436-INS-390 |
| chr9-27572644-INS-56 | chr9-27572644-INS-56 |
| chr9-27608960-INS-73 | chr9-27608960-INS-73 |
| chr9-27640262-INS-102 | chr9-27640262-INS-102 |
| chr9-27676991-INS-498 | chr9-27676991-INS-498 |
| chr9-27680595-INS-79 | chr9-27680595-INS-80 |
| chr9-27684104-INS-61 | chr9-27684104-INS-61 |
| chr9-27687112-INS-69 | chr9-27687122-INS-50 |
| chr9-27727410-INS-287 | chr9-27727410-INS-287 |
| chr9-27750363-INS-52 | chr9-27750363-INS-52 |

| | |
|---|---|
| chr9-27752512-INS-657 | chr9-27752512-INS-648 |
| chr9-27753785-INS-196 | chr9-27753785-INS-216 |
| chr9-27845219-INS-78 | chr9-27845216-INS-66 |
| chr9-27853344-INS-153 | chr9-27853344-INS-132 |
| chr9-27864002-INS-6277 | chr9-27864003-INS-6277 |
| chr9-27882565-INS-504 | chr9-27882565-INS-504 |
| chr9-27917210-INS-55 | chr9-27917211-INS-54 |
| chr9-27918736-INS-770 | chr9-27918736-INS-770 |
| chr9-27923421-INS-61 | chr9-27923421-INS-61 |
| chr9-27933662-INS-85 | chr9-27933668-INS-53 |
| chr9-27936942-INS-548 | chr9-27936942-INS-547 |
| chr9-27949449-INS-651 | chr9-27949449-INS-651 |
| chr9-27993131-INS-487 | chr9-27993131-INS-487 |
| chr9-28024996-INS-52 | chr9-28024996-INS-52 |
| chr9-28033108-INS-70 | chr9-28033108-INS-72 |
| chr9-28060145-INS-160 | chr9-28060145-INS-160 |
| chr9-28070867-INS-102 | chr9-28070882-INS-105 |
| chr9-28073740-INS-63 | chr9-28073740-INS-63 |
| chr9-28101408-INS-64 | chr9-28101408-INS-91 |
| chr9-28126906-INS-1246 | chr9-28126907-INS-1246 |
| chr9-28141601-INS-68 | chr9-28141602-INS-69 |
| chr9-28147430-INS-724 | chr9-28147430-INS-724 |
| chr9-28157809-INS-141 | chr9-28157809-INS-141 |
| chr9-28177287-INS-62 | chr9-28177287-INS-54 |
| chr9-28211834-INS-88 | chr9-28211834-INS-88 |
| chr9-28253557-INS-1015 | chr9-28253558-INS-1015 |
| chr9-28258963-INS-1893 | chr9-28258964-INS-1893 |
| chr9-28277221-INS-397 | chr9-28277221-INS-399 |
| chr9-28280489-INS-56 | chr9-28280486-INS-50 |
| chr9-28286073-INS-78 | chr9-28286073-INS-79 |
| chr9-28332589-INS-5548 | chr9-28332590-INS-5548 |
| chr9-28354612-INS-52 | chr9-28354612-INS-52 |
| chr9-28357038-INS-70 | chr9-28357038-INS-70 |
| chr9-28421236-INS-73 | chr9-28421226-INS-94 |
| chr9-28438721-INS-62 | chr9-28438721-INS-70 |
| chr9-28476789-INS-58 | chr9-28476789-INS-58 |
| chr9-28478064-INS-664 | chr9-28478064-INS-664 |
| chr9-28524608-INS-109 | chr9-28524608-INS-109 |
| chr9-28545915-INS-611 | chr9-28545915-INS-611 |
| chr9-28553202-INS-68 | chr9-28553209-INS-76 |
| chr9-28575856-INS-6464 | chr9-28575857-INS-6464 |
| chr9-28577964-INS-3186 | chr9-28577965-INS-3186 |
| chr9-28582420-INS-920 | chr9-28582420-INS-920 |
| chr9-28608171-INS-112 | chr9-28608171-INS-122 |
| chr9-28684101-INS-462 | chr9-28684101-INS-462 |
| chr9-28696959-INS-116 | chr9-28696959-INS-125 |
| chr9-28730411-INS-55 | chr9-28730411-INS-57 |
| chr9-28731961-INS-95 | chr9-28731963-INS-72 |
| chr9-28742716-INS-1751 | chr9-28742717-INS-1751 |
| chr9-28743538-INS-1892 | chr9-28743539-INS-1892 |
| chr9-28744085-INS-65 | chr9-28744085-INS-67 |
| chr9-28746818-INS-62 | chr9-28746834-INS-64 |
| chr9-28793476-INS-51 | chr9-28793476-INS-55 |
| chr9-28850597-INS-72 | chr9-28850597-INS-66 |
| chr9-28857008-INS-427 | chr9-28857008-INS-427 |
| chr9-28868132-INS-51 | chr9-28868132-INS-51 |
| chr9-28869595-INS-71 | chr9-28869595-INS-71 |
| chr9-28887229-INS-162 | chr9-28887229-INS-162 |
| chr9-28890242-INS-121 | chr9-28890242-INS-121 |
| chr9-28923101-INS-555 | chr9-28923101-INS-555 |
| chr9-28948811-INS-442 | chr9-28948811-INS-443 |
| chr9-28953832-INS-153 | chr9-28953832-INS-153 |
| chr9-28958878-INS-58 | chr9-28958878-INS-58 |
| chr9-28972992-INS-3342 | chr9-28972993-INS-3342 |
| chr9-28997357-INS-6631 | chr9-28997358-INS-6631 |
| chr9-29038049-INS-52 | chr9-29038049-INS-54 |
| chr9-29046287-INS-76 | chr9-29046287-INS-76 |
| chr9-29084800-INS-6649 | chr9-29084801-INS-6649 |
| chr9-29159039-INS-59 | chr9-29159039-INS-59 |
| chr9-29197577-INS-72 | chr9-29197617-INS-72 |
| chr9-29200823-INS-54 | chr9-29200823-INS-57 |
| chr9-29268444-INS-133 | chr9-29268439-INS-180 |
| chr9-29286590-INS-400 | chr9-29286590-INS-400 |
| chr9-29316681-INS-55 | chr9-29316673-INS-51 |
| chr9-29326794-INS-60 | chr9-29326794-INS-64 |
| chr9-29338584-INS-55 | chr9-29338584-INS-55 |
| chr9-29364991-INS-84 | chr9-29364987-INS-85 |

| | |
|---|---|
| chr9-29396790-INS-6463 | chr9-29396791-INS-6463 |
| chr9-29397331-INS-99 | chr9-29397331-INS-105 |
| chr9-29491197-INS-6380 | chr9-29491198-INS-6379 |
| chr9-29516839-INS-104 | chr9-29516839-INS-104 |
| chr9-29563784-INS-6749 | chr9-29563785-INS-6751 |
| chr9-29595244-INS-51 | chr9-29595244-INS-51 |
| chr9-29606502-INS-182 | chr9-29606502-INS-182 |
| chr9-29613445-INS-85 | chr9-29613445-INS-71 |
| chr9-29647035-INS-101 | chr9-29646995-INS-130 |
| chr9-29666521-INS-74 | chr9-29666517-INS-78 |
| chr9-29710889-INS-60 | chr9-29710889-INS-60 |
| chr9-29769003-INS-290 | chr9-29769003-INS-290 |
| chr9-29771249-INS-76 | chr9-29771249-INS-76 |
| chr9-29779848-INS-62 | chr9-29779848-INS-64 |
| chr9-29859652-INS-51 | chr9-29859652-INS-54 |
| chr9-29878163-INS-66 | chr9-29878163-INS-66 |
| chr9-29888192-INS-81 | chr9-29888192-INS-66 |
| chr9-29889289-INS-56 | chr9-29889289-INS-56 |
| chr9-29893987-INS-108 | chr9-29893992-INS-135 |
| chr9-29898961-INS-160 | chr9-29898961-INS-160 |
| chr9-29918075-INS-98 | chr9-29918074-INS-131 |
| chr9-29943193-INS-192 | chr9-29943193-INS-192 |
| chr9-29957202-INS-64 | chr9-29957216-INS-70 |
| chr9-29972524-INS-91 | chr9-29972522-INS-50 |
| chr9-29982290-INS-71 | chr9-29982290-INS-71 |
| chr9-29993391-INS-6415 | chr9-29993392-INS-6415 |
| chr9-30065308-INS-64 | chr9-30065308-INS-72 |
| chr9-30131322-INS-6255 | chr9-30131323-INS-6253 |
| chr9-30157356-INS-6384 | chr9-30157357-INS-6384 |
| chr9-30163583-INS-93 | chr9-30163557-INS-104 |
| chr9-30183149-INS-51 | chr9-30183149-INS-66 |
| chr9-30193464-INS-59 | chr9-30193464-INS-59 |
| chr9-30199624-INS-4199 | chr9-30199625-INS-4185 |
| chr9-30220419-INS-1863 | chr9-30220420-INS-1863 |
| chr9-30222542-INS-120 | chr9-30222542-INS-120 |
| chr9-30232766-INS-88 | chr9-30232766-INS-112 |
| chr9-30238312-INS-2061 | chr9-30238313-INS-2068 |
| chr9-30241303-INS-60 | chr9-30241327-INS-56 |
| chr9-30241788-INS-7113 | chr9-30241789-INS-7113 |
| chr9-30246413-INS-216 | chr9-30246414-INS-212 |
| chr9-30248213-INS-141 | chr9-30248209-INS-173 |
| chr9-30249862-INS-333 | chr9-30249862-INS-333 |
| chr9-30263967-INS-63 | chr9-30263967-INS-63 |
| chr9-30287153-INS-53 | chr9-30287153-INS-53 |
| chr9-30299016-INS-6639 | chr9-30299017-INS-6639 |
| chr9-30309263-INS-498 | chr9-30309263-INS-498 |
| chr9-30337790-INS-111 | chr9-30337790-INS-111 |
| chr9-30350222-INS-52 | chr9-30350222-INS-52 |
| chr9-30374146-INS-54 | chr9-30374146-INS-52 |
| chr9-30398736-INS-1753 | chr9-30398737-INS-1752 |
| chr9-30419099-INS-766 | chr9-30419099-INS-761 |
| chr9-30423939-INS-95 | chr9-30423939-INS-95 |
| chr9-30512042-INS-553 | chr9-30512042-INS-553 |
| chr9-30522510-INS-114 | chr9-30522510-INS-108 |
| chr9-30532832-INS-980 | chr9-30532832-INS-980 |
| chr9-30538733-INS-98 | chr9-30538767-INS-102 |
| chr9-30541250-INS-87 | chr9-30541240-INS-80 |
| chr9-30570064-INS-65 | chr9-30570064-INS-65 |
| chr9-30613298-INS-353 | chr9-30613298-INS-353 |
| chr9-30623362-INS-540 | chr9-30623357-INS-540 |
| chr9-30626352-INS-359 | chr9-30626352-INS-359 |
| chr9-30635373-INS-210 | chr9-30635381-INS-211 |
| chr9-30657981-INS-184 | chr9-30657956-INS-195 |
| chr9-30658564-INS-175 | chr9-30658564-INS-175 |
| chr9-30667903-INS-85 | chr9-30667902-INS-80 |
| chr9-30688936-INS-364 | chr9-30688936-INS-364 |
| chr9-30710313-INS-7816 | chr9-30710314-INS-7816 |
| chr9-30743297-INS-75 | chr9-30743297-INS-60 |
| chr9-30764101-INS-58 | chr9-30764101-INS-60 |
| chr9-30777120-INS-59 | chr9-30777120-INS-59 |
| chr9-30804567-INS-274 | chr9-30804567-INS-274 |
| chr9-30875479-INS-94 | chr9-30875477-INS-64 |
| chr9-30885975-INS-362 | chr9-30885975-INS-359 |
| chr9-30898584-INS-4234 | chr9-30898585-INS-4234 |
| chr9-30901949-INS-181 | chr9-30901949-INS-182 |
| chr9-30919707-INS-202 | chr9-30919707-INS-202 |
| chr9-30927067-INS-3507 | chr9-30927068-INS-3490 |

| | |
|---|---|
| chr9-30963572-INS-217 | chr9-30963572-INS-217 |
| chr9-30983090-INS-5545 | chr9-30983091-INS-5545 |
| chr9-31001340-INS-78 | chr9-31001331-INS-72 |
| chr9-31034018-INS-54 | chr9-31034017-INS-52 |
| chr9-31034024-INS-793 | chr9-31034024-INS-797 |
| chr9-31036206-INS-78 | chr9-31036206-INS-74 |
| chr9-31051765-INS-256 | chr9-31051765-INS-253 |
| chr9-31073268-INS-100 | chr9-31073268-INS-100 |
| chr9-31093495-INS-250 | chr9-31093500-INS-251 |
| chr9-31097077-INS-120 | chr9-31097077-INS-128 |
| chr9-31112997-INS-170 | chr9-31112997-INS-178 |
| chr9-31136986-INS-196 | chr9-31136986-INS-198 |
| chr9-31150577-INS-738 | chr9-31150577-INS-738 |
| chr9-31186179-INS-81 | chr9-31186179-INS-81 |
| chr9-31199762-INS-56 | chr9-31199762-INS-54 |
| chr9-31208631-INS-102 | chr9-31208599-INS-85 |
| chr9-31211911-INS-52 | chr9-31211911-INS-52 |
| chr9-31222854-INS-118 | chr9-31222854-INS-118 |
| chr9-31230225-INS-56 | chr9-31230225-INS-56 |
| chr9-31236214-INS-212 | chr9-31236214-INS-208 |
| chr9-31239641-INS-71 | chr9-31239609-INS-60 |
| chr9-31248977-INS-112 | chr9-31248977-INS-124 |
| chr9-31266455-INS-177 | chr9-31266455-INS-177 |
| chr9-31276924-INS-65 | chr9-31276924-INS-65 |
| chr9-31315210-INS-190 | chr9-31315210-INS-190 |
| chr9-31349554-INS-81 | chr9-31349554-INS-73 |
| chr9-31353094-INS-204 | chr9-31353094-INS-203 |
| chr9-31354063-INS-115 | chr9-31354063-INS-115 |
| chr9-31366569-INS-192 | chr9-31366569-INS-192 |
| chr9-31374345-INS-6355 | chr9-31374346-INS-6357 |
| chr9-31383142-INS-51 | chr9-31383147-INS-51 |
| chr9-31448895-INS-66 | chr9-31448895-INS-66 |
| chr9-31452705-INS-178 | chr9-31452705-INS-179 |
| chr9-31454742-INS-192 | chr9-31454749-INS-198 |
| chr9-31468852-INS-312 | chr9-31468852-INS-312 |
| chr9-31496723-INS-447 | chr9-31496723-INS-447 |
| chr9-31514765-INS-2876 | chr9-31514766-INS-2876 |
| chr9-31535175-INS-150 | chr9-31535175-INS-150 |
| chr9-31543513-INS-397 | chr9-31543515-INS-398 |
| chr9-31551288-INS-52 | chr9-31551288-INS-72 |
| chr9-31554400-INS-159 | chr9-31554400-INS-159 |
| chr9-31559189-INS-65 | chr9-31559189-INS-50 |
| chr9-31559864-INS-489 | chr9-31559864-INS-489 |
| chr9-31613128-INS-73 | chr9-31613128-INS-68 |
| chr9-31614201-INS-200 | chr9-31614201-INS-200 |
| chr9-31640290-INS-2924 | chr9-31640291-INS-2924 |
| chr9-31642821-INS-160 | chr9-31642819-INS-190 |
| chr9-31643780-INS-54 | chr9-31643788-INS-80 |
| chr9-31646282-INS-201 | chr9-31646282-INS-201 |
| chr9-31698344-INS-72 | chr9-31698344-INS-72 |
| chr9-31701233-INS-1386 | chr9-31701234-INS-1428 |
| chr9-31718083-INS-6460 | chr9-31718084-INS-6460 |
| chr9-31745209-INS-58 | chr9-31745209-INS-54 |
| chr9-31746429-INS-66 | chr9-31746429-INS-66 |
| chr9-31758485-INS-725 | chr9-31758485-INS-730 |
| chr9-31776488-INS-60 | chr9-31776488-INS-60 |
| chr9-31815826-INS-59 | chr9-31815826-INS-59 |
| chr9-31829359-INS-50 | chr9-31829359-INS-56 |
| chr9-31849118-INS-198 | chr9-31849118-INS-198 |
| chr9-31858415-INS-567 | chr9-31858415-INS-567 |
| chr9-31880427-INS-113 | chr9-31880427-INS-113 |
| chr9-31892971-INS-101 | chr9-31892971-INS-68 |
| chr9-31906867-INS-202 | chr9-31906867-INS-202 |
| chr9-31918956-INS-400 | chr9-31918952-INS-400 |
| chr9-31957051-INS-94 | chr9-31957051-INS-94 |
| chr9-31971139-INS-213 | chr9-31971139-INS-214 |
| chr9-31989369-INS-70 | chr9-31989368-INS-70 |
| chr9-31996117-INS-55 | chr9-31996117-INS-55 |
| chr9-32012321-INS-504 | chr9-32012321-INS-504 |
| chr9-32038751-INS-52 | chr9-32038751-INS-52 |
| chr9-32073712-INS-733 | chr9-32073712-INS-733 |
| chr9-32091795-INS-622 | chr9-32091798-INS-639 |
| chr9-32093033-INS-747 | chr9-32093033-INS-747 |
| chr9-32096109-INS-80 | chr9-32096109-INS-68 |
| chr9-32142607-INS-185 | chr9-32142607-INS-185 |
| chr9-32164693-INS-50 | chr9-32164693-INS-50 |
| chr9-32177234-INS-76 | chr9-32177234-INS-88 |

| | |
|---|---|
| chr9-32246901-INS-845 | chr9-32246901-INS-842 |
| chr9-32247684-INS-72 | chr9-32247684-INS-74 |
| chr9-32265798-INS-356 | chr9-32265814-INS-372 |
| chr9-32283510-INS-63 | chr9-32283510-INS-63 |
| chr9-32302641-INS-52 | chr9-32302641-INS-54 |
| chr9-32320209-INS-60 | chr9-32320209-INS-66 |
| chr9-32366385-INS-60 | chr9-32366385-INS-66 |
| chr9-32369937-INS-58 | chr9-32369937-INS-80 |
| chr9-32403548-INS-58 | chr9-32403558-INS-62 |
| chr9-32408358-INS-88 | chr9-32408358-INS-52 |
| chr9-32420559-INS-94 | chr9-32420559-INS-94 |
| chr9-32444450-INS-53 | chr9-32444446-INS-51 |
| chr9-32448815-INS-65 | chr9-32448815-INS-65 |
| chr9-32453375-INS-103 | chr9-32453375-INS-100 |
| chr9-32505117-INS-85 | chr9-32505140-INS-112 |
| chr9-32592943-INS-56 | chr9-32592944-INS-58 |
| chr9-32596213-INS-64 | chr9-32596213-INS-64 |
| chr9-32609482-INS-54 | chr9-32609482-INS-57 |
| chr9-32686578-INS-201 | chr9-32686578-INS-201 |
| chr9-32740309-INS-501 | chr9-32740309-INS-499 |
| chr9-32742872-INS-71 | chr9-32742872-INS-52 |
| chr9-32758121-INS-64 | chr9-32758121-INS-54 |
| chr9-32769092-INS-72 | chr9-32769092-INS-72 |
| chr9-32770991-INS-102 | chr9-32770991-INS-102 |
| chr9-32833696-INS-70 | chr9-32833696-INS-72 |
| chr9-32847235-INS-79 | chr9-32847235-INS-79 |
| chr9-32849250-INS-104 | chr9-32849250-INS-104 |
| chr9-32857172-INS-215 | chr9-32857172-INS-217 |
| chr9-32870412-INS-54 | chr9-32870412-INS-54 |
| chr9-32878133-INS-307 | chr9-32878133-INS-299 |
| chr9-32889009-INS-79 | chr9-32889009-INS-81 |
| chr9-32913571-INS-99 | chr9-32913571-INS-100 |
| chr9-32921396-INS-60 | chr9-32921413-INS-75 |
| chr9-32937645-INS-54 | chr9-32937645-INS-54 |
| chr9-32942468-INS-76 | chr9-32942466-INS-78 |
| chr9-32975698-INS-6620 | chr9-32975699-INS-6620 |
| chr9-33004524-INS-142 | chr9-33004569-INS-142 |
| chr9-33007672-INS-77 | chr9-33007672-INS-85 |
| chr9-33030194-INS-504 | chr9-33030194-INS-504 |
| chr9-33034700-INS-180 | chr9-33034700-INS-180 |
| chr9-33044012-INS-5171 | chr9-33044013-INS-5171 |
| chr9-33046597-INS-6600 | chr9-33046598-INS-6601 |
| chr9-33054203-INS-822 | chr9-33054203-INS-823 |
| chr9-33055007-INS-199 | chr9-33055007-INS-199 |
| chr9-33081499-INS-68 | chr9-33081499-INS-74 |
| chr9-33093489-INS-237 | chr9-33093489-INS-233 |
| chr9-33141271-INS-203 | chr9-33141271-INS-200 |
| chr9-33143851-INS-90 | chr9-33143833-INS-60 |
| chr9-33156453-INS-106 | chr9-33156445-INS-103 |
| chr9-33157716-INS-83 | chr9-33157716-INS-83 |
| chr9-33169383-INS-52 | chr9-33169383-INS-54 |
| chr9-33188412-INS-290 | chr9-33188412-INS-290 |
| chr9-33197324-INS-62 | chr9-33197324-INS-62 |
| chr9-33234931-INS-88 | chr9-33234931-INS-64 |
| chr9-33240824-INS-1132 | chr9-33240824-INS-1132 |
| chr9-33303179-INS-225 | chr9-33303179-INS-225 |
| chr9-33316808-INS-55 | chr9-33316809-INS-56 |
| chr9-33331412-INS-483 | chr9-33331412-INS-483 |
| chr9-33339870-INS-59 | chr9-33339870-INS-59 |
| chr9-33358916-INS-95 | chr9-33358916-INS-95 |
| chr9-33371659-INS-75 | chr9-33371659-INS-75 |
| chr9-33392733-INS-93 | chr9-33392739-INS-106 |
| chr9-33438946-INS-60 | chr9-33438946-INS-60 |
| chr9-33457164-INS-61 | chr9-33457164-INS-61 |
| chr9-33467530-INS-6133 | chr9-33467531-INS-6133 |
| chr9-33480946-INS-75 | chr9-33480946-INS-75 |
| chr9-33482410-INS-3188 | chr9-33482411-INS-3188 |
| chr9-33484440-INS-60 | chr9-33484441-INS-66 |
| chr9-33494488-INS-141 | chr9-33494495-INS-154 |
| chr9-33511612-INS-6351 | chr9-33511613-INS-6351 |
| chr9-33524416-INS-63 | chr9-33524411-INS-92 |
| chr9-33535299-INS-597 | chr9-33535299-INS-597 |
| chr9-33576753-INS-64 | chr9-33576753-INS-64 |
| chr9-33583901-INS-63 | chr9-33583901-INS-64 |
| chr9-33600022-INS-412 | chr9-33600022-INS-438 |
| chr9-33610584-INS-201 | chr9-33610584-INS-201 |
| chr9-33611871-INS-51 | chr9-33611871-INS-54 |

| | |
|---|---|
| chr9-33621088-INS-6389 | chr9-33621089-INS-6389 |
| chr9-33625439-INS-521 | chr9-33625439-INS-521 |
| chr9-33645757-INS-77 | chr9-33645757-INS-77 |
| chr9-33661804-INS-4977 | chr9-33661805-INS-4977 |
| chr9-33667323-INS-143 | chr9-33667323-INS-111 |
| chr9-33677324-INS-70 | chr9-33677324-INS-72 |
| chr9-33688750-INS-466 | chr9-33688750-INS-467 |
| chr9-33691494-INS-53 | chr9-33691494-INS-53 |
| chr9-33703391-INS-435 | chr9-33703391-INS-436 |
| chr9-33723954-INS-7093 | chr9-33723955-INS-7093 |
| chr9-33723955-INS-359 | chr9-33723955-INS-359 |
| chr9-33734849-INS-217 | chr9-33734849-INS-217 |
| chr9-33736499-INS-583 | chr9-33736499-INS-583 |
| chr9-33747166-INS-1467 | chr9-33747167-INS-1467 |
| chr9-33786437-INS-178 | chr9-33786437-INS-178 |
| chr9-33835262-INS-255 | chr9-33835249-INS-304 |
| chr9-33846609-INS-60 | chr9-33846609-INS-60 |
| chr9-33860188-INS-616 | chr9-33860188-INS-623 |
| chr9-33867108-INS-832 | chr9-33867108-INS-832 |
| chr9-33908278-INS-2717 | chr9-33908279-INS-2719 |
| chr9-33921089-INS-61 | chr9-33921089-INS-61 |
| chr9-33924105-INS-1704 | chr9-33924106-INS-1704 |
| chr9-33978135-INS-116 | chr9-33978135-INS-111 |
| chr9-34004321-INS-1146 | chr9-34004322-INS-1146 |
| chr9-34015471-INS-6351 | chr9-34015472-INS-6351 |
| chr9-34030181-INS-66 | chr9-34030181-INS-66 |
| chr9-34044587-INS-54 | chr9-34044608-INS-57 |
| chr9-34060159-INS-490 | chr9-34060159-INS-491 |
| chr9-34068880-INS-1189 | chr9-34068881-INS-1190 |
| chr9-34084711-INS-61 | chr9-34084724-INS-63 |
| chr9-34091495-INS-72 | chr9-34091495-INS-74 |
| chr9-34116040-INS-56 | chr9-34116040-INS-50 |
| chr9-34120985-INS-88 | chr9-34120985-INS-96 |
| chr9-34127679-INS-164 | chr9-34127679-INS-136 |
| chr9-34130665-INS-6296 | chr9-34130666-INS-6297 |
| chr9-34137028-INS-78 | chr9-34137028-INS-78 |
| chr9-34145900-INS-85 | chr9-34145900-INS-75 |
| chr9-34154060-INS-168 | chr9-34154060-INS-168 |
| chr9-34183405-INS-156 | chr9-34183405-INS-156 |
| chr9-34184576-INS-286 | chr9-34184579-INS-324 |
| chr9-34187334-INS-667 | chr9-34187334-INS-670 |
| chr9-34202974-INS-6297 | chr9-34202975-INS-6297 |
| chr9-34224504-INS-4931 | chr9-34224505-INS-4931 |
| chr9-34232703-INS-534 | chr9-34232703-INS-534 |
| chr9-34253580-INS-132 | chr9-34253580-INS-132 |
| chr9-34281846-INS-119 | chr9-34281849-INS-104 |
| chr9-34289834-INS-1888 | chr9-34289835-INS-1888 |
| chr9-34316084-INS-134 | chr9-34316084-INS-134 |
| chr9-34328001-INS-64 | chr9-34328001-INS-84 |
| chr9-34353978-INS-80 | chr9-34353978-INS-80 |
| chr9-34447518-INS-80 | chr9-34447521-INS-82 |
| chr9-34476592-INS-6677 | chr9-34476593-INS-6677 |
| chr9-34504577-INS-56 | chr9-34504577-INS-56 |
| chr9-34517998-INS-71 | chr9-34517998-INS-71 |
| chr9-34534493-INS-56 | chr9-34534493-INS-56 |
| chr9-34574330-INS-1297 | chr9-34574331-INS-1297 |
| chr9-34582674-INS-196 | chr9-34582674-INS-201 |
| chr9-34599259-INS-277 | chr9-34599259-INS-277 |
| chr9-34603347-INS-60 | chr9-34603352-INS-105 |
| chr9-34630565-INS-74 | chr9-34630576-INS-52 |
| chr9-34664898-INS-52 | chr9-34664898-INS-56 |
| chr9-34675818-INS-411 | chr9-34675818-INS-411 |
| chr9-34693666-INS-405 | chr9-34693666-INS-405 |
| chr9-34702300-INS-6440 | chr9-34702301-INS-6440 |
| chr9-34710363-INS-54 | chr9-34710363-INS-54 |
| chr9-34781833-INS-64 | chr9-34781833-INS-68 |
| chr9-34788298-INS-343 | chr9-34788298-INS-343 |
| chr9-34795466-INS-130 | chr9-34795466-INS-130 |
| chr9-34807466-INS-84 | chr9-34807466-INS-84 |
| chr9-34810982-INS-53 | chr9-34810982-INS-61 |
| chr9-34836715-INS-74 | chr9-34836715-INS-74 |
| chr9-34837769-INS-58 | chr9-34837769-INS-62 |
| chr9-34843094-INS-52 | chr9-34843094-INS-58 |
| chr9-34846525-INS-60 | chr9-34846525-INS-60 |
| chr9-34850048-INS-470 | chr9-34850048-INS-470 |
| chr9-34896935-INS-52 | chr9-34896935-INS-54 |
| chr9-34907774-INS-172 | chr9-34907774-INS-172 |

| | |
|---|---|
| chr9-34913761-INS-100 | chr9-34913761-INS-120 |
| chr9-34920816-INS-200 | chr9-34920816-INS-200 |
| chr9-34932111-INS-331 | chr9-34932111-INS-331 |
| chr9-34944707-INS-88 | chr9-34944707-INS-88 |
| chr9-34951174-INS-156 | chr9-34951174-INS-180 |
| chr9-34980156-INS-182 | chr9-34980157-INS-202 |
| chr9-35011284-INS-76 | chr9-35011284-INS-76 |
| chr9-35018786-INS-68 | chr9-35018786-INS-68 |
| chr9-35034380-INS-52 | chr9-35034380-INS-52 |
| chr9-35081148-INS-213 | chr9-35081148-INS-213 |
| chr9-35106763-INS-52 | chr9-35106763-INS-60 |
| chr9-35121794-INS-90 | chr9-35121794-INS-90 |
| chr9-35140636-INS-204 | chr9-35140636-INS-204 |
| chr9-35166649-INS-1799 | chr9-35166650-INS-1799 |
| chr9-35185075-INS-72 | chr9-35185056-INS-92 |
| chr9-35223076-INS-62 | chr9-35223076-INS-68 |
| chr9-35244137-INS-54 | chr9-35244137-INS-56 |
| chr9-35245788-INS-461 | chr9-35245788-INS-461 |
| chr9-35279529-INS-211 | chr9-35279529-INS-212 |
| chr9-35307420-INS-191 | chr9-35307420-INS-192 |
| chr9-35326411-INS-56 | chr9-35326423-INS-58 |
| chr9-35338409-INS-71 | chr9-35338409-INS-116 |
| chr9-35349561-INS-91 | chr9-35349561-INS-91 |
| chr9-35356934-INS-53 | chr9-35356934-INS-53 |
| chr9-35369173-INS-72 | chr9-35369174-INS-64 |
| chr9-35375597-INS-1155 | chr9-35375598-INS-1155 |
| chr9-35376718-INS-219 | chr9-35376718-INS-219 |
| chr9-35401779-INS-167 | chr9-35401779-INS-167 |
| chr9-35404609-INS-202 | chr9-35404609-INS-202 |
| chr9-35408542-INS-5535 | chr9-35408543-INS-5535 |
| chr9-35440878-INS-54 | chr9-35440878-INS-52 |
| chr9-35482298-INS-239 | chr9-35482300-INS-225 |
| chr9-35486215-INS-359 | chr9-35486215-INS-359 |
| chr9-35494985-INS-200 | chr9-35494985-INS-203 |
| chr9-35511778-INS-61 | chr9-35511778-INS-64 |
| chr9-35512453-INS-1162 | chr9-35512458-INS-1155 |
| chr9-35589981-INS-7098 | chr9-35589982-INS-7098 |
| chr9-35621853-INS-6240 | chr9-35621854-INS-6239 |
| chr9-35667247-INS-1956 | chr9-35667248-INS-1954 |
| chr9-35720450-INS-111 | chr9-35720450-INS-111 |
| chr9-35726075-INS-899 | chr9-35726075-INS-899 |
| chr9-35742729-INS-6934 | chr9-35742730-INS-6936 |
| chr9-35754672-INS-56 | chr9-35754684-INS-52 |
| chr9-35755652-INS-849 | chr9-35755652-INS-858 |
| chr9-35760420-INS-2905 | chr9-35760421-INS-2906 |
| chr9-35768336-INS-1204 | chr9-35768337-INS-1204 |
| chr9-35773085-INS-70 | chr9-35773085-INS-58 |
| chr9-35791200-INS-94 | chr9-35791200-INS-94 |
| chr9-35807138-INS-75 | chr9-35807138-INS-110 |
| chr9-35811545-INS-2576 | chr9-35811546-INS-2583 |
| chr9-35826447-INS-198 | chr9-35826447-INS-212 |
| chr9-35853126-INS-1892 | chr9-35853127-INS-1892 |
| chr9-35871314-INS-202 | chr9-35871314-INS-202 |
| chr9-35893465-INS-6634 | chr9-35893466-INS-6634 |
| chr9-35897541-INS-1770 | chr9-35897542-INS-1765 |
| chr9-35907876-INS-7113 | chr9-35907877-INS-7113 |
| chr9-35940773-INS-199 | chr9-35940773-INS-209 |
| chr9-35966277-INS-2965 | chr9-35966278-INS-2961 |
| chr9-35988543-INS-199 | chr9-35988543-INS-199 |
| chr9-36006815-INS-7211 | chr9-36006816-INS-7211 |
| chr9-36013474-INS-7074 | chr9-36013475-INS-7074 |
| chr9-36051838-INS-60 | chr9-36051838-INS-60 |
| chr9-36063235-INS-1047 | chr9-36063236-INS-1047 |
| chr9-36162163-INS-73 | chr9-36162199-INS-73 |
| chr9-36168153-INS-425 | chr9-36168153-INS-425 |
| chr9-36188764-INS-199 | chr9-36188764-INS-199 |
| chr9-36190789-INS-54 | chr9-36190790-INS-66 |
| chr9-36192461-INS-94 | chr9-36192462-INS-74 |
| chr9-36214350-INS-6192 | chr9-36214351-INS-6193 |
| chr9-36237085-INS-6803 | chr9-36237086-INS-6803 |
| chr9-36252042-INS-6336 | chr9-36252043-INS-6336 |
| chr9-36293594-INS-64 | chr9-36293594-INS-64 |
| chr9-36305364-INS-220 | chr9-36305364-INS-219 |
| chr9-36331559-INS-6715 | chr9-36331560-INS-6715 |
| chr9-36336763-INS-4023 | chr9-36336764-INS-4024 |
| chr9-36376271-INS-1271 | chr9-36376272-INS-1271 |
| chr9-36391719-INS-68 | chr9-36391719-INS-56 |

| | |
|---|---|
| chr9-36418612-INS-91 | chr9-36418612-INS-95 |
| chr9-36422716-INS-74 | chr9-36422716-INS-67 |
| chr9-36432481-INS-6450 | chr9-36432482-INS-6450 |
| chr9-36449952-INS-3037 | chr9-36449953-INS-3036 |
| chr9-36449953-INS-407 | chr9-36449953-INS-407 |
| chr9-36469547-INS-6731 | chr9-36469548-INS-6731 |
| chr9-36495245-INS-6694 | chr9-36495246-INS-6689 |
| chr9-36498610-INS-213 | chr9-36498610-INS-213 |
| chr9-36512446-INS-1283 | chr9-36512447-INS-1283 |
| chr9-36516029-INS-2254 | chr9-36516030-INS-2253 |
| chr9-36562520-INS-1044 | chr9-36562521-INS-1044 |
| chr9-36581744-INS-2799 | chr9-36581745-INS-2799 |
| chr9-36630706-INS-203 | chr9-36630706-INS-202 |
| chr9-36640891-INS-52 | chr9-36640891-INS-60 |
| chr9-36654994-INS-211 | chr9-36654994-INS-211 |
| chr9-36665479-INS-201 | chr9-36665479-INS-203 |
| chr9-36688712-INS-84 | chr9-36688712-INS-84 |
| chr9-36690417-INS-209 | chr9-36690416-INS-213 |
| chr9-36698386-INS-254 | chr9-36698386-INS-254 |
| chr9-36701437-INS-52 | chr9-36701437-INS-50 |
| chr9-36703926-INS-168 | chr9-36703926-INS-168 |
| chr9-36707189-INS-168 | chr9-36707189-INS-168 |
| chr9-36713054-INS-111 | chr9-36713057-INS-99 |
| chr9-36714333-INS-195 | chr9-36714333-INS-195 |
| chr9-36715616-INS-6452 | chr9-36715617-INS-6452 |
| chr9-36720800-INS-428 | chr9-36720800-INS-436 |
| chr9-36731410-INS-616 | chr9-36731410-INS-616 |
| chr9-36734578-INS-79 | chr9-36734578-INS-79 |
| chr9-36742388-INS-60 | chr9-36742388-INS-62 |
| chr9-36748192-INS-148 | chr9-36748187-INS-139 |
| chr9-36753584-INS-137 | chr9-36753584-INS-137 |
| chr9-36756266-INS-260 | chr9-36756266-INS-260 |
| chr9-36773600-INS-199 | chr9-36773600-INS-201 |
| chr9-36777369-INS-51 | chr9-36777322-INS-76 |
| chr9-36786897-INS-199 | chr9-36786897-INS-199 |
| chr9-36799630-INS-543 | chr9-36799606-INS-518 |
| chr9-36812959-INS-212 | chr9-36812959-INS-212 |
| chr9-36821996-INS-57 | chr9-36821996-INS-63 |
| chr9-36824870-INS-92 | chr9-36824870-INS-51 |
| chr9-36903392-INS-58 | chr9-36903392-INS-58 |
| chr9-36905386-INS-67 | chr9-36905396-INS-55 |
| chr9-36912263-INS-1891 | chr9-36912264-INS-1891 |
| chr9-36934245-INS-85 | chr9-36934245-INS-85 |
| chr9-36943433-INS-132 | chr9-36943439-INS-118 |
| chr9-36997019-INS-58 | chr9-36997019-INS-54 |
| chr9-37004032-INS-255 | chr9-37004032-INS-254 |
| chr9-37013880-INS-200 | chr9-37013880-INS-200 |
| chr9-37032692-INS-66 | chr9-37032692-INS-68 |
| chr9-37037180-INS-210 | chr9-37037180-INS-211 |
| chr9-37041421-INS-96 | chr9-37041421-INS-102 |
| chr9-37048112-INS-491 | chr9-37048112-INS-491 |
| chr9-37054078-INS-52 | chr9-37054078-INS-56 |
| chr9-37061117-INS-217 | chr9-37061123-INS-233 |
| chr9-37066323-INS-59 | chr9-37066323-INS-59 |
| chr9-37100449-INS-70 | chr9-37100447-INS-54 |
| chr9-37102991-INS-125 | chr9-37102987-INS-129 |
| chr9-37106141-INS-56 | chr9-37106141-INS-56 |
| chr9-37116002-INS-205 | chr9-37116002-INS-205 |
| chr9-37128055-INS-1190 | chr9-37128056-INS-1159 |
| chr9-37136093-INS-163 | chr9-37136075-INS-158 |
| chr9-37147511-INS-56 | chr9-37147511-INS-56 |
| chr9-37150050-INS-160 | chr9-37150050-INS-144 |
| chr9-37161469-INS-54 | chr9-37161469-INS-52 |
| chr9-37171797-INS-88 | chr9-37171797-INS-88 |
| chr9-37189272-INS-80 | chr9-37189272-INS-80 |
| chr9-37197790-INS-728 | chr9-37197790-INS-729 |
| chr9-37201339-INS-309 | chr9-37201339-INS-308 |
| chr9-37231146-INS-2300 | chr9-37231147-INS-2300 |
| chr9-37257258-INS-678 | chr9-37257258-INS-678 |
| chr9-37265349-INS-54 | chr9-37265317-INS-57 |
| chr9-37282464-INS-207 | chr9-37282464-INS-207 |
| chr9-37309787-INS-261 | chr9-37309775-INS-273 |
| chr9-37316699-INS-265 | chr9-37316699-INS-265 |
| chr9-37387904-INS-92 | chr9-37387904-INS-95 |
| chr9-37405138-INS-56 | chr9-37405129-INS-52 |
| chr9-37411614-INS-206 | chr9-37411614-INS-206 |
| chr9-37417968-INS-94 | chr9-37417968-INS-108 |

| | |
|---|---|
| chr9-37443851-INS-64 | chr9-37443853-INS-56 |
| chr9-37451997-INS-62 | chr9-37451997-INS-62 |
| chr9-37470199-INS-84 | chr9-37470199-INS-66 |
| chr9-37475747-INS-217 | chr9-37475747-INS-217 |
| chr9-37503893-INS-52 | chr9-37503893-INS-52 |
| chr9-37506392-INS-430 | chr9-37506392-INS-430 |
| chr9-37529353-INS-56 | chr9-37529353-INS-56 |
| chr9-37560397-INS-358 | chr9-37560397-INS-358 |
| chr9-37574583-INS-75 | chr9-37574583-INS-109 |
| chr9-37589639-INS-138 | chr9-37589639-INS-114 |
| chr9-37590341-INS-2975 | chr9-37590342-INS-2975 |
| chr9-37593178-INS-78 | chr9-37593182-INS-64 |
| chr9-37596090-INS-321 | chr9-37596090-INS-329 |
| chr9-37602532-INS-57 | chr9-37602532-INS-61 |
| chr9-37607729-INS-135 | chr9-37607729-INS-163 |
| chr9-37610205-INS-125 | chr9-37610208-INS-140 |
| chr9-37617479-INS-52 | chr9-37617479-INS-50 |
| chr9-37617902-INS-915 | chr9-37617902-INS-915 |
| chr9-37622244-INS-208 | chr9-37622244-INS-209 |
| chr9-37624462-INS-134 | chr9-37624462-INS-134 |
| chr9-37631261-INS-8428 | chr9-37631262-INS-8430 |
| chr9-37651956-INS-1232 | chr9-37651957-INS-1232 |
| chr9-37657038-INS-99 | chr9-37657038-INS-99 |
| chr9-37662412-INS-549 | chr9-37662412-INS-549 |
| chr9-37666589-INS-7247 | chr9-37666590-INS-7247 |
| chr9-37678575-INS-137 | chr9-37678575-INS-137 |
| chr9-37691107-INS-765 | chr9-37691107-INS-765 |
| chr9-37706783-INS-62 | chr9-37706783-INS-97 |
| chr9-37716993-INS-498 | chr9-37716993-INS-498 |
| chr9-37729409-INS-55 | chr9-37729409-INS-55 |
| chr9-37734718-INS-62 | chr9-37734718-INS-76 |
| chr9-37743261-INS-92 | chr9-37743261-INS-92 |
| chr9-37746974-INS-1051 | chr9-37746975-INS-1051 |
| chr9-37760949-INS-6369 | chr9-37760950-INS-6370 |
| chr9-37771155-INS-508 | chr9-37771155-INS-506 |
| chr9-37795311-INS-2158 | chr9-37795312-INS-2158 |
| chr9-37823624-INS-7123 | chr9-37823625-INS-7123 |
| chr9-37847866-INS-111 | chr9-37847866-INS-111 |
| chr9-37849465-INS-1030 | chr9-37849466-INS-1039 |
| chr9-37859808-INS-3136 | chr9-37859809-INS-3136 |
| chr9-37869706-INS-983 | chr9-37869706-INS-983 |
| chr9-37915344-INS-4399 | chr9-37915345-INS-4399 |
| chr9-37944351-INS-213 | chr9-37944350-INS-220 |
| chr9-37978097-INS-1096 | chr9-37978098-INS-1096 |
| chr9-37981028-INS-140 | chr9-37981045-INS-120 |
| chr9-37990206-INS-171 | chr9-37990206-INS-169 |
| chr9-37991110-INS-6282 | chr9-37991111-INS-6282 |
| chr9-38037534-INS-1506 | chr9-38037535-INS-1526 |
| chr9-38061756-INS-316 | chr9-38061756-INS-316 |
| chr9-38066922-INS-2145 | chr9-38066923-INS-2139 |
| chr9-38069866-INS-7090 | chr9-38069867-INS-7090 |
| chr9-38100572-INS-83 | chr9-38100579-INS-98 |
| chr9-38102488-INS-355 | chr9-38102488-INS-355 |
| chr9-38126803-INS-159 | chr9-38126803-INS-159 |
| chr9-38142387-INS-226 | chr9-38142387-INS-229 |
| chr9-38162898-INS-60 | chr9-38162898-INS-62 |
| chr9-38178181-INS-6180 | chr9-38178182-INS-6181 |
| chr9-38194709-INS-60 | chr9-38194709-INS-50 |
| chr9-38224346-INS-100 | chr9-38224345-INS-71 |
| chr9-38265789-INS-107 | chr9-38265789-INS-131 |
| chr9-38274227-INS-290 | chr9-38274227-INS-290 |
| chr9-38303472-INS-494 | chr9-38303472-INS-494 |
| chr9-38306618-INS-86 | chr9-38306618-INS-81 |
| chr9-38320988-INS-212 | chr9-38320988-INS-212 |
| chr9-38339737-INS-529 | chr9-38339737-INS-529 |
| chr9-38350411-INS-169 | chr9-38350411-INS-169 |
| chr9-38354418-INS-160 | chr9-38354462-INS-110 |
| chr9-38385755-INS-4601 | chr9-38385756-INS-4604 |
| chr9-38387316-INS-7729 | chr9-38387317-INS-7729 |
| chr9-38428827-INS-59 | chr9-38428827-INS-61 |
| chr9-38457256-INS-2980 | chr9-38457257-INS-2973 |
| chr9-38457934-INS-2709 | chr9-38457935-INS-2709 |
| chr9-38542180-INS-86 | chr9-38542173-INS-50 |
| chr9-38551198-INS-6193 | chr9-38551199-INS-6193 |
| chr9-38601022-INS-365 | chr9-38601022-INS-401 |
| chr9-38637678-INS-596 | chr9-38637678-INS-596 |
| chr9-38643535-INS-57 | chr9-38643535-INS-59 |

| | |
|---|---|
| chr9-38651197-INS-388 | chr9-38651197-INS-387 |
| chr9-38658073-INS-340 | chr9-38658073-INS-340 |
| chr9-38663242-INS-66 | chr9-38663242-INS-78 |
| chr9-38674861-INS-6448 | chr9-38674862-INS-6449 |
| chr9-38678266-INS-903 | chr9-38678266-INS-904 |
| chr9-38679327-INS-2703 | chr9-38679328-INS-2703 |
| chr9-38697578-INS-54 | chr9-38697578-INS-58 |
| chr9-38724022-INS-5369 | chr9-38724023-INS-5369 |
| chr9-38786754-INS-3209 | chr9-38786755-INS-3247 |
| chr9-38793006-INS-6455 | chr9-38793007-INS-6455 |
| chr9-38898407-INS-175 | chr9-38898407-INS-176 |
| chr9-38966655-INS-56 | chr9-38966655-INS-58 |
| chr9-38980935-INS-89 | chr9-38980935-INS-80 |
| chr9-39007594-INS-62 | chr9-39007594-INS-62 |
| chr9-39012778-INS-1891 | chr9-39012779-INS-1891 |
| chr9-39039580-INS-202 | chr9-39039580-INS-202 |
| chr9-39052159-INS-54 | chr9-39052157-INS-52 |
| chr9-39059966-INS-77 | chr9-39059966-INS-77 |
| chr9-39064116-INS-82 | chr9-39064116-INS-56 |
| chr9-39072045-INS-357 | chr9-39072045-INS-357 |
| chr9-39107125-INS-334 | chr9-39107125-INS-334 |
| chr9-39131650-INS-136 | chr9-39131640-INS-144 |
| chr9-39137284-INS-56 | chr9-39137284-INS-52 |
| chr9-39138454-INS-474 | chr9-39138454-INS-474 |
| chr9-39157805-INS-693 | chr9-39157805-INS-695 |
| chr9-39171789-INS-805 | chr9-39171789-INS-805 |
| chr9-39185356-INS-1692 | chr9-39185357-INS-1691 |
| chr9-39206180-INS-65 | chr9-39206179-INS-71 |
| chr9-39208054-INS-75 | chr9-39208054-INS-93 |
| chr9-39237572-INS-84 | chr9-39237572-INS-84 |
| chr9-39252340-INS-474 | chr9-39252340-INS-473 |
| chr9-39277753-INS-312 | chr9-39277753-INS-312 |
| chr9-39364492-INS-7269 | chr9-39364493-INS-7269 |
| chr9-39371658-INS-974 | chr9-39371658-INS-973 |
| chr9-39376155-INS-3281 | chr9-39376156-INS-3260 |
| chr9-39400087-INS-346 | chr9-39400087-INS-345 |
| chr9-39419561-INS-101 | chr9-39419572-INS-104 |
| chr9-39428192-INS-4675 | chr9-39428193-INS-4675 |
| chr9-39430351-INS-417 | chr9-39430351-INS-417 |
| chr9-39436307-INS-103 | chr9-39436310-INS-92 |
| chr9-39448659-INS-390 | chr9-39448659-INS-390 |
| chr9-39450779-INS-87 | chr9-39450779-INS-110 |
| chr9-39509772-INS-102 | chr9-39509767-INS-111 |
| chr9-39524176-INS-112 | chr9-39524176-INS-108 |
| chr9-39528386-INS-206 | chr9-39528372-INS-195 |
| chr9-39552416-INS-74 | chr9-39552416-INS-88 |
| chr9-39567958-INS-72 | chr9-39567958-INS-72 |
| chr9-39571452-INS-94 | chr9-39571452-INS-94 |
| chr9-39625963-INS-800 | chr9-39625963-INS-800 |
| chr9-39676859-INS-72 | chr9-39676859-INS-74 |
| chr9-39695719-INS-7358 | chr9-39695720-INS-7358 |
| chr9-39718492-INS-7130 | chr9-39718493-INS-7130 |
| chr9-39722739-INS-158 | chr9-39722739-INS-158 |
| chr9-39739911-INS-1104 | chr9-39739912-INS-1105 |
| chr9-39760711-INS-107 | chr9-39760717-INS-104 |
| chr9-39824666-INS-346 | chr9-39824666-INS-346 |
| chr9-39843422-INS-100 | chr9-39843422-INS-100 |
| chr9-39860196-INS-129 | chr9-39860196-INS-127 |
| chr9-39882725-INS-3250 | chr9-39882726-INS-3250 |
| chr9-40019322-INS-80 | chr9-40019323-INS-50 |
| chr9-40035392-INS-127 | chr9-40035397-INS-123 |
| chr9-40039651-INS-56 | chr9-40039643-INS-56 |
| chr9-40044082-INS-440 | chr9-40044082-INS-442 |
| chr9-40049477-INS-610 | chr9-40049478-INS-609 |
| chr9-40065178-INS-2631 | chr9-40065179-INS-2595 |
| chr9-40072431-INS-1341 | chr9-40072434-INS-1299 |
| chr9-40160059-INS-74 | chr9-40160059-INS-72 |
| chr9-40174424-INS-52 | chr9-40174424-INS-52 |
| chr9-40196415-INS-155 | chr9-40196403-INS-156 |
| chr9-40281432-INS-60 | chr9-40281432-INS-60 |
| chr9-40307077-INS-106 | chr9-40307078-INS-97 |
| chr9-40312491-INS-95 | chr9-40312491-INS-95 |
| chr9-40313412-INS-120 | chr9-40313412-INS-120 |
| chr9-40354827-INS-57 | chr9-40354827-INS-57 |
| chr9-40373157-INS-155 | chr9-40373157-INS-155 |
| chr9-40373741-INS-1834 | chr9-40373742-INS-1843 |
| chr9-40401174-INS-400 | chr9-40401174-INS-400 |

| | |
|---|---|
| chr9-40424955-INS-475 | chr9-40424956-INS-476 |
| chr9-40438631-INS-92 | chr9-40438631-INS-92 |
| chr9-40453171-INS-64 | chr9-40453171-INS-64 |
| chr9-40457345-INS-54 | chr9-40457323-INS-56 |
| chr9-40494266-INS-58 | chr9-40494261-INS-84 |
| chr9-40510920-INS-217 | chr9-40510920-INS-217 |
| chr9-40517238-INS-295 | chr9-40517238-INS-256 |
| chr9-40533972-INS-82 | chr9-40533972-INS-56 |
| chr9-40537195-INS-429 | chr9-40537195-INS-429 |
| chr9-40547726-INS-250 | chr9-40547726-INS-250 |
| chr9-40559624-INS-358 | chr9-40559624-INS-358 |
| chr9-40577254-INS-406 | chr9-40577254-INS-406 |
| chr9-40594800-INS-67 | chr9-40594800-INS-67 |
| chr9-40625487-INS-340 | chr9-40625487-INS-340 |
| chr9-40637782-INS-332 | chr9-40637782-INS-332 |
| chr9-40645270-INS-54 | chr9-40645295-INS-74 |
| chr9-40666586-INS-54 | chr9-40666586-INS-54 |
| chr9-40676287-INS-752 | chr9-40676287-INS-792 |
| chr9-40679704-INS-67 | chr9-40679704-INS-67 |
| chr9-40691297-INS-178 | chr9-40691297-INS-178 |
| chr9-40698709-INS-60 | chr9-40698709-INS-66 |
| chr9-40724407-INS-262 | chr9-40724407-INS-259 |
| chr9-40729642-INS-393 | chr9-40729642-INS-393 |
| chr9-40735275-INS-407 | chr9-40735275-INS-407 |
| chr9-40743558-INS-199 | chr9-40743558-INS-167 |
| chr9-40759242-INS-205 | chr9-40759242-INS-205 |
| chr9-40767197-INS-213 | chr9-40767197-INS-215 |
| chr9-40776101-INS-66 | chr9-40776085-INS-114 |
| chr9-40779624-INS-78 | chr9-40779624-INS-76 |
| chr9-40788562-INS-93 | chr9-40788562-INS-93 |
| chr9-40795299-INS-69 | chr9-40795299-INS-69 |
| chr9-40805271-INS-196 | chr9-40805271-INS-196 |
| chr9-40834281-INS-53 | chr9-40834281-INS-53 |
| chr9-40835645-INS-53 | chr9-40835644-INS-54 |
| chr9-40878421-INS-7130 | chr9-40878422-INS-7130 |
| chr9-40884141-INS-213 | chr9-40884141-INS-213 |
| chr9-40910928-INS-212 | chr9-40910928-INS-212 |
| chr9-40927309-INS-66 | chr9-40927317-INS-81 |
| chr9-40940277-INS-510 | chr9-40940277-INS-510 |
| chr9-40963717-INS-111 | chr9-40963711-INS-120 |
| chr9-41039292-INS-160 | chr9-41039292-INS-160 |
| chr9-41048716-INS-60 | chr9-41048716-INS-54 |
| chr9-41060628-INS-74 | chr9-41060643-INS-53 |
| chr9-41066753-INS-58 | chr9-41066753-INS-52 |
| chr9-41078729-INS-347 | chr9-41078734-INS-316 |
| chr9-41088730-INS-78 | chr9-41088711-INS-85 |
| chr9-41113417-INS-116 | chr9-41113411-INS-129 |
| chr9-41116835-INS-54 | chr9-41116835-INS-55 |
| chr9-41122888-INS-2841 | chr9-41122889-INS-2837 |
| chr9-41136330-INS-585 | chr9-41136330-INS-585 |
| chr9-41174648-INS-2333 | chr9-41174649-INS-2353 |
| chr9-41177741-INS-92 | chr9-41177745-INS-92 |
| chr9-41180105-INS-119 | chr9-41180105-INS-155 |
| chr9-41186468-INS-614 | chr9-41186468-INS-611 |
| chr9-41212979-INS-195 | chr9-41212979-INS-195 |
| chr9-41247708-INS-152 | chr9-41247708-INS-152 |
| chr9-41257904-INS-6837 | chr9-41257905-INS-6840 |
| chr9-41263137-INS-708 | chr9-41263137-INS-708 |
| chr9-41275081-INS-84 | chr9-41275081-INS-86 |
| chr9-41277241-INS-52 | chr9-41277241-INS-52 |
| chr9-41291642-INS-102 | chr9-41291643-INS-102 |
| chr9-41295450-INS-76 | chr9-41295450-INS-83 |
| chr9-41297775-INS-121 | chr9-41297775-INS-72 |
| chr9-41311248-INS-67 | chr9-41311249-INS-67 |
| chr9-41317830-INS-80 | chr9-41317831-INS-64 |
| chr9-41330182-INS-603 | chr9-41330182-INS-601 |
| chr9-41352327-INS-203 | chr9-41352327-INS-203 |
| chr9-41402345-INS-325 | chr9-41402345-INS-325 |
| chr9-41404261-INS-201 | chr9-41404261-INS-202 |
| chr9-41412206-INS-61 | chr9-41412206-INS-61 |
| chr9-41425308-INS-781 | chr9-41425308-INS-780 |
| chr9-41427166-INS-66 | chr9-41427166-INS-62 |
| chr9-41428932-INS-52 | chr9-41428932-INS-50 |
| chr9-41464247-INS-1036 | chr9-41464248-INS-1035 |
| chr9-41508431-INS-179 | chr9-41508431-INS-180 |
| chr9-41521007-INS-52 | chr9-41521007-INS-54 |
| chr9-41600065-INS-140 | chr9-41600046-INS-158 |

| | |
|---|---|
| chr9-41638266-INS-399 | chr9-41638266-INS-399 |
| chr9-41639408-INS-495 | chr9-41639408-INS-495 |
| chr9-41641477-INS-56 | chr9-41641477-INS-54 |
| chr9-41682980-INS-54 | chr9-41682978-INS-52 |
| chr9-41689521-INS-966 | chr9-41689521-INS-966 |
| chr9-41690874-INS-127 | chr9-41690874-INS-127 |
| chr9-41722387-INS-279 | chr9-41722387-INS-279 |
| chr9-41731453-INS-123 | chr9-41731453-INS-80 |
| chr9-41736484-INS-80 | chr9-41736487-INS-87 |
| chr9-41738499-INS-156 | chr9-41738499-INS-188 |
| chr9-41745820-INS-132 | chr9-41745820-INS-132 |
| chr9-41751411-INS-298 | chr9-41751411-INS-298 |
| chr9-41756447-INS-69 | chr9-41756427-INS-100 |
| chr9-41772360-INS-84 | chr9-41772370-INS-89 |
| chr9-41789570-INS-100 | chr9-41789570-INS-100 |
| chr9-41794463-INS-180 | chr9-41794463-INS-180 |
| chr9-41800523-INS-54 | chr9-41800523-INS-54 |
| chr9-41807547-INS-112 | chr9-41807547-INS-94 |
| chr9-41814380-INS-123 | chr9-41814380-INS-123 |
| chr9-41828513-INS-230 | chr9-41828513-INS-230 |
| chr9-41866022-INS-116 | chr9-41866006-INS-104 |
| chr9-41873908-INS-170 | chr9-41873908-INS-170 |
| chr9-41899244-INS-82 | chr9-41899223-INS-68 |
| chr9-41901733-INS-201 | chr9-41901733-INS-201 |
| chr9-41907709-INS-179 | chr9-41907709-INS-189 |
| chr9-41918787-INS-74 | chr9-41918791-INS-104 |
| chr9-41963016-INS-100 | chr9-41963016-INS-108 |
| chr9-41963989-INS-51 | chr9-41963990-INS-75 |
| chr9-41971498-INS-60 | chr9-41971498-INS-60 |
| chr9-41972284-INS-442 | chr9-41972284-INS-442 |
| chr9-42041996-INS-120 | chr9-42041996-INS-120 |
| chr9-42043796-INS-60 | chr9-42043788-INS-92 |
| chr9-42063720-INS-941 | chr9-42063719-INS-939 |
| chr9-42063835-INS-128 | chr9-42063836-INS-127 |
| chr9-42082947-INS-313 | chr9-42082950-INS-314 |
| chr9-42090198-INS-52 | chr9-42090198-INS-58 |
| chr9-42097146-INS-64 | chr9-42097146-INS-64 |
| chr9-42108718-INS-455 | chr9-42108718-INS-455 |
| chr9-42137965-INS-61 | chr9-42137965-INS-59 |
| chr9-42143535-INS-7098 | chr9-42143536-INS-7098 |
| chr9-42150486-INS-186 | chr9-42150486-INS-186 |
| chr9-42159000-INS-210 | chr9-42159000-INS-210 |
| chr9-42169180-INS-184 | chr9-42169180-INS-184 |
| chr9-42208877-INS-214 | chr9-42208877-INS-214 |
| chr9-42221039-INS-163 | chr9-42221028-INS-138 |
| chr9-42268349-INS-56 | chr9-42268349-INS-56 |
| chr9-42281216-INS-52 | chr9-42281216-INS-50 |
| chr9-42347420-INS-126 | chr9-42347423-INS-168 |
| chr9-42358060-INS-90 | chr9-42358060-INS-70 |
| chr9-42369997-INS-55 | chr9-42369997-INS-58 |
| chr9-42375744-INS-197 | chr9-42375744-INS-197 |
| chr9-42384107-INS-55 | chr9-42384105-INS-82 |
| chr9-42406227-INS-257 | chr9-42406227-INS-257 |
| chr9-42437873-INS-213 | chr9-42437873-INS-213 |
| chr9-42470478-INS-2906 | chr9-42470479-INS-2906 |
| chr9-42492374-INS-164 | chr9-42492374-INS-164 |
| chr9-42575450-INS-85 | chr9-42575446-INS-96 |
| chr9-42592878-INS-346 | chr9-42592878-INS-349 |
| chr9-42618142-INS-579 | chr9-42618133-INS-578 |
| chr9-42625862-INS-6743 | chr9-42625863-INS-6743 |
| chr9-42626254-INS-120 | chr9-42626254-INS-108 |
| chr9-42655800-INS-5174 | chr9-42655801-INS-5174 |
| chr9-42680065-INS-71 | chr9-42680047-INS-89 |
| chr9-42698056-INS-303 | chr9-42698056-INS-303 |
| chr9-42717753-INS-128 | chr9-42717753-INS-89 |
| chr9-42738023-INS-52 | chr9-42738023-INS-52 |
| chr9-42745629-INS-2427 | chr9-42745630-INS-2433 |
| chr9-42789275-INS-60 | chr9-42789275-INS-60 |
| chr9-42819858-INS-62 | chr9-42819858-INS-62 |
| chr9-42827298-INS-69 | chr9-42827298-INS-72 |
| chr9-42832005-INS-76 | chr9-42832005-INS-80 |
| chr9-42863405-INS-112 | chr9-42863406-INS-112 |
| chr9-42904626-INS-107 | chr9-42904659-INS-85 |
| chr9-42934469-INS-1052 | chr9-42934470-INS-1052 |
| chr9-42948904-INS-484 | chr9-42948904-INS-484 |
| chr9-42952527-INS-212 | chr9-42952527-INS-183 |
| chr9-42988700-INS-54 | chr9-42988700-INS-54 |

| | |
|---|---|
| chr9-43004744-INS-88 | chr9-43004744-INS-88 |
| chr9-43024311-INS-69 | chr9-43024315-INS-69 |
| chr9-43060276-INS-559 | chr9-43060276-INS-530 |
| chr9-43066091-INS-69 | chr9-43066086-INS-104 |
| chr9-43069561-INS-76 | chr9-43069561-INS-84 |
| chr9-43089554-INS-105 | chr9-43089554-INS-104 |
| chr9-43118168-INS-57 | chr9-43118168-INS-52 |
| chr9-43124766-INS-207 | chr9-43124766-INS-207 |
| chr9-43143118-INS-82 | chr9-43143100-INS-92 |
| chr9-43145434-INS-60 | chr9-43145434-INS-56 |
| chr9-43160734-INS-204 | chr9-43160734-INS-206 |
| chr9-43170714-INS-56 | chr9-43170714-INS-60 |
| chr9-43191750-INS-76 | chr9-43191750-INS-60 |
| chr9-43200722-INS-75 | chr9-43200722-INS-78 |
| chr9-43208751-INS-128 | chr9-43208751-INS-144 |
| chr9-43211029-INS-192 | chr9-43211029-INS-192 |
| chr9-43278967-INS-66 | chr9-43278967-INS-54 |
| chr9-43291083-INS-72 | chr9-43291089-INS-101 |
| chr9-43297727-INS-72 | chr9-43297727-INS-72 |
| chr9-43312633-INS-78 | chr9-43312670-INS-52 |
| chr9-43314376-INS-357 | chr9-43314376-INS-357 |
| chr9-43325776-INS-242 | chr9-43325776-INS-248 |
| chr9-43340237-INS-80 | chr9-43340237-INS-80 |
| chr9-43373250-INS-68 | chr9-43373250-INS-68 |
| chr9-43386145-INS-130 | chr9-43386145-INS-130 |
| chr9-43467647-INS-62 | chr9-43467647-INS-62 |
| chr9-43518606-INS-138 | chr9-43518606-INS-134 |
| chr9-43520810-INS-214 | chr9-43520811-INS-179 |
| chr9-43548994-INS-58 | chr9-43548994-INS-58 |
| chr9-43584267-INS-176 | chr9-43584276-INS-190 |
| chr9-43613724-INS-202 | chr9-43613724-INS-200 |
| chr9-43684079-INS-89 | chr9-43684076-INS-87 |
| chr9-43748461-INS-204 | chr9-43748461-INS-206 |
| chr9-43777522-INS-199 | chr9-43777528-INS-200 |
| chr9-43801470-INS-593 | chr9-43801470-INS-602 |
| chr9-43803392-INS-83 | chr9-43803392-INS-83 |
| chr9-43819752-INS-209 | chr9-43819752-INS-209 |
| chr9-43822245-INS-359 | chr9-43822245-INS-357 |
| chr9-43851124-INS-120 | chr9-43851132-INS-112 |
| chr9-43858187-INS-137 | chr9-43858144-INS-147 |
| chr9-43871802-INS-126 | chr9-43871802-INS-84 |
| chr9-43880344-INS-78 | chr9-43880350-INS-73 |
| chr9-43883947-INS-202 | chr9-43883947-INS-204 |
| chr9-43889093-INS-102 | chr9-43889100-INS-79 |
| chr9-43890906-INS-215 | chr9-43890906-INS-216 |
| chr9-43916025-INS-66 | chr9-43916025-INS-66 |
| chr9-43919308-INS-208 | chr9-43919308-INS-208 |
| chr9-43923885-INS-54 | chr9-43923885-INS-58 |
| chr9-43938959-INS-179 | chr9-43938959-INS-179 |
| chr9-43954018-INS-95 | chr9-43953983-INS-135 |
| chr9-43957560-INS-312 | chr9-43957560-INS-312 |
| chr9-43965206-INS-51 | chr9-43965206-INS-52 |
| chr9-43971605-INS-907 | chr9-43971615-INS-912 |
| chr9-43972839-INS-176 | chr9-43972839-INS-218 |
| chr9-43993570-INS-52 | chr9-43993570-INS-52 |
| chr9-44007256-INS-78 | chr9-44007256-INS-78 |
| chr9-44033456-INS-55 | chr9-44033456-INS-55 |
| chr9-44039129-INS-336 | chr9-44039129-INS-336 |
| chr9-44072884-INS-68 | chr9-44072884-INS-68 |
| chr9-44075645-INS-222 | chr9-44075645-INS-222 |
| chr9-44090090-INS-79 | chr9-44090090-INS-81 |
| chr9-44135538-INS-205 | chr9-44135538-INS-205 |
| chr9-44165932-INS-219 | chr9-44165932-INS-215 |
| chr9-44177780-INS-210 | chr9-44177780-INS-210 |
| chr9-44180896-INS-194 | chr9-44180896-INS-194 |
| chr9-44198200-INS-94 | chr9-44198200-INS-98 |
| chr9-44229008-INS-131 | chr9-44229008-INS-131 |
| chr9-44236718-INS-58 | chr9-44236718-INS-82 |
| chr9-44260652-INS-200 | chr9-44260652-INS-201 |
| chr9-44262829-INS-78 | chr9-44262829-INS-78 |
| chr9-44263831-INS-65 | chr9-44263831-INS-65 |
| chr9-44273709-INS-201 | chr9-44273709-INS-201 |
| chr9-44288615-INS-226 | chr9-44288615-INS-227 |
| chr9-44289726-INS-342 | chr9-44289726-INS-343 |
| chr9-44323576-INS-195 | chr9-44323576-INS-195 |
| chr9-44337008-INS-3057 | chr9-44337009-INS-3057 |
| chr9-44337191-INS-213 | chr9-44337191-INS-214 |

| | |
|---|---|
| chr9-44358257-INS-217 | chr9-44358257-INS-216 |
| chr9-44428097-INS-702 | chr9-44428097-INS-701 |
| chr9-44440343-INS-50 | chr9-44440343-INS-54 |
| chr9-44461954-INS-7077 | chr9-44461955-INS-7088 |
| chr9-44462293-INS-214 | chr9-44462293-INS-214 |
| chr9-44468972-INS-82 | chr9-44468972-INS-82 |
| chr9-44478292-INS-178 | chr9-44478292-INS-179 |
| chr9-44496707-INS-112 | chr9-44496707-INS-112 |
| chr9-44507327-INS-149 | chr9-44507325-INS-153 |
| chr9-44525027-INS-222 | chr9-44525027-INS-222 |
| chr9-44536629-INS-210 | chr9-44536629-INS-210 |
| chr9-44537366-INS-484 | chr9-44537366-INS-486 |
| chr9-44543419-INS-205 | chr9-44543419-INS-205 |
| chr9-44554853-INS-1006 | chr9-44554854-INS-1002 |
| chr9-44565451-INS-2234 | chr9-44565452-INS-2235 |
| chr9-44565874-INS-213 | chr9-44565874-INS-213 |
| chr9-44566372-INS-5500 | chr9-44566373-INS-5500 |
| chr9-44571173-INS-197 | chr9-44571173-INS-197 |
| chr9-44581160-INS-862 | chr9-44581160-INS-862 |
| chr9-44588984-INS-58 | chr9-44588984-INS-52 |
| chr9-44647737-INS-77 | chr9-44647737-INS-80 |
| chr9-44661992-INS-130 | chr9-44661992-INS-129 |
| chr9-44668084-INS-74 | chr9-44668084-INS-74 |
| chr9-44680906-INS-206 | chr9-44680906-INS-184 |
| chr9-44682753-INS-188 | chr9-44682753-INS-188 |
| chr9-44772631-INS-52 | chr9-44772631-INS-52 |
| chr9-44822029-INS-167 | chr9-44822029-INS-167 |
| chr9-44847267-INS-220 | chr9-44847267-INS-221 |
| chr9-44879101-INS-66 | chr9-44879101-INS-66 |
| chr9-44900996-INS-171 | chr9-44900994-INS-175 |
| chr9-44911662-INS-101 | chr9-44911662-INS-101 |
| chr9-44914909-INS-336 | chr9-44914909-INS-336 |
| chr9-44923974-INS-76 | chr9-44923974-INS-76 |
| chr9-44926849-INS-52 | chr9-44926849-INS-72 |
| chr9-44931858-INS-127 | chr9-44931862-INS-87 |
| chr9-44937489-INS-84 | chr9-44937489-INS-84 |
| chr9-44946689-INS-80 | chr9-44946684-INS-75 |
| chr9-44952681-INS-91 | chr9-44952681-INS-91 |
| chr9-44961496-INS-653 | chr9-44961466-INS-698 |
| chr9-44962247-INS-60 | chr9-44962256-INS-66 |
| chr9-44967731-INS-54 | chr9-44967731-INS-52 |
| chr9-44977136-INS-52 | chr9-44977136-INS-54 |
| chr9-45015184-INS-273 | chr9-45015184-INS-273 |
| chr9-45039966-INS-94 | chr9-45039966-INS-106 |
| chr9-45042001-INS-76 | chr9-45042001-INS-78 |
| chr9-45082695-INS-57 | chr9-45082695-INS-57 |
| chr9-45119961-INS-52 | chr9-45119961-INS-58 |
| chr9-45143148-INS-58 | chr9-45143148-INS-58 |
| chr9-45145430-INS-231 | chr9-45145430-INS-206 |
| chr9-45148742-INS-58 | chr9-45148742-INS-68 |
| chr9-45153720-INS-68 | chr9-45153704-INS-54 |
| chr9-45160103-INS-110 | chr9-45160103-INS-110 |
| chr9-45222505-INS-195 | chr9-45222464-INS-195 |
| chr9-45233788-INS-56 | chr9-45233789-INS-54 |
| chr9-45243297-INS-58 | chr9-45243325-INS-64 |
| chr9-45254944-INS-222 | chr9-45254944-INS-222 |
| chr9-45264145-INS-1283 | chr9-45264146-INS-1284 |
| chr9-45264517-INS-189 | chr9-45264517-INS-189 |
| chr9-45268391-INS-310 | chr9-45268391-INS-310 |
| chr9-45287109-INS-206 | chr9-45287109-INS-206 |
| chr9-45292174-INS-670 | chr9-45292174-INS-670 |
| chr9-45293237-INS-64 | chr9-45293237-INS-64 |
| chr9-45303594-INS-141 | chr9-45303594-INS-141 |
| chr9-45335544-INS-54 | chr9-45335544-INS-54 |
| chr9-45349774-INS-135 | chr9-45349774-INS-135 |
| chr9-45395210-INS-52 | chr9-45395210-INS-52 |
| chr9-45404757-INS-63 | chr9-45404753-INS-54 |
| chr9-45435135-INS-7086 | chr9-45435136-INS-7088 |
| chr9-45442927-INS-684 | chr9-45442927-INS-684 |
| chr9-45456148-INS-53 | chr9-45456121-INS-84 |
| chr9-45488497-INS-76 | chr9-45488497-INS-80 |
| chr9-45498420-INS-429 | chr9-45498420-INS-428 |
| chr9-45505908-INS-3086 | chr9-45505909-INS-3086 |
| chr9-45506190-INS-57 | chr9-45506190-INS-56 |
| chr9-45514625-INS-52 | chr9-45514625-INS-52 |
| chr9-45593009-INS-351 | chr9-45593009-INS-351 |
| chr9-45599991-INS-74 | chr9-45599989-INS-90 |

| | |
|---|---|
| chr9-45602355-INS-71 | chr9-45602355-INS-72 |
| chr9-45622054-INS-70 | chr9-45622054-INS-70 |
| chr9-45623853-INS-52 | chr9-45623853-INS-52 |
| chr9-45630642-INS-71 | chr9-45630642-INS-71 |
| chr9-45673363-INS-278 | chr9-45673363-INS-279 |
| chr9-45674867-INS-205 | chr9-45674867-INS-205 |
| chr9-45723345-INS-194 | chr9-45723345-INS-194 |
| chr9-45745941-INS-62 | chr9-45745970-INS-52 |
| chr9-45759548-INS-64 | chr9-45759548-INS-64 |
| chr9-45778608-INS-98 | chr9-45778608-INS-60 |
| chr9-45808060-INS-65 | chr9-45808060-INS-65 |
| chr9-45810088-INS-214 | chr9-45810088-INS-215 |
| chr9-45810720-INS-265 | chr9-45810725-INS-276 |
| chr9-45867689-INS-54 | chr9-45867689-INS-54 |
| chr9-45868249-INS-207 | chr9-45868249-INS-207 |
| chr9-45891135-INS-55 | chr9-45891135-INS-55 |
| chr9-45894756-INS-77 | chr9-45894754-INS-77 |
| chr9-45898541-INS-56 | chr9-45898541-INS-61 |
| chr9-45901498-INS-59 | chr9-45901498-INS-59 |
| chr9-45905605-INS-54 | chr9-45905605-INS-54 |
| chr9-45906291-INS-185 | chr9-45906291-INS-185 |
| chr9-45922293-INS-54 | chr9-45922282-INS-63 |
| chr9-45940711-INS-223 | chr9-45940711-INS-223 |
| chr9-45947304-INS-205 | chr9-45947304-INS-205 |
| chr9-45956816-INS-247 | chr9-45956816-INS-249 |
| chr9-45957316-INS-98 | chr9-45957316-INS-98 |
| chr9-45966340-INS-62 | chr9-45966340-INS-62 |
| chr9-45971218-INS-52 | chr9-45971218-INS-52 |
| chr9-45986402-INS-84 | chr9-45986402-INS-84 |
| chr9-45997807-INS-51 | chr9-45997807-INS-51 |
| chr9-45999684-INS-60 | chr9-45999684-INS-60 |
| chr9-46002772-INS-68 | chr9-46002772-INS-110 |
| chr9-46067152-INS-136 | chr9-46067153-INS-137 |
| chr9-46082034-INS-52 | chr9-46082034-INS-52 |
| chr9-46091041-INS-163 | chr9-46091041-INS-163 |
| chr9-46099543-INS-207 | chr9-46099543-INS-208 |
| chr9-46109351-INS-153 | chr9-46109351-INS-153 |
| chr9-46125541-INS-1165 | chr9-46125542-INS-1165 |
| chr9-46167412-INS-120 | chr9-46167412-INS-120 |
| chr9-46174040-INS-94 | chr9-46174052-INS-94 |
| chr9-46225856-INS-64 | chr9-46225856-INS-52 |
| chr9-46278817-INS-64 | chr9-46278817-INS-64 |
| chr9-46280846-INS-55 | chr9-46280846-INS-55 |
| chr9-46281782-INS-266 | chr9-46281782-INS-266 |
| chr9-46308568-INS-66 | chr9-46308568-INS-66 |
| chr9-46323682-INS-56 | chr9-46323682-INS-54 |
| chr9-46329287-INS-52 | chr9-46329287-INS-52 |
| chr9-46330575-INS-52 | chr9-46330575-INS-52 |
| chr9-46355488-INS-56 | chr9-46355484-INS-52 |
| chr9-46361227-INS-91 | chr9-46361227-INS-129 |
| chr9-46364606-INS-85 | chr9-46364606-INS-98 |
| chr9-46381252-INS-73 | chr9-46381252-INS-73 |
| chr9-46385180-INS-5413 | chr9-46385181-INS-5413 |
| chr9-46386648-INS-515 | chr9-46386648-INS-515 |
| chr9-46399301-INS-484 | chr9-46399302-INS-484 |
| chr9-46404029-INS-126 | chr9-46404029-INS-126 |
| chr9-46411143-INS-538 | chr9-46411143-INS-538 |
| chr9-46437637-INS-384 | chr9-46437637-INS-384 |
| chr9-46446942-INS-191 | chr9-46446942-INS-191 |
| chr9-46448627-INS-76 | chr9-46448594-INS-80 |
| chr9-46457404-INS-323 | chr9-46457402-INS-325 |
| chr9-46459593-INS-93 | chr9-46459593-INS-100 |
| chr9-46487869-INS-73 | chr9-46487869-INS-51 |
| chr9-46505516-INS-104 | chr9-46505531-INS-122 |
| chr9-46564052-INS-55 | chr9-46564053-INS-57 |
| chr9-46587698-INS-76 | chr9-46587698-INS-82 |
| chr9-46595144-INS-496 | chr9-46595144-INS-496 |
| chr9-46596141-INS-56 | chr9-46596141-INS-56 |
| chr9-46603691-INS-52 | chr9-46603691-INS-52 |
| chr9-46610272-INS-81 | chr9-46610272-INS-81 |
| chr9-46620497-INS-66 | chr9-46620472-INS-70 |
| chr9-46650201-INS-80 | chr9-46650201-INS-50 |
| chr9-46672201-INS-65 | chr9-46672184-INS-63 |
| chr9-46682995-INS-57 | chr9-46682956-INS-66 |
| chr9-46689788-INS-835 | chr9-46689788-INS-836 |
| chr9-46753779-INS-97 | chr9-46753777-INS-95 |
| chr9-46865252-INS-70 | chr9-46865252-INS-60 |

| | |
|---|---|
| chr9-46899893-INS-81 | chr9-46899893-INS-81 |
| chr9-46911658-INS-84 | chr9-46911690-INS-64 |
| chr9-46959804-INS-113 | chr9-46959809-INS-114 |
| chr9-46994440-INS-1888 | chr9-46994441-INS-1888 |
| chr9-47044181-INS-51 | chr9-47044182-INS-99 |
| chr9-47052705-INS-74 | chr9-47052705-INS-66 |
| chr9-47056661-INS-74 | chr9-47056652-INS-110 |
| chr9-47058718-INS-476 | chr9-47058718-INS-476 |
| chr9-47061329-INS-206 | chr9-47061351-INS-212 |
| chr9-47076154-INS-113 | chr9-47076154-INS-113 |
| chr9-47146465-INS-54 | chr9-47146465-INS-64 |
| chr9-47157840-INS-63 | chr9-47157840-INS-63 |
| chr9-47163065-INS-2018 | chr9-47163066-INS-2018 |
| chr9-47209807-INS-56 | chr9-47209807-INS-57 |
| chr9-47234739-INS-314 | chr9-47234739-INS-314 |
| chr9-47366985-INS-68 | chr9-47366975-INS-68 |
| chr9-47383744-INS-100 | chr9-47383746-INS-148 |
| chr9-47448502-INS-53 | chr9-47448502-INS-51 |
| chr9-47464286-INS-59 | chr9-47464286-INS-59 |
| chr9-47494637-INS-212 | chr9-47494637-INS-212 |
| chr9-47544049-INS-291 | chr9-47544047-INS-293 |
| chr9-47553231-INS-58 | chr9-47553231-INS-60 |
| chr9-47566802-INS-2193 | chr9-47566803-INS-2191 |
| chr9-47662597-INS-57 | chr9-47662597-INS-57 |
| chr9-47681033-INS-185 | chr9-47681033-INS-186 |
| chr9-47733951-INS-198 | chr9-47733951-INS-198 |
| chr9-47826296-INS-241 | chr9-47826296-INS-241 |
| chr9-47867255-INS-1830 | chr9-47867279-INS-1795 |
| chr9-47919818-INS-53 | chr9-47919819-INS-51 |
| chr9-47925365-INS-62 | chr9-47925348-INS-51 |
| chr9-47944545-INS-62 | chr9-47944545-INS-62 |
| chr9-48015131-INS-508 | chr9-48015131-INS-508 |
| chr9-48020071-INS-60 | chr9-48020071-INS-52 |
| chr9-48027831-INS-97 | chr9-48027825-INS-103 |
| chr9-48031445-INS-60 | chr9-48031445-INS-62 |
| chr9-48041508-INS-94 | chr9-48041508-INS-94 |
| chr9-48048240-INS-154 | chr9-48048240-INS-166 |
| chr9-48080662-INS-51 | chr9-48080656-INS-50 |
| chr9-48087809-INS-52 | chr9-48087809-INS-52 |
| chr9-48094738-INS-50 | chr9-48094738-INS-50 |
| chr9-48097863-INS-60 | chr9-48097864-INS-60 |
| chr9-48116809-INS-168 | chr9-48116810-INS-169 |
| chr9-48125640-INS-331 | chr9-48125640-INS-331 |
| chr9-48132029-INS-137 | chr9-48132065-INS-97 |
| chr9-48163255-INS-140 | chr9-48163255-INS-142 |
| chr9-48185401-INS-831 | chr9-48185401-INS-831 |
| chr9-48199960-INS-378 | chr9-48199960-INS-378 |
| chr9-48213358-INS-421 | chr9-48213358-INS-421 |
| chr9-48216096-INS-279 | chr9-48216096-INS-279 |
| chr9-48219096-INS-66 | chr9-48219096-INS-66 |
| chr9-48220311-INS-354 | chr9-48220311-INS-354 |
| chr9-48223107-INS-881 | chr9-48223107-INS-881 |
| chr9-48229121-INS-65 | chr9-48229121-INS-66 |
| chr9-48232137-INS-198 | chr9-48232126-INS-199 |
| chr9-48246628-INS-60 | chr9-48246628-INS-60 |
| chr9-48264510-INS-93 | chr9-48264510-INS-93 |
| chr9-48269969-INS-498 | chr9-48269969-INS-498 |
| chr9-48275084-INS-62 | chr9-48275084-INS-62 |
| chr9-48312668-INS-114 | chr9-48312668-INS-115 |
| chr9-48317290-INS-354 | chr9-48317290-INS-354 |
| chr9-48318465-INS-59 | chr9-48318422-INS-92 |
| chr9-48322225-INS-76 | chr9-48322225-INS-76 |
| chr9-48326599-INS-65 | chr9-48326599-INS-65 |
| chr9-48340526-INS-117 | chr9-48340526-INS-117 |
| chr9-48344326-INS-62 | chr9-48344326-INS-62 |
| chr9-48350391-INS-69 | chr9-48350373-INS-75 |
| chr9-48355980-INS-96 | chr9-48355980-INS-96 |
| chr9-48374678-INS-72 | chr9-48374678-INS-72 |
| chr9-48387519-INS-60 | chr9-48387519-INS-60 |
| chr9-48412839-INS-84 | chr9-48412839-INS-126 |
| chr9-48423602-INS-63 | chr9-48423602-INS-66 |
| chr9-48438450-INS-86 | chr9-48438450-INS-86 |
| chr9-48441242-INS-377 | chr9-48441242-INS-377 |
| chr9-48442251-INS-2786 | chr9-48442252-INS-2786 |
| chr9-48444468-INS-86 | chr9-48444468-INS-86 |
| chr9-48468780-INS-105 | chr9-48468781-INS-105 |
| chr9-48469410-INS-87 | chr9-48469412-INS-100 |

| | |
|---|---|
| chr9-48477712-INS-50 | chr9-48477712-INS-58 |
| chr9-48520732-INS-199 | chr9-48520732-INS-199 |
| chr9-48539222-INS-340 | chr9-48539222-INS-340 |
| chr9-48625758-INS-98 | chr9-48625758-INS-98 |
| chr9-48629720-INS-2972 | chr9-48629721-INS-2972 |
| chr9-48635911-INS-56 | chr9-48635911-INS-66 |
| chr9-48664779-INS-107 | chr9-48664779-INS-106 |
| chr9-48697675-INS-80 | chr9-48697631-INS-82 |
| chr9-48727875-INS-215 | chr9-48727875-INS-215 |
| chr9-48758434-INS-112 | chr9-48758434-INS-118 |
| chr9-48778989-INS-58 | chr9-48778989-INS-60 |
| chr9-48780036-INS-128 | chr9-48780036-INS-128 |
| chr9-48782465-INS-69 | chr9-48782465-INS-73 |
| chr9-48803499-INS-125 | chr9-48803497-INS-129 |
| chr9-48813649-INS-267 | chr9-48813649-INS-271 |
| chr9-48816845-INS-100 | chr9-48816845-INS-100 |
| chr9-48823699-INS-108 | chr9-48823699-INS-108 |
| chr9-48838663-INS-68 | chr9-48838668-INS-54 |
| chr9-48841852-INS-288 | chr9-48841852-INS-288 |
| chr9-48886367-INS-337 | chr9-48886367-INS-337 |
| chr9-48890364-INS-115 | chr9-48890364-INS-115 |
| chr9-48894773-INS-274 | chr9-48894773-INS-276 |
| chr9-48900632-INS-90 | chr9-48900632-INS-90 |
| chr9-48911872-INS-210 | chr9-48911872-INS-211 |
| chr9-48935561-INS-55 | chr9-48935561-INS-55 |
| chr9-48962694-INS-56 | chr9-48962690-INS-55 |
| chr9-48963831-INS-225 | chr9-48963831-INS-221 |
| chr9-48974224-INS-207 | chr9-48974224-INS-207 |
| chr9-48979561-INS-122 | chr9-48979561-INS-122 |
| chr9-48983388-INS-128 | chr9-48983388-INS-128 |
| chr9-48991596-INS-201 | chr9-48991596-INS-202 |
| chr9-49033919-INS-90 | chr9-49033919-INS-84 |
| chr9-49066914-INS-136 | chr9-49066913-INS-94 |
| chr9-49088120-INS-5040 | chr9-49088121-INS-5040 |
| chr9-49097123-INS-546 | chr9-49097123-INS-547 |
| chr9-49110739-INS-75 | chr9-49110739-INS-66 |
| chr9-49141665-INS-111 | chr9-49141665-INS-111 |
| chr9-49163003-INS-52 | chr9-49163003-INS-56 |
| chr9-49165588-INS-102 | chr9-49165588-INS-102 |
| chr9-49228329-INS-254 | chr9-49228308-INS-297 |
| chr9-49234808-INS-215 | chr9-49234808-INS-215 |
| chr9-49236908-INS-998 | chr9-49236908-INS-999 |
| chr9-49265790-INS-74 | chr9-49265790-INS-74 |
| chr9-49268865-INS-80 | chr9-49268865-INS-80 |
| chr9-49282059-INS-491 | chr9-49282059-INS-492 |
| chr9-49283778-INS-68 | chr9-49283792-INS-82 |
| chr9-49302579-INS-181 | chr9-49302579-INS-181 |
| chr9-49305222-INS-87 | chr9-49305222-INS-91 |
| chr9-49341286-INS-666 | chr9-49341286-INS-665 |
| chr9-49346992-INS-7201 | chr9-49346993-INS-7201 |
| chr9-49349040-INS-1017 | chr9-49349041-INS-1017 |
| chr9-49393409-INS-74 | chr9-49393409-INS-68 |
| chr9-49500455-INS-60 | chr9-49500455-INS-104 |
| chr9-49514883-INS-51 | chr9-49514883-INS-86 |
| chr9-49550035-INS-57 | chr9-49550035-INS-57 |
| chr9-49566398-INS-60 | chr9-49566398-INS-54 |
| chr9-49601394-INS-55 | chr9-49601394-INS-55 |
| chr9-49604905-INS-144 | chr9-49604905-INS-144 |
| chr9-49635205-INS-540 | chr9-49635205-INS-540 |
| chr9-49698455-INS-61 | chr9-49698463-INS-95 |
| chr9-49700602-INS-82 | chr9-49700602-INS-82 |
| chr9-49768562-INS-210 | chr9-49768562-INS-210 |
| chr9-49802826-INS-7766 | chr9-49802827-INS-7767 |
| chr9-49815941-INS-510 | chr9-49815941-INS-526 |
| chr9-49837064-INS-1105 | chr9-49837079-INS-1084 |
| chr9-49869767-INS-195 | chr9-49869767-INS-188 |
| chr9-49901994-INS-97 | chr9-49902015-INS-81 |
| chr9-49977905-INS-52 | chr9-49977861-INS-80 |
| chr9-50001902-INS-98 | chr9-50001902-INS-98 |
| chr9-50003261-INS-77 | chr9-50003288-INS-52 |
| chr9-50009842-INS-1894 | chr9-50009843-INS-1894 |
| chr9-50015076-INS-130 | chr9-50015076-INS-136 |
| chr9-50022829-INS-1292 | chr9-50022830-INS-1289 |
| chr9-50037076-INS-64 | chr9-50037076-INS-64 |
| chr9-50072552-INS-5339 | chr9-50072553-INS-5339 |
| chr9-50089837-INS-52 | chr9-50089838-INS-60 |
| chr9-50118360-INS-59 | chr9-50118360-INS-59 |

| | |
|---|---|
| chr9-50135936-INS-60 | chr9-50135936-INS-50 |
| chr9-50136925-INS-218 | chr9-50136925-INS-219 |
| chr9-50141143-INS-83 | chr9-50141130-INS-51 |
| chr9-50143154-INS-84 | chr9-50143154-INS-84 |
| chr9-50160732-INS-731 | chr9-50160732-INS-731 |
| chr9-50239673-INS-56 | chr9-50239673-INS-52 |
| chr9-50274691-INS-62 | chr9-50274691-INS-62 |
| chr9-50307045-INS-616 | chr9-50307045-INS-616 |
| chr9-50334195-INS-126 | chr9-50334217-INS-140 |
| chr9-50348316-INS-56 | chr9-50348316-INS-56 |
| chr9-50367543-INS-56 | chr9-50367533-INS-80 |
| chr9-50374354-INS-106 | chr9-50374357-INS-123 |
| chr9-50402213-INS-431 | chr9-50402213-INS-431 |
| chr9-50404972-INS-56 | chr9-50404972-INS-56 |
| chr9-50434468-INS-194 | chr9-50434468-INS-194 |
| chr9-50435349-INS-59 | chr9-50435349-INS-59 |
| chr9-50453885-INS-5130 | chr9-50453886-INS-5130 |
| chr9-50481886-INS-64 | chr9-50481886-INS-64 |
| chr9-50497979-INS-52 | chr9-50497979-INS-52 |
| chr9-50502102-INS-354 | chr9-50502102-INS-354 |
| chr9-50556745-INS-202 | chr9-50556745-INS-202 |
| chr9-50558540-INS-307 | chr9-50558540-INS-307 |
| chr9-50582942-INS-64 | chr9-50582946-INS-64 |
| chr9-50584489-INS-120 | chr9-50584489-INS-120 |
| chr9-50587959-INS-73 | chr9-50587959-INS-73 |
| chr9-50589872-INS-496 | chr9-50589872-INS-496 |
| chr9-50591367-INS-335 | chr9-50591367-INS-335 |
| chr9-50594152-INS-52 | chr9-50594152-INS-52 |
| chr9-50626256-INS-63 | chr9-50626256-INS-63 |
| chr9-50631429-INS-57 | chr9-50631429-INS-57 |
| chr9-50671307-INS-215 | chr9-50671307-INS-215 |
| chr9-50715932-INS-223 | chr9-50715932-INS-220 |
| chr9-50717246-INS-215 | chr9-50717246-INS-215 |
| chr9-50725953-INS-72 | chr9-50725953-INS-73 |
| chr9-50730323-INS-301 | chr9-50730323-INS-295 |
| chr9-50741377-INS-6978 | chr9-50741378-INS-6978 |
| chr9-50771344-INS-51 | chr9-50771344-INS-51 |
| chr9-50840195-INS-91 | chr9-50840195-INS-91 |
| chr9-50878952-INS-227 | chr9-50878952-INS-227 |
| chr9-50926818-INS-197 | chr9-50926818-INS-197 |
| chr9-50944553-INS-82 | chr9-50944553-INS-66 |
| chr9-50954762-INS-58 | chr9-50954762-INS-58 |
| chr9-50967629-INS-60 | chr9-50967629-INS-54 |
| chr9-50971858-INS-170 | chr9-50971858-INS-170 |
| chr9-50976645-INS-144 | chr9-50976645-INS-144 |
| chr9-50980696-INS-71 | chr9-50980696-INS-71 |
| chr9-50991636-INS-178 | chr9-50991636-INS-178 |
| chr9-51001064-INS-74 | chr9-51001064-INS-74 |
| chr9-51009797-INS-102 | chr9-51009812-INS-124 |
| chr9-51063275-INS-83 | chr9-51063276-INS-55 |
| chr9-51063579-INS-83 | chr9-51063579-INS-83 |
| chr9-51085838-INS-87 | chr9-51085839-INS-62 |
| chr9-51106312-INS-203 | chr9-51106312-INS-203 |
| chr9-51121945-INS-534 | chr9-51121945-INS-534 |
| chr9-51127881-INS-205 | chr9-51127881-INS-205 |
| chr9-51131316-INS-58 | chr9-51131316-INS-58 |
| chr9-51138970-INS-55 | chr9-51138970-INS-55 |
| chr9-51155826-INS-57 | chr9-51155826-INS-54 |
| chr9-51177185-INS-74 | chr9-51177177-INS-62 |
| chr9-51215463-INS-163 | chr9-51215463-INS-164 |
| chr9-51220752-INS-96 | chr9-51220735-INS-108 |
| chr9-51261519-INS-386 | chr9-51261519-INS-386 |
| chr9-51343205-INS-75 | chr9-51343205-INS-69 |
| chr9-51350624-INS-207 | chr9-51350624-INS-203 |
| chr9-51365239-INS-81 | chr9-51365239-INS-77 |
| chr9-51381129-INS-6232 | chr9-51381126-INS-6230 |
| chr9-51391082-INS-72 | chr9-51391068-INS-88 |
| chr9-51397223-INS-63 | chr9-51397223-INS-66 |
| chr9-51404669-INS-66 | chr9-51404659-INS-61 |
| chr9-51447355-INS-80 | chr9-51447331-INS-100 |
| chr9-51492534-INS-62 | chr9-51492534-INS-57 |
| chr9-51541542-INS-1871 | chr9-51541543-INS-1871 |
| chr9-51568006-INS-184 | chr9-51568006-INS-184 |
| chr9-51625272-INS-399 | chr9-51625272-INS-399 |
| chr9-51645349-INS-405 | chr9-51645349-INS-405 |
| chr9-51659716-INS-411 | chr9-51659716-INS-411 |
| chr9-51679153-INS-60 | chr9-51679153-INS-84 |

| | |
|---|---|
| chr9-51690431-INS-64 | chr9-51690431-INS-64 |
| chr9-51733870-INS-52 | chr9-51733870-INS-52 |
| chr9-51744970-INS-56 | chr9-51744970-INS-56 |
| chr9-51781171-INS-171 | chr9-51781171-INS-170 |
| chr9-51787105-INS-60 | chr9-51787105-INS-60 |
| chr9-51837370-INS-68 | chr9-51837370-INS-72 |
| chr9-51889733-INS-104 | chr9-51889752-INS-87 |
| chr9-51900473-INS-111 | chr9-51900473-INS-111 |
| chr9-51916677-INS-52 | chr9-51916677-INS-51 |
| chr9-51934290-INS-176 | chr9-51934290-INS-176 |
| chr9-51973731-INS-84 | chr9-51973731-INS-60 |
| chr9-51984020-INS-116 | chr9-51984020-INS-116 |
| chr9-52012949-INS-123 | chr9-52012991-INS-84 |
| chr9-52038735-INS-54 | chr9-52038736-INS-62 |
| chr9-52046981-INS-171 | chr9-52046981-INS-171 |
| chr9-52088005-INS-194 | chr9-52087988-INS-169 |
| chr9-52096152-INS-173 | chr9-52096152-INS-202 |
| chr9-52110950-INS-86 | chr9-52110950-INS-86 |
| chr9-52156567-INS-129 | chr9-52156567-INS-124 |
| chr9-52165905-INS-6353 | chr9-52165906-INS-6346 |
| chr9-52173830-INS-81 | chr9-52173831-INS-50 |
| chr9-52186757-INS-69 | chr9-52186757-INS-52 |
| chr9-52191473-INS-56 | chr9-52191473-INS-76 |
| chr9-52206798-INS-557 | chr9-52206798-INS-557 |
| chr9-52228597-INS-150 | chr9-52228597-INS-146 |
| chr9-52238897-INS-56 | chr9-52238934-INS-62 |
| chr9-52254853-INS-78 | chr9-52254853-INS-78 |
| chr9-52259059-INS-868 | chr9-52259059-INS-874 |
| chr9-52266628-INS-264 | chr9-52266628-INS-266 |
| chr9-52274881-INS-2916 | chr9-52274882-INS-2916 |
| chr9-52284924-INS-175 | chr9-52284924-INS-153 |
| chr9-52292703-INS-4250 | chr9-52292704-INS-4248 |
| chr9-52304117-INS-483 | chr9-52304117-INS-483 |
| chr9-52316300-INS-797 | chr9-52316300-INS-797 |
| chr9-52355191-INS-137 | chr9-52355191-INS-145 |
| chr9-52394234-INS-80 | chr9-52394234-INS-80 |
| chr9-52403894-INS-1333 | chr9-52403895-INS-1333 |
| chr9-52405378-INS-56 | chr9-52405378-INS-56 |
| chr9-52409587-INS-66 | chr9-52409587-INS-50 |
| chr9-52424206-INS-217 | chr9-52424206-INS-217 |
| chr9-52425691-INS-400 | chr9-52425691-INS-400 |
| chr9-52444286-INS-68 | chr9-52444286-INS-68 |
| chr9-52455085-INS-426 | chr9-52455085-INS-427 |
| chr9-52455085-INS-957 | chr9-52455085-INS-957 |
| chr9-52458085-INS-85 | chr9-52458068-INS-58 |
| chr9-52476152-INS-75 | chr9-52476152-INS-75 |
| chr9-52483113-INS-764 | chr9-52483113-INS-754 |
| chr9-52495918-INS-245 | chr9-52495918-INS-245 |
| chr9-52555928-INS-56 | chr9-52555928-INS-56 |
| chr9-52620530-INS-150 | chr9-52620531-INS-134 |
| chr9-52624683-INS-138 | chr9-52624683-INS-132 |
| chr9-52625366-INS-66 | chr9-52625366-INS-58 |
| chr9-52640744-INS-52 | chr9-52640744-INS-54 |
| chr9-52660277-INS-601 | chr9-52660274-INS-579 |
| chr9-52677161-INS-494 | chr9-52677161-INS-506 |
| chr9-52709217-INS-80 | chr9-52709213-INS-50 |
| chr9-52714916-INS-60 | chr9-52714916-INS-60 |
| chr9-52742447-INS-2618 | chr9-52742448-INS-2587 |
| chr9-52744938-INS-397 | chr9-52744938-INS-397 |
| chr9-52753215-INS-1463 | chr9-52753216-INS-1463 |
| chr9-52775054-INS-6389 | chr9-52775055-INS-6389 |
| chr9-52819891-INS-848 | chr9-52819899-INS-840 |
| chr9-52823628-INS-62 | chr9-52823627-INS-57 |
| chr9-52844425-INS-6186 | chr9-52844426-INS-6186 |
| chr9-52852117-INS-97 | chr9-52852107-INS-93 |
| chr9-52853963-INS-79 | chr9-52853963-INS-84 |
| chr9-52861566-INS-76 | chr9-52861566-INS-88 |
| chr9-52886271-INS-969 | chr9-52886271-INS-970 |
| chr9-52935280-INS-805 | chr9-52935280-INS-787 |
| chr9-52936896-INS-62 | chr9-52936896-INS-64 |
| chr9-52939511-INS-139 | chr9-52939511-INS-116 |
| chr9-52940782-INS-94 | chr9-52940782-INS-86 |
| chr9-52947875-INS-548 | chr9-52947875-INS-548 |
| chr9-52950903-INS-53 | chr9-52950903-INS-50 |
| chr9-52978950-INS-88 | chr9-52978950-INS-86 |
| chr9-52981729-INS-452 | chr9-52981729-INS-452 |
| chr9-52991897-INS-376 | chr9-52991897-INS-379 |

| | |
|---|---|
| chr9-53001780-INS-220 | chr9-53001780-INS-220 |
| chr9-53023289-INS-74 | chr9-53023289-INS-87 |
| chr9-53029318-INS-55 | chr9-53029318-INS-55 |
| chr9-53056528-INS-406 | chr9-53056528-INS-406 |
| chr9-53059332-INS-164 | chr9-53059332-INS-164 |
| chr9-53086485-INS-274 | chr9-53086485-INS-258 |
| chr9-53104715-INS-182 | chr9-53104715-INS-182 |
| chr9-53120840-INS-72 | chr9-53120840-INS-71 |
| chr9-53133708-INS-533 | chr9-53133703-INS-540 |
| chr9-53136090-INS-55 | chr9-53136090-INS-55 |
| chr9-53138071-INS-407 | chr9-53138071-INS-413 |
| chr9-53166148-INS-243 | chr9-53166148-INS-243 |
| chr9-53167157-INS-66 | chr9-53167157-INS-66 |
| chr9-53177292-INS-366 | chr9-53177292-INS-366 |
| chr9-53221578-INS-149 | chr9-53221578-INS-155 |
| chr9-53243936-INS-80 | chr9-53243921-INS-50 |
| chr9-53294539-INS-218 | chr9-53294539-INS-218 |
| chr9-53300334-INS-197 | chr9-53300334-INS-197 |
| chr9-53317560-INS-191 | chr9-53317560-INS-208 |
| chr9-53325446-INS-432 | chr9-53325446-INS-432 |
| chr9-53329319-INS-226 | chr9-53329319-INS-226 |
| chr9-53342143-INS-90 | chr9-53342143-INS-90 |
| chr9-53407037-INS-62 | chr9-53407029-INS-56 |
| chr9-53429993-INS-208 | chr9-53429993-INS-208 |
| chr9-53439407-INS-215 | chr9-53439407-INS-215 |
| chr9-53456981-INS-329 | chr9-53456981-INS-329 |
| chr9-53458475-INS-62 | chr9-53458429-INS-93 |
| chr9-53463832-INS-212 | chr9-53463832-INS-212 |
| chr9-53467489-INS-252 | chr9-53467489-INS-252 |
| chr9-53490656-INS-214 | chr9-53490656-INS-215 |
| chr9-53560069-INS-234 | chr9-53560069-INS-234 |
| chr9-53586510-INS-206 | chr9-53586511-INS-188 |
| chr9-53593756-INS-174 | chr9-53593756-INS-174 |
| chr9-53618555-INS-205 | chr9-53618555-INS-205 |
| chr9-53663893-INS-223 | chr9-53663893-INS-222 |
| chr9-53674465-INS-94 | chr9-53674465-INS-94 |
| chr9-53683342-INS-115 | chr9-53683342-INS-99 |
| chr9-53687106-INS-107 | chr9-53687106-INS-108 |
| chr9-53699244-INS-5214 | chr9-53699245-INS-5214 |
| chr9-53745708-INS-52 | chr9-53745708-INS-52 |
| chr9-53752825-INS-875 | chr9-53752825-INS-879 |
| chr9-53782102-INS-1069 | chr9-53782103-INS-1069 |
| chr9-53794520-INS-101 | chr9-53794520-INS-101 |
| chr9-53817520-INS-59 | chr9-53817520-INS-59 |
| chr9-53821179-INS-321 | chr9-53821179-INS-321 |
| chr9-53844959-INS-60 | chr9-53844959-INS-50 |
| chr9-53849267-INS-187 | chr9-53849267-INS-187 |
| chr9-53851960-INS-92 | chr9-53851965-INS-68 |
| chr9-53854908-INS-230 | chr9-53854908-INS-230 |
| chr9-53860708-INS-484 | chr9-53860708-INS-484 |
| chr9-53865241-INS-662 | chr9-53865241-INS-662 |
| chr9-53874453-INS-64 | chr9-53874453-INS-64 |
| chr9-53879673-INS-86 | chr9-53879673-INS-88 |
| chr9-53914948-INS-145 | chr9-53914960-INS-126 |
| chr9-53923419-INS-52 | chr9-53923413-INS-58 |
| chr9-53929823-INS-109 | chr9-53929823-INS-109 |
| chr9-53935754-INS-1314 | chr9-53935739-INS-1315 |
| chr9-53944134-INS-78 | chr9-53944100-INS-87 |
| chr9-53964718-INS-78 | chr9-53964718-INS-54 |
| chr9-53969829-INS-1892 | chr9-53969830-INS-1892 |
| chr9-53992570-INS-72 | chr9-53992568-INS-71 |
| chr9-54008863-INS-251 | chr9-54008870-INS-252 |
| chr9-54019942-INS-457 | chr9-54019942-INS-457 |
| chr9-54020959-INS-199 | chr9-54020959-INS-201 |
| chr9-54022257-INS-85 | chr9-54022257-INS-75 |
| chr9-54068861-INS-74 | chr9-54068835-INS-96 |
| chr9-54076070-INS-103 | chr9-54076070-INS-103 |
| chr9-54089702-INS-125 | chr9-54089703-INS-134 |
| chr9-54136778-INS-52 | chr9-54136778-INS-52 |
| chr9-54197103-INS-59 | chr9-54197103-INS-59 |
| chr9-54214349-INS-70 | chr9-54214349-INS-60 |
| chr9-54226637-INS-172 | chr9-54226637-INS-182 |
| chr9-54244495-INS-54 | chr9-54244495-INS-60 |
| chr9-54263960-INS-155 | chr9-54263960-INS-154 |
| chr9-54328489-INS-55 | chr9-54328489-INS-55 |
| chr9-54340556-INS-285 | chr9-54340556-INS-285 |
| chr9-54391558-INS-60 | chr9-54391558-INS-50 |

| | |
|---|---|
| chr9-54393968-INS-851 | chr9-54393968-INS-881 |
| chr9-54513703-INS-347 | chr9-54513703-INS-362 |
| chr9-54518130-INS-99 | chr9-54518141-INS-92 |
| chr9-54527859-INS-208 | chr9-54527859-INS-208 |
| chr9-54538132-INS-83 | chr9-54538132-INS-83 |
| chr9-54553704-INS-1889 | chr9-54553705-INS-1889 |
| chr9-54557679-INS-203 | chr9-54557679-INS-203 |
| chr9-54560746-INS-78 | chr9-54560746-INS-78 |
| chr9-54575655-INS-211 | chr9-54575655-INS-214 |
| chr9-54581934-INS-688 | chr9-54581934-INS-688 |
| chr9-54601529-INS-102 | chr9-54601549-INS-86 |
| chr9-54696104-INS-54 | chr9-54696099-INS-69 |
| chr9-54717941-INS-84 | chr9-54717942-INS-84 |
| chr9-54725394-INS-176 | chr9-54725389-INS-189 |
| chr9-54754430-INS-53 | chr9-54754430-INS-53 |
| chr9-54766159-INS-118 | chr9-54766159-INS-136 |
| chr9-54844157-INS-178 | chr9-54844153-INS-180 |
| chr9-54904409-INS-79 | chr9-54904409-INS-55 |
| chr9-54908670-INS-82 | chr9-54908670-INS-82 |
| chr9-54924953-INS-54 | chr9-54924953-INS-54 |
| chr9-54938427-INS-56 | chr9-54938427-INS-50 |
| chr9-54959843-INS-289 | chr9-54959843-INS-289 |
| chr9-54963153-INS-180 | chr9-54963153-INS-180 |
| chr9-55019343-INS-428 | chr9-55019343-INS-423 |
| chr9-55021052-INS-79 | chr9-55021052-INS-79 |
| chr9-55031343-INS-70 | chr9-55031343-INS-72 |
| chr9-55086873-INS-171 | chr9-55086850-INS-216 |
| chr9-55134403-INS-195 | chr9-55134403-INS-195 |
| chr9-55137473-INS-125 | chr9-55137462-INS-149 |
| chr9-55172395-INS-299 | chr9-55172395-INS-299 |
| chr9-55188674-INS-76 | chr9-55188674-INS-78 |
| chr9-55210780-INS-497 | chr9-55210780-INS-497 |
| chr9-55264889-INS-2702 | chr9-55264890-INS-2705 |
| chr9-55270427-INS-70 | chr9-55270398-INS-80 |
| chr9-55272641-INS-208 | chr9-55272641-INS-213 |
| chr9-55309130-INS-74 | chr9-55309130-INS-74 |
| chr9-55347881-INS-175 | chr9-55347881-INS-176 |
| chr9-55392382-INS-877 | chr9-55392382-INS-878 |
| chr9-55392911-INS-208 | chr9-55392911-INS-208 |
| chr9-55395771-INS-208 | chr9-55395771-INS-208 |
| chr9-55421750-INS-747 | chr9-55421750-INS-747 |
| chr9-55433335-INS-190 | chr9-55433335-INS-190 |
| chr9-55469609-INS-379 | chr9-55469609-INS-388 |
| chr9-55513928-INS-113 | chr9-55513928-INS-113 |
| chr9-55552266-INS-500 | chr9-55552266-INS-500 |
| chr9-55569877-INS-432 | chr9-55569877-INS-427 |
| chr9-55578250-INS-6314 | chr9-55578249-INS-6317 |
| chr9-55651831-INS-57 | chr9-55651831-INS-61 |
| chr9-55662583-INS-69 | chr9-55662583-INS-67 |
| chr9-55674388-INS-80 | chr9-55674388-INS-72 |
| chr9-55684153-INS-6599 | chr9-55684154-INS-6599 |
| chr9-55733624-INS-91 | chr9-55733624-INS-91 |
| chr9-55736328-INS-848 | chr9-55736328-INS-848 |
| chr9-55770669-INS-430 | chr9-55770669-INS-424 |
| chr9-55772126-INS-52 | chr9-55772126-INS-50 |
| chr9-55795712-INS-220 | chr9-55795712-INS-220 |
| chr9-55797835-INS-324 | chr9-55797835-INS-324 |
| chr9-55812007-INS-81 | chr9-55812025-INS-50 |
| chr9-55822163-INS-68 | chr9-55822163-INS-70 |
| chr9-55848482-INS-718 | chr9-55848482-INS-721 |
| chr9-55853824-INS-643 | chr9-55853824-INS-687 |
| chr9-55890336-INS-1074 | chr9-55890337-INS-1074 |
| chr9-55905343-INS-440 | chr9-55905343-INS-440 |
| chr9-55926068-INS-180 | chr9-55926068-INS-180 |
| chr9-55927376-INS-50 | chr9-55927369-INS-58 |
| chr9-55941406-INS-402 | chr9-55941406-INS-402 |
| chr9-55952194-INS-185 | chr9-55952194-INS-189 |
| chr9-55975655-INS-80 | chr9-55975651-INS-52 |
| chr9-56016960-INS-100 | chr9-56016960-INS-104 |
| chr9-56052283-INS-185 | chr9-56052283-INS-185 |
| chr9-56058696-INS-52 | chr9-56058700-INS-50 |
| chr9-56065958-INS-165 | chr9-56065958-INS-165 |
| chr9-56071709-INS-550 | chr9-56071709-INS-550 |
| chr9-56078771-INS-78 | chr9-56078771-INS-54 |
| chr9-56091841-INS-337 | chr9-56091841-INS-340 |
| chr9-56157475-INS-58 | chr9-56157475-INS-58 |
| chr9-56212121-INS-67 | chr9-56212121-INS-99 |

| | |
|---|---|
| chr9-56219702-INS-432 | chr9-56219702-INS-432 |
| chr9-56224209-INS-71 | chr9-56224201-INS-71 |
| chr9-56240767-INS-68 | chr9-56240767-INS-68 |
| chr9-56272648-INS-957 | chr9-56272648-INS-983 |
| chr9-56274683-INS-196 | chr9-56274683-INS-196 |
| chr9-56360908-INS-260 | chr9-56360908-INS-250 |
| chr9-56388171-INS-84 | chr9-56388171-INS-84 |
| chr9-56389952-INS-102 | chr9-56389952-INS-118 |
| chr9-56423156-INS-74 | chr9-56423195-INS-84 |
| chr9-56424978-INS-96 | chr9-56424963-INS-99 |
| chr9-56440961-INS-106 | chr9-56440961-INS-70 |
| chr9-56449584-INS-72 | chr9-56449584-INS-72 |
| chr9-56483858-INS-223 | chr9-56483858-INS-223 |
| chr9-56521877-INS-90 | chr9-56521877-INS-90 |
| chr9-56627227-INS-92 | chr9-56627227-INS-92 |
| chr9-56642002-INS-280 | chr9-56642002-INS-280 |
| chr9-56652141-INS-196 | chr9-56652141-INS-198 |
| chr9-56655715-INS-118 | chr9-56655716-INS-90 |
| chr9-56709079-INS-132 | chr9-56709100-INS-102 |
| chr9-56722994-INS-248 | chr9-56722994-INS-249 |
| chr9-56727990-INS-81 | chr9-56727990-INS-81 |
| chr9-56743953-INS-5221 | chr9-56743954-INS-5221 |
| chr9-56751774-INS-54 | chr9-56751774-INS-54 |
| chr9-56761069-INS-467 | chr9-56761069-INS-467 |
| chr9-56766630-INS-7119 | chr9-56766631-INS-7119 |
| chr9-56769523-INS-76 | chr9-56769523-INS-76 |
| chr9-56820619-INS-210 | chr9-56820619-INS-210 |
| chr9-56828755-INS-206 | chr9-56828755-INS-208 |
| chr9-56839558-INS-492 | chr9-56839557-INS-486 |
| chr9-56853758-INS-60 | chr9-56853758-INS-60 |
| chr9-56855559-INS-221 | chr9-56855559-INS-245 |
| chr9-56862844-INS-207 | chr9-56862844-INS-207 |
| chr9-56905462-INS-56 | chr9-56905462-INS-56 |
| chr9-56908286-INS-1021 | chr9-56908284-INS-1016 |
| chr9-56913488-INS-1087 | chr9-56913477-INS-1085 |
| chr9-56921549-INS-217 | chr9-56921549-INS-216 |
| chr9-56924216-INS-185 | chr9-56924216-INS-185 |
| chr9-56949023-INS-54 | chr9-56949017-INS-54 |
| chr9-56956482-INS-55 | chr9-56956482-INS-55 |
| chr9-56972182-INS-101 | chr9-56972182-INS-101 |
| chr9-56978589-INS-110 | chr9-56978589-INS-152 |
| chr9-57007419-INS-203 | chr9-57007419-INS-206 |
| chr9-57029332-INS-218 | chr9-57029332-INS-218 |
| chr9-57066994-INS-182 | chr9-57066996-INS-185 |
| chr9-57075024-INS-74 | chr9-57075000-INS-67 |
| chr9-57083075-INS-343 | chr9-57083075-INS-343 |
| chr9-57085297-INS-192 | chr9-57085297-INS-193 |
| chr9-57099357-INS-485 | chr9-57099357-INS-485 |
| chr9-57106436-INS-207 | chr9-57106436-INS-207 |
| chr9-57110606-INS-3764 | chr9-57110607-INS-3757 |
| chr9-57111809-INS-8831 | chr9-57111810-INS-8831 |
| chr9-57123336-INS-931 | chr9-57123336-INS-923 |
| chr9-57137981-INS-253 | chr9-57137981-INS-252 |
| chr9-57193485-INS-210 | chr9-57193485-INS-211 |
| chr9-57201353-INS-206 | chr9-57201353-INS-207 |
| chr9-57219668-INS-206 | chr9-57219668-INS-207 |
| chr9-57226385-INS-327 | chr9-57226385-INS-327 |
| chr9-57231606-INS-188 | chr9-57231606-INS-188 |
| chr9-57248205-INS-213 | chr9-57248205-INS-213 |
| chr9-57260736-INS-208 | chr9-57260752-INS-209 |
| chr9-57271377-INS-82 | chr9-57271377-INS-78 |
| chr9-57276826-INS-528 | chr9-57276834-INS-529 |
| chr9-57278373-INS-203 | chr9-57278373-INS-207 |
| chr9-57290496-INS-530 | chr9-57290496-INS-530 |
| chr9-57292062-INS-489 | chr9-57292062-INS-489 |
| chr9-57292983-INS-120 | chr9-57292983-INS-84 |
| chr9-57301997-INS-93 | chr9-57301997-INS-93 |
| chr9-57326868-INS-72 | chr9-57326868-INS-72 |
| chr9-57329519-INS-193 | chr9-57329519-INS-193 |
| chr9-57337985-INS-198 | chr9-57337985-INS-197 |
| chr9-57370958-INS-211 | chr9-57370958-INS-211 |
| chr9-57387249-INS-55 | chr9-57387249-INS-55 |
| chr9-57393058-INS-209 | chr9-57393058-INS-206 |
| chr9-57399151-INS-427 | chr9-57399149-INS-430 |
| chr9-57438349-INS-169 | chr9-57438349-INS-169 |
| chr9-57516123-INS-203 | chr9-57516123-INS-203 |
| chr9-57517096-INS-201 | chr9-57517096-INS-201 |

| | |
|---|---|
| chr9-57518793-INS-58 | chr9-57518782-INS-60 |
| chr9-57524203-INS-52 | chr9-57524212-INS-60 |
| chr9-57529663-INS-196 | chr9-57529663-INS-202 |
| chr9-57531000-INS-72 | chr9-57531000-INS-57 |
| chr9-57544247-INS-199 | chr9-57544247-INS-200 |
| chr9-57553251-INS-89 | chr9-57553251-INS-89 |
| chr9-57560300-INS-69 | chr9-57560312-INS-53 |
| chr9-57578082-INS-183 | chr9-57578082-INS-183 |
| chr9-57599501-INS-952 | chr9-57599501-INS-950 |
| chr9-57632284-INS-232 | chr9-57632284-INS-231 |
| chr9-57712255-INS-220 | chr9-57712255-INS-220 |
| chr9-57733661-INS-52 | chr9-57733661-INS-52 |
| chr9-57765497-INS-119 | chr9-57765497-INS-119 |
| chr9-57770473-INS-201 | chr9-57770473-INS-202 |
| chr9-57772861-INS-90 | chr9-57772862-INS-90 |
| chr9-57780233-INS-59 | chr9-57780233-INS-59 |
| chr9-57787596-INS-74 | chr9-57787592-INS-64 |
| chr9-57788927-INS-57 | chr9-57788934-INS-77 |
| chr9-57809624-INS-561 | chr9-57809624-INS-561 |
| chr9-57813281-INS-529 | chr9-57813281-INS-529 |
| chr9-57815907-INS-181 | chr9-57815907-INS-197 |
| chr9-57822188-INS-205 | chr9-57822188-INS-206 |
| chr9-57830155-INS-114 | chr9-57830155-INS-114 |
| chr9-57844047-INS-215 | chr9-57844047-INS-215 |
| chr9-57920916-INS-106 | chr9-57920916-INS-106 |
| chr9-57930268-INS-428 | chr9-57930263-INS-434 |
| chr9-57942680-INS-92 | chr9-57942680-INS-94 |
| chr9-57947241-INS-780 | chr9-57947241-INS-780 |
| chr9-57997876-INS-81 | chr9-57997876-INS-81 |
| chr9-58049694-INS-60 | chr9-58049694-INS-58 |
| chr9-58058978-INS-321 | chr9-58058978-INS-322 |
| chr9-58075101-INS-72 | chr9-58075101-INS-78 |
| chr9-58085649-INS-160 | chr9-58085649-INS-160 |
| chr9-58094873-INS-60 | chr9-58094873-INS-60 |
| chr9-58095668-INS-60 | chr9-58095668-INS-55 |
| chr9-58118780-INS-196 | chr9-58118780-INS-196 |
| chr9-58144297-INS-219 | chr9-58144297-INS-219 |
| chr9-58148338-INS-124 | chr9-58148338-INS-124 |
| chr9-58153490-INS-69 | chr9-58153490-INS-52 |
| chr9-58157488-INS-206 | chr9-58157488-INS-206 |
| chr9-58168880-INS-177 | chr9-58168880-INS-195 |
| chr9-58175086-INS-224 | chr9-58175086-INS-226 |
| chr9-58175643-INS-83 | chr9-58175647-INS-60 |
| chr9-58176165-INS-55 | chr9-58176165-INS-67 |
| chr9-58177728-INS-200 | chr9-58177728-INS-200 |
| chr9-58178975-INS-221 | chr9-58178975-INS-222 |
| chr9-58190706-INS-56 | chr9-58190707-INS-53 |
| chr9-58200337-INS-61 | chr9-58200335-INS-98 |
| chr9-58209096-INS-202 | chr9-58209096-INS-202 |
| chr9-58226265-INS-174 | chr9-58226266-INS-140 |
| chr9-58245241-INS-233 | chr9-58245241-INS-233 |
| chr9-58255818-INS-101 | chr9-58255818-INS-101 |
| chr9-58258098-INS-92 | chr9-58258099-INS-70 |
| chr9-58288532-INS-238 | chr9-58288533-INS-237 |
| chr9-58303718-INS-88 | chr9-58303718-INS-68 |
| chr9-58352189-INS-530 | chr9-58352189-INS-530 |
| chr9-58356331-INS-66 | chr9-58356331-INS-66 |
| chr9-58373530-INS-199 | chr9-58373530-INS-197 |
| chr9-58416498-INS-71 | chr9-58416498-INS-102 |
| chr9-58427317-INS-109 | chr9-58427317-INS-106 |
| chr9-58435545-INS-385 | chr9-58435545-INS-385 |
| chr9-58436208-INS-171 | chr9-58436208-INS-171 |
| chr9-58458175-INS-205 | chr9-58458190-INS-205 |
| chr9-58485611-INS-212 | chr9-58485611-INS-212 |
| chr9-58488477-INS-98 | chr9-58488476-INS-117 |
| chr9-58566510-INS-68 | chr9-58566510-INS-66 |
| chr9-58572064-INS-62 | chr9-58572064-INS-62 |
| chr9-58584472-INS-63 | chr9-58584472-INS-63 |
| chr9-58602395-INS-129 | chr9-58602395-INS-145 |
| chr9-58608440-INS-68 | chr9-58608440-INS-68 |
| chr9-58632766-INS-5362 | chr9-58632767-INS-5362 |
| chr9-58643715-INS-185 | chr9-58643715-INS-185 |
| chr9-58654428-INS-138 | chr9-58654428-INS-151 |
| chr9-58660066-INS-1890 | chr9-58660067-INS-1890 |
| chr9-58663772-INS-57 | chr9-58663772-INS-57 |
| chr9-58674781-INS-324 | chr9-58674759-INS-332 |
| chr9-58676417-INS-70 | chr9-58676417-INS-70 |

| | |
|---|---|
| chr9-58696179-INS-252 | chr9-58696188-INS-248 |
| chr9-58715489-INS-205 | chr9-58715489-INS-205 |
| chr9-58721485-INS-69 | chr9-58721485-INS-69 |
| chr9-58766112-INS-52 | chr9-58766112-INS-52 |
| chr9-58776916-INS-200 | chr9-58776909-INS-160 |
| chr9-58793785-INS-651 | chr9-58793785-INS-657 |
| chr9-58800617-INS-76 | chr9-58800619-INS-76 |
| chr9-58818121-INS-202 | chr9-58818121-INS-202 |
| chr9-58877629-INS-891 | chr9-58877629-INS-885 |
| chr9-58881677-INS-54 | chr9-58881677-INS-54 |
| chr9-58885449-INS-1575 | chr9-58885451-INS-1559 |
| chr9-58904503-INS-52 | chr9-58904503-INS-75 |
| chr9-58910673-INS-221 | chr9-58910673-INS-221 |
| chr9-58917421-INS-111 | chr9-58917421-INS-129 |
| chr9-58920267-INS-120 | chr9-58920267-INS-118 |
| chr9-58995144-INS-216 | chr9-58995144-INS-217 |
| chr9-59016844-INS-78 | chr9-59016844-INS-78 |
| chr9-59018673-INS-64 | chr9-59018673-INS-64 |
| chr9-59026039-INS-112 | chr9-59026039-INS-110 |
| chr9-59047640-INS-58 | chr9-59047640-INS-58 |
| chr9-59057576-INS-6401 | chr9-59057577-INS-6401 |
| chr9-59063859-INS-153 | chr9-59063859-INS-157 |
| chr9-59066761-INS-144 | chr9-59066761-INS-144 |
| chr9-59068680-INS-69 | chr9-59068680-INS-69 |
| chr9-59091138-INS-106 | chr9-59091139-INS-60 |
| chr9-59093289-INS-89 | chr9-59093293-INS-111 |
| chr9-59112887-INS-478 | chr9-59112887-INS-478 |
| chr9-59113582-INS-94 | chr9-59113582-INS-56 |
| chr9-59141259-INS-217 | chr9-59141259-INS-215 |
| chr9-59145339-INS-4725 | chr9-59145340-INS-4725 |
| chr9-59146489-INS-51 | chr9-59146489-INS-51 |
| chr9-59162586-INS-55 | chr9-59162586-INS-65 |
| chr9-59165901-INS-80 | chr9-59165901-INS-80 |
| chr9-59166214-INS-387 | chr9-59166214-INS-387 |
| chr9-59182459-INS-723 | chr9-59182459-INS-722 |
| chr9-59183374-INS-53 | chr9-59183374-INS-59 |
| chr9-59188666-INS-1886 | chr9-59188667-INS-1886 |
| chr9-59201974-INS-55 | chr9-59201974-INS-55 |
| chr9-59240498-INS-104 | chr9-59240498-INS-104 |
| chr9-59242813-INS-122 | chr9-59242813-INS-87 |
| chr9-59247435-INS-392 | chr9-59247435-INS-397 |
| chr9-59251693-INS-114 | chr9-59251693-INS-152 |
| chr9-59255137-INS-115 | chr9-59255137-INS-115 |
| chr9-59280169-INS-187 | chr9-59280169-INS-188 |
| chr9-59372797-INS-212 | chr9-59372797-INS-209 |
| chr9-59377505-INS-143 | chr9-59377506-INS-143 |
| chr9-59379605-INS-204 | chr9-59379605-INS-204 |
| chr9-59414030-INS-273 | chr9-59414030-INS-274 |
| chr9-59415707-INS-72 | chr9-59415705-INS-112 |
| chr9-59429168-INS-57 | chr9-59429168-INS-60 |
| chr9-59447517-INS-64 | chr9-59447517-INS-52 |
| chr9-59467192-INS-163 | chr9-59467192-INS-163 |
| chr9-59480550-INS-390 | chr9-59480550-INS-390 |
| chr9-59488929-INS-228 | chr9-59488929-INS-228 |
| chr9-59494217-INS-184 | chr9-59494217-INS-184 |
| chr9-59506176-INS-84 | chr9-59506167-INS-63 |
| chr9-59531087-INS-119 | chr9-59531087-INS-120 |
| chr9-59558624-INS-220 | chr9-59558624-INS-220 |
| chr9-59578444-INS-164 | chr9-59578461-INS-137 |
| chr9-59591668-INS-94 | chr9-59591693-INS-101 |
| chr9-59596861-INS-200 | chr9-59596861-INS-200 |
| chr9-59633917-INS-507 | chr9-59633917-INS-508 |
| chr9-59685896-INS-239 | chr9-59685896-INS-240 |
| chr9-59698879-INS-6436 | chr9-59698880-INS-6436 |
| chr9-59703221-INS-186 | chr9-59703221-INS-187 |
| chr9-59792749-INS-64 | chr9-59792749-INS-64 |
| chr9-59795074-INS-526 | chr9-59795074-INS-525 |
| chr9-59849742-INS-111 | chr9-59849746-INS-97 |
| chr9-59864530-INS-209 | chr9-59864530-INS-205 |
| chr9-59867214-INS-55 | chr9-59867214-INS-55 |
| chr9-59874853-INS-90 | chr9-59874853-INS-60 |
| chr9-59892977-INS-193 | chr9-59892980-INS-197 |
| chr9-59895810-INS-60 | chr9-59895810-INS-50 |
| chr9-59906025-INS-75 | chr9-59906025-INS-75 |
| chr9-59908009-INS-102 | chr9-59907996-INS-101 |
| chr9-59924956-INS-204 | chr9-59924956-INS-204 |
| chr9-59937760-INS-514 | chr9-59937760-INS-514 |

| | |
|---|---|
| chr9-59950575-INS-54 | chr9-59950575-INS-54 |
| chr9-59952839-INS-635 | chr9-59952839-INS-635 |
| chr9-59994553-INS-56 | chr9-59994553-INS-62 |
| chr9-60030287-INS-88 | chr9-60030287-INS-86 |
| chr9-60039364-INS-70 | chr9-60039364-INS-70 |
| chr9-60039933-INS-58 | chr9-60039933-INS-58 |
| chr9-60053235-INS-315 | chr9-60053245-INS-348 |
| chr9-60064476-INS-54 | chr9-60064476-INS-56 |
| chr9-60093060-INS-6111 | chr9-60093061-INS-6111 |
| chr9-60111427-INS-3426 | chr9-60111428-INS-3426 |
| chr9-60116616-INS-86 | chr9-60116616-INS-88 |
| chr9-60137404-INS-6777 | chr9-60137405-INS-6780 |
| chr9-60143242-INS-53 | chr9-60143242-INS-53 |
| chr9-60151433-INS-200 | chr9-60151433-INS-199 |
| chr9-60167569-INS-314 | chr9-60167569-INS-328 |
| chr9-60176491-INS-67 | chr9-60176491-INS-67 |
| chr9-60187661-INS-92 | chr9-60187657-INS-52 |
| chr9-60234644-INS-80 | chr9-60234644-INS-80 |
| chr9-60238563-INS-244 | chr9-60238563-INS-244 |
| chr9-60241896-INS-112 | chr9-60241896-INS-119 |
| chr9-60255923-INS-506 | chr9-60255924-INS-515 |
| chr9-60256434-INS-100 | chr9-60256411-INS-86 |
| chr9-60268593-INS-5627 | chr9-60268594-INS-5627 |
| chr9-60287287-INS-60 | chr9-60287287-INS-52 |
| chr9-60295856-INS-237 | chr9-60295856-INS-237 |
| chr9-60298281-INS-99 | chr9-60298281-INS-99 |
| chr9-60327058-INS-944 | chr9-60327058-INS-943 |
| chr9-60394379-INS-60 | chr9-60394393-INS-58 |
| chr9-60401641-INS-54 | chr9-60401641-INS-54 |
| chr9-60415174-INS-126 | chr9-60415174-INS-126 |
| chr9-60422967-INS-70 | chr9-60422967-INS-67 |
| chr9-60453941-INS-383 | chr9-60453941-INS-383 |
| chr9-60480712-INS-99 | chr9-60480712-INS-116 |
| chr9-60495829-INS-1033 | chr9-60495830-INS-1030 |
| chr9-60522617-INS-5219 | chr9-60522618-INS-5219 |
| chr9-60530833-INS-61 | chr9-60530840-INS-56 |
| chr9-60532689-INS-82 | chr9-60532689-INS-82 |
| chr9-60534640-INS-76 | chr9-60534643-INS-51 |
| chr9-60554519-INS-252 | chr9-60554519-INS-263 |
| chr9-60574559-INS-176 | chr9-60574559-INS-177 |
| chr9-60579505-INS-61 | chr9-60579505-INS-61 |
| chr9-60602269-INS-57 | chr9-60602269-INS-90 |
| chr9-60611786-INS-214 | chr9-60611786-INS-217 |
| chr9-60613732-INS-54 | chr9-60613732-INS-52 |
| chr9-60657805-INS-138 | chr9-60657811-INS-118 |
| chr9-60694673-INS-854 | chr9-60694673-INS-854 |
| chr9-60694677-INS-344 | chr9-60694677-INS-328 |
| chr9-60732573-INS-65 | chr9-60732573-INS-59 |
| chr9-60753990-INS-71 | chr9-60753990-INS-71 |
| chr9-60798047-INS-225 | chr9-60798047-INS-228 |
| chr9-60811631-INS-1266 | chr9-60811632-INS-1266 |
| chr9-60825499-INS-60 | chr9-60825499-INS-60 |
| chr9-60840476-INS-90 | chr9-60840476-INS-90 |
| chr9-60849298-INS-58 | chr9-60849298-INS-58 |
| chr9-60867994-INS-120 | chr9-60867994-INS-141 |
| chr9-60881299-INS-180 | chr9-60881299-INS-194 |
| chr9-60910867-INS-389 | chr9-60910867-INS-398 |
| chr9-60918414-INS-76 | chr9-60918414-INS-76 |
| chr9-60937207-INS-59 | chr9-60937186-INS-65 |
| chr9-60982183-INS-64 | chr9-60982175-INS-55 |
| chr9-61011031-INS-6466 | chr9-61011032-INS-6466 |
| chr9-61023327-INS-102 | chr9-61023327-INS-102 |
| chr9-61029879-INS-81 | chr9-61029879-INS-81 |
| chr9-61032034-INS-72 | chr9-61032034-INS-57 |
| chr9-61042564-INS-90 | chr9-61042564-INS-92 |
| chr9-61050942-INS-204 | chr9-61050942-INS-204 |
| chr9-61084025-INS-90 | chr9-61084025-INS-90 |
| chr9-61096770-INS-243 | chr9-61096770-INS-243 |
| chr9-61110079-INS-76 | chr9-61110041-INS-74 |
| chr9-61113075-INS-92 | chr9-61113075-INS-111 |
| chr9-61133180-INS-60 | chr9-61133180-INS-75 |
| chr9-61147996-INS-162 | chr9-61147996-INS-162 |
| chr9-61195395-INS-5556 | chr9-61195396-INS-5556 |
| chr9-61219656-INS-52 | chr9-61219697-INS-54 |
| chr9-61239050-INS-76 | chr9-61239050-INS-50 |
| chr9-61254267-INS-80 | chr9-61254267-INS-82 |
| chr9-61301240-INS-167 | chr9-61301240-INS-167 |

| | |
|---|---|
| chr9-61315170-INS-226 | chr9-61315170-INS-227 |
| chr9-61328433-INS-334 | chr9-61328433-INS-326 |
| chr9-61333195-INS-50 | chr9-61333195-INS-65 |
| chr9-61342299-INS-55 | chr9-61342299-INS-55 |
| chr9-61419196-INS-58 | chr9-61419196-INS-62 |
| chr9-61426065-INS-104 | chr9-61426065-INS-111 |
| chr9-61482322-INS-96 | chr9-61482322-INS-96 |
| chr9-61493334-INS-64 | chr9-61493337-INS-92 |
| chr9-61501861-INS-62 | chr9-61501856-INS-62 |
| chr9-61514383-INS-51 | chr9-61514383-INS-51 |
| chr9-61517661-INS-165 | chr9-61517661-INS-197 |
| chr9-61526845-INS-85 | chr9-61526845-INS-85 |
| chr9-61530869-INS-72 | chr9-61530866-INS-76 |
| chr9-61573920-INS-102 | chr9-61573888-INS-92 |
| chr9-61579151-INS-101 | chr9-61579151-INS-65 |
| chr9-61594044-INS-255 | chr9-61594044-INS-280 |
| chr9-61599423-INS-74 | chr9-61599423-INS-58 |
| chr9-61665461-INS-70 | chr9-61665456-INS-70 |
| chr9-61680011-INS-90 | chr9-61680011-INS-90 |
| chr9-61694023-INS-55 | chr9-61694023-INS-55 |
| chr9-61720134-INS-2917 | chr9-61720135-INS-2917 |
| chr9-61726183-INS-56 | chr9-61726186-INS-74 |
| chr9-61728808-INS-101 | chr9-61728808-INS-101 |
| chr9-61732998-INS-60 | chr9-61733004-INS-60 |
| chr9-61768931-INS-67 | chr9-61768931-INS-52 |
| chr9-61798552-INS-80 | chr9-61798552-INS-56 |
| chr9-61802855-INS-640 | chr9-61802856-INS-640 |
| chr9-61804298-INS-324 | chr9-61804301-INS-325 |
| chr9-61813968-INS-70 | chr9-61813968-INS-70 |
| chr9-61817408-INS-188 | chr9-61817408-INS-190 |
| chr9-61872270-INS-134 | chr9-61872270-INS-150 |
| chr9-61875333-INS-52 | chr9-61875356-INS-59 |
| chr9-61898383-INS-52 | chr9-61898383-INS-56 |
| chr9-61906173-INS-64 | chr9-61906157-INS-96 |
| chr9-61909764-INS-57 | chr9-61909768-INS-53 |
| chr9-61913908-INS-56 | chr9-61913920-INS-68 |
| chr9-61945346-INS-102 | chr9-61945346-INS-136 |
| chr9-61949150-INS-268 | chr9-61949150-INS-275 |
| chr9-61951549-INS-96 | chr9-61951549-INS-96 |
| chr9-61954375-INS-213 | chr9-61954375-INS-212 |
| chr9-61954818-INS-815 | chr9-61954818-INS-818 |
| chr9-61958198-INS-103 | chr9-61958198-INS-103 |
| chr9-62010170-INS-172 | chr9-62010170-INS-172 |
| chr9-62049086-INS-1314 | chr9-62049087-INS-1314 |
| chr9-62060185-INS-57 | chr9-62060185-INS-57 |
| chr9-62061206-INS-54 | chr9-62061206-INS-60 |
| chr9-62070681-INS-62 | chr9-62070681-INS-58 |
| chr9-62080967-INS-80 | chr9-62080967-INS-80 |
| chr9-62104004-INS-229 | chr9-62104004-INS-229 |
| chr9-62107270-INS-54 | chr9-62107270-INS-56 |
| chr9-62109114-INS-241 | chr9-62109100-INS-219 |
| chr9-62115440-INS-216 | chr9-62115440-INS-220 |
| chr9-62138066-INS-201 | chr9-62138066-INS-201 |
| chr9-62166507-INS-140 | chr9-62166507-INS-140 |
| chr9-62167287-INS-62 | chr9-62167287-INS-68 |
| chr9-62185094-INS-91 | chr9-62185094-INS-91 |
| chr9-62198695-INS-6390 | chr9-62198696-INS-6390 |
| chr9-62211405-INS-206 | chr9-62211405-INS-206 |
| chr9-62215420-INS-196 | chr9-62215420-INS-196 |
| chr9-62222062-INS-72 | chr9-62222063-INS-72 |
| chr9-62234680-INS-103 | chr9-62234680-INS-103 |
| chr9-62237100-INS-163 | chr9-62237100-INS-163 |
| chr9-62239349-INS-68 | chr9-62239352-INS-68 |
| chr9-62260691-INS-193 | chr9-62260691-INS-195 |
| chr9-62297980-INS-52 | chr9-62297980-INS-58 |
| chr9-62300943-INS-60 | chr9-62300940-INS-72 |
| chr9-62301886-INS-149 | chr9-62301882-INS-171 |
| chr9-62303825-INS-205 | chr9-62303825-INS-199 |
| chr9-62312132-INS-200 | chr9-62312132-INS-200 |
| chr9-62314771-INS-209 | chr9-62314771-INS-209 |
| chr9-62315880-INS-96 | chr9-62315880-INS-96 |
| chr9-62321201-INS-62 | chr9-62321201-INS-62 |
| chr9-62337176-INS-200 | chr9-62337176-INS-208 |
| chr9-62362002-INS-93 | chr9-62362002-INS-93 |
| chr9-62393825-INS-74 | chr9-62393825-INS-86 |
| chr9-62415024-INS-208 | chr9-62415024-INS-208 |
| chr9-62437074-INS-59 | chr9-62437074-INS-59 |

| | |
|---|---|
| chr9-62444917-INS-51 | chr9-62444953-INS-53 |
| chr9-62454538-INS-90 | chr9-62454514-INS-110 |
| chr9-62468274-INS-88 | chr9-62468274-INS-88 |
| chr9-62524819-INS-236 | chr9-62524819-INS-236 |
| chr9-62528855-INS-64 | chr9-62528855-INS-72 |
| chr9-62544916-INS-67 | chr9-62544912-INS-75 |
| chr9-62546417-INS-83 | chr9-62546413-INS-94 |
| chr9-62556983-INS-64 | chr9-62556983-INS-64 |
| chr9-62597160-INS-60 | chr9-62597160-INS-60 |
| chr9-62605288-INS-72 | chr9-62605288-INS-72 |
| chr9-62608626-INS-64 | chr9-62608626-INS-56 |
| chr9-62633067-INS-58 | chr9-62633067-INS-72 |
| chr9-62643578-INS-89 | chr9-62643576-INS-91 |
| chr9-62652086-INS-58 | chr9-62652086-INS-74 |
| chr9-62663577-INS-214 | chr9-62663577-INS-214 |
| chr9-62679359-INS-62 | chr9-62679373-INS-58 |
| chr9-62681065-INS-253 | chr9-62681065-INS-253 |
| chr9-62683434-INS-74 | chr9-62683434-INS-60 |
| chr9-62685729-INS-195 | chr9-62685729-INS-197 |
| chr9-62707669-INS-498 | chr9-62707669-INS-498 |
| chr9-62724049-INS-170 | chr9-62724030-INS-169 |
| chr9-62738758-INS-190 | chr9-62738758-INS-142 |
| chr9-62741764-INS-113 | chr9-62741764-INS-114 |
| chr9-62749837-INS-69 | chr9-62749837-INS-94 |
| chr9-62752689-INS-212 | chr9-62752689-INS-212 |
| chr9-62780761-INS-496 | chr9-62780761-INS-503 |
| chr9-62792490-INS-153 | chr9-62792490-INS-139 |
| chr9-62893363-INS-84 | chr9-62893363-INS-78 |
| chr9-62924780-INS-52 | chr9-62924780-INS-56 |
| chr9-62933229-INS-51 | chr9-62933221-INS-54 |
| chr9-62933708-INS-366 | chr9-62933709-INS-366 |
| chr9-62943235-INS-75 | chr9-62943235-INS-111 |
| chr9-62951997-INS-75 | chr9-62951997-INS-78 |
| chr9-62979421-INS-58 | chr9-62979421-INS-64 |
| chr9-62983307-INS-152 | chr9-62983300-INS-185 |
| chr9-63023901-INS-65 | chr9-63023902-INS-61 |
| chr9-63074376-INS-64 | chr9-63074376-INS-64 |
| chr9-63081571-INS-62 | chr9-63081568-INS-64 |
| chr9-63085709-INS-80 | chr9-63085709-INS-80 |
| chr9-63107854-INS-57 | chr9-63107850-INS-75 |
| chr9-63134612-INS-64 | chr9-63134612-INS-66 |
| chr9-63136844-INS-357 | chr9-63136844-INS-357 |
| chr9-63198567-INS-325 | chr9-63198567-INS-325 |
| chr9-63204043-INS-362 | chr9-63204043-INS-362 |
| chr9-63281203-INS-57 | chr9-63281203-INS-57 |
| chr9-63288462-INS-229 | chr9-63288462-INS-230 |
| chr9-63290301-INS-53 | chr9-63290301-INS-55 |
| chr9-63351369-INS-102 | chr9-63351369-INS-102 |
| chr9-63411406-INS-718 | chr9-63411406-INS-719 |
| chr9-63444859-INS-77 | chr9-63444859-INS-77 |
| chr9-63484810-INS-497 | chr9-63484811-INS-498 |
| chr9-63497328-INS-2033 | chr9-63497329-INS-2033 |
| chr9-63530874-INS-399 | chr9-63530874-INS-399 |
| chr9-63583493-INS-102 | chr9-63583497-INS-72 |
| chr9-63587901-INS-231 | chr9-63587902-INS-253 |
| chr9-63592376-INS-101 | chr9-63592376-INS-101 |
| chr9-63593578-INS-91 | chr9-63593587-INS-90 |
| chr9-63679632-INS-113 | chr9-63679632-INS-113 |
| chr9-63683917-INS-110 | chr9-63683917-INS-123 |
| chr9-63703514-INS-169 | chr9-63703514-INS-155 |
| chr9-63710257-INS-598 | chr9-63710257-INS-598 |
| chr9-63739145-INS-51 | chr9-63739145-INS-50 |
| chr9-63742673-INS-63 | chr9-63742673-INS-63 |
| chr9-63749361-INS-74 | chr9-63749361-INS-74 |
| chr9-63751168-INS-92 | chr9-63751168-INS-92 |
| chr9-63757223-INS-52 | chr9-63757223-INS-54 |
| chr9-63786711-INS-81 | chr9-63786711-INS-61 |
| chr9-63810619-INS-50 | chr9-63810619-INS-54 |
| chr9-63811843-INS-188 | chr9-63811843-INS-188 |
| chr9-63855337-INS-310 | chr9-63855337-INS-310 |
| chr9-63864622-INS-61 | chr9-63864653-INS-52 |
| chr9-63873900-INS-190 | chr9-63873900-INS-190 |
| chr9-63888888-INS-211 | chr9-63888888-INS-211 |
| chr9-63900774-INS-227 | chr9-63900774-INS-231 |
| chr9-63913413-INS-212 | chr9-63913413-INS-212 |
| chr9-63942326-INS-80 | chr9-63942326-INS-64 |
| chr9-63956133-INS-217 | chr9-63956133-INS-217 |

| | |
|---|---|
| chr9-63971454-INS-624 | chr9-63971458-INS-621 |
| chr9-63999951-INS-194 | chr9-63999951-INS-194 |
| chr9-64008645-INS-544 | chr9-64008676-INS-554 |
| chr9-64036067-INS-137 | chr9-64036034-INS-131 |
| chr9-64041104-INS-202 | chr9-64041104-INS-202 |
| chr9-64057950-INS-141 | chr9-64057950-INS-167 |
| chr9-64061753-INS-220 | chr9-64061753-INS-220 |
| chr9-64066696-INS-285 | chr9-64066696-INS-283 |
| chr9-64111821-INS-212 | chr9-64111821-INS-218 |
| chr9-64114818-INS-52 | chr9-64114841-INS-75 |
| chr9-64116997-INS-96 | chr9-64116997-INS-96 |
| chr9-64117907-INS-201 | chr9-64117861-INS-207 |
| chr9-64135386-INS-199 | chr9-64135386-INS-198 |
| chr9-64152609-INS-226 | chr9-64152609-INS-229 |
| chr9-64184145-INS-629 | chr9-64184145-INS-629 |
| chr9-64186324-INS-135 | chr9-64186324-INS-111 |
| chr9-64187606-INS-127 | chr9-64187606-INS-127 |
| chr9-64190686-INS-233 | chr9-64190686-INS-233 |
| chr9-64211295-INS-213 | chr9-64211295-INS-214 |
| chr9-64212431-INS-3964 | chr9-64212432-INS-3964 |
| chr9-64214093-INS-209 | chr9-64214093-INS-209 |
| chr9-64216779-INS-236 | chr9-64216779-INS-237 |
| chr9-64228237-INS-213 | chr9-64228237-INS-213 |
| chr9-64230109-INS-233 | chr9-64230109-INS-233 |
| chr9-64232394-INS-207 | chr9-64232406-INS-214 |
| chr9-64233525-INS-273 | chr9-64233510-INS-276 |
| chr9-64237016-INS-84 | chr9-64237014-INS-54 |
| chr9-64247294-INS-68 | chr9-64247294-INS-66 |
| chr9-64250174-INS-405 | chr9-64250174-INS-405 |
| chr9-64258755-INS-466 | chr9-64258755-INS-466 |
| chr9-64268099-INS-101 | chr9-64268060-INS-96 |
| chr9-64275371-INS-74 | chr9-64275371-INS-74 |
| chr9-64284573-INS-78 | chr9-64284573-INS-78 |
| chr9-64311958-INS-137 | chr9-64311958-INS-137 |
| chr9-64328373-INS-84 | chr9-64328373-INS-84 |
| chr9-64335896-INS-57 | chr9-64335896-INS-57 |
| chr9-64354832-INS-6403 | chr9-64354833-INS-6403 |
| chr9-64399379-INS-90 | chr9-64399379-INS-90 |
| chr9-64442747-INS-167 | chr9-64442745-INS-167 |
| chr9-64482485-INS-53 | chr9-64482485-INS-53 |
| chr9-64488034-INS-78 | chr9-64488034-INS-78 |
| chr9-64488773-INS-838 | chr9-64488773-INS-838 |
| chr9-64498585-INS-74 | chr9-64498569-INS-88 |
| chr9-64528692-INS-55 | chr9-64528692-INS-54 |
| chr9-64663838-INS-6387 | chr9-64663839-INS-6387 |
| chr9-64664322-INS-69 | chr9-64664342-INS-69 |
| chr9-64668598-INS-52 | chr9-64668598-INS-67 |
| chr9-64674924-INS-84 | chr9-64674924-INS-70 |
| chr9-64698982-INS-54 | chr9-64698978-INS-57 |
| chr9-64721036-INS-56 | chr9-64721036-INS-52 |
| chr9-64737221-INS-207 | chr9-64737221-INS-209 |
| chr9-64741073-INS-147 | chr9-64741066-INS-147 |
| chr9-64756046-INS-800 | chr9-64756048-INS-805 |
| chr9-64776976-INS-96 | chr9-64776976-INS-97 |
| chr9-64873087-INS-189 | chr9-64873091-INS-189 |
| chr9-64879164-INS-91 | chr9-64879164-INS-91 |
| chr9-64883338-INS-57 | chr9-64883338-INS-57 |
| chr9-64892358-INS-208 | chr9-64892358-INS-208 |
| chr9-64898797-INS-175 | chr9-64898800-INS-150 |
| chr9-64909557-INS-312 | chr9-64909557-INS-312 |
| chr9-64918337-INS-207 | chr9-64918337-INS-207 |
| chr9-64920920-INS-923 | chr9-64920920-INS-924 |
| chr9-64966667-INS-210 | chr9-64966667-INS-210 |
| chr9-64971432-INS-193 | chr9-64971432-INS-192 |
| chr9-64985720-INS-217 | chr9-64985720-INS-217 |
| chr9-64999012-INS-451 | chr9-64999013-INS-457 |
| chr9-65005729-INS-202 | chr9-65005729-INS-202 |
| chr9-65041252-INS-72 | chr9-65041248-INS-53 |
| chr9-65052904-INS-60 | chr9-65052902-INS-60 |
| chr9-65082235-INS-55 | chr9-65082238-INS-99 |
| chr9-65118218-INS-194 | chr9-65118218-INS-191 |
| chr9-65120756-INS-9047 | chr9-65120757-INS-9047 |
| chr9-65128919-INS-92 | chr9-65128915-INS-58 |
| chr9-65136853-INS-199 | chr9-65136853-INS-200 |
| chr9-65147394-INS-206 | chr9-65147394-INS-204 |
| chr9-65153638-INS-6676 | chr9-65153639-INS-6677 |
| chr9-65158202-INS-84 | chr9-65158202-INS-132 |

| | |
|---|---|
| chr9-65168795-INS-2382 | chr9-65168796-INS-2379 |
| chr9-65176366-INS-74 | chr9-65176372-INS-68 |
| chr9-65181945-INS-54 | chr9-65181945-INS-75 |
| chr9-65203411-INS-1361 | chr9-65203400-INS-1383 |
| chr9-65270129-INS-55 | chr9-65270129-INS-55 |
| chr9-65325331-INS-55 | chr9-65325331-INS-55 |
| chr9-65343782-INS-69 | chr9-65343782-INS-69 |
| chr9-65347242-INS-66 | chr9-65347223-INS-75 |
| chr9-65351357-INS-205 | chr9-65351357-INS-196 |
| chr9-65362210-INS-55 | chr9-65362210-INS-58 |
| chr9-65379768-INS-91 | chr9-65379770-INS-114 |
| chr9-65406147-INS-58 | chr9-65406147-INS-60 |
| chr9-65434142-INS-74 | chr9-65434142-INS-50 |
| chr9-65443111-INS-66 | chr9-65443074-INS-56 |
| chr9-65450179-INS-206 | chr9-65450179-INS-205 |
| chr9-65457870-INS-220 | chr9-65457870-INS-223 |
| chr9-65461316-INS-111 | chr9-65461288-INS-108 |
| chr9-65497789-INS-217 | chr9-65497789-INS-229 |
| chr9-65508343-INS-6966 | chr9-65508344-INS-6966 |
| chr9-65509032-INS-219 | chr9-65509032-INS-220 |
| chr9-65512540-INS-180 | chr9-65512538-INS-192 |
| chr9-65527343-INS-60 | chr9-65527344-INS-53 |
| chr9-65533941-INS-244 | chr9-65533941-INS-243 |
| chr9-65557635-INS-345 | chr9-65557636-INS-392 |
| chr9-65639745-INS-217 | chr9-65639745-INS-217 |
| chr9-65658053-INS-516 | chr9-65658053-INS-516 |
| chr9-65674131-INS-222 | chr9-65674131-INS-224 |
| chr9-65675028-INS-208 | chr9-65675028-INS-208 |
| chr9-65682139-INS-75 | chr9-65682139-INS-75 |
| chr9-65693697-INS-205 | chr9-65693697-INS-205 |
| chr9-65700800-INS-54 | chr9-65700800-INS-56 |
| chr9-65703469-INS-76 | chr9-65703469-INS-76 |
| chr9-65704421-INS-210 | chr9-65704422-INS-211 |
| chr9-65705402-INS-215 | chr9-65705402-INS-221 |
| chr9-65707067-INS-173 | chr9-65707067-INS-169 |
| chr9-65708564-INS-197 | chr9-65708564-INS-204 |
| chr9-65718956-INS-66 | chr9-65718956-INS-66 |
| chr9-65724201-INS-209 | chr9-65724201-INS-207 |
| chr9-65750608-INS-191 | chr9-65750608-INS-191 |
| chr9-65763893-INS-1864 | chr9-65763894-INS-1864 |
| chr9-65766352-INS-60 | chr9-65766349-INS-62 |
| chr9-65768160-INS-192 | chr9-65768160-INS-192 |
| chr9-65803535-INS-200 | chr9-65803535-INS-200 |
| chr9-65834019-INS-76 | chr9-65834019-INS-70 |
| chr9-65837226-INS-203 | chr9-65837226-INS-203 |
| chr9-65840671-INS-72 | chr9-65840671-INS-72 |
| chr9-65857047-INS-61 | chr9-65857047-INS-57 |
| chr9-65873744-INS-212 | chr9-65873743-INS-216 |
| chr9-65884762-INS-64 | chr9-65884762-INS-68 |
| chr9-65890640-INS-94 | chr9-65890641-INS-74 |
| chr9-65923401-INS-184 | chr9-65923401-INS-184 |
| chr9-65928249-INS-63 | chr9-65928232-INS-87 |
| chr9-65930517-INS-58 | chr9-65930517-INS-58 |
| chr9-65958929-INS-86 | chr9-65958916-INS-95 |
| chr9-65987780-INS-146 | chr9-65987780-INS-129 |
| chr9-66005027-INS-223 | chr9-66005027-INS-223 |
| chr9-66014833-INS-208 | chr9-66014833-INS-211 |
| chr9-66034734-INS-225 | chr9-66034734-INS-221 |
| chr9-66051607-INS-156 | chr9-66051607-INS-180 |
| chr9-66078580-INS-393 | chr9-66078580-INS-393 |
| chr9-66090397-INS-64 | chr9-66090383-INS-96 |
| chr9-66098199-INS-50 | chr9-66098197-INS-63 |
| chr9-66103758-INS-126 | chr9-66103758-INS-97 |
| chr9-66110347-INS-101 | chr9-66110347-INS-89 |
| chr9-66123296-INS-64 | chr9-66123296-INS-64 |
| chr9-66136612-INS-72 | chr9-66136612-INS-72 |
| chr9-66144776-INS-92 | chr9-66144805-INS-103 |
| chr9-66166069-INS-98 | chr9-66166069-INS-98 |
| chr9-66196003-INS-75 | chr9-66196003-INS-75 |
| chr9-66201807-INS-121 | chr9-66201827-INS-85 |
| chr9-66218631-INS-72 | chr9-66218649-INS-86 |
| chr9-66225216-INS-62 | chr9-66225216-INS-72 |
| chr9-66229880-INS-202 | chr9-66229880-INS-203 |
| chr9-66241041-INS-190 | chr9-66241041-INS-178 |
| chr9-66297949-INS-1272 | chr9-66297949-INS-1237 |
| chr9-66370836-INS-89 | chr9-66370836-INS-89 |
| chr9-66379605-INS-60 | chr9-66379605-INS-64 |

| | |
|---|---|
| chr9-66464306-INS-451 | chr9-66464306-INS-453 |
| chr9-66523058-INS-58 | chr9-66523058-INS-92 |
| chr9-66546334-INS-208 | chr9-66546334-INS-208 |
| chr9-66580155-INS-194 | chr9-66580155-INS-160 |
| chr9-66589983-INS-200 | chr9-66589983-INS-155 |
| chr9-66602690-INS-80 | chr9-66602690-INS-95 |
| chr9-66618540-INS-76 | chr9-66618526-INS-60 |
| chr9-66624375-INS-54 | chr9-66624419-INS-50 |
| chr9-66646841-INS-193 | chr9-66646841-INS-193 |
| chr9-66649895-INS-498 | chr9-66649895-INS-509 |
| chr9-66682027-INS-52 | chr9-66682027-INS-60 |
| chr9-66689401-INS-50 | chr9-66689387-INS-56 |
| chr9-66697495-INS-147 | chr9-66697490-INS-145 |
| chr9-66779026-INS-681 | chr9-66779026-INS-689 |
| chr9-66781679-INS-90 | chr9-66781673-INS-120 |
| chr9-66846654-INS-225 | chr9-66846654-INS-228 |
| chr9-66864111-INS-260 | chr9-66864111-INS-270 |
| chr9-66876072-INS-63 | chr9-66876053-INS-58 |
| chr9-66876925-INS-242 | chr9-66876925-INS-241 |
| chr9-66879994-INS-177 | chr9-66879994-INS-166 |
| chr9-66888254-INS-62 | chr9-66888254-INS-66 |
| chr9-66891319-INS-80 | chr9-66891319-INS-80 |
| chr9-66901448-INS-212 | chr9-66901448-INS-212 |
| chr9-66919811-INS-213 | chr9-66919811-INS-213 |
| chr9-66922222-INS-52 | chr9-66922222-INS-52 |
| chr9-66971512-INS-195 | chr9-66971512-INS-194 |
| chr9-66990218-INS-217 | chr9-66990218-INS-255 |
| chr9-66997736-INS-82 | chr9-66997736-INS-88 |
| chr9-67016620-INS-66 | chr9-67016620-INS-82 |
| chr9-67033295-INS-73 | chr9-67033297-INS-55 |
| chr9-67046935-INS-80 | chr9-67046935-INS-80 |
| chr9-67075463-INS-68 | chr9-67075454-INS-59 |
| chr9-67093234-INS-2377 | chr9-67093235-INS-2377 |
| chr9-67103923-INS-60 | chr9-67103923-INS-62 |
| chr9-67168021-INS-713 | chr9-67168021-INS-712 |
| chr9-67192182-INS-209 | chr9-67192182-INS-209 |
| chr9-67195756-INS-5579 | chr9-67195757-INS-5579 |
| chr9-67220041-INS-200 | chr9-67220041-INS-207 |
| chr9-67239459-INS-60 | chr9-67239459-INS-60 |
| chr9-67242599-INS-78 | chr9-67242605-INS-94 |
| chr9-67244158-INS-55 | chr9-67244176-INS-52 |
| chr9-67277660-INS-78 | chr9-67277660-INS-78 |
| chr9-67313364-INS-190 | chr9-67313364-INS-190 |
| chr9-67319285-INS-60 | chr9-67319285-INS-64 |
| chr9-67325110-INS-61 | chr9-67325110-INS-59 |
| chr9-67328921-INS-64 | chr9-67328921-INS-56 |
| chr9-67334934-INS-261 | chr9-67334930-INS-271 |
| chr9-67340541-INS-52 | chr9-67340541-INS-56 |
| chr9-67360779-INS-59 | chr9-67360779-INS-59 |
| chr9-67365491-INS-70 | chr9-67365491-INS-68 |
| chr9-67379303-INS-127 | chr9-67379308-INS-152 |
| chr9-67422679-INS-80 | chr9-67422679-INS-80 |
| chr9-67425692-INS-70 | chr9-67425692-INS-70 |
| chr9-67439961-INS-71 | chr9-67439961-INS-72 |
| chr9-67441144-INS-52 | chr9-67441104-INS-64 |
| chr9-67472991-INS-68 | chr9-67472986-INS-68 |
| chr9-67479829-INS-51 | chr9-67479825-INS-55 |
| chr9-67484683-INS-117 | chr9-67484683-INS-117 |
| chr9-67498493-INS-61 | chr9-67498525-INS-70 |
| chr9-67506006-INS-60 | chr9-67506006-INS-90 |
| chr9-67520343-INS-758 | chr9-67520343-INS-758 |
| chr9-67536007-INS-64 | chr9-67536007-INS-64 |
| chr9-67550409-INS-64 | chr9-67550409-INS-64 |
| chr9-67561212-INS-606 | chr9-67561212-INS-583 |
| chr9-67572926-INS-525 | chr9-67572926-INS-526 |
| chr9-67573717-INS-57 | chr9-67573717-INS-60 |
| chr9-67583628-INS-62 | chr9-67583638-INS-62 |
| chr9-67586604-INS-415 | chr9-67586604-INS-415 |
| chr9-67590899-INS-933 | chr9-67590899-INS-944 |
| chr9-67672752-INS-54 | chr9-67672755-INS-61 |
| chr9-67676956-INS-81 | chr9-67676956-INS-81 |
| chr9-67701809-INS-177 | chr9-67701809-INS-177 |
| chr9-67751186-INS-209 | chr9-67751186-INS-209 |
| chr9-67756361-INS-53 | chr9-67756361-INS-53 |
| chr9-67794637-INS-371 | chr9-67794637-INS-371 |
| chr9-67811661-INS-185 | chr9-67811661-INS-168 |
| chr9-67812883-INS-192 | chr9-67812883-INS-192 |

| | |
|---|---|
| chr9-67836995-INS-66 | chr9-67836977-INS-92 |
| chr9-67877372-INS-204 | chr9-67877372-INS-203 |
| chr9-67904280-INS-56 | chr9-67904280-INS-58 |
| chr9-67913618-INS-587 | chr9-67913618-INS-581 |
| chr9-67919212-INS-144 | chr9-67919200-INS-144 |
| chr9-67921483-INS-468 | chr9-67921483-INS-468 |
| chr9-67950044-INS-922 | chr9-67950044-INS-922 |
| chr9-67956583-INS-6372 | chr9-67956584-INS-6375 |
| chr9-67971909-INS-6422 | chr9-67971910-INS-6422 |
| chr9-67975852-INS-2066 | chr9-67975853-INS-2066 |
| chr9-68005110-INS-590 | chr9-68005110-INS-590 |
| chr9-68009815-INS-85 | chr9-68009819-INS-53 |
| chr9-68016730-INS-71 | chr9-68016730-INS-66 |
| chr9-68024958-INS-130 | chr9-68024929-INS-145 |
| chr9-68040734-INS-86 | chr9-68040734-INS-82 |
| chr9-68059156-INS-212 | chr9-68059156-INS-212 |
| chr9-68061587-INS-108 | chr9-68061619-INS-108 |
| chr9-68082479-INS-123 | chr9-68082479-INS-123 |
| chr9-68085065-INS-871 | chr9-68085065-INS-872 |
| chr9-68088739-INS-773 | chr9-68088739-INS-773 |
| chr9-68113231-INS-114 | chr9-68113231-INS-74 |
| chr9-68125676-INS-59 | chr9-68125676-INS-59 |
| chr9-68131566-INS-207 | chr9-68131566-INS-207 |
| chr9-68153717-INS-185 | chr9-68153717-INS-190 |
| chr9-68160326-INS-62 | chr9-68160326-INS-62 |
| chr9-68162887-INS-139 | chr9-68162887-INS-139 |
| chr9-68190106-INS-350 | chr9-68190106-INS-318 |
| chr9-68202758-INS-64 | chr9-68202774-INS-62 |
| chr9-68251220-INS-58 | chr9-68251220-INS-58 |
| chr9-68272174-INS-161 | chr9-68272176-INS-162 |
| chr9-68286013-INS-53 | chr9-68285979-INS-54 |
| chr9-68322700-INS-92 | chr9-68322700-INS-100 |
| chr9-68327755-INS-420 | chr9-68327755-INS-404 |
| chr9-68352176-INS-6658 | chr9-68352177-INS-6658 |
| chr9-68356662-INS-303 | chr9-68356662-INS-304 |
| chr9-68381609-INS-87 | chr9-68381609-INS-87 |
| chr9-68434618-INS-199 | chr9-68434618-INS-199 |
| chr9-68447245-INS-434 | chr9-68447245-INS-434 |
| chr9-68455425-INS-731 | chr9-68455425-INS-731 |
| chr9-68460407-INS-7085 | chr9-68460408-INS-7085 |
| chr9-68463921-INS-166 | chr9-68463921-INS-146 |
| chr9-68477105-INS-510 | chr9-68477105-INS-510 |
| chr9-68583125-INS-52 | chr9-68583125-INS-54 |
| chr9-68585646-INS-584 | chr9-68585646-INS-585 |
| chr9-68598056-INS-60 | chr9-68598056-INS-60 |
| chr9-68620309-INS-54 | chr9-68620303-INS-60 |
| chr9-68621509-INS-80 | chr9-68621509-INS-92 |
| chr9-68645355-INS-130 | chr9-68645355-INS-118 |
| chr9-68705525-INS-308 | chr9-68705525-INS-308 |
| chr9-68725323-INS-140 | chr9-68725323-INS-140 |
| chr9-68805614-INS-7385 | chr9-68805615-INS-7385 |
| chr9-68827897-INS-53 | chr9-68827897-INS-65 |
| chr9-68835439-INS-81 | chr9-68835478-INS-84 |
| chr9-68840667-INS-86 | chr9-68840667-INS-86 |
| chr9-68868104-INS-67 | chr9-68868104-INS-65 |
| chr9-68874644-INS-891 | chr9-68874644-INS-891 |
| chr9-68885409-INS-60 | chr9-68885409-INS-60 |
| chr9-68886468-INS-58 | chr9-68886468-INS-60 |
| chr9-68900860-INS-222 | chr9-68900861-INS-222 |
| chr9-68915718-INS-875 | chr9-68915718-INS-880 |
| chr9-68938643-INS-113 | chr9-68938643-INS-124 |
| chr9-68945519-INS-83 | chr9-68945516-INS-85 |
| chr9-68960525-INS-206 | chr9-68960525-INS-206 |
| chr9-68974318-INS-96 | chr9-68974318-INS-126 |
| chr9-69007637-INS-418 | chr9-69007637-INS-423 |
| chr9-69021311-INS-67 | chr9-69021311-INS-67 |
| chr9-69056098-INS-56 | chr9-69056098-INS-56 |
| chr9-69061259-INS-52 | chr9-69061271-INS-54 |
| chr9-69068117-INS-5251 | chr9-69068118-INS-5251 |
| chr9-69084968-INS-56 | chr9-69084968-INS-56 |
| chr9-69163279-INS-217 | chr9-69163279-INS-219 |
| chr9-69212644-INS-58 | chr9-69212644-INS-58 |
| chr9-69214896-INS-58 | chr9-69214896-INS-58 |
| chr9-69235327-INS-77 | chr9-69235327-INS-77 |
| chr9-69263679-INS-53 | chr9-69263683-INS-65 |
| chr9-69283093-INS-98 | chr9-69283089-INS-118 |
| chr9-69327146-INS-65 | chr9-69327132-INS-65 |

| | |
|---|---|
| chr9-69336608-INS-237 | chr9-69336608-INS-237 |
| chr9-69369776-INS-79 | chr9-69369778-INS-92 |
| chr9-69386183-INS-130 | chr9-69386183-INS-130 |
| chr9-69396816-INS-76 | chr9-69396810-INS-88 |
| chr9-69401417-INS-75 | chr9-69401418-INS-52 |
| chr9-69402007-INS-99 | chr9-69402007-INS-99 |
| chr9-69412220-INS-168 | chr9-69412221-INS-168 |
| chr9-69427716-INS-60 | chr9-69427716-INS-60 |
| chr9-69429892-INS-138 | chr9-69429892-INS-138 |
| chr9-69442937-INS-192 | chr9-69442937-INS-192 |
| chr9-69448285-INS-70 | chr9-69448285-INS-69 |
| chr9-69458833-INS-53 | chr9-69458821-INS-73 |
| chr9-69500685-INS-57 | chr9-69500685-INS-51 |
| chr9-69503257-INS-360 | chr9-69503257-INS-366 |
| chr9-69528716-INS-154 | chr9-69528716-INS-156 |
| chr9-69542399-INS-74 | chr9-69542399-INS-74 |
| chr9-69547304-INS-3139 | chr9-69547305-INS-3139 |
| chr9-69565261-INS-52 | chr9-69565261-INS-52 |
| chr9-69569059-INS-3124 | chr9-69569060-INS-3124 |
| chr9-69584065-INS-64 | chr9-69584065-INS-66 |
| chr9-69593884-INS-84 | chr9-69593884-INS-84 |
| chr9-69596963-INS-90 | chr9-69596963-INS-90 |
| chr9-69603893-INS-146 | chr9-69603893-INS-162 |
| chr9-69614603-INS-5548 | chr9-69614604-INS-5548 |
| chr9-69622027-INS-177 | chr9-69622027-INS-177 |
| chr9-69628880-INS-134 | chr9-69628880-INS-114 |
| chr9-69671571-INS-66 | chr9-69671571-INS-68 |
| chr9-69683013-INS-205 | chr9-69683013-INS-205 |
| chr9-69688814-INS-195 | chr9-69688814-INS-195 |
| chr9-69717433-INS-52 | chr9-69717433-INS-52 |
| chr9-69718612-INS-54 | chr9-69718612-INS-54 |
| chr9-69725011-INS-588 | chr9-69725011-INS-592 |
| chr9-69728197-INS-54 | chr9-69728197-INS-54 |
| chr9-69740012-INS-64 | chr9-69740012-INS-64 |
| chr9-69772451-INS-60 | chr9-69772451-INS-60 |
| chr9-69775708-INS-100 | chr9-69775708-INS-100 |
| chr9-69787828-INS-102 | chr9-69787828-INS-141 |
| chr9-69827050-INS-66 | chr9-69827050-INS-72 |
| chr9-69852997-INS-96 | chr9-69853001-INS-82 |
| chr9-69876479-INS-653 | chr9-69876479-INS-653 |
| chr9-69949581-INS-70 | chr9-69949581-INS-70 |
| chr9-69950895-INS-204 | chr9-69950895-INS-204 |
| chr9-69955042-INS-60 | chr9-69955042-INS-60 |
| chr9-70013959-INS-130 | chr9-70013959-INS-128 |
| chr9-70014961-INS-52 | chr9-70014961-INS-53 |
| chr9-70038247-INS-6226 | chr9-70038248-INS-6226 |
| chr9-70055340-INS-82 | chr9-70055341-INS-82 |
| chr9-70056045-INS-53 | chr9-70056045-INS-52 |
| chr9-70059798-INS-4555 | chr9-70059799-INS-4555 |
| chr9-70060555-INS-53 | chr9-70060555-INS-53 |
| chr9-70083299-INS-146 | chr9-70083299-INS-174 |
| chr9-70095650-INS-58 | chr9-70095650-INS-58 |
| chr9-70103210-INS-62 | chr9-70103210-INS-64 |
| chr9-70104766-INS-56 | chr9-70104766-INS-56 |
| chr9-70114001-INS-116 | chr9-70114001-INS-108 |
| chr9-70145747-INS-209 | chr9-70145747-INS-209 |
| chr9-70146445-INS-208 | chr9-70146445-INS-208 |
| chr9-70179517-INS-55 | chr9-70179517-INS-55 |
| chr9-70193619-INS-52 | chr9-70193619-INS-52 |
| chr9-70197447-INS-80 | chr9-70197447-INS-86 |
| chr9-70198531-INS-57 | chr9-70198531-INS-53 |
| chr9-70224909-INS-204 | chr9-70224909-INS-204 |
| chr9-70231695-INS-77 | chr9-70231695-INS-65 |
| chr9-70234760-INS-72 | chr9-70234760-INS-72 |
| chr9-70244924-INS-52 | chr9-70244921-INS-55 |
| chr9-70280778-INS-6443 | chr9-70280779-INS-6445 |
| chr9-70293985-INS-75 | chr9-70293985-INS-96 |
| chr9-70305495-INS-193 | chr9-70305495-INS-193 |
| chr9-70319336-INS-136 | chr9-70319316-INS-136 |
| chr9-70354452-INS-53 | chr9-70354452-INS-57 |
| chr9-70359440-INS-197 | chr9-70359440-INS-197 |
| chr9-70398994-INS-456 | chr9-70398994-INS-457 |
| chr9-70402834-INS-212 | chr9-70402834-INS-212 |
| chr9-70407462-INS-54 | chr9-70407462-INS-57 |
| chr9-70421533-INS-53 | chr9-70421568-INS-62 |
| chr9-70422567-INS-201 | chr9-70422567-INS-200 |
| chr9-70423431-INS-488 | chr9-70423431-INS-488 |

| | |
|---|---|
| chr9-70438864-INS-207 | chr9-70438864-INS-208 |
| chr9-70446649-INS-235 | chr9-70446649-INS-235 |
| chr9-70457931-INS-220 | chr9-70457931-INS-220 |
| chr9-70484800-INS-90 | chr9-70484800-INS-94 |
| chr9-70500274-INS-52 | chr9-70500274-INS-52 |
| chr9-70501476-INS-71 | chr9-70501476-INS-71 |
| chr9-70505471-INS-251 | chr9-70505471-INS-251 |
| chr9-70508246-INS-76 | chr9-70508246-INS-76 |
| chr9-70530581-INS-240 | chr9-70530601-INS-200 |
| chr9-70545543-INS-56 | chr9-70545543-INS-56 |
| chr9-70550948-INS-211 | chr9-70550948-INS-211 |
| chr9-70568045-INS-204 | chr9-70568045-INS-204 |
| chr9-70641245-INS-52 | chr9-70641245-INS-60 |
| chr9-70660385-INS-200 | chr9-70660385-INS-221 |
| chr9-70678603-INS-85 | chr9-70678603-INS-85 |
| chr9-70695948-INS-512 | chr9-70695965-INS-517 |
| chr9-70705303-INS-257 | chr9-70705303-INS-263 |
| chr9-70714038-INS-124 | chr9-70714038-INS-116 |
| chr9-70729201-INS-60 | chr9-70729201-INS-52 |
| chr9-70732322-INS-138 | chr9-70732322-INS-140 |
| chr9-70736197-INS-116 | chr9-70736201-INS-113 |
| chr9-70739395-INS-68 | chr9-70739395-INS-64 |
| chr9-70741419-INS-54 | chr9-70741420-INS-53 |
| chr9-70804010-INS-82 | chr9-70804010-INS-76 |
| chr9-70818966-INS-67 | chr9-70818966-INS-67 |
| chr9-70820168-INS-64 | chr9-70820169-INS-64 |
| chr9-70828956-INS-54 | chr9-70828956-INS-54 |
| chr9-70849781-INS-216 | chr9-70849781-INS-216 |
| chr9-70887840-INS-70 | chr9-70887840-INS-84 |
| chr9-70915312-INS-70 | chr9-70915312-INS-72 |
| chr9-70938295-INS-1174 | chr9-70938296-INS-1190 |
| chr9-70942557-INS-78 | chr9-70942545-INS-81 |
| chr9-70946460-INS-205 | chr9-70946460-INS-205 |
| chr9-71016761-INS-72 | chr9-71016761-INS-69 |
| chr9-71031141-INS-197 | chr9-71031141-INS-197 |
| chr9-71103725-INS-212 | chr9-71103725-INS-212 |
| chr9-71207641-INS-52 | chr9-71207641-INS-52 |
| chr9-71213335-INS-54 | chr9-71213335-INS-54 |
| chr9-71240483-INS-64 | chr9-71240483-INS-98 |
| chr9-71308062-INS-56 | chr9-71308062-INS-57 |
| chr9-71322665-INS-58 | chr9-71322665-INS-58 |
| chr9-71327601-INS-328 | chr9-71327601-INS-328 |
| chr9-71345864-INS-133 | chr9-71345874-INS-116 |
| chr9-71365133-INS-102 | chr9-71365133-INS-113 |
| chr9-71382755-INS-83 | chr9-71382755-INS-85 |
| chr9-71396407-INS-52 | chr9-71396410-INS-50 |
| chr9-71426701-INS-7083 | chr9-71426702-INS-7083 |
| chr9-71433388-INS-194 | chr9-71433388-INS-194 |
| chr9-71457447-INS-152 | chr9-71457447-INS-152 |
| chr9-71469242-INS-145 | chr9-71469242-INS-145 |
| chr9-71522208-INS-198 | chr9-71522208-INS-198 |
| chr9-71542111-INS-445 | chr9-71542111-INS-445 |
| chr9-71551931-INS-54 | chr9-71551932-INS-57 |
| chr9-71553296-INS-205 | chr9-71553296-INS-205 |
| chr9-71580778-INS-58 | chr9-71580778-INS-58 |
| chr9-71593250-INS-101 | chr9-71593252-INS-128 |
| chr9-71601043-INS-59 | chr9-71601040-INS-65 |
| chr9-71608973-INS-194 | chr9-71608973-INS-195 |
| chr9-71615870-INS-57 | chr9-71615870-INS-57 |
| chr9-71626673-INS-2426 | chr9-71626674-INS-2404 |
| chr9-71627238-INS-384 | chr9-71627235-INS-384 |
| chr9-71631527-INS-76 | chr9-71631527-INS-76 |
| chr9-71679894-INS-200 | chr9-71679894-INS-200 |
| chr9-71693440-INS-56 | chr9-71693440-INS-60 |
| chr9-71725982-INS-1887 | chr9-71725983-INS-1887 |
| chr9-71774884-INS-298 | chr9-71774884-INS-299 |
| chr9-71791407-INS-7181 | chr9-71791408-INS-7181 |
| chr9-71791936-INS-240 | chr9-71791936-INS-240 |
| chr9-71844271-INS-67 | chr9-71844271-INS-67 |
| chr9-71845584-INS-381 | chr9-71845584-INS-381 |
| chr9-71896032-INS-167 | chr9-71896032-INS-167 |
| chr9-71904294-INS-76 | chr9-71904294-INS-76 |
| chr9-71925987-INS-51 | chr9-71925987-INS-51 |
| chr9-71991140-INS-211 | chr9-71991140-INS-209 |
| chr9-72022653-INS-205 | chr9-72022653-INS-206 |
| chr9-72026166-INS-144 | chr9-72026166-INS-144 |
| chr9-72064946-INS-67 | chr9-72064946-INS-76 |

| | |
|---|---|
| chr9-72094567-INS-80 | chr9-72094567-INS-88 |
| chr9-72109060-INS-200 | chr9-72109060-INS-200 |
| chr9-72114932-INS-222 | chr9-72114932-INS-231 |
| chr9-72125360-INS-153 | chr9-72125358-INS-118 |
| chr9-72145679-INS-58 | chr9-72145679-INS-58 |
| chr9-72146453-INS-198 | chr9-72146453-INS-199 |
| chr9-72149969-INS-362 | chr9-72149969-INS-362 |
| chr9-72165418-INS-353 | chr9-72165418-INS-353 |
| chr9-72171166-INS-54 | chr9-72171166-INS-69 |
| chr9-72181602-INS-96 | chr9-72181602-INS-88 |
| chr9-72203918-INS-535 | chr9-72203918-INS-534 |
| chr9-72207527-INS-6440 | chr9-72207528-INS-6440 |
| chr9-72248755-INS-213 | chr9-72248755-INS-216 |
| chr9-72249176-INS-783 | chr9-72249182-INS-813 |
| chr9-72280356-INS-177 | chr9-72280357-INS-200 |
| chr9-72288751-INS-57 | chr9-72288751-INS-57 |
| chr9-72297494-INS-76 | chr9-72297494-INS-76 |
| chr9-72321980-INS-60 | chr9-72321980-INS-81 |
| chr9-72333660-INS-962 | chr9-72333660-INS-962 |
| chr9-72339528-INS-206 | chr9-72339528-INS-208 |
| chr9-72380778-INS-64 | chr9-72380778-INS-64 |
| chr9-72389882-INS-199 | chr9-72389882-INS-200 |
| chr9-72392397-INS-6412 | chr9-72392398-INS-6412 |
| chr9-72395005-INS-217 | chr9-72395006-INS-217 |
| chr9-72410710-INS-349 | chr9-72410708-INS-349 |
| chr9-72416528-INS-103 | chr9-72416504-INS-148 |
| chr9-72433263-INS-201 | chr9-72433251-INS-209 |
| chr9-72443703-INS-85 | chr9-72443703-INS-85 |
| chr9-72447229-INS-199 | chr9-72447229-INS-199 |
| chr9-72461428-INS-435 | chr9-72461428-INS-436 |
| chr9-72489086-INS-215 | chr9-72489086-INS-215 |
| chr9-72515607-INS-169 | chr9-72515607-INS-169 |
| chr9-72518203-INS-222 | chr9-72518203-INS-222 |
| chr9-72533658-INS-198 | chr9-72533658-INS-198 |
| chr9-72555389-INS-211 | chr9-72555389-INS-211 |
| chr9-72567287-INS-205 | chr9-72567287-INS-205 |
| chr9-72593804-INS-87 | chr9-72593804-INS-91 |
| chr9-72610887-INS-295 | chr9-72610887-INS-296 |
| chr9-72624167-INS-3063 | chr9-72624168-INS-3063 |
| chr9-72638349-INS-207 | chr9-72638349-INS-207 |
| chr9-72645339-INS-217 | chr9-72645339-INS-219 |
| chr9-72660777-INS-126 | chr9-72660777-INS-108 |
| chr9-72663345-INS-54 | chr9-72663345-INS-53 |
| chr9-72665699-INS-206 | chr9-72665699-INS-206 |
| chr9-72671400-INS-202 | chr9-72671400-INS-202 |
| chr9-72685614-INS-253 | chr9-72685614-INS-253 |
| chr9-72702529-INS-212 | chr9-72702529-INS-212 |
| chr9-72722104-INS-225 | chr9-72722104-INS-225 |
| chr9-72725933-INS-129 | chr9-72725933-INS-129 |
| chr9-72743410-INS-130 | chr9-72743410-INS-130 |
| chr9-72769964-INS-198 | chr9-72769964-INS-198 |
| chr9-72801664-INS-67 | chr9-72801664-INS-67 |
| chr9-72815515-INS-88 | chr9-72815515-INS-92 |
| chr9-72825262-INS-70 | chr9-72825263-INS-60 |
| chr9-72845132-INS-202 | chr9-72845132-INS-202 |
| chr9-72849267-INS-92 | chr9-72849267-INS-92 |
| chr9-72890187-INS-66 | chr9-72890187-INS-69 |
| chr9-72915703-INS-237 | chr9-72915703-INS-237 |
| chr9-72931634-INS-216 | chr9-72931634-INS-216 |
| chr9-72946323-INS-52 | chr9-72946323-INS-54 |
| chr9-72954394-INS-400 | chr9-72954394-INS-400 |
| chr9-72956720-INS-56 | chr9-72956720-INS-56 |
| chr9-72964905-INS-170 | chr9-72964905-INS-169 |
| chr9-72976641-INS-214 | chr9-72976641-INS-215 |
| chr9-72983348-INS-212 | chr9-72983348-INS-211 |
| chr9-72993499-INS-214 | chr9-72993499-INS-214 |
| chr9-72998345-INS-52 | chr9-72998345-INS-58 |
| chr9-73017469-INS-106 | chr9-73017469-INS-106 |
| chr9-73022911-INS-214 | chr9-73022911-INS-214 |
| chr9-73031660-INS-176 | chr9-73031661-INS-155 |
| chr9-73032966-INS-983 | chr9-73032966-INS-983 |
| chr9-73035000-INS-232 | chr9-73035000-INS-232 |
| chr9-73057186-INS-54 | chr9-73057161-INS-56 |
| chr9-73061838-INS-107 | chr9-73061838-INS-105 |
| chr9-73084938-INS-66 | chr9-73084938-INS-66 |
| chr9-73099445-INS-191 | chr9-73099445-INS-178 |
| chr9-73107181-INS-203 | chr9-73107181-INS-203 |

| | |
|---|---|
| chr9-73165260-INS-57 | chr9-73165260-INS-52 |
| chr9-73170277-INS-97 | chr9-73170277-INS-97 |
| chr9-73178344-INS-72 | chr9-73178330-INS-69 |
| chr9-73185120-INS-5298 | chr9-73185121-INS-5298 |
| chr9-73187277-INS-163 | chr9-73187277-INS-155 |
| chr9-73201938-INS-51 | chr9-73201929-INS-88 |
| chr9-73229118-INS-574 | chr9-73229118-INS-550 |
| chr9-73243031-INS-89 | chr9-73243031-INS-91 |
| chr9-73249283-INS-61 | chr9-73249283-INS-61 |
| chr9-73275795-INS-203 | chr9-73275795-INS-203 |
| chr9-73294073-INS-208 | chr9-73294073-INS-207 |
| chr9-73301201-INS-75 | chr9-73301175-INS-72 |
| chr9-73330436-INS-6269 | chr9-73330437-INS-6268 |
| chr9-73331204-INS-119 | chr9-73331204-INS-119 |
| chr9-73336408-INS-975 | chr9-73336408-INS-975 |
| chr9-73359298-INS-66 | chr9-73359298-INS-96 |
| chr9-73373050-INS-176 | chr9-73373050-INS-176 |
| chr9-73412902-INS-6467 | chr9-73412903-INS-6467 |
| chr9-73426202-INS-50 | chr9-73426202-INS-83 |
| chr9-73499587-INS-604 | chr9-73499587-INS-619 |
| chr9-73541540-INS-138 | chr9-73541540-INS-125 |
| chr9-73557244-INS-57 | chr9-73557242-INS-63 |
| chr9-73561242-INS-269 | chr9-73561232-INS-270 |
| chr9-73597667-INS-56 | chr9-73597654-INS-55 |
| chr9-73625811-INS-6407 | chr9-73625812-INS-6407 |
| chr9-73630786-INS-198 | chr9-73630786-INS-224 |
| chr9-73641630-INS-282 | chr9-73641638-INS-282 |
| chr9-73649926-INS-52 | chr9-73649926-INS-100 |
| chr9-73667215-INS-77 | chr9-73667215-INS-77 |
| chr9-73668097-INS-1867 | chr9-73668098-INS-1867 |
| chr9-73680788-INS-334 | chr9-73680788-INS-344 |
| chr9-73684075-INS-90 | chr9-73684084-INS-64 |
| chr9-73716298-INS-56 | chr9-73716345-INS-56 |
| chr9-73730218-INS-83 | chr9-73730218-INS-83 |
| chr9-73763290-INS-416 | chr9-73763290-INS-417 |
| chr9-73773285-INS-90 | chr9-73773255-INS-74 |
| chr9-73776687-INS-5227 | chr9-73776688-INS-5224 |
| chr9-73815475-INS-61 | chr9-73815475-INS-61 |
| chr9-73822723-INS-56 | chr9-73822723-INS-56 |
| chr9-73831110-INS-75 | chr9-73831110-INS-77 |
| chr9-73833151-INS-57 | chr9-73833151-INS-59 |
| chr9-73836330-INS-192 | chr9-73836304-INS-227 |
| chr9-73871294-INS-63 | chr9-73871294-INS-63 |
| chr9-73873871-INS-584 | chr9-73873871-INS-584 |
| chr9-73874915-INS-99 | chr9-73874915-INS-99 |
| chr9-73917234-INS-76 | chr9-73917234-INS-76 |
| chr9-73925790-INS-164 | chr9-73925790-INS-164 |
| chr9-73926832-INS-88 | chr9-73926832-INS-88 |
| chr9-73935878-INS-66 | chr9-73935878-INS-66 |
| chr9-73936720-INS-6500 | chr9-73936721-INS-6500 |
| chr9-73967530-INS-179 | chr9-73967530-INS-179 |
| chr9-73986104-INS-7109 | chr9-73986105-INS-7110 |
| chr9-74016162-INS-6503 | chr9-74016163-INS-6503 |
| chr9-74039543-INS-7184 | chr9-74039544-INS-7185 |
| chr9-74052818-INS-106 | chr9-74052826-INS-64 |
| chr9-74069639-INS-2572 | chr9-74069640-INS-2572 |
| chr9-74079788-INS-84 | chr9-74079788-INS-84 |
| chr9-74090272-INS-3150 | chr9-74090273-INS-3150 |
| chr9-74120181-INS-81 | chr9-74120181-INS-81 |
| chr9-74125117-INS-632 | chr9-74125117-INS-633 |
| chr9-74137186-INS-57 | chr9-74137186-INS-57 |
| chr9-74144947-INS-2784 | chr9-74144948-INS-2784 |
| chr9-74146849-INS-77 | chr9-74146849-INS-65 |
| chr9-74154252-INS-1074 | chr9-74154253-INS-1075 |
| chr9-74197718-INS-690 | chr9-74197726-INS-690 |
| chr9-74219254-INS-1752 | chr9-74219255-INS-1752 |
| chr9-74227975-INS-6374 | chr9-74227976-INS-6374 |
| chr9-74258466-INS-6832 | chr9-74258467-INS-6832 |
| chr9-74259924-INS-52 | chr9-74259925-INS-55 |
| chr9-74276426-INS-141 | chr9-74276418-INS-185 |
| chr9-74292113-INS-238 | chr9-74292113-INS-238 |
| chr9-74306622-INS-58 | chr9-74306622-INS-61 |
| chr9-74328113-INS-92 | chr9-74328114-INS-95 |
| chr9-74363900-INS-67 | chr9-74363918-INS-63 |
| chr9-74371387-INS-62 | chr9-74371373-INS-50 |
| chr9-74391706-INS-84 | chr9-74391722-INS-68 |
| chr9-74400644-INS-517 | chr9-74400644-INS-520 |

| | |
|---|---|
| chr9-74401477-INS-436 | chr9-74401477-INS-431 |
| chr9-74412511-INS-693 | chr9-74412511-INS-693 |
| chr9-74457073-INS-191 | chr9-74457073-INS-191 |
| chr9-74471529-INS-187 | chr9-74471521-INS-195 |
| chr9-74475978-INS-110 | chr9-74475978-INS-126 |
| chr9-74495943-INS-2035 | chr9-74495944-INS-2035 |
| chr9-74560501-INS-59 | chr9-74560501-INS-59 |
| chr9-74578016-INS-831 | chr9-74578016-INS-832 |
| chr9-74628248-INS-453 | chr9-74628248-INS-456 |
| chr9-74635125-INS-81 | chr9-74635125-INS-87 |
| chr9-74688940-INS-228 | chr9-74688940-INS-228 |
| chr9-74723654-INS-130 | chr9-74723654-INS-134 |
| chr9-74735569-INS-115 | chr9-74735569-INS-75 |
| chr9-74737608-INS-209 | chr9-74737608-INS-206 |
| chr9-74841394-INS-73 | chr9-74841394-INS-75 |
| chr9-74847672-INS-125 | chr9-74847694-INS-98 |
| chr9-74854949-INS-132 | chr9-74854949-INS-134 |
| chr9-74883608-INS-85 | chr9-74883608-INS-85 |
| chr9-74891415-INS-97 | chr9-74891425-INS-52 |
| chr9-74901547-INS-63 | chr9-74901547-INS-63 |
| chr9-74906994-INS-69 | chr9-74906994-INS-81 |
| chr9-74915186-INS-102 | chr9-74915186-INS-66 |
| chr9-74918139-INS-52 | chr9-74918139-INS-50 |
| chr9-74923678-INS-76 | chr9-74923678-INS-76 |
| chr9-74943982-INS-78 | chr9-74943982-INS-56 |
| chr9-74955042-INS-64 | chr9-74955042-INS-52 |
| chr9-74963573-INS-91 | chr9-74963573-INS-91 |
| chr9-74985207-INS-123 | chr9-74985207-INS-123 |
| chr9-74986712-INS-587 | chr9-74986724-INS-589 |
| chr9-74987861-INS-144 | chr9-74987861-INS-132 |
| chr9-74998218-INS-728 | chr9-74998218-INS-728 |
| chr9-75021719-INS-146 | chr9-75021682-INS-150 |
| chr9-75039097-INS-1440 | chr9-75039098-INS-1439 |
| chr9-75082055-INS-65 | chr9-75082067-INS-65 |
| chr9-75097804-INS-8732 | chr9-75097805-INS-8732 |
| chr9-75111847-INS-52 | chr9-75111847-INS-52 |
| chr9-75118863-INS-199 | chr9-75118863-INS-199 |
| chr9-75138683-INS-51 | chr9-75138683-INS-57 |
| chr9-75175281-INS-141 | chr9-75175281-INS-172 |
| chr9-75178165-INS-56 | chr9-75178177-INS-77 |
| chr9-75179000-INS-143 | chr9-75179000-INS-159 |
| chr9-75201753-INS-244 | chr9-75201753-INS-230 |
| chr9-75232531-INS-72 | chr9-75232531-INS-72 |
| chr9-75234039-INS-731 | chr9-75234039-INS-731 |
| chr9-75279726-INS-498 | chr9-75279726-INS-498 |
| chr9-75287236-INS-259 | chr9-75287233-INS-256 |
| chr9-75300218-INS-212 | chr9-75300218-INS-213 |
| chr9-75307014-INS-214 | chr9-75307014-INS-214 |
| chr9-75319045-INS-185 | chr9-75319045-INS-185 |
| chr9-75319626-INS-150 | chr9-75319626-INS-150 |
| chr9-75332785-INS-7156 | chr9-75332786-INS-7156 |
| chr9-75343171-INS-62 | chr9-75343171-INS-70 |
| chr9-75349718-INS-66 | chr9-75349720-INS-67 |
| chr9-75360406-INS-129 | chr9-75360406-INS-129 |
| chr9-75364315-INS-983 | chr9-75364315-INS-983 |
| chr9-75366519-INS-109 | chr9-75366519-INS-62 |
| chr9-75371442-INS-181 | chr9-75371442-INS-182 |
| chr9-75393618-INS-199 | chr9-75393618-INS-199 |
| chr9-75396135-INS-71 | chr9-75396135-INS-71 |
| chr9-75397225-INS-510 | chr9-75397225-INS-510 |
| chr9-75398208-INS-212 | chr9-75398208-INS-213 |
| chr9-75416403-INS-439 | chr9-75416403-INS-439 |
| chr9-75466542-INS-105 | chr9-75466542-INS-85 |
| chr9-75500325-INS-51 | chr9-75500325-INS-68 |
| chr9-75514550-INS-52 | chr9-75514550-INS-52 |
| chr9-75540035-INS-2011 | chr9-75540036-INS-2012 |
| chr9-75555251-INS-7135 | chr9-75555252-INS-7135 |
| chr9-75557092-INS-57 | chr9-75557092-INS-52 |
| chr9-75651031-INS-60 | chr9-75651031-INS-54 |
| chr9-75654307-INS-194 | chr9-75654307-INS-194 |
| chr9-75662043-INS-51 | chr9-75662043-INS-51 |
| chr9-75679086-INS-69 | chr9-75679086-INS-63 |
| chr9-75690161-INS-3510 | chr9-75690162-INS-3510 |
| chr9-75696394-INS-52 | chr9-75696394-INS-50 |
| chr9-75698969-INS-1888 | chr9-75698970-INS-1888 |
| chr9-75720317-INS-64 | chr9-75720317-INS-64 |
| chr9-75724390-INS-1152 | chr9-75724391-INS-1159 |

| | |
|---|---|
| chr9-75724851-INS-94 | chr9-75724851-INS-94 |
| chr9-75731097-INS-766 | chr9-75731097-INS-767 |
| chr9-75777083-INS-6893 | chr9-75777084-INS-6893 |
| chr9-75788024-INS-126 | chr9-75788024-INS-126 |
| chr9-75791873-INS-84 | chr9-75791876-INS-56 |
| chr9-75798460-INS-68 | chr9-75798460-INS-68 |
| chr9-75801901-INS-51 | chr9-75801901-INS-54 |
| chr9-75811866-INS-80 | chr9-75811866-INS-80 |
| chr9-75843019-INS-1191 | chr9-75843020-INS-1213 |
| chr9-75859352-INS-191 | chr9-75859352-INS-218 |
| chr9-75871463-INS-447 | chr9-75871463-INS-447 |
| chr9-75873771-INS-110 | chr9-75873771-INS-110 |
| chr9-75876291-INS-58 | chr9-75876291-INS-52 |
| chr9-75878143-INS-66 | chr9-75878151-INS-80 |
| chr9-75880116-INS-66 | chr9-75880093-INS-57 |
| chr9-75888184-INS-89 | chr9-75888184-INS-119 |
| chr9-75915783-INS-84 | chr9-75915783-INS-84 |
| chr9-75979010-INS-233 | chr9-75979014-INS-238 |
| chr9-75983041-INS-54 | chr9-75983041-INS-57 |
| chr9-75990538-INS-210 | chr9-75990538-INS-211 |
| chr9-75996602-INS-54 | chr9-75996602-INS-56 |
| chr9-76032029-INS-224 | chr9-76032029-INS-224 |
| chr9-76047218-INS-651 | chr9-76047218-INS-629 |
| chr9-76052214-INS-1460 | chr9-76052215-INS-1461 |
| chr9-76055297-INS-63 | chr9-76055297-INS-66 |
| chr9-76076593-INS-209 | chr9-76076609-INS-162 |
| chr9-76088003-INS-64 | chr9-76088003-INS-64 |
| chr9-76102640-INS-258 | chr9-76102640-INS-258 |
| chr9-76127693-INS-1272 | chr9-76127694-INS-1272 |
| chr9-76146988-INS-71 | chr9-76146988-INS-71 |
| chr9-76159113-INS-70 | chr9-76159113-INS-70 |
| chr9-76198935-INS-121 | chr9-76198935-INS-121 |
| chr9-76217228-INS-193 | chr9-76217229-INS-179 |
| chr9-76239173-INS-200 | chr9-76239173-INS-207 |
| chr9-76248321-INS-121 | chr9-76248321-INS-122 |
| chr9-76250229-INS-56 | chr9-76250261-INS-66 |
| chr9-76251143-INS-220 | chr9-76251143-INS-221 |
| chr9-76254625-INS-432 | chr9-76254625-INS-432 |
| chr9-76266780-INS-5541 | chr9-76266781-INS-5541 |
| chr9-76272774-INS-6789 | chr9-76272775-INS-6790 |
| chr9-76277122-INS-58 | chr9-76277122-INS-76 |
| chr9-76281933-INS-6540 | chr9-76281934-INS-6541 |
| chr9-76286612-INS-728 | chr9-76286612-INS-728 |
| chr9-76299227-INS-955 | chr9-76299227-INS-955 |
| chr9-76334706-INS-62 | chr9-76334698-INS-69 |
| chr9-76342796-INS-718 | chr9-76342796-INS-718 |
| chr9-76411229-INS-76 | chr9-76411229-INS-76 |
| chr9-76443205-INS-162 | chr9-76443205-INS-162 |
| chr9-76464872-INS-6879 | chr9-76464873-INS-6880 |
| chr9-76477107-INS-64 | chr9-76477107-INS-50 |
| chr9-76484314-INS-80 | chr9-76484314-INS-80 |
| chr9-76531295-INS-66 | chr9-76531287-INS-68 |
| chr9-76575245-INS-341 | chr9-76575245-INS-341 |
| chr9-76576032-INS-1767 | chr9-76576033-INS-1767 |
| chr9-76587177-INS-420 | chr9-76587177-INS-420 |
| chr9-76591968-INS-53 | chr9-76591968-INS-53 |
| chr9-76616197-INS-508 | chr9-76616197-INS-510 |
| chr9-76638101-INS-6596 | chr9-76638102-INS-6596 |
| chr9-76674893-INS-188 | chr9-76674893-INS-188 |
| chr9-76677911-INS-573 | chr9-76677911-INS-573 |
| chr9-76749016-INS-1510 | chr9-76749017-INS-1510 |
| chr9-76752177-INS-72 | chr9-76752177-INS-72 |
| chr9-76757906-INS-6656 | chr9-76757907-INS-6657 |
| chr9-76773964-INS-497 | chr9-76773964-INS-497 |
| chr9-76781493-INS-127 | chr9-76781493-INS-127 |
| chr9-76784582-INS-231 | chr9-76784582-INS-233 |
| chr9-76819618-INS-52 | chr9-76819617-INS-53 |
| chr9-76843993-INS-6176 | chr9-76843994-INS-6176 |
| chr9-76861875-INS-291 | chr9-76861875-INS-301 |
| chr9-76862848-INS-58 | chr9-76862859-INS-50 |
| chr9-76867257-INS-6159 | chr9-76867258-INS-6159 |
| chr9-76874403-INS-458 | chr9-76874402-INS-459 |
| chr9-76887924-INS-102 | chr9-76887924-INS-102 |
| chr9-76901828-INS-54 | chr9-76901828-INS-54 |
| chr9-76935942-INS-213 | chr9-76935945-INS-215 |
| chr9-77021333-INS-1339 | chr9-77021334-INS-1354 |
| chr9-77024478-INS-88 | chr9-77024478-INS-88 |

| | |
|---|---|
| chr9-77071137-INS-171 | chr9-77071137-INS-172 |
| chr9-77107949-INS-6429 | chr9-77107950-INS-6428 |
| chr9-77115981-INS-984 | chr9-77115981-INS-984 |
| chr9-77132885-INS-68 | chr9-77132869-INS-56 |
| chr9-77148566-INS-174 | chr9-77148566-INS-174 |
| chr9-77169685-INS-360 | chr9-77169685-INS-360 |
| chr9-77195935-INS-293 | chr9-77195935-INS-279 |
| chr9-77197665-INS-604 | chr9-77197665-INS-604 |
| chr9-77205323-INS-55 | chr9-77205324-INS-57 |
| chr9-77232131-INS-119 | chr9-77232131-INS-119 |
| chr9-77246829-INS-64 | chr9-77246829-INS-64 |
| chr9-77256984-INS-76 | chr9-77256976-INS-55 |
| chr9-77266880-INS-87 | chr9-77266880-INS-87 |
| chr9-77276625-INS-1860 | chr9-77276626-INS-1860 |
| chr9-77292547-INS-74 | chr9-77292548-INS-90 |
| chr9-77299505-INS-84 | chr9-77299505-INS-88 |
| chr9-77315349-INS-875 | chr9-77315349-INS-886 |
| chr9-77320605-INS-90 | chr9-77320606-INS-135 |
| chr9-77325705-INS-80 | chr9-77325705-INS-80 |
| chr9-77331678-INS-150 | chr9-77331678-INS-128 |
| chr9-77348547-INS-206 | chr9-77348547-INS-206 |
| chr9-77367988-INS-56 | chr9-77367988-INS-58 |
| chr9-77389602-INS-76 | chr9-77389602-INS-76 |
| chr9-77399275-INS-560 | chr9-77399275-INS-560 |
| chr9-77420227-INS-54 | chr9-77420263-INS-51 |
| chr9-77424452-INS-1893 | chr9-77424453-INS-1893 |
| chr9-77437696-INS-154 | chr9-77437687-INS-139 |
| chr9-77439126-INS-200 | chr9-77439126-INS-200 |
| chr9-77447878-INS-169 | chr9-77447878-INS-169 |
| chr9-77471146-INS-502 | chr9-77471146-INS-502 |
| chr9-77475137-INS-113 | chr9-77475137-INS-161 |
| chr9-77482125-INS-372 | chr9-77482125-INS-372 |
| chr9-77482957-INS-121 | chr9-77482957-INS-121 |
| chr9-77488209-INS-3222 | chr9-77488210-INS-3222 |
| chr9-77489917-INS-61 | chr9-77489921-INS-51 |
| chr9-77552021-INS-116 | chr9-77552059-INS-116 |
| chr9-77572745-INS-80 | chr9-77572745-INS-80 |
| chr9-77589743-INS-203 | chr9-77589743-INS-208 |
| chr9-77609736-INS-61 | chr9-77609736-INS-61 |
| chr9-77629324-INS-134 | chr9-77629324-INS-170 |
| chr9-77630571-INS-1246 | chr9-77630588-INS-1248 |
| chr9-77634038-INS-947 | chr9-77634038-INS-947 |
| chr9-77650381-INS-52 | chr9-77650381-INS-56 |
| chr9-77663688-INS-74 | chr9-77663688-INS-74 |
| chr9-77678867-INS-6646 | chr9-77678868-INS-6646 |
| chr9-77724817-INS-733 | chr9-77724817-INS-734 |
| chr9-77728855-INS-64 | chr9-77728855-INS-52 |
| chr9-77742908-INS-197 | chr9-77742908-INS-197 |
| chr9-77770202-INS-790 | chr9-77770202-INS-790 |
| chr9-77783815-INS-87 | chr9-77783799-INS-62 |
| chr9-77785461-INS-420 | chr9-77785461-INS-421 |
| chr9-77793583-INS-59 | chr9-77793583-INS-60 |
| chr9-77825448-INS-74 | chr9-77825448-INS-74 |
| chr9-77840477-INS-1071 | chr9-77840478-INS-1022 |
| chr9-77855331-INS-211 | chr9-77855331-INS-211 |
| chr9-77883639-INS-491 | chr9-77883639-INS-464 |
| chr9-77903535-INS-6440 | chr9-77903536-INS-6440 |
| chr9-77909555-INS-122 | chr9-77909555-INS-145 |
| chr9-77916765-INS-204 | chr9-77916765-INS-216 |
| chr9-77933413-INS-127 | chr9-77933413-INS-129 |
| chr9-77946323-INS-61 | chr9-77946324-INS-52 |
| chr9-77979134-INS-434 | chr9-77979134-INS-434 |
| chr9-78001182-INS-177 | chr9-78001182-INS-176 |
| chr9-78018379-INS-190 | chr9-78018379-INS-201 |
| chr9-78061493-INS-747 | chr9-78061493-INS-753 |
| chr9-78070313-INS-201 | chr9-78070313-INS-201 |
| chr9-78080907-INS-434 | chr9-78080907-INS-434 |
| chr9-78103292-INS-214 | chr9-78103292-INS-214 |
| chr9-78111081-INS-211 | chr9-78111081-INS-210 |
| chr9-78114942-INS-62 | chr9-78114990-INS-56 |
| chr9-78116103-INS-89 | chr9-78116103-INS-89 |
| chr9-78119236-INS-6457 | chr9-78119237-INS-6457 |
| chr9-78127239-INS-873 | chr9-78127239-INS-873 |
| chr9-78130513-INS-11203 | chr9-78130514-INS-11204 |
| chr9-78133543-INS-177 | chr9-78133544-INS-140 |
| chr9-78147945-INS-62 | chr9-78147945-INS-62 |
| chr9-78153969-INS-174 | chr9-78153969-INS-182 |

| | |
|---|---|
| chr9-78172314-INS-241 | chr9-78172315-INS-241 |
| chr9-78232186-INS-246 | chr9-78232186-INS-246 |
| chr9-78243479-INS-224 | chr9-78243479-INS-224 |
| chr9-78254773-INS-143 | chr9-78254773-INS-143 |
| chr9-78260671-INS-202 | chr9-78260671-INS-202 |
| chr9-78261963-INS-82 | chr9-78261962-INS-80 |
| chr9-78270679-INS-100 | chr9-78270679-INS-100 |
| chr9-78274237-INS-160 | chr9-78274237-INS-163 |
| chr9-78293800-INS-78 | chr9-78293822-INS-63 |
| chr9-78300077-INS-491 | chr9-78300077-INS-491 |
| chr9-78301429-INS-7214 | chr9-78301430-INS-7214 |
| chr9-78319179-INS-630 | chr9-78319179-INS-632 |
| chr9-78328806-INS-210 | chr9-78328806-INS-210 |
| chr9-78330177-INS-192 | chr9-78330177-INS-192 |
| chr9-78331105-INS-3028 | chr9-78331106-INS-3025 |
| chr9-78333759-INS-210 | chr9-78333759-INS-207 |
| chr9-78372369-INS-410 | chr9-78372369-INS-410 |
| chr9-78373861-INS-72 | chr9-78373861-INS-72 |
| chr9-78380172-INS-196 | chr9-78380172-INS-197 |
| chr9-78411639-INS-7346 | chr9-78411640-INS-7346 |
| chr9-78414634-INS-176 | chr9-78414645-INS-178 |
| chr9-78416343-INS-209 | chr9-78416344-INS-209 |
| chr9-78420461-INS-205 | chr9-78420461-INS-206 |
| chr9-78423238-INS-214 | chr9-78423238-INS-214 |
| chr9-78438049-INS-240 | chr9-78438049-INS-246 |
| chr9-78443330-INS-54 | chr9-78443359-INS-58 |
| chr9-78447112-INS-189 | chr9-78447112-INS-189 |
| chr9-78455043-INS-179 | chr9-78455043-INS-179 |
| chr9-78461085-INS-59 | chr9-78461110-INS-60 |
| chr9-78469379-INS-345 | chr9-78469379-INS-345 |
| chr9-78483463-INS-194 | chr9-78483463-INS-194 |
| chr9-78497264-INS-483 | chr9-78497264-INS-483 |
| chr9-78508428-INS-207 | chr9-78508428-INS-182 |
| chr9-78555796-INS-222 | chr9-78555796-INS-222 |
| chr9-78570285-INS-88 | chr9-78570285-INS-88 |
| chr9-78573406-INS-155 | chr9-78573406-INS-155 |
| chr9-78604269-INS-184 | chr9-78604269-INS-201 |
| chr9-78641554-INS-599 | chr9-78641554-INS-601 |
| chr9-78646275-INS-219 | chr9-78646278-INS-177 |
| chr9-78649275-INS-52 | chr9-78649275-INS-52 |
| chr9-78681334-INS-84 | chr9-78681334-INS-52 |
| chr9-78699372-INS-401 | chr9-78699372-INS-401 |
| chr9-78748463-INS-63 | chr9-78748463-INS-74 |
| chr9-78751548-INS-124 | chr9-78751594-INS-124 |
| chr9-78768807-INS-62 | chr9-78768807-INS-61 |
| chr9-78769337-INS-2828 | chr9-78769338-INS-2821 |
| chr9-78777650-INS-66 | chr9-78777651-INS-66 |
| chr9-78809067-INS-78 | chr9-78809067-INS-78 |
| chr9-78827213-INS-370 | chr9-78827213-INS-370 |
| chr9-78845348-INS-363 | chr9-78845348-INS-363 |
| chr9-78852841-INS-1310 | chr9-78852851-INS-1326 |
| chr9-78869491-INS-54 | chr9-78869491-INS-56 |
| chr9-78882940-INS-59 | chr9-78882940-INS-59 |
| chr9-78888867-INS-54 | chr9-78888837-INS-82 |
| chr9-78890541-INS-1855 | chr9-78890542-INS-1859 |
| chr9-78893537-INS-2808 | chr9-78893538-INS-2808 |
| chr9-78902926-INS-92 | chr9-78902927-INS-140 |
| chr9-78907253-INS-178 | chr9-78907253-INS-178 |
| chr9-78945936-INS-1057 | chr9-78945937-INS-1054 |
| chr9-78977787-INS-54 | chr9-78977787-INS-54 |
| chr9-78987153-INS-87 | chr9-78987131-INS-91 |
| chr9-78992406-INS-348 | chr9-78992406-INS-348 |
| chr9-79007091-INS-566 | chr9-79007091-INS-567 |
| chr9-79010216-INS-200 | chr9-79010216-INS-200 |
| chr9-79012557-INS-176 | chr9-79012557-INS-203 |
| chr9-79017532-INS-68 | chr9-79017537-INS-50 |
| chr9-79029482-INS-597 | chr9-79029482-INS-597 |
| chr9-79054355-INS-3219 | chr9-79054356-INS-3219 |
| chr9-79074279-INS-51 | chr9-79074279-INS-51 |
| chr9-79091085-INS-2389 | chr9-79091086-INS-2389 |
| chr9-79105295-INS-1451 | chr9-79105296-INS-1451 |
| chr9-79112555-INS-162 | chr9-79112547-INS-184 |
| chr9-79119049-INS-73 | chr9-79119049-INS-73 |
| chr9-79136071-INS-60 | chr9-79136071-INS-60 |
| chr9-79141770-INS-112 | chr9-79141770-INS-88 |
| chr9-79151388-INS-999 | chr9-79151388-INS-999 |
| chr9-79192945-INS-429 | chr9-79192945-INS-429 |

| | |
|---|---|
| chr9-79195801-INS-277 | chr9-79195801-INS-277 |
| chr9-79201733-INS-296 | chr9-79201736-INS-297 |
| chr9-79203777-INS-672 | chr9-79203777-INS-672 |
| chr9-79220711-INS-168 | chr9-79220711-INS-168 |
| chr9-79229235-INS-6299 | chr9-79229236-INS-6299 |
| chr9-79240601-INS-57 | chr9-79240601-INS-57 |
| chr9-79258284-INS-6292 | chr9-79258285-INS-6292 |
| chr9-79295118-INS-6593 | chr9-79295119-INS-6593 |
| chr9-79321547-INS-90 | chr9-79321547-INS-106 |
| chr9-79338888-INS-4889 | chr9-79338889-INS-4889 |
| chr9-79361284-INS-64 | chr9-79361284-INS-68 |
| chr9-79362906-INS-116 | chr9-79362906-INS-142 |
| chr9-79365050-INS-56 | chr9-79365050-INS-56 |
| chr9-79401142-INS-62 | chr9-79401142-INS-62 |
| chr9-79403855-INS-76 | chr9-79403855-INS-68 |
| chr9-79426659-INS-135 | chr9-79426665-INS-86 |
| chr9-79431797-INS-119 | chr9-79431843-INS-72 |
| chr9-79472398-INS-134 | chr9-79472398-INS-153 |
| chr9-79473896-INS-141 | chr9-79473896-INS-141 |
| chr9-79483788-INS-2073 | chr9-79483789-INS-2074 |
| chr9-79486674-INS-702 | chr9-79486674-INS-701 |
| chr9-79489190-INS-58 | chr9-79489190-INS-58 |
| chr9-79500306-INS-1428 | chr9-79500307-INS-1428 |
| chr9-79534496-INS-52 | chr9-79534496-INS-52 |
| chr9-79602486-INS-83 | chr9-79602473-INS-86 |
| chr9-79636235-INS-94 | chr9-79636233-INS-102 |
| chr9-79653135-INS-7848 | chr9-79653136-INS-7848 |
| chr9-79655834-INS-67 | chr9-79655834-INS-67 |
| chr9-79658330-INS-199 | chr9-79658330-INS-199 |
| chr9-79679945-INS-70 | chr9-79679946-INS-100 |
| chr9-79687991-INS-106 | chr9-79687991-INS-106 |
| chr9-79707929-INS-469 | chr9-79707929-INS-469 |
| chr9-79712207-INS-116 | chr9-79712210-INS-120 |
| chr9-79718650-INS-148 | chr9-79718648-INS-154 |
| chr9-79720647-INS-60 | chr9-79720647-INS-83 |
| chr9-79742965-INS-80 | chr9-79742965-INS-58 |
| chr9-79759031-INS-84 | chr9-79759020-INS-50 |
| chr9-79764582-INS-202 | chr9-79764582-INS-202 |
| chr9-79848145-INS-542 | chr9-79848145-INS-542 |
| chr9-79858204-INS-125 | chr9-79858204-INS-125 |
| chr9-79874015-INS-944 | chr9-79874015-INS-946 |
| chr9-79886352-INS-55 | chr9-79886352-INS-55 |
| chr9-79889531-INS-870 | chr9-79889531-INS-870 |
| chr9-79897453-INS-200 | chr9-79897453-INS-199 |
| chr9-79905926-INS-179 | chr9-79905926-INS-179 |
| chr9-79936146-INS-186 | chr9-79936132-INS-180 |
| chr9-79937165-INS-634 | chr9-79937165-INS-634 |
| chr9-79945820-INS-102 | chr9-79945820-INS-124 |
| chr9-79950131-INS-52 | chr9-79950131-INS-52 |
| chr9-79958102-INS-2144 | chr9-79958103-INS-2144 |
| chr9-79964106-INS-122 | chr9-79964094-INS-102 |
| chr9-79983106-INS-488 | chr9-79983106-INS-488 |
| chr9-79994591-INS-58 | chr9-79994591-INS-58 |
| chr9-80016316-INS-6440 | chr9-80016317-INS-6440 |
| chr9-80026646-INS-92 | chr9-80026646-INS-92 |
| chr9-80067678-INS-502 | chr9-80067678-INS-502 |
| chr9-80095180-INS-273 | chr9-80095180-INS-274 |
| chr9-80111569-INS-52 | chr9-80111569-INS-52 |
| chr9-80117323-INS-82 | chr9-80117323-INS-84 |
| chr9-80132214-INS-247 | chr9-80132214-INS-237 |
| chr9-80135165-INS-823 | chr9-80135165-INS-823 |
| chr9-80156769-INS-198 | chr9-80156769-INS-199 |
| chr9-80170043-INS-623 | chr9-80170043-INS-623 |
| chr9-80171445-INS-84 | chr9-80171445-INS-84 |
| chr9-80206029-INS-595 | chr9-80206029-INS-595 |
| chr9-80225144-INS-577 | chr9-80225144-INS-583 |
| chr9-80232594-INS-66 | chr9-80232594-INS-66 |
| chr9-80271724-INS-7060 | chr9-80271725-INS-7060 |
| chr9-80276943-INS-74 | chr9-80276943-INS-74 |
| chr9-80279174-INS-768 | chr9-80279175-INS-769 |
| chr9-80289299-INS-401 | chr9-80289299-INS-401 |
| chr9-80295542-INS-155 | chr9-80295542-INS-155 |
| chr9-80314502-INS-60 | chr9-80314502-INS-60 |
| chr9-80315728-INS-148 | chr9-80315728-INS-148 |
| chr9-80324164-INS-439 | chr9-80324164-INS-439 |
| chr9-80343998-INS-92 | chr9-80343991-INS-84 |
| chr9-80349467-INS-6731 | chr9-80349468-INS-6732 |

| | |
|---|---|
| chr9-80352869-INS-53 | chr9-80352875-INS-59 |
| chr9-80354864-INS-62 | chr9-80354844-INS-64 |
| chr9-80372384-INS-66 | chr9-80372381-INS-95 |
| chr9-80389261-INS-153 | chr9-80389261-INS-153 |
| chr9-80432799-INS-78 | chr9-80432799-INS-83 |
| chr9-80433841-INS-424 | chr9-80433841-INS-424 |
| chr9-80444734-INS-59 | chr9-80444734-INS-59 |
| chr9-80465639-INS-56 | chr9-80465639-INS-54 |
| chr9-80489507-INS-102 | chr9-80489507-INS-102 |
| chr9-80490753-INS-111 | chr9-80490753-INS-111 |
| chr9-80537453-INS-56 | chr9-80537453-INS-58 |
| chr9-80572956-INS-78 | chr9-80572950-INS-92 |
| chr9-80574979-INS-670 | chr9-80574979-INS-670 |
| chr9-80586640-INS-804 | chr9-80586634-INS-818 |
| chr9-80633158-INS-74 | chr9-80633158-INS-74 |
| chr9-80647874-INS-6530 | chr9-80647875-INS-6530 |
| chr9-80649277-INS-52 | chr9-80649277-INS-68 |
| chr9-80650305-INS-64 | chr9-80650305-INS-70 |
| chr9-80662099-INS-5951 | chr9-80662100-INS-5951 |
| chr9-80662780-INS-52 | chr9-80662780-INS-52 |
| chr9-80666775-INS-57 | chr9-80666775-INS-61 |
| chr9-80696197-INS-1993 | chr9-80696198-INS-1993 |
| chr9-80720996-INS-573 | chr9-80720996-INS-573 |
| chr9-80725117-INS-64 | chr9-80725117-INS-68 |
| chr9-80731510-INS-52 | chr9-80731524-INS-68 |
| chr9-80770023-INS-103 | chr9-80770028-INS-59 |
| chr9-80799184-INS-66 | chr9-80799184-INS-84 |
| chr9-80818689-INS-522 | chr9-80818689-INS-522 |
| chr9-80819339-INS-572 | chr9-80819339-INS-572 |
| chr9-80844450-INS-2858 | chr9-80844451-INS-2858 |
| chr9-80859801-INS-75 | chr9-80859801-INS-70 |
| chr9-80870196-INS-65 | chr9-80870190-INS-90 |
| chr9-80891683-INS-559 | chr9-80891683-INS-560 |
| chr9-80893920-INS-179 | chr9-80893920-INS-179 |
| chr9-80895697-INS-604 | chr9-80895693-INS-604 |
| chr9-80908456-INS-484 | chr9-80908456-INS-488 |
| chr9-80910756-INS-376 | chr9-80910756-INS-376 |
| chr9-80922414-INS-53 | chr9-80922414-INS-53 |
| chr9-80941881-INS-266 | chr9-80941881-INS-266 |
| chr9-80968669-INS-401 | chr9-80968669-INS-401 |
| chr9-80994369-INS-481 | chr9-80994369-INS-481 |
| chr9-80999068-INS-52 | chr9-80999068-INS-50 |
| chr9-81017120-INS-88 | chr9-81017120-INS-92 |
| chr9-81057193-INS-6248 | chr9-81057194-INS-6248 |
| chr9-81060346-INS-52 | chr9-81060346-INS-54 |
| chr9-81084456-INS-104 | chr9-81084456-INS-152 |
| chr9-81109601-INS-174 | chr9-81109601-INS-174 |
| chr9-81122270-INS-56 | chr9-81122270-INS-56 |
| chr9-81207805-INS-67 | chr9-81207805-INS-68 |
| chr9-81227418-INS-7016 | chr9-81227419-INS-7016 |
| chr9-81243426-INS-181 | chr9-81243426-INS-182 |
| chr9-81248776-INS-119 | chr9-81248820-INS-91 |
| chr9-81253868-INS-83 | chr9-81253868-INS-86 |
| chr9-81261224-INS-1113 | chr9-81261225-INS-1113 |
| chr9-81282001-INS-66 | chr9-81282001-INS-68 |
| chr9-81290591-INS-6462 | chr9-81290592-INS-6462 |
| chr9-81298914-INS-388 | chr9-81298914-INS-386 |
| chr9-81314982-INS-316 | chr9-81314983-INS-316 |
| chr9-81322351-INS-63 | chr9-81322341-INS-54 |
| chr9-81389750-INS-114 | chr9-81389747-INS-116 |
| chr9-81391869-INS-6380 | chr9-81391870-INS-6378 |
| chr9-81400329-INS-72 | chr9-81400329-INS-78 |
| chr9-81409286-INS-60 | chr9-81409286-INS-60 |
| chr9-81440589-INS-52 | chr9-81440589-INS-52 |
| chr9-81441633-INS-60 | chr9-81441633-INS-88 |
| chr9-81450528-INS-67 | chr9-81450526-INS-115 |
| chr9-81472981-INS-64 | chr9-81472981-INS-80 |
| chr9-81474698-INS-497 | chr9-81474698-INS-497 |
| chr9-81492136-INS-861 | chr9-81492136-INS-867 |
| chr9-81508734-INS-72 | chr9-81508734-INS-72 |
| chr9-81569778-INS-790 | chr9-81569778-INS-793 |
| chr9-81595416-INS-86 | chr9-81595419-INS-81 |
| chr9-81596144-INS-1893 | chr9-81596145-INS-1893 |
| chr9-81668093-INS-55 | chr9-81668093-INS-50 |
| chr9-81680752-INS-65 | chr9-81680752-INS-65 |
| chr9-81703107-INS-5239 | chr9-81703108-INS-5240 |
| chr9-81703360-INS-922 | chr9-81703360-INS-922 |

| | |
|---|---|
| chr9-81720753-INS-76 | chr9-81720755-INS-82 |
| chr9-81723638-INS-1204 | chr9-81723639-INS-1204 |
| chr9-81726736-INS-3882 | chr9-81726737-INS-3882 |
| chr9-81727956-INS-147 | chr9-81727956-INS-147 |
| chr9-81733585-INS-512 | chr9-81733585-INS-512 |
| chr9-81748066-INS-66 | chr9-81748066-INS-66 |
| chr9-81764152-INS-553 | chr9-81764153-INS-555 |
| chr9-81770148-INS-4515 | chr9-81770149-INS-4515 |
| chr9-81773662-INS-144 | chr9-81773663-INS-181 |
| chr9-81775083-INS-384 | chr9-81775086-INS-388 |
| chr9-81778727-INS-498 | chr9-81778727-INS-498 |
| chr9-81831321-INS-138 | chr9-81831321-INS-138 |
| chr9-81847639-INS-61 | chr9-81847639-INS-63 |
| chr9-81847809-INS-503 | chr9-81847809-INS-503 |
| chr9-81860840-INS-197 | chr9-81860840-INS-201 |
| chr9-81872782-INS-52 | chr9-81872782-INS-52 |
| chr9-81885810-INS-6367 | chr9-81885811-INS-6367 |
| chr9-81895465-INS-56 | chr9-81895469-INS-72 |
| chr9-81896394-INS-165 | chr9-81896394-INS-166 |
| chr9-81925400-INS-257 | chr9-81925400-INS-237 |
| chr9-81929597-INS-182 | chr9-81929598-INS-208 |
| chr9-81935092-INS-54 | chr9-81935069-INS-75 |
| chr9-81936475-INS-83 | chr9-81936475-INS-84 |
| chr9-81976285-INS-98 | chr9-81976286-INS-99 |
| chr9-81999289-INS-100 | chr9-81999254-INS-150 |
| chr9-82043051-INS-81 | chr9-82043072-INS-60 |
| chr9-82051203-INS-6659 | chr9-82051204-INS-6660 |
| chr9-82068455-INS-195 | chr9-82068455-INS-195 |
| chr9-82072850-INS-944 | chr9-82072850-INS-944 |
| chr9-82077244-INS-52 | chr9-82077244-INS-52 |
| chr9-82103415-INS-189 | chr9-82103418-INS-189 |
| chr9-82121316-INS-77 | chr9-82121316-INS-123 |
| chr9-82124150-INS-126 | chr9-82124192-INS-132 |
| chr9-82138846-INS-71 | chr9-82138846-INS-69 |
| chr9-82156599-INS-221 | chr9-82156599-INS-221 |
| chr9-82183365-INS-58 | chr9-82183365-INS-60 |
| chr9-82187286-INS-124 | chr9-82187286-INS-126 |
| chr9-82218015-INS-274 | chr9-82218015-INS-273 |
| chr9-82257506-INS-144 | chr9-82257506-INS-144 |
| chr9-82259295-INS-78 | chr9-82259295-INS-56 |
| chr9-82264611-INS-183 | chr9-82264611-INS-183 |
| chr9-82283959-INS-449 | chr9-82283959-INS-449 |
| chr9-82293728-INS-978 | chr9-82293728-INS-975 |
| chr9-82318016-INS-112 | chr9-82318016-INS-112 |
| chr9-82318978-INS-52 | chr9-82318963-INS-56 |
| chr9-82321254-INS-80 | chr9-82321254-INS-72 |
| chr9-82322333-INS-89 | chr9-82322344-INS-87 |
| chr9-82336265-INS-231 | chr9-82336265-INS-231 |
| chr9-82386122-INS-3205 | chr9-82386123-INS-3183 |
| chr9-82399357-INS-522 | chr9-82399357-INS-522 |
| chr9-82423476-INS-59 | chr9-82423436-INS-79 |
| chr9-82460448-INS-809 | chr9-82460448-INS-798 |
| chr9-82506368-INS-60 | chr9-82506368-INS-58 |
| chr9-82518564-INS-76 | chr9-82518564-INS-72 |
| chr9-82523392-INS-52 | chr9-82523344-INS-60 |
| chr9-82545414-INS-327 | chr9-82545414-INS-327 |
| chr9-82563265-INS-242 | chr9-82563265-INS-238 |
| chr9-82605982-INS-66 | chr9-82605990-INS-78 |
| chr9-82616410-INS-87 | chr9-82616406-INS-54 |
| chr9-82626462-INS-71 | chr9-82626462-INS-72 |
| chr9-82641579-INS-119 | chr9-82641587-INS-137 |
| chr9-82642712-INS-802 | chr9-82642712-INS-802 |
| chr9-82689714-INS-407 | chr9-82689714-INS-407 |
| chr9-82715645-INS-248 | chr9-82715645-INS-248 |
| chr9-82723949-INS-1023 | chr9-82723950-INS-1023 |
| chr9-82724850-INS-197 | chr9-82724851-INS-197 |
| chr9-82735398-INS-148 | chr9-82735398-INS-148 |
| chr9-82756921-INS-68 | chr9-82756921-INS-68 |
| chr9-82861381-INS-151 | chr9-82861381-INS-151 |
| chr9-82884968-INS-203 | chr9-82884968-INS-203 |
| chr9-82910226-INS-183 | chr9-82910226-INS-183 |
| chr9-82920824-INS-78 | chr9-82920830-INS-57 |
| chr9-82926569-INS-379 | chr9-82926569-INS-379 |
| chr9-82966435-INS-93 | chr9-82966435-INS-93 |
| chr9-82972771-INS-273 | chr9-82972735-INS-285 |
| chr9-83005981-INS-198 | chr9-83005981-INS-199 |
| chr9-83015881-INS-69 | chr9-83015881-INS-69 |

| | |
|---|---|
| chr9-83024991-INS-6487 | chr9-83024992-INS-6487 |
| chr9-83067439-INS-54 | chr9-83067439-INS-51 |
| chr9-83070569-INS-391 | chr9-83070569-INS-391 |
| chr9-83087511-INS-64 | chr9-83087511-INS-64 |
| chr9-83096657-INS-6420 | chr9-83096658-INS-6445 |
| chr9-83098378-INS-228 | chr9-83098370-INS-232 |
| chr9-83128372-INS-79 | chr9-83128401-INS-66 |
| chr9-83130918-INS-602 | chr9-83130918-INS-602 |
| chr9-83196970-INS-67 | chr9-83196970-INS-51 |
| chr9-83210348-INS-90 | chr9-83210353-INS-92 |
| chr9-83216485-INS-88 | chr9-83216485-INS-84 |
| chr9-83234936-INS-51 | chr9-83234936-INS-63 |
| chr9-83249050-INS-7207 | chr9-83249051-INS-7207 |
| chr9-83259047-INS-204 | chr9-83259047-INS-203 |
| chr9-83294809-INS-320 | chr9-83294809-INS-324 |
| chr9-83303654-INS-52 | chr9-83303654-INS-69 |
| chr9-83305948-INS-182 | chr9-83305948-INS-182 |
| chr9-83330782-INS-69 | chr9-83330787-INS-57 |
| chr9-83336411-INS-212 | chr9-83336411-INS-213 |
| chr9-83338347-INS-894 | chr9-83338347-INS-894 |
| chr9-83398436-INS-80 | chr9-83398406-INS-118 |
| chr9-83400357-INS-2747 | chr9-83400358-INS-2747 |
| chr9-83418393-INS-195 | chr9-83418393-INS-196 |
| chr9-83436520-INS-99 | chr9-83436520-INS-99 |
| chr9-83441265-INS-54 | chr9-83441265-INS-54 |
| chr9-83452841-INS-70 | chr9-83452841-INS-70 |
| chr9-83472369-INS-914 | chr9-83472370-INS-913 |
| chr9-83489032-INS-62 | chr9-83489032-INS-64 |
| chr9-83492044-INS-96 | chr9-83492042-INS-100 |
| chr9-83503467-INS-58 | chr9-83503467-INS-52 |
| chr9-83525260-INS-57 | chr9-83525260-INS-57 |
| chr9-83531264-INS-66 | chr9-83531264-INS-66 |
| chr9-83532210-INS-401 | chr9-83532210-INS-401 |
| chr9-83556668-INS-211 | chr9-83556668-INS-212 |
| chr9-83566054-INS-108 | chr9-83566054-INS-108 |
| chr9-83580076-INS-449 | chr9-83580076-INS-447 |
| chr9-83584325-INS-393 | chr9-83584325-INS-386 |
| chr9-83586339-INS-70 | chr9-83586310-INS-65 |
| chr9-83604363-INS-61 | chr9-83604364-INS-65 |
| chr9-83605659-INS-7146 | chr9-83605660-INS-7146 |
| chr9-83618073-INS-60 | chr9-83618073-INS-60 |
| chr9-83639967-INS-72 | chr9-83639967-INS-74 |
| chr9-83643098-INS-110 | chr9-83643098-INS-110 |
| chr9-83645660-INS-3836 | chr9-83645661-INS-3836 |
| chr9-83649064-INS-400 | chr9-83649064-INS-401 |
| chr9-83650797-INS-233 | chr9-83650797-INS-233 |
| chr9-83654160-INS-111 | chr9-83654160-INS-111 |
| chr9-83666059-INS-63 | chr9-83666059-INS-55 |
| chr9-83680138-INS-964 | chr9-83680138-INS-979 |
| chr9-83729803-INS-203 | chr9-83729803-INS-203 |
| chr9-83737583-INS-257 | chr9-83737583-INS-263 |
| chr9-83740172-INS-714 | chr9-83740172-INS-714 |
| chr9-83746615-INS-1512 | chr9-83746611-INS-1512 |
| chr9-83756212-INS-182 | chr9-83756212-INS-182 |
| chr9-83762166-INS-119 | chr9-83762166-INS-121 |
| chr9-83775744-INS-61 | chr9-83775744-INS-53 |
| chr9-83813994-INS-235 | chr9-83813994-INS-230 |
| chr9-83852600-INS-133 | chr9-83852600-INS-133 |
| chr9-83855685-INS-85 | chr9-83855661-INS-85 |
| chr9-83859408-INS-167 | chr9-83859408-INS-169 |
| chr9-83887043-INS-176 | chr9-83887036-INS-185 |
| chr9-83897661-INS-54 | chr9-83897659-INS-50 |
| chr9-83907083-INS-196 | chr9-83907083-INS-196 |
| chr9-83931884-INS-396 | chr9-83931884-INS-403 |
| chr9-83940638-INS-204 | chr9-83940621-INS-204 |
| chr9-83942567-INS-55 | chr9-83942567-INS-56 |
| chr9-83964248-INS-264 | chr9-83964248-INS-264 |
| chr9-83966741-INS-200 | chr9-83966741-INS-200 |
| chr9-83984360-INS-84 | chr9-83984360-INS-84 |
| chr9-83988717-INS-215 | chr9-83988717-INS-214 |
| chr9-83999986-INS-639 | chr9-83999986-INS-639 |
| chr9-84009035-INS-86 | chr9-84009035-INS-86 |
| chr9-84010980-INS-427 | chr9-84010997-INS-448 |
| chr9-84015715-INS-182 | chr9-84015715-INS-185 |
| chr9-84051692-INS-6163 | chr9-84051693-INS-6165 |
| chr9-84085928-INS-64 | chr9-84085929-INS-92 |
| chr9-84088671-INS-201 | chr9-84088671-INS-201 |

| | |
|---|---|
| chr9-84096576-INS-66 | chr9-84096576-INS-66 |
| chr9-84114171-INS-194 | chr9-84114171-INS-194 |
| chr9-84128968-INS-90 | chr9-84128968-INS-90 |
| chr9-84136409-INS-54 | chr9-84136409-INS-54 |
| chr9-84143454-INS-65 | chr9-84143461-INS-50 |
| chr9-84157452-INS-99 | chr9-84157452-INS-72 |
| chr9-84160718-INS-66 | chr9-84160718-INS-75 |
| chr9-84176648-INS-187 | chr9-84176647-INS-187 |
| chr9-84196012-INS-54 | chr9-84196019-INS-50 |
| chr9-84200232-INS-524 | chr9-84200232-INS-523 |
| chr9-84237921-INS-4940 | chr9-84237922-INS-4941 |
| chr9-84253644-INS-6425 | chr9-84253645-INS-6425 |
| chr9-84265217-INS-58 | chr9-84265217-INS-58 |
| chr9-84287635-INS-3685 | chr9-84287636-INS-3685 |
| chr9-84337793-INS-359 | chr9-84337793-INS-359 |
| chr9-84357779-INS-51 | chr9-84357765-INS-52 |
| chr9-84377282-INS-201 | chr9-84377282-INS-201 |
| chr9-84399460-INS-58 | chr9-84399460-INS-58 |
| chr9-84401939-INS-1108 | chr9-84401940-INS-1108 |
| chr9-84404149-INS-1574 | chr9-84404150-INS-1574 |
| chr9-84421828-INS-54 | chr9-84421837-INS-87 |
| chr9-84426372-INS-10829 | chr9-84426373-INS-10828 |
| chr9-84436741-INS-1402 | chr9-84436724-INS-1391 |
| chr9-84480407-INS-52 | chr9-84480407-INS-54 |
| chr9-84494271-INS-1592 | chr9-84494272-INS-1592 |
| chr9-84511612-INS-62 | chr9-84511612-INS-62 |
| chr9-84523064-INS-54 | chr9-84523064-INS-54 |
| chr9-84528227-INS-95 | chr9-84528223-INS-86 |
| chr9-84583270-INS-4585 | chr9-84583271-INS-4593 |
| chr9-84585279-INS-283 | chr9-84585279-INS-298 |
| chr9-84608237-INS-5061 | chr9-84608238-INS-5065 |
| chr9-84611460-INS-1891 | chr9-84611461-INS-1891 |
| chr9-84618710-INS-54 | chr9-84618710-INS-54 |
| chr9-84656439-INS-5574 | chr9-84656440-INS-5575 |
| chr9-84659875-INS-51 | chr9-84659870-INS-58 |
| chr9-84697756-INS-1676 | chr9-84697757-INS-1676 |
| chr9-84708323-INS-106 | chr9-84708331-INS-84 |
| chr9-84729286-INS-6440 | chr9-84729287-INS-6440 |
| chr9-84755624-INS-416 | chr9-84755620-INS-418 |
| chr9-84789193-INS-82 | chr9-84789193-INS-82 |
| chr9-84791042-INS-7244 | chr9-84791043-INS-7244 |
| chr9-84809775-INS-572 | chr9-84809775-INS-573 |
| chr9-84811046-INS-219 | chr9-84811046-INS-219 |
| chr9-84826188-INS-441 | chr9-84826188-INS-441 |
| chr9-84838350-INS-54 | chr9-84838350-INS-56 |
| chr9-84899224-INS-617 | chr9-84899224-INS-633 |
| chr9-84918680-INS-1878 | chr9-84918681-INS-1878 |
| chr9-84929931-INS-54 | chr9-84929956-INS-50 |
| chr9-84941523-INS-1703 | chr9-84941524-INS-1703 |
| chr9-84975066-INS-51 | chr9-84975066-INS-83 |
| chr9-84978847-INS-74 | chr9-84978847-INS-74 |
| chr9-85024949-INS-85 | chr9-85024949-INS-105 |
| chr9-85027093-INS-76 | chr9-85027082-INS-54 |
| chr9-85035794-INS-76 | chr9-85035801-INS-60 |
| chr9-85082578-INS-295 | chr9-85082578-INS-295 |
| chr9-85085358-INS-100 | chr9-85085358-INS-100 |
| chr9-85135469-INS-86 | chr9-85135469-INS-86 |
| chr9-85144333-INS-96 | chr9-85144333-INS-140 |
| chr9-85212769-INS-162 | chr9-85212763-INS-198 |
| chr9-85221985-INS-69 | chr9-85221985-INS-72 |
| chr9-85248965-INS-60 | chr9-85248965-INS-62 |
| chr9-85249646-INS-84 | chr9-85249646-INS-84 |
| chr9-85263694-INS-72 | chr9-85263688-INS-52 |
| chr9-85305341-INS-60 | chr9-85305348-INS-51 |
| chr9-85332582-INS-91 | chr9-85332582-INS-91 |
| chr9-85361109-INS-56 | chr9-85361109-INS-50 |
| chr9-85404409-INS-879 | chr9-85404409-INS-886 |
| chr9-85408797-INS-58 | chr9-85408797-INS-58 |
| chr9-85414184-INS-169 | chr9-85414184-INS-169 |
| chr9-85414869-INS-62 | chr9-85414873-INS-74 |
| chr9-85415973-INS-6242 | chr9-85415974-INS-6242 |
| chr9-85425695-INS-545 | chr9-85425693-INS-572 |
| chr9-85435310-INS-54 | chr9-85435310-INS-54 |
| chr9-85446624-INS-2106 | chr9-85446625-INS-2110 |
| chr9-85453616-INS-164 | chr9-85453616-INS-164 |
| chr9-85473598-INS-76 | chr9-85473598-INS-102 |
| chr9-85520284-INS-52 | chr9-85520279-INS-64 |

| | |
|---|---|
| chr9-85539569-INS-100 | chr9-85539569-INS-105 |
| chr9-85540866-INS-402 | chr9-85540866-INS-402 |
| chr9-85550121-INS-85 | chr9-85550144-INS-56 |
| chr9-85589193-INS-671 | chr9-85589193-INS-673 |
| chr9-85589913-INS-6455 | chr9-85589914-INS-6456 |
| chr9-85601272-INS-261 | chr9-85601272-INS-261 |
| chr9-85612527-INS-209 | chr9-85612527-INS-209 |
| chr9-85626653-INS-4864 | chr9-85626654-INS-4864 |
| chr9-85639288-INS-364 | chr9-85639288-INS-364 |
| chr9-85644280-INS-287 | chr9-85644280-INS-287 |
| chr9-85657677-INS-185 | chr9-85657677-INS-185 |
| chr9-85666495-INS-7065 | chr9-85666496-INS-7065 |
| chr9-85679084-INS-282 | chr9-85679084-INS-282 |
| chr9-85700604-INS-444 | chr9-85700604-INS-444 |
| chr9-85706914-INS-51 | chr9-85706914-INS-51 |
| chr9-85713243-INS-6765 | chr9-85713244-INS-6765 |
| chr9-85713575-INS-270 | chr9-85713575-INS-269 |
| chr9-85722833-INS-67 | chr9-85722835-INS-52 |
| chr9-85740511-INS-66 | chr9-85740538-INS-85 |
| chr9-85744233-INS-5652 | chr9-85744234-INS-5652 |
| chr9-85745211-INS-51 | chr9-85745255-INS-50 |
| chr9-85752254-INS-568 | chr9-85752254-INS-568 |
| chr9-85759275-INS-6859 | chr9-85759276-INS-6859 |
| chr9-85759425-INS-149 | chr9-85759425-INS-149 |
| chr9-85768050-INS-90 | chr9-85768050-INS-138 |
| chr9-85805201-INS-94 | chr9-85805203-INS-53 |
| chr9-85819079-INS-567 | chr9-85819079-INS-567 |
| chr9-85819961-INS-901 | chr9-85819961-INS-902 |
| chr9-85839951-INS-233 | chr9-85839951-INS-235 |
| chr9-85881477-INS-470 | chr9-85881477-INS-470 |
| chr9-85888629-INS-477 | chr9-85888629-INS-477 |
| chr9-85908037-INS-6306 | chr9-85908038-INS-6307 |
| chr9-85912470-INS-6499 | chr9-85912471-INS-6500 |
| chr9-85914091-INS-254 | chr9-85914091-INS-254 |
| chr9-85917142-INS-1894 | chr9-85917143-INS-1894 |
| chr9-85922011-INS-582 | chr9-85922011-INS-584 |
| chr9-85923740-INS-70 | chr9-85923740-INS-73 |
| chr9-85925778-INS-58 | chr9-85925778-INS-58 |
| chr9-85934999-INS-52 | chr9-85934999-INS-52 |
| chr9-85953388-INS-771 | chr9-85953388-INS-771 |
| chr9-85969047-INS-3224 | chr9-85969048-INS-3224 |
| chr9-85970309-INS-1385 | chr9-85970310-INS-1386 |
| chr9-85979489-INS-1667 | chr9-85979490-INS-1667 |
| chr9-86012161-INS-315 | chr9-86012161-INS-316 |
| chr9-86030783-INS-1602 | chr9-86030784-INS-1602 |
| chr9-86057646-INS-104 | chr9-86057646-INS-104 |
| chr9-86061316-INS-698 | chr9-86061316-INS-703 |
| chr9-86068340-INS-55 | chr9-86068345-INS-60 |
| chr9-86081302-INS-212 | chr9-86081302-INS-212 |
| chr9-86110921-INS-5326 | chr9-86110922-INS-5326 |
| chr9-86121925-INS-60 | chr9-86121925-INS-62 |
| chr9-86125436-INS-219 | chr9-86125436-INS-221 |
| chr9-86126737-INS-247 | chr9-86126737-INS-247 |
| chr9-86130806-INS-4966 | chr9-86130807-INS-4966 |
| chr9-86139036-INS-56 | chr9-86139036-INS-52 |
| chr9-86143238-INS-1985 | chr9-86143239-INS-1987 |
| chr9-86151348-INS-170 | chr9-86151348-INS-170 |
| chr9-86207984-INS-1522 | chr9-86207985-INS-1523 |
| chr9-86222628-INS-84 | chr9-86222628-INS-84 |
| chr9-86245239-INS-507 | chr9-86245239-INS-508 |
| chr9-86254837-INS-133 | chr9-86254837-INS-135 |
| chr9-86258536-INS-1358 | chr9-86258537-INS-1358 |
| chr9-86269757-INS-60 | chr9-86269757-INS-60 |
| chr9-86274858-INS-53 | chr9-86274859-INS-51 |
| chr9-86281509-INS-59 | chr9-86281509-INS-55 |
| chr9-86285481-INS-7249 | chr9-86285482-INS-7249 |
| chr9-86297154-INS-76 | chr9-86297154-INS-80 |
| chr9-86323229-INS-98 | chr9-86323229-INS-98 |
| chr9-86325335-INS-207 | chr9-86325359-INS-211 |
| chr9-86353624-INS-86 | chr9-86353624-INS-86 |
| chr9-86411941-INS-241 | chr9-86411941-INS-242 |
| chr9-86425291-INS-196 | chr9-86425291-INS-196 |
| chr9-86426240-INS-88 | chr9-86426240-INS-64 |
| chr9-86476658-INS-368 | chr9-86476658-INS-368 |
| chr9-86488633-INS-63 | chr9-86488633-INS-63 |
| chr9-86490417-INS-636 | chr9-86490417-INS-636 |
| chr9-86493423-INS-60 | chr9-86493428-INS-54 |

| | |
|---|---|
| chr9-86516430-INS-62 | chr9-86516430-INS-65 |
| chr9-86528977-INS-210 | chr9-86528977-INS-212 |
| chr9-86531496-INS-186 | chr9-86531496-INS-173 |
| chr9-86532601-INS-217 | chr9-86532601-INS-218 |
| chr9-86539701-INS-65 | chr9-86539706-INS-57 |
| chr9-86567683-INS-5173 | chr9-86567684-INS-5173 |
| chr9-86575409-INS-51 | chr9-86575408-INS-52 |
| chr9-86594468-INS-7122 | chr9-86594469-INS-7122 |
| chr9-86599935-INS-838 | chr9-86599935-INS-839 |
| chr9-86624458-INS-95 | chr9-86624454-INS-99 |
| chr9-86625143-INS-2954 | chr9-86625144-INS-2954 |
| chr9-86645576-INS-108 | chr9-86645576-INS-110 |
| chr9-86714397-INS-111 | chr9-86714397-INS-111 |
| chr9-86732458-INS-353 | chr9-86732458-INS-353 |
| chr9-86799448-INS-1553 | chr9-86799449-INS-1547 |
| chr9-86811325-INS-224 | chr9-86811325-INS-224 |
| chr9-86820954-INS-412 | chr9-86820954-INS-412 |
| chr9-86834563-INS-206 | chr9-86834551-INS-204 |
| chr9-86849230-INS-107 | chr9-86849230-INS-65 |
| chr9-86855009-INS-766 | chr9-86855009-INS-766 |
| chr9-86856288-INS-389 | chr9-86856286-INS-393 |
| chr9-86877399-INS-1491 | chr9-86877400-INS-1490 |
| chr9-86899456-INS-57 | chr9-86899456-INS-57 |
| chr9-86908643-INS-71 | chr9-86908643-INS-71 |
| chr9-86916050-INS-60 | chr9-86916050-INS-57 |
| chr9-86923585-INS-6705 | chr9-86923586-INS-6706 |
| chr9-86934053-INS-60 | chr9-86934053-INS-62 |
| chr9-86965691-INS-58 | chr9-86965691-INS-58 |
| chr9-86971644-INS-63 | chr9-86971644-INS-63 |
| chr9-86972557-INS-444 | chr9-86972556-INS-449 |
| chr9-87003496-INS-574 | chr9-87003496-INS-574 |
| chr9-87013451-INS-712 | chr9-87013451-INS-733 |
| chr9-87017399-INS-162 | chr9-87017399-INS-162 |
| chr9-87018902-INS-82 | chr9-87018902-INS-82 |
| chr9-87029344-INS-118 | chr9-87029335-INS-133 |
| chr9-87043776-INS-212 | chr9-87043776-INS-212 |
| chr9-87046282-INS-65 | chr9-87046282-INS-65 |
| chr9-87056181-INS-116 | chr9-87056181-INS-105 |
| chr9-87058163-INS-148 | chr9-87058163-INS-148 |
| chr9-87091159-INS-205 | chr9-87091159-INS-205 |
| chr9-87125376-INS-181 | chr9-87125376-INS-181 |
| chr9-87147318-INS-2868 | chr9-87147319-INS-2868 |
| chr9-87172495-INS-77 | chr9-87172495-INS-77 |
| chr9-87192153-INS-6290 | chr9-87192154-INS-6291 |
| chr9-87277302-INS-6485 | chr9-87277303-INS-6487 |
| chr9-87358968-INS-117 | chr9-87358968-INS-116 |
| chr9-87388908-INS-78 | chr9-87388908-INS-86 |
| chr9-87404066-INS-193 | chr9-87404066-INS-193 |
| chr9-87410889-INS-54 | chr9-87410889-INS-62 |
| chr9-87412550-INS-735 | chr9-87412550-INS-736 |
| chr9-87428466-INS-438 | chr9-87428466-INS-440 |
| chr9-87436231-INS-993 | chr9-87436231-INS-993 |
| chr9-87456313-INS-139 | chr9-87456313-INS-140 |
| chr9-87486113-INS-56 | chr9-87486113-INS-54 |
| chr9-87492162-INS-63 | chr9-87492162-INS-69 |
| chr9-87532688-INS-72 | chr9-87532688-INS-72 |
| chr9-87539841-INS-494 | chr9-87539841-INS-494 |
| chr9-87570633-INS-54 | chr9-87570633-INS-56 |
| chr9-87581884-INS-413 | chr9-87581884-INS-413 |
| chr9-87586253-INS-79 | chr9-87586253-INS-79 |
| chr9-87593473-INS-1274 | chr9-87593474-INS-1274 |
| chr9-87594399-INS-52 | chr9-87594411-INS-52 |
| chr9-87661021-INS-240 | chr9-87661022-INS-202 |
| chr9-87661883-INS-495 | chr9-87661883-INS-495 |
| chr9-87679127-INS-501 | chr9-87679127-INS-501 |
| chr9-87687108-INS-436 | chr9-87687108-INS-436 |
| chr9-87697521-INS-212 | chr9-87697521-INS-212 |
| chr9-87748813-INS-81 | chr9-87748813-INS-50 |
| chr9-87789054-INS-67 | chr9-87789054-INS-67 |
| chr9-87798641-INS-6656 | chr9-87798642-INS-6656 |
| chr9-87818854-INS-2581 | chr9-87818855-INS-2581 |
| chr9-87823277-INS-327 | chr9-87823277-INS-327 |
| chr9-87841979-INS-54 | chr9-87841979-INS-56 |
| chr9-87884477-INS-1367 | chr9-87884478-INS-1367 |
| chr9-87886362-INS-52 | chr9-87886377-INS-53 |
| chr9-87889154-INS-60 | chr9-87889152-INS-108 |
| chr9-87898777-INS-54 | chr9-87898777-INS-66 |

| | |
|---|---|
| chr9-87915891-INS-50 | chr9-87915891-INS-52 |
| chr9-87925370-INS-2743 | chr9-87925371-INS-2743 |
| chr9-87928962-INS-60 | chr9-87928962-INS-60 |
| chr9-87930866-INS-6377 | chr9-87930867-INS-6377 |
| chr9-87971805-INS-181 | chr9-87971806-INS-188 |
| chr9-88000659-INS-53 | chr9-88000659-INS-53 |
| chr9-88108883-INS-203 | chr9-88108883-INS-204 |
| chr9-88111818-INS-70 | chr9-88111818-INS-70 |
| chr9-88130079-INS-963 | chr9-88130079-INS-965 |
| chr9-88150842-INS-209 | chr9-88150842-INS-208 |
| chr9-88174623-INS-93 | chr9-88174645-INS-62 |
| chr9-88185748-INS-3535 | chr9-88185749-INS-3534 |
| chr9-88224294-INS-61 | chr9-88224281-INS-64 |
| chr9-88237338-INS-60 | chr9-88237299-INS-104 |
| chr9-88268658-INS-184 | chr9-88268654-INS-178 |
| chr9-88309420-INS-98 | chr9-88309405-INS-83 |
| chr9-88347374-INS-52 | chr9-88347374-INS-68 |
| chr9-88381572-INS-174 | chr9-88381572-INS-174 |
| chr9-88406753-INS-190 | chr9-88406753-INS-192 |
| chr9-88424803-INS-67 | chr9-88424803-INS-63 |
| chr9-88433946-INS-209 | chr9-88433946-INS-209 |
| chr9-88449207-INS-4940 | chr9-88449208-INS-4978 |
| chr9-88468440-INS-704 | chr9-88468440-INS-704 |
| chr9-88630378-INS-55 | chr9-88630378-INS-52 |
| chr9-88859480-INS-512 | chr9-88859480-INS-510 |
| chr9-88969308-INS-364 | chr9-88969308-INS-364 |
| chr9-89052858-INS-580 | chr9-89052860-INS-576 |
| chr9-89059700-INS-52 | chr9-89059700-INS-70 |
| chr9-89116764-INS-78 | chr9-89116770-INS-88 |
| chr9-89181124-INS-74 | chr9-89181124-INS-77 |
| chr9-89237572-INS-506 | chr9-89237565-INS-506 |
| chr9-89255840-INS-392 | chr9-89255840-INS-392 |
| chr9-89298139-INS-58 | chr9-89298139-INS-51 |
| chr9-89360615-INS-201 | chr9-89360615-INS-201 |
| chr9-89378246-INS-109 | chr9-89378256-INS-80 |
| chr9-89407896-INS-6313 | chr9-89407897-INS-6313 |
| chr9-89431059-INS-763 | chr9-89431059-INS-763 |
| chr9-89465247-INS-6830 | chr9-89465248-INS-6831 |
| chr9-89477239-INS-134 | chr9-89477239-INS-140 |
| chr9-89479051-INS-178 | chr9-89479051-INS-178 |
| chr9-89515061-INS-218 | chr9-89515083-INS-228 |
| chr9-89535380-INS-82 | chr9-89535380-INS-84 |
| chr9-89562945-INS-156 | chr9-89562945-INS-155 |
| chr9-89565686-INS-53 | chr9-89565684-INS-58 |
| chr9-89568142-INS-358 | chr9-89568142-INS-358 |
| chr9-89580667-INS-66 | chr9-89580667-INS-50 |
| chr9-89583491-INS-849 | chr9-89583491-INS-849 |
| chr9-89598418-INS-400 | chr9-89598418-INS-400 |
| chr9-89598811-INS-5040 | chr9-89598812-INS-5040 |
| chr9-89599641-INS-498 | chr9-89599641-INS-498 |
| chr9-89622176-INS-254 | chr9-89622176-INS-255 |
| chr9-89845918-INS-77 | chr9-89845918-INS-71 |
| chr9-89894498-INS-182 | chr9-89894498-INS-182 |
| chr9-89915533-INS-56 | chr9-89915533-INS-56 |
| chr9-89929280-INS-182 | chr9-89929280-INS-182 |
| chr9-89937268-INS-56 | chr9-89937268-INS-56 |
| chr9-89941731-INS-260 | chr9-89941731-INS-262 |
| chr9-89961126-INS-202 | chr9-89961126-INS-202 |
| chr9-89976400-INS-6472 | chr9-89976401-INS-6473 |
| chr9-89976768-INS-63 | chr9-89976768-INS-63 |
| chr9-90010673-INS-214 | chr9-90010669-INS-212 |
| chr9-90020498-INS-197 | chr9-90020498-INS-197 |
| chr9-90025365-INS-55 | chr9-90025365-INS-60 |
| chr9-90033895-INS-82 | chr9-90033895-INS-82 |
| chr9-90042965-INS-77 | chr9-90042996-INS-68 |
| chr9-90057387-INS-53 | chr9-90057382-INS-68 |
| chr9-90072659-INS-53 | chr9-90072659-INS-53 |
| chr9-90107687-INS-376 | chr9-90107687-INS-376 |
| chr9-90155757-INS-106 | chr9-90155757-INS-100 |
| chr9-90214936-INS-314 | chr9-90214932-INS-316 |
| chr9-90215723-INS-507 | chr9-90215724-INS-507 |
| chr9-90263748-INS-211 | chr9-90263748-INS-213 |
| chr9-90286166-INS-471 | chr9-90286166-INS-472 |
| chr9-90290603-INS-82 | chr9-90290603-INS-84 |
| chr9-90309490-INS-52 | chr9-90309490-INS-54 |
| chr9-90360253-INS-6211 | chr9-90360254-INS-6211 |
| chr9-90388237-INS-542 | chr9-90388244-INS-552 |

| | |
|---|---|
| chr9-90389737-INS-3536 | chr9-90389738-INS-3536 |
| chr9-90523835-INS-60 | chr9-90523843-INS-69 |
| chr9-90525791-INS-53 | chr9-90525791-INS-53 |
| chr9-90565310-INS-536 | chr9-90565310-INS-539 |
| chr9-90595757-INS-2504 | chr9-90595758-INS-2524 |
| chr9-90601947-INS-6009 | chr9-90601948-INS-6009 |
| chr9-90614835-INS-55 | chr9-90614835-INS-55 |
| chr9-90628826-INS-210 | chr9-90628826-INS-210 |
| chr9-90646043-INS-522 | chr9-90646043-INS-522 |
| chr9-90652272-INS-2806 | chr9-90652273-INS-2806 |
| chr9-90655231-INS-7137 | chr9-90655232-INS-7137 |
| chr9-90675015-INS-71 | chr9-90675015-INS-77 |
| chr9-90682009-INS-327 | chr9-90682009-INS-327 |
| chr9-90708434-INS-965 | chr9-90708434-INS-965 |
| chr9-90717300-INS-61 | chr9-90717300-INS-72 |
| chr9-90723980-INS-78 | chr9-90723980-INS-80 |
| chr9-90726227-INS-423 | chr9-90726227-INS-423 |
| chr9-90794148-INS-62 | chr9-90794148-INS-62 |
| chr9-90795237-INS-550 | chr9-90795237-INS-550 |
| chr9-90839740-INS-163 | chr9-90839740-INS-163 |
| chr9-90862047-INS-6173 | chr9-90862048-INS-6170 |
| chr9-90873116-INS-64 | chr9-90873116-INS-62 |
| chr9-90891112-INS-701 | chr9-90891112-INS-702 |
| chr9-90891992-INS-395 | chr9-90891992-INS-395 |
| chr9-90899377-INS-6982 | chr9-90899378-INS-6983 |
| chr9-90931772-INS-2264 | chr9-90931773-INS-2263 |
| chr9-90943071-INS-215 | chr9-90943071-INS-216 |
| chr9-91002785-INS-4859 | chr9-91002786-INS-4859 |
| chr9-91041737-INS-51 | chr9-91041737-INS-53 |
| chr9-91068638-INS-56 | chr9-91068638-INS-60 |
| chr9-91070471-INS-6849 | chr9-91070472-INS-6850 |
| chr9-91075664-INS-1021 | chr9-91075665-INS-1023 |
| chr9-91118276-INS-52 | chr9-91118289-INS-56 |
| chr9-91131866-INS-953 | chr9-91131866-INS-944 |
| chr9-91141788-INS-520 | chr9-91141788-INS-520 |
| chr9-91180823-INS-96 | chr9-91180823-INS-96 |
| chr9-91186843-INS-6665 | chr9-91186844-INS-6666 |
| chr9-91190000-INS-108 | chr9-91190000-INS-116 |
| chr9-91212617-INS-811 | chr9-91212617-INS-810 |
| chr9-91235551-INS-131 | chr9-91235549-INS-142 |
| chr9-91298941-INS-58 | chr9-91298941-INS-60 |
| chr9-91316010-INS-843 | chr9-91316010-INS-845 |
| chr9-91356433-INS-549 | chr9-91356433-INS-523 |
| chr9-91374813-INS-195 | chr9-91374813-INS-197 |
| chr9-91446537-INS-7857 | chr9-91446538-INS-7856 |
| chr9-91446589-INS-79 | chr9-91446598-INS-111 |
| chr9-91468376-INS-98 | chr9-91468392-INS-56 |
| chr9-91473048-INS-59 | chr9-91473048-INS-59 |
| chr9-91501327-INS-212 | chr9-91501327-INS-212 |
| chr9-91561818-INS-6378 | chr9-91561819-INS-6378 |
| chr9-91602121-INS-2357 | chr9-91602122-INS-2354 |
| chr9-91630339-INS-6915 | chr9-91630340-INS-6914 |
| chr9-91722799-INS-56 | chr9-91722801-INS-54 |
| chr9-91729469-INS-116 | chr9-91729469-INS-117 |
| chr9-91760643-INS-6448 | chr9-91760644-INS-6448 |
| chr9-91778120-INS-873 | chr9-91778120-INS-872 |
| chr9-91794518-INS-58 | chr9-91794513-INS-84 |
| chr9-91819970-INS-94 | chr9-91819970-INS-100 |
| chr9-91838621-INS-1875 | chr9-91838622-INS-1875 |
| chr9-91869179-INS-399 | chr9-91869179-INS-399 |
| chr9-91870619-INS-4817 | chr9-91870620-INS-4817 |
| chr9-91872538-INS-219 | chr9-91872538-INS-219 |
| chr9-91885063-INS-1683 | chr9-91885064-INS-1683 |
| chr9-91887661-INS-131 | chr9-91887664-INS-104 |
| chr9-91898980-INS-179 | chr9-91898980-INS-189 |
| chr9-91906036-INS-76 | chr9-91906036-INS-76 |
| chr9-91910205-INS-160 | chr9-91910205-INS-160 |
| chr9-91914619-INS-460 | chr9-91914619-INS-459 |
| chr9-91936766-INS-396 | chr9-91936766-INS-396 |
| chr9-91991447-INS-57 | chr9-91991447-INS-57 |
| chr9-92008890-INS-56 | chr9-92008890-INS-50 |
| chr9-92020341-INS-670 | chr9-92020341-INS-674 |
| chr9-92044217-INS-106 | chr9-92044217-INS-108 |
| chr9-92051970-INS-227 | chr9-92051970-INS-227 |
| chr9-92055067-INS-250 | chr9-92055067-INS-250 |
| chr9-92062751-INS-496 | chr9-92062751-INS-496 |
| chr9-92081069-INS-210 | chr9-92081069-INS-210 |

| | |
|---|---|
| chr9-92110233-INS-6698 | chr9-92110234-INS-6700 |
| chr9-92112236-INS-88 | chr9-92112237-INS-54 |
| chr9-92161354-INS-1776 | chr9-92161355-INS-1776 |
| chr9-92205975-INS-192 | chr9-92205975-INS-192 |
| chr9-92219095-INS-916 | chr9-92219095-INS-916 |
| chr9-92230304-INS-7144 | chr9-92230305-INS-7144 |
| chr9-92262347-INS-206 | chr9-92262347-INS-206 |
| chr9-92269740-INS-936 | chr9-92269740-INS-936 |
| chr9-92279792-INS-385 | chr9-92279792-INS-385 |
| chr9-92284383-INS-57 | chr9-92284383-INS-69 |
| chr9-92293853-INS-62 | chr9-92293841-INS-62 |
| chr9-92338839-INS-60 | chr9-92338839-INS-60 |
| chr9-92370174-INS-273 | chr9-92370195-INS-253 |
| chr9-92375521-INS-107 | chr9-92375521-INS-134 |
| chr9-92386006-INS-100 | chr9-92386006-INS-105 |
| chr9-92388047-INS-226 | chr9-92388047-INS-226 |
| chr9-92398229-INS-199 | chr9-92398229-INS-199 |
| chr9-92399570-INS-269 | chr9-92399570-INS-269 |
| chr9-92403223-INS-85 | chr9-92403223-INS-65 |
| chr9-92424818-INS-66 | chr9-92424856-INS-66 |
| chr9-92441626-INS-2489 | chr9-92441627-INS-2490 |
| chr9-92460919-INS-79 | chr9-92460919-INS-83 |
| chr9-92467950-INS-81 | chr9-92467950-INS-53 |
| chr9-92485461-INS-76 | chr9-92485461-INS-76 |
| chr9-92494468-INS-421 | chr9-92494468-INS-416 |
| chr9-92499302-INS-198 | chr9-92499302-INS-198 |
| chr9-92501718-INS-92 | chr9-92501714-INS-96 |
| chr9-92521154-INS-54 | chr9-92521154-INS-54 |
| chr9-92567038-INS-6164 | chr9-92567039-INS-6164 |
| chr9-92572082-INS-92 | chr9-92572082-INS-90 |
| chr9-92583033-INS-547 | chr9-92583033-INS-549 |
| chr9-92590622-INS-2308 | chr9-92590623-INS-2308 |
| chr9-92602737-INS-942 | chr9-92602737-INS-942 |
| chr9-92615105-INS-72 | chr9-92615105-INS-72 |
| chr9-92630872-INS-52 | chr9-92630872-INS-52 |
| chr9-92676197-INS-801 | chr9-92676197-INS-805 |
| chr9-92678146-INS-428 | chr9-92678146-INS-382 |
| chr9-92686515-INS-387 | chr9-92686516-INS-387 |
| chr9-92687367-INS-66 | chr9-92687367-INS-66 |
| chr9-92703634-INS-229 | chr9-92703634-INS-229 |
| chr9-92728481-INS-399 | chr9-92728482-INS-399 |
| chr9-92730300-INS-119 | chr9-92730300-INS-123 |
| chr9-92731421-INS-142 | chr9-92731421-INS-142 |
| chr9-92736045-INS-197 | chr9-92736045-INS-197 |
| chr9-92748275-INS-594 | chr9-92748275-INS-593 |
| chr9-92751250-INS-203 | chr9-92751250-INS-203 |
| chr9-92762719-INS-484 | chr9-92762719-INS-484 |
| chr9-92769844-INS-51 | chr9-92769845-INS-55 |
| chr9-92801493-INS-799 | chr9-92801493-INS-799 |
| chr9-92822748-INS-536 | chr9-92822748-INS-536 |
| chr9-92824473-INS-84 | chr9-92824473-INS-86 |
| chr9-92831656-INS-437 | chr9-92831656-INS-437 |
| chr9-92836115-INS-1893 | chr9-92836116-INS-1893 |
| chr9-92838943-INS-221 | chr9-92838943-INS-227 |
| chr9-92869826-INS-84 | chr9-92869816-INS-80 |
| chr9-92884628-INS-5665 | chr9-92884629-INS-5661 |
| chr9-92936230-INS-55 | chr9-92936230-INS-55 |
| chr9-92961274-INS-163 | chr9-92961274-INS-163 |
| chr9-92971682-INS-7796 | chr9-92971694-INS-7798 |
| chr9-93011842-INS-262 | chr9-93011842-INS-262 |
| chr9-93025583-INS-204 | chr9-93025583-INS-204 |
| chr9-93037991-INS-57 | chr9-93037991-INS-58 |
| chr9-93113917-INS-128 | chr9-93113917-INS-129 |
| chr9-93123925-INS-552 | chr9-93123925-INS-552 |
| chr9-93127434-INS-324 | chr9-93127434-INS-324 |
| chr9-93133059-INS-343 | chr9-93133059-INS-343 |
| chr9-93185046-INS-67 | chr9-93185046-INS-67 |
| chr9-93202083-INS-82 | chr9-93202083-INS-82 |
| chr9-93221486-INS-54 | chr9-93221486-INS-52 |
| chr9-93223236-INS-132 | chr9-93223266-INS-132 |
| chr9-93232747-INS-61 | chr9-93232747-INS-61 |
| chr9-93308146-INS-59 | chr9-93308146-INS-59 |
| chr9-93347261-INS-62 | chr9-93347258-INS-62 |
| chr9-93351019-INS-93 | chr9-93351016-INS-51 |
| chr9-93365315-INS-3130 | chr9-93365316-INS-3130 |
| chr9-93368666-INS-417 | chr9-93368666-INS-417 |
| chr9-93394736-INS-52 | chr9-93394736-INS-64 |

| | |
|---|---|
| chr9-93396958-INS-143 | chr9-93396958-INS-143 |
| chr9-93402668-INS-111 | chr9-93402668-INS-103 |
| chr9-93407186-INS-56 | chr9-93407186-INS-58 |
| chr9-93429705-INS-55 | chr9-93429705-INS-55 |
| chr9-93441789-INS-300 | chr9-93441789-INS-301 |
| chr9-93443345-INS-171 | chr9-93443345-INS-168 |
| chr9-93452859-INS-104 | chr9-93452867-INS-62 |
| chr9-93455470-INS-1387 | chr9-93455471-INS-1387 |
| chr9-93476058-INS-927 | chr9-93476058-INS-926 |
| chr9-93486730-INS-182 | chr9-93486730-INS-182 |
| chr9-93594561-INS-6767 | chr9-93594562-INS-6767 |
| chr9-93606356-INS-137 | chr9-93606312-INS-165 |
| chr9-93609050-INS-90 | chr9-93609050-INS-90 |
| chr9-93631592-INS-76 | chr9-93631572-INS-81 |
| chr9-93641824-INS-652 | chr9-93641829-INS-671 |
| chr9-93646221-INS-808 | chr9-93646221-INS-807 |
| chr9-93648654-INS-373 | chr9-93648654-INS-373 |
| chr9-93657767-INS-87 | chr9-93657767-INS-90 |
| chr9-93671407-INS-233 | chr9-93671407-INS-233 |
| chr9-93713452-INS-70 | chr9-93713452-INS-70 |
| chr9-93743318-INS-173 | chr9-93743318-INS-173 |
| chr9-93776319-INS-66 | chr9-93776319-INS-60 |
| chr9-93776442-INS-846 | chr9-93776442-INS-846 |
| chr9-93788077-INS-3691 | chr9-93788078-INS-3691 |
| chr9-93795759-INS-2631 | chr9-93795760-INS-2631 |
| chr9-93824416-INS-488 | chr9-93824416-INS-488 |
| chr9-93840518-INS-104 | chr9-93840518-INS-104 |
| chr9-93843553-INS-3709 | chr9-93843554-INS-3709 |
| chr9-93862224-INS-92 | chr9-93862224-INS-94 |
| chr9-93868137-INS-1259 | chr9-93868138-INS-1286 |
| chr9-93890682-INS-1084 | chr9-93890678-INS-1094 |
| chr9-93899046-INS-132 | chr9-93899064-INS-135 |
| chr9-93908901-INS-85 | chr9-93908901-INS-85 |
| chr9-93917881-INS-698 | chr9-93917881-INS-699 |
| chr9-93930981-INS-346 | chr9-93930981-INS-346 |
| chr9-93932401-INS-311 | chr9-93932401-INS-311 |
| chr9-93970264-INS-2168 | chr9-93970265-INS-2173 |
| chr9-93972151-INS-1880 | chr9-93972152-INS-1880 |
| chr9-93983252-INS-118 | chr9-93983252-INS-118 |
| chr9-93990892-INS-151 | chr9-93990892-INS-151 |
| chr9-94028240-INS-72 | chr9-94028240-INS-72 |
| chr9-94040476-INS-426 | chr9-94040476-INS-426 |
| chr9-94054374-INS-170 | chr9-94054374-INS-170 |
| chr9-94063950-INS-225 | chr9-94063950-INS-225 |
| chr9-94064575-INS-6691 | chr9-94064576-INS-6693 |
| chr9-94073961-INS-422 | chr9-94073961-INS-422 |
| chr9-94075007-INS-120 | chr9-94075007-INS-136 |
| chr9-94079962-INS-375 | chr9-94079962-INS-375 |
| chr9-94093990-INS-6905 | chr9-94093991-INS-6907 |
| chr9-94097991-INS-74 | chr9-94097991-INS-74 |
| chr9-94127947-INS-372 | chr9-94127947-INS-372 |
| chr9-94135247-INS-1648 | chr9-94135248-INS-1648 |
| chr9-94140896-INS-310 | chr9-94140896-INS-310 |
| chr9-94152928-INS-429 | chr9-94152928-INS-429 |
| chr9-94158166-INS-70 | chr9-94158166-INS-75 |
| chr9-94167750-INS-1497 | chr9-94167765-INS-1464 |
| chr9-94172337-INS-133 | chr9-94172337-INS-133 |
| chr9-94181574-INS-76 | chr9-94181574-INS-76 |
| chr9-94187294-INS-352 | chr9-94187294-INS-352 |
| chr9-94201425-INS-51 | chr9-94201425-INS-51 |
| chr9-94202911-INS-3235 | chr9-94202912-INS-3235 |
| chr9-94207427-INS-167 | chr9-94207427-INS-168 |
| chr9-94209120-INS-56 | chr9-94209121-INS-65 |
| chr9-94241012-INS-118 | chr9-94241012-INS-120 |
| chr9-94286957-INS-373 | chr9-94286957-INS-380 |
| chr9-94301989-INS-462 | chr9-94301989-INS-462 |
| chr9-94308398-INS-61 | chr9-94308398-INS-64 |
| chr9-94333035-INS-2411 | chr9-94333036-INS-2411 |
| chr9-94344736-INS-249 | chr9-94344736-INS-249 |
| chr9-94358066-INS-55 | chr9-94358066-INS-54 |
| chr9-94370272-INS-318 | chr9-94370272-INS-319 |
| chr9-94378759-INS-232 | chr9-94378753-INS-224 |
| chr9-94388877-INS-103 | chr9-94388877-INS-107 |
| chr9-94392718-INS-296 | chr9-94392718-INS-300 |
| chr9-94397602-INS-490 | chr9-94397602-INS-486 |
| chr9-94428875-INS-82 | chr9-94428875-INS-80 |
| chr9-94439765-INS-293 | chr9-94439765-INS-298 |

| | |
|---|---|
| chr9-94476413-INS-226 | chr9-94476413-INS-226 |
| chr9-94500128-INS-194 | chr9-94500128-INS-194 |
| chr9-94502314-INS-6226 | chr9-94502315-INS-6227 |
| chr9-94516676-INS-91 | chr9-94516701-INS-99 |
| chr9-94519210-INS-1208 | chr9-94519211-INS-1209 |
| chr9-94542312-INS-176 | chr9-94542312-INS-176 |
| chr9-94565905-INS-52 | chr9-94565898-INS-97 |
| chr9-94576249-INS-58 | chr9-94576249-INS-58 |
| chr9-94589359-INS-62 | chr9-94589355-INS-66 |
| chr9-94597477-INS-7024 | chr9-94597478-INS-7024 |
| chr9-94613689-INS-89 | chr9-94613689-INS-89 |
| chr9-94625671-INS-461 | chr9-94625671-INS-461 |
| chr9-94680905-INS-56 | chr9-94680905-INS-56 |
| chr9-94683527-INS-199 | chr9-94683527-INS-199 |
| chr9-94734278-INS-52 | chr9-94734278-INS-52 |
| chr9-94744979-INS-3529 | chr9-94744980-INS-3529 |
| chr9-94761948-INS-160 | chr9-94761948-INS-160 |
| chr9-94769839-INS-444 | chr9-94769839-INS-444 |
| chr9-94811193-INS-349 | chr9-94811180-INS-339 |
| chr9-94816575-INS-62 | chr9-94816575-INS-62 |
| chr9-94826754-INS-418 | chr9-94826754-INS-418 |
| chr9-94836508-INS-56 | chr9-94836508-INS-56 |
| chr9-94843908-INS-53 | chr9-94843905-INS-77 |
| chr9-94854733-INS-410 | chr9-94854733-INS-410 |
| chr9-94860641-INS-150 | chr9-94860641-INS-150 |
| chr9-94900758-INS-97 | chr9-94900757-INS-113 |
| chr9-94924247-INS-58 | chr9-94924247-INS-58 |
| chr9-94959431-INS-6238 | chr9-94959432-INS-6238 |
| chr9-94965938-INS-161 | chr9-94965938-INS-161 |
| chr9-94972092-INS-498 | chr9-94972092-INS-498 |
| chr9-94976019-INS-153 | chr9-94976029-INS-116 |
| chr9-95013042-INS-84 | chr9-95013031-INS-102 |
| chr9-95015194-INS-576 | chr9-95015194-INS-576 |
| chr9-95022463-INS-176 | chr9-95022463-INS-194 |
| chr9-95045137-INS-114 | chr9-95045137-INS-118 |
| chr9-95077165-INS-623 | chr9-95077165-INS-623 |
| chr9-95086757-INS-168 | chr9-95086757-INS-168 |
| chr9-95099730-INS-1223 | chr9-95099731-INS-1224 |
| chr9-95104947-INS-65 | chr9-95104947-INS-65 |
| chr9-95107173-INS-148 | chr9-95107175-INS-152 |
| chr9-95108736-INS-52 | chr9-95108736-INS-56 |
| chr9-95109929-INS-180 | chr9-95109929-INS-155 |
| chr9-95128799-INS-224 | chr9-95128799-INS-224 |
| chr9-95134003-INS-1284 | chr9-95134004-INS-1284 |
| chr9-95145933-INS-76 | chr9-95145933-INS-76 |
| chr9-95164373-INS-168 | chr9-95164373-INS-168 |
| chr9-95174728-INS-212 | chr9-95174728-INS-212 |
| chr9-95198143-INS-6322 | chr9-95198144-INS-6322 |
| chr9-95220756-INS-7118 | chr9-95220757-INS-7118 |
| chr9-95222199-INS-400 | chr9-95222204-INS-380 |
| chr9-95245177-INS-93 | chr9-95245169-INS-102 |
| chr9-95255621-INS-440 | chr9-95255621-INS-440 |
| chr9-95265594-INS-267 | chr9-95265594-INS-268 |
| chr9-95274056-INS-60 | chr9-95274042-INS-56 |
| chr9-95306510-INS-62 | chr9-95306484-INS-69 |
| chr9-95313414-INS-275 | chr9-95313414-INS-281 |
| chr9-95329957-INS-180 | chr9-95329959-INS-185 |
| chr9-95356880-INS-55 | chr9-95356880-INS-55 |
| chr9-95382562-INS-168 | chr9-95382562-INS-168 |
| chr9-95400981-INS-202 | chr9-95400981-INS-202 |
| chr9-95404469-INS-55 | chr9-95404469-INS-58 |
| chr9-95405771-INS-746 | chr9-95405771-INS-746 |
| chr9-95416266-INS-404 | chr9-95416266-INS-404 |
| chr9-95422073-INS-5419 | chr9-95422074-INS-5419 |
| chr9-95429659-INS-242 | chr9-95429659-INS-242 |
| chr9-95437971-INS-204 | chr9-95437971-INS-204 |
| chr9-95461345-INS-76 | chr9-95461345-INS-60 |
| chr9-95476346-INS-101 | chr9-95476346-INS-101 |
| chr9-95480116-INS-75 | chr9-95480116-INS-75 |
| chr9-95487661-INS-765 | chr9-95487662-INS-765 |
| chr9-95495592-INS-51 | chr9-95495592-INS-51 |
| chr9-95500302-INS-71 | chr9-95500290-INS-58 |
| chr9-95505766-INS-66 | chr9-95505766-INS-66 |
| chr9-95514345-INS-81 | chr9-95514351-INS-90 |
| chr9-95524427-INS-112 | chr9-95524427-INS-64 |
| chr9-95530750-INS-69 | chr9-95530749-INS-71 |
| chr9-95536120-INS-84 | chr9-95536120-INS-96 |

| | |
|---|---|
| chr9-95544507-INS-194 | chr9-95544517-INS-186 |
| chr9-95548898-INS-81 | chr9-95548902-INS-84 |
| chr9-95551334-INS-291 | chr9-95551362-INS-278 |
| chr9-95552671-INS-54 | chr9-95552671-INS-54 |
| chr9-95574347-INS-176 | chr9-95574345-INS-178 |
| chr9-95586059-INS-172 | chr9-95586059-INS-174 |
| chr9-95594384-INS-65 | chr9-95594385-INS-65 |
| chr9-95597188-INS-635 | chr9-95597188-INS-649 |
| chr9-95598646-INS-1507 | chr9-95598647-INS-1507 |
| chr9-95599464-INS-54 | chr9-95599465-INS-54 |
| chr9-95609942-INS-601 | chr9-95609942-INS-602 |
| chr9-95612470-INS-79 | chr9-95612470-INS-79 |
| chr9-95648957-INS-86 | chr9-95648957-INS-96 |
| chr9-95661707-INS-345 | chr9-95661707-INS-345 |
| chr9-95677690-INS-129 | chr9-95677690-INS-129 |
| chr9-95688095-INS-82 | chr9-95688095-INS-82 |
| chr9-95730118-INS-522 | chr9-95730118-INS-522 |
| chr9-95732799-INS-52 | chr9-95732797-INS-54 |
| chr9-95803593-INS-3204 | chr9-95803594-INS-3204 |
| chr9-95807563-INS-213 | chr9-95807563-INS-213 |
| chr9-95867917-INS-98 | chr9-95867917-INS-99 |
| chr9-95890884-INS-176 | chr9-95890884-INS-176 |
| chr9-95906912-INS-53 | chr9-95906912-INS-53 |
| chr9-95911128-INS-210 | chr9-95911128-INS-210 |
| chr9-95917962-INS-216 | chr9-95917962-INS-216 |
| chr9-95924511-INS-178 | chr9-95924511-INS-179 |
| chr9-95939144-INS-56 | chr9-95939144-INS-56 |
| chr9-95952190-INS-215 | chr9-95952190-INS-214 |
| chr9-95955326-INS-671 | chr9-95955326-INS-672 |
| chr9-95957840-INS-55 | chr9-95957841-INS-54 |
| chr9-95967002-INS-55 | chr9-95966987-INS-64 |
| chr9-95975601-INS-124 | chr9-95975601-INS-124 |
| chr9-95978015-INS-577 | chr9-95978015-INS-577 |
| chr9-95980573-INS-189 | chr9-95980573-INS-189 |
| chr9-96004386-INS-280 | chr9-96004386-INS-282 |
| chr9-96007835-INS-395 | chr9-96007835-INS-395 |
| chr9-96009672-INS-195 | chr9-96009672-INS-195 |
| chr9-96018171-INS-52 | chr9-96018168-INS-102 |
| chr9-96024950-INS-200 | chr9-96024950-INS-200 |
| chr9-96026150-INS-58 | chr9-96026150-INS-58 |
| chr9-96100838-INS-56 | chr9-96100838-INS-56 |
| chr9-96116707-INS-132 | chr9-96116705-INS-134 |
| chr9-96121483-INS-73 | chr9-96121484-INS-73 |
| chr9-96135535-INS-1410 | chr9-96135536-INS-1415 |
| chr9-96144831-INS-583 | chr9-96144831-INS-583 |
| chr9-96145379-INS-58 | chr9-96145380-INS-68 |
| chr9-96149702-INS-205 | chr9-96149702-INS-206 |
| chr9-96151039-INS-102 | chr9-96151033-INS-112 |
| chr9-96157263-INS-224 | chr9-96157263-INS-224 |
| chr9-96159781-INS-562 | chr9-96159781-INS-562 |
| chr9-96162101-INS-54 | chr9-96162101-INS-56 |
| chr9-96175403-INS-276 | chr9-96175403-INS-276 |
| chr9-96190362-INS-179 | chr9-96190362-INS-179 |
| chr9-96192452-INS-54 | chr9-96192430-INS-54 |
| chr9-96198063-INS-74 | chr9-96198061-INS-52 |
| chr9-96208386-INS-166 | chr9-96208386-INS-166 |
| chr9-96209465-INS-129 | chr9-96209465-INS-129 |
| chr9-96221253-INS-163 | chr9-96221253-INS-165 |
| chr9-96226336-INS-191 | chr9-96226336-INS-191 |
| chr9-96232000-INS-169 | chr9-96232000-INS-169 |
| chr9-96235330-INS-124 | chr9-96235330-INS-124 |
| chr9-96253478-INS-235 | chr9-96253479-INS-203 |
| chr9-96267262-INS-61 | chr9-96267262-INS-61 |
| chr9-96276049-INS-112 | chr9-96276049-INS-106 |
| chr9-96284311-INS-198 | chr9-96284311-INS-198 |
| chr9-96287342-INS-63 | chr9-96287342-INS-63 |
| chr9-96288261-INS-177 | chr9-96288261-INS-178 |
| chr9-96295895-INS-54 | chr9-96295895-INS-54 |
| chr9-96297741-INS-69 | chr9-96297737-INS-57 |
| chr9-96314393-INS-51 | chr9-96314398-INS-51 |
| chr9-96314625-INS-342 | chr9-96314625-INS-342 |
| chr9-96332500-INS-242 | chr9-96332500-INS-242 |
| chr9-96334193-INS-98 | chr9-96334193-INS-98 |
| chr9-96339225-INS-69 | chr9-96339225-INS-69 |
| chr9-96341998-INS-57 | chr9-96341998-INS-61 |
| chr9-96348369-INS-132 | chr9-96348369-INS-130 |
| chr9-96354296-INS-191 | chr9-96354296-INS-191 |

| | |
|---|---|
| chr9-96367957-INS-341 | chr9-96367957-INS-341 |
| chr9-96393701-INS-197 | chr9-96393701-INS-197 |
| chr9-96397376-INS-103 | chr9-96397376-INS-76 |
| chr9-96407938-INS-176 | chr9-96407938-INS-169 |
| chr9-96412198-INS-76 | chr9-96412198-INS-76 |
| chr9-96429846-INS-52 | chr9-96429828-INS-52 |
| chr9-96463999-INS-4321 | chr9-96464000-INS-4321 |
| chr9-96475270-INS-517 | chr9-96475270-INS-517 |
| chr9-96482935-INS-250 | chr9-96482935-INS-249 |
| chr9-96516764-INS-225 | chr9-96516764-INS-225 |
| chr9-96523891-INS-52 | chr9-96523891-INS-52 |
| chr9-96541212-INS-90 | chr9-96541212-INS-90 |
| chr9-96553915-INS-108 | chr9-96553915-INS-108 |
| chr9-96587394-INS-55 | chr9-96587394-INS-63 |
| chr9-96627088-INS-70 | chr9-96627088-INS-70 |
| chr9-96640111-INS-682 | chr9-96640111-INS-670 |
| chr9-96650147-INS-91 | chr9-96650146-INS-123 |
| chr9-96655175-INS-73 | chr9-96655191-INS-51 |
| chr9-96659295-INS-70 | chr9-96659295-INS-74 |
| chr9-96661406-INS-130 | chr9-96661406-INS-134 |
| chr9-96666707-INS-1813 | chr9-96666708-INS-1860 |
| chr9-96687813-INS-62 | chr9-96687806-INS-68 |
| chr9-96691422-INS-66 | chr9-96691422-INS-66 |
| chr9-96716024-INS-60 | chr9-96716024-INS-60 |
| chr9-96716666-INS-418 | chr9-96716666-INS-418 |
| chr9-96737156-INS-57 | chr9-96737156-INS-58 |
| chr9-96745315-INS-163 | chr9-96745315-INS-163 |
| chr9-96745356-INS-1429 | chr9-96745357-INS-1429 |
| chr9-96752431-INS-351 | chr9-96752431-INS-352 |
| chr9-96758982-INS-5272 | chr9-96758983-INS-5273 |
| chr9-96785932-INS-201 | chr9-96785932-INS-214 |
| chr9-96822627-INS-90 | chr9-96822622-INS-90 |
| chr9-96832418-INS-156 | chr9-96832418-INS-156 |
| chr9-96851422-INS-200 | chr9-96851422-INS-200 |
| chr9-96856448-INS-217 | chr9-96856448-INS-217 |
| chr9-96888288-INS-147 | chr9-96888288-INS-149 |
| chr9-96911614-INS-140 | chr9-96911614-INS-140 |
| chr9-96916981-INS-178 | chr9-96916982-INS-179 |
| chr9-96954260-INS-86 | chr9-96954260-INS-86 |
| chr9-96966702-INS-94 | chr9-96966702-INS-94 |
| chr9-97007375-INS-90 | chr9-97007375-INS-90 |
| chr9-97011660-INS-55 | chr9-97011672-INS-61 |
| chr9-97024908-INS-68 | chr9-97024908-INS-78 |
| chr9-97032631-INS-161 | chr9-97032631-INS-161 |
| chr9-97043841-INS-199 | chr9-97043841-INS-199 |
| chr9-97049204-INS-56 | chr9-97049241-INS-50 |
| chr9-97067287-INS-99 | chr9-97067287-INS-99 |
| chr9-97076207-INS-78 | chr9-97076207-INS-60 |
| chr9-97079221-INS-464 | chr9-97079221-INS-464 |
| chr9-97087626-INS-61 | chr9-97087626-INS-58 |
| chr9-97094195-INS-120 | chr9-97094195-INS-90 |
| chr9-97133048-INS-6260 | chr9-97133049-INS-6260 |
| chr9-97137893-INS-620 | chr9-97137893-INS-620 |
| chr9-97149827-INS-340 | chr9-97149827-INS-347 |
| chr9-97154807-INS-52 | chr9-97154807-INS-52 |
| chr9-97157241-INS-984 | chr9-97157241-INS-984 |
| chr9-97159570-INS-270 | chr9-97159570-INS-270 |
| chr9-97162079-INS-245 | chr9-97162079-INS-245 |
| chr9-97187759-INS-201 | chr9-97187759-INS-201 |
| chr9-97195658-INS-56 | chr9-97195658-INS-58 |
| chr9-97200346-INS-76 | chr9-97200339-INS-60 |
| chr9-97206278-INS-126 | chr9-97206278-INS-132 |
| chr9-97242249-INS-71 | chr9-97242228-INS-67 |
| chr9-97244071-INS-157 | chr9-97244072-INS-157 |
| chr9-97252977-INS-61 | chr9-97252977-INS-61 |
| chr9-97258323-INS-857 | chr9-97258323-INS-828 |
| chr9-97262854-INS-257 | chr9-97262854-INS-257 |
| chr9-97263910-INS-309 | chr9-97263910-INS-309 |
| chr9-97286092-INS-147 | chr9-97286092-INS-148 |
| chr9-97293046-INS-61 | chr9-97293046-INS-59 |
| chr9-97300838-INS-80 | chr9-97300838-INS-81 |
| chr9-97308989-INS-210 | chr9-97308989-INS-210 |
| chr9-97311658-INS-147 | chr9-97311658-INS-159 |
| chr9-97313000-INS-218 | chr9-97313000-INS-218 |
| chr9-97378177-INS-60 | chr9-97378177-INS-60 |
| chr9-97384287-INS-109 | chr9-97384288-INS-67 |
| chr9-97417924-INS-178 | chr9-97417924-INS-178 |

| | |
|---|---|
| chr9-97436241-INS-56 | chr9-97436241-INS-70 |
| chr9-97506849-INS-254 | chr9-97506849-INS-234 |
| chr9-97518190-INS-742 | chr9-97518190-INS-736 |
| chr9-97523927-INS-480 | chr9-97523927-INS-480 |
| chr9-97527734-INS-62 | chr9-97527726-INS-66 |
| chr9-97562600-INS-8391 | chr9-97562601-INS-8391 |
| chr9-97621760-INS-2709 | chr9-97621761-INS-2709 |
| chr9-97652758-INS-60 | chr9-97652760-INS-86 |
| chr9-97667847-INS-162 | chr9-97667847-INS-162 |
| chr9-97691380-INS-548 | chr9-97691380-INS-548 |
| chr9-97704297-INS-3082 | chr9-97704298-INS-3082 |
| chr9-97740154-INS-445 | chr9-97740154-INS-445 |
| chr9-97747436-INS-53 | chr9-97747436-INS-53 |
| chr9-97750621-INS-469 | chr9-97750621-INS-469 |
| chr9-97754410-INS-54 | chr9-97754410-INS-50 |
| chr9-97797414-INS-52 | chr9-97797414-INS-52 |
| chr9-97805157-INS-284 | chr9-97805157-INS-284 |
| chr9-97806353-INS-69 | chr9-97806353-INS-62 |
| chr9-97830750-INS-183 | chr9-97830750-INS-183 |
| chr9-97832326-INS-56 | chr9-97832326-INS-56 |
| chr9-97845521-INS-66 | chr9-97845521-INS-66 |
| chr9-97846438-INS-362 | chr9-97846438-INS-362 |
| chr9-97890561-INS-56 | chr9-97890561-INS-58 |
| chr9-97896520-INS-94 | chr9-97896530-INS-68 |
| chr9-97913550-INS-108 | chr9-97913550-INS-116 |
| chr9-97921457-INS-74 | chr9-97921457-INS-74 |
| chr9-97954738-INS-59 | chr9-97954738-INS-54 |
| chr9-97999560-INS-204 | chr9-97999560-INS-205 |
| chr9-98009935-INS-66 | chr9-98009935-INS-90 |
| chr9-98012291-INS-255 | chr9-98012288-INS-261 |
| chr9-98017784-INS-96 | chr9-98017745-INS-140 |
| chr9-98020034-INS-52 | chr9-98020034-INS-54 |
| chr9-98091973-INS-54 | chr9-98091973-INS-54 |
| chr9-98118638-INS-52 | chr9-98118635-INS-54 |
| chr9-98216260-INS-113 | chr9-98216261-INS-110 |
| chr9-98230083-INS-66 | chr9-98230083-INS-66 |
| chr9-98246336-INS-62 | chr9-98246336-INS-62 |
| chr9-98260563-INS-86 | chr9-98260563-INS-84 |
| chr9-98265878-INS-56 | chr9-98265878-INS-58 |
| chr9-98280266-INS-688 | chr9-98280266-INS-688 |
| chr9-98294618-INS-72 | chr9-98294618-INS-80 |
| chr9-98348249-INS-329 | chr9-98348250-INS-329 |
| chr9-98361987-INS-54 | chr9-98361987-INS-56 |
| chr9-98367983-INS-55 | chr9-98367983-INS-55 |
| chr9-98372065-INS-64 | chr9-98372065-INS-67 |
| chr9-98418605-INS-6898 | chr9-98418606-INS-6899 |
| chr9-98433375-INS-128 | chr9-98433375-INS-128 |
| chr9-98526806-INS-202 | chr9-98526806-INS-202 |
| chr9-98551718-INS-2013 | chr9-98551719-INS-2013 |
| chr9-98595145-INS-307 | chr9-98595146-INS-307 |
| chr9-98605653-INS-61 | chr9-98605653-INS-61 |
| chr9-98642538-INS-316 | chr9-98642538-INS-328 |
| chr9-98675352-INS-295 | chr9-98675352-INS-295 |
| chr9-98696989-INS-52 | chr9-98696989-INS-55 |
| chr9-98699352-INS-831 | chr9-98699352-INS-831 |
| chr9-98720104-INS-70 | chr9-98720104-INS-70 |
| chr9-98730942-INS-210 | chr9-98730942-INS-210 |
| chr9-98753242-INS-65 | chr9-98753242-INS-65 |
| chr9-98759691-INS-100 | chr9-98759691-INS-112 |
| chr9-98762808-INS-78 | chr9-98762808-INS-78 |
| chr9-98784405-INS-233 | chr9-98784405-INS-229 |
| chr9-98787033-INS-70 | chr9-98787033-INS-72 |
| chr9-98803651-INS-156 | chr9-98803651-INS-155 |
| chr9-98813825-INS-163 | chr9-98813824-INS-119 |
| chr9-98898246-INS-78 | chr9-98898246-INS-80 |
| chr9-98903892-INS-228 | chr9-98903892-INS-228 |
| chr9-98909648-INS-90 | chr9-98909648-INS-92 |
| chr9-98909861-INS-1944 | chr9-98909862-INS-1944 |
| chr9-98927706-INS-53 | chr9-98927706-INS-53 |
| chr9-98943538-INS-369 | chr9-98943538-INS-369 |
| chr9-98950053-INS-735 | chr9-98950053-INS-735 |
| chr9-98960023-INS-206 | chr9-98960023-INS-206 |
| chr9-98962807-INS-338 | chr9-98962807-INS-338 |
| chr9-98963055-INS-211 | chr9-98963055-INS-210 |
| chr9-98975000-INS-215 | chr9-98975000-INS-215 |
| chr9-98981945-INS-60 | chr9-98981953-INS-64 |
| chr9-98986751-INS-97 | chr9-98986751-INS-89 |

| | |
|---|---|
| chr9-98987991-INS-72 | chr9-98987991-INS-72 |
| chr9-98999985-INS-65 | chr9-98999986-INS-50 |
| chr9-99009237-INS-107 | chr9-99009237-INS-107 |
| chr9-99025856-INS-752 | chr9-99025859-INS-740 |
| chr9-99031536-INS-244 | chr9-99031536-INS-244 |
| chr9-99032651-INS-234 | chr9-99032651-INS-238 |
| chr9-99048790-INS-51 | chr9-99048784-INS-99 |
| chr9-99050800-INS-448 | chr9-99050801-INS-448 |
| chr9-99066065-INS-198 | chr9-99066065-INS-198 |
| chr9-99071001-INS-54 | chr9-99071000-INS-60 |
| chr9-99074442-INS-8701 | chr9-99074443-INS-8714 |
| chr9-99074637-INS-231 | chr9-99074637-INS-231 |
| chr9-99083019-INS-81 | chr9-99083019-INS-78 |
| chr9-99088287-INS-60 | chr9-99088287-INS-60 |
| chr9-99108233-INS-60 | chr9-99108233-INS-60 |
| chr9-99111299-INS-218 | chr9-99111299-INS-218 |
| chr9-99112172-INS-204 | chr9-99112172-INS-205 |
| chr9-99120091-INS-189 | chr9-99120091-INS-189 |
| chr9-99123274-INS-68 | chr9-99123274-INS-68 |
| chr9-99135204-INS-269 | chr9-99135204-INS-272 |
| chr9-99188459-INS-179 | chr9-99188468-INS-200 |
| chr9-99190243-INS-561 | chr9-99190243-INS-561 |
| chr9-99208834-INS-1864 | chr9-99208835-INS-1864 |
| chr9-99233133-INS-190 | chr9-99233133-INS-191 |
| chr9-99242935-INS-172 | chr9-99242935-INS-172 |
| chr9-99253744-INS-53 | chr9-99253744-INS-53 |
| chr9-99258144-INS-73 | chr9-99258140-INS-80 |
| chr9-99288114-INS-52 | chr9-99288114-INS-52 |
| chr9-99309526-INS-670 | chr9-99309526-INS-670 |
| chr9-99331254-INS-63 | chr9-99331281-INS-74 |
| chr9-99333737-INS-58 | chr9-99333737-INS-50 |
| chr9-99347429-INS-52 | chr9-99347429-INS-52 |
| chr9-99348525-INS-73 | chr9-99348526-INS-66 |
| chr9-99360097-INS-1755 | chr9-99360098-INS-1755 |
| chr9-99364546-INS-71 | chr9-99364546-INS-83 |
| chr9-99388823-INS-200 | chr9-99388822-INS-198 |
| chr9-99389562-INS-97 | chr9-99389557-INS-74 |
| chr9-99400619-INS-226 | chr9-99400619-INS-226 |
| chr9-99407639-INS-204 | chr9-99407639-INS-204 |
| chr9-99410226-INS-203 | chr9-99410226-INS-203 |
| chr9-99416004-INS-51 | chr9-99416004-INS-51 |
| chr9-99419014-INS-202 | chr9-99419014-INS-218 |
| chr9-99433070-INS-5255 | chr9-99433071-INS-5255 |
| chr9-99435573-INS-214 | chr9-99435573-INS-214 |
| chr9-99447788-INS-135 | chr9-99447788-INS-135 |
| chr9-99452981-INS-96 | chr9-99452981-INS-102 |
| chr9-99455301-INS-205 | chr9-99455301-INS-208 |
| chr9-99473157-INS-55 | chr9-99473157-INS-55 |
| chr9-99525856-INS-66 | chr9-99525856-INS-52 |
| chr9-99526327-INS-248 | chr9-99526327-INS-248 |
| chr9-99532447-INS-203 | chr9-99532447-INS-203 |
| chr9-99550276-INS-74 | chr9-99550276-INS-74 |
| chr9-99561351-INS-2075 | chr9-99561352-INS-2076 |
| chr9-99583277-INS-92 | chr9-99583277-INS-92 |
| chr9-99586391-INS-192 | chr9-99586391-INS-201 |
| chr9-99607012-INS-52 | chr9-99607012-INS-52 |
| chr9-99609314-INS-2474 | chr9-99609315-INS-2474 |
| chr9-99616139-INS-54 | chr9-99616139-INS-50 |
| chr9-99634294-INS-66 | chr9-99634292-INS-68 |
| chr9-99636053-INS-54 | chr9-99636053-INS-55 |
| chr9-99655133-INS-474 | chr9-99655133-INS-474 |
| chr9-99657600-INS-169 | chr9-99657600-INS-185 |
| chr9-99668226-INS-67 | chr9-99668226-INS-67 |
| chr9-99671486-INS-94 | chr9-99671485-INS-96 |
| chr9-99700551-INS-52 | chr9-99700551-INS-58 |
| chr9-99718511-INS-382 | chr9-99718511-INS-388 |
| chr9-99766750-INS-111 | chr9-99766756-INS-64 |
| chr9-99776678-INS-475 | chr9-99776678-INS-475 |
| chr9-99796350-INS-54 | chr9-99796350-INS-54 |
| chr9-99819432-INS-6573 | chr9-99819433-INS-6573 |
| chr9-99849665-INS-718 | chr9-99849665-INS-718 |
| chr9-99863417-INS-135 | chr9-99863436-INS-162 |
| chr9-99871280-INS-112 | chr9-99871287-INS-72 |
| chr9-99888376-INS-221 | chr9-99888374-INS-229 |
| chr9-99911143-INS-58 | chr9-99911143-INS-58 |
| chr9-99913513-INS-105 | chr9-99913513-INS-98 |
| chr9-99920748-INS-99 | chr9-99920748-INS-99 |

| | |
|---|---|
| chr9-99937017-INS-68 | chr9-99937017-INS-68 |
| chr9-99940775-INS-3566 | chr9-99940776-INS-3566 |
| chr9-99944354-INS-560 | chr9-99944354-INS-560 |
| chr9-99977511-INS-199 | chr9-99977511-INS-199 |
| chr9-99988141-INS-176 | chr9-99988141-INS-164 |
| chr9-99994000-INS-1342 | chr9-99993985-INS-1343 |
| chr9-100000394-INS-78 | chr9-100000394-INS-86 |
| chr9-100006306-INS-93 | chr9-100006306-INS-99 |
| chr9-100024087-INS-6176 | chr9-100024088-INS-6176 |
| chr9-100054236-INS-64 | chr9-100054236-INS-66 |
| chr9-100055694-INS-510 | chr9-100055694-INS-512 |
| chr9-100060955-INS-56 | chr9-100060923-INS-52 |
| chr9-100064074-INS-2858 | chr9-100064075-INS-2858 |
| chr9-100069587-INS-1116 | chr9-100069588-INS-1115 |
| chr9-100074768-INS-102 | chr9-100074808-INS-96 |
| chr9-100086452-INS-90 | chr9-100086452-INS-90 |
| chr9-100100994-INS-656 | chr9-100100995-INS-656 |
| chr9-100126018-INS-60 | chr9-100126018-INS-54 |
| chr9-100129929-INS-72 | chr9-100129929-INS-84 |
| chr9-100136298-INS-52 | chr9-100136298-INS-50 |
| chr9-100137828-INS-253 | chr9-100137828-INS-253 |
| chr9-100143917-INS-90 | chr9-100143904-INS-131 |
| chr9-100185435-INS-683 | chr9-100185435-INS-683 |
| chr9-100208707-INS-506 | chr9-100208707-INS-506 |
| chr9-100221838-INS-6440 | chr9-100221839-INS-6440 |
| chr9-100229125-INS-88 | chr9-100229125-INS-92 |
| chr9-100251929-INS-54 | chr9-100251930-INS-64 |
| chr9-100254285-INS-264 | chr9-100254285-INS-264 |
| chr9-100257489-INS-109 | chr9-100257489-INS-109 |
| chr9-100267593-INS-63 | chr9-100267594-INS-59 |
| chr9-100280403-INS-155 | chr9-100280403-INS-155 |
| chr9-100281839-INS-124 | chr9-100281857-INS-115 |
| chr9-100283584-INS-64 | chr9-100283584-INS-64 |
| chr9-100289273-INS-202 | chr9-100289273-INS-202 |
| chr9-100310735-INS-58 | chr9-100310735-INS-54 |
| chr9-100312973-INS-52 | chr9-100312973-INS-69 |
| chr9-100318398-INS-52 | chr9-100318398-INS-52 |
| chr9-100342824-INS-106 | chr9-100342824-INS-106 |
| chr9-100345336-INS-78 | chr9-100345305-INS-80 |
| chr9-100345794-INS-108 | chr9-100345794-INS-120 |
| chr9-100352786-INS-65 | chr9-100352808-INS-78 |
| chr9-100360891-INS-133 | chr9-100360889-INS-140 |
| chr9-100369509-INS-207 | chr9-100369509-INS-207 |
| chr9-100375611-INS-407 | chr9-100375611-INS-407 |
| chr9-100424952-INS-394 | chr9-100424952-INS-394 |
| chr9-100429647-INS-54 | chr9-100429670-INS-56 |
| chr9-100449848-INS-134 | chr9-100449848-INS-134 |
| chr9-100454750-INS-244 | chr9-100454750-INS-245 |
| chr9-100455872-INS-136 | chr9-100455872-INS-136 |
| chr9-100477771-INS-187 | chr9-100477771-INS-163 |
| chr9-100496681-INS-210 | chr9-100496682-INS-210 |
| chr9-100499653-INS-152 | chr9-100499653-INS-152 |
| chr9-100508107-INS-151 | chr9-100508107-INS-150 |
| chr9-100520367-INS-57 | chr9-100520350-INS-56 |
| chr9-100531345-INS-142 | chr9-100531345-INS-121 |
| chr9-100533063-INS-430 | chr9-100533067-INS-448 |
| chr9-100538876-INS-215 | chr9-100538876-INS-215 |
| chr9-100556480-INS-69 | chr9-100556470-INS-77 |
| chr9-100579830-INS-120 | chr9-100579830-INS-120 |
| chr9-100636976-INS-66 | chr9-100636976-INS-82 |
| chr9-100672416-INS-62 | chr9-100672416-INS-67 |
| chr9-100677012-INS-3972 | chr9-100677013-INS-3971 |
| chr9-100677950-INS-971 | chr9-100677951-INS-971 |
| chr9-100702211-INS-557 | chr9-100702211-INS-558 |
| chr9-100714883-INS-65 | chr9-100714889-INS-63 |
| chr9-100719644-INS-129 | chr9-100719644-INS-129 |
| chr9-100742526-INS-65 | chr9-100742525-INS-86 |
| chr9-100747619-INS-1900 | chr9-100747620-INS-1900 |
| chr9-100776208-INS-219 | chr9-100776208-INS-219 |
| chr9-100782680-INS-686 | chr9-100782707-INS-675 |
| chr9-100782805-INS-2484 | chr9-100782806-INS-2484 |
| chr9-100794992-INS-63 | chr9-100794992-INS-66 |
| chr9-100800992-INS-206 | chr9-100800992-INS-207 |
| chr9-100821751-INS-204 | chr9-100821751-INS-204 |
| chr9-100848281-INS-73 | chr9-100848318-INS-74 |
| chr9-100853083-INS-193 | chr9-100853083-INS-184 |
| chr9-100854047-INS-4525 | chr9-100854048-INS-4525 |

| | |
|---|---|
| chr9-100861595-INS-498 | chr9-100861595-INS-498 |
| chr9-100868551-INS-1696 | chr9-100868552-INS-1696 |
| chr9-100909290-INS-210 | chr9-100909289-INS-207 |
| chr9-100917266-INS-173 | chr9-100917266-INS-173 |
| chr9-100918976-INS-210 | chr9-100918976-INS-210 |
| chr9-100930963-INS-168 | chr9-100931009-INS-170 |
| chr9-100934804-INS-397 | chr9-100934804-INS-397 |
| chr9-100938449-INS-356 | chr9-100938449-INS-356 |
| chr9-100950018-INS-55 | chr9-100950028-INS-68 |
| chr9-100952558-INS-66 | chr9-100952558-INS-66 |
| chr9-100970165-INS-210 | chr9-100970165-INS-210 |
| chr9-100976480-INS-211 | chr9-100976480-INS-212 |
| chr9-100993254-INS-216 | chr9-100993254-INS-216 |
| chr9-101001989-INS-190 | chr9-101001989-INS-190 |
| chr9-101004289-INS-75 | chr9-101004289-INS-75 |
| chr9-101039538-INS-58 | chr9-101039538-INS-64 |
| chr9-101049690-INS-210 | chr9-101049690-INS-210 |
| chr9-101063671-INS-247 | chr9-101063671-INS-247 |
| chr9-101070757-INS-52 | chr9-101070757-INS-54 |
| chr9-101079157-INS-171 | chr9-101079181-INS-141 |
| chr9-101107254-INS-72 | chr9-101107254-INS-76 |
| chr9-101126143-INS-205 | chr9-101126143-INS-205 |
| chr9-101135486-INS-198 | chr9-101135486-INS-198 |
| chr9-101141664-INS-56 | chr9-101141664-INS-56 |
| chr9-101149672-INS-56 | chr9-101149672-INS-56 |
| chr9-101162383-INS-98 | chr9-101162383-INS-98 |
| chr9-101176480-INS-90 | chr9-101176514-INS-60 |
| chr9-101180405-INS-92 | chr9-101180405-INS-92 |
| chr9-101194382-INS-75 | chr9-101194382-INS-78 |
| chr9-101203267-INS-52 | chr9-101203267-INS-52 |
| chr9-101230407-INS-54 | chr9-101230407-INS-54 |
| chr9-101268076-INS-64 | chr9-101268077-INS-64 |
| chr9-101286301-INS-220 | chr9-101286301-INS-220 |
| chr9-101325492-INS-2532 | chr9-101325493-INS-2532 |
| chr9-101346113-INS-79 | chr9-101346113-INS-53 |
| chr9-101364254-INS-3050 | chr9-101364255-INS-3051 |
| chr9-101367228-INS-138 | chr9-101367248-INS-136 |
| chr9-101368805-INS-55 | chr9-101368803-INS-62 |
| chr9-101372264-INS-315 | chr9-101372266-INS-309 |
| chr9-101378935-INS-5313 | chr9-101378936-INS-5313 |
| chr9-101383607-INS-110 | chr9-101383609-INS-112 |
| chr9-101390356-INS-54 | chr9-101390356-INS-64 |
| chr9-101390758-INS-1427 | chr9-101390759-INS-1431 |
| chr9-101410401-INS-1072 | chr9-101410402-INS-1070 |
| chr9-101412726-INS-3206 | chr9-101412727-INS-3204 |
| chr9-101413138-INS-59 | chr9-101413138-INS-59 |
| chr9-101437411-INS-128 | chr9-101437411-INS-136 |
| chr9-101450838-INS-350 | chr9-101450838-INS-350 |
| chr9-101457774-INS-131 | chr9-101457774-INS-131 |
| chr9-101487461-INS-84 | chr9-101487458-INS-110 |
| chr9-101488809-INS-65 | chr9-101488809-INS-68 |
| chr9-101528646-INS-597 | chr9-101528646-INS-597 |
| chr9-101541943-INS-192 | chr9-101541943-INS-195 |
| chr9-101547471-INS-225 | chr9-101547471-INS-225 |
| chr9-101561247-INS-212 | chr9-101561247-INS-212 |
| chr9-101600559-INS-6657 | chr9-101600560-INS-6657 |
| chr9-101606383-INS-6219 | chr9-101606384-INS-6213 |
| chr9-101613485-INS-68 | chr9-101613485-INS-62 |
| chr9-101632224-INS-359 | chr9-101632224-INS-359 |
| chr9-101667218-INS-501 | chr9-101667218-INS-497 |
| chr9-101681563-INS-5100 | chr9-101681564-INS-5098 |
| chr9-101683537-INS-342 | chr9-101683538-INS-342 |
| chr9-101722687-INS-122 | chr9-101722687-INS-146 |
| chr9-101742671-INS-69 | chr9-101742671-INS-69 |
| chr9-101748317-INS-465 | chr9-101748317-INS-465 |
| chr9-101750940-INS-151 | chr9-101750940-INS-151 |
| chr9-101755657-INS-56 | chr9-101755657-INS-78 |
| chr9-101790385-INS-6285 | chr9-101790386-INS-6285 |
| chr9-101813198-INS-267 | chr9-101813198-INS-267 |
| chr9-101837481-INS-6055 | chr9-101837482-INS-6054 |
| chr9-101844096-INS-5992 | chr9-101844097-INS-5992 |
| chr9-101889004-INS-77 | chr9-101889004-INS-93 |
| chr9-101890662-INS-71 | chr9-101890662-INS-83 |
| chr9-101903410-INS-752 | chr9-101903410-INS-752 |
| chr9-101927469-INS-98 | chr9-101927469-INS-80 |
| chr9-101956819-INS-65 | chr9-101956819-INS-55 |
| chr9-101968410-INS-5114 | chr9-101968411-INS-5114 |

| | |
|---|---|
| chr9-101999194-INS-1582 | chr9-101999195-INS-1575 |
| chr9-102014310-INS-357 | chr9-102014310-INS-357 |
| chr9-102022413-INS-728 | chr9-102022413-INS-726 |
| chr9-102040525-INS-67 | chr9-102040525-INS-71 |
| chr9-102044106-INS-71 | chr9-102044106-INS-85 |
| chr9-102052998-INS-120 | chr9-102052999-INS-120 |
| chr9-102143468-INS-85 | chr9-102143468-INS-61 |
| chr9-102185353-INS-1448 | chr9-102185354-INS-1460 |
| chr9-102202645-INS-108 | chr9-102202645-INS-124 |
| chr9-102204341-INS-51 | chr9-102204341-INS-52 |
| chr9-102209614-INS-242 | chr9-102209614-INS-242 |
| chr9-102223838-INS-56 | chr9-102223838-INS-56 |
| chr9-102271144-INS-160 | chr9-102271149-INS-128 |
| chr9-102305919-INS-52 | chr9-102305919-INS-76 |
| chr9-102309219-INS-52 | chr9-102309192-INS-72 |
| chr9-102310962-INS-96 | chr9-102310962-INS-65 |
| chr9-102499610-INS-488 | chr9-102499610-INS-488 |
| chr9-102504513-INS-92 | chr9-102504513-INS-92 |
| chr9-102530965-INS-80 | chr9-102530965-INS-81 |
| chr9-102537458-INS-209 | chr9-102537458-INS-207 |
| chr9-102542332-INS-114 | chr9-102542332-INS-114 |
| chr9-102546549-INS-63 | chr9-102546550-INS-66 |
| chr9-102548309-INS-436 | chr9-102548309-INS-436 |
| chr9-102564040-INS-227 | chr9-102564040-INS-227 |
| chr9-102600597-INS-202 | chr9-102600597-INS-200 |
| chr9-102652098-INS-75 | chr9-102652098-INS-51 |
| chr9-102653913-INS-135 | chr9-102653913-INS-104 |
| chr9-102657847-INS-67 | chr9-102657847-INS-93 |
| chr9-102690987-INS-1242 | chr9-102690988-INS-1242 |
| chr9-102708428-INS-427 | chr9-102708428-INS-427 |
| chr9-102715102-INS-74 | chr9-102715102-INS-74 |
| chr9-102729217-INS-484 | chr9-102729217-INS-484 |
| chr9-102770996-INS-207 | chr9-102770996-INS-207 |
| chr9-102781925-INS-197 | chr9-102781925-INS-197 |
| chr9-102789340-INS-196 | chr9-102789340-INS-197 |
| chr9-102809876-INS-104 | chr9-102809876-INS-85 |
| chr9-102812898-INS-359 | chr9-102812898-INS-359 |
| chr9-102813461-INS-356 | chr9-102813461-INS-356 |
| chr9-102814786-INS-208 | chr9-102814786-INS-207 |
| chr9-102851797-INS-134 | chr9-102851797-INS-114 |
| chr9-102862238-INS-235 | chr9-102862238-INS-236 |
| chr9-102889095-INS-56 | chr9-102889072-INS-104 |
| chr9-102969013-INS-71 | chr9-102969013-INS-52 |
| chr9-102981066-INS-98 | chr9-102981072-INS-76 |
| chr9-103023446-INS-52 | chr9-103023446-INS-56 |
| chr9-103035357-INS-234 | chr9-103035357-INS-234 |
| chr9-103037310-INS-1060 | chr9-103037311-INS-1037 |
| chr9-103051740-INS-373 | chr9-103051740-INS-373 |
| chr9-103072618-INS-651 | chr9-103072620-INS-657 |
| chr9-103082530-INS-221 | chr9-103082530-INS-222 |
| chr9-103083400-INS-123 | chr9-103083400-INS-123 |
| chr9-103084688-INS-60 | chr9-103084688-INS-50 |
| chr9-103117809-INS-60 | chr9-103117809-INS-98 |
| chr9-103124794-INS-129 | chr9-103124794-INS-129 |
| chr9-103137447-INS-54 | chr9-103137485-INS-54 |
| chr9-103138778-INS-71 | chr9-103138782-INS-71 |
| chr9-103150975-INS-540 | chr9-103150975-INS-540 |
| chr9-103170851-INS-197 | chr9-103170851-INS-197 |
| chr9-103172955-INS-98 | chr9-103172955-INS-96 |
| chr9-103187211-INS-420 | chr9-103187246-INS-421 |
| chr9-103187217-INS-2704 | chr9-103187218-INS-2700 |
| chr9-103212771-INS-158 | chr9-103212771-INS-154 |
| chr9-103214454-INS-203 | chr9-103214454-INS-203 |
| chr9-103215596-INS-201 | chr9-103215596-INS-201 |
| chr9-103234492-INS-215 | chr9-103234492-INS-216 |
| chr9-103268404-INS-122 | chr9-103268404-INS-132 |
| chr9-103271219-INS-54 | chr9-103271235-INS-76 |
| chr9-103276244-INS-66 | chr9-103276244-INS-66 |
| chr9-103288811-INS-68 | chr9-103288811-INS-72 |
| chr9-103305563-INS-4780 | chr9-103305564-INS-4780 |
| chr9-103337916-INS-90 | chr9-103337916-INS-72 |
| chr9-103338996-INS-184 | chr9-103338996-INS-184 |
| chr9-103349258-INS-4413 | chr9-103349259-INS-4413 |
| chr9-103352525-INS-81 | chr9-103352525-INS-81 |
| chr9-103375071-INS-206 | chr9-103375072-INS-206 |
| chr9-103392186-INS-182 | chr9-103392186-INS-183 |
| chr9-103399803-INS-1890 | chr9-103399804-INS-1890 |

| | |
|---|---|
| chr9-103404854-INS-2681 | chr9-103404870-INS-2684 |
| chr9-103460179-INS-95 | chr9-103460179-INS-95 |
| chr9-103461202-INS-54 | chr9-103461202-INS-54 |
| chr9-103484919-INS-87 | chr9-103484919-INS-53 |
| chr9-103498623-INS-72 | chr9-103498623-INS-72 |
| chr9-103505491-INS-54 | chr9-103505491-INS-54 |
| chr9-103523628-INS-4419 | chr9-103523629-INS-4419 |
| chr9-103537934-INS-88 | chr9-103537934-INS-88 |
| chr9-103548002-INS-136 | chr9-103548002-INS-136 |
| chr9-103571407-INS-74 | chr9-103571431-INS-50 |
| chr9-103576690-INS-64 | chr9-103576690-INS-64 |
| chr9-103598125-INS-92 | chr9-103598125-INS-92 |
| chr9-103600691-INS-108 | chr9-103600701-INS-64 |
| chr9-103617911-INS-56 | chr9-103617911-INS-56 |
| chr9-103642226-INS-55 | chr9-103642242-INS-66 |
| chr9-103652670-INS-51 | chr9-103652670-INS-51 |
| chr9-103677236-INS-60 | chr9-103677236-INS-60 |
| chr9-103705796-INS-166 | chr9-103705796-INS-166 |
| chr9-103710263-INS-209 | chr9-103710263-INS-209 |
| chr9-103714870-INS-6355 | chr9-103714871-INS-6357 |
| chr9-103763290-INS-107 | chr9-103763290-INS-107 |
| chr9-103764672-INS-128 | chr9-103764673-INS-136 |
| chr9-103784829-INS-58 | chr9-103784829-INS-56 |
| chr9-103819204-INS-7205 | chr9-103819205-INS-7206 |
| chr9-103830895-INS-144 | chr9-103830895-INS-144 |
| chr9-103878636-INS-188 | chr9-103878636-INS-188 |
| chr9-103888562-INS-590 | chr9-103888560-INS-582 |
| chr9-103893397-INS-198 | chr9-103893397-INS-198 |
| chr9-103897032-INS-66 | chr9-103897032-INS-66 |
| chr9-103908015-INS-108 | chr9-103908015-INS-108 |
| chr9-103938727-INS-190 | chr9-103938727-INS-190 |
| chr9-103943530-INS-84 | chr9-103943530-INS-86 |
| chr9-103995036-INS-90 | chr9-103995023-INS-91 |
| chr9-104000605-INS-232 | chr9-104000605-INS-233 |
| chr9-104003269-INS-194 | chr9-104003269-INS-194 |
| chr9-104005583-INS-430 | chr9-104005583-INS-430 |
| chr9-104008213-INS-64 | chr9-104008213-INS-64 |
| chr9-104009569-INS-207 | chr9-104009525-INS-226 |
| chr9-104028092-INS-325 | chr9-104028092-INS-346 |
| chr9-104036643-INS-176 | chr9-104036643-INS-179 |
| chr9-104094887-INS-209 | chr9-104094887-INS-209 |
| chr9-104128126-INS-179 | chr9-104128126-INS-193 |
| chr9-104144224-INS-273 | chr9-104144224-INS-281 |
| chr9-104151355-INS-2822 | chr9-104151356-INS-2822 |
| chr9-104184578-INS-205 | chr9-104184578-INS-203 |
| chr9-104190216-INS-54 | chr9-104190216-INS-52 |
| chr9-104192690-INS-165 | chr9-104192690-INS-165 |
| chr9-104204236-INS-176 | chr9-104204236-INS-176 |
| chr9-104231134-INS-78 | chr9-104231134-INS-80 |
| chr9-104233825-INS-330 | chr9-104233825-INS-328 |
| chr9-104239404-INS-72 | chr9-104239417-INS-52 |
| chr9-104268701-INS-52 | chr9-104268701-INS-60 |
| chr9-104281831-INS-418 | chr9-104281836-INS-436 |
| chr9-104300146-INS-500 | chr9-104300146-INS-500 |
| chr9-104304153-INS-60 | chr9-104304153-INS-60 |
| chr9-104306557-INS-202 | chr9-104306556-INS-185 |
| chr9-104324314-INS-218 | chr9-104324314-INS-218 |
| chr9-104334594-INS-76 | chr9-104334594-INS-78 |
| chr9-104338359-INS-52 | chr9-104338362-INS-50 |
| chr9-104355283-INS-338 | chr9-104355283-INS-338 |
| chr9-104381369-INS-507 | chr9-104381369-INS-509 |
| chr9-104383596-INS-408 | chr9-104383596-INS-408 |
| chr9-104406225-INS-920 | chr9-104406224-INS-915 |
| chr9-104409075-INS-924 | chr9-104409075-INS-924 |
| chr9-104420929-INS-169 | chr9-104420929-INS-167 |
| chr9-104455190-INS-87 | chr9-104455190-INS-89 |
| chr9-104480053-INS-55 | chr9-104480053-INS-60 |
| chr9-104482413-INS-119 | chr9-104482413-INS-119 |
| chr9-104483781-INS-796 | chr9-104483781-INS-805 |
| chr9-104486081-INS-52 | chr9-104486081-INS-54 |
| chr9-104501711-INS-400 | chr9-104501711-INS-400 |
| chr9-104506405-INS-84 | chr9-104506405-INS-84 |
| chr9-104540564-INS-157 | chr9-104540564-INS-155 |
| chr9-104551505-INS-401 | chr9-104551505-INS-401 |
| chr9-104564276-INS-54 | chr9-104564276-INS-54 |
| chr9-104567524-INS-573 | chr9-104567524-INS-573 |
| chr9-104616690-INS-64 | chr9-104616690-INS-68 |

| | |
|---|---|
| chr9-104641333-INS-93 | chr9-104641333-INS-93 |
| chr9-104647943-INS-55 | chr9-104647970-INS-55 |
| chr9-104658142-INS-64 | chr9-104658138-INS-56 |
| chr9-104660215-INS-143 | chr9-104660237-INS-101 |
| chr9-104666815-INS-95 | chr9-104666815-INS-95 |
| chr9-104682989-INS-557 | chr9-104682989-INS-557 |
| chr9-104690388-INS-60 | chr9-104690388-INS-60 |
| chr9-104699367-INS-94 | chr9-104699369-INS-96 |
| chr9-104708104-INS-539 | chr9-104708104-INS-539 |
| chr9-104712769-INS-199 | chr9-104712769-INS-199 |
| chr9-104741465-INS-1710 | chr9-104741466-INS-1698 |
| chr9-104746447-INS-2255 | chr9-104746448-INS-2255 |
| chr9-104747732-INS-74 | chr9-104747732-INS-74 |
| chr9-104755836-INS-451 | chr9-104755836-INS-451 |
| chr9-104804407-INS-534 | chr9-104804407-INS-534 |
| chr9-104823358-INS-190 | chr9-104823358-INS-190 |
| chr9-104844414-INS-273 | chr9-104844417-INS-283 |
| chr9-104859753-INS-195 | chr9-104859753-INS-195 |
| chr9-104861472-INS-273 | chr9-104861472-INS-278 |
| chr9-104885116-INS-56 | chr9-104885131-INS-62 |
| chr9-104886780-INS-51 | chr9-104886771-INS-69 |
| chr9-104929501-INS-84 | chr9-104929474-INS-123 |
| chr9-104962621-INS-201 | chr9-104962621-INS-201 |
| chr9-104986137-INS-71 | chr9-104986137-INS-92 |
| chr9-105010293-INS-53 | chr9-105010305-INS-51 |
| chr9-105015560-INS-61 | chr9-105015560-INS-61 |
| chr9-105019453-INS-82 | chr9-105019453-INS-82 |
| chr9-105028034-INS-245 | chr9-105028034-INS-245 |
| chr9-105030501-INS-3538 | chr9-105030502-INS-3539 |
| chr9-105032425-INS-197 | chr9-105032425-INS-197 |
| chr9-105039659-INS-287 | chr9-105039659-INS-287 |
| chr9-105045416-INS-186 | chr9-105045416-INS-186 |
| chr9-105046525-INS-364 | chr9-105046525-INS-364 |
| chr9-105052374-INS-56 | chr9-105052374-INS-56 |
| chr9-105059288-INS-161 | chr9-105059288-INS-198 |
| chr9-105067073-INS-66 | chr9-105067084-INS-62 |
| chr9-105123847-INS-66 | chr9-105123860-INS-54 |
| chr9-105128039-INS-86 | chr9-105128039-INS-86 |
| chr9-105130359-INS-56 | chr9-105130359-INS-54 |
| chr9-105135866-INS-52 | chr9-105135866-INS-50 |
| chr9-105158055-INS-61 | chr9-105158055-INS-66 |
| chr9-105178982-INS-54 | chr9-105178982-INS-54 |
| chr9-105198524-INS-2768 | chr9-105198525-INS-2768 |
| chr9-105230487-INS-992 | chr9-105230487-INS-991 |
| chr9-105230911-INS-61 | chr9-105230911-INS-61 |
| chr9-105232201-INS-85 | chr9-105232201-INS-85 |
| chr9-105242104-INS-143 | chr9-105242144-INS-143 |
| chr9-105263240-INS-86 | chr9-105263240-INS-86 |
| chr9-105292994-INS-56 | chr9-105292994-INS-54 |
| chr9-105338074-INS-54 | chr9-105338074-INS-56 |
| chr9-105350067-INS-53 | chr9-105350067-INS-53 |
| chr9-105385895-INS-75 | chr9-105385895-INS-75 |
| chr9-105433438-INS-82 | chr9-105433438-INS-82 |
| chr9-105442528-INS-174 | chr9-105442528-INS-185 |
| chr9-105443855-INS-1861 | chr9-105443856-INS-1861 |
| chr9-105454006-INS-51 | chr9-105454006-INS-51 |
| chr9-105513567-INS-216 | chr9-105513567-INS-216 |
| chr9-105518513-INS-173 | chr9-105518513-INS-173 |
| chr9-105524026-INS-109 | chr9-105524026-INS-109 |
| chr9-105529202-INS-210 | chr9-105529202-INS-212 |
| chr9-105531844-INS-687 | chr9-105531844-INS-687 |
| chr9-105534371-INS-327 | chr9-105534371-INS-327 |
| chr9-105542621-INS-62 | chr9-105542621-INS-62 |
| chr9-105553382-INS-374 | chr9-105553382-INS-374 |
| chr9-105575484-INS-60 | chr9-105575484-INS-50 |
| chr9-105587460-INS-170 | chr9-105587460-INS-170 |
| chr9-105623228-INS-556 | chr9-105623228-INS-556 |
| chr9-105625253-INS-84 | chr9-105625253-INS-84 |
| chr9-105636934-INS-66 | chr9-105636934-INS-66 |
| chr9-105663537-INS-58 | chr9-105663537-INS-60 |
| chr9-105665363-INS-132 | chr9-105665363-INS-134 |
| chr9-105689523-INS-68 | chr9-105689523-INS-68 |
| chr9-105718148-INS-411 | chr9-105718148-INS-411 |
| chr9-105731100-INS-222 | chr9-105731100-INS-223 |
| chr9-105735785-INS-118 | chr9-105735785-INS-118 |
| chr9-105745234-INS-184 | chr9-105745242-INS-188 |
| chr9-105761245-INS-2652 | chr9-105761252-INS-2657 |

| | |
|---|---|
| chr9-105766713-INS-65 | chr9-105766713-INS-67 |
| chr9-105813086-INS-7495 | chr9-105813087-INS-7495 |
| chr9-105813802-INS-60 | chr9-105813802-INS-60 |
| chr9-105819660-INS-152 | chr9-105819661-INS-152 |
| chr9-105849870-INS-197 | chr9-105849870-INS-197 |
| chr9-105899883-INS-275 | chr9-105899883-INS-304 |
| chr9-105903425-INS-336 | chr9-105903425-INS-336 |
| chr9-105911200-INS-56 | chr9-105911200-INS-56 |
| chr9-105941568-INS-223 | chr9-105941568-INS-224 |
| chr9-105946084-INS-68 | chr9-105946084-INS-68 |
| chr9-105949015-INS-78 | chr9-105949015-INS-78 |
| chr9-105950557-INS-141 | chr9-105950557-INS-148 |
| chr9-105950557-INS-64 | chr9-105950539-INS-55 |
| chr9-105978202-INS-134 | chr9-105978202-INS-135 |
| chr9-105993677-INS-88 | chr9-105993708-INS-90 |
| chr9-105995324-INS-192 | chr9-105995324-INS-200 |
| chr9-105998625-INS-69 | chr9-105998625-INS-69 |
| chr9-106005432-INS-54 | chr9-106005432-INS-54 |
| chr9-106005820-INS-400 | chr9-106005820-INS-400 |
| chr9-106008998-INS-103 | chr9-106008998-INS-103 |
| chr9-106039500-INS-1103 | chr9-106039501-INS-1103 |
| chr9-106045003-INS-81 | chr9-106045003-INS-71 |
| chr9-106051154-INS-170 | chr9-106051154-INS-172 |
| chr9-106057411-INS-203 | chr9-106057409-INS-202 |
| chr9-106084969-INS-95 | chr9-106084969-INS-95 |
| chr9-106096703-INS-118 | chr9-106096703-INS-118 |
| chr9-106119917-INS-252 | chr9-106119917-INS-252 |
| chr9-106125767-INS-210 | chr9-106125767-INS-210 |
| chr9-106130461-INS-175 | chr9-106130461-INS-175 |
| chr9-106135496-INS-53 | chr9-106135496-INS-53 |
| chr9-106155050-INS-131 | chr9-106155050-INS-129 |
| chr9-106167029-INS-70 | chr9-106167029-INS-70 |
| chr9-106227284-INS-82 | chr9-106227280-INS-86 |
| chr9-106256334-INS-186 | chr9-106256334-INS-187 |
| chr9-106283200-INS-55 | chr9-106283200-INS-55 |
| chr9-106316693-INS-129 | chr9-106316693-INS-129 |
| chr9-106346007-INS-64 | chr9-106346007-INS-64 |
| chr9-106359371-INS-52 | chr9-106359371-INS-52 |
| chr9-106382098-INS-188 | chr9-106382098-INS-188 |
| chr9-106384773-INS-210 | chr9-106384773-INS-210 |
| chr9-106390958-INS-62 | chr9-106390958-INS-62 |
| chr9-106396892-INS-206 | chr9-106396892-INS-206 |
| chr9-106423962-INS-189 | chr9-106423962-INS-189 |
| chr9-106478883-INS-76 | chr9-106478883-INS-76 |
| chr9-106503059-INS-54 | chr9-106503059-INS-50 |
| chr9-106505886-INS-300 | chr9-106505886-INS-300 |
| chr9-106508620-INS-50 | chr9-106508619-INS-57 |
| chr9-106513917-INS-458 | chr9-106513965-INS-412 |
| chr9-106539277-INS-6442 | chr9-106539278-INS-6442 |
| chr9-106543025-INS-4519 | chr9-106543026-INS-4519 |
| chr9-106561012-INS-53 | chr9-106561012-INS-53 |
| chr9-106572061-INS-205 | chr9-106572061-INS-206 |
| chr9-106583850-INS-203 | chr9-106583850-INS-203 |
| chr9-106607861-INS-196 | chr9-106607861-INS-196 |
| chr9-106620405-INS-203 | chr9-106620405-INS-203 |
| chr9-106626430-INS-104 | chr9-106626430-INS-103 |
| chr9-106672610-INS-2761 | chr9-106672611-INS-2760 |
| chr9-106680262-INS-559 | chr9-106680262-INS-559 |
| chr9-106680869-INS-98 | chr9-106680870-INS-54 |
| chr9-106686873-INS-230 | chr9-106686873-INS-230 |
| chr9-106696815-INS-168 | chr9-106696815-INS-171 |
| chr9-106721279-INS-115 | chr9-106721279-INS-115 |
| chr9-106739725-INS-206 | chr9-106739725-INS-206 |
| chr9-106755792-INS-74 | chr9-106755792-INS-52 |
| chr9-106781381-INS-1787 | chr9-106781382-INS-1787 |
| chr9-106860606-INS-8641 | chr9-106860607-INS-8642 |
| chr9-106869786-INS-60 | chr9-106869786-INS-60 |
| chr9-106910446-INS-188 | chr9-106910446-INS-188 |
| chr9-106922058-INS-67 | chr9-106922058-INS-67 |
| chr9-106923421-INS-656 | chr9-106923421-INS-656 |
| chr9-106930967-INS-65 | chr9-106930967-INS-65 |
| chr9-106971745-INS-213 | chr9-106971745-INS-213 |
| chr9-106985310-INS-51 | chr9-106985310-INS-51 |
| chr9-106988761-INS-111 | chr9-106988761-INS-111 |
| chr9-107001704-INS-617 | chr9-107001691-INS-619 |
| chr9-107023712-INS-190 | chr9-107023712-INS-190 |
| chr9-107028330-INS-335 | chr9-107028330-INS-335 |

| | |
|---|---|
| chr9-107040300-INS-6686 | chr9-107040301-INS-6686 |
| chr9-107066569-INS-52 | chr9-107066569-INS-52 |
| chr9-107087095-INS-227 | chr9-107087095-INS-227 |
| chr9-107097285-INS-178 | chr9-107097287-INS-211 |
| chr9-107102512-INS-70 | chr9-107102512-INS-72 |
| chr9-107104388-INS-1546 | chr9-107104389-INS-1545 |
| chr9-107106870-INS-128 | chr9-107106870-INS-129 |
| chr9-107110077-INS-100 | chr9-107110077-INS-102 |
| chr9-107129790-INS-94 | chr9-107129790-INS-94 |
| chr9-107134965-INS-185 | chr9-107134965-INS-185 |
| chr9-107141793-INS-54 | chr9-107141793-INS-54 |
| chr9-107172413-INS-52 | chr9-107172413-INS-75 |
| chr9-107191555-INS-52 | chr9-107191555-INS-56 |
| chr9-107241239-INS-63 | chr9-107241240-INS-63 |
| chr9-107242450-INS-291 | chr9-107242450-INS-291 |
| chr9-107269882-INS-508 | chr9-107269882-INS-508 |
| chr9-107319647-INS-66 | chr9-107319639-INS-78 |
| chr9-107355956-INS-220 | chr9-107355956-INS-220 |
| chr9-107374075-INS-221 | chr9-107374075-INS-222 |
| chr9-107394939-INS-155 | chr9-107394939-INS-155 |
| chr9-107395928-INS-422 | chr9-107395928-INS-428 |
| chr9-107453697-INS-90 | chr9-107453697-INS-90 |
| chr9-107472780-INS-207 | chr9-107472780-INS-205 |
| chr9-107498839-INS-201 | chr9-107498839-INS-201 |
| chr9-107503203-INS-158 | chr9-107503203-INS-158 |
| chr9-107506289-INS-173 | chr9-107506334-INS-135 |
| chr9-107609345-INS-55 | chr9-107609312-INS-65 |
| chr9-107614119-INS-2207 | chr9-107614120-INS-2199 |
| chr9-107620868-INS-464 | chr9-107620868-INS-464 |
| chr9-107649720-INS-621 | chr9-107649720-INS-622 |
| chr9-107651944-INS-214 | chr9-107651944-INS-215 |
| chr9-107655694-INS-493 | chr9-107655694-INS-494 |
| chr9-107662871-INS-394 | chr9-107662871-INS-393 |
| chr9-107668774-INS-57 | chr9-107668774-INS-57 |
| chr9-107694683-INS-93 | chr9-107694687-INS-77 |
| chr9-107698556-INS-89 | chr9-107698567-INS-61 |
| chr9-107701082-INS-210 | chr9-107701082-INS-210 |
| chr9-107751955-INS-174 | chr9-107751955-INS-175 |
| chr9-107756353-INS-62 | chr9-107756353-INS-62 |
| chr9-107841363-INS-68 | chr9-107841363-INS-68 |
| chr9-107889881-INS-183 | chr9-107889881-INS-183 |
| chr9-107897702-INS-186 | chr9-107897702-INS-172 |
| chr9-107906003-INS-117 | chr9-107906003-INS-114 |
| chr9-107950283-INS-100 | chr9-107950283-INS-100 |
| chr9-108002296-INS-128 | chr9-108002296-INS-128 |
| chr9-108012902-INS-162 | chr9-108012868-INS-188 |
| chr9-108032836-INS-181 | chr9-108032836-INS-176 |
| chr9-108054150-INS-5079 | chr9-108054151-INS-5079 |
| chr9-108061884-INS-233 | chr9-108061884-INS-232 |
| chr9-108076839-INS-150 | chr9-108076839-INS-150 |
| chr9-108096819-INS-208 | chr9-108096819-INS-206 |
| chr9-108100650-INS-163 | chr9-108100650-INS-163 |
| chr9-108103926-INS-64 | chr9-108103926-INS-67 |
| chr9-108116641-INS-122 | chr9-108116641-INS-122 |
| chr9-108122469-INS-585 | chr9-108122469-INS-557 |
| chr9-108123357-INS-60 | chr9-108123357-INS-56 |
| chr9-108127266-INS-93 | chr9-108127266-INS-93 |
| chr9-108130792-INS-236 | chr9-108130792-INS-235 |
| chr9-108132082-INS-68 | chr9-108132082-INS-68 |
| chr9-108157116-INS-108 | chr9-108157116-INS-135 |
| chr9-108159644-INS-82 | chr9-108159640-INS-85 |
| chr9-108185176-INS-175 | chr9-108185176-INS-175 |
| chr9-108212169-INS-201 | chr9-108212169-INS-201 |
| chr9-108233040-INS-292 | chr9-108233040-INS-305 |
| chr9-108259510-INS-463 | chr9-108259510-INS-463 |
| chr9-108277670-INS-54 | chr9-108277670-INS-54 |
| chr9-108289256-INS-215 | chr9-108289256-INS-215 |
| chr9-108301981-INS-62 | chr9-108301980-INS-89 |
| chr9-108302950-INS-210 | chr9-108302957-INS-209 |
| chr9-108319574-INS-204 | chr9-108319574-INS-204 |
| chr9-108320506-INS-62 | chr9-108320506-INS-66 |
| chr9-108356911-INS-355 | chr9-108356911-INS-357 |
| chr9-108387876-INS-72 | chr9-108387876-INS-72 |
| chr9-108409890-INS-215 | chr9-108409890-INS-215 |
| chr9-108413920-INS-147 | chr9-108413924-INS-148 |
| chr9-108422034-INS-127 | chr9-108422033-INS-127 |
| chr9-108467955-INS-179 | chr9-108467955-INS-178 |

| | |
|---|---|
| chr9-108469887-INS-58 | chr9-108469887-INS-58 |
| chr9-108472123-INS-192 | chr9-108472123-INS-192 |
| chr9-108479442-INS-626 | chr9-108479442-INS-627 |
| chr9-108489946-INS-72 | chr9-108489946-INS-72 |
| chr9-108494732-INS-175 | chr9-108494732-INS-175 |
| chr9-108525289-INS-181 | chr9-108525289-INS-189 |
| chr9-108545375-INS-68 | chr9-108545399-INS-85 |
| chr9-108554942-INS-498 | chr9-108554942-INS-498 |
| chr9-108579139-INS-204 | chr9-108579139-INS-204 |
| chr9-108601362-INS-203 | chr9-108601362-INS-204 |
| chr9-108623694-INS-76 | chr9-108623694-INS-76 |
| chr9-108636646-INS-765 | chr9-108636646-INS-765 |
| chr9-108639344-INS-168 | chr9-108639349-INS-128 |
| chr9-108716752-INS-103 | chr9-108716715-INS-99 |
| chr9-108755862-INS-99 | chr9-108755863-INS-107 |
| chr9-108772918-INS-124 | chr9-108772918-INS-124 |
| chr9-108847732-INS-202 | chr9-108847732-INS-201 |
| chr9-108874034-INS-52 | chr9-108874034-INS-52 |
| chr9-108900160-INS-200 | chr9-108900141-INS-213 |
| chr9-108920392-INS-64 | chr9-108920392-INS-62 |
| chr9-108960954-INS-582 | chr9-108960954-INS-584 |
| chr9-108973550-INS-999 | chr9-108973550-INS-999 |
| chr9-108976012-INS-72 | chr9-108976012-INS-52 |
| chr9-108990831-INS-76 | chr9-108990831-INS-76 |
| chr9-109022022-INS-203 | chr9-109022035-INS-217 |
| chr9-109059881-INS-6487 | chr9-109059882-INS-6487 |
| chr9-109089786-INS-154 | chr9-109089791-INS-174 |
| chr9-109095515-INS-5905 | chr9-109095516-INS-5905 |
| chr9-109139448-INS-212 | chr9-109139449-INS-212 |
| chr9-109144524-INS-295 | chr9-109144524-INS-294 |
| chr9-109200399-INS-54 | chr9-109200399-INS-53 |
| chr9-109267059-INS-60 | chr9-109267059-INS-60 |
| chr9-109274025-INS-88 | chr9-109273987-INS-77 |
| chr9-109276202-INS-53 | chr9-109276199-INS-63 |
| chr9-109277876-INS-64 | chr9-109277876-INS-64 |
| chr9-109278763-INS-2197 | chr9-109278764-INS-2186 |
| chr9-109313839-INS-72 | chr9-109313839-INS-72 |
| chr9-109323074-INS-65 | chr9-109323075-INS-57 |
| chr9-109396813-INS-72 | chr9-109396813-INS-68 |
| chr9-109398472-INS-72 | chr9-109398472-INS-112 |
| chr9-109413775-INS-54 | chr9-109413775-INS-54 |
| chr9-109414617-INS-4223 | chr9-109414618-INS-4223 |
| chr9-109453151-INS-104 | chr9-109453141-INS-114 |
| chr9-109488645-INS-55 | chr9-109488645-INS-53 |
| chr9-109505195-INS-105 | chr9-109505195-INS-105 |
| chr9-109526708-INS-58 | chr9-109526708-INS-56 |
| chr9-109529642-INS-70 | chr9-109529661-INS-53 |
| chr9-109554238-INS-70 | chr9-109554238-INS-70 |
| chr9-109616567-INS-78 | chr9-109616567-INS-56 |
| chr9-109617598-INS-128 | chr9-109617634-INS-109 |
| chr9-109641735-INS-232 | chr9-109641735-INS-232 |
| chr9-109648760-INS-229 | chr9-109648760-INS-229 |
| chr9-109653446-INS-87 | chr9-109653446-INS-87 |
| chr9-109663938-INS-164 | chr9-109663938-INS-164 |
| chr9-109675558-INS-203 | chr9-109675558-INS-203 |
| chr9-109681111-INS-73 | chr9-109681111-INS-73 |
| chr9-109682736-INS-205 | chr9-109682736-INS-205 |
| chr9-109687810-INS-120 | chr9-109687810-INS-120 |
| chr9-109691366-INS-202 | chr9-109691366-INS-199 |
| chr9-109696005-INS-79 | chr9-109696005-INS-80 |
| chr9-109702659-INS-136 | chr9-109702659-INS-136 |
| chr9-109703771-INS-57 | chr9-109703763-INS-64 |
| chr9-109707396-INS-199 | chr9-109707396-INS-199 |
| chr9-109717538-INS-206 | chr9-109717538-INS-206 |
| chr9-109737938-INS-207 | chr9-109737938-INS-207 |
| chr9-109739624-INS-503 | chr9-109739624-INS-503 |
| chr9-109746974-INS-172 | chr9-109746974-INS-180 |
| chr9-109770112-INS-61 | chr9-109770112-INS-62 |
| chr9-109775112-INS-203 | chr9-109775112-INS-205 |
| chr9-109787260-INS-232 | chr9-109787262-INS-225 |
| chr9-109798005-INS-835 | chr9-109798007-INS-839 |
| chr9-109806087-INS-229 | chr9-109806087-INS-261 |
| chr9-109839186-INS-82 | chr9-109839186-INS-78 |
| chr9-109871934-INS-208 | chr9-109871934-INS-206 |
| chr9-109878861-INS-52 | chr9-109878839-INS-97 |
| chr9-109888375-INS-52 | chr9-109888403-INS-66 |
| chr9-109897267-INS-57 | chr9-109897267-INS-59 |

| | |
|---|---|
| chr9-109907458-INS-74 | chr9-109907458-INS-74 |
| chr9-109940380-INS-223 | chr9-109940380-INS-225 |
| chr9-109968167-INS-172 | chr9-109968167-INS-176 |
| chr9-109969399-INS-124 | chr9-109969396-INS-125 |
| chr9-109975397-INS-60 | chr9-109975351-INS-56 |
| chr9-110051465-INS-210 | chr9-110051465-INS-210 |
| chr9-110065290-INS-170 | chr9-110065290-INS-155 |
| chr9-110085847-INS-61 | chr9-110085847-INS-63 |
| chr9-110096737-INS-411 | chr9-110096752-INS-417 |
| chr9-110098913-INS-202 | chr9-110098913-INS-202 |
| chr9-110099601-INS-55 | chr9-110099606-INS-61 |
| chr9-110101943-INS-327 | chr9-110101943-INS-329 |
| chr9-110103318-INS-72 | chr9-110103318-INS-72 |
| chr9-110104565-INS-160 | chr9-110104563-INS-162 |
| chr9-110129812-INS-179 | chr9-110129812-INS-180 |
| chr9-110135563-INS-216 | chr9-110135563-INS-214 |
| chr9-110142280-INS-52 | chr9-110142280-INS-52 |
| chr9-110193078-INS-54 | chr9-110193079-INS-53 |
| chr9-110215209-INS-70 | chr9-110215209-INS-70 |
| chr9-110241248-INS-125 | chr9-110241248-INS-133 |
| chr9-110259707-INS-56 | chr9-110259707-INS-80 |
| chr9-110284877-INS-222 | chr9-110284877-INS-222 |
| chr9-110286140-INS-360 | chr9-110286142-INS-353 |
| chr9-110309765-INS-197 | chr9-110309765-INS-197 |
| chr9-110313655-INS-55 | chr9-110313655-INS-55 |
| chr9-110321132-INS-110 | chr9-110321132-INS-110 |
| chr9-110326699-INS-181 | chr9-110326699-INS-181 |
| chr9-110329094-INS-216 | chr9-110329094-INS-216 |
| chr9-110380860-INS-209 | chr9-110380861-INS-208 |
| chr9-110381724-INS-167 | chr9-110381724-INS-167 |
| chr9-110402219-INS-192 | chr9-110402219-INS-192 |
| chr9-110414525-INS-205 | chr9-110414525-INS-205 |
| chr9-110460270-INS-94 | chr9-110460271-INS-95 |
| chr9-110496226-INS-78 | chr9-110496229-INS-81 |
| chr9-110500810-INS-6439 | chr9-110500811-INS-6439 |
| chr9-110503501-INS-125 | chr9-110503479-INS-112 |
| chr9-110519074-INS-210 | chr9-110519074-INS-210 |
| chr9-110545821-INS-54 | chr9-110545821-INS-62 |
| chr9-110596697-INS-99 | chr9-110596697-INS-125 |
| chr9-110613256-INS-4413 | chr9-110613257-INS-4413 |
| chr9-110623895-INS-52 | chr9-110623895-INS-52 |
| chr9-110636289-INS-64 | chr9-110636289-INS-60 |
| chr9-110637541-INS-214 | chr9-110637541-INS-214 |
| chr9-110640814-INS-908 | chr9-110640814-INS-908 |
| chr9-110650255-INS-141 | chr9-110650255-INS-165 |
| chr9-110658716-INS-599 | chr9-110658716-INS-622 |
| chr9-110679690-INS-74 | chr9-110679690-INS-74 |
| chr9-110694079-INS-65 | chr9-110694079-INS-65 |
| chr9-110696129-INS-54 | chr9-110696129-INS-54 |
| chr9-110756151-INS-161 | chr9-110756151-INS-161 |
| chr9-110759683-INS-417 | chr9-110759683-INS-417 |
| chr9-110772076-INS-169 | chr9-110772076-INS-169 |
| chr9-110779026-INS-91 | chr9-110779026-INS-91 |
| chr9-110786761-INS-54 | chr9-110786747-INS-57 |
| chr9-110788614-INS-205 | chr9-110788614-INS-205 |
| chr9-110790963-INS-733 | chr9-110790963-INS-733 |
| chr9-110794914-INS-211 | chr9-110794914-INS-211 |
| chr9-110796486-INS-127 | chr9-110796487-INS-131 |
| chr9-110797078-INS-52 | chr9-110797082-INS-53 |
| chr9-110798436-INS-208 | chr9-110798436-INS-208 |
| chr9-110807259-INS-221 | chr9-110807259-INS-214 |
| chr9-110813640-INS-416 | chr9-110813640-INS-416 |
| chr9-110844300-INS-53 | chr9-110844301-INS-53 |
| chr9-110849500-INS-214 | chr9-110849500-INS-214 |
| chr9-110866077-INS-76 | chr9-110866055-INS-98 |
| chr9-110898754-INS-99 | chr9-110898755-INS-63 |
| chr9-110901915-INS-439 | chr9-110901915-INS-438 |
| chr9-110904935-INS-54 | chr9-110904914-INS-76 |
| chr9-110906103-INS-75 | chr9-110906103-INS-117 |
| chr9-110908912-INS-165 | chr9-110908912-INS-165 |
| chr9-110918479-INS-87 | chr9-110918485-INS-92 |
| chr9-110919322-INS-208 | chr9-110919322-INS-208 |
| chr9-110924685-INS-82 | chr9-110924690-INS-90 |
| chr9-110928383-INS-193 | chr9-110928395-INS-206 |
| chr9-110930703-INS-75 | chr9-110930703-INS-76 |
| chr9-110933533-INS-196 | chr9-110933529-INS-196 |
| chr9-110953283-INS-211 | chr9-110953283-INS-211 |

| | |
|---|---|
| chr9-110954055-INS-148 | chr9-110954055-INS-148 |
| chr9-110964747-INS-145 | chr9-110964747-INS-145 |
| chr9-110965905-INS-199 | chr9-110965905-INS-199 |
| chr9-110990942-INS-6432 | chr9-110990943-INS-6432 |
| chr9-110997948-INS-135 | chr9-110997948-INS-132 |
| chr9-111020896-INS-83 | chr9-111020896-INS-83 |
| chr9-111031143-INS-1191 | chr9-111031144-INS-1191 |
| chr9-111039872-INS-119 | chr9-111039872-INS-124 |
| chr9-111065752-INS-113 | chr9-111065751-INS-113 |
| chr9-111071499-INS-77 | chr9-111071499-INS-77 |
| chr9-111087686-INS-478 | chr9-111087686-INS-478 |
| chr9-111091866-INS-6454 | chr9-111091867-INS-6454 |
| chr9-111095065-INS-55 | chr9-111095065-INS-55 |
| chr9-111112301-INS-110 | chr9-111112301-INS-122 |
| chr9-111124161-INS-88 | chr9-111124161-INS-88 |
| chr9-111131597-INS-52 | chr9-111131597-INS-53 |
| chr9-111145108-INS-51 | chr9-111145108-INS-53 |
| chr9-111162212-INS-186 | chr9-111162212-INS-186 |
| chr9-111165070-INS-175 | chr9-111165070-INS-175 |
| chr9-111185186-INS-84 | chr9-111185186-INS-84 |
| chr9-111248214-INS-77 | chr9-111248214-INS-77 |
| chr9-111249625-INS-222 | chr9-111249616-INS-186 |
| chr9-111260021-INS-184 | chr9-111260021-INS-184 |
| chr9-111262589-INS-67 | chr9-111262589-INS-71 |
| chr9-111276695-INS-63 | chr9-111276695-INS-66 |
| chr9-111281864-INS-178 | chr9-111281864-INS-208 |
| chr9-111288453-INS-163 | chr9-111288453-INS-149 |
| chr9-111295407-INS-398 | chr9-111295407-INS-402 |
| chr9-111302370-INS-65 | chr9-111302370-INS-59 |
| chr9-111309682-INS-63 | chr9-111309682-INS-63 |
| chr9-111334129-INS-322 | chr9-111334129-INS-339 |
| chr9-111336224-INS-71 | chr9-111336196-INS-60 |
| chr9-111349382-INS-1236 | chr9-111349383-INS-1236 |
| chr9-111367137-INS-493 | chr9-111367137-INS-493 |
| chr9-111378460-INS-138 | chr9-111378460-INS-138 |
| chr9-111400269-INS-82 | chr9-111400269-INS-82 |
| chr9-111416462-INS-66 | chr9-111416462-INS-66 |
| chr9-111424026-INS-280 | chr9-111424026-INS-280 |
| chr9-111427459-INS-52 | chr9-111427459-INS-52 |
| chr9-111464710-INS-6211 | chr9-111464711-INS-6211 |
| chr9-111483077-INS-55 | chr9-111483072-INS-50 |
| chr9-111505674-INS-243 | chr9-111505680-INS-234 |
| chr9-111522074-INS-144 | chr9-111522102-INS-160 |
| chr9-111538901-INS-88 | chr9-111538901-INS-90 |
| chr9-111540786-INS-84 | chr9-111540786-INS-88 |
| chr9-111550954-INS-356 | chr9-111550943-INS-357 |
| chr9-111558819-INS-70 | chr9-111558819-INS-78 |
| chr9-111603158-INS-114 | chr9-111603158-INS-114 |
| chr9-111621868-INS-98 | chr9-111621868-INS-98 |
| chr9-111683737-INS-177 | chr9-111683737-INS-177 |
| chr9-111695792-INS-83 | chr9-111695792-INS-83 |
| chr9-111701289-INS-60 | chr9-111701289-INS-60 |
| chr9-111721983-INS-118 | chr9-111721983-INS-110 |
| chr9-111729212-INS-57 | chr9-111729208-INS-59 |
| chr9-111740193-INS-811 | chr9-111740193-INS-811 |
| chr9-111747197-INS-124 | chr9-111747197-INS-124 |
| chr9-111748069-INS-95 | chr9-111748070-INS-73 |
| chr9-111764459-INS-75 | chr9-111764459-INS-66 |
| chr9-111774060-INS-553 | chr9-111774060-INS-553 |
| chr9-111803123-INS-226 | chr9-111803141-INS-196 |
| chr9-111811017-INS-522 | chr9-111811017-INS-522 |
| chr9-111825826-INS-72 | chr9-111825843-INS-88 |
| chr9-111828439-INS-54 | chr9-111828439-INS-54 |
| chr9-111839502-INS-52 | chr9-111839502-INS-52 |
| chr9-111845907-INS-67 | chr9-111845907-INS-67 |
| chr9-111850978-INS-145 | chr9-111850978-INS-145 |
| chr9-111858396-INS-107 | chr9-111858366-INS-96 |
| chr9-111889381-INS-358 | chr9-111889381-INS-358 |
| chr9-111891119-INS-295 | chr9-111891119-INS-311 |
| chr9-111916006-INS-78 | chr9-111915991-INS-57 |
| chr9-111917279-INS-64 | chr9-111917279-INS-68 |
| chr9-111921590-INS-55 | chr9-111921558-INS-54 |
| chr9-111925651-INS-98 | chr9-111925651-INS-102 |
| chr9-111936959-INS-195 | chr9-111937005-INS-184 |
| chr9-111942753-INS-85 | chr9-111942753-INS-85 |
| chr9-111962856-INS-367 | chr9-111962856-INS-367 |
| chr9-111967041-INS-66 | chr9-111967041-INS-66 |

| | |
|---|---|
| chr9-112024983-INS-148 | chr9-112024983-INS-148 |
| chr9-112040738-INS-104 | chr9-112040738-INS-110 |
| chr9-112093815-INS-51 | chr9-112093815-INS-66 |
| chr9-112114297-INS-225 | chr9-112114284-INS-226 |
| chr9-112115277-INS-97 | chr9-112115277-INS-119 |
| chr9-112118651-INS-114 | chr9-112118651-INS-120 |
| chr9-112126433-INS-555 | chr9-112126434-INS-554 |
| chr9-112158284-INS-52 | chr9-112158284-INS-54 |
| chr9-112181248-INS-68 | chr9-112181248-INS-82 |
| chr9-112212322-INS-6241 | chr9-112212323-INS-6242 |
| chr9-112222399-INS-66 | chr9-112222399-INS-69 |
| chr9-112228761-INS-78 | chr9-112228761-INS-86 |
| chr9-112251264-INS-498 | chr9-112251264-INS-498 |
| chr9-112270156-INS-273 | chr9-112270156-INS-273 |
| chr9-112276932-INS-132 | chr9-112276932-INS-132 |
| chr9-112281661-INS-92 | chr9-112281661-INS-70 |
| chr9-112295759-INS-58 | chr9-112295759-INS-58 |
| chr9-112298274-INS-76 | chr9-112298293-INS-76 |
| chr9-112355969-INS-349 | chr9-112355969-INS-349 |
| chr9-112372340-INS-335 | chr9-112372340-INS-321 |
| chr9-112373135-INS-82 | chr9-112373135-INS-52 |
| chr9-112381650-INS-61 | chr9-112381632-INS-64 |
| chr9-112391565-INS-52 | chr9-112391565-INS-54 |
| chr9-112409893-INS-171 | chr9-112409890-INS-175 |
| chr9-112429007-INS-1099 | chr9-112429008-INS-1098 |
| chr9-112434446-INS-2573 | chr9-112434447-INS-2573 |
| chr9-112465037-INS-2543 | chr9-112465038-INS-2524 |
| chr9-112486274-INS-437 | chr9-112486274-INS-437 |
| chr9-112517014-INS-76 | chr9-112517014-INS-78 |
| chr9-112529605-INS-377 | chr9-112529605-INS-377 |
| chr9-112553808-INS-56 | chr9-112553808-INS-56 |
| chr9-112582832-INS-51 | chr9-112582832-INS-51 |
| chr9-112594778-INS-6158 | chr9-112594779-INS-6158 |
| chr9-112624132-INS-86 | chr9-112624129-INS-70 |
| chr9-112650986-INS-70 | chr9-112650986-INS-70 |
| chr9-112677381-INS-80 | chr9-112677381-INS-60 |
| chr9-112683923-INS-98 | chr9-112683917-INS-96 |
| chr9-112700300-INS-951 | chr9-112700300-INS-951 |
| chr9-112708754-INS-332 | chr9-112708754-INS-332 |
| chr9-112712859-INS-60 | chr9-112712859-INS-60 |
| chr9-112725432-INS-2542 | chr9-112725433-INS-2542 |
| chr9-112730011-INS-51 | chr9-112730012-INS-66 |
| chr9-112740216-INS-2866 | chr9-112740217-INS-2866 |
| chr9-112758910-INS-72 | chr9-112758910-INS-66 |
| chr9-112768591-INS-233 | chr9-112768591-INS-233 |
| chr9-112769904-INS-74 | chr9-112769905-INS-58 |
| chr9-112777390-INS-100 | chr9-112777390-INS-72 |
| chr9-112785290-INS-211 | chr9-112785290-INS-211 |
| chr9-112790108-INS-5313 | chr9-112790108-INS-5313 |
| chr9-112810780-INS-94 | chr9-112810780-INS-94 |
| chr9-112834488-INS-767 | chr9-112834488-INS-770 |
| chr9-112841914-INS-318 | chr9-112841914-INS-318 |
| chr9-112884741-INS-74 | chr9-112884731-INS-51 |
| chr9-112900120-INS-59 | chr9-112900120-INS-59 |
| chr9-112906228-INS-138 | chr9-112906228-INS-138 |
| chr9-112909309-INS-87 | chr9-112909343-INS-86 |
| chr9-112910480-INS-6423 | chr9-112910481-INS-6424 |
| chr9-112916697-INS-52 | chr9-112916697-INS-56 |
| chr9-112920638-INS-66 | chr9-112920638-INS-110 |
| chr9-112921633-INS-426 | chr9-112921633-INS-426 |
| chr9-112930693-INS-56 | chr9-112930693-INS-56 |
| chr9-112931703-INS-160 | chr9-112931703-INS-160 |
| chr9-112932929-INS-1511 | chr9-112932930-INS-1511 |
| chr9-112933603-INS-57 | chr9-112933622-INS-76 |
| chr9-112947436-INS-902 | chr9-112947435-INS-901 |
| chr9-112951090-INS-66 | chr9-112951090-INS-66 |
| chr9-112955518-INS-52 | chr9-112955518-INS-52 |
| chr9-112958326-INS-847 | chr9-112958326-INS-847 |
| chr9-112972070-INS-763 | chr9-112972070-INS-763 |
| chr9-112972911-INS-186 | chr9-112972911-INS-190 |
| chr9-112975113-INS-63 | chr9-112975101-INS-51 |
| chr9-112975526-INS-64 | chr9-112975526-INS-64 |
| chr9-112987968-INS-73 | chr9-112987944-INS-109 |
| chr9-113025531-INS-61 | chr9-113025531-INS-64 |
| chr9-113035710-INS-121 | chr9-113035710-INS-121 |
| chr9-113043758-INS-56 | chr9-113043757-INS-54 |
| chr9-113057322-INS-82 | chr9-113057322-INS-82 |

| | |
|---|---|
| chr9-113059067-INS-235 | chr9-113059067-INS-235 |
| chr9-113060222-INS-104 | chr9-113060222-INS-104 |
| chr9-113064241-INS-398 | chr9-113064241-INS-398 |
| chr9-113067886-INS-109 | chr9-113067886-INS-109 |
| chr9-113100151-INS-52 | chr9-113100151-INS-55 |
| chr9-113113109-INS-52 | chr9-113113109-INS-50 |
| chr9-113116454-INS-64 | chr9-113116454-INS-68 |
| chr9-113118655-INS-68 | chr9-113118655-INS-68 |
| chr9-113130919-INS-7689 | chr9-113130920-INS-7737 |
| chr9-113132349-INS-82 | chr9-113132349-INS-82 |
| chr9-113141133-INS-418 | chr9-113141133-INS-418 |
| chr9-113155811-INS-538 | chr9-113155811-INS-535 |
| chr9-113159243-INS-258 | chr9-113159243-INS-258 |
| chr9-113159845-INS-92 | chr9-113159845-INS-54 |
| chr9-113166922-INS-185 | chr9-113166924-INS-185 |
| chr9-113197366-INS-54 | chr9-113197366-INS-54 |
| chr9-113199574-INS-325 | chr9-113199574-INS-325 |
| chr9-113201912-INS-76 | chr9-113201915-INS-60 |
| chr9-113208915-INS-77 | chr9-113208916-INS-77 |
| chr9-113212469-INS-396 | chr9-113212469-INS-392 |
| chr9-113215683-INS-99 | chr9-113215683-INS-99 |
| chr9-113222250-INS-196 | chr9-113222250-INS-197 |
| chr9-113225781-INS-72 | chr9-113225781-INS-72 |
| chr9-113229793-INS-142 | chr9-113229793-INS-144 |
| chr9-113239992-INS-50 | chr9-113240001-INS-66 |
| chr9-113242012-INS-744 | chr9-113241962-INS-756 |
| chr9-113242013-INS-213 | chr9-113242013-INS-213 |
| chr9-113250013-INS-76 | chr9-113250013-INS-97 |
| chr9-113265601-INS-467 | chr9-113265601-INS-483 |
| chr9-113267286-INS-84 | chr9-113267286-INS-92 |
| chr9-113269670-INS-102 | chr9-113269670-INS-102 |
| chr9-113271576-INS-139 | chr9-113271576-INS-139 |
| chr9-113284082-INS-400 | chr9-113284082-INS-400 |
| chr9-113286990-INS-427 | chr9-113286990-INS-427 |
| chr9-113310341-INS-78 | chr9-113310341-INS-70 |
| chr9-113312380-INS-116 | chr9-113312380-INS-116 |
| chr9-113343089-INS-52 | chr9-113343089-INS-52 |
| chr9-113372554-INS-553 | chr9-113372554-INS-553 |
| chr9-113388787-INS-54 | chr9-113388823-INS-62 |
| chr9-113396983-INS-6251 | chr9-113396984-INS-6251 |
| chr9-113418808-INS-440 | chr9-113418808-INS-440 |
| chr9-113443898-INS-78 | chr9-113443898-INS-70 |
| chr9-113448659-INS-54 | chr9-113448659-INS-54 |
| chr9-113464051-INS-164 | chr9-113464056-INS-168 |
| chr9-113469590-INS-53 | chr9-113469590-INS-63 |
| chr9-113491985-INS-96 | chr9-113491985-INS-96 |
| chr9-113494732-INS-623 | chr9-113494732-INS-624 |
| chr9-113519424-INS-227 | chr9-113519424-INS-227 |
| chr9-113538637-INS-217 | chr9-113538637-INS-217 |
| chr9-113554127-INS-88 | chr9-113554127-INS-88 |
| chr9-113559459-INS-5482 | chr9-113559460-INS-5482 |
| chr9-113565025-INS-70 | chr9-113565025-INS-74 |
| chr9-113576571-INS-223 | chr9-113576571-INS-223 |
| chr9-113591866-INS-1288 | chr9-113591867-INS-1288 |
| chr9-113599098-INS-206 | chr9-113599098-INS-206 |
| chr9-113619877-INS-7049 | chr9-113619878-INS-7049 |
| chr9-113625282-INS-4780 | chr9-113625283-INS-4780 |
| chr9-113632089-INS-84 | chr9-113632089-INS-56 |
| chr9-113635154-INS-106 | chr9-113635143-INS-71 |
| chr9-113638323-INS-312 | chr9-113638323-INS-312 |
| chr9-113653984-INS-67 | chr9-113653984-INS-75 |
| chr9-113666158-INS-116 | chr9-113666137-INS-147 |
| chr9-113720530-INS-355 | chr9-113720530-INS-355 |
| chr9-113726087-INS-386 | chr9-113726087-INS-386 |
| chr9-113731789-INS-399 | chr9-113731789-INS-399 |
| chr9-113771624-INS-68 | chr9-113771624-INS-76 |
| chr9-113794144-INS-194 | chr9-113794144-INS-194 |
| chr9-113817971-INS-198 | chr9-113817990-INS-221 |
| chr9-113824442-INS-223 | chr9-113824442-INS-225 |
| chr9-113838787-INS-110 | chr9-113838787-INS-110 |
| chr9-113853620-INS-74 | chr9-113853633-INS-81 |
| chr9-113858324-INS-7165 | chr9-113858325-INS-7165 |
| chr9-113881200-INS-6452 | chr9-113881201-INS-6452 |
| chr9-113914213-INS-119 | chr9-113914213-INS-154 |
| chr9-113928924-INS-57 | chr9-113928925-INS-54 |
| chr9-113969304-INS-130 | chr9-113969304-INS-130 |
| chr9-114012458-INS-114 | chr9-114012458-INS-84 |

| | |
|---|---|
| chr9-114039990-INS-71 | chr9-114039990-INS-69 |
| chr9-114046143-INS-148 | chr9-114046143-INS-148 |
| chr9-114135979-INS-600 | chr9-114135974-INS-603 |
| chr9-114205139-INS-205 | chr9-114205139-INS-204 |
| chr9-114209840-INS-84 | chr9-114209834-INS-82 |
| chr9-114212761-INS-812 | chr9-114212761-INS-812 |
| chr9-114221760-INS-237 | chr9-114221760-INS-237 |
| chr9-114226326-INS-213 | chr9-114226326-INS-215 |
| chr9-114228918-INS-68 | chr9-114228918-INS-78 |
| chr9-114245698-INS-5428 | chr9-114245699-INS-5428 |
| chr9-114247852-INS-212 | chr9-114247852-INS-211 |
| chr9-114252179-INS-67 | chr9-114252216-INS-67 |
| chr9-114261078-INS-60 | chr9-114261078-INS-60 |
| chr9-114279215-INS-212 | chr9-114279215-INS-213 |
| chr9-114285039-INS-204 | chr9-114285039-INS-215 |
| chr9-114330807-INS-97 | chr9-114330807-INS-97 |
| chr9-114346297-INS-199 | chr9-114346297-INS-199 |
| chr9-114359583-INS-58 | chr9-114359583-INS-58 |
| chr9-114375189-INS-226 | chr9-114375186-INS-212 |
| chr9-114401028-INS-157 | chr9-114401028-INS-156 |
| chr9-114412196-INS-209 | chr9-114412197-INS-202 |
| chr9-114418562-INS-60 | chr9-114418562-INS-60 |
| chr9-114421826-INS-251 | chr9-114421826-INS-251 |
| chr9-114430769-INS-194 | chr9-114430769-INS-195 |
| chr9-114437286-INS-222 | chr9-114437286-INS-222 |
| chr9-114439543-INS-94 | chr9-114439561-INS-50 |
| chr9-114452796-INS-217 | chr9-114452796-INS-217 |
| chr9-114475206-INS-195 | chr9-114475206-INS-195 |
| chr9-114476493-INS-197 | chr9-114476493-INS-197 |
| chr9-114477122-INS-206 | chr9-114477121-INS-221 |
| chr9-114479873-INS-215 | chr9-114479873-INS-215 |
| chr9-114482669-INS-84 | chr9-114482669-INS-78 |
| chr9-114483715-INS-166 | chr9-114483718-INS-184 |
| chr9-114484888-INS-60 | chr9-114484888-INS-60 |
| chr9-114503343-INS-403 | chr9-114503333-INS-406 |
| chr9-114504558-INS-62 | chr9-114504579-INS-72 |
| chr9-114514533-INS-96 | chr9-114514533-INS-82 |
| chr9-114526734-INS-60 | chr9-114526734-INS-60 |
| chr9-114534849-INS-285 | chr9-114534849-INS-285 |
| chr9-114550296-INS-195 | chr9-114550296-INS-195 |
| chr9-114572658-INS-227 | chr9-114572658-INS-227 |
| chr9-114573152-INS-1888 | chr9-114573153-INS-1888 |
| chr9-114585184-INS-112 | chr9-114585184-INS-112 |
| chr9-114597125-INS-220 | chr9-114597125-INS-220 |
| chr9-114598569-INS-56 | chr9-114598569-INS-56 |
| chr9-114602962-INS-54 | chr9-114602962-INS-58 |
| chr9-114616949-INS-170 | chr9-114616949-INS-170 |
| chr9-114618294-INS-90 | chr9-114618294-INS-90 |
| chr9-114620269-INS-301 | chr9-114620269-INS-313 |
| chr9-114621853-INS-70 | chr9-114621856-INS-87 |
| chr9-114623791-INS-225 | chr9-114623791-INS-226 |
| chr9-114639353-INS-57 | chr9-114639353-INS-57 |
| chr9-114644505-INS-223 | chr9-114644505-INS-223 |
| chr9-114658037-INS-56 | chr9-114658037-INS-58 |
| chr9-114661605-INS-342 | chr9-114661606-INS-342 |
| chr9-114675255-INS-134 | chr9-114675255-INS-132 |
| chr9-114679263-INS-2768 | chr9-114679262-INS-2767 |
| chr9-114700289-INS-120 | chr9-114700289-INS-90 |
| chr9-114745561-INS-121 | chr9-114745561-INS-121 |
| chr9-114748014-INS-74 | chr9-114748014-INS-78 |
| chr9-114752480-INS-166 | chr9-114752480-INS-167 |
| chr9-114775073-INS-214 | chr9-114775074-INS-213 |
| chr9-114779643-INS-52 | chr9-114779643-INS-50 |
| chr9-114782527-INS-298 | chr9-114782527-INS-298 |
| chr9-114783577-INS-81 | chr9-114783595-INS-81 |
| chr9-114816254-INS-471 | chr9-114816254-INS-470 |
| chr9-114839641-INS-240 | chr9-114839641-INS-242 |
| chr9-114847866-INS-468 | chr9-114847866-INS-468 |
| chr9-114860980-INS-66 | chr9-114860980-INS-66 |
| chr9-114868732-INS-118 | chr9-114868732-INS-126 |
| chr9-114878525-INS-203 | chr9-114878525-INS-203 |
| chr9-114880467-INS-216 | chr9-114880467-INS-176 |
| chr9-114882888-INS-52 | chr9-114882888-INS-65 |
| chr9-114885465-INS-54 | chr9-114885465-INS-56 |
| chr9-114898052-INS-54 | chr9-114898057-INS-63 |
| chr9-114899479-INS-112 | chr9-114899479-INS-112 |
| chr9-114903363-INS-56 | chr9-114903363-INS-56 |

| | |
|---|---|
| chr9-114915739-INS-80 | chr9-114915743-INS-72 |
| chr9-114920580-INS-64 | chr9-114920580-INS-64 |
| chr9-114937766-INS-59 | chr9-114937766-INS-59 |
| chr9-114944473-INS-162 | chr9-114944473-INS-162 |
| chr9-114945375-INS-705 | chr9-114945375-INS-705 |
| chr9-114962987-INS-280 | chr9-114962987-INS-280 |
| chr9-114966860-INS-77 | chr9-114966862-INS-56 |
| chr9-115008383-INS-72 | chr9-115008383-INS-72 |
| chr9-115024043-INS-141 | chr9-115024038-INS-151 |
| chr9-115031210-INS-66 | chr9-115031210-INS-60 |
| chr9-115034633-INS-58 | chr9-115034633-INS-75 |
| chr9-115037296-INS-191 | chr9-115037296-INS-191 |
| chr9-115052426-INS-230 | chr9-115052426-INS-230 |
| chr9-115096359-INS-126 | chr9-115096359-INS-125 |
| chr9-115127304-INS-211 | chr9-115127304-INS-214 |
| chr9-115169362-INS-56 | chr9-115169362-INS-86 |
| chr9-115173432-INS-788 | chr9-115173432-INS-789 |
| chr9-115175530-INS-70 | chr9-115175530-INS-70 |
| chr9-115181257-INS-401 | chr9-115181257-INS-401 |
| chr9-115188292-INS-444 | chr9-115188292-INS-443 |
| chr9-115191513-INS-313 | chr9-115191513-INS-328 |
| chr9-115193944-INS-2389 | chr9-115193945-INS-2417 |
| chr9-115196774-INS-105 | chr9-115196799-INS-70 |
| chr9-115208550-INS-524 | chr9-115208550-INS-524 |
| chr9-115222333-INS-56 | chr9-115222333-INS-52 |
| chr9-115232799-INS-459 | chr9-115232799-INS-458 |
| chr9-115239620-INS-76 | chr9-115239620-INS-76 |
| chr9-115242828-INS-59 | chr9-115242808-INS-77 |
| chr9-115272995-INS-65 | chr9-115272995-INS-67 |
| chr9-115318021-INS-63 | chr9-115318021-INS-81 |
| chr9-115320404-INS-504 | chr9-115320404-INS-504 |
| chr9-115329027-INS-99 | chr9-115329027-INS-99 |
| chr9-115359833-INS-206 | chr9-115359833-INS-206 |
| chr9-115362665-INS-64 | chr9-115362665-INS-64 |
| chr9-115375820-INS-51 | chr9-115375783-INS-56 |
| chr9-115393855-INS-62 | chr9-115393855-INS-62 |
| chr9-115414875-INS-119 | chr9-115414875-INS-109 |
| chr9-115418762-INS-62 | chr9-115418770-INS-62 |
| chr9-115473140-INS-151 | chr9-115473140-INS-151 |
| chr9-115495056-INS-60 | chr9-115495056-INS-60 |
| chr9-115502303-INS-78 | chr9-115502303-INS-78 |
| chr9-115513502-INS-56 | chr9-115513502-INS-70 |
| chr9-115522853-INS-72 | chr9-115522853-INS-80 |
| chr9-115536944-INS-260 | chr9-115536944-INS-260 |
| chr9-115541035-INS-184 | chr9-115541034-INS-183 |
| chr9-115562410-INS-433 | chr9-115562410-INS-434 |
| chr9-115571321-INS-86 | chr9-115571321-INS-88 |
| chr9-115581587-INS-128 | chr9-115581584-INS-100 |
| chr9-115584573-INS-58 | chr9-115584573-INS-58 |
| chr9-115614643-INS-106 | chr9-115614643-INS-106 |
| chr9-115629441-INS-210 | chr9-115629441-INS-210 |
| chr9-115630182-INS-6452 | chr9-115630183-INS-6452 |
| chr9-115645801-INS-400 | chr9-115645801-INS-403 |
| chr9-115654040-INS-902 | chr9-115654040-INS-890 |
| chr9-115665705-INS-109 | chr9-115665705-INS-109 |
| chr9-115692046-INS-86 | chr9-115692094-INS-50 |
| chr9-115693361-INS-261 | chr9-115693361-INS-268 |
| chr9-115701730-INS-62 | chr9-115701720-INS-52 |
| chr9-115712267-INS-159 | chr9-115712267-INS-159 |
| chr9-115716552-INS-145 | chr9-115716544-INS-150 |
| chr9-115717949-INS-143 | chr9-115717949-INS-143 |
| chr9-115795679-INS-77 | chr9-115795679-INS-55 |
| chr9-115804468-INS-53 | chr9-115804481-INS-65 |
| chr9-115810620-INS-91 | chr9-115810607-INS-103 |
| chr9-115830374-INS-252 | chr9-115830371-INS-268 |
| chr9-115851442-INS-72 | chr9-115851442-INS-76 |
| chr9-115853567-INS-70 | chr9-115853567-INS-52 |
| chr9-115859518-INS-52 | chr9-115859518-INS-60 |
| chr9-115872529-INS-94 | chr9-115872529-INS-78 |
| chr9-115882085-INS-144 | chr9-115882085-INS-144 |
| chr9-115888345-INS-1259 | chr9-115888346-INS-1228 |
| chr9-115924067-INS-739 | chr9-115924067-INS-743 |
| chr9-115924997-INS-68 | chr9-115924997-INS-68 |
| chr9-115938179-INS-641 | chr9-115938179-INS-640 |
| chr9-115968411-INS-7216 | chr9-115968412-INS-7217 |
| chr9-115970266-INS-54 | chr9-115970266-INS-54 |
| chr9-115985195-INS-112 | chr9-115985195-INS-148 |

| | |
|---|---|
| chr9-116006128-INS-939 | chr9-116006128-INS-939 |
| chr9-116026612-INS-70 | chr9-116026592-INS-106 |
| chr9-116036686-INS-60 | chr9-116036686-INS-60 |
| chr9-116051530-INS-56 | chr9-116051530-INS-50 |
| chr9-116069057-INS-3311 | chr9-116069058-INS-3311 |
| chr9-116069433-INS-616 | chr9-116069433-INS-616 |
| chr9-116110843-INS-7483 | chr9-116110844-INS-7483 |
| chr9-116113544-INS-65 | chr9-116113543-INS-75 |
| chr9-116119677-INS-76 | chr9-116119677-INS-76 |
| chr9-116123608-INS-102 | chr9-116123594-INS-95 |
| chr9-116132212-INS-64 | chr9-116132212-INS-84 |
| chr9-116237750-INS-225 | chr9-116237750-INS-225 |
| chr9-116303680-INS-56 | chr9-116303669-INS-92 |
| chr9-116339333-INS-91 | chr9-116339333-INS-91 |
| chr9-116352121-INS-67 | chr9-116352121-INS-78 |
| chr9-116360085-INS-58 | chr9-116360085-INS-68 |
| chr9-116367435-INS-56 | chr9-116367435-INS-52 |
| chr9-116416414-INS-54 | chr9-116416414-INS-72 |
| chr9-116436192-INS-150 | chr9-116436188-INS-151 |
| chr9-116467112-INS-69 | chr9-116467112-INS-59 |
| chr9-116478082-INS-58 | chr9-116478072-INS-66 |
| chr9-116493255-INS-322 | chr9-116493255-INS-324 |
| chr9-116512998-INS-128 | chr9-116512998-INS-114 |
| chr9-116523769-INS-57 | chr9-116523769-INS-52 |
| chr9-116544396-INS-137 | chr9-116544396-INS-137 |
| chr9-116560240-INS-54 | chr9-116560240-INS-54 |
| chr9-116577136-INS-729 | chr9-116577136-INS-729 |
| chr9-116590185-INS-125 | chr9-116590185-INS-102 |
| chr9-116599933-INS-51 | chr9-116599933-INS-51 |
| chr9-116609655-INS-234 | chr9-116609655-INS-234 |
| chr9-116645567-INS-71 | chr9-116645564-INS-74 |
| chr9-116655899-INS-52 | chr9-116655899-INS-52 |
| chr9-116660060-INS-126 | chr9-116660060-INS-126 |
| chr9-116695492-INS-1893 | chr9-116695493-INS-1893 |
| chr9-116713170-INS-388 | chr9-116713170-INS-388 |
| chr9-116735120-INS-402 | chr9-116735114-INS-404 |
| chr9-116736419-INS-63 | chr9-116736429-INS-88 |
| chr9-116750226-INS-56 | chr9-116750222-INS-60 |
| chr9-116759710-INS-65 | chr9-116759710-INS-65 |
| chr9-116763047-INS-57 | chr9-116763047-INS-57 |
| chr9-116768003-INS-114 | chr9-116768003-INS-115 |
| chr9-116773478-INS-71 | chr9-116773478-INS-71 |
| chr9-116794169-INS-89 | chr9-116794169-INS-92 |
| chr9-116799900-INS-72 | chr9-116799900-INS-52 |
| chr9-116819971-INS-188 | chr9-116819971-INS-188 |
| chr9-116844831-INS-111 | chr9-116844831-INS-111 |
| chr9-116862071-INS-6401 | chr9-116862072-INS-6401 |
| chr9-116866700-INS-62 | chr9-116866700-INS-62 |
| chr9-116870525-INS-66 | chr9-116870525-INS-66 |
| chr9-116873461-INS-56 | chr9-116873461-INS-56 |
| chr9-116901363-INS-54 | chr9-116901363-INS-55 |
| chr9-116924663-INS-399 | chr9-116924661-INS-400 |
| chr9-116934080-INS-7114 | chr9-116934081-INS-7114 |
| chr9-116936009-INS-101 | chr9-116935987-INS-148 |
| chr9-116967364-INS-64 | chr9-116967364-INS-66 |
| chr9-116969941-INS-822 | chr9-116969941-INS-822 |
| chr9-116979510-INS-6675 | chr9-116979511-INS-6675 |
| chr9-117000845-INS-58 | chr9-117000845-INS-52 |
| chr9-117002485-INS-51 | chr9-117002485-INS-51 |
| chr9-117033829-INS-2535 | chr9-117033830-INS-2535 |
| chr9-117085971-INS-7204 | chr9-117085972-INS-7204 |
| chr9-117101424-INS-51 | chr9-117101442-INS-69 |
| chr9-117122178-INS-101 | chr9-117122185-INS-72 |
| chr9-117133045-INS-59 | chr9-117133045-INS-63 |
| chr9-117135804-INS-85 | chr9-117135804-INS-54 |
| chr9-117149370-INS-215 | chr9-117149370-INS-217 |
| chr9-117152773-INS-1864 | chr9-117152774-INS-1864 |
| chr9-117165935-INS-201 | chr9-117165935-INS-201 |
| chr9-117186741-INS-92 | chr9-117186741-INS-92 |
| chr9-117200060-INS-51 | chr9-117200060-INS-69 |
| chr9-117211873-INS-233 | chr9-117211873-INS-232 |
| chr9-117221788-INS-4583 | chr9-117221789-INS-4583 |
| chr9-117256688-INS-1098 | chr9-117256689-INS-1097 |
| chr9-117259814-INS-150 | chr9-117259814-INS-138 |
| chr9-117275897-INS-270 | chr9-117275898-INS-273 |
| chr9-117295833-INS-131 | chr9-117295852-INS-108 |
| chr9-117308512-INS-86 | chr9-117308512-INS-52 |

| | |
|---|---|
| chr9-117322683-INS-3540 | chr9-117322691-INS-3504 |
| chr9-117336369-INS-66 | chr9-117336369-INS-66 |
| chr9-117341880-INS-54 | chr9-117341864-INS-76 |
| chr9-117353965-INS-93 | chr9-117353965-INS-93 |
| chr9-117362760-INS-72 | chr9-117362726-INS-76 |
| chr9-117388537-INS-102 | chr9-117388537-INS-102 |
| chr9-117417711-INS-69 | chr9-117417717-INS-51 |
| chr9-117464368-INS-106 | chr9-117464364-INS-108 |
| chr9-117516097-INS-303 | chr9-117516097-INS-305 |
| chr9-117540765-INS-54 | chr9-117540765-INS-54 |
| chr9-117566577-INS-115 | chr9-117566585-INS-70 |
| chr9-117583307-INS-297 | chr9-117583307-INS-298 |
| chr9-117589016-INS-198 | chr9-117589016-INS-200 |
| chr9-117607080-INS-53 | chr9-117607080-INS-65 |
| chr9-117615431-INS-61 | chr9-117615460-INS-56 |
| chr9-117617478-INS-75 | chr9-117617478-INS-78 |
| chr9-117626949-INS-86 | chr9-117626949-INS-86 |
| chr9-117629740-INS-173 | chr9-117629740-INS-173 |
| chr9-117633583-INS-227 | chr9-117633583-INS-229 |
| chr9-117638215-INS-56 | chr9-117638249-INS-54 |
| chr9-117643133-INS-324 | chr9-117643133-INS-324 |
| chr9-117658573-INS-52 | chr9-117658573-INS-52 |
| chr9-117661353-INS-80 | chr9-117661353-INS-75 |
| chr9-117676528-INS-214 | chr9-117676528-INS-214 |
| chr9-117682788-INS-53 | chr9-117682788-INS-57 |
| chr9-117685216-INS-286 | chr9-117685216-INS-286 |
| chr9-117687113-INS-92 | chr9-117687113-INS-92 |
| chr9-117692343-INS-124 | chr9-117692343-INS-124 |
| chr9-117712895-INS-74 | chr9-117712895-INS-78 |
| chr9-117744002-INS-65 | chr9-117744023-INS-50 |
| chr9-117746603-INS-159 | chr9-117746612-INS-156 |
| chr9-117754117-INS-128 | chr9-117754117-INS-128 |
| chr9-117760561-INS-207 | chr9-117760561-INS-207 |
| chr9-117798105-INS-121 | chr9-117798099-INS-127 |
| chr9-117805142-INS-1450 | chr9-117805143-INS-1422 |
| chr9-117820705-INS-166 | chr9-117820715-INS-150 |
| chr9-117822654-INS-62 | chr9-117822654-INS-66 |
| chr9-117825183-INS-156 | chr9-117825183-INS-158 |
| chr9-117929955-INS-64 | chr9-117929955-INS-78 |
| chr9-117931367-INS-54 | chr9-117931367-INS-56 |
| chr9-117939876-INS-52 | chr9-117939876-INS-52 |
| chr9-117955094-INS-62 | chr9-117955094-INS-62 |
| chr9-117965869-INS-277 | chr9-117965869-INS-277 |
| chr9-117974873-INS-164 | chr9-117974873-INS-164 |
| chr9-117981235-INS-859 | chr9-117981235-INS-855 |
| chr9-118115377-INS-518 | chr9-118115377-INS-486 |
| chr9-118124412-INS-400 | chr9-118124424-INS-438 |
| chr9-118133980-INS-64 | chr9-118133980-INS-64 |
| chr9-118140609-INS-80 | chr9-118140620-INS-80 |
| chr9-118143742-INS-73 | chr9-118143742-INS-73 |
| chr9-118149145-INS-231 | chr9-118149145-INS-231 |
| chr9-118172093-INS-431 | chr9-118172093-INS-431 |
| chr9-118215353-INS-191 | chr9-118215328-INS-226 |
| chr9-118223210-INS-71 | chr9-118223210-INS-71 |
| chr9-118249308-INS-307 | chr9-118249302-INS-330 |
| chr9-118264441-INS-138 | chr9-118264443-INS-94 |
| chr9-118277088-INS-97 | chr9-118277088-INS-99 |
| chr9-118330788-INS-60 | chr9-118330788-INS-60 |
| chr9-118364128-INS-133 | chr9-118364128-INS-96 |
| chr9-118399903-INS-58 | chr9-118399904-INS-57 |
| chr9-118440132-INS-130 | chr9-118440123-INS-92 |
| chr9-118456725-INS-79 | chr9-118456725-INS-79 |
| chr9-118460770-INS-150 | chr9-118460770-INS-135 |
| chr9-118469943-INS-214 | chr9-118469943-INS-214 |
| chr9-118476484-INS-613 | chr9-118476484-INS-634 |
| chr9-118503534-INS-51 | chr9-118503534-INS-51 |
| chr9-118539577-INS-6577 | chr9-118539578-INS-6577 |
| chr9-118555885-INS-60 | chr9-118555883-INS-64 |
| chr9-118583150-INS-51 | chr9-118583150-INS-51 |
| chr9-118590532-INS-128 | chr9-118590532-INS-133 |
| chr9-118598654-INS-60 | chr9-118598654-INS-58 |
| chr9-118599815-INS-146 | chr9-118599815-INS-146 |
| chr9-118603281-INS-56 | chr9-118603281-INS-58 |
| chr9-118610444-INS-894 | chr9-118610444-INS-894 |
| chr9-118649571-INS-64 | chr9-118649572-INS-96 |
| chr9-118653463-INS-51 | chr9-118653463-INS-51 |
| chr9-118655820-INS-108 | chr9-118655820-INS-108 |

| | |
|---|---|
| chr9-118660640-INS-51 | chr9-118660640-INS-51 |
| chr9-118661522-INS-445 | chr9-118661522-INS-445 |
| chr9-118715444-INS-375 | chr9-118715444-INS-375 |
| chr9-118725622-INS-74 | chr9-118725622-INS-74 |
| chr9-118744897-INS-194 | chr9-118744914-INS-198 |
| chr9-118761098-INS-144 | chr9-118761098-INS-144 |
| chr9-118767711-INS-200 | chr9-118767711-INS-200 |
| chr9-118785978-INS-175 | chr9-118785978-INS-198 |
| chr9-118805562-INS-6422 | chr9-118805563-INS-6457 |
| chr9-118890228-INS-204 | chr9-118890228-INS-204 |
| chr9-119006205-INS-66 | chr9-119006201-INS-56 |
| chr9-119009253-INS-356 | chr9-119009253-INS-356 |
| chr9-119019820-INS-4531 | chr9-119019821-INS-4531 |
| chr9-119020593-INS-607 | chr9-119020593-INS-607 |
| chr9-119045827-INS-434 | chr9-119045827-INS-434 |
| chr9-119089052-INS-70 | chr9-119089052-INS-70 |
| chr9-119111977-INS-6089 | chr9-119111978-INS-6089 |
| chr9-119114890-INS-145 | chr9-119114890-INS-145 |
| chr9-119117520-INS-205 | chr9-119117526-INS-204 |
| chr9-119119232-INS-199 | chr9-119119232-INS-199 |
| chr9-119129396-INS-208 | chr9-119129396-INS-206 |
| chr9-119182844-INS-240 | chr9-119182845-INS-234 |
| chr9-119193124-INS-56 | chr9-119193124-INS-60 |
| chr9-119204061-INS-202 | chr9-119204052-INS-185 |
| chr9-119205352-INS-207 | chr9-119205353-INS-203 |
| chr9-119214983-INS-201 | chr9-119214983-INS-201 |
| chr9-119218307-INS-180 | chr9-119218291-INS-224 |
| chr9-119225014-INS-203 | chr9-119225014-INS-199 |
| chr9-119251129-INS-89 | chr9-119251128-INS-79 |
| chr9-119271437-INS-63 | chr9-119271437-INS-64 |
| chr9-119281174-INS-185 | chr9-119281174-INS-185 |
| chr9-119288556-INS-93 | chr9-119288556-INS-96 |
| chr9-119295110-INS-395 | chr9-119295110-INS-395 |
| chr9-119301476-INS-69 | chr9-119301471-INS-74 |
| chr9-119337128-INS-116 | chr9-119337136-INS-118 |
| chr9-119364799-INS-124 | chr9-119364799-INS-156 |
| chr9-119377870-INS-77 | chr9-119377870-INS-64 |
| chr9-119393504-INS-59 | chr9-119393507-INS-87 |
| chr9-119399108-INS-63 | chr9-119399108-INS-57 |
| chr9-119423745-INS-216 | chr9-119423745-INS-246 |
| chr9-119428045-INS-82 | chr9-119428045-INS-86 |
| chr9-119430651-INS-128 | chr9-119430651-INS-134 |
| chr9-119437617-INS-57 | chr9-119437617-INS-57 |
| chr9-119479431-INS-308 | chr9-119479431-INS-308 |
| chr9-119484596-INS-79 | chr9-119484596-INS-52 |
| chr9-119503595-INS-232 | chr9-119503596-INS-231 |
| chr9-119522368-INS-66 | chr9-119522368-INS-66 |
| chr9-119524155-INS-81 | chr9-119524156-INS-51 |
| chr9-119526888-INS-62 | chr9-119526888-INS-63 |
| chr9-119535547-INS-136 | chr9-119535525-INS-168 |
| chr9-119552786-INS-72 | chr9-119552786-INS-72 |
| chr9-119565945-INS-100 | chr9-119565943-INS-92 |
| chr9-119582152-INS-60 | chr9-119582152-INS-60 |
| chr9-119605395-INS-86 | chr9-119605395-INS-84 |
| chr9-119607984-INS-364 | chr9-119607984-INS-364 |
| chr9-119610364-INS-66 | chr9-119610364-INS-60 |
| chr9-119623861-INS-1864 | chr9-119623862-INS-1864 |
| chr9-119629014-INS-256 | chr9-119629014-INS-256 |
| chr9-119635231-INS-1256 | chr9-119635232-INS-1256 |
| chr9-119658968-INS-1087 | chr9-119658969-INS-1087 |
| chr9-119668379-INS-59 | chr9-119668388-INS-85 |
| chr9-119682376-INS-97 | chr9-119682393-INS-67 |
| chr9-119688584-INS-56 | chr9-119688584-INS-60 |
| chr9-119722576-INS-439 | chr9-119722576-INS-439 |
| chr9-119744240-INS-60 | chr9-119744240-INS-62 |
| chr9-119761332-INS-53 | chr9-119761332-INS-53 |
| chr9-119784715-INS-57 | chr9-119784716-INS-72 |
| chr9-119788992-INS-101 | chr9-119788992-INS-101 |
| chr9-119814747-INS-85 | chr9-119814750-INS-91 |
| chr9-119821547-INS-60 | chr9-119821528-INS-106 |
| chr9-119826110-INS-248 | chr9-119826110-INS-251 |
| chr9-119858941-INS-57 | chr9-119858942-INS-57 |
| chr9-119862084-INS-700 | chr9-119862084-INS-700 |
| chr9-119865525-INS-52 | chr9-119865525-INS-52 |
| chr9-119868265-INS-207 | chr9-119868265-INS-207 |
| chr9-119878210-INS-84 | chr9-119878210-INS-84 |
| chr9-119890125-INS-414 | chr9-119890125-INS-414 |

| | |
|---|---|
| chr9-119910287-INS-194 | chr9-119910288-INS-193 |
| chr9-119912672-INS-158 | chr9-119912672-INS-162 |
| chr9-119919584-INS-250 | chr9-119919584-INS-250 |
| chr9-119925671-INS-497 | chr9-119925671-INS-497 |
| chr9-119931097-INS-102 | chr9-119931097-INS-84 |
| chr9-119943621-INS-57 | chr9-119943621-INS-57 |
| chr9-119952632-INS-73 | chr9-119952637-INS-111 |
| chr9-119956585-INS-146 | chr9-119956585-INS-146 |
| chr9-119960737-INS-365 | chr9-119960737-INS-365 |
| chr9-119966385-INS-70 | chr9-119966398-INS-52 |
| chr9-119967118-INS-78 | chr9-119967130-INS-64 |
| chr9-119971051-INS-198 | chr9-119971051-INS-199 |
| chr9-119992390-INS-196 | chr9-119992390-INS-196 |
| chr9-120012365-INS-60 | chr9-120012365-INS-60 |
| chr9-120015664-INS-63 | chr9-120015664-INS-63 |
| chr9-120017874-INS-51 | chr9-120017874-INS-51 |
| chr9-120018888-INS-73 | chr9-120018888-INS-73 |
| chr9-120024042-INS-157 | chr9-120024042-INS-156 |
| chr9-120030476-INS-56 | chr9-120030478-INS-100 |
| chr9-120046724-INS-72 | chr9-120046724-INS-75 |
| chr9-120056044-INS-58 | chr9-120056045-INS-56 |
| chr9-120058576-INS-53 | chr9-120058573-INS-61 |
| chr9-120071031-INS-60 | chr9-120071031-INS-66 |
| chr9-120077324-INS-358 | chr9-120077324-INS-358 |
| chr9-120085530-INS-130 | chr9-120085530-INS-130 |
| chr9-120087825-INS-53 | chr9-120087825-INS-53 |
| chr9-120101793-INS-57 | chr9-120101801-INS-56 |
| chr9-120116833-INS-1460 | chr9-120116834-INS-1460 |
| chr9-120117676-INS-95 | chr9-120117676-INS-56 |
| chr9-120129081-INS-82 | chr9-120129040-INS-82 |
| chr9-120157201-INS-59 | chr9-120157203-INS-70 |
| chr9-120163165-INS-66 | chr9-120163168-INS-54 |
| chr9-120168352-INS-55 | chr9-120168352-INS-55 |
| chr9-120171272-INS-80 | chr9-120171272-INS-56 |
| chr9-120208110-INS-73 | chr9-120208110-INS-78 |
| chr9-120214425-INS-306 | chr9-120214425-INS-306 |
| chr9-120242818-INS-205 | chr9-120242818-INS-205 |
| chr9-120254091-INS-58 | chr9-120254083-INS-52 |
| chr9-120272395-INS-136 | chr9-120272408-INS-150 |
| chr9-120334154-INS-189 | chr9-120334154-INS-209 |
| chr9-120336969-INS-306 | chr9-120336951-INS-306 |
| chr9-120344568-INS-199 | chr9-120344568-INS-188 |
| chr9-120347749-INS-58 | chr9-120347749-INS-93 |
| chr9-120351523-INS-73 | chr9-120351523-INS-73 |
| chr9-120352718-INS-68 | chr9-120352718-INS-51 |
| chr9-120391543-INS-60 | chr9-120391543-INS-64 |
| chr9-120405743-INS-106 | chr9-120405743-INS-107 |
| chr9-120412386-INS-71 | chr9-120412386-INS-53 |
| chr9-120413551-INS-229 | chr9-120413551-INS-229 |
| chr9-120427143-INS-412 | chr9-120427143-INS-412 |
| chr9-120440631-INS-56 | chr9-120440631-INS-56 |
| chr9-120445144-INS-78 | chr9-120445165-INS-78 |
| chr9-120492106-INS-204 | chr9-120492106-INS-205 |
| chr9-120497857-INS-181 | chr9-120497857-INS-191 |
| chr9-120500373-INS-76 | chr9-120500373-INS-76 |
| chr9-120523089-INS-68 | chr9-120523089-INS-68 |
| chr9-120524048-INS-401 | chr9-120524048-INS-401 |
| chr9-120528469-INS-208 | chr9-120528469-INS-209 |
| chr9-120541884-INS-219 | chr9-120541884-INS-217 |
| chr9-120555631-INS-383 | chr9-120555625-INS-384 |
| chr9-120564936-INS-254 | chr9-120564936-INS-254 |
| chr9-120574686-INS-102 | chr9-120574686-INS-143 |
| chr9-120576472-INS-207 | chr9-120576472-INS-207 |
| chr9-120600760-INS-209 | chr9-120600760-INS-209 |
| chr9-120606475-INS-3230 | chr9-120606470-INS-3230 |
| chr9-120606596-INS-120 | chr9-120606596-INS-120 |
| chr9-120610062-INS-234 | chr9-120610062-INS-235 |
| chr9-120636596-INS-52 | chr9-120636594-INS-54 |
| chr9-120671410-INS-55 | chr9-120671410-INS-55 |
| chr9-120682917-INS-437 | chr9-120682917-INS-437 |
| chr9-120692269-INS-78 | chr9-120692244-INS-84 |
| chr9-120696265-INS-85 | chr9-120696263-INS-85 |
| chr9-120715245-INS-174 | chr9-120715284-INS-173 |
| chr9-120726899-INS-74 | chr9-120726899-INS-74 |
| chr9-120739427-INS-58 | chr9-120739398-INS-50 |
| chr9-120741445-INS-498 | chr9-120741445-INS-498 |
| chr9-120745096-INS-207 | chr9-120745096-INS-208 |

| | |
|---|---|
| chr9-120754979-INS-56 | chr9-120755006-INS-76 |
| chr9-120798897-INS-231 | chr9-120798897-INS-231 |
| chr9-120801940-INS-179 | chr9-120801940-INS-179 |
| chr9-120818864-INS-58 | chr9-120818864-INS-58 |
| chr9-120824582-INS-120 | chr9-120824582-INS-120 |
| chr9-120839060-INS-328 | chr9-120839060-INS-328 |
| chr9-120862096-INS-55 | chr9-120862096-INS-55 |
| chr9-120862538-INS-1008 | chr9-120862539-INS-1008 |
| chr9-120883025-INS-193 | chr9-120883024-INS-212 |
| chr9-120884779-INS-128 | chr9-120884779-INS-128 |
| chr9-120892568-INS-498 | chr9-120892568-INS-498 |
| chr9-120914754-INS-2785 | chr9-120914755-INS-2785 |
| chr9-120937167-INS-454 | chr9-120937167-INS-429 |
| chr9-120944899-INS-76 | chr9-120944859-INS-86 |
| chr9-120946470-INS-3902 | chr9-120946471-INS-3904 |
| chr9-120957108-INS-64 | chr9-120957108-INS-64 |
| chr9-120958578-INS-98 | chr9-120958578-INS-98 |
| chr9-120965447-INS-90 | chr9-120965447-INS-90 |
| chr9-120987126-INS-52 | chr9-120987126-INS-52 |
| chr9-120990142-INS-92 | chr9-120990142-INS-92 |
| chr9-120991493-INS-63 | chr9-120991493-INS-84 |
| chr9-120993734-INS-522 | chr9-120993734-INS-522 |
| chr9-120994664-INS-440 | chr9-120994664-INS-440 |
| chr9-121003839-INS-539 | chr9-121003839-INS-539 |
| chr9-121032275-INS-102 | chr9-121032305-INS-66 |
| chr9-121078422-INS-106 | chr9-121078419-INS-88 |
| chr9-121081787-INS-177 | chr9-121081787-INS-177 |
| chr9-121082277-INS-229 | chr9-121082277-INS-230 |
| chr9-121084434-INS-58 | chr9-121084434-INS-58 |
| chr9-121107098-INS-227 | chr9-121107098-INS-227 |
| chr9-121121552-INS-254 | chr9-121121552-INS-254 |
| chr9-121123893-INS-399 | chr9-121123893-INS-400 |
| chr9-121149650-INS-54 | chr9-121149650-INS-54 |
| chr9-121164016-INS-58 | chr9-121164016-INS-50 |
| chr9-121203657-INS-56 | chr9-121203657-INS-56 |
| chr9-121209034-INS-71 | chr9-121209034-INS-80 |
| chr9-121221404-INS-61 | chr9-121221404-INS-61 |
| chr9-121222580-INS-219 | chr9-121222586-INS-245 |
| chr9-121226425-INS-64 | chr9-121226435-INS-67 |
| chr9-121276718-INS-133 | chr9-121276719-INS-118 |
| chr9-121291103-INS-413 | chr9-121291103-INS-414 |
| chr9-121294751-INS-74 | chr9-121294749-INS-74 |
| chr9-121314779-INS-138 | chr9-121314826-INS-132 |
| chr9-121334452-INS-54 | chr9-121334452-INS-64 |
| chr9-121347354-INS-1301 | chr9-121347355-INS-1274 |
| chr9-121357436-INS-174 | chr9-121357430-INS-173 |
| chr9-121365004-INS-80 | chr9-121364972-INS-80 |
| chr9-121375007-INS-55 | chr9-121375007-INS-55 |
| chr9-121390696-INS-189 | chr9-121390696-INS-189 |
| chr9-121399809-INS-138 | chr9-121399809-INS-138 |
| chr9-121403461-INS-59 | chr9-121403473-INS-59 |
| chr9-121413835-INS-336 | chr9-121413840-INS-336 |
| chr9-121420529-INS-7089 | chr9-121420530-INS-7089 |
| chr9-121420582-INS-146 | chr9-121420582-INS-146 |
| chr9-121469334-INS-100 | chr9-121469352-INS-100 |
| chr9-121528112-INS-213 | chr9-121528112-INS-213 |
| chr9-121633466-INS-777 | chr9-121633466-INS-758 |
| chr9-121641721-INS-252 | chr9-121641721-INS-252 |
| chr9-121662598-INS-312 | chr9-121662611-INS-312 |
| chr9-121665244-INS-1073 | chr9-121665245-INS-1073 |
| chr9-121674469-INS-5957 | chr9-121674470-INS-5957 |
| chr9-121676116-INS-124 | chr9-121676117-INS-141 |
| chr9-121680793-INS-315 | chr9-121680793-INS-315 |
| chr9-121689533-INS-210 | chr9-121689533-INS-210 |
| chr9-121695266-INS-81 | chr9-121695266-INS-81 |
| chr9-121702202-INS-354 | chr9-121702246-INS-359 |
| chr9-121725568-INS-76 | chr9-121725595-INS-80 |
| chr9-121757964-INS-94 | chr9-121757964-INS-108 |
| chr9-121761741-INS-72 | chr9-121761709-INS-87 |
| chr9-121778153-INS-440 | chr9-121778153-INS-440 |
| chr9-121789927-INS-197 | chr9-121789925-INS-241 |
| chr9-121798184-INS-80 | chr9-121798184-INS-80 |
| chr9-121810105-INS-1889 | chr9-121810106-INS-1889 |
| chr9-121868208-INS-117 | chr9-121868208-INS-117 |
| chr9-121872328-INS-77 | chr9-121872328-INS-69 |
| chr9-121900843-INS-339 | chr9-121900843-INS-339 |
| chr9-121948829-INS-52 | chr9-121948829-INS-54 |

| | |
|---|---|
| chr9-121967999-INS-54 | chr9-121967999-INS-54 |
| chr9-121978331-INS-135 | chr9-121978350-INS-98 |
| chr9-122010962-INS-482 | chr9-122010962-INS-457 |
| chr9-122015704-INS-222 | chr9-122015704-INS-222 |
| chr9-122044745-INS-253 | chr9-122044739-INS-258 |
| chr9-122059892-INS-76 | chr9-122059893-INS-50 |
| chr9-122071787-INS-72 | chr9-122071787-INS-80 |
| chr9-122074997-INS-448 | chr9-122074997-INS-448 |
| chr9-122079482-INS-121 | chr9-122079482-INS-121 |
| chr9-122123953-INS-135 | chr9-122123953-INS-135 |
| chr9-122146445-INS-189 | chr9-122146445-INS-181 |
| chr9-122179859-INS-57 | chr9-122179859-INS-57 |
| chr9-122189105-INS-66 | chr9-122189105-INS-66 |
| chr9-122213770-INS-794 | chr9-122213770-INS-794 |
| chr9-122250994-INS-58 | chr9-122250994-INS-56 |
| chr9-122284167-INS-2837 | chr9-122284173-INS-2831 |
| chr9-122291659-INS-5237 | chr9-122291660-INS-5237 |
| chr9-122304776-INS-420 | chr9-122304776-INS-420 |
| chr9-122375300-INS-421 | chr9-122375300-INS-421 |
| chr9-122394043-INS-73 | chr9-122394043-INS-73 |
| chr9-122404983-INS-76 | chr9-122404983-INS-73 |
| chr9-122405883-INS-77 | chr9-122405883-INS-121 |
| chr9-122423800-INS-92 | chr9-122423804-INS-52 |
| chr9-122432502-INS-70 | chr9-122432502-INS-70 |
| chr9-122448899-INS-590 | chr9-122448899-INS-590 |
| chr9-122479753-INS-152 | chr9-122479776-INS-133 |
| chr9-122485582-INS-178 | chr9-122485582-INS-180 |
| chr9-122512159-INS-93 | chr9-122512191-INS-79 |
| chr9-122525826-INS-65 | chr9-122525808-INS-51 |
| chr9-122541532-INS-184 | chr9-122541532-INS-184 |
| chr9-122546385-INS-422 | chr9-122546385-INS-422 |
| chr9-122565183-INS-486 | chr9-122565183-INS-486 |
| chr9-122581339-INS-158 | chr9-122581339-INS-158 |
| chr9-122593517-INS-79 | chr9-122593517-INS-79 |
| chr9-122595725-INS-52 | chr9-122595725-INS-52 |
| chr9-122630375-INS-98 | chr9-122630375-INS-98 |
| chr9-122631298-INS-198 | chr9-122631298-INS-198 |
| chr9-122645748-INS-411 | chr9-122645748-INS-411 |
| chr9-122651102-INS-200 | chr9-122651102-INS-200 |
| chr9-122691254-INS-104 | chr9-122691254-INS-104 |
| chr9-122709404-INS-4667 | chr9-122709405-INS-4667 |
| chr9-122709405-INS-440 | chr9-122709405-INS-440 |
| chr9-122729710-INS-465 | chr9-122729710-INS-468 |
| chr9-122739891-INS-184 | chr9-122739891-INS-184 |
| chr9-122761212-INS-662 | chr9-122761212-INS-662 |
| chr9-122785321-INS-60 | chr9-122785321-INS-64 |
| chr9-122785776-INS-340 | chr9-122785776-INS-340 |
| chr9-122788279-INS-64 | chr9-122788286-INS-102 |
| chr9-122791319-INS-155 | chr9-122791319-INS-155 |
| chr9-122793101-INS-138 | chr9-122793101-INS-130 |
| chr9-122798362-INS-56 | chr9-122798362-INS-56 |
| chr9-122800870-INS-61 | chr9-122800870-INS-61 |
| chr9-122816505-INS-56 | chr9-122816510-INS-61 |
| chr9-122827147-INS-66 | chr9-122827147-INS-60 |
| chr9-122839312-INS-80 | chr9-122839312-INS-80 |
| chr9-122857296-INS-236 | chr9-122857296-INS-258 |
| chr9-122858892-INS-423 | chr9-122858892-INS-423 |
| chr9-122860305-INS-202 | chr9-122860305-INS-200 |
| chr9-122872251-INS-967 | chr9-122872251-INS-969 |
| chr9-122891929-INS-74 | chr9-122891929-INS-75 |
| chr9-122894575-INS-220 | chr9-122894589-INS-215 |
| chr9-122903408-INS-68 | chr9-122903408-INS-68 |
| chr9-122904409-INS-310 | chr9-122904409-INS-310 |
| chr9-122910465-INS-99 | chr9-122910465-INS-79 |
| chr9-122952579-INS-515 | chr9-122952579-INS-515 |
| chr9-122953583-INS-52 | chr9-122953591-INS-52 |
| chr9-122958008-INS-79 | chr9-122958008-INS-79 |
| chr9-122963360-INS-92 | chr9-122963362-INS-82 |
| chr9-122969688-INS-86 | chr9-122969688-INS-66 |
| chr9-122987837-INS-61 | chr9-122987838-INS-56 |
| chr9-123010739-INS-54 | chr9-123010739-INS-95 |
| chr9-123027859-INS-173 | chr9-123027859-INS-173 |
| chr9-123040033-INS-71 | chr9-123040033-INS-92 |
| chr9-123081734-INS-141 | chr9-123081734-INS-142 |
| chr9-123085788-INS-54 | chr9-123085788-INS-56 |
| chr9-123121117-INS-82 | chr9-123121071-INS-96 |
| chr9-123128202-INS-74 | chr9-123128202-INS-74 |

| | |
|---|---|
| chr9-123141084-INS-64 | chr9-123141084-INS-64 |
| chr9-123144359-INS-231 | chr9-123144359-INS-231 |
| chr9-123161748-INS-875 | chr9-123161749-INS-875 |
| chr9-123186238-INS-401 | chr9-123186238-INS-400 |
| chr9-123209598-INS-80 | chr9-123209598-INS-84 |
| chr9-123211121-INS-55 | chr9-123211117-INS-52 |
| chr9-123219612-INS-215 | chr9-123219612-INS-215 |
| chr9-123223633-INS-193 | chr9-123223633-INS-195 |
| chr9-123246718-INS-62 | chr9-123246718-INS-66 |
| chr9-123253571-INS-213 | chr9-123253571-INS-212 |
| chr9-123261928-INS-314 | chr9-123261928-INS-321 |
| chr9-123274562-INS-180 | chr9-123274562-INS-180 |
| chr9-123277817-INS-148 | chr9-123277817-INS-148 |
| chr9-123283950-INS-858 | chr9-123283953-INS-869 |
| chr9-123292742-INS-56 | chr9-123292742-INS-56 |
| chr9-123318053-INS-111 | chr9-123318053-INS-113 |
| chr9-123356725-INS-54 | chr9-123356725-INS-56 |
| chr9-123392256-INS-96 | chr9-123392256-INS-100 |
| chr9-123423249-INS-72 | chr9-123423249-INS-72 |
| chr9-123451367-INS-88 | chr9-123451371-INS-102 |
| chr9-123478514-INS-434 | chr9-123478514-INS-434 |
| chr9-123500947-INS-99 | chr9-123500947-INS-99 |
| chr9-123516147-INS-168 | chr9-123516147-INS-168 |
| chr9-123549310-INS-208 | chr9-123549310-INS-208 |
| chr9-123553813-INS-205 | chr9-123553813-INS-205 |
| chr9-123554586-INS-377 | chr9-123554586-INS-377 |
| chr9-123570604-INS-58 | chr9-123570590-INS-106 |
| chr9-123584513-INS-385 | chr9-123584513-INS-386 |
| chr9-123602677-INS-168 | chr9-123602673-INS-163 |
| chr9-123630996-INS-209 | chr9-123630996-INS-209 |
| chr9-123652765-INS-96 | chr9-123652765-INS-69 |
| chr9-123693383-INS-186 | chr9-123693396-INS-201 |
| chr9-123694565-INS-135 | chr9-123694565-INS-135 |
| chr9-123730011-INS-69 | chr9-123730012-INS-69 |
| chr9-123749853-INS-7237 | chr9-123749854-INS-7237 |
| chr9-123782415-INS-1889 | chr9-123782416-INS-1889 |
| chr9-123825689-INS-6963 | chr9-123825690-INS-6965 |
| chr9-123892535-INS-54 | chr9-123892535-INS-54 |
| chr9-124063718-INS-413 | chr9-124063718-INS-412 |
| chr9-124141074-INS-108 | chr9-124141074-INS-96 |
| chr9-124147726-INS-1297 | chr9-124147727-INS-1297 |
| chr9-124196885-INS-58 | chr9-124196885-INS-58 |
| chr9-124244610-INS-1270 | chr9-124244611-INS-1270 |
| chrX-3372321-INS-95 | chrX-3372321-INS-95 |
| chrX-4257679-INS-67 | chrX-4257679-INS-66 |
| chrX-5249740-INS-100 | chrX-5249740-INS-108 |
| chrX-5292950-INS-1232 | chrX-5292951-INS-1231 |
| chrX-5338126-INS-201 | chrX-5338126-INS-203 |
| chrX-5422563-INS-2125 | chrX-5422564-INS-2126 |
| chrX-5503401-INS-1211 | chrX-5503402-INS-1211 |
| chrX-5507466-INS-56 | chrX-5507466-INS-56 |
| chrX-5519540-INS-6682 | chrX-5519541-INS-6682 |
| chrX-5531427-INS-167 | chrX-5531427-INS-167 |
| chrX-5536977-INS-3187 | chrX-5536978-INS-3187 |
| chrX-5624397-INS-64 | chrX-5624394-INS-68 |
| chrX-5646523-INS-174 | chrX-5646523-INS-160 |
| chrX-5678143-INS-2948 | chrX-5678144-INS-2948 |
| chrX-5716439-INS-6329 | chrX-5716440-INS-6326 |
| chrX-5747481-INS-81 | chrX-5747481-INS-93 |
| chrX-5802809-INS-6546 | chrX-5802810-INS-6547 |
| chrX-5807355-INS-55 | chrX-5807355-INS-51 |
| chrX-5841380-INS-59 | chrX-5841380-INS-59 |
| chrX-5848851-INS-1890 | chrX-5848852-INS-1890 |
| chrX-5887101-INS-58 | chrX-5887101-INS-89 |
| chrX-5893767-INS-79 | chrX-5893767-INS-85 |
| chrX-5923992-INS-3508 | chrX-5923993-INS-3510 |
| chrX-5933731-INS-197 | chrX-5933731-INS-197 |
| chrX-5935582-INS-113 | chrX-5935610-INS-131 |
| chrX-5954585-INS-2412 | chrX-5954586-INS-2417 |
| chrX-5969957-INS-204 | chrX-5969957-INS-204 |
| chrX-6000387-INS-6443 | chrX-6000388-INS-6443 |
| chrX-6074559-INS-184 | chrX-6074559-INS-184 |
| chrX-6096192-INS-56 | chrX-6096216-INS-56 |
| chrX-6124676-INS-176 | chrX-6124676-INS-176 |
| chrX-6134272-INS-187 | chrX-6134272-INS-186 |
| chrX-6135152-INS-1892 | chrX-6135153-INS-1892 |
| chrX-6143413-INS-114 | chrX-6143413-INS-101 |

| | |
|---|---|
| chrX-6152621-INS-85 | chrX-6152621-INS-67 |
| chrX-6154231-INS-94 | chrX-6154237-INS-74 |
| chrX-6176964-INS-171 | chrX-6176964-INS-171 |
| chrX-6187071-INS-8934 | chrX-6187072-INS-8937 |
| chrX-6195110-INS-56 | chrX-6195110-INS-56 |
| chrX-6204027-INS-58 | chrX-6204027-INS-50 |
| chrX-6218280-INS-1116 | chrX-6218281-INS-1116 |
| chrX-6225338-INS-142 | chrX-6225384-INS-138 |
| chrX-6227278-INS-81 | chrX-6227278-INS-81 |
| chrX-6251889-INS-200 | chrX-6251880-INS-200 |
| chrX-6256419-INS-390 | chrX-6256419-INS-390 |
| chrX-6271478-INS-238 | chrX-6271478-INS-238 |
| chrX-6299001-INS-394 | chrX-6299001-INS-394 |
| chrX-6339147-INS-6684 | chrX-6339148-INS-6684 |
| chrX-6353168-INS-96 | chrX-6353168-INS-96 |
| chrX-6362427-INS-101 | chrX-6362427-INS-60 |
| chrX-6385795-INS-124 | chrX-6385795-INS-122 |
| chrX-6427524-INS-633 | chrX-6427524-INS-633 |
| chrX-6432543-INS-5313 | chrX-6432544-INS-5313 |
| chrX-6560751-INS-203 | chrX-6560751-INS-203 |
| chrX-6578242-INS-55 | chrX-6578242-INS-65 |
| chrX-6597547-INS-121 | chrX-6597535-INS-92 |
| chrX-6612480-INS-576 | chrX-6612480-INS-577 |
| chrX-6623508-INS-70 | chrX-6623508-INS-69 |
| chrX-6636814-INS-354 | chrX-6636814-INS-389 |
| chrX-6650509-INS-78 | chrX-6650509-INS-78 |
| chrX-6690671-INS-72 | chrX-6690672-INS-60 |
| chrX-6695117-INS-185 | chrX-6695117-INS-185 |
| chrX-6700566-INS-55 | chrX-6700566-INS-55 |
| chrX-6732771-INS-205 | chrX-6732772-INS-206 |
| chrX-6778440-INS-70 | chrX-6778440-INS-70 |
| chrX-6782829-INS-122 | chrX-6782859-INS-96 |
| chrX-6850294-INS-55 | chrX-6850249-INS-55 |
| chrX-6855837-INS-96 | chrX-6855837-INS-102 |
| chrX-6869358-INS-77 | chrX-6869359-INS-77 |
| chrX-6908966-INS-198 | chrX-6908966-INS-179 |
| chrX-6937156-INS-102 | chrX-6937156-INS-102 |
| chrX-6954495-INS-75 | chrX-6954453-INS-100 |
| chrX-6955611-INS-1671 | chrX-6955612-INS-1672 |
| chrX-6972316-INS-111 | chrX-6972331-INS-67 |
| chrX-6974524-INS-2216 | chrX-6974525-INS-2218 |
| chrX-7049151-INS-86 | chrX-7049107-INS-95 |
| chrX-7074632-INS-61 | chrX-7074632-INS-61 |
| chrX-7108795-INS-58 | chrX-7108795-INS-70 |
| chrX-7131132-INS-262 | chrX-7131132-INS-262 |
| chrX-7141062-INS-714 | chrX-7141062-INS-713 |
| chrX-7249831-INS-84 | chrX-7249835-INS-65 |
| chrX-7267633-INS-211 | chrX-7267633-INS-210 |
| chrX-7302372-INS-199 | chrX-7302372-INS-199 |
| chrX-7363508-INS-69 | chrX-7363508-INS-69 |
| chrX-7382157-INS-65 | chrX-7382157-INS-60 |
| chrX-7386485-INS-6621 | chrX-7386486-INS-6621 |
| chrX-7415962-INS-202 | chrX-7415962-INS-201 |
| chrX-7428859-INS-96 | chrX-7428863-INS-90 |
| chrX-7470785-INS-206 | chrX-7470785-INS-206 |
| chrX-7528243-INS-51 | chrX-7528197-INS-60 |
| chrX-7544195-INS-5226 | chrX-7544196-INS-5226 |
| chrX-7550597-INS-195 | chrX-7550597-INS-195 |
| chrX-7555356-INS-205 | chrX-7555360-INS-201 |
| chrX-7558617-INS-84 | chrX-7558622-INS-80 |
| chrX-7563891-INS-178 | chrX-7563891-INS-184 |
| chrX-7584400-INS-211 | chrX-7584400-INS-211 |
| chrX-7609935-INS-57 | chrX-7609935-INS-57 |
| chrX-7610934-INS-120 | chrX-7610934-INS-168 |
| chrX-7650202-INS-203 | chrX-7650202-INS-203 |
| chrX-7655264-INS-187 | chrX-7655264-INS-187 |
| chrX-7692448-INS-214 | chrX-7692449-INS-214 |
| chrX-7696953-INS-112 | chrX-7696953-INS-100 |
| chrX-7699279-INS-81 | chrX-7699280-INS-81 |
| chrX-7764955-INS-164 | chrX-7764955-INS-166 |
| chrX-7780771-INS-218 | chrX-7780771-INS-216 |
| chrX-7819288-INS-78 | chrX-7819272-INS-58 |
| chrX-7854108-INS-62 | chrX-7854108-INS-62 |
| chrX-7860046-INS-106 | chrX-7860046-INS-96 |
| chrX-7867944-INS-61 | chrX-7867944-INS-62 |
| chrX-7873738-INS-198 | chrX-7873738-INS-199 |
| chrX-7874771-INS-209 | chrX-7874771-INS-208 |

| | |
|---|---|
| chrX-7877525-INS-214 | chrX-7877525-INS-214 |
| chrX-7887010-INS-197 | chrX-7887010-INS-197 |
| chrX-7905377-INS-216 | chrX-7905377-INS-216 |
| chrX-7909266-INS-60 | chrX-7909266-INS-60 |
| chrX-7910347-INS-101 | chrX-7910347-INS-101 |
| chrX-7912349-INS-56 | chrX-7912343-INS-60 |
| chrX-7960472-INS-201 | chrX-7960472-INS-203 |
| chrX-7961439-INS-77 | chrX-7961431-INS-95 |
| chrX-7973852-INS-68 | chrX-7973852-INS-80 |
| chrX-7979775-INS-616 | chrX-7979775-INS-615 |
| chrX-7986496-INS-150 | chrX-7986496-INS-150 |
| chrX-8003204-INS-192 | chrX-8003204-INS-189 |
| chrX-8033604-INS-94 | chrX-8033604-INS-96 |
| chrX-8035790-INS-174 | chrX-8035790-INS-171 |
| chrX-8037629-INS-200 | chrX-8037629-INS-200 |
| chrX-8044534-INS-208 | chrX-8044534-INS-212 |
| chrX-8045478-INS-281 | chrX-8045478-INS-281 |
| chrX-8081671-INS-120 | chrX-8081681-INS-98 |
| chrX-8101591-INS-194 | chrX-8101595-INS-195 |
| chrX-8102647-INS-87 | chrX-8102651-INS-60 |
| chrX-8165187-INS-127 | chrX-8165187-INS-123 |
| chrX-8166559-INS-597 | chrX-8166559-INS-597 |
| chrX-8180149-INS-59 | chrX-8180149-INS-77 |
| chrX-8218755-INS-305 | chrX-8218755-INS-308 |
| chrX-8399025-INS-199 | chrX-8399025-INS-199 |
| chrX-8573708-INS-286 | chrX-8573708-INS-286 |
| chrX-8596665-INS-741 | chrX-8596665-INS-741 |
| chrX-8623056-INS-427 | chrX-8623056-INS-427 |
| chrX-8726428-INS-573 | chrX-8726428-INS-574 |
| chrX-8825397-INS-88 | chrX-8825408-INS-90 |
| chrX-8839656-INS-52 | chrX-8839656-INS-56 |
| chrX-8902880-INS-108 | chrX-8902880-INS-138 |
| chrX-8909413-INS-71 | chrX-8909413-INS-71 |
| chrX-8949548-INS-51 | chrX-8949549-INS-51 |
| chrX-8975082-INS-5413 | chrX-8975083-INS-5413 |
| chrX-8976450-INS-79 | chrX-8976450-INS-79 |
| chrX-8982106-INS-253 | chrX-8982106-INS-254 |
| chrX-8983516-INS-52 | chrX-8983516-INS-72 |
| chrX-8990445-INS-162 | chrX-8990445-INS-142 |
| chrX-8995656-INS-623 | chrX-8995656-INS-622 |
| chrX-9010730-INS-713 | chrX-9010730-INS-712 |
| chrX-9016494-INS-872 | chrX-9016494-INS-872 |
| chrX-9024159-INS-2438 | chrX-9024160-INS-2438 |
| chrX-9027177-INS-215 | chrX-9027177-INS-215 |
| chrX-9041664-INS-377 | chrX-9041664-INS-377 |
| chrX-9049132-INS-978 | chrX-9049132-INS-979 |
| chrX-9077087-INS-657 | chrX-9077087-INS-652 |
| chrX-9129048-INS-96 | chrX-9129048-INS-96 |
| chrX-9183167-INS-7358 | chrX-9183168-INS-7358 |
| chrX-9208916-INS-108 | chrX-9208916-INS-108 |
| chrX-9218465-INS-209 | chrX-9218465-INS-209 |
| chrX-9230123-INS-4868 | chrX-9230124-INS-4868 |
| chrX-9244517-INS-108 | chrX-9244517-INS-108 |
| chrX-9264110-INS-160 | chrX-9264110-INS-160 |
| chrX-9275347-INS-873 | chrX-9275336-INS-862 |
| chrX-9276552-INS-54 | chrX-9276552-INS-54 |
| chrX-9282047-INS-179 | chrX-9282047-INS-167 |
| chrX-9283166-INS-104 | chrX-9283166-INS-104 |
| chrX-9286023-INS-262 | chrX-9286023-INS-262 |
| chrX-9337238-INS-102 | chrX-9337238-INS-102 |
| chrX-9353050-INS-5476 | chrX-9353051-INS-5476 |
| chrX-9364695-INS-6397 | chrX-9364696-INS-6397 |
| chrX-9382498-INS-80 | chrX-9382498-INS-80 |
| chrX-9390274-INS-367 | chrX-9390274-INS-367 |
| chrX-9433197-INS-940 | chrX-9433197-INS-940 |
| chrX-9450199-INS-160 | chrX-9450199-INS-160 |
| chrX-9460664-INS-6573 | chrX-9460665-INS-6572 |
| chrX-9473151-INS-871 | chrX-9473151-INS-874 |
| chrX-9488785-INS-53 | chrX-9488785-INS-53 |
| chrX-9495307-INS-201 | chrX-9495307-INS-201 |
| chrX-9499544-INS-56 | chrX-9499544-INS-56 |
| chrX-9500523-INS-108 | chrX-9500524-INS-108 |
| chrX-9506430-INS-251 | chrX-9506430-INS-254 |
| chrX-9515578-INS-104 | chrX-9515578-INS-104 |
| chrX-9565086-INS-2953 | chrX-9565087-INS-2953 |
| chrX-9584914-INS-811 | chrX-9584915-INS-811 |
| chrX-9615730-INS-332 | chrX-9615730-INS-332 |

| | |
|---|---|
| chrX-9651121-INS-71 | chrX-9651121-INS-55 |
| chrX-9676274-INS-58 | chrX-9676274-INS-58 |
| chrX-9736701-INS-4232 | chrX-9736702-INS-4232 |
| chrX-9784630-INS-51 | chrX-9784601-INS-79 |
| chrX-9818635-INS-178 | chrX-9818635-INS-178 |
| chrX-9838865-INS-398 | chrX-9838865-INS-398 |
| chrX-9852198-INS-59 | chrX-9852198-INS-59 |
| chrX-9862452-INS-348 | chrX-9862452-INS-348 |
| chrX-9888092-INS-106 | chrX-9888092-INS-106 |
| chrX-9888768-INS-52 | chrX-9888768-INS-76 |
| chrX-9922503-INS-378 | chrX-9922503-INS-378 |
| chrX-9937886-INS-80 | chrX-9937886-INS-80 |
| chrX-9947069-INS-255 | chrX-9947069-INS-255 |
| chrX-9950580-INS-358 | chrX-9950580-INS-358 |
| chrX-9975415-INS-71 | chrX-9975415-INS-78 |
| chrX-9978443-INS-60 | chrX-9978443-INS-60 |
| chrX-9981287-INS-535 | chrX-9981287-INS-535 |
| chrX-10012031-INS-1169 | chrX-10012032-INS-1168 |
| chrX-10014937-INS-56 | chrX-10014937-INS-56 |
| chrX-10030725-INS-62 | chrX-10030725-INS-62 |
| chrX-10041390-INS-121 | chrX-10041391-INS-85 |
| chrX-10068081-INS-226 | chrX-10068081-INS-226 |
| chrX-10077301-INS-548 | chrX-10077302-INS-548 |
| chrX-10108534-INS-537 | chrX-10108534-INS-536 |
| chrX-10149386-INS-162 | chrX-10149386-INS-186 |
| chrX-10172767-INS-3000 | chrX-10172768-INS-3000 |
| chrX-10173147-INS-607 | chrX-10173147-INS-608 |
| chrX-10181215-INS-563 | chrX-10181215-INS-563 |
| chrX-10202427-INS-396 | chrX-10202427-INS-394 |
| chrX-10213589-INS-52 | chrX-10213589-INS-52 |
| chrX-10224717-INS-215 | chrX-10224715-INS-218 |
| chrX-10228608-INS-1718 | chrX-10228609-INS-1718 |
| chrX-10229772-INS-531 | chrX-10229772-INS-485 |
| chrX-10247015-INS-594 | chrX-10247015-INS-594 |
| chrX-10285479-INS-95 | chrX-10285454-INS-95 |
| chrX-10309541-INS-1444 | chrX-10309542-INS-1444 |
| chrX-10348926-INS-56 | chrX-10348926-INS-74 |
| chrX-10421607-INS-68 | chrX-10421607-INS-70 |
| chrX-10472010-INS-54 | chrX-10472010-INS-54 |
| chrX-10495882-INS-54 | chrX-10495883-INS-54 |
| chrX-10527957-INS-155 | chrX-10527957-INS-155 |
| chrX-10538903-INS-70 | chrX-10538903-INS-70 |
| chrX-10566017-INS-161 | chrX-10566017-INS-161 |
| chrX-10579044-INS-53 | chrX-10579037-INS-54 |
| chrX-10591851-INS-72 | chrX-10591851-INS-72 |
| chrX-10615249-INS-690 | chrX-10615249-INS-690 |
| chrX-10619266-INS-72 | chrX-10619254-INS-60 |
| chrX-10664608-INS-206 | chrX-10664608-INS-208 |
| chrX-10692428-INS-454 | chrX-10692428-INS-453 |
| chrX-10703307-INS-775 | chrX-10703307-INS-781 |
| chrX-10713880-INS-163 | chrX-10713881-INS-168 |
| chrX-10755102-INS-594 | chrX-10755108-INS-598 |
| chrX-10756677-INS-227 | chrX-10756677-INS-231 |
| chrX-10760762-INS-78 | chrX-10760770-INS-64 |
| chrX-10763229-INS-563 | chrX-10763229-INS-563 |
| chrX-10791031-INS-786 | chrX-10791031-INS-783 |
| chrX-10816471-INS-199 | chrX-10816471-INS-199 |
| chrX-10828299-INS-1298 | chrX-10828300-INS-1298 |
| chrX-10900158-INS-153 | chrX-10900158-INS-152 |
| chrX-10930564-INS-113 | chrX-10930565-INS-112 |
| chrX-10952863-INS-56 | chrX-10952863-INS-52 |
| chrX-10972258-INS-530 | chrX-10972252-INS-530 |
| chrX-11037609-INS-1227 | chrX-11037610-INS-1227 |
| chrX-11042520-INS-52 | chrX-11042520-INS-52 |
| chrX-11048959-INS-378 | chrX-11048959-INS-381 |
| chrX-11053565-INS-62 | chrX-11053565-INS-86 |
| chrX-11072229-INS-2931 | chrX-11072230-INS-2931 |
| chrX-11094926-INS-51 | chrX-11094926-INS-51 |
| chrX-11108632-INS-76 | chrX-11108632-INS-76 |
| chrX-11130575-INS-124 | chrX-11130576-INS-105 |
| chrX-11133488-INS-120 | chrX-11133488-INS-120 |
| chrX-11136289-INS-66 | chrX-11136287-INS-52 |
| chrX-11157543-INS-302 | chrX-11157543-INS-302 |
| chrX-11203050-INS-63 | chrX-11203056-INS-74 |
| chrX-11233605-INS-6476 | chrX-11233606-INS-6474 |
| chrX-11241303-INS-1303 | chrX-11241304-INS-1274 |
| chrX-11245220-INS-60 | chrX-11245220-INS-60 |

| | |
|---|---|
| chrX-11318913-INS-99 | chrX-11318956-INS-88 |
| chrX-11373420-INS-100 | chrX-11373418-INS-104 |
| chrX-11377749-INS-51 | chrX-11377729-INS-101 |
| chrX-11398946-INS-89 | chrX-11398958-INS-88 |
| chrX-11524951-INS-66 | chrX-11524951-INS-68 |
| chrX-11528772-INS-1890 | chrX-11528773-INS-1890 |
| chrX-11542449-INS-95 | chrX-11542475-INS-61 |
| chrX-11584788-INS-67 | chrX-11584788-INS-69 |
| chrX-11604560-INS-414 | chrX-11604558-INS-450 |
| chrX-11616228-INS-124 | chrX-11616228-INS-124 |
| chrX-11635333-INS-196 | chrX-11635333-INS-196 |
| chrX-11654544-INS-554 | chrX-11654544-INS-559 |
| chrX-11664576-INS-54 | chrX-11664588-INS-76 |
| chrX-11678553-INS-332 | chrX-11678553-INS-332 |
| chrX-11723106-INS-7267 | chrX-11723107-INS-7267 |
| chrX-11746477-INS-113 | chrX-11746477-INS-113 |
| chrX-11771500-INS-83 | chrX-11771482-INS-95 |
| chrX-11775199-INS-60 | chrX-11775199-INS-63 |
| chrX-11798084-INS-78 | chrX-11798084-INS-83 |
| chrX-11816476-INS-343 | chrX-11816453-INS-342 |
| chrX-11830313-INS-232 | chrX-11830313-INS-256 |
| chrX-11836292-INS-72 | chrX-11836292-INS-72 |
| chrX-11850034-INS-324 | chrX-11849990-INS-327 |
| chrX-11868053-INS-58 | chrX-11868053-INS-62 |
| chrX-11888525-INS-115 | chrX-11888562-INS-66 |
| chrX-11894348-INS-147 | chrX-11894324-INS-121 |
| chrX-11897040-INS-166 | chrX-11897040-INS-166 |
| chrX-11907609-INS-84 | chrX-11907611-INS-90 |
| chrX-11989895-INS-82 | chrX-11989897-INS-73 |
| chrX-12058217-INS-74 | chrX-12058217-INS-64 |
| chrX-12063084-INS-56 | chrX-12063084-INS-56 |
| chrX-12075851-INS-256 | chrX-12075851-INS-256 |
| chrX-12079504-INS-150 | chrX-12079491-INS-124 |
| chrX-12093252-INS-390 | chrX-12093252-INS-389 |
| chrX-12107295-INS-72 | chrX-12107295-INS-64 |
| chrX-12119876-INS-165 | chrX-12119876-INS-166 |
| chrX-12144254-INS-770 | chrX-12144254-INS-770 |
| chrX-12151953-INS-78 | chrX-12151956-INS-78 |
| chrX-12158945-INS-6401 | chrX-12158946-INS-6401 |
| chrX-12168332-INS-65 | chrX-12168332-INS-68 |
| chrX-12170550-INS-363 | chrX-12170550-INS-364 |
| chrX-12176400-INS-75 | chrX-12176400-INS-75 |
| chrX-12179138-INS-490 | chrX-12179136-INS-499 |
| chrX-12230986-INS-57 | chrX-12230986-INS-56 |
| chrX-12278242-INS-64 | chrX-12278242-INS-64 |
| chrX-12303893-INS-57 | chrX-12303902-INS-71 |
| chrX-12349661-INS-104 | chrX-12349661-INS-104 |
| chrX-12386086-INS-201 | chrX-12386086-INS-201 |
| chrX-12405283-INS-213 | chrX-12405283-INS-214 |
| chrX-12408089-INS-208 | chrX-12408090-INS-204 |
| chrX-12433764-INS-65 | chrX-12433795-INS-55 |
| chrX-12440744-INS-182 | chrX-12440744-INS-180 |
| chrX-12447126-INS-78 | chrX-12447144-INS-76 |
| chrX-12467832-INS-150 | chrX-12467832-INS-170 |
| chrX-12476036-INS-315 | chrX-12476040-INS-321 |
| chrX-12503420-INS-196 | chrX-12503420-INS-196 |
| chrX-12601329-INS-417 | chrX-12601329-INS-417 |
| chrX-12638906-INS-214 | chrX-12638906-INS-213 |
| chrX-12653219-INS-64 | chrX-12653219-INS-64 |
| chrX-12654183-INS-231 | chrX-12654162-INS-211 |
| chrX-12656747-INS-402 | chrX-12656747-INS-402 |
| chrX-12664666-INS-198 | chrX-12664666-INS-198 |
| chrX-12670248-INS-181 | chrX-12670248-INS-181 |
| chrX-12680956-INS-54 | chrX-12680952-INS-74 |
| chrX-12684297-INS-204 | chrX-12684297-INS-204 |
| chrX-12686193-INS-200 | chrX-12686193-INS-200 |
| chrX-12687524-INS-174 | chrX-12687524-INS-181 |
| chrX-12697491-INS-200 | chrX-12697491-INS-201 |
| chrX-12700937-INS-68 | chrX-12700937-INS-74 |
| chrX-12709919-INS-240 | chrX-12709919-INS-251 |
| chrX-12734400-INS-259 | chrX-12734400-INS-259 |
| chrX-12741080-INS-65 | chrX-12741080-INS-65 |
| chrX-12747856-INS-191 | chrX-12747855-INS-194 |
| chrX-12771748-INS-205 | chrX-12771748-INS-205 |
| chrX-12775675-INS-212 | chrX-12775675-INS-212 |
| chrX-12778679-INS-56 | chrX-12778655-INS-76 |
| chrX-12779803-INS-71 | chrX-12779806-INS-73 |

| | |
|---|---|
| chrX-12835025-INS-210 | chrX-12835025-INS-210 |
| chrX-12849895-INS-61 | chrX-12849895-INS-67 |
| chrX-12867418-INS-83 | chrX-12867418-INS-83 |
| chrX-12883212-INS-270 | chrX-12883212-INS-270 |
| chrX-12883991-INS-205 | chrX-12883991-INS-208 |
| chrX-12931712-INS-68 | chrX-12931739-INS-61 |
| chrX-12993965-INS-122 | chrX-12993965-INS-122 |
| chrX-13001770-INS-83 | chrX-13001770-INS-59 |
| chrX-13005400-INS-2004 | chrX-13005401-INS-2004 |
| chrX-13012818-INS-214 | chrX-13012818-INS-210 |
| chrX-13019458-INS-214 | chrX-13019458-INS-214 |
| chrX-13064813-INS-202 | chrX-13064813-INS-202 |
| chrX-13071152-INS-70 | chrX-13071152-INS-63 |
| chrX-13085575-INS-72 | chrX-13085575-INS-72 |
| chrX-13089093-INS-594 | chrX-13089093-INS-594 |
| chrX-13096106-INS-243 | chrX-13096106-INS-243 |
| chrX-13117240-INS-64 | chrX-13117240-INS-60 |
| chrX-13127031-INS-197 | chrX-13127031-INS-197 |
| chrX-13214430-INS-76 | chrX-13214430-INS-64 |
| chrX-13223964-INS-634 | chrX-13223964-INS-634 |
| chrX-13228226-INS-261 | chrX-13228226-INS-261 |
| chrX-13233884-INS-194 | chrX-13233884-INS-195 |
| chrX-13235527-INS-3578 | chrX-13235528-INS-3581 |
| chrX-13237246-INS-199 | chrX-13237246-INS-199 |
| chrX-13263420-INS-90 | chrX-13263418-INS-62 |
| chrX-13274224-INS-345 | chrX-13274224-INS-345 |
| chrX-13280842-INS-473 | chrX-13280842-INS-474 |
| chrX-13292699-INS-75 | chrX-13292699-INS-75 |
| chrX-13293224-INS-880 | chrX-13293224-INS-899 |
| chrX-13297082-INS-82 | chrX-13297082-INS-81 |
| chrX-13300260-INS-66 | chrX-13300260-INS-66 |
| chrX-13302613-INS-109 | chrX-13302613-INS-109 |
| chrX-13307701-INS-223 | chrX-13307701-INS-223 |
| chrX-13320106-INS-668 | chrX-13320106-INS-668 |
| chrX-13320870-INS-57 | chrX-13320870-INS-56 |
| chrX-13340128-INS-52 | chrX-13340134-INS-67 |
| chrX-13343265-INS-62 | chrX-13343280-INS-70 |
| chrX-13346328-INS-70 | chrX-13346328-INS-52 |
| chrX-13376300-INS-135 | chrX-13376290-INS-130 |
| chrX-13419909-INS-67 | chrX-13419909-INS-67 |
| chrX-13426727-INS-68 | chrX-13426727-INS-68 |
| chrX-13434211-INS-173 | chrX-13434211-INS-173 |
| chrX-13448655-INS-60 | chrX-13448655-INS-55 |
| chrX-13494608-INS-739 | chrX-13494608-INS-737 |
| chrX-13500166-INS-90 | chrX-13500166-INS-90 |
| chrX-13522225-INS-76 | chrX-13522225-INS-76 |
| chrX-13529446-INS-119 | chrX-13529446-INS-150 |
| chrX-13554244-INS-373 | chrX-13554244-INS-373 |
| chrX-13562316-INS-7134 | chrX-13562317-INS-7134 |
| chrX-13564751-INS-454 | chrX-13564751-INS-454 |
| chrX-13572939-INS-52 | chrX-13572940-INS-52 |
| chrX-13655960-INS-7325 | chrX-13655946-INS-7311 |
| chrX-13674120-INS-51 | chrX-13674120-INS-51 |
| chrX-13727188-INS-87 | chrX-13727188-INS-88 |
| chrX-13750439-INS-172 | chrX-13750447-INS-183 |
| chrX-13767583-INS-362 | chrX-13767583-INS-371 |
| chrX-13774719-INS-235 | chrX-13774719-INS-217 |
| chrX-13871646-INS-790 | chrX-13871646-INS-790 |
| chrX-13874812-INS-60 | chrX-13874810-INS-60 |
| chrX-13913087-INS-6387 | chrX-13913088-INS-6386 |
| chrX-13960216-INS-363 | chrX-13960216-INS-363 |
| chrX-13961853-INS-6223 | chrX-13961854-INS-6224 |
| chrX-13973981-INS-507 | chrX-13973981-INS-507 |
| chrX-13993669-INS-519 | chrX-13993669-INS-519 |
| chrX-14000547-INS-2027 | chrX-14000548-INS-2036 |
| chrX-14001301-INS-909 | chrX-14001301-INS-909 |
| chrX-14022913-INS-810 | chrX-14022913-INS-810 |
| chrX-14048578-INS-324 | chrX-14048578-INS-324 |
| chrX-14050254-INS-63 | chrX-14050254-INS-64 |
| chrX-14068945-INS-480 | chrX-14068945-INS-480 |
| chrX-14081902-INS-900 | chrX-14081902-INS-900 |
| chrX-14114858-INS-442 | chrX-14114858-INS-442 |
| chrX-14137139-INS-7104 | chrX-14137140-INS-7104 |
| chrX-14186751-INS-111 | chrX-14186751-INS-110 |
| chrX-14244472-INS-58 | chrX-14244472-INS-58 |
| chrX-14258652-INS-54 | chrX-14258652-INS-54 |
| chrX-14304852-INS-7080 | chrX-14304853-INS-7080 |

| | |
|---|---|
| chrX-14334093-INS-70 | chrX-14334093-INS-65 |
| chrX-14353572-INS-355 | chrX-14353568-INS-314 |
| chrX-14362748-INS-5188 | chrX-14362749-INS-5196 |
| chrX-14368138-INS-118 | chrX-14368138-INS-115 |
| chrX-14414360-INS-451 | chrX-14414360-INS-451 |
| chrX-14422600-INS-479 | chrX-14422600-INS-479 |
| chrX-14515108-INS-212 | chrX-14515108-INS-214 |
| chrX-14551042-INS-381 | chrX-14551042-INS-381 |
| chrX-14564165-INS-64 | chrX-14564207-INS-64 |
| chrX-14581959-INS-433 | chrX-14581959-INS-433 |
| chrX-14585879-INS-409 | chrX-14585879-INS-409 |
| chrX-14594188-INS-2329 | chrX-14594189-INS-2329 |
| chrX-14627139-INS-1217 | chrX-14627137-INS-1206 |
| chrX-14627409-INS-332 | chrX-14627409-INS-332 |
| chrX-14668181-INS-142 | chrX-14668181-INS-167 |
| chrX-14742377-INS-70 | chrX-14742377-INS-70 |
| chrX-14757088-INS-260 | chrX-14757088-INS-260 |
| chrX-14757292-INS-1620 | chrX-14757293-INS-1620 |
| chrX-14787725-INS-1131 | chrX-14787726-INS-1131 |
| chrX-14791699-INS-419 | chrX-14791699-INS-416 |
| chrX-14803198-INS-364 | chrX-14803198-INS-364 |
| chrX-14820163-INS-61 | chrX-14820143-INS-60 |
| chrX-14889278-INS-53 | chrX-14889278-INS-88 |
| chrX-14916227-INS-1053 | chrX-14916228-INS-1055 |
| chrX-14968055-INS-77 | chrX-14968062-INS-78 |
| chrX-14993843-INS-127 | chrX-14993841-INS-127 |
| chrX-15020521-INS-126 | chrX-15020521-INS-128 |
| chrX-15036957-INS-58 | chrX-15036918-INS-50 |
| chrX-15044818-INS-109 | chrX-15044818-INS-157 |
| chrX-15072302-INS-84 | chrX-15072302-INS-102 |
| chrX-15099493-INS-2235 | chrX-15099497-INS-2230 |
| chrX-15121302-INS-226 | chrX-15121281-INS-214 |
| chrX-15187395-INS-939 | chrX-15187395-INS-938 |
| chrX-15196347-INS-2342 | chrX-15196348-INS-2352 |
| chrX-15256827-INS-196 | chrX-15256827-INS-196 |
| chrX-15268212-INS-68 | chrX-15268212-INS-68 |
| chrX-15283170-INS-383 | chrX-15283170-INS-383 |
| chrX-15325317-INS-88 | chrX-15325317-INS-90 |
| chrX-15336137-INS-939 | chrX-15336141-INS-939 |
| chrX-15368593-INS-527 | chrX-15368593-INS-530 |
| chrX-15382009-INS-56 | chrX-15382009-INS-60 |
| chrX-15401930-INS-407 | chrX-15401930-INS-407 |
| chrX-15444332-INS-52 | chrX-15444332-INS-52 |
| chrX-15498528-INS-423 | chrX-15498533-INS-428 |
| chrX-15517163-INS-205 | chrX-15517163-INS-203 |
| chrX-15528224-INS-1826 | chrX-15528225-INS-1841 |
| chrX-15531903-INS-559 | chrX-15531903-INS-559 |
| chrX-15557245-INS-812 | chrX-15557245-INS-815 |
| chrX-15638381-INS-138 | chrX-15638381-INS-138 |
| chrX-15649315-INS-592 | chrX-15649315-INS-592 |
| chrX-15692857-INS-1956 | chrX-15692858-INS-1956 |
| chrX-15697217-INS-8413 | chrX-15697218-INS-8413 |
| chrX-15729122-INS-124 | chrX-15729122-INS-122 |
| chrX-15748907-INS-6362 | chrX-15748908-INS-6363 |
| chrX-15809690-INS-84 | chrX-15809690-INS-66 |
| chrX-15857075-INS-119 | chrX-15857076-INS-122 |
| chrX-15882984-INS-60 | chrX-15882984-INS-60 |
| chrX-15916609-INS-210 | chrX-15916609-INS-210 |
| chrX-15987020-INS-380 | chrX-15987020-INS-380 |
| chrX-15995971-INS-60 | chrX-15995971-INS-60 |
| chrX-16045762-INS-101 | chrX-16045762-INS-101 |
| chrX-16054398-INS-407 | chrX-16054398-INS-411 |
| chrX-16058304-INS-1943 | chrX-16058305-INS-1947 |
| chrX-16085894-INS-2899 | chrX-16085895-INS-2901 |
| chrX-16119584-INS-81 | chrX-16119579-INS-103 |
| chrX-16149809-INS-337 | chrX-16149809-INS-337 |
| chrX-16152658-INS-137 | chrX-16152658-INS-132 |
| chrX-16184675-INS-6417 | chrX-16184676-INS-6417 |
| chrX-16240869-INS-1993 | chrX-16240870-INS-1993 |
| chrX-16260736-INS-70 | chrX-16260724-INS-60 |
| chrX-16274338-INS-6695 | chrX-16274339-INS-6695 |
| chrX-16293731-INS-1493 | chrX-16293732-INS-1492 |
| chrX-16315498-INS-228 | chrX-16315498-INS-229 |
| chrX-16366309-INS-1347 | chrX-16366310-INS-1346 |
| chrX-16378341-INS-55 | chrX-16378349-INS-80 |
| chrX-16482408-INS-52 | chrX-16482408-INS-60 |
| chrX-16503778-INS-399 | chrX-16503782-INS-398 |

| | |
|---|---|
| chrX-16520586-INS-593 | chrX-16520586-INS-594 |
| chrX-16541863-INS-201 | chrX-16541863-INS-209 |
| chrX-16563892-INS-8532 | chrX-16563895-INS-8540 |
| chrX-16566549-INS-424 | chrX-16566549-INS-424 |
| chrX-16578335-INS-180 | chrX-16578335-INS-180 |
| chrX-16597742-INS-205 | chrX-16597742-INS-205 |
| chrX-16659301-INS-269 | chrX-16659301-INS-285 |
| chrX-16687712-INS-114 | chrX-16687719-INS-126 |
| chrX-16692696-INS-52 | chrX-16692696-INS-50 |
| chrX-16712373-INS-140 | chrX-16712373-INS-141 |
| chrX-16785729-INS-140 | chrX-16785713-INS-140 |
| chrX-16799523-INS-98 | chrX-16799523-INS-98 |
| chrX-16815994-INS-2381 | chrX-16815995-INS-2380 |
| chrX-16823002-INS-249 | chrX-16823002-INS-245 |
| chrX-16854275-INS-1164 | chrX-16854276-INS-1164 |
| chrX-16914599-INS-79 | chrX-16914599-INS-79 |
| chrX-16925987-INS-125 | chrX-16925987-INS-124 |
| chrX-16947155-INS-1453 | chrX-16947158-INS-1441 |
| chrX-17001021-INS-333 | chrX-17001021-INS-337 |
| chrX-17012140-INS-789 | chrX-17012140-INS-790 |
| chrX-17050558-INS-111 | chrX-17050558-INS-111 |
| chrX-17056434-INS-174 | chrX-17056434-INS-174 |
| chrX-17058735-INS-1236 | chrX-17058736-INS-1236 |
| chrX-17067481-INS-63 | chrX-17067481-INS-63 |
| chrX-17113952-INS-52 | chrX-17113953-INS-52 |
| chrX-17152552-INS-2782 | chrX-17152553-INS-2783 |
| chrX-17180197-INS-844 | chrX-17180197-INS-845 |
| chrX-17196064-INS-69 | chrX-17196064-INS-69 |
| chrX-17215636-INS-357 | chrX-17215636-INS-356 |
| chrX-17223836-INS-1517 | chrX-17223837-INS-1518 |
| chrX-17234459-INS-52 | chrX-17234459-INS-52 |
| chrX-17255125-INS-4244 | chrX-17255126-INS-4244 |
| chrX-17264782-INS-2527 | chrX-17264783-INS-2527 |
| chrX-17275028-INS-500 | chrX-17275028-INS-504 |
| chrX-17276164-INS-906 | chrX-17276164-INS-902 |
| chrX-17284172-INS-552 | chrX-17284172-INS-552 |
| chrX-17304294-INS-207 | chrX-17304294-INS-207 |
| chrX-17323809-INS-58 | chrX-17323809-INS-58 |
| chrX-17332200-INS-328 | chrX-17332200-INS-328 |
| chrX-17341399-INS-59 | chrX-17341400-INS-64 |
| chrX-17345684-INS-107 | chrX-17345693-INS-58 |
| chrX-17398910-INS-62 | chrX-17398910-INS-56 |
| chrX-17404307-INS-64 | chrX-17404307-INS-66 |
| chrX-17415709-INS-68 | chrX-17415709-INS-68 |
| chrX-17426259-INS-375 | chrX-17426259-INS-375 |
| chrX-17435182-INS-212 | chrX-17435182-INS-212 |
| chrX-17523154-INS-171 | chrX-17523191-INS-161 |
| chrX-17604887-INS-3317 | chrX-17604888-INS-3317 |
| chrX-17691261-INS-1886 | chrX-17691262-INS-1886 |
| chrX-17698462-INS-6630 | chrX-17698463-INS-6630 |
| chrX-17740173-INS-1734 | chrX-17740174-INS-1734 |
| chrX-17771650-INS-77 | chrX-17771641-INS-77 |
| chrX-17776827-INS-87 | chrX-17776827-INS-126 |
| chrX-17784239-INS-689 | chrX-17784239-INS-699 |
| chrX-17809627-INS-52 | chrX-17809632-INS-56 |
| chrX-17819073-INS-6854 | chrX-17819074-INS-6855 |
| chrX-17828524-INS-124 | chrX-17828524-INS-124 |
| chrX-17839073-INS-3326 | chrX-17839074-INS-3328 |
| chrX-17843189-INS-113 | chrX-17843229-INS-75 |
| chrX-17893674-INS-127 | chrX-17893674-INS-127 |
| chrX-17921107-INS-3294 | chrX-17921108-INS-3290 |
| chrX-17936395-INS-214 | chrX-17936395-INS-214 |
| chrX-17960560-INS-6468 | chrX-17960561-INS-6460 |
| chrX-17966425-INS-324 | chrX-17966425-INS-329 |
| chrX-17988421-INS-886 | chrX-17988421-INS-886 |
| chrX-18053092-INS-247 | chrX-18053092-INS-247 |
| chrX-18072188-INS-185 | chrX-18072188-INS-185 |
| chrX-18139607-INS-1570 | chrX-18139608-INS-1570 |
| chrX-18140673-INS-54 | chrX-18140673-INS-52 |
| chrX-18148773-INS-172 | chrX-18148773-INS-166 |
| chrX-18221457-INS-203 | chrX-18221457-INS-203 |
| chrX-18267102-INS-241 | chrX-18267102-INS-242 |
| chrX-18274107-INS-2861 | chrX-18274108-INS-2858 |
| chrX-18278545-INS-69 | chrX-18278545-INS-89 |
| chrX-18279484-INS-708 | chrX-18279484-INS-707 |
| chrX-18287695-INS-124 | chrX-18287695-INS-124 |
| chrX-18290401-INS-78 | chrX-18290402-INS-66 |

| | |
|---|---|
| chrX-18307723-INS-470 | chrX-18307723-INS-471 |
| chrX-18319644-INS-51 | chrX-18319644-INS-59 |
| chrX-18327971-INS-368 | chrX-18327971-INS-370 |
| chrX-18359266-INS-118 | chrX-18359256-INS-120 |
| chrX-18390648-INS-109 | chrX-18390636-INS-154 |
| chrX-18405262-INS-213 | chrX-18405262-INS-212 |
| chrX-18430791-INS-76 | chrX-18430797-INS-93 |
| chrX-18440935-INS-64 | chrX-18440932-INS-84 |
| chrX-18441231-INS-4575 | chrX-18441232-INS-4575 |
| chrX-18448681-INS-60 | chrX-18448681-INS-60 |
| chrX-18526619-INS-54 | chrX-18526622-INS-50 |
| chrX-18541808-INS-978 | chrX-18541808-INS-981 |
| chrX-18546885-INS-58 | chrX-18546885-INS-62 |
| chrX-18549239-INS-6253 | chrX-18549240-INS-6257 |
| chrX-18566261-INS-3189 | chrX-18566262-INS-3197 |
| chrX-18613616-INS-6208 | chrX-18613617-INS-6208 |
| chrX-18616366-INS-2159 | chrX-18616367-INS-2158 |
| chrX-18639956-INS-4750 | chrX-18639957-INS-4750 |
| chrX-18657617-INS-427 | chrX-18657617-INS-427 |
| chrX-18684519-INS-524 | chrX-18684519-INS-525 |
| chrX-18702004-INS-657 | chrX-18702004-INS-657 |
| chrX-18710584-INS-6380 | chrX-18710585-INS-6380 |
| chrX-18767627-INS-530 | chrX-18767627-INS-530 |
| chrX-18786025-INS-690 | chrX-18786025-INS-692 |
| chrX-18791750-INS-6459 | chrX-18791751-INS-6460 |
| chrX-18795231-INS-64 | chrX-18795231-INS-70 |
| chrX-18810883-INS-55 | chrX-18810883-INS-100 |
| chrX-18850621-INS-1200 | chrX-18850622-INS-1186 |
| chrX-18855855-INS-3252 | chrX-18855856-INS-3253 |
| chrX-18868329-INS-7002 | chrX-18868330-INS-7001 |
| chrX-18889492-INS-701 | chrX-18889492-INS-694 |
| chrX-18892374-INS-66 | chrX-18892374-INS-66 |
| chrX-18894235-INS-60 | chrX-18894235-INS-58 |
| chrX-18908903-INS-60 | chrX-18908903-INS-72 |
| chrX-18918518-INS-6327 | chrX-18918519-INS-6327 |
| chrX-18970878-INS-805 | chrX-18970878-INS-805 |
| chrX-18995406-INS-199 | chrX-18995420-INS-199 |
| chrX-19004973-INS-195 | chrX-19004973-INS-195 |
| chrX-19007059-INS-245 | chrX-19007059-INS-245 |
| chrX-19032952-INS-78 | chrX-19032952-INS-80 |
| chrX-19051540-INS-822 | chrX-19051556-INS-817 |
| chrX-19077619-INS-6344 | chrX-19077620-INS-6344 |
| chrX-19114660-INS-78 | chrX-19114660-INS-73 |
| chrX-19126780-INS-77 | chrX-19126780-INS-79 |
| chrX-19184424-INS-75 | chrX-19184424-INS-111 |
| chrX-19213933-INS-558 | chrX-19213934-INS-553 |
| chrX-19275566-INS-126 | chrX-19275566-INS-126 |
| chrX-19280437-INS-385 | chrX-19280437-INS-385 |
| chrX-19280788-INS-1869 | chrX-19280789-INS-1869 |
| chrX-19286178-INS-1801 | chrX-19286179-INS-1801 |
| chrX-19310202-INS-199 | chrX-19310202-INS-197 |
| chrX-19327825-INS-855 | chrX-19327825-INS-853 |
| chrX-19339723-INS-388 | chrX-19339723-INS-411 |
| chrX-19358784-INS-226 | chrX-19358784-INS-226 |
| chrX-19400725-INS-220 | chrX-19400725-INS-219 |
| chrX-19550631-INS-55 | chrX-19550631-INS-51 |
| chrX-19565888-INS-252 | chrX-19565919-INS-253 |
| chrX-19592044-INS-4228 | chrX-19592045-INS-4227 |
| chrX-19598365-INS-843 | chrX-19598365-INS-843 |
| chrX-19602920-INS-6466 | chrX-19602921-INS-6464 |
| chrX-19628084-INS-482 | chrX-19628101-INS-481 |
| chrX-19634087-INS-486 | chrX-19634087-INS-486 |
| chrX-19641508-INS-300 | chrX-19641509-INS-300 |
| chrX-19654647-INS-828 | chrX-19654647-INS-828 |
| chrX-19658976-INS-187 | chrX-19658976-INS-188 |
| chrX-19696444-INS-102 | chrX-19696444-INS-108 |
| chrX-19712352-INS-1867 | chrX-19712353-INS-1867 |
| chrX-19742160-INS-700 | chrX-19742160-INS-700 |
| chrX-19759310-INS-820 | chrX-19759310-INS-820 |
| chrX-19774700-INS-399 | chrX-19774700-INS-399 |
| chrX-19805628-INS-7018 | chrX-19805629-INS-7024 |
| chrX-19817735-INS-58 | chrX-19817735-INS-58 |
| chrX-19821850-INS-1173 | chrX-19821851-INS-1172 |
| chrX-19879275-INS-51 | chrX-19879275-INS-51 |
| chrX-19899981-INS-76 | chrX-19899981-INS-76 |
| chrX-19901087-INS-5542 | chrX-19901088-INS-5542 |
| chrX-19959234-INS-336 | chrX-19959234-INS-336 |

| | |
|---|---|
| chrX-19975214-INS-197 | chrX-19975214-INS-197 |
| chrX-19978411-INS-759 | chrX-19978411-INS-759 |
| chrX-19979540-INS-194 | chrX-19979540-INS-194 |
| chrX-19987260-INS-105 | chrX-19987260-INS-84 |
| chrX-19988804-INS-212 | chrX-19988804-INS-212 |
| chrX-20020079-INS-454 | chrX-20020079-INS-454 |
| chrX-20031851-INS-315 | chrX-20031851-INS-315 |
| chrX-20050554-INS-830 | chrX-20050554-INS-830 |
| chrX-20081600-INS-145 | chrX-20081600-INS-115 |
| chrX-20083564-INS-1821 | chrX-20083565-INS-1821 |
| chrX-20084138-INS-268 | chrX-20084138-INS-268 |
| chrX-20120390-INS-127 | chrX-20120390-INS-127 |
| chrX-20160111-INS-59 | chrX-20160111-INS-61 |
| chrX-20165724-INS-88 | chrX-20165724-INS-90 |
| chrX-20170217-INS-69 | chrX-20170219-INS-52 |
| chrX-20182767-INS-58 | chrX-20182767-INS-66 |
| chrX-20198385-INS-936 | chrX-20198367-INS-936 |
| chrX-20208654-INS-140 | chrX-20208615-INS-131 |
| chrX-20211749-INS-586 | chrX-20211749-INS-586 |
| chrX-20238607-INS-148 | chrX-20238607-INS-147 |
| chrX-20256558-INS-162 | chrX-20256558-INS-162 |
| chrX-20260871-INS-211 | chrX-20260871-INS-203 |
| chrX-20303377-INS-219 | chrX-20303377-INS-220 |
| chrX-20319823-INS-55 | chrX-20319823-INS-55 |
| chrX-20330317-INS-209 | chrX-20330317-INS-207 |
| chrX-20346902-INS-134 | chrX-20346905-INS-130 |
| chrX-20354732-INS-144 | chrX-20354732-INS-144 |
| chrX-20399737-INS-429 | chrX-20399737-INS-434 |
| chrX-20428369-INS-134 | chrX-20428365-INS-132 |
| chrX-20429068-INS-1298 | chrX-20429069-INS-1298 |
| chrX-20453318-INS-348 | chrX-20453318-INS-348 |
| chrX-20466423-INS-201 | chrX-20466423-INS-201 |
| chrX-20480353-INS-206 | chrX-20480353-INS-206 |
| chrX-20491228-INS-171 | chrX-20491228-INS-171 |
| chrX-20521533-INS-198 | chrX-20521533-INS-198 |
| chrX-20523201-INS-212 | chrX-20523201-INS-212 |
| chrX-20551015-INS-181 | chrX-20551015-INS-177 |
| chrX-20552905-INS-237 | chrX-20552905-INS-240 |
| chrX-20583871-INS-91 | chrX-20583876-INS-75 |
| chrX-20592891-INS-161 | chrX-20592891-INS-161 |
| chrX-20601253-INS-66 | chrX-20601253-INS-66 |
| chrX-20605493-INS-194 | chrX-20605493-INS-194 |
| chrX-20605959-INS-236 | chrX-20605959-INS-228 |
| chrX-20607147-INS-59 | chrX-20607147-INS-59 |
| chrX-20639701-INS-249 | chrX-20639701-INS-249 |
| chrX-20645706-INS-75 | chrX-20645706-INS-75 |
| chrX-20646850-INS-68 | chrX-20646850-INS-80 |
| chrX-20650856-INS-174 | chrX-20650856-INS-174 |
| chrX-20652086-INS-182 | chrX-20652086-INS-182 |
| chrX-20654373-INS-314 | chrX-20654373-INS-313 |
| chrX-20661824-INS-84 | chrX-20661827-INS-95 |
| chrX-20669312-INS-207 | chrX-20669312-INS-207 |
| chrX-20672790-INS-232 | chrX-20672790-INS-229 |
| chrX-20692076-INS-116 | chrX-20692088-INS-119 |
| chrX-20707011-INS-92 | chrX-20707011-INS-92 |
| chrX-20742906-INS-108 | chrX-20742906-INS-82 |
| chrX-20749875-INS-443 | chrX-20749875-INS-443 |
| chrX-20751849-INS-202 | chrX-20751849-INS-198 |
| chrX-20757403-INS-384 | chrX-20757403-INS-384 |
| chrX-20758154-INS-943 | chrX-20758172-INS-932 |
| chrX-20760964-INS-169 | chrX-20760964-INS-170 |
| chrX-20763614-INS-177 | chrX-20763614-INS-185 |
| chrX-20773812-INS-169 | chrX-20773812-INS-169 |
| chrX-20809083-INS-80 | chrX-20809083-INS-86 |
| chrX-20844546-INS-920 | chrX-20844546-INS-919 |
| chrX-20845339-INS-445 | chrX-20845339-INS-445 |
| chrX-20854964-INS-3241 | chrX-20854965-INS-3255 |
| chrX-20859535-INS-1274 | chrX-20859539-INS-1275 |
| chrX-20862161-INS-5348 | chrX-20862162-INS-5348 |
| chrX-20882210-INS-86 | chrX-20882210-INS-93 |
| chrX-20903300-INS-1058 | chrX-20903301-INS-1058 |
| chrX-20907229-INS-6415 | chrX-20907230-INS-6416 |
| chrX-20911252-INS-362 | chrX-20911252-INS-362 |
| chrX-20960277-INS-8888 | chrX-20960278-INS-8889 |
| chrX-20964614-INS-2085 | chrX-20964615-INS-2087 |
| chrX-20987512-INS-719 | chrX-20987512-INS-726 |
| chrX-21008423-INS-498 | chrX-21008423-INS-498 |

| | |
|---|---|
| chrX-21014337-INS-950 | chrX-21014335-INS-923 |
| chrX-21040347-INS-568 | chrX-21040347-INS-571 |
| chrX-21043009-INS-296 | chrX-21043009-INS-296 |
| chrX-21054456-INS-107 | chrX-21054456-INS-107 |
| chrX-21137224-INS-2480 | chrX-21137225-INS-2480 |
| chrX-21165852-INS-668 | chrX-21165858-INS-675 |
| chrX-21179805-INS-146 | chrX-21179805-INS-146 |
| chrX-21182392-INS-71 | chrX-21182392-INS-74 |
| chrX-21195226-INS-200 | chrX-21195226-INS-198 |
| chrX-21214197-INS-194 | chrX-21214197-INS-194 |
| chrX-21221850-INS-917 | chrX-21221850-INS-918 |
| chrX-21222123-INS-8327 | chrX-21222124-INS-8327 |
| chrX-21227868-INS-51 | chrX-21227868-INS-51 |
| chrX-21257436-INS-56 | chrX-21257436-INS-56 |
| chrX-21321752-INS-309 | chrX-21321752-INS-310 |
| chrX-21332484-INS-80 | chrX-21332484-INS-80 |
| chrX-21369195-INS-197 | chrX-21369192-INS-208 |
| chrX-21404564-INS-81 | chrX-21404545-INS-125 |
| chrX-21423604-INS-4040 | chrX-21423605-INS-4040 |
| chrX-21439642-INS-198 | chrX-21439642-INS-203 |
| chrX-21449741-INS-56 | chrX-21449742-INS-103 |
| chrX-21458153-INS-2101 | chrX-21458162-INS-2083 |
| chrX-21458315-INS-84 | chrX-21458315-INS-76 |
| chrX-21483918-INS-2820 | chrX-21483919-INS-2820 |
| chrX-21528716-INS-776 | chrX-21528716-INS-776 |
| chrX-21534627-INS-54 | chrX-21534614-INS-60 |
| chrX-21548589-INS-102 | chrX-21548589-INS-135 |
| chrX-21555775-INS-1511 | chrX-21555776-INS-1511 |
| chrX-21570654-INS-185 | chrX-21570654-INS-186 |
| chrX-21573453-INS-64 | chrX-21573453-INS-54 |
| chrX-21596343-INS-67 | chrX-21596343-INS-68 |
| chrX-21633159-INS-76 | chrX-21633130-INS-68 |
| chrX-21634561-INS-181 | chrX-21634561-INS-178 |
| chrX-21660784-INS-149 | chrX-21660784-INS-149 |
| chrX-21661176-INS-6415 | chrX-21661177-INS-6415 |
| chrX-21671440-INS-55 | chrX-21671440-INS-55 |
| chrX-21702535-INS-647 | chrX-21702535-INS-647 |
| chrX-21719753-INS-934 | chrX-21719753-INS-934 |
| chrX-21748319-INS-54 | chrX-21748319-INS-52 |
| chrX-21768610-INS-57 | chrX-21768610-INS-57 |
| chrX-21789351-INS-3069 | chrX-21789352-INS-3069 |
| chrX-21835726-INS-60 | chrX-21835726-INS-60 |
| chrX-21871198-INS-579 | chrX-21871198-INS-580 |
| chrX-21874403-INS-6489 | chrX-21874404-INS-6489 |
| chrX-21936027-INS-138 | chrX-21936027-INS-138 |
| chrX-22091584-INS-7285 | chrX-22091585-INS-7285 |
| chrX-22125324-INS-54 | chrX-22125324-INS-54 |
| chrX-22148277-INS-66 | chrX-22148277-INS-66 |
| chrX-22151136-INS-382 | chrX-22151137-INS-381 |
| chrX-22175144-INS-111 | chrX-22175144-INS-138 |
| chrX-22183529-INS-6219 | chrX-22183530-INS-6216 |
| chrX-22272789-INS-3796 | chrX-22272790-INS-3796 |
| chrX-22323168-INS-1049 | chrX-22323169-INS-1049 |
| chrX-22357469-INS-1699 | chrX-22357470-INS-1699 |
| chrX-22365730-INS-98 | chrX-22365724-INS-116 |
| chrX-22455591-INS-374 | chrX-22455591-INS-363 |
| chrX-22473441-INS-333 | chrX-22473442-INS-333 |
| chrX-22488046-INS-6523 | chrX-22488047-INS-6524 |
| chrX-22498239-INS-64 | chrX-22498239-INS-64 |
| chrX-22523534-INS-51 | chrX-22523535-INS-51 |
| chrX-22535893-INS-3158 | chrX-22535894-INS-3158 |
| chrX-22569929-INS-56 | chrX-22569929-INS-60 |
| chrX-22571372-INS-921 | chrX-22571372-INS-921 |
| chrX-22575723-INS-120 | chrX-22575723-INS-121 |
| chrX-22593065-INS-65 | chrX-22593065-INS-66 |
| chrX-22594460-INS-3491 | chrX-22594461-INS-3491 |
| chrX-22604044-INS-2753 | chrX-22604045-INS-2755 |
| chrX-22631677-INS-6853 | chrX-22631678-INS-6853 |
| chrX-22634539-INS-198 | chrX-22634539-INS-198 |
| chrX-22649453-INS-2445 | chrX-22649454-INS-2445 |
| chrX-22693022-INS-4190 | chrX-22693023-INS-4191 |
| chrX-22776891-INS-52 | chrX-22776891-INS-50 |
| chrX-22798302-INS-2448 | chrX-22798303-INS-2448 |
| chrX-22800586-INS-467 | chrX-22800577-INS-473 |
| chrX-22816697-INS-56 | chrX-22816705-INS-58 |
| chrX-22830888-INS-1523 | chrX-22830889-INS-1523 |
| chrX-22839744-INS-6524 | chrX-22839745-INS-6512 |

| | |
|---|---|
| chrX-22857909-INS-64 | chrX-22857909-INS-64 |
| chrX-22884399-INS-60 | chrX-22884399-INS-60 |
| chrX-22891357-INS-1890 | chrX-22891358-INS-1890 |
| chrX-22914602-INS-58 | chrX-22914598-INS-50 |
| chrX-22921512-INS-200 | chrX-22921512-INS-200 |
| chrX-22961363-INS-80 | chrX-22961351-INS-104 |
| chrX-22995670-INS-176 | chrX-22995670-INS-178 |
| chrX-22996667-INS-6710 | chrX-22996668-INS-6710 |
| chrX-23029661-INS-90 | chrX-23029661-INS-90 |
| chrX-23041694-INS-325 | chrX-23041694-INS-325 |
| chrX-23057397-INS-1402 | chrX-23057398-INS-1404 |
| chrX-23067277-INS-177 | chrX-23067271-INS-180 |
| chrX-23076613-INS-510 | chrX-23076613-INS-506 |
| chrX-23128306-INS-90 | chrX-23128290-INS-65 |
| chrX-23140286-INS-103 | chrX-23140284-INS-63 |
| chrX-23167256-INS-154 | chrX-23167254-INS-146 |
| chrX-23197155-INS-245 | chrX-23197155-INS-245 |
| chrX-23271415-INS-715 | chrX-23271415-INS-715 |
| chrX-23286587-INS-110 | chrX-23286587-INS-108 |
| chrX-23293522-INS-417 | chrX-23293522-INS-417 |
| chrX-23346870-INS-67 | chrX-23346870-INS-67 |
| chrX-23492899-INS-492 | chrX-23492899-INS-498 |
| chrX-23497222-INS-390 | chrX-23497222-INS-390 |
| chrX-23531519-INS-6768 | chrX-23531520-INS-6769 |
| chrX-23597124-INS-3488 | chrX-23597125-INS-3488 |
| chrX-23689463-INS-57 | chrX-23689449-INS-62 |
| chrX-24596821-INS-66 | chrX-24596821-INS-66 |
| chrX-24691950-INS-1887 | chrX-24691951-INS-1887 |
| chrX-25909785-INS-60 | chrX-25909785-INS-60 |
| chrX-35014800-INS-175 | chrX-35014800-INS-176 |
| chrX-35024096-INS-385 | chrX-35024096-INS-385 |
| chrX-35042090-INS-54 | chrX-35042058-INS-52 |
| chrX-35063442-INS-1890 | chrX-35063443-INS-1890 |
| chrX-35086040-INS-522 | chrX-35086040-INS-522 |
| chrX-35117704-INS-52 | chrX-35117698-INS-54 |
| chrX-35140623-INS-512 | chrX-35140623-INS-512 |
| chrX-35197276-INS-64 | chrX-35197276-INS-90 |
| chrX-35220283-INS-70 | chrX-35220283-INS-58 |
| chrX-35224511-INS-418 | chrX-35224511-INS-418 |
| chrX-35227716-INS-59 | chrX-35227716-INS-92 |
| chrX-35230538-INS-429 | chrX-35230538-INS-428 |
| chrX-35248407-INS-177 | chrX-35248407-INS-175 |
| chrX-35267518-INS-547 | chrX-35267518-INS-547 |
| chrX-35342242-INS-854 | chrX-35342242-INS-854 |
| chrX-35373835-INS-216 | chrX-35373835-INS-216 |
| chrX-35456648-INS-150 | chrX-35456648-INS-150 |
| chrX-35502537-INS-60 | chrX-35502537-INS-62 |
| chrX-35504262-INS-58 | chrX-35504262-INS-74 |
| chrX-35534928-INS-51 | chrX-35534930-INS-82 |
| chrX-35557904-INS-478 | chrX-35557904-INS-478 |
| chrX-35623185-INS-51 | chrX-35623192-INS-62 |
| chrX-35643227-INS-498 | chrX-35643227-INS-498 |
| chrX-35671455-INS-1884 | chrX-35671456-INS-1884 |
| chrX-35745110-INS-656 | chrX-35745110-INS-656 |
| chrX-35856573-INS-6965 | chrX-35856574-INS-6967 |
| chrX-35927694-INS-2106 | chrX-35927695-INS-2106 |
| chrX-35977042-INS-194 | chrX-35977042-INS-193 |
| chrX-35982406-INS-914 | chrX-35982406-INS-912 |
| chrX-36009545-INS-197 | chrX-36009545-INS-197 |
| chrX-36016152-INS-769 | chrX-36016152-INS-770 |
| chrX-36051782-INS-696 | chrX-36051782-INS-691 |
| chrX-36077659-INS-165 | chrX-36077659-INS-165 |
| chrX-36095145-INS-247 | chrX-36095146-INS-221 |
| chrX-36099370-INS-172 | chrX-36099370-INS-177 |
| chrX-36198588-INS-1223 | chrX-36198589-INS-1193 |
| chrX-36212423-INS-119 | chrX-36212424-INS-110 |
| chrX-36265841-INS-200 | chrX-36265841-INS-200 |
| chrX-36336829-INS-1284 | chrX-36336830-INS-1285 |
| chrX-36338311-INS-172 | chrX-36338311-INS-198 |
| chrX-36341603-INS-396 | chrX-36341603-INS-397 |
| chrX-36343284-INS-217 | chrX-36343284-INS-217 |
| chrX-36350015-INS-309 | chrX-36350015-INS-310 |
| chrX-36377440-INS-64 | chrX-36377440-INS-64 |
| chrX-36379508-INS-91 | chrX-36379497-INS-81 |
| chrX-36391766-INS-202 | chrX-36391766-INS-202 |
| chrX-36396026-INS-351 | chrX-36396036-INS-346 |
| chrX-36398378-INS-1166 | chrX-36398379-INS-1166 |

| | |
|---|---|
| chrX-36402121-INS-163 | chrX-36402121-INS-165 |
| chrX-36405208-INS-193 | chrX-36405208-INS-191 |
| chrX-36442801-INS-111 | chrX-36442801-INS-111 |
| chrX-36446775-INS-86 | chrX-36446775-INS-74 |
| chrX-36461039-INS-2925 | chrX-36461040-INS-2925 |
| chrX-36471738-INS-68 | chrX-36471738-INS-56 |
| chrX-36866670-INS-210 | chrX-36866670-INS-210 |
| chrX-36903467-INS-52 | chrX-36903453-INS-52 |
| chrX-36928700-INS-1206 | chrX-36928702-INS-1203 |
| chrX-36978791-INS-60 | chrX-36978761-INS-90 |
| chrX-37048760-INS-211 | chrX-37048760-INS-211 |
| chrX-37185135-INS-52 | chrX-37185145-INS-62 |
| chrX-37250933-INS-129 | chrX-37250933-INS-129 |
| chrX-37256593-INS-78 | chrX-37256593-INS-78 |
| chrX-37300172-INS-153 | chrX-37300172-INS-167 |
| chrX-37336335-INS-100 | chrX-37336335-INS-102 |
| chrX-37350576-INS-68 | chrX-37350573-INS-71 |
| chrX-37456225-INS-1571 | chrX-37456226-INS-1569 |
| chrX-37462417-INS-78 | chrX-37462417-INS-78 |
| chrX-37477069-INS-6440 | chrX-37477070-INS-6440 |
| chrX-37488268-INS-454 | chrX-37488267-INS-431 |
| chrX-37522468-INS-112 | chrX-37522486-INS-134 |
| chrX-37533965-INS-92 | chrX-37533951-INS-107 |
| chrX-37546881-INS-385 | chrX-37546882-INS-388 |
| chrX-37548806-INS-166 | chrX-37548806-INS-166 |
| chrX-37578693-INS-580 | chrX-37578693-INS-585 |
| chrX-37695296-INS-65 | chrX-37695299-INS-72 |
| chrX-37763224-INS-134 | chrX-37763229-INS-170 |
| chrX-37768902-INS-362 | chrX-37768902-INS-363 |
| chrX-37792704-INS-167 | chrX-37792704-INS-167 |
| chrX-37847212-INS-94 | chrX-37847216-INS-118 |
| chrX-37848355-INS-3821 | chrX-37848356-INS-3822 |
| chrX-37873104-INS-424 | chrX-37873104-INS-424 |
| chrX-37952051-INS-52 | chrX-37952051-INS-54 |
| chrX-37970017-INS-7115 | chrX-37970018-INS-7117 |
| chrX-38001929-INS-6623 | chrX-38001930-INS-6624 |
| chrX-38034188-INS-1980 | chrX-38034189-INS-1981 |
| chrX-38048977-INS-6687 | chrX-38048978-INS-6688 |
| chrX-38082495-INS-96 | chrX-38082495-INS-70 |
| chrX-38088250-INS-87 | chrX-38088250-INS-87 |
| chrX-38112434-INS-399 | chrX-38112434-INS-399 |
| chrX-38203020-INS-72 | chrX-38203020-INS-60 |
| chrX-38271212-INS-1183 | chrX-38271213-INS-1179 |
| chrX-38303244-INS-419 | chrX-38303244-INS-407 |
| chrX-38330400-INS-66 | chrX-38330400-INS-62 |
| chrX-38358054-INS-497 | chrX-38358054-INS-497 |
| chrX-38374584-INS-106 | chrX-38374584-INS-107 |
| chrX-38538588-INS-229 | chrX-38538588-INS-230 |
| chrX-38574711-INS-6335 | chrX-38574712-INS-6341 |
| chrX-38575284-INS-80 | chrX-38575284-INS-80 |
| chrX-38633340-INS-3215 | chrX-38633323-INS-3213 |
| chrX-38634386-INS-222 | chrX-38634384-INS-236 |
| chrX-38641123-INS-7234 | chrX-38641124-INS-7234 |
| chrX-38726116-INS-6178 | chrX-38726117-INS-6178 |
| chrX-38739795-INS-56 | chrX-38739795-INS-56 |
| chrX-38881395-INS-421 | chrX-38881395-INS-421 |
| chrX-38894196-INS-65 | chrX-38894168-INS-56 |
| chrX-38901702-INS-6442 | chrX-38901703-INS-6442 |
| chrX-38926842-INS-7081 | chrX-38926843-INS-7081 |
| chrX-39033402-INS-122 | chrX-39033402-INS-110 |
| chrX-39075942-INS-389 | chrX-39075942-INS-389 |
| chrX-39080454-INS-55 | chrX-39080468-INS-56 |
| chrX-39105237-INS-70 | chrX-39105237-INS-70 |
| chrX-39148975-INS-56 | chrX-39148982-INS-56 |
| chrX-39187332-INS-92 | chrX-39187332-INS-60 |
| chrX-39229375-INS-544 | chrX-39229351-INS-545 |
| chrX-39246857-INS-280 | chrX-39246857-INS-280 |
| chrX-39249803-INS-6608 | chrX-39249804-INS-6608 |
| chrX-39271417-INS-208 | chrX-39271417-INS-208 |
| chrX-39306802-INS-93 | chrX-39306771-INS-93 |
| chrX-39445519-INS-519 | chrX-39445519-INS-521 |
| chrX-39505953-INS-6380 | chrX-39505954-INS-6380 |
| chrX-39521352-INS-314 | chrX-39521352-INS-323 |
| chrX-39527526-INS-6272 | chrX-39527527-INS-6273 |
| chrX-39552390-INS-211 | chrX-39552390-INS-211 |
| chrX-39583348-INS-1822 | chrX-39583349-INS-1824 |
| chrX-39597284-INS-506 | chrX-39597284-INS-506 |

| | |
|---|---|
| chrX-39632399-INS-2697 | chrX-39632400-INS-2698 |
| chrX-39772265-INS-3322 | chrX-39772266-INS-3322 |
| chrX-39794446-INS-6523 | chrX-39794447-INS-6526 |
| chrX-39811359-INS-92 | chrX-39811393-INS-50 |
| chrX-39820771-INS-63 | chrX-39820771-INS-60 |
| chrX-39823911-INS-2397 | chrX-39823912-INS-2395 |
| chrX-39896139-INS-4563 | chrX-39896140-INS-4560 |
| chrX-39921383-INS-920 | chrX-39921383-INS-922 |
| chrX-39975963-INS-1376 | chrX-39975964-INS-1380 |
| chrX-39976003-INS-61 | chrX-39976027-INS-56 |
| chrX-40001880-INS-1550 | chrX-40001896-INS-1553 |
| chrX-40021426-INS-58 | chrX-40021444-INS-50 |
| chrX-40059650-INS-74 | chrX-40059654-INS-74 |
| chrX-40075534-INS-84 | chrX-40075536-INS-66 |
| chrX-40078706-INS-3000 | chrX-40078707-INS-3000 |
| chrX-40153462-INS-3838 | chrX-40153463-INS-3839 |
| chrX-40200320-INS-7157 | chrX-40200321-INS-7158 |
| chrX-40200893-INS-148 | chrX-40200893-INS-129 |
| chrX-40208305-INS-6203 | chrX-40208306-INS-6203 |
| chrX-40289593-INS-146 | chrX-40289593-INS-146 |
| chrX-40334109-INS-4320 | chrX-40334110-INS-4320 |
| chrX-40370687-INS-60 | chrX-40370687-INS-50 |
| chrX-40385084-INS-356 | chrX-40385084-INS-356 |
| chrX-40392381-INS-146 | chrX-40392381-INS-146 |
| chrX-40462233-INS-1538 | chrX-40462234-INS-1538 |
| chrX-40480999-INS-69 | chrX-40480999-INS-63 |
| chrX-40486052-INS-68 | chrX-40486052-INS-81 |
| chrX-40539426-INS-7114 | chrX-40539427-INS-7114 |
| chrX-40569377-INS-54 | chrX-40569377-INS-52 |
| chrX-40593006-INS-1896 | chrX-40593007-INS-1907 |
| chrX-40597346-INS-158 | chrX-40597346-INS-142 |
| chrX-40606174-INS-2523 | chrX-40606175-INS-2523 |
| chrX-40618890-INS-6690 | chrX-40618891-INS-6690 |
| chrX-40825883-INS-382 | chrX-40825883-INS-384 |
| chrX-40869443-INS-6646 | chrX-40869444-INS-6646 |
| chrX-40972439-INS-51 | chrX-40972437-INS-53 |
| chrX-40974620-INS-485 | chrX-40974618-INS-489 |
| chrX-40994006-INS-7082 | chrX-40994007-INS-7082 |
| chrX-41021736-INS-2769 | chrX-41021737-INS-2769 |
| chrX-41074528-INS-448 | chrX-41074497-INS-452 |
| chrX-41119664-INS-51 | chrX-41119664-INS-55 |
| chrX-41125075-INS-176 | chrX-41125075-INS-178 |
| chrX-41129333-INS-417 | chrX-41129337-INS-426 |
| chrX-41198010-INS-51 | chrX-41198010-INS-51 |
| chrX-41248948-INS-626 | chrX-41248948-INS-626 |
| chrX-41256172-INS-187 | chrX-41256170-INS-186 |
| chrX-41344501-INS-696 | chrX-41344501-INS-697 |
| chrX-41400955-INS-564 | chrX-41400955-INS-563 |
| chrX-41414402-INS-54 | chrX-41414400-INS-58 |
| chrX-41508710-INS-234 | chrX-41508710-INS-232 |
| chrX-41557340-INS-586 | chrX-41557340-INS-586 |
| chrX-41581351-INS-124 | chrX-41581384-INS-76 |
| chrX-41641798-INS-122 | chrX-41641808-INS-99 |
| chrX-41662984-INS-60 | chrX-41662984-INS-72 |
| chrX-41668611-INS-625 | chrX-41668611-INS-626 |
| chrX-41670929-INS-273 | chrX-41670929-INS-288 |
| chrX-41695730-INS-54 | chrX-41695730-INS-53 |
| chrX-41800929-INS-321 | chrX-41800929-INS-321 |
| chrX-41813100-INS-73 | chrX-41813100-INS-57 |
| chrX-41821127-INS-1555 | chrX-41821128-INS-1556 |
| chrX-41871801-INS-6460 | chrX-41871802-INS-6460 |
| chrX-41890989-INS-927 | chrX-41890989-INS-927 |
| chrX-41910921-INS-160 | chrX-41910918-INS-162 |
| chrX-41921158-INS-683 | chrX-41921158-INS-683 |
| chrX-41936701-INS-6651 | chrX-41936702-INS-6654 |
| chrX-42015611-INS-212 | chrX-42015611-INS-212 |
| chrX-42025535-INS-390 | chrX-42025535-INS-390 |
| chrX-42065268-INS-6774 | chrX-42065269-INS-6773 |
| chrX-42129291-INS-1868 | chrX-42129292-INS-1868 |
| chrX-42228164-INS-6925 | chrX-42228165-INS-6939 |
| chrX-42241019-INS-177 | chrX-42241019-INS-179 |
| chrX-42272930-INS-90 | chrX-42272930-INS-90 |
| chrX-42282780-INS-1862 | chrX-42282781-INS-1862 |
| chrX-42347435-INS-6680 | chrX-42347436-INS-6673 |
| chrX-42434992-INS-580 | chrX-42434992-INS-580 |
| chrX-42501845-INS-172 | chrX-42501845-INS-170 |
| chrX-42514032-INS-73 | chrX-42514038-INS-73 |

| | |
|---|---|
| chrX-42539494-INS-890 | chrX-42539494-INS-869 |
| chrX-42568311-INS-692 | chrX-42568311-INS-692 |
| chrX-42603388-INS-133 | chrX-42603388-INS-133 |
| chrX-42604639-INS-171 | chrX-42604639-INS-171 |
| chrX-42606972-INS-6165 | chrX-42606973-INS-6164 |
| chrX-42612424-INS-210 | chrX-42612424-INS-209 |
| chrX-42617307-INS-1241 | chrX-42617308-INS-1241 |
| chrX-42686499-INS-196 | chrX-42686499-INS-194 |
| chrX-42792645-INS-6464 | chrX-42792646-INS-6459 |
| chrX-42795157-INS-2860 | chrX-42795158-INS-2860 |
| chrX-42806017-INS-184 | chrX-42806017-INS-186 |
| chrX-42816445-INS-68 | chrX-42816445-INS-56 |
| chrX-42826859-INS-108 | chrX-42826859-INS-108 |
| chrX-42834495-INS-604 | chrX-42834495-INS-605 |
| chrX-42850169-INS-3209 | chrX-42850170-INS-3230 |
| chrX-42923644-INS-4735 | chrX-42923645-INS-4735 |
| chrX-42932452-INS-64 | chrX-42932455-INS-53 |
| chrX-43057436-INS-2586 | chrX-43057437-INS-2586 |
| chrX-43150421-INS-3324 | chrX-43150422-INS-3317 |
| chrX-43155168-INS-6294 | chrX-43155169-INS-6294 |
| chrX-43158445-INS-371 | chrX-43158446-INS-377 |
| chrX-43175805-INS-81 | chrX-43175805-INS-81 |
| chrX-43210819-INS-198 | chrX-43210819-INS-199 |
| chrX-43228919-INS-166 | chrX-43228922-INS-162 |
| chrX-43233431-INS-63 | chrX-43233431-INS-66 |
| chrX-43241968-INS-1007 | chrX-43241969-INS-1008 |
| chrX-43268587-INS-101 | chrX-43268587-INS-51 |
| chrX-43274246-INS-94 | chrX-43274246-INS-102 |
| chrX-43276119-INS-772 | chrX-43276119-INS-774 |
| chrX-43293209-INS-208 | chrX-43293209-INS-208 |
| chrX-43310321-INS-61 | chrX-43310326-INS-71 |
| chrX-43369190-INS-965 | chrX-43369190-INS-965 |
| chrX-43371480-INS-6654 | chrX-43371481-INS-6654 |
| chrX-43487473-INS-76 | chrX-43487473-INS-72 |
| chrX-43561956-INS-118 | chrX-43561953-INS-122 |
| chrX-43594720-INS-646 | chrX-43594733-INS-655 |
| chrX-43651780-INS-349 | chrX-43651780-INS-349 |
| chrX-43657415-INS-524 | chrX-43657415-INS-524 |
| chrX-43707228-INS-71 | chrX-43707233-INS-63 |
| chrX-43751060-INS-203 | chrX-43751060-INS-203 |
| chrX-43765177-INS-1124 | chrX-43765178-INS-1126 |
| chrX-43937762-INS-3312 | chrX-43937763-INS-3312 |
| chrX-43979694-INS-571 | chrX-43979694-INS-570 |
| chrX-44047648-INS-334 | chrX-44047648-INS-346 |
| chrX-44170331-INS-72 | chrX-44170331-INS-72 |
| chrX-44195947-INS-3002 | chrX-44195948-INS-3002 |
| chrX-44198469-INS-2713 | chrX-44198470-INS-2713 |
| chrX-44233043-INS-239 | chrX-44233073-INS-252 |
| chrX-44241073-INS-454 | chrX-44241073-INS-458 |
| chrX-44296804-INS-981 | chrX-44296804-INS-981 |
| chrX-44308796-INS-4945 | chrX-44308797-INS-4947 |
| chrX-44349768-INS-7656 | chrX-44349769-INS-7660 |
| chrX-44351620-INS-195 | chrX-44351620-INS-195 |
| chrX-44428669-INS-100 | chrX-44428663-INS-108 |
| chrX-44446605-INS-66 | chrX-44446599-INS-72 |
| chrX-44477898-INS-6684 | chrX-44477899-INS-6685 |
| chrX-44600240-INS-55 | chrX-44600229-INS-55 |
| chrX-44621611-INS-251 | chrX-44621611-INS-251 |
| chrX-44676090-INS-170 | chrX-44676090-INS-170 |
| chrX-44677421-INS-2765 | chrX-44677422-INS-2765 |
| chrX-44682189-INS-6383 | chrX-44682190-INS-6378 |
| chrX-44711975-INS-7080 | chrX-44711976-INS-7079 |
| chrX-44727156-INS-174 | chrX-44727137-INS-193 |
| chrX-44745746-INS-62 | chrX-44745752-INS-64 |
| chrX-44748171-INS-359 | chrX-44748171-INS-359 |
| chrX-44776622-INS-1149 | chrX-44776623-INS-1131 |
| chrX-44782970-INS-87 | chrX-44782970-INS-51 |
| chrX-44789782-INS-664 | chrX-44789782-INS-664 |
| chrX-44868175-INS-505 | chrX-44868175-INS-505 |
| chrX-44875600-INS-654 | chrX-44875600-INS-656 |
| chrX-44910881-INS-51 | chrX-44910881-INS-69 |
| chrX-44936061-INS-1435 | chrX-44936062-INS-1435 |
| chrX-44951392-INS-1862 | chrX-44951393-INS-1862 |
| chrX-44969531-INS-197 | chrX-44969531-INS-198 |
| chrX-45025376-INS-3645 | chrX-45025377-INS-3641 |
| chrX-45093286-INS-1107 | chrX-45093287-INS-1107 |
| chrX-45113080-INS-576 | chrX-45113080-INS-576 |

| | |
|---|---|
| chrX-45118566-INS-6447 | chrX-45118567-INS-6445 |
| chrX-45130624-INS-1279 | chrX-45130625-INS-1279 |
| chrX-45149733-INS-381 | chrX-45149733-INS-382 |
| chrX-45212611-INS-51 | chrX-45212611-INS-51 |
| chrX-45230380-INS-2140 | chrX-45230381-INS-2140 |
| chrX-45232014-INS-513 | chrX-45232014-INS-513 |
| chrX-45240222-INS-64 | chrX-45240264-INS-60 |
| chrX-45267109-INS-96 | chrX-45267109-INS-84 |
| chrX-45284527-INS-6295 | chrX-45284528-INS-6296 |
| chrX-45295873-INS-108 | chrX-45295873-INS-85 |
| chrX-45405421-INS-1483 | chrX-45405422-INS-1483 |
| chrX-45415183-INS-3104 | chrX-45415184-INS-3108 |
| chrX-45430905-INS-64 | chrX-45430905-INS-80 |
| chrX-45478292-INS-675 | chrX-45478292-INS-675 |
| chrX-45485796-INS-56 | chrX-45485792-INS-92 |
| chrX-45546253-INS-74 | chrX-45546253-INS-72 |
| chrX-45569095-INS-7188 | chrX-45569096-INS-7202 |
| chrX-45664468-INS-102 | chrX-45664479-INS-135 |
| chrX-45697569-INS-216 | chrX-45697569-INS-218 |
| chrX-45707296-INS-162 | chrX-45707296-INS-162 |
| chrX-45718508-INS-2936 | chrX-45718509-INS-2936 |
| chrX-45732176-INS-1208 | chrX-45732177-INS-1209 |
| chrX-45790232-INS-56 | chrX-45790232-INS-56 |
| chrX-45805563-INS-2845 | chrX-45805564-INS-2845 |
| chrX-45805949-INS-119 | chrX-45805949-INS-119 |
| chrX-45831819-INS-2904 | chrX-45831820-INS-2904 |
| chrX-45851591-INS-397 | chrX-45851591-INS-397 |
| chrX-45915305-INS-3032 | chrX-45915306-INS-3032 |
| chrX-45916738-INS-86 | chrX-45916738-INS-78 |
| chrX-45995475-INS-565 | chrX-45995472-INS-565 |
| chrX-46034889-INS-72 | chrX-46034862-INS-56 |
| chrX-46149610-INS-119 | chrX-46149610-INS-152 |
| chrX-46210322-INS-60 | chrX-46210322-INS-52 |
| chrX-46283691-INS-211 | chrX-46283691-INS-212 |
| chrX-46291717-INS-1891 | chrX-46291718-INS-1891 |
| chrX-46407798-INS-739 | chrX-46407798-INS-738 |
| chrX-46473550-INS-117 | chrX-46473594-INS-120 |
| chrX-46532229-INS-172 | chrX-46532229-INS-176 |
| chrX-46592134-INS-115 | chrX-46592131-INS-75 |
| chrX-46650659-INS-107 | chrX-46650659-INS-97 |
| chrX-46664698-INS-80 | chrX-46664698-INS-50 |
| chrX-46805495-INS-1886 | chrX-46805496-INS-1886 |
| chrX-46953627-INS-1893 | chrX-46953628-INS-1893 |
| chrX-46971625-INS-216 | chrX-46971625-INS-217 |
| chrX-47031217-INS-2127 | chrX-47031229-INS-2128 |
| chrX-47066925-INS-7180 | chrX-47066926-INS-7180 |
| chrX-47082958-INS-141 | chrX-47082958-INS-142 |
| chrX-47096343-INS-208 | chrX-47096343-INS-210 |
| chrX-47104222-INS-60 | chrX-47104222-INS-60 |
| chrX-47129945-INS-197 | chrX-47129945-INS-197 |
| chrX-47148851-INS-7245 | chrX-47148852-INS-7246 |
| chrX-47160147-INS-100 | chrX-47160147-INS-94 |
| chrX-47173928-INS-64 | chrX-47173928-INS-64 |
| chrX-47176212-INS-6304 | chrX-47176213-INS-6307 |
| chrX-47192190-INS-118 | chrX-47192165-INS-93 |
| chrX-47200195-INS-205 | chrX-47200195-INS-203 |
| chrX-47252315-INS-81 | chrX-47252315-INS-83 |
| chrX-47281917-INS-197 | chrX-47281917-INS-197 |
| chrX-47303431-INS-83 | chrX-47303447-INS-64 |
| chrX-47308143-INS-211 | chrX-47308143-INS-211 |
| chrX-47315723-INS-68 | chrX-47315723-INS-68 |
| chrX-47318198-INS-200 | chrX-47318198-INS-200 |
| chrX-47321673-INS-185 | chrX-47321673-INS-185 |
| chrX-47329325-INS-60 | chrX-47329325-INS-72 |
| chrX-47334123-INS-191 | chrX-47334123-INS-186 |
| chrX-47340305-INS-1867 | chrX-47340306-INS-1867 |
| chrX-47342368-INS-5546 | chrX-47342369-INS-5546 |
| chrX-47381549-INS-50 | chrX-47381549-INS-57 |
| chrX-47386062-INS-523 | chrX-47386062-INS-523 |
| chrX-47411342-INS-60 | chrX-47411342-INS-60 |
| chrX-47416070-INS-59 | chrX-47416055-INS-76 |
| chrX-47529316-INS-163 | chrX-47529316-INS-164 |
| chrX-47561505-INS-52 | chrX-47561505-INS-52 |
| chrX-47589964-INS-83 | chrX-47589988-INS-58 |
| chrX-47604498-INS-68 | chrX-47604516-INS-55 |
| chrX-47675536-INS-76 | chrX-47675536-INS-88 |
| chrX-47676990-INS-118 | chrX-47676990-INS-118 |

| | |
|---|---|
| chrX-47690461-INS-6401 | chrX-47690462-INS-6401 |
| chrX-47734322-INS-371 | chrX-47734322-INS-371 |
| chrX-47787015-INS-378 | chrX-47786992-INS-351 |
| chrX-47803215-INS-1136 | chrX-47803216-INS-1136 |
| chrX-47868295-INS-54 | chrX-47868295-INS-56 |
| chrX-47890135-INS-1194 | chrX-47890136-INS-1194 |
| chrX-47891172-INS-63 | chrX-47891172-INS-63 |
| chrX-47894946-INS-3333 | chrX-47894947-INS-3333 |
| chrX-47945211-INS-55 | chrX-47945211-INS-55 |
| chrX-47956463-INS-68 | chrX-47956487-INS-52 |
| chrX-47967480-INS-75 | chrX-47967480-INS-75 |
| chrX-47978578-INS-724 | chrX-47978578-INS-724 |
| chrX-47980841-INS-615 | chrX-47980841-INS-614 |
| chrX-47983914-INS-830 | chrX-47983914-INS-809 |
| chrX-47995270-INS-103 | chrX-47995277-INS-127 |
| chrX-48001655-INS-5224 | chrX-48001656-INS-5224 |
| chrX-48034491-INS-550 | chrX-48034491-INS-550 |
| chrX-48049827-INS-60 | chrX-48049827-INS-60 |
| chrX-48084887-INS-60 | chrX-48084890-INS-50 |
| chrX-48121638-INS-128 | chrX-48121638-INS-128 |
| chrX-48133534-INS-110 | chrX-48133529-INS-105 |
| chrX-48145616-INS-475 | chrX-48145616-INS-472 |
| chrX-48150621-INS-129 | chrX-48150621-INS-129 |
| chrX-48154841-INS-345 | chrX-48154841-INS-344 |
| chrX-48170733-INS-566 | chrX-48170733-INS-567 |
| chrX-48172385-INS-687 | chrX-48172385-INS-687 |
| chrX-48210753-INS-6522 | chrX-48210754-INS-6523 |
| chrX-48219963-INS-1204 | chrX-48219964-INS-1204 |
| chrX-48227298-INS-121 | chrX-48227298-INS-121 |
| chrX-48242339-INS-7128 | chrX-48242340-INS-7130 |
| chrX-48281001-INS-56 | chrX-48281001-INS-58 |
| chrX-48288308-INS-6926 | chrX-48288309-INS-6927 |
| chrX-48298507-INS-68 | chrX-48298507-INS-68 |
| chrX-48316996-INS-569 | chrX-48316996-INS-570 |
| chrX-48324269-INS-391 | chrX-48324269-INS-391 |
| chrX-48340624-INS-54 | chrX-48340624-INS-98 |
| chrX-48351044-INS-54 | chrX-48351044-INS-54 |
| chrX-48360101-INS-559 | chrX-48360101-INS-561 |
| chrX-48393607-INS-294 | chrX-48393607-INS-294 |
| chrX-48397392-INS-114 | chrX-48397392-INS-76 |
| chrX-48408579-INS-293 | chrX-48408579-INS-293 |
| chrX-48428118-INS-88 | chrX-48428131-INS-50 |
| chrX-48482074-INS-423 | chrX-48482074-INS-434 |
| chrX-48496353-INS-226 | chrX-48496353-INS-206 |
| chrX-48506894-INS-201 | chrX-48506894-INS-201 |
| chrX-48528240-INS-574 | chrX-48528240-INS-574 |
| chrX-48607897-INS-3032 | chrX-48607898-INS-3032 |
| chrX-48611994-INS-7052 | chrX-48611995-INS-7052 |
| chrX-48686738-INS-55 | chrX-48686738-INS-54 |
| chrX-48741620-INS-75 | chrX-48741620-INS-81 |
| chrX-48787330-INS-429 | chrX-48787330-INS-428 |
| chrX-48801296-INS-460 | chrX-48801296-INS-460 |
| chrX-48915096-INS-701 | chrX-48915096-INS-705 |
| chrX-49071655-INS-6576 | chrX-49071656-INS-6576 |
| chrX-49078060-INS-2149 | chrX-49078061-INS-2153 |
| chrX-49082819-INS-1512 | chrX-49082820-INS-1511 |
| chrX-49131496-INS-6862 | chrX-49131497-INS-6862 |
| chrX-49166385-INS-2088 | chrX-49166386-INS-2088 |
| chrX-49212557-INS-88 | chrX-49212557-INS-52 |
| chrX-49225061-INS-150 | chrX-49225061-INS-150 |
| chrX-49294858-INS-98 | chrX-49294858-INS-94 |
| chrX-49337678-INS-7469 | chrX-49337679-INS-7470 |
| chrX-49359177-INS-6264 | chrX-49359178-INS-6264 |
| chrX-49404297-INS-794 | chrX-49404297-INS-794 |
| chrX-49422780-INS-651 | chrX-49422780-INS-651 |
| chrX-49431100-INS-838 | chrX-49431100-INS-854 |
| chrX-49436978-INS-163 | chrX-49436978-INS-163 |
| chrX-49451530-INS-177 | chrX-49451530-INS-170 |
| chrX-49604497-INS-1206 | chrX-49604498-INS-1206 |
| chrX-49642171-INS-97 | chrX-49642171-INS-97 |
| chrX-49645336-INS-1151 | chrX-49645337-INS-1151 |
| chrX-49725147-INS-212 | chrX-49725147-INS-212 |
| chrX-49726934-INS-720 | chrX-49726934-INS-721 |
| chrX-49740585-INS-629 | chrX-49740559-INS-645 |
| chrX-49792056-INS-173 | chrX-49792056-INS-173 |
| chrX-49798275-INS-983 | chrX-49798275-INS-983 |
| chrX-49828330-INS-109 | chrX-49828330-INS-109 |

| | |
|---|---|
| chrX-49828806-INS-156 | chrX-49828806-INS-156 |
| chrX-49854913-INS-9234 | chrX-49854914-INS-9234 |
| chrX-49879958-INS-561 | chrX-49879958-INS-561 |
| chrX-49961383-INS-83 | chrX-49961341-INS-76 |
| chrX-50014560-INS-54 | chrX-50014560-INS-54 |
| chrX-50034694-INS-7451 | chrX-50034695-INS-7452 |
| chrX-50076245-INS-190 | chrX-50076245-INS-186 |
| chrX-50082633-INS-405 | chrX-50082633-INS-450 |
| chrX-50085492-INS-6436 | chrX-50085493-INS-6436 |
| chrX-50217188-INS-6443 | chrX-50217189-INS-6443 |
| chrX-50298821-INS-872 | chrX-50298820-INS-873 |
| chrX-50301838-INS-207 | chrX-50301838-INS-210 |
| chrX-50341007-INS-60 | chrX-50341007-INS-50 |
| chrX-50384418-INS-850 | chrX-50384418-INS-848 |
| chrX-50401921-INS-2663 | chrX-50401922-INS-2663 |
| chrX-50486134-INS-6753 | chrX-50486135-INS-6757 |
| chrX-50498060-INS-1890 | chrX-50498061-INS-1889 |
| chrX-50518654-INS-222 | chrX-50518654-INS-224 |
| chrX-50543419-INS-195 | chrX-50543419-INS-188 |
| chrX-50600584-INS-51 | chrX-50600584-INS-51 |
| chrX-50605468-INS-128 | chrX-50605468-INS-163 |
| chrX-50633716-INS-3293 | chrX-50633717-INS-3288 |
| chrX-50743254-INS-91 | chrX-50743266-INS-90 |
| chrX-50792904-INS-72 | chrX-50792904-INS-72 |
| chrX-50793926-INS-70 | chrX-50793926-INS-71 |
| chrX-50828646-INS-212 | chrX-50828646-INS-212 |
| chrX-50832737-INS-120 | chrX-50832737-INS-84 |
| chrX-50866441-INS-6710 | chrX-50866442-INS-6711 |
| chrX-50877161-INS-194 | chrX-50877161-INS-194 |
| chrX-50890495-INS-478 | chrX-50890490-INS-434 |
| chrX-50930982-INS-60 | chrX-50930982-INS-61 |
| chrX-50955454-INS-53 | chrX-50955454-INS-53 |
| chrX-50968564-INS-182 | chrX-50968564-INS-221 |
| chrX-50975428-INS-66 | chrX-50975428-INS-66 |
| chrX-50979005-INS-455 | chrX-50979005-INS-455 |
| chrX-51001110-INS-145 | chrX-51001110-INS-145 |
| chrX-51009363-INS-249 | chrX-51009363-INS-246 |
| chrX-51013356-INS-277 | chrX-51013356-INS-277 |
| chrX-51034768-INS-109 | chrX-51034768-INS-109 |
| chrX-51090898-INS-381 | chrX-51090898-INS-381 |
| chrX-51095953-INS-140 | chrX-51095953-INS-128 |
| chrX-51107467-INS-100 | chrX-51107467-INS-99 |
| chrX-51133153-INS-72 | chrX-51133171-INS-100 |
| chrX-51156476-INS-68 | chrX-51156476-INS-66 |
| chrX-51159226-INS-78 | chrX-51159226-INS-78 |
| chrX-51168142-INS-701 | chrX-51168142-INS-697 |
| chrX-51187366-INS-370 | chrX-51187366-INS-370 |
| chrX-51265233-INS-390 | chrX-51265233-INS-390 |
| chrX-51280097-INS-61 | chrX-51280097-INS-87 |
| chrX-51290975-INS-63 | chrX-51290975-INS-57 |
| chrX-51294761-INS-211 | chrX-51294761-INS-211 |
| chrX-51336974-INS-190 | chrX-51336974-INS-190 |
| chrX-51343153-INS-56 | chrX-51343151-INS-58 |
| chrX-51343760-INS-93 | chrX-51343760-INS-93 |
| chrX-51349153-INS-164 | chrX-51349153-INS-164 |
| chrX-51353915-INS-264 | chrX-51353915-INS-267 |
| chrX-51395155-INS-62 | chrX-51395155-INS-64 |
| chrX-51418874-INS-213 | chrX-51418874-INS-208 |
| chrX-51479702-INS-453 | chrX-51479702-INS-454 |
| chrX-51553223-INS-60 | chrX-51553223-INS-62 |
| chrX-51555111-INS-58 | chrX-51555111-INS-58 |
| chrX-51623086-INS-58 | chrX-51623087-INS-67 |
| chrX-51631068-INS-974 | chrX-51631068-INS-976 |
| chrX-51669597-INS-93 | chrX-51669597-INS-93 |
| chrX-51692577-INS-180 | chrX-51692573-INS-169 |
| chrX-51739573-INS-75 | chrX-51739573-INS-84 |
| chrX-51796598-INS-204 | chrX-51796598-INS-204 |
| chrX-51799914-INS-1891 | chrX-51799915-INS-1891 |
| chrX-51804691-INS-101 | chrX-51804691-INS-86 |
| chrX-51822396-INS-202 | chrX-51822396-INS-202 |
| chrX-51824432-INS-7327 | chrX-51824433-INS-7342 |
| chrX-51858721-INS-52 | chrX-51858721-INS-52 |
| chrX-51864433-INS-188 | chrX-51864433-INS-188 |
| chrX-51865491-INS-91 | chrX-51865491-INS-91 |
| chrX-51866247-INS-418 | chrX-51866247-INS-418 |
| chrX-51871948-INS-7113 | chrX-51871949-INS-7113 |
| chrX-51877442-INS-116 | chrX-51877442-INS-116 |

| | |
|---|---|
| chrX-51878628-INS-53 | chrX-51878628-INS-59 |
| chrX-51889477-INS-192 | chrX-51889477-INS-194 |
| chrX-51906046-INS-74 | chrX-51906040-INS-116 |
| chrX-51912029-INS-5630 | chrX-51912030-INS-5630 |
| chrX-51915720-INS-747 | chrX-51915721-INS-747 |
| chrX-51915788-INS-177 | chrX-51915801-INS-202 |
| chrX-51922266-INS-56 | chrX-51922266-INS-83 |
| chrX-51927453-INS-925 | chrX-51927453-INS-926 |
| chrX-51954350-INS-64 | chrX-51954350-INS-60 |
| chrX-51965035-INS-62 | chrX-51965035-INS-62 |
| chrX-51975458-INS-100 | chrX-51975482-INS-52 |
| chrX-51994216-INS-83 | chrX-51994216-INS-76 |
| chrX-52038486-INS-229 | chrX-52038486-INS-229 |
| chrX-52039538-INS-180 | chrX-52039538-INS-181 |
| chrX-52059605-INS-329 | chrX-52059605-INS-327 |
| chrX-52060829-INS-141 | chrX-52060829-INS-117 |
| chrX-52062535-INS-189 | chrX-52062547-INS-196 |
| chrX-52062990-INS-5319 | chrX-52062991-INS-5319 |
| chrX-52102111-INS-200 | chrX-52102111-INS-197 |
| chrX-52107057-INS-216 | chrX-52107057-INS-216 |
| chrX-52129546-INS-150 | chrX-52129548-INS-150 |
| chrX-52157272-INS-96 | chrX-52157272-INS-112 |
| chrX-52245162-INS-85 | chrX-52245162-INS-111 |
| chrX-52249632-INS-213 | chrX-52249632-INS-223 |
| chrX-52273874-INS-84 | chrX-52273874-INS-84 |
| chrX-52299973-INS-363 | chrX-52299973-INS-369 |
| chrX-52360699-INS-61 | chrX-52360699-INS-61 |
| chrX-52429746-INS-4339 | chrX-52429747-INS-4339 |
| chrX-52462563-INS-120 | chrX-52462563-INS-137 |
| chrX-52616865-INS-80 | chrX-52616865-INS-80 |
| chrX-52624130-INS-58 | chrX-52624130-INS-63 |
| chrX-52658830-INS-80 | chrX-52658830-INS-80 |
| chrX-52670158-INS-5425 | chrX-52670159-INS-5425 |
| chrX-52706281-INS-1886 | chrX-52706282-INS-1886 |
| chrX-52711532-INS-142 | chrX-52711532-INS-144 |
| chrX-52722459-INS-1862 | chrX-52722460-INS-1862 |
| chrX-52758813-INS-194 | chrX-52758813-INS-194 |
| chrX-52765829-INS-111 | chrX-52765829-INS-114 |
| chrX-52842375-INS-209 | chrX-52842375-INS-209 |
| chrX-52980869-INS-911 | chrX-52980869-INS-911 |
| chrX-52986591-INS-140 | chrX-52986591-INS-96 |
| chrX-52987278-INS-1062 | chrX-52987279-INS-1058 |
| chrX-53115360-INS-213 | chrX-53115360-INS-213 |
| chrX-53147387-INS-902 | chrX-53147387-INS-898 |
| chrX-53162899-INS-53 | chrX-53162899-INS-53 |
| chrX-54646312-INS-208 | chrX-54646312-INS-208 |
| chrX-54885130-INS-3762 | chrX-54885131-INS-3769 |
| chrX-54921290-INS-6467 | chrX-54921291-INS-6469 |
| chrX-55157833-INS-2261 | chrX-55157840-INS-2251 |
| chrX-55170158-INS-748 | chrX-55170158-INS-747 |
| chrX-55172807-INS-68 | chrX-55172807-INS-68 |
| chrX-55180838-INS-6393 | chrX-55180839-INS-6394 |
| chrX-55215351-INS-162 | chrX-55215354-INS-140 |
| chrX-55257162-INS-266 | chrX-55257162-INS-266 |
| chrX-55299646-INS-231 | chrX-55299646-INS-233 |
| chrX-55375457-INS-326 | chrX-55375457-INS-326 |
| chrX-55380185-INS-1014 | chrX-55380186-INS-1012 |
| chrX-55426133-INS-65 | chrX-55426133-INS-65 |
| chrX-55470218-INS-63 | chrX-55470218-INS-63 |
| chrX-55473796-INS-107 | chrX-55473771-INS-108 |
| chrX-55490988-INS-91 | chrX-55490988-INS-91 |
| chrX-55562082-INS-6435 | chrX-55562083-INS-6435 |
| chrX-55578231-INS-135 | chrX-55578231-INS-135 |
| chrX-55582779-INS-1348 | chrX-55582780-INS-1348 |
| chrX-55672433-INS-484 | chrX-55672433-INS-484 |
| chrX-55673090-INS-118 | chrX-55673090-INS-118 |
| chrX-55729589-INS-57 | chrX-55729589-INS-57 |
| chrX-55742320-INS-104 | chrX-55742339-INS-59 |
| chrX-55752901-INS-315 | chrX-55752901-INS-315 |
| chrX-55761294-INS-100 | chrX-55761294-INS-80 |
| chrX-55766230-INS-102 | chrX-55766270-INS-75 |
| chrX-55802777-INS-109 | chrX-55802783-INS-97 |
| chrX-55847879-INS-51 | chrX-55847879-INS-53 |
| chrX-55869954-INS-92 | chrX-55869998-INS-88 |
| chrX-55877812-INS-312 | chrX-55877849-INS-312 |
| chrX-55895532-INS-399 | chrX-55895532-INS-400 |
| chrX-55938872-INS-196 | chrX-55938872-INS-196 |

| | |
|---|---|
| chrX-55951375-INS-60 | chrX-55951375-INS-66 |
| chrX-55968849-INS-53 | chrX-55968849-INS-53 |
| chrX-56036370-INS-132 | chrX-56036370-INS-132 |
| chrX-56051806-INS-155 | chrX-56051806-INS-155 |
| chrX-56060720-INS-1062 | chrX-56060721-INS-1062 |
| chrX-56072632-INS-66 | chrX-56072632-INS-66 |
| chrX-56075322-INS-829 | chrX-56075322-INS-829 |
| chrX-56168513-INS-93 | chrX-56168514-INS-84 |
| chrX-56169858-INS-2536 | chrX-56169859-INS-2536 |
| chrX-56235109-INS-7525 | chrX-56235110-INS-7526 |
| chrX-56304494-INS-71 | chrX-56304494-INS-71 |
| chrX-56322722-INS-56 | chrX-56322724-INS-50 |
| chrX-56326397-INS-474 | chrX-56326397-INS-474 |
| chrX-56327611-INS-76 | chrX-56327611-INS-76 |
| chrX-56381915-INS-173 | chrX-56381915-INS-173 |
| chrX-56408940-INS-3556 | chrX-56408941-INS-3557 |
| chrX-56450952-INS-4510 | chrX-56450953-INS-4510 |
| chrX-56478215-INS-253 | chrX-56478215-INS-253 |
| chrX-56551250-INS-80 | chrX-56551250-INS-80 |
| chrX-56680593-INS-132 | chrX-56680593-INS-84 |
| chrX-56709507-INS-644 | chrX-56709507-INS-644 |
| chrX-56721077-INS-55 | chrX-56721077-INS-57 |
| chrX-56782381-INS-5726 | chrX-56782382-INS-5726 |
| chrX-56787020-INS-2674 | chrX-56787021-INS-2674 |
| chrX-56787337-INS-68 | chrX-56787293-INS-78 |
| chrX-56788358-INS-463 | chrX-56788358-INS-463 |
| chrX-56793191-INS-83 | chrX-56793189-INS-91 |
| chrX-56822823-INS-6550 | chrX-56822824-INS-6550 |
| chrX-56869927-INS-113 | chrX-56869927-INS-113 |
| chrX-56881958-INS-136 | chrX-56881958-INS-136 |
| chrX-56900571-INS-441 | chrX-56900571-INS-441 |
| chrX-56946878-INS-54 | chrX-56946878-INS-54 |
| chrX-56991825-INS-53 | chrX-56991825-INS-53 |
| chrX-56995441-INS-137 | chrX-56995441-INS-137 |
| chrX-57035945-INS-59 | chrX-57035942-INS-82 |
| chrX-57049750-INS-5533 | chrX-57049751-INS-5533 |
| chrX-57057008-INS-182 | chrX-57057009-INS-170 |
| chrX-57088895-INS-71 | chrX-57088888-INS-58 |
| chrX-57092954-INS-403 | chrX-57092954-INS-403 |
| chrX-57118024-INS-86 | chrX-57118024-INS-86 |
| chrX-57123225-INS-84 | chrX-57123225-INS-124 |
| chrX-57145393-INS-54 | chrX-57145394-INS-50 |
| chrX-57156824-INS-198 | chrX-57156824-INS-190 |
| chrX-57159304-INS-74 | chrX-57159289-INS-66 |
| chrX-57169527-INS-199 | chrX-57169527-INS-200 |
| chrX-57205694-INS-93 | chrX-57205688-INS-51 |
| chrX-57216249-INS-135 | chrX-57216249-INS-138 |
| chrX-57247051-INS-2398 | chrX-57247052-INS-2398 |
| chrX-57278153-INS-766 | chrX-57278153-INS-767 |
| chrX-57298070-INS-272 | chrX-57298070-INS-272 |
| chrX-57369208-INS-380 | chrX-57369208-INS-380 |
| chrX-57454315-INS-78 | chrX-57454315-INS-62 |
| chrX-57470851-INS-66 | chrX-57470821-INS-84 |
| chrX-57493746-INS-174 | chrX-57493746-INS-174 |
| chrX-57551684-INS-106 | chrX-57551684-INS-107 |
| chrX-57587261-INS-6888 | chrX-57587262-INS-6888 |
| chrX-57610445-INS-7138 | chrX-57610446-INS-7138 |
| chrX-57618660-INS-3015 | chrX-57618661-INS-3015 |
| chrX-57659035-INS-210 | chrX-57659035-INS-210 |
| chrX-57668855-INS-128 | chrX-57668855-INS-128 |
| chrX-57693240-INS-64 | chrX-57693240-INS-60 |
| chrX-57694701-INS-6846 | chrX-57694702-INS-6846 |
| chrX-57776206-INS-6369 | chrX-57776207-INS-6369 |
| chrX-57812978-INS-6212 | chrX-57812979-INS-6212 |
| chrX-57831813-INS-214 | chrX-57831813-INS-214 |
| chrX-57837295-INS-75 | chrX-57837295-INS-75 |
| chrX-57863624-INS-72 | chrX-57863624-INS-72 |
| chrX-57881324-INS-84 | chrX-57881324-INS-84 |
| chrX-57905848-INS-122 | chrX-57905848-INS-139 |
| chrX-57940214-INS-1455 | chrX-57940215-INS-1455 |
| chrX-57969921-INS-270 | chrX-57969921-INS-270 |
| chrX-57972808-INS-7628 | chrX-57972809-INS-7630 |
| chrX-57975904-INS-1128 | chrX-57975905-INS-1128 |
| chrX-57977125-INS-277 | chrX-57977125-INS-277 |
| chrX-57996930-INS-119 | chrX-57996930-INS-109 |
| chrX-58013002-INS-6598 | chrX-58013003-INS-6599 |
| chrX-58023769-INS-1690 | chrX-58023770-INS-1690 |

| | |
|---|---|
| chrX-58112371-INS-549 | chrX-58112371-INS-544 |
| chrX-58156919-INS-113 | chrX-58156919-INS-113 |
| chrX-58196545-INS-221 | chrX-58196545-INS-221 |
| chrX-58240494-INS-503 | chrX-58240494-INS-506 |
| chrX-58260670-INS-96 | chrX-58260670-INS-96 |
| chrX-58271141-INS-6544 | chrX-58271142-INS-6545 |
| chrX-58305053-INS-1894 | chrX-58305054-INS-1900 |
| chrX-58429278-INS-6135 | chrX-58429279-INS-6151 |
| chrX-58453363-INS-117 | chrX-58453363-INS-117 |
| chrX-58466490-INS-1323 | chrX-58466491-INS-1323 |
| chrX-58499384-INS-221 | chrX-58499384-INS-220 |
| chrX-58534169-INS-51 | chrX-58534169-INS-52 |
| chrX-58535204-INS-53 | chrX-58535198-INS-56 |
| chrX-58538173-INS-123 | chrX-58538173-INS-120 |
| chrX-58550190-INS-157 | chrX-58550193-INS-175 |
| chrX-58586280-INS-344 | chrX-58586280-INS-344 |
| chrX-58618272-INS-203 | chrX-58618272-INS-203 |
| chrX-58642919-INS-1183 | chrX-58642938-INS-1184 |
| chrX-58654676-INS-1085 | chrX-58654677-INS-1081 |
| chrX-58655224-INS-1659 | chrX-58655225-INS-1659 |
| chrX-58736952-INS-60 | chrX-58736952-INS-60 |
| chrX-58739994-INS-294 | chrX-58739994-INS-294 |
| chrX-58766124-INS-67 | chrX-58766124-INS-67 |
| chrX-58766761-INS-351 | chrX-58766761-INS-357 |
| chrX-58774124-INS-204 | chrX-58774124-INS-204 |
| chrX-58776619-INS-6180 | chrX-58776620-INS-6180 |
| chrX-58807040-INS-55 | chrX-58807040-INS-65 |
| chrX-58830061-INS-457 | chrX-58830061-INS-457 |
| chrX-58833611-INS-212 | chrX-58833611-INS-212 |
| chrX-58887252-INS-2309 | chrX-58887253-INS-2309 |
| chrX-58891981-INS-600 | chrX-58891981-INS-600 |
| chrX-58895013-INS-214 | chrX-58895013-INS-214 |
| chrX-58911495-INS-1062 | chrX-58911496-INS-1063 |
| chrX-58921846-INS-111 | chrX-58921846-INS-111 |
| chrX-58937758-INS-68 | chrX-58937758-INS-69 |
| chrX-58963431-INS-247 | chrX-58963431-INS-247 |
| chrX-59031761-INS-116 | chrX-59031761-INS-116 |
| chrX-59057023-INS-168 | chrX-59057023-INS-168 |
| chrX-59066882-INS-2458 | chrX-59066883-INS-2458 |
| chrX-59092546-INS-202 | chrX-59092546-INS-202 |
| chrX-59102961-INS-317 | chrX-59102961-INS-329 |
| chrX-59112376-INS-204 | chrX-59112369-INS-205 |
| chrX-59146562-INS-134 | chrX-59146562-INS-134 |
| chrX-59160743-INS-467 | chrX-59160743-INS-467 |
| chrX-59162675-INS-201 | chrX-59162675-INS-202 |
| chrX-59200832-INS-201 | chrX-59200832-INS-199 |
| chrX-59203847-INS-61 | chrX-59203848-INS-60 |
| chrX-59226741-INS-66 | chrX-59226741-INS-58 |
| chrX-59297547-INS-160 | chrX-59297547-INS-160 |
| chrX-59312585-INS-158 | chrX-59312585-INS-158 |
| chrX-59351553-INS-232 | chrX-59351552-INS-224 |
| chrX-59353699-INS-96 | chrX-59353703-INS-112 |
| chrX-59355129-INS-163 | chrX-59355129-INS-163 |
| chrX-59356132-INS-7088 | chrX-59356133-INS-7086 |
| chrX-59417383-INS-2889 | chrX-59417384-INS-2899 |
| chrX-59423146-INS-806 | chrX-59423145-INS-809 |
| chrX-59492756-INS-52 | chrX-59492756-INS-52 |
| chrX-59504205-INS-101 | chrX-59504205-INS-101 |
| chrX-59529238-INS-63 | chrX-59529238-INS-65 |
| chrX-59545502-INS-76 | chrX-59545540-INS-54 |
| chrX-59548303-INS-60 | chrX-59548303-INS-54 |
| chrX-59576372-INS-644 | chrX-59576372-INS-646 |
| chrX-59649244-INS-58 | chrX-59649244-INS-58 |
| chrX-59657198-INS-114 | chrX-59657198-INS-114 |
| chrX-59681655-INS-2249 | chrX-59681656-INS-2249 |
| chrX-59716385-INS-72 | chrX-59716370-INS-86 |
| chrX-59756797-INS-1647 | chrX-59756798-INS-1647 |
| chrX-59776349-INS-476 | chrX-59776350-INS-476 |
| chrX-59815250-INS-176 | chrX-59815250-INS-224 |
| chrX-59835004-INS-327 | chrX-59835004-INS-327 |
| chrX-59837626-INS-51 | chrX-59837626-INS-51 |
| chrX-59900154-INS-101 | chrX-59900154-INS-99 |
| chrX-59906661-INS-1016 | chrX-59906662-INS-1016 |
| chrX-59950193-INS-52 | chrX-59950193-INS-51 |
| chrX-60064078-INS-78 | chrX-60064078-INS-78 |
| chrX-60066424-INS-1203 | chrX-60066425-INS-1216 |
| chrX-60095310-INS-102 | chrX-60095319-INS-140 |

| | |
|---|---|
| chrX-60113670-INS-354 | chrX-60113670-INS-354 |
| chrX-60115552-INS-172 | chrX-60115552-INS-171 |
| chrX-60131065-INS-7646 | chrX-60131066-INS-7652 |
| chrX-60211963-INS-163 | chrX-60211963-INS-164 |
| chrX-60233441-INS-170 | chrX-60233473-INS-175 |
| chrX-60264876-INS-62 | chrX-60264876-INS-62 |
| chrX-60271521-INS-418 | chrX-60271521-INS-418 |
| chrX-60275310-INS-378 | chrX-60275310-INS-379 |
| chrX-60285126-INS-958 | chrX-60285126-INS-961 |
| chrX-60285706-INS-100 | chrX-60285705-INS-96 |
| chrX-60327977-INS-84 | chrX-60327977-INS-72 |
| chrX-60346083-INS-101 | chrX-60346087-INS-74 |
| chrX-60367696-INS-209 | chrX-60367696-INS-211 |
| chrX-60408092-INS-114 | chrX-60408085-INS-74 |
| chrX-60480185-INS-124 | chrX-60480184-INS-128 |
| chrX-60549684-INS-240 | chrX-60549684-INS-241 |
| chrX-60552695-INS-58 | chrX-60552695-INS-58 |
| chrX-60560298-INS-527 | chrX-60560298-INS-512 |
| chrX-60564625-INS-6608 | chrX-60564626-INS-6615 |
| chrX-60582636-INS-124 | chrX-60582637-INS-124 |
| chrX-60632811-INS-123 | chrX-60632857-INS-74 |
| chrX-60641093-INS-2800 | chrX-60641094-INS-2770 |
| chrX-60647286-INS-52 | chrX-60647286-INS-52 |
| chrX-60665687-INS-1297 | chrX-60665688-INS-1297 |
| chrX-60695746-INS-60 | chrX-60695746-INS-66 |
| chrX-60724809-INS-250 | chrX-60724809-INS-250 |
| chrX-60750082-INS-324 | chrX-60750082-INS-316 |
| chrX-60823240-INS-1611 | chrX-60823241-INS-1601 |
| chrX-60850990-INS-1490 | chrX-60850991-INS-1491 |
| chrX-60904573-INS-3812 | chrX-60904574-INS-3812 |
| chrX-60921395-INS-93 | chrX-60921395-INS-54 |
| chrX-60939685-INS-52 | chrX-60939685-INS-50 |
| chrX-60955552-INS-3032 | chrX-60955553-INS-3028 |
| chrX-60965270-INS-56 | chrX-60965270-INS-56 |
| chrX-60977888-INS-110 | chrX-60977888-INS-110 |
| chrX-60980004-INS-205 | chrX-60979963-INS-205 |
| chrX-60995128-INS-342 | chrX-60995128-INS-342 |
| chrX-61039839-INS-6212 | chrX-61039840-INS-6227 |
| chrX-61072360-INS-153 | chrX-61072360-INS-153 |
| chrX-61075026-INS-651 | chrX-61075025-INS-635 |
| chrX-61131199-INS-740 | chrX-61131199-INS-740 |
| chrX-61150550-INS-61 | chrX-61150550-INS-61 |
| chrX-61172963-INS-334 | chrX-61172969-INS-321 |
| chrX-61215426-INS-410 | chrX-61215426-INS-400 |
| chrX-61229006-INS-211 | chrX-61229006-INS-211 |
| chrX-61241612-INS-203 | chrX-61241612-INS-203 |
| chrX-61301492-INS-55 | chrX-61301492-INS-85 |
| chrX-61324112-INS-60 | chrX-61324112-INS-62 |
| chrX-61373010-INS-220 | chrX-61373010-INS-221 |
| chrX-61399810-INS-392 | chrX-61399801-INS-407 |
| chrX-61459820-INS-112 | chrX-61459820-INS-116 |
| chrX-61474700-INS-60 | chrX-61474700-INS-58 |
| chrX-61517970-INS-522 | chrX-61517970-INS-522 |
| chrX-61563364-INS-56 | chrX-61563393-INS-56 |
| chrX-61619210-INS-1067 | chrX-61619211-INS-1067 |
| chrX-61641578-INS-615 | chrX-61641578-INS-615 |
| chrX-61659219-INS-566 | chrX-61659219-INS-566 |
| chrX-61715967-INS-52 | chrX-61715967-INS-52 |
| chrX-61760218-INS-51 | chrX-61760217-INS-64 |
| chrX-61788927-INS-72 | chrX-61788927-INS-60 |
| chrX-61840205-INS-68 | chrX-61840205-INS-63 |
| chrX-61890858-INS-231 | chrX-61890858-INS-208 |
| chrX-61899965-INS-56 | chrX-61899961-INS-60 |
| chrX-61904411-INS-282 | chrX-61904409-INS-280 |
| chrX-62030421-INS-90 | chrX-62030421-INS-86 |
| chrX-62039797-INS-60 | chrX-62039784-INS-70 |
| chrX-62092079-INS-56 | chrX-62092123-INS-56 |
| chrX-62093155-INS-970 | chrX-62093155-INS-1002 |
| chrX-62094606-INS-151 | chrX-62094606-INS-151 |
| chrX-62100047-INS-281 | chrX-62100047-INS-281 |
| chrX-62104117-INS-1166 | chrX-62104102-INS-1145 |
| chrX-62124150-INS-6332 | chrX-62124151-INS-6332 |
| chrX-62152393-INS-618 | chrX-62152393-INS-618 |
| chrX-62171227-INS-69 | chrX-62171227-INS-50 |
| chrX-62193966-INS-70 | chrX-62193966-INS-88 |
| chrX-62208830-INS-7117 | chrX-62208830-INS-7097 |
| chrX-62236898-INS-6967 | chrX-62236899-INS-6969 |

| | |
|---|---|
| chrX-62247231-INS-70 | chrX-62247231-INS-52 |
| chrX-62254547-INS-1862 | chrX-62254548-INS-1862 |
| chrX-62260929-INS-56 | chrX-62260929-INS-56 |
| chrX-62267395-INS-1018 | chrX-62267396-INS-1018 |
| chrX-62275117-INS-58 | chrX-62275117-INS-58 |
| chrX-62286401-INS-6446 | chrX-62286402-INS-6446 |
| chrX-62306552-INS-7177 | chrX-62306553-INS-7178 |
| chrX-62334732-INS-859 | chrX-62334732-INS-860 |
| chrX-62336954-INS-52 | chrX-62336970-INS-50 |
| chrX-62341749-INS-53 | chrX-62341749-INS-53 |
| chrX-62365795-INS-68 | chrX-62365795-INS-64 |
| chrX-62366556-INS-8429 | chrX-62366557-INS-8431 |
| chrX-62427364-INS-155 | chrX-62427364-INS-155 |
| chrX-62436856-INS-3395 | chrX-62436857-INS-3395 |
| chrX-62461982-INS-203 | chrX-62461982-INS-203 |
| chrX-62467434-INS-668 | chrX-62467434-INS-668 |
| chrX-62475239-INS-6181 | chrX-62475240-INS-6181 |
| chrX-62497671-INS-196 | chrX-62497671-INS-193 |
| chrX-62575079-INS-70 | chrX-62575079-INS-75 |
| chrX-62576520-INS-2557 | chrX-62576521-INS-2561 |
| chrX-62607374-INS-1722 | chrX-62607375-INS-1722 |
| chrX-62610867-INS-2408 | chrX-62610868-INS-2406 |
| chrX-62699614-INS-533 | chrX-62699614-INS-533 |
| chrX-62731206-INS-5852 | chrX-62731207-INS-5852 |
| chrX-62773762-INS-54 | chrX-62773762-INS-69 |
| chrX-62790328-INS-58 | chrX-62790328-INS-58 |
| chrX-62795626-INS-492 | chrX-62795626-INS-493 |
| chrX-62807835-INS-83 | chrX-62807835-INS-60 |
| chrX-62856241-INS-498 | chrX-62856241-INS-498 |
| chrX-62911839-INS-101 | chrX-62911841-INS-56 |
| chrX-63031151-INS-2981 | chrX-63031152-INS-2981 |
| chrX-63088598-INS-2350 | chrX-63088599-INS-2350 |
| chrX-63107625-INS-341 | chrX-63107625-INS-341 |
| chrX-63159080-INS-1130 | chrX-63159081-INS-1128 |
| chrX-63185293-INS-2445 | chrX-63185294-INS-2445 |
| chrX-63202386-INS-56 | chrX-63202365-INS-84 |
| chrX-63293512-INS-73 | chrX-63293512-INS-75 |
| chrX-63337688-INS-410 | chrX-63337688-INS-411 |
| chrX-63395841-INS-6330 | chrX-63395842-INS-6327 |
| chrX-63413482-INS-54 | chrX-63413482-INS-54 |
| chrX-63447429-INS-3390 | chrX-63447430-INS-3392 |
| chrX-63486495-INS-1475 | chrX-63486496-INS-1475 |
| chrX-63491261-INS-68 | chrX-63491261-INS-68 |
| chrX-63500641-INS-3684 | chrX-63500642-INS-3684 |
| chrX-63514792-INS-57 | chrX-63514837-INS-60 |
| chrX-63583004-INS-214 | chrX-63583004-INS-214 |
| chrX-63651612-INS-2109 | chrX-63651613-INS-2113 |
| chrX-63722703-INS-400 | chrX-63722703-INS-400 |
| chrX-63741911-INS-62 | chrX-63741911-INS-60 |
| chrX-63761567-INS-2810 | chrX-63761568-INS-2810 |
| chrX-63765875-INS-6668 | chrX-63765876-INS-6668 |
| chrX-63801122-INS-55 | chrX-63801122-INS-55 |
| chrX-63821275-INS-201 | chrX-63821275-INS-206 |
| chrX-63854465-INS-2018 | chrX-63854466-INS-2018 |
| chrX-63906102-INS-6419 | chrX-63906103-INS-6421 |
| chrX-63938673-INS-1752 | chrX-63938674-INS-1754 |
| chrX-63944662-INS-463 | chrX-63944662-INS-463 |
| chrX-63954720-INS-5553 | chrX-63954721-INS-5553 |
| chrX-63961582-INS-4141 | chrX-63961583-INS-4115 |
| chrX-63998843-INS-743 | chrX-63998843-INS-743 |
| chrX-64024407-INS-526 | chrX-64024390-INS-557 |
| chrX-64103421-INS-6907 | chrX-64103422-INS-6907 |
| chrX-64113926-INS-834 | chrX-64113927-INS-833 |
| chrX-64122300-INS-57 | chrX-64122300-INS-57 |
| chrX-64164205-INS-76 | chrX-64164203-INS-76 |
| chrX-64179494-INS-7302 | chrX-64179495-INS-7302 |
| chrX-64208547-INS-211 | chrX-64208547-INS-212 |
| chrX-64213741-INS-293 | chrX-64213741-INS-294 |
| chrX-64221945-INS-67 | chrX-64221916-INS-58 |
| chrX-64258333-INS-514 | chrX-64258333-INS-514 |
| chrX-64293149-INS-54 | chrX-64293150-INS-81 |
| chrX-64301243-INS-569 | chrX-64301243-INS-569 |
| chrX-64322154-INS-1337 | chrX-64322155-INS-1338 |
| chrX-64343558-INS-6606 | chrX-64343559-INS-6607 |
| chrX-64365502-INS-973 | chrX-64365502-INS-973 |
| chrX-64385686-INS-1501 | chrX-64385687-INS-1501 |
| chrX-64404526-INS-4928 | chrX-64404527-INS-4928 |

| | |
|---|---|
| chrX-64412240-INS-201 | chrX-64412240-INS-201 |
| chrX-64447739-INS-486 | chrX-64447739-INS-487 |
| chrX-64466295-INS-108 | chrX-64466295-INS-72 |
| chrX-64481363-INS-432 | chrX-64481363-INS-432 |
| chrX-64482392-INS-4107 | chrX-64482393-INS-4106 |
| chrX-64483675-INS-3639 | chrX-64483676-INS-3639 |
| chrX-64505955-INS-74 | chrX-64505955-INS-62 |
| chrX-64522373-INS-51 | chrX-64522374-INS-54 |
| chrX-64526124-INS-756 | chrX-64526124-INS-753 |
| chrX-64560390-INS-203 | chrX-64560390-INS-203 |
| chrX-64562498-INS-60 | chrX-64562498-INS-60 |
| chrX-64568611-INS-56 | chrX-64568611-INS-60 |
| chrX-64606203-INS-2472 | chrX-64606204-INS-2472 |
| chrX-64625870-INS-2746 | chrX-64625871-INS-2747 |
| chrX-64626744-INS-53 | chrX-64626744-INS-55 |
| chrX-64650819-INS-78 | chrX-64650844-INS-58 |
| chrX-64653446-INS-278 | chrX-64653445-INS-279 |
| chrX-64666829-INS-3116 | chrX-64666830-INS-3116 |
| chrX-64677735-INS-1083 | chrX-64677736-INS-1085 |
| chrX-64745264-INS-79 | chrX-64745264-INS-75 |
| chrX-64745734-INS-614 | chrX-64745734-INS-614 |
| chrX-64772079-INS-331 | chrX-64772079-INS-331 |
| chrX-64818014-INS-363 | chrX-64818014-INS-363 |
| chrX-64902569-INS-224 | chrX-64902569-INS-222 |
| chrX-64919684-INS-607 | chrX-64919684-INS-600 |
| chrX-64941596-INS-52 | chrX-64941596-INS-52 |
| chrX-64949425-INS-55 | chrX-64949430-INS-94 |
| chrX-64953312-INS-210 | chrX-64953303-INS-204 |
| chrX-64954918-INS-1493 | chrX-64954928-INS-1495 |
| chrX-65004870-INS-782 | chrX-65004870-INS-783 |
| chrX-65031212-INS-214 | chrX-65031212-INS-214 |
| chrX-65068415-INS-143 | chrX-65068415-INS-109 |
| chrX-65097926-INS-527 | chrX-65097926-INS-527 |
| chrX-65109968-INS-837 | chrX-65109968-INS-837 |
| chrX-65142844-INS-60 | chrX-65142844-INS-70 |
| chrX-65165451-INS-133 | chrX-65165451-INS-140 |
| chrX-65192911-INS-2701 | chrX-65192912-INS-2701 |
| chrX-65288733-INS-78 | chrX-65288733-INS-78 |
| chrX-65334315-INS-358 | chrX-65334315-INS-358 |
| chrX-65349721-INS-973 | chrX-65349713-INS-972 |
| chrX-65363183-INS-839 | chrX-65363183-INS-839 |
| chrX-65385002-INS-4166 | chrX-65385003-INS-4166 |
| chrX-65438577-INS-3906 | chrX-65438578-INS-3906 |
| chrX-65467358-INS-2760 | chrX-65467359-INS-2760 |
| chrX-65472222-INS-388 | chrX-65472222-INS-386 |
| chrX-65490678-INS-740 | chrX-65490678-INS-718 |
| chrX-65491459-INS-60 | chrX-65491460-INS-52 |
| chrX-65493548-INS-71 | chrX-65493548-INS-55 |
| chrX-65524559-INS-186 | chrX-65524559-INS-180 |
| chrX-65525498-INS-6687 | chrX-65525499-INS-6688 |
| chrX-65529002-INS-54 | chrX-65529000-INS-50 |
| chrX-65566142-INS-56 | chrX-65566142-INS-55 |
| chrX-65603881-INS-55 | chrX-65603881-INS-55 |
| chrX-65621354-INS-66 | chrX-65621354-INS-66 |
| chrX-65661569-INS-154 | chrX-65661569-INS-154 |
| chrX-65685547-INS-210 | chrX-65685547-INS-209 |
| chrX-65740009-INS-164 | chrX-65740009-INS-164 |
| chrX-65808636-INS-497 | chrX-65808636-INS-497 |
| chrX-65834938-INS-53 | chrX-65834938-INS-53 |
| chrX-65865948-INS-446 | chrX-65865948-INS-446 |
| chrX-65886154-INS-308 | chrX-65886154-INS-308 |
| chrX-65986388-INS-51 | chrX-65986389-INS-88 |
| chrX-66009835-INS-6442 | chrX-66009836-INS-6442 |
| chrX-66026806-INS-183 | chrX-66026806-INS-192 |
| chrX-66103543-INS-59 | chrX-66103543-INS-54 |
| chrX-66105548-INS-74 | chrX-66105548-INS-72 |
| chrX-66157634-INS-180 | chrX-66157634-INS-181 |
| chrX-66185855-INS-53 | chrX-66185855-INS-79 |
| chrX-66198472-INS-199 | chrX-66198472-INS-199 |
| chrX-66203620-INS-1285 | chrX-66203621-INS-1285 |
| chrX-66207762-INS-6390 | chrX-66207763-INS-6390 |
| chrX-66235113-INS-1474 | chrX-66235114-INS-1474 |
| chrX-66245164-INS-3401 | chrX-66245165-INS-3401 |
| chrX-66250887-INS-106 | chrX-66250887-INS-106 |
| chrX-66252713-INS-202 | chrX-66252713-INS-203 |
| chrX-66323127-INS-76 | chrX-66323116-INS-60 |
| chrX-66338735-INS-61 | chrX-66338735-INS-61 |

| | |
|---|---|
| chrX-66341199-INS-66 | chrX-66341200-INS-72 |
| chrX-66361286-INS-59 | chrX-66361286-INS-79 |
| chrX-66361404-INS-4421 | chrX-66361405-INS-4424 |
| chrX-66376100-INS-398 | chrX-66376100-INS-399 |
| chrX-66386163-INS-7010 | chrX-66386164-INS-7010 |
| chrX-66403167-INS-468 | chrX-66403167-INS-468 |
| chrX-66409138-INS-63 | chrX-66409138-INS-63 |
| chrX-66410596-INS-4598 | chrX-66410597-INS-4597 |
| chrX-66445247-INS-370 | chrX-66445247-INS-370 |
| chrX-66448510-INS-389 | chrX-66448510-INS-389 |
| chrX-66449336-INS-140 | chrX-66449336-INS-140 |
| chrX-66464950-INS-6442 | chrX-66464951-INS-6441 |
| chrX-66487985-INS-75 | chrX-66487985-INS-75 |
| chrX-66506952-INS-72 | chrX-66506930-INS-76 |
| chrX-66551952-INS-58 | chrX-66551952-INS-58 |
| chrX-66573229-INS-68 | chrX-66573229-INS-72 |
| chrX-66575004-INS-68 | chrX-66575004-INS-68 |
| chrX-66696034-INS-183 | chrX-66696034-INS-183 |
| chrX-66696742-INS-1495 | chrX-66696743-INS-1495 |
| chrX-66724306-INS-68 | chrX-66724306-INS-66 |
| chrX-66734874-INS-64 | chrX-66734874-INS-69 |
| chrX-66741541-INS-54 | chrX-66741567-INS-69 |
| chrX-66896900-INS-1612 | chrX-66896901-INS-1612 |
| chrX-66935531-INS-113 | chrX-66935533-INS-111 |
| chrX-66943470-INS-322 | chrX-66943470-INS-322 |
| chrX-66988564-INS-390 | chrX-66988564-INS-390 |
| chrX-67022411-INS-732 | chrX-67022411-INS-732 |
| chrX-67036852-INS-7157 | chrX-67036853-INS-7158 |
| chrX-67056575-INS-84 | chrX-67056575-INS-88 |
| chrX-67065354-INS-104 | chrX-67065354-INS-104 |
| chrX-67085892-INS-76 | chrX-67085892-INS-118 |
| chrX-67232540-INS-585 | chrX-67232540-INS-596 |
| chrX-67256787-INS-320 | chrX-67256787-INS-322 |
| chrX-67310765-INS-459 | chrX-67310765-INS-466 |
| chrX-67323309-INS-97 | chrX-67323296-INS-136 |
| chrX-67352190-INS-827 | chrX-67352190-INS-827 |
| chrX-67364864-INS-56 | chrX-67364864-INS-56 |
| chrX-67418001-INS-142 | chrX-67418001-INS-142 |
| chrX-67432933-INS-52 | chrX-67432933-INS-52 |
| chrX-67528791-INS-91 | chrX-67528791-INS-99 |
| chrX-67556628-INS-58 | chrX-67556628-INS-58 |
| chrX-67579045-INS-278 | chrX-67579055-INS-275 |
| chrX-67593375-INS-193 | chrX-67593375-INS-193 |
| chrX-67617103-INS-57 | chrX-67617099-INS-52 |
| chrX-67632569-INS-396 | chrX-67632569-INS-396 |
| chrX-67638185-INS-474 | chrX-67638185-INS-474 |
| chrX-67645414-INS-523 | chrX-67645414-INS-522 |
| chrX-67661465-INS-166 | chrX-67661465-INS-166 |
| chrX-67735689-INS-76 | chrX-67735689-INS-76 |
| chrX-67781569-INS-2242 | chrX-67781570-INS-2263 |
| chrX-67790373-INS-103 | chrX-67790373-INS-102 |
| chrX-67800329-INS-3202 | chrX-67800330-INS-3202 |
| chrX-67870193-INS-129 | chrX-67870193-INS-129 |
| chrX-67876335-INS-6862 | chrX-67876336-INS-6863 |
| chrX-67891162-INS-74 | chrX-67891162-INS-74 |
| chrX-67925047-INS-74 | chrX-67925047-INS-74 |
| chrX-67931589-INS-689 | chrX-67931589-INS-689 |
| chrX-67950954-INS-945 | chrX-67950954-INS-957 |
| chrX-67967066-INS-4469 | chrX-67967067-INS-4469 |
| chrX-67973135-INS-480 | chrX-67973135-INS-480 |
| chrX-67981662-INS-496 | chrX-67981662-INS-495 |
| chrX-67996096-INS-503 | chrX-67996096-INS-505 |
| chrX-68031143-INS-6274 | chrX-68031144-INS-6274 |
| chrX-68038710-INS-169 | chrX-68038710-INS-169 |
| chrX-68071430-INS-217 | chrX-68071430-INS-217 |
| chrX-68080612-INS-6467 | chrX-68080613-INS-6467 |
| chrX-68095393-INS-2176 | chrX-68095394-INS-2176 |
| chrX-68112083-INS-2281 | chrX-68112084-INS-2281 |
| chrX-68132643-INS-435 | chrX-68132643-INS-435 |
| chrX-68139499-INS-78 | chrX-68139499-INS-78 |
| chrX-68158810-INS-9908 | chrX-68158811-INS-9908 |
| chrX-68166371-INS-78 | chrX-68166371-INS-78 |
| chrX-68201208-INS-6865 | chrX-68201209-INS-6866 |
| chrX-68280097-INS-68 | chrX-68280097-INS-66 |
| chrX-68288862-INS-1966 | chrX-68288863-INS-1966 |
| chrX-68300514-INS-409 | chrX-68300514-INS-409 |
| chrX-68333901-INS-275 | chrX-68333901-INS-275 |

| | |
|---|---|
| chrX-68337621-INS-557 | chrX-68337621-INS-557 |
| chrX-68374591-INS-469 | chrX-68374591-INS-469 |
| chrX-68381816-INS-124 | chrX-68381816-INS-120 |
| chrX-68382846-INS-251 | chrX-68382846-INS-251 |
| chrX-68389448-INS-93 | chrX-68389421-INS-96 |
| chrX-68563697-INS-64 | chrX-68563697-INS-64 |
| chrX-68632551-INS-1176 | chrX-68632552-INS-1167 |
| chrX-68635389-INS-112 | chrX-68635389-INS-112 |
| chrX-68682835-INS-207 | chrX-68682835-INS-210 |
| chrX-68711695-INS-5249 | chrX-68711696-INS-5249 |
| chrX-68789392-INS-78 | chrX-68789392-INS-66 |
| chrX-68839990-INS-1759 | chrX-68839991-INS-1760 |
| chrX-68954379-INS-1857 | chrX-68954380-INS-1857 |
| chrX-68958936-INS-209 | chrX-68958936-INS-209 |
| chrX-68971816-INS-1896 | chrX-68971817-INS-1896 |
| chrX-68973572-INS-210 | chrX-68973573-INS-210 |
| chrX-68984433-INS-795 | chrX-68984434-INS-794 |
| chrX-69012704-INS-73 | chrX-69012707-INS-77 |
| chrX-69082886-INS-469 | chrX-69082886-INS-469 |
| chrX-69095566-INS-219 | chrX-69095566-INS-219 |
| chrX-69096482-INS-911 | chrX-69096482-INS-912 |
| chrX-69099092-INS-212 | chrX-69099092-INS-212 |
| chrX-69157758-INS-9042 | chrX-69157759-INS-9047 |
| chrX-69170620-INS-489 | chrX-69170620-INS-489 |
| chrX-69179621-INS-89 | chrX-69179608-INS-88 |
| chrX-69195628-INS-455 | chrX-69195628-INS-455 |
| chrX-69197963-INS-146 | chrX-69197964-INS-146 |
| chrX-69201676-INS-6578 | chrX-69201677-INS-6577 |
| chrX-69221941-INS-1371 | chrX-69221942-INS-1371 |
| chrX-69257295-INS-160 | chrX-69257295-INS-178 |
| chrX-69269576-INS-1419 | chrX-69269577-INS-1407 |
| chrX-69270991-INS-527 | chrX-69270992-INS-527 |
| chrX-69323261-INS-153 | chrX-69323261-INS-153 |
| chrX-69391377-INS-197 | chrX-69391377-INS-197 |
| chrX-69420664-INS-421 | chrX-69420658-INS-421 |
| chrX-69423845-INS-486 | chrX-69423843-INS-487 |
| chrX-69490000-INS-653 | chrX-69490000-INS-652 |
| chrX-69511758-INS-181 | chrX-69511758-INS-181 |
| chrX-69516569-INS-402 | chrX-69516569-INS-401 |
| chrX-69565825-INS-124 | chrX-69565825-INS-124 |
| chrX-69568736-INS-6228 | chrX-69568737-INS-6224 |
| chrX-69577902-INS-91 | chrX-69577902-INS-91 |
| chrX-69589687-INS-60 | chrX-69589687-INS-60 |
| chrX-69597551-INS-5549 | chrX-69597552-INS-5549 |
| chrX-69601147-INS-1575 | chrX-69601148-INS-1575 |
| chrX-69631573-INS-2800 | chrX-69631574-INS-2800 |
| chrX-69636497-INS-6932 | chrX-69636498-INS-6934 |
| chrX-69645505-INS-6399 | chrX-69645506-INS-6402 |
| chrX-69652890-INS-58 | chrX-69652890-INS-60 |
| chrX-69666161-INS-56 | chrX-69666159-INS-52 |
| chrX-69688763-INS-2610 | chrX-69688764-INS-2594 |
| chrX-69717077-INS-1303 | chrX-69717078-INS-1304 |
| chrX-69729844-INS-54 | chrX-69729855-INS-67 |
| chrX-69778362-INS-427 | chrX-69778362-INS-427 |
| chrX-69811573-INS-56 | chrX-69811573-INS-56 |
| chrX-69819838-INS-1636 | chrX-69819839-INS-1636 |
| chrX-69819847-INS-3409 | chrX-69819853-INS-3403 |
| chrX-69831314-INS-6487 | chrX-69831315-INS-6488 |
| chrX-69857796-INS-56 | chrX-69857796-INS-52 |
| chrX-69871367-INS-219 | chrX-69871367-INS-218 |
| chrX-69872046-INS-53 | chrX-69872046-INS-53 |
| chrX-69910416-INS-852 | chrX-69910416-INS-852 |
| chrX-69924057-INS-59 | chrX-69924057-INS-67 |
| chrX-69948049-INS-211 | chrX-69948049-INS-211 |
| chrX-70013631-INS-293 | chrX-70013631-INS-278 |
| chrX-70015805-INS-1076 | chrX-70015806-INS-1080 |
| chrX-70030579-INS-7180 | chrX-70030580-INS-7180 |
| chrX-70053335-INS-58 | chrX-70053335-INS-60 |
| chrX-70110410-INS-534 | chrX-70110410-INS-534 |
| chrX-70148575-INS-53 | chrX-70148575-INS-53 |
| chrX-70201040-INS-60 | chrX-70201040-INS-60 |
| chrX-70216773-INS-63 | chrX-70216773-INS-60 |
| chrX-70333844-INS-3320 | chrX-70333845-INS-3320 |
| chrX-70335126-INS-5544 | chrX-70335127-INS-5544 |
| chrX-70347287-INS-65 | chrX-70347287-INS-65 |
| chrX-70362520-INS-548 | chrX-70362520-INS-548 |
| chrX-70435033-INS-271 | chrX-70435033-INS-271 |

| | |
|---|---|
| chrX-70471510-INS-736 | chrX-70471510-INS-736 |
| chrX-70499475-INS-69 | chrX-70499475-INS-69 |
| chrX-70500246-INS-340 | chrX-70500246-INS-340 |
| chrX-70506644-INS-238 | chrX-70506644-INS-234 |
| chrX-70548319-INS-477 | chrX-70548319-INS-475 |
| chrX-70552728-INS-122 | chrX-70552733-INS-134 |
| chrX-70558140-INS-949 | chrX-70558140-INS-955 |
| chrX-70566143-INS-51 | chrX-70566142-INS-82 |
| chrX-70590241-INS-434 | chrX-70590241-INS-416 |
| chrX-70610567-INS-71 | chrX-70610567-INS-71 |
| chrX-70626791-INS-311 | chrX-70626791-INS-311 |
| chrX-70627714-INS-217 | chrX-70627714-INS-217 |
| chrX-70641296-INS-662 | chrX-70641296-INS-663 |
| chrX-70659266-INS-400 | chrX-70659266-INS-400 |
| chrX-70667546-INS-425 | chrX-70667546-INS-425 |
| chrX-70670459-INS-705 | chrX-70670459-INS-706 |
| chrX-70776752-INS-224 | chrX-70776752-INS-224 |
| chrX-70793089-INS-212 | chrX-70793089-INS-212 |
| chrX-70814728-INS-54 | chrX-70814728-INS-60 |
| chrX-70817957-INS-68 | chrX-70817957-INS-68 |
| chrX-70853692-INS-60 | chrX-70853693-INS-63 |
| chrX-70870225-INS-6804 | chrX-70870226-INS-6805 |
| chrX-70892958-INS-137 | chrX-70892958-INS-137 |
| chrX-70894095-INS-507 | chrX-70894095-INS-509 |
| chrX-70907222-INS-56 | chrX-70907211-INS-60 |
| chrX-70935373-INS-64 | chrX-70935361-INS-68 |
| chrX-70977785-INS-78 | chrX-70977798-INS-78 |
| chrX-71000678-INS-62 | chrX-71000699-INS-67 |
| chrX-71023099-INS-684 | chrX-71023092-INS-689 |
| chrX-71031178-INS-56 | chrX-71031178-INS-56 |
| chrX-71038298-INS-171 | chrX-71038298-INS-171 |
| chrX-71069346-INS-3208 | chrX-71069347-INS-3208 |
| chrX-71102417-INS-394 | chrX-71102417-INS-392 |
| chrX-71134721-INS-2054 | chrX-71134722-INS-2054 |
| chrX-71183588-INS-2949 | chrX-71183589-INS-2957 |
| chrX-71227302-INS-200 | chrX-71227302-INS-205 |
| chrX-71234640-INS-4395 | chrX-71234641-INS-4395 |
| chrX-71234641-INS-340 | chrX-71234641-INS-340 |
| chrX-71241889-INS-411 | chrX-71241889-INS-408 |
| chrX-71255920-INS-567 | chrX-71255920-INS-567 |
| chrX-71290397-INS-56 | chrX-71290356-INS-92 |
| chrX-71296516-INS-71 | chrX-71296516-INS-63 |
| chrX-71339691-INS-802 | chrX-71339691-INS-801 |
| chrX-71355231-INS-625 | chrX-71355232-INS-625 |
| chrX-71371103-INS-472 | chrX-71371103-INS-472 |
| chrX-71375422-INS-66 | chrX-71375422-INS-61 |
| chrX-71408763-INS-628 | chrX-71408763-INS-628 |
| chrX-71423055-INS-645 | chrX-71423055-INS-645 |
| chrX-71452146-INS-100 | chrX-71452146-INS-100 |
| chrX-71523439-INS-83 | chrX-71523439-INS-68 |
| chrX-71531664-INS-112 | chrX-71531664-INS-116 |
| chrX-71575534-INS-52 | chrX-71575535-INS-50 |
| chrX-71579031-INS-742 | chrX-71579031-INS-743 |
| chrX-71580379-INS-77 | chrX-71580396-INS-85 |
| chrX-71604800-INS-61 | chrX-71604832-INS-64 |
| chrX-71643009-INS-89 | chrX-71643009-INS-53 |
| chrX-71704332-INS-3333 | chrX-71704333-INS-3330 |
| chrX-71738015-INS-177 | chrX-71738015-INS-177 |
| chrX-71803839-INS-202 | chrX-71803839-INS-202 |
| chrX-71847928-INS-84 | chrX-71847928-INS-84 |
| chrX-71864198-INS-86 | chrX-71864198-INS-75 |
| chrX-71866554-INS-596 | chrX-71866554-INS-596 |
| chrX-71895167-INS-65 | chrX-71895126-INS-66 |
| chrX-71897162-INS-243 | chrX-71897162-INS-243 |
| chrX-71899609-INS-6495 | chrX-71899610-INS-6496 |
| chrX-71901605-INS-60 | chrX-71901605-INS-60 |
| chrX-71929901-INS-182 | chrX-71929901-INS-182 |
| chrX-71932602-INS-94 | chrX-71932602-INS-85 |
| chrX-71932860-INS-1009 | chrX-71932861-INS-1010 |
| chrX-71971247-INS-198 | chrX-71971247-INS-198 |
| chrX-72002095-INS-77 | chrX-72002095-INS-77 |
| chrX-72011992-INS-57 | chrX-72011999-INS-57 |
| chrX-72033859-INS-144 | chrX-72033859-INS-144 |
| chrX-72061292-INS-280 | chrX-72061292-INS-280 |
| chrX-72069144-INS-207 | chrX-72069144-INS-207 |
| chrX-72084091-INS-56 | chrX-72084091-INS-64 |
| chrX-72126679-INS-199 | chrX-72126679-INS-199 |

| | |
|---|---|
| chrX-72238373-INS-91 | chrX-72238389-INS-57 |
| chrX-72275949-INS-83 | chrX-72275949-INS-83 |
| chrX-72336637-INS-139 | chrX-72336637-INS-140 |
| chrX-72377692-INS-55 | chrX-72377692-INS-66 |
| chrX-72438546-INS-92 | chrX-72438542-INS-50 |
| chrX-72487399-INS-66 | chrX-72487399-INS-66 |
| chrX-72490236-INS-204 | chrX-72490236-INS-205 |
| chrX-72491222-INS-129 | chrX-72491222-INS-121 |
| chrX-72653255-INS-94 | chrX-72653255-INS-93 |
| chrX-72680415-INS-63 | chrX-72680415-INS-67 |
| chrX-72739180-INS-902 | chrX-72739180-INS-899 |
| chrX-72744993-INS-82 | chrX-72744993-INS-82 |
| chrX-72746172-INS-68 | chrX-72746172-INS-68 |
| chrX-72746642-INS-218 | chrX-72746642-INS-219 |
| chrX-72749507-INS-434 | chrX-72749507-INS-435 |
| chrX-72752101-INS-456 | chrX-72752101-INS-456 |
| chrX-72764964-INS-202 | chrX-72764964-INS-202 |
| chrX-72770101-INS-207 | chrX-72770101-INS-207 |
| chrX-72785778-INS-70 | chrX-72785779-INS-92 |
| chrX-72790919-INS-203 | chrX-72790920-INS-202 |
| chrX-72799480-INS-89 | chrX-72799480-INS-89 |
| chrX-72799542-INS-1888 | chrX-72799543-INS-1888 |
| chrX-72816851-INS-206 | chrX-72816851-INS-206 |
| chrX-72819003-INS-210 | chrX-72819003-INS-207 |
| chrX-72852169-INS-198 | chrX-72852169-INS-198 |
| chrX-72861763-INS-60 | chrX-72861763-INS-60 |
| chrX-72932368-INS-179 | chrX-72932368-INS-182 |
| chrX-72946931-INS-102 | chrX-72946931-INS-110 |
| chrX-72966394-INS-196 | chrX-72966394-INS-196 |
| chrX-73013726-INS-173 | chrX-73013726-INS-173 |
| chrX-73031071-INS-1012 | chrX-73031072-INS-1012 |
| chrX-73033502-INS-183 | chrX-73033502-INS-183 |
| chrX-73149502-INS-385 | chrX-73149502-INS-386 |
| chrX-73157533-INS-55 | chrX-73157533-INS-50 |
| chrX-73249841-INS-139 | chrX-73249841-INS-139 |
| chrX-73262064-INS-192 | chrX-73262064-INS-192 |
| chrX-73265347-INS-193 | chrX-73265347-INS-193 |
| chrX-73280275-INS-187 | chrX-73280275-INS-155 |
| chrX-73289135-INS-57 | chrX-73289135-INS-61 |
| chrX-73304158-INS-127 | chrX-73304166-INS-166 |
| chrX-73307168-INS-186 | chrX-73307168-INS-186 |
| chrX-73319403-INS-159 | chrX-73319403-INS-155 |
| chrX-73329651-INS-222 | chrX-73329651-INS-223 |
| chrX-73334234-INS-65 | chrX-73334234-INS-65 |
| chrX-73353132-INS-259 | chrX-73353132-INS-259 |
| chrX-73356004-INS-179 | chrX-73356004-INS-175 |
| chrX-73370302-INS-80 | chrX-73370302-INS-80 |
| chrX-73405023-INS-62 | chrX-73405023-INS-58 |
| chrX-73417088-INS-208 | chrX-73417088-INS-208 |
| chrX-73422226-INS-64 | chrX-73422220-INS-52 |
| chrX-73423255-INS-56 | chrX-73423211-INS-60 |
| chrX-73438756-INS-1051 | chrX-73438757-INS-1050 |
| chrX-73441697-INS-201 | chrX-73441697-INS-201 |
| chrX-73461698-INS-91 | chrX-73461698-INS-91 |
| chrX-73477213-INS-188 | chrX-73477213-INS-188 |
| chrX-73490611-INS-215 | chrX-73490611-INS-215 |
| chrX-73516017-INS-5150 | chrX-73516018-INS-5150 |
| chrX-73519975-INS-180 | chrX-73519975-INS-181 |
| chrX-73521243-INS-4795 | chrX-73521244-INS-4795 |
| chrX-73624979-INS-1893 | chrX-73624980-INS-1893 |
| chrX-73641279-INS-143 | chrX-73641265-INS-160 |
| chrX-73772100-INS-60 | chrX-73772100-INS-60 |
| chrX-73779775-INS-77 | chrX-73779775-INS-77 |
| chrX-73787403-INS-162 | chrX-73787403-INS-162 |
| chrX-73788455-INS-78 | chrX-73788455-INS-78 |
| chrX-73794720-INS-198 | chrX-73794720-INS-199 |
| chrX-73795739-INS-118 | chrX-73795739-INS-118 |
| chrX-73805796-INS-401 | chrX-73805796-INS-401 |
| chrX-73809676-INS-6475 | chrX-73809677-INS-6475 |
| chrX-73824268-INS-92 | chrX-73824269-INS-92 |
| chrX-73877841-INS-151 | chrX-73877841-INS-127 |
| chrX-73896054-INS-92 | chrX-73896054-INS-92 |
| chrX-73925982-INS-75 | chrX-73925982-INS-125 |
| chrX-73944018-INS-64 | chrX-73944018-INS-63 |
| chrX-74003011-INS-882 | chrX-74003011-INS-882 |
| chrX-74010443-INS-3125 | chrX-74010444-INS-3125 |
| chrX-74093161-INS-207 | chrX-74093164-INS-212 |

| | |
|---|---|
| chrX-74135530-INS-206 | chrX-74135530-INS-207 |
| chrX-74190326-INS-138 | chrX-74190326-INS-168 |
| chrX-74237123-INS-81 | chrX-74237123-INS-70 |
| chrX-74258582-INS-6466 | chrX-74258583-INS-6466 |
| chrX-74267307-INS-91 | chrX-74267307-INS-92 |
| chrX-74268121-INS-122 | chrX-74268121-INS-122 |
| chrX-74269973-INS-107 | chrX-74269973-INS-107 |
| chrX-74303392-INS-51 | chrX-74303392-INS-51 |
| chrX-74306942-INS-3607 | chrX-74306943-INS-3607 |
| chrX-74319545-INS-6364 | chrX-74319546-INS-6364 |
| chrX-74319569-INS-2252 | chrX-74319570-INS-2250 |
| chrX-74343308-INS-498 | chrX-74343308-INS-498 |
| chrX-74360900-INS-56 | chrX-74360900-INS-56 |
| chrX-74373664-INS-6224 | chrX-74373665-INS-6224 |
| chrX-74407587-INS-202 | chrX-74407587-INS-204 |
| chrX-74410617-INS-562 | chrX-74410617-INS-562 |
| chrX-74423514-INS-59 | chrX-74423514-INS-59 |
| chrX-74433124-INS-1037 | chrX-74433125-INS-1037 |
| chrX-74434085-INS-70 | chrX-74434085-INS-70 |
| chrX-74435600-INS-6643 | chrX-74435601-INS-6644 |
| chrX-74439103-INS-757 | chrX-74439103-INS-758 |
| chrX-74472776-INS-86 | chrX-74472776-INS-86 |
| chrX-74484178-INS-115 | chrX-74484178-INS-115 |
| chrX-74518134-INS-942 | chrX-74518134-INS-942 |
| chrX-74546013-INS-1374 | chrX-74546014-INS-1330 |
| chrX-74563091-INS-142 | chrX-74563091-INS-142 |
| chrX-74625634-INS-130 | chrX-74625650-INS-133 |
| chrX-74642200-INS-1566 | chrX-74642201-INS-1566 |
| chrX-74658422-INS-205 | chrX-74658422-INS-208 |
| chrX-74686958-INS-204 | chrX-74686958-INS-204 |
| chrX-74712106-INS-174 | chrX-74712106-INS-174 |
| chrX-74723516-INS-138 | chrX-74723516-INS-156 |
| chrX-74732505-INS-7143 | chrX-74732506-INS-7143 |
| chrX-74762925-INS-203 | chrX-74762925-INS-203 |
| chrX-74769014-INS-54 | chrX-74769014-INS-52 |
| chrX-74788859-INS-962 | chrX-74788860-INS-962 |
| chrX-74810071-INS-210 | chrX-74810071-INS-210 |
| chrX-74814933-INS-224 | chrX-74814933-INS-229 |
| chrX-74816317-INS-59 | chrX-74816317-INS-59 |
| chrX-74842774-INS-200 | chrX-74842774-INS-200 |
| chrX-74853224-INS-207 | chrX-74853224-INS-207 |
| chrX-74901641-INS-219 | chrX-74901641-INS-219 |
| chrX-74909038-INS-6676 | chrX-74909039-INS-6679 |
| chrX-74914601-INS-68 | chrX-74914601-INS-72 |
| chrX-74922351-INS-80 | chrX-74922388-INS-58 |
| chrX-74941904-INS-5298 | chrX-74941905-INS-5298 |
| chrX-74994750-INS-214 | chrX-74994750-INS-214 |
| chrX-75048206-INS-185 | chrX-75048206-INS-185 |
| chrX-75101882-INS-577 | chrX-75101882-INS-577 |
| chrX-75113611-INS-65 | chrX-75113636-INS-57 |
| chrX-75115254-INS-4850 | chrX-75115255-INS-4850 |
| chrX-75120032-INS-59 | chrX-75120032-INS-62 |
| chrX-75128541-INS-126 | chrX-75128541-INS-106 |
| chrX-75155268-INS-68 | chrX-75155268-INS-68 |
| chrX-75156486-INS-63 | chrX-75156486-INS-75 |
| chrX-75170642-INS-3261 | chrX-75170643-INS-3261 |
| chrX-75180737-INS-161 | chrX-75180737-INS-161 |
| chrX-75196657-INS-198 | chrX-75196657-INS-198 |
| chrX-75206552-INS-430 | chrX-75206552-INS-430 |
| chrX-75212990-INS-364 | chrX-75212990-INS-363 |
| chrX-75293565-INS-73 | chrX-75293565-INS-73 |
| chrX-75345612-INS-166 | chrX-75345612-INS-166 |
| chrX-75359787-INS-56 | chrX-75359785-INS-52 |
| chrX-75371457-INS-1061 | chrX-75371458-INS-1061 |
| chrX-75421242-INS-7145 | chrX-75421243-INS-7149 |
| chrX-75422327-INS-58 | chrX-75422327-INS-105 |
| chrX-75429780-INS-277 | chrX-75429780-INS-277 |
| chrX-75456447-INS-97 | chrX-75456447-INS-92 |
| chrX-75464511-INS-2006 | chrX-75464512-INS-2006 |
| chrX-75478567-INS-1072 | chrX-75478568-INS-1072 |
| chrX-75533306-INS-2337 | chrX-75533307-INS-2336 |
| chrX-75560957-INS-8861 | chrX-75560958-INS-8835 |
| chrX-75587822-INS-70 | chrX-75587822-INS-70 |
| chrX-75596626-INS-3294 | chrX-75596627-INS-3279 |
| chrX-75596926-INS-792 | chrX-75596927-INS-792 |
| chrX-75611101-INS-6817 | chrX-75611102-INS-6819 |
| chrX-75628560-INS-85 | chrX-75628560-INS-90 |

| | |
|---|---|
| chrX-75631877-INS-6633 | chrX-75631878-INS-6633 |
| chrX-75637897-INS-211 | chrX-75637897-INS-211 |
| chrX-75639511-INS-6395 | chrX-75639512-INS-6396 |
| chrX-75656196-INS-164 | chrX-75656196-INS-165 |
| chrX-75665990-INS-332 | chrX-75665990-INS-332 |
| chrX-75691480-INS-1595 | chrX-75691481-INS-1596 |
| chrX-75696153-INS-2899 | chrX-75696154-INS-2899 |
| chrX-75709197-INS-516 | chrX-75709197-INS-516 |
| chrX-75755081-INS-399 | chrX-75755081-INS-399 |
| chrX-75761591-INS-857 | chrX-75761591-INS-853 |
| chrX-75776307-INS-6224 | chrX-75776308-INS-6224 |
| chrX-75799496-INS-175 | chrX-75799496-INS-175 |
| chrX-75822768-INS-4595 | chrX-75822769-INS-4594 |
| chrX-75823555-INS-518 | chrX-75823555-INS-518 |
| chrX-75826133-INS-91 | chrX-75826133-INS-91 |
| chrX-75834557-INS-2740 | chrX-75834558-INS-2740 |
| chrX-75846467-INS-156 | chrX-75846467-INS-183 |
| chrX-75860944-INS-6672 | chrX-75860945-INS-6672 |
| chrX-75866063-INS-156 | chrX-75866063-INS-144 |
| chrX-75889371-INS-61 | chrX-75889371-INS-59 |
| chrX-75906883-INS-385 | chrX-75906883-INS-386 |
| chrX-75980575-INS-232 | chrX-75980575-INS-233 |
| chrX-75989879-INS-578 | chrX-75989879-INS-578 |
| chrX-76009642-INS-55 | chrX-76009642-INS-55 |
| chrX-76028246-INS-6466 | chrX-76028247-INS-6466 |
| chrX-76053930-INS-173 | chrX-76053902-INS-159 |
| chrX-76089229-INS-1201 | chrX-76089230-INS-1201 |
| chrX-76099545-INS-64 | chrX-76099570-INS-72 |
| chrX-76128077-INS-52 | chrX-76128077-INS-52 |
| chrX-76136995-INS-204 | chrX-76136995-INS-204 |
| chrX-76167878-INS-2944 | chrX-76167879-INS-2943 |
| chrX-76174250-INS-6494 | chrX-76174251-INS-6493 |
| chrX-76194940-INS-87 | chrX-76194940-INS-93 |
| chrX-76205593-INS-72 | chrX-76205595-INS-66 |
| chrX-76210532-INS-199 | chrX-76210532-INS-199 |
| chrX-76224535-INS-197 | chrX-76224535-INS-197 |
| chrX-76228489-INS-755 | chrX-76228489-INS-776 |
| chrX-76235914-INS-402 | chrX-76235914-INS-402 |
| chrX-76274887-INS-1719 | chrX-76274888-INS-1721 |
| chrX-76277258-INS-129 | chrX-76277256-INS-107 |
| chrX-76411234-INS-58 | chrX-76411265-INS-53 |
| chrX-76412829-INS-52 | chrX-76412829-INS-53 |
| chrX-76418838-INS-111 | chrX-76418838-INS-111 |
| chrX-76423895-INS-164 | chrX-76423904-INS-122 |
| chrX-76456697-INS-60 | chrX-76456717-INS-60 |
| chrX-76460266-INS-1892 | chrX-76460267-INS-1892 |
| chrX-76485199-INS-70 | chrX-76485199-INS-70 |
| chrX-76499429-INS-97 | chrX-76499429-INS-104 |
| chrX-76503684-INS-131 | chrX-76503692-INS-117 |
| chrX-76527434-INS-1890 | chrX-76527435-INS-1890 |
| chrX-76615337-INS-222 | chrX-76615337-INS-222 |
| chrX-76630112-INS-7195 | chrX-76630113-INS-7195 |
| chrX-76641973-INS-57 | chrX-76642001-INS-60 |
| chrX-76657789-INS-108 | chrX-76657789-INS-108 |
| chrX-76722320-INS-95 | chrX-76722321-INS-70 |
| chrX-76748587-INS-220 | chrX-76748587-INS-220 |
| chrX-76752323-INS-197 | chrX-76752323-INS-199 |
| chrX-76760074-INS-53 | chrX-76760074-INS-54 |
| chrX-76760721-INS-447 | chrX-76760730-INS-405 |
| chrX-76760732-INS-1299 | chrX-76760733-INS-1287 |
| chrX-76793103-INS-97 | chrX-76793102-INS-60 |
| chrX-76809436-INS-92 | chrX-76809436-INS-92 |
| chrX-76812602-INS-208 | chrX-76812602-INS-208 |
| chrX-76827037-INS-72 | chrX-76827037-INS-56 |
| chrX-76886301-INS-652 | chrX-76886301-INS-653 |
| chrX-76888106-INS-421 | chrX-76888106-INS-421 |
| chrX-76927204-INS-7113 | chrX-76927205-INS-7113 |
| chrX-76953439-INS-204 | chrX-76953450-INS-204 |
| chrX-76976535-INS-2984 | chrX-76976536-INS-2984 |
| chrX-76980739-INS-51 | chrX-76980739-INS-51 |
| chrX-77005235-INS-124 | chrX-77005235-INS-124 |
| chrX-77063028-INS-398 | chrX-77063028-INS-398 |
| chrX-77074916-INS-200 | chrX-77074916-INS-200 |
| chrX-77080722-INS-255 | chrX-77080722-INS-255 |
| chrX-77102818-INS-774 | chrX-77102818-INS-772 |
| chrX-77120689-INS-2271 | chrX-77120690-INS-2271 |
| chrX-77125907-INS-199 | chrX-77125907-INS-199 |

| | |
|---|---|
| chrX-77128242-INS-1049 | chrX-77128243-INS-1049 |
| chrX-77132457-INS-86 | chrX-77132457-INS-87 |
| chrX-77138608-INS-257 | chrX-77138608-INS-257 |
| chrX-77166530-INS-2738 | chrX-77166530-INS-2735 |
| chrX-77174979-INS-74 | chrX-77174977-INS-84 |
| chrX-77179056-INS-1668 | chrX-77179057-INS-1672 |
| chrX-77192312-INS-96 | chrX-77192280-INS-118 |
| chrX-77207294-INS-509 | chrX-77207294-INS-509 |
| chrX-77216355-INS-6442 | chrX-77216356-INS-6442 |
| chrX-77255582-INS-158 | chrX-77255582-INS-158 |
| chrX-77258950-INS-174 | chrX-77258950-INS-174 |
| chrX-77260965-INS-1627 | chrX-77260966-INS-1627 |
| chrX-77286908-INS-80 | chrX-77286951-INS-56 |
| chrX-77299839-INS-2777 | chrX-77299840-INS-2777 |
| chrX-77310952-INS-1473 | chrX-77310953-INS-1474 |
| chrX-77317209-INS-384 | chrX-77317209-INS-384 |
| chrX-77318555-INS-65 | chrX-77318555-INS-66 |
| chrX-77333840-INS-498 | chrX-77333840-INS-498 |
| chrX-77334672-INS-498 | chrX-77334672-INS-498 |
| chrX-77335717-INS-1134 | chrX-77335718-INS-1134 |
| chrX-77369609-INS-1095 | chrX-77369610-INS-1081 |
| chrX-77370133-INS-62 | chrX-77370133-INS-70 |
| chrX-77435433-INS-53 | chrX-77435433-INS-58 |
| chrX-77454156-INS-120 | chrX-77454156-INS-119 |
| chrX-77462370-INS-564 | chrX-77462370-INS-564 |
| chrX-77539632-INS-398 | chrX-77539632-INS-384 |
| chrX-77586481-INS-164 | chrX-77586481-INS-163 |
| chrX-77649370-INS-838 | chrX-77649370-INS-838 |
| chrX-77651425-INS-498 | chrX-77651425-INS-498 |
| chrX-77682716-INS-2380 | chrX-77682717-INS-2380 |
| chrX-77714559-INS-901 | chrX-77714559-INS-901 |
| chrX-77745642-INS-407 | chrX-77745642-INS-407 |
| chrX-77745839-INS-4416 | chrX-77745840-INS-4415 |
| chrX-77800495-INS-6390 | chrX-77800496-INS-6390 |
| chrX-77852152-INS-409 | chrX-77852152-INS-409 |
| chrX-77871843-INS-2471 | chrX-77871844-INS-2467 |
| chrX-77888761-INS-84 | chrX-77888761-INS-84 |
| chrX-77892111-INS-3472 | chrX-77892112-INS-3472 |
| chrX-77893229-INS-3325 | chrX-77893230-INS-3323 |
| chrX-77897475-INS-9938 | chrX-77897476-INS-9939 |
| chrX-78044820-INS-3044 | chrX-78044821-INS-3044 |
| chrX-78086123-INS-528 | chrX-78086123-INS-529 |
| chrX-78123963-INS-2815 | chrX-78123964-INS-2815 |
| chrX-78141945-INS-129 | chrX-78141948-INS-132 |
| chrX-78150303-INS-98 | chrX-78150303-INS-58 |
| chrX-78176049-INS-87 | chrX-78176088-INS-87 |
| chrX-78177106-INS-481 | chrX-78177106-INS-481 |
| chrX-78196047-INS-892 | chrX-78196047-INS-893 |
| chrX-78197312-INS-2887 | chrX-78197313-INS-2887 |
| chrX-78216701-INS-759 | chrX-78216701-INS-748 |
| chrX-78230645-INS-60 | chrX-78230645-INS-60 |
| chrX-78253161-INS-88 | chrX-78253161-INS-88 |
| chrX-78370751-INS-1239 | chrX-78370752-INS-1243 |
| chrX-78435921-INS-6410 | chrX-78435922-INS-6410 |
| chrX-78464773-INS-772 | chrX-78464773-INS-749 |
| chrX-78478525-INS-74 | chrX-78478525-INS-84 |
| chrX-78479337-INS-72 | chrX-78479383-INS-50 |
| chrX-78490287-INS-166 | chrX-78490287-INS-166 |
| chrX-78607924-INS-1887 | chrX-78607925-INS-1887 |
| chrX-78623164-INS-2471 | chrX-78623165-INS-2471 |
| chrX-78660251-INS-202 | chrX-78660251-INS-202 |
| chrX-78661298-INS-164 | chrX-78661298-INS-161 |
| chrX-78711657-INS-51 | chrX-78711657-INS-51 |
| chrX-78727248-INS-427 | chrX-78727248-INS-428 |
| chrX-78780908-INS-826 | chrX-78780908-INS-826 |
| chrX-78811359-INS-81 | chrX-78811364-INS-75 |
| chrX-78820785-INS-58 | chrX-78820785-INS-60 |
| chrX-78922122-INS-8308 | chrX-78922123-INS-8308 |
| chrX-78922235-INS-62 | chrX-78922236-INS-70 |
| chrX-78931534-INS-72 | chrX-78931540-INS-50 |
| chrX-78958943-INS-166 | chrX-78958943-INS-166 |
| chrX-78997269-INS-197 | chrX-78997269-INS-199 |
| chrX-79054931-INS-377 | chrX-79054931-INS-378 |
| chrX-79138465-INS-4726 | chrX-79138466-INS-4726 |
| chrX-79141001-INS-52 | chrX-79141010-INS-56 |
| chrX-79168862-INS-239 | chrX-79168818-INS-239 |
| chrX-79197763-INS-4888 | chrX-79197764-INS-4888 |

| | |
|---|---|
| chrX-79198522-INS-169 | chrX-79198522-INS-171 |
| chrX-79202573-INS-163 | chrX-79202573-INS-167 |
| chrX-79210491-INS-50 | chrX-79210493-INS-64 |
| chrX-79224337-INS-876 | chrX-79224337-INS-886 |
| chrX-79293097-INS-3500 | chrX-79293097-INS-3480 |
| chrX-79334391-INS-1906 | chrX-79334392-INS-1905 |
| chrX-79368415-INS-4526 | chrX-79368416-INS-4526 |
| chrX-79386466-INS-7103 | chrX-79386467-INS-7103 |
| chrX-79387351-INS-3266 | chrX-79387352-INS-3267 |
| chrX-79422560-INS-603 | chrX-79422560-INS-603 |
| chrX-79463735-INS-55 | chrX-79463761-INS-61 |
| chrX-79478744-INS-6495 | chrX-79478745-INS-6496 |
| chrX-79503288-INS-4636 | chrX-79503289-INS-4639 |
| chrX-79505116-INS-75 | chrX-79505126-INS-60 |
| chrX-79515710-INS-914 | chrX-79515710-INS-914 |
| chrX-79522605-INS-370 | chrX-79522605-INS-370 |
| chrX-79551560-INS-3086 | chrX-79551561-INS-3051 |
| chrX-79567130-INS-545 | chrX-79567130-INS-545 |
| chrX-79572081-INS-106 | chrX-79572081-INS-56 |
| chrX-79616597-INS-3048 | chrX-79616598-INS-3054 |
| chrX-79673276-INS-96 | chrX-79673276-INS-96 |
| chrX-79679221-INS-126 | chrX-79679221-INS-126 |
| chrX-79686888-INS-1586 | chrX-79686889-INS-1586 |
| chrX-79732797-INS-767 | chrX-79732797-INS-768 |
| chrX-79744463-INS-4501 | chrX-79744464-INS-4501 |
| chrX-79750278-INS-52 | chrX-79750278-INS-56 |
| chrX-79771803-INS-89 | chrX-79771803-INS-91 |
| chrX-79788718-INS-52 | chrX-79788718-INS-50 |
| chrX-79832039-INS-781 | chrX-79832039-INS-782 |
| chrX-79886622-INS-488 | chrX-79886622-INS-501 |
| chrX-79919626-INS-63 | chrX-79919626-INS-63 |
| chrX-79930621-INS-470 | chrX-79930621-INS-470 |
| chrX-79932400-INS-60 | chrX-79932400-INS-60 |
| chrX-79940374-INS-7226 | chrX-79940375-INS-7226 |
| chrX-80034882-INS-152 | chrX-80034882-INS-152 |
| chrX-80048853-INS-319 | chrX-80048866-INS-318 |
| chrX-80072331-INS-216 | chrX-80072331-INS-217 |
| chrX-80158122-INS-66 | chrX-80158122-INS-66 |
| chrX-80171913-INS-355 | chrX-80171913-INS-355 |
| chrX-80219831-INS-63 | chrX-80219799-INS-55 |
| chrX-80227093-INS-78 | chrX-80227094-INS-79 |
| chrX-80231692-INS-149 | chrX-80231692-INS-153 |
| chrX-80232588-INS-4268 | chrX-80232589-INS-4268 |
| chrX-80235543-INS-104 | chrX-80235529-INS-100 |
| chrX-80257188-INS-52 | chrX-80257188-INS-50 |
| chrX-80316149-INS-6369 | chrX-80316150-INS-6369 |
| chrX-80353741-INS-194 | chrX-80353744-INS-195 |
| chrX-80360104-INS-184 | chrX-80360104-INS-187 |
| chrX-80369130-INS-411 | chrX-80369130-INS-411 |
| chrX-80372642-INS-95 | chrX-80372642-INS-110 |
| chrX-80375367-INS-6330 | chrX-80375368-INS-6334 |
| chrX-80387791-INS-561 | chrX-80387791-INS-562 |
| chrX-80423517-INS-1202 | chrX-80423518-INS-1197 |
| chrX-80426193-INS-480 | chrX-80426193-INS-488 |
| chrX-80450405-INS-53 | chrX-80450405-INS-54 |
| chrX-80456057-INS-56 | chrX-80456044-INS-60 |
| chrX-80485912-INS-5154 | chrX-80485913-INS-5154 |
| chrX-80486644-INS-83 | chrX-80486644-INS-52 |
| chrX-80499366-INS-228 | chrX-80499366-INS-252 |
| chrX-80505792-INS-77 | chrX-80505792-INS-89 |
| chrX-80557075-INS-709 | chrX-80557075-INS-706 |
| chrX-80598447-INS-4710 | chrX-80598448-INS-4710 |
| chrX-80657877-INS-2581 | chrX-80657878-INS-2582 |
| chrX-80662361-INS-122 | chrX-80662361-INS-85 |
| chrX-80830910-INS-192 | chrX-80830910-INS-192 |
| chrX-80906732-INS-208 | chrX-80906733-INS-208 |
| chrX-80945757-INS-463 | chrX-80945757-INS-462 |
| chrX-80953058-INS-316 | chrX-80953058-INS-316 |
| chrX-80955049-INS-73 | chrX-80955049-INS-72 |
| chrX-80986576-INS-3168 | chrX-80986577-INS-3168 |
| chrX-80996109-INS-54 | chrX-80996109-INS-54 |
| chrX-80998537-INS-54 | chrX-80998537-INS-54 |
| chrX-81019322-INS-209 | chrX-81019322-INS-209 |
| chrX-81089319-INS-93 | chrX-81089315-INS-102 |
| chrX-81117376-INS-4180 | chrX-81117377-INS-4180 |
| chrX-81160333-INS-459 | chrX-81160330-INS-462 |
| chrX-81179108-INS-7324 | chrX-81179109-INS-7324 |

| | |
|---|---|
| chrX-81184594-INS-1608 | chrX-81184595-INS-1608 |
| chrX-81185014-INS-52 | chrX-81185014-INS-50 |
| chrX-81209542-INS-160 | chrX-81209542-INS-160 |
| chrX-81325857-INS-6272 | chrX-81325858-INS-6272 |
| chrX-81446647-INS-456 | chrX-81446647-INS-456 |
| chrX-81478155-INS-2807 | chrX-81478156-INS-2808 |
| chrX-81496117-INS-58 | chrX-81496117-INS-58 |
| chrX-81512439-INS-218 | chrX-81512439-INS-218 |
| chrX-81553056-INS-497 | chrX-81553056-INS-497 |
| chrX-81581358-INS-728 | chrX-81581358-INS-728 |
| chrX-81583351-INS-52 | chrX-81583351-INS-54 |
| chrX-81645724-INS-168 | chrX-81645724-INS-174 |
| chrX-81664098-INS-4454 | chrX-81664099-INS-4454 |
| chrX-81690597-INS-216 | chrX-81690597-INS-216 |
| chrX-81730940-INS-2473 | chrX-81730941-INS-2457 |
| chrX-81744757-INS-241 | chrX-81744757-INS-242 |
| chrX-81774783-INS-373 | chrX-81774783-INS-374 |
| chrX-81807808-INS-133 | chrX-81807808-INS-137 |
| chrX-81894244-INS-73 | chrX-81894244-INS-55 |
| chrX-81918457-INS-67 | chrX-81918461-INS-61 |
| chrX-81938639-INS-76 | chrX-81938639-INS-81 |
| chrX-81962525-INS-212 | chrX-81962525-INS-212 |
| chrX-82005498-INS-65 | chrX-82005499-INS-74 |
| chrX-82045325-INS-52 | chrX-82045325-INS-52 |
| chrX-82058852-INS-882 | chrX-82058852-INS-882 |
| chrX-82059495-INS-158 | chrX-82059495-INS-158 |
| chrX-82096171-INS-3339 | chrX-82096172-INS-3339 |
| chrX-82134016-INS-66 | chrX-82134024-INS-66 |
| chrX-82143726-INS-67 | chrX-82143711-INS-64 |
| chrX-82169274-INS-180 | chrX-82169274-INS-198 |
| chrX-82173172-INS-55 | chrX-82173172-INS-55 |
| chrX-82176757-INS-7538 | chrX-82176758-INS-7539 |
| chrX-82248595-INS-616 | chrX-82248595-INS-616 |
| chrX-82271504-INS-82 | chrX-82271504-INS-106 |
| chrX-82382550-INS-173 | chrX-82382549-INS-198 |
| chrX-82400303-INS-60 | chrX-82400298-INS-61 |
| chrX-82413572-INS-51 | chrX-82413572-INS-51 |
| chrX-82483132-INS-335 | chrX-82483132-INS-336 |
| chrX-82485345-INS-183 | chrX-82485345-INS-183 |
| chrX-82488568-INS-3180 | chrX-82488569-INS-3180 |
| chrX-82502727-INS-810 | chrX-82502727-INS-810 |
| chrX-82518617-INS-92 | chrX-82518613-INS-94 |
| chrX-82532542-INS-147 | chrX-82532542-INS-147 |
| chrX-82604744-INS-60 | chrX-82604744-INS-60 |
| chrX-82638476-INS-66 | chrX-82638476-INS-66 |
| chrX-82640409-INS-68 | chrX-82640409-INS-68 |
| chrX-82650644-INS-68 | chrX-82650644-INS-68 |
| chrX-82666274-INS-65 | chrX-82666274-INS-66 |
| chrX-82667019-INS-5177 | chrX-82667020-INS-5177 |
| chrX-82668180-INS-65 | chrX-82668180-INS-66 |
| chrX-82675868-INS-68 | chrX-82675868-INS-76 |
| chrX-82691223-INS-63 | chrX-82691223-INS-63 |
| chrX-82730270-INS-58 | chrX-82730270-INS-64 |
| chrX-82743444-INS-59 | chrX-82743444-INS-59 |
| chrX-82818371-INS-290 | chrX-82818371-INS-291 |
| chrX-82854921-INS-146 | chrX-82854921-INS-146 |
| chrX-82860348-INS-52 | chrX-82860348-INS-50 |
| chrX-82885652-INS-1639 | chrX-82885653-INS-1640 |
| chrX-82921462-INS-739 | chrX-82921462-INS-739 |
| chrX-82948980-INS-2522 | chrX-82948981-INS-2521 |
| chrX-82975063-INS-598 | chrX-82975063-INS-598 |
| chrX-82990418-INS-245 | chrX-82990418-INS-246 |
| chrX-83118650-INS-6575 | chrX-83118651-INS-6575 |
| chrX-83136948-INS-2872 | chrX-83136949-INS-2873 |
| chrX-83139608-INS-762 | chrX-83139608-INS-765 |
| chrX-83166009-INS-101 | chrX-83166009-INS-101 |
| chrX-83186998-INS-314 | chrX-83186998-INS-318 |
| chrX-83194322-INS-323 | chrX-83194322-INS-307 |
| chrX-83222270-INS-424 | chrX-83222270-INS-429 |
| chrX-83259636-INS-6667 | chrX-83259637-INS-6667 |
| chrX-83266519-INS-58 | chrX-83266519-INS-58 |
| chrX-83271372-INS-52 | chrX-83271372-INS-68 |
| chrX-83284082-INS-262 | chrX-83284082-INS-262 |
| chrX-83284439-INS-1583 | chrX-83284440-INS-1600 |
| chrX-83357917-INS-85 | chrX-83357917-INS-87 |
| chrX-83410380-INS-654 | chrX-83410380-INS-654 |
| chrX-83443738-INS-212 | chrX-83443738-INS-212 |

| | |
|---|---|
| chrX-83466955-INS-553 | chrX-83466955-INS-553 |
| chrX-83470556-INS-61 | chrX-83470556-INS-52 |
| chrX-83494810-INS-5850 | chrX-83494811-INS-5850 |
| chrX-83525565-INS-55 | chrX-83525565-INS-55 |
| chrX-83568308-INS-107 | chrX-83568310-INS-108 |
| chrX-83602169-INS-629 | chrX-83602169-INS-631 |
| chrX-83605534-INS-2033 | chrX-83605535-INS-2031 |
| chrX-83615803-INS-52 | chrX-83615804-INS-52 |
| chrX-83627151-INS-55 | chrX-83627151-INS-55 |
| chrX-83628345-INS-4529 | chrX-83628346-INS-4529 |
| chrX-83644128-INS-342 | chrX-83644128-INS-342 |
| chrX-83711097-INS-56 | chrX-83711137-INS-64 |
| chrX-83723792-INS-1150 | chrX-83723793-INS-1150 |
| chrX-83724959-INS-481 | chrX-83724959-INS-491 |
| chrX-83735110-INS-211 | chrX-83735110-INS-211 |
| chrX-83740615-INS-401 | chrX-83740615-INS-401 |
| chrX-83741469-INS-2499 | chrX-83741470-INS-2499 |
| chrX-83756201-INS-755 | chrX-83756201-INS-755 |
| chrX-83776284-INS-149 | chrX-83776284-INS-149 |
| chrX-83814941-INS-80 | chrX-83814941-INS-90 |
| chrX-83873699-INS-103 | chrX-83873699-INS-103 |
| chrX-83885490-INS-455 | chrX-83885489-INS-458 |
| chrX-83939006-INS-68 | chrX-83939006-INS-62 |
| chrX-83945582-INS-82 | chrX-83945582-INS-82 |
| chrX-83985384-INS-52 | chrX-83985384-INS-52 |
| chrX-84015750-INS-958 | chrX-84015750-INS-957 |
| chrX-84019381-INS-1701 | chrX-84019382-INS-1701 |
| chrX-84037519-INS-335 | chrX-84037519-INS-335 |
| chrX-84068535-INS-972 | chrX-84068535-INS-972 |
| chrX-84138951-INS-333 | chrX-84138954-INS-297 |
| chrX-84170050-INS-147 | chrX-84170050-INS-147 |
| chrX-84177484-INS-147 | chrX-84177484-INS-147 |
| chrX-84231354-INS-55 | chrX-84231354-INS-55 |
| chrX-84276144-INS-902 | chrX-84276144-INS-903 |
| chrX-84328221-INS-6502 | chrX-84328222-INS-6504 |
| chrX-84332814-INS-148 | chrX-84332814-INS-148 |
| chrX-84423215-INS-1893 | chrX-84423216-INS-1893 |
| chrX-84427320-INS-1893 | chrX-84427321-INS-1893 |
| chrX-84437257-INS-202 | chrX-84437257-INS-202 |
| chrX-84465561-INS-620 | chrX-84465561-INS-622 |
| chrX-84495408-INS-6551 | chrX-84495409-INS-6553 |
| chrX-84501955-INS-6271 | chrX-84501956-INS-6271 |
| chrX-84504543-INS-1125 | chrX-84504544-INS-1125 |
| chrX-84513701-INS-525 | chrX-84513701-INS-525 |
| chrX-84530663-INS-6495 | chrX-84530664-INS-6495 |
| chrX-84558806-INS-2031 | chrX-84558807-INS-2031 |
| chrX-84569101-INS-100 | chrX-84569103-INS-50 |
| chrX-84585467-INS-874 | chrX-84585467-INS-874 |
| chrX-84625893-INS-1605 | chrX-84625894-INS-1605 |
| chrX-84641770-INS-1248 | chrX-84641771-INS-1247 |
| chrX-84707083-INS-364 | chrX-84707083-INS-364 |
| chrX-84714484-INS-1506 | chrX-84714485-INS-1506 |
| chrX-84716759-INS-62 | chrX-84716759-INS-65 |
| chrX-84725470-INS-214 | chrX-84725466-INS-248 |
| chrX-84781937-INS-60 | chrX-84781937-INS-62 |
| chrX-84786595-INS-60 | chrX-84786595-INS-60 |
| chrX-84824545-INS-1674 | chrX-84824546-INS-1675 |
| chrX-84828525-INS-86 | chrX-84828525-INS-78 |
| chrX-84849701-INS-100 | chrX-84849697-INS-105 |
| chrX-84868390-INS-98 | chrX-84868390-INS-98 |
| chrX-84886670-INS-1894 | chrX-84886671-INS-1894 |
| chrX-84955707-INS-101 | chrX-84955736-INS-99 |
| chrX-84981866-INS-80 | chrX-84981843-INS-108 |
| chrX-85033723-INS-230 | chrX-85033723-INS-215 |
| chrX-85039776-INS-1051 | chrX-85039777-INS-1052 |
| chrX-85070758-INS-1704 | chrX-85070759-INS-1704 |
| chrX-85073652-INS-74 | chrX-85073652-INS-74 |
| chrX-85129413-INS-2054 | chrX-85129414-INS-2054 |
| chrX-85135297-INS-6201 | chrX-85135298-INS-6201 |
| chrX-85157345-INS-610 | chrX-85157345-INS-610 |
| chrX-85177351-INS-76 | chrX-85177363-INS-95 |
| chrX-85180863-INS-307 | chrX-85180863-INS-307 |
| chrX-85185477-INS-3054 | chrX-85185478-INS-3054 |
| chrX-85194152-INS-443 | chrX-85194152-INS-443 |
| chrX-85195224-INS-169 | chrX-85195224-INS-169 |
| chrX-85209493-INS-181 | chrX-85209493-INS-181 |
| chrX-85247926-INS-4317 | chrX-85247927-INS-4317 |

| | |
|---|---|
| chrX-85257656-INS-450 | chrX-85257647-INS-454 |
| chrX-85316171-INS-70 | chrX-85316151-INS-50 |
| chrX-85318598-INS-814 | chrX-85318598-INS-810 |
| chrX-85345264-INS-68 | chrX-85345264-INS-68 |
| chrX-85366415-INS-180 | chrX-85366415-INS-180 |
| chrX-85393174-INS-733 | chrX-85393168-INS-762 |
| chrX-85404244-INS-772 | chrX-85404244-INS-772 |
| chrX-85418730-INS-199 | chrX-85418730-INS-199 |
| chrX-85448519-INS-4557 | chrX-85448520-INS-4557 |
| chrX-85466478-INS-75 | chrX-85466478-INS-75 |
| chrX-85492840-INS-7084 | chrX-85492841-INS-7084 |
| chrX-85504971-INS-52 | chrX-85504971-INS-78 |
| chrX-85540155-INS-202 | chrX-85540155-INS-201 |
| chrX-85582466-INS-98 | chrX-85582458-INS-75 |
| chrX-85590126-INS-564 | chrX-85590126-INS-565 |
| chrX-85602724-INS-120 | chrX-85602724-INS-138 |
| chrX-85626654-INS-416 | chrX-85626654-INS-416 |
| chrX-85652144-INS-6691 | chrX-85652142-INS-6678 |
| chrX-85664343-INS-1300 | chrX-85664344-INS-1300 |
| chrX-85675391-INS-78 | chrX-85675391-INS-78 |
| chrX-85682436-INS-400 | chrX-85682441-INS-400 |
| chrX-85749804-INS-2948 | chrX-85749805-INS-2948 |
| chrX-85820648-INS-597 | chrX-85820648-INS-597 |
| chrX-85825275-INS-885 | chrX-85825275-INS-882 |
| chrX-85833916-INS-152 | chrX-85833916-INS-152 |
| chrX-85853347-INS-690 | chrX-85853347-INS-692 |
| chrX-85862391-INS-56 | chrX-85862391-INS-58 |
| chrX-85884310-INS-183 | chrX-85884310-INS-183 |
| chrX-85948515-INS-52 | chrX-85948515-INS-51 |
| chrX-85950504-INS-1107 | chrX-85950505-INS-1106 |
| chrX-85952096-INS-207 | chrX-85952096-INS-207 |
| chrX-85969713-INS-55 | chrX-85969718-INS-85 |
| chrX-86008896-INS-64 | chrX-86008882-INS-51 |
| chrX-86014064-INS-94 | chrX-86014063-INS-96 |
| chrX-86081109-INS-6983 | chrX-86081110-INS-6983 |
| chrX-86117310-INS-115 | chrX-86117310-INS-116 |
| chrX-86133566-INS-52 | chrX-86133566-INS-64 |
| chrX-86148518-INS-210 | chrX-86148518-INS-210 |
| chrX-86164827-INS-4564 | chrX-86164828-INS-4564 |
| chrX-86219131-INS-4807 | chrX-86219132-INS-4807 |
| chrX-86328846-INS-6926 | chrX-86328847-INS-6921 |
| chrX-86334501-INS-6500 | chrX-86334502-INS-6501 |
| chrX-86338186-INS-669 | chrX-86338186-INS-669 |
| chrX-86357552-INS-6368 | chrX-86357553-INS-6368 |
| chrX-86371258-INS-6877 | chrX-86371259-INS-6878 |
| chrX-86416001-INS-432 | chrX-86416001-INS-432 |
| chrX-86416251-INS-3720 | chrX-86416252-INS-3719 |
| chrX-86419034-INS-1041 | chrX-86419035-INS-1041 |
| chrX-86437869-INS-134 | chrX-86437869-INS-135 |
| chrX-86459677-INS-1034 | chrX-86459678-INS-1034 |
| chrX-86465032-INS-6384 | chrX-86465033-INS-6384 |
| chrX-86477520-INS-140 | chrX-86477520-INS-140 |
| chrX-86487151-INS-649 | chrX-86487151-INS-649 |
| chrX-86492777-INS-176 | chrX-86492777-INS-175 |
| chrX-86525438-INS-6428 | chrX-86525439-INS-6429 |
| chrX-86541920-INS-3718 | chrX-86541913-INS-3737 |
| chrX-86551315-INS-224 | chrX-86551315-INS-223 |
| chrX-86570154-INS-386 | chrX-86570154-INS-386 |
| chrX-86580900-INS-100 | chrX-86580900-INS-100 |
| chrX-86583202-INS-568 | chrX-86583202-INS-568 |
| chrX-86658940-INS-429 | chrX-86658940-INS-429 |
| chrX-86664285-INS-6210 | chrX-86664286-INS-6210 |
| chrX-86667097-INS-2111 | chrX-86667098-INS-2111 |
| chrX-86688773-INS-60 | chrX-86688773-INS-69 |
| chrX-86695913-INS-1485 | chrX-86695914-INS-1484 |
| chrX-86712286-INS-6591 | chrX-86712287-INS-6591 |
| chrX-86717692-INS-79 | chrX-86717693-INS-79 |
| chrX-86721775-INS-54 | chrX-86721775-INS-56 |
| chrX-86724951-INS-905 | chrX-86724951-INS-904 |
| chrX-86729534-INS-574 | chrX-86729534-INS-572 |
| chrX-86743800-INS-4813 | chrX-86743801-INS-4813 |
| chrX-86757185-INS-107 | chrX-86757185-INS-107 |
| chrX-86767787-INS-177 | chrX-86767787-INS-178 |
| chrX-86800373-INS-72 | chrX-86800373-INS-56 |
| chrX-86810751-INS-1363 | chrX-86810752-INS-1363 |
| chrX-86823719-INS-434 | chrX-86823719-INS-434 |
| chrX-86834676-INS-2298 | chrX-86834677-INS-2299 |

| | |
|---|---|
| chrX-86841501-INS-1888 | chrX-86841502-INS-1888 |
| chrX-86900334-INS-564 | chrX-86900334-INS-564 |
| chrX-86982749-INS-80 | chrX-86982749-INS-82 |
| chrX-87026311-INS-137 | chrX-87026311-INS-137 |
| chrX-87030634-INS-60 | chrX-87030634-INS-59 |
| chrX-87047872-INS-363 | chrX-87047872-INS-363 |
| chrX-87063830-INS-574 | chrX-87063847-INS-607 |
| chrX-87092950-INS-788 | chrX-87092951-INS-787 |
| chrX-87127045-INS-3104 | chrX-87127046-INS-3097 |
| chrX-87170393-INS-56 | chrX-87170393-INS-68 |
| chrX-87222704-INS-6374 | chrX-87222705-INS-6368 |
| chrX-87302839-INS-371 | chrX-87302839-INS-371 |
| chrX-87381975-INS-664 | chrX-87381975-INS-664 |
| chrX-87398035-INS-54 | chrX-87398035-INS-58 |
| chrX-87404358-INS-84 | chrX-87404365-INS-78 |
| chrX-87416163-INS-96 | chrX-87416148-INS-63 |
| chrX-87451224-INS-793 | chrX-87451224-INS-793 |
| chrX-87498454-INS-55 | chrX-87498454-INS-55 |
| chrX-87533127-INS-5412 | chrX-87533128-INS-5412 |
| chrX-87564625-INS-393 | chrX-87564625-INS-393 |
| chrX-87615881-INS-2775 | chrX-87615882-INS-2775 |
| chrX-87682952-INS-2809 | chrX-87682953-INS-2827 |
| chrX-87709836-INS-74 | chrX-87709836-INS-76 |
| chrX-87719882-INS-129 | chrX-87719882-INS-129 |
| chrX-87734272-INS-1306 | chrX-87734273-INS-1307 |
| chrX-87739124-INS-6453 | chrX-87739125-INS-6453 |
| chrX-87822400-INS-396 | chrX-87822400-INS-396 |
| chrX-87831044-INS-2470 | chrX-87831045-INS-2473 |
| chrX-87859597-INS-838 | chrX-87859597-INS-838 |
| chrX-87863926-INS-191 | chrX-87863935-INS-197 |
| chrX-87870582-INS-1024 | chrX-87870583-INS-1024 |
| chrX-87891582-INS-4286 | chrX-87891583-INS-4286 |
| chrX-87942244-INS-52 | chrX-87942244-INS-52 |
| chrX-87950375-INS-6369 | chrX-87950376-INS-6368 |
| chrX-87955055-INS-180 | chrX-87955055-INS-180 |
| chrX-87976295-INS-6797 | chrX-87976296-INS-6800 |
| chrX-88027604-INS-137 | chrX-88027604-INS-145 |
| chrX-88029973-INS-1323 | chrX-88029969-INS-1319 |
| chrX-88068155-INS-54 | chrX-88068155-INS-52 |
| chrX-88070534-INS-4172 | chrX-88070535-INS-4183 |
| chrX-88121301-INS-108 | chrX-88121301-INS-113 |
| chrX-88153124-INS-445 | chrX-88153124-INS-445 |
| chrX-88177672-INS-1893 | chrX-88177673-INS-1893 |
| chrX-88204864-INS-52 | chrX-88204864-INS-56 |
| chrX-88230219-INS-180 | chrX-88230212-INS-162 |
| chrX-88234673-INS-98 | chrX-88234673-INS-97 |
| chrX-88272545-INS-51 | chrX-88272545-INS-51 |
| chrX-88309247-INS-58 | chrX-88309247-INS-58 |
| chrX-88311889-INS-2516 | chrX-88311890-INS-2518 |
| chrX-88332963-INS-771 | chrX-88332963-INS-787 |
| chrX-88369908-INS-199 | chrX-88369908-INS-199 |
| chrX-88378514-INS-7183 | chrX-88378515-INS-7183 |
| chrX-88385022-INS-3096 | chrX-88385023-INS-3099 |
| chrX-88389467-INS-69 | chrX-88389467-INS-69 |
| chrX-88391119-INS-1482 | chrX-88391120-INS-1482 |
| chrX-88421907-INS-78 | chrX-88421907-INS-103 |
| chrX-88438858-INS-609 | chrX-88438858-INS-609 |
| chrX-88478527-INS-70 | chrX-88478502-INS-66 |
| chrX-88485524-INS-1895 | chrX-88485525-INS-1895 |
| chrX-88493440-INS-315 | chrX-88493440-INS-315 |
| chrX-88495232-INS-463 | chrX-88495232-INS-463 |
| chrX-88498659-INS-494 | chrX-88498659-INS-494 |
| chrX-88519072-INS-487 | chrX-88519072-INS-487 |
| chrX-88529640-INS-120 | chrX-88529640-INS-145 |
| chrX-88591784-INS-1765 | chrX-88591785-INS-1765 |
| chrX-88640457-INS-1484 | chrX-88640458-INS-1484 |
| chrX-88666780-INS-972 | chrX-88666780-INS-972 |
| chrX-88674388-INS-198 | chrX-88674388-INS-198 |
| chrX-88713994-INS-173 | chrX-88713994-INS-173 |
| chrX-88781261-INS-1676 | chrX-88781262-INS-1680 |
| chrX-88821089-INS-148 | chrX-88821089-INS-149 |
| chrX-88843588-INS-2576 | chrX-88843589-INS-2572 |
| chrX-88895260-INS-105 | chrX-88895260-INS-135 |
| chrX-88948619-INS-5553 | chrX-88948620-INS-5554 |
| chrX-88972774-INS-7942 | chrX-88972775-INS-7942 |
| chrX-89021062-INS-56 | chrX-89021062-INS-56 |
| chrX-89025976-INS-59 | chrX-89025978-INS-57 |

| | |
|---|---|
| chrX-89038158-INS-204 | chrX-89038158-INS-204 |
| chrX-89039592-INS-395 | chrX-89039592-INS-395 |
| chrX-89065421-INS-6489 | chrX-89065422-INS-6490 |
| chrX-89089458-INS-68 | chrX-89089458-INS-68 |
| chrX-89090907-INS-57 | chrX-89090919-INS-63 |
| chrX-89130400-INS-52 | chrX-89130410-INS-60 |
| chrX-89130966-INS-685 | chrX-89130966-INS-685 |
| chrX-89132432-INS-1783 | chrX-89132433-INS-1781 |
| chrX-89149912-INS-566 | chrX-89149912-INS-567 |
| chrX-89236510-INS-1702 | chrX-89236511-INS-1702 |
| chrX-89241455-INS-1296 | chrX-89241465-INS-1285 |
| chrX-89345874-INS-227 | chrX-89345874-INS-227 |
| chrX-89350973-INS-480 | chrX-89350973-INS-480 |
| chrX-89413165-INS-5549 | chrX-89413166-INS-5549 |
| chrX-89492340-INS-6316 | chrX-89492341-INS-6316 |
| chrX-89494557-INS-78 | chrX-89494557-INS-80 |
| chrX-89509427-INS-54 | chrX-89509427-INS-54 |
| chrX-89525739-INS-6392 | chrX-89525740-INS-6394 |
| chrX-89525779-INS-1559 | chrX-89525780-INS-1559 |
| chrX-89534179-INS-135 | chrX-89534179-INS-135 |
| chrX-89583662-INS-51 | chrX-89583662-INS-50 |
| chrX-89604336-INS-535 | chrX-89604336-INS-535 |
| chrX-89662597-INS-90 | chrX-89662597-INS-95 |
| chrX-89679209-INS-487 | chrX-89679209-INS-487 |
| chrX-89707315-INS-71 | chrX-89707334-INS-70 |
| chrX-89715590-INS-208 | chrX-89715590-INS-208 |
| chrX-89775978-INS-7094 | chrX-89775979-INS-7094 |
| chrX-89778934-INS-58 | chrX-89778934-INS-58 |
| chrX-89786247-INS-589 | chrX-89786247-INS-589 |
| chrX-89818970-INS-212 | chrX-89818970-INS-212 |
| chrX-89823305-INS-2928 | chrX-89823306-INS-2928 |
| chrX-89829851-INS-2139 | chrX-89829852-INS-2139 |
| chrX-89833968-INS-6410 | chrX-89833969-INS-6410 |
| chrX-89892735-INS-54 | chrX-89892735-INS-68 |
| chrX-89895123-INS-51 | chrX-89895123-INS-51 |
| chrX-89944923-INS-51 | chrX-89944923-INS-51 |
| chrX-89963910-INS-173 | chrX-89963910-INS-175 |
| chrX-89985183-INS-4714 | chrX-89985184-INS-4714 |
| chrX-89987488-INS-2343 | chrX-89987489-INS-2343 |
| chrX-90009654-INS-87 | chrX-90009656-INS-60 |
| chrX-90020514-INS-60 | chrX-90020539-INS-50 |
| chrX-90030674-INS-58 | chrX-90030674-INS-60 |
| chrX-90038245-INS-6450 | chrX-90038246-INS-6450 |
| chrX-90070506-INS-130 | chrX-90070506-INS-130 |
| chrX-90090006-INS-623 | chrX-90090006-INS-601 |
| chrX-90115563-INS-2566 | chrX-90115564-INS-2566 |
| chrX-90134643-INS-101 | chrX-90134642-INS-122 |
| chrX-90155277-INS-64 | chrX-90155277-INS-64 |
| chrX-90161907-INS-101 | chrX-90161904-INS-106 |
| chrX-90212435-INS-449 | chrX-90212435-INS-449 |
| chrX-90241004-INS-203 | chrX-90241005-INS-203 |
| chrX-90243974-INS-1773 | chrX-90243975-INS-1774 |
| chrX-90252707-INS-477 | chrX-90252707-INS-478 |
| chrX-90270467-INS-203 | chrX-90270467-INS-200 |
| chrX-90278748-INS-77 | chrX-90278744-INS-81 |
| chrX-90290977-INS-4149 | chrX-90290978-INS-4149 |
| chrX-90319329-INS-56 | chrX-90319329-INS-52 |
| chrX-90330805-INS-55 | chrX-90330784-INS-64 |
| chrX-90340285-INS-687 | chrX-90340285-INS-687 |
| chrX-90356286-INS-274 | chrX-90356277-INS-272 |
| chrX-90377402-INS-2936 | chrX-90377403-INS-2936 |
| chrX-90412283-INS-3010 | chrX-90412284-INS-3010 |
| chrX-90420777-INS-4770 | chrX-90420778-INS-4770 |
| chrX-90538058-INS-2364 | chrX-90538059-INS-2369 |
| chrX-90553608-INS-52 | chrX-90553608-INS-52 |
| chrX-90591536-INS-153 | chrX-90591536-INS-162 |
| chrX-90600594-INS-210 | chrX-90600594-INS-210 |
| chrX-90614955-INS-170 | chrX-90614955-INS-170 |
| chrX-90623521-INS-270 | chrX-90623521-INS-271 |
| chrX-90664050-INS-1511 | chrX-90664051-INS-1511 |
| chrX-90716365-INS-176 | chrX-90716365-INS-176 |
| chrX-90744710-INS-348 | chrX-90744710-INS-348 |
| chrX-90759309-INS-205 | chrX-90759309-INS-205 |
| chrX-90836510-INS-499 | chrX-90836510-INS-498 |
| chrX-90853461-INS-86 | chrX-90853461-INS-64 |
| chrX-90901933-INS-199 | chrX-90901933-INS-202 |
| chrX-90922163-INS-747 | chrX-90922164-INS-754 |

| | |
|---|---|
| chrX-90954132-INS-339 | chrX-90954132-INS-339 |
| chrX-91003479-INS-63 | chrX-91003483-INS-59 |
| chrX-91028939-INS-454 | chrX-91028939-INS-454 |
| chrX-91045118-INS-510 | chrX-91045118-INS-510 |
| chrX-91047548-INS-206 | chrX-91047548-INS-206 |
| chrX-91085422-INS-270 | chrX-91085422-INS-271 |
| chrX-91116065-INS-2345 | chrX-91116066-INS-2321 |
| chrX-91172176-INS-80 | chrX-91172190-INS-70 |
| chrX-91195852-INS-155 | chrX-91195848-INS-170 |
| chrX-91199437-INS-6584 | chrX-91199437-INS-6583 |
| chrX-91224760-INS-60 | chrX-91224760-INS-85 |
| chrX-91237721-INS-1330 | chrX-91237722-INS-1317 |
| chrX-91271055-INS-86 | chrX-91271055-INS-86 |
| chrX-91284853-INS-76 | chrX-91284853-INS-76 |
| chrX-91322319-INS-396 | chrX-91322319-INS-396 |
| chrX-91330948-INS-106 | chrX-91330948-INS-58 |
| chrX-91428564-INS-110 | chrX-91428564-INS-108 |
| chrX-91493289-INS-1382 | chrX-91493290-INS-1382 |
| chrX-91654715-INS-2640 | chrX-91654716-INS-2641 |
| chrX-91698070-INS-279 | chrX-91698070-INS-279 |
| chrX-91704176-INS-853 | chrX-91704176-INS-830 |
| chrX-91706188-INS-539 | chrX-91706188-INS-539 |
| chrX-91711678-INS-116 | chrX-91711678-INS-110 |
| chrX-91732006-INS-6724 | chrX-91732007-INS-6725 |
| chrX-91759988-INS-6658 | chrX-91759989-INS-6660 |
| chrX-91776442-INS-311 | chrX-91776442-INS-312 |
| chrX-91850900-INS-145 | chrX-91850900-INS-159 |
| chrX-91875010-INS-6423 | chrX-91875011-INS-6423 |
| chrX-91880418-INS-52 | chrX-91880418-INS-52 |
| chrX-91896535-INS-52 | chrX-91896535-INS-52 |
| chrX-91898785-INS-852 | chrX-91898781-INS-853 |
| chrX-91923429-INS-1415 | chrX-91923430-INS-1415 |
| chrX-91941499-INS-88 | chrX-91941514-INS-68 |
| chrX-92014283-INS-176 | chrX-92014281-INS-175 |
| chrX-92018667-INS-745 | chrX-92018667-INS-745 |
| chrX-92031619-INS-209 | chrX-92031619-INS-210 |
| chrX-92037406-INS-84 | chrX-92037406-INS-86 |
| chrX-92070708-INS-578 | chrX-92070708-INS-578 |
| chrX-92153862-INS-618 | chrX-92153862-INS-617 |
| chrX-92187060-INS-6182 | chrX-92187061-INS-6180 |
| chrX-92211732-INS-54 | chrX-92211732-INS-54 |
| chrX-92243115-INS-772 | chrX-92243115-INS-770 |
| chrX-92248498-INS-314 | chrX-92248498-INS-314 |
| chrX-92310570-INS-8229 | chrX-92310571-INS-8229 |
| chrX-92324507-INS-104 | chrX-92324497-INS-112 |
| chrX-92543049-INS-162 | chrX-92543008-INS-154 |
| chrX-92562527-INS-207 | chrX-92562527-INS-209 |
| chrX-92595446-INS-52 | chrX-92595431-INS-68 |
| chrX-92611649-INS-209 | chrX-92611649-INS-208 |
| chrX-92612215-INS-508 | chrX-92612215-INS-508 |
| chrX-92622028-INS-195 | chrX-92622026-INS-205 |
| chrX-92635863-INS-69 | chrX-92635863-INS-66 |
| chrX-92652958-INS-193 | chrX-92652958-INS-191 |
| chrX-92685629-INS-101 | chrX-92685650-INS-110 |
| chrX-92715785-INS-6425 | chrX-92715786-INS-6426 |
| chrX-92717194-INS-169 | chrX-92717194-INS-187 |
| chrX-92720259-INS-201 | chrX-92720259-INS-200 |
| chrX-92735220-INS-185 | chrX-92735220-INS-185 |
| chrX-92758930-INS-230 | chrX-92758930-INS-229 |
| chrX-92780693-INS-215 | chrX-92780693-INS-215 |
| chrX-92793740-INS-198 | chrX-92793740-INS-198 |
| chrX-92796794-INS-103 | chrX-92796794-INS-103 |
| chrX-92830644-INS-150 | chrX-92830649-INS-158 |
| chrX-92848596-INS-468 | chrX-92848596-INS-468 |
| chrX-92852392-INS-85 | chrX-92852392-INS-85 |
| chrX-92868969-INS-87 | chrX-92868969-INS-87 |
| chrX-92873184-INS-207 | chrX-92873184-INS-207 |
| chrX-92910772-INS-73 | chrX-92910772-INS-73 |
| chrX-92927089-INS-1889 | chrX-92927090-INS-1889 |
| chrX-92941722-INS-1893 | chrX-92941723-INS-1893 |
| chrX-92952030-INS-180 | chrX-92952030-INS-180 |
| chrX-92954388-INS-86 | chrX-92954388-INS-86 |
| chrX-93004194-INS-132 | chrX-93004164-INS-133 |
| chrX-93052678-INS-6215 | chrX-93052679-INS-6214 |
| chrX-93059222-INS-497 | chrX-93059222-INS-497 |
| chrX-93089276-INS-114 | chrX-93089277-INS-109 |
| chrX-93133520-INS-147 | chrX-93133520-INS-141 |

| | |
|---|---|
| chrX-93224826-INS-221 | chrX-93224826-INS-221 |
| chrX-93245363-INS-82 | chrX-93245363-INS-82 |
| chrX-93246019-INS-12455 | chrX-93246020-INS-12455 |
| chrX-93262742-INS-195 | chrX-93262742-INS-195 |
| chrX-93286038-INS-176 | chrX-93286038-INS-184 |
| chrX-93304481-INS-321 | chrX-93304481-INS-322 |
| chrX-93342126-INS-60 | chrX-93342126-INS-66 |
| chrX-93464673-INS-908 | chrX-93464673-INS-908 |
| chrX-93470202-INS-54 | chrX-93470202-INS-54 |
| chrX-93478570-INS-105 | chrX-93478570-INS-105 |
| chrX-93513783-INS-71 | chrX-93513783-INS-71 |
| chrX-93522451-INS-121 | chrX-93522451-INS-124 |
| chrX-93538461-INS-84 | chrX-93538462-INS-84 |
| chrX-93538921-INS-195 | chrX-93538921-INS-195 |
| chrX-93542056-INS-107 | chrX-93542030-INS-114 |
| chrX-93549489-INS-509 | chrX-93549489-INS-510 |
| chrX-93550685-INS-54 | chrX-93550685-INS-60 |
| chrX-93562052-INS-52 | chrX-93562048-INS-56 |
| chrX-93573735-INS-144 | chrX-93573740-INS-143 |
| chrX-93644560-INS-182 | chrX-93644560-INS-174 |
| chrX-93654793-INS-50 | chrX-93654793-INS-64 |
| chrX-93660901-INS-2535 | chrX-93660902-INS-2541 |
| chrX-93661365-INS-1610 | chrX-93661366-INS-1610 |
| chrX-93664513-INS-4884 | chrX-93664514-INS-4883 |
| chrX-93686025-INS-51 | chrX-93686025-INS-51 |
| chrX-93697836-INS-6240 | chrX-93697837-INS-6231 |
| chrX-93735258-INS-94 | chrX-93735258-INS-94 |
| chrX-93761979-INS-123 | chrX-93761979-INS-123 |
| chrX-93825520-INS-325 | chrX-93825520-INS-324 |
| chrX-93848116-INS-123 | chrX-93848116-INS-123 |
| chrX-93896445-INS-433 | chrX-93896445-INS-433 |
| chrX-93906569-INS-473 | chrX-93906569-INS-473 |
| chrX-93916532-INS-57 | chrX-93916529-INS-56 |
| chrX-94030427-INS-113 | chrX-94030427-INS-113 |
| chrX-94133449-INS-931 | chrX-94133449-INS-931 |
| chrX-94195920-INS-104 | chrX-94195921-INS-136 |
| chrX-94213687-INS-70 | chrX-94213687-INS-118 |
| chrX-94248317-INS-2736 | chrX-94248318-INS-2737 |
| chrX-94252150-INS-1476 | chrX-94252151-INS-1476 |
| chrX-94285706-INS-90 | chrX-94285706-INS-90 |
| chrX-94335138-INS-416 | chrX-94335139-INS-416 |
| chrX-94336317-INS-2162 | chrX-94336318-INS-2162 |
| chrX-94342920-INS-72 | chrX-94342920-INS-63 |
| chrX-94353647-INS-214 | chrX-94353647-INS-214 |
| chrX-94428693-INS-52 | chrX-94428699-INS-54 |
| chrX-94470890-INS-60 | chrX-94470890-INS-66 |
| chrX-94485585-INS-376 | chrX-94485585-INS-379 |
| chrX-94517243-INS-136 | chrX-94517243-INS-136 |
| chrX-94557397-INS-1010 | chrX-94557392-INS-1025 |
| chrX-94566492-INS-192 | chrX-94566492-INS-196 |
| chrX-94568521-INS-3091 | chrX-94568522-INS-3091 |
| chrX-94577416-INS-5224 | chrX-94577417-INS-5224 |
| chrX-94608869-INS-78 | chrX-94608871-INS-68 |
| chrX-94618828-INS-1823 | chrX-94618829-INS-1823 |
| chrX-94620001-INS-62 | chrX-94620001-INS-60 |
| chrX-94636029-INS-52 | chrX-94636052-INS-78 |
| chrX-94641165-INS-51 | chrX-94641165-INS-54 |
| chrX-94649382-INS-61 | chrX-94649382-INS-61 |
| chrX-94665130-INS-440 | chrX-94665130-INS-440 |
| chrX-94693584-INS-1705 | chrX-94693585-INS-1705 |
| chrX-94747840-INS-2192 | chrX-94747841-INS-2195 |
| chrX-94773131-INS-107 | chrX-94773131-INS-107 |
| chrX-94785350-INS-125 | chrX-94785350-INS-125 |
| chrX-94787377-INS-208 | chrX-94787377-INS-208 |
| chrX-94820505-INS-85 | chrX-94820505-INS-52 |
| chrX-94844582-INS-3761 | chrX-94844583-INS-3757 |
| chrX-94903805-INS-201 | chrX-94903805-INS-201 |
| chrX-94934756-INS-52 | chrX-94934756-INS-52 |
| chrX-94967414-INS-50 | chrX-94967408-INS-50 |
| chrX-94983261-INS-208 | chrX-94983261-INS-208 |
| chrX-95016270-INS-498 | chrX-95016270-INS-498 |
| chrX-95020115-INS-4400 | chrX-95020116-INS-4400 |
| chrX-95035525-INS-120 | chrX-95035525-INS-120 |
| chrX-95039766-INS-486 | chrX-95039766-INS-486 |
| chrX-95042628-INS-137 | chrX-95042628-INS-141 |
| chrX-95062911-INS-1172 | chrX-95062912-INS-1172 |
| chrX-95078638-INS-394 | chrX-95078638-INS-378 |

| | |
|---|---|
| chrX-95091974-INS-54 | chrX-95091974-INS-52 |
| chrX-95114438-INS-76 | chrX-95114438-INS-80 |
| chrX-95123070-INS-908 | chrX-95123070-INS-907 |
| chrX-95207875-INS-61 | chrX-95207875-INS-60 |
| chrX-95237098-INS-1890 | chrX-95237099-INS-1890 |
| chrX-95307547-INS-200 | chrX-95307547-INS-200 |
| chrX-95319621-INS-2735 | chrX-95319622-INS-2732 |
| chrX-95359428-INS-139 | chrX-95359428-INS-138 |
| chrX-95383687-INS-52 | chrX-95383687-INS-60 |
| chrX-95459017-INS-1820 | chrX-95459018-INS-1820 |
| chrX-95485255-INS-959 | chrX-95485255-INS-958 |
| chrX-95514958-INS-313 | chrX-95514958-INS-312 |
| chrX-95606057-INS-84 | chrX-95606057-INS-84 |
| chrX-95610149-INS-470 | chrX-95610149-INS-470 |
| chrX-95642293-INS-6177 | chrX-95642294-INS-6177 |
| chrX-95664385-INS-465 | chrX-95664385-INS-465 |
| chrX-95664674-INS-7133 | chrX-95664675-INS-7135 |
| chrX-95678431-INS-66 | chrX-95678431-INS-66 |
| chrX-95697653-INS-787 | chrX-95697653-INS-788 |
| chrX-95741088-INS-188 | chrX-95741088-INS-213 |
| chrX-95800147-INS-1916 | chrX-95800148-INS-1917 |
| chrX-95812283-INS-54 | chrX-95812281-INS-52 |
| chrX-95947497-INS-52 | chrX-95947497-INS-52 |
| chrX-95961875-INS-659 | chrX-95961875-INS-659 |
| chrX-95993933-INS-69 | chrX-95993933-INS-50 |
| chrX-96136856-INS-367 | chrX-96136856-INS-367 |
| chrX-96151027-INS-1842 | chrX-96151028-INS-1842 |
| chrX-96161008-INS-954 | chrX-96161008-INS-955 |
| chrX-96208803-INS-1354 | chrX-96208804-INS-1355 |
| chrX-96226349-INS-76 | chrX-96226349-INS-68 |
| chrX-96245378-INS-1610 | chrX-96245379-INS-1610 |
| chrX-96287176-INS-399 | chrX-96287176-INS-399 |
| chrX-96295959-INS-115 | chrX-96296001-INS-67 |
| chrX-96342294-INS-421 | chrX-96342305-INS-451 |
| chrX-96400025-INS-102 | chrX-96400025-INS-102 |
| chrX-96442220-INS-399 | chrX-96442220-INS-399 |
| chrX-96450804-INS-2749 | chrX-96450805-INS-2749 |
| chrX-96472873-INS-730 | chrX-96472873-INS-730 |
| chrX-96486583-INS-157 | chrX-96486583-INS-157 |
| chrX-96515336-INS-456 | chrX-96515336-INS-464 |
| chrX-96518718-INS-537 | chrX-96518718-INS-536 |
| chrX-96538261-INS-95 | chrX-96538261-INS-95 |
| chrX-96592792-INS-661 | chrX-96592792-INS-666 |
| chrX-96611780-INS-390 | chrX-96611780-INS-390 |
| chrX-96674306-INS-77 | chrX-96674306-INS-75 |
| chrX-96682916-INS-203 | chrX-96682916-INS-203 |
| chrX-96687892-INS-77 | chrX-96687892-INS-77 |
| chrX-96710017-INS-443 | chrX-96710017-INS-421 |
| chrX-96753693-INS-292 | chrX-96753693-INS-292 |
| chrX-96770862-INS-70 | chrX-96770862-INS-70 |
| chrX-96819962-INS-6248 | chrX-96819963-INS-6248 |
| chrX-96845168-INS-1289 | chrX-96845169-INS-1288 |
| chrX-96877235-INS-214 | chrX-96877235-INS-214 |
| chrX-96893604-INS-100 | chrX-96893604-INS-100 |
| chrX-97042338-INS-69 | chrX-97042348-INS-61 |
| chrX-97113781-INS-85 | chrX-97113781-INS-84 |
| chrX-97224398-INS-1916 | chrX-97224399-INS-1914 |
| chrX-97289029-INS-65 | chrX-97289029-INS-65 |
| chrX-97290851-INS-2228 | chrX-97290852-INS-2228 |
| chrX-97317953-INS-6662 | chrX-97317954-INS-6661 |
| chrX-97474146-INS-536 | chrX-97474146-INS-536 |
| chrX-97494558-INS-6437 | chrX-97494559-INS-6437 |
| chrX-97540759-INS-187 | chrX-97540759-INS-187 |
| chrX-97580409-INS-430 | chrX-97580409-INS-430 |
| chrX-97588514-INS-78 | chrX-97588514-INS-78 |
| chrX-97595109-INS-55 | chrX-97595109-INS-57 |
| chrX-97645491-INS-349 | chrX-97645491-INS-349 |
| chrX-97680471-INS-96 | chrX-97680471-INS-103 |
| chrX-97715527-INS-319 | chrX-97715527-INS-317 |
| chrX-97768206-INS-53 | chrX-97768204-INS-78 |
| chrX-97785241-INS-268 | chrX-97785242-INS-268 |
| chrX-97857314-INS-2459 | chrX-97857315-INS-2460 |
| chrX-97863581-INS-231 | chrX-97863581-INS-231 |
| chrX-97866520-INS-200 | chrX-97866520-INS-204 |
| chrX-97877165-INS-448 | chrX-97877165-INS-448 |
| chrX-97887362-INS-54 | chrX-97887362-INS-54 |
| chrX-97888482-INS-164 | chrX-97888482-INS-180 |

| | |
|---|---|
| chrX-97892032-INS-1668 | chrX-97892033-INS-1668 |
| chrX-97898727-INS-327 | chrX-97898767-INS-324 |
| chrX-97941103-INS-233 | chrX-97941103-INS-234 |
| chrX-97946371-INS-75 | chrX-97946400-INS-84 |
| chrX-97947367-INS-4697 | chrX-97947368-INS-4697 |
| chrX-97959199-INS-479 | chrX-97959209-INS-462 |
| chrX-97976587-INS-52 | chrX-97976587-INS-52 |
| chrX-97981135-INS-250 | chrX-97981135-INS-250 |
| chrX-98000876-INS-60 | chrX-98000877-INS-59 |
| chrX-98006383-INS-198 | chrX-98006383-INS-198 |
| chrX-98026968-INS-138 | chrX-98026968-INS-126 |
| chrX-98127854-INS-627 | chrX-98127854-INS-625 |
| chrX-98128406-INS-98 | chrX-98128408-INS-113 |
| chrX-98129845-INS-71 | chrX-98129845-INS-96 |
| chrX-98237885-INS-77 | chrX-98237886-INS-90 |
| chrX-98358251-INS-90 | chrX-98358251-INS-90 |
| chrX-98395226-INS-4731 | chrX-98395227-INS-4731 |
| chrX-98436074-INS-66 | chrX-98436074-INS-68 |
| chrX-98483008-INS-59 | chrX-98483008-INS-81 |
| chrX-98537269-INS-92 | chrX-98537269-INS-122 |
| chrX-98566810-INS-55 | chrX-98566810-INS-55 |
| chrX-98573964-INS-206 | chrX-98573964-INS-206 |
| chrX-98604661-INS-88 | chrX-98604661-INS-88 |
| chrX-98647233-INS-54 | chrX-98647233-INS-54 |
| chrX-98655215-INS-83 | chrX-98655215-INS-85 |
| chrX-98701144-INS-75 | chrX-98701144-INS-51 |
| chrX-98705143-INS-79 | chrX-98705167-INS-65 |
| chrX-98730426-INS-70 | chrX-98730426-INS-52 |
| chrX-98760651-INS-209 | chrX-98760651-INS-209 |
| chrX-98770762-INS-59 | chrX-98770761-INS-100 |
| chrX-98787090-INS-59 | chrX-98787090-INS-59 |
| chrX-98821904-INS-2841 | chrX-98821905-INS-2843 |
| chrX-98903577-INS-1575 | chrX-98903578-INS-1599 |
| chrX-98922707-INS-1866 | chrX-98922708-INS-1866 |
| chrX-98924955-INS-63 | chrX-98924957-INS-60 |
| chrX-98969099-INS-115 | chrX-98969099-INS-93 |
| chrX-98969695-INS-201 | chrX-98969695-INS-201 |
| chrX-98976116-INS-7357 | chrX-98976117-INS-7346 |
| chrX-99015591-INS-2944 | chrX-99015592-INS-2944 |
| chrX-99052949-INS-533 | chrX-99052949-INS-533 |
| chrX-99083724-INS-475 | chrX-99083724-INS-475 |
| chrX-99094162-INS-216 | chrX-99094162-INS-216 |
| chrX-99096069-INS-1146 | chrX-99096070-INS-1146 |
| chrX-99102416-INS-531 | chrX-99102416-INS-531 |
| chrX-99181645-INS-197 | chrX-99181645-INS-198 |
| chrX-99202284-INS-582 | chrX-99202284-INS-583 |
| chrX-99255168-INS-470 | chrX-99255168-INS-470 |
| chrX-99259844-INS-52 | chrX-99259844-INS-52 |
| chrX-99268687-INS-77 | chrX-99268687-INS-79 |
| chrX-99270462-INS-81 | chrX-99270462-INS-72 |
| chrX-99329521-INS-789 | chrX-99329521-INS-789 |
| chrX-99425436-INS-214 | chrX-99425436-INS-215 |
| chrX-99500673-INS-1827 | chrX-99500674-INS-1827 |
| chrX-99505848-INS-179 | chrX-99505848-INS-182 |
| chrX-99548410-INS-67 | chrX-99548459-INS-67 |
| chrX-99574275-INS-207 | chrX-99574275-INS-212 |
| chrX-99640811-INS-93 | chrX-99640811-INS-93 |
| chrX-99651611-INS-104 | chrX-99651616-INS-72 |
| chrX-99657814-INS-60 | chrX-99657814-INS-60 |
| chrX-99688772-INS-6890 | chrX-99688773-INS-6891 |
| chrX-99720008-INS-66 | chrX-99720008-INS-66 |
| chrX-99724607-INS-198 | chrX-99724607-INS-198 |
| chrX-99726244-INS-284 | chrX-99726244-INS-284 |
| chrX-99726694-INS-875 | chrX-99726694-INS-881 |
| chrX-99735532-INS-194 | chrX-99735532-INS-195 |
| chrX-99761312-INS-219 | chrX-99761312-INS-219 |
| chrX-99844733-INS-124 | chrX-99844758-INS-88 |
| chrX-99891021-INS-853 | chrX-99891021-INS-853 |
| chrX-99932169-INS-504 | chrX-99932169-INS-504 |
| chrX-99984464-INS-785 | chrX-99984464-INS-785 |
| chrX-100013062-INS-3315 | chrX-100013063-INS-3314 |
| chrX-100027531-INS-491 | chrX-100027531-INS-491 |
| chrX-100035834-INS-52 | chrX-100035857-INS-54 |
| chrX-100038768-INS-66 | chrX-100038768-INS-64 |
| chrX-100043779-INS-122 | chrX-100043779-INS-121 |
| chrX-100053650-INS-70 | chrX-100053627-INS-72 |
| chrX-100104822-INS-136 | chrX-100104829-INS-95 |

| | |
|---|---|
| chrX-100112795-INS-757 | chrX-100112795-INS-757 |
| chrX-100155185-INS-5217 | chrX-100155186-INS-5217 |
| chrX-100157965-INS-52 | chrX-100157966-INS-56 |
| chrX-100169193-INS-170 | chrX-100169197-INS-174 |
| chrX-100178603-INS-112 | chrX-100178603-INS-112 |
| chrX-100195748-INS-195 | chrX-100195748-INS-195 |
| chrX-100203731-INS-1921 | chrX-100203732-INS-1922 |
| chrX-100314472-INS-98 | chrX-100314472-INS-98 |
| chrX-100322548-INS-173 | chrX-100322548-INS-173 |
| chrX-100323422-INS-79 | chrX-100323422-INS-88 |
| chrX-100330973-INS-206 | chrX-100330973-INS-206 |
| chrX-100374306-INS-202 | chrX-100374306-INS-200 |
| chrX-100378393-INS-163 | chrX-100378393-INS-163 |
| chrX-100382294-INS-52 | chrX-100382294-INS-56 |
| chrX-100395081-INS-97 | chrX-100395081-INS-97 |
| chrX-100399233-INS-94 | chrX-100399233-INS-52 |
| chrX-100406766-INS-107 | chrX-100406775-INS-80 |
| chrX-100446759-INS-168 | chrX-100446759-INS-170 |
| chrX-100493932-INS-227 | chrX-100493932-INS-227 |
| chrX-100506170-INS-438 | chrX-100506170-INS-437 |
| chrX-100515363-INS-200 | chrX-100515363-INS-201 |
| chrX-100522824-INS-204 | chrX-100522824-INS-204 |
| chrX-100531722-INS-58 | chrX-100531722-INS-58 |
| chrX-100545890-INS-68 | chrX-100545890-INS-68 |
| chrX-100553531-INS-194 | chrX-100553531-INS-194 |
| chrX-100555993-INS-425 | chrX-100555993-INS-425 |
| chrX-100587002-INS-839 | chrX-100587002-INS-839 |
| chrX-100596528-INS-215 | chrX-100596528-INS-216 |
| chrX-100629196-INS-310 | chrX-100629191-INS-300 |
| chrX-100629769-INS-93 | chrX-100629769-INS-93 |
| chrX-100654785-INS-81 | chrX-100654785-INS-82 |
| chrX-100657327-INS-838 | chrX-100657327-INS-838 |
| chrX-100665989-INS-87 | chrX-100665995-INS-58 |
| chrX-100677717-INS-206 | chrX-100677717-INS-212 |
| chrX-100729131-INS-493 | chrX-100729131-INS-497 |
| chrX-100736077-INS-208 | chrX-100736077-INS-209 |
| chrX-100743150-INS-247 | chrX-100743150-INS-235 |
| chrX-100748331-INS-96 | chrX-100748331-INS-96 |
| chrX-100858706-INS-244 | chrX-100858706-INS-244 |
| chrX-100866589-INS-78 | chrX-100866593-INS-60 |
| chrX-100870800-INS-79 | chrX-100870800-INS-68 |
| chrX-100892848-INS-59 | chrX-100892832-INS-107 |
| chrX-100979005-INS-105 | chrX-100979005-INS-104 |
| chrX-100982459-INS-64 | chrX-100982459-INS-68 |
| chrX-100985601-INS-205 | chrX-100985601-INS-203 |
| chrX-101020386-INS-219 | chrX-101020386-INS-219 |
| chrX-101036839-INS-90 | chrX-101036839-INS-50 |
| chrX-101043925-INS-86 | chrX-101043925-INS-87 |
| chrX-101085832-INS-57 | chrX-101085832-INS-55 |
| chrX-101148759-INS-1477 | chrX-101148760-INS-1477 |
| chrX-101171423-INS-232 | chrX-101171423-INS-227 |
| chrX-101174113-INS-208 | chrX-101174113-INS-213 |
| chrX-101180109-INS-155 | chrX-101180109-INS-155 |
| chrX-101215973-INS-212 | chrX-101215973-INS-212 |
| chrX-101220707-INS-4800 | chrX-101220708-INS-4800 |
| chrX-101224324-INS-277 | chrX-101224324-INS-277 |
| chrX-101252262-INS-4414 | chrX-101252263-INS-4414 |
| chrX-101268260-INS-198 | chrX-101268260-INS-198 |
| chrX-101308552-INS-196 | chrX-101308547-INS-216 |
| chrX-101338548-INS-506 | chrX-101338548-INS-509 |
| chrX-101339956-INS-301 | chrX-101339956-INS-301 |
| chrX-101340813-INS-58 | chrX-101340813-INS-54 |
| chrX-101342153-INS-98 | chrX-101342153-INS-68 |
| chrX-101373695-INS-131 | chrX-101373695-INS-131 |
| chrX-101419070-INS-78 | chrX-101419068-INS-56 |
| chrX-101439451-INS-1458 | chrX-101439452-INS-1462 |
| chrX-101496210-INS-77 | chrX-101496210-INS-77 |
| chrX-101534888-INS-54 | chrX-101534888-INS-54 |
| chrX-101555955-INS-120 | chrX-101555955-INS-120 |
| chrX-101583757-INS-3220 | chrX-101583758-INS-3222 |
| chrX-101592049-INS-158 | chrX-101592049-INS-171 |
| chrX-101604861-INS-624 | chrX-101604862-INS-624 |
| chrX-101620177-INS-217 | chrX-101620177-INS-198 |
| chrX-101638183-INS-923 | chrX-101638183-INS-908 |
| chrX-101638937-INS-193 | chrX-101638937-INS-192 |
| chrX-101641339-INS-319 | chrX-101641339-INS-316 |
| chrX-101795269-INS-80 | chrX-101795269-INS-130 |

| | |
|---|---|
| chrX-101832563-INS-447 | chrX-101832563-INS-447 |
| chrX-101834478-INS-320 | chrX-101834478-INS-320 |
| chrX-101944109-INS-59 | chrX-101944109-INS-59 |
| chrX-101994618-INS-462 | chrX-101994618-INS-462 |
| chrX-102021809-INS-1227 | chrX-102021810-INS-1227 |
| chrX-102030499-INS-2612 | chrX-102030500-INS-2612 |
| chrX-102046876-INS-238 | chrX-102046876-INS-267 |
| chrX-102051465-INS-112 | chrX-102051465-INS-111 |
| chrX-102066405-INS-86 | chrX-102066417-INS-58 |
| chrX-102116660-INS-111 | chrX-102116660-INS-111 |
| chrX-102141531-INS-2820 | chrX-102141532-INS-2820 |
| chrX-102219260-INS-201 | chrX-102219260-INS-203 |
| chrX-102231726-INS-72 | chrX-102231726-INS-76 |
| chrX-102242129-INS-245 | chrX-102242173-INS-219 |
| chrX-102271276-INS-218 | chrX-102271276-INS-218 |
| chrX-102277088-INS-597 | chrX-102277088-INS-597 |
| chrX-102278714-INS-55 | chrX-102278699-INS-75 |
| chrX-102284795-INS-187 | chrX-102284795-INS-186 |
| chrX-102291757-INS-402 | chrX-102291757-INS-405 |
| chrX-102293357-INS-198 | chrX-102293357-INS-198 |
| chrX-102304714-INS-66 | chrX-102304714-INS-66 |
| chrX-102316118-INS-153 | chrX-102316118-INS-147 |
| chrX-102320227-INS-70 | chrX-102320193-INS-99 |
| chrX-102340394-INS-228 | chrX-102340394-INS-229 |
| chrX-102343247-INS-7116 | chrX-102343248-INS-7116 |
| chrX-102357250-INS-231 | chrX-102357250-INS-232 |
| chrX-102381514-INS-58 | chrX-102381514-INS-59 |
| chrX-102398178-INS-190 | chrX-102398178-INS-190 |
| chrX-102404416-INS-208 | chrX-102404416-INS-205 |
| chrX-102410995-INS-203 | chrX-102410995-INS-201 |
| chrX-102431460-INS-239 | chrX-102431460-INS-238 |
| chrX-102457802-INS-209 | chrX-102457802-INS-209 |
| chrX-102472533-INS-202 | chrX-102472533-INS-203 |
| chrX-102477308-INS-207 | chrX-102477308-INS-231 |
| chrX-102492303-INS-203 | chrX-102492303-INS-203 |
| chrX-102551240-INS-100 | chrX-102551240-INS-90 |
| chrX-102578227-INS-204 | chrX-102578227-INS-204 |
| chrX-102588487-INS-191 | chrX-102588487-INS-193 |
| chrX-102601685-INS-112 | chrX-102601684-INS-115 |
| chrX-102602885-INS-198 | chrX-102602885-INS-197 |
| chrX-102609336-INS-62 | chrX-102609336-INS-62 |
| chrX-102638724-INS-623 | chrX-102638724-INS-617 |
| chrX-102651741-INS-5555 | chrX-102651742-INS-5555 |
| chrX-102697420-INS-64 | chrX-102697420-INS-64 |
| chrX-102708358-INS-485 | chrX-102708358-INS-491 |
| chrX-102715115-INS-5143 | chrX-102715116-INS-5143 |
| chrX-102721029-INS-92 | chrX-102721029-INS-83 |
| chrX-102729782-INS-546 | chrX-102729782-INS-546 |
| chrX-102733521-INS-64 | chrX-102733521-INS-68 |
| chrX-102733681-INS-470 | chrX-102733681-INS-470 |
| chrX-102767187-INS-525 | chrX-102767187-INS-525 |
| chrX-102776749-INS-208 | chrX-102776749-INS-208 |
| chrX-102804723-INS-536 | chrX-102804723-INS-543 |
| chrX-102827244-INS-141 | chrX-102827244-INS-141 |
| chrX-102842993-INS-85 | chrX-102843001-INS-70 |
| chrX-102954072-INS-2140 | chrX-102954073-INS-2145 |
| chrX-102955718-INS-651 | chrX-102955719-INS-651 |
| chrX-102963933-INS-259 | chrX-102963933-INS-259 |
| chrX-102971344-INS-200 | chrX-102971344-INS-200 |
| chrX-102976334-INS-112 | chrX-102976334-INS-112 |
| chrX-102984064-INS-341 | chrX-102984064-INS-341 |
| chrX-102992431-INS-5129 | chrX-102992432-INS-5129 |
| chrX-102992985-INS-204 | chrX-102992985-INS-201 |
| chrX-102995992-INS-516 | chrX-102995992-INS-516 |
| chrX-102998935-INS-204 | chrX-102998935-INS-204 |
| chrX-103015407-INS-66 | chrX-103015407-INS-66 |
| chrX-103034561-INS-88 | chrX-103034561-INS-88 |
| chrX-103075607-INS-56 | chrX-103075607-INS-54 |
| chrX-103114655-INS-114 | chrX-103114655-INS-116 |
| chrX-103119317-INS-175 | chrX-103119317-INS-174 |
| chrX-103142755-INS-220 | chrX-103142755-INS-220 |
| chrX-103199441-INS-202 | chrX-103199441-INS-202 |
| chrX-103203023-INS-226 | chrX-103203023-INS-225 |
| chrX-103204073-INS-209 | chrX-103204073-INS-209 |
| chrX-103247152-INS-202 | chrX-103247152-INS-202 |
| chrX-103263975-INS-1810 | chrX-103263976-INS-1794 |
| chrX-103279159-INS-86 | chrX-103279130-INS-70 |

| | |
|---|---|
| chrX-103282172-INS-226 | chrX-103282172-INS-226 |
| chrX-103290300-INS-208 | chrX-103290300-INS-208 |
| chrX-103295051-INS-63 | chrX-103295016-INS-104 |
| chrX-103319373-INS-255 | chrX-103319373-INS-255 |
| chrX-103327971-INS-181 | chrX-103327971-INS-181 |
| chrX-103336829-INS-70 | chrX-103336829-INS-78 |
| chrX-103339168-INS-6215 | chrX-103339169-INS-6216 |
| chrX-103341241-INS-209 | chrX-103341241-INS-208 |
| chrX-103401460-INS-213 | chrX-103401460-INS-213 |
| chrX-103402472-INS-216 | chrX-103402472-INS-218 |
| chrX-103430057-INS-78 | chrX-103430057-INS-78 |
| chrX-103455077-INS-640 | chrX-103455077-INS-641 |
| chrX-103490973-INS-489 | chrX-103490973-INS-474 |
| chrX-103567429-INS-864 | chrX-103567429-INS-865 |
| chrX-103753528-INS-435 | chrX-103753528-INS-434 |
| chrX-103775354-INS-74 | chrX-103775354-INS-74 |
| chrX-103817148-INS-6243 | chrX-103817149-INS-6244 |
| chrX-103818313-INS-394 | chrX-103818313-INS-394 |
| chrX-103820999-INS-2857 | chrX-103821000-INS-2858 |
| chrX-103830579-INS-1530 | chrX-103830580-INS-1530 |
| chrX-103840520-INS-498 | chrX-103840520-INS-498 |
| chrX-103870858-INS-400 | chrX-103870858-INS-400 |
| chrX-103885709-INS-316 | chrX-103885709-INS-316 |
| chrX-103892063-INS-392 | chrX-103892063-INS-392 |
| chrX-103903337-INS-204 | chrX-103903338-INS-204 |
| chrX-103926815-INS-75 | chrX-103926815-INS-76 |
| chrX-103949979-INS-3079 | chrX-103949980-INS-3089 |
| chrX-103963991-INS-55 | chrX-103963991-INS-50 |
| chrX-103989987-INS-93 | chrX-103989987-INS-93 |
| chrX-104001049-INS-60 | chrX-104001049-INS-72 |
| chrX-104052290-INS-54 | chrX-104052292-INS-69 |
| chrX-104062283-INS-260 | chrX-104062282-INS-259 |
| chrX-104071750-INS-6754 | chrX-104071751-INS-6754 |
| chrX-104081437-INS-88 | chrX-104081437-INS-86 |
| chrX-104093983-INS-588 | chrX-104093983-INS-572 |
| chrX-104111680-INS-977 | chrX-104111680-INS-977 |
| chrX-104116895-INS-3271 | chrX-104116896-INS-3271 |
| chrX-104128288-INS-202 | chrX-104128288-INS-202 |
| chrX-104131763-INS-171 | chrX-104131763-INS-171 |
| chrX-104134541-INS-277 | chrX-104134548-INS-262 |
| chrX-104149018-INS-56 | chrX-104149018-INS-60 |
| chrX-104151204-INS-206 | chrX-104151204-INS-203 |
| chrX-104156244-INS-489 | chrX-104156244-INS-489 |
| chrX-104182593-INS-69 | chrX-104182593-INS-101 |
| chrX-104197238-INS-80 | chrX-104197238-INS-80 |
| chrX-104202699-INS-332 | chrX-104202699-INS-332 |
| chrX-104212645-INS-7164 | chrX-104212646-INS-7164 |
| chrX-104259664-INS-6489 | chrX-104259665-INS-6489 |
| chrX-104290543-INS-764 | chrX-104290544-INS-765 |
| chrX-104305875-INS-801 | chrX-104305875-INS-794 |
| chrX-104313296-INS-284 | chrX-104313312-INS-255 |
| chrX-104323782-INS-67 | chrX-104323782-INS-67 |
| chrX-104462169-INS-2217 | chrX-104462170-INS-2216 |
| chrX-104471540-INS-140 | chrX-104471541-INS-134 |
| chrX-104498157-INS-2834 | chrX-104498158-INS-2832 |
| chrX-104521440-INS-7762 | chrX-104521441-INS-7763 |
| chrX-104575450-INS-197 | chrX-104575450-INS-197 |
| chrX-104607652-INS-72 | chrX-104607652-INS-72 |
| chrX-104609366-INS-6828 | chrX-104609367-INS-6828 |
| chrX-104625412-INS-395 | chrX-104625412-INS-395 |
| chrX-104641666-INS-208 | chrX-104641666-INS-208 |
| chrX-104668420-INS-146 | chrX-104668421-INS-104 |
| chrX-104707277-INS-450 | chrX-104707277-INS-446 |
| chrX-104711355-INS-65 | chrX-104711355-INS-50 |
| chrX-104738738-INS-214 | chrX-104738738-INS-214 |
| chrX-104760368-INS-198 | chrX-104760368-INS-204 |
| chrX-104771438-INS-147 | chrX-104771487-INS-116 |
| chrX-104834473-INS-64 | chrX-104834473-INS-64 |
| chrX-104893306-INS-213 | chrX-104893306-INS-213 |
| chrX-104899851-INS-52 | chrX-104899851-INS-52 |
| chrX-104901666-INS-52 | chrX-104901666-INS-52 |
| chrX-104908487-INS-76 | chrX-104908487-INS-58 |
| chrX-104955799-INS-272 | chrX-104955799-INS-272 |
| chrX-104968994-INS-216 | chrX-104968989-INS-204 |
| chrX-104971085-INS-198 | chrX-104971085-INS-197 |
| chrX-104977666-INS-58 | chrX-104977666-INS-58 |
| chrX-104986567-INS-53 | chrX-104986551-INS-75 |

| | |
|---|---|
| chrX-104996505-INS-59 | chrX-104996505-INS-50 |
| chrX-104998209-INS-5093 | chrX-104998210-INS-5093 |
| chrX-104998425-INS-203 | chrX-104998425-INS-203 |
| chrX-104998722-INS-2038 | chrX-104998723-INS-2038 |
| chrX-105000785-INS-5388 | chrX-105000786-INS-5388 |
| chrX-105001185-INS-253 | chrX-105001185-INS-253 |
| chrX-105012665-INS-130 | chrX-105012665-INS-159 |
| chrX-105039473-INS-593 | chrX-105039473-INS-593 |
| chrX-105069003-INS-180 | chrX-105069003-INS-183 |
| chrX-105111877-INS-498 | chrX-105111877-INS-498 |
| chrX-105140548-INS-57 | chrX-105140548-INS-57 |
| chrX-105152354-INS-191 | chrX-105152354-INS-192 |
| chrX-105162056-INS-347 | chrX-105162056-INS-347 |
| chrX-105235142-INS-316 | chrX-105235142-INS-317 |
| chrX-105256152-INS-741 | chrX-105256152-INS-741 |
| chrX-105270684-INS-98 | chrX-105270702-INS-115 |
| chrX-105282408-INS-6998 | chrX-105282409-INS-6998 |
| chrX-105304628-INS-6557 | chrX-105304629-INS-6557 |
| chrX-105307768-INS-776 | chrX-105307768-INS-776 |
| chrX-105312147-INS-53 | chrX-105312174-INS-53 |
| chrX-105314962-INS-6519 | chrX-105314963-INS-6520 |
| chrX-105317353-INS-1542 | chrX-105317354-INS-1542 |
| chrX-105374209-INS-402 | chrX-105374209-INS-402 |
| chrX-105383699-INS-91 | chrX-105383671-INS-80 |
| chrX-105391802-INS-9706 | chrX-105391803-INS-9706 |
| chrX-105395178-INS-4657 | chrX-105395179-INS-4657 |
| chrX-105398848-INS-2035 | chrX-105398849-INS-2036 |
| chrX-105401320-INS-64 | chrX-105401311-INS-64 |
| chrX-105409828-INS-9140 | chrX-105409829-INS-9141 |
| chrX-105415528-INS-188 | chrX-105415524-INS-191 |
| chrX-105418486-INS-363 | chrX-105418486-INS-363 |
| chrX-105492162-INS-65 | chrX-105492162-INS-65 |
| chrX-105509044-INS-7522 | chrX-105509045-INS-7522 |
| chrX-105534930-INS-176 | chrX-105534930-INS-181 |
| chrX-105561464-INS-88 | chrX-105561464-INS-126 |
| chrX-105633714-INS-386 | chrX-105633714-INS-386 |
| chrX-105669785-INS-347 | chrX-105669785-INS-347 |
| chrX-105693496-INS-161 | chrX-105693496-INS-164 |
| chrX-105696304-INS-6434 | chrX-105696305-INS-6434 |
| chrX-105698828-INS-411 | chrX-105698828-INS-411 |
| chrX-105704329-INS-2811 | chrX-105704330-INS-2811 |
| chrX-105715147-INS-6314 | chrX-105715148-INS-6311 |
| chrX-105729732-INS-997 | chrX-105729732-INS-998 |
| chrX-105742422-INS-165 | chrX-105742422-INS-163 |
| chrX-105765842-INS-80 | chrX-105765842-INS-80 |
| chrX-105766853-INS-810 | chrX-105766853-INS-792 |
| chrX-105769875-INS-968 | chrX-105769919-INS-973 |
| chrX-105791370-INS-2962 | chrX-105791371-INS-2944 |
| chrX-105811911-INS-66 | chrX-105811911-INS-72 |
| chrX-105838221-INS-72 | chrX-105838221-INS-72 |
| chrX-105871379-INS-112 | chrX-105871379-INS-112 |
| chrX-105889136-INS-52 | chrX-105889136-INS-68 |
| chrX-105902330-INS-6427 | chrX-105902331-INS-6445 |
| chrX-105914162-INS-300 | chrX-105914162-INS-308 |
| chrX-105928021-INS-602 | chrX-105928021-INS-602 |
| chrX-105981557-INS-7095 | chrX-105981558-INS-7095 |
| chrX-105986450-INS-83 | chrX-105986450-INS-83 |
| chrX-105996766-INS-389 | chrX-105996766-INS-389 |
| chrX-106001933-INS-570 | chrX-106001933-INS-568 |
| chrX-106021468-INS-7092 | chrX-106021469-INS-7093 |
| chrX-106074980-INS-9806 | chrX-106074981-INS-9807 |
| chrX-106086874-INS-2281 | chrX-106086875-INS-2281 |
| chrX-106108369-INS-702 | chrX-106108369-INS-702 |
| chrX-106116876-INS-86 | chrX-106116876-INS-84 |
| chrX-106172889-INS-396 | chrX-106172889-INS-395 |
| chrX-106179326-INS-6885 | chrX-106179327-INS-6885 |
| chrX-106242543-INS-144 | chrX-106242543-INS-144 |
| chrX-106255863-INS-1593 | chrX-106255864-INS-1593 |
| chrX-106257967-INS-255 | chrX-106257973-INS-256 |
| chrX-106317717-INS-718 | chrX-106317720-INS-720 |
| chrX-106370072-INS-55 | chrX-106370072-INS-65 |
| chrX-106384378-INS-68 | chrX-106384378-INS-68 |
| chrX-106400981-INS-6307 | chrX-106400982-INS-6267 |
| chrX-106407338-INS-157 | chrX-106407338-INS-157 |
| chrX-106412461-INS-62 | chrX-106412461-INS-63 |
| chrX-106435132-INS-68 | chrX-106435132-INS-68 |
| chrX-106454538-INS-6249 | chrX-106454539-INS-6249 |

| | |
|---|---|
| chrX-106480544-INS-58 | chrX-106480544-INS-58 |
| chrX-106498471-INS-72 | chrX-106498471-INS-76 |
| chrX-106523747-INS-2213 | chrX-106523748-INS-2213 |
| chrX-106549786-INS-52 | chrX-106549786-INS-54 |
| chrX-106590506-INS-151 | chrX-106590506-INS-151 |
| chrX-106606456-INS-344 | chrX-106606456-INS-344 |
| chrX-106607667-INS-3687 | chrX-106607668-INS-3684 |
| chrX-106630782-INS-2672 | chrX-106630783-INS-2696 |
| chrX-106645357-INS-1016 | chrX-106645358-INS-1020 |
| chrX-106687600-INS-828 | chrX-106687600-INS-828 |
| chrX-106693222-INS-6203 | chrX-106693223-INS-6203 |
| chrX-106742652-INS-3838 | chrX-106742653-INS-3838 |
| chrX-106745665-INS-68 | chrX-106745665-INS-68 |
| chrX-106760839-INS-155 | chrX-106760839-INS-155 |
| chrX-106766918-INS-204 | chrX-106766918-INS-204 |
| chrX-106833770-INS-482 | chrX-106833770-INS-482 |
| chrX-106845694-INS-501 | chrX-106845694-INS-501 |
| chrX-106871677-INS-55 | chrX-106871677-INS-68 |
| chrX-106878437-INS-745 | chrX-106878437-INS-745 |
| chrX-106889160-INS-566 | chrX-106889160-INS-566 |
| chrX-106925497-INS-75 | chrX-106925497-INS-75 |
| chrX-106933754-INS-6500 | chrX-106933755-INS-6501 |
| chrX-106954706-INS-6714 | chrX-106954707-INS-6715 |
| chrX-106961989-INS-70 | chrX-106961989-INS-74 |
| chrX-107040975-INS-2953 | chrX-107040976-INS-2953 |
| chrX-107057142-INS-4349 | chrX-107057143-INS-4351 |
| chrX-107065101-INS-233 | chrX-107065101-INS-233 |
| chrX-107081137-INS-92 | chrX-107081137-INS-72 |
| chrX-107084000-INS-663 | chrX-107084000-INS-665 |
| chrX-107122022-INS-4127 | chrX-107122023-INS-4127 |
| chrX-107125127-INS-188 | chrX-107125127-INS-188 |
| chrX-107144411-INS-1661 | chrX-107144412-INS-1675 |
| chrX-107171946-INS-54 | chrX-107171944-INS-71 |
| chrX-107173760-INS-3654 | chrX-107173761-INS-3655 |
| chrX-107190001-INS-86 | chrX-107189995-INS-62 |
| chrX-107193154-INS-125 | chrX-107193139-INS-140 |
| chrX-107205929-INS-1136 | chrX-107205930-INS-1136 |
| chrX-107208805-INS-3258 | chrX-107208806-INS-3253 |
| chrX-107242846-INS-361 | chrX-107242846-INS-362 |
| chrX-107276567-INS-627 | chrX-107276567-INS-626 |
| chrX-107281924-INS-479 | chrX-107281917-INS-478 |
| chrX-107285727-INS-52 | chrX-107285727-INS-96 |
| chrX-107329980-INS-59 | chrX-107329980-INS-59 |
| chrX-107333287-INS-2819 | chrX-107333288-INS-2819 |
| chrX-107365078-INS-2875 | chrX-107365079-INS-2875 |
| chrX-107375056-INS-54 | chrX-107375056-INS-54 |
| chrX-107491698-INS-254 | chrX-107491698-INS-254 |
| chrX-107495739-INS-81 | chrX-107495739-INS-81 |
| chrX-107630540-INS-199 | chrX-107630543-INS-199 |
| chrX-107676848-INS-1358 | chrX-107676849-INS-1358 |
| chrX-107729948-INS-3984 | chrX-107729949-INS-3966 |
| chrX-107748592-INS-6541 | chrX-107748593-INS-6547 |
| chrX-107754231-INS-7014 | chrX-107754232-INS-7014 |
| chrX-107757902-INS-60 | chrX-107757913-INS-87 |
| chrX-107758002-INS-1292 | chrX-107758003-INS-1292 |
| chrX-107770734-INS-66 | chrX-107770734-INS-93 |
| chrX-107788344-INS-54 | chrX-107788344-INS-56 |
| chrX-107806274-INS-58 | chrX-107806274-INS-56 |
| chrX-107817684-INS-56 | chrX-107817684-INS-58 |
| chrX-107821519-INS-211 | chrX-107821519-INS-211 |
| chrX-107849568-INS-64 | chrX-107849568-INS-56 |
| chrX-107874309-INS-120 | chrX-107874309-INS-112 |
| chrX-107934934-INS-3569 | chrX-107934935-INS-3569 |
| chrX-107994845-INS-5755 | chrX-107994846-INS-5755 |
| chrX-108008512-INS-74 | chrX-108008512-INS-74 |
| chrX-108032474-INS-6649 | chrX-108032475-INS-6650 |
| chrX-108043085-INS-121 | chrX-108043085-INS-121 |
| chrX-108059706-INS-57 | chrX-108059706-INS-69 |
| chrX-108071216-INS-52 | chrX-108071216-INS-52 |
| chrX-108118958-INS-1363 | chrX-108118959-INS-1363 |
| chrX-108126455-INS-7097 | chrX-108126456-INS-7097 |
| chrX-108129151-INS-999 | chrX-108129151-INS-994 |
| chrX-108156903-INS-181 | chrX-108156903-INS-171 |
| chrX-108175022-INS-61 | chrX-108175023-INS-58 |
| chrX-108191264-INS-55 | chrX-108191264-INS-71 |
| chrX-108197621-INS-175 | chrX-108197621-INS-167 |
| chrX-108203533-INS-305 | chrX-108203533-INS-306 |

| | |
|---|---|
| chrX-108249557-INS-59 | chrX-108249557-INS-59 |
| chrX-108338221-INS-4952 | chrX-108338222-INS-4952 |
| chrX-108343191-INS-561 | chrX-108343191-INS-561 |
| chrX-108346450-INS-5380 | chrX-108346451-INS-5380 |
| chrX-108380203-INS-297 | chrX-108380202-INS-303 |
| chrX-108391594-INS-474 | chrX-108391594-INS-474 |
| chrX-108402068-INS-3175 | chrX-108402069-INS-3175 |
| chrX-108450189-INS-1890 | chrX-108450190-INS-1890 |
| chrX-108463924-INS-6698 | chrX-108463925-INS-6698 |
| chrX-108471872-INS-6719 | chrX-108471873-INS-6721 |
| chrX-108495921-INS-1228 | chrX-108495922-INS-1228 |
| chrX-108503671-INS-106 | chrX-108503660-INS-139 |
| chrX-108517595-INS-214 | chrX-108517595-INS-214 |
| chrX-108529038-INS-876 | chrX-108529038-INS-876 |
| chrX-108537736-INS-6402 | chrX-108537737-INS-6401 |
| chrX-108550083-INS-126 | chrX-108550083-INS-156 |
| chrX-108591640-INS-582 | chrX-108591640-INS-582 |
| chrX-108597191-INS-7092 | chrX-108597192-INS-7092 |
| chrX-108668466-INS-2545 | chrX-108668467-INS-2541 |
| chrX-108762410-INS-112 | chrX-108762410-INS-112 |
| chrX-108787679-INS-104 | chrX-108787679-INS-104 |
| chrX-108822750-INS-7088 | chrX-108822751-INS-7088 |
| chrX-108841742-INS-6358 | chrX-108841743-INS-6359 |
| chrX-108844125-INS-56 | chrX-108844147-INS-57 |
| chrX-108846673-INS-1460 | chrX-108846668-INS-1460 |
| chrX-108885385-INS-202 | chrX-108885385-INS-203 |
| chrX-108912465-INS-2157 | chrX-108912466-INS-2157 |
| chrX-108968404-INS-410 | chrX-108968404-INS-410 |
| chrX-108975794-INS-95 | chrX-108975794-INS-55 |
| chrX-109009417-INS-428 | chrX-109009417-INS-428 |
| chrX-109014586-INS-62 | chrX-109014586-INS-54 |
| chrX-109033472-INS-3307 | chrX-109033473-INS-3307 |
| chrX-109058217-INS-1052 | chrX-109058218-INS-1052 |
| chrX-109068987-INS-51 | chrX-109068963-INS-58 |
| chrX-109073525-INS-346 | chrX-109073525-INS-346 |
| chrX-109090663-INS-4500 | chrX-109090664-INS-4500 |
| chrX-109141677-INS-121 | chrX-109141677-INS-141 |
| chrX-109149317-INS-63 | chrX-109149317-INS-63 |
| chrX-109241997-INS-3240 | chrX-109241998-INS-3218 |
| chrX-109259510-INS-91 | chrX-109259492-INS-84 |
| chrX-109278607-INS-6465 | chrX-109278608-INS-6465 |
| chrX-109282961-INS-2799 | chrX-109282962-INS-2779 |
| chrX-109282962-INS-62 | chrX-109282962-INS-60 |
| chrX-109284841-INS-1903 | chrX-109284842-INS-1903 |
| chrX-109317544-INS-5313 | chrX-109317545-INS-5313 |
| chrX-109324541-INS-139 | chrX-109324541-INS-139 |
| chrX-109407345-INS-6385 | chrX-109407346-INS-6385 |
| chrX-109420916-INS-169 | chrX-109420916-INS-169 |
| chrX-109422893-INS-381 | chrX-109422893-INS-381 |
| chrX-109431927-INS-9857 | chrX-109431928-INS-9857 |
| chrX-109454765-INS-77 | chrX-109454765-INS-72 |
| chrX-109514772-INS-69 | chrX-109514772-INS-62 |
| chrX-109538011-INS-477 | chrX-109538011-INS-477 |
| chrX-109582643-INS-557 | chrX-109582643-INS-557 |
| chrX-109590266-INS-2885 | chrX-109590267-INS-2886 |
| chrX-109608793-INS-221 | chrX-109608793-INS-237 |
| chrX-109635858-INS-2871 | chrX-109635859-INS-2917 |
| chrX-109652136-INS-87 | chrX-109652120-INS-92 |
| chrX-109653246-INS-1128 | chrX-109653247-INS-1129 |
| chrX-109664298-INS-6233 | chrX-109664299-INS-6233 |
| chrX-109676307-INS-570 | chrX-109676305-INS-612 |
| chrX-109700392-INS-1289 | chrX-109700393-INS-1289 |
| chrX-109724203-INS-1501 | chrX-109724204-INS-1503 |
| chrX-109724702-INS-3029 | chrX-109724703-INS-3029 |
| chrX-109793073-INS-184 | chrX-109793073-INS-184 |
| chrX-109802660-INS-1942 | chrX-109802673-INS-1935 |
| chrX-109819473-INS-2104 | chrX-109819474-INS-2104 |
| chrX-109885957-INS-1135 | chrX-109885958-INS-1135 |
| chrX-109907920-INS-5313 | chrX-109907921-INS-5313 |
| chrX-109941095-INS-1893 | chrX-109941096-INS-1893 |
| chrX-110017148-INS-658 | chrX-110017148-INS-654 |
| chrX-110027364-INS-1440 | chrX-110027365-INS-1440 |
| chrX-110110110-INS-7227 | chrX-110110111-INS-7227 |
| chrX-110183254-INS-1316 | chrX-110183255-INS-1321 |
| chrX-110188400-INS-68 | chrX-110188400-INS-68 |
| chrX-110232539-INS-55 | chrX-110232547-INS-57 |
| chrX-110241566-INS-203 | chrX-110241566-INS-203 |

| | |
|---|---|
| chrX-110248422-INS-2290 | chrX-110248423-INS-2294 |
| chrX-110260552-INS-77 | chrX-110260552-INS-81 |
| chrX-110267264-INS-1543 | chrX-110267265-INS-1542 |
| chrX-110275516-INS-55 | chrX-110275516-INS-55 |
| chrX-110285087-INS-56 | chrX-110285087-INS-56 |
| chrX-110289430-INS-114 | chrX-110289430-INS-114 |
| chrX-110292852-INS-61 | chrX-110292852-INS-65 |
| chrX-110296645-INS-381 | chrX-110296645-INS-381 |
| chrX-110300123-INS-60 | chrX-110300123-INS-60 |
| chrX-110347324-INS-395 | chrX-110347324-INS-395 |
| chrX-110386810-INS-54 | chrX-110386810-INS-56 |
| chrX-110406365-INS-208 | chrX-110406365-INS-208 |
| chrX-110461643-INS-107 | chrX-110461642-INS-85 |
| chrX-110462328-INS-1679 | chrX-110462329-INS-1679 |
| chrX-110499988-INS-50 | chrX-110499988-INS-50 |
| chrX-110575751-INS-6304 | chrX-110575752-INS-6305 |
| chrX-110578008-INS-932 | chrX-110578009-INS-931 |
| chrX-110605068-INS-1118 | chrX-110605069-INS-1118 |
| chrX-110641121-INS-2971 | chrX-110641122-INS-2971 |
| chrX-110673917-INS-100 | chrX-110673917-INS-111 |
| chrX-110676728-INS-3545 | chrX-110676729-INS-3546 |
| chrX-110689951-INS-2521 | chrX-110689952-INS-2521 |
| chrX-110698836-INS-56 | chrX-110698836-INS-66 |
| chrX-110714794-INS-6753 | chrX-110714795-INS-6756 |
| chrX-110750352-INS-538 | chrX-110750352-INS-538 |
| chrX-110924626-INS-1184 | chrX-110924627-INS-1184 |
| chrX-110933312-INS-68 | chrX-110933312-INS-68 |
| chrX-110940711-INS-2798 | chrX-110940712-INS-2798 |
| chrX-110975157-INS-52 | chrX-110975157-INS-50 |
| chrX-111027720-INS-56 | chrX-111027720-INS-56 |
| chrX-111032726-INS-1101 | chrX-111032727-INS-1101 |
| chrX-111067870-INS-592 | chrX-111067870-INS-592 |
| chrX-111319225-INS-56 | chrX-111319225-INS-54 |
| chrX-111326256-INS-72 | chrX-111326256-INS-72 |
| chrX-111345016-INS-199 | chrX-111345016-INS-199 |
| chrX-111355879-INS-726 | chrX-111355879-INS-726 |
| chrX-111373122-INS-390 | chrX-111373122-INS-390 |
| chrX-111447460-INS-141 | chrX-111447460-INS-141 |
| chrX-111565562-INS-990 | chrX-111565562-INS-990 |
| chrX-111580137-INS-1040 | chrX-111580138-INS-1041 |
| chrX-111599630-INS-79 | chrX-111599630-INS-78 |
| chrX-111622137-INS-59 | chrX-111622137-INS-59 |
| chrX-111688530-INS-7104 | chrX-111688531-INS-7104 |
| chrX-111697679-INS-6282 | chrX-111697680-INS-6280 |
| chrX-111706986-INS-56 | chrX-111706986-INS-56 |
| chrX-111734006-INS-100 | chrX-111734006-INS-88 |
| chrX-111817793-INS-803 | chrX-111817793-INS-803 |
| chrX-111822270-INS-353 | chrX-111822270-INS-353 |
| chrX-111854854-INS-6457 | chrX-111854855-INS-6477 |
| chrX-111868705-INS-56 | chrX-111868705-INS-56 |
| chrX-111879143-INS-6236 | chrX-111879144-INS-6236 |
| chrX-111882013-INS-397 | chrX-111882013-INS-397 |
| chrX-111907044-INS-9784 | chrX-111907056-INS-9771 |
| chrX-111925822-INS-4672 | chrX-111925823-INS-4672 |
| chrX-111943338-INS-198 | chrX-111943338-INS-198 |
| chrX-111958682-INS-2742 | chrX-111958683-INS-2743 |
| chrX-112013817-INS-7198 | chrX-112013818-INS-7198 |
| chrX-112019539-INS-5152 | chrX-112019540-INS-5152 |
| chrX-112113285-INS-1426 | chrX-112113286-INS-1426 |
| chrX-112127822-INS-54 | chrX-112127826-INS-50 |
| chrX-112176974-INS-8819 | chrX-112176975-INS-8819 |
| chrX-112230960-INS-7157 | chrX-112230961-INS-7157 |
| chrX-112297400-INS-118 | chrX-112297400-INS-132 |
| chrX-112314653-INS-449 | chrX-112314653-INS-449 |
| chrX-112327318-INS-112 | chrX-112327305-INS-157 |
| chrX-112334405-INS-68 | chrX-112334405-INS-68 |
| chrX-112347154-INS-109 | chrX-112347149-INS-106 |
| chrX-112349396-INS-132 | chrX-112349396-INS-132 |
| chrX-112372380-INS-99 | chrX-112372380-INS-128 |
| chrX-112377012-INS-60 | chrX-112377026-INS-63 |
| chrX-112381832-INS-199 | chrX-112381832-INS-199 |
| chrX-112436786-INS-78 | chrX-112436786-INS-74 |
| chrX-112487298-INS-60 | chrX-112487298-INS-78 |
| chrX-112495376-INS-701 | chrX-112495376-INS-701 |
| chrX-112497782-INS-187 | chrX-112497782-INS-187 |
| chrX-112503531-INS-6499 | chrX-112503531-INS-6505 |
| chrX-112519861-INS-1890 | chrX-112519862-INS-1890 |

| | |
|---|---|
| chrX-112554722-INS-3068 | chrX-112554723-INS-3068 |
| chrX-112564442-INS-70 | chrX-112564442-INS-72 |
| chrX-112570643-INS-242 | chrX-112570643-INS-242 |
| chrX-112606909-INS-575 | chrX-112606909-INS-575 |
| chrX-112621280-INS-884 | chrX-112621280-INS-884 |
| chrX-112655948-INS-183 | chrX-112655949-INS-184 |
| chrX-112700075-INS-6688 | chrX-112700076-INS-6688 |
| chrX-112746619-INS-496 | chrX-112746619-INS-496 |
| chrX-112750140-INS-7416 | chrX-112750141-INS-7415 |
| chrX-112751634-INS-273 | chrX-112751634-INS-273 |
| chrX-112761747-INS-200 | chrX-112761747-INS-200 |
| chrX-112765113-INS-64 | chrX-112765113-INS-54 |
| chrX-112773926-INS-117 | chrX-112773926-INS-117 |
| chrX-112799842-INS-88 | chrX-112799844-INS-58 |
| chrX-112860284-INS-280 | chrX-112860284-INS-278 |
| chrX-112883311-INS-565 | chrX-112883311-INS-565 |
| chrX-112900898-INS-682 | chrX-112900908-INS-653 |
| chrX-112915572-INS-135 | chrX-112915572-INS-135 |
| chrX-112992219-INS-6442 | chrX-112992220-INS-6442 |
| chrX-113150814-INS-56 | chrX-113150842-INS-60 |
| chrX-113235273-INS-52 | chrX-113235273-INS-52 |
| chrX-113365171-INS-7122 | chrX-113365172-INS-7122 |
| chrX-113365172-INS-375 | chrX-113365172-INS-375 |
| chrX-113373272-INS-72 | chrX-113373279-INS-73 |
| chrX-113416858-INS-343 | chrX-113416858-INS-343 |
| chrX-113448988-INS-201 | chrX-113448988-INS-201 |
| chrX-113479310-INS-7086 | chrX-113479311-INS-7087 |
| chrX-113534222-INS-188 | chrX-113534222-INS-188 |
| chrX-113539543-INS-944 | chrX-113539543-INS-944 |
| chrX-113548920-INS-332 | chrX-113548920-INS-332 |
| chrX-113677209-INS-411 | chrX-113677209-INS-409 |
| chrX-113683684-INS-130 | chrX-113683684-INS-142 |
| chrX-113752922-INS-4907 | chrX-113752923-INS-4908 |
| chrX-113771431-INS-337 | chrX-113771431-INS-336 |
| chrX-113825449-INS-159 | chrX-113825449-INS-159 |
| chrX-113853591-INS-8293 | chrX-113853592-INS-8296 |
| chrX-113860627-INS-136 | chrX-113860627-INS-137 |
| chrX-113864458-INS-235 | chrX-113864459-INS-235 |
| chrX-113865168-INS-155 | chrX-113865169-INS-165 |
| chrX-113879750-INS-482 | chrX-113879750-INS-482 |
| chrX-113882493-INS-230 | chrX-113882493-INS-231 |
| chrX-113887195-INS-473 | chrX-113887195-INS-473 |
| chrX-113895601-INS-434 | chrX-113895601-INS-434 |
| chrX-113931499-INS-57 | chrX-113931499-INS-76 |
| chrX-113963714-INS-162 | chrX-113963714-INS-162 |
| chrX-113981109-INS-413 | chrX-113981109-INS-413 |
| chrX-114009889-INS-175 | chrX-114009894-INS-180 |
| chrX-114012186-INS-95 | chrX-114012187-INS-95 |
| chrX-114061291-INS-54 | chrX-114061291-INS-60 |
| chrX-114081974-INS-7080 | chrX-114081975-INS-7080 |
| chrX-114101075-INS-1957 | chrX-114101076-INS-1958 |
| chrX-114115667-INS-6810 | chrX-114115668-INS-6810 |
| chrX-114134259-INS-2645 | chrX-114134260-INS-2645 |
| chrX-114156011-INS-364 | chrX-114156011-INS-364 |
| chrX-114170802-INS-74 | chrX-114170802-INS-78 |
| chrX-114177269-INS-4270 | chrX-114177270-INS-4266 |
| chrX-114191343-INS-58 | chrX-114191357-INS-62 |
| chrX-114193671-INS-6426 | chrX-114193672-INS-6426 |
| chrX-114211471-INS-484 | chrX-114211471-INS-484 |
| chrX-114225653-INS-132 | chrX-114225653-INS-132 |
| chrX-114254072-INS-6436 | chrX-114254073-INS-6438 |
| chrX-114257160-INS-64 | chrX-114257160-INS-64 |
| chrX-114304283-INS-3187 | chrX-114304284-INS-3187 |
| chrX-114334322-INS-891 | chrX-114334322-INS-891 |
| chrX-114349303-INS-55 | chrX-114349303-INS-55 |
| chrX-114370676-INS-126 | chrX-114370679-INS-137 |
| chrX-114394827-INS-4500 | chrX-114394828-INS-4500 |
| chrX-114506327-INS-145 | chrX-114506327-INS-145 |
| chrX-114538994-INS-4137 | chrX-114538995-INS-4137 |
| chrX-114544474-INS-482 | chrX-114544474-INS-492 |
| chrX-114563617-INS-385 | chrX-114563617-INS-385 |
| chrX-114610531-INS-60 | chrX-114610531-INS-60 |
| chrX-114639353-INS-51 | chrX-114639353-INS-54 |
| chrX-114725056-INS-98 | chrX-114725054-INS-97 |
| chrX-114782583-INS-64 | chrX-114782583-INS-64 |
| chrX-114922668-INS-825 | chrX-114922668-INS-825 |
| chrX-114925974-INS-395 | chrX-114925974-INS-395 |

| | |
|---|---|
| chrX-114932782-INS-53 | chrX-114932778-INS-56 |
| chrX-114940835-INS-2040 | chrX-114940836-INS-2040 |
| chrX-114941158-INS-61 | chrX-114941176-INS-88 |
| chrX-114967501-INS-347 | chrX-114967501-INS-348 |
| chrX-114971815-INS-554 | chrX-114971815-INS-554 |
| chrX-114996494-INS-331 | chrX-114996494-INS-323 |
| chrX-115034418-INS-66 | chrX-115034418-INS-50 |
| chrX-115060259-INS-92 | chrX-115060229-INS-79 |
| chrX-115110697-INS-1353 | chrX-115110698-INS-1352 |
| chrX-115193632-INS-68 | chrX-115193632-INS-74 |
| chrX-115218987-INS-1886 | chrX-115218988-INS-1886 |
| chrX-115229393-INS-6171 | chrX-115229394-INS-6171 |
| chrX-115264207-INS-65 | chrX-115264226-INS-80 |
| chrX-115298131-INS-358 | chrX-115298131-INS-358 |
| chrX-115403176-INS-6401 | chrX-115403177-INS-6401 |
| chrX-115435571-INS-494 | chrX-115435571-INS-494 |
| chrX-115497386-INS-396 | chrX-115497386-INS-396 |
| chrX-115536152-INS-92 | chrX-115536149-INS-113 |
| chrX-115573517-INS-3450 | chrX-115573518-INS-3450 |
| chrX-115580854-INS-5311 | chrX-115580855-INS-5311 |
| chrX-115630928-INS-135 | chrX-115630928-INS-93 |
| chrX-115659325-INS-162 | chrX-115659325-INS-161 |
| chrX-115662534-INS-6595 | chrX-115662535-INS-6595 |
| chrX-115686150-INS-440 | chrX-115686150-INS-440 |
| chrX-115718136-INS-3658 | chrX-115718137-INS-3658 |
| chrX-115768301-INS-4593 | chrX-115768302-INS-4593 |
| chrX-115820611-INS-3077 | chrX-115820612-INS-3077 |
| chrX-115842097-INS-331 | chrX-115842097-INS-330 |
| chrX-115846091-INS-58 | chrX-115846092-INS-66 |
| chrX-115855534-INS-566 | chrX-115855534-INS-566 |
| chrX-115877503-INS-118 | chrX-115877504-INS-119 |
| chrX-115933203-INS-371 | chrX-115933203-INS-372 |
| chrX-115986436-INS-4553 | chrX-115986437-INS-4553 |
| chrX-116010002-INS-56 | chrX-116010002-INS-60 |
| chrX-116012161-INS-341 | chrX-116012161-INS-341 |
| chrX-116017813-INS-161 | chrX-116017813-INS-125 |
| chrX-116093547-INS-2536 | chrX-116093548-INS-2536 |
| chrX-116097324-INS-6396 | chrX-116097325-INS-6396 |
| chrX-116200937-INS-151 | chrX-116200937-INS-151 |
| chrX-116215745-INS-161 | chrX-116215789-INS-111 |
| chrX-116215782-INS-699 | chrX-116215782-INS-687 |
| chrX-116219137-INS-64 | chrX-116219137-INS-64 |
| chrX-116243004-INS-1090 | chrX-116243005-INS-1070 |
| chrX-116334824-INS-74 | chrX-116334824-INS-74 |
| chrX-116339726-INS-2786 | chrX-116339727-INS-2786 |
| chrX-116342650-INS-1673 | chrX-116342651-INS-1673 |
| chrX-116389345-INS-358 | chrX-116389346-INS-358 |
| chrX-116397533-INS-64 | chrX-116397533-INS-66 |
| chrX-116399890-INS-576 | chrX-116399890-INS-577 |
| chrX-116400928-INS-62 | chrX-116400916-INS-50 |
| chrX-116423602-INS-947 | chrX-116423602-INS-947 |
| chrX-116424272-INS-90 | chrX-116424265-INS-95 |
| chrX-116425371-INS-149 | chrX-116425371-INS-151 |
| chrX-116482759-INS-66 | chrX-116482759-INS-66 |
| chrX-116568660-INS-422 | chrX-116568660-INS-422 |
| chrX-116586302-INS-211 | chrX-116586302-INS-211 |
| chrX-116587506-INS-6464 | chrX-116587507-INS-6466 |
| chrX-116629264-INS-2572 | chrX-116629265-INS-2572 |
| chrX-116693962-INS-76 | chrX-116693962-INS-74 |
| chrX-116697682-INS-495 | chrX-116697682-INS-495 |
| chrX-116717300-INS-9384 | chrX-116717301-INS-9384 |
| chrX-116779932-INS-3107 | chrX-116779933-INS-3107 |
| chrX-116803470-INS-7069 | chrX-116803498-INS-7054 |
| chrX-116805220-INS-85 | chrX-116805220-INS-68 |
| chrX-116818437-INS-7121 | chrX-116818438-INS-7121 |
| chrX-116827780-INS-6591 | chrX-116827781-INS-6591 |
| chrX-116843876-INS-960 | chrX-116843876-INS-960 |
| chrX-116898164-INS-2454 | chrX-116898165-INS-2454 |
| chrX-116921483-INS-446 | chrX-116921483-INS-444 |
| chrX-116957864-INS-217 | chrX-116957864-INS-217 |
| chrX-116988828-INS-1479 | chrX-116988829-INS-1479 |
| chrX-117044586-INS-5480 | chrX-117044587-INS-5480 |
| chrX-117059447-INS-255 | chrX-117059448-INS-221 |
| chrX-117107673-INS-58 | chrX-117107673-INS-58 |
| chrX-117125879-INS-1887 | chrX-117125880-INS-1887 |
| chrX-117129475-INS-6926 | chrX-117129476-INS-6927 |
| chrX-117169166-INS-87 | chrX-117169166-INS-87 |

| | |
|---|---|
| chrX-117176977-INS-85 | chrX-117176977-INS-55 |
| chrX-117210565-INS-87 | chrX-117210565-INS-81 |
| chrX-117219188-INS-55 | chrX-117219188-INS-55 |
| chrX-117222290-INS-1307 | chrX-117222291-INS-1307 |
| chrX-117244188-INS-637 | chrX-117244188-INS-637 |
| chrX-117338797-INS-598 | chrX-117338797-INS-598 |
| chrX-117388500-INS-598 | chrX-117388500-INS-598 |
| chrX-117440103-INS-6499 | chrX-117440104-INS-6500 |
| chrX-117508331-INS-60 | chrX-117508331-INS-55 |
| chrX-117515565-INS-750 | chrX-117515562-INS-759 |
| chrX-117558286-INS-3167 | chrX-117558287-INS-3167 |
| chrX-117593028-INS-72 | chrX-117593025-INS-75 |
| chrX-117651263-INS-6392 | chrX-117651264-INS-6392 |
| chrX-117680165-INS-1471 | chrX-117680166-INS-1469 |
| chrX-117735464-INS-2949 | chrX-117735465-INS-2949 |
| chrX-117742349-INS-497 | chrX-117742350-INS-497 |
| chrX-117756496-INS-62 | chrX-117756477-INS-94 |
| chrX-117767816-INS-152 | chrX-117767816-INS-152 |
| chrX-117820582-INS-53 | chrX-117820582-INS-83 |
| chrX-117846911-INS-81 | chrX-117846911-INS-81 |
| chrX-117863120-INS-54 | chrX-117863120-INS-54 |
| chrX-117960488-INS-6418 | chrX-117960489-INS-6417 |
| chrX-118027455-INS-557 | chrX-118027455-INS-557 |
| chrX-118036062-INS-280 | chrX-118036062-INS-325 |
| chrX-118075455-INS-232 | chrX-118075455-INS-232 |
| chrX-118082617-INS-162 | chrX-118082617-INS-162 |
| chrX-118094057-INS-97 | chrX-118094057-INS-97 |
| chrX-118096140-INS-104 | chrX-118096120-INS-92 |
| chrX-118122755-INS-53 | chrX-118122755-INS-53 |
| chrX-118125944-INS-53 | chrX-118125944-INS-53 |
| chrX-118188723-INS-903 | chrX-118188723-INS-903 |
| chrX-118216218-INS-6463 | chrX-118216219-INS-6463 |
| chrX-118228091-INS-78 | chrX-118228091-INS-74 |
| chrX-118238291-INS-138 | chrX-118238291-INS-120 |
| chrX-118245743-INS-51 | chrX-118245743-INS-51 |
| chrX-118268179-INS-1530 | chrX-118268180-INS-1530 |
| chrX-118306024-INS-630 | chrX-118306024-INS-630 |
| chrX-118312553-INS-183 | chrX-118312553-INS-183 |
| chrX-118324581-INS-83 | chrX-118324581-INS-119 |
| chrX-118333217-INS-64 | chrX-118333217-INS-72 |
| chrX-118358743-INS-991 | chrX-118358743-INS-992 |
| chrX-118364651-INS-606 | chrX-118364651-INS-605 |
| chrX-118405768-INS-477 | chrX-118405768-INS-477 |
| chrX-118448537-INS-208 | chrX-118448537-INS-226 |
| chrX-118501449-INS-3557 | chrX-118501450-INS-3557 |
| chrX-118518012-INS-9344 | chrX-118518013-INS-9344 |
| chrX-118539992-INS-6670 | chrX-118539993-INS-6670 |
| chrX-118545410-INS-6450 | chrX-118545411-INS-6450 |
| chrX-118604081-INS-1699 | chrX-118604082-INS-1699 |
| chrX-118608765-INS-2975 | chrX-118608766-INS-2975 |
| chrX-118610778-INS-1238 | chrX-118610779-INS-1239 |
| chrX-118635804-INS-106 | chrX-118635804-INS-106 |
| chrX-118641631-INS-641 | chrX-118641631-INS-641 |
| chrX-118642556-INS-5601 | chrX-118642557-INS-5601 |
| chrX-118655181-INS-1803 | chrX-118655182-INS-1803 |
| chrX-118674855-INS-636 | chrX-118674855-INS-636 |
| chrX-118737511-INS-87 | chrX-118737529-INS-88 |
| chrX-118742859-INS-501 | chrX-118742859-INS-501 |
| chrX-118790938-INS-51 | chrX-118790938-INS-51 |
| chrX-118833734-INS-6381 | chrX-118833735-INS-6381 |
| chrX-118836630-INS-111 | chrX-118836630-INS-112 |
| chrX-118871671-INS-220 | chrX-118871671-INS-220 |
| chrX-118916101-INS-491 | chrX-118916101-INS-491 |
| chrX-118924194-INS-282 | chrX-118924178-INS-284 |
| chrX-118925216-INS-516 | chrX-118925216-INS-516 |
| chrX-118976912-INS-78 | chrX-118976913-INS-74 |
| chrX-118985044-INS-6600 | chrX-118985045-INS-6600 |
| chrX-119019629-INS-6398 | chrX-119019630-INS-6398 |
| chrX-119059585-INS-168 | chrX-119059585-INS-168 |
| chrX-119066059-INS-683 | chrX-119066059-INS-683 |
| chrX-119106098-INS-70 | chrX-119106098-INS-70 |
| chrX-119110974-INS-2333 | chrX-119110975-INS-2334 |
| chrX-119126505-INS-80 | chrX-119126505-INS-82 |
| chrX-119130417-INS-970 | chrX-119130417-INS-970 |
| chrX-119140489-INS-200 | chrX-119140489-INS-200 |
| chrX-119146530-INS-948 | chrX-119146530-INS-948 |
| chrX-119213699-INS-73 | chrX-119213699-INS-78 |

| | |
|---|---|
| chrX-119240251-INS-151 | chrX-119240251-INS-150 |
| chrX-119250520-INS-1006 | chrX-119250521-INS-1000 |
| chrX-119253619-INS-58 | chrX-119253634-INS-68 |
| chrX-119254122-INS-6376 | chrX-119254123-INS-6376 |
| chrX-119258846-INS-6563 | chrX-119258847-INS-6563 |
| chrX-119278969-INS-710 | chrX-119278969-INS-731 |
| chrX-119291310-INS-1143 | chrX-119291311-INS-1143 |
| chrX-119325939-INS-118 | chrX-119325939-INS-112 |
| chrX-119329483-INS-549 | chrX-119329483-INS-549 |
| chrX-119341405-INS-4103 | chrX-119341406-INS-4103 |
| chrX-119347503-INS-58 | chrX-119347503-INS-60 |
| chrX-119349785-INS-2159 | chrX-119349786-INS-2159 |
| chrX-119353538-INS-1523 | chrX-119353539-INS-1523 |
| chrX-119366457-INS-6826 | chrX-119366458-INS-6828 |
| chrX-119431075-INS-7102 | chrX-119431076-INS-7104 |
| chrX-119436252-INS-54 | chrX-119436219-INS-85 |
| chrX-119440793-INS-58 | chrX-119440793-INS-90 |
| chrX-119452236-INS-3133 | chrX-119452237-INS-3133 |
| chrX-119477702-INS-63 | chrX-119477702-INS-63 |
| chrX-119493148-INS-251 | chrX-119493148-INS-251 |
| chrX-119502890-INS-119 | chrX-119502890-INS-119 |
| chrX-119508124-INS-581 | chrX-119508124-INS-581 |
| chrX-119511651-INS-53 | chrX-119511651-INS-58 |
| chrX-119513790-INS-6588 | chrX-119513791-INS-6590 |
| chrX-119539319-INS-6618 | chrX-119539320-INS-6620 |
| chrX-119559622-INS-2042 | chrX-119559623-INS-2039 |
| chrX-119614462-INS-9766 | chrX-119614463-INS-9766 |
| chrX-119629782-INS-1221 | chrX-119629783-INS-1221 |
| chrX-119645485-INS-216 | chrX-119645485-INS-216 |
| chrX-119666504-INS-198 | chrX-119666504-INS-198 |
| chrX-119681970-INS-63 | chrX-119681970-INS-63 |
| chrX-119684323-INS-176 | chrX-119684323-INS-176 |
| chrX-119708454-INS-6439 | chrX-119708455-INS-6439 |
| chrX-119715018-INS-347 | chrX-119715016-INS-345 |
| chrX-119793342-INS-6330 | chrX-119793343-INS-6331 |
| chrX-119804855-INS-54 | chrX-119804855-INS-50 |
| chrX-119835012-INS-64 | chrX-119835012-INS-64 |
| chrX-119844955-INS-264 | chrX-119844955-INS-264 |
| chrX-119852129-INS-6531 | chrX-119852130-INS-6533 |
| chrX-119913151-INS-990 | chrX-119913151-INS-983 |
| chrX-119957465-INS-438 | chrX-119957465-INS-438 |
| chrX-119968011-INS-66 | chrX-119968011-INS-66 |
| chrX-120006605-INS-86 | chrX-120006605-INS-86 |
| chrX-120034211-INS-6800 | chrX-120034212-INS-6800 |
| chrX-120039276-INS-342 | chrX-120039276-INS-342 |
| chrX-120049546-INS-2699 | chrX-120049547-INS-2699 |
| chrX-120132132-INS-51 | chrX-120132132-INS-52 |
| chrX-120133381-INS-6410 | chrX-120133382-INS-6410 |
| chrX-120135089-INS-259 | chrX-120135089-INS-259 |
| chrX-120137320-INS-154 | chrX-120137320-INS-154 |
| chrX-120140924-INS-1397 | chrX-120140925-INS-1397 |
| chrX-120157671-INS-92 | chrX-120157671-INS-92 |
| chrX-120162287-INS-59 | chrX-120162270-INS-63 |
| chrX-120163383-INS-8847 | chrX-120163384-INS-8847 |
| chrX-120211172-INS-748 | chrX-120211172-INS-748 |
| chrX-120240178-INS-111 | chrX-120240178-INS-115 |
| chrX-120248112-INS-2135 | chrX-120248113-INS-2135 |
| chrX-120250997-INS-192 | chrX-120250997-INS-192 |
| chrX-120256959-INS-518 | chrX-120256959-INS-518 |
| chrX-120346861-INS-136 | chrX-120346861-INS-136 |
| chrX-120362318-INS-848 | chrX-120362318-INS-848 |
| chrX-120383065-INS-851 | chrX-120383065-INS-851 |
| chrX-120384266-INS-1524 | chrX-120384267-INS-1524 |
| chrX-120387841-INS-92 | chrX-120387841-INS-92 |
| chrX-120390410-INS-6189 | chrX-120390411-INS-6189 |
| chrX-120397248-INS-66 | chrX-120397258-INS-64 |
| chrX-120411298-INS-57 | chrX-120411298-INS-57 |
| chrX-120411860-INS-6570 | chrX-120411861-INS-6570 |
| chrX-120440413-INS-51 | chrX-120440414-INS-54 |
| chrX-120472160-INS-2923 | chrX-120472161-INS-2923 |
| chrX-120478264-INS-5515 | chrX-120478265-INS-5518 |
| chrX-120478332-INS-456 | chrX-120478332-INS-467 |
| chrX-120494601-INS-68 | chrX-120494601-INS-68 |
| chrX-120509586-INS-2567 | chrX-120509587-INS-2567 |
| chrX-120528376-INS-6395 | chrX-120528377-INS-6397 |
| chrX-120545722-INS-52 | chrX-120545722-INS-52 |
| chrX-120566717-INS-68 | chrX-120566670-INS-74 |

| | |
|---|---|
| chrX-120573711-INS-2769 | chrX-120573712-INS-2769 |
| chrX-120576614-INS-14623 | chrX-120576615-INS-14623 |
| chrX-120601136-INS-52 | chrX-120601136-INS-50 |
| chrX-120612248-INS-749 | chrX-120612248-INS-748 |
| chrX-120658895-INS-243 | chrX-120658895-INS-243 |
| chrX-120689871-INS-1006 | chrX-120689872-INS-1006 |
| chrX-120705012-INS-610 | chrX-120705012-INS-610 |
| chrX-120706998-INS-83 | chrX-120706999-INS-83 |
| chrX-120709346-INS-516 | chrX-120709346-INS-517 |
| chrX-120716240-INS-516 | chrX-120716240-INS-516 |
| chrX-120721167-INS-328 | chrX-120721167-INS-328 |
| chrX-120728520-INS-127 | chrX-120728512-INS-104 |
| chrX-120744038-INS-7132 | chrX-120744039-INS-7132 |
| chrX-120751208-INS-497 | chrX-120751208-INS-497 |
| chrX-120780619-INS-533 | chrX-120780619-INS-537 |
| chrX-120787691-INS-2671 | chrX-120787692-INS-2672 |
| chrX-120816143-INS-95 | chrX-120816144-INS-117 |
| chrX-120829861-INS-2747 | chrX-120829861-INS-2741 |
| chrX-120893310-INS-501 | chrX-120893310-INS-502 |
| chrX-120942487-INS-7937 | chrX-120942488-INS-7937 |
| chrX-120947645-INS-2831 | chrX-120947646-INS-2831 |
| chrX-120996904-INS-1059 | chrX-120996905-INS-1059 |
| chrX-121006819-INS-95 | chrX-121006827-INS-86 |
| chrX-121021535-INS-184 | chrX-121021535-INS-184 |
| chrX-121044061-INS-438 | chrX-121044061-INS-438 |
| chrX-121059798-INS-2492 | chrX-121059799-INS-2492 |
| chrX-121068864-INS-575 | chrX-121068864-INS-576 |
| chrX-121073106-INS-201 | chrX-121073104-INS-201 |
| chrX-121088353-INS-722 | chrX-121088353-INS-722 |
| chrX-121111068-INS-104 | chrX-121111068-INS-110 |
| chrX-121205862-INS-174 | chrX-121205862-INS-174 |
| chrX-121215964-INS-86 | chrX-121215964-INS-86 |
| chrX-121283405-INS-5336 | chrX-121283405-INS-5336 |
| chrX-121301846-INS-571 | chrX-121301846-INS-573 |
| chrX-121303414-INS-94 | chrX-121303414-INS-94 |
| chrX-121310157-INS-79 | chrX-121310157-INS-92 |
| chrX-121315026-INS-60 | chrX-121315026-INS-60 |
| chrX-121321166-INS-3712 | chrX-121321167-INS-3712 |
| chrX-121362692-INS-6397 | chrX-121362693-INS-6396 |
| chrX-121367007-INS-448 | chrX-121367007-INS-447 |
| chrX-121367948-INS-993 | chrX-121367948-INS-993 |
| chrX-121384923-INS-219 | chrX-121384923-INS-219 |
| chrX-121398946-INS-196 | chrX-121398946-INS-196 |
| chrX-121480258-INS-6014 | chrX-121480259-INS-6014 |
| chrX-121507013-INS-7094 | chrX-121507014-INS-7094 |
| chrX-121562708-INS-78 | chrX-121562708-INS-78 |
| chrX-121579612-INS-70 | chrX-121579612-INS-98 |
| chrX-121601522-INS-62 | chrX-121601522-INS-62 |
| chrX-121635445-INS-5262 | chrX-121635446-INS-5262 |
| chrX-121718692-INS-498 | chrX-121718692-INS-498 |
| chrX-121738513-INS-51 | chrX-121738513-INS-53 |
| chrX-121751965-INS-105 | chrX-121751965-INS-105 |
| chrX-121769436-INS-53 | chrX-121769436-INS-52 |
| chrX-121772211-INS-402 | chrX-121772211-INS-402 |
| chrX-121805437-INS-52 | chrX-121805438-INS-60 |
| chrX-121830591-INS-62 | chrX-121830590-INS-76 |
| chrX-121875412-INS-144 | chrX-121875412-INS-143 |
| chrX-121877158-INS-6488 | chrX-121877159-INS-6488 |
| chrX-121923239-INS-377 | chrX-121923239-INS-377 |
| chrX-121942799-INS-60 | chrX-121942800-INS-62 |
| chrX-121943696-INS-85 | chrX-121943686-INS-73 |
| chrX-122004268-INS-84 | chrX-122004268-INS-56 |
| chrX-122396586-INS-55 | chrX-122396568-INS-50 |
| chrX-122944501-INS-75 | chrX-122944501-INS-68 |
| chrX-123120514-INS-190 | chrX-123120514-INS-190 |
| chrX-123127019-INS-120 | chrX-123127020-INS-120 |
| chrX-123128591-INS-51 | chrX-123128592-INS-59 |
| chrX-123148656-INS-59 | chrX-123148656-INS-62 |
| chrX-123241335-INS-60 | chrX-123241356-INS-64 |
| chrX-125008747-INS-133 | chrX-125008747-INS-125 |
| chrX-125076881-INS-58 | chrX-125076881-INS-60 |
| chrX-125120955-INS-455 | chrX-125120955-INS-455 |
| chrX-125125248-INS-949 | chrX-125125248-INS-949 |
| chrX-125281635-INS-254 | chrX-125281635-INS-254 |
| chrX-125290336-INS-8325 | chrX-125290337-INS-8325 |
| chrX-125354724-INS-1891 | chrX-125354725-INS-1891 |
| chrX-125363037-INS-57 | chrX-125363037-INS-57 |

| | |
|---|---|
| chrX-125366586-INS-120 | chrX-125366576-INS-125 |
| chrX-125439493-INS-388 | chrX-125439493-INS-388 |
| chrX-125476493-INS-6402 | chrX-125476494-INS-6402 |
| chrX-125489078-INS-226 | chrX-125489078-INS-226 |
| chrX-125516718-INS-2829 | chrX-125516719-INS-2829 |
| chrX-125518596-INS-4598 | chrX-125518597-INS-4598 |
| chrX-125536752-INS-154 | chrX-125536752-INS-154 |
| chrX-125587334-INS-2954 | chrX-125587335-INS-2955 |
| chrX-125721041-INS-71 | chrX-125721041-INS-53 |
| chrX-125734222-INS-162 | chrX-125734222-INS-162 |
| chrX-125775996-INS-57 | chrX-125775996-INS-77 |
| chrX-125803354-INS-51 | chrX-125803354-INS-63 |
| chrX-125849104-INS-5657 | chrX-125849105-INS-5657 |
| chrX-125854436-INS-707 | chrX-125854436-INS-708 |
| chrX-125884314-INS-561 | chrX-125884314-INS-561 |
| chrX-125939520-INS-153 | chrX-125939520-INS-153 |
| chrX-125941524-INS-436 | chrX-125941524-INS-436 |
| chrX-125943345-INS-251 | chrX-125943345-INS-251 |
| chrX-125955075-INS-57 | chrX-125955075-INS-57 |
| chrX-125976079-INS-576 | chrX-125976079-INS-573 |
| chrX-125981164-INS-232 | chrX-125981164-INS-232 |
| chrX-125982809-INS-4084 | chrX-125982810-INS-4082 |
| chrX-125999221-INS-227 | chrX-125999221-INS-227 |
| chrX-126007098-INS-169 | chrX-126007098-INS-169 |
| chrX-126065678-INS-93 | chrX-126065677-INS-105 |
| chrX-126133924-INS-98 | chrX-126133924-INS-96 |
| chrX-126157899-INS-2732 | chrX-126157900-INS-2724 |
| chrX-126162424-INS-95 | chrX-126162424-INS-85 |
| chrX-126218642-INS-66 | chrX-126218642-INS-72 |
| chrX-126223160-INS-67 | chrX-126223160-INS-69 |
| chrX-126270758-INS-56 | chrX-126270758-INS-64 |
| chrX-126297571-INS-6383 | chrX-126297572-INS-6383 |
| chrX-126329967-INS-204 | chrX-126329967-INS-204 |
| chrX-126345500-INS-56 | chrX-126345500-INS-56 |
| chrX-126379991-INS-607 | chrX-126379991-INS-609 |
| chrX-126410836-INS-90 | chrX-126410836-INS-100 |
| chrX-126460317-INS-1001 | chrX-126460318-INS-999 |
| chrX-126462487-INS-149 | chrX-126462489-INS-150 |
| chrX-126504879-INS-781 | chrX-126504879-INS-781 |
| chrX-126522431-INS-84 | chrX-126522431-INS-84 |
| chrX-126529563-INS-96 | chrX-126529560-INS-94 |
| chrX-126537573-INS-85 | chrX-126537573-INS-66 |
| chrX-126592006-INS-66 | chrX-126592006-INS-66 |
| chrX-126621986-INS-101 | chrX-126621986-INS-101 |
| chrX-126625723-INS-137 | chrX-126625723-INS-117 |
| chrX-126644261-INS-70 | chrX-126644261-INS-70 |
| chrX-126682572-INS-160 | chrX-126682572-INS-172 |
| chrX-126691042-INS-103 | chrX-126691042-INS-97 |
| chrX-126698388-INS-222 | chrX-126698388-INS-222 |
| chrX-126777571-INS-1388 | chrX-126777572-INS-1388 |
| chrX-126793086-INS-57 | chrX-126793103-INS-68 |
| chrX-126801425-INS-6214 | chrX-126801426-INS-6214 |
| chrX-126805066-INS-118 | chrX-126805066-INS-112 |
| chrX-126806068-INS-54 | chrX-126806068-INS-54 |
| chrX-126807134-INS-3186 | chrX-126807135-INS-3186 |
| chrX-126807362-INS-151 | chrX-126807362-INS-155 |
| chrX-126825986-INS-777 | chrX-126825986-INS-777 |
| chrX-126838306-INS-66 | chrX-126838306-INS-66 |
| chrX-126878742-INS-3373 | chrX-126878743-INS-3373 |
| chrX-126894711-INS-1116 | chrX-126894709-INS-1120 |
| chrX-126907413-INS-135 | chrX-126907413-INS-135 |
| chrX-126909390-INS-206 | chrX-126909348-INS-250 |
| chrX-126936102-INS-215 | chrX-126936102-INS-201 |
| chrX-126957044-INS-5169 | chrX-126957045-INS-5169 |
| chrX-127014660-INS-7142 | chrX-127014661-INS-7142 |
| chrX-127020156-INS-3079 | chrX-127020157-INS-3079 |
| chrX-127021797-INS-51 | chrX-127021797-INS-57 |
| chrX-127054843-INS-447 | chrX-127054843-INS-447 |
| chrX-127078923-INS-1681 | chrX-127078924-INS-1681 |
| chrX-127082743-INS-60 | chrX-127082743-INS-55 |
| chrX-127088845-INS-54 | chrX-127088845-INS-54 |
| chrX-127097678-INS-704 | chrX-127097678-INS-704 |
| chrX-127104329-INS-96 | chrX-127104329-INS-100 |
| chrX-127184692-INS-4311 | chrX-127184693-INS-4311 |
| chrX-127195878-INS-65 | chrX-127195875-INS-61 |
| chrX-127202320-INS-6569 | chrX-127202321-INS-6569 |
| chrX-127229910-INS-6380 | chrX-127229911-INS-6380 |

| | |
|---|---|
| chrX-127260483-INS-62 | chrX-127260483-INS-66 |
| chrX-127329136-INS-6363 | chrX-127329137-INS-6362 |
| chrX-127362151-INS-218 | chrX-127362151-INS-218 |
| chrX-127376132-INS-792 | chrX-127376132-INS-793 |
| chrX-127407436-INS-414 | chrX-127407436-INS-414 |
| chrX-127438036-INS-655 | chrX-127438036-INS-655 |
| chrX-127443635-INS-444 | chrX-127443635-INS-460 |
| chrX-127488111-INS-5549 | chrX-127488112-INS-5550 |
| chrX-127536115-INS-51 | chrX-127536115-INS-51 |
| chrX-127643386-INS-4199 | chrX-127643387-INS-4199 |
| chrX-127648798-INS-3955 | chrX-127648799-INS-3956 |
| chrX-127668704-INS-91 | chrX-127668704-INS-66 |
| chrX-127728305-INS-55 | chrX-127728305-INS-55 |
| chrX-127745711-INS-55 | chrX-127745726-INS-86 |
| chrX-127872298-INS-151 | chrX-127872263-INS-142 |
| chrX-127931322-INS-235 | chrX-127931351-INS-204 |
| chrX-128081562-INS-149 | chrX-128081562-INS-150 |
| chrX-128137205-INS-6605 | chrX-128137206-INS-6595 |
| chrX-128221324-INS-214 | chrX-128221324-INS-214 |
| chrX-128234200-INS-60 | chrX-128234200-INS-72 |
| chrX-128316095-INS-55 | chrX-128316122-INS-58 |
| chrX-128348280-INS-208 | chrX-128348280-INS-208 |
| chrX-128373366-INS-56 | chrX-128373407-INS-56 |
| chrX-128379165-INS-1889 | chrX-128379166-INS-1889 |
| chrX-128395233-INS-6385 | chrX-128395234-INS-6378 |
| chrX-128533553-INS-169 | chrX-128533553-INS-167 |
| chrX-128534996-INS-225 | chrX-128534996-INS-235 |
| chrX-128537342-INS-964 | chrX-128537342-INS-965 |
| chrX-128576440-INS-580 | chrX-128576440-INS-581 |
| chrX-128631421-INS-1394 | chrX-128631422-INS-1397 |
| chrX-128641094-INS-179 | chrX-128641094-INS-180 |
| chrX-128648491-INS-52 | chrX-128648491-INS-52 |
| chrX-128654262-INS-231 | chrX-128654262-INS-231 |
| chrX-128676574-INS-197 | chrX-128676574-INS-197 |
| chrX-128682001-INS-52 | chrX-128681973-INS-50 |
| chrX-128691082-INS-3650 | chrX-128691083-INS-3650 |
| chrX-128720198-INS-6933 | chrX-128720199-INS-6935 |
| chrX-128825889-INS-209 | chrX-128825889-INS-210 |
| chrX-128842754-INS-760 | chrX-128842754-INS-756 |
| chrX-128873039-INS-494 | chrX-128873039-INS-494 |
| chrX-128885114-INS-64 | chrX-128885114-INS-64 |
| chrX-129017684-INS-3372 | chrX-129017685-INS-3364 |
| chrX-129142435-INS-133 | chrX-129142435-INS-133 |
| chrX-129272071-INS-78 | chrX-129272076-INS-50 |
| chrX-129283365-INS-114 | chrX-129283365-INS-90 |
| chrX-129335008-INS-58 | chrX-129335008-INS-50 |
| chrX-129400953-INS-1251 | chrX-129400954-INS-1251 |
| chrX-129466261-INS-5324 | chrX-129466262-INS-5324 |
| chrX-129493254-INS-284 | chrX-129493254-INS-284 |
| chrX-129532812-INS-4785 | chrX-129532813-INS-4785 |
| chrX-129535578-INS-221 | chrX-129535578-INS-221 |
| chrX-129602955-INS-4995 | chrX-129602955-INS-4995 |
| chrX-129619468-INS-6435 | chrX-129619469-INS-6435 |
| chrX-129627575-INS-82 | chrX-129627575-INS-88 |
| chrX-129638955-INS-92 | chrX-129638955-INS-86 |
| chrX-129711080-INS-54 | chrX-129711080-INS-56 |
| chrX-129711976-INS-6539 | chrX-129711977-INS-6541 |
| chrX-129786700-INS-369 | chrX-129786700-INS-367 |
| chrX-129796836-INS-6353 | chrX-129796837-INS-6353 |
| chrX-129825371-INS-4412 | chrX-129825372-INS-4412 |
| chrX-129837970-INS-56 | chrX-129837971-INS-66 |
| chrX-130019760-INS-82 | chrX-130019747-INS-113 |
| chrX-130035079-INS-141 | chrX-130035099-INS-100 |
| chrX-130076681-INS-82 | chrX-130076681-INS-82 |
| chrX-130135514-INS-459 | chrX-130135514-INS-459 |
| chrX-130192744-INS-5332 | chrX-130192745-INS-5332 |
| chrX-130200336-INS-1622 | chrX-130200337-INS-1616 |
| chrX-130206821-INS-404 | chrX-130206821-INS-406 |
| chrX-130230642-INS-4359 | chrX-130230643-INS-4359 |
| chrX-130230643-INS-340 | chrX-130230643-INS-340 |
| chrX-130272944-INS-51 | chrX-130272944-INS-54 |
| chrX-130311586-INS-1580 | chrX-130311587-INS-1580 |
| chrX-130421063-INS-137 | chrX-130421062-INS-163 |
| chrX-130485662-INS-7077 | chrX-130485663-INS-7078 |
| chrX-130581388-INS-635 | chrX-130581388-INS-634 |
| chrX-130623699-INS-116 | chrX-130623701-INS-107 |
| chrX-130733566-INS-1111 | chrX-130733575-INS-1100 |

| | |
|---|---|
| chrX-130742383-INS-161 | chrX-130742383-INS-161 |
| chrX-130746688-INS-75 | chrX-130746689-INS-90 |
| chrX-130802541-INS-5370 | chrX-130802542-INS-5370 |
| chrX-130821568-INS-1866 | chrX-130821569-INS-1866 |
| chrX-130916436-INS-112 | chrX-130916436-INS-112 |
| chrX-130953816-INS-811 | chrX-130953816-INS-775 |
| chrX-130955278-INS-4291 | chrX-130955279-INS-4291 |
| chrX-130970963-INS-2349 | chrX-130970964-INS-2349 |
| chrX-131013338-INS-5180 | chrX-131013339-INS-5180 |
| chrX-131019744-INS-55 | chrX-131019744-INS-65 |
| chrX-131020787-INS-72 | chrX-131020787-INS-72 |
| chrX-131023871-INS-118 | chrX-131023871-INS-118 |
| chrX-131032294-INS-674 | chrX-131032294-INS-674 |
| chrX-131052652-INS-53 | chrX-131052652-INS-53 |
| chrX-131065704-INS-73 | chrX-131065704-INS-73 |
| chrX-131083314-INS-1542 | chrX-131083315-INS-1541 |
| chrX-131137766-INS-1282 | chrX-131137767-INS-1269 |
| chrX-131178605-INS-61 | chrX-131178605-INS-61 |
| chrX-131205666-INS-73 | chrX-131205666-INS-56 |
| chrX-131242065-INS-70 | chrX-131242063-INS-72 |
| chrX-131284727-INS-1875 | chrX-131284728-INS-1875 |
| chrX-131294015-INS-2867 | chrX-131294016-INS-2867 |
| chrX-131312045-INS-51 | chrX-131312045-INS-51 |
| chrX-131321613-INS-1732 | chrX-131321614-INS-1732 |
| chrX-131321796-INS-498 | chrX-131321796-INS-498 |
| chrX-131343800-INS-508 | chrX-131343800-INS-508 |
| chrX-131355922-INS-53 | chrX-131355928-INS-51 |
| chrX-131363901-INS-569 | chrX-131363901-INS-570 |
| chrX-131371575-INS-493 | chrX-131371575-INS-493 |
| chrX-131375304-INS-6464 | chrX-131375305-INS-6464 |
| chrX-131383531-INS-3428 | chrX-131383532-INS-3428 |
| chrX-131398070-INS-493 | chrX-131398070-INS-494 |
| chrX-131429171-INS-848 | chrX-131429171-INS-843 |
| chrX-131447822-INS-51 | chrX-131447822-INS-51 |
| chrX-131520907-INS-6919 | chrX-131520908-INS-6922 |
| chrX-131566995-INS-1867 | chrX-131566996-INS-1867 |
| chrX-131570571-INS-7184 | chrX-131570572-INS-7184 |
| chrX-131624536-INS-169 | chrX-131624536-INS-169 |
| chrX-131657091-INS-3370 | chrX-131657092-INS-3370 |
| chrX-131691106-INS-289 | chrX-131691106-INS-289 |
| chrX-131692534-INS-67 | chrX-131692534-INS-67 |
| chrX-131694297-INS-7151 | chrX-131694298-INS-7152 |
| chrX-131730176-INS-553 | chrX-131730176-INS-553 |
| chrX-131754426-INS-6199 | chrX-131754440-INS-6200 |
| chrX-131769558-INS-5173 | chrX-131769559-INS-5174 |
| chrX-131780851-INS-1379 | chrX-131780852-INS-1380 |
| chrX-131811307-INS-6862 | chrX-131811308-INS-6863 |
| chrX-131882213-INS-614 | chrX-131882213-INS-636 |
| chrX-131901299-INS-828 | chrX-131901299-INS-828 |
| chrX-131905547-INS-1100 | chrX-131905548-INS-1100 |
| chrX-131923490-INS-243 | chrX-131923490-INS-243 |
| chrX-131950129-INS-1085 | chrX-131950130-INS-1086 |
| chrX-131997169-INS-4287 | chrX-131997170-INS-4283 |
| chrX-132000102-INS-6082 | chrX-132000103-INS-6082 |
| chrX-132043707-INS-3820 | chrX-132043708-INS-3819 |
| chrX-132058078-INS-809 | chrX-132058050-INS-847 |
| chrX-132127929-INS-531 | chrX-132127929-INS-531 |
| chrX-132169257-INS-76 | chrX-132169257-INS-76 |
| chrX-132178511-INS-83 | chrX-132178535-INS-58 |
| chrX-132190024-INS-104 | chrX-132190024-INS-101 |
| chrX-132246537-INS-212 | chrX-132246537-INS-212 |
| chrX-132294054-INS-6473 | chrX-132294055-INS-6473 |
| chrX-132339251-INS-240 | chrX-132339251-INS-240 |
| chrX-132360526-INS-3857 | chrX-132360527-INS-3857 |
| chrX-132373613-INS-2455 | chrX-132373614-INS-2457 |
| chrX-132457941-INS-6905 | chrX-132457942-INS-6905 |
| chrX-132459924-INS-472 | chrX-132459924-INS-474 |
| chrX-132467663-INS-726 | chrX-132467663-INS-727 |
| chrX-132480672-INS-99 | chrX-132480672-INS-94 |
| chrX-132482464-INS-70 | chrX-132482464-INS-70 |
| chrX-132595736-INS-7106 | chrX-132595737-INS-7106 |
| chrX-132623717-INS-92 | chrX-132623679-INS-128 |
| chrX-132679284-INS-864 | chrX-132679284-INS-864 |
| chrX-132688606-INS-6873 | chrX-132688607-INS-6873 |
| chrX-132814816-INS-54 | chrX-132814824-INS-68 |
| chrX-132833879-INS-207 | chrX-132833893-INS-176 |
| chrX-132857892-INS-543 | chrX-132857892-INS-515 |

| | |
|---|---|
| chrX-132909037-INS-654 | chrX-132909037-INS-655 |
| chrX-132932053-INS-694 | chrX-132932053-INS-694 |
| chrX-132934875-INS-483 | chrX-132934875-INS-479 |
| chrX-132938415-INS-111 | chrX-132938445-INS-111 |
| chrX-132948967-INS-1892 | chrX-132948963-INS-1893 |
| chrX-132995685-INS-52 | chrX-132995686-INS-71 |
| chrX-133009925-INS-57 | chrX-133009925-INS-60 |
| chrX-133064650-INS-141 | chrX-133064650-INS-141 |
| chrX-133070003-INS-175 | chrX-133070003-INS-175 |
| chrX-133071923-INS-7521 | chrX-133071924-INS-7524 |
| chrX-133095372-INS-77 | chrX-133095372-INS-77 |
| chrX-133109705-INS-480 | chrX-133109705-INS-482 |
| chrX-133140687-INS-440 | chrX-133140687-INS-440 |
| chrX-133246883-INS-494 | chrX-133246868-INS-495 |
| chrX-133250541-INS-208 | chrX-133250541-INS-208 |
| chrX-133288529-INS-63 | chrX-133288529-INS-63 |
| chrX-133289501-INS-917 | chrX-133289501-INS-920 |
| chrX-133311625-INS-206 | chrX-133311625-INS-206 |
| chrX-133362234-INS-61 | chrX-133362234-INS-61 |
| chrX-133385063-INS-327 | chrX-133385063-INS-345 |
| chrX-133393971-INS-139 | chrX-133393971-INS-138 |
| chrX-133429127-INS-1899 | chrX-133429128-INS-1899 |
| chrX-133462513-INS-80 | chrX-133462513-INS-80 |
| chrX-133463122-INS-71 | chrX-133463122-INS-71 |
| chrX-133479575-INS-70 | chrX-133479575-INS-50 |
| chrX-133494940-INS-407 | chrX-133494940-INS-399 |
| chrX-133540322-INS-1890 | chrX-133540323-INS-1890 |
| chrX-133627344-INS-577 | chrX-133627344-INS-557 |
| chrX-133667014-INS-56 | chrX-133667014-INS-60 |
| chrX-133671793-INS-74 | chrX-133671793-INS-74 |
| chrX-133685323-INS-199 | chrX-133685323-INS-199 |
| chrX-133692787-INS-458 | chrX-133692787-INS-467 |
| chrX-133694075-INS-2929 | chrX-133694076-INS-2929 |
| chrX-133723211-INS-57 | chrX-133723211-INS-57 |
| chrX-133729918-INS-219 | chrX-133729918-INS-218 |
| chrX-133734261-INS-1890 | chrX-133734262-INS-1890 |
| chrX-133744674-INS-2159 | chrX-133744675-INS-2158 |
| chrX-133746690-INS-55 | chrX-133746690-INS-50 |
| chrX-133755542-INS-55 | chrX-133755542-INS-55 |
| chrX-133772554-INS-6422 | chrX-133772555-INS-6422 |
| chrX-133807379-INS-573 | chrX-133807379-INS-569 |
| chrX-133833204-INS-440 | chrX-133833204-INS-440 |
| chrX-133861213-INS-60 | chrX-133861213-INS-60 |
| chrX-133899431-INS-6210 | chrX-133899432-INS-6210 |
| chrX-133972120-INS-394 | chrX-133972120-INS-393 |
| chrX-133975829-INS-83 | chrX-133975830-INS-83 |
| chrX-133988496-INS-962 | chrX-133988496-INS-971 |
| chrX-133990283-INS-64 | chrX-133990283-INS-64 |
| chrX-133993235-INS-378 | chrX-133993235-INS-375 |
| chrX-134000491-INS-234 | chrX-134000491-INS-234 |
| chrX-134001232-INS-78 | chrX-134001232-INS-78 |
| chrX-134081141-INS-76 | chrX-134081141-INS-76 |
| chrX-134092100-INS-86 | chrX-134092100-INS-86 |
| chrX-134104496-INS-485 | chrX-134104496-INS-484 |
| chrX-134114333-INS-198 | chrX-134114333-INS-186 |
| chrX-134124670-INS-95 | chrX-134124672-INS-96 |
| chrX-134127133-INS-578 | chrX-134127133-INS-578 |
| chrX-134128535-INS-203 | chrX-134128535-INS-204 |
| chrX-134164663-INS-148 | chrX-134164663-INS-140 |
| chrX-134192799-INS-280 | chrX-134192799-INS-280 |
| chrX-134245568-INS-797 | chrX-134245568-INS-798 |
| chrX-134252016-INS-109 | chrX-134252039-INS-63 |
| chrX-134265170-INS-72 | chrX-134265170-INS-72 |
| chrX-134455942-INS-200 | chrX-134455950-INS-200 |
| chrX-134457424-INS-876 | chrX-134457424-INS-875 |
| chrX-134461478-INS-783 | chrX-134461478-INS-784 |
| chrX-134464099-INS-66 | chrX-134464099-INS-68 |
| chrX-134472122-INS-64 | chrX-134472122-INS-64 |
| chrX-134477209-INS-121 | chrX-134477209-INS-121 |
| chrX-134507291-INS-7873 | chrX-134507292-INS-7873 |
| chrX-134538782-INS-214 | chrX-134538782-INS-214 |
| chrX-134572453-INS-68 | chrX-134572453-INS-62 |
| chrX-134588425-INS-76 | chrX-134588413-INS-84 |
| chrX-134607218-INS-1446 | chrX-134607219-INS-1447 |
| chrX-134695760-INS-1888 | chrX-134695761-INS-1888 |
| chrX-134719439-INS-421 | chrX-134719439-INS-419 |
| chrX-134761763-INS-72 | chrX-134761763-INS-72 |

| | |
|---|---|
| chrX-134769344-INS-261 | chrX-134769344-INS-266 |
| chrX-134809705-INS-189 | chrX-134809705-INS-190 |
| chrX-134839661-INS-170 | chrX-134839661-INS-170 |
| chrX-134851126-INS-201 | chrX-134851122-INS-200 |
| chrX-134855420-INS-6523 | chrX-134855421-INS-6523 |
| chrX-134862964-INS-69 | chrX-134862964-INS-59 |
| chrX-134865009-INS-5172 | chrX-134865010-INS-5172 |
| chrX-134869357-INS-121 | chrX-134869357-INS-122 |
| chrX-134887791-INS-103 | chrX-134887791-INS-104 |
| chrX-134890008-INS-105 | chrX-134890008-INS-105 |
| chrX-134891711-INS-82 | chrX-134891711-INS-88 |
| chrX-134906243-INS-79 | chrX-134906239-INS-91 |
| chrX-134948905-INS-1247 | chrX-134948906-INS-1247 |
| chrX-135009077-INS-194 | chrX-135009077-INS-190 |
| chrX-135020353-INS-116 | chrX-135020353-INS-116 |
| chrX-135021469-INS-303 | chrX-135021469-INS-303 |
| chrX-135049941-INS-209 | chrX-135049941-INS-209 |
| chrX-135060485-INS-861 | chrX-135060477-INS-855 |
| chrX-135066111-INS-60 | chrX-135066106-INS-52 |
| chrX-135070290-INS-204 | chrX-135070290-INS-205 |
| chrX-135107279-INS-52 | chrX-135107282-INS-84 |
| chrX-135134118-INS-187 | chrX-135134118-INS-187 |
| chrX-135138431-INS-247 | chrX-135138431-INS-247 |
| chrX-135182474-INS-145 | chrX-135182474-INS-145 |
| chrX-135243006-INS-4817 | chrX-135243007-INS-4817 |
| chrX-135253459-INS-6456 | chrX-135253460-INS-6456 |
| chrX-135284535-INS-54 | chrX-135284535-INS-54 |
| chrX-135297900-INS-74 | chrX-135297900-INS-56 |
| chrX-135323931-INS-618 | chrX-135323931-INS-618 |
| chrX-135332063-INS-5523 | chrX-135332064-INS-5523 |
| chrX-135353294-INS-608 | chrX-135353294-INS-608 |
| chrX-135366553-INS-6665 | chrX-135366554-INS-6665 |
| chrX-135389786-INS-80 | chrX-135389786-INS-80 |
| chrX-135421854-INS-113 | chrX-135421854-INS-113 |
| chrX-135460418-INS-77 | chrX-135460418-INS-54 |
| chrX-135475194-INS-215 | chrX-135475194-INS-215 |
| chrX-135511495-INS-6605 | chrX-135511496-INS-6605 |
| chrX-135534114-INS-1893 | chrX-135534115-INS-1893 |
| chrX-135539829-INS-75 | chrX-135539829-INS-77 |
| chrX-135590003-INS-400 | chrX-135590003-INS-399 |
| chrX-135595123-INS-283 | chrX-135595123-INS-271 |
| chrX-135598279-INS-89 | chrX-135598273-INS-85 |
| chrX-135617901-INS-868 | chrX-135617901-INS-868 |
| chrX-135659573-INS-62 | chrX-135659573-INS-72 |
| chrX-135677551-INS-56 | chrX-135677524-INS-94 |
| chrX-135699289-INS-6298 | chrX-135699290-INS-6298 |
| chrX-135705774-INS-1312 | chrX-135705775-INS-1312 |
| chrX-135707374-INS-109 | chrX-135707391-INS-122 |
| chrX-135711883-INS-1890 | chrX-135711884-INS-1890 |
| chrX-135742508-INS-735 | chrX-135742508-INS-735 |
| chrX-135822380-INS-63 | chrX-135822350-INS-88 |
| chrX-135825947-INS-241 | chrX-135825947-INS-241 |
| chrX-135874162-INS-508 | chrX-135874162-INS-508 |
| chrX-135881077-INS-212 | chrX-135881077-INS-226 |
| chrX-135905053-INS-214 | chrX-135905053-INS-215 |
| chrX-135969148-INS-114 | chrX-135969148-INS-108 |
| chrX-135984584-INS-208 | chrX-135984584-INS-209 |
| chrX-135987938-INS-87 | chrX-135987939-INS-55 |
| chrX-136053277-INS-227 | chrX-136053277-INS-235 |
| chrX-136075662-INS-258 | chrX-136075662-INS-258 |
| chrX-136077666-INS-1887 | chrX-136077667-INS-1890 |
| chrX-136077707-INS-6452 | chrX-136077708-INS-6452 |
| chrX-136093562-INS-207 | chrX-136093562-INS-207 |
| chrX-136123443-INS-82 | chrX-136123443-INS-82 |
| chrX-136127839-INS-342 | chrX-136127839-INS-341 |
| chrX-136140657-INS-118 | chrX-136140647-INS-110 |
| chrX-136150322-INS-4174 | chrX-136150323-INS-4174 |
| chrX-136155043-INS-87 | chrX-136155043-INS-87 |
| chrX-136171077-INS-7156 | chrX-136171078-INS-7158 |
| chrX-136223549-INS-603 | chrX-136223549-INS-605 |
| chrX-136295301-INS-512 | chrX-136295301-INS-511 |
| chrX-136344590-INS-1153 | chrX-136344591-INS-1153 |
| chrX-136375230-INS-57 | chrX-136375230-INS-72 |
| chrX-136378907-INS-155 | chrX-136378907-INS-154 |
| chrX-136466541-INS-413 | chrX-136466541-INS-414 |
| chrX-136467487-INS-56 | chrX-136467487-INS-56 |
| chrX-136481273-INS-6439 | chrX-136481274-INS-6440 |

| | |
|---|---|
| chrX-136488588-INS-268 | chrX-136488588-INS-268 |
| chrX-136490054-INS-462 | chrX-136490054-INS-462 |
| chrX-136578923-INS-51 | chrX-136578921-INS-50 |
| chrX-136656870-INS-4355 | chrX-136656871-INS-4355 |
| chrX-136666703-INS-649 | chrX-136666703-INS-650 |
| chrX-136693685-INS-528 | chrX-136693685-INS-529 |
| chrX-136719739-INS-6880 | chrX-136719740-INS-6881 |
| chrX-136747549-INS-58 | chrX-136747549-INS-60 |
| chrX-136771103-INS-184 | chrX-136771087-INS-187 |
| chrX-136882463-INS-790 | chrX-136882463-INS-798 |
| chrX-136892082-INS-771 | chrX-136892082-INS-772 |
| chrX-136902397-INS-316 | chrX-136902397-INS-314 |
| chrX-136989483-INS-54 | chrX-136989483-INS-53 |
| chrX-136991256-INS-193 | chrX-136991221-INS-214 |
| chrX-136992859-INS-52 | chrX-136992859-INS-52 |
| chrX-136994004-INS-90 | chrX-136994012-INS-104 |
| chrX-137001294-INS-54 | chrX-137001334-INS-50 |
| chrX-137041549-INS-1594 | chrX-137041550-INS-1596 |
| chrX-137045256-INS-97 | chrX-137045292-INS-69 |
| chrX-137093963-INS-322 | chrX-137093963-INS-322 |
| chrX-137097398-INS-1005 | chrX-137097399-INS-999 |
| chrX-137108681-INS-56 | chrX-137108651-INS-60 |
| chrX-137145233-INS-65 | chrX-137145218-INS-80 |
| chrX-137162162-INS-54 | chrX-137162162-INS-54 |
| chrX-137204554-INS-285 | chrX-137204554-INS-285 |
| chrX-137273925-INS-132 | chrX-137273925-INS-132 |
| chrX-137387974-INS-775 | chrX-137387974-INS-775 |
| chrX-137402035-INS-340 | chrX-137402032-INS-346 |
| chrX-137435581-INS-3192 | chrX-137435582-INS-3193 |
| chrX-137438553-INS-7344 | chrX-137438554-INS-7344 |
| chrX-137454918-INS-80 | chrX-137454905-INS-114 |
| chrX-137456169-INS-194 | chrX-137456169-INS-195 |
| chrX-137500676-INS-85 | chrX-137500676-INS-85 |
| chrX-137509297-INS-62 | chrX-137509297-INS-62 |
| chrX-137528401-INS-53 | chrX-137528407-INS-80 |
| chrX-137620618-INS-822 | chrX-137620618-INS-827 |
| chrX-137629422-INS-70 | chrX-137629422-INS-72 |
| chrX-137666863-INS-77 | chrX-137666861-INS-50 |
| chrX-137685358-INS-72 | chrX-137685358-INS-92 |
| chrX-137693400-INS-492 | chrX-137693400-INS-492 |
| chrX-137696316-INS-72 | chrX-137696316-INS-72 |
| chrX-137750678-INS-66 | chrX-137750678-INS-99 |
| chrX-137763339-INS-1169 | chrX-137763340-INS-1169 |
| chrX-137764245-INS-129 | chrX-137764259-INS-102 |
| chrX-137781514-INS-6615 | chrX-137781515-INS-6615 |
| chrX-137798820-INS-951 | chrX-137798820-INS-953 |
| chrX-137911831-INS-280 | chrX-137911831-INS-281 |
| chrX-137915914-INS-88 | chrX-137915958-INS-90 |
| chrX-137930634-INS-77 | chrX-137930634-INS-77 |
| chrX-137933180-INS-117 | chrX-137933182-INS-112 |
| chrX-137934221-INS-6224 | chrX-137934222-INS-6226 |
| chrX-137938696-INS-64 | chrX-137938696-INS-64 |
| chrX-137940391-INS-52 | chrX-137940391-INS-54 |
| chrX-137941394-INS-736 | chrX-137941394-INS-736 |
| chrX-137947555-INS-1198 | chrX-137947555-INS-1200 |
| chrX-137958679-INS-82 | chrX-137958672-INS-62 |
| chrX-137963209-INS-80 | chrX-137963190-INS-78 |
| chrX-137999517-INS-1817 | chrX-137999518-INS-1817 |
| chrX-138001719-INS-54 | chrX-138001719-INS-59 |
| chrX-138010833-INS-131 | chrX-138010833-INS-159 |
| chrX-138029601-INS-205 | chrX-138029601-INS-204 |
| chrX-138044038-INS-485 | chrX-138044038-INS-485 |
| chrX-138176474-INS-772 | chrX-138176474-INS-772 |
| chrX-138179161-INS-400 | chrX-138179161-INS-400 |
| chrX-138267859-INS-703 | chrX-138267859-INS-703 |
| chrX-138306879-INS-727 | chrX-138306879-INS-727 |
| chrX-138314561-INS-846 | chrX-138314561-INS-848 |
| chrX-138335859-INS-94 | chrX-138335859-INS-100 |
| chrX-138341349-INS-181 | chrX-138341349-INS-181 |
| chrX-138366457-INS-54 | chrX-138366457-INS-54 |
| chrX-138430285-INS-122 | chrX-138430285-INS-121 |
| chrX-138439691-INS-475 | chrX-138439691-INS-475 |
| chrX-138443244-INS-206 | chrX-138443244-INS-206 |
| chrX-138457339-INS-63 | chrX-138457339-INS-65 |
| chrX-138469030-INS-52 | chrX-138469030-INS-55 |
| chrX-138476757-INS-57 | chrX-138476757-INS-69 |
| chrX-138488695-INS-58 | chrX-138488695-INS-56 |

| | |
|---|---|
| chrX-138520275-INS-64 | chrX-138520275-INS-66 |
| chrX-138529079-INS-809 | chrX-138529079-INS-810 |
| chrX-138545020-INS-60 | chrX-138545020-INS-64 |
| chrX-138571133-INS-2702 | chrX-138571134-INS-2702 |
| chrX-138600264-INS-54 | chrX-138600264-INS-86 |
| chrX-138606432-INS-78 | chrX-138606432-INS-78 |
| chrX-138615454-INS-467 | chrX-138615454-INS-468 |
| chrX-138619256-INS-100 | chrX-138619299-INS-56 |
| chrX-138620457-INS-6386 | chrX-138620458-INS-6386 |
| chrX-138651098-INS-440 | chrX-138651098-INS-440 |
| chrX-138662424-INS-512 | chrX-138662424-INS-512 |
| chrX-138668976-INS-901 | chrX-138668976-INS-901 |
| chrX-138694458-INS-71 | chrX-138694414-INS-72 |
| chrX-138708293-INS-54 | chrX-138708293-INS-56 |
| chrX-138728418-INS-1886 | chrX-138728419-INS-1886 |
| chrX-138812007-INS-100 | chrX-138812007-INS-100 |
| chrX-138859075-INS-80 | chrX-138859075-INS-80 |
| chrX-138966160-INS-141 | chrX-138966160-INS-141 |
| chrX-139017136-INS-342 | chrX-139017136-INS-335 |
| chrX-139024634-INS-128 | chrX-139024634-INS-128 |
| chrX-139042823-INS-4173 | chrX-139042824-INS-4173 |
| chrX-139099434-INS-1718 | chrX-139099435-INS-1718 |
| chrX-139101217-INS-423 | chrX-139101217-INS-423 |
| chrX-139180582-INS-51 | chrX-139180574-INS-52 |
| chrX-139227237-INS-715 | chrX-139227237-INS-716 |
| chrX-139249849-INS-58 | chrX-139249849-INS-58 |
| chrX-139282699-INS-74 | chrX-139282683-INS-59 |
| chrX-139307493-INS-76 | chrX-139307493-INS-76 |
| chrX-139339040-INS-217 | chrX-139339040-INS-217 |
| chrX-139341607-INS-109 | chrX-139341607-INS-107 |
| chrX-139397490-INS-54 | chrX-139397491-INS-52 |
| chrX-139429518-INS-72 | chrX-139429518-INS-95 |
| chrX-139433500-INS-52 | chrX-139433500-INS-87 |
| chrX-139507619-INS-250 | chrX-139507619-INS-242 |
| chrX-139521738-INS-84 | chrX-139521699-INS-80 |
| chrX-139530224-INS-5320 | chrX-139530225-INS-5320 |
| chrX-139609021-INS-1058 | chrX-139609022-INS-1058 |
| chrX-139610916-INS-198 | chrX-139610908-INS-162 |
| chrX-139639668-INS-74 | chrX-139639669-INS-53 |
| chrX-139657651-INS-173 | chrX-139657651-INS-137 |
| chrX-139680124-INS-97 | chrX-139680124-INS-97 |
| chrX-139716286-INS-61 | chrX-139716286-INS-72 |
| chrX-139722595-INS-81 | chrX-139722559-INS-117 |
| chrX-139780871-INS-109 | chrX-139780871-INS-109 |
| chrX-139782844-INS-198 | chrX-139782844-INS-198 |
| chrX-139823229-INS-327 | chrX-139823229-INS-327 |
| chrX-139848624-INS-6494 | chrX-139848625-INS-6497 |
| chrX-139859070-INS-78 | chrX-139859070-INS-82 |
| chrX-139898097-INS-225 | chrX-139898097-INS-225 |
| chrX-139899625-INS-90 | chrX-139899625-INS-90 |
| chrX-139906115-INS-78 | chrX-139906079-INS-80 |
| chrX-139908433-INS-1116 | chrX-139908434-INS-1115 |
| chrX-139914537-INS-110 | chrX-139914531-INS-78 |
| chrX-139926051-INS-75 | chrX-139926028-INS-82 |
| chrX-139995420-INS-6513 | chrX-139995421-INS-6514 |
| chrX-139997680-INS-268 | chrX-139997680-INS-269 |
| chrX-140035116-INS-52 | chrX-140035116-INS-52 |
| chrX-140036903-INS-94 | chrX-140036903-INS-94 |
| chrX-140039700-INS-6652 | chrX-140039701-INS-6653 |
| chrX-140073209-INS-159 | chrX-140073209-INS-182 |
| chrX-140078297-INS-104 | chrX-140078297-INS-104 |
| chrX-140091717-INS-63 | chrX-140091717-INS-55 |
| chrX-140097601-INS-143 | chrX-140097601-INS-98 |
| chrX-140098662-INS-117 | chrX-140098662-INS-121 |
| chrX-140118689-INS-144 | chrX-140118689-INS-138 |
| chrX-140122350-INS-166 | chrX-140122350-INS-165 |
| chrX-140131554-INS-534 | chrX-140131554-INS-534 |
| chrX-140147253-INS-216 | chrX-140147253-INS-216 |
| chrX-140154789-INS-5547 | chrX-140154790-INS-5547 |
| chrX-140197884-INS-2570 | chrX-140197885-INS-2570 |
| chrX-140212111-INS-222 | chrX-140212111-INS-222 |
| chrX-140216432-INS-58 | chrX-140216426-INS-56 |
| chrX-140224468-INS-3578 | chrX-140224469-INS-3578 |
| chrX-140282137-INS-6507 | chrX-140282138-INS-6507 |
| chrX-140294216-INS-744 | chrX-140294216-INS-744 |
| chrX-140309684-INS-56 | chrX-140309684-INS-56 |
| chrX-140388644-INS-217 | chrX-140388644-INS-217 |

| | |
|---|---|
| chrX-140417167-INS-84 | chrX-140417167-INS-82 |
| chrX-140459495-INS-89 | chrX-140459522-INS-85 |
| chrX-140530481-INS-424 | chrX-140530481-INS-424 |
| chrX-140531523-INS-56 | chrX-140531523-INS-63 |
| chrX-140560782-INS-130 | chrX-140560782-INS-131 |
| chrX-140562208-INS-60 | chrX-140562226-INS-60 |
| chrX-140621340-INS-680 | chrX-140621340-INS-681 |
| chrX-140625747-INS-224 | chrX-140625748-INS-224 |
| chrX-140629882-INS-1890 | chrX-140629883-INS-1890 |
| chrX-140642557-INS-58 | chrX-140642527-INS-89 |
| chrX-140656639-INS-72 | chrX-140656639-INS-72 |
| chrX-140689230-INS-1254 | chrX-140689231-INS-1253 |
| chrX-140704000-INS-146 | chrX-140704000-INS-142 |
| chrX-140709045-INS-1164 | chrX-140709046-INS-1167 |
| chrX-140711335-INS-124 | chrX-140711335-INS-174 |
| chrX-140729934-INS-52 | chrX-140729934-INS-94 |
| chrX-140748143-INS-498 | chrX-140748143-INS-498 |
| chrX-140770558-INS-60 | chrX-140770558-INS-60 |
| chrX-140799759-INS-5277 | chrX-140799760-INS-5277 |
| chrX-140811058-INS-145 | chrX-140811058-INS-145 |
| chrX-140825279-INS-357 | chrX-140825279-INS-357 |
| chrX-140833201-INS-181 | chrX-140833201-INS-175 |
| chrX-140916859-INS-131 | chrX-140916859-INS-128 |
| chrX-140927417-INS-201 | chrX-140927417-INS-201 |
| chrX-140960898-INS-53 | chrX-140960895-INS-78 |
| chrX-141002545-INS-108 | chrX-141002545-INS-114 |
| chrX-141011556-INS-51 | chrX-141011557-INS-66 |
| chrX-141033008-INS-237 | chrX-141033008-INS-237 |
| chrX-141072897-INS-6351 | chrX-141072898-INS-6350 |
| chrX-141109033-INS-56 | chrX-141109036-INS-100 |
| chrX-141122486-INS-3035 | chrX-141122487-INS-3035 |
| chrX-141129589-INS-54 | chrX-141129603-INS-82 |
| chrX-141148056-INS-682 | chrX-141148071-INS-681 |
| chrX-141167680-INS-100 | chrX-141167680-INS-108 |
| chrX-141175939-INS-53 | chrX-141175940-INS-52 |
| chrX-141188920-INS-194 | chrX-141188920-INS-195 |
| chrX-141191243-INS-172 | chrX-141191243-INS-174 |
| chrX-141211915-INS-180 | chrX-141211915-INS-181 |
| chrX-141225629-INS-92 | chrX-141225629-INS-96 |
| chrX-141227604-INS-55 | chrX-141227605-INS-87 |
| chrX-141233777-INS-227 | chrX-141233781-INS-217 |
| chrX-141234956-INS-1069 | chrX-141234957-INS-1071 |
| chrX-141239863-INS-457 | chrX-141239863-INS-457 |
| chrX-141272572-INS-69 | chrX-141272568-INS-69 |
| chrX-141295992-INS-72 | chrX-141295992-INS-68 |
| chrX-141312198-INS-235 | chrX-141312198-INS-235 |
| chrX-141363539-INS-55 | chrX-141363539-INS-65 |
| chrX-141415912-INS-54 | chrX-141415912-INS-54 |
| chrX-141428929-INS-72 | chrX-141428917-INS-105 |
| chrX-141432552-INS-202 | chrX-141432552-INS-203 |
| chrX-141438486-INS-113 | chrX-141438486-INS-113 |
| chrX-141476601-INS-62 | chrX-141476568-INS-57 |
| chrX-141498051-INS-214 | chrX-141498051-INS-214 |
| chrX-141503562-INS-187 | chrX-141503562-INS-159 |
| chrX-141522316-INS-121 | chrX-141522316-INS-121 |
| chrX-141594441-INS-694 | chrX-141594441-INS-692 |
| chrX-141610653-INS-80 | chrX-141610673-INS-52 |
| chrX-141628442-INS-578 | chrX-141628442-INS-577 |
| chrX-141693573-INS-66 | chrX-141693573-INS-66 |
| chrX-141704347-INS-833 | chrX-141704347-INS-833 |
| chrX-141721250-INS-128 | chrX-141721250-INS-128 |
| chrX-141778464-INS-237 | chrX-141778464-INS-237 |
| chrX-141814867-INS-215 | chrX-141814867-INS-214 |
| chrX-141836084-INS-52 | chrX-141836084-INS-52 |
| chrX-141841108-INS-184 | chrX-141841108-INS-184 |
| chrX-141844463-INS-84 | chrX-141844463-INS-84 |
| chrX-141856011-INS-105 | chrX-141856011-INS-70 |
| chrX-141900772-INS-62 | chrX-141900772-INS-61 |
| chrX-141920435-INS-140 | chrX-141920435-INS-140 |
| chrX-141968171-INS-77 | chrX-141968171-INS-77 |
| chrX-141989566-INS-1625 | chrX-141989567-INS-1625 |
| chrX-141996138-INS-458 | chrX-141996138-INS-457 |
| chrX-142020495-INS-84 | chrX-142020513-INS-84 |
| chrX-142155493-INS-321 | chrX-142155497-INS-327 |
| chrX-142157126-INS-6451 | chrX-142157127-INS-6451 |
| chrX-142232923-INS-88 | chrX-142232923-INS-64 |
| chrX-142256510-INS-56 | chrX-142256510-INS-56 |

| | |
|---|---|
| chrX-142263839-INS-289 | chrX-142263838-INS-293 |
| chrX-142343458-INS-98 | chrX-142343458-INS-113 |
| chrX-142399331-INS-547 | chrX-142399331-INS-547 |
| chrX-142418613-INS-64 | chrX-142418613-INS-64 |
| chrX-142420972-INS-71 | chrX-142420972-INS-73 |
| chrX-142443418-INS-1890 | chrX-142443419-INS-1890 |
| chrX-142445269-INS-172 | chrX-142445269-INS-173 |
| chrX-142463213-INS-65 | chrX-142463213-INS-61 |
| chrX-142488267-INS-59 | chrX-142488265-INS-53 |
| chrX-142538858-INS-6318 | chrX-142538859-INS-6323 |
| chrX-142544612-INS-181 | chrX-142544612-INS-231 |
| chrX-142556987-INS-72 | chrX-142556987-INS-72 |
| chrX-142587957-INS-80 | chrX-142587957-INS-70 |
| chrX-142633334-INS-464 | chrX-142633334-INS-465 |
| chrX-142684579-INS-113 | chrX-142684579-INS-113 |
| chrX-142690308-INS-65 | chrX-142690308-INS-50 |
| chrX-142705955-INS-82 | chrX-142705943-INS-81 |
| chrX-142733862-INS-343 | chrX-142733862-INS-343 |
| chrX-142763617-INS-582 | chrX-142763617-INS-582 |
| chrX-142797575-INS-213 | chrX-142797580-INS-214 |
| chrX-142863165-INS-1104 | chrX-142863166-INS-1104 |
| chrX-142883006-INS-53 | chrX-142882956-INS-54 |
| chrX-142934628-INS-8986 | chrX-142934629-INS-8986 |
| chrX-142967749-INS-2618 | chrX-142967750-INS-2617 |
| chrX-142968543-INS-774 | chrX-142968543-INS-774 |
| chrX-142970677-INS-6557 | chrX-142970678-INS-6558 |
| chrX-142974760-INS-96 | chrX-142974760-INS-100 |
| chrX-142982689-INS-141 | chrX-142982688-INS-186 |
| chrX-142987451-INS-2248 | chrX-142987452-INS-2248 |
| chrX-143020035-INS-179 | chrX-143020035-INS-178 |
| chrX-143036535-INS-164 | chrX-143036555-INS-155 |
| chrX-143076526-INS-184 | chrX-143076526-INS-185 |
| chrX-143080574-INS-4763 | chrX-143080575-INS-4765 |
| chrX-143082675-INS-53 | chrX-143082675-INS-90 |
| chrX-143089838-INS-609 | chrX-143089838-INS-605 |
| chrX-143116854-INS-82 | chrX-143116854-INS-82 |
| chrX-143129710-INS-143 | chrX-143129710-INS-143 |
| chrX-143132542-INS-66 | chrX-143132542-INS-66 |
| chrX-143171885-INS-6403 | chrX-143171886-INS-6403 |
| chrX-143181406-INS-104 | chrX-143181406-INS-68 |
| chrX-143198607-INS-401 | chrX-143198607-INS-401 |
| chrX-143265646-INS-59 | chrX-143265647-INS-59 |
| chrX-143268041-INS-202 | chrX-143268041-INS-201 |
| chrX-143272694-INS-778 | chrX-143272694-INS-778 |
| chrX-143285761-INS-6442 | chrX-143285762-INS-6442 |
| chrX-143295508-INS-263 | chrX-143295508-INS-263 |
| chrX-143317761-INS-4224 | chrX-143317770-INS-4214 |
| chrX-143321640-INS-6535 | chrX-143321641-INS-6536 |
| chrX-143321650-INS-2512 | chrX-143321651-INS-2513 |
| chrX-143328087-INS-2170 | chrX-143328088-INS-2170 |
| chrX-143357086-INS-667 | chrX-143357086-INS-667 |
| chrX-143360357-INS-123 | chrX-143360357-INS-122 |
| chrX-143363527-INS-212 | chrX-143363527-INS-212 |
| chrX-143377269-INS-197 | chrX-143377269-INS-197 |
| chrX-143395227-INS-3426 | chrX-143395228-INS-3427 |
| chrX-143432118-INS-249 | chrX-143432118-INS-249 |
| chrX-143450231-INS-536 | chrX-143450231-INS-536 |
| chrX-143560305-INS-388 | chrX-143560305-INS-387 |
| chrX-143565647-INS-52 | chrX-143565648-INS-52 |
| chrX-143634599-INS-142 | chrX-143634610-INS-136 |
| chrX-143675465-INS-82 | chrX-143675465-INS-70 |
| chrX-143704428-INS-65 | chrX-143704428-INS-55 |
| chrX-143719195-INS-488 | chrX-143719195-INS-488 |
| chrX-143731407-INS-2362 | chrX-143731408-INS-2361 |
| chrX-143733190-INS-6195 | chrX-143733191-INS-6199 |
| chrX-143785433-INS-56 | chrX-143785433-INS-58 |
| chrX-143793611-INS-356 | chrX-143793611-INS-354 |
| chrX-143795009-INS-764 | chrX-143795009-INS-763 |
| chrX-143796285-INS-59 | chrX-143796285-INS-77 |
| chrX-143817506-INS-130 | chrX-143817506-INS-130 |
| chrX-143870645-INS-376 | chrX-143870645-INS-376 |
| chrX-143906555-INS-1315 | chrX-143906556-INS-1317 |
| chrX-143916272-INS-825 | chrX-143916272-INS-824 |
| chrX-143943654-INS-87 | chrX-143943654-INS-71 |
| chrX-143969247-INS-6563 | chrX-143969248-INS-6563 |
| chrX-143982039-INS-1163 | chrX-143982041-INS-1155 |
| chrX-144039905-INS-1121 | chrX-144039906-INS-1121 |

| | |
|---|---|
| chrX-144051516-INS-649 | chrX-144051516-INS-649 |
| chrX-144053162-INS-946 | chrX-144053162-INS-947 |
| chrX-144056858-INS-1452 | chrX-144056859-INS-1457 |
| chrX-144097745-INS-214 | chrX-144097745-INS-214 |
| chrX-144115145-INS-71 | chrX-144115145-INS-71 |
| chrX-144136648-INS-590 | chrX-144136648-INS-588 |
| chrX-144143507-INS-152 | chrX-144143507-INS-152 |
| chrX-144164328-INS-939 | chrX-144164329-INS-939 |
| chrX-144179669-INS-524 | chrX-144179669-INS-524 |
| chrX-144206751-INS-106 | chrX-144206751-INS-112 |
| chrX-144315015-INS-322 | chrX-144315015-INS-322 |
| chrX-144335977-INS-387 | chrX-144335977-INS-388 |
| chrX-144379822-INS-80 | chrX-144379822-INS-80 |
| chrX-144380984-INS-1548 | chrX-144380985-INS-1550 |
| chrX-144395016-INS-727 | chrX-144395016-INS-727 |
| chrX-144401450-INS-4631 | chrX-144401451-INS-4631 |
| chrX-144415216-INS-6387 | chrX-144415217-INS-6388 |
| chrX-144419855-INS-4761 | chrX-144419856-INS-4761 |
| chrX-144463592-INS-72 | chrX-144463592-INS-72 |
| chrX-144554488-INS-1219 | chrX-144554489-INS-1211 |
| chrX-144557795-INS-65 | chrX-144557795-INS-59 |
| chrX-144563478-INS-208 | chrX-144563478-INS-208 |
| chrX-144604905-INS-1245 | chrX-144604906-INS-1249 |
| chrX-144633995-INS-426 | chrX-144633995-INS-426 |
| chrX-144646973-INS-2061 | chrX-144646974-INS-2062 |
| chrX-144725515-INS-67 | chrX-144725515-INS-67 |
| chrX-144750553-INS-5576 | chrX-144750554-INS-5576 |
| chrX-144753376-INS-425 | chrX-144753376-INS-425 |
| chrX-144754495-INS-651 | chrX-144754495-INS-651 |
| chrX-144758512-INS-4515 | chrX-144758513-INS-4515 |
| chrX-144785060-INS-92 | chrX-144785060-INS-92 |
| chrX-144838579-INS-7430 | chrX-144838580-INS-7430 |
| chrX-144871548-INS-64 | chrX-144871548-INS-64 |
| chrX-144890114-INS-85 | chrX-144890103-INS-78 |
| chrX-144950155-INS-60 | chrX-144950155-INS-60 |
| chrX-144965362-INS-400 | chrX-144965362-INS-400 |
| chrX-144972768-INS-7240 | chrX-144972769-INS-7240 |
| chrX-144975914-INS-2600 | chrX-144975915-INS-2600 |
| chrX-144978713-INS-6290 | chrX-144978714-INS-6290 |
| chrX-144989424-INS-5000 | chrX-144989425-INS-5000 |
| chrX-144989480-INS-197 | chrX-144989480-INS-197 |
| chrX-144998954-INS-66 | chrX-144998954-INS-66 |
| chrX-145002157-INS-62 | chrX-145002157-INS-62 |
| chrX-145016711-INS-58 | chrX-145016711-INS-57 |
| chrX-145041913-INS-52 | chrX-145041914-INS-52 |
| chrX-145056255-INS-60 | chrX-145056255-INS-60 |
| chrX-145063705-INS-146 | chrX-145063705-INS-146 |
| chrX-145128756-INS-547 | chrX-145128756-INS-547 |
| chrX-145155552-INS-292 | chrX-145155552-INS-293 |
| chrX-145156474-INS-67 | chrX-145156474-INS-52 |
| chrX-145165781-INS-198 | chrX-145165781-INS-198 |
| chrX-145220218-INS-81 | chrX-145220220-INS-84 |
| chrX-145245502-INS-1302 | chrX-145245503-INS-1302 |
| chrX-145307070-INS-66 | chrX-145307070-INS-51 |
| chrX-145309918-INS-205 | chrX-145309918-INS-227 |
| chrX-145327712-INS-55 | chrX-145327712-INS-55 |
| chrX-145342746-INS-7824 | chrX-145342747-INS-7824 |
| chrX-145406930-INS-3209 | chrX-145406931-INS-3209 |
| chrX-145409053-INS-6379 | chrX-145409069-INS-6380 |
| chrX-145464567-INS-102 | chrX-145464551-INS-119 |
| chrX-145465859-INS-1701 | chrX-145465860-INS-1701 |
| chrX-145472985-INS-98 | chrX-145472985-INS-50 |
| chrX-145493027-INS-7843 | chrX-145493028-INS-7846 |
| chrX-145515737-INS-506 | chrX-145515737-INS-506 |
| chrX-145530935-INS-500 | chrX-145530935-INS-500 |
| chrX-145568999-INS-884 | chrX-145568999-INS-883 |
| chrX-145571570-INS-6180 | chrX-145571571-INS-6179 |
| chrX-145588078-INS-7335 | chrX-145588079-INS-7336 |
| chrX-145597997-INS-4117 | chrX-145597998-INS-4119 |
| chrX-145633688-INS-593 | chrX-145633688-INS-593 |
| chrX-145634463-INS-1489 | chrX-145634464-INS-1488 |
| chrX-145637198-INS-110 | chrX-145637198-INS-131 |
| chrX-145647615-INS-6516 | chrX-145647616-INS-6516 |
| chrX-145654117-INS-110 | chrX-145654114-INS-116 |
| chrX-145695860-INS-57 | chrX-145695860-INS-93 |
| chrX-145698031-INS-571 | chrX-145698031-INS-572 |
| chrX-145698821-INS-116 | chrX-145698821-INS-116 |

| | |
|---|---|
| chrX-145710137-INS-60 | chrX-145710185-INS-54 |
| chrX-145725457-INS-1441 | chrX-145725458-INS-1441 |
| chrX-145728900-INS-5411 | chrX-145728901-INS-5410 |
| chrX-145734385-INS-234 | chrX-145734385-INS-270 |
| chrX-145765582-INS-152 | chrX-145765582-INS-164 |
| chrX-145778882-INS-604 | chrX-145778882-INS-604 |
| chrX-145820467-INS-7355 | chrX-145820468-INS-7355 |
| chrX-145883877-INS-83 | chrX-145883877-INS-83 |
| chrX-145889565-INS-423 | chrX-145889565-INS-419 |
| chrX-145893200-INS-212 | chrX-145893200-INS-212 |
| chrX-145920220-INS-82 | chrX-145920220-INS-84 |
| chrX-145962757-INS-53 | chrX-145962765-INS-93 |
| chrX-145989055-INS-882 | chrX-145989055-INS-882 |
| chrX-146020524-INS-53 | chrX-146020524-INS-53 |
| chrX-146031558-INS-70 | chrX-146031558-INS-74 |
| chrX-146032211-INS-218 | chrX-146032211-INS-217 |
| chrX-146034184-INS-308 | chrX-146034184-INS-308 |
| chrX-146040483-INS-357 | chrX-146040483-INS-357 |
| chrX-146062006-INS-1967 | chrX-146062007-INS-1967 |
| chrX-146123753-INS-63 | chrX-146123756-INS-63 |
| chrX-146130043-INS-889 | chrX-146130043-INS-889 |
| chrX-146137444-INS-146 | chrX-146137444-INS-158 |
| chrX-146158329-INS-5969 | chrX-146158330-INS-5971 |
| chrX-146243453-INS-231 | chrX-146243453-INS-231 |
| chrX-146303129-INS-2624 | chrX-146303130-INS-2623 |
| chrX-146306293-INS-69 | chrX-146306293-INS-69 |
| chrX-146307505-INS-225 | chrX-146307505-INS-225 |
| chrX-146317136-INS-305 | chrX-146317136-INS-305 |
| chrX-146372890-INS-1890 | chrX-146372891-INS-1890 |
| chrX-146389851-INS-1642 | chrX-146389852-INS-1637 |
| chrX-146444870-INS-6731 | chrX-146444871-INS-6732 |
| chrX-146450013-INS-639 | chrX-146450013-INS-639 |
| chrX-146471423-INS-3037 | chrX-146471424-INS-3036 |
| chrX-146527365-INS-56 | chrX-146527365-INS-62 |
| chrX-146529402-INS-7140 | chrX-146529403-INS-7144 |
| chrX-146559118-INS-65 | chrX-146559123-INS-52 |
| chrX-146723564-INS-57 | chrX-146723564-INS-57 |
| chrX-148503992-INS-4346 | chrX-148503993-INS-4346 |
| chrX-148886307-INS-1414 | chrX-148886308-INS-1426 |
| chrX-148896268-INS-4604 | chrX-148896269-INS-4604 |
| chrX-148896521-INS-352 | chrX-148896521-INS-349 |
| chrX-148936879-INS-6415 | chrX-148936880-INS-6416 |
| chrX-148939056-INS-138 | chrX-148939056-INS-138 |
| chrX-148960681-INS-83 | chrX-148960681-INS-97 |
| chrX-148961794-INS-218 | chrX-148961794-INS-226 |
| chrX-148999807-INS-964 | chrX-148999807-INS-961 |
| chrX-149001118-INS-3000 | chrX-149001119-INS-2985 |
| chrX-149020736-INS-1386 | chrX-149020737-INS-1386 |
| chrX-149041174-INS-6497 | chrX-149041175-INS-6496 |
| chrX-149075611-INS-1071 | chrX-149075612-INS-1073 |
| chrX-149076377-INS-5486 | chrX-149076377-INS-5491 |
| chrX-149080502-INS-342 | chrX-149080502-INS-343 |
| chrX-149082681-INS-353 | chrX-149082681-INS-353 |
| chrX-149090711-INS-6440 | chrX-149090712-INS-6440 |
| chrX-149127674-INS-51 | chrX-149127674-INS-51 |
| chrX-149129712-INS-94 | chrX-149129710-INS-52 |
| chrX-149131877-INS-101 | chrX-149131876-INS-110 |
| chrX-149158544-INS-3172 | chrX-149158545-INS-3172 |
| chrX-149162735-INS-236 | chrX-149162735-INS-236 |
| chrX-149175610-INS-125 | chrX-149175608-INS-119 |
| chrX-149199473-INS-211 | chrX-149199473-INS-207 |
| chrX-149205272-INS-239 | chrX-149205272-INS-239 |
| chrX-149224732-INS-79 | chrX-149224732-INS-79 |
| chrX-149230407-INS-761 | chrX-149230407-INS-761 |
| chrX-149246214-INS-259 | chrX-149246214-INS-261 |
| chrX-149259035-INS-600 | chrX-149259035-INS-600 |
| chrX-149261830-INS-60 | chrX-149261830-INS-59 |
| chrX-149273972-INS-6655 | chrX-149273973-INS-6658 |
| chrX-149288969-INS-627 | chrX-149288969-INS-625 |
| chrX-149298448-INS-53 | chrX-149298449-INS-53 |
| chrX-149328405-INS-399 | chrX-149328405-INS-399 |
| chrX-149342016-INS-72 | chrX-149342016-INS-78 |
| chrX-149348386-INS-66 | chrX-149348386-INS-64 |
| chrX-149355859-INS-52 | chrX-149355901-INS-52 |
| chrX-149360072-INS-427 | chrX-149360072-INS-428 |
| chrX-149424673-INS-1165 | chrX-149424674-INS-1127 |
| chrX-149491594-INS-202 | chrX-149491594-INS-202 |

| | |
|---|---|
| chrX-149575313-INS-1010 | chrX-149575314-INS-1010 |
| chrX-149577442-INS-52 | chrX-149577442-INS-52 |
| chrX-149608384-INS-89 | chrX-149608384-INS-90 |
| chrX-149652844-INS-74 | chrX-149652844-INS-76 |
| chrX-149683482-INS-381 | chrX-149683482-INS-381 |
| chrX-149687542-INS-68 | chrX-149687542-INS-68 |
| chrX-149716251-INS-84 | chrX-149716231-INS-64 |
| chrX-149743658-INS-58 | chrX-149743655-INS-68 |
| chrX-149760826-INS-354 | chrX-149760826-INS-354 |
| chrX-149795906-INS-202 | chrX-149795908-INS-189 |
| chrX-149802087-INS-496 | chrX-149802088-INS-465 |
| chrX-149806517-INS-2385 | chrX-149806518-INS-2385 |
| chrX-149815244-INS-51 | chrX-149815245-INS-54 |
| chrX-149817014-INS-89 | chrX-149817014-INS-89 |
| chrX-149828247-INS-202 | chrX-149828247-INS-202 |
| chrX-149905803-INS-645 | chrX-149905803-INS-645 |
| chrX-149917913-INS-66 | chrX-149917913-INS-66 |
| chrX-149936807-INS-825 | chrX-149936807-INS-825 |
| chrX-149943077-INS-203 | chrX-149943062-INS-204 |
| chrX-149945813-INS-220 | chrX-149945813-INS-218 |
| chrX-149957936-INS-196 | chrX-149957936-INS-197 |
| chrX-149958999-INS-206 | chrX-149958999-INS-206 |
| chrX-149969387-INS-229 | chrX-149969387-INS-229 |
| chrX-149984057-INS-83 | chrX-149984057-INS-83 |
| chrX-149994242-INS-212 | chrX-149994242-INS-212 |
| chrX-149995270-INS-205 | chrX-149995270-INS-205 |
| chrX-150002519-INS-60 | chrX-150002517-INS-76 |
| chrX-150012670-INS-53 | chrX-150012670-INS-53 |
| chrX-150030330-INS-561 | chrX-150030330-INS-561 |
| chrX-150035436-INS-750 | chrX-150035436-INS-750 |
| chrX-150044075-INS-231 | chrX-150044075-INS-231 |
| chrX-150049751-INS-76 | chrX-150049751-INS-52 |
| chrX-150112922-INS-5718 | chrX-150112923-INS-5718 |
| chrX-150119614-INS-243 | chrX-150119614-INS-243 |
| chrX-150125801-INS-198 | chrX-150125801-INS-199 |
| chrX-150141399-INS-520 | chrX-150141399-INS-518 |
| chrX-150157459-INS-66 | chrX-150157459-INS-66 |
| chrX-150169317-INS-4203 | chrX-150169318-INS-4203 |
| chrX-150173920-INS-1340 | chrX-150173921-INS-1340 |
| chrX-150194652-INS-7211 | chrX-150194653-INS-7213 |
| chrX-150216822-INS-64 | chrX-150216822-INS-73 |
| chrX-150243047-INS-1954 | chrX-150243048-INS-1954 |
| chrX-150258752-INS-1372 | chrX-150258753-INS-1373 |
| chrX-150264595-INS-4785 | chrX-150264596-INS-4785 |
| chrX-150271036-INS-1205 | chrX-150271037-INS-1207 |
| chrX-150332804-INS-370 | chrX-150332804-INS-369 |
| chrX-150417198-INS-319 | chrX-150417198-INS-321 |
| chrX-150430800-INS-69 | chrX-150430800-INS-52 |
| chrX-150432603-INS-1890 | chrX-150432604-INS-1890 |
| chrX-150449366-INS-2620 | chrX-150449367-INS-2595 |
| chrX-150455923-INS-54 | chrX-150455923-INS-58 |
| chrX-150496402-INS-3083 | chrX-150496403-INS-3085 |
| chrX-150503385-INS-239 | chrX-150503385-INS-214 |
| chrX-150505094-INS-103 | chrX-150505094-INS-103 |
| chrX-150510290-INS-212 | chrX-150510290-INS-211 |
| chrX-150515008-INS-481 | chrX-150515008-INS-481 |
| chrX-150543025-INS-566 | chrX-150543025-INS-566 |
| chrX-150553605-INS-175 | chrX-150553605-INS-177 |
| chrX-150556687-INS-268 | chrX-150556687-INS-268 |
| chrX-150601799-INS-225 | chrX-150601799-INS-245 |
| chrX-150644747-INS-2413 | chrX-150644748-INS-2413 |
| chrX-150655345-INS-229 | chrX-150655345-INS-229 |
| chrX-150660174-INS-60 | chrX-150660174-INS-52 |
| chrX-150710118-INS-56 | chrX-150710118-INS-64 |
| chrX-150728012-INS-113 | chrX-150728008-INS-117 |
| chrX-150731851-INS-207 | chrX-150731851-INS-205 |
| chrX-150742994-INS-59 | chrX-150742994-INS-54 |
| chrX-150748957-INS-87 | chrX-150748957-INS-111 |
| chrX-150755863-INS-54 | chrX-150755863-INS-54 |
| chrX-150769612-INS-52 | chrX-150769612-INS-58 |
| chrX-150796497-INS-208 | chrX-150796497-INS-208 |
| chrX-150813286-INS-205 | chrX-150813286-INS-205 |
| chrX-150823823-INS-213 | chrX-150823823-INS-214 |
| chrX-150853685-INS-417 | chrX-150853685-INS-417 |
| chrX-150882379-INS-1993 | chrX-150882348-INS-1980 |
| chrX-150889054-INS-212 | chrX-150889054-INS-212 |
| chrX-150891491-INS-657 | chrX-150891492-INS-647 |

| | |
|---|---|
| chrX-150895389-INS-208 | chrX-150895389-INS-205 |
| chrX-150910556-INS-61 | chrX-150910559-INS-69 |
| chrX-150968484-INS-203 | chrX-150968484-INS-203 |
| chrX-150994577-INS-59 | chrX-150994577-INS-59 |
| chrX-151003499-INS-682 | chrX-151003499-INS-697 |
| chrX-151018061-INS-205 | chrX-151018061-INS-207 |
| chrX-151048375-INS-70 | chrX-151048375-INS-70 |
| chrX-151091660-INS-5095 | chrX-151091661-INS-5095 |
| chrX-151139427-INS-83 | chrX-151139427-INS-56 |
| chrX-151228744-INS-5451 | chrX-151228745-INS-5451 |
| chrX-151251885-INS-551 | chrX-151251886-INS-549 |
| chrX-151280317-INS-7076 | chrX-151280318-INS-7076 |
| chrX-151336283-INS-1444 | chrX-151336284-INS-1444 |
| chrX-151361829-INS-106 | chrX-151361829-INS-106 |
| chrX-151365480-INS-440 | chrX-151365480-INS-435 |
| chrX-151366748-INS-126 | chrX-151366748-INS-81 |
| chrX-151371347-INS-75 | chrX-151371347-INS-53 |
| chrX-151374778-INS-74 | chrX-151374787-INS-75 |
| chrX-151391652-INS-80 | chrX-151391652-INS-82 |
| chrX-151407918-INS-497 | chrX-151407918-INS-497 |
| chrX-151411474-INS-631 | chrX-151411474-INS-630 |
| chrX-151435909-INS-1008 | chrX-151435910-INS-1008 |
| chrX-151471042-INS-245 | chrX-151471042-INS-245 |
| chrX-151494590-INS-199 | chrX-151494590-INS-198 |
| chrX-151533454-INS-497 | chrX-151533454-INS-489 |
| chrX-151541258-INS-216 | chrX-151541258-INS-216 |
| chrX-151561978-INS-246 | chrX-151561977-INS-246 |
| chrX-151563486-INS-3596 | chrX-151563487-INS-3596 |
| chrX-151569499-INS-103 | chrX-151569499-INS-103 |
| chrX-151575493-INS-210 | chrX-151575493-INS-213 |
| chrX-151578948-INS-68 | chrX-151578960-INS-68 |
| chrX-151610666-INS-480 | chrX-151610666-INS-479 |
| chrX-151703729-INS-86 | chrX-151703736-INS-87 |
| chrX-151709953-INS-472 | chrX-151709953-INS-472 |
| chrX-151782070-INS-5767 | chrX-151782071-INS-5767 |
| chrX-151811850-INS-51 | chrX-151811850-INS-55 |
| chrX-151855215-INS-76 | chrX-151855215-INS-74 |
| chrX-151870301-INS-1192 | chrX-151870302-INS-1193 |
| chrX-151876976-INS-539 | chrX-151876976-INS-540 |
| chrX-151892423-INS-185 | chrX-151892423-INS-185 |
| chrX-151918646-INS-201 | chrX-151918647-INS-201 |
| chrX-151935058-INS-55 | chrX-151935058-INS-55 |
| chrX-151947163-INS-873 | chrX-151947163-INS-873 |
| chrX-151948487-INS-54 | chrX-151948497-INS-50 |
| chrX-151952422-INS-72 | chrX-151952422-INS-51 |
| chrX-151960056-INS-200 | chrX-151960056-INS-201 |
| chrX-151972280-INS-73 | chrX-151972280-INS-73 |
| chrX-151985649-INS-5312 | chrX-151985650-INS-5312 |
| chrX-151993851-INS-52 | chrX-151993851-INS-76 |
| chrX-152006353-INS-5253 | chrX-152006354-INS-5254 |
| chrX-152016943-INS-55 | chrX-152016943-INS-55 |
| chrX-152023657-INS-341 | chrX-152023657-INS-338 |
| chrX-152054793-INS-5185 | chrX-152054794-INS-5185 |
| chrX-152140407-INS-231 | chrX-152140407-INS-231 |
| chrX-152156420-INS-711 | chrX-152156434-INS-711 |
| chrX-152177707-INS-125 | chrX-152177707-INS-125 |
| chrX-152189475-INS-203 | chrX-152189475-INS-203 |
| chrX-152230942-INS-459 | chrX-152230942-INS-459 |
| chrX-152242900-INS-1345 | chrX-152242901-INS-1345 |
| chrX-152254133-INS-56 | chrX-152254133-INS-56 |
| chrX-152297436-INS-77 | chrX-152297436-INS-65 |
| chrX-152299327-INS-253 | chrX-152299327-INS-251 |
| chrX-152316251-INS-11413 | chrX-152316252-INS-11413 |
| chrX-152381485-INS-770 | chrX-152381485-INS-771 |
| chrX-152382832-INS-250 | chrX-152382832-INS-250 |
| chrX-152400362-INS-2265 | chrX-152400363-INS-2265 |
| chrX-152402551-INS-739 | chrX-152402551-INS-740 |
| chrX-152411154-INS-6716 | chrX-152411155-INS-6716 |
| chrX-152414410-INS-60 | chrX-152414410-INS-60 |
| chrX-152420364-INS-2821 | chrX-152420365-INS-2821 |
| chrX-152441276-INS-101 | chrX-152441276-INS-53 |
| chrX-152500878-INS-76 | chrX-152500878-INS-76 |
| chrX-152545540-INS-56 | chrX-152545540-INS-56 |
| chrX-152603648-INS-119 | chrX-152603648-INS-119 |
| chrX-152625249-INS-89 | chrX-152625249-INS-91 |
| chrX-152683250-INS-54 | chrX-152683250-INS-56 |
| chrX-152705636-INS-209 | chrX-152705636-INS-209 |

| | |
|---|---|
| chrX-152728151-INS-68 | chrX-152728151-INS-68 |
| chrX-152761367-INS-93 | chrX-152761367-INS-97 |
| chrX-152997474-INS-204 | chrX-152997474-INS-204 |
| chrX-153035912-INS-8818 | chrX-153035913-INS-8820 |
| chrX-153051750-INS-55 | chrX-153051766-INS-64 |
| chrX-153084885-INS-72 | chrX-153084885-INS-67 |
| chrX-153094128-INS-6684 | chrX-153094129-INS-6684 |
| chrX-153243034-INS-872 | chrX-153243034-INS-872 |
| chrX-153295024-INS-387 | chrX-153295024-INS-387 |
| chrX-153362980-INS-58 | chrX-153362980-INS-52 |
| chrX-153405831-INS-207 | chrX-153405831-INS-206 |
| chrX-153409653-INS-81 | chrX-153409653-INS-81 |
| chrX-153411606-INS-60 | chrX-153411606-INS-60 |
| chrX-153486712-INS-112 | chrX-153486712-INS-107 |
| chrX-153545490-INS-6181 | chrX-153545491-INS-6181 |
| chrX-153577985-INS-1276 | chrX-153577986-INS-1276 |
| chrX-153684723-INS-284 | chrX-153684723-INS-283 |
| chrX-153830150-INS-8535 | chrX-153830150-INS-8542 |
| chrX-153868341-INS-498 | chrX-153868341-INS-500 |
| chrX-153894278-INS-213 | chrX-153894278-INS-213 |
| chrX-153921050-INS-5246 | chrX-153921051-INS-5246 |
| chrX-153930502-INS-7626 | chrX-153930503-INS-7626 |
| chrX-153963009-INS-87 | chrX-153963009-INS-58 |
| chrX-153965900-INS-100 | chrX-153965900-INS-100 |
| chrX-153967667-INS-62 | chrX-153967667-INS-64 |
| chrX-153976106-INS-112 | chrX-153976102-INS-116 |
| chrX-153983450-INS-212 | chrX-153983450-INS-212 |
| chrX-153991008-INS-182 | chrX-153991008-INS-182 |
| chrX-154020631-INS-177 | chrX-154020631-INS-177 |
| chrX-154049967-INS-50 | chrX-154049971-INS-54 |
| chrX-154053211-INS-242 | chrX-154053211-INS-242 |
| chrX-154056980-INS-61 | chrX-154056980-INS-61 |
| chrX-154063109-INS-906 | chrX-154063109-INS-911 |
| chrX-154068732-INS-102 | chrX-154068716-INS-131 |
| chrX-154079535-INS-449 | chrX-154079535-INS-445 |
| chrX-154126895-INS-7083 | chrX-154126896-INS-7083 |
| chrX-154133940-INS-73 | chrX-154133940-INS-93 |
| chrX-154144238-INS-64 | chrX-154144238-INS-56 |
| chrX-154194675-INS-442 | chrX-154194675-INS-443 |
| chrX-154199588-INS-6682 | chrX-154199589-INS-6681 |
| chrX-154227697-INS-58 | chrX-154227697-INS-56 |
| chrX-154249619-INS-82 | chrX-154249619-INS-86 |
| chrX-154255299-INS-635 | chrX-154255299-INS-635 |
| chrX-154271801-INS-113 | chrX-154271801-INS-90 |
| chrX-154319469-INS-93 | chrX-154319469-INS-113 |
| chrX-154332260-INS-56 | chrX-154332260-INS-56 |
| chrX-154352440-INS-191 | chrX-154352440-INS-191 |
| chrX-154372271-INS-756 | chrX-154372271-INS-755 |
| chrX-154430850-INS-6804 | chrX-154430851-INS-6804 |
| chrX-154462299-INS-64 | chrX-154462299-INS-64 |
| chrX-154475553-INS-52 | chrX-154475553-INS-52 |
| chrX-154498755-INS-6326 | chrX-154498756-INS-6326 |
| chrX-154502657-INS-392 | chrX-154502657-INS-391 |
| chrX-154503810-INS-54 | chrX-154503810-INS-52 |
| chrX-154522135-INS-67 | chrX-154522135-INS-67 |
| chrX-154582929-INS-70 | chrX-154582929-INS-90 |
| chrX-154612068-INS-7579 | chrX-154612069-INS-7575 |
| chrX-154631854-INS-58 | chrX-154631850-INS-74 |
| chrX-154734068-INS-4756 | chrX-154734069-INS-4756 |
| chrX-154751228-INS-6906 | chrX-154751229-INS-6906 |
| chrX-154771050-INS-90 | chrX-154771050-INS-96 |
| chrX-154823396-INS-130 | chrX-154823396-INS-130 |
| chrX-154877631-INS-197 | chrX-154877631-INS-197 |
| chrX-154936634-INS-359 | chrX-154936634-INS-359 |
| chrX-154949997-INS-521 | chrX-154949997-INS-521 |
| chrX-154955484-INS-492 | chrX-154955484-INS-498 |
| chrX-154993872-INS-150 | chrX-154993870-INS-181 |
| chrX-155027434-INS-320 | chrX-155027433-INS-275 |
| chrX-155047732-INS-241 | chrX-155047732-INS-241 |
| chrX-155047951-INS-1431 | chrX-155047952-INS-1430 |
| chrX-155084689-INS-4498 | chrX-155084702-INS-4485 |
| chrX-155100303-INS-2923 | chrX-155100304-INS-2923 |
| chrX-155143868-INS-199 | chrX-155143868-INS-199 |
| chrX-155150043-INS-1592 | chrX-155150044-INS-1593 |
| chrX-155162970-INS-218 | chrX-155162970-INS-218 |
| chrX-155174345-INS-1367 | chrX-155174346-INS-1367 |
| chrX-155181161-INS-59 | chrX-155181112-INS-82 |

| | |
|---|---|
| chrX-155188849-INS-65 | chrX-155188849-INS-65 |
| chrX-155192349-INS-767 | chrX-155192349-INS-767 |
| chrX-155194027-INS-6376 | chrX-155194028-INS-6375 |
| chrX-155200273-INS-6450 | chrX-155200274-INS-6450 |
| chrX-155213386-INS-58 | chrX-155213386-INS-54 |
| chrX-155214686-INS-429 | chrX-155214686-INS-426 |
| chrX-155264873-INS-3915 | chrX-155264874-INS-3915 |
| chrX-155279059-INS-452 | chrX-155279059-INS-452 |
| chrX-155301870-INS-64 | chrX-155301870-INS-60 |
| chrX-155311206-INS-400 | chrX-155311206-INS-408 |
| chrX-155356625-INS-56 | chrX-155356625-INS-52 |
| chrX-155376190-INS-112 | chrX-155376190-INS-115 |
| chrX-155476951-INS-204 | chrX-155476951-INS-204 |
| chrX-155497267-INS-1939 | chrX-155497268-INS-1944 |
| chrX-155574400-INS-1002 | chrX-155574401-INS-1002 |
| chrX-155591399-INS-199 | chrX-155591399-INS-199 |
| chrX-155627789-INS-89 | chrX-155627789-INS-89 |
| chrX-155642115-INS-65 | chrX-155642115-INS-65 |
| chrX-155644739-INS-781 | chrX-155644739-INS-781 |
| chrX-155673338-INS-5146 | chrX-155673339-INS-5146 |
| chrX-155679868-INS-430 | chrX-155679851-INS-441 |
| chrX-155709016-INS-53 | chrX-155709012-INS-61 |
| chrX-155746600-INS-56 | chrX-155746600-INS-58 |
| chrX-155762478-INS-90 | chrX-155762478-INS-90 |
| chrX-155784197-INS-229 | chrX-155784197-INS-228 |
| chrX-155792391-INS-198 | chrX-155792391-INS-198 |
| chrX-155798745-INS-891 | chrX-155798745-INS-891 |
| chrX-155814586-INS-551 | chrX-155814586-INS-552 |
| chrX-155827255-INS-161 | chrX-155827255-INS-162 |
| chrX-155828764-INS-187 | chrX-155828764-INS-187 |
| chrX-155833934-INS-2189 | chrX-155833935-INS-2190 |
| chrX-155917522-INS-52 | chrX-155917522-INS-53 |
| chrX-155934566-INS-68 | chrX-155934566-INS-50 |
| chrX-155981479-INS-7238 | chrX-155981480-INS-7239 |
| chrX-155989602-INS-5218 | chrX-155989603-INS-5218 |
| chrX-155993608-INS-91 | chrX-155993608-INS-91 |
| chrX-155995423-INS-60 | chrX-155995423-INS-70 |
| chrX-156002837-INS-131 | chrX-156002837-INS-134 |
| chrX-156043659-INS-1974 | chrX-156043660-INS-1950 |
| chrX-156074061-INS-94 | chrX-156074061-INS-94 |
| chrX-156092024-INS-599 | chrX-156092024-INS-594 |
| chrX-156124561-INS-989 | chrX-156124561-INS-989 |
| chrX-156142019-INS-125 | chrX-156142018-INS-120 |
| chrX-156149183-INS-217 | chrX-156149183-INS-217 |
| chrX-156163929-INS-119 | chrX-156163929-INS-119 |
| chrX-156164566-INS-73 | chrX-156164563-INS-77 |
| chrX-156195945-INS-64 | chrX-156195945-INS-57 |
| chrX-156200296-INS-74 | chrX-156200303-INS-75 |
| chrX-156215502-INS-289 | chrX-156215502-INS-289 |
| chrX-156233170-INS-101 | chrX-156233170-INS-97 |
| chrX-156239630-INS-72 | chrX-156239635-INS-66 |
| chrX-156240817-INS-241 | chrX-156240817-INS-241 |
| chrX-156245731-INS-313 | chrX-156245731-INS-295 |
| chrX-156257556-INS-72 | chrX-156257556-INS-68 |
| chrX-156263539-INS-175 | chrX-156263539-INS-155 |
| chrX-156265491-INS-216 | chrX-156265491-INS-215 |
| chrX-156280994-INS-390 | chrX-156280994-INS-390 |
| chrX-156282630-INS-67 | chrX-156282628-INS-70 |
| chrX-156287450-INS-226 | chrX-156287450-INS-212 |
| chrX-156294085-INS-133 | chrX-156294085-INS-133 |
| chrX-156303243-INS-52 | chrX-156303218-INS-51 |
| chrX-156304916-INS-4290 | chrX-156304920-INS-4289 |
| chrX-156306227-INS-206 | chrX-156306227-INS-206 |
| chrX-156309249-INS-226 | chrX-156309249-INS-226 |
| chrX-156323762-INS-72 | chrX-156323772-INS-72 |
| chrX-156327760-INS-69 | chrX-156327760-INS-69 |
| chrX-156353129-INS-482 | chrX-156353129-INS-481 |
| chrX-156356440-INS-201 | chrX-156356440-INS-202 |
| chrX-156386974-INS-200 | chrX-156386974-INS-200 |
| chrX-156411550-INS-390 | chrX-156411550-INS-389 |
| chrX-156422419-INS-555 | chrX-156422419-INS-556 |
| chrX-156445079-INS-171 | chrX-156445079-INS-171 |
| chrX-156452174-INS-4718 | chrX-156452175-INS-4718 |
| chrX-156452926-INS-70 | chrX-156452926-INS-70 |
| chrX-156544421-INS-461 | chrX-156544421-INS-461 |
| chrX-156563220-INS-386 | chrX-156563220-INS-386 |
| chrX-156567265-INS-6684 | chrX-156567266-INS-6684 |

| | |
|---|---|
| chrX-156573610-INS-58 | chrX-156573610-INS-58 |
| chrX-156630092-INS-74 | chrX-156630092-INS-74 |
| chrX-156643925-INS-75 | chrX-156643925-INS-87 |
| chrX-156656943-INS-317 | chrX-156656943-INS-318 |
| chrX-156666738-INS-135 | chrX-156666738-INS-134 |
| chrX-156725952-INS-6910 | chrX-156725953-INS-6911 |
| chrX-156731740-INS-98 | chrX-156731730-INS-100 |
| chrX-156764086-INS-2850 | chrX-156764087-INS-2851 |
| chrX-156773211-INS-694 | chrX-156773211-INS-694 |
| chrX-156786991-INS-74 | chrX-156786991-INS-74 |
| chrX-156792183-INS-1374 | chrX-156792184-INS-1374 |
| chrX-156832899-INS-608 | chrX-156832899-INS-608 |
| chrX-156869389-INS-176 | chrX-156869387-INS-197 |
| chrX-156874506-INS-7133 | chrX-156874507-INS-7133 |
| chrX-156914878-INS-1885 | chrX-156914879-INS-1885 |
| chrX-156923747-INS-57 | chrX-156923719-INS-87 |
| chrX-156925192-INS-208 | chrX-156925192-INS-208 |
| chrX-156987705-INS-6386 | chrX-156987706-INS-6418 |
| chrX-156992903-INS-100 | chrX-156992903-INS-100 |
| chrX-157025352-INS-6807 | chrX-157025366-INS-6824 |
| chrX-157044284-INS-77 | chrX-157044283-INS-75 |
| chrX-157071793-INS-76 | chrX-157071793-INS-76 |
| chrX-157084125-INS-2699 | chrX-157084126-INS-2699 |
| chrX-157132821-INS-231 | chrX-157132821-INS-231 |
| chrX-157147240-INS-52 | chrX-157147240-INS-52 |
| chrX-157222586-INS-6481 | chrX-157222587-INS-6477 |
| chrX-157253878-INS-68 | chrX-157253874-INS-68 |
| chrX-157261110-INS-3374 | chrX-157261111-INS-3374 |
| chrX-157282596-INS-903 | chrX-157282596-INS-903 |
| chrX-157311049-INS-181 | chrX-157311049-INS-182 |
| chrX-157365869-INS-64 | chrX-157365869-INS-59 |
| chrX-157423479-INS-99 | chrX-157423479-INS-99 |
| chrX-157504583-INS-50 | chrX-157504583-INS-84 |
| chrX-157506642-INS-67 | chrX-157506642-INS-105 |
| chrX-157530083-INS-60 | chrX-157530112-INS-74 |
| chrX-157536093-INS-4805 | chrX-157536094-INS-4805 |
| chrX-157548499-INS-72 | chrX-157548499-INS-72 |
| chrX-157567576-INS-514 | chrX-157567576-INS-542 |
| chrX-157568266-INS-6610 | chrX-157568267-INS-6610 |
| chrX-157572212-INS-123 | chrX-157572212-INS-80 |
| chrX-157573129-INS-111 | chrX-157573129-INS-111 |
| chrX-157623106-INS-73 | chrX-157623106-INS-96 |
| chrX-157625746-INS-691 | chrX-157625746-INS-691 |
| chrX-157630314-INS-2954 | chrX-157630315-INS-2954 |
| chrX-157633406-INS-195 | chrX-157633406-INS-195 |
| chrX-157639268-INS-67 | chrX-157639268-INS-66 |
| chrX-157677248-INS-3028 | chrX-157677249-INS-3028 |
| chrX-157702662-INS-64 | chrX-157702662-INS-64 |
| chrX-157710492-INS-1272 | chrX-157710493-INS-1272 |
| chrX-157727405-INS-76 | chrX-157727405-INS-76 |
| chrX-157730135-INS-261 | chrX-157730135-INS-261 |
| chrX-157762361-INS-7150 | chrX-157762362-INS-7150 |
| chrX-157772255-INS-220 | chrX-157772255-INS-220 |
| chrX-157809432-INS-1477 | chrX-157809433-INS-1477 |
| chrX-157831322-INS-524 | chrX-157831322-INS-524 |
| chrX-157838693-INS-6438 | chrX-157838694-INS-6438 |
| chrX-157845642-INS-777 | chrX-157845642-INS-777 |
| chrX-157882985-INS-148 | chrX-157882985-INS-148 |
| chrX-157929833-INS-82 | chrX-157929833-INS-82 |
| chrX-157939493-INS-59 | chrX-157939495-INS-60 |
| chrX-157955979-INS-653 | chrX-157955979-INS-654 |
| chrX-157988873-INS-62 | chrX-157988839-INS-66 |
| chrX-157991637-INS-122 | chrX-157991637-INS-128 |
| chrX-158014963-INS-3333 | chrX-158014964-INS-3318 |
| chrX-158022917-INS-162 | chrX-158022917-INS-159 |
| chrX-158037395-INS-69 | chrX-158037415-INS-71 |
| chrX-158071343-INS-115 | chrX-158071343-INS-115 |
| chrX-158114711-INS-224 | chrX-158114711-INS-213 |
| chrX-158116832-INS-1046 | chrX-158116833-INS-1024 |
| chrX-158120381-INS-74 | chrX-158120381-INS-74 |
| chrX-158123758-INS-3656 | chrX-158123759-INS-3656 |
| chrX-158130962-INS-3110 | chrX-158130963-INS-3110 |
| chrX-158162633-INS-133 | chrX-158162633-INS-133 |
| chrX-158172162-INS-113 | chrX-158172162-INS-113 |
| chrX-158195035-INS-60 | chrX-158195035-INS-60 |
| chrX-158196344-INS-227 | chrX-158196344-INS-227 |
| chrX-158204446-INS-1153 | chrX-158204447-INS-1153 |

| | |
|---|---|
| chrX-158205663-INS-57 | chrX-158205663-INS-57 |
| chrX-158230555-INS-118 | chrX-158230555-INS-118 |
| chrX-158245646-INS-205 | chrX-158245646-INS-207 |
| chrX-158253721-INS-54 | chrX-158253746-INS-63 |
| chrX-158300031-INS-260 | chrX-158300031-INS-260 |
| chrX-158311498-INS-73 | chrX-158311498-INS-73 |
| chrX-158345972-INS-138 | chrX-158345972-INS-138 |
| chrX-158359430-INS-887 | chrX-158359430-INS-872 |
| chrX-158362396-INS-267 | chrX-158362396-INS-267 |
| chrX-158372355-INS-207 | chrX-158372355-INS-201 |
| chrX-158388884-INS-178 | chrX-158388884-INS-178 |
| chrX-158395273-INS-72 | chrX-158395273-INS-72 |
| chrX-158415940-INS-206 | chrX-158415940-INS-203 |
| chrX-158442471-INS-168 | chrX-158442462-INS-164 |
| chrX-158444931-INS-505 | chrX-158444931-INS-505 |
| chrX-158458750-INS-58 | chrX-158458750-INS-58 |
| chrX-158465238-INS-104 | chrX-158465214-INS-79 |
| chrX-158515315-INS-90 | chrX-158515315-INS-138 |
| chrX-158566537-INS-392 | chrX-158566537-INS-392 |
| chrX-158587418-INS-130 | chrX-158587418-INS-170 |
| chrX-158603191-INS-208 | chrX-158603191-INS-208 |
| chrX-158613057-INS-2579 | chrX-158613058-INS-2579 |
| chrX-158615622-INS-72 | chrX-158615622-INS-72 |
| chrX-158644606-INS-76 | chrX-158644606-INS-76 |
| chrX-158646590-INS-495 | chrX-158646590-INS-459 |
| chrX-158656900-INS-1904 | chrX-158656901-INS-1904 |
| chrX-158692041-INS-56 | chrX-158692042-INS-61 |
| chrX-158723574-INS-1190 | chrX-158723575-INS-1199 |
| chrX-158723634-INS-143 | chrX-158723634-INS-152 |
| chrX-158770391-INS-262 | chrX-158770391-INS-264 |
| chrX-158786924-INS-3157 | chrX-158786925-INS-3107 |
| chrX-158797390-INS-7437 | chrX-158797391-INS-7438 |
| chrX-158823435-INS-687 | chrX-158823435-INS-687 |
| chrX-158831732-INS-200 | chrX-158831752-INS-206 |
| chrX-158832235-INS-52 | chrX-158832235-INS-54 |
| chrX-158866390-INS-185 | chrX-158866390-INS-187 |
| chrX-158902757-INS-67 | chrX-158902757-INS-58 |
| chrX-158913308-INS-190 | chrX-158913308-INS-190 |
| chrX-158933916-INS-1894 | chrX-158933917-INS-1894 |
| chrX-158934278-INS-492 | chrX-158934278-INS-492 |
| chrX-158936797-INS-827 | chrX-158936797-INS-827 |
| chrX-158955561-INS-120 | chrX-158955566-INS-77 |
| chrX-158969280-INS-4551 | chrX-158969281-INS-4552 |
| chrX-158981350-INS-201 | chrX-158981350-INS-201 |
| chrX-159007103-INS-318 | chrX-159007109-INS-311 |
| chrX-159014546-INS-529 | chrX-159014546-INS-530 |
| chrX-159021289-INS-72 | chrX-159021289-INS-72 |
| chrX-159037784-INS-572 | chrX-159037783-INS-572 |
| chrX-159040248-INS-1269 | chrX-159040249-INS-1270 |
| chrX-159053270-INS-2078 | chrX-159053271-INS-2079 |
| chrX-159097309-INS-517 | chrX-159097309-INS-517 |
| chrX-159169194-INS-842 | chrX-159169194-INS-842 |
| chrX-159194394-INS-848 | chrX-159194394-INS-847 |
| chrX-159231142-INS-944 | chrX-159231142-INS-933 |
| chrX-159239190-INS-73 | chrX-159239190-INS-73 |
| chrX-159257932-INS-64 | chrX-159257932-INS-64 |
| chrX-159330480-INS-56 | chrX-159330480-INS-58 |
| chrX-159424639-INS-81 | chrX-159424639-INS-84 |
| chrX-159425253-INS-517 | chrX-159425253-INS-517 |
| chrX-159431011-INS-970 | chrX-159431011-INS-970 |
| chrX-159466889-INS-736 | chrX-159466889-INS-736 |
| chrX-159484600-INS-7127 | chrX-159484601-INS-7126 |
| chrX-159506734-INS-4558 | chrX-159506735-INS-4558 |
| chrX-159534550-INS-54 | chrX-159534550-INS-54 |
| chrX-159544346-INS-81 | chrX-159544346-INS-84 |
| chrX-159561419-INS-177 | chrX-159561419-INS-177 |
| chrX-159563775-INS-222 | chrX-159563775-INS-222 |
| chrX-159571519-INS-7158 | chrX-159571520-INS-7158 |
| chrX-159600742-INS-139 | chrX-159600754-INS-134 |
| chrX-159633072-INS-186 | chrX-159633072-INS-190 |
| chrX-159672249-INS-179 | chrX-159672249-INS-184 |
| chrX-159720404-INS-79 | chrX-159720404-INS-83 |
| chrX-159759963-INS-61 | chrX-159759929-INS-60 |
| chrX-159813958-INS-196 | chrX-159813958-INS-198 |
| chrX-159879300-INS-7162 | chrX-159879301-INS-7162 |
| chrX-159882096-INS-222 | chrX-159882096-INS-221 |
| chrX-159893954-INS-71 | chrX-159893954-INS-71 |

| | |
|---|---|
| chrX-159897329-INS-5225 | chrX-159897330-INS-5225 |
| chrX-159903026-INS-78 | chrX-159903026-INS-78 |
| chrX-159928639-INS-75 | chrX-159928649-INS-58 |
| chrX-159983646-INS-1493 | chrX-159983647-INS-1493 |
| chrX-160021641-INS-625 | chrX-160021641-INS-625 |
| chrX-160026598-INS-111 | chrX-160026598-INS-102 |
| chrX-160057822-INS-1713 | chrX-160057823-INS-1715 |
| chrX-160080014-INS-72 | chrX-160080014-INS-90 |
| chrX-160087018-INS-199 | chrX-160087018-INS-199 |
| chrX-160099514-INS-235 | chrX-160099514-INS-236 |
| chrX-160121793-INS-84 | chrX-160121793-INS-52 |
| chrX-160166229-INS-60 | chrX-160166229-INS-60 |
| chrX-160179976-INS-152 | chrX-160179976-INS-147 |
| chrX-160195699-INS-423 | chrX-160195699-INS-422 |
| chrX-160242227-INS-649 | chrX-160242227-INS-649 |
| chrX-160282297-INS-2292 | chrX-160282298-INS-2292 |
| chrX-160329868-INS-213 | chrX-160329868-INS-212 |
| chrX-160383122-INS-3033 | chrX-160383123-INS-3034 |
| chrX-160424879-INS-3257 | chrX-160424880-INS-3257 |
| chrX-160465306-INS-54 | chrX-160465306-INS-54 |
| chrX-160522735-INS-52 | chrX-160522735-INS-52 |
| chrX-160595897-INS-96 | chrX-160595897-INS-96 |
| chrX-160671753-INS-8991 | chrX-160671754-INS-8992 |
| chrX-160684168-INS-5127 | chrX-160684169-INS-5132 |
| chrX-160711458-INS-323 | chrX-160711458-INS-323 |
| chrX-160729019-INS-196 | chrX-160729016-INS-197 |
| chrX-160738189-INS-6903 | chrX-160738190-INS-6904 |
| chrX-160769066-INS-57 | chrX-160769066-INS-57 |
| chrX-160791453-INS-218 | chrX-160791453-INS-218 |
| chrX-160871875-INS-52 | chrX-160871875-INS-56 |
| chrX-160887665-INS-54 | chrX-160887665-INS-56 |
| chrX-160897888-INS-6342 | chrX-160897889-INS-6341 |
| chrX-160994492-INS-213 | chrX-160994492-INS-208 |
| chrX-161034856-INS-1701 | chrX-161034857-INS-1701 |
| chrX-161039347-INS-213 | chrX-161039347-INS-213 |
| chrX-161089558-INS-1830 | chrX-161089559-INS-1831 |
| chrX-161130053-INS-96 | chrX-161130053-INS-90 |
| chrX-161130609-INS-102 | chrX-161130609-INS-102 |
| chrX-161136868-INS-2187 | chrX-161136869-INS-2208 |
| chrX-161137911-INS-74 | chrX-161137911-INS-58 |
| chrX-161155233-INS-6674 | chrX-161155234-INS-6674 |
| chrX-161166450-INS-7363 | chrX-161166451-INS-7363 |
| chrX-161179022-INS-809 | chrX-161179023-INS-806 |
| chrX-161222072-INS-85 | chrX-161222072-INS-88 |
| chrX-161262714-INS-194 | chrX-161262714-INS-177 |
| chrX-161369826-INS-729 | chrX-161369826-INS-730 |
| chrX-161371266-INS-441 | chrX-161371266-INS-474 |
| chrX-161371668-INS-2916 | chrX-161371669-INS-2917 |
| chrX-161393905-INS-326 | chrX-161393905-INS-326 |
| chrX-161412898-INS-1118 | chrX-161412899-INS-1118 |
| chrX-161414632-INS-115 | chrX-161414632-INS-115 |
| chrX-161464213-INS-400 | chrX-161464213-INS-400 |
| chrX-161468807-INS-81 | chrX-161468831-INS-68 |
| chrX-161475586-INS-201 | chrX-161475586-INS-201 |
| chrX-161493114-INS-199 | chrX-161493114-INS-199 |
| chrX-161503276-INS-185 | chrX-161503283-INS-182 |
| chrX-161521584-INS-207 | chrX-161521584-INS-208 |
| chrX-161532549-INS-332 | chrX-161532551-INS-334 |
| chrX-161537095-INS-85 | chrX-161537100-INS-119 |
| chrX-161537969-INS-546 | chrX-161537982-INS-553 |
| chrX-161576224-INS-165 | chrX-161576224-INS-165 |
| chrX-161578643-INS-5005 | chrX-161578644-INS-5007 |
| chrX-161608389-INS-220 | chrX-161608389-INS-221 |
| chrX-161618467-INS-186 | chrX-161618467-INS-186 |
| chrX-161628059-INS-214 | chrX-161628059-INS-214 |
| chrX-161633396-INS-155 | chrX-161633397-INS-155 |
| chrX-161636584-INS-68 | chrX-161636582-INS-71 |
| chrX-161637852-INS-176 | chrX-161637852-INS-177 |
| chrX-161639540-INS-611 | chrX-161639540-INS-611 |
| chrX-161644901-INS-168 | chrX-161644901-INS-168 |
| chrX-161646916-INS-80 | chrX-161646916-INS-58 |
| chrX-161650873-INS-200 | chrX-161650873-INS-200 |
| chrX-161663766-INS-210 | chrX-161663766-INS-211 |
| chrX-161675808-INS-213 | chrX-161675825-INS-217 |
| chrX-161689500-INS-285 | chrX-161689500-INS-285 |
| chrX-161690996-INS-55 | chrX-161690996-INS-51 |
| chrX-161692445-INS-62 | chrX-161692445-INS-62 |

| | |
|---|---|
| chrX-161693686-INS-169 | chrX-161693686-INS-170 |
| chrX-161695964-INS-175 | chrX-161695964-INS-176 |
| chrX-161700620-INS-5219 | chrX-161700621-INS-5219 |
| chrX-161701498-INS-352 | chrX-161701498-INS-352 |
| chrX-161704801-INS-282 | chrX-161704801-INS-282 |
| chrX-161728178-INS-387 | chrX-161728178-INS-387 |
| chrX-161739267-INS-331 | chrX-161739267-INS-331 |
| chrX-161740943-INS-68 | chrX-161740943-INS-68 |
| chrX-161751019-INS-854 | chrX-161751019-INS-854 |
| chrX-161771201-INS-97 | chrX-161771201-INS-95 |
| chrX-161772962-INS-206 | chrX-161772962-INS-203 |
| chrX-161791760-INS-62 | chrX-161791760-INS-62 |
| chrX-161824831-INS-460 | chrX-161824831-INS-460 |
| chrX-161827412-INS-766 | chrX-161827412-INS-765 |
| chrX-161844310-INS-194 | chrX-161844303-INS-198 |
| chrX-161854016-INS-67 | chrX-161854016-INS-67 |
| chrX-161878033-INS-54 | chrX-161878029-INS-62 |
| chrX-161937239-INS-69 | chrX-161937227-INS-52 |
| chrX-161969034-INS-162 | chrX-161969034-INS-160 |
| chrX-161994648-INS-144 | chrX-161994644-INS-143 |
| chrX-161995950-INS-221 | chrX-161995950-INS-218 |
| chrX-162012842-INS-1293 | chrX-162012843-INS-1293 |
| chrX-162052207-INS-68 | chrX-162052207-INS-64 |
| chrX-162057576-INS-54 | chrX-162057580-INS-52 |
| chrX-162104002-INS-56 | chrX-162104002-INS-56 |
| chrX-162106562-INS-1905 | chrX-162106563-INS-1905 |
| chrX-162144933-INS-825 | chrX-162144933-INS-797 |
| chrX-162145816-INS-56 | chrX-162145816-INS-81 |
| chrX-162202224-INS-126 | chrX-162202262-INS-134 |
| chrX-162204795-INS-114 | chrX-162204795-INS-120 |
| chrX-162289733-INS-199 | chrX-162289733-INS-199 |
| chrX-162300734-INS-207 | chrX-162300734-INS-207 |
| chrX-162302651-INS-4149 | chrX-162302634-INS-4150 |
| chrX-162310414-INS-132 | chrX-162310414-INS-158 |
| chrX-162405644-INS-989 | chrX-162405644-INS-990 |
| chrX-162408471-INS-56 | chrX-162408439-INS-65 |
| chrX-162427827-INS-150 | chrX-162427827-INS-150 |
| chrX-162442556-INS-226 | chrX-162442556-INS-226 |
| chrX-162492941-INS-566 | chrX-162492941-INS-566 |
| chrX-162503990-INS-100 | chrX-162503990-INS-60 |
| chrX-162508623-INS-64 | chrX-162508623-INS-56 |
| chrX-162552616-INS-72 | chrX-162552616-INS-64 |
| chrX-162588873-INS-89 | chrX-162588866-INS-92 |
| chrX-162591326-INS-7150 | chrX-162591327-INS-7150 |
| chrX-162597546-INS-977 | chrX-162597546-INS-977 |
| chrX-162603618-INS-60 | chrX-162603595-INS-105 |
| chrX-162611818-INS-55 | chrX-162611818-INS-50 |
| chrX-162639174-INS-207 | chrX-162639174-INS-207 |
| chrX-162641870-INS-96 | chrX-162641910-INS-102 |
| chrX-162650099-INS-126 | chrX-162650097-INS-83 |
| chrX-162678443-INS-54 | chrX-162678443-INS-54 |
| chrX-162704929-INS-233 | chrX-162704929-INS-232 |
| chrX-162747242-INS-510 | chrX-162747242-INS-521 |
| chrX-162751322-INS-78 | chrX-162751322-INS-86 |
| chrX-162767971-INS-177 | chrX-162767971-INS-177 |
| chrX-162779399-INS-201 | chrX-162779399-INS-201 |
| chrX-162817790-INS-60 | chrX-162817790-INS-52 |
| chrX-162868086-INS-66 | chrX-162868086-INS-66 |
| chrX-162871156-INS-51 | chrX-162871156-INS-58 |
| chrX-162889391-INS-109 | chrX-162889391-INS-109 |
| chrX-162891757-INS-224 | chrX-162891757-INS-227 |
| chrX-162914188-INS-84 | chrX-162914188-INS-72 |
| chrX-162954070-INS-86 | chrX-162954070-INS-86 |
| chrX-162965515-INS-52 | chrX-162965515-INS-60 |
| chrX-162988502-INS-87 | chrX-162988526-INS-92 |
| chrX-162999298-INS-300 | chrX-162999298-INS-300 |
| chrX-163000834-INS-378 | chrX-163000834-INS-382 |
| chrX-163020604-INS-109 | chrX-163020604-INS-139 |
| chrX-163028055-INS-1463 | chrX-163028056-INS-1445 |
| chrX-163051224-INS-51 | chrX-163051224-INS-51 |
| chrX-163087986-INS-198 | chrX-163087986-INS-198 |
| chrX-163111000-INS-4181 | chrX-163111001-INS-4182 |
| chrX-163121295-INS-66 | chrX-163121295-INS-62 |
| chrX-163141278-INS-52 | chrX-163141259-INS-91 |
| chrX-163184530-INS-99 | chrX-163184530-INS-99 |
| chrX-163237749-INS-1127 | chrX-163237750-INS-1126 |
| chrX-163240184-INS-60 | chrX-163240184-INS-60 |

| | |
|---|---|
| chrX-163246614-INS-96 | chrX-163246629-INS-98 |
| chrX-163251551-INS-80 | chrX-163251551-INS-79 |
| chrX-163320041-INS-123 | chrX-163320041-INS-126 |
| chrX-163453962-INS-50 | chrX-163453975-INS-63 |
| chrX-163470388-INS-64 | chrX-163470388-INS-66 |
| chrX-163477707-INS-326 | chrX-163477707-INS-321 |
| chrX-163518650-INS-332 | chrX-163518650-INS-332 |
| chrX-163533444-INS-201 | chrX-163533469-INS-201 |
| chrX-163587948-INS-409 | chrX-163587948-INS-409 |
| chrX-163600135-INS-56 | chrX-163600135-INS-61 |
| chrX-163612098-INS-69 | chrX-163612090-INS-68 |
| chrX-163622117-INS-53 | chrX-163622117-INS-51 |
| chrX-163637566-INS-1225 | chrX-163637567-INS-1226 |
| chrX-163637570-INS-452 | chrX-163637606-INS-464 |
| chrX-163643716-INS-62 | chrX-163643700-INS-70 |
| chrX-163689603-INS-100 | chrX-163689603-INS-100 |
| chrX-163789837-INS-60 | chrX-163789837-INS-58 |
| chrX-163795935-INS-73 | chrX-163795935-INS-56 |
| chrX-163837425-INS-78 | chrX-163837425-INS-78 |
| chrX-163854788-INS-608 | chrX-163854788-INS-609 |
| chrX-163879192-INS-419 | chrX-163879192-INS-419 |
| chrX-163897338-INS-492 | chrX-163897338-INS-492 |
| chrX-163902548-INS-55 | chrX-163902548-INS-60 |
| chrX-163933900-INS-266 | chrX-163933900-INS-270 |
| chrX-163941563-INS-208 | chrX-163941563-INS-208 |
| chrX-163993602-INS-202 | chrX-163993602-INS-202 |
| chrX-164041929-INS-52 | chrX-164041914-INS-50 |
| chrX-164044485-INS-1455 | chrX-164044486-INS-1456 |
| chrX-164047950-INS-512 | chrX-164047950-INS-512 |
| chrX-164090993-INS-201 | chrX-164090993-INS-201 |
| chrX-164159023-INS-1891 | chrX-164159024-INS-1891 |
| chrX-164164076-INS-813 | chrX-164164076-INS-813 |
| chrX-164181959-INS-7393 | chrX-164181960-INS-7393 |
| chrX-164240119-INS-824 | chrX-164240119-INS-824 |
| chrX-164277756-INS-56 | chrX-164277756-INS-52 |
| chrX-164296766-INS-60 | chrX-164296766-INS-60 |
| chrX-164314995-INS-70 | chrX-164314995-INS-52 |
| chrX-164397600-INS-467 | chrX-164397600-INS-467 |
| chrX-164454039-INS-4373 | chrX-164454040-INS-4373 |
| chrX-164467745-INS-91 | chrX-164467745-INS-91 |
| chrX-164543014-INS-150 | chrX-164543014-INS-150 |
| chrX-164546993-INS-1866 | chrX-164546994-INS-1866 |
| chrX-164550916-INS-240 | chrX-164550916-INS-217 |
| chrX-164572212-INS-79 | chrX-164572212-INS-81 |
| chrX-164607662-INS-3353 | chrX-164607663-INS-3352 |
| chrX-164613557-INS-528 | chrX-164613557-INS-527 |
| chrX-164631104-INS-414 | chrX-164631104-INS-413 |
| chrX-164633252-INS-78 | chrX-164633252-INS-78 |
| chrX-164646815-INS-235 | chrX-164646815-INS-235 |
| chrX-164680363-INS-71 | chrX-164680359-INS-54 |
| chrX-164686136-INS-7125 | chrX-164686137-INS-7125 |
| chrX-164690740-INS-80 | chrX-164690740-INS-78 |
| chrX-164716198-INS-152 | chrX-164716198-INS-152 |
| chrX-164742962-INS-7082 | chrX-164742963-INS-7082 |
| chrX-164745097-INS-87 | chrX-164745097-INS-82 |
| chrX-164750294-INS-504 | chrX-164750294-INS-504 |
| chrX-164821532-INS-185 | chrX-164821532-INS-188 |
| chrX-164823744-INS-84 | chrX-164823744-INS-79 |
| chrX-164836381-INS-64 | chrX-164836391-INS-54 |
| chrX-164837348-INS-461 | chrX-164837348-INS-473 |
| chrX-164845994-INS-199 | chrX-164845994-INS-199 |
| chrX-164863253-INS-1723 | chrX-164863254-INS-1724 |
| chrX-164863703-INS-6454 | chrX-164863704-INS-6454 |
| chrX-164881066-INS-187 | chrX-164881066-INS-187 |
| chrX-164883934-INS-684 | chrX-164883934-INS-684 |
| chrX-164887890-INS-60 | chrX-164887912-INS-60 |
| chrX-164891343-INS-6604 | chrX-164891344-INS-6604 |
| chrX-164892971-INS-524 | chrX-164892971-INS-524 |
| chrX-164910113-INS-1703 | chrX-164910114-INS-1703 |
| chrX-164934238-INS-373 | chrX-164934238-INS-373 |
| chrX-164984231-INS-4801 | chrX-164984232-INS-4801 |
| chrX-164985768-INS-326 | chrX-164985768-INS-326 |
| chrX-164993258-INS-384 | chrX-164993258-INS-384 |
| chrX-164997200-INS-105 | chrX-164997200-INS-108 |
| chrX-165056100-INS-185 | chrX-165056100-INS-185 |
| chrX-165059948-INS-637 | chrX-165059948-INS-659 |
| chrX-165103249-INS-62 | chrX-165103246-INS-66 |

| | |
|---|---|
| chrX-165114196-INS-54 | chrX-165114196-INS-62 |
| chrX-165130357-INS-53 | chrX-165130357-INS-53 |
| chrX-165153008-INS-56 | chrX-165153008-INS-56 |
| chrX-165162332-INS-61 | chrX-165162332-INS-62 |
| chrX-165213261-INS-1893 | chrX-165213262-INS-1893 |
| chrX-165250388-INS-99 | chrX-165250388-INS-99 |
| chrX-165273447-INS-6756 | chrX-165273448-INS-6756 |
| chrX-165278722-INS-68 | chrX-165278736-INS-91 |
| chrX-165282499-INS-1125 | chrX-165282500-INS-1169 |
| chrX-165287481-INS-284 | chrX-165287481-INS-285 |
| chrX-165295871-INS-68 | chrX-165295871-INS-78 |
| chrX-165304502-INS-196 | chrX-165304502-INS-197 |
| chrX-165305726-INS-67 | chrX-165305726-INS-69 |
| chrX-165307952-INS-170 | chrX-165307952-INS-170 |
| chrX-165318798-INS-360 | chrX-165318798-INS-360 |
| chrX-165325869-INS-2965 | chrX-165325870-INS-2968 |
| chrX-165333042-INS-164 | chrX-165333042-INS-175 |
| chrX-165337087-INS-209 | chrX-165337087-INS-212 |
| chrX-165340233-INS-401 | chrX-165340233-INS-401 |
| chrX-165341103-INS-6477 | chrX-165341104-INS-6477 |
| chrX-165341571-INS-107 | chrX-165341589-INS-85 |
| chrX-165378877-INS-59 | chrX-165378877-INS-59 |
| chrX-165412859-INS-551 | chrX-165412861-INS-551 |
| chrX-165424173-INS-322 | chrX-165424173-INS-322 |
| chrX-165430699-INS-89 | chrX-165430694-INS-76 |
| chrX-165433393-INS-425 | chrX-165433393-INS-425 |
| chrX-165460914-INS-1104 | chrX-165460915-INS-1103 |
| chrX-165475111-INS-8267 | chrX-165475112-INS-8268 |
| chrX-165495854-INS-2367 | chrX-165495855-INS-2368 |
| chrX-165517139-INS-489 | chrX-165517139-INS-487 |
| chrX-165530593-INS-120 | chrX-165530593-INS-122 |
| chrX-165540934-INS-147 | chrX-165540934-INS-148 |
| chrX-165572778-INS-58 | chrX-165572772-INS-62 |
| chrX-165645501-INS-563 | chrX-165645501-INS-519 |
| chrX-165646966-INS-1886 | chrX-165646967-INS-1886 |
| chrX-165652443-INS-6244 | chrX-165652444-INS-6244 |
| chrX-165770214-INS-66 | chrX-165770214-INS-66 |
| chrX-165781914-INS-241 | chrX-165781914-INS-241 |
| chrX-165791502-INS-71 | chrX-165791502-INS-81 |
| chrX-165831661-INS-155 | chrX-165831682-INS-165 |
| chrX-165863149-INS-107 | chrX-165863150-INS-107 |
| chrX-165879343-INS-105 | chrX-165879343-INS-66 |
| chrX-165896214-INS-59 | chrX-165896214-INS-59 |
| chrX-165911163-INS-1133 | chrX-165911164-INS-1133 |
| chrX-165911795-INS-52 | chrX-165911795-INS-52 |
| chrX-165916862-INS-66 | chrX-165916862-INS-57 |
| chrX-165971326-INS-72 | chrX-165971308-INS-98 |
| chrX-165978143-INS-120 | chrX-165978143-INS-120 |
| chrX-165984785-INS-54 | chrX-165984805-INS-60 |
| chrX-166030374-INS-7076 | chrX-166030375-INS-7076 |
| chrX-166053587-INS-58 | chrX-166053587-INS-58 |
| chrX-166058673-INS-54 | chrX-166058673-INS-54 |
| chrX-166065130-INS-198 | chrX-166065130-INS-198 |
| chrX-166106743-INS-239 | chrX-166106743-INS-236 |
| chrX-166120221-INS-54 | chrX-166120221-INS-54 |
| chrX-166162997-INS-6475 | chrX-166162998-INS-6475 |
| chrX-166179590-INS-998 | chrX-166179590-INS-999 |
| chrX-166184920-INS-283 | chrX-166184920-INS-283 |
| chrX-166197792-INS-56 | chrX-166197792-INS-56 |
| chrX-166223396-INS-91 | chrX-166223396-INS-91 |
| chrX-166226662-INS-54 | chrX-166226662-INS-63 |
| chrX-166237166-INS-508 | chrX-166237166-INS-508 |
| chrX-166277300-INS-52 | chrX-166277300-INS-52 |
| chrX-166307481-INS-63 | chrX-166307481-INS-59 |
| chrX-166321101-INS-60 | chrX-166321101-INS-60 |
| chrX-166324820-INS-6237 | chrX-166324821-INS-6237 |
| chrX-166331115-INS-55 | chrX-166331115-INS-52 |
| chrX-166340939-INS-493 | chrX-166340939-INS-493 |
| chrX-166366553-INS-147 | chrX-166366551-INS-187 |
| chrX-166369261-INS-64 | chrX-166369261-INS-58 |
| chrX-166384310-INS-56 | chrX-166384321-INS-57 |
| chrX-166575142-INS-97 | chrX-166575142-INS-102 |
| chrX-166576033-INS-385 | chrX-166576033-INS-385 |
| chrX-166626361-INS-418 | chrX-166626361-INS-418 |
| chrX-166638713-INS-50 | chrX-166638713-INS-52 |
| chrX-166653057-INS-54 | chrX-166653086-INS-52 |
| chrX-166687147-INS-3292 | chrX-166687148-INS-3292 |

| | |
|---|---|
| chrX-166704824-INS-465 | chrX-166704824-INS-465 |
| chrX-166723530-INS-440 | chrX-166723530-INS-440 |
| chrX-166740566-INS-1200 | chrX-166740567-INS-1201 |
| chrX-166812472-INS-56 | chrX-166812472-INS-72 |
| chrX-166882079-INS-68 | chrX-166882079-INS-72 |
| chrX-166907402-INS-57 | chrX-166907367-INS-76 |
| chrX-166917604-INS-754 | chrX-166917604-INS-754 |
| chrX-166922204-INS-116 | chrX-166922204-INS-104 |
| chrX-166957817-INS-115 | chrX-166957817-INS-115 |
| chrX-167003419-INS-979 | chrX-167003419-INS-979 |
| chrX-167031820-INS-60 | chrX-167031820-INS-60 |
| chrX-167035882-INS-6475 | chrX-167035883-INS-6474 |
| chrX-167091526-INS-1035 | chrX-167091527-INS-1035 |
| chrX-167120665-INS-62 | chrX-167120665-INS-62 |
| chrX-167134328-INS-77 | chrX-167134325-INS-119 |
| chrX-167184351-INS-513 | chrX-167184351-INS-513 |
| chrX-167205781-INS-52 | chrX-167205781-INS-52 |
| chrX-167241549-INS-92 | chrX-167241549-INS-92 |
| chrX-167268510-INS-5318 | chrX-167268511-INS-5318 |
| chrX-167316756-INS-840 | chrX-167316756-INS-840 |
| chrX-167325447-INS-72 | chrX-167325447-INS-60 |
| chrX-167471486-INS-5749 | chrX-167471487-INS-5749 |
| chrX-167522040-INS-233 | chrX-167522040-INS-233 |
| chrX-167525204-INS-1272 | chrX-167525205-INS-1272 |
| chrX-167539984-INS-7325 | chrX-167539985-INS-7325 |
| chrX-167551452-INS-4099 | chrX-167551453-INS-4100 |
| chrX-167563672-INS-144 | chrX-167563672-INS-154 |
| chrX-167581087-INS-52 | chrX-167581087-INS-52 |
| chrX-167623319-INS-5595 | chrX-167623320-INS-5595 |
| chrX-167641840-INS-579 | chrX-167641840-INS-579 |
| chrX-167675377-INS-213 | chrX-167675377-INS-213 |
| chrX-167681427-INS-58 | chrX-167681427-INS-58 |
| chrX-167736659-INS-6851 | chrX-167736660-INS-6851 |
| chrX-167753879-INS-292 | chrX-167753879-INS-292 |
| chrX-167758292-INS-70 | chrX-167758251-INS-90 |
| chrX-167805949-INS-3177 | chrX-167805950-INS-3177 |
| chrX-167835676-INS-90 | chrX-167835676-INS-90 |
| chrX-167862628-INS-102 | chrX-167862628-INS-102 |
| chrX-167875364-INS-58 | chrX-167875364-INS-58 |
| chrX-167892379-INS-60 | chrX-167892379-INS-58 |
| chrX-167896537-INS-68 | chrX-167896537-INS-66 |
| chrX-167943226-INS-212 | chrX-167943226-INS-212 |
| chrX-167957155-INS-60 | chrX-167957155-INS-62 |
| chrX-167972597-INS-61 | chrX-167972598-INS-63 |
| chrX-168087229-INS-60 | chrX-168087237-INS-50 |
| chrX-168209343-INS-3074 | chrX-168209344-INS-3074 |
| chrX-168217385-INS-62 | chrX-168217385-INS-64 |
| chrX-168219807-INS-78 | chrX-168219807-INS-78 |
| chrX-168381523-INS-56 | chrX-168381523-INS-56 |
| chrX-168410623-INS-2227 | chrX-168410607-INS-2233 |
| chrX-168442869-INS-960 | chrX-168442869-INS-960 |
| chrX-168449018-INS-515 | chrX-168449018-INS-515 |
| chrX-168454552-INS-459 | chrX-168454552-INS-459 |
| chrX-168473872-INS-2625 | chrX-168473873-INS-2612 |
| chrX-168480049-INS-75 | chrX-168480049-INS-75 |
| chrX-168618286-INS-602 | chrX-168618286-INS-618 |
| chrX-168642485-INS-351 | chrX-168642485-INS-351 |
| chrX-168695691-INS-161 | chrX-168695691-INS-161 |
| chrX-168755547-INS-202 | chrX-168755537-INS-208 |
| chrX-168759179-INS-86 | chrX-168759179-INS-84 |
| chrX-168775107-INS-311 | chrX-168775107-INS-310 |
| chrX-168803252-INS-176 | chrX-168803252-INS-176 |
| chrX-169069157-INS-90 | chrX-169069152-INS-107 |
| chrX-169081066-INS-185 | chrX-169081097-INS-154 |
| chrX-169082141-INS-62 | chrX-169082141-INS-62 |
| chrX-169085595-INS-92 | chrX-169085595-INS-92 |
| chrX-169091723-INS-216 | chrX-169091723-INS-215 |
| chrX-169100942-INS-72 | chrX-169100942-INS-72 |
| chrX-169103584-INS-106 | chrX-169103584-INS-106 |
| chrX-169104313-INS-237 | chrX-169104286-INS-247 |
| chrX-169113429-INS-61 | chrX-169113429-INS-61 |
| chrX-169309362-INS-62 | chrX-169309351-INS-63 |

**Identified_By_Us**
chr1-3068908-INS-88
chr1-3084405-INS-506
chr1-3093692-INS-380
chr1-3099484-INS-119
chr1-3174972-INS-75
chr1-3185346-INS-522
chr1-3235671-INS-630
chr1-3242619-INS-324
chr1-3247086-INS-8220
chr1-3282171-INS-134
chr1-3292141-INS-63
chr1-3357791-INS-51
chr1-3369968-INS-385
chr1-3370762-INS-98
chr1-3385272-INS-358
chr1-3472915-INS-122
chr1-3478551-INS-216
chr1-3497983-INS-561
chr1-3600501-INS-150
chr1-3601185-INS-83
chr1-3641130-INS-56
chr1-3656535-INS-326
chr1-3668361-INS-94
chr1-3680253-INS-219
chr1-3720486-INS-330
chr1-3779778-INS-426
chr1-3871663-INS-6636
chr1-3909233-INS-90
chr1-3955322-INS-179
chr1-3956948-INS-3171
chr1-3965262-INS-6442
chr1-3984765-INS-6213
chr1-3998473-INS-72
chr1-4030154-INS-460
chr1-4037480-INS-6377
chr1-4041461-INS-6411
chr1-4050528-INS-7264
chr1-4113636-INS-68
chr1-4116443-INS-407
chr1-4120438-INS-591
chr1-4155628-INS-208
chr1-4172829-INS-178
chr1-4179899-INS-329
chr1-4189407-INS-79
chr1-4220326-INS-198
chr1-4227442-INS-6062
chr1-4230376-INS-498
chr1-4233123-INS-6817
chr1-4258165-INS-324
chr1-4298375-INS-56
chr1-4326912-INS-65
chr1-4328032-INS-68
chr1-4355441-INS-53
chr1-4368896-INS-544
chr1-4400120-INS-62
chr1-4423362-INS-572
chr1-4452073-INS-65
chr1-4459783-INS-601
chr1-4464969-INS-78
chr1-4469913-INS-425
chr1-4512059-INS-498
chr1-4520353-INS-248
chr1-4525156-INS-133
chr1-4531641-INS-230
chr1-4546610-INS-87
chr1-4565240-INS-64
chr1-4586404-INS-151
chr1-4587457-INS-64
chr1-4597420-INS-72
chr1-4601171-INS-128
chr1-4608253-INS-146
chr1-4621080-INS-6402
chr1-4630141-INS-424
chr1-4644257-INS-135
chr1-4649269-INS-400
chr1-4690127-INS-56

chr1-4695876-INS-275
chr1-4698888-INS-80
chr1-4699756-INS-5443
chr1-4703819-INS-4103
chr1-4707824-INS-583
chr1-4720962-INS-7215
chr1-4750477-INS-52
chr1-4752613-INS-78
chr1-4769286-INS-5547
chr1-4778602-INS-501
chr1-4818866-INS-94
chr1-4823362-INS-167
chr1-4826104-INS-116
chr1-4851852-INS-136
chr1-4859378-INS-9047
chr1-4865772-INS-106
chr1-4876652-INS-63
chr1-4888290-INS-6473
chr1-4894653-INS-290
chr1-4905400-INS-111
chr1-4930538-INS-107
chr1-4971884-INS-748
chr1-4988618-INS-199
chr1-4992654-INS-102
chr1-5001717-INS-172
chr1-5011978-INS-4058
chr1-5014143-INS-55
chr1-5034353-INS-972
chr1-5074416-INS-272
chr1-5082230-INS-58
chr1-5086840-INS-102
chr1-5092630-INS-138
chr1-5099057-INS-6215
chr1-5136746-INS-103
chr1-5140898-INS-404
chr1-5142654-INS-147
chr1-5142764-INS-1491
chr1-5158050-INS-62
chr1-5243743-INS-5619
chr1-5257494-INS-232
chr1-5288760-INS-332
chr1-5290206-INS-182
chr1-5296349-INS-120
chr1-5309374-INS-798
chr1-5332034-INS-90
chr1-5334116-INS-138
chr1-5370172-INS-245
chr1-5386691-INS-518
chr1-5398807-INS-779
chr1-5405392-INS-328
chr1-5409307-INS-485
chr1-5431330-INS-7101
chr1-5432381-INS-63
chr1-5467405-INS-118
chr1-5473404-INS-98
chr1-5489559-INS-476
chr1-5512647-INS-74
chr1-5527405-INS-70
chr1-5540186-INS-203
chr1-5552988-INS-1888
chr1-5579755-INS-290
chr1-5601564-INS-431
chr1-5628852-INS-354
chr1-5630291-INS-565
chr1-5637093-INS-81
chr1-5652048-INS-104
chr1-5664002-INS-130
chr1-5671761-INS-52
chr1-5685098-INS-743
chr1-5689987-INS-831
chr1-5722813-INS-211
chr1-5792429-INS-57
chr1-5806386-INS-215
chr1-5815729-INS-215
chr1-5844820-INS-65
chr1-5849294-INS-343
chr1-5866940-INS-110

chr1-5872078-INS-711
chr1-5883769-INS-8757
chr1-5884112-INS-331
chr1-5885938-INS-1893
chr1-5886911-INS-116
chr1-5900964-INS-234
chr1-5901169-INS-307
chr1-5930054-INS-164
chr1-5993131-INS-169
chr1-6004584-INS-233
chr1-6032984-INS-118
chr1-6087250-INS-241
chr1-6097228-INS-198
chr1-6106160-INS-331
chr1-6143146-INS-51
chr1-6185599-INS-55
chr1-6186841-INS-81
chr1-6246598-INS-238
chr1-6257047-INS-9531
chr1-6257426-INS-2366
chr1-6261301-INS-212
chr1-6278639-INS-171
chr1-6309378-INS-53
chr1-6310614-INS-142
chr1-6327805-INS-100
chr1-6334614-INS-499
chr1-6373552-INS-131
chr1-6387728-INS-316
chr1-6398337-INS-117
chr1-6421652-INS-125
chr1-6442013-INS-326
chr1-6454042-INS-6921
chr1-6455001-INS-142
chr1-6485395-INS-74
chr1-6489419-INS-72
chr1-6499294-INS-166
chr1-6515581-INS-7484
chr1-6529236-INS-56
chr1-6587156-INS-90
chr1-6648278-INS-89
chr1-6652121-INS-76
chr1-6660939-INS-96
chr1-6662037-INS-56
chr1-6671345-INS-125
chr1-6680633-INS-505
chr1-6732219-INS-380
chr1-6736204-INS-906
chr1-6834420-INS-579
chr1-6851458-INS-209
chr1-6861113-INS-172
chr1-6891932-INS-6906
chr1-6935547-INS-130
chr1-6949038-INS-66
chr1-6961952-INS-169
chr1-6983318-INS-377
chr1-7008592-INS-368
chr1-7012089-INS-348
chr1-7050945-INS-511
chr1-7051695-INS-66
chr1-7073732-INS-65
chr1-7101103-INS-104
chr1-7105131-INS-83
chr1-7111706-INS-196
chr1-7146528-INS-92
chr1-7148572-INS-115
chr1-7158618-INS-5545
chr1-7162161-INS-102
chr1-7177345-INS-63
chr1-7182902-INS-88
chr1-7203065-INS-233
chr1-7215414-INS-54
chr1-7220785-INS-129
chr1-7234509-INS-244
chr1-7277069-INS-75
chr1-7296185-INS-624
chr1-7325567-INS-343
chr1-7361820-INS-54

chr1-7380554-INS-439
chr1-7392855-INS-205
chr1-7428346-INS-341
chr1-7467134-INS-361
chr1-7485631-INS-66
chr1-7519159-INS-416
chr1-7538330-INS-76
chr1-7538979-INS-498
chr1-7562471-INS-94
chr1-7579537-INS-93
chr1-7593211-INS-126
chr1-7594317-INS-5542
chr1-7621945-INS-99
chr1-7661773-INS-937
chr1-7731216-INS-298
chr1-7749120-INS-111
chr1-7750848-INS-1894
chr1-7822520-INS-387
chr1-7843629-INS-348
chr1-7847661-INS-144
chr1-7953858-INS-111
chr1-7975093-INS-1990
chr1-8034989-INS-85
chr1-8105393-INS-69
chr1-8133583-INS-76
chr1-8166121-INS-52
chr1-8166688-INS-81
chr1-8188098-INS-70
chr1-8218264-INS-443
chr1-8220071-INS-6468
chr1-8227572-INS-72
chr1-8276371-INS-530
chr1-8279239-INS-168
chr1-8279848-INS-4412
chr1-8294366-INS-59
chr1-8301887-INS-274
chr1-8343174-INS-637
chr1-8360713-INS-160
chr1-8369867-INS-128
chr1-8387781-INS-787
chr1-8409455-INS-6723
chr1-8448505-INS-64
chr1-8563387-INS-79
chr1-8565503-INS-6269
chr1-8598357-INS-256
chr1-8600804-INS-55
chr1-8614045-INS-142
chr1-8622981-INS-358
chr1-8650127-INS-467
chr1-8676268-INS-120
chr1-8701573-INS-94
chr1-8713212-INS-183
chr1-8721602-INS-1080
chr1-8772507-INS-57
chr1-8773675-INS-242
chr1-8800534-INS-6460
chr1-8815776-INS-81
chr1-8833270-INS-545
chr1-8971386-INS-154
chr1-9015036-INS-519
chr1-9020431-INS-98
chr1-9023694-INS-492
chr1-9033498-INS-681
chr1-9060881-INS-839
chr1-9099604-INS-72
chr1-9109173-INS-664
chr1-9124574-INS-57
chr1-9136841-INS-468
chr1-9172101-INS-198
chr1-9181840-INS-204
chr1-9245884-INS-310
chr1-9255214-INS-342
chr1-9263083-INS-181
chr1-9279576-INS-6500
chr1-9287748-INS-390
chr1-9290848-INS-7318
chr1-9294843-INS-300

chr1-9387582-INS-527
chr1-9406079-INS-469
chr1-9407444-INS-437
chr1-9441416-INS-190
chr1-9512591-INS-52
chr1-9539146-INS-131
chr1-9578672-INS-295
chr1-9589292-INS-59
chr1-9589343-INS-6440
chr1-9590705-INS-79
chr1-9614923-INS-138
chr1-9653419-INS-277
chr1-9662601-INS-55
chr1-9670735-INS-626
chr1-9705761-INS-133
chr1-9708239-INS-66
chr1-9716872-INS-120
chr1-9724040-INS-89
chr1-9727643-INS-54
chr1-9773594-INS-181
chr1-9782730-INS-406
chr1-9783530-INS-196
chr1-9813380-INS-202
chr1-9819237-INS-609
chr1-9820693-INS-198
chr1-9824790-INS-51
chr1-9827392-INS-118
chr1-9854770-INS-546
chr1-9859065-INS-60
chr1-9896814-INS-316
chr1-9900705-INS-56
chr1-9951878-INS-512
chr1-9980142-INS-59
chr1-9989510-INS-1144
chr1-10004417-INS-900
chr1-10009734-INS-56
chr1-10010784-INS-137
chr1-10019094-INS-210
chr1-10042935-INS-352
chr1-10059672-INS-244
chr1-10071661-INS-95
chr1-10090225-INS-230
chr1-10095676-INS-112
chr1-10112757-INS-53
chr1-10132221-INS-60
chr1-10133482-INS-209
chr1-10170795-INS-479
chr1-10172311-INS-207
chr1-10182325-INS-251
chr1-10197959-INS-56
chr1-10229100-INS-213
chr1-10247758-INS-78
chr1-10259804-INS-77
chr1-10298578-INS-173
chr1-10325260-INS-211
chr1-10332337-INS-196
chr1-10339383-INS-154
chr1-10341932-INS-60
chr1-10343301-INS-7273
chr1-10344813-INS-72
chr1-10360590-INS-431
chr1-10364711-INS-64
chr1-10378351-INS-182
chr1-10383689-INS-54
chr1-10396810-INS-107
chr1-10397985-INS-67
chr1-10446712-INS-94
chr1-10463487-INS-204
chr1-10470947-INS-234
chr1-10560764-INS-127
chr1-10576879-INS-1160
chr1-10577068-INS-171
chr1-10587398-INS-471
chr1-10590767-INS-6411
chr1-10596260-INS-67
chr1-10596897-INS-90
chr1-10598308-INS-58

chr1-10619234-INS-334
chr1-10620622-INS-267
chr1-10696626-INS-6418
chr1-10705316-INS-558
chr1-10720752-INS-62
chr1-10747034-INS-705
chr1-10823226-INS-7207
chr1-10829488-INS-52
chr1-10865699-INS-620
chr1-10873037-INS-58
chr1-10915088-INS-810
chr1-10923627-INS-86
chr1-10938254-INS-109
chr1-10957019-INS-2149
chr1-10968410-INS-62
chr1-10974356-INS-199
chr1-10975785-INS-62
chr1-10997678-INS-144
chr1-10998572-INS-457
chr1-11016023-INS-147
chr1-11026755-INS-534
chr1-11059297-INS-2089
chr1-11062906-INS-8154
chr1-11078326-INS-186
chr1-11128121-INS-67
chr1-11163462-INS-194
chr1-11216727-INS-70
chr1-11233251-INS-456
chr1-11262344-INS-80
chr1-11269077-INS-56
chr1-11318113-INS-150
chr1-11322189-INS-6253
chr1-11326771-INS-52
chr1-11337551-INS-4521
chr1-11344607-INS-6481
chr1-11384651-INS-127
chr1-11406574-INS-113
chr1-11407663-INS-326
chr1-11412990-INS-73
chr1-11413783-INS-5140
chr1-11425367-INS-56
chr1-11428894-INS-6627
chr1-11452949-INS-6544
chr1-11454103-INS-498
chr1-11462523-INS-515
chr1-11464281-INS-8418
chr1-11469873-INS-9090
chr1-11470370-INS-230
chr1-11474335-INS-5543
chr1-11487780-INS-298
chr1-11516309-INS-118
chr1-11520138-INS-449
chr1-11543182-INS-52
chr1-11554825-INS-1889
chr1-11573025-INS-469
chr1-11587744-INS-131
chr1-11652524-INS-72
chr1-11676360-INS-6467
chr1-11680375-INS-60
chr1-11696976-INS-424
chr1-11720494-INS-224
chr1-11722872-INS-387
chr1-11805629-INS-63
chr1-11823709-INS-525
chr1-11861493-INS-114
chr1-11872528-INS-4978
chr1-11964620-INS-6801
chr1-11987306-INS-6404
chr1-11987996-INS-129
chr1-12056139-INS-341
chr1-12087826-INS-426
chr1-12136357-INS-54
chr1-12153114-INS-74
chr1-12193947-INS-400
chr1-12196107-INS-85
chr1-12197414-INS-68
chr1-12246547-INS-53

chr1-12267350-INS-242
chr1-12268771-INS-250
chr1-12317408-INS-363
chr1-12331686-INS-54
chr1-12355146-INS-101
chr1-12382024-INS-452
chr1-12383320-INS-216
chr1-12424276-INS-497
chr1-12436226-INS-126
chr1-12443428-INS-400
chr1-12467897-INS-7107
chr1-12478412-INS-91
chr1-12484036-INS-55
chr1-12500338-INS-175
chr1-12501737-INS-763
chr1-12507142-INS-60
chr1-12543662-INS-58
chr1-12557209-INS-385
chr1-12572283-INS-83
chr1-12575614-INS-134
chr1-12590736-INS-52
chr1-12592495-INS-224
chr1-12610088-INS-59
chr1-12697248-INS-837
chr1-12708898-INS-424
chr1-12722436-INS-511
chr1-12728026-INS-541
chr1-12729923-INS-52
chr1-12757909-INS-68
chr1-12807701-INS-95
chr1-12828921-INS-64
chr1-12831931-INS-76
chr1-12879209-INS-216
chr1-12890462-INS-197
chr1-12891888-INS-78
chr1-12910016-INS-89
chr1-12911533-INS-203
chr1-12985732-INS-5222
chr1-13022284-INS-341
chr1-13028660-INS-157
chr1-13033703-INS-144
chr1-13043149-INS-190
chr1-13064493-INS-974
chr1-13066636-INS-87
chr1-13077244-INS-528
chr1-13080072-INS-68
chr1-13085500-INS-53
chr1-13101336-INS-61
chr1-13109827-INS-184
chr1-13114180-INS-433
chr1-13118796-INS-230
chr1-13122103-INS-151
chr1-13193274-INS-132
chr1-13214613-INS-5510
chr1-13230855-INS-79
chr1-13247982-INS-200
chr1-13273773-INS-148
chr1-13321302-INS-68
chr1-13324310-INS-104
chr1-13330390-INS-498
chr1-13332045-INS-140
chr1-13343266-INS-104
chr1-13364376-INS-120
chr1-13388457-INS-718
chr1-13472699-INS-202
chr1-13478504-INS-62
chr1-13489605-INS-347
chr1-13502791-INS-5766
chr1-13502815-INS-72
chr1-13518518-INS-442
chr1-13521957-INS-183
chr1-13532703-INS-6403
chr1-13557182-INS-6437
chr1-13592006-INS-72
chr1-13629959-INS-583
chr1-13641224-INS-177
chr1-13663994-INS-102

chr1-13704541-INS-201
chr1-13721098-INS-68
chr1-13732881-INS-119
chr1-13734530-INS-66
chr1-13746568-INS-5365
chr1-13779276-INS-69
chr1-13807552-INS-566
chr1-13818690-INS-327
chr1-13821878-INS-6765
chr1-13824908-INS-60
chr1-13832549-INS-164
chr1-13841465-INS-193
chr1-13845886-INS-76
chr1-13846244-INS-6517
chr1-13903000-INS-60
chr1-13919331-INS-252
chr1-13938722-INS-320
chr1-13995552-INS-52
chr1-14018044-INS-1397
chr1-14032866-INS-171
chr1-14037233-INS-7148
chr1-14045962-INS-6397
chr1-14064956-INS-73
chr1-14067116-INS-294
chr1-14075777-INS-203
chr1-14077647-INS-562
chr1-14078964-INS-84
chr1-14083822-INS-220
chr1-14104719-INS-55
chr1-14113369-INS-307
chr1-14119971-INS-98
chr1-14122066-INS-1746
chr1-14127482-INS-578
chr1-14130341-INS-202
chr1-14141604-INS-153
chr1-14152324-INS-7344
chr1-14165698-INS-7206
chr1-14172167-INS-54
chr1-14193686-INS-84
chr1-14201348-INS-6375
chr1-14267598-INS-320
chr1-14336271-INS-116
chr1-14339942-INS-291
chr1-14368836-INS-52
chr1-14401971-INS-82
chr1-14434966-INS-120
chr1-14446465-INS-172
chr1-14470806-INS-423
chr1-14479736-INS-8281
chr1-14505179-INS-441
chr1-14507380-INS-395
chr1-14517922-INS-125
chr1-14525894-INS-82
chr1-14590300-INS-101
chr1-14614197-INS-144
chr1-14638049-INS-766
chr1-14643106-INS-116
chr1-14656054-INS-329
chr1-14663391-INS-431
chr1-14701968-INS-226
chr1-14736101-INS-495
chr1-14755529-INS-105
chr1-14764170-INS-339
chr1-14800214-INS-153
chr1-14816582-INS-271
chr1-14830655-INS-399
chr1-14868145-INS-58
chr1-14883924-INS-52
chr1-14893199-INS-58
chr1-14898586-INS-227
chr1-14934159-INS-58
chr1-14952664-INS-245
chr1-14969857-INS-162
chr1-14985992-INS-761
chr1-15046006-INS-408
chr1-15050981-INS-56
chr1-15052934-INS-65

chr1-15105619-INS-6664
chr1-15119372-INS-127
chr1-15140090-INS-319
chr1-15146226-INS-73
chr1-15159511-INS-7104
chr1-15173849-INS-181
chr1-15189201-INS-69
chr1-15197916-INS-617
chr1-15211727-INS-342
chr1-15214140-INS-2999
chr1-15243999-INS-63
chr1-15265926-INS-8827
chr1-15278705-INS-82
chr1-15281676-INS-881
chr1-15307819-INS-6223
chr1-15310519-INS-56
chr1-15327254-INS-610
chr1-15335782-INS-7420
chr1-15395425-INS-495
chr1-15406401-INS-6404
chr1-15414880-INS-182
chr1-15450220-INS-512
chr1-15459591-INS-96
chr1-15490516-INS-54
chr1-15514027-INS-4439
chr1-15522603-INS-69
chr1-15523823-INS-195
chr1-15578827-INS-105
chr1-15630023-INS-3210
chr1-15636921-INS-1265
chr1-15650438-INS-1254
chr1-15702048-INS-68
chr1-15704451-INS-80
chr1-15708092-INS-55
chr1-15712127-INS-288
chr1-15716486-INS-209
chr1-15725421-INS-138
chr1-15728320-INS-195
chr1-15742314-INS-135
chr1-15744321-INS-52
chr1-15746493-INS-933
chr1-15758080-INS-594
chr1-15771219-INS-221
chr1-15777838-INS-249
chr1-15793503-INS-237
chr1-15800411-INS-239
chr1-15827849-INS-72
chr1-15836878-INS-5486
chr1-15843436-INS-129
chr1-15855918-INS-604
chr1-15856151-INS-151
chr1-15860832-INS-59
chr1-15914133-INS-193
chr1-15923778-INS-354
chr1-15940650-INS-200
chr1-15946828-INS-108
chr1-15958883-INS-132
chr1-15974671-INS-175
chr1-15980849-INS-359
chr1-16011846-INS-59
chr1-16032613-INS-344
chr1-16037329-INS-93
chr1-16046213-INS-858
chr1-16049080-INS-150
chr1-16058505-INS-331
chr1-16075663-INS-52
chr1-16089957-INS-491
chr1-16091013-INS-54
chr1-16142664-INS-900
chr1-16144611-INS-74
chr1-16146646-INS-58
chr1-16155128-INS-72
chr1-16160225-INS-529
chr1-16221541-INS-994
chr1-16256749-INS-68
chr1-16266028-INS-998
chr1-16274312-INS-343

chr1-16395734-INS-197
chr1-16403486-INS-641
chr1-16412730-INS-81
chr1-16413122-INS-1016
chr1-16413227-INS-90
chr1-16429441-INS-247
chr1-16459539-INS-320
chr1-16474964-INS-4038
chr1-16475496-INS-956
chr1-16480639-INS-102
chr1-16485919-INS-410
chr1-16510022-INS-81
chr1-16536762-INS-6459
chr1-16538602-INS-66
chr1-16550163-INS-197
chr1-16551378-INS-340
chr1-16553473-INS-556
chr1-16554669-INS-74
chr1-16555677-INS-104
chr1-16557715-INS-832
chr1-16602745-INS-274
chr1-16608200-INS-73
chr1-16617700-INS-146
chr1-16663833-INS-57
chr1-16668662-INS-60
chr1-16700353-INS-69
chr1-16743704-INS-203
chr1-16778111-INS-960
chr1-16787194-INS-135
chr1-16841799-INS-321
chr1-16850982-INS-64
chr1-16856976-INS-968
chr1-16859834-INS-67
chr1-16897050-INS-54
chr1-16918131-INS-68
chr1-16931238-INS-5604
chr1-16975760-INS-540
chr1-16996664-INS-6375
chr1-17016237-INS-100
chr1-17034807-INS-256
chr1-17038140-INS-6525
chr1-17112217-INS-470
chr1-17160040-INS-526
chr1-17199666-INS-500
chr1-17229448-INS-68
chr1-17259555-INS-4401
chr1-17270235-INS-56
chr1-17270635-INS-781
chr1-17280185-INS-75
chr1-17290402-INS-217
chr1-17302105-INS-130
chr1-17347226-INS-961
chr1-17369603-INS-151
chr1-17375866-INS-69
chr1-17406618-INS-68
chr1-17419837-INS-52
chr1-17430510-INS-54
chr1-17434127-INS-130
chr1-17436671-INS-819
chr1-17467994-INS-53
chr1-17492209-INS-120
chr1-17498766-INS-114
chr1-17512304-INS-4881
chr1-17517492-INS-366
chr1-17586628-INS-2961
chr1-17598178-INS-112
chr1-17622429-INS-209
chr1-17632328-INS-385
chr1-17656490-INS-6518
chr1-17704832-INS-53
chr1-17718680-INS-514
chr1-17745424-INS-874
chr1-17746134-INS-4432
chr1-17760441-INS-465
chr1-17768778-INS-92
chr1-17775582-INS-206
chr1-17783635-INS-55

chr1-17786140-INS-391
chr1-17819835-INS-91
chr1-17925703-INS-2841
chr1-17980151-INS-6611
chr1-18007567-INS-59
chr1-18039615-INS-122
chr1-18073439-INS-3332
chr1-18075930-INS-7594
chr1-18079736-INS-5476
chr1-18105273-INS-53
chr1-18119918-INS-5531
chr1-18182076-INS-2547
chr1-18203894-INS-199
chr1-18203984-INS-6946
chr1-18217449-INS-7466
chr1-18252663-INS-51
chr1-18268402-INS-384
chr1-18277111-INS-311
chr1-18286285-INS-204
chr1-18295335-INS-5286
chr1-18313116-INS-6440
chr1-18318948-INS-3116
chr1-18386777-INS-7111
chr1-18424625-INS-359
chr1-18425346-INS-54
chr1-18430963-INS-199
chr1-18434504-INS-79
chr1-18439759-INS-3743
chr1-18441163-INS-102
chr1-18458810-INS-8021
chr1-18477279-INS-227
chr1-18478864-INS-7114
chr1-18503660-INS-5551
chr1-18512124-INS-521
chr1-18525501-INS-539
chr1-18562133-INS-6689
chr1-18567145-INS-312
chr1-18572686-INS-356
chr1-18576705-INS-64
chr1-18668243-INS-52
chr1-18671831-INS-189
chr1-18682495-INS-8560
chr1-18686083-INS-447
chr1-18695780-INS-81
chr1-18724466-INS-166
chr1-18766007-INS-56
chr1-18774681-INS-95
chr1-18775633-INS-800
chr1-18779225-INS-70
chr1-18782448-INS-198
chr1-18797318-INS-863
chr1-18810632-INS-8351
chr1-18849377-INS-208
chr1-18910469-INS-6301
chr1-18927048-INS-3324
chr1-18934861-INS-64
chr1-18935805-INS-6360
chr1-18979972-INS-324
chr1-19018874-INS-54
chr1-19042552-INS-59
chr1-19050739-INS-4437
chr1-19052536-INS-6926
chr1-19054378-INS-94
chr1-19055673-INS-449
chr1-19096105-INS-897
chr1-19098686-INS-94
chr1-19103201-INS-448
chr1-19104392-INS-80
chr1-19114362-INS-84
chr1-19123033-INS-6410
chr1-19136457-INS-60
chr1-19147434-INS-374
chr1-19150820-INS-103
chr1-19209215-INS-1893
chr1-19213617-INS-58
chr1-19221187-INS-170
chr1-19226604-INS-61

chr1-19227400-INS-618
chr1-19251166-INS-207
chr1-19251959-INS-82
chr1-19253709-INS-56
chr1-19256044-INS-310
chr1-19272784-INS-220
chr1-19273560-INS-223
chr1-19275134-INS-57
chr1-19285786-INS-72
chr1-19311951-INS-871
chr1-19322959-INS-74
chr1-19323631-INS-2971
chr1-19330969-INS-72
chr1-19339327-INS-205
chr1-19366883-INS-6655
chr1-19368550-INS-6428
chr1-19389860-INS-119
chr1-19392214-INS-175
chr1-19403468-INS-289
chr1-19409443-INS-350
chr1-19416023-INS-83
chr1-19421432-INS-58
chr1-19426718-INS-6572
chr1-19437258-INS-6238
chr1-19483525-INS-166
chr1-19493214-INS-1392
chr1-19496872-INS-6906
chr1-19526377-INS-182
chr1-19551310-INS-89
chr1-19566789-INS-6919
chr1-19610590-INS-205
chr1-19655922-INS-1211
chr1-19662796-INS-92
chr1-19677373-INS-6474
chr1-19679379-INS-835
chr1-19688586-INS-58
chr1-19712678-INS-465
chr1-19741318-INS-964
chr1-19755366-INS-6389
chr1-19764969-INS-167
chr1-19801745-INS-77
chr1-19805946-INS-130
chr1-19820515-INS-193
chr1-19824937-INS-64
chr1-19841119-INS-894
chr1-19860563-INS-112
chr1-19867533-INS-55
chr1-19906961-INS-597
chr1-19921711-INS-282
chr1-19925197-INS-106
chr1-19942441-INS-115
chr1-19977267-INS-80
chr1-20001702-INS-106
chr1-20003966-INS-1306
chr1-20046351-INS-172
chr1-20073803-INS-222
chr1-20085229-INS-506
chr1-20090271-INS-340
chr1-20102120-INS-63
chr1-20113809-INS-1686
chr1-20122855-INS-6215
chr1-20126677-INS-1657
chr1-20133784-INS-693
chr1-20142235-INS-62
chr1-20182008-INS-101
chr1-20187021-INS-674
chr1-20190992-INS-7076
chr1-20199646-INS-58
chr1-20234414-INS-6387
chr1-20245955-INS-950
chr1-20248178-INS-52
chr1-20262127-INS-230
chr1-20272658-INS-192
chr1-20280740-INS-57
chr1-20314331-INS-53
chr1-20328827-INS-1890
chr1-20340123-INS-52

chr1-20342770-INS-109
chr1-20343781-INS-1864
chr1-20367403-INS-54
chr1-20370953-INS-6566
chr1-20374728-INS-228
chr1-20391721-INS-207
chr1-20440279-INS-177
chr1-20440389-INS-609
chr1-20442371-INS-585
chr1-20450088-INS-150
chr1-20471783-INS-112
chr1-20518687-INS-81
chr1-20523984-INS-76
chr1-20532660-INS-312
chr1-20534221-INS-52
chr1-20534922-INS-371
chr1-20546060-INS-108
chr1-20547219-INS-90
chr1-20554272-INS-347
chr1-20556602-INS-7324
chr1-20566486-INS-71
chr1-20569253-INS-54
chr1-20570654-INS-70
chr1-20579502-INS-87
chr1-20590898-INS-64
chr1-20612000-INS-80
chr1-20636274-INS-62
chr1-20658278-INS-174
chr1-20672026-INS-6830
chr1-20674576-INS-276
chr1-20679343-INS-85
chr1-20724940-INS-90
chr1-20740496-INS-89
chr1-20746009-INS-101
chr1-20748686-INS-119
chr1-20758140-INS-81
chr1-20765365-INS-449
chr1-20768075-INS-866
chr1-20785464-INS-56
chr1-20789129-INS-244
chr1-20794070-INS-79
chr1-20839149-INS-263
chr1-20860896-INS-96
chr1-20870421-INS-139
chr1-20890259-INS-94
chr1-20897302-INS-78
chr1-20917976-INS-92
chr1-20919407-INS-60
chr1-20927565-INS-379
chr1-20929781-INS-345
chr1-20940427-INS-376
chr1-20985241-INS-83
chr1-21014126-INS-114
chr1-21027015-INS-50
chr1-21035721-INS-390
chr1-21049424-INS-80
chr1-21055875-INS-72
chr1-21060820-INS-156
chr1-21078967-INS-66
chr1-21094052-INS-399
chr1-21095000-INS-442
chr1-21095916-INS-80
chr1-21151010-INS-114
chr1-21157580-INS-198
chr1-21228404-INS-140
chr1-21235250-INS-221
chr1-21236162-INS-54
chr1-21240131-INS-5146
chr1-21253431-INS-132
chr1-21254572-INS-6413
chr1-21255342-INS-53
chr1-21256842-INS-82
chr1-21260678-INS-54
chr1-21290964-INS-64
chr1-21295917-INS-63
chr1-21301394-INS-103
chr1-21316191-INS-128

chr1-21339556-INS-56
chr1-21381849-INS-90
chr1-21395586-INS-401
chr1-21398484-INS-210
chr1-21409033-INS-59
chr1-21411772-INS-73
chr1-21414401-INS-8674
chr1-21416618-INS-152
chr1-21474207-INS-206
chr1-21483049-INS-124
chr1-21486827-INS-267
chr1-21489130-INS-56
chr1-21490222-INS-192
chr1-21504965-INS-87
chr1-21510463-INS-218
chr1-21519125-INS-391
chr1-21527253-INS-64
chr1-21541040-INS-127
chr1-21565692-INS-269
chr1-21572030-INS-6694
chr1-21577923-INS-7851
chr1-21588191-INS-136
chr1-21592919-INS-54
chr1-21599795-INS-399
chr1-21632445-INS-80
chr1-21636565-INS-231
chr1-21643532-INS-6775
chr1-21658924-INS-401
chr1-21661549-INS-68
chr1-21665555-INS-321
chr1-21672706-INS-96
chr1-21687319-INS-147
chr1-21690264-INS-52
chr1-21695699-INS-200
chr1-21702957-INS-1008
chr1-21703146-INS-384
chr1-21706937-INS-264
chr1-21753572-INS-109
chr1-21756313-INS-164
chr1-21758460-INS-7336
chr1-21758885-INS-88
chr1-21767862-INS-467
chr1-21773093-INS-51
chr1-21780396-INS-68
chr1-21810388-INS-753
chr1-21816050-INS-134
chr1-21843736-INS-130
chr1-21844749-INS-525
chr1-21886878-INS-53
chr1-21902876-INS-652
chr1-21922101-INS-115
chr1-21930302-INS-957
chr1-21933124-INS-106
chr1-21937207-INS-6949
chr1-21955874-INS-222
chr1-21968885-INS-129
chr1-21988483-INS-152
chr1-22000908-INS-134
chr1-22003759-INS-165
chr1-22020583-INS-66
chr1-22042182-INS-68
chr1-22046690-INS-59
chr1-22053269-INS-146
chr1-22055502-INS-7204
chr1-22061543-INS-66
chr1-22074935-INS-104
chr1-22076345-INS-445
chr1-22090733-INS-99
chr1-22090913-INS-567
chr1-22102447-INS-113
chr1-22104213-INS-88
chr1-22106557-INS-249
chr1-22131616-INS-416
chr1-22144803-INS-113
chr1-22154216-INS-474
chr1-22175093-INS-88
chr1-22184908-INS-364

chr1-22206194-INS-492
chr1-22208415-INS-104
chr1-22212928-INS-386
chr1-22219557-INS-193
chr1-22260499-INS-6869
chr1-22270189-INS-122
chr1-22273481-INS-51
chr1-22285806-INS-54
chr1-22295432-INS-132
chr1-22307097-INS-193
chr1-22308097-INS-98
chr1-22317750-INS-75
chr1-22321433-INS-942
chr1-22333497-INS-54
chr1-22344354-INS-7185
chr1-22344556-INS-697
chr1-22351538-INS-176
chr1-22359840-INS-116
chr1-22373900-INS-70
chr1-22379977-INS-225
chr1-22391628-INS-68
chr1-22394640-INS-721
chr1-22422332-INS-107
chr1-22431761-INS-174
chr1-22434665-INS-186
chr1-22442527-INS-427
chr1-22444841-INS-216
chr1-22447595-INS-100
chr1-22486539-INS-51
chr1-22498363-INS-80
chr1-22506435-INS-123
chr1-22508699-INS-102
chr1-22543045-INS-53
chr1-22561785-INS-55
chr1-22589140-INS-112
chr1-22619601-INS-67
chr1-22659238-INS-79
chr1-22668916-INS-60
chr1-22672081-INS-148
chr1-22677354-INS-83
chr1-22684328-INS-76
chr1-22755605-INS-268
chr1-22758507-INS-548
chr1-22776530-INS-187
chr1-22781828-INS-238
chr1-22787178-INS-81
chr1-22788442-INS-128
chr1-22807278-INS-68
chr1-22814727-INS-114
chr1-22846569-INS-4265
chr1-22847897-INS-6704
chr1-22873076-INS-400
chr1-22878170-INS-368
chr1-22880589-INS-57
chr1-22893143-INS-1864
chr1-22916122-INS-83
chr1-22933506-INS-68
chr1-22935107-INS-64
chr1-22936910-INS-56
chr1-22939601-INS-56
chr1-22942075-INS-793
chr1-22947689-INS-124
chr1-22975061-INS-214
chr1-22982340-INS-262
chr1-22991714-INS-525
chr1-23061869-INS-851
chr1-23062957-INS-210
chr1-23067930-INS-59
chr1-23090442-INS-128
chr1-23122448-INS-676
chr1-23127732-INS-153
chr1-23178377-INS-54
chr1-23179884-INS-150
chr1-23187491-INS-121
chr1-23189768-INS-250
chr1-23197574-INS-834
chr1-23224971-INS-130

chr1-23260509-INS-76
chr1-23266392-INS-55
chr1-23284207-INS-1731
chr1-23314136-INS-55
chr1-23322779-INS-8993
chr1-23332521-INS-306
chr1-23380614-INS-226
chr1-23399367-INS-60
chr1-23439703-INS-2044
chr1-23449756-INS-51
chr1-23463481-INS-95
chr1-23470922-INS-118
chr1-23478187-INS-3386
chr1-23481429-INS-60
chr1-23505539-INS-144
chr1-23513476-INS-54
chr1-23549084-INS-661
chr1-23566748-INS-411
chr1-23570060-INS-65
chr1-23576169-INS-80
chr1-23597251-INS-68
chr1-23612594-INS-132
chr1-23615961-INS-182
chr1-23631251-INS-66
chr1-23638913-INS-373
chr1-23661382-INS-99
chr1-23668406-INS-1021
chr1-23712362-INS-6862
chr1-23720186-INS-367
chr1-23751459-INS-71
chr1-23760743-INS-101
chr1-23776735-INS-2868
chr1-23782693-INS-52
chr1-23805656-INS-79
chr1-23814327-INS-201
chr1-23829823-INS-90
chr1-23838669-INS-52
chr1-23848021-INS-257
chr1-23860157-INS-66
chr1-23870229-INS-620
chr1-23877519-INS-51
chr1-23905401-INS-646
chr1-23947226-INS-284
chr1-23957445-INS-131
chr1-23963167-INS-120
chr1-23975671-INS-157
chr1-23983763-INS-458
chr1-24002244-INS-108
chr1-24018167-INS-59
chr1-24061250-INS-198
chr1-24065241-INS-448
chr1-24081365-INS-68
chr1-24090446-INS-554
chr1-24128014-INS-1242
chr1-24129310-INS-122
chr1-24150093-INS-78
chr1-24151910-INS-53
chr1-24156666-INS-343
chr1-24167955-INS-5377
chr1-24168237-INS-82
chr1-24171531-INS-6494
chr1-24181013-INS-60
chr1-24220133-INS-533
chr1-24224807-INS-378
chr1-24247972-INS-63
chr1-24264335-INS-340
chr1-24306153-INS-165
chr1-24307500-INS-75
chr1-24322213-INS-224
chr1-24323381-INS-85
chr1-24347425-INS-955
chr1-24358036-INS-132
chr1-24363933-INS-138
chr1-24370557-INS-327
chr1-24370616-INS-196
chr1-24372257-INS-56
chr1-24376308-INS-64

chr1-24398770-INS-223
chr1-24404189-INS-58
chr1-24405129-INS-148
chr1-24442783-INS-90
chr1-24445320-INS-136
chr1-24463225-INS-291
chr1-24475170-INS-58
chr1-24475851-INS-257
chr1-24492776-INS-82
chr1-24497923-INS-54
chr1-24526783-INS-219
chr1-24542344-INS-207
chr1-24557970-INS-76
chr1-24559249-INS-507
chr1-24565805-INS-764
chr1-24576835-INS-181
chr1-24601535-INS-58
chr1-24604666-INS-160
chr1-24607700-INS-6401
chr1-24632991-INS-111
chr1-24635588-INS-389
chr1-24665705-INS-51
chr1-24672603-INS-244
chr1-24701950-INS-61
chr1-24702564-INS-53
chr1-24714296-INS-182
chr1-24715631-INS-52
chr1-24718668-INS-96
chr1-24720440-INS-560
chr1-24723118-INS-218
chr1-24723132-INS-1204
chr1-24728380-INS-348
chr1-24732880-INS-129
chr1-24734816-INS-197
chr1-24749821-INS-287
chr1-24760427-INS-408
chr1-24800324-INS-196
chr1-24816257-INS-4678
chr1-24821119-INS-100
chr1-24867930-INS-82
chr1-24868537-INS-302
chr1-24877080-INS-224
chr1-24903094-INS-56
chr1-24909108-INS-233
chr1-24916798-INS-119
chr1-24928522-INS-274
chr1-24937707-INS-1892
chr1-24939269-INS-73
chr1-24944269-INS-54
chr1-24962614-INS-6440
chr1-24963444-INS-60
chr1-24968904-INS-93
chr1-24970272-INS-439
chr1-24995043-INS-497
chr1-24998124-INS-352
chr1-25013992-INS-4132
chr1-25015988-INS-6738
chr1-25026271-INS-4354
chr1-25026613-INS-102
chr1-25046247-INS-287
chr1-25091382-INS-64
chr1-25144195-INS-108
chr1-25144900-INS-53
chr1-25175813-INS-59
chr1-25204124-INS-74
chr1-25209218-INS-142
chr1-25227107-INS-864
chr1-25227718-INS-231
chr1-25238252-INS-65
chr1-25242497-INS-266
chr1-25243963-INS-496
chr1-25245410-INS-463
chr1-25256876-INS-126
chr1-25272733-INS-7087
chr1-25291982-INS-173
chr1-25315565-INS-93
chr1-25370133-INS-334

chr1-25371441-INS-455
chr1-25410032-INS-522
chr1-25410458-INS-51
chr1-25415315-INS-595
chr1-25464694-INS-54
chr1-25476592-INS-96
chr1-25481556-INS-6674
chr1-25487693-INS-6858
chr1-25488689-INS-59
chr1-25491259-INS-61
chr1-25497680-INS-290
chr1-25499612-INS-6403
chr1-25506583-INS-87
chr1-25514085-INS-60
chr1-25541013-INS-87
chr1-25556637-INS-992
chr1-25588346-INS-74
chr1-25588439-INS-657
chr1-25607142-INS-58
chr1-25643716-INS-1530
chr1-25657452-INS-74
chr1-25668640-INS-121
chr1-25673185-INS-400
chr1-25685809-INS-489
chr1-25689323-INS-197
chr1-25692251-INS-80
chr1-25697608-INS-63
chr1-25724354-INS-58
chr1-25751234-INS-56
chr1-25757117-INS-2143
chr1-25773660-INS-301
chr1-25780402-INS-76
chr1-25804088-INS-318
chr1-25807738-INS-72
chr1-25813042-INS-580
chr1-25821353-INS-7061
chr1-25827361-INS-6353
chr1-25838950-INS-343
chr1-25843144-INS-51
chr1-25874825-INS-291
chr1-25896633-INS-240
chr1-25899083-INS-411
chr1-25907168-INS-559
chr1-25912880-INS-76
chr1-25936577-INS-305
chr1-25940544-INS-118
chr1-25945228-INS-65
chr1-25983072-INS-56
chr1-25995487-INS-205
chr1-26012167-INS-559
chr1-26022852-INS-251
chr1-26023397-INS-398
chr1-26033839-INS-172
chr1-26035349-INS-6475
chr1-26042985-INS-400
chr1-26056041-INS-141
chr1-26058527-INS-72
chr1-26074531-INS-6641
chr1-26076452-INS-65
chr1-26095197-INS-52
chr1-26096615-INS-160
chr1-26097861-INS-4883
chr1-26118397-INS-105
chr1-26150853-INS-132
chr1-26170048-INS-81
chr1-26225027-INS-515
chr1-26231627-INS-409
chr1-26240100-INS-56
chr1-26244643-INS-6462
chr1-26245607-INS-5871
chr1-26249719-INS-576
chr1-26263748-INS-822
chr1-26276997-INS-823
chr1-26278930-INS-53
chr1-26280510-INS-57
chr1-26285607-INS-72
chr1-26291122-INS-144

chr1-26313693-INS-64
chr1-26313810-INS-285
chr1-26341735-INS-6689
chr1-26365635-INS-327
chr1-26370005-INS-287
chr1-26374022-INS-60
chr1-26380483-INS-66
chr1-26399026-INS-93
chr1-26436841-INS-93
chr1-26440707-INS-6903
chr1-26496147-INS-276
chr1-26498255-INS-459
chr1-26507642-INS-6924
chr1-26620531-INS-50
chr1-26621297-INS-935
chr1-26627301-INS-2077
chr1-26665330-INS-70
chr1-26675912-INS-1883
chr1-26688410-INS-117
chr1-26710201-INS-71
chr1-26713121-INS-61
chr1-26725895-INS-15310
chr1-26757322-INS-7846
chr1-26764847-INS-52
chr1-26783015-INS-600
chr1-26786057-INS-135
chr1-26807056-INS-983
chr1-26808464-INS-292
chr1-26808841-INS-196
chr1-26818286-INS-240
chr1-26845823-INS-314
chr1-26873435-INS-7121
chr1-26875221-INS-199
chr1-26925149-INS-479
chr1-26933895-INS-243
chr1-26945648-INS-120
chr1-26945809-INS-1249
chr1-26976308-INS-344
chr1-26999824-INS-271
chr1-27004304-INS-538
chr1-27043054-INS-64
chr1-27118051-INS-599
chr1-27131316-INS-996
chr1-27137456-INS-7242
chr1-27142801-INS-406
chr1-27148005-INS-90
chr1-27179201-INS-214
chr1-27197332-INS-4859
chr1-27210396-INS-64
chr1-27218105-INS-480
chr1-27218105-INS-498
chr1-27233328-INS-208
chr1-27269782-INS-63
chr1-27287239-INS-173
chr1-27301090-INS-700
chr1-27420910-INS-67
chr1-27421981-INS-58
chr1-27466951-INS-116
chr1-27470278-INS-52
chr1-27475427-INS-4540
chr1-27497047-INS-5863
chr1-27508581-INS-658
chr1-27510604-INS-7145
chr1-27514196-INS-2123
chr1-27521025-INS-1271
chr1-27532425-INS-2739
chr1-27549864-INS-106
chr1-27579946-INS-122
chr1-27586557-INS-56
chr1-27587646-INS-534
chr1-27612698-INS-220
chr1-27626305-INS-948
chr1-27635951-INS-120
chr1-27641741-INS-62
chr1-27683767-INS-82
chr1-27689540-INS-352
chr1-27725254-INS-99

chr1-27741973-INS-230
chr1-27751652-INS-1637
chr1-27760090-INS-56
chr1-27765909-INS-96
chr1-27780270-INS-5415
chr1-27808093-INS-127
chr1-27869910-INS-952
chr1-27881562-INS-353
chr1-27953138-INS-193
chr1-27971426-INS-71
chr1-27989432-INS-956
chr1-27990158-INS-164
chr1-28011407-INS-1720
chr1-28033951-INS-222
chr1-28036403-INS-216
chr1-28039901-INS-180
chr1-28047249-INS-135
chr1-28056593-INS-196
chr1-28066222-INS-436
chr1-28089198-INS-80
chr1-28137706-INS-61
chr1-28214850-INS-70
chr1-28221022-INS-69
chr1-28237882-INS-437
chr1-28256065-INS-102
chr1-28261013-INS-505
chr1-28295604-INS-54
chr1-28314124-INS-635
chr1-28316526-INS-458
chr1-28335851-INS-52
chr1-28367596-INS-592
chr1-28376219-INS-681
chr1-28378598-INS-428
chr1-28406932-INS-69
chr1-28462540-INS-69
chr1-28464550-INS-53
chr1-28508830-INS-310
chr1-28513209-INS-185
chr1-28517165-INS-6011
chr1-28527440-INS-6799
chr1-28527766-INS-56
chr1-28568108-INS-506
chr1-28575102-INS-308
chr1-28618558-INS-67
chr1-28626033-INS-533
chr1-28631398-INS-674
chr1-28645884-INS-360
chr1-28653622-INS-332
chr1-28666838-INS-92
chr1-28676075-INS-6846
chr1-28693597-INS-772
chr1-28722892-INS-58
chr1-28726824-INS-79
chr1-28747505-INS-60
chr1-28750947-INS-216
chr1-28759982-INS-145
chr1-28825186-INS-371
chr1-28846583-INS-68
chr1-28848231-INS-552
chr1-28848843-INS-55
chr1-28852860-INS-778
chr1-28855656-INS-60
chr1-28869189-INS-63
chr1-28880077-INS-88
chr1-28890234-INS-78
chr1-28894496-INS-160
chr1-28894559-INS-1444
chr1-28895135-INS-785
chr1-28918516-INS-605
chr1-28919435-INS-84
chr1-28931545-INS-7095
chr1-28955517-INS-3036
chr1-28961052-INS-54
chr1-29021561-INS-54
chr1-29035969-INS-52
chr1-29046637-INS-131
chr1-29058036-INS-828

chr1-29125121-INS-208
chr1-29157820-INS-52
chr1-29164871-INS-621
chr1-29169989-INS-820
chr1-29173134-INS-573
chr1-29219068-INS-482
chr1-29276817-INS-192
chr1-29302102-INS-156
chr1-29329037-INS-123
chr1-29334859-INS-103
chr1-29346843-INS-70
chr1-29371941-INS-199
chr1-29397499-INS-264
chr1-29404710-INS-184
chr1-29415227-INS-398
chr1-29436257-INS-1257
chr1-29440297-INS-108
chr1-29445188-INS-6261
chr1-29491987-INS-6475
chr1-29509528-INS-1888
chr1-29513621-INS-465
chr1-29518031-INS-113
chr1-29539101-INS-260
chr1-29551229-INS-6551
chr1-29571849-INS-6475
chr1-29603168-INS-6431
chr1-29626196-INS-95
chr1-29713154-INS-68
chr1-29725623-INS-360
chr1-29741739-INS-688
chr1-29755841-INS-5157
chr1-29782160-INS-570
chr1-29787365-INS-2622
chr1-29799076-INS-82
chr1-29811410-INS-132
chr1-29823620-INS-51
chr1-29856150-INS-179
chr1-29861391-INS-6845
chr1-29862486-INS-562
chr1-29865389-INS-178
chr1-29872123-INS-54
chr1-29880862-INS-5186
chr1-29893243-INS-3473
chr1-29895617-INS-70
chr1-29899259-INS-143
chr1-29921823-INS-359
chr1-29939936-INS-74
chr1-29951541-INS-98
chr1-29955149-INS-52
chr1-29958129-INS-75
chr1-29960329-INS-1428
chr1-29968642-INS-75
chr1-29992232-INS-4420
chr1-30000201-INS-239
chr1-30046120-INS-159
chr1-30066803-INS-7150
chr1-30075223-INS-426
chr1-30095208-INS-54
chr1-30143083-INS-639
chr1-30153292-INS-7206
chr1-30159219-INS-82
chr1-30171590-INS-399
chr1-30172746-INS-327
chr1-30179153-INS-128
chr1-30209925-INS-52
chr1-30224463-INS-326
chr1-30226852-INS-320
chr1-30243715-INS-185
chr1-30269693-INS-64
chr1-30279896-INS-452
chr1-30284282-INS-60
chr1-30290156-INS-307
chr1-30294946-INS-73
chr1-30301534-INS-1972
chr1-30325351-INS-472
chr1-30339140-INS-6444
chr1-30341340-INS-80

chr1-30354806-INS-973
chr1-30355083-INS-6482
chr1-30379182-INS-162
chr1-30389292-INS-6967
chr1-30395702-INS-177
chr1-30403643-INS-92
chr1-30410544-INS-603
chr1-30424261-INS-303
chr1-30450475-INS-135
chr1-30455429-INS-70
chr1-30458916-INS-6225
chr1-30471825-INS-223
chr1-30478721-INS-464
chr1-30491147-INS-60
chr1-30508667-INS-262
chr1-30514088-INS-6689
chr1-30518730-INS-897
chr1-30544135-INS-51
chr1-30554143-INS-105
chr1-30561520-INS-4543
chr1-30563766-INS-2311
chr1-30581330-INS-203
chr1-30595637-INS-761
chr1-30618833-INS-60
chr1-30620461-INS-53
chr1-30628869-INS-546
chr1-30635967-INS-6214
chr1-30636574-INS-56
chr1-30638755-INS-670
chr1-30639276-INS-53
chr1-30692635-INS-53
chr1-30733599-INS-6385
chr1-30736775-INS-204
chr1-30739548-INS-56
chr1-30753627-INS-199
chr1-30780335-INS-156
chr1-30785801-INS-5259
chr1-30786237-INS-148
chr1-30816423-INS-160
chr1-30839338-INS-439
chr1-30840518-INS-207
chr1-30860585-INS-66
chr1-30861771-INS-232
chr1-30866597-INS-129
chr1-30882330-INS-515
chr1-30890587-INS-6666
chr1-30892763-INS-425
chr1-30914375-INS-602
chr1-30916027-INS-372
chr1-30926774-INS-477
chr1-30947532-INS-977
chr1-30956336-INS-74
chr1-30957943-INS-124
chr1-30980183-INS-54
chr1-30992213-INS-215
chr1-31028972-INS-86
chr1-31062202-INS-7335
chr1-31064946-INS-598
chr1-31066776-INS-98
chr1-31078790-INS-74
chr1-31106221-INS-7077
chr1-31108438-INS-4824
chr1-31112665-INS-268
chr1-31168324-INS-51
chr1-31171768-INS-2373
chr1-31194938-INS-688
chr1-31199955-INS-915
chr1-31237795-INS-893
chr1-31244055-INS-947
chr1-31248886-INS-52
chr1-31262931-INS-208
chr1-31283864-INS-58
chr1-31301233-INS-58
chr1-31302987-INS-58
chr1-31332667-INS-2866
chr1-31338245-INS-4076
chr1-31359940-INS-694

chr1-31365261-INS-54
chr1-31377057-INS-340
chr1-31391578-INS-154
chr1-31400382-INS-116
chr1-31408348-INS-52
chr1-31413409-INS-73
chr1-31445066-INS-299
chr1-31486652-INS-72
chr1-31519337-INS-245
chr1-31535779-INS-56
chr1-31553540-INS-171
chr1-31554476-INS-7117
chr1-31558742-INS-52
chr1-31574685-INS-745
chr1-31609322-INS-126
chr1-31612608-INS-57
chr1-31644266-INS-445
chr1-31658631-INS-209
chr1-31682350-INS-440
chr1-31689968-INS-57
chr1-31721390-INS-3618
chr1-31749110-INS-1864
chr1-31764015-INS-68
chr1-31765931-INS-6044
chr1-31767987-INS-77
chr1-31775008-INS-342
chr1-31781359-INS-309
chr1-31785599-INS-104
chr1-31800000-INS-120
chr1-31801034-INS-569
chr1-31809979-INS-52
chr1-31831396-INS-194
chr1-31834859-INS-301
chr1-31842861-INS-6660
chr1-31874290-INS-870
chr1-31913366-INS-610
chr1-31914659-INS-144
chr1-31917835-INS-97
chr1-31922734-INS-143
chr1-31946137-INS-140
chr1-31954474-INS-108
chr1-31988551-INS-124
chr1-31991859-INS-58
chr1-31996035-INS-80
chr1-32004685-INS-5967
chr1-32021143-INS-57
chr1-32083184-INS-5096
chr1-32088735-INS-84
chr1-32089924-INS-128
chr1-32136962-INS-538
chr1-32141031-INS-58
chr1-32143555-INS-132
chr1-32214973-INS-51
chr1-32216278-INS-57
chr1-32306672-INS-144
chr1-32317487-INS-492
chr1-32339630-INS-250
chr1-32364536-INS-52
chr1-32383290-INS-69
chr1-32390367-INS-209
chr1-32394983-INS-74
chr1-32402193-INS-265
chr1-32408982-INS-54
chr1-32419590-INS-664
chr1-32439514-INS-882
chr1-32444250-INS-581
chr1-32450146-INS-81
chr1-32450807-INS-54
chr1-32452215-INS-66
chr1-32457078-INS-160
chr1-32462557-INS-446
chr1-32464618-INS-3198
chr1-32487527-INS-145
chr1-32488742-INS-247
chr1-32501302-INS-160
chr1-32507296-INS-372
chr1-32514701-INS-64

chr1-32546429-INS-368
chr1-32557991-INS-196
chr1-32581844-INS-60
chr1-32608228-INS-54
chr1-32623985-INS-66
chr1-32660983-INS-64
chr1-32664269-INS-221
chr1-32675470-INS-72
chr1-32682942-INS-184
chr1-32684274-INS-156
chr1-32702586-INS-678
chr1-32717449-INS-74
chr1-32736892-INS-58
chr1-32737511-INS-270
chr1-32808029-INS-87
chr1-32809622-INS-56
chr1-32811075-INS-424
chr1-32818010-INS-96
chr1-32838993-INS-558
chr1-32859227-INS-56
chr1-32859813-INS-57
chr1-32860564-INS-707
chr1-32864196-INS-4550
chr1-32879911-INS-369
chr1-32888463-INS-56
chr1-32913086-INS-462
chr1-32923540-INS-54
chr1-32937046-INS-6206
chr1-32971389-INS-285
chr1-32977115-INS-56
chr1-32988792-INS-52
chr1-32993059-INS-98
chr1-32996328-INS-498
chr1-32997967-INS-7243
chr1-33004168-INS-3096
chr1-33020037-INS-7012
chr1-33030375-INS-446
chr1-33045418-INS-153
chr1-33048867-INS-103
chr1-33058728-INS-850
chr1-33097866-INS-7003
chr1-33136093-INS-114
chr1-33138378-INS-434
chr1-33139387-INS-209
chr1-33173142-INS-737
chr1-33173904-INS-489
chr1-33177970-INS-301
chr1-33215666-INS-80
chr1-33221559-INS-6101
chr1-33227062-INS-109
chr1-33229203-INS-138
chr1-33234765-INS-144
chr1-33235937-INS-6703
chr1-33245788-INS-565
chr1-33258529-INS-7234
chr1-33260132-INS-305
chr1-33276047-INS-351
chr1-33295858-INS-413
chr1-33298623-INS-381
chr1-33316853-INS-6458
chr1-33317403-INS-408
chr1-33325071-INS-210
chr1-33328602-INS-94
chr1-33334323-INS-60
chr1-33348026-INS-202
chr1-33351608-INS-52
chr1-33369903-INS-6241
chr1-33389270-INS-114
chr1-33408295-INS-367
chr1-33422067-INS-1544
chr1-33439439-INS-512
chr1-33459140-INS-3182
chr1-33491776-INS-2575
chr1-33505657-INS-346
chr1-33525421-INS-294
chr1-33530835-INS-57
chr1-33532168-INS-117

chr1-33541547-INS-75
chr1-33544026-INS-73
chr1-33566088-INS-218
chr1-33594525-INS-391
chr1-33614394-INS-55
chr1-33641053-INS-64
chr1-33652021-INS-339
chr1-33664408-INS-116
chr1-33676269-INS-52
chr1-33685541-INS-897
chr1-33703414-INS-308
chr1-33706510-INS-85
chr1-33715859-INS-67
chr1-33720652-INS-123
chr1-33721548-INS-762
chr1-33725339-INS-223
chr1-33727318-INS-59
chr1-33729998-INS-204
chr1-33741008-INS-52
chr1-33744766-INS-526
chr1-33755489-INS-62
chr1-33767882-INS-166
chr1-33781891-INS-288
chr1-33811054-INS-62
chr1-33820854-INS-215
chr1-33881953-INS-615
chr1-33885631-INS-5671
chr1-33906570-INS-658
chr1-33938959-INS-72
chr1-33940799-INS-62
chr1-33950254-INS-196
chr1-33963013-INS-395
chr1-33970380-INS-60
chr1-33972958-INS-53
chr1-33998658-INS-125
chr1-34000187-INS-460
chr1-34011919-INS-52
chr1-34020420-INS-151
chr1-34040156-INS-656
chr1-34041176-INS-186
chr1-34046496-INS-51
chr1-34094483-INS-205
chr1-34096162-INS-2528
chr1-34111822-INS-101
chr1-34149360-INS-341
chr1-34175238-INS-206
chr1-34183361-INS-67
chr1-34190739-INS-58
chr1-34269538-INS-225
chr1-34286401-INS-55
chr1-34301449-INS-61
chr1-34303931-INS-140
chr1-34321270-INS-207
chr1-34325878-INS-104
chr1-34335735-INS-52
chr1-34376042-INS-194
chr1-34382535-INS-7459
chr1-34395448-INS-169
chr1-34406492-INS-79
chr1-34420015-INS-176
chr1-34423700-INS-88
chr1-34424926-INS-400
chr1-34437947-INS-711
chr1-34439824-INS-222
chr1-34448039-INS-220
chr1-34449819-INS-66
chr1-34451289-INS-54
chr1-34494477-INS-52
chr1-34536569-INS-57
chr1-34550143-INS-68
chr1-34554337-INS-93
chr1-34569163-INS-434
chr1-34570810-INS-61
chr1-34582808-INS-52
chr1-34589584-INS-221
chr1-34590021-INS-84
chr1-34593795-INS-219

chr1-34594444-INS-351
chr1-34616574-INS-100
chr1-34624940-INS-56
chr1-34647326-INS-74
chr1-34663918-INS-556
chr1-34668743-INS-112
chr1-34682421-INS-71
chr1-34691699-INS-200
chr1-34713060-INS-90
chr1-34718163-INS-52
chr1-34753755-INS-208
chr1-34755375-INS-76
chr1-34760306-INS-79
chr1-34768416-INS-82
chr1-34769309-INS-63
chr1-34773081-INS-52
chr1-34778167-INS-56
chr1-34791129-INS-105
chr1-34821505-INS-401
chr1-34886963-INS-360
chr1-34898366-INS-120
chr1-34899533-INS-6697
chr1-34905831-INS-90
chr1-34919487-INS-116
chr1-34933868-INS-116
chr1-34936299-INS-70
chr1-34951854-INS-72
chr1-34968645-INS-278
chr1-34977430-INS-164
chr1-35000197-INS-649
chr1-35023010-INS-70
chr1-35033331-INS-52
chr1-35038852-INS-52
chr1-35066284-INS-203
chr1-35068709-INS-6217
chr1-35079897-INS-174
chr1-35112454-INS-52
chr1-35143999-INS-4747
chr1-35145778-INS-268
chr1-35145834-INS-577
chr1-35153629-INS-377
chr1-35163477-INS-60
chr1-35166015-INS-227
chr1-35250903-INS-92
chr1-35251909-INS-53
chr1-35260185-INS-116
chr1-35276878-INS-6799
chr1-35290129-INS-207
chr1-35306086-INS-58
chr1-35308445-INS-96
chr1-35309854-INS-127
chr1-35317934-INS-94
chr1-35347400-INS-409
chr1-35360718-INS-94
chr1-35397004-INS-99
chr1-35402329-INS-763
chr1-35433766-INS-134
chr1-35445097-INS-7111
chr1-35454443-INS-308
chr1-35459105-INS-66
chr1-35464662-INS-6212
chr1-35472931-INS-344
chr1-35502295-INS-5131
chr1-35507917-INS-158
chr1-35517113-INS-6472
chr1-35536638-INS-428
chr1-35538584-INS-99
chr1-35539650-INS-113
chr1-35557899-INS-574
chr1-35605106-INS-156
chr1-35624366-INS-71
chr1-35630446-INS-443
chr1-35631956-INS-442
chr1-35637726-INS-63
chr1-35644900-INS-383
chr1-35651724-INS-72
chr1-35656018-INS-69

chr1-35669824-INS-92
chr1-35679090-INS-249
chr1-35685216-INS-422
chr1-35685968-INS-52
chr1-35695892-INS-120
chr1-35744667-INS-130
chr1-35748059-INS-160
chr1-35763843-INS-68
chr1-35767765-INS-6687
chr1-35774778-INS-205
chr1-35785791-INS-394
chr1-35788432-INS-1346
chr1-35809997-INS-88
chr1-35818123-INS-384
chr1-35837521-INS-56
chr1-35838271-INS-676
chr1-35838352-INS-132
chr1-35844306-INS-134
chr1-35845764-INS-147
chr1-35855072-INS-660
chr1-35875638-INS-868
chr1-35878225-INS-116
chr1-35881163-INS-204
chr1-35891481-INS-174
chr1-35909534-INS-75
chr1-35922052-INS-167
chr1-35926077-INS-6355
chr1-35939096-INS-198
chr1-35961961-INS-376
chr1-36005619-INS-60
chr1-36048936-INS-742
chr1-36061189-INS-337
chr1-36062957-INS-66
chr1-36064745-INS-218
chr1-36068027-INS-60
chr1-36068546-INS-387
chr1-36080283-INS-56
chr1-36081145-INS-332
chr1-36123570-INS-56
chr1-36136643-INS-113
chr1-36163625-INS-148
chr1-36168882-INS-96
chr1-36173194-INS-281
chr1-36196739-INS-69
chr1-36225258-INS-151
chr1-36243887-INS-204
chr1-36256021-INS-168
chr1-36289763-INS-53
chr1-36291726-INS-208
chr1-36293034-INS-399
chr1-36301259-INS-180
chr1-36313314-INS-94
chr1-36326495-INS-264
chr1-36330237-INS-71
chr1-36331659-INS-437
chr1-36394322-INS-208
chr1-36401685-INS-216
chr1-36429831-INS-75
chr1-36455141-INS-64
chr1-36465153-INS-248
chr1-36487111-INS-415
chr1-36500756-INS-332
chr1-36502043-INS-60
chr1-36530448-INS-96
chr1-36536485-INS-175
chr1-36553702-INS-63
chr1-36583729-INS-203
chr1-36590414-INS-222
chr1-36605905-INS-175
chr1-36635581-INS-244
chr1-36658808-INS-150
chr1-36661601-INS-215
chr1-36676906-INS-109
chr1-36701087-INS-65
chr1-36702483-INS-73
chr1-36720452-INS-86
chr1-36732507-INS-55

chr1-36766171-INS-311
chr1-36772643-INS-198
chr1-36776002-INS-51
chr1-36782920-INS-66
chr1-36783971-INS-164
chr1-36796567-INS-130
chr1-36812291-INS-470
chr1-36829115-INS-79
chr1-36835479-INS-91
chr1-36839661-INS-138
chr1-36841902-INS-136
chr1-36842105-INS-1526
chr1-36848785-INS-300
chr1-36914450-INS-140
chr1-36949352-INS-950
chr1-36970458-INS-437
chr1-37003215-INS-278
chr1-37006059-INS-199
chr1-37012413-INS-186
chr1-37015285-INS-206
chr1-37019754-INS-187
chr1-37023064-INS-191
chr1-37024010-INS-60
chr1-37042031-INS-70
chr1-37048318-INS-94
chr1-37073423-INS-186
chr1-37075164-INS-97
chr1-37076066-INS-55
chr1-37078066-INS-237
chr1-37107715-INS-332
chr1-37111211-INS-294
chr1-37145969-INS-353
chr1-37150780-INS-427
chr1-37155667-INS-54
chr1-37162840-INS-150
chr1-37170597-INS-139
chr1-37181188-INS-111
chr1-37223923-INS-81
chr1-37235363-INS-55
chr1-37245674-INS-188
chr1-37249744-INS-58
chr1-37250889-INS-146
chr1-37256265-INS-124
chr1-37262508-INS-287
chr1-37271958-INS-1012
chr1-37276158-INS-201
chr1-37289288-INS-97
chr1-37309187-INS-95
chr1-37334839-INS-57
chr1-37351888-INS-398
chr1-37352516-INS-320
chr1-37352822-INS-407
chr1-37358591-INS-394
chr1-37364017-INS-252
chr1-37430826-INS-51
chr1-37435470-INS-165
chr1-37451155-INS-74
chr1-37457066-INS-64
chr1-37459831-INS-472
chr1-37462101-INS-145
chr1-37467264-INS-92
chr1-37473832-INS-419
chr1-37526451-INS-58
chr1-37533181-INS-114
chr1-37554371-INS-229
chr1-37563336-INS-270
chr1-37587676-INS-398
chr1-37606668-INS-426
chr1-37611668-INS-57
chr1-37621670-INS-398
chr1-37622983-INS-1467
chr1-37629663-INS-79
chr1-37657120-INS-58
chr1-37672134-INS-186
chr1-37703189-INS-54
chr1-37705145-INS-340
chr1-37768181-INS-66

chr1-37769516-INS-83
chr1-37775371-INS-91
chr1-37793032-INS-1100
chr1-37800801-INS-257
chr1-37856672-INS-86
chr1-37867375-INS-58
chr1-37868783-INS-340
chr1-37883286-INS-252
chr1-37891824-INS-62
chr1-37892311-INS-148
chr1-37902055-INS-564
chr1-37953208-INS-168
chr1-37954514-INS-101
chr1-37972561-INS-224
chr1-37973061-INS-1158
chr1-37999191-INS-126
chr1-38003031-INS-71
chr1-38069507-INS-360
chr1-38087362-INS-147
chr1-38109155-INS-484
chr1-38111477-INS-192
chr1-38129245-INS-79
chr1-38140944-INS-3658
chr1-38141169-INS-76
chr1-38174269-INS-61
chr1-38210415-INS-188
chr1-38214250-INS-60
chr1-38223649-INS-56
chr1-38226705-INS-953
chr1-38232283-INS-194
chr1-38279351-INS-600
chr1-38311175-INS-62
chr1-38315905-INS-434
chr1-38321243-INS-118
chr1-38394859-INS-185
chr1-38426006-INS-180
chr1-38443422-INS-168
chr1-38448014-INS-80
chr1-38469487-INS-63
chr1-38495381-INS-54
chr1-38499723-INS-64
chr1-38507642-INS-53
chr1-38528271-INS-117
chr1-38575525-INS-63
chr1-38605487-INS-316
chr1-38616105-INS-62
chr1-38618393-INS-179
chr1-38625663-INS-154
chr1-38649889-INS-73
chr1-38676325-INS-104
chr1-38678080-INS-80
chr1-38683275-INS-76
chr1-38692737-INS-230
chr1-38696291-INS-126
chr1-38706671-INS-70
chr1-38741783-INS-86
chr1-38755499-INS-159
chr1-38755876-INS-512
chr1-38770485-INS-160
chr1-38777361-INS-138
chr1-38784112-INS-161
chr1-38817695-INS-72
chr1-38833629-INS-175
chr1-38839235-INS-66
chr1-38840873-INS-873
chr1-38848486-INS-331
chr1-38860203-INS-522
chr1-38877483-INS-399
chr1-38906630-INS-78
chr1-38907828-INS-51
chr1-38911917-INS-52
chr1-38924921-INS-216
chr1-38943761-INS-64
chr1-38955072-INS-80
chr1-38955868-INS-6545
chr1-38970424-INS-6401
chr1-38970938-INS-1018

chr1-38971055-INS-62
chr1-38997867-INS-93
chr1-39031713-INS-64
chr1-39033729-INS-59
chr1-39082959-INS-796
chr1-39094922-INS-645
chr1-39132054-INS-64
chr1-39146790-INS-258
chr1-39163819-INS-358
chr1-39201243-INS-6383
chr1-39203949-INS-3709
chr1-39215248-INS-72
chr1-39227237-INS-52
chr1-39270836-INS-197
chr1-39296444-INS-69
chr1-39321530-INS-170
chr1-39322836-INS-64
chr1-39323549-INS-94
chr1-39334013-INS-247
chr1-39343454-INS-312
chr1-39357363-INS-360
chr1-39367118-INS-200
chr1-39370896-INS-78
chr1-39377403-INS-611
chr1-39381190-INS-106
chr1-39424040-INS-76
chr1-39461114-INS-280
chr1-39465364-INS-528
chr1-39476301-INS-462
chr1-39482580-INS-78
chr1-39521802-INS-56
chr1-39524818-INS-115
chr1-39538698-INS-81
chr1-39556512-INS-207
chr1-39563696-INS-8723
chr1-39572716-INS-166
chr1-39585134-INS-205
chr1-39589225-INS-92
chr1-39621962-INS-674
chr1-39626414-INS-107
chr1-39636115-INS-58
chr1-39642573-INS-56
chr1-39648694-INS-231
chr1-39650888-INS-54
chr1-39653612-INS-79
chr1-39665200-INS-78
chr1-39672592-INS-196
chr1-39705736-INS-52
chr1-39714434-INS-166
chr1-39716065-INS-62
chr1-39724233-INS-260
chr1-39727985-INS-280
chr1-39737472-INS-65
chr1-39754754-INS-56
chr1-39813567-INS-62
chr1-39814884-INS-289
chr1-39831231-INS-353
chr1-39832978-INS-61
chr1-39848012-INS-86
chr1-39853861-INS-1891
chr1-39854554-INS-208
chr1-39887461-INS-429
chr1-39898214-INS-77
chr1-39947709-INS-392
chr1-40012208-INS-52
chr1-40014809-INS-201
chr1-40025589-INS-364
chr1-40088759-INS-507
chr1-40095334-INS-643
chr1-40122237-INS-101
chr1-40127440-INS-6401
chr1-40143827-INS-543
chr1-40144856-INS-279
chr1-40162567-INS-58
chr1-40174727-INS-162
chr1-40222650-INS-371
chr1-40226260-INS-78

chr1-40233230-INS-62
chr1-40249410-INS-93
chr1-40254915-INS-213
chr1-40257669-INS-236
chr1-40268791-INS-142
chr1-40271343-INS-7164
chr1-40290264-INS-50
chr1-40297710-INS-101
chr1-40316930-INS-83
chr1-40319476-INS-655
chr1-40371077-INS-123
chr1-40379760-INS-64
chr1-40383801-INS-255
chr1-40399727-INS-84
chr1-40424230-INS-96
chr1-40434428-INS-178
chr1-40435705-INS-124
chr1-40444616-INS-199
chr1-40446479-INS-6428
chr1-40447744-INS-4347
chr1-40448936-INS-554
chr1-40467779-INS-300
chr1-40521599-INS-177
chr1-40601558-INS-132
chr1-40602823-INS-284
chr1-40688492-INS-59
chr1-40695805-INS-88
chr1-40705531-INS-83
chr1-40707172-INS-59
chr1-40710135-INS-161
chr1-40716844-INS-68
chr1-40730213-INS-75
chr1-40750597-INS-199
chr1-40764060-INS-810
chr1-40770183-INS-412
chr1-40789305-INS-82
chr1-40792183-INS-133
chr1-40831554-INS-650
chr1-40833695-INS-201
chr1-40869266-INS-123
chr1-40912877-INS-165
chr1-40921468-INS-88
chr1-40942961-INS-803
chr1-40950907-INS-792
chr1-40951572-INS-66
chr1-40959266-INS-52
chr1-40966897-INS-200
chr1-40983631-INS-110
chr1-40985330-INS-128
chr1-40986535-INS-100
chr1-40991518-INS-2557
chr1-40992587-INS-84
chr1-40999274-INS-117
chr1-41009903-INS-2674
chr1-41014277-INS-655
chr1-41015272-INS-120
chr1-41017446-INS-6265
chr1-41023179-INS-79
chr1-41033306-INS-141
chr1-41054378-INS-53
chr1-41058246-INS-1862
chr1-41069811-INS-112
chr1-41075264-INS-84
chr1-41138011-INS-188
chr1-41138421-INS-6422
chr1-41151528-INS-97
chr1-41180731-INS-162
chr1-41212672-INS-123
chr1-41215165-INS-55
chr1-41224311-INS-66
chr1-41230679-INS-116
chr1-41235280-INS-484
chr1-41237305-INS-55
chr1-41242646-INS-480
chr1-41309418-INS-571
chr1-41317739-INS-57
chr1-41322886-INS-155

chr1-41335485-INS-176
chr1-41372300-INS-56
chr1-41375631-INS-107
chr1-41395254-INS-61
chr1-41406710-INS-62
chr1-41425887-INS-64
chr1-41443522-INS-90
chr1-41454876-INS-147
chr1-41492054-INS-162
chr1-41494671-INS-640
chr1-41523141-INS-258
chr1-41528095-INS-82
chr1-41537388-INS-255
chr1-41546242-INS-125
chr1-41575917-INS-51
chr1-41585391-INS-195
chr1-41601714-INS-54
chr1-41602964-INS-6249
chr1-41621639-INS-201
chr1-41631546-INS-944
chr1-41632285-INS-96
chr1-41657077-INS-81
chr1-41673556-INS-60
chr1-41685837-INS-57
chr1-41689939-INS-300
chr1-41732859-INS-344
chr1-41740586-INS-115
chr1-41753427-INS-60
chr1-41767967-INS-103
chr1-41779033-INS-210
chr1-41796183-INS-54
chr1-41810654-INS-92
chr1-41820175-INS-55
chr1-41821595-INS-887
chr1-41825235-INS-56
chr1-41835562-INS-320
chr1-41857575-INS-115
chr1-41861354-INS-68
chr1-41873825-INS-77
chr1-41877826-INS-123
chr1-41878147-INS-161
chr1-41893908-INS-203
chr1-41905812-INS-59
chr1-41907170-INS-58
chr1-41912738-INS-597
chr1-41916026-INS-251
chr1-41920933-INS-128
chr1-41923996-INS-193
chr1-41950215-INS-224
chr1-41965793-INS-517
chr1-41971766-INS-6403
chr1-41974633-INS-54
chr1-41985222-INS-199
chr1-42011151-INS-51
chr1-42015160-INS-187
chr1-42017402-INS-126
chr1-42024297-INS-96
chr1-42057673-INS-77
chr1-42060523-INS-217
chr1-42074560-INS-132
chr1-42077150-INS-459
chr1-42107711-INS-78
chr1-42119006-INS-343
chr1-42121150-INS-124
chr1-42129553-INS-79
chr1-42141913-INS-63
chr1-42155098-INS-57
chr1-42157809-INS-796
chr1-42163906-INS-5844
chr1-42169228-INS-84
chr1-42174202-INS-84
chr1-42212468-INS-92
chr1-42216836-INS-53
chr1-42226735-INS-401
chr1-42244045-INS-56
chr1-42261207-INS-53
chr1-42290247-INS-67

chr1-42318807-INS-60
chr1-42328800-INS-80
chr1-42336567-INS-331
chr1-42340215-INS-76
chr1-42345935-INS-63
chr1-42353847-INS-6435
chr1-42354793-INS-52
chr1-42385603-INS-422
chr1-42409361-INS-142
chr1-42425375-INS-973
chr1-42429489-INS-350
chr1-42440241-INS-55
chr1-42459329-INS-113
chr1-42460630-INS-5993
chr1-42461659-INS-72
chr1-42471685-INS-264
chr1-42472848-INS-82
chr1-42476876-INS-98
chr1-42482241-INS-486
chr1-42495367-INS-83
chr1-42513940-INS-5210
chr1-42517655-INS-118
chr1-42521321-INS-6536
chr1-42546077-INS-82
chr1-42611858-INS-6664
chr1-42616319-INS-756
chr1-42627159-INS-471
chr1-42640744-INS-2988
chr1-42680453-INS-76
chr1-42707917-INS-75
chr1-42709988-INS-74
chr1-42755973-INS-229
chr1-42764868-INS-108
chr1-42777682-INS-125
chr1-42787072-INS-58
chr1-42794695-INS-7179
chr1-42796042-INS-387
chr1-42802890-INS-414
chr1-42813996-INS-95
chr1-42824700-INS-54
chr1-42827222-INS-140
chr1-42836499-INS-304
chr1-42837862-INS-108
chr1-42845385-INS-208
chr1-42860812-INS-400
chr1-42869414-INS-77
chr1-42889535-INS-53
chr1-42902145-INS-52
chr1-42907738-INS-155
chr1-42968565-INS-105
chr1-42976699-INS-390
chr1-42982404-INS-175
chr1-42986711-INS-339
chr1-42986849-INS-7413
chr1-42997208-INS-140
chr1-42997779-INS-326
chr1-42998657-INS-80
chr1-43003372-INS-731
chr1-43007809-INS-410
chr1-43024105-INS-256
chr1-43053482-INS-113
chr1-43074963-INS-68
chr1-43077835-INS-100
chr1-43080602-INS-212
chr1-43117246-INS-722
chr1-43164961-INS-70
chr1-43185681-INS-61
chr1-43217276-INS-238
chr1-43237062-INS-61
chr1-43247299-INS-57
chr1-43300911-INS-557
chr1-43305469-INS-374
chr1-43316503-INS-966
chr1-43321360-INS-78
chr1-43329880-INS-1686
chr1-43334713-INS-396
chr1-43338978-INS-64

chr1-43356984-INS-157
chr1-43361285-INS-400
chr1-43410895-INS-76
chr1-43424997-INS-152
chr1-43463245-INS-232
chr1-43469720-INS-122
chr1-43479093-INS-106
chr1-43482110-INS-268
chr1-43490470-INS-64
chr1-43502347-INS-220
chr1-43511869-INS-853
chr1-43530243-INS-171
chr1-43564698-INS-1142
chr1-43568886-INS-64
chr1-43573036-INS-79
chr1-43584131-INS-129
chr1-43598647-INS-84
chr1-43613428-INS-120
chr1-43614993-INS-281
chr1-43616538-INS-7883
chr1-43619263-INS-202
chr1-43629468-INS-111
chr1-43638487-INS-194
chr1-43640231-INS-60
chr1-43644937-INS-338
chr1-43646985-INS-7202
chr1-43680139-INS-53
chr1-43697619-INS-73
chr1-43724500-INS-58
chr1-43739396-INS-104
chr1-43745352-INS-92
chr1-43745537-INS-2287
chr1-43793690-INS-145
chr1-43811437-INS-52
chr1-43817606-INS-4034
chr1-43817643-INS-122
chr1-43854585-INS-204
chr1-43904608-INS-194
chr1-43917221-INS-378
chr1-43923047-INS-78
chr1-43925008-INS-197
chr1-43935090-INS-62
chr1-43939628-INS-101
chr1-43942498-INS-62
chr1-43952557-INS-507
chr1-43955618-INS-76
chr1-43981946-INS-6241
chr1-43982311-INS-2947
chr1-43987766-INS-112
chr1-44006353-INS-3653
chr1-44046525-INS-6550
chr1-44072026-INS-742
chr1-44074593-INS-2133
chr1-44075610-INS-197
chr1-44121262-INS-565
chr1-44122392-INS-165
chr1-44172607-INS-508
chr1-44178986-INS-64
chr1-44181612-INS-678
chr1-44193963-INS-256
chr1-44197937-INS-84
chr1-44208318-INS-55
chr1-44209836-INS-518
chr1-44249644-INS-588
chr1-44261552-INS-6367
chr1-44278445-INS-61
chr1-44283392-INS-399
chr1-44316466-INS-7135
chr1-44321442-INS-6907
chr1-44324639-INS-416
chr1-44335229-INS-72
chr1-44340650-INS-203
chr1-44357813-INS-57
chr1-44365842-INS-369
chr1-44368217-INS-517
chr1-44371107-INS-194
chr1-44477949-INS-643

chr1-44478280-INS-67
chr1-44489138-INS-167
chr1-44490616-INS-288
chr1-44494115-INS-7126
chr1-44506825-INS-7120
chr1-44532560-INS-96
chr1-44555697-INS-572
chr1-44590321-INS-56
chr1-44601936-INS-157
chr1-44615138-INS-6907
chr1-44637127-INS-487
chr1-44638829-INS-243
chr1-44662842-INS-365
chr1-44666029-INS-52
chr1-44667638-INS-110
chr1-44723688-INS-6191
chr1-44724801-INS-231
chr1-44768029-INS-191
chr1-44786090-INS-177
chr1-44793210-INS-118
chr1-44801936-INS-77
chr1-44831241-INS-438
chr1-44850269-INS-91
chr1-44878888-INS-176
chr1-44914037-INS-134
chr1-44920982-INS-259
chr1-44935549-INS-7283
chr1-44953899-INS-4816
chr1-44954889-INS-6203
chr1-44964904-INS-183
chr1-44993043-INS-3990
chr1-45028914-INS-4497
chr1-45040000-INS-54
chr1-45066338-INS-226
chr1-45068243-INS-130
chr1-45069530-INS-6413
chr1-45081548-INS-165
chr1-45092692-INS-64
chr1-45109980-INS-6882
chr1-45153623-INS-227
chr1-45159967-INS-208
chr1-45179468-INS-78
chr1-45187877-INS-807
chr1-45196864-INS-6572
chr1-45226549-INS-99
chr1-45236313-INS-68
chr1-45263604-INS-57
chr1-45281005-INS-285
chr1-45293231-INS-55
chr1-45353454-INS-66
chr1-45393424-INS-400
chr1-45405639-INS-6456
chr1-45416670-INS-82
chr1-45418167-INS-54
chr1-45435714-INS-413
chr1-45446610-INS-52
chr1-45460328-INS-6178
chr1-45516374-INS-184
chr1-45537989-INS-144
chr1-45549075-INS-90
chr1-45558744-INS-84
chr1-45562399-INS-430
chr1-45573526-INS-533
chr1-45600915-INS-204
chr1-45661404-INS-64
chr1-45689967-INS-6408
chr1-45705684-INS-67
chr1-45720844-INS-6367
chr1-45762528-INS-439
chr1-45803716-INS-58
chr1-45807373-INS-6389
chr1-45833692-INS-72
chr1-45846966-INS-96
chr1-45880553-INS-59
chr1-45908899-INS-584
chr1-45921287-INS-168
chr1-45929875-INS-138

chr1-45935458-INS-60
chr1-45936559-INS-272
chr1-45994940-INS-164
chr1-46001279-INS-51
chr1-46047892-INS-432
chr1-46128135-INS-1101
chr1-46128161-INS-265
chr1-46158681-INS-123
chr1-46192284-INS-4468
chr1-46254917-INS-834
chr1-46265602-INS-136
chr1-46277403-INS-106
chr1-46286954-INS-745
chr1-46291409-INS-55
chr1-46314174-INS-107
chr1-46316895-INS-6928
chr1-46378880-INS-7320
chr1-46399291-INS-507
chr1-46400267-INS-2837
chr1-46431383-INS-6332
chr1-46431440-INS-6922
chr1-46437050-INS-7085
chr1-46450328-INS-619
chr1-46495315-INS-114
chr1-46519846-INS-86
chr1-46525065-INS-401
chr1-46533546-INS-6265
chr1-46565225-INS-121
chr1-46580429-INS-7021
chr1-46610917-INS-1034
chr1-46643572-INS-6818
chr1-46648037-INS-115
chr1-46691054-INS-6369
chr1-46710203-INS-6708
chr1-46722188-INS-339
chr1-46734531-INS-220
chr1-46735556-INS-540
chr1-46804713-INS-6485
chr1-46808724-INS-6433
chr1-46819724-INS-66
chr1-46824669-INS-114
chr1-46833969-INS-1574
chr1-46876615-INS-4765
chr1-46921161-INS-67
chr1-46925710-INS-82
chr1-46949066-INS-210
chr1-46957059-INS-243
chr1-46962123-INS-200
chr1-46978929-INS-64
chr1-47006852-INS-4396
chr1-47022896-INS-400
chr1-47026530-INS-103
chr1-47060136-INS-892
chr1-47061382-INS-645
chr1-47061382-INS-78
chr1-47062022-INS-471
chr1-47067241-INS-6520
chr1-47072827-INS-792
chr1-47078647-INS-78
chr1-47087306-INS-456
chr1-47089697-INS-122
chr1-47125934-INS-636
chr1-47126877-INS-501
chr1-47128108-INS-5523
chr1-47154342-INS-580
chr1-47159978-INS-2987
chr1-47166741-INS-175
chr1-47176396-INS-642
chr1-47179948-INS-7090
chr1-47196834-INS-734
chr1-47209430-INS-6406
chr1-47227465-INS-51
chr1-47241699-INS-60
chr1-47253928-INS-6420
chr1-47256850-INS-437
chr1-47274883-INS-52
chr1-47292941-INS-93

chr1-47312476-INS-1946
chr1-47335666-INS-504
chr1-47366068-INS-1914
chr1-47370948-INS-6504
chr1-47407269-INS-610
chr1-47409182-INS-400
chr1-47423466-INS-2397
chr1-47432428-INS-75
chr1-47453190-INS-54
chr1-47454309-INS-839
chr1-47456718-INS-320
chr1-47462484-INS-70
chr1-47466267-INS-895
chr1-47489187-INS-330
chr1-47506498-INS-58
chr1-47507757-INS-64
chr1-47513966-INS-393
chr1-47519975-INS-66
chr1-47526403-INS-396
chr1-47610489-INS-55
chr1-47614065-INS-59
chr1-47643637-INS-515
chr1-47684840-INS-278
chr1-47689278-INS-670
chr1-47703652-INS-498
chr1-47711572-INS-96
chr1-47713723-INS-246
chr1-47720494-INS-201
chr1-47733845-INS-291
chr1-47741196-INS-185
chr1-47756355-INS-2621
chr1-47763700-INS-7113
chr1-47771786-INS-393
chr1-47781227-INS-7166
chr1-47787571-INS-160
chr1-47790224-INS-6440
chr1-47803020-INS-7445
chr1-47805120-INS-1504
chr1-47811947-INS-1011
chr1-47855595-INS-844
chr1-47887135-INS-88
chr1-47892717-INS-520
chr1-47980871-INS-957
chr1-47996762-INS-75
chr1-48002583-INS-329
chr1-48013067-INS-385
chr1-48027065-INS-124
chr1-48028709-INS-56
chr1-48030775-INS-208
chr1-48043707-INS-58
chr1-48057443-INS-2066
chr1-48058997-INS-52
chr1-48067119-INS-256
chr1-48069559-INS-259
chr1-48079407-INS-430
chr1-48111488-INS-661
chr1-48123623-INS-2927
chr1-48125902-INS-60
chr1-48189339-INS-209
chr1-48248538-INS-200
chr1-48259572-INS-486
chr1-48274673-INS-1752
chr1-48293480-INS-106
chr1-48294369-INS-156
chr1-48296033-INS-125
chr1-48297206-INS-154
chr1-48300835-INS-99
chr1-48314862-INS-52
chr1-48380695-INS-2916
chr1-48420615-INS-705
chr1-48424425-INS-655
chr1-48431232-INS-78
chr1-48434657-INS-211
chr1-48448519-INS-259
chr1-48463350-INS-207
chr1-48464819-INS-241
chr1-48472447-INS-84

chr1-48500969-INS-7162
chr1-48518812-INS-246
chr1-48522193-INS-56
chr1-48529041-INS-327
chr1-48531559-INS-62
chr1-48533087-INS-56
chr1-48559735-INS-199
chr1-48567421-INS-968
chr1-48570741-INS-6905
chr1-48578818-INS-384
chr1-48583447-INS-2199
chr1-48599854-INS-6185
chr1-48605120-INS-54
chr1-48674902-INS-91
chr1-48703166-INS-118
chr1-48716820-INS-60
chr1-48733894-INS-5477
chr1-48738650-INS-54
chr1-48741559-INS-153
chr1-48744706-INS-72
chr1-48745707-INS-366
chr1-48758285-INS-774
chr1-48763589-INS-176
chr1-48772673-INS-694
chr1-48783958-INS-597
chr1-48793384-INS-359
chr1-48820005-INS-123
chr1-48868490-INS-128
chr1-48888876-INS-420
chr1-48912949-INS-199
chr1-48934581-INS-58
chr1-48937405-INS-132
chr1-48949575-INS-156
chr1-48972471-INS-143
chr1-48977955-INS-382
chr1-48985426-INS-76
chr1-49003872-INS-116
chr1-49005428-INS-419
chr1-49059443-INS-127
chr1-49064790-INS-58
chr1-49070229-INS-56
chr1-49083090-INS-79
chr1-49094616-INS-177
chr1-49096699-INS-653
chr1-49136744-INS-7330
chr1-49141485-INS-6215
chr1-49158683-INS-211
chr1-49164443-INS-953
chr1-49169183-INS-110
chr1-49170948-INS-83
chr1-49202663-INS-161
chr1-49264622-INS-839
chr1-49274805-INS-699
chr1-49288495-INS-390
chr1-49299612-INS-60
chr1-49346686-INS-102
chr1-49350004-INS-279
chr1-49401928-INS-68
chr1-49412763-INS-56
chr1-49419744-INS-489
chr1-49435030-INS-72
chr1-49436989-INS-67
chr1-49522801-INS-71
chr1-49540018-INS-239
chr1-49540985-INS-56
chr1-49543097-INS-453
chr1-49548595-INS-387
chr1-49555050-INS-176
chr1-49566638-INS-456
chr1-49592532-INS-78
chr1-49603204-INS-70
chr1-49618032-INS-134
chr1-49665143-INS-107
chr1-49696382-INS-527
chr1-49698589-INS-51
chr1-49712036-INS-77
chr1-49721345-INS-109

chr1-49727035-INS-748
chr1-49735453-INS-116
chr1-49741346-INS-98
chr1-49774956-INS-175
chr1-49785180-INS-1485
chr1-49790813-INS-74
chr1-49814485-INS-72
chr1-49816815-INS-725
chr1-49854132-INS-690
chr1-49870844-INS-6811
chr1-49885439-INS-322
chr1-49892633-INS-238
chr1-49895640-INS-146
chr1-49905198-INS-123
chr1-49935639-INS-114
chr1-49954440-INS-379
chr1-49960419-INS-58
chr1-49972304-INS-176
chr1-49983302-INS-651
chr1-50037946-INS-135
chr1-50038909-INS-174
chr1-50086332-INS-240
chr1-50093808-INS-5191
chr1-50110705-INS-166
chr1-50117829-INS-115
chr1-50123928-INS-332
chr1-50130328-INS-191
chr1-50133241-INS-557
chr1-50137220-INS-141
chr1-50139401-INS-704
chr1-50197112-INS-97
chr1-50214673-INS-170
chr1-50216787-INS-51
chr1-50241348-INS-168
chr1-50269181-INS-238
chr1-50272916-INS-90
chr1-50280701-INS-752
chr1-50309235-INS-6287
chr1-50313880-INS-7151
chr1-50322132-INS-53
chr1-50340829-INS-263
chr1-50362816-INS-6593
chr1-50386844-INS-6253
chr1-50433906-INS-52
chr1-50501354-INS-3573
chr1-50539859-INS-94
chr1-50568962-INS-85
chr1-50573191-INS-124
chr1-50603650-INS-6641
chr1-50637947-INS-261
chr1-50665975-INS-348
chr1-50688167-INS-77
chr1-50706145-INS-63
chr1-50711555-INS-248
chr1-50735863-INS-58
chr1-50742419-INS-51
chr1-50767595-INS-367
chr1-50786888-INS-108
chr1-50795209-INS-271
chr1-50839303-INS-54
chr1-50857136-INS-54
chr1-50869769-INS-484
chr1-50875950-INS-5244
chr1-50920787-INS-199
chr1-50923337-INS-394
chr1-50931085-INS-390
chr1-50935082-INS-108
chr1-50963311-INS-57
chr1-50974836-INS-523
chr1-50993347-INS-233
chr1-51000328-INS-7321
chr1-51007147-INS-155
chr1-51012848-INS-186
chr1-51013501-INS-6382
chr1-51045349-INS-358
chr1-51054791-INS-91
chr1-51060613-INS-153

chr1-51087546-INS-2026
chr1-51094874-INS-156
chr1-51102801-INS-94
chr1-51110198-INS-238
chr1-51119268-INS-525
chr1-51130039-INS-84
chr1-51133710-INS-96
chr1-51151331-INS-124
chr1-51161709-INS-103
chr1-51169461-INS-60
chr1-51195051-INS-167
chr1-51204172-INS-1342
chr1-51207018-INS-68
chr1-51269434-INS-54
chr1-51279436-INS-126
chr1-51297093-INS-5195
chr1-51313184-INS-395
chr1-51316986-INS-109
chr1-51318756-INS-149
chr1-51320561-INS-229
chr1-51322874-INS-1890
chr1-51324077-INS-70
chr1-51343084-INS-92
chr1-51350983-INS-182
chr1-51353271-INS-6411
chr1-51380806-INS-77
chr1-51381289-INS-147
chr1-51383182-INS-56
chr1-51402063-INS-127
chr1-51431291-INS-134
chr1-51445169-INS-726
chr1-51456815-INS-63
chr1-51520994-INS-138
chr1-51564461-INS-89
chr1-51609330-INS-5544
chr1-51610131-INS-75
chr1-51643990-INS-63
chr1-51684389-INS-186
chr1-51701503-INS-165
chr1-51714999-INS-70
chr1-51716674-INS-162
chr1-51724878-INS-54
chr1-51736063-INS-248
chr1-51739652-INS-542
chr1-51757737-INS-58
chr1-51810945-INS-316
chr1-51844224-INS-83
chr1-51856793-INS-247
chr1-51870058-INS-370
chr1-51921169-INS-75
chr1-51938009-INS-131
chr1-51945992-INS-136
chr1-51970433-INS-165
chr1-51991449-INS-401
chr1-51999131-INS-58
chr1-52004282-INS-5421
chr1-52008124-INS-104
chr1-52023612-INS-150
chr1-52028112-INS-56
chr1-52040658-INS-96
chr1-52064828-INS-261
chr1-52089302-INS-329
chr1-52092633-INS-198
chr1-52114939-INS-68
chr1-52118634-INS-87
chr1-52154498-INS-509
chr1-52172978-INS-122
chr1-52197291-INS-188
chr1-52208009-INS-368
chr1-52209605-INS-1166
chr1-52223569-INS-85
chr1-52289027-INS-54
chr1-52290608-INS-559
chr1-52302853-INS-81
chr1-52316326-INS-243
chr1-52327422-INS-146
chr1-52350373-INS-56

chr1-52362897-INS-109
chr1-52383404-INS-98
chr1-52398712-INS-6656
chr1-52403313-INS-60
chr1-52415367-INS-514
chr1-52419919-INS-189
chr1-52421335-INS-3767
chr1-52435688-INS-987
chr1-52451367-INS-7370
chr1-52499735-INS-773
chr1-52543515-INS-134
chr1-52559890-INS-144
chr1-52571054-INS-142
chr1-52572782-INS-180
chr1-52617162-INS-2783
chr1-52618286-INS-272
chr1-52634984-INS-6439
chr1-52642800-INS-60
chr1-52713566-INS-198
chr1-52719911-INS-494
chr1-52733280-INS-800
chr1-52734867-INS-310
chr1-52741585-INS-1294
chr1-52779655-INS-507
chr1-52781374-INS-336
chr1-52791648-INS-268
chr1-52794890-INS-81
chr1-52810207-INS-106
chr1-52813537-INS-5157
chr1-52825479-INS-138
chr1-52839356-INS-70
chr1-52878152-INS-107
chr1-52878828-INS-59
chr1-52891453-INS-187
chr1-52891571-INS-910
chr1-52893451-INS-742
chr1-52895299-INS-66
chr1-52906116-INS-62
chr1-52909116-INS-1154
chr1-52936554-INS-81
chr1-52946837-INS-706
chr1-52947847-INS-400
chr1-52956357-INS-267
chr1-52969708-INS-78
chr1-52972675-INS-120
chr1-52997472-INS-245
chr1-53002833-INS-67
chr1-53006492-INS-404
chr1-53042831-INS-269
chr1-53047840-INS-62
chr1-53052607-INS-684
chr1-53060298-INS-871
chr1-53060392-INS-473
chr1-53075069-INS-568
chr1-53096333-INS-509
chr1-53110968-INS-312
chr1-53117141-INS-212
chr1-53123103-INS-191
chr1-53128601-INS-59
chr1-53139009-INS-60
chr1-53140909-INS-6664
chr1-53186005-INS-226
chr1-53191263-INS-442
chr1-53208216-INS-783
chr1-53218180-INS-465
chr1-53219261-INS-152
chr1-53222442-INS-85
chr1-53247974-INS-444
chr1-53270494-INS-606
chr1-53273368-INS-7428
chr1-53284893-INS-399
chr1-53289409-INS-189
chr1-53304320-INS-69
chr1-53338544-INS-897
chr1-53345231-INS-456
chr1-53359744-INS-1863
chr1-53368276-INS-203

chr1-53369715-INS-78
chr1-53370720-INS-66
chr1-53377996-INS-100
chr1-53425496-INS-534
chr1-53431596-INS-95
chr1-53434030-INS-468
chr1-53438888-INS-201
chr1-53490339-INS-145
chr1-53520333-INS-548
chr1-53531846-INS-4558
chr1-53567066-INS-6284
chr1-53596407-INS-144
chr1-53603978-INS-230
chr1-53606388-INS-5542
chr1-53608169-INS-7075
chr1-53660601-INS-5381
chr1-53683225-INS-1894
chr1-53684610-INS-97
chr1-53691845-INS-307
chr1-53712662-INS-2254
chr1-53723055-INS-58
chr1-53725194-INS-109
chr1-53743252-INS-434
chr1-53744303-INS-274
chr1-53824963-INS-54
chr1-53842593-INS-123
chr1-53863738-INS-59
chr1-53872091-INS-147
chr1-53892892-INS-5402
chr1-53899267-INS-115
chr1-53920105-INS-172
chr1-53925142-INS-498
chr1-53954658-INS-183
chr1-53972957-INS-56
chr1-53978242-INS-55
chr1-53985693-INS-756
chr1-53988425-INS-6442
chr1-54019964-INS-314
chr1-54023763-INS-51
chr1-54027633-INS-423
chr1-54060026-INS-114
chr1-54103602-INS-6282
chr1-54121059-INS-554
chr1-54126194-INS-79
chr1-54142619-INS-52
chr1-54174376-INS-81
chr1-54194102-INS-81
chr1-54201233-INS-6575
chr1-54217317-INS-122
chr1-54220561-INS-131
chr1-54260553-INS-2318
chr1-54278971-INS-4620
chr1-54314453-INS-401
chr1-54317390-INS-213
chr1-54406040-INS-476
chr1-54443231-INS-565
chr1-54474368-INS-75
chr1-54491740-INS-217
chr1-54495705-INS-80
chr1-54502496-INS-113
chr1-54531616-INS-60
chr1-54558553-INS-191
chr1-54570428-INS-839
chr1-54581038-INS-7309
chr1-54597835-INS-324
chr1-54603667-INS-322
chr1-54606704-INS-946
chr1-54611884-INS-92
chr1-54614628-INS-623
chr1-54625030-INS-72
chr1-54633325-INS-117
chr1-54655854-INS-298
chr1-54656801-INS-6201
chr1-54661727-INS-6966
chr1-54661881-INS-60
chr1-54662689-INS-5267
chr1-54671873-INS-401

chr1-54728654-INS-98
chr1-54758612-INS-946
chr1-54771226-INS-156
chr1-54783898-INS-86
chr1-54790980-INS-88
chr1-54799235-INS-60
chr1-54801358-INS-134
chr1-54822897-INS-234
chr1-54825026-INS-97
chr1-54829710-INS-82
chr1-54840871-INS-171
chr1-54847388-INS-117
chr1-54858191-INS-60
chr1-54863697-INS-265
chr1-54874691-INS-60
chr1-54876061-INS-199
chr1-54897785-INS-6730
chr1-54898335-INS-90
chr1-54921610-INS-125
chr1-54925673-INS-266
chr1-54970905-INS-89
chr1-54996472-INS-52
chr1-55008091-INS-202
chr1-55010068-INS-84
chr1-55012272-INS-343
chr1-55015500-INS-183
chr1-55017057-INS-213
chr1-55038930-INS-197
chr1-55058228-INS-93
chr1-55059008-INS-223
chr1-55069706-INS-190
chr1-55087046-INS-600
chr1-55090117-INS-64
chr1-55093672-INS-204
chr1-55094381-INS-207
chr1-55102049-INS-190
chr1-55107712-INS-52
chr1-55125275-INS-133
chr1-55129000-INS-193
chr1-55139446-INS-206
chr1-55141106-INS-240
chr1-55156808-INS-185
chr1-55169412-INS-99
chr1-55178209-INS-61
chr1-55186371-INS-432
chr1-55186371-INS-864
chr1-55190599-INS-268
chr1-55194433-INS-80
chr1-55194867-INS-148
chr1-55201147-INS-93
chr1-55204123-INS-62
chr1-55208135-INS-403
chr1-55209800-INS-4159
chr1-55218190-INS-209
chr1-55218888-INS-202
chr1-55220360-INS-2562
chr1-55237557-INS-501
chr1-55239457-INS-161
chr1-55243588-INS-202
chr1-55253031-INS-293
chr1-55274935-INS-100
chr1-55288833-INS-211
chr1-55329729-INS-7092
chr1-55335101-INS-1068
chr1-55335859-INS-57
chr1-55339009-INS-111
chr1-55342283-INS-165
chr1-55362387-INS-6204
chr1-55362948-INS-498
chr1-55363430-INS-6645
chr1-55372301-INS-6676
chr1-55373196-INS-54
chr1-55462584-INS-401
chr1-55493641-INS-748
chr1-55533348-INS-126
chr1-55551963-INS-303
chr1-55555209-INS-52

chr1-55555888-INS-6454
chr1-55563322-INS-64
chr1-55567778-INS-58
chr1-55579876-INS-1175
chr1-55582808-INS-6718
chr1-55603413-INS-66
chr1-55613944-INS-160
chr1-55641529-INS-52
chr1-55652096-INS-82
chr1-55663149-INS-330
chr1-55666127-INS-52
chr1-55674388-INS-191
chr1-55707242-INS-161
chr1-55720079-INS-5550
chr1-55722595-INS-382
chr1-55726985-INS-60
chr1-55729368-INS-92
chr1-55750678-INS-54
chr1-55751038-INS-719
chr1-55805918-INS-6309
chr1-55824632-INS-80
chr1-55825290-INS-8689
chr1-55832684-INS-171
chr1-55837482-INS-336
chr1-55863131-INS-82
chr1-55867030-INS-51
chr1-55883952-INS-53
chr1-55909421-INS-414
chr1-55911921-INS-62
chr1-55921008-INS-123
chr1-55928397-INS-194
chr1-55941054-INS-58
chr1-55956062-INS-66
chr1-55962038-INS-52
chr1-55991516-INS-168
chr1-55992408-INS-348
chr1-56003893-INS-1111
chr1-56020029-INS-629
chr1-56051039-INS-72
chr1-56089333-INS-442
chr1-56094261-INS-580
chr1-56098185-INS-96
chr1-56109887-INS-701
chr1-56113562-INS-1453
chr1-56150497-INS-69
chr1-56156174-INS-88
chr1-56160427-INS-92
chr1-56161174-INS-6930
chr1-56186170-INS-88
chr1-56229595-INS-111
chr1-56237331-INS-143
chr1-56242241-INS-155
chr1-56244205-INS-88
chr1-56276292-INS-618
chr1-56280802-INS-66
chr1-56336884-INS-488
chr1-56357549-INS-55
chr1-56362667-INS-90
chr1-56375094-INS-379
chr1-56377799-INS-390
chr1-56393766-INS-2888
chr1-56415428-INS-7447
chr1-56422081-INS-1894
chr1-56454978-INS-184
chr1-56463264-INS-740
chr1-56482724-INS-60
chr1-56487541-INS-62
chr1-56501407-INS-7036
chr1-56544669-INS-102
chr1-56565510-INS-195
chr1-56575879-INS-1361
chr1-56606062-INS-286
chr1-56623238-INS-80
chr1-56631710-INS-6675
chr1-56635983-INS-6449
chr1-56637392-INS-252
chr1-56645524-INS-730

chr1-56647701-INS-52
chr1-56664347-INS-383
chr1-56698499-INS-56
chr1-56701991-INS-228
chr1-56709463-INS-209
chr1-56712687-INS-6690
chr1-56713146-INS-345
chr1-56717514-INS-114
chr1-56723009-INS-7356
chr1-56743474-INS-152
chr1-56751004-INS-414
chr1-56754143-INS-212
chr1-56775288-INS-86
chr1-56785490-INS-60
chr1-56828655-INS-475
chr1-56842625-INS-134
chr1-56907024-INS-62
chr1-56909869-INS-185
chr1-56917141-INS-70
chr1-56919991-INS-67
chr1-56955437-INS-163
chr1-56962783-INS-52
chr1-56966709-INS-200
chr1-56977065-INS-333
chr1-57015161-INS-58
chr1-57022240-INS-227
chr1-57052138-INS-56
chr1-57061848-INS-86
chr1-57074437-INS-113
chr1-57077512-INS-64
chr1-57083095-INS-132
chr1-57085650-INS-262
chr1-57116353-INS-57
chr1-57130711-INS-79
chr1-57164745-INS-137
chr1-57185428-INS-53
chr1-57204208-INS-84
chr1-57209048-INS-99
chr1-57210544-INS-6653
chr1-57233075-INS-479
chr1-57261774-INS-4512
chr1-57261788-INS-56
chr1-57268739-INS-58
chr1-57294293-INS-127
chr1-57310311-INS-70
chr1-57315473-INS-203
chr1-57316975-INS-56
chr1-57318512-INS-186
chr1-57350822-INS-7370
chr1-57359580-INS-105
chr1-57373695-INS-774
chr1-57380438-INS-60
chr1-57382224-INS-136
chr1-57386942-INS-57
chr1-57393297-INS-64
chr1-57419895-INS-160
chr1-57455110-INS-70
chr1-57458924-INS-76
chr1-57462224-INS-154
chr1-57511581-INS-6564
chr1-57520009-INS-372
chr1-57535063-INS-324
chr1-57553611-INS-60
chr1-57567689-INS-181
chr1-57569015-INS-66
chr1-57575162-INS-355
chr1-57581392-INS-598
chr1-57587765-INS-130
chr1-57592578-INS-385
chr1-57606208-INS-151
chr1-57621524-INS-57
chr1-57638676-INS-265
chr1-57672089-INS-3426
chr1-57685296-INS-161
chr1-57691566-INS-114
chr1-57694172-INS-52
chr1-57695749-INS-171

chr1-57697240-INS-56
chr1-57703652-INS-831
chr1-57744654-INS-86
chr1-57753541-INS-92
chr1-57768645-INS-187
chr1-57792147-INS-343
chr1-57848643-INS-54
chr1-57891538-INS-72
chr1-57906927-INS-4142
chr1-57908546-INS-51
chr1-57909732-INS-105
chr1-57923766-INS-206
chr1-57944248-INS-598
chr1-57952676-INS-63
chr1-57984709-INS-72
chr1-57998655-INS-6936
chr1-58038222-INS-206
chr1-58054216-INS-99
chr1-58065060-INS-119
chr1-58070839-INS-84
chr1-58071943-INS-56
chr1-58080940-INS-67
chr1-58096868-INS-55
chr1-58099920-INS-107
chr1-58125776-INS-63
chr1-58129246-INS-153
chr1-58175531-INS-79
chr1-58202209-INS-181
chr1-58207187-INS-6438
chr1-58213945-INS-59
chr1-58232920-INS-413
chr1-58246684-INS-105
chr1-58269800-INS-54
chr1-58288377-INS-99
chr1-58301568-INS-78
chr1-58303957-INS-79
chr1-58308928-INS-101
chr1-58333773-INS-3134
chr1-58333937-INS-146
chr1-58376141-INS-84
chr1-58378584-INS-176
chr1-58383425-INS-75
chr1-58389010-INS-56
chr1-58414135-INS-270
chr1-58414452-INS-445
chr1-58417259-INS-223
chr1-58420812-INS-176
chr1-58423924-INS-203
chr1-58425172-INS-211
chr1-58433065-INS-242
chr1-58454798-INS-77
chr1-58466005-INS-211
chr1-58469635-INS-5239
chr1-58474549-INS-202
chr1-58482394-INS-213
chr1-58489483-INS-406
chr1-58494256-INS-171
chr1-58525751-INS-56
chr1-58541406-INS-196
chr1-58550258-INS-356
chr1-58575248-INS-103
chr1-58578889-INS-803
chr1-58602845-INS-982
chr1-58605358-INS-92
chr1-58635172-INS-204
chr1-58644336-INS-58
chr1-58647372-INS-52
chr1-58675955-INS-887
chr1-58678235-INS-60
chr1-58692544-INS-225
chr1-58751294-INS-214
chr1-58783253-INS-826
chr1-58798547-INS-54
chr1-58803903-INS-194
chr1-58809306-INS-731
chr1-58819838-INS-198
chr1-58820611-INS-1045

chr1-58826483-INS-57
chr1-58843402-INS-6185
chr1-58847539-INS-499
chr1-58878582-INS-492
chr1-58879624-INS-52
chr1-58883009-INS-1518
chr1-58935126-INS-599
chr1-58935926-INS-135
chr1-59046000-INS-895
chr1-59046179-INS-1122
chr1-59048504-INS-99
chr1-59054664-INS-70
chr1-59090712-INS-211
chr1-59113109-INS-70
chr1-59123141-INS-508
chr1-59137296-INS-160
chr1-59140749-INS-118
chr1-59161742-INS-80
chr1-59180581-INS-213
chr1-59188386-INS-196
chr1-59201179-INS-496
chr1-59213803-INS-242
chr1-59216411-INS-268
chr1-59251309-INS-5549
chr1-59252472-INS-136
chr1-59264145-INS-78
chr1-59286963-INS-75
chr1-59288334-INS-56
chr1-59290909-INS-57
chr1-59308599-INS-348
chr1-59331344-INS-168
chr1-59333041-INS-83
chr1-59377097-INS-604
chr1-59394034-INS-88
chr1-59400096-INS-194
chr1-59406029-INS-946
chr1-59413227-INS-203
chr1-59430205-INS-56
chr1-59447221-INS-252
chr1-59454822-INS-341
chr1-59502465-INS-184
chr1-59503593-INS-193
chr1-59507527-INS-120
chr1-59518756-INS-1096
chr1-59519770-INS-81
chr1-59526518-INS-212
chr1-59538071-INS-207
chr1-59545143-INS-7180
chr1-59577345-INS-69
chr1-59586496-INS-57
chr1-59588809-INS-393
chr1-59596214-INS-216
chr1-59601255-INS-52
chr1-59605693-INS-55
chr1-59617235-INS-207
chr1-59640253-INS-120
chr1-59644161-INS-54
chr1-59653513-INS-570
chr1-59659446-INS-278
chr1-59683442-INS-132
chr1-59686118-INS-166
chr1-59692676-INS-248
chr1-59702251-INS-680
chr1-59707504-INS-238
chr1-59711061-INS-210
chr1-59716307-INS-74
chr1-59745613-INS-144
chr1-59757121-INS-59
chr1-59780218-INS-173
chr1-59785548-INS-199
chr1-59786637-INS-201
chr1-59796362-INS-150
chr1-59825845-INS-229
chr1-59827757-INS-267
chr1-59871441-INS-200
chr1-59877368-INS-56
chr1-59882449-INS-66

chr1-59885185-INS-206
chr1-59889080-INS-62
chr1-59892336-INS-195
chr1-59911861-INS-207
chr1-59927160-INS-94
chr1-59930523-INS-136
chr1-59950009-INS-51
chr1-59973252-INS-207
chr1-59975178-INS-185
chr1-59991618-INS-113
chr1-59993258-INS-140
chr1-59997233-INS-382
chr1-60037531-INS-51
chr1-60051432-INS-227
chr1-60055648-INS-71
chr1-60091665-INS-6686
chr1-60100997-INS-364
chr1-60102743-INS-221
chr1-60103822-INS-607
chr1-60107293-INS-205
chr1-60141982-INS-396
chr1-60156410-INS-159
chr1-60170951-INS-943
chr1-60171833-INS-268
chr1-60175557-INS-7432
chr1-60180766-INS-721
chr1-60182355-INS-117
chr1-60183855-INS-53
chr1-60215173-INS-216
chr1-60222890-INS-84
chr1-60226269-INS-153
chr1-60227965-INS-149
chr1-60234437-INS-399
chr1-60243413-INS-426
chr1-60245045-INS-107
chr1-60246269-INS-60
chr1-60251714-INS-54
chr1-60271947-INS-241
chr1-60277167-INS-800
chr1-60289337-INS-67
chr1-60299026-INS-1876
chr1-60310049-INS-156
chr1-60375404-INS-193
chr1-60393436-INS-205
chr1-60394419-INS-152
chr1-60400794-INS-429
chr1-60436853-INS-602
chr1-60460001-INS-1593
chr1-60466342-INS-162
chr1-60474347-INS-3215
chr1-60493264-INS-178
chr1-60508274-INS-195
chr1-60513146-INS-205
chr1-60530826-INS-106
chr1-60542250-INS-126
chr1-60557982-INS-3256
chr1-60578546-INS-76
chr1-60602127-INS-209
chr1-60616165-INS-56
chr1-60619721-INS-60
chr1-60649408-INS-498
chr1-60668314-INS-344
chr1-60686002-INS-168
chr1-60693804-INS-87
chr1-60701417-INS-6579
chr1-60702503-INS-823
chr1-60716088-INS-58
chr1-60739972-INS-440
chr1-60751956-INS-4865
chr1-60759313-INS-74
chr1-60784138-INS-857
chr1-60819560-INS-58
chr1-60819574-INS-6452
chr1-60845302-INS-531
chr1-60868095-INS-253
chr1-60910389-INS-140
chr1-60917242-INS-54

chr1-60923823-INS-62
chr1-60929279-INS-136
chr1-60959003-INS-7434
chr1-60983143-INS-7187
chr1-60997049-INS-84
chr1-61030869-INS-1296
chr1-61048881-INS-82
chr1-61071381-INS-148
chr1-61111632-INS-187
chr1-61122482-INS-80
chr1-61138809-INS-60
chr1-61152216-INS-1383
chr1-61186949-INS-84
chr1-61194589-INS-6670
chr1-61196097-INS-454
chr1-61199488-INS-134
chr1-61232574-INS-174
chr1-61257324-INS-72
chr1-61257992-INS-3022
chr1-61267039-INS-548
chr1-61272013-INS-81
chr1-61278533-INS-497
chr1-61291061-INS-258
chr1-61306964-INS-77
chr1-61333296-INS-82
chr1-61338864-INS-55
chr1-61361282-INS-66
chr1-61400142-INS-401
chr1-61407550-INS-402
chr1-61407609-INS-1206
chr1-61426416-INS-62
chr1-61453161-INS-55
chr1-61482636-INS-70
chr1-61494611-INS-1256
chr1-61506623-INS-135
chr1-61507589-INS-434
chr1-61527097-INS-489
chr1-61528960-INS-426
chr1-61541666-INS-120
chr1-61548535-INS-493
chr1-61570787-INS-52
chr1-61572119-INS-73
chr1-61574677-INS-84
chr1-61589884-INS-510
chr1-61595261-INS-2820
chr1-61596019-INS-6190
chr1-61616491-INS-6370
chr1-61621019-INS-408
chr1-61631725-INS-6195
chr1-61640617-INS-997
chr1-61661792-INS-60
chr1-61662180-INS-7562
chr1-61698513-INS-166
chr1-61702567-INS-1363
chr1-61733135-INS-66
chr1-61739331-INS-58
chr1-61763416-INS-198
chr1-61767722-INS-99
chr1-61774660-INS-402
chr1-61779328-INS-149
chr1-61797079-INS-164
chr1-61812702-INS-54
chr1-61813494-INS-236
chr1-61838161-INS-64
chr1-61841127-INS-224
chr1-61863840-INS-183
chr1-61880543-INS-614
chr1-61882621-INS-230
chr1-61901038-INS-69
chr1-61908196-INS-200
chr1-61928253-INS-64
chr1-61946763-INS-401
chr1-61982467-INS-195
chr1-62005774-INS-139
chr1-62010730-INS-6431
chr1-62018760-INS-62
chr1-62054551-INS-57

chr1-62073431-INS-2905
chr1-62147704-INS-6548
chr1-62184695-INS-566
chr1-62195723-INS-6741
chr1-62202361-INS-6313
chr1-62235803-INS-330
chr1-62243679-INS-756
chr1-62307689-INS-65
chr1-62317405-INS-72
chr1-62345090-INS-63
chr1-62416167-INS-294
chr1-62436630-INS-59
chr1-62503974-INS-4769
chr1-62507931-INS-86
chr1-62537706-INS-115
chr1-62553964-INS-231
chr1-62629634-INS-6684
chr1-62782942-INS-54
chr1-62794608-INS-170
chr1-62817589-INS-56
chr1-62874805-INS-58
chr1-62892460-INS-96
chr1-62896632-INS-120
chr1-62915911-INS-71
chr1-62963454-INS-158
chr1-63004051-INS-56
chr1-63059919-INS-104
chr1-63123103-INS-174
chr1-63127879-INS-197
chr1-63134612-INS-70
chr1-63177789-INS-52
chr1-63190082-INS-451
chr1-63205841-INS-183
chr1-63231220-INS-70
chr1-63246275-INS-197
chr1-63248359-INS-70
chr1-63254417-INS-199
chr1-63257858-INS-205
chr1-63278891-INS-90
chr1-63283422-INS-543
chr1-63285879-INS-197
chr1-63289377-INS-64
chr1-63324468-INS-498
chr1-63331988-INS-68
chr1-63338323-INS-201
chr1-63350797-INS-197
chr1-63368982-INS-1170
chr1-63377145-INS-5554
chr1-63379529-INS-116
chr1-63383180-INS-393
chr1-63394611-INS-343
chr1-63448194-INS-115
chr1-63482229-INS-108
chr1-63501282-INS-179
chr1-63501283-INS-156
chr1-63506343-INS-84
chr1-63514351-INS-90
chr1-63515900-INS-459
chr1-63544592-INS-86
chr1-63581283-INS-6666
chr1-63597260-INS-105
chr1-63599422-INS-188
chr1-63629156-INS-62
chr1-63630718-INS-253
chr1-63647466-INS-870
chr1-63669164-INS-1403
chr1-63669382-INS-103
chr1-63669553-INS-3749
chr1-63674790-INS-286
chr1-63678816-INS-128
chr1-63681300-INS-516
chr1-63718816-INS-207
chr1-63727873-INS-551
chr1-63746347-INS-932
chr1-63747235-INS-194
chr1-63767056-INS-251
chr1-63824397-INS-408

chr1-63836701-INS-57
chr1-63843311-INS-204
chr1-63864832-INS-54
chr1-63958376-INS-69
chr1-63967973-INS-69
chr1-63968527-INS-1120
chr1-63978295-INS-162
chr1-63987413-INS-339
chr1-63995887-INS-225
chr1-63996848-INS-55
chr1-64001084-INS-70
chr1-64020442-INS-94
chr1-64027572-INS-216
chr1-64041649-INS-6433
chr1-64076718-INS-276
chr1-64090166-INS-203
chr1-64145889-INS-612
chr1-64147735-INS-5567
chr1-64158874-INS-121
chr1-64208910-INS-56
chr1-64215192-INS-105
chr1-64228039-INS-74
chr1-64255324-INS-380
chr1-64272371-INS-72
chr1-64278282-INS-227
chr1-64283239-INS-76
chr1-64291263-INS-75
chr1-64295472-INS-58
chr1-64311555-INS-1888
chr1-64313494-INS-57
chr1-64327215-INS-72
chr1-64335726-INS-242
chr1-64352538-INS-184
chr1-64360887-INS-54
chr1-64364659-INS-118
chr1-64402555-INS-1888
chr1-64434822-INS-643
chr1-64437885-INS-340
chr1-64448132-INS-1863
chr1-64456303-INS-199
chr1-64486497-INS-100
chr1-64490489-INS-108
chr1-64501185-INS-52
chr1-64505852-INS-59
chr1-64524034-INS-60
chr1-64541745-INS-60
chr1-64551824-INS-156
chr1-64568361-INS-56
chr1-64583172-INS-344
chr1-64587379-INS-7277
chr1-64594084-INS-68
chr1-64626219-INS-111
chr1-64653751-INS-113
chr1-64666074-INS-6690
chr1-64668607-INS-146
chr1-64670888-INS-204
chr1-64675221-INS-82
chr1-64696971-INS-56
chr1-64702289-INS-120
chr1-64792714-INS-60
chr1-64873796-INS-203
chr1-64877230-INS-71
chr1-64887543-INS-164
chr1-64893985-INS-68
chr1-64908572-INS-66
chr1-64923044-INS-346
chr1-64937588-INS-2893
chr1-64940041-INS-203
chr1-64980002-INS-120
chr1-65008114-INS-81
chr1-65011378-INS-2931
chr1-65013221-INS-400
chr1-65015544-INS-98
chr1-65028292-INS-124
chr1-65029971-INS-8985
chr1-65030506-INS-72
chr1-65040659-INS-168

chr1-65043319-INS-250
chr1-65083391-INS-64
chr1-65090939-INS-523
chr1-65094139-INS-78
chr1-65098729-INS-56
chr1-65099743-INS-773
chr1-65114285-INS-64
chr1-65117848-INS-73
chr1-65155114-INS-88
chr1-65183999-INS-185
chr1-65199847-INS-113
chr1-65233328-INS-179
chr1-65240277-INS-54
chr1-65258093-INS-95
chr1-65298180-INS-6253
chr1-65317295-INS-52
chr1-65319399-INS-241
chr1-65338120-INS-6730
chr1-65344373-INS-934
chr1-65367881-INS-172
chr1-65384065-INS-54
chr1-65391041-INS-976
chr1-65394166-INS-52
chr1-65418116-INS-134
chr1-65424487-INS-196
chr1-65426868-INS-110
chr1-65478318-INS-4877
chr1-65481946-INS-786
chr1-65483832-INS-2888
chr1-65507422-INS-177
chr1-65514351-INS-461
chr1-65533410-INS-102
chr1-65554240-INS-63
chr1-65577787-INS-1147
chr1-65600901-INS-6675
chr1-65601968-INS-254
chr1-65609449-INS-6277
chr1-65630943-INS-795
chr1-65632554-INS-527
chr1-65633894-INS-6566
chr1-65644937-INS-573
chr1-65645652-INS-6573
chr1-65666821-INS-275
chr1-65699449-INS-6310
chr1-65709486-INS-316
chr1-65725365-INS-56
chr1-65728397-INS-10443
chr1-65747584-INS-64
chr1-65785397-INS-82
chr1-65810652-INS-144
chr1-65847597-INS-60
chr1-65880073-INS-515
chr1-65885815-INS-69
chr1-65897794-INS-139
chr1-65902039-INS-327
chr1-65919787-INS-789
chr1-65927546-INS-57
chr1-65964946-INS-608
chr1-65983967-INS-5037
chr1-66005880-INS-69
chr1-66027169-INS-54
chr1-66028272-INS-6731
chr1-66036929-INS-6461
chr1-66069793-INS-198
chr1-66099777-INS-52
chr1-66103238-INS-507
chr1-66112954-INS-204
chr1-66133797-INS-83
chr1-66202869-INS-7219
chr1-66209960-INS-54
chr1-66243222-INS-66
chr1-66245110-INS-664
chr1-66260489-INS-73
chr1-66285240-INS-750
chr1-66287667-INS-958
chr1-66293377-INS-54
chr1-66305032-INS-294

chr1-66307977-INS-385
chr1-66364943-INS-440
chr1-66384643-INS-60
chr1-66413308-INS-7151
chr1-66417408-INS-136
chr1-66487706-INS-280
chr1-66498496-INS-109
chr1-66518749-INS-135
chr1-66521632-INS-454
chr1-66527727-INS-590
chr1-66530053-INS-64
chr1-66531531-INS-128
chr1-66536088-INS-167
chr1-66536961-INS-7206
chr1-66538572-INS-211
chr1-66567344-INS-63
chr1-66568535-INS-184
chr1-66590880-INS-60
chr1-66606795-INS-2930
chr1-66622267-INS-6573
chr1-66679039-INS-329
chr1-66711226-INS-55
chr1-66732973-INS-84
chr1-66786259-INS-151
chr1-66787515-INS-2182
chr1-66796839-INS-426
chr1-66811793-INS-153
chr1-66844620-INS-428
chr1-66879443-INS-54
chr1-66898826-INS-118
chr1-66924062-INS-182
chr1-66967498-INS-142
chr1-66991208-INS-195
chr1-67009776-INS-6475
chr1-67012824-INS-610
chr1-67020778-INS-159
chr1-67022228-INS-66
chr1-67024337-INS-59
chr1-67046963-INS-65
chr1-67050919-INS-145
chr1-67083901-INS-52
chr1-67095749-INS-256
chr1-67102549-INS-405
chr1-67108725-INS-184
chr1-67120187-INS-376
chr1-67181258-INS-56
chr1-67198887-INS-103
chr1-67232912-INS-653
chr1-67242292-INS-168
chr1-67275726-INS-156
chr1-67277578-INS-110
chr1-67302027-INS-515
chr1-67304801-INS-728
chr1-67309398-INS-332
chr1-67310664-INS-64
chr1-67328329-INS-55
chr1-67377124-INS-53
chr1-67398195-INS-3465
chr1-67403207-INS-6656
chr1-67404292-INS-7741
chr1-67407418-INS-56
chr1-67409205-INS-6284
chr1-67425481-INS-503
chr1-67427920-INS-379
chr1-67444442-INS-51
chr1-67451389-INS-54
chr1-67456154-INS-821
chr1-67467759-INS-57
chr1-67482832-INS-98
chr1-67515236-INS-220
chr1-67519771-INS-165
chr1-67527749-INS-167
chr1-67601589-INS-205
chr1-67602852-INS-938
chr1-67654322-INS-297
chr1-67677254-INS-769
chr1-67707531-INS-60

chr1-67728694-INS-56
chr1-67749320-INS-6487
chr1-67766708-INS-157
chr1-67798182-INS-95
chr1-67803673-INS-339
chr1-67862708-INS-564
chr1-67879626-INS-77
chr1-67889332-INS-476
chr1-67905260-INS-314
chr1-67906091-INS-79
chr1-67935476-INS-170
chr1-67962098-INS-69
chr1-67972834-INS-398
chr1-67980905-INS-106
chr1-67986770-INS-56
chr1-68020789-INS-106
chr1-68039040-INS-51
chr1-68130042-INS-257
chr1-68163743-INS-3387
chr1-68184681-INS-57
chr1-68192115-INS-381
chr1-68220873-INS-393
chr1-68256238-INS-581
chr1-68262369-INS-821
chr1-68262924-INS-160
chr1-68273037-INS-421
chr1-68278998-INS-63
chr1-68284778-INS-54
chr1-68294752-INS-101
chr1-68302683-INS-96
chr1-68328452-INS-6975
chr1-68333415-INS-88
chr1-68365702-INS-630
chr1-68376563-INS-52
chr1-68403538-INS-194
chr1-68413131-INS-136
chr1-68425861-INS-6878
chr1-68481207-INS-272
chr1-68490446-INS-57
chr1-68499895-INS-56
chr1-68501464-INS-480
chr1-68504032-INS-52
chr1-68518840-INS-358
chr1-68532993-INS-761
chr1-68547204-INS-94
chr1-68548621-INS-56
chr1-68580386-INS-237
chr1-68592576-INS-81
chr1-68597599-INS-128
chr1-68606199-INS-75
chr1-68616457-INS-693
chr1-68656825-INS-199
chr1-68657891-INS-74
chr1-68700259-INS-278
chr1-68724921-INS-64
chr1-68725771-INS-308
chr1-68740169-INS-3471
chr1-68771464-INS-90
chr1-68781955-INS-104
chr1-68797122-INS-105
chr1-68805597-INS-133
chr1-68818790-INS-441
chr1-68830879-INS-150
chr1-68831934-INS-546
chr1-68872256-INS-200
chr1-68899513-INS-58
chr1-68936290-INS-69
chr1-68947902-INS-58
chr1-68988982-INS-202
chr1-68997902-INS-792
chr1-69005614-INS-661
chr1-69024367-INS-86
chr1-69058275-INS-921
chr1-69058376-INS-214
chr1-69071078-INS-168
chr1-69089090-INS-212
chr1-69092221-INS-233

chr1-69163119-INS-178
chr1-69172777-INS-102
chr1-69177618-INS-230
chr1-69182110-INS-60
chr1-69192025-INS-682
chr1-69193451-INS-65
chr1-69193955-INS-71
chr1-69205117-INS-479
chr1-69211404-INS-178
chr1-69245561-INS-61
chr1-69293571-INS-641
chr1-69347104-INS-57
chr1-69380978-INS-70
chr1-69381888-INS-6200
chr1-69389658-INS-6374
chr1-69425571-INS-79
chr1-69441016-INS-613
chr1-69493057-INS-463
chr1-69525278-INS-135
chr1-69526794-INS-632
chr1-69570228-INS-70
chr1-69579573-INS-3752
chr1-69579850-INS-1030
chr1-69626097-INS-161
chr1-69641569-INS-567
chr1-69654131-INS-54
chr1-69655882-INS-240
chr1-69699263-INS-52
chr1-69709970-INS-83
chr1-69712663-INS-214
chr1-69735846-INS-96
chr1-69761886-INS-297
chr1-69775994-INS-106
chr1-69792835-INS-266
chr1-69803337-INS-309
chr1-69805490-INS-397
chr1-69819262-INS-131
chr1-69857564-INS-6228
chr1-69858913-INS-62
chr1-69864269-INS-5316
chr1-69870155-INS-55
chr1-69871306-INS-312
chr1-69874154-INS-565
chr1-69883865-INS-108
chr1-69928968-INS-6651
chr1-69933467-INS-6403
chr1-69953640-INS-655
chr1-69991506-INS-150
chr1-69992544-INS-393
chr1-70024267-INS-987
chr1-70032364-INS-267
chr1-70038859-INS-354
chr1-70045145-INS-6473
chr1-70064727-INS-5046
chr1-70071310-INS-61
chr1-70077625-INS-327
chr1-70092656-INS-149
chr1-70093207-INS-85
chr1-70096493-INS-222
chr1-70108699-INS-3183
chr1-70109957-INS-6654
chr1-70118951-INS-228
chr1-70123914-INS-6510
chr1-70131246-INS-124
chr1-70159751-INS-60
chr1-70164653-INS-2492
chr1-70184115-INS-6334
chr1-70201764-INS-6215
chr1-70242128-INS-142
chr1-70246701-INS-412
chr1-70256409-INS-62
chr1-70292440-INS-70
chr1-70309291-INS-92
chr1-70327255-INS-7449
chr1-70335629-INS-128
chr1-70338729-INS-2423
chr1-70350299-INS-14381

chr1-70366055-INS-549
chr1-70371748-INS-7106
chr1-70375748-INS-7107
chr1-70382885-INS-6474
chr1-70391363-INS-3147
chr1-70395824-INS-1000
chr1-70396034-INS-131
chr1-70403037-INS-194
chr1-70412244-INS-513
chr1-70433572-INS-53
chr1-70437295-INS-186
chr1-70449811-INS-651
chr1-70458972-INS-1858
chr1-70464616-INS-77
chr1-70477530-INS-73
chr1-70503772-INS-198
chr1-70512728-INS-147
chr1-70528230-INS-318
chr1-70543117-INS-562
chr1-70553038-INS-183
chr1-70626822-INS-184
chr1-70643468-INS-6870
chr1-70643508-INS-382
chr1-70649390-INS-83
chr1-70678177-INS-90
chr1-70684497-INS-152
chr1-70704082-INS-342
chr1-70713565-INS-61
chr1-70737664-INS-66
chr1-70748422-INS-7960
chr1-70762013-INS-197
chr1-70788669-INS-61
chr1-70791145-INS-64
chr1-70812523-INS-630
chr1-70820040-INS-343
chr1-70821647-INS-132
chr1-70822592-INS-319
chr1-70865620-INS-305
chr1-70876207-INS-3087
chr1-70881259-INS-624
chr1-70893639-INS-6446
chr1-70909823-INS-74
chr1-70920368-INS-57
chr1-70930539-INS-365
chr1-70938494-INS-552
chr1-70942013-INS-425
chr1-70943337-INS-418
chr1-70945183-INS-100
chr1-70962257-INS-401
chr1-70972298-INS-502
chr1-70994413-INS-535
chr1-71004415-INS-6194
chr1-71015991-INS-117
chr1-71029260-INS-78
chr1-71050079-INS-816
chr1-71091852-INS-144
chr1-71092991-INS-224
chr1-71096123-INS-133
chr1-71107209-INS-1893
chr1-71134258-INS-235
chr1-71150672-INS-96
chr1-71167985-INS-6285
chr1-71187631-INS-570
chr1-71203025-INS-93
chr1-71203472-INS-7585
chr1-71208534-INS-120
chr1-71213839-INS-6927
chr1-71213989-INS-111
chr1-71224505-INS-571
chr1-71230880-INS-134
chr1-71265790-INS-55
chr1-71289719-INS-212
chr1-71297687-INS-6478
chr1-71375514-INS-299
chr1-71400363-INS-168
chr1-71403537-INS-70
chr1-71407399-INS-88

chr1-71436324-INS-1965
chr1-71437946-INS-57
chr1-71475330-INS-3661
chr1-71501710-INS-52
chr1-71511318-INS-104
chr1-71528363-INS-145
chr1-71580117-INS-502
chr1-71591775-INS-87
chr1-71612921-INS-96
chr1-71613869-INS-130
chr1-71619741-INS-125
chr1-71621716-INS-112
chr1-71624372-INS-60
chr1-71627498-INS-198
chr1-71642352-INS-227
chr1-71649148-INS-199
chr1-71673141-INS-137
chr1-71674177-INS-194
chr1-71707455-INS-2340
chr1-71713121-INS-68
chr1-71733330-INS-207
chr1-71749802-INS-97
chr1-71751603-INS-199
chr1-71765735-INS-180
chr1-71773838-INS-191
chr1-71776025-INS-189
chr1-71783230-INS-156
chr1-71806953-INS-82
chr1-71814007-INS-144
chr1-71817737-INS-6232
chr1-71831248-INS-222
chr1-71848965-INS-178
chr1-71849960-INS-6676
chr1-71874609-INS-467
chr1-71895737-INS-265
chr1-71897818-INS-388
chr1-71910651-INS-908
chr1-71914807-INS-120
chr1-71949269-INS-361
chr1-71954658-INS-1108
chr1-71972299-INS-76
chr1-71978181-INS-108
chr1-72011970-INS-90
chr1-72014033-INS-492
chr1-72020394-INS-56
chr1-72049849-INS-62
chr1-72093601-INS-448
chr1-72094379-INS-203
chr1-72109651-INS-6374
chr1-72116345-INS-72
chr1-72130342-INS-358
chr1-72143113-INS-110
chr1-72180486-INS-52
chr1-72211902-INS-64
chr1-72217788-INS-56
chr1-72231634-INS-372
chr1-72231741-INS-1250
chr1-72233949-INS-206
chr1-72240960-INS-63
chr1-72273249-INS-52
chr1-72285176-INS-328
chr1-72293025-INS-435
chr1-72314444-INS-241
chr1-72318442-INS-213
chr1-72320063-INS-205
chr1-72325440-INS-194
chr1-72338208-INS-51
chr1-72352920-INS-96
chr1-72357224-INS-399
chr1-72361006-INS-60
chr1-72392236-INS-52
chr1-72404050-INS-183
chr1-72420620-INS-90
chr1-72429005-INS-85
chr1-72441218-INS-58
chr1-72467364-INS-80
chr1-72477786-INS-51

chr1-72490675-INS-52
chr1-72520121-INS-61
chr1-72551588-INS-438
chr1-72555065-INS-393
chr1-72580653-INS-124
chr1-72588819-INS-340
chr1-72633305-INS-6454
chr1-72661829-INS-370
chr1-72681808-INS-486
chr1-72707370-INS-364
chr1-72719962-INS-76
chr1-72740933-INS-51
chr1-72746965-INS-231
chr1-72760939-INS-60
chr1-72764040-INS-53
chr1-72793559-INS-79
chr1-72810518-INS-68
chr1-72832813-INS-56
chr1-72833396-INS-581
chr1-72840270-INS-184
chr1-72846572-INS-401
chr1-72848222-INS-180
chr1-72850140-INS-71
chr1-72858881-INS-194
chr1-72874087-INS-56
chr1-72884820-INS-60
chr1-72885745-INS-87
chr1-72886882-INS-205
chr1-72914757-INS-154
chr1-72922648-INS-57
chr1-72937047-INS-56
chr1-72945312-INS-201
chr1-72950095-INS-377
chr1-72998046-INS-521
chr1-73049645-INS-126
chr1-73066559-INS-59
chr1-73079738-INS-63
chr1-73082185-INS-192
chr1-73094951-INS-70
chr1-73109059-INS-219
chr1-73130360-INS-51
chr1-73144139-INS-267
chr1-73158649-INS-208
chr1-73160326-INS-129
chr1-73163368-INS-328
chr1-73166146-INS-446
chr1-73228867-INS-977
chr1-73269281-INS-80
chr1-73286100-INS-325
chr1-73288360-INS-95
chr1-73313433-INS-536
chr1-73336132-INS-7423
chr1-73339124-INS-54
chr1-73339851-INS-176
chr1-73353607-INS-56
chr1-73364966-INS-6398
chr1-73391949-INS-92
chr1-73394900-INS-831
chr1-73401833-INS-201
chr1-73408116-INS-92
chr1-73424797-INS-6589
chr1-73430427-INS-110
chr1-73444935-INS-244
chr1-73452235-INS-57
chr1-73517783-INS-64
chr1-73533837-INS-134
chr1-73562971-INS-336
chr1-73567471-INS-6514
chr1-73570089-INS-52
chr1-73614499-INS-94
chr1-73618470-INS-5485
chr1-73626568-INS-56
chr1-73661258-INS-537
chr1-73670253-INS-57
chr1-73671474-INS-726
chr1-73677102-INS-77
chr1-73679550-INS-60

chr1-73694021-INS-226
chr1-73728366-INS-260
chr1-73744738-INS-6440
chr1-73760532-INS-1117
chr1-73766212-INS-212
chr1-73772494-INS-76
chr1-73779050-INS-123
chr1-73783431-INS-65
chr1-73789500-INS-102
chr1-73810055-INS-7217
chr1-73821925-INS-888
chr1-73821933-INS-270
chr1-73872057-INS-108
chr1-73873846-INS-401
chr1-73874608-INS-70
chr1-73877473-INS-146
chr1-73905483-INS-4671
chr1-73905594-INS-6224
chr1-73915240-INS-202
chr1-73915989-INS-60
chr1-73916604-INS-66
chr1-73916961-INS-7136
chr1-73941583-INS-190
chr1-73942926-INS-788
chr1-73963096-INS-230
chr1-73984190-INS-54
chr1-74004345-INS-125
chr1-74020450-INS-166
chr1-74043200-INS-60
chr1-74047492-INS-3411
chr1-74084322-INS-57
chr1-74099867-INS-52
chr1-74119911-INS-76
chr1-74149505-INS-434
chr1-74151472-INS-116
chr1-74175542-INS-56
chr1-74181311-INS-5306
chr1-74194644-INS-884
chr1-74211446-INS-7188
chr1-74230696-INS-182
chr1-74242083-INS-207
chr1-74247372-INS-106
chr1-74256430-INS-208
chr1-74262232-INS-222
chr1-74270945-INS-174
chr1-74280631-INS-197
chr1-74285967-INS-5656
chr1-74289204-INS-738
chr1-74311352-INS-157
chr1-74353122-INS-60
chr1-74383920-INS-184
chr1-74384161-INS-1647
chr1-74393219-INS-59
chr1-74394594-INS-205
chr1-74400271-INS-581
chr1-74415501-INS-217
chr1-74422000-INS-156
chr1-74438701-INS-218
chr1-74447448-INS-420
chr1-74471085-INS-238
chr1-74481952-INS-204
chr1-74483305-INS-59
chr1-74551225-INS-5316
chr1-74578975-INS-57
chr1-74591817-INS-78
chr1-74595348-INS-208
chr1-74598129-INS-64
chr1-74634658-INS-57
chr1-74656883-INS-228
chr1-74657898-INS-133
chr1-74661005-INS-128
chr1-74670328-INS-186
chr1-74683408-INS-53
chr1-74683625-INS-131
chr1-74687531-INS-211
chr1-74689934-INS-77
chr1-74714538-INS-204

chr1-74716807-INS-114
chr1-74763997-INS-424
chr1-74766050-INS-219
chr1-74768397-INS-202
chr1-74778567-INS-52
chr1-74788820-INS-227
chr1-74789652-INS-184
chr1-74808183-INS-76
chr1-74849756-INS-218
chr1-74852283-INS-180
chr1-74870511-INS-205
chr1-74874573-INS-188
chr1-74884075-INS-141
chr1-74901002-INS-196
chr1-74917038-INS-62
chr1-74928146-INS-207
chr1-74929036-INS-56
chr1-74934400-INS-119
chr1-74952999-INS-99
chr1-74963102-INS-52
chr1-74966662-INS-84
chr1-74995280-INS-79
chr1-75007887-INS-244
chr1-75024824-INS-1916
chr1-75034945-INS-86
chr1-75042483-INS-186
chr1-75058622-INS-209
chr1-75067843-INS-67
chr1-75073552-INS-55
chr1-75096517-INS-185
chr1-75130781-INS-88
chr1-75143152-INS-7130
chr1-75143655-INS-162
chr1-75171057-INS-179
chr1-75198742-INS-81
chr1-75201451-INS-74
chr1-75204006-INS-69
chr1-75225527-INS-118
chr1-75233420-INS-140
chr1-75244651-INS-569
chr1-75266485-INS-1070
chr1-75281831-INS-138
chr1-75296794-INS-113
chr1-75347556-INS-204
chr1-75349571-INS-240
chr1-75357704-INS-205
chr1-75367272-INS-149
chr1-75391412-INS-54
chr1-75435513-INS-69
chr1-75438264-INS-415
chr1-75480637-INS-108
chr1-75494670-INS-2460
chr1-75500876-INS-132
chr1-75516166-INS-613
chr1-75517601-INS-6299
chr1-75549266-INS-156
chr1-75556805-INS-3096
chr1-75561001-INS-133
chr1-75565153-INS-60
chr1-75584496-INS-549
chr1-75599377-INS-213
chr1-75612843-INS-486
chr1-75631750-INS-114
chr1-75640333-INS-69
chr1-75651440-INS-434
chr1-75678059-INS-92
chr1-75684187-INS-73
chr1-75709736-INS-82
chr1-75738347-INS-1554
chr1-75814230-INS-132
chr1-75831560-INS-53
chr1-75855952-INS-733
chr1-75863300-INS-341
chr1-75865959-INS-163
chr1-75867122-INS-5273
chr1-75906784-INS-59
chr1-75911694-INS-5294

chr1-75919333-INS-7212
chr1-75921175-INS-93
chr1-75923911-INS-72
chr1-75967885-INS-1184
chr1-75993185-INS-11790
chr1-75993186-INS-129
chr1-76010317-INS-6442
chr1-76029525-INS-143
chr1-76044762-INS-69
chr1-76047469-INS-293
chr1-76048084-INS-895
chr1-76054476-INS-195
chr1-76064220-INS-326
chr1-76072143-INS-57
chr1-76077637-INS-64
chr1-76108091-INS-81
chr1-76115206-INS-160
chr1-76125000-INS-232
chr1-76138101-INS-68
chr1-76146665-INS-75
chr1-76147920-INS-399
chr1-76157038-INS-459
chr1-76170834-INS-291
chr1-76179995-INS-62
chr1-76199285-INS-61
chr1-76206381-INS-74
chr1-76216520-INS-187
chr1-76235723-INS-263
chr1-76250065-INS-6989
chr1-76276497-INS-334
chr1-76312023-INS-71
chr1-76318272-INS-1335
chr1-76322036-INS-2289
chr1-76324769-INS-124
chr1-76344963-INS-6611
chr1-76355402-INS-2184
chr1-76360777-INS-498
chr1-76381222-INS-60
chr1-76385084-INS-7972
chr1-76388116-INS-80
chr1-76394913-INS-51
chr1-76412644-INS-51
chr1-76418994-INS-256
chr1-76462793-INS-126
chr1-76470305-INS-180
chr1-76490983-INS-71
chr1-76496688-INS-86
chr1-76546983-INS-725
chr1-76556621-INS-717
chr1-76577491-INS-2125
chr1-76584753-INS-95
chr1-76601423-INS-88
chr1-76640819-INS-52
chr1-76643853-INS-163
chr1-76648863-INS-440
chr1-76662549-INS-65
chr1-76668424-INS-66
chr1-76745217-INS-90
chr1-76761962-INS-267
chr1-76779357-INS-2082
chr1-76790210-INS-289
chr1-76807224-INS-274
chr1-76810257-INS-202
chr1-76825114-INS-57
chr1-76868486-INS-695
chr1-76876992-INS-53
chr1-76887475-INS-498
chr1-76901164-INS-67
chr1-76906713-INS-406
chr1-76906727-INS-211
chr1-76915417-INS-88
chr1-76968703-INS-87
chr1-77002705-INS-1514
chr1-77019443-INS-440
chr1-77020516-INS-213
chr1-77025313-INS-205
chr1-77028977-INS-106

chr1-77053860-INS-7179
chr1-77055232-INS-54
chr1-77058005-INS-88
chr1-77083471-INS-195
chr1-77084599-INS-877
chr1-77085895-INS-193
chr1-77138437-INS-557
chr1-77139852-INS-193
chr1-77147902-INS-7266
chr1-77156482-INS-3224
chr1-77165569-INS-54
chr1-77167150-INS-760
chr1-77175677-INS-476
chr1-77182514-INS-117
chr1-77183959-INS-210
chr1-77193769-INS-83
chr1-77211453-INS-53
chr1-77215223-INS-7299
chr1-77283105-INS-86
chr1-77299279-INS-180
chr1-77328392-INS-111
chr1-77331101-INS-161
chr1-77349462-INS-52
chr1-77436820-INS-107
chr1-77444499-INS-6371
chr1-77448857-INS-219
chr1-77451059-INS-424
chr1-77481584-INS-81
chr1-77513869-INS-86
chr1-77514816-INS-167
chr1-77517534-INS-144
chr1-77550608-INS-6792
chr1-77561957-INS-105
chr1-77567654-INS-80
chr1-77580157-INS-56
chr1-77582765-INS-70
chr1-77583912-INS-144
chr1-77587177-INS-224
chr1-77658653-INS-56
chr1-77671086-INS-191
chr1-77691336-INS-59
chr1-77693060-INS-64
chr1-77706923-INS-168
chr1-77748301-INS-234
chr1-77782909-INS-210
chr1-77796311-INS-71
chr1-77798548-INS-128
chr1-77808151-INS-6607
chr1-77811140-INS-3781
chr1-77814155-INS-675
chr1-77814912-INS-6243
chr1-77836944-INS-261
chr1-77840828-INS-256
chr1-77845452-INS-283
chr1-77850471-INS-802
chr1-77854508-INS-4589
chr1-77854509-INS-440
chr1-77854931-INS-636
chr1-77855489-INS-174
chr1-77868300-INS-110
chr1-77869288-INS-52
chr1-77872155-INS-151
chr1-77872557-INS-6985
chr1-77875271-INS-117
chr1-77896141-INS-387
chr1-77897663-INS-638
chr1-77921724-INS-56
chr1-77934778-INS-133
chr1-77950261-INS-63
chr1-77952698-INS-94
chr1-77965703-INS-76
chr1-77976668-INS-198
chr1-77989845-INS-80
chr1-78008812-INS-7118
chr1-78017576-INS-564
chr1-78044804-INS-68
chr1-78065913-INS-156

chr1-78110177-INS-90
chr1-78124213-INS-339
chr1-78124966-INS-60
chr1-78131284-INS-96
chr1-78137647-INS-1018
chr1-78151758-INS-56
chr1-78226956-INS-72
chr1-78237117-INS-399
chr1-78243027-INS-68
chr1-78278040-INS-1487
chr1-78311748-INS-468
chr1-78315944-INS-120
chr1-78318998-INS-157
chr1-78321224-INS-142
chr1-78322334-INS-432
chr1-78339115-INS-103
chr1-78360290-INS-201
chr1-78375013-INS-101
chr1-78390220-INS-224
chr1-78432722-INS-62
chr1-78460679-INS-56
chr1-78473019-INS-57
chr1-78481058-INS-470
chr1-78532196-INS-390
chr1-78540926-INS-115
chr1-78548545-INS-86
chr1-78550857-INS-2342
chr1-78554825-INS-194
chr1-78557438-INS-156
chr1-78559030-INS-209
chr1-78559842-INS-96
chr1-78567571-INS-144
chr1-78568784-INS-65
chr1-78571082-INS-94
chr1-78574046-INS-274
chr1-78576735-INS-135
chr1-78584207-INS-97
chr1-78585002-INS-52
chr1-78593618-INS-551
chr1-78594336-INS-104
chr1-78604357-INS-60
chr1-78607871-INS-124
chr1-78609613-INS-52
chr1-78621137-INS-92
chr1-78665837-INS-91
chr1-78669037-INS-84
chr1-78670658-INS-637
chr1-78696615-INS-138
chr1-78703563-INS-281
chr1-78718853-INS-6692
chr1-78721664-INS-254
chr1-78733312-INS-76
chr1-78735213-INS-149
chr1-78757362-INS-414
chr1-78786037-INS-61
chr1-78786482-INS-149
chr1-78810110-INS-54
chr1-78835411-INS-55
chr1-78854549-INS-82
chr1-78857971-INS-271
chr1-78875722-INS-6754
chr1-78891423-INS-1064
chr1-78900777-INS-6273
chr1-78908505-INS-133
chr1-78918383-INS-6758
chr1-78936236-INS-116
chr1-78957334-INS-260
chr1-78969365-INS-128
chr1-78971952-INS-946
chr1-78984057-INS-60
chr1-79005944-INS-74
chr1-79006825-INS-1424
chr1-79018250-INS-687
chr1-79031876-INS-8926
chr1-79083347-INS-133
chr1-79111693-INS-1573
chr1-79115704-INS-6327

chr1-79141565-INS-360
chr1-79144700-INS-163
chr1-79146673-INS-992
chr1-79151470-INS-938
chr1-79162383-INS-79
chr1-79173079-INS-421
chr1-79187672-INS-68
chr1-79216145-INS-284
chr1-79218434-INS-7092
chr1-79252043-INS-982
chr1-79262289-INS-62
chr1-79270483-INS-93
chr1-79272906-INS-6223
chr1-79282766-INS-186
chr1-79291407-INS-71
chr1-79298792-INS-321
chr1-79300217-INS-873
chr1-79315555-INS-212
chr1-79324211-INS-251
chr1-79333295-INS-153
chr1-79351375-INS-240
chr1-79355760-INS-447
chr1-79393009-INS-313
chr1-79419009-INS-6681
chr1-79424906-INS-52
chr1-79431537-INS-54
chr1-79435317-INS-1894
chr1-79441984-INS-63
chr1-79492506-INS-112
chr1-79497871-INS-196
chr1-79504003-INS-95
chr1-79505275-INS-54
chr1-79505480-INS-260
chr1-79513415-INS-69
chr1-79540780-INS-70
chr1-79548311-INS-399
chr1-79562854-INS-146
chr1-79570205-INS-96
chr1-79574499-INS-752
chr1-79577634-INS-171
chr1-79614565-INS-76
chr1-79638021-INS-72
chr1-79652405-INS-125
chr1-79687216-INS-667
chr1-79692890-INS-118
chr1-79710232-INS-54
chr1-79722824-INS-1490
chr1-79727789-INS-800
chr1-79727969-INS-875
chr1-79766145-INS-4637
chr1-79773939-INS-51
chr1-79783471-INS-451
chr1-79814225-INS-138
chr1-79820559-INS-6601
chr1-79854169-INS-341
chr1-79859129-INS-73
chr1-79882887-INS-55
chr1-79891419-INS-1741
chr1-79895053-INS-131
chr1-79895055-INS-639
chr1-79927824-INS-52
chr1-79967235-INS-121
chr1-79968964-INS-86
chr1-79993928-INS-64
chr1-79998095-INS-140
chr1-80070920-INS-6918
chr1-80133091-INS-342
chr1-80146413-INS-6481
chr1-80147958-INS-76
chr1-80149576-INS-199
chr1-80156269-INS-832
chr1-80173477-INS-56
chr1-80175026-INS-167
chr1-80186088-INS-59
chr1-80293198-INS-740
chr1-80323458-INS-72
chr1-80325826-INS-76

chr1-80327771-INS-58
chr1-80329212-INS-52
chr1-80363754-INS-54
chr1-80378736-INS-80
chr1-80394015-INS-910
chr1-80401163-INS-1044
chr1-80401168-INS-932
chr1-80404641-INS-54
chr1-80419842-INS-224
chr1-80430572-INS-114
chr1-80435404-INS-59
chr1-80447382-INS-113
chr1-80448883-INS-6176
chr1-80456057-INS-252
chr1-80465896-INS-396
chr1-80487097-INS-6578
chr1-80491892-INS-399
chr1-80507261-INS-54
chr1-80514965-INS-295
chr1-80523472-INS-223
chr1-80525922-INS-66
chr1-80582047-INS-139
chr1-80583254-INS-93
chr1-80595427-INS-50
chr1-80638706-INS-113
chr1-80650301-INS-83
chr1-80660366-INS-118
chr1-80683042-INS-156
chr1-80684763-INS-71
chr1-80692926-INS-77
chr1-80728456-INS-50
chr1-80739627-INS-156
chr1-80746985-INS-120
chr1-80751961-INS-52
chr1-80802407-INS-62
chr1-80805321-INS-52
chr1-80817771-INS-611
chr1-80825512-INS-6492
chr1-80827086-INS-342
chr1-80828339-INS-1641
chr1-80851021-INS-488
chr1-80875254-INS-165
chr1-80878174-INS-112
chr1-80883845-INS-6777
chr1-80884919-INS-56
chr1-80890596-INS-52
chr1-80903633-INS-110
chr1-80906506-INS-6806
chr1-80912989-INS-201
chr1-80933890-INS-351
chr1-80956920-INS-166
chr1-80973560-INS-79
chr1-80975894-INS-196
chr1-80994713-INS-421
chr1-80995383-INS-722
chr1-81011557-INS-876
chr1-81018399-INS-546
chr1-81026381-INS-144
chr1-81033245-INS-70
chr1-81048929-INS-80
chr1-81101939-INS-461
chr1-81103907-INS-66
chr1-81106228-INS-59
chr1-81111054-INS-97
chr1-81114819-INS-58
chr1-81118746-INS-98
chr1-81124002-INS-51
chr1-81129829-INS-304
chr1-81131302-INS-55
chr1-81132712-INS-102
chr1-81149960-INS-499
chr1-81150647-INS-3663
chr1-81153724-INS-52
chr1-81180453-INS-827
chr1-81198667-INS-3132
chr1-81206387-INS-74
chr1-81265139-INS-3254

chr1-81286950-INS-66
chr1-81288982-INS-54
chr1-81291131-INS-199
chr1-81299346-INS-65
chr1-81304624-INS-170
chr1-81305244-INS-351
chr1-81312015-INS-168
chr1-81329980-INS-81
chr1-81375270-INS-7029
chr1-81387806-INS-70
chr1-81389417-INS-115
chr1-81412928-INS-737
chr1-81413984-INS-427
chr1-81457985-INS-482
chr1-81475401-INS-1893
chr1-81522708-INS-99
chr1-81524930-INS-6498
chr1-81528353-INS-133
chr1-81533742-INS-192
chr1-81562042-INS-76
chr1-81569782-INS-6924
chr1-81572245-INS-66
chr1-81580697-INS-750
chr1-81582618-INS-52
chr1-81609945-INS-135
chr1-81647744-INS-721
chr1-81652792-INS-140
chr1-81683228-INS-76
chr1-81685273-INS-1867
chr1-81692756-INS-189
chr1-81697358-INS-504
chr1-81703090-INS-106
chr1-81703243-INS-6455
chr1-81704461-INS-412
chr1-81714374-INS-3728
chr1-81717475-INS-199
chr1-81744064-INS-69
chr1-81751817-INS-6464
chr1-81760215-INS-69
chr1-81861180-INS-56
chr1-81879469-INS-246
chr1-81885485-INS-54
chr1-81907195-INS-80
chr1-81930296-INS-58
chr1-81937121-INS-710
chr1-81957279-INS-54
chr1-81978263-INS-78
chr1-81994066-INS-52
chr1-82009793-INS-75
chr1-82015132-INS-60
chr1-82025953-INS-87
chr1-82029529-INS-6183
chr1-82031288-INS-70
chr1-82042026-INS-55
chr1-82052776-INS-1131
chr1-82052833-INS-203
chr1-82056506-INS-198
chr1-82059722-INS-498
chr1-82073744-INS-260
chr1-82076827-INS-159
chr1-82078703-INS-54
chr1-82090687-INS-60
chr1-82091338-INS-1061
chr1-82104426-INS-80
chr1-82111916-INS-110
chr1-82116793-INS-339
chr1-82131252-INS-87
chr1-82145286-INS-587
chr1-82174192-INS-120
chr1-82193740-INS-1045
chr1-82205853-INS-86
chr1-82236953-INS-54
chr1-82258554-INS-161
chr1-82322310-INS-6355
chr1-82345312-INS-1892
chr1-82370306-INS-58
chr1-82394167-INS-85

chr1-82420196-INS-51
chr1-82438345-INS-55
chr1-82455319-INS-91
chr1-82502377-INS-56
chr1-82515254-INS-180
chr1-82548117-INS-94
chr1-82560837-INS-239
chr1-82563282-INS-58
chr1-82572473-INS-220
chr1-82580425-INS-437
chr1-82592033-INS-58
chr1-82593797-INS-82
chr1-82597074-INS-66
chr1-82610085-INS-51
chr1-82619860-INS-57
chr1-82625381-INS-119
chr1-82629160-INS-51
chr1-82630658-INS-876
chr1-82639640-INS-76
chr1-82646475-INS-55
chr1-82651004-INS-72
chr1-82668526-INS-84
chr1-82712946-INS-52
chr1-82746417-INS-2343
chr1-82748860-INS-75
chr1-82751826-INS-1888
chr1-82753932-INS-62
chr1-82756063-INS-385
chr1-82781041-INS-54
chr1-82785277-INS-60
chr1-82790814-INS-399
chr1-82794103-INS-56
chr1-82796389-INS-78
chr1-82808778-INS-55
chr1-82814260-INS-56
chr1-82822258-INS-64
chr1-82827805-INS-2393
chr1-82829600-INS-319
chr1-82835238-INS-203
chr1-82853176-INS-521
chr1-82886330-INS-51
chr1-82904792-INS-247
chr1-82922015-INS-56
chr1-82933542-INS-61
chr1-82976760-INS-104
chr1-82980159-INS-84
chr1-82997797-INS-64
chr1-83003674-INS-68
chr1-83011076-INS-59
chr1-83021620-INS-86
chr1-83032487-INS-88
chr1-83035037-INS-444
chr1-83040556-INS-327
chr1-83060986-INS-241
chr1-83091375-INS-62
chr1-83106527-INS-323
chr1-83115066-INS-71
chr1-83131109-INS-162
chr1-83139685-INS-61
chr1-83147620-INS-595
chr1-83165221-INS-134
chr1-83174700-INS-132
chr1-83217793-INS-76
chr1-83219051-INS-176
chr1-83237138-INS-13264
chr1-83252460-INS-52
chr1-83276964-INS-5290
chr1-83296694-INS-401
chr1-83322677-INS-250
chr1-83342400-INS-93
chr1-83368752-INS-64
chr1-83377339-INS-70
chr1-83379142-INS-400
chr1-83410866-INS-6208
chr1-83417091-INS-111
chr1-83450368-INS-349
chr1-83454613-INS-73

chr1-83465770-INS-69
chr1-83468171-INS-860
chr1-83471752-INS-64
chr1-83498540-INS-1886
chr1-83518531-INS-6572
chr1-83549984-INS-886
chr1-83552173-INS-80
chr1-83557772-INS-534
chr1-83561522-INS-62
chr1-83574701-INS-56
chr1-83579143-INS-994
chr1-83589267-INS-5024
chr1-83615958-INS-774
chr1-83619029-INS-481
chr1-83642720-INS-219
chr1-83668707-INS-131
chr1-83678175-INS-73
chr1-83678603-INS-204
chr1-83689980-INS-198
chr1-83704927-INS-72
chr1-83711259-INS-54
chr1-83722387-INS-58
chr1-83757864-INS-1236
chr1-83788101-INS-54
chr1-83813108-INS-7308
chr1-83823250-INS-56
chr1-83841062-INS-82
chr1-83879541-INS-57
chr1-83938361-INS-52
chr1-83945251-INS-6449
chr1-83949915-INS-3555
chr1-83994425-INS-330
chr1-83996410-INS-166
chr1-84024414-INS-65
chr1-84067958-INS-85
chr1-84074215-INS-52
chr1-84114153-INS-76
chr1-84131782-INS-578
chr1-84133355-INS-81
chr1-84168658-INS-480
chr1-84187628-INS-65
chr1-84192978-INS-205
chr1-84202530-INS-412
chr1-84205033-INS-190
chr1-84205829-INS-7149
chr1-84207381-INS-52
chr1-84208846-INS-52
chr1-84224592-INS-108
chr1-84228258-INS-150
chr1-84252940-INS-66
chr1-84263805-INS-60
chr1-84269783-INS-525
chr1-84275002-INS-799
chr1-84277006-INS-237
chr1-84278790-INS-283
chr1-84280215-INS-399
chr1-84287292-INS-79
chr1-84301200-INS-394
chr1-84315996-INS-91
chr1-84319153-INS-571
chr1-84350199-INS-65
chr1-84372573-INS-183
chr1-84405035-INS-56
chr1-84408141-INS-191
chr1-84436160-INS-62
chr1-84447694-INS-96
chr1-84447861-INS-7119
chr1-84480112-INS-66
chr1-84481749-INS-69
chr1-84486209-INS-141
chr1-84492237-INS-498
chr1-84497723-INS-122
chr1-84498163-INS-743
chr1-84504917-INS-59
chr1-84530377-INS-3118
chr1-84561468-INS-295
chr1-84569772-INS-82

chr1-84581495-INS-277
chr1-84621822-INS-105
chr1-84653851-INS-338
chr1-84655522-INS-150
chr1-84735402-INS-73
chr1-84756395-INS-137
chr1-84766463-INS-169
chr1-84772949-INS-207
chr1-84783493-INS-6440
chr1-84789384-INS-299
chr1-84791192-INS-218
chr1-84800440-INS-141
chr1-84806486-INS-170
chr1-84811989-INS-193
chr1-84819581-INS-339
chr1-84823822-INS-174
chr1-84837199-INS-204
chr1-84842595-INS-122
chr1-84843510-INS-262
chr1-84859617-INS-65
chr1-84863143-INS-58
chr1-84866884-INS-55
chr1-84869015-INS-177
chr1-84883420-INS-65
chr1-84894189-INS-197
chr1-84904370-INS-214
chr1-84909836-INS-159
chr1-84921743-INS-178
chr1-84926225-INS-144
chr1-84936285-INS-242
chr1-84942542-INS-3065
chr1-85002575-INS-6442
chr1-85078461-INS-5344
chr1-85082441-INS-5481
chr1-85197676-INS-1306
chr1-85222370-INS-118
chr1-85436476-INS-710
chr1-85526172-INS-75
chr1-85527132-INS-498
chr1-85534803-INS-231
chr1-85578050-INS-132
chr1-85579275-INS-88
chr1-85594927-INS-90
chr1-85604355-INS-139
chr1-85619362-INS-212
chr1-85637380-INS-149
chr1-85673443-INS-2433
chr1-85687351-INS-61
chr1-85696147-INS-185
chr1-85710110-INS-129
chr1-85711956-INS-52
chr1-85727101-INS-194
chr1-85730586-INS-185
chr1-85746902-INS-213
chr1-85762084-INS-127
chr1-85788478-INS-98
chr1-85798659-INS-60
chr1-85802265-INS-58
chr1-85826293-INS-70
chr1-85849373-INS-551
chr1-85853582-INS-76
chr1-85874550-INS-66
chr1-85879364-INS-147
chr1-85899403-INS-170
chr1-85910158-INS-225
chr1-85931316-INS-302
chr1-85960910-INS-229
chr1-85965551-INS-291
chr1-85977630-INS-111
chr1-85985244-INS-542
chr1-85990042-INS-548
chr1-86061850-INS-1306
chr1-86073407-INS-72
chr1-86081438-INS-60
chr1-86102832-INS-92
chr1-86114518-INS-52
chr1-86118497-INS-120

chr1-86122837-INS-1890
chr1-86129389-INS-1892
chr1-86160049-INS-1802
chr1-86168127-INS-248
chr1-86198410-INS-58
chr1-86204807-INS-76
chr1-86205808-INS-74
chr1-86218230-INS-208
chr1-86221578-INS-112
chr1-86227671-INS-217
chr1-86255459-INS-1889
chr1-86257931-INS-69
chr1-86261012-INS-90
chr1-86261679-INS-231
chr1-86266524-INS-169
chr1-86281436-INS-188
chr1-86292691-INS-201
chr1-86302023-INS-1156
chr1-86305645-INS-72
chr1-86312026-INS-67
chr1-86321868-INS-100
chr1-86335813-INS-228
chr1-86337606-INS-163
chr1-86343890-INS-208
chr1-86364981-INS-198
chr1-86365471-INS-226
chr1-86366582-INS-201
chr1-86383727-INS-257
chr1-86399392-INS-339
chr1-86409180-INS-201
chr1-86411265-INS-217
chr1-86412339-INS-208
chr1-86416541-INS-61
chr1-86419794-INS-126
chr1-86443581-INS-179
chr1-86448860-INS-996
chr1-86451320-INS-91
chr1-86489715-INS-73
chr1-86513924-INS-208
chr1-86531247-INS-146
chr1-86561126-INS-360
chr1-86570219-INS-203
chr1-86579827-INS-208
chr1-86614201-INS-201
chr1-86617809-INS-174
chr1-86669517-INS-167
chr1-86685779-INS-201
chr1-86688332-INS-368
chr1-86690884-INS-263
chr1-86698455-INS-6987
chr1-86714327-INS-6709
chr1-86717130-INS-96
chr1-86724493-INS-101
chr1-86743687-INS-52
chr1-86752165-INS-58
chr1-86758325-INS-956
chr1-86759903-INS-425
chr1-86762390-INS-2395
chr1-86769321-INS-88
chr1-86783936-INS-6632
chr1-86809833-INS-92
chr1-86810385-INS-66
chr1-86812682-INS-141
chr1-86833591-INS-84
chr1-86866673-INS-176
chr1-86904755-INS-53
chr1-86910549-INS-1892
chr1-86967554-INS-196
chr1-87011870-INS-161
chr1-87049076-INS-98
chr1-87109391-INS-379
chr1-87141003-INS-189
chr1-87145156-INS-70
chr1-87178756-INS-244
chr1-87191430-INS-239
chr1-87196887-INS-78
chr1-87207133-INS-472

chr1-87242166-INS-64
chr1-87261865-INS-180
chr1-87286978-INS-411
chr1-87292406-INS-217
chr1-87294745-INS-75
chr1-87318154-INS-89
chr1-87351044-INS-113
chr1-87362158-INS-121
chr1-87372357-INS-65
chr1-87382596-INS-74
chr1-87397621-INS-115
chr1-87418620-INS-208
chr1-87446992-INS-51
chr1-87455037-INS-217
chr1-87482501-INS-52
chr1-87521552-INS-7348
chr1-87529023-INS-233
chr1-87567111-INS-187
chr1-87603585-INS-170
chr1-87606657-INS-175
chr1-87633936-INS-64
chr1-87636048-INS-184
chr1-87637570-INS-182
chr1-87654475-INS-66
chr1-87658783-INS-213
chr1-87668355-INS-168
chr1-87709843-INS-400
chr1-87711890-INS-75
chr1-87716658-INS-201
chr1-87801492-INS-266
chr1-87823418-INS-408
chr1-87838724-INS-73
chr1-87860191-INS-928
chr1-87884665-INS-88
chr1-87890020-INS-428
chr1-87912687-INS-90
chr1-87952880-INS-7196
chr1-87973728-INS-132
chr1-88017686-INS-91
chr1-88025628-INS-164
chr1-88028125-INS-52
chr1-88028294-INS-869
chr1-88045194-INS-181
chr1-88070989-INS-94
chr1-88071800-INS-169
chr1-88090637-INS-264
chr1-88099889-INS-64
chr1-88115477-INS-52
chr1-88136431-INS-72
chr1-88149218-INS-61
chr1-88180951-INS-255
chr1-88218705-INS-1282
chr1-88254743-INS-760
chr1-88260358-INS-388
chr1-88291452-INS-116
chr1-88295345-INS-52
chr1-88297945-INS-286
chr1-88313772-INS-150
chr1-88325516-INS-62
chr1-88346803-INS-120
chr1-88371919-INS-282
chr1-88385279-INS-114
chr1-88386543-INS-355
chr1-88388408-INS-608
chr1-88389511-INS-53
chr1-88397455-INS-55
chr1-88431142-INS-57
chr1-88441142-INS-72
chr1-88498169-INS-502
chr1-88515543-INS-76
chr1-88517133-INS-148
chr1-88519125-INS-200
chr1-88521011-INS-3231
chr1-88530334-INS-508
chr1-88556637-INS-213
chr1-88564834-INS-92
chr1-88568478-INS-482

chr1-88570418-INS-214
chr1-88607698-INS-797
chr1-88662964-INS-397
chr1-88675092-INS-368
chr1-88681276-INS-58
chr1-88688307-INS-69
chr1-88702386-INS-83
chr1-88712642-INS-111
chr1-88716519-INS-62
chr1-88717548-INS-193
chr1-88739680-INS-97
chr1-88776841-INS-161
chr1-88798644-INS-328
chr1-88857915-INS-151
chr1-88859785-INS-75
chr1-88917919-INS-399
chr1-88939801-INS-60
chr1-88968851-INS-224
chr1-88973010-INS-167
chr1-88983914-INS-172
chr1-88987078-INS-55
chr1-88994886-INS-208
chr1-89016914-INS-69
chr1-89039644-INS-433
chr1-89041023-INS-5473
chr1-89062557-INS-107
chr1-89131483-INS-1894
chr1-89131533-INS-2793
chr1-89211873-INS-325
chr1-89222700-INS-1449
chr1-89229836-INS-148
chr1-89259139-INS-162
chr1-89270232-INS-491
chr1-89287923-INS-158
chr1-89310749-INS-124
chr1-89317835-INS-94
chr1-89347422-INS-571
chr1-89349002-INS-213
chr1-89397403-INS-140
chr1-89429745-INS-209
chr1-89439962-INS-357
chr1-89462509-INS-138
chr1-89473255-INS-176
chr1-89549092-INS-62
chr1-89606342-INS-62
chr1-89639079-INS-81
chr1-89688600-INS-63
chr1-89692931-INS-52
chr1-89742679-INS-63
chr1-89818413-INS-220
chr1-89831828-INS-82
chr1-89866141-INS-56
chr1-89880000-INS-190
chr1-89911723-INS-168
chr1-89960534-INS-393
chr1-89974786-INS-130
chr1-90005690-INS-457
chr1-90022043-INS-86
chr1-90036495-INS-577
chr1-90056125-INS-742
chr1-90056130-INS-57
chr1-90061069-INS-457
chr1-90068605-INS-299
chr1-90079761-INS-105
chr1-90102751-INS-96
chr1-90112476-INS-72
chr1-90114775-INS-147
chr1-90128398-INS-188
chr1-90145173-INS-82
chr1-90154795-INS-56
chr1-90169426-INS-68
chr1-90221877-INS-435
chr1-90222567-INS-84
chr1-90227546-INS-51
chr1-90233527-INS-166
chr1-90240814-INS-401
chr1-90253418-INS-262

chr1-90284792-INS-190
chr1-90303507-INS-96
chr1-90334987-INS-195
chr1-90345097-INS-75
chr1-90364570-INS-345
chr1-90378568-INS-72
chr1-90388359-INS-996
chr1-90398005-INS-2188
chr1-90416869-INS-54
chr1-90418617-INS-54
chr1-90422366-INS-2959
chr1-90444220-INS-544
chr1-90483447-INS-94
chr1-90506868-INS-114
chr1-90565570-INS-60
chr1-90571239-INS-147
chr1-90585929-INS-65
chr1-90612293-INS-63
chr1-90632457-INS-52
chr1-90637305-INS-67
chr1-90656095-INS-202
chr1-90664972-INS-225
chr1-90674679-INS-52
chr1-90684163-INS-107
chr1-90687851-INS-56
chr1-90735085-INS-61
chr1-90736557-INS-55
chr1-90747975-INS-110
chr1-90772556-INS-60
chr1-90780855-INS-408
chr1-90802124-INS-971
chr1-90821675-INS-54
chr1-90841638-INS-692
chr1-90858442-INS-324
chr1-90867337-INS-128
chr1-90887586-INS-146
chr1-90892489-INS-102
chr1-90899313-INS-98
chr1-90961381-INS-100
chr1-90994949-INS-873
chr1-91008538-INS-92
chr1-91052846-INS-312
chr1-91066307-INS-103
chr1-91087923-INS-52
chr1-91091032-INS-851
chr1-91096940-INS-462
chr1-91108640-INS-59
chr1-91128728-INS-62
chr1-91142460-INS-306
chr1-91156735-INS-535
chr1-91165917-INS-64
chr1-91172219-INS-156
chr1-91199860-INS-296
chr1-91207102-INS-277
chr1-91221361-INS-412
chr1-91232035-INS-1875
chr1-91248850-INS-98
chr1-91350392-INS-90
chr1-91369928-INS-363
chr1-91412746-INS-212
chr1-91417703-INS-93
chr1-91446269-INS-86
chr1-91452957-INS-204
chr1-91457567-INS-68
chr1-91466180-INS-244
chr1-91471232-INS-211
chr1-91503271-INS-56
chr1-91504501-INS-160
chr1-91507018-INS-63
chr1-91536974-INS-76
chr1-91555038-INS-64
chr1-91566384-INS-558
chr1-91579250-INS-3098
chr1-91599855-INS-6401
chr1-91600382-INS-52
chr1-91615312-INS-86
chr1-91619917-INS-412

chr1-91633805-INS-472
chr1-91646880-INS-120
chr1-91658092-INS-294
chr1-91662083-INS-77
chr1-91666908-INS-64
chr1-91672849-INS-53
chr1-91675247-INS-75
chr1-91680433-INS-64
chr1-91691291-INS-52
chr1-91724878-INS-60
chr1-91751765-INS-60
chr1-91770927-INS-87
chr1-91809691-INS-102
chr1-91836853-INS-104
chr1-91863476-INS-108
chr1-91957292-INS-172
chr1-91993332-INS-2082
chr1-92000980-INS-54
chr1-92055728-INS-341
chr1-92117839-INS-84
chr1-92142316-INS-111
chr1-92162204-INS-5343
chr1-92164452-INS-66
chr1-92190974-INS-1883
chr1-92208136-INS-374
chr1-92217001-INS-92
chr1-92219535-INS-231
chr1-92238455-INS-72
chr1-92249856-INS-119
chr1-92255995-INS-100
chr1-92259065-INS-2628
chr1-92265461-INS-203
chr1-92286024-INS-80
chr1-92307631-INS-153
chr1-92311854-INS-93
chr1-92312357-INS-1862
chr1-92352853-INS-224
chr1-92417891-INS-5057
chr1-92436382-INS-200
chr1-92442438-INS-168
chr1-92463186-INS-82
chr1-92467553-INS-55
chr1-92484618-INS-71
chr1-92497286-INS-264
chr1-92521402-INS-779
chr1-92529502-INS-123
chr1-92540499-INS-113
chr1-92544109-INS-866
chr1-92552554-INS-3629
chr1-92562366-INS-154
chr1-92584555-INS-1342
chr1-92584556-INS-822
chr1-92594048-INS-498
chr1-92607273-INS-3549
chr1-92610574-INS-224
chr1-92630222-INS-70
chr1-92641645-INS-85
chr1-92666431-INS-53
chr1-92670871-INS-88
chr1-92687912-INS-104
chr1-92696702-INS-62
chr1-92728851-INS-5313
chr1-92733570-INS-54
chr1-92742002-INS-196
chr1-92765191-INS-6761
chr1-92796487-INS-84
chr1-92801703-INS-184
chr1-92807861-INS-153
chr1-92814272-INS-57
chr1-92818980-INS-191
chr1-92822251-INS-117
chr1-92828149-INS-358
chr1-92853293-INS-308
chr1-92860599-INS-305
chr1-92907020-INS-159
chr1-92914224-INS-160
chr1-92942957-INS-131

chr1-92972939-INS-52
chr1-92976368-INS-215
chr1-92990752-INS-146
chr1-93018829-INS-249
chr1-93020556-INS-56
chr1-93021854-INS-59
chr1-93047156-INS-51
chr1-93061527-INS-458
chr1-93115034-INS-110
chr1-93168604-INS-63
chr1-93217504-INS-439
chr1-93220124-INS-6435
chr1-93243221-INS-1884
chr1-93270065-INS-827
chr1-93276312-INS-223
chr1-93304905-INS-539
chr1-93309377-INS-169
chr1-93321019-INS-56
chr1-93363198-INS-119
chr1-93375193-INS-106
chr1-93444703-INS-184
chr1-93447082-INS-52
chr1-93451077-INS-219
chr1-93458833-INS-74
chr1-93480304-INS-56
chr1-93525864-INS-671
chr1-93534657-INS-166
chr1-93538215-INS-135
chr1-93539413-INS-63
chr1-93564948-INS-99
chr1-93567551-INS-119
chr1-93577375-INS-54
chr1-93580152-INS-377
chr1-93585897-INS-267
chr1-93595343-INS-53
chr1-93597183-INS-460
chr1-93600382-INS-331
chr1-93603886-INS-157
chr1-93605678-INS-52
chr1-93610606-INS-188
chr1-93622404-INS-7453
chr1-93630119-INS-57
chr1-93659961-INS-72
chr1-93702073-INS-185
chr1-93705100-INS-826
chr1-93707164-INS-208
chr1-93723648-INS-226
chr1-93725583-INS-689
chr1-93730100-INS-191
chr1-93737044-INS-221
chr1-93742381-INS-162
chr1-93745023-INS-200
chr1-93753389-INS-70
chr1-93764561-INS-99
chr1-93781744-INS-840
chr1-93784572-INS-127
chr1-93784825-INS-5978
chr1-93805821-INS-241
chr1-93821126-INS-72
chr1-93837080-INS-63
chr1-93880840-INS-158
chr1-93891206-INS-217
chr1-93907096-INS-118
chr1-93945601-INS-70
chr1-93958770-INS-160
chr1-93988395-INS-104
chr1-94012627-INS-204
chr1-94022041-INS-654
chr1-94025124-INS-3693
chr1-94026970-INS-2468
chr1-94049364-INS-1678
chr1-94052867-INS-7167
chr1-94056573-INS-1287
chr1-94057954-INS-2154
chr1-94063727-INS-6271
chr1-94065499-INS-84
chr1-94072028-INS-8734

chr1-94073347-INS-75
chr1-94100972-INS-103
chr1-94104415-INS-6543
chr1-94106396-INS-7817
chr1-94110240-INS-5574
chr1-94150646-INS-100
chr1-94153638-INS-357
chr1-94154395-INS-6396
chr1-94173236-INS-7014
chr1-94188824-INS-208
chr1-94210973-INS-232
chr1-94214254-INS-80
chr1-94229894-INS-850
chr1-94236582-INS-166
chr1-94239022-INS-116
chr1-94246565-INS-58
chr1-94259970-INS-65
chr1-94264657-INS-66
chr1-94267526-INS-6681
chr1-94280424-INS-79
chr1-94291595-INS-5979
chr1-94297244-INS-6792
chr1-94303587-INS-501
chr1-94323392-INS-223
chr1-94325444-INS-7197
chr1-94334782-INS-112
chr1-94367690-INS-415
chr1-94373852-INS-764
chr1-94374284-INS-7098
chr1-94382398-INS-52
chr1-94382774-INS-189
chr1-94425651-INS-6609
chr1-94431007-INS-123
chr1-94433581-INS-6231
chr1-94448443-INS-58
chr1-94451073-INS-2962
chr1-94452262-INS-52
chr1-94454623-INS-56
chr1-94468918-INS-413
chr1-94474202-INS-395
chr1-94477147-INS-72
chr1-94502156-INS-582
chr1-94507166-INS-451
chr1-94514739-INS-515
chr1-94533908-INS-77
chr1-94534804-INS-487
chr1-94555887-INS-284
chr1-94561220-INS-112
chr1-94566853-INS-1198
chr1-94568965-INS-128
chr1-94590893-INS-626
chr1-94595638-INS-105
chr1-94608028-INS-473
chr1-94624299-INS-190
chr1-94661541-INS-863
chr1-94661998-INS-6800
chr1-94694325-INS-85
chr1-94696571-INS-222
chr1-94701897-INS-74
chr1-94715213-INS-445
chr1-94765402-INS-58
chr1-94767264-INS-224
chr1-94776348-INS-615
chr1-94778648-INS-157
chr1-94804300-INS-83
chr1-94817367-INS-76
chr1-94828227-INS-110
chr1-94860095-INS-708
chr1-94873044-INS-7195
chr1-94889241-INS-135
chr1-94903164-INS-56
chr1-94911258-INS-222
chr1-94919029-INS-66
chr1-94934298-INS-104
chr1-94944021-INS-142
chr1-94974697-INS-5549
chr1-94996343-INS-788

chr1-95002294-INS-358
chr1-95014211-INS-434
chr1-95020524-INS-3002
chr1-95024621-INS-180
chr1-95046110-INS-199
chr1-95107689-INS-6442
chr1-95141193-INS-6491
chr1-95145998-INS-54
chr1-95168441-INS-104
chr1-95176683-INS-226
chr1-95179648-INS-159
chr1-95192119-INS-50
chr1-95216943-INS-112
chr1-95230544-INS-88
chr1-95249734-INS-78
chr1-95253264-INS-531
chr1-95254107-INS-80
chr1-95267960-INS-724
chr1-95273644-INS-611
chr1-95280961-INS-52
chr1-95293561-INS-4533
chr1-95294789-INS-7173
chr1-95306572-INS-6481
chr1-95317384-INS-56
chr1-95320745-INS-80
chr1-95334058-INS-432
chr1-95348798-INS-5136
chr1-95349667-INS-64
chr1-95355915-INS-707
chr1-95365844-INS-56
chr1-95369210-INS-5329
chr1-95370662-INS-76
chr1-95391170-INS-64
chr1-95393607-INS-291
chr1-95402101-INS-449
chr1-95402103-INS-454
chr1-95415688-INS-174
chr1-95422890-INS-147
chr1-95445435-INS-207
chr1-95448382-INS-56
chr1-95454166-INS-6478
chr1-95455166-INS-327
chr1-95470631-INS-124
chr1-95533956-INS-173
chr1-95537404-INS-477
chr1-95557130-INS-1319
chr1-95559553-INS-281
chr1-95601845-INS-502
chr1-95608949-INS-324
chr1-95645943-INS-7027
chr1-95657571-INS-65
chr1-95660247-INS-539
chr1-95662133-INS-87
chr1-95672249-INS-6468
chr1-95677757-INS-1474
chr1-95694265-INS-6255
chr1-95700563-INS-6686
chr1-95703131-INS-396
chr1-95709485-INS-506
chr1-95776587-INS-7081
chr1-95785717-INS-5414
chr1-95786757-INS-410
chr1-95793081-INS-10245
chr1-95798645-INS-380
chr1-95799605-INS-6605
chr1-95811643-INS-140
chr1-95839138-INS-324
chr1-95847345-INS-4940
chr1-95849905-INS-764
chr1-95902354-INS-196
chr1-95927010-INS-6534
chr1-95938904-INS-53
chr1-95952207-INS-937
chr1-95956918-INS-7009
chr1-95979035-INS-492
chr1-95994252-INS-366
chr1-96036087-INS-328

chr1-96038184-INS-371
chr1-96043558-INS-129
chr1-96079042-INS-142
chr1-96096260-INS-78
chr1-96119016-INS-208
chr1-96130092-INS-1099
chr1-96136561-INS-4681
chr1-96160713-INS-938
chr1-96180773-INS-76
chr1-96197899-INS-443
chr1-96206778-INS-997
chr1-96213776-INS-6226
chr1-96215417-INS-331
chr1-96221862-INS-7928
chr1-96235433-INS-51
chr1-96235906-INS-3573
chr1-96238129-INS-492
chr1-96248894-INS-283
chr1-96249611-INS-54
chr1-96257545-INS-185
chr1-96269810-INS-78
chr1-96275611-INS-212
chr1-96277606-INS-392
chr1-96294200-INS-1226
chr1-96295435-INS-63
chr1-96298477-INS-233
chr1-96310354-INS-917
chr1-96311487-INS-495
chr1-96313936-INS-51
chr1-96338079-INS-56
chr1-96347394-INS-89
chr1-96352294-INS-339
chr1-96365111-INS-64
chr1-96384600-INS-4100
chr1-96385856-INS-205
chr1-96394160-INS-202
chr1-96409202-INS-113
chr1-96416689-INS-6284
chr1-96422359-INS-88
chr1-96442529-INS-51
chr1-96454113-INS-575
chr1-96485103-INS-6601
chr1-96529146-INS-131
chr1-96537522-INS-295
chr1-96550870-INS-2483
chr1-96557245-INS-95
chr1-96560061-INS-5906
chr1-96561962-INS-214
chr1-96565667-INS-1345
chr1-96566349-INS-60
chr1-96586873-INS-247
chr1-96596300-INS-3102
chr1-96604497-INS-76
chr1-96666328-INS-133
chr1-96675368-INS-94
chr1-96682340-INS-599
chr1-96685412-INS-125
chr1-96691540-INS-479
chr1-96703785-INS-509
chr1-96713188-INS-51
chr1-96723523-INS-53
chr1-96777563-INS-52
chr1-96777688-INS-510
chr1-96798337-INS-126
chr1-96813112-INS-7067
chr1-96833052-INS-6702
chr1-96835978-INS-497
chr1-96838937-INS-78
chr1-96839486-INS-6605
chr1-96840262-INS-91
chr1-96848228-INS-314
chr1-96854609-INS-186
chr1-96860695-INS-6401
chr1-96861087-INS-589
chr1-96885930-INS-212
chr1-96900906-INS-295
chr1-96918240-INS-75

chr1-96957767-INS-348
chr1-96981349-INS-99
chr1-97005713-INS-77
chr1-97031648-INS-276
chr1-97035177-INS-231
chr1-97041495-INS-6536
chr1-97055191-INS-410
chr1-97085706-INS-6774
chr1-97094092-INS-54
chr1-97096730-INS-91
chr1-97149537-INS-6563
chr1-97183106-INS-736
chr1-97216162-INS-91
chr1-97238311-INS-355
chr1-97263263-INS-6658
chr1-97263412-INS-52
chr1-97266884-INS-310
chr1-97271137-INS-1458
chr1-97272625-INS-200
chr1-97283128-INS-772
chr1-97283896-INS-7635
chr1-97292162-INS-56
chr1-97302824-INS-6295
chr1-97340275-INS-124
chr1-97397360-INS-538
chr1-97405234-INS-834
chr1-97408755-INS-1077
chr1-97433736-INS-447
chr1-97440598-INS-401
chr1-97480100-INS-819
chr1-97485017-INS-84
chr1-97496230-INS-609
chr1-97517349-INS-2920
chr1-97540138-INS-6482
chr1-97548040-INS-6592
chr1-97551636-INS-84
chr1-97565164-INS-149
chr1-97626053-INS-58
chr1-97628437-INS-450
chr1-97652688-INS-53
chr1-97661161-INS-55
chr1-97676434-INS-197
chr1-97681312-INS-88
chr1-97682132-INS-372
chr1-97714985-INS-273
chr1-97724142-INS-440
chr1-97729794-INS-449
chr1-97740979-INS-77
chr1-97776884-INS-6201
chr1-97790526-INS-6641
chr1-97809020-INS-64
chr1-97810549-INS-719
chr1-97814028-INS-298
chr1-97833852-INS-277
chr1-97846476-INS-180
chr1-97847295-INS-155
chr1-97874221-INS-61
chr1-97890912-INS-53
chr1-97892959-INS-690
chr1-97907602-INS-95
chr1-97909537-INS-58
chr1-97923480-INS-68
chr1-97934031-INS-132
chr1-97935798-INS-223
chr1-97937626-INS-104
chr1-97972779-INS-3411
chr1-98006003-INS-96
chr1-98019220-INS-52
chr1-98055664-INS-243
chr1-98070484-INS-88
chr1-98086956-INS-7087
chr1-98087714-INS-3690
chr1-98091858-INS-79
chr1-98096261-INS-484
chr1-98108643-INS-75
chr1-98114040-INS-206
chr1-98117393-INS-6532

chr1-98122873-INS-210
chr1-98128613-INS-510
chr1-98136849-INS-195
chr1-98146035-INS-346
chr1-98147077-INS-89
chr1-98149933-INS-85
chr1-98159312-INS-6442
chr1-98162358-INS-100
chr1-98173943-INS-126
chr1-98178902-INS-2364
chr1-98181706-INS-5276
chr1-98195851-INS-166
chr1-98199199-INS-173
chr1-98225873-INS-498
chr1-98227244-INS-54
chr1-98235137-INS-1257
chr1-98240369-INS-341
chr1-98241172-INS-6686
chr1-98243782-INS-1546
chr1-98274099-INS-306
chr1-98275775-INS-470
chr1-98289439-INS-191
chr1-98293015-INS-4511
chr1-98294177-INS-212
chr1-98303008-INS-77
chr1-98325235-INS-498
chr1-98330000-INS-203
chr1-98333350-INS-7629
chr1-98346000-INS-5088
chr1-98360229-INS-954
chr1-98375409-INS-59
chr1-98399315-INS-6668
chr1-98402775-INS-53
chr1-98405645-INS-189
chr1-98414436-INS-88
chr1-98422756-INS-2169
chr1-98437980-INS-6189
chr1-98445167-INS-225
chr1-98448317-INS-70
chr1-98452320-INS-334
chr1-98485440-INS-4017
chr1-98491625-INS-719
chr1-98496243-INS-6496
chr1-98505337-INS-461
chr1-98531934-INS-292
chr1-98556957-INS-6407
chr1-98561513-INS-85
chr1-98597564-INS-419
chr1-98599402-INS-352
chr1-98600211-INS-386
chr1-98607815-INS-79
chr1-98614029-INS-53
chr1-98640859-INS-132
chr1-98644534-INS-120
chr1-98655940-INS-368
chr1-98662000-INS-1519
chr1-98667688-INS-1073
chr1-98667973-INS-51
chr1-98671288-INS-403
chr1-98680500-INS-6283
chr1-98687318-INS-1071
chr1-98735578-INS-7637
chr1-98743414-INS-619
chr1-98758863-INS-498
chr1-98775351-INS-6479
chr1-98779404-INS-333
chr1-98795362-INS-71
chr1-98808096-INS-112
chr1-98819514-INS-1107
chr1-98834234-INS-57
chr1-98886949-INS-2528
chr1-98913875-INS-135
chr1-98920948-INS-146
chr1-98926049-INS-2514
chr1-98947083-INS-69
chr1-98960674-INS-96
chr1-98963529-INS-200

chr1-98993121-INS-505
chr1-98998850-INS-304
chr1-99000183-INS-132
chr1-99008181-INS-108
chr1-99051499-INS-154
chr1-99070921-INS-7188
chr1-99103371-INS-97
chr1-99109443-INS-188
chr1-99110228-INS-198
chr1-99118943-INS-71
chr1-99122077-INS-115
chr1-99139288-INS-96
chr1-99154923-INS-6279
chr1-99200115-INS-145
chr1-99213542-INS-241
chr1-99214825-INS-118
chr1-99228225-INS-106
chr1-99234968-INS-75
chr1-99244328-INS-64
chr1-99268688-INS-121
chr1-99269741-INS-212
chr1-99289424-INS-112
chr1-99301222-INS-59
chr1-99306939-INS-133
chr1-99309306-INS-52
chr1-99310650-INS-124
chr1-99317805-INS-2976
chr1-99321348-INS-72
chr1-99351919-INS-68
chr1-99371677-INS-941
chr1-99373473-INS-633
chr1-99391182-INS-84
chr1-99397316-INS-64
chr1-99428218-INS-70
chr1-99449000-INS-739
chr1-99480282-INS-144
chr1-99499924-INS-90
chr1-99503725-INS-332
chr1-99512090-INS-163
chr1-99550423-INS-474
chr1-99558729-INS-91
chr1-99570983-INS-68
chr1-99580862-INS-399
chr1-99592571-INS-365
chr1-99599680-INS-7349
chr1-99603591-INS-665
chr1-99628588-INS-900
chr1-99632517-INS-875
chr1-99633915-INS-65
chr1-99650287-INS-1858
chr1-99653801-INS-6216
chr1-99685867-INS-51
chr1-99689628-INS-60
chr1-99694443-INS-51
chr1-99697462-INS-150
chr1-99699741-INS-80
chr1-99706149-INS-162
chr1-99708887-INS-75
chr1-99735912-INS-5307
chr1-99789097-INS-81
chr1-99792283-INS-581
chr1-99793288-INS-437
chr1-99800508-INS-114
chr1-99813010-INS-182
chr1-99852824-INS-54
chr1-99859175-INS-69
chr1-99872338-INS-317
chr1-99881154-INS-81
chr1-99881286-INS-619
chr1-99884607-INS-379
chr1-99890248-INS-2933
chr1-99893091-INS-259
chr1-99910152-INS-129
chr1-99910670-INS-4311
chr1-99911278-INS-6567
chr1-99926802-INS-116
chr1-99950535-INS-54

chr1-99960779-INS-100
chr1-99966635-INS-6694
chr1-99980318-INS-3036
chr1-99984033-INS-6254
chr1-99989520-INS-193
chr1-100005147-INS-158
chr1-100011180-INS-7108
chr1-100035171-INS-72
chr1-100037774-INS-989
chr1-100038844-INS-459
chr1-100049295-INS-541
chr1-100064444-INS-87
chr1-100067522-INS-170
chr1-100069237-INS-72
chr1-100084096-INS-474
chr1-100097022-INS-62
chr1-100111438-INS-106
chr1-100166287-INS-86
chr1-100168487-INS-154
chr1-100235868-INS-60
chr1-100242021-INS-381
chr1-100248975-INS-54
chr1-100262490-INS-527
chr1-100288161-INS-137
chr1-100290411-INS-441
chr1-100293327-INS-357
chr1-100296461-INS-1452
chr1-100301819-INS-76
chr1-100324729-INS-410
chr1-100354074-INS-84
chr1-100379515-INS-53
chr1-100385246-INS-225
chr1-100389539-INS-586
chr1-100395189-INS-363
chr1-100436459-INS-56
chr1-100452248-INS-966
chr1-100466394-INS-58
chr1-100480898-INS-196
chr1-100487359-INS-201
chr1-100507786-INS-222
chr1-100507786-INS-365
chr1-100535392-INS-2793
chr1-100543324-INS-8756
chr1-100550985-INS-54
chr1-100553429-INS-6758
chr1-100561607-INS-342
chr1-100584849-INS-6175
chr1-100592209-INS-7247
chr1-100604120-INS-6826
chr1-100604554-INS-3951
chr1-100612500-INS-498
chr1-100618435-INS-78
chr1-100727029-INS-56
chr1-100804802-INS-216
chr1-100852678-INS-142
chr1-100854325-INS-739
chr1-100859094-INS-791
chr1-100859912-INS-74
chr1-100873996-INS-69
chr1-100877362-INS-72
chr1-100885292-INS-173
chr1-100905084-INS-64
chr1-100915113-INS-72
chr1-100992244-INS-62
chr1-100999717-INS-77
chr1-101013322-INS-740
chr1-101029665-INS-112
chr1-101200443-INS-114
chr1-101203650-INS-480
chr1-101205054-INS-6835
chr1-101220276-INS-117
chr1-101221648-INS-71
chr1-101227244-INS-72
chr1-101229403-INS-1468
chr1-101243974-INS-300
chr1-101264452-INS-204
chr1-101268385-INS-3941

chr1-101278443-INS-135
chr1-101280080-INS-613
chr1-101285103-INS-203
chr1-101286948-INS-6578
chr1-101326275-INS-78
chr1-101332432-INS-467
chr1-101336654-INS-177
chr1-101337764-INS-198
chr1-101341460-INS-3050
chr1-101353093-INS-162
chr1-101366269-INS-73
chr1-101370659-INS-1808
chr1-101392768-INS-326
chr1-101451177-INS-6769
chr1-101475170-INS-79
chr1-101481310-INS-482
chr1-101494025-INS-104
chr1-101495286-INS-69
chr1-101513876-INS-150
chr1-101550967-INS-6471
chr1-101574354-INS-320
chr1-101574358-INS-6389
chr1-101606874-INS-884
chr1-101608679-INS-157
chr1-101609882-INS-52
chr1-101610989-INS-608
chr1-101613545-INS-169
chr1-101624094-INS-6612
chr1-101658014-INS-62
chr1-101731867-INS-552
chr1-101735322-INS-70
chr1-101761969-INS-7181
chr1-101813988-INS-3950
chr1-101876972-INS-462
chr1-101879652-INS-244
chr1-101900092-INS-1941
chr1-101902966-INS-75
chr1-101905034-INS-6389
chr1-101924892-INS-116
chr1-101943366-INS-6667
chr1-101958773-INS-60
chr1-101962517-INS-303
chr1-102033915-INS-201
chr1-102085779-INS-6393
chr1-102102760-INS-150
chr1-102106273-INS-86
chr1-102161879-INS-100
chr1-102183279-INS-201
chr1-102201564-INS-119
chr1-102244813-INS-244
chr1-102259122-INS-72
chr1-102260072-INS-250
chr1-102279452-INS-6232
chr1-102279705-INS-66
chr1-102293980-INS-65
chr1-102316611-INS-391
chr1-102328721-INS-6688
chr1-102347302-INS-68
chr1-102349525-INS-6452
chr1-102357211-INS-6923
chr1-102358435-INS-71
chr1-102368252-INS-140
chr1-102373638-INS-398
chr1-102375303-INS-309
chr1-102384267-INS-539
chr1-102425104-INS-70
chr1-102431063-INS-76
chr1-102467708-INS-80
chr1-102485332-INS-147
chr1-102514986-INS-172
chr1-102534013-INS-245
chr1-102603678-INS-66
chr1-102616585-INS-204
chr1-102624976-INS-498
chr1-102629109-INS-52
chr1-102635112-INS-84
chr1-102662171-INS-345

chr1-102665238-INS-86
chr1-102720339-INS-105
chr1-102767026-INS-190
chr1-102773282-INS-88
chr1-102777495-INS-98
chr1-102784626-INS-6592
chr1-102789998-INS-228
chr1-102873009-INS-119
chr1-102882189-INS-334
chr1-102887941-INS-52
chr1-102928502-INS-214
chr1-102974167-INS-637
chr1-102986046-INS-601
chr1-103019133-INS-386
chr1-103032530-INS-969
chr1-103061909-INS-63
chr1-103093099-INS-314
chr1-103122170-INS-180
chr1-103143895-INS-51
chr1-103161238-INS-62
chr1-103166850-INS-138
chr1-103188420-INS-235
chr1-103229845-INS-60
chr1-103293395-INS-166
chr1-103347763-INS-60
chr1-103390351-INS-116
chr1-103411561-INS-105
chr1-103455969-INS-113
chr1-103458801-INS-52
chr1-103499236-INS-501
chr1-103509748-INS-6439
chr1-103552533-INS-56
chr1-103585963-INS-67
chr1-103604988-INS-5648
chr1-103614571-INS-1485
chr1-103634626-INS-162
chr1-103646106-INS-574
chr1-103654214-INS-54
chr1-103676526-INS-72
chr1-103677696-INS-64
chr1-103678777-INS-346
chr1-103686343-INS-445
chr1-103708108-INS-6246
chr1-103719083-INS-130
chr1-103727144-INS-72
chr1-103747340-INS-329
chr1-103754611-INS-328
chr1-103765152-INS-6590
chr1-103771388-INS-176
chr1-103787845-INS-159
chr1-103824724-INS-13977
chr1-103835085-INS-210
chr1-103838145-INS-658
chr1-103862732-INS-6309
chr1-103880105-INS-123
chr1-103885581-INS-7067
chr1-103894119-INS-74
chr1-103897511-INS-75
chr1-103914881-INS-96
chr1-103918543-INS-51
chr1-103931999-INS-111
chr1-103955266-INS-355
chr1-103979781-INS-142
chr1-103989211-INS-3272
chr1-103996646-INS-13817
chr1-104003428-INS-126
chr1-104009256-INS-204
chr1-104018011-INS-84
chr1-104060426-INS-58
chr1-104097718-INS-56
chr1-104103005-INS-100
chr1-104115018-INS-6264
chr1-104160316-INS-6818
chr1-104210128-INS-543
chr1-104234703-INS-51
chr1-104239046-INS-60
chr1-104240385-INS-116

chr1-104244279-INS-477
chr1-104247300-INS-67
chr1-104265429-INS-88
chr1-104280110-INS-3615
chr1-104296231-INS-204
chr1-104299213-INS-58
chr1-104319123-INS-404
chr1-104326123-INS-95
chr1-104341115-INS-84
chr1-104372522-INS-68
chr1-104403670-INS-5023
chr1-104404186-INS-340
chr1-104410060-INS-52
chr1-104415392-INS-7080
chr1-104457390-INS-190
chr1-104458144-INS-52
chr1-104465668-INS-210
chr1-104486280-INS-56
chr1-104492873-INS-445
chr1-104496702-INS-78
chr1-104524252-INS-7224
chr1-104527655-INS-588
chr1-104530867-INS-140
chr1-104531246-INS-877
chr1-104531475-INS-72
chr1-104534508-INS-60
chr1-104536954-INS-71
chr1-104551424-INS-75
chr1-104577821-INS-6440
chr1-104599128-INS-325
chr1-104599429-INS-967
chr1-104612156-INS-67
chr1-104621991-INS-94
chr1-104639712-INS-143
chr1-104656771-INS-52
chr1-104681739-INS-6474
chr1-104694368-INS-844
chr1-104709494-INS-61
chr1-104717827-INS-107
chr1-104735979-INS-52
chr1-104753912-INS-109
chr1-104776861-INS-4951
chr1-104795284-INS-6790
chr1-104816813-INS-477
chr1-104827023-INS-352
chr1-104850053-INS-332
chr1-104869561-INS-55
chr1-104874117-INS-78
chr1-104902875-INS-398
chr1-104904024-INS-352
chr1-104915502-INS-195
chr1-104930786-INS-266
chr1-104959301-INS-393
chr1-104965483-INS-111
chr1-104990672-INS-83
chr1-105006318-INS-72
chr1-105010384-INS-199
chr1-105023209-INS-194
chr1-105027778-INS-434
chr1-105040905-INS-964
chr1-105057782-INS-59
chr1-105062305-INS-671
chr1-105076917-INS-206
chr1-105152791-INS-268
chr1-105176577-INS-56
chr1-105189939-INS-386
chr1-105241796-INS-98
chr1-105254777-INS-220
chr1-105271510-INS-6858
chr1-105275415-INS-6436
chr1-105277187-INS-297
chr1-105289418-INS-56
chr1-105310206-INS-65
chr1-105312725-INS-126
chr1-105328697-INS-146
chr1-105342036-INS-6466
chr1-105348679-INS-265

chr1-105352582-INS-6488
chr1-105352688-INS-7172
chr1-105353597-INS-68
chr1-105356489-INS-387
chr1-105359997-INS-56
chr1-105362545-INS-392
chr1-105367705-INS-675
chr1-105371456-INS-428
chr1-105373058-INS-603
chr1-105418167-INS-454
chr1-105428874-INS-596
chr1-105437260-INS-254
chr1-105479050-INS-466
chr1-105489808-INS-6234
chr1-105549339-INS-75
chr1-105552856-INS-506
chr1-105580229-INS-726
chr1-105609092-INS-163
chr1-105624853-INS-190
chr1-105628952-INS-108
chr1-105652453-INS-59
chr1-105666993-INS-441
chr1-105698060-INS-216
chr1-105717743-INS-87
chr1-105739792-INS-63
chr1-105747470-INS-626
chr1-105748121-INS-56
chr1-105750451-INS-208
chr1-105767208-INS-211
chr1-105786290-INS-618
chr1-105791076-INS-108
chr1-105795414-INS-65
chr1-105810802-INS-69
chr1-105818321-INS-85
chr1-105821695-INS-354
chr1-105835013-INS-6440
chr1-105887964-INS-141
chr1-105900285-INS-67
chr1-105972362-INS-87
chr1-105975787-INS-76
chr1-106011138-INS-762
chr1-106012624-INS-984
chr1-106016790-INS-275
chr1-106020670-INS-53
chr1-106033765-INS-52
chr1-106036572-INS-103
chr1-106081530-INS-93
chr1-106096822-INS-7372
chr1-106125586-INS-106
chr1-106174995-INS-163
chr1-106190258-INS-553
chr1-106204911-INS-67
chr1-106209317-INS-807
chr1-106212050-INS-4490
chr1-106219237-INS-175
chr1-106229197-INS-56
chr1-106231410-INS-548
chr1-106258138-INS-3136
chr1-106260174-INS-6888
chr1-106278087-INS-116
chr1-106278912-INS-184
chr1-106285260-INS-133
chr1-106289859-INS-55
chr1-106294647-INS-76
chr1-106332669-INS-85
chr1-106338315-INS-181
chr1-106358314-INS-671
chr1-106365265-INS-6338
chr1-106370343-INS-264
chr1-106397417-INS-60
chr1-106409210-INS-76
chr1-106446390-INS-6413
chr1-106452158-INS-66
chr1-106475087-INS-58
chr1-106484698-INS-88
chr1-106518009-INS-71
chr1-106520956-INS-56

chr1-106549754-INS-56
chr1-106619101-INS-165
chr1-106645576-INS-1538
chr1-106645726-INS-274
chr1-106672381-INS-446
chr1-106756559-INS-100
chr1-106760180-INS-255
chr1-106762497-INS-104
chr1-106785175-INS-90
chr1-106790006-INS-55
chr1-106796327-INS-60
chr1-106856980-INS-504
chr1-106868380-INS-119
chr1-106871271-INS-404
chr1-106913376-INS-205
chr1-106918193-INS-613
chr1-106962075-INS-174
chr1-106979882-INS-72
chr1-106994277-INS-251
chr1-107006556-INS-208
chr1-107017984-INS-496
chr1-107034968-INS-3838
chr1-107039107-INS-568
chr1-107045593-INS-6379
chr1-107057800-INS-3893
chr1-107058076-INS-80
chr1-107112387-INS-57
chr1-107113406-INS-855
chr1-107113415-INS-856
chr1-107163833-INS-585
chr1-107173877-INS-98
chr1-107198209-INS-7399
chr1-107295920-INS-146
chr1-107334619-INS-57
chr1-107335478-INS-528
chr1-107347144-INS-228
chr1-107369872-INS-64
chr1-107418545-INS-83
chr1-107421423-INS-312
chr1-107422855-INS-234
chr1-107433249-INS-103
chr1-107448256-INS-56
chr1-107465864-INS-72
chr1-107477683-INS-518
chr1-107501893-INS-7347
chr1-107538993-INS-686
chr1-107547996-INS-103
chr1-107577432-INS-89
chr1-107593636-INS-371
chr1-107615217-INS-71
chr1-107618560-INS-197
chr1-107624052-INS-328
chr1-107625927-INS-134
chr1-107668331-INS-180
chr1-107701348-INS-404
chr1-107704431-INS-52
chr1-107761480-INS-6221
chr1-107770128-INS-62
chr1-107786422-INS-52
chr1-107824701-INS-60
chr1-107872359-INS-59
chr1-107876534-INS-119
chr1-107891184-INS-282
chr1-107898055-INS-330
chr1-107933705-INS-101
chr1-107952421-INS-337
chr1-107969588-INS-2245
chr1-108002945-INS-143
chr1-108006617-INS-56
chr1-108040481-INS-629
chr1-108053861-INS-181
chr1-108075260-INS-400
chr1-108085959-INS-155
chr1-108105026-INS-63
chr1-108106750-INS-64
chr1-108129583-INS-647
chr1-108134478-INS-158

chr1-108137446-INS-327
chr1-108155620-INS-211
chr1-108174036-INS-58
chr1-108178521-INS-6946
chr1-108188169-INS-13934
chr1-108189680-INS-142
chr1-108199078-INS-199
chr1-108230417-INS-659
chr1-108236803-INS-451
chr1-108262880-INS-7486
chr1-108264555-INS-510
chr1-108273132-INS-60
chr1-108316641-INS-74
chr1-108319399-INS-5630
chr1-108388557-INS-6401
chr1-108391447-INS-2673
chr1-108401944-INS-6732
chr1-108408478-INS-565
chr1-108411802-INS-66
chr1-108455202-INS-63
chr1-108457367-INS-63
chr1-108462017-INS-64
chr1-108462504-INS-2899
chr1-108470117-INS-7231
chr1-108484603-INS-475
chr1-108494505-INS-6170
chr1-108495285-INS-69
chr1-108534247-INS-201
chr1-108608974-INS-345
chr1-108620777-INS-127
chr1-108640660-INS-59
chr1-108712936-INS-55
chr1-108723611-INS-599
chr1-108729691-INS-7979
chr1-108747548-INS-200
chr1-108759425-INS-151
chr1-108762056-INS-83
chr1-108796298-INS-373
chr1-108798880-INS-63
chr1-108803763-INS-642
chr1-108815393-INS-158
chr1-108816737-INS-462
chr1-108834945-INS-424
chr1-108868789-INS-268
chr1-108871459-INS-51
chr1-108884757-INS-6582
chr1-108892324-INS-5032
chr1-108894368-INS-69
chr1-108906131-INS-54
chr1-108961955-INS-2231
chr1-109007378-INS-6804
chr1-109020980-INS-6252
chr1-109023745-INS-6769
chr1-109041455-INS-89
chr1-109054972-INS-293
chr1-109055074-INS-6536
chr1-109170180-INS-131
chr1-109183935-INS-2585
chr1-109213120-INS-54
chr1-109215365-INS-5141
chr1-109215402-INS-7138
chr1-109231446-INS-7135
chr1-109231891-INS-52
chr1-109256166-INS-975
chr1-109265607-INS-626
chr1-109286096-INS-236
chr1-109287358-INS-171
chr1-109288562-INS-341
chr1-109299197-INS-296
chr1-109304875-INS-120
chr1-109403120-INS-6480
chr1-109427730-INS-53
chr1-109526903-INS-59
chr1-109528633-INS-501
chr1-109536998-INS-88
chr1-109559089-INS-6489
chr1-109560634-INS-816

chr1-109564074-INS-213
chr1-109574520-INS-102
chr1-109579863-INS-339
chr1-109587579-INS-92
chr1-109594561-INS-90
chr1-109598915-INS-76
chr1-109606879-INS-208
chr1-109607098-INS-7161
chr1-109622143-INS-193
chr1-109623242-INS-498
chr1-109672713-INS-400
chr1-109695048-INS-158
chr1-109715859-INS-441
chr1-109746691-INS-792
chr1-109749584-INS-180
chr1-109749710-INS-598
chr1-109758976-INS-130
chr1-109764479-INS-63
chr1-109768682-INS-6442
chr1-109769347-INS-6372
chr1-109769699-INS-437
chr1-109819481-INS-52
chr1-109822487-INS-60
chr1-109824887-INS-439
chr1-109841738-INS-411
chr1-109862423-INS-76
chr1-109870682-INS-186
chr1-109872969-INS-362
chr1-109873780-INS-170
chr1-109884257-INS-54
chr1-109896904-INS-73
chr1-109908017-INS-72
chr1-109919441-INS-70
chr1-109927747-INS-6511
chr1-109933638-INS-52
chr1-109944504-INS-406
chr1-109989549-INS-60
chr1-110004891-INS-141
chr1-110034452-INS-120
chr1-110041096-INS-58
chr1-110045100-INS-65
chr1-110048141-INS-532
chr1-110075933-INS-499
chr1-110111733-INS-418
chr1-110112230-INS-52
chr1-110146859-INS-92
chr1-110189648-INS-60
chr1-110281917-INS-167
chr1-110306993-INS-748
chr1-110321934-INS-366
chr1-110377123-INS-60
chr1-110401485-INS-102
chr1-110412717-INS-674
chr1-110420216-INS-257
chr1-110464217-INS-58
chr1-110467225-INS-577
chr1-110493039-INS-98
chr1-110501503-INS-86
chr1-110508396-INS-753
chr1-110509584-INS-6882
chr1-110511202-INS-6663
chr1-110529562-INS-80
chr1-110530123-INS-703
chr1-110537236-INS-211
chr1-110541002-INS-4683
chr1-110554301-INS-6691
chr1-110559437-INS-1661
chr1-110623877-INS-6270
chr1-110624480-INS-205
chr1-110648005-INS-56
chr1-110648549-INS-55
chr1-110672130-INS-59
chr1-110698464-INS-73
chr1-110726652-INS-56
chr1-110742406-INS-77
chr1-110764662-INS-212
chr1-110785671-INS-94

chr1-110829219-INS-243
chr1-110843941-INS-254
chr1-110874387-INS-62
chr1-110922879-INS-3573
chr1-110934185-INS-58
chr1-110940247-INS-574
chr1-110968342-INS-5251
chr1-110977621-INS-6530
chr1-110997795-INS-959
chr1-111001361-INS-60
chr1-111013179-INS-2395
chr1-111055857-INS-218
chr1-111080340-INS-69
chr1-111101641-INS-74
chr1-111107387-INS-59
chr1-111111320-INS-100
chr1-111176653-INS-58
chr1-111228265-INS-209
chr1-111230498-INS-4434
chr1-111248714-INS-76
chr1-111263818-INS-475
chr1-111269873-INS-84
chr1-111271784-INS-138
chr1-111288646-INS-491
chr1-111298124-INS-131
chr1-111302157-INS-844
chr1-111318368-INS-160
chr1-111331969-INS-130
chr1-111349404-INS-150
chr1-111402699-INS-487
chr1-111406193-INS-397
chr1-111411879-INS-68
chr1-111482532-INS-6191
chr1-111488283-INS-4372
chr1-111489317-INS-796
chr1-111496884-INS-265
chr1-111531983-INS-105
chr1-111539140-INS-746
chr1-111542825-INS-66
chr1-111591653-INS-134
chr1-111598544-INS-113
chr1-111609494-INS-54
chr1-111651909-INS-52
chr1-111655630-INS-54
chr1-111671685-INS-343
chr1-111691686-INS-592
chr1-111696943-INS-385
chr1-111719269-INS-624
chr1-111720763-INS-6984
chr1-111737410-INS-140
chr1-111776149-INS-227
chr1-111803070-INS-256
chr1-111845023-INS-171
chr1-111853484-INS-58
chr1-111867134-INS-127
chr1-111870693-INS-508
chr1-111880629-INS-6058
chr1-111900848-INS-116
chr1-111965794-INS-5551
chr1-111969934-INS-7322
chr1-111976779-INS-457
chr1-112018922-INS-54
chr1-112024166-INS-5505
chr1-112032440-INS-77
chr1-112041326-INS-3374
chr1-112046720-INS-6136
chr1-112104157-INS-593
chr1-112119787-INS-79
chr1-112148103-INS-212
chr1-112180002-INS-224
chr1-112205113-INS-54
chr1-112215746-INS-6405
chr1-112224751-INS-8844
chr1-112241168-INS-54
chr1-112247837-INS-6154
chr1-112265967-INS-100
chr1-112273819-INS-58

chr1-112280124-INS-929
chr1-112289942-INS-6787
chr1-112303774-INS-54
chr1-112320936-INS-56
chr1-112334367-INS-119
chr1-112362946-INS-74
chr1-112429269-INS-7179
chr1-112444652-INS-1889
chr1-112454531-INS-2390
chr1-112463162-INS-3703
chr1-112473577-INS-6390
chr1-112490561-INS-1014
chr1-112501540-INS-548
chr1-112512707-INS-139
chr1-112517624-INS-498
chr1-112530281-INS-843
chr1-112565016-INS-226
chr1-112840120-INS-88
chr1-112853888-INS-452
chr1-112874828-INS-540
chr1-112938165-INS-196
chr1-112953377-INS-74
chr1-112958931-INS-6652
chr1-112972484-INS-557
chr1-112991279-INS-6132
chr1-112994683-INS-7220
chr1-113007882-INS-125
chr1-113032320-INS-62
chr1-113034171-INS-57
chr1-113038360-INS-329
chr1-113071711-INS-138
chr1-113072087-INS-120
chr1-113077352-INS-57
chr1-113140396-INS-7007
chr1-113164142-INS-62
chr1-113193436-INS-620
chr1-113224148-INS-6403
chr1-113234721-INS-6381
chr1-113235368-INS-84
chr1-113253336-INS-133
chr1-113285241-INS-140
chr1-113301093-INS-333
chr1-113307047-INS-6394
chr1-113308869-INS-397
chr1-113332631-INS-86
chr1-113374451-INS-175
chr1-113386898-INS-527
chr1-113428753-INS-69
chr1-113439646-INS-7004
chr1-113484302-INS-163
chr1-113524757-INS-66
chr1-113589410-INS-52
chr1-113612711-INS-75
chr1-113628748-INS-6909
chr1-113637968-INS-276
chr1-113677458-INS-6401
chr1-113688608-INS-6401
chr1-113692630-INS-255
chr1-113693915-INS-719
chr1-113709471-INS-94
chr1-113721751-INS-170
chr1-113743777-INS-412
chr1-113808859-INS-7053
chr1-113830183-INS-7100
chr1-113841314-INS-6386
chr1-113841315-INS-224
chr1-113850012-INS-60
chr1-113857060-INS-115
chr1-113861489-INS-2593
chr1-113871608-INS-85
chr1-113889836-INS-76
chr1-113892749-INS-98
chr1-113903142-INS-91
chr1-113916495-INS-639
chr1-113950159-INS-221
chr1-113974757-INS-196
chr1-114000528-INS-1892

chr1-114019722-INS-59
chr1-114057302-INS-395
chr1-114069433-INS-144
chr1-114073405-INS-70
chr1-114091748-INS-3129
chr1-114138302-INS-68
chr1-114148840-INS-72
chr1-114177578-INS-1888
chr1-114185521-INS-140
chr1-114197387-INS-219
chr1-114264702-INS-52
chr1-114288835-INS-65
chr1-114327463-INS-92
chr1-114345970-INS-172
chr1-114365689-INS-1079
chr1-114367736-INS-395
chr1-114398581-INS-1791
chr1-114452992-INS-67
chr1-114462884-INS-651
chr1-114473390-INS-5242
chr1-114479978-INS-202
chr1-114485270-INS-212
chr1-114520848-INS-6391
chr1-114538988-INS-5314
chr1-114614153-INS-105
chr1-114621895-INS-466
chr1-114653307-INS-6532
chr1-114682139-INS-6207
chr1-114701047-INS-105
chr1-114720304-INS-6475
chr1-114724791-INS-594
chr1-114724883-INS-122
chr1-114726888-INS-1253
chr1-114740086-INS-273
chr1-114742462-INS-732
chr1-114746727-INS-60
chr1-114756490-INS-135
chr1-114771142-INS-206
chr1-114775216-INS-56
chr1-114777813-INS-669
chr1-114778840-INS-202
chr1-114813379-INS-179
chr1-114831163-INS-6541
chr1-114847598-INS-140
chr1-114857436-INS-168
chr1-114865890-INS-731
chr1-114883388-INS-900
chr1-114895307-INS-8734
chr1-114905829-INS-56
chr1-114906494-INS-428
chr1-114931762-INS-619
chr1-114937070-INS-170
chr1-114937070-INS-957
chr1-114956734-INS-52
chr1-114982348-INS-6378
chr1-114991598-INS-2525
chr1-114997964-INS-257
chr1-115010945-INS-54
chr1-115064166-INS-416
chr1-115077756-INS-312
chr1-115082619-INS-56
chr1-115092350-INS-329
chr1-115171377-INS-5039
chr1-115182122-INS-51
chr1-115187473-INS-180
chr1-115192375-INS-237
chr1-115196232-INS-64
chr1-115197958-INS-71
chr1-115201692-INS-99
chr1-115218971-INS-138
chr1-115223415-INS-90
chr1-115244273-INS-107
chr1-115248627-INS-8734
chr1-115255535-INS-54
chr1-115264815-INS-58
chr1-115368412-INS-6601
chr1-115399704-INS-6476

chr1-115474858-INS-117
chr1-115485642-INS-328
chr1-115535290-INS-57
chr1-115562140-INS-3342
chr1-115591282-INS-212
chr1-115656089-INS-212
chr1-115663272-INS-3827
chr1-115665321-INS-71
chr1-115674521-INS-146
chr1-115699568-INS-198
chr1-115706439-INS-3309
chr1-115711045-INS-685
chr1-115711066-INS-56
chr1-115714543-INS-302
chr1-115748359-INS-1661
chr1-115768688-INS-6330
chr1-115772795-INS-973
chr1-115773011-INS-126
chr1-115789065-INS-435
chr1-115803048-INS-6241
chr1-115803952-INS-94
chr1-115829196-INS-6375
chr1-115837564-INS-92
chr1-115855076-INS-159
chr1-115856913-INS-156
chr1-115858006-INS-521
chr1-115866845-INS-322
chr1-115869671-INS-74
chr1-115874765-INS-62
chr1-115881777-INS-233
chr1-115927720-INS-107
chr1-115939187-INS-698
chr1-115953409-INS-56
chr1-115968177-INS-193
chr1-115969321-INS-71
chr1-115988407-INS-72
chr1-115991077-INS-324
chr1-115994334-INS-1892
chr1-115996987-INS-1376
chr1-116000998-INS-124
chr1-116002723-INS-201
chr1-116005646-INS-8192
chr1-116010495-INS-52
chr1-116018000-INS-59
chr1-116024368-INS-60
chr1-116036729-INS-673
chr1-116050209-INS-190
chr1-116052673-INS-91
chr1-116085458-INS-317
chr1-116086908-INS-59
chr1-116093667-INS-426
chr1-116095465-INS-55
chr1-116098285-INS-814
chr1-116104612-INS-120
chr1-116108601-INS-282
chr1-116145391-INS-112
chr1-116160608-INS-620
chr1-116188174-INS-54
chr1-116207871-INS-6360
chr1-116217110-INS-6381
chr1-116240706-INS-3469
chr1-116245449-INS-1712
chr1-116255443-INS-932
chr1-116300155-INS-359
chr1-116305892-INS-51
chr1-116328868-INS-6800
chr1-116340141-INS-102
chr1-116379463-INS-804
chr1-116394713-INS-86
chr1-116418339-INS-262
chr1-116427238-INS-398
chr1-116429642-INS-364
chr1-116440606-INS-3065
chr1-116450616-INS-197
chr1-116464981-INS-348
chr1-116489470-INS-70
chr1-116496938-INS-55

chr1-116515957-INS-70
chr1-116564472-INS-485
chr1-116565231-INS-6701
chr1-116567264-INS-4350
chr1-116585655-INS-51
chr1-116595914-INS-57
chr1-116676773-INS-52
chr1-116679645-INS-6253
chr1-116681065-INS-380
chr1-116716343-INS-64
chr1-116740699-INS-58
chr1-116780160-INS-76
chr1-116801555-INS-737
chr1-116836030-INS-59
chr1-116838805-INS-106
chr1-116839948-INS-350
chr1-116842087-INS-2350
chr1-116844046-INS-6059
chr1-116851915-INS-1136
chr1-116857378-INS-6761
chr1-116868440-INS-58
chr1-116877345-INS-464
chr1-116885641-INS-72
chr1-116909499-INS-546
chr1-116923193-INS-168
chr1-116929550-INS-6826
chr1-116937294-INS-70
chr1-116941373-INS-776
chr1-117027711-INS-103
chr1-117032178-INS-6415
chr1-117063180-INS-6363
chr1-117097351-INS-6939
chr1-117157354-INS-5169
chr1-117163719-INS-295
chr1-117169299-INS-87
chr1-117223054-INS-144
chr1-117230432-INS-335
chr1-117249773-INS-5426
chr1-117269955-INS-758
chr1-117270589-INS-373
chr1-117278133-INS-121
chr1-117296076-INS-699
chr1-117324370-INS-108
chr1-117340370-INS-84
chr1-117343650-INS-56
chr1-117350042-INS-6154
chr1-117372427-INS-549
chr1-117373076-INS-450
chr1-117389943-INS-522
chr1-117447505-INS-305
chr1-117455072-INS-76
chr1-117456764-INS-51
chr1-117462454-INS-61
chr1-117467275-INS-6656
chr1-117490123-INS-315
chr1-117508357-INS-194
chr1-117523780-INS-94
chr1-117527053-INS-6682
chr1-117533662-INS-6601
chr1-117600817-INS-888
chr1-117645350-INS-181
chr1-117647245-INS-535
chr1-117683228-INS-2891
chr1-117707045-INS-6795
chr1-117734122-INS-122
chr1-117838733-INS-4564
chr1-117870433-INS-153
chr1-117870758-INS-6665
chr1-117876915-INS-6434
chr1-117910084-INS-221
chr1-117914188-INS-336
chr1-117975273-INS-160
chr1-118024051-INS-8626
chr1-118030637-INS-7072
chr1-118036509-INS-8626
chr1-118047792-INS-5397
chr1-118061972-INS-1202

chr1-118090342-INS-97
chr1-118128515-INS-168
chr1-118131660-INS-696
chr1-118139873-INS-212
chr1-118144882-INS-3838
chr1-118156651-INS-4522
chr1-118170979-INS-6381
chr1-118188467-INS-5449
chr1-118195178-INS-6488
chr1-118198865-INS-1866
chr1-118203140-INS-321
chr1-118231570-INS-242
chr1-118247028-INS-69
chr1-118268372-INS-9047
chr1-118278243-INS-396
chr1-118288943-INS-1888
chr1-118315720-INS-52
chr1-118320258-INS-96
chr1-118329583-INS-69
chr1-118332115-INS-215
chr1-118334929-INS-54
chr1-118348547-INS-153
chr1-118372548-INS-103
chr1-118388581-INS-8948
chr1-118422769-INS-56
chr1-118424340-INS-303
chr1-118430424-INS-78
chr1-118461847-INS-80
chr1-118478222-INS-174
chr1-118480393-INS-5227
chr1-118486999-INS-160
chr1-118506198-INS-54
chr1-118563028-INS-211
chr1-118582029-INS-58
chr1-118590354-INS-69
chr1-118596622-INS-408
chr1-118616478-INS-69
chr1-118624958-INS-186
chr1-118639188-INS-177
chr1-118640170-INS-62
chr1-118655974-INS-53
chr1-118663512-INS-52
chr1-118665887-INS-70
chr1-118683039-INS-95
chr1-118712090-INS-209
chr1-118721469-INS-700
chr1-118725197-INS-250
chr1-118750369-INS-1892
chr1-118790846-INS-80
chr1-118809249-INS-88
chr1-118856093-INS-204
chr1-118884600-INS-60
chr1-118896476-INS-92
chr1-118931496-INS-75
chr1-118970239-INS-202
chr1-118972330-INS-88
chr1-118977098-INS-64
chr1-118991345-INS-184
chr1-118995308-INS-99
chr1-118999396-INS-894
chr1-119000788-INS-179
chr1-119014444-INS-158
chr1-119016613-INS-196
chr1-119024389-INS-185
chr1-119033542-INS-2667
chr1-119041647-INS-54
chr1-119042329-INS-64
chr1-119045992-INS-115
chr1-119070274-INS-56
chr1-119087309-INS-145
chr1-119102097-INS-53
chr1-119156184-INS-109
chr1-119206923-INS-55
chr1-119209214-INS-273
chr1-119217228-INS-51
chr1-119269122-INS-211
chr1-119296467-INS-72

chr1-119304862-INS-978
chr1-119320106-INS-147
chr1-119333238-INS-140
chr1-119347824-INS-78
chr1-119352219-INS-62
chr1-119373851-INS-54
chr1-119384260-INS-82
chr1-119393883-INS-187
chr1-119403285-INS-381
chr1-119406000-INS-104
chr1-119413182-INS-298
chr1-119415857-INS-55
chr1-119418985-INS-174
chr1-119440035-INS-60
chr1-119443989-INS-121
chr1-119458259-INS-492
chr1-119459517-INS-2416
chr1-119478638-INS-95
chr1-119481607-INS-500
chr1-119514932-INS-203
chr1-119526904-INS-393
chr1-119555026-INS-96
chr1-119567533-INS-168
chr1-119567977-INS-682
chr1-119576892-INS-66
chr1-119604262-INS-140
chr1-119637572-INS-5086
chr1-119663651-INS-1433
chr1-119686156-INS-220
chr1-119712005-INS-296
chr1-119728215-INS-177
chr1-119732630-INS-73
chr1-119738763-INS-173
chr1-119780348-INS-64
chr1-119808029-INS-140
chr1-119811671-INS-96
chr1-119817725-INS-242
chr1-119821333-INS-56
chr1-119832662-INS-216
chr1-119860479-INS-173
chr1-119878489-INS-392
chr1-119908095-INS-110
chr1-119918128-INS-56
chr1-119926416-INS-790
chr1-119927779-INS-228
chr1-119929527-INS-55
chr1-119945838-INS-433
chr1-119968637-INS-195
chr1-119973490-INS-138
chr1-119975047-INS-240
chr1-119976412-INS-179
chr1-120008765-INS-59
chr1-120061890-INS-84
chr1-120087565-INS-109
chr1-120103354-INS-203
chr1-120117624-INS-183
chr1-120133493-INS-146
chr1-120138788-INS-90
chr1-120141465-INS-55
chr1-120173054-INS-246
chr1-120185666-INS-194
chr1-120207687-INS-65
chr1-120212881-INS-96
chr1-120214525-INS-60
chr1-120237995-INS-98
chr1-120245554-INS-440
chr1-120263240-INS-98
chr1-120265245-INS-176
chr1-120267937-INS-395
chr1-120285342-INS-1058
chr1-120295267-INS-65
chr1-120340378-INS-137
chr1-120345240-INS-114
chr1-120441523-INS-57
chr1-120446944-INS-56
chr1-120450700-INS-178
chr1-120459909-INS-160

chr1-120470218-INS-133
chr1-120476745-INS-84
chr1-120506935-INS-144
chr1-120548590-INS-158
chr1-120557687-INS-263
chr1-120562980-INS-122
chr1-120584203-INS-58
chr1-120585836-INS-160
chr1-120592162-INS-93
chr1-120602411-INS-66
chr1-120608445-INS-392
chr1-120609319-INS-52
chr1-120611583-INS-513
chr1-120627574-INS-105
chr1-120679465-INS-153
chr1-120695697-INS-364
chr1-120702025-INS-73
chr1-120703659-INS-92
chr1-120713301-INS-59
chr1-120728613-INS-58
chr1-120749008-INS-6483
chr1-120749165-INS-928
chr1-120762020-INS-146
chr1-120814660-INS-53
chr1-120816104-INS-90
chr1-120835078-INS-80
chr1-120849195-INS-104
chr1-120855623-INS-121
chr1-120875430-INS-54
chr1-120878365-INS-926
chr1-120881395-INS-109
chr1-120886211-INS-484
chr1-120909107-INS-91
chr1-120937119-INS-54
chr1-120937455-INS-400
chr1-120999856-INS-503
chr1-121003424-INS-2634
chr1-121013703-INS-56
chr1-121015457-INS-565
chr1-121022440-INS-104
chr1-121046807-INS-121
chr1-121076827-INS-6144
chr1-121093752-INS-105
chr1-121111374-INS-70
chr1-121118190-INS-508
chr1-121119720-INS-54
chr1-121125951-INS-401
chr1-121126600-INS-154
chr1-121144122-INS-208
chr1-121148183-INS-102
chr1-121158265-INS-204
chr1-121158600-INS-6442
chr1-121210302-INS-413
chr1-121214400-INS-66
chr1-121224139-INS-72
chr1-121228185-INS-156
chr1-121240798-INS-721
chr1-121246399-INS-1888
chr1-121246499-INS-715
chr1-121251692-INS-170
chr1-121290958-INS-54
chr1-121294054-INS-72
chr1-121297385-INS-56
chr1-121313260-INS-311
chr1-121331777-INS-58
chr1-121362691-INS-53
chr1-121384371-INS-63
chr1-121389898-INS-384
chr1-121396592-INS-56
chr1-121423818-INS-168
chr1-121447857-INS-56
chr1-121471645-INS-571
chr1-121489951-INS-231
chr1-121496304-INS-148
chr1-121506119-INS-240
chr1-121522235-INS-335
chr1-121537281-INS-200

chr1-121561739-INS-790
chr1-121582894-INS-498
chr1-121584712-INS-77
chr1-121627312-INS-76
chr1-121658236-INS-6362
chr1-121663475-INS-108
chr1-121688541-INS-6579
chr1-121735623-INS-6784
chr1-121741445-INS-99
chr1-121760396-INS-52
chr1-121761625-INS-328
chr1-121773836-INS-52
chr1-121792230-INS-861
chr1-121794006-INS-262
chr1-121796024-INS-617
chr1-121800245-INS-66
chr1-121813520-INS-552
chr1-121831677-INS-179
chr1-121842171-INS-98
chr1-121845194-INS-59
chr1-121850312-INS-112
chr1-121892484-INS-192
chr1-121895533-INS-84
chr1-121897404-INS-201
chr1-121899152-INS-60
chr1-121900916-INS-170
chr1-121906480-INS-70
chr1-121907237-INS-57
chr1-121924036-INS-126
chr1-121927583-INS-263
chr1-121931682-INS-548
chr1-121940112-INS-54
chr1-121952493-INS-106
chr1-121955988-INS-76
chr1-121977900-INS-66
chr1-121999301-INS-104
chr1-122011011-INS-56
chr1-122013628-INS-90
chr1-122029769-INS-4154
chr1-122032486-INS-53
chr1-122042495-INS-6401
chr1-122068985-INS-152
chr1-122072813-INS-5939
chr1-122108904-INS-116
chr1-122121261-INS-116
chr1-122126531-INS-60
chr1-122131622-INS-438
chr1-122142968-INS-467
chr1-122212274-INS-222
chr1-122216196-INS-932
chr1-122230212-INS-234
chr1-122232011-INS-1032
chr1-122237944-INS-77
chr1-122241862-INS-184
chr1-122276089-INS-395
chr1-122300599-INS-70
chr1-122326791-INS-87
chr1-122334215-INS-238
chr1-122340064-INS-58
chr1-122363963-INS-356
chr1-122384127-INS-454
chr1-122384632-INS-993
chr1-122386866-INS-365
chr1-122388872-INS-6658
chr1-122403368-INS-56
chr1-122406362-INS-96
chr1-122416694-INS-52
chr1-122423481-INS-72
chr1-122459902-INS-52
chr1-122465971-INS-367
chr1-122492818-INS-326
chr1-122497124-INS-645
chr1-122514364-INS-54
chr1-122515820-INS-112
chr1-122527332-INS-120
chr1-122533424-INS-51
chr1-122535923-INS-53

chr1-122538171-INS-76
chr1-122543277-INS-111
chr1-122548673-INS-1912
chr1-122560463-INS-68
chr1-122573218-INS-7908
chr1-122577631-INS-5227
chr1-122625547-INS-70
chr1-122634098-INS-85
chr1-122658456-INS-7360
chr1-122664967-INS-70
chr1-122668135-INS-7092
chr1-122674342-INS-356
chr1-122684427-INS-6703
chr1-122704694-INS-6521
chr1-122728965-INS-470
chr1-122729111-INS-2675
chr1-122737420-INS-72
chr1-122755611-INS-64
chr1-122760052-INS-276
chr1-122760659-INS-7126
chr1-122763394-INS-92
chr1-122782697-INS-88
chr1-122814640-INS-65
chr1-122834873-INS-278
chr1-122836723-INS-169
chr1-122851262-INS-76
chr1-122905740-INS-6625
chr1-122913326-INS-62
chr1-122955841-INS-255
chr1-122972335-INS-165
chr1-122989307-INS-132
chr1-122994170-INS-201
chr1-123024689-INS-57
chr1-123026156-INS-839
chr1-123032060-INS-6147
chr1-123055139-INS-116
chr1-123058408-INS-93
chr1-123104483-INS-82
chr1-123110722-INS-704
chr1-123114548-INS-108
chr1-123120813-INS-208
chr1-123159729-INS-79
chr1-123177866-INS-142
chr1-123194757-INS-54
chr1-123196597-INS-131
chr1-123213210-INS-381
chr1-123231442-INS-1434
chr1-123231610-INS-300
chr1-123231647-INS-3311
chr1-123231648-INS-2032
chr1-123231649-INS-1518
chr1-123237284-INS-60
chr1-123254401-INS-951
chr1-123279887-INS-69
chr1-123283555-INS-1766
chr1-123306147-INS-6193
chr1-123341387-INS-358
chr1-123343882-INS-156
chr1-123371986-INS-199
chr1-123376673-INS-166
chr1-123383515-INS-225
chr1-123398941-INS-62
chr1-123401057-INS-53
chr1-123419447-INS-2811
chr1-123448967-INS-6516
chr1-123452293-INS-70
chr1-123458712-INS-2021
chr1-123485225-INS-134
chr1-123514915-INS-856
chr1-123529810-INS-783
chr1-123542471-INS-970
chr1-123546537-INS-92
chr1-123555611-INS-6483
chr1-123557018-INS-261
chr1-123559587-INS-6679
chr1-123600655-INS-56
chr1-123619757-INS-528

chr1-123631065-INS-307
chr1-123647197-INS-6442
chr1-123652138-INS-66
chr1-123657091-INS-815
chr1-123659160-INS-1819
chr1-123659519-INS-77
chr1-123677491-INS-6431
chr1-123698367-INS-257
chr1-123700958-INS-66
chr1-123709395-INS-1190
chr1-123709408-INS-144
chr1-123709456-INS-149
chr1-123719916-INS-202
chr1-123728999-INS-336
chr1-123729627-INS-7590
chr1-123739851-INS-51
chr1-123746356-INS-63
chr1-123751155-INS-115
chr1-123762968-INS-1757
chr1-123774812-INS-1209
chr1-123778306-INS-965
chr1-123795108-INS-72
chr1-123814410-INS-145
chr1-123829566-INS-53
chr1-123838271-INS-180
chr1-123843679-INS-212
chr1-123852594-INS-1176
chr1-123871359-INS-166
chr1-123875991-INS-198
chr1-123889697-INS-108
chr1-123895498-INS-433
chr1-123912015-INS-54
chr1-123915501-INS-376
chr1-123917156-INS-205
chr1-123936936-INS-189
chr1-123944300-INS-406
chr1-123949639-INS-55
chr1-123951405-INS-163
chr1-123957548-INS-839
chr1-123960348-INS-60
chr1-123980318-INS-170
chr1-123982668-INS-137
chr1-123983393-INS-52
chr1-123990424-INS-84
chr1-124001478-INS-78
chr1-124009048-INS-54
chr1-124020109-INS-65
chr1-124032142-INS-60
chr1-124041145-INS-68
chr1-124057129-INS-51
chr1-124104342-INS-73
chr1-124105865-INS-52
chr1-124140549-INS-659
chr1-124143674-INS-79
chr1-124152937-INS-52
chr1-124154211-INS-203
chr1-124173860-INS-92
chr1-124178840-INS-1058
chr1-124193707-INS-701
chr1-124203202-INS-6570
chr1-124206479-INS-54
chr1-124208478-INS-1311
chr1-124243800-INS-70
chr1-124284241-INS-260
chr1-124297147-INS-504
chr1-124299277-INS-1154
chr1-124319138-INS-387
chr1-124324848-INS-852
chr1-124333390-INS-888
chr1-124346244-INS-118
chr1-124347570-INS-70
chr1-124350278-INS-359
chr1-124373082-INS-68
chr1-124383044-INS-927
chr1-124390677-INS-316
chr1-124391751-INS-227
chr1-124396339-INS-390

chr1-124413900-INS-455
chr1-124424741-INS-181
chr1-124431348-INS-90
chr1-124466485-INS-108
chr1-124483339-INS-997
chr1-124527327-INS-290
chr1-124551801-INS-74
chr1-124576642-INS-260
chr1-124602646-INS-162
chr1-124610281-INS-6440
chr1-124613584-INS-53
chr1-124621164-INS-285
chr1-124622232-INS-331
chr1-124627097-INS-214
chr1-124649735-INS-59
chr1-124654307-INS-51
chr1-124666402-INS-685
chr1-124673482-INS-154
chr1-124683940-INS-140
chr1-124757217-INS-547
chr1-124759769-INS-80
chr1-124764720-INS-4247
chr1-124793246-INS-64
chr1-124800070-INS-868
chr1-124813360-INS-58
chr1-124821823-INS-450
chr1-124829254-INS-876
chr1-124833128-INS-74
chr1-124867532-INS-62
chr1-124891833-INS-81
chr1-124892342-INS-99
chr1-124960460-INS-68
chr1-124965130-INS-350
chr1-124975816-INS-52
chr1-125072984-INS-66
chr1-125081313-INS-97
chr1-125110319-INS-61
chr1-125177637-INS-201
chr1-125193329-INS-198
chr1-125228275-INS-72
chr1-125236775-INS-193
chr1-125253164-INS-68
chr1-125280052-INS-986
chr1-125317785-INS-57
chr1-125342206-INS-111
chr1-125342649-INS-172
chr1-125345189-INS-395
chr1-125352941-INS-159
chr1-125364894-INS-80
chr1-125401640-INS-8720
chr1-125438262-INS-62
chr1-125443923-INS-224
chr1-125446462-INS-70
chr1-125458829-INS-196
chr1-125468702-INS-60
chr1-125471193-INS-79
chr1-125475844-INS-405
chr1-125531405-INS-72
chr1-125544628-INS-58
chr1-125550354-INS-70
chr1-125568287-INS-8284
chr1-125573119-INS-72
chr1-125594873-INS-322
chr1-125596569-INS-58
chr1-125602550-INS-591
chr1-125619332-INS-66
chr1-125635915-INS-168
chr1-125640785-INS-7111
chr1-125674835-INS-68
chr1-125675789-INS-64
chr1-125686412-INS-6733
chr1-125689098-INS-62
chr1-125698355-INS-118
chr1-125700335-INS-66
chr1-125707534-INS-104
chr1-125756086-INS-340
chr1-125760032-INS-66

chr1-125767415-INS-112
chr1-125770844-INS-115
chr1-125805488-INS-439
chr1-125809901-INS-124
chr1-125849993-INS-56
chr1-125857108-INS-52
chr1-125918786-INS-54
chr1-125925957-INS-610
chr1-125959766-INS-196
chr1-125973295-INS-60
chr1-125988344-INS-111
chr1-126006526-INS-69
chr1-126009726-INS-188
chr1-126012878-INS-72
chr1-126019604-INS-69
chr1-126020222-INS-238
chr1-126034681-INS-52
chr1-126051216-INS-117
chr1-126089830-INS-403
chr1-126094811-INS-87
chr1-126097461-INS-72
chr1-126115864-INS-55
chr1-126129299-INS-167
chr1-126158141-INS-436
chr1-126177707-INS-66
chr1-126182313-INS-172
chr1-126193631-INS-211
chr1-126218495-INS-51
chr1-126221822-INS-7308
chr1-126223835-INS-51
chr1-126229610-INS-6574
chr1-126239379-INS-137
chr1-126281495-INS-543
chr1-126284513-INS-413
chr1-126308544-INS-424
chr1-126369748-INS-52
chr1-126407445-INS-60
chr1-126421096-INS-356
chr1-126446863-INS-136
chr1-126449304-INS-58
chr1-126476692-INS-147
chr1-126483725-INS-121
chr1-126488185-INS-183
chr1-126504106-INS-6888
chr1-126517135-INS-172
chr1-126548608-INS-60
chr1-126550653-INS-133
chr1-126558903-INS-400
chr1-126563759-INS-131
chr1-126577109-INS-126
chr1-126583612-INS-6589
chr1-126585515-INS-138
chr1-126603422-INS-508
chr1-126604283-INS-78
chr1-126627495-INS-146
chr1-126643374-INS-541
chr1-126653101-INS-365
chr1-126673779-INS-87
chr1-126674536-INS-221
chr1-126684785-INS-64
chr1-126688064-INS-83
chr1-126706594-INS-72
chr1-126713827-INS-157
chr1-126748182-INS-72
chr1-126753819-INS-76
chr1-126761493-INS-167
chr1-126765185-INS-450
chr1-126780045-INS-116
chr1-126798779-INS-942
chr1-126809340-INS-68
chr1-126819358-INS-245
chr1-126822309-INS-187
chr1-126843280-INS-199
chr1-126857868-INS-78
chr1-126879230-INS-1888
chr1-126906085-INS-55
chr1-126924434-INS-86

chr1-126926713-INS-193
chr1-126928879-INS-402
chr1-126934841-INS-205
chr1-126938163-INS-199
chr1-126939444-INS-285
chr1-126954637-INS-131
chr1-126964999-INS-85
chr1-126968898-INS-144
chr1-126971982-INS-252
chr1-126975900-INS-56
chr1-126980414-INS-483
chr1-126991812-INS-60
chr1-126999073-INS-170
chr1-127025632-INS-498
chr1-127028525-INS-1465
chr1-127046803-INS-61
chr1-127049203-INS-548
chr1-127053853-INS-219
chr1-127056805-INS-81
chr1-127070374-INS-209
chr1-127071492-INS-398
chr1-127072005-INS-433
chr1-127080301-INS-67
chr1-127085001-INS-120
chr1-127091087-INS-126
chr1-127138432-INS-121
chr1-127143863-INS-171
chr1-127160485-INS-78
chr1-127160745-INS-7310
chr1-127166615-INS-52
chr1-127172949-INS-304
chr1-127183287-INS-215
chr1-127189548-INS-85
chr1-127191160-INS-198
chr1-127200792-INS-197
chr1-127205234-INS-341
chr1-127243888-INS-700
chr1-127259802-INS-127
chr1-127274292-INS-285
chr1-127311837-INS-422
chr1-127320251-INS-194
chr1-127322922-INS-70
chr1-127333605-INS-84
chr1-127339178-INS-68
chr1-127377868-INS-57
chr1-127391328-INS-106
chr1-127409153-INS-51
chr1-127415541-INS-56
chr1-127421794-INS-160
chr1-127422193-INS-1339
chr1-127422557-INS-60
chr1-127437917-INS-94
chr1-127460360-INS-62
chr1-127471403-INS-754
chr1-127496544-INS-6677
chr1-127502281-INS-146
chr1-127504256-INS-121
chr1-127505431-INS-267
chr1-127506898-INS-854
chr1-127528273-INS-62
chr1-127531170-INS-288
chr1-127557103-INS-84
chr1-127571859-INS-62
chr1-127612026-INS-70
chr1-127625257-INS-6537
chr1-127632830-INS-89
chr1-127643370-INS-211
chr1-127648596-INS-58
chr1-127665475-INS-54
chr1-127684188-INS-814
chr1-127688310-INS-108
chr1-127689742-INS-61
chr1-127695241-INS-468
chr1-127705108-INS-74
chr1-127707312-INS-313
chr1-127709618-INS-436
chr1-127740418-INS-57

chr1-127740712-INS-289
chr1-127746026-INS-74
chr1-127746386-INS-229
chr1-127755711-INS-123
chr1-127761828-INS-81
chr1-127763172-INS-208
chr1-127768314-INS-611
chr1-127817282-INS-423
chr1-127828397-INS-205
chr1-127833983-INS-101
chr1-127834580-INS-174
chr1-127853353-INS-108
chr1-127885785-INS-88
chr1-127893364-INS-183
chr1-127904917-INS-208
chr1-127919424-INS-188
chr1-127920849-INS-94
chr1-127935243-INS-56
chr1-127936680-INS-196
chr1-127948413-INS-6442
chr1-127954873-INS-537
chr1-127989943-INS-64
chr1-127991548-INS-75
chr1-127992969-INS-258
chr1-127993296-INS-215
chr1-127993425-INS-817
chr1-127996169-INS-210
chr1-127997793-INS-213
chr1-128007334-INS-62
chr1-128038138-INS-179
chr1-128041032-INS-172
chr1-128066751-INS-5167
chr1-128070319-INS-4835
chr1-128071736-INS-217
chr1-128084471-INS-1223
chr1-128141207-INS-6444
chr1-128142105-INS-66
chr1-128158691-INS-104
chr1-128189075-INS-166
chr1-128194794-INS-5197
chr1-128200334-INS-212
chr1-128217456-INS-240
chr1-128224047-INS-146
chr1-128227212-INS-232
chr1-128236238-INS-172
chr1-128238847-INS-251
chr1-128239668-INS-188
chr1-128270406-INS-202
chr1-128278366-INS-176
chr1-128295148-INS-55
chr1-128333823-INS-513
chr1-128376955-INS-382
chr1-128376957-INS-83
chr1-128427207-INS-236
chr1-128444128-INS-334
chr1-128444803-INS-267
chr1-128462150-INS-367
chr1-128480208-INS-70
chr1-128482126-INS-176
chr1-128497575-INS-198
chr1-128531927-INS-136
chr1-128561507-INS-56
chr1-128604123-INS-52
chr1-128610100-INS-120
chr1-128615734-INS-64
chr1-128637068-INS-390
chr1-128666925-INS-64
chr1-128670919-INS-65
chr1-128697649-INS-76
chr1-128710773-INS-54
chr1-128711167-INS-54
chr1-128712893-INS-210
chr1-128716627-INS-51
chr1-128730614-INS-132
chr1-128737474-INS-56
chr1-128752525-INS-88
chr1-128761942-INS-212

chr1-128765941-INS-60
chr1-128768586-INS-99
chr1-128773929-INS-268
chr1-128786666-INS-80
chr1-128798164-INS-127
chr1-128801650-INS-146
chr1-128802068-INS-758
chr1-128823500-INS-110
chr1-128830591-INS-380
chr1-128843384-INS-463
chr1-128844100-INS-1197
chr1-128851130-INS-54
chr1-128859387-INS-308
chr1-128877468-INS-74
chr1-128885805-INS-72
chr1-128889636-INS-957
chr1-128894060-INS-1193
chr1-128906787-INS-6289
chr1-128910194-INS-6288
chr1-128950876-INS-164
chr1-128952930-INS-473
chr1-128976996-INS-64
chr1-128989142-INS-75
chr1-129027927-INS-61
chr1-129028143-INS-6899
chr1-129035267-INS-54
chr1-129040525-INS-103
chr1-129053151-INS-338
chr1-129066636-INS-7358
chr1-129082923-INS-189
chr1-129097108-INS-219
chr1-129137455-INS-6694
chr1-129144203-INS-234
chr1-129155662-INS-6614
chr1-129178088-INS-52
chr1-129180257-INS-331
chr1-129186485-INS-182
chr1-129189391-INS-62
chr1-129196832-INS-376
chr1-129199615-INS-66
chr1-129214429-INS-77
chr1-129216774-INS-70
chr1-129218505-INS-96
chr1-129223613-INS-70
chr1-129224642-INS-681
chr1-129224695-INS-3415
chr1-129227831-INS-176
chr1-129239225-INS-278
chr1-129264834-INS-747
chr1-129274710-INS-109
chr1-129286277-INS-65
chr1-129288832-INS-779
chr1-129293302-INS-74
chr1-129309169-INS-954
chr1-129312220-INS-994
chr1-129321015-INS-398
chr1-129372229-INS-310
chr1-129375367-INS-444
chr1-129383433-INS-263
chr1-129395954-INS-743
chr1-129426682-INS-172
chr1-129447948-INS-130
chr1-129476428-INS-92
chr1-129481273-INS-124
chr1-129514225-INS-87
chr1-129602064-INS-52
chr1-129635795-INS-6678
chr1-129640345-INS-65
chr1-129652273-INS-591
chr1-129656835-INS-6276
chr1-129689186-INS-318
chr1-129707643-INS-2453
chr1-129711818-INS-59
chr1-129716757-INS-850
chr1-129731747-INS-248
chr1-129752984-INS-87
chr1-129758134-INS-64

chr1-129770832-INS-6794
chr1-129773084-INS-248
chr1-129783596-INS-384
chr1-129784430-INS-92
chr1-129787991-INS-579
chr1-129809625-INS-430
chr1-129811195-INS-6330
chr1-129812615-INS-195
chr1-129815588-INS-101
chr1-129816798-INS-7107
chr1-129821415-INS-10063
chr1-129822626-INS-55
chr1-129832523-INS-100
chr1-129864714-INS-6559
chr1-129865376-INS-2867
chr1-129874225-INS-6873
chr1-129907283-INS-177
chr1-129921082-INS-436
chr1-129939596-INS-51
chr1-129970259-INS-7141
chr1-129974277-INS-52
chr1-129996558-INS-511
chr1-130000163-INS-875
chr1-130004062-INS-89
chr1-130016684-INS-196
chr1-130023760-INS-6916
chr1-130069967-INS-202
chr1-130074479-INS-6782
chr1-130084491-INS-60
chr1-130086571-INS-52
chr1-130115884-INS-120
chr1-130116526-INS-68
chr1-130134544-INS-132
chr1-130137568-INS-88
chr1-130145476-INS-61
chr1-130149293-INS-6520
chr1-130160892-INS-6368
chr1-130170265-INS-7463
chr1-130170874-INS-178
chr1-130187391-INS-327
chr1-130220022-INS-58
chr1-130223555-INS-237
chr1-130241437-INS-270
chr1-130245138-INS-6226
chr1-130256255-INS-85
chr1-130262266-INS-59
chr1-130264473-INS-60
chr1-130272349-INS-111
chr1-130305583-INS-112
chr1-130323896-INS-110
chr1-130336715-INS-234
chr1-130371529-INS-1626
chr1-130388450-INS-1935
chr1-130426197-INS-166
chr1-130435083-INS-56
chr1-130440267-INS-75
chr1-130441036-INS-716
chr1-130443470-INS-60
chr1-130453441-INS-149
chr1-130457594-INS-6442
chr1-130460731-INS-472
chr1-130461394-INS-7347
chr1-130479070-INS-130
chr1-130496578-INS-88
chr1-130516789-INS-81
chr1-130538770-INS-59
chr1-130540442-INS-722
chr1-130540443-INS-186
chr1-130555610-INS-172
chr1-130582165-INS-559
chr1-130587518-INS-495
chr1-130596266-INS-117
chr1-130622402-INS-66
chr1-130654354-INS-6381
chr1-130671871-INS-147
chr1-130697108-INS-117
chr1-130701840-INS-144

chr1-130712226-INS-585
chr1-130747799-INS-5394
chr1-130762820-INS-148
chr1-130780768-INS-330
chr1-130783088-INS-794
chr1-130783917-INS-270
chr1-130832423-INS-284
chr1-130835429-INS-70
chr1-130838244-INS-51
chr1-130844163-INS-64
chr1-130875494-INS-233
chr1-130879442-INS-962
chr1-130888007-INS-111
chr1-130914499-INS-52
chr1-130976225-INS-98
chr1-131030308-INS-210
chr1-131030926-INS-66
chr1-131035277-INS-201
chr1-131045660-INS-136
chr1-131064700-INS-154
chr1-131106911-INS-152
chr1-131143937-INS-205
chr1-131154378-INS-80
chr1-131159550-INS-76
chr1-131218319-INS-198
chr1-131254038-INS-66
chr1-131288904-INS-51
chr1-131318647-INS-1986
chr1-131346395-INS-150
chr1-131349295-INS-199
chr1-131354099-INS-126
chr1-131356184-INS-211
chr1-131394871-INS-6118
chr1-131398490-INS-1561
chr1-131417046-INS-251
chr1-131439259-INS-71
chr1-131441974-INS-68
chr1-131481527-INS-224
chr1-131483621-INS-594
chr1-131495111-INS-89
chr1-131540650-INS-135
chr1-131545525-INS-98
chr1-131546628-INS-60
chr1-131554577-INS-72
chr1-131559231-INS-65
chr1-131566541-INS-59
chr1-131567529-INS-75
chr1-131580267-INS-100
chr1-131582584-INS-94
chr1-131595258-INS-59
chr1-131602029-INS-110
chr1-131645138-INS-71
chr1-131687322-INS-97
chr1-131694282-INS-548
chr1-131735880-INS-836
chr1-131753699-INS-266
chr1-131803561-INS-136
chr1-131813399-INS-138
chr1-131817461-INS-52
chr1-131827639-INS-91
chr1-131836078-INS-1892
chr1-131840679-INS-189
chr1-131850271-INS-190
chr1-131852823-INS-293
chr1-131855204-INS-326
chr1-131876853-INS-57
chr1-131881677-INS-51
chr1-131893301-INS-141
chr1-131912656-INS-101
chr1-131920895-INS-198
chr1-131925970-INS-390
chr1-131929491-INS-404
chr1-131955135-INS-217
chr1-131958620-INS-122
chr1-131982580-INS-196
chr1-132006439-INS-121
chr1-132015153-INS-301

chr1-132028189-INS-432
chr1-132032051-INS-98
chr1-132033516-INS-128
chr1-132036021-INS-207
chr1-132054063-INS-78
chr1-132058952-INS-696
chr1-132059213-INS-1350
chr1-132107593-INS-83
chr1-132109149-INS-208
chr1-132121224-INS-171
chr1-132131145-INS-498
chr1-132147492-INS-108
chr1-132151070-INS-168
chr1-132152145-INS-195
chr1-132183489-INS-464
chr1-132188113-INS-392
chr1-132201551-INS-224
chr1-132201973-INS-344
chr1-132214952-INS-81
chr1-132222991-INS-52
chr1-132225442-INS-52
chr1-132239639-INS-716
chr1-132253029-INS-206
chr1-132256977-INS-127
chr1-132275377-INS-393
chr1-132279080-INS-93
chr1-132299433-INS-991
chr1-132310462-INS-234
chr1-132312145-INS-74
chr1-132325629-INS-228
chr1-132326758-INS-54
chr1-132335106-INS-649
chr1-132347134-INS-213
chr1-132358607-INS-5259
chr1-132381421-INS-7296
chr1-132399710-INS-199
chr1-132409942-INS-226
chr1-132419290-INS-52
chr1-132455100-INS-52
chr1-132534070-INS-211
chr1-132539477-INS-116
chr1-132560931-INS-331
chr1-132585105-INS-333
chr1-132604524-INS-504
chr1-132607335-INS-82
chr1-132617656-INS-603
chr1-132628791-INS-67
chr1-132633961-INS-80
chr1-132635765-INS-198
chr1-132641081-INS-130
chr1-132653900-INS-114
chr1-132700390-INS-79
chr1-132733314-INS-84
chr1-132734680-INS-152
chr1-132752588-INS-138
chr1-132764281-INS-227
chr1-132770238-INS-56
chr1-132786604-INS-58
chr1-132850464-INS-876
chr1-132856090-INS-142
chr1-132871237-INS-4699
chr1-132875162-INS-87
chr1-132877570-INS-64
chr1-132879448-INS-53
chr1-132897874-INS-220
chr1-132903342-INS-55
chr1-132904507-INS-93
chr1-132908008-INS-100
chr1-132921006-INS-211
chr1-132929496-INS-282
chr1-132934215-INS-180
chr1-132938070-INS-68
chr1-132940559-INS-170
chr1-132950879-INS-232
chr1-132968295-INS-205
chr1-132969556-INS-223
chr1-132970656-INS-344

chr1-132998205-INS-102
chr1-133012512-INS-52
chr1-133044931-INS-203
chr1-133045680-INS-598
chr1-133069319-INS-58
chr1-133072914-INS-209
chr1-133077384-INS-1157
chr1-133093991-INS-159
chr1-133104383-INS-175
chr1-133126419-INS-201
chr1-133130138-INS-541
chr1-133194842-INS-186
chr1-133217719-INS-215
chr1-133236353-INS-196
chr1-133240233-INS-55
chr1-133250985-INS-53
chr1-133266179-INS-118
chr1-133320192-INS-50
chr1-133332190-INS-239
chr1-133357195-INS-342
chr1-133375053-INS-53
chr1-133390136-INS-113
chr1-133394257-INS-3620
chr1-133400449-INS-126
chr1-133425756-INS-119
chr1-133432244-INS-246
chr1-133451366-INS-186
chr1-133455054-INS-99
chr1-133485730-INS-434
chr1-133490661-INS-215
chr1-133493707-INS-143
chr1-133496892-INS-62
chr1-133511390-INS-51
chr1-133523392-INS-199
chr1-133523596-INS-2486
chr1-133523838-INS-199
chr1-133555497-INS-382
chr1-133573896-INS-127
chr1-133589962-INS-81
chr1-133624284-INS-52
chr1-133627625-INS-87
chr1-133672180-INS-78
chr1-133691842-INS-56
chr1-133701964-INS-56
chr1-133733629-INS-71
chr1-133783318-INS-73
chr1-133808096-INS-196
chr1-133816071-INS-104
chr1-133842059-INS-63
chr1-133853672-INS-157
chr1-133858141-INS-58
chr1-133868207-INS-64
chr1-133874770-INS-346
chr1-133888966-INS-158
chr1-133894579-INS-340
chr1-133926366-INS-155
chr1-133939078-INS-78
chr1-133977469-INS-302
chr1-133994084-INS-57
chr1-134040100-INS-880
chr1-134042226-INS-82
chr1-134047214-INS-132
chr1-134050122-INS-1886
chr1-134077468-INS-64
chr1-134091016-INS-268
chr1-134096794-INS-511
chr1-134109031-INS-51
chr1-134148631-INS-321
chr1-134190865-INS-331
chr1-134192403-INS-250
chr1-134205851-INS-61
chr1-134231276-INS-263
chr1-134231285-INS-772
chr1-134241565-INS-111
chr1-134282170-INS-85
chr1-134318900-INS-180
chr1-134322023-INS-497

chr1-134323713-INS-80
chr1-134339327-INS-70
chr1-134371365-INS-70
chr1-134374585-INS-141
chr1-134381095-INS-225
chr1-134401762-INS-202
chr1-134422126-INS-85
chr1-134440730-INS-59
chr1-134442207-INS-211
chr1-134462256-INS-124
chr1-134463219-INS-336
chr1-134512319-INS-418
chr1-134515003-INS-61
chr1-134530901-INS-53
chr1-134534019-INS-78
chr1-134580301-INS-400
chr1-134595702-INS-90
chr1-134600538-INS-235
chr1-134606016-INS-76
chr1-134622372-INS-108
chr1-134659759-INS-319
chr1-134663748-INS-175
chr1-134663886-INS-222
chr1-134692859-INS-76
chr1-134708615-INS-337
chr1-134727167-INS-417
chr1-134752524-INS-178
chr1-134757878-INS-61
chr1-134784519-INS-873
chr1-134815687-INS-74
chr1-134818067-INS-111
chr1-134828766-INS-236
chr1-134873591-INS-7207
chr1-134886834-INS-314
chr1-134931504-INS-74
chr1-134974739-INS-195
chr1-134983851-INS-407
chr1-134987339-INS-158
chr1-135041641-INS-52
chr1-135065463-INS-139
chr1-135112425-INS-54
chr1-135114695-INS-110
chr1-135129422-INS-311
chr1-135141515-INS-461
chr1-135158165-INS-252
chr1-135201621-INS-249
chr1-135205379-INS-209
chr1-135219244-INS-5546
chr1-135220840-INS-250
chr1-135234833-INS-185
chr1-135235328-INS-490
chr1-135277819-INS-57
chr1-135278923-INS-141
chr1-135287313-INS-316
chr1-135288494-INS-69
chr1-135293677-INS-736
chr1-135321004-INS-178
chr1-135329029-INS-219
chr1-135344598-INS-156
chr1-135355171-INS-365
chr1-135363118-INS-135
chr1-135399196-INS-190
chr1-135401813-INS-237
chr1-135407675-INS-85
chr1-135463591-INS-131
chr1-135464614-INS-254
chr1-135479286-INS-80
chr1-135504475-INS-401
chr1-135523776-INS-64
chr1-135539555-INS-343
chr1-135545534-INS-64
chr1-135548294-INS-192
chr1-135570456-INS-141
chr1-135574130-INS-214
chr1-135582536-INS-279
chr1-135607461-INS-210
chr1-135620759-INS-201

chr1-135641347-INS-211
chr1-135644960-INS-53
chr1-135666943-INS-100
chr1-135690294-INS-402
chr1-135715990-INS-178
chr1-135774725-INS-108
chr1-135784412-INS-53
chr1-135816615-INS-54
chr1-135849348-INS-249
chr1-135869151-INS-56
chr1-135869865-INS-92
chr1-135877046-INS-58
chr1-135880248-INS-225
chr1-135888489-INS-96
chr1-135928515-INS-203
chr1-135960905-INS-81
chr1-135974339-INS-100
chr1-135987838-INS-771
chr1-136000702-INS-143
chr1-136001682-INS-154
chr1-136011679-INS-90
chr1-136018191-INS-363
chr1-136041492-INS-116
chr1-136066979-INS-91
chr1-136075195-INS-89
chr1-136115639-INS-209
chr1-136120591-INS-62
chr1-136139444-INS-70
chr1-136180234-INS-61
chr1-136193710-INS-61
chr1-136255144-INS-90
chr1-136291461-INS-55
chr1-136300535-INS-125
chr1-136311979-INS-66
chr1-136321622-INS-324
chr1-136324966-INS-496
chr1-136327561-INS-68
chr1-136333011-INS-52
chr1-136334811-INS-79
chr1-136366870-INS-243
chr1-136387687-INS-141
chr1-136402118-INS-137
chr1-136405918-INS-601
chr1-136413865-INS-58
chr1-136438294-INS-89
chr1-136462021-INS-120
chr1-136462661-INS-759
chr1-136488323-INS-167
chr1-136493406-INS-196
chr1-136519826-INS-397
chr1-136525303-INS-291
chr1-136532012-INS-200
chr1-136534463-INS-117
chr1-136537013-INS-653
chr1-136564457-INS-582
chr1-136591061-INS-198
chr1-136609512-INS-171
chr1-136612864-INS-217
chr1-136618504-INS-175
chr1-136619772-INS-198
chr1-136629691-INS-178
chr1-136630897-INS-229
chr1-136638218-INS-194
chr1-136644253-INS-204
chr1-136660125-INS-63
chr1-136666518-INS-134
chr1-136686230-INS-245
chr1-136690830-INS-412
chr1-136720148-INS-95
chr1-136727068-INS-63
chr1-136730272-INS-204
chr1-136755974-INS-190
chr1-136757076-INS-202
chr1-136759435-INS-67
chr1-136799527-INS-233
chr1-136844366-INS-146
chr1-136847514-INS-52

chr1-136854464-INS-520
chr1-136857332-INS-187
chr1-136868903-INS-56
chr1-136879438-INS-211
chr1-136886432-INS-51
chr1-136902198-INS-301
chr1-136905710-INS-99
chr1-136907135-INS-54
chr1-136912328-INS-169
chr1-136913375-INS-225
chr1-136919333-INS-56
chr1-136925190-INS-384
chr1-136933143-INS-5461
chr1-136952600-INS-179
chr1-136957944-INS-213
chr1-136958041-INS-809
chr1-136965222-INS-204
chr1-136973321-INS-378
chr1-136987013-INS-55
chr1-137015396-INS-170
chr1-137019060-INS-108
chr1-137026069-INS-54
chr1-137029137-INS-198
chr1-137039424-INS-548
chr1-137072141-INS-6452
chr1-137094041-INS-301
chr1-137099558-INS-84
chr1-137119417-INS-278
chr1-137126384-INS-60
chr1-137143897-INS-447
chr1-137147266-INS-57
chr1-137160181-INS-870
chr1-137211939-INS-75
chr1-137217241-INS-572
chr1-137225742-INS-340
chr1-137226827-INS-68
chr1-137251486-INS-69
chr1-137262777-INS-181
chr1-137272754-INS-424
chr1-137290305-INS-822
chr1-137294695-INS-216
chr1-137305593-INS-51
chr1-137320562-INS-275
chr1-137321274-INS-7128
chr1-137323251-INS-412
chr1-137325619-INS-51
chr1-137326718-INS-439
chr1-137332540-INS-498
chr1-137343789-INS-103
chr1-137377512-INS-242
chr1-137401133-INS-180
chr1-137433639-INS-2424
chr1-137445117-INS-224
chr1-137450703-INS-532
chr1-137460088-INS-255
chr1-137467201-INS-309
chr1-137481208-INS-199
chr1-137504490-INS-61
chr1-137524231-INS-5247
chr1-137528718-INS-2722
chr1-137533909-INS-644
chr1-137538021-INS-351
chr1-137543334-INS-6442
chr1-137565580-INS-238
chr1-137568365-INS-497
chr1-137579314-INS-99
chr1-137596256-INS-505
chr1-137654708-INS-65
chr1-137682576-INS-342
chr1-137712667-INS-552
chr1-137726464-INS-6573
chr1-137727894-INS-1292
chr1-137803710-INS-3601
chr1-137805601-INS-89
chr1-137809291-INS-673
chr1-137829898-INS-6660
chr1-137879667-INS-393

chr1-137881718-INS-1748
chr1-137896136-INS-56
chr1-137934004-INS-155
chr1-137935325-INS-75
chr1-137945311-INS-167
chr1-137956675-INS-111
chr1-137968726-INS-175
chr1-137994542-INS-6810
chr1-137997165-INS-921
chr1-137997698-INS-64
chr1-138024857-INS-168
chr1-138035186-INS-174
chr1-138040927-INS-66
chr1-138050424-INS-70
chr1-138075544-INS-132
chr1-138091379-INS-60
chr1-138132956-INS-6412
chr1-138149295-INS-711
chr1-138164834-INS-113
chr1-138166862-INS-115
chr1-138175316-INS-102
chr1-138181991-INS-111
chr1-138182767-INS-2320
chr1-138202039-INS-491
chr1-138211100-INS-441
chr1-138225400-INS-56
chr1-138227762-INS-459
chr1-138248285-INS-68
chr1-138263590-INS-74
chr1-138284586-INS-92
chr1-138296312-INS-60
chr1-138348168-INS-52
chr1-138370996-INS-63
chr1-138377595-INS-122
chr1-138448833-INS-222
chr1-138449724-INS-204
chr1-138452952-INS-71
chr1-138456737-INS-66
chr1-138472668-INS-508
chr1-138513714-INS-567
chr1-138525785-INS-294
chr1-138526706-INS-64
chr1-138534202-INS-369
chr1-138541958-INS-358
chr1-138558751-INS-184
chr1-138581232-INS-172
chr1-138587367-INS-128
chr1-138609425-INS-552
chr1-138611260-INS-214
chr1-138630403-INS-78
chr1-138633337-INS-98
chr1-138637602-INS-558
chr1-138646541-INS-839
chr1-138654547-INS-211
chr1-138681272-INS-116
chr1-138686386-INS-192
chr1-138704335-INS-81
chr1-138710372-INS-6699
chr1-138715115-INS-56
chr1-138725856-INS-135
chr1-138749966-INS-56
chr1-138754651-INS-84
chr1-138774831-INS-105
chr1-138777257-INS-220
chr1-138786783-INS-339
chr1-138808225-INS-213
chr1-138812089-INS-102
chr1-138832807-INS-74
chr1-138846348-INS-206
chr1-138856192-INS-52
chr1-138862376-INS-68
chr1-138912482-INS-211
chr1-138914626-INS-65
chr1-138978052-INS-51
chr1-139068339-INS-599
chr1-139075408-INS-52
chr1-139084209-INS-474

chr1-139119063-INS-67
chr1-139142274-INS-79
chr1-139163358-INS-164
chr1-139166673-INS-5711
chr1-139192421-INS-6698
chr1-139213401-INS-69
chr1-139232179-INS-98
chr1-139234275-INS-233
chr1-139247414-INS-156
chr1-139267576-INS-96
chr1-139283885-INS-64
chr1-139332208-INS-499
chr1-139339610-INS-103
chr1-139342966-INS-167
chr1-139362445-INS-214
chr1-139380950-INS-204
chr1-139432225-INS-58
chr1-139459754-INS-124
chr1-139461997-INS-203
chr1-139482282-INS-2600
chr1-139489267-INS-55
chr1-139502629-INS-199
chr1-139520244-INS-498
chr1-139520985-INS-160
chr1-139562624-INS-903
chr1-139701765-INS-170
chr1-139776354-INS-6501
chr1-139807744-INS-6539
chr1-139820256-INS-5355
chr1-139824029-INS-54
chr1-139859497-INS-7354
chr1-139949410-INS-312
chr1-139962545-INS-608
chr1-139990419-INS-410
chr1-140066525-INS-52
chr1-140070659-INS-504
chr1-140098304-INS-176
chr1-140102848-INS-51
chr1-140128836-INS-792
chr1-140152510-INS-67
chr1-140198043-INS-75
chr1-140208677-INS-278
chr1-140210639-INS-88
chr1-140226969-INS-265
chr1-140255444-INS-112
chr1-140272634-INS-52
chr1-140292588-INS-201
chr1-140341853-INS-57
chr1-140343880-INS-53
chr1-140374438-INS-490
chr1-140375886-INS-216
chr1-140376958-INS-851
chr1-140383785-INS-81
chr1-140405733-INS-458
chr1-140430674-INS-77
chr1-140463323-INS-285
chr1-140468589-INS-120
chr1-140471681-INS-93
chr1-140472514-INS-64
chr1-140484681-INS-579
chr1-140494719-INS-7153
chr1-140495503-INS-86
chr1-140498833-INS-172
chr1-140506656-INS-2207
chr1-140506854-INS-427
chr1-140513364-INS-64
chr1-140515892-INS-6387
chr1-140518540-INS-137
chr1-140552906-INS-315
chr1-140596955-INS-401
chr1-140614052-INS-409
chr1-140616967-INS-452
chr1-140630702-INS-61
chr1-140635200-INS-4550
chr1-140637416-INS-6651
chr1-140645672-INS-6317
chr1-140646349-INS-72

chr1-140648777-INS-5972
chr1-140658807-INS-248
chr1-140666042-INS-337
chr1-140667472-INS-68
chr1-140686098-INS-55
chr1-140710349-INS-6672
chr1-140711367-INS-62
chr1-140732903-INS-5293
chr1-140737114-INS-113
chr1-140743686-INS-102
chr1-140767595-INS-6409
chr1-140771751-INS-72
chr1-140772311-INS-151
chr1-140774106-INS-64
chr1-140781760-INS-51
chr1-140808889-INS-60
chr1-140811894-INS-52
chr1-140816677-INS-248
chr1-140821011-INS-304
chr1-140839293-INS-68
chr1-140841150-INS-60
chr1-140849617-INS-641
chr1-140868911-INS-60
chr1-140881181-INS-64
chr1-140911188-INS-56
chr1-140913536-INS-87
chr1-140961319-INS-6408
chr1-140981582-INS-102
chr1-140982651-INS-66
chr1-141034722-INS-147
chr1-141067853-INS-6173
chr1-141084538-INS-384
chr1-141090727-INS-354
chr1-141092379-INS-801
chr1-141094000-INS-74
chr1-141094797-INS-6468
chr1-141110968-INS-54
chr1-141185093-INS-484
chr1-141188459-INS-62
chr1-141202584-INS-5199
chr1-141203156-INS-58
chr1-141219114-INS-59
chr1-141220621-INS-447
chr1-141225649-INS-237
chr1-141240740-INS-244
chr1-141276793-INS-647
chr1-141280824-INS-6417
chr1-141308959-INS-248
chr1-141312866-INS-197
chr1-141314028-INS-73
chr1-141320176-INS-386
chr1-141339502-INS-116
chr1-141349454-INS-63
chr1-141353964-INS-61
chr1-141358597-INS-9043
chr1-141375566-INS-413
chr1-141402634-INS-331
chr1-141411022-INS-133
chr1-141417385-INS-72
chr1-141419996-INS-66
chr1-141431570-INS-5373
chr1-141450658-INS-52
chr1-141478603-INS-7138
chr1-141536305-INS-240
chr1-141552718-INS-55
chr1-141564967-INS-328
chr1-141570846-INS-72
chr1-141586121-INS-76
chr1-141595698-INS-59
chr1-141631133-INS-212
chr1-141634029-INS-531
chr1-141657294-INS-104
chr1-141661168-INS-2423
chr1-141662210-INS-59
chr1-141725598-INS-64
chr1-141749224-INS-578
chr1-141770991-INS-68

chr1-141776099-INS-1530
chr1-141777492-INS-5499
chr1-141888093-INS-52
chr1-141916765-INS-442
chr1-141944462-INS-6943
chr1-141967705-INS-58
chr1-141979495-INS-133
chr1-142009131-INS-56
chr1-142050086-INS-105
chr1-142056996-INS-128
chr1-142061572-INS-92
chr1-142067490-INS-1889
chr1-142068040-INS-109
chr1-142101637-INS-5240
chr1-142115724-INS-321
chr1-142117663-INS-60
chr1-142118912-INS-157
chr1-142132070-INS-107
chr1-142139006-INS-80
chr1-142152738-INS-1209
chr1-142153854-INS-526
chr1-142163600-INS-97
chr1-142173040-INS-379
chr1-142185047-INS-540
chr1-142227528-INS-105
chr1-142247800-INS-6564
chr1-142252693-INS-768
chr1-142263487-INS-60
chr1-142269619-INS-8084
chr1-142276789-INS-773
chr1-142303059-INS-314
chr1-142315403-INS-6717
chr1-142316596-INS-65
chr1-142330159-INS-6200
chr1-142334418-INS-129
chr1-142366095-INS-507
chr1-142383665-INS-56
chr1-142397734-INS-65
chr1-142407528-INS-80
chr1-142417637-INS-60
chr1-142439089-INS-54
chr1-142440139-INS-52
chr1-142449317-INS-207
chr1-142500098-INS-615
chr1-142545774-INS-212
chr1-142554777-INS-6747
chr1-142555738-INS-6185
chr1-142561512-INS-7003
chr1-142563646-INS-398
chr1-142571607-INS-69
chr1-142578337-INS-98
chr1-142580411-INS-102
chr1-142608079-INS-597
chr1-142609374-INS-6212
chr1-142646889-INS-269
chr1-142674731-INS-169
chr1-142678802-INS-655
chr1-142692661-INS-493
chr1-142701998-INS-84
chr1-142721349-INS-3427
chr1-142731382-INS-277
chr1-142737133-INS-2196
chr1-142765470-INS-57
chr1-142774144-INS-117
chr1-142775725-INS-99
chr1-142779062-INS-615
chr1-142784382-INS-150
chr1-142786340-INS-58
chr1-142812066-INS-303
chr1-142842074-INS-6442
chr1-142846658-INS-964
chr1-142872091-INS-67
chr1-142878324-INS-203
chr1-142881608-INS-497
chr1-142894362-INS-561
chr1-142925587-INS-60
chr1-142928410-INS-203

chr1-142937340-INS-689
chr1-142971607-INS-5326
chr1-142975355-INS-203
chr1-143019727-INS-219
chr1-143022892-INS-569
chr1-143043453-INS-814
chr1-143056944-INS-6200
chr1-143078387-INS-5983
chr1-143102292-INS-126
chr1-143105744-INS-6900
chr1-143112443-INS-6692
chr1-143125810-INS-114
chr1-143145405-INS-84
chr1-143148510-INS-6438
chr1-143149613-INS-879
chr1-143171706-INS-237
chr1-143172883-INS-59
chr1-143181403-INS-322
chr1-143192011-INS-51
chr1-143195511-INS-110
chr1-143203444-INS-5122
chr1-143206297-INS-649
chr1-143208972-INS-399
chr1-143226206-INS-74
chr1-143235214-INS-50
chr1-143245963-INS-7123
chr1-143262520-INS-139
chr1-143265585-INS-302
chr1-143279126-INS-864
chr1-143289948-INS-166
chr1-143291991-INS-6633
chr1-143292643-INS-276
chr1-143302927-INS-7008
chr1-143304033-INS-66
chr1-143306693-INS-190
chr1-143308354-INS-416
chr1-143326767-INS-419
chr1-143333009-INS-8215
chr1-143342496-INS-56
chr1-143352223-INS-402
chr1-143359938-INS-708
chr1-143360787-INS-6983
chr1-143363545-INS-164
chr1-143383705-INS-2333
chr1-143406385-INS-130
chr1-143414534-INS-179
chr1-143421451-INS-6438
chr1-143428042-INS-209
chr1-143435318-INS-333
chr1-143445402-INS-533
chr1-143448723-INS-7727
chr1-143492197-INS-214
chr1-143547953-INS-61
chr1-143554410-INS-207
chr1-143600196-INS-188
chr1-143601349-INS-114
chr1-143605350-INS-5182
chr1-143609044-INS-7142
chr1-143621249-INS-5019
chr1-143626407-INS-190
chr1-143680062-INS-210
chr1-143701401-INS-778
chr1-143708171-INS-502
chr1-143715694-INS-110
chr1-143779299-INS-986
chr1-143781992-INS-62
chr1-143791268-INS-9500
chr1-143809395-INS-54
chr1-143811426-INS-560
chr1-143830161-INS-7482
chr1-143831155-INS-164
chr1-143833707-INS-6613
chr1-143859109-INS-58
chr1-143865387-INS-801
chr1-143880517-INS-126
chr1-143885852-INS-6956
chr1-143886257-INS-861

chr1-143919205-INS-52
chr1-143937447-INS-2395
chr1-143955387-INS-434
chr1-143958722-INS-484
chr1-143992633-INS-209
chr1-144021493-INS-560
chr1-144039183-INS-60
chr1-144062257-INS-140
chr1-144074084-INS-282
chr1-144086277-INS-169
chr1-144091734-INS-6046
chr1-144103548-INS-937
chr1-144133440-INS-6369
chr1-144142417-INS-51
chr1-144144148-INS-7086
chr1-144205215-INS-67
chr1-144268129-INS-174
chr1-144282204-INS-123
chr1-144293847-INS-169
chr1-144313781-INS-78
chr1-144315230-INS-182
chr1-144322947-INS-6801
chr1-144323267-INS-187
chr1-144343120-INS-52
chr1-144428437-INS-57
chr1-144438519-INS-88
chr1-144454939-INS-785
chr1-144458622-INS-6452
chr1-144482552-INS-5543
chr1-144488510-INS-138
chr1-144500228-INS-212
chr1-144543792-INS-65
chr1-144558427-INS-177
chr1-144561463-INS-80
chr1-144578773-INS-2598
chr1-144580099-INS-55
chr1-144586207-INS-57
chr1-144596799-INS-63
chr1-144605863-INS-115
chr1-144625958-INS-54
chr1-144627700-INS-716
chr1-144655928-INS-369
chr1-144664030-INS-57
chr1-144665277-INS-105
chr1-144667309-INS-64
chr1-144712061-INS-106
chr1-144753719-INS-60
chr1-144767463-INS-507
chr1-144775354-INS-62
chr1-144807431-INS-6715
chr1-144808188-INS-103
chr1-144847179-INS-197
chr1-144848290-INS-68
chr1-144880995-INS-112
chr1-144951857-INS-132
chr1-144962447-INS-64
chr1-144996733-INS-55
chr1-145011883-INS-310
chr1-145046527-INS-307
chr1-145092206-INS-52
chr1-145130880-INS-424
chr1-145169913-INS-52
chr1-145171141-INS-75
chr1-145172880-INS-701
chr1-145214849-INS-481
chr1-145230405-INS-476
chr1-145243776-INS-327
chr1-145290542-INS-3085
chr1-145290999-INS-78
chr1-145297115-INS-7128
chr1-145330190-INS-188
chr1-145334927-INS-52
chr1-145342034-INS-58
chr1-145342201-INS-5974
chr1-145374643-INS-953
chr1-145394260-INS-151
chr1-145423240-INS-3030

chr1-145438820-INS-67
chr1-145443527-INS-56
chr1-145454665-INS-183
chr1-145500436-INS-93
chr1-145505591-INS-6638
chr1-145528120-INS-273
chr1-145530679-INS-158
chr1-145544171-INS-358
chr1-145545413-INS-65
chr1-145552799-INS-75
chr1-145588088-INS-327
chr1-145592709-INS-493
chr1-145594840-INS-608
chr1-145634672-INS-7118
chr1-145684230-INS-7154
chr1-145702891-INS-194
chr1-145712040-INS-6587
chr1-145724777-INS-470
chr1-145733082-INS-125
chr1-145772982-INS-526
chr1-145778348-INS-906
chr1-145780393-INS-58
chr1-145792838-INS-1805
chr1-145839023-INS-293
chr1-145851877-INS-7104
chr1-145856655-INS-172
chr1-145860259-INS-446
chr1-145876591-INS-57
chr1-145889743-INS-175
chr1-145891941-INS-342
chr1-145893035-INS-3378
chr1-145910794-INS-72
chr1-145911802-INS-85
chr1-145915168-INS-6360
chr1-145922741-INS-243
chr1-145923664-INS-327
chr1-145925370-INS-669
chr1-145931255-INS-153
chr1-145932945-INS-73
chr1-145944708-INS-162
chr1-145978457-INS-111
chr1-146018754-INS-236
chr1-146119742-INS-866
chr1-146137584-INS-196
chr1-146141726-INS-214
chr1-146143115-INS-239
chr1-146171414-INS-3630
chr1-146193433-INS-53
chr1-146203683-INS-196
chr1-146264113-INS-286
chr1-146281274-INS-2892
chr1-146298518-INS-123
chr1-146312254-INS-54
chr1-146331718-INS-76
chr1-146379532-INS-118
chr1-146438329-INS-569
chr1-146442034-INS-373
chr1-146501370-INS-81
chr1-146505241-INS-501
chr1-146509573-INS-6443
chr1-146521449-INS-306
chr1-146545140-INS-6821
chr1-146570727-INS-552
chr1-146582124-INS-54
chr1-146599755-INS-3841
chr1-146611264-INS-64
chr1-146613548-INS-744
chr1-146626591-INS-52
chr1-146641701-INS-52
chr1-146671489-INS-54
chr1-146689064-INS-55
chr1-146699426-INS-64
chr1-146703087-INS-213
chr1-146713663-INS-80
chr1-146720959-INS-5203
chr1-146725564-INS-6458
chr1-146736794-INS-57

chr1-146772835-INS-66
chr1-146775957-INS-6261
chr1-146828340-INS-1776
chr1-146864120-INS-56
chr1-146872212-INS-5276
chr1-146877367-INS-199
chr1-146877384-INS-854
chr1-146901386-INS-180
chr1-146902437-INS-5329
chr1-146903477-INS-206
chr1-146927654-INS-66
chr1-146937574-INS-84
chr1-146938369-INS-7983
chr1-146963962-INS-66
chr1-146993032-INS-482
chr1-146996624-INS-3002
chr1-147007442-INS-81
chr1-147017537-INS-64
chr1-147053930-INS-2890
chr1-147057940-INS-367
chr1-147073115-INS-97
chr1-147079120-INS-219
chr1-147085904-INS-5547
chr1-147103100-INS-52
chr1-147104652-INS-548
chr1-147108008-INS-210
chr1-147114505-INS-107
chr1-147134003-INS-423
chr1-147140402-INS-483
chr1-147149244-INS-6452
chr1-147149247-INS-213
chr1-147150796-INS-491
chr1-147157046-INS-810
chr1-147169852-INS-130
chr1-147220273-INS-66
chr1-147283932-INS-62
chr1-147298954-INS-712
chr1-147309543-INS-54
chr1-147315287-INS-64
chr1-147341233-INS-246
chr1-147364685-INS-72
chr1-147386825-INS-286
chr1-147389001-INS-843
chr1-147406940-INS-125
chr1-147414122-INS-72
chr1-147452060-INS-257
chr1-147497502-INS-54
chr1-147521350-INS-80
chr1-147584402-INS-6183
chr1-147640430-INS-485
chr1-147670712-INS-291
chr1-147679859-INS-784
chr1-147772089-INS-60
chr1-147799146-INS-374
chr1-147848107-INS-396
chr1-147851633-INS-378
chr1-147855816-INS-1339
chr1-147896109-INS-52
chr1-147901027-INS-243
chr1-147959280-INS-78
chr1-147966461-INS-150
chr1-147983511-INS-363
chr1-147994932-INS-131
chr1-148032167-INS-358
chr1-148057451-INS-488
chr1-148134790-INS-925
chr1-148162225-INS-88
chr1-148170110-INS-56
chr1-148247165-INS-58
chr1-148316762-INS-81
chr1-148327551-INS-57
chr1-148341236-INS-597
chr1-148341709-INS-976
chr1-148346068-INS-957
chr1-148347743-INS-60
chr1-148355568-INS-972
chr1-148415274-INS-454

chr1-148467059-INS-358
chr1-148506338-INS-6890
chr1-148537463-INS-7383
chr1-148576049-INS-7090
chr1-148590647-INS-5167
chr1-148598549-INS-1092
chr1-148598910-INS-565
chr1-148613695-INS-433
chr1-148617297-INS-2883
chr1-148637603-INS-1490
chr1-148649798-INS-7062
chr1-148654296-INS-53
chr1-148661276-INS-64
chr1-148663599-INS-69
chr1-148682699-INS-350
chr1-148721684-INS-63
chr1-148728653-INS-226
chr1-148736168-INS-55
chr1-148736816-INS-6403
chr1-148740434-INS-126
chr1-148766136-INS-965
chr1-148818355-INS-219
chr1-148835794-INS-162
chr1-148843790-INS-6377
chr1-148844901-INS-635
chr1-148866715-INS-4699
chr1-148875396-INS-60
chr1-148885212-INS-56
chr1-148885230-INS-654
chr1-148898433-INS-219
chr1-148917673-INS-54
chr1-148930399-INS-52
chr1-148932127-INS-51
chr1-148958266-INS-412
chr1-148958271-INS-4050
chr1-148973346-INS-65
chr1-148977525-INS-5613
chr1-148982462-INS-390
chr1-148985364-INS-138
chr1-148996657-INS-3149
chr1-149004576-INS-6363
chr1-149016789-INS-490
chr1-149035452-INS-419
chr1-149076859-INS-63
chr1-149084156-INS-5347
chr1-149107188-INS-58
chr1-149111561-INS-88
chr1-149127241-INS-100
chr1-149143982-INS-735
chr1-149207741-INS-193
chr1-149218262-INS-52
chr1-149219497-INS-1750
chr1-149228962-INS-59
chr1-149245820-INS-56
chr1-149246840-INS-2483
chr1-149265378-INS-55
chr1-149267980-INS-60
chr1-149286312-INS-578
chr1-149310890-INS-6395
chr1-149312662-INS-1597
chr1-149315091-INS-860
chr1-149317588-INS-83
chr1-149321368-INS-356
chr1-149337568-INS-6274
chr1-149357690-INS-6633
chr1-149358766-INS-159
chr1-149361351-INS-84
chr1-149375369-INS-6636
chr1-149378433-INS-66
chr1-149400456-INS-6564
chr1-149403219-INS-99
chr1-149411136-INS-159
chr1-149418315-INS-255
chr1-149430497-INS-132
chr1-149441330-INS-1059
chr1-149448628-INS-267
chr1-149469799-INS-7153

chr1-149473002-INS-301
chr1-149492430-INS-200
chr1-149494976-INS-56
chr1-149496481-INS-331
chr1-149516280-INS-357
chr1-149538018-INS-333
chr1-149550668-INS-753
chr1-149562583-INS-84
chr1-149569381-INS-55
chr1-149583135-INS-6265
chr1-149595487-INS-68
chr1-149602188-INS-53
chr1-149619917-INS-64
chr1-149632714-INS-56
chr1-149639261-INS-850
chr1-149641556-INS-496
chr1-149646690-INS-665
chr1-149654938-INS-85
chr1-149681127-INS-60
chr1-149681978-INS-7118
chr1-149693810-INS-238
chr1-149705308-INS-390
chr1-149713024-INS-54
chr1-149714682-INS-768
chr1-149735926-INS-55
chr1-149762690-INS-97
chr1-149764780-INS-92
chr1-149798681-INS-175
chr1-149800366-INS-157
chr1-149803942-INS-302
chr1-149806334-INS-514
chr1-149884106-INS-241
chr1-149902538-INS-206
chr1-149911280-INS-114
chr1-149944448-INS-53
chr1-149955567-INS-127
chr1-149957519-INS-112
chr1-149959107-INS-585
chr1-149965005-INS-63
chr1-149984944-INS-191
chr1-149995735-INS-61
chr1-150028219-INS-58
chr1-150057307-INS-6413
chr1-150077978-INS-581
chr1-150138124-INS-55
chr1-150162335-INS-475
chr1-150181556-INS-1886
chr1-150199788-INS-200
chr1-150233155-INS-114
chr1-150240168-INS-203
chr1-150283653-INS-512
chr1-150307131-INS-51
chr1-150323405-INS-1012
chr1-150344403-INS-115
chr1-150349447-INS-56
chr1-150372318-INS-65
chr1-150373299-INS-6283
chr1-150396159-INS-409
chr1-150482997-INS-120
chr1-150484365-INS-101
chr1-150485211-INS-3250
chr1-150501350-INS-54
chr1-150505347-INS-203
chr1-150512914-INS-710
chr1-150538620-INS-52
chr1-150555469-INS-129
chr1-150558737-INS-54
chr1-150563400-INS-63
chr1-150565872-INS-116
chr1-150576876-INS-56
chr1-150622079-INS-130
chr1-150622423-INS-607
chr1-150657594-INS-984
chr1-150716463-INS-615
chr1-150765881-INS-94
chr1-150773169-INS-6883
chr1-150808928-INS-1890

chr1-150813656-INS-58
chr1-150843977-INS-886
chr1-150846915-INS-81
chr1-150869935-INS-60
chr1-150902663-INS-384
chr1-150924151-INS-96
chr1-150931054-INS-62
chr1-150934181-INS-60
chr1-150972973-INS-99
chr1-150998713-INS-293
chr1-151005587-INS-268
chr1-151008255-INS-58
chr1-151008815-INS-211
chr1-151011510-INS-64
chr1-151016387-INS-68
chr1-151021545-INS-8371
chr1-151026715-INS-56
chr1-151060193-INS-353
chr1-151087725-INS-732
chr1-151110679-INS-134
chr1-151112008-INS-75
chr1-151127114-INS-86
chr1-151143705-INS-80
chr1-151158777-INS-122
chr1-151179890-INS-116
chr1-151218304-INS-59
chr1-151258051-INS-52
chr1-151272779-INS-60
chr1-151321724-INS-171
chr1-151340484-INS-79
chr1-151361530-INS-157
chr1-151384602-INS-63
chr1-151394151-INS-66
chr1-151400646-INS-222
chr1-151407753-INS-73
chr1-151408297-INS-240
chr1-151438805-INS-71
chr1-151440583-INS-132
chr1-151443323-INS-125
chr1-151450542-INS-464
chr1-151455864-INS-112
chr1-151463249-INS-112
chr1-151464803-INS-126
chr1-151497395-INS-96
chr1-151523559-INS-90
chr1-151527434-INS-99
chr1-151533439-INS-167
chr1-151555294-INS-262
chr1-151564038-INS-98
chr1-151598277-INS-248
chr1-151602667-INS-152
chr1-151673813-INS-55
chr1-151677251-INS-110
chr1-151677251-INS-86
chr1-151683444-INS-90
chr1-151689348-INS-210
chr1-151710512-INS-70
chr1-151716262-INS-215
chr1-151780444-INS-218
chr1-151789686-INS-76
chr1-151802766-INS-84
chr1-151861863-INS-606
chr1-151880170-INS-160
chr1-151892454-INS-68
chr1-151897991-INS-206
chr1-151916562-INS-732
chr1-151918583-INS-156
chr1-151928959-INS-399
chr1-151941983-INS-5313
chr1-151942507-INS-59
chr1-151947686-INS-223
chr1-151952751-INS-84
chr1-151986908-INS-68
chr1-151991241-INS-52
chr1-151995011-INS-122
chr1-152012267-INS-677
chr1-152014449-INS-61

chr1-152022240-INS-70
chr1-152023329-INS-53
chr1-152030001-INS-155
chr1-152032342-INS-59
chr1-152047680-INS-891
chr1-152053871-INS-52
chr1-152066022-INS-713
chr1-152091137-INS-189
chr1-152114481-INS-83
chr1-152143464-INS-102
chr1-152154519-INS-58
chr1-152155000-INS-90
chr1-152156008-INS-122
chr1-152159635-INS-102
chr1-152184809-INS-52
chr1-152255550-INS-98
chr1-152286099-INS-51
chr1-152302916-INS-401
chr1-152387150-INS-96
chr1-152396171-INS-213
chr1-152435877-INS-63
chr1-152437489-INS-78
chr1-152481541-INS-189
chr1-152485832-INS-58
chr1-152494123-INS-167
chr1-152506913-INS-114
chr1-152511880-INS-277
chr1-152545186-INS-272
chr1-152549446-INS-225
chr1-152552259-INS-54
chr1-152555815-INS-160
chr1-152578342-INS-72
chr1-152592686-INS-204
chr1-152600790-INS-116
chr1-152603732-INS-383
chr1-152607754-INS-189
chr1-152612279-INS-85
chr1-152627576-INS-74
chr1-152634215-INS-118
chr1-152638996-INS-610
chr1-152682288-INS-1893
chr1-152688323-INS-143
chr1-152690014-INS-309
chr1-152693222-INS-121
chr1-152727092-INS-210
chr1-152740618-INS-54
chr1-152755855-INS-154
chr1-152762607-INS-207
chr1-152770650-INS-214
chr1-152792648-INS-202
chr1-152798502-INS-599
chr1-152867865-INS-113
chr1-152884461-INS-52
chr1-152900440-INS-201
chr1-152915313-INS-201
chr1-152944278-INS-51
chr1-152979400-INS-266
chr1-152995290-INS-85
chr1-153027569-INS-498
chr1-153089068-INS-274
chr1-153090649-INS-260
chr1-153094375-INS-53
chr1-153111758-INS-55
chr1-153142346-INS-173
chr1-153153674-INS-86
chr1-153165258-INS-198
chr1-153169646-INS-200
chr1-153175547-INS-68
chr1-153177652-INS-110
chr1-153221329-INS-233
chr1-153225803-INS-110
chr1-153229855-INS-192
chr1-153245230-INS-428
chr1-153248155-INS-77
chr1-153289612-INS-51
chr1-153315812-INS-178
chr1-153322904-INS-51

chr1-153357226-INS-168
chr1-153358931-INS-124
chr1-153369559-INS-70
chr1-153385691-INS-208
chr1-153393880-INS-84
chr1-153397703-INS-229
chr1-153398942-INS-212
chr1-153408246-INS-253
chr1-153425862-INS-298
chr1-153429476-INS-186
chr1-153444394-INS-117
chr1-153451150-INS-100
chr1-153457869-INS-58
chr1-153462259-INS-611
chr1-153464528-INS-498
chr1-153466408-INS-506
chr1-153468712-INS-944
chr1-153479964-INS-180
chr1-153482322-INS-78
chr1-153501029-INS-98
chr1-153533082-INS-52
chr1-153550414-INS-155
chr1-153560413-INS-129
chr1-153574089-INS-7102
chr1-153586237-INS-56
chr1-153598288-INS-108
chr1-153606043-INS-144
chr1-153610839-INS-105
chr1-153615811-INS-339
chr1-153663234-INS-379
chr1-153670841-INS-730
chr1-153670930-INS-247
chr1-153694099-INS-112
chr1-153698954-INS-149
chr1-153702592-INS-94
chr1-153705047-INS-97
chr1-153709035-INS-70
chr1-153716296-INS-77
chr1-153757097-INS-86
chr1-153758077-INS-339
chr1-153780000-INS-69
chr1-153801429-INS-129
chr1-153810341-INS-358
chr1-153837295-INS-486
chr1-153849864-INS-54
chr1-153863529-INS-112
chr1-153866795-INS-66
chr1-153875376-INS-54
chr1-153892255-INS-1154
chr1-153897358-INS-282
chr1-153916104-INS-51
chr1-153919940-INS-176
chr1-153929133-INS-496
chr1-153937921-INS-60
chr1-153952712-INS-226
chr1-154034216-INS-416
chr1-154048851-INS-123
chr1-154055837-INS-419
chr1-154063792-INS-164
chr1-154065384-INS-558
chr1-154065472-INS-190
chr1-154118083-INS-944
chr1-154118867-INS-201
chr1-154144928-INS-52
chr1-154146294-INS-240
chr1-154167174-INS-1860
chr1-154178704-INS-164
chr1-154207066-INS-107
chr1-154227947-INS-423
chr1-154239089-INS-147
chr1-154255791-INS-253
chr1-154264681-INS-372
chr1-154302478-INS-76
chr1-154322823-INS-167
chr1-154324995-INS-99
chr1-154336559-INS-76
chr1-154366925-INS-66

chr1-154401417-INS-168
chr1-154416681-INS-128
chr1-154420174-INS-445
chr1-154469019-INS-342
chr1-154471897-INS-60
chr1-154485016-INS-205
chr1-154489235-INS-6696
chr1-154512711-INS-51
chr1-154535358-INS-145
chr1-154589471-INS-7811
chr1-154607328-INS-106
chr1-154612845-INS-52
chr1-154616753-INS-3444
chr1-154657234-INS-196
chr1-154665248-INS-965
chr1-154728542-INS-357
chr1-154747977-INS-949
chr1-154757363-INS-52
chr1-154787045-INS-176
chr1-154825490-INS-56
chr1-154834843-INS-341
chr1-154853840-INS-186
chr1-154855497-INS-148
chr1-154864362-INS-66
chr1-154884229-INS-164
chr1-154895815-INS-94
chr1-154898414-INS-52
chr1-154916941-INS-5549
chr1-154942160-INS-210
chr1-154946256-INS-52
chr1-154949949-INS-213
chr1-154953360-INS-182
chr1-154955750-INS-116
chr1-154968206-INS-127
chr1-154992173-INS-112
chr1-155002110-INS-201
chr1-155003932-INS-220
chr1-155006789-INS-116
chr1-155009439-INS-3060
chr1-155029066-INS-203
chr1-155038182-INS-299
chr1-155049110-INS-69
chr1-155059320-INS-80
chr1-155086765-INS-206
chr1-155096872-INS-58
chr1-155107965-INS-77
chr1-155150727-INS-117
chr1-155167750-INS-233
chr1-155176232-INS-796
chr1-155179468-INS-343
chr1-155181462-INS-122
chr1-155192178-INS-55
chr1-155254314-INS-6393
chr1-155256398-INS-202
chr1-155263488-INS-222
chr1-155276954-INS-59
chr1-155312724-INS-1706
chr1-155402064-INS-129
chr1-155410891-INS-76
chr1-155447162-INS-197
chr1-155473643-INS-180
chr1-155477474-INS-112
chr1-155479456-INS-236
chr1-155484910-INS-57
chr1-155497168-INS-476
chr1-155523595-INS-1039
chr1-155534146-INS-91
chr1-155540009-INS-418
chr1-155540769-INS-160
chr1-155598453-INS-71
chr1-155636578-INS-12114
chr1-155693828-INS-163
chr1-155694676-INS-162
chr1-155749814-INS-78
chr1-155823591-INS-220
chr1-155844212-INS-257
chr1-155855547-INS-60

chr1-155867820-INS-186
chr1-155916411-INS-99
chr1-155925057-INS-198
chr1-155962140-INS-407
chr1-155968590-INS-161
chr1-155974592-INS-61
chr1-155990073-INS-237
chr1-155996729-INS-190
chr1-156008453-INS-58
chr1-156011647-INS-52
chr1-156033215-INS-64
chr1-156071202-INS-209
chr1-156072949-INS-52
chr1-156076696-INS-182
chr1-156086253-INS-113
chr1-156089397-INS-343
chr1-156092199-INS-491
chr1-156093699-INS-218
chr1-156108298-INS-198
chr1-156111267-INS-329
chr1-156157875-INS-56
chr1-156180425-INS-400
chr1-156192409-INS-620
chr1-156217573-INS-94
chr1-156229300-INS-144
chr1-156254652-INS-199
chr1-156259391-INS-124
chr1-156269154-INS-131
chr1-156277418-INS-265
chr1-156295147-INS-94
chr1-156303078-INS-53
chr1-156305877-INS-72
chr1-156308939-INS-205
chr1-156316827-INS-191
chr1-156323626-INS-176
chr1-156329386-INS-72
chr1-156345233-INS-104
chr1-156364761-INS-138
chr1-156368237-INS-108
chr1-156374219-INS-324
chr1-156379579-INS-216
chr1-156407418-INS-598
chr1-156413569-INS-167
chr1-156426121-INS-401
chr1-156430505-INS-70
chr1-156434455-INS-130
chr1-156448172-INS-1622
chr1-156448173-INS-158
chr1-156448173-INS-940
chr1-156494949-INS-70
chr1-156497595-INS-178
chr1-156502780-INS-178
chr1-156504225-INS-91
chr1-156537187-INS-130
chr1-156551502-INS-274
chr1-156553995-INS-66
chr1-156558206-INS-210
chr1-156566910-INS-269
chr1-156570224-INS-426
chr1-156579297-INS-58
chr1-156584866-INS-177
chr1-156592685-INS-899
chr1-156599642-INS-551
chr1-156623640-INS-131
chr1-156629489-INS-93
chr1-156654767-INS-907
chr1-156683167-INS-72
chr1-156734913-INS-53
chr1-156739584-INS-66
chr1-156745707-INS-238
chr1-156761435-INS-62
chr1-156770901-INS-57
chr1-156773384-INS-169
chr1-156782223-INS-5630
chr1-156793865-INS-514
chr1-156812115-INS-150
chr1-156818446-INS-56

chr1-156860000-INS-123
chr1-156868754-INS-220
chr1-156878107-INS-386
chr1-156899804-INS-452
chr1-156911745-INS-117
chr1-156940690-INS-144
chr1-156962572-INS-210
chr1-156965251-INS-137
chr1-157018064-INS-172
chr1-157022616-INS-60
chr1-157060990-INS-56
chr1-157105421-INS-60
chr1-157116265-INS-215
chr1-157122017-INS-124
chr1-157136996-INS-100
chr1-157141918-INS-755
chr1-157144891-INS-208
chr1-157165272-INS-715
chr1-157167890-INS-54
chr1-157178167-INS-793
chr1-157193817-INS-71
chr1-157195285-INS-310
chr1-157198640-INS-217
chr1-157218198-INS-207
chr1-157248802-INS-176
chr1-157265249-INS-81
chr1-157278744-INS-100
chr1-157289170-INS-240
chr1-157291534-INS-892
chr1-157291541-INS-58
chr1-157308552-INS-221
chr1-157320415-INS-213
chr1-157335156-INS-54
chr1-157356246-INS-402
chr1-157357176-INS-697
chr1-157369438-INS-160
chr1-157386379-INS-214
chr1-157389198-INS-274
chr1-157390934-INS-102
chr1-157416225-INS-52
chr1-157433693-INS-105
chr1-157449533-INS-70
chr1-157451785-INS-52
chr1-157517614-INS-51
chr1-157565022-INS-90
chr1-157575559-INS-368
chr1-157587058-INS-54
chr1-157610733-INS-2792
chr1-157614419-INS-148
chr1-157615359-INS-87
chr1-157645421-INS-104
chr1-157647160-INS-111
chr1-157650231-INS-68
chr1-157659910-INS-285
chr1-157663551-INS-6408
chr1-157670017-INS-13259
chr1-157692818-INS-54
chr1-157702957-INS-462
chr1-157753180-INS-296
chr1-157756978-INS-218
chr1-157769297-INS-80
chr1-157772453-INS-169
chr1-157832342-INS-113
chr1-157835134-INS-52
chr1-157835210-INS-7784
chr1-157873203-INS-149
chr1-157923321-INS-78
chr1-157928839-INS-328
chr1-157944592-INS-7242
chr1-157956274-INS-149
chr1-157966043-INS-51
chr1-157981853-INS-145
chr1-158002494-INS-290
chr1-158030490-INS-400
chr1-158089392-INS-234
chr1-158101632-INS-547
chr1-158104210-INS-56

chr1-158109207-INS-70
chr1-158130596-INS-72
chr1-158139986-INS-56
chr1-158147094-INS-92
chr1-158149621-INS-370
chr1-158151072-INS-5377
chr1-158158134-INS-51
chr1-158210774-INS-82
chr1-158233208-INS-60
chr1-158239059-INS-200
chr1-158250322-INS-94
chr1-158288782-INS-89
chr1-158306153-INS-60
chr1-158317628-INS-79
chr1-158319123-INS-54
chr1-158323407-INS-4739
chr1-158324915-INS-51
chr1-158345976-INS-412
chr1-158354178-INS-525
chr1-158358521-INS-112
chr1-158443964-INS-5209
chr1-158455795-INS-6474
chr1-158501725-INS-56
chr1-158512977-INS-57
chr1-158531890-INS-79
chr1-158546894-INS-8349
chr1-158562274-INS-168
chr1-158570044-INS-60
chr1-158574164-INS-70
chr1-158597182-INS-590
chr1-158602577-INS-312
chr1-158609327-INS-826
chr1-158648650-INS-58
chr1-158679816-INS-138
chr1-158707351-INS-212
chr1-158729732-INS-171
chr1-158735571-INS-466
chr1-158740607-INS-53
chr1-158745261-INS-152
chr1-158798658-INS-6326
chr1-158800786-INS-61
chr1-158803542-INS-130
chr1-158815989-INS-7366
chr1-158867255-INS-454
chr1-158890589-INS-83
chr1-158905843-INS-174
chr1-158923916-INS-339
chr1-158925526-INS-129
chr1-158935483-INS-216
chr1-158989699-INS-132
chr1-158997679-INS-76
chr1-159009206-INS-94
chr1-159078453-INS-164
chr1-159171367-INS-160
chr1-159174439-INS-782
chr1-159204832-INS-504
chr1-159207209-INS-200
chr1-159212243-INS-79
chr1-159254318-INS-178
chr1-159271501-INS-381
chr1-159276118-INS-66
chr1-159277773-INS-68
chr1-159302136-INS-161
chr1-159329184-INS-54
chr1-159350528-INS-61
chr1-159355250-INS-66
chr1-159389686-INS-92
chr1-159397238-INS-175
chr1-159401046-INS-5274
chr1-159401450-INS-124
chr1-159427133-INS-80
chr1-159480122-INS-432
chr1-159484903-INS-53
chr1-159487621-INS-226
chr1-159490680-INS-417
chr1-159534255-INS-652
chr1-159560421-INS-128

chr1-159573212-INS-185
chr1-159588394-INS-85
chr1-159595434-INS-99
chr1-159615484-INS-63
chr1-159654871-INS-80
chr1-159659495-INS-66
chr1-159668521-INS-66
chr1-159669698-INS-64
chr1-159672251-INS-112
chr1-159713581-INS-195
chr1-159729200-INS-74
chr1-159745962-INS-80
chr1-159783031-INS-70
chr1-159807587-INS-6897
chr1-159818800-INS-54
chr1-159831472-INS-6633
chr1-159842227-INS-313
chr1-159852215-INS-170
chr1-159858543-INS-401
chr1-159862721-INS-52
chr1-159881107-INS-60
chr1-159939004-INS-260
chr1-159967750-INS-126
chr1-159970355-INS-6436
chr1-159974841-INS-344
chr1-159986606-INS-61
chr1-159992852-INS-189
chr1-159994209-INS-75
chr1-160028902-INS-373
chr1-160227906-INS-153
chr1-160234240-INS-65
chr1-160241328-INS-224
chr1-160263366-INS-143
chr1-160299914-INS-73
chr1-160323877-INS-76
chr1-160327946-INS-358
chr1-160329488-INS-60
chr1-160332744-INS-206
chr1-160357212-INS-676
chr1-160376487-INS-66
chr1-160385692-INS-111
chr1-160390900-INS-84
chr1-160418932-INS-627
chr1-160430929-INS-213
chr1-160445585-INS-219
chr1-160450506-INS-57
chr1-160476934-INS-307
chr1-160485997-INS-806
chr1-160493888-INS-351
chr1-160524939-INS-269
chr1-160545431-INS-189
chr1-160556318-INS-821
chr1-160558492-INS-218
chr1-160570465-INS-76
chr1-160572238-INS-965
chr1-160585744-INS-220
chr1-160598714-INS-152
chr1-160633400-INS-192
chr1-160667251-INS-329
chr1-160676606-INS-215
chr1-160685933-INS-76
chr1-160690223-INS-208
chr1-160705774-INS-52
chr1-160705951-INS-6195
chr1-160715796-INS-638
chr1-160716674-INS-164
chr1-160721023-INS-172
chr1-160725070-INS-180
chr1-160728633-INS-213
chr1-160774604-INS-63
chr1-160776304-INS-189
chr1-160788796-INS-81
chr1-160841070-INS-170
chr1-160902037-INS-154
chr1-160904350-INS-68
chr1-160919270-INS-195
chr1-160956806-INS-64

chr1-160980995-INS-120
chr1-161031548-INS-55
chr1-161043835-INS-68
chr1-161081405-INS-66
chr1-161086457-INS-192
chr1-161102780-INS-55
chr1-161115414-INS-122
chr1-161207624-INS-78
chr1-161211695-INS-64
chr1-161217992-INS-546
chr1-161240266-INS-72
chr1-161242792-INS-67
chr1-161248785-INS-125
chr1-161261254-INS-125
chr1-161275702-INS-92
chr1-161282429-INS-935
chr1-161308445-INS-189
chr1-161333085-INS-82
chr1-161346405-INS-484
chr1-161348709-INS-170
chr1-161357744-INS-7778
chr1-161363489-INS-3520
chr1-161379406-INS-6504
chr1-161382319-INS-75
chr1-161414951-INS-54
chr1-161421266-INS-196
chr1-161422663-INS-58
chr1-161426875-INS-104
chr1-161442409-INS-94
chr1-161445924-INS-78
chr1-161450209-INS-80
chr1-161473164-INS-6509
chr1-161539866-INS-194
chr1-161566020-INS-6477
chr1-161569882-INS-158
chr1-161573832-INS-7226
chr1-161576523-INS-59
chr1-161577345-INS-151
chr1-161590578-INS-188
chr1-161592610-INS-2680
chr1-161635841-INS-72
chr1-161641980-INS-100
chr1-161676814-INS-425
chr1-161676915-INS-55
chr1-161719925-INS-72
chr1-161756581-INS-1763
chr1-161780500-INS-83
chr1-161825457-INS-288
chr1-161840041-INS-52
chr1-161890759-INS-165
chr1-161917333-INS-116
chr1-161933531-INS-7207
chr1-161957228-INS-230
chr1-161967482-INS-100
chr1-161974540-INS-54
chr1-161978126-INS-106
chr1-161995159-INS-214
chr1-162022693-INS-75
chr1-162048767-INS-179
chr1-162063340-INS-7443
chr1-162064627-INS-56
chr1-162124353-INS-86
chr1-162165148-INS-6459
chr1-162194700-INS-363
chr1-162212023-INS-68
chr1-162213013-INS-56
chr1-162251800-INS-109
chr1-162292343-INS-86
chr1-162328704-INS-549
chr1-162340973-INS-78
chr1-162348760-INS-167
chr1-162354080-INS-180
chr1-162391864-INS-272
chr1-162396917-INS-7971
chr1-162419261-INS-122
chr1-162431154-INS-193
chr1-162434590-INS-1102

chr1-162446120-INS-168
chr1-162478680-INS-190
chr1-162479623-INS-423
chr1-162491250-INS-67
chr1-162494807-INS-180
chr1-162512002-INS-204
chr1-162614927-INS-334
chr1-162639393-INS-965
chr1-162679526-INS-51
chr1-162684691-INS-104
chr1-162695573-INS-222
chr1-162696514-INS-95
chr1-162709073-INS-4617
chr1-162735302-INS-418
chr1-162750794-INS-5217
chr1-162752845-INS-288
chr1-162796898-INS-200
chr1-162805393-INS-220
chr1-162851898-INS-203
chr1-162869727-INS-4335
chr1-162902380-INS-260
chr1-162915351-INS-135
chr1-162961595-INS-158
chr1-162964464-INS-62
chr1-162971232-INS-590
chr1-163003167-INS-71
chr1-163056842-INS-67
chr1-163084650-INS-69
chr1-163085957-INS-399
chr1-163087919-INS-69
chr1-163167085-INS-111
chr1-163333677-INS-277
chr1-163337338-INS-7106
chr1-163340230-INS-154
chr1-163360775-INS-83
chr1-163430933-INS-836
chr1-163452497-INS-63
chr1-163471320-INS-3336
chr1-163578351-INS-56
chr1-163657484-INS-108
chr1-163660349-INS-53
chr1-163692691-INS-110
chr1-163699738-INS-65
chr1-163700677-INS-102
chr1-163701622-INS-58
chr1-163710235-INS-211
chr1-163719152-INS-528
chr1-163748390-INS-51
chr1-163764767-INS-93
chr1-163831478-INS-534
chr1-163840200-INS-218
chr1-163842329-INS-122
chr1-163858221-INS-196
chr1-163880746-INS-64
chr1-163883551-INS-61
chr1-163892693-INS-170
chr1-163896394-INS-737
chr1-163900500-INS-69
chr1-163920844-INS-93
chr1-163922178-INS-506
chr1-163966923-INS-139
chr1-164010131-INS-71
chr1-164018936-INS-271
chr1-164021215-INS-160
chr1-164023924-INS-449
chr1-164030053-INS-105
chr1-164068593-INS-60
chr1-164099462-INS-74
chr1-164113315-INS-242
chr1-164131654-INS-196
chr1-164205762-INS-145
chr1-164215434-INS-216
chr1-164287825-INS-53
chr1-164302727-INS-168
chr1-164306066-INS-53
chr1-164315010-INS-96
chr1-164318292-INS-745

chr1-164332229-INS-100
chr1-164335761-INS-207
chr1-164338208-INS-316
chr1-164348066-INS-64
chr1-164367384-INS-63
chr1-164367787-INS-191
chr1-164388889-INS-71
chr1-164428609-INS-71
chr1-164440522-INS-98
chr1-164470775-INS-164
chr1-164498132-INS-340
chr1-164503478-INS-58
chr1-164507563-INS-1126
chr1-164507779-INS-193
chr1-164558350-INS-91
chr1-164562351-INS-114
chr1-164583064-INS-304
chr1-164589716-INS-360
chr1-164601651-INS-429
chr1-164603499-INS-2577
chr1-164605848-INS-78
chr1-164606854-INS-626
chr1-164616364-INS-96
chr1-164705977-INS-50
chr1-164708546-INS-496
chr1-164738649-INS-2046
chr1-164740727-INS-104
chr1-164780697-INS-187
chr1-164799984-INS-455
chr1-164822238-INS-51
chr1-164824445-INS-126
chr1-164841821-INS-54
chr1-164850608-INS-144
chr1-164862592-INS-5515
chr1-164882945-INS-7145
chr1-164889740-INS-75
chr1-164899107-INS-67
chr1-164937730-INS-136
chr1-164946733-INS-262
chr1-164974224-INS-80
chr1-164982804-INS-82
chr1-164996858-INS-70
chr1-164998294-INS-166
chr1-165026624-INS-192
chr1-165039901-INS-56
chr1-165045947-INS-293
chr1-165068653-INS-55
chr1-165078901-INS-105
chr1-165093477-INS-157
chr1-165112229-INS-61
chr1-165156494-INS-135
chr1-165156607-INS-446
chr1-165161850-INS-460
chr1-165172644-INS-66
chr1-165217499-INS-564
chr1-165259975-INS-824
chr1-165271425-INS-664
chr1-165278622-INS-214
chr1-165300025-INS-57
chr1-165311367-INS-893
chr1-165319404-INS-242
chr1-165339380-INS-99
chr1-165373201-INS-60
chr1-165377724-INS-76
chr1-165402845-INS-245
chr1-165407512-INS-66
chr1-165413892-INS-837
chr1-165438969-INS-54
chr1-165463442-INS-186
chr1-165465035-INS-58
chr1-165500276-INS-99
chr1-165500928-INS-109
chr1-165504166-INS-63
chr1-165506287-INS-127
chr1-165531504-INS-116
chr1-165548413-INS-55
chr1-165560150-INS-51

chr1-165567034-INS-241
chr1-165569411-INS-86
chr1-165584770-INS-57
chr1-165632883-INS-1431
chr1-165639376-INS-100
chr1-165645277-INS-80
chr1-165691167-INS-268
chr1-165725066-INS-101
chr1-165743353-INS-246
chr1-165753748-INS-2957
chr1-165756082-INS-56
chr1-165758264-INS-201
chr1-165767389-INS-81
chr1-165771283-INS-728
chr1-165806507-INS-7339
chr1-165809280-INS-204
chr1-165824276-INS-624
chr1-165845329-INS-259
chr1-165859298-INS-109
chr1-165860303-INS-504
chr1-165861653-INS-114
chr1-165875089-INS-806
chr1-165888232-INS-81
chr1-165891197-INS-144
chr1-165898948-INS-528
chr1-165905511-INS-162
chr1-165933099-INS-79
chr1-165949231-INS-52
chr1-165975827-INS-803
chr1-165976046-INS-60
chr1-165989406-INS-176
chr1-165998244-INS-124
chr1-166016368-INS-366
chr1-166017864-INS-6325
chr1-166025718-INS-439
chr1-166046587-INS-253
chr1-166108028-INS-151
chr1-166128312-INS-77
chr1-166155423-INS-58
chr1-166174332-INS-961
chr1-166175957-INS-52
chr1-166179066-INS-53
chr1-166203602-INS-55
chr1-166252324-INS-214
chr1-166269104-INS-1888
chr1-166272114-INS-1470
chr1-166276995-INS-179
chr1-166305516-INS-756
chr1-166353399-INS-63
chr1-166354755-INS-5730
chr1-166472752-INS-779
chr1-166484164-INS-1302
chr1-166484894-INS-58
chr1-166494181-INS-54
chr1-166509893-INS-391
chr1-166518413-INS-6827
chr1-166546562-INS-87
chr1-166563781-INS-67
chr1-166572567-INS-86
chr1-166574119-INS-132
chr1-166577278-INS-63
chr1-166598848-INS-189
chr1-166599378-INS-85
chr1-166617663-INS-400
chr1-166638283-INS-87
chr1-166649402-INS-143
chr1-166659021-INS-251
chr1-166664751-INS-89
chr1-166664848-INS-60
chr1-166684968-INS-4492
chr1-166705451-INS-62
chr1-166712910-INS-200
chr1-166724755-INS-130
chr1-166759078-INS-1120
chr1-166759085-INS-854
chr1-166772087-INS-222
chr1-166812224-INS-84

chr1-166813798-INS-79
chr1-166815566-INS-612
chr1-166839325-INS-54
chr1-166842544-INS-109
chr1-166853122-INS-577
chr1-166870527-INS-76
chr1-166876396-INS-6721
chr1-166890341-INS-6292
chr1-166931842-INS-66
chr1-166942216-INS-57
chr1-166959774-INS-120
chr1-166962747-INS-82
chr1-166976679-INS-560
chr1-167039417-INS-65
chr1-167047686-INS-526
chr1-167056083-INS-57
chr1-167082204-INS-642
chr1-167092685-INS-156
chr1-167098634-INS-304
chr1-167099297-INS-207
chr1-167118101-INS-231
chr1-167122861-INS-117
chr1-167125217-INS-175
chr1-167168113-INS-220
chr1-167213199-INS-69
chr1-167255871-INS-176
chr1-167260326-INS-52
chr1-167280413-INS-178
chr1-167303978-INS-1609
chr1-167308807-INS-497
chr1-167317394-INS-74
chr1-167331438-INS-158
chr1-167356001-INS-53
chr1-167406030-INS-131
chr1-167414861-INS-62
chr1-167423814-INS-58
chr1-167426225-INS-56
chr1-167450716-INS-51
chr1-167469952-INS-714
chr1-167498892-INS-469
chr1-167513111-INS-218
chr1-167540904-INS-69
chr1-167591151-INS-279
chr1-167616051-INS-168
chr1-167636728-INS-187
chr1-167640723-INS-82
chr1-167648674-INS-74
chr1-167669696-INS-231
chr1-167729282-INS-52
chr1-167736626-INS-206
chr1-167818289-INS-726
chr1-167830316-INS-6401
chr1-167834168-INS-68
chr1-167837407-INS-1020
chr1-167839190-INS-80
chr1-167870045-INS-58
chr1-167895542-INS-53
chr1-167897875-INS-1960
chr1-167897876-INS-58
chr1-167907903-INS-77
chr1-167922504-INS-3006
chr1-167953482-INS-4095
chr1-167954888-INS-94
chr1-167956299-INS-57
chr1-167973252-INS-821
chr1-168030268-INS-148
chr1-168079171-INS-423
chr1-168088736-INS-379
chr1-168115070-INS-164
chr1-168121201-INS-157
chr1-168146296-INS-293
chr1-168157150-INS-52
chr1-168224871-INS-777
chr1-168226020-INS-165
chr1-168274090-INS-203
chr1-168291372-INS-122
chr1-168295613-INS-52

chr1-168328463-INS-57
chr1-168330903-INS-52
chr1-168352474-INS-680
chr1-168363999-INS-469
chr1-168382261-INS-232
chr1-168400841-INS-3956
chr1-168411682-INS-6463
chr1-168417170-INS-288
chr1-168437688-INS-2102
chr1-168445947-INS-199
chr1-168465225-INS-7669
chr1-168485544-INS-123
chr1-168515231-INS-71
chr1-168530610-INS-72
chr1-168556858-INS-162
chr1-168558227-INS-108
chr1-168590864-INS-65
chr1-168610954-INS-4431
chr1-168654315-INS-52
chr1-168667869-INS-82
chr1-168678033-INS-51
chr1-168772707-INS-52
chr1-168808076-INS-169
chr1-168822813-INS-76
chr1-168848336-INS-80
chr1-168878994-INS-139
chr1-168892197-INS-64
chr1-168893095-INS-6712
chr1-168905246-INS-112
chr1-168952061-INS-96
chr1-168964281-INS-55
chr1-169000391-INS-102
chr1-169001862-INS-421
chr1-169026176-INS-854
chr1-169105696-INS-4715
chr1-169123563-INS-180
chr1-169135488-INS-136
chr1-169176201-INS-271
chr1-169176589-INS-1337
chr1-169211581-INS-51
chr1-169214962-INS-55
chr1-169217133-INS-54
chr1-169217887-INS-72
chr1-169225836-INS-144
chr1-169253467-INS-214
chr1-169259753-INS-92
chr1-169279390-INS-115
chr1-169347906-INS-58
chr1-169349683-INS-60
chr1-169388143-INS-52
chr1-169427301-INS-230
chr1-169431775-INS-166
chr1-169440091-INS-290
chr1-169442454-INS-247
chr1-169452092-INS-175
chr1-169530369-INS-6235
chr1-169536833-INS-178
chr1-169554117-INS-146
chr1-169707229-INS-179
chr1-169722761-INS-68
chr1-169735131-INS-64
chr1-169757371-INS-69
chr1-169785624-INS-279
chr1-169808469-INS-103
chr1-169810704-INS-396
chr1-169819891-INS-387
chr1-169850841-INS-121
chr1-169878094-INS-51
chr1-169890628-INS-65
chr1-169919778-INS-60
chr1-169930242-INS-52
chr1-169993026-INS-644
chr1-170004013-INS-984
chr1-170028802-INS-208
chr1-170062532-INS-740
chr1-170109280-INS-64
chr1-170135213-INS-54

chr1-170144430-INS-62
chr1-170147267-INS-72
chr1-170154145-INS-90
chr1-170206654-INS-75
chr1-170217733-INS-169
chr1-170220305-INS-358
chr1-170249057-INS-169
chr1-170264107-INS-164
chr1-170275528-INS-53
chr1-170281558-INS-64
chr1-170296732-INS-107
chr1-170304800-INS-915
chr1-170316134-INS-97
chr1-170345526-INS-135
chr1-170351123-INS-68
chr1-170352478-INS-146
chr1-170357931-INS-92
chr1-170365306-INS-446
chr1-170374591-INS-62
chr1-170377144-INS-79
chr1-170401854-INS-250
chr1-170407009-INS-120
chr1-170410858-INS-139
chr1-170420132-INS-497
chr1-170445329-INS-120
chr1-170446959-INS-192
chr1-170518141-INS-94
chr1-170519949-INS-6799
chr1-170521276-INS-6684
chr1-170522069-INS-142
chr1-170526264-INS-195
chr1-170632436-INS-53
chr1-170644593-INS-206
chr1-170671228-INS-110
chr1-170674772-INS-102
chr1-170695579-INS-71
chr1-170718467-INS-718
chr1-170718994-INS-7080
chr1-170739289-INS-84
chr1-170765299-INS-176
chr1-170811108-INS-302
chr1-170825017-INS-507
chr1-170833482-INS-149
chr1-170839665-INS-209
chr1-170876123-INS-83
chr1-170883712-INS-158
chr1-170884605-INS-230
chr1-170891253-INS-579
chr1-170919400-INS-223
chr1-170922485-INS-538
chr1-170936339-INS-3051
chr1-170945860-INS-56
chr1-170990139-INS-211
chr1-171015431-INS-7482
chr1-171025024-INS-81
chr1-171029633-INS-217
chr1-171031708-INS-205
chr1-171071795-INS-214
chr1-171092946-INS-160
chr1-171094484-INS-917
chr1-171123611-INS-52
chr1-171125615-INS-184
chr1-171132552-INS-184
chr1-171136644-INS-209
chr1-171152416-INS-219
chr1-171153400-INS-62
chr1-171155818-INS-181
chr1-171164486-INS-216
chr1-171182985-INS-178
chr1-171190242-INS-138
chr1-171192472-INS-58
chr1-171209009-INS-130
chr1-171227959-INS-63
chr1-171248893-INS-245
chr1-171254350-INS-213
chr1-171269877-INS-659
chr1-171295978-INS-72

chr1-171299738-INS-245
chr1-171304080-INS-202
chr1-171318028-INS-64
chr1-171324039-INS-143
chr1-171325992-INS-117
chr1-171327116-INS-543
chr1-171328602-INS-85
chr1-171329567-INS-935
chr1-171338981-INS-77
chr1-171371713-INS-132
chr1-171375316-INS-59
chr1-171418357-INS-53
chr1-171450640-INS-60
chr1-171455777-INS-401
chr1-171458026-INS-72
chr1-171505101-INS-136
chr1-171611871-INS-202
chr1-171625129-INS-435
chr1-171628587-INS-382
chr1-171664840-INS-7151
chr1-171666894-INS-215
chr1-171724801-INS-241
chr1-171765717-INS-227
chr1-171769902-INS-86
chr1-171774247-INS-842
chr1-171776737-INS-60
chr1-171781651-INS-477
chr1-171795048-INS-113
chr1-171805468-INS-132
chr1-171861207-INS-2884
chr1-171861771-INS-105
chr1-171869322-INS-58
chr1-171890093-INS-193
chr1-171891741-INS-51
chr1-171902020-INS-64
chr1-171907684-INS-209
chr1-171933505-INS-223
chr1-171936705-INS-111
chr1-171958021-INS-214
chr1-171965780-INS-199
chr1-171986113-INS-211
chr1-171990716-INS-198
chr1-172044227-INS-72
chr1-172054186-INS-1123
chr1-172061149-INS-106
chr1-172083165-INS-198
chr1-172165324-INS-54
chr1-172167613-INS-59
chr1-172211990-INS-7438
chr1-172213924-INS-564
chr1-172233496-INS-98
chr1-172239654-INS-227
chr1-172240778-INS-213
chr1-172243779-INS-52
chr1-172246549-INS-201
chr1-172247042-INS-265
chr1-172266527-INS-60
chr1-172279082-INS-52
chr1-172300691-INS-785
chr1-172346514-INS-222
chr1-172419234-INS-2355
chr1-172438089-INS-72
chr1-172473852-INS-52
chr1-172489088-INS-117
chr1-172498886-INS-101
chr1-172520848-INS-56
chr1-172524070-INS-61
chr1-172537306-INS-644
chr1-172621966-INS-506
chr1-172623078-INS-62
chr1-172624036-INS-86
chr1-172653888-INS-110
chr1-172688160-INS-7200
chr1-172690734-INS-53
chr1-172703632-INS-1656
chr1-172755467-INS-241
chr1-172791339-INS-145

chr1-172808626-INS-292
chr1-172822860-INS-278
chr1-172868853-INS-64
chr1-172869831-INS-558
chr1-172904439-INS-68
chr1-172910828-INS-65
chr1-172930667-INS-52
chr1-173072805-INS-429
chr1-173088165-INS-207
chr1-173099728-INS-601
chr1-173104011-INS-112
chr1-173118357-INS-2653
chr1-173124086-INS-209
chr1-173134931-INS-96
chr1-173144599-INS-222
chr1-173210043-INS-5065
chr1-173227610-INS-847
chr1-173246126-INS-54
chr1-173261350-INS-70
chr1-173293184-INS-4080
chr1-173296708-INS-228
chr1-173302826-INS-117
chr1-173326983-INS-94
chr1-173433988-INS-181
chr1-173439529-INS-54
chr1-173443287-INS-58
chr1-173483075-INS-496
chr1-173568295-INS-81
chr1-173581269-INS-393
chr1-173597648-INS-641
chr1-173603188-INS-67
chr1-173626466-INS-73
chr1-173648761-INS-51
chr1-173649017-INS-820
chr1-173668846-INS-62
chr1-173671886-INS-62
chr1-173704033-INS-55
chr1-173716272-INS-203
chr1-173743618-INS-641
chr1-173768589-INS-6252
chr1-173815775-INS-52
chr1-173845523-INS-199
chr1-173858750-INS-265
chr1-173913254-INS-69
chr1-173933631-INS-258
chr1-174031121-INS-65
chr1-174041901-INS-183
chr1-174064155-INS-604
chr1-174089545-INS-940
chr1-174130656-INS-99
chr1-174145931-INS-137
chr1-174151966-INS-66
chr1-174162837-INS-162
chr1-174213937-INS-56
chr1-174239225-INS-190
chr1-174244224-INS-440
chr1-174274637-INS-181
chr1-174279520-INS-216
chr1-174302572-INS-469
chr1-174315508-INS-524
chr1-174318764-INS-106
chr1-174328917-INS-107
chr1-174343193-INS-206
chr1-174345044-INS-79
chr1-174349085-INS-72
chr1-174359043-INS-122
chr1-174367502-INS-90
chr1-174417307-INS-201
chr1-174420238-INS-2932
chr1-174445971-INS-91
chr1-174476769-INS-237
chr1-174517445-INS-6361
chr1-174532900-INS-232
chr1-174551893-INS-82
chr1-174595546-INS-79
chr1-174618458-INS-109
chr1-174620023-INS-264

chr1-174643639-INS-315
chr1-174643797-INS-321
chr1-174670323-INS-903
chr1-174682714-INS-86
chr1-174688904-INS-55
chr1-174699544-INS-123
chr1-174755834-INS-56
chr1-174829957-INS-51
chr1-174831437-INS-6203
chr1-174854102-INS-659
chr1-174858092-INS-51
chr1-174867956-INS-78
chr1-174981353-INS-504
chr1-174993312-INS-6488
chr1-174994186-INS-106
chr1-175024712-INS-759
chr1-175036618-INS-141
chr1-175059835-INS-211
chr1-175094580-INS-526
chr1-175099657-INS-514
chr1-175122242-INS-62
chr1-175126343-INS-150
chr1-175132570-INS-51
chr1-175136544-INS-164
chr1-175141017-INS-453
chr1-175192941-INS-876
chr1-175269164-INS-84
chr1-175272995-INS-66
chr1-175274479-INS-90
chr1-175280129-INS-787
chr1-175280185-INS-68
chr1-175294794-INS-62
chr1-175313804-INS-5369
chr1-175330646-INS-60
chr1-175358310-INS-516
chr1-175374919-INS-68
chr1-175409750-INS-233
chr1-175416014-INS-75
chr1-175430643-INS-84
chr1-175454694-INS-611
chr1-175463300-INS-57
chr1-175503621-INS-54
chr1-175522807-INS-507
chr1-175525732-INS-147
chr1-175535230-INS-356
chr1-175538519-INS-50
chr1-175542863-INS-487
chr1-175546445-INS-106
chr1-175556503-INS-163
chr1-175571882-INS-497
chr1-175581572-INS-90
chr1-175583350-INS-52
chr1-175608738-INS-417
chr1-175612751-INS-54
chr1-175631462-INS-54
chr1-175661910-INS-104
chr1-175664651-INS-52
chr1-175714789-INS-96
chr1-175723349-INS-423
chr1-175749812-INS-56
chr1-175757651-INS-145
chr1-175840811-INS-64
chr1-175845393-INS-469
chr1-175865098-INS-511
chr1-175870081-INS-56
chr1-175880664-INS-4314
chr1-175885234-INS-66
chr1-175891386-INS-144
chr1-175898608-INS-97
chr1-175918271-INS-204
chr1-175921109-INS-6415
chr1-175928120-INS-56
chr1-175942918-INS-75
chr1-175953159-INS-120
chr1-175960369-INS-865
chr1-175973223-INS-716
chr1-175977444-INS-87

chr1-175981350-INS-593
chr1-175992400-INS-491
chr1-176000423-INS-165
chr1-176016998-INS-135
chr1-176023724-INS-52
chr1-176039402-INS-401
chr1-176042724-INS-195
chr1-176048994-INS-62
chr1-176072298-INS-52
chr1-176123243-INS-6127
chr1-176123737-INS-346
chr1-176134584-INS-61
chr1-176145638-INS-60
chr1-176146330-INS-95
chr1-176164179-INS-52
chr1-176168389-INS-63
chr1-176178474-INS-130
chr1-176191700-INS-464
chr1-176212115-INS-54
chr1-176235303-INS-2804
chr1-176245813-INS-9978
chr1-176253416-INS-60
chr1-176258727-INS-384
chr1-176273327-INS-133
chr1-176274393-INS-5457
chr1-176287536-INS-332
chr1-176295602-INS-412
chr1-176301662-INS-134
chr1-176344187-INS-56
chr1-176351864-INS-505
chr1-176354925-INS-3406
chr1-176392437-INS-6680
chr1-176395184-INS-6260
chr1-176409121-INS-57
chr1-176419642-INS-493
chr1-176426204-INS-84
chr1-176428227-INS-6684
chr1-176433724-INS-62
chr1-176452981-INS-5547
chr1-176461351-INS-70
chr1-176463769-INS-7172
chr1-176480999-INS-89
chr1-176489318-INS-64
chr1-176498645-INS-7313
chr1-176508317-INS-7694
chr1-176528078-INS-287
chr1-176533209-INS-1873
chr1-176538424-INS-519
chr1-176549081-INS-256
chr1-176549765-INS-108
chr1-176566066-INS-450
chr1-176569174-INS-262
chr1-176611222-INS-64
chr1-176647214-INS-252
chr1-176654564-INS-400
chr1-176671460-INS-6374
chr1-176690495-INS-116
chr1-176694343-INS-137
chr1-176698368-INS-52
chr1-176705021-INS-6417
chr1-176711558-INS-832
chr1-176752987-INS-872
chr1-176761896-INS-55
chr1-176786256-INS-61
chr1-176787695-INS-53
chr1-176805099-INS-54
chr1-176828032-INS-84
chr1-176829732-INS-72
chr1-176836623-INS-843
chr1-176850818-INS-194
chr1-176868065-INS-243
chr1-176882855-INS-517
chr1-176891685-INS-182
chr1-176919103-INS-6411
chr1-176924224-INS-183
chr1-176934745-INS-210
chr1-176939927-INS-64

chr1-176940970-INS-1037
chr1-176966849-INS-57
chr1-176994702-INS-662
chr1-177009038-INS-60
chr1-177035590-INS-1071
chr1-177064511-INS-286
chr1-177067603-INS-150
chr1-177069952-INS-570
chr1-177071641-INS-52
chr1-177111445-INS-66
chr1-177119010-INS-205
chr1-177129110-INS-444
chr1-177129957-INS-154
chr1-177130605-INS-52
chr1-177141210-INS-247
chr1-177149154-INS-1475
chr1-177157327-INS-179
chr1-177186809-INS-363
chr1-177198258-INS-236
chr1-177202158-INS-68
chr1-177228460-INS-452
chr1-177238120-INS-52
chr1-177242087-INS-412
chr1-177248566-INS-64
chr1-177289718-INS-65
chr1-177305439-INS-52
chr1-177339742-INS-102
chr1-177406343-INS-158
chr1-177407424-INS-169
chr1-177411772-INS-432
chr1-177416302-INS-6489
chr1-177430782-INS-7957
chr1-177498309-INS-89
chr1-177522362-INS-95
chr1-177523726-INS-84
chr1-177525542-INS-58
chr1-177530894-INS-220
chr1-177535596-INS-2473
chr1-177537216-INS-210
chr1-177539596-INS-91
chr1-177544199-INS-448
chr1-177549982-INS-89
chr1-177566173-INS-120
chr1-177567332-INS-1894
chr1-177576752-INS-93
chr1-177594271-INS-391
chr1-177615092-INS-56
chr1-177639712-INS-60
chr1-177643357-INS-287
chr1-177643478-INS-387
chr1-177685603-INS-150
chr1-177704418-INS-68
chr1-177709668-INS-1892
chr1-177727700-INS-6499
chr1-177733013-INS-57
chr1-177785065-INS-368
chr1-177875630-INS-83
chr1-177934714-INS-312
chr1-177940311-INS-53
chr1-178006455-INS-3854
chr1-178036853-INS-1119
chr1-178106957-INS-104
chr1-178107070-INS-911
chr1-178113084-INS-332
chr1-178124058-INS-328
chr1-178125263-INS-539
chr1-178131740-INS-622
chr1-178141014-INS-64
chr1-178170484-INS-618
chr1-178183079-INS-58
chr1-178189711-INS-624
chr1-178204133-INS-52
chr1-178206621-INS-162
chr1-178211839-INS-66
chr1-178214910-INS-238
chr1-178243569-INS-267
chr1-178247754-INS-85

chr1-178248677-INS-57
chr1-178260796-INS-204
chr1-178271144-INS-64
chr1-178275127-INS-506
chr1-178287396-INS-67
chr1-178293959-INS-152
chr1-178323523-INS-64
chr1-178346560-INS-132
chr1-178359286-INS-109
chr1-178376702-INS-308
chr1-178400652-INS-180
chr1-178400697-INS-6571
chr1-178427043-INS-122
chr1-178429890-INS-56
chr1-178464320-INS-188
chr1-178477143-INS-186
chr1-178484241-INS-461
chr1-178487379-INS-74
chr1-178496123-INS-138
chr1-178505197-INS-52
chr1-178513954-INS-89
chr1-178539365-INS-179
chr1-178552110-INS-2166
chr1-178554412-INS-228
chr1-178580962-INS-1228
chr1-178598398-INS-53
chr1-178600982-INS-186
chr1-178604308-INS-59
chr1-178627341-INS-129
chr1-178646107-INS-229
chr1-178656279-INS-52
chr1-178678549-INS-903
chr1-178684860-INS-59
chr1-178702506-INS-167
chr1-178707097-INS-115
chr1-178733914-INS-67
chr1-178759059-INS-62
chr1-178818105-INS-54
chr1-178820778-INS-80
chr1-178896262-INS-70
chr1-178896970-INS-6701
chr1-178935607-INS-53
chr1-178967557-INS-56
chr1-178975903-INS-56
chr1-178976857-INS-615
chr1-178987705-INS-204
chr1-179012968-INS-333
chr1-179030478-INS-53
chr1-179062467-INS-58
chr1-179107798-INS-53
chr1-179124460-INS-80
chr1-179125420-INS-56
chr1-179156837-INS-68
chr1-179167458-INS-228
chr1-179222366-INS-463
chr1-179238832-INS-57
chr1-179239790-INS-343
chr1-179249385-INS-64
chr1-179253113-INS-212
chr1-179284795-INS-67
chr1-179289073-INS-81
chr1-179294397-INS-84
chr1-179298351-INS-51
chr1-179311469-INS-55
chr1-179335851-INS-216
chr1-179337660-INS-56
chr1-179348813-INS-5024
chr1-179372670-INS-68
chr1-179381621-INS-509
chr1-179385431-INS-182
chr1-179400630-INS-94
chr1-179404650-INS-54
chr1-179417585-INS-211
chr1-179425143-INS-60
chr1-179438451-INS-140
chr1-179439930-INS-109
chr1-179449598-INS-56

chr1-179471719-INS-508
chr1-179474276-INS-90
chr1-179491533-INS-1188
chr1-179491699-INS-199
chr1-179498269-INS-214
chr1-179503012-INS-161
chr1-179511663-INS-206
chr1-179515674-INS-609
chr1-179522479-INS-455
chr1-179535290-INS-192
chr1-179539372-INS-593
chr1-179545490-INS-209
chr1-179547266-INS-548
chr1-179555067-INS-86
chr1-179566131-INS-382
chr1-179587443-INS-205
chr1-179649994-INS-158
chr1-179667780-INS-231
chr1-179668775-INS-88
chr1-179673159-INS-72
chr1-179700177-INS-193
chr1-179702742-INS-61
chr1-179705306-INS-59
chr1-179710120-INS-6868
chr1-179715328-INS-257
chr1-179715328-INS-700
chr1-179734709-INS-202
chr1-179739184-INS-58
chr1-179743731-INS-201
chr1-179751447-INS-6536
chr1-179752493-INS-6449
chr1-179857378-INS-195
chr1-179859501-INS-585
chr1-179860713-INS-188
chr1-179868509-INS-361
chr1-179869442-INS-203
chr1-179882940-INS-56
chr1-179888254-INS-603
chr1-179928080-INS-2995
chr1-179968099-INS-232
chr1-179976084-INS-6224
chr1-179982433-INS-56
chr1-180008708-INS-116
chr1-180032284-INS-211
chr1-180051500-INS-4848
chr1-180088160-INS-92
chr1-180110848-INS-51
chr1-180114421-INS-210
chr1-180166375-INS-2001
chr1-180178842-INS-227
chr1-180180307-INS-56
chr1-180189680-INS-60
chr1-180192568-INS-159
chr1-180198710-INS-113
chr1-180206722-INS-69
chr1-180219689-INS-71
chr1-180246957-INS-100
chr1-180278640-INS-204
chr1-180287955-INS-132
chr1-180296911-INS-61
chr1-180298803-INS-360
chr1-180300874-INS-73
chr1-180316407-INS-560
chr1-180323342-INS-104
chr1-180325018-INS-52
chr1-180336293-INS-113
chr1-180342806-INS-181
chr1-180349860-INS-93
chr1-180363267-INS-124
chr1-180370427-INS-210
chr1-180385412-INS-52
chr1-180390255-INS-171
chr1-180392090-INS-180
chr1-180402367-INS-132
chr1-180405772-INS-288
chr1-180438043-INS-332
chr1-180450940-INS-267

chr1-180462908-INS-633
chr1-180476960-INS-263
chr1-180489791-INS-214
chr1-180502097-INS-123
chr1-180512521-INS-231
chr1-180519729-INS-92
chr1-180523888-INS-92
chr1-180545175-INS-168
chr1-180577478-INS-207
chr1-180594014-INS-213
chr1-180609907-INS-85
chr1-180612425-INS-363
chr1-180619040-INS-96
chr1-180621095-INS-136
chr1-180622561-INS-569
chr1-180649232-INS-233
chr1-180656507-INS-181
chr1-180662672-INS-137
chr1-180665740-INS-51
chr1-180666551-INS-56
chr1-180668838-INS-115
chr1-180675189-INS-226
chr1-180679409-INS-7344
chr1-180683869-INS-173
chr1-180686894-INS-196
chr1-180689670-INS-208
chr1-180691926-INS-54
chr1-180700346-INS-394
chr1-180704464-INS-59
chr1-180708311-INS-433
chr1-180714667-INS-207
chr1-180727648-INS-71
chr1-180728687-INS-167
chr1-180736039-INS-206
chr1-180747695-INS-96
chr1-180756796-INS-425
chr1-180763362-INS-60
chr1-180768064-INS-129
chr1-180782243-INS-245
chr1-180792655-INS-206
chr1-180793940-INS-52
chr1-180808488-INS-258
chr1-180815045-INS-292
chr1-180821742-INS-125
chr1-180825446-INS-849
chr1-180858402-INS-196
chr1-180889001-INS-400
chr1-180898094-INS-149
chr1-180901491-INS-208
chr1-180906569-INS-210
chr1-180910070-INS-352
chr1-180912582-INS-51
chr1-180916439-INS-66
chr1-180925656-INS-62
chr1-180949815-INS-338
chr1-180952602-INS-211
chr1-180964227-INS-105
chr1-180966020-INS-61
chr1-180986654-INS-187
chr1-180988312-INS-181
chr1-181018544-INS-324
chr1-181021650-INS-209
chr1-181028772-INS-58
chr1-181030280-INS-214
chr1-181032695-INS-141
chr1-181060655-INS-231
chr1-181064921-INS-207
chr1-181068426-INS-80
chr1-181070553-INS-99
chr1-181075443-INS-2370
chr1-181077441-INS-75
chr1-181102248-INS-56
chr1-181122733-INS-166
chr1-181134238-INS-148
chr1-181141840-INS-524
chr1-181159006-INS-237
chr1-181182743-INS-64

chr1-181188667-INS-207
chr1-181199845-INS-92
chr1-181205936-INS-63
chr1-181227899-INS-163
chr1-181229861-INS-446
chr1-181233161-INS-52
chr1-181240050-INS-121
chr1-181273985-INS-210
chr1-181280453-INS-1475
chr1-181282961-INS-134
chr1-181317357-INS-79
chr1-181328509-INS-68
chr1-181336136-INS-151
chr1-181348221-INS-445
chr1-181360710-INS-54
chr1-181364410-INS-91
chr1-181366780-INS-148
chr1-181390032-INS-55
chr1-181390900-INS-211
chr1-181391641-INS-136
chr1-181398954-INS-66
chr1-181399918-INS-171
chr1-181409156-INS-669
chr1-181419170-INS-567
chr1-181423754-INS-246
chr1-181426664-INS-498
chr1-181431721-INS-6160
chr1-181461065-INS-140
chr1-181467114-INS-70
chr1-181502729-INS-189
chr1-181518843-INS-194
chr1-181525216-INS-118
chr1-181537037-INS-75
chr1-181555327-INS-205
chr1-181556575-INS-286
chr1-181561601-INS-401
chr1-181568209-INS-52
chr1-181571196-INS-180
chr1-181571306-INS-1280
chr1-181596231-INS-154
chr1-181610055-INS-441
chr1-181619777-INS-300
chr1-181622486-INS-145
chr1-181626688-INS-268
chr1-181637748-INS-383
chr1-181653962-INS-64
chr1-181655523-INS-102
chr1-181656792-INS-216
chr1-181678425-INS-196
chr1-181685877-INS-84
chr1-181691084-INS-431
chr1-181703153-INS-174
chr1-181716488-INS-186
chr1-181719025-INS-63
chr1-181739867-INS-746
chr1-181743550-INS-140
chr1-181759461-INS-114
chr1-181761732-INS-68
chr1-181783296-INS-85
chr1-181810507-INS-141
chr1-181812109-INS-523
chr1-181836380-INS-193
chr1-181851637-INS-203
chr1-181858507-INS-56
chr1-181862458-INS-127
chr1-181886507-INS-207
chr1-181889364-INS-162
chr1-181891885-INS-408
chr1-181907839-INS-7088
chr1-181916462-INS-7296
chr1-181923796-INS-103
chr1-181925165-INS-68
chr1-181928292-INS-52
chr1-181933274-INS-518
chr1-181946663-INS-530
chr1-181971865-INS-63
chr1-181974914-INS-70

chr1-181980971-INS-526
chr1-181985860-INS-82
chr1-182002742-INS-5619
chr1-182006588-INS-6401
chr1-182017146-INS-248
chr1-182023181-INS-62
chr1-182025644-INS-934
chr1-182025782-INS-128
chr1-182038741-INS-377
chr1-182046359-INS-60
chr1-182049313-INS-208
chr1-182059410-INS-62
chr1-182113380-INS-83
chr1-182124180-INS-2157
chr1-182151368-INS-210
chr1-182154238-INS-84
chr1-182161889-INS-1040
chr1-182181999-INS-77
chr1-182193226-INS-756
chr1-182201287-INS-202
chr1-182223173-INS-208
chr1-182230370-INS-86
chr1-182231474-INS-78
chr1-182255300-INS-166
chr1-182265169-INS-148
chr1-182294121-INS-440
chr1-182314669-INS-86
chr1-182332695-INS-220
chr1-182334681-INS-437
chr1-182337143-INS-69
chr1-182339219-INS-7484
chr1-182359390-INS-201
chr1-182364924-INS-6199
chr1-182366481-INS-535
chr1-182367600-INS-99
chr1-182374733-INS-6458
chr1-182376395-INS-162
chr1-182381437-INS-121
chr1-182381752-INS-125
chr1-182386266-INS-84
chr1-182388350-INS-63
chr1-182403165-INS-105
chr1-182422699-INS-73
chr1-182423849-INS-56
chr1-182430561-INS-75
chr1-182450371-INS-88
chr1-182454181-INS-70
chr1-182458688-INS-558
chr1-182465886-INS-60
chr1-182490132-INS-70
chr1-182492142-INS-58
chr1-182503268-INS-208
chr1-182505908-INS-65
chr1-182526496-INS-526
chr1-182564282-INS-54
chr1-182564806-INS-72
chr1-182581226-INS-313
chr1-182618745-INS-116
chr1-182624064-INS-80
chr1-182639192-INS-217
chr1-182643921-INS-71
chr1-182649868-INS-80
chr1-182658469-INS-174
chr1-182705615-INS-588
chr1-182710928-INS-53
chr1-182712014-INS-180
chr1-182719389-INS-68
chr1-182727605-INS-1340
chr1-182727606-INS-603
chr1-182728755-INS-58
chr1-182741077-INS-52
chr1-182765591-INS-460
chr1-182769230-INS-325
chr1-182790069-INS-58
chr1-182808050-INS-199
chr1-182812641-INS-75
chr1-182839000-INS-184

chr1-182863084-INS-56
chr1-182876369-INS-114
chr1-182877419-INS-105
chr1-182880087-INS-96
chr1-182896614-INS-289
chr1-182934083-INS-56
chr1-182938546-INS-52
chr1-182943639-INS-136
chr1-182951799-INS-62
chr1-182962222-INS-89
chr1-182963376-INS-54
chr1-182980557-INS-51
chr1-182995099-INS-103
chr1-183011618-INS-130
chr1-183024975-INS-4419
chr1-183071055-INS-98
chr1-183081016-INS-77
chr1-183084845-INS-90
chr1-183091501-INS-1891
chr1-183106796-INS-217
chr1-183116966-INS-200
chr1-183123877-INS-55
chr1-183130372-INS-80
chr1-183163962-INS-84
chr1-183172568-INS-160
chr1-183177794-INS-420
chr1-183179948-INS-396
chr1-183182831-INS-205
chr1-183185918-INS-222
chr1-183206826-INS-63
chr1-183222779-INS-176
chr1-183235837-INS-75
chr1-183250806-INS-288
chr1-183257413-INS-112
chr1-183260474-INS-64
chr1-183263772-INS-111
chr1-183266293-INS-250
chr1-183267859-INS-110
chr1-183270193-INS-76
chr1-183286373-INS-6448
chr1-183294935-INS-88
chr1-183304485-INS-226
chr1-183305553-INS-55
chr1-183311047-INS-645
chr1-183316602-INS-144
chr1-183322658-INS-86
chr1-183349485-INS-165
chr1-183350108-INS-151
chr1-183355535-INS-481
chr1-183357789-INS-138
chr1-183369597-INS-66
chr1-183373246-INS-82
chr1-183384727-INS-74
chr1-183392116-INS-848
chr1-183398540-INS-56
chr1-183400022-INS-89
chr1-183415510-INS-6793
chr1-183438901-INS-86
chr1-183443460-INS-6401
chr1-183458062-INS-70
chr1-183471897-INS-141
chr1-183481967-INS-64
chr1-183493369-INS-56
chr1-183541596-INS-367
chr1-183572940-INS-7186
chr1-183600933-INS-303
chr1-183603473-INS-148
chr1-183619550-INS-725
chr1-183621026-INS-84
chr1-183622429-INS-52
chr1-183624966-INS-122
chr1-183629132-INS-6460
chr1-183635405-INS-59
chr1-183654721-INS-55
chr1-183663062-INS-7304
chr1-183667402-INS-97
chr1-183670020-INS-112

chr1-183671263-INS-371
chr1-183687489-INS-69
chr1-183692406-INS-95
chr1-183694613-INS-58
chr1-183701400-INS-62
chr1-183724673-INS-377
chr1-183739058-INS-258
chr1-183742604-INS-120
chr1-183745467-INS-292
chr1-183823819-INS-60
chr1-183835781-INS-85
chr1-183838460-INS-399
chr1-183845828-INS-52
chr1-183847738-INS-113
chr1-183851158-INS-52
chr1-183851807-INS-245
chr1-183855528-INS-56
chr1-183874488-INS-291
chr1-183875945-INS-103
chr1-183906561-INS-64
chr1-183911846-INS-187
chr1-183923159-INS-241
chr1-183929659-INS-514
chr1-183942146-INS-938
chr1-183964292-INS-53
chr1-183970591-INS-396
chr1-183975878-INS-69
chr1-184004222-INS-61
chr1-184041609-INS-399
chr1-184046366-INS-139
chr1-184062827-INS-53
chr1-184065043-INS-97
chr1-184068530-INS-140
chr1-184069584-INS-94
chr1-184087369-INS-54
chr1-184106963-INS-132
chr1-184111346-INS-53
chr1-184120138-INS-92
chr1-184122125-INS-68
chr1-184140528-INS-798
chr1-184143356-INS-70
chr1-184148170-INS-112
chr1-184150054-INS-243
chr1-184152344-INS-179
chr1-184153785-INS-509
chr1-184161213-INS-56
chr1-184179539-INS-55
chr1-184190881-INS-204
chr1-184217788-INS-160
chr1-184224096-INS-141
chr1-184229597-INS-152
chr1-184230747-INS-95
chr1-184240475-INS-61
chr1-184241555-INS-56
chr1-184303873-INS-837
chr1-184311280-INS-592
chr1-184319475-INS-1045
chr1-184319565-INS-615
chr1-184357941-INS-7254
chr1-184364527-INS-58
chr1-184383886-INS-72
chr1-184390276-INS-64
chr1-184401351-INS-112
chr1-184415501-INS-159
chr1-184426194-INS-691
chr1-184440582-INS-67
chr1-184446342-INS-175
chr1-184449803-INS-55
chr1-184474689-INS-205
chr1-184479738-INS-135
chr1-184486337-INS-513
chr1-184488996-INS-84
chr1-184493598-INS-60
chr1-184495540-INS-126
chr1-184518818-INS-184
chr1-184525379-INS-160
chr1-184526610-INS-195

chr1-184546895-INS-129
chr1-184547504-INS-770
chr1-184562812-INS-485
chr1-184580004-INS-555
chr1-184591259-INS-399
chr1-184594860-INS-81
chr1-184612190-INS-68
chr1-184626292-INS-975
chr1-184631819-INS-210
chr1-184644904-INS-2086
chr1-184644942-INS-548
chr1-184658756-INS-297
chr1-184661030-INS-251
chr1-184665738-INS-124
chr1-184676385-INS-234
chr1-184684705-INS-257
chr1-184687309-INS-130
chr1-184690317-INS-356
chr1-184708113-INS-196
chr1-184712151-INS-56
chr1-184713468-INS-4956
chr1-184722710-INS-151
chr1-184726755-INS-514
chr1-184738936-INS-52
chr1-184740029-INS-94
chr1-184751849-INS-218
chr1-184753107-INS-53
chr1-184756941-INS-378
chr1-184766351-INS-67
chr1-184788651-INS-57
chr1-184795264-INS-64
chr1-184799752-INS-57
chr1-184810842-INS-189
chr1-184815969-INS-100
chr1-184859009-INS-82
chr1-184861696-INS-63
chr1-184861706-INS-1263
chr1-184861707-INS-842
chr1-184874080-INS-78
chr1-184896145-INS-56
chr1-184901505-INS-324
chr1-184905373-INS-180
chr1-184922540-INS-53
chr1-184926029-INS-637
chr1-184933074-INS-69
chr1-184934107-INS-178
chr1-184943538-INS-504
chr1-184946862-INS-540
chr1-184947151-INS-675
chr1-184949254-INS-1502
chr1-184951032-INS-179
chr1-184951555-INS-176
chr1-184980421-INS-51
chr1-184987256-INS-53
chr1-184993369-INS-64
chr1-185014221-INS-214
chr1-185021147-INS-518
chr1-185026306-INS-84
chr1-185040339-INS-273
chr1-185053333-INS-211
chr1-185062307-INS-273
chr1-185064048-INS-179
chr1-185084802-INS-229
chr1-185103457-INS-207
chr1-185118113-INS-500
chr1-185170292-INS-60
chr1-185193899-INS-150
chr1-185201253-INS-76
chr1-185214590-INS-242
chr1-185216992-INS-99
chr1-185228949-INS-99
chr1-185254197-INS-174
chr1-185257161-INS-210
chr1-185272736-INS-97
chr1-185274711-INS-52
chr1-185321497-INS-72
chr1-185324903-INS-86

chr1-185327821-INS-70
chr1-185389327-INS-52
chr1-185410510-INS-129
chr1-185415944-INS-95
chr1-185432646-INS-5315
chr1-185451272-INS-170
chr1-185452899-INS-78
chr1-185470361-INS-577
chr1-185480584-INS-170
chr1-185495257-INS-52
chr1-185504640-INS-345
chr1-185524061-INS-7139
chr1-185567257-INS-81
chr1-185585104-INS-198
chr1-185590217-INS-444
chr1-185604244-INS-99
chr1-185607299-INS-291
chr1-185616216-INS-400
chr1-185639318-INS-201
chr1-185645634-INS-150
chr1-185658648-INS-52
chr1-185681903-INS-361
chr1-185711119-INS-173
chr1-185721577-INS-533
chr1-185742150-INS-679
chr1-185746384-INS-74
chr1-185750165-INS-168
chr1-185762956-INS-292
chr1-185770554-INS-69
chr1-185791886-INS-100
chr1-185793138-INS-357
chr1-185834024-INS-395
chr1-185894971-INS-172
chr1-185913883-INS-718
chr1-185917194-INS-121
chr1-185934550-INS-200
chr1-185938802-INS-66
chr1-185955375-INS-53
chr1-185977158-INS-100
chr1-186006059-INS-161
chr1-186045599-INS-154
chr1-186067891-INS-148
chr1-186068755-INS-7556
chr1-186094069-INS-678
chr1-186102679-INS-253
chr1-186105205-INS-56
chr1-186121650-INS-489
chr1-186123684-INS-98
chr1-186124892-INS-257
chr1-186132728-INS-149
chr1-186147398-INS-119
chr1-186153937-INS-60
chr1-186159789-INS-65
chr1-186164975-INS-143
chr1-186191123-INS-325
chr1-186193447-INS-94
chr1-186203207-INS-58
chr1-186212145-INS-52
chr1-186216545-INS-96
chr1-186227726-INS-5543
chr1-186230340-INS-5270
chr1-186230574-INS-116
chr1-186235880-INS-125
chr1-186240193-INS-78
chr1-186267162-INS-136
chr1-186269397-INS-6778
chr1-186278626-INS-64
chr1-186285430-INS-119
chr1-186296250-INS-557
chr1-186338358-INS-127
chr1-186351273-INS-126
chr1-186353940-INS-201
chr1-186366015-INS-118
chr1-186382678-INS-51
chr1-186421466-INS-68
chr1-186427069-INS-283
chr1-186440857-INS-212

chr1-186452640-INS-90
chr1-186464415-INS-1798
chr1-186467153-INS-646
chr1-186473136-INS-51
chr1-186479034-INS-201
chr1-186527800-INS-69
chr1-186532331-INS-112
chr1-186540275-INS-331
chr1-186553658-INS-183
chr1-186571730-INS-390
chr1-186592637-INS-105
chr1-186595947-INS-138
chr1-186611236-INS-89
chr1-186658712-INS-71
chr1-186678244-INS-78
chr1-186684498-INS-229
chr1-186705195-INS-84
chr1-186707982-INS-184
chr1-186709973-INS-150
chr1-186731220-INS-69
chr1-186737082-INS-60
chr1-186767601-INS-216
chr1-186788865-INS-79
chr1-186819915-INS-170
chr1-186820944-INS-77
chr1-186834322-INS-102
chr1-186880592-INS-143
chr1-186888903-INS-348
chr1-186896901-INS-581
chr1-186922512-INS-155
chr1-186932338-INS-486
chr1-186937332-INS-155
chr1-186943525-INS-203
chr1-186988590-INS-437
chr1-186997931-INS-55
chr1-187092536-INS-314
chr1-187094958-INS-306
chr1-187111308-INS-133
chr1-187143118-INS-72
chr1-187149068-INS-247
chr1-187178041-INS-68
chr1-187200483-INS-107
chr1-187231159-INS-6100
chr1-187282130-INS-222
chr1-187320961-INS-60
chr1-187323493-INS-273
chr1-187334210-INS-56
chr1-187352032-INS-75
chr1-187361795-INS-52
chr1-187430421-INS-796
chr1-187448752-INS-450
chr1-187482089-INS-254
chr1-187484525-INS-86
chr1-187491194-INS-62
chr1-187500504-INS-66
chr1-187520808-INS-56
chr1-187522685-INS-74
chr1-187539755-INS-157
chr1-187548160-INS-105
chr1-187600099-INS-83
chr1-187611953-INS-937
chr1-187626671-INS-108
chr1-187643955-INS-80
chr1-187646625-INS-52
chr1-187671376-INS-93
chr1-187686847-INS-209
chr1-187692305-INS-116
chr1-187701193-INS-79
chr1-187733137-INS-54
chr1-187756742-INS-155
chr1-187774917-INS-84
chr1-187781749-INS-61
chr1-187783049-INS-102
chr1-187788912-INS-58
chr1-187816525-INS-111
chr1-187862973-INS-130
chr1-187875312-INS-77

chr1-187886718-INS-52
chr1-187890206-INS-300
chr1-187895380-INS-62
chr1-187903828-INS-144
chr1-187927925-INS-111
chr1-187929665-INS-136
chr1-187934159-INS-837
chr1-187934408-INS-70
chr1-187950385-INS-74
chr1-187991383-INS-55
chr1-188033981-INS-292
chr1-188060210-INS-62
chr1-188078170-INS-176
chr1-188080276-INS-115
chr1-188130941-INS-192
chr1-188136148-INS-628
chr1-188178159-INS-187
chr1-188183371-INS-62
chr1-188204304-INS-499
chr1-188266543-INS-220
chr1-188279382-INS-70
chr1-188296884-INS-106
chr1-188310525-INS-239
chr1-188311508-INS-102
chr1-188316115-INS-112
chr1-188331438-INS-68
chr1-188334270-INS-183
chr1-188349733-INS-99
chr1-188364765-INS-150
chr1-188370643-INS-62
chr1-188381653-INS-103
chr1-188386048-INS-207
chr1-188395801-INS-341
chr1-188438241-INS-201
chr1-188465477-INS-195
chr1-188470146-INS-56
chr1-188492914-INS-402
chr1-188512153-INS-563
chr1-188522385-INS-112
chr1-188535140-INS-651
chr1-188537369-INS-62
chr1-188540711-INS-443
chr1-188553693-INS-58
chr1-188563842-INS-76
chr1-188579031-INS-148
chr1-188587274-INS-208
chr1-188606019-INS-88
chr1-188633442-INS-398
chr1-188663253-INS-5632
chr1-188676600-INS-372
chr1-188682482-INS-213
chr1-188696328-INS-6347
chr1-188721224-INS-116
chr1-188747173-INS-212
chr1-188789740-INS-124
chr1-188804474-INS-154
chr1-188805624-INS-7442
chr1-188810224-INS-6455
chr1-188816734-INS-921
chr1-188820504-INS-567
chr1-188830486-INS-416
chr1-188837215-INS-8508
chr1-188846509-INS-328
chr1-188851512-INS-88
chr1-188862925-INS-187
chr1-188867732-INS-301
chr1-188878230-INS-228
chr1-188887671-INS-791
chr1-188893877-INS-70
chr1-188908374-INS-279
chr1-188910540-INS-74
chr1-188914266-INS-504
chr1-188915530-INS-752
chr1-188921602-INS-703
chr1-188933225-INS-142
chr1-188983147-INS-64
chr1-189010367-INS-77

chr1-189012627-INS-170
chr1-189013547-INS-136
chr1-189037128-INS-91
chr1-189052055-INS-189
chr1-189061139-INS-200
chr1-189095554-INS-388
chr1-189102980-INS-204
chr1-189118637-INS-58
chr1-189136406-INS-56
chr1-189141126-INS-56
chr1-189147312-INS-7143
chr1-189214778-INS-110
chr1-189216976-INS-4088
chr1-189218761-INS-70
chr1-189221757-INS-881
chr1-189233948-INS-234
chr1-189234806-INS-78
chr1-189240327-INS-90
chr1-189254558-INS-199
chr1-189269295-INS-177
chr1-189270280-INS-534
chr1-189271964-INS-592
chr1-189273120-INS-6385
chr1-189301108-INS-54
chr1-189306688-INS-92
chr1-189317975-INS-203
chr1-189330254-INS-87
chr1-189330887-INS-148
chr1-189335936-INS-210
chr1-189337128-INS-57
chr1-189343889-INS-72
chr1-189364035-INS-149
chr1-189367559-INS-851
chr1-189427638-INS-320
chr1-189439352-INS-408
chr1-189457336-INS-83
chr1-189462527-INS-80
chr1-189468220-INS-147
chr1-189468999-INS-55
chr1-189471704-INS-54
chr1-189483768-INS-80
chr1-189529611-INS-2012
chr1-189546044-INS-339
chr1-189557794-INS-69
chr1-189601740-INS-82
chr1-189625450-INS-308
chr1-189638186-INS-787
chr1-189638310-INS-78
chr1-189639170-INS-73
chr1-189658385-INS-473
chr1-189658980-INS-63
chr1-189673929-INS-85
chr1-189713054-INS-59
chr1-189715908-INS-202
chr1-189716931-INS-58
chr1-189723518-INS-86
chr1-189728896-INS-121
chr1-189730536-INS-57
chr1-189761659-INS-335
chr1-189763422-INS-184
chr1-189764306-INS-462
chr1-189780359-INS-168
chr1-189790529-INS-355
chr1-189791012-INS-457
chr1-189800093-INS-199
chr1-189821063-INS-80
chr1-189866479-INS-177
chr1-189910511-INS-72
chr1-189916435-INS-355
chr1-189919624-INS-5268
chr1-189920305-INS-104
chr1-189926166-INS-204
chr1-189929082-INS-76
chr1-189935331-INS-166
chr1-189955567-INS-210
chr1-189959739-INS-4581
chr1-189967663-INS-333

chr1-189999062-INS-551
chr1-190019918-INS-75
chr1-190020980-INS-100
chr1-190035452-INS-100
chr1-190036010-INS-75
chr1-190036260-INS-1748
chr1-190046194-INS-304
chr1-190064093-INS-398
chr1-190093794-INS-125
chr1-190111539-INS-91
chr1-190120639-INS-51
chr1-190121035-INS-1558
chr1-190121053-INS-54
chr1-190124796-INS-102
chr1-190147399-INS-140
chr1-190150280-INS-60
chr1-190165294-INS-78
chr1-190170337-INS-82
chr1-190179142-INS-246
chr1-190182991-INS-60
chr1-190184427-INS-214
chr1-190185500-INS-56
chr1-190191013-INS-52
chr1-190193736-INS-75
chr1-190211849-INS-505
chr1-190231151-INS-130
chr1-190234199-INS-330
chr1-190254325-INS-938
chr1-190254494-INS-138
chr1-190259497-INS-189
chr1-190284984-INS-58
chr1-190289759-INS-124
chr1-190289864-INS-810
chr1-190290965-INS-70
chr1-190309000-INS-78
chr1-190322731-INS-75
chr1-190333385-INS-138
chr1-190339690-INS-66
chr1-190345263-INS-454
chr1-190353788-INS-70
chr1-190355263-INS-83
chr1-190361104-INS-51
chr1-190362456-INS-54
chr1-190379462-INS-380
chr1-190389747-INS-84
chr1-190406668-INS-70
chr1-190481276-INS-335
chr1-190487413-INS-56
chr1-190492538-INS-58
chr1-190499752-INS-64
chr1-190507758-INS-400
chr1-190517102-INS-649
chr1-190519953-INS-510
chr1-190535334-INS-120
chr1-190548005-INS-263
chr1-190557587-INS-61
chr1-190563619-INS-67
chr1-190568556-INS-58
chr1-190572459-INS-323
chr1-190577562-INS-304
chr1-190583729-INS-92
chr1-190589779-INS-386
chr1-190604806-INS-146
chr1-190610791-INS-52
chr1-190612737-INS-132
chr1-190624858-INS-514
chr1-190630551-INS-60
chr1-190633325-INS-358
chr1-190674617-INS-199
chr1-190688251-INS-206
chr1-190689384-INS-137
chr1-190697961-INS-156
chr1-190712208-INS-315
chr1-190719918-INS-68
chr1-190722150-INS-360
chr1-190727091-INS-171
chr1-190729793-INS-119

chr1-190736764-INS-335
chr1-190738816-INS-176
chr1-190741237-INS-498
chr1-190763094-INS-95
chr1-190770185-INS-125
chr1-190783820-INS-61
chr1-190783847-INS-353
chr1-190785947-INS-153
chr1-190802888-INS-758
chr1-190805272-INS-8279
chr1-190811275-INS-228
chr1-190819100-INS-219
chr1-190844527-INS-181
chr1-190882833-INS-3159
chr1-190884314-INS-130
chr1-190887747-INS-182
chr1-190916263-INS-80
chr1-190934344-INS-386
chr1-190943076-INS-52
chr1-190944663-INS-788
chr1-190951636-INS-202
chr1-190954988-INS-91
chr1-190987845-INS-173
chr1-191005699-INS-763
chr1-191006716-INS-95
chr1-191012181-INS-122
chr1-191024249-INS-58
chr1-191025569-INS-269
chr1-191027786-INS-109
chr1-191032878-INS-187
chr1-191039732-INS-58
chr1-191041755-INS-206
chr1-191046083-INS-53
chr1-191048508-INS-54
chr1-191050799-INS-173
chr1-191065199-INS-476
chr1-191067255-INS-1891
chr1-191071344-INS-104
chr1-191100374-INS-211
chr1-191132623-INS-136
chr1-191151964-INS-73
chr1-191165725-INS-202
chr1-191183591-INS-78
chr1-191188928-INS-189
chr1-191205545-INS-138
chr1-191220377-INS-204
chr1-191227767-INS-241
chr1-191240465-INS-228
chr1-191258287-INS-196
chr1-191261042-INS-230
chr1-191269046-INS-60
chr1-191275310-INS-59
chr1-191288108-INS-782
chr1-191348605-INS-53
chr1-191349212-INS-426
chr1-191351120-INS-212
chr1-191359715-INS-58
chr1-191384393-INS-64
chr1-191405017-INS-6876
chr1-191413465-INS-76
chr1-191432223-INS-166
chr1-191444846-INS-108
chr1-191461798-INS-61
chr1-191463132-INS-60
chr1-191468248-INS-110
chr1-191469324-INS-74
chr1-191482530-INS-71
chr1-191486127-INS-198
chr1-191501863-INS-152
chr1-191517042-INS-198
chr1-191522966-INS-175
chr1-191532517-INS-6452
chr1-191532667-INS-516
chr1-191535192-INS-4547
chr1-191536157-INS-183
chr1-191546661-INS-711
chr1-191547511-INS-155

chr1-191555912-INS-173
chr1-191588511-INS-186
chr1-191614651-INS-71
chr1-191618669-INS-87
chr1-191621140-INS-76
chr1-191628427-INS-73
chr1-191630971-INS-834
chr1-191636600-INS-52
chr1-191650122-INS-62
chr1-191660791-INS-1274
chr1-191684631-INS-335
chr1-191692947-INS-60
chr1-191707377-INS-302
chr1-191726583-INS-442
chr1-191727739-INS-161
chr1-191730033-INS-84
chr1-191731387-INS-187
chr1-191744014-INS-52
chr1-191752649-INS-63
chr1-191782068-INS-58
chr1-191845876-INS-7183
chr1-191850770-INS-102
chr1-191855401-INS-933
chr1-191865847-INS-298
chr1-191889563-INS-53
chr1-191900907-INS-179
chr1-191911135-INS-76
chr1-191934125-INS-68
chr1-191956615-INS-171
chr1-191962120-INS-6605
chr1-191973122-INS-68
chr1-191978503-INS-54
chr1-191981334-INS-456
chr1-191984858-INS-179
chr1-191996875-INS-64
chr1-192033412-INS-89
chr1-192048552-INS-90
chr1-192070782-INS-92
chr1-192089545-INS-5399
chr1-192100360-INS-200
chr1-192102839-INS-469
chr1-192108373-INS-52
chr1-192110023-INS-62
chr1-192110956-INS-62
chr1-192111904-INS-3871
chr1-192139090-INS-71
chr1-192167834-INS-73
chr1-192173533-INS-66
chr1-192196105-INS-96
chr1-192200852-INS-1863
chr1-192206116-INS-57
chr1-192215549-INS-72
chr1-192229576-INS-401
chr1-192230257-INS-144
chr1-192236541-INS-87
chr1-192244466-INS-66
chr1-192246648-INS-537
chr1-192263922-INS-92
chr1-192264717-INS-117
chr1-192268863-INS-120
chr1-192305240-INS-132
chr1-192347522-INS-81
chr1-192348452-INS-984
chr1-192348565-INS-1162
chr1-192359931-INS-380
chr1-192399524-INS-636
chr1-192399999-INS-67
chr1-192411104-INS-193
chr1-192415683-INS-87
chr1-192431578-INS-204
chr1-192440509-INS-200
chr1-192452835-INS-60
chr1-192471989-INS-172
chr1-192475731-INS-414
chr1-192490024-INS-104
chr1-192527962-INS-118
chr1-192541278-INS-207

chr1-192561001-INS-69
chr1-192584700-INS-80
chr1-192589648-INS-260
chr1-192592995-INS-81
chr1-192614456-INS-313
chr1-192631833-INS-751
chr1-192650796-INS-219
chr1-192676016-INS-210
chr1-192688095-INS-51
chr1-192715040-INS-91
chr1-192719975-INS-55
chr1-192728301-INS-80
chr1-192742623-INS-86
chr1-192780637-INS-112
chr1-192797343-INS-71
chr1-192798683-INS-70
chr1-192821981-INS-88
chr1-192874786-INS-166
chr1-192884678-INS-88
chr1-192895127-INS-60
chr1-192899804-INS-227
chr1-192916073-INS-65
chr1-193027008-INS-90
chr1-193034586-INS-58
chr1-193067886-INS-418
chr1-193079898-INS-3190
chr1-193086414-INS-89
chr1-193093680-INS-60
chr1-193106349-INS-7087
chr1-193107226-INS-55
chr1-193115356-INS-2903
chr1-193118015-INS-917
chr1-193131053-INS-84
chr1-193133338-INS-62
chr1-193154309-INS-81
chr1-193170001-INS-409
chr1-193180347-INS-65
chr1-193245668-INS-139
chr1-193248898-INS-64
chr1-193261876-INS-82
chr1-193315679-INS-64
chr1-193334292-INS-599
chr1-193354465-INS-5218
chr1-193358087-INS-639
chr1-193386160-INS-80
chr1-193389202-INS-163
chr1-193448862-INS-390
chr1-193457079-INS-391
chr1-193471256-INS-90
chr1-193492303-INS-201
chr1-193498664-INS-88
chr1-193501435-INS-252
chr1-193535075-INS-684
chr1-193535909-INS-5181
chr1-193538021-INS-999
chr1-193557706-INS-285
chr1-193562690-INS-947
chr1-193593023-INS-214
chr1-193594197-INS-54
chr1-193602737-INS-156
chr1-193607772-INS-59
chr1-193643204-INS-200
chr1-193657941-INS-738
chr1-193659864-INS-7331
chr1-193667292-INS-1133
chr1-193669823-INS-51
chr1-193677430-INS-60
chr1-193690756-INS-95
chr1-193697733-INS-408
chr1-193715636-INS-56
chr1-193726975-INS-111
chr1-193824976-INS-1228
chr1-193832166-INS-288
chr1-193846432-INS-400
chr1-193859910-INS-76
chr1-193869021-INS-128
chr1-193882477-INS-466

chr1-193885657-INS-71
chr1-193896701-INS-400
chr1-193912136-INS-59
chr1-193917981-INS-164
chr1-193929305-INS-976
chr1-193986064-INS-58
chr1-194007508-INS-221
chr1-194022660-INS-81
chr1-194039754-INS-6902
chr1-194049184-INS-180
chr1-194074959-INS-7295
chr1-194101376-INS-833
chr1-194112698-INS-54
chr1-194123270-INS-60
chr1-194123435-INS-6190
chr1-194123494-INS-262
chr1-194130580-INS-4165
chr1-194155009-INS-6291
chr1-194193692-INS-398
chr1-194212600-INS-6605
chr1-194234263-INS-77
chr1-194281753-INS-516
chr1-194331118-INS-275
chr1-194331661-INS-86
chr1-194334051-INS-66
chr1-194351095-INS-66
chr1-194368379-INS-80
chr1-194405050-INS-57
chr1-194440876-INS-77
chr1-194443343-INS-168
chr1-194500242-INS-245
chr1-194527175-INS-82
chr1-194534576-INS-62
chr1-194570044-INS-94
chr1-194577180-INS-58
chr1-194605214-INS-1763
chr1-194606914-INS-51
chr1-194617042-INS-56
chr1-194646984-INS-445
chr1-194653727-INS-509
chr1-194785561-INS-96
chr1-194790443-INS-540
chr1-194862335-INS-86
chr1-194874581-INS-62
chr1-194877493-INS-57
chr1-194880008-INS-128
chr1-194885087-INS-548
chr1-194907251-INS-242
chr1-194915016-INS-221
chr1-194952343-INS-1446
chr1-194988900-INS-11771
chr1-194993883-INS-6220
chr1-194997822-INS-2566
chr1-195019804-INS-517
chr10-3100156-INS-132
chr10-3110460-INS-153
chr10-3111670-INS-233
chr10-3126153-INS-120
chr10-3215115-INS-58
chr10-3259349-INS-6344
chr10-3280332-INS-86
chr10-3286604-INS-492
chr10-3294390-INS-610
chr10-3297188-INS-62
chr10-3307089-INS-132
chr10-3323418-INS-52
chr10-3359894-INS-1504
chr10-3359894-INS-158
chr10-3360470-INS-357
chr10-3428629-INS-170
chr10-3429955-INS-146
chr10-3444536-INS-402
chr10-3456406-INS-188
chr10-3465145-INS-52
chr10-3469138-INS-228
chr10-3470934-INS-197
chr10-3503200-INS-5788

chr10-3504512-INS-288
chr10-3505994-INS-365
chr10-3514520-INS-1609
chr10-3514954-INS-424
chr10-3518657-INS-3610
chr10-3521307-INS-2658
chr10-3521421-INS-62
chr10-3542435-INS-6519
chr10-3542946-INS-6423
chr10-3592104-INS-86
chr10-3601928-INS-60
chr10-3638299-INS-108
chr10-3644571-INS-56
chr10-3658843-INS-71
chr10-3661058-INS-130
chr10-3666435-INS-325
chr10-3683770-INS-522
chr10-3697340-INS-8980
chr10-3703425-INS-99
chr10-3709250-INS-328
chr10-3714055-INS-65
chr10-3719915-INS-113
chr10-3725305-INS-330
chr10-3742663-INS-54
chr10-3746311-INS-52
chr10-3753501-INS-53
chr10-3756147-INS-64
chr10-3764501-INS-64
chr10-3805280-INS-55
chr10-3808288-INS-60
chr10-3811536-INS-76
chr10-3817322-INS-224
chr10-3836535-INS-80
chr10-3849364-INS-479
chr10-3861666-INS-173
chr10-3866406-INS-162
chr10-3897937-INS-158
chr10-3920859-INS-103
chr10-3962488-INS-148
chr10-3994420-INS-485
chr10-3997995-INS-333
chr10-4012116-INS-64
chr10-4015034-INS-59
chr10-4036277-INS-401
chr10-4040328-INS-54
chr10-4063239-INS-212
chr10-4096700-INS-202
chr10-4106347-INS-54
chr10-4108469-INS-54
chr10-4114479-INS-78
chr10-4119085-INS-191
chr10-4151513-INS-205
chr10-4173630-INS-201
chr10-4175360-INS-112
chr10-4180477-INS-167
chr10-4187891-INS-276
chr10-4189370-INS-5283
chr10-4191325-INS-123
chr10-4191670-INS-302
chr10-4195787-INS-62
chr10-4199404-INS-132
chr10-4211705-INS-142
chr10-4228535-INS-68
chr10-4230690-INS-60
chr10-4238162-INS-112
chr10-4240653-INS-128
chr10-4241787-INS-64
chr10-4249344-INS-99
chr10-4264967-INS-73
chr10-4280109-INS-142
chr10-4298601-INS-226
chr10-4313776-INS-170
chr10-4329456-INS-99
chr10-4344506-INS-134
chr10-4351166-INS-85
chr10-4394339-INS-112
chr10-4399281-INS-51

chr10-4429319-INS-64
chr10-4431446-INS-202
chr10-4437634-INS-76
chr10-4448819-INS-5148
chr10-4474160-INS-6972
chr10-4480067-INS-78
chr10-4491187-INS-209
chr10-4494990-INS-72
chr10-4500948-INS-244
chr10-4504493-INS-925
chr10-4505231-INS-84
chr10-4505599-INS-542
chr10-4527647-INS-68
chr10-4551516-INS-64
chr10-4578707-INS-56
chr10-4581943-INS-364
chr10-4594911-INS-661
chr10-4594993-INS-910
chr10-4600737-INS-182
chr10-4604243-INS-400
chr10-4606403-INS-55
chr10-4608501-INS-211
chr10-4615000-INS-91
chr10-4619040-INS-359
chr10-4622933-INS-59
chr10-4658734-INS-78
chr10-4668747-INS-483
chr10-4680344-INS-57
chr10-4702937-INS-118
chr10-4750143-INS-78
chr10-4756007-INS-172
chr10-4771070-INS-301
chr10-4776631-INS-66
chr10-4802323-INS-58
chr10-4802966-INS-81
chr10-4805115-INS-6464
chr10-4813877-INS-57
chr10-4822395-INS-7807
chr10-4827794-INS-183
chr10-4829958-INS-6412
chr10-4870355-INS-639
chr10-4885531-INS-6442
chr10-4888424-INS-99
chr10-4907333-INS-447
chr10-4933183-INS-60
chr10-4937591-INS-323
chr10-4943254-INS-55
chr10-4956189-INS-72
chr10-4959947-INS-56
chr10-4960923-INS-227
chr10-4985125-INS-68
chr10-5010006-INS-567
chr10-5012018-INS-258
chr10-5019330-INS-114
chr10-5022603-INS-74
chr10-5036553-INS-396
chr10-5039107-INS-144
chr10-5077444-INS-60
chr10-5097512-INS-137
chr10-5161532-INS-88
chr10-5177307-INS-62
chr10-5205272-INS-125
chr10-5210831-INS-52
chr10-5216274-INS-56
chr10-5238334-INS-155
chr10-5254095-INS-260
chr10-5256105-INS-68
chr10-5256826-INS-148
chr10-5281320-INS-256
chr10-5286412-INS-87
chr10-5289976-INS-54
chr10-5291168-INS-247
chr10-5298902-INS-399
chr10-5305442-INS-520
chr10-5312991-INS-76
chr10-5327133-INS-1058
chr10-5333733-INS-62

chr10-5364346-INS-58
chr10-5387417-INS-268
chr10-5395428-INS-376
chr10-5426705-INS-5741
chr10-5439120-INS-988
chr10-5441599-INS-57
chr10-5443154-INS-6537
chr10-5465205-INS-54
chr10-5473911-INS-214
chr10-5489580-INS-54
chr10-5496044-INS-89
chr10-5515246-INS-506
chr10-5554117-INS-409
chr10-5557413-INS-88
chr10-5562252-INS-258
chr10-5601861-INS-55
chr10-5619834-INS-78
chr10-5704343-INS-207
chr10-5740145-INS-52
chr10-5746274-INS-154
chr10-5791675-INS-90
chr10-5830842-INS-79
chr10-5835429-INS-1816
chr10-5857966-INS-54
chr10-5887639-INS-198
chr10-5901624-INS-179
chr10-5964946-INS-493
chr10-5967148-INS-67
chr10-5971218-INS-149
chr10-5982080-INS-252
chr10-6012057-INS-7076
chr10-6021159-INS-62
chr10-6023974-INS-343
chr10-6025695-INS-98
chr10-6045548-INS-85
chr10-6048434-INS-75
chr10-6059563-INS-337
chr10-6065732-INS-302
chr10-6076826-INS-561
chr10-6088475-INS-52
chr10-6103135-INS-7260
chr10-6112276-INS-6426
chr10-6155827-INS-375
chr10-6166092-INS-179
chr10-6168071-INS-54
chr10-6169335-INS-85
chr10-6171588-INS-2970
chr10-6195630-INS-1664
chr10-6207741-INS-879
chr10-6214300-INS-3095
chr10-6214461-INS-60
chr10-6216980-INS-134
chr10-6220882-INS-2751
chr10-6257848-INS-7275
chr10-6264302-INS-67
chr10-6275518-INS-1860
chr10-6284880-INS-65
chr10-6289681-INS-96
chr10-6318697-INS-7344
chr10-6336770-INS-60
chr10-6344925-INS-70
chr10-6360840-INS-267
chr10-6391906-INS-201
chr10-6395394-INS-77
chr10-6396399-INS-321
chr10-6401353-INS-1890
chr10-6406285-INS-64
chr10-6419651-INS-91
chr10-6422341-INS-75
chr10-6435033-INS-8032
chr10-6500698-INS-430
chr10-6515162-INS-100
chr10-6531523-INS-186
chr10-6537164-INS-53
chr10-6549534-INS-62
chr10-6558984-INS-61
chr10-6566023-INS-55

chr10-6580628-INS-154
chr10-6580742-INS-225
chr10-6583300-INS-162
chr10-6587688-INS-53
chr10-6604729-INS-208
chr10-6618038-INS-861
chr10-6629325-INS-162
chr10-6632188-INS-85
chr10-6660261-INS-112
chr10-6672517-INS-424
chr10-6677387-INS-409
chr10-6679644-INS-77
chr10-6691034-INS-429
chr10-6692248-INS-52
chr10-6699065-INS-398
chr10-6706993-INS-144
chr10-6720665-INS-233
chr10-6728782-INS-2872
chr10-6743521-INS-370
chr10-6751814-INS-64
chr10-6757910-INS-140
chr10-6759790-INS-76
chr10-6781413-INS-320
chr10-6792094-INS-56
chr10-6796088-INS-592
chr10-6811382-INS-98
chr10-6832578-INS-183
chr10-6837057-INS-64
chr10-6838622-INS-128
chr10-6846943-INS-610
chr10-6849700-INS-60
chr10-6868599-INS-60
chr10-6880037-INS-55
chr10-6917218-INS-327
chr10-6920086-INS-54
chr10-6934429-INS-819
chr10-6936925-INS-6214
chr10-6954142-INS-280
chr10-6961995-INS-6646
chr10-6973722-INS-200
chr10-6995184-INS-105
chr10-6999145-INS-260
chr10-7012558-INS-190
chr10-7015971-INS-52
chr10-7020980-INS-90
chr10-7053026-INS-129
chr10-7062463-INS-70
chr10-7093670-INS-66
chr10-7098010-INS-88
chr10-7119244-INS-452
chr10-7134107-INS-204
chr10-7143703-INS-187
chr10-7158117-INS-68
chr10-7167126-INS-53
chr10-7175640-INS-259
chr10-7189451-INS-206
chr10-7198098-INS-207
chr10-7219391-INS-222
chr10-7224340-INS-143
chr10-7227184-INS-69
chr10-7232187-INS-146
chr10-7260891-INS-322
chr10-7307794-INS-57
chr10-7314734-INS-94
chr10-7321591-INS-512
chr10-7358354-INS-182
chr10-7369725-INS-52
chr10-7423116-INS-55
chr10-7434031-INS-87
chr10-7441233-INS-400
chr10-7445065-INS-65
chr10-7479868-INS-150
chr10-7483138-INS-108
chr10-7496577-INS-100
chr10-7509585-INS-240
chr10-7518923-INS-5293
chr10-7519499-INS-406

chr10-7532249-INS-183
chr10-7534117-INS-2932
chr10-7546277-INS-92
chr10-7574055-INS-66
chr10-7580888-INS-136
chr10-7582273-INS-368
chr10-7596901-INS-69
chr10-7599477-INS-195
chr10-7602817-INS-54
chr10-7608935-INS-190
chr10-7617466-INS-7146
chr10-7680092-INS-154
chr10-7680903-INS-203
chr10-7685003-INS-203
chr10-7695202-INS-90
chr10-7716200-INS-202
chr10-7732809-INS-1885
chr10-7732810-INS-400
chr10-7750012-INS-64
chr10-7756357-INS-320
chr10-7778765-INS-260
chr10-7783802-INS-64
chr10-7789010-INS-119
chr10-7790076-INS-950
chr10-7817649-INS-68
chr10-7826470-INS-58
chr10-7834501-INS-173
chr10-7855596-INS-91
chr10-7862857-INS-422
chr10-7874136-INS-64
chr10-7889198-INS-518
chr10-7896908-INS-52
chr10-7899057-INS-483
chr10-7934714-INS-942
chr10-7988276-INS-90
chr10-7992175-INS-517
chr10-8007758-INS-112
chr10-8020335-INS-55
chr10-8027476-INS-458
chr10-8036842-INS-267
chr10-8060096-INS-119
chr10-8067884-INS-65
chr10-8070350-INS-168
chr10-8073850-INS-230
chr10-8091129-INS-299
chr10-8135309-INS-51
chr10-8150211-INS-385
chr10-8168859-INS-78
chr10-8168937-INS-1197
chr10-8169153-INS-391
chr10-8175930-INS-82
chr10-8180865-INS-198
chr10-8181531-INS-59
chr10-8198501-INS-127
chr10-8224549-INS-148
chr10-8232189-INS-56
chr10-8242430-INS-102
chr10-8243897-INS-175
chr10-8252037-INS-93
chr10-8265345-INS-51
chr10-8291716-INS-58
chr10-8299463-INS-68
chr10-8328430-INS-124
chr10-8329926-INS-262
chr10-8330079-INS-1239
chr10-8330366-INS-286
chr10-8330431-INS-1909
chr10-8331572-INS-61
chr10-8337419-INS-118
chr10-8353367-INS-141
chr10-8358316-INS-190
chr10-8366193-INS-127
chr10-8385354-INS-130
chr10-8442658-INS-93
chr10-8451062-INS-508
chr10-8453318-INS-197
chr10-8462140-INS-75

chr10-8471471-INS-78
chr10-8473660-INS-81
chr10-8494949-INS-400
chr10-8509299-INS-62
chr10-8515668-INS-177
chr10-8521391-INS-55
chr10-8531512-INS-52
chr10-8544073-INS-133
chr10-8554082-INS-4587
chr10-8554812-INS-56
chr10-8555726-INS-317
chr10-8571492-INS-244
chr10-8576882-INS-297
chr10-8608919-INS-247
chr10-8651522-INS-60
chr10-8652355-INS-299
chr10-8666334-INS-207
chr10-8701654-INS-117
chr10-8715616-INS-58
chr10-8717294-INS-7088
chr10-8717763-INS-112
chr10-8718840-INS-64
chr10-8725680-INS-189
chr10-8725809-INS-752
chr10-8730525-INS-65
chr10-8735763-INS-193
chr10-8748408-INS-167
chr10-8757937-INS-213
chr10-8760740-INS-56
chr10-8765288-INS-135
chr10-8766145-INS-5406
chr10-8767643-INS-84
chr10-8773062-INS-138
chr10-8781292-INS-51
chr10-8789656-INS-56
chr10-8790700-INS-229
chr10-8792427-INS-215
chr10-8795111-INS-80
chr10-8819059-INS-149
chr10-8822603-INS-214
chr10-8838004-INS-123
chr10-8862732-INS-58
chr10-8865922-INS-333
chr10-8867816-INS-516
chr10-8882075-INS-132
chr10-8893398-INS-152
chr10-8919504-INS-245
chr10-8952111-INS-70
chr10-8977771-INS-99
chr10-8983873-INS-8594
chr10-8988137-INS-233
chr10-8994618-INS-53
chr10-9019698-INS-4034
chr10-9029684-INS-176
chr10-9031228-INS-368
chr10-9032915-INS-222
chr10-9056851-INS-4276
chr10-9061277-INS-801
chr10-9071000-INS-127
chr10-9091298-INS-112
chr10-9098414-INS-76
chr10-9143213-INS-161
chr10-9187589-INS-125
chr10-9190852-INS-53
chr10-9220052-INS-130
chr10-9253824-INS-6666
chr10-9257616-INS-70
chr10-9258412-INS-235
chr10-9271418-INS-145
chr10-9271493-INS-969
chr10-9301662-INS-449
chr10-9306620-INS-274
chr10-9322883-INS-6449
chr10-9339311-INS-65
chr10-9348366-INS-92
chr10-9358430-INS-144
chr10-9365304-INS-70

chr10-9368636-INS-3993
chr10-9373386-INS-68
chr10-9389866-INS-767
chr10-9391378-INS-119
chr10-9403116-INS-52
chr10-9405852-INS-65
chr10-9407195-INS-698
chr10-9447401-INS-401
chr10-9473180-INS-135
chr10-9477362-INS-254
chr10-9491082-INS-352
chr10-9493031-INS-252
chr10-9498002-INS-313
chr10-9537293-INS-57
chr10-9541415-INS-409
chr10-9562502-INS-60
chr10-9564745-INS-76
chr10-9565701-INS-253
chr10-9615336-INS-98
chr10-9628218-INS-73
chr10-9637806-INS-79
chr10-9653702-INS-116
chr10-9667763-INS-58
chr10-9668073-INS-830
chr10-9675820-INS-56
chr10-9699876-INS-60
chr10-9707465-INS-64
chr10-9729332-INS-180
chr10-9735463-INS-138
chr10-9748236-INS-154
chr10-9752980-INS-142
chr10-9760621-INS-94
chr10-9766003-INS-52
chr10-9784877-INS-73
chr10-9810252-INS-53
chr10-9813363-INS-75
chr10-9832045-INS-72
chr10-9842922-INS-339
chr10-9854506-INS-73
chr10-9858423-INS-188
chr10-9875272-INS-138
chr10-9883926-INS-77
chr10-9928901-INS-234
chr10-9942378-INS-90
chr10-9988311-INS-68
chr10-10029593-INS-146
chr10-10082662-INS-5291
chr10-10162437-INS-621
chr10-10178314-INS-70
chr10-10186308-INS-72
chr10-10197764-INS-60
chr10-10232174-INS-56
chr10-10271179-INS-113
chr10-10288493-INS-70
chr10-10306889-INS-63
chr10-10344201-INS-63
chr10-10347175-INS-60
chr10-10355364-INS-188
chr10-10419756-INS-129
chr10-10423706-INS-162
chr10-10439448-INS-500
chr10-10458018-INS-65
chr10-10474316-INS-52
chr10-10516488-INS-211
chr10-10527243-INS-81
chr10-10581022-INS-7084
chr10-10658972-INS-104
chr10-10660124-INS-3020
chr10-10682679-INS-84
chr10-10700782-INS-302
chr10-10703358-INS-152
chr10-10715345-INS-327
chr10-10736118-INS-94
chr10-10769748-INS-62
chr10-10771092-INS-6396
chr10-10773654-INS-6486
chr10-10775088-INS-884

chr10-10781089-INS-82
chr10-10790489-INS-83
chr10-10809398-INS-295
chr10-10816643-INS-76
chr10-10827094-INS-68
chr10-10829694-INS-1683
chr10-10843665-INS-58
chr10-10849942-INS-6712
chr10-10859481-INS-6673
chr10-10887184-INS-879
chr10-10888399-INS-79
chr10-10893463-INS-58
chr10-10899561-INS-258
chr10-10900607-INS-141
chr10-10913646-INS-56
chr10-10934665-INS-225
chr10-10947846-INS-104
chr10-10955022-INS-52
chr10-10960873-INS-61
chr10-10964816-INS-90
chr10-10985810-INS-76
chr10-11033719-INS-198
chr10-11041263-INS-212
chr10-11060617-INS-512
chr10-11068713-INS-320
chr10-11076719-INS-212
chr10-11085507-INS-84
chr10-11107447-INS-395
chr10-11122790-INS-559
chr10-11133170-INS-521
chr10-11139158-INS-540
chr10-11145615-INS-575
chr10-11149736-INS-327
chr10-11151752-INS-224
chr10-11159382-INS-196
chr10-11163923-INS-207
chr10-11163970-INS-7172
chr10-11213312-INS-574
chr10-11222094-INS-3373
chr10-11230793-INS-606
chr10-11284085-INS-1895
chr10-11294496-INS-187
chr10-11305720-INS-954
chr10-11309411-INS-108
chr10-11314694-INS-208
chr10-11323036-INS-108
chr10-11349459-INS-54
chr10-11371272-INS-67
chr10-11394978-INS-207
chr10-11396589-INS-534
chr10-11417631-INS-139
chr10-11429679-INS-157
chr10-11434324-INS-7024
chr10-11438181-INS-398
chr10-11448963-INS-54
chr10-11455816-INS-214
chr10-11466335-INS-50
chr10-11479479-INS-433
chr10-11493656-INS-217
chr10-11510553-INS-101
chr10-11521695-INS-210
chr10-11539106-INS-205
chr10-11540611-INS-463
chr10-11546947-INS-232
chr10-11589143-INS-7085
chr10-11600867-INS-56
chr10-11613738-INS-6378
chr10-11617875-INS-107
chr10-11627542-INS-160
chr10-11638392-INS-94
chr10-11639603-INS-90
chr10-11644753-INS-126
chr10-11657287-INS-272
chr10-11659184-INS-224
chr10-11670052-INS-64
chr10-11672020-INS-202
chr10-11672577-INS-55

chr10-11683807-INS-122
chr10-11696806-INS-513
chr10-11709308-INS-1274
chr10-11712710-INS-183
chr10-11739170-INS-419
chr10-11748570-INS-837
chr10-11765983-INS-250
chr10-11770176-INS-53
chr10-11774122-INS-109
chr10-11799158-INS-64
chr10-11805175-INS-511
chr10-11839069-INS-66
chr10-11839445-INS-511
chr10-11864711-INS-4859
chr10-11866073-INS-1800
chr10-11870380-INS-63
chr10-11872276-INS-70
chr10-11884636-INS-55
chr10-11894830-INS-78
chr10-11898692-INS-129
chr10-11952542-INS-398
chr10-11956209-INS-268
chr10-11978857-INS-300
chr10-11979548-INS-64
chr10-11994682-INS-87
chr10-11997155-INS-132
chr10-12002275-INS-83
chr10-12013130-INS-52
chr10-12046976-INS-2945
chr10-12075251-INS-72
chr10-12102432-INS-329
chr10-12127505-INS-227
chr10-12128063-INS-3769
chr10-12130703-INS-7984
chr10-12159278-INS-6304
chr10-12162793-INS-6768
chr10-12167073-INS-831
chr10-12220444-INS-1831
chr10-12229113-INS-356
chr10-12236502-INS-824
chr10-12240973-INS-400
chr10-12244765-INS-129
chr10-12250894-INS-70
chr10-12253059-INS-6438
chr10-12254683-INS-53
chr10-12280336-INS-54
chr10-12298313-INS-98
chr10-12318932-INS-518
chr10-12332929-INS-90
chr10-12336310-INS-208
chr10-12347319-INS-68
chr10-12366372-INS-180
chr10-12388225-INS-124
chr10-12389133-INS-925
chr10-12423895-INS-141
chr10-12433741-INS-147
chr10-12449583-INS-203
chr10-12454288-INS-6507
chr10-12463092-INS-53
chr10-12493843-INS-66
chr10-12502565-INS-173
chr10-12514819-INS-91
chr10-12520995-INS-184
chr10-12529332-INS-62
chr10-12538443-INS-114
chr10-12547920-INS-210
chr10-12554014-INS-648
chr10-12555254-INS-130
chr10-12559747-INS-73
chr10-12570429-INS-112
chr10-12573116-INS-60
chr10-12593317-INS-212
chr10-12597439-INS-514
chr10-12598660-INS-216
chr10-12614631-INS-998
chr10-12637827-INS-94
chr10-12675852-INS-222

chr10-12702141-INS-5542
chr10-12707557-INS-4549
chr10-12710493-INS-53
chr10-12746920-INS-261
chr10-12747457-INS-705
chr10-12750266-INS-91
chr10-12753044-INS-326
chr10-12777887-INS-72
chr10-12807423-INS-102
chr10-12808752-INS-359
chr10-12812018-INS-230
chr10-12812536-INS-147
chr10-12829135-INS-62
chr10-12858748-INS-63
chr10-12868660-INS-207
chr10-12872713-INS-66
chr10-12883625-INS-63
chr10-12887751-INS-175
chr10-12906280-INS-86
chr10-12959443-INS-227
chr10-13018239-INS-53
chr10-13025128-INS-236
chr10-13034986-INS-239
chr10-13040869-INS-224
chr10-13063616-INS-52
chr10-13071611-INS-140
chr10-13086458-INS-60
chr10-13094083-INS-148
chr10-13105639-INS-5358
chr10-13128760-INS-416
chr10-13141865-INS-118
chr10-13155841-INS-242
chr10-13178830-INS-412
chr10-13183105-INS-54
chr10-13194046-INS-57
chr10-13196098-INS-186
chr10-13226779-INS-721
chr10-13235172-INS-157
chr10-13246301-INS-6397
chr10-13271132-INS-149
chr10-13274445-INS-118
chr10-13275335-INS-366
chr10-13278964-INS-57
chr10-13282676-INS-338
chr10-13284623-INS-519
chr10-13294048-INS-54
chr10-13306497-INS-73
chr10-13322598-INS-68
chr10-13340629-INS-86
chr10-13360229-INS-7076
chr10-13368424-INS-51
chr10-13376004-INS-453
chr10-13410230-INS-4869
chr10-13422066-INS-290
chr10-13430519-INS-52
chr10-13434194-INS-3183
chr10-13461753-INS-98
chr10-13489504-INS-69
chr10-13491433-INS-52
chr10-13494852-INS-129
chr10-13518232-INS-400
chr10-13527683-INS-344
chr10-13544288-INS-208
chr10-13607425-INS-59
chr10-13611137-INS-3005
chr10-13611330-INS-825
chr10-13620943-INS-72
chr10-13642560-INS-56
chr10-13662181-INS-72
chr10-13667950-INS-265
chr10-13690746-INS-581
chr10-13724077-INS-162
chr10-13748506-INS-206
chr10-13755987-INS-52
chr10-13760026-INS-64
chr10-13770713-INS-424
chr10-13783950-INS-326

chr10-13801091-INS-56
chr10-13816947-INS-217
chr10-13844409-INS-103
chr10-13886251-INS-60
chr10-13888667-INS-96
chr10-13921046-INS-52
chr10-13923671-INS-74
chr10-13927924-INS-560
chr10-13937195-INS-756
chr10-13959129-INS-72
chr10-13983210-INS-927
chr10-14001089-INS-57
chr10-14023776-INS-74
chr10-14049659-INS-1361
chr10-14095238-INS-678
chr10-14116620-INS-425
chr10-14119087-INS-6257
chr10-14132216-INS-180
chr10-14204511-INS-5219
chr10-14212026-INS-3791
chr10-14212085-INS-60
chr10-14221653-INS-136
chr10-14234891-INS-2655
chr10-14247877-INS-228
chr10-14248671-INS-668
chr10-14252162-INS-158
chr10-14256707-INS-218
chr10-14271493-INS-127
chr10-14274031-INS-6872
chr10-14293676-INS-58
chr10-14413508-INS-68
chr10-14421531-INS-110
chr10-14470019-INS-518
chr10-14485746-INS-1894
chr10-14491776-INS-854
chr10-14494465-INS-112
chr10-14495979-INS-6419
chr10-14525797-INS-68
chr10-14528812-INS-483
chr10-14536935-INS-68
chr10-14564514-INS-6442
chr10-14590933-INS-5186
chr10-14608499-INS-144
chr10-14610124-INS-647
chr10-14627032-INS-53
chr10-14672451-INS-417
chr10-14684255-INS-255
chr10-14686628-INS-61
chr10-14694222-INS-505
chr10-14748063-INS-248
chr10-14774563-INS-64
chr10-14803283-INS-6378
chr10-14809558-INS-400
chr10-14831484-INS-85
chr10-14833469-INS-132
chr10-14839120-INS-7199
chr10-14840721-INS-450
chr10-14844794-INS-8390
chr10-14859755-INS-121
chr10-14861895-INS-548
chr10-14876693-INS-2470
chr10-14883543-INS-58
chr10-14948788-INS-789
chr10-14953194-INS-6601
chr10-14971765-INS-564
chr10-14975729-INS-1416
chr10-14976917-INS-361
chr10-15012999-INS-66
chr10-15018058-INS-55
chr10-15063363-INS-530
chr10-15065790-INS-2196
chr10-15088090-INS-6234
chr10-15096234-INS-98
chr10-15098139-INS-82
chr10-15119847-INS-525
chr10-15130279-INS-76
chr10-15175962-INS-56

chr10-15177058-INS-55
chr10-15182059-INS-7046
chr10-15190923-INS-2116
chr10-15196327-INS-1335
chr10-15200375-INS-202
chr10-15244456-INS-274
chr10-15250167-INS-432
chr10-15289878-INS-59
chr10-15297172-INS-102
chr10-15309857-INS-633
chr10-15335930-INS-194
chr10-15362206-INS-249
chr10-15371285-INS-85
chr10-15389379-INS-60
chr10-15390631-INS-4408
chr10-15400471-INS-60
chr10-15417464-INS-58
chr10-15450511-INS-3024
chr10-15460043-INS-3111
chr10-15489564-INS-173
chr10-15508517-INS-5311
chr10-15520272-INS-89
chr10-15527834-INS-58
chr10-15535435-INS-147
chr10-15544518-INS-269
chr10-15545598-INS-677
chr10-15599026-INS-243
chr10-15625663-INS-58
chr10-15626573-INS-60
chr10-15637501-INS-108
chr10-15656465-INS-208
chr10-15665252-INS-540
chr10-15666944-INS-550
chr10-15696361-INS-667
chr10-15730611-INS-52
chr10-15731903-INS-435
chr10-15779137-INS-52
chr10-15780626-INS-436
chr10-15828747-INS-54
chr10-15835167-INS-491
chr10-15839449-INS-60
chr10-15855477-INS-153
chr10-15890169-INS-6281
chr10-15892105-INS-229
chr10-15904473-INS-110
chr10-15916268-INS-293
chr10-15920359-INS-738
chr10-15950059-INS-76
chr10-15954322-INS-378
chr10-15955477-INS-152
chr10-15978453-INS-5297
chr10-15993974-INS-54
chr10-16019973-INS-75
chr10-16026203-INS-173
chr10-16044179-INS-2636
chr10-16068773-INS-240
chr10-16083755-INS-201
chr10-16088428-INS-134
chr10-16090061-INS-541
chr10-16102470-INS-70
chr10-16112508-INS-183
chr10-16175009-INS-52
chr10-16182940-INS-148
chr10-16190944-INS-6438
chr10-16216548-INS-150
chr10-16233458-INS-235
chr10-16237107-INS-321
chr10-16243852-INS-258
chr10-16258017-INS-75
chr10-16277042-INS-482
chr10-16321120-INS-281
chr10-16360261-INS-187
chr10-16366524-INS-52
chr10-16369375-INS-152
chr10-16375732-INS-6391
chr10-16378453-INS-81
chr10-16403260-INS-63

chr10-16424731-INS-582
chr10-16464034-INS-1991
chr10-16471632-INS-7111
chr10-16481394-INS-110
chr10-16517197-INS-485
chr10-16518808-INS-350
chr10-16529259-INS-340
chr10-16547138-INS-6718
chr10-16550791-INS-615
chr10-16565637-INS-72
chr10-16593669-INS-62
chr10-16611458-INS-6435
chr10-16635606-INS-62
chr10-16672237-INS-494
chr10-16677090-INS-56
chr10-16705631-INS-54
chr10-16718300-INS-562
chr10-16735283-INS-227
chr10-16738351-INS-193
chr10-16742104-INS-203
chr10-16743902-INS-211
chr10-16758797-INS-111
chr10-16778367-INS-52
chr10-16793896-INS-2952
chr10-16827746-INS-401
chr10-16835787-INS-64
chr10-16839734-INS-388
chr10-16852427-INS-52
chr10-16864700-INS-510
chr10-16881251-INS-11702
chr10-16904698-INS-2908
chr10-16961036-INS-84
chr10-16964829-INS-292
chr10-16968972-INS-6455
chr10-16975166-INS-165
chr10-16980871-INS-203
chr10-17003306-INS-106
chr10-17006349-INS-6398
chr10-17019394-INS-424
chr10-17041897-INS-65
chr10-17044720-INS-96
chr10-17074171-INS-78
chr10-17083912-INS-127
chr10-17104508-INS-6250
chr10-17120237-INS-62
chr10-17133502-INS-470
chr10-17164526-INS-131
chr10-17188057-INS-60
chr10-17226717-INS-82
chr10-17242497-INS-101
chr10-17245845-INS-183
chr10-17249127-INS-83
chr10-17268602-INS-104
chr10-17281436-INS-171
chr10-17283473-INS-99
chr10-17329837-INS-60
chr10-17370226-INS-95
chr10-17380071-INS-7181
chr10-17398077-INS-59
chr10-17434104-INS-499
chr10-17434826-INS-124
chr10-17466707-INS-514
chr10-17471606-INS-6417
chr10-17479604-INS-199
chr10-17482351-INS-72
chr10-17486058-INS-115
chr10-17507989-INS-63
chr10-17565023-INS-315
chr10-17590394-INS-84
chr10-17593386-INS-68
chr10-17606475-INS-120
chr10-17623745-INS-496
chr10-17640409-INS-313
chr10-17656543-INS-51
chr10-17678393-INS-138
chr10-17687938-INS-3416
chr10-17692161-INS-124

chr10-17695886-INS-6485
chr10-17697598-INS-147
chr10-17748265-INS-3534
chr10-17750697-INS-7114
chr10-17752844-INS-56
chr10-17753643-INS-317
chr10-17760472-INS-7402
chr10-17822608-INS-60
chr10-17835935-INS-435
chr10-17854793-INS-80
chr10-17856952-INS-6915
chr10-17866304-INS-242
chr10-17869183-INS-312
chr10-17881345-INS-60
chr10-17902529-INS-795
chr10-17921120-INS-410
chr10-17947336-INS-292
chr10-17947676-INS-190
chr10-17948438-INS-433
chr10-17956693-INS-906
chr10-17959288-INS-93
chr10-17963073-INS-52
chr10-17972501-INS-61
chr10-17976708-INS-64
chr10-17999546-INS-237
chr10-18008281-INS-115
chr10-18021261-INS-207
chr10-18051319-INS-59
chr10-18063532-INS-566
chr10-18066122-INS-71
chr10-18076994-INS-208
chr10-18095588-INS-118
chr10-18096782-INS-501
chr10-18111572-INS-116
chr10-18134367-INS-60
chr10-18159297-INS-7521
chr10-18172028-INS-102
chr10-18234990-INS-117
chr10-18260584-INS-658
chr10-18263867-INS-656
chr10-18279594-INS-201
chr10-18288495-INS-146
chr10-18303153-INS-201
chr10-18304072-INS-74
chr10-18328500-INS-66
chr10-18343871-INS-376
chr10-18355884-INS-83
chr10-18388900-INS-211
chr10-18411791-INS-5983
chr10-18414736-INS-135
chr10-18414962-INS-3203
chr10-18421329-INS-62
chr10-18444743-INS-52
chr10-18452633-INS-324
chr10-18458599-INS-64
chr10-18469784-INS-272
chr10-18504597-INS-547
chr10-18512770-INS-710
chr10-18528875-INS-154
chr10-18542828-INS-72
chr10-18545122-INS-2131
chr10-18557701-INS-122
chr10-18566223-INS-89
chr10-18567142-INS-78
chr10-18574189-INS-98
chr10-18581296-INS-65
chr10-18585668-INS-236
chr10-18666951-INS-590
chr10-18669577-INS-549
chr10-18680073-INS-842
chr10-18688734-INS-204
chr10-18758437-INS-85
chr10-18764185-INS-2492
chr10-18784911-INS-197
chr10-18792069-INS-73
chr10-18793571-INS-294
chr10-18799773-INS-712

chr10-18804404-INS-163
chr10-18814091-INS-55
chr10-18820836-INS-52
chr10-18823240-INS-647
chr10-18826474-INS-73
chr10-18832830-INS-925
chr10-18846815-INS-7156
chr10-18864765-INS-586
chr10-18870667-INS-78
chr10-18894409-INS-69
chr10-18903315-INS-442
chr10-18911418-INS-866
chr10-18957868-INS-73
chr10-18961504-INS-831
chr10-18963239-INS-66
chr10-18990653-INS-73
chr10-19006520-INS-60
chr10-19007245-INS-519
chr10-19008545-INS-529
chr10-19010641-INS-280
chr10-19014771-INS-60
chr10-19065883-INS-50
chr10-19073100-INS-169
chr10-19074312-INS-151
chr10-19077830-INS-66
chr10-19104465-INS-132
chr10-19113978-INS-172
chr10-19136350-INS-185
chr10-19151473-INS-208
chr10-19163879-INS-6706
chr10-19167087-INS-50
chr10-19206414-INS-764
chr10-19210420-INS-183
chr10-19220145-INS-174
chr10-19235738-INS-67
chr10-19242604-INS-422
chr10-19268977-INS-61
chr10-19312098-INS-401
chr10-19342146-INS-108
chr10-19377012-INS-51
chr10-19381491-INS-346
chr10-19390510-INS-322
chr10-19397807-INS-207
chr10-19402989-INS-54
chr10-19405183-INS-68
chr10-19462724-INS-58
chr10-19481194-INS-548
chr10-19507745-INS-387
chr10-19512677-INS-75
chr10-19516230-INS-2825
chr10-19517926-INS-494
chr10-19531117-INS-653
chr10-19544018-INS-352
chr10-19554810-INS-72
chr10-19557301-INS-319
chr10-19586004-INS-72
chr10-19621783-INS-3508
chr10-19627813-INS-316
chr10-19637926-INS-153
chr10-19647490-INS-218
chr10-19656263-INS-693
chr10-19669310-INS-452
chr10-19677704-INS-63
chr10-19691962-INS-785
chr10-19692923-INS-7483
chr10-19705657-INS-262
chr10-19706808-INS-74
chr10-19710085-INS-53
chr10-19733982-INS-57
chr10-19794193-INS-117
chr10-19797342-INS-84
chr10-19800011-INS-173
chr10-19823762-INS-404
chr10-19883259-INS-218
chr10-19895027-INS-101
chr10-19905557-INS-156
chr10-19909313-INS-7139

chr10-19916397-INS-229
chr10-19921644-INS-132
chr10-19928755-INS-89
chr10-19961809-INS-4866
chr10-20020197-INS-70
chr10-20026699-INS-99
chr10-20038048-INS-64
chr10-20039664-INS-163
chr10-20052846-INS-97
chr10-20060569-INS-196
chr10-20062726-INS-159
chr10-20065564-INS-80
chr10-20084135-INS-191
chr10-20112112-INS-72
chr10-20113487-INS-167
chr10-20121784-INS-257
chr10-20138228-INS-102
chr10-20162944-INS-52
chr10-20168114-INS-153
chr10-20183072-INS-633
chr10-20192393-INS-471
chr10-20222556-INS-58
chr10-20232567-INS-525
chr10-20233607-INS-112
chr10-20280033-INS-133
chr10-20281895-INS-101
chr10-20289943-INS-77
chr10-20314636-INS-126
chr10-20322868-INS-184
chr10-20331996-INS-54
chr10-20337016-INS-126
chr10-20346775-INS-56
chr10-20396440-INS-341
chr10-20403051-INS-123
chr10-20411120-INS-60
chr10-20442635-INS-120
chr10-20450291-INS-86
chr10-20468252-INS-54
chr10-20470871-INS-60
chr10-20478847-INS-57
chr10-20480337-INS-74
chr10-20499526-INS-86
chr10-20536318-INS-112
chr10-20551976-INS-80
chr10-20557106-INS-173
chr10-20572990-INS-78
chr10-20580708-INS-52
chr10-20588083-INS-104
chr10-20589526-INS-411
chr10-20602285-INS-79
chr10-20621556-INS-82
chr10-20639122-INS-401
chr10-20658019-INS-358
chr10-20666192-INS-79
chr10-20672819-INS-58
chr10-20680727-INS-104
chr10-20715941-INS-240
chr10-20728156-INS-53
chr10-20729632-INS-927
chr10-20751544-INS-177
chr10-20784163-INS-148
chr10-20790499-INS-247
chr10-20794384-INS-90
chr10-20805414-INS-121
chr10-20830332-INS-62
chr10-20836297-INS-126
chr10-20850800-INS-178
chr10-20857129-INS-182
chr10-20879335-INS-219
chr10-20881780-INS-696
chr10-20890269-INS-187
chr10-20911688-INS-93
chr10-20913577-INS-645
chr10-20920277-INS-207
chr10-20922213-INS-169
chr10-20922316-INS-410
chr10-20927479-INS-87

chr10-20927521-INS-173
chr10-20944031-INS-839
chr10-20952845-INS-61
chr10-21025824-INS-144
chr10-21041887-INS-1358
chr10-21045049-INS-637
chr10-21073480-INS-66
chr10-21134259-INS-78
chr10-21137294-INS-361
chr10-21138738-INS-281
chr10-21154397-INS-204
chr10-21200238-INS-737
chr10-21238502-INS-93
chr10-21251741-INS-347
chr10-21255248-INS-66
chr10-21273822-INS-932
chr10-21277104-INS-78
chr10-21286279-INS-420
chr10-21288937-INS-252
chr10-21290118-INS-608
chr10-21305074-INS-428
chr10-21308514-INS-177
chr10-21311285-INS-56
chr10-21323503-INS-440
chr10-21330859-INS-176
chr10-21343124-INS-155
chr10-21369659-INS-434
chr10-21409279-INS-269
chr10-21419302-INS-51
chr10-21427712-INS-122
chr10-21448842-INS-96
chr10-21497197-INS-6970
chr10-21526007-INS-121
chr10-21531230-INS-201
chr10-21537259-INS-333
chr10-21553315-INS-135
chr10-21567840-INS-53
chr10-21636974-INS-50
chr10-21645649-INS-51
chr10-21648478-INS-7428
chr10-21671088-INS-52
chr10-21676969-INS-52
chr10-21679673-INS-143
chr10-21696723-INS-107
chr10-21702667-INS-80
chr10-21719996-INS-494
chr10-21724816-INS-614
chr10-21739541-INS-136
chr10-21746717-INS-296
chr10-21762897-INS-234
chr10-21778087-INS-256
chr10-21788623-INS-69
chr10-21805986-INS-52
chr10-21837510-INS-54
chr10-21840316-INS-100
chr10-21844039-INS-213
chr10-21845912-INS-97
chr10-21854203-INS-276
chr10-21891584-INS-195
chr10-21894826-INS-197
chr10-21906734-INS-220
chr10-21988348-INS-724
chr10-21990627-INS-169
chr10-22007948-INS-343
chr10-22020021-INS-151
chr10-22023408-INS-572
chr10-22048684-INS-952
chr10-22048870-INS-66
chr10-22073901-INS-191
chr10-22094052-INS-327
chr10-22109124-INS-121
chr10-22114156-INS-4083
chr10-22133994-INS-57
chr10-22151303-INS-667
chr10-22237190-INS-231
chr10-22284281-INS-335
chr10-22299351-INS-111

chr10-22337594-INS-64
chr10-22346640-INS-1221
chr10-22346707-INS-428
chr10-22347181-INS-165
chr10-22371534-INS-61
chr10-22448112-INS-67
chr10-22472175-INS-7081
chr10-22497074-INS-56
chr10-22523779-INS-131
chr10-22533785-INS-132
chr10-22538297-INS-351
chr10-22567249-INS-6536
chr10-22569663-INS-1411
chr10-22571166-INS-989
chr10-22575904-INS-141
chr10-22599483-INS-60
chr10-22649654-INS-195
chr10-22662650-INS-96
chr10-22681793-INS-897
chr10-22689116-INS-363
chr10-22711791-INS-802
chr10-22712475-INS-173
chr10-22726286-INS-78
chr10-22753912-INS-135
chr10-22785542-INS-775
chr10-22800939-INS-927
chr10-22836393-INS-6941
chr10-22837024-INS-96
chr10-22859363-INS-53
chr10-22866947-INS-59
chr10-22877832-INS-108
chr10-22902407-INS-66
chr10-22920215-INS-88
chr10-22934804-INS-110
chr10-22938676-INS-52
chr10-22944319-INS-200
chr10-22961073-INS-368
chr10-22970969-INS-133
chr10-22979363-INS-397
chr10-22994874-INS-159
chr10-23004172-INS-232
chr10-23011074-INS-68
chr10-23060548-INS-546
chr10-23072626-INS-134
chr10-23087754-INS-327
chr10-23106975-INS-132
chr10-23150547-INS-93
chr10-23172979-INS-287
chr10-23175375-INS-244
chr10-23211268-INS-933
chr10-23219110-INS-52
chr10-23242133-INS-275
chr10-23250899-INS-149
chr10-23341865-INS-616
chr10-23351500-INS-318
chr10-23352289-INS-51
chr10-23354393-INS-367
chr10-23364025-INS-52
chr10-23399883-INS-67
chr10-23401999-INS-301
chr10-23406577-INS-72
chr10-23415805-INS-189
chr10-23444774-INS-411
chr10-23475545-INS-78
chr10-23480142-INS-222
chr10-23502990-INS-92
chr10-23518028-INS-177
chr10-23540091-INS-268
chr10-23544693-INS-915
chr10-23546019-INS-53
chr10-23560050-INS-609
chr10-23576574-INS-490
chr10-23588839-INS-459
chr10-23609322-INS-163
chr10-23620250-INS-973
chr10-23622490-INS-679
chr10-23639911-INS-143

chr10-23646174-INS-164
chr10-23648768-INS-277
chr10-23666486-INS-58
chr10-23669291-INS-153
chr10-23676551-INS-71
chr10-23684382-INS-56
chr10-23685370-INS-133
chr10-23698219-INS-88
chr10-23704628-INS-109
chr10-23719507-INS-224
chr10-23739546-INS-79
chr10-23741279-INS-623
chr10-23767363-INS-66
chr10-23774906-INS-205
chr10-23798461-INS-52
chr10-23801406-INS-62
chr10-23818255-INS-76
chr10-23824940-INS-82
chr10-23829660-INS-529
chr10-23835875-INS-297
chr10-23864508-INS-99
chr10-23905584-INS-201
chr10-23908528-INS-300
chr10-23932185-INS-77
chr10-23939613-INS-86
chr10-23951781-INS-186
chr10-23953745-INS-793
chr10-23961045-INS-56
chr10-23964692-INS-329
chr10-23971005-INS-98
chr10-23975372-INS-198
chr10-23976410-INS-129
chr10-23995481-INS-1892
chr10-24004984-INS-52
chr10-24018415-INS-937
chr10-24020865-INS-69
chr10-24037405-INS-110
chr10-24087891-INS-139
chr10-24100709-INS-101
chr10-24113315-INS-600
chr10-24156127-INS-581
chr10-24174919-INS-59
chr10-24177083-INS-95
chr10-24179230-INS-767
chr10-24187406-INS-54
chr10-24188693-INS-51
chr10-24243769-INS-98
chr10-24248802-INS-141
chr10-24261011-INS-125
chr10-24324911-INS-148
chr10-24376141-INS-7492
chr10-24460270-INS-395
chr10-24476862-INS-319
chr10-24484765-INS-334
chr10-24497940-INS-199
chr10-24502315-INS-155
chr10-24528323-INS-419
chr10-24548082-INS-68
chr10-24581901-INS-175
chr10-24589179-INS-86
chr10-24591045-INS-188
chr10-24599595-INS-67
chr10-24643466-INS-180
chr10-24658092-INS-62
chr10-24661719-INS-218
chr10-24665313-INS-201
chr10-24666789-INS-472
chr10-24672519-INS-203
chr10-24676641-INS-79
chr10-24697548-INS-175
chr10-24708661-INS-98
chr10-24713226-INS-60
chr10-24724483-INS-76
chr10-24733505-INS-6401
chr10-24755143-INS-164
chr10-24760516-INS-108
chr10-24796129-INS-53

chr10-24800211-INS-120
chr10-24818118-INS-145
chr10-24823827-INS-60
chr10-24838868-INS-99
chr10-24862727-INS-84
chr10-24877849-INS-790
chr10-24883942-INS-156
chr10-24901442-INS-659
chr10-24905838-INS-445
chr10-24916806-INS-292
chr10-24933047-INS-171
chr10-24940086-INS-343
chr10-24950937-INS-56
chr10-24953092-INS-95
chr10-24968092-INS-261
chr10-25010099-INS-58
chr10-25061281-INS-54
chr10-25066251-INS-72
chr10-25084842-INS-95
chr10-25098678-INS-210
chr10-25106040-INS-68
chr10-25116436-INS-492
chr10-25128101-INS-132
chr10-25134278-INS-128
chr10-25137868-INS-205
chr10-25158265-INS-128
chr10-25168447-INS-70
chr10-25204756-INS-108
chr10-25223013-INS-100
chr10-25228729-INS-132
chr10-25232666-INS-1844
chr10-25232667-INS-966
chr10-25232686-INS-1283
chr10-25235072-INS-122
chr10-25236928-INS-433
chr10-25287775-INS-57
chr10-25289789-INS-222
chr10-25308227-INS-5563
chr10-25312198-INS-144
chr10-25316828-INS-401
chr10-25334615-INS-140
chr10-25358107-INS-234
chr10-25364361-INS-959
chr10-25395014-INS-453
chr10-25403791-INS-96
chr10-25405089-INS-128
chr10-25415183-INS-408
chr10-25436798-INS-117
chr10-25494808-INS-6569
chr10-25523664-INS-192
chr10-25599275-INS-148
chr10-25623802-INS-273
chr10-25651901-INS-569
chr10-25725389-INS-6278
chr10-25745165-INS-930
chr10-25766856-INS-56
chr10-25785736-INS-161
chr10-25822886-INS-416
chr10-25825778-INS-6008
chr10-25852254-INS-358
chr10-25859343-INS-523
chr10-25871657-INS-949
chr10-25881025-INS-487
chr10-25886209-INS-60
chr10-25890275-INS-102
chr10-25914500-INS-281
chr10-25920010-INS-5545
chr10-25933951-INS-96
chr10-25955682-INS-175
chr10-25961535-INS-121
chr10-25962326-INS-386
chr10-25972383-INS-70
chr10-25995272-INS-68
chr10-26031165-INS-224
chr10-26079504-INS-124
chr10-26103444-INS-64
chr10-26130407-INS-60

chr10-26142459-INS-163
chr10-26169857-INS-230
chr10-26206308-INS-199
chr10-26238541-INS-99
chr10-26242696-INS-61
chr10-26256664-INS-287
chr10-26267548-INS-207
chr10-26307338-INS-53
chr10-26309036-INS-209
chr10-26313754-INS-90
chr10-26324774-INS-191
chr10-26341229-INS-184
chr10-26343063-INS-56
chr10-26347255-INS-124
chr10-26348314-INS-116
chr10-26349559-INS-455
chr10-26358817-INS-575
chr10-26389816-INS-368
chr10-26400230-INS-55
chr10-26417876-INS-108
chr10-26446905-INS-95
chr10-26485496-INS-216
chr10-26509063-INS-60
chr10-26554428-INS-60
chr10-26565638-INS-184
chr10-26586040-INS-56
chr10-26614857-INS-53
chr10-26627001-INS-389
chr10-26648316-INS-58
chr10-26673658-INS-5550
chr10-26676288-INS-232
chr10-26677739-INS-272
chr10-26706216-INS-398
chr10-26710456-INS-410
chr10-26723881-INS-105
chr10-26729391-INS-3488
chr10-26734959-INS-201
chr10-26766157-INS-98
chr10-26767968-INS-107
chr10-26769019-INS-65
chr10-26770680-INS-80
chr10-26771826-INS-433
chr10-26774830-INS-223
chr10-26795422-INS-204
chr10-26823841-INS-52
chr10-26838284-INS-210
chr10-26966808-INS-1208
chr10-26969301-INS-66
chr10-26986110-INS-139
chr10-26995395-INS-105
chr10-26996998-INS-2711
chr10-27008219-INS-86
chr10-27055471-INS-533
chr10-27073222-INS-268
chr10-27105105-INS-52
chr10-27106778-INS-51
chr10-27127223-INS-54
chr10-27144409-INS-1119
chr10-27167214-INS-76
chr10-27177814-INS-658
chr10-27246694-INS-821
chr10-27273312-INS-2144
chr10-27273810-INS-2137
chr10-27296275-INS-6237
chr10-27296825-INS-1473
chr10-27347899-INS-205
chr10-27371751-INS-505
chr10-27381325-INS-446
chr10-27387865-INS-168
chr10-27389374-INS-530
chr10-27412678-INS-123
chr10-27446542-INS-303
chr10-27451443-INS-143
chr10-27454221-INS-8149
chr10-27457802-INS-3012
chr10-27462787-INS-77
chr10-27480488-INS-126

chr10-27483212-INS-1230
chr10-27494422-INS-74
chr10-27521011-INS-404
chr10-27524469-INS-63
chr10-27547743-INS-78
chr10-27550938-INS-205
chr10-27574868-INS-58
chr10-27579310-INS-7026
chr10-27585814-INS-339
chr10-27606030-INS-192
chr10-27606847-INS-6235
chr10-27615968-INS-56
chr10-27616942-INS-466
chr10-27656662-INS-524
chr10-27686862-INS-618
chr10-27701008-INS-3536
chr10-27721216-INS-1909
chr10-27743977-INS-4121
chr10-27760942-INS-5975
chr10-27763557-INS-6698
chr10-27765394-INS-217
chr10-27767896-INS-384
chr10-27772227-INS-417
chr10-27826138-INS-6427
chr10-27847831-INS-134
chr10-27877131-INS-6615
chr10-27887500-INS-136
chr10-27890807-INS-92
chr10-27895184-INS-536
chr10-27897227-INS-264
chr10-27931716-INS-108
chr10-27945819-INS-60
chr10-27970595-INS-2488
chr10-28001955-INS-56
chr10-28033616-INS-225
chr10-28052811-INS-885
chr10-28061278-INS-81
chr10-28073704-INS-120
chr10-28080319-INS-219
chr10-28088784-INS-96
chr10-28117535-INS-56
chr10-28145887-INS-51
chr10-28154794-INS-321
chr10-28160094-INS-223
chr10-28186926-INS-52
chr10-28194669-INS-257
chr10-28212544-INS-245
chr10-28216483-INS-55
chr10-28223013-INS-65
chr10-28229155-INS-208
chr10-28247860-INS-88
chr10-28301422-INS-320
chr10-28310200-INS-680
chr10-28329711-INS-230
chr10-28350922-INS-52
chr10-28420045-INS-71
chr10-28427330-INS-300
chr10-28454197-INS-140
chr10-28455716-INS-203
chr10-28469904-INS-54
chr10-28492880-INS-94
chr10-28536368-INS-68
chr10-28540679-INS-117
chr10-28543770-INS-54
chr10-28544959-INS-81
chr10-28549362-INS-112
chr10-28559368-INS-171
chr10-28564562-INS-283
chr10-28576472-INS-536
chr10-28587988-INS-59
chr10-28606635-INS-120
chr10-28617255-INS-233
chr10-28626095-INS-63
chr10-28629453-INS-6832
chr10-28652858-INS-6462
chr10-28666499-INS-6484
chr10-28667174-INS-164

chr10-28676992-INS-181
chr10-28685227-INS-68
chr10-28694843-INS-632
chr10-28709878-INS-292
chr10-28712310-INS-399
chr10-28720923-INS-146
chr10-28723222-INS-80
chr10-28751383-INS-58
chr10-28763971-INS-793
chr10-28782276-INS-2587
chr10-28835716-INS-110
chr10-28839582-INS-6591
chr10-28883114-INS-102
chr10-28923193-INS-95
chr10-28928899-INS-84
chr10-28930337-INS-132
chr10-28931784-INS-197
chr10-28988732-INS-92
chr10-29012039-INS-330
chr10-29051927-INS-8472
chr10-29054008-INS-6588
chr10-29054160-INS-341
chr10-29071526-INS-243
chr10-29081411-INS-197
chr10-29084071-INS-63
chr10-29089069-INS-58
chr10-29101861-INS-96
chr10-29128715-INS-206
chr10-29163189-INS-89
chr10-29167572-INS-357
chr10-29187217-INS-8940
chr10-29197184-INS-218
chr10-29200084-INS-351
chr10-29230949-INS-65
chr10-29253672-INS-6642
chr10-29287711-INS-399
chr10-29297845-INS-193
chr10-29308343-INS-139
chr10-29310733-INS-126
chr10-29321929-INS-68
chr10-29332966-INS-89
chr10-29350953-INS-850
chr10-29361002-INS-111
chr10-29367657-INS-209
chr10-29377009-INS-60
chr10-29386038-INS-492
chr10-29449698-INS-874
chr10-29464509-INS-174
chr10-29476433-INS-57
chr10-29513823-INS-76
chr10-29543446-INS-88
chr10-29545957-INS-52
chr10-29570431-INS-56
chr10-29583857-INS-80
chr10-29585247-INS-76
chr10-29592011-INS-52
chr10-29600371-INS-122
chr10-29614834-INS-343
chr10-29624665-INS-60
chr10-29630210-INS-2574
chr10-29633214-INS-1045
chr10-29658901-INS-87
chr10-29739064-INS-493
chr10-29746087-INS-83
chr10-29798767-INS-54
chr10-29810543-INS-1685
chr10-29829642-INS-88
chr10-29844790-INS-6294
chr10-29852660-INS-352
chr10-29864154-INS-68
chr10-29866336-INS-228
chr10-29870619-INS-841
chr10-29889388-INS-487
chr10-29929025-INS-80
chr10-29954716-INS-4628
chr10-29967991-INS-133
chr10-29976491-INS-3617

chr10-29977569-INS-2333
chr10-29984650-INS-2643
chr10-30005556-INS-201
chr10-30018940-INS-91
chr10-30046272-INS-7134
chr10-30054869-INS-5477
chr10-30057619-INS-89
chr10-30066840-INS-113
chr10-30075457-INS-165
chr10-30090115-INS-64
chr10-30112250-INS-78
chr10-30129747-INS-6477
chr10-30142308-INS-69
chr10-30193749-INS-4528
chr10-30202953-INS-121
chr10-30252813-INS-451
chr10-30285567-INS-178
chr10-30296009-INS-61
chr10-30306277-INS-327
chr10-30374252-INS-66
chr10-30382710-INS-54
chr10-30395007-INS-169
chr10-30401540-INS-85
chr10-30461471-INS-55
chr10-30463181-INS-156
chr10-30473101-INS-504
chr10-30512949-INS-6560
chr10-30522449-INS-6622
chr10-30543037-INS-128
chr10-30556060-INS-901
chr10-30638504-INS-81
chr10-30640593-INS-52
chr10-30681589-INS-149
chr10-30723248-INS-132
chr10-30746599-INS-321
chr10-30764806-INS-229
chr10-30766446-INS-402
chr10-30788291-INS-932
chr10-30791595-INS-6363
chr10-30799300-INS-154
chr10-30802178-INS-65
chr10-30809646-INS-231
chr10-30813115-INS-99
chr10-30889949-INS-81
chr10-30920503-INS-198
chr10-30927255-INS-82
chr10-30965310-INS-75
chr10-30976340-INS-893
chr10-30984785-INS-599
chr10-30991306-INS-138
chr10-31016858-INS-183
chr10-31031382-INS-412
chr10-31057674-INS-5547
chr10-31058347-INS-607
chr10-31082959-INS-52
chr10-31106774-INS-691
chr10-31174145-INS-349
chr10-31175024-INS-280
chr10-31181472-INS-375
chr10-31184979-INS-586
chr10-31213298-INS-980
chr10-31225679-INS-640
chr10-31236699-INS-1888
chr10-31255875-INS-694
chr10-31267373-INS-62
chr10-31279414-INS-650
chr10-31287438-INS-165
chr10-31292759-INS-5637
chr10-31301973-INS-124
chr10-31310628-INS-65
chr10-31318730-INS-185
chr10-31320918-INS-54
chr10-31331257-INS-390
chr10-31333455-INS-326
chr10-31340477-INS-122
chr10-31342194-INS-7288
chr10-31354156-INS-1019

chr10-31354157-INS-919
chr10-31357274-INS-609
chr10-31379022-INS-466
chr10-31383996-INS-166
chr10-31391761-INS-72
chr10-31446474-INS-541
chr10-31479761-INS-662
chr10-31487170-INS-133
chr10-31505203-INS-181
chr10-31508794-INS-7009
chr10-31536899-INS-5072
chr10-31540173-INS-2623
chr10-31548844-INS-211
chr10-31552449-INS-197
chr10-31561209-INS-189
chr10-31571365-INS-436
chr10-31572669-INS-123
chr10-31610123-INS-74
chr10-31621395-INS-6460
chr10-31639708-INS-300
chr10-31664098-INS-68
chr10-31750831-INS-201
chr10-31763157-INS-85
chr10-31769522-INS-55
chr10-31772910-INS-842
chr10-31786542-INS-54
chr10-31789498-INS-86
chr10-31805691-INS-880
chr10-31831008-INS-154
chr10-31831407-INS-6391
chr10-31837248-INS-66
chr10-31842007-INS-247
chr10-31871444-INS-772
chr10-31929533-INS-193
chr10-31945234-INS-101
chr10-31950540-INS-164
chr10-31993998-INS-117
chr10-31996228-INS-195
chr10-31999810-INS-2546
chr10-32012499-INS-188
chr10-32035589-INS-118
chr10-32042780-INS-6899
chr10-32047649-INS-202
chr10-32050149-INS-88
chr10-32063724-INS-1500
chr10-32077556-INS-6061
chr10-32083586-INS-7417
chr10-32120435-INS-205
chr10-32137846-INS-60
chr10-32172709-INS-240
chr10-32185937-INS-6395
chr10-32265783-INS-542
chr10-32336087-INS-621
chr10-32336823-INS-119
chr10-32344303-INS-60
chr10-32399605-INS-55
chr10-32455904-INS-286
chr10-32463819-INS-399
chr10-32573842-INS-441
chr10-32604944-INS-62
chr10-32613681-INS-108
chr10-32623543-INS-5373
chr10-32632562-INS-74
chr10-32675757-INS-647
chr10-32686275-INS-168
chr10-32708066-INS-68
chr10-32711874-INS-202
chr10-32723201-INS-177
chr10-32724626-INS-5549
chr10-32731944-INS-6494
chr10-32754370-INS-64
chr10-32766481-INS-60
chr10-32777053-INS-2665
chr10-32788389-INS-6220
chr10-32816782-INS-11931
chr10-32822785-INS-4862
chr10-32838716-INS-889

chr10-32853565-INS-6145
chr10-32865089-INS-149
chr10-32873095-INS-366
chr10-32879787-INS-827
chr10-32924294-INS-187
chr10-32933941-INS-52
chr10-32950040-INS-172
chr10-32954098-INS-69
chr10-32973365-INS-256
chr10-32988469-INS-57
chr10-33012256-INS-209
chr10-33023478-INS-70
chr10-33023746-INS-3532
chr10-33036692-INS-7409
chr10-33059925-INS-546
chr10-33071842-INS-112
chr10-33085542-INS-155
chr10-33110572-INS-210
chr10-33119723-INS-94
chr10-33180342-INS-337
chr10-33205450-INS-111
chr10-33214935-INS-83
chr10-33304847-INS-181
chr10-33318349-INS-70
chr10-33374448-INS-77
chr10-33389399-INS-51
chr10-33420895-INS-56
chr10-33444280-INS-135
chr10-33497087-INS-871
chr10-33499296-INS-109
chr10-33506225-INS-717
chr10-33525849-INS-93
chr10-33539918-INS-4470
chr10-33578791-INS-218
chr10-33588333-INS-78
chr10-33601340-INS-535
chr10-33657284-INS-60
chr10-33663226-INS-496
chr10-33708616-INS-211
chr10-33718147-INS-120
chr10-33760116-INS-108
chr10-33770992-INS-536
chr10-33809025-INS-134
chr10-33822748-INS-100
chr10-33829218-INS-500
chr10-33831141-INS-4837
chr10-33831233-INS-327
chr10-33851549-INS-447
chr10-33869726-INS-171
chr10-33872929-INS-85
chr10-33880309-INS-54
chr10-33887864-INS-307
chr10-33893715-INS-324
chr10-33903896-INS-994
chr10-33905089-INS-366
chr10-33909522-INS-88
chr10-33925810-INS-291
chr10-33929415-INS-223
chr10-33930591-INS-397
chr10-33935290-INS-137
chr10-33950869-INS-61
chr10-34010899-INS-96
chr10-34017524-INS-207
chr10-34018282-INS-403
chr10-34031244-INS-51
chr10-34033631-INS-140
chr10-34040212-INS-63
chr10-34041473-INS-110
chr10-34056073-INS-205
chr10-34059428-INS-67
chr10-34082022-INS-205
chr10-34116014-INS-392
chr10-34125517-INS-437
chr10-34168826-INS-357
chr10-34181324-INS-62
chr10-34191021-INS-184
chr10-34192259-INS-131

chr10-34222511-INS-208
chr10-34226790-INS-328
chr10-34323013-INS-52
chr10-34344989-INS-74
chr10-34348375-INS-56
chr10-34373999-INS-62
chr10-34385155-INS-6376
chr10-34392413-INS-248
chr10-34408240-INS-88
chr10-34419313-INS-483
chr10-34427035-INS-2737
chr10-34441963-INS-367
chr10-34455738-INS-6939
chr10-34457135-INS-95
chr10-34461549-INS-695
chr10-34474038-INS-143
chr10-34493349-INS-206
chr10-34535110-INS-129
chr10-34573164-INS-66
chr10-34634711-INS-176
chr10-34675819-INS-89
chr10-34707140-INS-896
chr10-34708717-INS-148
chr10-34757359-INS-301
chr10-34795098-INS-277
chr10-34837899-INS-64
chr10-34844521-INS-386
chr10-34956037-INS-214
chr10-34959605-INS-829
chr10-34965976-INS-53
chr10-34985019-INS-3805
chr10-35006382-INS-96
chr10-35033392-INS-327
chr10-35038945-INS-81
chr10-35067337-INS-90
chr10-35076683-INS-67
chr10-35099761-INS-75
chr10-35116373-INS-126
chr10-35120384-INS-86
chr10-35124183-INS-5547
chr10-35283220-INS-298
chr10-35286876-INS-205
chr10-35293607-INS-53
chr10-35334220-INS-7141
chr10-35336407-INS-6128
chr10-35340965-INS-64
chr10-35355076-INS-537
chr10-35357479-INS-480
chr10-35463934-INS-179
chr10-35467769-INS-72
chr10-35489337-INS-56
chr10-35493940-INS-561
chr10-35575487-INS-56
chr10-35592583-INS-78
chr10-35594968-INS-53
chr10-35634073-INS-4309
chr10-35634232-INS-597
chr10-35636237-INS-3552
chr10-35665136-INS-512
chr10-35672477-INS-103
chr10-35780698-INS-372
chr10-35820004-INS-62
chr10-35822431-INS-6732
chr10-35840600-INS-1475
chr10-35875111-INS-53
chr10-35878766-INS-107
chr10-35924752-INS-151
chr10-35940058-INS-75
chr10-35946364-INS-64
chr10-35955951-INS-58
chr10-35964103-INS-7524
chr10-35977551-INS-110
chr10-36003776-INS-71
chr10-36005854-INS-155
chr10-36022354-INS-122
chr10-36031737-INS-330
chr10-36066013-INS-51

chr10-36105456-INS-7100
chr10-36135781-INS-6419
chr10-36176960-INS-146
chr10-36194080-INS-322
chr10-36198752-INS-53
chr10-36202652-INS-92
chr10-36214061-INS-70
chr10-36229404-INS-269
chr10-36251460-INS-55
chr10-36253630-INS-756
chr10-36274535-INS-55
chr10-36314039-INS-485
chr10-36379578-INS-136
chr10-36422880-INS-249
chr10-36487256-INS-533
chr10-36490459-INS-713
chr10-36523499-INS-574
chr10-36625347-INS-235
chr10-36657505-INS-67
chr10-36695673-INS-79
chr10-36719018-INS-184
chr10-36753657-INS-464
chr10-36780450-INS-165
chr10-36794104-INS-92
chr10-36806033-INS-118
chr10-36809733-INS-183
chr10-36826205-INS-139
chr10-36846975-INS-51
chr10-36856291-INS-82
chr10-36889395-INS-131
chr10-36904540-INS-59
chr10-36931117-INS-221
chr10-36978169-INS-102
chr10-36988202-INS-99
chr10-37003668-INS-96
chr10-37018084-INS-110
chr10-37026223-INS-75
chr10-37088972-INS-81
chr10-37103030-INS-109
chr10-37113838-INS-98
chr10-37115989-INS-54
chr10-37121645-INS-148
chr10-37123672-INS-60
chr10-37145052-INS-74
chr10-37157397-INS-139
chr10-37181114-INS-66
chr10-37183036-INS-469
chr10-37215607-INS-62
chr10-37217768-INS-100
chr10-37233214-INS-98
chr10-37247333-INS-58
chr10-37282723-INS-513
chr10-37284954-INS-185
chr10-37324994-INS-81
chr10-37335315-INS-147
chr10-37339866-INS-6910
chr10-37342814-INS-177
chr10-37347358-INS-102
chr10-37353166-INS-317
chr10-37361913-INS-756
chr10-37367130-INS-96
chr10-37374204-INS-97
chr10-37388037-INS-74
chr10-37401734-INS-61
chr10-37420178-INS-64
chr10-37451271-INS-742
chr10-37463972-INS-54
chr10-37498534-INS-758
chr10-37551329-INS-295
chr10-37573228-INS-6662
chr10-37573693-INS-6172
chr10-37583798-INS-212
chr10-37587098-INS-6484
chr10-37589266-INS-52
chr10-37599567-INS-106
chr10-37613342-INS-360
chr10-37627548-INS-56

chr10-37653254-INS-58
chr10-37662690-INS-799
chr10-37685412-INS-140
chr10-37692973-INS-160
chr10-37698047-INS-56
chr10-37736042-INS-495
chr10-37820334-INS-178
chr10-37829294-INS-351
chr10-37841900-INS-1420
chr10-37850692-INS-61
chr10-37943844-INS-222
chr10-37953255-INS-66
chr10-37983084-INS-87
chr10-37985661-INS-719
chr10-38002954-INS-54
chr10-38009459-INS-7476
chr10-38011689-INS-114
chr10-38045265-INS-210
chr10-38045868-INS-57
chr10-38063415-INS-107
chr10-38078739-INS-70
chr10-38086592-INS-52
chr10-38091030-INS-957
chr10-38161586-INS-89
chr10-38166236-INS-452
chr10-38181351-INS-166
chr10-38189342-INS-61
chr10-38217202-INS-73
chr10-38228563-INS-325
chr10-38294071-INS-115
chr10-38312726-INS-7733
chr10-38314139-INS-146
chr10-38321332-INS-167
chr10-38322429-INS-62
chr10-38324810-INS-6409
chr10-38327882-INS-239
chr10-38338530-INS-5302
chr10-38381996-INS-172
chr10-38421639-INS-713
chr10-38454492-INS-51
chr10-38463028-INS-156
chr10-38497508-INS-263
chr10-38507598-INS-224
chr10-38508803-INS-77
chr10-38512486-INS-281
chr10-38540736-INS-385
chr10-38574757-INS-54
chr10-38579241-INS-165
chr10-38581624-INS-935
chr10-38585861-INS-65
chr10-38603074-INS-64
chr10-38656317-INS-93
chr10-38684191-INS-811
chr10-38691329-INS-75
chr10-38703615-INS-401
chr10-38718078-INS-1342
chr10-38730081-INS-66
chr10-38733980-INS-6283
chr10-38739046-INS-822
chr10-38757871-INS-68
chr10-38761185-INS-7645
chr10-38774851-INS-147
chr10-38797063-INS-7327
chr10-38803068-INS-375
chr10-38822431-INS-62
chr10-38825533-INS-72
chr10-38852519-INS-71
chr10-38865689-INS-924
chr10-38899220-INS-76
chr10-38909255-INS-716
chr10-38913645-INS-52
chr10-38920225-INS-70
chr10-38967065-INS-210
chr10-38992771-INS-91
chr10-38995544-INS-81
chr10-39000894-INS-131
chr10-39016207-INS-219

chr10-39021051-INS-677
chr10-39023451-INS-214
chr10-39032342-INS-928
chr10-39035126-INS-111
chr10-39036682-INS-84
chr10-39043147-INS-816
chr10-39044290-INS-124
chr10-39077324-INS-76
chr10-39082354-INS-253
chr10-39084520-INS-66
chr10-39093923-INS-331
chr10-39119027-INS-201
chr10-39132666-INS-109
chr10-39139166-INS-62
chr10-39142874-INS-140
chr10-39147879-INS-149
chr10-39193822-INS-202
chr10-39195430-INS-664
chr10-39260795-INS-116
chr10-39266126-INS-356
chr10-39272009-INS-97
chr10-39281372-INS-220
chr10-39283390-INS-6493
chr10-39302098-INS-584
chr10-39308052-INS-181
chr10-39313624-INS-1886
chr10-39325047-INS-73
chr10-39350871-INS-292
chr10-39358997-INS-5743
chr10-39359237-INS-548
chr10-39386955-INS-96
chr10-39391515-INS-223
chr10-39421871-INS-209
chr10-39429017-INS-142
chr10-39433189-INS-60
chr10-39437514-INS-51
chr10-39443968-INS-75
chr10-39458856-INS-204
chr10-39474650-INS-259
chr10-39512924-INS-202
chr10-39561934-INS-147
chr10-39571221-INS-54
chr10-39588202-INS-52
chr10-39667450-INS-154
chr10-39677814-INS-304
chr10-39690599-INS-204
chr10-39707215-INS-51
chr10-39715828-INS-396
chr10-39769278-INS-154
chr10-39781021-INS-56
chr10-39784320-INS-276
chr10-39807432-INS-946
chr10-39814366-INS-341
chr10-39819616-INS-55
chr10-39852485-INS-66
chr10-39874944-INS-213
chr10-39884381-INS-168
chr10-39903969-INS-91
chr10-39905582-INS-84
chr10-39918507-INS-71
chr10-39941334-INS-460
chr10-39961691-INS-5566
chr10-40010145-INS-474
chr10-40027741-INS-390
chr10-40035133-INS-88
chr10-40046724-INS-147
chr10-40048780-INS-78
chr10-40053249-INS-82
chr10-40063846-INS-173
chr10-40068301-INS-511
chr10-40073506-INS-588
chr10-40094349-INS-98
chr10-40112641-INS-80
chr10-40117278-INS-270
chr10-40131393-INS-471
chr10-40143527-INS-374
chr10-40161377-INS-194

chr10-40166283-INS-81
chr10-40173817-INS-202
chr10-40191435-INS-205
chr10-40207430-INS-174
chr10-40219405-INS-330
chr10-40222333-INS-137
chr10-40236102-INS-123
chr10-40242034-INS-7155
chr10-40271970-INS-915
chr10-40295980-INS-123
chr10-40328753-INS-165
chr10-40341721-INS-221
chr10-40347468-INS-60
chr10-40358788-INS-61
chr10-40384133-INS-80
chr10-40398456-INS-52
chr10-40408944-INS-94
chr10-40423307-INS-54
chr10-40426780-INS-172
chr10-40451904-INS-184
chr10-40455315-INS-52
chr10-40463282-INS-100
chr10-40471443-INS-158
chr10-40479848-INS-72
chr10-40485009-INS-85
chr10-40517748-INS-91
chr10-40522449-INS-718
chr10-40522997-INS-1773
chr10-40527203-INS-493
chr10-40529716-INS-69
chr10-40586380-INS-256
chr10-40588806-INS-54
chr10-40617922-INS-58
chr10-40618637-INS-103
chr10-40653398-INS-140
chr10-40660310-INS-509
chr10-40670347-INS-68
chr10-40682908-INS-333
chr10-40687625-INS-73
chr10-40691997-INS-59
chr10-40693151-INS-374
chr10-40695672-INS-183
chr10-40708505-INS-130
chr10-40712702-INS-162
chr10-40744558-INS-147
chr10-40749467-INS-94
chr10-40829997-INS-338
chr10-40864298-INS-804
chr10-40872559-INS-5904
chr10-40880965-INS-152
chr10-40886124-INS-52
chr10-40908773-INS-323
chr10-40916814-INS-85
chr10-40936353-INS-6493
chr10-40948711-INS-63
chr10-40955795-INS-72
chr10-40970409-INS-60
chr10-40971733-INS-55
chr10-40982956-INS-438
chr10-40987950-INS-66
chr10-41002831-INS-202
chr10-41005260-INS-409
chr10-41007279-INS-177
chr10-41024227-INS-92
chr10-41040021-INS-166
chr10-41070220-INS-78
chr10-41096394-INS-132
chr10-41102164-INS-323
chr10-41109703-INS-135
chr10-41180517-INS-95
chr10-41197519-INS-161
chr10-41209841-INS-564
chr10-41216719-INS-58
chr10-41237506-INS-195
chr10-41241410-INS-188
chr10-41249826-INS-222
chr10-41253370-INS-328

chr10-41257571-INS-202
chr10-41262113-INS-351
chr10-41273776-INS-86
chr10-41277886-INS-64
chr10-41288098-INS-399
chr10-41295092-INS-133
chr10-41310941-INS-396
chr10-41311940-INS-51
chr10-41330084-INS-218
chr10-41358676-INS-86
chr10-41389048-INS-80
chr10-41441477-INS-198
chr10-41458067-INS-5543
chr10-41461052-INS-120
chr10-41475825-INS-132
chr10-41503011-INS-57
chr10-41514062-INS-634
chr10-41522816-INS-88
chr10-41523784-INS-325
chr10-41525459-INS-405
chr10-41531325-INS-54
chr10-41534184-INS-492
chr10-41541073-INS-151
chr10-41553116-INS-249
chr10-41582394-INS-105
chr10-41587219-INS-211
chr10-41597062-INS-85
chr10-41605877-INS-89
chr10-41609445-INS-66
chr10-41623840-INS-529
chr10-41628223-INS-59
chr10-41634951-INS-149
chr10-41637815-INS-8057
chr10-41639673-INS-153
chr10-41641443-INS-6955
chr10-41643330-INS-570
chr10-41646165-INS-84
chr10-41667690-INS-302
chr10-41693442-INS-284
chr10-41708846-INS-321
chr10-41728827-INS-67
chr10-41743183-INS-144
chr10-41750974-INS-93
chr10-41754077-INS-169
chr10-41755961-INS-145
chr10-41759553-INS-67
chr10-41771486-INS-228
chr10-41781555-INS-98
chr10-41806017-INS-81
chr10-41822463-INS-104
chr10-41848418-INS-138
chr10-41856439-INS-72
chr10-41935722-INS-117
chr10-41951592-INS-98
chr10-41960456-INS-69
chr10-42013414-INS-265
chr10-42014257-INS-69
chr10-42049621-INS-360
chr10-42052119-INS-201
chr10-42056670-INS-55
chr10-42074244-INS-60
chr10-42087763-INS-230
chr10-42096344-INS-143
chr10-42102854-INS-176
chr10-42111381-INS-215
chr10-42129034-INS-532
chr10-42158034-INS-65
chr10-42167065-INS-400
chr10-42175822-INS-179
chr10-42186398-INS-90
chr10-42187775-INS-72
chr10-42200046-INS-107
chr10-42212485-INS-337
chr10-42219970-INS-69
chr10-42234002-INS-60
chr10-42283715-INS-196
chr10-42294845-INS-76

chr10-42305485-INS-103
chr10-42320269-INS-51
chr10-42321267-INS-400
chr10-42326062-INS-122
chr10-42373289-INS-54
chr10-42374295-INS-57
chr10-42412741-INS-54
chr10-42424093-INS-122
chr10-42428116-INS-117
chr10-42430187-INS-60
chr10-42443739-INS-226
chr10-42451018-INS-53
chr10-42473166-INS-205
chr10-42475021-INS-54
chr10-42476664-INS-224
chr10-42502072-INS-84
chr10-42532864-INS-204
chr10-42551043-INS-84
chr10-42557513-INS-191
chr10-42581988-INS-809
chr10-42610254-INS-230
chr10-42615108-INS-59
chr10-42673356-INS-62
chr10-42689667-INS-82
chr10-42716105-INS-8335
chr10-42722008-INS-481
chr10-42728861-INS-291
chr10-42749856-INS-66
chr10-42757358-INS-74
chr10-42768175-INS-132
chr10-42769667-INS-393
chr10-42784950-INS-315
chr10-42793864-INS-62
chr10-42816988-INS-106
chr10-42824678-INS-100
chr10-42842927-INS-623
chr10-42849836-INS-52
chr10-42874784-INS-288
chr10-42884851-INS-52
chr10-42906927-INS-80
chr10-42911494-INS-78
chr10-42915258-INS-85
chr10-42917492-INS-683
chr10-42920163-INS-67
chr10-42938546-INS-54
chr10-42948293-INS-185
chr10-42949556-INS-1892
chr10-42953374-INS-52
chr10-42984311-INS-340
chr10-42991397-INS-56
chr10-43059234-INS-67
chr10-43069911-INS-143
chr10-43113254-INS-134
chr10-43122509-INS-54
chr10-43140625-INS-59
chr10-43157101-INS-132
chr10-43183468-INS-137
chr10-43191602-INS-56
chr10-43195107-INS-331
chr10-43205925-INS-55
chr10-43243929-INS-408
chr10-43254091-INS-86
chr10-43262901-INS-74
chr10-43264429-INS-56
chr10-43278326-INS-184
chr10-43291154-INS-614
chr10-43307208-INS-56
chr10-43309725-INS-56
chr10-43321223-INS-80
chr10-43323524-INS-191
chr10-43327549-INS-62
chr10-43329830-INS-203
chr10-43331308-INS-183
chr10-43335339-INS-212
chr10-43336255-INS-70
chr10-43369941-INS-652
chr10-43371551-INS-207

chr10-43374215-INS-479
chr10-43389955-INS-54
chr10-43422536-INS-124
chr10-43424778-INS-227
chr10-43438577-INS-1398
chr10-43439244-INS-286
chr10-43464721-INS-120
chr10-43479465-INS-5369
chr10-43487474-INS-111
chr10-43493955-INS-138
chr10-43496052-INS-56
chr10-43501787-INS-66
chr10-43513156-INS-122
chr10-43516477-INS-84
chr10-43526280-INS-78
chr10-43551564-INS-196
chr10-43564869-INS-66
chr10-43580151-INS-177
chr10-43596033-INS-68
chr10-43632752-INS-68
chr10-43653083-INS-160
chr10-43663831-INS-355
chr10-43669696-INS-176
chr10-43677320-INS-65
chr10-43696180-INS-547
chr10-43721323-INS-82
chr10-43763565-INS-422
chr10-43772330-INS-440
chr10-43775380-INS-74
chr10-43782036-INS-137
chr10-43814379-INS-224
chr10-43830681-INS-52
chr10-43844218-INS-135
chr10-43861382-INS-52
chr10-43865016-INS-249
chr10-43876932-INS-137
chr10-43910544-INS-56
chr10-43935760-INS-136
chr10-43960904-INS-540
chr10-43990638-INS-312
chr10-43996428-INS-163
chr10-43998097-INS-138
chr10-44053204-INS-62
chr10-44095755-INS-422
chr10-44106717-INS-608
chr10-44143440-INS-70
chr10-44183241-INS-171
chr10-44189561-INS-287
chr10-44200892-INS-202
chr10-44235307-INS-339
chr10-44244106-INS-170
chr10-44245594-INS-71
chr10-44258507-INS-58
chr10-44309994-INS-92
chr10-44339647-INS-88
chr10-44344097-INS-174
chr10-44346926-INS-168
chr10-44352177-INS-100
chr10-44358370-INS-53
chr10-44374159-INS-189
chr10-44410987-INS-55
chr10-44428639-INS-966
chr10-44434844-INS-627
chr10-44440122-INS-7483
chr10-44455891-INS-890
chr10-44463564-INS-436
chr10-44471440-INS-270
chr10-44505231-INS-126
chr10-44524712-INS-202
chr10-44536991-INS-375
chr10-44548895-INS-2575
chr10-44571149-INS-107
chr10-44577413-INS-1180
chr10-44654432-INS-53
chr10-44666142-INS-88
chr10-44689384-INS-199
chr10-44696803-INS-655

chr10-44711014-INS-390
chr10-44771371-INS-74
chr10-44791480-INS-61
chr10-44802174-INS-56
chr10-44813770-INS-3421
chr10-44814891-INS-171
chr10-44840428-INS-803
chr10-44849619-INS-60
chr10-44859414-INS-80
chr10-44875233-INS-54
chr10-44884235-INS-7485
chr10-44886096-INS-52
chr10-44910253-INS-100
chr10-44910785-INS-152
chr10-44916072-INS-59
chr10-44925839-INS-51
chr10-44934466-INS-3219
chr10-44934712-INS-477
chr10-44976860-INS-72
chr10-44995182-INS-128
chr10-45035402-INS-61
chr10-45060694-INS-52
chr10-45096882-INS-59
chr10-45110568-INS-206
chr10-45142291-INS-608
chr10-45147288-INS-440
chr10-45156014-INS-122
chr10-45167325-INS-575
chr10-45169756-INS-6429
chr10-45225467-INS-70
chr10-45229527-INS-439
chr10-45234740-INS-95
chr10-45236717-INS-275
chr10-45241632-INS-78
chr10-45251870-INS-64
chr10-45272013-INS-204
chr10-45319512-INS-217
chr10-45320905-INS-100
chr10-45336379-INS-232
chr10-45374670-INS-294
chr10-45377412-INS-172
chr10-45403677-INS-231
chr10-45423423-INS-458
chr10-45423464-INS-3969
chr10-45440409-INS-109
chr10-45468984-INS-536
chr10-45472784-INS-476
chr10-45502653-INS-62
chr10-45513464-INS-583
chr10-45537212-INS-52
chr10-45539003-INS-1029
chr10-45540329-INS-183
chr10-45572624-INS-219
chr10-45575202-INS-474
chr10-45579597-INS-144
chr10-45581551-INS-127
chr10-45581808-INS-6428
chr10-45590980-INS-7204
chr10-45646509-INS-52
chr10-45660611-INS-240
chr10-45702632-INS-656
chr10-45722096-INS-128
chr10-45739668-INS-53
chr10-45742292-INS-898
chr10-45744779-INS-52
chr10-45768306-INS-124
chr10-45772019-INS-52
chr10-45795577-INS-144
chr10-45801804-INS-223
chr10-45809623-INS-52
chr10-45811825-INS-452
chr10-45863837-INS-64
chr10-45878031-INS-892
chr10-45894573-INS-4050
chr10-45897840-INS-72
chr10-45907329-INS-343
chr10-45931456-INS-292

chr10-45935520-INS-217
chr10-45974911-INS-950
chr10-45979909-INS-52
chr10-45989759-INS-223
chr10-45993565-INS-220
chr10-46006572-INS-80
chr10-46009012-INS-818
chr10-46021613-INS-605
chr10-46031929-INS-7309
chr10-46044974-INS-385
chr10-46053074-INS-89
chr10-46056488-INS-432
chr10-46069369-INS-60
chr10-46087839-INS-76
chr10-46098069-INS-143
chr10-46117681-INS-616
chr10-46119715-INS-167
chr10-46130908-INS-52
chr10-46142407-INS-161
chr10-46154690-INS-174
chr10-46173250-INS-84
chr10-46192161-INS-605
chr10-46204684-INS-64
chr10-46238809-INS-108
chr10-46270529-INS-409
chr10-46296516-INS-707
chr10-46321142-INS-139
chr10-46336929-INS-206
chr10-46368874-INS-118
chr10-46388475-INS-183
chr10-46408678-INS-600
chr10-46416351-INS-377
chr10-46428662-INS-236
chr10-46465408-INS-400
chr10-46470411-INS-184
chr10-46483226-INS-635
chr10-46499182-INS-58
chr10-46506170-INS-951
chr10-46540145-INS-80
chr10-46552776-INS-498
chr10-46574772-INS-6593
chr10-46613920-INS-514
chr10-46614562-INS-346
chr10-46622288-INS-236
chr10-46639040-INS-53
chr10-46649636-INS-6372
chr10-46653050-INS-2945
chr10-46660945-INS-184
chr10-46691251-INS-177
chr10-46726794-INS-139
chr10-46746174-INS-163
chr10-46771194-INS-60
chr10-46773032-INS-225
chr10-46801872-INS-5361
chr10-46808468-INS-553
chr10-46830450-INS-440
chr10-46888449-INS-6908
chr10-46899066-INS-67
chr10-46915806-INS-975
chr10-46921631-INS-190
chr10-46937233-INS-6648
chr10-46959034-INS-64
chr10-46987505-INS-60
chr10-46989684-INS-223
chr10-46999580-INS-56
chr10-47044183-INS-117
chr10-47046765-INS-273
chr10-47054434-INS-199
chr10-47057491-INS-189
chr10-47113797-INS-56
chr10-47114662-INS-406
chr10-47116541-INS-96
chr10-47118354-INS-157
chr10-47127923-INS-500
chr10-47139007-INS-383
chr10-47149009-INS-6803
chr10-47179182-INS-52

chr10-47190262-INS-62
chr10-47198591-INS-6180
chr10-47261503-INS-6543
chr10-47279926-INS-55
chr10-47290790-INS-61
chr10-47309381-INS-123
chr10-47310240-INS-421
chr10-47340093-INS-74
chr10-47421876-INS-200
chr10-47458177-INS-475
chr10-47528496-INS-95
chr10-47538793-INS-56
chr10-47606028-INS-502
chr10-47669046-INS-2949
chr10-47685411-INS-3172
chr10-47690895-INS-157
chr10-47696379-INS-62
chr10-47718589-INS-6575
chr10-47750619-INS-6193
chr10-47761362-INS-1050
chr10-47800479-INS-165
chr10-47823000-INS-4868
chr10-47847987-INS-92
chr10-47913385-INS-393
chr10-47924291-INS-68
chr10-47990865-INS-209
chr10-48002790-INS-70
chr10-48010194-INS-299
chr10-48122012-INS-53
chr10-48125132-INS-59
chr10-48137957-INS-58
chr10-48155646-INS-122
chr10-48164766-INS-3123
chr10-48166690-INS-254
chr10-48169625-INS-2832
chr10-48169927-INS-3634
chr10-48177543-INS-65
chr10-48198130-INS-84
chr10-48218022-INS-396
chr10-48236333-INS-6478
chr10-48274062-INS-113
chr10-48288865-INS-58
chr10-48302676-INS-72
chr10-48318848-INS-6380
chr10-48345830-INS-419
chr10-48350358-INS-88
chr10-48353686-INS-94
chr10-48389463-INS-75
chr10-48402407-INS-5159
chr10-48449301-INS-53
chr10-48459991-INS-62
chr10-48531519-INS-56
chr10-48571801-INS-121
chr10-48607412-INS-621
chr10-48607814-INS-953
chr10-48609677-INS-136
chr10-48625565-INS-1050
chr10-48671533-INS-62
chr10-48684385-INS-3592
chr10-48700368-INS-333
chr10-48718833-INS-6884
chr10-48725877-INS-2215
chr10-48746175-INS-835
chr10-48755866-INS-53
chr10-48767138-INS-94
chr10-48771805-INS-253
chr10-48828705-INS-107
chr10-48875749-INS-555
chr10-48917779-INS-108
chr10-48937615-INS-571
chr10-48953461-INS-179
chr10-48955207-INS-70
chr10-48962479-INS-324
chr10-48994892-INS-558
chr10-49000728-INS-55
chr10-49069522-INS-91
chr10-49083936-INS-1541

chr10-49088352-INS-252
chr10-49095018-INS-91
chr10-49111662-INS-60
chr10-49115776-INS-69
chr10-49190566-INS-58
chr10-49219331-INS-149
chr10-49227656-INS-6364
chr10-49228165-INS-333
chr10-49229079-INS-460
chr10-49247188-INS-73
chr10-49253263-INS-371
chr10-49294068-INS-472
chr10-49311906-INS-868
chr10-49335515-INS-508
chr10-49350376-INS-913
chr10-49355161-INS-874
chr10-49406650-INS-100
chr10-49412069-INS-95
chr10-49420562-INS-328
chr10-49468176-INS-260
chr10-49471375-INS-62
chr10-49543553-INS-198
chr10-49555962-INS-74
chr10-49557946-INS-959
chr10-49573483-INS-940
chr10-49582201-INS-54
chr10-49592108-INS-241
chr10-49604111-INS-54
chr10-49623967-INS-59
chr10-49624763-INS-1900
chr10-49645576-INS-118
chr10-49654543-INS-75
chr10-49656007-INS-72
chr10-49678807-INS-342
chr10-49695717-INS-318
chr10-49708162-INS-684
chr10-49720134-INS-329
chr10-49723974-INS-191
chr10-49772769-INS-353
chr10-49785145-INS-205
chr10-49848124-INS-51
chr10-49853297-INS-546
chr10-49856350-INS-2855
chr10-49863537-INS-9950
chr10-49867083-INS-515
chr10-49896680-INS-68
chr10-49903047-INS-752
chr10-49925345-INS-60
chr10-49936051-INS-539
chr10-49943449-INS-59
chr10-49956182-INS-295
chr10-49973147-INS-71
chr10-49981378-INS-388
chr10-50007699-INS-530
chr10-50015468-INS-52
chr10-50053436-INS-6521
chr10-50079532-INS-64
chr10-50108383-INS-344
chr10-50111185-INS-266
chr10-50115475-INS-86
chr10-50129783-INS-303
chr10-50156296-INS-79
chr10-50159353-INS-66
chr10-50183408-INS-5721
chr10-50197661-INS-132
chr10-50201375-INS-64
chr10-50218503-INS-70
chr10-50235571-INS-262
chr10-50251460-INS-151
chr10-50253122-INS-118
chr10-50260177-INS-6985
chr10-50286554-INS-986
chr10-50290772-INS-59
chr10-50308383-INS-99
chr10-50309266-INS-75
chr10-50315471-INS-764
chr10-50322940-INS-6928

chr10-50348409-INS-65
chr10-50355518-INS-7213
chr10-50357257-INS-180
chr10-50362287-INS-5122
chr10-50366984-INS-120
chr10-50381338-INS-5542
chr10-50395166-INS-64
chr10-50405306-INS-448
chr10-50422078-INS-64
chr10-50430931-INS-199
chr10-50437289-INS-6650
chr10-50457287-INS-209
chr10-50460200-INS-955
chr10-50476170-INS-857
chr10-50505197-INS-317
chr10-50535497-INS-5064
chr10-50575855-INS-389
chr10-50600459-INS-1731
chr10-50603708-INS-1396
chr10-50612468-INS-97
chr10-50630276-INS-58
chr10-50684188-INS-70
chr10-50691642-INS-6480
chr10-50693819-INS-487
chr10-50698151-INS-11159
chr10-50704548-INS-560
chr10-50705337-INS-135
chr10-50718511-INS-1932
chr10-50728797-INS-6664
chr10-50732602-INS-199
chr10-50735974-INS-73
chr10-50743805-INS-7909
chr10-50751426-INS-728
chr10-50757305-INS-147
chr10-50810144-INS-170
chr10-50811188-INS-87
chr10-50811820-INS-112
chr10-50813815-INS-55
chr10-50816466-INS-718
chr10-50825103-INS-276
chr10-50848683-INS-60
chr10-50855108-INS-7298
chr10-50874224-INS-83
chr10-50882435-INS-358
chr10-50904238-INS-60
chr10-50909645-INS-6178
chr10-50910853-INS-207
chr10-50919034-INS-51
chr10-50926401-INS-11996
chr10-50929121-INS-110
chr10-50934295-INS-218
chr10-50936213-INS-64
chr10-50942389-INS-84
chr10-50950426-INS-63
chr10-50951766-INS-90
chr10-50967842-INS-394
chr10-50981134-INS-967
chr10-50981860-INS-565
chr10-51018386-INS-132
chr10-51023045-INS-70
chr10-51050138-INS-60
chr10-51105017-INS-198
chr10-51136353-INS-6565
chr10-51141008-INS-3086
chr10-51142511-INS-137
chr10-51169185-INS-420
chr10-51181081-INS-6488
chr10-51188244-INS-56
chr10-51191735-INS-93
chr10-51194951-INS-7530
chr10-51199737-INS-226
chr10-51207146-INS-72
chr10-51219112-INS-1157
chr10-51219113-INS-755
chr10-51241353-INS-3247
chr10-51242894-INS-57
chr10-51248812-INS-388

chr10-51255780-INS-62
chr10-51258391-INS-104
chr10-51281378-INS-629
chr10-51283758-INS-51
chr10-51291260-INS-3940
chr10-51294140-INS-2834
chr10-51294698-INS-328
chr10-51304253-INS-4272
chr10-51365219-INS-52
chr10-51394695-INS-91
chr10-51402191-INS-59
chr10-51407723-INS-159
chr10-51410199-INS-59
chr10-51415510-INS-55
chr10-51443807-INS-256
chr10-51468714-INS-56
chr10-51476166-INS-62
chr10-51489887-INS-87
chr10-51512951-INS-662
chr10-51535469-INS-84
chr10-51538147-INS-68
chr10-51541658-INS-737
chr10-51571541-INS-58
chr10-51580749-INS-107
chr10-51582000-INS-104
chr10-51590685-INS-7125
chr10-51593343-INS-55
chr10-51604585-INS-145
chr10-51607816-INS-7425
chr10-51616449-INS-254
chr10-51647259-INS-904
chr10-51655316-INS-112
chr10-51660700-INS-92
chr10-51707126-INS-681
chr10-51721302-INS-967
chr10-51723855-INS-51
chr10-51743463-INS-231
chr10-51754280-INS-228
chr10-51783282-INS-340
chr10-51785208-INS-60
chr10-51794166-INS-74
chr10-51795780-INS-66
chr10-51801414-INS-59
chr10-51808987-INS-6496
chr10-51809466-INS-1864
chr10-51829634-INS-479
chr10-51833407-INS-603
chr10-51844102-INS-1893
chr10-51848109-INS-6585
chr10-51862575-INS-1598
chr10-51874265-INS-210
chr10-51877245-INS-64
chr10-51893114-INS-6502
chr10-51895464-INS-112
chr10-51908200-INS-390
chr10-51941892-INS-73
chr10-51945492-INS-171
chr10-51967474-INS-66
chr10-51999722-INS-63
chr10-52008434-INS-60
chr10-52027010-INS-66
chr10-52027950-INS-197
chr10-52032756-INS-4231
chr10-52035587-INS-54
chr10-52044657-INS-214
chr10-52046902-INS-368
chr10-52048503-INS-136
chr10-52074990-INS-306
chr10-52106369-INS-158
chr10-52126832-INS-58
chr10-52133400-INS-202
chr10-52146890-INS-92
chr10-52158535-INS-96
chr10-52161300-INS-100
chr10-52171282-INS-129
chr10-52179320-INS-61
chr10-52194700-INS-52

chr10-52201536-INS-954
chr10-52202521-INS-142
chr10-52219133-INS-78
chr10-52233722-INS-205
chr10-52288415-INS-3085
chr10-52304698-INS-210
chr10-52310981-INS-317
chr10-52337288-INS-60
chr10-52354950-INS-400
chr10-52360073-INS-561
chr10-52364078-INS-5971
chr10-52382132-INS-1857
chr10-52415089-INS-6213
chr10-52415952-INS-6206
chr10-52454699-INS-364
chr10-52467247-INS-90
chr10-52467572-INS-461
chr10-52496520-INS-68
chr10-52498590-INS-123
chr10-52520890-INS-112
chr10-52593073-INS-113
chr10-52622130-INS-868
chr10-52625607-INS-196
chr10-52634603-INS-649
chr10-52646214-INS-62
chr10-52654257-INS-183
chr10-52664472-INS-305
chr10-52680463-INS-73
chr10-52724049-INS-421
chr10-52732339-INS-6546
chr10-52742719-INS-99
chr10-52744333-INS-201
chr10-52767476-INS-295
chr10-52773668-INS-80
chr10-52774274-INS-553
chr10-52784913-INS-396
chr10-52787786-INS-53
chr10-52789915-INS-182
chr10-52802906-INS-401
chr10-52818127-INS-67
chr10-52833718-INS-809
chr10-52835398-INS-56
chr10-52844790-INS-56
chr10-52878102-INS-6546
chr10-52887682-INS-84
chr10-52913716-INS-63
chr10-52918116-INS-181
chr10-52940625-INS-1866
chr10-52969030-INS-203
chr10-52987478-INS-123
chr10-52990719-INS-52
chr10-53038006-INS-887
chr10-53042339-INS-876
chr10-53091682-INS-60
chr10-53094989-INS-341
chr10-53102339-INS-6771
chr10-53106786-INS-54
chr10-53109822-INS-130
chr10-53147532-INS-449
chr10-53163153-INS-79
chr10-53168851-INS-783
chr10-53178710-INS-6705
chr10-53183722-INS-696
chr10-53185912-INS-6905
chr10-53186360-INS-6302
chr10-53188134-INS-75
chr10-53190510-INS-7415
chr10-53193320-INS-962
chr10-53196564-INS-176
chr10-53205197-INS-72
chr10-53233346-INS-7239
chr10-53262447-INS-365
chr10-53266119-INS-6668
chr10-53290021-INS-78
chr10-53293669-INS-53
chr10-53315068-INS-499
chr10-53328052-INS-147

chr10-53334156-INS-310
chr10-53339166-INS-1891
chr10-53348592-INS-3383
chr10-53353278-INS-201
chr10-53356477-INS-278
chr10-53361858-INS-54
chr10-53362096-INS-964
chr10-53365157-INS-386
chr10-53371223-INS-6436
chr10-53383087-INS-6122
chr10-53396396-INS-185
chr10-53398160-INS-489
chr10-53409858-INS-102
chr10-53413249-INS-60
chr10-53423398-INS-72
chr10-53426879-INS-646
chr10-53437731-INS-6901
chr10-53440145-INS-516
chr10-53464055-INS-156
chr10-53481857-INS-229
chr10-53507866-INS-65
chr10-53537378-INS-108
chr10-53549560-INS-221
chr10-53559776-INS-62
chr10-53563302-INS-402
chr10-53570099-INS-56
chr10-53580473-INS-1889
chr10-53597780-INS-55
chr10-53602203-INS-52
chr10-53604721-INS-61
chr10-53622227-INS-654
chr10-53641484-INS-553
chr10-53649016-INS-51
chr10-53655262-INS-769
chr10-53687529-INS-7256
chr10-53701285-INS-7188
chr10-53713211-INS-7253
chr10-53719662-INS-132
chr10-53720338-INS-55
chr10-53724975-INS-554
chr10-53727272-INS-665
chr10-53729507-INS-247
chr10-53731496-INS-227
chr10-53732532-INS-93
chr10-53738506-INS-382
chr10-53745980-INS-76
chr10-53747906-INS-56
chr10-53756972-INS-56
chr10-53760971-INS-68
chr10-53788837-INS-364
chr10-53790112-INS-6413
chr10-53793396-INS-68
chr10-53819451-INS-301
chr10-53857198-INS-1545
chr10-53888306-INS-190
chr10-53901364-INS-617
chr10-53906466-INS-125
chr10-53921066-INS-85
chr10-53965396-INS-133
chr10-53976188-INS-69
chr10-53986873-INS-259
chr10-54007872-INS-108
chr10-54026895-INS-877
chr10-54048844-INS-68
chr10-54061954-INS-6372
chr10-54109099-INS-112
chr10-54114061-INS-325
chr10-54119312-INS-60
chr10-54148233-INS-301
chr10-54153185-INS-6468
chr10-54165855-INS-6979
chr10-54180673-INS-466
chr10-54189539-INS-791
chr10-54213117-INS-687
chr10-54218441-INS-125
chr10-54258638-INS-358
chr10-54270414-INS-103

chr10-54287769-INS-51
chr10-54296398-INS-102
chr10-54306225-INS-126
chr10-54309149-INS-67
chr10-54312025-INS-74
chr10-54316078-INS-112
chr10-54322561-INS-400
chr10-54331836-INS-127
chr10-54340935-INS-497
chr10-54350898-INS-74
chr10-54368790-INS-430
chr10-54369877-INS-282
chr10-54385865-INS-515
chr10-54396451-INS-70
chr10-54416862-INS-1893
chr10-54422485-INS-101
chr10-54424215-INS-68
chr10-54459453-INS-328
chr10-54504398-INS-140
chr10-54505532-INS-652
chr10-54508962-INS-324
chr10-54526857-INS-667
chr10-54536372-INS-95
chr10-54545273-INS-441
chr10-54553024-INS-6047
chr10-54599915-INS-200
chr10-54605212-INS-72
chr10-54615808-INS-934
chr10-54625320-INS-371
chr10-54641092-INS-466
chr10-54662870-INS-262
chr10-54666024-INS-388
chr10-54674513-INS-71
chr10-54716866-INS-54
chr10-54722333-INS-474
chr10-54751008-INS-76
chr10-54761911-INS-4123
chr10-54808870-INS-478
chr10-54828495-INS-469
chr10-54829434-INS-131
chr10-54853321-INS-506
chr10-54881398-INS-76
chr10-54968494-INS-417
chr10-54981224-INS-67
chr10-54994919-INS-59
chr10-55021701-INS-59
chr10-55025930-INS-105
chr10-55029435-INS-316
chr10-55032185-INS-1789
chr10-55047707-INS-54
chr10-55077739-INS-162
chr10-55123252-INS-249
chr10-55142874-INS-961
chr10-55144545-INS-359
chr10-55168483-INS-126
chr10-55169351-INS-544
chr10-55195438-INS-58
chr10-55199714-INS-84
chr10-55202352-INS-8331
chr10-55231645-INS-6311
chr10-55246740-INS-645
chr10-55283833-INS-90
chr10-55308555-INS-210
chr10-55313355-INS-4036
chr10-55328951-INS-56
chr10-55331620-INS-56
chr10-55332800-INS-78
chr10-55335068-INS-283
chr10-55342495-INS-62
chr10-55387929-INS-2520
chr10-55392942-INS-217
chr10-55400954-INS-142
chr10-55435929-INS-519
chr10-55449445-INS-178
chr10-55469207-INS-906
chr10-55486234-INS-72
chr10-55490438-INS-216

chr10-55534317-INS-4661
chr10-55538870-INS-152
chr10-55563920-INS-142
chr10-55569655-INS-69
chr10-55570791-INS-417
chr10-55584106-INS-55
chr10-55605452-INS-192
chr10-55641537-INS-441
chr10-55714121-INS-5068
chr10-55717874-INS-730
chr10-55743297-INS-67
chr10-55749549-INS-486
chr10-55765365-INS-76
chr10-55768848-INS-296
chr10-55770821-INS-214
chr10-55811678-INS-60
chr10-55818221-INS-5983
chr10-55821195-INS-738
chr10-55823113-INS-676
chr10-55847210-INS-5292
chr10-55862207-INS-353
chr10-55868302-INS-243
chr10-55868389-INS-699
chr10-55869530-INS-62
chr10-55876007-INS-6908
chr10-55882829-INS-63
chr10-55893463-INS-162
chr10-55901299-INS-214
chr10-55917729-INS-531
chr10-55918705-INS-72
chr10-55929825-INS-401
chr10-55930976-INS-6599
chr10-55934400-INS-1100
chr10-55934513-INS-152
chr10-55990036-INS-51
chr10-56013233-INS-401
chr10-56032755-INS-67
chr10-56071030-INS-362
chr10-56089684-INS-79
chr10-56092593-INS-167
chr10-56123384-INS-62
chr10-56137891-INS-116
chr10-56138858-INS-62
chr10-56141306-INS-155
chr10-56143916-INS-91
chr10-56154452-INS-214
chr10-56154975-INS-69
chr10-56156657-INS-52
chr10-56190908-INS-108
chr10-56205436-INS-125
chr10-56220822-INS-967
chr10-56228798-INS-56
chr10-56247060-INS-276
chr10-56263055-INS-105
chr10-56266797-INS-93
chr10-56268201-INS-233
chr10-56273605-INS-51
chr10-56278812-INS-75
chr10-56289952-INS-222
chr10-56293017-INS-325
chr10-56296168-INS-220
chr10-56312148-INS-234
chr10-56333335-INS-233
chr10-56344118-INS-253
chr10-56345976-INS-646
chr10-56352026-INS-81
chr10-56355293-INS-99
chr10-56382992-INS-380
chr10-56400342-INS-106
chr10-56403759-INS-258
chr10-56421218-INS-53
chr10-56427635-INS-56
chr10-56431249-INS-178
chr10-56434175-INS-466
chr10-56463059-INS-420
chr10-56466342-INS-6318
chr10-56470219-INS-149

chr10-56506616-INS-741
chr10-56517348-INS-214
chr10-56535459-INS-694
chr10-56586069-INS-65
chr10-56587092-INS-842
chr10-56609964-INS-6529
chr10-56618455-INS-114
chr10-56640218-INS-53
chr10-56674362-INS-95
chr10-56678977-INS-171
chr10-56680555-INS-116
chr10-56684732-INS-110
chr10-56712033-INS-6695
chr10-56758567-INS-186
chr10-56774585-INS-213
chr10-56800027-INS-2513
chr10-56808833-INS-6280
chr10-56812830-INS-82
chr10-56841657-INS-1865
chr10-56843476-INS-201
chr10-56845572-INS-387
chr10-56848919-INS-262
chr10-56860779-INS-138
chr10-56862734-INS-182
chr10-56868686-INS-108
chr10-56873840-INS-2218
chr10-56875607-INS-54
chr10-56911260-INS-86
chr10-56926568-INS-243
chr10-56927284-INS-70
chr10-56935004-INS-52
chr10-56936756-INS-54
chr10-56942625-INS-331
chr10-56972511-INS-6405
chr10-56998248-INS-6161
chr10-57012248-INS-259
chr10-57059986-INS-71
chr10-57076431-INS-487
chr10-57087076-INS-203
chr10-57091836-INS-96
chr10-57095932-INS-62
chr10-57138153-INS-330
chr10-57156049-INS-56
chr10-57169908-INS-731
chr10-57170977-INS-672
chr10-57172898-INS-6320
chr10-57174840-INS-892
chr10-57189160-INS-88
chr10-57253163-INS-51
chr10-57254393-INS-397
chr10-57269946-INS-187
chr10-57270240-INS-3811
chr10-57270429-INS-6595
chr10-57296884-INS-317
chr10-57351606-INS-52
chr10-57368098-INS-170
chr10-57400565-INS-144
chr10-57434924-INS-394
chr10-57438385-INS-505
chr10-57444424-INS-66
chr10-57461423-INS-370
chr10-57468030-INS-628
chr10-57472506-INS-270
chr10-57507704-INS-68
chr10-57511584-INS-208
chr10-57551690-INS-185
chr10-57572826-INS-60
chr10-57591724-INS-67
chr10-57606309-INS-56
chr10-57629358-INS-400
chr10-57632474-INS-59
chr10-57635832-INS-322
chr10-57691422-INS-1890
chr10-57693056-INS-76
chr10-57700798-INS-141
chr10-57728047-INS-422
chr10-57742475-INS-484

chr10-57745238-INS-55
chr10-57978967-INS-54
chr10-57983883-INS-54
chr10-58056224-INS-88
chr10-58061047-INS-54
chr10-58077163-INS-57
chr10-58082914-INS-198
chr10-58116513-INS-167
chr10-58146552-INS-236
chr10-58192729-INS-206
chr10-58195522-INS-61
chr10-58208150-INS-105
chr10-58210738-INS-110
chr10-58221724-INS-57
chr10-58223751-INS-62
chr10-58228568-INS-202
chr10-58232903-INS-89
chr10-58242386-INS-201
chr10-58242646-INS-957
chr10-58247446-INS-93
chr10-58252112-INS-723
chr10-58255883-INS-170
chr10-58277871-INS-87
chr10-58287045-INS-181
chr10-58289398-INS-557
chr10-58296257-INS-508
chr10-58346163-INS-197
chr10-58350230-INS-112
chr10-58366219-INS-53
chr10-58384531-INS-63
chr10-58385973-INS-133
chr10-58414348-INS-211
chr10-58425545-INS-629
chr10-58429441-INS-7082
chr10-58439432-INS-115
chr10-58455651-INS-1890
chr10-58457478-INS-111
chr10-58469619-INS-165
chr10-58502157-INS-116
chr10-58520062-INS-7484
chr10-58538168-INS-285
chr10-58542228-INS-458
chr10-58547038-INS-118
chr10-58563769-INS-2131
chr10-58616691-INS-214
chr10-58620355-INS-75
chr10-58639660-INS-199
chr10-58641159-INS-67
chr10-58663927-INS-817
chr10-58667203-INS-84
chr10-58668812-INS-169
chr10-58676207-INS-59
chr10-58699376-INS-6801
chr10-58728544-INS-576
chr10-58736409-INS-65
chr10-58737880-INS-201
chr10-58741882-INS-60
chr10-58809496-INS-5258
chr10-58821887-INS-97
chr10-58842023-INS-124
chr10-58870086-INS-201
chr10-58887278-INS-92
chr10-58931662-INS-6295
chr10-58962912-INS-83
chr10-58964240-INS-69
chr10-58972893-INS-482
chr10-58975028-INS-78
chr10-58998757-INS-254
chr10-59079365-INS-150
chr10-59097705-INS-70
chr10-59107781-INS-197
chr10-59120053-INS-58
chr10-59121248-INS-65
chr10-59122810-INS-601
chr10-59129208-INS-162
chr10-59133140-INS-178
chr10-59135114-INS-128

chr10-59157926-INS-656
chr10-59182218-INS-1838
chr10-59227726-INS-745
chr10-59285191-INS-102
chr10-59295627-INS-268
chr10-59300007-INS-685
chr10-59431553-INS-608
chr10-59454526-INS-163
chr10-59461258-INS-176
chr10-59465778-INS-56
chr10-59521213-INS-284
chr10-59530304-INS-4950
chr10-59530497-INS-189
chr10-59556561-INS-354
chr10-59559712-INS-54
chr10-59578175-INS-69
chr10-59583519-INS-60
chr10-59620024-INS-226
chr10-59636820-INS-67
chr10-59659848-INS-219
chr10-59666134-INS-184
chr10-59672906-INS-248
chr10-59675463-INS-207
chr10-59684338-INS-61
chr10-59706548-INS-60
chr10-59713279-INS-202
chr10-59731790-INS-91
chr10-59742356-INS-62
chr10-59745793-INS-203
chr10-59753741-INS-190
chr10-59759232-INS-199
chr10-59761059-INS-217
chr10-59772841-INS-83
chr10-59777419-INS-209
chr10-59780917-INS-128
chr10-59802714-INS-936
chr10-59828344-INS-217
chr10-59837285-INS-349
chr10-59860054-INS-6447
chr10-59865437-INS-7099
chr10-59873172-INS-68
chr10-59886360-INS-204
chr10-59913997-INS-210
chr10-59950725-INS-66
chr10-59963145-INS-5358
chr10-59972104-INS-657
chr10-59986476-INS-122
chr10-60003966-INS-551
chr10-60006964-INS-526
chr10-60010998-INS-195
chr10-60025056-INS-170
chr10-60032480-INS-55
chr10-60045843-INS-207
chr10-60071024-INS-54
chr10-60073100-INS-146
chr10-60117840-INS-199
chr10-60119451-INS-103
chr10-60127168-INS-53
chr10-60143121-INS-54
chr10-60164606-INS-692
chr10-60168611-INS-81
chr10-60178707-INS-74
chr10-60224088-INS-196
chr10-60225735-INS-1890
chr10-60260450-INS-845
chr10-60273808-INS-53
chr10-60312810-INS-211
chr10-60337511-INS-802
chr10-60343332-INS-88
chr10-60350782-INS-74
chr10-60362738-INS-150
chr10-60391752-INS-82
chr10-60399482-INS-99
chr10-60402875-INS-72
chr10-60425725-INS-83
chr10-60469598-INS-212
chr10-60473866-INS-5262

chr10-60487635-INS-938
chr10-60490631-INS-367
chr10-60508401-INS-52
chr10-60516828-INS-208
chr10-60517514-INS-179
chr10-60537769-INS-68
chr10-60538070-INS-377
chr10-60543116-INS-124
chr10-60548742-INS-64
chr10-60566959-INS-56
chr10-60581993-INS-104
chr10-60585074-INS-235
chr10-60592574-INS-53
chr10-60611634-INS-208
chr10-60641131-INS-56
chr10-60673187-INS-130
chr10-60679007-INS-176
chr10-60689845-INS-111
chr10-60691266-INS-138
chr10-60695610-INS-54
chr10-60720818-INS-60
chr10-60730277-INS-5342
chr10-60755810-INS-51
chr10-60757618-INS-3068
chr10-60769901-INS-53
chr10-60785741-INS-68
chr10-60799652-INS-78
chr10-60801012-INS-52
chr10-60819229-INS-79
chr10-60823344-INS-79
chr10-60826285-INS-5218
chr10-60836276-INS-285
chr10-60841443-INS-574
chr10-60842646-INS-105
chr10-60877392-INS-399
chr10-60885722-INS-190
chr10-60902946-INS-63
chr10-60913731-INS-387
chr10-60920029-INS-231
chr10-60920588-INS-144
chr10-60933320-INS-778
chr10-60943008-INS-179
chr10-60952918-INS-225
chr10-60990009-INS-54
chr10-61012995-INS-99
chr10-61036118-INS-58
chr10-61045398-INS-197
chr10-61045825-INS-358
chr10-61057925-INS-54
chr10-61083969-INS-360
chr10-61087503-INS-152
chr10-61093914-INS-55
chr10-61147506-INS-131
chr10-61154337-INS-174
chr10-61155007-INS-82
chr10-61172610-INS-52
chr10-61175989-INS-194
chr10-61185569-INS-194
chr10-61199968-INS-136
chr10-61228690-INS-51
chr10-61231370-INS-186
chr10-61233225-INS-361
chr10-61234327-INS-212
chr10-61287341-INS-137
chr10-61297229-INS-182
chr10-61305735-INS-215
chr10-61316370-INS-230
chr10-61322327-INS-61
chr10-61344095-INS-5261
chr10-61354433-INS-261
chr10-61366318-INS-529
chr10-61383265-INS-52
chr10-61413290-INS-860
chr10-61421855-INS-653
chr10-61448085-INS-145
chr10-61464353-INS-131
chr10-61469200-INS-85

chr10-61474441-INS-437
chr10-61478186-INS-198
chr10-61485369-INS-139
chr10-61582081-INS-54
chr10-61590969-INS-195
chr10-61596479-INS-85
chr10-61614891-INS-106
chr10-61633171-INS-829
chr10-61665050-INS-103
chr10-61670851-INS-57
chr10-61694251-INS-88
chr10-61715421-INS-51
chr10-61740233-INS-78
chr10-61744009-INS-151
chr10-61757378-INS-135
chr10-61768222-INS-55
chr10-61788041-INS-82
chr10-61808956-INS-106
chr10-61835852-INS-245
chr10-61859265-INS-94
chr10-61871424-INS-84
chr10-61881528-INS-58
chr10-61913436-INS-102
chr10-61926855-INS-222
chr10-61931593-INS-53
chr10-61943520-INS-63
chr10-61952830-INS-147
chr10-61967546-INS-87
chr10-61973040-INS-54
chr10-61974218-INS-199
chr10-61979336-INS-263
chr10-61984527-INS-173
chr10-62002316-INS-440
chr10-62019900-INS-127
chr10-62050739-INS-70
chr10-62073203-INS-366
chr10-62094286-INS-80
chr10-62101814-INS-54
chr10-62133327-INS-263
chr10-62136529-INS-114
chr10-62148856-INS-84
chr10-62155483-INS-230
chr10-62164624-INS-111
chr10-62168966-INS-52
chr10-62196287-INS-105
chr10-62213291-INS-51
chr10-62221315-INS-106
chr10-62232118-INS-117
chr10-62233726-INS-555
chr10-62239123-INS-63
chr10-62262024-INS-52
chr10-62276618-INS-91
chr10-62290954-INS-212
chr10-62293322-INS-218
chr10-62297256-INS-170
chr10-62308837-INS-218
chr10-62315242-INS-196
chr10-62318341-INS-167
chr10-62324441-INS-190
chr10-62327237-INS-122
chr10-62328131-INS-170
chr10-62329487-INS-260
chr10-62331319-INS-316
chr10-62346415-INS-56
chr10-62350599-INS-160
chr10-62352793-INS-691
chr10-62361105-INS-130
chr10-62383604-INS-68
chr10-62399974-INS-234
chr10-62411702-INS-3035
chr10-62427848-INS-205
chr10-62467242-INS-202
chr10-62486076-INS-117
chr10-62493972-INS-58
chr10-62535395-INS-66
chr10-62536661-INS-398
chr10-62555559-INS-152

chr10-62556251-INS-905
chr10-62559958-INS-335
chr10-62566993-INS-155
chr10-62575608-INS-52
chr10-62583507-INS-106
chr10-62594661-INS-183
chr10-62612828-INS-187
chr10-62650356-INS-214
chr10-62660849-INS-210
chr10-62670494-INS-58
chr10-62686794-INS-80
chr10-62689608-INS-51
chr10-62705547-INS-142
chr10-62709861-INS-229
chr10-62716053-INS-132
chr10-62724700-INS-202
chr10-62729473-INS-214
chr10-62740717-INS-68
chr10-62741418-INS-68
chr10-62747964-INS-130
chr10-62751528-INS-93
chr10-62762545-INS-57
chr10-62778087-INS-435
chr10-62783600-INS-51
chr10-62790502-INS-118
chr10-62792936-INS-85
chr10-62795848-INS-80
chr10-62800152-INS-377
chr10-62806387-INS-234
chr10-62808928-INS-54
chr10-62811588-INS-4277
chr10-62815456-INS-658
chr10-62824558-INS-276
chr10-62882016-INS-87
chr10-62886197-INS-103
chr10-62890678-INS-268
chr10-62894735-INS-91
chr10-62896859-INS-211
chr10-62901402-INS-212
chr10-62903060-INS-98
chr10-62918815-INS-72
chr10-62932280-INS-204
chr10-62932810-INS-921
chr10-62946981-INS-51
chr10-62951285-INS-97
chr10-62953293-INS-100
chr10-62956828-INS-792
chr10-62964816-INS-54
chr10-62968641-INS-58
chr10-62989255-INS-169
chr10-63002691-INS-120
chr10-63019444-INS-377
chr10-63023458-INS-74
chr10-63035246-INS-52
chr10-63041314-INS-57
chr10-63049810-INS-60
chr10-63076001-INS-152
chr10-63083305-INS-73
chr10-63088900-INS-206
chr10-63095929-INS-163
chr10-63108470-INS-80
chr10-63114613-INS-362
chr10-63116730-INS-70
chr10-63131046-INS-197
chr10-63139072-INS-725
chr10-63168099-INS-344
chr10-63176269-INS-82
chr10-63177896-INS-225
chr10-63194574-INS-164
chr10-63198015-INS-169
chr10-63202993-INS-106
chr10-63207488-INS-66
chr10-63213143-INS-60
chr10-63218743-INS-194
chr10-63223651-INS-194
chr10-63225007-INS-56
chr10-63234545-INS-203

chr10-63247944-INS-52
chr10-63250555-INS-154
chr10-63258456-INS-72
chr10-63259421-INS-81
chr10-63264203-INS-56
chr10-63269720-INS-196
chr10-63278438-INS-185
chr10-63300804-INS-388
chr10-63302223-INS-79
chr10-63302484-INS-435
chr10-63315761-INS-181
chr10-63322572-INS-6176
chr10-63322624-INS-338
chr10-63328832-INS-56
chr10-63340582-INS-94
chr10-63342605-INS-267
chr10-63347400-INS-85
chr10-63363603-INS-68
chr10-63382423-INS-145
chr10-63407500-INS-734
chr10-63412559-INS-51
chr10-63413498-INS-82
chr10-63414275-INS-140
chr10-63424424-INS-52
chr10-63430129-INS-174
chr10-63433501-INS-85
chr10-63445769-INS-88
chr10-63455703-INS-180
chr10-63465455-INS-123
chr10-63488495-INS-704
chr10-63511143-INS-52
chr10-63525239-INS-147
chr10-63527183-INS-3794
chr10-63534886-INS-122
chr10-63572256-INS-52
chr10-63610667-INS-1147
chr10-63618143-INS-80
chr10-63627046-INS-112
chr10-63637282-INS-895
chr10-63647912-INS-472
chr10-63655871-INS-143
chr10-63665213-INS-85
chr10-63680826-INS-52
chr10-63691396-INS-50
chr10-63692399-INS-58
chr10-63740625-INS-145
chr10-63776460-INS-54
chr10-63790171-INS-86
chr10-63790753-INS-125
chr10-63804925-INS-55
chr10-63807617-INS-280
chr10-63812626-INS-168
chr10-63817879-INS-930
chr10-63832944-INS-92
chr10-63880812-INS-355
chr10-63909911-INS-7189
chr10-63936093-INS-58
chr10-63979902-INS-6807
chr10-63981745-INS-710
chr10-63989698-INS-112
chr10-64023163-INS-636
chr10-64028762-INS-58
chr10-64032836-INS-53
chr10-64036421-INS-59
chr10-64052326-INS-80
chr10-64054892-INS-52
chr10-64059626-INS-68
chr10-64108710-INS-847
chr10-64111082-INS-628
chr10-64120271-INS-484
chr10-64125318-INS-82
chr10-64163016-INS-62
chr10-64191671-INS-112
chr10-64212230-INS-109
chr10-64242577-INS-89
chr10-64265341-INS-325
chr10-64290724-INS-155

chr10-64291093-INS-344
chr10-64300913-INS-218
chr10-64304500-INS-64
chr10-64306922-INS-51
chr10-64343880-INS-55
chr10-64386105-INS-253
chr10-64387152-INS-58
chr10-64410657-INS-6387
chr10-64441370-INS-3148
chr10-64446289-INS-70
chr10-64448463-INS-536
chr10-64465544-INS-360
chr10-64482945-INS-81
chr10-64501084-INS-678
chr10-64509872-INS-70
chr10-64548270-INS-103
chr10-64575658-INS-810
chr10-64603595-INS-6505
chr10-64608388-INS-2609
chr10-64625179-INS-187
chr10-64640515-INS-130
chr10-64646813-INS-76
chr10-64661032-INS-695
chr10-64684882-INS-227
chr10-64691146-INS-728
chr10-64692654-INS-155
chr10-64711559-INS-102
chr10-64713726-INS-88
chr10-64728824-INS-110
chr10-64753520-INS-522
chr10-64754607-INS-174
chr10-64796932-INS-87
chr10-64802168-INS-195
chr10-64805561-INS-52
chr10-64815837-INS-96
chr10-64842771-INS-190
chr10-64844320-INS-65
chr10-64895700-INS-400
chr10-64899124-INS-423
chr10-64904988-INS-71
chr10-64907699-INS-313
chr10-64917305-INS-113
chr10-64922032-INS-51
chr10-64928267-INS-67
chr10-64938193-INS-96
chr10-64952479-INS-81
chr10-64959626-INS-58
chr10-64960203-INS-134
chr10-64975929-INS-64
chr10-64992488-INS-62
chr10-64996965-INS-539
chr10-65012773-INS-260
chr10-65021622-INS-113
chr10-65053725-INS-121
chr10-65068148-INS-69
chr10-65077493-INS-594
chr10-65083450-INS-284
chr10-65109945-INS-60
chr10-65115253-INS-1913
chr10-65131899-INS-492
chr10-65136232-INS-202
chr10-65144195-INS-459
chr10-65185317-INS-523
chr10-65196950-INS-86
chr10-65207658-INS-154
chr10-65243469-INS-123
chr10-65246511-INS-68
chr10-65253612-INS-795
chr10-65257111-INS-316
chr10-65258702-INS-102
chr10-65273172-INS-56
chr10-65273184-INS-928
chr10-65280492-INS-57
chr10-65297955-INS-604
chr10-65311045-INS-54
chr10-65315611-INS-331
chr10-65319501-INS-395

chr10-65333493-INS-752
chr10-65362164-INS-54
chr10-65383354-INS-591
chr10-65418631-INS-52
chr10-65421552-INS-721
chr10-65459601-INS-280
chr10-65483799-INS-132
chr10-65487635-INS-75
chr10-65490722-INS-196
chr10-65492099-INS-465
chr10-65537764-INS-569
chr10-65564885-INS-992
chr10-65573850-INS-356
chr10-65597838-INS-182
chr10-65606442-INS-364
chr10-65616198-INS-2272
chr10-65618043-INS-85
chr10-65620805-INS-56
chr10-65624956-INS-3031
chr10-65625809-INS-1027
chr10-65637873-INS-84
chr10-65659089-INS-217
chr10-65672567-INS-91
chr10-65677190-INS-872
chr10-65684504-INS-448
chr10-65686619-INS-299
chr10-65750195-INS-5844
chr10-65765049-INS-65
chr10-65770927-INS-530
chr10-65785141-INS-7237
chr10-65802016-INS-51
chr10-65816755-INS-926
chr10-65821877-INS-243
chr10-65852517-INS-69
chr10-65865983-INS-6125
chr10-65870859-INS-194
chr10-65871118-INS-6469
chr10-65874403-INS-57
chr10-65877740-INS-352
chr10-65878853-INS-246
chr10-65918843-INS-53
chr10-65936647-INS-984
chr10-65939424-INS-66
chr10-65970689-INS-6196
chr10-65975834-INS-1151
chr10-65987750-INS-737
chr10-65992154-INS-52
chr10-66007312-INS-62
chr10-66015270-INS-433
chr10-66028252-INS-4875
chr10-66030568-INS-149
chr10-66053524-INS-164
chr10-66053526-INS-520
chr10-66059036-INS-272
chr10-66072317-INS-790
chr10-66093296-INS-7751
chr10-66094008-INS-246
chr10-66105500-INS-312
chr10-66124458-INS-58
chr10-66130297-INS-60
chr10-66164597-INS-60
chr10-66172031-INS-104
chr10-66183786-INS-153
chr10-66194283-INS-100
chr10-66204048-INS-944
chr10-66210456-INS-381
chr10-66220994-INS-211
chr10-66226543-INS-55
chr10-66233727-INS-6389
chr10-66241576-INS-567
chr10-66245055-INS-117
chr10-66266407-INS-1502
chr10-66274298-INS-123
chr10-66276468-INS-60
chr10-66297828-INS-53
chr10-66309575-INS-6252
chr10-66325724-INS-124

chr10-66354107-INS-80
chr10-66366302-INS-240
chr10-66370495-INS-57
chr10-66377927-INS-107
chr10-66390480-INS-265
chr10-66428003-INS-80
chr10-66433105-INS-75
chr10-66437694-INS-51
chr10-66442420-INS-81
chr10-66455710-INS-5284
chr10-66462636-INS-337
chr10-66467693-INS-52
chr10-66485319-INS-148
chr10-66519747-INS-6940
chr10-66530687-INS-86
chr10-66546971-INS-94
chr10-66550616-INS-83
chr10-66552048-INS-6303
chr10-66576410-INS-168
chr10-66582618-INS-81
chr10-66585265-INS-143
chr10-66587011-INS-97
chr10-66607275-INS-58
chr10-66607798-INS-59
chr10-66622131-INS-52
chr10-66637950-INS-141
chr10-66638945-INS-246
chr10-66642174-INS-204
chr10-66651135-INS-72
chr10-66654394-INS-596
chr10-66665421-INS-372
chr10-66680292-INS-216
chr10-66683197-INS-208
chr10-66706076-INS-52
chr10-66724231-INS-171
chr10-66733914-INS-175
chr10-66745801-INS-177
chr10-66761345-INS-51
chr10-66778094-INS-206
chr10-66780796-INS-158
chr10-66785079-INS-122
chr10-66793584-INS-223
chr10-66797327-INS-211
chr10-66802071-INS-54
chr10-66805520-INS-200
chr10-66824207-INS-220
chr10-66832594-INS-52
chr10-66836301-INS-142
chr10-66840843-INS-204
chr10-66842320-INS-544
chr10-66844334-INS-237
chr10-66848321-INS-81
chr10-66873952-INS-92
chr10-66878496-INS-181
chr10-66916086-INS-212
chr10-66921700-INS-243
chr10-66929568-INS-152
chr10-66966099-INS-70
chr10-66997301-INS-210
chr10-67006793-INS-240
chr10-67011500-INS-207
chr10-67045926-INS-224
chr10-67078330-INS-120
chr10-67095411-INS-186
chr10-67099557-INS-86
chr10-67108808-INS-145
chr10-67110244-INS-66
chr10-67134155-INS-216
chr10-67146526-INS-242
chr10-67158039-INS-113
chr10-67160054-INS-116
chr10-67164216-INS-170
chr10-67166222-INS-276
chr10-67175081-INS-402
chr10-67175731-INS-64
chr10-67183193-INS-60
chr10-67183850-INS-79

chr10-67185461-INS-401
chr10-67187549-INS-112
chr10-67202346-INS-56
chr10-67204525-INS-605
chr10-67210813-INS-66
chr10-67233826-INS-112
chr10-67251994-INS-234
chr10-67278173-INS-624
chr10-67296327-INS-500
chr10-67310644-INS-58
chr10-67313940-INS-108
chr10-67329302-INS-69
chr10-67331021-INS-54
chr10-67336339-INS-114
chr10-67337598-INS-52
chr10-67342477-INS-58
chr10-67372691-INS-124
chr10-67403979-INS-126
chr10-67407199-INS-65
chr10-67433334-INS-146
chr10-67439534-INS-439
chr10-67443162-INS-112
chr10-67460038-INS-66
chr10-67495970-INS-105
chr10-67582711-INS-137
chr10-67605977-INS-53
chr10-67607764-INS-7110
chr10-67614726-INS-165
chr10-67626366-INS-135
chr10-67633030-INS-342
chr10-67638769-INS-56
chr10-67671385-INS-52
chr10-67688030-INS-364
chr10-67703915-INS-1253
chr10-67705497-INS-61
chr10-67707080-INS-80
chr10-67709413-INS-127
chr10-67716417-INS-127
chr10-67718423-INS-523
chr10-67755134-INS-58
chr10-67763458-INS-56
chr10-67772248-INS-221
chr10-67778962-INS-76
chr10-67779381-INS-99
chr10-67788103-INS-71
chr10-67789993-INS-60
chr10-67802862-INS-66
chr10-67821989-INS-137
chr10-67824281-INS-248
chr10-67829980-INS-60
chr10-67869546-INS-296
chr10-67873582-INS-147
chr10-67906656-INS-58
chr10-67909035-INS-210
chr10-67947858-INS-70
chr10-67950378-INS-128
chr10-67979760-INS-157
chr10-68091808-INS-52
chr10-68118069-INS-82
chr10-68127641-INS-52
chr10-68133077-INS-213
chr10-68141505-INS-915
chr10-68143195-INS-7317
chr10-68189104-INS-378
chr10-68202516-INS-70
chr10-68211413-INS-69
chr10-68305208-INS-114
chr10-68315059-INS-6559
chr10-68331970-INS-238
chr10-68337690-INS-6363
chr10-68397701-INS-509
chr10-68401133-INS-642
chr10-68407159-INS-68
chr10-68454707-INS-165
chr10-68474310-INS-66
chr10-68481492-INS-62
chr10-68482763-INS-132

chr10-68531666-INS-151
chr10-68539931-INS-138
chr10-68616690-INS-53
chr10-68624929-INS-56
chr10-68626629-INS-202
chr10-68642803-INS-87
chr10-68644639-INS-396
chr10-68650127-INS-74
chr10-68651286-INS-72
chr10-68668781-INS-112
chr10-68682962-INS-51
chr10-68687985-INS-2151
chr10-68688070-INS-151
chr10-68695295-INS-52
chr10-68697269-INS-1617
chr10-68705102-INS-188
chr10-68719065-INS-172
chr10-68719087-INS-889
chr10-68723144-INS-337
chr10-68757298-INS-56
chr10-68761810-INS-61
chr10-68805687-INS-388
chr10-68810132-INS-94
chr10-68845919-INS-153
chr10-68893728-INS-54
chr10-68917990-INS-128
chr10-68938627-INS-296
chr10-68948101-INS-397
chr10-68985114-INS-74
chr10-69008484-INS-72
chr10-69029302-INS-329
chr10-69042304-INS-538
chr10-69055661-INS-53
chr10-69058838-INS-288
chr10-69083685-INS-60
chr10-69094294-INS-157
chr10-69105378-INS-68
chr10-69110016-INS-109
chr10-69128841-INS-7264
chr10-69134181-INS-236
chr10-69171773-INS-91
chr10-69198240-INS-163
chr10-69217492-INS-78
chr10-69231932-INS-121
chr10-69233091-INS-170
chr10-69233497-INS-142
chr10-69248246-INS-105
chr10-69251292-INS-197
chr10-69252479-INS-848
chr10-69262691-INS-51
chr10-69271799-INS-55
chr10-69277167-INS-70
chr10-69308049-INS-62
chr10-69315138-INS-139
chr10-69351948-INS-98
chr10-69355858-INS-125
chr10-69361237-INS-108
chr10-69375363-INS-148
chr10-69387149-INS-148
chr10-69390260-INS-65
chr10-69401187-INS-88
chr10-69411791-INS-190
chr10-69422692-INS-317
chr10-69445577-INS-218
chr10-69456120-INS-78
chr10-69464234-INS-53
chr10-69478619-INS-119
chr10-69488158-INS-6576
chr10-69493734-INS-363
chr10-69503066-INS-116
chr10-69506053-INS-288
chr10-69509072-INS-623
chr10-69524245-INS-526
chr10-69533651-INS-82
chr10-69545210-INS-118
chr10-69552580-INS-59
chr10-69584303-INS-316

chr10-69590782-INS-168
chr10-69593609-INS-115
chr10-69598725-INS-98
chr10-69599502-INS-72
chr10-69602158-INS-118
chr10-69609997-INS-79
chr10-69627511-INS-56
chr10-69641622-INS-1175
chr10-69666703-INS-152
chr10-69670976-INS-92
chr10-69683555-INS-90
chr10-69690616-INS-121
chr10-69699484-INS-309
chr10-69713303-INS-192
chr10-69714160-INS-76
chr10-69720317-INS-148
chr10-69721504-INS-74
chr10-69730082-INS-140
chr10-69745787-INS-138
chr10-69750354-INS-276
chr10-69759539-INS-177
chr10-69768342-INS-102
chr10-69769680-INS-100
chr10-69771172-INS-213
chr10-69776360-INS-68
chr10-69783910-INS-641
chr10-69799658-INS-108
chr10-69854460-INS-220
chr10-69855268-INS-80
chr10-69858684-INS-124
chr10-69868303-INS-111
chr10-69875306-INS-79
chr10-69884044-INS-65
chr10-69900654-INS-793
chr10-69910459-INS-142
chr10-69922514-INS-345
chr10-69925755-INS-206
chr10-69929156-INS-143
chr10-69935410-INS-221
chr10-69958376-INS-53
chr10-69998603-INS-72
chr10-70000431-INS-58
chr10-70016247-INS-55
chr10-70051841-INS-57
chr10-70056621-INS-865
chr10-70082041-INS-105
chr10-70082702-INS-117
chr10-70110052-INS-116
chr10-70135237-INS-355
chr10-70142541-INS-399
chr10-70143629-INS-56
chr10-70163599-INS-413
chr10-70165778-INS-548
chr10-70189697-INS-167
chr10-70202162-INS-72
chr10-70222943-INS-51
chr10-70228802-INS-471
chr10-70236334-INS-56
chr10-70237772-INS-361
chr10-70244256-INS-95
chr10-70249430-INS-395
chr10-70252413-INS-66
chr10-70254841-INS-285
chr10-70262048-INS-118
chr10-70265460-INS-184
chr10-70270291-INS-208
chr10-70297282-INS-413
chr10-70302846-INS-1777
chr10-70334266-INS-81
chr10-70343988-INS-67
chr10-70363238-INS-497
chr10-70463969-INS-270
chr10-70470622-INS-115
chr10-70524279-INS-51
chr10-70525330-INS-163
chr10-70552759-INS-52
chr10-70564387-INS-400

chr10-70609009-INS-71
chr10-70613589-INS-730
chr10-70643012-INS-52
chr10-70649464-INS-60
chr10-70659095-INS-78
chr10-70697910-INS-64
chr10-70733162-INS-122
chr10-70754026-INS-95
chr10-70790280-INS-137
chr10-70792248-INS-55
chr10-70793901-INS-255
chr10-70805746-INS-55
chr10-70858543-INS-55
chr10-70861940-INS-972
chr10-70893840-INS-6487
chr10-70894316-INS-677
chr10-70936264-INS-666
chr10-70974628-INS-99
chr10-71000422-INS-159
chr10-71028547-INS-168
chr10-71032332-INS-121
chr10-71039654-INS-67
chr10-71059450-INS-481
chr10-71061694-INS-112
chr10-71062118-INS-308
chr10-71077275-INS-225
chr10-71095046-INS-195
chr10-71105724-INS-473
chr10-71111710-INS-66
chr10-71119009-INS-59
chr10-71129951-INS-191
chr10-71133653-INS-66
chr10-71139086-INS-109
chr10-71153985-INS-194
chr10-71155805-INS-468
chr10-71203771-INS-105
chr10-71206366-INS-5351
chr10-71211786-INS-77
chr10-71216203-INS-106
chr10-71239073-INS-64
chr10-71252959-INS-130
chr10-71261550-INS-424
chr10-71281011-INS-66
chr10-71291194-INS-66
chr10-71308795-INS-663
chr10-71328205-INS-401
chr10-71330617-INS-498
chr10-71337836-INS-66
chr10-71341217-INS-90
chr10-71355097-INS-400
chr10-71364317-INS-434
chr10-71367545-INS-958
chr10-71379393-INS-676
chr10-71413719-INS-639
chr10-71415133-INS-616
chr10-71434404-INS-57
chr10-71438924-INS-78
chr10-71450721-INS-62
chr10-71477144-INS-229
chr10-71490231-INS-53
chr10-71494219-INS-124
chr10-71501791-INS-66
chr10-71525551-INS-100
chr10-71526991-INS-118
chr10-71560968-INS-52
chr10-71571262-INS-357
chr10-71585987-INS-801
chr10-71614295-INS-439
chr10-71633298-INS-913
chr10-71649879-INS-445
chr10-71664708-INS-6710
chr10-71670998-INS-569
chr10-71715630-INS-81
chr10-71734854-INS-682
chr10-71738887-INS-6588
chr10-71741197-INS-6272
chr10-71762995-INS-218

chr10-71779282-INS-225
chr10-71781297-INS-497
chr10-71858408-INS-293
chr10-71860177-INS-400
chr10-71873963-INS-231
chr10-71888185-INS-66
chr10-71893077-INS-7441
chr10-71916309-INS-54
chr10-71941932-INS-63
chr10-71944624-INS-88
chr10-71947973-INS-56
chr10-71958318-INS-82
chr10-71968935-INS-170
chr10-71983231-INS-513
chr10-72003697-INS-5007
chr10-72013663-INS-6319
chr10-72039325-INS-1168
chr10-72081354-INS-174
chr10-72092494-INS-56
chr10-72098463-INS-59
chr10-72119047-INS-211
chr10-72125162-INS-235
chr10-72126202-INS-266
chr10-72140084-INS-5988
chr10-72144688-INS-132
chr10-72157420-INS-54
chr10-72170411-INS-209
chr10-72172129-INS-314
chr10-72180928-INS-495
chr10-72185961-INS-72
chr10-72203114-INS-60
chr10-72210930-INS-54
chr10-72217894-INS-88
chr10-72220392-INS-72
chr10-72225225-INS-100
chr10-72236716-INS-52
chr10-72256161-INS-446
chr10-72269411-INS-598
chr10-72272139-INS-77
chr10-72305463-INS-64
chr10-72351872-INS-751
chr10-72384287-INS-2030
chr10-72385862-INS-4934
chr10-72398305-INS-6382
chr10-72404764-INS-3019
chr10-72409777-INS-7125
chr10-72412559-INS-6931
chr10-72436952-INS-114
chr10-72442798-INS-6214
chr10-72456362-INS-74
chr10-72472280-INS-68
chr10-72496062-INS-66
chr10-72519888-INS-84
chr10-72520592-INS-457
chr10-72526033-INS-291
chr10-72526757-INS-206
chr10-72536433-INS-57
chr10-72577662-INS-54
chr10-72582620-INS-860
chr10-72614321-INS-353
chr10-72650829-INS-99
chr10-72653286-INS-65
chr10-72661464-INS-72
chr10-72662866-INS-261
chr10-72682761-INS-156
chr10-72685778-INS-226
chr10-72703687-INS-95
chr10-72763132-INS-200
chr10-72809215-INS-147
chr10-72813608-INS-563
chr10-72827809-INS-498
chr10-72845142-INS-516
chr10-72845426-INS-4136
chr10-72852908-INS-276
chr10-72873845-INS-66
chr10-72879966-INS-98
chr10-72893030-INS-147

chr10-72894249-INS-6529
chr10-72895522-INS-1157
chr10-72904139-INS-115
chr10-72926497-INS-950
chr10-72976553-INS-152
chr10-72991714-INS-794
chr10-73007070-INS-6804
chr10-73007220-INS-83
chr10-73010365-INS-136
chr10-73015160-INS-6303
chr10-73023311-INS-220
chr10-73024477-INS-6261
chr10-73046515-INS-260
chr10-73047403-INS-66
chr10-73068851-INS-66
chr10-73072527-INS-67
chr10-73081829-INS-54
chr10-73174117-INS-463
chr10-73193595-INS-60
chr10-73205847-INS-201
chr10-73231929-INS-328
chr10-73234709-INS-70
chr10-73263449-INS-68
chr10-73266753-INS-57
chr10-73303728-INS-720
chr10-73313094-INS-1243
chr10-73332360-INS-321
chr10-73347999-INS-5662
chr10-73353289-INS-162
chr10-73356417-INS-68
chr10-73382890-INS-369
chr10-73388718-INS-64
chr10-73394243-INS-170
chr10-73409039-INS-53
chr10-73428784-INS-710
chr10-73447037-INS-192
chr10-73455212-INS-288
chr10-73508119-INS-85
chr10-73513524-INS-65
chr10-73550502-INS-122
chr10-73555901-INS-183
chr10-73593503-INS-419
chr10-73594986-INS-52
chr10-73632888-INS-64
chr10-73637422-INS-156
chr10-73639532-INS-95
chr10-73648498-INS-573
chr10-73674920-INS-76
chr10-73679971-INS-73
chr10-73688360-INS-172
chr10-73704277-INS-212
chr10-73713171-INS-104
chr10-73743717-INS-582
chr10-73752316-INS-68
chr10-73808370-INS-573
chr10-73835670-INS-54
chr10-73878236-INS-839
chr10-73879914-INS-75
chr10-73898207-INS-353
chr10-73913746-INS-111
chr10-73917912-INS-75
chr10-73952312-INS-732
chr10-73969317-INS-266
chr10-73979709-INS-6616
chr10-73985114-INS-64
chr10-74010908-INS-64
chr10-74046714-INS-218
chr10-74067136-INS-304
chr10-74087431-INS-734
chr10-74090335-INS-6795
chr10-74121555-INS-278
chr10-74124476-INS-81
chr10-74127553-INS-593
chr10-74142342-INS-64
chr10-74153821-INS-469
chr10-74196449-INS-117
chr10-74205951-INS-62

chr10-74206307-INS-85
chr10-74210639-INS-60
chr10-74228652-INS-744
chr10-74229381-INS-193
chr10-74248654-INS-7118
chr10-74262842-INS-506
chr10-74263154-INS-6699
chr10-74266063-INS-58
chr10-74291957-INS-82
chr10-74295436-INS-176
chr10-74296697-INS-70
chr10-74327904-INS-55
chr10-74341511-INS-56
chr10-74345460-INS-82
chr10-74366027-INS-194
chr10-74399917-INS-82
chr10-74416725-INS-624
chr10-74421755-INS-180
chr10-74432463-INS-914
chr10-74432687-INS-132
chr10-74445935-INS-187
chr10-74446260-INS-6385
chr10-74470884-INS-67
chr10-74476527-INS-82
chr10-74517188-INS-90
chr10-74547490-INS-513
chr10-74562792-INS-241
chr10-74573598-INS-1144
chr10-74595167-INS-143
chr10-74600325-INS-6192
chr10-74618058-INS-1279
chr10-74638834-INS-5229
chr10-74645243-INS-8520
chr10-74660092-INS-132
chr10-74665617-INS-8334
chr10-74674195-INS-120
chr10-74681223-INS-59
chr10-74690127-INS-71
chr10-74692956-INS-219
chr10-74704128-INS-84
chr10-74746018-INS-62
chr10-74813209-INS-60
chr10-74839755-INS-498
chr10-74841197-INS-204
chr10-74869960-INS-94
chr10-74872428-INS-85
chr10-74876092-INS-110
chr10-74891209-INS-99
chr10-74930819-INS-62
chr10-74951355-INS-123
chr10-74957324-INS-199
chr10-74967269-INS-186
chr10-75028322-INS-88
chr10-75048633-INS-67
chr10-75069281-INS-51
chr10-75069281-INS-900
chr10-75119665-INS-256
chr10-75129187-INS-189
chr10-75218117-INS-91
chr10-75269856-INS-219
chr10-75289046-INS-66
chr10-75313231-INS-64
chr10-75328151-INS-321
chr10-75333839-INS-2792
chr10-75346929-INS-91
chr10-75364000-INS-172
chr10-75409430-INS-75
chr10-75426379-INS-65
chr10-75430123-INS-206
chr10-75432601-INS-6399
chr10-75455809-INS-95
chr10-75461618-INS-211
chr10-75543842-INS-179
chr10-75596607-INS-608
chr10-75596649-INS-416
chr10-75626473-INS-339
chr10-75627377-INS-57

chr10-75636625-INS-211
chr10-75643476-INS-78
chr10-75652716-INS-104
chr10-75654848-INS-378
chr10-75661383-INS-159
chr10-75662813-INS-112
chr10-75666157-INS-1163
chr10-75672110-INS-213
chr10-75683731-INS-375
chr10-75687635-INS-70
chr10-75708189-INS-176
chr10-75711749-INS-8736
chr10-75712378-INS-180
chr10-75766522-INS-203
chr10-75777311-INS-51
chr10-75781601-INS-128
chr10-75789382-INS-208
chr10-75799929-INS-93
chr10-75800319-INS-195
chr10-75802817-INS-74
chr10-75813415-INS-57
chr10-75850159-INS-888
chr10-75852900-INS-579
chr10-75872080-INS-53
chr10-75880938-INS-96
chr10-75886936-INS-77
chr10-75892041-INS-398
chr10-75893634-INS-210
chr10-75903946-INS-64
chr10-75929394-INS-87
chr10-75930439-INS-77
chr10-75955565-INS-225
chr10-75961023-INS-92
chr10-75976124-INS-141
chr10-75981731-INS-53
chr10-75984376-INS-57
chr10-76003116-INS-128
chr10-76014265-INS-89
chr10-76047214-INS-395
chr10-76082493-INS-103
chr10-76142238-INS-102
chr10-76166747-INS-199
chr10-76196918-INS-100
chr10-76208140-INS-173
chr10-76209336-INS-201
chr10-76221362-INS-854
chr10-76226462-INS-58
chr10-76238448-INS-213
chr10-76255012-INS-121
chr10-76263247-INS-271
chr10-76271281-INS-205
chr10-76279494-INS-5687
chr10-76280377-INS-162
chr10-76281683-INS-202
chr10-76290945-INS-53
chr10-76302532-INS-331
chr10-76317084-INS-166
chr10-76320439-INS-56
chr10-76336725-INS-202
chr10-76337745-INS-214
chr10-76346044-INS-91
chr10-76354779-INS-213
chr10-76365132-INS-231
chr10-76387017-INS-101
chr10-76393609-INS-69
chr10-76401849-INS-285
chr10-76421900-INS-57
chr10-76429235-INS-114
chr10-76434963-INS-127
chr10-76456547-INS-240
chr10-76488340-INS-51
chr10-76501999-INS-1926
chr10-76505818-INS-431
chr10-76508747-INS-62
chr10-76516294-INS-128
chr10-76519806-INS-82
chr10-76558199-INS-198

chr10-76581380-INS-353
chr10-76588139-INS-63
chr10-76662878-INS-255
chr10-76663836-INS-53
chr10-76671245-INS-217
chr10-76678884-INS-66
chr10-76687500-INS-498
chr10-76716917-INS-142
chr10-76737185-INS-78
chr10-76753805-INS-371
chr10-76780211-INS-56
chr10-76786256-INS-170
chr10-76880957-INS-224
chr10-76927018-INS-64
chr10-76935932-INS-208
chr10-76937702-INS-216
chr10-76938733-INS-259
chr10-76940695-INS-222
chr10-76957586-INS-92
chr10-76965524-INS-58
chr10-76988172-INS-65
chr10-77005665-INS-252
chr10-77008298-INS-56
chr10-77014818-INS-62
chr10-77016887-INS-441
chr10-77019112-INS-176
chr10-77019763-INS-1587
chr10-77021775-INS-199
chr10-77031130-INS-61
chr10-77037491-INS-80
chr10-77039964-INS-88
chr10-77065229-INS-70
chr10-77069297-INS-71
chr10-77074941-INS-202
chr10-77090393-INS-5546
chr10-77117661-INS-251
chr10-77120237-INS-56
chr10-77124243-INS-476
chr10-77132707-INS-2579
chr10-77133556-INS-145
chr10-77149163-INS-5000
chr10-77156610-INS-217
chr10-77185277-INS-310
chr10-77211626-INS-56
chr10-77227568-INS-6498
chr10-77233688-INS-136
chr10-77235680-INS-55
chr10-77264264-INS-105
chr10-77282290-INS-250
chr10-77286973-INS-78
chr10-77313060-INS-526
chr10-77319465-INS-847
chr10-77357039-INS-129
chr10-77421262-INS-698
chr10-77456960-INS-216
chr10-77491550-INS-7240
chr10-77508228-INS-325
chr10-77529410-INS-120
chr10-77530937-INS-1486
chr10-77535927-INS-416
chr10-77551738-INS-60
chr10-77564213-INS-134
chr10-77639408-INS-303
chr10-77657384-INS-85
chr10-77686221-INS-60
chr10-77691866-INS-124
chr10-77693292-INS-411
chr10-77697982-INS-61
chr10-77714474-INS-831
chr10-77725546-INS-53
chr10-77743840-INS-56
chr10-77755871-INS-125
chr10-77756900-INS-60
chr10-77776240-INS-301
chr10-77779687-INS-117
chr10-77786322-INS-63
chr10-77799108-INS-52

chr10-77828459-INS-165
chr10-77834446-INS-84
chr10-77855138-INS-79
chr10-77862495-INS-122
chr10-77862775-INS-3522
chr10-77866483-INS-64
chr10-77869880-INS-175
chr10-77892871-INS-199
chr10-77901276-INS-72
chr10-77902107-INS-1906
chr10-77902601-INS-1051
chr10-77904155-INS-270
chr10-77916040-INS-102
chr10-77945907-INS-177
chr10-77949602-INS-87
chr10-77964227-INS-249
chr10-77968543-INS-81
chr10-77992221-INS-51
chr10-77993831-INS-308
chr10-78045619-INS-144
chr10-78068153-INS-56
chr10-78074155-INS-66
chr10-78093255-INS-55
chr10-78115333-INS-240
chr10-78128325-INS-77
chr10-78168602-INS-226
chr10-78175604-INS-93
chr10-78202005-INS-512
chr10-78220437-INS-81
chr10-78220985-INS-201
chr10-78221802-INS-262
chr10-78234510-INS-575
chr10-78258118-INS-155
chr10-78274594-INS-648
chr10-78317125-INS-82
chr10-78322170-INS-352
chr10-78331223-INS-602
chr10-78334804-INS-727
chr10-78342825-INS-406
chr10-78350609-INS-90
chr10-78359496-INS-205
chr10-78371544-INS-7191
chr10-78381079-INS-7707
chr10-78438432-INS-594
chr10-78441205-INS-105
chr10-78464548-INS-133
chr10-78466683-INS-54
chr10-78469688-INS-160
chr10-78494124-INS-286
chr10-78499926-INS-54
chr10-78501787-INS-206
chr10-78506959-INS-52
chr10-78513364-INS-169
chr10-78524518-INS-58
chr10-78542960-INS-58
chr10-78552690-INS-170
chr10-78563380-INS-150
chr10-78573071-INS-144
chr10-78576165-INS-297
chr10-78583350-INS-6251
chr10-78594930-INS-80
chr10-78628526-INS-142
chr10-78638269-INS-250
chr10-78662662-INS-187
chr10-78672702-INS-603
chr10-78679021-INS-8405
chr10-78684445-INS-56
chr10-78703448-INS-2858
chr10-78704734-INS-184
chr10-78752119-INS-201
chr10-78753852-INS-189
chr10-78818307-INS-5282
chr10-78823387-INS-7225
chr10-78827037-INS-6403
chr10-78828850-INS-59
chr10-78855517-INS-362
chr10-78982159-INS-198

chr10-78986062-INS-80
chr10-79014882-INS-2965
chr10-79024689-INS-7325
chr10-79052678-INS-2439
chr10-79062609-INS-7922
chr10-79144647-INS-439
chr10-79146791-INS-231
chr10-79194574-INS-392
chr10-79226627-INS-52
chr10-79237364-INS-2657
chr10-79265367-INS-6173
chr10-79296498-INS-329
chr10-79331652-INS-249
chr10-79343353-INS-635
chr10-79345178-INS-784
chr10-79360900-INS-93
chr10-79383928-INS-105
chr10-79409945-INS-98
chr10-79413515-INS-60
chr10-79434516-INS-264
chr10-79527062-INS-209
chr10-79536167-INS-201
chr10-79539740-INS-189
chr10-79583159-INS-201
chr10-79584106-INS-104
chr10-79585104-INS-222
chr10-79589571-INS-198
chr10-79595722-INS-202
chr10-79598194-INS-235
chr10-79605493-INS-251
chr10-79608233-INS-256
chr10-79646628-INS-219
chr10-79661056-INS-172
chr10-79681383-INS-185
chr10-79708144-INS-492
chr10-79711968-INS-180
chr10-79725647-INS-168
chr10-79775926-INS-51
chr10-79785698-INS-227
chr10-79793348-INS-106
chr10-79848354-INS-132
chr10-79873904-INS-210
chr10-79877945-INS-221
chr10-79891659-INS-119
chr10-79904861-INS-262
chr10-79924161-INS-212
chr10-79928086-INS-873
chr10-79946657-INS-399
chr10-80002129-INS-204
chr10-80036098-INS-128
chr10-80052680-INS-166
chr10-80059584-INS-204
chr10-80111598-INS-213
chr10-80126741-INS-63
chr10-80132274-INS-86
chr10-80135241-INS-52
chr10-80137189-INS-215
chr10-80144365-INS-51
chr10-80198773-INS-113
chr10-80204172-INS-2672
chr10-80209359-INS-202
chr10-80210285-INS-71
chr10-80241844-INS-586
chr10-80244294-INS-112
chr10-80248135-INS-144
chr10-80257869-INS-96
chr10-80260518-INS-115
chr10-80273271-INS-81
chr10-80289021-INS-239
chr10-80303590-INS-180
chr10-80309642-INS-92
chr10-80319022-INS-55
chr10-80322573-INS-206
chr10-80324675-INS-83
chr10-80344568-INS-279
chr10-80366532-INS-556
chr10-80388606-INS-55

chr10-80394650-INS-148
chr10-80399036-INS-558
chr10-80416029-INS-213
chr10-80481577-INS-56
chr10-80490117-INS-239
chr10-80496122-INS-204
chr10-80498524-INS-250
chr10-80520801-INS-52
chr10-80523566-INS-209
chr10-80527313-INS-293
chr10-80548922-INS-208
chr10-80550398-INS-65
chr10-80571536-INS-301
chr10-80582871-INS-65
chr10-80585176-INS-106
chr10-80593466-INS-207
chr10-80625714-INS-204
chr10-80638307-INS-142
chr10-80647891-INS-143
chr10-80649683-INS-214
chr10-80656069-INS-56
chr10-80666386-INS-177
chr10-80690190-INS-255
chr10-80705276-INS-55
chr10-80713454-INS-202
chr10-80718900-INS-92
chr10-80730304-INS-53
chr10-80786478-INS-51
chr10-80810793-INS-57
chr10-80817964-INS-184
chr10-80820215-INS-238
chr10-80824761-INS-191
chr10-80826151-INS-81
chr10-80829484-INS-209
chr10-80841043-INS-157
chr10-80871465-INS-157
chr10-80886430-INS-184
chr10-80910356-INS-123
chr10-80923625-INS-178
chr10-80927925-INS-134
chr10-80938805-INS-213
chr10-81000621-INS-52
chr10-81032204-INS-206
chr10-81083677-INS-271
chr10-81092122-INS-170
chr10-81122002-INS-219
chr10-81132812-INS-97
chr10-81167186-INS-102
chr10-81188709-INS-243
chr10-81190054-INS-207
chr10-81200565-INS-52
chr10-81211354-INS-755
chr10-81219520-INS-470
chr10-81259182-INS-94
chr10-81267974-INS-84
chr10-81305640-INS-67
chr10-81315364-INS-120
chr10-81358925-INS-219
chr10-81361786-INS-183
chr10-81372831-INS-5512
chr10-81374111-INS-240
chr10-81436457-INS-198
chr10-81439084-INS-202
chr10-81447545-INS-257
chr10-81453337-INS-5678
chr10-81454885-INS-201
chr10-81455253-INS-431
chr10-81466421-INS-71
chr10-81483752-INS-58
chr10-81491253-INS-164
chr10-81493526-INS-247
chr10-81548130-INS-5558
chr10-81617284-INS-5120
chr10-81632745-INS-85
chr10-81643381-INS-139
chr10-81792093-INS-86
chr10-81802248-INS-174

chr10-81816592-INS-61
chr10-81845438-INS-201
chr10-81875550-INS-184
chr10-81887032-INS-57
chr10-81927370-INS-265
chr10-81932775-INS-196
chr10-81938200-INS-13743
chr10-81955704-INS-338
chr10-81961654-INS-573
chr10-81979708-INS-298
chr10-81992430-INS-300
chr10-81997277-INS-507
chr10-82030643-INS-185
chr10-82093433-INS-400
chr10-82116328-INS-3821
chr10-82135907-INS-208
chr10-82173766-INS-230
chr10-82185406-INS-59
chr10-82202531-INS-62
chr10-82214042-INS-251
chr10-82246604-INS-60
chr10-82259777-INS-84
chr10-82261747-INS-59
chr10-82264501-INS-3739
chr10-82303830-INS-506
chr10-82318714-INS-66
chr10-82340255-INS-6063
chr10-82341475-INS-104
chr10-82361174-INS-6194
chr10-82440417-INS-212
chr10-82442977-INS-144
chr10-82451379-INS-368
chr10-82473319-INS-164
chr10-82492462-INS-52
chr10-82501428-INS-514
chr10-82504007-INS-81
chr10-82515838-INS-69
chr10-82519784-INS-109
chr10-82522785-INS-381
chr10-82528402-INS-253
chr10-82546035-INS-760
chr10-82571670-INS-279
chr10-82610611-INS-64
chr10-82632087-INS-57
chr10-82640186-INS-214
chr10-82649312-INS-176
chr10-82651293-INS-102
chr10-82655396-INS-181
chr10-82663757-INS-167
chr10-82669532-INS-53
chr10-82676282-INS-52
chr10-82703774-INS-201
chr10-82709471-INS-197
chr10-82729667-INS-54
chr10-82748749-INS-57
chr10-82756597-INS-105
chr10-82801193-INS-55
chr10-82848010-INS-107
chr10-82849220-INS-62
chr10-82859120-INS-128
chr10-82871927-INS-212
chr10-82947595-INS-95
chr10-82955287-INS-72
chr10-82964779-INS-112
chr10-83003425-INS-242
chr10-83010054-INS-64
chr10-83019241-INS-116
chr10-83043844-INS-498
chr10-83053980-INS-78
chr10-83080064-INS-58
chr10-83082781-INS-104
chr10-83120947-INS-86
chr10-83121518-INS-51
chr10-83144162-INS-140
chr10-83150361-INS-73
chr10-83155948-INS-778
chr10-83185772-INS-62

chr10-83213067-INS-487
chr10-83234636-INS-5591
chr10-83260968-INS-410
chr10-83316129-INS-371
chr10-83320368-INS-1832
chr10-83333792-INS-532
chr10-83343340-INS-156
chr10-83389064-INS-162
chr10-83408370-INS-76
chr10-83439841-INS-735
chr10-83447715-INS-5225
chr10-83450512-INS-504
chr10-83451835-INS-75
chr10-83472965-INS-189
chr10-83492232-INS-98
chr10-83494413-INS-56
chr10-83503074-INS-185
chr10-83505107-INS-541
chr10-83505956-INS-291
chr10-83515955-INS-204
chr10-83535767-INS-488
chr10-83566370-INS-112
chr10-83568648-INS-109
chr10-83571447-INS-59
chr10-83573091-INS-116
chr10-83575894-INS-51
chr10-83580788-INS-193
chr10-83632455-INS-400
chr10-83637886-INS-81
chr10-83640525-INS-66
chr10-83657911-INS-96
chr10-83691177-INS-1432
chr10-83691315-INS-56
chr10-83696414-INS-54
chr10-83699302-INS-93
chr10-83709233-INS-336
chr10-83735599-INS-70
chr10-83741030-INS-149
chr10-83743582-INS-58
chr10-83770490-INS-441
chr10-83773646-INS-430
chr10-83815719-INS-68
chr10-83816694-INS-6619
chr10-83819762-INS-76
chr10-83825347-INS-77
chr10-83867627-INS-56
chr10-83916791-INS-84
chr10-83918229-INS-224
chr10-83961393-INS-545
chr10-83990760-INS-137
chr10-84019978-INS-68
chr10-84022115-INS-52
chr10-84023667-INS-6380
chr10-84027854-INS-80
chr10-84076584-INS-295
chr10-84107948-INS-154
chr10-84119357-INS-119
chr10-84122899-INS-124
chr10-84127052-INS-274
chr10-84143638-INS-102
chr10-84190504-INS-60
chr10-84212665-INS-58
chr10-84214353-INS-55
chr10-84284169-INS-52
chr10-84290166-INS-7087
chr10-84300889-INS-110
chr10-84307022-INS-101
chr10-84321388-INS-100
chr10-84328214-INS-59
chr10-84328972-INS-55
chr10-84347524-INS-86
chr10-84367874-INS-492
chr10-84390826-INS-263
chr10-84404231-INS-350
chr10-84417090-INS-98
chr10-84426487-INS-66
chr10-84430412-INS-367

chr10-84434618-INS-193
chr10-84448898-INS-73
chr10-84462593-INS-251
chr10-84479990-INS-217
chr10-84500028-INS-489
chr10-84532324-INS-2907
chr10-84551572-INS-300
chr10-84551893-INS-7139
chr10-84574452-INS-179
chr10-84600766-INS-212
chr10-84603336-INS-181
chr10-84615551-INS-52
chr10-84621087-INS-52
chr10-84641310-INS-129
chr10-84643059-INS-56
chr10-84644772-INS-302
chr10-84665710-INS-51
chr10-84691207-INS-55
chr10-84695065-INS-687
chr10-84716439-INS-200
chr10-84725855-INS-60
chr10-84728693-INS-597
chr10-84756399-INS-52
chr10-84757571-INS-72
chr10-84762402-INS-230
chr10-84770558-INS-70
chr10-84791095-INS-55
chr10-84803342-INS-52
chr10-84816668-INS-52
chr10-84844046-INS-206
chr10-84866352-INS-212
chr10-84868484-INS-453
chr10-84876551-INS-134
chr10-84884194-INS-196
chr10-84885414-INS-753
chr10-84885415-INS-253
chr10-84886521-INS-58
chr10-84899409-INS-216
chr10-84901230-INS-198
chr10-84905122-INS-134
chr10-84909619-INS-7766
chr10-84916259-INS-209
chr10-84919534-INS-68
chr10-84921453-INS-160
chr10-84925509-INS-210
chr10-84927383-INS-220
chr10-84928321-INS-209
chr10-84931930-INS-6816
chr10-84933152-INS-4762
chr10-84947288-INS-77
chr10-84974489-INS-58
chr10-84978145-INS-64
chr10-84984151-INS-508
chr10-85010484-INS-208
chr10-85029846-INS-58
chr10-85047525-INS-164
chr10-85055430-INS-544
chr10-85099449-INS-58
chr10-85108759-INS-75
chr10-85116960-INS-132
chr10-85153022-INS-806
chr10-85186602-INS-590
chr10-85262281-INS-492
chr10-85267078-INS-56
chr10-85308370-INS-96
chr10-85323719-INS-105
chr10-85331853-INS-258
chr10-85343118-INS-202
chr10-85351987-INS-127
chr10-85379750-INS-210
chr10-85383396-INS-185
chr10-85387255-INS-90
chr10-85423589-INS-586
chr10-85491371-INS-76
chr10-85506724-INS-72
chr10-85510989-INS-147
chr10-85539086-INS-180

chr10-85579580-INS-6605
chr10-85673595-INS-6715
chr10-85679026-INS-92
chr10-85696595-INS-5198
chr10-85725441-INS-176
chr10-85748186-INS-214
chr10-85794141-INS-108
chr10-85809279-INS-62
chr10-85811436-INS-210
chr10-85812338-INS-222
chr10-85814516-INS-404
chr10-85836101-INS-7095
chr10-85836772-INS-80
chr10-85872572-INS-152
chr10-85878418-INS-112
chr10-85887193-INS-294
chr10-85928776-INS-83
chr10-85967712-INS-200
chr10-85969137-INS-84
chr10-85979895-INS-147
chr10-86012947-INS-500
chr10-86017188-INS-77
chr10-86038989-INS-6276
chr10-86047635-INS-100
chr10-86051280-INS-87
chr10-86060500-INS-104
chr10-86060923-INS-7282
chr10-86065048-INS-967
chr10-86066993-INS-396
chr10-86101555-INS-98
chr10-86103097-INS-198
chr10-86103223-INS-6205
chr10-86123199-INS-72
chr10-86136367-INS-5281
chr10-86173084-INS-72
chr10-86197098-INS-390
chr10-86216065-INS-237
chr10-86249114-INS-127
chr10-86254496-INS-675
chr10-86275999-INS-214
chr10-86285485-INS-224
chr10-86286924-INS-235
chr10-86288032-INS-79
chr10-86305798-INS-73
chr10-86318113-INS-76
chr10-86339709-INS-684
chr10-86344476-INS-548
chr10-86372402-INS-114
chr10-86393381-INS-449
chr10-86413627-INS-6442
chr10-86419486-INS-54
chr10-86475658-INS-3646
chr10-86504590-INS-242
chr10-86519636-INS-6965
chr10-86543078-INS-58
chr10-86559345-INS-64
chr10-86586375-INS-583
chr10-86595138-INS-168
chr10-86603383-INS-228
chr10-86613931-INS-76
chr10-86628304-INS-60
chr10-86641910-INS-57
chr10-86643503-INS-52
chr10-86648428-INS-634
chr10-86681266-INS-336
chr10-86700576-INS-86
chr10-86741080-INS-1123
chr10-86742743-INS-239
chr10-86765097-INS-205
chr10-86768586-INS-52
chr10-86772713-INS-64
chr10-86793910-INS-64
chr10-86798413-INS-165
chr10-86808064-INS-80
chr10-86824860-INS-62
chr10-86836596-INS-465
chr10-86839258-INS-65

chr10-86857658-INS-126
chr10-86862926-INS-182
chr10-86915085-INS-144
chr10-86935584-INS-60
chr10-86964155-INS-164
chr10-86986429-INS-224
chr10-86995314-INS-88
chr10-87037894-INS-59
chr10-87081788-INS-113
chr10-87089407-INS-75
chr10-87090497-INS-72
chr10-87097561-INS-7200
chr10-87117578-INS-110
chr10-87126623-INS-62
chr10-87162682-INS-136
chr10-87182666-INS-80
chr10-87196140-INS-326
chr10-87202693-INS-56
chr10-87218515-INS-247
chr10-87222564-INS-6271
chr10-87223951-INS-85
chr10-87255732-INS-66
chr10-87269067-INS-750
chr10-87284624-INS-91
chr10-87303476-INS-378
chr10-87395873-INS-461
chr10-87398728-INS-54
chr10-87428965-INS-81
chr10-87440722-INS-6504
chr10-87453012-INS-80
chr10-87453594-INS-92
chr10-87471966-INS-96
chr10-87506392-INS-56
chr10-87517866-INS-105
chr10-87533016-INS-886
chr10-87559652-INS-150
chr10-87592596-INS-273
chr10-87627137-INS-186
chr10-87645839-INS-58
chr10-87647116-INS-105
chr10-87660946-INS-139
chr10-87675939-INS-774
chr10-87679644-INS-490
chr10-87716699-INS-178
chr10-87755560-INS-569
chr10-87765870-INS-65
chr10-87794999-INS-57
chr10-87795093-INS-634
chr10-87798713-INS-241
chr10-87805668-INS-93
chr10-87806763-INS-100
chr10-87847148-INS-962
chr10-87866975-INS-105
chr10-87882379-INS-116
chr10-87884869-INS-71
chr10-87888839-INS-72
chr10-87891346-INS-355
chr10-87920574-INS-4075
chr10-87939894-INS-144
chr10-87948504-INS-96
chr10-87969457-INS-1067
chr10-87977139-INS-1860
chr10-87993015-INS-171
chr10-88028104-INS-228
chr10-88033287-INS-172
chr10-88046472-INS-227
chr10-88051628-INS-157
chr10-88063856-INS-193
chr10-88077052-INS-54
chr10-88081953-INS-160
chr10-88088258-INS-109
chr10-88124116-INS-213
chr10-88155439-INS-51
chr10-88168229-INS-440
chr10-88169016-INS-295
chr10-88184210-INS-737
chr10-88214446-INS-64

chr10-88223115-INS-197
chr10-88230158-INS-227
chr10-88232417-INS-124
chr10-88240486-INS-58
chr10-88242119-INS-75
chr10-88275680-INS-952
chr10-88287842-INS-109
chr10-88294925-INS-193
chr10-88299340-INS-54
chr10-88323755-INS-320
chr10-88334644-INS-6444
chr10-88335829-INS-177
chr10-88342215-INS-398
chr10-88358067-INS-160
chr10-88368277-INS-72
chr10-88368744-INS-99
chr10-88371721-INS-236
chr10-88375798-INS-65
chr10-88405077-INS-95
chr10-88413290-INS-230
chr10-88443991-INS-337
chr10-88446599-INS-70
chr10-88454737-INS-7143
chr10-88455313-INS-52
chr10-88473006-INS-449
chr10-88479111-INS-93
chr10-88497293-INS-236
chr10-88503501-INS-90
chr10-88516581-INS-616
chr10-88532470-INS-60
chr10-88547258-INS-78
chr10-88552331-INS-76
chr10-88557932-INS-51
chr10-88571073-INS-80
chr10-88573839-INS-72
chr10-88577004-INS-220
chr10-88595409-INS-52
chr10-88607141-INS-575
chr10-88610258-INS-96
chr10-88617049-INS-100
chr10-88617745-INS-798
chr10-88636553-INS-215
chr10-88642581-INS-92
chr10-88647524-INS-319
chr10-88655618-INS-233
chr10-88658957-INS-608
chr10-88663074-INS-88
chr10-88676726-INS-6575
chr10-88679705-INS-60
chr10-88681587-INS-239
chr10-88697358-INS-505
chr10-88706050-INS-651
chr10-88706059-INS-197
chr10-88713265-INS-400
chr10-88716308-INS-153
chr10-88727093-INS-127
chr10-88734906-INS-522
chr10-88746280-INS-59
chr10-88755873-INS-144
chr10-88778084-INS-204
chr10-88779020-INS-52
chr10-88780088-INS-54
chr10-88783198-INS-112
chr10-88790830-INS-55
chr10-88797114-INS-87
chr10-88813484-INS-111
chr10-88824173-INS-207
chr10-88832589-INS-78
chr10-88875348-INS-176
chr10-88902464-INS-59
chr10-88908976-INS-55
chr10-88919970-INS-4955
chr10-88920230-INS-55
chr10-88921302-INS-130
chr10-88926621-INS-86
chr10-88937730-INS-80
chr10-88944250-INS-60

chr10-88968397-INS-476
chr10-88987837-INS-56
chr10-89000349-INS-227
chr10-89009784-INS-95
chr10-89012698-INS-72
chr10-89028247-INS-576
chr10-89042312-INS-76
chr10-89042651-INS-61
chr10-89057530-INS-6409
chr10-89063169-INS-74
chr10-89082908-INS-90
chr10-89083015-INS-7192
chr10-89109495-INS-67
chr10-89113275-INS-97
chr10-89137509-INS-938
chr10-89142795-INS-82
chr10-89145371-INS-7173
chr10-89147434-INS-2132
chr10-89148827-INS-2453
chr10-89156723-INS-102
chr10-89159876-INS-58
chr10-89168904-INS-57
chr10-89209017-INS-425
chr10-89217931-INS-191
chr10-89224604-INS-577
chr10-89231207-INS-62
chr10-89240928-INS-507
chr10-89268386-INS-66
chr10-89288199-INS-462
chr10-89316186-INS-113
chr10-89336734-INS-64
chr10-89350557-INS-68
chr10-89382429-INS-966
chr10-89389323-INS-1445
chr10-89405272-INS-351
chr10-89411558-INS-52
chr10-89415071-INS-3390
chr10-89424462-INS-136
chr10-89448010-INS-767
chr10-89448046-INS-53
chr10-89467510-INS-94
chr10-89472912-INS-53
chr10-89482032-INS-58
chr10-89483894-INS-902
chr10-89484687-INS-57
chr10-89495233-INS-73
chr10-89506512-INS-159
chr10-89509277-INS-199
chr10-89518236-INS-132
chr10-89527554-INS-122
chr10-89551612-INS-211
chr10-89553171-INS-260
chr10-89577566-INS-70
chr10-89592018-INS-206
chr10-89593008-INS-68
chr10-89602328-INS-74
chr10-89609788-INS-7070
chr10-89624410-INS-52
chr10-89634869-INS-643
chr10-89642816-INS-148
chr10-89650789-INS-56
chr10-89684412-INS-100
chr10-89689947-INS-55
chr10-89714579-INS-204
chr10-89717200-INS-52
chr10-89748668-INS-106
chr10-89772989-INS-545
chr10-89801946-INS-151
chr10-89854605-INS-62
chr10-89862935-INS-88
chr10-89873047-INS-206
chr10-89874568-INS-299
chr10-89959138-INS-329
chr10-89973585-INS-585
chr10-89980110-INS-217
chr10-90014789-INS-7285
chr10-90017378-INS-117

chr10-90029838-INS-72
chr10-90040310-INS-297
chr10-90067587-INS-59
chr10-90075489-INS-111
chr10-90076856-INS-4489
chr10-90081221-INS-60
chr10-90081411-INS-462
chr10-90084560-INS-811
chr10-90096390-INS-730
chr10-90100901-INS-205
chr10-90122381-INS-171
chr10-90126788-INS-6391
chr10-90130991-INS-72
chr10-90139239-INS-78
chr10-90158448-INS-151
chr10-90229977-INS-95
chr10-90238289-INS-52
chr10-90250046-INS-70
chr10-90300007-INS-658
chr10-90306704-INS-204
chr10-90338251-INS-76
chr10-90376080-INS-82
chr10-90388685-INS-429
chr10-90392698-INS-730
chr10-90401916-INS-195
chr10-90414428-INS-225
chr10-90430276-INS-80
chr10-90439619-INS-453
chr10-90441904-INS-7735
chr10-90466477-INS-258
chr10-90496856-INS-270
chr10-90498144-INS-96
chr10-90515010-INS-51
chr10-90531768-INS-55
chr10-90556581-INS-551
chr10-90565456-INS-6503
chr10-90607705-INS-87
chr10-90640490-INS-114
chr10-90641776-INS-56
chr10-90659022-INS-6685
chr10-90699108-INS-220
chr10-90714817-INS-387
chr10-90729629-INS-251
chr10-90751545-INS-88
chr10-90799429-INS-57
chr10-90812076-INS-1475
chr10-90814207-INS-8675
chr10-90817240-INS-5227
chr10-90820028-INS-72
chr10-90871431-INS-109
chr10-90879716-INS-68
chr10-90890756-INS-822
chr10-90898887-INS-190
chr10-90906249-INS-162
chr10-90922109-INS-262
chr10-90926629-INS-164
chr10-90951549-INS-168
chr10-90971112-INS-7123
chr10-90972838-INS-210
chr10-90976638-INS-7126
chr10-90977467-INS-174
chr10-90980480-INS-251
chr10-90986036-INS-212
chr10-91012231-INS-98
chr10-91056167-INS-73
chr10-91056787-INS-982
chr10-91066814-INS-7230
chr10-91067776-INS-123
chr10-91085325-INS-513
chr10-91134777-INS-315
chr10-91153369-INS-283
chr10-91200625-INS-80
chr10-91243372-INS-968
chr10-91247748-INS-215
chr10-91269800-INS-192
chr10-91275524-INS-101
chr10-91282345-INS-523

chr10-91284753-INS-589
chr10-91308348-INS-63
chr10-91334043-INS-52
chr10-91347401-INS-936
chr10-91366726-INS-64
chr10-91380430-INS-150
chr10-91401847-INS-587
chr10-91428859-INS-5344
chr10-91429504-INS-6349
chr10-91430797-INS-133
chr10-91435535-INS-853
chr10-91449899-INS-1858
chr10-91456995-INS-366
chr10-91469522-INS-54
chr10-91470697-INS-141
chr10-91480045-INS-60
chr10-91487631-INS-365
chr10-91495046-INS-64
chr10-91497928-INS-89
chr10-91501285-INS-352
chr10-91543203-INS-54
chr10-91565745-INS-60
chr10-91572944-INS-146
chr10-91577114-INS-956
chr10-91583438-INS-753
chr10-91615585-INS-6470
chr10-91643185-INS-107
chr10-91680387-INS-160
chr10-91685111-INS-124
chr10-91695701-INS-438
chr10-91711100-INS-93
chr10-91716581-INS-120
chr10-91799545-INS-163
chr10-91809533-INS-986
chr10-91844305-INS-75
chr10-91863541-INS-72
chr10-91872502-INS-7079
chr10-91877607-INS-51
chr10-91894913-INS-288
chr10-91910712-INS-51
chr10-91917928-INS-345
chr10-91942721-INS-98
chr10-91950345-INS-61
chr10-91957625-INS-108
chr10-91963742-INS-451
chr10-91966474-INS-54
chr10-92017894-INS-183
chr10-92027433-INS-433
chr10-92040682-INS-75
chr10-92074278-INS-576
chr10-92077369-INS-492
chr10-92104359-INS-779
chr10-92143448-INS-62
chr10-92183697-INS-6791
chr10-92187595-INS-87
chr10-92208586-INS-6380
chr10-92214251-INS-606
chr10-92224195-INS-163
chr10-92226476-INS-122
chr10-92229929-INS-64
chr10-92271791-INS-704
chr10-92298756-INS-78
chr10-92348833-INS-370
chr10-92377669-INS-72
chr10-92390440-INS-592
chr10-92396074-INS-115
chr10-92422667-INS-52
chr10-92425167-INS-90
chr10-92429879-INS-1858
chr10-92434260-INS-652
chr10-92447874-INS-55
chr10-92474028-INS-120
chr10-92528443-INS-222
chr10-92547306-INS-3232
chr10-92583911-INS-78
chr10-92593395-INS-202
chr10-92613377-INS-62

chr10-92645226-INS-214
chr10-92669491-INS-400
chr10-92682640-INS-6440
chr10-92689633-INS-66
chr10-92697209-INS-636
chr10-92713552-INS-956
chr10-92713895-INS-60
chr10-92721800-INS-54
chr10-92726806-INS-64
chr10-92752387-INS-146
chr10-92755833-INS-66
chr10-92776600-INS-2756
chr10-92842059-INS-163
chr10-92858957-INS-445
chr10-92867194-INS-60
chr10-92887819-INS-139
chr10-92900170-INS-65
chr10-92910596-INS-217
chr10-92937158-INS-52
chr10-92942085-INS-399
chr10-92969356-INS-51
chr10-92977014-INS-100
chr10-92985845-INS-74
chr10-93031156-INS-102
chr10-93034319-INS-78
chr10-93058620-INS-74
chr10-93059036-INS-182
chr10-93065263-INS-388
chr10-93070297-INS-141
chr10-93076134-INS-63
chr10-93082903-INS-107
chr10-93098147-INS-56
chr10-93099640-INS-434
chr10-93104270-INS-191
chr10-93108856-INS-111
chr10-93125903-INS-156
chr10-93159916-INS-989
chr10-93170317-INS-73
chr10-93171141-INS-93
chr10-93173965-INS-55
chr10-93176994-INS-53
chr10-93180907-INS-401
chr10-93186164-INS-126
chr10-93210944-INS-114
chr10-93236054-INS-408
chr10-93251412-INS-198
chr10-93253717-INS-135
chr10-93304187-INS-193
chr10-93306060-INS-276
chr10-93326233-INS-197
chr10-93353197-INS-135
chr10-93376928-INS-61
chr10-93396838-INS-146
chr10-93408194-INS-198
chr10-93443688-INS-144
chr10-93445948-INS-497
chr10-93467202-INS-278
chr10-93472260-INS-179
chr10-93491469-INS-112
chr10-93499510-INS-195
chr10-93502356-INS-128
chr10-93506681-INS-71
chr10-93511082-INS-64
chr10-93513518-INS-106
chr10-93516989-INS-711
chr10-93523022-INS-66
chr10-93533037-INS-97
chr10-93561267-INS-230
chr10-93568381-INS-445
chr10-93580509-INS-203
chr10-93583842-INS-399
chr10-93593687-INS-66
chr10-93648078-INS-456
chr10-93656928-INS-256
chr10-93664580-INS-135
chr10-93691042-INS-153
chr10-93700320-INS-56

chr10-93702715-INS-90
chr10-93727012-INS-56
chr10-93766945-INS-58
chr10-93796976-INS-52
chr10-93804648-INS-5544
chr10-93814319-INS-61
chr10-93844761-INS-54
chr10-93888745-INS-409
chr10-93906929-INS-900
chr10-93909425-INS-77
chr10-93916801-INS-133
chr10-93931652-INS-61
chr10-93979486-INS-824
chr10-93996038-INS-69
chr10-93997499-INS-241
chr10-94023098-INS-201
chr10-94026888-INS-569
chr10-94028369-INS-61
chr10-94075288-INS-362
chr10-94140261-INS-693
chr10-94149678-INS-54
chr10-94161280-INS-3011
chr10-94167883-INS-55
chr10-94172739-INS-56
chr10-94206328-INS-96
chr10-94210819-INS-5693
chr10-94223266-INS-7109
chr10-94225740-INS-126
chr10-94233050-INS-75
chr10-94243558-INS-94
chr10-94249229-INS-162
chr10-94256125-INS-52
chr10-94281829-INS-424
chr10-94320109-INS-92
chr10-94330473-INS-115
chr10-94333353-INS-2911
chr10-94349049-INS-57
chr10-94359468-INS-204
chr10-94374617-INS-80
chr10-94392575-INS-342
chr10-94424087-INS-92
chr10-94502399-INS-225
chr10-94513194-INS-141
chr10-94516577-INS-252
chr10-94536414-INS-78
chr10-94564018-INS-56
chr10-94577715-INS-6231
chr10-94584627-INS-221
chr10-94598606-INS-207
chr10-94605166-INS-100
chr10-94614050-INS-58
chr10-94664970-INS-99
chr10-94670265-INS-164
chr10-94722272-INS-64
chr10-94744727-INS-63
chr10-94749053-INS-413
chr10-94751044-INS-116
chr10-94788930-INS-131
chr10-94808248-INS-69
chr10-94817214-INS-229
chr10-94842628-INS-84
chr10-94850374-INS-7188
chr10-94853360-INS-498
chr10-94880561-INS-79
chr10-94886655-INS-6442
chr10-94887805-INS-184
chr10-94889629-INS-69
chr10-94932886-INS-392
chr10-94959495-INS-52
chr10-94964869-INS-62
chr10-94990119-INS-104
chr10-94992786-INS-58
chr10-95000237-INS-402
chr10-95004771-INS-68
chr10-95016525-INS-102
chr10-95019670-INS-52
chr10-95038580-INS-953

chr10-95056777-INS-99
chr10-95061968-INS-413
chr10-95065078-INS-56
chr10-95077536-INS-132
chr10-95083707-INS-53
chr10-95085788-INS-74
chr10-95106917-INS-149
chr10-95110011-INS-394
chr10-95121021-INS-332
chr10-95129322-INS-279
chr10-95133178-INS-143
chr10-95141806-INS-58
chr10-95145846-INS-56
chr10-95147989-INS-70
chr10-95151255-INS-56
chr10-95174614-INS-290
chr10-95180515-INS-178
chr10-95204414-INS-70
chr10-95229924-INS-85
chr10-95237923-INS-130
chr10-95268908-INS-313
chr10-95277419-INS-115
chr10-95289931-INS-86
chr10-95295646-INS-206
chr10-95302636-INS-51
chr10-95304798-INS-350
chr10-95307600-INS-124
chr10-95309283-INS-390
chr10-95323334-INS-193
chr10-95324373-INS-192
chr10-95335968-INS-88
chr10-95342674-INS-507
chr10-95345332-INS-188
chr10-95347613-INS-63
chr10-95361647-INS-53
chr10-95394340-INS-68
chr10-95402540-INS-60
chr10-95405372-INS-57
chr10-95413111-INS-62
chr10-95420467-INS-316
chr10-95426591-INS-1880
chr10-95430818-INS-127
chr10-95434027-INS-80
chr10-95451431-INS-78
chr10-95462904-INS-152
chr10-95490860-INS-197
chr10-95504416-INS-188
chr10-95519698-INS-343
chr10-95537996-INS-100
chr10-95545316-INS-189
chr10-95545853-INS-85
chr10-95565154-INS-175
chr10-95566861-INS-220
chr10-95579299-INS-124
chr10-95581026-INS-199
chr10-95585042-INS-147
chr10-95595903-INS-222
chr10-95614649-INS-222
chr10-95630418-INS-164
chr10-95652587-INS-167
chr10-95697031-INS-55
chr10-95700303-INS-80
chr10-95708039-INS-55
chr10-95748891-INS-6516
chr10-95764336-INS-1341
chr10-95770033-INS-311
chr10-95827772-INS-53
chr10-95848938-INS-111
chr10-95856492-INS-301
chr10-95862133-INS-789
chr10-95921184-INS-70
chr10-95930384-INS-106
chr10-95939200-INS-54
chr10-95953551-INS-72
chr10-95962891-INS-185
chr10-96001438-INS-81
chr10-96002002-INS-3537

chr10-96007473-INS-587
chr10-96038018-INS-61
chr10-96048635-INS-110
chr10-96055505-INS-2882
chr10-96061467-INS-136
chr10-96096161-INS-68
chr10-96124211-INS-200
chr10-96149106-INS-262
chr10-96172748-INS-88
chr10-96175605-INS-140
chr10-96176464-INS-235
chr10-96183201-INS-146
chr10-96190234-INS-275
chr10-96196142-INS-74
chr10-96198856-INS-64
chr10-96219653-INS-151
chr10-96237532-INS-70
chr10-96278708-INS-337
chr10-96339733-INS-62
chr10-96355871-INS-713
chr10-96355906-INS-57
chr10-96391504-INS-76
chr10-96399575-INS-149
chr10-96404660-INS-172
chr10-96432054-INS-98
chr10-96462102-INS-410
chr10-96483649-INS-54
chr10-96487535-INS-69
chr10-96488538-INS-78
chr10-96511372-INS-53
chr10-96525308-INS-110
chr10-96527878-INS-288
chr10-96549396-INS-189
chr10-96550272-INS-1483
chr10-96551587-INS-52
chr10-96593881-INS-5229
chr10-96622968-INS-53
chr10-96676652-INS-440
chr10-96701233-INS-152
chr10-96711959-INS-869
chr10-96769454-INS-78
chr10-96814297-INS-52
chr10-96826552-INS-56
chr10-96866563-INS-7108
chr10-96870728-INS-81
chr10-96872827-INS-136
chr10-96877342-INS-7086
chr10-96921408-INS-289
chr10-96936590-INS-226
chr10-96943436-INS-83
chr10-96953745-INS-65
chr10-96968293-INS-874
chr10-96976262-INS-103
chr10-96979290-INS-94
chr10-97021905-INS-212
chr10-97041824-INS-133
chr10-97050641-INS-60
chr10-97055064-INS-99
chr10-97069840-INS-592
chr10-97087886-INS-68
chr10-97121772-INS-172
chr10-97143880-INS-345
chr10-97151500-INS-4446
chr10-97186251-INS-140
chr10-97188329-INS-445
chr10-97192647-INS-124
chr10-97195732-INS-538
chr10-97207183-INS-93
chr10-97208941-INS-89
chr10-97211748-INS-138
chr10-97214063-INS-74
chr10-97221427-INS-510
chr10-97248209-INS-4153
chr10-97254989-INS-210
chr10-97264554-INS-586
chr10-97284412-INS-230
chr10-97285869-INS-113

chr10-97288617-INS-213
chr10-97301231-INS-672
chr10-97317141-INS-79
chr10-97321739-INS-226
chr10-97326928-INS-85
chr10-97331438-INS-59
chr10-97338006-INS-218
chr10-97372661-INS-116
chr10-97380569-INS-72
chr10-97411028-INS-82
chr10-97430580-INS-61
chr10-97455242-INS-89
chr10-97464678-INS-111
chr10-97475452-INS-132
chr10-97503027-INS-894
chr10-97521995-INS-186
chr10-97528315-INS-61
chr10-97535310-INS-52
chr10-97545318-INS-92
chr10-97546340-INS-112
chr10-97556332-INS-453
chr10-97587217-INS-208
chr10-97596993-INS-1272
chr10-97605638-INS-552
chr10-97618772-INS-197
chr10-97640564-INS-54
chr10-97650215-INS-62
chr10-97659090-INS-292
chr10-97698832-INS-64
chr10-97723296-INS-55
chr10-97733807-INS-97
chr10-97736354-INS-75
chr10-97758167-INS-591
chr10-97766632-INS-113
chr10-97769515-INS-5029
chr10-97771095-INS-197
chr10-97815337-INS-75
chr10-97826634-INS-63
chr10-97841063-INS-7038
chr10-97853419-INS-211
chr10-97855841-INS-143
chr10-97856222-INS-102
chr10-97861463-INS-7119
chr10-97864079-INS-177
chr10-97871937-INS-814
chr10-97900702-INS-134
chr10-97913696-INS-496
chr10-97921445-INS-75
chr10-97926297-INS-144
chr10-97930579-INS-1476
chr10-97953884-INS-100
chr10-97961890-INS-109
chr10-97985409-INS-191
chr10-97987362-INS-128
chr10-97989724-INS-482
chr10-97990269-INS-90
chr10-97994950-INS-69
chr10-97998992-INS-2632
chr10-98014021-INS-259
chr10-98019699-INS-99
chr10-98026068-INS-198
chr10-98047039-INS-116
chr10-98078715-INS-324
chr10-98105220-INS-753
chr10-98125340-INS-66
chr10-98127067-INS-136
chr10-98139248-INS-51
chr10-98157690-INS-546
chr10-98214362-INS-70
chr10-98216733-INS-66
chr10-98218481-INS-964
chr10-98225571-INS-126
chr10-98229848-INS-75
chr10-98254113-INS-58
chr10-98261669-INS-674
chr10-98296161-INS-98
chr10-98309695-INS-209

chr10-98312210-INS-92
chr10-98324724-INS-148
chr10-98357820-INS-129
chr10-98412009-INS-189
chr10-98455018-INS-722
chr10-98461189-INS-63
chr10-98483120-INS-91
chr10-98485516-INS-113
chr10-98503521-INS-163
chr10-98516635-INS-200
chr10-98525218-INS-81
chr10-98586392-INS-572
chr10-98600842-INS-66
chr10-98608198-INS-116
chr10-98609240-INS-53
chr10-98612208-INS-186
chr10-98632495-INS-703
chr10-98644995-INS-75
chr10-98659340-INS-61
chr10-98667261-INS-78
chr10-98670422-INS-709
chr10-98687391-INS-459
chr10-98811135-INS-363
chr10-98879114-INS-88
chr10-98884486-INS-156
chr10-98888242-INS-66
chr10-98901502-INS-59
chr10-98940695-INS-401
chr10-98953522-INS-53
chr10-98961447-INS-89
chr10-98966411-INS-98
chr10-98983346-INS-252
chr10-98992240-INS-191
chr10-99000221-INS-249
chr10-99024573-INS-440
chr10-99042853-INS-119
chr10-99049498-INS-71
chr10-99068272-INS-218
chr10-99086312-INS-227
chr10-99162018-INS-180
chr10-99183263-INS-6691
chr10-99184984-INS-432
chr10-99200983-INS-206
chr10-99244383-INS-76
chr10-99246230-INS-59
chr10-99263693-INS-638
chr10-99266323-INS-73
chr10-99277122-INS-586
chr10-99297968-INS-113
chr10-99302931-INS-122
chr10-99306354-INS-60
chr10-99316719-INS-5245
chr10-99322581-INS-700
chr10-99325175-INS-440
chr10-99330547-INS-188
chr10-99336148-INS-711
chr10-99351560-INS-439
chr10-99353260-INS-5188
chr10-99394349-INS-352
chr10-99413549-INS-646
chr10-99420830-INS-714
chr10-99460617-INS-280
chr10-99464138-INS-119
chr10-99496211-INS-104
chr10-99497566-INS-356
chr10-99507736-INS-58
chr10-99535475-INS-907
chr10-99551016-INS-3049
chr10-99564855-INS-75
chr10-99566067-INS-220
chr10-99567873-INS-72
chr10-99639275-INS-749
chr10-99728341-INS-132
chr10-99745824-INS-177
chr10-99773007-INS-52
chr10-99773700-INS-500
chr10-99812606-INS-100

chr10-99862985-INS-52
chr10-99880259-INS-596
chr10-99883487-INS-94
chr10-99883945-INS-981
chr10-99922929-INS-210
chr10-99982685-INS-98
chr10-99984921-INS-74
chr10-99997024-INS-193
chr10-100003037-INS-768
chr10-100009618-INS-55
chr10-100011380-INS-162
chr10-100058388-INS-125
chr10-100106105-INS-52
chr10-100263927-INS-101
chr10-100359847-INS-426
chr10-100438990-INS-422
chr10-100448578-INS-212
chr10-100491542-INS-6198
chr10-100497843-INS-72
chr10-100519474-INS-355
chr10-100522472-INS-498
chr10-100527666-INS-88
chr10-100537264-INS-112
chr10-100561241-INS-433
chr10-100593575-INS-540
chr10-100627372-INS-541
chr10-100658913-INS-628
chr10-100661470-INS-84
chr10-100675385-INS-5303
chr10-100697809-INS-96
chr10-100704495-INS-322
chr10-100710969-INS-238
chr10-100730977-INS-85
chr10-100743763-INS-96
chr10-100783840-INS-139
chr10-100823565-INS-523
chr10-100838709-INS-345
chr10-100852821-INS-100
chr10-100854536-INS-192
chr10-100857968-INS-120
chr10-100888466-INS-336
chr10-100919488-INS-160
chr10-100936145-INS-139
chr10-100938492-INS-114
chr10-100973200-INS-112
chr10-100978360-INS-103
chr10-100994612-INS-71
chr10-101023328-INS-53
chr10-101033601-INS-51
chr10-101047103-INS-70
chr10-101057793-INS-584
chr10-101097179-INS-60
chr10-101116829-INS-121
chr10-101127693-INS-128
chr10-101160599-INS-516
chr10-101164337-INS-166
chr10-101167220-INS-357
chr10-101171484-INS-52
chr10-101199338-INS-62
chr10-101233645-INS-163
chr10-101256225-INS-53
chr10-101281177-INS-52
chr10-101293009-INS-429
chr10-101300775-INS-256
chr10-101351804-INS-69
chr10-101383078-INS-294
chr10-101384594-INS-199
chr10-101418961-INS-79
chr10-101428057-INS-78
chr10-101477281-INS-78
chr10-101478256-INS-115
chr10-101495978-INS-52
chr10-101503307-INS-314
chr10-101539926-INS-517
chr10-101553477-INS-181
chr10-101587907-INS-52
chr10-101601221-INS-126

chr10-101612539-INS-79
chr10-101651295-INS-132
chr10-101653744-INS-52
chr10-101663227-INS-199
chr10-101668554-INS-66
chr10-101675646-INS-173
chr10-101789668-INS-75
chr10-101798930-INS-211
chr10-101818859-INS-278
chr10-101846711-INS-70
chr10-101865522-INS-130
chr10-101880133-INS-601
chr10-101880987-INS-54
chr10-101882977-INS-56
chr10-101896327-INS-230
chr10-101967542-INS-64
chr10-101968903-INS-87
chr10-101973978-INS-254
chr10-101986431-INS-79
chr10-102007156-INS-512
chr10-102040807-INS-92
chr10-102059699-INS-614
chr10-102082978-INS-159
chr10-102087961-INS-296
chr10-102092482-INS-111
chr10-102195130-INS-97
chr10-102196540-INS-376
chr10-102216404-INS-378
chr10-102227901-INS-371
chr10-102233383-INS-221
chr10-102245947-INS-647
chr10-102247981-INS-102
chr10-102254737-INS-390
chr10-102313616-INS-205
chr10-102323122-INS-563
chr10-102336705-INS-101
chr10-102345163-INS-165
chr10-102358939-INS-74
chr10-102361898-INS-60
chr10-102401720-INS-59
chr10-102404577-INS-520
chr10-102405495-INS-72
chr10-102447196-INS-53
chr10-102451139-INS-314
chr10-102451815-INS-3203
chr10-102452912-INS-508
chr10-102458265-INS-103
chr10-102500355-INS-162
chr10-102535342-INS-97
chr10-102566923-INS-77
chr10-102570981-INS-100
chr10-102584576-INS-869
chr10-102612042-INS-179
chr10-102620308-INS-938
chr10-102640125-INS-333
chr10-102641690-INS-870
chr10-102651004-INS-64
chr10-102671950-INS-451
chr10-102674908-INS-392
chr10-102691851-INS-116
chr10-102713170-INS-427
chr10-102721624-INS-99
chr10-102733018-INS-114
chr10-102739019-INS-52
chr10-102743392-INS-54
chr10-102745945-INS-119
chr10-102747683-INS-152
chr10-102748862-INS-84
chr10-102770231-INS-578
chr10-102786379-INS-509
chr10-102799139-INS-3216
chr10-102802595-INS-137
chr10-102846167-INS-52
chr10-102849253-INS-171
chr10-102868550-INS-56
chr10-102880155-INS-54
chr10-102889021-INS-312

chr10-102909530-INS-55
chr10-102912096-INS-580
chr10-102931633-INS-423
chr10-102942355-INS-60
chr10-102968502-INS-122
chr10-103001559-INS-90
chr10-103007720-INS-114
chr10-103027080-INS-53
chr10-103097670-INS-121
chr10-103111710-INS-138
chr10-103128687-INS-153
chr10-103144706-INS-684
chr10-103146226-INS-149
chr10-103173830-INS-66
chr10-103217089-INS-60
chr10-103256052-INS-90
chr10-103333277-INS-60
chr10-103358566-INS-7309
chr10-103382896-INS-7099
chr10-103383923-INS-1288
chr10-103387216-INS-73
chr10-103394289-INS-88
chr10-103404283-INS-252
chr10-103411301-INS-207
chr10-103444785-INS-388
chr10-103489381-INS-53
chr10-103522317-INS-199
chr10-103590009-INS-212
chr10-103615614-INS-622
chr10-103652828-INS-51
chr10-103675719-INS-244
chr10-103736898-INS-54
chr10-103784173-INS-384
chr10-103821757-INS-859
chr10-103835050-INS-67
chr10-103844550-INS-344
chr10-103846092-INS-289
chr10-103904170-INS-52
chr10-103938528-INS-929
chr10-103959208-INS-78
chr10-104047814-INS-52
chr10-104052362-INS-70
chr10-104054834-INS-80
chr10-104056189-INS-166
chr10-104129308-INS-51
chr10-104203930-INS-65
chr10-104228110-INS-56
chr10-104233805-INS-497
chr10-104235902-INS-6401
chr10-104245755-INS-337
chr10-104268084-INS-63
chr10-104351198-INS-551
chr10-104368044-INS-388
chr10-104382130-INS-86
chr10-104518455-INS-511
chr10-104548838-INS-52
chr10-104577469-INS-904
chr10-104600247-INS-73
chr10-104611529-INS-171
chr10-104619863-INS-134
chr10-104661132-INS-674
chr10-104662409-INS-800
chr10-104665048-INS-54
chr10-104669098-INS-594
chr10-104711837-INS-98
chr10-104714497-INS-57
chr10-104738566-INS-88
chr10-104748191-INS-52
chr10-104764416-INS-456
chr10-104766589-INS-63
chr10-104768758-INS-56
chr10-104783620-INS-701
chr10-104788312-INS-92
chr10-104801986-INS-52
chr10-104843626-INS-383
chr10-104844348-INS-51
chr10-104844416-INS-246

chr10-104869616-INS-887
chr10-104889208-INS-122
chr10-104890247-INS-300
chr10-104895227-INS-329
chr10-104911691-INS-722
chr10-104918385-INS-80
chr10-104975296-INS-711
chr10-104976576-INS-325
chr10-105018263-INS-112
chr10-105022166-INS-58
chr10-105027756-INS-875
chr10-105032047-INS-580
chr10-105052439-INS-741
chr10-105063713-INS-6233
chr10-105126265-INS-62
chr10-105134302-INS-793
chr10-105146824-INS-378
chr10-105149821-INS-66
chr10-105173389-INS-61
chr10-105181329-INS-790
chr10-105218072-INS-1545
chr10-105256426-INS-863
chr10-105256498-INS-50
chr10-105277606-INS-56
chr10-105292375-INS-151
chr10-105300982-INS-293
chr10-105343285-INS-386
chr10-105357567-INS-144
chr10-105366464-INS-86
chr10-105394634-INS-204
chr10-105396036-INS-55
chr10-105418421-INS-62
chr10-105446623-INS-331
chr10-105449376-INS-410
chr10-105464693-INS-80
chr10-105468143-INS-111
chr10-105491866-INS-154
chr10-105507964-INS-534
chr10-105538896-INS-73
chr10-105539895-INS-69
chr10-105544960-INS-6654
chr10-105562628-INS-3640
chr10-105567165-INS-416
chr10-105606268-INS-60
chr10-105607676-INS-108
chr10-105609923-INS-378
chr10-105614916-INS-714
chr10-105627175-INS-69
chr10-105633708-INS-222
chr10-105637739-INS-521
chr10-105651890-INS-103
chr10-105664858-INS-149
chr10-105684366-INS-74
chr10-105689962-INS-459
chr10-105702386-INS-72
chr10-105708090-INS-517
chr10-105775172-INS-305
chr10-105776146-INS-7283
chr10-105782237-INS-66
chr10-105793211-INS-358
chr10-105795625-INS-6797
chr10-105801303-INS-7349
chr10-105802068-INS-488
chr10-105812223-INS-58
chr10-105840614-INS-76
chr10-105867376-INS-400
chr10-105877968-INS-71
chr10-105895729-INS-7543
chr10-105895871-INS-127
chr10-105898209-INS-90
chr10-105908585-INS-7109
chr10-105908963-INS-83
chr10-105923695-INS-57
chr10-105931542-INS-54
chr10-105954769-INS-584
chr10-105966897-INS-153
chr10-105978234-INS-4999

chr10-105979116-INS-677
chr10-105995276-INS-504
chr10-106017950-INS-829
chr10-106034275-INS-648
chr10-106040894-INS-111
chr10-106042381-INS-848
chr10-106053378-INS-389
chr10-106062112-INS-446
chr10-106077035-INS-328
chr10-106093561-INS-140
chr10-106095212-INS-92
chr10-106096001-INS-7103
chr10-106099147-INS-1531
chr10-106116839-INS-60
chr10-106133016-INS-72
chr10-106140742-INS-263
chr10-106180760-INS-95
chr10-106197033-INS-919
chr10-106218598-INS-201
chr10-106278186-INS-207
chr10-106280621-INS-248
chr10-106281254-INS-52
chr10-106285336-INS-484
chr10-106287686-INS-7172
chr10-106288402-INS-7164
chr10-106291735-INS-92
chr10-106319828-INS-56
chr10-106335056-INS-243
chr10-106336621-INS-61
chr10-106405526-INS-4739
chr10-106419665-INS-149
chr10-106427380-INS-101
chr10-106478791-INS-57
chr10-106507218-INS-948
chr10-106510824-INS-196
chr10-106531656-INS-758
chr10-106535778-INS-1888
chr10-106548564-INS-109
chr10-106582937-INS-79
chr10-106585543-INS-183
chr10-106610819-INS-333
chr10-106611405-INS-838
chr10-106624285-INS-104
chr10-106639290-INS-62
chr10-106643142-INS-75
chr10-106647663-INS-311
chr10-106648527-INS-466
chr10-106652071-INS-474
chr10-106723008-INS-58
chr10-106742558-INS-106
chr10-106749026-INS-66
chr10-106813918-INS-390
chr10-106825834-INS-1605
chr10-106841943-INS-447
chr10-106861197-INS-53
chr10-106891363-INS-92
chr10-106892712-INS-899
chr10-106898033-INS-178
chr10-106906265-INS-52
chr10-106922749-INS-153
chr10-106946621-INS-215
chr10-106979322-INS-56
chr10-107036559-INS-112
chr10-107043762-INS-654
chr10-107053243-INS-648
chr10-107054667-INS-531
chr10-107057899-INS-62
chr10-107098338-INS-72
chr10-107122745-INS-59
chr10-107129561-INS-117
chr10-107171847-INS-54
chr10-107189762-INS-98
chr10-107263104-INS-56
chr10-107290146-INS-150
chr10-107411416-INS-97
chr10-107423848-INS-145
chr10-107426374-INS-60

chr10-107436287-INS-196
chr10-107490885-INS-80
chr10-107503694-INS-130
chr10-107515645-INS-61
chr10-107517017-INS-62
chr10-107530452-INS-302
chr10-107532155-INS-466
chr10-107555923-INS-60
chr10-107580849-INS-496
chr10-107587362-INS-183
chr10-107589383-INS-53
chr10-107592622-INS-756
chr10-107604581-INS-52
chr10-107610698-INS-402
chr10-107678579-INS-91
chr10-107700680-INS-72
chr10-107702944-INS-103
chr10-107712329-INS-65
chr10-107753991-INS-70
chr10-107787241-INS-593
chr10-107833795-INS-6385
chr10-107877402-INS-176
chr10-107900581-INS-323
chr10-107910566-INS-94
chr10-107911886-INS-79
chr10-107913556-INS-154
chr10-107950888-INS-152
chr10-107953907-INS-120
chr10-107960613-INS-819
chr10-107961666-INS-67
chr10-107975613-INS-6134
chr10-108033125-INS-76
chr10-108059064-INS-908
chr10-108067590-INS-113
chr10-108101972-INS-206
chr10-108116677-INS-52
chr10-108124609-INS-187
chr10-108125727-INS-205
chr10-108136746-INS-57
chr10-108148590-INS-86
chr10-108153022-INS-128
chr10-108154684-INS-94
chr10-108164700-INS-56
chr10-108172546-INS-187
chr10-108182407-INS-217
chr10-108185627-INS-196
chr10-108189479-INS-54
chr10-108197001-INS-86
chr10-108226812-INS-261
chr10-108310613-INS-383
chr10-108339079-INS-188
chr10-108349860-INS-6695
chr10-108364359-INS-131
chr10-108370654-INS-197
chr10-108428205-INS-786
chr10-108445270-INS-144
chr10-108453500-INS-65
chr10-108461599-INS-270
chr10-108476736-INS-69
chr10-108495086-INS-429
chr10-108517985-INS-96
chr10-108544879-INS-53
chr10-108547906-INS-7955
chr10-108556008-INS-170
chr10-108575444-INS-295
chr10-108586607-INS-674
chr10-108594100-INS-71
chr10-108598444-INS-74
chr10-108632443-INS-7092
chr10-108679889-INS-65
chr10-108685855-INS-230
chr10-108688168-INS-457
chr10-108690771-INS-499
chr10-108691406-INS-656
chr10-108693998-INS-328
chr10-108700837-INS-1860
chr10-108719986-INS-8076

chr10-108721162-INS-207
chr10-108726305-INS-84
chr10-108730326-INS-96
chr10-108740878-INS-82
chr10-108744206-INS-6715
chr10-108767725-INS-348
chr10-108776884-INS-251
chr10-108792222-INS-892
chr10-108850797-INS-56
chr10-108881756-INS-76
chr10-108897275-INS-402
chr10-108935775-INS-291
chr10-108954865-INS-137
chr10-109029035-INS-89
chr10-109069835-INS-5129
chr10-109086413-INS-55
chr10-109088849-INS-211
chr10-109104378-INS-369
chr10-109105671-INS-52
chr10-109140455-INS-55
chr10-109160105-INS-626
chr10-109162621-INS-51
chr10-109167442-INS-438
chr10-109176258-INS-325
chr10-109178912-INS-878
chr10-109200762-INS-65
chr10-109226727-INS-237
chr10-109251249-INS-299
chr10-109267982-INS-1191
chr10-109268524-INS-56
chr10-109298067-INS-139
chr10-109300709-INS-55
chr10-109309753-INS-204
chr10-109326225-INS-5539
chr10-109349399-INS-96
chr10-109354775-INS-461
chr10-109370559-INS-54
chr10-109379716-INS-986
chr10-109412753-INS-5309
chr10-109423943-INS-151
chr10-109511567-INS-87
chr10-109535247-INS-117
chr10-109552565-INS-201
chr10-109558859-INS-53
chr10-109562646-INS-390
chr10-109587026-INS-222
chr10-109614251-INS-56
chr10-109618721-INS-120
chr10-109622613-INS-125
chr10-109626055-INS-51
chr10-109634709-INS-132
chr10-109653642-INS-69
chr10-109658976-INS-205
chr10-109663659-INS-90
chr10-109667281-INS-64
chr10-109754104-INS-57
chr10-109773842-INS-6741
chr10-109784145-INS-55
chr10-109815398-INS-767
chr10-109817337-INS-94
chr10-109819855-INS-468
chr10-109826546-INS-203
chr10-109835220-INS-139
chr10-109837268-INS-59
chr10-109860145-INS-774
chr10-109876207-INS-385
chr10-109916328-INS-261
chr10-109919838-INS-71
chr10-109920646-INS-147
chr10-109949015-INS-100
chr10-109959300-INS-60
chr10-109985264-INS-257
chr10-110001529-INS-227
chr10-110022199-INS-125
chr10-110024656-INS-212
chr10-110029451-INS-196
chr10-110035739-INS-141

chr10-110048364-INS-65
chr10-110070488-INS-64
chr10-110081927-INS-54
chr10-110088522-INS-214
chr10-110094407-INS-51
chr10-110109053-INS-81
chr10-110109816-INS-142
chr10-110114036-INS-253
chr10-110117909-INS-175
chr10-110172099-INS-71
chr10-110193564-INS-80
chr10-110197967-INS-311
chr10-110224567-INS-79
chr10-110268609-INS-270
chr10-110270630-INS-80
chr10-110282185-INS-55
chr10-110287128-INS-201
chr10-110312795-INS-112
chr10-110328170-INS-74
chr10-110332112-INS-627
chr10-110352354-INS-197
chr10-110364879-INS-126
chr10-110385819-INS-284
chr10-110396481-INS-6868
chr10-110423966-INS-64
chr10-110439965-INS-55
chr10-110480036-INS-188
chr10-110489871-INS-124
chr10-110495009-INS-982
chr10-110502824-INS-6930
chr10-110508609-INS-65
chr10-110514543-INS-150
chr10-110531718-INS-129
chr10-110538862-INS-132
chr10-110541689-INS-58
chr10-110545076-INS-97
chr10-110562425-INS-63
chr10-110564292-INS-60
chr10-110572120-INS-407
chr10-110579456-INS-74
chr10-110594949-INS-440
chr10-110607148-INS-140
chr10-110637240-INS-748
chr10-110646264-INS-208
chr10-110647979-INS-468
chr10-110655317-INS-209
chr10-110665118-INS-111
chr10-110666961-INS-66
chr10-110670829-INS-116
chr10-110678711-INS-1011
chr10-110678730-INS-611
chr10-110703107-INS-103
chr10-110736738-INS-99
chr10-110738900-INS-400
chr10-110771848-INS-119
chr10-110773788-INS-304
chr10-110787968-INS-112
chr10-110791677-INS-203
chr10-110803108-INS-101
chr10-110831427-INS-197
chr10-110854249-INS-197
chr10-110880262-INS-416
chr10-110892876-INS-179
chr10-110896219-INS-7260
chr10-110904488-INS-278
chr10-110909944-INS-208
chr10-110912012-INS-60
chr10-110916474-INS-225
chr10-110953937-INS-56
chr10-110962808-INS-382
chr10-110967741-INS-5314
chr10-111023460-INS-67
chr10-111030579-INS-214
chr10-111032577-INS-6478
chr10-111040767-INS-801
chr10-111068479-INS-75
chr10-111083262-INS-215

chr10-111088476-INS-242
chr10-111105793-INS-213
chr10-111109131-INS-626
chr10-111110214-INS-205
chr10-111131285-INS-62
chr10-111141288-INS-97
chr10-111151679-INS-100
chr10-111170990-INS-68
chr10-111181347-INS-108
chr10-111191117-INS-77
chr10-111194387-INS-201
chr10-111213707-INS-132
chr10-111229162-INS-188
chr10-111242286-INS-4946
chr10-111268826-INS-222
chr10-111271263-INS-246
chr10-111278951-INS-52
chr10-111281756-INS-78
chr10-111283205-INS-210
chr10-111283844-INS-60
chr10-111287331-INS-91
chr10-111304625-INS-52
chr10-111363315-INS-81
chr10-111382101-INS-264
chr10-111383330-INS-5554
chr10-111390559-INS-243
chr10-111394937-INS-72
chr10-111398321-INS-176
chr10-111432190-INS-98
chr10-111433195-INS-5144
chr10-111437460-INS-140
chr10-111445694-INS-291
chr10-111456652-INS-102
chr10-111471251-INS-105
chr10-111481784-INS-124
chr10-111501012-INS-550
chr10-111504500-INS-75
chr10-111523183-INS-62
chr10-111550346-INS-190
chr10-111557688-INS-78
chr10-111594020-INS-704
chr10-111596836-INS-117
chr10-111620490-INS-5321
chr10-111627992-INS-6401
chr10-111637740-INS-59
chr10-111653814-INS-979
chr10-111655496-INS-143
chr10-111681489-INS-91
chr10-111688336-INS-107
chr10-111730797-INS-121
chr10-111738521-INS-309
chr10-111744467-INS-230
chr10-111766816-INS-431
chr10-111773466-INS-52
chr10-111775414-INS-98
chr10-111794660-INS-463
chr10-111799276-INS-199
chr10-111799469-INS-566
chr10-111801466-INS-978
chr10-111806346-INS-317
chr10-111814401-INS-160
chr10-111815533-INS-142
chr10-111821118-INS-164
chr10-111844272-INS-52
chr10-111859982-INS-86
chr10-111863748-INS-51
chr10-111884027-INS-64
chr10-111980691-INS-157
chr10-111996322-INS-911
chr10-112044161-INS-77
chr10-112046375-INS-78
chr10-112055440-INS-134
chr10-112060639-INS-153
chr10-112088670-INS-119
chr10-112109618-INS-1889
chr10-112147615-INS-453
chr10-112157378-INS-6362

chr10-112217862-INS-52
chr10-112231987-INS-152
chr10-112239400-INS-53
chr10-112252821-INS-519
chr10-112267253-INS-697
chr10-112279515-INS-163
chr10-112302402-INS-122
chr10-112318585-INS-76
chr10-112322031-INS-94
chr10-112328425-INS-144
chr10-112367927-INS-71
chr10-112368268-INS-76
chr10-112394514-INS-399
chr10-112419615-INS-148
chr10-112442798-INS-4458
chr10-112443254-INS-619
chr10-112449252-INS-216
chr10-112453126-INS-749
chr10-112460316-INS-86
chr10-112467134-INS-55
chr10-112481986-INS-4207
chr10-112484904-INS-1459
chr10-112498762-INS-60
chr10-112503516-INS-861
chr10-112508278-INS-66
chr10-112512735-INS-629
chr10-112517376-INS-716
chr10-112525705-INS-329
chr10-112530850-INS-171
chr10-112545401-INS-128
chr10-112576150-INS-280
chr10-112586728-INS-145
chr10-112595075-INS-1090
chr10-112595535-INS-105
chr10-112598742-INS-174
chr10-112599410-INS-56
chr10-112599669-INS-701
chr10-112606097-INS-105
chr10-112615826-INS-6382
chr10-112643211-INS-188
chr10-112658995-INS-153
chr10-112676320-INS-864
chr10-112698839-INS-56
chr10-112718942-INS-2633
chr10-112748288-INS-174
chr10-112751746-INS-55
chr10-112759516-INS-208
chr10-112767243-INS-461
chr10-112770115-INS-80
chr10-112775754-INS-864
chr10-112781615-INS-201
chr10-112788989-INS-107
chr10-112791319-INS-629
chr10-112794036-INS-170
chr10-112814479-INS-6342
chr10-112835941-INS-128
chr10-112863311-INS-401
chr10-112892036-INS-102
chr10-112911844-INS-105
chr10-112920564-INS-100
chr10-112924688-INS-347
chr10-112925898-INS-310
chr10-112929539-INS-84
chr10-112930916-INS-1146
chr10-112955922-INS-341
chr10-112972477-INS-115
chr10-112979671-INS-52
chr10-112982587-INS-62
chr10-112984556-INS-537
chr10-113014122-INS-126
chr10-113032732-INS-72
chr10-113044834-INS-3554
chr10-113055461-INS-72
chr10-113060543-INS-57
chr10-113062625-INS-54
chr10-113075095-INS-116
chr10-113078423-INS-118

chr10-113094573-INS-347
chr10-113098597-INS-142
chr10-113102845-INS-749
chr10-113104043-INS-77
chr10-113109437-INS-72
chr10-113110622-INS-517
chr10-113113134-INS-965
chr10-113126216-INS-6401
chr10-113129013-INS-64
chr10-113181972-INS-120
chr10-113186352-INS-148
chr10-113202180-INS-205
chr10-113209855-INS-251
chr10-113252646-INS-503
chr10-113259494-INS-543
chr10-113265686-INS-205
chr10-113276326-INS-52
chr10-113288813-INS-397
chr10-113301193-INS-78
chr10-113311690-INS-94
chr10-113314344-INS-52
chr10-113320540-INS-976
chr10-113330827-INS-135
chr10-113377757-INS-180
chr10-113386682-INS-401
chr10-113403970-INS-98
chr10-113407837-INS-559
chr10-113429595-INS-844
chr10-113433660-INS-493
chr10-113443904-INS-468
chr10-113468373-INS-51
chr10-113470156-INS-156
chr10-113476411-INS-226
chr10-113477741-INS-175
chr10-113491893-INS-64
chr10-113498078-INS-62
chr10-113507737-INS-66
chr10-113533131-INS-102
chr10-113536027-INS-77
chr10-113551567-INS-326
chr10-113566964-INS-696
chr10-113577141-INS-68
chr10-113594195-INS-74
chr10-113598316-INS-740
chr10-113608927-INS-55
chr10-113632951-INS-68
chr10-113634878-INS-3383
chr10-113650712-INS-854
chr10-113661705-INS-166
chr10-113682612-INS-55
chr10-113689098-INS-411
chr10-113691443-INS-60
chr10-113709749-INS-52
chr10-113756250-INS-60
chr10-113758080-INS-70
chr10-113765583-INS-103
chr10-113767241-INS-215
chr10-113803318-INS-345
chr10-113817500-INS-122
chr10-113828751-INS-471
chr10-113870726-INS-512
chr10-113936245-INS-82
chr10-113944292-INS-156
chr10-113949819-INS-302
chr10-113963383-INS-640
chr10-113978667-INS-94
chr10-114022578-INS-268
chr10-114025970-INS-55
chr10-114028094-INS-67
chr10-114035474-INS-131
chr10-114043990-INS-110
chr10-114063882-INS-436
chr10-114068284-INS-107
chr10-114103691-INS-64
chr10-114118010-INS-555
chr10-114125118-INS-522
chr10-114149452-INS-128

chr10-114153750-INS-214
chr10-114153816-INS-4250
chr10-114159698-INS-397
chr10-114196164-INS-528
chr10-114220469-INS-53
chr10-114239597-INS-423
chr10-114263915-INS-206
chr10-114265007-INS-54
chr10-114281424-INS-83
chr10-114289455-INS-54
chr10-114291574-INS-360
chr10-114300351-INS-89
chr10-114311710-INS-62
chr10-114313184-INS-176
chr10-114329357-INS-63
chr10-114350320-INS-57
chr10-114392971-INS-171
chr10-114395004-INS-611
chr10-114397236-INS-75
chr10-114425715-INS-72
chr10-114472513-INS-54
chr10-114474952-INS-60
chr10-114528020-INS-55
chr10-114532448-INS-62
chr10-114563111-INS-251
chr10-114563973-INS-99
chr10-114582678-INS-55
chr10-114586843-INS-179
chr10-114599466-INS-57
chr10-114603730-INS-591
chr10-114619855-INS-378
chr10-114651788-INS-189
chr10-114682056-INS-61
chr10-114717087-INS-130
chr10-114720359-INS-87
chr10-114784218-INS-137
chr10-114800481-INS-365
chr10-114830107-INS-53
chr10-114854676-INS-330
chr10-114863622-INS-58
chr10-114871496-INS-54
chr10-114880540-INS-110
chr10-114886472-INS-54
chr10-114887786-INS-7445
chr10-114892019-INS-57
chr10-114901732-INS-56
chr10-114940146-INS-740
chr10-114961332-INS-78
chr10-114976225-INS-51
chr10-114983614-INS-55
chr10-114988446-INS-693
chr10-114990824-INS-397
chr10-115004579-INS-80
chr10-115022016-INS-212
chr10-115028474-INS-78
chr10-115029849-INS-134
chr10-115042465-INS-583
chr10-115060669-INS-329
chr10-115079424-INS-685
chr10-115095033-INS-399
chr10-115098762-INS-319
chr10-115102018-INS-66
chr10-115112934-INS-126
chr10-115123744-INS-56
chr10-115163677-INS-253
chr10-115166901-INS-208
chr10-115171739-INS-210
chr10-115185474-INS-200
chr10-115227546-INS-190
chr10-115233458-INS-129
chr10-115237028-INS-91
chr10-115256181-INS-916
chr10-115271525-INS-172
chr10-115296159-INS-305
chr10-115319013-INS-104
chr10-115333740-INS-483
chr10-115336983-INS-169

chr10-115343702-INS-399
chr10-115355161-INS-70
chr10-115368057-INS-99
chr10-115381228-INS-121
chr10-115385021-INS-56
chr10-115391910-INS-82
chr10-115408833-INS-170
chr10-115432656-INS-7117
chr10-115456633-INS-358
chr10-115466075-INS-135
chr10-115484532-INS-84
chr10-115497712-INS-247
chr10-115539186-INS-69
chr10-115572768-INS-150
chr10-115583410-INS-822
chr10-115588872-INS-457
chr10-115607375-INS-223
chr10-115616678-INS-191
chr10-115632304-INS-86
chr10-115649711-INS-104
chr10-115674393-INS-75
chr10-115692855-INS-339
chr10-115696775-INS-54
chr10-115700581-INS-161
chr10-115703409-INS-167
chr10-115712690-INS-222
chr10-115718310-INS-60
chr10-115749865-INS-52
chr10-115758758-INS-66
chr10-115773432-INS-75
chr10-115804143-INS-94
chr10-115807451-INS-175
chr10-115813555-INS-52
chr10-115836702-INS-64
chr10-115847327-INS-68
chr10-115853042-INS-60
chr10-115859848-INS-55
chr10-115878831-INS-165
chr10-115882107-INS-58
chr10-115885874-INS-470
chr10-115896944-INS-72
chr10-115901208-INS-106
chr10-115926042-INS-149
chr10-115953746-INS-54
chr10-115971341-INS-75
chr10-115981250-INS-638
chr10-116009694-INS-82
chr10-116031389-INS-533
chr10-116045594-INS-506
chr10-116047243-INS-163
chr10-116078960-INS-161
chr10-116089121-INS-158
chr10-116094198-INS-386
chr10-116117363-INS-66
chr10-116127015-INS-195
chr10-116142888-INS-74
chr10-116148026-INS-70
chr10-116160235-INS-310
chr10-116251609-INS-359
chr10-116299332-INS-68
chr10-116316508-INS-533
chr10-116325136-INS-113
chr10-116360957-INS-208
chr10-116384434-INS-210
chr10-116414947-INS-101
chr10-116425457-INS-66
chr10-116453060-INS-81
chr10-116454062-INS-351
chr10-116469892-INS-52
chr10-116480784-INS-398
chr10-116484424-INS-128
chr10-116485776-INS-175
chr10-116497530-INS-204
chr10-116516829-INS-190
chr10-116523991-INS-942
chr10-116527077-INS-102
chr10-116534053-INS-112

chr10-116538576-INS-53
chr10-116548247-INS-80
chr10-116565962-INS-53
chr10-116570863-INS-635
chr10-116572584-INS-52
chr10-116576374-INS-63
chr10-116588120-INS-668
chr10-116606222-INS-84
chr10-116611243-INS-832
chr10-116619248-INS-336
chr10-116623758-INS-74
chr10-116625478-INS-191
chr10-116635859-INS-105
chr10-116637639-INS-76
chr10-116647737-INS-162
chr10-116650630-INS-81
chr10-116654649-INS-216
chr10-116669431-INS-171
chr10-116691472-INS-616
chr10-116697027-INS-769
chr10-116699655-INS-97
chr10-116702752-INS-120
chr10-116712668-INS-70
chr10-116714504-INS-114
chr10-116714889-INS-142
chr10-116716543-INS-56
chr10-116722960-INS-3092
chr10-116725544-INS-159
chr10-116738937-INS-52
chr10-116743159-INS-80
chr10-116760000-INS-62
chr10-116777406-INS-160
chr10-116801785-INS-70
chr10-116806995-INS-184
chr10-116816036-INS-211
chr10-116832626-INS-174
chr10-116841083-INS-98
chr10-116846105-INS-118
chr10-116853510-INS-158
chr10-116863947-INS-90
chr10-116876689-INS-66
chr10-116877615-INS-74
chr10-116885935-INS-212
chr10-116892659-INS-90
chr10-116894161-INS-216
chr10-116895398-INS-96
chr10-116897927-INS-231
chr10-116921211-INS-315
chr10-116933851-INS-1202
chr10-116955120-INS-207
chr10-116961138-INS-197
chr10-116971120-INS-54
chr10-116991053-INS-52
chr10-116996883-INS-190
chr10-117000172-INS-184
chr10-117014631-INS-197
chr10-117032568-INS-224
chr10-117037363-INS-112
chr10-117045767-INS-102
chr10-117049475-INS-80
chr10-117050568-INS-63
chr10-117068378-INS-288
chr10-117071710-INS-238
chr10-117081426-INS-58
chr10-117129153-INS-738
chr10-117134387-INS-51
chr10-117137332-INS-62
chr10-117164804-INS-84
chr10-117205654-INS-56
chr10-117213825-INS-5086
chr10-117221376-INS-693
chr10-117235293-INS-85
chr10-117240025-INS-61
chr10-117246511-INS-136
chr10-117260673-INS-221
chr10-117272640-INS-56
chr10-117276848-INS-344

chr10-117284148-INS-64
chr10-117289010-INS-63
chr10-117305087-INS-458
chr10-117315394-INS-271
chr10-117332861-INS-219
chr10-117347872-INS-60
chr10-117361288-INS-138
chr10-117377260-INS-175
chr10-117428617-INS-97
chr10-117446858-INS-210
chr10-117453370-INS-330
chr10-117470276-INS-208
chr10-117472922-INS-168
chr10-117485059-INS-202
chr10-117490865-INS-172
chr10-117492960-INS-80
chr10-117496863-INS-156
chr10-117498187-INS-175
chr10-117518720-INS-60
chr10-117524173-INS-5059
chr10-117529224-INS-54
chr10-117535107-INS-75
chr10-117553290-INS-51
chr10-117558489-INS-135
chr10-117560090-INS-194
chr10-117563202-INS-102
chr10-117589355-INS-177
chr10-117599401-INS-448
chr10-117605007-INS-3498
chr10-117605162-INS-2406
chr10-117605224-INS-280
chr10-117605284-INS-2694
chr10-117616406-INS-198
chr10-117617833-INS-59
chr10-117647047-INS-156
chr10-117691084-INS-86
chr10-117697824-INS-192
chr10-117699905-INS-322
chr10-117705483-INS-176
chr10-117723809-INS-165
chr10-117725446-INS-72
chr10-117727927-INS-80
chr10-117728741-INS-51
chr10-117736538-INS-178
chr10-117740306-INS-152
chr10-117747441-INS-406
chr10-117748288-INS-567
chr10-117761346-INS-354
chr10-117771670-INS-172
chr10-117782361-INS-52
chr10-117785655-INS-171
chr10-117794697-INS-429
chr10-117848187-INS-96
chr10-117851333-INS-56
chr10-117852394-INS-129
chr10-117894834-INS-72
chr10-117923072-INS-161
chr10-117989296-INS-136
chr10-118012103-INS-575
chr10-118026615-INS-6228
chr10-118033931-INS-276
chr10-118048055-INS-135
chr10-118088005-INS-173
chr10-118130999-INS-682
chr10-118143262-INS-55
chr10-118205668-INS-57
chr10-118209348-INS-67
chr10-118212822-INS-76
chr10-118227632-INS-74
chr10-118237232-INS-144
chr10-118251216-INS-73
chr10-118254917-INS-56
chr10-118256450-INS-122
chr10-118257524-INS-431
chr10-118272562-INS-52
chr10-118274291-INS-70
chr10-118288667-INS-176

chr10-118293076-INS-154
chr10-118297312-INS-1401
chr10-118297546-INS-59
chr10-118302223-INS-90
chr10-118303412-INS-7239
chr10-118306018-INS-56
chr10-118314260-INS-330
chr10-118322038-INS-2798
chr10-118322298-INS-494
chr10-118329972-INS-5031
chr10-118333173-INS-142
chr10-118345226-INS-116
chr10-118350412-INS-65
chr10-118421333-INS-389
chr10-118429296-INS-52
chr10-118447468-INS-126
chr10-118461423-INS-106
chr10-118486951-INS-66
chr10-118493564-INS-88
chr10-118495217-INS-159
chr10-118497958-INS-51
chr10-118502743-INS-145
chr10-118534743-INS-485
chr10-118552950-INS-60
chr10-118571973-INS-86
chr10-118574178-INS-206
chr10-118615182-INS-629
chr10-118648061-INS-96
chr10-118649763-INS-99
chr10-118651022-INS-63
chr10-118654972-INS-350
chr10-118669504-INS-70
chr10-118710567-INS-141
chr10-118734124-INS-798
chr10-118742154-INS-62
chr10-118743601-INS-78
chr10-118745160-INS-74
chr10-118814824-INS-57
chr10-118821825-INS-124
chr10-118828093-INS-50
chr10-118843187-INS-131
chr10-118886762-INS-66
chr10-118915827-INS-74
chr10-118919192-INS-84
chr10-118940409-INS-141
chr10-118942054-INS-56
chr10-118975163-INS-70
chr10-118991053-INS-125
chr10-118997745-INS-54
chr10-119001689-INS-65
chr10-119029756-INS-674
chr10-119035086-INS-938
chr10-119055820-INS-117
chr10-119081487-INS-215
chr10-119087389-INS-80
chr10-119099969-INS-99
chr10-119102723-INS-63
chr10-119103949-INS-66
chr10-119107028-INS-399
chr10-119144647-INS-890
chr10-119171751-INS-482
chr10-119226898-INS-68
chr10-119229528-INS-66
chr10-119253277-INS-222
chr10-119258769-INS-59
chr10-119259025-INS-338
chr10-119262981-INS-65
chr10-119264702-INS-401
chr10-119265275-INS-196
chr10-119271706-INS-722
chr10-119282479-INS-75
chr10-119318348-INS-68
chr10-119325607-INS-54
chr10-119334880-INS-66
chr10-119336443-INS-317
chr10-119340773-INS-55
chr10-119343475-INS-155

chr10-119380149-INS-387
chr10-119431362-INS-85
chr10-119465954-INS-730
chr10-119485713-INS-800
chr10-119488728-INS-82
chr10-119505830-INS-221
chr10-119515657-INS-304
chr10-119567712-INS-111
chr10-119576366-INS-72
chr10-119588775-INS-473
chr10-119591372-INS-116
chr10-119611166-INS-81
chr10-119628778-INS-6752
chr10-119658832-INS-134
chr10-119662336-INS-144
chr10-119672614-INS-221
chr10-119681454-INS-74
chr10-119696281-INS-95
chr10-119713744-INS-68
chr10-119722749-INS-150
chr10-119742587-INS-116
chr10-119746931-INS-7142
chr10-119795470-INS-481
chr10-119806377-INS-119
chr10-119822545-INS-137
chr10-119847796-INS-93
chr10-119877403-INS-399
chr10-119884920-INS-510
chr10-119914930-INS-134
chr10-119923702-INS-85
chr10-119934590-INS-124
chr10-119956806-INS-7128
chr10-119959902-INS-60
chr10-119971428-INS-53
chr10-119978598-INS-61
chr10-119980336-INS-202
chr10-119989276-INS-5671
chr10-119989758-INS-221
chr10-120011528-INS-81
chr10-120036248-INS-57
chr10-120041741-INS-192
chr10-120047577-INS-744
chr10-120051622-INS-166
chr10-120052721-INS-60
chr10-120053590-INS-71
chr10-120058525-INS-252
chr10-120058954-INS-213
chr10-120063677-INS-341
chr10-120074601-INS-70
chr10-120109867-INS-221
chr10-120113917-INS-52
chr10-120129091-INS-118
chr10-120135616-INS-57
chr10-120144055-INS-111
chr10-120146235-INS-118
chr10-120148952-INS-64
chr10-120150355-INS-146
chr10-120169804-INS-357
chr10-120170932-INS-80
chr10-120201654-INS-250
chr10-120234205-INS-51
chr10-120237278-INS-203
chr10-120254914-INS-168
chr10-120256377-INS-152
chr10-120283648-INS-128
chr10-120285441-INS-61
chr10-120291929-INS-205
chr10-120300225-INS-217
chr10-120326702-INS-118
chr10-120330422-INS-60
chr10-120350311-INS-320
chr10-120351334-INS-87
chr10-120390091-INS-62
chr10-120392489-INS-503
chr10-120397471-INS-227
chr10-120402821-INS-61
chr10-120404795-INS-272

chr10-120406182-INS-120
chr10-120411241-INS-541
chr10-120417023-INS-204
chr10-120417806-INS-155
chr10-120429250-INS-82
chr10-120462249-INS-161
chr10-120468190-INS-113
chr10-120480220-INS-112
chr10-120483388-INS-108
chr10-120502006-INS-273
chr10-120521521-INS-95
chr10-120526765-INS-120
chr10-120535993-INS-206
chr10-120559680-INS-852
chr10-120590186-INS-84
chr10-120609625-INS-140
chr10-120638579-INS-64
chr10-120641227-INS-58
chr10-120656227-INS-66
chr10-120674161-INS-93
chr10-120683376-INS-324
chr10-120687735-INS-258
chr10-120701691-INS-283
chr10-120714981-INS-117
chr10-120728049-INS-52
chr10-120729019-INS-88
chr10-120731000-INS-5671
chr10-120733214-INS-154
chr10-120739938-INS-53
chr10-120744296-INS-227
chr10-120751155-INS-97
chr10-120784459-INS-206
chr10-120801616-INS-180
chr10-120804355-INS-351
chr10-120805763-INS-205
chr10-120811428-INS-442
chr10-120832222-INS-1092
chr10-120832416-INS-809
chr10-120836318-INS-143
chr10-120837426-INS-57
chr10-120869075-INS-67
chr10-120872506-INS-80
chr10-120881002-INS-60
chr10-120893824-INS-156
chr10-120901110-INS-252
chr10-120934760-INS-62
chr10-120964813-INS-529
chr10-120971765-INS-652
chr10-120982210-INS-94
chr10-120984072-INS-143
chr10-120998486-INS-750
chr10-121040609-INS-79
chr10-121085604-INS-68
chr10-121114806-INS-142
chr10-121120908-INS-409
chr10-121190096-INS-220
chr10-121193837-INS-4784
chr10-121208488-INS-195
chr10-121209337-INS-174
chr10-121218007-INS-250
chr10-121286064-INS-289
chr10-121321154-INS-116
chr10-121326695-INS-205
chr10-121354657-INS-137
chr10-121355556-INS-91
chr10-121356696-INS-139
chr10-121358367-INS-196
chr10-121368938-INS-199
chr10-121375246-INS-606
chr10-121398480-INS-61
chr10-121398920-INS-186
chr10-121403056-INS-8326
chr10-121415182-INS-166
chr10-121439717-INS-282
chr10-121448866-INS-144
chr10-121467701-INS-64
chr10-121482929-INS-475

chr10-121484522-INS-59
chr10-121491102-INS-130
chr10-121504134-INS-401
chr10-121532092-INS-204
chr10-121553239-INS-130
chr10-121554104-INS-139
chr10-121565380-INS-221
chr10-121579133-INS-217
chr10-121594806-INS-623
chr10-121610476-INS-64
chr10-121656349-INS-56
chr10-121665710-INS-203
chr10-121674354-INS-205
chr10-121704408-INS-114
chr10-121713753-INS-174
chr10-121733244-INS-494
chr10-121737896-INS-653
chr10-121749987-INS-359
chr10-121760224-INS-261
chr10-121761691-INS-216
chr10-121771657-INS-51
chr10-121802816-INS-60
chr10-121812377-INS-163
chr10-121825265-INS-100
chr10-121830228-INS-965
chr10-121838403-INS-53
chr10-121838433-INS-1425
chr10-121868593-INS-65
chr10-121889357-INS-64
chr10-121909846-INS-899
chr10-121928828-INS-54
chr10-121933364-INS-64
chr10-121941393-INS-58
chr10-121963167-INS-122
chr10-121980352-INS-400
chr10-121981487-INS-141
chr10-121983492-INS-267
chr10-122028337-INS-98
chr10-122080725-INS-77
chr10-122098464-INS-64
chr10-122103931-INS-51
chr10-122124370-INS-834
chr10-122125410-INS-58
chr10-122147425-INS-170
chr10-122155439-INS-79
chr10-122160003-INS-56
chr10-122170750-INS-6888
chr10-122226832-INS-872
chr10-122235827-INS-60
chr10-122236180-INS-687
chr10-122237572-INS-79
chr10-122241442-INS-65
chr10-122269281-INS-56
chr10-122279229-INS-747
chr10-122280385-INS-202
chr10-122304883-INS-51
chr10-122322870-INS-75
chr10-122324753-INS-208
chr10-122336166-INS-6741
chr10-122363668-INS-119
chr10-122368466-INS-69
chr10-122373747-INS-53
chr10-122382488-INS-60
chr10-122395299-INS-509
chr10-122424480-INS-77
chr10-122442681-INS-677
chr10-122447713-INS-109
chr10-122467588-INS-201
chr10-122492265-INS-2701
chr10-122494609-INS-80
chr10-122504068-INS-444
chr10-122531646-INS-58
chr10-122533429-INS-84
chr10-122535148-INS-190
chr10-122549072-INS-54
chr10-122568164-INS-104
chr10-122577504-INS-66

chr10-122586854-INS-113
chr10-122599444-INS-66
chr10-122608003-INS-360
chr10-122612400-INS-63
chr10-122620497-INS-52
chr10-122621990-INS-354
chr10-122627657-INS-84
chr10-122629644-INS-57
chr10-122632258-INS-62
chr10-122663329-INS-76
chr10-122688185-INS-60
chr10-122689438-INS-70
chr10-122719013-INS-100
chr10-122757661-INS-201
chr10-122766017-INS-96
chr10-122771478-INS-360
chr10-122773166-INS-928
chr10-122788833-INS-84
chr10-122798966-INS-82
chr10-122803122-INS-4825
chr10-122825459-INS-279
chr10-122828785-INS-216
chr10-122855796-INS-71
chr10-122862653-INS-204
chr10-122864744-INS-218
chr10-122897728-INS-202
chr10-122910341-INS-436
chr10-122914135-INS-51
chr10-122916377-INS-466
chr10-122937376-INS-52
chr10-122940260-INS-296
chr10-122954798-INS-72
chr10-122971525-INS-90
chr10-123054461-INS-63
chr10-123058543-INS-748
chr10-123068359-INS-122
chr10-123069623-INS-310
chr10-123078042-INS-506
chr10-123079943-INS-416
chr10-123087208-INS-63
chr10-123132280-INS-96
chr10-123137512-INS-61
chr10-123158717-INS-56
chr10-123170697-INS-95
chr10-123179658-INS-137
chr10-123180728-INS-144
chr10-123207683-INS-407
chr10-123229104-INS-303
chr10-123240090-INS-73
chr10-123247755-INS-461
chr10-123255958-INS-92
chr10-123281843-INS-4037
chr10-123288392-INS-5372
chr10-123288868-INS-76
chr10-123298229-INS-929
chr10-123312828-INS-88
chr10-123324263-INS-167
chr10-123372252-INS-1014
chr10-123385028-INS-327
chr10-123420654-INS-56
chr10-123426107-INS-190
chr10-123428796-INS-88
chr10-123433310-INS-106
chr10-123445900-INS-352
chr10-123464630-INS-69
chr10-123465095-INS-1847
chr10-123501498-INS-228
chr10-123504446-INS-63
chr10-123511735-INS-110
chr10-123552699-INS-80
chr10-123554486-INS-56
chr10-123567107-INS-123
chr10-123601925-INS-60
chr10-123611561-INS-54
chr10-123628260-INS-98
chr10-123645779-INS-100
chr10-123683204-INS-265

chr10-123725177-INS-86
chr10-123750968-INS-392
chr10-123808967-INS-133
chr10-123812035-INS-72
chr10-123826157-INS-6564
chr10-123843819-INS-245
chr10-123859222-INS-99
chr10-123874008-INS-67
chr10-123874730-INS-412
chr10-123880183-INS-73
chr10-123905021-INS-104
chr10-123907879-INS-52
chr10-123911427-INS-145
chr10-123936571-INS-91
chr10-123939219-INS-606
chr10-123953207-INS-5313
chr10-123957745-INS-85
chr10-123959218-INS-199
chr10-123960534-INS-128
chr10-123971134-INS-50
chr10-123991051-INS-129
chr10-124007074-INS-224
chr10-124016986-INS-643
chr10-124023030-INS-66
chr10-124043662-INS-108
chr10-124073673-INS-326
chr10-124123723-INS-66
chr10-124128768-INS-129
chr10-124130757-INS-54
chr10-124170226-INS-153
chr10-124175559-INS-81
chr10-124203384-INS-184
chr10-124209544-INS-340
chr10-124209821-INS-6223
chr10-124222978-INS-196
chr10-124225098-INS-84
chr10-124243956-INS-116
chr10-124256368-INS-7125
chr10-124258871-INS-3020
chr10-124284781-INS-104
chr10-124285848-INS-218
chr10-124314824-INS-730
chr10-124320833-INS-104
chr10-124321710-INS-971
chr10-124326350-INS-68
chr10-124327720-INS-72
chr10-124328025-INS-916
chr10-124365749-INS-52
chr10-124375242-INS-252
chr10-124424700-INS-94
chr10-124452710-INS-77
chr10-124468165-INS-78
chr10-124481227-INS-5174
chr10-124512275-INS-1700
chr10-124524961-INS-6431
chr10-124563085-INS-185
chr10-124617611-INS-92
chr10-124634887-INS-394
chr10-124643790-INS-67
chr10-124658049-INS-175
chr10-124658666-INS-54
chr10-124703335-INS-357
chr10-124705411-INS-61
chr10-124711534-INS-84
chr10-124712314-INS-529
chr10-124718583-INS-74
chr10-124728551-INS-64
chr10-124754729-INS-53
chr10-124760216-INS-98
chr10-124772786-INS-96
chr10-124809496-INS-107
chr10-124809505-INS-604
chr10-124858168-INS-85
chr10-124866348-INS-411
chr10-124868456-INS-365
chr10-124902843-INS-77
chr10-124904113-INS-405

chr10-124906010-INS-638
chr10-124918526-INS-61
chr10-124959615-INS-329
chr10-124962373-INS-92
chr10-124963589-INS-476
chr10-124967084-INS-121
chr10-124997536-INS-829
chr10-124997537-INS-220
chr10-125006259-INS-78
chr10-125012783-INS-241
chr10-125035103-INS-810
chr10-125062208-INS-66
chr10-125078886-INS-753
chr10-125093739-INS-221
chr10-125110571-INS-359
chr10-125114674-INS-425
chr10-125116683-INS-318
chr10-125157035-INS-427
chr10-125172287-INS-419
chr10-125177575-INS-866
chr10-125182479-INS-357
chr10-125222824-INS-52
chr10-125223332-INS-57
chr10-125230814-INS-597
chr10-125282619-INS-56
chr10-125315773-INS-226
chr10-125318374-INS-55
chr10-125330179-INS-65
chr10-125354352-INS-6791
chr10-125359988-INS-140
chr10-125378254-INS-52
chr10-125388949-INS-189
chr10-125406971-INS-156
chr10-125433718-INS-950
chr10-125434626-INS-301
chr10-125435737-INS-277
chr10-125442681-INS-168
chr10-125444444-INS-332
chr10-125444929-INS-708
chr10-125449644-INS-146
chr10-125456852-INS-76
chr10-125468097-INS-144
chr10-125471783-INS-99
chr10-125480065-INS-70
chr10-125498883-INS-98
chr10-125499710-INS-219
chr10-125513613-INS-120
chr10-125528456-INS-124
chr10-125535524-INS-66
chr10-125537560-INS-121
chr10-125567440-INS-658
chr10-125571570-INS-380
chr10-125580298-INS-89
chr10-125594742-INS-255
chr10-125628712-INS-204
chr10-125640798-INS-343
chr10-125645145-INS-66
chr10-125666874-INS-55
chr10-125670996-INS-66
chr10-125680695-INS-114
chr10-125686185-INS-52
chr10-125696090-INS-506
chr10-125701256-INS-59
chr10-125716714-INS-66
chr10-125739745-INS-65
chr10-125755604-INS-360
chr10-125773092-INS-53
chr10-125777007-INS-86
chr10-125812087-INS-309
chr10-125813623-INS-66
chr10-125815493-INS-77
chr10-125820202-INS-58
chr10-125825473-INS-6454
chr10-125834273-INS-53
chr10-125837490-INS-70
chr10-125844236-INS-147
chr10-125893590-INS-213

chr10-125901137-INS-116
chr10-125902170-INS-6868
chr10-125912883-INS-114
chr10-125940566-INS-93
chr10-125941916-INS-72
chr10-125946911-INS-1777
chr10-125971538-INS-180
chr10-125976321-INS-567
chr10-125981755-INS-99
chr10-126006456-INS-173
chr10-126020923-INS-66
chr10-126024843-INS-6419
chr10-126035332-INS-178
chr10-126045881-INS-468
chr10-126069807-INS-201
chr10-126084634-INS-333
chr10-126104861-INS-209
chr10-126115803-INS-66
chr10-126121460-INS-537
chr10-126160418-INS-54
chr10-126189727-INS-490
chr10-126200807-INS-94
chr10-126206660-INS-258
chr10-126214136-INS-331
chr10-126232062-INS-96
chr10-126232983-INS-292
chr10-126265167-INS-62
chr10-126268369-INS-86
chr10-126300169-INS-59
chr10-126308151-INS-102
chr10-126316979-INS-328
chr10-126333486-INS-119
chr10-126343931-INS-68
chr10-126350151-INS-66
chr10-126352770-INS-209
chr10-126364105-INS-188
chr10-126385516-INS-72
chr10-126386011-INS-108
chr10-126395912-INS-119
chr10-126412597-INS-64
chr10-126428077-INS-352
chr10-126447180-INS-85
chr10-126450718-INS-68
chr10-126476988-INS-162
chr10-126495231-INS-138
chr10-126499244-INS-96
chr10-126501075-INS-208
chr10-126503673-INS-430
chr10-126549948-INS-139
chr10-126559605-INS-341
chr10-126581400-INS-97
chr10-126613297-INS-7034
chr10-126616787-INS-77
chr10-126616975-INS-63
chr10-126626426-INS-163
chr10-126632869-INS-171
chr10-126653191-INS-155
chr10-126678699-INS-76
chr10-126697389-INS-372
chr10-126707394-INS-108
chr10-126708530-INS-183
chr10-126744732-INS-159
chr10-126752773-INS-83
chr10-126757678-INS-137
chr10-126761263-INS-438
chr10-126765154-INS-65
chr10-126776066-INS-60
chr10-126804195-INS-63
chr10-126819646-INS-324
chr10-126841300-INS-210
chr10-126862395-INS-198
chr10-126871758-INS-208
chr10-126937051-INS-6485
chr10-126948773-INS-69
chr10-126964486-INS-507
chr10-127044099-INS-216
chr10-127066169-INS-92

chr10-127068380-INS-69
chr10-127070502-INS-460
chr10-127078120-INS-186
chr10-127089330-INS-189
chr10-127096254-INS-64
chr10-127099720-INS-85
chr10-127107522-INS-141
chr10-127109648-INS-96
chr10-127127790-INS-207
chr10-127135644-INS-104
chr10-127156518-INS-211
chr10-127164556-INS-204
chr10-127182969-INS-1933
chr10-127194508-INS-575
chr10-127199916-INS-205
chr10-127213063-INS-193
chr10-127214122-INS-53
chr10-127239664-INS-81
chr10-127242064-INS-545
chr10-127254320-INS-117
chr10-127259293-INS-110
chr10-127291876-INS-659
chr10-127329246-INS-164
chr10-127398389-INS-92
chr10-127400850-INS-256
chr10-127403484-INS-196
chr10-127446523-INS-207
chr10-127461218-INS-62
chr10-127466517-INS-241
chr10-127471495-INS-406
chr10-127472933-INS-56
chr10-127481939-INS-84
chr10-127485318-INS-106
chr10-127516483-INS-203
chr10-127520399-INS-58
chr10-127521468-INS-199
chr10-127537180-INS-267
chr10-127552301-INS-224
chr10-127588291-INS-6425
chr10-127602382-INS-211
chr10-127619307-INS-235
chr10-127642797-INS-311
chr10-127649579-INS-469
chr10-127661890-INS-142
chr10-127666615-INS-338
chr10-127689067-INS-400
chr10-127702392-INS-52
chr10-127715264-INS-258
chr10-127744157-INS-185
chr10-127753178-INS-78
chr10-127770169-INS-556
chr10-127773582-INS-132
chr10-127774532-INS-180
chr10-127776474-INS-474
chr10-127777420-INS-241
chr10-127782561-INS-120
chr10-127794526-INS-264
chr10-127799776-INS-371
chr10-127803744-INS-184
chr10-127822133-INS-76
chr10-127825594-INS-191
chr10-127836708-INS-58
chr10-127838256-INS-316
chr10-127846111-INS-233
chr10-127847276-INS-67
chr10-127860319-INS-96
chr10-127877994-INS-288
chr10-127888451-INS-261
chr10-127891334-INS-299
chr10-127896537-INS-200
chr10-127908886-INS-199
chr10-127911336-INS-205
chr10-127923796-INS-211
chr10-127934413-INS-52
chr10-127952961-INS-241
chr10-127987661-INS-54
chr10-127998406-INS-466

chr10-128001945-INS-410
chr10-128029775-INS-201
chr10-128033871-INS-817
chr10-128033880-INS-563
chr10-128035869-INS-263
chr10-128059189-INS-873
chr10-128072956-INS-72
chr10-128076618-INS-83
chr10-128093964-INS-212
chr10-128100212-INS-205
chr10-128101892-INS-400
chr10-128104228-INS-198
chr10-128110412-INS-191
chr10-128154998-INS-213
chr10-128159905-INS-172
chr10-128166203-INS-194
chr10-128202619-INS-205
chr10-128205933-INS-248
chr10-128210111-INS-211
chr10-128252975-INS-208
chr10-128254850-INS-212
chr10-128256447-INS-221
chr10-128257698-INS-235
chr10-128263668-INS-208
chr10-128265384-INS-65
chr10-128267943-INS-72
chr10-128277818-INS-1426
chr10-128286076-INS-98
chr10-128300838-INS-103
chr10-128314068-INS-72
chr10-128317824-INS-57
chr10-128324400-INS-106
chr10-128335896-INS-58
chr10-128355873-INS-208
chr10-128369082-INS-52
chr10-128370239-INS-186
chr10-128451395-INS-447
chr10-128457496-INS-110
chr10-128463144-INS-163
chr10-128464488-INS-179
chr10-128499023-INS-56
chr10-128501170-INS-92
chr10-128517522-INS-67
chr10-128544178-INS-76
chr10-128546371-INS-59
chr10-128558750-INS-165
chr10-128565714-INS-147
chr10-128575333-INS-52
chr10-128588105-INS-99
chr10-128595416-INS-62
chr10-128621960-INS-183
chr10-128669844-INS-56
chr10-128703135-INS-644
chr10-128712154-INS-172
chr10-128729479-INS-195
chr10-128740295-INS-52
chr10-128743280-INS-125
chr10-128750435-INS-124
chr10-128760433-INS-62
chr10-128760946-INS-173
chr10-128781722-INS-67
chr10-128790645-INS-196
chr10-128798364-INS-71
chr10-128798863-INS-154
chr10-128807409-INS-332
chr10-128811035-INS-199
chr10-128820507-INS-4982
chr10-128828083-INS-110
chr10-128831905-INS-1862
chr10-128838716-INS-394
chr10-128841677-INS-340
chr10-128860307-INS-84
chr10-128878475-INS-117
chr10-128883894-INS-137
chr10-128888642-INS-6018
chr10-128889903-INS-9902
chr10-128920530-INS-159

chr10-128921071-INS-60
chr10-128955114-INS-404
chr10-128956449-INS-52
chr10-128988744-INS-6510
chr10-129000924-INS-74
chr10-129002591-INS-6858
chr10-129008085-INS-6401
chr10-129038624-INS-80
chr10-129106249-INS-759
chr10-129130315-INS-965
chr10-129134337-INS-395
chr10-129141361-INS-1132
chr10-129164413-INS-92
chr10-129189533-INS-66
chr10-129258210-INS-6421
chr10-129297172-INS-6440
chr10-129329286-INS-233
chr10-129347216-INS-390
chr10-129364810-INS-683
chr10-129365627-INS-66
chr10-129390080-INS-90
chr10-129409786-INS-5479
chr10-129411153-INS-275
chr10-129411263-INS-3169
chr10-129439558-INS-62
chr10-129465478-INS-78
chr10-129467994-INS-556
chr10-129472582-INS-974
chr10-129479750-INS-6175
chr10-129484720-INS-817
chr10-129515126-INS-500
chr10-129526044-INS-959
chr10-129531105-INS-641
chr10-129539364-INS-342
chr10-129566751-INS-399
chr10-129577892-INS-313
chr10-129589675-INS-363
chr10-129625345-INS-66
chr10-129647372-INS-445
chr10-129659316-INS-390
chr10-129663576-INS-72
chr10-129665023-INS-9709
chr10-129669880-INS-155
chr10-129670492-INS-6211
chr10-129687372-INS-72
chr10-129723680-INS-615
chr10-129749372-INS-595
chr10-129768059-INS-919
chr10-129780211-INS-112
chr10-129780579-INS-6504
chr10-129786323-INS-4413
chr10-129788540-INS-7333
chr10-129806350-INS-323
chr10-129815092-INS-146
chr10-129829079-INS-154
chr10-129838784-INS-7657
chr10-129841122-INS-619
chr10-129851978-INS-58
chr10-129863415-INS-55
chr10-129880803-INS-64
chr10-129897923-INS-999
chr10-129909472-INS-109
chr10-129940618-INS-97
chr10-129963093-INS-64
chr10-129974890-INS-62
chr10-130028527-INS-6184
chr10-130043117-INS-325
chr10-130061379-INS-52
chr10-130062013-INS-6385
chr10-130071309-INS-95
chr10-130115117-INS-58
chr10-130123369-INS-414
chr10-130134682-INS-152
chr10-130149640-INS-64
chr10-130201876-INS-4599
chr10-130208029-INS-5492
chr10-130209152-INS-113

chr10-130213361-INS-249
chr10-130225317-INS-378
chr10-130231632-INS-10194
chr10-130241616-INS-7183
chr10-130271657-INS-6906
chr10-130272498-INS-1495
chr10-130287679-INS-358
chr10-130289118-INS-2447
chr10-130295946-INS-74
chr10-130296661-INS-7034
chr10-130302497-INS-6678
chr10-130302816-INS-53
chr10-130305131-INS-6242
chr10-130326556-INS-6705
chr10-130332354-INS-721
chr10-130345096-INS-7109
chr10-130345768-INS-60
chr10-130375885-INS-599
chr10-130403996-INS-7114
chr10-130425393-INS-316
chr11-3070295-INS-241
chr11-3070628-INS-272
chr11-3071726-INS-69
chr11-3073164-INS-62
chr11-3088172-INS-147
chr11-3107192-INS-199
chr11-3114361-INS-64
chr11-3141229-INS-390
chr11-3144461-INS-88
chr11-3147275-INS-110
chr11-3151152-INS-536
chr11-3161780-INS-193
chr11-3171488-INS-218
chr11-3204193-INS-358
chr11-3218869-INS-206
chr11-3228392-INS-151
chr11-3233711-INS-5756
chr11-3264916-INS-91
chr11-3270373-INS-498
chr11-3277347-INS-638
chr11-3292245-INS-62
chr11-3300012-INS-90
chr11-3311854-INS-197
chr11-3340689-INS-324
chr11-3345130-INS-199
chr11-3350309-INS-59
chr11-3377698-INS-368
chr11-3395175-INS-324
chr11-3417909-INS-468
chr11-3421938-INS-139
chr11-3435728-INS-60
chr11-3473608-INS-504
chr11-3499169-INS-126
chr11-3508416-INS-92
chr11-3518446-INS-107
chr11-3536669-INS-59
chr11-3537422-INS-115
chr11-3539871-INS-72
chr11-3541145-INS-238
chr11-3543966-INS-228
chr11-3546634-INS-5257
chr11-3546733-INS-153
chr11-3548693-INS-179
chr11-3564150-INS-128
chr11-3588482-INS-195
chr11-3600890-INS-220
chr11-3614302-INS-190
chr11-3650764-INS-500
chr11-3656821-INS-52
chr11-3665822-INS-120
chr11-3672659-INS-240
chr11-3675697-INS-57
chr11-3679129-INS-231
chr11-3690858-INS-57
chr11-3691225-INS-201
chr11-3703185-INS-60
chr11-3711230-INS-62

chr11-3737899-INS-280
chr11-3738341-INS-394
chr11-3739208-INS-92
chr11-3779720-INS-532
chr11-3782218-INS-185
chr11-3793562-INS-445
chr11-3802554-INS-606
chr11-3811775-INS-778
chr11-3826113-INS-230
chr11-3828063-INS-58
chr11-3849068-INS-167
chr11-3854172-INS-525
chr11-3876929-INS-67
chr11-3893256-INS-198
chr11-3896251-INS-598
chr11-3909486-INS-187
chr11-3916943-INS-231
chr11-3922318-INS-116
chr11-3931040-INS-235
chr11-3936298-INS-94
chr11-3941265-INS-96
chr11-3945245-INS-69
chr11-3958095-INS-134
chr11-3961294-INS-441
chr11-3973124-INS-2837
chr11-3978178-INS-59
chr11-3979301-INS-142
chr11-4006244-INS-545
chr11-4015534-INS-92
chr11-4019041-INS-75
chr11-4027718-INS-127
chr11-4047704-INS-64
chr11-4050915-INS-66
chr11-4070176-INS-157
chr11-4076125-INS-221
chr11-4080066-INS-218
chr11-4095665-INS-214
chr11-4113620-INS-212
chr11-4118959-INS-153
chr11-4127006-INS-455
chr11-4172107-INS-53
chr11-4174225-INS-68
chr11-4181835-INS-1236
chr11-4182162-INS-346
chr11-4188680-INS-106
chr11-4194122-INS-68
chr11-4196660-INS-69
chr11-4198813-INS-149
chr11-4199743-INS-76
chr11-4203426-INS-312
chr11-4239954-INS-94
chr11-4241322-INS-253
chr11-4264849-INS-688
chr11-4268484-INS-102
chr11-4272673-INS-63
chr11-4276599-INS-110
chr11-4282289-INS-115
chr11-4297764-INS-56
chr11-4298748-INS-505
chr11-4306191-INS-809
chr11-4306221-INS-1169
chr11-4306736-INS-57
chr11-4307925-INS-225
chr11-4321451-INS-61
chr11-4327261-INS-3081
chr11-4330977-INS-215
chr11-4339879-INS-147
chr11-4358039-INS-1886
chr11-4372224-INS-213
chr11-4373003-INS-64
chr11-4374904-INS-63
chr11-4387085-INS-295
chr11-4406201-INS-55
chr11-4406454-INS-2152
chr11-4407548-INS-164
chr11-4418594-INS-498
chr11-4439181-INS-211

chr11-4442314-INS-201
chr11-4468247-INS-52
chr11-4469421-INS-388
chr11-4482617-INS-667
chr11-4502194-INS-112
chr11-4505063-INS-203
chr11-4510516-INS-91
chr11-4523401-INS-331
chr11-4523880-INS-395
chr11-4529892-INS-239
chr11-4538824-INS-65
chr11-4547359-INS-558
chr11-4556266-INS-66
chr11-4566995-INS-401
chr11-4568171-INS-102
chr11-4574729-INS-52
chr11-4578355-INS-208
chr11-4580444-INS-211
chr11-4584018-INS-128
chr11-4597698-INS-299
chr11-4602389-INS-204
chr11-4604535-INS-59
chr11-4606036-INS-172
chr11-4609795-INS-50
chr11-4639909-INS-58
chr11-4641777-INS-156
chr11-4643351-INS-124
chr11-4652618-INS-202
chr11-4665766-INS-6436
chr11-4672873-INS-216
chr11-4678970-INS-241
chr11-4717995-INS-52
chr11-4743972-INS-62
chr11-4750982-INS-64
chr11-4752828-INS-193
chr11-4756790-INS-189
chr11-4767952-INS-317
chr11-4772530-INS-500
chr11-4778397-INS-167
chr11-4786910-INS-207
chr11-4788233-INS-386
chr11-4789916-INS-188
chr11-4803397-INS-118
chr11-4804692-INS-141
chr11-4806648-INS-179
chr11-4808917-INS-201
chr11-4814668-INS-203
chr11-4818695-INS-107
chr11-4820524-INS-377
chr11-4831077-INS-201
chr11-4848834-INS-58
chr11-4851708-INS-353
chr11-4854761-INS-71
chr11-4858923-INS-57
chr11-4863069-INS-186
chr11-4869520-INS-224
chr11-4875732-INS-54
chr11-4881170-INS-231
chr11-4884072-INS-358
chr11-4885243-INS-2704
chr11-4885378-INS-76
chr11-4901195-INS-76
chr11-4902028-INS-248
chr11-4906127-INS-144
chr11-4910321-INS-213
chr11-4911201-INS-231
chr11-4920323-INS-222
chr11-4927608-INS-201
chr11-4931946-INS-90
chr11-4933499-INS-128
chr11-4947854-INS-201
chr11-4949405-INS-606
chr11-4964190-INS-56
chr11-4969236-INS-54
chr11-4971813-INS-795
chr11-4977464-INS-60
chr11-5002017-INS-221

chr11-5004830-INS-116
chr11-5016225-INS-261
chr11-5022158-INS-212
chr11-5052348-INS-105
chr11-5066829-INS-193
chr11-5069987-INS-243
chr11-5070830-INS-70
chr11-5075323-INS-295
chr11-5103332-INS-54
chr11-5118887-INS-391
chr11-5119702-INS-653
chr11-5128642-INS-328
chr11-5132914-INS-65
chr11-5134715-INS-210
chr11-5136166-INS-1170
chr11-5136446-INS-314
chr11-5138918-INS-169
chr11-5144064-INS-247
chr11-5153268-INS-180
chr11-5155156-INS-56
chr11-5159647-INS-136
chr11-5161967-INS-62
chr11-5172191-INS-80
chr11-5179630-INS-194
chr11-5182381-INS-498
chr11-5209502-INS-105
chr11-5213939-INS-8734
chr11-5239547-INS-73
chr11-5249391-INS-388
chr11-5251578-INS-65
chr11-5278021-INS-52
chr11-5298605-INS-59
chr11-5318648-INS-51
chr11-5334565-INS-472
chr11-5345004-INS-205
chr11-5346340-INS-112
chr11-5351232-INS-153
chr11-5356344-INS-230
chr11-5380765-INS-370
chr11-5405091-INS-172
chr11-5411540-INS-196
chr11-5416061-INS-306
chr11-5416969-INS-158
chr11-5421007-INS-55
chr11-5442984-INS-100
chr11-5443635-INS-133
chr11-5445873-INS-51
chr11-5446483-INS-7051
chr11-5451200-INS-120
chr11-5453048-INS-236
chr11-5462837-INS-201
chr11-5467582-INS-375
chr11-5487836-INS-907
chr11-5493986-INS-8985
chr11-5495306-INS-209
chr11-5496363-INS-5528
chr11-5498397-INS-233
chr11-5513334-INS-368
chr11-5526904-INS-510
chr11-5541270-INS-300
chr11-5544285-INS-188
chr11-5650854-INS-56
chr11-5671179-INS-88
chr11-5685814-INS-152
chr11-5700839-INS-257
chr11-5704573-INS-183
chr11-5725055-INS-122
chr11-5730964-INS-123
chr11-5742898-INS-200
chr11-5743966-INS-141
chr11-5756981-INS-963
chr11-5784241-INS-350
chr11-5795187-INS-115
chr11-5814669-INS-196
chr11-5827381-INS-214
chr11-5830316-INS-216
chr11-5835739-INS-435

chr11-5841735-INS-82
chr11-5863032-INS-132
chr11-5894572-INS-96
chr11-5902091-INS-409
chr11-5904750-INS-96
chr11-5909757-INS-124
chr11-5919231-INS-82
chr11-5962136-INS-177
chr11-5973218-INS-68
chr11-5977037-INS-230
chr11-5978474-INS-203
chr11-6022059-INS-58
chr11-6023176-INS-150
chr11-6025076-INS-227
chr11-6032021-INS-64
chr11-6083617-INS-490
chr11-6087871-INS-74
chr11-6111450-INS-194
chr11-6141457-INS-278
chr11-6154092-INS-400
chr11-6156211-INS-57
chr11-6159646-INS-85
chr11-6166944-INS-65
chr11-6184084-INS-776
chr11-6198898-INS-57
chr11-6220547-INS-205
chr11-6228918-INS-185
chr11-6232824-INS-338
chr11-6254542-INS-205
chr11-6265105-INS-76
chr11-6277599-INS-172
chr11-6281629-INS-207
chr11-6291414-INS-759
chr11-6302566-INS-218
chr11-6322322-INS-222
chr11-6323609-INS-104
chr11-6330206-INS-235
chr11-6332161-INS-204
chr11-6342964-INS-240
chr11-6358235-INS-498
chr11-6361091-INS-138
chr11-6369367-INS-204
chr11-6383074-INS-281
chr11-6385002-INS-653
chr11-6396637-INS-191
chr11-6407815-INS-195
chr11-6433836-INS-316
chr11-6440614-INS-215
chr11-6441775-INS-234
chr11-6444854-INS-193
chr11-6454337-INS-554
chr11-6455173-INS-114
chr11-6465507-INS-163
chr11-6491400-INS-75
chr11-6503781-INS-462
chr11-6511349-INS-187
chr11-6522912-INS-79
chr11-6525224-INS-205
chr11-6526258-INS-206
chr11-6532031-INS-236
chr11-6557396-INS-221
chr11-6560546-INS-219
chr11-6562713-INS-443
chr11-6572641-INS-57
chr11-6583284-INS-198
chr11-6589789-INS-152
chr11-6592397-INS-260
chr11-6594973-INS-421
chr11-6615184-INS-338
chr11-6625560-INS-98
chr11-6678795-INS-8834
chr11-6685783-INS-203
chr11-6714600-INS-122
chr11-6722884-INS-80
chr11-6724055-INS-599
chr11-6737310-INS-106
chr11-6758946-INS-60

chr11-6761583-INS-7239
chr11-6764234-INS-498
chr11-6780180-INS-54
chr11-6788159-INS-140
chr11-6822625-INS-157
chr11-6827225-INS-58
chr11-6837589-INS-173
chr11-6838084-INS-239
chr11-6847512-INS-328
chr11-6858263-INS-196
chr11-6888670-INS-54
chr11-6900952-INS-345
chr11-6913245-INS-63
chr11-6935959-INS-129
chr11-6937719-INS-96
chr11-6942883-INS-61
chr11-6958853-INS-481
chr11-6960519-INS-506
chr11-6988703-INS-2406
chr11-7003912-INS-1890
chr11-7004499-INS-498
chr11-7031239-INS-128
chr11-7043538-INS-5346
chr11-7048203-INS-76
chr11-7051743-INS-512
chr11-7179260-INS-504
chr11-7187363-INS-167
chr11-7194273-INS-82
chr11-7196014-INS-343
chr11-7199827-INS-298
chr11-7223148-INS-52
chr11-7226481-INS-7292
chr11-7241203-INS-60
chr11-7246423-INS-53
chr11-7249002-INS-2922
chr11-7270206-INS-468
chr11-7279173-INS-232
chr11-7288785-INS-54
chr11-7294527-INS-80
chr11-7302891-INS-412
chr11-7311555-INS-266
chr11-7329261-INS-175
chr11-7339242-INS-498
chr11-7364348-INS-54
chr11-7430206-INS-74
chr11-7459161-INS-7159
chr11-7484376-INS-2982
chr11-7484634-INS-164
chr11-7499630-INS-231
chr11-7506167-INS-1387
chr11-7551150-INS-61
chr11-7558707-INS-144
chr11-7566414-INS-147
chr11-7578828-INS-190
chr11-7584476-INS-149
chr11-7589637-INS-558
chr11-7590900-INS-90
chr11-7596577-INS-340
chr11-7613231-INS-329
chr11-7641680-INS-270
chr11-7644009-INS-399
chr11-7646974-INS-5539
chr11-7657203-INS-51
chr11-7658130-INS-340
chr11-7663884-INS-141
chr11-7680834-INS-155
chr11-7690263-INS-191
chr11-7695514-INS-60
chr11-7736940-INS-124
chr11-7840029-INS-272
chr11-7849483-INS-515
chr11-7863900-INS-234
chr11-7867813-INS-58
chr11-7896973-INS-4186
chr11-7921564-INS-208
chr11-7932772-INS-5170
chr11-7941467-INS-111

chr11-7980448-INS-62
chr11-7997180-INS-247
chr11-8000553-INS-60
chr11-8002740-INS-2115
chr11-8014785-INS-93
chr11-8038090-INS-6208
chr11-8059941-INS-134
chr11-8064724-INS-103
chr11-8077352-INS-52
chr11-8080223-INS-62
chr11-8082134-INS-324
chr11-8109660-INS-335
chr11-8115767-INS-662
chr11-8162681-INS-249
chr11-8184746-INS-74
chr11-8185985-INS-420
chr11-8190484-INS-58
chr11-8211734-INS-5267
chr11-8232521-INS-83
chr11-8249037-INS-446
chr11-8249827-INS-130
chr11-8307468-INS-118
chr11-8317713-INS-905
chr11-8324801-INS-173
chr11-8331704-INS-99
chr11-8339233-INS-192
chr11-8345301-INS-303
chr11-8369285-INS-225
chr11-8490275-INS-92
chr11-8501681-INS-163
chr11-8556505-INS-62
chr11-8558473-INS-123
chr11-8581316-INS-275
chr11-8584509-INS-55
chr11-8598094-INS-88
chr11-8616133-INS-120
chr11-8627292-INS-202
chr11-8630455-INS-64
chr11-8681288-INS-574
chr11-8697454-INS-897
chr11-8707453-INS-205
chr11-8731243-INS-66
chr11-8752083-INS-201
chr11-8756701-INS-1027
chr11-8758377-INS-200
chr11-8760042-INS-184
chr11-8765414-INS-85
chr11-8866923-INS-246
chr11-8879737-INS-167
chr11-8881758-INS-65
chr11-8882652-INS-68
chr11-8886447-INS-111
chr11-8918043-INS-63
chr11-8921159-INS-190
chr11-8939111-INS-73
chr11-8971355-INS-70
chr11-8981770-INS-62
chr11-9009468-INS-526
chr11-9012750-INS-160
chr11-9016868-INS-53
chr11-9046125-INS-612
chr11-9068596-INS-57
chr11-9080247-INS-178
chr11-9093796-INS-182
chr11-9139713-INS-172
chr11-9139734-INS-415
chr11-9175279-INS-230
chr11-9194458-INS-68
chr11-9199369-INS-127
chr11-9228959-INS-129
chr11-9254403-INS-98
chr11-9265749-INS-255
chr11-9302337-INS-6469
chr11-9423578-INS-156
chr11-9444318-INS-65
chr11-9492843-INS-55
chr11-9513511-INS-51

chr11-9540619-INS-52
chr11-9570991-INS-52
chr11-9587350-INS-53
chr11-9654564-INS-352
chr11-9666990-INS-349
chr11-9674318-INS-1147
chr11-9687982-INS-52
chr11-9725143-INS-506
chr11-9742801-INS-52
chr11-9747867-INS-250
chr11-9768125-INS-58
chr11-9812898-INS-89
chr11-9813007-INS-521
chr11-9823798-INS-138
chr11-9827435-INS-187
chr11-9834508-INS-95
chr11-9847195-INS-167
chr11-9848936-INS-345
chr11-9883953-INS-137
chr11-9899555-INS-91
chr11-9910139-INS-203
chr11-9919187-INS-7117
chr11-9936069-INS-2161
chr11-9954556-INS-56
chr11-9989718-INS-401
chr11-10001342-INS-223
chr11-10003724-INS-8419
chr11-10046388-INS-120
chr11-10079605-INS-73
chr11-10088217-INS-273
chr11-10124261-INS-1384
chr11-10137534-INS-194
chr11-10147184-INS-234
chr11-10181141-INS-99
chr11-10199112-INS-67
chr11-10201464-INS-862
chr11-10238045-INS-52
chr11-10251330-INS-117
chr11-10353339-INS-939
chr11-10358023-INS-1367
chr11-10375153-INS-113
chr11-10397040-INS-60
chr11-10408678-INS-289
chr11-10420455-INS-2080
chr11-10436689-INS-6880
chr11-10456978-INS-606
chr11-10469931-INS-53
chr11-10478189-INS-59
chr11-10514294-INS-493
chr11-10528703-INS-120
chr11-10553634-INS-463
chr11-10571005-INS-209
chr11-10601567-INS-59
chr11-10602038-INS-752
chr11-10609445-INS-355
chr11-10631212-INS-817
chr11-10634061-INS-6413
chr11-10647642-INS-63
chr11-10690963-INS-5017
chr11-10691073-INS-349
chr11-10699297-INS-263
chr11-10702538-INS-6297
chr11-10703122-INS-843
chr11-10742829-INS-76
chr11-10747983-INS-680
chr11-10807985-INS-98
chr11-10811356-INS-54
chr11-10816725-INS-216
chr11-10820699-INS-225
chr11-10826172-INS-243
chr11-10837284-INS-522
chr11-10855008-INS-133
chr11-10876936-INS-3353
chr11-10902636-INS-128
chr11-10922597-INS-67
chr11-10933897-INS-407
chr11-10957169-INS-354

chr11-10960602-INS-58
chr11-10985214-INS-68
chr11-10998944-INS-212
chr11-11023848-INS-6447
chr11-11042158-INS-5798
chr11-11103382-INS-56
chr11-11105715-INS-234
chr11-11113673-INS-96
chr11-11151119-INS-70
chr11-11166787-INS-313
chr11-11176646-INS-191
chr11-11180120-INS-5549
chr11-11186999-INS-94
chr11-11204406-INS-72
chr11-11219601-INS-130
chr11-11225981-INS-574
chr11-11247032-INS-95
chr11-11320328-INS-243
chr11-11357009-INS-59
chr11-11370231-INS-58
chr11-11386748-INS-78
chr11-11402975-INS-399
chr11-11406257-INS-188
chr11-11409309-INS-54
chr11-11418327-INS-138
chr11-11463138-INS-54
chr11-11467126-INS-204
chr11-11468947-INS-56
chr11-11473587-INS-230
chr11-11479518-INS-76
chr11-11523128-INS-572
chr11-11538147-INS-60
chr11-11568833-INS-83
chr11-11577641-INS-66
chr11-11603849-INS-90
chr11-11632577-INS-128
chr11-11638002-INS-290
chr11-11644024-INS-53
chr11-11717954-INS-223
chr11-11731288-INS-70
chr11-11732516-INS-167
chr11-11734277-INS-74
chr11-11735088-INS-120
chr11-11753596-INS-6937
chr11-11777815-INS-218
chr11-11800490-INS-61
chr11-11825784-INS-87
chr11-11836294-INS-327
chr11-11844240-INS-203
chr11-11855114-INS-99
chr11-11856588-INS-84
chr11-11861277-INS-162
chr11-11871379-INS-136
chr11-11876296-INS-62
chr11-11886093-INS-209
chr11-11899090-INS-223
chr11-11933858-INS-291
chr11-11974465-INS-147
chr11-11997886-INS-57
chr11-12012415-INS-7538
chr11-12018188-INS-635
chr11-12020788-INS-313
chr11-12022212-INS-143
chr11-12030710-INS-185
chr11-12035881-INS-198
chr11-12075181-INS-989
chr11-12096473-INS-208
chr11-12106229-INS-7652
chr11-12122979-INS-64
chr11-12131235-INS-256
chr11-12133515-INS-169
chr11-12162456-INS-296
chr11-12166514-INS-1451
chr11-12178474-INS-151
chr11-12188758-INS-92
chr11-12218371-INS-66
chr11-12256090-INS-7085

chr11-12262009-INS-171
chr11-12283950-INS-1892
chr11-12302070-INS-1642
chr11-12302960-INS-102
chr11-12360859-INS-225
chr11-12365510-INS-85
chr11-12431111-INS-93
chr11-12449456-INS-55
chr11-12451746-INS-399
chr11-12475812-INS-131
chr11-12485833-INS-438
chr11-12526672-INS-73
chr11-12534667-INS-829
chr11-12558500-INS-335
chr11-12643008-INS-98
chr11-12643806-INS-590
chr11-12693100-INS-610
chr11-12769986-INS-460
chr11-12784726-INS-11474
chr11-12794703-INS-6440
chr11-12870053-INS-53
chr11-12872351-INS-78
chr11-12881440-INS-60
chr11-12922698-INS-479
chr11-12932874-INS-60
chr11-12944028-INS-530
chr11-12973880-INS-507
chr11-12976506-INS-491
chr11-13018532-INS-401
chr11-13034580-INS-71
chr11-13049283-INS-312
chr11-13066460-INS-8003
chr11-13076844-INS-856
chr11-13078517-INS-200
chr11-13084264-INS-297
chr11-13097683-INS-530
chr11-13107063-INS-187
chr11-13132541-INS-62
chr11-13133028-INS-204
chr11-13145165-INS-53
chr11-13184878-INS-7178
chr11-13225535-INS-59
chr11-13307019-INS-51
chr11-13309869-INS-404
chr11-13316322-INS-436
chr11-13323196-INS-558
chr11-13329110-INS-212
chr11-13357500-INS-686
chr11-13367432-INS-544
chr11-13375966-INS-89
chr11-13401471-INS-666
chr11-13404737-INS-861
chr11-13416092-INS-82
chr11-13431429-INS-902
chr11-13431558-INS-96
chr11-13435254-INS-57
chr11-13440759-INS-1815
chr11-13447335-INS-57
chr11-13492121-INS-225
chr11-13516152-INS-200
chr11-13537014-INS-1186
chr11-13544906-INS-52
chr11-13545672-INS-51
chr11-13557486-INS-92
chr11-13580498-INS-70
chr11-13627654-INS-110
chr11-13646810-INS-3030
chr11-13652450-INS-166
chr11-13712772-INS-135
chr11-13718152-INS-106
chr11-13738073-INS-6225
chr11-13750955-INS-6200
chr11-13761295-INS-214
chr11-13775473-INS-58
chr11-13822266-INS-82
chr11-13854126-INS-55
chr11-13869895-INS-897

chr11-13888085-INS-68
chr11-13897190-INS-203
chr11-13940144-INS-388
chr11-13943145-INS-528
chr11-13949616-INS-6386
chr11-14007771-INS-105
chr11-14110875-INS-590
chr11-14130596-INS-222
chr11-14181247-INS-85
chr11-14185216-INS-280
chr11-14198424-INS-219
chr11-14224402-INS-76
chr11-14236162-INS-995
chr11-14237425-INS-88
chr11-14240489-INS-143
chr11-14280910-INS-123
chr11-14282878-INS-154
chr11-14298087-INS-52
chr11-14299105-INS-120
chr11-14328297-INS-358
chr11-14334671-INS-85
chr11-14338131-INS-54
chr11-14349484-INS-496
chr11-14369645-INS-145
chr11-14370841-INS-63
chr11-14396416-INS-386
chr11-14420177-INS-342
chr11-14430616-INS-57
chr11-14433464-INS-56
chr11-14460658-INS-235
chr11-14505777-INS-72
chr11-14536977-INS-193
chr11-14539732-INS-890
chr11-14547125-INS-1888
chr11-14584280-INS-70
chr11-14592188-INS-6509
chr11-14616994-INS-576
chr11-14647936-INS-62
chr11-14671169-INS-398
chr11-14684795-INS-448
chr11-14698288-INS-52
chr11-14721539-INS-106
chr11-14743598-INS-215
chr11-14750091-INS-61
chr11-14756646-INS-325
chr11-14765233-INS-563
chr11-14780972-INS-482
chr11-14785707-INS-1891
chr11-14790150-INS-515
chr11-14820064-INS-220
chr11-14827953-INS-214
chr11-14842718-INS-6689
chr11-14843085-INS-156
chr11-14864148-INS-400
chr11-14864942-INS-6395
chr11-14892370-INS-6659
chr11-14945683-INS-60
chr11-14955165-INS-75
chr11-14995610-INS-123
chr11-15012772-INS-148
chr11-15062099-INS-121
chr11-15072483-INS-63
chr11-15081949-INS-80
chr11-15103449-INS-446
chr11-15147895-INS-75
chr11-15154707-INS-435
chr11-15192436-INS-147
chr11-15202837-INS-197
chr11-15271273-INS-56
chr11-15315204-INS-5350
chr11-15316443-INS-4352
chr11-15329883-INS-6337
chr11-15368642-INS-70
chr11-15375855-INS-245
chr11-15378542-INS-309
chr11-15379502-INS-88
chr11-15382668-INS-218

chr11-15404912-INS-285
chr11-15437448-INS-56
chr11-15441533-INS-233
chr11-15444463-INS-234
chr11-15470473-INS-149
chr11-15473030-INS-52
chr11-15473920-INS-1399
chr11-15475824-INS-52
chr11-15540893-INS-6597
chr11-15540988-INS-236
chr11-15573030-INS-193
chr11-15576884-INS-86
chr11-15587481-INS-81
chr11-15592662-INS-892
chr11-15601847-INS-725
chr11-15616012-INS-906
chr11-15641008-INS-782
chr11-15705449-INS-171
chr11-15711725-INS-328
chr11-15714179-INS-479
chr11-15723818-INS-6480
chr11-15724510-INS-114
chr11-15740912-INS-188
chr11-15815073-INS-139
chr11-15828292-INS-671
chr11-15851254-INS-62
chr11-15852312-INS-662
chr11-15925481-INS-72
chr11-15946140-INS-63
chr11-15988556-INS-54
chr11-16032491-INS-278
chr11-16036906-INS-226
chr11-16039961-INS-290
chr11-16069301-INS-630
chr11-16076639-INS-454
chr11-16083688-INS-172
chr11-16095837-INS-51
chr11-16133951-INS-199
chr11-16156134-INS-75
chr11-16163479-INS-203
chr11-16193468-INS-132
chr11-16208330-INS-66
chr11-16210385-INS-82
chr11-16222991-INS-54
chr11-16236194-INS-102
chr11-16270681-INS-450
chr11-16330163-INS-165
chr11-16348278-INS-180
chr11-16373068-INS-939
chr11-16402472-INS-825
chr11-16437568-INS-749
chr11-16454387-INS-52
chr11-16460763-INS-60
chr11-16601320-INS-313
chr11-16611189-INS-124
chr11-16631973-INS-6389
chr11-16637423-INS-94
chr11-16654600-INS-114
chr11-16660428-INS-200
chr11-16661731-INS-499
chr11-16739907-INS-86
chr11-16755694-INS-809
chr11-16829349-INS-79
chr11-16842084-INS-6732
chr11-16842374-INS-105
chr11-16883451-INS-110
chr11-16901192-INS-81
chr11-16904529-INS-6877
chr11-16914004-INS-237
chr11-16928537-INS-278
chr11-16930267-INS-147
chr11-16946351-INS-54
chr11-16956840-INS-96
chr11-16974213-INS-440
chr11-16977509-INS-823
chr11-16985078-INS-94
chr11-16986098-INS-108

chr11-16989829-INS-85
chr11-16993536-INS-179
chr11-17000903-INS-56
chr11-17013406-INS-198
chr11-17040578-INS-59
chr11-17068434-INS-53
chr11-17075652-INS-6684
chr11-17091755-INS-80
chr11-17102254-INS-124
chr11-17108621-INS-225
chr11-17112451-INS-54
chr11-17115530-INS-733
chr11-17119054-INS-2728
chr11-17123265-INS-79
chr11-17142595-INS-388
chr11-17150490-INS-588
chr11-17179216-INS-205
chr11-17206034-INS-142
chr11-17207408-INS-178
chr11-17208747-INS-6456
chr11-17244612-INS-1894
chr11-17249987-INS-5548
chr11-17250214-INS-655
chr11-17273657-INS-74
chr11-17276413-INS-60
chr11-17279103-INS-7078
chr11-17284196-INS-108
chr11-17284521-INS-445
chr11-17298379-INS-59
chr11-17300284-INS-178
chr11-17309613-INS-172
chr11-17316786-INS-575
chr11-17361522-INS-56
chr11-17365222-INS-52
chr11-17371726-INS-82
chr11-17374470-INS-354
chr11-17403033-INS-52
chr11-17429489-INS-72
chr11-17433977-INS-69
chr11-17436522-INS-361
chr11-17442324-INS-273
chr11-17502109-INS-316
chr11-17524674-INS-141
chr11-17533159-INS-745
chr11-17545086-INS-6438
chr11-17558953-INS-61
chr11-17570457-INS-6370
chr11-17633277-INS-56
chr11-17637326-INS-52
chr11-17641912-INS-6384
chr11-17668715-INS-283
chr11-17708881-INS-106
chr11-17733468-INS-1356
chr11-17774181-INS-6114
chr11-17776715-INS-6554
chr11-17782504-INS-347
chr11-17802801-INS-393
chr11-17808299-INS-51
chr11-17828652-INS-71
chr11-17831928-INS-66
chr11-17833976-INS-205
chr11-17839395-INS-179
chr11-17841311-INS-203
chr11-17855491-INS-400
chr11-17861807-INS-447
chr11-17873023-INS-150
chr11-17906983-INS-105
chr11-17907232-INS-5959
chr11-17912348-INS-7103
chr11-17950226-INS-117
chr11-17982664-INS-775
chr11-17988273-INS-762
chr11-18009931-INS-69
chr11-18017252-INS-52
chr11-18022619-INS-756
chr11-18049288-INS-376
chr11-18090136-INS-58

chr11-18110683-INS-211
chr11-18113112-INS-186
chr11-18146234-INS-148
chr11-18161718-INS-6646
chr11-18173704-INS-6628
chr11-18180377-INS-52
chr11-18191086-INS-468
chr11-18194888-INS-186
chr11-18196823-INS-228
chr11-18202838-INS-63
chr11-18232672-INS-83
chr11-18241259-INS-187
chr11-18257004-INS-2691
chr11-18268138-INS-113
chr11-18281754-INS-179
chr11-18285965-INS-1592
chr11-18328238-INS-205
chr11-18345027-INS-314
chr11-18359101-INS-257
chr11-18393068-INS-338
chr11-18416377-INS-6555
chr11-18433360-INS-200
chr11-18461693-INS-639
chr11-18470573-INS-57
chr11-18471093-INS-59
chr11-18475498-INS-1457
chr11-18477406-INS-94
chr11-18490390-INS-6672
chr11-18500098-INS-202
chr11-18501015-INS-6254
chr11-18503868-INS-619
chr11-18511479-INS-163
chr11-18534019-INS-131
chr11-18546238-INS-51
chr11-18553149-INS-816
chr11-18569342-INS-96
chr11-18609301-INS-6208
chr11-18613828-INS-6140
chr11-18633882-INS-57
chr11-18653835-INS-104
chr11-18658447-INS-178
chr11-18695081-INS-78
chr11-18716990-INS-396
chr11-18741560-INS-408
chr11-18748395-INS-84
chr11-18751694-INS-56
chr11-18808001-INS-315
chr11-18808959-INS-2598
chr11-18877123-INS-290
chr11-18903523-INS-833
chr11-18903593-INS-203
chr11-18948971-INS-105
chr11-19012702-INS-295
chr11-19042303-INS-1893
chr11-19071277-INS-7040
chr11-19082506-INS-485
chr11-19086108-INS-65
chr11-19088911-INS-52
chr11-19093160-INS-81
chr11-19101765-INS-67
chr11-19113914-INS-226
chr11-19129564-INS-144
chr11-19131546-INS-75
chr11-19185843-INS-56
chr11-19208566-INS-171
chr11-19235275-INS-70
chr11-19258836-INS-52
chr11-19275786-INS-687
chr11-19295508-INS-51
chr11-19336919-INS-52
chr11-19338104-INS-122
chr11-19341621-INS-108
chr11-19347544-INS-56
chr11-19351675-INS-719
chr11-19354171-INS-646
chr11-19372951-INS-7194
chr11-19382270-INS-81

chr11-19382738-INS-288
chr11-19414791-INS-505
chr11-19433229-INS-68
chr11-19445397-INS-56
chr11-19475649-INS-141
chr11-19485191-INS-223
chr11-19505951-INS-699
chr11-19521469-INS-7073
chr11-19531575-INS-156
chr11-19544373-INS-186
chr11-19561527-INS-62
chr11-19570766-INS-210
chr11-19596442-INS-73
chr11-19606447-INS-506
chr11-19615132-INS-115
chr11-19620952-INS-120
chr11-19635096-INS-103
chr11-19651723-INS-383
chr11-19685860-INS-103
chr11-19693091-INS-68
chr11-19702002-INS-258
chr11-19762646-INS-112
chr11-19782258-INS-94
chr11-19790800-INS-102
chr11-19800711-INS-444
chr11-19822640-INS-144
chr11-19834000-INS-84
chr11-19845450-INS-57
chr11-19846752-INS-309
chr11-19855710-INS-211
chr11-19863833-INS-210
chr11-19873452-INS-202
chr11-19884693-INS-148
chr11-19890278-INS-51
chr11-19903917-INS-231
chr11-19912966-INS-127
chr11-19913382-INS-89
chr11-19927268-INS-106
chr11-19938096-INS-219
chr11-19940817-INS-107
chr11-19944947-INS-207
chr11-19948031-INS-211
chr11-19954717-INS-163
chr11-19959811-INS-387
chr11-19960353-INS-108
chr11-19965926-INS-201
chr11-19969012-INS-146
chr11-19979184-INS-133
chr11-19983476-INS-60
chr11-20004356-INS-201
chr11-20021911-INS-186
chr11-20031905-INS-76
chr11-20057018-INS-196
chr11-20064349-INS-210
chr11-20069672-INS-193
chr11-20070223-INS-136
chr11-20081971-INS-199
chr11-20082664-INS-72
chr11-20084279-INS-529
chr11-20088188-INS-216
chr11-20098442-INS-214
chr11-20099842-INS-70
chr11-20101224-INS-184
chr11-20109542-INS-59
chr11-20109920-INS-183
chr11-20112528-INS-116
chr11-20113635-INS-221
chr11-20117222-INS-239
chr11-20119865-INS-186
chr11-20128627-INS-402
chr11-20130165-INS-201
chr11-20145579-INS-5198
chr11-20166987-INS-201
chr11-20185837-INS-195
chr11-20186518-INS-1887
chr11-20186692-INS-223
chr11-20206951-INS-403

chr11-20209135-INS-56
chr11-20212751-INS-92
chr11-20222473-INS-194
chr11-20224290-INS-102
chr11-20281673-INS-79
chr11-20296858-INS-51
chr11-20306916-INS-588
chr11-20309755-INS-101
chr11-20315309-INS-2621
chr11-20315526-INS-296
chr11-20329056-INS-882
chr11-20329130-INS-76
chr11-20329478-INS-773
chr11-20335602-INS-7426
chr11-20338426-INS-7095
chr11-20360908-INS-240
chr11-20365095-INS-180
chr11-20374191-INS-223
chr11-20386149-INS-66
chr11-20390412-INS-101
chr11-20402600-INS-360
chr11-20408446-INS-59
chr11-20414994-INS-59
chr11-20436641-INS-430
chr11-20440273-INS-265
chr11-20447763-INS-122
chr11-20474135-INS-66
chr11-20525394-INS-60
chr11-20527190-INS-340
chr11-20534479-INS-201
chr11-20536206-INS-211
chr11-20541232-INS-55
chr11-20549001-INS-152
chr11-20560437-INS-92
chr11-20577746-INS-69
chr11-20596824-INS-217
chr11-20654055-INS-342
chr11-20680570-INS-471
chr11-20696660-INS-121
chr11-20701346-INS-801
chr11-20721921-INS-6111
chr11-20756629-INS-51
chr11-20784458-INS-71
chr11-20802896-INS-953
chr11-20818712-INS-71
chr11-20821704-INS-69
chr11-20838391-INS-4515
chr11-20879202-INS-123
chr11-20886541-INS-66
chr11-20890145-INS-56
chr11-20933195-INS-462
chr11-20964157-INS-66
chr11-21009515-INS-458
chr11-21010758-INS-142
chr11-21031384-INS-86
chr11-21033503-INS-356
chr11-21047362-INS-74
chr11-21089832-INS-76
chr11-21103755-INS-72
chr11-21112670-INS-1888
chr11-21153147-INS-56
chr11-21191378-INS-70
chr11-21203930-INS-162
chr11-21220883-INS-232
chr11-21300680-INS-1328
chr11-21303944-INS-477
chr11-21347496-INS-286
chr11-21353559-INS-66
chr11-21396369-INS-365
chr11-21426666-INS-199
chr11-21469519-INS-547
chr11-21491882-INS-355
chr11-21496852-INS-313
chr11-21498001-INS-53
chr11-21503910-INS-193
chr11-21504080-INS-4405
chr11-21534631-INS-93

chr11-21538667-INS-69
chr11-21566939-INS-503
chr11-21584871-INS-53
chr11-21620153-INS-675
chr11-21622676-INS-149
chr11-21660617-INS-128
chr11-21689938-INS-80
chr11-21704818-INS-730
chr11-21704898-INS-112
chr11-21714895-INS-98
chr11-21744892-INS-7276
chr11-21750841-INS-409
chr11-21751834-INS-6719
chr11-21766043-INS-278
chr11-21799390-INS-70
chr11-21822719-INS-142
chr11-21836304-INS-160
chr11-21837958-INS-60
chr11-21866293-INS-505
chr11-21869327-INS-123
chr11-21902528-INS-358
chr11-21909746-INS-136
chr11-21917293-INS-183
chr11-21939155-INS-220
chr11-21942539-INS-104
chr11-21952421-INS-99
chr11-21963312-INS-354
chr11-21969789-INS-983
chr11-21974905-INS-198
chr11-22122381-INS-581
chr11-22154308-INS-56
chr11-22177154-INS-364
chr11-22180062-INS-74
chr11-22201084-INS-511
chr11-22223325-INS-859
chr11-22245095-INS-358
chr11-22334403-INS-7119
chr11-22344148-INS-234
chr11-22357885-INS-919
chr11-22363981-INS-394
chr11-22387405-INS-54
chr11-22399528-INS-61
chr11-22402185-INS-54
chr11-22462177-INS-8988
chr11-22470740-INS-85
chr11-22488859-INS-435
chr11-22498443-INS-400
chr11-22504520-INS-392
chr11-22506175-INS-222
chr11-22516770-INS-126
chr11-22540761-INS-68
chr11-22543347-INS-60
chr11-22555935-INS-216
chr11-22596670-INS-128
chr11-22646778-INS-94
chr11-22663828-INS-329
chr11-22664914-INS-53
chr11-22676184-INS-92
chr11-22707127-INS-92
chr11-22711283-INS-59
chr11-22730608-INS-85
chr11-22861531-INS-155
chr11-22873117-INS-121
chr11-22876107-INS-57
chr11-22892983-INS-212
chr11-22907622-INS-842
chr11-22925332-INS-232
chr11-22930865-INS-1860
chr11-22937374-INS-632
chr11-22953016-INS-210
chr11-22961328-INS-82
chr11-22971804-INS-629
chr11-22986311-INS-55
chr11-23008400-INS-62
chr11-23023549-INS-73
chr11-23039063-INS-468
chr11-23065151-INS-198

chr11-23079279-INS-178
chr11-23083538-INS-323
chr11-23093577-INS-52
chr11-23095374-INS-55
chr11-23100871-INS-520
chr11-23113763-INS-473
chr11-23126936-INS-111
chr11-23129097-INS-4781
chr11-23155797-INS-444
chr11-23187024-INS-637
chr11-23199851-INS-5203
chr11-23248977-INS-168
chr11-23254596-INS-120
chr11-23259517-INS-206
chr11-23287415-INS-205
chr11-23319639-INS-197
chr11-23328318-INS-52
chr11-23339505-INS-182
chr11-23401624-INS-225
chr11-23429982-INS-52
chr11-23473574-INS-187
chr11-23486567-INS-147
chr11-23521048-INS-984
chr11-23521058-INS-122
chr11-23522314-INS-386
chr11-23533408-INS-68
chr11-23555772-INS-206
chr11-23574225-INS-57
chr11-23603656-INS-220
chr11-23606958-INS-81
chr11-23628205-INS-613
chr11-23640645-INS-60
chr11-23679444-INS-200
chr11-23711974-INS-172
chr11-23727724-INS-936
chr11-23768608-INS-135
chr11-23791057-INS-212
chr11-23832738-INS-205
chr11-23871765-INS-640
chr11-23886978-INS-90
chr11-23898072-INS-342
chr11-23902795-INS-212
chr11-23906457-INS-7043
chr11-23907813-INS-113
chr11-23912033-INS-107
chr11-23918553-INS-56
chr11-23932860-INS-61
chr11-23977329-INS-277
chr11-23985447-INS-70
chr11-24011858-INS-59
chr11-24017347-INS-50
chr11-24050423-INS-83
chr11-24054110-INS-61
chr11-24064741-INS-108
chr11-24108525-INS-128
chr11-24150137-INS-60
chr11-24174009-INS-87
chr11-24185754-INS-98
chr11-24196145-INS-108
chr11-24202031-INS-304
chr11-24202600-INS-50
chr11-24229155-INS-202
chr11-24239906-INS-168
chr11-24270455-INS-221
chr11-24286152-INS-951
chr11-24287174-INS-83
chr11-24292517-INS-158
chr11-24307450-INS-158
chr11-24345142-INS-428
chr11-24346792-INS-1888
chr11-24450998-INS-60
chr11-24487397-INS-176
chr11-24533257-INS-154
chr11-24571444-INS-96
chr11-24583558-INS-99
chr11-24584215-INS-1946
chr11-24592220-INS-120

chr11-24595891-INS-469
chr11-24625635-INS-64
chr11-24667483-INS-154
chr11-24685597-INS-155
chr11-24728223-INS-58
chr11-24731074-INS-66
chr11-24748538-INS-59
chr11-24750587-INS-6112
chr11-24790280-INS-58
chr11-24821233-INS-60
chr11-24927114-INS-51
chr11-24939979-INS-56
chr11-24948504-INS-82
chr11-24951615-INS-73
chr11-24953162-INS-2064
chr11-24989667-INS-66
chr11-24998987-INS-98
chr11-25033839-INS-66
chr11-25044782-INS-116
chr11-25066672-INS-853
chr11-25081133-INS-53
chr11-25084104-INS-62
chr11-25206741-INS-68
chr11-25243292-INS-404
chr11-25263049-INS-56
chr11-25265734-INS-8855
chr11-25343105-INS-95
chr11-25386956-INS-81
chr11-25449603-INS-62
chr11-25451913-INS-93
chr11-25466912-INS-164
chr11-25489152-INS-126
chr11-25492442-INS-62
chr11-25520361-INS-6442
chr11-25532857-INS-147
chr11-25538274-INS-597
chr11-25549131-INS-58
chr11-25559658-INS-361
chr11-25574655-INS-52
chr11-25575703-INS-169
chr11-25577917-INS-6284
chr11-25583395-INS-148
chr11-25589160-INS-72
chr11-25624193-INS-1883
chr11-25648462-INS-202
chr11-25664246-INS-111
chr11-25674111-INS-58
chr11-25676261-INS-137
chr11-25723737-INS-51
chr11-25733956-INS-59
chr11-25781607-INS-139
chr11-25784354-INS-54
chr11-25813800-INS-60
chr11-25844402-INS-204
chr11-25844572-INS-466
chr11-25859635-INS-908
chr11-25860799-INS-6480
chr11-25884781-INS-152
chr11-25889383-INS-1229
chr11-25976249-INS-52
chr11-26014619-INS-64
chr11-26037390-INS-224
chr11-26062691-INS-64
chr11-26095054-INS-260
chr11-26113143-INS-76
chr11-26164626-INS-409
chr11-26178517-INS-61
chr11-26204951-INS-207
chr11-26212671-INS-392
chr11-26226006-INS-63
chr11-26241410-INS-99
chr11-26255285-INS-123
chr11-26281063-INS-58
chr11-26286331-INS-506
chr11-26304543-INS-231
chr11-26328525-INS-82
chr11-26333474-INS-389

chr11-26349089-INS-511
chr11-26361994-INS-214
chr11-26363154-INS-154
chr11-26391146-INS-54
chr11-26395357-INS-8787
chr11-26453052-INS-73
chr11-26461534-INS-358
chr11-26472210-INS-358
chr11-26478829-INS-83
chr11-26479277-INS-865
chr11-26494903-INS-82
chr11-26538477-INS-208
chr11-26556408-INS-132
chr11-26561509-INS-384
chr11-26579430-INS-3394
chr11-26593896-INS-77
chr11-26596216-INS-843
chr11-26599425-INS-59
chr11-26617638-INS-56
chr11-26628110-INS-56
chr11-26632676-INS-6558
chr11-26636275-INS-222
chr11-26636649-INS-5841
chr11-26645754-INS-6663
chr11-26647054-INS-196
chr11-26654833-INS-65
chr11-26658099-INS-76
chr11-26660774-INS-58
chr11-26664384-INS-7049
chr11-26671987-INS-6452
chr11-26682862-INS-212
chr11-26711574-INS-6512
chr11-26729056-INS-202
chr11-26729943-INS-322
chr11-26771442-INS-158
chr11-26778778-INS-172
chr11-26783421-INS-358
chr11-26793834-INS-209
chr11-26804544-INS-90
chr11-26838478-INS-112
chr11-26863373-INS-1181
chr11-26883777-INS-461
chr11-26890680-INS-575
chr11-26891483-INS-186
chr11-26903771-INS-52
chr11-26963779-INS-472
chr11-27001567-INS-832
chr11-27033833-INS-93
chr11-27039239-INS-253
chr11-27051223-INS-455
chr11-27199041-INS-144
chr11-27216290-INS-178
chr11-27226350-INS-8578
chr11-27240310-INS-304
chr11-27269784-INS-134
chr11-27283276-INS-191
chr11-27286216-INS-184
chr11-27294400-INS-402
chr11-27299594-INS-619
chr11-27369266-INS-293
chr11-27371051-INS-52
chr11-27374612-INS-6667
chr11-27380538-INS-51
chr11-27417863-INS-520
chr11-27427805-INS-76
chr11-27449353-INS-69
chr11-27459494-INS-632
chr11-27507189-INS-60
chr11-27533282-INS-54
chr11-27545503-INS-228
chr11-27556767-INS-55
chr11-27557778-INS-746
chr11-27562615-INS-57
chr11-27569683-INS-6359
chr11-27601555-INS-400
chr11-27621509-INS-119
chr11-27633273-INS-5986

chr11-27662619-INS-4911
chr11-27664308-INS-7781
chr11-27668398-INS-343
chr11-27684507-INS-140
chr11-27698906-INS-6386
chr11-27707085-INS-393
chr11-27710560-INS-185
chr11-27730130-INS-343
chr11-27744966-INS-257
chr11-27791916-INS-56
chr11-27814464-INS-155
chr11-27815940-INS-70
chr11-27837387-INS-1626
chr11-27848706-INS-497
chr11-27852053-INS-65
chr11-27856616-INS-56
chr11-27876629-INS-7131
chr11-27889786-INS-136
chr11-27894223-INS-71
chr11-27897318-INS-548
chr11-27944101-INS-853
chr11-28017769-INS-2990
chr11-28025459-INS-497
chr11-28078094-INS-931
chr11-28143965-INS-148
chr11-28159101-INS-549
chr11-28207891-INS-445
chr11-28229559-INS-592
chr11-28233755-INS-362
chr11-28239938-INS-1892
chr11-28258863-INS-70
chr11-28315948-INS-72
chr11-28321304-INS-1622
chr11-28357331-INS-7376
chr11-28377473-INS-940
chr11-28383245-INS-270
chr11-28383250-INS-599
chr11-28405546-INS-513
chr11-28431903-INS-96
chr11-28447795-INS-107
chr11-28462116-INS-67
chr11-28465499-INS-139
chr11-28474649-INS-4344
chr11-28508001-INS-118
chr11-28564538-INS-4671
chr11-28600979-INS-89
chr11-28620152-INS-54
chr11-28627890-INS-305
chr11-28639046-INS-171
chr11-28656483-INS-4562
chr11-28690426-INS-345
chr11-28693598-INS-587
chr11-28700087-INS-801
chr11-28749466-INS-6703
chr11-28766515-INS-6634
chr11-28767582-INS-349
chr11-28776331-INS-209
chr11-28777140-INS-6442
chr11-28787767-INS-88
chr11-28808898-INS-123
chr11-28832871-INS-317
chr11-28845120-INS-52
chr11-28851762-INS-58
chr11-28896157-INS-211
chr11-28906074-INS-54
chr11-28938802-INS-1892
chr11-28941703-INS-6360
chr11-28971006-INS-837
chr11-28984742-INS-113
chr11-28989645-INS-205
chr11-28991393-INS-198
chr11-28998649-INS-1893
chr11-29011710-INS-542
chr11-29020993-INS-87
chr11-29029908-INS-112
chr11-29038819-INS-2211
chr11-29040107-INS-6421

chr11-29063881-INS-81
chr11-29070587-INS-97
chr11-29109831-INS-151
chr11-29126732-INS-76
chr11-29136583-INS-52
chr11-29142987-INS-51
chr11-29180314-INS-1191
chr11-29191384-INS-408
chr11-29217095-INS-486
chr11-29219115-INS-263
chr11-29224223-INS-265
chr11-29242061-INS-67
chr11-29245352-INS-493
chr11-29296963-INS-205
chr11-29351989-INS-467
chr11-29361403-INS-455
chr11-29396129-INS-54
chr11-29423908-INS-137
chr11-29429427-INS-193
chr11-29486994-INS-188
chr11-29502109-INS-6440
chr11-29505762-INS-207
chr11-29508186-INS-82
chr11-29509824-INS-51
chr11-29515767-INS-51
chr11-29551738-INS-121
chr11-29553177-INS-189
chr11-29560598-INS-69
chr11-29565699-INS-68
chr11-29579516-INS-826
chr11-29597525-INS-195
chr11-29599171-INS-183
chr11-29600447-INS-202
chr11-29602459-INS-202
chr11-29603025-INS-351
chr11-29619454-INS-145
chr11-29673569-INS-202
chr11-29731278-INS-212
chr11-29744111-INS-63
chr11-29750003-INS-207
chr11-29750180-INS-5336
chr11-29753914-INS-202
chr11-29796158-INS-6999
chr11-29797909-INS-278
chr11-29800035-INS-152
chr11-29810287-INS-963
chr11-29825595-INS-188
chr11-29885333-INS-274
chr11-29897539-INS-258
chr11-29905049-INS-67
chr11-29923013-INS-145
chr11-29948830-INS-55
chr11-29962915-INS-54
chr11-29973937-INS-116
chr11-30002031-INS-271
chr11-30052526-INS-512
chr11-30057849-INS-117
chr11-30076461-INS-82
chr11-30078207-INS-95
chr11-30164233-INS-66
chr11-30183375-INS-1025
chr11-30187044-INS-286
chr11-30211189-INS-200
chr11-30257350-INS-1864
chr11-30278150-INS-51
chr11-30330192-INS-399
chr11-30350225-INS-560
chr11-30378436-INS-108
chr11-30382842-INS-122
chr11-30409901-INS-213
chr11-30424036-INS-75
chr11-30442532-INS-512
chr11-30443058-INS-211
chr11-30444232-INS-341
chr11-30472277-INS-889
chr11-30495436-INS-144
chr11-30523526-INS-52

chr11-30543286-INS-408
chr11-30570907-INS-976
chr11-30579292-INS-92
chr11-30593569-INS-138
chr11-30602922-INS-841
chr11-30612173-INS-110
chr11-30624020-INS-225
chr11-30626040-INS-784
chr11-30641719-INS-203
chr11-30643439-INS-406
chr11-30687258-INS-889
chr11-30710076-INS-283
chr11-30711084-INS-206
chr11-30732794-INS-56
chr11-30741861-INS-1892
chr11-30755145-INS-525
chr11-30758741-INS-54
chr11-30792661-INS-84
chr11-30813844-INS-1893
chr11-30840211-INS-60
chr11-30855733-INS-51
chr11-30883506-INS-156
chr11-30900363-INS-269
chr11-30931487-INS-117
chr11-30973174-INS-62
chr11-30984946-INS-120
chr11-31015048-INS-90
chr11-31071173-INS-572
chr11-31120049-INS-52
chr11-31192398-INS-428
chr11-31220133-INS-214
chr11-31254769-INS-94
chr11-31305328-INS-74
chr11-31337444-INS-163
chr11-31342932-INS-488
chr11-31344517-INS-370
chr11-31366756-INS-485
chr11-31425735-INS-104
chr11-31441168-INS-635
chr11-31443801-INS-128
chr11-31450418-INS-52
chr11-31466152-INS-54
chr11-31505724-INS-340
chr11-31525728-INS-52
chr11-31529596-INS-71
chr11-31536782-INS-278
chr11-31678429-INS-53
chr11-31698318-INS-85
chr11-31720147-INS-100
chr11-31724114-INS-201
chr11-31849275-INS-114
chr11-31934114-INS-342
chr11-31940027-INS-404
chr11-31948312-INS-854
chr11-31956191-INS-99
chr11-31989359-INS-255
chr11-32013186-INS-153
chr11-32053045-INS-141
chr11-32053587-INS-55
chr11-32077945-INS-81
chr11-32089976-INS-53
chr11-32113215-INS-347
chr11-32142348-INS-223
chr11-32150886-INS-81
chr11-32167262-INS-130
chr11-32190794-INS-134
chr11-32221162-INS-163
chr11-32242301-INS-5436
chr11-32243504-INS-80
chr11-32301554-INS-61
chr11-32319823-INS-472
chr11-32325831-INS-465
chr11-32335222-INS-70
chr11-32339318-INS-198
chr11-32397299-INS-327
chr11-32399356-INS-197
chr11-32427875-INS-5218

chr11-32447284-INS-51
chr11-32448284-INS-167
chr11-32453108-INS-115
chr11-32465486-INS-82
chr11-32471897-INS-213
chr11-32476898-INS-441
chr11-32491358-INS-104
chr11-32508523-INS-52
chr11-32523990-INS-70
chr11-32528247-INS-64
chr11-32529746-INS-106
chr11-32538242-INS-69
chr11-32552039-INS-70
chr11-32557077-INS-227
chr11-32565359-INS-70
chr11-32571428-INS-91
chr11-32578711-INS-64
chr11-32584429-INS-205
chr11-32586988-INS-206
chr11-32647323-INS-148
chr11-32647375-INS-888
chr11-32674281-INS-225
chr11-32679252-INS-210
chr11-32703192-INS-120
chr11-32714217-INS-264
chr11-32716715-INS-464
chr11-32742789-INS-76
chr11-32747878-INS-105
chr11-32763595-INS-52
chr11-32764101-INS-141
chr11-32769634-INS-489
chr11-32771285-INS-104
chr11-32789656-INS-87
chr11-32792722-INS-885
chr11-32820564-INS-335
chr11-32842591-INS-267
chr11-32851137-INS-94
chr11-32854132-INS-340
chr11-32876815-INS-240
chr11-32886619-INS-282
chr11-32896339-INS-616
chr11-32913274-INS-60
chr11-32926266-INS-88
chr11-32933721-INS-212
chr11-32941786-INS-1892
chr11-32953414-INS-702
chr11-32956253-INS-120
chr11-32971886-INS-60
chr11-32974894-INS-210
chr11-32976865-INS-358
chr11-32988263-INS-3377
chr11-32996338-INS-6369
chr11-33063817-INS-93
chr11-33075059-INS-52
chr11-33077942-INS-136
chr11-33088335-INS-75
chr11-33096162-INS-51
chr11-33118868-INS-204
chr11-33194093-INS-7082
chr11-33197098-INS-231
chr11-33203722-INS-400
chr11-33278272-INS-52
chr11-33337533-INS-399
chr11-33394422-INS-53
chr11-33410789-INS-128
chr11-33430871-INS-198
chr11-33451217-INS-79
chr11-33459564-INS-573
chr11-33487105-INS-112
chr11-33488851-INS-186
chr11-33519480-INS-92
chr11-33533978-INS-86
chr11-33556031-INS-499
chr11-33557425-INS-136
chr11-33592387-INS-68
chr11-33605801-INS-165
chr11-33613089-INS-56

chr11-33617012-INS-166
chr11-33643080-INS-96
chr11-33643415-INS-199
chr11-33644281-INS-146
chr11-33668715-INS-74
chr11-33674739-INS-321
chr11-33684326-INS-399
chr11-33691059-INS-70
chr11-33719693-INS-183
chr11-33733708-INS-288
chr11-33743117-INS-206
chr11-33764050-INS-265
chr11-33778620-INS-103
chr11-33860463-INS-143
chr11-33870858-INS-177
chr11-33876732-INS-75
chr11-33901370-INS-686
chr11-33923106-INS-441
chr11-33928348-INS-600
chr11-33959382-INS-198
chr11-33980524-INS-516
chr11-33988413-INS-507
chr11-34023274-INS-67
chr11-34040511-INS-61
chr11-34046686-INS-111
chr11-34102201-INS-54
chr11-34123772-INS-117
chr11-34195136-INS-89
chr11-34201621-INS-73
chr11-34261603-INS-64
chr11-34263818-INS-58
chr11-34313495-INS-133
chr11-34332854-INS-284
chr11-34341808-INS-64
chr11-34365244-INS-68
chr11-34378384-INS-51
chr11-34447099-INS-771
chr11-34457799-INS-136
chr11-34462569-INS-251
chr11-34475750-INS-661
chr11-34498720-INS-404
chr11-34500129-INS-72
chr11-34503342-INS-84
chr11-34577888-INS-195
chr11-34589286-INS-79
chr11-34619985-INS-419
chr11-34630926-INS-383
chr11-34632887-INS-165
chr11-34651973-INS-64
chr11-34658069-INS-88
chr11-34660068-INS-120
chr11-34665788-INS-51
chr11-34673873-INS-54
chr11-34677880-INS-69
chr11-34679129-INS-60
chr11-34689329-INS-970
chr11-34703836-INS-158
chr11-34730362-INS-73
chr11-34735687-INS-401
chr11-34775568-INS-84
chr11-34790074-INS-108
chr11-34791640-INS-62
chr11-34839069-INS-703
chr11-34846231-INS-392
chr11-34868319-INS-197
chr11-34875178-INS-66
chr11-34897091-INS-164
chr11-34901233-INS-607
chr11-34901290-INS-167
chr11-34914966-INS-148
chr11-34959213-INS-90
chr11-34988128-INS-103
chr11-35090734-INS-138
chr11-35099763-INS-397
chr11-35110783-INS-388
chr11-35114112-INS-52
chr11-35121909-INS-1204

chr11-35124983-INS-203
chr11-35135142-INS-96
chr11-35202251-INS-103
chr11-35221246-INS-105
chr11-35224738-INS-283
chr11-35232344-INS-6591
chr11-35245685-INS-131
chr11-35247809-INS-210
chr11-35250146-INS-506
chr11-35294564-INS-389
chr11-35305124-INS-445
chr11-35316099-INS-117
chr11-35319717-INS-55
chr11-35349503-INS-204
chr11-35372995-INS-456
chr11-35376373-INS-6401
chr11-35422724-INS-65
chr11-35424400-INS-116
chr11-35440082-INS-76
chr11-35445453-INS-152
chr11-35455612-INS-86
chr11-35502051-INS-172
chr11-35551307-INS-144
chr11-35558476-INS-224
chr11-35563030-INS-76
chr11-35564922-INS-241
chr11-35581333-INS-64
chr11-35586167-INS-258
chr11-35612195-INS-292
chr11-35619637-INS-278
chr11-35622673-INS-1054
chr11-35637303-INS-72
chr11-35638575-INS-74
chr11-35646785-INS-67
chr11-35662744-INS-68
chr11-35664527-INS-172
chr11-35718469-INS-3524
chr11-35719083-INS-183
chr11-35734007-INS-743
chr11-35736167-INS-400
chr11-35753631-INS-92
chr11-35792758-INS-248
chr11-35849895-INS-96
chr11-35851318-INS-79
chr11-35858506-INS-218
chr11-35880776-INS-401
chr11-35885737-INS-171
chr11-35886773-INS-2469
chr11-35896467-INS-141
chr11-35903140-INS-74
chr11-35913027-INS-498
chr11-35916954-INS-74
chr11-35948462-INS-163
chr11-35958856-INS-432
chr11-35961664-INS-85
chr11-35963480-INS-6450
chr11-35998294-INS-53
chr11-36035125-INS-81
chr11-36036808-INS-58
chr11-36063049-INS-6708
chr11-36084308-INS-96
chr11-36111978-INS-101
chr11-36231514-INS-61
chr11-36256265-INS-58
chr11-36259736-INS-64
chr11-36269027-INS-890
chr11-36271821-INS-75
chr11-36317400-INS-112
chr11-36336367-INS-215
chr11-36339460-INS-54
chr11-36415783-INS-401
chr11-36456151-INS-388
chr11-36484839-INS-296
chr11-36485706-INS-118
chr11-36493407-INS-3115
chr11-36525135-INS-329
chr11-36552096-INS-93

chr11-36564690-INS-76
chr11-36635217-INS-183
chr11-36644152-INS-80
chr11-36692467-INS-438
chr11-36724938-INS-197
chr11-36770138-INS-144
chr11-36771894-INS-117
chr11-36800317-INS-108
chr11-36812169-INS-266
chr11-36818831-INS-116
chr11-36822194-INS-230
chr11-36866257-INS-52
chr11-36883141-INS-71
chr11-36903001-INS-56
chr11-36925459-INS-138
chr11-36946276-INS-132
chr11-36964602-INS-52
chr11-36977197-INS-6612
chr11-36983173-INS-672
chr11-36995866-INS-6440
chr11-37003475-INS-72
chr11-37031209-INS-52
chr11-37039209-INS-67
chr11-37082455-INS-151
chr11-37099714-INS-265
chr11-37145049-INS-69
chr11-37149556-INS-54
chr11-37152806-INS-5645
chr11-37159149-INS-885
chr11-37174331-INS-106
chr11-37190744-INS-60
chr11-37236541-INS-56
chr11-37246842-INS-394
chr11-37273873-INS-511
chr11-37334673-INS-3618
chr11-37335490-INS-70
chr11-37389715-INS-52
chr11-37436478-INS-288
chr11-37450286-INS-56
chr11-37451239-INS-215
chr11-37456580-INS-316
chr11-37487942-INS-177
chr11-37495855-INS-52
chr11-37522973-INS-5386
chr11-37538828-INS-6817
chr11-37587778-INS-54
chr11-37678567-INS-59
chr11-37730706-INS-151
chr11-37738444-INS-558
chr11-37744617-INS-244
chr11-37745791-INS-476
chr11-37786520-INS-58
chr11-37808575-INS-130
chr11-37842913-INS-56
chr11-37851712-INS-108
chr11-37859886-INS-269
chr11-37861841-INS-136
chr11-37901253-INS-56
chr11-37931477-INS-100
chr11-37935831-INS-6536
chr11-37971879-INS-331
chr11-38013467-INS-190
chr11-38014043-INS-267
chr11-38044604-INS-458
chr11-38057567-INS-63
chr11-38066816-INS-64
chr11-38085500-INS-100
chr11-38112447-INS-131
chr11-38131966-INS-6537
chr11-38135263-INS-144
chr11-38149247-INS-106
chr11-38176058-INS-84
chr11-38197648-INS-67
chr11-38201551-INS-58
chr11-38206436-INS-4370
chr11-38211045-INS-94
chr11-38259998-INS-55

chr11-38277982-INS-54
chr11-38290550-INS-359
chr11-38367702-INS-67
chr11-38394927-INS-57
chr11-38430363-INS-196
chr11-38488886-INS-102
chr11-38582688-INS-78
chr11-38594673-INS-148
chr11-38660525-INS-60
chr11-38700361-INS-70
chr11-38725230-INS-205
chr11-38770458-INS-7286
chr11-38770961-INS-108
chr11-38774558-INS-53
chr11-38787679-INS-58
chr11-38788841-INS-7904
chr11-38800050-INS-416
chr11-38836500-INS-447
chr11-38857556-INS-104
chr11-38915264-INS-179
chr11-38916900-INS-60
chr11-38927270-INS-489
chr11-38935485-INS-398
chr11-38938020-INS-172
chr11-38947969-INS-619
chr11-38975133-INS-74
chr11-38977239-INS-69
chr11-38981613-INS-51
chr11-38983293-INS-882
chr11-39049909-INS-7299
chr11-39053572-INS-89
chr11-39075300-INS-56
chr11-39112355-INS-68
chr11-39168132-INS-141
chr11-39168931-INS-915
chr11-39170353-INS-78
chr11-39188376-INS-4388
chr11-39193353-INS-206
chr11-39193828-INS-1037
chr11-39202460-INS-72
chr11-39210563-INS-351
chr11-39221505-INS-90
chr11-39222252-INS-64
chr11-39230046-INS-280
chr11-39245425-INS-116
chr11-39261643-INS-6157
chr11-39284109-INS-127
chr11-39299410-INS-714
chr11-39303408-INS-488
chr11-39310040-INS-868
chr11-39361231-INS-263
chr11-39401299-INS-202
chr11-39410348-INS-220
chr11-39413425-INS-104
chr11-39430468-INS-65
chr11-39435518-INS-122
chr11-39446874-INS-56
chr11-39450279-INS-57
chr11-39452629-INS-52
chr11-39455127-INS-513
chr11-39457750-INS-546
chr11-39477297-INS-195
chr11-39478206-INS-58
chr11-39483676-INS-86
chr11-39492106-INS-174
chr11-39496987-INS-505
chr11-39509312-INS-612
chr11-39513708-INS-3164
chr11-39521096-INS-80
chr11-39555297-INS-6515
chr11-39555838-INS-78
chr11-39561216-INS-58
chr11-39573710-INS-59
chr11-39606554-INS-60
chr11-39620705-INS-78
chr11-39621754-INS-131
chr11-39623530-INS-60

chr11-39628286-INS-72
chr11-39648329-INS-7097
chr11-39649899-INS-93
chr11-39668579-INS-6687
chr11-39681262-INS-60
chr11-39695415-INS-6406
chr11-39702336-INS-271
chr11-39725268-INS-279
chr11-39738281-INS-219
chr11-39750460-INS-117
chr11-39770157-INS-500
chr11-39777005-INS-4900
chr11-39791333-INS-164
chr11-39846952-INS-55
chr11-39858646-INS-87
chr11-39861177-INS-68
chr11-39876815-INS-121
chr11-39876979-INS-5796
chr11-39894101-INS-777
chr11-39945493-INS-252
chr11-39964318-INS-62
chr11-39975268-INS-61
chr11-39990609-INS-55
chr11-40008001-INS-51
chr11-40017901-INS-219
chr11-40031684-INS-351
chr11-40034403-INS-70
chr11-40058134-INS-65
chr11-40072242-INS-891
chr11-40077717-INS-1494
chr11-40078490-INS-190
chr11-40090555-INS-6489
chr11-40095608-INS-873
chr11-40107721-INS-52
chr11-40119372-INS-7058
chr11-40192627-INS-696
chr11-40197353-INS-139
chr11-40203046-INS-773
chr11-40204821-INS-250
chr11-40211695-INS-70
chr11-40220438-INS-56
chr11-40222344-INS-6391
chr11-40230056-INS-62
chr11-40239386-INS-202
chr11-40262540-INS-413
chr11-40266702-INS-128
chr11-40288066-INS-551
chr11-40313505-INS-6215
chr11-40338002-INS-77
chr11-40388994-INS-57
chr11-40394043-INS-128
chr11-40405029-INS-473
chr11-40409845-INS-142
chr11-40414647-INS-105
chr11-40428439-INS-59
chr11-40438776-INS-170
chr11-40454752-INS-1802
chr11-40476481-INS-52
chr11-40479174-INS-515
chr11-40485642-INS-53
chr11-40494026-INS-63
chr11-40499915-INS-400
chr11-40537128-INS-486
chr11-40541257-INS-97
chr11-40543452-INS-506
chr11-40579132-INS-127
chr11-40586227-INS-826
chr11-40596534-INS-314
chr11-40614439-INS-55
chr11-40636960-INS-206
chr11-40649124-INS-400
chr11-40654207-INS-58
chr11-40670221-INS-54
chr11-40679822-INS-207
chr11-40682509-INS-156
chr11-40706843-INS-54
chr11-40764886-INS-140

chr11-40776608-INS-55
chr11-40780251-INS-159
chr11-40781659-INS-54
chr11-40795281-INS-571
chr11-40817393-INS-216
chr11-40831177-INS-597
chr11-40841887-INS-69
chr11-40890842-INS-956
chr11-40893903-INS-941
chr11-40915767-INS-6785
chr11-40928055-INS-1861
chr11-40945116-INS-442
chr11-40976487-INS-393
chr11-41009762-INS-3686
chr11-41011998-INS-6656
chr11-41018759-INS-324
chr11-41039908-INS-378
chr11-41057779-INS-483
chr11-41061822-INS-52
chr11-41074023-INS-6198
chr11-41075035-INS-161
chr11-41158243-INS-66
chr11-41177966-INS-1094
chr11-41182751-INS-67
chr11-41193184-INS-88
chr11-41194695-INS-286
chr11-41204630-INS-952
chr11-41211652-INS-91
chr11-41220868-INS-783
chr11-41236055-INS-66
chr11-41252405-INS-6226
chr11-41285918-INS-333
chr11-41320037-INS-190
chr11-41320345-INS-347
chr11-41321877-INS-123
chr11-41330778-INS-120
chr11-41333608-INS-67
chr11-41379434-INS-80
chr11-41394986-INS-54
chr11-41435492-INS-121
chr11-41460809-INS-786
chr11-41500867-INS-523
chr11-41502832-INS-54
chr11-41521034-INS-457
chr11-41528337-INS-404
chr11-41557360-INS-181
chr11-41574313-INS-288
chr11-41612102-INS-6620
chr11-41619109-INS-70
chr11-41683656-INS-260
chr11-41694139-INS-307
chr11-41695613-INS-57
chr11-41736802-INS-82
chr11-41792198-INS-456
chr11-41795208-INS-112
chr11-41819731-INS-6554
chr11-41832021-INS-72
chr11-41840381-INS-129
chr11-41851692-INS-87
chr11-41904121-INS-278
chr11-41909064-INS-72
chr11-41930931-INS-72
chr11-41946970-INS-4053
chr11-41956905-INS-216
chr11-41958431-INS-166
chr11-41987453-INS-52
chr11-41995804-INS-561
chr11-42085236-INS-246
chr11-42096163-INS-702
chr11-42109532-INS-497
chr11-42134052-INS-338
chr11-42143616-INS-60
chr11-42188585-INS-54
chr11-42196184-INS-56
chr11-42198766-INS-65
chr11-42218497-INS-112
chr11-42222754-INS-60

chr11-42244835-INS-7019
chr11-42247028-INS-400
chr11-42256183-INS-108
chr11-42261683-INS-450
chr11-42282320-INS-58
chr11-42314501-INS-100
chr11-42327116-INS-61
chr11-42364190-INS-628
chr11-42368335-INS-63
chr11-42384837-INS-669
chr11-42389053-INS-860
chr11-42459747-INS-276
chr11-42468532-INS-555
chr11-42481454-INS-469
chr11-42525730-INS-60
chr11-42539003-INS-60
chr11-42607021-INS-61
chr11-42609964-INS-213
chr11-42612200-INS-659
chr11-42623255-INS-438
chr11-42647373-INS-7091
chr11-42648996-INS-257
chr11-42682193-INS-56
chr11-42686772-INS-1002
chr11-42686798-INS-981
chr11-42686800-INS-1466
chr11-42693357-INS-54
chr11-42720951-INS-461
chr11-42739718-INS-56
chr11-42783705-INS-83
chr11-42793326-INS-87
chr11-42822232-INS-116
chr11-42869061-INS-507
chr11-42872254-INS-630
chr11-42919368-INS-64
chr11-42920656-INS-328
chr11-42936687-INS-77
chr11-42947947-INS-132
chr11-42975886-INS-904
chr11-42979252-INS-358
chr11-42992065-INS-95
chr11-43016553-INS-70
chr11-43028835-INS-132
chr11-43035633-INS-86
chr11-43048572-INS-624
chr11-43050299-INS-145
chr11-43064493-INS-53
chr11-43075117-INS-406
chr11-43082569-INS-798
chr11-43096264-INS-56
chr11-43099764-INS-92
chr11-43123663-INS-312
chr11-43185945-INS-72
chr11-43214229-INS-155
chr11-43236630-INS-52
chr11-43248581-INS-57
chr11-43252014-INS-54
chr11-43257200-INS-55
chr11-43269823-INS-96
chr11-43271965-INS-166
chr11-43276550-INS-116
chr11-43282705-INS-291
chr11-43293221-INS-139
chr11-43297689-INS-5658
chr11-43308723-INS-65
chr11-43330787-INS-176
chr11-43331637-INS-897
chr11-43339179-INS-880
chr11-43341788-INS-163
chr11-43343531-INS-4882
chr11-43360145-INS-60
chr11-43368690-INS-56
chr11-43380416-INS-981
chr11-43391685-INS-108
chr11-43398338-INS-52
chr11-43417607-INS-182
chr11-43432581-INS-55

chr11-43438553-INS-55
chr11-43467921-INS-53
chr11-43473517-INS-163
chr11-43488799-INS-279
chr11-43490471-INS-202
chr11-43493251-INS-58
chr11-43525991-INS-213
chr11-43547789-INS-95
chr11-43548844-INS-423
chr11-43575891-INS-88
chr11-43594316-INS-168
chr11-43607399-INS-225
chr11-43626622-INS-96
chr11-43630378-INS-124
chr11-43649972-INS-51
chr11-43656183-INS-65
chr11-43668807-INS-111
chr11-43669975-INS-324
chr11-43672316-INS-90
chr11-43685984-INS-6442
chr11-43701727-INS-77
chr11-43720777-INS-135
chr11-43743128-INS-128
chr11-43757401-INS-98
chr11-43765822-INS-93
chr11-43769485-INS-126
chr11-43781789-INS-245
chr11-43803868-INS-53
chr11-43809553-INS-93
chr11-43816937-INS-117
chr11-43825118-INS-341
chr11-43829973-INS-135
chr11-43847845-INS-403
chr11-43850643-INS-153
chr11-43861608-INS-52
chr11-43870141-INS-75
chr11-43889789-INS-139
chr11-43895161-INS-2349
chr11-43913216-INS-82
chr11-43915013-INS-68
chr11-43916929-INS-79
chr11-43922926-INS-5417
chr11-43934477-INS-318
chr11-43935156-INS-96
chr11-43938414-INS-95
chr11-43942353-INS-6786
chr11-43989421-INS-1390
chr11-44013247-INS-208
chr11-44072572-INS-53
chr11-44123038-INS-124
chr11-44157742-INS-295
chr11-44159990-INS-76
chr11-44170352-INS-322
chr11-44179691-INS-400
chr11-44183514-INS-94
chr11-44183539-INS-682
chr11-44208560-INS-982
chr11-44211313-INS-136
chr11-44212136-INS-6504
chr11-44212769-INS-7640
chr11-44234186-INS-441
chr11-44236082-INS-65
chr11-44239598-INS-241
chr11-44261893-INS-52
chr11-44281590-INS-572
chr11-44284268-INS-90
chr11-44288973-INS-55
chr11-44315582-INS-88
chr11-44331300-INS-988
chr11-44335343-INS-509
chr11-44338554-INS-52
chr11-44339884-INS-145
chr11-44340943-INS-632
chr11-44349442-INS-160
chr11-44353499-INS-212
chr11-44353638-INS-723
chr11-44356140-INS-92

chr11-44365183-INS-56
chr11-44373376-INS-51
chr11-44391062-INS-131
chr11-44400901-INS-169
chr11-44427248-INS-156
chr11-44431293-INS-64
chr11-44434933-INS-64
chr11-44498376-INS-117
chr11-44526641-INS-68
chr11-44527673-INS-225
chr11-44551325-INS-222
chr11-44558200-INS-84
chr11-44608334-INS-56
chr11-44616872-INS-106
chr11-44630095-INS-61
chr11-44633162-INS-236
chr11-44642832-INS-84
chr11-44656840-INS-102
chr11-44670391-INS-55
chr11-44677643-INS-91
chr11-44699563-INS-101
chr11-44713491-INS-104
chr11-44720252-INS-80
chr11-44732968-INS-67
chr11-44737968-INS-74
chr11-44776153-INS-112
chr11-44803582-INS-281
chr11-44849061-INS-436
chr11-44858771-INS-134
chr11-44891030-INS-59
chr11-44893476-INS-104
chr11-44910569-INS-108
chr11-44929609-INS-62
chr11-44953828-INS-728
chr11-44958999-INS-54
chr11-44960502-INS-56
chr11-44994020-INS-401
chr11-44995151-INS-66
chr11-45002747-INS-60
chr11-45021988-INS-124
chr11-45040449-INS-526
chr11-45096605-INS-60
chr11-45099834-INS-6397
chr11-45100242-INS-51
chr11-45107757-INS-80
chr11-45108211-INS-104
chr11-45116097-INS-61
chr11-45124277-INS-336
chr11-45133715-INS-612
chr11-45193908-INS-56
chr11-45238850-INS-398
chr11-45245416-INS-112
chr11-45252585-INS-83
chr11-45255045-INS-246
chr11-45269352-INS-56
chr11-45329639-INS-6281
chr11-45359922-INS-6931
chr11-45373478-INS-5714
chr11-45383600-INS-65
chr11-45400056-INS-160
chr11-45408658-INS-97
chr11-45409386-INS-85
chr11-45420522-INS-6479
chr11-45427944-INS-54
chr11-45445610-INS-52
chr11-45469291-INS-56
chr11-45473942-INS-402
chr11-45487685-INS-345
chr11-45489958-INS-53
chr11-45505832-INS-401
chr11-45518933-INS-89
chr11-45524784-INS-149
chr11-45536996-INS-201
chr11-45567751-INS-1891
chr11-45567986-INS-394
chr11-45574799-INS-963
chr11-45606219-INS-77

chr11-45614828-INS-624
chr11-45636473-INS-12424
chr11-45645100-INS-889
chr11-45646109-INS-237
chr11-45677118-INS-6403
chr11-45679746-INS-357
chr11-45694386-INS-392
chr11-45703345-INS-124
chr11-45723707-INS-124
chr11-45725808-INS-60
chr11-45740416-INS-200
chr11-45758354-INS-213
chr11-45779345-INS-422
chr11-45781840-INS-215
chr11-45791516-INS-6401
chr11-45814253-INS-120
chr11-45829193-INS-198
chr11-45830893-INS-202
chr11-45905506-INS-54
chr11-45915548-INS-184
chr11-45923210-INS-197
chr11-45925985-INS-407
chr11-45931012-INS-181
chr11-45934243-INS-68
chr11-45935606-INS-334
chr11-45952290-INS-83
chr11-45966613-INS-206
chr11-45997163-INS-514
chr11-46003566-INS-138
chr11-46012073-INS-205
chr11-46027666-INS-75
chr11-46028945-INS-200
chr11-46039757-INS-62
chr11-46046562-INS-1891
chr11-46051928-INS-272
chr11-46058059-INS-202
chr11-46060581-INS-52
chr11-46066478-INS-68
chr11-46072849-INS-214
chr11-46083236-INS-358
chr11-46093501-INS-211
chr11-46101645-INS-345
chr11-46111828-INS-116
chr11-46145824-INS-55
chr11-46153377-INS-306
chr11-46175917-INS-72
chr11-46203651-INS-322
chr11-46214894-INS-56
chr11-46219768-INS-56
chr11-46223956-INS-56
chr11-46224552-INS-56
chr11-46228358-INS-61
chr11-46230451-INS-119
chr11-46230954-INS-56
chr11-46276626-INS-958
chr11-46291206-INS-191
chr11-46293833-INS-6443
chr11-46310891-INS-180
chr11-46328888-INS-99
chr11-46337794-INS-1526
chr11-46352211-INS-969
chr11-46362853-INS-87
chr11-46378867-INS-205
chr11-46407574-INS-181
chr11-46429799-INS-95
chr11-46436160-INS-70
chr11-46454288-INS-207
chr11-46455576-INS-84
chr11-46456647-INS-226
chr11-46459919-INS-85
chr11-46468943-INS-76
chr11-46528431-INS-54
chr11-46553393-INS-181
chr11-46556067-INS-233
chr11-46582628-INS-54
chr11-46587060-INS-53
chr11-46598637-INS-53

chr11-46612241-INS-438
chr11-46621242-INS-6590
chr11-46635917-INS-61
chr11-46661760-INS-7327
chr11-46686299-INS-234
chr11-46690400-INS-66
chr11-46692432-INS-70
chr11-46725406-INS-572
chr11-46751309-INS-210
chr11-46756954-INS-951
chr11-46769875-INS-170
chr11-46787039-INS-502
chr11-46790850-INS-66
chr11-46823962-INS-2911
chr11-46827940-INS-344
chr11-46870901-INS-61
chr11-46895488-INS-60
chr11-46901278-INS-134
chr11-46922293-INS-60
chr11-46933069-INS-76
chr11-46963711-INS-166
chr11-46966200-INS-502
chr11-47013899-INS-446
chr11-47022569-INS-195
chr11-47031722-INS-457
chr11-47046429-INS-90
chr11-47095085-INS-55
chr11-47099233-INS-89
chr11-47114026-INS-81
chr11-47166019-INS-488
chr11-47189789-INS-240
chr11-47199509-INS-56
chr11-47220545-INS-102
chr11-47250132-INS-68
chr11-47251678-INS-6182
chr11-47256746-INS-5401
chr11-47291320-INS-201
chr11-47299711-INS-1231
chr11-47307304-INS-110
chr11-47327364-INS-141
chr11-47332362-INS-53
chr11-47349124-INS-175
chr11-47368805-INS-7464
chr11-47440819-INS-548
chr11-47447120-INS-75
chr11-47494084-INS-547
chr11-47500762-INS-399
chr11-47503309-INS-962
chr11-47531963-INS-101
chr11-47538441-INS-461
chr11-47556474-INS-81
chr11-47564146-INS-147
chr11-47567793-INS-120
chr11-47591249-INS-374
chr11-47615638-INS-62
chr11-47623040-INS-3821
chr11-47627411-INS-445
chr11-47648299-INS-94
chr11-47653633-INS-55
chr11-47695896-INS-192
chr11-47700437-INS-1239
chr11-47708360-INS-7119
chr11-47741224-INS-212
chr11-47752546-INS-258
chr11-47768712-INS-51
chr11-47779096-INS-240
chr11-47787236-INS-60
chr11-47821920-INS-178
chr11-47827080-INS-6953
chr11-47827388-INS-133
chr11-47849455-INS-1218
chr11-47850288-INS-164
chr11-47877923-INS-195
chr11-47899475-INS-3935
chr11-47906085-INS-6740
chr11-47916934-INS-146
chr11-47943102-INS-152

chr11-47955741-INS-8199
chr11-47962780-INS-6026
chr11-47963242-INS-608
chr11-47968717-INS-6440
chr11-47968718-INS-497
chr11-47970017-INS-2638
chr11-47991834-INS-51
chr11-47998731-INS-618
chr11-48032951-INS-309
chr11-48039360-INS-401
chr11-48073917-INS-794
chr11-48082535-INS-76
chr11-48088276-INS-132
chr11-48088608-INS-436
chr11-48089694-INS-443
chr11-48099138-INS-88
chr11-48100278-INS-74
chr11-48132676-INS-271
chr11-48178952-INS-56
chr11-48192601-INS-80
chr11-48198239-INS-388
chr11-48201542-INS-77
chr11-48204130-INS-359
chr11-48215397-INS-170
chr11-48232328-INS-52
chr11-48245742-INS-110
chr11-48252100-INS-177
chr11-48286224-INS-68
chr11-48332393-INS-175
chr11-48355469-INS-117
chr11-48384049-INS-692
chr11-48404091-INS-2948
chr11-48404733-INS-6376
chr11-48408768-INS-3862
chr11-48447678-INS-6796
chr11-48459957-INS-819
chr11-48477701-INS-97
chr11-48478001-INS-1524
chr11-48479510-INS-52
chr11-48495688-INS-85
chr11-48500100-INS-6793
chr11-48505413-INS-6593
chr11-48525759-INS-105
chr11-48533018-INS-6114
chr11-48535761-INS-8836
chr11-48541358-INS-58
chr11-48570934-INS-205
chr11-48575024-INS-927
chr11-48613094-INS-214
chr11-48615529-INS-6686
chr11-48618548-INS-512
chr11-48639044-INS-95
chr11-48645133-INS-153
chr11-48658345-INS-2542
chr11-48661027-INS-54
chr11-48665982-INS-497
chr11-48671728-INS-371
chr11-48672819-INS-208
chr11-48679533-INS-60
chr11-48681677-INS-65
chr11-48687564-INS-200
chr11-48690122-INS-8419
chr11-48710581-INS-214
chr11-48719443-INS-130
chr11-48724281-INS-54
chr11-48729850-INS-132
chr11-48751494-INS-302
chr11-48751738-INS-965
chr11-48755945-INS-128
chr11-48771530-INS-185
chr11-48796639-INS-196
chr11-48818500-INS-101
chr11-48835772-INS-6401
chr11-48854692-INS-111
chr11-48871031-INS-92
chr11-48898160-INS-62
chr11-48909649-INS-200

chr11-48955566-INS-75
chr11-48964401-INS-6382
chr11-48988451-INS-449
chr11-48996569-INS-118
chr11-49000247-INS-56
chr11-49014463-INS-11118
chr11-49024462-INS-82
chr11-49036705-INS-74
chr11-49037795-INS-72
chr11-49039003-INS-71
chr11-49042536-INS-529
chr11-49073287-INS-98
chr11-49093603-INS-51
chr11-49095150-INS-205
chr11-49101729-INS-673
chr11-49117853-INS-396
chr11-49143617-INS-3190
chr11-49149964-INS-684
chr11-49158432-INS-72
chr11-49203318-INS-68
chr11-49247915-INS-202
chr11-49255516-INS-94
chr11-49300487-INS-876
chr11-49318171-INS-92
chr11-49318357-INS-494
chr11-49345828-INS-6999
chr11-49375105-INS-2548
chr11-49391385-INS-3873
chr11-49404164-INS-5757
chr11-49428001-INS-3901
chr11-49438837-INS-64
chr11-49454134-INS-310
chr11-49463612-INS-136
chr11-49521824-INS-123
chr11-49552836-INS-118
chr11-49567394-INS-61
chr11-49584352-INS-57
chr11-49605161-INS-60
chr11-49619494-INS-94
chr11-49643125-INS-56
chr11-49651568-INS-56
chr11-49655961-INS-182
chr11-49661474-INS-83
chr11-49666287-INS-102
chr11-49667163-INS-358
chr11-49668645-INS-473
chr11-49670452-INS-290
chr11-49703075-INS-52
chr11-49712261-INS-74
chr11-49725642-INS-92
chr11-49726717-INS-143
chr11-49748182-INS-70
chr11-49781709-INS-439
chr11-49814101-INS-120
chr11-49819521-INS-77
chr11-49851093-INS-66
chr11-49864324-INS-840
chr11-49864372-INS-136
chr11-49865769-INS-209
chr11-49868229-INS-163
chr11-49878107-INS-202
chr11-49879523-INS-648
chr11-49880431-INS-113
chr11-49900992-INS-171
chr11-49921294-INS-175
chr11-49939530-INS-168
chr11-49955530-INS-62
chr11-49958662-INS-203
chr11-49959861-INS-220
chr11-49961094-INS-216
chr11-49969414-INS-171
chr11-49983588-INS-202
chr11-50019005-INS-54
chr11-50024983-INS-204
chr11-50046182-INS-95
chr11-50047169-INS-80
chr11-50057436-INS-99

chr11-50064878-INS-778
chr11-50080508-INS-213
chr11-50084006-INS-6438
chr11-50094786-INS-176
chr11-50096744-INS-206
chr11-50154773-INS-204
chr11-50155668-INS-81
chr11-50158811-INS-469
chr11-50167063-INS-52
chr11-50178896-INS-67
chr11-50187838-INS-445
chr11-50194457-INS-205
chr11-50214666-INS-106
chr11-50224383-INS-60
chr11-50247065-INS-196
chr11-50252611-INS-62
chr11-50259438-INS-160
chr11-50314502-INS-409
chr11-50340419-INS-4062
chr11-50347776-INS-66
chr11-50368956-INS-141
chr11-50376280-INS-104
chr11-50377616-INS-233
chr11-50397426-INS-67
chr11-50404781-INS-255
chr11-50409389-INS-341
chr11-50413319-INS-52
chr11-50428002-INS-488
chr11-50473138-INS-669
chr11-50486148-INS-54
chr11-50510567-INS-237
chr11-50557393-INS-576
chr11-50588684-INS-365
chr11-50607993-INS-114
chr11-50616697-INS-177
chr11-50652421-INS-64
chr11-50662988-INS-59
chr11-50664822-INS-368
chr11-50690857-INS-61
chr11-50721952-INS-59
chr11-50739397-INS-53
chr11-50771991-INS-268
chr11-50784717-INS-890
chr11-50818322-INS-212
chr11-50830415-INS-1926
chr11-50832285-INS-7181
chr11-50832449-INS-4707
chr11-50866985-INS-400
chr11-50878086-INS-4420
chr11-50881780-INS-735
chr11-50934210-INS-22972
chr11-50945852-INS-873
chr11-50948339-INS-3755
chr11-50965311-INS-443
chr11-50966163-INS-214
chr11-50970666-INS-79
chr11-50990539-INS-66
chr11-51019497-INS-486
chr11-51023215-INS-60
chr11-51057442-INS-96
chr11-51074063-INS-538
chr11-51080567-INS-6231
chr11-51094395-INS-399
chr11-51109163-INS-130
chr11-51138969-INS-54
chr11-51141117-INS-217
chr11-51160035-INS-57
chr11-51198056-INS-6243
chr11-51203619-INS-126
chr11-51205094-INS-62
chr11-51212809-INS-56
chr11-51215306-INS-93
chr11-51224380-INS-69
chr11-51235598-INS-68
chr11-51241728-INS-111
chr11-51261053-INS-2948
chr11-51271520-INS-67

chr11-51279533-INS-89
chr11-51291024-INS-873
chr11-51307888-INS-52
chr11-51311771-INS-236
chr11-51313036-INS-222
chr11-51321489-INS-110
chr11-51326358-INS-68
chr11-51390288-INS-99
chr11-51401470-INS-64
chr11-51404817-INS-52
chr11-51423611-INS-391
chr11-51428412-INS-339
chr11-51441624-INS-93
chr11-51464351-INS-212
chr11-51465806-INS-345
chr11-51479518-INS-164
chr11-51482672-INS-213
chr11-51505729-INS-151
chr11-51524037-INS-150
chr11-51540431-INS-251
chr11-51544122-INS-184
chr11-51551064-INS-140
chr11-51566970-INS-66
chr11-51568147-INS-206
chr11-51569842-INS-88
chr11-51573354-INS-253
chr11-51600978-INS-52
chr11-51615368-INS-360
chr11-51625140-INS-129
chr11-51649153-INS-53
chr11-51653195-INS-57
chr11-51657273-INS-627
chr11-51658170-INS-197
chr11-51667144-INS-8985
chr11-51676209-INS-190
chr11-51680621-INS-181
chr11-51687684-INS-52
chr11-51690095-INS-214
chr11-51690283-INS-2872
chr11-51696727-INS-162
chr11-51701510-INS-52
chr11-51701964-INS-197
chr11-51782336-INS-105
chr11-51789508-INS-104
chr11-51802160-INS-221
chr11-51808193-INS-71
chr11-51814690-INS-62
chr11-51824476-INS-131
chr11-51834783-INS-52
chr11-51846316-INS-126
chr11-51856886-INS-2775
chr11-51883063-INS-509
chr11-51899685-INS-167
chr11-51906083-INS-109
chr11-51906676-INS-67
chr11-51921349-INS-125
chr11-51934936-INS-278
chr11-51939816-INS-162
chr11-51951843-INS-106
chr11-51984969-INS-5219
chr11-52000995-INS-89
chr11-52022159-INS-52
chr11-52235358-INS-180
chr11-52253518-INS-54
chr11-52294470-INS-213
chr11-52340784-INS-337
chr11-52350413-INS-332
chr11-52366482-INS-209
chr11-52389663-INS-192
chr11-52398193-INS-340
chr11-52405699-INS-203
chr11-52409758-INS-140
chr11-52427580-INS-55
chr11-52428581-INS-874
chr11-52449189-INS-209
chr11-52460582-INS-54
chr11-52474150-INS-112

chr11-52479882-INS-56
chr11-52497566-INS-347
chr11-52515206-INS-393
chr11-52554049-INS-198
chr11-52558995-INS-54
chr11-52559271-INS-464
chr11-52577476-INS-681
chr11-52602933-INS-178
chr11-52605120-INS-54
chr11-52606267-INS-733
chr11-52612906-INS-78
chr11-52613938-INS-61
chr11-52621683-INS-104
chr11-52630223-INS-90
chr11-52635022-INS-1044
chr11-52645654-INS-5293
chr11-52654888-INS-66
chr11-52663909-INS-93
chr11-52678520-INS-470
chr11-52727429-INS-449
chr11-52784208-INS-109
chr11-52796798-INS-66
chr11-52799022-INS-65
chr11-52843870-INS-100
chr11-52855346-INS-6425
chr11-52876606-INS-56
chr11-52902828-INS-60
chr11-52933415-INS-124
chr11-52944957-INS-54
chr11-52982231-INS-94
chr11-52995157-INS-62
chr11-53022859-INS-419
chr11-53045734-INS-68
chr11-53056782-INS-54
chr11-53093705-INS-3279
chr11-53102452-INS-468
chr11-53104740-INS-144
chr11-53122041-INS-205
chr11-53133137-INS-179
chr11-53137826-INS-95
chr11-53144618-INS-169
chr11-53170197-INS-263
chr11-53203894-INS-53
chr11-53206381-INS-72
chr11-53225784-INS-230
chr11-53230840-INS-646
chr11-53237720-INS-211
chr11-53241077-INS-63
chr11-53247043-INS-191
chr11-53286836-INS-132
chr11-53317603-INS-239
chr11-53323246-INS-173
chr11-53329424-INS-176
chr11-53338300-INS-201
chr11-53359869-INS-210
chr11-53372760-INS-52
chr11-53376824-INS-51
chr11-53389760-INS-56
chr11-53391096-INS-225
chr11-53403398-INS-106
chr11-53404644-INS-757
chr11-53409684-INS-104
chr11-53441846-INS-199
chr11-53469116-INS-150
chr11-53501079-INS-198
chr11-53510748-INS-220
chr11-53516901-INS-78
chr11-53528614-INS-116
chr11-53530485-INS-83
chr11-53532918-INS-552
chr11-53552241-INS-293
chr11-53573320-INS-196
chr11-53617165-INS-170
chr11-53618829-INS-354
chr11-53625059-INS-211
chr11-53628721-INS-53
chr11-53636020-INS-143

chr11-53640017-INS-434
chr11-53642346-INS-72
chr11-53649301-INS-91
chr11-53652595-INS-59
chr11-53669564-INS-135
chr11-53671605-INS-75
chr11-53677557-INS-105
chr11-53687344-INS-52
chr11-53698016-INS-196
chr11-53726220-INS-197
chr11-53759413-INS-400
chr11-53776962-INS-195
chr11-53788380-INS-203
chr11-53800947-INS-142
chr11-53823755-INS-153
chr11-53844440-INS-98
chr11-53852600-INS-219
chr11-53857215-INS-5712
chr11-53873363-INS-82
chr11-53907703-INS-60
chr11-53921973-INS-274
chr11-53925881-INS-51
chr11-53983355-INS-257
chr11-53985578-INS-187
chr11-53986232-INS-80
chr11-53987820-INS-194
chr11-54007489-INS-51
chr11-54018886-INS-94
chr11-54060634-INS-94
chr11-54073964-INS-393
chr11-54077666-INS-81
chr11-54089694-INS-5584
chr11-54099924-INS-5344
chr11-54109153-INS-6439
chr11-54114299-INS-425
chr11-54125815-INS-109
chr11-54205519-INS-96
chr11-54237758-INS-418
chr11-54240476-INS-279
chr11-54258437-INS-105
chr11-54262005-INS-75
chr11-54280417-INS-54
chr11-54281227-INS-381
chr11-54305827-INS-90
chr11-54317840-INS-61
chr11-54326871-INS-73
chr11-54364282-INS-109
chr11-54375845-INS-443
chr11-54380983-INS-60
chr11-54386012-INS-75
chr11-54402492-INS-222
chr11-54404829-INS-92
chr11-54413211-INS-162
chr11-54421155-INS-66
chr11-54434836-INS-194
chr11-54444431-INS-54
chr11-54465859-INS-211
chr11-54466711-INS-189
chr11-54499563-INS-347
chr11-54501247-INS-216
chr11-54517619-INS-6291
chr11-54553117-INS-216
chr11-54573707-INS-237
chr11-54582673-INS-52
chr11-54591266-INS-180
chr11-54592835-INS-142
chr11-54593868-INS-788
chr11-54599038-INS-198
chr11-54604201-INS-130
chr11-54631900-INS-6454
chr11-54649846-INS-80
chr11-54692410-INS-123
chr11-54702623-INS-74
chr11-54705795-INS-256
chr11-54719478-INS-221
chr11-54727125-INS-839
chr11-54730080-INS-191

chr11-54748193-INS-201
chr11-54768367-INS-507
chr11-54782136-INS-205
chr11-54791195-INS-61
chr11-54792438-INS-211
chr11-54824259-INS-200
chr11-54835599-INS-264
chr11-54836958-INS-54
chr11-54880333-INS-64
chr11-54883603-INS-76
chr11-54891940-INS-118
chr11-54925786-INS-69
chr11-54957358-INS-306
chr11-54959511-INS-176
chr11-54962980-INS-95
chr11-55042967-INS-52
chr11-55047137-INS-81
chr11-55086001-INS-64
chr11-55091886-INS-64
chr11-55095898-INS-167
chr11-55118180-INS-78
chr11-55121839-INS-200
chr11-55177511-INS-56
chr11-55185968-INS-55
chr11-55214647-INS-478
chr11-55219665-INS-51
chr11-55241621-INS-56
chr11-55265043-INS-785
chr11-55268805-INS-686
chr11-55281234-INS-189
chr11-55286553-INS-430
chr11-55294915-INS-460
chr11-55295750-INS-174
chr11-55302324-INS-57
chr11-55308450-INS-56
chr11-55325021-INS-230
chr11-55326611-INS-207
chr11-55333668-INS-66
chr11-55354324-INS-71
chr11-55356486-INS-68
chr11-55370774-INS-581
chr11-55384865-INS-169
chr11-55406791-INS-56
chr11-55420901-INS-299
chr11-55427804-INS-270
chr11-55454289-INS-388
chr11-55458672-INS-136
chr11-55470026-INS-81
chr11-55507512-INS-638
chr11-55509735-INS-50
chr11-55534425-INS-60
chr11-55538098-INS-66
chr11-55580043-INS-72
chr11-55609548-INS-140
chr11-55624913-INS-54
chr11-55632794-INS-834
chr11-55638589-INS-334
chr11-55678005-INS-189
chr11-55714048-INS-107
chr11-55720554-INS-72
chr11-55737734-INS-58
chr11-55761368-INS-318
chr11-55772934-INS-57
chr11-55810806-INS-240
chr11-55829782-INS-153
chr11-55867261-INS-226
chr11-55880361-INS-439
chr11-55882574-INS-6439
chr11-55892313-INS-405
chr11-55903178-INS-651
chr11-55923354-INS-58
chr11-55928160-INS-68
chr11-55945776-INS-134
chr11-55954101-INS-78
chr11-55998038-INS-52
chr11-56016377-INS-361
chr11-56067130-INS-6541

chr11-56078250-INS-641
chr11-56086684-INS-67
chr11-56104090-INS-77
chr11-56110797-INS-172
chr11-56126864-INS-308
chr11-56157216-INS-114
chr11-56171685-INS-147
chr11-56177448-INS-113
chr11-56180831-INS-338
chr11-56192126-INS-62
chr11-56200160-INS-68
chr11-56210111-INS-451
chr11-56219072-INS-6188
chr11-56221165-INS-7778
chr11-56258849-INS-61
chr11-56269895-INS-313
chr11-56274143-INS-60
chr11-56302891-INS-294
chr11-56358338-INS-224
chr11-56391732-INS-84
chr11-56398432-INS-174
chr11-56420499-INS-52
chr11-56423785-INS-71
chr11-56436304-INS-400
chr11-56436629-INS-5535
chr11-56442507-INS-53
chr11-56447820-INS-348
chr11-56478548-INS-199
chr11-56490339-INS-943
chr11-56505384-INS-56
chr11-56512585-INS-72
chr11-56532280-INS-309
chr11-56552156-INS-125
chr11-56596525-INS-481
chr11-56603225-INS-92
chr11-56612382-INS-69
chr11-56647223-INS-124
chr11-56663639-INS-61
chr11-56684628-INS-271
chr11-56730545-INS-72
chr11-56732582-INS-60
chr11-56734806-INS-53
chr11-56755937-INS-102
chr11-56758210-INS-57
chr11-56798725-INS-54
chr11-56820607-INS-399
chr11-56843520-INS-52
chr11-56860899-INS-245
chr11-56865690-INS-114
chr11-56891411-INS-58
chr11-56905582-INS-56
chr11-56975162-INS-78
chr11-56989425-INS-584
chr11-57004713-INS-106
chr11-57017176-INS-524
chr11-57032864-INS-357
chr11-57055472-INS-60
chr11-57077380-INS-6924
chr11-57127029-INS-52
chr11-57150689-INS-679
chr11-57174734-INS-59
chr11-57186571-INS-76
chr11-57193534-INS-102
chr11-57205984-INS-176
chr11-57219741-INS-53
chr11-57227585-INS-112
chr11-57266631-INS-105
chr11-57275032-INS-420
chr11-57279489-INS-133
chr11-57289155-INS-52
chr11-57292981-INS-658
chr11-57313266-INS-207
chr11-57327300-INS-675
chr11-57348454-INS-88
chr11-57361777-INS-153
chr11-57397188-INS-333
chr11-57419408-INS-146

chr11-57425161-INS-60
chr11-57484999-INS-56
chr11-57493701-INS-423
chr11-57495564-INS-84
chr11-57497928-INS-54
chr11-57522003-INS-212
chr11-57547451-INS-3412
chr11-57575816-INS-78
chr11-57579955-INS-114
chr11-57587560-INS-339
chr11-57646983-INS-498
chr11-57663218-INS-204
chr11-57683154-INS-87
chr11-57703619-INS-201
chr11-57707812-INS-526
chr11-57712885-INS-218
chr11-57717446-INS-251
chr11-57743832-INS-360
chr11-57775878-INS-64
chr11-57778204-INS-418
chr11-57783555-INS-266
chr11-57797131-INS-172
chr11-57824491-INS-54
chr11-57826815-INS-344
chr11-57842669-INS-292
chr11-57860637-INS-877
chr11-57865346-INS-180
chr11-57867976-INS-130
chr11-57876633-INS-351
chr11-57878985-INS-190
chr11-57892863-INS-58
chr11-57902835-INS-69
chr11-57909978-INS-107
chr11-57923057-INS-259
chr11-57923695-INS-228
chr11-57927204-INS-207
chr11-57958672-INS-112
chr11-57962272-INS-393
chr11-57971266-INS-62
chr11-57979340-INS-57
chr11-57985549-INS-60
chr11-58002477-INS-147
chr11-58046264-INS-103
chr11-58048666-INS-209
chr11-58050322-INS-74
chr11-58058847-INS-428
chr11-58061388-INS-162
chr11-58082045-INS-76
chr11-58089863-INS-67
chr11-58091989-INS-56
chr11-58095874-INS-176
chr11-58106371-INS-213
chr11-58113816-INS-65
chr11-58121005-INS-201
chr11-58122745-INS-96
chr11-58136133-INS-67
chr11-58156215-INS-195
chr11-58158548-INS-208
chr11-58173214-INS-133
chr11-58180478-INS-62
chr11-58188210-INS-190
chr11-58196350-INS-206
chr11-58203022-INS-217
chr11-58235190-INS-153
chr11-58245342-INS-65
chr11-58247113-INS-3889
chr11-58263273-INS-458
chr11-58269178-INS-79
chr11-58272281-INS-78
chr11-58279613-INS-52
chr11-58284469-INS-212
chr11-58290620-INS-137
chr11-58339519-INS-812
chr11-58349599-INS-52
chr11-58414526-INS-366
chr11-58430150-INS-527
chr11-58435440-INS-845

chr11-58461742-INS-64
chr11-58492929-INS-60
chr11-58502348-INS-57
chr11-58519484-INS-84
chr11-58534380-INS-482
chr11-58624338-INS-89
chr11-58628484-INS-57
chr11-58635724-INS-56
chr11-58671764-INS-213
chr11-58681694-INS-344
chr11-58688419-INS-607
chr11-58689171-INS-80
chr11-58704641-INS-80
chr11-58757364-INS-71
chr11-58780350-INS-148
chr11-58784764-INS-199
chr11-58785793-INS-55
chr11-58787639-INS-674
chr11-58806451-INS-312
chr11-58810904-INS-52
chr11-58817488-INS-100
chr11-58833690-INS-354
chr11-58853329-INS-99
chr11-58860074-INS-220
chr11-58877064-INS-105
chr11-58949294-INS-51
chr11-58950483-INS-60
chr11-58957410-INS-136
chr11-58970036-INS-164
chr11-58991959-INS-56
chr11-59000428-INS-132
chr11-59010320-INS-96
chr11-59011717-INS-866
chr11-59047601-INS-710
chr11-59051262-INS-327
chr11-59093568-INS-130
chr11-59128976-INS-55
chr11-59143159-INS-212
chr11-59172039-INS-179
chr11-59192858-INS-51
chr11-59211889-INS-1888
chr11-59228815-INS-186
chr11-59230999-INS-79
chr11-59270150-INS-5549
chr11-59276810-INS-736
chr11-59300965-INS-224
chr11-59322027-INS-70
chr11-59332378-INS-88
chr11-59357585-INS-256
chr11-59386307-INS-105
chr11-59390059-INS-68
chr11-59415098-INS-57
chr11-59424067-INS-134
chr11-59426332-INS-64
chr11-59427659-INS-60
chr11-59441670-INS-105
chr11-59448355-INS-89
chr11-59471873-INS-373
chr11-59476104-INS-274
chr11-59501363-INS-8735
chr11-59507484-INS-6440
chr11-59516972-INS-186
chr11-59521750-INS-161
chr11-59538464-INS-150
chr11-59540732-INS-52
chr11-59543761-INS-56
chr11-59611356-INS-53
chr11-59613624-INS-68
chr11-59642304-INS-436
chr11-59676666-INS-66
chr11-59693814-INS-94
chr11-59738411-INS-9092
chr11-59763051-INS-72
chr11-59764892-INS-128
chr11-59790513-INS-195
chr11-59796596-INS-112
chr11-59800491-INS-51

chr11-59820637-INS-420
chr11-59835607-INS-1287
chr11-59843797-INS-197
chr11-59846549-INS-157
chr11-59885869-INS-328
chr11-59918232-INS-186
chr11-59933737-INS-1889
chr11-59949881-INS-168
chr11-59954645-INS-61
chr11-59957159-INS-202
chr11-59966910-INS-69
chr11-59986460-INS-239
chr11-60059008-INS-81
chr11-60144337-INS-65
chr11-60200594-INS-397
chr11-60203924-INS-306
chr11-60215352-INS-231
chr11-60247657-INS-486
chr11-60260123-INS-216
chr11-60273485-INS-55
chr11-60274420-INS-84
chr11-60302323-INS-307
chr11-60332491-INS-87
chr11-60342113-INS-1692
chr11-60343382-INS-208
chr11-60404585-INS-67
chr11-60406563-INS-86
chr11-60424862-INS-415
chr11-60459028-INS-139
chr11-60466425-INS-66
chr11-60562001-INS-66
chr11-60567138-INS-56
chr11-60634553-INS-207
chr11-60652332-INS-104
chr11-60686700-INS-290
chr11-60692185-INS-301
chr11-60692697-INS-109
chr11-60699821-INS-197
chr11-60707713-INS-129
chr11-60713167-INS-457
chr11-60714303-INS-92
chr11-60724748-INS-197
chr11-60729386-INS-181
chr11-60731036-INS-189
chr11-60736069-INS-54
chr11-60741332-INS-197
chr11-60744648-INS-68
chr11-60750131-INS-138
chr11-60760280-INS-54
chr11-60776875-INS-199
chr11-60789346-INS-191
chr11-60793081-INS-205
chr11-60811451-INS-55
chr11-60814786-INS-182
chr11-60819031-INS-238
chr11-60819868-INS-82
chr11-60828156-INS-51
chr11-60836574-INS-57
chr11-60841182-INS-224
chr11-60847066-INS-434
chr11-60852142-INS-211
chr11-60855453-INS-106
chr11-60856692-INS-51
chr11-60865186-INS-101
chr11-60885707-INS-5548
chr11-60885901-INS-115
chr11-60906694-INS-3775
chr11-60918796-INS-197
chr11-60930094-INS-108
chr11-60931426-INS-105
chr11-60949874-INS-98
chr11-60959467-INS-234
chr11-60968976-INS-179
chr11-60970438-INS-51
chr11-60979023-INS-219
chr11-60983499-INS-80
chr11-61014558-INS-92

chr11-61015196-INS-92
chr11-61027822-INS-118
chr11-61030904-INS-153
chr11-61033084-INS-163
chr11-61036008-INS-61
chr11-61056155-INS-220
chr11-61069641-INS-497
chr11-61074375-INS-311
chr11-61078520-INS-211
chr11-61148226-INS-113
chr11-61151904-INS-316
chr11-61163578-INS-109
chr11-61167020-INS-66
chr11-61171169-INS-60
chr11-61184024-INS-185
chr11-61188481-INS-78
chr11-61198793-INS-360
chr11-61203779-INS-107
chr11-61209342-INS-117
chr11-61217201-INS-60
chr11-61226727-INS-198
chr11-61236340-INS-124
chr11-61237407-INS-401
chr11-61264875-INS-84
chr11-61299293-INS-54
chr11-61450341-INS-212
chr11-61476222-INS-60
chr11-61487306-INS-115
chr11-61492946-INS-226
chr11-61505876-INS-54
chr11-61524289-INS-81
chr11-61531223-INS-137
chr11-61631648-INS-58
chr11-61643368-INS-217
chr11-61644715-INS-339
chr11-61651987-INS-174
chr11-61654834-INS-56
chr11-61658777-INS-53
chr11-61673187-INS-314
chr11-61682622-INS-916
chr11-61693749-INS-54
chr11-61696053-INS-54
chr11-61753477-INS-4908
chr11-61778717-INS-295
chr11-61804970-INS-224
chr11-61832732-INS-991
chr11-61860553-INS-4564
chr11-61864551-INS-132
chr11-61869308-INS-806
chr11-61871341-INS-590
chr11-61875658-INS-74
chr11-61885633-INS-81
chr11-61887344-INS-87
chr11-61911594-INS-216
chr11-61938893-INS-207
chr11-61943835-INS-176
chr11-61958302-INS-65
chr11-61961652-INS-639
chr11-61986983-INS-177
chr11-62004344-INS-58
chr11-62021807-INS-70
chr11-62041021-INS-76
chr11-62060536-INS-201
chr11-62072155-INS-86
chr11-62090801-INS-52
chr11-62095745-INS-324
chr11-62097941-INS-233
chr11-62118598-INS-121
chr11-62138971-INS-64
chr11-62166442-INS-53
chr11-62216599-INS-61
chr11-62235128-INS-96
chr11-62244337-INS-411
chr11-62248955-INS-187
chr11-62252366-INS-123
chr11-62254837-INS-52
chr11-62258709-INS-56

chr11-62271164-INS-157
chr11-62273971-INS-110
chr11-62283734-INS-125
chr11-62287296-INS-54
chr11-62301047-INS-141
chr11-62301154-INS-992
chr11-62307399-INS-217
chr11-62309515-INS-203
chr11-62311173-INS-167
chr11-62313098-INS-216
chr11-62319658-INS-214
chr11-62333294-INS-216
chr11-62352887-INS-216
chr11-62373836-INS-199
chr11-62378219-INS-178
chr11-62379950-INS-463
chr11-62389057-INS-79
chr11-62398367-INS-219
chr11-62404965-INS-205
chr11-62413801-INS-198
chr11-62427599-INS-137
chr11-62435466-INS-225
chr11-62435960-INS-213
chr11-62438262-INS-60
chr11-62438266-INS-953
chr11-62444794-INS-129
chr11-62447369-INS-198
chr11-62453127-INS-170
chr11-62456669-INS-179
chr11-62489218-INS-732
chr11-62492468-INS-206
chr11-62499550-INS-211
chr11-62521565-INS-214
chr11-62523002-INS-144
chr11-62533785-INS-636
chr11-62574220-INS-964
chr11-62584444-INS-193
chr11-62625096-INS-121
chr11-62634047-INS-218
chr11-62637972-INS-155
chr11-62653473-INS-299
chr11-62655326-INS-59
chr11-62656025-INS-111
chr11-62663817-INS-218
chr11-62695258-INS-252
chr11-62704115-INS-120
chr11-62704654-INS-126
chr11-62709194-INS-184
chr11-62728914-INS-175
chr11-62734082-INS-236
chr11-62742332-INS-117
chr11-62762009-INS-179
chr11-62775609-INS-5287
chr11-62806163-INS-111
chr11-62808382-INS-492
chr11-62811859-INS-57
chr11-62818079-INS-286
chr11-62827793-INS-72
chr11-62835744-INS-86
chr11-62851535-INS-69
chr11-62865362-INS-626
chr11-62872408-INS-66
chr11-62879029-INS-68
chr11-62883156-INS-72
chr11-62899894-INS-114
chr11-62906153-INS-113
chr11-62912199-INS-115
chr11-62924890-INS-80
chr11-62960746-INS-68
chr11-62971356-INS-242
chr11-62974240-INS-58
chr11-62977070-INS-98
chr11-62978525-INS-522
chr11-63013394-INS-156
chr11-63016387-INS-51
chr11-63029604-INS-110
chr11-63106799-INS-62

chr11-63139943-INS-346
chr11-63164782-INS-74
chr11-63167181-INS-288
chr11-63179905-INS-127
chr11-63197111-INS-4805
chr11-63223523-INS-773
chr11-63272210-INS-110
chr11-63285678-INS-774
chr11-63298011-INS-108
chr11-63346930-INS-864
chr11-63376416-INS-97
chr11-63385244-INS-76
chr11-63409038-INS-184
chr11-63449842-INS-51
chr11-63455845-INS-72
chr11-63465144-INS-132
chr11-63491518-INS-62
chr11-63526636-INS-121
chr11-63530584-INS-371
chr11-63560815-INS-241
chr11-63567200-INS-264
chr11-63612539-INS-115
chr11-63625046-INS-173
chr11-63643391-INS-64
chr11-63653390-INS-6666
chr11-63654753-INS-75
chr11-63665554-INS-142
chr11-63684560-INS-6443
chr11-63686969-INS-656
chr11-63708130-INS-203
chr11-63722283-INS-55
chr11-63732372-INS-6716
chr11-63744464-INS-60
chr11-63757755-INS-108
chr11-63765262-INS-857
chr11-63825315-INS-51
chr11-63828688-INS-52
chr11-63835625-INS-86
chr11-63845210-INS-84
chr11-63847166-INS-236
chr11-63860377-INS-303
chr11-63865267-INS-77
chr11-63897350-INS-52
chr11-63927805-INS-80
chr11-63933505-INS-112
chr11-63948424-INS-55
chr11-63975588-INS-5700
chr11-63997010-INS-498
chr11-64007827-INS-564
chr11-64009033-INS-56
chr11-64014663-INS-111
chr11-64021740-INS-187
chr11-64030764-INS-55
chr11-64037960-INS-212
chr11-64054335-INS-104
chr11-64086112-INS-411
chr11-64096896-INS-63
chr11-64106362-INS-56
chr11-64139069-INS-65
chr11-64141597-INS-147
chr11-64237012-INS-7536
chr11-64259979-INS-56
chr11-64286560-INS-178
chr11-64299418-INS-152
chr11-64304172-INS-170
chr11-64315319-INS-70
chr11-64335608-INS-166
chr11-64346129-INS-90
chr11-64354279-INS-75
chr11-64382974-INS-6694
chr11-64389111-INS-56
chr11-64431228-INS-178
chr11-64442755-INS-122
chr11-64443545-INS-218
chr11-64471493-INS-106
chr11-64510858-INS-440
chr11-64510924-INS-216

chr11-64568825-INS-6434
chr11-64572703-INS-344
chr11-64596918-INS-879
chr11-64604432-INS-150
chr11-64608907-INS-219
chr11-64644032-INS-7114
chr11-64659194-INS-728
chr11-64676875-INS-58
chr11-64677619-INS-61
chr11-64689787-INS-60
chr11-64694644-INS-3343
chr11-64718917-INS-192
chr11-64726665-INS-67
chr11-64732875-INS-127
chr11-64739699-INS-52
chr11-64759712-INS-55
chr11-64765262-INS-75
chr11-64781135-INS-341
chr11-64785929-INS-58
chr11-64789365-INS-86
chr11-64799042-INS-81
chr11-64802161-INS-428
chr11-64805716-INS-88
chr11-64811756-INS-76
chr11-64819407-INS-787
chr11-64832823-INS-6477
chr11-64837905-INS-141
chr11-64892697-INS-75
chr11-64923834-INS-205
chr11-64926963-INS-52
chr11-64940776-INS-6400
chr11-64945801-INS-145
chr11-64967916-INS-3971
chr11-64971205-INS-6439
chr11-64976052-INS-1893
chr11-64985158-INS-58
chr11-65005825-INS-218
chr11-65037911-INS-2270
chr11-65062745-INS-1565
chr11-65072664-INS-192
chr11-65082634-INS-77
chr11-65094275-INS-102
chr11-65108880-INS-121
chr11-65116045-INS-199
chr11-65127651-INS-189
chr11-65134223-INS-128
chr11-65134301-INS-150
chr11-65163125-INS-167
chr11-65192973-INS-63
chr11-65202833-INS-94
chr11-65209283-INS-52
chr11-65222034-INS-116
chr11-65237811-INS-71
chr11-65258505-INS-145
chr11-65315763-INS-72
chr11-65320194-INS-81
chr11-65333270-INS-130
chr11-65363629-INS-68
chr11-65364820-INS-170
chr11-65380434-INS-66
chr11-65381317-INS-78
chr11-65397075-INS-110
chr11-65406135-INS-78
chr11-65408451-INS-67
chr11-65412340-INS-493
chr11-65422720-INS-58
chr11-65442824-INS-154
chr11-65466338-INS-183
chr11-65474595-INS-622
chr11-65500841-INS-110
chr11-65509176-INS-6432
chr11-65516938-INS-51
chr11-65540190-INS-191
chr11-65560700-INS-178
chr11-65601507-INS-112
chr11-65608147-INS-295
chr11-65629702-INS-70

chr11-65637824-INS-148
chr11-65644041-INS-413
chr11-65662263-INS-106
chr11-65677341-INS-72
chr11-65707959-INS-102
chr11-65715356-INS-58
chr11-65726267-INS-101
chr11-65731588-INS-62
chr11-65736599-INS-735
chr11-65737272-INS-1579
chr11-65775804-INS-454
chr11-65814477-INS-212
chr11-65838242-INS-72
chr11-65841518-INS-3928
chr11-65869897-INS-6550
chr11-65871543-INS-73
chr11-65908894-INS-60
chr11-65935845-INS-68
chr11-65948906-INS-583
chr11-65974768-INS-777
chr11-65991238-INS-96
chr11-66018683-INS-55
chr11-66019739-INS-899
chr11-66042124-INS-305
chr11-66062974-INS-56
chr11-66069415-INS-402
chr11-66081428-INS-910
chr11-66120094-INS-93
chr11-66121978-INS-528
chr11-66167046-INS-7772
chr11-66171998-INS-88
chr11-66187047-INS-143
chr11-66229741-INS-125
chr11-66230775-INS-621
chr11-66244040-INS-135
chr11-66246377-INS-127
chr11-66260967-INS-63
chr11-66299076-INS-93
chr11-66317500-INS-127
chr11-66331995-INS-417
chr11-66334480-INS-361
chr11-66341333-INS-60
chr11-66358195-INS-556
chr11-66358967-INS-69
chr11-66361589-INS-1274
chr11-66361974-INS-56
chr11-66371505-INS-159
chr11-66373794-INS-106
chr11-66398687-INS-274
chr11-66467272-INS-66
chr11-66481716-INS-7114
chr11-66481871-INS-177
chr11-66488532-INS-56
chr11-66537605-INS-319
chr11-66539782-INS-6281
chr11-66560770-INS-6689
chr11-66568966-INS-64
chr11-66587117-INS-68
chr11-66593526-INS-56
chr11-66595202-INS-889
chr11-66608488-INS-64
chr11-66616580-INS-163
chr11-66631418-INS-68
chr11-66655661-INS-204
chr11-66657656-INS-1059
chr11-66657963-INS-108
chr11-66678634-INS-189
chr11-66699399-INS-59
chr11-66708052-INS-56
chr11-66728319-INS-52
chr11-66737766-INS-415
chr11-66748270-INS-231
chr11-66762485-INS-64
chr11-66769727-INS-296
chr11-66774833-INS-60
chr11-66795628-INS-200
chr11-66824384-INS-218

chr11-66831012-INS-337
chr11-66832115-INS-604
chr11-66857128-INS-112
chr11-66859045-INS-211
chr11-66861830-INS-212
chr11-66866699-INS-56
chr11-66877188-INS-196
chr11-66902076-INS-1620
chr11-66902725-INS-83
chr11-66910321-INS-82
chr11-66935364-INS-57
chr11-66940386-INS-51
chr11-66982205-INS-72
chr11-66983575-INS-289
chr11-66983810-INS-224
chr11-66999132-INS-216
chr11-67002974-INS-63
chr11-67004760-INS-222
chr11-67006242-INS-124
chr11-67008666-INS-92
chr11-67020435-INS-87
chr11-67043834-INS-61
chr11-67051747-INS-88
chr11-67054021-INS-4276
chr11-67058159-INS-156
chr11-67149422-INS-85
chr11-67176247-INS-51
chr11-67178530-INS-137
chr11-67189979-INS-73
chr11-67191171-INS-128
chr11-67206212-INS-54
chr11-67228023-INS-371
chr11-67243749-INS-55
chr11-67244079-INS-947
chr11-67260569-INS-60
chr11-67296337-INS-53
chr11-67299180-INS-105
chr11-67338972-INS-371
chr11-67349039-INS-216
chr11-67351740-INS-160
chr11-67354875-INS-449
chr11-67362511-INS-174
chr11-67377821-INS-134
chr11-67383911-INS-109
chr11-67405388-INS-82
chr11-67420158-INS-265
chr11-67428173-INS-354
chr11-67436998-INS-294
chr11-67451503-INS-55
chr11-67471342-INS-54
chr11-67483806-INS-59
chr11-67485073-INS-72
chr11-67508807-INS-72
chr11-67516747-INS-107
chr11-67528368-INS-69
chr11-67549307-INS-57
chr11-67564785-INS-102
chr11-67564860-INS-660
chr11-67593126-INS-142
chr11-67634161-INS-108
chr11-67656773-INS-72
chr11-67658822-INS-117
chr11-67672892-INS-86
chr11-67675596-INS-60
chr11-67709483-INS-345
chr11-67754951-INS-52
chr11-67788466-INS-55
chr11-67800362-INS-76
chr11-67833673-INS-60
chr11-67840900-INS-234
chr11-67844116-INS-52
chr11-67870930-INS-343
chr11-67895882-INS-103
chr11-67899226-INS-57
chr11-67936740-INS-86
chr11-67952398-INS-6439
chr11-67969713-INS-444

chr11-67982606-INS-84
chr11-67985023-INS-206
chr11-67996146-INS-202
chr11-68030636-INS-325
chr11-68051818-INS-62
chr11-68089194-INS-6447
chr11-68121959-INS-6440
chr11-68227096-INS-243
chr11-68248829-INS-53
chr11-68317398-INS-80
chr11-68338634-INS-112
chr11-68397166-INS-111
chr11-68426566-INS-110
chr11-68445328-INS-59
chr11-68455082-INS-156
chr11-68459567-INS-139
chr11-68464361-INS-54
chr11-68464596-INS-544
chr11-68482178-INS-84
chr11-68496932-INS-101
chr11-68499142-INS-60
chr11-68523587-INS-1523
chr11-68524842-INS-203
chr11-68547141-INS-85
chr11-68609571-INS-6554
chr11-68617319-INS-208
chr11-68621250-INS-96
chr11-68641039-INS-104
chr11-68646127-INS-54
chr11-68656331-INS-6444
chr11-68677835-INS-139
chr11-68726616-INS-153
chr11-68729209-INS-192
chr11-68765986-INS-184
chr11-68771073-INS-216
chr11-68772458-INS-210
chr11-68776092-INS-196
chr11-68795896-INS-56
chr11-68797090-INS-80
chr11-68799279-INS-626
chr11-68818166-INS-132
chr11-68829813-INS-363
chr11-68833606-INS-199
chr11-68838660-INS-240
chr11-68841746-INS-253
chr11-68850015-INS-476
chr11-68853506-INS-257
chr11-68867922-INS-238
chr11-68871682-INS-436
chr11-68876076-INS-70
chr11-68877637-INS-133
chr11-68886803-INS-171
chr11-68893936-INS-65
chr11-68905111-INS-396
chr11-68914565-INS-210
chr11-68923404-INS-242
chr11-68928249-INS-166
chr11-68936296-INS-265
chr11-68948969-INS-196
chr11-68958932-INS-124
chr11-68967987-INS-151
chr11-68977637-INS-56
chr11-69004885-INS-282
chr11-69041228-INS-203
chr11-69044009-INS-54
chr11-69049890-INS-108
chr11-69052841-INS-155
chr11-69068636-INS-296
chr11-69069674-INS-257
chr11-69070928-INS-716
chr11-69072128-INS-222
chr11-69082710-INS-311
chr11-69104129-INS-167
chr11-69104376-INS-882
chr11-69119183-INS-55
chr11-69126037-INS-65
chr11-69160281-INS-102

chr11-69164772-INS-52
chr11-69166064-INS-71
chr11-69178524-INS-207
chr11-69186455-INS-93
chr11-69188053-INS-410
chr11-69192080-INS-158
chr11-69210420-INS-99
chr11-69221686-INS-58
chr11-69265544-INS-211
chr11-69267785-INS-210
chr11-69293755-INS-117
chr11-69314206-INS-52
chr11-69342607-INS-321
chr11-69353994-INS-194
chr11-69372973-INS-80
chr11-69426595-INS-176
chr11-69436164-INS-54
chr11-69441486-INS-747
chr11-69470409-INS-207
chr11-69487879-INS-88
chr11-69500033-INS-51
chr11-69510899-INS-64
chr11-69511843-INS-192
chr11-69515561-INS-249
chr11-69530044-INS-193
chr11-69552356-INS-196
chr11-69554030-INS-106
chr11-69604977-INS-573
chr11-69613221-INS-220
chr11-69621187-INS-508
chr11-69681545-INS-205
chr11-69701661-INS-93
chr11-69713189-INS-213
chr11-69732526-INS-354
chr11-69756280-INS-126
chr11-69765031-INS-154
chr11-69803052-INS-53
chr11-69819737-INS-59
chr11-69822434-INS-297
chr11-69823863-INS-184
chr11-69844740-INS-348
chr11-69846429-INS-76
chr11-69850147-INS-208
chr11-69853232-INS-194
chr11-69860491-INS-443
chr11-69862517-INS-196
chr11-69897505-INS-206
chr11-69957023-INS-4750
chr11-69962588-INS-52
chr11-69970293-INS-173
chr11-69989393-INS-199
chr11-70000330-INS-182
chr11-70008648-INS-194
chr11-70024668-INS-60
chr11-70065782-INS-63
chr11-70083701-INS-399
chr11-70093372-INS-134
chr11-70100815-INS-6440
chr11-70146690-INS-52
chr11-70148545-INS-126
chr11-70152307-INS-70
chr11-70159305-INS-234
chr11-70164745-INS-62
chr11-70174453-INS-479
chr11-70188544-INS-211
chr11-70214705-INS-54
chr11-70226446-INS-5506
chr11-70233586-INS-63
chr11-70248217-INS-74
chr11-70261813-INS-85
chr11-70274071-INS-480
chr11-70305908-INS-311
chr11-70317104-INS-223
chr11-70333826-INS-63
chr11-70338533-INS-54
chr11-70345386-INS-169
chr11-70358019-INS-212

chr11-70359089-INS-54
chr11-70370934-INS-241
chr11-70380427-INS-214
chr11-70386317-INS-5561
chr11-70387921-INS-200
chr11-70392459-INS-406
chr11-70427964-INS-358
chr11-70430625-INS-169
chr11-70461707-INS-1975
chr11-70488240-INS-189
chr11-70493325-INS-242
chr11-70495178-INS-444
chr11-70500650-INS-471
chr11-70500650-INS-978
chr11-70528941-INS-200
chr11-70554410-INS-83
chr11-70583114-INS-191
chr11-70648932-INS-5167
chr11-70653418-INS-234
chr11-70665268-INS-973
chr11-70697352-INS-204
chr11-70728153-INS-56
chr11-70733387-INS-212
chr11-70745449-INS-263
chr11-70751550-INS-989
chr11-70754819-INS-627
chr11-70756414-INS-644
chr11-70789008-INS-77
chr11-70791585-INS-201
chr11-70804617-INS-205
chr11-70807160-INS-75
chr11-70815783-INS-223
chr11-70822080-INS-52
chr11-70856867-INS-509
chr11-70858056-INS-262
chr11-70867126-INS-73
chr11-70883243-INS-198
chr11-70887475-INS-200
chr11-70927854-INS-2990
chr11-70930406-INS-301
chr11-71081001-INS-330
chr11-71085823-INS-356
chr11-71104385-INS-1288
chr11-71111808-INS-427
chr11-71117212-INS-496
chr11-71156091-INS-164
chr11-71187589-INS-132
chr11-71221904-INS-8813
chr11-71265022-INS-6560
chr11-71274969-INS-207
chr11-71291185-INS-9423
chr11-71312188-INS-7165
chr11-71323477-INS-144
chr11-71349698-INS-64
chr11-71355596-INS-96
chr11-71356462-INS-59
chr11-71357121-INS-127
chr11-71377734-INS-88
chr11-71404025-INS-288
chr11-71411188-INS-341
chr11-71427096-INS-199
chr11-71428140-INS-63
chr11-71439341-INS-225
chr11-71441445-INS-175
chr11-71464876-INS-108
chr11-71505276-INS-54
chr11-71518862-INS-72
chr11-71522131-INS-577
chr11-71541903-INS-82
chr11-71545549-INS-52
chr11-71558548-INS-58
chr11-71565820-INS-58
chr11-71580574-INS-63
chr11-71585944-INS-249
chr11-71605006-INS-421
chr11-71616180-INS-509
chr11-71702246-INS-59

chr11-71720231-INS-4363
chr11-71724585-INS-156
chr11-71773306-INS-146
chr11-71783173-INS-6960
chr11-71783199-INS-52
chr11-71796432-INS-124
chr11-71844458-INS-57
chr11-71861992-INS-92
chr11-71865966-INS-358
chr11-71880366-INS-306
chr11-71898400-INS-107
chr11-71909482-INS-54
chr11-71911374-INS-137
chr11-71976885-INS-94
chr11-72000512-INS-182
chr11-72008561-INS-290
chr11-72036694-INS-106
chr11-72054912-INS-297
chr11-72073347-INS-132
chr11-72109303-INS-64
chr11-72121717-INS-93
chr11-72123670-INS-56
chr11-72128103-INS-96
chr11-72137392-INS-170
chr11-72140911-INS-142
chr11-72162558-INS-80
chr11-72164967-INS-60
chr11-72166605-INS-204
chr11-72201055-INS-205
chr11-72201951-INS-564
chr11-72204137-INS-183
chr11-72213804-INS-190
chr11-72215638-INS-429
chr11-72239424-INS-87
chr11-72244046-INS-207
chr11-72263188-INS-209
chr11-72264414-INS-128
chr11-72322182-INS-62
chr11-72325238-INS-321
chr11-72381633-INS-243
chr11-72412738-INS-64
chr11-72423333-INS-110
chr11-72427642-INS-1871
chr11-72434725-INS-207
chr11-72445742-INS-55
chr11-72451099-INS-176
chr11-72452528-INS-92
chr11-72481519-INS-7117
chr11-72490072-INS-207
chr11-72548948-INS-61
chr11-72569672-INS-917
chr11-72594554-INS-57
chr11-72628292-INS-168
chr11-72632250-INS-64
chr11-72650156-INS-134
chr11-72654939-INS-64
chr11-72656929-INS-714
chr11-72657045-INS-1036
chr11-72672279-INS-145
chr11-72709605-INS-192
chr11-72748008-INS-329
chr11-72752194-INS-364
chr11-72807900-INS-67
chr11-72841250-INS-54
chr11-72881156-INS-62
chr11-72913361-INS-51
chr11-72922324-INS-215
chr11-72923554-INS-63
chr11-72937136-INS-63
chr11-72949501-INS-218
chr11-72957005-INS-73
chr11-72963202-INS-70
chr11-72972930-INS-52
chr11-72990156-INS-99
chr11-72994150-INS-64
chr11-73004855-INS-560
chr11-73006267-INS-51

chr11-73032935-INS-81
chr11-73065397-INS-60
chr11-73103775-INS-55
chr11-73108080-INS-155
chr11-73113587-INS-862
chr11-73123276-INS-127
chr11-73147157-INS-6693
chr11-73167861-INS-206
chr11-73172912-INS-82
chr11-73182226-INS-1958
chr11-73183191-INS-138
chr11-73194786-INS-423
chr11-73201435-INS-121
chr11-73206746-INS-56
chr11-73236541-INS-844
chr11-73238333-INS-52
chr11-73283496-INS-588
chr11-73299068-INS-667
chr11-73347127-INS-52
chr11-73375727-INS-56
chr11-73381685-INS-6461
chr11-73391079-INS-196
chr11-73399550-INS-1573
chr11-73405183-INS-341
chr11-73412219-INS-3233
chr11-73459116-INS-6250
chr11-73480617-INS-1747
chr11-73494686-INS-1745
chr11-73496315-INS-76
chr11-73500817-INS-159
chr11-73523596-INS-6582
chr11-73552234-INS-338
chr11-73563238-INS-5089
chr11-73609034-INS-330
chr11-73637767-INS-63
chr11-73649627-INS-6361
chr11-73679985-INS-299
chr11-73682767-INS-407
chr11-73687895-INS-117
chr11-73708123-INS-130
chr11-73711164-INS-471
chr11-73731626-INS-120
chr11-73733202-INS-980
chr11-73738005-INS-581
chr11-73750084-INS-327
chr11-73767378-INS-327
chr11-73777191-INS-4479
chr11-73782966-INS-60
chr11-73802618-INS-6928
chr11-73820258-INS-6937
chr11-73821613-INS-3599
chr11-73823991-INS-63
chr11-73825519-INS-374
chr11-73827240-INS-6673
chr11-73832246-INS-70
chr11-73848470-INS-74
chr11-73903542-INS-54
chr11-73938733-INS-153
chr11-73940848-INS-52
chr11-73956070-INS-158
chr11-73962964-INS-182
chr11-73972137-INS-59
chr11-74003415-INS-808
chr11-74066666-INS-482
chr11-74073079-INS-2931
chr11-74109650-INS-5374
chr11-74148250-INS-225
chr11-74182484-INS-398
chr11-74186314-INS-229
chr11-74261231-INS-199
chr11-74290103-INS-56
chr11-74318967-INS-98
chr11-74334912-INS-160
chr11-74338298-INS-698
chr11-74351224-INS-80
chr11-74367237-INS-221
chr11-74386043-INS-59

chr11-74402620-INS-72
chr11-74415604-INS-60
chr11-74417259-INS-232
chr11-74425023-INS-276
chr11-74430423-INS-134
chr11-74437432-INS-126
chr11-74452907-INS-192
chr11-74458381-INS-93
chr11-74473672-INS-58
chr11-74491676-INS-56
chr11-74496908-INS-199
chr11-74507582-INS-60
chr11-74516269-INS-186
chr11-74517658-INS-214
chr11-74528683-INS-131
chr11-74536249-INS-196
chr11-74538071-INS-266
chr11-74553767-INS-137
chr11-74579367-INS-66
chr11-74594963-INS-187
chr11-74596526-INS-229
chr11-74597690-INS-97
chr11-74606346-INS-110
chr11-74617272-INS-192
chr11-74622881-INS-3255
chr11-74641017-INS-203
chr11-74643127-INS-71
chr11-74650193-INS-239
chr11-74657903-INS-361
chr11-74658930-INS-65
chr11-74668750-INS-149
chr11-74670567-INS-231
chr11-74680017-INS-229
chr11-74687291-INS-364
chr11-74689517-INS-211
chr11-74692229-INS-219
chr11-74697812-INS-240
chr11-74746709-INS-66
chr11-74755319-INS-213
chr11-74756918-INS-58
chr11-74801754-INS-68
chr11-74830283-INS-197
chr11-74841303-INS-118
chr11-74849627-INS-312
chr11-74891230-INS-725
chr11-74902135-INS-160
chr11-74921386-INS-173
chr11-74930853-INS-207
chr11-74977264-INS-194
chr11-74984045-INS-390
chr11-74998170-INS-223
chr11-75006158-INS-230
chr11-75032574-INS-121
chr11-75035847-INS-52
chr11-75070207-INS-117
chr11-75072660-INS-127
chr11-75090076-INS-56
chr11-75097175-INS-88
chr11-75117300-INS-324
chr11-75129206-INS-185
chr11-75131028-INS-89
chr11-75134435-INS-202
chr11-75140334-INS-196
chr11-75147890-INS-155
chr11-75165938-INS-619
chr11-75166908-INS-622
chr11-75218211-INS-63
chr11-75220508-INS-228
chr11-75245724-INS-400
chr11-75263454-INS-261
chr11-75274049-INS-940
chr11-75282511-INS-201
chr11-75286391-INS-65
chr11-75296946-INS-494
chr11-75301381-INS-417
chr11-75302987-INS-273
chr11-75304600-INS-71

chr11-75309745-INS-230
chr11-75317449-INS-331
chr11-75321992-INS-5345
chr11-75348189-INS-51
chr11-75365006-INS-223
chr11-75378196-INS-52
chr11-75387660-INS-208
chr11-75397774-INS-241
chr11-75398338-INS-123
chr11-75424492-INS-179
chr11-75437409-INS-80
chr11-75526771-INS-148
chr11-75542280-INS-195
chr11-75550180-INS-200
chr11-75567457-INS-1142
chr11-75567632-INS-237
chr11-75581355-INS-116
chr11-75587776-INS-573
chr11-75601942-INS-71
chr11-75614842-INS-167
chr11-75623052-INS-117
chr11-75628889-INS-224
chr11-75630126-INS-67
chr11-75631419-INS-75
chr11-75633823-INS-205
chr11-75635579-INS-58
chr11-75638048-INS-180
chr11-75640608-INS-204
chr11-75650655-INS-192
chr11-75669825-INS-51
chr11-75697102-INS-175
chr11-75707652-INS-54
chr11-75710317-INS-101
chr11-75736552-INS-64
chr11-75738140-INS-75
chr11-75754155-INS-736
chr11-75756352-INS-152
chr11-75758078-INS-61
chr11-75772712-INS-186
chr11-75777676-INS-315
chr11-75809189-INS-360
chr11-75813987-INS-54
chr11-75823089-INS-356
chr11-75833797-INS-110
chr11-75840855-INS-134
chr11-75842629-INS-146
chr11-75851978-INS-220
chr11-75863457-INS-611
chr11-75888314-INS-462
chr11-75890287-INS-184
chr11-75896459-INS-213
chr11-75898842-INS-183
chr11-75905201-INS-184
chr11-75907622-INS-59
chr11-75914833-INS-93
chr11-75919516-INS-213
chr11-75922949-INS-66
chr11-75929242-INS-151
chr11-75968855-INS-275
chr11-75981772-INS-52
chr11-75999016-INS-419
chr11-76008136-INS-112
chr11-76010472-INS-68
chr11-76047121-INS-199
chr11-76078982-INS-105
chr11-76084404-INS-70
chr11-76118707-INS-174
chr11-76126520-INS-66
chr11-76128730-INS-200
chr11-76160012-INS-54
chr11-76161529-INS-200
chr11-76165852-INS-51
chr11-76166963-INS-243
chr11-76170552-INS-209
chr11-76187588-INS-174
chr11-76196675-INS-213
chr11-76206310-INS-210

chr11-76211553-INS-80
chr11-76214682-INS-71
chr11-76219986-INS-160
chr11-76222851-INS-51
chr11-76226576-INS-96
chr11-76242065-INS-50
chr11-76244409-INS-104
chr11-76279050-INS-205
chr11-76303957-INS-140
chr11-76310812-INS-62
chr11-76311676-INS-212
chr11-76312706-INS-203
chr11-76337528-INS-429
chr11-76352969-INS-153
chr11-76409917-INS-55
chr11-76419896-INS-215
chr11-76481805-INS-507
chr11-76486870-INS-204
chr11-76493136-INS-181
chr11-76508127-INS-52
chr11-76526697-INS-84
chr11-76539028-INS-74
chr11-76544411-INS-5290
chr11-76559581-INS-212
chr11-76579764-INS-80
chr11-76591030-INS-56
chr11-76610775-INS-58
chr11-76613621-INS-235
chr11-76644239-INS-118
chr11-76669478-INS-97
chr11-76673643-INS-323
chr11-76690038-INS-52
chr11-76744697-INS-85
chr11-76772461-INS-79
chr11-76785472-INS-627
chr11-76809803-INS-719
chr11-76851403-INS-218
chr11-76880689-INS-158
chr11-76919434-INS-175
chr11-76925216-INS-681
chr11-76925918-INS-81
chr11-76932537-INS-207
chr11-76936391-INS-111
chr11-76942095-INS-70
chr11-76951013-INS-59
chr11-76973963-INS-120
chr11-76983535-INS-58
chr11-76985812-INS-130
chr11-76991957-INS-215
chr11-76993348-INS-581
chr11-77002952-INS-122
chr11-77010391-INS-252
chr11-77018208-INS-214
chr11-77037361-INS-109
chr11-77040453-INS-51
chr11-77042116-INS-52
chr11-77051890-INS-52
chr11-77059416-INS-511
chr11-77065878-INS-65
chr11-77069881-INS-727
chr11-77079102-INS-88
chr11-77084093-INS-70
chr11-77093239-INS-702
chr11-77094580-INS-210
chr11-77170507-INS-70
chr11-77183667-INS-54
chr11-77197719-INS-116
chr11-77221143-INS-138
chr11-77258289-INS-2838
chr11-77259480-INS-62
chr11-77265321-INS-114
chr11-77268445-INS-52
chr11-77268894-INS-6601
chr11-77286680-INS-209
chr11-77288164-INS-141
chr11-77318384-INS-246
chr11-77351276-INS-66

chr11-77447192-INS-213
chr11-77477444-INS-61
chr11-77481312-INS-217
chr11-77483961-INS-226
chr11-77500731-INS-61
chr11-77503265-INS-278
chr11-77524466-INS-374
chr11-77527127-INS-250
chr11-77528910-INS-298
chr11-77547414-INS-203
chr11-77556049-INS-91
chr11-77563034-INS-5814
chr11-77566565-INS-52
chr11-77566980-INS-227
chr11-77571904-INS-218
chr11-77588855-INS-192
chr11-77596818-INS-210
chr11-77599786-INS-384
chr11-77616673-INS-81
chr11-77620563-INS-193
chr11-77633454-INS-111
chr11-77636212-INS-145
chr11-77637082-INS-54
chr11-77640432-INS-70
chr11-77642350-INS-53
chr11-77646277-INS-181
chr11-77648077-INS-216
chr11-77650456-INS-52
chr11-77652468-INS-286
chr11-77663603-INS-199
chr11-77669574-INS-54
chr11-77673906-INS-191
chr11-77675883-INS-83
chr11-77697376-INS-207
chr11-77701157-INS-209
chr11-77714725-INS-200
chr11-77717448-INS-214
chr11-77725049-INS-217
chr11-77796573-INS-202
chr11-77799171-INS-81
chr11-77799836-INS-221
chr11-77804452-INS-253
chr11-77818474-INS-87
chr11-77820014-INS-5562
chr11-77832591-INS-51
chr11-77833712-INS-74
chr11-77858458-INS-73
chr11-77871859-INS-84
chr11-77881294-INS-176
chr11-77887841-INS-659
chr11-77888687-INS-94
chr11-77907070-INS-64
chr11-77931784-INS-221
chr11-77936369-INS-218
chr11-77937550-INS-174
chr11-77938021-INS-214
chr11-77947826-INS-212
chr11-77951843-INS-205
chr11-77954689-INS-221
chr11-77959810-INS-189
chr11-77967252-INS-199
chr11-77976371-INS-356
chr11-77978181-INS-176
chr11-77991053-INS-68
chr11-78003157-INS-241
chr11-78031344-INS-306
chr11-78041661-INS-177
chr11-78060659-INS-380
chr11-78081784-INS-53
chr11-78089283-INS-329
chr11-78091713-INS-470
chr11-78105489-INS-184
chr11-78114063-INS-221
chr11-78189566-INS-204
chr11-78209913-INS-60
chr11-78231597-INS-184
chr11-78304164-INS-206

chr11-78305987-INS-70
chr11-78318082-INS-391
chr11-78320516-INS-5313
chr11-78320580-INS-8766
chr11-78323764-INS-51
chr11-78328437-INS-323
chr11-78332097-INS-55
chr11-78337459-INS-181
chr11-78351037-INS-211
chr11-78384824-INS-168
chr11-78435203-INS-283
chr11-78437091-INS-128
chr11-78454672-INS-57
chr11-78460092-INS-53
chr11-78482374-INS-131
chr11-78503060-INS-201
chr11-78511569-INS-208
chr11-78524975-INS-1051
chr11-78550100-INS-60
chr11-78563385-INS-167
chr11-78588975-INS-200
chr11-78615646-INS-502
chr11-78631601-INS-113
chr11-78646140-INS-82
chr11-78647527-INS-55
chr11-78650790-INS-130
chr11-78651844-INS-225
chr11-78653910-INS-155
chr11-78658809-INS-498
chr11-78667950-INS-7131
chr11-78697797-INS-178
chr11-78700274-INS-57
chr11-78710117-INS-52
chr11-78722004-INS-206
chr11-78723682-INS-197
chr11-78727915-INS-352
chr11-78735298-INS-228
chr11-78748975-INS-52
chr11-78753774-INS-67
chr11-78757012-INS-77
chr11-78759687-INS-236
chr11-78761957-INS-212
chr11-78763071-INS-142
chr11-78766041-INS-256
chr11-78769440-INS-65
chr11-78771550-INS-54
chr11-78785550-INS-122
chr11-78817864-INS-68
chr11-78821677-INS-94
chr11-78847179-INS-379
chr11-78852996-INS-359
chr11-78856087-INS-51
chr11-78879932-INS-55
chr11-78887650-INS-68
chr11-78890011-INS-68
chr11-78896434-INS-207
chr11-78899348-INS-110
chr11-78913138-INS-67
chr11-78933120-INS-54
chr11-78941033-INS-62
chr11-78943222-INS-515
chr11-78981430-INS-88
chr11-78990912-INS-50
chr11-79022237-INS-114
chr11-79022733-INS-50
chr11-79043277-INS-70
chr11-79091846-INS-97
chr11-79093908-INS-173
chr11-79096740-INS-70
chr11-79097925-INS-78
chr11-79099009-INS-54
chr11-79100670-INS-201
chr11-79101967-INS-200
chr11-79118553-INS-210
chr11-79141038-INS-56
chr11-79145801-INS-231
chr11-79154649-INS-345

chr11-79160116-INS-53
chr11-79165636-INS-116
chr11-79166856-INS-71
chr11-79171775-INS-52
chr11-79188864-INS-189
chr11-79189703-INS-398
chr11-79190536-INS-145
chr11-79203359-INS-354
chr11-79222140-INS-6844
chr11-79243966-INS-109
chr11-79264982-INS-60
chr11-79295224-INS-60
chr11-79313895-INS-216
chr11-79321632-INS-75
chr11-79322428-INS-220
chr11-79342556-INS-174
chr11-79387463-INS-215
chr11-79411366-INS-5548
chr11-79430190-INS-166
chr11-79474307-INS-146
chr11-79518613-INS-198
chr11-79519635-INS-96
chr11-79521156-INS-106
chr11-79527537-INS-100
chr11-79530369-INS-212
chr11-79533807-INS-90
chr11-79536666-INS-182
chr11-79540563-INS-127
chr11-79558469-INS-71
chr11-79562383-INS-132
chr11-79579944-INS-74
chr11-79590827-INS-58
chr11-79593170-INS-84
chr11-79593469-INS-427
chr11-79607170-INS-65
chr11-79610974-INS-551
chr11-79643131-INS-229
chr11-79658138-INS-125
chr11-79665842-INS-53
chr11-79690260-INS-116
chr11-79710046-INS-84
chr11-79725018-INS-70
chr11-79728821-INS-336
chr11-79742919-INS-149
chr11-79754741-INS-209
chr11-79772505-INS-159
chr11-79803945-INS-202
chr11-79804501-INS-63
chr11-79816796-INS-54
chr11-79830563-INS-52
chr11-79849132-INS-217
chr11-79853843-INS-277
chr11-79862059-INS-206
chr11-79872256-INS-193
chr11-79878142-INS-175
chr11-79887164-INS-147
chr11-79911380-INS-198
chr11-79914139-INS-212
chr11-79915108-INS-244
chr11-79927895-INS-194
chr11-79930611-INS-56
chr11-79932219-INS-197
chr11-79934791-INS-166
chr11-79947673-INS-185
chr11-79954047-INS-188
chr11-79954687-INS-204
chr11-79960016-INS-179
chr11-79961977-INS-259
chr11-79975777-INS-57
chr11-79977406-INS-174
chr11-79977984-INS-223
chr11-79979023-INS-339
chr11-79990118-INS-225
chr11-79996069-INS-214
chr11-80002617-INS-56
chr11-80006969-INS-271
chr11-80024517-INS-201

chr11-80051912-INS-201
chr11-80066126-INS-242
chr11-80070715-INS-52
chr11-80072991-INS-202
chr11-80081067-INS-328
chr11-80088310-INS-848
chr11-80088329-INS-86
chr11-80094763-INS-209
chr11-80097198-INS-733
chr11-80099020-INS-365
chr11-80112194-INS-55
chr11-80127346-INS-217
chr11-80130429-INS-208
chr11-80159956-INS-109
chr11-80165012-INS-53
chr11-80172589-INS-392
chr11-80182436-INS-5000
chr11-80198238-INS-129
chr11-80204492-INS-64
chr11-80216609-INS-256
chr11-80229412-INS-102
chr11-80237912-INS-182
chr11-80248833-INS-70
chr11-80250232-INS-2102
chr11-80258315-INS-201
chr11-80271097-INS-202
chr11-80276663-INS-309
chr11-80283159-INS-212
chr11-80308947-INS-217
chr11-80341496-INS-416
chr11-80348112-INS-74
chr11-80356853-INS-157
chr11-80376136-INS-56
chr11-80398617-INS-299
chr11-80465957-INS-743
chr11-80467733-INS-52
chr11-80488049-INS-123
chr11-80490637-INS-228
chr11-80491643-INS-85
chr11-80511440-INS-279
chr11-80518694-INS-56
chr11-80522327-INS-70
chr11-80524196-INS-7493
chr11-80564516-INS-248
chr11-80574748-INS-122
chr11-80585918-INS-6637
chr11-80608641-INS-56
chr11-80632803-INS-181
chr11-80637382-INS-65
chr11-80641730-INS-432
chr11-80663369-INS-60
chr11-80673759-INS-167
chr11-80688195-INS-7090
chr11-80696382-INS-141
chr11-80699720-INS-127
chr11-80718111-INS-66
chr11-80732063-INS-68
chr11-80733030-INS-54
chr11-80737612-INS-112
chr11-80746231-INS-86
chr11-80752772-INS-155
chr11-80755177-INS-73
chr11-80772077-INS-126
chr11-80775717-INS-59
chr11-80779775-INS-83
chr11-80808409-INS-123
chr11-80811198-INS-639
chr11-80822348-INS-80
chr11-80834999-INS-533
chr11-80836304-INS-68
chr11-80844328-INS-82
chr11-80845616-INS-143
chr11-80858545-INS-70
chr11-80887477-INS-54
chr11-80936009-INS-304
chr11-80957357-INS-492
chr11-80972167-INS-52

chr11-80972716-INS-337
chr11-80974392-INS-6485
chr11-80996439-INS-5931
chr11-81007344-INS-336
chr11-81016417-INS-100
chr11-81040200-INS-95
chr11-81073663-INS-312
chr11-81126845-INS-186
chr11-81134643-INS-55
chr11-81169435-INS-97
chr11-81191970-INS-68
chr11-81195585-INS-72
chr11-81201389-INS-7165
chr11-81251379-INS-1891
chr11-81283400-INS-87
chr11-81289657-INS-5281
chr11-81329517-INS-59
chr11-81351343-INS-213
chr11-81354030-INS-551
chr11-81362840-INS-157
chr11-81373114-INS-55
chr11-81379180-INS-71
chr11-81380996-INS-63
chr11-81382673-INS-733
chr11-81390529-INS-58
chr11-81417327-INS-73
chr11-81427559-INS-69
chr11-81458603-INS-211
chr11-81464492-INS-111
chr11-81506849-INS-110
chr11-81508939-INS-224
chr11-81512319-INS-92
chr11-81512985-INS-73
chr11-81544810-INS-93
chr11-81590493-INS-56
chr11-81596902-INS-88
chr11-81619434-INS-212
chr11-81621915-INS-54
chr11-81640515-INS-92
chr11-81647240-INS-57
chr11-81661718-INS-184
chr11-81666721-INS-110
chr11-81689866-INS-56
chr11-81695312-INS-53
chr11-81723315-INS-54
chr11-81744242-INS-92
chr11-81773604-INS-52
chr11-81782863-INS-450
chr11-81804942-INS-57
chr11-81805955-INS-62
chr11-81819545-INS-204
chr11-81824081-INS-57
chr11-81828064-INS-67
chr11-81830414-INS-133
chr11-81834214-INS-6685
chr11-81843265-INS-85
chr11-81896621-INS-100
chr11-81900425-INS-97
chr11-81916488-INS-56
chr11-81917484-INS-4589
chr11-81940645-INS-52
chr11-81943775-INS-725
chr11-81972351-INS-84
chr11-81999146-INS-445
chr11-82010251-INS-261
chr11-82019390-INS-199
chr11-82022320-INS-445
chr11-82087288-INS-61
chr11-82092090-INS-662
chr11-82107853-INS-314
chr11-82161505-INS-54
chr11-82179442-INS-51
chr11-82179993-INS-144
chr11-82186738-INS-500
chr11-82195125-INS-207
chr11-82386941-INS-7116
chr11-82413387-INS-158

chr11-82440978-INS-649
chr11-82441963-INS-7103
chr11-82462396-INS-87
chr11-82499548-INS-70
chr11-82514414-INS-62
chr11-82532382-INS-56
chr11-82544431-INS-281
chr11-82578043-INS-101
chr11-82623879-INS-236
chr11-82638998-INS-205
chr11-82652239-INS-54
chr11-82663808-INS-324
chr11-82664809-INS-219
chr11-82670251-INS-308
chr11-82675970-INS-60
chr11-82683307-INS-226
chr11-82706671-INS-187
chr11-82707576-INS-56
chr11-82708244-INS-53
chr11-82758921-INS-86
chr11-82765926-INS-397
chr11-82786324-INS-208
chr11-82797089-INS-104
chr11-82821629-INS-272
chr11-82822221-INS-354
chr11-82823058-INS-442
chr11-82823292-INS-1324
chr11-82826289-INS-70
chr11-82863506-INS-90
chr11-82866424-INS-3214
chr11-82867856-INS-229
chr11-82885874-INS-55
chr11-82888679-INS-69
chr11-82889138-INS-178
chr11-82901976-INS-87
chr11-82914066-INS-181
chr11-82981721-INS-6401
chr11-82998774-INS-7127
chr11-83016714-INS-96
chr11-83028404-INS-55
chr11-83063399-INS-846
chr11-83175786-INS-139
chr11-83191061-INS-105
chr11-83193427-INS-291
chr11-83207507-INS-175
chr11-83208579-INS-58
chr11-83235988-INS-406
chr11-83268556-INS-367
chr11-83275160-INS-66
chr11-83347757-INS-244
chr11-83383721-INS-235
chr11-83385869-INS-255
chr11-83415334-INS-241
chr11-83415996-INS-157
chr11-83446560-INS-151
chr11-83450162-INS-62
chr11-83452793-INS-202
chr11-83456447-INS-92
chr11-83460624-INS-58
chr11-83470998-INS-57
chr11-83489210-INS-58
chr11-83533908-INS-137
chr11-83565063-INS-499
chr11-83600188-INS-208
chr11-83613107-INS-400
chr11-83675094-INS-60
chr11-83677727-INS-723
chr11-83730639-INS-66
chr11-83748252-INS-63
chr11-83750573-INS-99
chr11-83766722-INS-58
chr11-83767879-INS-214
chr11-83774992-INS-84
chr11-83776968-INS-75
chr11-83779139-INS-146
chr11-83811626-INS-459
chr11-83822667-INS-206

chr11-83835304-INS-269
chr11-83836569-INS-181
chr11-83857086-INS-181
chr11-83868274-INS-227
chr11-83874562-INS-286
chr11-83888464-INS-76
chr11-83898529-INS-207
chr11-83905666-INS-75
chr11-83913707-INS-220
chr11-83936331-INS-179
chr11-83950886-INS-188
chr11-83968370-INS-72
chr11-83998633-INS-171
chr11-84001837-INS-218
chr11-84031794-INS-53
chr11-84049154-INS-308
chr11-84061085-INS-334
chr11-84077927-INS-55
chr11-84083006-INS-54
chr11-84088194-INS-60
chr11-84097250-INS-176
chr11-84098466-INS-97
chr11-84123400-INS-58
chr11-84124947-INS-158
chr11-84223254-INS-115
chr11-84226518-INS-65
chr11-84236020-INS-74
chr11-84255924-INS-622
chr11-84277289-INS-54
chr11-84278475-INS-280
chr11-84302141-INS-208
chr11-84321556-INS-127
chr11-84356110-INS-223
chr11-84358197-INS-58
chr11-84367487-INS-212
chr11-84368965-INS-58
chr11-84375978-INS-61
chr11-84394538-INS-198
chr11-84419902-INS-56
chr11-84436771-INS-63
chr11-84438381-INS-118
chr11-84463243-INS-126
chr11-84483606-INS-62
chr11-84519847-INS-118
chr11-84522822-INS-149
chr11-84579603-INS-589
chr11-84595233-INS-510
chr11-84601163-INS-105
chr11-84606870-INS-87
chr11-84644570-INS-69
chr11-84675974-INS-72
chr11-84696209-INS-85
chr11-84715349-INS-257
chr11-84717389-INS-205
chr11-84733188-INS-310
chr11-84758456-INS-210
chr11-84803373-INS-350
chr11-84814624-INS-235
chr11-84815977-INS-357
chr11-84827087-INS-80
chr11-84837371-INS-510
chr11-84849696-INS-70
chr11-84868851-INS-140
chr11-84914166-INS-60
chr11-84919207-INS-195
chr11-84922381-INS-101
chr11-84965291-INS-162
chr11-84976319-INS-55
chr11-84986520-INS-200
chr11-85001897-INS-64
chr11-85020833-INS-237
chr11-85024800-INS-64
chr11-85034240-INS-77
chr11-85035708-INS-182
chr11-85041606-INS-153
chr11-85046322-INS-94
chr11-85057892-INS-270

chr11-85060089-INS-326
chr11-85072995-INS-127
chr11-85141117-INS-85
chr11-85141483-INS-74
chr11-85150140-INS-7033
chr11-85157022-INS-56
chr11-85160837-INS-190
chr11-85183635-INS-62
chr11-85184641-INS-2145
chr11-85186998-INS-195
chr11-85196943-INS-169
chr11-85199241-INS-74
chr11-85216019-INS-7100
chr11-85226346-INS-695
chr11-85229361-INS-148
chr11-85239919-INS-584
chr11-85288669-INS-90
chr11-85301754-INS-547
chr11-85319495-INS-54
chr11-85347357-INS-229
chr11-85418541-INS-206
chr11-85459940-INS-56
chr11-85466168-INS-206
chr11-85493259-INS-72
chr11-85525817-INS-182
chr11-85539460-INS-867
chr11-85550783-INS-229
chr11-85565035-INS-1029
chr11-85565112-INS-176
chr11-85565193-INS-186
chr11-85571448-INS-54
chr11-85624360-INS-811
chr11-85655189-INS-400
chr11-85671796-INS-4854
chr11-85714613-INS-93
chr11-85745144-INS-109
chr11-85761816-INS-199
chr11-85769405-INS-76
chr11-85810975-INS-131
chr11-85820844-INS-399
chr11-85844444-INS-76
chr11-85866182-INS-709
chr11-85872940-INS-83
chr11-85879713-INS-96
chr11-85928511-INS-53
chr11-85961844-INS-52
chr11-86000940-INS-334
chr11-86006456-INS-108
chr11-86024165-INS-213
chr11-86028532-INS-591
chr11-86057061-INS-144
chr11-86063476-INS-156
chr11-86065701-INS-204
chr11-86077747-INS-205
chr11-86081254-INS-168
chr11-86093214-INS-63
chr11-86119882-INS-220
chr11-86122705-INS-146
chr11-86125144-INS-62
chr11-86126962-INS-78
chr11-86135560-INS-208
chr11-86143769-INS-221
chr11-86150059-INS-58
chr11-86183428-INS-79
chr11-86195235-INS-722
chr11-86205136-INS-271
chr11-86227532-INS-229
chr11-86230440-INS-148
chr11-86235985-INS-62
chr11-86255394-INS-52
chr11-86264459-INS-56
chr11-86266503-INS-193
chr11-86280883-INS-97
chr11-86326753-INS-88
chr11-86329901-INS-70
chr11-86332075-INS-212
chr11-86360991-INS-108

chr11-86364895-INS-816
chr11-86369771-INS-76
chr11-86379683-INS-268
chr11-86388627-INS-282
chr11-86431486-INS-113
chr11-86436959-INS-193
chr11-86438326-INS-200
chr11-86460781-INS-116
chr11-86499280-INS-62
chr11-86538792-INS-53
chr11-86538972-INS-5935
chr11-86567164-INS-874
chr11-86568116-INS-56
chr11-86623617-INS-52
chr11-86671837-INS-98
chr11-86676306-INS-99
chr11-86708148-INS-79
chr11-86722873-INS-294
chr11-86728893-INS-54
chr11-86735486-INS-202
chr11-86750268-INS-182
chr11-86772689-INS-222
chr11-86797167-INS-252
chr11-86817646-INS-51
chr11-86824742-INS-51
chr11-86861624-INS-271
chr11-86867284-INS-131
chr11-86895757-INS-169
chr11-86906749-INS-64
chr11-86910242-INS-208
chr11-86911730-INS-87
chr11-86917897-INS-67
chr11-86919917-INS-384
chr11-86919922-INS-950
chr11-86922791-INS-100
chr11-86939472-INS-625
chr11-86963609-INS-245
chr11-86983112-INS-76
chr11-86989712-INS-207
chr11-86991236-INS-168
chr11-87032742-INS-217
chr11-87045571-INS-58
chr11-87047361-INS-488
chr11-87053368-INS-5091
chr11-87055248-INS-204
chr11-87075292-INS-66
chr11-87077811-INS-68
chr11-87090921-INS-102
chr11-87093550-INS-53
chr11-87117334-INS-78
chr11-87135897-INS-72
chr11-87153905-INS-106
chr11-87154542-INS-52
chr11-87158155-INS-90
chr11-87196257-INS-581
chr11-87213526-INS-133
chr11-87266908-INS-197
chr11-87289596-INS-205
chr11-87304343-INS-254
chr11-87328916-INS-184
chr11-87333220-INS-598
chr11-87339726-INS-469
chr11-87343549-INS-213
chr11-87353683-INS-83
chr11-87374687-INS-410
chr11-87375965-INS-83
chr11-87380518-INS-54
chr11-87382783-INS-488
chr11-87389596-INS-177
chr11-87393606-INS-176
chr11-87395862-INS-829
chr11-87399854-INS-153
chr11-87403718-INS-240
chr11-87408388-INS-208
chr11-87413380-INS-72
chr11-87447479-INS-220
chr11-87471444-INS-211

chr11-87523483-INS-62
chr11-87524614-INS-81
chr11-87525679-INS-66
chr11-87532103-INS-69
chr11-87533628-INS-106
chr11-87535543-INS-77
chr11-87558433-INS-52
chr11-87564978-INS-215
chr11-87577540-INS-63
chr11-87582660-INS-60
chr11-87633752-INS-56
chr11-87641736-INS-55
chr11-87648874-INS-155
chr11-87658384-INS-70
chr11-87717049-INS-168
chr11-87724198-INS-451
chr11-87733932-INS-5849
chr11-87736046-INS-56
chr11-87738971-INS-62
chr11-87772138-INS-153
chr11-87780987-INS-106
chr11-87807707-INS-63
chr11-87822397-INS-127
chr11-87859551-INS-185
chr11-87860709-INS-53
chr11-87865071-INS-204
chr11-87889386-INS-196
chr11-87901579-INS-219
chr11-87925839-INS-106
chr11-87949078-INS-109
chr11-87955537-INS-185
chr11-88012186-INS-207
chr11-88015558-INS-61
chr11-88022218-INS-219
chr11-88035471-INS-482
chr11-88059089-INS-601
chr11-88064044-INS-58
chr11-88072844-INS-103
chr11-88075881-INS-208
chr11-88096553-INS-133
chr11-88109404-INS-175
chr11-88120298-INS-222
chr11-88147667-INS-110
chr11-88158572-INS-170
chr11-88178547-INS-114
chr11-88197412-INS-252
chr11-88204990-INS-58
chr11-88226979-INS-6966
chr11-88252048-INS-80
chr11-88269055-INS-212
chr11-88292014-INS-225
chr11-88330647-INS-64
chr11-88335354-INS-189
chr11-88364985-INS-136
chr11-88440264-INS-216
chr11-88454470-INS-153
chr11-88482713-INS-56
chr11-88485189-INS-135
chr11-88497273-INS-201
chr11-88565432-INS-78
chr11-88583856-INS-58
chr11-88641142-INS-218
chr11-88649864-INS-62
chr11-88660213-INS-54
chr11-88670914-INS-175
chr11-88695797-INS-56
chr11-88718776-INS-79
chr11-88724625-INS-190
chr11-88733382-INS-219
chr11-88744494-INS-80
chr11-88745679-INS-233
chr11-88761500-INS-206
chr11-88769904-INS-534
chr11-88771083-INS-54
chr11-88771717-INS-463
chr11-88775145-INS-190
chr11-88780938-INS-197

chr11-88789089-INS-400
chr11-88822165-INS-222
chr11-88837265-INS-119
chr11-88847081-INS-359
chr11-88860734-INS-57
chr11-88863027-INS-477
chr11-88896850-INS-358
chr11-88914610-INS-119
chr11-88922513-INS-462
chr11-88924384-INS-207
chr11-88940218-INS-66
chr11-88942875-INS-206
chr11-88953487-INS-196
chr11-88955899-INS-166
chr11-88956909-INS-182
chr11-88957920-INS-54
chr11-88970818-INS-229
chr11-88978605-INS-117
chr11-88987025-INS-84
chr11-89005867-INS-54
chr11-89030417-INS-679
chr11-89031793-INS-228
chr11-89041858-INS-188
chr11-89067939-INS-54
chr11-89081369-INS-140
chr11-89085094-INS-72
chr11-89120009-INS-135
chr11-89123622-INS-6442
chr11-89127756-INS-3417
chr11-89138665-INS-54
chr11-89146195-INS-497
chr11-89165980-INS-233
chr11-89183772-INS-167
chr11-89206622-INS-51
chr11-89210317-INS-68
chr11-89221101-INS-3770
chr11-89230554-INS-58
chr11-89234811-INS-176
chr11-89237248-INS-636
chr11-89260614-INS-83
chr11-89266429-INS-392
chr11-89270952-INS-322
chr11-89294926-INS-1084
chr11-89305064-INS-65
chr11-89331160-INS-54
chr11-89340343-INS-53
chr11-89374159-INS-57
chr11-89392555-INS-85
chr11-89408473-INS-151
chr11-89426515-INS-66
chr11-89438483-INS-160
chr11-89453469-INS-7171
chr11-89476228-INS-66
chr11-89491323-INS-355
chr11-89508720-INS-123
chr11-89520746-INS-148
chr11-89532519-INS-369
chr11-89534134-INS-70
chr11-89593428-INS-430
chr11-89599857-INS-401
chr11-89699723-INS-60
chr11-89701621-INS-72
chr11-89731449-INS-74
chr11-89745095-INS-977
chr11-89755127-INS-98
chr11-89781836-INS-61
chr11-89810219-INS-60
chr11-89830122-INS-906
chr11-89847117-INS-73
chr11-89907864-INS-70
chr11-89912932-INS-217
chr11-89914713-INS-328
chr11-89959883-INS-197
chr11-89981884-INS-70
chr11-89995148-INS-231
chr11-90065050-INS-87
chr11-90096236-INS-54

chr11-90099653-INS-401
chr11-90108081-INS-78
chr11-90110340-INS-95
chr11-90113892-INS-120
chr11-90165917-INS-155
chr11-90177205-INS-2382
chr11-90206990-INS-59
chr11-90241642-INS-56
chr11-90270289-INS-72
chr11-90294387-INS-8294
chr11-90312616-INS-7963
chr11-90334168-INS-7070
chr11-90346413-INS-92
chr11-90356198-INS-404
chr11-90367211-INS-174
chr11-90375405-INS-51
chr11-90426462-INS-143
chr11-90428292-INS-56
chr11-90430124-INS-1873
chr11-90433558-INS-210
chr11-90466785-INS-276
chr11-90562977-INS-132
chr11-90564319-INS-161
chr11-90565379-INS-131
chr11-90583937-INS-102
chr11-90583960-INS-658
chr11-90599130-INS-93
chr11-90610846-INS-122
chr11-90621209-INS-169
chr11-90631476-INS-4405
chr11-90675317-INS-251
chr11-90683946-INS-195
chr11-90704809-INS-117
chr11-90720586-INS-51
chr11-90748372-INS-2834
chr11-90759134-INS-188
chr11-90764427-INS-347
chr11-90782268-INS-807
chr11-90785984-INS-323
chr11-90805220-INS-277
chr11-90812882-INS-60
chr11-90823822-INS-7100
chr11-90833698-INS-499
chr11-90836243-INS-224
chr11-90839624-INS-545
chr11-90846874-INS-78
chr11-90848591-INS-215
chr11-90857678-INS-111
chr11-90880540-INS-62
chr11-90887083-INS-634
chr11-90918435-INS-68
chr11-90934090-INS-71
chr11-90955324-INS-342
chr11-90963330-INS-469
chr11-90972485-INS-140
chr11-90976886-INS-154
chr11-90983149-INS-5420
chr11-90986509-INS-189
chr11-90988168-INS-55
chr11-91002806-INS-62
chr11-91004385-INS-201
chr11-91006964-INS-52
chr11-91029744-INS-205
chr11-91036152-INS-261
chr11-91054492-INS-964
chr11-91057234-INS-210
chr11-91066402-INS-91
chr11-91068036-INS-199
chr11-91069794-INS-85
chr11-91084121-INS-209
chr11-91085402-INS-53
chr11-91089534-INS-316
chr11-91101641-INS-5630
chr11-91122966-INS-206
chr11-91153754-INS-909
chr11-91178652-INS-532
chr11-91179968-INS-127

chr11-91182363-INS-6717
chr11-91203043-INS-52
chr11-91226119-INS-197
chr11-91245220-INS-78
chr11-91260274-INS-60
chr11-91277146-INS-56
chr11-91279351-INS-566
chr11-91296819-INS-174
chr11-91327367-INS-159
chr11-91332339-INS-56
chr11-91346766-INS-171
chr11-91354949-INS-7110
chr11-91374450-INS-139
chr11-91375469-INS-139
chr11-91376294-INS-235
chr11-91413632-INS-72
chr11-91437903-INS-527
chr11-91458685-INS-59
chr11-91469720-INS-86
chr11-91523777-INS-724
chr11-91545356-INS-60
chr11-91564809-INS-66
chr11-91579832-INS-67
chr11-91604434-INS-423
chr11-91645791-INS-435
chr11-91645899-INS-6217
chr11-91648814-INS-56
chr11-91659315-INS-114
chr11-91662715-INS-979
chr11-91674712-INS-224
chr11-91695335-INS-125
chr11-91730216-INS-106
chr11-91766693-INS-6474
chr11-91789076-INS-525
chr11-91798247-INS-644
chr11-91799717-INS-133
chr11-91807671-INS-216
chr11-91823500-INS-625
chr11-91846395-INS-123
chr11-91850104-INS-86
chr11-91860734-INS-54
chr11-91863365-INS-132
chr11-91866954-INS-6376
chr11-91869984-INS-60
chr11-91877745-INS-497
chr11-91890595-INS-71
chr11-91911009-INS-62
chr11-91919893-INS-7086
chr11-91941494-INS-673
chr11-91959862-INS-196
chr11-91963919-INS-98
chr11-91977049-INS-531
chr11-91983276-INS-208
chr11-91989330-INS-59
chr11-92000531-INS-52
chr11-92006770-INS-167
chr11-92013418-INS-52
chr11-92025359-INS-897
chr11-92025371-INS-315
chr11-92028935-INS-58
chr11-92080038-INS-840
chr11-92082329-INS-104
chr11-92124563-INS-456
chr11-92164651-INS-5331
chr11-92178925-INS-6401
chr11-92200245-INS-6605
chr11-92217528-INS-73
chr11-92234351-INS-123
chr11-92237813-INS-123
chr11-92240120-INS-831
chr11-92247967-INS-52
chr11-92252244-INS-52
chr11-92254117-INS-58
chr11-92262745-INS-201
chr11-92275348-INS-596
chr11-92280192-INS-351
chr11-92284914-INS-59

chr11-92306897-INS-75
chr11-92324274-INS-72
chr11-92364573-INS-182
chr11-92374435-INS-54
chr11-92381120-INS-2825
chr11-92432018-INS-512
chr11-92442814-INS-163
chr11-92454772-INS-73
chr11-92459536-INS-298
chr11-92504899-INS-203
chr11-92531403-INS-84
chr11-92537949-INS-60
chr11-92542665-INS-581
chr11-92544632-INS-67
chr11-92558357-INS-90
chr11-92573100-INS-137
chr11-92613690-INS-704
chr11-92616139-INS-7186
chr11-92628407-INS-60
chr11-92631582-INS-6593
chr11-92639345-INS-60
chr11-92644061-INS-134
chr11-92660757-INS-54
chr11-92667167-INS-660
chr11-92686377-INS-7243
chr11-92688494-INS-118
chr11-92696480-INS-52
chr11-92711910-INS-219
chr11-92714206-INS-355
chr11-92722444-INS-8413
chr11-92729161-INS-1886
chr11-92732177-INS-4057
chr11-92744480-INS-80
chr11-92760532-INS-6958
chr11-92772841-INS-507
chr11-92777263-INS-353
chr11-92784164-INS-526
chr11-92788336-INS-74
chr11-92792129-INS-3038
chr11-92799655-INS-75
chr11-92810722-INS-4778
chr11-92820336-INS-3196
chr11-92836243-INS-101
chr11-92864602-INS-128
chr11-92875417-INS-115
chr11-92895891-INS-424
chr11-92902332-INS-665
chr11-92916016-INS-58
chr11-92923775-INS-134
chr11-92925516-INS-136
chr11-92928761-INS-575
chr11-92973346-INS-373
chr11-92974802-INS-322
chr11-92982012-INS-131
chr11-93009794-INS-1887
chr11-93049325-INS-599
chr11-93064214-INS-164
chr11-93068387-INS-86
chr11-93069858-INS-239
chr11-93112359-INS-57
chr11-93148429-INS-6941
chr11-93162243-INS-1722
chr11-93168097-INS-72
chr11-93201318-INS-101
chr11-93206188-INS-60
chr11-93207391-INS-529
chr11-93222206-INS-443
chr11-93233986-INS-382
chr11-93262108-INS-519
chr11-93268725-INS-158
chr11-93276264-INS-62
chr11-93280417-INS-57
chr11-93298793-INS-304
chr11-93300455-INS-62
chr11-93346338-INS-578
chr11-93349410-INS-69
chr11-93358369-INS-704

chr11-93415074-INS-58
chr11-93473156-INS-370
chr11-93489561-INS-52
chr11-93491476-INS-92
chr11-93493725-INS-313
chr11-93495570-INS-175
chr11-93499430-INS-518
chr11-93503622-INS-102
chr11-93511483-INS-202
chr11-93514889-INS-172
chr11-93520930-INS-51
chr11-93527013-INS-400
chr11-93577988-INS-70
chr11-93579035-INS-56
chr11-93581239-INS-126
chr11-93593201-INS-60
chr11-93595273-INS-231
chr11-93597355-INS-125
chr11-93669422-INS-200
chr11-93680546-INS-54
chr11-93726233-INS-163
chr11-93729969-INS-60
chr11-93744679-INS-82
chr11-93748729-INS-75
chr11-93762331-INS-471
chr11-93780047-INS-180
chr11-93797819-INS-217
chr11-93808574-INS-120
chr11-93845496-INS-163
chr11-93851340-INS-223
chr11-93852362-INS-211
chr11-93871013-INS-383
chr11-93897976-INS-270
chr11-93944641-INS-102
chr11-93945734-INS-130
chr11-93959689-INS-55
chr11-93975283-INS-97
chr11-94001304-INS-202
chr11-94033796-INS-68
chr11-94035598-INS-169
chr11-94053434-INS-113
chr11-94092130-INS-52
chr11-94111568-INS-63
chr11-94117128-INS-67
chr11-94141406-INS-77
chr11-94146534-INS-57
chr11-94152233-INS-734
chr11-94153971-INS-159
chr11-94190982-INS-198
chr11-94224899-INS-207
chr11-94225000-INS-6442
chr11-94229083-INS-245
chr11-94247495-INS-52
chr11-94260326-INS-56
chr11-94298150-INS-182
chr11-94308801-INS-166
chr11-94326400-INS-58
chr11-94344761-INS-107
chr11-94362604-INS-52
chr11-94390670-INS-60
chr11-94406920-INS-204
chr11-94408349-INS-90
chr11-94409428-INS-196
chr11-94447042-INS-123
chr11-94448825-INS-135
chr11-94474816-INS-138
chr11-94478340-INS-638
chr11-94490805-INS-54
chr11-94498006-INS-60
chr11-94500472-INS-52
chr11-94506017-INS-192
chr11-94510886-INS-78
chr11-94531376-INS-115
chr11-94552629-INS-164
chr11-94577144-INS-59
chr11-94582030-INS-534
chr11-94587881-INS-210

chr11-94601900-INS-242
chr11-94605665-INS-494
chr11-94615516-INS-191
chr11-94623730-INS-115
chr11-94627275-INS-152
chr11-94630324-INS-249
chr11-94631048-INS-197
chr11-94638180-INS-99
chr11-94638740-INS-56
chr11-94652896-INS-56
chr11-94655255-INS-1866
chr11-94670927-INS-177
chr11-94681053-INS-384
chr11-94681054-INS-269
chr11-94721218-INS-82
chr11-94742949-INS-72
chr11-94754584-INS-109
chr11-94758141-INS-266
chr11-94781188-INS-64
chr11-94856698-INS-477
chr11-94857670-INS-223
chr11-94874553-INS-490
chr11-94904379-INS-767
chr11-94904404-INS-301
chr11-94914881-INS-232
chr11-94920392-INS-80
chr11-94940752-INS-78
chr11-94949725-INS-176
chr11-94973696-INS-185
chr11-94975656-INS-222
chr11-94981143-INS-71
chr11-94985034-INS-202
chr11-95004780-INS-66
chr11-95055227-INS-52
chr11-95090192-INS-224
chr11-95108679-INS-151
chr11-95116712-INS-56
chr11-95117963-INS-88
chr11-95126186-INS-52
chr11-95137819-INS-76
chr11-95155574-INS-82
chr11-95198016-INS-134
chr11-95202247-INS-101
chr11-95214095-INS-202
chr11-95216071-INS-173
chr11-95239669-INS-204
chr11-95263639-INS-180
chr11-95286451-INS-56
chr11-95317084-INS-203
chr11-95324129-INS-209
chr11-95354627-INS-202
chr11-95360166-INS-72
chr11-95369771-INS-193
chr11-95397277-INS-196
chr11-95428392-INS-132
chr11-95451998-INS-160
chr11-95461995-INS-67
chr11-95498685-INS-180
chr11-95526421-INS-59
chr11-95528701-INS-140
chr11-95551479-INS-192
chr11-95555290-INS-220
chr11-95590999-INS-271
chr11-95598007-INS-116
chr11-95611547-INS-112
chr11-95613478-INS-204
chr11-95676392-INS-228
chr11-95678830-INS-193
chr11-95707938-INS-54
chr11-95761551-INS-60
chr11-95771160-INS-198
chr11-95796333-INS-200
chr11-95801384-INS-206
chr11-95806536-INS-88
chr11-95811579-INS-159
chr11-95811579-INS-203
chr11-95812564-INS-164

chr11-95816678-INS-65
chr11-95828104-INS-260
chr11-95829591-INS-193
chr11-95829876-INS-125
chr11-95837627-INS-143
chr11-95838159-INS-113
chr11-95838465-INS-346
chr11-95846228-INS-177
chr11-95863222-INS-125
chr11-95877869-INS-83
chr11-95885990-INS-214
chr11-95894293-INS-94
chr11-95905861-INS-182
chr11-95913488-INS-69
chr11-95913941-INS-78
chr11-95917159-INS-56
chr11-95967695-INS-1118
chr11-95969340-INS-246
chr11-95972389-INS-51
chr11-95976795-INS-144
chr11-95982666-INS-104
chr11-95990891-INS-126
chr11-95994945-INS-212
chr11-96003094-INS-237
chr11-96022125-INS-60
chr11-96023768-INS-328
chr11-96025593-INS-69
chr11-96035874-INS-213
chr11-96044759-INS-82
chr11-96070364-INS-113
chr11-96084827-INS-54
chr11-96126586-INS-78
chr11-96128725-INS-262
chr11-96130282-INS-483
chr11-96131036-INS-66
chr11-96131559-INS-52
chr11-96134898-INS-188
chr11-96137338-INS-112
chr11-96142222-INS-68
chr11-96148401-INS-70
chr11-96156754-INS-64
chr11-96189711-INS-191
chr11-96215574-INS-179
chr11-96268891-INS-471
chr11-96270293-INS-87
chr11-96278171-INS-58
chr11-96280502-INS-78
chr11-96280924-INS-4538
chr11-96289270-INS-53
chr11-96326462-INS-336
chr11-96331880-INS-128
chr11-96341597-INS-57
chr11-96357312-INS-199
chr11-96363945-INS-71
chr11-96367692-INS-69
chr11-96370935-INS-507
chr11-96378027-INS-219
chr11-96380962-INS-88
chr11-96392755-INS-72
chr11-96398380-INS-196
chr11-96403687-INS-52
chr11-96407047-INS-75
chr11-96420451-INS-54
chr11-96452951-INS-81
chr11-96478120-INS-84
chr11-96494033-INS-52
chr11-96509616-INS-82
chr11-96539681-INS-73
chr11-96550273-INS-6442
chr11-96550591-INS-56
chr11-96557111-INS-158
chr11-96583010-INS-261
chr11-96593814-INS-111
chr11-96596552-INS-210
chr11-96610660-INS-96
chr11-96614562-INS-409
chr11-96639889-INS-76

chr11-96642814-INS-64
chr11-96660774-INS-199
chr11-96664224-INS-8682
chr11-96672711-INS-177
chr11-96679687-INS-207
chr11-96682467-INS-280
chr11-96697257-INS-214
chr11-96745171-INS-210
chr11-96770749-INS-295
chr11-96784901-INS-201
chr11-96786822-INS-56
chr11-96795393-INS-167
chr11-96809378-INS-68
chr11-96811149-INS-106
chr11-96817799-INS-204
chr11-96822202-INS-249
chr11-96823645-INS-138
chr11-96888550-INS-56
chr11-96903298-INS-67
chr11-96918585-INS-291
chr11-96925362-INS-95
chr11-96942241-INS-66
chr11-96953877-INS-187
chr11-96957321-INS-69
chr11-96964960-INS-77
chr11-96979198-INS-174
chr11-96998662-INS-207
chr11-97001433-INS-63
chr11-97070572-INS-255
chr11-97080011-INS-180
chr11-97086525-INS-54
chr11-97095646-INS-203
chr11-97107675-INS-192
chr11-97109614-INS-307
chr11-97119743-INS-214
chr11-97126503-INS-211
chr11-97138972-INS-211
chr11-97156561-INS-210
chr11-97160327-INS-346
chr11-97172010-INS-217
chr11-97191230-INS-142
chr11-97197960-INS-123
chr11-97207032-INS-199
chr11-97225126-INS-233
chr11-97230366-INS-103
chr11-97240876-INS-132
chr11-97242933-INS-60
chr11-97246034-INS-171
chr11-97279405-INS-204
chr11-97299467-INS-227
chr11-97310709-INS-51
chr11-97323714-INS-199
chr11-97345735-INS-66
chr11-97388648-INS-927
chr11-97405920-INS-219
chr11-97408859-INS-214
chr11-97418093-INS-60
chr11-97420076-INS-307
chr11-97424881-INS-52
chr11-97450773-INS-134
chr11-97472376-INS-395
chr11-97475248-INS-62
chr11-97483702-INS-382
chr11-97492917-INS-64
chr11-97504166-INS-348
chr11-97506815-INS-112
chr11-97510544-INS-352
chr11-97511481-INS-113
chr11-97512227-INS-193
chr11-97522258-INS-58
chr11-97524929-INS-56
chr11-97546170-INS-208
chr11-97606053-INS-222
chr11-97620982-INS-130
chr11-97646278-INS-199
chr11-97648857-INS-54
chr11-97661318-INS-697

chr11-97665906-INS-168
chr11-97670628-INS-184
chr11-97707620-INS-187
chr11-97735560-INS-311
chr11-97736668-INS-210
chr11-97749439-INS-52
chr11-97750204-INS-56
chr11-97751105-INS-494
chr11-97755281-INS-243
chr11-97758311-INS-179
chr11-97767314-INS-78
chr11-97777083-INS-369
chr11-97781596-INS-122
chr11-97784404-INS-146
chr11-97785345-INS-255
chr11-97799643-INS-3737
chr11-97803626-INS-241
chr11-97825789-INS-81
chr11-97827194-INS-182
chr11-97842844-INS-193
chr11-97849551-INS-197
chr11-97851155-INS-196
chr11-97858787-INS-224
chr11-97864591-INS-118
chr11-97882425-INS-70
chr11-97889659-INS-68
chr11-97907043-INS-54
chr11-97937204-INS-200
chr11-97972075-INS-200
chr11-97997604-INS-241
chr11-98011539-INS-96
chr11-98032053-INS-205
chr11-98033322-INS-240
chr11-98064789-INS-839
chr11-98069864-INS-131
chr11-98079835-INS-206
chr11-98085113-INS-79
chr11-98087617-INS-177
chr11-98088603-INS-232
chr11-98109981-INS-181
chr11-98126594-INS-144
chr11-98153329-INS-204
chr11-98167697-INS-5315
chr11-98175962-INS-54
chr11-98181622-INS-218
chr11-98183046-INS-390
chr11-98193516-INS-241
chr11-98196904-INS-55
chr11-98205723-INS-70
chr11-98236754-INS-53
chr11-98254558-INS-201
chr11-98272198-INS-252
chr11-98292452-INS-68
chr11-98304217-INS-58
chr11-98308695-INS-147
chr11-98385924-INS-249
chr11-98389203-INS-156
chr11-98395191-INS-475
chr11-98398438-INS-61
chr11-98420151-INS-206
chr11-98424413-INS-59
chr11-98431063-INS-80
chr11-98463138-INS-96
chr11-98464518-INS-464
chr11-98469512-INS-880
chr11-98480799-INS-83
chr11-98484140-INS-463
chr11-98496245-INS-204
chr11-98498013-INS-108
chr11-98518324-INS-5103
chr11-98570072-INS-417
chr11-98594827-INS-205
chr11-98617936-INS-64
chr11-98627599-INS-167
chr11-98630561-INS-188
chr11-98651062-INS-57
chr11-98667434-INS-108

chr11-98699290-INS-190
chr11-98711414-INS-217
chr11-98723431-INS-222
chr11-98734358-INS-56
chr11-98739602-INS-86
chr11-98749619-INS-237
chr11-98754599-INS-90
chr11-98765340-INS-175
chr11-98770141-INS-205
chr11-98788422-INS-67
chr11-98790142-INS-163
chr11-98823776-INS-56
chr11-98825293-INS-207
chr11-98830385-INS-60
chr11-98839073-INS-82
chr11-98868618-INS-237
chr11-98870134-INS-52
chr11-98881167-INS-165
chr11-98886327-INS-160
chr11-98901058-INS-187
chr11-98923254-INS-146
chr11-98956249-INS-102
chr11-98981807-INS-213
chr11-98982997-INS-164
chr11-98984734-INS-55
chr11-99004418-INS-149
chr11-99095964-INS-157
chr11-99108232-INS-88
chr11-99132539-INS-58
chr11-99132704-INS-923
chr11-99138622-INS-56
chr11-99160984-INS-408
chr11-99171219-INS-148
chr11-99172550-INS-56
chr11-99186780-INS-107
chr11-99189728-INS-52
chr11-99195115-INS-68
chr11-99249202-INS-487
chr11-99262454-INS-68
chr11-99276823-INS-60
chr11-99284042-INS-81
chr11-99318234-INS-66
chr11-99332755-INS-56
chr11-99343236-INS-112
chr11-99404219-INS-55
chr11-99411495-INS-175
chr11-99412912-INS-159
chr11-99414738-INS-196
chr11-99418124-INS-144
chr11-99423922-INS-108
chr11-99440157-INS-118
chr11-99444125-INS-75
chr11-99452575-INS-74
chr11-99461818-INS-435
chr11-99477359-INS-65
chr11-99500214-INS-440
chr11-99503841-INS-6585
chr11-99546273-INS-423
chr11-99549559-INS-150
chr11-99572989-INS-271
chr11-99583235-INS-52
chr11-99611736-INS-94
chr11-99628893-INS-51
chr11-99667945-INS-59
chr11-99679013-INS-6312
chr11-99679151-INS-220
chr11-99682415-INS-4056
chr11-99698697-INS-120
chr11-99705987-INS-105
chr11-99715210-INS-5293
chr11-99797088-INS-66
chr11-99839625-INS-204
chr11-99844538-INS-6227
chr11-99844539-INS-256
chr11-99856147-INS-163
chr11-99927158-INS-72
chr11-99936233-INS-69

chr11-99939871-INS-96
chr11-99977626-INS-437
chr11-100029285-INS-60
chr11-100043018-INS-56
chr11-100050250-INS-53
chr11-100052938-INS-322
chr11-100062282-INS-66
chr11-100063479-INS-510
chr11-100067071-INS-108
chr11-100072178-INS-223
chr11-100073266-INS-82
chr11-100103274-INS-266
chr11-100145715-INS-53
chr11-100147031-INS-60
chr11-100161555-INS-57
chr11-100166290-INS-96
chr11-100166595-INS-5552
chr11-100168382-INS-82
chr11-100190439-INS-135
chr11-100200397-INS-52
chr11-100202404-INS-72
chr11-100205025-INS-247
chr11-100226829-INS-89
chr11-100230264-INS-54
chr11-100237756-INS-54
chr11-100255405-INS-209
chr11-100293920-INS-66
chr11-100322612-INS-201
chr11-100323384-INS-117
chr11-100342713-INS-123
chr11-100359531-INS-201
chr11-100365393-INS-285
chr11-100371207-INS-229
chr11-100378832-INS-238
chr11-100398185-INS-297
chr11-100425452-INS-206
chr11-100428032-INS-69
chr11-100432061-INS-199
chr11-100433961-INS-225
chr11-100445161-INS-160
chr11-100448580-INS-192
chr11-100449720-INS-251
chr11-100457185-INS-225
chr11-100458473-INS-192
chr11-100498808-INS-61
chr11-100512933-INS-204
chr11-100527184-INS-516
chr11-100533998-INS-98
chr11-100540751-INS-113
chr11-100545065-INS-54
chr11-100567807-INS-76
chr11-100570927-INS-193
chr11-100573761-INS-633
chr11-100573779-INS-82
chr11-100619980-INS-194
chr11-100651075-INS-184
chr11-100658527-INS-214
chr11-100664468-INS-201
chr11-100669327-INS-6524
chr11-100671168-INS-188
chr11-100682233-INS-406
chr11-100705081-INS-197
chr11-100714347-INS-222
chr11-100715226-INS-61
chr11-100726798-INS-212
chr11-100727889-INS-80
chr11-100748361-INS-81
chr11-100795446-INS-60
chr11-100806342-INS-143
chr11-100817614-INS-203
chr11-100826424-INS-62
chr11-100833754-INS-130
chr11-100835440-INS-52
chr11-100851010-INS-178
chr11-100867618-INS-398
chr11-100876267-INS-139
chr11-100877036-INS-185

chr11-100888975-INS-93
chr11-100896313-INS-178
chr11-100898742-INS-55
chr11-100910575-INS-178
chr11-100930204-INS-51
chr11-100932087-INS-53
chr11-100946787-INS-215
chr11-100986919-INS-224
chr11-100990809-INS-238
chr11-101004524-INS-374
chr11-101016821-INS-60
chr11-101044222-INS-202
chr11-101049619-INS-170
chr11-101091164-INS-199
chr11-101103390-INS-133
chr11-101105009-INS-208
chr11-101128289-INS-506
chr11-101145681-INS-213
chr11-101148289-INS-191
chr11-101195568-INS-61
chr11-101199138-INS-146
chr11-101235762-INS-100
chr11-101247795-INS-58
chr11-101250672-INS-212
chr11-101252037-INS-203
chr11-101274632-INS-204
chr11-101284154-INS-873
chr11-101300209-INS-106
chr11-101303926-INS-416
chr11-101308983-INS-125
chr11-101340748-INS-204
chr11-101344188-INS-157
chr11-101351642-INS-183
chr11-101389261-INS-125
chr11-101390488-INS-172
chr11-101408841-INS-233
chr11-101411713-INS-152
chr11-101429329-INS-112
chr11-101437058-INS-120
chr11-101441153-INS-246
chr11-101475244-INS-128
chr11-101479238-INS-75
chr11-101493715-INS-204
chr11-101495211-INS-210
chr11-101495944-INS-414
chr11-101519754-INS-196
chr11-101528155-INS-5448
chr11-101529356-INS-203
chr11-101533445-INS-199
chr11-101549454-INS-84
chr11-101566637-INS-139
chr11-101572116-INS-207
chr11-101584548-INS-143
chr11-101603564-INS-89
chr11-101606176-INS-136
chr11-101610042-INS-212
chr11-101631237-INS-120
chr11-101631256-INS-1540
chr11-101632946-INS-439
chr11-101637062-INS-644
chr11-101650573-INS-62
chr11-101651877-INS-81
chr11-101653511-INS-202
chr11-101680709-INS-124
chr11-101715050-INS-230
chr11-101739873-INS-170
chr11-101756830-INS-80
chr11-101790667-INS-636
chr11-101804683-INS-89
chr11-101808457-INS-51
chr11-101811610-INS-191
chr11-101831685-INS-64
chr11-101841962-INS-54
chr11-101847219-INS-61
chr11-101867331-INS-318
chr11-101871045-INS-204
chr11-101885040-INS-145

chr11-101903108-INS-189
chr11-101910296-INS-72
chr11-101913736-INS-52
chr11-101922375-INS-56
chr11-101925287-INS-201
chr11-101926241-INS-135
chr11-101953356-INS-62
chr11-101955295-INS-114
chr11-101969575-INS-255
chr11-101984284-INS-90
chr11-101990563-INS-927
chr11-102005033-INS-327
chr11-102005821-INS-192
chr11-102007425-INS-125
chr11-102027854-INS-74
chr11-102031274-INS-63
chr11-102034596-INS-279
chr11-102067529-INS-200
chr11-102079168-INS-104
chr11-102094796-INS-205
chr11-102103997-INS-155
chr11-102143470-INS-60
chr11-102156097-INS-82
chr11-102161461-INS-128
chr11-102170036-INS-72
chr11-102173164-INS-244
chr11-102191682-INS-57
chr11-102213566-INS-75
chr11-102227673-INS-213
chr11-102241386-INS-206
chr11-102262416-INS-56
chr11-102275676-INS-62
chr11-102311251-INS-206
chr11-102314527-INS-289
chr11-102354425-INS-113
chr11-102357463-INS-317
chr11-102374168-INS-52
chr11-102375989-INS-180
chr11-102384588-INS-170
chr11-102415045-INS-63
chr11-102416848-INS-200
chr11-102423076-INS-277
chr11-102457442-INS-155
chr11-102482030-INS-73
chr11-102489964-INS-244
chr11-102504631-INS-67
chr11-102512974-INS-736
chr11-102525685-INS-209
chr11-102532639-INS-389
chr11-102538391-INS-232
chr11-102541685-INS-242
chr11-102547111-INS-399
chr11-102551035-INS-117
chr11-102557856-INS-229
chr11-102599050-INS-289
chr11-102604204-INS-175
chr11-102620767-INS-74
chr11-102637544-INS-210
chr11-102639752-INS-210
chr11-102673025-INS-60
chr11-102680330-INS-414
chr11-102685666-INS-195
chr11-102696204-INS-220
chr11-102699369-INS-112
chr11-102704811-INS-179
chr11-102714590-INS-175
chr11-102719488-INS-117
chr11-102724810-INS-935
chr11-102754575-INS-181
chr11-102766508-INS-227
chr11-102767193-INS-202
chr11-102781900-INS-81
chr11-102785790-INS-64
chr11-102824941-INS-187
chr11-102840549-INS-62
chr11-102862079-INS-93
chr11-102867061-INS-118

chr11-102876656-INS-465
chr11-102902722-INS-189
chr11-102904860-INS-196
chr11-102914812-INS-56
chr11-102932570-INS-52
chr11-102943870-INS-194
chr11-102950395-INS-337
chr11-102953415-INS-210
chr11-102961025-INS-58
chr11-102970356-INS-54
chr11-102987884-INS-102
chr11-103021986-INS-175
chr11-103028579-INS-58
chr11-103041912-INS-54
chr11-103043488-INS-195
chr11-103044484-INS-157
chr11-103046120-INS-147
chr11-103048262-INS-98
chr11-103052818-INS-193
chr11-103055943-INS-237
chr11-103140137-INS-213
chr11-103142201-INS-140
chr11-103152750-INS-62
chr11-103164838-INS-52
chr11-103165653-INS-229
chr11-103178729-INS-641
chr11-103203462-INS-55
chr11-103233644-INS-52
chr11-103279554-INS-156
chr11-103294042-INS-72
chr11-103304359-INS-102
chr11-103306194-INS-59
chr11-103310879-INS-474
chr11-103311874-INS-783
chr11-103312058-INS-1034
chr11-103314534-INS-498
chr11-103338021-INS-197
chr11-103344259-INS-293
chr11-103379521-INS-487
chr11-103398513-INS-6369
chr11-103400514-INS-176
chr11-103427085-INS-110
chr11-103437400-INS-95
chr11-103467764-INS-97
chr11-103497416-INS-94
chr11-103545514-INS-60
chr11-103554758-INS-757
chr11-103556377-INS-864
chr11-103558607-INS-59
chr11-103560820-INS-206
chr11-103565972-INS-145
chr11-103572661-INS-289
chr11-103573756-INS-59
chr11-103597100-INS-173
chr11-103631783-INS-68
chr11-103648062-INS-55
chr11-103652311-INS-94
chr11-103657046-INS-109
chr11-103674863-INS-60
chr11-103744018-INS-58
chr11-103759419-INS-56
chr11-103764199-INS-206
chr11-103772714-INS-207
chr11-103805031-INS-914
chr11-103821446-INS-220
chr11-103830374-INS-207
chr11-103853703-INS-5373
chr11-103853880-INS-464
chr11-103895712-INS-149
chr11-103905437-INS-53
chr11-103912445-INS-383
chr11-103932150-INS-160
chr11-103950095-INS-126
chr11-103960410-INS-235
chr11-104012213-INS-51
chr11-104036585-INS-56
chr11-104070219-INS-65

chr11-104071306-INS-5556
chr11-104095491-INS-105
chr11-104112208-INS-116
chr11-104125798-INS-51
chr11-104148158-INS-51
chr11-104179352-INS-208
chr11-104187857-INS-512
chr11-104205497-INS-202
chr11-104214558-INS-249
chr11-104231798-INS-487
chr11-104238736-INS-204
chr11-104242510-INS-259
chr11-104272769-INS-759
chr11-104280959-INS-66
chr11-104319977-INS-237
chr11-104334648-INS-200
chr11-104337415-INS-197
chr11-104347588-INS-84
chr11-104356057-INS-219
chr11-104367840-INS-151
chr11-104369567-INS-172
chr11-104372227-INS-421
chr11-104374437-INS-161
chr11-104380292-INS-58
chr11-104405210-INS-5319
chr11-104410805-INS-208
chr11-104417625-INS-399
chr11-104422755-INS-794
chr11-104459776-INS-121
chr11-104479289-INS-339
chr11-104498116-INS-92
chr11-104503695-INS-56
chr11-104527666-INS-106
chr11-104538988-INS-56
chr11-104558841-INS-66
chr11-104561457-INS-506
chr11-104562536-INS-119
chr11-104594227-INS-52
chr11-104608772-INS-608
chr11-104611644-INS-99
chr11-104616175-INS-80
chr11-104618290-INS-72
chr11-104633067-INS-54
chr11-104639108-INS-194
chr11-104664794-INS-833
chr11-104676149-INS-65
chr11-104697780-INS-87
chr11-104705569-INS-367
chr11-104728735-INS-118
chr11-104735252-INS-506
chr11-104740907-INS-608
chr11-104763556-INS-148
chr11-104764794-INS-60
chr11-104770384-INS-433
chr11-104783331-INS-201
chr11-104795675-INS-109
chr11-104825820-INS-82
chr11-104844198-INS-140
chr11-104849352-INS-102
chr11-104865183-INS-557
chr11-104888845-INS-6511
chr11-104910001-INS-324
chr11-104929688-INS-158
chr11-104935229-INS-64
chr11-104954749-INS-99
chr11-104961658-INS-66
chr11-104967778-INS-161
chr11-104973525-INS-90
chr11-104983650-INS-55
chr11-104992844-INS-55
chr11-104998229-INS-52
chr11-105036169-INS-339
chr11-105037950-INS-174
chr11-105039747-INS-177
chr11-105055685-INS-206
chr11-105058882-INS-94
chr11-105061619-INS-169

chr11-105073872-INS-198
chr11-105075595-INS-80
chr11-105079399-INS-6440
chr11-105088467-INS-212
chr11-105095043-INS-53
chr11-105226440-INS-262
chr11-105236708-INS-114
chr11-105239919-INS-99
chr11-105253587-INS-99
chr11-105277072-INS-171
chr11-105285458-INS-79
chr11-105310861-INS-243
chr11-105312690-INS-124
chr11-105350711-INS-51
chr11-105356310-INS-73
chr11-105370928-INS-52
chr11-105391641-INS-55
chr11-105406273-INS-116
chr11-105420749-INS-88
chr11-105450985-INS-60
chr11-105470889-INS-484
chr11-105496082-INS-174
chr11-105505838-INS-123
chr11-105569259-INS-75
chr11-105571380-INS-7189
chr11-105582821-INS-120
chr11-105604116-INS-104
chr11-105630127-INS-94
chr11-105646051-INS-365
chr11-105672545-INS-6174
chr11-105684934-INS-1067
chr11-105696471-INS-6802
chr11-105700298-INS-6522
chr11-105714072-INS-207
chr11-105715041-INS-171
chr11-105735049-INS-57
chr11-105743116-INS-8969
chr11-105750297-INS-6433
chr11-105756910-INS-184
chr11-105764467-INS-64
chr11-105778611-INS-448
chr11-105796168-INS-64
chr11-105810154-INS-5217
chr11-105841185-INS-87
chr11-105873151-INS-292
chr11-105893308-INS-70
chr11-105908622-INS-207
chr11-105926709-INS-176
chr11-105934141-INS-80
chr11-105940928-INS-196
chr11-105953905-INS-228
chr11-106007281-INS-494
chr11-106057408-INS-128
chr11-106075885-INS-196
chr11-106079334-INS-505
chr11-106082570-INS-212
chr11-106088956-INS-506
chr11-106095498-INS-124
chr11-106111989-INS-90
chr11-106131252-INS-139
chr11-106196274-INS-96
chr11-106213200-INS-80
chr11-106214171-INS-64
chr11-106218685-INS-104
chr11-106260028-INS-206
chr11-106262046-INS-76
chr11-106262464-INS-234
chr11-106264153-INS-102
chr11-106270289-INS-72
chr11-106293167-INS-327
chr11-106303096-INS-155
chr11-106304869-INS-221
chr11-106315309-INS-259
chr11-106327160-INS-202
chr11-106346910-INS-216
chr11-106388033-INS-222
chr11-106391475-INS-89

chr11-106399631-INS-215
chr11-106406298-INS-208
chr11-106429827-INS-211
chr11-106438767-INS-268
chr11-106440031-INS-119
chr11-106452909-INS-166
chr11-106471905-INS-170
chr11-106475167-INS-512
chr11-106482294-INS-68
chr11-106504900-INS-77
chr11-106511551-INS-81
chr11-106518382-INS-165
chr11-106538642-INS-120
chr11-106552054-INS-60
chr11-106561193-INS-399
chr11-106567126-INS-56
chr11-106582176-INS-103
chr11-106583360-INS-246
chr11-106587349-INS-185
chr11-106602897-INS-68
chr11-106606749-INS-190
chr11-106608135-INS-203
chr11-106608701-INS-66
chr11-106622943-INS-396
chr11-106634402-INS-225
chr11-106635111-INS-79
chr11-106656335-INS-6444
chr11-106663194-INS-212
chr11-106667896-INS-93
chr11-106682420-INS-131
chr11-106697428-INS-140
chr11-106724892-INS-77
chr11-106741759-INS-209
chr11-106744999-INS-157
chr11-106756599-INS-319
chr11-106763360-INS-83
chr11-106767399-INS-494
chr11-106790506-INS-83
chr11-106791798-INS-209
chr11-106797342-INS-193
chr11-106806275-INS-109
chr11-106827576-INS-69
chr11-106846236-INS-51
chr11-106852058-INS-193
chr11-106854882-INS-206
chr11-106864080-INS-221
chr11-106868261-INS-163
chr11-106872237-INS-62
chr11-106874839-INS-199
chr11-106895399-INS-58
chr11-106898445-INS-236
chr11-106905136-INS-236
chr11-106907615-INS-84
chr11-106911856-INS-1860
chr11-106915597-INS-6401
chr11-106930930-INS-220
chr11-106939994-INS-78
chr11-106947556-INS-215
chr11-106950712-INS-390
chr11-106970320-INS-136
chr11-106979537-INS-325
chr11-106987008-INS-217
chr11-106993869-INS-264
chr11-107000046-INS-86
chr11-107007991-INS-206
chr11-107034258-INS-175
chr11-107039228-INS-199
chr11-107043094-INS-213
chr11-107044863-INS-214
chr11-107045867-INS-545
chr11-107053345-INS-429
chr11-107063668-INS-221
chr11-107066702-INS-52
chr11-107083922-INS-77
chr11-107085050-INS-63
chr11-107090018-INS-60
chr11-107097028-INS-56

chr11-107111830-INS-216
chr11-107122167-INS-427
chr11-107134015-INS-62
chr11-107156975-INS-192
chr11-107159288-INS-6452
chr11-107164194-INS-303
chr11-107169014-INS-537
chr11-107182624-INS-216
chr11-107185871-INS-80
chr11-107196944-INS-207
chr11-107201011-INS-222
chr11-107214866-INS-235
chr11-107222152-INS-261
chr11-107225024-INS-385
chr11-107232877-INS-58
chr11-107240902-INS-213
chr11-107249385-INS-210
chr11-107280032-INS-202
chr11-107284450-INS-366
chr11-107292506-INS-244
chr11-107303820-INS-207
chr11-107311228-INS-228
chr11-107312089-INS-683
chr11-107336256-INS-3988
chr11-107343992-INS-181
chr11-107351316-INS-211
chr11-107358672-INS-65
chr11-107362856-INS-96
chr11-107363381-INS-189
chr11-107366748-INS-198
chr11-107381760-INS-69
chr11-107390499-INS-56
chr11-107394759-INS-140
chr11-107396547-INS-172
chr11-107398674-INS-68
chr11-107400162-INS-245
chr11-107410280-INS-380
chr11-107425733-INS-53
chr11-107428326-INS-127
chr11-107432457-INS-213
chr11-107450862-INS-126
chr11-107453109-INS-167
chr11-107456039-INS-166
chr11-107491692-INS-210
chr11-107513401-INS-130
chr11-107514448-INS-289
chr11-107549238-INS-156
chr11-107552288-INS-196
chr11-107557975-INS-69
chr11-107570438-INS-135
chr11-107591225-INS-111
chr11-107612127-INS-193
chr11-107643731-INS-100
chr11-107660587-INS-195
chr11-107682465-INS-98
chr11-107706716-INS-141
chr11-107731958-INS-700
chr11-107741565-INS-85
chr11-107743248-INS-84
chr11-107744846-INS-3340
chr11-107747282-INS-66
chr11-107747926-INS-327
chr11-107754138-INS-118
chr11-107762038-INS-144
chr11-107775692-INS-132
chr11-107779872-INS-68
chr11-107781488-INS-197
chr11-107820027-INS-186
chr11-107833055-INS-52
chr11-107845551-INS-251
chr11-107845580-INS-1200
chr11-107881525-INS-265
chr11-107893724-INS-118
chr11-107902374-INS-7290
chr11-107902566-INS-159
chr11-107904442-INS-167
chr11-107920990-INS-399

chr11-107930411-INS-84
chr11-107961331-INS-59
chr11-107967307-INS-7150
chr11-107974384-INS-205
chr11-108009402-INS-5514
chr11-108043174-INS-68
chr11-108087650-INS-69
chr11-108140704-INS-72
chr11-108150612-INS-55
chr11-108156154-INS-470
chr11-108157705-INS-126
chr11-108159431-INS-66
chr11-108161057-INS-101
chr11-108184798-INS-60
chr11-108193987-INS-52
chr11-108210294-INS-72
chr11-108254132-INS-185
chr11-108259856-INS-435
chr11-108262611-INS-590
chr11-108309404-INS-78
chr11-108325395-INS-282
chr11-108333494-INS-255
chr11-108339044-INS-55
chr11-108348838-INS-378
chr11-108351449-INS-168
chr11-108363619-INS-220
chr11-108365021-INS-76
chr11-108394918-INS-252
chr11-108395857-INS-82
chr11-108401589-INS-98
chr11-108406579-INS-896
chr11-108430228-INS-530
chr11-108442620-INS-217
chr11-108470952-INS-120
chr11-108480649-INS-127
chr11-108484318-INS-99
chr11-108487525-INS-66
chr11-108495646-INS-80
chr11-108497746-INS-92
chr11-108506989-INS-132
chr11-108516242-INS-7770
chr11-108520137-INS-107
chr11-108556234-INS-100
chr11-108566740-INS-6655
chr11-108581074-INS-52
chr11-108587643-INS-937
chr11-108589876-INS-56
chr11-108619479-INS-60
chr11-108621127-INS-7106
chr11-108621796-INS-93
chr11-108676477-INS-158
chr11-108708400-INS-186
chr11-108709973-INS-367
chr11-108726093-INS-878
chr11-108757513-INS-203
chr11-108766035-INS-68
chr11-108770070-INS-240
chr11-108800054-INS-61
chr11-108809288-INS-142
chr11-108824226-INS-432
chr11-108843583-INS-202
chr11-108847121-INS-674
chr11-108859622-INS-106
chr11-108879405-INS-60
chr11-108898606-INS-198
chr11-108907523-INS-60
chr11-108908623-INS-7086
chr11-108982390-INS-168
chr11-109000045-INS-83
chr11-109004874-INS-100
chr11-109029785-INS-63
chr11-109032559-INS-219
chr11-109041537-INS-96
chr11-109047525-INS-464
chr11-109050440-INS-459
chr11-109053739-INS-96
chr11-109061453-INS-141

chr11-109068365-INS-54
chr11-109070332-INS-58
chr11-109088557-INS-182
chr11-109089024-INS-158
chr11-109092012-INS-99
chr11-109102082-INS-91
chr11-109108697-INS-202
chr11-109111529-INS-106
chr11-109140679-INS-88
chr11-109163984-INS-109
chr11-109178654-INS-101
chr11-109179533-INS-82
chr11-109190380-INS-208
chr11-109195978-INS-169
chr11-109202349-INS-72
chr11-109225380-INS-53
chr11-109236189-INS-225
chr11-109248742-INS-84
chr11-109285019-INS-61
chr11-109289650-INS-198
chr11-109307313-INS-186
chr11-109332879-INS-59
chr11-109344987-INS-185
chr11-109352464-INS-62
chr11-109392361-INS-112
chr11-109395291-INS-115
chr11-109396343-INS-62
chr11-109406147-INS-120
chr11-109406827-INS-64
chr11-109407149-INS-76
chr11-109442434-INS-56
chr11-109471618-INS-124
chr11-109484316-INS-66
chr11-109495929-INS-83
chr11-109527902-INS-149
chr11-109582372-INS-88
chr11-109594416-INS-53
chr11-109602295-INS-74
chr11-109633185-INS-63
chr11-109635805-INS-540
chr11-109640417-INS-56
chr11-109654349-INS-52
chr11-109669378-INS-758
chr11-109672774-INS-548
chr11-109676226-INS-199
chr11-109705530-INS-67
chr11-109748844-INS-89
chr11-109768837-INS-166
chr11-109793252-INS-80
chr11-109805669-INS-138
chr11-109819194-INS-441
chr11-109846989-INS-77
chr11-109877268-INS-164
chr11-109879297-INS-135
chr11-109883893-INS-783
chr11-109888712-INS-241
chr11-109934374-INS-64
chr11-109961566-INS-85
chr11-109964569-INS-201
chr11-109965447-INS-60
chr11-110008837-INS-104
chr11-110035787-INS-72
chr11-110036634-INS-283
chr11-110052473-INS-81
chr11-110058461-INS-56
chr11-110070777-INS-440
chr11-110083043-INS-70
chr11-110092224-INS-68
chr11-110106914-INS-496
chr11-110144087-INS-200
chr11-110145612-INS-349
chr11-110150079-INS-73
chr11-110157954-INS-71
chr11-110203676-INS-960
chr11-110204660-INS-300
chr11-110229366-INS-414
chr11-110245066-INS-109

chr11-110256359-INS-130
chr11-110270001-INS-63
chr11-110274520-INS-135
chr11-110276873-INS-197
chr11-110289609-INS-62
chr11-110319840-INS-71
chr11-110324811-INS-5329
chr11-110332281-INS-60
chr11-110344646-INS-71
chr11-110347269-INS-60
chr11-110349250-INS-212
chr11-110354757-INS-234
chr11-110358413-INS-62
chr11-110359495-INS-108
chr11-110384627-INS-53
chr11-110389292-INS-58
chr11-110408068-INS-52
chr11-110418692-INS-203
chr11-110438296-INS-53
chr11-110448606-INS-164
chr11-110449936-INS-152
chr11-110475167-INS-150
chr11-110495356-INS-257
chr11-110503348-INS-203
chr11-110504600-INS-253
chr11-110536417-INS-406
chr11-110543980-INS-60
chr11-110559743-INS-227
chr11-110590556-INS-55
chr11-110605213-INS-82
chr11-110610952-INS-694
chr11-110628512-INS-796
chr11-110644116-INS-68
chr11-110650779-INS-54
chr11-110669136-INS-6931
chr11-110688932-INS-7519
chr11-110691787-INS-60
chr11-110694414-INS-3932
chr11-110731734-INS-589
chr11-110734766-INS-537
chr11-110791412-INS-80
chr11-110793029-INS-422
chr11-110803152-INS-117
chr11-110807864-INS-96
chr11-110840132-INS-156
chr11-110850544-INS-63
chr11-110859676-INS-472
chr11-110882631-INS-52
chr11-110887454-INS-72
chr11-110929112-INS-54
chr11-110937352-INS-106
chr11-110944469-INS-290
chr11-110948592-INS-64
chr11-110954490-INS-90
chr11-110964534-INS-56
chr11-110988779-INS-56
chr11-110997331-INS-54
chr11-111098257-INS-173
chr11-111101102-INS-61
chr11-111158619-INS-351
chr11-111159785-INS-137
chr11-111196890-INS-50
chr11-111213457-INS-504
chr11-111214981-INS-341
chr11-111235911-INS-85
chr11-111243645-INS-239
chr11-111273353-INS-71
chr11-111321395-INS-60
chr11-111327643-INS-84
chr11-111330364-INS-62
chr11-111332118-INS-309
chr11-111333847-INS-68
chr11-111343429-INS-426
chr11-111452821-INS-931
chr11-111467021-INS-68
chr11-111513639-INS-174
chr11-111527489-INS-156

chr11-111529409-INS-95
chr11-111567932-INS-67
chr11-111610224-INS-244
chr11-111613225-INS-76
chr11-111621810-INS-510
chr11-111641131-INS-607
chr11-111653293-INS-112
chr11-111656258-INS-51
chr11-111675235-INS-330
chr11-111695042-INS-534
chr11-111702829-INS-409
chr11-111748701-INS-74
chr11-111760288-INS-110
chr11-111776975-INS-115
chr11-111792444-INS-497
chr11-111802678-INS-70
chr11-111811775-INS-57
chr11-111823633-INS-290
chr11-111841027-INS-5269
chr11-111869914-INS-64
chr11-111899784-INS-70
chr11-111921902-INS-94
chr11-111984537-INS-293
chr11-111993113-INS-130
chr11-112030548-INS-1892
chr11-112030857-INS-56
chr11-112037848-INS-64
chr11-112042268-INS-59
chr11-112044056-INS-60
chr11-112055412-INS-64
chr11-112070094-INS-70
chr11-112082807-INS-62
chr11-112087649-INS-55
chr11-112089978-INS-60
chr11-112111284-INS-144
chr11-112114694-INS-66
chr11-112117190-INS-63
chr11-112144494-INS-64
chr11-112145389-INS-164
chr11-112172878-INS-87
chr11-112180166-INS-103
chr11-112192479-INS-56
chr11-112206017-INS-52
chr11-112213114-INS-185
chr11-112245508-INS-211
chr11-112270939-INS-125
chr11-112293826-INS-591
chr11-112295804-INS-185
chr11-112336508-INS-55
chr11-112342351-INS-140
chr11-112360080-INS-60
chr11-112362119-INS-78
chr11-112366688-INS-135
chr11-112372675-INS-56
chr11-112406088-INS-249
chr11-112412916-INS-56
chr11-112419284-INS-69
chr11-112438971-INS-124
chr11-112450485-INS-170
chr11-112456356-INS-69
chr11-112461981-INS-252
chr11-112490770-INS-152
chr11-112525253-INS-81
chr11-112528661-INS-105
chr11-112532585-INS-265
chr11-112538240-INS-52
chr11-112540959-INS-147
chr11-112542232-INS-53
chr11-112559840-INS-59
chr11-112585232-INS-130
chr11-112625797-INS-64
chr11-112631946-INS-147
chr11-112635937-INS-75
chr11-112648661-INS-109
chr11-112652732-INS-529
chr11-112660042-INS-97
chr11-112682587-INS-77

chr11-112685283-INS-68
chr11-112748162-INS-355
chr11-112753151-INS-60
chr11-112758649-INS-56
chr11-112762941-INS-60
chr11-112773738-INS-96
chr11-112776809-INS-255
chr11-112785100-INS-60
chr11-112790990-INS-60
chr11-112791849-INS-697
chr11-112795626-INS-117
chr11-112802773-INS-111
chr11-112803869-INS-5549
chr11-112808945-INS-96
chr11-112810082-INS-64
chr11-112820416-INS-127
chr11-112834667-INS-142
chr11-112835214-INS-912
chr11-112893772-INS-291
chr11-112898574-INS-151
chr11-112905576-INS-79
chr11-112926496-INS-68
chr11-112933238-INS-1883
chr11-112942742-INS-52
chr11-112952758-INS-454
chr11-113025147-INS-98
chr11-113055638-INS-360
chr11-113060461-INS-98
chr11-113065535-INS-93
chr11-113066925-INS-72
chr11-113070233-INS-68
chr11-113078340-INS-74
chr11-113084220-INS-216
chr11-113094220-INS-172
chr11-113128608-INS-80
chr11-113166782-INS-132
chr11-113168719-INS-54
chr11-113173090-INS-134
chr11-113197114-INS-97
chr11-113197227-INS-539
chr11-113218200-INS-209
chr11-113231282-INS-360
chr11-113239722-INS-168
chr11-113254933-INS-53
chr11-113275209-INS-58
chr11-113296549-INS-100
chr11-113300549-INS-52
chr11-113316512-INS-106
chr11-113372120-INS-497
chr11-113382302-INS-71
chr11-113393978-INS-213
chr11-113414539-INS-55
chr11-113429854-INS-64
chr11-113436070-INS-77
chr11-113444177-INS-60
chr11-113455774-INS-150
chr11-113489785-INS-136
chr11-113507271-INS-144
chr11-113525242-INS-246
chr11-113537530-INS-7483
chr11-113542187-INS-525
chr11-113560742-INS-178
chr11-113576994-INS-105
chr11-113591658-INS-178
chr11-113602404-INS-63
chr11-113613208-INS-203
chr11-113614757-INS-207
chr11-113615633-INS-5543
chr11-113665907-INS-81
chr11-113674584-INS-54
chr11-113678959-INS-66
chr11-113681123-INS-329
chr11-113681702-INS-56
chr11-113683223-INS-91
chr11-113695191-INS-65
chr11-113705265-INS-59
chr11-113706486-INS-71

chr11-113717529-INS-66
chr11-113727060-INS-60
chr11-113728549-INS-401
chr11-113734439-INS-75
chr11-113752273-INS-56
chr11-113812311-INS-166
chr11-113828405-INS-191
chr11-113836422-INS-66
chr11-113941436-INS-57
chr11-113943820-INS-52
chr11-113974899-INS-98
chr11-113979343-INS-171
chr11-113988636-INS-78
chr11-113989748-INS-55
chr11-114011700-INS-62
chr11-114060659-INS-223
chr11-114070707-INS-427
chr11-114085682-INS-84
chr11-114090573-INS-58
chr11-114096887-INS-152
chr11-114109877-INS-138
chr11-114111426-INS-75
chr11-114156529-INS-90
chr11-114187254-INS-80
chr11-114190499-INS-84
chr11-114210292-INS-980
chr11-114248688-INS-68
chr11-114262583-INS-183
chr11-114288496-INS-92
chr11-114302775-INS-400
chr11-114303297-INS-181
chr11-114308148-INS-116
chr11-114325176-INS-239
chr11-114331534-INS-269
chr11-114334056-INS-166
chr11-114359892-INS-5342
chr11-114395414-INS-120
chr11-114404612-INS-134
chr11-114418599-INS-52
chr11-114422048-INS-368
chr11-114432638-INS-209
chr11-114462620-INS-176
chr11-114485204-INS-530
chr11-114498792-INS-145
chr11-114511661-INS-89
chr11-114518237-INS-358
chr11-114522694-INS-682
chr11-114525460-INS-52
chr11-114528029-INS-70
chr11-114529646-INS-51
chr11-114531483-INS-192
chr11-114556243-INS-52
chr11-114557494-INS-108
chr11-114588328-INS-100
chr11-114590228-INS-194
chr11-114598733-INS-203
chr11-114600306-INS-66
chr11-114606945-INS-204
chr11-114616633-INS-561
chr11-114617296-INS-211
chr11-114644813-INS-399
chr11-114645793-INS-78
chr11-114653780-INS-208
chr11-114655227-INS-146
chr11-114669594-INS-201
chr11-114674866-INS-180
chr11-114688638-INS-354
chr11-114707853-INS-396
chr11-114731589-INS-52
chr11-114734305-INS-53
chr11-114736481-INS-183
chr11-114737033-INS-204
chr11-114751195-INS-145
chr11-114756185-INS-64
chr11-114777477-INS-120
chr11-114778443-INS-287
chr11-114779184-INS-91

chr11-114799472-INS-56
chr11-114806876-INS-457
chr11-114808712-INS-264
chr11-114818968-INS-52
chr11-114821025-INS-153
chr11-114829792-INS-348
chr11-114850812-INS-1128
chr11-114879211-INS-643
chr11-114879262-INS-88
chr11-114889972-INS-248
chr11-114909719-INS-57
chr11-114914254-INS-68
chr11-114922214-INS-191
chr11-114931183-INS-62
chr11-114935004-INS-180
chr11-114941168-INS-62
chr11-114963486-INS-64
chr11-114971285-INS-182
chr11-114978856-INS-125
chr11-114988508-INS-55
chr11-114993366-INS-137
chr11-115004996-INS-198
chr11-115021061-INS-270
chr11-115032915-INS-107
chr11-115033963-INS-192
chr11-115037272-INS-87
chr11-115049898-INS-245
chr11-115076728-INS-95
chr11-115081542-INS-212
chr11-115091387-INS-178
chr11-115142413-INS-53
chr11-115145509-INS-218
chr11-115179501-INS-167
chr11-115185727-INS-60
chr11-115198788-INS-129
chr11-115201898-INS-126
chr11-115268628-INS-228
chr11-115283261-INS-656
chr11-115285235-INS-1888
chr11-115287706-INS-123
chr11-115290419-INS-108
chr11-115320364-INS-214
chr11-115326094-INS-52
chr11-115339668-INS-487
chr11-115342601-INS-100
chr11-115357253-INS-183
chr11-115361803-INS-185
chr11-115365782-INS-188
chr11-115368625-INS-213
chr11-115373714-INS-211
chr11-115384237-INS-210
chr11-115391311-INS-451
chr11-115407008-INS-113
chr11-115412828-INS-155
chr11-115416413-INS-262
chr11-115419048-INS-218
chr11-115433391-INS-206
chr11-115434460-INS-378
chr11-115438323-INS-643
chr11-115440791-INS-319
chr11-115442364-INS-923
chr11-115444385-INS-120
chr11-115448278-INS-230
chr11-115451146-INS-2864
chr11-115454521-INS-217
chr11-115463625-INS-74
chr11-115470685-INS-75
chr11-115474773-INS-2398
chr11-115482691-INS-60
chr11-115493472-INS-214
chr11-115496243-INS-106
chr11-115520680-INS-218
chr11-115543960-INS-176
chr11-115545572-INS-411
chr11-115549722-INS-199
chr11-115553354-INS-117
chr11-115558366-INS-210

chr11-115566963-INS-122
chr11-115569099-INS-64
chr11-115582021-INS-114
chr11-115593839-INS-54
chr11-115596890-INS-167
chr11-115606532-INS-8735
chr11-115609222-INS-326
chr11-115610149-INS-533
chr11-115619523-INS-160
chr11-115624145-INS-364
chr11-115641325-INS-150
chr11-115642496-INS-57
chr11-115661651-INS-144
chr11-115665834-INS-237
chr11-115677935-INS-238
chr11-115725408-INS-65
chr11-115753495-INS-208
chr11-115760561-INS-170
chr11-115762905-INS-186
chr11-115766675-INS-200
chr11-115814910-INS-54
chr11-115816393-INS-98
chr11-115832235-INS-166
chr11-115835355-INS-210
chr11-115854950-INS-52
chr11-115859063-INS-190
chr11-115883716-INS-53
chr11-115887087-INS-88
chr11-115906103-INS-198
chr11-115911991-INS-202
chr11-115918935-INS-197
chr11-115922889-INS-165
chr11-115927370-INS-207
chr11-115932485-INS-53
chr11-115942933-INS-253
chr11-115988321-INS-209
chr11-115995746-INS-176
chr11-115999972-INS-76
chr11-116043978-INS-967
chr11-116046611-INS-201
chr11-116051424-INS-72
chr11-116053547-INS-244
chr11-116055023-INS-155
chr11-116060428-INS-363
chr11-116062571-INS-348
chr11-116077858-INS-188
chr11-116079257-INS-56
chr11-116086661-INS-170
chr11-116126648-INS-211
chr11-116153877-INS-199
chr11-116155058-INS-195
chr11-116156388-INS-176
chr11-116160513-INS-104
chr11-116167278-INS-140
chr11-116177852-INS-132
chr11-116182242-INS-67
chr11-116185914-INS-234
chr11-116190415-INS-264
chr11-116204939-INS-212
chr11-116214331-INS-143
chr11-116243715-INS-317
chr11-116248018-INS-241
chr11-116252060-INS-155
chr11-116257514-INS-1674
chr11-116265244-INS-209
chr11-116313942-INS-188
chr11-116322164-INS-74
chr11-116348263-INS-136
chr11-116357827-INS-56
chr11-116364641-INS-165
chr11-116367283-INS-55
chr11-116369814-INS-202
chr11-116373137-INS-188
chr11-116376279-INS-203
chr11-116377915-INS-336
chr11-116385903-INS-205
chr11-116389294-INS-6122

chr11-116390921-INS-100
chr11-116393539-INS-166
chr11-116412365-INS-209
chr11-116416338-INS-280
chr11-116421288-INS-51
chr11-116450928-INS-206
chr11-116486379-INS-113
chr11-116504396-INS-81
chr11-116507944-INS-86
chr11-116516855-INS-185
chr11-116523546-INS-203
chr11-116550453-INS-220
chr11-116558248-INS-52
chr11-116569468-INS-182
chr11-116573024-INS-197
chr11-116588160-INS-230
chr11-116596221-INS-9047
chr11-116596551-INS-122
chr11-116606819-INS-356
chr11-116611914-INS-200
chr11-116613321-INS-515
chr11-116624285-INS-502
chr11-116636520-INS-280
chr11-116639404-INS-52
chr11-116642992-INS-9853
chr11-116661402-INS-108
chr11-116665798-INS-60
chr11-116672197-INS-332
chr11-116698327-INS-74
chr11-116778454-INS-233
chr11-116779551-INS-90
chr11-116781816-INS-348
chr11-116783676-INS-242
chr11-116805043-INS-54
chr11-116815908-INS-56
chr11-116847894-INS-57
chr11-116888363-INS-84
chr11-116893823-INS-570
chr11-116905154-INS-757
chr11-116907390-INS-242
chr11-116907400-INS-6401
chr11-116923599-INS-186
chr11-116939073-INS-59
chr11-116958515-INS-92
chr11-116960596-INS-133
chr11-116963663-INS-206
chr11-116976608-INS-56
chr11-116986831-INS-300
chr11-116990255-INS-216
chr11-117012753-INS-497
chr11-117017328-INS-1111
chr11-117062319-INS-102
chr11-117073728-INS-782
chr11-117087955-INS-66
chr11-117094994-INS-201
chr11-117112656-INS-70
chr11-117128454-INS-191
chr11-117144025-INS-5337
chr11-117172986-INS-111
chr11-117183748-INS-104
chr11-117183971-INS-251
chr11-117192149-INS-207
chr11-117197548-INS-203
chr11-117198293-INS-66
chr11-117213049-INS-130
chr11-117239127-INS-122
chr11-117313879-INS-127
chr11-117315825-INS-55
chr11-117321014-INS-168
chr11-117332575-INS-122
chr11-117341324-INS-55
chr11-117343035-INS-222
chr11-117351922-INS-60
chr11-117354688-INS-162
chr11-117369428-INS-208
chr11-117376147-INS-72
chr11-117404483-INS-202

chr11-117405554-INS-107
chr11-117409717-INS-203
chr11-117425484-INS-114
chr11-117433185-INS-232
chr11-117438865-INS-52
chr11-117449396-INS-244
chr11-117464299-INS-394
chr11-117469285-INS-70
chr11-117470901-INS-209
chr11-117476890-INS-71
chr11-117482247-INS-183
chr11-117494419-INS-52
chr11-117496453-INS-700
chr11-117555392-INS-70
chr11-117589768-INS-90
chr11-117596073-INS-94
chr11-117601370-INS-52
chr11-117606222-INS-205
chr11-117607919-INS-130
chr11-117638903-INS-123
chr11-117659334-INS-243
chr11-117680315-INS-193
chr11-117686274-INS-164
chr11-117691740-INS-105
chr11-117707036-INS-198
chr11-117717721-INS-235
chr11-117719755-INS-901
chr11-117735371-INS-195
chr11-117736530-INS-191
chr11-117773472-INS-204
chr11-117795489-INS-175
chr11-117834551-INS-161
chr11-117835104-INS-738
chr11-117841133-INS-651
chr11-117841648-INS-55
chr11-117845476-INS-134
chr11-117868362-INS-360
chr11-117879570-INS-206
chr11-117893686-INS-204
chr11-117911216-INS-202
chr11-117923618-INS-732
chr11-117944461-INS-335
chr11-117959536-INS-149
chr11-117960547-INS-171
chr11-117965337-INS-187
chr11-117968442-INS-754
chr11-117968452-INS-80
chr11-117970708-INS-77
chr11-117986775-INS-227
chr11-117995006-INS-55
chr11-118023659-INS-363
chr11-118030742-INS-176
chr11-118038267-INS-1026
chr11-118038738-INS-104
chr11-118069156-INS-52
chr11-118108744-INS-148
chr11-118141233-INS-998
chr11-118148810-INS-440
chr11-118166565-INS-67
chr11-118216742-INS-75
chr11-118226364-INS-70
chr11-118237284-INS-498
chr11-118267457-INS-319
chr11-118278766-INS-80
chr11-118325421-INS-7280
chr11-118347240-INS-177
chr11-118354904-INS-54
chr11-118358790-INS-93
chr11-118361484-INS-510
chr11-118451584-INS-145
chr11-118471087-INS-166
chr11-118487620-INS-83
chr11-118540829-INS-57
chr11-118551256-INS-68
chr11-118574391-INS-387
chr11-118578786-INS-259
chr11-118615359-INS-72

chr11-118637425-INS-173
chr11-118676786-INS-205
chr11-118688895-INS-70
chr11-118697927-INS-82
chr11-118707981-INS-66
chr11-118747752-INS-70
chr11-118760977-INS-132
chr11-118763748-INS-149
chr11-118770016-INS-66
chr11-118804529-INS-68
chr11-118805050-INS-128
chr11-118816860-INS-56
chr11-118844264-INS-84
chr11-118850864-INS-70
chr11-118940454-INS-69
chr11-118952193-INS-54
chr11-118965909-INS-262
chr11-118995721-INS-440
chr11-119002020-INS-205
chr11-119007858-INS-54
chr11-119009629-INS-159
chr11-119012602-INS-53
chr11-119030344-INS-171
chr11-119044272-INS-115
chr11-119059219-INS-175
chr11-119070391-INS-59
chr11-119082166-INS-346
chr11-119117499-INS-197
chr11-119128817-INS-105
chr11-119135272-INS-61
chr11-119137673-INS-58
chr11-119153398-INS-58
chr11-119188554-INS-146
chr11-119197368-INS-252
chr11-119200073-INS-203
chr11-119202309-INS-200
chr11-119203085-INS-127
chr11-119253914-INS-181
chr11-119261984-INS-170
chr11-119266153-INS-68
chr11-119270030-INS-214
chr11-119281696-INS-106
chr11-119288362-INS-193
chr11-119301953-INS-204
chr11-119319710-INS-68
chr11-119353412-INS-506
chr11-119356552-INS-537
chr11-119384817-INS-387
chr11-119389214-INS-159
chr11-119412163-INS-4773
chr11-119418583-INS-193
chr11-119459419-INS-68
chr11-119462402-INS-434
chr11-119474130-INS-76
chr11-119483335-INS-147
chr11-119486463-INS-127
chr11-119489144-INS-106
chr11-119507445-INS-67
chr11-119509922-INS-80
chr11-119519843-INS-166
chr11-119537683-INS-204
chr11-119538977-INS-260
chr11-119541121-INS-78
chr11-119545230-INS-60
chr11-119556210-INS-56
chr11-119561745-INS-64
chr11-119606762-INS-53
chr11-119620164-INS-2116
chr11-119626572-INS-116
chr11-119636717-INS-93
chr11-119651873-INS-56
chr11-119656219-INS-62
chr11-119710488-INS-92
chr11-119778288-INS-55
chr11-119779853-INS-213
chr11-119837057-INS-96
chr11-119860402-INS-306

chr11-119864334-INS-56
chr11-119933965-INS-143
chr11-119939807-INS-136
chr11-119944201-INS-95
chr11-119969043-INS-209
chr11-119970103-INS-187
chr11-119978431-INS-167
chr11-120005578-INS-154
chr11-120017399-INS-58
chr11-120023792-INS-72
chr11-120053448-INS-57
chr11-120064994-INS-66
chr11-120075846-INS-68
chr11-120094486-INS-441
chr11-120105815-INS-153
chr11-120111361-INS-54
chr11-120133044-INS-210
chr11-120137192-INS-189
chr11-120149211-INS-75
chr11-120187865-INS-211
chr11-120191781-INS-90
chr11-120192806-INS-4775
chr11-120199405-INS-56
chr11-120227049-INS-218
chr11-120307859-INS-209
chr11-120322816-INS-710
chr11-120322819-INS-536
chr11-120326592-INS-83
chr11-120330824-INS-362
chr11-120333771-INS-78
chr11-120339835-INS-1888
chr11-120340448-INS-201
chr11-120341026-INS-107
chr11-120359978-INS-188
chr11-120365648-INS-225
chr11-120371882-INS-238
chr11-120377977-INS-207
chr11-120396438-INS-86
chr11-120397536-INS-66
chr11-120407911-INS-396
chr11-120410346-INS-74
chr11-120412771-INS-64
chr11-120455494-INS-189
chr11-120475491-INS-373
chr11-120492704-INS-244
chr11-120504883-INS-222
chr11-120512506-INS-495
chr11-120513669-INS-215
chr11-120528544-INS-200
chr11-120529769-INS-161
chr11-120531481-INS-221
chr11-120544000-INS-379
chr11-120546428-INS-66
chr11-120553433-INS-180
chr11-120571921-INS-543
chr11-120589632-INS-211
chr11-120598445-INS-62
chr11-120629680-INS-170
chr11-120638255-INS-198
chr11-120655817-INS-136
chr11-120666472-INS-88
chr11-120730346-INS-54
chr11-120746766-INS-437
chr11-120754754-INS-53
chr11-120757024-INS-203
chr11-120768487-INS-150
chr11-120793663-INS-211
chr11-120804952-INS-177
chr11-120824229-INS-194
chr11-120829524-INS-66
chr11-120906556-INS-1892
chr11-120906966-INS-2840
chr11-120907364-INS-400
chr11-120911133-INS-52
chr11-120912251-INS-290
chr11-120918952-INS-3004
chr11-120919274-INS-212

chr11-120920509-INS-591
chr11-120921035-INS-318
chr11-120923728-INS-4948
chr11-120923729-INS-683
chr11-120977901-INS-213
chr11-120987346-INS-228
chr11-120989635-INS-79
chr11-121006550-INS-55
chr11-121023938-INS-5349
chr11-121040391-INS-175
chr11-121070024-INS-95
chr11-121072604-INS-247
chr11-121075592-INS-233
chr11-121081914-INS-178
chr11-121085435-INS-142
chr11-121087355-INS-538
chr11-121090149-INS-80
chr11-121125249-INS-249
chr11-121131915-INS-198
chr11-121170395-INS-236
chr11-121180625-INS-209
chr11-121196266-INS-73
chr11-121255379-INS-127
chr11-121255625-INS-1194
chr11-121284102-INS-6401
chr11-121296123-INS-213
chr11-121321701-INS-71
chr11-121336713-INS-60
chr11-121381889-INS-259
chr11-121393672-INS-72
chr11-121453724-INS-51
chr11-121536742-INS-68
chr11-121545210-INS-89
chr11-121554385-INS-152
chr11-121655792-INS-939
chr11-121655999-INS-72
chr11-121704996-INS-6502
chr11-121723534-INS-408
chr11-121724096-INS-678
chr11-121726819-INS-63
chr11-121727465-INS-210
chr11-121743514-INS-443
chr11-121811471-INS-203
chr11-121842379-INS-1398
chr11-121863014-INS-441
chr12-3060602-INS-60
chr12-3080262-INS-57
chr12-3091367-INS-189
chr12-3141066-INS-228
chr12-3142933-INS-59
chr12-3146398-INS-9791
chr12-3179378-INS-549
chr12-3194500-INS-130
chr12-3220876-INS-8988
chr12-3242556-INS-6445
chr12-3247501-INS-1830
chr12-3284571-INS-619
chr12-3325067-INS-55
chr12-3350255-INS-82
chr12-3369273-INS-100
chr12-3369650-INS-77
chr12-3379605-INS-60
chr12-3405438-INS-88
chr12-3428370-INS-62
chr12-3443714-INS-633
chr12-3446090-INS-262
chr12-3462169-INS-91
chr12-3468357-INS-74
chr12-3482457-INS-6442
chr12-3493523-INS-276
chr12-3499476-INS-343
chr12-3543919-INS-94
chr12-3565048-INS-218
chr12-3579194-INS-200
chr12-3586446-INS-224
chr12-3592980-INS-119
chr12-3606785-INS-558

chr12-3611965-INS-142
chr12-3628840-INS-438
chr12-3639609-INS-159
chr12-3648482-INS-60
chr12-3658016-INS-52
chr12-3665971-INS-206
chr12-3671938-INS-6003
chr12-3698577-INS-208
chr12-3711665-INS-72
chr12-3730082-INS-80
chr12-3738470-INS-663
chr12-3775620-INS-155
chr12-3780047-INS-248
chr12-3783617-INS-225
chr12-3788518-INS-56
chr12-3814305-INS-205
chr12-3816709-INS-720
chr12-3821791-INS-220
chr12-3840786-INS-51
chr12-3850389-INS-195
chr12-3892804-INS-79
chr12-3902434-INS-209
chr12-3908436-INS-2724
chr12-3919133-INS-74
chr12-3948532-INS-239
chr12-3966357-INS-119
chr12-3975793-INS-64
chr12-3979841-INS-78
chr12-3980327-INS-70
chr12-3990845-INS-197
chr12-4002528-INS-53
chr12-4007898-INS-210
chr12-4013159-INS-200
chr12-4016730-INS-211
chr12-4019302-INS-90
chr12-4023264-INS-257
chr12-4043734-INS-294
chr12-4062335-INS-6404
chr12-4063417-INS-146
chr12-4073404-INS-66
chr12-4076498-INS-100
chr12-4082367-INS-69
chr12-4098847-INS-209
chr12-4101920-INS-130
chr12-4106229-INS-76
chr12-4108223-INS-69
chr12-4112328-INS-76
chr12-4134074-INS-666
chr12-4139475-INS-190
chr12-4143675-INS-300
chr12-4149934-INS-202
chr12-4159420-INS-76
chr12-4161813-INS-1500
chr12-4161868-INS-224
chr12-4161927-INS-2288
chr12-4163370-INS-303
chr12-4174482-INS-66
chr12-4176217-INS-61
chr12-4179435-INS-60
chr12-4180446-INS-666
chr12-4190513-INS-194
chr12-4204236-INS-191
chr12-4206393-INS-206
chr12-4212455-INS-251
chr12-4213363-INS-132
chr12-4227152-INS-299
chr12-4233193-INS-171
chr12-4254508-INS-105
chr12-4271647-INS-3626
chr12-4271731-INS-3194
chr12-4318976-INS-222
chr12-4330708-INS-58
chr12-4334790-INS-188
chr12-4356374-INS-110
chr12-4358483-INS-5436
chr12-4360537-INS-69
chr12-4388396-INS-894

chr12-4389333-INS-7150
chr12-4398075-INS-6475
chr12-4398633-INS-102
chr12-4418411-INS-206
chr12-4431146-INS-140
chr12-4444064-INS-58
chr12-4463826-INS-216
chr12-4468998-INS-170
chr12-4505258-INS-205
chr12-4513649-INS-143
chr12-4517255-INS-170
chr12-4517337-INS-3876
chr12-4538168-INS-85
chr12-4559128-INS-1392
chr12-4559235-INS-445
chr12-4562903-INS-158
chr12-4567298-INS-6303
chr12-4571239-INS-55
chr12-4580983-INS-120
chr12-4595200-INS-68
chr12-4596044-INS-209
chr12-4597115-INS-218
chr12-4603351-INS-493
chr12-4607866-INS-210
chr12-4616915-INS-75
chr12-4620983-INS-209
chr12-4628535-INS-193
chr12-4646870-INS-212
chr12-4652037-INS-63
chr12-4696872-INS-132
chr12-4711564-INS-238
chr12-4720804-INS-6497
chr12-4737975-INS-111
chr12-4743306-INS-68
chr12-4749385-INS-733
chr12-4753412-INS-498
chr12-4760921-INS-69
chr12-4767049-INS-60
chr12-4784144-INS-58
chr12-4786591-INS-51
chr12-4815728-INS-108
chr12-4819152-INS-102
chr12-4847873-INS-391
chr12-4849331-INS-212
chr12-4862781-INS-213
chr12-4869480-INS-56
chr12-4875594-INS-96
chr12-4878192-INS-621
chr12-4893461-INS-154
chr12-4895949-INS-72
chr12-4905534-INS-56
chr12-4911159-INS-4934
chr12-4943494-INS-226
chr12-4949204-INS-71
chr12-4972853-INS-175
chr12-4985652-INS-264
chr12-4988446-INS-178
chr12-5005763-INS-294
chr12-5010185-INS-155
chr12-5013725-INS-201
chr12-5016626-INS-260
chr12-5030696-INS-5508
chr12-5033196-INS-6849
chr12-5068453-INS-376
chr12-5091390-INS-678
chr12-5098359-INS-70
chr12-5101836-INS-1827
chr12-5113609-INS-2121
chr12-5115872-INS-92
chr12-5160253-INS-55
chr12-5173801-INS-68
chr12-5174318-INS-123
chr12-5190821-INS-74
chr12-5191925-INS-59
chr12-5220031-INS-199
chr12-5236066-INS-205
chr12-5241234-INS-78

chr12-5259277-INS-62
chr12-5330728-INS-99
chr12-5336809-INS-253
chr12-5341579-INS-157
chr12-5357206-INS-93
chr12-5367256-INS-52
chr12-5401365-INS-114
chr12-5445084-INS-164
chr12-5458072-INS-54
chr12-5483166-INS-206
chr12-5505040-INS-296
chr12-5505977-INS-6457
chr12-5506796-INS-295
chr12-5509448-INS-6373
chr12-5515408-INS-284
chr12-5531101-INS-2152
chr12-5556807-INS-246
chr12-5579985-INS-75
chr12-5587619-INS-90
chr12-5590559-INS-51
chr12-5642419-INS-444
chr12-5648844-INS-70
chr12-5654004-INS-121
chr12-5655722-INS-400
chr12-5676961-INS-447
chr12-5690496-INS-119
chr12-5692747-INS-7042
chr12-5729046-INS-6388
chr12-5771752-INS-160
chr12-5774670-INS-722
chr12-5794859-INS-332
chr12-5801711-INS-219
chr12-5825611-INS-8190
chr12-5865946-INS-108
chr12-5893309-INS-475
chr12-5911192-INS-6452
chr12-5913241-INS-634
chr12-5915523-INS-468
chr12-5925660-INS-901
chr12-5934860-INS-405
chr12-5935724-INS-71
chr12-5937219-INS-6241
chr12-5963565-INS-348
chr12-5966866-INS-736
chr12-5972630-INS-64
chr12-5985572-INS-102
chr12-5990572-INS-169
chr12-6017856-INS-352
chr12-6063252-INS-212
chr12-6072260-INS-2044
chr12-6104895-INS-72
chr12-6110113-INS-111
chr12-6150037-INS-76
chr12-6154710-INS-2873
chr12-6162102-INS-52
chr12-6173105-INS-198
chr12-6178744-INS-84
chr12-6185055-INS-144
chr12-6191367-INS-2875
chr12-6214967-INS-56
chr12-6227695-INS-58
chr12-6269796-INS-322
chr12-6271703-INS-5182
chr12-6301970-INS-513
chr12-6304257-INS-7346
chr12-6304315-INS-6510
chr12-6337392-INS-632
chr12-6338029-INS-7094
chr12-6339612-INS-67
chr12-6343207-INS-66
chr12-6347031-INS-944
chr12-6349729-INS-1655
chr12-6359846-INS-97
chr12-6369595-INS-373
chr12-6370212-INS-6506
chr12-6373466-INS-58
chr12-6401247-INS-52

chr12-6403785-INS-87
chr12-6414009-INS-52
chr12-6444899-INS-665
chr12-6479311-INS-120
chr12-6481385-INS-355
chr12-6489654-INS-83
chr12-6497849-INS-129
chr12-6499122-INS-186
chr12-6509447-INS-220
chr12-6516086-INS-66
chr12-6522978-INS-72
chr12-6525147-INS-6271
chr12-6556346-INS-294
chr12-6589271-INS-144
chr12-6619183-INS-209
chr12-6676895-INS-55
chr12-6705612-INS-204
chr12-6736798-INS-64
chr12-6744765-INS-84
chr12-6755311-INS-6442
chr12-6766371-INS-64
chr12-6784530-INS-59
chr12-6794385-INS-249
chr12-6797152-INS-460
chr12-6800860-INS-716
chr12-6804854-INS-55
chr12-6813282-INS-117
chr12-6813363-INS-764
chr12-6834868-INS-173
chr12-6847198-INS-145
chr12-6881218-INS-148
chr12-6888552-INS-7338
chr12-6890115-INS-201
chr12-6894215-INS-410
chr12-6898103-INS-156
chr12-6902449-INS-230
chr12-6915042-INS-73
chr12-6924229-INS-148
chr12-6929570-INS-1495
chr12-6930059-INS-350
chr12-6936433-INS-328
chr12-6947690-INS-70
chr12-6950262-INS-386
chr12-6956554-INS-6089
chr12-6962022-INS-52
chr12-6964004-INS-6890
chr12-6978455-INS-182
chr12-6978848-INS-221
chr12-7074794-INS-141
chr12-7083734-INS-54
chr12-7092692-INS-78
chr12-7113799-INS-1841
chr12-7119214-INS-60
chr12-7138469-INS-52
chr12-7142151-INS-58
chr12-7157775-INS-7165
chr12-7181306-INS-2855
chr12-7182347-INS-52
chr12-7188225-INS-81
chr12-7217749-INS-7111
chr12-7220427-INS-113
chr12-7223046-INS-197
chr12-7250252-INS-326
chr12-7255406-INS-66
chr12-7304727-INS-6247
chr12-7313575-INS-79
chr12-7315356-INS-54
chr12-7335083-INS-201
chr12-7347567-INS-315
chr12-7355545-INS-89
chr12-7374365-INS-164
chr12-7383579-INS-62
chr12-7401364-INS-176
chr12-7403503-INS-554
chr12-7405545-INS-248
chr12-7425496-INS-140
chr12-7434295-INS-72

chr12-7439656-INS-103
chr12-7459862-INS-87
chr12-7489746-INS-578
chr12-7497423-INS-131
chr12-7509018-INS-153
chr12-7511983-INS-68
chr12-7513503-INS-64
chr12-7545559-INS-6815
chr12-7570324-INS-92
chr12-7584262-INS-946
chr12-7631385-INS-6194
chr12-7631586-INS-400
chr12-7635289-INS-2650
chr12-7635808-INS-1886
chr12-7645819-INS-4706
chr12-7649952-INS-6449
chr12-7654316-INS-408
chr12-7656974-INS-56
chr12-7659523-INS-62
chr12-7670852-INS-56
chr12-7674499-INS-2912
chr12-7688268-INS-64
chr12-7711916-INS-56
chr12-7715147-INS-112
chr12-7716367-INS-376
chr12-7721614-INS-85
chr12-7736354-INS-161
chr12-7753311-INS-6243
chr12-7769958-INS-92
chr12-7772369-INS-193
chr12-7787666-INS-499
chr12-7799527-INS-126
chr12-7828704-INS-67
chr12-7837683-INS-5850
chr12-7843537-INS-688
chr12-7850011-INS-115
chr12-7851204-INS-6230
chr12-7870335-INS-207
chr12-7877790-INS-386
chr12-7885326-INS-61
chr12-7886549-INS-76
chr12-7887587-INS-1126
chr12-7920828-INS-6440
chr12-7938211-INS-69
chr12-7939351-INS-559
chr12-7959308-INS-73
chr12-7968272-INS-56
chr12-7999481-INS-116
chr12-8026387-INS-109
chr12-8034135-INS-503
chr12-8047750-INS-194
chr12-8073133-INS-7129
chr12-8079157-INS-54
chr12-8082695-INS-1403
chr12-8104106-INS-382
chr12-8120691-INS-6399
chr12-8123357-INS-8749
chr12-8126679-INS-745
chr12-8126992-INS-7128
chr12-8159835-INS-97
chr12-8195487-INS-765
chr12-8199768-INS-444
chr12-8224660-INS-797
chr12-8224692-INS-52
chr12-8273177-INS-66
chr12-8280703-INS-229
chr12-8292562-INS-6197
chr12-8307935-INS-213
chr12-8312078-INS-175
chr12-8342822-INS-90
chr12-8357223-INS-89
chr12-8357586-INS-134
chr12-8406307-INS-66
chr12-8417508-INS-55
chr12-8419049-INS-198
chr12-8424328-INS-193
chr12-8438068-INS-356

chr12-8443693-INS-52
chr12-8444402-INS-375
chr12-8480092-INS-3549
chr12-8481446-INS-62
chr12-8483676-INS-117
chr12-8513595-INS-914
chr12-8522538-INS-120
chr12-8528322-INS-449
chr12-8531621-INS-151
chr12-8540813-INS-132
chr12-8554072-INS-128
chr12-8568119-INS-228
chr12-8606995-INS-422
chr12-8648833-INS-8837
chr12-8692633-INS-704
chr12-8743685-INS-61
chr12-8813448-INS-386
chr12-8817165-INS-76
chr12-8826549-INS-200
chr12-8891042-INS-96
chr12-8914793-INS-268
chr12-8917555-INS-198
chr12-8922484-INS-1888
chr12-8941137-INS-5505
chr12-8941315-INS-206
chr12-8942421-INS-72
chr12-8943549-INS-2700
chr12-8990445-INS-102
chr12-9005926-INS-416
chr12-9033348-INS-112
chr12-9040921-INS-60
chr12-9043296-INS-889
chr12-9054881-INS-269
chr12-9056598-INS-4965
chr12-9057239-INS-808
chr12-9087370-INS-961
chr12-9091343-INS-968
chr12-9131187-INS-526
chr12-9136646-INS-120
chr12-9146955-INS-114
chr12-9152383-INS-60
chr12-9161628-INS-116
chr12-9185958-INS-202
chr12-9188212-INS-163
chr12-9191586-INS-56
chr12-9224836-INS-52
chr12-9252402-INS-55
chr12-9269503-INS-140
chr12-9280509-INS-167
chr12-9285301-INS-56
chr12-9298158-INS-90
chr12-9337623-INS-965
chr12-9342114-INS-396
chr12-9346265-INS-286
chr12-9354515-INS-198
chr12-9355591-INS-854
chr12-9399377-INS-190
chr12-9405491-INS-51
chr12-9414871-INS-104
chr12-9428213-INS-587
chr12-9454263-INS-4009
chr12-9454815-INS-81
chr12-9460156-INS-408
chr12-9493937-INS-168
chr12-9499552-INS-243
chr12-9510183-INS-327
chr12-9524607-INS-66
chr12-9528497-INS-54
chr12-9536077-INS-149
chr12-9560851-INS-403
chr12-9575539-INS-146
chr12-9684076-INS-63
chr12-9688596-INS-75
chr12-9700535-INS-126
chr12-9750298-INS-1816
chr12-9752880-INS-6728
chr12-9755090-INS-5564

chr12-9780596-INS-96
chr12-9801954-INS-75
chr12-9803097-INS-126
chr12-9814738-INS-78
chr12-9828854-INS-258
chr12-9837068-INS-258
chr12-9837182-INS-54
chr12-9842356-INS-914
chr12-9844657-INS-125
chr12-9858096-INS-114
chr12-9884629-INS-3456
chr12-9920180-INS-60
chr12-9925259-INS-80
chr12-9928970-INS-60
chr12-9946435-INS-217
chr12-9962126-INS-164
chr12-9964850-INS-69
chr12-9984965-INS-645
chr12-9985697-INS-66
chr12-9987889-INS-10058
chr12-10014455-INS-7126
chr12-10056324-INS-59
chr12-10066235-INS-2261
chr12-10096610-INS-188
chr12-10112999-INS-6564
chr12-10113134-INS-52
chr12-10117316-INS-7003
chr12-10126867-INS-565
chr12-10127238-INS-4200
chr12-10131760-INS-59
chr12-10132455-INS-88
chr12-10134147-INS-118
chr12-10154299-INS-508
chr12-10155264-INS-322
chr12-10164755-INS-56
chr12-10183857-INS-52
chr12-10196729-INS-58
chr12-10220357-INS-6583
chr12-10228977-INS-88
chr12-10267351-INS-64
chr12-10276029-INS-185
chr12-10287815-INS-122
chr12-10289476-INS-66
chr12-10290832-INS-227
chr12-10349956-INS-114
chr12-10365706-INS-325
chr12-10374751-INS-199
chr12-10378794-INS-51
chr12-10404284-INS-208
chr12-10406902-INS-3592
chr12-10408193-INS-455
chr12-10409316-INS-336
chr12-10458861-INS-2847
chr12-10458862-INS-410
chr12-10465562-INS-1634
chr12-10478895-INS-142
chr12-10503529-INS-241
chr12-10518841-INS-68
chr12-10528627-INS-6657
chr12-10534118-INS-554
chr12-10543080-INS-73
chr12-10544112-INS-6442
chr12-10566585-INS-313
chr12-10605072-INS-129
chr12-10607290-INS-202
chr12-10649019-INS-519
chr12-10651152-INS-4739
chr12-10661372-INS-594
chr12-10662647-INS-291
chr12-10666576-INS-397
chr12-10672200-INS-213
chr12-10693485-INS-6224
chr12-10695874-INS-6010
chr12-10701662-INS-64
chr12-10703489-INS-831
chr12-10706655-INS-6508
chr12-10719537-INS-6488

chr12-10730431-INS-68
chr12-10764215-INS-6659
chr12-10770307-INS-120
chr12-10774804-INS-882
chr12-10785961-INS-100
chr12-10791302-INS-74
chr12-10793263-INS-243
chr12-10805662-INS-149
chr12-10815288-INS-399
chr12-10830485-INS-295
chr12-10847152-INS-88
chr12-10860234-INS-101
chr12-10882867-INS-56
chr12-10889282-INS-76
chr12-10903555-INS-258
chr12-10905515-INS-63
chr12-10908498-INS-53
chr12-10928711-INS-68
chr12-10944972-INS-8285
chr12-10945147-INS-58
chr12-10955409-INS-72
chr12-10957849-INS-6649
chr12-10962100-INS-201
chr12-10989207-INS-83
chr12-11000059-INS-226
chr12-11011559-INS-428
chr12-11011875-INS-312
chr12-11012211-INS-3215
chr12-11012510-INS-168
chr12-11012530-INS-2222
chr12-11013406-INS-107
chr12-11014936-INS-110
chr12-11057789-INS-52
chr12-11059173-INS-851
chr12-11061699-INS-106
chr12-11063705-INS-6639
chr12-11064912-INS-87
chr12-11068558-INS-308
chr12-11075610-INS-6196
chr12-11077681-INS-89
chr12-11090565-INS-78
chr12-11099786-INS-236
chr12-11103580-INS-7621
chr12-11106066-INS-6480
chr12-11112917-INS-219
chr12-11120950-INS-498
chr12-11136769-INS-72
chr12-11164710-INS-80
chr12-11169112-INS-109
chr12-11185259-INS-77
chr12-11267263-INS-56
chr12-11277379-INS-220
chr12-11293485-INS-504
chr12-11308969-INS-54
chr12-11328347-INS-75
chr12-11334479-INS-113
chr12-11352411-INS-5311
chr12-11357698-INS-998
chr12-11361012-INS-68
chr12-11378550-INS-176
chr12-11384167-INS-55
chr12-11395422-INS-59
chr12-11397739-INS-70
chr12-11400684-INS-410
chr12-11411512-INS-110
chr12-11417935-INS-132
chr12-11447391-INS-475
chr12-11454229-INS-67
chr12-11475074-INS-118
chr12-11492230-INS-131
chr12-11502814-INS-216
chr12-11555376-INS-6064
chr12-11568262-INS-1193
chr12-11572160-INS-6499
chr12-11585225-INS-280
chr12-11592325-INS-629
chr12-11625629-INS-55

chr12-11636837-INS-61
chr12-11642134-INS-6985
chr12-11656952-INS-61
chr12-11662217-INS-52
chr12-11668083-INS-538
chr12-11684910-INS-116
chr12-11702087-INS-301
chr12-11704877-INS-674
chr12-11712001-INS-153
chr12-11718493-INS-936
chr12-11730721-INS-1492
chr12-11734620-INS-238
chr12-11753843-INS-152
chr12-11765633-INS-64
chr12-11776436-INS-120
chr12-11791681-INS-54
chr12-11796948-INS-65
chr12-11798153-INS-57
chr12-11833475-INS-115
chr12-11852777-INS-907
chr12-11863919-INS-184
chr12-11867013-INS-345
chr12-11868180-INS-401
chr12-11877200-INS-292
chr12-11880391-INS-6800
chr12-11880913-INS-671
chr12-11888996-INS-6401
chr12-11897957-INS-724
chr12-11897979-INS-78
chr12-11900877-INS-114
chr12-11915888-INS-209
chr12-11922544-INS-320
chr12-11930765-INS-72
chr12-11960006-INS-56
chr12-11982777-INS-602
chr12-11984080-INS-577
chr12-11990017-INS-423
chr12-12045211-INS-217
chr12-12046879-INS-88
chr12-12048822-INS-202
chr12-12069576-INS-56
chr12-12077540-INS-967
chr12-12081640-INS-263
chr12-12097296-INS-185
chr12-12100713-INS-9106
chr12-12104079-INS-374
chr12-12154101-INS-68
chr12-12168834-INS-767
chr12-12175484-INS-6546
chr12-12179736-INS-193
chr12-12188669-INS-96
chr12-12189390-INS-52
chr12-12200916-INS-80
chr12-12212975-INS-322
chr12-12237440-INS-114
chr12-12243930-INS-106
chr12-12250155-INS-509
chr12-12260013-INS-52
chr12-12288183-INS-189
chr12-12295870-INS-249
chr12-12300704-INS-678
chr12-12325764-INS-116
chr12-12336277-INS-872
chr12-12362140-INS-410
chr12-12388287-INS-553
chr12-12403121-INS-54
chr12-12413341-INS-259
chr12-12505240-INS-772
chr12-12515864-INS-398
chr12-12531148-INS-121
chr12-12558902-INS-92
chr12-12611016-INS-6401
chr12-12623394-INS-866
chr12-12668778-INS-484
chr12-12673035-INS-120
chr12-12691995-INS-56
chr12-12742116-INS-184

chr12-12746161-INS-123
chr12-12758197-INS-176
chr12-12769356-INS-138
chr12-12779080-INS-87
chr12-12790978-INS-439
chr12-12800393-INS-62
chr12-12807986-INS-212
chr12-12832370-INS-106
chr12-12836041-INS-72
chr12-12857148-INS-196
chr12-12879507-INS-289
chr12-12880690-INS-169
chr12-12887715-INS-177
chr12-12894176-INS-221
chr12-12903732-INS-183
chr12-12945807-INS-109
chr12-12971831-INS-216
chr12-12979589-INS-264
chr12-13003192-INS-75
chr12-13006648-INS-116
chr12-13012258-INS-75
chr12-13014922-INS-201
chr12-13032700-INS-210
chr12-13033727-INS-63
chr12-13049833-INS-58
chr12-13062976-INS-213
chr12-13065383-INS-137
chr12-13121765-INS-52
chr12-13124545-INS-139
chr12-13148625-INS-172
chr12-13153030-INS-206
chr12-13154816-INS-105
chr12-13180420-INS-400
chr12-13185890-INS-75
chr12-13187153-INS-100
chr12-13194480-INS-487
chr12-13199418-INS-196
chr12-13227514-INS-459
chr12-13236553-INS-56
chr12-13240100-INS-127
chr12-13251324-INS-1893
chr12-13280153-INS-219
chr12-13290531-INS-80
chr12-13294413-INS-154
chr12-13308135-INS-68
chr12-13314885-INS-156
chr12-13332950-INS-113
chr12-13384548-INS-67
chr12-13396991-INS-60
chr12-13397408-INS-652
chr12-13409881-INS-110
chr12-13411504-INS-157
chr12-13440306-INS-199
chr12-13440349-INS-3603
chr12-13479266-INS-90
chr12-13527796-INS-546
chr12-13536010-INS-72
chr12-13553792-INS-136
chr12-13555537-INS-180
chr12-13561250-INS-3982
chr12-13578353-INS-612
chr12-13579545-INS-310
chr12-13585133-INS-80
chr12-13592799-INS-666
chr12-13617571-INS-244
chr12-13650696-INS-130
chr12-13662924-INS-452
chr12-13665518-INS-58
chr12-13669098-INS-51
chr12-13671108-INS-149
chr12-13685968-INS-175
chr12-13724329-INS-54
chr12-13725702-INS-59
chr12-13726797-INS-68
chr12-13740016-INS-276
chr12-13741579-INS-68
chr12-13781620-INS-311

chr12-13784907-INS-1353
chr12-13788537-INS-60
chr12-13814506-INS-54
chr12-13815521-INS-331
chr12-13819771-INS-113
chr12-13824497-INS-6909
chr12-13827443-INS-68
chr12-13834076-INS-413
chr12-13835665-INS-6434
chr12-13840622-INS-65
chr12-13840900-INS-399
chr12-13909758-INS-6356
chr12-13941566-INS-6807
chr12-13948101-INS-395
chr12-13984603-INS-56
chr12-14007108-INS-5055
chr12-14010792-INS-313
chr12-14029107-INS-180
chr12-14056676-INS-361
chr12-14086651-INS-196
chr12-14109104-INS-77
chr12-14125666-INS-1888
chr12-14138702-INS-82
chr12-14144165-INS-417
chr12-14152079-INS-115
chr12-14160738-INS-315
chr12-14171393-INS-57
chr12-14171947-INS-111
chr12-14178029-INS-548
chr12-14182539-INS-58
chr12-14184092-INS-386
chr12-14209253-INS-487
chr12-14216264-INS-68
chr12-14224835-INS-528
chr12-14232613-INS-51
chr12-14232750-INS-6453
chr12-14249367-INS-58
chr12-14259093-INS-128
chr12-14261908-INS-56
chr12-14364180-INS-60
chr12-14485704-INS-6403
chr12-14494780-INS-496
chr12-14510092-INS-226
chr12-14520411-INS-4944
chr12-14531179-INS-2139
chr12-14572459-INS-51
chr12-14576594-INS-52
chr12-14595064-INS-177
chr12-14596075-INS-194
chr12-14621421-INS-623
chr12-14635066-INS-74
chr12-14638230-INS-62
chr12-14670139-INS-6670
chr12-14689297-INS-183
chr12-14690374-INS-66
chr12-14702520-INS-6250
chr12-14718494-INS-103
chr12-14734256-INS-1704
chr12-14735349-INS-280
chr12-14738263-INS-520
chr12-14742040-INS-4351
chr12-14767099-INS-317
chr12-14786637-INS-74
chr12-14816220-INS-1912
chr12-14871396-INS-7162
chr12-14880251-INS-192
chr12-14888893-INS-1072
chr12-14889647-INS-73
chr12-14909422-INS-108
chr12-14921997-INS-808
chr12-14925112-INS-62
chr12-14940937-INS-70
chr12-14943036-INS-102
chr12-14957379-INS-401
chr12-14981926-INS-55
chr12-15004573-INS-6376
chr12-15021452-INS-157

chr12-15038863-INS-715
chr12-15056576-INS-749
chr12-15062330-INS-342
chr12-15072951-INS-949
chr12-15074835-INS-79
chr12-15076469-INS-64
chr12-15095453-INS-58
chr12-15120929-INS-107
chr12-15149863-INS-54
chr12-15180050-INS-56
chr12-15192417-INS-103
chr12-15195100-INS-54
chr12-15199145-INS-802
chr12-15208805-INS-497
chr12-15211945-INS-5338
chr12-15213320-INS-825
chr12-15221275-INS-119
chr12-15225186-INS-52
chr12-15239253-INS-294
chr12-15257335-INS-149
chr12-15277150-INS-159
chr12-15286750-INS-527
chr12-15300158-INS-906
chr12-15300409-INS-135
chr12-15330947-INS-197
chr12-15334086-INS-356
chr12-15339114-INS-64
chr12-15347099-INS-158
chr12-15369865-INS-576
chr12-15378384-INS-67
chr12-15380498-INS-194
chr12-15397163-INS-234
chr12-15455975-INS-497
chr12-15465327-INS-63
chr12-15469263-INS-64
chr12-15480137-INS-122
chr12-15487612-INS-505
chr12-15496490-INS-276
chr12-15612101-INS-54
chr12-15624868-INS-69
chr12-15630476-INS-144
chr12-15643695-INS-56
chr12-15650510-INS-506
chr12-15662687-INS-6178
chr12-15685364-INS-141
chr12-15735721-INS-109
chr12-15742939-INS-341
chr12-15753432-INS-100
chr12-15754985-INS-51
chr12-15768452-INS-200
chr12-15772017-INS-90
chr12-15798088-INS-118
chr12-15806505-INS-310
chr12-15831725-INS-155
chr12-15833999-INS-87
chr12-15846154-INS-88
chr12-15857774-INS-60
chr12-15862132-INS-136
chr12-15875632-INS-72
chr12-15876794-INS-88
chr12-15878169-INS-58
chr12-15880496-INS-117
chr12-15912136-INS-56
chr12-15913598-INS-6407
chr12-15914931-INS-519
chr12-15922084-INS-307
chr12-15945260-INS-174
chr12-15972563-INS-342
chr12-15988249-INS-56
chr12-15998271-INS-54
chr12-16002699-INS-616
chr12-16018966-INS-246
chr12-16063083-INS-85
chr12-16110476-INS-114
chr12-16113146-INS-6284
chr12-16115278-INS-70
chr12-16124920-INS-75

chr12-16130250-INS-606
chr12-16145554-INS-6464
chr12-16165810-INS-195
chr12-16175117-INS-887
chr12-16176710-INS-55
chr12-16185424-INS-131
chr12-16217088-INS-56
chr12-16275313-INS-277
chr12-16282067-INS-52
chr12-16286694-INS-64
chr12-16303544-INS-57
chr12-16306187-INS-304
chr12-16309651-INS-63
chr12-16318234-INS-805
chr12-16345103-INS-90
chr12-16349548-INS-154
chr12-16365295-INS-468
chr12-16408684-INS-105
chr12-16415829-INS-7213
chr12-16431977-INS-769
chr12-16438544-INS-67
chr12-16467806-INS-72
chr12-16524151-INS-78
chr12-16528825-INS-70
chr12-16554249-INS-473
chr12-16571555-INS-65
chr12-16624201-INS-88
chr12-16626902-INS-112
chr12-16630064-INS-72
chr12-16635307-INS-89
chr12-16649609-INS-502
chr12-16652665-INS-52
chr12-16682558-INS-192
chr12-16686138-INS-374
chr12-16686943-INS-134
chr12-16692946-INS-82
chr12-16750508-INS-54
chr12-16829483-INS-1864
chr12-16830034-INS-68
chr12-16847170-INS-199
chr12-16870991-INS-212
chr12-16883058-INS-53
chr12-16885844-INS-2435
chr12-16892399-INS-278
chr12-16912170-INS-52
chr12-16922553-INS-201
chr12-16931441-INS-209
chr12-16973566-INS-2000
chr12-16975303-INS-288
chr12-16983607-INS-239
chr12-16997569-INS-223
chr12-16999243-INS-63
chr12-17016236-INS-179
chr12-17077743-INS-5237
chr12-17098308-INS-6662
chr12-17103282-INS-325
chr12-17105796-INS-67
chr12-17115316-INS-272
chr12-17117358-INS-77
chr12-17128751-INS-174
chr12-17144496-INS-86
chr12-17159073-INS-985
chr12-17159968-INS-133
chr12-17172377-INS-120
chr12-17172839-INS-82
chr12-17182245-INS-54
chr12-17191884-INS-575
chr12-17197297-INS-76
chr12-17205462-INS-3432
chr12-17207598-INS-142
chr12-17233002-INS-462
chr12-17237740-INS-207
chr12-17238635-INS-77
chr12-17248919-INS-56
chr12-17268972-INS-58
chr12-17282053-INS-208
chr12-17289552-INS-6440

chr12-17289845-INS-99
chr12-17302471-INS-226
chr12-17328725-INS-82
chr12-17346651-INS-92
chr12-17406245-INS-185
chr12-17412021-INS-217
chr12-17416457-INS-245
chr12-17436946-INS-496
chr12-17445125-INS-161
chr12-17446813-INS-112
chr12-17473873-INS-52
chr12-17492598-INS-63
chr12-17504514-INS-94
chr12-17508175-INS-365
chr12-17519518-INS-178
chr12-17537788-INS-57
chr12-17544120-INS-54
chr12-17573305-INS-52
chr12-17581274-INS-82
chr12-17583489-INS-87
chr12-17612678-INS-60
chr12-17631495-INS-213
chr12-17631727-INS-5148
chr12-17655515-INS-249
chr12-17660506-INS-60
chr12-17668823-INS-114
chr12-17684881-INS-99
chr12-17687293-INS-214
chr12-17724767-INS-59
chr12-17821463-INS-740
chr12-17840952-INS-61
chr12-17861099-INS-53
chr12-17869230-INS-394
chr12-17961919-INS-5319
chr12-17963964-INS-76
chr12-17996034-INS-73
chr12-17997893-INS-85
chr12-18042041-INS-85
chr12-18122225-INS-59
chr12-18132391-INS-53
chr12-18139226-INS-3469
chr12-18166525-INS-610
chr12-18222593-INS-85
chr12-18259026-INS-185
chr12-18314518-INS-134
chr12-18318806-INS-504
chr12-18370482-INS-2711
chr12-18435797-INS-134
chr12-18455439-INS-169
chr12-18534989-INS-66
chr12-18599490-INS-93
chr12-18618344-INS-96
chr12-18625793-INS-95
chr12-18630009-INS-527
chr12-18637409-INS-201
chr12-18640146-INS-56
chr12-18652394-INS-2020
chr12-18664709-INS-514
chr12-18688929-INS-327
chr12-18698689-INS-84
chr12-18699948-INS-84
chr12-18725310-INS-179
chr12-18745607-INS-60
chr12-18746927-INS-459
chr12-18770488-INS-6452
chr12-18771710-INS-1000
chr12-18778153-INS-327
chr12-18838053-INS-254
chr12-18893010-INS-605
chr12-19013839-INS-53
chr12-19020823-INS-281
chr12-19052225-INS-6442
chr12-19058798-INS-57
chr12-19091437-INS-209
chr12-19116242-INS-3081
chr12-19223450-INS-62
chr12-19225577-INS-233

chr12-19271838-INS-422
chr12-19296210-INS-843
chr12-19344458-INS-475
chr12-19351979-INS-179
chr12-19353282-INS-84
chr12-19362514-INS-64
chr12-19411135-INS-212
chr12-19435747-INS-70
chr12-19436869-INS-313
chr12-19446168-INS-422
chr12-19466849-INS-70
chr12-19472843-INS-56
chr12-19485450-INS-121
chr12-19522399-INS-7089
chr12-19547437-INS-3533
chr12-19550969-INS-168
chr12-19647377-INS-811
chr12-19843388-INS-6079
chr12-19971828-INS-72
chr12-19972796-INS-191
chr12-19990580-INS-84
chr12-20017680-INS-129
chr12-20026900-INS-84
chr12-20061635-INS-64
chr12-20064947-INS-282
chr12-20066025-INS-325
chr12-20072401-INS-1876
chr12-20079003-INS-9004
chr12-20135243-INS-251
chr12-20145051-INS-2860
chr12-20191992-INS-60
chr12-20213451-INS-51
chr12-20251288-INS-114
chr12-20298129-INS-359
chr12-20299895-INS-6879
chr12-20321200-INS-7451
chr12-20334172-INS-102
chr12-20372167-INS-84
chr12-20399948-INS-52
chr12-20409697-INS-393
chr12-20436811-INS-448
chr12-20441060-INS-63
chr12-20461143-INS-250
chr12-20471279-INS-401
chr12-20499738-INS-100
chr12-20610522-INS-96
chr12-20624455-INS-841
chr12-20627324-INS-52
chr12-20633210-INS-287
chr12-20636039-INS-137
chr12-20648359-INS-172
chr12-20656305-INS-73
chr12-20669815-INS-314
chr12-20685996-INS-249
chr12-20721349-INS-154
chr12-20727506-INS-8822
chr12-20738675-INS-199
chr12-20805902-INS-287
chr12-20824935-INS-8362
chr12-20853633-INS-186
chr12-20892053-INS-120
chr12-20909692-INS-62
chr12-20910077-INS-106
chr12-20934142-INS-563
chr12-20986435-INS-91
chr12-20990451-INS-91
chr12-20999463-INS-230
chr12-21007943-INS-59
chr12-21022696-INS-6771
chr12-21045592-INS-114
chr12-21070922-INS-1478
chr12-21085512-INS-76
chr12-21098189-INS-186
chr12-21105429-INS-52
chr12-21166276-INS-174
chr12-21199088-INS-100
chr12-21265202-INS-120

chr12-21276099-INS-221
chr12-21282785-INS-199
chr12-21311411-INS-195
chr12-21325579-INS-316
chr12-21332204-INS-970
chr12-21340941-INS-190
chr12-21365609-INS-313
chr12-21365609-INS-422
chr12-21420524-INS-193
chr12-21438015-INS-217
chr12-21442881-INS-70
chr12-21472982-INS-176
chr12-21478414-INS-456
chr12-21488997-INS-52
chr12-21496396-INS-201
chr12-21515603-INS-321
chr12-21530788-INS-72
chr12-21538222-INS-108
chr12-21545358-INS-57
chr12-21559802-INS-92
chr12-21567569-INS-403
chr12-21568384-INS-816
chr12-21648756-INS-61
chr12-21774117-INS-725
chr12-21776488-INS-126
chr12-21778612-INS-220
chr12-21795742-INS-95
chr12-21809212-INS-420
chr12-21818278-INS-65
chr12-21829261-INS-642
chr12-21842451-INS-68
chr12-21862399-INS-114
chr12-21868342-INS-4831
chr12-21930640-INS-508
chr12-22010297-INS-212
chr12-22011500-INS-184
chr12-22017050-INS-160
chr12-22043073-INS-168
chr12-22128399-INS-7471
chr12-22143889-INS-215
chr12-22344260-INS-1708
chr12-22395833-INS-4604
chr12-22444968-INS-7165
chr12-22497683-INS-84
chr12-22498152-INS-423
chr12-22508698-INS-422
chr12-22570338-INS-53
chr12-22590484-INS-85
chr12-22630890-INS-118
chr12-22682852-INS-813
chr12-22710271-INS-715
chr12-22775125-INS-84
chr12-22790822-INS-58
chr12-22797503-INS-537
chr12-22803402-INS-91
chr12-22830423-INS-59
chr12-22837709-INS-51
chr12-22859296-INS-95
chr12-22887150-INS-76
chr12-22968937-INS-211
chr12-23055441-INS-288
chr12-23118971-INS-574
chr12-23125141-INS-64
chr12-23131581-INS-75
chr12-23179384-INS-1023
chr12-23198003-INS-84
chr12-23234948-INS-229
chr12-23242361-INS-176
chr12-23319329-INS-3807
chr12-23337216-INS-209
chr12-23460618-INS-116
chr12-23589421-INS-9134
chr12-23603991-INS-7168
chr12-23613786-INS-7098
chr12-23687430-INS-7185
chr12-23727918-INS-4951
chr12-23772638-INS-67

chr12-23774886-INS-159
chr12-23801291-INS-57
chr12-23803313-INS-116
chr12-23831658-INS-252
chr12-23837967-INS-2286
chr12-23846080-INS-343
chr12-23909537-INS-62
chr12-23975348-INS-175
chr12-23979516-INS-591
chr12-24022617-INS-219
chr12-24069550-INS-4756
chr12-24082083-INS-56
chr12-24111802-INS-5922
chr12-24121785-INS-168
chr12-24142298-INS-172
chr12-24165844-INS-72
chr12-24205303-INS-118
chr12-24260569-INS-78
chr12-24297657-INS-114
chr12-24345124-INS-123
chr12-24353022-INS-56
chr12-24387994-INS-126
chr12-24405682-INS-58
chr12-24408037-INS-5639
chr12-24471526-INS-176
chr12-24497832-INS-231
chr12-24501110-INS-4788
chr12-24515743-INS-453
chr12-24527786-INS-416
chr12-24532186-INS-416
chr12-24535658-INS-145
chr12-24538335-INS-76
chr12-24556584-INS-1219
chr12-24566332-INS-210
chr12-24595284-INS-90
chr12-24597358-INS-444
chr12-24669792-INS-183
chr12-24680892-INS-51
chr12-24725124-INS-158
chr12-24728225-INS-152
chr12-24755032-INS-206
chr12-24769783-INS-430
chr12-24770656-INS-201
chr12-24772704-INS-424
chr12-24778544-INS-128
chr12-24781005-INS-200
chr12-24785660-INS-115
chr12-24796028-INS-161
chr12-24802948-INS-54
chr12-24818629-INS-214
chr12-24828631-INS-205
chr12-24843783-INS-197
chr12-24848871-INS-209
chr12-24850799-INS-70
chr12-24859654-INS-58
chr12-24864812-INS-201
chr12-24868234-INS-57
chr12-24871370-INS-222
chr12-24875501-INS-72
chr12-24928874-INS-73
chr12-24930031-INS-256
chr12-24941957-INS-173
chr12-24947891-INS-156
chr12-24957431-INS-60
chr12-24972795-INS-69
chr12-24984417-INS-202
chr12-25004591-INS-286
chr12-25035740-INS-978
chr12-25071733-INS-176
chr12-25076410-INS-417
chr12-25105483-INS-198
chr12-25123276-INS-1440
chr12-25123355-INS-590
chr12-25125799-INS-61
chr12-25130983-INS-116
chr12-25135170-INS-438
chr12-25153401-INS-128

chr12-25156450-INS-205
chr12-25167458-INS-168
chr12-25171121-INS-192
chr12-25179938-INS-204
chr12-25198496-INS-58
chr12-25205841-INS-126
chr12-25254823-INS-74
chr12-25280054-INS-142
chr12-25287449-INS-385
chr12-25293207-INS-144
chr12-25297770-INS-542
chr12-25326370-INS-389
chr12-25328048-INS-92
chr12-25381015-INS-6399
chr12-25387493-INS-306
chr12-25401354-INS-357
chr12-25418596-INS-250
chr12-25434292-INS-276
chr12-25464672-INS-51
chr12-25467126-INS-147
chr12-25476362-INS-87
chr12-25548255-INS-52
chr12-25551862-INS-128
chr12-25600870-INS-1888
chr12-25605575-INS-134
chr12-25649065-INS-327
chr12-25665943-INS-295
chr12-25755116-INS-7327
chr12-25757126-INS-6446
chr12-25812691-INS-60
chr12-25848457-INS-82
chr12-25869515-INS-57
chr12-25884767-INS-7102
chr12-25946993-INS-60
chr12-25956277-INS-52
chr12-25985768-INS-432
chr12-26007994-INS-197
chr12-26009517-INS-56
chr12-26048194-INS-83
chr12-26097365-INS-431
chr12-26106479-INS-171
chr12-26114641-INS-6464
chr12-26160681-INS-119
chr12-26207449-INS-6461
chr12-26264050-INS-400
chr12-26267634-INS-218
chr12-26286946-INS-320
chr12-26393192-INS-55
chr12-26405190-INS-219
chr12-26412191-INS-52
chr12-26422938-INS-114
chr12-26433220-INS-60
chr12-26457603-INS-100
chr12-26473378-INS-72
chr12-26508968-INS-216
chr12-26510068-INS-100
chr12-26522025-INS-79
chr12-26525075-INS-99
chr12-26533842-INS-75
chr12-26542179-INS-54
chr12-26546224-INS-96
chr12-26577788-INS-157
chr12-26594429-INS-52
chr12-26596025-INS-74
chr12-26620027-INS-127
chr12-26648676-INS-356
chr12-26648676-INS-724
chr12-26666073-INS-305
chr12-26672573-INS-202
chr12-26688796-INS-59
chr12-26690856-INS-56
chr12-26757465-INS-870
chr12-26771952-INS-72
chr12-26811599-INS-174
chr12-26816503-INS-108
chr12-26818892-INS-6279
chr12-26822846-INS-52

chr12-26837384-INS-53
chr12-26852242-INS-225
chr12-26856811-INS-910
chr12-26871006-INS-56
chr12-26878946-INS-7085
chr12-26883287-INS-54
chr12-26892343-INS-52
chr12-26903165-INS-961
chr12-26919043-INS-145
chr12-26932759-INS-549
chr12-26958981-INS-247
chr12-26978364-INS-60
chr12-26986376-INS-66
chr12-26988389-INS-68
chr12-26994858-INS-126
chr12-27024591-INS-62
chr12-27035553-INS-408
chr12-27038601-INS-86
chr12-27069531-INS-66
chr12-27145572-INS-102
chr12-27186758-INS-430
chr12-27201950-INS-60
chr12-27215142-INS-105
chr12-27220729-INS-69
chr12-27253935-INS-6264
chr12-27257736-INS-113
chr12-27265325-INS-199
chr12-27301252-INS-52
chr12-27317124-INS-62
chr12-27322389-INS-785
chr12-27338117-INS-173
chr12-27339252-INS-86
chr12-27367014-INS-92
chr12-27401648-INS-71
chr12-27420114-INS-108
chr12-27420743-INS-61
chr12-27448366-INS-1430
chr12-27459191-INS-107
chr12-27484809-INS-366
chr12-27489101-INS-52
chr12-27491204-INS-192
chr12-27527485-INS-51
chr12-27528592-INS-814
chr12-27533962-INS-843
chr12-27535299-INS-4380
chr12-27536423-INS-58
chr12-27538037-INS-708
chr12-27538078-INS-56
chr12-27568460-INS-95
chr12-27577284-INS-155
chr12-27594452-INS-4728
chr12-27611874-INS-690
chr12-27621274-INS-97
chr12-27638503-INS-64
chr12-27656145-INS-341
chr12-27657427-INS-94
chr12-27670988-INS-7081
chr12-27759802-INS-8706
chr12-27761837-INS-161
chr12-27766380-INS-1888
chr12-27774678-INS-57
chr12-27782983-INS-935
chr12-27801043-INS-75
chr12-27810876-INS-341
chr12-27815834-INS-604
chr12-27902839-INS-421
chr12-27916966-INS-597
chr12-27927656-INS-489
chr12-27933367-INS-86
chr12-27961429-INS-100
chr12-27972311-INS-338
chr12-28001365-INS-54
chr12-28004588-INS-58
chr12-28009189-INS-280
chr12-28037451-INS-401
chr12-28046102-INS-204
chr12-28059959-INS-841

chr12-28106253-INS-55
chr12-28107744-INS-58
chr12-28117168-INS-514
chr12-28139302-INS-54
chr12-28164408-INS-502
chr12-28167538-INS-122
chr12-28171852-INS-156
chr12-28241542-INS-201
chr12-28291493-INS-129
chr12-28301412-INS-58
chr12-28304258-INS-327
chr12-28305386-INS-7141
chr12-28316611-INS-99
chr12-28367149-INS-581
chr12-28418787-INS-66
chr12-28422300-INS-110
chr12-28428842-INS-567
chr12-28433392-INS-67
chr12-28440789-INS-177
chr12-28446436-INS-68
chr12-28478633-INS-6200
chr12-28492143-INS-303
chr12-28514205-INS-54
chr12-28519153-INS-65
chr12-28522219-INS-154
chr12-28544557-INS-70
chr12-28558816-INS-73
chr12-28575395-INS-183
chr12-28586675-INS-55
chr12-28595747-INS-686
chr12-28603025-INS-58
chr12-28605146-INS-89
chr12-28644043-INS-52
chr12-28646736-INS-60
chr12-28654235-INS-140
chr12-28679996-INS-168
chr12-28692074-INS-627
chr12-28735114-INS-73
chr12-28820126-INS-535
chr12-28823324-INS-109
chr12-28841547-INS-423
chr12-28844833-INS-55
chr12-28849636-INS-61
chr12-28871502-INS-152
chr12-28917555-INS-251
chr12-28929868-INS-350
chr12-28954690-INS-139
chr12-28976764-INS-75
chr12-29010494-INS-62
chr12-29026618-INS-651
chr12-29081691-INS-400
chr12-29085024-INS-95
chr12-29098349-INS-71
chr12-29104988-INS-134
chr12-29106055-INS-6766
chr12-29108832-INS-492
chr12-29113325-INS-52
chr12-29156567-INS-218
chr12-29163652-INS-51
chr12-29194447-INS-91
chr12-29202328-INS-191
chr12-29250107-INS-2854
chr12-29255219-INS-342
chr12-29276681-INS-105
chr12-29280364-INS-58
chr12-29312665-INS-155
chr12-29364462-INS-87
chr12-29366857-INS-55
chr12-29401634-INS-190
chr12-29417799-INS-163
chr12-29444227-INS-52
chr12-29493549-INS-61
chr12-29507144-INS-523
chr12-29529395-INS-684
chr12-29531350-INS-121
chr12-29555471-INS-66
chr12-29568579-INS-68

chr12-29600153-INS-194
chr12-29612314-INS-370
chr12-29638094-INS-147
chr12-29673620-INS-6874
chr12-29741884-INS-202
chr12-29899533-INS-71
chr12-29909660-INS-186
chr12-29914530-INS-56
chr12-29965833-INS-82
chr12-30045553-INS-64
chr12-30048263-INS-112
chr12-30120543-INS-60
chr12-30150166-INS-354
chr12-30168341-INS-214
chr12-30177082-INS-56
chr12-30193593-INS-6919
chr12-30230162-INS-169
chr12-30233585-INS-6790
chr12-30256736-INS-78
chr12-30259759-INS-152
chr12-30269536-INS-910
chr12-30270894-INS-113
chr12-30275101-INS-54
chr12-30289757-INS-56
chr12-30296660-INS-53
chr12-30308975-INS-130
chr12-30313443-INS-66
chr12-30372353-INS-6135
chr12-30404183-INS-68
chr12-30412339-INS-79
chr12-30413287-INS-342
chr12-30432852-INS-113
chr12-30441171-INS-51
chr12-30455028-INS-52
chr12-30469901-INS-190
chr12-30472924-INS-277
chr12-30489699-INS-328
chr12-30502079-INS-203
chr12-30524180-INS-252
chr12-30527662-INS-7086
chr12-30539401-INS-132
chr12-30543587-INS-63
chr12-30568337-INS-58
chr12-30575779-INS-138
chr12-30584480-INS-72
chr12-30593171-INS-943
chr12-30598497-INS-202
chr12-30622058-INS-1726
chr12-30628558-INS-1170
chr12-30628986-INS-7617
chr12-30629038-INS-6967
chr12-30641368-INS-107
chr12-30658563-INS-422
chr12-30686104-INS-60
chr12-30692313-INS-362
chr12-30697411-INS-506
chr12-30704273-INS-3964
chr12-30705950-INS-358
chr12-30712904-INS-103
chr12-30742975-INS-4294
chr12-30768937-INS-6663
chr12-30785091-INS-338
chr12-30788448-INS-58
chr12-30792309-INS-56
chr12-30812854-INS-1479
chr12-30838519-INS-3102
chr12-30882426-INS-5555
chr12-30899538-INS-266
chr12-30906333-INS-455
chr12-30906871-INS-1712
chr12-30951093-INS-326
chr12-30973359-INS-278
chr12-30998784-INS-54
chr12-31034891-INS-251
chr12-31052137-INS-116
chr12-31078124-INS-116
chr12-31136561-INS-150

chr12-31138551-INS-86
chr12-31146084-INS-200
chr12-31183017-INS-211
chr12-31185270-INS-61
chr12-31197007-INS-128
chr12-31207204-INS-78
chr12-31210843-INS-267
chr12-31224256-INS-598
chr12-31253392-INS-392
chr12-31257123-INS-474
chr12-31273465-INS-209
chr12-31277012-INS-62
chr12-31279346-INS-102
chr12-31291172-INS-167
chr12-31293319-INS-196
chr12-31299958-INS-54
chr12-31311835-INS-400
chr12-31322529-INS-90
chr12-31338502-INS-72
chr12-31343077-INS-64
chr12-31395938-INS-211
chr12-31418991-INS-324
chr12-31437412-INS-70
chr12-31453184-INS-74
chr12-31466317-INS-175
chr12-31491162-INS-156
chr12-31510239-INS-142
chr12-31516768-INS-106
chr12-31519486-INS-302
chr12-31523251-INS-100
chr12-31545973-INS-180
chr12-31557109-INS-86
chr12-31558457-INS-4824
chr12-31622013-INS-163
chr12-31641043-INS-471
chr12-31653093-INS-799
chr12-31691392-INS-121
chr12-31697180-INS-223
chr12-31708093-INS-821
chr12-31712523-INS-151
chr12-31714598-INS-551
chr12-31737384-INS-241
chr12-31756468-INS-98
chr12-31765727-INS-188
chr12-31775412-INS-511
chr12-31812042-INS-76
chr12-31820643-INS-114
chr12-31832855-INS-60
chr12-31843578-INS-226
chr12-31847648-INS-241
chr12-31863504-INS-74
chr12-31865045-INS-2013
chr12-31881254-INS-68
chr12-31921981-INS-119
chr12-31933019-INS-161
chr12-31937605-INS-432
chr12-32058640-INS-384
chr12-32059194-INS-165
chr12-32060075-INS-150
chr12-32082645-INS-222
chr12-32100629-INS-701
chr12-32143830-INS-259
chr12-32171436-INS-411
chr12-32191427-INS-140
chr12-32193517-INS-204
chr12-32219907-INS-203
chr12-32239239-INS-181
chr12-32256572-INS-65
chr12-32275250-INS-338
chr12-32276055-INS-139
chr12-32283343-INS-62
chr12-32297578-INS-73
chr12-32315097-INS-417
chr12-32333224-INS-92
chr12-32348294-INS-399
chr12-32364686-INS-136
chr12-32410649-INS-67

chr12-32460038-INS-202
chr12-32461109-INS-493
chr12-32468039-INS-212
chr12-32535122-INS-147
chr12-32547195-INS-125
chr12-32557582-INS-52
chr12-32562392-INS-68
chr12-32568296-INS-209
chr12-32569886-INS-53
chr12-32576009-INS-61
chr12-32600678-INS-1792
chr12-32605456-INS-5476
chr12-32636315-INS-119
chr12-32673335-INS-100
chr12-32682227-INS-533
chr12-32699172-INS-56
chr12-32774116-INS-581
chr12-32812743-INS-276
chr12-32835863-INS-112
chr12-32891689-INS-58
chr12-32893450-INS-115
chr12-32898467-INS-442
chr12-32908949-INS-102
chr12-32914736-INS-144
chr12-32928486-INS-372
chr12-32929997-INS-319
chr12-32933657-INS-1893
chr12-32940472-INS-101
chr12-32966746-INS-63
chr12-32971697-INS-582
chr12-32972231-INS-985
chr12-32981474-INS-551
chr12-32985312-INS-298
chr12-33035622-INS-590
chr12-33036716-INS-2963
chr12-33036896-INS-273
chr12-33047929-INS-213
chr12-33049738-INS-506
chr12-33054084-INS-80
chr12-33065612-INS-54
chr12-33110851-INS-62
chr12-33125206-INS-52
chr12-33128730-INS-63
chr12-33134598-INS-99
chr12-33156280-INS-86
chr12-33195282-INS-3420
chr12-33243096-INS-59
chr12-33246917-INS-515
chr12-33251061-INS-127
chr12-33253354-INS-109
chr12-33270002-INS-105
chr12-33298156-INS-58
chr12-33303606-INS-78
chr12-33307547-INS-84
chr12-33310084-INS-250
chr12-33325705-INS-188
chr12-33349040-INS-96
chr12-33370099-INS-51
chr12-33371172-INS-174
chr12-33377260-INS-760
chr12-33398103-INS-69
chr12-33427741-INS-224
chr12-33431167-INS-80
chr12-33456906-INS-203
chr12-33460870-INS-110
chr12-33480398-INS-55
chr12-33485851-INS-201
chr12-33492181-INS-721
chr12-33506826-INS-303
chr12-33510675-INS-103
chr12-33520037-INS-130
chr12-33522344-INS-80
chr12-33611794-INS-8096
chr12-33612558-INS-70
chr12-33613167-INS-416
chr12-33646155-INS-60
chr12-33649265-INS-76

chr12-33690353-INS-59
chr12-33703254-INS-63
chr12-33703775-INS-703
chr12-33736686-INS-491
chr12-33749094-INS-58
chr12-33757984-INS-198
chr12-33775569-INS-57
chr12-33776546-INS-3681
chr12-33808965-INS-81
chr12-33809110-INS-1878
chr12-33810370-INS-58
chr12-33838981-INS-55
chr12-33845607-INS-435
chr12-33869155-INS-60
chr12-33875330-INS-134
chr12-33884787-INS-99
chr12-33911583-INS-129
chr12-33933250-INS-314
chr12-33937785-INS-294
chr12-33937785-INS-545
chr12-33952652-INS-64
chr12-33956397-INS-101
chr12-33999851-INS-58
chr12-34025901-INS-197
chr12-34050226-INS-207
chr12-34058501-INS-6233
chr12-34074378-INS-293
chr12-34081592-INS-475
chr12-34087212-INS-395
chr12-34090619-INS-124
chr12-34094961-INS-55
chr12-34133599-INS-405
chr12-34150106-INS-181
chr12-34154364-INS-75
chr12-34167366-INS-113
chr12-34171331-INS-497
chr12-34192491-INS-835
chr12-34197681-INS-143
chr12-34245741-INS-2291
chr12-34280844-INS-6611
chr12-34282321-INS-74
chr12-34288712-INS-165
chr12-34296208-INS-157
chr12-34310141-INS-294
chr12-34322966-INS-304
chr12-34370328-INS-337
chr12-34373591-INS-371
chr12-34374597-INS-215
chr12-34387192-INS-79
chr12-34390441-INS-98
chr12-34395272-INS-114
chr12-34453975-INS-56
chr12-34468707-INS-155
chr12-34489947-INS-580
chr12-34493477-INS-152
chr12-34519937-INS-107
chr12-34522908-INS-632
chr12-34530022-INS-135
chr12-34588856-INS-147
chr12-34609564-INS-102
chr12-34620721-INS-351
chr12-34635903-INS-72
chr12-34649432-INS-50
chr12-34656174-INS-226
chr12-34659286-INS-104
chr12-34680250-INS-280
chr12-34703255-INS-57
chr12-34706978-INS-174
chr12-34716634-INS-717
chr12-34718425-INS-206
chr12-34729909-INS-102
chr12-34745224-INS-81
chr12-34749062-INS-624
chr12-34753991-INS-463
chr12-34755221-INS-186
chr12-34769924-INS-72
chr12-34775848-INS-66

chr12-34783692-INS-206
chr12-34795431-INS-486
chr12-34796251-INS-53
chr12-34820687-INS-63
chr12-34835887-INS-100
chr12-34864144-INS-54
chr12-34900522-INS-126
chr12-34908816-INS-256
chr12-34936601-INS-116
chr12-34940666-INS-74
chr12-34948720-INS-214
chr12-34953227-INS-55
chr12-34955484-INS-283
chr12-34969297-INS-295
chr12-34987860-INS-85
chr12-34993033-INS-57
chr12-35022162-INS-88
chr12-35046670-INS-268
chr12-35062503-INS-201
chr12-35075074-INS-62
chr12-35077333-INS-211
chr12-35093002-INS-159
chr12-35111935-INS-64
chr12-35136716-INS-481
chr12-35165679-INS-106
chr12-35186281-INS-274
chr12-35186281-INS-826
chr12-35225466-INS-171
chr12-35263776-INS-87
chr12-35268709-INS-770
chr12-35270633-INS-477
chr12-35301203-INS-189
chr12-35375760-INS-7508
chr12-35388044-INS-6280
chr12-35392332-INS-6396
chr12-35420183-INS-56
chr12-35422122-INS-56
chr12-35480900-INS-205
chr12-35501481-INS-100
chr12-35502827-INS-191
chr12-35541078-INS-55
chr12-35576126-INS-56
chr12-35598994-INS-66
chr12-35602646-INS-467
chr12-35622572-INS-633
chr12-35648026-INS-60
chr12-35667706-INS-63
chr12-35668876-INS-175
chr12-35704293-INS-52
chr12-35719505-INS-52
chr12-35730534-INS-53
chr12-35731424-INS-607
chr12-35746352-INS-129
chr12-35772163-INS-515
chr12-35796770-INS-787
chr12-35804202-INS-60
chr12-35812660-INS-179
chr12-35872556-INS-6596
chr12-35891778-INS-88
chr12-35894130-INS-932
chr12-35904486-INS-636
chr12-35936550-INS-209
chr12-35937660-INS-329
chr12-35943329-INS-975
chr12-35965957-INS-6441
chr12-35967340-INS-62
chr12-35972187-INS-191
chr12-36002824-INS-2564
chr12-36014127-INS-6402
chr12-36022007-INS-1077
chr12-36028227-INS-6164
chr12-36032370-INS-58
chr12-36049259-INS-71
chr12-36050051-INS-168
chr12-36059773-INS-535
chr12-36068846-INS-123
chr12-36072188-INS-245

chr12-36094481-INS-58
chr12-36099418-INS-340
chr12-36132461-INS-52
chr12-36142372-INS-128
chr12-36192663-INS-124
chr12-36230410-INS-651
chr12-36245447-INS-494
chr12-36249572-INS-208
chr12-36263626-INS-109
chr12-36264204-INS-177
chr12-36281290-INS-53
chr12-36282417-INS-567
chr12-36294044-INS-797
chr12-36313453-INS-154
chr12-36322265-INS-54
chr12-36377255-INS-940
chr12-36380372-INS-7007
chr12-36400217-INS-862
chr12-36403715-INS-84
chr12-36484398-INS-399
chr12-36500134-INS-184
chr12-36503918-INS-110
chr12-36529339-INS-240
chr12-36534010-INS-114
chr12-36550713-INS-565
chr12-36563557-INS-68
chr12-36622432-INS-351
chr12-36632514-INS-262
chr12-36647337-INS-130
chr12-36656664-INS-341
chr12-36699059-INS-103
chr12-36705264-INS-8460
chr12-36742982-INS-101
chr12-36750529-INS-6373
chr12-36801666-INS-98
chr12-36802759-INS-151
chr12-36808531-INS-266
chr12-36818330-INS-6761
chr12-36818758-INS-7093
chr12-36832320-INS-132
chr12-36851003-INS-1888
chr12-36858170-INS-542
chr12-36888468-INS-56
chr12-36935551-INS-72
chr12-36936928-INS-374
chr12-36955477-INS-270
chr12-36956068-INS-60
chr12-36958537-INS-326
chr12-37025344-INS-404
chr12-37030342-INS-214
chr12-37034327-INS-4095
chr12-37038865-INS-190
chr12-37040532-INS-6718
chr12-37048060-INS-102
chr12-37051935-INS-67
chr12-37056457-INS-53
chr12-37076582-INS-430
chr12-37090739-INS-224
chr12-37131156-INS-216
chr12-37140521-INS-302
chr12-37160546-INS-66
chr12-37197542-INS-173
chr12-37231319-INS-528
chr12-37236640-INS-339
chr12-37248600-INS-56
chr12-37294218-INS-225
chr12-37296862-INS-50
chr12-37298334-INS-994
chr12-37336345-INS-72
chr12-37340401-INS-138
chr12-37363407-INS-99
chr12-37364670-INS-885
chr12-37383313-INS-80
chr12-37405271-INS-135
chr12-37433087-INS-6152
chr12-37436029-INS-833
chr12-37439494-INS-750

chr12-37451627-INS-130
chr12-37457485-INS-164
chr12-37470212-INS-118
chr12-37480726-INS-478
chr12-37481203-INS-206
chr12-37489320-INS-396
chr12-37501301-INS-127
chr12-37501895-INS-2755
chr12-37504103-INS-52
chr12-37540313-INS-64
chr12-37565418-INS-6786
chr12-37571676-INS-123
chr12-37606121-INS-54
chr12-37623239-INS-52
chr12-37637648-INS-154
chr12-37640278-INS-235
chr12-37644637-INS-4607
chr12-37670781-INS-633
chr12-37746269-INS-276
chr12-37747255-INS-243
chr12-37753894-INS-6436
chr12-37758373-INS-217
chr12-37812065-INS-360
chr12-37835529-INS-6435
chr12-37853730-INS-507
chr12-37860446-INS-58
chr12-37883694-INS-60
chr12-37884193-INS-64
chr12-37916966-INS-111
chr12-37919205-INS-546
chr12-37924322-INS-60
chr12-37925144-INS-72
chr12-37980302-INS-58
chr12-38013165-INS-123
chr12-38035981-INS-202
chr12-38051127-INS-148
chr12-38072627-INS-567
chr12-38079368-INS-6559
chr12-38086718-INS-79
chr12-38113154-INS-64
chr12-38120829-INS-397
chr12-38156800-INS-53
chr12-38212264-INS-6707
chr12-38230369-INS-2368
chr12-38235829-INS-246
chr12-38241896-INS-593
chr12-38259590-INS-61
chr12-38265258-INS-58
chr12-38286727-INS-98
chr12-38289714-INS-56
chr12-38290959-INS-157
chr12-38306787-INS-799
chr12-38322783-INS-148
chr12-38329107-INS-138
chr12-38363454-INS-992
chr12-38385176-INS-66
chr12-38389253-INS-194
chr12-38391435-INS-121
chr12-38417492-INS-58
chr12-38422496-INS-169
chr12-38423093-INS-6656
chr12-38475446-INS-6499
chr12-38478608-INS-84
chr12-38487721-INS-55
chr12-38495540-INS-62
chr12-38527108-INS-3647
chr12-38535506-INS-6247
chr12-38542745-INS-142
chr12-38589004-INS-87
chr12-38612316-INS-556
chr12-38642882-INS-57
chr12-38651476-INS-169
chr12-38671654-INS-60
chr12-38671815-INS-772
chr12-38672910-INS-7350
chr12-38677366-INS-134
chr12-38726832-INS-118

chr12-38736656-INS-204
chr12-38741371-INS-437
chr12-38754692-INS-108
chr12-38757628-INS-266
chr12-38759052-INS-70
chr12-38765802-INS-809
chr12-38802272-INS-328
chr12-38803819-INS-55
chr12-38880709-INS-59
chr12-38900032-INS-5548
chr12-38926156-INS-53
chr12-38933582-INS-126
chr12-38972878-INS-171
chr12-38975296-INS-441
chr12-39000844-INS-363
chr12-39005903-INS-203
chr12-39013239-INS-443
chr12-39017640-INS-348
chr12-39026875-INS-54
chr12-39034927-INS-4258
chr12-39051396-INS-315
chr12-39074816-INS-447
chr12-39143530-INS-138
chr12-39144138-INS-2745
chr12-39146337-INS-52
chr12-39149423-INS-91
chr12-39154355-INS-69
chr12-39174806-INS-114
chr12-39207962-INS-112
chr12-39298182-INS-355
chr12-39312170-INS-768
chr12-39316421-INS-52
chr12-39320932-INS-62
chr12-39330534-INS-8034
chr12-39366173-INS-868
chr12-39366258-INS-55
chr12-39385040-INS-54
chr12-39445922-INS-358
chr12-39462318-INS-56
chr12-39481872-INS-100
chr12-39488056-INS-54
chr12-39492271-INS-70
chr12-39503920-INS-261
chr12-39559843-INS-130
chr12-39566830-INS-214
chr12-39587096-INS-181
chr12-39606777-INS-6490
chr12-39645907-INS-54
chr12-39647393-INS-6870
chr12-39669820-INS-51
chr12-39689732-INS-302
chr12-39729094-INS-100
chr12-39732803-INS-335
chr12-39738842-INS-296
chr12-39776262-INS-60
chr12-39809234-INS-121
chr12-39821325-INS-1889
chr12-39853950-INS-53
chr12-39890040-INS-200
chr12-39895505-INS-432
chr12-39898833-INS-88
chr12-39903526-INS-377
chr12-39925305-INS-7185
chr12-39930123-INS-845
chr12-39930141-INS-94
chr12-39937485-INS-95
chr12-39940739-INS-514
chr12-39960312-INS-119
chr12-39967982-INS-199
chr12-39969556-INS-538
chr12-39971884-INS-7635
chr12-39977547-INS-64
chr12-39978163-INS-58
chr12-39982007-INS-75
chr12-39983053-INS-276
chr12-39996535-INS-60
chr12-39999119-INS-58

chr12-40026594-INS-805
chr12-40036237-INS-140
chr12-40040405-INS-491
chr12-40047960-INS-234
chr12-40069450-INS-145
chr12-40080335-INS-84
chr12-40091024-INS-433
chr12-40098379-INS-143
chr12-40105981-INS-199
chr12-40108742-INS-6411
chr12-40130105-INS-58
chr12-40130820-INS-60
chr12-40133176-INS-230
chr12-40163687-INS-1399
chr12-40169898-INS-170
chr12-40207596-INS-6514
chr12-40216819-INS-6618
chr12-40223094-INS-60
chr12-40232422-INS-75
chr12-40234446-INS-522
chr12-40235586-INS-71
chr12-40247677-INS-210
chr12-40258576-INS-469
chr12-40307000-INS-132
chr12-40322066-INS-289
chr12-40322631-INS-333
chr12-40324097-INS-72
chr12-40331938-INS-126
chr12-40334760-INS-65
chr12-40356087-INS-483
chr12-40360002-INS-82
chr12-40373454-INS-60
chr12-40385749-INS-310
chr12-40396731-INS-206
chr12-40447081-INS-750
chr12-40451554-INS-6601
chr12-40470651-INS-62
chr12-40482943-INS-582
chr12-40500039-INS-120
chr12-40505940-INS-82
chr12-40515288-INS-90
chr12-40545644-INS-60
chr12-40567318-INS-191
chr12-40569775-INS-498
chr12-40580415-INS-7532
chr12-40582137-INS-4128
chr12-40677373-INS-345
chr12-40695131-INS-180
chr12-40697743-INS-308
chr12-40705105-INS-67
chr12-40719297-INS-86
chr12-40724571-INS-6199
chr12-40729084-INS-6475
chr12-40731928-INS-92
chr12-40746680-INS-64
chr12-40747922-INS-62
chr12-40753327-INS-96
chr12-40755537-INS-252
chr12-40801187-INS-53
chr12-40828149-INS-75
chr12-40837536-INS-68
chr12-40841197-INS-400
chr12-40845908-INS-70
chr12-40871026-INS-252
chr12-40921033-INS-131
chr12-40922659-INS-244
chr12-40924291-INS-60
chr12-40938502-INS-152
chr12-40961733-INS-104
chr12-40980700-INS-657
chr12-40983051-INS-80
chr12-40993461-INS-180
chr12-41016769-INS-113
chr12-41027982-INS-130
chr12-41031389-INS-6854
chr12-41039875-INS-186
chr12-41043005-INS-155

chr12-41064826-INS-66
chr12-41066249-INS-60
chr12-41076365-INS-54
chr12-41078657-INS-53
chr12-41091463-INS-86
chr12-41098946-INS-663
chr12-41107162-INS-52
chr12-41117243-INS-751
chr12-41121850-INS-90
chr12-41143543-INS-63
chr12-41145032-INS-80
chr12-41165210-INS-200
chr12-41174133-INS-52
chr12-41182067-INS-1218
chr12-41186374-INS-579
chr12-41221384-INS-63
chr12-41225081-INS-54
chr12-41225662-INS-6861
chr12-41261249-INS-161
chr12-41265427-INS-83
chr12-41271361-INS-63
chr12-41276206-INS-86
chr12-41284837-INS-6459
chr12-41285723-INS-6791
chr12-41303621-INS-934
chr12-41305530-INS-64
chr12-41309584-INS-1442
chr12-41324762-INS-56
chr12-41364037-INS-56
chr12-41379237-INS-52
chr12-41403097-INS-598
chr12-41409484-INS-363
chr12-41425040-INS-767
chr12-41437229-INS-53
chr12-41445407-INS-586
chr12-41468619-INS-344
chr12-41473500-INS-93
chr12-41513692-INS-103
chr12-41520249-INS-60
chr12-41550182-INS-6996
chr12-41569745-INS-68
chr12-41577811-INS-106
chr12-41584649-INS-52
chr12-41594385-INS-58
chr12-41622558-INS-56
chr12-41623839-INS-3022
chr12-41644312-INS-540
chr12-41663572-INS-6902
chr12-41670428-INS-7128
chr12-41723415-INS-128
chr12-41724421-INS-1799
chr12-41737482-INS-6413
chr12-41754524-INS-203
chr12-41775336-INS-530
chr12-41788571-INS-85
chr12-41812104-INS-56
chr12-41826804-INS-96
chr12-41856082-INS-67
chr12-41872584-INS-51
chr12-41893156-INS-76
chr12-41918266-INS-6369
chr12-41925217-INS-6274
chr12-41933390-INS-4525
chr12-41946993-INS-168
chr12-41968263-INS-133
chr12-41969623-INS-121
chr12-41983714-INS-72
chr12-42003245-INS-7171
chr12-42009153-INS-6900
chr12-42009987-INS-147
chr12-42066980-INS-126
chr12-42076161-INS-445
chr12-42077132-INS-637
chr12-42091052-INS-6719
chr12-42097068-INS-94
chr12-42149885-INS-2816
chr12-42163858-INS-116

chr12-42174613-INS-66
chr12-42204400-INS-68
chr12-42208019-INS-65
chr12-42214284-INS-449
chr12-42257523-INS-508
chr12-42276615-INS-92
chr12-42302625-INS-96
chr12-42303873-INS-751
chr12-42305253-INS-6592
chr12-42328599-INS-66
chr12-42340019-INS-209
chr12-42349274-INS-52
chr12-42360608-INS-7042
chr12-42390837-INS-7092
chr12-42392646-INS-874
chr12-42453441-INS-58
chr12-42456089-INS-381
chr12-42461695-INS-52
chr12-42471298-INS-769
chr12-42510842-INS-168
chr12-42520996-INS-727
chr12-42524702-INS-6942
chr12-42532619-INS-2948
chr12-42538873-INS-54
chr12-42543247-INS-123
chr12-42548916-INS-903
chr12-42555671-INS-136
chr12-42565519-INS-184
chr12-42582880-INS-337
chr12-42597715-INS-666
chr12-42626332-INS-148
chr12-42679000-INS-246
chr12-42680643-INS-559
chr12-42689585-INS-1188
chr12-42723010-INS-766
chr12-42726823-INS-98
chr12-42741146-INS-72
chr12-42755641-INS-89
chr12-42771694-INS-1891
chr12-42772523-INS-365
chr12-42784317-INS-379
chr12-42796294-INS-200
chr12-42813490-INS-769
chr12-42847546-INS-2427
chr12-42852351-INS-7229
chr12-42862268-INS-816
chr12-42868661-INS-244
chr12-42898579-INS-7333
chr12-42909265-INS-61
chr12-42912094-INS-106
chr12-42927002-INS-167
chr12-42944729-INS-169
chr12-42950881-INS-2953
chr12-42983826-INS-711
chr12-42986548-INS-545
chr12-43006911-INS-64
chr12-43007458-INS-6647
chr12-43013087-INS-1891
chr12-43064929-INS-193
chr12-43076558-INS-448
chr12-43078319-INS-6732
chr12-43094486-INS-90
chr12-43097124-INS-239
chr12-43102333-INS-875
chr12-43140099-INS-6584
chr12-43142570-INS-154
chr12-43145094-INS-115
chr12-43155253-INS-6489
chr12-43160947-INS-518
chr12-43164214-INS-195
chr12-43167359-INS-85
chr12-43171630-INS-59
chr12-43194365-INS-87
chr12-43221264-INS-3158
chr12-43221613-INS-1387
chr12-43265713-INS-140
chr12-43278238-INS-57

chr12-43282400-INS-90
chr12-43286827-INS-168
chr12-43418804-INS-628
chr12-43427952-INS-3198
chr12-43428296-INS-72
chr12-43452374-INS-88
chr12-43464262-INS-656
chr12-43470311-INS-6507
chr12-43503867-INS-1684
chr12-43507552-INS-119
chr12-43522761-INS-308
chr12-43533109-INS-6572
chr12-43537986-INS-387
chr12-43539813-INS-830
chr12-43554914-INS-80
chr12-43555174-INS-471
chr12-43557523-INS-4395
chr12-43559626-INS-6995
chr12-43576625-INS-65
chr12-43577602-INS-165
chr12-43594852-INS-839
chr12-43597861-INS-285
chr12-43608923-INS-136
chr12-43640484-INS-68
chr12-43646548-INS-7745
chr12-43671511-INS-937
chr12-43675545-INS-585
chr12-43687014-INS-580
chr12-43702050-INS-9092
chr12-43703665-INS-3627
chr12-43730924-INS-861
chr12-43735986-INS-67
chr12-43795537-INS-89
chr12-43802081-INS-578
chr12-43808299-INS-439
chr12-43815236-INS-914
chr12-43831253-INS-251
chr12-43866281-INS-222
chr12-43914762-INS-3244
chr12-44016159-INS-1339
chr12-44020992-INS-78
chr12-44024795-INS-6466
chr12-44029982-INS-53
chr12-44062680-INS-975
chr12-44084062-INS-57
chr12-44086529-INS-105
chr12-44108257-INS-2315
chr12-44120892-INS-250
chr12-44127843-INS-184
chr12-44136246-INS-180
chr12-44184336-INS-52
chr12-44212973-INS-60
chr12-44215423-INS-927
chr12-44226669-INS-6209
chr12-44229171-INS-777
chr12-44239921-INS-144
chr12-44244150-INS-52
chr12-44252269-INS-496
chr12-44263841-INS-137
chr12-44274974-INS-52
chr12-44277620-INS-75
chr12-44328629-INS-779
chr12-44401382-INS-2838
chr12-44426739-INS-50
chr12-44439329-INS-139
chr12-44447706-INS-350
chr12-44457775-INS-160
chr12-44460720-INS-189
chr12-44468382-INS-196
chr12-44480747-INS-227
chr12-44522990-INS-89
chr12-44528676-INS-1627
chr12-44577743-INS-593
chr12-44591418-INS-78
chr12-44611548-INS-101
chr12-44675411-INS-212
chr12-44678143-INS-6233

chr12-44686983-INS-6307
chr12-44752728-INS-7172
chr12-44758983-INS-61
chr12-44772317-INS-400
chr12-44773605-INS-358
chr12-44776228-INS-341
chr12-44795398-INS-1449
chr12-44805043-INS-57
chr12-44819656-INS-98
chr12-44869299-INS-58
chr12-44876656-INS-234
chr12-44883751-INS-229
chr12-44892535-INS-318
chr12-44929067-INS-524
chr12-44932742-INS-104
chr12-44957375-INS-60
chr12-44973683-INS-115
chr12-44981769-INS-71
chr12-44989328-INS-69
chr12-45012930-INS-109
chr12-45017838-INS-173
chr12-45024434-INS-6338
chr12-45099442-INS-78
chr12-45110681-INS-52
chr12-45125999-INS-69
chr12-45132910-INS-2487
chr12-45140039-INS-252
chr12-45158297-INS-6894
chr12-45165331-INS-66
chr12-45181997-INS-73
chr12-45190283-INS-940
chr12-45204049-INS-848
chr12-45205326-INS-110
chr12-45209215-INS-178
chr12-45241878-INS-6801
chr12-45262083-INS-358
chr12-45352931-INS-96
chr12-45353026-INS-873
chr12-45368508-INS-60
chr12-45394780-INS-421
chr12-45446820-INS-76
chr12-45447149-INS-116
chr12-45449566-INS-516
chr12-45456669-INS-68
chr12-45477993-INS-144
chr12-45495962-INS-6640
chr12-45508507-INS-4806
chr12-45511030-INS-6445
chr12-45517150-INS-62
chr12-45551572-INS-179
chr12-45568252-INS-6832
chr12-45574734-INS-68
chr12-45586500-INS-159
chr12-45624468-INS-226
chr12-45658767-INS-67
chr12-45679268-INS-199
chr12-45685777-INS-6743
chr12-45686727-INS-235
chr12-45705920-INS-216
chr12-45723453-INS-1128
chr12-45744076-INS-186
chr12-45745873-INS-2474
chr12-45751514-INS-6621
chr12-45762517-INS-10242
chr12-45769352-INS-67
chr12-45782833-INS-438
chr12-45783182-INS-6816
chr12-45798587-INS-1743
chr12-45807680-INS-6489
chr12-45809558-INS-173
chr12-45822587-INS-6250
chr12-45822908-INS-143
chr12-45833357-INS-356
chr12-45843094-INS-248
chr12-45846276-INS-626
chr12-45858292-INS-351
chr12-45859921-INS-397

chr12-45863938-INS-139
chr12-45893157-INS-205
chr12-45919776-INS-465
chr12-45930169-INS-63
chr12-45936695-INS-7083
chr12-45939377-INS-118
chr12-45939452-INS-6011
chr12-45958884-INS-84
chr12-45987383-INS-69
chr12-46012760-INS-6670
chr12-46020821-INS-174
chr12-46048904-INS-72
chr12-46062531-INS-7281
chr12-46066335-INS-104
chr12-46070196-INS-90
chr12-46070252-INS-1888
chr12-46080698-INS-112
chr12-46091124-INS-80
chr12-46127795-INS-306
chr12-46143122-INS-64
chr12-46177504-INS-70
chr12-46183764-INS-992
chr12-46215980-INS-163
chr12-46231828-INS-142
chr12-46243649-INS-136
chr12-46264750-INS-445
chr12-46266773-INS-77
chr12-46272147-INS-182
chr12-46274934-INS-592
chr12-46281088-INS-137
chr12-46287005-INS-202
chr12-46293421-INS-64
chr12-46307127-INS-65
chr12-46325678-INS-3150
chr12-46336625-INS-54
chr12-46362820-INS-220
chr12-46375337-INS-59
chr12-46389419-INS-187
chr12-46409914-INS-100
chr12-46413132-INS-223
chr12-46417717-INS-52
chr12-46509141-INS-6244
chr12-46586597-INS-276
chr12-46601103-INS-487
chr12-46617322-INS-87
chr12-46633299-INS-6776
chr12-46728212-INS-187
chr12-46732850-INS-6681
chr12-46754657-INS-354
chr12-46755503-INS-65
chr12-46772769-INS-398
chr12-46782542-INS-64
chr12-46880339-INS-818
chr12-46899962-INS-53
chr12-46909620-INS-62
chr12-46919225-INS-61
chr12-46925374-INS-113
chr12-46927642-INS-1698
chr12-46933144-INS-74
chr12-46950561-INS-52
chr12-46956589-INS-898
chr12-46958666-INS-52
chr12-46966503-INS-146
chr12-46982180-INS-6399
chr12-46988953-INS-6646
chr12-46994645-INS-73
chr12-47009286-INS-2722
chr12-47043894-INS-282
chr12-47044223-INS-249
chr12-47048891-INS-6983
chr12-47051874-INS-165
chr12-47093555-INS-187
chr12-47109809-INS-136
chr12-47134532-INS-60
chr12-47139135-INS-60
chr12-47179826-INS-195
chr12-47203811-INS-96

chr12-47217373-INS-7375
chr12-47219701-INS-61
chr12-47230840-INS-194
chr12-47232951-INS-3241
chr12-47260585-INS-232
chr12-47339552-INS-508
chr12-47365063-INS-73
chr12-47372731-INS-124
chr12-47373500-INS-56
chr12-47386183-INS-333
chr12-47404349-INS-4786
chr12-47419303-INS-484
chr12-47419724-INS-6509
chr12-47437604-INS-129
chr12-47444138-INS-68
chr12-47471038-INS-125
chr12-47497572-INS-1893
chr12-47506353-INS-60
chr12-47518669-INS-2945
chr12-47526712-INS-210
chr12-47548089-INS-195
chr12-47548570-INS-322
chr12-47560668-INS-6199
chr12-47561777-INS-65
chr12-47595841-INS-102
chr12-47597696-INS-241
chr12-47603665-INS-2748
chr12-47615113-INS-6695
chr12-47620094-INS-3801
chr12-47623556-INS-6252
chr12-47633745-INS-173
chr12-47636487-INS-237
chr12-47661867-INS-74
chr12-47684363-INS-177
chr12-47688042-INS-2954
chr12-47690705-INS-361
chr12-47702383-INS-443
chr12-47705055-INS-535
chr12-47706439-INS-1882
chr12-47745693-INS-10492
chr12-47746242-INS-53
chr12-47747515-INS-1221
chr12-47751667-INS-94
chr12-47797269-INS-63
chr12-47808613-INS-105
chr12-47820690-INS-2618
chr12-47820740-INS-1380
chr12-47820772-INS-1176
chr12-47831827-INS-126
chr12-47854095-INS-52
chr12-47883968-INS-392
chr12-47886799-INS-817
chr12-47887900-INS-344
chr12-47897249-INS-4770
chr12-47905097-INS-5721
chr12-47913895-INS-328
chr12-47922284-INS-486
chr12-47975642-INS-6640
chr12-47976971-INS-68
chr12-47999721-INS-4534
chr12-48016538-INS-385
chr12-48048230-INS-60
chr12-48060984-INS-54
chr12-48064191-INS-1290
chr12-48064202-INS-3192
chr12-48075381-INS-128
chr12-48081489-INS-178
chr12-48088087-INS-244
chr12-48103706-INS-141
chr12-48124884-INS-54
chr12-48175961-INS-329
chr12-48190526-INS-985
chr12-48202343-INS-8016
chr12-48214232-INS-72
chr12-48243552-INS-268
chr12-48258715-INS-87
chr12-48260766-INS-192

chr12-48263983-INS-66
chr12-48279493-INS-302
chr12-48374321-INS-582
chr12-48387319-INS-213
chr12-48396512-INS-73
chr12-48452655-INS-184
chr12-48459467-INS-515
chr12-48472639-INS-197
chr12-48482298-INS-4313
chr12-48534831-INS-58
chr12-48538827-INS-138
chr12-48541996-INS-4200
chr12-48557749-INS-82
chr12-48558408-INS-6185
chr12-48566915-INS-5655
chr12-48629231-INS-181
chr12-48632376-INS-1007
chr12-48644340-INS-7218
chr12-48652808-INS-1319
chr12-48660384-INS-125
chr12-48662862-INS-6398
chr12-48666102-INS-53
chr12-48689740-INS-52
chr12-48703621-INS-6836
chr12-48721416-INS-62
chr12-48746964-INS-1624
chr12-48750438-INS-497
chr12-48782001-INS-439
chr12-48802551-INS-327
chr12-48804488-INS-168
chr12-48807924-INS-114
chr12-48810425-INS-52
chr12-48819412-INS-69
chr12-48829068-INS-126
chr12-48831523-INS-482
chr12-48842685-INS-70
chr12-48847615-INS-201
chr12-48850249-INS-404
chr12-48869042-INS-64
chr12-48869579-INS-3485
chr12-48877642-INS-58
chr12-48884446-INS-93
chr12-48906613-INS-92
chr12-48916097-INS-867
chr12-48935100-INS-53
chr12-48946867-INS-59
chr12-48976820-INS-52
chr12-48997342-INS-70
chr12-49001327-INS-78
chr12-49016824-INS-7093
chr12-49019809-INS-982
chr12-49024961-INS-4875
chr12-49026237-INS-544
chr12-49027786-INS-59
chr12-49030623-INS-112
chr12-49032125-INS-685
chr12-49049611-INS-134
chr12-49096239-INS-731
chr12-49105700-INS-6648
chr12-49111825-INS-462
chr12-49140712-INS-933
chr12-49155045-INS-6675
chr12-49162532-INS-7171
chr12-49171663-INS-214
chr12-49184781-INS-201
chr12-49192477-INS-451
chr12-49204531-INS-6961
chr12-49213308-INS-2821
chr12-49218305-INS-510
chr12-49226920-INS-328
chr12-49246804-INS-6392
chr12-49259895-INS-6382
chr12-49286457-INS-2694
chr12-49301815-INS-5268
chr12-49316753-INS-906
chr12-49327735-INS-55
chr12-49365358-INS-56

chr12-49406194-INS-63
chr12-49464041-INS-724
chr12-49477460-INS-488
chr12-49490849-INS-158
chr12-49494650-INS-542
chr12-49522717-INS-6926
chr12-49523889-INS-391
chr12-49535419-INS-512
chr12-49545576-INS-65
chr12-49548519-INS-1293
chr12-49564078-INS-484
chr12-49605182-INS-494
chr12-49611754-INS-81
chr12-49612914-INS-74
chr12-49631447-INS-338
chr12-49633449-INS-325
chr12-49641022-INS-124
chr12-49646908-INS-84
chr12-49663869-INS-391
chr12-49670031-INS-56
chr12-49687848-INS-81
chr12-49702689-INS-57
chr12-49702856-INS-479
chr12-49718770-INS-125
chr12-49722519-INS-178
chr12-49740399-INS-859
chr12-49757265-INS-153
chr12-49766504-INS-85
chr12-49769225-INS-58
chr12-49789511-INS-684
chr12-49803602-INS-84
chr12-49814319-INS-915
chr12-49817791-INS-56
chr12-49828609-INS-100
chr12-49834674-INS-62
chr12-49847355-INS-1870
chr12-49849881-INS-762
chr12-49857624-INS-53
chr12-49870818-INS-101
chr12-49886097-INS-196
chr12-49887334-INS-180
chr12-49902542-INS-2727
chr12-49915418-INS-6283
chr12-49917414-INS-101
chr12-49921410-INS-689
chr12-49937856-INS-395
chr12-49944950-INS-679
chr12-49955161-INS-143
chr12-49961120-INS-193
chr12-49985685-INS-62
chr12-50006781-INS-669
chr12-50017755-INS-256
chr12-50026072-INS-70
chr12-50043589-INS-55
chr12-50050438-INS-56
chr12-50056225-INS-100
chr12-50060976-INS-114
chr12-50061032-INS-668
chr12-50077297-INS-73
chr12-50081878-INS-175
chr12-50095545-INS-775
chr12-50106340-INS-51
chr12-50111973-INS-812
chr12-50140039-INS-6394
chr12-50145552-INS-6049
chr12-50156076-INS-52
chr12-50184954-INS-6890
chr12-50222986-INS-145
chr12-50240258-INS-82
chr12-50244108-INS-60
chr12-50288568-INS-106
chr12-50298760-INS-452
chr12-50327861-INS-221
chr12-50352504-INS-51
chr12-50368138-INS-2004
chr12-50381721-INS-745
chr12-50405761-INS-666

chr12-50418882-INS-58
chr12-50421158-INS-133
chr12-50443535-INS-6671
chr12-50463911-INS-150
chr12-50526215-INS-215
chr12-50529717-INS-382
chr12-50535953-INS-441
chr12-50591389-INS-245
chr12-50641146-INS-500
chr12-50709776-INS-5538
chr12-50714461-INS-6519
chr12-50720886-INS-119
chr12-50734523-INS-1798
chr12-50749743-INS-64
chr12-50769422-INS-54
chr12-50781128-INS-88
chr12-50782365-INS-1808
chr12-50805245-INS-292
chr12-50816704-INS-359
chr12-50856466-INS-317
chr12-50860436-INS-174
chr12-50860919-INS-6401
chr12-50873227-INS-55
chr12-50880178-INS-881
chr12-50884496-INS-528
chr12-50899730-INS-92
chr12-50906045-INS-56
chr12-50921800-INS-768
chr12-50927870-INS-951
chr12-50930255-INS-82
chr12-50938873-INS-6829
chr12-50965154-INS-5330
chr12-50994279-INS-69
chr12-51031442-INS-6920
chr12-51042087-INS-354
chr12-51050729-INS-198
chr12-51057586-INS-790
chr12-51070958-INS-67
chr12-51071895-INS-198
chr12-51105935-INS-1365
chr12-51115977-INS-75
chr12-51135229-INS-97
chr12-51138483-INS-166
chr12-51156671-INS-87
chr12-51163252-INS-401
chr12-51163800-INS-7167
chr12-51179336-INS-551
chr12-51210362-INS-7096
chr12-51214733-INS-131
chr12-51222722-INS-6712
chr12-51223144-INS-102
chr12-51232326-INS-312
chr12-51266085-INS-6488
chr12-51269894-INS-275
chr12-51279039-INS-57
chr12-51292842-INS-188
chr12-51295346-INS-52
chr12-51296043-INS-51
chr12-51306179-INS-58
chr12-51319356-INS-113
chr12-51326880-INS-70
chr12-51334141-INS-172
chr12-51345398-INS-6689
chr12-51349325-INS-7151
chr12-51367589-INS-233
chr12-51383414-INS-123
chr12-51386647-INS-317
chr12-51434912-INS-51
chr12-51437956-INS-161
chr12-51442754-INS-54
chr12-51447529-INS-102
chr12-51449825-INS-52
chr12-51451659-INS-772
chr12-51458306-INS-496
chr12-51493662-INS-80
chr12-51496953-INS-58
chr12-51497963-INS-283

chr12-51501730-INS-210
chr12-51530690-INS-203
chr12-51553257-INS-57
chr12-51555256-INS-188
chr12-51570283-INS-168
chr12-51590929-INS-51
chr12-51603231-INS-376
chr12-51607590-INS-76
chr12-51611960-INS-280
chr12-51621681-INS-422
chr12-51623159-INS-214
chr12-51625470-INS-61
chr12-51635913-INS-466
chr12-51668314-INS-1283
chr12-51668554-INS-213
chr12-51671234-INS-105
chr12-51685455-INS-104
chr12-51693329-INS-58
chr12-51698319-INS-51
chr12-51712872-INS-226
chr12-51717641-INS-265
chr12-51727016-INS-56
chr12-51746854-INS-264
chr12-51758524-INS-261
chr12-51762680-INS-152
chr12-51767315-INS-222
chr12-51770533-INS-622
chr12-51771584-INS-64
chr12-51772950-INS-186
chr12-51779989-INS-68
chr12-51791765-INS-470
chr12-51801522-INS-76
chr12-51807011-INS-215
chr12-51847385-INS-216
chr12-51870689-INS-197
chr12-51871327-INS-109
chr12-51886159-INS-86
chr12-51889858-INS-171
chr12-51892322-INS-181
chr12-51897724-INS-208
chr12-51898158-INS-7547
chr12-51903942-INS-54
chr12-51925399-INS-52
chr12-51927165-INS-200
chr12-51934632-INS-396
chr12-51938722-INS-63
chr12-51975470-INS-213
chr12-51982138-INS-60
chr12-51989318-INS-208
chr12-51998183-INS-5549
chr12-52001795-INS-141
chr12-52006899-INS-212
chr12-52011839-INS-168
chr12-52021530-INS-560
chr12-52024092-INS-55
chr12-52025476-INS-191
chr12-52031143-INS-329
chr12-52066214-INS-230
chr12-52082733-INS-400
chr12-52083809-INS-621
chr12-52089088-INS-56
chr12-52122268-INS-76
chr12-52126879-INS-498
chr12-52141348-INS-6442
chr12-52156362-INS-781
chr12-52164117-INS-6160
chr12-52177602-INS-54
chr12-52178223-INS-100
chr12-52181616-INS-56
chr12-52189911-INS-2692
chr12-52193247-INS-6535
chr12-52197711-INS-76
chr12-52200296-INS-205
chr12-52206925-INS-2334
chr12-52237286-INS-328
chr12-52258696-INS-57
chr12-52288811-INS-647

chr12-52290732-INS-294
chr12-52316811-INS-82
chr12-52330864-INS-1864
chr12-52351590-INS-889
chr12-52380531-INS-66
chr12-52403449-INS-5277
chr12-52409980-INS-570
chr12-52419441-INS-52
chr12-52427788-INS-59
chr12-52438410-INS-132
chr12-52440081-INS-64
chr12-52458276-INS-51
chr12-52474800-INS-54
chr12-52500489-INS-68
chr12-52528745-INS-60
chr12-52570827-INS-51
chr12-52610927-INS-210
chr12-52634162-INS-217
chr12-52635343-INS-205
chr12-52642838-INS-434
chr12-52654439-INS-93
chr12-52655726-INS-181
chr12-52665838-INS-125
chr12-52687877-INS-229
chr12-52702541-INS-91
chr12-52705052-INS-90
chr12-52710386-INS-201
chr12-52743656-INS-431
chr12-52752410-INS-96
chr12-52771554-INS-6252
chr12-52802373-INS-6698
chr12-52804050-INS-392
chr12-52833881-INS-260
chr12-52873526-INS-68
chr12-52907807-INS-312
chr12-52943300-INS-471
chr12-52953130-INS-85
chr12-52953842-INS-935
chr12-52953843-INS-181
chr12-52956364-INS-777
chr12-52957666-INS-83
chr12-52958878-INS-1893
chr12-52976286-INS-514
chr12-52976839-INS-458
chr12-52979096-INS-141
chr12-52980468-INS-56
chr12-52997304-INS-875
chr12-53007295-INS-564
chr12-53051601-INS-91
chr12-53056588-INS-68
chr12-53077483-INS-232
chr12-53102929-INS-80
chr12-53128686-INS-403
chr12-53135220-INS-212
chr12-53137642-INS-450
chr12-53144564-INS-79
chr12-53178507-INS-65
chr12-53182454-INS-304
chr12-53259716-INS-172
chr12-53266906-INS-71
chr12-53279040-INS-6580
chr12-53317207-INS-175
chr12-53319169-INS-56
chr12-53326075-INS-92
chr12-53352165-INS-52
chr12-53373029-INS-56
chr12-53377952-INS-51
chr12-53392386-INS-616
chr12-53411552-INS-4043
chr12-53414490-INS-110
chr12-53434370-INS-104
chr12-53437517-INS-624
chr12-53456815-INS-52
chr12-53458794-INS-54
chr12-53479086-INS-225
chr12-53500317-INS-86
chr12-53502066-INS-6708

chr12-53527420-INS-329
chr12-53532611-INS-96
chr12-53557346-INS-91
chr12-53563711-INS-74
chr12-53631762-INS-386
chr12-53650643-INS-646
chr12-53728043-INS-636
chr12-53740733-INS-77
chr12-53763456-INS-317
chr12-53768081-INS-240
chr12-53770700-INS-66
chr12-53820195-INS-6493
chr12-53827654-INS-223
chr12-53834298-INS-428
chr12-53847586-INS-1688
chr12-53881681-INS-119
chr12-53941391-INS-51
chr12-53953548-INS-72
chr12-53975648-INS-82
chr12-53991530-INS-64
chr12-54046712-INS-225
chr12-54078165-INS-90
chr12-54110264-INS-222
chr12-54128046-INS-51
chr12-54169925-INS-7256
chr12-54187492-INS-5540
chr12-54189487-INS-8985
chr12-54203299-INS-761
chr12-54208329-INS-94
chr12-54211866-INS-67
chr12-54220470-INS-321
chr12-54224014-INS-93
chr12-54233327-INS-74
chr12-54236382-INS-106
chr12-54248693-INS-183
chr12-54271000-INS-220
chr12-54292318-INS-504
chr12-54306609-INS-206
chr12-54308176-INS-81
chr12-54312718-INS-210
chr12-54320359-INS-140
chr12-54348461-INS-166
chr12-54352450-INS-84
chr12-54373513-INS-60
chr12-54375044-INS-93
chr12-54376021-INS-60
chr12-54384907-INS-168
chr12-54394277-INS-65
chr12-54398021-INS-51
chr12-54420219-INS-60
chr12-54462069-INS-430
chr12-54470514-INS-52
chr12-54475618-INS-5296
chr12-54480286-INS-400
chr12-54486394-INS-64
chr12-54490406-INS-78
chr12-54502377-INS-58
chr12-54512040-INS-217
chr12-54530040-INS-96
chr12-54546241-INS-116
chr12-54547965-INS-103
chr12-54563142-INS-63
chr12-54565696-INS-5411
chr12-54568065-INS-58
chr12-54591527-INS-62
chr12-54607922-INS-142
chr12-54612017-INS-101
chr12-54612540-INS-70
chr12-54622089-INS-70
chr12-54624103-INS-226
chr12-54637282-INS-80
chr12-54638401-INS-52
chr12-54641975-INS-83
chr12-54643464-INS-231
chr12-54646215-INS-293
chr12-54647677-INS-530
chr12-54668402-INS-61

chr12-54670891-INS-99
chr12-54687370-INS-104
chr12-54689274-INS-110
chr12-54691234-INS-611
chr12-54704140-INS-202
chr12-54716227-INS-193
chr12-54725677-INS-124
chr12-54728130-INS-168
chr12-54732220-INS-173
chr12-54738337-INS-579
chr12-54764144-INS-112
chr12-54766776-INS-194
chr12-54769008-INS-3046
chr12-54772144-INS-197
chr12-54775711-INS-104
chr12-54778342-INS-97
chr12-54779888-INS-76
chr12-54805030-INS-53
chr12-54812824-INS-52
chr12-54813399-INS-213
chr12-54815870-INS-227
chr12-54820255-INS-137
chr12-54838088-INS-114
chr12-54847772-INS-213
chr12-54848153-INS-53
chr12-54850460-INS-156
chr12-54852182-INS-175
chr12-54856880-INS-123
chr12-54863976-INS-194
chr12-54878779-INS-252
chr12-54880378-INS-58
chr12-54891054-INS-63
chr12-54892025-INS-188
chr12-54893329-INS-184
chr12-54934544-INS-175
chr12-54936444-INS-165
chr12-54942104-INS-659
chr12-54943905-INS-771
chr12-54957862-INS-222
chr12-54966061-INS-481
chr12-54967708-INS-130
chr12-54981187-INS-210
chr12-55000469-INS-121
chr12-55014014-INS-171
chr12-55025199-INS-81
chr12-55030479-INS-232
chr12-55041059-INS-1864
chr12-55056771-INS-6440
chr12-55058294-INS-211
chr12-55075347-INS-259
chr12-55080571-INS-55
chr12-55113948-INS-58
chr12-55156336-INS-239
chr12-55298701-INS-210
chr12-55309193-INS-207
chr12-55317693-INS-193
chr12-55320532-INS-58
chr12-55324772-INS-77
chr12-55327273-INS-236
chr12-55331410-INS-208
chr12-55332630-INS-734
chr12-55339389-INS-206
chr12-55354709-INS-276
chr12-55363478-INS-196
chr12-55376161-INS-676
chr12-55385170-INS-162
chr12-55402545-INS-252
chr12-55403678-INS-211
chr12-55409125-INS-155
chr12-55418194-INS-385
chr12-55433586-INS-240
chr12-55439218-INS-65
chr12-55448146-INS-173
chr12-55452657-INS-574
chr12-55468997-INS-538
chr12-55477663-INS-100
chr12-55514396-INS-206

chr12-55522229-INS-66
chr12-55525046-INS-52
chr12-55551338-INS-68
chr12-55559551-INS-61
chr12-55561255-INS-99
chr12-55572719-INS-265
chr12-55577325-INS-510
chr12-55590281-INS-6873
chr12-55594686-INS-200
chr12-55596044-INS-213
chr12-55603391-INS-385
chr12-55607062-INS-200
chr12-55626970-INS-151
chr12-55633862-INS-2980
chr12-55635323-INS-60
chr12-55706695-INS-60
chr12-55713171-INS-107
chr12-55735025-INS-7303
chr12-55756758-INS-54
chr12-55763720-INS-180
chr12-55789168-INS-55
chr12-55789622-INS-244
chr12-55791688-INS-223
chr12-55805151-INS-65
chr12-55820506-INS-145
chr12-55824284-INS-356
chr12-55828075-INS-238
chr12-55856497-INS-78
chr12-55862356-INS-434
chr12-55879383-INS-231
chr12-55886646-INS-61
chr12-55893101-INS-231
chr12-55893569-INS-397
chr12-55897041-INS-99
chr12-55902728-INS-52
chr12-55911479-INS-126
chr12-55943788-INS-370
chr12-55955972-INS-376
chr12-55960730-INS-324
chr12-55961262-INS-52
chr12-55964931-INS-52
chr12-55972337-INS-270
chr12-55976343-INS-99
chr12-55984381-INS-83
chr12-55993223-INS-89
chr12-56003687-INS-60
chr12-56004305-INS-194
chr12-56004335-INS-8799
chr12-56019189-INS-106
chr12-56070339-INS-240
chr12-56086479-INS-7131
chr12-56109202-INS-560
chr12-56135875-INS-66
chr12-56138981-INS-92
chr12-56176617-INS-62
chr12-56186129-INS-539
chr12-56188420-INS-96
chr12-56248903-INS-85
chr12-56271784-INS-107
chr12-56274652-INS-494
chr12-56285788-INS-6856
chr12-56310002-INS-159
chr12-56311357-INS-50
chr12-56335149-INS-546
chr12-56337441-INS-98
chr12-56376105-INS-418
chr12-56388049-INS-56
chr12-56393435-INS-60
chr12-56400721-INS-595
chr12-56425237-INS-75
chr12-56445549-INS-51
chr12-56450155-INS-64
chr12-56456308-INS-73
chr12-56469424-INS-94
chr12-56479738-INS-591
chr12-56492374-INS-936
chr12-56510356-INS-80

chr12-56527763-INS-56
chr12-56541469-INS-260
chr12-56543635-INS-72
chr12-56591461-INS-5213
chr12-56627913-INS-214
chr12-56657314-INS-99
chr12-56686676-INS-92
chr12-56702766-INS-430
chr12-56707039-INS-74
chr12-56712307-INS-64
chr12-56718409-INS-62
chr12-56739000-INS-63
chr12-56754617-INS-64
chr12-56762015-INS-443
chr12-56774508-INS-51
chr12-56807212-INS-213
chr12-56808717-INS-711
chr12-56812664-INS-581
chr12-56824596-INS-80
chr12-56840104-INS-66
chr12-56843533-INS-56
chr12-56849910-INS-148
chr12-56893580-INS-51
chr12-56906117-INS-51
chr12-56914502-INS-2050
chr12-56922721-INS-116
chr12-56938999-INS-418
chr12-56945941-INS-4600
chr12-56948957-INS-74
chr12-56956902-INS-345
chr12-56972568-INS-3027
chr12-56980694-INS-60
chr12-57012666-INS-951
chr12-57061922-INS-170
chr12-57064032-INS-614
chr12-57088220-INS-169
chr12-57091511-INS-64
chr12-57144705-INS-62
chr12-57153714-INS-64
chr12-57179534-INS-118
chr12-57226009-INS-87
chr12-57230029-INS-180
chr12-57233544-INS-151
chr12-57244573-INS-189
chr12-57257276-INS-311
chr12-57259615-INS-114
chr12-57266580-INS-51
chr12-57291615-INS-105
chr12-57295100-INS-559
chr12-57300252-INS-376
chr12-57306098-INS-52
chr12-57310708-INS-176
chr12-57312457-INS-82
chr12-57313500-INS-109
chr12-57344499-INS-90
chr12-57400106-INS-614
chr12-57405785-INS-58
chr12-57431666-INS-7019
chr12-57450167-INS-1469
chr12-57474652-INS-130
chr12-57479016-INS-758
chr12-57485693-INS-108
chr12-57488795-INS-275
chr12-57495110-INS-731
chr12-57500841-INS-709
chr12-57511587-INS-184
chr12-57524981-INS-62
chr12-57541511-INS-214
chr12-57545507-INS-344
chr12-57560904-INS-144
chr12-57568446-INS-65
chr12-57571900-INS-685
chr12-57601155-INS-78
chr12-57644375-INS-495
chr12-57661476-INS-85
chr12-57690019-INS-132
chr12-57700467-INS-341

chr12-57716685-INS-54
chr12-57734019-INS-78
chr12-57744856-INS-111
chr12-57748115-INS-245
chr12-57775673-INS-199
chr12-57800505-INS-480
chr12-57815300-INS-110
chr12-57820402-INS-465
chr12-57826640-INS-173
chr12-57827794-INS-5165
chr12-57837208-INS-6574
chr12-57847513-INS-420
chr12-57861043-INS-1894
chr12-57890702-INS-710
chr12-57891751-INS-1061
chr12-57895989-INS-295
chr12-57909001-INS-5869
chr12-57911198-INS-56
chr12-57912298-INS-917
chr12-57912578-INS-6690
chr12-57916889-INS-350
chr12-57926391-INS-5019
chr12-57940706-INS-229
chr12-57964398-INS-194
chr12-58011104-INS-453
chr12-58017371-INS-329
chr12-58022478-INS-84
chr12-58027783-INS-56
chr12-58028195-INS-153
chr12-58034937-INS-232
chr12-58052153-INS-55
chr12-58069333-INS-241
chr12-58072662-INS-1702
chr12-58077215-INS-417
chr12-58091670-INS-54
chr12-58119419-INS-7075
chr12-58121415-INS-53
chr12-58124210-INS-7790
chr12-58134937-INS-237
chr12-58160123-INS-120
chr12-58166574-INS-207
chr12-58173079-INS-610
chr12-58190673-INS-6452
chr12-58203603-INS-208
chr12-58241321-INS-64
chr12-58266053-INS-108
chr12-58266061-INS-505
chr12-58280501-INS-493
chr12-58286569-INS-169
chr12-58309287-INS-391
chr12-58320087-INS-73
chr12-58331304-INS-660
chr12-58344185-INS-258
chr12-58377563-INS-5793
chr12-58384419-INS-232
chr12-58421711-INS-136
chr12-58444702-INS-116
chr12-58480216-INS-424
chr12-58498570-INS-279
chr12-58546522-INS-200
chr12-58550183-INS-221
chr12-58582444-INS-52
chr12-58588336-INS-76
chr12-58588509-INS-262
chr12-58606068-INS-435
chr12-58609356-INS-59
chr12-58635768-INS-122
chr12-58705175-INS-105
chr12-58706729-INS-200
chr12-58719466-INS-342
chr12-58724215-INS-324
chr12-58731338-INS-858
chr12-58748934-INS-504
chr12-58760050-INS-104
chr12-58792123-INS-74
chr12-58809430-INS-126
chr12-58847139-INS-237

chr12-58858238-INS-157
chr12-58895809-INS-60
chr12-58971293-INS-452
chr12-58981979-INS-260
chr12-58988061-INS-160
chr12-59021249-INS-215
chr12-59028255-INS-154
chr12-59032576-INS-1748
chr12-59037290-INS-501
chr12-59043718-INS-115
chr12-59045756-INS-173
chr12-59048443-INS-6297
chr12-59049580-INS-121
chr12-59054506-INS-200
chr12-59061385-INS-230
chr12-59069153-INS-203
chr12-59072700-INS-220
chr12-59078363-INS-201
chr12-59081029-INS-183
chr12-59093717-INS-173
chr12-59098885-INS-4379
chr12-59104571-INS-192
chr12-59112162-INS-219
chr12-59117410-INS-70
chr12-59136531-INS-358
chr12-59144422-INS-149
chr12-59167706-INS-178
chr12-59197995-INS-74
chr12-59203171-INS-210
chr12-59211165-INS-529
chr12-59213990-INS-194
chr12-59214610-INS-66
chr12-59230074-INS-71
chr12-59232632-INS-56
chr12-59247182-INS-321
chr12-59253547-INS-101
chr12-59259283-INS-7093
chr12-59260280-INS-214
chr12-59268166-INS-349
chr12-59281469-INS-6804
chr12-59296527-INS-178
chr12-59309883-INS-400
chr12-59313337-INS-73
chr12-59314383-INS-966
chr12-59337406-INS-354
chr12-59361837-INS-5228
chr12-59375116-INS-54
chr12-59378198-INS-271
chr12-59379330-INS-78
chr12-59432407-INS-186
chr12-59437593-INS-259
chr12-59450143-INS-627
chr12-59458840-INS-6618
chr12-59468975-INS-2811
chr12-59472901-INS-64
chr12-59484224-INS-140
chr12-59523867-INS-140
chr12-59529519-INS-560
chr12-59591962-INS-639
chr12-59594855-INS-144
chr12-59598111-INS-82
chr12-59607763-INS-271
chr12-59625432-INS-6607
chr12-59639370-INS-184
chr12-59667924-INS-173
chr12-59671881-INS-403
chr12-59709157-INS-258
chr12-59717420-INS-751
chr12-59717443-INS-554
chr12-59728658-INS-98
chr12-59735311-INS-52
chr12-59743775-INS-390
chr12-59763673-INS-1310
chr12-59780263-INS-250
chr12-59815067-INS-80
chr12-59817006-INS-171
chr12-59842156-INS-56

chr12-59855864-INS-60
chr12-59861290-INS-66
chr12-59882341-INS-6917
chr12-59883099-INS-55
chr12-59896738-INS-209
chr12-59921459-INS-263
chr12-59931857-INS-343
chr12-59994308-INS-194
chr12-60029436-INS-198
chr12-60034338-INS-6585
chr12-60036228-INS-85
chr12-60074856-INS-151
chr12-60098973-INS-573
chr12-60106295-INS-272
chr12-60107663-INS-540
chr12-60111835-INS-229
chr12-60136589-INS-362
chr12-60142734-INS-76
chr12-60163308-INS-92
chr12-60201505-INS-582
chr12-60225320-INS-53
chr12-60226489-INS-169
chr12-60229523-INS-194
chr12-60231816-INS-734
chr12-60268147-INS-140
chr12-60306655-INS-57
chr12-60365029-INS-72
chr12-60380687-INS-420
chr12-60387171-INS-354
chr12-60397718-INS-110
chr12-60414756-INS-451
chr12-60436194-INS-5727
chr12-60455015-INS-53
chr12-60501556-INS-405
chr12-60502363-INS-56
chr12-60521855-INS-413
chr12-60535671-INS-268
chr12-60545395-INS-745
chr12-60559427-INS-166
chr12-60564189-INS-6994
chr12-60572989-INS-171
chr12-60585343-INS-142
chr12-60589461-INS-54
chr12-60599707-INS-765
chr12-60606299-INS-275
chr12-60624170-INS-78
chr12-60633096-INS-378
chr12-60717377-INS-325
chr12-60722474-INS-5600
chr12-60735315-INS-74
chr12-60769770-INS-3054
chr12-60814026-INS-989
chr12-60820974-INS-52
chr12-60827186-INS-6277
chr12-60850140-INS-72
chr12-60855660-INS-4024
chr12-60856495-INS-711
chr12-60875009-INS-5194
chr12-60880007-INS-276
chr12-60899316-INS-382
chr12-60910199-INS-1763
chr12-60913145-INS-6433
chr12-60928037-INS-64
chr12-60972359-INS-1281
chr12-60978138-INS-279
chr12-61003487-INS-140
chr12-61027013-INS-52
chr12-61037161-INS-257
chr12-61039108-INS-92
chr12-61082523-INS-134
chr12-61093314-INS-75
chr12-61095205-INS-78
chr12-61134224-INS-185
chr12-61139853-INS-403
chr12-61201477-INS-214
chr12-61218459-INS-746
chr12-61227464-INS-2314

chr12-61231982-INS-104
chr12-61261716-INS-57
chr12-61284267-INS-135
chr12-61290816-INS-66
chr12-61308430-INS-90
chr12-61357863-INS-130
chr12-61377190-INS-6265
chr12-61392470-INS-6394
chr12-61407564-INS-221
chr12-61427206-INS-154
chr12-61427264-INS-652
chr12-61430776-INS-3010
chr12-61435070-INS-353
chr12-61442841-INS-447
chr12-61447469-INS-58
chr12-61451275-INS-222
chr12-61456460-INS-56
chr12-61460499-INS-181
chr12-61474977-INS-92
chr12-61479834-INS-564
chr12-61481863-INS-5144
chr12-61543135-INS-829
chr12-61560894-INS-6908
chr12-61563420-INS-265
chr12-61591531-INS-413
chr12-61602172-INS-6810
chr12-61607158-INS-375
chr12-61614990-INS-109
chr12-61619895-INS-96
chr12-61632556-INS-72
chr12-61654528-INS-246
chr12-61663154-INS-54
chr12-61679799-INS-212
chr12-61723165-INS-60
chr12-61732732-INS-214
chr12-61739959-INS-472
chr12-61816485-INS-566
chr12-61832594-INS-85
chr12-61839916-INS-797
chr12-61854834-INS-6410
chr12-61871026-INS-65
chr12-61883740-INS-556
chr12-61898589-INS-459
chr12-61904070-INS-1364
chr12-61915493-INS-633
chr12-61925365-INS-523
chr12-61940256-INS-2487
chr12-61977223-INS-8010
chr12-61983843-INS-98
chr12-61987193-INS-1989
chr12-62050662-INS-2781
chr12-62065799-INS-63
chr12-62072237-INS-769
chr12-62075771-INS-69
chr12-62112633-INS-56
chr12-62133306-INS-157
chr12-62135019-INS-2755
chr12-62146035-INS-1890
chr12-62147642-INS-7096
chr12-62157648-INS-7234
chr12-62159324-INS-100
chr12-62160204-INS-1094
chr12-62205429-INS-296
chr12-62222578-INS-62
chr12-62237332-INS-251
chr12-62249406-INS-5017
chr12-62251596-INS-129
chr12-62259245-INS-3180
chr12-62269362-INS-6592
chr12-62269489-INS-339
chr12-62285220-INS-95
chr12-62289587-INS-51
chr12-62302287-INS-52
chr12-62311559-INS-6399
chr12-62396718-INS-84
chr12-62400967-INS-138
chr12-62407709-INS-104

chr12-62409580-INS-68
chr12-62413413-INS-96
chr12-62433940-INS-660
chr12-62437385-INS-524
chr12-62461942-INS-6721
chr12-62472039-INS-78
chr12-62486111-INS-73
chr12-62489958-INS-7166
chr12-62522458-INS-152
chr12-62531624-INS-718
chr12-62539865-INS-7115
chr12-62542186-INS-103
chr12-62544759-INS-7615
chr12-62545249-INS-170
chr12-62562939-INS-900
chr12-62567158-INS-752
chr12-62568043-INS-178
chr12-62595902-INS-212
chr12-62598907-INS-138
chr12-62604656-INS-552
chr12-62623282-INS-7031
chr12-62625099-INS-7082
chr12-62627321-INS-103
chr12-62675020-INS-375
chr12-62694261-INS-6717
chr12-62725838-INS-51
chr12-62748708-INS-253
chr12-62766666-INS-183
chr12-62772313-INS-397
chr12-62825835-INS-54
chr12-62841612-INS-56
chr12-62852106-INS-62
chr12-62858044-INS-69
chr12-62876806-INS-99
chr12-62877140-INS-7683
chr12-62881297-INS-825
chr12-62886400-INS-382
chr12-62887987-INS-6686
chr12-62898265-INS-171
chr12-62925561-INS-219
chr12-62950825-INS-427
chr12-62977342-INS-51
chr12-62982350-INS-690
chr12-62987074-INS-71
chr12-62991072-INS-1864
chr12-62997431-INS-52
chr12-63006949-INS-424
chr12-63030858-INS-55
chr12-63063172-INS-107
chr12-63073355-INS-81
chr12-63155991-INS-1033
chr12-63177371-INS-86
chr12-63188204-INS-466
chr12-63188225-INS-1730
chr12-63188240-INS-1249
chr12-63247530-INS-6863
chr12-63265030-INS-359
chr12-63276965-INS-87
chr12-63294077-INS-66
chr12-63305523-INS-73
chr12-63344122-INS-200
chr12-63363369-INS-244
chr12-63381208-INS-56
chr12-63395884-INS-59
chr12-63449799-INS-4865
chr12-63475751-INS-62
chr12-63485229-INS-6428
chr12-63521834-INS-59
chr12-63528540-INS-84
chr12-63546605-INS-225
chr12-63566419-INS-5188
chr12-63595160-INS-6317
chr12-63631443-INS-327
chr12-63670786-INS-422
chr12-63674549-INS-622
chr12-63701338-INS-357
chr12-63769004-INS-785

chr12-63784574-INS-399
chr12-63807008-INS-503
chr12-63815623-INS-159
chr12-63823314-INS-55
chr12-63825645-INS-56
chr12-63836398-INS-129
chr12-63872825-INS-56
chr12-63901054-INS-5174
chr12-63909668-INS-326
chr12-63913534-INS-84
chr12-63936985-INS-54
chr12-63968086-INS-319
chr12-63974696-INS-4356
chr12-63984332-INS-144
chr12-64019534-INS-437
chr12-64043060-INS-52
chr12-64055687-INS-327
chr12-64084211-INS-406
chr12-64096192-INS-54
chr12-64096506-INS-7108
chr12-64121433-INS-324
chr12-64130978-INS-706
chr12-64153651-INS-273
chr12-64162447-INS-535
chr12-64172654-INS-68
chr12-64212279-INS-1686
chr12-64215110-INS-200
chr12-64224801-INS-104
chr12-64226968-INS-6928
chr12-64241564-INS-6361
chr12-64249776-INS-4400
chr12-64277459-INS-65
chr12-64315332-INS-493
chr12-64339052-INS-74
chr12-64377019-INS-66
chr12-64421954-INS-263
chr12-64434041-INS-77
chr12-64472036-INS-312
chr12-64480430-INS-57
chr12-64519538-INS-144
chr12-64526555-INS-6428
chr12-64540684-INS-55
chr12-64551606-INS-147
chr12-64560253-INS-110
chr12-64596442-INS-6412
chr12-64597643-INS-291
chr12-64605782-INS-646
chr12-64609562-INS-392
chr12-64645601-INS-118
chr12-64665803-INS-235
chr12-64677932-INS-583
chr12-64699413-INS-451
chr12-64703051-INS-629
chr12-64710875-INS-62
chr12-64735278-INS-6379
chr12-64740289-INS-102
chr12-64741440-INS-545
chr12-64743328-INS-103
chr12-64759731-INS-230
chr12-64762588-INS-805
chr12-64796457-INS-124
chr12-64797334-INS-3261
chr12-64827762-INS-1214
chr12-64829947-INS-92
chr12-64837673-INS-71
chr12-64852419-INS-882
chr12-64855202-INS-640
chr12-64882731-INS-54
chr12-64883600-INS-282
chr12-64892998-INS-9548
chr12-64906093-INS-269
chr12-64907487-INS-5106
chr12-64909276-INS-290
chr12-64911435-INS-738
chr12-64917940-INS-621
chr12-64921954-INS-7203
chr12-64925357-INS-2844

chr12-64930365-INS-6235
chr12-64933629-INS-208
chr12-64954976-INS-7086
chr12-64961434-INS-57
chr12-64976263-INS-73
chr12-64982586-INS-193
chr12-64986678-INS-332
chr12-64993343-INS-56
chr12-64995354-INS-60
chr12-65020078-INS-6550
chr12-65021127-INS-62
chr12-65023642-INS-183
chr12-65034429-INS-51
chr12-65034924-INS-401
chr12-65040247-INS-200
chr12-65046175-INS-221
chr12-65049559-INS-356
chr12-65052597-INS-211
chr12-65062481-INS-484
chr12-65064009-INS-56
chr12-65085491-INS-131
chr12-65110238-INS-79
chr12-65131993-INS-178
chr12-65136877-INS-204
chr12-65158873-INS-94
chr12-65178769-INS-96
chr12-65220630-INS-275
chr12-65223439-INS-7065
chr12-65240509-INS-51
chr12-65245961-INS-950
chr12-65257974-INS-4345
chr12-65264263-INS-63
chr12-65304399-INS-231
chr12-65317837-INS-62
chr12-65340145-INS-64
chr12-65348865-INS-78
chr12-65349185-INS-177
chr12-65352334-INS-7181
chr12-65369079-INS-93
chr12-65369664-INS-3367
chr12-65406931-INS-122
chr12-65411607-INS-60
chr12-65443643-INS-523
chr12-65444551-INS-143
chr12-65446764-INS-224
chr12-65449662-INS-7002
chr12-65469782-INS-100
chr12-65582965-INS-118
chr12-65596942-INS-72
chr12-65606410-INS-555
chr12-65612516-INS-179
chr12-65616504-INS-62
chr12-65617035-INS-801
chr12-65638886-INS-111
chr12-65661396-INS-64
chr12-65716573-INS-52
chr12-65730697-INS-76
chr12-65734699-INS-200
chr12-65736708-INS-313
chr12-65759603-INS-174
chr12-65767974-INS-316
chr12-65772240-INS-64
chr12-65777033-INS-58
chr12-65778141-INS-840
chr12-65798685-INS-74
chr12-65824672-INS-6378
chr12-65857491-INS-75
chr12-65863179-INS-6439
chr12-65867987-INS-63
chr12-65913538-INS-68
chr12-65918965-INS-314
chr12-65931336-INS-166
chr12-65953554-INS-6802
chr12-65970777-INS-702
chr12-65971606-INS-137
chr12-65984551-INS-63
chr12-65989798-INS-696

chr12-65994357-INS-233
chr12-65997080-INS-185
chr12-66068667-INS-54
chr12-66074305-INS-56
chr12-66087111-INS-5707
chr12-66099645-INS-57
chr12-66124862-INS-66
chr12-66133693-INS-56
chr12-66140156-INS-180
chr12-66192728-INS-77
chr12-66199307-INS-58
chr12-66219056-INS-60
chr12-66220219-INS-56
chr12-66222808-INS-545
chr12-66261720-INS-610
chr12-66270080-INS-197
chr12-66273071-INS-52
chr12-66287770-INS-123
chr12-66288565-INS-6156
chr12-66290625-INS-624
chr12-66320103-INS-89
chr12-66335115-INS-490
chr12-66337681-INS-60
chr12-66339060-INS-54
chr12-66408447-INS-366
chr12-66411913-INS-52
chr12-66416491-INS-134
chr12-66439419-INS-269
chr12-66441004-INS-316
chr12-66485067-INS-191
chr12-66512486-INS-66
chr12-66531303-INS-169
chr12-66583461-INS-482
chr12-66603643-INS-120
chr12-66614108-INS-56
chr12-66627625-INS-68
chr12-66635205-INS-132
chr12-66644919-INS-344
chr12-66652324-INS-344
chr12-66696405-INS-68
chr12-66697853-INS-116
chr12-66713793-INS-203
chr12-66731034-INS-64
chr12-66754250-INS-189
chr12-66758635-INS-108
chr12-66769272-INS-279
chr12-66777382-INS-137
chr12-66796858-INS-56
chr12-66804631-INS-2844
chr12-66861210-INS-6363
chr12-66873170-INS-4494
chr12-66919073-INS-202
chr12-66925364-INS-384
chr12-66957383-INS-512
chr12-66998309-INS-81
chr12-66998314-INS-6838
chr12-66998614-INS-108
chr12-67039038-INS-138
chr12-67044614-INS-420
chr12-67054997-INS-71
chr12-67064577-INS-58
chr12-67079424-INS-220
chr12-67084755-INS-100
chr12-67096113-INS-647
chr12-67108199-INS-73
chr12-67129784-INS-52
chr12-67152856-INS-7171
chr12-67159047-INS-731
chr12-67160237-INS-64
chr12-67174070-INS-58
chr12-67207987-INS-86
chr12-67217679-INS-517
chr12-67223664-INS-80
chr12-67227282-INS-89
chr12-67271632-INS-411
chr12-67279214-INS-76
chr12-67292601-INS-209

chr12-67304892-INS-198
chr12-67322008-INS-199
chr12-67328697-INS-624
chr12-67357442-INS-101
chr12-67374702-INS-290
chr12-67378687-INS-486
chr12-67394904-INS-180
chr12-67402610-INS-76
chr12-67421613-INS-864
chr12-67459528-INS-63
chr12-67463777-INS-520
chr12-67499268-INS-66
chr12-67524463-INS-68
chr12-67531352-INS-74
chr12-67538264-INS-67
chr12-67540783-INS-447
chr12-67542467-INS-56
chr12-67543599-INS-181
chr12-67546212-INS-2764
chr12-67549362-INS-138
chr12-67570316-INS-84
chr12-67591760-INS-6461
chr12-67594823-INS-2704
chr12-67596484-INS-177
chr12-67598846-INS-381
chr12-67607907-INS-207
chr12-67609205-INS-96
chr12-67713587-INS-120
chr12-67731020-INS-271
chr12-67740365-INS-410
chr12-67770050-INS-97
chr12-67790023-INS-732
chr12-67808723-INS-89
chr12-67836892-INS-6452
chr12-67840074-INS-92
chr12-67841165-INS-6363
chr12-67848467-INS-51
chr12-67850607-INS-462
chr12-67884338-INS-6103
chr12-67884742-INS-58
chr12-67890320-INS-7225
chr12-67898393-INS-471
chr12-67899529-INS-5437
chr12-67936813-INS-193
chr12-67962093-INS-284
chr12-67982853-INS-55
chr12-67991957-INS-5425
chr12-67993968-INS-94
chr12-68008418-INS-223
chr12-68055529-INS-130
chr12-68074150-INS-6703
chr12-68075383-INS-378
chr12-68077070-INS-511
chr12-68080731-INS-249
chr12-68108527-INS-88
chr12-68109093-INS-573
chr12-68120726-INS-127
chr12-68130894-INS-365
chr12-68133854-INS-320
chr12-68139987-INS-670
chr12-68145202-INS-609
chr12-68159482-INS-56
chr12-68180049-INS-7740
chr12-68183196-INS-108
chr12-68186077-INS-6211
chr12-68225776-INS-138
chr12-68242224-INS-917
chr12-68250820-INS-60
chr12-68251592-INS-386
chr12-68264878-INS-52
chr12-68279912-INS-431
chr12-68305542-INS-770
chr12-68309681-INS-75
chr12-68330254-INS-116
chr12-68332532-INS-693
chr12-68341743-INS-320
chr12-68366311-INS-612

chr12-68370389-INS-310
chr12-68388702-INS-7370
chr12-68390212-INS-520
chr12-68396813-INS-56
chr12-68398161-INS-51
chr12-68415128-INS-1056
chr12-68429072-INS-104
chr12-68432181-INS-256
chr12-68440675-INS-69
chr12-68505882-INS-360
chr12-68549294-INS-106
chr12-68568434-INS-85
chr12-68584865-INS-84
chr12-68602032-INS-204
chr12-68633952-INS-56
chr12-68650669-INS-989
chr12-68654870-INS-200
chr12-68658355-INS-462
chr12-68659945-INS-139
chr12-68681287-INS-398
chr12-68695877-INS-405
chr12-68701208-INS-132
chr12-68711445-INS-79
chr12-68714083-INS-115
chr12-68732743-INS-60
chr12-68745496-INS-57
chr12-68771612-INS-74
chr12-68776914-INS-173
chr12-68779054-INS-296
chr12-68790137-INS-144
chr12-68804285-INS-6609
chr12-68811299-INS-94
chr12-68817749-INS-494
chr12-68818992-INS-96
chr12-68835709-INS-56
chr12-68860947-INS-270
chr12-68863093-INS-615
chr12-68882817-INS-171
chr12-68909921-INS-895
chr12-68910839-INS-64
chr12-68914400-INS-85
chr12-68918847-INS-666
chr12-68932133-INS-72
chr12-68935360-INS-96
chr12-68937649-INS-6182
chr12-68939858-INS-8412
chr12-68948229-INS-6604
chr12-68971129-INS-6698
chr12-68982462-INS-210
chr12-69015528-INS-241
chr12-69047422-INS-417
chr12-69100023-INS-221
chr12-69112225-INS-588
chr12-69119556-INS-330
chr12-69126828-INS-7261
chr12-69133634-INS-56
chr12-69140754-INS-55
chr12-69146732-INS-114
chr12-69153973-INS-475
chr12-69164947-INS-182
chr12-69167941-INS-146
chr12-69169428-INS-220
chr12-69170827-INS-56
chr12-69191941-INS-157
chr12-69219646-INS-473
chr12-69220860-INS-69
chr12-69241478-INS-78
chr12-69245729-INS-202
chr12-69252300-INS-180
chr12-69252720-INS-5011
chr12-69269433-INS-214
chr12-69277096-INS-382
chr12-69283631-INS-182
chr12-69285114-INS-198
chr12-69292755-INS-293
chr12-69301402-INS-262
chr12-69307608-INS-60

chr12-69311152-INS-202
chr12-69319063-INS-139
chr12-69324186-INS-270
chr12-69327966-INS-69
chr12-69332345-INS-197
chr12-69333494-INS-70
chr12-69335959-INS-213
chr12-69340501-INS-119
chr12-69350141-INS-523
chr12-69358598-INS-185
chr12-69372369-INS-66
chr12-69374833-INS-128
chr12-69396440-INS-238
chr12-69401058-INS-71
chr12-69434201-INS-210
chr12-69452023-INS-162
chr12-69453215-INS-205
chr12-69478006-INS-74
chr12-69490378-INS-140
chr12-69497632-INS-65
chr12-69525812-INS-52
chr12-69535572-INS-51
chr12-69551900-INS-219
chr12-69556247-INS-953
chr12-69556715-INS-163
chr12-69564085-INS-1630
chr12-69567853-INS-53
chr12-69612838-INS-66
chr12-69634364-INS-62
chr12-69645824-INS-208
chr12-69653317-INS-183
chr12-69659415-INS-108
chr12-69671016-INS-117
chr12-69671300-INS-408
chr12-69687966-INS-54
chr12-69694417-INS-52
chr12-69717393-INS-187
chr12-69720562-INS-173
chr12-69722281-INS-194
chr12-69761065-INS-83
chr12-69778482-INS-5461
chr12-69779239-INS-5520
chr12-69781382-INS-208
chr12-69790853-INS-54
chr12-69811920-INS-187
chr12-69817404-INS-3319
chr12-69817631-INS-204
chr12-69827084-INS-210
chr12-69834377-INS-312
chr12-69842007-INS-173
chr12-69848837-INS-4570
chr12-69848849-INS-343
chr12-69874895-INS-63
chr12-69887307-INS-149
chr12-69892685-INS-76
chr12-69900370-INS-206
chr12-69916383-INS-117
chr12-69961372-INS-487
chr12-70037493-INS-404
chr12-70039156-INS-422
chr12-70055241-INS-6194
chr12-70064272-INS-98
chr12-70072122-INS-6399
chr12-70123098-INS-74
chr12-70136746-INS-101
chr12-70168988-INS-159
chr12-70183830-INS-372
chr12-70188927-INS-55
chr12-70203711-INS-549
chr12-70221831-INS-97
chr12-70255212-INS-88
chr12-70286286-INS-56
chr12-70297351-INS-54
chr12-70319408-INS-66
chr12-70358159-INS-827
chr12-70368782-INS-864
chr12-70400378-INS-108

chr12-70423552-INS-955
chr12-70424244-INS-60
chr12-70434826-INS-51
chr12-70437105-INS-201
chr12-70448728-INS-72
chr12-70455796-INS-78
chr12-70476039-INS-113
chr12-70478264-INS-82
chr12-70492569-INS-441
chr12-70559596-INS-989
chr12-70569484-INS-316
chr12-70588084-INS-3091
chr12-70599130-INS-96
chr12-70615448-INS-228
chr12-70616741-INS-681
chr12-70618402-INS-6849
chr12-70663861-INS-86
chr12-70667150-INS-180
chr12-70677220-INS-81
chr12-70686009-INS-1885
chr12-70690308-INS-76
chr12-70692196-INS-243
chr12-70693720-INS-354
chr12-70721399-INS-60
chr12-70730576-INS-103
chr12-70744040-INS-60
chr12-70764135-INS-60
chr12-70777452-INS-185
chr12-70787122-INS-80
chr12-70792537-INS-81
chr12-70811666-INS-204
chr12-70819174-INS-259
chr12-70841798-INS-51
chr12-70842701-INS-184
chr12-70892480-INS-174
chr12-70913209-INS-170
chr12-70924695-INS-333
chr12-70944808-INS-182
chr12-70957341-INS-443
chr12-70979323-INS-207
chr12-70981773-INS-159
chr12-70995839-INS-169
chr12-70998858-INS-192
chr12-71007140-INS-75
chr12-71010766-INS-1890
chr12-71019311-INS-62
chr12-71025853-INS-167
chr12-71041697-INS-495
chr12-71043265-INS-237
chr12-71058751-INS-230
chr12-71065214-INS-164
chr12-71075232-INS-122
chr12-71116429-INS-137
chr12-71118476-INS-82
chr12-71146644-INS-222
chr12-71152919-INS-60
chr12-71163984-INS-845
chr12-71165860-INS-208
chr12-71167022-INS-199
chr12-71179525-INS-596
chr12-71184855-INS-373
chr12-71200442-INS-58
chr12-71207401-INS-52
chr12-71215060-INS-94
chr12-71233005-INS-60
chr12-71243278-INS-429
chr12-71246813-INS-54
chr12-71248324-INS-266
chr12-71264203-INS-2929
chr12-71287850-INS-102
chr12-71291071-INS-100
chr12-71293427-INS-372
chr12-71306256-INS-183
chr12-71310571-INS-392
chr12-71314347-INS-72
chr12-71342013-INS-211
chr12-71367072-INS-225

chr12-71373995-INS-54
chr12-71375653-INS-71
chr12-71392263-INS-199
chr12-71422675-INS-135
chr12-71440794-INS-67
chr12-71458652-INS-373
chr12-71462233-INS-106
chr12-71468057-INS-65
chr12-71468826-INS-376
chr12-71471391-INS-71
chr12-71472395-INS-352
chr12-71513605-INS-90
chr12-71546133-INS-203
chr12-71566863-INS-63
chr12-71570768-INS-115
chr12-71572035-INS-205
chr12-71591058-INS-126
chr12-71592574-INS-119
chr12-71603832-INS-2130
chr12-71654030-INS-64
chr12-71676936-INS-90
chr12-71681397-INS-847
chr12-71689289-INS-65
chr12-71716324-INS-63
chr12-71726046-INS-953
chr12-71729390-INS-102
chr12-71736028-INS-52
chr12-71761173-INS-79
chr12-71762200-INS-78
chr12-71766267-INS-260
chr12-71771875-INS-67
chr12-71775271-INS-173
chr12-71780204-INS-84
chr12-71785522-INS-75
chr12-71798378-INS-63
chr12-71806192-INS-340
chr12-71809149-INS-308
chr12-71820293-INS-96
chr12-71864606-INS-157
chr12-71934208-INS-175
chr12-71941798-INS-135
chr12-71952769-INS-56
chr12-71961268-INS-75
chr12-71963034-INS-122
chr12-71970455-INS-65
chr12-71978606-INS-66
chr12-72048977-INS-4596
chr12-72051055-INS-7323
chr12-72055429-INS-132
chr12-72058579-INS-506
chr12-72060808-INS-64
chr12-72090358-INS-190
chr12-72106246-INS-146
chr12-72138830-INS-5427
chr12-72140353-INS-52
chr12-72142850-INS-125
chr12-72159907-INS-187
chr12-72161122-INS-52
chr12-72162393-INS-677
chr12-72209717-INS-6450
chr12-72230701-INS-177
chr12-72239732-INS-6688
chr12-72246570-INS-333
chr12-72249843-INS-424
chr12-72268943-INS-292
chr12-72278746-INS-85
chr12-72312656-INS-64
chr12-72316089-INS-186
chr12-72325884-INS-72
chr12-72347418-INS-216
chr12-72354139-INS-99
chr12-72362777-INS-3512
chr12-72368519-INS-64
chr12-72373856-INS-218
chr12-72380501-INS-276
chr12-72466586-INS-329
chr12-72473949-INS-247

chr12-72479793-INS-122
chr12-72490349-INS-53
chr12-72491531-INS-182
chr12-72501604-INS-56
chr12-72503607-INS-497
chr12-72505107-INS-503
chr12-72515630-INS-58
chr12-72524175-INS-277
chr12-72580195-INS-557
chr12-72582640-INS-197
chr12-72612698-INS-680
chr12-72654926-INS-292
chr12-72680878-INS-51
chr12-72682691-INS-57
chr12-72694544-INS-164
chr12-72696158-INS-524
chr12-72700784-INS-405
chr12-72722815-INS-5512
chr12-72723328-INS-74
chr12-72725476-INS-65
chr12-72728736-INS-519
chr12-72738723-INS-209
chr12-72740968-INS-433
chr12-72741612-INS-69
chr12-72751148-INS-76
chr12-72755635-INS-121
chr12-72759131-INS-73
chr12-72762645-INS-365
chr12-72765561-INS-507
chr12-72771643-INS-217
chr12-72772945-INS-194
chr12-72777882-INS-93
chr12-72781557-INS-353
chr12-72783860-INS-191
chr12-72803003-INS-369
chr12-72808314-INS-52
chr12-72814021-INS-205
chr12-72820993-INS-57
chr12-72823684-INS-202
chr12-72826601-INS-205
chr12-72868668-INS-5349
chr12-72871357-INS-116
chr12-72892521-INS-88
chr12-72895723-INS-198
chr12-72898338-INS-150
chr12-72904585-INS-66
chr12-72925578-INS-285
chr12-72937391-INS-124
chr12-72945361-INS-83
chr12-72973189-INS-206
chr12-72985995-INS-54
chr12-72997506-INS-441
chr12-73006150-INS-53
chr12-73039124-INS-60
chr12-73056219-INS-97
chr12-73056496-INS-90
chr12-73074123-INS-894
chr12-73113241-INS-63
chr12-73114678-INS-62
chr12-73118383-INS-797
chr12-73135107-INS-60
chr12-73140612-INS-200
chr12-73149707-INS-96
chr12-73154413-INS-184
chr12-73162462-INS-152
chr12-73164001-INS-233
chr12-73165306-INS-64
chr12-73166668-INS-63
chr12-73169398-INS-77
chr12-73172770-INS-581
chr12-73174304-INS-210
chr12-73186806-INS-192
chr12-73230198-INS-440
chr12-73240552-INS-232
chr12-73244852-INS-56
chr12-73284058-INS-556
chr12-73301548-INS-117

chr12-73333091-INS-166
chr12-73354328-INS-204
chr12-73377122-INS-215
chr12-73383567-INS-69
chr12-73434474-INS-547
chr12-73447103-INS-167
chr12-73458906-INS-101
chr12-73470424-INS-206
chr12-73473079-INS-91
chr12-73501726-INS-167
chr12-73514413-INS-310
chr12-73528345-INS-102
chr12-73532452-INS-396
chr12-73541512-INS-259
chr12-73559723-INS-116
chr12-73576807-INS-66
chr12-73580556-INS-104
chr12-73585280-INS-64
chr12-73597841-INS-57
chr12-73608048-INS-228
chr12-73609861-INS-56
chr12-73614993-INS-203
chr12-73621245-INS-217
chr12-73622623-INS-155
chr12-73729397-INS-94
chr12-73767066-INS-56
chr12-73771718-INS-57
chr12-73782454-INS-134
chr12-73791841-INS-288
chr12-73804970-INS-64
chr12-73812973-INS-110
chr12-73819094-INS-70
chr12-73820030-INS-663
chr12-73831642-INS-52
chr12-73866521-INS-70
chr12-73870736-INS-400
chr12-73911304-INS-6388
chr12-73916878-INS-6278
chr12-73931265-INS-6170
chr12-73939718-INS-113
chr12-73947462-INS-64
chr12-73979014-INS-92
chr12-74020158-INS-56
chr12-74026513-INS-58
chr12-74040653-INS-99
chr12-74041902-INS-217
chr12-74059016-INS-161
chr12-74061052-INS-65
chr12-74071265-INS-567
chr12-74074228-INS-6886
chr12-74103019-INS-266
chr12-74121394-INS-282
chr12-74124745-INS-991
chr12-74127735-INS-118
chr12-74165816-INS-72
chr12-74166099-INS-89
chr12-74173720-INS-84
chr12-74210848-INS-258
chr12-74230225-INS-238
chr12-74289394-INS-56
chr12-74298775-INS-6409
chr12-74321804-INS-114
chr12-74360945-INS-266
chr12-74362827-INS-55
chr12-74367891-INS-54
chr12-74378234-INS-62
chr12-74393597-INS-527
chr12-74395050-INS-56
chr12-74398995-INS-401
chr12-74468745-INS-57
chr12-74473097-INS-86
chr12-74480260-INS-6375
chr12-74494004-INS-52
chr12-74512680-INS-10663
chr12-74520096-INS-2975
chr12-74539594-INS-68
chr12-74540829-INS-56

chr12-74559396-INS-58
chr12-74575688-INS-167
chr12-74580952-INS-60
chr12-74594326-INS-64
chr12-74596326-INS-670
chr12-74600234-INS-7376
chr12-74609461-INS-6678
chr12-74619030-INS-903
chr12-74656837-INS-66
chr12-74661749-INS-55
chr12-74676263-INS-70
chr12-74680719-INS-6445
chr12-74689604-INS-346
chr12-74715239-INS-228
chr12-74729666-INS-488
chr12-74737538-INS-738
chr12-74745015-INS-77
chr12-74748306-INS-7494
chr12-74766633-INS-55
chr12-74772949-INS-160
chr12-74788376-INS-55
chr12-74791187-INS-152
chr12-74793944-INS-256
chr12-74827718-INS-212
chr12-74830476-INS-106
chr12-74861383-INS-91
chr12-74869795-INS-205
chr12-74874780-INS-160
chr12-74926709-INS-74
chr12-74929091-INS-121
chr12-75023895-INS-82
chr12-75025218-INS-876
chr12-75035183-INS-92
chr12-75046803-INS-229
chr12-75066826-INS-171
chr12-75070430-INS-135
chr12-75074669-INS-76
chr12-75086776-INS-60
chr12-75095981-INS-54
chr12-75106558-INS-264
chr12-75116346-INS-840
chr12-75118073-INS-103
chr12-75145752-INS-201
chr12-75153437-INS-172
chr12-75159597-INS-64
chr12-75164639-INS-342
chr12-75182124-INS-259
chr12-75186910-INS-730
chr12-75190565-INS-123
chr12-75194143-INS-170
chr12-75196640-INS-63
chr12-75232583-INS-53
chr12-75242764-INS-2249
chr12-75247021-INS-70
chr12-75248216-INS-1860
chr12-75258025-INS-347
chr12-75263260-INS-158
chr12-75284215-INS-105
chr12-75288593-INS-132
chr12-75291952-INS-6575
chr12-75302574-INS-566
chr12-75313398-INS-263
chr12-75316662-INS-358
chr12-75322557-INS-9400
chr12-75325527-INS-396
chr12-75335029-INS-71
chr12-75343882-INS-113
chr12-75345623-INS-95
chr12-75373185-INS-132
chr12-75405982-INS-417
chr12-75413788-INS-56
chr12-75447810-INS-502
chr12-75453365-INS-498
chr12-75464214-INS-200
chr12-75469985-INS-215
chr12-75473319-INS-79
chr12-75487258-INS-2597

chr12-75502820-INS-78
chr12-75510592-INS-208
chr12-75553477-INS-60
chr12-75573259-INS-84
chr12-75581345-INS-60
chr12-75601750-INS-226
chr12-75606688-INS-125
chr12-75613093-INS-493
chr12-75614963-INS-408
chr12-75632873-INS-236
chr12-75656686-INS-148
chr12-75669994-INS-184
chr12-75671454-INS-226
chr12-75677401-INS-223
chr12-75692504-INS-260
chr12-75707107-INS-198
chr12-75708063-INS-173
chr12-75709708-INS-204
chr12-75727737-INS-64
chr12-75732530-INS-70
chr12-75738474-INS-981
chr12-75741660-INS-190
chr12-75746527-INS-72
chr12-75747371-INS-6413
chr12-75747587-INS-146
chr12-75754578-INS-57
chr12-75764574-INS-202
chr12-75788034-INS-222
chr12-75788538-INS-194
chr12-75791202-INS-83
chr12-75809813-INS-5412
chr12-75810565-INS-2928
chr12-75812387-INS-5159
chr12-75820547-INS-67
chr12-75832083-INS-70
chr12-75845392-INS-69
chr12-75848500-INS-70
chr12-75851827-INS-2986
chr12-75856176-INS-401
chr12-75893264-INS-70
chr12-75896466-INS-124
chr12-75902492-INS-198
chr12-75903867-INS-218
chr12-75905314-INS-124
chr12-75908397-INS-61
chr12-75917029-INS-66
chr12-75919405-INS-80
chr12-75940032-INS-54
chr12-75970799-INS-111
chr12-75974656-INS-60
chr12-75976348-INS-56
chr12-75995335-INS-110
chr12-75999236-INS-206
chr12-76007547-INS-160
chr12-76019414-INS-252
chr12-76034777-INS-195
chr12-76040402-INS-73
chr12-76041082-INS-179
chr12-76086427-INS-67
chr12-76092757-INS-195
chr12-76102154-INS-216
chr12-76109435-INS-399
chr12-76109675-INS-6400
chr12-76125057-INS-56
chr12-76140086-INS-166
chr12-76171898-INS-151
chr12-76177942-INS-203
chr12-76185137-INS-116
chr12-76202515-INS-749
chr12-76206338-INS-75
chr12-76211290-INS-117
chr12-76215216-INS-213
chr12-76230093-INS-81
chr12-76238295-INS-131
chr12-76238636-INS-83
chr12-76244684-INS-201
chr12-76249270-INS-468

chr12-76251752-INS-200
chr12-76280730-INS-257
chr12-76287757-INS-119
chr12-76308057-INS-55
chr12-76340292-INS-58
chr12-76348614-INS-241
chr12-76365220-INS-53
chr12-76383756-INS-195
chr12-76402326-INS-114
chr12-76405070-INS-226
chr12-76428013-INS-222
chr12-76431342-INS-57
chr12-76438766-INS-198
chr12-76446105-INS-56
chr12-76473425-INS-97
chr12-76486248-INS-215
chr12-76502805-INS-164
chr12-76506525-INS-176
chr12-76510530-INS-70
chr12-76514705-INS-119
chr12-76518027-INS-78
chr12-76538616-INS-174
chr12-76560851-INS-719
chr12-76572011-INS-196
chr12-76586274-INS-180
chr12-76589648-INS-52
chr12-76603557-INS-72
chr12-76630413-INS-138
chr12-76683247-INS-77
chr12-76710308-INS-52
chr12-76715801-INS-56
chr12-76725276-INS-197
chr12-76735738-INS-132
chr12-76774576-INS-79
chr12-76796074-INS-111
chr12-76797194-INS-214
chr12-76825979-INS-202
chr12-76828088-INS-150
chr12-76834616-INS-134
chr12-76858521-INS-90
chr12-76859103-INS-58
chr12-76872844-INS-593
chr12-76873419-INS-181
chr12-76874989-INS-164
chr12-76875065-INS-6402
chr12-76876862-INS-338
chr12-76878109-INS-208
chr12-76883096-INS-264
chr12-76892986-INS-80
chr12-76895666-INS-201
chr12-76899606-INS-408
chr12-76900257-INS-99
chr12-76916391-INS-387
chr12-76937048-INS-54
chr12-76940977-INS-61
chr12-76949109-INS-122
chr12-76965797-INS-218
chr12-76979541-INS-204
chr12-77015720-INS-86
chr12-77021549-INS-181
chr12-77027862-INS-61
chr12-77037955-INS-56
chr12-77039608-INS-209
chr12-77043787-INS-53
chr12-77057964-INS-205
chr12-77065948-INS-53
chr12-77074835-INS-871
chr12-77078562-INS-167
chr12-77079549-INS-189
chr12-77124464-INS-63
chr12-77153905-INS-105
chr12-77182582-INS-488
chr12-77190453-INS-79
chr12-77195537-INS-82
chr12-77203964-INS-905
chr12-77218718-INS-104
chr12-77229440-INS-155

chr12-77240168-INS-82
chr12-77247188-INS-165
chr12-77278398-INS-55
chr12-77298001-INS-434
chr12-77305406-INS-58
chr12-77315467-INS-55
chr12-77346596-INS-1892
chr12-77349590-INS-402
chr12-77351383-INS-918
chr12-77356201-INS-390
chr12-77358545-INS-7214
chr12-77379589-INS-166
chr12-77393626-INS-6522
chr12-77396298-INS-60
chr12-77397563-INS-310
chr12-77423588-INS-6669
chr12-77423795-INS-401
chr12-77446576-INS-983
chr12-77457704-INS-212
chr12-77474826-INS-358
chr12-77524323-INS-4344
chr12-77554577-INS-195
chr12-77558020-INS-84
chr12-77559534-INS-112
chr12-77570651-INS-400
chr12-77604450-INS-108
chr12-77614558-INS-54
chr12-77624389-INS-144
chr12-77638864-INS-954
chr12-77649624-INS-120
chr12-77652171-INS-221
chr12-77690411-INS-83
chr12-77698049-INS-358
chr12-77700145-INS-77
chr12-77707831-INS-55
chr12-77709662-INS-64
chr12-77757741-INS-8691
chr12-77782558-INS-7111
chr12-77783286-INS-399
chr12-77785214-INS-84
chr12-77859919-INS-473
chr12-77883511-INS-86
chr12-77940004-INS-60
chr12-77955129-INS-55
chr12-77971593-INS-537
chr12-77997900-INS-452
chr12-78012209-INS-149
chr12-78037379-INS-322
chr12-78038291-INS-79
chr12-78043052-INS-2778
chr12-78050339-INS-6074
chr12-78063641-INS-132
chr12-78072823-INS-959
chr12-78077857-INS-1885
chr12-78093468-INS-127
chr12-78103526-INS-6496
chr12-78109540-INS-223
chr12-78134393-INS-232
chr12-78138197-INS-106
chr12-78144478-INS-58
chr12-78161422-INS-64
chr12-78187620-INS-358
chr12-78194592-INS-6731
chr12-78199740-INS-70
chr12-78261003-INS-72
chr12-78315076-INS-387
chr12-78319314-INS-6901
chr12-78325796-INS-255
chr12-78398219-INS-6215
chr12-78409293-INS-200
chr12-78425592-INS-139
chr12-78443213-INS-322
chr12-78492461-INS-108
chr12-78497966-INS-118
chr12-78498211-INS-1921
chr12-78526301-INS-55
chr12-78545001-INS-80

chr12-78559214-INS-357
chr12-78599463-INS-531
chr12-78616754-INS-721
chr12-78622258-INS-274
chr12-78645232-INS-60
chr12-78650780-INS-225
chr12-78669398-INS-622
chr12-78702289-INS-199
chr12-78731921-INS-145
chr12-78739029-INS-78
chr12-78739458-INS-6469
chr12-78754218-INS-653
chr12-78783862-INS-60
chr12-78789626-INS-1407
chr12-78945857-INS-59
chr12-78952721-INS-68
chr12-78964833-INS-210
chr12-78979928-INS-68
chr12-78983209-INS-499
chr12-78988119-INS-112
chr12-78993545-INS-490
chr12-78999932-INS-84
chr12-79028552-INS-58
chr12-79059573-INS-201
chr12-79067388-INS-99
chr12-79099613-INS-80
chr12-79106344-INS-55
chr12-79132275-INS-208
chr12-79184129-INS-217
chr12-79195463-INS-72
chr12-79208642-INS-84
chr12-79228980-INS-58
chr12-79297422-INS-419
chr12-79306425-INS-127
chr12-79310286-INS-90
chr12-79331111-INS-326
chr12-79340401-INS-613
chr12-79352036-INS-536
chr12-79356126-INS-58
chr12-79359104-INS-96
chr12-79392659-INS-510
chr12-79397685-INS-66
chr12-79408246-INS-72
chr12-79454000-INS-92
chr12-79492470-INS-54
chr12-79504960-INS-52
chr12-79537535-INS-174
chr12-79543564-INS-86
chr12-79563660-INS-78
chr12-79568186-INS-497
chr12-79603465-INS-151
chr12-79608194-INS-88
chr12-79617964-INS-58
chr12-79641408-INS-8447
chr12-79641702-INS-160
chr12-79650081-INS-488
chr12-79669906-INS-68
chr12-79671933-INS-104
chr12-79700440-INS-199
chr12-79715152-INS-116
chr12-79724417-INS-143
chr12-79726853-INS-215
chr12-79735429-INS-54
chr12-79783362-INS-202
chr12-79785170-INS-63
chr12-79865386-INS-58
chr12-79920546-INS-72
chr12-79968544-INS-60
chr12-79977902-INS-877
chr12-79985892-INS-94
chr12-79988679-INS-117
chr12-79991343-INS-351
chr12-80011096-INS-66
chr12-80027182-INS-205
chr12-80037039-INS-200
chr12-80088123-INS-73
chr12-80100164-INS-54

chr12-80114065-INS-56
chr12-80117813-INS-241
chr12-80151672-INS-61
chr12-80160306-INS-79
chr12-80168551-INS-90
chr12-80175802-INS-70
chr12-80203665-INS-149
chr12-80205644-INS-262
chr12-80286949-INS-1889
chr12-80312162-INS-196
chr12-80314143-INS-419
chr12-80321945-INS-119
chr12-80334824-INS-156
chr12-80344836-INS-105
chr12-80354015-INS-134
chr12-80358738-INS-172
chr12-80359963-INS-54
chr12-80377402-INS-56
chr12-80382211-INS-248
chr12-80407988-INS-179
chr12-80454405-INS-293
chr12-80460597-INS-908
chr12-80461804-INS-211
chr12-80477138-INS-223
chr12-80479031-INS-5679
chr12-80484560-INS-204
chr12-80516211-INS-207
chr12-80560399-INS-510
chr12-80604840-INS-187
chr12-80606367-INS-400
chr12-80611742-INS-73
chr12-80672447-INS-75
chr12-80687439-INS-217
chr12-80718265-INS-448
chr12-80725651-INS-291
chr12-80744966-INS-191
chr12-80749231-INS-243
chr12-80749236-INS-7149
chr12-80757929-INS-199
chr12-80764303-INS-92
chr12-80792348-INS-145
chr12-80828498-INS-260
chr12-80841546-INS-76
chr12-80847295-INS-75
chr12-80879868-INS-144
chr12-80881917-INS-324
chr12-80889088-INS-623
chr12-80930288-INS-52
chr12-80938231-INS-750
chr12-80939783-INS-139
chr12-80964055-INS-126
chr12-80982880-INS-355
chr12-80985788-INS-67
chr12-81007757-INS-81
chr12-81031266-INS-52
chr12-81033537-INS-630
chr12-81041211-INS-73
chr12-81043120-INS-252
chr12-81053462-INS-207
chr12-81060218-INS-103
chr12-81087107-INS-201
chr12-81095770-INS-54
chr12-81097370-INS-185
chr12-81113260-INS-505
chr12-81114541-INS-298
chr12-81118661-INS-80
chr12-81145238-INS-331
chr12-81169960-INS-586
chr12-81174034-INS-56
chr12-81194184-INS-64
chr12-81197299-INS-64
chr12-81204140-INS-75
chr12-81225607-INS-106
chr12-81240218-INS-168
chr12-81263219-INS-64
chr12-81275240-INS-70
chr12-81332006-INS-6754

chr12-81334641-INS-522
chr12-81342413-INS-252
chr12-81344082-INS-449
chr12-81345890-INS-509
chr12-81346822-INS-639
chr12-81363802-INS-66
chr12-81370290-INS-1112
chr12-81374452-INS-82
chr12-81376815-INS-56
chr12-81452598-INS-128
chr12-81455508-INS-242
chr12-81477078-INS-472
chr12-81480917-INS-401
chr12-81501580-INS-325
chr12-81509849-INS-203
chr12-81534555-INS-130
chr12-81550894-INS-151
chr12-81560767-INS-68
chr12-81564138-INS-200
chr12-81608277-INS-52
chr12-81620169-INS-587
chr12-81644328-INS-994
chr12-81644550-INS-200
chr12-81652846-INS-67
chr12-81653573-INS-6627
chr12-81658305-INS-5630
chr12-81738363-INS-122
chr12-81769717-INS-52
chr12-81776774-INS-113
chr12-81799694-INS-128
chr12-81815779-INS-260
chr12-81818134-INS-367
chr12-81845493-INS-115
chr12-81855787-INS-378
chr12-81860387-INS-52
chr12-81866637-INS-1864
chr12-81869025-INS-84
chr12-81878544-INS-193
chr12-81881603-INS-486
chr12-81921957-INS-69
chr12-81931974-INS-289
chr12-81932203-INS-291
chr12-81954676-INS-515
chr12-81977219-INS-52
chr12-81983539-INS-169
chr12-81987644-INS-388
chr12-81998135-INS-624
chr12-82012245-INS-112
chr12-82022334-INS-210
chr12-82032412-INS-240
chr12-82040724-INS-68
chr12-82058442-INS-572
chr12-82061267-INS-347
chr12-82092093-INS-347
chr12-82100914-INS-108
chr12-82118057-INS-82
chr12-82121925-INS-76
chr12-82130911-INS-199
chr12-82159419-INS-63
chr12-82160351-INS-640
chr12-82196355-INS-54
chr12-82208553-INS-400
chr12-82210873-INS-59
chr12-82241055-INS-78
chr12-82246861-INS-320
chr12-82250039-INS-6134
chr12-82250410-INS-129
chr12-82313016-INS-506
chr12-82320022-INS-255
chr12-82328403-INS-936
chr12-82344802-INS-70
chr12-82345553-INS-66
chr12-82364695-INS-109
chr12-82365457-INS-3292
chr12-82365496-INS-709
chr12-82365674-INS-3509
chr12-82366006-INS-1202

chr12-82400872-INS-53
chr12-82415615-INS-179
chr12-82496141-INS-52
chr12-82524315-INS-475
chr12-82528772-INS-358
chr12-82530427-INS-62
chr12-82534381-INS-62
chr12-82539160-INS-172
chr12-82540316-INS-52
chr12-82562073-INS-84
chr12-82564421-INS-3327
chr12-82578992-INS-422
chr12-82583646-INS-8245
chr12-82601068-INS-183
chr12-82606834-INS-421
chr12-82612006-INS-179
chr12-82654425-INS-129
chr12-82713736-INS-1894
chr12-82718440-INS-55
chr12-82753926-INS-310
chr12-82773831-INS-5549
chr12-82817655-INS-888
chr12-82821118-INS-317
chr12-82828879-INS-83
chr12-82841126-INS-6700
chr12-82859568-INS-201
chr12-82886371-INS-62
chr12-82889370-INS-78
chr12-82892466-INS-279
chr12-82894177-INS-141
chr12-82901753-INS-66
chr12-82904924-INS-670
chr12-82918567-INS-117
chr12-82941112-INS-58
chr12-82953906-INS-343
chr12-82970209-INS-108
chr12-82992234-INS-214
chr12-83015520-INS-87
chr12-83017909-INS-74
chr12-83032672-INS-205
chr12-83060016-INS-88
chr12-83067214-INS-60
chr12-83069084-INS-75
chr12-83090440-INS-116
chr12-83123728-INS-112
chr12-83126201-INS-828
chr12-83153806-INS-163
chr12-83196390-INS-883
chr12-83209945-INS-62
chr12-83223771-INS-196
chr12-83226174-INS-68
chr12-83242930-INS-1056
chr12-83242998-INS-676
chr12-83244913-INS-186
chr12-83277324-INS-431
chr12-83292436-INS-75
chr12-83315438-INS-5221
chr12-83380964-INS-163
chr12-83385954-INS-104
chr12-83390833-INS-56
chr12-83418882-INS-52
chr12-83466081-INS-184
chr12-83475657-INS-226
chr12-83481060-INS-246
chr12-83485160-INS-208
chr12-83488730-INS-122
chr12-83517572-INS-233
chr12-83533441-INS-68
chr12-83546576-INS-171
chr12-83548476-INS-604
chr12-83548485-INS-71
chr12-83555396-INS-193
chr12-83572608-INS-202
chr12-83587243-INS-255
chr12-83589456-INS-184
chr12-83602352-INS-93
chr12-83612316-INS-96

chr12-83616506-INS-194
chr12-83620578-INS-191
chr12-83623882-INS-198
chr12-83631226-INS-70
chr12-83631549-INS-71
chr12-83639670-INS-224
chr12-83651045-INS-68
chr12-83659575-INS-56
chr12-83660186-INS-136
chr12-83666609-INS-155
chr12-83669661-INS-124
chr12-83697544-INS-186
chr12-83705313-INS-174
chr12-83712722-INS-188
chr12-83720779-INS-202
chr12-83726337-INS-347
chr12-83754097-INS-181
chr12-83776388-INS-455
chr12-83783344-INS-215
chr12-83793888-INS-434
chr12-83796906-INS-166
chr12-83797653-INS-191
chr12-83817996-INS-75
chr12-83819629-INS-203
chr12-83820442-INS-100
chr12-83821803-INS-82
chr12-83833832-INS-255
chr12-83852887-INS-54
chr12-83853777-INS-205
chr12-83861286-INS-200
chr12-83863360-INS-200
chr12-83865378-INS-839
chr12-83892102-INS-212
chr12-83893776-INS-400
chr12-83898229-INS-176
chr12-83907183-INS-199
chr12-83913740-INS-8879
chr12-83917333-INS-154
chr12-83919119-INS-71
chr12-83930645-INS-51
chr12-83938885-INS-72
chr12-83946449-INS-253
chr12-83952644-INS-212
chr12-83989749-INS-194
chr12-83991181-INS-227
chr12-83992895-INS-111
chr12-83996671-INS-111
chr12-84012306-INS-176
chr12-84027377-INS-66
chr12-84032469-INS-345
chr12-84036159-INS-195
chr12-84050203-INS-2498
chr12-84051728-INS-54
chr12-84091656-INS-68
chr12-84096226-INS-154
chr12-84098726-INS-199
chr12-84114262-INS-90
chr12-84153130-INS-75
chr12-84163151-INS-70
chr12-84163528-INS-55
chr12-84169697-INS-66
chr12-84187047-INS-71
chr12-84218224-INS-88
chr12-84241188-INS-62
chr12-84274575-INS-186
chr12-84277319-INS-116
chr12-84283942-INS-147
chr12-84297388-INS-736
chr12-84298852-INS-112
chr12-84306422-INS-184
chr12-84308774-INS-91
chr12-84310879-INS-100
chr12-84314739-INS-498
chr12-84321528-INS-425
chr12-84323356-INS-191
chr12-84327267-INS-133
chr12-84329537-INS-1548

chr12-84329617-INS-189
chr12-84337549-INS-124
chr12-84340107-INS-272
chr12-84346073-INS-177
chr12-84349243-INS-244
chr12-84350493-INS-74
chr12-84356999-INS-57
chr12-84382057-INS-904
chr12-84383483-INS-266
chr12-84398897-INS-197
chr12-84399468-INS-75
chr12-84403787-INS-62
chr12-84462454-INS-154
chr12-84472112-INS-68
chr12-84482593-INS-305
chr12-84486888-INS-210
chr12-84495316-INS-181
chr12-84535395-INS-213
chr12-84541815-INS-89
chr12-84562438-INS-390
chr12-84572600-INS-54
chr12-84587371-INS-481
chr12-84590054-INS-66
chr12-84592073-INS-52
chr12-84594187-INS-94
chr12-84601239-INS-1111
chr12-84603075-INS-58
chr12-84609619-INS-102
chr12-84615159-INS-223
chr12-84621237-INS-68
chr12-84631493-INS-63
chr12-84642576-INS-232
chr12-84669434-INS-96
chr12-84677512-INS-57
chr12-84680316-INS-807
chr12-84684300-INS-237
chr12-84716519-INS-195
chr12-84723287-INS-292
chr12-84733667-INS-132
chr12-84747682-INS-5180
chr12-84766492-INS-204
chr12-84783820-INS-5938
chr12-84787463-INS-7069
chr12-84789234-INS-402
chr12-84815447-INS-881
chr12-84860238-INS-511
chr12-84865724-INS-54
chr12-84886179-INS-2956
chr12-84902267-INS-320
chr12-84919939-INS-84
chr12-84930023-INS-159
chr12-84946127-INS-90
chr12-84952489-INS-70
chr12-84971396-INS-77
chr12-84972712-INS-672
chr12-84978456-INS-64
chr12-84988879-INS-67
chr12-84993916-INS-72
chr12-84997228-INS-222
chr12-85001536-INS-176
chr12-85005847-INS-696
chr12-85008425-INS-216
chr12-85013016-INS-55
chr12-85046611-INS-175
chr12-85097733-INS-55
chr12-85135606-INS-62
chr12-85152813-INS-1855
chr12-85175926-INS-54
chr12-85178950-INS-132
chr12-85206831-INS-215
chr12-85210943-INS-201
chr12-85229967-INS-55
chr12-85233814-INS-104
chr12-85245500-INS-159
chr12-85286126-INS-146
chr12-85295491-INS-88
chr12-85321325-INS-371

chr12-85333562-INS-74
chr12-85371249-INS-498
chr12-85386524-INS-214
chr12-85396352-INS-71
chr12-85398412-INS-203
chr12-85527277-INS-51
chr12-85543181-INS-77
chr12-85552798-INS-343
chr12-85556100-INS-393
chr12-85570437-INS-416
chr12-85587241-INS-56
chr12-85591402-INS-200
chr12-85597540-INS-258
chr12-85624870-INS-204
chr12-85653274-INS-177
chr12-85659928-INS-422
chr12-85708553-INS-52
chr12-85715249-INS-72
chr12-85717730-INS-5233
chr12-85718400-INS-88
chr12-85719044-INS-52
chr12-85721410-INS-370
chr12-85739570-INS-58
chr12-85760030-INS-250
chr12-85764070-INS-90
chr12-85769967-INS-90
chr12-85779679-INS-186
chr12-85783588-INS-147
chr12-85785724-INS-221
chr12-85810474-INS-160
chr12-85849397-INS-209
chr12-85880721-INS-1493
chr12-85896733-INS-72
chr12-85920256-INS-232
chr12-85922612-INS-124
chr12-85952816-INS-51
chr12-86004613-INS-209
chr12-86014261-INS-67
chr12-86054964-INS-1219
chr12-86057828-INS-112
chr12-86063775-INS-195
chr12-86108275-INS-52
chr12-86126991-INS-64
chr12-86137537-INS-191
chr12-86160368-INS-230
chr12-86164280-INS-140
chr12-86165125-INS-144
chr12-86195735-INS-106
chr12-86201926-INS-58
chr12-86211654-INS-106
chr12-86214686-INS-212
chr12-86225031-INS-57
chr12-86230905-INS-204
chr12-86256349-INS-52
chr12-86261580-INS-77
chr12-86262615-INS-66
chr12-86283515-INS-111
chr12-86285967-INS-89
chr12-86288401-INS-53
chr12-86293134-INS-85
chr12-86347158-INS-1978
chr12-86394180-INS-54
chr12-86404941-INS-131
chr12-86415014-INS-52
chr12-86430277-INS-340
chr12-86440040-INS-85
chr12-86459912-INS-73
chr12-86489931-INS-89
chr12-86494492-INS-318
chr12-86525739-INS-52
chr12-86528442-INS-98
chr12-86549787-INS-58
chr12-86557150-INS-64
chr12-86568994-INS-122
chr12-86569062-INS-5315
chr12-86581951-INS-66
chr12-86608621-INS-129

chr12-86614210-INS-56
chr12-86619851-INS-210
chr12-86654633-INS-210
chr12-86678822-INS-393
chr12-86680400-INS-182
chr12-86707354-INS-112
chr12-86713499-INS-74
chr12-86755897-INS-1367
chr12-86786672-INS-568
chr12-86791042-INS-200
chr12-86799833-INS-172
chr12-86816056-INS-70
chr12-86832346-INS-385
chr12-86835403-INS-64
chr12-86847953-INS-64
chr12-86850495-INS-107
chr12-86859459-INS-157
chr12-86866701-INS-57
chr12-86869417-INS-174
chr12-86890544-INS-366
chr12-86911859-INS-174
chr12-86915066-INS-189
chr12-86949790-INS-68
chr12-86956207-INS-237
chr12-86968534-INS-209
chr12-86970376-INS-212
chr12-86971736-INS-208
chr12-86973375-INS-130
chr12-86981754-INS-54
chr12-86983328-INS-450
chr12-86989589-INS-206
chr12-86991164-INS-169
chr12-87016399-INS-52
chr12-87026271-INS-57
chr12-87034447-INS-6454
chr12-87044701-INS-54
chr12-87047115-INS-208
chr12-87056890-INS-56
chr12-87058114-INS-220
chr12-87097703-INS-52
chr12-87126157-INS-64
chr12-87132929-INS-228
chr12-87211724-INS-6452
chr12-87227807-INS-117
chr12-87235458-INS-154
chr12-87242404-INS-203
chr12-87256983-INS-211
chr12-87270114-INS-210
chr12-87292034-INS-57
chr12-87310996-INS-204
chr12-87315911-INS-203
chr12-87328069-INS-214
chr12-87330816-INS-445
chr12-87332650-INS-195
chr12-87344447-INS-125
chr12-87383704-INS-2390
chr12-87390909-INS-512
chr12-87410240-INS-165
chr12-87414275-INS-217
chr12-87427090-INS-64
chr12-87431868-INS-52
chr12-87442103-INS-130
chr12-87444222-INS-352
chr12-87444898-INS-160
chr12-87447275-INS-189
chr12-87449871-INS-72
chr12-87467618-INS-426
chr12-87489228-INS-114
chr12-87492253-INS-324
chr12-87494406-INS-163
chr12-87502275-INS-108
chr12-87535013-INS-57
chr12-87630839-INS-235
chr12-87637294-INS-69
chr12-87650654-INS-58
chr12-87728467-INS-60
chr12-87777034-INS-320

chr12-87777933-INS-142
chr12-87780577-INS-60
chr12-87801322-INS-405
chr12-87805611-INS-655
chr12-87897265-INS-73
chr12-88012922-INS-7080
chr12-88020069-INS-4970
chr12-88045642-INS-4503
chr12-88046324-INS-63
chr12-88051105-INS-185
chr12-88077382-INS-58
chr12-88154000-INS-55
chr12-88184368-INS-5260
chr12-88185705-INS-3024
chr12-88186230-INS-2725
chr12-88233621-INS-2872
chr12-88252918-INS-291
chr12-88255389-INS-64
chr12-88257134-INS-5555
chr12-88321387-INS-57
chr12-88325008-INS-84
chr12-88336201-INS-181
chr12-88347178-INS-96
chr12-88368601-INS-256
chr12-88377719-INS-71
chr12-88380637-INS-207
chr12-88385594-INS-568
chr12-88386273-INS-58
chr12-88464444-INS-84
chr12-88471094-INS-72
chr12-88484773-INS-156
chr12-88529644-INS-54
chr12-88534160-INS-664
chr12-88541860-INS-97
chr12-88556381-INS-105
chr12-88583410-INS-3769
chr12-88587849-INS-63
chr12-88600745-INS-79
chr12-88609040-INS-62
chr12-88647725-INS-142
chr12-88654100-INS-62
chr12-88655186-INS-6273
chr12-88680826-INS-316
chr12-88713273-INS-118
chr12-88721457-INS-53
chr12-88728575-INS-113
chr12-88747268-INS-74
chr12-88755504-INS-104
chr12-88763736-INS-108
chr12-88785257-INS-617
chr12-88811306-INS-54
chr12-88836168-INS-52
chr12-88842745-INS-116
chr12-88852006-INS-202
chr12-88894848-INS-155
chr12-88932194-INS-111
chr12-88938412-INS-51
chr12-88961221-INS-1312
chr12-88975204-INS-224
chr12-88995196-INS-490
chr12-89033174-INS-52
chr12-89061239-INS-76
chr12-89099884-INS-74
chr12-89107277-INS-65
chr12-89120790-INS-68
chr12-89124819-INS-210
chr12-89133262-INS-64
chr12-89134042-INS-120
chr12-89142065-INS-546
chr12-89149456-INS-192
chr12-89163255-INS-6179
chr12-89168852-INS-51
chr12-89176773-INS-64
chr12-89209464-INS-168
chr12-89224608-INS-246
chr12-89246053-INS-6463
chr12-89251813-INS-156

chr12-89270692-INS-72
chr12-89274469-INS-288
chr12-89278267-INS-104
chr12-89299651-INS-51
chr12-89344405-INS-75
chr12-89360030-INS-134
chr12-89362756-INS-2812
chr12-89391630-INS-806
chr12-89400762-INS-92
chr12-89417333-INS-297
chr12-89419695-INS-175
chr12-89431092-INS-72
chr12-89489908-INS-55
chr12-89546621-INS-66
chr12-89577801-INS-70
chr12-89597276-INS-217
chr12-89631311-INS-761
chr12-89643529-INS-291
chr12-89676810-INS-58
chr12-89688216-INS-7170
chr12-89766340-INS-5361
chr12-89815423-INS-495
chr12-89833950-INS-113
chr12-89868938-INS-68
chr12-89877821-INS-60
chr12-89882270-INS-6376
chr12-89926900-INS-52
chr12-89942429-INS-54
chr12-89944981-INS-186
chr12-89956427-INS-64
chr12-89964245-INS-84
chr12-89967278-INS-387
chr12-89972626-INS-6652
chr12-90001732-INS-85
chr12-90002826-INS-60
chr12-90015481-INS-399
chr12-90028835-INS-162
chr12-90046664-INS-6914
chr12-90061029-INS-100
chr12-90074691-INS-78
chr12-90093980-INS-228
chr12-90101127-INS-977
chr12-90104245-INS-259
chr12-90121514-INS-62
chr12-90130343-INS-55
chr12-90133690-INS-728
chr12-90148194-INS-100
chr12-90151258-INS-76
chr12-90161927-INS-167
chr12-90180185-INS-200
chr12-90180893-INS-60
chr12-90193723-INS-171
chr12-90197885-INS-401
chr12-90219578-INS-679
chr12-90226958-INS-69
chr12-90238601-INS-6273
chr12-90240352-INS-72
chr12-90259505-INS-80
chr12-90260764-INS-1091
chr12-90261617-INS-510
chr12-90266932-INS-57
chr12-90268719-INS-111
chr12-90269860-INS-66
chr12-90271921-INS-644
chr12-90295339-INS-142
chr12-90317397-INS-396
chr12-90337838-INS-112
chr12-90342567-INS-7191
chr12-90348530-INS-568
chr12-90376766-INS-59
chr12-90394229-INS-102
chr12-90397308-INS-265
chr12-90427297-INS-471
chr12-90435904-INS-151
chr12-90476026-INS-102
chr12-90481407-INS-7218
chr12-90482885-INS-101

chr12-90494966-INS-54
chr12-90496887-INS-195
chr12-90530992-INS-8506
chr12-90537545-INS-538
chr12-90545061-INS-536
chr12-90549979-INS-517
chr12-90567061-INS-106
chr12-90598029-INS-58
chr12-90607163-INS-980
chr12-90616206-INS-105
chr12-90656391-INS-8881
chr12-90670791-INS-72
chr12-90688808-INS-609
chr12-90707661-INS-199
chr12-90713287-INS-169
chr12-90727154-INS-902
chr12-90741586-INS-59
chr12-90757958-INS-549
chr12-90758469-INS-52
chr12-90769871-INS-202
chr12-90770022-INS-4544
chr12-90782670-INS-141
chr12-90787227-INS-648
chr12-90795534-INS-627
chr12-90819487-INS-53
chr12-90825557-INS-133
chr12-90830929-INS-101
chr12-90833851-INS-1359
chr12-90845730-INS-301
chr12-90854593-INS-401
chr12-90872172-INS-1890
chr12-90874341-INS-261
chr12-90894613-INS-89
chr12-90901228-INS-96
chr12-90902468-INS-3861
chr12-90903962-INS-52
chr12-90928578-INS-176
chr12-90936808-INS-142
chr12-90945264-INS-54
chr12-90953785-INS-6190
chr12-90962467-INS-2129
chr12-90971119-INS-125
chr12-90991832-INS-102
chr12-91025808-INS-246
chr12-91039598-INS-515
chr12-91103740-INS-55
chr12-91114573-INS-222
chr12-91121927-INS-6863
chr12-91128184-INS-210
chr12-91129899-INS-120
chr12-91149977-INS-90
chr12-91155340-INS-535
chr12-91175665-INS-407
chr12-91177641-INS-146
chr12-91186472-INS-354
chr12-91202201-INS-174
chr12-91221749-INS-90
chr12-91235273-INS-94
chr12-91237769-INS-600
chr12-91238751-INS-159
chr12-91241027-INS-594
chr12-91251827-INS-1445
chr12-91260152-INS-175
chr12-91271484-INS-405
chr12-91278898-INS-165
chr12-91282228-INS-68
chr12-91292966-INS-997
chr12-91297265-INS-51
chr12-91303298-INS-712
chr12-91305657-INS-216
chr12-91315328-INS-68
chr12-91351862-INS-68
chr12-91365582-INS-110
chr12-91374510-INS-67
chr12-91395071-INS-72
chr12-91397535-INS-734
chr12-91467670-INS-74

chr12-91471059-INS-972
chr12-91473414-INS-203
chr12-91476601-INS-195
chr12-91503933-INS-8740
chr12-91505427-INS-55
chr12-91513036-INS-587
chr12-91513061-INS-114
chr12-91518199-INS-59
chr12-91529142-INS-6315
chr12-91531798-INS-439
chr12-91536811-INS-410
chr12-91560298-INS-770
chr12-91575978-INS-681
chr12-91575978-INS-71
chr12-91577231-INS-75
chr12-91588180-INS-80
chr12-91591273-INS-83
chr12-91592606-INS-139
chr12-91596488-INS-373
chr12-91607269-INS-163
chr12-91613310-INS-214
chr12-91644468-INS-53
chr12-91684078-INS-110
chr12-91697348-INS-63
chr12-91723121-INS-52
chr12-91736165-INS-56
chr12-91742229-INS-196
chr12-91750072-INS-56
chr12-91760012-INS-202
chr12-91761171-INS-6529
chr12-91790102-INS-140
chr12-91827786-INS-795
chr12-91831272-INS-993
chr12-91857917-INS-3841
chr12-91861667-INS-53
chr12-91868103-INS-2896
chr12-91876827-INS-69
chr12-91880376-INS-351
chr12-91883588-INS-52
chr12-91893204-INS-73
chr12-91894870-INS-7140
chr12-91896721-INS-7304
chr12-91899930-INS-138
chr12-91900576-INS-4253
chr12-91922958-INS-371
chr12-91933178-INS-51
chr12-91936110-INS-1021
chr12-91964719-INS-117
chr12-91973764-INS-52
chr12-91994120-INS-850
chr12-91995312-INS-163
chr12-92009763-INS-72
chr12-92016776-INS-61
chr12-92032767-INS-83
chr12-92056982-INS-6239
chr12-92059481-INS-7082
chr12-92070022-INS-382
chr12-92088035-INS-442
chr12-92101174-INS-147
chr12-92122502-INS-240
chr12-92128048-INS-219
chr12-92132349-INS-1020
chr12-92140415-INS-222
chr12-92143873-INS-56
chr12-92147889-INS-196
chr12-92158021-INS-682
chr12-92172279-INS-60
chr12-92221400-INS-1886
chr12-92248431-INS-80
chr12-92251412-INS-155
chr12-92257519-INS-733
chr12-92264391-INS-173
chr12-92268046-INS-58
chr12-92278079-INS-63
chr12-92281795-INS-7190
chr12-92293186-INS-52
chr12-92305739-INS-767

chr12-92337046-INS-95
chr12-92343869-INS-147
chr12-92344252-INS-304
chr12-92370887-INS-52
chr12-92375366-INS-81
chr12-92381907-INS-1894
chr12-92393754-INS-569
chr12-92405093-INS-5211
chr12-92416186-INS-281
chr12-92416245-INS-990
chr12-92416875-INS-947
chr12-92433295-INS-51
chr12-92456069-INS-548
chr12-92480902-INS-426
chr12-92505326-INS-7112
chr12-92523350-INS-171
chr12-92549562-INS-754
chr12-92555253-INS-105
chr12-92566354-INS-462
chr12-92577223-INS-857
chr12-92597096-INS-354
chr12-92601289-INS-1055
chr12-92637899-INS-68
chr12-92647164-INS-430
chr12-92652117-INS-455
chr12-92656819-INS-400
chr12-92679136-INS-61
chr12-92698369-INS-7024
chr12-92699668-INS-461
chr12-92704380-INS-54
chr12-92715850-INS-59
chr12-92719669-INS-68
chr12-92733868-INS-217
chr12-92741891-INS-6615
chr12-92744663-INS-102
chr12-92757559-INS-62
chr12-92764410-INS-7123
chr12-92836239-INS-6704
chr12-92847254-INS-245
chr12-92898903-INS-205
chr12-92903817-INS-95
chr12-92913242-INS-307
chr12-92919428-INS-2966
chr12-92938671-INS-772
chr12-92956675-INS-167
chr12-92963165-INS-361
chr12-93024085-INS-963
chr12-93026524-INS-331
chr12-93033101-INS-58
chr12-93037747-INS-6463
chr12-93051589-INS-118
chr12-93056890-INS-162
chr12-93064323-INS-2931
chr12-93066290-INS-191
chr12-93079795-INS-452
chr12-93080902-INS-6218
chr12-93105186-INS-635
chr12-93150011-INS-382
chr12-93170839-INS-55
chr12-93178188-INS-205
chr12-93179253-INS-202
chr12-93184265-INS-64
chr12-93248483-INS-268
chr12-93270787-INS-243
chr12-93278787-INS-6699
chr12-93280165-INS-76
chr12-93281871-INS-88
chr12-93302185-INS-58
chr12-93310674-INS-52
chr12-93323483-INS-218
chr12-93324560-INS-88
chr12-93336139-INS-215
chr12-93359965-INS-79
chr12-93402026-INS-2761
chr12-93420367-INS-55
chr12-93425523-INS-4879
chr12-93433519-INS-78

chr12-93454367-INS-56
chr12-93519018-INS-68
chr12-93566888-INS-111
chr12-93580776-INS-826
chr12-93584847-INS-5316
chr12-93619400-INS-128
chr12-93635438-INS-106
chr12-93653504-INS-58
chr12-93684969-INS-169
chr12-93687213-INS-318
chr12-93719732-INS-1864
chr12-93746172-INS-90
chr12-93753188-INS-363
chr12-93757410-INS-55
chr12-93761404-INS-7740
chr12-93771138-INS-60
chr12-93783909-INS-62
chr12-93790167-INS-54
chr12-93802245-INS-339
chr12-93809057-INS-191
chr12-93819715-INS-88
chr12-93827645-INS-124
chr12-93830083-INS-114
chr12-93843354-INS-796
chr12-93848417-INS-63
chr12-93862769-INS-212
chr12-93904778-INS-69
chr12-93913946-INS-60
chr12-93922913-INS-90
chr12-93934368-INS-56
chr12-93961384-INS-64
chr12-93971284-INS-95
chr12-93979586-INS-429
chr12-93981106-INS-54
chr12-94042891-INS-2854
chr12-94068059-INS-5262
chr12-94088449-INS-5984
chr12-94092848-INS-230
chr12-94102666-INS-124
chr12-94126729-INS-6215
chr12-94151215-INS-133
chr12-94196686-INS-309
chr12-94201546-INS-292
chr12-94257164-INS-349
chr12-94325251-INS-88
chr12-94334735-INS-8057
chr12-94335386-INS-54
chr12-94344498-INS-381
chr12-94360754-INS-66
chr12-94366440-INS-978
chr12-94374325-INS-6940
chr12-94376935-INS-639
chr12-94376946-INS-5433
chr12-94413846-INS-6648
chr12-94426334-INS-5344
chr12-94434750-INS-112
chr12-94436045-INS-6675
chr12-94438641-INS-82
chr12-94444695-INS-324
chr12-94463812-INS-73
chr12-94505200-INS-184
chr12-94524106-INS-7101
chr12-94546260-INS-322
chr12-94566548-INS-360
chr12-94571983-INS-110
chr12-94596476-INS-6328
chr12-94711788-INS-226
chr12-94724216-INS-801
chr12-94750068-INS-576
chr12-94821678-INS-424
chr12-94825113-INS-600
chr12-94834388-INS-76
chr12-94841837-INS-53
chr12-94858477-INS-172
chr12-94861464-INS-628
chr12-94884119-INS-1811
chr12-94912248-INS-6375

chr12-94917748-INS-120
chr12-94975634-INS-51
chr12-94984672-INS-438
chr12-95001741-INS-172
chr12-95053089-INS-85
chr12-95073406-INS-6561
chr12-95074951-INS-393
chr12-95099230-INS-54
chr12-95136291-INS-4949
chr12-95223312-INS-74
chr12-95248241-INS-549
chr12-95267363-INS-61
chr12-95268997-INS-1055
chr12-95274935-INS-6881
chr12-95293957-INS-92
chr12-95307552-INS-636
chr12-95352415-INS-102
chr12-95378557-INS-862
chr12-95394889-INS-6483
chr12-95423272-INS-56
chr12-95426232-INS-56
chr12-95456638-INS-540
chr12-95458689-INS-56
chr12-95500748-INS-52
chr12-95504936-INS-54
chr12-95521199-INS-248
chr12-95530129-INS-52
chr12-95545024-INS-72
chr12-95627895-INS-58
chr12-95631692-INS-216
chr12-95656590-INS-144
chr12-95707888-INS-76
chr12-95713420-INS-403
chr12-95827688-INS-4458
chr12-95839467-INS-840
chr12-95842319-INS-69
chr12-95869121-INS-55
chr12-95874925-INS-70
chr12-95878606-INS-93
chr12-95928188-INS-386
chr12-95942423-INS-54
chr12-95957036-INS-126
chr12-95980391-INS-87
chr12-95994593-INS-460
chr12-95997681-INS-86
chr12-96009853-INS-71
chr12-96043777-INS-75
chr12-96087618-INS-93
chr12-96159037-INS-99
chr12-96185291-INS-385
chr12-96232416-INS-52
chr12-96240443-INS-63
chr12-96269807-INS-292
chr12-96273312-INS-1582
chr12-96305898-INS-54
chr12-96309121-INS-58
chr12-96312687-INS-208
chr12-96340949-INS-120
chr12-96354887-INS-53
chr12-96357242-INS-6696
chr12-96357926-INS-1786
chr12-96362089-INS-453
chr12-96362847-INS-77
chr12-96370125-INS-52
chr12-96373293-INS-6303
chr12-96374602-INS-403
chr12-96385381-INS-78
chr12-96404313-INS-6726
chr12-96405260-INS-2776
chr12-96417256-INS-489
chr12-96426471-INS-202
chr12-96447696-INS-1132
chr12-96465608-INS-218
chr12-96469061-INS-207
chr12-96471011-INS-58
chr12-96472267-INS-73
chr12-96497821-INS-122

chr12-96502027-INS-60
chr12-96573437-INS-56
chr12-96591467-INS-59
chr12-96599020-INS-72
chr12-96600158-INS-153
chr12-96639966-INS-505
chr12-96666976-INS-184
chr12-96685737-INS-166
chr12-96693991-INS-62
chr12-96707577-INS-376
chr12-96733284-INS-358
chr12-96742011-INS-142
chr12-96764722-INS-1252
chr12-96770803-INS-56
chr12-96773689-INS-444
chr12-96799378-INS-62
chr12-96809436-INS-5549
chr12-96827032-INS-59
chr12-96830132-INS-291
chr12-96858525-INS-57
chr12-96865393-INS-54
chr12-96881097-INS-168
chr12-96917699-INS-56
chr12-96962261-INS-604
chr12-97013702-INS-106
chr12-97021834-INS-127
chr12-97024201-INS-71
chr12-97052491-INS-137
chr12-97055814-INS-167
chr12-97068409-INS-498
chr12-97069500-INS-51
chr12-97071753-INS-92
chr12-97076292-INS-57
chr12-97085445-INS-104
chr12-97090153-INS-6660
chr12-97092605-INS-160
chr12-97104324-INS-6401
chr12-97108566-INS-71
chr12-97112006-INS-158
chr12-97121301-INS-62
chr12-97154403-INS-5363
chr12-97158246-INS-75
chr12-97160658-INS-550
chr12-97163903-INS-191
chr12-97169846-INS-347
chr12-97187432-INS-504
chr12-97188556-INS-99
chr12-97192643-INS-94
chr12-97208665-INS-985
chr12-97211823-INS-594
chr12-97231954-INS-60
chr12-97261330-INS-57
chr12-97266629-INS-74
chr12-97280893-INS-60
chr12-97300102-INS-314
chr12-97338496-INS-90
chr12-97356964-INS-196
chr12-97365011-INS-60
chr12-97395555-INS-56
chr12-97404288-INS-381
chr12-97435111-INS-263
chr12-97437751-INS-53
chr12-97451102-INS-60
chr12-97469121-INS-130
chr12-97500212-INS-64
chr12-97504745-INS-90
chr12-97508369-INS-361
chr12-97532401-INS-285
chr12-97560957-INS-3017
chr12-97560958-INS-632
chr12-97561749-INS-75
chr12-97578650-INS-334
chr12-97580074-INS-196
chr12-97609544-INS-70
chr12-97612036-INS-892
chr12-97628311-INS-68
chr12-97642509-INS-199

chr12-97657386-INS-6504
chr12-97692772-INS-6628
chr12-97708004-INS-257
chr12-97709347-INS-198
chr12-97728691-INS-75
chr12-97736659-INS-274
chr12-97805120-INS-57
chr12-97835567-INS-128
chr12-97837203-INS-64
chr12-97842661-INS-400
chr12-97867144-INS-61
chr12-97872082-INS-399
chr12-97880340-INS-52
chr12-97896824-INS-54
chr12-97907718-INS-471
chr12-97910324-INS-140
chr12-97942446-INS-114
chr12-97953409-INS-225
chr12-97970139-INS-132
chr12-97979071-INS-203
chr12-97992341-INS-52
chr12-97993692-INS-110
chr12-98027069-INS-5383
chr12-98037731-INS-195
chr12-98058810-INS-55
chr12-98070113-INS-53
chr12-98077415-INS-1378
chr12-98080847-INS-60
chr12-98087861-INS-199
chr12-98089488-INS-7435
chr12-98095615-INS-7392
chr12-98110559-INS-1177
chr12-98118324-INS-4944
chr12-98135367-INS-400
chr12-98149791-INS-3099
chr12-98152406-INS-7194
chr12-98157145-INS-55
chr12-98172911-INS-370
chr12-98198179-INS-219
chr12-98200768-INS-211
chr12-98226735-INS-55
chr12-98239373-INS-88
chr12-98246502-INS-130
chr12-98252618-INS-66
chr12-98256428-INS-1094
chr12-98260029-INS-944
chr12-98280093-INS-118
chr12-98283125-INS-62
chr12-98293647-INS-166
chr12-98296364-INS-62
chr12-98317976-INS-147
chr12-98347971-INS-160
chr12-98356763-INS-332
chr12-98358413-INS-881
chr12-98360602-INS-300
chr12-98379052-INS-86
chr12-98384039-INS-566
chr12-98440260-INS-71
chr12-98508953-INS-199
chr12-98521405-INS-106
chr12-98525880-INS-52
chr12-98543888-INS-83
chr12-98551831-INS-997
chr12-98565361-INS-69
chr12-98585288-INS-117
chr12-98610320-INS-75
chr12-98616214-INS-196
chr12-98618612-INS-141
chr12-98638717-INS-55
chr12-98646108-INS-68
chr12-98648740-INS-58
chr12-98664449-INS-82
chr12-98672652-INS-55
chr12-98674560-INS-222
chr12-98677646-INS-6458
chr12-98741262-INS-98
chr12-98759555-INS-85

chr12-98784272-INS-5192
chr12-98794650-INS-59
chr12-98795928-INS-105
chr12-98807238-INS-222
chr12-98809624-INS-58
chr12-98843204-INS-691
chr12-98864009-INS-240
chr12-98874444-INS-68
chr12-98896672-INS-56
chr12-98928495-INS-550
chr12-98934551-INS-204
chr12-98946123-INS-67
chr12-98955269-INS-5217
chr12-98964717-INS-4990
chr12-98986630-INS-84
chr12-99009316-INS-174
chr12-99012283-INS-51
chr12-99018106-INS-447
chr12-99029147-INS-70
chr12-99049163-INS-88
chr12-99081725-INS-461
chr12-99092402-INS-97
chr12-99131472-INS-77
chr12-99140984-INS-64
chr12-99221586-INS-255
chr12-99225978-INS-166
chr12-99227483-INS-58
chr12-99326438-INS-53
chr12-99339813-INS-640
chr12-99350094-INS-194
chr12-99355960-INS-58
chr12-99362277-INS-222
chr12-99371016-INS-183
chr12-99387548-INS-168
chr12-99394201-INS-163
chr12-99404174-INS-137
chr12-99407949-INS-65
chr12-99428465-INS-58
chr12-99439165-INS-83
chr12-99440537-INS-225
chr12-99457873-INS-92
chr12-99486819-INS-65
chr12-99507790-INS-87
chr12-99512259-INS-54
chr12-99516234-INS-54
chr12-99517159-INS-173
chr12-99526742-INS-1889
chr12-99544472-INS-113
chr12-99599056-INS-58
chr12-99615406-INS-53
chr12-99633610-INS-1835
chr12-99639789-INS-5532
chr12-99665573-INS-427
chr12-99671723-INS-221
chr12-99680272-INS-212
chr12-99748633-INS-406
chr12-99760551-INS-102
chr12-99763235-INS-102
chr12-99773621-INS-65
chr12-99790573-INS-225
chr12-99791798-INS-93
chr12-99794085-INS-558
chr12-99835590-INS-71
chr12-99838028-INS-146
chr12-99882813-INS-232
chr12-99892858-INS-127
chr12-99905808-INS-56
chr12-99928072-INS-80
chr12-99934704-INS-143
chr12-99938477-INS-66
chr12-99965561-INS-112
chr12-99970108-INS-59
chr12-99988942-INS-54
chr12-100005529-INS-128
chr12-100034864-INS-79
chr12-100040540-INS-70
chr12-100065399-INS-138

chr12-100074034-INS-149
chr12-100084241-INS-88
chr12-100105142-INS-189
chr12-100116881-INS-277
chr12-100119575-INS-231
chr12-100124681-INS-58
chr12-100137120-INS-2116
chr12-100139890-INS-203
chr12-100160525-INS-166
chr12-100182702-INS-102
chr12-100186595-INS-51
chr12-100187683-INS-80
chr12-100221672-INS-357
chr12-100226055-INS-71
chr12-100251190-INS-300
chr12-100263755-INS-397
chr12-100308902-INS-167
chr12-100344000-INS-159
chr12-100347586-INS-350
chr12-100352852-INS-73
chr12-100436056-INS-540
chr12-100440086-INS-808
chr12-100443056-INS-646
chr12-100448400-INS-601
chr12-100453787-INS-64
chr12-100456966-INS-2343
chr12-100457955-INS-2442
chr12-100457988-INS-1617
chr12-100460000-INS-60
chr12-100464482-INS-167
chr12-100465933-INS-64
chr12-100468295-INS-208
chr12-100472160-INS-147
chr12-100483986-INS-197
chr12-100504263-INS-75
chr12-100532729-INS-408
chr12-100555537-INS-172
chr12-100579929-INS-90
chr12-100597023-INS-64
chr12-100610996-INS-3004
chr12-100631470-INS-461
chr12-100631993-INS-65
chr12-100655379-INS-1887
chr12-100667997-INS-108
chr12-100690025-INS-206
chr12-100694632-INS-64
chr12-100704285-INS-80
chr12-100717896-INS-66
chr12-100718674-INS-51
chr12-100721024-INS-435
chr12-100724037-INS-85
chr12-100727751-INS-88
chr12-100737231-INS-56
chr12-100748307-INS-212
chr12-100785040-INS-116
chr12-100795527-INS-507
chr12-100800715-INS-187
chr12-100804732-INS-209
chr12-100808163-INS-66
chr12-100812507-INS-510
chr12-100845800-INS-228
chr12-100846311-INS-250
chr12-100863574-INS-588
chr12-100871169-INS-52
chr12-100871795-INS-528
chr12-100903371-INS-217
chr12-100915845-INS-142
chr12-100931552-INS-78
chr12-100934091-INS-193
chr12-100939409-INS-315
chr12-100943737-INS-172
chr12-100954274-INS-57
chr12-100987357-INS-718
chr12-100988186-INS-79
chr12-101001016-INS-126
chr12-101076895-INS-209
chr12-101079370-INS-72

chr12-101082453-INS-197
chr12-101095277-INS-6503
chr12-101095436-INS-57
chr12-101100647-INS-6684
chr12-101111541-INS-65
chr12-101117814-INS-1873
chr12-101117815-INS-399
chr12-101119421-INS-6442
chr12-101150675-INS-319
chr12-101160665-INS-401
chr12-101172751-INS-259
chr12-101249998-INS-529
chr12-101258632-INS-739
chr12-101285822-INS-2919
chr12-101285833-INS-876
chr12-101303792-INS-6612
chr12-101327007-INS-649
chr12-101328689-INS-73
chr12-101340650-INS-113
chr12-101342302-INS-483
chr12-101345091-INS-388
chr12-101355291-INS-400
chr12-101359273-INS-53
chr12-101360364-INS-734
chr12-101364776-INS-746
chr12-101365562-INS-7158
chr12-101369271-INS-345
chr12-101373365-INS-55
chr12-101408562-INS-325
chr12-101426832-INS-111
chr12-101427276-INS-352
chr12-101439052-INS-89
chr12-101446800-INS-142
chr12-101466969-INS-197
chr12-101472955-INS-691
chr12-101479868-INS-758
chr12-101516146-INS-370
chr12-101552981-INS-690
chr12-101560956-INS-249
chr12-101566876-INS-70
chr12-101589564-INS-96
chr12-101622114-INS-654
chr12-101631837-INS-372
chr12-101647483-INS-76
chr12-101658189-INS-967
chr12-101705273-INS-213
chr12-101709018-INS-57
chr12-101710517-INS-97
chr12-101730843-INS-230
chr12-101752312-INS-219
chr12-101775633-INS-68
chr12-101780177-INS-582
chr12-101790832-INS-288
chr12-101792991-INS-66
chr12-101806204-INS-354
chr12-101807428-INS-186
chr12-101820642-INS-80
chr12-101823978-INS-129
chr12-101824200-INS-387
chr12-101857650-INS-79
chr12-101866585-INS-66
chr12-101887119-INS-858
chr12-101904255-INS-205
chr12-101910072-INS-172
chr12-101916122-INS-235
chr12-101930847-INS-104
chr12-101958247-INS-144
chr12-101964288-INS-204
chr12-101978268-INS-400
chr12-101985890-INS-7084
chr12-102007195-INS-165
chr12-102019464-INS-94
chr12-102027451-INS-55
chr12-102035696-INS-97
chr12-102037281-INS-533
chr12-102052270-INS-69
chr12-102053529-INS-70

chr12-102077453-INS-7053
chr12-102079741-INS-6026
chr12-102083668-INS-811
chr12-102083677-INS-1101
chr12-102122660-INS-322
chr12-102124979-INS-73
chr12-102125216-INS-349
chr12-102155587-INS-356
chr12-102158482-INS-311
chr12-102179083-INS-141
chr12-102191878-INS-206
chr12-102204717-INS-182
chr12-102217337-INS-52
chr12-102227618-INS-224
chr12-102251324-INS-371
chr12-102257970-INS-80
chr12-102261291-INS-109
chr12-102267101-INS-92
chr12-102272052-INS-85
chr12-102285366-INS-7250
chr12-102297629-INS-55
chr12-102318746-INS-84
chr12-102321962-INS-188
chr12-102326851-INS-329
chr12-102336690-INS-111
chr12-102342738-INS-60
chr12-102343879-INS-144
chr12-102366705-INS-143
chr12-102369264-INS-71
chr12-102391399-INS-150
chr12-102397369-INS-217
chr12-102412413-INS-79
chr12-102418271-INS-151
chr12-102441977-INS-160
chr12-102463634-INS-90
chr12-102467649-INS-59
chr12-102485870-INS-148
chr12-102489449-INS-170
chr12-102502889-INS-117
chr12-102514319-INS-86
chr12-102514498-INS-8940
chr12-102523304-INS-214
chr12-102530597-INS-205
chr12-102533867-INS-54
chr12-102554247-INS-116
chr12-102556671-INS-67
chr12-102584077-INS-94
chr12-102604740-INS-80
chr12-102624055-INS-192
chr12-102626204-INS-112
chr12-102637571-INS-83
chr12-102661996-INS-445
chr12-102679805-INS-191
chr12-102680998-INS-108
chr12-102689859-INS-160
chr12-102702447-INS-55
chr12-102734016-INS-71
chr12-102735786-INS-327
chr12-102755993-INS-217
chr12-102760260-INS-60
chr12-102764337-INS-56
chr12-102785294-INS-184
chr12-102794871-INS-375
chr12-102822419-INS-77
chr12-102849941-INS-336
chr12-102854952-INS-237
chr12-102856283-INS-504
chr12-102864521-INS-60
chr12-102866913-INS-63
chr12-102870866-INS-155
chr12-102871364-INS-801
chr12-102873879-INS-512
chr12-102887101-INS-53
chr12-102890794-INS-581
chr12-102892121-INS-205
chr12-102895466-INS-56
chr12-102898540-INS-92

chr12-102904046-INS-410
chr12-102904953-INS-155
chr12-102910557-INS-65
chr12-102924480-INS-53
chr12-102931102-INS-273
chr12-102938594-INS-141
chr12-102940745-INS-602
chr12-102976842-INS-315
chr12-102986440-INS-52
chr12-103005861-INS-74
chr12-103007692-INS-750
chr12-103023091-INS-51
chr12-103045517-INS-118
chr12-103117783-INS-210
chr12-103123267-INS-80
chr12-103141246-INS-88
chr12-103144422-INS-493
chr12-103190643-INS-55
chr12-103192035-INS-442
chr12-103238951-INS-187
chr12-103274343-INS-208
chr12-103288654-INS-78
chr12-103300603-INS-156
chr12-103315966-INS-66
chr12-103321681-INS-52
chr12-103330513-INS-208
chr12-103338912-INS-52
chr12-103351800-INS-180
chr12-103400429-INS-91
chr12-103402398-INS-95
chr12-103409865-INS-921
chr12-103411020-INS-156
chr12-103467844-INS-68
chr12-103477527-INS-72
chr12-103510333-INS-191
chr12-103511863-INS-87
chr12-103515523-INS-441
chr12-103562212-INS-86
chr12-103568172-INS-918
chr12-103603603-INS-68
chr12-103612253-INS-70
chr12-103629535-INS-94
chr12-103630582-INS-106
chr12-103633435-INS-107
chr12-103707716-INS-62
chr12-103709786-INS-400
chr12-103710661-INS-144
chr12-103715163-INS-63
chr12-103967081-INS-278
chr12-103985913-INS-210
chr12-104024541-INS-60
chr12-104025730-INS-60
chr12-104030090-INS-6894
chr12-104062867-INS-362
chr12-104075160-INS-227
chr12-104078591-INS-4550
chr12-104078610-INS-5259
chr12-104094422-INS-51
chr12-104119120-INS-56
chr12-104166493-INS-82
chr12-104176099-INS-65
chr12-104188186-INS-372
chr12-104197837-INS-628
chr12-104204055-INS-431
chr12-104208788-INS-983
chr12-104281508-INS-57
chr12-104292452-INS-194
chr12-104335705-INS-234
chr12-104371984-INS-136
chr12-104388036-INS-80
chr12-104408704-INS-232
chr12-104408983-INS-1050
chr12-104415293-INS-4413
chr12-104419706-INS-56
chr12-104424598-INS-70
chr12-104448190-INS-114
chr12-104469505-INS-128

chr12-104478384-INS-341
chr12-104501014-INS-82
chr12-104505075-INS-105
chr12-104506959-INS-58
chr12-104516892-INS-414
chr12-104527097-INS-64
chr12-104532270-INS-72
chr12-104542041-INS-86
chr12-104611142-INS-130
chr12-104630062-INS-56
chr12-104633529-INS-201
chr12-104644992-INS-61
chr12-104649125-INS-204
chr12-104668260-INS-206
chr12-104701987-INS-328
chr12-104714494-INS-363
chr12-104752217-INS-265
chr12-104786492-INS-118
chr12-104793905-INS-306
chr12-104801313-INS-128
chr12-104859877-INS-66
chr12-104866523-INS-203
chr12-104877518-INS-210
chr12-104879036-INS-2982
chr12-104881600-INS-130
chr12-104888692-INS-208
chr12-104903949-INS-76
chr12-104943632-INS-205
chr12-104946886-INS-460
chr12-104952344-INS-184
chr12-104960335-INS-99
chr12-104962216-INS-64
chr12-104972646-INS-84
chr12-104985561-INS-192
chr12-104992453-INS-202
chr12-104998106-INS-88
chr12-105016900-INS-68
chr12-105039374-INS-189
chr12-105054806-INS-559
chr12-105061369-INS-202
chr12-105074606-INS-261
chr12-105075449-INS-62
chr12-105090259-INS-213
chr12-105098144-INS-66
chr12-105173606-INS-117
chr12-105212772-INS-64
chr12-105236619-INS-73
chr12-105245642-INS-648
chr12-105251696-INS-504
chr12-105310587-INS-58
chr12-105352808-INS-117
chr12-105355265-INS-78
chr12-105356274-INS-75
chr12-105365768-INS-77
chr12-105378652-INS-577
chr12-105382990-INS-220
chr12-105384020-INS-52
chr12-105385379-INS-51
chr12-105402481-INS-767
chr12-105404229-INS-581
chr12-105435204-INS-595
chr12-105468529-INS-198
chr12-105471277-INS-61
chr12-105491001-INS-90
chr12-105506659-INS-78
chr12-105516330-INS-353
chr12-105577928-INS-3029
chr12-105578543-INS-241
chr12-105597271-INS-216
chr12-105621902-INS-120
chr12-105672080-INS-168
chr12-105678007-INS-459
chr12-105685538-INS-177
chr12-105687779-INS-144
chr12-105691410-INS-535
chr12-105696820-INS-132
chr12-105699933-INS-51

chr12-105702979-INS-366
chr12-105713361-INS-59
chr12-105718543-INS-76
chr12-105727074-INS-97
chr12-105733947-INS-165
chr12-105760046-INS-101
chr12-105808447-INS-197
chr12-105813663-INS-68
chr12-105817840-INS-63
chr12-105835240-INS-79
chr12-105839616-INS-221
chr12-105849468-INS-188
chr12-105849490-INS-183
chr12-105851768-INS-136
chr12-105853629-INS-341
chr12-105874506-INS-110
chr12-105875664-INS-63
chr12-105882396-INS-108
chr12-105888726-INS-96
chr12-105893169-INS-52
chr12-105906172-INS-55
chr12-105908638-INS-5412
chr12-105914548-INS-100
chr12-105923150-INS-248
chr12-105941867-INS-213
chr12-105952069-INS-188
chr12-105966974-INS-95
chr12-105973728-INS-261
chr12-105979368-INS-58
chr12-105984139-INS-83
chr12-105993422-INS-126
chr12-106011589-INS-58
chr12-106033170-INS-59
chr12-106060597-INS-80
chr12-106072610-INS-156
chr12-106088687-INS-199
chr12-106093607-INS-209
chr12-106096766-INS-61
chr12-106098527-INS-69
chr12-106108565-INS-101
chr12-106112399-INS-112
chr12-106124982-INS-105
chr12-106132047-INS-368
chr12-106143248-INS-147
chr12-106147997-INS-394
chr12-106184385-INS-64
chr12-106194357-INS-494
chr12-106246972-INS-52
chr12-106251435-INS-134
chr12-106259247-INS-598
chr12-106285921-INS-75
chr12-106292428-INS-52
chr12-106304583-INS-114
chr12-106321491-INS-164
chr12-106328672-INS-94
chr12-106350144-INS-52
chr12-106358352-INS-548
chr12-106360377-INS-81
chr12-106375809-INS-52
chr12-106389838-INS-78
chr12-106395009-INS-288
chr12-106398482-INS-70
chr12-106401980-INS-78
chr12-106430391-INS-60
chr12-106432391-INS-145
chr12-106456699-INS-68
chr12-106472885-INS-166
chr12-106476138-INS-51
chr12-106479676-INS-176
chr12-106479753-INS-1370
chr12-106485811-INS-58
chr12-106489238-INS-53
chr12-106504749-INS-58
chr12-106507999-INS-327
chr12-106519766-INS-341
chr12-106533135-INS-186
chr12-106543194-INS-122

chr12-106562127-INS-130
chr12-106569103-INS-127
chr12-106604174-INS-76
chr12-106612708-INS-194
chr12-106630618-INS-60
chr12-106651113-INS-273
chr12-106655511-INS-52
chr12-106660947-INS-111
chr12-106692602-INS-57
chr12-106706378-INS-72
chr12-106707613-INS-58
chr12-106745078-INS-290
chr12-106762389-INS-247
chr12-106765906-INS-479
chr12-106826056-INS-100
chr12-106844238-INS-53
chr12-106847572-INS-194
chr12-106867749-INS-548
chr12-106873252-INS-60
chr12-106875062-INS-3016
chr12-106886051-INS-125
chr12-106930649-INS-50
chr12-106938744-INS-72
chr12-106945709-INS-95
chr12-106952030-INS-237
chr12-106953461-INS-452
chr12-106959281-INS-55
chr12-106960866-INS-107
chr12-106966017-INS-54
chr12-106971205-INS-60
chr12-107012575-INS-6442
chr12-107014727-INS-143
chr12-107036644-INS-95
chr12-107041600-INS-98
chr12-107069796-INS-69
chr12-107081692-INS-68
chr12-107090322-INS-64
chr12-107124391-INS-55
chr12-107128483-INS-112
chr12-107135503-INS-96
chr12-107174281-INS-105
chr12-107221287-INS-52
chr12-107229911-INS-57
chr12-107264613-INS-122
chr12-107270306-INS-229
chr12-107285035-INS-222
chr12-107303244-INS-65
chr12-107307222-INS-91
chr12-107332458-INS-92
chr12-107343920-INS-112
chr12-107384333-INS-54
chr12-107409638-INS-110
chr12-107421217-INS-705
chr12-107422519-INS-77
chr12-107428603-INS-58
chr12-107456415-INS-102
chr12-107473103-INS-366
chr12-107485495-INS-56
chr12-107489133-INS-186
chr12-107499686-INS-150
chr12-107506853-INS-174
chr12-107527112-INS-140
chr12-107529333-INS-89
chr12-107546800-INS-64
chr12-107556292-INS-71
chr12-107573162-INS-118
chr12-107575080-INS-85
chr12-107575115-INS-558
chr12-107585564-INS-131
chr12-107598347-INS-54
chr12-107601654-INS-110
chr12-107664448-INS-66
chr12-107692553-INS-77
chr12-107697677-INS-108
chr12-107711951-INS-139
chr12-107734158-INS-54
chr12-107737978-INS-180

chr12-107745556-INS-66
chr12-107769933-INS-71
chr12-107786269-INS-142
chr12-107792708-INS-73
chr12-107800551-INS-115
chr12-107803556-INS-294
chr12-107805865-INS-188
chr12-107855986-INS-64
chr12-107860740-INS-112
chr12-107861446-INS-54
chr12-107873806-INS-154
chr12-107889869-INS-359
chr12-107898109-INS-68
chr12-107909288-INS-192
chr12-107915257-INS-54
chr12-107933055-INS-57
chr12-107990080-INS-186
chr12-108003239-INS-56
chr12-108011007-INS-218
chr12-108035578-INS-72
chr12-108074446-INS-60
chr12-108105529-INS-366
chr12-108111260-INS-51
chr12-108148210-INS-167
chr12-108173333-INS-83
chr12-108176271-INS-58
chr12-108200293-INS-216
chr12-108214860-INS-148
chr12-108218896-INS-69
chr12-108226648-INS-233
chr12-108234669-INS-168
chr12-108246098-INS-240
chr12-108249309-INS-198
chr12-108263578-INS-84
chr12-108287668-INS-63
chr12-108290835-INS-100
chr12-108298498-INS-381
chr12-108304029-INS-216
chr12-108315201-INS-59
chr12-108316079-INS-172
chr12-108328748-INS-142
chr12-108335639-INS-336
chr12-108345380-INS-217
chr12-108385898-INS-111
chr12-108392169-INS-113
chr12-108434834-INS-424
chr12-108437662-INS-464
chr12-108439442-INS-134
chr12-108451791-INS-258
chr12-108508086-INS-270
chr12-108508860-INS-225
chr12-108549112-INS-192
chr12-108585444-INS-5549
chr12-108622331-INS-60
chr12-108624058-INS-365
chr12-108645494-INS-523
chr12-108656190-INS-59
chr12-108664289-INS-172
chr12-108692841-INS-170
chr12-108701551-INS-318
chr12-108717299-INS-98
chr12-108728554-INS-119
chr12-108729853-INS-505
chr12-108733526-INS-51
chr12-108747933-INS-399
chr12-108748859-INS-56
chr12-108750230-INS-206
chr12-108797348-INS-204
chr12-108806094-INS-86
chr12-108810573-INS-207
chr12-108839627-INS-116
chr12-108842152-INS-71
chr12-108899000-INS-145
chr12-108911782-INS-229
chr12-108932376-INS-230
chr12-108967177-INS-74
chr12-108968403-INS-110

chr12-108968879-INS-80
chr12-108984089-INS-118
chr12-108985715-INS-82
chr12-109003256-INS-52
chr12-109028604-INS-98
chr12-109038875-INS-106
chr12-109047443-INS-67
chr12-109064726-INS-53
chr12-109071422-INS-84
chr12-109079750-INS-192
chr12-109096859-INS-697
chr12-109100579-INS-73
chr12-109106124-INS-280
chr12-109116742-INS-418
chr12-109127802-INS-620
chr12-109157753-INS-295
chr12-109160275-INS-6125
chr12-109259604-INS-52
chr12-109261057-INS-389
chr12-109263485-INS-68
chr12-109314643-INS-679
chr12-109328765-INS-211
chr12-109333192-INS-232
chr12-109344219-INS-105
chr12-109352371-INS-126
chr12-109396995-INS-100
chr12-109410692-INS-64
chr12-109440694-INS-201
chr12-109441523-INS-116
chr12-109446557-INS-76
chr12-109487111-INS-74
chr12-109592850-INS-72
chr12-109642234-INS-56
chr12-109645360-INS-54
chr12-109717170-INS-56
chr12-109719275-INS-59
chr12-109730490-INS-219
chr12-109731883-INS-514
chr12-109757034-INS-155
chr12-109793311-INS-52
chr12-109816549-INS-208
chr12-109823621-INS-456
chr12-109825532-INS-491
chr12-109826557-INS-138
chr12-109843868-INS-6365
chr12-109857865-INS-59
chr12-109897766-INS-162
chr12-109905718-INS-515
chr12-109913544-INS-62
chr12-109931809-INS-81
chr12-109954012-INS-125
chr12-109955482-INS-633
chr12-109966578-INS-67
chr12-109968096-INS-171
chr12-109973123-INS-62
chr12-109977305-INS-195
chr12-109979892-INS-443
chr12-109989018-INS-68
chr12-110019747-INS-166
chr12-110030158-INS-99
chr12-110043055-INS-441
chr12-110045777-INS-318
chr12-110066934-INS-60
chr12-110076101-INS-202
chr12-110076118-INS-782
chr12-110081832-INS-87
chr12-110100698-INS-83
chr12-110132872-INS-68
chr12-110141583-INS-377
chr12-110159633-INS-134
chr12-110184256-INS-51
chr12-110213516-INS-89
chr12-110227156-INS-417
chr12-110264513-INS-150
chr12-110281455-INS-104
chr12-110293564-INS-898
chr12-110295339-INS-114

chr12-110298115-INS-69
chr12-110299266-INS-211
chr12-110335022-INS-653
chr12-110336232-INS-270
chr12-110337663-INS-102
chr12-110337911-INS-230
chr12-110349402-INS-875
chr12-110361966-INS-209
chr12-110366183-INS-83
chr12-110381000-INS-182
chr12-110387213-INS-214
chr12-110409230-INS-248
chr12-110418761-INS-71
chr12-110437115-INS-127
chr12-110457603-INS-214
chr12-110461633-INS-504
chr12-110467719-INS-57
chr12-110469882-INS-217
chr12-110476960-INS-138
chr12-110479047-INS-52
chr12-110484692-INS-197
chr12-110542848-INS-64
chr12-110556393-INS-217
chr12-110573910-INS-427
chr12-110582207-INS-195
chr12-110589139-INS-396
chr12-110591255-INS-54
chr12-110605807-INS-315
chr12-110617454-INS-245
chr12-110641859-INS-206
chr12-110646241-INS-204
chr12-110646842-INS-57
chr12-110665547-INS-124
chr12-110667979-INS-76
chr12-110672258-INS-217
chr12-110686149-INS-1154
chr12-110686335-INS-99
chr12-110687471-INS-198
chr12-110689131-INS-418
chr12-110690301-INS-194
chr12-110703425-INS-68
chr12-110716763-INS-780
chr12-110717317-INS-219
chr12-110733200-INS-76
chr12-110733709-INS-194
chr12-110775331-INS-198
chr12-110792578-INS-52
chr12-110807504-INS-120
chr12-110843121-INS-52
chr12-110847946-INS-556
chr12-110848873-INS-211
chr12-110850205-INS-75
chr12-110866912-INS-199
chr12-110880888-INS-192
chr12-110891373-INS-488
chr12-110917761-INS-182
chr12-110922156-INS-217
chr12-110923462-INS-134
chr12-110925218-INS-472
chr12-110928915-INS-60
chr12-110955279-INS-58
chr12-110956925-INS-79
chr12-110977163-INS-342
chr12-110978245-INS-183
chr12-110992277-INS-53
chr12-111031208-INS-1134
chr12-111031240-INS-222
chr12-111031304-INS-3283
chr12-111039531-INS-134
chr12-111039616-INS-314
chr12-111047351-INS-200
chr12-111088402-INS-96
chr12-111093435-INS-106
chr12-111107138-INS-58
chr12-111127060-INS-85
chr12-111128446-INS-164
chr12-111129831-INS-226

chr12-111172867-INS-147
chr12-111178289-INS-116
chr12-111184655-INS-225
chr12-111185853-INS-274
chr12-111252590-INS-540
chr12-111279215-INS-228
chr12-111285411-INS-76
chr12-111289217-INS-68
chr12-111307616-INS-169
chr12-111308168-INS-113
chr12-111315642-INS-162
chr12-111330030-INS-89
chr12-111345748-INS-120
chr12-111370244-INS-76
chr12-111377231-INS-51
chr12-111400179-INS-156
chr12-111433129-INS-60
chr12-111437514-INS-86
chr12-111445444-INS-423
chr12-111447040-INS-215
chr12-111448050-INS-108
chr12-111457159-INS-68
chr12-111473160-INS-259
chr12-111474479-INS-171
chr12-111517838-INS-198
chr12-111531403-INS-358
chr12-111547420-INS-198
chr12-111552759-INS-193
chr12-111561872-INS-224
chr12-111570180-INS-167
chr12-111571278-INS-231
chr12-111575197-INS-222
chr12-111580232-INS-282
chr12-111585037-INS-243
chr12-111610137-INS-163
chr12-111618145-INS-202
chr12-111628348-INS-145
chr12-111639496-INS-236
chr12-111645941-INS-370
chr12-111646073-INS-525
chr12-111668367-INS-82
chr12-111690481-INS-201
chr12-111695160-INS-220
chr12-111702982-INS-168
chr12-111708405-INS-66
chr12-111718978-INS-207
chr12-111731110-INS-205
chr12-111732684-INS-52
chr12-111737595-INS-52
chr12-111751766-INS-329
chr12-111768933-INS-124
chr12-111773731-INS-86
chr12-111782298-INS-68
chr12-111816568-INS-152
chr12-111823104-INS-208
chr12-111828584-INS-62
chr12-111834752-INS-1627
chr12-111857791-INS-58
chr12-111884561-INS-4751
chr12-111888055-INS-80
chr12-111915299-INS-123
chr12-111923916-INS-251
chr12-111964101-INS-486
chr12-112002154-INS-452
chr12-112014155-INS-898
chr12-112048432-INS-52
chr12-112049522-INS-182
chr12-112055857-INS-225
chr12-112103579-INS-220
chr12-112109885-INS-5544
chr12-112146422-INS-160
chr12-112160711-INS-84
chr12-112225836-INS-478
chr12-112228429-INS-91
chr12-112254137-INS-66
chr12-112263935-INS-124
chr12-112267386-INS-400

chr12-112280366-INS-59
chr12-112284345-INS-84
chr12-112292289-INS-252
chr12-112295722-INS-62
chr12-112320390-INS-281
chr12-112347417-INS-107
chr12-112377666-INS-71
chr12-112390320-INS-60
chr12-112446737-INS-112
chr12-112451167-INS-361
chr12-112479321-INS-183
chr12-112480764-INS-131
chr12-112509827-INS-664
chr12-112511370-INS-156
chr12-112516468-INS-80
chr12-112521303-INS-208
chr12-112523614-INS-208
chr12-112528409-INS-198
chr12-112548114-INS-90
chr12-112548631-INS-330
chr12-112598155-INS-199
chr12-112654835-INS-196
chr12-112658747-INS-964
chr12-112666096-INS-208
chr12-112675102-INS-53
chr12-112679026-INS-236
chr12-112685122-INS-206
chr12-112690364-INS-212
chr12-112697041-INS-71
chr12-112718015-INS-201
chr12-112725455-INS-225
chr12-112729290-INS-96
chr12-112730687-INS-175
chr12-112736956-INS-102
chr12-112753727-INS-135
chr12-112757204-INS-82
chr12-112764524-INS-254
chr12-112786366-INS-200
chr12-112800614-INS-209
chr12-112801923-INS-147
chr12-112808308-INS-50
chr12-112819376-INS-166
chr12-112886831-INS-214
chr12-112900969-INS-84
chr12-112942829-INS-95
chr12-112968310-INS-210
chr12-112970985-INS-78
chr12-112973359-INS-103
chr12-112985635-INS-201
chr12-113007704-INS-214
chr12-113012423-INS-120
chr12-113019899-INS-207
chr12-113023257-INS-52
chr12-113077650-INS-54
chr12-113079483-INS-455
chr12-113087417-INS-53
chr12-113129654-INS-198
chr12-113134890-INS-292
chr12-113138884-INS-200
chr12-113183986-INS-96
chr12-113193016-INS-356
chr12-113202186-INS-166
chr12-113226377-INS-157
chr12-113226577-INS-628
chr12-113234172-INS-58
chr12-113247681-INS-151
chr12-113258015-INS-363
chr12-113274801-INS-196
chr12-113276224-INS-537
chr12-113298955-INS-162
chr12-113299265-INS-833
chr12-113319120-INS-137
chr12-113327868-INS-117
chr12-113328469-INS-138
chr12-113381947-INS-52
chr12-113387979-INS-1037
chr12-113388142-INS-258

chr12-113388746-INS-157
chr12-113406988-INS-6474
chr12-113442734-INS-143
chr12-113457088-INS-204
chr12-113457922-INS-379
chr12-113481117-INS-88
chr12-113485885-INS-5316
chr12-113497022-INS-64
chr12-113517209-INS-594
chr12-113540125-INS-15406
chr12-113694064-INS-168
chr12-113781995-INS-68
chr12-113840907-INS-689
chr12-113907124-INS-238
chr12-113913173-INS-276
chr12-113941693-INS-203
chr12-113949662-INS-312
chr12-113956023-INS-387
chr12-113972158-INS-426
chr12-113978162-INS-196
chr12-113978207-INS-194
chr12-113980948-INS-76
chr12-114022365-INS-196
chr12-114039893-INS-95
chr12-114054469-INS-5510
chr12-114134113-INS-64
chr12-114134950-INS-120
chr12-114145930-INS-56
chr12-114175055-INS-204
chr12-114187742-INS-457
chr12-114205434-INS-362
chr12-114240121-INS-56
chr12-114247847-INS-206
chr12-114268793-INS-564
chr12-114298379-INS-161
chr12-114302344-INS-60
chr12-114303313-INS-52
chr12-114336379-INS-180
chr12-114351711-INS-54
chr12-114423748-INS-68
chr12-114456009-INS-52
chr12-114478269-INS-915
chr12-114483821-INS-3317
chr12-114533144-INS-158
chr12-114543916-INS-560
chr12-114553098-INS-60
chr12-114574784-INS-453
chr12-114669836-INS-72
chr12-114723711-INS-59
chr12-114729417-INS-806
chr12-114835185-INS-229
chr12-114905209-INS-162
chr12-114907407-INS-208
chr12-114914353-INS-155
chr12-115019890-INS-61
chr12-115062966-INS-93
chr12-115119090-INS-69
chr12-115138512-INS-455
chr12-115312727-INS-101
chr12-115351491-INS-75
chr12-115372431-INS-7270
chr12-115438509-INS-438
chr12-115514985-INS-136
chr12-115572138-INS-54
chr12-115606005-INS-168
chr12-115660415-INS-168
chr12-115747784-INS-1660
chr12-115820143-INS-74
chr12-115848351-INS-54
chr12-115855683-INS-150
chr12-115887586-INS-278
chr12-115914208-INS-2928
chr12-115925001-INS-76
chr12-115960818-INS-244
chr12-115966247-INS-184
chr12-115973887-INS-55
chr12-116001483-INS-519

chr12-116001590-INS-866
chr12-116038197-INS-128
chr12-116044346-INS-7068
chr12-116096393-INS-101
chr12-116170513-INS-224
chr12-116172050-INS-584
chr12-116199836-INS-856
chr12-116205676-INS-595
chr12-116222478-INS-69
chr12-116252508-INS-388
chr12-116298077-INS-486
chr12-116354584-INS-179
chr12-116375685-INS-66
chr12-116418059-INS-55
chr12-116441756-INS-8755
chr12-116445005-INS-72
chr12-116497597-INS-6545
chr12-116551881-INS-632
chr12-116594796-INS-62
chr12-116622750-INS-242
chr12-116639023-INS-385
chr12-116642103-INS-205
chr12-116648547-INS-6563
chr12-116662839-INS-80
chr12-116726191-INS-144
chr12-116729728-INS-6273
chr12-116730511-INS-984
chr12-116739599-INS-59
chr12-116745645-INS-677
chr12-116781250-INS-660
chr12-116822296-INS-1141
chr12-116862092-INS-76
chr12-116869327-INS-81
chr12-116870541-INS-375
chr12-116881266-INS-76
chr12-116886228-INS-612
chr12-116897255-INS-264
chr12-116917471-INS-6475
chr12-116922814-INS-64
chr12-116945205-INS-54
chr12-116947807-INS-52
chr12-116971190-INS-116
chr12-116984040-INS-87
chr12-116984680-INS-162
chr12-116996943-INS-94
chr12-117012474-INS-359
chr12-117015844-INS-136
chr12-117028143-INS-84
chr12-117044781-INS-80
chr12-117127064-INS-809
chr12-117135335-INS-64
chr12-117143937-INS-216
chr12-117147778-INS-791
chr12-117149104-INS-220
chr12-117157695-INS-70
chr12-117173647-INS-72
chr12-117183091-INS-58
chr12-117203113-INS-107
chr12-117251064-INS-56
chr12-117258155-INS-66
chr12-117262659-INS-182
chr12-117303076-INS-279
chr12-117316885-INS-293
chr12-117318357-INS-146
chr12-117367087-INS-96
chr12-117367380-INS-349
chr12-117379469-INS-104
chr12-117397054-INS-195
chr12-117401926-INS-400
chr12-117403687-INS-495
chr12-117409353-INS-64
chr12-117414548-INS-57
chr12-117434180-INS-76
chr12-117442239-INS-167
chr12-117452703-INS-772
chr12-117459911-INS-207
chr12-117489422-INS-165

chr12-117517487-INS-52
chr12-117534487-INS-52
chr12-117552065-INS-278
chr12-117552154-INS-1200
chr12-117552986-INS-155
chr12-117561481-INS-52
chr12-117578134-INS-68
chr12-117581646-INS-131
chr12-117582973-INS-203
chr12-117603937-INS-84
chr12-117613576-INS-276
chr12-117615369-INS-322
chr12-117637688-INS-64
chr12-117638956-INS-60
chr12-117648927-INS-251
chr12-117676121-INS-203
chr12-117680438-INS-97
chr12-117687101-INS-121
chr12-117711278-INS-84
chr12-117714277-INS-156
chr12-117724114-INS-328
chr12-117725984-INS-221
chr12-117752169-INS-169
chr12-117786831-INS-6754
chr12-117804390-INS-7167
chr12-117813132-INS-105
chr12-117825568-INS-334
chr12-117825915-INS-131
chr12-117829573-INS-55
chr12-117848920-INS-5244
chr12-117869708-INS-528
chr12-117877062-INS-89
chr12-117895636-INS-535
chr12-117913150-INS-171
chr12-117920390-INS-126
chr12-117927713-INS-111
chr12-117935575-INS-849
chr12-117950649-INS-81
chr12-117965671-INS-52
chr12-117970776-INS-54
chr12-118004652-INS-135
chr12-118044581-INS-401
chr12-118062863-INS-79
chr12-118066690-INS-63
chr12-118070959-INS-103
chr12-118143034-INS-116
chr12-118273105-INS-54
chr12-118274901-INS-549
chr12-118295018-INS-55
chr12-118349500-INS-1689
chr12-118361208-INS-559
chr12-118385398-INS-158
chr12-118437297-INS-991
chr12-118466508-INS-395
chr12-118474372-INS-145
chr12-118483331-INS-683
chr12-118498602-INS-234
chr12-118513142-INS-102
chr12-118536430-INS-66
chr12-118605117-INS-108
chr12-118721491-INS-99
chr12-118756745-INS-337
chr12-118758970-INS-62
chr12-118767166-INS-129
chr12-118774783-INS-133
chr12-118781755-INS-62
chr12-118788215-INS-6442
chr12-118791568-INS-5549
chr12-118811585-INS-62
chr12-118825436-INS-207
chr12-118826516-INS-177
chr12-118827967-INS-2181
chr12-118864035-INS-74
chr12-118865377-INS-60
chr12-118878654-INS-77
chr12-118928749-INS-5843
chr12-118945819-INS-334

chr12-118953036-INS-195
chr12-118968081-INS-2018
chr12-118973273-INS-6178
chr12-119010655-INS-65
chr12-119014489-INS-256
chr12-119045890-INS-7078
chr12-119068981-INS-109
chr12-119070189-INS-466
chr12-119072240-INS-420
chr12-119083578-INS-301
chr12-119085334-INS-6389
chr12-119098385-INS-60
chr12-119107564-INS-399
chr12-119235347-INS-61
chr12-119239463-INS-475
chr12-119269883-INS-55
chr12-119272070-INS-7288
chr12-119274903-INS-6211
chr12-119276279-INS-52
chr12-119281009-INS-82
chr12-119293869-INS-2932
chr12-119295793-INS-6235
chr12-119306762-INS-51
chr12-119350918-INS-90
chr12-119388531-INS-575
chr12-119391505-INS-339
chr12-119401311-INS-6440
chr12-119457945-INS-708
chr12-119462809-INS-82
chr12-119492543-INS-52
chr12-119505402-INS-7091
chr12-119509998-INS-6539
chr12-119513883-INS-56
chr12-119541017-INS-121
chr12-119541947-INS-53
chr12-119549773-INS-203
chr12-119575630-INS-521
chr12-119601179-INS-879
chr12-119613186-INS-6426
chr12-119616450-INS-90
chr12-119681670-INS-3024
chr12-119714558-INS-4446
chr12-119743213-INS-6197
chr12-119751693-INS-1922
chr12-119757320-INS-7092
chr12-119759629-INS-5846
chr12-119785063-INS-5962
chr12-119786788-INS-333
chr12-119799502-INS-623
chr12-119800665-INS-6949
chr12-119803918-INS-834
chr12-119807998-INS-6210
chr12-119822825-INS-133
chr12-119824695-INS-72
chr12-119831111-INS-305
chr12-119865427-INS-6513
chr12-119877173-INS-6864
chr12-119888671-INS-1041
chr12-119893290-INS-6215
chr12-119923343-INS-373
chr12-119940682-INS-121
chr12-119967431-INS-5237
chr13-3099100-INS-208
chr13-3237246-INS-96
chr13-3357989-INS-879
chr13-3367662-INS-53
chr13-3371544-INS-197
chr13-3377471-INS-695
chr13-3449584-INS-498
chr13-3456430-INS-342
chr13-3465761-INS-52
chr13-3476422-INS-960
chr13-3514989-INS-7083
chr13-3523740-INS-8128
chr13-3526055-INS-5434
chr13-3542643-INS-119
chr13-3549684-INS-3472

chr13-3563682-INS-2640
chr13-3570061-INS-397
chr13-3576134-INS-497
chr13-3578372-INS-603
chr13-3595267-INS-167
chr13-3637777-INS-178
chr13-3643195-INS-6475
chr13-3647753-INS-504
chr13-3683551-INS-58
chr13-3698313-INS-231
chr13-3701161-INS-59
chr13-3707798-INS-64
chr13-3708691-INS-201
chr13-3712034-INS-74
chr13-3744210-INS-400
chr13-3747528-INS-54
chr13-3751538-INS-186
chr13-3777364-INS-160
chr13-3845640-INS-125
chr13-3914318-INS-672
chr13-3929586-INS-3010
chr13-3931809-INS-72
chr13-3948253-INS-195
chr13-3950188-INS-739
chr13-3951765-INS-2041
chr13-3952659-INS-3565
chr13-3961260-INS-1889
chr13-3964125-INS-92
chr13-3971274-INS-529
chr13-3995747-INS-74
chr13-4022343-INS-60
chr13-4031147-INS-54
chr13-4074053-INS-6276
chr13-4080383-INS-84
chr13-4086365-INS-138
chr13-4086711-INS-191
chr13-4128095-INS-52
chr13-4186436-INS-5393
chr13-4186821-INS-74
chr13-4214903-INS-407
chr13-4217842-INS-86
chr13-4298723-INS-332
chr13-4344571-INS-325
chr13-4413970-INS-1890
chr13-4415755-INS-188
chr13-4418970-INS-332
chr13-4425547-INS-63
chr13-4437934-INS-148
chr13-4438607-INS-361
chr13-4440289-INS-356
chr13-4450433-INS-498
chr13-4489564-INS-66
chr13-4498384-INS-6964
chr13-4500350-INS-289
chr13-4500532-INS-60
chr13-4504848-INS-978
chr13-4519701-INS-52
chr13-4550283-INS-3689
chr13-4556978-INS-4337
chr13-4576643-INS-5480
chr13-4577481-INS-1278
chr13-4601833-INS-868
chr13-4613534-INS-7049
chr13-4656123-INS-506
chr13-4656833-INS-122
chr13-4663692-INS-117
chr13-4688878-INS-112
chr13-4693115-INS-151
chr13-4701309-INS-164
chr13-4713274-INS-495
chr13-4730802-INS-197
chr13-4737070-INS-60
chr13-4766542-INS-69
chr13-4779736-INS-127
chr13-4794447-INS-618
chr13-4801331-INS-595
chr13-4802739-INS-162

chr13-4805031-INS-1860
chr13-4823082-INS-63
chr13-4826432-INS-139
chr13-4828419-INS-144
chr13-4873190-INS-66
chr13-4876416-INS-500
chr13-4887797-INS-396
chr13-4909078-INS-7124
chr13-4927768-INS-120
chr13-4937763-INS-191
chr13-4940066-INS-631
chr13-5009712-INS-6205
chr13-5042029-INS-470
chr13-5087499-INS-7113
chr13-5089453-INS-196
chr13-5091720-INS-7642
chr13-5094098-INS-76
chr13-5096664-INS-880
chr13-5123772-INS-6699
chr13-5134305-INS-769
chr13-5156608-INS-7047
chr13-5203548-INS-375
chr13-5210806-INS-399
chr13-5244705-INS-65
chr13-5254231-INS-5869
chr13-5261415-INS-2239
chr13-5283431-INS-1565
chr13-5312147-INS-392
chr13-5316317-INS-6818
chr13-5371353-INS-56
chr13-5373181-INS-100
chr13-5396670-INS-147
chr13-5397920-INS-58
chr13-5417854-INS-299
chr13-5449014-INS-211
chr13-5461786-INS-105
chr13-5535816-INS-146
chr13-5549462-INS-80
chr13-5579774-INS-61
chr13-5590288-INS-52
chr13-5617589-INS-529
chr13-5619615-INS-90
chr13-5630226-INS-782
chr13-5648021-INS-505
chr13-5661778-INS-68
chr13-5714998-INS-906
chr13-5749891-INS-57
chr13-5750532-INS-522
chr13-5754650-INS-72
chr13-5777908-INS-78
chr13-5789319-INS-66
chr13-5830618-INS-58
chr13-5844324-INS-108
chr13-5849381-INS-432
chr13-5870381-INS-231
chr13-5948245-INS-55
chr13-5981321-INS-112
chr13-5995699-INS-414
chr13-5999633-INS-407
chr13-6005241-INS-609
chr13-6022125-INS-62
chr13-6047350-INS-396
chr13-6058680-INS-68
chr13-6065669-INS-202
chr13-6075985-INS-68
chr13-6078753-INS-66
chr13-6089011-INS-166
chr13-6099879-INS-72
chr13-6133824-INS-176
chr13-6137770-INS-94
chr13-6155834-INS-2268
chr13-6159769-INS-2252
chr13-6170021-INS-6381
chr13-6185436-INS-263
chr13-6189181-INS-81
chr13-6229576-INS-82
chr13-6245010-INS-59

chr13-6260061-INS-68
chr13-6267578-INS-151
chr13-6302078-INS-74
chr13-6307692-INS-56
chr13-6326442-INS-52
chr13-6327013-INS-64
chr13-6328423-INS-6694
chr13-6336985-INS-175
chr13-6364067-INS-162
chr13-6365846-INS-145
chr13-6377681-INS-107
chr13-6392235-INS-169
chr13-6396846-INS-497
chr13-6416823-INS-250
chr13-6423037-INS-76
chr13-6446451-INS-78
chr13-6473829-INS-6270
chr13-6479797-INS-505
chr13-6481359-INS-373
chr13-6482536-INS-110
chr13-6500619-INS-390
chr13-6502294-INS-68
chr13-6503145-INS-126
chr13-6506124-INS-421
chr13-6515014-INS-492
chr13-6526694-INS-206
chr13-6526752-INS-2381
chr13-6531166-INS-2408
chr13-6550477-INS-308
chr13-6550975-INS-166
chr13-6554506-INS-6535
chr13-6564719-INS-69
chr13-6581920-INS-66
chr13-6591045-INS-86
chr13-6632243-INS-52
chr13-6633791-INS-173
chr13-6656497-INS-240
chr13-6703471-INS-215
chr13-6706224-INS-152
chr13-6707124-INS-215
chr13-6726132-INS-336
chr13-6734495-INS-5490
chr13-6737156-INS-105
chr13-6748355-INS-1290
chr13-6754868-INS-161
chr13-6768772-INS-142
chr13-6783634-INS-4265
chr13-6798275-INS-58
chr13-6805039-INS-490
chr13-6819510-INS-54
chr13-6839981-INS-107
chr13-6860161-INS-324
chr13-6865018-INS-6663
chr13-6867396-INS-6216
chr13-6890242-INS-387
chr13-6904383-INS-5087
chr13-6908211-INS-6309
chr13-6919119-INS-134
chr13-6921935-INS-493
chr13-6937499-INS-518
chr13-6943587-INS-258
chr13-6946138-INS-52
chr13-6950712-INS-54
chr13-6974573-INS-572
chr13-7000679-INS-4396
chr13-7004900-INS-51
chr13-7064855-INS-196
chr13-7078254-INS-58
chr13-7099050-INS-551
chr13-7118690-INS-53
chr13-7127399-INS-68
chr13-7137896-INS-55
chr13-7149640-INS-6389
chr13-7154309-INS-163
chr13-7154621-INS-9045
chr13-7165939-INS-88
chr13-7181171-INS-72

chr13-7230038-INS-120
chr13-7233862-INS-6999
chr13-7255808-INS-161
chr13-7258641-INS-8414
chr13-7262509-INS-150
chr13-7308974-INS-68
chr13-7314137-INS-54
chr13-7317420-INS-296
chr13-7343783-INS-58
chr13-7347400-INS-127
chr13-7350957-INS-7208
chr13-7362871-INS-73
chr13-7371460-INS-75
chr13-7379777-INS-51
chr13-7387776-INS-52
chr13-7400433-INS-68
chr13-7441319-INS-271
chr13-7445592-INS-317
chr13-7465962-INS-719
chr13-7475773-INS-140
chr13-7484393-INS-6573
chr13-7499871-INS-400
chr13-7526462-INS-121
chr13-7534682-INS-86
chr13-7545151-INS-237
chr13-7565468-INS-400
chr13-7578458-INS-52
chr13-7590391-INS-60
chr13-7603321-INS-7529
chr13-7646546-INS-271
chr13-7658515-INS-234
chr13-7685563-INS-680
chr13-7709758-INS-72
chr13-7723202-INS-7212
chr13-7754813-INS-155
chr13-7767612-INS-142
chr13-7812599-INS-538
chr13-7816315-INS-3039
chr13-7857153-INS-1132
chr13-7899488-INS-683
chr13-7956731-INS-52
chr13-7976723-INS-435
chr13-7984736-INS-59
chr13-7992752-INS-8822
chr13-7998939-INS-58
chr13-8011372-INS-225
chr13-8011623-INS-135
chr13-8012424-INS-331
chr13-8075446-INS-773
chr13-8129019-INS-131
chr13-8146910-INS-643
chr13-8151442-INS-168
chr13-8205346-INS-6275
chr13-8242966-INS-75
chr13-8265408-INS-53
chr13-8272499-INS-416
chr13-8284894-INS-116
chr13-8314053-INS-204
chr13-8341617-INS-237
chr13-8375622-INS-110
chr13-8381317-INS-52
chr13-8396411-INS-196
chr13-8400266-INS-161
chr13-8404273-INS-68
chr13-8408936-INS-260
chr13-8414233-INS-99
chr13-8429931-INS-293
chr13-8433993-INS-6418
chr13-8441778-INS-2021
chr13-8462418-INS-67
chr13-8470937-INS-844
chr13-8475252-INS-160
chr13-8501176-INS-60
chr13-8525577-INS-133
chr13-8531814-INS-205
chr13-8545534-INS-306
chr13-8560490-INS-86

chr13-8576772-INS-219
chr13-8602720-INS-124
chr13-8634050-INS-58
chr13-8666430-INS-82
chr13-8695102-INS-215
chr13-8713634-INS-474
chr13-8740259-INS-105
chr13-8741210-INS-345
chr13-8781463-INS-60
chr13-8782643-INS-56
chr13-8802171-INS-75
chr13-8804631-INS-255
chr13-8851698-INS-126
chr13-8871506-INS-129
chr13-8877779-INS-168
chr13-8886440-INS-102
chr13-8901609-INS-302
chr13-8958660-INS-502
chr13-8961548-INS-206
chr13-9013069-INS-193
chr13-9040321-INS-212
chr13-9054191-INS-438
chr13-9054999-INS-188
chr13-9062100-INS-196
chr13-9070019-INS-237
chr13-9141016-INS-518
chr13-9153029-INS-213
chr13-9182441-INS-307
chr13-9188052-INS-280
chr13-9203107-INS-302
chr13-9210993-INS-211
chr13-9217614-INS-59
chr13-9218267-INS-1895
chr13-9227336-INS-184
chr13-9265593-INS-370
chr13-9267552-INS-221
chr13-9268602-INS-184
chr13-9272214-INS-235
chr13-9300617-INS-52
chr13-9335915-INS-142
chr13-9367135-INS-491
chr13-9369138-INS-89
chr13-9371569-INS-222
chr13-9414917-INS-52
chr13-9443930-INS-202
chr13-9491550-INS-768
chr13-9522778-INS-7292
chr13-9530776-INS-54
chr13-9583553-INS-81
chr13-9601972-INS-868
chr13-9676946-INS-121
chr13-9693767-INS-151
chr13-9714010-INS-320
chr13-9743812-INS-70
chr13-9783193-INS-162
chr13-9787572-INS-158
chr13-9801460-INS-228
chr13-9852393-INS-82
chr13-9859594-INS-113
chr13-9863418-INS-58
chr13-9875857-INS-94
chr13-9895718-INS-88
chr13-9920022-INS-108
chr13-9933592-INS-238
chr13-9950636-INS-232
chr13-9957001-INS-548
chr13-9966129-INS-159
chr13-9975261-INS-228
chr13-9983671-INS-69
chr13-10021325-INS-372
chr13-10034841-INS-75
chr13-10064619-INS-165
chr13-10080788-INS-135
chr13-10082287-INS-6369
chr13-10098865-INS-401
chr13-10129923-INS-78
chr13-10148444-INS-54

chr13-10149439-INS-364
chr13-10158876-INS-390
chr13-10163314-INS-54
chr13-10169509-INS-85
chr13-10176135-INS-133
chr13-10183235-INS-484
chr13-10232210-INS-119
chr13-10236949-INS-58
chr13-10239199-INS-53
chr13-10258972-INS-66
chr13-10264077-INS-64
chr13-10271270-INS-93
chr13-10275808-INS-385
chr13-10287213-INS-203
chr13-10327594-INS-114
chr13-10331964-INS-221
chr13-10346244-INS-74
chr13-10381467-INS-735
chr13-10400368-INS-656
chr13-10431125-INS-56
chr13-10431714-INS-61
chr13-10464629-INS-72
chr13-10474931-INS-131
chr13-10493232-INS-52
chr13-10532588-INS-241
chr13-10553640-INS-94
chr13-10558850-INS-81
chr13-10563450-INS-95
chr13-10577616-INS-58
chr13-10586346-INS-70
chr13-10589013-INS-827
chr13-10593014-INS-364
chr13-10615313-INS-297
chr13-10620636-INS-55
chr13-10622024-INS-163
chr13-10648005-INS-64
chr13-10670151-INS-507
chr13-10679754-INS-113
chr13-10682544-INS-74
chr13-10684897-INS-115
chr13-10690502-INS-6471
chr13-10781967-INS-62
chr13-10795145-INS-520
chr13-10862751-INS-310
chr13-10879938-INS-58
chr13-10917396-INS-98
chr13-10962671-INS-53
chr13-10984381-INS-62
chr13-11004609-INS-3499
chr13-11039720-INS-293
chr13-11050613-INS-60
chr13-11068643-INS-330
chr13-11077775-INS-538
chr13-11081561-INS-8420
chr13-11089605-INS-6645
chr13-11095457-INS-51
chr13-11095505-INS-2734
chr13-11097307-INS-54
chr13-11098734-INS-6936
chr13-11104325-INS-628
chr13-11105567-INS-121
chr13-11115403-INS-591
chr13-11138508-INS-145
chr13-11139726-INS-338
chr13-11141904-INS-83
chr13-11173627-INS-52
chr13-11199759-INS-110
chr13-11209192-INS-93
chr13-11223619-INS-74
chr13-11238228-INS-70
chr13-11251078-INS-401
chr13-11257038-INS-429
chr13-11298836-INS-90
chr13-11299080-INS-212
chr13-11306576-INS-173
chr13-11326011-INS-70
chr13-11339340-INS-110

chr13-11361481-INS-76
chr13-11362398-INS-62
chr13-11384884-INS-703
chr13-11408099-INS-5543
chr13-11422131-INS-57
chr13-11427424-INS-340
chr13-11442242-INS-158
chr13-11463512-INS-57
chr13-11470129-INS-406
chr13-11500242-INS-770
chr13-11506506-INS-6232
chr13-11527535-INS-6233
chr13-11531574-INS-140
chr13-11593577-INS-74
chr13-11597875-INS-61
chr13-11618052-INS-126
chr13-11629472-INS-432
chr13-11643756-INS-136
chr13-11658025-INS-105
chr13-11666655-INS-100
chr13-11672810-INS-613
chr13-11680258-INS-580
chr13-11686979-INS-390
chr13-11717541-INS-215
chr13-11737097-INS-448
chr13-11748208-INS-254
chr13-11768682-INS-623
chr13-11799682-INS-60
chr13-11816269-INS-608
chr13-11817699-INS-54
chr13-11842129-INS-320
chr13-11895967-INS-70
chr13-11900299-INS-862
chr13-11910456-INS-52
chr13-11919659-INS-70
chr13-11927741-INS-191
chr13-11935441-INS-171
chr13-11948240-INS-69
chr13-12009249-INS-189
chr13-12061709-INS-136
chr13-12073222-INS-89
chr13-12090846-INS-875
chr13-12093744-INS-931
chr13-12114729-INS-274
chr13-12123991-INS-138
chr13-12131611-INS-207
chr13-12166583-INS-134
chr13-12177788-INS-875
chr13-12182158-INS-106
chr13-12183255-INS-375
chr13-12220442-INS-224
chr13-12242870-INS-161
chr13-12252459-INS-143
chr13-12263757-INS-293
chr13-12281235-INS-56
chr13-12300517-INS-98
chr13-12317962-INS-155
chr13-12320385-INS-53
chr13-12322453-INS-467
chr13-12328399-INS-72
chr13-12336776-INS-56
chr13-12344018-INS-68
chr13-12370480-INS-61
chr13-12375415-INS-207
chr13-12381081-INS-127
chr13-12394327-INS-864
chr13-12420248-INS-8739
chr13-12433370-INS-100
chr13-12434700-INS-232
chr13-12451703-INS-132
chr13-12456614-INS-219
chr13-12507692-INS-255
chr13-12521100-INS-63
chr13-12524347-INS-194
chr13-12538865-INS-76
chr13-12577770-INS-53
chr13-12586240-INS-225

chr13-12612367-INS-79
chr13-12640529-INS-55
chr13-12647993-INS-123
chr13-12649655-INS-307
chr13-12652811-INS-200
chr13-12662041-INS-4039
chr13-12701217-INS-172
chr13-12704586-INS-157
chr13-12709958-INS-6454
chr13-12710542-INS-167
chr13-12712077-INS-107
chr13-12727372-INS-67
chr13-12728169-INS-60
chr13-12772834-INS-7163
chr13-12780588-INS-137
chr13-12985456-INS-67
chr13-13236451-INS-54
chr13-13376053-INS-401
chr13-13537139-INS-232
chr13-13549022-INS-501
chr13-13550884-INS-91
chr13-13552617-INS-288
chr13-13560820-INS-5557
chr13-13584721-INS-3329
chr13-13584722-INS-578
chr13-13589709-INS-269
chr13-13596070-INS-172
chr13-13599449-INS-184
chr13-13607206-INS-168
chr13-13622225-INS-75
chr13-13622740-INS-53
chr13-13639137-INS-571
chr13-13658093-INS-59
chr13-13669086-INS-190
chr13-13687156-INS-86
chr13-13696794-INS-78
chr13-13702227-INS-74
chr13-13723948-INS-92
chr13-13727574-INS-400
chr13-13769797-INS-378
chr13-13790599-INS-54
chr13-13793760-INS-480
chr13-13795399-INS-132
chr13-13796802-INS-126
chr13-13806630-INS-112
chr13-13834581-INS-448
chr13-13852932-INS-226
chr13-13867600-INS-993
chr13-13873260-INS-70
chr13-13876838-INS-210
chr13-13893787-INS-408
chr13-13928552-INS-1742
chr13-13933655-INS-187
chr13-13952677-INS-53
chr13-13968798-INS-85
chr13-13977732-INS-56
chr13-13997480-INS-183
chr13-14014458-INS-374
chr13-14029233-INS-54
chr13-14031906-INS-181
chr13-14035643-INS-173
chr13-14037404-INS-101
chr13-14041463-INS-64
chr13-14043093-INS-64
chr13-14056120-INS-712
chr13-14068085-INS-6454
chr13-14070393-INS-607
chr13-14072145-INS-80
chr13-14074287-INS-233
chr13-14095485-INS-51
chr13-14120801-INS-810
chr13-14128317-INS-90
chr13-14156827-INS-73
chr13-14171381-INS-1053
chr13-14171381-INS-284
chr13-14173341-INS-56
chr13-14187128-INS-94

chr13-14208947-INS-138
chr13-14213308-INS-265
chr13-14223893-INS-529
chr13-14228184-INS-54
chr13-14249649-INS-292
chr13-14267389-INS-105
chr13-14268676-INS-401
chr13-14299243-INS-115
chr13-14319224-INS-96
chr13-14332511-INS-74
chr13-14400588-INS-84
chr13-14454267-INS-249
chr13-14540779-INS-2978
chr13-14547470-INS-143
chr13-14548247-INS-516
chr13-14557106-INS-169
chr13-14562129-INS-758
chr13-14595118-INS-793
chr13-14616334-INS-63
chr13-14632714-INS-420
chr13-14642996-INS-210
chr13-14679911-INS-7305
chr13-14726125-INS-54
chr13-14733047-INS-6345
chr13-14757995-INS-1013
chr13-14759728-INS-125
chr13-14760125-INS-6214
chr13-14761178-INS-504
chr13-14762815-INS-132
chr13-14763958-INS-6574
chr13-14764766-INS-503
chr13-14816661-INS-237
chr13-14826505-INS-59
chr13-14830776-INS-76
chr13-14844249-INS-213
chr13-14847309-INS-6180
chr13-14876930-INS-492
chr13-14884771-INS-366
chr13-14901386-INS-520
chr13-14913541-INS-175
chr13-14922798-INS-7066
chr13-14930889-INS-170
chr13-14931729-INS-5499
chr13-14950192-INS-1990
chr13-14950991-INS-5533
chr13-14950994-INS-4724
chr13-14953984-INS-598
chr13-14958653-INS-106
chr13-14958983-INS-6416
chr13-15010647-INS-108
chr13-15053343-INS-58
chr13-15077064-INS-75
chr13-15139898-INS-56
chr13-15141796-INS-75
chr13-15166486-INS-53
chr13-15223313-INS-2754
chr13-15266525-INS-54
chr13-15281778-INS-664
chr13-15308771-INS-2957
chr13-15322396-INS-226
chr13-15330814-INS-52
chr13-15383014-INS-6675
chr13-15399648-INS-132
chr13-15400412-INS-312
chr13-15421132-INS-1885
chr13-15436549-INS-1858
chr13-15437092-INS-74
chr13-15441067-INS-439
chr13-15464782-INS-51
chr13-15468665-INS-516
chr13-15484731-INS-89
chr13-15485765-INS-204
chr13-15494414-INS-7223
chr13-15502366-INS-672
chr13-15504523-INS-120
chr13-15523394-INS-764
chr13-15529353-INS-293

chr13-15542739-INS-130
chr13-15551955-INS-52
chr13-15556418-INS-146
chr13-15557179-INS-358
chr13-15559886-INS-6754
chr13-15561793-INS-148
chr13-15584124-INS-89
chr13-15597167-INS-623
chr13-15597553-INS-7015
chr13-15628184-INS-147
chr13-15640051-INS-56
chr13-15672662-INS-77
chr13-15678620-INS-75
chr13-15689369-INS-140
chr13-15719194-INS-72
chr13-15735629-INS-149
chr13-15744449-INS-276
chr13-15750705-INS-179
chr13-15760114-INS-142
chr13-15783443-INS-183
chr13-15832818-INS-240
chr13-15854037-INS-7039
chr13-15864378-INS-6302
chr13-15884020-INS-61
chr13-15897104-INS-312
chr13-15898993-INS-95
chr13-15936447-INS-63
chr13-15941170-INS-481
chr13-15947829-INS-68
chr13-15958957-INS-5440
chr13-15972857-INS-6363
chr13-15996470-INS-214
chr13-16001071-INS-86
chr13-16002500-INS-84
chr13-16008077-INS-455
chr13-16034569-INS-512
chr13-16202288-INS-242
chr13-16216573-INS-144
chr13-16232881-INS-809
chr13-16238119-INS-817
chr13-16269722-INS-842
chr13-16309418-INS-375
chr13-16318748-INS-352
chr13-16324461-INS-7209
chr13-16346990-INS-137
chr13-16389909-INS-213
chr13-16394042-INS-51
chr13-16400579-INS-54
chr13-16420329-INS-54
chr13-16437353-INS-6369
chr13-16525493-INS-99
chr13-16540913-INS-6245
chr13-16550240-INS-358
chr13-16578515-INS-62
chr13-16616378-INS-3902
chr13-16634887-INS-154
chr13-16643692-INS-154
chr13-16658572-INS-6267
chr13-16668366-INS-7113
chr13-16668564-INS-1299
chr13-16669893-INS-289
chr13-16688409-INS-5500
chr13-16697141-INS-64
chr13-16703027-INS-61
chr13-16720379-INS-163
chr13-16730985-INS-407
chr13-16766353-INS-77
chr13-16768336-INS-134
chr13-16815231-INS-81
chr13-16824696-INS-3822
chr13-16831948-INS-93
chr13-16873325-INS-225
chr13-16880773-INS-109
chr13-16890793-INS-52
chr13-16895196-INS-321
chr13-16912468-INS-7179
chr13-16921545-INS-429

chr13-16927750-INS-990
chr13-16940547-INS-51
chr13-16954015-INS-167
chr13-16966057-INS-177
chr13-16971551-INS-751
chr13-16973808-INS-594
chr13-16980975-INS-549
chr13-16982284-INS-180
chr13-16983127-INS-69
chr13-16992615-INS-55
chr13-17018584-INS-75
chr13-17026242-INS-229
chr13-17039310-INS-133
chr13-17040445-INS-484
chr13-17047432-INS-6440
chr13-17074834-INS-7089
chr13-17082654-INS-66
chr13-17126297-INS-6662
chr13-17127655-INS-434
chr13-17137136-INS-55
chr13-17155716-INS-554
chr13-17164403-INS-66
chr13-17169635-INS-284
chr13-17175788-INS-101
chr13-17185557-INS-882
chr13-17245819-INS-363
chr13-17255294-INS-216
chr13-17268313-INS-76
chr13-17274924-INS-74
chr13-17286130-INS-400
chr13-17307107-INS-98
chr13-17312617-INS-52
chr13-17341342-INS-223
chr13-17348077-INS-56
chr13-17388287-INS-52
chr13-17404828-INS-2555
chr13-17439367-INS-521
chr13-17494360-INS-7013
chr13-17522426-INS-596
chr13-17524611-INS-335
chr13-17529300-INS-497
chr13-17532619-INS-53
chr13-17540826-INS-148
chr13-17556759-INS-6359
chr13-17581976-INS-6116
chr13-17623377-INS-187
chr13-17663586-INS-340
chr13-17672754-INS-6490
chr13-17716109-INS-55
chr13-17734491-INS-174
chr13-17736112-INS-194
chr13-17763356-INS-966
chr13-17772222-INS-340
chr13-17819826-INS-7113
chr13-17838012-INS-269
chr13-17851784-INS-204
chr13-17878047-INS-553
chr13-17927175-INS-72
chr13-17985880-INS-436
chr13-18006649-INS-56
chr13-18021508-INS-359
chr13-18035742-INS-51
chr13-18038621-INS-156
chr13-18042205-INS-82
chr13-18057907-INS-52
chr13-18095949-INS-5864
chr13-18134214-INS-221
chr13-18147822-INS-399
chr13-18186057-INS-78
chr13-18191745-INS-120
chr13-18194005-INS-480
chr13-18201217-INS-87
chr13-18225315-INS-6592
chr13-18230479-INS-70
chr13-18247183-INS-506
chr13-18252279-INS-164
chr13-18274959-INS-541

chr13-18284009-INS-81
chr13-18338906-INS-136
chr13-18351170-INS-539
chr13-18372948-INS-73
chr13-18377212-INS-206
chr13-18397275-INS-213
chr13-18434220-INS-81
chr13-18444950-INS-157
chr13-18446376-INS-56
chr13-18447362-INS-216
chr13-18467425-INS-52
chr13-18470114-INS-67
chr13-18476107-INS-57
chr13-18482779-INS-92
chr13-18491844-INS-63
chr13-18496956-INS-58
chr13-18497924-INS-58
chr13-18518491-INS-162
chr13-18541128-INS-68
chr13-18589034-INS-226
chr13-18602392-INS-60
chr13-18615463-INS-173
chr13-18625449-INS-76
chr13-18633622-INS-553
chr13-18644443-INS-102
chr13-18653044-INS-62
chr13-18655599-INS-431
chr13-18693970-INS-56
chr13-18744209-INS-51
chr13-18758341-INS-54
chr13-18762436-INS-58
chr13-18770566-INS-69
chr13-18803909-INS-6989
chr13-18808315-INS-3055
chr13-18835074-INS-66
chr13-18855926-INS-52
chr13-18868916-INS-2010
chr13-18885841-INS-4446
chr13-18896223-INS-397
chr13-18909006-INS-289
chr13-18910396-INS-177
chr13-18926503-INS-761
chr13-18931930-INS-201
chr13-18952792-INS-120
chr13-18957902-INS-54
chr13-18959476-INS-462
chr13-18960724-INS-53
chr13-18985941-INS-117
chr13-19000593-INS-54
chr13-19001926-INS-55
chr13-19008909-INS-189
chr13-19063149-INS-308
chr13-19070186-INS-192
chr13-19072731-INS-375
chr13-19087659-INS-396
chr13-19097217-INS-377
chr13-19211784-INS-74
chr13-19221916-INS-222
chr13-19222849-INS-340
chr13-19253025-INS-5732
chr13-19260410-INS-6808
chr13-19265643-INS-13386
chr13-19277006-INS-52
chr13-19280357-INS-73
chr13-19291235-INS-111
chr13-19324917-INS-924
chr13-19368718-INS-151
chr13-19400920-INS-57
chr13-19406801-INS-83
chr13-19468301-INS-284
chr13-19478713-INS-62
chr13-19483773-INS-354
chr13-19492066-INS-7008
chr13-19509928-INS-67
chr13-19514776-INS-6387
chr13-19522171-INS-55
chr13-19530964-INS-193

chr13-19544211-INS-304
chr13-19546205-INS-102
chr13-19565408-INS-203
chr13-19584856-INS-328
chr13-19634754-INS-160
chr13-19651552-INS-60
chr13-19658177-INS-532
chr13-19660407-INS-275
chr13-19666245-INS-93
chr13-19683010-INS-66
chr13-19697400-INS-120
chr13-19707571-INS-88
chr13-19712344-INS-5545
chr13-19716310-INS-6906
chr13-19784425-INS-508
chr13-19830827-INS-656
chr13-19834588-INS-162
chr13-19849020-INS-198
chr13-19886739-INS-70
chr13-19892564-INS-72
chr13-19898901-INS-522
chr13-19900627-INS-888
chr13-19908589-INS-6796
chr13-19931773-INS-118
chr13-19962359-INS-205
chr13-19986082-INS-327
chr13-20000979-INS-1864
chr13-20026752-INS-802
chr13-20031612-INS-612
chr13-20057495-INS-6387
chr13-20092927-INS-127
chr13-20102135-INS-75
chr13-20103871-INS-407
chr13-20106000-INS-113
chr13-20123985-INS-947
chr13-20157166-INS-9568
chr13-20161005-INS-324
chr13-20174802-INS-60
chr13-20182518-INS-338
chr13-20197893-INS-370
chr13-20208599-INS-95
chr13-20209494-INS-204
chr13-20210495-INS-53
chr13-20262801-INS-52
chr13-20273280-INS-79
chr13-20278371-INS-200
chr13-20280005-INS-84
chr13-20292965-INS-371
chr13-20311239-INS-69
chr13-20314368-INS-56
chr13-20326260-INS-236
chr13-20329558-INS-132
chr13-20353630-INS-122
chr13-20358719-INS-901
chr13-20363365-INS-144
chr13-20383381-INS-197
chr13-20385290-INS-171
chr13-20394669-INS-358
chr13-20420897-INS-91
chr13-20432295-INS-123
chr13-20438638-INS-178
chr13-20448175-INS-133
chr13-20464788-INS-68
chr13-20484116-INS-64
chr13-20486245-INS-75
chr13-20492599-INS-190
chr13-20514802-INS-90
chr13-20535340-INS-615
chr13-20615928-INS-104
chr13-20621785-INS-64
chr13-20632317-INS-56
chr13-20651178-INS-166
chr13-20685736-INS-228
chr13-20698219-INS-1039
chr13-20700782-INS-369
chr13-20737857-INS-60
chr13-20801242-INS-52

chr13-20813991-INS-57
chr13-20827750-INS-7096
chr13-20828584-INS-192
chr13-20833125-INS-6403
chr13-20883329-INS-55
chr13-20887581-INS-116
chr13-20894835-INS-51
chr13-20913741-INS-493
chr13-20916296-INS-6268
chr13-20925350-INS-108
chr13-20945092-INS-247
chr13-20954862-INS-407
chr13-20971530-INS-79
chr13-20971593-INS-7510
chr13-20975175-INS-91
chr13-20989531-INS-196
chr13-20990618-INS-6546
chr13-21004356-INS-66
chr13-21010780-INS-73
chr13-21026653-INS-7055
chr13-21040376-INS-93
chr13-21042395-INS-152
chr13-21074737-INS-396
chr13-21089940-INS-52
chr13-21097974-INS-92
chr13-21103856-INS-80
chr13-21108695-INS-120
chr13-21119101-INS-53
chr13-21122655-INS-6859
chr13-21128988-INS-344
chr13-21136852-INS-272
chr13-21146652-INS-916
chr13-21174812-INS-180
chr13-21183791-INS-631
chr13-21188199-INS-118
chr13-21202804-INS-420
chr13-21206532-INS-7178
chr13-21220911-INS-7290
chr13-21277221-INS-502
chr13-21279788-INS-413
chr13-21286184-INS-59
chr13-21295219-INS-6203
chr13-21309247-INS-232
chr13-21310011-INS-1360
chr13-21320508-INS-206
chr13-21337194-INS-167
chr13-21340004-INS-3094
chr13-21366380-INS-135
chr13-21368485-INS-189
chr13-21396563-INS-52
chr13-21413516-INS-497
chr13-21458776-INS-60
chr13-21462997-INS-179
chr13-21465662-INS-527
chr13-21477549-INS-186
chr13-21478867-INS-448
chr13-21494649-INS-362
chr13-21532831-INS-944
chr13-21538858-INS-83
chr13-21611430-INS-247
chr13-21624419-INS-61
chr13-21631459-INS-96
chr13-21646165-INS-52
chr13-21671116-INS-709
chr13-21694643-INS-6722
chr13-21733179-INS-177
chr13-21739708-INS-191
chr13-21747310-INS-6287
chr13-21762259-INS-391
chr13-21794887-INS-60
chr13-21807577-INS-554
chr13-21876368-INS-54
chr13-21885712-INS-76
chr13-21896410-INS-92
chr13-21910940-INS-71
chr13-21911721-INS-53
chr13-21928978-INS-180

chr13-21936055-INS-58
chr13-21945942-INS-649
chr13-21960562-INS-779
chr13-21962171-INS-118
chr13-21965345-INS-422
chr13-21990919-INS-595
chr13-22009749-INS-577
chr13-22013718-INS-195
chr13-22015055-INS-467
chr13-22021324-INS-595
chr13-22027450-INS-255
chr13-22037868-INS-698
chr13-22037990-INS-54
chr13-22041321-INS-59
chr13-22054394-INS-68
chr13-22062041-INS-283
chr13-22066069-INS-691
chr13-22068386-INS-97
chr13-22079431-INS-65
chr13-22089192-INS-511
chr13-22099463-INS-60
chr13-22106337-INS-83
chr13-22149034-INS-60
chr13-22171250-INS-209
chr13-22194308-INS-879
chr13-22195264-INS-246
chr13-22206326-INS-199
chr13-22209781-INS-110
chr13-22212567-INS-765
chr13-22214496-INS-151
chr13-22218000-INS-120
chr13-22222558-INS-180
chr13-22237964-INS-117
chr13-22277102-INS-60
chr13-22280550-INS-159
chr13-22344192-INS-177
chr13-22350801-INS-4803
chr13-22376503-INS-2702
chr13-22420435-INS-4564
chr13-22424510-INS-183
chr13-22429590-INS-578
chr13-22432941-INS-110
chr13-22443146-INS-498
chr13-22444789-INS-401
chr13-22472308-INS-274
chr13-22486700-INS-58
chr13-22502521-INS-127
chr13-22505921-INS-64
chr13-22513486-INS-6294
chr13-22517529-INS-6597
chr13-22522910-INS-6382
chr13-22552829-INS-61
chr13-22558296-INS-489
chr13-22559442-INS-71
chr13-22586934-INS-6615
chr13-22624496-INS-3764
chr13-22627822-INS-447
chr13-22647197-INS-393
chr13-22670841-INS-686
chr13-22672541-INS-348
chr13-22674425-INS-109
chr13-22770341-INS-151
chr13-22781882-INS-236
chr13-22803420-INS-189
chr13-22849176-INS-58
chr13-22861194-INS-72
chr13-22872697-INS-5549
chr13-22939002-INS-954
chr13-22950585-INS-52
chr13-22951237-INS-7067
chr13-23021409-INS-109
chr13-23023496-INS-6010
chr13-23031521-INS-3802
chr13-23046497-INS-54
chr13-23064726-INS-96
chr13-23076883-INS-151
chr13-23083388-INS-62

chr13-23124862-INS-62
chr13-23171106-INS-52
chr13-23201709-INS-58
chr13-23217806-INS-104
chr13-23225393-INS-268
chr13-23225942-INS-148
chr13-23234050-INS-96
chr13-23256877-INS-367
chr13-23271088-INS-7248
chr13-23278616-INS-3077
chr13-23295677-INS-5311
chr13-23314511-INS-2449
chr13-23340383-INS-136
chr13-23381915-INS-6643
chr13-23408265-INS-445
chr13-23408839-INS-127
chr13-23410487-INS-69
chr13-23425139-INS-592
chr13-23427968-INS-224
chr13-23429146-INS-207
chr13-23432338-INS-213
chr13-23462005-INS-394
chr13-23467124-INS-488
chr13-23475269-INS-63
chr13-23484560-INS-106
chr13-23486109-INS-83
chr13-23489095-INS-6210
chr13-23503798-INS-58
chr13-23507818-INS-2795
chr13-23514449-INS-132
chr13-23521543-INS-95
chr13-23598520-INS-74
chr13-23601585-INS-173
chr13-23601673-INS-342
chr13-23614821-INS-115
chr13-23651416-INS-212
chr13-23654214-INS-74
chr13-23660209-INS-231
chr13-23674656-INS-6602
chr13-23675666-INS-679
chr13-23678350-INS-7010
chr13-23681732-INS-156
chr13-23700331-INS-93
chr13-23718891-INS-284
chr13-23721731-INS-52
chr13-23765971-INS-782
chr13-23791335-INS-54
chr13-23847597-INS-163
chr13-23849095-INS-4359
chr13-23849523-INS-477
chr13-23858539-INS-71
chr13-23875993-INS-7069
chr13-23885609-INS-58
chr13-23886935-INS-1475
chr13-23890723-INS-302
chr13-23901039-INS-134
chr13-23901680-INS-528
chr13-23915268-INS-53
chr13-23924750-INS-199
chr13-23951525-INS-54
chr13-23976297-INS-168
chr13-23984552-INS-140
chr13-24017104-INS-108
chr13-24030528-INS-181
chr13-24047423-INS-439
chr13-24117013-INS-90
chr13-24135199-INS-118
chr13-24137260-INS-72
chr13-24138338-INS-5544
chr13-24139203-INS-115
chr13-24144669-INS-75
chr13-24151053-INS-561
chr13-24153509-INS-937
chr13-24168796-INS-206
chr13-24170196-INS-274
chr13-24176012-INS-110
chr13-24191237-INS-329

chr13-24288722-INS-966
chr13-24288944-INS-289
chr13-24340084-INS-137
chr13-24348651-INS-344
chr13-24352572-INS-72
chr13-24355142-INS-57
chr13-24370432-INS-400
chr13-24381083-INS-508
chr13-24437904-INS-108
chr13-24444730-INS-51
chr13-24451101-INS-52
chr13-24468957-INS-66
chr13-24484197-INS-51
chr13-24525803-INS-86
chr13-24534062-INS-6366
chr13-24535192-INS-640
chr13-24569785-INS-640
chr13-24579156-INS-84
chr13-24591828-INS-204
chr13-24605782-INS-58
chr13-24628137-INS-64
chr13-24630711-INS-56
chr13-24682321-INS-67
chr13-24686091-INS-54
chr13-24755086-INS-192
chr13-24794013-INS-400
chr13-24819971-INS-81
chr13-24827594-INS-191
chr13-24842992-INS-58
chr13-24884669-INS-243
chr13-24891690-INS-67
chr13-24910311-INS-105
chr13-24911632-INS-53
chr13-24922922-INS-91
chr13-24940171-INS-76
chr13-24955598-INS-146
chr13-24969420-INS-126
chr13-24978589-INS-268
chr13-25023030-INS-208
chr13-25056755-INS-122
chr13-25058031-INS-525
chr13-25061110-INS-399
chr13-25063307-INS-196
chr13-25081247-INS-150
chr13-25088833-INS-86
chr13-25098987-INS-172
chr13-25111031-INS-202
chr13-25116274-INS-508
chr13-25162975-INS-62
chr13-25167669-INS-51
chr13-25183624-INS-85
chr13-25205392-INS-100
chr13-25221418-INS-833
chr13-25234656-INS-75
chr13-25247530-INS-150
chr13-25257157-INS-722
chr13-25275266-INS-78
chr13-25279680-INS-202
chr13-25285359-INS-215
chr13-25304158-INS-400
chr13-25327456-INS-60
chr13-25330554-INS-302
chr13-25335829-INS-132
chr13-25342348-INS-653
chr13-25364699-INS-93
chr13-25370449-INS-135
chr13-25385413-INS-54
chr13-25394525-INS-54
chr13-25395400-INS-56
chr13-25408132-INS-100
chr13-25413010-INS-8698
chr13-25466186-INS-528
chr13-25495681-INS-70
chr13-25530906-INS-158
chr13-25591798-INS-6440
chr13-25612254-INS-69
chr13-25625354-INS-66

chr13-25634393-INS-167
chr13-25649723-INS-153
chr13-25688080-INS-6526
chr13-25690384-INS-182
chr13-25696210-INS-6378
chr13-25704370-INS-138
chr13-25708685-INS-53
chr13-25717212-INS-165
chr13-25726901-INS-208
chr13-25767708-INS-6410
chr13-25785331-INS-71
chr13-25789997-INS-673
chr13-25827620-INS-130
chr13-25861096-INS-210
chr13-25885238-INS-174
chr13-25888217-INS-1950
chr13-25895952-INS-6301
chr13-25896524-INS-90
chr13-25901405-INS-80
chr13-25907741-INS-52
chr13-25921093-INS-204
chr13-25938411-INS-96
chr13-25953786-INS-565
chr13-25960668-INS-94
chr13-26015441-INS-459
chr13-26038477-INS-243
chr13-26039963-INS-61
chr13-26043107-INS-739
chr13-26056873-INS-831
chr13-26077110-INS-62
chr13-26104353-INS-52
chr13-26120723-INS-227
chr13-26145121-INS-904
chr13-26146832-INS-132
chr13-26167193-INS-166
chr13-26169910-INS-157
chr13-26181107-INS-141
chr13-26210827-INS-101
chr13-26229069-INS-62
chr13-26250033-INS-553
chr13-26251335-INS-156
chr13-26267462-INS-76
chr13-26273524-INS-528
chr13-26278386-INS-173
chr13-26303704-INS-60
chr13-26308634-INS-476
chr13-26313485-INS-52
chr13-26322141-INS-6605
chr13-26325544-INS-198
chr13-26337120-INS-65
chr13-26344700-INS-55
chr13-26351504-INS-6312
chr13-26358287-INS-426
chr13-26372809-INS-190
chr13-26374204-INS-199
chr13-26401360-INS-1225
chr13-26403408-INS-407
chr13-26418553-INS-829
chr13-26434193-INS-302
chr13-26438724-INS-629
chr13-26444096-INS-6258
chr13-26447103-INS-7483
chr13-26451973-INS-4278
chr13-26468558-INS-142
chr13-26474933-INS-529
chr13-26485986-INS-4957
chr13-26506796-INS-319
chr13-26514201-INS-400
chr13-26515343-INS-54
chr13-26534377-INS-178
chr13-26539632-INS-52
chr13-26567997-INS-237
chr13-26587870-INS-255
chr13-26589437-INS-62
chr13-26595261-INS-64
chr13-26600479-INS-135
chr13-26604307-INS-77

chr13-26625252-INS-212
chr13-26640037-INS-444
chr13-26640083-INS-259
chr13-26644226-INS-536
chr13-26645732-INS-52
chr13-26653902-INS-108
chr13-26682065-INS-64
chr13-26692945-INS-113
chr13-26718074-INS-6426
chr13-26719801-INS-64
chr13-26733661-INS-214
chr13-26736722-INS-82
chr13-26739433-INS-62
chr13-26764693-INS-62
chr13-26767077-INS-135
chr13-26782464-INS-331
chr13-26786643-INS-573
chr13-26826547-INS-123
chr13-26829437-INS-57
chr13-26852836-INS-119
chr13-26862750-INS-6471
chr13-26880418-INS-6741
chr13-26897864-INS-182
chr13-26900708-INS-60
chr13-26916567-INS-638
chr13-26954951-INS-6175
chr13-26957074-INS-170
chr13-26973996-INS-85
chr13-26981144-INS-52
chr13-26999954-INS-1675
chr13-27000332-INS-742
chr13-27058501-INS-59
chr13-27088389-INS-128
chr13-27090080-INS-197
chr13-27109975-INS-547
chr13-27113052-INS-64
chr13-27117540-INS-105
chr13-27179369-INS-54
chr13-27181247-INS-52
chr13-27208965-INS-78
chr13-27219066-INS-110
chr13-27228760-INS-1357
chr13-27240356-INS-59
chr13-27243347-INS-485
chr13-27275124-INS-252
chr13-27318127-INS-178
chr13-27354163-INS-249
chr13-27361625-INS-64
chr13-27363981-INS-75
chr13-27386061-INS-432
chr13-27386210-INS-7076
chr13-27387374-INS-65
chr13-27389039-INS-74
chr13-27390100-INS-160
chr13-27409603-INS-64
chr13-27427904-INS-52
chr13-27432438-INS-380
chr13-27440213-INS-55
chr13-27452798-INS-192
chr13-27477695-INS-182
chr13-27483519-INS-104
chr13-27517874-INS-97
chr13-27519685-INS-208
chr13-27533299-INS-56
chr13-27537792-INS-104
chr13-27569761-INS-55
chr13-27577954-INS-134
chr13-27579696-INS-846
chr13-27593933-INS-63
chr13-27595785-INS-330
chr13-27620528-INS-53
chr13-27645434-INS-6271
chr13-27654159-INS-2859
chr13-27677771-INS-121
chr13-27695268-INS-79
chr13-27708896-INS-140
chr13-27735784-INS-563

chr13-27738797-INS-6726
chr13-27743556-INS-284
chr13-27753706-INS-64
chr13-27779837-INS-365
chr13-27789546-INS-388
chr13-27835200-INS-53
chr13-27841814-INS-50
chr13-27844185-INS-53
chr13-27854818-INS-230
chr13-27887105-INS-963
chr13-27928344-INS-528
chr13-27935975-INS-139
chr13-27943833-INS-57
chr13-27955921-INS-109
chr13-27995952-INS-56
chr13-27996832-INS-6581
chr13-28016984-INS-106
chr13-28090015-INS-125
chr13-28092302-INS-218
chr13-28095557-INS-471
chr13-28140416-INS-104
chr13-28157386-INS-463
chr13-28161806-INS-540
chr13-28165544-INS-52
chr13-28176561-INS-632
chr13-28187768-INS-1156
chr13-28196242-INS-88
chr13-28212923-INS-232
chr13-28319912-INS-168
chr13-28334134-INS-192
chr13-28353552-INS-98
chr13-28365384-INS-63
chr13-28407718-INS-56
chr13-28440806-INS-60
chr13-28443854-INS-485
chr13-28454283-INS-3117
chr13-28467344-INS-581
chr13-28477494-INS-52
chr13-28491232-INS-4154
chr13-28492379-INS-358
chr13-28535932-INS-156
chr13-28548060-INS-6495
chr13-28548061-INS-168
chr13-28550273-INS-665
chr13-28555177-INS-599
chr13-28562130-INS-196
chr13-28572549-INS-545
chr13-28580513-INS-55
chr13-28585734-INS-79
chr13-28627496-INS-6282
chr13-28750449-INS-264
chr13-28754050-INS-807
chr13-28758121-INS-55
chr13-28800577-INS-454
chr13-28834286-INS-118
chr13-28852911-INS-92
chr13-28860098-INS-101
chr13-28861476-INS-133
chr13-28878303-INS-56
chr13-28907629-INS-53
chr13-28917674-INS-351
chr13-28945349-INS-102
chr13-28960776-INS-52
chr13-29002760-INS-83
chr13-29004222-INS-76
chr13-29022054-INS-141
chr13-29030623-INS-62
chr13-29034221-INS-75
chr13-29037431-INS-92
chr13-29078748-INS-78
chr13-29079218-INS-203
chr13-29120504-INS-79
chr13-29146437-INS-394
chr13-29172622-INS-1778
chr13-29177310-INS-101
chr13-29180909-INS-326
chr13-29194384-INS-3787

chr13-29204391-INS-468
chr13-29210341-INS-202
chr13-29213554-INS-67
chr13-29219890-INS-763
chr13-29222794-INS-51
chr13-29227390-INS-54
chr13-29263450-INS-485
chr13-29276445-INS-56
chr13-29309241-INS-106
chr13-29332196-INS-549
chr13-29335253-INS-55
chr13-29339960-INS-78
chr13-29352023-INS-135
chr13-29362936-INS-81
chr13-29367917-INS-496
chr13-29429199-INS-80
chr13-29447348-INS-121
chr13-29450262-INS-219
chr13-29468133-INS-109
chr13-29503848-INS-117
chr13-29513549-INS-355
chr13-29514124-INS-74
chr13-29582207-INS-121
chr13-29662211-INS-450
chr13-29662454-INS-4134
chr13-29678480-INS-185
chr13-29702036-INS-97
chr13-29708500-INS-203
chr13-29711710-INS-145
chr13-29718692-INS-183
chr13-29727677-INS-117
chr13-29760135-INS-288
chr13-29761719-INS-3937
chr13-29762379-INS-115
chr13-29782061-INS-64
chr13-29788589-INS-60
chr13-29809283-INS-458
chr13-29811293-INS-95
chr13-29813124-INS-65
chr13-29829420-INS-104
chr13-29857247-INS-212
chr13-29872710-INS-84
chr13-29884403-INS-400
chr13-29886813-INS-136
chr13-29901509-INS-60
chr13-29927590-INS-405
chr13-29934626-INS-158
chr13-29936771-INS-52
chr13-29957195-INS-58
chr13-29986484-INS-103
chr13-29990649-INS-6805
chr13-30032645-INS-58
chr13-30043296-INS-289
chr13-30046354-INS-309
chr13-30047823-INS-129
chr13-30057646-INS-293
chr13-30127574-INS-60
chr13-30144899-INS-56
chr13-30187993-INS-54
chr13-30205309-INS-52
chr13-30222089-INS-54
chr13-30264638-INS-222
chr13-30291813-INS-202
chr13-30297228-INS-72
chr13-30298862-INS-54
chr13-30327896-INS-366
chr13-30329630-INS-52
chr13-30340758-INS-84
chr13-30348536-INS-136
chr13-30383608-INS-52
chr13-30395992-INS-341
chr13-30400533-INS-346
chr13-30443661-INS-68
chr13-30462430-INS-391
chr13-30471332-INS-294
chr13-30475692-INS-374
chr13-30524502-INS-78

chr13-30529952-INS-267
chr13-30534455-INS-923
chr13-30538727-INS-56
chr13-30549778-INS-214
chr13-30640382-INS-6215
chr13-30659502-INS-58
chr13-30677314-INS-149
chr13-30680724-INS-447
chr13-30692464-INS-97
chr13-30694347-INS-498
chr13-30737894-INS-365
chr13-30752621-INS-85
chr13-30759945-INS-150
chr13-30762077-INS-124
chr13-30768793-INS-55
chr13-30781959-INS-185
chr13-30804180-INS-179
chr13-30806806-INS-185
chr13-30808121-INS-53
chr13-30829303-INS-57
chr13-30884035-INS-105
chr13-30891142-INS-53
chr13-30902831-INS-156
chr13-30917723-INS-106
chr13-30927721-INS-103
chr13-30937718-INS-58
chr13-30956841-INS-87
chr13-30962694-INS-81
chr13-30978487-INS-391
chr13-30997643-INS-112
chr13-31008577-INS-54
chr13-31018696-INS-158
chr13-31023533-INS-76
chr13-31036493-INS-538
chr13-31050300-INS-432
chr13-31068230-INS-66
chr13-31071873-INS-120
chr13-31161673-INS-158
chr13-31192457-INS-64
chr13-31230150-INS-237
chr13-31242026-INS-54
chr13-31250940-INS-174
chr13-31276390-INS-6791
chr13-31288584-INS-80
chr13-31316758-INS-6368
chr13-31334488-INS-2850
chr13-31346548-INS-651
chr13-31354269-INS-124
chr13-31358873-INS-109
chr13-31362281-INS-51
chr13-31369995-INS-82
chr13-31384879-INS-54
chr13-31386375-INS-253
chr13-31406094-INS-51
chr13-31429624-INS-7675
chr13-31429808-INS-57
chr13-31438560-INS-383
chr13-31452593-INS-136
chr13-31471204-INS-408
chr13-31478402-INS-66
chr13-31512237-INS-167
chr13-31538543-INS-534
chr13-31573653-INS-203
chr13-31575772-INS-86
chr13-31576829-INS-80
chr13-31584638-INS-188
chr13-31614678-INS-58
chr13-31616943-INS-58
chr13-31618400-INS-321
chr13-31621966-INS-53
chr13-31637338-INS-1726
chr13-31640368-INS-100
chr13-31667086-INS-54
chr13-31675356-INS-90
chr13-31716498-INS-197
chr13-31726141-INS-130
chr13-31727519-INS-93

chr13-31737089-INS-364
chr13-31758929-INS-85
chr13-31760464-INS-85
chr13-31768711-INS-247
chr13-31795520-INS-74
chr13-31803504-INS-69
chr13-31820127-INS-162
chr13-31822109-INS-195
chr13-31839381-INS-102
chr13-31869637-INS-439
chr13-31875591-INS-62
chr13-31876332-INS-130
chr13-31879676-INS-452
chr13-31893267-INS-100
chr13-31947423-INS-104
chr13-31958168-INS-57
chr13-31962078-INS-132
chr13-31967069-INS-231
chr13-32011229-INS-474
chr13-32031042-INS-218
chr13-32039832-INS-96
chr13-32078352-INS-172
chr13-32081907-INS-97
chr13-32085448-INS-72
chr13-32115924-INS-109
chr13-32118985-INS-1176
chr13-32119631-INS-250
chr13-32182230-INS-143
chr13-32206488-INS-435
chr13-32217675-INS-75
chr13-32219058-INS-53
chr13-32236998-INS-1191
chr13-32238559-INS-80
chr13-32245325-INS-101
chr13-32280728-INS-80
chr13-32281017-INS-430
chr13-32301458-INS-507
chr13-32309651-INS-59
chr13-32332750-INS-6275
chr13-32349555-INS-6518
chr13-32388013-INS-54
chr13-32416428-INS-100
chr13-32419090-INS-296
chr13-32423247-INS-60
chr13-32443138-INS-54
chr13-32459319-INS-351
chr13-32466830-INS-55
chr13-32479873-INS-662
chr13-32482171-INS-72
chr13-32484270-INS-94
chr13-32484772-INS-2572
chr13-32493149-INS-153
chr13-32495130-INS-6405
chr13-32511113-INS-171
chr13-32547999-INS-440
chr13-32550522-INS-183
chr13-32552163-INS-214
chr13-32558444-INS-92
chr13-32564022-INS-163
chr13-32580016-INS-943
chr13-32581906-INS-150
chr13-32587696-INS-64
chr13-32599116-INS-156
chr13-32608345-INS-505
chr13-32616295-INS-207
chr13-32618533-INS-321
chr13-32620174-INS-121
chr13-32628039-INS-466
chr13-32637764-INS-438
chr13-32640637-INS-245
chr13-32660896-INS-945
chr13-32661736-INS-52
chr13-32682951-INS-77
chr13-32687280-INS-75
chr13-32690268-INS-218
chr13-32695106-INS-76
chr13-32700155-INS-211

chr13-32710093-INS-413
chr13-32714294-INS-58
chr13-32716793-INS-58
chr13-32720802-INS-56
chr13-32721813-INS-293
chr13-32726744-INS-72
chr13-32741677-INS-204
chr13-32752998-INS-324
chr13-32770324-INS-241
chr13-32783283-INS-55
chr13-32808531-INS-852
chr13-32819917-INS-52
chr13-32828321-INS-58
chr13-32830249-INS-187
chr13-32835387-INS-61
chr13-32837256-INS-63
chr13-32874921-INS-51
chr13-32900378-INS-446
chr13-32908253-INS-69
chr13-32926774-INS-177
chr13-32937766-INS-376
chr13-32980680-INS-52
chr13-32993922-INS-56
chr13-33023118-INS-64
chr13-33051002-INS-608
chr13-33062783-INS-166
chr13-33075513-INS-5225
chr13-33085969-INS-497
chr13-33088092-INS-241
chr13-33105295-INS-60
chr13-33107777-INS-206
chr13-33111435-INS-202
chr13-33150681-INS-5284
chr13-33167576-INS-6710
chr13-33173351-INS-60
chr13-33178136-INS-503
chr13-33197247-INS-60
chr13-33200855-INS-358
chr13-33211847-INS-503
chr13-33230509-INS-180
chr13-33253509-INS-66
chr13-33257985-INS-114
chr13-33273147-INS-358
chr13-33281497-INS-81
chr13-33301858-INS-401
chr13-33368533-INS-52
chr13-33473216-INS-59
chr13-33505861-INS-114
chr13-33507004-INS-133
chr13-33547852-INS-59
chr13-33551263-INS-394
chr13-33618303-INS-57
chr13-33629016-INS-1129
chr13-33652655-INS-998
chr13-33694891-INS-329
chr13-33749362-INS-84
chr13-33756761-INS-1239
chr13-33756762-INS-851
chr13-33798523-INS-191
chr13-33843936-INS-129
chr13-33860946-INS-103
chr13-33920301-INS-8701
chr13-33968866-INS-183
chr13-33971881-INS-82
chr13-34014855-INS-109
chr13-34040031-INS-6440
chr13-34041269-INS-114
chr13-34082989-INS-51
chr13-34085477-INS-3863
chr13-34091368-INS-100
chr13-34095684-INS-291
chr13-34123589-INS-121
chr13-34131498-INS-66
chr13-34178140-INS-52
chr13-34179811-INS-250
chr13-34182820-INS-64
chr13-34220869-INS-328

chr13-34222168-INS-62
chr13-34227564-INS-148
chr13-34279921-INS-107
chr13-34287299-INS-4400
chr13-34287827-INS-4268
chr13-34327709-INS-898
chr13-34343611-INS-72
chr13-34393153-INS-161
chr13-34424877-INS-380
chr13-34502360-INS-56
chr13-34504131-INS-90
chr13-34552040-INS-128
chr13-34557419-INS-967
chr13-34594186-INS-110
chr13-34613876-INS-77
chr13-34664711-INS-321
chr13-34677429-INS-6303
chr13-34688004-INS-6666
chr13-34708253-INS-105
chr13-34714876-INS-242
chr13-34721123-INS-166
chr13-34754845-INS-68
chr13-34756805-INS-70
chr13-34759188-INS-3554
chr13-34774236-INS-408
chr13-34775190-INS-59
chr13-34777146-INS-949
chr13-34794776-INS-167
chr13-34804261-INS-1773
chr13-34808095-INS-225
chr13-34851958-INS-56
chr13-34871366-INS-315
chr13-34874705-INS-275
chr13-34882199-INS-200
chr13-34884057-INS-60
chr13-34896268-INS-58
chr13-34932539-INS-343
chr13-34939986-INS-92
chr13-34944165-INS-220
chr13-34945058-INS-357
chr13-34969346-INS-594
chr13-34994235-INS-508
chr13-35011013-INS-56
chr13-35013417-INS-90
chr13-35028668-INS-56
chr13-35045871-INS-940
chr13-35068526-INS-64
chr13-35098870-INS-205
chr13-35125309-INS-510
chr13-35132336-INS-665
chr13-35134424-INS-90
chr13-35137934-INS-53
chr13-35168408-INS-167
chr13-35261633-INS-255
chr13-35263745-INS-1309
chr13-35264444-INS-184
chr13-35295000-INS-63
chr13-35299190-INS-909
chr13-35314719-INS-84
chr13-35348587-INS-586
chr13-35351989-INS-983
chr13-35371581-INS-61
chr13-35379513-INS-83
chr13-35383225-INS-559
chr13-35387363-INS-81
chr13-35392204-INS-6273
chr13-35392697-INS-408
chr13-35399132-INS-66
chr13-35416288-INS-6773
chr13-35485240-INS-6193
chr13-35487006-INS-144
chr13-35492914-INS-57
chr13-35550138-INS-97
chr13-35555505-INS-179
chr13-35578790-INS-230
chr13-35596663-INS-197
chr13-35638804-INS-492

chr13-35653944-INS-96
chr13-35706148-INS-62
chr13-35720017-INS-6212
chr13-35797892-INS-83
chr13-35805085-INS-659
chr13-35825166-INS-66
chr13-35840920-INS-71
chr13-35844663-INS-358
chr13-35847097-INS-404
chr13-35874598-INS-350
chr13-35883796-INS-56
chr13-35889608-INS-70
chr13-35889982-INS-126
chr13-35897881-INS-54
chr13-35901257-INS-7327
chr13-35919746-INS-117
chr13-35921452-INS-251
chr13-35956941-INS-68
chr13-35959800-INS-187
chr13-35962257-INS-231
chr13-35978691-INS-52
chr13-35999771-INS-264
chr13-36006016-INS-132
chr13-36018802-INS-402
chr13-36025972-INS-401
chr13-36033272-INS-208
chr13-36052324-INS-56
chr13-36053316-INS-56
chr13-36067957-INS-128
chr13-36074377-INS-124
chr13-36079965-INS-213
chr13-36099421-INS-173
chr13-36118787-INS-68
chr13-36130439-INS-58
chr13-36139731-INS-590
chr13-36140944-INS-55
chr13-36144653-INS-175
chr13-36146938-INS-141
chr13-36159779-INS-808
chr13-36160473-INS-204
chr13-36163130-INS-201
chr13-36164013-INS-7274
chr13-36165102-INS-614
chr13-36167855-INS-258
chr13-36182086-INS-542
chr13-36196284-INS-302
chr13-36198907-INS-233
chr13-36217371-INS-69
chr13-36268173-INS-355
chr13-36273148-INS-1296
chr13-36295104-INS-199
chr13-36296164-INS-121
chr13-36309819-INS-792
chr13-36315791-INS-158
chr13-36327798-INS-243
chr13-36343726-INS-93
chr13-36344678-INS-6385
chr13-36359135-INS-200
chr13-36362251-INS-228
chr13-36363801-INS-1262
chr13-36366133-INS-475
chr13-36390139-INS-168
chr13-36407700-INS-343
chr13-36439994-INS-120
chr13-36454590-INS-96
chr13-36456574-INS-100
chr13-36460561-INS-801
chr13-36483334-INS-52
chr13-36491955-INS-52
chr13-36494418-INS-448
chr13-36546633-INS-76
chr13-36552563-INS-136
chr13-36583393-INS-378
chr13-36607642-INS-65
chr13-36634785-INS-327
chr13-36653297-INS-84
chr13-36659272-INS-252

chr13-36687263-INS-167
chr13-36718274-INS-371
chr13-36719026-INS-142
chr13-36732538-INS-6523
chr13-36751076-INS-214
chr13-36789049-INS-68
chr13-36794439-INS-52
chr13-36805418-INS-134
chr13-36811128-INS-140
chr13-36815328-INS-12215
chr13-36820981-INS-1895
chr13-36822082-INS-104
chr13-36829024-INS-75
chr13-36856951-INS-122
chr13-36880901-INS-104
chr13-36894130-INS-164
chr13-36895202-INS-505
chr13-36902761-INS-744
chr13-36925707-INS-439
chr13-36926438-INS-251
chr13-36941609-INS-52
chr13-36958094-INS-72
chr13-36959487-INS-53
chr13-36985480-INS-808
chr13-37007870-INS-805
chr13-37019444-INS-399
chr13-37026477-INS-68
chr13-37046376-INS-135
chr13-37055649-INS-93
chr13-37059635-INS-60
chr13-37067453-INS-60
chr13-37067606-INS-1910
chr13-37080346-INS-261
chr13-37092179-INS-92
chr13-37105184-INS-89
chr13-37114476-INS-59
chr13-37115940-INS-83
chr13-37118104-INS-70
chr13-37144911-INS-99
chr13-37153890-INS-125
chr13-37162530-INS-252
chr13-37171053-INS-60
chr13-37189814-INS-141
chr13-37199825-INS-526
chr13-37280904-INS-154
chr13-37342156-INS-179
chr13-37346685-INS-296
chr13-37349405-INS-6270
chr13-37350727-INS-104
chr13-37370019-INS-7643
chr13-37374617-INS-84
chr13-37391823-INS-598
chr13-37421904-INS-211
chr13-37427847-INS-93
chr13-37432336-INS-1888
chr13-37507714-INS-54
chr13-37516550-INS-518
chr13-37518372-INS-406
chr13-37532046-INS-193
chr13-37543021-INS-219
chr13-37586400-INS-51
chr13-37599705-INS-51
chr13-37602923-INS-55
chr13-37611738-INS-329
chr13-37617633-INS-56
chr13-37621587-INS-118
chr13-37666517-INS-133
chr13-37675083-INS-242
chr13-37679918-INS-50
chr13-37680510-INS-132
chr13-37690062-INS-99
chr13-37697567-INS-62
chr13-37698030-INS-54
chr13-37705078-INS-199
chr13-37729783-INS-204
chr13-37760864-INS-206
chr13-37768800-INS-54

chr13-37790582-INS-58
chr13-37791080-INS-85
chr13-37798117-INS-71
chr13-37824934-INS-123
chr13-37839189-INS-215
chr13-37851324-INS-172
chr13-37855143-INS-91
chr13-37855907-INS-498
chr13-37895445-INS-98
chr13-37928884-INS-151
chr13-37942200-INS-226
chr13-37960122-INS-73
chr13-37961204-INS-120
chr13-37968642-INS-208
chr13-37998199-INS-228
chr13-38006192-INS-60
chr13-38020293-INS-51
chr13-38043374-INS-264
chr13-38047943-INS-106
chr13-38095784-INS-62
chr13-38111442-INS-154
chr13-38112615-INS-67
chr13-38117453-INS-71
chr13-38140321-INS-199
chr13-38145427-INS-202
chr13-38148453-INS-64
chr13-38173718-INS-1240
chr13-38178438-INS-52
chr13-38183148-INS-486
chr13-38241280-INS-89
chr13-38282026-INS-86
chr13-38313006-INS-8940
chr13-38318823-INS-87
chr13-38319328-INS-176
chr13-38323016-INS-56
chr13-38347481-INS-461
chr13-38356502-INS-384
chr13-38371121-INS-54
chr13-38375058-INS-184
chr13-38388251-INS-56
chr13-38393443-INS-71
chr13-38439117-INS-166
chr13-38447317-INS-52
chr13-38454614-INS-176
chr13-38461744-INS-76
chr13-38465297-INS-75
chr13-38477046-INS-135
chr13-38506486-INS-303
chr13-38518202-INS-67
chr13-38522102-INS-144
chr13-38522374-INS-196
chr13-38522511-INS-2865
chr13-38554968-INS-285
chr13-38557161-INS-60
chr13-38581550-INS-762
chr13-38588188-INS-92
chr13-38611926-INS-52
chr13-38660323-INS-85
chr13-38662707-INS-51
chr13-38679058-INS-177
chr13-38715047-INS-72
chr13-38727729-INS-100
chr13-38735486-INS-128
chr13-38766003-INS-1965
chr13-38773095-INS-111
chr13-38773617-INS-143
chr13-38802081-INS-203
chr13-38806342-INS-62
chr13-38819995-INS-206
chr13-38826872-INS-67
chr13-38837800-INS-56
chr13-38868032-INS-82
chr13-38869257-INS-139
chr13-38883141-INS-330
chr13-38888021-INS-199
chr13-38892163-INS-87
chr13-38896208-INS-87

chr13-38946721-INS-204
chr13-38956006-INS-70
chr13-38960100-INS-434
chr13-38962848-INS-303
chr13-38969895-INS-187
chr13-38978793-INS-72
chr13-38994449-INS-62
chr13-38996223-INS-117
chr13-39014002-INS-63
chr13-39021391-INS-236
chr13-39055244-INS-451
chr13-39056694-INS-400
chr13-39081303-INS-371
chr13-39093724-INS-530
chr13-39133951-INS-135
chr13-39169103-INS-168
chr13-39199059-INS-66
chr13-39200468-INS-436
chr13-39210118-INS-92
chr13-39213066-INS-6873
chr13-39228354-INS-62
chr13-39243047-INS-53
chr13-39245015-INS-55
chr13-39295576-INS-58
chr13-39338597-INS-149
chr13-39342109-INS-222
chr13-39343043-INS-397
chr13-39347520-INS-426
chr13-39370696-INS-1092
chr13-39373475-INS-86
chr13-39376303-INS-292
chr13-39390277-INS-52
chr13-39395522-INS-98
chr13-39401977-INS-70
chr13-39403705-INS-61
chr13-39410556-INS-568
chr13-39426607-INS-395
chr13-39467137-INS-94
chr13-39495221-INS-111
chr13-39503721-INS-74
chr13-39506853-INS-354
chr13-39517730-INS-70
chr13-39540941-INS-158
chr13-39551397-INS-54
chr13-39585594-INS-863
chr13-39590686-INS-478
chr13-39610691-INS-60
chr13-39633062-INS-63
chr13-39640067-INS-97
chr13-39720793-INS-240
chr13-39727548-INS-116
chr13-39753809-INS-163
chr13-39763445-INS-328
chr13-39768674-INS-4211
chr13-39787001-INS-84
chr13-39801265-INS-109
chr13-39837258-INS-70
chr13-39852372-INS-78
chr13-39868259-INS-71
chr13-39874761-INS-186
chr13-39884115-INS-369
chr13-39911869-INS-60
chr13-39918402-INS-6361
chr13-39927066-INS-1943
chr13-39951148-INS-56
chr13-39974578-INS-3286
chr13-39975926-INS-118
chr13-39988313-INS-92
chr13-40003251-INS-58
chr13-40012317-INS-163
chr13-40032268-INS-52
chr13-40037104-INS-90
chr13-40066805-INS-150
chr13-40088679-INS-78
chr13-40137769-INS-76
chr13-40151898-INS-92
chr13-40158936-INS-403

chr13-40166498-INS-785
chr13-40170831-INS-80
chr13-40183498-INS-62
chr13-40190140-INS-224
chr13-40194571-INS-12449
chr13-40229729-INS-52
chr13-40243044-INS-585
chr13-40248551-INS-128
chr13-40261396-INS-199
chr13-40284030-INS-138
chr13-40305711-INS-66
chr13-40324587-INS-5548
chr13-40327082-INS-161
chr13-40329626-INS-169
chr13-40331795-INS-104
chr13-40334903-INS-173
chr13-40336462-INS-401
chr13-40337448-INS-126
chr13-40339665-INS-400
chr13-40359356-INS-305
chr13-40381405-INS-78
chr13-40416238-INS-56
chr13-40419246-INS-207
chr13-40461260-INS-110
chr13-40471928-INS-518
chr13-40472429-INS-874
chr13-40495499-INS-52
chr13-40520669-INS-70
chr13-40529569-INS-53
chr13-40534141-INS-88
chr13-40598635-INS-275
chr13-40615646-INS-54
chr13-40635517-INS-159
chr13-40641243-INS-435
chr13-40655991-INS-79
chr13-40673644-INS-109
chr13-40685941-INS-71
chr13-40692260-INS-57
chr13-40701625-INS-207
chr13-40704364-INS-63
chr13-40709640-INS-54
chr13-40720817-INS-1142
chr13-40730038-INS-108
chr13-40749443-INS-6691
chr13-40754139-INS-633
chr13-40762898-INS-3148
chr13-40767921-INS-171
chr13-40775427-INS-215
chr13-40776537-INS-153
chr13-40809432-INS-223
chr13-40852832-INS-90
chr13-40860993-INS-60
chr13-40886022-INS-66
chr13-40909373-INS-186
chr13-40912016-INS-59
chr13-40938923-INS-260
chr13-40948529-INS-417
chr13-40951773-INS-591
chr13-40978796-INS-72
chr13-40981198-INS-325
chr13-40993586-INS-210
chr13-41006808-INS-253
chr13-41008817-INS-101
chr13-41032442-INS-7109
chr13-41036733-INS-101
chr13-41044741-INS-453
chr13-41049899-INS-182
chr13-41063052-INS-52
chr13-41068574-INS-54
chr13-41099029-INS-55
chr13-41144292-INS-69
chr13-41147406-INS-60
chr13-41156407-INS-55
chr13-41182638-INS-51
chr13-41184434-INS-198
chr13-41192823-INS-405
chr13-41194924-INS-69

chr13-41218860-INS-498
chr13-41229524-INS-93
chr13-41237669-INS-92
chr13-41240989-INS-60
chr13-41249450-INS-52
chr13-41250819-INS-67
chr13-41252664-INS-55
chr13-41276097-INS-65
chr13-41288867-INS-107
chr13-41303027-INS-56
chr13-41310551-INS-175
chr13-41332408-INS-609
chr13-41346747-INS-214
chr13-41352701-INS-7664
chr13-41365560-INS-159
chr13-41377848-INS-441
chr13-41382393-INS-72
chr13-41382394-INS-1278
chr13-41385635-INS-6126
chr13-41399461-INS-53
chr13-41406620-INS-74
chr13-41410452-INS-821
chr13-41420629-INS-142
chr13-41434690-INS-108
chr13-41444186-INS-74
chr13-41454576-INS-558
chr13-41472122-INS-78
chr13-41479066-INS-735
chr13-41509049-INS-160
chr13-41536113-INS-172
chr13-41567067-INS-401
chr13-41583950-INS-80
chr13-41643891-INS-180
chr13-41775580-INS-200
chr13-41780787-INS-341
chr13-41785944-INS-83
chr13-41805740-INS-104
chr13-41811418-INS-64
chr13-41812472-INS-91
chr13-41816270-INS-156
chr13-41825616-INS-180
chr13-41860252-INS-68
chr13-41868477-INS-62
chr13-41883697-INS-120
chr13-41908052-INS-1445
chr13-41909781-INS-1152
chr13-41916623-INS-6360
chr13-41934556-INS-94
chr13-41956971-INS-869
chr13-41959424-INS-57
chr13-41975994-INS-80
chr13-41979461-INS-59
chr13-41981033-INS-564
chr13-41987044-INS-589
chr13-42063825-INS-79
chr13-42090430-INS-203
chr13-42099682-INS-83
chr13-42105210-INS-69
chr13-42111117-INS-69
chr13-42112899-INS-517
chr13-42117402-INS-5265
chr13-42163569-INS-51
chr13-42174299-INS-401
chr13-42177742-INS-239
chr13-42184242-INS-99
chr13-42248467-INS-51
chr13-42255405-INS-402
chr13-42327790-INS-236
chr13-42383805-INS-175
chr13-42400648-INS-58
chr13-42402309-INS-54
chr13-42404555-INS-113
chr13-42405060-INS-300
chr13-42406610-INS-90
chr13-42417417-INS-402
chr13-42422597-INS-81
chr13-42445573-INS-64

chr13-42480015-INS-208
chr13-42480785-INS-364
chr13-42495642-INS-129
chr13-42496225-INS-65
chr13-42501124-INS-218
chr13-42503555-INS-51
chr13-42531135-INS-82
chr13-42533113-INS-132
chr13-42549526-INS-550
chr13-42565772-INS-253
chr13-42580886-INS-695
chr13-42616259-INS-99
chr13-42618056-INS-398
chr13-42625523-INS-463
chr13-42657299-INS-307
chr13-42705462-INS-56
chr13-42728170-INS-68
chr13-42735792-INS-1890
chr13-42757662-INS-70
chr13-42759723-INS-74
chr13-42769382-INS-52
chr13-42782612-INS-70
chr13-42795226-INS-91
chr13-42801132-INS-493
chr13-42823899-INS-668
chr13-42866885-INS-69
chr13-42892511-INS-129
chr13-42925079-INS-67
chr13-42934135-INS-60
chr13-42938313-INS-109
chr13-42940343-INS-85
chr13-42953706-INS-258
chr13-43003653-INS-6657
chr13-43012222-INS-170
chr13-43013399-INS-52
chr13-43026710-INS-175
chr13-43051064-INS-78
chr13-43063114-INS-267
chr13-43072229-INS-152
chr13-43078322-INS-52
chr13-43089009-INS-56
chr13-43099177-INS-55
chr13-43104204-INS-56
chr13-43126519-INS-75
chr13-43127525-INS-52
chr13-43133209-INS-201
chr13-43152457-INS-57
chr13-43168093-INS-136
chr13-43189515-INS-66
chr13-43198669-INS-72
chr13-43201856-INS-912
chr13-43204948-INS-93
chr13-43206777-INS-552
chr13-43243466-INS-257
chr13-43256080-INS-76
chr13-43258721-INS-324
chr13-43261341-INS-68
chr13-43286855-INS-88
chr13-43288845-INS-52
chr13-43289862-INS-75
chr13-43313705-INS-88
chr13-43319448-INS-76
chr13-43343563-INS-207
chr13-43371505-INS-96
chr13-43382185-INS-206
chr13-43401725-INS-78
chr13-43409820-INS-64
chr13-43423567-INS-142
chr13-43481658-INS-52
chr13-43502053-INS-289
chr13-43509069-INS-114
chr13-43515158-INS-120
chr13-43522757-INS-53
chr13-43522922-INS-222
chr13-43525919-INS-84
chr13-43528970-INS-519
chr13-43539727-INS-439

chr13-43616517-INS-229
chr13-43618947-INS-158
chr13-43661509-INS-197
chr13-43662865-INS-314
chr13-43675134-INS-52
chr13-43679474-INS-8985
chr13-43682891-INS-396
chr13-43698969-INS-215
chr13-43722768-INS-52
chr13-43724514-INS-669
chr13-43726397-INS-72
chr13-43727166-INS-434
chr13-43746442-INS-61
chr13-43764983-INS-55
chr13-43765876-INS-206
chr13-43808242-INS-70
chr13-43816086-INS-449
chr13-43842649-INS-207
chr13-43867633-INS-99
chr13-43881386-INS-130
chr13-43911110-INS-175
chr13-43914681-INS-409
chr13-43915645-INS-152
chr13-43917797-INS-7205
chr13-43922844-INS-102
chr13-43924658-INS-56
chr13-43940772-INS-98
chr13-43987227-INS-94
chr13-43991264-INS-60
chr13-44029216-INS-58
chr13-44030927-INS-359
chr13-44034789-INS-169
chr13-44038185-INS-121
chr13-44040821-INS-58
chr13-44066694-INS-107
chr13-44078178-INS-125
chr13-44102759-INS-226
chr13-44129769-INS-448
chr13-44139462-INS-116
chr13-44149623-INS-54
chr13-44152519-INS-64
chr13-44167052-INS-97
chr13-44175130-INS-58
chr13-44194414-INS-55
chr13-44202388-INS-124
chr13-44205706-INS-160
chr13-44207638-INS-65
chr13-44218100-INS-86
chr13-44223921-INS-191
chr13-44245822-INS-108
chr13-44286789-INS-144
chr13-44288076-INS-192
chr13-44291640-INS-60
chr13-44310559-INS-54
chr13-44325992-INS-104
chr13-44364319-INS-95
chr13-44392775-INS-111
chr13-44398082-INS-197
chr13-44403141-INS-96
chr13-44425303-INS-80
chr13-44464177-INS-504
chr13-44467640-INS-63
chr13-44480759-INS-97
chr13-44487247-INS-399
chr13-44494787-INS-88
chr13-44508740-INS-472
chr13-44525653-INS-82
chr13-44528010-INS-67
chr13-44557210-INS-95
chr13-44584634-INS-2865
chr13-44603773-INS-57
chr13-44607544-INS-189
chr13-44615707-INS-290
chr13-44636889-INS-141
chr13-44644776-INS-684
chr13-44654114-INS-53
chr13-44658100-INS-180

chr13-44679825-INS-325
chr13-44685528-INS-93
chr13-44687026-INS-424
chr13-44700274-INS-95
chr13-44710213-INS-85
chr13-44711148-INS-58
chr13-44712466-INS-544
chr13-44715603-INS-56
chr13-44728696-INS-6441
chr13-44734583-INS-211
chr13-44739623-INS-52
chr13-44743260-INS-221
chr13-44744231-INS-55
chr13-44758214-INS-210
chr13-44769917-INS-237
chr13-44786249-INS-86
chr13-44787318-INS-201
chr13-44794589-INS-104
chr13-44803504-INS-67
chr13-44805830-INS-56
chr13-44808828-INS-224
chr13-44821564-INS-54
chr13-44829159-INS-253
chr13-44836266-INS-75
chr13-44851108-INS-55
chr13-44851720-INS-201
chr13-44855979-INS-223
chr13-44857267-INS-7368
chr13-44871185-INS-201
chr13-44878345-INS-210
chr13-44880259-INS-84
chr13-44881297-INS-64
chr13-44894830-INS-7485
chr13-44895895-INS-173
chr13-44898747-INS-251
chr13-44912242-INS-55
chr13-44913935-INS-76
chr13-44918054-INS-105
chr13-44919346-INS-106
chr13-44920094-INS-124
chr13-44924572-INS-170
chr13-44936187-INS-197
chr13-44961546-INS-59
chr13-44973932-INS-73
chr13-44979571-INS-118
chr13-44982398-INS-204
chr13-45004104-INS-193
chr13-45019336-INS-106
chr13-45021903-INS-56
chr13-45023575-INS-242
chr13-45036325-INS-55
chr13-45049500-INS-192
chr13-45065239-INS-95
chr13-45082252-INS-575
chr13-45085941-INS-102
chr13-45128720-INS-543
chr13-45142263-INS-248
chr13-45160968-INS-52
chr13-45177597-INS-57
chr13-45191569-INS-536
chr13-45192133-INS-54
chr13-45200042-INS-216
chr13-45200768-INS-61
chr13-45222236-INS-206
chr13-45238431-INS-972
chr13-45248055-INS-56
chr13-45283871-INS-58
chr13-45284678-INS-344
chr13-45295231-INS-66
chr13-45301499-INS-501
chr13-45307779-INS-65
chr13-45317255-INS-199
chr13-45347698-INS-203
chr13-45363005-INS-264
chr13-45377588-INS-202
chr13-45385699-INS-5562
chr13-45399419-INS-78

chr13-45457056-INS-79
chr13-45480005-INS-438
chr13-45518088-INS-54
chr13-45525160-INS-60
chr13-45528594-INS-60
chr13-45529165-INS-56
chr13-45534911-INS-94
chr13-45540928-INS-116
chr13-45555191-INS-114
chr13-45568331-INS-76
chr13-45593736-INS-59
chr13-45624717-INS-210
chr13-45631513-INS-206
chr13-45633155-INS-445
chr13-45645844-INS-100
chr13-45649963-INS-337
chr13-45653685-INS-214
chr13-45661719-INS-112
chr13-45671158-INS-199
chr13-45719913-INS-73
chr13-45731952-INS-120
chr13-45772437-INS-401
chr13-45791801-INS-176
chr13-45797789-INS-57
chr13-45804878-INS-85
chr13-45812498-INS-64
chr13-45828063-INS-65
chr13-45855868-INS-370
chr13-45858769-INS-206
chr13-45868730-INS-100
chr13-45886545-INS-229
chr13-45891612-INS-70
chr13-45893760-INS-70
chr13-45916576-INS-252
chr13-45930662-INS-63
chr13-45948242-INS-367
chr13-45988975-INS-97
chr13-45992575-INS-64
chr13-45994855-INS-71
chr13-45999025-INS-964
chr13-46006145-INS-81
chr13-46009352-INS-69
chr13-46031937-INS-162
chr13-46038188-INS-2485
chr13-46051462-INS-265
chr13-46053470-INS-145
chr13-46073152-INS-68
chr13-46079407-INS-98
chr13-46101371-INS-96
chr13-46103072-INS-77
chr13-46128045-INS-1891
chr13-46129192-INS-148
chr13-46157274-INS-148
chr13-46185549-INS-56
chr13-46188203-INS-158
chr13-46189259-INS-554
chr13-46194858-INS-168
chr13-46203698-INS-7188
chr13-46211446-INS-72
chr13-46212935-INS-590
chr13-46216458-INS-370
chr13-46219823-INS-64
chr13-46232133-INS-242
chr13-46247151-INS-119
chr13-46268867-INS-51
chr13-46270563-INS-54
chr13-46289174-INS-217
chr13-46295415-INS-136
chr13-46301283-INS-242
chr13-46312447-INS-53
chr13-46320236-INS-171
chr13-46325342-INS-758
chr13-46328120-INS-184
chr13-46360905-INS-63
chr13-46364969-INS-60
chr13-46371474-INS-75
chr13-46396616-INS-139

chr13-46404990-INS-445
chr13-46406479-INS-979
chr13-46413979-INS-256
chr13-46420393-INS-97
chr13-46423580-INS-60
chr13-46427557-INS-68
chr13-46434414-INS-99
chr13-46464141-INS-208
chr13-46510456-INS-142
chr13-46538344-INS-66
chr13-46580880-INS-83
chr13-46591736-INS-72
chr13-46615666-INS-215
chr13-46618322-INS-110
chr13-46643651-INS-112
chr13-46666452-INS-146
chr13-46672380-INS-93
chr13-46689800-INS-57
chr13-46692932-INS-225
chr13-46695045-INS-100
chr13-46700781-INS-56
chr13-46727962-INS-52
chr13-46729641-INS-201
chr13-46755103-INS-52
chr13-46755992-INS-484
chr13-46782341-INS-60
chr13-46814148-INS-216
chr13-46841526-INS-57
chr13-46883627-INS-211
chr13-46892465-INS-59
chr13-46916103-INS-158
chr13-46920726-INS-76
chr13-46927746-INS-65
chr13-46929475-INS-506
chr13-46956833-INS-266
chr13-47005357-INS-205
chr13-47040822-INS-91
chr13-47047835-INS-62
chr13-47050824-INS-95
chr13-47072020-INS-83
chr13-47074098-INS-58
chr13-47097495-INS-199
chr13-47110449-INS-111
chr13-47114389-INS-230
chr13-47121491-INS-72
chr13-47122942-INS-55
chr13-47124760-INS-242
chr13-47143949-INS-158
chr13-47148149-INS-736
chr13-47153732-INS-105
chr13-47165037-INS-97
chr13-47189228-INS-111
chr13-47220035-INS-744
chr13-47233226-INS-277
chr13-47235873-INS-122
chr13-47282064-INS-56
chr13-47290753-INS-114
chr13-47297139-INS-55
chr13-47336060-INS-66
chr13-47411062-INS-65
chr13-47416065-INS-56
chr13-47423566-INS-54
chr13-47436360-INS-212
chr13-47440414-INS-52
chr13-47467081-INS-60
chr13-47470754-INS-358
chr13-47473938-INS-59
chr13-47473975-INS-872
chr13-47481616-INS-95
chr13-47494199-INS-64
chr13-47503967-INS-224
chr13-47519741-INS-436
chr13-47526334-INS-344
chr13-47530718-INS-78
chr13-47546808-INS-76
chr13-47577618-INS-6595
chr13-47580364-INS-143

chr13-47592791-INS-655
chr13-47619886-INS-151
chr13-47639796-INS-200
chr13-47655240-INS-93
chr13-47734797-INS-67
chr13-47737832-INS-88
chr13-47738923-INS-232
chr13-47745670-INS-174
chr13-47749384-INS-781
chr13-47756113-INS-52
chr13-47762938-INS-333
chr13-47769772-INS-52
chr13-47778958-INS-182
chr13-47781403-INS-358
chr13-47786894-INS-125
chr13-47843379-INS-155
chr13-47850575-INS-60
chr13-47881765-INS-6864
chr13-47884668-INS-604
chr13-47886576-INS-65
chr13-47893831-INS-208
chr13-47903501-INS-79
chr13-47948437-INS-63
chr13-47959028-INS-63
chr13-47969859-INS-94
chr13-47973597-INS-168
chr13-47993937-INS-85
chr13-48010335-INS-223
chr13-48017242-INS-74
chr13-48023024-INS-60
chr13-48058967-INS-58
chr13-48061979-INS-340
chr13-48081762-INS-206
chr13-48102571-INS-313
chr13-48120696-INS-138
chr13-48156248-INS-75
chr13-48161238-INS-122
chr13-48163459-INS-702
chr13-48232752-INS-53
chr13-48234203-INS-890
chr13-48235975-INS-61
chr13-48242168-INS-241
chr13-48253418-INS-84
chr13-48278880-INS-262
chr13-48287867-INS-386
chr13-48295730-INS-62
chr13-48317193-INS-284
chr13-48332132-INS-58
chr13-48337443-INS-57
chr13-48351403-INS-54
chr13-48363013-INS-1870
chr13-48381040-INS-66
chr13-48427326-INS-66
chr13-48450974-INS-318
chr13-48486704-INS-84
chr13-48488150-INS-55
chr13-48491547-INS-52
chr13-48519703-INS-175
chr13-48526257-INS-110
chr13-48529713-INS-62
chr13-48564741-INS-173
chr13-48571783-INS-882
chr13-48572360-INS-77
chr13-48587519-INS-165
chr13-48607114-INS-189
chr13-48611338-INS-56
chr13-48620887-INS-63
chr13-48625437-INS-212
chr13-48680864-INS-160
chr13-48705462-INS-189
chr13-48710894-INS-60
chr13-48724426-INS-56
chr13-48754856-INS-271
chr13-48756923-INS-200
chr13-48758342-INS-595
chr13-48760272-INS-105
chr13-48768761-INS-98

chr13-48786689-INS-149
chr13-48788295-INS-67
chr13-48791896-INS-203
chr13-48794118-INS-54
chr13-48824145-INS-138
chr13-48827576-INS-105
chr13-48830500-INS-150
chr13-48844856-INS-327
chr13-48864405-INS-106
chr13-48887661-INS-55
chr13-48895383-INS-53
chr13-48899076-INS-58
chr13-48905956-INS-512
chr13-48918058-INS-58
chr13-48932825-INS-102
chr13-48942692-INS-87
chr13-48958333-INS-319
chr13-48963456-INS-52
chr13-49009321-INS-209
chr13-49028188-INS-52
chr13-49097737-INS-197
chr13-49108103-INS-89
chr13-49115800-INS-98
chr13-49129842-INS-188
chr13-49141398-INS-497
chr13-49167817-INS-519
chr13-49168588-INS-51
chr13-49179838-INS-50
chr13-49230591-INS-56
chr13-49232362-INS-280
chr13-49250227-INS-209
chr13-49271333-INS-90
chr13-49274691-INS-93
chr13-49276758-INS-71
chr13-49280581-INS-1897
chr13-49293340-INS-233
chr13-49313033-INS-399
chr13-49326961-INS-157
chr13-49342545-INS-52
chr13-49350932-INS-138
chr13-49374355-INS-74
chr13-49415975-INS-484
chr13-49423507-INS-98
chr13-49431188-INS-275
chr13-49440080-INS-59
chr13-49449507-INS-734
chr13-49461425-INS-60
chr13-49476832-INS-62
chr13-49524341-INS-126
chr13-49529416-INS-89
chr13-49535530-INS-433
chr13-49560201-INS-76
chr13-49560843-INS-843
chr13-49573511-INS-74
chr13-49583182-INS-223
chr13-49653737-INS-81
chr13-49753201-INS-1163
chr13-49753573-INS-180
chr13-49773574-INS-119
chr13-49863712-INS-202
chr13-49893608-INS-249
chr13-49907229-INS-216
chr13-49910575-INS-69
chr13-49935179-INS-97
chr13-49969453-INS-242
chr13-49999464-INS-144
chr13-50011795-INS-122
chr13-50038663-INS-89
chr13-50046640-INS-320
chr13-50048186-INS-206
chr13-50063062-INS-5933
chr13-50086549-INS-852
chr13-50108035-INS-51
chr13-50114513-INS-4458
chr13-50119893-INS-78
chr13-50158948-INS-3545
chr13-50168788-INS-473

chr13-50177767-INS-7287
chr13-50183809-INS-3833
chr13-50226083-INS-5510
chr13-50231380-INS-3016
chr13-50424878-INS-104
chr13-50470780-INS-7484
chr13-50579539-INS-52
chr13-50593023-INS-88
chr13-50604394-INS-52
chr13-50612271-INS-228
chr13-50627437-INS-66
chr13-50725668-INS-78
chr13-50927411-INS-473
chr13-50940151-INS-109
chr13-50966342-INS-127
chr13-50968556-INS-429
chr13-50972463-INS-120
chr13-50976611-INS-86
chr13-50977680-INS-498
chr13-50988594-INS-54
chr13-51004410-INS-320
chr13-51016406-INS-142
chr13-51046304-INS-97
chr13-51102195-INS-267
chr13-51115586-INS-63
chr13-51117687-INS-88
chr13-51121734-INS-393
chr13-51132471-INS-580
chr13-51134698-INS-944
chr13-51136407-INS-54
chr13-51138513-INS-219
chr13-51147727-INS-122
chr13-51151405-INS-395
chr13-51156844-INS-60
chr13-51169153-INS-104
chr13-51191679-INS-965
chr13-51196831-INS-55
chr13-51220774-INS-52
chr13-51227848-INS-5849
chr13-51232545-INS-233
chr13-51236497-INS-67
chr13-51246043-INS-201
chr13-51250518-INS-53
chr13-51258466-INS-5384
chr13-51260817-INS-108
chr13-51279660-INS-51
chr13-51293202-INS-53
chr13-51330266-INS-186
chr13-51342965-INS-615
chr13-51344599-INS-133
chr13-51368081-INS-162
chr13-51396159-INS-60
chr13-51398525-INS-192
chr13-51405849-INS-489
chr13-51407758-INS-60
chr13-51428694-INS-51
chr13-51437957-INS-156
chr13-51444574-INS-123
chr13-51446388-INS-221
chr13-51448142-INS-907
chr13-51483616-INS-105
chr13-51509557-INS-66
chr13-51527443-INS-277
chr13-51546029-INS-62
chr13-51554273-INS-150
chr13-51587435-INS-56
chr13-51635660-INS-298
chr13-51693495-INS-347
chr13-51700580-INS-54
chr13-51709280-INS-204
chr13-51727260-INS-69
chr13-51744525-INS-212
chr13-51761014-INS-79
chr13-51763784-INS-160
chr13-51786647-INS-628
chr13-51818193-INS-319
chr13-51832177-INS-215

chr13-51833349-INS-60
chr13-51860761-INS-90
chr13-51882190-INS-77
chr13-51887481-INS-60
chr13-51928772-INS-219
chr13-51940762-INS-236
chr13-51973390-INS-89
chr13-51983703-INS-3497
chr13-51984054-INS-403
chr13-51998592-INS-204
chr13-52028755-INS-102
chr13-52029941-INS-178
chr13-52051286-INS-241
chr13-52064872-INS-153
chr13-52082363-INS-124
chr13-52095299-INS-555
chr13-52128038-INS-82
chr13-52146153-INS-216
chr13-52167052-INS-77
chr13-52178747-INS-211
chr13-52183408-INS-66
chr13-52187561-INS-125
chr13-52193152-INS-54
chr13-52236290-INS-54
chr13-52237346-INS-313
chr13-52243155-INS-7079
chr13-52260680-INS-60
chr13-52278216-INS-461
chr13-52336016-INS-60
chr13-52339795-INS-76
chr13-52340071-INS-895
chr13-52345864-INS-239
chr13-52352068-INS-51
chr13-52364904-INS-4326
chr13-52380940-INS-74
chr13-52391908-INS-427
chr13-52397932-INS-75
chr13-52399185-INS-120
chr13-52413031-INS-580
chr13-52421990-INS-3212
chr13-52432071-INS-371
chr13-52468121-INS-324
chr13-52469359-INS-156
chr13-52478579-INS-146
chr13-52495868-INS-135
chr13-52545472-INS-322
chr13-52551315-INS-278
chr13-52579028-INS-60
chr13-52591411-INS-58
chr13-52610557-INS-212
chr13-52637509-INS-60
chr13-52640871-INS-60
chr13-52644900-INS-58
chr13-52646059-INS-79
chr13-52649177-INS-100
chr13-52658366-INS-52
chr13-52662606-INS-56
chr13-52663181-INS-5548
chr13-52669418-INS-412
chr13-52695101-INS-82
chr13-52697442-INS-76
chr13-52706162-INS-242
chr13-52739621-INS-82
chr13-52742460-INS-150
chr13-52745943-INS-125
chr13-52752244-INS-72
chr13-52771702-INS-222
chr13-52772869-INS-63
chr13-52809566-INS-75
chr13-52823780-INS-56
chr13-52828777-INS-52
chr13-52836772-INS-201
chr13-52856950-INS-79
chr13-52896492-INS-56
chr13-52899939-INS-163
chr13-52918812-INS-1497
chr13-52939723-INS-60

chr13-52946423-INS-130
chr13-52968138-INS-216
chr13-52977463-INS-59
chr13-52996576-INS-206
chr13-53007686-INS-400
chr13-53036577-INS-383
chr13-53052509-INS-183
chr13-53091450-INS-207
chr13-53111374-INS-93
chr13-53120832-INS-135
chr13-53137854-INS-206
chr13-53140582-INS-206
chr13-53149932-INS-52
chr13-53173419-INS-98
chr13-53183271-INS-6440
chr13-53194938-INS-66
chr13-53203791-INS-73
chr13-53204232-INS-167
chr13-53213653-INS-77
chr13-53240651-INS-52
chr13-53275480-INS-352
chr13-53283438-INS-142
chr13-53354147-INS-270
chr13-53365560-INS-208
chr13-53380037-INS-60
chr13-53394037-INS-111
chr13-53401095-INS-52
chr13-53424892-INS-379
chr13-53434468-INS-149
chr13-53460595-INS-233
chr13-53473574-INS-59
chr13-53476795-INS-59
chr13-53483964-INS-56
chr13-53484881-INS-64
chr13-53515790-INS-944
chr13-53515846-INS-117
chr13-53535627-INS-207
chr13-53555209-INS-394
chr13-53584427-INS-149
chr13-53632979-INS-73
chr13-53637915-INS-95
chr13-53642472-INS-108
chr13-53653024-INS-97
chr13-53654481-INS-87
chr13-53691467-INS-422
chr13-53722765-INS-330
chr13-53732876-INS-213
chr13-53753964-INS-242
chr13-53758829-INS-78
chr13-53777510-INS-332
chr13-53787720-INS-52
chr13-53798299-INS-234
chr13-53844801-INS-87
chr13-53882768-INS-119
chr13-53887537-INS-289
chr13-53891340-INS-6415
chr13-53896883-INS-262
chr13-53920740-INS-292
chr13-53986548-INS-115
chr13-53990229-INS-54
chr13-53990544-INS-63
chr13-53996269-INS-157
chr13-54003236-INS-53
chr13-54006722-INS-58
chr13-54025810-INS-221
chr13-54027264-INS-544
chr13-54035669-INS-68
chr13-54067725-INS-5249
chr13-54081458-INS-59
chr13-54084525-INS-90
chr13-54086148-INS-126
chr13-54093307-INS-202
chr13-54133909-INS-109
chr13-54146477-INS-177
chr13-54149847-INS-149
chr13-54173945-INS-344
chr13-54200976-INS-144

chr13-54236762-INS-108
chr13-54262823-INS-82
chr13-54272140-INS-5550
chr13-54301693-INS-64
chr13-54320578-INS-239
chr13-54327481-INS-71
chr13-54409735-INS-55
chr13-54432941-INS-56
chr13-54450277-INS-107
chr13-54472968-INS-233
chr13-54514036-INS-175
chr13-54516453-INS-90
chr13-54527195-INS-205
chr13-54530818-INS-58
chr13-54554443-INS-5451
chr13-54566284-INS-80
chr13-54614944-INS-76
chr13-54617431-INS-382
chr13-54636425-INS-320
chr13-54650071-INS-293
chr13-54656313-INS-86
chr13-54696756-INS-103
chr13-54697738-INS-280
chr13-54734695-INS-159
chr13-54756287-INS-202
chr13-54766613-INS-121
chr13-54767386-INS-54
chr13-54801151-INS-64
chr13-54805230-INS-128
chr13-54806542-INS-208
chr13-54846665-INS-78
chr13-54857834-INS-97
chr13-54930672-INS-598
chr13-54931437-INS-8671
chr13-54935993-INS-81
chr13-54958475-INS-222
chr13-54961499-INS-56
chr13-54967172-INS-120
chr13-54978765-INS-159
chr13-55004686-INS-208
chr13-55053458-INS-198
chr13-55076666-INS-56
chr13-55188503-INS-231
chr13-55203023-INS-362
chr13-55219169-INS-54
chr13-55234980-INS-105
chr13-55260134-INS-220
chr13-55264941-INS-910
chr13-55265648-INS-199
chr13-55273561-INS-6456
chr13-55278789-INS-206
chr13-55284879-INS-142
chr13-55295198-INS-64
chr13-55305421-INS-52
chr13-55306422-INS-209
chr13-55309615-INS-72
chr13-55325961-INS-491
chr13-55338867-INS-106
chr13-55343019-INS-169
chr13-55346150-INS-203
chr13-55348631-INS-166
chr13-55372187-INS-203
chr13-55385070-INS-121
chr13-55406368-INS-877
chr13-55438593-INS-96
chr13-55454487-INS-195
chr13-55480074-INS-125
chr13-55490765-INS-449
chr13-55524127-INS-51
chr13-55529920-INS-211
chr13-55533057-INS-191
chr13-55533959-INS-390
chr13-55535384-INS-62
chr13-55579597-INS-293
chr13-55605770-INS-499
chr13-55637050-INS-199
chr13-55678866-INS-471

chr13-55678866-INS-684
chr13-55712934-INS-286
chr13-55738506-INS-77
chr13-55738776-INS-614
chr13-55740528-INS-235
chr13-55781358-INS-199
chr13-55785804-INS-92
chr13-55790061-INS-207
chr13-55795901-INS-55
chr13-55797005-INS-152
chr13-55808839-INS-56
chr13-55813182-INS-206
chr13-55833325-INS-183
chr13-55844705-INS-234
chr13-55855633-INS-203
chr13-55856261-INS-51
chr13-55866138-INS-199
chr13-55870577-INS-100
chr13-55870935-INS-79
chr13-55878012-INS-170
chr13-55917789-INS-99
chr13-55921357-INS-6415
chr13-55933261-INS-7161
chr13-55962218-INS-158
chr13-55967534-INS-206
chr13-55991567-INS-204
chr13-56004700-INS-140
chr13-56016734-INS-1205
chr13-56044431-INS-398
chr13-56068835-INS-67
chr13-56079696-INS-240
chr13-56086806-INS-60
chr13-56129574-INS-62
chr13-56141993-INS-498
chr13-56163870-INS-115
chr13-56190356-INS-898
chr13-56196760-INS-53
chr13-56224338-INS-6442
chr13-56247215-INS-1841
chr13-56247216-INS-348
chr13-56269860-INS-66
chr13-56290308-INS-52
chr13-56306608-INS-122
chr13-56319416-INS-64
chr13-56346228-INS-169
chr13-56378681-INS-73
chr13-56382843-INS-55
chr13-56387277-INS-300
chr13-56391879-INS-132
chr13-56423752-INS-961
chr13-56426917-INS-76
chr13-56428790-INS-67
chr13-56430393-INS-334
chr13-56446935-INS-80
chr13-56517278-INS-429
chr13-56526910-INS-95
chr13-56598059-INS-401
chr13-56608394-INS-53
chr13-56624363-INS-62
chr13-56643035-INS-68
chr13-56712600-INS-68
chr13-56714819-INS-606
chr13-56732887-INS-90
chr13-56777167-INS-922
chr13-56794122-INS-60
chr13-56799508-INS-152
chr13-56810778-INS-54
chr13-56821020-INS-96
chr13-56827210-INS-175
chr13-56834986-INS-210
chr13-56878473-INS-78
chr13-56892866-INS-150
chr13-56915950-INS-159
chr13-56919176-INS-58
chr13-56948060-INS-143
chr13-57001807-INS-73
chr13-57012974-INS-52

chr13-57013836-INS-621
chr13-57022593-INS-314
chr13-57022812-INS-636
chr13-57069955-INS-71
chr13-57083530-INS-252
chr13-57090759-INS-2197
chr13-57096120-INS-198
chr13-57117896-INS-95
chr13-57121819-INS-67
chr13-57131961-INS-56
chr13-57152761-INS-169
chr13-57186982-INS-259
chr13-57190314-INS-196
chr13-57191904-INS-1721
chr13-57206583-INS-59
chr13-57227108-INS-90
chr13-57245013-INS-6669
chr13-57250631-INS-267
chr13-57251636-INS-58
chr13-57251939-INS-6403
chr13-57259825-INS-275
chr13-57261376-INS-52
chr13-57264743-INS-110
chr13-57269244-INS-68
chr13-57292225-INS-581
chr13-57309793-INS-167
chr13-57316015-INS-139
chr13-57323465-INS-972
chr13-57341236-INS-172
chr13-57343246-INS-111
chr13-57374353-INS-401
chr13-57375529-INS-132
chr13-57376288-INS-3007
chr13-57376336-INS-2508
chr13-57376336-INS-304
chr13-57381898-INS-7102
chr13-57389941-INS-68
chr13-57401261-INS-52
chr13-57408854-INS-62
chr13-57428155-INS-88
chr13-57431387-INS-60
chr13-57479351-INS-5542
chr13-57492890-INS-509
chr13-57499758-INS-63
chr13-57513526-INS-165
chr13-57524879-INS-62
chr13-57526074-INS-6907
chr13-57535216-INS-156
chr13-57547767-INS-90
chr13-57615753-INS-60
chr13-57635071-INS-83
chr13-57658779-INS-68
chr13-57663085-INS-170
chr13-57696580-INS-59
chr13-57748287-INS-86
chr13-57806307-INS-266
chr13-57806620-INS-2344
chr13-57806771-INS-985
chr13-57810156-INS-241
chr13-57895257-INS-58
chr13-57900942-INS-83
chr13-57911863-INS-124
chr13-57947442-INS-164
chr13-57971030-INS-128
chr13-57992625-INS-57
chr13-58001384-INS-198
chr13-58007541-INS-104
chr13-58039468-INS-4520
chr13-58047863-INS-132
chr13-58058068-INS-1092
chr13-58058426-INS-527
chr13-58076191-INS-52
chr13-58080518-INS-90
chr13-58101978-INS-360
chr13-58105018-INS-57
chr13-58113420-INS-336
chr13-58118933-INS-345

chr13-58118933-INS-570
chr13-58125970-INS-54
chr13-58135399-INS-52
chr13-58146001-INS-140
chr13-58155092-INS-205
chr13-58189697-INS-66
chr13-58207408-INS-225
chr13-58228548-INS-51
chr13-58233838-INS-62
chr13-58238490-INS-56
chr13-58241283-INS-401
chr13-58244724-INS-75
chr13-58256730-INS-196
chr13-58257515-INS-55
chr13-58268551-INS-190
chr13-58279672-INS-7135
chr13-58293497-INS-5255
chr13-58297025-INS-174
chr13-58308759-INS-88
chr13-58319475-INS-214
chr13-58351997-INS-121
chr13-58360062-INS-283
chr13-58370432-INS-241
chr13-58373675-INS-91
chr13-58388475-INS-259
chr13-58391457-INS-394
chr13-58399152-INS-179
chr13-58409581-INS-369
chr13-58410414-INS-1856
chr13-58425145-INS-162
chr13-58432896-INS-122
chr13-58451527-INS-207
chr13-58457111-INS-129
chr13-58490792-INS-58
chr13-58499489-INS-408
chr13-58504065-INS-199
chr13-58508666-INS-179
chr13-58518438-INS-489
chr13-58583825-INS-95
chr13-58609072-INS-88
chr13-58632750-INS-219
chr13-58640420-INS-88
chr13-58692737-INS-258
chr13-58703031-INS-58
chr13-58731534-INS-182
chr13-58734506-INS-67
chr13-58738609-INS-54
chr13-58774599-INS-88
chr13-58783440-INS-64
chr13-58787943-INS-462
chr13-58790261-INS-133
chr13-58797751-INS-53
chr13-58798424-INS-52
chr13-58808360-INS-89
chr13-58826077-INS-212
chr13-58857825-INS-257
chr13-58859406-INS-180
chr13-58879261-INS-113
chr13-58888168-INS-144
chr13-58888283-INS-684
chr13-58892671-INS-77
chr13-58900415-INS-83
chr13-58912824-INS-93
chr13-58937857-INS-213
chr13-58952092-INS-116
chr13-58952789-INS-460
chr13-58961032-INS-227
chr13-58969448-INS-400
chr13-59017292-INS-318
chr13-59021325-INS-76
chr13-59035623-INS-163
chr13-59054722-INS-74
chr13-59060451-INS-256
chr13-59080500-INS-64
chr13-59100850-INS-56
chr13-59102635-INS-54
chr13-59120868-INS-74

chr13-59146451-INS-60
chr13-59179941-INS-253
chr13-59189579-INS-55
chr13-59198101-INS-86
chr13-59237378-INS-849
chr13-59286168-INS-76
chr13-59300457-INS-100
chr13-59317146-INS-92
chr13-59318873-INS-277
chr13-59345214-INS-1661
chr13-59369059-INS-1369
chr13-59381071-INS-124
chr13-59387956-INS-64
chr13-59404522-INS-95
chr13-59418866-INS-477
chr13-59448638-INS-215
chr13-59468601-INS-71
chr13-59482261-INS-149
chr13-59485703-INS-94
chr13-59511525-INS-54
chr13-59522486-INS-565
chr13-59557711-INS-67
chr13-59559120-INS-199
chr13-59575520-INS-55
chr13-59611487-INS-288
chr13-59625752-INS-432
chr13-59638027-INS-66
chr13-59642788-INS-189
chr13-59670558-INS-363
chr13-59674088-INS-336
chr13-59682894-INS-272
chr13-59689675-INS-66
chr13-59693176-INS-375
chr13-59716539-INS-152
chr13-59719683-INS-234
chr13-59728923-INS-854
chr13-59744349-INS-108
chr13-59753206-INS-94
chr13-59767709-INS-150
chr13-59768965-INS-5382
chr13-59786051-INS-209
chr13-59788052-INS-196
chr13-59792527-INS-63
chr13-59794156-INS-236
chr13-59821937-INS-86
chr13-59847408-INS-123
chr13-59857913-INS-358
chr13-59868821-INS-56
chr13-59884451-INS-211
chr13-59902244-INS-4123
chr13-59913017-INS-117
chr13-59919667-INS-229
chr13-59932140-INS-108
chr13-59934772-INS-200
chr13-59941482-INS-205
chr13-59945593-INS-678
chr13-59953137-INS-123
chr13-59963952-INS-66
chr13-59971889-INS-85
chr13-59995709-INS-410
chr13-60004131-INS-271
chr13-60025224-INS-262
chr13-60046842-INS-164
chr13-60048776-INS-5243
chr13-60052285-INS-3760
chr13-60056965-INS-8326
chr13-60058407-INS-8881
chr13-60061827-INS-156
chr13-60074485-INS-68
chr13-60084305-INS-148
chr13-60086253-INS-748
chr13-60099478-INS-268
chr13-60106581-INS-193
chr13-60111841-INS-64
chr13-60115461-INS-204
chr13-60119069-INS-422
chr13-60128878-INS-427

chr13-60131188-INS-267
chr13-60147326-INS-143
chr13-60157663-INS-122
chr13-60168294-INS-416
chr13-60181140-INS-7108
chr13-60222435-INS-174
chr13-60244773-INS-890
chr13-60262700-INS-149
chr13-60271532-INS-64
chr13-60302604-INS-54
chr13-60307650-INS-118
chr13-60307965-INS-368
chr13-60312687-INS-333
chr13-60318641-INS-122
chr13-60327364-INS-90
chr13-60338383-INS-128
chr13-60377814-INS-62
chr13-60394669-INS-167
chr13-60415225-INS-52
chr13-60427832-INS-91
chr13-60435175-INS-209
chr13-60441630-INS-236
chr13-60443084-INS-223
chr13-60445979-INS-279
chr13-60452509-INS-73
chr13-60460087-INS-68
chr13-60494534-INS-600
chr13-60516240-INS-54
chr13-60546326-INS-53
chr13-60548363-INS-273
chr13-60560040-INS-209
chr13-60590309-INS-455
chr13-60590329-INS-1890
chr13-60591538-INS-717
chr13-60611435-INS-205
chr13-60617382-INS-364
chr13-60629630-INS-52
chr13-60648994-INS-282
chr13-60652575-INS-976
chr13-60665905-INS-5639
chr13-60671597-INS-66
chr13-60679643-INS-76
chr13-60681040-INS-60
chr13-60696572-INS-229
chr13-60698068-INS-6229
chr13-60734012-INS-400
chr13-60740052-INS-387
chr13-60747927-INS-164
chr13-60758862-INS-130
chr13-60772759-INS-227
chr13-60803501-INS-96
chr13-60829449-INS-102
chr13-60850902-INS-452
chr13-60860927-INS-104
chr13-60872054-INS-52
chr13-60928062-INS-76
chr13-60928907-INS-6652
chr13-60936311-INS-109
chr13-60957876-INS-416
chr13-60958803-INS-248
chr13-60970546-INS-98
chr13-60985913-INS-115
chr13-60991460-INS-71
chr13-61014654-INS-119
chr13-61023802-INS-439
chr13-61039802-INS-139
chr13-61053719-INS-66
chr13-61055181-INS-62
chr13-61060720-INS-511
chr13-61066030-INS-81
chr13-61067402-INS-890
chr13-61090188-INS-284
chr13-61092495-INS-98
chr13-61110116-INS-115
chr13-61112901-INS-197
chr13-61128221-INS-416
chr13-61135789-INS-64

chr13-61252394-INS-259
chr13-61271997-INS-68
chr13-61314240-INS-1667
chr13-61314352-INS-404
chr13-61322189-INS-422
chr13-61363171-INS-56
chr13-61394611-INS-1305
chr13-61426893-INS-85
chr13-61489342-INS-347
chr13-61504911-INS-52
chr13-61527546-INS-412
chr13-61558850-INS-113
chr13-61561930-INS-236
chr13-61565252-INS-166
chr13-61567556-INS-222
chr13-61581670-INS-6213
chr13-61593388-INS-8456
chr13-61637964-INS-243
chr13-61638825-INS-118
chr13-61663050-INS-169
chr13-61710449-INS-1377
chr13-61732310-INS-1016
chr13-61733123-INS-3120
chr13-61738518-INS-113
chr13-61790719-INS-63
chr13-61795309-INS-208
chr13-61805303-INS-129
chr13-61829406-INS-7166
chr13-61841097-INS-260
chr13-61892663-INS-252
chr13-61899073-INS-856
chr13-61904833-INS-5448
chr13-61907977-INS-173
chr13-61954916-INS-100
chr13-62066276-INS-113
chr13-62069401-INS-767
chr13-62070307-INS-72
chr13-62141198-INS-351
chr13-62143067-INS-358
chr13-62144506-INS-200
chr13-62150837-INS-81
chr13-62182491-INS-242
chr13-62194772-INS-191
chr13-62197201-INS-588
chr13-62200847-INS-92
chr13-62206321-INS-722
chr13-62206585-INS-194
chr13-62217017-INS-90
chr13-62218367-INS-337
chr13-62347979-INS-3327
chr13-62353533-INS-1893
chr13-62425764-INS-56
chr13-62427356-INS-5282
chr13-62486143-INS-84
chr13-62502048-INS-116
chr13-62519276-INS-213
chr13-62553603-INS-384
chr13-62567220-INS-424
chr13-62574346-INS-98
chr13-62590622-INS-281
chr13-62596677-INS-196
chr13-62601933-INS-84
chr13-62605668-INS-524
chr13-62608377-INS-4020
chr13-62611964-INS-427
chr13-62636907-INS-250
chr13-62749573-INS-391
chr13-62804884-INS-138
chr13-62832483-INS-976
chr13-62846853-INS-195
chr13-62848408-INS-90
chr13-62854546-INS-290
chr13-62857403-INS-5313
chr13-62857749-INS-164
chr13-62899728-INS-527
chr13-62954247-INS-628
chr13-62958606-INS-205

chr13-62966884-INS-65
chr13-62978552-INS-200
chr13-62993629-INS-63
chr13-63003845-INS-258
chr13-63004346-INS-421
chr13-63017482-INS-78
chr13-63020426-INS-229
chr13-63037525-INS-203
chr13-63042887-INS-259
chr13-63046500-INS-401
chr13-63053531-INS-246
chr13-63062833-INS-210
chr13-63074594-INS-54
chr13-63076214-INS-78
chr13-63093925-INS-495
chr13-63109674-INS-216
chr13-63113987-INS-68
chr13-63122111-INS-1890
chr13-63151396-INS-360
chr13-63180587-INS-68
chr13-63188923-INS-66
chr13-63194303-INS-192
chr13-63204429-INS-230
chr13-63230233-INS-53
chr13-63245229-INS-144
chr13-63262307-INS-70
chr13-63278725-INS-236
chr13-63296990-INS-208
chr13-63312357-INS-83
chr13-63313100-INS-142
chr13-63320716-INS-194
chr13-63330044-INS-166
chr13-63344374-INS-101
chr13-63348838-INS-54
chr13-63348853-INS-757
chr13-63392033-INS-98
chr13-63396845-INS-238
chr13-63443478-INS-247
chr13-63459105-INS-57
chr13-63473669-INS-200
chr13-63486491-INS-357
chr13-63500956-INS-55
chr13-63502392-INS-59
chr13-63512360-INS-354
chr13-63522611-INS-2829
chr13-63527297-INS-190
chr13-63532785-INS-88
chr13-63532944-INS-5085
chr13-63536407-INS-239
chr13-63539537-INS-115
chr13-63570186-INS-80
chr13-63614216-INS-176
chr13-63615102-INS-84
chr13-63617934-INS-55
chr13-63705197-INS-54
chr13-63726927-INS-107
chr13-63729941-INS-54
chr13-63738207-INS-7232
chr13-63739195-INS-141
chr13-63748379-INS-502
chr13-63759092-INS-75
chr13-63772638-INS-2958
chr13-63785738-INS-411
chr13-63790226-INS-68
chr13-63834840-INS-157
chr13-63870310-INS-112
chr13-63876972-INS-153
chr13-63880394-INS-346
chr13-63928182-INS-53
chr13-63930256-INS-88
chr13-63950444-INS-176
chr13-63961281-INS-123
chr13-63970333-INS-91
chr13-63980688-INS-840
chr13-63991312-INS-204
chr13-64022890-INS-241
chr13-64052131-INS-363

chr13-64057874-INS-96
chr13-64061499-INS-52
chr13-64091201-INS-496
chr13-64096749-INS-305
chr13-64184226-INS-242
chr13-64195186-INS-398
chr13-64221383-INS-54
chr13-64234357-INS-435
chr13-64242069-INS-50
chr13-64247339-INS-188
chr13-64261996-INS-200
chr13-64271871-INS-52
chr13-64274809-INS-211
chr13-64288041-INS-152
chr13-64289079-INS-52
chr13-64291145-INS-229
chr13-64297761-INS-204
chr13-64315005-INS-78
chr13-64317240-INS-138
chr13-64322172-INS-70
chr13-64339205-INS-196
chr13-64367948-INS-168
chr13-64388040-INS-202
chr13-64414931-INS-152
chr13-64417866-INS-162
chr13-64424278-INS-64
chr13-64433415-INS-127
chr13-64462744-INS-115
chr13-64468715-INS-207
chr13-64473025-INS-176
chr13-64477557-INS-155
chr13-64478598-INS-190
chr13-64493576-INS-85
chr13-64506480-INS-2322
chr13-64508650-INS-220
chr13-64519912-INS-233
chr13-64521793-INS-202
chr13-64534975-INS-170
chr13-64542013-INS-175
chr13-64543394-INS-73
chr13-64547215-INS-103
chr13-64577228-INS-114
chr13-64611140-INS-321
chr13-64611967-INS-576
chr13-64625019-INS-105
chr13-64643860-INS-414
chr13-64683140-INS-330
chr13-64695591-INS-633
chr13-64701544-INS-55
chr13-64721395-INS-58
chr13-64748394-INS-91
chr13-64765379-INS-6252
chr13-64766552-INS-8536
chr13-64799459-INS-52
chr13-64840794-INS-1486
chr13-64857735-INS-6479
chr13-64865078-INS-7794
chr13-64893640-INS-53
chr13-64912641-INS-7330
chr13-64915989-INS-365
chr13-64922694-INS-891
chr13-64925805-INS-6395
chr13-64938967-INS-60
chr13-64948064-INS-218
chr13-64965103-INS-224
chr13-64991698-INS-117
chr13-65002348-INS-73
chr13-65058100-INS-828
chr13-65089802-INS-241
chr13-65199894-INS-786
chr13-65210173-INS-88
chr13-65269067-INS-8358
chr13-65276927-INS-52
chr13-65305843-INS-212
chr13-65306316-INS-882
chr13-65310203-INS-88
chr13-65360103-INS-63

chr13-65387048-INS-52
chr13-65431122-INS-1892
chr13-65476322-INS-135
chr13-65510725-INS-2743
chr13-65538757-INS-6434
chr13-65545527-INS-84
chr13-65622584-INS-485
chr13-65659890-INS-66
chr13-65660298-INS-168
chr13-65722040-INS-361
chr13-65741766-INS-203
chr13-65754553-INS-60
chr13-65757950-INS-490
chr13-65763547-INS-54
chr13-65766852-INS-91
chr13-65839073-INS-252
chr13-65857576-INS-51
chr13-65944376-INS-5939
chr13-65948945-INS-60
chr13-65989225-INS-200
chr13-65994787-INS-208
chr13-66002060-INS-223
chr13-66047143-INS-72
chr13-66055879-INS-125
chr13-66080457-INS-207
chr13-66132020-INS-218
chr13-66141863-INS-215
chr13-66188296-INS-504
chr13-66191408-INS-69
chr13-66311194-INS-128
chr13-66311706-INS-84
chr13-66314603-INS-61
chr13-66348332-INS-204
chr13-66395152-INS-204
chr13-66413777-INS-209
chr13-66420540-INS-3150
chr13-66543464-INS-658
chr13-66599865-INS-65
chr13-66606826-INS-52
chr13-66616124-INS-480
chr13-66631723-INS-412
chr13-66658542-INS-201
chr13-66690574-INS-258
chr13-66720706-INS-195
chr13-66746215-INS-250
chr13-66802751-INS-4725
chr13-66818635-INS-208
chr13-66879469-INS-63
chr13-66882871-INS-482
chr13-66907689-INS-646
chr13-67010679-INS-564
chr13-67020973-INS-7378
chr13-67027225-INS-8819
chr13-67056673-INS-191
chr13-67058833-INS-415
chr13-67083352-INS-72
chr13-67106983-INS-737
chr13-67111291-INS-222
chr13-67152629-INS-354
chr13-67158414-INS-199
chr13-67177436-INS-6251
chr13-67178866-INS-2951
chr13-67214286-INS-64
chr13-67220190-INS-56
chr13-67251808-INS-74
chr13-67265478-INS-95
chr13-67289978-INS-226
chr13-67314019-INS-1977
chr13-67319997-INS-202
chr13-67324089-INS-195
chr13-67346276-INS-156
chr13-67348271-INS-5306
chr13-67365745-INS-53
chr13-67402908-INS-6519
chr13-67447329-INS-56
chr13-67469984-INS-210
chr13-67476230-INS-184

chr13-67499036-INS-58
chr13-67502546-INS-72
chr13-67502655-INS-369
chr13-67503009-INS-97
chr13-67511110-INS-201
chr13-67550167-INS-7277
chr13-67551901-INS-7291
chr13-67553474-INS-56
chr13-67563432-INS-224
chr13-67564498-INS-316
chr13-67572521-INS-598
chr13-67574468-INS-806
chr13-67582112-INS-94
chr13-67583922-INS-229
chr13-67588647-INS-169
chr13-67610825-INS-223
chr13-67622508-INS-242
chr13-67626548-INS-229
chr13-67628085-INS-62
chr13-67636608-INS-207
chr13-67642607-INS-421
chr13-67679665-INS-867
chr13-67715374-INS-68
chr13-67730564-INS-180
chr13-67764195-INS-686
chr13-67773519-INS-597
chr13-67774861-INS-202
chr13-67799988-INS-173
chr13-67800876-INS-261
chr13-67814761-INS-54
chr13-67822981-INS-283
chr13-67826595-INS-222
chr13-67828508-INS-180
chr13-67842563-INS-188
chr13-67844371-INS-861
chr13-67858113-INS-203
chr13-67902093-INS-329
chr13-67947963-INS-437
chr13-67996191-INS-69
chr13-68006094-INS-82
chr13-68014150-INS-7212
chr13-68016643-INS-280
chr13-68020773-INS-148
chr13-68021852-INS-894
chr13-68023123-INS-419
chr13-68028749-INS-84
chr13-68048053-INS-203
chr13-68054241-INS-127
chr13-68056652-INS-104
chr13-68093385-INS-308
chr13-68147614-INS-135
chr13-68155405-INS-7080
chr13-68287805-INS-78
chr13-68385326-INS-91
chr13-68418497-INS-59
chr13-68435520-INS-235
chr13-68445156-INS-111
chr13-68449108-INS-140
chr13-68450689-INS-75
chr13-68461080-INS-64
chr13-68585191-INS-64
chr13-68588693-INS-6436
chr13-68588868-INS-1741
chr13-68637226-INS-121
chr13-68643913-INS-308
chr13-68753659-INS-114
chr13-68770851-INS-766
chr13-68806903-INS-351
chr13-68847668-INS-387
chr13-68848984-INS-490
chr13-68896655-INS-354
chr13-68916368-INS-54
chr13-68953886-INS-63
chr13-68993659-INS-6420
chr13-69002631-INS-439
chr13-69006024-INS-193
chr13-69013577-INS-97

chr13-69028593-INS-62
chr13-69052960-INS-51
chr13-69059399-INS-147
chr13-69061692-INS-209
chr13-69066368-INS-212
chr13-69070630-INS-1302
chr13-69075242-INS-85
chr13-69076776-INS-66
chr13-69095023-INS-703
chr13-69098296-INS-131
chr13-69103364-INS-282
chr13-69116621-INS-133
chr13-69131580-INS-64
chr13-69160212-INS-57
chr13-69166263-INS-477
chr13-69206541-INS-324
chr13-69222842-INS-6394
chr13-69224821-INS-60
chr13-69287714-INS-68
chr13-69310903-INS-6272
chr13-69314895-INS-925
chr13-69339684-INS-93
chr13-69344479-INS-99
chr13-69403632-INS-330
chr13-69417833-INS-92
chr13-69446215-INS-489
chr13-69449809-INS-927
chr13-69467719-INS-490
chr13-69479396-INS-53
chr13-69485214-INS-317
chr13-69517747-INS-135
chr13-69518671-INS-265
chr13-69525481-INS-212
chr13-69541905-INS-55
chr13-69547454-INS-90
chr13-69548516-INS-7837
chr13-69571966-INS-504
chr13-69575056-INS-80
chr13-69575345-INS-396
chr13-69579913-INS-105
chr13-69588875-INS-852
chr13-69623213-INS-979
chr13-69625139-INS-310
chr13-69626887-INS-88
chr13-69632751-INS-358
chr13-69653235-INS-177
chr13-69659908-INS-70
chr13-69664304-INS-184
chr13-69691630-INS-221
chr13-69709000-INS-90
chr13-69710509-INS-86
chr13-69712138-INS-328
chr13-69723558-INS-78
chr13-69729034-INS-72
chr13-69730630-INS-498
chr13-69748214-INS-220
chr13-69748534-INS-8940
chr13-69769058-INS-207
chr13-69785881-INS-126
chr13-69805164-INS-131
chr13-69805430-INS-373
chr13-69809636-INS-6452
chr13-69813857-INS-400
chr13-69822224-INS-386
chr13-69822976-INS-1891
chr13-69825522-INS-127
chr13-69828266-INS-325
chr13-69843050-INS-1288
chr13-69846388-INS-212
chr13-69885984-INS-66
chr13-69890337-INS-417
chr13-69908017-INS-492
chr13-69908910-INS-88
chr13-69921148-INS-192
chr13-69951292-INS-806
chr13-69960954-INS-51
chr13-69976042-INS-58

chr13-69990728-INS-334
chr13-70012676-INS-54
chr13-70025500-INS-600
chr13-70048299-INS-76
chr13-70065171-INS-350
chr13-70073753-INS-915
chr13-70086954-INS-126
chr13-70115954-INS-6652
chr13-70146650-INS-103
chr13-70153805-INS-495
chr13-70156975-INS-549
chr13-70167584-INS-358
chr13-70169530-INS-94
chr13-70189455-INS-68
chr13-70224991-INS-229
chr13-70328673-INS-118
chr13-70340048-INS-153
chr13-70341947-INS-267
chr13-70357281-INS-112
chr13-70366419-INS-68
chr13-70366647-INS-6596
chr13-70387104-INS-102
chr13-70418197-INS-112
chr13-70455296-INS-6803
chr13-70471569-INS-6434
chr13-70472907-INS-53
chr13-70485184-INS-89
chr13-70487895-INS-6372
chr13-70501538-INS-56
chr13-70530089-INS-100
chr13-70531756-INS-70
chr13-70533190-INS-1087
chr13-70546950-INS-54
chr13-70559768-INS-1219
chr13-70582628-INS-57
chr13-70678320-INS-205
chr13-70678426-INS-553
chr13-70689604-INS-80
chr13-70690556-INS-3313
chr13-70717977-INS-189
chr13-70723453-INS-400
chr13-70743086-INS-161
chr13-70771709-INS-592
chr13-70787111-INS-120
chr13-70806754-INS-82
chr13-70807897-INS-362
chr13-70819672-INS-68
chr13-70820957-INS-981
chr13-70850487-INS-101
chr13-70860804-INS-99
chr13-70870128-INS-349
chr13-70885652-INS-336
chr13-70911978-INS-435
chr13-70934317-INS-162
chr13-70935189-INS-355
chr13-70977242-INS-218
chr13-71005165-INS-145
chr13-71008308-INS-150
chr13-71013960-INS-59
chr13-71015930-INS-66
chr13-71018446-INS-60
chr13-71041644-INS-80
chr13-71063983-INS-541
chr13-71103450-INS-53
chr13-71112119-INS-60
chr13-71121006-INS-707
chr13-71127699-INS-67
chr13-71147363-INS-57
chr13-71168438-INS-182
chr13-71185570-INS-113
chr13-71229926-INS-6307
chr13-71251430-INS-103
chr13-71260704-INS-54
chr13-71301190-INS-282
chr13-71304935-INS-64
chr13-71327421-INS-56
chr13-71331226-INS-64

chr13-71352902-INS-54
chr13-71355013-INS-58
chr13-71359853-INS-54
chr13-71386680-INS-397
chr13-71397453-INS-1866
chr13-71406943-INS-59
chr13-71432067-INS-359
chr13-71439003-INS-80
chr13-71471282-INS-110
chr13-71473884-INS-52
chr13-71483337-INS-358
chr13-71517487-INS-51
chr13-71540699-INS-132
chr13-71545492-INS-422
chr13-71557330-INS-81
chr13-71569573-INS-389
chr13-71575367-INS-54
chr13-71612466-INS-107
chr13-71619923-INS-334
chr13-71619933-INS-670
chr13-71655473-INS-6436
chr13-71682682-INS-358
chr13-71695340-INS-87
chr13-71715544-INS-156
chr13-71717838-INS-758
chr13-71729924-INS-7027
chr13-71744962-INS-181
chr13-71748327-INS-70
chr13-71756378-INS-60
chr13-71803242-INS-54
chr13-71860291-INS-4368
chr13-71903533-INS-92
chr13-71927039-INS-74
chr13-71934968-INS-330
chr13-71982142-INS-116
chr13-72010274-INS-196
chr13-72027905-INS-213
chr13-72051708-INS-52
chr13-72051950-INS-3179
chr13-72092635-INS-142
chr13-72104928-INS-198
chr13-72145737-INS-88
chr13-72151043-INS-304
chr13-72158834-INS-56
chr13-72160679-INS-204
chr13-72217574-INS-60
chr13-72233289-INS-592
chr13-72240257-INS-52
chr13-72243202-INS-108
chr13-72252446-INS-79
chr13-72264623-INS-52
chr13-72322295-INS-388
chr13-72327341-INS-902
chr13-72362208-INS-114
chr13-72378062-INS-55
chr13-72429844-INS-331
chr13-72455417-INS-201
chr13-72457051-INS-481
chr13-72459434-INS-457
chr13-72474511-INS-83
chr13-72505340-INS-305
chr13-72515337-INS-56
chr13-72525932-INS-135
chr13-72542953-INS-390
chr13-72551198-INS-6484
chr13-72581393-INS-521
chr13-72582076-INS-79
chr13-72590758-INS-383
chr13-72598295-INS-6444
chr13-72606983-INS-55
chr13-72611782-INS-56
chr13-72695199-INS-62
chr13-72704347-INS-7890
chr13-72704906-INS-71
chr13-72727293-INS-181
chr13-72728950-INS-66
chr13-72742830-INS-65

chr13-72758443-INS-54
chr13-72760846-INS-132
chr13-72783423-INS-95
chr13-72810719-INS-84
chr13-72834056-INS-8580
chr13-72849842-INS-640
chr13-72867599-INS-132
chr13-72894533-INS-121
chr13-72902266-INS-140
chr13-72935688-INS-190
chr13-72940363-INS-755
chr13-72952207-INS-68
chr13-72958523-INS-108
chr13-72961336-INS-51
chr13-72969315-INS-99
chr13-72977475-INS-400
chr13-73003898-INS-70
chr13-73027117-INS-120
chr13-73042141-INS-7198
chr13-73048758-INS-1040
chr13-73122159-INS-68
chr13-73123523-INS-59
chr13-73132608-INS-81
chr13-73158856-INS-184
chr13-73164868-INS-54
chr13-73178021-INS-68
chr13-73179681-INS-698
chr13-73193881-INS-51
chr13-73206589-INS-119
chr13-73208907-INS-110
chr13-73244813-INS-132
chr13-73259429-INS-169
chr13-73263706-INS-303
chr13-73266600-INS-52
chr13-73271011-INS-112
chr13-73296196-INS-354
chr13-73311863-INS-57
chr13-73313885-INS-110
chr13-73317120-INS-110
chr13-73320579-INS-391
chr13-73344853-INS-209
chr13-73348367-INS-805
chr13-73352810-INS-64
chr13-73357755-INS-4936
chr13-73364871-INS-2399
chr13-73373814-INS-6246
chr13-73401451-INS-201
chr13-73414217-INS-157
chr13-73463143-INS-115
chr13-73466263-INS-183
chr13-73496977-INS-447
chr13-73498993-INS-5195
chr13-73513189-INS-131
chr13-73517543-INS-196
chr13-73537089-INS-354
chr13-73570205-INS-226
chr13-73604681-INS-97
chr13-73627974-INS-53
chr13-73628858-INS-64
chr13-73653353-INS-66
chr13-73661382-INS-537
chr13-73687823-INS-100
chr13-73700299-INS-76
chr13-73711593-INS-86
chr13-73727423-INS-530
chr13-73730500-INS-512
chr13-73732994-INS-61
chr13-73744842-INS-7058
chr13-73749115-INS-391
chr13-73770396-INS-213
chr13-73777728-INS-489
chr13-73781167-INS-191
chr13-73796231-INS-81
chr13-73796724-INS-156
chr13-73798364-INS-84
chr13-73807953-INS-74
chr13-73810256-INS-69

chr13-73818565-INS-211
chr13-73821264-INS-233
chr13-73824899-INS-105
chr13-73827834-INS-6412
chr13-73846512-INS-66
chr13-73861931-INS-54
chr13-73873397-INS-258
chr13-73886390-INS-186
chr13-73892348-INS-110
chr13-73894172-INS-245
chr13-73899038-INS-227
chr13-73902390-INS-204
chr13-73906579-INS-263
chr13-73949869-INS-222
chr13-73978659-INS-52
chr13-73979269-INS-147
chr13-73982874-INS-53
chr13-73989587-INS-62
chr13-73996746-INS-87
chr13-73998924-INS-67
chr13-74043702-INS-58
chr13-74046607-INS-194
chr13-74050849-INS-186
chr13-74053575-INS-269
chr13-74062093-INS-295
chr13-74069113-INS-191
chr13-74141337-INS-145
chr13-74156885-INS-66
chr13-74166225-INS-199
chr13-74187912-INS-1062
chr13-74187917-INS-459
chr13-74195568-INS-81
chr13-74221396-INS-52
chr13-74223282-INS-596
chr13-74232487-INS-103
chr13-74245166-INS-711
chr13-74343531-INS-194
chr13-74344628-INS-60
chr13-74345563-INS-2739
chr13-74351050-INS-64
chr13-74372100-INS-75
chr13-74374705-INS-99
chr13-74411806-INS-758
chr13-74430467-INS-54
chr13-74446835-INS-56
chr13-74454924-INS-76
chr13-74456933-INS-57
chr13-74481700-INS-113
chr13-74492034-INS-2886
chr13-74504354-INS-85
chr13-74534984-INS-205
chr13-74541569-INS-831
chr13-74543510-INS-329
chr13-74547992-INS-52
chr13-74557661-INS-596
chr13-74559786-INS-194
chr13-74563621-INS-333
chr13-74573443-INS-182
chr13-74635281-INS-68
chr13-74660130-INS-80
chr13-74674766-INS-390
chr13-74697929-INS-6378
chr13-74721447-INS-467
chr13-74778285-INS-53
chr13-74794183-INS-290
chr13-74809763-INS-180
chr13-74876606-INS-7322
chr13-74892521-INS-200
chr13-74936361-INS-89
chr13-74971750-INS-101
chr13-74987915-INS-6449
chr13-74992268-INS-172
chr13-74993268-INS-88
chr13-74994200-INS-6605
chr13-75019129-INS-822
chr13-75054410-INS-66
chr13-75061818-INS-64

chr13-75062919-INS-138
chr13-75074703-INS-963
chr13-75127737-INS-66
chr13-75132919-INS-476
chr13-75143693-INS-162
chr13-75146832-INS-403
chr13-75149210-INS-62
chr13-75156833-INS-242
chr13-75202018-INS-6508
chr13-75363961-INS-123
chr13-75367368-INS-95
chr13-75373237-INS-53
chr13-75411121-INS-138
chr13-75416632-INS-52
chr13-75417230-INS-58
chr13-75431941-INS-77
chr13-75485455-INS-396
chr13-75489608-INS-201
chr13-75492970-INS-394
chr13-75535856-INS-544
chr13-75537529-INS-623
chr13-75586519-INS-59
chr13-75614729-INS-204
chr13-75620303-INS-128
chr13-75624828-INS-58
chr13-75626347-INS-121
chr13-75633887-INS-198
chr13-75661518-INS-1893
chr13-75663323-INS-418
chr13-75667889-INS-66
chr13-75670450-INS-72
chr13-75694193-INS-145
chr13-75697816-INS-79
chr13-75702578-INS-354
chr13-75707386-INS-66
chr13-75736611-INS-263
chr13-75748426-INS-5551
chr13-75754412-INS-417
chr13-75756059-INS-55
chr13-75758576-INS-209
chr13-75762810-INS-468
chr13-75776302-INS-398
chr13-75802846-INS-476
chr13-75807210-INS-202
chr13-75811882-INS-182
chr13-75841743-INS-187
chr13-75843826-INS-549
chr13-75849778-INS-262
chr13-75850889-INS-87
chr13-75851644-INS-81
chr13-75906570-INS-523
chr13-75907593-INS-105
chr13-75911516-INS-52
chr13-75953466-INS-630
chr13-75954458-INS-7213
chr13-75991057-INS-316
chr13-76036748-INS-982
chr13-76041511-INS-58
chr13-76065697-INS-447
chr13-76132521-INS-420
chr13-76133635-INS-7171
chr13-76141082-INS-99
chr13-76142067-INS-78
chr13-76142067-INS-780
chr13-76155394-INS-393
chr13-76199116-INS-189
chr13-76218397-INS-62
chr13-76302854-INS-220
chr13-76326771-INS-978
chr13-76329147-INS-546
chr13-76360153-INS-62
chr13-76382359-INS-745
chr13-76394581-INS-112
chr13-76432958-INS-342
chr13-76483669-INS-421
chr13-76526380-INS-539
chr13-76560051-INS-138

chr13-76561093-INS-275
chr13-76591689-INS-163
chr13-76614087-INS-60
chr13-76685059-INS-528
chr13-76725734-INS-257
chr13-76738279-INS-112
chr13-76739176-INS-1034
chr13-76766138-INS-243
chr13-76776333-INS-76
chr13-76781093-INS-210
chr13-76786362-INS-7178
chr13-76829841-INS-126
chr13-76844068-INS-218
chr13-76886804-INS-145
chr13-76893678-INS-68
chr13-76907117-INS-63
chr13-76962810-INS-179
chr13-77010844-INS-149
chr13-77024355-INS-72
chr13-77024379-INS-2181
chr13-77028188-INS-6106
chr13-77084033-INS-159
chr13-77088438-INS-430
chr13-77154677-INS-54
chr13-77164843-INS-179
chr13-77209466-INS-60
chr13-77242743-INS-87
chr13-77257779-INS-202
chr13-77259666-INS-902
chr13-77296352-INS-262
chr13-77308135-INS-54
chr13-77342552-INS-60
chr13-77346805-INS-157
chr13-77385634-INS-737
chr13-77425177-INS-6553
chr13-77428124-INS-91
chr13-77442964-INS-400
chr13-77446213-INS-111
chr13-77453168-INS-72
chr13-77466463-INS-60
chr13-77541076-INS-91
chr13-77553004-INS-270
chr13-77599687-INS-1843
chr13-77645905-INS-1397
chr13-77646915-INS-504
chr13-77662559-INS-6408
chr13-77673853-INS-374
chr13-77675699-INS-591
chr13-77682245-INS-6839
chr13-77694184-INS-195
chr13-77700612-INS-66
chr13-77710544-INS-56
chr13-77727806-INS-51
chr13-77742327-INS-76
chr13-77744681-INS-181
chr13-77746209-INS-103
chr13-77791431-INS-76
chr13-77837087-INS-653
chr13-77842727-INS-68
chr13-77851936-INS-5319
chr13-77852666-INS-6929
chr13-77861830-INS-2911
chr13-77868752-INS-241
chr13-77885067-INS-79
chr13-77888670-INS-706
chr13-77911376-INS-68
chr13-77912163-INS-411
chr13-77930255-INS-77
chr13-77952514-INS-1895
chr13-77996498-INS-1558
chr13-78009545-INS-6163
chr13-78014744-INS-164
chr13-78024197-INS-2108
chr13-78035604-INS-627
chr13-78056093-INS-351
chr13-78087721-INS-647
chr13-78099473-INS-199

chr13-78130283-INS-96
chr13-78144501-INS-516
chr13-78165450-INS-84
chr13-78194801-INS-72
chr13-78213490-INS-449
chr13-78219168-INS-1152
chr13-78234625-INS-54
chr13-78260616-INS-7085
chr13-78264483-INS-68
chr13-78269483-INS-172
chr13-78271609-INS-868
chr13-78285767-INS-1540
chr13-78286990-INS-203
chr13-78297158-INS-143
chr13-78298518-INS-1715
chr13-78369570-INS-271
chr13-78372925-INS-452
chr13-78382540-INS-399
chr13-78388185-INS-533
chr13-78396115-INS-360
chr13-78399217-INS-7722
chr13-78401101-INS-432
chr13-78401725-INS-54
chr13-78407548-INS-75
chr13-78422486-INS-115
chr13-78422969-INS-146
chr13-78424215-INS-56
chr13-78426167-INS-3799
chr13-78456324-INS-224
chr13-78480187-INS-115
chr13-78487002-INS-202
chr13-78495649-INS-90
chr13-78499308-INS-58
chr13-78505192-INS-1893
chr13-78513288-INS-99
chr13-78514929-INS-7362
chr13-78518718-INS-8066
chr13-78521094-INS-6241
chr13-78534435-INS-7154
chr13-78557371-INS-71
chr13-78576715-INS-58
chr13-78578125-INS-126
chr13-78600648-INS-53
chr13-78623939-INS-3997
chr13-78624818-INS-6600
chr13-78647892-INS-428
chr13-78652245-INS-705
chr13-78653631-INS-7700
chr13-78657054-INS-260
chr13-78668049-INS-7390
chr13-78670570-INS-111
chr13-78676820-INS-121
chr13-78686736-INS-474
chr13-78687044-INS-7003
chr13-78701277-INS-62
chr13-78762921-INS-236
chr13-78783283-INS-70
chr13-78831569-INS-4392
chr13-78839012-INS-96
chr13-78858905-INS-101
chr13-78859965-INS-197
chr13-78871882-INS-57
chr13-78875611-INS-710
chr13-78914726-INS-454
chr13-78928445-INS-2661
chr13-78944948-INS-376
chr13-78974824-INS-71
chr13-78974885-INS-447
chr13-78984747-INS-150
chr13-78993588-INS-5547
chr13-78994871-INS-6440
chr13-79023626-INS-104
chr13-79035346-INS-90
chr13-79037046-INS-649
chr13-79040890-INS-721
chr13-79052185-INS-6853
chr13-79072312-INS-215

chr13-79072589-INS-6264
chr13-79110252-INS-6285
chr13-79113714-INS-6583
chr13-79126496-INS-107
chr13-79130785-INS-1012
chr13-79140623-INS-608
chr13-79160180-INS-1582
chr13-79167515-INS-6126
chr13-79172111-INS-119
chr13-79209374-INS-1886
chr13-79211985-INS-1553
chr13-79215798-INS-64
chr13-79221939-INS-68
chr13-79228909-INS-298
chr13-79235478-INS-7124
chr13-79259823-INS-3251
chr13-79262290-INS-65
chr13-79274802-INS-4960
chr13-79292569-INS-5102
chr13-79302799-INS-118
chr13-79310548-INS-3126
chr13-79317929-INS-7341
chr13-79402575-INS-56
chr13-79407616-INS-88
chr13-79428588-INS-54
chr13-79438213-INS-166
chr13-79441019-INS-444
chr13-79441710-INS-76
chr13-79476475-INS-52
chr13-79502545-INS-5062
chr13-79509713-INS-2669
chr13-79515242-INS-123
chr13-79531431-INS-64
chr13-79551158-INS-2099
chr13-79552749-INS-56
chr13-79555130-INS-352
chr13-79561567-INS-128
chr13-79585276-INS-52
chr13-79646561-INS-7081
chr13-79648741-INS-478
chr13-79683923-INS-102
chr13-79727950-INS-63
chr13-79728990-INS-69
chr13-79752471-INS-1895
chr13-79757320-INS-6344
chr13-79774455-INS-284
chr13-79783261-INS-992
chr13-79785945-INS-51
chr13-79806490-INS-68
chr13-79808823-INS-69
chr13-79822207-INS-52
chr13-79823683-INS-53
chr13-79833340-INS-140
chr13-79834374-INS-135
chr13-79843808-INS-522
chr13-79892317-INS-400
chr13-79920119-INS-202
chr13-79924809-INS-192
chr13-79935537-INS-99
chr13-79983258-INS-388
chr13-79989289-INS-6454
chr13-80012968-INS-252
chr13-80018572-INS-7793
chr13-80020061-INS-54
chr13-80035414-INS-5903
chr13-80081452-INS-1110
chr13-80084955-INS-3220
chr13-80106758-INS-107
chr13-80118043-INS-7080
chr13-80121433-INS-952
chr13-80127657-INS-74
chr13-80132148-INS-359
chr13-80142450-INS-109
chr13-80159743-INS-92
chr13-80162414-INS-65
chr13-80163843-INS-421
chr13-80170854-INS-7108

chr13-80194614-INS-95
chr13-80197062-INS-69
chr13-80223923-INS-7078
chr13-80238587-INS-105
chr13-80257006-INS-118
chr13-80281339-INS-113
chr13-80281358-INS-272
chr13-80332669-INS-498
chr13-80335358-INS-74
chr13-80378040-INS-219
chr13-80393393-INS-5313
chr13-80411144-INS-988
chr13-80428552-INS-55
chr13-80450510-INS-6399
chr13-80479050-INS-3128
chr13-80489633-INS-3842
chr13-80496207-INS-87
chr13-80534175-INS-6537
chr13-80563175-INS-91
chr13-80563862-INS-3698
chr13-80598249-INS-227
chr13-80600231-INS-757
chr13-80624155-INS-7043
chr13-80645277-INS-3676
chr13-80645577-INS-528
chr13-80666726-INS-212
chr13-80712450-INS-401
chr13-80738894-INS-223
chr13-80772956-INS-496
chr13-80777171-INS-64
chr13-80777697-INS-2571
chr13-80784406-INS-610
chr13-80802615-INS-52
chr13-80809024-INS-560
chr13-80822887-INS-58
chr13-80831758-INS-208
chr13-80846344-INS-103
chr13-80856503-INS-129
chr13-80861938-INS-109
chr13-80866000-INS-51
chr13-80927099-INS-989
chr13-80963264-INS-363
chr13-81013291-INS-64
chr13-81050541-INS-1286
chr13-81074641-INS-292
chr13-81087934-INS-53
chr13-81096219-INS-211
chr13-81100635-INS-473
chr13-81102947-INS-54
chr13-81145041-INS-55
chr13-81156050-INS-132
chr13-81174601-INS-114
chr13-81178490-INS-343
chr13-81220490-INS-487
chr13-81267704-INS-60
chr13-81275371-INS-66
chr13-81279784-INS-628
chr13-81296012-INS-7590
chr13-81316849-INS-80
chr13-81333850-INS-208
chr13-81407613-INS-123
chr13-81426026-INS-6710
chr13-81426728-INS-6660
chr13-81449370-INS-6403
chr13-81459336-INS-52
chr13-81483678-INS-58
chr13-81498277-INS-58
chr13-81531585-INS-98
chr13-81552069-INS-176
chr13-81561971-INS-156
chr13-81575609-INS-87
chr13-81580020-INS-92
chr13-81617696-INS-744
chr13-81621777-INS-610
chr13-81664043-INS-80
chr13-81672155-INS-62
chr13-81679399-INS-73

chr13-81688627-INS-68
chr13-81696042-INS-64
chr13-81714676-INS-152
chr13-81721132-INS-94
chr13-81768175-INS-57
chr13-81792532-INS-7153
chr13-81807958-INS-219
chr13-81814429-INS-152
chr13-81832117-INS-224
chr13-81836530-INS-215
chr13-81849937-INS-385
chr13-81865243-INS-253
chr13-81874553-INS-70
chr13-81905984-INS-705
chr13-81933783-INS-445
chr13-81961880-INS-87
chr13-81974982-INS-359
chr13-81994728-INS-55
chr13-82005891-INS-694
chr13-82028679-INS-179
chr13-82052088-INS-125
chr13-82059583-INS-140
chr13-82066368-INS-163
chr13-82076349-INS-74
chr13-82079507-INS-193
chr13-82086662-INS-196
chr13-82092055-INS-390
chr13-82096434-INS-128
chr13-82113048-INS-194
chr13-82146686-INS-55
chr13-82192674-INS-119
chr13-82196610-INS-288
chr13-82208732-INS-163
chr13-82279970-INS-166
chr13-82298004-INS-648
chr13-82304211-INS-158
chr13-82305322-INS-102
chr13-82324400-INS-114
chr13-82333963-INS-130
chr13-82363041-INS-61
chr13-82364331-INS-328
chr13-82370664-INS-378
chr13-82375495-INS-382
chr13-82388776-INS-6283
chr13-82392791-INS-729
chr13-82403494-INS-6673
chr13-82444381-INS-117
chr13-82454469-INS-190
chr13-82461050-INS-1817
chr13-82471912-INS-4358
chr13-82473367-INS-6645
chr13-82511866-INS-6947
chr13-82521309-INS-7242
chr13-82527965-INS-435
chr13-82553801-INS-182
chr13-82579086-INS-1534
chr13-82593922-INS-68
chr13-82610088-INS-252
chr13-82618417-INS-5433
chr13-82618845-INS-385
chr13-82659959-INS-51
chr13-82700796-INS-54
chr13-82757492-INS-2539
chr13-82780106-INS-895
chr13-82784379-INS-455
chr13-82813618-INS-108
chr13-82814898-INS-6424
chr13-82827603-INS-61
chr13-82834448-INS-7116
chr13-82879097-INS-114
chr13-82884956-INS-84
chr13-82905740-INS-60
chr13-82912533-INS-245
chr13-82913347-INS-834
chr13-82939815-INS-444
chr13-82957885-INS-210
chr13-82960974-INS-138

chr13-82978465-INS-184
chr13-83002359-INS-58
chr13-83006904-INS-317
chr13-83025502-INS-58
chr13-83028768-INS-94
chr13-83042137-INS-151
chr13-83047156-INS-167
chr13-83069734-INS-192
chr13-83074870-INS-426
chr13-83100321-INS-819
chr13-83141234-INS-70
chr13-83205451-INS-60
chr13-83206828-INS-335
chr13-83219612-INS-207
chr13-83262513-INS-224
chr13-83317877-INS-685
chr13-83319103-INS-142
chr13-83338066-INS-84
chr13-83360530-INS-377
chr13-83368171-INS-72
chr13-83390764-INS-61
chr13-83397831-INS-64
chr13-83423716-INS-6377
chr13-83429840-INS-511
chr13-83432534-INS-51
chr13-83434808-INS-51
chr13-83460532-INS-1933
chr13-83473086-INS-55
chr13-83474183-INS-63
chr13-83487427-INS-60
chr13-83507634-INS-52
chr13-83516833-INS-6412
chr13-83541610-INS-135
chr13-83565203-INS-130
chr13-83566596-INS-202
chr13-83583777-INS-51
chr13-83608417-INS-160
chr13-83611699-INS-322
chr13-83617240-INS-260
chr13-83634284-INS-206
chr13-83635479-INS-109
chr13-83646788-INS-884
chr13-83701560-INS-927
chr13-83709288-INS-324
chr13-83739091-INS-1209
chr13-83739092-INS-579
chr13-83754158-INS-291
chr13-83833578-INS-6246
chr13-83860292-INS-78
chr13-83863001-INS-94
chr13-83869265-INS-86
chr13-83885860-INS-506
chr13-83940425-INS-71
chr13-83941719-INS-272
chr13-83975844-INS-318
chr13-83978791-INS-94
chr13-83990734-INS-8670
chr13-84009850-INS-296
chr13-84051105-INS-7669
chr13-84051789-INS-250
chr13-84060698-INS-258
chr13-84063321-INS-353
chr13-84082664-INS-59
chr13-84190222-INS-330
chr13-84195710-INS-4132
chr13-84207453-INS-60
chr13-84226221-INS-787
chr13-84250436-INS-540
chr13-84276582-INS-298
chr13-84293579-INS-62
chr13-84313086-INS-54
chr13-84405053-INS-69
chr13-84426029-INS-180
chr13-84454744-INS-189
chr13-84466348-INS-6841
chr13-84468501-INS-6159
chr13-84478131-INS-3542

chr13-84479258-INS-655
chr13-84488228-INS-180
chr13-84511823-INS-1891
chr13-84512133-INS-75
chr13-84521006-INS-92
chr13-84523549-INS-491
chr13-84542089-INS-120
chr13-84544475-INS-7308
chr13-84546028-INS-63
chr13-84558318-INS-76
chr13-84577598-INS-850
chr13-84580935-INS-7320
chr13-84581394-INS-6296
chr13-84596678-INS-54
chr13-84600671-INS-7558
chr13-84607668-INS-1303
chr13-84610750-INS-1588
chr13-84622489-INS-68
chr13-84648502-INS-370
chr13-84665065-INS-359
chr13-84775310-INS-111
chr13-84797043-INS-120
chr13-84802400-INS-97
chr13-84806545-INS-65
chr13-84815970-INS-236
chr13-84825279-INS-61
chr13-84843571-INS-152
chr13-84848898-INS-95
chr13-84868329-INS-6698
chr13-84882102-INS-6468
chr13-84882367-INS-6997
chr13-84894015-INS-128
chr13-84895612-INS-62
chr13-84992338-INS-205
chr13-84993377-INS-178
chr13-85012268-INS-167
chr13-85017921-INS-98
chr13-85021901-INS-508
chr13-85034810-INS-218
chr13-85037933-INS-80
chr13-85042762-INS-258
chr13-85067577-INS-52
chr13-85075496-INS-52
chr13-85076766-INS-608
chr13-85078480-INS-327
chr13-85093389-INS-108
chr13-85116347-INS-84
chr13-85151880-INS-287
chr13-85166940-INS-358
chr13-85169218-INS-62
chr13-85187669-INS-811
chr13-85279607-INS-524
chr13-85358336-INS-415
chr13-85396432-INS-225
chr13-85403142-INS-887
chr13-85499545-INS-140
chr13-85568753-INS-874
chr13-85664269-INS-108
chr13-85665885-INS-255
chr13-85680308-INS-51
chr13-85693683-INS-429
chr13-85702838-INS-52
chr13-85746264-INS-223
chr13-85760968-INS-6242
chr13-85770023-INS-101
chr13-85805271-INS-201
chr13-85812145-INS-403
chr13-85818707-INS-6718
chr13-85820549-INS-208
chr13-85864175-INS-54
chr13-85882137-INS-71
chr13-85900493-INS-58
chr13-85905861-INS-6880
chr13-85911795-INS-132
chr13-85913440-INS-6416
chr13-85915194-INS-166
chr13-85927592-INS-81

chr13-85957576-INS-51
chr13-85968705-INS-288
chr13-85979322-INS-99
chr13-85996287-INS-61
chr13-86032773-INS-6621
chr13-86035989-INS-57
chr13-86043011-INS-53
chr13-86045889-INS-3752
chr13-86047582-INS-1323
chr13-86085043-INS-6204
chr13-86090819-INS-58
chr13-86118351-INS-262
chr13-86139774-INS-100
chr13-86158091-INS-228
chr13-86177043-INS-104
chr13-86196109-INS-100
chr13-86200769-INS-428
chr13-86233770-INS-494
chr13-86238220-INS-93
chr13-86245883-INS-486
chr13-86247467-INS-208
chr13-86248579-INS-153
chr13-86251768-INS-55
chr13-86294604-INS-331
chr13-86294930-INS-6599
chr13-86331226-INS-128
chr13-86356169-INS-300
chr13-86361325-INS-78
chr13-86377170-INS-212
chr13-86403918-INS-194
chr13-86434841-INS-104
chr13-86477652-INS-679
chr13-86524704-INS-55
chr13-86527703-INS-530
chr13-86530127-INS-99
chr13-86564062-INS-149
chr13-86567777-INS-6439
chr13-86595659-INS-78
chr13-86611407-INS-212
chr13-86615870-INS-59
chr13-86622323-INS-115
chr13-86686396-INS-198
chr13-86760408-INS-486
chr13-86793555-INS-108
chr13-86820531-INS-529
chr13-86840669-INS-160
chr13-86896858-INS-163
chr13-86900463-INS-6237
chr13-86915673-INS-311
chr13-86920275-INS-333
chr13-86944481-INS-6386
chr13-86948758-INS-130
chr13-86952535-INS-496
chr13-86956360-INS-90
chr13-86961886-INS-163
chr13-86973986-INS-57
chr13-86989696-INS-138
chr13-87040240-INS-101
chr13-87056036-INS-1866
chr13-87065187-INS-94
chr13-87145423-INS-6383
chr13-87222003-INS-5913
chr13-87244694-INS-354
chr13-87265454-INS-118
chr13-87314985-INS-2060
chr13-87374899-INS-302
chr13-87400920-INS-131
chr13-87409773-INS-172
chr13-87421222-INS-104
chr13-87446199-INS-575
chr13-87451050-INS-199
chr13-87481294-INS-78
chr13-87489970-INS-54
chr13-87503873-INS-57
chr13-87516643-INS-6715
chr13-87533902-INS-150
chr13-87554607-INS-365

chr13-87563420-INS-69
chr13-87572690-INS-64
chr13-87599059-INS-64
chr13-87602254-INS-6157
chr13-87663654-INS-219
chr13-87677926-INS-207
chr13-87693393-INS-240
chr13-87739112-INS-529
chr13-87751203-INS-504
chr13-87763126-INS-649
chr13-87783241-INS-363
chr13-87787937-INS-5316
chr13-87792838-INS-184
chr13-87793388-INS-5369
chr13-87793909-INS-80
chr13-87804467-INS-247
chr13-87813090-INS-64
chr13-87822705-INS-118
chr13-87862569-INS-3891
chr13-87864736-INS-3205
chr13-87946945-INS-1192
chr13-87975960-INS-366
chr13-88041608-INS-6984
chr13-88055049-INS-76
chr13-88066460-INS-884
chr13-88087891-INS-214
chr13-88150449-INS-56
chr13-88152274-INS-5401
chr13-88179868-INS-476
chr13-88247192-INS-487
chr13-88256356-INS-189
chr13-88297507-INS-627
chr13-88301594-INS-972
chr13-88302585-INS-2417
chr13-88413937-INS-66
chr13-88428604-INS-432
chr13-88440962-INS-73
chr13-88531277-INS-330
chr13-88572564-INS-67
chr13-88584899-INS-454
chr13-88616761-INS-52
chr13-88637831-INS-75
chr13-88657130-INS-92
chr13-88793987-INS-91
chr13-88796745-INS-11501
chr13-88797562-INS-5383
chr13-88820838-INS-69
chr13-88826298-INS-393
chr13-88844638-INS-149
chr13-88931523-INS-138
chr13-88932196-INS-6480
chr13-88933465-INS-416
chr13-88960035-INS-52
chr13-88982285-INS-162
chr13-88988145-INS-489
chr13-89034357-INS-447
chr13-89045133-INS-220
chr13-89052338-INS-100
chr13-89057513-INS-6506
chr13-89102452-INS-391
chr13-89158503-INS-157
chr13-89160437-INS-85
chr13-89180469-INS-80
chr13-89227467-INS-90
chr13-89247468-INS-62
chr13-89248393-INS-62
chr13-89277896-INS-54
chr13-89299439-INS-275
chr13-89312919-INS-703
chr13-89326512-INS-162
chr13-89351371-INS-54
chr13-89373196-INS-165
chr13-89387066-INS-178
chr13-89392027-INS-90
chr13-89421929-INS-110
chr13-89454900-INS-476
chr13-89461462-INS-78

chr13-89462564-INS-109
chr13-89479974-INS-1622
chr13-89480559-INS-638
chr13-89489819-INS-629
chr13-89500706-INS-504
chr13-89505384-INS-213
chr13-89547333-INS-308
chr13-89559449-INS-6452
chr13-89566936-INS-333
chr13-89595729-INS-106
chr13-89601005-INS-2980
chr13-89625025-INS-380
chr13-89628877-INS-94
chr13-89629348-INS-219
chr13-89645861-INS-52
chr13-89653619-INS-244
chr13-89687164-INS-84
chr13-89699074-INS-5160
chr13-89699090-INS-429
chr13-89709070-INS-80
chr13-89710427-INS-328
chr13-89721497-INS-60
chr13-89729728-INS-537
chr13-89745044-INS-68
chr13-89769939-INS-75
chr13-89772138-INS-320
chr13-89781942-INS-166
chr13-89790123-INS-5549
chr13-89790555-INS-6226
chr13-89796210-INS-107
chr13-89815402-INS-452
chr13-89885071-INS-163
chr13-89895738-INS-487
chr13-89897987-INS-62
chr13-89898784-INS-261
chr13-89934929-INS-6683
chr13-89939304-INS-494
chr13-89952621-INS-633
chr13-89973021-INS-134
chr13-89974580-INS-224
chr13-89985180-INS-75
chr13-90005735-INS-6435
chr13-90023416-INS-82
chr13-90055546-INS-488
chr13-90070574-INS-321
chr13-90092884-INS-120
chr13-90098907-INS-84
chr13-90101494-INS-365
chr13-90103121-INS-7389
chr13-90109269-INS-89
chr13-90150384-INS-504
chr13-90162353-INS-57
chr13-90202224-INS-108
chr13-90221311-INS-51
chr13-90224050-INS-275
chr13-90244545-INS-90
chr13-90256752-INS-173
chr13-90260294-INS-1865
chr13-90266978-INS-393
chr13-90267627-INS-5289
chr13-90277756-INS-295
chr13-90288814-INS-7249
chr13-90305481-INS-197
chr13-90311158-INS-70
chr13-90339607-INS-56
chr13-90343269-INS-554
chr13-90348878-INS-765
chr13-90355716-INS-150
chr13-90377212-INS-327
chr13-90389184-INS-297
chr13-90451202-INS-52
chr13-90489504-INS-323
chr13-90533085-INS-72
chr13-90567078-INS-72
chr13-90578130-INS-94
chr13-90581910-INS-457
chr13-90625269-INS-7234

chr13-90696756-INS-53
chr13-90709341-INS-482
chr13-90714518-INS-214
chr13-90735628-INS-70
chr13-90735977-INS-7094
chr13-90760892-INS-62
chr13-90762954-INS-2703
chr13-90765535-INS-8256
chr13-90767806-INS-60
chr13-90783061-INS-56
chr13-90790806-INS-52
chr13-90807764-INS-72
chr13-90810510-INS-83
chr13-90838254-INS-367
chr13-90840979-INS-11790
chr13-90842778-INS-729
chr13-90844851-INS-517
chr13-90847204-INS-1093
chr13-90939445-INS-376
chr13-90956401-INS-60
chr13-90957479-INS-75
chr13-90968500-INS-116
chr13-90975291-INS-483
chr13-91017920-INS-7073
chr13-91018925-INS-598
chr13-91041217-INS-243
chr13-91048772-INS-138
chr13-91066203-INS-399
chr13-91072951-INS-207
chr13-91112323-INS-73
chr13-91118237-INS-704
chr13-91127745-INS-587
chr13-91132733-INS-502
chr13-91143575-INS-85
chr13-91144739-INS-7640
chr13-91152898-INS-260
chr13-91165583-INS-92
chr13-91172380-INS-126
chr13-91214004-INS-58
chr13-91219426-INS-824
chr13-91227378-INS-297
chr13-91229687-INS-61
chr13-91233601-INS-412
chr13-91316850-INS-300
chr13-91321953-INS-208
chr13-91322475-INS-453
chr13-91325685-INS-172
chr13-91344035-INS-178
chr13-91364746-INS-72
chr13-91391371-INS-241
chr13-91440429-INS-192
chr13-91475394-INS-58
chr13-91488362-INS-774
chr13-91497807-INS-557
chr13-91551261-INS-458
chr13-91582736-INS-496
chr13-91590111-INS-59
chr13-91615735-INS-80
chr13-91623439-INS-1889
chr13-91633079-INS-65
chr13-91644797-INS-1063
chr13-91654453-INS-1946
chr13-91671076-INS-388
chr13-91678616-INS-97
chr13-91691768-INS-155
chr13-91692904-INS-373
chr13-91714126-INS-68
chr13-91742836-INS-165
chr13-91783162-INS-199
chr13-91787051-INS-306
chr13-91788867-INS-61
chr13-91896372-INS-419
chr13-91908510-INS-95
chr13-91935826-INS-56
chr13-91943630-INS-1041
chr13-91947858-INS-59
chr13-91956073-INS-66

chr13-91981669-INS-54
chr13-92002997-INS-244
chr13-92018636-INS-82
chr13-92040494-INS-96
chr13-92047348-INS-203
chr13-92055754-INS-91
chr13-92059903-INS-76
chr13-92068523-INS-760
chr13-92102545-INS-63
chr13-92103066-INS-200
chr13-92108211-INS-60
chr13-92124848-INS-962
chr13-92132158-INS-55
chr13-92143029-INS-412
chr13-92146374-INS-258
chr13-92147043-INS-144
chr13-92153449-INS-101
chr13-92166538-INS-384
chr13-92176184-INS-400
chr13-92195734-INS-76
chr13-92203076-INS-632
chr13-92231925-INS-1444
chr13-92278149-INS-6484
chr13-92324757-INS-290
chr13-92329796-INS-61
chr13-92331406-INS-70
chr13-92338234-INS-110
chr13-92343196-INS-6773
chr13-92352910-INS-267
chr13-92366425-INS-185
chr13-92385322-INS-220
chr13-92396640-INS-207
chr13-92435820-INS-62
chr13-92448558-INS-134
chr13-92462054-INS-72
chr13-92469330-INS-989
chr13-92478267-INS-169
chr13-92493578-INS-363
chr13-92514426-INS-163
chr13-92549543-INS-354
chr13-92558852-INS-194
chr13-92566865-INS-148
chr13-92601310-INS-187
chr13-92625999-INS-606
chr13-92638296-INS-51
chr13-92661577-INS-1889
chr13-92662009-INS-462
chr13-92665253-INS-706
chr13-92680902-INS-227
chr13-92701277-INS-153
chr13-92720495-INS-68
chr13-92743188-INS-350
chr13-92750374-INS-7218
chr13-92763513-INS-171
chr13-92768488-INS-156
chr13-92774149-INS-90
chr13-92778393-INS-72
chr13-92779496-INS-155
chr13-92802100-INS-500
chr13-92803994-INS-196
chr13-92847135-INS-52
chr13-92851837-INS-167
chr13-92872578-INS-64
chr13-92878204-INS-285
chr13-92906946-INS-84
chr13-92910303-INS-172
chr13-92927794-INS-255
chr13-92935771-INS-5411
chr13-92996681-INS-138
chr13-93032811-INS-197
chr13-93073478-INS-330
chr13-93078111-INS-818
chr13-93122609-INS-411
chr13-93148690-INS-224
chr13-93159538-INS-84
chr13-93175921-INS-191
chr13-93184906-INS-2766

chr13-93192039-INS-96
chr13-93217769-INS-80
chr13-93219107-INS-100
chr13-93233253-INS-145
chr13-93298321-INS-210
chr13-93302586-INS-221
chr13-93335028-INS-234
chr13-93341649-INS-52
chr13-93348122-INS-207
chr13-93365269-INS-249
chr13-93374936-INS-55
chr13-93376439-INS-544
chr13-93395453-INS-329
chr13-93426423-INS-162
chr13-93433967-INS-139
chr13-93466993-INS-54
chr13-93476964-INS-74
chr13-93486694-INS-112
chr13-93527941-INS-324
chr13-93556789-INS-173
chr13-93563635-INS-150
chr13-93564302-INS-194
chr13-93576858-INS-519
chr13-93576888-INS-1337
chr13-93578894-INS-212
chr13-93600061-INS-168
chr13-93619601-INS-107
chr13-93626036-INS-56
chr13-93637880-INS-84
chr13-93649737-INS-106
chr13-93686385-INS-375
chr13-93699699-INS-197
chr13-93705307-INS-170
chr13-93717846-INS-119
chr13-93735138-INS-69
chr13-93736152-INS-171
chr13-93764009-INS-246
chr13-93766754-INS-113
chr13-93772521-INS-84
chr13-93797944-INS-193
chr13-93813047-INS-5548
chr13-93841674-INS-405
chr13-93850989-INS-259
chr13-93864630-INS-603
chr13-93865414-INS-240
chr13-93885150-INS-53
chr13-93894084-INS-503
chr13-93900156-INS-204
chr13-93902579-INS-198
chr13-93905594-INS-172
chr13-93909769-INS-52
chr13-93920264-INS-247
chr13-93920779-INS-226
chr13-93923113-INS-60
chr13-93926328-INS-528
chr13-93935874-INS-111
chr13-93945222-INS-77
chr13-93953971-INS-304
chr13-93963208-INS-328
chr13-93969058-INS-192
chr13-94000315-INS-54
chr13-94022901-INS-60
chr13-94046123-INS-84
chr13-94052980-INS-107
chr13-94054061-INS-154
chr13-94062745-INS-555
chr13-94064598-INS-167
chr13-94085369-INS-98
chr13-94120832-INS-75
chr13-94144223-INS-6296
chr13-94144392-INS-106
chr13-94150376-INS-88
chr13-94174776-INS-185
chr13-94187404-INS-911
chr13-94205119-INS-490
chr13-94205734-INS-206
chr13-94210638-INS-223

chr13-94241863-INS-54
chr13-94254204-INS-519
chr13-94268194-INS-197
chr13-94287907-INS-52
chr13-94297228-INS-196
chr13-94312314-INS-63
chr13-94322088-INS-51
chr13-94327122-INS-210
chr13-94341756-INS-177
chr13-94347283-INS-114
chr13-94359312-INS-117
chr13-94365341-INS-440
chr13-94377282-INS-387
chr13-94377736-INS-502
chr13-94415094-INS-60
chr13-94425172-INS-204
chr13-94426323-INS-80
chr13-94427280-INS-241
chr13-94434265-INS-143
chr13-94437240-INS-84
chr13-94447331-INS-331
chr13-94448093-INS-507
chr13-94449263-INS-184
chr13-94462453-INS-439
chr13-94470610-INS-328
chr13-94473158-INS-3540
chr13-94477583-INS-116
chr13-94483268-INS-7738
chr13-94488850-INS-354
chr13-94500118-INS-154
chr13-94506019-INS-60
chr13-94522772-INS-102
chr13-94525575-INS-69
chr13-94528036-INS-622
chr13-94561204-INS-185
chr13-94568619-INS-156
chr13-94572786-INS-65
chr13-94573996-INS-197
chr13-94574011-INS-1794
chr13-94597812-INS-222
chr13-94668806-INS-385
chr13-94706360-INS-210
chr13-94739747-INS-317
chr13-94750114-INS-80
chr13-94752169-INS-71
chr13-94752611-INS-171
chr13-94754184-INS-78
chr13-94787102-INS-160
chr13-94793060-INS-119
chr13-94799134-INS-53
chr13-94807355-INS-135
chr13-94810336-INS-180
chr13-94825782-INS-135
chr13-94829280-INS-90
chr13-94834612-INS-388
chr13-94852171-INS-70
chr13-94855740-INS-80
chr13-94867646-INS-457
chr13-94869027-INS-88
chr13-94895798-INS-254
chr13-94924326-INS-118
chr13-94956386-INS-214
chr13-94971972-INS-64
chr13-94979804-INS-502
chr13-94988532-INS-96
chr13-94989494-INS-204
chr13-95000245-INS-101
chr13-95001639-INS-76
chr13-95004411-INS-476
chr13-95037504-INS-121
chr13-95053519-INS-163
chr13-95093354-INS-163
chr13-95096391-INS-662
chr13-95103521-INS-355
chr13-95114839-INS-879
chr13-95134880-INS-100
chr13-95138590-INS-84

chr13-95146260-INS-5547
chr13-95148989-INS-180
chr13-95174306-INS-72
chr13-95228708-INS-205
chr13-95246826-INS-428
chr13-95252425-INS-186
chr13-95265440-INS-235
chr13-95286400-INS-147
chr13-95287345-INS-58
chr13-95299992-INS-135
chr13-95306421-INS-51
chr13-95322524-INS-64
chr13-95325457-INS-87
chr13-95337348-INS-88
chr13-95372479-INS-54
chr13-95378580-INS-1424
chr13-95391043-INS-86
chr13-95401800-INS-54
chr13-95415506-INS-122
chr13-95437326-INS-59
chr13-95437887-INS-60
chr13-95450804-INS-86
chr13-95452975-INS-51
chr13-95465327-INS-58
chr13-95488289-INS-61
chr13-95490130-INS-89
chr13-95565342-INS-85
chr13-95588644-INS-101
chr13-95613177-INS-102
chr13-95625801-INS-64
chr13-95632846-INS-237
chr13-95664033-INS-64
chr13-95682044-INS-157
chr13-95685807-INS-68
chr13-95697006-INS-175
chr13-95701868-INS-58
chr13-95721038-INS-56
chr13-95732398-INS-687
chr13-95734196-INS-53
chr13-95756223-INS-160
chr13-95757711-INS-426
chr13-95801596-INS-136
chr13-95824974-INS-106
chr13-95830549-INS-230
chr13-95931397-INS-221
chr13-95957867-INS-136
chr13-95965654-INS-108
chr13-95967475-INS-417
chr13-95974839-INS-60
chr13-95991448-INS-60
chr13-96005051-INS-378
chr13-96016048-INS-216
chr13-96019418-INS-53
chr13-96058653-INS-193
chr13-96073179-INS-228
chr13-96126079-INS-118
chr13-96127188-INS-642
chr13-96146711-INS-400
chr13-96182532-INS-172
chr13-96186072-INS-144
chr13-96193487-INS-273
chr13-96201554-INS-1535
chr13-96205705-INS-58
chr13-96259156-INS-99
chr13-96261091-INS-135
chr13-96277164-INS-399
chr13-96282281-INS-185
chr13-96285691-INS-111
chr13-96305871-INS-293
chr13-96319038-INS-51
chr13-96344734-INS-545
chr13-96348733-INS-60
chr13-96351925-INS-54
chr13-96354923-INS-92
chr13-96362628-INS-313
chr13-96392887-INS-54
chr13-96442004-INS-7086

chr13-96463359-INS-573
chr13-96473793-INS-62
chr13-96485352-INS-78
chr13-96500467-INS-208
chr13-96506486-INS-99
chr13-96513295-INS-86
chr13-96518585-INS-50
chr13-96528405-INS-203
chr13-96529443-INS-732
chr13-96548358-INS-76
chr13-96559414-INS-297
chr13-96585578-INS-575
chr13-96597479-INS-73
chr13-96602538-INS-316
chr13-96640835-INS-569
chr13-96642943-INS-138
chr13-96661811-INS-400
chr13-96707837-INS-161
chr13-96708821-INS-196
chr13-96723383-INS-100
chr13-96758412-INS-373
chr13-96795068-INS-203
chr13-96802978-INS-292
chr13-96810945-INS-63
chr13-96833191-INS-195
chr13-96835009-INS-519
chr13-96855853-INS-530
chr13-96881725-INS-142
chr13-96890298-INS-121
chr13-96898293-INS-119
chr13-96908183-INS-151
chr13-96935670-INS-66
chr13-96971481-INS-69
chr13-96987580-INS-579
chr13-96994370-INS-55
chr13-97050734-INS-50
chr13-97052229-INS-53
chr13-97053282-INS-617
chr13-97089223-INS-1765
chr13-97093732-INS-5446
chr13-97113353-INS-124
chr13-97117265-INS-63
chr13-97145577-INS-526
chr13-97152370-INS-69
chr13-97153578-INS-204
chr13-97157165-INS-201
chr13-97170738-INS-84
chr13-97192311-INS-188
chr13-97194868-INS-196
chr13-97195743-INS-209
chr13-97198989-INS-93
chr13-97201103-INS-53
chr13-97214679-INS-83
chr13-97223005-INS-75
chr13-97228766-INS-264
chr13-97230867-INS-127
chr13-97239283-INS-407
chr13-97268578-INS-204
chr13-97271109-INS-206
chr13-97303673-INS-86
chr13-97308561-INS-982
chr13-97331622-INS-55
chr13-97336219-INS-5542
chr13-97414616-INS-564
chr13-97453493-INS-461
chr13-97457468-INS-555
chr13-97496381-INS-401
chr13-97505604-INS-96
chr13-97506196-INS-51
chr13-97508852-INS-117
chr13-97527511-INS-64
chr13-97529350-INS-72
chr13-97536976-INS-137
chr13-97543257-INS-56
chr13-97590092-INS-177
chr13-97613448-INS-442
chr13-97661879-INS-327

chr13-97663889-INS-107
chr13-97701310-INS-718
chr13-97712740-INS-201
chr13-97715517-INS-66
chr13-97719435-INS-60
chr13-97721850-INS-339
chr13-97753398-INS-64
chr13-97776766-INS-99
chr13-97783505-INS-56
chr13-97810193-INS-52
chr13-97821191-INS-87
chr13-97899178-INS-56
chr13-97918128-INS-132
chr13-97943177-INS-53
chr13-97947693-INS-57
chr13-97995260-INS-116
chr13-98013767-INS-86
chr13-98022360-INS-217
chr13-98081875-INS-80
chr13-98111002-INS-55
chr13-98116861-INS-156
chr13-98131581-INS-109
chr13-98136471-INS-88
chr13-98148010-INS-285
chr13-98186664-INS-66
chr13-98192700-INS-200
chr13-98200780-INS-651
chr13-98208174-INS-127
chr13-98240624-INS-238
chr13-98280075-INS-53
chr13-98302589-INS-93
chr13-98313445-INS-56
chr13-98335411-INS-165
chr13-98351893-INS-99
chr13-98358184-INS-169
chr13-98360111-INS-57
chr13-98366249-INS-75
chr13-98374830-INS-248
chr13-98376455-INS-52
chr13-98382224-INS-321
chr13-98394902-INS-120
chr13-98416749-INS-60
chr13-98419647-INS-514
chr13-98426754-INS-80
chr13-98429219-INS-76
chr13-98433548-INS-51
chr13-98435923-INS-157
chr13-98441752-INS-76
chr13-98468362-INS-65
chr13-98469648-INS-96
chr13-98470990-INS-68
chr13-98483468-INS-113
chr13-98498346-INS-54
chr13-98517770-INS-281
chr13-98524734-INS-68
chr13-98534526-INS-344
chr13-98547472-INS-138
chr13-98561269-INS-83
chr13-98562716-INS-88
chr13-98570310-INS-204
chr13-98594859-INS-146
chr13-98611927-INS-155
chr13-98631874-INS-96
chr13-98632277-INS-335
chr13-98647024-INS-93
chr13-98667959-INS-58
chr13-98673777-INS-179
chr13-98689326-INS-100
chr13-98689669-INS-775
chr13-98702012-INS-5547
chr13-98711330-INS-74
chr13-98743028-INS-254
chr13-98762293-INS-52
chr13-98796168-INS-179
chr13-98813144-INS-527
chr13-98850293-INS-100
chr13-98853762-INS-194

chr13-98857765-INS-92
chr13-98871973-INS-84
chr13-98882183-INS-204
chr13-98890728-INS-135
chr13-98922852-INS-67
chr13-98952811-INS-198
chr13-98959810-INS-184
chr13-98969607-INS-610
chr13-98994042-INS-208
chr13-98995678-INS-206
chr13-99019041-INS-53
chr13-99030245-INS-89
chr13-99067612-INS-480
chr13-99070441-INS-1905
chr13-99087520-INS-172
chr13-99112757-INS-249
chr13-99117527-INS-193
chr13-99119114-INS-158
chr13-99132706-INS-4104
chr13-99147640-INS-190
chr13-99186595-INS-53
chr13-99189689-INS-96
chr13-99192825-INS-58
chr13-99209066-INS-90
chr13-99221471-INS-132
chr13-99222236-INS-62
chr13-99222236-INS-99
chr13-99227586-INS-443
chr13-99256133-INS-257
chr13-99281760-INS-52
chr13-99300406-INS-53
chr13-99307935-INS-93
chr13-99331491-INS-87
chr13-99351383-INS-94
chr13-99353450-INS-57
chr13-99354813-INS-162
chr13-99363960-INS-85
chr13-99401505-INS-468
chr13-99404925-INS-449
chr13-99424834-INS-60
chr13-99428481-INS-399
chr13-99429978-INS-320
chr13-99438485-INS-164
chr13-99485775-INS-220
chr13-99492143-INS-255
chr13-99507590-INS-56
chr13-99508942-INS-103
chr13-99524734-INS-231
chr13-99660399-INS-104
chr13-99701037-INS-440
chr13-99707341-INS-129
chr13-99787622-INS-688
chr13-99808185-INS-66
chr13-99818594-INS-51
chr13-99822445-INS-116
chr13-99844988-INS-156
chr13-99864058-INS-161
chr13-99902935-INS-76
chr13-99912458-INS-477
chr13-99912568-INS-1534
chr13-99931406-INS-5543
chr13-99964740-INS-506
chr13-99968171-INS-342
chr13-99982268-INS-51
chr13-99984872-INS-850
chr13-99986871-INS-56
chr13-100017727-INS-623
chr13-100018326-INS-124
chr13-100033136-INS-96
chr13-100045898-INS-155
chr13-100049539-INS-166
chr13-100069707-INS-459
chr13-100075195-INS-532
chr13-100084777-INS-78
chr13-100098145-INS-146
chr13-100130648-INS-188
chr13-100157698-INS-168

chr13-100181714-INS-129
chr13-100183999-INS-65
chr13-100194525-INS-800
chr13-100199028-INS-243
chr13-100201450-INS-201
chr13-100213002-INS-65
chr13-100219472-INS-401
chr13-100230321-INS-163
chr13-100247419-INS-530
chr13-100248718-INS-449
chr13-100261091-INS-231
chr13-100263934-INS-70
chr13-100269907-INS-78
chr13-100280273-INS-566
chr13-100293968-INS-375
chr13-100306282-INS-157
chr13-100331638-INS-439
chr13-100339690-INS-76
chr13-100357058-INS-64
chr13-100370570-INS-416
chr13-100401423-INS-2620
chr13-100401945-INS-203
chr13-100421348-INS-344
chr13-100468226-INS-226
chr13-100528463-INS-616
chr13-100546064-INS-405
chr13-100572897-INS-78
chr13-100582627-INS-178
chr13-100598467-INS-8457
chr13-100617867-INS-170
chr13-100619005-INS-521
chr13-100636347-INS-192
chr13-100647189-INS-128
chr13-100661292-INS-125
chr13-100662459-INS-167
chr13-100674258-INS-173
chr13-100678658-INS-207
chr13-100681742-INS-154
chr13-100695512-INS-226
chr13-100700499-INS-197
chr13-100705547-INS-614
chr13-100706469-INS-198
chr13-100761366-INS-213
chr13-100762708-INS-109
chr13-100773949-INS-94
chr13-100780157-INS-208
chr13-100784643-INS-166
chr13-100789434-INS-55
chr13-100793714-INS-173
chr13-100799152-INS-225
chr13-100820395-INS-178
chr13-100822954-INS-288
chr13-100829027-INS-928
chr13-100843211-INS-202
chr13-100857944-INS-138
chr13-100860198-INS-211
chr13-100865022-INS-185
chr13-100873955-INS-230
chr13-100879712-INS-316
chr13-100910524-INS-90
chr13-100926327-INS-54
chr13-100927886-INS-193
chr13-100932867-INS-304
chr13-100943600-INS-302
chr13-100949345-INS-65
chr13-100959059-INS-94
chr13-100962916-INS-122
chr13-100974838-INS-84
chr13-100989015-INS-507
chr13-101002196-INS-346
chr13-101007205-INS-206
chr13-101010620-INS-457
chr13-101014979-INS-54
chr13-101025418-INS-63
chr13-101032020-INS-530
chr13-101032957-INS-124
chr13-101046627-INS-52

chr13-101056180-INS-52
chr13-101075438-INS-6667
chr13-101090323-INS-180
chr13-101099252-INS-77
chr13-101116082-INS-579
chr13-101127390-INS-351
chr13-101131572-INS-75
chr13-101141339-INS-144
chr13-101171225-INS-75
chr13-101235498-INS-824
chr13-101246629-INS-1241
chr13-101262887-INS-6257
chr13-101306683-INS-444
chr13-101314609-INS-212
chr13-101399942-INS-139
chr13-101403101-INS-232
chr13-101514616-INS-396
chr13-101531628-INS-326
chr13-101623175-INS-63
chr13-101676863-INS-59
chr13-101692354-INS-918
chr13-101695761-INS-211
chr13-101698682-INS-54
chr13-101711492-INS-160
chr13-101721948-INS-52
chr13-101747200-INS-3043
chr13-101795181-INS-59
chr13-101804565-INS-108
chr13-101813789-INS-52
chr13-101821104-INS-61
chr13-101847963-INS-193
chr13-101868881-INS-52
chr13-101881776-INS-70
chr13-101912937-INS-71
chr13-101912975-INS-545
chr13-101942806-INS-422
chr13-101945024-INS-537
chr13-101998223-INS-114
chr13-102019714-INS-82
chr13-102061823-INS-132
chr13-102081624-INS-52
chr13-102111422-INS-51
chr13-102120371-INS-61
chr13-102137050-INS-199
chr13-102140687-INS-3222
chr13-102143855-INS-102
chr13-102161218-INS-67
chr13-102172162-INS-72
chr13-102182855-INS-150
chr13-102189994-INS-338
chr13-102202898-INS-58
chr13-102209518-INS-92
chr13-102211894-INS-146
chr13-102215160-INS-58
chr13-102220996-INS-341
chr13-102223332-INS-55
chr13-102233009-INS-98
chr13-102242027-INS-97
chr13-102248869-INS-988
chr13-102249008-INS-84
chr13-102282662-INS-316
chr13-102289992-INS-124
chr13-102299902-INS-63
chr13-102312734-INS-76
chr13-102347386-INS-128
chr13-102351836-INS-71
chr13-102372791-INS-88
chr13-102384926-INS-733
chr13-102398684-INS-59
chr13-102420428-INS-59
chr13-102440343-INS-167
chr13-102450776-INS-309
chr13-102471307-INS-66
chr13-102472327-INS-52
chr13-102487454-INS-496
chr13-102493700-INS-61
chr13-102506025-INS-409

chr13-102526564-INS-84
chr13-102549211-INS-60
chr13-102550844-INS-151
chr13-102555480-INS-6401
chr13-102555831-INS-5610
chr13-102591275-INS-84
chr13-102602161-INS-6433
chr13-102605664-INS-135
chr13-102659448-INS-88
chr13-102672120-INS-961
chr13-102676012-INS-68
chr13-102678414-INS-57
chr13-102696618-INS-920
chr13-102707856-INS-147
chr13-102715920-INS-301
chr13-102717849-INS-55
chr13-102736022-INS-60
chr13-102750166-INS-69
chr13-102768046-INS-140
chr13-102776722-INS-65
chr13-102787533-INS-56
chr13-102799226-INS-198
chr13-102816699-INS-59
chr13-102831967-INS-124
chr13-102835740-INS-61
chr13-102851006-INS-272
chr13-102852865-INS-645
chr13-102886718-INS-88
chr13-102922038-INS-401
chr13-102987203-INS-121
chr13-103008821-INS-77
chr13-103031715-INS-58
chr13-103035862-INS-65
chr13-103057610-INS-379
chr13-103061295-INS-110
chr13-103074545-INS-90
chr13-103132834-INS-995
chr13-103178790-INS-54
chr13-103219381-INS-66
chr13-103220944-INS-201
chr13-103252919-INS-347
chr13-103253956-INS-364
chr13-103255769-INS-98
chr13-103293129-INS-108
chr13-103312968-INS-211
chr13-103319145-INS-212
chr13-103321177-INS-141
chr13-103337394-INS-52
chr13-103358506-INS-72
chr13-103364643-INS-941
chr13-103373104-INS-85
chr13-103416844-INS-211
chr13-103423244-INS-70
chr13-103435245-INS-72
chr13-103475032-INS-183
chr13-103508427-INS-218
chr13-103528527-INS-2768
chr13-103532283-INS-199
chr13-103548778-INS-1455
chr13-103573161-INS-60
chr13-103592640-INS-176
chr13-103622920-INS-181
chr13-103649751-INS-314
chr13-103656818-INS-67
chr13-103656824-INS-1416
chr13-103656854-INS-732
chr13-103656883-INS-1129
chr13-103658690-INS-4531
chr13-103681876-INS-198
chr13-103697785-INS-131
chr13-103711791-INS-202
chr13-103714251-INS-329
chr13-103729926-INS-80
chr13-103776074-INS-54
chr13-103778848-INS-60
chr13-103798222-INS-64
chr13-103802270-INS-89

chr13-103861026-INS-7284
chr13-103871763-INS-172
chr13-103884403-INS-212
chr13-103892857-INS-118
chr13-103951085-INS-148
chr13-103969163-INS-54
chr13-103972062-INS-205
chr13-104005097-INS-1125
chr13-104006750-INS-1349
chr13-104012297-INS-6383
chr13-104031226-INS-6485
chr13-104078356-INS-65
chr13-104081563-INS-216
chr13-104107305-INS-52
chr13-104109206-INS-160
chr13-104120597-INS-135
chr13-104127753-INS-222
chr13-104148797-INS-88
chr13-104151086-INS-277
chr13-104166443-INS-58
chr13-104180115-INS-418
chr13-104193576-INS-210
chr13-104220835-INS-242
chr13-104225333-INS-57
chr13-104235309-INS-618
chr13-104236924-INS-105
chr13-104244363-INS-6319
chr13-104274117-INS-136
chr13-104282863-INS-106
chr13-104295583-INS-194
chr13-104312679-INS-204
chr13-104345301-INS-598
chr13-104441173-INS-213
chr13-104456194-INS-1201
chr13-104471342-INS-4670
chr13-104473992-INS-162
chr13-104485902-INS-403
chr13-104514538-INS-105
chr13-104542383-INS-86
chr13-104559227-INS-61
chr13-104597922-INS-354
chr13-104647193-INS-57
chr13-104680111-INS-56
chr13-104690603-INS-408
chr13-104692288-INS-88
chr13-104712631-INS-98
chr13-104714158-INS-82
chr13-104745043-INS-119
chr13-104827511-INS-87
chr13-104879522-INS-1092
chr13-104879548-INS-1638
chr13-104879821-INS-222
chr13-104885069-INS-3248
chr13-104889941-INS-1006
chr13-104890146-INS-69
chr13-104919172-INS-1311
chr13-104939276-INS-60
chr13-104968272-INS-182
chr13-104992163-INS-262
chr13-105027881-INS-231
chr13-105030863-INS-56
chr13-105050966-INS-87
chr13-105087715-INS-64
chr13-105091898-INS-150
chr13-105111558-INS-116
chr13-105115698-INS-96
chr13-105124135-INS-392
chr13-105142497-INS-139
chr13-105169956-INS-296
chr13-105203913-INS-6438
chr13-105273408-INS-73
chr13-105287001-INS-400
chr13-105301410-INS-76
chr13-105306379-INS-428
chr13-105308059-INS-51
chr13-105317525-INS-481
chr13-105323401-INS-79

chr13-105389916-INS-6269
chr13-105397477-INS-804
chr13-105418520-INS-111
chr13-105436878-INS-52
chr13-105469724-INS-53
chr13-105477815-INS-108
chr13-105486124-INS-4462
chr13-105561813-INS-63
chr13-105567824-INS-677
chr13-105576810-INS-188
chr13-105585125-INS-160
chr13-105603425-INS-388
chr13-105613093-INS-595
chr13-105617796-INS-447
chr13-105657304-INS-114
chr13-105665249-INS-221
chr13-105691740-INS-211
chr13-105708572-INS-66
chr13-105734745-INS-440
chr13-105774598-INS-76
chr13-105780045-INS-7023
chr13-105784719-INS-7124
chr13-105793849-INS-309
chr13-105807895-INS-6695
chr13-105832830-INS-391
chr13-105839740-INS-839
chr13-105868416-INS-265
chr13-105906373-INS-64
chr13-105929071-INS-213
chr13-105975007-INS-299
chr13-105980510-INS-1888
chr13-105988350-INS-52
chr13-106017710-INS-772
chr13-106020815-INS-235
chr13-106065779-INS-175
chr13-106096476-INS-540
chr13-106177012-INS-687
chr13-106197578-INS-7922
chr13-106251262-INS-231
chr13-106262380-INS-96
chr13-106288747-INS-64
chr13-106297659-INS-131
chr13-106324087-INS-126
chr13-106343410-INS-247
chr13-106353451-INS-117
chr13-106355580-INS-96
chr13-106381318-INS-201
chr13-106391505-INS-172
chr13-106397736-INS-415
chr13-106416432-INS-775
chr13-106438375-INS-174
chr13-106456960-INS-145
chr13-106465562-INS-239
chr13-106469756-INS-592
chr13-106471460-INS-221
chr13-106477592-INS-474
chr13-106499246-INS-70
chr13-106500634-INS-7150
chr13-106526355-INS-6226
chr13-106533335-INS-63
chr13-106537769-INS-7247
chr13-106539705-INS-113
chr13-106541302-INS-64
chr13-106549205-INS-298
chr13-106561390-INS-60
chr13-106573814-INS-385
chr13-106601725-INS-144
chr13-106603196-INS-54
chr13-106623871-INS-196
chr13-106637089-INS-161
chr13-106649471-INS-176
chr13-106658035-INS-234
chr13-106671604-INS-326
chr13-106685877-INS-3350
chr13-106703826-INS-178
chr13-106707930-INS-104
chr13-106739150-INS-7088

chr13-106805696-INS-6305
chr13-106902396-INS-112
chr13-106911768-INS-1416
chr13-106912132-INS-85
chr13-106912704-INS-575
chr13-106918069-INS-60
chr13-106919814-INS-6921
chr13-106932689-INS-213
chr13-106935576-INS-58
chr13-106967049-INS-55
chr13-106967822-INS-867
chr13-106973861-INS-123
chr13-106977647-INS-292
chr13-106983797-INS-74
chr13-106986666-INS-81
chr13-106989694-INS-98
chr13-107003208-INS-70
chr13-107010037-INS-64
chr13-107014057-INS-197
chr13-107019307-INS-66
chr13-107031899-INS-51
chr13-107039324-INS-400
chr13-107044814-INS-146
chr13-107047883-INS-83
chr13-107061310-INS-251
chr13-107064427-INS-90
chr13-107070152-INS-617
chr13-107078956-INS-579
chr13-107090267-INS-368
chr13-107113906-INS-160
chr13-107114965-INS-5375
chr13-107135511-INS-257
chr13-107141678-INS-108
chr13-107197478-INS-184
chr13-107206658-INS-233
chr13-107212980-INS-102
chr13-107215311-INS-58
chr13-107220593-INS-179
chr13-107238568-INS-230
chr13-107254486-INS-363
chr13-107258949-INS-585
chr13-107265783-INS-53
chr13-107290582-INS-191
chr13-107312973-INS-76
chr13-107321402-INS-357
chr13-107346545-INS-100
chr13-107378037-INS-72
chr13-107424479-INS-76
chr13-107450818-INS-109
chr13-107464472-INS-78
chr13-107500938-INS-81
chr13-107511676-INS-313
chr13-107533272-INS-383
chr13-107562030-INS-100
chr13-107563591-INS-54
chr13-107597523-INS-67
chr13-107603836-INS-83
chr13-107617787-INS-961
chr13-107619434-INS-64
chr13-107630716-INS-59
chr13-107635264-INS-54
chr13-107653029-INS-72
chr13-107666089-INS-103
chr13-107686499-INS-95
chr13-107709997-INS-61
chr13-107742588-INS-176
chr13-107767465-INS-52
chr13-107769068-INS-90
chr13-107782199-INS-55
chr13-107791625-INS-230
chr13-107805435-INS-64
chr13-107824360-INS-60
chr13-107826737-INS-117
chr13-107845927-INS-928
chr13-107851531-INS-708
chr13-107923196-INS-58
chr13-107927278-INS-593

chr13-107930899-INS-60
chr13-107935102-INS-7142
chr13-107956136-INS-150
chr13-107994789-INS-2925
chr13-107994998-INS-76
chr13-108019469-INS-64
chr13-108055612-INS-138
chr13-108074053-INS-72
chr13-108117431-INS-280
chr13-108152979-INS-183
chr13-108189864-INS-150
chr13-108201166-INS-83
chr13-108223364-INS-61
chr13-108248385-INS-95
chr13-108249396-INS-248
chr13-108257945-INS-78
chr13-108343090-INS-78
chr13-108347113-INS-57
chr13-108353396-INS-198
chr13-108370671-INS-79
chr13-108425099-INS-170
chr13-108458597-INS-52
chr13-108465388-INS-132
chr13-108466825-INS-141
chr13-108488171-INS-61
chr13-108492328-INS-74
chr13-108496771-INS-204
chr13-108497839-INS-57
chr13-108506133-INS-132
chr13-108526516-INS-6248
chr13-108527662-INS-1729
chr13-108528989-INS-3140
chr13-108555996-INS-197
chr13-108576593-INS-614
chr13-108587267-INS-51
chr13-108590986-INS-243
chr13-108596619-INS-809
chr13-108621074-INS-76
chr13-108647637-INS-202
chr13-108648689-INS-98
chr13-108696347-INS-474
chr13-108701992-INS-94
chr13-108730664-INS-60
chr13-108735211-INS-79
chr13-108739485-INS-6401
chr13-108764049-INS-369
chr13-108783338-INS-273
chr13-108786869-INS-434
chr13-108802359-INS-394
chr13-108827935-INS-328
chr13-108832275-INS-115
chr13-108854383-INS-851
chr13-108919675-INS-6534
chr13-108938650-INS-242
chr13-108948158-INS-141
chr13-108971887-INS-131
chr13-108972833-INS-563
chr13-108989279-INS-86
chr13-109027723-INS-538
chr13-109037031-INS-57
chr13-109045175-INS-144
chr13-109069197-INS-204
chr13-109086973-INS-60
chr13-109089253-INS-54
chr13-109110164-INS-103
chr13-109122522-INS-354
chr13-109122582-INS-877
chr13-109131631-INS-205
chr13-109164503-INS-111
chr13-109187883-INS-94
chr13-109235120-INS-553
chr13-109259836-INS-526
chr13-109264535-INS-125
chr13-109323397-INS-129
chr13-109329821-INS-72
chr13-109335346-INS-102
chr13-109340720-INS-210

chr13-109349646-INS-79
chr13-109371168-INS-86
chr13-109380857-INS-53
chr13-109391552-INS-55
chr13-109392954-INS-76
chr13-109403066-INS-69
chr13-109442886-INS-399
chr13-109459580-INS-126
chr13-109508815-INS-636
chr13-109539820-INS-98
chr13-109553595-INS-6754
chr13-109565680-INS-60
chr13-109589110-INS-91
chr13-109629251-INS-116
chr13-109630226-INS-52
chr13-109635504-INS-154
chr13-109665189-INS-50
chr13-109687387-INS-361
chr13-109688449-INS-420
chr13-109708046-INS-121
chr13-109722120-INS-146
chr13-109724678-INS-128
chr13-109730554-INS-71
chr13-109746070-INS-235
chr13-109764964-INS-198
chr13-109774917-INS-200
chr13-109786107-INS-51
chr13-109800899-INS-58
chr13-109806890-INS-404
chr13-109850233-INS-108
chr13-109874484-INS-58
chr13-109882927-INS-133
chr13-109920680-INS-78
chr13-109924399-INS-59
chr13-109937070-INS-95
chr13-109967925-INS-310
chr13-109978814-INS-111
chr13-110016967-INS-141
chr13-110021012-INS-61
chr13-110022744-INS-58
chr13-110024307-INS-166
chr13-110033969-INS-52
chr13-110036026-INS-161
chr13-110046152-INS-68
chr13-110052989-INS-61
chr13-110068241-INS-258
chr13-110079431-INS-711
chr13-110083641-INS-106
chr13-110099024-INS-519
chr13-110139095-INS-69
chr13-110159211-INS-146
chr13-110188302-INS-328
chr13-110213591-INS-66
chr13-110238396-INS-116
chr13-110245265-INS-116
chr13-110264703-INS-60
chr13-110270581-INS-208
chr13-110277135-INS-646
chr13-110280543-INS-51
chr13-110351114-INS-65
chr13-110353493-INS-326
chr13-110392951-INS-60
chr13-110400529-INS-57
chr13-110412061-INS-221
chr13-110423296-INS-85
chr13-110434092-INS-52
chr13-110456324-INS-795
chr13-110485626-INS-158
chr13-110495824-INS-57
chr13-110505251-INS-200
chr13-110519896-INS-747
chr13-110547890-INS-125
chr13-110566088-INS-54
chr13-110578986-INS-577
chr13-110582140-INS-62
chr13-110596247-INS-224
chr13-110598993-INS-157

chr13-110625890-INS-71
chr13-110634651-INS-369
chr13-110645157-INS-50
chr13-110653491-INS-151
chr13-110680864-INS-84
chr13-110693821-INS-396
chr13-110706513-INS-196
chr13-110727382-INS-99
chr13-110747546-INS-384
chr13-110807370-INS-65
chr13-110843351-INS-350
chr13-110846867-INS-161
chr13-110862846-INS-316
chr13-110875319-INS-75
chr13-110885371-INS-132
chr13-110898152-INS-433
chr13-110948826-INS-569
chr13-110959386-INS-57
chr13-111034983-INS-76
chr13-111055943-INS-180
chr13-111058916-INS-264
chr13-111077307-INS-127
chr13-111096309-INS-262
chr13-111101264-INS-55
chr13-111192838-INS-309
chr13-111195899-INS-364
chr13-111210633-INS-212
chr13-111232261-INS-72
chr13-111272341-INS-657
chr13-111276847-INS-50
chr13-111291868-INS-312
chr13-111343373-INS-97
chr13-111362673-INS-350
chr13-111362828-INS-745
chr13-111364720-INS-578
chr13-111367365-INS-108
chr13-111416107-INS-211
chr13-111429787-INS-321
chr13-111433659-INS-910
chr13-111442790-INS-84
chr13-111463356-INS-182
chr13-111465441-INS-67
chr13-111477455-INS-140
chr13-111489840-INS-331
chr13-111494777-INS-60
chr13-111499057-INS-556
chr13-111513276-INS-292
chr13-111535704-INS-415
chr13-111559710-INS-137
chr13-111566198-INS-208
chr13-111567956-INS-64
chr13-111576490-INS-310
chr13-111622386-INS-62
chr13-111656498-INS-116
chr13-111666476-INS-648
chr13-111666526-INS-793
chr13-111667813-INS-519
chr13-111687031-INS-79
chr13-111693083-INS-192
chr13-111704198-INS-69
chr13-111709641-INS-456
chr13-111727931-INS-434
chr13-111729383-INS-84
chr13-111770078-INS-71
chr13-111859048-INS-210
chr13-111879560-INS-110
chr13-111897027-INS-51
chr13-111906994-INS-90
chr13-111911308-INS-86
chr13-111934949-INS-131
chr13-111952887-INS-142
chr13-111955329-INS-66
chr13-111991616-INS-189
chr13-111999376-INS-66
chr13-112003979-INS-96
chr13-112029485-INS-226
chr13-112030482-INS-286

chr13-112031152-INS-1917
chr13-112031525-INS-54
chr13-112055697-INS-1134
chr13-112055927-INS-141
chr13-112083168-INS-741
chr13-112154666-INS-72
chr13-112176861-INS-91
chr13-112213470-INS-420
chr13-112233122-INS-278
chr13-112247336-INS-469
chr13-112247430-INS-941
chr13-112250304-INS-95
chr13-112266172-INS-304
chr13-112270036-INS-98
chr13-112271627-INS-492
chr13-112285801-INS-267
chr13-112287916-INS-219
chr13-112303513-INS-62
chr13-112317934-INS-211
chr13-112356455-INS-270
chr13-112386836-INS-320
chr13-112391803-INS-107
chr13-112391831-INS-83
chr13-112410142-INS-903
chr13-112413908-INS-56
chr13-112421406-INS-63
chr13-112421705-INS-63
chr13-112439993-INS-65
chr13-112506981-INS-155
chr13-112521993-INS-52
chr13-112535023-INS-128
chr13-112546132-INS-192
chr13-112550915-INS-316
chr13-112565410-INS-174
chr13-112567136-INS-124
chr13-112580675-INS-115
chr13-112583072-INS-122
chr13-112628991-INS-480
chr13-112659052-INS-66
chr13-112676472-INS-138
chr13-112684857-INS-127
chr13-112700932-INS-168
chr13-112726000-INS-842
chr13-112764486-INS-60
chr13-112768738-INS-254
chr13-112776304-INS-60
chr13-112783437-INS-53
chr13-112824517-INS-82
chr13-112832792-INS-586
chr13-112835198-INS-102
chr13-112842483-INS-612
chr13-112854443-INS-106
chr13-112861276-INS-110
chr13-112861604-INS-167
chr13-112862029-INS-244
chr13-112866492-INS-108
chr13-112867151-INS-185
chr13-112884303-INS-6416
chr13-112885278-INS-51
chr13-112907958-INS-56
chr13-112913682-INS-175
chr13-112914589-INS-6475
chr13-112915724-INS-212
chr13-112917503-INS-346
chr13-112920199-INS-175
chr13-112928289-INS-114
chr13-112932449-INS-360
chr13-112949300-INS-184
chr13-112956679-INS-304
chr13-112958319-INS-283
chr13-112966610-INS-280
chr13-113024669-INS-54
chr13-113044279-INS-170
chr13-113053123-INS-486
chr13-113055927-INS-595
chr13-113073277-INS-191
chr13-113097135-INS-57

chr13-113109318-INS-205
chr13-113112340-INS-203
chr13-113122171-INS-104
chr13-113129462-INS-590
chr13-113139966-INS-179
chr13-113140498-INS-498
chr13-113148325-INS-382
chr13-113155401-INS-207
chr13-113189269-INS-568
chr13-113192768-INS-209
chr13-113193374-INS-90
chr13-113203527-INS-266
chr13-113222950-INS-65
chr13-113242176-INS-592
chr13-113252931-INS-219
chr13-113255522-INS-106
chr13-113267120-INS-116
chr13-113289742-INS-60
chr13-113312556-INS-55
chr13-113323098-INS-95
chr13-113328946-INS-65
chr13-113345534-INS-136
chr13-113349553-INS-52
chr13-113355057-INS-88
chr13-113361387-INS-497
chr13-113369654-INS-276
chr13-113372821-INS-69
chr13-113386904-INS-342
chr13-113398610-INS-506
chr13-113400775-INS-205
chr13-113401560-INS-65
chr13-113466002-INS-51
chr13-113476911-INS-496
chr13-113481878-INS-408
chr13-113501842-INS-355
chr13-113517583-INS-579
chr13-113523261-INS-150
chr13-113541212-INS-69
chr13-113543019-INS-52
chr13-113568927-INS-92
chr13-113592660-INS-54
chr13-113593583-INS-252
chr13-113615234-INS-184
chr13-113648066-INS-157
chr13-113653850-INS-201
chr13-113660040-INS-90
chr13-113666207-INS-58
chr13-113669679-INS-117
chr13-113715174-INS-216
chr13-113719396-INS-203
chr13-113745877-INS-110
chr13-113747360-INS-62
chr13-113763385-INS-75
chr13-113769215-INS-1892
chr13-113793249-INS-399
chr13-113842659-INS-449
chr13-113850189-INS-7073
chr13-113908559-INS-268
chr13-113921678-INS-80
chr13-113926251-INS-112
chr13-113927975-INS-76
chr13-113942989-INS-132
chr13-113952269-INS-72
chr13-113955375-INS-160
chr13-113973379-INS-381
chr13-113986702-INS-308
chr13-113988382-INS-603
chr13-114015950-INS-228
chr13-114069091-INS-75
chr13-114092537-INS-58
chr13-114114666-INS-220
chr13-114117818-INS-162
chr13-114125031-INS-141
chr13-114144923-INS-903
chr13-114154111-INS-88
chr13-114162222-INS-194
chr13-114164366-INS-459

chr13-114204363-INS-226
chr13-114258249-INS-156
chr13-114270669-INS-184
chr13-114283747-INS-815
chr13-114299216-INS-56
chr13-114322416-INS-81
chr13-114338798-INS-81
chr13-114366720-INS-234
chr13-114386401-INS-145
chr13-114388561-INS-498
chr13-114413883-INS-168
chr13-114425008-INS-280
chr13-114426018-INS-204
chr13-114437652-INS-142
chr13-114454409-INS-51
chr13-114458092-INS-421
chr13-114510519-INS-779
chr13-114511174-INS-133
chr13-114532427-INS-53
chr13-114540628-INS-519
chr13-114554974-INS-498
chr13-114569753-INS-54
chr13-114577993-INS-57
chr13-114611422-INS-209
chr13-114623508-INS-420
chr13-114634739-INS-295
chr13-114644542-INS-151
chr13-114647287-INS-64
chr13-114662696-INS-54
chr13-114672261-INS-280
chr13-114734228-INS-798
chr13-114741519-INS-142
chr13-114747126-INS-59
chr13-114774720-INS-138
chr13-114801138-INS-937
chr13-114802010-INS-85
chr13-114806044-INS-60
chr13-114808423-INS-106
chr13-114833203-INS-52
chr13-114892090-INS-993
chr13-114895212-INS-105
chr13-114907464-INS-147
chr13-114920935-INS-350
chr13-114935521-INS-51
chr13-114943378-INS-945
chr13-114945094-INS-76
chr13-114953541-INS-68
chr13-114964264-INS-94
chr13-114965308-INS-65
chr13-114988697-INS-72
chr13-114999209-INS-637
chr13-115000834-INS-165
chr13-115018513-INS-402
chr13-115033007-INS-119
chr13-115048581-INS-206
chr13-115092561-INS-72
chr13-115125434-INS-173
chr13-115137486-INS-180
chr13-115148007-INS-56
chr13-115151198-INS-84
chr13-115175341-INS-701
chr13-115183625-INS-86
chr13-115189491-INS-58
chr13-115194823-INS-62
chr13-115202837-INS-353
chr13-115210977-INS-185
chr13-115234035-INS-70
chr13-115251737-INS-192
chr13-115265211-INS-1266
chr13-115272758-INS-454
chr13-115283656-INS-456
chr13-115317281-INS-445
chr13-115353581-INS-52
chr13-115384455-INS-164
chr13-115488163-INS-3805
chr13-115489427-INS-208
chr13-115520997-INS-51

chr13-115529692-INS-620
chr13-115579385-INS-93
chr13-115594195-INS-60
chr13-115595074-INS-222
chr13-115606180-INS-62
chr13-115609511-INS-7085
chr13-115617067-INS-73
chr13-115619979-INS-7225
chr13-115643663-INS-79
chr13-115666215-INS-6370
chr13-115684537-INS-54
chr13-115702220-INS-131
chr13-115753330-INS-55
chr13-115759945-INS-53
chr13-115779586-INS-193
chr13-115780307-INS-62
chr13-115799475-INS-512
chr13-115808794-INS-184
chr13-115870074-INS-212
chr13-115878739-INS-54
chr13-115883488-INS-808
chr13-115886606-INS-74
chr13-115901119-INS-93
chr13-115958532-INS-817
chr13-115978024-INS-509
chr13-115994074-INS-424
chr13-116049302-INS-140
chr13-116060749-INS-56
chr13-116062439-INS-2473
chr13-116075593-INS-470
chr13-116079658-INS-361
chr13-116119478-INS-813
chr13-116139833-INS-539
chr13-116143142-INS-58
chr13-116190663-INS-7102
chr13-116221451-INS-52
chr13-116225358-INS-687
chr13-116233501-INS-212
chr13-116253891-INS-6985
chr13-116256276-INS-6442
chr13-116278124-INS-192
chr13-116295916-INS-197
chr13-116301016-INS-58
chr13-116339718-INS-578
chr13-116341106-INS-386
chr13-116353604-INS-339
chr13-116368292-INS-459
chr13-116382949-INS-108
chr13-116389752-INS-209
chr13-116408933-INS-260
chr13-116441280-INS-127
chr13-116462943-INS-508
chr13-116476488-INS-369
chr13-116496007-INS-52
chr13-116518411-INS-64
chr13-116536642-INS-66
chr13-116546416-INS-98
chr13-116569541-INS-55
chr13-116590879-INS-6507
chr13-116594419-INS-6662
chr13-116609322-INS-942
chr13-116621700-INS-207
chr13-116630991-INS-354
chr13-116685238-INS-6770
chr13-116692893-INS-51
chr13-116749396-INS-414
chr13-116752123-INS-152
chr13-116780979-INS-267
chr13-116794078-INS-373
chr13-116854400-INS-112
chr13-116860394-INS-98
chr13-116870090-INS-62
chr13-116908959-INS-635
chr13-116938173-INS-238
chr13-116967964-INS-7289
chr13-116970749-INS-68
chr13-117008999-INS-52

chr13-117009387-INS-380
chr13-117012843-INS-5608
chr13-117043006-INS-52
chr13-117080015-INS-427
chr13-117089060-INS-53
chr13-117101353-INS-170
chr13-117144494-INS-68
chr13-117156122-INS-203
chr13-117183189-INS-57
chr13-117206052-INS-86
chr13-117207726-INS-623
chr13-117215071-INS-68
chr13-117220858-INS-238
chr13-117221581-INS-7276
chr13-117227639-INS-2317
chr13-117229410-INS-83
chr13-117246294-INS-67
chr13-117278792-INS-354
chr13-117287500-INS-78
chr13-117296019-INS-400
chr13-117297788-INS-80
chr13-117315459-INS-135
chr13-117334985-INS-184
chr13-117354991-INS-53
chr13-117362908-INS-109
chr13-117398571-INS-57
chr13-117412909-INS-132
chr13-117444630-INS-6939
chr13-117452686-INS-434
chr13-117489495-INS-126
chr13-117498439-INS-5392
chr13-117502761-INS-96
chr13-117508148-INS-6728
chr13-117508939-INS-486
chr13-117511506-INS-7081
chr13-117526674-INS-92
chr13-117533173-INS-810
chr13-117541404-INS-939
chr13-117556395-INS-377
chr13-117565134-INS-69
chr13-117571399-INS-536
chr13-117594102-INS-72
chr13-117609725-INS-101
chr13-117630531-INS-192
chr13-117631863-INS-941
chr13-117635716-INS-622
chr13-117636327-INS-85
chr13-117637928-INS-275
chr13-117657165-INS-513
chr13-117661579-INS-862
chr13-117701329-INS-51
chr13-117728085-INS-440
chr13-117766945-INS-177
chr13-117781770-INS-52
chr13-117815004-INS-101
chr13-117825243-INS-268
chr13-117828476-INS-700
chr13-117832021-INS-75
chr13-117879920-INS-83
chr13-117908074-INS-70
chr13-117922812-INS-603
chr13-117981017-INS-70
chr13-117984778-INS-68
chr13-117990292-INS-52
chr13-118038366-INS-687
chr13-118042016-INS-214
chr13-118050054-INS-167
chr13-118058011-INS-626
chr13-118060650-INS-100
chr13-118082654-INS-52
chr13-118083137-INS-869
chr13-118092518-INS-220
chr13-118129963-INS-66
chr13-118164704-INS-54
chr13-118182093-INS-610
chr13-118183802-INS-70
chr13-118224328-INS-354

chr13-118253366-INS-234
chr13-118312267-INS-52
chr13-118341879-INS-56
chr13-118349456-INS-136
chr13-118351970-INS-185
chr13-118367662-INS-69
chr13-118389419-INS-87
chr13-118400660-INS-196
chr13-118408611-INS-60
chr13-118438381-INS-137
chr13-118455675-INS-1197
chr13-118493290-INS-188
chr13-118530564-INS-633
chr13-118534084-INS-201
chr13-118542121-INS-455
chr13-118577504-INS-4058
chr13-118625943-INS-143
chr13-118649962-INS-365
chr13-118689090-INS-192
chr13-118738253-INS-55
chr13-118753251-INS-52
chr13-118757238-INS-119
chr13-118764076-INS-737
chr13-118770219-INS-151
chr13-118778950-INS-102
chr13-118796686-INS-134
chr13-118831301-INS-294
chr13-118833928-INS-194
chr13-118841025-INS-148
chr13-118895232-INS-91
chr13-118941568-INS-165
chr13-118952592-INS-90
chr13-118975822-INS-498
chr13-118997854-INS-192
chr13-119003606-INS-79
chr13-119020305-INS-8133
chr13-119058391-INS-6123
chr13-119063945-INS-3066
chr13-119069651-INS-230
chr13-119072449-INS-219
chr13-119076344-INS-52
chr13-119076879-INS-61
chr13-119094820-INS-51
chr13-119097740-INS-358
chr13-119113703-INS-56
chr13-119114911-INS-120
chr13-119160211-INS-63
chr13-119211292-INS-123
chr13-119244438-INS-103
chr13-119253252-INS-51
chr13-119255431-INS-339
chr13-119276799-INS-6721
chr13-119289798-INS-200
chr13-119302819-INS-192
chr13-119310785-INS-114
chr13-119332536-INS-194
chr13-119374608-INS-433
chr13-119377730-INS-9300
chr13-119380210-INS-117
chr13-119386841-INS-6212
chr13-119392674-INS-665
chr13-119471215-INS-475
chr13-119476320-INS-58
chr13-119477591-INS-987
chr13-119504079-INS-193
chr13-119511984-INS-16400
chr13-119532132-INS-210
chr13-119537707-INS-444
chr13-119539693-INS-112
chr13-119541833-INS-60
chr13-119564659-INS-458
chr13-119566313-INS-106
chr13-119586195-INS-180
chr13-119597729-INS-215
chr13-119609587-INS-429
chr13-119616329-INS-219
chr13-119619351-INS-894

chr13-120059018-INS-60
chr13-120063656-INS-649
chr13-120065060-INS-1985
chr13-120089117-INS-55
chr13-120146903-INS-66
chr13-120152268-INS-378
chr13-120175146-INS-212
chr13-120182826-INS-841
chr13-120194266-INS-440
chr13-120217106-INS-103
chr13-120225303-INS-232
chr13-120256432-INS-201
chr13-120268493-INS-54
chr13-120269894-INS-487
chr13-120272926-INS-271
chr13-120273778-INS-8323
chr13-120292682-INS-78
chr13-120310200-INS-353
chr13-120344183-INS-2993
chr13-120344576-INS-602
chr13-120345975-INS-800
chr13-120397353-INS-53
chr13-120429466-INS-2447
chr13-120436199-INS-795
chr13-120483030-INS-371
chr13-120497993-INS-2708
chr13-120503063-INS-371
chr13-120549399-INS-805
chr13-120556014-INS-2394
chr13-120566957-INS-66
chr13-120591622-INS-65
chr13-120622233-INS-262
chr13-120622775-INS-83
chr13-120626967-INS-250
chr13-120654501-INS-199
chr13-120656474-INS-129
chr13-120671062-INS-5523
chr13-120674136-INS-354
chr13-120711319-INS-330
chr13-120722232-INS-118
chr13-120736219-INS-51
chr14-3060075-INS-52
chr14-3072473-INS-4285
chr14-3114621-INS-389
chr14-3129882-INS-6559
chr14-3137764-INS-537
chr14-3155581-INS-54
chr14-3220722-INS-235
chr14-3326610-INS-861
chr14-3332291-INS-85
chr14-3367245-INS-52
chr14-3375001-INS-460
chr14-3411586-INS-178
chr14-3412037-INS-751
chr14-3413775-INS-860
chr14-3415744-INS-54
chr14-3417364-INS-54
chr14-3420123-INS-108
chr14-3424015-INS-61
chr14-3424860-INS-401
chr14-3425618-INS-430
chr14-3443855-INS-3736
chr14-3464656-INS-5530
chr14-3500470-INS-54
chr14-3552533-INS-205
chr14-3575137-INS-67
chr14-3631996-INS-438
chr14-3663056-INS-1324
chr14-3717997-INS-7450
chr14-3789828-INS-308
chr14-3800085-INS-144
chr14-3801485-INS-310
chr14-3826005-INS-52
chr14-3845768-INS-105
chr14-3903489-INS-62
chr14-3912327-INS-598
chr14-3927155-INS-88

chr14-3928181-INS-64
chr14-3930285-INS-986
chr14-3946695-INS-4892
chr14-3991657-INS-370
chr14-4002867-INS-513
chr14-4020094-INS-498
chr14-4091448-INS-56
chr14-4110697-INS-135
chr14-4121150-INS-98
chr14-4130676-INS-56
chr14-4159644-INS-1892
chr14-4169690-INS-475
chr14-4171282-INS-224
chr14-4175806-INS-87
chr14-4192263-INS-117
chr14-4194146-INS-5553
chr14-4223258-INS-52
chr14-4248782-INS-291
chr14-4273899-INS-52
chr14-4359788-INS-1984
chr14-4361098-INS-60
chr14-4379328-INS-675
chr14-4460251-INS-378
chr14-4465177-INS-59
chr14-4481180-INS-79
chr14-4512744-INS-80
chr14-4545252-INS-136
chr14-4552241-INS-292
chr14-4559211-INS-112
chr14-4573969-INS-211
chr14-4597415-INS-64
chr14-4599552-INS-58
chr14-4603246-INS-248
chr14-4604398-INS-54
chr14-4611775-INS-99
chr14-4650068-INS-179
chr14-4666152-INS-249
chr14-4687938-INS-136
chr14-4694259-INS-103
chr14-4753682-INS-126
chr14-4784328-INS-104
chr14-4793264-INS-949
chr14-4795584-INS-72
chr14-4819810-INS-52
chr14-4833256-INS-54
chr14-4838186-INS-53
chr14-4844275-INS-79
chr14-4895928-INS-256
chr14-4904400-INS-62
chr14-4913538-INS-57
chr14-4943975-INS-379
chr14-4957858-INS-383
chr14-4961516-INS-706
chr14-4995879-INS-230
chr14-5002090-INS-6954
chr14-5016874-INS-58
chr14-5122538-INS-115
chr14-5140144-INS-73
chr14-5177183-INS-257
chr14-5209361-INS-189
chr14-5243019-INS-680
chr14-5247172-INS-94
chr14-5276892-INS-72
chr14-5291438-INS-335
chr14-5316179-INS-56
chr14-5325690-INS-492
chr14-5384410-INS-216
chr14-5384541-INS-488
chr14-5391194-INS-71
chr14-5408158-INS-4491
chr14-5431666-INS-581
chr14-5478007-INS-280
chr14-5482940-INS-60
chr14-5551798-INS-85
chr14-5554507-INS-105
chr14-5555126-INS-1004
chr14-5571457-INS-5207

chr14-5579523-INS-111
chr14-5586225-INS-421
chr14-5592679-INS-854
chr14-5604413-INS-66
chr14-5616015-INS-90
chr14-5620532-INS-146
chr14-5652011-INS-60
chr14-5675072-INS-523
chr14-5700398-INS-142
chr14-5705580-INS-151
chr14-5714623-INS-517
chr14-5728146-INS-88
chr14-5729797-INS-548
chr14-5742941-INS-53
chr14-5785449-INS-68
chr14-5789498-INS-382
chr14-5841137-INS-62
chr14-5844152-INS-218
chr14-5890908-INS-159
chr14-5915894-INS-7483
chr14-5926541-INS-241
chr14-5944609-INS-272
chr14-5966012-INS-366
chr14-5967889-INS-137
chr14-5978950-INS-120
chr14-5994093-INS-509
chr14-6020972-INS-241
chr14-6022937-INS-368
chr14-6036784-INS-155
chr14-6082185-INS-201
chr14-6101797-INS-107
chr14-6124340-INS-129
chr14-6136620-INS-79
chr14-6174168-INS-71
chr14-6186451-INS-189
chr14-6230449-INS-81
chr14-6232060-INS-94
chr14-6261095-INS-638
chr14-6269783-INS-113
chr14-6283229-INS-62
chr14-6301560-INS-62
chr14-6338808-INS-5363
chr14-6343726-INS-461
chr14-6351636-INS-7131
chr14-6360517-INS-7095
chr14-6361487-INS-305
chr14-6368661-INS-361
chr14-6373393-INS-51
chr14-6405888-INS-1255
chr14-6418033-INS-5015
chr14-6419681-INS-76
chr14-6423583-INS-6359
chr14-6445348-INS-117
chr14-6463301-INS-65
chr14-6481005-INS-498
chr14-6500022-INS-63
chr14-6511458-INS-192
chr14-6515033-INS-56
chr14-6524542-INS-62
chr14-6560416-INS-57
chr14-6583188-INS-830
chr14-6583219-INS-271
chr14-6622019-INS-6387
chr14-6625077-INS-1017
chr14-6625078-INS-998
chr14-6638614-INS-507
chr14-6654910-INS-338
chr14-6668437-INS-431
chr14-6674926-INS-544
chr14-6688195-INS-435
chr14-6706673-INS-56
chr14-6710823-INS-140
chr14-6720468-INS-562
chr14-6760710-INS-124
chr14-6822014-INS-74
chr14-6822851-INS-230
chr14-6824258-INS-69

chr14-6849791-INS-56
chr14-6855048-INS-892
chr14-6877214-INS-1589
chr14-6890843-INS-6502
chr14-6898801-INS-97
chr14-6903061-INS-5211
chr14-6914468-INS-3370
chr14-6942594-INS-6662
chr14-6973095-INS-6168
chr14-6982098-INS-6726
chr14-6989168-INS-525
chr14-7041110-INS-52
chr14-7042891-INS-1827
chr14-7060864-INS-57
chr14-7067640-INS-279
chr14-7071344-INS-90
chr14-7089270-INS-258
chr14-7144793-INS-76
chr14-7157157-INS-197
chr14-7182933-INS-54
chr14-7186590-INS-329
chr14-7203384-INS-402
chr14-7211451-INS-6882
chr14-7232346-INS-70
chr14-7266333-INS-380
chr14-7269681-INS-6205
chr14-7271346-INS-61
chr14-7311066-INS-354
chr14-7321763-INS-68
chr14-7455320-INS-50
chr14-7477991-INS-612
chr14-7485691-INS-238
chr14-7515062-INS-106
chr14-7523808-INS-205
chr14-7555157-INS-353
chr14-7560734-INS-283
chr14-7584229-INS-54
chr14-7595823-INS-174
chr14-7598547-INS-76
chr14-7603400-INS-54
chr14-7605640-INS-305
chr14-7611681-INS-55
chr14-7663417-INS-145
chr14-7666674-INS-97
chr14-7674479-INS-369
chr14-7708033-INS-158
chr14-7760273-INS-156
chr14-7764290-INS-126
chr14-7803018-INS-54
chr14-7806364-INS-6902
chr14-7838084-INS-206
chr14-7849258-INS-83
chr14-7850564-INS-4754
chr14-7853995-INS-85
chr14-7872967-INS-736
chr14-7880176-INS-526
chr14-7887804-INS-713
chr14-7947452-INS-82
chr14-7983709-INS-85
chr14-8067658-INS-203
chr14-8073854-INS-235
chr14-8097247-INS-6213
chr14-8101073-INS-5404
chr14-8115242-INS-69
chr14-8136433-INS-109
chr14-8167168-INS-653
chr14-8167246-INS-989
chr14-8174859-INS-72
chr14-8230941-INS-71
chr14-8273835-INS-62
chr14-8282109-INS-51
chr14-8288269-INS-100
chr14-8291290-INS-52
chr14-8365514-INS-70
chr14-8370767-INS-7171
chr14-8402865-INS-66
chr14-8428907-INS-115

chr14-8445740-INS-220
chr14-8464583-INS-75
chr14-8472761-INS-121
chr14-8479057-INS-145
chr14-8491433-INS-582
chr14-8494111-INS-120
chr14-8497084-INS-284
chr14-8513807-INS-165
chr14-8554132-INS-989
chr14-8563682-INS-190
chr14-8622908-INS-405
chr14-8703236-INS-344
chr14-8704547-INS-345
chr14-8726312-INS-202
chr14-8736825-INS-689
chr14-8748330-INS-6487
chr14-8753584-INS-5545
chr14-8758699-INS-52
chr14-8764049-INS-91
chr14-8768802-INS-78
chr14-8785291-INS-52
chr14-8785360-INS-7476
chr14-8805567-INS-439
chr14-8807499-INS-52
chr14-8811298-INS-230
chr14-8858289-INS-572
chr14-8867104-INS-75
chr14-8868848-INS-1797
chr14-8869742-INS-486
chr14-8876573-INS-58
chr14-8901010-INS-120
chr14-8906098-INS-5554
chr14-8919682-INS-845
chr14-8960209-INS-5283
chr14-8980653-INS-51
chr14-9001276-INS-119
chr14-9044390-INS-658
chr14-9053997-INS-6680
chr14-9055179-INS-382
chr14-9058753-INS-6183
chr14-9069329-INS-214
chr14-9090819-INS-6174
chr14-9113842-INS-73
chr14-9138434-INS-304
chr14-9155168-INS-348
chr14-9180528-INS-70
chr14-9183406-INS-120
chr14-9188731-INS-179
chr14-9194915-INS-52
chr14-9195885-INS-87
chr14-9276015-INS-10908
chr14-9333589-INS-138
chr14-9353590-INS-53
chr14-9424000-INS-740
chr14-9424584-INS-76
chr14-9435112-INS-68
chr14-9474667-INS-329
chr14-9524993-INS-182
chr14-9539498-INS-492
chr14-9548720-INS-300
chr14-9558781-INS-7851
chr14-9652467-INS-512
chr14-9661898-INS-136
chr14-9676299-INS-860
chr14-9691496-INS-144
chr14-9702403-INS-71
chr14-9706130-INS-128
chr14-9707446-INS-77
chr14-9720840-INS-203
chr14-9721954-INS-240
chr14-9748133-INS-101
chr14-9814663-INS-57
chr14-9823094-INS-71
chr14-9870496-INS-511
chr14-9881640-INS-104
chr14-9886568-INS-829
chr14-9888550-INS-58

chr14-9900755-INS-165
chr14-9915494-INS-85
chr14-9937002-INS-116
chr14-9974393-INS-314
chr14-10027393-INS-65
chr14-10032402-INS-100
chr14-10088246-INS-440
chr14-10133848-INS-662
chr14-10188392-INS-90
chr14-10194623-INS-6401
chr14-10205088-INS-109
chr14-10257884-INS-66
chr14-10265857-INS-56
chr14-10286559-INS-74
chr14-10298128-INS-61
chr14-10319820-INS-52
chr14-10331505-INS-117
chr14-10366276-INS-126
chr14-10394013-INS-63
chr14-10395481-INS-334
chr14-10445684-INS-176
chr14-10467929-INS-80
chr14-10525304-INS-122
chr14-10526882-INS-75
chr14-10535373-INS-221
chr14-10558953-INS-111
chr14-10579917-INS-590
chr14-10608074-INS-59
chr14-10615241-INS-120
chr14-10671437-INS-224
chr14-10714089-INS-279
chr14-10732862-INS-65
chr14-10757971-INS-152
chr14-10783807-INS-214
chr14-10810737-INS-116
chr14-10815103-INS-103
chr14-10822424-INS-53
chr14-10884209-INS-58
chr14-10905315-INS-76
chr14-10923611-INS-264
chr14-10928753-INS-238
chr14-10989263-INS-65
chr14-11044382-INS-198
chr14-11061302-INS-54
chr14-11153869-INS-102
chr14-11167342-INS-399
chr14-11170745-INS-395
chr14-11209939-INS-900
chr14-11217611-INS-349
chr14-11220175-INS-215
chr14-11245737-INS-318
chr14-11252548-INS-60
chr14-11283630-INS-328
chr14-11290039-INS-296
chr14-11358850-INS-65
chr14-11381098-INS-294
chr14-11400748-INS-96
chr14-11410242-INS-251
chr14-11411748-INS-51
chr14-11421584-INS-720
chr14-11422668-INS-252
chr14-11423389-INS-360
chr14-11449756-INS-218
chr14-11492425-INS-495
chr14-11503981-INS-149
chr14-11512930-INS-102
chr14-11578848-INS-52
chr14-11580241-INS-544
chr14-11586296-INS-552
chr14-11608876-INS-62
chr14-11617467-INS-274
chr14-11643679-INS-72
chr14-11654952-INS-700
chr14-11672585-INS-120
chr14-11683829-INS-6705
chr14-11692390-INS-5548
chr14-11697615-INS-933

chr14-11705636-INS-1875
chr14-11711817-INS-74
chr14-11770243-INS-100
chr14-11776944-INS-6202
chr14-11779570-INS-1296
chr14-11825420-INS-768
chr14-11837972-INS-115
chr14-11879514-INS-675
chr14-11882255-INS-64
chr14-11884777-INS-174
chr14-11916311-INS-126
chr14-11938168-INS-52
chr14-11939612-INS-438
chr14-11959258-INS-62
chr14-12016044-INS-74
chr14-12034464-INS-61
chr14-12068428-INS-355
chr14-12071879-INS-123
chr14-12138152-INS-55
chr14-12170091-INS-792
chr14-12173611-INS-310
chr14-12178897-INS-970
chr14-12240210-INS-54
chr14-12241334-INS-87
chr14-12250258-INS-71
chr14-12265388-INS-227
chr14-12276250-INS-568
chr14-12338238-INS-70
chr14-12388908-INS-113
chr14-12389468-INS-52
chr14-12473248-INS-102
chr14-12489065-INS-486
chr14-12489571-INS-455
chr14-12509564-INS-158
chr14-12516970-INS-104
chr14-12543099-INS-739
chr14-12560369-INS-56
chr14-12627385-INS-64
chr14-12633990-INS-126
chr14-12635599-INS-60
chr14-12695719-INS-57
chr14-12782776-INS-222
chr14-12783976-INS-123
chr14-12787061-INS-365
chr14-12794524-INS-144
chr14-12828336-INS-132
chr14-12844542-INS-514
chr14-12858830-INS-63
chr14-12906241-INS-279
chr14-12909104-INS-54
chr14-12920023-INS-339
chr14-12932248-INS-59
chr14-12935138-INS-173
chr14-12953477-INS-2419
chr14-12973284-INS-88
chr14-12999527-INS-52
chr14-13010817-INS-64
chr14-13014263-INS-328
chr14-13017637-INS-588
chr14-13071895-INS-874
chr14-13075099-INS-64
chr14-13081719-INS-95
chr14-13149829-INS-5020
chr14-13171301-INS-66
chr14-13200601-INS-56
chr14-13231153-INS-5283
chr14-13268092-INS-166
chr14-13310548-INS-59
chr14-13312547-INS-1403
chr14-13397374-INS-117
chr14-13431975-INS-179
chr14-13459186-INS-58
chr14-13471415-INS-119
chr14-13472827-INS-7876
chr14-13475472-INS-7370
chr14-13485755-INS-68
chr14-13487742-INS-62

chr14-13513233-INS-73
chr14-13514735-INS-60
chr14-13529809-INS-1383
chr14-13531252-INS-992
chr14-13568247-INS-204
chr14-13568735-INS-405
chr14-13572876-INS-411
chr14-13589141-INS-214
chr14-13607150-INS-1323
chr14-13617891-INS-379
chr14-13627139-INS-292
chr14-13649290-INS-51
chr14-13697642-INS-6253
chr14-13699938-INS-351
chr14-13706384-INS-837
chr14-13710152-INS-72
chr14-13731668-INS-6768
chr14-13738375-INS-72
chr14-13748936-INS-515
chr14-13758938-INS-359
chr14-13761530-INS-79
chr14-13768979-INS-207
chr14-13776885-INS-243
chr14-13778882-INS-363
chr14-13838892-INS-52
chr14-13851632-INS-6170
chr14-13855186-INS-145
chr14-13863123-INS-303
chr14-13894720-INS-119
chr14-13932263-INS-2122
chr14-13961036-INS-62
chr14-14013221-INS-109
chr14-14014356-INS-6454
chr14-14025698-INS-98
chr14-14027169-INS-1022
chr14-14032940-INS-262
chr14-14071406-INS-892
chr14-14076842-INS-126
chr14-14087065-INS-54
chr14-14089123-INS-53
chr14-14090623-INS-67
chr14-14110302-INS-158
chr14-14147789-INS-96
chr14-14178106-INS-719
chr14-14180146-INS-207
chr14-14185871-INS-158
chr14-14189248-INS-660
chr14-14216652-INS-62
chr14-14270619-INS-160
chr14-14285051-INS-204
chr14-14293037-INS-167
chr14-14295454-INS-163
chr14-14319796-INS-267
chr14-14356644-INS-199
chr14-14363240-INS-155
chr14-14365591-INS-94
chr14-14377646-INS-142
chr14-14402348-INS-1521
chr14-14409695-INS-7304
chr14-14428366-INS-66
chr14-14430764-INS-732
chr14-14442688-INS-74
chr14-14462627-INS-140
chr14-14473784-INS-909
chr14-14480825-INS-122
chr14-14538178-INS-84
chr14-14541574-INS-94
chr14-14546185-INS-214
chr14-14567198-INS-206
chr14-14598889-INS-2622
chr14-14630820-INS-83
chr14-14657910-INS-231
chr14-14659332-INS-331
chr14-14662809-INS-172
chr14-14670941-INS-155
chr14-14671920-INS-311
chr14-14704138-INS-72

chr14-14716714-INS-225
chr14-14719268-INS-448
chr14-14720891-INS-81
chr14-14725519-INS-468
chr14-14731756-INS-479
chr14-14741372-INS-200
chr14-14780587-INS-1533
chr14-14793502-INS-11184
chr14-14793651-INS-1948
chr14-14861820-INS-54
chr14-14896802-INS-98
chr14-14933095-INS-86
chr14-14955426-INS-98
chr14-15008566-INS-110
chr14-15012405-INS-310
chr14-15087428-INS-2483
chr14-15095844-INS-134
chr14-15096200-INS-179
chr14-15104080-INS-58
chr14-15361777-INS-88
chr14-15417970-INS-194
chr14-15420616-INS-136
chr14-15472900-INS-61
chr14-15528836-INS-57
chr14-15559067-INS-417
chr14-15575746-INS-303
chr14-15576772-INS-57
chr14-15610525-INS-247
chr14-15832060-INS-1348
chr14-15832091-INS-225
chr14-15878606-INS-86
chr14-16035376-INS-377
chr14-16039801-INS-96
chr14-16056743-INS-65
chr14-16100668-INS-1452
chr14-16141042-INS-518
chr14-16153110-INS-7202
chr14-16156641-INS-1215
chr14-16182939-INS-1094
chr14-16381155-INS-955
chr14-16442979-INS-658
chr14-16617068-INS-202
chr14-16762512-INS-2054
chr14-17085623-INS-107
chr14-17115059-INS-2054
chr14-17430735-INS-95
chr14-17500489-INS-270
chr14-17509222-INS-2468
chr14-17553807-INS-1890
chr14-17554320-INS-3046
chr14-17677329-INS-180
chr14-17697564-INS-87
chr14-17713091-INS-183
chr14-17736323-INS-610
chr14-17793712-INS-203
chr14-17845412-INS-989
chr14-17856306-INS-64
chr14-17905756-INS-1891
chr14-17922530-INS-946
chr14-17966619-INS-97
chr14-18022348-INS-482
chr14-18056028-INS-99
chr14-18116488-INS-953
chr14-18132694-INS-70
chr14-18159245-INS-3803
chr14-18168771-INS-350
chr14-18234729-INS-132
chr14-18259431-INS-212
chr14-18371738-INS-208
chr14-18455177-INS-100
chr14-18551752-INS-284
chr14-18622891-INS-93
chr14-18685187-INS-1079
chr14-19019682-INS-94
chr14-19123145-INS-146
chr14-19161477-INS-1234
chr14-19162886-INS-449

chr14-19176212-INS-64
chr14-19247278-INS-183
chr14-19367661-INS-489
chr14-19387699-INS-171
chr14-19417112-INS-61
chr14-19422442-INS-148
chr14-19442306-INS-235
chr14-19451490-INS-2987
chr14-19464851-INS-575
chr14-19574869-INS-5559
chr14-19660396-INS-225
chr14-19660552-INS-271
chr14-19712648-INS-515
chr14-19717247-INS-400
chr14-19774390-INS-128
chr14-19787065-INS-82
chr14-19807321-INS-147
chr14-19816252-INS-78
chr14-19831946-INS-78
chr14-19832274-INS-750
chr14-19859569-INS-67
chr14-19860501-INS-147
chr14-19863627-INS-72
chr14-19902508-INS-127
chr14-19909282-INS-53
chr14-19926979-INS-94
chr14-19962385-INS-269
chr14-19974557-INS-60
chr14-19976134-INS-99
chr14-19983239-INS-60
chr14-19990600-INS-192
chr14-19997923-INS-182
chr14-20004609-INS-203
chr14-20007068-INS-220
chr14-20015340-INS-211
chr14-20020166-INS-82
chr14-20022822-INS-705
chr14-20035702-INS-90
chr14-20054944-INS-122
chr14-20080041-INS-68
chr14-20083636-INS-197
chr14-20090052-INS-67
chr14-20092534-INS-93
chr14-20092850-INS-90
chr14-20106725-INS-100
chr14-20114963-INS-186
chr14-20118458-INS-406
chr14-20146155-INS-192
chr14-20147410-INS-74
chr14-20166965-INS-60
chr14-20179903-INS-172
chr14-20197171-INS-80
chr14-20212137-INS-168
chr14-20255745-INS-194
chr14-20259139-INS-63
chr14-20261108-INS-94
chr14-20266989-INS-111
chr14-20269323-INS-6420
chr14-20270025-INS-215
chr14-20270517-INS-531
chr14-20272367-INS-129
chr14-20316028-INS-98
chr14-20342855-INS-83
chr14-20343617-INS-271
chr14-20346191-INS-76
chr14-20349577-INS-121
chr14-20353201-INS-50
chr14-20357177-INS-53
chr14-20361357-INS-374
chr14-20362927-INS-99
chr14-20373077-INS-67
chr14-20377015-INS-177
chr14-20381349-INS-218
chr14-20383239-INS-114
chr14-20393410-INS-227
chr14-20424638-INS-224
chr14-20469813-INS-158

chr14-20470347-INS-273
chr14-20478614-INS-197
chr14-20507520-INS-136
chr14-20525301-INS-440
chr14-20525519-INS-549
chr14-20532535-INS-183
chr14-20534350-INS-201
chr14-20562249-INS-199
chr14-20618842-INS-669
chr14-20619915-INS-199
chr14-20622536-INS-942
chr14-20650942-INS-309
chr14-20662014-INS-62
chr14-20664095-INS-439
chr14-20671989-INS-198
chr14-20676726-INS-181
chr14-20707689-INS-158
chr14-20722861-INS-66
chr14-20728880-INS-5245
chr14-20758382-INS-68
chr14-20762520-INS-384
chr14-20782977-INS-237
chr14-20794380-INS-103
chr14-20860777-INS-342
chr14-20872753-INS-155
chr14-20895750-INS-233
chr14-20904643-INS-118
chr14-20927678-INS-366
chr14-20947722-INS-87
chr14-20955490-INS-64
chr14-20957381-INS-69
chr14-20993041-INS-118
chr14-20997246-INS-1893
chr14-21012402-INS-51
chr14-21053821-INS-54
chr14-21054511-INS-280
chr14-21107702-INS-59
chr14-21114040-INS-53
chr14-21115223-INS-78
chr14-21130084-INS-60
chr14-21154001-INS-178
chr14-21163313-INS-221
chr14-21229766-INS-68
chr14-21236739-INS-87
chr14-21261157-INS-129
chr14-21286421-INS-128
chr14-21292469-INS-58
chr14-21296779-INS-71
chr14-21330402-INS-637
chr14-21375983-INS-968
chr14-21387958-INS-259
chr14-21402350-INS-212
chr14-21443858-INS-1856
chr14-21451198-INS-68
chr14-21477136-INS-95
chr14-21494680-INS-184
chr14-21503449-INS-197
chr14-21524319-INS-72
chr14-21530332-INS-214
chr14-21538514-INS-566
chr14-21558807-INS-140
chr14-21565607-INS-164
chr14-21573220-INS-219
chr14-21583636-INS-53
chr14-21598626-INS-332
chr14-21600276-INS-232
chr14-21612281-INS-209
chr14-21637314-INS-87
chr14-21668509-INS-54
chr14-21676999-INS-204
chr14-21680136-INS-189
chr14-21685984-INS-137
chr14-21692691-INS-222
chr14-21711280-INS-75
chr14-21732608-INS-65
chr14-21744840-INS-922
chr14-21767961-INS-173

chr14-21772094-INS-63
chr14-21781604-INS-109
chr14-21788346-INS-265
chr14-21806027-INS-98
chr14-21807243-INS-364
chr14-21827557-INS-99
chr14-21833415-INS-244
chr14-21869511-INS-381
chr14-21875725-INS-220
chr14-21889882-INS-74
chr14-21904968-INS-64
chr14-21936297-INS-56
chr14-21940936-INS-146
chr14-21948850-INS-82
chr14-21949470-INS-80
chr14-21950505-INS-67
chr14-21970937-INS-60
chr14-21978826-INS-102
chr14-21983136-INS-73
chr14-21993058-INS-83
chr14-22002648-INS-266
chr14-22014381-INS-152
chr14-22059057-INS-124
chr14-22062376-INS-78
chr14-22080271-INS-163
chr14-22082519-INS-144
chr14-22103516-INS-73
chr14-22130282-INS-122
chr14-22159100-INS-60
chr14-22167234-INS-183
chr14-22178217-INS-60
chr14-22199017-INS-69
chr14-22209300-INS-387
chr14-22214546-INS-100
chr14-22252521-INS-104
chr14-22256811-INS-401
chr14-22264972-INS-150
chr14-22275289-INS-68
chr14-22288684-INS-117
chr14-22322356-INS-96
chr14-22327327-INS-399
chr14-22346452-INS-129
chr14-22349278-INS-82
chr14-22350020-INS-124
chr14-22375757-INS-256
chr14-22399006-INS-80
chr14-22400016-INS-737
chr14-22406848-INS-146
chr14-22474611-INS-103
chr14-22524059-INS-135
chr14-22533925-INS-1123
chr14-22540937-INS-2514
chr14-22590902-INS-57
chr14-22592541-INS-127
chr14-22610036-INS-52
chr14-22627288-INS-53
chr14-22633033-INS-55
chr14-22660530-INS-55
chr14-22666597-INS-114
chr14-22667117-INS-132
chr14-22670089-INS-52
chr14-22688419-INS-73
chr14-22695601-INS-400
chr14-22704667-INS-222
chr14-22709641-INS-57
chr14-22713004-INS-59
chr14-22747138-INS-111
chr14-22762845-INS-66
chr14-22818160-INS-453
chr14-22820789-INS-89
chr14-22855653-INS-65
chr14-22891354-INS-6917
chr14-22906346-INS-194
chr14-22939732-INS-185
chr14-22950207-INS-62
chr14-22954911-INS-298
chr14-22962730-INS-58

chr14-22979608-INS-617
chr14-22981160-INS-64
chr14-22988102-INS-95
chr14-22991487-INS-59
chr14-23037315-INS-178
chr14-23182989-INS-7123
chr14-23204678-INS-129
chr14-23214447-INS-390
chr14-23245602-INS-199
chr14-23253894-INS-6189
chr14-23277860-INS-312
chr14-23323688-INS-54
chr14-23333369-INS-52
chr14-23368613-INS-909
chr14-23376571-INS-76
chr14-23383305-INS-68
chr14-23394518-INS-551
chr14-23403706-INS-153
chr14-23410185-INS-113
chr14-23419498-INS-106
chr14-23434711-INS-77
chr14-23458708-INS-81
chr14-23561014-INS-103
chr14-23566649-INS-64
chr14-23571992-INS-70
chr14-23578332-INS-200
chr14-23622250-INS-68
chr14-23633652-INS-154
chr14-23719002-INS-1884
chr14-23728186-INS-223
chr14-23777626-INS-67
chr14-23829798-INS-7835
chr14-23901770-INS-164
chr14-23932063-INS-56
chr14-23962677-INS-128
chr14-23980537-INS-96
chr14-23996910-INS-60
chr14-24027707-INS-66
chr14-24057689-INS-58
chr14-24069390-INS-6233
chr14-24079987-INS-288
chr14-24121565-INS-220
chr14-24126358-INS-7215
chr14-24158794-INS-100
chr14-24218112-INS-62
chr14-24237077-INS-55
chr14-24254525-INS-66
chr14-24263048-INS-242
chr14-24322040-INS-85
chr14-24375431-INS-392
chr14-24405915-INS-134
chr14-24442605-INS-92
chr14-24453873-INS-141
chr14-24473634-INS-53
chr14-24486490-INS-96
chr14-24508152-INS-207
chr14-24529694-INS-303
chr14-24532226-INS-190
chr14-24570943-INS-133
chr14-24590683-INS-151
chr14-24630654-INS-598
chr14-24632482-INS-172
chr14-24658945-INS-147
chr14-24664933-INS-196
chr14-24674973-INS-143
chr14-24675413-INS-145
chr14-24688244-INS-67
chr14-24725341-INS-68
chr14-24744459-INS-81
chr14-24758937-INS-105
chr14-24892886-INS-268
chr14-24899158-INS-54
chr14-24902916-INS-72
chr14-24912996-INS-271
chr14-24921390-INS-354
chr14-24930084-INS-68
chr14-24932983-INS-386

chr14-24946674-INS-547
chr14-24958631-INS-122
chr14-24967960-INS-51
chr14-24988395-INS-83
chr14-24989397-INS-136
chr14-25043836-INS-155
chr14-25050647-INS-119
chr14-25077854-INS-173
chr14-25081950-INS-72
chr14-25118293-INS-52
chr14-25148500-INS-121
chr14-25153925-INS-118
chr14-25171188-INS-96
chr14-25184784-INS-82
chr14-25190092-INS-60
chr14-25198677-INS-158
chr14-25233119-INS-159
chr14-25307796-INS-51
chr14-25325791-INS-59
chr14-25356211-INS-196
chr14-25359867-INS-72
chr14-25380254-INS-271
chr14-25396252-INS-156
chr14-25421240-INS-109
chr14-25502344-INS-85
chr14-25515934-INS-54
chr14-25587360-INS-79
chr14-25601549-INS-56
chr14-25612076-INS-71
chr14-25635196-INS-121
chr14-25648763-INS-207
chr14-25685341-INS-62
chr14-25702964-INS-316
chr14-25744457-INS-350
chr14-25745468-INS-115
chr14-25762147-INS-298
chr14-25763718-INS-54
chr14-25791996-INS-281
chr14-25792518-INS-205
chr14-25834473-INS-288
chr14-25838658-INS-94
chr14-25890602-INS-327
chr14-25901607-INS-103
chr14-25908647-INS-218
chr14-25909965-INS-400
chr14-25921226-INS-202
chr14-25926990-INS-727
chr14-25932283-INS-159
chr14-25943261-INS-5280
chr14-25952262-INS-116
chr14-25971960-INS-81
chr14-25988362-INS-297
chr14-25993044-INS-53
chr14-26003607-INS-464
chr14-26005005-INS-324
chr14-26026613-INS-398
chr14-26119211-INS-207
chr14-26120041-INS-496
chr14-26147971-INS-330
chr14-26153869-INS-91
chr14-26167876-INS-155
chr14-26205788-INS-54
chr14-26208463-INS-225
chr14-26211275-INS-51
chr14-26232946-INS-52
chr14-26258701-INS-2754
chr14-26279900-INS-1887
chr14-26281226-INS-331
chr14-26289832-INS-124
chr14-26304545-INS-81
chr14-26307037-INS-60
chr14-26311756-INS-387
chr14-26314296-INS-60
chr14-26341197-INS-881
chr14-26387409-INS-211
chr14-26398408-INS-251
chr14-26408509-INS-1716

chr14-26408564-INS-265
chr14-26408564-INS-819
chr14-26408626-INS-2166
chr14-26408698-INS-2795
chr14-26408764-INS-2394
chr14-26433276-INS-885
chr14-26458372-INS-116
chr14-26461260-INS-108
chr14-26464146-INS-587
chr14-26472303-INS-266
chr14-26474455-INS-162
chr14-26493408-INS-145
chr14-26517762-INS-200
chr14-26518431-INS-57
chr14-26521689-INS-114
chr14-26532196-INS-61
chr14-26543313-INS-83
chr14-26570374-INS-638
chr14-26573846-INS-224
chr14-26587668-INS-168
chr14-26598996-INS-120
chr14-26609774-INS-60
chr14-26610344-INS-62
chr14-26628771-INS-2874
chr14-26629810-INS-53
chr14-26661635-INS-87
chr14-26667154-INS-216
chr14-26676100-INS-536
chr14-26694068-INS-5105
chr14-26709274-INS-84
chr14-26740111-INS-164
chr14-26768735-INS-199
chr14-26806173-INS-194
chr14-26808957-INS-400
chr14-26825568-INS-56
chr14-26830511-INS-86
chr14-26832966-INS-147
chr14-26849510-INS-200
chr14-26854808-INS-217
chr14-26893117-INS-83
chr14-26894487-INS-56
chr14-26935854-INS-193
chr14-26942868-INS-94
chr14-26947128-INS-328
chr14-26970493-INS-602
chr14-26977601-INS-179
chr14-26993708-INS-218
chr14-27004526-INS-395
chr14-27027313-INS-52
chr14-27043066-INS-400
chr14-27063028-INS-76
chr14-27094494-INS-533
chr14-27108908-INS-192
chr14-27123316-INS-224
chr14-27155392-INS-312
chr14-27225619-INS-57
chr14-27227116-INS-52
chr14-27231375-INS-54
chr14-27273997-INS-338
chr14-27310100-INS-399
chr14-27314625-INS-501
chr14-27319880-INS-328
chr14-27331780-INS-128
chr14-27353355-INS-111
chr14-27363764-INS-64
chr14-27381036-INS-157
chr14-27386678-INS-76
chr14-27387170-INS-270
chr14-27400165-INS-71
chr14-27414526-INS-109
chr14-27449143-INS-488
chr14-27460902-INS-95
chr14-27473372-INS-120
chr14-27482989-INS-118
chr14-27483449-INS-85
chr14-27505781-INS-79
chr14-27515344-INS-985

chr14-27529408-INS-352
chr14-27537149-INS-147
chr14-27562749-INS-284
chr14-27578519-INS-98
chr14-27583499-INS-196
chr14-27600052-INS-78
chr14-27608001-INS-1867
chr14-27609264-INS-52
chr14-27637404-INS-6170
chr14-27640688-INS-1181
chr14-27672072-INS-329
chr14-27721098-INS-328
chr14-27731909-INS-135
chr14-27737194-INS-185
chr14-27753957-INS-51
chr14-27768834-INS-180
chr14-27777106-INS-382
chr14-27793912-INS-96
chr14-27797389-INS-75
chr14-27797997-INS-66
chr14-27858520-INS-179
chr14-27883946-INS-257
chr14-27910952-INS-206
chr14-27935188-INS-66
chr14-27952648-INS-376
chr14-27962309-INS-76
chr14-27978029-INS-168
chr14-27981534-INS-62
chr14-27999880-INS-143
chr14-28029566-INS-52
chr14-28070676-INS-122
chr14-28081326-INS-51
chr14-28142945-INS-498
chr14-28152402-INS-67
chr14-28174669-INS-339
chr14-28196946-INS-155
chr14-28200608-INS-1888
chr14-28211088-INS-107
chr14-28238621-INS-216
chr14-28269022-INS-412
chr14-28292163-INS-344
chr14-28296822-INS-84
chr14-28299007-INS-55
chr14-28300864-INS-6045
chr14-28311244-INS-1893
chr14-28335959-INS-89
chr14-28374828-INS-476
chr14-28409525-INS-68
chr14-28419441-INS-182
chr14-28462513-INS-138
chr14-28464032-INS-416
chr14-28505346-INS-70
chr14-28508745-INS-200
chr14-28522980-INS-177
chr14-28539961-INS-160
chr14-28561870-INS-54
chr14-28578033-INS-138
chr14-28594861-INS-131
chr14-28635549-INS-51
chr14-28653596-INS-70
chr14-28654188-INS-115
chr14-28662159-INS-693
chr14-28687118-INS-58
chr14-28693559-INS-168
chr14-28719283-INS-966
chr14-28722625-INS-259
chr14-28739725-INS-356
chr14-28753176-INS-116
chr14-28800290-INS-60
chr14-28825897-INS-103
chr14-28831918-INS-67
chr14-28868519-INS-830
chr14-28883483-INS-74
chr14-28884159-INS-6167
chr14-28933554-INS-88
chr14-28955787-INS-68
chr14-28980299-INS-2884

chr14-28981839-INS-5555
chr14-28985797-INS-62
chr14-29043645-INS-77
chr14-29070686-INS-62
chr14-29084827-INS-56
chr14-29099757-INS-78
chr14-29102402-INS-406
chr14-29149415-INS-56
chr14-29202140-INS-1296
chr14-29228529-INS-437
chr14-29249917-INS-251
chr14-29284672-INS-76
chr14-29306313-INS-214
chr14-29336508-INS-89
chr14-29352868-INS-314
chr14-29358110-INS-160
chr14-29360931-INS-217
chr14-29389635-INS-475
chr14-29401835-INS-75
chr14-29423462-INS-63
chr14-29456812-INS-439
chr14-29464012-INS-229
chr14-29584094-INS-77
chr14-29605327-INS-358
chr14-29612583-INS-142
chr14-29633047-INS-57
chr14-29639561-INS-443
chr14-29656739-INS-2432
chr14-29661259-INS-72
chr14-29680156-INS-300
chr14-29709322-INS-201
chr14-29749388-INS-307
chr14-29776378-INS-66
chr14-29781833-INS-140
chr14-29783834-INS-240
chr14-29812794-INS-84
chr14-29867284-INS-192
chr14-29867765-INS-6442
chr14-29867921-INS-159
chr14-29882816-INS-116
chr14-29891778-INS-281
chr14-29898691-INS-310
chr14-29925188-INS-58
chr14-29959398-INS-54
chr14-29979111-INS-78
chr14-29989091-INS-74
chr14-30002814-INS-348
chr14-30010040-INS-65
chr14-30015313-INS-60
chr14-30050264-INS-92
chr14-30078989-INS-68
chr14-30103713-INS-182
chr14-30104705-INS-102
chr14-30112291-INS-74
chr14-30113224-INS-57
chr14-30114744-INS-143
chr14-30115359-INS-102
chr14-30124492-INS-110
chr14-30136992-INS-441
chr14-30156968-INS-660
chr14-30160508-INS-136
chr14-30168601-INS-221
chr14-30173860-INS-128
chr14-30243037-INS-313
chr14-30261602-INS-116
chr14-30263285-INS-92
chr14-30268095-INS-183
chr14-30290693-INS-233
chr14-30296584-INS-209
chr14-30304234-INS-163
chr14-30304574-INS-85
chr14-30308059-INS-169
chr14-30375599-INS-531
chr14-30385607-INS-456
chr14-30411409-INS-115
chr14-30423551-INS-266
chr14-30430273-INS-204

chr14-30433620-INS-186
chr14-30484142-INS-56
chr14-30490701-INS-58
chr14-30514338-INS-116
chr14-30571871-INS-199
chr14-30582699-INS-253
chr14-30607239-INS-52
chr14-30610675-INS-53
chr14-30615528-INS-210
chr14-30621063-INS-212
chr14-30628675-INS-258
chr14-30654138-INS-5543
chr14-30681697-INS-193
chr14-30685129-INS-85
chr14-30686466-INS-200
chr14-30693959-INS-453
chr14-30713459-INS-297
chr14-30717643-INS-192
chr14-30720166-INS-207
chr14-30721135-INS-127
chr14-30727144-INS-72
chr14-30771573-INS-63
chr14-30837809-INS-199
chr14-30844719-INS-165
chr14-30849914-INS-96
chr14-30851421-INS-200
chr14-30897555-INS-55
chr14-30919648-INS-198
chr14-30926090-INS-128
chr14-30967724-INS-336
chr14-31002858-INS-195
chr14-31025144-INS-206
chr14-31030556-INS-54
chr14-31035060-INS-6204
chr14-31066307-INS-165
chr14-31071188-INS-449
chr14-31074107-INS-208
chr14-31092852-INS-240
chr14-31108801-INS-84
chr14-31170024-INS-5438
chr14-31186292-INS-537
chr14-31195147-INS-153
chr14-31205475-INS-212
chr14-31240028-INS-400
chr14-31248808-INS-168
chr14-31264785-INS-81
chr14-31272813-INS-109
chr14-31275137-INS-60
chr14-31292277-INS-95
chr14-31296952-INS-6405
chr14-31299344-INS-53
chr14-31312258-INS-230
chr14-31334206-INS-239
chr14-31336325-INS-432
chr14-31369035-INS-98
chr14-31370388-INS-106
chr14-31371301-INS-198
chr14-31380918-INS-278
chr14-31396788-INS-208
chr14-31404833-INS-223
chr14-31414749-INS-288
chr14-31420145-INS-419
chr14-31431755-INS-403
chr14-31445258-INS-187
chr14-31458254-INS-144
chr14-31482749-INS-668
chr14-31529496-INS-233
chr14-31543913-INS-486
chr14-31589546-INS-197
chr14-31592787-INS-561
chr14-31595827-INS-249
chr14-31597364-INS-52
chr14-31606763-INS-89
chr14-31617461-INS-75
chr14-31635077-INS-89
chr14-31668032-INS-99
chr14-31674796-INS-274

chr14-31681747-INS-55
chr14-31692734-INS-169
chr14-31696103-INS-522
chr14-31705305-INS-53
chr14-31717080-INS-979
chr14-31719125-INS-64
chr14-31733760-INS-687
chr14-31744725-INS-381
chr14-31762647-INS-76
chr14-31768755-INS-405
chr14-31785243-INS-60
chr14-31789877-INS-327
chr14-31793392-INS-390
chr14-31801658-INS-310
chr14-31802338-INS-87
chr14-31812511-INS-120
chr14-31814895-INS-377
chr14-31816774-INS-210
chr14-31818983-INS-211
chr14-31829169-INS-114
chr14-31836312-INS-202
chr14-31844867-INS-159
chr14-31853521-INS-100
chr14-31855293-INS-75
chr14-31865094-INS-61
chr14-31866854-INS-208
chr14-31876829-INS-126
chr14-31899196-INS-261
chr14-31902870-INS-204
chr14-31905302-INS-193
chr14-31909227-INS-59
chr14-31917202-INS-153
chr14-31920138-INS-51
chr14-31924632-INS-5577
chr14-31926047-INS-291
chr14-31939059-INS-92
chr14-31945419-INS-446
chr14-31950993-INS-315
chr14-31963111-INS-170
chr14-31985141-INS-1864
chr14-32017671-INS-64
chr14-32025905-INS-224
chr14-32037525-INS-774
chr14-32070746-INS-124
chr14-32073426-INS-59
chr14-32082338-INS-83
chr14-32084705-INS-150
chr14-32089861-INS-193
chr14-32090947-INS-982
chr14-32124572-INS-391
chr14-32128674-INS-148
chr14-32129083-INS-107
chr14-32172609-INS-60
chr14-32201253-INS-176
chr14-32205603-INS-179
chr14-32214242-INS-448
chr14-32216177-INS-5635
chr14-32222215-INS-209
chr14-32234046-INS-193
chr14-32245085-INS-6440
chr14-32250564-INS-182
chr14-32280089-INS-492
chr14-32296604-INS-60
chr14-32314483-INS-58
chr14-32326768-INS-143
chr14-32331493-INS-57
chr14-32340110-INS-76
chr14-32340544-INS-767
chr14-32364474-INS-88
chr14-32365962-INS-76
chr14-32369261-INS-121
chr14-32380772-INS-6442
chr14-32387789-INS-65
chr14-32432295-INS-97
chr14-32470055-INS-310
chr14-32481336-INS-399
chr14-32485639-INS-216

chr14-32491855-INS-62
chr14-32494133-INS-726
chr14-32497231-INS-107
chr14-32497657-INS-831
chr14-32565941-INS-75
chr14-32604801-INS-6649
chr14-32629544-INS-60
chr14-32644080-INS-70
chr14-32655074-INS-2346
chr14-32682045-INS-254
chr14-32752775-INS-192
chr14-32770412-INS-100
chr14-32771747-INS-160
chr14-32772609-INS-356
chr14-32799453-INS-56
chr14-32807450-INS-117
chr14-32808198-INS-400
chr14-32813400-INS-176
chr14-32828355-INS-64
chr14-32833207-INS-116
chr14-32843015-INS-68
chr14-32879549-INS-114
chr14-32881242-INS-51
chr14-32898567-INS-117
chr14-32902992-INS-181
chr14-32913608-INS-235
chr14-32916720-INS-144
chr14-32932247-INS-89
chr14-32963269-INS-295
chr14-32965997-INS-114
chr14-32981019-INS-58
chr14-32983702-INS-138
chr14-32998800-INS-72
chr14-33009328-INS-113
chr14-33021775-INS-56
chr14-33034995-INS-222
chr14-33061775-INS-54
chr14-33076688-INS-97
chr14-33081579-INS-2877
chr14-33086126-INS-137
chr14-33096809-INS-76
chr14-33153312-INS-7419
chr14-33167412-INS-7055
chr14-33187970-INS-66
chr14-33196803-INS-64
chr14-33201494-INS-58
chr14-33216094-INS-62
chr14-33219799-INS-67
chr14-33274828-INS-66
chr14-33278743-INS-386
chr14-33282280-INS-105
chr14-33289637-INS-54
chr14-33302668-INS-479
chr14-33313406-INS-93
chr14-33321939-INS-88
chr14-33333375-INS-159
chr14-33349672-INS-201
chr14-33404665-INS-63
chr14-33411556-INS-90
chr14-33457060-INS-59
chr14-33460791-INS-54
chr14-33466259-INS-54
chr14-33505072-INS-67
chr14-33525437-INS-280
chr14-33528659-INS-60
chr14-33540669-INS-65
chr14-33545592-INS-60
chr14-33573765-INS-52
chr14-33585634-INS-358
chr14-33594411-INS-89
chr14-33606687-INS-192
chr14-33626843-INS-184
chr14-33641147-INS-199
chr14-33656466-INS-64
chr14-33659319-INS-276
chr14-33661894-INS-189
chr14-33672063-INS-121

chr14-33701193-INS-54
chr14-33755216-INS-145
chr14-33764362-INS-127
chr14-33786556-INS-60
chr14-33803149-INS-52
chr14-33807259-INS-214
chr14-33830845-INS-140
chr14-33862028-INS-7138
chr14-33901046-INS-918
chr14-33921447-INS-215
chr14-33932686-INS-218
chr14-33936250-INS-396
chr14-33955773-INS-184
chr14-33967827-INS-204
chr14-33971902-INS-86
chr14-33974277-INS-68
chr14-33998662-INS-53
chr14-33999603-INS-86
chr14-34024540-INS-209
chr14-34062739-INS-182
chr14-34076433-INS-209
chr14-34091052-INS-189
chr14-34102354-INS-66
chr14-34122916-INS-84
chr14-34142659-INS-118
chr14-34170421-INS-88
chr14-34172904-INS-199
chr14-34179253-INS-72
chr14-34183664-INS-146
chr14-34188738-INS-211
chr14-34208886-INS-60
chr14-34216526-INS-425
chr14-34222470-INS-60
chr14-34236928-INS-132
chr14-34254941-INS-243
chr14-34273160-INS-159
chr14-34291220-INS-100
chr14-34302451-INS-168
chr14-34323358-INS-388
chr14-34328931-INS-282
chr14-34343774-INS-694
chr14-34351356-INS-200
chr14-34368132-INS-109
chr14-34382181-INS-117
chr14-34400053-INS-58
chr14-34416773-INS-199
chr14-34435119-INS-3183
chr14-34437489-INS-69
chr14-34454973-INS-580
chr14-34456249-INS-197
chr14-34480457-INS-198
chr14-34488001-INS-192
chr14-34532430-INS-72
chr14-34562614-INS-156
chr14-34574671-INS-359
chr14-34587989-INS-55
chr14-34600895-INS-99
chr14-34603583-INS-190
chr14-34621556-INS-58
chr14-34634946-INS-380
chr14-34658316-INS-614
chr14-34694092-INS-82
chr14-34697770-INS-224
chr14-34700778-INS-62
chr14-34761757-INS-5569
chr14-34779599-INS-115
chr14-34783253-INS-7136
chr14-34796011-INS-52
chr14-34800419-INS-80
chr14-34825564-INS-84
chr14-34836680-INS-268
chr14-34843180-INS-58
chr14-34855076-INS-100
chr14-34916894-INS-70
chr14-34920274-INS-196
chr14-34952893-INS-2186
chr14-34954940-INS-1215

chr14-34957577-INS-455
chr14-34964837-INS-56
chr14-35009483-INS-6617
chr14-35023276-INS-55
chr14-35068732-INS-275
chr14-35090402-INS-64
chr14-35108678-INS-1894
chr14-35109636-INS-2590
chr14-35135791-INS-8812
chr14-35143716-INS-774
chr14-35162867-INS-7247
chr14-35176040-INS-57
chr14-35201908-INS-6937
chr14-35217161-INS-261
chr14-35228089-INS-52
chr14-35236221-INS-62
chr14-35241748-INS-506
chr14-35271192-INS-88
chr14-35288173-INS-55
chr14-35310497-INS-50
chr14-35320519-INS-210
chr14-35347969-INS-82
chr14-35355382-INS-62
chr14-35369915-INS-210
chr14-35425652-INS-75
chr14-35437816-INS-62
chr14-35448523-INS-114
chr14-35454254-INS-387
chr14-35456114-INS-3030
chr14-35456482-INS-81
chr14-35457919-INS-185
chr14-35460959-INS-1864
chr14-35499848-INS-172
chr14-35502016-INS-62
chr14-35566740-INS-787
chr14-35592899-INS-408
chr14-35631570-INS-341
chr14-35634953-INS-72
chr14-35651754-INS-52
chr14-35683444-INS-68
chr14-35687534-INS-102
chr14-35708414-INS-100
chr14-35710612-INS-759
chr14-35731295-INS-64
chr14-35736572-INS-64
chr14-35767252-INS-7089
chr14-35828124-INS-503
chr14-35870406-INS-471
chr14-35907277-INS-257
chr14-35918334-INS-4708
chr14-35926544-INS-51
chr14-35942165-INS-179
chr14-35962348-INS-1561
chr14-35978455-INS-6566
chr14-36000937-INS-164
chr14-36019562-INS-215
chr14-36035241-INS-106
chr14-36040139-INS-6790
chr14-36085604-INS-947
chr14-36178068-INS-219
chr14-36183519-INS-177
chr14-36183963-INS-470
chr14-36188511-INS-6645
chr14-36215609-INS-6282
chr14-36216240-INS-1885
chr14-36226366-INS-884
chr14-36241918-INS-206
chr14-36261161-INS-75
chr14-36283549-INS-224
chr14-36283885-INS-84
chr14-36299004-INS-153
chr14-36301354-INS-59
chr14-36342461-INS-6554
chr14-36346955-INS-4144
chr14-36369722-INS-531
chr14-36427184-INS-479
chr14-36444469-INS-64

chr14-36452293-INS-54
chr14-36455376-INS-216
chr14-36465502-INS-355
chr14-36477945-INS-219
chr14-36498676-INS-59
chr14-36513896-INS-76
chr14-36544647-INS-72
chr14-36546676-INS-276
chr14-36551077-INS-206
chr14-36583182-INS-124
chr14-36603405-INS-153
chr14-36625521-INS-54
chr14-36627491-INS-70
chr14-36632795-INS-384
chr14-36641168-INS-88
chr14-36644940-INS-130
chr14-36655852-INS-119
chr14-36655885-INS-528
chr14-36666925-INS-85
chr14-36699866-INS-7024
chr14-36714219-INS-99
chr14-36721872-INS-2948
chr14-36730372-INS-681
chr14-36730764-INS-5541
chr14-36745791-INS-60
chr14-36747453-INS-155
chr14-36748368-INS-146
chr14-36795820-INS-867
chr14-36850483-INS-158
chr14-36866009-INS-59
chr14-36888759-INS-59
chr14-36908780-INS-524
chr14-36919713-INS-212
chr14-37065463-INS-461
chr14-37105715-INS-100
chr14-37151192-INS-82
chr14-37173970-INS-86
chr14-37178475-INS-51
chr14-37184861-INS-6809
chr14-37200961-INS-108
chr14-37207168-INS-52
chr14-37256397-INS-903
chr14-37267757-INS-102
chr14-37282934-INS-433
chr14-37304601-INS-6529
chr14-37308095-INS-551
chr14-37322470-INS-6187
chr14-37326107-INS-97
chr14-37328378-INS-6990
chr14-37368211-INS-6412
chr14-37371194-INS-158
chr14-37385264-INS-400
chr14-37397235-INS-51
chr14-37401105-INS-112
chr14-37423465-INS-60
chr14-37433276-INS-123
chr14-37471087-INS-6403
chr14-37498451-INS-520
chr14-37517098-INS-52
chr14-37517267-INS-6672
chr14-37544121-INS-54
chr14-37577842-INS-60
chr14-37641668-INS-173
chr14-37677812-INS-453
chr14-37686975-INS-341
chr14-37708232-INS-120
chr14-37714205-INS-57
chr14-37716942-INS-6331
chr14-37727126-INS-710
chr14-37784720-INS-194
chr14-37800626-INS-237
chr14-37839718-INS-4084
chr14-37849879-INS-127
chr14-37863586-INS-66
chr14-37941828-INS-344
chr14-37952213-INS-555
chr14-37958487-INS-1399

chr14-37963435-INS-169
chr14-37965100-INS-7183
chr14-37971265-INS-6399
chr14-37974413-INS-6545
chr14-38024485-INS-130
chr14-38058066-INS-268
chr14-38061597-INS-591
chr14-38075770-INS-192
chr14-38082509-INS-6380
chr14-38091720-INS-106
chr14-38141522-INS-358
chr14-38155669-INS-110
chr14-38160878-INS-4495
chr14-38161632-INS-70
chr14-38212265-INS-167
chr14-38222376-INS-296
chr14-38235268-INS-72
chr14-38236578-INS-98
chr14-38243474-INS-351
chr14-38253490-INS-1774
chr14-38264662-INS-412
chr14-38270851-INS-182
chr14-38276118-INS-88
chr14-38282871-INS-893
chr14-38312674-INS-348
chr14-38333877-INS-51
chr14-38356596-INS-164
chr14-38358173-INS-69
chr14-38360767-INS-77
chr14-38396066-INS-231
chr14-38443897-INS-417
chr14-38453554-INS-69
chr14-38470505-INS-88
chr14-38483293-INS-86
chr14-38523176-INS-789
chr14-38526248-INS-681
chr14-38566181-INS-677
chr14-38582071-INS-52
chr14-38595505-INS-80
chr14-38613141-INS-194
chr14-38617328-INS-1223
chr14-38621235-INS-155
chr14-38660411-INS-201
chr14-38693546-INS-103
chr14-38723639-INS-138
chr14-38736955-INS-975
chr14-38737765-INS-51
chr14-38740610-INS-650
chr14-38749006-INS-251
chr14-38750457-INS-112
chr14-38760622-INS-94
chr14-38767717-INS-52
chr14-38780672-INS-112
chr14-38781053-INS-160
chr14-38785682-INS-475
chr14-38809617-INS-7302
chr14-38831115-INS-68
chr14-38844476-INS-231
chr14-38878260-INS-276
chr14-38894257-INS-59
chr14-38936698-INS-58
chr14-38939728-INS-130
chr14-38959148-INS-184
chr14-38961731-INS-56
chr14-38964720-INS-58
chr14-38967099-INS-85
chr14-38971416-INS-524
chr14-38981007-INS-684
chr14-39056565-INS-98
chr14-39062897-INS-400
chr14-39068909-INS-309
chr14-39073546-INS-116
chr14-39086835-INS-53
chr14-39089100-INS-69
chr14-39102491-INS-109
chr14-39195800-INS-58
chr14-39198081-INS-6044

chr14-39203378-INS-665
chr14-39241999-INS-78
chr14-39244520-INS-487
chr14-39257653-INS-270
chr14-39260884-INS-235
chr14-39285202-INS-150
chr14-39360302-INS-329
chr14-39366049-INS-871
chr14-39367969-INS-400
chr14-39382007-INS-2550
chr14-39413765-INS-6419
chr14-39418847-INS-6394
chr14-39418943-INS-741
chr14-39421841-INS-61
chr14-39471354-INS-6599
chr14-39497030-INS-117
chr14-39509919-INS-52
chr14-39559790-INS-78
chr14-39583667-INS-126
chr14-39596919-INS-301
chr14-39608915-INS-132
chr14-39619389-INS-260
chr14-39694231-INS-283
chr14-39695427-INS-139
chr14-39748740-INS-96
chr14-39784128-INS-488
chr14-39797002-INS-60
chr14-39819229-INS-424
chr14-39828151-INS-6435
chr14-39831691-INS-258
chr14-39894736-INS-358
chr14-39914804-INS-72
chr14-39977261-INS-69
chr14-39987145-INS-286
chr14-39994473-INS-212
chr14-40053777-INS-6684
chr14-40084841-INS-66
chr14-40086934-INS-65
chr14-40091434-INS-133
chr14-40091600-INS-7659
chr14-40095143-INS-61
chr14-40096787-INS-245
chr14-40110318-INS-115
chr14-40114694-INS-358
chr14-40138883-INS-269
chr14-40144143-INS-900
chr14-40182113-INS-519
chr14-40226030-INS-6633
chr14-40265853-INS-86
chr14-40286242-INS-4745
chr14-40286575-INS-72
chr14-40305300-INS-371
chr14-40311930-INS-217
chr14-40318391-INS-212
chr14-40343657-INS-322
chr14-40363504-INS-6327
chr14-40395730-INS-208
chr14-40401257-INS-52
chr14-40406877-INS-65
chr14-40410022-INS-84
chr14-40433750-INS-57
chr14-40490313-INS-8606
chr14-40532532-INS-66
chr14-40551455-INS-60
chr14-40575827-INS-256
chr14-40583117-INS-401
chr14-40602758-INS-193
chr14-40605928-INS-207
chr14-40608335-INS-212
chr14-40634398-INS-62
chr14-40634491-INS-941
chr14-40733144-INS-211
chr14-40734814-INS-82
chr14-40747542-INS-191
chr14-40754147-INS-183
chr14-40792234-INS-166
chr14-40803565-INS-93

chr14-40807154-INS-63
chr14-40810744-INS-154
chr14-40816658-INS-86
chr14-40830971-INS-157
chr14-40858294-INS-211
chr14-40886162-INS-142
chr14-40893671-INS-76
chr14-40899473-INS-110
chr14-40904493-INS-155
chr14-40910703-INS-273
chr14-40913963-INS-58
chr14-40936283-INS-144
chr14-40960331-INS-7313
chr14-41044449-INS-64
chr14-41067596-INS-71
chr14-41143716-INS-109
chr14-41207945-INS-156
chr14-41229763-INS-294
chr14-41259086-INS-58
chr14-41264496-INS-79
chr14-41329911-INS-52
chr14-41439475-INS-4758
chr14-41490916-INS-154
chr14-41772220-INS-88
chr14-41801614-INS-72
chr14-42069064-INS-56
chr14-42089827-INS-55
chr14-42232103-INS-171
chr14-42266428-INS-188
chr14-42548924-INS-172
chr14-42556520-INS-79
chr14-42557891-INS-502
chr14-42674041-INS-363
chr14-42680205-INS-188
chr14-42906939-INS-63
chr14-42907932-INS-502
chr14-42928217-INS-560
chr14-42968727-INS-511
chr14-43068459-INS-78
chr14-43073222-INS-76
chr14-43075919-INS-424
chr14-43167719-INS-6250
chr14-43766283-INS-100
chr14-43842004-INS-126
chr14-43916341-INS-139
chr14-43918231-INS-5313
chr14-43985188-INS-89
chr14-44009928-INS-1580
chr14-44032616-INS-5065
chr14-44069516-INS-68
chr14-44073706-INS-521
chr14-44276238-INS-383
chr14-44537036-INS-145
chr14-44805093-INS-6401
chr14-44830798-INS-1495
chr14-44865885-INS-441
chr14-44985152-INS-454
chr14-45023993-INS-401
chr14-45031811-INS-76
chr14-45068223-INS-216
chr14-45072572-INS-612
chr14-45075095-INS-93
chr14-45088945-INS-7167
chr14-45093926-INS-55
chr14-45117570-INS-90
chr14-45145182-INS-116
chr14-45163709-INS-418
chr14-45232507-INS-61
chr14-45236959-INS-6131
chr14-45240799-INS-69
chr14-45250017-INS-166
chr14-45257204-INS-206
chr14-45271823-INS-57
chr14-45297323-INS-180
chr14-45343367-INS-70
chr14-45359076-INS-190
chr14-45392873-INS-94

chr14-45404265-INS-909
chr14-45414685-INS-68
chr14-45431667-INS-116
chr14-45478517-INS-821
chr14-45532697-INS-120
chr14-45536627-INS-183
chr14-45552329-INS-284
chr14-45555271-INS-2870
chr14-45560777-INS-404
chr14-45562261-INS-386
chr14-45565254-INS-206
chr14-45573904-INS-54
chr14-45585621-INS-65
chr14-45587198-INS-106
chr14-45590083-INS-91
chr14-45597551-INS-596
chr14-45630316-INS-491
chr14-45633758-INS-65
chr14-45649231-INS-500
chr14-45653659-INS-2286
chr14-45685827-INS-499
chr14-45689145-INS-56
chr14-45704354-INS-65
chr14-45721061-INS-65
chr14-45773030-INS-199
chr14-45774244-INS-66
chr14-45841536-INS-56
chr14-45846653-INS-144
chr14-45890757-INS-196
chr14-45906783-INS-202
chr14-45910491-INS-69
chr14-45924564-INS-270
chr14-45930840-INS-822
chr14-45934884-INS-78
chr14-45938141-INS-100
chr14-45949230-INS-413
chr14-45969659-INS-487
chr14-45996178-INS-190
chr14-46007352-INS-179
chr14-46008297-INS-116
chr14-46009118-INS-496
chr14-46024728-INS-79
chr14-46053473-INS-233
chr14-46056596-INS-553
chr14-46110121-INS-52
chr14-46135923-INS-751
chr14-46150374-INS-399
chr14-46170397-INS-880
chr14-46173674-INS-6401
chr14-46177573-INS-101
chr14-46178108-INS-6308
chr14-46194742-INS-992
chr14-46195004-INS-75
chr14-46207113-INS-990
chr14-46231944-INS-6643
chr14-46238388-INS-108
chr14-46241307-INS-339
chr14-46273944-INS-68
chr14-46298216-INS-94
chr14-46300683-INS-52
chr14-46330574-INS-119
chr14-46347192-INS-52
chr14-46351037-INS-202
chr14-46353122-INS-265
chr14-46409445-INS-70
chr14-46493377-INS-51
chr14-46498906-INS-159
chr14-46522491-INS-139
chr14-46531194-INS-6994
chr14-46532811-INS-490
chr14-46536015-INS-393
chr14-46545949-INS-229
chr14-46547793-INS-354
chr14-46549279-INS-53
chr14-46599610-INS-184
chr14-46663548-INS-169
chr14-46665013-INS-209

chr14-46672305-INS-445
chr14-46678489-INS-202
chr14-46691041-INS-72
chr14-46696770-INS-447
chr14-46702717-INS-2842
chr14-46707315-INS-184
chr14-46739730-INS-90
chr14-46791258-INS-404
chr14-46794045-INS-54
chr14-46851097-INS-156
chr14-46853426-INS-102
chr14-46874615-INS-51
chr14-46886113-INS-402
chr14-46894468-INS-60
chr14-46926937-INS-152
chr14-46928384-INS-84
chr14-46931931-INS-60
chr14-46971246-INS-73
chr14-46976218-INS-199
chr14-46992993-INS-201
chr14-47024133-INS-620
chr14-47038256-INS-405
chr14-47039911-INS-153
chr14-47049353-INS-99
chr14-47067794-INS-184
chr14-47092252-INS-104
chr14-47109194-INS-132
chr14-47137122-INS-148
chr14-47144321-INS-399
chr14-47156928-INS-424
chr14-47160743-INS-123
chr14-47175027-INS-112
chr14-47198202-INS-229
chr14-47201408-INS-73
chr14-47202333-INS-66
chr14-47223359-INS-56
chr14-47258078-INS-58
chr14-47270908-INS-498
chr14-47276209-INS-80
chr14-47323348-INS-465
chr14-47351616-INS-58
chr14-47354785-INS-140
chr14-47355806-INS-62
chr14-47385451-INS-279
chr14-47393472-INS-491
chr14-47411879-INS-55
chr14-47412845-INS-94
chr14-47457950-INS-192
chr14-47466350-INS-137
chr14-47480562-INS-175
chr14-47510843-INS-51
chr14-47533031-INS-263
chr14-47538784-INS-159
chr14-47547665-INS-160
chr14-47550830-INS-244
chr14-47552591-INS-57
chr14-47556424-INS-148
chr14-47564018-INS-74
chr14-47574126-INS-202
chr14-47577637-INS-5635
chr14-47624610-INS-652
chr14-47656922-INS-192
chr14-47674864-INS-201
chr14-47688800-INS-93
chr14-47714578-INS-142
chr14-47770114-INS-173
chr14-47791306-INS-51
chr14-47812360-INS-66
chr14-47828984-INS-72
chr14-47837082-INS-289
chr14-47849579-INS-82
chr14-47867176-INS-200
chr14-47882937-INS-79
chr14-47890205-INS-132
chr14-47925372-INS-99
chr14-47959512-INS-801
chr14-47974917-INS-214

chr14-47984563-INS-280
chr14-48032051-INS-470
chr14-48042756-INS-279
chr14-48045038-INS-204
chr14-48055302-INS-51
chr14-48072457-INS-159
chr14-48074461-INS-67
chr14-48118438-INS-968
chr14-48121555-INS-227
chr14-48126430-INS-130
chr14-48128495-INS-60
chr14-48145059-INS-58
chr14-48153061-INS-138
chr14-48167319-INS-63
chr14-48184680-INS-52
chr14-48201364-INS-165
chr14-48250579-INS-61
chr14-48282022-INS-181
chr14-48284788-INS-124
chr14-48288594-INS-298
chr14-48301383-INS-68
chr14-48321262-INS-120
chr14-48322999-INS-68
chr14-48341462-INS-227
chr14-48364204-INS-280
chr14-48372907-INS-736
chr14-48377498-INS-66
chr14-48424387-INS-60
chr14-48433910-INS-158
chr14-48481816-INS-54
chr14-48504524-INS-128
chr14-48541167-INS-130
chr14-48557894-INS-60
chr14-48568772-INS-76
chr14-48604074-INS-491
chr14-48653127-INS-182
chr14-48664375-INS-116
chr14-48687355-INS-141
chr14-48735398-INS-84
chr14-48739029-INS-402
chr14-48754949-INS-156
chr14-48756014-INS-222
chr14-48784064-INS-63
chr14-48815726-INS-70
chr14-48817748-INS-85
chr14-48818929-INS-56
chr14-48822686-INS-60
chr14-48833098-INS-65
chr14-48836541-INS-57
chr14-48875350-INS-62
chr14-48878340-INS-90
chr14-48922030-INS-118
chr14-48931460-INS-218
chr14-48943596-INS-84
chr14-48985312-INS-88
chr14-48986378-INS-55
chr14-49019682-INS-209
chr14-49045765-INS-81
chr14-49079407-INS-54
chr14-49094569-INS-1740
chr14-49094787-INS-1350
chr14-49094787-INS-538
chr14-49120722-INS-438
chr14-49122455-INS-1508
chr14-49144224-INS-52
chr14-49167722-INS-73
chr14-49174626-INS-7514
chr14-49176634-INS-400
chr14-49186915-INS-65
chr14-49214551-INS-76
chr14-49216113-INS-6150
chr14-49245870-INS-162
chr14-49247026-INS-208
chr14-49262580-INS-395
chr14-49290998-INS-198
chr14-49317953-INS-175
chr14-49362094-INS-72

chr14-49394766-INS-120
chr14-49400407-INS-171
chr14-49443412-INS-101
chr14-49451727-INS-7430
chr14-49453056-INS-774
chr14-49453179-INS-1368
chr14-49453208-INS-1076
chr14-49467409-INS-54
chr14-49471711-INS-612
chr14-49498168-INS-134
chr14-49544847-INS-153
chr14-49560223-INS-6015
chr14-49564916-INS-6744
chr14-49566305-INS-73
chr14-49572938-INS-498
chr14-49578671-INS-78
chr14-49580672-INS-3254
chr14-49594572-INS-98
chr14-49608528-INS-62
chr14-49609026-INS-86
chr14-49625558-INS-570
chr14-49629794-INS-491
chr14-49634031-INS-141
chr14-49651431-INS-56
chr14-49673729-INS-64
chr14-49679751-INS-6396
chr14-49722001-INS-88
chr14-49762334-INS-58
chr14-49763906-INS-343
chr14-49819707-INS-400
chr14-49820703-INS-71
chr14-49829759-INS-54
chr14-49837558-INS-56
chr14-49864419-INS-6792
chr14-49885849-INS-530
chr14-49890423-INS-404
chr14-49902293-INS-111
chr14-49905073-INS-211
chr14-49979771-INS-443
chr14-50000161-INS-673
chr14-50004428-INS-79
chr14-50026610-INS-356
chr14-50030727-INS-70
chr14-50064376-INS-69
chr14-50098315-INS-72
chr14-50126461-INS-594
chr14-50171277-INS-55
chr14-50171909-INS-270
chr14-50185564-INS-319
chr14-50189787-INS-5226
chr14-50190371-INS-1719
chr14-50192010-INS-104
chr14-50247273-INS-439
chr14-50262320-INS-4980
chr14-50276138-INS-55
chr14-50283359-INS-65
chr14-50287657-INS-250
chr14-50335583-INS-7324
chr14-50345591-INS-1770
chr14-50349191-INS-142
chr14-50349606-INS-7022
chr14-50366325-INS-120
chr14-50387793-INS-247
chr14-50398468-INS-145
chr14-50404114-INS-52
chr14-50405010-INS-73
chr14-50405834-INS-296
chr14-50421901-INS-463
chr14-50438483-INS-4372
chr14-50466193-INS-149
chr14-50470103-INS-207
chr14-50569336-INS-452
chr14-50570935-INS-120
chr14-50579552-INS-195
chr14-50600974-INS-434
chr14-50605070-INS-6206
chr14-50613720-INS-1400

chr14-50642077-INS-596
chr14-50648482-INS-52
chr14-50670023-INS-7318
chr14-50696675-INS-5941
chr14-50705914-INS-193
chr14-50708302-INS-362
chr14-50743953-INS-84
chr14-50755332-INS-6228
chr14-50756292-INS-243
chr14-50763139-INS-846
chr14-50766304-INS-94
chr14-50784763-INS-95
chr14-50789479-INS-791
chr14-50795613-INS-64
chr14-50814815-INS-68
chr14-50818397-INS-405
chr14-50828073-INS-138
chr14-50857462-INS-402
chr14-50864471-INS-1089
chr14-50879009-INS-866
chr14-50895304-INS-432
chr14-50972627-INS-210
chr14-50978515-INS-52
chr14-50994107-INS-73
chr14-51009114-INS-6410
chr14-51011381-INS-101
chr14-51038086-INS-61
chr14-51040646-INS-57
chr14-51062692-INS-56
chr14-51072629-INS-169
chr14-51114568-INS-62
chr14-51117112-INS-209
chr14-51123456-INS-200
chr14-51139365-INS-203
chr14-51143220-INS-206
chr14-51210173-INS-52
chr14-51214861-INS-72
chr14-51240246-INS-189
chr14-51253616-INS-206
chr14-51265221-INS-701
chr14-51267601-INS-200
chr14-51269669-INS-137
chr14-51272955-INS-180
chr14-51313923-INS-199
chr14-51317827-INS-815
chr14-51341753-INS-126
chr14-51345967-INS-123
chr14-51360168-INS-129
chr14-51375638-INS-571
chr14-51394423-INS-194
chr14-51439654-INS-89
chr14-51441881-INS-58
chr14-51465545-INS-156
chr14-51480779-INS-324
chr14-51511004-INS-400
chr14-51517930-INS-58
chr14-51522692-INS-222
chr14-51524543-INS-125
chr14-51525647-INS-294
chr14-51601148-INS-7771
chr14-51624365-INS-82
chr14-51667089-INS-572
chr14-51723496-INS-7282
chr14-51726118-INS-5385
chr14-51759277-INS-428
chr14-51808996-INS-84
chr14-51809432-INS-74
chr14-51842596-INS-6869
chr14-51890143-INS-6438
chr14-51930021-INS-418
chr14-51950049-INS-384
chr14-51958059-INS-64
chr14-52106680-INS-66
chr14-52109198-INS-431
chr14-52137917-INS-72
chr14-52177155-INS-196
chr14-52181490-INS-344

chr14-52212180-INS-135
chr14-52225223-INS-186
chr14-52230417-INS-204
chr14-52232160-INS-206
chr14-52245489-INS-52
chr14-52261376-INS-210
chr14-52265582-INS-407
chr14-52266910-INS-133
chr14-52268862-INS-206
chr14-52284755-INS-235
chr14-52286701-INS-123
chr14-52295364-INS-206
chr14-52302078-INS-206
chr14-52305025-INS-420
chr14-52306973-INS-175
chr14-52310262-INS-218
chr14-52314143-INS-193
chr14-52315671-INS-273
chr14-52325667-INS-77
chr14-52327492-INS-56
chr14-52374581-INS-130
chr14-52379948-INS-56
chr14-52391980-INS-328
chr14-52402512-INS-247
chr14-52403974-INS-369
chr14-52410827-INS-238
chr14-52424963-INS-223
chr14-52428357-INS-75
chr14-52444037-INS-66
chr14-52471607-INS-73
chr14-52515759-INS-52
chr14-52527238-INS-55
chr14-52541787-INS-166
chr14-52546838-INS-156
chr14-52549564-INS-236
chr14-52574317-INS-288
chr14-52582328-INS-108
chr14-52583009-INS-200
chr14-52607477-INS-112
chr14-52649530-INS-564
chr14-52654730-INS-654
chr14-52661521-INS-74
chr14-52663080-INS-291
chr14-52675594-INS-326
chr14-52697396-INS-199
chr14-52702539-INS-800
chr14-52727084-INS-126
chr14-52730815-INS-72
chr14-52731924-INS-7518
chr14-52737423-INS-4770
chr14-52755795-INS-1893
chr14-52757096-INS-472
chr14-52757593-INS-6921
chr14-52761604-INS-6613
chr14-52773257-INS-929
chr14-52842848-INS-6552
chr14-52901937-INS-394
chr14-52946645-INS-125
chr14-52969442-INS-3362
chr14-52991399-INS-5485
chr14-53020324-INS-97
chr14-53036493-INS-136
chr14-53079706-INS-858
chr14-53081037-INS-57
chr14-53089417-INS-163
chr14-53171680-INS-55
chr14-53179568-INS-74
chr14-53193127-INS-60
chr14-53248989-INS-118
chr14-53257650-INS-52
chr14-53257664-INS-540
chr14-53263872-INS-54
chr14-53287520-INS-319
chr14-53289166-INS-108
chr14-53309313-INS-53
chr14-53346485-INS-50
chr14-53362397-INS-543

chr14-53373646-INS-339
chr14-53376722-INS-157
chr14-53377751-INS-54
chr14-53379059-INS-78
chr14-53382122-INS-128
chr14-53387894-INS-163
chr14-53390753-INS-858
chr14-53392085-INS-55
chr14-53400461-INS-163
chr14-53490597-INS-78
chr14-53522892-INS-171
chr14-53547786-INS-588
chr14-53561875-INS-102
chr14-53563246-INS-314
chr14-53566312-INS-131
chr14-53568335-INS-328
chr14-53657612-INS-538
chr14-53677770-INS-605
chr14-53711664-INS-52
chr14-53714309-INS-213
chr14-53720304-INS-549
chr14-53732495-INS-142
chr14-53738634-INS-375
chr14-53758122-INS-202
chr14-53760913-INS-422
chr14-53774684-INS-54
chr14-53779194-INS-76
chr14-53793845-INS-112
chr14-53844878-INS-210
chr14-53871599-INS-130
chr14-53894058-INS-88
chr14-53905222-INS-171
chr14-53920205-INS-222
chr14-53940782-INS-212
chr14-53944621-INS-176
chr14-53953946-INS-549
chr14-53955817-INS-61
chr14-53960537-INS-54
chr14-53965814-INS-445
chr14-54036338-INS-3749
chr14-54075642-INS-60
chr14-54080549-INS-52
chr14-54082301-INS-381
chr14-54105359-INS-400
chr14-54141952-INS-96
chr14-54152462-INS-77
chr14-54157392-INS-363
chr14-54163653-INS-133
chr14-54180544-INS-70
chr14-54195207-INS-480
chr14-54210526-INS-61
chr14-54214429-INS-140
chr14-54240691-INS-301
chr14-54293438-INS-143
chr14-54311039-INS-152
chr14-54322559-INS-6692
chr14-54329376-INS-76
chr14-54329399-INS-4827
chr14-54338824-INS-111
chr14-54342626-INS-70
chr14-54350367-INS-868
chr14-54365644-INS-52
chr14-54380883-INS-54
chr14-54415983-INS-134
chr14-54426283-INS-6446
chr14-54448224-INS-88
chr14-54452286-INS-78
chr14-54454338-INS-72
chr14-54469114-INS-96
chr14-54501086-INS-71
chr14-54504173-INS-701
chr14-54513452-INS-749
chr14-54551456-INS-105
chr14-54557122-INS-223
chr14-54566463-INS-851
chr14-54571606-INS-201
chr14-54579742-INS-431

chr14-54589010-INS-91
chr14-54589969-INS-3349
chr14-54591426-INS-58
chr14-54603384-INS-145
chr14-54623883-INS-151
chr14-54627043-INS-217
chr14-54633820-INS-52
chr14-54639147-INS-212
chr14-54658585-INS-99
chr14-54659021-INS-201
chr14-54662143-INS-54
chr14-54672701-INS-60
chr14-54696106-INS-116
chr14-54723488-INS-69
chr14-54757397-INS-178
chr14-54758541-INS-403
chr14-54761317-INS-154
chr14-54780630-INS-235
chr14-54781043-INS-213
chr14-54793459-INS-57
chr14-54797130-INS-192
chr14-54819175-INS-195
chr14-54844470-INS-105
chr14-54866850-INS-493
chr14-54871713-INS-110
chr14-54908515-INS-191
chr14-54925274-INS-223
chr14-54939101-INS-153
chr14-54946177-INS-66
chr14-54951103-INS-52
chr14-54980816-INS-208
chr14-54988118-INS-200
chr14-55002776-INS-66
chr14-55021361-INS-167
chr14-55102400-INS-103
chr14-55117438-INS-201
chr14-55160453-INS-322
chr14-55165645-INS-150
chr14-55192362-INS-186
chr14-55200009-INS-96
chr14-55219261-INS-55
chr14-55257489-INS-198
chr14-55269247-INS-136
chr14-55275707-INS-203
chr14-55350512-INS-210
chr14-55356328-INS-80
chr14-55367265-INS-773
chr14-55375501-INS-58
chr14-55391985-INS-231
chr14-55402894-INS-280
chr14-55403414-INS-207
chr14-55421574-INS-88
chr14-55435014-INS-182
chr14-55449765-INS-101
chr14-55461072-INS-154
chr14-55468416-INS-343
chr14-55481584-INS-6440
chr14-55500257-INS-506
chr14-55503389-INS-58
chr14-55522435-INS-958
chr14-55534222-INS-205
chr14-55547791-INS-354
chr14-55572594-INS-88
chr14-55583832-INS-105
chr14-55620022-INS-287
chr14-55631561-INS-68
chr14-55634355-INS-827
chr14-55635273-INS-58
chr14-55670663-INS-58
chr14-55672230-INS-338
chr14-55673516-INS-329
chr14-55674319-INS-218
chr14-55682983-INS-52
chr14-55692227-INS-51
chr14-55693617-INS-228
chr14-55700120-INS-52
chr14-55705591-INS-166

chr14-55714016-INS-84
chr14-55715707-INS-151
chr14-55722736-INS-95
chr14-55749858-INS-89
chr14-55759223-INS-236
chr14-55772824-INS-386
chr14-55776388-INS-60
chr14-55780709-INS-63
chr14-55785023-INS-54
chr14-55793534-INS-56
chr14-55796788-INS-205
chr14-55817703-INS-144
chr14-55825235-INS-152
chr14-55844590-INS-117
chr14-55852670-INS-273
chr14-55925142-INS-90
chr14-55929782-INS-129
chr14-55934896-INS-5543
chr14-55961656-INS-596
chr14-55963444-INS-64
chr14-55966714-INS-228
chr14-55996959-INS-54
chr14-56010518-INS-149
chr14-56114296-INS-60
chr14-56128004-INS-67
chr14-56131202-INS-96
chr14-56140612-INS-58
chr14-56141260-INS-5118
chr14-56142869-INS-5366
chr14-56150464-INS-246
chr14-56152196-INS-168
chr14-56165831-INS-56
chr14-56173761-INS-497
chr14-56177742-INS-213
chr14-56185237-INS-228
chr14-56198354-INS-843
chr14-56211866-INS-4926
chr14-56234732-INS-685
chr14-56247541-INS-2999
chr14-56273169-INS-4491
chr14-56311525-INS-6436
chr14-56315903-INS-844
chr14-56333746-INS-260
chr14-56372505-INS-667
chr14-56398163-INS-137
chr14-56414376-INS-414
chr14-56447980-INS-674
chr14-56449312-INS-6270
chr14-56452156-INS-2866
chr14-56463965-INS-421
chr14-56496344-INS-166
chr14-56517515-INS-694
chr14-56540267-INS-138
chr14-56542735-INS-423
chr14-56555506-INS-145
chr14-56561390-INS-55
chr14-56562746-INS-95
chr14-56576292-INS-78
chr14-56585247-INS-107
chr14-56613910-INS-54
chr14-56616565-INS-487
chr14-56627394-INS-65
chr14-56629097-INS-52
chr14-56642912-INS-54
chr14-56656408-INS-336
chr14-56669427-INS-96
chr14-56699747-INS-153
chr14-56707797-INS-181
chr14-56709298-INS-96
chr14-56732058-INS-84
chr14-56736451-INS-77
chr14-56749518-INS-54
chr14-56762993-INS-242
chr14-56778818-INS-97
chr14-56787599-INS-104
chr14-56793049-INS-71
chr14-56797374-INS-70

chr14-56799684-INS-277
chr14-56805687-INS-54
chr14-56808620-INS-8985
chr14-56819667-INS-175
chr14-56827667-INS-599
chr14-56833508-INS-700
chr14-56834631-INS-443
chr14-56840448-INS-54
chr14-56850322-INS-54
chr14-56857691-INS-68
chr14-56860668-INS-92
chr14-56863203-INS-4862
chr14-56864383-INS-181
chr14-56865034-INS-672
chr14-56866243-INS-176
chr14-56871597-INS-112
chr14-56874948-INS-180
chr14-56878890-INS-86
chr14-56880478-INS-92
chr14-56893784-INS-316
chr14-56899751-INS-133
chr14-56918600-INS-181
chr14-56921992-INS-115
chr14-56969709-INS-204
chr14-56980275-INS-61
chr14-57000867-INS-151
chr14-57005436-INS-161
chr14-57021854-INS-182
chr14-57024424-INS-63
chr14-57036048-INS-228
chr14-57041423-INS-219
chr14-57055649-INS-65
chr14-57056206-INS-398
chr14-57061917-INS-98
chr14-57062705-INS-220
chr14-57068796-INS-205
chr14-57086958-INS-448
chr14-57093708-INS-194
chr14-57105031-INS-172
chr14-57111893-INS-212
chr14-57113479-INS-210
chr14-57115117-INS-3384
chr14-57128024-INS-66
chr14-57129824-INS-198
chr14-57133097-INS-168
chr14-57143320-INS-100
chr14-57146233-INS-212
chr14-57159716-INS-583
chr14-57160647-INS-107
chr14-57181304-INS-288
chr14-57188809-INS-319
chr14-57218220-INS-68
chr14-57248386-INS-3163
chr14-57254850-INS-7276
chr14-57269292-INS-60
chr14-57298503-INS-53
chr14-57307185-INS-64
chr14-57316360-INS-59
chr14-57328584-INS-85
chr14-57334100-INS-78
chr14-57340608-INS-64
chr14-57383903-INS-110
chr14-57409471-INS-183
chr14-57502516-INS-52
chr14-57515509-INS-52
chr14-57536492-INS-381
chr14-57578093-INS-58
chr14-57586492-INS-52
chr14-57586492-INS-64
chr14-57590989-INS-53
chr14-57595461-INS-55
chr14-57653726-INS-101
chr14-57667650-INS-180
chr14-57669814-INS-78
chr14-57719140-INS-956
chr14-57720030-INS-106
chr14-57758453-INS-112

chr14-57760518-INS-133
chr14-57768249-INS-193
chr14-57781612-INS-180
chr14-57790800-INS-70
chr14-57800461-INS-51
chr14-57874391-INS-1349
chr14-57876271-INS-126
chr14-57903652-INS-54
chr14-57917738-INS-183
chr14-57921316-INS-244
chr14-57939038-INS-52
chr14-57988379-INS-232
chr14-57989097-INS-69
chr14-57993311-INS-207
chr14-58017982-INS-235
chr14-58018586-INS-195
chr14-58026378-INS-629
chr14-58032153-INS-530
chr14-58045023-INS-99
chr14-58053303-INS-678
chr14-58072440-INS-203
chr14-58107259-INS-204
chr14-58109918-INS-252
chr14-58112029-INS-69
chr14-58122932-INS-201
chr14-58143535-INS-177
chr14-58149657-INS-59
chr14-58153848-INS-5552
chr14-58160133-INS-889
chr14-58162062-INS-172
chr14-58165310-INS-317
chr14-58168390-INS-70
chr14-58184410-INS-114
chr14-58189413-INS-498
chr14-58224850-INS-120
chr14-58253244-INS-349
chr14-58256880-INS-165
chr14-58266882-INS-113
chr14-58283146-INS-52
chr14-58303599-INS-368
chr14-58319765-INS-151
chr14-58333904-INS-64
chr14-58343122-INS-52
chr14-58361333-INS-65
chr14-58375163-INS-5931
chr14-58380325-INS-99
chr14-58381789-INS-68
chr14-58399159-INS-124
chr14-58402646-INS-742
chr14-58410756-INS-90
chr14-58445091-INS-440
chr14-58456087-INS-305
chr14-58458709-INS-573
chr14-58461506-INS-316
chr14-58481334-INS-270
chr14-58498960-INS-1786
chr14-58505738-INS-65
chr14-58513142-INS-86
chr14-58518624-INS-52
chr14-58526382-INS-70
chr14-58541828-INS-78
chr14-58560496-INS-87
chr14-58594309-INS-72
chr14-58597397-INS-54
chr14-58614099-INS-516
chr14-58642684-INS-94
chr14-58669112-INS-898
chr14-58674361-INS-63
chr14-58714195-INS-5648
chr14-58714821-INS-199
chr14-58723500-INS-415
chr14-58729212-INS-5207
chr14-58735282-INS-56
chr14-58741769-INS-52
chr14-58752603-INS-169
chr14-58800009-INS-56
chr14-58803222-INS-131

chr14-58808922-INS-53
chr14-58815125-INS-228
chr14-58819248-INS-236
chr14-58821871-INS-76
chr14-58831302-INS-675
chr14-58850566-INS-52
chr14-58882844-INS-52
chr14-58917285-INS-84
chr14-58926733-INS-492
chr14-58932175-INS-168
chr14-58945772-INS-289
chr14-58953152-INS-106
chr14-58957580-INS-68
chr14-58960681-INS-56
chr14-58961459-INS-507
chr14-58979590-INS-899
chr14-58983384-INS-178
chr14-58984850-INS-63
chr14-58990135-INS-245
chr14-58996133-INS-68
chr14-59032311-INS-510
chr14-59047507-INS-365
chr14-59052219-INS-69
chr14-59079158-INS-52
chr14-59095062-INS-109
chr14-59100075-INS-66
chr14-59112397-INS-413
chr14-59143513-INS-1343
chr14-59145930-INS-114
chr14-59161126-INS-6381
chr14-59189615-INS-454
chr14-59195954-INS-228
chr14-59197040-INS-54
chr14-59219889-INS-56
chr14-59223343-INS-54
chr14-59227688-INS-74
chr14-59229826-INS-97
chr14-59246498-INS-143
chr14-59264612-INS-65
chr14-59290527-INS-74
chr14-59315443-INS-96
chr14-59316837-INS-200
chr14-59318269-INS-838
chr14-59345149-INS-1886
chr14-59349514-INS-132
chr14-59354373-INS-349
chr14-59356308-INS-163
chr14-59389682-INS-140
chr14-59409401-INS-1912
chr14-59425340-INS-7437
chr14-59425465-INS-900
chr14-59430194-INS-54
chr14-59430929-INS-7366
chr14-59442727-INS-131
chr14-59453377-INS-707
chr14-59473853-INS-90
chr14-59483683-INS-227
chr14-59530328-INS-60
chr14-59595121-INS-527
chr14-59610171-INS-3156
chr14-59619528-INS-1181
chr14-59623302-INS-162
chr14-59624708-INS-102
chr14-59640191-INS-64
chr14-59642253-INS-731
chr14-59656100-INS-434
chr14-59683202-INS-630
chr14-59694975-INS-455
chr14-59698617-INS-3713
chr14-59700807-INS-155
chr14-59709124-INS-131
chr14-59713873-INS-7130
chr14-59721061-INS-57
chr14-59722710-INS-2410
chr14-59814434-INS-344
chr14-59845095-INS-70
chr14-59846810-INS-63

chr14-59861951-INS-1207
chr14-59892641-INS-88
chr14-59895933-INS-162
chr14-59947240-INS-6223
chr14-59956411-INS-111
chr14-59977566-INS-68
chr14-59983018-INS-54
chr14-59984448-INS-232
chr14-59987184-INS-65
chr14-60009377-INS-71
chr14-60048555-INS-201
chr14-60051836-INS-66
chr14-60076278-INS-68
chr14-60087556-INS-82
chr14-60090370-INS-687
chr14-60097298-INS-63
chr14-60112177-INS-110
chr14-60117435-INS-266
chr14-60129153-INS-66
chr14-60142074-INS-6779
chr14-60149100-INS-161
chr14-60162910-INS-70
chr14-60188071-INS-64
chr14-60190354-INS-69
chr14-60192676-INS-110
chr14-60196898-INS-644
chr14-60215916-INS-59
chr14-60240324-INS-79
chr14-60259847-INS-99
chr14-60261517-INS-428
chr14-60264266-INS-521
chr14-60276301-INS-203
chr14-60278669-INS-1063
chr14-60299338-INS-486
chr14-60307226-INS-257
chr14-60313813-INS-50
chr14-60318202-INS-57
chr14-60334844-INS-432
chr14-60342833-INS-2068
chr14-60383334-INS-155
chr14-60389979-INS-192
chr14-60439465-INS-140
chr14-60440609-INS-126
chr14-60449886-INS-105
chr14-60475082-INS-169
chr14-60512635-INS-81
chr14-60585690-INS-105
chr14-60587155-INS-414
chr14-60598124-INS-120
chr14-60608063-INS-168
chr14-60622162-INS-127
chr14-60625842-INS-60
chr14-60628002-INS-374
chr14-60630048-INS-111
chr14-60647018-INS-52
chr14-60658091-INS-6451
chr14-60662974-INS-703
chr14-60689166-INS-72
chr14-60706258-INS-196
chr14-60721012-INS-83
chr14-60728771-INS-526
chr14-60731970-INS-50
chr14-60739898-INS-62
chr14-60741360-INS-325
chr14-60761693-INS-78
chr14-60764587-INS-126
chr14-60766407-INS-129
chr14-60767379-INS-71
chr14-60784489-INS-54
chr14-60815656-INS-92
chr14-60823719-INS-137
chr14-60831405-INS-95
chr14-60841721-INS-54
chr14-60843005-INS-76
chr14-60844676-INS-446
chr14-60942003-INS-7420
chr14-60962682-INS-199

chr14-60989449-INS-130
chr14-61013647-INS-252
chr14-61014713-INS-52
chr14-61021532-INS-155
chr14-61024329-INS-438
chr14-61025078-INS-78
chr14-61029333-INS-121
chr14-61047528-INS-473
chr14-61055577-INS-120
chr14-61056736-INS-84
chr14-61073641-INS-572
chr14-61082332-INS-412
chr14-61086861-INS-55
chr14-61140723-INS-1285
chr14-61156509-INS-74
chr14-61162626-INS-203
chr14-61171297-INS-53
chr14-61217970-INS-697
chr14-61219427-INS-579
chr14-61226997-INS-607
chr14-61231559-INS-229
chr14-61232201-INS-177
chr14-61236245-INS-120
chr14-61240666-INS-54
chr14-61258775-INS-64
chr14-61265311-INS-173
chr14-61290256-INS-216
chr14-61334167-INS-217
chr14-61337766-INS-57
chr14-61348754-INS-115
chr14-61368035-INS-222
chr14-61377818-INS-1888
chr14-61407285-INS-56
chr14-61453428-INS-504
chr14-61456878-INS-318
chr14-61459442-INS-84
chr14-61461229-INS-80
chr14-61463063-INS-408
chr14-61477931-INS-118
chr14-61483696-INS-51
chr14-61484957-INS-480
chr14-61515051-INS-93
chr14-61519075-INS-123
chr14-61536627-INS-153
chr14-61543749-INS-5283
chr14-61553979-INS-208
chr14-61567325-INS-209
chr14-61591022-INS-422
chr14-61627458-INS-176
chr14-61640341-INS-84
chr14-61640829-INS-204
chr14-61674655-INS-71
chr14-61685040-INS-384
chr14-61723016-INS-71
chr14-61729362-INS-198
chr14-61738189-INS-52
chr14-61740497-INS-60
chr14-61743731-INS-54
chr14-61758019-INS-598
chr14-61763641-INS-72
chr14-61796093-INS-220
chr14-61801155-INS-7001
chr14-61804275-INS-178
chr14-61829422-INS-201
chr14-61831664-INS-224
chr14-61838730-INS-410
chr14-61847279-INS-216
chr14-61853033-INS-312
chr14-61856869-INS-82
chr14-61861737-INS-269
chr14-61865296-INS-72
chr14-61891023-INS-180
chr14-61945047-INS-146
chr14-61947341-INS-66
chr14-61998378-INS-135
chr14-62002100-INS-66
chr14-62035542-INS-213

chr14-62036519-INS-720
chr14-62040372-INS-129
chr14-62043448-INS-206
chr14-62059249-INS-103
chr14-62060729-INS-202
chr14-62096673-INS-229
chr14-62139854-INS-144
chr14-62165029-INS-255
chr14-62177730-INS-77
chr14-62189868-INS-109
chr14-62207784-INS-228
chr14-62208081-INS-612
chr14-62225480-INS-502
chr14-62237390-INS-247
chr14-62241017-INS-5542
chr14-62242576-INS-653
chr14-62264359-INS-79
chr14-62272647-INS-148
chr14-62293692-INS-123
chr14-62296590-INS-149
chr14-62309185-INS-342
chr14-62311546-INS-928
chr14-62318175-INS-73
chr14-62319505-INS-315
chr14-62322793-INS-449
chr14-62328247-INS-118
chr14-62331007-INS-6486
chr14-62334173-INS-114
chr14-62337859-INS-144
chr14-62344723-INS-67
chr14-62417869-INS-120
chr14-62424610-INS-122
chr14-62427241-INS-181
chr14-62456939-INS-245
chr14-62459633-INS-1249
chr14-62465971-INS-118
chr14-62468700-INS-200
chr14-62469802-INS-54
chr14-62472512-INS-53
chr14-62481742-INS-60
chr14-62489316-INS-55
chr14-62498217-INS-267
chr14-62519755-INS-123
chr14-62523103-INS-146
chr14-62524313-INS-372
chr14-62541772-INS-6429
chr14-62582835-INS-134
chr14-62590501-INS-52
chr14-62596321-INS-3819
chr14-62605457-INS-52
chr14-62612067-INS-182
chr14-62613838-INS-903
chr14-62617936-INS-140
chr14-62629027-INS-74
chr14-62636847-INS-152
chr14-62668579-INS-154
chr14-62670679-INS-375
chr14-62691371-INS-124
chr14-62699152-INS-118
chr14-62703425-INS-177
chr14-62706312-INS-114
chr14-62714979-INS-52
chr14-62765573-INS-79
chr14-62776707-INS-112
chr14-62799273-INS-6543
chr14-62803309-INS-60
chr14-62833940-INS-163
chr14-62835787-INS-389
chr14-62838737-INS-78
chr14-62868005-INS-443
chr14-62879328-INS-68
chr14-62906390-INS-153
chr14-62977755-INS-84
chr14-63006917-INS-66
chr14-63024124-INS-196
chr14-63030726-INS-54
chr14-63034717-INS-201

chr14-63037090-INS-52
chr14-63048675-INS-66
chr14-63052901-INS-8419
chr14-63058891-INS-57
chr14-63104548-INS-616
chr14-63108418-INS-152
chr14-63116123-INS-138
chr14-63124313-INS-353
chr14-63131738-INS-316
chr14-63170914-INS-329
chr14-63178852-INS-363
chr14-63204755-INS-511
chr14-63213418-INS-406
chr14-63214264-INS-94
chr14-63216046-INS-281
chr14-63225559-INS-254
chr14-63242573-INS-887
chr14-63257451-INS-116
chr14-63294224-INS-106
chr14-63308936-INS-101
chr14-63314881-INS-68
chr14-63322968-INS-109
chr14-63324153-INS-57
chr14-63329739-INS-79
chr14-63337082-INS-64
chr14-63354509-INS-204
chr14-63385580-INS-334
chr14-63430762-INS-75
chr14-63453111-INS-87
chr14-63514069-INS-934
chr14-63517753-INS-1374
chr14-63518034-INS-11772
chr14-63518050-INS-3678
chr14-63518056-INS-4842
chr14-63520866-INS-96
chr14-63532324-INS-126
chr14-63536510-INS-113
chr14-63550906-INS-54
chr14-63552456-INS-218
chr14-63565586-INS-78
chr14-63569267-INS-56
chr14-63578312-INS-210
chr14-63583084-INS-52
chr14-63590121-INS-406
chr14-63592160-INS-51
chr14-63639360-INS-141
chr14-63660724-INS-493
chr14-63662926-INS-199
chr14-63674409-INS-109
chr14-63705821-INS-195
chr14-63712984-INS-68
chr14-63719159-INS-92
chr14-63732290-INS-689
chr14-63755829-INS-185
chr14-63789165-INS-322
chr14-63800217-INS-54
chr14-63817863-INS-62
chr14-63831874-INS-174
chr14-63834809-INS-52
chr14-63930971-INS-196
chr14-63933126-INS-78
chr14-63986847-INS-56
chr14-64002725-INS-176
chr14-64007608-INS-52
chr14-64100889-INS-68
chr14-64104850-INS-119
chr14-64242767-INS-101
chr14-64244584-INS-57
chr14-64252446-INS-71
chr14-64289278-INS-62
chr14-64309558-INS-242
chr14-64338415-INS-106
chr14-64419464-INS-52
chr14-64455596-INS-210
chr14-64461571-INS-354
chr14-64478360-INS-68
chr14-64499934-INS-239

chr14-64524359-INS-128
chr14-64537087-INS-70
chr14-64587008-INS-57
chr14-64596330-INS-810
chr14-64615200-INS-5564
chr14-64628582-INS-92
chr14-64646534-INS-190
chr14-64648451-INS-64
chr14-64659159-INS-271
chr14-64671003-INS-1522
chr14-64671190-INS-174
chr14-64713615-INS-932
chr14-64717546-INS-104
chr14-64766304-INS-7368
chr14-64785500-INS-70
chr14-64803203-INS-59
chr14-64843161-INS-96
chr14-64844183-INS-173
chr14-64852056-INS-174
chr14-64855047-INS-162
chr14-64859911-INS-207
chr14-64869410-INS-56
chr14-64874229-INS-2036
chr14-64878109-INS-69
chr14-64878318-INS-235
chr14-64879732-INS-113
chr14-64884885-INS-205
chr14-64885302-INS-218
chr14-64910171-INS-84
chr14-64921237-INS-401
chr14-64924532-INS-52
chr14-64926825-INS-199
chr14-64927974-INS-482
chr14-64977408-INS-251
chr14-64996433-INS-202
chr14-65164656-INS-1941
chr14-65204721-INS-64
chr14-65207737-INS-228
chr14-65243012-INS-400
chr14-65296066-INS-360
chr14-65306110-INS-64
chr14-65319275-INS-82
chr14-65351520-INS-75
chr14-65356497-INS-242
chr14-65362871-INS-66
chr14-65365121-INS-198
chr14-65384503-INS-207
chr14-65391737-INS-195
chr14-65419726-INS-51
chr14-65422005-INS-247
chr14-65447799-INS-64
chr14-65465767-INS-120
chr14-65470608-INS-88
chr14-65485712-INS-178
chr14-65487582-INS-263
chr14-65507091-INS-379
chr14-65523786-INS-58
chr14-65534689-INS-202
chr14-65540181-INS-200
chr14-65541459-INS-63
chr14-65565795-INS-201
chr14-65567362-INS-72
chr14-65585237-INS-100
chr14-65590919-INS-86
chr14-65600119-INS-62
chr14-65602965-INS-226
chr14-65611356-INS-55
chr14-65622778-INS-60
chr14-65644980-INS-424
chr14-65664738-INS-111
chr14-65669431-INS-124
chr14-65672290-INS-326
chr14-65678086-INS-629
chr14-65691514-INS-345
chr14-65704474-INS-2854
chr14-65708249-INS-89
chr14-65717402-INS-80

chr14-65741865-INS-216
chr14-65756212-INS-803
chr14-65757685-INS-64
chr14-65772505-INS-59
chr14-65791957-INS-290
chr14-65795405-INS-374
chr14-65812671-INS-55
chr14-65861723-INS-293
chr14-65866920-INS-214
chr14-65867576-INS-96
chr14-65904716-INS-190
chr14-65925014-INS-53
chr14-65936274-INS-96
chr14-65942308-INS-106
chr14-65945619-INS-201
chr14-65958661-INS-53
chr14-65988475-INS-592
chr14-65998566-INS-4981
chr14-66007128-INS-85
chr14-66017709-INS-69
chr14-66020292-INS-5548
chr14-66041586-INS-54
chr14-66041866-INS-599
chr14-66045418-INS-56
chr14-66052119-INS-206
chr14-66064358-INS-772
chr14-66073444-INS-6350
chr14-66083620-INS-52
chr14-66086393-INS-112
chr14-66138248-INS-66
chr14-66143350-INS-109
chr14-66163002-INS-56
chr14-66165443-INS-111
chr14-66192321-INS-213
chr14-66258418-INS-183
chr14-66264616-INS-60
chr14-66320715-INS-90
chr14-66323569-INS-165
chr14-66333004-INS-124
chr14-66363075-INS-52
chr14-66372216-INS-86
chr14-66377977-INS-88
chr14-66422261-INS-168
chr14-66432807-INS-200
chr14-66438153-INS-67
chr14-66462089-INS-202
chr14-66463873-INS-54
chr14-66467998-INS-70
chr14-66488972-INS-90
chr14-66567075-INS-322
chr14-66632678-INS-73
chr14-66634821-INS-172
chr14-66636107-INS-248
chr14-66643298-INS-6904
chr14-66682814-INS-156
chr14-66696409-INS-100
chr14-66711182-INS-269
chr14-66732404-INS-64
chr14-66735705-INS-60
chr14-66753499-INS-130
chr14-66772980-INS-84
chr14-66774114-INS-84
chr14-66802985-INS-96
chr14-66862450-INS-327
chr14-66871522-INS-52
chr14-66927268-INS-74
chr14-66936729-INS-70
chr14-66944058-INS-52
chr14-66947854-INS-69
chr14-66948939-INS-442
chr14-66956360-INS-102
chr14-66971260-INS-1331
chr14-66994539-INS-7808
chr14-67013084-INS-55
chr14-67015885-INS-104
chr14-67068831-INS-141
chr14-67082580-INS-374

chr14-67117799-INS-132
chr14-67146053-INS-124
chr14-67149228-INS-185
chr14-67187024-INS-67
chr14-67193287-INS-107
chr14-67197709-INS-197
chr14-67209417-INS-165
chr14-67217436-INS-66
chr14-67218850-INS-518
chr14-67220992-INS-550
chr14-67260122-INS-247
chr14-67264137-INS-66
chr14-67266165-INS-126
chr14-67280158-INS-201
chr14-67325356-INS-83
chr14-67330772-INS-111
chr14-67347258-INS-61
chr14-67371463-INS-183
chr14-67374757-INS-169
chr14-67376157-INS-58
chr14-67383627-INS-60
chr14-67400341-INS-94
chr14-67402902-INS-108
chr14-67409217-INS-129
chr14-67418896-INS-94
chr14-67444729-INS-289
chr14-67456311-INS-140
chr14-67502453-INS-54
chr14-67523361-INS-118
chr14-67530052-INS-52
chr14-67538210-INS-52
chr14-67597250-INS-1435
chr14-67617748-INS-60
chr14-67632976-INS-97
chr14-67676492-INS-78
chr14-67719357-INS-7285
chr14-67749518-INS-115
chr14-67763528-INS-144
chr14-67777294-INS-190
chr14-67794150-INS-223
chr14-67821049-INS-192
chr14-67828262-INS-51
chr14-67833332-INS-492
chr14-67840881-INS-119
chr14-67868200-INS-78
chr14-67871425-INS-92
chr14-67903977-INS-57
chr14-67913127-INS-82
chr14-67941183-INS-535
chr14-67950245-INS-641
chr14-67968680-INS-608
chr14-67982041-INS-191
chr14-67986534-INS-127
chr14-67991625-INS-228
chr14-67994185-INS-163
chr14-68035390-INS-105
chr14-68038811-INS-1408
chr14-68042735-INS-266
chr14-68046782-INS-124
chr14-68055164-INS-657
chr14-68057668-INS-205
chr14-68074515-INS-227
chr14-68088724-INS-304
chr14-68091666-INS-86
chr14-68099209-INS-220
chr14-68108183-INS-392
chr14-68111943-INS-256
chr14-68134408-INS-104
chr14-68174835-INS-141
chr14-68176635-INS-100
chr14-68192089-INS-179
chr14-68228544-INS-519
chr14-68231840-INS-103
chr14-68237702-INS-615
chr14-68252970-INS-514
chr14-68270098-INS-208
chr14-68274392-INS-231

chr14-68290464-INS-72
chr14-68293162-INS-426
chr14-68308107-INS-814
chr14-68309883-INS-70
chr14-68328888-INS-360
chr14-68359886-INS-69
chr14-68387625-INS-56
chr14-68389703-INS-54
chr14-68410360-INS-57
chr14-68423584-INS-6576
chr14-68444449-INS-60
chr14-68479070-INS-76
chr14-68507705-INS-74
chr14-68518463-INS-106
chr14-68530010-INS-198
chr14-68545206-INS-346
chr14-68568819-INS-6506
chr14-68572439-INS-56
chr14-68630152-INS-5546
chr14-68665674-INS-335
chr14-68696668-INS-55
chr14-68703807-INS-1330
chr14-68706829-INS-385
chr14-68728550-INS-54
chr14-68736656-INS-61
chr14-68741662-INS-99
chr14-68746397-INS-502
chr14-68765866-INS-116
chr14-68769779-INS-247
chr14-68780601-INS-415
chr14-68813473-INS-55
chr14-68826663-INS-386
chr14-68839105-INS-247
chr14-68855730-INS-607
chr14-68856492-INS-1739
chr14-68864025-INS-927
chr14-68887167-INS-74
chr14-68894755-INS-54
chr14-68913022-INS-122
chr14-68925259-INS-502
chr14-68945903-INS-211
chr14-68980658-INS-6888
chr14-68995806-INS-6222
chr14-69034832-INS-418
chr14-69079542-INS-56
chr14-69096671-INS-506
chr14-69121941-INS-53
chr14-69152110-INS-3745
chr14-69171194-INS-67
chr14-69190100-INS-509
chr14-69207686-INS-139
chr14-69208996-INS-134
chr14-69213737-INS-6876
chr14-69226742-INS-5370
chr14-69287387-INS-2738
chr14-69289193-INS-6292
chr14-69289699-INS-182
chr14-69309455-INS-51
chr14-69317934-INS-6006
chr14-69320034-INS-4785
chr14-69320172-INS-6181
chr14-69362091-INS-189
chr14-69368252-INS-6440
chr14-69375530-INS-75
chr14-69384625-INS-58
chr14-69387348-INS-144
chr14-69424444-INS-68
chr14-69427408-INS-107
chr14-69449115-INS-119
chr14-69452521-INS-60
chr14-69466662-INS-66
chr14-69468408-INS-1636
chr14-69531502-INS-1888
chr14-69782508-INS-4167
chr14-69912952-INS-95
chr14-69930749-INS-78
chr14-69935379-INS-171

chr14-69962850-INS-498
chr14-69964123-INS-5338
chr14-70045047-INS-4400
chr14-70064757-INS-80
chr14-70101929-INS-188
chr14-70102042-INS-204
chr14-70166917-INS-145
chr14-70174490-INS-115
chr14-70192637-INS-471
chr14-70197235-INS-181
chr14-70208073-INS-61
chr14-70210677-INS-59
chr14-70212561-INS-165
chr14-70220173-INS-206
chr14-70222708-INS-54
chr14-70266494-INS-92
chr14-70273140-INS-56
chr14-70277739-INS-71
chr14-70288527-INS-76
chr14-70290766-INS-57
chr14-70298840-INS-56
chr14-70366593-INS-78
chr14-70399512-INS-218
chr14-70410089-INS-54
chr14-70433926-INS-89
chr14-70437406-INS-242
chr14-70444862-INS-131
chr14-70448531-INS-168
chr14-70456944-INS-72
chr14-70458309-INS-72
chr14-70460692-INS-60
chr14-70471225-INS-5268
chr14-70488217-INS-190
chr14-70556838-INS-62
chr14-70606798-INS-112
chr14-70608566-INS-80
chr14-70620152-INS-76
chr14-70626096-INS-56
chr14-70627648-INS-134
chr14-70633346-INS-203
chr14-70652636-INS-260
chr14-70660725-INS-182
chr14-70670171-INS-54
chr14-70672260-INS-203
chr14-70679712-INS-215
chr14-70700185-INS-57
chr14-70731817-INS-81
chr14-70740465-INS-137
chr14-70746623-INS-205
chr14-70751127-INS-68
chr14-70765370-INS-202
chr14-70779942-INS-119
chr14-70782129-INS-776
chr14-70804171-INS-250
chr14-70847325-INS-255
chr14-70848270-INS-114
chr14-70849667-INS-65
chr14-70850973-INS-107
chr14-70854244-INS-54
chr14-70887632-INS-213
chr14-70891542-INS-147
chr14-70893780-INS-199
chr14-70914038-INS-166
chr14-70916808-INS-111
chr14-70952114-INS-213
chr14-70957657-INS-125
chr14-70958890-INS-88
chr14-70994176-INS-56
chr14-70999245-INS-213
chr14-71007474-INS-262
chr14-71043877-INS-51
chr14-71047047-INS-204
chr14-71078377-INS-108
chr14-71140759-INS-61
chr14-71148322-INS-90
chr14-71148852-INS-80
chr14-71205530-INS-54

chr14-71210100-INS-103
chr14-71216059-INS-76
chr14-71245744-INS-6756
chr14-71263236-INS-101
chr14-71284042-INS-644
chr14-71287428-INS-148
chr14-71303600-INS-69
chr14-71320981-INS-7239
chr14-71322124-INS-7800
chr14-71328667-INS-106
chr14-71339206-INS-286
chr14-71343513-INS-4362
chr14-71376309-INS-53
chr14-71378336-INS-6413
chr14-71386724-INS-75
chr14-71402469-INS-56
chr14-71405763-INS-67
chr14-71407278-INS-400
chr14-71417463-INS-5304
chr14-71422770-INS-55
chr14-71428029-INS-60
chr14-71429908-INS-6440
chr14-71441988-INS-68
chr14-71474038-INS-2010
chr14-71488300-INS-95
chr14-71508632-INS-7076
chr14-71553452-INS-97
chr14-71562065-INS-163
chr14-71563519-INS-263
chr14-71566803-INS-1864
chr14-71572577-INS-142
chr14-71603457-INS-168
chr14-71613406-INS-433
chr14-71648197-INS-52
chr14-71658201-INS-395
chr14-71659850-INS-83
chr14-71671401-INS-54
chr14-71702179-INS-7494
chr14-71713236-INS-586
chr14-71715246-INS-6808
chr14-71754267-INS-1390
chr14-71775517-INS-713
chr14-71782800-INS-398
chr14-71806191-INS-154
chr14-71812106-INS-86
chr14-71844515-INS-397
chr14-71847367-INS-462
chr14-71852091-INS-52
chr14-71865893-INS-236
chr14-71867025-INS-276
chr14-71895324-INS-89
chr14-71900983-INS-2388
chr14-71907359-INS-69
chr14-71916439-INS-58
chr14-71921381-INS-1016
chr14-71921460-INS-1688
chr14-71957306-INS-310
chr14-71958700-INS-474
chr14-71971343-INS-163
chr14-71975932-INS-591
chr14-71988917-INS-7924
chr14-72015000-INS-76
chr14-72017134-INS-108
chr14-72019037-INS-7379
chr14-72032865-INS-6181
chr14-72044973-INS-318
chr14-72052548-INS-213
chr14-72052635-INS-6643
chr14-72070099-INS-80
chr14-72079395-INS-198
chr14-72082274-INS-59
chr14-72100817-INS-105
chr14-72105253-INS-409
chr14-72105425-INS-7299
chr14-72110664-INS-84
chr14-72153937-INS-137
chr14-72156344-INS-225

chr14-72220203-INS-100
chr14-72230517-INS-102
chr14-72233456-INS-51
chr14-72235834-INS-87
chr14-72258698-INS-165
chr14-72266616-INS-56
chr14-72293304-INS-7121
chr14-72301701-INS-676
chr14-72329430-INS-6561
chr14-72353267-INS-66
chr14-72384619-INS-64
chr14-72396711-INS-63
chr14-72428935-INS-526
chr14-72435796-INS-228
chr14-72440100-INS-57
chr14-72460173-INS-58
chr14-72474768-INS-52
chr14-72523889-INS-228
chr14-72534442-INS-54
chr14-72540643-INS-6707
chr14-72543096-INS-141
chr14-72560494-INS-84
chr14-72596710-INS-84
chr14-72615610-INS-6586
chr14-72618036-INS-6646
chr14-72623448-INS-82
chr14-72637338-INS-942
chr14-72640653-INS-673
chr14-72640737-INS-53
chr14-72642051-INS-199
chr14-72643042-INS-557
chr14-72647678-INS-199
chr14-72664713-INS-68
chr14-72666542-INS-412
chr14-72686624-INS-76
chr14-72736341-INS-492
chr14-72740780-INS-60
chr14-72746514-INS-74
chr14-72748437-INS-53
chr14-72755772-INS-105
chr14-72805888-INS-249
chr14-72829832-INS-2129
chr14-72852377-INS-340
chr14-72876883-INS-415
chr14-72909968-INS-199
chr14-72912167-INS-825
chr14-72912433-INS-941
chr14-72955956-INS-124
chr14-72957900-INS-164
chr14-72962699-INS-525
chr14-72965299-INS-224
chr14-72990888-INS-961
chr14-72991910-INS-56
chr14-73003124-INS-137
chr14-73012781-INS-62
chr14-73029591-INS-2178
chr14-73042120-INS-565
chr14-73045817-INS-54
chr14-73047325-INS-594
chr14-73052450-INS-102
chr14-73061677-INS-5215
chr14-73069474-INS-154
chr14-73074986-INS-131
chr14-73080362-INS-55
chr14-73094330-INS-70
chr14-73096827-INS-83
chr14-73099849-INS-85
chr14-73128555-INS-314
chr14-73135347-INS-90
chr14-73145895-INS-7112
chr14-73153579-INS-212
chr14-73164088-INS-143
chr14-73165249-INS-64
chr14-73178070-INS-7945
chr14-73199853-INS-76
chr14-73200028-INS-6371
chr14-73205267-INS-52

chr14-73211951-INS-401
chr14-73252447-INS-64
chr14-73257508-INS-6524
chr14-73300637-INS-5858
chr14-73317851-INS-410
chr14-73384322-INS-70
chr14-73392999-INS-135
chr14-73396008-INS-79
chr14-73450032-INS-383
chr14-73454592-INS-99
chr14-73467380-INS-515
chr14-73468431-INS-71
chr14-73480879-INS-100
chr14-73482877-INS-350
chr14-73483771-INS-547
chr14-73492966-INS-445
chr14-73499034-INS-519
chr14-73552898-INS-83
chr14-73602682-INS-74
chr14-73655352-INS-834
chr14-73664044-INS-54
chr14-73686483-INS-54
chr14-73718895-INS-6744
chr14-73719455-INS-79
chr14-73732456-INS-792
chr14-73733406-INS-52
chr14-73737830-INS-591
chr14-73745082-INS-89
chr14-73746505-INS-66
chr14-73750403-INS-206
chr14-73776509-INS-215
chr14-73782440-INS-225
chr14-73814314-INS-8796
chr14-73822326-INS-75
chr14-73824877-INS-6727
chr14-73825242-INS-783
chr14-73830266-INS-572
chr14-73845315-INS-163
chr14-73860373-INS-400
chr14-73876678-INS-56
chr14-73877542-INS-60
chr14-73881922-INS-329
chr14-73898874-INS-400
chr14-73936572-INS-145
chr14-73943470-INS-174
chr14-73974976-INS-106
chr14-73984117-INS-7018
chr14-73989150-INS-234
chr14-73989309-INS-417
chr14-73991736-INS-443
chr14-74036366-INS-826
chr14-74100059-INS-178
chr14-74126958-INS-124
chr14-74152915-INS-903
chr14-74155624-INS-73
chr14-74158170-INS-1212
chr14-74159504-INS-6491
chr14-74162667-INS-864
chr14-74171165-INS-7132
chr14-74187493-INS-816
chr14-74193567-INS-442
chr14-74221686-INS-82
chr14-74228989-INS-55
chr14-74237241-INS-78
chr14-74238748-INS-68
chr14-74240288-INS-330
chr14-74258473-INS-71
chr14-74265921-INS-92
chr14-74280615-INS-77
chr14-74303896-INS-1891
chr14-74313763-INS-56
chr14-74325066-INS-408
chr14-74325841-INS-639
chr14-74345592-INS-6167
chr14-74350835-INS-7173
chr14-74380275-INS-60
chr14-74384588-INS-54

chr14-74416096-INS-95
chr14-74453724-INS-58
chr14-74460896-INS-54
chr14-74465022-INS-52
chr14-74470089-INS-7367
chr14-74477515-INS-6261
chr14-74488710-INS-225
chr14-74494660-INS-430
chr14-74502738-INS-481
chr14-74514304-INS-136
chr14-74530176-INS-63
chr14-74544170-INS-116
chr14-74564665-INS-572
chr14-74571575-INS-797
chr14-74579815-INS-292
chr14-74585370-INS-465
chr14-74600513-INS-7129
chr14-74610188-INS-495
chr14-74624784-INS-53
chr14-74630777-INS-402
chr14-74636393-INS-57
chr14-74654482-INS-499
chr14-74667461-INS-232
chr14-74671985-INS-441
chr14-74678597-INS-416
chr14-74691922-INS-301
chr14-74697771-INS-60
chr14-74715066-INS-58
chr14-74728242-INS-167
chr14-74746169-INS-64
chr14-74752814-INS-327
chr14-74770033-INS-134
chr14-74778824-INS-296
chr14-74797424-INS-89
chr14-74798440-INS-76
chr14-74798619-INS-630
chr14-74815882-INS-763
chr14-74853899-INS-217
chr14-74893986-INS-169
chr14-74898359-INS-81
chr14-74909515-INS-186
chr14-74922885-INS-72
chr14-74926780-INS-114
chr14-74992492-INS-255
chr14-75003984-INS-188
chr14-75044977-INS-1144
chr14-75044978-INS-988
chr14-75053577-INS-182
chr14-75060720-INS-203
chr14-75068350-INS-54
chr14-75086902-INS-131
chr14-75095752-INS-58
chr14-75097470-INS-327
chr14-75105732-INS-112
chr14-75142875-INS-69
chr14-75188699-INS-4682
chr14-75238146-INS-102
chr14-75245435-INS-154
chr14-75247320-INS-121
chr14-75253369-INS-482
chr14-75257886-INS-365
chr14-75291464-INS-66
chr14-75294921-INS-57
chr14-75296707-INS-55
chr14-75309949-INS-670
chr14-75311768-INS-56
chr14-75335311-INS-56
chr14-75379380-INS-207
chr14-75400642-INS-115
chr14-75441745-INS-147
chr14-75443100-INS-165
chr14-75444741-INS-85
chr14-75446007-INS-680
chr14-75454472-INS-541
chr14-75492166-INS-498
chr14-75494513-INS-66
chr14-75504764-INS-1572

chr14-75506248-INS-385
chr14-75515459-INS-65
chr14-75519984-INS-563
chr14-75547506-INS-107
chr14-75555514-INS-513
chr14-75601856-INS-475
chr14-75623872-INS-85
chr14-75672142-INS-528
chr14-75679897-INS-63
chr14-75682209-INS-132
chr14-75692430-INS-51
chr14-75704446-INS-366
chr14-75706555-INS-70
chr14-75735439-INS-497
chr14-75745229-INS-52
chr14-75780762-INS-60
chr14-75786684-INS-463
chr14-75794849-INS-477
chr14-75798767-INS-56
chr14-75807080-INS-60
chr14-75810561-INS-58
chr14-75812460-INS-53
chr14-75824486-INS-120
chr14-75833283-INS-6714
chr14-75836092-INS-193
chr14-75837805-INS-116
chr14-75858606-INS-576
chr14-75862153-INS-491
chr14-75896616-INS-134
chr14-75911544-INS-102
chr14-75932278-INS-211
chr14-75961509-INS-197
chr14-75969560-INS-503
chr14-76003658-INS-128
chr14-76005922-INS-213
chr14-76020889-INS-150
chr14-76034188-INS-76
chr14-76035685-INS-221
chr14-76041547-INS-76
chr14-76042463-INS-68
chr14-76060565-INS-5031
chr14-76061939-INS-225
chr14-76064201-INS-60
chr14-76070087-INS-100
chr14-76089753-INS-186
chr14-76114755-INS-58
chr14-76116572-INS-194
chr14-76118241-INS-54
chr14-76139214-INS-695
chr14-76140350-INS-960
chr14-76147087-INS-345
chr14-76191712-INS-54
chr14-76207152-INS-721
chr14-76211334-INS-110
chr14-76215473-INS-76
chr14-76218590-INS-61
chr14-76254127-INS-666
chr14-76256070-INS-189
chr14-76260347-INS-58
chr14-76268640-INS-150
chr14-76284051-INS-304
chr14-76292468-INS-88
chr14-76295484-INS-439
chr14-76299930-INS-360
chr14-76305650-INS-51
chr14-76308340-INS-56
chr14-76320816-INS-244
chr14-76336925-INS-52
chr14-76341867-INS-487
chr14-76351466-INS-416
chr14-76356471-INS-52
chr14-76367453-INS-6440
chr14-76370272-INS-312
chr14-76374974-INS-202
chr14-76377112-INS-51
chr14-76383866-INS-7264
chr14-76397003-INS-159

chr14-76405556-INS-63
chr14-76413609-INS-52
chr14-76430545-INS-304
chr14-76431601-INS-66
chr14-76437067-INS-5544
chr14-76452710-INS-100
chr14-76457754-INS-471
chr14-76462545-INS-104
chr14-76474603-INS-7190
chr14-76484030-INS-483
chr14-76485418-INS-103
chr14-76488710-INS-390
chr14-76494790-INS-663
chr14-76522925-INS-101
chr14-76526778-INS-78
chr14-76530195-INS-473
chr14-76533160-INS-269
chr14-76535198-INS-97
chr14-76552262-INS-397
chr14-76558552-INS-55
chr14-76577213-INS-208
chr14-76597980-INS-703
chr14-76599458-INS-2265
chr14-76606432-INS-312
chr14-76620178-INS-579
chr14-76631745-INS-57
chr14-76631871-INS-786
chr14-76632498-INS-487
chr14-76650232-INS-103
chr14-76697536-INS-207
chr14-76718822-INS-103
chr14-76721774-INS-73
chr14-76736862-INS-76
chr14-76745829-INS-131
chr14-76749130-INS-52
chr14-76754974-INS-56
chr14-76760946-INS-210
chr14-76765214-INS-54
chr14-76798338-INS-239
chr14-76818946-INS-80
chr14-76880710-INS-1167
chr14-76882219-INS-52
chr14-76884310-INS-323
chr14-76901660-INS-131
chr14-76920428-INS-138
chr14-76933202-INS-52
chr14-76958086-INS-516
chr14-76974084-INS-74
chr14-76991942-INS-817
chr14-76992645-INS-66
chr14-77004811-INS-69
chr14-77012049-INS-101
chr14-77085960-INS-326
chr14-77090785-INS-69
chr14-77096972-INS-60
chr14-77134641-INS-331
chr14-77145564-INS-1988
chr14-77149295-INS-101
chr14-77168350-INS-5546
chr14-77199198-INS-133
chr14-77204814-INS-244
chr14-77210922-INS-300
chr14-77214877-INS-323
chr14-77260329-INS-350
chr14-77269918-INS-434
chr14-77273511-INS-63
chr14-77314566-INS-208
chr14-77315561-INS-203
chr14-77321621-INS-6475
chr14-77331134-INS-99
chr14-77344419-INS-50
chr14-77350290-INS-6194
chr14-77365018-INS-52
chr14-77398238-INS-58
chr14-77417397-INS-137
chr14-77441204-INS-180
chr14-77444190-INS-129

chr14-77449536-INS-70
chr14-77451507-INS-60
chr14-77457755-INS-63
chr14-77461536-INS-105
chr14-77482451-INS-108
chr14-77494803-INS-114
chr14-77526961-INS-4848
chr14-77530255-INS-2627
chr14-77533469-INS-304
chr14-77542879-INS-1634
chr14-77552171-INS-101
chr14-77556715-INS-72
chr14-77618573-INS-57
chr14-77620109-INS-6280
chr14-77624147-INS-54
chr14-77631428-INS-56
chr14-77639958-INS-58
chr14-77676901-INS-272
chr14-77682030-INS-67
chr14-77683145-INS-289
chr14-77704073-INS-55
chr14-77732694-INS-83
chr14-77755898-INS-52
chr14-77807101-INS-210
chr14-77858170-INS-7301
chr14-77867381-INS-62
chr14-77873691-INS-214
chr14-77899168-INS-215
chr14-77935315-INS-64
chr14-77937847-INS-130
chr14-78017905-INS-138
chr14-78056894-INS-158
chr14-78079737-INS-65
chr14-78173590-INS-327
chr14-78182842-INS-115
chr14-78190020-INS-138
chr14-78192571-INS-6947
chr14-78197331-INS-645
chr14-78210328-INS-59
chr14-78225157-INS-54
chr14-78241621-INS-288
chr14-78249079-INS-82
chr14-78283168-INS-177
chr14-78301273-INS-62
chr14-78321884-INS-4824
chr14-78331767-INS-72
chr14-78345533-INS-77
chr14-78366017-INS-281
chr14-78371925-INS-322
chr14-78397899-INS-7693
chr14-78402312-INS-4841
chr14-78439651-INS-82
chr14-78459532-INS-73
chr14-78465112-INS-58
chr14-78476150-INS-110
chr14-78494625-INS-56
chr14-78504693-INS-146
chr14-78517983-INS-145
chr14-78534385-INS-62
chr14-78561016-INS-111
chr14-78588115-INS-1504
chr14-78602866-INS-72
chr14-78610434-INS-1038
chr14-78623637-INS-634
chr14-78626100-INS-54
chr14-78638462-INS-56
chr14-78692742-INS-67
chr14-78698350-INS-106
chr14-78751813-INS-248
chr14-78780692-INS-618
chr14-78785536-INS-324
chr14-78787175-INS-160
chr14-78832311-INS-191
chr14-78834905-INS-196
chr14-78849945-INS-63
chr14-78853441-INS-828
chr14-78853448-INS-1622

chr14-78888296-INS-644
chr14-78930456-INS-52
chr14-78940595-INS-151
chr14-78977114-INS-68
chr14-79000988-INS-197
chr14-79019264-INS-353
chr14-79046633-INS-58
chr14-79050379-INS-523
chr14-79066140-INS-80
chr14-79098630-INS-331
chr14-79098811-INS-6455
chr14-79101164-INS-557
chr14-79132547-INS-70
chr14-79209532-INS-176
chr14-79212299-INS-68
chr14-79231509-INS-381
chr14-79233172-INS-114
chr14-79261815-INS-307
chr14-79288068-INS-109
chr14-79294681-INS-466
chr14-79295979-INS-141
chr14-79296554-INS-144
chr14-79355176-INS-51
chr14-79394323-INS-452
chr14-79407324-INS-963
chr14-79413561-INS-445
chr14-79416121-INS-312
chr14-79419504-INS-507
chr14-79488862-INS-52
chr14-79499786-INS-64
chr14-79502884-INS-504
chr14-79512547-INS-82
chr14-79526889-INS-301
chr14-79557412-INS-123
chr14-79624109-INS-186
chr14-79649748-INS-114
chr14-79652650-INS-1492
chr14-79656297-INS-186
chr14-79685851-INS-70
chr14-79687010-INS-203
chr14-79694399-INS-5545
chr14-79703946-INS-350
chr14-79708754-INS-247
chr14-79732491-INS-81
chr14-79747373-INS-91
chr14-79752333-INS-61
chr14-79763528-INS-130
chr14-79766027-INS-136
chr14-79790825-INS-205
chr14-79792235-INS-304
chr14-79815439-INS-143
chr14-79820147-INS-52
chr14-79830774-INS-59
chr14-79833012-INS-91
chr14-79841590-INS-176
chr14-79844272-INS-102
chr14-79889138-INS-64
chr14-79903112-INS-104
chr14-79904640-INS-389
chr14-79925847-INS-180
chr14-79943893-INS-206
chr14-79946659-INS-218
chr14-79951786-INS-211
chr14-79968482-INS-66
chr14-79989436-INS-109
chr14-80010586-INS-76
chr14-80012937-INS-92
chr14-80030561-INS-114
chr14-80054449-INS-206
chr14-80055512-INS-961
chr14-80092931-INS-62
chr14-80097652-INS-54
chr14-80106799-INS-110
chr14-80123877-INS-66
chr14-80143558-INS-195
chr14-80144751-INS-82
chr14-80148690-INS-71

chr14-80168470-INS-538
chr14-80169915-INS-121
chr14-80183355-INS-91
chr14-80185712-INS-300
chr14-80190942-INS-453
chr14-80197882-INS-79
chr14-80203601-INS-6195
chr14-80203603-INS-73
chr14-80205671-INS-54
chr14-80228889-INS-356
chr14-80232249-INS-145
chr14-80234348-INS-68
chr14-80246845-INS-61
chr14-80257967-INS-336
chr14-80264215-INS-150
chr14-80268684-INS-2175
chr14-80279925-INS-1009
chr14-80287908-INS-6631
chr14-80292298-INS-59
chr14-80296198-INS-52
chr14-80304069-INS-727
chr14-80316208-INS-1302
chr14-80335103-INS-122
chr14-80359670-INS-643
chr14-80368300-INS-66
chr14-80370335-INS-58
chr14-80371672-INS-153
chr14-80395598-INS-6648
chr14-80397902-INS-99
chr14-80404390-INS-307
chr14-80413547-INS-157
chr14-80433411-INS-95
chr14-80437863-INS-56
chr14-80485368-INS-130
chr14-80505067-INS-411
chr14-80530305-INS-3663
chr14-80531334-INS-692
chr14-80540383-INS-6892
chr14-80553157-INS-58
chr14-80554078-INS-6611
chr14-80559211-INS-6187
chr14-80579107-INS-400
chr14-80604861-INS-61
chr14-80605816-INS-1787
chr14-80608945-INS-52
chr14-80616996-INS-106
chr14-80620900-INS-7254
chr14-80631468-INS-80
chr14-80637496-INS-652
chr14-80640705-INS-135
chr14-80646952-INS-60
chr14-80651578-INS-145
chr14-80652395-INS-207
chr14-80665739-INS-51
chr14-80685541-INS-255
chr14-80701109-INS-524
chr14-80745949-INS-271
chr14-80759267-INS-64
chr14-80764753-INS-59
chr14-80780539-INS-72
chr14-80785043-INS-630
chr14-80790293-INS-52
chr14-80823513-INS-299
chr14-80846362-INS-240
chr14-80857034-INS-326
chr14-80857034-INS-525
chr14-80884949-INS-72
chr14-80888982-INS-8659
chr14-80888997-INS-425
chr14-80904731-INS-53
chr14-80919384-INS-71
chr14-80922891-INS-733
chr14-80924138-INS-219
chr14-80935583-INS-8748
chr14-80935584-INS-578
chr14-80948239-INS-586
chr14-80958703-INS-4046

chr14-80964092-INS-1889
chr14-80964527-INS-2206
chr14-81008865-INS-218
chr14-81012150-INS-471
chr14-81020631-INS-115
chr14-81031869-INS-444
chr14-81037687-INS-120
chr14-81041926-INS-51
chr14-81063898-INS-7164
chr14-81085642-INS-72
chr14-81109207-INS-66
chr14-81111102-INS-830
chr14-81111815-INS-480
chr14-81118958-INS-235
chr14-81131982-INS-6301
chr14-81145383-INS-842
chr14-81169801-INS-124
chr14-81173798-INS-401
chr14-81192496-INS-117
chr14-81196070-INS-58
chr14-81246656-INS-125
chr14-81285851-INS-342
chr14-81290383-INS-52
chr14-81299984-INS-106
chr14-81312336-INS-570
chr14-81360866-INS-876
chr14-81367760-INS-112
chr14-81370161-INS-159
chr14-81370794-INS-88
chr14-81372634-INS-576
chr14-81394284-INS-320
chr14-81404648-INS-67
chr14-81405622-INS-152
chr14-81413097-INS-7080
chr14-81447291-INS-61
chr14-81449415-INS-321
chr14-81450981-INS-54
chr14-81452365-INS-56
chr14-81517658-INS-344
chr14-81533442-INS-78
chr14-81536754-INS-311
chr14-81540410-INS-7395
chr14-81544375-INS-591
chr14-81549367-INS-198
chr14-81549618-INS-739
chr14-81551153-INS-69
chr14-81569178-INS-200
chr14-81580683-INS-188
chr14-81585522-INS-6172
chr14-81616453-INS-151
chr14-81622685-INS-91
chr14-81625360-INS-634
chr14-81627156-INS-91
chr14-81628809-INS-235
chr14-81636684-INS-137
chr14-81639256-INS-6940
chr14-81647649-INS-115
chr14-81658110-INS-79
chr14-81691116-INS-275
chr14-81705151-INS-118
chr14-81707287-INS-551
chr14-81716175-INS-68
chr14-81729348-INS-309
chr14-81742397-INS-6493
chr14-81745133-INS-1593
chr14-81748350-INS-72
chr14-81759451-INS-54
chr14-81768828-INS-112
chr14-81812816-INS-606
chr14-81861560-INS-814
chr14-81869897-INS-3673
chr14-81875137-INS-607
chr14-81884305-INS-90
chr14-81912355-INS-54
chr14-81920201-INS-2391
chr14-81939545-INS-317
chr14-81943870-INS-196

chr14-81946569-INS-1377
chr14-81958867-INS-80
chr14-81964540-INS-188
chr14-81975918-INS-53
chr14-81989229-INS-226
chr14-81991529-INS-120
chr14-81995312-INS-57
chr14-82012025-INS-105
chr14-82020833-INS-241
chr14-82040792-INS-55
chr14-82042358-INS-418
chr14-82044736-INS-179
chr14-82054692-INS-64
chr14-82057257-INS-156
chr14-82066167-INS-93
chr14-82066963-INS-6222
chr14-82067360-INS-82
chr14-82078424-INS-478
chr14-82127497-INS-162
chr14-82137967-INS-414
chr14-82149164-INS-6237
chr14-82197164-INS-60
chr14-82228171-INS-180
chr14-82240006-INS-112
chr14-82281941-INS-178
chr14-82286788-INS-6097
chr14-82297121-INS-79
chr14-82346937-INS-126
chr14-82354190-INS-63
chr14-82359673-INS-327
chr14-82368927-INS-168
chr14-82384927-INS-153
chr14-82395750-INS-222
chr14-82412093-INS-146
chr14-82447146-INS-74
chr14-82470349-INS-68
chr14-82472287-INS-56
chr14-82481980-INS-52
chr14-82486837-INS-150
chr14-82495145-INS-162
chr14-82536747-INS-608
chr14-82547550-INS-120
chr14-82578267-INS-189
chr14-82587864-INS-52
chr14-82604087-INS-56
chr14-82609683-INS-6180
chr14-82627380-INS-407
chr14-82642555-INS-263
chr14-82643739-INS-52
chr14-82659558-INS-60
chr14-82673277-INS-197
chr14-82676482-INS-738
chr14-82679498-INS-400
chr14-82710088-INS-1581
chr14-82727573-INS-59
chr14-82739222-INS-656
chr14-82767301-INS-120
chr14-82772630-INS-1597
chr14-82843845-INS-190
chr14-82877618-INS-975
chr14-82919096-INS-1130
chr14-82955588-INS-8584
chr14-82960912-INS-86
chr14-82968260-INS-765
chr14-82983381-INS-61
chr14-82992618-INS-507
chr14-82997280-INS-585
chr14-83005215-INS-271
chr14-83015573-INS-52
chr14-83025604-INS-248
chr14-83074532-INS-5430
chr14-83084068-INS-7031
chr14-83091726-INS-57
chr14-83093906-INS-64
chr14-83123662-INS-161
chr14-83133520-INS-608
chr14-83134967-INS-64

chr14-83155504-INS-127
chr14-83188229-INS-90
chr14-83257271-INS-323
chr14-83259003-INS-350
chr14-83294678-INS-57
chr14-83301160-INS-462
chr14-83330125-INS-60
chr14-83338894-INS-426
chr14-83344994-INS-400
chr14-83347245-INS-259
chr14-83354453-INS-62
chr14-83374884-INS-106
chr14-83382427-INS-52
chr14-83470420-INS-75
chr14-83480128-INS-95
chr14-83498828-INS-530
chr14-83505166-INS-396
chr14-83516998-INS-6201
chr14-83520895-INS-408
chr14-83524165-INS-2986
chr14-83528928-INS-377
chr14-83547879-INS-6160
chr14-83558064-INS-379
chr14-83566090-INS-64
chr14-83589457-INS-254
chr14-83597375-INS-113
chr14-83638262-INS-83
chr14-83686013-INS-478
chr14-83697127-INS-6716
chr14-83702057-INS-320
chr14-83809745-INS-547
chr14-83843037-INS-537
chr14-83863758-INS-448
chr14-83886843-INS-153
chr14-83894991-INS-65
chr14-83898279-INS-4567
chr14-83924361-INS-102
chr14-83957952-INS-56
chr14-83985808-INS-52
chr14-84011622-INS-572
chr14-84036366-INS-81
chr14-84054633-INS-70
chr14-84115006-INS-6403
chr14-84116356-INS-191
chr14-84124790-INS-98
chr14-84130054-INS-159
chr14-84192319-INS-6976
chr14-84193724-INS-252
chr14-84200397-INS-7087
chr14-84226251-INS-492
chr14-84247959-INS-7099
chr14-84259771-INS-449
chr14-84270136-INS-323
chr14-84290849-INS-209
chr14-84292906-INS-56
chr14-84335684-INS-78
chr14-84336457-INS-6447
chr14-84342067-INS-2449
chr14-84343952-INS-6391
chr14-84372496-INS-801
chr14-84375580-INS-205
chr14-84377322-INS-6862
chr14-84410162-INS-969
chr14-84419096-INS-112
chr14-84442625-INS-887
chr14-84450161-INS-116
chr14-84474000-INS-6392
chr14-84481532-INS-348
chr14-84530255-INS-7234
chr14-84531904-INS-652
chr14-84547464-INS-150
chr14-84563632-INS-189
chr14-84600960-INS-6161
chr14-84612163-INS-302
chr14-84616346-INS-434
chr14-84617314-INS-182
chr14-84617958-INS-628

chr14-84621147-INS-95
chr14-84632705-INS-6480
chr14-84634564-INS-1575
chr14-84637666-INS-239
chr14-84639673-INS-405
chr14-84647804-INS-3009
chr14-84660358-INS-216
chr14-84663617-INS-72
chr14-84700821-INS-60
chr14-84718702-INS-507
chr14-84740249-INS-135
chr14-84786065-INS-6332
chr14-84793852-INS-5280
chr14-84797989-INS-2176
chr14-84803604-INS-346
chr14-84811244-INS-78
chr14-84820173-INS-53
chr14-84863242-INS-581
chr14-84878810-INS-423
chr14-84891084-INS-711
chr14-84917661-INS-6295
chr14-84920544-INS-118
chr14-84975435-INS-410
chr14-84976026-INS-73
chr14-84992123-INS-558
chr14-85012624-INS-56
chr14-85042870-INS-504
chr14-85049554-INS-207
chr14-85122357-INS-163
chr14-85141571-INS-154
chr14-85143254-INS-3113
chr14-85185017-INS-291
chr14-85205751-INS-588
chr14-85257838-INS-52
chr14-85288925-INS-75
chr14-85291431-INS-120
chr14-85305043-INS-1117
chr14-85340286-INS-102
chr14-85404075-INS-715
chr14-85425496-INS-115
chr14-85435531-INS-6673
chr14-85445743-INS-531
chr14-85449640-INS-6447
chr14-85493518-INS-59
chr14-85534177-INS-296
chr14-85537889-INS-305
chr14-85550668-INS-68
chr14-85551399-INS-64
chr14-85615069-INS-129
chr14-85637993-INS-631
chr14-85642264-INS-117
chr14-85664250-INS-560
chr14-85672473-INS-160
chr14-85714767-INS-7324
chr14-85717630-INS-128
chr14-85739712-INS-82
chr14-85812303-INS-303
chr14-85833461-INS-537
chr14-85844822-INS-101
chr14-85887333-INS-738
chr14-85893496-INS-2070
chr14-85915888-INS-93
chr14-85929135-INS-516
chr14-85946052-INS-2762
chr14-85965834-INS-58
chr14-85997089-INS-160
chr14-85999132-INS-58
chr14-86001360-INS-248
chr14-86002685-INS-89
chr14-86009344-INS-57
chr14-86023242-INS-212
chr14-86036914-INS-382
chr14-86068453-INS-222
chr14-86087301-INS-77
chr14-86108634-INS-168
chr14-86121804-INS-7130
chr14-86165261-INS-521

chr14-86229656-INS-116
chr14-86238106-INS-288
chr14-86289107-INS-146
chr14-86315990-INS-149
chr14-86341340-INS-540
chr14-86349464-INS-2899
chr14-86382676-INS-265
chr14-86388626-INS-345
chr14-86401050-INS-306
chr14-86405495-INS-558
chr14-86406743-INS-268
chr14-86410479-INS-746
chr14-86443266-INS-300
chr14-86452541-INS-55
chr14-86459633-INS-56
chr14-86501496-INS-53
chr14-86504299-INS-349
chr14-86507632-INS-67
chr14-86534944-INS-96
chr14-86536879-INS-86
chr14-86557168-INS-313
chr14-86560518-INS-106
chr14-86645019-INS-365
chr14-86658715-INS-568
chr14-86719186-INS-194
chr14-86721143-INS-6458
chr14-86750635-INS-6728
chr14-86765252-INS-57
chr14-86771724-INS-400
chr14-86778444-INS-6419
chr14-86780951-INS-112
chr14-86807781-INS-348
chr14-86825863-INS-148
chr14-86838682-INS-271
chr14-86871522-INS-400
chr14-86901279-INS-6626
chr14-86960432-INS-107
chr14-86963329-INS-124
chr14-87002297-INS-54
chr14-87030257-INS-209
chr14-87033819-INS-6879
chr14-87069184-INS-97
chr14-87077069-INS-961
chr14-87124889-INS-119
chr14-87135747-INS-124
chr14-87166269-INS-738
chr14-87200578-INS-54
chr14-87217223-INS-186
chr14-87226544-INS-64
chr14-87228463-INS-400
chr14-87232170-INS-53
chr14-87251765-INS-285
chr14-87305348-INS-96
chr14-87319322-INS-85
chr14-87325827-INS-54
chr14-87337207-INS-52
chr14-87341368-INS-82
chr14-87353922-INS-717
chr14-87399628-INS-183
chr14-87408543-INS-122
chr14-87410681-INS-147
chr14-87423734-INS-124
chr14-87451235-INS-6711
chr14-87454143-INS-281
chr14-87477881-INS-6600
chr14-87481130-INS-145
chr14-87487418-INS-192
chr14-87498631-INS-56
chr14-87502035-INS-193
chr14-87508655-INS-58
chr14-87534108-INS-102
chr14-87576608-INS-69
chr14-87606052-INS-324
chr14-87606069-INS-173
chr14-87621268-INS-7150
chr14-87677441-INS-367
chr14-87697490-INS-228

chr14-87700492-INS-180
chr14-87701021-INS-640
chr14-87708089-INS-277
chr14-87718648-INS-238
chr14-87721671-INS-334
chr14-87727907-INS-425
chr14-87753025-INS-202
chr14-87767273-INS-201
chr14-87778170-INS-108
chr14-87807722-INS-58
chr14-87812438-INS-470
chr14-87817667-INS-6694
chr14-87850816-INS-198
chr14-87859527-INS-138
chr14-87859974-INS-687
chr14-87866184-INS-112
chr14-87887028-INS-63
chr14-87908628-INS-70
chr14-87915805-INS-1903
chr14-87945308-INS-407
chr14-87946319-INS-144
chr14-87963734-INS-268
chr14-87968806-INS-6380
chr14-87970897-INS-3684
chr14-87999397-INS-5852
chr14-88015042-INS-64
chr14-88022929-INS-58
chr14-88040206-INS-99
chr14-88123321-INS-177
chr14-88129412-INS-52
chr14-88144831-INS-404
chr14-88180451-INS-835
chr14-88216327-INS-691
chr14-88236772-INS-196
chr14-88242215-INS-169
chr14-88245753-INS-386
chr14-88256993-INS-255
chr14-88335945-INS-60
chr14-88353900-INS-498
chr14-88356196-INS-114
chr14-88406139-INS-616
chr14-88420434-INS-76
chr14-88442338-INS-6391
chr14-88463929-INS-359
chr14-88467157-INS-80
chr14-88467777-INS-842
chr14-88481862-INS-60
chr14-88498744-INS-862
chr14-88504764-INS-133
chr14-88528005-INS-1313
chr14-88540114-INS-275
chr14-88547146-INS-718
chr14-88562342-INS-58
chr14-88568397-INS-82
chr14-88577564-INS-6436
chr14-88583522-INS-6273
chr14-88611947-INS-206
chr14-88618185-INS-5378
chr14-88622540-INS-6357
chr14-88662154-INS-53
chr14-88712720-INS-106
chr14-88720169-INS-6227
chr14-88733472-INS-203
chr14-88745467-INS-67
chr14-88746894-INS-54
chr14-88783260-INS-628
chr14-88790123-INS-530
chr14-88806924-INS-158
chr14-88810922-INS-1406
chr14-88859777-INS-56
chr14-88878166-INS-7562
chr14-88935172-INS-6638
chr14-88940063-INS-251
chr14-88940253-INS-225
chr14-88956195-INS-136
chr14-89019892-INS-239
chr14-89096613-INS-54

chr14-89101243-INS-3733
chr14-89117263-INS-77
chr14-89214911-INS-6942
chr14-89226777-INS-77
chr14-89259940-INS-955
chr14-89277019-INS-81
chr14-89349035-INS-55
chr14-89468270-INS-97
chr14-89480928-INS-884
chr14-89536212-INS-152
chr14-89553730-INS-144
chr14-89580287-INS-164
chr14-89591955-INS-68
chr14-89626348-INS-5215
chr14-89710817-INS-320
chr14-89712034-INS-192
chr14-89766549-INS-221
chr14-89808550-INS-251
chr14-89811835-INS-222
chr14-89824875-INS-181
chr14-89881030-INS-348
chr14-89953660-INS-202
chr14-89982796-INS-53
chr14-89995884-INS-443
chr14-90007257-INS-878
chr14-90040606-INS-55
chr14-90054270-INS-517
chr14-90072448-INS-6216
chr14-90099174-INS-862
chr14-90103007-INS-55
chr14-90104618-INS-815
chr14-90167729-INS-88
chr14-90178293-INS-468
chr14-90186645-INS-403
chr14-90199817-INS-1041
chr14-90218213-INS-106
chr14-90221806-INS-55
chr14-90229427-INS-240
chr14-90266118-INS-60
chr14-90269790-INS-177
chr14-90339281-INS-103
chr14-90348860-INS-73
chr14-90406560-INS-84
chr14-90422961-INS-339
chr14-90468120-INS-110
chr14-90511060-INS-2285
chr14-90563762-INS-390
chr14-90568766-INS-90
chr14-90592961-INS-72
chr14-90594748-INS-818
chr14-90614045-INS-50
chr14-90614844-INS-759
chr14-90702389-INS-113
chr14-90729363-INS-469
chr14-90737418-INS-109
chr14-90792166-INS-1693
chr14-90794181-INS-145
chr14-90796148-INS-1529
chr14-90830453-INS-58
chr14-90847654-INS-128
chr14-90853221-INS-6894
chr14-90893018-INS-325
chr14-90929893-INS-206
chr14-91007697-INS-212
chr14-91012429-INS-57
chr14-91019683-INS-425
chr14-91021273-INS-197
chr14-91042021-INS-377
chr14-91056703-INS-58
chr14-91057675-INS-91
chr14-91153584-INS-199
chr14-91155669-INS-61
chr14-91179879-INS-65
chr14-91181265-INS-2121
chr14-91182280-INS-51
chr14-91227110-INS-7105
chr14-91243142-INS-95

chr14-91247092-INS-193
chr14-91254544-INS-611
chr14-91262107-INS-54
chr14-91280052-INS-90
chr14-91332931-INS-202
chr14-91339136-INS-496
chr14-91347892-INS-204
chr14-91349439-INS-6461
chr14-91362398-INS-349
chr14-91420292-INS-6631
chr14-91435243-INS-118
chr14-91450868-INS-169
chr14-91453415-INS-88
chr14-91490021-INS-178
chr14-91506234-INS-6229
chr14-91527427-INS-133
chr14-91599451-INS-79
chr14-91609116-INS-213
chr14-91615030-INS-52
chr14-91615576-INS-113
chr14-91645777-INS-77
chr14-91647652-INS-6177
chr14-91686844-INS-6755
chr14-91699639-INS-13331
chr14-91703579-INS-6379
chr14-91714264-INS-63
chr14-91723232-INS-212
chr14-91725611-INS-414
chr14-91743962-INS-81
chr14-91744189-INS-6402
chr14-91775996-INS-122
chr14-91779563-INS-65
chr14-91780686-INS-363
chr14-91784475-INS-616
chr14-91805314-INS-451
chr14-91809018-INS-160
chr14-91813253-INS-357
chr14-91825991-INS-7800
chr14-91827312-INS-375
chr14-91832875-INS-577
chr14-91885937-INS-53
chr14-91890572-INS-104
chr14-91917826-INS-181
chr14-91925124-INS-85
chr14-91967253-INS-65
chr14-91975681-INS-6476
chr14-91981075-INS-64
chr14-92007053-INS-568
chr14-92048874-INS-64
chr14-92052903-INS-294
chr14-92072051-INS-791
chr14-92073129-INS-358
chr14-92094160-INS-71
chr14-92120895-INS-88
chr14-92141644-INS-78
chr14-92154486-INS-334
chr14-92249354-INS-234
chr14-92262762-INS-54
chr14-92364385-INS-505
chr14-92366407-INS-882
chr14-92386759-INS-56
chr14-92390858-INS-52
chr14-92400386-INS-52
chr14-92401259-INS-847
chr14-92432356-INS-215
chr14-92442219-INS-334
chr14-92497256-INS-67
chr14-92502583-INS-51
chr14-92534761-INS-7618
chr14-92550025-INS-127
chr14-92551292-INS-546
chr14-92566007-INS-4362
chr14-92588822-INS-181
chr14-92599766-INS-111
chr14-92612433-INS-238
chr14-92628301-INS-120
chr14-92680882-INS-96

chr14-92696000-INS-92
chr14-92746244-INS-63
chr14-92746905-INS-64
chr14-92810891-INS-276
chr14-92812527-INS-502
chr14-92819645-INS-117
chr14-92844936-INS-6381
chr14-92847587-INS-300
chr14-92848722-INS-51
chr14-92857355-INS-52
chr14-92861498-INS-64
chr14-92894666-INS-52
chr14-92905864-INS-980
chr14-92906961-INS-51
chr14-92919176-INS-74
chr14-92939649-INS-67
chr14-92949016-INS-206
chr14-92954807-INS-94
chr14-92959412-INS-210
chr14-92972974-INS-321
chr14-93009413-INS-115
chr14-93013077-INS-207
chr14-93014343-INS-649
chr14-93016870-INS-2107
chr14-93021051-INS-56
chr14-93031700-INS-55
chr14-93053185-INS-52
chr14-93059983-INS-811
chr14-93075917-INS-543
chr14-93077741-INS-53
chr14-93148704-INS-823
chr14-93187305-INS-518
chr14-93253547-INS-6537
chr14-93266729-INS-140
chr14-93269706-INS-3166
chr14-93292651-INS-6311
chr14-93321129-INS-450
chr14-93337863-INS-90
chr14-93340746-INS-112
chr14-93350882-INS-221
chr14-93354268-INS-938
chr14-93359734-INS-138
chr14-93378207-INS-494
chr14-93387669-INS-66
chr14-93388483-INS-6383
chr14-93394169-INS-660
chr14-93400574-INS-92
chr14-93419815-INS-237
chr14-93440731-INS-65
chr14-93463532-INS-103
chr14-93514396-INS-344
chr14-93551117-INS-61
chr14-93558270-INS-66
chr14-93590024-INS-69
chr14-93590633-INS-55
chr14-93647566-INS-351
chr14-93686641-INS-208
chr14-93762011-INS-67
chr14-93797617-INS-123
chr14-93834977-INS-663
chr14-93838541-INS-128
chr14-93865471-INS-68
chr14-93914177-INS-447
chr14-93942645-INS-125
chr14-93956597-INS-56
chr14-93960426-INS-58
chr14-93960842-INS-558
chr14-94018598-INS-210
chr14-94037550-INS-159
chr14-94049563-INS-118
chr14-94053070-INS-776
chr14-94073132-INS-89
chr14-94085707-INS-179
chr14-94088529-INS-140
chr14-94109256-INS-92
chr14-94110635-INS-53
chr14-94136059-INS-60

chr14-94194646-INS-108
chr14-94196328-INS-54
chr14-94214079-INS-136
chr14-94219956-INS-52
chr14-94229901-INS-124
chr14-94247549-INS-59
chr14-94298898-INS-58
chr14-94320347-INS-3208
chr14-94322526-INS-279
chr14-94328402-INS-306
chr14-94352411-INS-823
chr14-94352660-INS-1405
chr14-94371506-INS-384
chr14-94388226-INS-443
chr14-94414166-INS-824
chr14-94448257-INS-6463
chr14-94459130-INS-52
chr14-94531420-INS-64
chr14-94538586-INS-167
chr14-94543078-INS-649
chr14-94604841-INS-214
chr14-94609968-INS-92
chr14-94649734-INS-730
chr14-94658536-INS-562
chr14-94664238-INS-56
chr14-94674990-INS-72
chr14-94747440-INS-548
chr14-94748245-INS-52
chr14-94752489-INS-115
chr14-94763456-INS-474
chr14-94806883-INS-138
chr14-94835712-INS-69
chr14-94838965-INS-513
chr14-94842195-INS-140
chr14-94899454-INS-71
chr14-94920185-INS-271
chr14-94934574-INS-531
chr14-95052097-INS-166
chr14-95056915-INS-56
chr14-95065638-INS-161
chr14-95120983-INS-81
chr14-95132635-INS-52
chr14-95135604-INS-223
chr14-95136657-INS-152
chr14-95163518-INS-1069
chr14-95166723-INS-473
chr14-95172707-INS-288
chr14-95175309-INS-708
chr14-95181391-INS-439
chr14-95214331-INS-66
chr14-95225614-INS-84
chr14-95226989-INS-585
chr14-95229391-INS-56
chr14-95273297-INS-58
chr14-95322403-INS-84
chr14-95360687-INS-144
chr14-95446323-INS-132
chr14-95495754-INS-108
chr14-95507006-INS-142
chr14-95520756-INS-883
chr14-95521293-INS-359
chr14-95536136-INS-70
chr14-95549766-INS-395
chr14-95569786-INS-3408
chr14-95584995-INS-7080
chr14-95645360-INS-784
chr14-95667859-INS-6511
chr14-95690060-INS-283
chr14-95726820-INS-286
chr14-95729538-INS-55
chr14-95735486-INS-59
chr14-95774994-INS-6891
chr14-95788786-INS-58
chr14-95817307-INS-54
chr14-95836091-INS-2140
chr14-95873719-INS-794
chr14-95880302-INS-894

chr14-95892898-INS-1653
chr14-95905411-INS-178
chr14-95951031-INS-114
chr14-95954903-INS-245
chr14-95960303-INS-105
chr14-95971812-INS-222
chr14-95983104-INS-205
chr14-96011097-INS-60
chr14-96022905-INS-431
chr14-96034688-INS-61
chr14-96048788-INS-65
chr14-96052782-INS-60
chr14-96101982-INS-96
chr14-96116224-INS-56
chr14-96143848-INS-199
chr14-96160625-INS-74
chr14-96164190-INS-68
chr14-96167196-INS-69
chr14-96171884-INS-482
chr14-96181019-INS-781
chr14-96198568-INS-216
chr14-96244467-INS-67
chr14-96253637-INS-131
chr14-96306854-INS-498
chr14-96317677-INS-263
chr14-96366322-INS-54
chr14-96388917-INS-55
chr14-96390639-INS-383
chr14-96462017-INS-116
chr14-96475518-INS-106
chr14-96486419-INS-116
chr14-96490010-INS-291
chr14-96587694-INS-547
chr14-96687453-INS-6142
chr14-96687745-INS-119
chr14-96693331-INS-340
chr14-96769520-INS-224
chr14-96774992-INS-137
chr14-96792990-INS-1478
chr14-96799526-INS-54
chr14-96846037-INS-139
chr14-96920441-INS-54
chr14-96966668-INS-911
chr14-96989670-INS-734
chr14-97004791-INS-6987
chr14-97007459-INS-4713
chr14-97008625-INS-940
chr14-97012551-INS-7203
chr14-97022149-INS-1571
chr14-97042790-INS-1735
chr14-97071818-INS-120
chr14-97078192-INS-6386
chr14-97084081-INS-598
chr14-97100159-INS-65
chr14-97102375-INS-7155
chr14-97129173-INS-343
chr14-97143990-INS-93
chr14-97166708-INS-53
chr14-97202912-INS-271
chr14-97213287-INS-203
chr14-97215016-INS-51
chr14-97241780-INS-59
chr14-97292720-INS-336
chr14-97297932-INS-72
chr14-97303995-INS-377
chr14-97318964-INS-192
chr14-97322537-INS-56
chr14-97324333-INS-630
chr14-97326211-INS-405
chr14-97335177-INS-666
chr14-97418578-INS-238
chr14-97452397-INS-314
chr14-97453559-INS-58
chr14-97483306-INS-136
chr14-97497211-INS-52
chr14-97533210-INS-530
chr14-97540367-INS-2431

chr14-97555157-INS-419
chr14-97559417-INS-81
chr14-97574350-INS-137
chr14-97597894-INS-6570
chr14-97631258-INS-56
chr14-97664042-INS-128
chr14-97665254-INS-315
chr14-97680265-INS-100
chr14-97698903-INS-276
chr14-97735884-INS-7485
chr14-97736264-INS-106
chr14-97762485-INS-667
chr14-97813419-INS-6887
chr14-97820347-INS-295
chr14-97835286-INS-78
chr14-97842833-INS-58
chr14-97850249-INS-59
chr14-97883863-INS-67
chr14-97909344-INS-120
chr14-97915431-INS-98
chr14-97923236-INS-429
chr14-97930642-INS-544
chr14-97938743-INS-61
chr14-97951854-INS-80
chr14-97979397-INS-592
chr14-97983232-INS-54
chr14-98048239-INS-331
chr14-98049926-INS-692
chr14-98058946-INS-115
chr14-98069638-INS-66
chr14-98090340-INS-269
chr14-98098689-INS-101
chr14-98100985-INS-6439
chr14-98113429-INS-102
chr14-98116209-INS-7162
chr14-98136303-INS-1114
chr14-98154546-INS-65
chr14-98158851-INS-55
chr14-98189676-INS-6300
chr14-98231919-INS-59
chr14-98249538-INS-65
chr14-98252306-INS-560
chr14-98329184-INS-60
chr14-98363634-INS-139
chr14-98367167-INS-674
chr14-98378027-INS-789
chr14-98407728-INS-70
chr14-98430373-INS-292
chr14-98438814-INS-53
chr14-98483720-INS-88
chr14-98526829-INS-52
chr14-98568063-INS-132
chr14-98575675-INS-1352
chr14-98584376-INS-136
chr14-98588770-INS-124
chr14-98622490-INS-221
chr14-98656748-INS-81
chr14-98662083-INS-52
chr14-98666392-INS-74
chr14-98671766-INS-717
chr14-98714097-INS-80
chr14-98717760-INS-225
chr14-98727055-INS-66
chr14-98743963-INS-990
chr14-98746828-INS-132
chr14-98762506-INS-502
chr14-98764022-INS-56
chr14-98781486-INS-58
chr14-98785345-INS-167
chr14-98816861-INS-57
chr14-98831623-INS-94
chr14-98892653-INS-6343
chr14-98914391-INS-294
chr14-98914523-INS-693
chr14-98916023-INS-593
chr14-98938865-INS-90
chr14-98951943-INS-65

chr14-98953846-INS-170
chr14-98958251-INS-134
chr14-98962150-INS-941
chr14-98978788-INS-130
chr14-98991450-INS-59
chr14-99008233-INS-196
chr14-99061539-INS-162
chr14-99068968-INS-1974
chr14-99105348-INS-77
chr14-99142628-INS-64
chr14-99146394-INS-168
chr14-99204745-INS-121
chr14-99213479-INS-103
chr14-99216962-INS-186
chr14-99226287-INS-53
chr14-99235936-INS-139
chr14-99249429-INS-410
chr14-99293731-INS-191
chr14-99308192-INS-71
chr14-99376800-INS-130
chr14-99389292-INS-401
chr14-99393315-INS-201
chr14-99470027-INS-75
chr14-99471641-INS-73
chr14-99510022-INS-139
chr14-99525631-INS-53
chr14-99530513-INS-114
chr14-99532661-INS-200
chr14-99538505-INS-83
chr14-99540223-INS-134
chr14-99558348-INS-245
chr14-99563087-INS-54
chr14-99568236-INS-113
chr14-99576051-INS-60
chr14-99589615-INS-201
chr14-99617738-INS-200
chr14-99629867-INS-54
chr14-99648011-INS-63
chr14-99667271-INS-110
chr14-99676987-INS-81
chr14-99699115-INS-339
chr14-99699624-INS-72
chr14-99705664-INS-280
chr14-99714098-INS-123
chr14-99742382-INS-63
chr14-99764015-INS-65
chr14-99776330-INS-140
chr14-99855506-INS-88
chr14-99877066-INS-76
chr14-99919486-INS-157
chr14-99932842-INS-385
chr14-99940498-INS-193
chr14-99950910-INS-6451
chr14-100036372-INS-475
chr14-100060113-INS-498
chr14-100098932-INS-7799
chr14-100154161-INS-280
chr14-100186607-INS-110
chr14-100207385-INS-59
chr14-100217148-INS-100
chr14-100240456-INS-2549
chr14-100311753-INS-279
chr14-100318851-INS-71
chr14-100348563-INS-172
chr14-100356478-INS-245
chr14-100360739-INS-130
chr14-100418040-INS-164
chr14-100425971-INS-62
chr14-100455779-INS-114
chr14-100503866-INS-56
chr14-100514063-INS-97
chr14-100554163-INS-58
chr14-100554250-INS-6361
chr14-100568652-INS-587
chr14-100572171-INS-6520
chr14-100589805-INS-224
chr14-100613534-INS-71

chr14-100648228-INS-95
chr14-100674343-INS-60
chr14-100677795-INS-108
chr14-100735460-INS-140
chr14-100740284-INS-353
chr14-100741188-INS-53
chr14-100765044-INS-6680
chr14-100769611-INS-214
chr14-100779066-INS-350
chr14-100812732-INS-3316
chr14-100819558-INS-6381
chr14-100827830-INS-151
chr14-100832826-INS-114
chr14-100861056-INS-52
chr14-100864753-INS-127
chr14-100865794-INS-402
chr14-100873979-INS-6280
chr14-100892495-INS-416
chr14-100894284-INS-963
chr14-100901073-INS-69
chr14-100906897-INS-199
chr14-100908219-INS-402
chr14-100910048-INS-6361
chr14-100934709-INS-205
chr14-100967220-INS-254
chr14-100975034-INS-209
chr14-100986574-INS-96
chr14-101005181-INS-364
chr14-101011479-INS-1947
chr14-101014785-INS-877
chr14-101042254-INS-54
chr14-101047992-INS-847
chr14-101072400-INS-5196
chr14-101077298-INS-662
chr14-101086016-INS-372
chr14-101134773-INS-330
chr14-101176387-INS-171
chr14-101214127-INS-563
chr14-101214547-INS-6678
chr14-101217202-INS-161
chr14-101252607-INS-374
chr14-101264879-INS-756
chr14-101266975-INS-108
chr14-101280852-INS-57
chr14-101301220-INS-383
chr14-101325160-INS-126
chr14-101359002-INS-145
chr14-101382224-INS-55
chr14-101384288-INS-7121
chr14-101427440-INS-56
chr14-101434656-INS-110
chr14-101449041-INS-403
chr14-101455665-INS-293
chr14-101468705-INS-95
chr14-101500523-INS-127
chr14-101515954-INS-914
chr14-101532354-INS-830
chr14-101567099-INS-56
chr14-101571967-INS-218
chr14-101587090-INS-248
chr14-101697015-INS-52
chr14-101704566-INS-115
chr14-101723163-INS-94
chr14-101744135-INS-206
chr14-101753642-INS-54
chr14-101758506-INS-77
chr14-101765955-INS-168
chr14-101770839-INS-90
chr14-101806916-INS-227
chr14-101808604-INS-382
chr14-101812556-INS-68
chr14-101813111-INS-485
chr14-101818429-INS-87
chr14-101830233-INS-74
chr14-101836365-INS-170
chr14-101887162-INS-183
chr14-101897635-INS-101

chr14-101920179-INS-202
chr14-101925709-INS-160
chr14-101928665-INS-139
chr14-101948147-INS-170
chr14-101961290-INS-924
chr14-101982480-INS-203
chr14-101994103-INS-140
chr14-101998050-INS-70
chr14-101999057-INS-56
chr14-102004232-INS-72
chr14-102005677-INS-193
chr14-102027018-INS-55
chr14-102074064-INS-938
chr14-102082494-INS-188
chr14-102117935-INS-72
chr14-102133180-INS-228
chr14-102146912-INS-175
chr14-102180410-INS-760
chr14-102184133-INS-5087
chr14-102187045-INS-109
chr14-102191001-INS-198
chr14-102208049-INS-152
chr14-102235892-INS-54
chr14-102250343-INS-181
chr14-102264418-INS-68
chr14-102277718-INS-123
chr14-102282503-INS-1022
chr14-102287823-INS-491
chr14-102314748-INS-111
chr14-102319382-INS-519
chr14-102326777-INS-136
chr14-102369141-INS-150
chr14-102413890-INS-172
chr14-102418348-INS-72
chr14-102438743-INS-211
chr14-102442330-INS-247
chr14-102461881-INS-675
chr14-102500807-INS-474
chr14-102512847-INS-61
chr14-102636055-INS-4864
chr14-102661745-INS-1153
chr14-102678302-INS-328
chr14-102693760-INS-377
chr14-102716390-INS-78
chr14-102717064-INS-277
chr14-102727841-INS-227
chr14-102743851-INS-144
chr14-102753836-INS-104
chr14-102761195-INS-298
chr14-102766888-INS-168
chr14-102772113-INS-229
chr14-102777341-INS-7134
chr14-102781901-INS-104
chr14-102791603-INS-52
chr14-102796879-INS-214
chr14-102815293-INS-55
chr14-102837049-INS-99
chr14-102844355-INS-56
chr14-102861159-INS-66
chr14-102863181-INS-365
chr14-102913588-INS-164
chr14-102917354-INS-239
chr14-102931967-INS-145
chr14-102965511-INS-310
chr14-102991500-INS-125
chr14-103005702-INS-70
chr14-103034292-INS-441
chr14-103034316-INS-5254
chr14-103039462-INS-225
chr14-103056966-INS-52
chr14-103058389-INS-82
chr14-103072031-INS-354
chr14-103078337-INS-1267
chr14-103093892-INS-1998
chr14-103096180-INS-89
chr14-103099653-INS-6262
chr14-103128419-INS-51

chr14-103139557-INS-112
chr14-103148379-INS-598
chr14-103149494-INS-68
chr14-103186760-INS-106
chr14-103187945-INS-178
chr14-103190532-INS-1437
chr14-103192043-INS-76
chr14-103204044-INS-113
chr14-103226045-INS-54
chr14-103233258-INS-708
chr14-103233297-INS-206
chr14-103241630-INS-210
chr14-103242089-INS-4025
chr14-103247311-INS-112
chr14-103248316-INS-6206
chr14-103265367-INS-52
chr14-103283108-INS-56
chr14-103283132-INS-701
chr14-103297422-INS-1236
chr14-103297423-INS-969
chr14-103344117-INS-183
chr14-103350149-INS-61
chr14-103385842-INS-68
chr14-103404999-INS-70
chr14-103430107-INS-51
chr14-103445897-INS-804
chr14-103486258-INS-69
chr14-103499238-INS-57
chr14-103504461-INS-110
chr14-103521174-INS-4368
chr14-103575357-INS-435
chr14-103586608-INS-60
chr14-103599307-INS-120
chr14-103613352-INS-64
chr14-103624253-INS-697
chr14-103671191-INS-65
chr14-103673046-INS-5276
chr14-103702094-INS-326
chr14-103703905-INS-80
chr14-103734303-INS-166
chr14-103743143-INS-250
chr14-103744119-INS-385
chr14-103758451-INS-61
chr14-103787462-INS-292
chr14-103796208-INS-56
chr14-103812052-INS-59
chr14-103819555-INS-7152
chr14-103823798-INS-850
chr14-103828031-INS-275
chr14-103847127-INS-118
chr14-103857081-INS-54
chr14-103866115-INS-71
chr14-103873081-INS-105
chr14-103907934-INS-55
chr14-103910622-INS-745
chr14-103910663-INS-100
chr14-103911953-INS-827
chr14-103914578-INS-3202
chr14-103914944-INS-265
chr14-103915154-INS-1179
chr14-103947641-INS-200
chr14-103950501-INS-193
chr14-103970855-INS-110
chr14-103990748-INS-117
chr14-104008659-INS-472
chr14-104029466-INS-216
chr14-104037507-INS-498
chr14-104044577-INS-117
chr14-104069424-INS-113
chr14-104077265-INS-99
chr14-104078913-INS-6447
chr14-104079112-INS-54
chr14-104089669-INS-200
chr14-104093807-INS-72
chr14-104095697-INS-269
chr14-104102175-INS-59
chr14-104123423-INS-197

chr14-104127206-INS-204
chr14-104131660-INS-96
chr14-104135226-INS-6436
chr14-104135559-INS-87
chr14-104147225-INS-132
chr14-104152112-INS-7796
chr14-104160351-INS-301
chr14-104166726-INS-88
chr14-104169519-INS-70
chr14-104176884-INS-56
chr14-104177621-INS-239
chr14-104185114-INS-100
chr14-104200397-INS-60
chr14-104209240-INS-389
chr14-104214571-INS-561
chr14-104220072-INS-379
chr14-104221258-INS-63
chr14-104228519-INS-160
chr14-104239663-INS-223
chr14-104241251-INS-226
chr14-104282226-INS-51
chr14-104308819-INS-212
chr14-104309407-INS-424
chr14-104321422-INS-72
chr14-104400501-INS-7048
chr14-104405443-INS-118
chr14-104414861-INS-98
chr14-104415806-INS-154
chr14-104452989-INS-322
chr14-104471518-INS-484
chr14-104472849-INS-69
chr14-104547065-INS-526
chr14-104549093-INS-104
chr14-104563086-INS-6643
chr14-104564147-INS-52
chr14-104570187-INS-273
chr14-104578399-INS-891
chr14-104605902-INS-936
chr14-104609816-INS-81
chr14-104621879-INS-139
chr14-104629753-INS-950
chr14-104668113-INS-256
chr14-104691178-INS-66
chr14-104730840-INS-490
chr14-104732582-INS-62
chr14-104737847-INS-917
chr14-104750513-INS-90
chr14-104762119-INS-1894
chr14-104776171-INS-62
chr14-104778739-INS-286
chr14-104783037-INS-60
chr14-104792430-INS-61
chr14-104881585-INS-76
chr14-104885214-INS-334
chr14-104891931-INS-55
chr14-104899430-INS-520
chr14-104901004-INS-108
chr14-104904458-INS-56
chr14-104932837-INS-188
chr14-104955667-INS-7204
chr14-104958829-INS-7240
chr14-104959559-INS-5256
chr14-104960510-INS-304
chr14-104960722-INS-3255
chr14-104982143-INS-6095
chr14-104998611-INS-54
chr14-105004377-INS-735
chr14-105010015-INS-82
chr14-105013442-INS-62
chr14-105036621-INS-54
chr14-105046055-INS-6447
chr14-105055881-INS-1891
chr14-105077319-INS-102
chr14-105083040-INS-166
chr14-105091105-INS-80
chr14-105097763-INS-81
chr14-105106662-INS-77

chr14-105110325-INS-498
chr14-105112012-INS-71
chr14-105126609-INS-3085
chr14-105133200-INS-144
chr14-105167923-INS-161
chr14-105190002-INS-6428
chr14-105192457-INS-54
chr14-105199065-INS-56
chr14-105201214-INS-52
chr14-105206732-INS-831
chr14-105208859-INS-215
chr14-105214173-INS-6793
chr14-105223262-INS-67
chr14-105232323-INS-99
chr14-105256388-INS-7158
chr14-105293640-INS-198
chr14-105298471-INS-95
chr14-105301444-INS-146
chr14-105311323-INS-60
chr14-105317989-INS-716
chr14-105318027-INS-162
chr14-105330380-INS-85
chr14-105334236-INS-5002
chr14-105341489-INS-114
chr14-105394969-INS-225
chr14-105400824-INS-173
chr14-105403803-INS-194
chr14-105418435-INS-392
chr14-105422256-INS-126
chr14-105455988-INS-168
chr14-105460779-INS-446
chr14-105467455-INS-201
chr14-105474769-INS-68
chr14-105501783-INS-228
chr14-105527255-INS-68
chr14-105527745-INS-59
chr14-105529210-INS-118
chr14-105530244-INS-211
chr14-105552810-INS-231
chr14-105557457-INS-135
chr14-105559650-INS-316
chr14-105577187-INS-534
chr14-105585943-INS-61
chr14-105592168-INS-400
chr14-105595062-INS-114
chr14-105595903-INS-289
chr14-105606089-INS-52
chr14-105609887-INS-65
chr14-105612926-INS-116
chr14-105622392-INS-693
chr14-105632918-INS-70
chr14-105644108-INS-248
chr14-105646084-INS-816
chr14-105646816-INS-74
chr14-105653532-INS-292
chr14-105658825-INS-602
chr14-105670927-INS-144
chr14-105673794-INS-56
chr14-105691763-INS-391
chr14-105714391-INS-64
chr14-105715593-INS-145
chr14-105720108-INS-88
chr14-105723437-INS-97
chr14-105725452-INS-935
chr14-105733632-INS-60
chr14-105737672-INS-178
chr14-105752483-INS-788
chr14-105768375-INS-530
chr14-105775356-INS-52
chr14-105776839-INS-120
chr14-105784923-INS-335
chr14-105786257-INS-60
chr14-105800387-INS-159
chr14-105811902-INS-126
chr14-105815893-INS-55
chr14-105828970-INS-663
chr14-105861476-INS-281

chr14-105876663-INS-167
chr14-105891209-INS-113
chr14-105895487-INS-90
chr14-105899270-INS-61
chr14-105923060-INS-52
chr14-105957228-INS-482
chr14-105972870-INS-196
chr14-105975194-INS-316
chr14-106016594-INS-61
chr14-106031990-INS-86
chr14-106050676-INS-926
chr14-106054395-INS-84
chr14-106058477-INS-166
chr14-106070569-INS-52
chr14-106072708-INS-216
chr14-106081164-INS-92
chr14-106091722-INS-211
chr14-106096204-INS-86
chr14-106101834-INS-169
chr14-106112442-INS-140
chr14-106117884-INS-124
chr14-106122096-INS-136
chr14-106126569-INS-223
chr14-106127218-INS-5556
chr14-106139781-INS-67
chr14-106156076-INS-217
chr14-106158772-INS-56
chr14-106172226-INS-214
chr14-106187831-INS-52
chr14-106189252-INS-381
chr14-106189944-INS-51
chr14-106191819-INS-6596
chr14-106192091-INS-76
chr14-106193856-INS-153
chr14-106223507-INS-64
chr14-106237229-INS-7558
chr14-106243035-INS-74
chr14-106246067-INS-205
chr14-106250471-INS-575
chr14-106255930-INS-51
chr14-106276513-INS-96
chr14-106286299-INS-999
chr14-106286722-INS-67
chr14-106292479-INS-80
chr14-106297992-INS-99
chr14-106314186-INS-6875
chr14-106326750-INS-57
chr14-106388333-INS-400
chr14-106396027-INS-2871
chr14-106405285-INS-827
chr14-106406394-INS-61
chr14-106416677-INS-77
chr14-106419325-INS-139
chr14-106422712-INS-302
chr14-106427913-INS-116
chr14-106450438-INS-72
chr14-106454399-INS-210
chr14-106466321-INS-448
chr14-106468971-INS-52
chr14-106478158-INS-54
chr14-106485296-INS-408
chr14-106487929-INS-88
chr14-106490321-INS-445
chr14-106490387-INS-7184
chr14-106504448-INS-307
chr14-106510581-INS-294
chr14-106586372-INS-52
chr14-106594376-INS-278
chr14-106612902-INS-58
chr14-106627768-INS-86
chr14-106632590-INS-88
chr14-106633700-INS-105
chr14-106641224-INS-112
chr14-106644148-INS-84
chr14-106655648-INS-64
chr14-106657134-INS-78
chr14-106723129-INS-6452

chr14-106733212-INS-716
chr14-106735764-INS-863
chr14-106762612-INS-750
chr14-106765225-INS-83
chr14-106769194-INS-122
chr14-106790123-INS-80
chr14-106815216-INS-568
chr14-106867822-INS-321
chr14-106885933-INS-1821
chr14-106890630-INS-445
chr14-106905381-INS-74
chr14-106914711-INS-52
chr14-106919032-INS-577
chr14-106921154-INS-63
chr14-106954815-INS-164
chr14-106959190-INS-208
chr14-106967328-INS-66
chr14-106980975-INS-594
chr14-107000859-INS-133
chr14-107066194-INS-161
chr14-107086792-INS-95
chr14-107100285-INS-66
chr14-107110492-INS-115
chr14-107111901-INS-132
chr14-107116099-INS-246
chr14-107130574-INS-156
chr14-107139173-INS-1078
chr14-107149034-INS-508
chr14-107156921-INS-235
chr14-107188833-INS-6675
chr14-107193212-INS-643
chr14-107211939-INS-67
chr14-107222845-INS-69
chr14-107225629-INS-457
chr14-107229641-INS-165
chr14-107242387-INS-88
chr14-107242776-INS-447
chr14-107243207-INS-1606
chr14-107244442-INS-86
chr14-107273081-INS-60
chr14-107282502-INS-6915
chr14-107295392-INS-407
chr14-107301294-INS-52
chr14-107302152-INS-13146
chr14-107311113-INS-6454
chr14-107312824-INS-301
chr14-107336689-INS-2804
chr14-107341333-INS-455
chr14-107369491-INS-247
chr14-107387523-INS-6381
chr14-107389475-INS-169
chr14-107411255-INS-191
chr14-107415767-INS-6142
chr14-107418680-INS-303
chr14-107429151-INS-270
chr14-107459615-INS-72
chr14-107462123-INS-6589
chr14-107464897-INS-6185
chr14-107468624-INS-56
chr14-107478078-INS-6538
chr14-107478079-INS-448
chr14-107490609-INS-498
chr14-107493036-INS-80
chr14-107509993-INS-146
chr14-107520311-INS-1516
chr14-107528471-INS-400
chr14-107530260-INS-52
chr14-107543034-INS-270
chr14-107577567-INS-76
chr14-107589125-INS-173
chr14-107595879-INS-549
chr14-107611200-INS-179
chr14-107641244-INS-878
chr14-107656999-INS-182
chr14-107673213-INS-58
chr14-107682725-INS-95
chr14-107692165-INS-984

chr14-107699890-INS-674
chr14-107709879-INS-209
chr14-107718902-INS-339
chr14-107730827-INS-59
chr14-107748337-INS-245
chr14-107754163-INS-68
chr14-107779956-INS-148
chr14-107794938-INS-97
chr14-107804218-INS-53
chr14-107811127-INS-208
chr14-107826878-INS-5282
chr14-107876972-INS-53
chr14-107883494-INS-218
chr14-107888852-INS-297
chr14-107888854-INS-2728
chr14-107893489-INS-58
chr14-107914840-INS-2370
chr14-107927933-INS-129
chr14-107943264-INS-61
chr14-107945984-INS-6171
chr14-107965227-INS-503
chr14-107974493-INS-119
chr14-107985410-INS-56
chr14-107993667-INS-176
chr14-107994492-INS-212
chr14-108008364-INS-6437
chr14-108030781-INS-439
chr14-108044450-INS-6359
chr14-108046302-INS-253
chr14-108059716-INS-58
chr14-108076974-INS-1891
chr14-108103470-INS-510
chr14-108127818-INS-536
chr14-108146232-INS-196
chr14-108151585-INS-98
chr14-108159198-INS-68
chr14-108175433-INS-141
chr14-108188391-INS-170
chr14-108193396-INS-55
chr14-108199083-INS-426
chr14-108201310-INS-7157
chr14-108203921-INS-361
chr14-108213425-INS-390
chr14-108216486-INS-183
chr14-108217827-INS-106
chr14-108241192-INS-96
chr14-108242611-INS-702
chr14-108269459-INS-90
chr14-108284963-INS-253
chr14-108337737-INS-332
chr14-108347539-INS-60
chr14-108349040-INS-262
chr14-108360971-INS-966
chr14-108381084-INS-175
chr14-108382756-INS-540
chr14-108397637-INS-7094
chr14-108422115-INS-430
chr14-108438218-INS-2947
chr14-108447553-INS-742
chr14-108461997-INS-5736
chr14-108464033-INS-55
chr14-108467837-INS-940
chr14-108480875-INS-60
chr14-108485756-INS-196
chr14-108510413-INS-344
chr14-108530893-INS-64
chr14-108548043-INS-156
chr14-108560860-INS-119
chr14-108572741-INS-6459
chr14-108601078-INS-1931
chr14-108601596-INS-1895
chr14-108609709-INS-250
chr14-108614872-INS-510
chr14-108624031-INS-56
chr14-108632662-INS-53
chr14-108639367-INS-52
chr14-108648350-INS-72

chr14-108663682-INS-272
chr14-108667001-INS-68
chr14-108706660-INS-206
chr14-108736621-INS-962
chr14-108749396-INS-7186
chr14-108753357-INS-164
chr14-108776352-INS-6582
chr14-108780627-INS-144
chr14-108819924-INS-112
chr14-108827527-INS-63
chr14-108844579-INS-271
chr14-108874715-INS-141
chr14-108884392-INS-206
chr14-108979337-INS-477
chr14-108988516-INS-452
chr14-108990821-INS-183
chr14-109024131-INS-68
chr14-109034436-INS-542
chr14-109036190-INS-104
chr14-109046838-INS-416
chr14-109061547-INS-201
chr14-109081356-INS-6374
chr14-109097719-INS-56
chr14-109109374-INS-768
chr14-109121329-INS-51
chr14-109143414-INS-55
chr14-109155921-INS-86
chr14-109178907-INS-121
chr14-109183764-INS-348
chr14-109225603-INS-98
chr14-109226254-INS-58
chr14-109263597-INS-2282
chr14-109281611-INS-418
chr14-109291431-INS-701
chr14-109293632-INS-5195
chr14-109305582-INS-433
chr14-109328856-INS-55
chr14-109371804-INS-6487
chr14-109388122-INS-649
chr14-109441166-INS-875
chr14-109441965-INS-94
chr14-109462056-INS-181
chr14-109519403-INS-252
chr14-109542724-INS-52
chr14-109559062-INS-125
chr14-109588497-INS-78
chr14-109591498-INS-402
chr14-109595415-INS-957
chr14-109624651-INS-7103
chr14-109626292-INS-143
chr14-109658544-INS-81
chr14-109659416-INS-515
chr14-109669290-INS-51
chr14-109682156-INS-56
chr14-109690686-INS-158
chr14-109698332-INS-52
chr14-109704334-INS-203
chr14-109715243-INS-216
chr14-109715872-INS-253
chr14-109740037-INS-796
chr14-109745388-INS-1025
chr14-109755816-INS-7068
chr14-109838202-INS-54
chr14-109871900-INS-5038
chr14-109872935-INS-809
chr14-109899559-INS-342
chr14-109913528-INS-63
chr14-109915587-INS-86
chr14-109920335-INS-58
chr14-109923639-INS-335
chr14-109934601-INS-218
chr14-109938783-INS-97
chr14-109941185-INS-218
chr14-109952537-INS-127
chr14-109992911-INS-184
chr14-109997429-INS-64
chr14-110004706-INS-85

chr14-110021590-INS-54
chr14-110056343-INS-55
chr14-110062298-INS-52
chr14-110083275-INS-216
chr14-110144875-INS-319
chr14-110163833-INS-6424
chr14-110203032-INS-7317
chr14-110259896-INS-161
chr14-110285189-INS-951
chr14-110336876-INS-72
chr14-110350371-INS-165
chr14-110363104-INS-58
chr14-110371585-INS-999
chr14-110387345-INS-179
chr14-110463859-INS-907
chr14-110466085-INS-66
chr14-110468943-INS-56
chr14-110475908-INS-52
chr14-110475962-INS-502
chr14-110520699-INS-70
chr14-110537091-INS-733
chr14-110545524-INS-138
chr14-110551657-INS-63
chr14-110592528-INS-4426
chr14-110614436-INS-421
chr14-110619205-INS-5300
chr14-110621949-INS-6576
chr14-110647574-INS-5052
chr14-110671090-INS-64
chr14-110708223-INS-7123
chr14-110713203-INS-81
chr14-110737534-INS-89
chr14-110770995-INS-6528
chr14-110820894-INS-200
chr14-110888693-INS-237
chr14-110903770-INS-80
chr14-110951778-INS-79
chr14-110954396-INS-158
chr14-110959624-INS-156
chr14-110976437-INS-95
chr14-110994703-INS-65
chr14-111002986-INS-220
chr14-111045882-INS-490
chr14-111057040-INS-82
chr14-111064936-INS-1888
chr14-111134841-INS-58
chr14-111158176-INS-80
chr14-111218729-INS-312
chr14-111220024-INS-217
chr14-111269009-INS-151
chr14-111285033-INS-263
chr14-111295659-INS-84
chr14-111305074-INS-63
chr14-111306670-INS-54
chr14-111313203-INS-898
chr14-111385394-INS-103
chr14-111388727-INS-344
chr14-111405387-INS-765
chr14-111414596-INS-539
chr14-111415542-INS-95
chr14-111420729-INS-74
chr14-111435496-INS-483
chr14-111447340-INS-88
chr14-111493747-INS-932
chr14-111498577-INS-78
chr14-111508047-INS-67
chr14-111517255-INS-139
chr14-111526136-INS-209
chr14-111546528-INS-102
chr14-111569552-INS-1895
chr14-111598149-INS-103
chr14-111612549-INS-71
chr14-111612954-INS-4155
chr14-111629964-INS-69
chr14-111634193-INS-464
chr14-111639850-INS-65
chr14-111645667-INS-398

chr14-111649744-INS-135
chr14-111666432-INS-56
chr14-111681613-INS-325
chr14-111706138-INS-531
chr14-111708104-INS-4799
chr14-111718851-INS-56
chr14-111750846-INS-54
chr14-111808180-INS-123
chr14-111811109-INS-394
chr14-111859399-INS-137
chr14-111869508-INS-299
chr14-111897028-INS-52
chr14-111979916-INS-74
chr14-111985237-INS-6658
chr14-112002931-INS-63
chr14-112041391-INS-61
chr14-112056034-INS-93
chr14-112059033-INS-7333
chr14-112105558-INS-440
chr14-112131195-INS-7292
chr14-112140287-INS-3449
chr14-112152701-INS-6698
chr14-112155422-INS-53
chr14-112188089-INS-74
chr14-112193491-INS-6726
chr14-112203085-INS-76
chr14-112207389-INS-128
chr14-112230058-INS-135
chr14-112235786-INS-67
chr14-112250440-INS-350
chr14-112250758-INS-603
chr14-112278292-INS-92
chr14-112285342-INS-60
chr14-112285850-INS-69
chr14-112300523-INS-58
chr14-112316778-INS-54
chr14-112322940-INS-63
chr14-112350108-INS-62
chr14-112354137-INS-476
chr14-112356221-INS-379
chr14-112374908-INS-51
chr14-112380628-INS-88
chr14-112395274-INS-621
chr14-112425031-INS-584
chr14-112428893-INS-61
chr14-112433011-INS-820
chr14-112438867-INS-123
chr14-112440923-INS-111
chr14-112441907-INS-619
chr14-112453853-INS-201
chr14-112465209-INS-80
chr14-112474704-INS-54
chr14-112502902-INS-64
chr14-112540620-INS-6710
chr14-112603141-INS-1864
chr14-112616265-INS-365
chr14-112624716-INS-315
chr14-112646509-INS-6215
chr14-112652588-INS-492
chr14-112663817-INS-63
chr14-112701081-INS-105
chr14-112704734-INS-288
chr14-112712992-INS-104
chr14-112724680-INS-216
chr14-112732307-INS-275
chr14-112774460-INS-53
chr14-112777819-INS-104
chr14-112795369-INS-197
chr14-112843563-INS-542
chr14-112849655-INS-67
chr14-112860563-INS-62
chr14-112862224-INS-54
chr14-112881782-INS-180
chr14-112894477-INS-111
chr14-112903465-INS-482
chr14-112905132-INS-334
chr14-112927547-INS-502

chr14-112930801-INS-7292
chr14-112959395-INS-211
chr14-113001219-INS-74
chr14-113042572-INS-76
chr14-113058102-INS-53
chr14-113060164-INS-482
chr14-113067700-INS-872
chr14-113072523-INS-480
chr14-113101236-INS-52
chr14-113110489-INS-6709
chr14-113115908-INS-83
chr14-113116843-INS-475
chr14-113129585-INS-54
chr14-113136515-INS-80
chr14-113151870-INS-185
chr14-113166230-INS-75
chr14-113191212-INS-60
chr14-113192967-INS-339
chr14-113208539-INS-545
chr14-113220345-INS-239
chr14-113231180-INS-301
chr14-113239978-INS-449
chr14-113252854-INS-7091
chr14-113288030-INS-5449
chr14-113294651-INS-125
chr14-113335548-INS-4732
chr14-113335933-INS-60
chr14-113350169-INS-516
chr14-113365210-INS-131
chr14-113366854-INS-60
chr14-113382365-INS-158
chr14-113386687-INS-105
chr14-113422171-INS-76
chr14-113498823-INS-70
chr14-113508529-INS-6887
chr14-113516019-INS-69
chr14-113518877-INS-6435
chr14-113539899-INS-304
chr14-113544085-INS-5048
chr14-113576279-INS-82
chr14-113595540-INS-108
chr14-113693781-INS-176
chr14-113710539-INS-351
chr14-113712136-INS-53
chr14-113713503-INS-203
chr14-113760474-INS-548
chr14-113762229-INS-430
chr14-113766688-INS-63
chr14-113773679-INS-7905
chr14-113791740-INS-302
chr14-113799621-INS-547
chr14-113806588-INS-401
chr14-113814208-INS-504
chr14-113849046-INS-136
chr14-113869558-INS-6419
chr14-113896924-INS-764
chr14-113939972-INS-203
chr14-113952804-INS-63
chr14-113982041-INS-7122
chr14-113995622-INS-108
chr14-114049435-INS-1540
chr14-114059906-INS-150
chr14-114077710-INS-1890
chr14-114087446-INS-279
chr14-114110396-INS-6468
chr14-114129183-INS-6670
chr14-114157639-INS-80
chr14-114161016-INS-55
chr14-114193521-INS-76
chr14-114207047-INS-8095
chr14-114235385-INS-3344
chr14-114256998-INS-170
chr14-114274207-INS-255
chr14-114290020-INS-328
chr14-114299929-INS-333
chr14-114344464-INS-53
chr14-114378156-INS-99

chr14-114431740-INS-54
chr14-114466638-INS-419
chr14-114471340-INS-749
chr14-114487046-INS-357
chr14-114507801-INS-608
chr14-114511456-INS-6470
chr14-114548793-INS-67
chr14-114578924-INS-310
chr14-114586052-INS-106
chr14-114598552-INS-649
chr14-114599758-INS-228
chr14-114601838-INS-94
chr14-114606688-INS-199
chr14-114609983-INS-62
chr14-114616716-INS-4496
chr14-114632011-INS-7095
chr14-114643509-INS-954
chr14-114654955-INS-580
chr14-114664487-INS-834
chr14-114686622-INS-219
chr14-114688637-INS-845
chr14-114697831-INS-397
chr14-114708126-INS-64
chr14-114736976-INS-61
chr14-114753110-INS-51
chr14-114760404-INS-226
chr14-114770929-INS-830
chr14-114773037-INS-4414
chr14-114786523-INS-108
chr14-114787647-INS-78
chr14-114794281-INS-207
chr14-114800472-INS-375
chr14-114802607-INS-212
chr14-114815326-INS-61
chr14-114826386-INS-7017
chr14-114826895-INS-176
chr14-114838216-INS-63
chr14-114858207-INS-80
chr14-114864831-INS-96
chr14-114877432-INS-56
chr14-114886746-INS-54
chr14-114935681-INS-107
chr14-114939143-INS-588
chr14-114943332-INS-3061
chr14-114956723-INS-75
chr14-114986230-INS-7841
chr14-114986471-INS-133
chr14-114991600-INS-439
chr14-114993330-INS-1888
chr14-114995589-INS-7105
chr14-114999674-INS-252
chr14-115007405-INS-144
chr14-115010279-INS-160
chr14-115022155-INS-201
chr14-115032963-INS-1926
chr14-115035771-INS-400
chr14-115040518-INS-3287
chr14-115048055-INS-171
chr14-115050044-INS-57
chr14-115097052-INS-293
chr14-115114425-INS-117
chr14-115124892-INS-62
chr14-115133005-INS-179
chr14-115153645-INS-56
chr14-115155708-INS-640
chr14-115159833-INS-56
chr14-115162042-INS-3172
chr14-115174462-INS-6818
chr14-115182689-INS-413
chr14-115183901-INS-7566
chr14-115196727-INS-328
chr14-115197294-INS-105
chr14-115244128-INS-6864
chr14-115250129-INS-218
chr14-115250484-INS-268
chr14-115253964-INS-438
chr14-115271006-INS-382

chr14-115276644-INS-52
chr14-115288963-INS-144
chr14-115298687-INS-544
chr14-115316584-INS-824
chr14-115317145-INS-60
chr14-115326236-INS-60
chr14-115331987-INS-139
chr14-115342642-INS-52
chr14-115363987-INS-150
chr14-115365919-INS-6462
chr14-115405153-INS-193
chr14-115410639-INS-186
chr14-115412651-INS-1281
chr14-115421674-INS-441
chr14-115426836-INS-72
chr14-115435593-INS-384
chr14-115437718-INS-7057
chr14-115444184-INS-100
chr14-115444396-INS-699
chr14-115445662-INS-96
chr14-115490125-INS-78
chr14-115491208-INS-84
chr14-115500680-INS-6472
chr14-115568297-INS-231
chr14-115571111-INS-8309
chr14-115583161-INS-4494
chr14-115597166-INS-201
chr14-115616500-INS-582
chr14-115620705-INS-8337
chr14-115637489-INS-235
chr14-115643036-INS-300
chr14-115649749-INS-5319
chr14-115665253-INS-74
chr14-115680516-INS-64
chr14-115711531-INS-7965
chr14-115718747-INS-177
chr14-115741124-INS-1893
chr14-115742025-INS-117
chr14-115761719-INS-88
chr14-115778610-INS-2948
chr14-115782563-INS-187
chr14-115805511-INS-296
chr14-115833764-INS-55
chr14-115857595-INS-156
chr14-115859720-INS-246
chr14-115864447-INS-994
chr14-115880493-INS-60
chr14-115903681-INS-2918
chr14-115909106-INS-60
chr14-115912784-INS-125
chr14-115917069-INS-58
chr14-115932476-INS-557
chr14-115939709-INS-74
chr14-115953968-INS-826
chr14-115963847-INS-6706
chr14-115973451-INS-152
chr14-115993839-INS-295
chr14-115997067-INS-923
chr14-115999552-INS-81
chr14-116004715-INS-52
chr14-116024019-INS-138
chr14-116025945-INS-165
chr14-116031670-INS-68
chr14-116033209-INS-60
chr14-116051421-INS-182
chr14-116057718-INS-78
chr14-116058775-INS-60
chr14-116072192-INS-112
chr14-116081086-INS-248
chr14-116098721-INS-98
chr14-116109362-INS-7515
chr14-116120725-INS-60
chr14-116137821-INS-385
chr14-116140442-INS-3619
chr14-116158221-INS-1894
chr14-116179315-INS-175
chr14-116185967-INS-6157

chr14-116186016-INS-368
chr14-116193586-INS-329
chr14-116247675-INS-59
chr14-116251525-INS-64
chr14-116343913-INS-166
chr14-116347978-INS-6570
chr14-116387931-INS-150
chr14-116389788-INS-77
chr14-116425675-INS-60
chr14-116449873-INS-117
chr14-116452398-INS-464
chr14-116453079-INS-51
chr14-116481809-INS-123
chr14-116484500-INS-109
chr14-116498597-INS-78
chr14-116500967-INS-174
chr14-116504652-INS-105
chr14-116509004-INS-762
chr14-116514323-INS-765
chr14-116516042-INS-130
chr14-116535486-INS-84
chr14-116540907-INS-184
chr14-116550507-INS-332
chr14-116554235-INS-5309
chr14-116571411-INS-63
chr14-116571528-INS-796
chr14-116580572-INS-108
chr14-116588150-INS-6380
chr14-116589924-INS-6764
chr14-116599985-INS-100
chr14-116645411-INS-112
chr14-116657618-INS-368
chr14-116663099-INS-324
chr14-116742342-INS-94
chr14-116752475-INS-6202
chr14-116781724-INS-885
chr14-116787206-INS-10984
chr14-116792870-INS-361
chr14-116794822-INS-6805
chr14-116799981-INS-87
chr14-116821016-INS-134
chr14-116831326-INS-8070
chr14-116840952-INS-66
chr14-116842835-INS-594
chr14-116902629-INS-57
chr14-116927526-INS-195
chr14-116933827-INS-199
chr14-116933896-INS-770
chr14-116953240-INS-520
chr14-116982500-INS-95
chr14-116986735-INS-100
chr14-116991908-INS-6399
chr14-117033872-INS-52
chr14-117052950-INS-109
chr14-117064855-INS-72
chr14-117111264-INS-194
chr14-117122097-INS-75
chr14-117144256-INS-91
chr14-117159360-INS-52
chr14-117170288-INS-69
chr14-117172243-INS-90
chr14-117177623-INS-78
chr14-117199664-INS-137
chr14-117208122-INS-60
chr14-117211517-INS-129
chr14-117223005-INS-212
chr14-117228928-INS-74
chr14-117253997-INS-56
chr14-117255557-INS-83
chr14-117283848-INS-639
chr14-117312945-INS-606
chr14-117321257-INS-93
chr14-117322696-INS-332
chr14-117358598-INS-819
chr14-117363545-INS-2151
chr14-117365690-INS-119
chr14-117391222-INS-593

chr14-117393281-INS-70
chr14-117397100-INS-202
chr14-117405510-INS-507
chr14-117426078-INS-63
chr14-117435037-INS-405
chr14-117440556-INS-59
chr14-117452436-INS-54
chr14-117460316-INS-210
chr14-117489094-INS-196
chr14-117491139-INS-57
chr14-117506616-INS-531
chr14-117549991-INS-92
chr14-117554172-INS-158
chr14-117578853-INS-801
chr14-117585713-INS-119
chr14-117591967-INS-104
chr14-117604284-INS-98
chr14-117608389-INS-62
chr14-117619067-INS-96
chr14-117620204-INS-70
chr14-117642407-INS-72
chr14-117645632-INS-714
chr14-117650103-INS-135
chr14-117667256-INS-236
chr14-117669060-INS-6451
chr14-117678938-INS-180
chr14-117682139-INS-1275
chr14-117692616-INS-786
chr14-117696363-INS-585
chr14-117706480-INS-58
chr14-117708175-INS-105
chr14-117716426-INS-81
chr14-117724025-INS-7058
chr14-117732754-INS-627
chr14-117744168-INS-327
chr14-117759615-INS-60
chr14-117765406-INS-298
chr14-117766181-INS-54
chr14-117772170-INS-2602
chr14-117808041-INS-471
chr14-117811185-INS-182
chr14-117817891-INS-1337
chr14-117835738-INS-6592
chr14-117843250-INS-59
chr14-117844131-INS-6889
chr14-117845350-INS-321
chr14-117850020-INS-155
chr14-117894536-INS-156
chr14-117901891-INS-161
chr14-117903549-INS-180
chr14-117906342-INS-62
chr14-117907652-INS-60
chr14-117958024-INS-126
chr14-117959533-INS-92
chr14-117976735-INS-192
chr14-117985692-INS-142
chr14-117988402-INS-169
chr14-117997795-INS-125
chr14-118017588-INS-167
chr14-118025478-INS-158
chr14-118027972-INS-544
chr14-118049290-INS-271
chr14-118051902-INS-72
chr14-118061537-INS-7483
chr14-118069722-INS-105
chr14-118077709-INS-692
chr14-118078807-INS-132
chr14-118081443-INS-58
chr14-118082373-INS-59
chr14-118107283-INS-54
chr14-118138765-INS-172
chr14-118148687-INS-74
chr14-118167000-INS-94
chr14-118183297-INS-58
chr14-118184263-INS-109
chr14-118190314-INS-86
chr14-118214322-INS-57

chr14-118223583-INS-134
chr14-118234282-INS-180
chr14-118237431-INS-100
chr14-118249255-INS-221
chr14-118252902-INS-6434
chr14-118255462-INS-51
chr14-118261174-INS-510
chr14-118263484-INS-6081
chr14-118266069-INS-108
chr14-118272134-INS-67
chr14-118279098-INS-364
chr14-118331539-INS-76
chr14-118345654-INS-106
chr14-118356747-INS-181
chr14-118363212-INS-193
chr14-118383130-INS-77
chr14-118389334-INS-323
chr14-118390591-INS-192
chr14-118393674-INS-223
chr14-118403076-INS-6452
chr14-118408009-INS-278
chr14-118427845-INS-64
chr14-118436454-INS-127
chr14-118443713-INS-132
chr14-118445442-INS-957
chr14-118458082-INS-204
chr14-118466613-INS-115
chr14-118483548-INS-104
chr14-118488031-INS-54
chr14-118530745-INS-117
chr14-118554362-INS-52
chr14-118571703-INS-54
chr14-118573023-INS-97
chr14-118596003-INS-6347
chr14-118619698-INS-203
chr14-118627379-INS-265
chr14-118679355-INS-63
chr14-118689362-INS-573
chr14-118693894-INS-82
chr14-118705686-INS-426
chr14-118708930-INS-134
chr14-118732101-INS-57
chr14-118738586-INS-130
chr14-118746009-INS-304
chr14-118764073-INS-111
chr14-118781041-INS-261
chr14-118827876-INS-112
chr14-118863260-INS-173
chr14-118874817-INS-201
chr14-118878315-INS-168
chr14-118896796-INS-60
chr14-118903115-INS-5253
chr14-118907219-INS-122
chr14-118909091-INS-53
chr14-118958275-INS-211
chr14-118959247-INS-84
chr14-118972579-INS-103
chr14-118972893-INS-162
chr14-118989575-INS-123
chr14-119006424-INS-633
chr14-119006435-INS-107
chr14-119008459-INS-254
chr14-119015814-INS-198
chr14-119022273-INS-220
chr14-119038803-INS-290
chr14-119039746-INS-342
chr14-119066034-INS-75
chr14-119072312-INS-1890
chr14-119073040-INS-78
chr14-119091545-INS-58
chr14-119111415-INS-667
chr14-119116052-INS-755
chr14-119118975-INS-161
chr14-119210263-INS-54
chr14-119251056-INS-397
chr14-119257747-INS-94
chr14-119263808-INS-580

chr14-119265635-INS-178
chr14-119267392-INS-408
chr14-119271169-INS-137
chr14-119271450-INS-224
chr14-119293331-INS-6852
chr14-119298619-INS-199
chr14-119304686-INS-101
chr14-119315961-INS-228
chr14-119318457-INS-173
chr14-119323895-INS-662
chr14-119341821-INS-123
chr14-119347567-INS-411
chr14-119357030-INS-6531
chr14-119382446-INS-249
chr14-119384634-INS-61
chr14-119388289-INS-7045
chr14-119406906-INS-76
chr14-119416093-INS-6462
chr14-119418273-INS-187
chr14-119437659-INS-2854
chr14-119461503-INS-543
chr14-119469908-INS-505
chr14-119513571-INS-191
chr14-119536926-INS-124
chr14-119596754-INS-193
chr14-119611167-INS-54
chr14-119645236-INS-59
chr14-119664089-INS-331
chr14-119669149-INS-246
chr14-119713049-INS-85
chr14-119722463-INS-114
chr14-119759318-INS-162
chr14-119767499-INS-327
chr14-119825509-INS-100
chr14-119826823-INS-548
chr14-119905657-INS-66
chr14-119956320-INS-226
chr14-119964329-INS-308
chr14-119986178-INS-65
chr14-120013006-INS-360
chr14-120017268-INS-233
chr14-120049739-INS-741
chr14-120062042-INS-71
chr14-120066102-INS-176
chr14-120076915-INS-169
chr14-120110872-INS-63
chr14-120123196-INS-384
chr14-120129009-INS-317
chr14-120155819-INS-72
chr14-120187410-INS-84
chr14-120191169-INS-60
chr14-120246164-INS-329
chr14-120261778-INS-80
chr14-120263353-INS-63
chr14-120269725-INS-1770
chr14-120271797-INS-52
chr14-120295037-INS-56
chr14-120346700-INS-134
chr14-120366174-INS-86
chr14-120370937-INS-98
chr14-120371791-INS-167
chr14-120390505-INS-118
chr14-120416092-INS-54
chr14-120464375-INS-109
chr14-120480453-INS-434
chr14-120485950-INS-54
chr14-120496990-INS-177
chr14-120534816-INS-60
chr14-120549603-INS-87
chr14-120588299-INS-64
chr14-120601501-INS-92
chr14-120617462-INS-204
chr14-120659684-INS-76
chr14-120691545-INS-399
chr14-120694500-INS-77
chr14-120721843-INS-55
chr14-120740142-INS-263

chr14-120762755-INS-86
chr14-120782773-INS-316
chr14-120783642-INS-61
chr14-120820075-INS-754
chr14-120834104-INS-142
chr14-120863876-INS-62
chr14-120894716-INS-106
chr14-120921973-INS-202
chr14-120923963-INS-177
chr14-120928376-INS-529
chr14-120953706-INS-147
chr14-120966956-INS-55
chr14-120997566-INS-460
chr14-121014117-INS-524
chr14-121045114-INS-511
chr14-121055331-INS-498
chr14-121068115-INS-411
chr14-121095343-INS-146
chr14-121109252-INS-590
chr14-121141284-INS-244
chr14-121158949-INS-223
chr14-121169782-INS-98
chr14-121194357-INS-2260
chr14-121201344-INS-154
chr14-121207972-INS-430
chr14-121210562-INS-105
chr14-121235738-INS-621
chr14-121253127-INS-72
chr14-121261685-INS-75
chr14-121276088-INS-213
chr14-121300255-INS-59
chr14-121302930-INS-354
chr14-121332495-INS-56
chr14-121341729-INS-210
chr14-121344347-INS-55
chr14-121350855-INS-194
chr14-121369055-INS-251
chr14-121403750-INS-72
chr14-121416924-INS-119
chr14-121417466-INS-222
chr14-121426259-INS-56
chr14-121439330-INS-128
chr14-121449687-INS-88
chr14-121463714-INS-138
chr14-121471461-INS-201
chr14-121478672-INS-190
chr14-121495141-INS-92
chr14-121543141-INS-84
chr14-121544488-INS-155
chr14-121607224-INS-182
chr14-121618601-INS-104
chr14-121632066-INS-58
chr14-121653963-INS-61
chr14-121662521-INS-467
chr14-121671763-INS-116
chr14-121676380-INS-135
chr14-121678323-INS-60
chr14-121680587-INS-205
chr14-121689483-INS-515
chr14-121693635-INS-509
chr14-121700713-INS-70
chr14-121701056-INS-70
chr14-121708872-INS-190
chr14-121718961-INS-534
chr14-121720600-INS-79
chr14-121756173-INS-168
chr14-121767734-INS-124
chr14-121803805-INS-256
chr14-121812642-INS-350
chr14-121818312-INS-234
chr14-121820561-INS-69
chr14-121822772-INS-163
chr14-121841533-INS-318
chr14-121847065-INS-113
chr14-121852695-INS-60
chr14-121858357-INS-60
chr14-121899006-INS-148

chr14-121910447-INS-451
chr14-121952074-INS-190
chr14-121955512-INS-174
chr14-121959673-INS-359
chr14-121972298-INS-142
chr14-121985583-INS-498
chr14-121998123-INS-71
chr14-122005512-INS-234
chr14-122010520-INS-228
chr14-122044184-INS-507
chr14-122046587-INS-316
chr14-122049277-INS-220
chr14-122061081-INS-178
chr14-122068864-INS-222
chr14-122079226-INS-56
chr14-122084726-INS-528
chr14-122092041-INS-411
chr14-122134803-INS-149
chr14-122148484-INS-333
chr14-122158070-INS-100
chr14-122188777-INS-329
chr14-122205365-INS-58
chr14-122214910-INS-185
chr14-122240008-INS-895
chr14-122245671-INS-60
chr14-122250786-INS-140
chr14-122269191-INS-54
chr14-122280546-INS-7139
chr14-122281337-INS-65
chr14-122307258-INS-272
chr14-122311658-INS-326
chr14-122332699-INS-981
chr14-122346754-INS-190
chr14-122350153-INS-170
chr14-122370107-INS-61
chr14-122392260-INS-207
chr14-122415271-INS-53
chr14-122428666-INS-202
chr14-122448616-INS-54
chr14-122492656-INS-201
chr14-122497457-INS-205
chr14-122498964-INS-196
chr14-122513048-INS-243
chr14-122522979-INS-52
chr14-122533919-INS-250
chr14-122534717-INS-137
chr14-122554414-INS-182
chr14-122559577-INS-191
chr14-122580233-INS-60
chr14-122584684-INS-51
chr14-122586699-INS-54
chr14-122589637-INS-214
chr14-122595330-INS-57
chr14-122636947-INS-228
chr14-122646519-INS-223
chr14-122656741-INS-72
chr14-122668164-INS-197
chr14-122669955-INS-72
chr14-122672900-INS-279
chr14-122677367-INS-81
chr14-122678640-INS-64
chr14-122680377-INS-84
chr14-122681957-INS-157
chr14-122682406-INS-370
chr14-122738355-INS-4354
chr14-122753202-INS-206
chr14-122754675-INS-134
chr14-122768716-INS-441
chr14-122801743-INS-236
chr14-122821479-INS-60
chr14-122823336-INS-85
chr14-122836033-INS-180
chr14-122837988-INS-204
chr14-122843275-INS-137
chr14-122849457-INS-765
chr14-122853662-INS-81
chr14-122855486-INS-102

chr14-122878962-INS-159
chr14-122910389-INS-106
chr14-122917525-INS-96
chr14-122925671-INS-473
chr14-122929101-INS-140
chr14-122931853-INS-60
chr14-122939947-INS-132
chr14-122962763-INS-63
chr14-122965223-INS-141
chr14-122992582-INS-849
chr14-123045632-INS-358
chr14-123048652-INS-400
chr14-123060985-INS-108
chr14-123068138-INS-122
chr14-123090817-INS-285
chr14-123097032-INS-392
chr14-123136391-INS-496
chr14-123167514-INS-205
chr14-123231967-INS-119
chr14-123256863-INS-265
chr14-123266106-INS-72
chr14-123313593-INS-58
chr14-123317167-INS-6794
chr14-123318411-INS-315
chr14-123323389-INS-6192
chr14-123351341-INS-59
chr14-123380673-INS-6655
chr14-123402829-INS-95
chr14-123406057-INS-59
chr14-123413793-INS-197
chr14-123423145-INS-772
chr14-123432904-INS-873
chr14-123434444-INS-167
chr14-123450930-INS-76
chr14-123454013-INS-6190
chr14-123471086-INS-6496
chr14-123484397-INS-80
chr14-123525417-INS-73
chr14-123531522-INS-274
chr14-123536738-INS-63
chr14-123552892-INS-674
chr14-123562945-INS-154
chr14-123593307-INS-72
chr14-123625352-INS-710
chr14-123646040-INS-386
chr14-123673816-INS-129
chr14-123689034-INS-54
chr14-123717797-INS-126
chr14-123743247-INS-385
chr14-123746128-INS-87
chr14-123753620-INS-60
chr14-123765334-INS-159
chr14-123793927-INS-6534
chr14-123810162-INS-51
chr14-123849703-INS-5128
chr14-123854177-INS-64
chr14-123856251-INS-6923
chr14-123871453-INS-153
chr14-123874718-INS-336
chr14-123914864-INS-76
chr14-123917829-INS-199
chr14-123951343-INS-386
chr14-123987988-INS-93
chr14-124003933-INS-440
chr14-124005361-INS-99
chr14-124010369-INS-209
chr14-124054284-INS-72
chr14-124067966-INS-199
chr14-124099488-INS-6894
chr14-124103861-INS-1541
chr14-124109407-INS-172
chr14-124123042-INS-60
chr14-124139723-INS-198
chr14-124202261-INS-62
chr14-124219157-INS-54
chr14-124223364-INS-9247
chr14-124230617-INS-738

chr14-124241213-INS-175
chr14-124270663-INS-548
chr14-124294858-INS-54
chr14-124309906-INS-278
chr14-124326626-INS-539
chr14-124342663-INS-92
chr14-124346547-INS-448
chr14-124357021-INS-73
chr14-124361614-INS-56
chr14-124369049-INS-91
chr14-124390485-INS-688
chr14-124433620-INS-241
chr14-124440301-INS-6178
chr14-124448787-INS-440
chr14-124489830-INS-443
chr14-124497083-INS-6563
chr14-124499292-INS-6610
chr14-124504662-INS-615
chr14-124531688-INS-94
chr14-124556429-INS-197
chr14-124581250-INS-117
chr14-124587598-INS-153
chr14-124618224-INS-110
chr14-124627520-INS-149
chr14-124653126-INS-408
chr14-124669619-INS-190
chr14-124698231-INS-113
chr14-124715715-INS-6231
chr14-124719053-INS-440
chr14-124736406-INS-85
chr14-124742514-INS-99
chr14-124769099-INS-2679
chr14-124773408-INS-56
chr14-124776701-INS-152
chr14-124789466-INS-7950
chr14-124834712-INS-10338
chr14-124854631-INS-682
chr14-124860171-INS-52
chr14-124866977-INS-6227
chr14-124881648-INS-236
chr14-124882572-INS-5527
chr14-124898703-INS-821
chr14-124905365-INS-188
chr14-124919420-INS-651
chr14-124919960-INS-6329
chr14-124950618-INS-321
chr14-124962183-INS-6924
chr14-124968781-INS-120
chr14-124980736-INS-96
chr14-125008464-INS-12892
chr14-125012137-INS-280
chr14-125014382-INS-6413
chr14-125035687-INS-486
chr15-3055524-INS-151
chr15-3058295-INS-70
chr15-3087480-INS-6442
chr15-3099030-INS-63
chr15-3124015-INS-60
chr15-3158974-INS-265
chr15-3169988-INS-60
chr15-3194504-INS-56
chr15-3201871-INS-68
chr15-3212312-INS-57
chr15-3216309-INS-6208
chr15-3217429-INS-52
chr15-3243954-INS-52
chr15-3244282-INS-201
chr15-3245580-INS-61
chr15-3263583-INS-108
chr15-3271282-INS-497
chr15-3278857-INS-332
chr15-3320469-INS-52
chr15-3325029-INS-80
chr15-3355687-INS-127
chr15-3379771-INS-63
chr15-3397696-INS-116
chr15-3430481-INS-7300

chr15-3432325-INS-150
chr15-3433947-INS-81
chr15-3442867-INS-5046
chr15-3442944-INS-69
chr15-3462123-INS-80
chr15-3488549-INS-60
chr15-3497678-INS-84
chr15-3519280-INS-70
chr15-3527905-INS-182
chr15-3535142-INS-364
chr15-3553192-INS-265
chr15-3555839-INS-83
chr15-3561866-INS-97
chr15-3590441-INS-435
chr15-3592203-INS-872
chr15-3592783-INS-52
chr15-3595097-INS-909
chr15-3605860-INS-103
chr15-3606758-INS-378
chr15-3609664-INS-414
chr15-3625548-INS-119
chr15-3635334-INS-64
chr15-3663139-INS-4061
chr15-3713655-INS-6660
chr15-3732020-INS-343
chr15-3811378-INS-229
chr15-3814811-INS-87
chr15-3834540-INS-52
chr15-3842032-INS-6585
chr15-3918162-INS-52
chr15-3966771-INS-127
chr15-3991186-INS-199
chr15-3996819-INS-149
chr15-4018782-INS-203
chr15-4036396-INS-526
chr15-4052205-INS-70
chr15-4133074-INS-456
chr15-4161193-INS-60
chr15-4164058-INS-105
chr15-4168884-INS-667
chr15-4177247-INS-77
chr15-4202744-INS-82
chr15-4207736-INS-121
chr15-4214257-INS-109
chr15-4221926-INS-579
chr15-4226344-INS-329
chr15-4234796-INS-552
chr15-4243742-INS-87
chr15-4248479-INS-92
chr15-4249789-INS-90
chr15-4268090-INS-400
chr15-4273495-INS-6573
chr15-4274782-INS-54
chr15-4296049-INS-62
chr15-4303739-INS-171
chr15-4311992-INS-6890
chr15-4329211-INS-2896
chr15-4337822-INS-7085
chr15-4343591-INS-58
chr15-4380524-INS-275
chr15-4385612-INS-323
chr15-4401331-INS-170
chr15-4436453-INS-1318
chr15-4453539-INS-6280
chr15-4462601-INS-164
chr15-4467399-INS-83
chr15-4545558-INS-63
chr15-4574451-INS-1886
chr15-4578358-INS-2336
chr15-4579029-INS-64
chr15-4581673-INS-236
chr15-4603210-INS-6514
chr15-4603368-INS-145
chr15-4612861-INS-1547
chr15-4621355-INS-55
chr15-4634040-INS-99
chr15-4647766-INS-128

chr15-4763498-INS-63
chr15-4780823-INS-550
chr15-4816939-INS-871
chr15-4842194-INS-264
chr15-4850277-INS-328
chr15-4853982-INS-84
chr15-4859662-INS-121
chr15-4860653-INS-63
chr15-4862843-INS-7186
chr15-4881342-INS-58
chr15-4905685-INS-293
chr15-4906869-INS-994
chr15-4959802-INS-129
chr15-4961064-INS-165
chr15-4966536-INS-68
chr15-4969655-INS-58
chr15-4971576-INS-181
chr15-4977246-INS-185
chr15-4979344-INS-574
chr15-4989452-INS-989
chr15-4992507-INS-102
chr15-5005457-INS-2441
chr15-5007147-INS-53
chr15-5045559-INS-196
chr15-5050039-INS-56
chr15-5055382-INS-188
chr15-5063114-INS-72
chr15-5070429-INS-84
chr15-5083330-INS-305
chr15-5108965-INS-431
chr15-5115513-INS-64
chr15-5120436-INS-109
chr15-5128246-INS-60
chr15-5131780-INS-816
chr15-5132762-INS-303
chr15-5138048-INS-5170
chr15-5139029-INS-182
chr15-5143551-INS-97
chr15-5155448-INS-74
chr15-5162653-INS-408
chr15-5170678-INS-309
chr15-5193924-INS-195
chr15-5197613-INS-408
chr15-5239168-INS-64
chr15-5239928-INS-182
chr15-5252748-INS-53
chr15-5286127-INS-52
chr15-5309343-INS-145
chr15-5312026-INS-128
chr15-5340621-INS-92
chr15-5348279-INS-190
chr15-5349758-INS-65
chr15-5360639-INS-147
chr15-5368812-INS-329
chr15-5370949-INS-101
chr15-5384780-INS-95
chr15-5454741-INS-1884
chr15-5462478-INS-1785
chr15-5493132-INS-408
chr15-5495621-INS-860
chr15-5512633-INS-55
chr15-5512948-INS-7629
chr15-5513570-INS-1889
chr15-5515901-INS-108
chr15-5548568-INS-1416
chr15-5597525-INS-245
chr15-5614672-INS-251
chr15-5616994-INS-205
chr15-5618183-INS-488
chr15-5633602-INS-102
chr15-5646351-INS-74
chr15-5664903-INS-63
chr15-5688568-INS-718
chr15-5690274-INS-6216
chr15-5702668-INS-111
chr15-5720493-INS-221
chr15-5723637-INS-94

chr15-5727828-INS-152
chr15-5750741-INS-867
chr15-5758426-INS-294
chr15-5779422-INS-509
chr15-5790873-INS-86
chr15-5798462-INS-205
chr15-5838805-INS-148
chr15-5840300-INS-6501
chr15-5845216-INS-64
chr15-5879068-INS-76
chr15-5887986-INS-184
chr15-5897810-INS-4893
chr15-5913134-INS-4365
chr15-5929864-INS-81
chr15-5960636-INS-6584
chr15-5962510-INS-3886
chr15-5976318-INS-6043
chr15-5980677-INS-152
chr15-5980981-INS-4950
chr15-5987247-INS-6050
chr15-5998176-INS-128
chr15-6022237-INS-208
chr15-6030789-INS-78
chr15-6032453-INS-174
chr15-6072812-INS-60
chr15-6073398-INS-275
chr15-6074515-INS-204
chr15-6085967-INS-254
chr15-6095360-INS-217
chr15-6133301-INS-228
chr15-6176682-INS-372
chr15-6216486-INS-78
chr15-6222678-INS-66
chr15-6240076-INS-420
chr15-6243995-INS-110
chr15-6254218-INS-51
chr15-6284975-INS-94
chr15-6296110-INS-180
chr15-6321980-INS-204
chr15-6342825-INS-124
chr15-6372616-INS-210
chr15-6397464-INS-60
chr15-6404271-INS-54
chr15-6405502-INS-6572
chr15-6410092-INS-2245
chr15-6410093-INS-883
chr15-6412933-INS-336
chr15-6421269-INS-102
chr15-6426766-INS-138
chr15-6434383-INS-1590
chr15-6435164-INS-67
chr15-6439920-INS-1113
chr15-6439921-INS-369
chr15-6455161-INS-52
chr15-6462831-INS-74
chr15-6485758-INS-147
chr15-6488816-INS-54
chr15-6524136-INS-104
chr15-6528674-INS-88
chr15-6574687-INS-451
chr15-6581038-INS-68
chr15-6586379-INS-60
chr15-6621984-INS-195
chr15-6627061-INS-252
chr15-6634147-INS-497
chr15-6655055-INS-575
chr15-6660313-INS-369
chr15-6662638-INS-173
chr15-6669251-INS-279
chr15-6677178-INS-52
chr15-6682361-INS-160
chr15-6700658-INS-165
chr15-6719402-INS-56
chr15-6720761-INS-51
chr15-6784780-INS-51
chr15-6878247-INS-65
chr15-6892018-INS-81

chr15-6896807-INS-76
chr15-6908473-INS-113
chr15-6922110-INS-208
chr15-6932764-INS-6440
chr15-6955298-INS-1862
chr15-6962450-INS-155
chr15-6968657-INS-446
chr15-7006133-INS-88
chr15-7015212-INS-155
chr15-7015573-INS-80
chr15-7019166-INS-133
chr15-7036564-INS-62
chr15-7082159-INS-192
chr15-7090511-INS-196
chr15-7095528-INS-57
chr15-7104692-INS-208
chr15-7106158-INS-5887
chr15-7109690-INS-6452
chr15-7121183-INS-959
chr15-7154187-INS-81
chr15-7183079-INS-74
chr15-7190058-INS-66
chr15-7198246-INS-207
chr15-7209124-INS-66
chr15-7218220-INS-122
chr15-7232262-INS-64
chr15-7238100-INS-207
chr15-7240768-INS-54
chr15-7243904-INS-104
chr15-7245018-INS-108
chr15-7251015-INS-776
chr15-7301656-INS-129
chr15-7313292-INS-121
chr15-7328885-INS-300
chr15-7362469-INS-51
chr15-7366144-INS-153
chr15-7383902-INS-180
chr15-7388706-INS-54
chr15-7418645-INS-132
chr15-7419870-INS-5352
chr15-7437622-INS-52
chr15-7443418-INS-90
chr15-7447595-INS-6594
chr15-7447975-INS-202
chr15-7460968-INS-148
chr15-7475240-INS-359
chr15-7482412-INS-7064
chr15-7502130-INS-129
chr15-7528873-INS-143
chr15-7530980-INS-167
chr15-7532090-INS-391
chr15-7543244-INS-60
chr15-7581847-INS-144
chr15-7583235-INS-6403
chr15-7586130-INS-425
chr15-7603419-INS-277
chr15-7614048-INS-133
chr15-7649027-INS-7075
chr15-7677649-INS-54
chr15-7687322-INS-621
chr15-7694913-INS-130
chr15-7704390-INS-148
chr15-7707008-INS-665
chr15-7732825-INS-326
chr15-7756269-INS-66
chr15-7764621-INS-216
chr15-7768551-INS-633
chr15-7776069-INS-176
chr15-7782613-INS-69
chr15-7790723-INS-55
chr15-7793556-INS-63
chr15-7805767-INS-153
chr15-7834757-INS-114
chr15-7861397-INS-88
chr15-7875128-INS-216
chr15-7875622-INS-96
chr15-7878156-INS-62

chr15-7919645-INS-600
chr15-7929774-INS-71
chr15-7932686-INS-55
chr15-7947950-INS-98
chr15-7948537-INS-336
chr15-7965045-INS-4057
chr15-7978043-INS-168
chr15-7997603-INS-219
chr15-8007568-INS-194
chr15-8013680-INS-59
chr15-8019223-INS-2907
chr15-8021521-INS-446
chr15-8025281-INS-2109
chr15-8025328-INS-470
chr15-8027482-INS-60
chr15-8038527-INS-6398
chr15-8043743-INS-275
chr15-8058986-INS-238
chr15-8061982-INS-59
chr15-8077494-INS-905
chr15-8089429-INS-153
chr15-8096786-INS-208
chr15-8107123-INS-2722
chr15-8115221-INS-219
chr15-8116557-INS-73
chr15-8120556-INS-195
chr15-8129160-INS-187
chr15-8149816-INS-104
chr15-8157022-INS-114
chr15-8174244-INS-152
chr15-8189758-INS-52
chr15-8196816-INS-214
chr15-8204633-INS-270
chr15-8206135-INS-225
chr15-8212264-INS-203
chr15-8213590-INS-192
chr15-8252697-INS-204
chr15-8257477-INS-538
chr15-8262124-INS-3317
chr15-8263423-INS-1864
chr15-8278862-INS-98
chr15-8289501-INS-725
chr15-8293164-INS-1151
chr15-8293254-INS-253
chr15-8333780-INS-132
chr15-8337930-INS-202
chr15-8387528-INS-224
chr15-8394989-INS-220
chr15-8398656-INS-190
chr15-8416581-INS-72
chr15-8427753-INS-89
chr15-8436240-INS-106
chr15-8462806-INS-66
chr15-8508938-INS-104
chr15-8521289-INS-66
chr15-8533360-INS-307
chr15-8536613-INS-341
chr15-8552941-INS-194
chr15-8562196-INS-204
chr15-8563782-INS-229
chr15-8572483-INS-862
chr15-8584652-INS-2921
chr15-8587199-INS-66
chr15-8606123-INS-56
chr15-8607502-INS-90
chr15-8610237-INS-69
chr15-8621406-INS-174
chr15-8635323-INS-62
chr15-8641429-INS-143
chr15-8672625-INS-83
chr15-8674742-INS-80
chr15-8688149-INS-66
chr15-8705593-INS-123
chr15-8706072-INS-424
chr15-8717677-INS-56
chr15-8730018-INS-321
chr15-8736632-INS-181

chr15-8742755-INS-97
chr15-8750802-INS-135
chr15-8767383-INS-67
chr15-8768873-INS-111
chr15-8770080-INS-187
chr15-8774563-INS-180
chr15-8779299-INS-66
chr15-8782970-INS-422
chr15-8791836-INS-51
chr15-8794026-INS-92
chr15-8808653-INS-182
chr15-8814259-INS-386
chr15-8824426-INS-140
chr15-8841588-INS-158
chr15-8853970-INS-265
chr15-8868345-INS-628
chr15-8871330-INS-222
chr15-8888998-INS-212
chr15-8911163-INS-57
chr15-8915433-INS-54
chr15-8920588-INS-278
chr15-8929249-INS-193
chr15-8976679-INS-60
chr15-9011325-INS-296
chr15-9028833-INS-52
chr15-9036030-INS-66
chr15-9042061-INS-7171
chr15-9059536-INS-129
chr15-9075836-INS-226
chr15-9078802-INS-52
chr15-9084468-INS-7066
chr15-9085461-INS-369
chr15-9085499-INS-152
chr15-9086358-INS-70
chr15-9092562-INS-89
chr15-9095382-INS-65
chr15-9099736-INS-308
chr15-9122172-INS-211
chr15-9126603-INS-91
chr15-9128776-INS-165
chr15-9171884-INS-204
chr15-9199833-INS-386
chr15-9229420-INS-3784
chr15-9229580-INS-65
chr15-9231339-INS-540
chr15-9244766-INS-6497
chr15-9262294-INS-925
chr15-9264775-INS-210
chr15-9289145-INS-58
chr15-9293012-INS-290
chr15-9303400-INS-174
chr15-9304800-INS-64
chr15-9330129-INS-80
chr15-9366688-INS-8954
chr15-9368899-INS-64
chr15-9378092-INS-6491
chr15-9415835-INS-6601
chr15-9433371-INS-76
chr15-9447039-INS-58
chr15-9454164-INS-549
chr15-9458160-INS-68
chr15-9461776-INS-162
chr15-9470269-INS-290
chr15-9474539-INS-56
chr15-9515868-INS-120
chr15-9525076-INS-56
chr15-9579914-INS-243
chr15-9620413-INS-171
chr15-9623527-INS-913
chr15-9645405-INS-108
chr15-9665856-INS-6370
chr15-9672525-INS-424
chr15-9703127-INS-219
chr15-9717378-INS-105
chr15-9726449-INS-77
chr15-9740380-INS-53
chr15-9741141-INS-813

chr15-9765014-INS-7508
chr15-9767015-INS-6773
chr15-9770351-INS-230
chr15-9786823-INS-6586
chr15-9821373-INS-670
chr15-9839312-INS-481
chr15-9845798-INS-6201
chr15-9848790-INS-78
chr15-9853109-INS-872
chr15-9863981-INS-430
chr15-9866942-INS-464
chr15-9873430-INS-312
chr15-9887205-INS-4305
chr15-9888296-INS-3193
chr15-9902317-INS-6236
chr15-9909878-INS-447
chr15-9920628-INS-2400
chr15-9925753-INS-118
chr15-9927781-INS-6196
chr15-9937535-INS-6135
chr15-9942048-INS-7204
chr15-9947650-INS-67
chr15-9954666-INS-341
chr15-9977895-INS-289
chr15-10002964-INS-376
chr15-10004603-INS-731
chr15-10009351-INS-74
chr15-10026590-INS-57
chr15-10052975-INS-241
chr15-10073171-INS-4101
chr15-10075048-INS-356
chr15-10086364-INS-95
chr15-10086678-INS-323
chr15-10113704-INS-331
chr15-10118938-INS-77
chr15-10165063-INS-104
chr15-10168269-INS-54
chr15-10169889-INS-1464
chr15-10171671-INS-6582
chr15-10174824-INS-104
chr15-10197726-INS-51
chr15-10202432-INS-153
chr15-10221263-INS-63
chr15-10242524-INS-89
chr15-10244610-INS-128
chr15-10256673-INS-518
chr15-10273725-INS-383
chr15-10312640-INS-165
chr15-10370923-INS-134
chr15-10378124-INS-53
chr15-10395995-INS-100
chr15-10397108-INS-424
chr15-10445626-INS-261
chr15-10456838-INS-153
chr15-10497499-INS-5389
chr15-10521956-INS-95
chr15-10534721-INS-186
chr15-10538176-INS-119
chr15-10568871-INS-80
chr15-10570618-INS-209
chr15-10572347-INS-65
chr15-10581510-INS-84
chr15-10583617-INS-59
chr15-10590441-INS-60
chr15-10591269-INS-327
chr15-10685538-INS-204
chr15-10720195-INS-77
chr15-10767561-INS-480
chr15-10789436-INS-71
chr15-10792763-INS-212
chr15-10793525-INS-7182
chr15-10803399-INS-128
chr15-10822455-INS-104
chr15-10826640-INS-1883
chr15-10831880-INS-693
chr15-10846812-INS-6805
chr15-10851884-INS-94

chr15-10874489-INS-1475
chr15-10884079-INS-5100
chr15-10894484-INS-196
chr15-10922242-INS-671
chr15-10929797-INS-59
chr15-11055478-INS-630
chr15-11063807-INS-80
chr15-11066353-INS-56
chr15-11073372-INS-234
chr15-11105287-INS-121
chr15-11112417-INS-54
chr15-11127005-INS-60
chr15-11128446-INS-56
chr15-11133973-INS-83
chr15-11136570-INS-1723
chr15-11176032-INS-54
chr15-11185484-INS-86
chr15-11196198-INS-58
chr15-11201384-INS-83
chr15-11233962-INS-286
chr15-11270884-INS-2810
chr15-11273742-INS-328
chr15-11293989-INS-108
chr15-11334777-INS-124
chr15-11341558-INS-156
chr15-11350405-INS-86
chr15-11368962-INS-174
chr15-11389219-INS-365
chr15-11389369-INS-912
chr15-11413740-INS-68
chr15-11429210-INS-770
chr15-11429236-INS-365
chr15-11429321-INS-1088
chr15-11438210-INS-77
chr15-11439768-INS-784
chr15-11453215-INS-126
chr15-11519745-INS-54
chr15-11528033-INS-329
chr15-11535897-INS-325
chr15-11538552-INS-53
chr15-11550803-INS-58
chr15-11563611-INS-405
chr15-11570025-INS-112
chr15-11589370-INS-71
chr15-11593176-INS-62
chr15-11594582-INS-61
chr15-11619311-INS-1339
chr15-11629992-INS-96
chr15-11644782-INS-196
chr15-11666069-INS-142
chr15-11671773-INS-58
chr15-11681106-INS-54
chr15-11691230-INS-96
chr15-11703979-INS-438
chr15-11705849-INS-1493
chr15-11747013-INS-182
chr15-11751494-INS-56
chr15-11804697-INS-471
chr15-11814955-INS-353
chr15-11816875-INS-84
chr15-11820340-INS-106
chr15-11832741-INS-99
chr15-11853530-INS-456
chr15-11868713-INS-6448
chr15-11872927-INS-164
chr15-11873906-INS-60
chr15-11876292-INS-2187
chr15-11884333-INS-64
chr15-11885579-INS-297
chr15-11907672-INS-138
chr15-11912128-INS-706
chr15-11974363-INS-140
chr15-11975796-INS-52
chr15-11999981-INS-118
chr15-12007443-INS-212
chr15-12013096-INS-54
chr15-12017348-INS-7061

chr15-12051172-INS-209
chr15-12065455-INS-53
chr15-12074398-INS-430
chr15-12095745-INS-66
chr15-12099326-INS-204
chr15-12103782-INS-235
chr15-12112063-INS-3420
chr15-12116535-INS-61
chr15-12123567-INS-138
chr15-12134713-INS-68
chr15-12145876-INS-77
chr15-12157142-INS-167
chr15-12160388-INS-656
chr15-12202461-INS-52
chr15-12217398-INS-229
chr15-12219408-INS-5410
chr15-12229287-INS-134
chr15-12237442-INS-74
chr15-12239036-INS-100
chr15-12249929-INS-929
chr15-12253981-INS-766
chr15-12265241-INS-54
chr15-12278702-INS-361
chr15-12288471-INS-77
chr15-12290933-INS-254
chr15-12310318-INS-499
chr15-12317619-INS-52
chr15-12341037-INS-181
chr15-12352011-INS-78
chr15-12359273-INS-60
chr15-12410965-INS-115
chr15-12418383-INS-52
chr15-12421173-INS-201
chr15-12470100-INS-60
chr15-12471371-INS-1890
chr15-12471372-INS-400
chr15-12483137-INS-6450
chr15-12524246-INS-245
chr15-12546748-INS-215
chr15-12547716-INS-285
chr15-12583070-INS-263
chr15-12594778-INS-60
chr15-12614860-INS-128
chr15-12621819-INS-121
chr15-12648237-INS-132
chr15-12662365-INS-70
chr15-12663767-INS-80
chr15-12690865-INS-301
chr15-12708880-INS-111
chr15-12735257-INS-307
chr15-12812642-INS-53
chr15-12813424-INS-289
chr15-12820855-INS-322
chr15-12873096-INS-795
chr15-12873304-INS-758
chr15-12877993-INS-252
chr15-12889510-INS-157
chr15-12931812-INS-197
chr15-12976254-INS-52
chr15-13020050-INS-563
chr15-13068947-INS-56
chr15-13080242-INS-53
chr15-13143704-INS-143
chr15-13147427-INS-246
chr15-13196186-INS-114
chr15-13215141-INS-266
chr15-13220560-INS-50
chr15-13247600-INS-218
chr15-13250716-INS-147
chr15-13263035-INS-336
chr15-13269360-INS-6454
chr15-13278246-INS-60
chr15-13299714-INS-458
chr15-13370824-INS-61
chr15-13383397-INS-58
chr15-13388382-INS-6222
chr15-13402503-INS-492

chr15-13425400-INS-88
chr15-13448725-INS-180
chr15-13451584-INS-58
chr15-13510289-INS-102
chr15-13524462-INS-2040
chr15-13543886-INS-54
chr15-13548710-INS-179
chr15-13553858-INS-193
chr15-13574868-INS-324
chr15-13590649-INS-140
chr15-13594517-INS-119
chr15-13602285-INS-66
chr15-13603019-INS-259
chr15-13619116-INS-81
chr15-13644601-INS-61
chr15-13647873-INS-62
chr15-13649746-INS-77
chr15-13680068-INS-104
chr15-13681984-INS-109
chr15-13696286-INS-491
chr15-13699413-INS-266
chr15-13711632-INS-342
chr15-13714973-INS-6995
chr15-13737192-INS-535
chr15-13775846-INS-962
chr15-13784085-INS-84
chr15-13787522-INS-262
chr15-13804149-INS-52
chr15-13829417-INS-92
chr15-13853582-INS-922
chr15-13859415-INS-91
chr15-13910196-INS-98
chr15-13939764-INS-470
chr15-13957323-INS-108
chr15-14029486-INS-241
chr15-14049249-INS-975
chr15-14058078-INS-1449
chr15-14076133-INS-5101
chr15-14086665-INS-937
chr15-14090453-INS-155
chr15-14093951-INS-57
chr15-14103474-INS-69
chr15-14167933-INS-583
chr15-14193085-INS-371
chr15-14195366-INS-56
chr15-14197101-INS-88
chr15-14211363-INS-136
chr15-14214208-INS-99
chr15-14235440-INS-288
chr15-14252344-INS-127
chr15-14252571-INS-195
chr15-14260028-INS-400
chr15-14292248-INS-123
chr15-14297970-INS-209
chr15-14304681-INS-66
chr15-14321189-INS-83
chr15-14324440-INS-77
chr15-14330694-INS-3442
chr15-14335470-INS-56
chr15-14336821-INS-76
chr15-14382544-INS-211
chr15-14384961-INS-434
chr15-14414268-INS-6563
chr15-14422025-INS-6583
chr15-14431382-INS-120
chr15-14439745-INS-120
chr15-14444753-INS-98
chr15-14445964-INS-184
chr15-14449902-INS-502
chr15-14452332-INS-53
chr15-14463582-INS-54
chr15-14465602-INS-189
chr15-14467310-INS-352
chr15-14524526-INS-246
chr15-14524910-INS-136
chr15-14526192-INS-153
chr15-14541925-INS-56

chr15-14561386-INS-250
chr15-14583937-INS-6725
chr15-14587775-INS-5353
chr15-14598025-INS-55
chr15-14602035-INS-52
chr15-14612964-INS-96
chr15-14652553-INS-91
chr15-14674819-INS-136
chr15-14675559-INS-520
chr15-14702956-INS-62
chr15-14735366-INS-81
chr15-14750661-INS-125
chr15-14753148-INS-6200
chr15-14764891-INS-74
chr15-14767400-INS-59
chr15-14770919-INS-99
chr15-14785338-INS-87
chr15-14792814-INS-6575
chr15-14799323-INS-60
chr15-14817863-INS-327
chr15-14825207-INS-2492
chr15-14837830-INS-142
chr15-14886706-INS-8809
chr15-14891007-INS-376
chr15-14912420-INS-2346
chr15-14914670-INS-201
chr15-14916455-INS-57
chr15-14954700-INS-1177
chr15-14957366-INS-92
chr15-14963437-INS-7957
chr15-14986080-INS-213
chr15-14986391-INS-8578
chr15-15000622-INS-52
chr15-15044785-INS-85
chr15-15047988-INS-59
chr15-15091315-INS-797
chr15-15102233-INS-55
chr15-15108833-INS-82
chr15-15118995-INS-95
chr15-15131589-INS-202
chr15-15177040-INS-361
chr15-15190211-INS-54
chr15-15208250-INS-384
chr15-15232024-INS-67
chr15-15241917-INS-483
chr15-15247320-INS-6375
chr15-15249285-INS-194
chr15-15254324-INS-403
chr15-15265506-INS-214
chr15-15286773-INS-68
chr15-15316299-INS-635
chr15-15321985-INS-248
chr15-15332083-INS-99
chr15-15344123-INS-505
chr15-15349310-INS-127
chr15-15355255-INS-358
chr15-15358256-INS-57
chr15-15365385-INS-58
chr15-15376491-INS-491
chr15-15397059-INS-5542
chr15-15397889-INS-171
chr15-15414449-INS-537
chr15-15456250-INS-370
chr15-15463001-INS-55
chr15-15472459-INS-963
chr15-15541788-INS-63
chr15-15570089-INS-124
chr15-15570733-INS-68
chr15-15580921-INS-66
chr15-15596405-INS-340
chr15-15610097-INS-52
chr15-15610683-INS-56
chr15-15615602-INS-53
chr15-15618269-INS-884
chr15-15649319-INS-4686
chr15-15671968-INS-55
chr15-15675948-INS-69

chr15-15699786-INS-69
chr15-15701144-INS-272
chr15-15725418-INS-76
chr15-15743141-INS-118
chr15-15775460-INS-796
chr15-15847306-INS-236
chr15-15864442-INS-6249
chr15-15870622-INS-70
chr15-15876695-INS-94
chr15-15887137-INS-984
chr15-15887760-INS-1685
chr15-15901961-INS-1837
chr15-15908419-INS-212
chr15-15912710-INS-2490
chr15-15934280-INS-386
chr15-15951233-INS-580
chr15-15959440-INS-445
chr15-16041515-INS-136
chr15-16054923-INS-890
chr15-16069310-INS-58
chr15-16071612-INS-6129
chr15-16108567-INS-126
chr15-16124754-INS-412
chr15-16129720-INS-70
chr15-16133344-INS-63
chr15-16136167-INS-62
chr15-16137762-INS-101
chr15-16156037-INS-364
chr15-16191271-INS-55
chr15-16210897-INS-154
chr15-16218230-INS-50
chr15-16220510-INS-197
chr15-16226925-INS-206
chr15-16241040-INS-246
chr15-16262144-INS-312
chr15-16271138-INS-141
chr15-16298576-INS-92
chr15-16341498-INS-744
chr15-16371440-INS-70
chr15-16380199-INS-676
chr15-16381040-INS-480
chr15-16397741-INS-212
chr15-16416104-INS-84
chr15-16417908-INS-60
chr15-16425655-INS-426
chr15-16440376-INS-447
chr15-16447519-INS-59
chr15-16469466-INS-600
chr15-16482805-INS-95
chr15-16487226-INS-376
chr15-16496167-INS-6498
chr15-16519156-INS-139
chr15-16536515-INS-100
chr15-16539703-INS-66
chr15-16545495-INS-324
chr15-16546906-INS-187
chr15-16671213-INS-6422
chr15-16734115-INS-87
chr15-16757800-INS-406
chr15-16774820-INS-640
chr15-16776563-INS-6100
chr15-16781886-INS-93
chr15-16783441-INS-105
chr15-16788557-INS-80
chr15-16796197-INS-67
chr15-16797976-INS-140
chr15-16806966-INS-71
chr15-16843450-INS-84
chr15-16860313-INS-202
chr15-16874471-INS-67
chr15-16884736-INS-206
chr15-16922505-INS-51
chr15-17010360-INS-68
chr15-17053122-INS-748
chr15-17084962-INS-56
chr15-17112860-INS-297
chr15-17115286-INS-96

chr15-17185361-INS-468
chr15-17215682-INS-60
chr15-17257327-INS-5547
chr15-17260130-INS-479
chr15-17264072-INS-235
chr15-17279647-INS-796
chr15-17294366-INS-92
chr15-17307609-INS-73
chr15-17337365-INS-199
chr15-17357559-INS-52
chr15-17365549-INS-130
chr15-17392449-INS-390
chr15-17406801-INS-142
chr15-17409304-INS-542
chr15-17414105-INS-70
chr15-17415284-INS-52
chr15-17417775-INS-88
chr15-17425095-INS-228
chr15-17474426-INS-754
chr15-17494798-INS-68
chr15-17515403-INS-3342
chr15-17584243-INS-449
chr15-17584248-INS-2818
chr15-17589547-INS-312
chr15-17622093-INS-58
chr15-17632751-INS-679
chr15-17727907-INS-2725
chr15-17728249-INS-181
chr15-17734013-INS-231
chr15-17789303-INS-208
chr15-17852928-INS-58
chr15-17858218-INS-762
chr15-17890564-INS-522
chr15-17915015-INS-56
chr15-17918191-INS-4341
chr15-17933893-INS-52
chr15-17937866-INS-115
chr15-17987494-INS-56
chr15-17997653-INS-403
chr15-18009117-INS-6460
chr15-18022150-INS-4779
chr15-18041477-INS-493
chr15-18058139-INS-109
chr15-18067288-INS-354
chr15-18069617-INS-387
chr15-18084656-INS-52
chr15-18098757-INS-843
chr15-18099519-INS-347
chr15-18116686-INS-86
chr15-18123607-INS-57
chr15-18131505-INS-67
chr15-18158450-INS-52
chr15-18169944-INS-2800
chr15-18209002-INS-883
chr15-18209880-INS-746
chr15-18210530-INS-334
chr15-18223409-INS-88
chr15-18228664-INS-60
chr15-18258954-INS-324
chr15-18275448-INS-60
chr15-18311699-INS-6680
chr15-18329375-INS-63
chr15-18338534-INS-67
chr15-18358877-INS-258
chr15-18375222-INS-127
chr15-18400122-INS-207
chr15-18401331-INS-121
chr15-18404421-INS-355
chr15-18408757-INS-180
chr15-18418941-INS-6745
chr15-18483676-INS-432
chr15-18484888-INS-68
chr15-18501144-INS-58
chr15-18512443-INS-132
chr15-18513622-INS-54
chr15-18517581-INS-338
chr15-18530425-INS-234

chr15-18534621-INS-57
chr15-18544067-INS-62
chr15-18550184-INS-7397
chr15-18553088-INS-7113
chr15-18555339-INS-105
chr15-18558421-INS-4678
chr15-18560849-INS-68
chr15-18563865-INS-222
chr15-18587392-INS-7166
chr15-18593621-INS-2269
chr15-18612911-INS-2229
chr15-18613146-INS-63
chr15-18619806-INS-50
chr15-18626856-INS-81
chr15-18675500-INS-172
chr15-18683972-INS-72
chr15-18708958-INS-3030
chr15-18735925-INS-96
chr15-18746918-INS-209
chr15-18759674-INS-88
chr15-18765481-INS-989
chr15-18815050-INS-61
chr15-18854838-INS-7592
chr15-18854914-INS-925
chr15-18860989-INS-328
chr15-18864001-INS-138
chr15-18868933-INS-6222
chr15-18869372-INS-6675
chr15-18876805-INS-5655
chr15-18884150-INS-516
chr15-18895292-INS-80
chr15-18981016-INS-59
chr15-18982766-INS-7077
chr15-18999009-INS-84
chr15-19056426-INS-330
chr15-19067048-INS-1951
chr15-19071564-INS-904
chr15-19073159-INS-104
chr15-19084847-INS-63
chr15-19118484-INS-60
chr15-19129672-INS-358
chr15-19130231-INS-68
chr15-19143855-INS-6669
chr15-19166999-INS-60
chr15-19182874-INS-1455
chr15-19196716-INS-193
chr15-19210573-INS-682
chr15-19226981-INS-612
chr15-19236336-INS-75
chr15-19248428-INS-434
chr15-19251232-INS-4692
chr15-19259358-INS-206
chr15-19274186-INS-54
chr15-19302709-INS-10648
chr15-19303908-INS-54
chr15-19315312-INS-84
chr15-19325847-INS-299
chr15-19349332-INS-98
chr15-19355707-INS-60
chr15-19363808-INS-413
chr15-19416546-INS-313
chr15-19427286-INS-516
chr15-19469274-INS-2930
chr15-19492376-INS-6264
chr15-19520908-INS-55
chr15-19543634-INS-91
chr15-19548603-INS-52
chr15-19614986-INS-400
chr15-19662292-INS-76
chr15-19664270-INS-5251
chr15-19665438-INS-111
chr15-19667108-INS-120
chr15-19749227-INS-58
chr15-19783685-INS-74
chr15-19797972-INS-5259
chr15-19817472-INS-385
chr15-19822578-INS-55

chr15-19830201-INS-74
chr15-19838322-INS-2482
chr15-19852967-INS-114
chr15-19868059-INS-79
chr15-19869036-INS-971
chr15-19872640-INS-198
chr15-19880661-INS-57
chr15-19881710-INS-894
chr15-19882231-INS-134
chr15-19893630-INS-211
chr15-19960744-INS-103
chr15-19961624-INS-55
chr15-20032926-INS-427
chr15-20055172-INS-834
chr15-20058128-INS-229
chr15-20058786-INS-499
chr15-20067990-INS-76
chr15-20082068-INS-60
chr15-20089255-INS-82
chr15-20108495-INS-463
chr15-20110341-INS-60
chr15-20151209-INS-6171
chr15-20198416-INS-149
chr15-20199946-INS-108
chr15-20211800-INS-496
chr15-20214402-INS-175
chr15-20220102-INS-84
chr15-20238412-INS-112
chr15-20260017-INS-5293
chr15-20260845-INS-63
chr15-20264263-INS-555
chr15-20265989-INS-462
chr15-20282049-INS-102
chr15-20310665-INS-708
chr15-20362955-INS-106
chr15-20370444-INS-1951
chr15-20376850-INS-51
chr15-20395050-INS-151
chr15-20403551-INS-148
chr15-20403946-INS-785
chr15-20429141-INS-62
chr15-20440226-INS-52
chr15-20445116-INS-53
chr15-20482919-INS-6528
chr15-20488321-INS-668
chr15-20567753-INS-115
chr15-20585745-INS-71
chr15-20611614-INS-541
chr15-20625653-INS-275
chr15-20632936-INS-474
chr15-20641151-INS-6433
chr15-20654582-INS-7131
chr15-20663179-INS-94
chr15-20687641-INS-58
chr15-20698116-INS-74
chr15-20702162-INS-319
chr15-20704222-INS-54
chr15-20705981-INS-92
chr15-20721904-INS-633
chr15-20724261-INS-614
chr15-20744257-INS-72
chr15-20748987-INS-72
chr15-20765506-INS-73
chr15-20770313-INS-7714
chr15-20771142-INS-186
chr15-20802012-INS-448
chr15-20804975-INS-79
chr15-20808567-INS-108
chr15-20858939-INS-88
chr15-20861304-INS-70
chr15-20864426-INS-377
chr15-20870403-INS-6739
chr15-20875507-INS-60
chr15-20879106-INS-7090
chr15-20879569-INS-54
chr15-20904070-INS-640
chr15-20904891-INS-66

chr15-20930270-INS-886
chr15-20937086-INS-354
chr15-20942142-INS-248
chr15-20946142-INS-60
chr15-20953671-INS-243
chr15-20956338-INS-224
chr15-20964979-INS-529
chr15-20970561-INS-6258
chr15-21029519-INS-1407
chr15-21033442-INS-1284
chr15-21071005-INS-97
chr15-21072861-INS-673
chr15-21101836-INS-132
chr15-21119907-INS-1891
chr15-21138217-INS-507
chr15-21145928-INS-349
chr15-21195659-INS-70
chr15-21200065-INS-9045
chr15-21220327-INS-184
chr15-21222716-INS-78
chr15-21230228-INS-152
chr15-21242682-INS-548
chr15-21247836-INS-412
chr15-21258868-INS-67
chr15-21295982-INS-255
chr15-21299676-INS-538
chr15-21302717-INS-100
chr15-21315960-INS-6208
chr15-21316452-INS-84
chr15-21321569-INS-155
chr15-21342224-INS-212
chr15-21358442-INS-60
chr15-21361520-INS-74
chr15-21367436-INS-63
chr15-21384054-INS-5182
chr15-21389873-INS-75
chr15-21403275-INS-146
chr15-21407099-INS-202
chr15-21428846-INS-324
chr15-21430709-INS-88
chr15-21438603-INS-6933
chr15-21438692-INS-116
chr15-21443128-INS-298
chr15-21480625-INS-175
chr15-21489685-INS-119
chr15-21494395-INS-590
chr15-21541748-INS-126
chr15-21543870-INS-64
chr15-21554922-INS-74
chr15-21599679-INS-63
chr15-21609089-INS-459
chr15-21610115-INS-83
chr15-21614592-INS-5005
chr15-21636581-INS-60
chr15-21638978-INS-686
chr15-21646564-INS-172
chr15-21667714-INS-85
chr15-21700504-INS-904
chr15-21701054-INS-82
chr15-21727333-INS-94
chr15-21736556-INS-80
chr15-21738773-INS-191
chr15-21780089-INS-598
chr15-21823994-INS-409
chr15-21829635-INS-5930
chr15-21831791-INS-4396
chr15-21833862-INS-3599
chr15-21841167-INS-70
chr15-21849687-INS-6469
chr15-21885379-INS-2880
chr15-21886155-INS-211
chr15-21893388-INS-422
chr15-21898199-INS-66
chr15-21910056-INS-66
chr15-21930278-INS-56
chr15-21932874-INS-112
chr15-21971096-INS-84

chr15-21977287-INS-63
chr15-21986338-INS-246
chr15-22022468-INS-761
chr15-22027770-INS-90
chr15-22045779-INS-114
chr15-22082550-INS-52
chr15-22091604-INS-54
chr15-22116205-INS-58
chr15-22126146-INS-206
chr15-22159325-INS-56
chr15-22170754-INS-190
chr15-22174209-INS-360
chr15-22175017-INS-54
chr15-22177513-INS-121
chr15-22191483-INS-4329
chr15-22201758-INS-52
chr15-22227249-INS-357
chr15-22239121-INS-603
chr15-22255292-INS-184
chr15-22309930-INS-314
chr15-22315229-INS-106
chr15-22318647-INS-2959
chr15-22329693-INS-6862
chr15-22341136-INS-813
chr15-22350148-INS-333
chr15-22354517-INS-86
chr15-22357857-INS-780
chr15-22428809-INS-276
chr15-22445212-INS-56
chr15-22452923-INS-6395
chr15-22461995-INS-305
chr15-22474613-INS-1543
chr15-22478229-INS-202
chr15-22479804-INS-2621
chr15-22486647-INS-135
chr15-22500928-INS-432
chr15-22512298-INS-592
chr15-22517680-INS-413
chr15-22518696-INS-66
chr15-22520746-INS-571
chr15-22578505-INS-209
chr15-22582314-INS-54
chr15-22615141-INS-194
chr15-22619453-INS-7157
chr15-22629336-INS-58
chr15-22672755-INS-94
chr15-22680280-INS-102
chr15-22692844-INS-62
chr15-22707453-INS-157
chr15-22738829-INS-88
chr15-22780444-INS-53
chr15-22791550-INS-6028
chr15-22799717-INS-550
chr15-22799717-INS-935
chr15-22829470-INS-58
chr15-22835337-INS-247
chr15-22840929-INS-441
chr15-22848857-INS-70
chr15-22859542-INS-221
chr15-22878511-INS-69
chr15-22887124-INS-60
chr15-22915268-INS-58
chr15-22927020-INS-71
chr15-22943501-INS-106
chr15-22957603-INS-221
chr15-23001117-INS-162
chr15-23015856-INS-85
chr15-23020479-INS-161
chr15-23021762-INS-198
chr15-23025762-INS-406
chr15-23031376-INS-106
chr15-23044900-INS-358
chr15-23094201-INS-162
chr15-23118685-INS-71
chr15-23132477-INS-152
chr15-23145257-INS-386
chr15-23147468-INS-76

chr15-23154329-INS-77
chr15-23159048-INS-98
chr15-23172103-INS-65
chr15-23184049-INS-979
chr15-23214676-INS-64
chr15-23219102-INS-197
chr15-23318553-INS-60
chr15-23323490-INS-193
chr15-23326370-INS-433
chr15-23393538-INS-67
chr15-23401532-INS-527
chr15-23411978-INS-235
chr15-23423884-INS-450
chr15-23425905-INS-250
chr15-23432000-INS-78
chr15-23439368-INS-405
chr15-23445227-INS-54
chr15-23452973-INS-54
chr15-23455162-INS-169
chr15-23486391-INS-59
chr15-23487481-INS-51
chr15-23543769-INS-5810
chr15-23547887-INS-158
chr15-23550669-INS-52
chr15-23577794-INS-416
chr15-23582781-INS-52
chr15-23611053-INS-52
chr15-23635301-INS-122
chr15-23641643-INS-52
chr15-23642793-INS-108
chr15-23686259-INS-204
chr15-23690802-INS-94
chr15-23792844-INS-57
chr15-23804287-INS-3166
chr15-23812534-INS-66
chr15-23818201-INS-80
chr15-23880960-INS-315
chr15-23896403-INS-503
chr15-23971419-INS-1888
chr15-23988764-INS-107
chr15-24003357-INS-64
chr15-24007062-INS-79
chr15-24023955-INS-64
chr15-24050193-INS-9487
chr15-24050614-INS-105
chr15-24055353-INS-52
chr15-24092093-INS-74
chr15-24093886-INS-90
chr15-24106026-INS-60
chr15-24145863-INS-390
chr15-24147509-INS-330
chr15-24152536-INS-52
chr15-24161037-INS-153
chr15-24186707-INS-358
chr15-24202512-INS-242
chr15-24242030-INS-56
chr15-24248539-INS-78
chr15-24289205-INS-59
chr15-24302584-INS-6399
chr15-24337078-INS-7957
chr15-24398823-INS-76
chr15-24404799-INS-66
chr15-24479230-INS-56
chr15-24490456-INS-668
chr15-24499339-INS-67
chr15-24511639-INS-2557
chr15-24531258-INS-157
chr15-24544172-INS-561
chr15-24553806-INS-119
chr15-24556792-INS-3064
chr15-24577568-INS-130
chr15-24614690-INS-273
chr15-24624678-INS-769
chr15-24632959-INS-63
chr15-24687935-INS-307
chr15-24699471-INS-214
chr15-24717448-INS-830

chr15-24724469-INS-854
chr15-24759603-INS-57
chr15-24771718-INS-476
chr15-24790961-INS-560
chr15-24809878-INS-54
chr15-24839044-INS-6797
chr15-24842243-INS-64
chr15-24843519-INS-426
chr15-24849344-INS-6695
chr15-24889904-INS-446
chr15-24905758-INS-116
chr15-24922730-INS-53
chr15-24924486-INS-170
chr15-24931802-INS-107
chr15-24978506-INS-124
chr15-24981886-INS-827
chr15-25013143-INS-443
chr15-25020955-INS-77
chr15-25025389-INS-118
chr15-25060204-INS-65
chr15-25066616-INS-6282
chr15-25072302-INS-88
chr15-25094661-INS-51
chr15-25127714-INS-59
chr15-25150595-INS-84
chr15-25152198-INS-443
chr15-25156353-INS-162
chr15-25182730-INS-351
chr15-25202109-INS-66
chr15-25218001-INS-345
chr15-25271328-INS-7303
chr15-25313952-INS-88
chr15-25315309-INS-73
chr15-25328088-INS-54
chr15-25328643-INS-84
chr15-25337200-INS-66
chr15-25338293-INS-380
chr15-25374449-INS-128
chr15-25436207-INS-550
chr15-25436706-INS-148
chr15-25439156-INS-325
chr15-25440958-INS-234
chr15-25447133-INS-62
chr15-25455913-INS-53
chr15-25485921-INS-116
chr15-25524305-INS-81
chr15-25538644-INS-132
chr15-25548016-INS-157
chr15-25553586-INS-150
chr15-25564581-INS-58
chr15-25570361-INS-270
chr15-25574554-INS-150
chr15-25586026-INS-56
chr15-25587443-INS-80
chr15-25615652-INS-58
chr15-25621239-INS-204
chr15-25644121-INS-122
chr15-25662043-INS-361
chr15-25677393-INS-220
chr15-25681548-INS-750
chr15-25684585-INS-164
chr15-25694782-INS-497
chr15-25705221-INS-51
chr15-25710534-INS-256
chr15-25725475-INS-192
chr15-25739058-INS-380
chr15-25746879-INS-51
chr15-25751895-INS-128
chr15-25755967-INS-196
chr15-25769072-INS-354
chr15-25813819-INS-199
chr15-25827287-INS-173
chr15-25847980-INS-220
chr15-25848962-INS-68
chr15-25894420-INS-144
chr15-25900478-INS-142
chr15-25901649-INS-128

chr15-25928827-INS-7084
chr15-25940049-INS-66
chr15-25981802-INS-7110
chr15-25982827-INS-149
chr15-25986811-INS-79
chr15-25991818-INS-744
chr15-26001199-INS-475
chr15-26002171-INS-464
chr15-26020962-INS-51
chr15-26023044-INS-59
chr15-26024309-INS-485
chr15-26042212-INS-56
chr15-26057085-INS-524
chr15-26062819-INS-289
chr15-26071227-INS-59
chr15-26078139-INS-325
chr15-26078797-INS-6452
chr15-26081567-INS-2137
chr15-26088990-INS-279
chr15-26090730-INS-453
chr15-26110280-INS-79
chr15-26122732-INS-774
chr15-26139302-INS-453
chr15-26141611-INS-5164
chr15-26147985-INS-52
chr15-26157808-INS-590
chr15-26167448-INS-64
chr15-26177919-INS-303
chr15-26203780-INS-695
chr15-26204746-INS-340
chr15-26216385-INS-310
chr15-26239660-INS-1829
chr15-26251169-INS-6470
chr15-26251171-INS-498
chr15-26264220-INS-64
chr15-26269713-INS-165
chr15-26274740-INS-64
chr15-26278717-INS-996
chr15-26291280-INS-497
chr15-26302699-INS-132
chr15-26307526-INS-54
chr15-26319529-INS-54
chr15-26341457-INS-54
chr15-26373276-INS-835
chr15-26375331-INS-165
chr15-26405375-INS-84
chr15-26423942-INS-76
chr15-26449335-INS-93
chr15-26454563-INS-295
chr15-26458393-INS-202
chr15-26460882-INS-195
chr15-26486356-INS-184
chr15-26524606-INS-129
chr15-26553994-INS-654
chr15-26557583-INS-100
chr15-26561111-INS-155
chr15-26563551-INS-99
chr15-26574328-INS-114
chr15-26595420-INS-5016
chr15-26604818-INS-1065
chr15-26606398-INS-65
chr15-26607027-INS-65
chr15-26617404-INS-6808
chr15-26660745-INS-262
chr15-26662497-INS-3475
chr15-26665146-INS-294
chr15-26667380-INS-70
chr15-26670669-INS-498
chr15-26672770-INS-184
chr15-26690077-INS-495
chr15-26703026-INS-207
chr15-26707140-INS-256
chr15-26722235-INS-60
chr15-26726109-INS-756
chr15-26728375-INS-113
chr15-26735588-INS-73
chr15-26760138-INS-52

chr15-26803574-INS-5518
chr15-26806069-INS-120
chr15-26831898-INS-389
chr15-26852834-INS-243
chr15-26855478-INS-426
chr15-26866823-INS-302
chr15-26880136-INS-102
chr15-26889662-INS-51
chr15-26901081-INS-435
chr15-26918447-INS-2110
chr15-26918478-INS-788
chr15-26920980-INS-345
chr15-26923918-INS-75
chr15-26929022-INS-58
chr15-26930520-INS-51
chr15-26934220-INS-61
chr15-26971353-INS-67
chr15-26974659-INS-64
chr15-26978557-INS-6451
chr15-26991281-INS-197
chr15-26994610-INS-97
chr15-27015967-INS-105
chr15-27033869-INS-160
chr15-27072206-INS-340
chr15-27081092-INS-128
chr15-27083120-INS-4528
chr15-27098934-INS-221
chr15-27106393-INS-313
chr15-27156548-INS-477
chr15-27158107-INS-6801
chr15-27168280-INS-670
chr15-27179718-INS-146
chr15-27222835-INS-92
chr15-27224949-INS-204
chr15-27228772-INS-108
chr15-27242407-INS-197
chr15-27254302-INS-418
chr15-27285102-INS-797
chr15-27285305-INS-5049
chr15-27314622-INS-322
chr15-27356810-INS-187
chr15-27380338-INS-181
chr15-27428406-INS-900
chr15-27431287-INS-59
chr15-27464635-INS-547
chr15-27482232-INS-297
chr15-27485355-INS-230
chr15-27528750-INS-355
chr15-27561140-INS-100
chr15-27580156-INS-68
chr15-27611185-INS-93
chr15-27632999-INS-769
chr15-27645051-INS-60
chr15-27654975-INS-79
chr15-27663324-INS-222
chr15-27674965-INS-52
chr15-27677807-INS-211
chr15-27699164-INS-177
chr15-27700772-INS-122
chr15-27704069-INS-501
chr15-27742025-INS-68
chr15-27764996-INS-184
chr15-27766119-INS-52
chr15-27812088-INS-78
chr15-27828114-INS-408
chr15-27873248-INS-229
chr15-27874519-INS-205
chr15-27901820-INS-616
chr15-27917726-INS-135
chr15-27935681-INS-171
chr15-27938295-INS-65
chr15-27973728-INS-52
chr15-27980860-INS-115
chr15-27986189-INS-215
chr15-28011620-INS-89
chr15-28016588-INS-105
chr15-28047771-INS-209

chr15-28049897-INS-178
chr15-28050979-INS-234
chr15-28051797-INS-193
chr15-28054525-INS-58
chr15-28109913-INS-82
chr15-28122940-INS-159
chr15-28134764-INS-53
chr15-28137272-INS-56
chr15-28140002-INS-173
chr15-28144124-INS-65
chr15-28147296-INS-142
chr15-28153920-INS-107
chr15-28175010-INS-63
chr15-28200518-INS-136
chr15-28214318-INS-397
chr15-28231136-INS-354
chr15-28236290-INS-128
chr15-28243828-INS-439
chr15-28249949-INS-556
chr15-28272843-INS-975
chr15-28325355-INS-574
chr15-28333613-INS-286
chr15-28359471-INS-330
chr15-28385084-INS-72
chr15-28397608-INS-124
chr15-28426024-INS-5197
chr15-28457004-INS-65
chr15-28460482-INS-222
chr15-28463895-INS-211
chr15-28476865-INS-588
chr15-28482406-INS-7074
chr15-28492948-INS-77
chr15-28500659-INS-243
chr15-28504618-INS-201
chr15-28518198-INS-140
chr15-28525574-INS-98
chr15-28527684-INS-166
chr15-28546598-INS-1891
chr15-28553043-INS-397
chr15-28559876-INS-660
chr15-28563393-INS-131
chr15-28570365-INS-200
chr15-28586268-INS-307
chr15-28606420-INS-187
chr15-28609317-INS-105
chr15-28614057-INS-82
chr15-28619324-INS-498
chr15-28628987-INS-108
chr15-28632689-INS-259
chr15-28658693-INS-400
chr15-28659592-INS-56
chr15-28694505-INS-560
chr15-28700709-INS-266
chr15-28702286-INS-540
chr15-28712427-INS-171
chr15-28712443-INS-1206
chr15-28718443-INS-959
chr15-28721041-INS-311
chr15-28734173-INS-445
chr15-28734996-INS-83
chr15-28760568-INS-6862
chr15-28768183-INS-141
chr15-28772409-INS-167
chr15-28790797-INS-371
chr15-28800771-INS-2785
chr15-28813701-INS-449
chr15-28818152-INS-93
chr15-28841380-INS-263
chr15-28873061-INS-55
chr15-28887880-INS-106
chr15-28889468-INS-129
chr15-28898608-INS-4250
chr15-28993995-INS-53
chr15-28996508-INS-6411
chr15-28996962-INS-203
chr15-28999394-INS-92
chr15-29028930-INS-3726

chr15-29029700-INS-53
chr15-29030257-INS-1313
chr15-29039108-INS-1189
chr15-29041833-INS-60
chr15-29048571-INS-6664
chr15-29050991-INS-503
chr15-29053385-INS-52
chr15-29053671-INS-1129
chr15-29103295-INS-76
chr15-29111451-INS-2308
chr15-29114580-INS-134
chr15-29123835-INS-64
chr15-29149154-INS-172
chr15-29161248-INS-133
chr15-29180801-INS-90
chr15-29212842-INS-175
chr15-29227260-INS-74
chr15-29231213-INS-7329
chr15-29251669-INS-62
chr15-29264686-INS-113
chr15-29317241-INS-90
chr15-29332445-INS-2961
chr15-29351093-INS-148
chr15-29355595-INS-6902
chr15-29359882-INS-125
chr15-29378253-INS-583
chr15-29402490-INS-852
chr15-29407342-INS-82
chr15-29441001-INS-103
chr15-29447989-INS-431
chr15-29458481-INS-58
chr15-29460238-INS-969
chr15-29495927-INS-398
chr15-29500887-INS-55
chr15-29506999-INS-189
chr15-29512588-INS-62
chr15-29514709-INS-674
chr15-29532769-INS-1013
chr15-29541764-INS-1107
chr15-29543840-INS-6317
chr15-29585025-INS-267
chr15-29606479-INS-398
chr15-29609392-INS-6484
chr15-29610221-INS-73
chr15-29614209-INS-585
chr15-29629206-INS-178
chr15-29654706-INS-66
chr15-29659638-INS-75
chr15-29664549-INS-6198
chr15-29688795-INS-154
chr15-29689437-INS-532
chr15-29727430-INS-78
chr15-29744426-INS-134
chr15-29772309-INS-512
chr15-29778489-INS-398
chr15-29780516-INS-70
chr15-29792949-INS-285
chr15-29793882-INS-118
chr15-29822177-INS-52
chr15-29830138-INS-173
chr15-29832771-INS-668
chr15-29838017-INS-72
chr15-29858628-INS-91
chr15-29875016-INS-102
chr15-29911091-INS-72
chr15-29919476-INS-572
chr15-29921000-INS-66
chr15-29932368-INS-55
chr15-29948327-INS-74
chr15-29969945-INS-355
chr15-29984114-INS-208
chr15-30006265-INS-78
chr15-30007436-INS-6717
chr15-30060986-INS-111
chr15-30061759-INS-224
chr15-30085809-INS-2397
chr15-30114783-INS-114

chr15-30114977-INS-574
chr15-30117079-INS-209
chr15-30117699-INS-156
chr15-30150528-INS-504
chr15-30158697-INS-400
chr15-30164377-INS-116
chr15-30175262-INS-174
chr15-30191994-INS-5529
chr15-30195620-INS-52
chr15-30197980-INS-58
chr15-30199733-INS-57
chr15-30204975-INS-336
chr15-30208789-INS-340
chr15-30244813-INS-70
chr15-30251333-INS-700
chr15-30264721-INS-588
chr15-30272903-INS-6753
chr15-30275072-INS-395
chr15-30279039-INS-3385
chr15-30279793-INS-51
chr15-30292139-INS-60
chr15-30299543-INS-90
chr15-30304760-INS-2478
chr15-30334049-INS-71
chr15-30342004-INS-60
chr15-30359644-INS-108
chr15-30363297-INS-124
chr15-30453502-INS-6476
chr15-30454191-INS-6979
chr15-30480178-INS-436
chr15-30484038-INS-359
chr15-30504555-INS-390
chr15-30511210-INS-502
chr15-30537914-INS-92
chr15-30571485-INS-320
chr15-30592519-INS-3777
chr15-30638064-INS-219
chr15-30641419-INS-56
chr15-30649251-INS-180
chr15-30657759-INS-54
chr15-30677742-INS-779
chr15-30677806-INS-186
chr15-30689048-INS-319
chr15-30724775-INS-6269
chr15-30728543-INS-208
chr15-30730635-INS-54
chr15-30739657-INS-913
chr15-30744848-INS-227
chr15-30780303-INS-125
chr15-30783415-INS-90
chr15-30798924-INS-66
chr15-30841149-INS-67
chr15-30847536-INS-110
chr15-30850308-INS-162
chr15-30894878-INS-455
chr15-30924592-INS-229
chr15-30930721-INS-56
chr15-30944205-INS-86
chr15-30949141-INS-110
chr15-30959617-INS-54
chr15-30978261-INS-218
chr15-30994305-INS-78
chr15-30995095-INS-108
chr15-31036330-INS-62
chr15-31043858-INS-198
chr15-31044513-INS-978
chr15-31051003-INS-177
chr15-31087501-INS-63
chr15-31138046-INS-317
chr15-31147862-INS-60
chr15-31157296-INS-51
chr15-31158205-INS-696
chr15-31192874-INS-70
chr15-31208553-INS-57
chr15-31213333-INS-64
chr15-31230115-INS-119
chr15-31232102-INS-208

chr15-31236966-INS-111
chr15-31319367-INS-102
chr15-31327834-INS-58
chr15-31339061-INS-117
chr15-31343143-INS-219
chr15-31350765-INS-347
chr15-31355102-INS-137
chr15-31373596-INS-548
chr15-31378788-INS-400
chr15-31398903-INS-679
chr15-31413650-INS-198
chr15-31431499-INS-51
chr15-31440790-INS-147
chr15-31447376-INS-83
chr15-31513500-INS-63
chr15-31515052-INS-309
chr15-31518583-INS-202
chr15-31570199-INS-58
chr15-31571460-INS-214
chr15-31573516-INS-127
chr15-31597289-INS-93
chr15-31605460-INS-720
chr15-31606478-INS-203
chr15-31607229-INS-180
chr15-31626221-INS-80
chr15-31634179-INS-110
chr15-31657713-INS-935
chr15-31663762-INS-328
chr15-31698419-INS-64
chr15-31703160-INS-416
chr15-31712544-INS-70
chr15-31745000-INS-76
chr15-31785658-INS-71
chr15-31792424-INS-6292
chr15-31805487-INS-455
chr15-31819127-INS-65
chr15-31820478-INS-943
chr15-31871868-INS-125
chr15-31903097-INS-6453
chr15-31905308-INS-463
chr15-31915294-INS-445
chr15-31940748-INS-55
chr15-31968923-INS-146
chr15-31986778-INS-80
chr15-32007716-INS-202
chr15-32017863-INS-1891
chr15-32020765-INS-493
chr15-32086901-INS-930
chr15-32109993-INS-716
chr15-32110897-INS-75
chr15-32126598-INS-6240
chr15-32148135-INS-4390
chr15-32152992-INS-6650
chr15-32155281-INS-240
chr15-32159550-INS-6222
chr15-32171280-INS-60
chr15-32196665-INS-157
chr15-32235180-INS-165
chr15-32254167-INS-56
chr15-32288585-INS-60
chr15-32289527-INS-208
chr15-32293871-INS-138
chr15-32305910-INS-113
chr15-32339176-INS-56
chr15-32384855-INS-69
chr15-32412325-INS-89
chr15-32414737-INS-54
chr15-32415889-INS-211
chr15-32419636-INS-189
chr15-32491913-INS-267
chr15-32499169-INS-517
chr15-32532115-INS-722
chr15-32542835-INS-7135
chr15-32553076-INS-401
chr15-32599135-INS-68
chr15-32610417-INS-576
chr15-32628233-INS-70

chr15-32632293-INS-60
chr15-32653505-INS-793
chr15-32680511-INS-821
chr15-32683900-INS-68
chr15-32705011-INS-491
chr15-32709445-INS-452
chr15-32716731-INS-718
chr15-32747311-INS-210
chr15-32764161-INS-60
chr15-32853188-INS-435
chr15-32854290-INS-55
chr15-32880842-INS-129
chr15-32892559-INS-103
chr15-32929657-INS-77
chr15-32997019-INS-702
chr15-33093016-INS-6413
chr15-33161332-INS-625
chr15-33167828-INS-108
chr15-33264550-INS-710
chr15-33285939-INS-128
chr15-33290712-INS-438
chr15-33293029-INS-240
chr15-33300364-INS-98
chr15-33354351-INS-4322
chr15-33367270-INS-5275
chr15-33428478-INS-517
chr15-33432371-INS-1325
chr15-33438764-INS-1109
chr15-33461161-INS-79
chr15-33470916-INS-339
chr15-33474781-INS-6208
chr15-33484414-INS-2033
chr15-33512143-INS-73
chr15-33557453-INS-199
chr15-33572551-INS-170
chr15-33582137-INS-95
chr15-33592539-INS-690
chr15-33639373-INS-1865
chr15-33655757-INS-186
chr15-33673739-INS-96
chr15-33786167-INS-155
chr15-33820399-INS-6578
chr15-33822647-INS-3745
chr15-33862816-INS-3575
chr15-33876567-INS-6438
chr15-33910844-INS-115
chr15-33932785-INS-519
chr15-33976070-INS-64
chr15-33990376-INS-60
chr15-33995607-INS-127
chr15-33996131-INS-67
chr15-33997959-INS-75
chr15-34042726-INS-85
chr15-34057811-INS-316
chr15-34148383-INS-290
chr15-34174581-INS-198
chr15-34176245-INS-78
chr15-34182233-INS-73
chr15-34196970-INS-352
chr15-34212080-INS-67
chr15-34216655-INS-501
chr15-34221127-INS-892
chr15-34252216-INS-52
chr15-34268735-INS-70
chr15-34290529-INS-122
chr15-34293652-INS-60
chr15-34298293-INS-128
chr15-34302320-INS-208
chr15-34317640-INS-400
chr15-34333296-INS-5573
chr15-34361488-INS-592
chr15-34406729-INS-52
chr15-34408698-INS-80
chr15-34413773-INS-226
chr15-34414849-INS-1851
chr15-34420919-INS-212
chr15-34421926-INS-78

chr15-34422314-INS-195
chr15-34452017-INS-241
chr15-34455218-INS-232
chr15-34460830-INS-202
chr15-34486857-INS-58
chr15-34505544-INS-82
chr15-34509285-INS-208
chr15-34514993-INS-348
chr15-34516774-INS-134
chr15-34528838-INS-808
chr15-34534194-INS-55
chr15-34562074-INS-510
chr15-34581138-INS-68
chr15-34610542-INS-486
chr15-34615223-INS-196
chr15-34616342-INS-56
chr15-34630401-INS-140
chr15-34635201-INS-255
chr15-34654974-INS-51
chr15-34672323-INS-197
chr15-34672896-INS-1004
chr15-34682819-INS-429
chr15-34697579-INS-114
chr15-34776806-INS-52
chr15-34902544-INS-68
chr15-34928216-INS-150
chr15-34956329-INS-488
chr15-34981673-INS-2436
chr15-35000329-INS-1445
chr15-35008250-INS-69
chr15-35015449-INS-87
chr15-35050536-INS-533
chr15-35052679-INS-199
chr15-35075936-INS-162
chr15-35084462-INS-52
chr15-35095662-INS-221
chr15-35099172-INS-116
chr15-35102342-INS-70
chr15-35130039-INS-200
chr15-35232492-INS-199
chr15-35247266-INS-52
chr15-35290991-INS-201
chr15-35344359-INS-51
chr15-35350261-INS-55
chr15-35353565-INS-1892
chr15-35355483-INS-552
chr15-35360189-INS-1893
chr15-35410057-INS-701
chr15-35450036-INS-55
chr15-35456180-INS-186
chr15-35465030-INS-6223
chr15-35469070-INS-7096
chr15-35491010-INS-405
chr15-35502220-INS-97
chr15-35504437-INS-670
chr15-35547430-INS-400
chr15-35565580-INS-841
chr15-35590102-INS-319
chr15-35598656-INS-306
chr15-35602945-INS-195
chr15-35630217-INS-539
chr15-35653714-INS-204
chr15-35701109-INS-5540
chr15-35717914-INS-517
chr15-35719592-INS-200
chr15-35735860-INS-1866
chr15-35747620-INS-159
chr15-35753654-INS-144
chr15-35768369-INS-57
chr15-35795318-INS-90
chr15-35827965-INS-68
chr15-35858332-INS-154
chr15-35892250-INS-56
chr15-35894946-INS-52
chr15-35896053-INS-394
chr15-35903917-INS-150
chr15-35906248-INS-215

chr15-35909321-INS-130
chr15-35917103-INS-130
chr15-35943036-INS-7484
chr15-35944892-INS-688
chr15-35953128-INS-198
chr15-35953450-INS-6162
chr15-35959693-INS-479
chr15-35980490-INS-56
chr15-36012274-INS-136
chr15-36029188-INS-349
chr15-36077997-INS-3277
chr15-36104880-INS-832
chr15-36106493-INS-216
chr15-36127087-INS-129
chr15-36152633-INS-52
chr15-36188373-INS-599
chr15-36190093-INS-173
chr15-36194007-INS-64
chr15-36203029-INS-964
chr15-36219325-INS-58
chr15-36223476-INS-62
chr15-36249654-INS-94
chr15-36275972-INS-340
chr15-36287050-INS-106
chr15-36312989-INS-304
chr15-36324231-INS-577
chr15-36331048-INS-500
chr15-36332994-INS-62
chr15-36348422-INS-219
chr15-36351096-INS-299
chr15-36385369-INS-654
chr15-36392339-INS-120
chr15-36407267-INS-182
chr15-36428390-INS-779
chr15-36440186-INS-5149
chr15-36444271-INS-95
chr15-36445686-INS-214
chr15-36455001-INS-276
chr15-36457824-INS-70
chr15-36466639-INS-228
chr15-36480350-INS-863
chr15-36531020-INS-760
chr15-36537156-INS-82
chr15-36572872-INS-1893
chr15-36583746-INS-122
chr15-36591571-INS-362
chr15-36611475-INS-203
chr15-36616856-INS-68
chr15-36622275-INS-147
chr15-36624443-INS-72
chr15-36627211-INS-204
chr15-36643737-INS-117
chr15-36656660-INS-110
chr15-36679764-INS-199
chr15-36681714-INS-203
chr15-36689768-INS-926
chr15-36690068-INS-88
chr15-36693068-INS-188
chr15-36702078-INS-187
chr15-36710686-INS-143
chr15-36713745-INS-59
chr15-36723434-INS-175
chr15-36727235-INS-197
chr15-36733160-INS-169
chr15-36737588-INS-135
chr15-36740296-INS-2975
chr15-36744953-INS-102
chr15-36746419-INS-65
chr15-36750270-INS-7189
chr15-36758451-INS-471
chr15-36767556-INS-130
chr15-36784570-INS-212
chr15-36798078-INS-57
chr15-36848719-INS-124
chr15-36863468-INS-122
chr15-36887691-INS-132
chr15-36909300-INS-153

chr15-36916256-INS-187
chr15-36920970-INS-240
chr15-36923860-INS-148
chr15-36954035-INS-880
chr15-36972498-INS-109
chr15-36972777-INS-4605
chr15-36976724-INS-151
chr15-36977668-INS-104
chr15-37001461-INS-229
chr15-37023810-INS-969
chr15-37048109-INS-629
chr15-37051880-INS-168
chr15-37057146-INS-454
chr15-37062106-INS-66
chr15-37084778-INS-93
chr15-37086713-INS-183
chr15-37106632-INS-231
chr15-37130604-INS-336
chr15-37147355-INS-110
chr15-37170791-INS-250
chr15-37191982-INS-200
chr15-37194117-INS-83
chr15-37203679-INS-920
chr15-37212610-INS-400
chr15-37216964-INS-253
chr15-37225961-INS-130
chr15-37231340-INS-206
chr15-37265689-INS-819
chr15-37277313-INS-177
chr15-37279375-INS-67
chr15-37309643-INS-193
chr15-37316295-INS-1904
chr15-37335734-INS-51
chr15-37349105-INS-51
chr15-37385616-INS-63
chr15-37394724-INS-348
chr15-37403334-INS-467
chr15-37404322-INS-117
chr15-37416393-INS-114
chr15-37474210-INS-94
chr15-37490146-INS-198
chr15-37509000-INS-63
chr15-37518646-INS-57
chr15-37530768-INS-58
chr15-37538058-INS-162
chr15-37538938-INS-180
chr15-37542159-INS-140
chr15-37545006-INS-60
chr15-37553042-INS-85
chr15-37566469-INS-237
chr15-37567685-INS-574
chr15-37587521-INS-174
chr15-37596478-INS-540
chr15-37631992-INS-400
chr15-37635201-INS-215
chr15-37639222-INS-57
chr15-37647145-INS-78
chr15-37655716-INS-6561
chr15-37668933-INS-463
chr15-37674493-INS-68
chr15-37674799-INS-4173
chr15-37699171-INS-68
chr15-37707097-INS-57
chr15-37709583-INS-110
chr15-37725585-INS-62
chr15-37751354-INS-54
chr15-37786806-INS-223
chr15-37799869-INS-102
chr15-37815642-INS-452
chr15-37816659-INS-101
chr15-37852679-INS-164
chr15-37859848-INS-206
chr15-37904866-INS-64
chr15-37907226-INS-2466
chr15-37908323-INS-99
chr15-37925874-INS-5318
chr15-37936995-INS-347

chr15-37982003-INS-152
chr15-37986806-INS-228
chr15-37988160-INS-57
chr15-38014151-INS-207
chr15-38018497-INS-218
chr15-38024482-INS-54
chr15-38026029-INS-63
chr15-38047239-INS-67
chr15-38077026-INS-254
chr15-38090343-INS-154
chr15-38107595-INS-7076
chr15-38119771-INS-476
chr15-38121214-INS-69
chr15-38130250-INS-387
chr15-38136902-INS-422
chr15-38138657-INS-120
chr15-38146985-INS-68
chr15-38163232-INS-153
chr15-38172858-INS-254
chr15-38183144-INS-58
chr15-38187141-INS-532
chr15-38206204-INS-52
chr15-38211014-INS-526
chr15-38241530-INS-567
chr15-38250539-INS-88
chr15-38259343-INS-200
chr15-38279236-INS-100
chr15-38281661-INS-237
chr15-38285421-INS-242
chr15-38331415-INS-54
chr15-38331845-INS-123
chr15-38345690-INS-55
chr15-38370265-INS-243
chr15-38383403-INS-240
chr15-38394757-INS-68
chr15-38396949-INS-204
chr15-38407012-INS-54
chr15-38427214-INS-57
chr15-38429298-INS-223
chr15-38469224-INS-70
chr15-38492530-INS-840
chr15-38500287-INS-193
chr15-38505490-INS-210
chr15-38527658-INS-348
chr15-38535352-INS-215
chr15-38540466-INS-196
chr15-38553425-INS-463
chr15-38554437-INS-404
chr15-38592850-INS-65
chr15-38595577-INS-54
chr15-38624504-INS-52
chr15-38692037-INS-54
chr15-38709099-INS-115
chr15-38712597-INS-145
chr15-38734721-INS-255
chr15-38738684-INS-51
chr15-38751775-INS-479
chr15-38756845-INS-68
chr15-38759214-INS-200
chr15-38761246-INS-65
chr15-38803979-INS-63
chr15-38847375-INS-94
chr15-38856733-INS-682
chr15-38858288-INS-98
chr15-38885911-INS-60
chr15-38937505-INS-495
chr15-38949446-INS-52
chr15-38952586-INS-60
chr15-38981614-INS-428
chr15-38998728-INS-737
chr15-39012707-INS-7114
chr15-39088139-INS-55
chr15-39091283-INS-6462
chr15-39100406-INS-65
chr15-39102847-INS-126
chr15-39119248-INS-3176
chr15-39166472-INS-56

chr15-39168408-INS-6387
chr15-39175150-INS-150
chr15-39188779-INS-172
chr15-39207624-INS-65
chr15-39219939-INS-393
chr15-39221545-INS-150
chr15-39227614-INS-538
chr15-39238228-INS-724
chr15-39252018-INS-192
chr15-39269477-INS-6736
chr15-39270806-INS-52
chr15-39286627-INS-650
chr15-39337809-INS-6718
chr15-39340685-INS-194
chr15-39344464-INS-187
chr15-39364879-INS-82
chr15-39384644-INS-64
chr15-39399961-INS-575
chr15-39422124-INS-180
chr15-39463330-INS-220
chr15-39463863-INS-4518
chr15-39476150-INS-55
chr15-39488794-INS-169
chr15-39510218-INS-91
chr15-39558692-INS-377
chr15-39605979-INS-347
chr15-39611934-INS-335
chr15-39613498-INS-58
chr15-39617396-INS-55
chr15-39625085-INS-403
chr15-39653271-INS-73
chr15-39658104-INS-99
chr15-39666299-INS-118
chr15-39678763-INS-220
chr15-39692523-INS-101
chr15-39727411-INS-179
chr15-39760352-INS-723
chr15-39777920-INS-347
chr15-39782030-INS-117
chr15-39787940-INS-478
chr15-39800643-INS-4357
chr15-39820588-INS-69
chr15-39821899-INS-61
chr15-39828012-INS-200
chr15-39841485-INS-495
chr15-39859979-INS-83
chr15-39880114-INS-67
chr15-39925560-INS-84
chr15-39941467-INS-359
chr15-39949589-INS-58
chr15-39982983-INS-79
chr15-40018158-INS-68
chr15-40022807-INS-52
chr15-40068583-INS-208
chr15-40081519-INS-315
chr15-40135790-INS-825
chr15-40137022-INS-203
chr15-40146403-INS-81
chr15-40154559-INS-6838
chr15-40196346-INS-5988
chr15-40213310-INS-52
chr15-40239296-INS-101
chr15-40272112-INS-130
chr15-40304528-INS-551
chr15-40319898-INS-60
chr15-40331660-INS-106
chr15-40337592-INS-66
chr15-40359305-INS-196
chr15-40360235-INS-66
chr15-40391830-INS-267
chr15-40396533-INS-68
chr15-40399993-INS-167
chr15-40403761-INS-6207
chr15-40403765-INS-331
chr15-40407362-INS-298
chr15-40412536-INS-161
chr15-40424227-INS-1337

chr15-40425091-INS-6482
chr15-40432499-INS-581
chr15-40435273-INS-379
chr15-40435956-INS-4963
chr15-40449667-INS-6022
chr15-40455170-INS-89
chr15-40459697-INS-436
chr15-40463927-INS-82
chr15-40474901-INS-54
chr15-40504839-INS-188
chr15-40528350-INS-96
chr15-40535087-INS-5344
chr15-40535877-INS-601
chr15-40555348-INS-74
chr15-40591454-INS-323
chr15-40620812-INS-514
chr15-40637873-INS-334
chr15-40645708-INS-53
chr15-40673049-INS-142
chr15-40704262-INS-96
chr15-40705415-INS-64
chr15-40772832-INS-101
chr15-40778641-INS-690
chr15-40792871-INS-252
chr15-40795304-INS-83
chr15-40812653-INS-51
chr15-40837301-INS-254
chr15-40843445-INS-225
chr15-40871499-INS-6229
chr15-40908138-INS-253
chr15-40928675-INS-86
chr15-40931195-INS-89
chr15-40938500-INS-82
chr15-40946863-INS-76
chr15-40968410-INS-195
chr15-40987771-INS-125
chr15-41021644-INS-129
chr15-41031107-INS-105
chr15-41059661-INS-1358
chr15-41064341-INS-59
chr15-41064872-INS-6521
chr15-41132844-INS-418
chr15-41193969-INS-89
chr15-41202497-INS-6945
chr15-41233153-INS-180
chr15-41246621-INS-568
chr15-41259359-INS-10858
chr15-41266845-INS-52
chr15-41273315-INS-96
chr15-41319017-INS-60
chr15-41328802-INS-308
chr15-41328897-INS-525
chr15-41345783-INS-902
chr15-41351045-INS-741
chr15-41357843-INS-74
chr15-41412039-INS-3668
chr15-41424246-INS-73
chr15-41429413-INS-75
chr15-41441043-INS-112
chr15-41464147-INS-112
chr15-41511275-INS-77
chr15-41521327-INS-101
chr15-41523319-INS-630
chr15-41532317-INS-59
chr15-41552361-INS-400
chr15-41556093-INS-58
chr15-41567623-INS-64
chr15-41590783-INS-306
chr15-41603810-INS-182
chr15-41627050-INS-2920
chr15-41646293-INS-211
chr15-41681481-INS-116
chr15-41697330-INS-464
chr15-41725187-INS-52
chr15-41731693-INS-60
chr15-41747167-INS-7166
chr15-41750935-INS-110

chr15-41766297-INS-208
chr15-41767803-INS-53
chr15-41770775-INS-201
chr15-41772815-INS-202
chr15-41779396-INS-73
chr15-41782761-INS-61
chr15-41788229-INS-232
chr15-41789883-INS-410
chr15-41810235-INS-325
chr15-41818892-INS-7469
chr15-41843560-INS-1116
chr15-41853664-INS-113
chr15-41862898-INS-91
chr15-41892581-INS-56
chr15-41898764-INS-55
chr15-41907540-INS-282
chr15-41912970-INS-871
chr15-41914691-INS-6255
chr15-41919231-INS-492
chr15-41921782-INS-882
chr15-41946494-INS-375
chr15-41950125-INS-3343
chr15-41961149-INS-498
chr15-41979588-INS-52
chr15-41989496-INS-288
chr15-42014006-INS-963
chr15-42016790-INS-56
chr15-42026558-INS-52
chr15-42062885-INS-321
chr15-42070343-INS-67
chr15-42093976-INS-319
chr15-42127224-INS-2862
chr15-42174620-INS-84
chr15-42175685-INS-721
chr15-42195094-INS-827
chr15-42208283-INS-117
chr15-42218265-INS-115
chr15-42229623-INS-71
chr15-42276717-INS-76
chr15-42324878-INS-139
chr15-42334128-INS-7074
chr15-42350357-INS-134
chr15-42352796-INS-236
chr15-42356705-INS-6267
chr15-42373362-INS-200
chr15-42395990-INS-358
chr15-42409941-INS-92
chr15-42425773-INS-212
chr15-42426949-INS-90
chr15-42431239-INS-56
chr15-42435307-INS-70
chr15-42483446-INS-358
chr15-42521330-INS-6539
chr15-42522318-INS-313
chr15-42597984-INS-94
chr15-42619983-INS-59
chr15-42627008-INS-76
chr15-42638046-INS-465
chr15-42646060-INS-201
chr15-42647960-INS-643
chr15-42706615-INS-3797
chr15-42731856-INS-410
chr15-42737206-INS-92
chr15-42753008-INS-284
chr15-42767211-INS-468
chr15-42768827-INS-916
chr15-42769140-INS-63
chr15-42779992-INS-328
chr15-42781869-INS-651
chr15-42791523-INS-163
chr15-42817410-INS-554
chr15-42828156-INS-80
chr15-42860727-INS-142
chr15-42868876-INS-2289
chr15-42869466-INS-277
chr15-42877609-INS-7260
chr15-42895170-INS-76

chr15-42926426-INS-804
chr15-43032659-INS-134
chr15-43071880-INS-6438
chr15-43101068-INS-194
chr15-43106152-INS-66
chr15-43204513-INS-7014
chr15-43208232-INS-6354
chr15-43217235-INS-82
chr15-43225336-INS-6612
chr15-43237513-INS-184
chr15-43242304-INS-172
chr15-43253841-INS-134
chr15-43261188-INS-4125
chr15-43265841-INS-609
chr15-43270126-INS-1170
chr15-43274363-INS-6445
chr15-43280033-INS-3963
chr15-43286147-INS-169
chr15-43287553-INS-212
chr15-43289549-INS-483
chr15-43327391-INS-70
chr15-43331539-INS-139
chr15-43397365-INS-71
chr15-43403309-INS-622
chr15-43405107-INS-526
chr15-43429537-INS-998
chr15-43483302-INS-76
chr15-43486955-INS-363
chr15-43507951-INS-436
chr15-43542150-INS-421
chr15-43547555-INS-682
chr15-43550640-INS-73
chr15-43557533-INS-877
chr15-43568938-INS-243
chr15-43619062-INS-50
chr15-43635884-INS-234
chr15-43638069-INS-150
chr15-43653961-INS-6454
chr15-43665445-INS-62
chr15-43687705-INS-60
chr15-43708340-INS-204
chr15-43710157-INS-315
chr15-43710984-INS-56
chr15-43723412-INS-112
chr15-43724281-INS-6255
chr15-43732722-INS-51
chr15-43740377-INS-5167
chr15-43752182-INS-6655
chr15-43756072-INS-887
chr15-43763657-INS-6389
chr15-43808425-INS-217
chr15-43815237-INS-7340
chr15-43819305-INS-6318
chr15-43820839-INS-437
chr15-43831325-INS-216
chr15-43845013-INS-355
chr15-43852604-INS-946
chr15-43854295-INS-991
chr15-43858150-INS-64
chr15-43880870-INS-210
chr15-43881989-INS-280
chr15-43910117-INS-128
chr15-43918748-INS-508
chr15-43933622-INS-54
chr15-43968440-INS-348
chr15-43986535-INS-798
chr15-43989510-INS-382
chr15-43993582-INS-3980
chr15-43995379-INS-130
chr15-43998143-INS-120
chr15-44068587-INS-78
chr15-44118474-INS-211
chr15-44132891-INS-70
chr15-44139586-INS-60
chr15-44140685-INS-180
chr15-44169059-INS-226
chr15-44213260-INS-56

chr15-44229475-INS-220
chr15-44260600-INS-861
chr15-44265068-INS-54
chr15-44312278-INS-6483
chr15-44319493-INS-84
chr15-44327259-INS-6701
chr15-44330177-INS-110
chr15-44332519-INS-1042
chr15-44355832-INS-288
chr15-44411674-INS-64
chr15-44457065-INS-181
chr15-44497355-INS-974
chr15-44497618-INS-72
chr15-44502297-INS-55
chr15-44524890-INS-65
chr15-44543546-INS-80
chr15-44545814-INS-156
chr15-44556961-INS-206
chr15-44599886-INS-170
chr15-44630017-INS-55
chr15-44632874-INS-353
chr15-44636545-INS-8954
chr15-44659129-INS-428
chr15-44669857-INS-148
chr15-44688062-INS-60
chr15-44707099-INS-61
chr15-44711226-INS-63
chr15-44722610-INS-951
chr15-44735844-INS-224
chr15-44759063-INS-280
chr15-44784341-INS-184
chr15-44785477-INS-56
chr15-44796121-INS-6433
chr15-44804003-INS-507
chr15-44826780-INS-59
chr15-44832841-INS-94
chr15-44849098-INS-681
chr15-44850844-INS-67
chr15-44860693-INS-335
chr15-44873934-INS-561
chr15-44882405-INS-5547
chr15-44899449-INS-740
chr15-44901528-INS-1984
chr15-44935556-INS-484
chr15-44937903-INS-6862
chr15-44952909-INS-131
chr15-44958679-INS-560
chr15-44982936-INS-607
chr15-44983609-INS-98
chr15-45006807-INS-66
chr15-45015853-INS-114
chr15-45021471-INS-93
chr15-45032273-INS-388
chr15-45036837-INS-236
chr15-45043098-INS-52
chr15-45061525-INS-60
chr15-45068535-INS-7102
chr15-45078913-INS-107
chr15-45081900-INS-154
chr15-45132491-INS-109
chr15-45146252-INS-529
chr15-45156077-INS-7257
chr15-45169612-INS-7141
chr15-45190377-INS-566
chr15-45194998-INS-127
chr15-45237582-INS-78
chr15-45261403-INS-58
chr15-45262772-INS-7082
chr15-45291771-INS-689
chr15-45323684-INS-70
chr15-45357297-INS-184
chr15-45367119-INS-59
chr15-45386044-INS-130
chr15-45402963-INS-357
chr15-45403756-INS-4703
chr15-45423045-INS-92
chr15-45444677-INS-6297

chr15-45453552-INS-846
chr15-45457331-INS-7412
chr15-45466725-INS-371
chr15-45477359-INS-354
chr15-45487815-INS-193
chr15-45488974-INS-72
chr15-45507402-INS-53
chr15-45520300-INS-73
chr15-45527649-INS-387
chr15-45543209-INS-261
chr15-45549959-INS-287
chr15-45557767-INS-64
chr15-45567308-INS-64
chr15-45582912-INS-53
chr15-45613661-INS-83
chr15-45617929-INS-107
chr15-45650325-INS-6694
chr15-45712168-INS-183
chr15-45732959-INS-84
chr15-45768195-INS-70
chr15-45778235-INS-477
chr15-45872040-INS-498
chr15-45882590-INS-409
chr15-45950382-INS-882
chr15-45959139-INS-168
chr15-45960344-INS-292
chr15-46015086-INS-778
chr15-46044440-INS-54
chr15-46046662-INS-192
chr15-46060603-INS-103
chr15-46061853-INS-60
chr15-46113447-INS-188
chr15-46126402-INS-53
chr15-46170012-INS-339
chr15-46174062-INS-75
chr15-46193143-INS-718
chr15-46222847-INS-61
chr15-46226995-INS-214
chr15-46294529-INS-57
chr15-46308400-INS-64
chr15-46377672-INS-3386
chr15-46460012-INS-6203
chr15-46464197-INS-226
chr15-46480206-INS-263
chr15-46495563-INS-52
chr15-46500113-INS-498
chr15-46533717-INS-658
chr15-46561744-INS-56
chr15-46585669-INS-55
chr15-46587722-INS-54
chr15-46609333-INS-135
chr15-46624210-INS-2145
chr15-46633122-INS-235
chr15-46660312-INS-62
chr15-46793750-INS-6402
chr15-46803961-INS-6239
chr15-46815936-INS-8457
chr15-46852612-INS-6887
chr15-46882474-INS-80
chr15-46889580-INS-727
chr15-46897527-INS-56
chr15-46901026-INS-270
chr15-46943919-INS-458
chr15-46946470-INS-138
chr15-46949555-INS-52
chr15-46959511-INS-97
chr15-46984746-INS-202
chr15-47022450-INS-62
chr15-47027910-INS-870
chr15-47040689-INS-152
chr15-47058805-INS-486
chr15-47141340-INS-443
chr15-47178475-INS-6294
chr15-47180299-INS-67
chr15-47205315-INS-146
chr15-47230062-INS-450
chr15-47272019-INS-68

chr15-47288572-INS-62
chr15-47310547-INS-108
chr15-47397434-INS-553
chr15-47441352-INS-668
chr15-47446667-INS-180
chr15-47489909-INS-868
chr15-47499162-INS-413
chr15-47578738-INS-52
chr15-47592243-INS-60
chr15-47606522-INS-89
chr15-47655356-INS-102
chr15-47666999-INS-7183
chr15-47674625-INS-74
chr15-47677957-INS-64
chr15-47704163-INS-64
chr15-47707388-INS-4922
chr15-47756692-INS-333
chr15-47778613-INS-178
chr15-47794178-INS-82
chr15-47798660-INS-438
chr15-47806428-INS-970
chr15-47824908-INS-107
chr15-47846854-INS-159
chr15-47861997-INS-465
chr15-47878399-INS-524
chr15-47878500-INS-1864
chr15-47890764-INS-56
chr15-47955728-INS-604
chr15-47963265-INS-588
chr15-47970159-INS-399
chr15-47975702-INS-351
chr15-47977553-INS-428
chr15-47995678-INS-709
chr15-48006938-INS-411
chr15-48012975-INS-84
chr15-48017390-INS-179
chr15-48039938-INS-71
chr15-48044403-INS-369
chr15-48046992-INS-80
chr15-48064632-INS-212
chr15-48082560-INS-194
chr15-48085042-INS-9356
chr15-48117886-INS-129
chr15-48123041-INS-51
chr15-48141436-INS-121
chr15-48145303-INS-396
chr15-48149377-INS-64
chr15-48199210-INS-1654
chr15-48218533-INS-62
chr15-48222964-INS-2136
chr15-48299395-INS-53
chr15-48304751-INS-500
chr15-48328719-INS-360
chr15-48331564-INS-121
chr15-48370131-INS-198
chr15-48376026-INS-58
chr15-48390181-INS-58
chr15-48392510-INS-189
chr15-48406101-INS-187
chr15-48420735-INS-508
chr15-48428499-INS-70
chr15-48449062-INS-120
chr15-48465387-INS-87
chr15-48477518-INS-68
chr15-48478639-INS-309
chr15-48520086-INS-290
chr15-48540139-INS-188
chr15-48584077-INS-68
chr15-48599889-INS-611
chr15-48611691-INS-56
chr15-48624345-INS-205
chr15-48629796-INS-340
chr15-48642972-INS-582
chr15-48674439-INS-6632
chr15-48713014-INS-332
chr15-48727450-INS-60
chr15-48754406-INS-911

chr15-48756443-INS-93
chr15-48763480-INS-549
chr15-48810947-INS-127
chr15-48821735-INS-51
chr15-48828221-INS-831
chr15-48838980-INS-948
chr15-48918497-INS-6584
chr15-48938901-INS-60
chr15-48941524-INS-5553
chr15-48949905-INS-54
chr15-48970813-INS-8255
chr15-48981695-INS-6650
chr15-49016135-INS-68
chr15-49083292-INS-4198
chr15-49147541-INS-4913
chr15-49170836-INS-151
chr15-49174064-INS-196
chr15-49191310-INS-6127
chr15-49217367-INS-2876
chr15-49221271-INS-413
chr15-49234326-INS-51
chr15-49264349-INS-825
chr15-49270752-INS-134
chr15-49284491-INS-52
chr15-49288073-INS-105
chr15-49322105-INS-54
chr15-49326369-INS-1887
chr15-49348766-INS-629
chr15-49399540-INS-895
chr15-49411365-INS-451
chr15-49430518-INS-701
chr15-49443511-INS-138
chr15-49446426-INS-68
chr15-49460857-INS-6378
chr15-49480842-INS-90
chr15-49492994-INS-209
chr15-49513559-INS-651
chr15-49525556-INS-69
chr15-49546522-INS-56
chr15-49548580-INS-58
chr15-49567534-INS-58
chr15-49587666-INS-79
chr15-49617047-INS-589
chr15-49621240-INS-131
chr15-49642686-INS-2463
chr15-49647000-INS-6864
chr15-49681316-INS-173
chr15-49730652-INS-2439
chr15-49739936-INS-119
chr15-49741056-INS-864
chr15-49753425-INS-3192
chr15-49759354-INS-52
chr15-49776574-INS-1888
chr15-49788277-INS-6232
chr15-49790472-INS-53
chr15-49803162-INS-777
chr15-49804843-INS-156
chr15-49808092-INS-332
chr15-49876183-INS-208
chr15-49878483-INS-6452
chr15-49895150-INS-61
chr15-49906921-INS-145
chr15-49913184-INS-64
chr15-49916920-INS-5219
chr15-49919617-INS-217
chr15-49919717-INS-662
chr15-49920781-INS-200
chr15-49930830-INS-55
chr15-49945686-INS-78
chr15-49967035-INS-67
chr15-49974824-INS-109
chr15-49982015-INS-88
chr15-50014492-INS-66
chr15-50015538-INS-112
chr15-50019033-INS-58
chr15-50027704-INS-401
chr15-50051852-INS-315

chr15-50080699-INS-81
chr15-50130295-INS-281
chr15-50133667-INS-55
chr15-50165247-INS-87
chr15-50171541-INS-131
chr15-50209004-INS-308
chr15-50210026-INS-320
chr15-50224895-INS-247
chr15-50236302-INS-629
chr15-50239155-INS-185
chr15-50258379-INS-234
chr15-50259278-INS-179
chr15-50273436-INS-7187
chr15-50283860-INS-147
chr15-50292207-INS-89
chr15-50333228-INS-468
chr15-50339741-INS-91
chr15-50346439-INS-633
chr15-50363957-INS-329
chr15-50375044-INS-72
chr15-50384419-INS-180
chr15-50387697-INS-80
chr15-50415702-INS-6955
chr15-50433860-INS-525
chr15-50454386-INS-612
chr15-50478324-INS-1889
chr15-50480994-INS-4846
chr15-50533280-INS-193
chr15-50539598-INS-6270
chr15-50542664-INS-126
chr15-50550492-INS-175
chr15-50580517-INS-230
chr15-50583971-INS-137
chr15-50596010-INS-132
chr15-50602770-INS-100
chr15-50611381-INS-969
chr15-50659029-INS-555
chr15-50672664-INS-82
chr15-50720930-INS-74
chr15-50726761-INS-64
chr15-50739502-INS-56
chr15-50761724-INS-443
chr15-50781581-INS-161
chr15-50783824-INS-115
chr15-50795519-INS-74
chr15-50806519-INS-310
chr15-50822047-INS-52
chr15-50859079-INS-66
chr15-50870153-INS-2489
chr15-50891628-INS-198
chr15-50892989-INS-80
chr15-50927486-INS-53
chr15-50943311-INS-176
chr15-50984725-INS-54
chr15-51015936-INS-329
chr15-51030288-INS-106
chr15-51032419-INS-7381
chr15-51057260-INS-75
chr15-51058781-INS-289
chr15-51061648-INS-181
chr15-51089899-INS-568
chr15-51106272-INS-80
chr15-51111727-INS-72
chr15-51160800-INS-60
chr15-51176698-INS-421
chr15-51184892-INS-65
chr15-51215963-INS-906
chr15-51220617-INS-590
chr15-51244642-INS-58
chr15-51270544-INS-51
chr15-51283767-INS-69
chr15-51284690-INS-203
chr15-51313241-INS-6352
chr15-51317360-INS-60
chr15-51317773-INS-8199
chr15-51345893-INS-248
chr15-51353177-INS-60

chr15-51367819-INS-498
chr15-51400657-INS-79
chr15-51449621-INS-257
chr15-51476425-INS-1338
chr15-51488706-INS-81
chr15-51490555-INS-904
chr15-51500709-INS-537
chr15-51508149-INS-105
chr15-51526788-INS-113
chr15-51545100-INS-194
chr15-51592296-INS-51
chr15-51597850-INS-3230
chr15-51609648-INS-6517
chr15-51650825-INS-249
chr15-51710978-INS-156
chr15-51742186-INS-203
chr15-51743844-INS-348
chr15-51748196-INS-400
chr15-51762340-INS-7306
chr15-51763780-INS-797
chr15-51795814-INS-186
chr15-51806657-INS-710
chr15-51817943-INS-938
chr15-51839854-INS-215
chr15-51851347-INS-164
chr15-51858699-INS-99
chr15-51867270-INS-439
chr15-51885952-INS-324
chr15-51894972-INS-515
chr15-51905489-INS-210
chr15-51925819-INS-63
chr15-51929228-INS-69
chr15-51936081-INS-1036
chr15-51955619-INS-388
chr15-51960452-INS-51
chr15-51982126-INS-667
chr15-51987623-INS-66
chr15-51996577-INS-66
chr15-51998758-INS-206
chr15-52004671-INS-905
chr15-52037915-INS-122
chr15-52038005-INS-224
chr15-52073062-INS-530
chr15-52081274-INS-78
chr15-52108700-INS-489
chr15-52115104-INS-52
chr15-52120094-INS-7041
chr15-52124247-INS-110
chr15-52130529-INS-200
chr15-52133875-INS-6380
chr15-52182519-INS-512
chr15-52186662-INS-181
chr15-52201235-INS-127
chr15-52203811-INS-74
chr15-52208539-INS-557
chr15-52210445-INS-186
chr15-52219753-INS-234
chr15-52229349-INS-57
chr15-52235776-INS-188
chr15-52246039-INS-358
chr15-52248189-INS-51
chr15-52253169-INS-504
chr15-52283600-INS-406
chr15-52316692-INS-697
chr15-52355059-INS-400
chr15-52359398-INS-213
chr15-52383390-INS-61
chr15-52385308-INS-984
chr15-52385430-INS-359
chr15-52394584-INS-60
chr15-52395361-INS-87
chr15-52397401-INS-407
chr15-52433436-INS-769
chr15-52461104-INS-2150
chr15-52479348-INS-60
chr15-52533151-INS-90
chr15-52546634-INS-430

chr15-52632843-INS-180
chr15-52663963-INS-192
chr15-52683067-INS-3100
chr15-52697638-INS-6608
chr15-52709745-INS-243
chr15-52732087-INS-104
chr15-52764262-INS-268
chr15-52767644-INS-94
chr15-52821585-INS-90
chr15-52824159-INS-137
chr15-52856487-INS-855
chr15-52874916-INS-141
chr15-52883543-INS-122
chr15-52915439-INS-148
chr15-52916018-INS-660
chr15-52918614-INS-51
chr15-52932296-INS-77
chr15-52944720-INS-243
chr15-53011969-INS-198
chr15-53036079-INS-163
chr15-53047319-INS-212
chr15-53067138-INS-359
chr15-53081195-INS-77
chr15-53088063-INS-188
chr15-53101638-INS-6401
chr15-53108143-INS-53
chr15-53118672-INS-235
chr15-53138546-INS-557
chr15-53142297-INS-64
chr15-53150975-INS-193
chr15-53201225-INS-56
chr15-53223905-INS-76
chr15-53230136-INS-98
chr15-53231550-INS-229
chr15-53269566-INS-164
chr15-53282076-INS-109
chr15-53284609-INS-304
chr15-53285828-INS-70
chr15-53291683-INS-443
chr15-53305422-INS-947
chr15-53309742-INS-305
chr15-53344643-INS-55
chr15-53346357-INS-53
chr15-53349862-INS-81
chr15-53358497-INS-91
chr15-53372263-INS-112
chr15-53374352-INS-6498
chr15-53383360-INS-63
chr15-53435171-INS-142
chr15-53454496-INS-6481
chr15-53474341-INS-55
chr15-53483876-INS-80
chr15-53495497-INS-53
chr15-53527025-INS-6641
chr15-53544102-INS-401
chr15-53548124-INS-58
chr15-53558837-INS-76
chr15-53567248-INS-577
chr15-53589361-INS-70
chr15-53617061-INS-954
chr15-53636839-INS-195
chr15-53649005-INS-770
chr15-53656870-INS-6907
chr15-53658237-INS-79
chr15-53659436-INS-58
chr15-53662993-INS-219
chr15-53676631-INS-69
chr15-53738468-INS-726
chr15-53758561-INS-215
chr15-53777593-INS-55
chr15-53780783-INS-108
chr15-53801752-INS-56
chr15-53804845-INS-456
chr15-53835034-INS-52
chr15-53860648-INS-146
chr15-53885454-INS-226
chr15-53887214-INS-201

chr15-53889122-INS-188
chr15-53949184-INS-51
chr15-53959006-INS-217
chr15-53987323-INS-56
chr15-54008467-INS-302
chr15-54017868-INS-620
chr15-54040352-INS-196
chr15-54042482-INS-100
chr15-54048604-INS-1888
chr15-54064323-INS-1209
chr15-54074692-INS-212
chr15-54085544-INS-92
chr15-54089546-INS-60
chr15-54127863-INS-938
chr15-54129362-INS-53
chr15-54148232-INS-62
chr15-54155738-INS-194
chr15-54159356-INS-942
chr15-54172119-INS-499
chr15-54190484-INS-193
chr15-54191017-INS-67
chr15-54199900-INS-376
chr15-54205040-INS-65
chr15-54239036-INS-386
chr15-54247556-INS-52
chr15-54261601-INS-54
chr15-54267932-INS-1265
chr15-54296383-INS-58
chr15-54344082-INS-411
chr15-54345007-INS-336
chr15-54347025-INS-333
chr15-54349382-INS-176
chr15-54351682-INS-88
chr15-54353645-INS-695
chr15-54356040-INS-6878
chr15-54359195-INS-66
chr15-54370036-INS-69
chr15-54374612-INS-92
chr15-54393651-INS-6339
chr15-54403123-INS-64
chr15-54416616-INS-533
chr15-54426201-INS-76
chr15-54428092-INS-327
chr15-54432314-INS-52
chr15-54478282-INS-6424
chr15-54478651-INS-393
chr15-54486081-INS-68
chr15-54487273-INS-109
chr15-54505356-INS-609
chr15-54513411-INS-160
chr15-54521980-INS-8066
chr15-54526424-INS-65
chr15-54543545-INS-6445
chr15-54550860-INS-101
chr15-54553975-INS-66
chr15-54581200-INS-55
chr15-54593912-INS-389
chr15-54598684-INS-72
chr15-54603009-INS-104
chr15-54604377-INS-60
chr15-54605457-INS-58
chr15-54611852-INS-78
chr15-54623248-INS-568
chr15-54632854-INS-282
chr15-54651755-INS-114
chr15-54672161-INS-161
chr15-54686125-INS-258
chr15-54694160-INS-6333
chr15-54707572-INS-52
chr15-54748816-INS-117
chr15-54788835-INS-309
chr15-54806614-INS-63
chr15-54812752-INS-128
chr15-54842131-INS-71
chr15-54843553-INS-118
chr15-54849916-INS-583
chr15-54854834-INS-205

chr15-54864437-INS-53
chr15-54867805-INS-53
chr15-54895792-INS-195
chr15-54925190-INS-880
chr15-54931510-INS-116
chr15-54939823-INS-185
chr15-54951870-INS-66
chr15-54956769-INS-167
chr15-54958798-INS-181
chr15-54961595-INS-76
chr15-54973447-INS-134
chr15-54974205-INS-712
chr15-54974721-INS-73
chr15-55001264-INS-86
chr15-55036143-INS-81
chr15-55037523-INS-66
chr15-55038869-INS-161
chr15-55077809-INS-165
chr15-55087442-INS-67
chr15-55088673-INS-762
chr15-55097489-INS-54
chr15-55101819-INS-510
chr15-55104680-INS-429
chr15-55130290-INS-77
chr15-55159595-INS-104
chr15-55161932-INS-270
chr15-55163611-INS-196
chr15-55191869-INS-351
chr15-55216916-INS-79
chr15-55222065-INS-419
chr15-55250094-INS-134
chr15-55252968-INS-201
chr15-55254281-INS-67
chr15-55277569-INS-65
chr15-55285878-INS-296
chr15-55299370-INS-547
chr15-55316527-INS-68
chr15-55319312-INS-86
chr15-55321697-INS-55
chr15-55322706-INS-212
chr15-55332460-INS-202
chr15-55350332-INS-516
chr15-55369749-INS-221
chr15-55387473-INS-391
chr15-55397215-INS-731
chr15-55433395-INS-80
chr15-55504436-INS-134
chr15-55508785-INS-297
chr15-55516674-INS-72
chr15-55526180-INS-311
chr15-55540864-INS-415
chr15-55547318-INS-211
chr15-55550736-INS-93
chr15-55551827-INS-56
chr15-55566143-INS-105
chr15-55568292-INS-65
chr15-55568336-INS-590
chr15-55587587-INS-7465
chr15-55591548-INS-53
chr15-55593031-INS-193
chr15-55604272-INS-1391
chr15-55611939-INS-52
chr15-55619047-INS-74
chr15-55643390-INS-594
chr15-55646943-INS-6439
chr15-55670219-INS-353
chr15-55676862-INS-173
chr15-55686157-INS-113
chr15-55688008-INS-101
chr15-55718462-INS-1054
chr15-55727039-INS-71
chr15-55747370-INS-60
chr15-55751526-INS-446
chr15-55763021-INS-260
chr15-55772034-INS-211
chr15-55810331-INS-55
chr15-55825216-INS-86

chr15-55826016-INS-692
chr15-55836971-INS-233
chr15-55847754-INS-6781
chr15-55849180-INS-1892
chr15-55859890-INS-174
chr15-55860176-INS-5067
chr15-55863559-INS-405
chr15-55880326-INS-78
chr15-55888399-INS-161
chr15-55892283-INS-398
chr15-55907319-INS-5828
chr15-55914530-INS-6516
chr15-55917739-INS-85
chr15-55931074-INS-128
chr15-55960105-INS-506
chr15-55976868-INS-60
chr15-55991226-INS-197
chr15-56022046-INS-6394
chr15-56068063-INS-240
chr15-56088131-INS-269
chr15-56106711-INS-58
chr15-56146013-INS-772
chr15-56150212-INS-1529
chr15-56161941-INS-630
chr15-56176522-INS-52
chr15-56178522-INS-54
chr15-56199339-INS-103
chr15-56203636-INS-66
chr15-56211571-INS-55
chr15-56232756-INS-58
chr15-56242889-INS-400
chr15-56244745-INS-94
chr15-56277889-INS-682
chr15-56287407-INS-12706
chr15-56296020-INS-256
chr15-56309403-INS-66
chr15-56325303-INS-84
chr15-56357184-INS-52
chr15-56405012-INS-445
chr15-56422403-INS-805
chr15-56430666-INS-66
chr15-56443657-INS-52
chr15-56447323-INS-197
chr15-56455372-INS-2146
chr15-56468177-INS-924
chr15-56478046-INS-63
chr15-56481097-INS-195
chr15-56484259-INS-159
chr15-56490440-INS-209
chr15-56496580-INS-58
chr15-56520191-INS-172
chr15-56532157-INS-205
chr15-56539066-INS-747
chr15-56550032-INS-470
chr15-56558394-INS-625
chr15-56568989-INS-158
chr15-56594876-INS-62
chr15-56596772-INS-56
chr15-56602161-INS-66
chr15-56612252-INS-389
chr15-56622051-INS-61
chr15-56625587-INS-64
chr15-56629172-INS-212
chr15-56661862-INS-818
chr15-56663279-INS-473
chr15-56669947-INS-177
chr15-56688909-INS-6211
chr15-56709604-INS-490
chr15-56710689-INS-197
chr15-56716886-INS-80
chr15-56728210-INS-248
chr15-56730120-INS-131
chr15-56758982-INS-54
chr15-56762588-INS-524
chr15-56778731-INS-6309
chr15-56785906-INS-6654
chr15-56786513-INS-6266

chr15-56788305-INS-6452
chr15-56827409-INS-208
chr15-56828241-INS-272
chr15-56830753-INS-7009
chr15-56837547-INS-120
chr15-56840282-INS-167
chr15-56858613-INS-7192
chr15-56900098-INS-227
chr15-56901554-INS-344
chr15-56905127-INS-105
chr15-56907505-INS-6379
chr15-56911765-INS-564
chr15-56911787-INS-658
chr15-56959980-INS-148
chr15-56971851-INS-136
chr15-56973569-INS-189
chr15-56982947-INS-77
chr15-56987753-INS-187
chr15-57001614-INS-89
chr15-57023549-INS-68
chr15-57033792-INS-70
chr15-57039206-INS-55
chr15-57066786-INS-166
chr15-57091611-INS-68
chr15-57104767-INS-239
chr15-57109194-INS-107
chr15-57119255-INS-169
chr15-57150895-INS-53
chr15-57179559-INS-175
chr15-57192131-INS-112
chr15-57194056-INS-101
chr15-57197941-INS-881
chr15-57206309-INS-55
chr15-57213979-INS-78
chr15-57223469-INS-431
chr15-57224038-INS-6400
chr15-57233213-INS-84
chr15-57234584-INS-162
chr15-57242722-INS-436
chr15-57244663-INS-449
chr15-57254063-INS-54
chr15-57269218-INS-77
chr15-57272574-INS-120
chr15-57297209-INS-88
chr15-57309135-INS-53
chr15-57317410-INS-6858
chr15-57319061-INS-5938
chr15-57325167-INS-6393
chr15-57336548-INS-50
chr15-57364511-INS-103
chr15-57374646-INS-1883
chr15-57378714-INS-68
chr15-57405461-INS-60
chr15-57422748-INS-259
chr15-57435694-INS-6688
chr15-57436432-INS-400
chr15-57476864-INS-54
chr15-57481569-INS-91
chr15-57488675-INS-84
chr15-57499918-INS-74
chr15-57514396-INS-60
chr15-57522442-INS-523
chr15-57530762-INS-214
chr15-57595559-INS-857
chr15-57611780-INS-65
chr15-57619384-INS-55
chr15-57634065-INS-68
chr15-57636651-INS-56
chr15-57648957-INS-96
chr15-57651391-INS-320
chr15-57667810-INS-222
chr15-57691403-INS-285
chr15-57703417-INS-493
chr15-57704390-INS-216
chr15-57707095-INS-210
chr15-57716490-INS-81
chr15-57718888-INS-210

chr15-57723265-INS-58
chr15-57727291-INS-3888
chr15-57760564-INS-52
chr15-57776972-INS-244
chr15-57781769-INS-786
chr15-57789391-INS-56
chr15-57793766-INS-78
chr15-57820762-INS-243
chr15-57823899-INS-108
chr15-57830678-INS-59
chr15-57848577-INS-56
chr15-57855818-INS-190
chr15-57857039-INS-186
chr15-57859653-INS-65
chr15-57898544-INS-129
chr15-57926621-INS-76
chr15-57941754-INS-145
chr15-57943812-INS-640
chr15-57956945-INS-72
chr15-57960352-INS-53
chr15-57973330-INS-231
chr15-57999502-INS-161
chr15-58011011-INS-95
chr15-58014661-INS-54
chr15-58036457-INS-224
chr15-58049432-INS-197
chr15-58051303-INS-224
chr15-58055910-INS-84
chr15-58056797-INS-888
chr15-58059829-INS-254
chr15-58065764-INS-54
chr15-58072741-INS-98
chr15-58091946-INS-51
chr15-58099361-INS-62
chr15-58108043-INS-125
chr15-58110474-INS-498
chr15-58112611-INS-139
chr15-58117312-INS-67
chr15-58171970-INS-505
chr15-58173584-INS-120
chr15-58197826-INS-70
chr15-58211324-INS-78
chr15-58218824-INS-188
chr15-58221670-INS-302
chr15-58223104-INS-217
chr15-58239063-INS-129
chr15-58246572-INS-70
chr15-58248287-INS-92
chr15-58294332-INS-72
chr15-58311232-INS-3488
chr15-58324797-INS-6843
chr15-58334367-INS-161
chr15-58347534-INS-77
chr15-58351843-INS-62
chr15-58361464-INS-64
chr15-58376050-INS-88
chr15-58379979-INS-174
chr15-58386421-INS-125
chr15-58400766-INS-128
chr15-58412975-INS-344
chr15-58422033-INS-309
chr15-58424051-INS-257
chr15-58474847-INS-255
chr15-58494803-INS-388
chr15-58500477-INS-310
chr15-58512794-INS-671
chr15-58517943-INS-52
chr15-58533669-INS-168
chr15-58556868-INS-325
chr15-58562483-INS-60
chr15-58563806-INS-90
chr15-58574824-INS-91
chr15-58590643-INS-125
chr15-58596076-INS-180
chr15-58601489-INS-389
chr15-58609088-INS-86
chr15-58659315-INS-159

chr15-58665648-INS-955
chr15-58688598-INS-121
chr15-58693353-INS-97
chr15-58712951-INS-6499
chr15-58714268-INS-327
chr15-58725915-INS-268
chr15-58742229-INS-193
chr15-58744138-INS-598
chr15-58753022-INS-230
chr15-58759582-INS-190
chr15-58764071-INS-58
chr15-58785483-INS-52
chr15-58786518-INS-74
chr15-58810378-INS-596
chr15-58849446-INS-622
chr15-58865107-INS-64
chr15-58911127-INS-401
chr15-58914644-INS-56
chr15-58918987-INS-50
chr15-58934931-INS-4840
chr15-58941593-INS-415
chr15-58949426-INS-68
chr15-58959473-INS-57
chr15-58975552-INS-125
chr15-58984136-INS-87
chr15-58988926-INS-82
chr15-59005707-INS-96
chr15-59015922-INS-82
chr15-59024445-INS-8575
chr15-59032558-INS-72
chr15-59052488-INS-1062
chr15-59052488-INS-942
chr15-59052589-INS-268
chr15-59057544-INS-62
chr15-59062073-INS-60
chr15-59064340-INS-123
chr15-59065727-INS-99
chr15-59080377-INS-98
chr15-59083873-INS-60
chr15-59085989-INS-68
chr15-59099003-INS-212
chr15-59107762-INS-52
chr15-59114484-INS-156
chr15-59137083-INS-150
chr15-59157560-INS-52
chr15-59164446-INS-112
chr15-59176979-INS-149
chr15-59198113-INS-116
chr15-59207478-INS-210
chr15-59216046-INS-243
chr15-59251918-INS-182
chr15-59259148-INS-434
chr15-59261370-INS-625
chr15-59282718-INS-74
chr15-59287555-INS-982
chr15-59306454-INS-117
chr15-59318516-INS-201
chr15-59319352-INS-205
chr15-59358495-INS-80
chr15-59367011-INS-78
chr15-59374031-INS-466
chr15-59418448-INS-112
chr15-59432282-INS-92
chr15-59451215-INS-704
chr15-59485270-INS-257
chr15-59509488-INS-157
chr15-59512377-INS-110
chr15-59536488-INS-54
chr15-59540164-INS-758
chr15-59543397-INS-90
chr15-59555649-INS-100
chr15-59557796-INS-758
chr15-59562709-INS-112
chr15-59563867-INS-219
chr15-59572664-INS-86
chr15-59575665-INS-1866
chr15-59577552-INS-51

chr15-59597077-INS-166
chr15-59601804-INS-225
chr15-59623101-INS-981
chr15-59624523-INS-155
chr15-59626692-INS-860
chr15-59628841-INS-169
chr15-59643326-INS-96
chr15-59648385-INS-6497
chr15-59648823-INS-72
chr15-59661685-INS-141
chr15-59684723-INS-87
chr15-59689553-INS-52
chr15-59711822-INS-112
chr15-59740962-INS-107
chr15-59752002-INS-137
chr15-59756651-INS-80
chr15-59760710-INS-109
chr15-59762224-INS-97
chr15-59774988-INS-542
chr15-59777693-INS-2819
chr15-59784755-INS-54
chr15-59790503-INS-143
chr15-59823711-INS-60
chr15-59831528-INS-109
chr15-59838990-INS-2904
chr15-59841411-INS-738
chr15-59851492-INS-68
chr15-59859611-INS-69
chr15-59878107-INS-75
chr15-59895266-INS-97
chr15-59914338-INS-82
chr15-59928563-INS-52
chr15-59938478-INS-52
chr15-59941569-INS-319
chr15-59946764-INS-6421
chr15-59953142-INS-73
chr15-59982925-INS-535
chr15-59997730-INS-72
chr15-60014085-INS-210
chr15-60038501-INS-349
chr15-60063900-INS-55
chr15-60067745-INS-59
chr15-60091379-INS-945
chr15-60099562-INS-377
chr15-60112951-INS-6406
chr15-60124329-INS-61
chr15-60135886-INS-191
chr15-60151055-INS-844
chr15-60157639-INS-199
chr15-60184788-INS-90
chr15-60185113-INS-86
chr15-60290291-INS-57
chr15-60293286-INS-489
chr15-60367716-INS-401
chr15-60377485-INS-327
chr15-60385866-INS-73
chr15-60393327-INS-152
chr15-60412090-INS-818
chr15-60425656-INS-869
chr15-60431199-INS-85
chr15-60439325-INS-72
chr15-60440365-INS-152
chr15-60471004-INS-157
chr15-60477680-INS-186
chr15-60479509-INS-141
chr15-60482757-INS-93
chr15-60490305-INS-58
chr15-60497547-INS-150
chr15-60508027-INS-439
chr15-60511496-INS-478
chr15-60539924-INS-5319
chr15-60541814-INS-573
chr15-60546292-INS-56
chr15-60551733-INS-58
chr15-60552214-INS-154
chr15-60558273-INS-201
chr15-60568164-INS-54

chr15-60572862-INS-80
chr15-60583032-INS-455
chr15-60601084-INS-176
chr15-60602062-INS-101
chr15-60608589-INS-82
chr15-60616656-INS-100
chr15-60629701-INS-69
chr15-60640621-INS-224
chr15-60641939-INS-395
chr15-60643619-INS-386
chr15-60667974-INS-183
chr15-60668941-INS-5420
chr15-60685198-INS-59
chr15-60693155-INS-353
chr15-60708561-INS-53
chr15-60712992-INS-548
chr15-60715893-INS-124
chr15-60722572-INS-77
chr15-60723560-INS-82
chr15-60727767-INS-52
chr15-60738591-INS-6435
chr15-60739813-INS-69
chr15-60743295-INS-167
chr15-60745000-INS-60
chr15-60782124-INS-56
chr15-60813180-INS-912
chr15-60828197-INS-87
chr15-60853075-INS-1024
chr15-60861438-INS-795
chr15-60867679-INS-76
chr15-60869230-INS-66
chr15-60877781-INS-295
chr15-60894417-INS-498
chr15-60904982-INS-1237
chr15-60906838-INS-62
chr15-60912501-INS-75
chr15-60929493-INS-155
chr15-60930697-INS-56
chr15-60940477-INS-103
chr15-60960703-INS-199
chr15-60973455-INS-3848
chr15-60980498-INS-60
chr15-61009850-INS-267
chr15-61028388-INS-87
chr15-61060173-INS-54
chr15-61068801-INS-259
chr15-61084209-INS-454
chr15-61103459-INS-544
chr15-61109560-INS-52
chr15-61147636-INS-6253
chr15-61168598-INS-986
chr15-61187882-INS-128
chr15-61188757-INS-3125
chr15-61192452-INS-2663
chr15-61192687-INS-56
chr15-61197795-INS-390
chr15-61201787-INS-670
chr15-61231123-INS-170
chr15-61241130-INS-499
chr15-61254711-INS-300
chr15-61260762-INS-6683
chr15-61275713-INS-432
chr15-61311212-INS-56
chr15-61336936-INS-450
chr15-61396848-INS-54
chr15-61446904-INS-8018
chr15-61448002-INS-398
chr15-61449250-INS-54
chr15-61455497-INS-698
chr15-61457384-INS-73
chr15-61458108-INS-398
chr15-61486850-INS-78
chr15-61498238-INS-55
chr15-61512341-INS-590
chr15-61516731-INS-77
chr15-61523731-INS-209
chr15-61524642-INS-60

chr15-61539776-INS-5541
chr15-61558446-INS-303
chr15-61570232-INS-387
chr15-61591987-INS-7000
chr15-61592624-INS-286
chr15-61601083-INS-98
chr15-61613020-INS-330
chr15-61613127-INS-833
chr15-61634454-INS-278
chr15-61636628-INS-949
chr15-61650743-INS-53
chr15-61674508-INS-124
chr15-61685419-INS-711
chr15-61685419-INS-98
chr15-61694026-INS-257
chr15-61701493-INS-74
chr15-61711928-INS-79
chr15-61720593-INS-53
chr15-61729906-INS-204
chr15-61735490-INS-186
chr15-61767002-INS-6964
chr15-61801665-INS-650
chr15-61841977-INS-199
chr15-61845137-INS-326
chr15-61848734-INS-120
chr15-61852170-INS-770
chr15-61878995-INS-202
chr15-61899728-INS-54
chr15-61911962-INS-73
chr15-61916723-INS-79
chr15-61922048-INS-103
chr15-61956477-INS-256
chr15-61958502-INS-56
chr15-62014622-INS-6413
chr15-62023055-INS-174
chr15-62065667-INS-61
chr15-62090016-INS-53
chr15-62105891-INS-453
chr15-62124763-INS-590
chr15-62149090-INS-78
chr15-62166262-INS-791
chr15-62171669-INS-61
chr15-62181223-INS-212
chr15-62190300-INS-296
chr15-62200361-INS-69
chr15-62202238-INS-232
chr15-62204766-INS-51
chr15-62271993-INS-136
chr15-62279420-INS-158
chr15-62282554-INS-756
chr15-62284688-INS-361
chr15-62297371-INS-64
chr15-62306309-INS-7741
chr15-62328657-INS-201
chr15-62339533-INS-319
chr15-62348195-INS-2957
chr15-62348723-INS-83
chr15-62363135-INS-7298
chr15-62370165-INS-53
chr15-62402141-INS-1679
chr15-62410875-INS-935
chr15-62412691-INS-6158
chr15-62415734-INS-384
chr15-62431767-INS-92
chr15-62445728-INS-6608
chr15-62456087-INS-234
chr15-62458186-INS-54
chr15-62483371-INS-216
chr15-62487188-INS-52
chr15-62498007-INS-203
chr15-62519638-INS-94
chr15-62528330-INS-6465
chr15-62533451-INS-91
chr15-62556327-INS-208
chr15-62571475-INS-55
chr15-62574212-INS-7086
chr15-62588613-INS-3036

chr15-62590157-INS-212
chr15-62605018-INS-148
chr15-62620756-INS-299
chr15-62666847-INS-147
chr15-62668295-INS-611
chr15-62678960-INS-125
chr15-62683744-INS-129
chr15-62725296-INS-54
chr15-62734036-INS-53
chr15-62744155-INS-136
chr15-62748958-INS-655
chr15-62751349-INS-303
chr15-62758754-INS-142
chr15-62762341-INS-268
chr15-62809174-INS-494
chr15-62831110-INS-57
chr15-62846581-INS-212
chr15-62851048-INS-6793
chr15-62882808-INS-97
chr15-62890687-INS-125
chr15-62891645-INS-58
chr15-62926020-INS-59
chr15-62947504-INS-88
chr15-62978265-INS-60
chr15-63016487-INS-74
chr15-63020897-INS-811
chr15-63023157-INS-257
chr15-63040295-INS-67
chr15-63057593-INS-6758
chr15-63057757-INS-157
chr15-63074224-INS-63
chr15-63084664-INS-152
chr15-63096125-INS-63
chr15-63101306-INS-475
chr15-63121853-INS-464
chr15-63129327-INS-242
chr15-63152599-INS-693
chr15-63162569-INS-76
chr15-63185868-INS-439
chr15-63191346-INS-161
chr15-63196265-INS-627
chr15-63214970-INS-831
chr15-63245287-INS-77
chr15-63246293-INS-153
chr15-63269441-INS-58
chr15-63281632-INS-143
chr15-63281710-INS-672
chr15-63307436-INS-730
chr15-63315704-INS-55
chr15-63331059-INS-474
chr15-63357673-INS-54
chr15-63358886-INS-576
chr15-63369851-INS-73
chr15-63374201-INS-54
chr15-63444617-INS-168
chr15-63454967-INS-181
chr15-63458522-INS-174
chr15-63481908-INS-260
chr15-63517004-INS-932
chr15-63538157-INS-1089
chr15-63540319-INS-281
chr15-63541373-INS-164
chr15-63545448-INS-796
chr15-63559895-INS-358
chr15-63567350-INS-120
chr15-63573894-INS-382
chr15-63588435-INS-134
chr15-63603612-INS-108
chr15-63624462-INS-240
chr15-63633069-INS-67
chr15-63658845-INS-2262
chr15-63734343-INS-68
chr15-63740015-INS-159
chr15-63762193-INS-250
chr15-63784608-INS-67
chr15-63839501-INS-683
chr15-63860215-INS-286

chr15-63885882-INS-245
chr15-63898234-INS-80
chr15-63903490-INS-80
chr15-63908722-INS-60
chr15-63918692-INS-185
chr15-63927972-INS-331
chr15-63949173-INS-56
chr15-64004809-INS-172
chr15-64020944-INS-107
chr15-64062085-INS-1584
chr15-64068322-INS-784
chr15-64098439-INS-408
chr15-64145819-INS-236
chr15-64146360-INS-61
chr15-64168715-INS-750
chr15-64191722-INS-114
chr15-64194152-INS-61
chr15-64207825-INS-80
chr15-64218317-INS-349
chr15-64223295-INS-907
chr15-64271098-INS-165
chr15-64284997-INS-461
chr15-64294151-INS-60
chr15-64294895-INS-195
chr15-64298999-INS-539
chr15-64305657-INS-57
chr15-64318580-INS-502
chr15-64330107-INS-6363
chr15-64333863-INS-64
chr15-64344830-INS-2996
chr15-64359634-INS-55
chr15-64380680-INS-51
chr15-64390467-INS-799
chr15-64417544-INS-218
chr15-64421562-INS-8165
chr15-64433407-INS-394
chr15-64474923-INS-2960
chr15-64506200-INS-1492
chr15-64535144-INS-272
chr15-64538086-INS-52
chr15-64556823-INS-68
chr15-64561043-INS-346
chr15-64562719-INS-78
chr15-64564179-INS-59
chr15-64574996-INS-1085
chr15-64607388-INS-277
chr15-64619168-INS-902
chr15-64630072-INS-6665
chr15-64632726-INS-285
chr15-64648412-INS-113
chr15-64655161-INS-204
chr15-64667141-INS-52
chr15-64669892-INS-131
chr15-64681549-INS-172
chr15-64692534-INS-7314
chr15-64692723-INS-900
chr15-64693822-INS-224
chr15-64701332-INS-72
chr15-64712278-INS-111
chr15-64721372-INS-76
chr15-64732146-INS-86
chr15-64736509-INS-118
chr15-64761302-INS-52
chr15-64784976-INS-717
chr15-64811852-INS-51
chr15-64817327-INS-90
chr15-64842255-INS-78
chr15-64873616-INS-60
chr15-64879706-INS-72
chr15-64909084-INS-359
chr15-64910863-INS-88
chr15-64933260-INS-112
chr15-64943787-INS-67
chr15-64945549-INS-51
chr15-64980760-INS-143
chr15-64982053-INS-6600
chr15-64982223-INS-62

chr15-64983326-INS-1666
chr15-64984332-INS-265
chr15-64999258-INS-82
chr15-65007173-INS-65
chr15-65010236-INS-3052
chr15-65018727-INS-2440
chr15-65049641-INS-73
chr15-65065566-INS-68
chr15-65066603-INS-497
chr15-65068966-INS-157
chr15-65081573-INS-4921
chr15-65112602-INS-79
chr15-65114062-INS-93
chr15-65132832-INS-141
chr15-65133072-INS-462
chr15-65134571-INS-6442
chr15-65138218-INS-678
chr15-65140808-INS-7105
chr15-65167161-INS-117
chr15-65170588-INS-5402
chr15-65171629-INS-165
chr15-65185632-INS-65
chr15-65190656-INS-600
chr15-65195565-INS-231
chr15-65210579-INS-56
chr15-65220697-INS-431
chr15-65229472-INS-441
chr15-65251780-INS-60
chr15-65253065-INS-516
chr15-65307687-INS-496
chr15-65308479-INS-6452
chr15-65316283-INS-5371
chr15-65355050-INS-1890
chr15-65366902-INS-1257
chr15-65408816-INS-125
chr15-65410431-INS-52
chr15-65441622-INS-925
chr15-65455767-INS-335
chr15-65461347-INS-308
chr15-65474040-INS-721
chr15-65480392-INS-941
chr15-65494448-INS-219
chr15-65509969-INS-184
chr15-65511071-INS-86
chr15-65512578-INS-52
chr15-65525643-INS-129
chr15-65526685-INS-411
chr15-65528235-INS-875
chr15-65535114-INS-75
chr15-65551908-INS-79
chr15-65561000-INS-6254
chr15-65570357-INS-281
chr15-65575996-INS-82
chr15-65578063-INS-63
chr15-65580935-INS-114
chr15-65581872-INS-331
chr15-65608385-INS-54
chr15-65616931-INS-5884
chr15-65626622-INS-430
chr15-65647193-INS-562
chr15-65649916-INS-165
chr15-65655162-INS-230
chr15-65664541-INS-216
chr15-65679888-INS-537
chr15-65687838-INS-197
chr15-65710220-INS-220
chr15-65721055-INS-112
chr15-65726304-INS-70
chr15-65739954-INS-72
chr15-65749419-INS-100
chr15-65799513-INS-1073
chr15-65807747-INS-158
chr15-65838401-INS-65
chr15-65839143-INS-296
chr15-65873148-INS-341
chr15-65879439-INS-191
chr15-65915327-INS-879

chr15-65951285-INS-934
chr15-65974415-INS-53
chr15-65990211-INS-849
chr15-65995988-INS-429
chr15-65998426-INS-946
chr15-66013233-INS-89
chr15-66028687-INS-68
chr15-66043587-INS-332
chr15-66049290-INS-202
chr15-66053691-INS-62
chr15-66058269-INS-6384
chr15-66105704-INS-6685
chr15-66114744-INS-60
chr15-66119195-INS-79
chr15-66126934-INS-54
chr15-66139752-INS-75
chr15-66145776-INS-991
chr15-66172088-INS-6910
chr15-66195467-INS-791
chr15-66198963-INS-59
chr15-66199043-INS-448
chr15-66201919-INS-7100
chr15-66224520-INS-74
chr15-66230580-INS-872
chr15-66232875-INS-1892
chr15-66234215-INS-787
chr15-66246371-INS-471
chr15-66262152-INS-1303
chr15-66284715-INS-144
chr15-66288949-INS-98
chr15-66297464-INS-316
chr15-66306848-INS-447
chr15-66316310-INS-201
chr15-66336614-INS-75
chr15-66343848-INS-138
chr15-66402531-INS-7138
chr15-66438974-INS-70
chr15-66443446-INS-358
chr15-66451303-INS-177
chr15-66520491-INS-6075
chr15-66526824-INS-2468
chr15-66526937-INS-199
chr15-66547505-INS-102
chr15-66548713-INS-54
chr15-66583721-INS-84
chr15-66631137-INS-64
chr15-66648829-INS-65
chr15-66650561-INS-329
chr15-66667561-INS-96
chr15-66682309-INS-97
chr15-66693772-INS-435
chr15-66704690-INS-103
chr15-66715851-INS-143
chr15-66731190-INS-76
chr15-66743481-INS-206
chr15-66744623-INS-2519
chr15-66757097-INS-735
chr15-66774998-INS-646
chr15-66781311-INS-198
chr15-66785283-INS-7291
chr15-66786561-INS-100
chr15-66805396-INS-157
chr15-66820766-INS-88
chr15-66861768-INS-6545
chr15-66879198-INS-211
chr15-66882069-INS-385
chr15-66896756-INS-226
chr15-66899135-INS-3853
chr15-66914757-INS-140
chr15-66929228-INS-617
chr15-66932808-INS-66
chr15-67034111-INS-70
chr15-67037025-INS-709
chr15-67066831-INS-113
chr15-67070679-INS-56
chr15-67090223-INS-138
chr15-67095677-INS-874

chr15-67125233-INS-94
chr15-67125307-INS-6876
chr15-67146363-INS-6439
chr15-67164801-INS-3002
chr15-67212400-INS-100
chr15-67218230-INS-120
chr15-67234293-INS-244
chr15-67240464-INS-59
chr15-67247894-INS-1892
chr15-67272910-INS-61
chr15-67284280-INS-1050
chr15-67292157-INS-12491
chr15-67310531-INS-52
chr15-67344705-INS-102
chr15-67358180-INS-94
chr15-67359067-INS-52
chr15-67365620-INS-93
chr15-67370256-INS-603
chr15-67382713-INS-165
chr15-67389222-INS-190
chr15-67402775-INS-56
chr15-67404085-INS-211
chr15-67405625-INS-416
chr15-67435139-INS-6603
chr15-67446281-INS-6161
chr15-67462888-INS-6952
chr15-67496261-INS-133
chr15-67548175-INS-6233
chr15-67558232-INS-690
chr15-67596610-INS-84
chr15-67638000-INS-178
chr15-67680749-INS-715
chr15-67690009-INS-6291
chr15-67695404-INS-4515
chr15-67699247-INS-573
chr15-67728069-INS-59
chr15-67735694-INS-92
chr15-67738305-INS-128
chr15-67749621-INS-115
chr15-67769233-INS-62
chr15-67772815-INS-362
chr15-67790165-INS-8295
chr15-67828926-INS-146
chr15-67852781-INS-187
chr15-67874515-INS-521
chr15-67964741-INS-60
chr15-67968116-INS-220
chr15-67971366-INS-194
chr15-68008485-INS-77
chr15-68026796-INS-468
chr15-68032496-INS-76
chr15-68071028-INS-514
chr15-68086383-INS-168
chr15-68088632-INS-782
chr15-68094601-INS-623
chr15-68095707-INS-90
chr15-68103582-INS-120
chr15-68106510-INS-61
chr15-68150223-INS-541
chr15-68151805-INS-412
chr15-68160740-INS-60
chr15-68167901-INS-220
chr15-68175604-INS-7489
chr15-68200248-INS-568
chr15-68212643-INS-323
chr15-68265053-INS-78
chr15-68280661-INS-52
chr15-68293208-INS-56
chr15-68310891-INS-213
chr15-68316136-INS-75
chr15-68331199-INS-111
chr15-68362081-INS-104
chr15-68398180-INS-163
chr15-68412971-INS-908
chr15-68423365-INS-103
chr15-68433609-INS-6666
chr15-68438344-INS-5222

chr15-68450099-INS-441
chr15-68457039-INS-6399
chr15-68457246-INS-386
chr15-68481779-INS-112
chr15-68534026-INS-65
chr15-68561150-INS-52
chr15-68569229-INS-6401
chr15-68569768-INS-58
chr15-68582558-INS-83
chr15-68592738-INS-528
chr15-68597850-INS-914
chr15-68613728-INS-164
chr15-68633146-INS-412
chr15-68638444-INS-453
chr15-68645722-INS-196
chr15-68648870-INS-70
chr15-68657382-INS-159
chr15-68659738-INS-128
chr15-68725327-INS-3826
chr15-68731240-INS-152
chr15-68733330-INS-6237
chr15-68737855-INS-358
chr15-68742150-INS-202
chr15-68747031-INS-6216
chr15-68771326-INS-139
chr15-68772972-INS-53
chr15-68776476-INS-52
chr15-68822509-INS-1658
chr15-68832604-INS-109
chr15-68847275-INS-58
chr15-68861577-INS-100
chr15-68871458-INS-96
chr15-68872794-INS-73
chr15-68886362-INS-56
chr15-68897253-INS-1061
chr15-68897448-INS-192
chr15-68897449-INS-1538
chr15-68941315-INS-193
chr15-68961542-INS-60
chr15-68972477-INS-303
chr15-68977334-INS-114
chr15-68984717-INS-3122
chr15-69005861-INS-6149
chr15-69028829-INS-66
chr15-69040132-INS-109
chr15-69057446-INS-61
chr15-69077733-INS-61
chr15-69078438-INS-59
chr15-69095668-INS-6905
chr15-69100044-INS-6656
chr15-69174631-INS-590
chr15-69189540-INS-58
chr15-69215839-INS-6467
chr15-69224671-INS-416
chr15-69249807-INS-52
chr15-69276256-INS-64
chr15-69277785-INS-59
chr15-69312056-INS-274
chr15-69319121-INS-70
chr15-69323548-INS-551
chr15-69325731-INS-480
chr15-69329148-INS-58
chr15-69330856-INS-62
chr15-69345857-INS-94
chr15-69379076-INS-67
chr15-69415242-INS-207
chr15-69420672-INS-75
chr15-69426669-INS-307
chr15-69428519-INS-178
chr15-69467202-INS-290
chr15-69486192-INS-108
chr15-69494995-INS-83
chr15-69502691-INS-7191
chr15-69509655-INS-112
chr15-69521827-INS-60
chr15-69522269-INS-625
chr15-69533003-INS-939

chr15-69535485-INS-94
chr15-69541969-INS-6204
chr15-69551352-INS-146
chr15-69595214-INS-329
chr15-69601329-INS-60
chr15-69649757-INS-881
chr15-69660996-INS-74
chr15-69702280-INS-82
chr15-69704048-INS-614
chr15-69719551-INS-227
chr15-69739507-INS-129
chr15-69746174-INS-378
chr15-69751571-INS-88
chr15-69780823-INS-502
chr15-69792475-INS-203
chr15-69799297-INS-56
chr15-69818517-INS-272
chr15-69837598-INS-561
chr15-69848147-INS-150
chr15-69889456-INS-58
chr15-69899119-INS-76
chr15-69901761-INS-538
chr15-69922863-INS-65
chr15-69931678-INS-91
chr15-69934336-INS-112
chr15-69941157-INS-308
chr15-69990435-INS-54
chr15-70003038-INS-5245
chr15-70006282-INS-1578
chr15-70015975-INS-202
chr15-70049317-INS-60
chr15-70096538-INS-513
chr15-70102258-INS-341
chr15-70126119-INS-308
chr15-70132718-INS-82
chr15-70153619-INS-685
chr15-70154616-INS-54
chr15-70201825-INS-173
chr15-70212873-INS-66
chr15-70215694-INS-168
chr15-70229744-INS-75
chr15-70234852-INS-52
chr15-70259413-INS-62
chr15-70277726-INS-52
chr15-70304028-INS-56
chr15-70328738-INS-274
chr15-70333045-INS-137
chr15-70362619-INS-82
chr15-70365722-INS-142
chr15-70375695-INS-198
chr15-70420713-INS-626
chr15-70423854-INS-6590
chr15-70429416-INS-102
chr15-70467211-INS-193
chr15-70508429-INS-450
chr15-70520012-INS-185
chr15-70521980-INS-82
chr15-70524468-INS-67
chr15-70542538-INS-323
chr15-70548984-INS-2753
chr15-70562302-INS-106
chr15-70574774-INS-166
chr15-70576041-INS-7126
chr15-70605777-INS-431
chr15-70634939-INS-975
chr15-70639358-INS-56
chr15-70644363-INS-302
chr15-70659997-INS-343
chr15-70673739-INS-1385
chr15-70699103-INS-272
chr15-70717291-INS-284
chr15-70730954-INS-468
chr15-70743403-INS-7067
chr15-70743652-INS-95
chr15-70751604-INS-54
chr15-70763961-INS-56
chr15-70791700-INS-718

chr15-70794634-INS-536
chr15-70798000-INS-1605
chr15-70815161-INS-92
chr15-70822139-INS-62
chr15-70837513-INS-212
chr15-70856582-INS-7110
chr15-70859020-INS-625
chr15-70864786-INS-932
chr15-70906424-INS-54
chr15-70920632-INS-369
chr15-70939073-INS-108
chr15-70971445-INS-128
chr15-71009292-INS-462
chr15-71013213-INS-8007
chr15-71027513-INS-60
chr15-71088151-INS-4818
chr15-71110239-INS-337
chr15-71134676-INS-56
chr15-71158515-INS-6595
chr15-71170150-INS-70
chr15-71178762-INS-139
chr15-71198065-INS-107
chr15-71198656-INS-54
chr15-71208001-INS-4720
chr15-71218781-INS-214
chr15-71227497-INS-1886
chr15-71240003-INS-540
chr15-71243550-INS-185
chr15-71253210-INS-83
chr15-71258476-INS-185
chr15-71265071-INS-422
chr15-71287066-INS-716
chr15-71289173-INS-482
chr15-71310029-INS-86
chr15-71313280-INS-400
chr15-71323580-INS-74
chr15-71323866-INS-2698
chr15-71339351-INS-319
chr15-71344341-INS-60
chr15-71398350-INS-120
chr15-71427663-INS-58
chr15-71440000-INS-304
chr15-71442342-INS-154
chr15-71479262-INS-6692
chr15-71480051-INS-400
chr15-71485349-INS-6396
chr15-71505990-INS-6776
chr15-71505993-INS-89
chr15-71506931-INS-365
chr15-71554501-INS-69
chr15-71560504-INS-104
chr15-71596477-INS-84
chr15-71609688-INS-106
chr15-71613132-INS-122
chr15-71630214-INS-405
chr15-71646378-INS-353
chr15-71648594-INS-121
chr15-71654095-INS-839
chr15-71674691-INS-7721
chr15-71686483-INS-199
chr15-71714328-INS-73
chr15-71746314-INS-55
chr15-71752273-INS-322
chr15-71766219-INS-60
chr15-71774548-INS-393
chr15-71801141-INS-66
chr15-71841428-INS-60
chr15-71861886-INS-1862
chr15-71862584-INS-232
chr15-71893859-INS-82
chr15-71917051-INS-96
chr15-71918115-INS-118
chr15-71928927-INS-128
chr15-71943296-INS-80
chr15-71952328-INS-6267
chr15-71966691-INS-317
chr15-71983811-INS-76

chr15-71986933-INS-2139
chr15-71995917-INS-740
chr15-71997804-INS-301
chr15-72005661-INS-689
chr15-72032198-INS-324
chr15-72036047-INS-183
chr15-72081383-INS-148
chr15-72099751-INS-191
chr15-72128404-INS-3906
chr15-72164344-INS-201
chr15-72187733-INS-471
chr15-72198297-INS-364
chr15-72207353-INS-60
chr15-72242195-INS-80
chr15-72244677-INS-6648
chr15-72267387-INS-7131
chr15-72280428-INS-6488
chr15-72286160-INS-225
chr15-72323054-INS-437
chr15-72329318-INS-6204
chr15-72342504-INS-137
chr15-72453447-INS-80
chr15-72463457-INS-78
chr15-72494631-INS-130
chr15-72502353-INS-70
chr15-72506776-INS-66
chr15-72542736-INS-6487
chr15-72568246-INS-68
chr15-72588630-INS-64
chr15-72597853-INS-114
chr15-72633360-INS-55
chr15-72645999-INS-127
chr15-72663708-INS-68
chr15-72716016-INS-282
chr15-72719787-INS-204
chr15-72722369-INS-1410
chr15-72722370-INS-470
chr15-72732347-INS-123
chr15-72732976-INS-767
chr15-72738134-INS-443
chr15-72758581-INS-6182
chr15-72767174-INS-76
chr15-72777974-INS-51
chr15-72803677-INS-206
chr15-72809320-INS-6801
chr15-72816728-INS-60
chr15-72835468-INS-291
chr15-72849977-INS-94
chr15-72862693-INS-68
chr15-72868980-INS-128
chr15-72901760-INS-70
chr15-72911635-INS-54
chr15-72918432-INS-201
chr15-72920383-INS-109
chr15-72927033-INS-81
chr15-72929422-INS-203
chr15-72933284-INS-66
chr15-72937432-INS-55
chr15-72941411-INS-204
chr15-72947196-INS-303
chr15-72955770-INS-5215
chr15-72983357-INS-92
chr15-72984779-INS-52
chr15-73009593-INS-52
chr15-73038232-INS-54
chr15-73069028-INS-199
chr15-73092430-INS-198
chr15-73099692-INS-64
chr15-73124944-INS-199
chr15-73137191-INS-1796
chr15-73148838-INS-155
chr15-73210009-INS-113
chr15-73222464-INS-108
chr15-73257109-INS-52
chr15-73257920-INS-60
chr15-73277233-INS-112
chr15-73319452-INS-133

chr15-73327474-INS-227
chr15-73344961-INS-204
chr15-73352180-INS-100
chr15-73375209-INS-204
chr15-73423187-INS-204
chr15-73430824-INS-52
chr15-73460869-INS-247
chr15-73470577-INS-182
chr15-73484255-INS-308
chr15-73491232-INS-111
chr15-73508017-INS-54
chr15-73518940-INS-536
chr15-73539842-INS-62
chr15-73589573-INS-214
chr15-73631200-INS-77
chr15-73646751-INS-85
chr15-73655631-INS-74
chr15-73656419-INS-1391
chr15-73712960-INS-95
chr15-73717193-INS-190
chr15-73753839-INS-332
chr15-73763598-INS-69
chr15-73777733-INS-157
chr15-73792756-INS-160
chr15-73826279-INS-5551
chr15-73889098-INS-56
chr15-73896623-INS-52
chr15-73904365-INS-76
chr15-73977527-INS-263
chr15-73998238-INS-58
chr15-74015088-INS-173
chr15-74019879-INS-116
chr15-74073515-INS-80
chr15-74084249-INS-90
chr15-74096629-INS-399
chr15-74108132-INS-572
chr15-74110127-INS-62
chr15-74155457-INS-55
chr15-74157434-INS-99
chr15-74176015-INS-831
chr15-74212222-INS-121
chr15-74212443-INS-1206
chr15-74216435-INS-90
chr15-74231122-INS-76
chr15-74319788-INS-66
chr15-74423129-INS-51
chr15-74495983-INS-64
chr15-74501016-INS-57
chr15-74549943-INS-108
chr15-74613956-INS-118
chr15-74629396-INS-365
chr15-74630923-INS-381
chr15-74657115-INS-52
chr15-74657949-INS-200
chr15-74676198-INS-73
chr15-74691406-INS-373
chr15-74695706-INS-139
chr15-74703059-INS-769
chr15-74768724-INS-328
chr15-74776586-INS-65
chr15-74780385-INS-60
chr15-74781069-INS-130
chr15-74785798-INS-257
chr15-74789722-INS-125
chr15-74792275-INS-506
chr15-74801937-INS-275
chr15-74807649-INS-58
chr15-74825374-INS-71
chr15-74837069-INS-100
chr15-74892552-INS-51
chr15-74894641-INS-521
chr15-74919378-INS-256
chr15-74922280-INS-3398
chr15-74926065-INS-61
chr15-75012469-INS-99
chr15-75014970-INS-65
chr15-75018926-INS-134

chr15-75022557-INS-53
chr15-75034269-INS-54
chr15-75037545-INS-352
chr15-75074480-INS-57
chr15-75095247-INS-340
chr15-75120327-INS-2344
chr15-75134193-INS-103
chr15-75137346-INS-60
chr15-75163799-INS-157
chr15-75179335-INS-203
chr15-75195500-INS-7371
chr15-75248567-INS-5950
chr15-75297010-INS-989
chr15-75313974-INS-85
chr15-75355782-INS-708
chr15-75375266-INS-587
chr15-75410937-INS-54
chr15-75422134-INS-73
chr15-75449850-INS-58
chr15-75462965-INS-227
chr15-75473822-INS-138
chr15-75481351-INS-514
chr15-75492592-INS-386
chr15-75516618-INS-400
chr15-75519874-INS-59
chr15-75539110-INS-72
chr15-75545169-INS-61
chr15-75546517-INS-85
chr15-75567020-INS-198
chr15-75571047-INS-90
chr15-75596012-INS-54
chr15-75598935-INS-110
chr15-75605035-INS-58
chr15-75652676-INS-147
chr15-75657873-INS-689
chr15-75664649-INS-167
chr15-75665907-INS-378
chr15-75687356-INS-92
chr15-75698556-INS-55
chr15-75699617-INS-199
chr15-75732409-INS-99
chr15-75747273-INS-392
chr15-75751012-INS-204
chr15-75785506-INS-205
chr15-75794854-INS-55
chr15-75838749-INS-353
chr15-75841729-INS-78
chr15-75850159-INS-94
chr15-75872780-INS-60
chr15-75896136-INS-187
chr15-75899726-INS-62
chr15-75908536-INS-56
chr15-75999320-INS-101
chr15-76014333-INS-58
chr15-76018402-INS-72
chr15-76049606-INS-63
chr15-76103710-INS-218
chr15-76123307-INS-248
chr15-76170413-INS-245
chr15-76193389-INS-97
chr15-76208437-INS-283
chr15-76251846-INS-200
chr15-76260634-INS-64
chr15-76271043-INS-686
chr15-76275862-INS-61
chr15-76278929-INS-218
chr15-76299186-INS-183
chr15-76302457-INS-942
chr15-76304360-INS-90
chr15-76314143-INS-52
chr15-76375762-INS-214
chr15-76379864-INS-222
chr15-76385843-INS-119
chr15-76391938-INS-113
chr15-76432097-INS-97
chr15-76454844-INS-358
chr15-76461433-INS-8945

chr15-76468633-INS-106
chr15-76494296-INS-61
chr15-76499062-INS-54
chr15-76525084-INS-183
chr15-76547304-INS-271
chr15-76559541-INS-54
chr15-76561798-INS-196
chr15-76609930-INS-107
chr15-76625895-INS-169
chr15-76628347-INS-213
chr15-76629750-INS-243
chr15-76630589-INS-399
chr15-76637347-INS-152
chr15-76640355-INS-124
chr15-76663096-INS-62
chr15-76668547-INS-66
chr15-76675260-INS-705
chr15-76677170-INS-215
chr15-76697334-INS-192
chr15-76714135-INS-58
chr15-76739305-INS-228
chr15-76744053-INS-301
chr15-76745858-INS-181
chr15-76748147-INS-524
chr15-76750775-INS-433
chr15-76758574-INS-207
chr15-76794459-INS-77
chr15-76796513-INS-211
chr15-76797785-INS-51
chr15-76798619-INS-107
chr15-76826839-INS-243
chr15-76841009-INS-74
chr15-76873233-INS-372
chr15-76878306-INS-436
chr15-76879615-INS-67
chr15-76886877-INS-202
chr15-76890076-INS-127
chr15-76919916-INS-435
chr15-76934633-INS-82
chr15-76937419-INS-73
chr15-76957593-INS-66
chr15-76959259-INS-216
chr15-76984586-INS-86
chr15-76985407-INS-92
chr15-77029309-INS-219
chr15-77055922-INS-90
chr15-77063586-INS-697
chr15-77088500-INS-166
chr15-77090868-INS-76
chr15-77097785-INS-102
chr15-77113211-INS-90
chr15-77117516-INS-68
chr15-77122655-INS-498
chr15-77149842-INS-59
chr15-77155412-INS-460
chr15-77179818-INS-58
chr15-77219573-INS-228
chr15-77233271-INS-110
chr15-77241372-INS-189
chr15-77241983-INS-202
chr15-77246544-INS-64
chr15-77285353-INS-52
chr15-77295751-INS-4810
chr15-77328048-INS-426
chr15-77333757-INS-99
chr15-77344384-INS-6180
chr15-77354720-INS-66
chr15-77360349-INS-2857
chr15-77430993-INS-199
chr15-77462066-INS-57
chr15-77470735-INS-89
chr15-77550579-INS-145
chr15-77559729-INS-528
chr15-77578747-INS-197
chr15-77580648-INS-297
chr15-77638375-INS-55
chr15-77639668-INS-62

chr15-77677550-INS-386
chr15-77703512-INS-52
chr15-77713503-INS-66
chr15-77719170-INS-214
chr15-77737541-INS-70
chr15-77739002-INS-94
chr15-77753083-INS-197
chr15-77777941-INS-128
chr15-77795707-INS-302
chr15-77810487-INS-95
chr15-77814890-INS-335
chr15-77839119-INS-98
chr15-77865980-INS-179
chr15-77887481-INS-96
chr15-77898653-INS-132
chr15-77929866-INS-54
chr15-77955702-INS-75
chr15-77961738-INS-195
chr15-77998971-INS-133
chr15-78010723-INS-114
chr15-78028215-INS-65
chr15-78062304-INS-80
chr15-78136114-INS-114
chr15-78140912-INS-1041
chr15-78140923-INS-1287
chr15-78140942-INS-1558
chr15-78140994-INS-725
chr15-78141261-INS-896
chr15-78151345-INS-66
chr15-78155169-INS-78
chr15-78163474-INS-197
chr15-78164357-INS-55
chr15-78193930-INS-65
chr15-78201335-INS-52
chr15-78238154-INS-90
chr15-78239612-INS-52
chr15-78246365-INS-56
chr15-78249987-INS-432
chr15-78257023-INS-117
chr15-78267258-INS-96
chr15-78275661-INS-56
chr15-78277410-INS-183
chr15-78282568-INS-401
chr15-78288554-INS-86
chr15-78314123-INS-53
chr15-78315461-INS-108
chr15-78338743-INS-201
chr15-78370929-INS-112
chr15-78383883-INS-206
chr15-78394990-INS-61
chr15-78399506-INS-116
chr15-78401349-INS-81
chr15-78418298-INS-102
chr15-78436640-INS-195
chr15-78447508-INS-928
chr15-78447513-INS-1429
chr15-78456375-INS-88
chr15-78457703-INS-765
chr15-78467312-INS-593
chr15-78477776-INS-55
chr15-78486856-INS-53
chr15-78548490-INS-52
chr15-78593830-INS-79
chr15-78631669-INS-57
chr15-78650535-INS-5678
chr15-78652846-INS-308
chr15-78666440-INS-201
chr15-78673461-INS-245
chr15-78693252-INS-162
chr15-78710297-INS-52
chr15-78714710-INS-242
chr15-78726404-INS-51
chr15-78741853-INS-263
chr15-78745338-INS-208
chr15-78747297-INS-200
chr15-78803734-INS-54
chr15-78805039-INS-130

chr15-78809969-INS-93
chr15-78813406-INS-221
chr15-78816828-INS-191
chr15-78837435-INS-55
chr15-78882679-INS-182
chr15-78892487-INS-64
chr15-78894025-INS-248
chr15-78904563-INS-121
chr15-78905467-INS-193
chr15-78917771-INS-124
chr15-78931317-INS-354
chr15-78933215-INS-174
chr15-78935205-INS-4355
chr15-78938822-INS-72
chr15-78950173-INS-196
chr15-78953359-INS-117
chr15-78957021-INS-179
chr15-78977341-INS-227
chr15-78979648-INS-205
chr15-78982915-INS-60
chr15-79000588-INS-167
chr15-79009676-INS-135
chr15-79026611-INS-207
chr15-79061496-INS-135
chr15-79066678-INS-99
chr15-79068911-INS-132
chr15-79083712-INS-54
chr15-79093918-INS-71
chr15-79117678-INS-223
chr15-79121311-INS-182
chr15-79160892-INS-84
chr15-79172136-INS-438
chr15-79172343-INS-784
chr15-79176629-INS-97
chr15-79206771-INS-56
chr15-79222766-INS-199
chr15-79251990-INS-84
chr15-79292370-INS-174
chr15-79294265-INS-521
chr15-79329653-INS-193
chr15-79346746-INS-65
chr15-79349207-INS-74
chr15-79429947-INS-348
chr15-79436208-INS-250
chr15-79480040-INS-97
chr15-79491096-INS-107
chr15-79494697-INS-64
chr15-79505745-INS-607
chr15-79518698-INS-198
chr15-79529402-INS-81
chr15-79547830-INS-487
chr15-79553044-INS-505
chr15-79642868-INS-207
chr15-79663610-INS-285
chr15-79665548-INS-67
chr15-79667132-INS-52
chr15-79667233-INS-91
chr15-79705346-INS-167
chr15-79733123-INS-169
chr15-79735705-INS-259
chr15-79749173-INS-175
chr15-79750823-INS-225
chr15-79776114-INS-535
chr15-79783862-INS-191
chr15-79826570-INS-61
chr15-79829889-INS-186
chr15-79849711-INS-198
chr15-79871582-INS-54
chr15-80000503-INS-56
chr15-80006797-INS-578
chr15-80011456-INS-205
chr15-80013937-INS-195
chr15-80035323-INS-199
chr15-80041086-INS-146
chr15-80051727-INS-53
chr15-80068119-INS-237
chr15-80079117-INS-999

chr15-80124949-INS-120
chr15-80147727-INS-230
chr15-80159382-INS-111
chr15-80186690-INS-554
chr15-80215291-INS-192
chr15-80311841-INS-52
chr15-80328692-INS-111
chr15-80351461-INS-57
chr15-80354388-INS-84
chr15-80399590-INS-77
chr15-80408423-INS-64
chr15-80411504-INS-350
chr15-80412501-INS-72
chr15-80413686-INS-52
chr15-80417273-INS-64
chr15-80426633-INS-235
chr15-80474615-INS-166
chr15-80481083-INS-238
chr15-80482321-INS-93
chr15-80496304-INS-63
chr15-80496813-INS-281
chr15-80507629-INS-59
chr15-80522409-INS-173
chr15-80535258-INS-81
chr15-80579148-INS-89
chr15-80587695-INS-70
chr15-80589652-INS-123
chr15-80617264-INS-52
chr15-80636085-INS-264
chr15-80655172-INS-173
chr15-80667260-INS-170
chr15-80677204-INS-296
chr15-80685873-INS-447
chr15-80691232-INS-208
chr15-80727029-INS-245
chr15-80744869-INS-154
chr15-80747195-INS-70
chr15-80750301-INS-144
chr15-80794942-INS-206
chr15-80815026-INS-55
chr15-80827467-INS-99
chr15-80866037-INS-179
chr15-80869390-INS-315
chr15-80912244-INS-529
chr15-80915280-INS-60
chr15-80935844-INS-334
chr15-80986969-INS-92
chr15-81012985-INS-173
chr15-81014960-INS-273
chr15-81023869-INS-988
chr15-81041562-INS-140
chr15-81053436-INS-205
chr15-81069012-INS-192
chr15-81100573-INS-3090
chr15-81100591-INS-2468
chr15-81129874-INS-65
chr15-81155077-INS-260
chr15-81160750-INS-270
chr15-81174484-INS-200
chr15-81183289-INS-83
chr15-81190707-INS-77
chr15-81191595-INS-125
chr15-81193181-INS-241
chr15-81216415-INS-644
chr15-81267891-INS-96
chr15-81300965-INS-181
chr15-81302522-INS-117
chr15-81316573-INS-214
chr15-81329342-INS-68
chr15-81331746-INS-91
chr15-81343167-INS-204
chr15-81345588-INS-234
chr15-81363270-INS-146
chr15-81371690-INS-194
chr15-81372974-INS-191
chr15-81384935-INS-215
chr15-81401715-INS-73

chr15-81403608-INS-145
chr15-81405892-INS-448
chr15-81413556-INS-432
chr15-81423050-INS-506
chr15-81428622-INS-5152
chr15-81433744-INS-212
chr15-81434384-INS-235
chr15-81435198-INS-132
chr15-81441216-INS-586
chr15-81448118-INS-213
chr15-81452037-INS-163
chr15-81456643-INS-104
chr15-81463124-INS-139
chr15-81465129-INS-85
chr15-81475247-INS-140
chr15-81475722-INS-245
chr15-81484381-INS-210
chr15-81487991-INS-202
chr15-81496960-INS-218
chr15-81526923-INS-82
chr15-81546101-INS-952
chr15-81570062-INS-172
chr15-81574271-INS-506
chr15-81576920-INS-481
chr15-81618171-INS-143
chr15-81624253-INS-427
chr15-81636686-INS-81
chr15-81641761-INS-156
chr15-81653467-INS-78
chr15-81659564-INS-210
chr15-81700542-INS-69
chr15-81708389-INS-60
chr15-81730775-INS-51
chr15-81744748-INS-5462
chr15-81792333-INS-200
chr15-81807228-INS-108
chr15-81812326-INS-112
chr15-81815577-INS-93
chr15-81827173-INS-76
chr15-81847214-INS-178
chr15-81848843-INS-822
chr15-81866144-INS-161
chr15-81875056-INS-198
chr15-81881069-INS-57
chr15-81888858-INS-193
chr15-81904571-INS-273
chr15-81905703-INS-76
chr15-81913034-INS-228
chr15-81924977-INS-215
chr15-81944881-INS-188
chr15-81951681-INS-252
chr15-81981633-INS-132
chr15-81992350-INS-134
chr15-82001267-INS-6442
chr15-82007357-INS-201
chr15-82018430-INS-175
chr15-82047478-INS-139
chr15-82049079-INS-240
chr15-82077687-INS-213
chr15-82080920-INS-86
chr15-82085045-INS-203
chr15-82101265-INS-5554
chr15-82111399-INS-56
chr15-82115448-INS-385
chr15-82121881-INS-72
chr15-82129621-INS-204
chr15-82132420-INS-218
chr15-82147178-INS-457
chr15-82152761-INS-188
chr15-82157381-INS-89
chr15-82162910-INS-6473
chr15-82168280-INS-205
chr15-82169323-INS-392
chr15-82188633-INS-330
chr15-82194013-INS-58
chr15-82201521-INS-110
chr15-82243277-INS-4649

chr15-82246195-INS-93
chr15-82300196-INS-268
chr15-82330025-INS-84
chr15-82347067-INS-1919
chr15-82377900-INS-5419
chr15-82444119-INS-521
chr15-82447535-INS-6334
chr15-82452887-INS-498
chr15-82459655-INS-476
chr15-82467761-INS-207
chr15-82500736-INS-93
chr15-82508033-INS-55
chr15-82525326-INS-205
chr15-82554181-INS-938
chr15-82557368-INS-1269
chr15-82569224-INS-1757
chr15-82583210-INS-853
chr15-82594163-INS-350
chr15-82597346-INS-201
chr15-82606461-INS-522
chr15-82624483-INS-105
chr15-82629662-INS-232
chr15-82657849-INS-6683
chr15-82660914-INS-1099
chr15-82660945-INS-3718
chr15-82701189-INS-56
chr15-82771256-INS-275
chr15-82854818-INS-195
chr15-82863308-INS-282
chr15-82902169-INS-222
chr15-82920685-INS-230
chr15-82922808-INS-54
chr15-82949791-INS-146
chr15-82951087-INS-79
chr15-82955132-INS-56
chr15-82957761-INS-52
chr15-82964104-INS-78
chr15-82968176-INS-113
chr15-82990311-INS-84
chr15-82994033-INS-210
chr15-82995874-INS-367
chr15-83013322-INS-419
chr15-83015011-INS-101
chr15-83030767-INS-450
chr15-83048380-INS-108
chr15-83058951-INS-6413
chr15-83060220-INS-99
chr15-83066025-INS-71
chr15-83069303-INS-219
chr15-83121308-INS-109
chr15-83122980-INS-83
chr15-83132979-INS-310
chr15-83144271-INS-202
chr15-83149394-INS-1893
chr15-83151916-INS-6682
chr15-83154918-INS-95
chr15-83155586-INS-76
chr15-83164877-INS-188
chr15-83167498-INS-7281
chr15-83171709-INS-131
chr15-83174278-INS-105
chr15-83193628-INS-88
chr15-83212911-INS-100
chr15-83220823-INS-245
chr15-83221743-INS-141
chr15-83223772-INS-109
chr15-83236602-INS-203
chr15-83252545-INS-65
chr15-83266875-INS-455
chr15-83297989-INS-172
chr15-83309619-INS-7398
chr15-83322575-INS-68
chr15-83336964-INS-54
chr15-83342556-INS-198
chr15-83360958-INS-206
chr15-83366712-INS-99
chr15-83386222-INS-59

chr15-83389146-INS-145
chr15-83401176-INS-84
chr15-83405132-INS-786
chr15-83432218-INS-56
chr15-83439075-INS-256
chr15-83441313-INS-430
chr15-83449069-INS-115
chr15-83450774-INS-120
chr15-83457077-INS-159
chr15-83477071-INS-4418
chr15-83481712-INS-219
chr15-83503327-INS-99
chr15-83507096-INS-163
chr15-83568805-INS-755
chr15-83601457-INS-80
chr15-83617450-INS-56
chr15-83624396-INS-100
chr15-83626758-INS-281
chr15-83659876-INS-73
chr15-83737149-INS-52
chr15-83744417-INS-193
chr15-83765076-INS-126
chr15-83769085-INS-81
chr15-83783383-INS-137
chr15-83785010-INS-109
chr15-83802185-INS-1893
chr15-83804552-INS-392
chr15-83806649-INS-53
chr15-83833355-INS-374
chr15-83848373-INS-432
chr15-83853627-INS-103
chr15-83863289-INS-661
chr15-83878614-INS-67
chr15-83905723-INS-395
chr15-83924822-INS-718
chr15-83937277-INS-81
chr15-83946381-INS-74
chr15-83952513-INS-133
chr15-83979901-INS-60
chr15-83981736-INS-58
chr15-84021285-INS-125
chr15-84027659-INS-190
chr15-84034419-INS-227
chr15-84051079-INS-218
chr15-84055948-INS-154
chr15-84064066-INS-59
chr15-84065350-INS-451
chr15-84085753-INS-360
chr15-84088114-INS-175
chr15-84105275-INS-62
chr15-84106535-INS-62
chr15-84118782-INS-100
chr15-84143365-INS-208
chr15-84147872-INS-82
chr15-84151712-INS-2490
chr15-84179695-INS-62
chr15-84192654-INS-78
chr15-84211567-INS-68
chr15-84214395-INS-54
chr15-84242325-INS-205
chr15-84268762-INS-329
chr15-84281234-INS-256
chr15-84315040-INS-196
chr15-84330825-INS-128
chr15-84334082-INS-65
chr15-84364780-INS-64
chr15-84386796-INS-176
chr15-84398995-INS-271
chr15-84408850-INS-53
chr15-84417706-INS-782
chr15-84426469-INS-158
chr15-84428707-INS-368
chr15-84456444-INS-173
chr15-84490244-INS-51
chr15-84518961-INS-172
chr15-84524550-INS-72
chr15-84581368-INS-62

chr15-84659565-INS-52
chr15-84662764-INS-60
chr15-84674941-INS-127
chr15-84698062-INS-890
chr15-84723289-INS-466
chr15-84730422-INS-63
chr15-84748975-INS-208
chr15-84800168-INS-118
chr15-84839130-INS-4812
chr15-84854832-INS-60
chr15-84862296-INS-125
chr15-84863491-INS-51
chr15-84877390-INS-252
chr15-84882846-INS-79
chr15-84884709-INS-188
chr15-84913001-INS-203
chr15-84947818-INS-206
chr15-84971640-INS-6484
chr15-84972710-INS-81
chr15-84978053-INS-304
chr15-84980425-INS-728
chr15-84982657-INS-358
chr15-84983661-INS-51
chr15-85002549-INS-80
chr15-85013753-INS-167
chr15-85033886-INS-343
chr15-85064001-INS-386
chr15-85064555-INS-64
chr15-85070633-INS-812
chr15-85082961-INS-80
chr15-85119828-INS-517
chr15-85127242-INS-52
chr15-85131995-INS-54
chr15-85133878-INS-302
chr15-85145084-INS-405
chr15-85163963-INS-130
chr15-85167828-INS-296
chr15-85179132-INS-219
chr15-85184350-INS-126
chr15-85186780-INS-63
chr15-85187390-INS-466
chr15-85191154-INS-112
chr15-85213029-INS-59
chr15-85216935-INS-82
chr15-85241673-INS-153
chr15-85251979-INS-187
chr15-85256077-INS-855
chr15-85283947-INS-84
chr15-85310417-INS-243
chr15-85327725-INS-131
chr15-85338382-INS-149
chr15-85355839-INS-73
chr15-85380789-INS-181
chr15-85382452-INS-147
chr15-85386371-INS-165
chr15-85423173-INS-90
chr15-85426022-INS-206
chr15-85426895-INS-58
chr15-85477657-INS-54
chr15-85493444-INS-312
chr15-85511051-INS-54
chr15-85557600-INS-56
chr15-85582168-INS-84
chr15-85588169-INS-52
chr15-85598808-INS-110
chr15-85601431-INS-222
chr15-85611471-INS-152
chr15-85614929-INS-117
chr15-85618364-INS-57
chr15-85625803-INS-92
chr15-85662452-INS-56
chr15-85664745-INS-72
chr15-85666302-INS-55
chr15-85677481-INS-60
chr15-85684382-INS-84
chr15-85724141-INS-102
chr15-85728233-INS-122

chr15-85771374-INS-496
chr15-85773129-INS-199
chr15-85792474-INS-445
chr15-85800266-INS-165
chr15-85818130-INS-549
chr15-85833928-INS-61
chr15-85849546-INS-105
chr15-85856335-INS-54
chr15-85874843-INS-209
chr15-85889221-INS-59
chr15-85903978-INS-206
chr15-85928650-INS-53
chr15-85949071-INS-240
chr15-85950309-INS-62
chr15-85969731-INS-151
chr15-85972965-INS-409
chr15-86007289-INS-81
chr15-86033204-INS-184
chr15-86045128-INS-212
chr15-86046245-INS-95
chr15-86067244-INS-6461
chr15-86078937-INS-214
chr15-86135677-INS-90
chr15-86150353-INS-611
chr15-86151749-INS-224
chr15-86183493-INS-4771
chr15-86192110-INS-57
chr15-86208002-INS-611
chr15-86242111-INS-5417
chr15-86274871-INS-135
chr15-86276031-INS-62
chr15-86285159-INS-202
chr15-86338136-INS-420
chr15-86344828-INS-600
chr15-86362604-INS-589
chr15-86479439-INS-83
chr15-86488534-INS-416
chr15-86500167-INS-72
chr15-86515284-INS-61
chr15-86517708-INS-299
chr15-86545365-INS-119
chr15-86589764-INS-108
chr15-86621187-INS-67
chr15-86632894-INS-288
chr15-86639033-INS-274
chr15-86648999-INS-116
chr15-86677719-INS-319
chr15-86693429-INS-78
chr15-86699626-INS-56
chr15-86705695-INS-398
chr15-86715471-INS-56
chr15-86724957-INS-63
chr15-86737753-INS-65
chr15-86757237-INS-153
chr15-86758331-INS-98
chr15-86760301-INS-565
chr15-86761739-INS-51
chr15-86770513-INS-102
chr15-86788812-INS-62
chr15-86799948-INS-239
chr15-86806463-INS-86
chr15-86809052-INS-477
chr15-86812370-INS-94
chr15-86824234-INS-542
chr15-86830422-INS-188
chr15-86842895-INS-76
chr15-86847339-INS-53
chr15-86891783-INS-191
chr15-86896109-INS-91
chr15-86898262-INS-116
chr15-86911537-INS-101
chr15-86916040-INS-624
chr15-86918965-INS-148
chr15-86943718-INS-52
chr15-86967793-INS-56
chr15-86975061-INS-125
chr15-86977474-INS-67

chr15-86977488-INS-613
chr15-87066039-INS-82
chr15-87070468-INS-171
chr15-87082788-INS-106
chr15-87090171-INS-264
chr15-87102355-INS-198
chr15-87122662-INS-175
chr15-87128446-INS-67
chr15-87131618-INS-56
chr15-87151281-INS-73
chr15-87174982-INS-55
chr15-87186335-INS-94
chr15-87198195-INS-6634
chr15-87201434-INS-51
chr15-87203061-INS-66
chr15-87208839-INS-160
chr15-87212685-INS-137
chr15-87230765-INS-60
chr15-87240369-INS-119
chr15-87242557-INS-124
chr15-87280801-INS-168
chr15-87331856-INS-385
chr15-87342270-INS-297
chr15-87343331-INS-328
chr15-87354138-INS-932
chr15-87363657-INS-108
chr15-87384462-INS-61
chr15-87397043-INS-54
chr15-87436846-INS-56
chr15-87493271-INS-218
chr15-87549026-INS-66
chr15-87564968-INS-6451
chr15-87582466-INS-76
chr15-87599136-INS-146
chr15-87657330-INS-237
chr15-87670741-INS-6708
chr15-87679469-INS-631
chr15-87683006-INS-70
chr15-87684068-INS-216
chr15-87705680-INS-60
chr15-87710678-INS-66
chr15-87735176-INS-180
chr15-87750446-INS-138
chr15-87752588-INS-343
chr15-87766644-INS-131
chr15-87779735-INS-66
chr15-87793059-INS-327
chr15-87823975-INS-190
chr15-87837256-INS-113
chr15-87848391-INS-155
chr15-87857654-INS-208
chr15-87887913-INS-6246
chr15-87889265-INS-1711
chr15-87891049-INS-1750
chr15-87905464-INS-58
chr15-87911514-INS-111
chr15-87915296-INS-55
chr15-87917674-INS-401
chr15-87924199-INS-79
chr15-87945798-INS-345
chr15-87988108-INS-171
chr15-87995169-INS-72
chr15-87996175-INS-6259
chr15-88031178-INS-393
chr15-88046546-INS-52
chr15-88068458-INS-159
chr15-88084132-INS-644
chr15-88108091-INS-901
chr15-88137373-INS-78
chr15-88148554-INS-89
chr15-88169343-INS-54
chr15-88171480-INS-793
chr15-88180329-INS-146
chr15-88188146-INS-95
chr15-88193824-INS-132
chr15-88219146-INS-6470
chr15-88226832-INS-308

chr15-88240697-INS-58
chr15-88257509-INS-70
chr15-88262264-INS-1891
chr15-88265594-INS-71
chr15-88286880-INS-255
chr15-88329168-INS-60
chr15-88346814-INS-270
chr15-88351695-INS-99
chr15-88355722-INS-154
chr15-88387055-INS-54
chr15-88391194-INS-208
chr15-88397134-INS-101
chr15-88435696-INS-227
chr15-88488396-INS-61
chr15-88508043-INS-200
chr15-88511473-INS-60
chr15-88513597-INS-291
chr15-88515809-INS-90
chr15-88605794-INS-214
chr15-88616151-INS-60
chr15-88625017-INS-228
chr15-88626940-INS-57
chr15-88629048-INS-199
chr15-88637946-INS-135
chr15-88655067-INS-192
chr15-88672499-INS-54
chr15-88685912-INS-486
chr15-88687724-INS-51
chr15-88691668-INS-84
chr15-88717530-INS-120
chr15-88722749-INS-157
chr15-88766855-INS-7093
chr15-88778475-INS-748
chr15-88783120-INS-75
chr15-88795436-INS-66
chr15-88796453-INS-161
chr15-88815448-INS-305
chr15-88859664-INS-251
chr15-88904219-INS-192
chr15-88930316-INS-199
chr15-88966105-INS-378
chr15-88975595-INS-195
chr15-89001672-INS-62
chr15-89008530-INS-213
chr15-89018124-INS-63
chr15-89022808-INS-64
chr15-89050419-INS-220
chr15-89085690-INS-168
chr15-89090861-INS-173
chr15-89093840-INS-120
chr15-89103455-INS-104
chr15-89103557-INS-161
chr15-89112685-INS-206
chr15-89117787-INS-235
chr15-89125539-INS-119
chr15-89128309-INS-217
chr15-89132958-INS-171
chr15-89136219-INS-194
chr15-89136363-INS-1891
chr15-89145025-INS-230
chr15-89146858-INS-69
chr15-89183016-INS-214
chr15-89202091-INS-892
chr15-89215074-INS-922
chr15-89224231-INS-60
chr15-89226671-INS-73
chr15-89228080-INS-194
chr15-89229662-INS-82
chr15-89233618-INS-136
chr15-89296500-INS-232
chr15-89321747-INS-234
chr15-89326522-INS-150
chr15-89331554-INS-195
chr15-89365386-INS-102
chr15-89366676-INS-54
chr15-89373751-INS-947
chr15-89391954-INS-210

chr15-89404957-INS-271
chr15-89511146-INS-68
chr15-89527756-INS-390
chr15-89543172-INS-5587
chr15-89568386-INS-6892
chr15-89583058-INS-355
chr15-89637592-INS-154
chr15-89643989-INS-74
chr15-89648918-INS-2607
chr15-89653977-INS-129
chr15-89659104-INS-243
chr15-89672359-INS-256
chr15-89703731-INS-201
chr15-89709774-INS-183
chr15-89711889-INS-900
chr15-89734949-INS-255
chr15-89765847-INS-59
chr15-89777004-INS-83
chr15-89785352-INS-400
chr15-89799931-INS-54
chr15-89801941-INS-102
chr15-89857269-INS-67
chr15-89876939-INS-56
chr15-89883448-INS-875
chr15-89887057-INS-292
chr15-89888014-INS-113
chr15-89895519-INS-55
chr15-89902950-INS-72
chr15-89930503-INS-51
chr15-89943326-INS-731
chr15-89950337-INS-329
chr15-89967113-INS-934
chr15-90016241-INS-327
chr15-90100737-INS-51
chr15-90107501-INS-73
chr15-90134356-INS-178
chr15-90137136-INS-492
chr15-90147646-INS-64
chr15-90149845-INS-798
chr15-90166909-INS-151
chr15-90171205-INS-284
chr15-90208112-INS-60
chr15-90210505-INS-375
chr15-90212992-INS-129
chr15-90214597-INS-290
chr15-90215589-INS-321
chr15-90234313-INS-394
chr15-90247909-INS-51
chr15-90253259-INS-183
chr15-90255288-INS-92
chr15-90262024-INS-56
chr15-90275189-INS-60
chr15-90320935-INS-173
chr15-90326715-INS-51
chr15-90344162-INS-426
chr15-90354045-INS-98
chr15-90373462-INS-90
chr15-90375589-INS-105
chr15-90416899-INS-250
chr15-90433170-INS-170
chr15-90450423-INS-95
chr15-90471524-INS-58
chr15-90485196-INS-72
chr15-90489953-INS-87
chr15-90519777-INS-54
chr15-90528970-INS-52
chr15-90597852-INS-65
chr15-90600037-INS-169
chr15-90602442-INS-104
chr15-90602773-INS-2733
chr15-90621200-INS-252
chr15-90628461-INS-529
chr15-90630084-INS-57
chr15-90637432-INS-63
chr15-90660620-INS-92
chr15-90694398-INS-141
chr15-90717937-INS-142

chr15-90727896-INS-58
chr15-90731162-INS-509
chr15-90752277-INS-52
chr15-90759683-INS-74
chr15-90768202-INS-77
chr15-90779629-INS-74
chr15-90788104-INS-64
chr15-90807930-INS-366
chr15-90810020-INS-351
chr15-90816079-INS-107
chr15-90827758-INS-142
chr15-90832379-INS-223
chr15-90856155-INS-102
chr15-90882520-INS-167
chr15-90976191-INS-354
chr15-90990910-INS-66
chr15-91047130-INS-360
chr15-91048079-INS-64
chr15-91053445-INS-58
chr15-91123931-INS-64
chr15-91124702-INS-63
chr15-91157546-INS-88
chr15-91186742-INS-122
chr15-91190462-INS-60
chr15-91209402-INS-532
chr15-91219202-INS-82
chr15-91221019-INS-340
chr15-91231465-INS-7505
chr15-91231753-INS-77
chr15-91252537-INS-724
chr15-91291759-INS-400
chr15-91295126-INS-125
chr15-91302537-INS-416
chr15-91328556-INS-107
chr15-91344667-INS-125
chr15-91349109-INS-489
chr15-91360426-INS-183
chr15-91377413-INS-54
chr15-91382002-INS-55
chr15-91395196-INS-497
chr15-91412200-INS-51
chr15-91423395-INS-85
chr15-91462699-INS-92
chr15-91465262-INS-66
chr15-91480897-INS-96
chr15-91511532-INS-93
chr15-91524596-INS-127
chr15-91549108-INS-362
chr15-91577511-INS-68
chr15-91592933-INS-58
chr15-91624622-INS-106
chr15-91625837-INS-190
chr15-91642784-INS-232
chr15-91677236-INS-568
chr15-91687960-INS-436
chr15-91689214-INS-132
chr15-91690709-INS-78
chr15-91712775-INS-440
chr15-91733813-INS-116
chr15-91741097-INS-99
chr15-91749017-INS-64
chr15-91751936-INS-56
chr15-91752793-INS-66
chr15-91753224-INS-385
chr15-91773434-INS-228
chr15-91782071-INS-272
chr15-91833182-INS-63
chr15-91858519-INS-513
chr15-91880745-INS-228
chr15-91881073-INS-768
chr15-91891280-INS-1184
chr15-91891772-INS-415
chr15-91892949-INS-53
chr15-91939352-INS-79
chr15-91967025-INS-107
chr15-91976704-INS-138
chr15-91989032-INS-225

chr15-92036983-INS-64
chr15-92046322-INS-778
chr15-92052578-INS-6487
chr15-92067740-INS-147
chr15-92077652-INS-197
chr15-92103028-INS-86
chr15-92103854-INS-62
chr15-92115072-INS-265
chr15-92116795-INS-56
chr15-92119967-INS-297
chr15-92152168-INS-62
chr15-92162431-INS-60
chr15-92208973-INS-400
chr15-92215744-INS-490
chr15-92219667-INS-114
chr15-92249771-INS-52
chr15-92265677-INS-289
chr15-92275594-INS-58
chr15-92291738-INS-542
chr15-92319481-INS-140
chr15-92434535-INS-60
chr15-92454888-INS-52
chr15-92467871-INS-313
chr15-92471432-INS-75
chr15-92477902-INS-68
chr15-92480574-INS-81
chr15-92491819-INS-109
chr15-92523139-INS-5192
chr15-92525481-INS-339
chr15-92558689-INS-269
chr15-92606553-INS-60
chr15-92618758-INS-7905
chr15-92665611-INS-193
chr15-92671715-INS-155
chr15-92699417-INS-56
chr15-92725003-INS-57
chr15-92748537-INS-365
chr15-92751248-INS-356
chr15-92777087-INS-117
chr15-92803369-INS-60
chr15-92814750-INS-58
chr15-92864678-INS-261
chr15-92868017-INS-180
chr15-92894843-INS-329
chr15-92950340-INS-74
chr15-92974729-INS-6734
chr15-92985080-INS-64
chr15-92991386-INS-514
chr15-93021896-INS-340
chr15-93033822-INS-864
chr15-93039841-INS-220
chr15-93045304-INS-156
chr15-93047276-INS-447
chr15-93067625-INS-636
chr15-93079998-INS-150
chr15-93102264-INS-72
chr15-93112798-INS-146
chr15-93115152-INS-467
chr15-93117673-INS-434
chr15-93120771-INS-591
chr15-93131320-INS-406
chr15-93134144-INS-78
chr15-93148263-INS-5354
chr15-93161222-INS-174
chr15-93185412-INS-108
chr15-93204996-INS-2812
chr15-93206976-INS-419
chr15-93214038-INS-57
chr15-93218610-INS-212
chr15-93221609-INS-220
chr15-93222858-INS-364
chr15-93228202-INS-322
chr15-93246948-INS-208
chr15-93263814-INS-199
chr15-93272308-INS-683
chr15-93276658-INS-994
chr15-93284995-INS-197

chr15-93298188-INS-231
chr15-93303092-INS-268
chr15-93307424-INS-234
chr15-93319757-INS-68
chr15-93351665-INS-65
chr15-93359610-INS-294
chr15-93366376-INS-199
chr15-93366969-INS-2046
chr15-93368446-INS-96
chr15-93380895-INS-204
chr15-93382946-INS-93
chr15-93424940-INS-262
chr15-93436160-INS-73
chr15-93454838-INS-53
chr15-93462588-INS-609
chr15-93465974-INS-95
chr15-93467986-INS-213
chr15-93505605-INS-196
chr15-93555470-INS-206
chr15-93565766-INS-91
chr15-93575252-INS-61
chr15-93577302-INS-377
chr15-93586630-INS-217
chr15-93602339-INS-211
chr15-93605436-INS-74
chr15-93634755-INS-140
chr15-93661898-INS-2216
chr15-93676078-INS-60
chr15-93706846-INS-590
chr15-93797293-INS-358
chr15-93811868-INS-243
chr15-93811931-INS-285
chr15-93820357-INS-98
chr15-93829118-INS-223
chr15-93832243-INS-140
chr15-93835076-INS-52
chr15-93838534-INS-215
chr15-93839910-INS-66
chr15-93899925-INS-138
chr15-93926749-INS-133
chr15-93932870-INS-222
chr15-93963500-INS-398
chr15-93990660-INS-97
chr15-93994510-INS-246
chr15-94007303-INS-59
chr15-94029523-INS-117
chr15-94042842-INS-372
chr15-94059783-INS-187
chr15-94074992-INS-72
chr15-94134771-INS-75
chr15-94142442-INS-147
chr15-94154731-INS-125
chr15-94184616-INS-62
chr15-94191916-INS-208
chr15-94222990-INS-359
chr15-94246428-INS-69
chr15-94260807-INS-179
chr15-94265806-INS-55
chr15-94289055-INS-126
chr15-94291653-INS-146
chr15-94294484-INS-228
chr15-94296165-INS-63
chr15-94324408-INS-163
chr15-94425232-INS-260
chr15-94438560-INS-60
chr15-94485756-INS-60
chr15-94515909-INS-255
chr15-94521740-INS-59
chr15-94535176-INS-60
chr15-94558210-INS-170
chr15-94612715-INS-86
chr15-94629429-INS-1288
chr15-94644313-INS-66
chr15-94650946-INS-430
chr15-94674698-INS-100
chr15-94693138-INS-88
chr15-94717552-INS-55

chr15-94728728-INS-155
chr15-94743140-INS-59
chr15-94761648-INS-760
chr15-94762741-INS-65
chr15-94803718-INS-180
chr15-94818101-INS-249
chr15-94829066-INS-244
chr15-94845828-INS-93
chr15-94847448-INS-84
chr15-94851209-INS-801
chr15-94852693-INS-336
chr15-94920930-INS-55
chr15-94924873-INS-107
chr15-94929162-INS-63
chr15-94935398-INS-71
chr15-94937584-INS-358
chr15-94941368-INS-71
chr15-94950309-INS-182
chr15-94959460-INS-148
chr15-94969411-INS-128
chr15-95008065-INS-54
chr15-95060758-INS-257
chr15-95062728-INS-761
chr15-95077124-INS-1133
chr15-95088239-INS-118
chr15-95095895-INS-60
chr15-95105925-INS-201
chr15-95107725-INS-123
chr15-95121402-INS-64
chr15-95140401-INS-61
chr15-95157511-INS-68
chr15-95163915-INS-203
chr15-95166661-INS-311
chr15-95170316-INS-396
chr15-95184550-INS-6442
chr15-95223970-INS-589
chr15-95226035-INS-61
chr15-95244890-INS-136
chr15-95254636-INS-159
chr15-95256050-INS-57
chr15-95270021-INS-56
chr15-95273022-INS-124
chr15-95276096-INS-815
chr15-95284323-INS-55
chr15-95284851-INS-5083
chr15-95290316-INS-431
chr15-95321146-INS-92
chr15-95335214-INS-76
chr15-95350417-INS-197
chr15-95355211-INS-398
chr15-95357405-INS-69
chr15-95409644-INS-100
chr15-95419583-INS-657
chr15-95420337-INS-105
chr15-95456036-INS-72
chr15-95496914-INS-508
chr15-95497836-INS-162
chr15-95528875-INS-561
chr15-95553600-INS-174
chr15-95553879-INS-656
chr15-95556085-INS-184
chr15-95559240-INS-192
chr15-95579417-INS-389
chr15-95614610-INS-324
chr15-95616275-INS-55
chr15-95649629-INS-86
chr15-95661668-INS-58
chr15-95664486-INS-527
chr15-95665192-INS-55
chr15-95681294-INS-233
chr15-95729257-INS-168
chr15-95752599-INS-82
chr15-95783512-INS-86
chr15-95789297-INS-112
chr15-95915596-INS-93
chr15-95975281-INS-296
chr15-95983698-INS-6459

chr15-95984058-INS-485
chr15-95986058-INS-168
chr15-95997189-INS-255
chr15-96030267-INS-517
chr15-96038710-INS-81
chr15-96043427-INS-59
chr15-96049092-INS-158
chr15-96077947-INS-55
chr15-96091077-INS-334
chr15-96095956-INS-61
chr15-96112171-INS-229
chr15-96113161-INS-449
chr15-96147921-INS-430
chr15-96153905-INS-74
chr15-96157018-INS-252
chr15-96157652-INS-196
chr15-96175470-INS-478
chr15-96210142-INS-96
chr15-96229263-INS-294
chr15-96230431-INS-129
chr15-96232304-INS-260
chr15-96239141-INS-97
chr15-96247107-INS-201
chr15-96255147-INS-202
chr15-96292407-INS-103
chr15-96314214-INS-225
chr15-96317779-INS-96
chr15-96325316-INS-52
chr15-96368937-INS-754
chr15-96378493-INS-385
chr15-96386726-INS-70
chr15-96390403-INS-105
chr15-96391725-INS-52
chr15-96413624-INS-528
chr15-96417464-INS-289
chr15-96419522-INS-71
chr15-96433467-INS-194
chr15-96457606-INS-84
chr15-96460902-INS-208
chr15-96494017-INS-391
chr15-96495382-INS-1407
chr15-96512847-INS-549
chr15-96516681-INS-216
chr15-96529567-INS-66
chr15-96540250-INS-61
chr15-96555940-INS-55
chr15-96562088-INS-187
chr15-96562594-INS-194
chr15-96570946-INS-62
chr15-96574839-INS-67
chr15-96578266-INS-202
chr15-96604549-INS-90
chr15-96612720-INS-165
chr15-96614752-INS-204
chr15-96691133-INS-70
chr15-96692682-INS-163
chr15-96705494-INS-109
chr15-96716143-INS-144
chr15-96747510-INS-59
chr15-96764584-INS-197
chr15-96782030-INS-72
chr15-96801078-INS-58
chr15-96808711-INS-111
chr15-96828965-INS-66
chr15-96842853-INS-320
chr15-96848211-INS-199
chr15-96854226-INS-72
chr15-96903346-INS-113
chr15-96912792-INS-194
chr15-96933167-INS-52
chr15-96944321-INS-109
chr15-96952397-INS-74
chr15-96976903-INS-241
chr15-96999113-INS-62
chr15-97015859-INS-51
chr15-97030721-INS-64
chr15-97036272-INS-61

chr15-97045228-INS-66
chr15-97052751-INS-251
chr15-97062999-INS-197
chr15-97080084-INS-143
chr15-97081687-INS-127
chr15-97084953-INS-334
chr15-97153685-INS-218
chr15-97166613-INS-65
chr15-97171662-INS-82
chr15-97180060-INS-178
chr15-97192122-INS-171
chr15-97194559-INS-223
chr15-97202407-INS-266
chr15-97221119-INS-70
chr15-97224143-INS-117
chr15-97227014-INS-459
chr15-97227773-INS-536
chr15-97233296-INS-135
chr15-97234885-INS-144
chr15-97237944-INS-80
chr15-97253165-INS-147
chr15-97257506-INS-51
chr15-97260354-INS-53
chr15-97266696-INS-336
chr15-97270888-INS-225
chr15-97293185-INS-161
chr15-97311707-INS-671
chr15-97331587-INS-639
chr15-97336839-INS-95
chr15-97346087-INS-300
chr15-97355175-INS-470
chr15-97363455-INS-52
chr15-97419548-INS-129
chr15-97464758-INS-120
chr15-97467552-INS-338
chr15-97480596-INS-548
chr15-97496877-INS-208
chr15-97502573-INS-116
chr15-97509460-INS-196
chr15-97511237-INS-140
chr15-97515631-INS-63
chr15-97515769-INS-275
chr15-97519543-INS-159
chr15-97535999-INS-655
chr15-97550281-INS-104
chr15-97552529-INS-294
chr15-97566917-INS-303
chr15-97577436-INS-98
chr15-97580369-INS-140
chr15-97584011-INS-231
chr15-97586492-INS-73
chr15-97590065-INS-131
chr15-97592810-INS-181
chr15-97598632-INS-130
chr15-97613251-INS-135
chr15-97614406-INS-142
chr15-97661160-INS-173
chr15-97676222-INS-51
chr15-97703952-INS-123
chr15-97713583-INS-53
chr15-97716955-INS-227
chr15-97719018-INS-102
chr15-97720552-INS-56
chr15-97725840-INS-73
chr15-97746664-INS-260
chr15-97747966-INS-232
chr15-97800070-INS-176
chr15-97810193-INS-53
chr15-97816712-INS-744
chr15-97818586-INS-179
chr15-97819000-INS-261
chr15-97831680-INS-59
chr15-97840937-INS-150
chr15-97843458-INS-116
chr15-97884975-INS-136
chr15-97887876-INS-76
chr15-97922447-INS-222

chr15-97956439-INS-64
chr15-97977594-INS-57
chr15-97989585-INS-458
chr15-97996761-INS-94
chr15-98018995-INS-59
chr15-98027425-INS-126
chr15-98032004-INS-280
chr15-98039329-INS-139
chr15-98044575-INS-220
chr15-98058317-INS-116
chr15-98104066-INS-303
chr15-98112496-INS-376
chr15-98113724-INS-65
chr15-98121116-INS-219
chr15-98121131-INS-56
chr15-98132205-INS-72
chr15-98138300-INS-215
chr15-98145783-INS-165
chr15-98146004-INS-55
chr15-98150157-INS-201
chr15-98158706-INS-149
chr15-98160384-INS-134
chr15-98187307-INS-108
chr15-98197514-INS-220
chr15-98245250-INS-155
chr15-98293020-INS-333
chr15-98314790-INS-202
chr15-98322193-INS-54
chr15-98333272-INS-148
chr15-98355643-INS-537
chr15-98372527-INS-132
chr15-98394751-INS-309
chr15-98400567-INS-64
chr15-98401932-INS-5512
chr15-98414261-INS-215
chr15-98414820-INS-295
chr15-98415411-INS-152
chr15-98419119-INS-70
chr15-98421704-INS-134
chr15-98434193-INS-109
chr15-98484589-INS-77
chr15-98491728-INS-75
chr15-98509460-INS-182
chr15-98516406-INS-199
chr15-98517339-INS-144
chr15-98519091-INS-51
chr15-98523488-INS-72
chr15-98536595-INS-51
chr15-98563423-INS-170
chr15-98569585-INS-207
chr15-98583050-INS-172
chr15-98585767-INS-72
chr15-98591378-INS-66
chr15-98593105-INS-144
chr15-98598773-INS-72
chr15-98599792-INS-78
chr15-98612882-INS-210
chr15-98650074-INS-177
chr15-98684248-INS-216
chr15-98695416-INS-57
chr15-98698940-INS-117
chr15-98717212-INS-210
chr15-98723646-INS-204
chr15-98748903-INS-87
chr15-98783049-INS-148
chr15-98818036-INS-233
chr15-98820283-INS-241
chr15-98828086-INS-200
chr15-98834771-INS-165
chr15-98855889-INS-694
chr15-98865470-INS-91
chr15-98866640-INS-63
chr15-98871580-INS-201
chr15-98873141-INS-196
chr15-98893455-INS-347
chr15-98904116-INS-174
chr15-98914920-INS-177

chr15-98917873-INS-70
chr15-98919098-INS-213
chr15-98925622-INS-149
chr15-98946102-INS-605
chr15-98946604-INS-60
chr15-98956969-INS-206
chr15-98961224-INS-219
chr15-98969556-INS-203
chr15-99000127-INS-146
chr15-99003241-INS-152
chr15-99022891-INS-260
chr15-99033535-INS-98
chr15-99050082-INS-55
chr15-99056307-INS-169
chr15-99063000-INS-85
chr15-99066645-INS-230
chr15-99077859-INS-52
chr15-99083122-INS-72
chr15-99095764-INS-117
chr15-99101040-INS-245
chr15-99113915-INS-199
chr15-99117194-INS-184
chr15-99120874-INS-68
chr15-99129401-INS-58
chr15-99138134-INS-212
chr15-99156196-INS-57
chr15-99161819-INS-116
chr15-99163413-INS-219
chr15-99223920-INS-180
chr15-99224800-INS-116
chr15-99246831-INS-101
chr15-99281470-INS-60
chr15-99313548-INS-156
chr15-99340757-INS-156
chr15-99344809-INS-195
chr15-99345979-INS-96
chr15-99357054-INS-90
chr15-99374504-INS-564
chr15-99379009-INS-361
chr15-99387880-INS-337
chr15-99434247-INS-91
chr15-99436752-INS-129
chr15-99438204-INS-320
chr15-99449436-INS-64
chr15-99466995-INS-96
chr15-99468173-INS-103
chr15-99469778-INS-6405
chr15-99473163-INS-205
chr15-99507704-INS-344
chr15-99516450-INS-178
chr15-99531243-INS-626
chr15-99539064-INS-155
chr15-99548853-INS-76
chr15-99565480-INS-212
chr15-99576095-INS-133
chr15-99579367-INS-122
chr15-99583511-INS-168
chr15-99595142-INS-206
chr15-99611895-INS-116
chr15-99613133-INS-489
chr15-99624462-INS-66
chr15-99630659-INS-171
chr15-99656994-INS-322
chr15-99659361-INS-5244
chr15-99682365-INS-7134
chr15-99725621-INS-302
chr15-99730802-INS-847
chr15-99732285-INS-190
chr15-99743767-INS-152
chr15-99763246-INS-123
chr15-99771763-INS-73
chr15-99801607-INS-58
chr15-99809620-INS-52
chr15-99810044-INS-649
chr15-99827455-INS-63
chr15-99829022-INS-89
chr15-99831137-INS-883

chr15-99835474-INS-65
chr15-99841182-INS-223
chr15-99864266-INS-84
chr15-99874263-INS-249
chr15-99876854-INS-87
chr15-99886145-INS-296
chr15-99900688-INS-373
chr15-99914897-INS-204
chr15-99921021-INS-203
chr15-99924881-INS-96
chr15-99926037-INS-869
chr15-99933205-INS-479
chr15-99935063-INS-74
chr15-99935552-INS-139
chr15-99945510-INS-64
chr15-99972328-INS-94
chr15-99973088-INS-65
chr15-99983134-INS-205
chr15-99992166-INS-106
chr15-99995607-INS-242
chr15-100002893-INS-66
chr15-100014375-INS-96
chr15-100020850-INS-69
chr15-100034717-INS-203
chr15-100035594-INS-56
chr15-100042724-INS-54
chr15-100053664-INS-122
chr15-100063990-INS-2078
chr15-100068870-INS-201
chr15-100071901-INS-65
chr15-100089547-INS-90
chr15-100090695-INS-136
chr15-100093540-INS-98
chr15-100096329-INS-142
chr15-100109410-INS-220
chr15-100123118-INS-77
chr15-100134375-INS-218
chr15-100138167-INS-316
chr15-100160042-INS-333
chr15-100163935-INS-96
chr15-100169917-INS-175
chr15-100171720-INS-1894
chr15-100174982-INS-6508
chr15-100176141-INS-54
chr15-100176938-INS-106
chr15-100182082-INS-182
chr15-100189645-INS-123
chr15-100200158-INS-360
chr15-100217404-INS-385
chr15-100221983-INS-208
chr15-100228607-INS-62
chr15-100234960-INS-109
chr15-100254910-INS-54
chr15-100268518-INS-112
chr15-100273829-INS-51
chr15-100280690-INS-146
chr15-100283637-INS-58
chr15-100301026-INS-182
chr15-100317713-INS-6440
chr15-100325032-INS-84
chr15-100341459-INS-97
chr15-100342994-INS-199
chr15-100345329-INS-128
chr15-100352554-INS-273
chr15-100355762-INS-63
chr15-100372497-INS-58
chr15-100375036-INS-52
chr15-100385887-INS-810
chr15-100395100-INS-173
chr15-100404177-INS-204
chr15-100409551-INS-323
chr15-100412455-INS-73
chr15-100425262-INS-354
chr15-100434342-INS-311
chr15-100440237-INS-309
chr15-100443622-INS-76
chr15-100443952-INS-100

chr15-100451985-INS-84
chr15-100453625-INS-481
chr15-100465561-INS-217
chr15-100483089-INS-84
chr15-100495894-INS-92
chr15-100500834-INS-97
chr15-100513001-INS-55
chr15-100522608-INS-207
chr15-100544089-INS-175
chr15-100546349-INS-4320
chr15-100546357-INS-1029
chr15-100569383-INS-53
chr15-100570549-INS-52
chr15-100580258-INS-125
chr15-100597299-INS-321
chr15-100604874-INS-710
chr15-100607696-INS-118
chr15-100610132-INS-52
chr15-100617616-INS-323
chr15-100623377-INS-195
chr15-100624427-INS-345
chr15-100636033-INS-52
chr15-100642562-INS-310
chr15-100647636-INS-170
chr15-100649698-INS-238
chr15-100690589-INS-266
chr15-100710746-INS-64
chr15-100726299-INS-175
chr15-100729815-INS-235
chr15-100731935-INS-69
chr15-100734851-INS-192
chr15-100772521-INS-183
chr15-100774543-INS-386
chr15-100790992-INS-95
chr15-100800120-INS-451
chr15-100800865-INS-115
chr15-100814913-INS-828
chr15-100818371-INS-230
chr15-100822570-INS-56
chr15-100828281-INS-202
chr15-100855300-INS-121
chr15-100860371-INS-54
chr15-100863360-INS-446
chr15-100864849-INS-240
chr15-100885604-INS-99
chr15-100908285-INS-123
chr15-100956419-INS-133
chr15-100988657-INS-148
chr15-101010850-INS-128
chr15-101016105-INS-130
chr15-101039276-INS-52
chr15-101040380-INS-60
chr15-101049899-INS-109
chr15-101054023-INS-98
chr15-101069180-INS-245
chr15-101085640-INS-88
chr15-101091089-INS-212
chr15-101120645-INS-67
chr15-101124510-INS-140
chr15-101147728-INS-207
chr15-101149593-INS-215
chr15-101174863-INS-88
chr15-101184191-INS-98
chr15-101185811-INS-152
chr15-101189373-INS-364
chr15-101192638-INS-418
chr15-101196999-INS-136
chr15-101198676-INS-51
chr15-101212989-INS-223
chr15-101214204-INS-92
chr15-101215362-INS-149
chr15-101219874-INS-372
chr15-101222402-INS-90
chr15-101223501-INS-137
chr15-101256923-INS-404
chr15-101258155-INS-512
chr15-101278470-INS-66

chr15-101283380-INS-58
chr15-101295028-INS-52
chr15-101299976-INS-103
chr15-101325329-INS-149
chr15-101335993-INS-375
chr15-101337342-INS-223
chr15-101346294-INS-136
chr15-101351722-INS-52
chr15-101362473-INS-567
chr15-101381120-INS-133
chr15-101382304-INS-70
chr15-101402893-INS-55
chr15-101417204-INS-68
chr15-101425573-INS-56
chr15-101429657-INS-67
chr15-101432328-INS-1525
chr15-101489822-INS-131
chr15-101502537-INS-7203
chr15-101502740-INS-359
chr15-101503641-INS-72
chr15-101510377-INS-241
chr15-101523073-INS-75
chr15-101534777-INS-191
chr15-101546874-INS-55
chr15-101564260-INS-423
chr15-101607005-INS-261
chr15-101607911-INS-249
chr15-101615151-INS-323
chr15-101655482-INS-111
chr15-101659193-INS-53
chr15-101685445-INS-402
chr15-101744118-INS-55
chr15-101745912-INS-84
chr15-101745916-INS-750
chr15-101753264-INS-93
chr15-101803871-INS-58
chr15-101811287-INS-107
chr15-101815579-INS-183
chr15-101825231-INS-67
chr15-101868430-INS-104
chr15-101871864-INS-100
chr15-101890949-INS-104
chr15-101893750-INS-197
chr15-101918192-INS-68
chr15-101929010-INS-86
chr15-101935882-INS-90
chr15-101946052-INS-68
chr15-101961924-INS-400
chr15-101962634-INS-202
chr15-101966098-INS-60
chr15-101970571-INS-399
chr15-101977434-INS-226
chr15-101985755-INS-214
chr15-101987179-INS-74
chr15-102039048-INS-69
chr15-102048840-INS-186
chr15-102077181-INS-416
chr15-102082835-INS-51
chr15-102092657-INS-108
chr15-102099561-INS-200
chr15-102124166-INS-169
chr15-102161882-INS-169
chr15-102177180-INS-73
chr15-102183493-INS-206
chr15-102191954-INS-187
chr15-102195337-INS-216
chr15-102201372-INS-125
chr15-102202981-INS-201
chr15-102209025-INS-204
chr15-102211369-INS-199
chr15-102222754-INS-150
chr15-102233557-INS-306
chr15-102270374-INS-160
chr15-102278266-INS-104
chr15-102288457-INS-139
chr15-102292246-INS-137
chr15-102294309-INS-187

chr15-102305981-INS-56
chr15-102311404-INS-197
chr15-102313368-INS-193
chr15-102323513-INS-204
chr15-102333993-INS-309
chr15-102349547-INS-124
chr15-102361893-INS-54
chr15-102364173-INS-99
chr15-102404209-INS-102
chr15-102417300-INS-197
chr15-102447912-INS-211
chr15-102478463-INS-3328
chr15-102483821-INS-135
chr15-102485916-INS-181
chr15-102522743-INS-235
chr15-102538729-INS-176
chr15-102546461-INS-124
chr15-102547647-INS-286
chr15-102549979-INS-164
chr15-102567899-INS-202
chr15-102570003-INS-193
chr15-102609525-INS-70
chr15-102630030-INS-60
chr15-102649236-INS-83
chr15-102665697-INS-151
chr15-102681072-INS-70
chr15-102683910-INS-666
chr15-102696349-INS-80
chr15-102725743-INS-180
chr15-102747307-INS-87
chr15-102775483-INS-892
chr15-102779863-INS-786
chr15-102802956-INS-57
chr15-102969428-INS-153
chr15-102981200-INS-62
chr15-102993185-INS-204
chr15-103027162-INS-271
chr15-103072405-INS-260
chr15-103083333-INS-101
chr15-103084904-INS-55
chr15-103116318-INS-234
chr15-103141462-INS-206
chr15-103143394-INS-63
chr15-103162814-INS-60
chr15-103169085-INS-206
chr15-103172093-INS-100
chr15-103173319-INS-197
chr15-103176221-INS-76
chr15-103182997-INS-203
chr15-103193068-INS-206
chr15-103221842-INS-205
chr15-103271936-INS-123
chr15-103280939-INS-69
chr15-103281490-INS-329
chr15-103314857-INS-198
chr15-103319700-INS-424
chr15-103332289-INS-256
chr15-103352586-INS-206
chr15-103392708-INS-198
chr15-103423034-INS-185
chr15-103465471-INS-110
chr15-103474392-INS-284
chr15-103491990-INS-68
chr15-103493775-INS-358
chr15-103504458-INS-5995
chr15-103572013-INS-53
chr15-103573044-INS-76
chr15-103636140-INS-211
chr15-103667208-INS-412
chr15-103764819-INS-4093
chr15-103768691-INS-125
chr15-103771187-INS-797
chr15-103809537-INS-51
chr15-103909162-INS-130
chr15-103909615-INS-246
chr15-103944855-INS-108
chr15-103955624-INS-130

chr16-3244459-INS-53
chr16-3247545-INS-52
chr16-3263464-INS-261
chr16-3286023-INS-1074
chr16-3315200-INS-202
chr16-3391743-INS-4726
chr16-3395691-INS-5543
chr16-3466672-INS-81
chr16-3469493-INS-345
chr16-3475518-INS-100
chr16-3476637-INS-60
chr16-3479489-INS-570
chr16-3480245-INS-239
chr16-3507660-INS-981
chr16-3523865-INS-876
chr16-3543387-INS-390
chr16-3577404-INS-836
chr16-3598651-INS-217
chr16-3636015-INS-1893
chr16-3731550-INS-109
chr16-3739832-INS-52
chr16-3745796-INS-726
chr16-3762415-INS-60
chr16-3784670-INS-204
chr16-3820764-INS-206
chr16-3860426-INS-214
chr16-3941954-INS-280
chr16-3960650-INS-819
chr16-3969804-INS-399
chr16-3989906-INS-181
chr16-4018161-INS-95
chr16-4054108-INS-81
chr16-4059387-INS-52
chr16-4068090-INS-51
chr16-4070649-INS-198
chr16-4096363-INS-239
chr16-4119080-INS-171
chr16-4122700-INS-440
chr16-4152650-INS-456
chr16-4182330-INS-76
chr16-4185862-INS-418
chr16-4190332-INS-93
chr16-4203055-INS-82
chr16-4207476-INS-60
chr16-4210182-INS-89
chr16-4220518-INS-111
chr16-4278751-INS-215
chr16-4305077-INS-3191
chr16-4324767-INS-215
chr16-4340128-INS-232
chr16-4364529-INS-283
chr16-4366652-INS-256
chr16-4372770-INS-230
chr16-4385097-INS-266
chr16-4405139-INS-5549
chr16-4463007-INS-457
chr16-4471112-INS-249
chr16-4480249-INS-52
chr16-4491826-INS-313
chr16-4505860-INS-179
chr16-4519620-INS-76
chr16-4527435-INS-419
chr16-4539400-INS-79
chr16-4549418-INS-4359
chr16-4553044-INS-57
chr16-4569655-INS-52
chr16-4590091-INS-5547
chr16-4613175-INS-151
chr16-4614432-INS-579
chr16-4615172-INS-571
chr16-4623263-INS-198
chr16-4660822-INS-400
chr16-4679861-INS-365
chr16-4699703-INS-185
chr16-4711502-INS-59
chr16-4724008-INS-982
chr16-4735523-INS-52

chr16-4753480-INS-265
chr16-4772955-INS-184
chr16-4790954-INS-233
chr16-4791605-INS-52
chr16-4799901-INS-52
chr16-4801020-INS-192
chr16-4809204-INS-83
chr16-4813192-INS-117
chr16-4860289-INS-148
chr16-4879232-INS-555
chr16-4945095-INS-54
chr16-4948389-INS-73
chr16-4980819-INS-70
chr16-4984078-INS-344
chr16-4997394-INS-79
chr16-4998934-INS-519
chr16-5005594-INS-199
chr16-5006483-INS-54
chr16-5022389-INS-80
chr16-5059837-INS-86
chr16-5070638-INS-208
chr16-5074360-INS-84
chr16-5087150-INS-150
chr16-5099162-INS-157
chr16-5124094-INS-52
chr16-5138304-INS-130
chr16-5152109-INS-97
chr16-5152730-INS-60
chr16-5163975-INS-72
chr16-5185242-INS-60
chr16-5221402-INS-151
chr16-5223798-INS-95
chr16-5235937-INS-63
chr16-5251679-INS-107
chr16-5253229-INS-741
chr16-5262145-INS-60
chr16-5287561-INS-74
chr16-5298813-INS-217
chr16-5300474-INS-72
chr16-5307404-INS-145
chr16-5332320-INS-90
chr16-5338503-INS-644
chr16-5340525-INS-63
chr16-5347115-INS-52
chr16-5361732-INS-114
chr16-5362823-INS-222
chr16-5369100-INS-260
chr16-5372493-INS-57
chr16-5402491-INS-70
chr16-5404572-INS-6165
chr16-5407940-INS-140
chr16-5477475-INS-55
chr16-5483738-INS-112
chr16-5567229-INS-150
chr16-5607532-INS-161
chr16-5614190-INS-76
chr16-5622950-INS-133
chr16-5625946-INS-242
chr16-5640266-INS-496
chr16-5646185-INS-226
chr16-5677094-INS-55
chr16-5687913-INS-58
chr16-5700137-INS-75
chr16-5733734-INS-56
chr16-5769790-INS-399
chr16-5815585-INS-114
chr16-5829352-INS-74
chr16-5833560-INS-123
chr16-5836218-INS-232
chr16-5847773-INS-69
chr16-5872167-INS-76
chr16-5891271-INS-127
chr16-5905262-INS-142
chr16-5921110-INS-323
chr16-5925856-INS-174
chr16-5983265-INS-56
chr16-5992682-INS-78

chr16-6048142-INS-80
chr16-6051227-INS-74
chr16-6065066-INS-158
chr16-6076470-INS-865
chr16-6080394-INS-85
chr16-6089173-INS-50
chr16-6134156-INS-109
chr16-6181471-INS-357
chr16-6261325-INS-51
chr16-6292326-INS-154
chr16-6297313-INS-84
chr16-6322664-INS-296
chr16-6331168-INS-124
chr16-6368705-INS-66
chr16-6378019-INS-793
chr16-6395350-INS-245
chr16-6410511-INS-60
chr16-6431100-INS-111
chr16-6448785-INS-106
chr16-6459455-INS-62
chr16-6463166-INS-298
chr16-6513829-INS-218
chr16-6516932-INS-52
chr16-6521649-INS-691
chr16-6535609-INS-69
chr16-6543626-INS-334
chr16-6614743-INS-468
chr16-6630021-INS-57
chr16-6637757-INS-102
chr16-6686423-INS-56
chr16-6692431-INS-69
chr16-6708468-INS-76
chr16-6714684-INS-68
chr16-6732029-INS-63
chr16-6778461-INS-120
chr16-6828592-INS-400
chr16-6853309-INS-211
chr16-6895175-INS-84
chr16-6938265-INS-587
chr16-6967025-INS-93
chr16-6991891-INS-457
chr16-7005606-INS-56
chr16-7086240-INS-58
chr16-7102383-INS-94
chr16-7123365-INS-76
chr16-7190319-INS-55
chr16-7251910-INS-94
chr16-7356478-INS-349
chr16-7387129-INS-102
chr16-7440645-INS-106
chr16-7444389-INS-611
chr16-7482464-INS-572
chr16-7497955-INS-56
chr16-7505259-INS-56
chr16-7506372-INS-583
chr16-7508855-INS-110
chr16-7509499-INS-60
chr16-7511011-INS-65
chr16-7513173-INS-196
chr16-7518935-INS-117
chr16-7534479-INS-52
chr16-7542620-INS-108
chr16-7544031-INS-53
chr16-7552274-INS-60
chr16-7590647-INS-258
chr16-7602137-INS-180
chr16-7606731-INS-483
chr16-7613417-INS-166
chr16-7644832-INS-57
chr16-7652295-INS-97
chr16-7700742-INS-236
chr16-7711994-INS-102
chr16-7722623-INS-238
chr16-7730762-INS-238
chr16-7755839-INS-376
chr16-7813778-INS-429
chr16-7823192-INS-76

chr16-7835444-INS-171
chr16-7837073-INS-830
chr16-7840304-INS-82
chr16-7887635-INS-125
chr16-7894726-INS-588
chr16-7910305-INS-116
chr16-7921883-INS-147
chr16-7928940-INS-273
chr16-7934448-INS-8452
chr16-7946094-INS-64
chr16-7978418-INS-122
chr16-8006181-INS-88
chr16-8019938-INS-125
chr16-8030273-INS-314
chr16-8053244-INS-168
chr16-8060301-INS-397
chr16-8072643-INS-5512
chr16-8115214-INS-796
chr16-8149139-INS-60
chr16-8173834-INS-84
chr16-8179648-INS-56
chr16-8181930-INS-358
chr16-8193568-INS-108
chr16-8218885-INS-333
chr16-8236458-INS-938
chr16-8237512-INS-55
chr16-8261509-INS-199
chr16-8266032-INS-72
chr16-8270050-INS-94
chr16-8278325-INS-2650
chr16-8278495-INS-187
chr16-8322082-INS-61
chr16-8334235-INS-54
chr16-8346207-INS-483
chr16-8349004-INS-74
chr16-8382240-INS-192
chr16-8393817-INS-67
chr16-8412020-INS-562
chr16-8419811-INS-100
chr16-8422752-INS-78
chr16-8430107-INS-94
chr16-8453280-INS-532
chr16-8459110-INS-215
chr16-8498009-INS-121
chr16-8501584-INS-201
chr16-8522422-INS-163
chr16-8542493-INS-120
chr16-8545653-INS-543
chr16-8560317-INS-52
chr16-8571721-INS-128
chr16-8590624-INS-60
chr16-8607915-INS-52
chr16-8610468-INS-205
chr16-8630498-INS-58
chr16-8637571-INS-76
chr16-8639065-INS-54
chr16-8691129-INS-233
chr16-8696606-INS-296
chr16-8707347-INS-72
chr16-8717212-INS-687
chr16-8720070-INS-104
chr16-8721420-INS-567
chr16-8724371-INS-256
chr16-8726681-INS-831
chr16-8734039-INS-132
chr16-8738461-INS-312
chr16-8782194-INS-107
chr16-8783459-INS-192
chr16-8790836-INS-196
chr16-8801039-INS-523
chr16-8820874-INS-396
chr16-8829105-INS-99
chr16-8839955-INS-52
chr16-8841031-INS-56
chr16-8842901-INS-69
chr16-8850086-INS-205
chr16-8855208-INS-175

chr16-8863387-INS-831
chr16-8874109-INS-140
chr16-8887964-INS-139
chr16-8897882-INS-831
chr16-8908511-INS-1481
chr16-8919683-INS-105
chr16-8920300-INS-171
chr16-8930592-INS-470
chr16-8968515-INS-99
chr16-8982191-INS-263
chr16-9025700-INS-332
chr16-9039585-INS-71
chr16-9082266-INS-294
chr16-9084459-INS-182
chr16-9105026-INS-143
chr16-9108363-INS-5142
chr16-9187447-INS-213
chr16-9204631-INS-97
chr16-9237690-INS-269
chr16-9257640-INS-69
chr16-9274388-INS-52
chr16-9278126-INS-96
chr16-9283720-INS-192
chr16-9298627-INS-726
chr16-9317455-INS-118
chr16-9377662-INS-1886
chr16-9404231-INS-128
chr16-9408520-INS-108
chr16-9431257-INS-54
chr16-9432764-INS-71
chr16-9435774-INS-129
chr16-9451020-INS-52
chr16-9456383-INS-135
chr16-9471547-INS-58
chr16-9486144-INS-54
chr16-9494497-INS-84
chr16-9541196-INS-125
chr16-9638630-INS-132
chr16-9663616-INS-69
chr16-9680746-INS-89
chr16-9704994-INS-54
chr16-9711172-INS-56
chr16-9737233-INS-97
chr16-9738750-INS-185
chr16-9747125-INS-95
chr16-9777157-INS-56
chr16-9778220-INS-477
chr16-9785853-INS-297
chr16-9802605-INS-52
chr16-9817763-INS-142
chr16-9819350-INS-68
chr16-9845004-INS-58
chr16-9894038-INS-129
chr16-9894277-INS-416
chr16-9899945-INS-5549
chr16-9903978-INS-212
chr16-9911610-INS-130
chr16-9913009-INS-199
chr16-9938408-INS-212
chr16-9968673-INS-113
chr16-9971769-INS-338
chr16-9984647-INS-86
chr16-10046899-INS-532
chr16-10066550-INS-75
chr16-10083662-INS-204
chr16-10103599-INS-222
chr16-10107960-INS-747
chr16-10117364-INS-110
chr16-10147323-INS-71
chr16-10151336-INS-589
chr16-10157339-INS-151
chr16-10158831-INS-206
chr16-10176902-INS-428
chr16-10181441-INS-335
chr16-10188696-INS-234
chr16-10204321-INS-66
chr16-10216261-INS-342

chr16-10261156-INS-1885
chr16-10278014-INS-96
chr16-10281431-INS-275
chr16-10287530-INS-340
chr16-10292708-INS-75
chr16-10312136-INS-780
chr16-10332743-INS-552
chr16-10333358-INS-68
chr16-10341450-INS-113
chr16-10368092-INS-118
chr16-10421751-INS-2122
chr16-10426350-INS-318
chr16-10454951-INS-52
chr16-10485697-INS-66
chr16-10536171-INS-330
chr16-10578419-INS-4086
chr16-10587944-INS-399
chr16-10604548-INS-65
chr16-10658800-INS-240
chr16-10674068-INS-94
chr16-10674310-INS-361
chr16-10679294-INS-76
chr16-10688226-INS-550
chr16-10704244-INS-264
chr16-10712752-INS-168
chr16-10717569-INS-85
chr16-10741495-INS-156
chr16-10749520-INS-294
chr16-10767406-INS-162
chr16-10773601-INS-78
chr16-10790633-INS-72
chr16-10809775-INS-97
chr16-10813509-INS-74
chr16-10841583-INS-164
chr16-10844101-INS-132
chr16-10855870-INS-101
chr16-10861435-INS-148
chr16-10866002-INS-101
chr16-10878664-INS-180
chr16-10921628-INS-220
chr16-10928765-INS-72
chr16-10935995-INS-202
chr16-10961325-INS-73
chr16-10973594-INS-124
chr16-10978761-INS-353
chr16-10979981-INS-126
chr16-10981543-INS-122
chr16-10981578-INS-274
chr16-10987613-INS-106
chr16-10997029-INS-51
chr16-10999046-INS-350
chr16-11026597-INS-54
chr16-11043562-INS-412
chr16-11066923-INS-161
chr16-11079899-INS-141
chr16-11083634-INS-87
chr16-11093164-INS-458
chr16-11104961-INS-201
chr16-11109966-INS-400
chr16-11113774-INS-194
chr16-11129210-INS-54
chr16-11139017-INS-97
chr16-11158288-INS-200
chr16-11163409-INS-71
chr16-11176734-INS-390
chr16-11179204-INS-709
chr16-11199849-INS-100
chr16-11200958-INS-279
chr16-11220805-INS-65
chr16-11224780-INS-96
chr16-11233643-INS-692
chr16-11241879-INS-114
chr16-11294702-INS-79
chr16-11309823-INS-578
chr16-11318091-INS-192
chr16-11368143-INS-54
chr16-11372977-INS-282

chr16-11408544-INS-62
chr16-11424885-INS-84
chr16-11453042-INS-409
chr16-11493075-INS-182
chr16-11505219-INS-90
chr16-11521713-INS-384
chr16-11524668-INS-721
chr16-11576194-INS-5543
chr16-11600589-INS-74
chr16-11624772-INS-900
chr16-11660093-INS-66
chr16-11669860-INS-200
chr16-11669930-INS-681
chr16-11690143-INS-68
chr16-11692543-INS-147
chr16-11742228-INS-104
chr16-11744430-INS-197
chr16-11755225-INS-257
chr16-11760782-INS-71
chr16-11790523-INS-560
chr16-11854128-INS-222
chr16-11857067-INS-144
chr16-11866848-INS-150
chr16-11879292-INS-516
chr16-11885051-INS-142
chr16-11888483-INS-112
chr16-11893335-INS-142
chr16-11909179-INS-97
chr16-11918424-INS-71
chr16-11924862-INS-56
chr16-11940605-INS-52
chr16-11948050-INS-66
chr16-11962714-INS-57
chr16-11968387-INS-66
chr16-11973548-INS-164
chr16-12014241-INS-461
chr16-12022770-INS-574
chr16-12027326-INS-96
chr16-12031501-INS-173
chr16-12037059-INS-62
chr16-12040615-INS-88
chr16-12055471-INS-700
chr16-12079567-INS-100
chr16-12094968-INS-61
chr16-12113835-INS-132
chr16-12146613-INS-6462
chr16-12148931-INS-201
chr16-12170688-INS-174
chr16-12183060-INS-81
chr16-12265493-INS-78
chr16-12265842-INS-3614
chr16-12297728-INS-179
chr16-12307770-INS-57
chr16-12357998-INS-948
chr16-12407631-INS-68
chr16-12423852-INS-66
chr16-12427094-INS-451
chr16-12455060-INS-1882
chr16-12474293-INS-265
chr16-12494916-INS-54
chr16-12498019-INS-336
chr16-12509260-INS-549
chr16-12527654-INS-54
chr16-12544791-INS-69
chr16-12549059-INS-74
chr16-12564118-INS-574
chr16-12582006-INS-54
chr16-12607875-INS-153
chr16-12624204-INS-6742
chr16-12638769-INS-78
chr16-12647950-INS-265
chr16-12648052-INS-319
chr16-12700211-INS-113
chr16-12707876-INS-71
chr16-12708916-INS-232
chr16-12712734-INS-86
chr16-12725244-INS-82

chr16-12749382-INS-201
chr16-12750957-INS-510
chr16-12807920-INS-3238
chr16-12812630-INS-84
chr16-12820524-INS-56
chr16-12840808-INS-60
chr16-12846416-INS-121
chr16-12851542-INS-60
chr16-12855731-INS-86
chr16-12863032-INS-68
chr16-12870498-INS-848
chr16-12878591-INS-95
chr16-12913786-INS-289
chr16-12914528-INS-90
chr16-12917284-INS-130
chr16-12938292-INS-81
chr16-12976711-INS-84
chr16-12991781-INS-232
chr16-13081609-INS-192
chr16-13113429-INS-6476
chr16-13127906-INS-391
chr16-13130265-INS-829
chr16-13181526-INS-69
chr16-13203423-INS-89
chr16-13216587-INS-220
chr16-13223301-INS-1900
chr16-13237506-INS-57
chr16-13259630-INS-89
chr16-13272926-INS-56
chr16-13309489-INS-391
chr16-13310543-INS-52
chr16-13315226-INS-239
chr16-13321420-INS-70
chr16-13381042-INS-60
chr16-13385420-INS-414
chr16-13397685-INS-70
chr16-13404457-INS-323
chr16-13409354-INS-501
chr16-13436291-INS-483
chr16-13458728-INS-302
chr16-13460639-INS-155
chr16-13463329-INS-147
chr16-13481276-INS-206
chr16-13486261-INS-5249
chr16-13496965-INS-990
chr16-13498214-INS-172
chr16-13504187-INS-278
chr16-13510684-INS-84
chr16-13512456-INS-131
chr16-13542285-INS-209
chr16-13552635-INS-174
chr16-13559766-INS-199
chr16-13561186-INS-56
chr16-13570236-INS-97
chr16-13577607-INS-124
chr16-13578138-INS-148
chr16-13584825-INS-82
chr16-13592337-INS-209
chr16-13594862-INS-65
chr16-13607985-INS-68
chr16-13620984-INS-72
chr16-13632243-INS-220
chr16-13634961-INS-190
chr16-13642809-INS-306
chr16-13653456-INS-981
chr16-13721279-INS-149
chr16-13726736-INS-6468
chr16-13746710-INS-206
chr16-13779619-INS-188
chr16-13796293-INS-223
chr16-13814485-INS-52
chr16-13854829-INS-682
chr16-13855768-INS-118
chr16-13860864-INS-97
chr16-13866687-INS-74
chr16-13890739-INS-333
chr16-13955112-INS-65

chr16-13957723-INS-213
chr16-13964214-INS-183
chr16-13965204-INS-106
chr16-13983599-INS-184
chr16-14002141-INS-269
chr16-14006911-INS-1157
chr16-14010976-INS-263
chr16-14032310-INS-389
chr16-14050194-INS-152
chr16-14051072-INS-375
chr16-14077524-INS-116
chr16-14092885-INS-239
chr16-14103421-INS-146
chr16-14114106-INS-76
chr16-14116821-INS-403
chr16-14164411-INS-176
chr16-14187616-INS-208
chr16-14225286-INS-195
chr16-14236924-INS-527
chr16-14238482-INS-210
chr16-14256341-INS-243
chr16-14272024-INS-77
chr16-14280295-INS-70
chr16-14294425-INS-70
chr16-14320730-INS-219
chr16-14333278-INS-58
chr16-14340078-INS-233
chr16-14340377-INS-7542
chr16-14341381-INS-234
chr16-14375505-INS-207
chr16-14379317-INS-170
chr16-14380974-INS-133
chr16-14383206-INS-328
chr16-14400197-INS-706
chr16-14401577-INS-166
chr16-14406255-INS-6160
chr16-14413736-INS-8951
chr16-14416846-INS-604
chr16-14425813-INS-321
chr16-14428139-INS-172
chr16-14437568-INS-116
chr16-14449296-INS-77
chr16-14474577-INS-110
chr16-14485615-INS-238
chr16-14516480-INS-68
chr16-14552915-INS-713
chr16-14585289-INS-125
chr16-14600079-INS-53
chr16-14607780-INS-417
chr16-14642321-INS-54
chr16-14678545-INS-354
chr16-14684357-INS-325
chr16-14690596-INS-406
chr16-14772415-INS-77
chr16-14777522-INS-325
chr16-14779500-INS-54
chr16-14781070-INS-188
chr16-14789300-INS-70
chr16-14816746-INS-6945
chr16-14838373-INS-418
chr16-14857885-INS-167
chr16-14884444-INS-470
chr16-14895720-INS-1837
chr16-14897848-INS-1580
chr16-14905954-INS-51
chr16-14919226-INS-6003
chr16-14921702-INS-147
chr16-14928907-INS-518
chr16-14950798-INS-6274
chr16-14951257-INS-229
chr16-14967348-INS-115
chr16-14974031-INS-6178
chr16-15032094-INS-98
chr16-15054823-INS-162
chr16-15120239-INS-498
chr16-15127384-INS-400
chr16-15128598-INS-458

chr16-15179303-INS-813
chr16-15195857-INS-65
chr16-15221287-INS-6430
chr16-15348964-INS-253
chr16-15350575-INS-212
chr16-15363064-INS-1456
chr16-15417859-INS-76
chr16-15421953-INS-501
chr16-15442746-INS-75
chr16-15444539-INS-166
chr16-15448763-INS-824
chr16-15489060-INS-1782
chr16-15505368-INS-189
chr16-15579678-INS-361
chr16-15586748-INS-167
chr16-15588340-INS-231
chr16-15615945-INS-89
chr16-15617196-INS-52
chr16-15625354-INS-215
chr16-15627976-INS-229
chr16-15635825-INS-873
chr16-15667167-INS-58
chr16-15670778-INS-225
chr16-15675348-INS-138
chr16-15676968-INS-400
chr16-15716991-INS-180
chr16-15732158-INS-81
chr16-15738527-INS-106
chr16-15748447-INS-150
chr16-15754731-INS-100
chr16-15758788-INS-108
chr16-15763110-INS-515
chr16-15804341-INS-7728
chr16-15809704-INS-82
chr16-15821243-INS-506
chr16-15874956-INS-172
chr16-15881906-INS-137
chr16-15902526-INS-576
chr16-15924594-INS-69
chr16-15929073-INS-736
chr16-15933088-INS-5783
chr16-15935948-INS-7487
chr16-15966153-INS-191
chr16-15981593-INS-208
chr16-16004907-INS-159
chr16-16086097-INS-60
chr16-16088544-INS-224
chr16-16102474-INS-56
chr16-16111847-INS-7204
chr16-16114734-INS-3994
chr16-16116644-INS-480
chr16-16139788-INS-2314
chr16-16148540-INS-579
chr16-16179417-INS-2828
chr16-16272454-INS-304
chr16-16286528-INS-2205
chr16-16299518-INS-54
chr16-16311308-INS-206
chr16-16314240-INS-466
chr16-16316089-INS-968
chr16-16318960-INS-319
chr16-16332146-INS-370
chr16-16358320-INS-136
chr16-16364872-INS-54
chr16-16376917-INS-74
chr16-16379009-INS-96
chr16-16405985-INS-61
chr16-16430574-INS-6798
chr16-16432050-INS-7945
chr16-16460588-INS-6956
chr16-16461369-INS-113
chr16-16463704-INS-56
chr16-16466421-INS-556
chr16-16468453-INS-463
chr16-16489763-INS-6598
chr16-16498251-INS-6403
chr16-16524410-INS-327

chr16-16531126-INS-194
chr16-16546636-INS-7305
chr16-16547120-INS-515
chr16-16547891-INS-3022
chr16-16556168-INS-8304
chr16-16556940-INS-188
chr16-16560378-INS-426
chr16-16563901-INS-582
chr16-16572395-INS-546
chr16-16579041-INS-206
chr16-16585321-INS-719
chr16-16585927-INS-80
chr16-16588667-INS-209
chr16-16592719-INS-102
chr16-16595799-INS-66
chr16-16604216-INS-189
chr16-16626211-INS-138
chr16-16632651-INS-280
chr16-16682351-INS-135
chr16-16687633-INS-199
chr16-16693269-INS-56
chr16-16695153-INS-211
chr16-16711351-INS-54
chr16-16761702-INS-71
chr16-16789197-INS-201
chr16-16796664-INS-52
chr16-16809541-INS-201
chr16-16823973-INS-189
chr16-16871790-INS-195
chr16-16902013-INS-216
chr16-16912001-INS-214
chr16-16929715-INS-298
chr16-16932713-INS-237
chr16-16945447-INS-73
chr16-16950824-INS-56
chr16-16955120-INS-238
chr16-16956175-INS-160
chr16-16958680-INS-794
chr16-16961009-INS-192
chr16-16982832-INS-298
chr16-16988534-INS-620
chr16-16991585-INS-76
chr16-17002387-INS-184
chr16-17012977-INS-179
chr16-17020914-INS-52
chr16-17021284-INS-195
chr16-17036759-INS-61
chr16-17041129-INS-219
chr16-17068997-INS-82
chr16-17072422-INS-216
chr16-17082992-INS-318
chr16-17090190-INS-219
chr16-17123631-INS-200
chr16-17192207-INS-83
chr16-17237545-INS-206
chr16-17244541-INS-84
chr16-17266916-INS-52
chr16-17315150-INS-118
chr16-17316282-INS-205
chr16-17334637-INS-55
chr16-17352478-INS-102
chr16-17363250-INS-57
chr16-17403946-INS-151
chr16-17439850-INS-348
chr16-17458537-INS-69
chr16-17497275-INS-58
chr16-17503988-INS-180
chr16-17510932-INS-137
chr16-17561016-INS-332
chr16-17566276-INS-959
chr16-17581114-INS-198
chr16-17587593-INS-350
chr16-17590872-INS-99
chr16-17600904-INS-201
chr16-17607765-INS-54
chr16-17629843-INS-108
chr16-17644861-INS-411

chr16-17657484-INS-7240
chr16-17673428-INS-80
chr16-17679729-INS-146
chr16-17731780-INS-54
chr16-17762168-INS-120
chr16-17764332-INS-195
chr16-17767769-INS-52
chr16-17785791-INS-349
chr16-17788843-INS-61
chr16-17791938-INS-205
chr16-17807046-INS-721
chr16-17819449-INS-438
chr16-17837962-INS-82
chr16-17856721-INS-257
chr16-17862425-INS-210
chr16-17864537-INS-104
chr16-17899225-INS-344
chr16-17920235-INS-598
chr16-17940091-INS-139
chr16-17954908-INS-188
chr16-17995265-INS-163
chr16-18007115-INS-673
chr16-18009601-INS-62
chr16-18092205-INS-53
chr16-18108561-INS-208
chr16-18112064-INS-101
chr16-18121917-INS-354
chr16-18130313-INS-331
chr16-18156969-INS-103
chr16-18182546-INS-205
chr16-18184794-INS-172
chr16-18188177-INS-217
chr16-18193032-INS-211
chr16-18204796-INS-195
chr16-18226056-INS-201
chr16-18236518-INS-69
chr16-18241649-INS-188
chr16-18243319-INS-205
chr16-18250849-INS-57
chr16-18257768-INS-72
chr16-18260833-INS-240
chr16-18269967-INS-326
chr16-18273471-INS-177
chr16-18285747-INS-183
chr16-18288096-INS-196
chr16-18329051-INS-96
chr16-18331577-INS-56
chr16-18346927-INS-463
chr16-18348552-INS-121
chr16-18353436-INS-194
chr16-18360804-INS-124
chr16-18372364-INS-72
chr16-18375878-INS-205
chr16-18381193-INS-62
chr16-18389748-INS-63
chr16-18411436-INS-205
chr16-18421735-INS-270
chr16-18426228-INS-734
chr16-18443246-INS-54
chr16-18449374-INS-381
chr16-18489698-INS-78
chr16-18504156-INS-701
chr16-18507864-INS-589
chr16-18512604-INS-203
chr16-18517765-INS-119
chr16-18519443-INS-498
chr16-18520730-INS-215
chr16-18545901-INS-210
chr16-18555682-INS-183
chr16-18556120-INS-258
chr16-18556918-INS-70
chr16-18563018-INS-60
chr16-18566015-INS-2310
chr16-18568641-INS-5237
chr16-18577740-INS-360
chr16-18582204-INS-169
chr16-18617964-INS-72

chr16-18642384-INS-80
chr16-18647244-INS-208
chr16-18656916-INS-209
chr16-18660602-INS-169
chr16-18665521-INS-87
chr16-18682144-INS-205
chr16-18703501-INS-436
chr16-18721134-INS-264
chr16-18727162-INS-239
chr16-18731761-INS-69
chr16-18733624-INS-203
chr16-18733951-INS-130
chr16-18749355-INS-206
chr16-18760885-INS-294
chr16-18762224-INS-500
chr16-18763702-INS-57
chr16-18921972-INS-56
chr16-18973052-INS-88
chr16-18974893-INS-395
chr16-19056923-INS-671
chr16-19081654-INS-60
chr16-19083561-INS-2707
chr16-19092074-INS-7323
chr16-19097928-INS-2334
chr16-19098748-INS-6208
chr16-19148758-INS-6153
chr16-19154211-INS-200
chr16-19156042-INS-88
chr16-19167120-INS-212
chr16-19179768-INS-210
chr16-19229434-INS-432
chr16-19246563-INS-102
chr16-19254959-INS-70
chr16-19261060-INS-401
chr16-19326261-INS-362
chr16-19338382-INS-592
chr16-19388768-INS-104
chr16-19441391-INS-219
chr16-19444839-INS-202
chr16-19446466-INS-261
chr16-19484395-INS-51
chr16-19521879-INS-126
chr16-19539939-INS-166
chr16-19542931-INS-84
chr16-19577639-INS-100
chr16-19598277-INS-75
chr16-19600009-INS-190
chr16-19604503-INS-281
chr16-19645382-INS-1266
chr16-19651926-INS-204
chr16-19668459-INS-294
chr16-19675208-INS-61
chr16-19701555-INS-51
chr16-19711656-INS-164
chr16-19712641-INS-573
chr16-19721365-INS-578
chr16-19737515-INS-57
chr16-19773192-INS-66
chr16-19780967-INS-90
chr16-19784391-INS-58
chr16-19787399-INS-66
chr16-19789142-INS-90
chr16-19806827-INS-275
chr16-19818553-INS-430
chr16-19838741-INS-132
chr16-19890606-INS-113
chr16-19891265-INS-8940
chr16-19924928-INS-89
chr16-19929138-INS-65
chr16-19959300-INS-142
chr16-19965007-INS-222
chr16-19975427-INS-204
chr16-19999820-INS-253
chr16-20007427-INS-522
chr16-20013913-INS-116
chr16-20018900-INS-219
chr16-20061333-INS-70

chr16-20062521-INS-227
chr16-20066586-INS-481
chr16-20068060-INS-187
chr16-20073127-INS-395
chr16-20096539-INS-167
chr16-20097414-INS-386
chr16-20110494-INS-201
chr16-20127711-INS-141
chr16-20145980-INS-60
chr16-20147904-INS-149
chr16-20188087-INS-62
chr16-20190383-INS-105
chr16-20217215-INS-109
chr16-20225348-INS-82
chr16-20228176-INS-205
chr16-20231756-INS-178
chr16-20251982-INS-164
chr16-20262375-INS-396
chr16-20266065-INS-212
chr16-20267098-INS-175
chr16-20272312-INS-231
chr16-20284901-INS-369
chr16-20297228-INS-353
chr16-20316493-INS-5432
chr16-20333263-INS-89
chr16-20338143-INS-56
chr16-20358379-INS-104
chr16-20380787-INS-176
chr16-20391206-INS-109
chr16-20399960-INS-66
chr16-20404865-INS-385
chr16-20406380-INS-407
chr16-20489210-INS-221
chr16-20538016-INS-198
chr16-20541856-INS-306
chr16-20542364-INS-200
chr16-20546562-INS-145
chr16-20565060-INS-179
chr16-20567628-INS-389
chr16-20574562-INS-383
chr16-20580771-INS-389
chr16-20602334-INS-194
chr16-20612536-INS-173
chr16-20628635-INS-169
chr16-20682887-INS-484
chr16-20782258-INS-209
chr16-20814555-INS-165
chr16-20821676-INS-342
chr16-20839589-INS-366
chr16-20880037-INS-617
chr16-20895352-INS-50
chr16-20906846-INS-66
chr16-20913845-INS-779
chr16-20955105-INS-210
chr16-20956195-INS-196
chr16-20963516-INS-266
chr16-20967830-INS-215
chr16-20989117-INS-462
chr16-21003863-INS-199
chr16-21016336-INS-132
chr16-21061469-INS-1884
chr16-21064069-INS-116
chr16-21070833-INS-99
chr16-21094061-INS-210
chr16-21108807-INS-217
chr16-21112777-INS-66
chr16-21164580-INS-196
chr16-21166370-INS-300
chr16-21173089-INS-83
chr16-21173728-INS-229
chr16-21187530-INS-215
chr16-21190848-INS-81
chr16-21199929-INS-147
chr16-21202792-INS-438
chr16-21249623-INS-206
chr16-21255433-INS-144
chr16-21256316-INS-346

chr16-21292779-INS-63
chr16-21300883-INS-714
chr16-21325946-INS-76
chr16-21330465-INS-72
chr16-21365893-INS-76
chr16-21370207-INS-70
chr16-21371327-INS-68
chr16-21394495-INS-205
chr16-21410836-INS-340
chr16-21413539-INS-343
chr16-21431908-INS-57
chr16-21447363-INS-440
chr16-21447873-INS-332
chr16-21487477-INS-150
chr16-21495082-INS-59
chr16-21525006-INS-150
chr16-21536123-INS-94
chr16-21545293-INS-291
chr16-21583809-INS-68
chr16-21585967-INS-364
chr16-21615658-INS-70
chr16-21622225-INS-177
chr16-21625162-INS-755
chr16-21626816-INS-66
chr16-21632794-INS-591
chr16-21637920-INS-108
chr16-21643171-INS-198
chr16-21661576-INS-295
chr16-21695316-INS-78
chr16-21698040-INS-442
chr16-21703818-INS-111
chr16-21705327-INS-201
chr16-21723413-INS-145
chr16-21726985-INS-53
chr16-21737945-INS-202
chr16-21779138-INS-115
chr16-21797132-INS-205
chr16-21811361-INS-54
chr16-21821153-INS-207
chr16-21827306-INS-222
chr16-21831400-INS-174
chr16-21849044-INS-67
chr16-21856907-INS-60
chr16-21861835-INS-60
chr16-21874135-INS-60
chr16-21876939-INS-187
chr16-21881031-INS-487
chr16-21889662-INS-123
chr16-21894115-INS-114
chr16-21896202-INS-549
chr16-21914084-INS-72
chr16-21955746-INS-128
chr16-21963771-INS-73
chr16-21965700-INS-209
chr16-21973740-INS-225
chr16-21996065-INS-106
chr16-21998309-INS-163
chr16-22012166-INS-179
chr16-22016433-INS-65
chr16-22030736-INS-133
chr16-22032911-INS-223
chr16-22043047-INS-132
chr16-22044880-INS-115
chr16-22046751-INS-280
chr16-22087708-INS-5543
chr16-22102756-INS-64
chr16-22137044-INS-241
chr16-22147350-INS-135
chr16-22162814-INS-302
chr16-22165994-INS-97
chr16-22182733-INS-59
chr16-22194306-INS-200
chr16-22195814-INS-262
chr16-22225242-INS-163
chr16-22263169-INS-60
chr16-22269730-INS-63
chr16-22269851-INS-54

chr16-22277302-INS-210
chr16-22304338-INS-400
chr16-22330480-INS-130
chr16-22342394-INS-200
chr16-22348303-INS-104
chr16-22379307-INS-101
chr16-22394936-INS-233
chr16-22408898-INS-67
chr16-22439959-INS-198
chr16-22485235-INS-58
chr16-22516588-INS-111
chr16-22527421-INS-919
chr16-22531384-INS-198
chr16-22546069-INS-62
chr16-22554280-INS-54
chr16-22595682-INS-80
chr16-22607175-INS-69
chr16-22612567-INS-130
chr16-22617990-INS-145
chr16-22620417-INS-113
chr16-22635655-INS-145
chr16-22640379-INS-90
chr16-22647794-INS-405
chr16-22706993-INS-282
chr16-22735217-INS-364
chr16-22784963-INS-86
chr16-22835922-INS-667
chr16-22865168-INS-190
chr16-22883839-INS-90
chr16-22924870-INS-91
chr16-22935138-INS-210
chr16-22937885-INS-210
chr16-22937885-INS-211
chr16-22940791-INS-319
chr16-22962894-INS-344
chr16-22970111-INS-60
chr16-22973084-INS-180
chr16-22976702-INS-200
chr16-22977611-INS-67
chr16-22982570-INS-148
chr16-23008076-INS-214
chr16-23012412-INS-57
chr16-23013315-INS-94
chr16-23024138-INS-67
chr16-23030563-INS-60
chr16-23037550-INS-100
chr16-23039473-INS-250
chr16-23047013-INS-71
chr16-23047787-INS-538
chr16-23051974-INS-655
chr16-23059116-INS-200
chr16-23072853-INS-442
chr16-23104740-INS-63
chr16-23143991-INS-152
chr16-23169864-INS-52
chr16-23180227-INS-275
chr16-23181436-INS-51
chr16-23182840-INS-151
chr16-23184981-INS-272
chr16-23200057-INS-434
chr16-23223202-INS-60
chr16-23230598-INS-72
chr16-23255350-INS-115
chr16-23262701-INS-75
chr16-23296412-INS-70
chr16-23296907-INS-80
chr16-23297989-INS-54
chr16-23308567-INS-55
chr16-23324139-INS-55
chr16-23352727-INS-56
chr16-23354285-INS-163
chr16-23367028-INS-152
chr16-23370497-INS-79
chr16-23379776-INS-144
chr16-23384960-INS-679
chr16-23391869-INS-140
chr16-23402913-INS-96

chr16-23456183-INS-60
chr16-23457101-INS-92
chr16-23485396-INS-138
chr16-23508228-INS-128
chr16-23516775-INS-256
chr16-23544706-INS-82
chr16-23549634-INS-668
chr16-23551257-INS-64
chr16-23553793-INS-77
chr16-23604949-INS-3267
chr16-23612199-INS-86
chr16-23641729-INS-60
chr16-23650995-INS-106
chr16-23673792-INS-336
chr16-23689767-INS-469
chr16-23701105-INS-392
chr16-23710212-INS-60
chr16-23748801-INS-557
chr16-23762403-INS-88
chr16-23768479-INS-704
chr16-23768481-INS-110
chr16-23804134-INS-59
chr16-23811834-INS-56
chr16-23812344-INS-71
chr16-23828731-INS-347
chr16-23886879-INS-162
chr16-23927476-INS-285
chr16-23940865-INS-224
chr16-23944397-INS-64
chr16-23957404-INS-390
chr16-23965382-INS-69
chr16-24000269-INS-898
chr16-24036908-INS-134
chr16-24071270-INS-1059
chr16-24084772-INS-60
chr16-24118428-INS-72
chr16-24121855-INS-1012
chr16-24124701-INS-2789
chr16-24127487-INS-199
chr16-24144701-INS-6371
chr16-24165884-INS-1483
chr16-24189506-INS-650
chr16-24196928-INS-81
chr16-24203218-INS-396
chr16-24205667-INS-179
chr16-24224688-INS-135
chr16-24299238-INS-157
chr16-24303053-INS-105
chr16-24308026-INS-57
chr16-24311415-INS-126
chr16-24394076-INS-55
chr16-24415957-INS-187
chr16-24423161-INS-1775
chr16-24439891-INS-3022
chr16-24452276-INS-94
chr16-24456130-INS-1821
chr16-24463620-INS-408
chr16-24473990-INS-53
chr16-24480814-INS-63
chr16-24538452-INS-336
chr16-24551285-INS-72
chr16-24552995-INS-82
chr16-24575704-INS-56
chr16-24593079-INS-175
chr16-24603700-INS-55
chr16-24605183-INS-63
chr16-24617405-INS-219
chr16-24627016-INS-400
chr16-24629576-INS-211
chr16-24651584-INS-59
chr16-24653228-INS-847
chr16-24685965-INS-57
chr16-24696010-INS-83
chr16-24707348-INS-399
chr16-24715859-INS-650
chr16-24749133-INS-66
chr16-24828690-INS-68

chr16-24838183-INS-77
chr16-24844923-INS-5357
chr16-24846527-INS-264
chr16-24849915-INS-250
chr16-24850855-INS-229
chr16-24855404-INS-90
chr16-24900150-INS-935
chr16-24912123-INS-54
chr16-24934281-INS-341
chr16-24936004-INS-53
chr16-24945843-INS-499
chr16-24951445-INS-943
chr16-24959359-INS-6547
chr16-24965838-INS-154
chr16-24967254-INS-305
chr16-24974859-INS-109
chr16-25004080-INS-114
chr16-25018017-INS-167
chr16-25025596-INS-144
chr16-25034089-INS-397
chr16-25051183-INS-116
chr16-25056277-INS-58
chr16-25059467-INS-100
chr16-25089146-INS-281
chr16-25094164-INS-121
chr16-25098020-INS-85
chr16-25124670-INS-154
chr16-25153874-INS-6461
chr16-25174955-INS-53
chr16-25175504-INS-51
chr16-25203231-INS-6696
chr16-25229601-INS-288
chr16-25235429-INS-587
chr16-25237659-INS-62
chr16-25271576-INS-87
chr16-25274353-INS-51
chr16-25277345-INS-206
chr16-25283192-INS-60
chr16-25292715-INS-54
chr16-25312855-INS-617
chr16-25315548-INS-176
chr16-25350414-INS-7223
chr16-25382522-INS-128
chr16-25383891-INS-500
chr16-25398575-INS-627
chr16-25400556-INS-880
chr16-25409272-INS-451
chr16-25416735-INS-431
chr16-25419440-INS-96
chr16-25429411-INS-164
chr16-25446840-INS-194
chr16-25467090-INS-54
chr16-25483146-INS-398
chr16-25491377-INS-123
chr16-25507085-INS-811
chr16-25585985-INS-172
chr16-25638608-INS-72
chr16-25667614-INS-65
chr16-25691063-INS-141
chr16-25705382-INS-54
chr16-25756657-INS-400
chr16-25778718-INS-107
chr16-25779687-INS-54
chr16-25798325-INS-5267
chr16-25808830-INS-7408
chr16-25834227-INS-94
chr16-25854980-INS-152
chr16-25877612-INS-53
chr16-25880478-INS-378
chr16-25901551-INS-57
chr16-25906685-INS-76
chr16-25913480-INS-76
chr16-25920145-INS-55
chr16-25939090-INS-7871
chr16-25949308-INS-3026
chr16-25961330-INS-122
chr16-25964415-INS-430

chr16-25967794-INS-328
chr16-26003603-INS-54
chr16-26019341-INS-56
chr16-26084075-INS-54
chr16-26098625-INS-82
chr16-26111410-INS-104
chr16-26124975-INS-198
chr16-26182673-INS-334
chr16-26191348-INS-64
chr16-26195287-INS-111
chr16-26203538-INS-195
chr16-26204581-INS-7161
chr16-26217930-INS-135
chr16-26280983-INS-94
chr16-26283514-INS-57
chr16-26312394-INS-59
chr16-26353700-INS-400
chr16-26359199-INS-6628
chr16-26368684-INS-315
chr16-26397897-INS-431
chr16-26417862-INS-466
chr16-26417967-INS-206
chr16-26449442-INS-735
chr16-26450603-INS-65
chr16-26451506-INS-634
chr16-26460819-INS-62
chr16-26473196-INS-62
chr16-26480384-INS-208
chr16-26491286-INS-1006
chr16-26491287-INS-981
chr16-26502755-INS-105
chr16-26515372-INS-68
chr16-26517198-INS-137
chr16-26518328-INS-6478
chr16-26520888-INS-329
chr16-26552768-INS-147
chr16-26557592-INS-314
chr16-26567926-INS-542
chr16-26574687-INS-175
chr16-26595792-INS-93
chr16-26601764-INS-204
chr16-26605537-INS-79
chr16-26624215-INS-460
chr16-26663785-INS-56
chr16-26703145-INS-6940
chr16-26739220-INS-52
chr16-26755372-INS-246
chr16-26765826-INS-157
chr16-26769574-INS-76
chr16-26792802-INS-70
chr16-26814314-INS-55
chr16-26830820-INS-55
chr16-26842171-INS-217
chr16-26844681-INS-70
chr16-26857929-INS-460
chr16-26881674-INS-86
chr16-26887648-INS-6278
chr16-26922319-INS-71
chr16-26924855-INS-388
chr16-26929472-INS-201
chr16-26940336-INS-54
chr16-26979829-INS-68
chr16-27003765-INS-219
chr16-27057380-INS-56
chr16-27058410-INS-498
chr16-27105523-INS-152
chr16-27122853-INS-358
chr16-27138698-INS-192
chr16-27179902-INS-82
chr16-27194683-INS-111
chr16-27207686-INS-123
chr16-27241921-INS-100
chr16-27312388-INS-868
chr16-27314615-INS-290
chr16-27315426-INS-7108
chr16-27326449-INS-79
chr16-27340866-INS-64

chr16-27344749-INS-72
chr16-27346680-INS-135
chr16-27360782-INS-120
chr16-27376895-INS-121
chr16-27390487-INS-100
chr16-27398266-INS-351
chr16-27452042-INS-172
chr16-27464068-INS-389
chr16-27489093-INS-6082
chr16-27496725-INS-62
chr16-27505825-INS-200
chr16-27511709-INS-52
chr16-27520130-INS-2724
chr16-27522950-INS-703
chr16-27582153-INS-297
chr16-27600296-INS-54
chr16-27634857-INS-193
chr16-27741309-INS-101
chr16-27752272-INS-6401
chr16-27770869-INS-54
chr16-27811340-INS-980
chr16-27827033-INS-60
chr16-27853511-INS-81
chr16-27855340-INS-205
chr16-27859038-INS-259
chr16-27874391-INS-5217
chr16-27876111-INS-51
chr16-27881226-INS-131
chr16-27911471-INS-195
chr16-27932190-INS-351
chr16-27958587-INS-94
chr16-27959858-INS-395
chr16-27975928-INS-199
chr16-27987359-INS-54
chr16-28009398-INS-917
chr16-28009431-INS-265
chr16-28019215-INS-350
chr16-28040476-INS-464
chr16-28044208-INS-743
chr16-28069971-INS-776
chr16-28079750-INS-258
chr16-28082983-INS-70
chr16-28092633-INS-58
chr16-28109891-INS-448
chr16-28128493-INS-56
chr16-28130031-INS-168
chr16-28132344-INS-3294
chr16-28132382-INS-1242
chr16-28149745-INS-544
chr16-28158750-INS-6459
chr16-28160849-INS-174
chr16-28166047-INS-324
chr16-28180703-INS-870
chr16-28187138-INS-677
chr16-28215174-INS-66
chr16-28218954-INS-56
chr16-28240870-INS-56
chr16-28244861-INS-278
chr16-28279377-INS-6888
chr16-28294013-INS-222
chr16-28311047-INS-143
chr16-28397084-INS-66
chr16-28404414-INS-321
chr16-28412367-INS-326
chr16-28418408-INS-73
chr16-28458965-INS-6271
chr16-28459550-INS-74
chr16-28488390-INS-81
chr16-28497781-INS-70
chr16-28500222-INS-88
chr16-28506534-INS-63
chr16-28546100-INS-716
chr16-28560061-INS-102
chr16-28584575-INS-196
chr16-28609070-INS-164
chr16-28619317-INS-867
chr16-28668296-INS-498

chr16-28686334-INS-7148
chr16-28697024-INS-82
chr16-28698057-INS-58
chr16-28701375-INS-176
chr16-28732166-INS-7138
chr16-28753855-INS-52
chr16-28773725-INS-105
chr16-28784944-INS-130
chr16-28826919-INS-259
chr16-28829693-INS-62
chr16-28856826-INS-176
chr16-28868508-INS-104
chr16-28893534-INS-102
chr16-28900756-INS-201
chr16-28906059-INS-6451
chr16-28956293-INS-102
chr16-28962418-INS-535
chr16-29016967-INS-5629
chr16-29021736-INS-6250
chr16-29026015-INS-6605
chr16-29039521-INS-53
chr16-29072230-INS-56
chr16-29119627-INS-455
chr16-29122639-INS-110
chr16-29149381-INS-85
chr16-29157401-INS-114
chr16-29164283-INS-72
chr16-29164932-INS-505
chr16-29170816-INS-6144
chr16-29199070-INS-57
chr16-29220755-INS-136
chr16-29234758-INS-58
chr16-29236927-INS-170
chr16-29237713-INS-6532
chr16-29250403-INS-6898
chr16-29252012-INS-197
chr16-29257235-INS-121
chr16-29271948-INS-142
chr16-29278792-INS-52
chr16-29316838-INS-148
chr16-29358414-INS-296
chr16-29368566-INS-74
chr16-29374246-INS-231
chr16-29414159-INS-212
chr16-29429946-INS-185
chr16-29484133-INS-80
chr16-29503978-INS-252
chr16-29504716-INS-106
chr16-29511102-INS-58
chr16-29532115-INS-55
chr16-29543025-INS-116
chr16-29547147-INS-55
chr16-29551206-INS-137
chr16-29590393-INS-69
chr16-29592506-INS-115
chr16-29612262-INS-195
chr16-29614943-INS-99
chr16-29624383-INS-170
chr16-29625947-INS-130
chr16-29635366-INS-284
chr16-29643925-INS-79
chr16-29646229-INS-451
chr16-29658781-INS-148
chr16-29661411-INS-595
chr16-29664225-INS-56
chr16-29710231-INS-199
chr16-29711905-INS-219
chr16-29717669-INS-186
chr16-29724945-INS-282
chr16-29731737-INS-131
chr16-29736794-INS-150
chr16-29746785-INS-459
chr16-29760358-INS-175
chr16-29820518-INS-62
chr16-29821933-INS-104
chr16-29850956-INS-122
chr16-29859853-INS-159

chr16-29880932-INS-60
chr16-29917356-INS-119
chr16-29934478-INS-66
chr16-29936282-INS-64
chr16-29940383-INS-712
chr16-29943906-INS-507
chr16-29954319-INS-55
chr16-29956484-INS-274
chr16-29963345-INS-188
chr16-29972174-INS-100
chr16-29975254-INS-52
chr16-30004428-INS-210
chr16-30005450-INS-181
chr16-30037095-INS-442
chr16-30037823-INS-62
chr16-30052119-INS-84
chr16-30066342-INS-58
chr16-30081496-INS-158
chr16-30118155-INS-95
chr16-30158034-INS-54
chr16-30165693-INS-151
chr16-30188757-INS-231
chr16-30216898-INS-200
chr16-30220642-INS-128
chr16-30230290-INS-183
chr16-30238562-INS-422
chr16-30241029-INS-57
chr16-30251124-INS-54
chr16-30261034-INS-173
chr16-30273457-INS-76
chr16-30300065-INS-114
chr16-30342554-INS-90
chr16-30344561-INS-84
chr16-30355656-INS-171
chr16-30379362-INS-208
chr16-30410703-INS-206
chr16-30417771-INS-570
chr16-30430787-INS-188
chr16-30454874-INS-128
chr16-30460036-INS-136
chr16-30462685-INS-84
chr16-30465648-INS-323
chr16-30466341-INS-326
chr16-30485365-INS-75
chr16-30494465-INS-54
chr16-30499719-INS-125
chr16-30513574-INS-189
chr16-30528057-INS-67
chr16-30536602-INS-272
chr16-30537179-INS-306
chr16-30537766-INS-837
chr16-30544740-INS-73
chr16-30574954-INS-109
chr16-30604235-INS-109
chr16-30638087-INS-100
chr16-30644295-INS-56
chr16-30646051-INS-276
chr16-30693688-INS-189
chr16-30700776-INS-70
chr16-30704060-INS-345
chr16-30717453-INS-120
chr16-30721485-INS-60
chr16-30745713-INS-295
chr16-30747983-INS-53
chr16-30754556-INS-398
chr16-30768873-INS-189
chr16-30779966-INS-64
chr16-30795474-INS-87
chr16-30807550-INS-205
chr16-30842075-INS-65
chr16-30845156-INS-155
chr16-30848304-INS-136
chr16-30850064-INS-7142
chr16-30860402-INS-185
chr16-30861454-INS-450
chr16-30867355-INS-118
chr16-30891858-INS-201

chr16-30924149-INS-184
chr16-30957636-INS-56
chr16-30969325-INS-54
chr16-30974700-INS-76
chr16-30981603-INS-261
chr16-30987707-INS-201
chr16-30990907-INS-7224
chr16-30993092-INS-137
chr16-30997924-INS-54
chr16-31000794-INS-223
chr16-31010521-INS-775
chr16-31050860-INS-56
chr16-31082650-INS-59
chr16-31084042-INS-1163
chr16-31098560-INS-196
chr16-31142704-INS-326
chr16-31166795-INS-112
chr16-31186571-INS-319
chr16-31188420-INS-110
chr16-31210091-INS-1891
chr16-31212063-INS-211
chr16-31230787-INS-112
chr16-31239066-INS-68
chr16-31249715-INS-196
chr16-31258717-INS-285
chr16-31263593-INS-125
chr16-31266680-INS-84
chr16-31285926-INS-5330
chr16-31291019-INS-92
chr16-31323841-INS-157
chr16-31325872-INS-327
chr16-31328344-INS-224
chr16-31331228-INS-70
chr16-31331536-INS-204
chr16-31338301-INS-86
chr16-31361538-INS-209
chr16-31365662-INS-134
chr16-31367483-INS-5253
chr16-31387812-INS-486
chr16-31393638-INS-574
chr16-31408878-INS-5466
chr16-31408949-INS-400
chr16-31417895-INS-6436
chr16-31454550-INS-161
chr16-31463966-INS-112
chr16-31477478-INS-233
chr16-31486616-INS-150
chr16-31493765-INS-199
chr16-31509786-INS-161
chr16-31510877-INS-77
chr16-31522798-INS-973
chr16-31529676-INS-54
chr16-31539765-INS-87
chr16-31548138-INS-572
chr16-31586301-INS-5420
chr16-31630197-INS-63
chr16-31637039-INS-431
chr16-31644641-INS-174
chr16-31647633-INS-195
chr16-31658645-INS-216
chr16-31663818-INS-182
chr16-31678275-INS-53
chr16-31692479-INS-195
chr16-31741793-INS-453
chr16-31742922-INS-279
chr16-31743329-INS-486
chr16-31744747-INS-196
chr16-31756417-INS-754
chr16-31760467-INS-118
chr16-31791981-INS-62
chr16-31800620-INS-193
chr16-31812644-INS-134
chr16-31826703-INS-195
chr16-31845307-INS-238
chr16-31859610-INS-202
chr16-31865510-INS-659
chr16-31878505-INS-235

chr16-31892988-INS-68
chr16-31900742-INS-183
chr16-31901765-INS-916
chr16-31904727-INS-205
chr16-31905012-INS-5515
chr16-31913450-INS-173
chr16-31915457-INS-228
chr16-31917914-INS-59
chr16-31922046-INS-69
chr16-31922927-INS-208
chr16-31925432-INS-243
chr16-31927846-INS-144
chr16-31928880-INS-197
chr16-31930290-INS-58
chr16-31936124-INS-73
chr16-31939211-INS-73
chr16-31941415-INS-57
chr16-31959109-INS-104
chr16-31964506-INS-180
chr16-31975080-INS-73
chr16-31981012-INS-88
chr16-31999122-INS-96
chr16-31999685-INS-76
chr16-32001403-INS-190
chr16-32012014-INS-170
chr16-32012750-INS-636
chr16-32030985-INS-66
chr16-32036492-INS-54
chr16-32062035-INS-7080
chr16-32072027-INS-156
chr16-32074943-INS-56
chr16-32077442-INS-167
chr16-32079442-INS-56
chr16-32080982-INS-56
chr16-32081634-INS-55
chr16-32083016-INS-179
chr16-32105311-INS-67
chr16-32127396-INS-122
chr16-32134639-INS-408
chr16-32137738-INS-187
chr16-32157354-INS-231
chr16-32158995-INS-129
chr16-32228577-INS-167
chr16-32242232-INS-173
chr16-32251566-INS-203
chr16-32253635-INS-52
chr16-32262767-INS-55
chr16-32267678-INS-53
chr16-32285670-INS-104
chr16-32302753-INS-211
chr16-32305888-INS-299
chr16-32309493-INS-81
chr16-32316622-INS-64
chr16-32318277-INS-56
chr16-32325521-INS-54
chr16-32336221-INS-60
chr16-32351210-INS-856
chr16-32355485-INS-235
chr16-32359834-INS-215
chr16-32366729-INS-205
chr16-32372374-INS-237
chr16-32381661-INS-186
chr16-32392245-INS-57
chr16-32393689-INS-155
chr16-32396224-INS-684
chr16-32411307-INS-85
chr16-32417464-INS-130
chr16-32418384-INS-222
chr16-32418840-INS-5348
chr16-32420957-INS-59
chr16-32428677-INS-65
chr16-32445015-INS-173
chr16-32462574-INS-126
chr16-32470914-INS-248
chr16-32512535-INS-190
chr16-32521481-INS-164
chr16-32553938-INS-82

chr16-32555361-INS-61
chr16-32568285-INS-425
chr16-32572401-INS-372
chr16-32573704-INS-744
chr16-32574492-INS-744
chr16-32576214-INS-372
chr16-32595370-INS-70
chr16-32616695-INS-56
chr16-32644634-INS-121
chr16-32646404-INS-67
chr16-32660686-INS-154
chr16-32666730-INS-573
chr16-32686475-INS-154
chr16-32696373-INS-96
chr16-32704322-INS-1210
chr16-32716604-INS-66
chr16-32730321-INS-450
chr16-32766584-INS-63
chr16-32772684-INS-185
chr16-32779545-INS-565
chr16-32809946-INS-104
chr16-32824375-INS-213
chr16-32831890-INS-173
chr16-32838965-INS-98
chr16-32841268-INS-5259
chr16-32851830-INS-194
chr16-32868700-INS-3269
chr16-32910114-INS-221
chr16-32913654-INS-197
chr16-32940614-INS-181
chr16-32942154-INS-203
chr16-32952040-INS-72
chr16-32967181-INS-54
chr16-32975275-INS-208
chr16-32978647-INS-354
chr16-32980442-INS-209
chr16-32982834-INS-177
chr16-32985912-INS-369
chr16-32991535-INS-267
chr16-32994958-INS-58
chr16-33008969-INS-52
chr16-33010216-INS-212
chr16-33020371-INS-223
chr16-33057922-INS-109
chr16-33066171-INS-56
chr16-33082274-INS-135
chr16-33091681-INS-164
chr16-33129999-INS-138
chr16-33180846-INS-94
chr16-33183434-INS-3138
chr16-33227090-INS-145
chr16-33239927-INS-114
chr16-33250241-INS-596
chr16-33290712-INS-85
chr16-33347982-INS-52
chr16-33351659-INS-1845
chr16-33372041-INS-62
chr16-33385242-INS-244
chr16-33402281-INS-76
chr16-33414366-INS-278
chr16-33417680-INS-170
chr16-33420310-INS-106
chr16-33431296-INS-311
chr16-33438074-INS-109
chr16-33440980-INS-56
chr16-33470902-INS-201
chr16-33472065-INS-202
chr16-33481999-INS-78
chr16-33509604-INS-72
chr16-33524727-INS-52
chr16-33532116-INS-164
chr16-33550528-INS-207
chr16-33569863-INS-66
chr16-33587537-INS-137
chr16-33594244-INS-177
chr16-33596448-INS-57
chr16-33599039-INS-61

chr16-33615691-INS-63
chr16-33619609-INS-72
chr16-33628828-INS-168
chr16-33658458-INS-332
chr16-33662363-INS-173
chr16-33674216-INS-734
chr16-33674262-INS-160
chr16-33677516-INS-192
chr16-33678641-INS-80
chr16-33684155-INS-214
chr16-33685339-INS-69
chr16-33701375-INS-66
chr16-33702836-INS-57
chr16-33706108-INS-319
chr16-33726600-INS-95
chr16-33730153-INS-121
chr16-33734720-INS-515
chr16-33739106-INS-90
chr16-33745478-INS-88
chr16-33747562-INS-244
chr16-33798715-INS-51
chr16-33800877-INS-234
chr16-33804636-INS-103
chr16-33811855-INS-193
chr16-33837479-INS-228
chr16-33861724-INS-57
chr16-33865669-INS-99
chr16-33907503-INS-182
chr16-33915560-INS-86
chr16-33919449-INS-59
chr16-33934565-INS-62
chr16-33935653-INS-172
chr16-33953850-INS-125
chr16-33956011-INS-189
chr16-33959929-INS-378
chr16-33995694-INS-232
chr16-34011289-INS-388
chr16-34019259-INS-65
chr16-34028055-INS-100
chr16-34054963-INS-136
chr16-34065165-INS-91
chr16-34068393-INS-60
chr16-34107523-INS-73
chr16-34121048-INS-62
chr16-34154025-INS-133
chr16-34167878-INS-54
chr16-34172375-INS-464
chr16-34173483-INS-84
chr16-34175784-INS-58
chr16-34205136-INS-6441
chr16-34211231-INS-200
chr16-34260266-INS-50
chr16-34270464-INS-90
chr16-34355148-INS-71
chr16-34396027-INS-86
chr16-34444023-INS-369
chr16-34452153-INS-440
chr16-34461302-INS-205
chr16-34472716-INS-57
chr16-34475561-INS-181
chr16-34483229-INS-237
chr16-34495521-INS-68
chr16-34501172-INS-612
chr16-34570989-INS-203
chr16-34573074-INS-83
chr16-34582507-INS-143
chr16-34594703-INS-227
chr16-34651101-INS-112
chr16-34684382-INS-106
chr16-34697253-INS-61
chr16-34723027-INS-78
chr16-34737785-INS-104
chr16-34743981-INS-83
chr16-34757758-INS-54
chr16-34816966-INS-68
chr16-34818388-INS-54
chr16-34834205-INS-538

chr16-34861697-INS-211
chr16-34862696-INS-206
chr16-34874258-INS-57
chr16-34890168-INS-56
chr16-34894383-INS-202
chr16-34935869-INS-92
chr16-34957083-INS-556
chr16-34995172-INS-52
chr16-34998081-INS-106
chr16-35029946-INS-199
chr16-35092781-INS-234
chr16-35104657-INS-78
chr16-35107627-INS-206
chr16-35126754-INS-214
chr16-35153859-INS-425
chr16-35157246-INS-119
chr16-35162834-INS-107
chr16-35172206-INS-195
chr16-35191142-INS-68
chr16-35202662-INS-65
chr16-35206394-INS-216
chr16-35209521-INS-204
chr16-35210056-INS-64
chr16-35272618-INS-93
chr16-35273781-INS-136
chr16-35278427-INS-99
chr16-35288750-INS-53
chr16-35289593-INS-76
chr16-35299137-INS-91
chr16-35311809-INS-112
chr16-35342531-INS-1483
chr16-35411770-INS-178
chr16-35428547-INS-477
chr16-35435568-INS-419
chr16-35440180-INS-99
chr16-35456803-INS-85
chr16-35464634-INS-343
chr16-35521277-INS-460
chr16-35539524-INS-103
chr16-35548359-INS-180
chr16-35575702-INS-208
chr16-35583086-INS-512
chr16-35583938-INS-75
chr16-35585745-INS-256
chr16-35600669-INS-204
chr16-35635944-INS-60
chr16-35638082-INS-300
chr16-35644468-INS-317
chr16-35658232-INS-753
chr16-35665447-INS-70
chr16-35694462-INS-201
chr16-35700418-INS-82
chr16-35704385-INS-61
chr16-35704942-INS-62
chr16-35711104-INS-155
chr16-35720587-INS-213
chr16-35724490-INS-88
chr16-35728141-INS-497
chr16-35731375-INS-54
chr16-35738244-INS-100
chr16-35739290-INS-216
chr16-35744234-INS-262
chr16-35758687-INS-239
chr16-35786421-INS-309
chr16-35797134-INS-169
chr16-35798285-INS-390
chr16-35839095-INS-263
chr16-35861293-INS-554
chr16-35900783-INS-53
chr16-35901559-INS-303
chr16-35903723-INS-207
chr16-35925183-INS-67
chr16-35939337-INS-244
chr16-35940189-INS-202
chr16-35989632-INS-189
chr16-35991611-INS-235
chr16-36029425-INS-216

chr16-36033408-INS-53
chr16-36128190-INS-519
chr16-36133125-INS-548
chr16-36137974-INS-523
chr16-36140621-INS-892
chr16-36157200-INS-483
chr16-36163364-INS-127
chr16-36181691-INS-6523
chr16-36197500-INS-480
chr16-36216125-INS-168
chr16-36224659-INS-57
chr16-36258420-INS-68
chr16-36271226-INS-98
chr16-36295517-INS-106
chr16-36298867-INS-105
chr16-36303230-INS-56
chr16-36306785-INS-108
chr16-36340490-INS-51
chr16-36345666-INS-257
chr16-36348824-INS-68
chr16-36379620-INS-675
chr16-36384578-INS-70
chr16-36402004-INS-84
chr16-36408072-INS-513
chr16-36409049-INS-62
chr16-36416206-INS-117
chr16-36422691-INS-51
chr16-36431683-INS-4807
chr16-36491707-INS-179
chr16-36551387-INS-5297
chr16-36574823-INS-94
chr16-36602474-INS-415
chr16-36630123-INS-252
chr16-36639781-INS-82
chr16-36649197-INS-64
chr16-36653829-INS-209
chr16-36671077-INS-1332
chr16-36772167-INS-66
chr16-36776194-INS-71
chr16-36797912-INS-91
chr16-36817333-INS-389
chr16-36854955-INS-100
chr16-36907119-INS-289
chr16-36918470-INS-1568
chr16-36928937-INS-325
chr16-36929460-INS-242
chr16-36937153-INS-591
chr16-36947016-INS-286
chr16-36952870-INS-68
chr16-36957780-INS-216
chr16-36995882-INS-900
chr16-36997808-INS-185
chr16-37005960-INS-165
chr16-37019331-INS-695
chr16-37021428-INS-871
chr16-37025582-INS-538
chr16-37031952-INS-658
chr16-37032518-INS-120
chr16-37039057-INS-297
chr16-37051311-INS-442
chr16-37102288-INS-202
chr16-37104507-INS-5730
chr16-37196202-INS-213
chr16-37214022-INS-118
chr16-37237621-INS-54
chr16-37248915-INS-112
chr16-37286325-INS-548
chr16-37318310-INS-6457
chr16-37363567-INS-99
chr16-37425976-INS-310
chr16-37453836-INS-231
chr16-37484940-INS-64
chr16-37494265-INS-54
chr16-37539855-INS-74
chr16-37565838-INS-182
chr16-37575706-INS-121
chr16-37579883-INS-356

chr16-37594160-INS-115
chr16-37609275-INS-87
chr16-37630681-INS-61
chr16-37632043-INS-80
chr16-37689865-INS-58
chr16-37719041-INS-62
chr16-37734499-INS-246
chr16-37744770-INS-54
chr16-37749661-INS-66
chr16-37755940-INS-54
chr16-37766183-INS-128
chr16-37781218-INS-316
chr16-37803426-INS-216
chr16-37815226-INS-52
chr16-37831230-INS-81
chr16-37839068-INS-80
chr16-37844958-INS-503
chr16-37848376-INS-147
chr16-37862449-INS-135
chr16-37889056-INS-81
chr16-37905689-INS-204
chr16-37927730-INS-171
chr16-37936612-INS-60
chr16-37946675-INS-91
chr16-37956980-INS-52
chr16-38000289-INS-111
chr16-38034034-INS-79
chr16-38042641-INS-70
chr16-38052656-INS-387
chr16-38068533-INS-831
chr16-38080207-INS-56
chr16-38085883-INS-5549
chr16-38099344-INS-57
chr16-38106015-INS-157
chr16-38107263-INS-224
chr16-38137562-INS-193
chr16-38145646-INS-5198
chr16-38160484-INS-220
chr16-38213649-INS-82
chr16-38224492-INS-173
chr16-38225461-INS-100
chr16-38229818-INS-303
chr16-38260859-INS-168
chr16-38298133-INS-117
chr16-38339799-INS-100
chr16-38341421-INS-215
chr16-38362399-INS-170
chr16-38364572-INS-106
chr16-38373245-INS-188
chr16-38375222-INS-60
chr16-38383804-INS-235
chr16-38397696-INS-293
chr16-38406317-INS-238
chr16-38410897-INS-237
chr16-38471222-INS-56
chr16-38589353-INS-116
chr16-38606616-INS-161
chr16-38640374-INS-366
chr16-38640539-INS-366
chr16-38640545-INS-914
chr16-38643404-INS-170
chr16-38658372-INS-923
chr16-38732113-INS-276
chr16-38746786-INS-108
chr16-38788413-INS-80
chr16-38868129-INS-62
chr16-38873048-INS-56
chr16-38914105-INS-116
chr16-38916242-INS-94
chr16-38964721-INS-58
chr16-38968518-INS-135
chr16-38993332-INS-1602
chr16-38998812-INS-60
chr16-39001713-INS-560
chr16-39018053-INS-70
chr16-39023569-INS-822
chr16-39037895-INS-52

chr16-39046110-INS-209
chr16-39052755-INS-230
chr16-39057073-INS-167
chr16-39058398-INS-96
chr16-39081586-INS-96
chr16-39086667-INS-121
chr16-39096600-INS-71
chr16-39118788-INS-3274
chr16-39134077-INS-72
chr16-39153213-INS-767
chr16-39161381-INS-435
chr16-39173782-INS-124
chr16-39193306-INS-238
chr16-39195487-INS-78
chr16-39214095-INS-65
chr16-39224891-INS-603
chr16-39244772-INS-7652
chr16-39306589-INS-63
chr16-39307622-INS-79
chr16-39329099-INS-1277
chr16-39329842-INS-6905
chr16-39343801-INS-569
chr16-39353064-INS-57
chr16-39376321-INS-62
chr16-39380380-INS-328
chr16-39391759-INS-64
chr16-39402424-INS-200
chr16-39426368-INS-201
chr16-39463450-INS-2996
chr16-39466764-INS-734
chr16-39470005-INS-481
chr16-39471548-INS-305
chr16-39502536-INS-236
chr16-39518330-INS-52
chr16-39524814-INS-401
chr16-39529801-INS-933
chr16-39533466-INS-6227
chr16-39536804-INS-90
chr16-39550965-INS-3688
chr16-39554065-INS-110
chr16-39557877-INS-889
chr16-39632435-INS-79
chr16-39648929-INS-105
chr16-39650869-INS-196
chr16-39661515-INS-107
chr16-39708916-INS-71
chr16-39761365-INS-959
chr16-39784742-INS-262
chr16-39829629-INS-57
chr16-39845876-INS-61
chr16-39848589-INS-68
chr16-39877189-INS-56
chr16-39889430-INS-62
chr16-39892992-INS-98
chr16-39903668-INS-54
chr16-39905900-INS-64
chr16-39909475-INS-330
chr16-39958884-INS-6401
chr16-40005334-INS-58
chr16-40008256-INS-51
chr16-40026799-INS-83
chr16-40029295-INS-52
chr16-40044801-INS-667
chr16-40059448-INS-59
chr16-40084318-INS-64
chr16-40106631-INS-76
chr16-40108346-INS-890
chr16-40120664-INS-605
chr16-40131754-INS-76
chr16-40156788-INS-51
chr16-40161694-INS-522
chr16-40181713-INS-6434
chr16-40195317-INS-211
chr16-40196428-INS-844
chr16-40200852-INS-434
chr16-40246774-INS-253
chr16-40256777-INS-697

chr16-40324384-INS-598
chr16-40325487-INS-116
chr16-40333776-INS-1120
chr16-40339150-INS-6707
chr16-40353630-INS-598
chr16-40359948-INS-94
chr16-40384608-INS-6229
chr16-40403610-INS-73
chr16-40461159-INS-176
chr16-40470848-INS-6673
chr16-40493077-INS-326
chr16-40549449-INS-457
chr16-40569058-INS-217
chr16-40592929-INS-57
chr16-40608414-INS-7581
chr16-40615446-INS-381
chr16-40634671-INS-214
chr16-40646298-INS-6678
chr16-40663644-INS-58
chr16-40686886-INS-106
chr16-40690627-INS-761
chr16-40709007-INS-76
chr16-40712446-INS-542
chr16-40754607-INS-66
chr16-40763966-INS-339
chr16-40779461-INS-106
chr16-40793555-INS-200
chr16-40796285-INS-445
chr16-40808905-INS-114
chr16-40840069-INS-114
chr16-40874408-INS-96
chr16-40891222-INS-2929
chr16-40895639-INS-92
chr16-40929188-INS-66
chr16-40962101-INS-142
chr16-40974749-INS-267
chr16-40977027-INS-55
chr16-40991867-INS-117
chr16-40995107-INS-118
chr16-41002544-INS-944
chr16-41004403-INS-109
chr16-41009548-INS-88
chr16-41015036-INS-614
chr16-41027815-INS-84
chr16-41031898-INS-203
chr16-41061452-INS-6504
chr16-41102179-INS-63
chr16-41105485-INS-102
chr16-41107788-INS-377
chr16-41134204-INS-162
chr16-41143788-INS-258
chr16-41164321-INS-99
chr16-41166047-INS-178
chr16-41180284-INS-93
chr16-41190324-INS-83
chr16-41197659-INS-8843
chr16-41205824-INS-417
chr16-41212172-INS-215
chr16-41223592-INS-6374
chr16-41269347-INS-501
chr16-41275951-INS-396
chr16-41279944-INS-164
chr16-41284700-INS-177
chr16-41292021-INS-768
chr16-41295784-INS-239
chr16-41303303-INS-155
chr16-41330974-INS-52
chr16-41342768-INS-106
chr16-41345407-INS-208
chr16-41350602-INS-56
chr16-41372151-INS-170
chr16-41382718-INS-107
chr16-41404718-INS-135
chr16-41412883-INS-55
chr16-41415769-INS-7535
chr16-41443029-INS-208
chr16-41452347-INS-58

chr16-41455007-INS-405
chr16-41471534-INS-141
chr16-41477275-INS-226
chr16-41507469-INS-60
chr16-41509067-INS-59
chr16-41516391-INS-56
chr16-41547009-INS-227
chr16-41553673-INS-307
chr16-41638513-INS-52
chr16-41653522-INS-330
chr16-41657657-INS-1808
chr16-41669680-INS-84
chr16-41682980-INS-1888
chr16-41686145-INS-52
chr16-41695027-INS-94
chr16-41705463-INS-846
chr16-41735920-INS-65
chr16-41739722-INS-99
chr16-41741150-INS-68
chr16-41754189-INS-71
chr16-41760935-INS-236
chr16-41795671-INS-158
chr16-41809135-INS-113
chr16-41809278-INS-137
chr16-41815342-INS-68
chr16-41842568-INS-64
chr16-41845814-INS-6456
chr16-41850503-INS-257
chr16-41851374-INS-7076
chr16-41871166-INS-56
chr16-41896529-INS-7316
chr16-41896970-INS-400
chr16-41936746-INS-174
chr16-41976152-INS-86
chr16-42013875-INS-57
chr16-42044076-INS-57
chr16-42052508-INS-233
chr16-42080200-INS-146
chr16-42105868-INS-199
chr16-42134611-INS-182
chr16-42144661-INS-442
chr16-42152995-INS-65
chr16-42164227-INS-68
chr16-42173647-INS-6401
chr16-42192344-INS-76
chr16-42197271-INS-5839
chr16-42203023-INS-327
chr16-42207258-INS-395
chr16-42229572-INS-6632
chr16-42233264-INS-180
chr16-42259217-INS-70
chr16-42269893-INS-200
chr16-42295554-INS-62
chr16-42310625-INS-65
chr16-42326523-INS-84
chr16-42374974-INS-941
chr16-42390680-INS-58
chr16-42413156-INS-131
chr16-42421248-INS-6503
chr16-42454606-INS-72
chr16-42457115-INS-5632
chr16-42477752-INS-240
chr16-42498001-INS-136
chr16-42500837-INS-6678
chr16-42506602-INS-2821
chr16-42507823-INS-275
chr16-42542755-INS-182
chr16-42574918-INS-51
chr16-42585092-INS-108
chr16-42599832-INS-208
chr16-42603528-INS-55
chr16-42608729-INS-62
chr16-42610147-INS-264
chr16-42614763-INS-120
chr16-42621627-INS-154
chr16-42651594-INS-7108
chr16-42709027-INS-54

chr16-42714481-INS-410
chr16-42716544-INS-233
chr16-42726022-INS-61
chr16-42774327-INS-61
chr16-42777236-INS-63
chr16-42810571-INS-60
chr16-42839349-INS-92
chr16-42881372-INS-52
chr16-42920776-INS-259
chr16-42928069-INS-399
chr16-42928426-INS-1967
chr16-42977509-INS-513
chr16-43040812-INS-1893
chr16-43084768-INS-205
chr16-43086920-INS-69
chr16-43138363-INS-117
chr16-43149890-INS-1862
chr16-43300795-INS-62
chr16-43309118-INS-5365
chr16-43315749-INS-685
chr16-43377685-INS-67
chr16-43409899-INS-214
chr16-43480627-INS-240
chr16-43483665-INS-103
chr16-43489625-INS-337
chr16-43501475-INS-657
chr16-43517993-INS-69
chr16-43536129-INS-107
chr16-43555016-INS-206
chr16-43564163-INS-189
chr16-43593536-INS-216
chr16-43615013-INS-602
chr16-43630339-INS-190
chr16-43650372-INS-183
chr16-43653290-INS-336
chr16-43657936-INS-6217
chr16-43672671-INS-56
chr16-43689273-INS-206
chr16-43702225-INS-200
chr16-43721249-INS-6224
chr16-43731897-INS-59
chr16-43735110-INS-1192
chr16-43741523-INS-54
chr16-43743534-INS-52
chr16-43765400-INS-60
chr16-43807250-INS-447
chr16-43817970-INS-916
chr16-43824391-INS-54
chr16-43825740-INS-70
chr16-43884428-INS-451
chr16-43890240-INS-60
chr16-43906509-INS-156
chr16-43924443-INS-79
chr16-43954635-INS-139
chr16-43958325-INS-1888
chr16-43975765-INS-2436
chr16-44014101-INS-51
chr16-44015398-INS-190
chr16-44016678-INS-234
chr16-44017330-INS-75
chr16-44031046-INS-545
chr16-44091367-INS-103
chr16-44116477-INS-114
chr16-44123015-INS-296
chr16-44133042-INS-144
chr16-44135823-INS-51
chr16-44150638-INS-195
chr16-44178452-INS-186
chr16-44198286-INS-165
chr16-44202102-INS-185
chr16-44209241-INS-532
chr16-44223953-INS-62
chr16-44228187-INS-96
chr16-44231287-INS-127
chr16-44241464-INS-56
chr16-44250111-INS-808
chr16-44309464-INS-525

chr16-44310795-INS-289
chr16-44317130-INS-148
chr16-44343329-INS-73
chr16-44345554-INS-57
chr16-44375579-INS-58
chr16-44381896-INS-58
chr16-44395177-INS-4402
chr16-44397851-INS-54
chr16-44410228-INS-204
chr16-44419161-INS-53
chr16-44439046-INS-157
chr16-44447864-INS-783
chr16-44471797-INS-132
chr16-44480264-INS-63
chr16-44491079-INS-161
chr16-44504410-INS-423
chr16-44512932-INS-414
chr16-44522457-INS-219
chr16-44525947-INS-148
chr16-44537711-INS-664
chr16-44540618-INS-72
chr16-44547023-INS-595
chr16-44551008-INS-282
chr16-44592221-INS-56
chr16-44595494-INS-66
chr16-44611183-INS-60
chr16-44617865-INS-7171
chr16-44626952-INS-242
chr16-44636613-INS-251
chr16-44641402-INS-216
chr16-44740659-INS-955
chr16-44758844-INS-447
chr16-44760427-INS-398
chr16-44774225-INS-70
chr16-44797134-INS-52
chr16-44803045-INS-74
chr16-44810386-INS-51
chr16-44851944-INS-80
chr16-44860067-INS-151
chr16-44878190-INS-181
chr16-44905752-INS-182
chr16-44907142-INS-123
chr16-44938271-INS-1000
chr16-45016544-INS-965
chr16-45019845-INS-400
chr16-45021914-INS-140
chr16-45035880-INS-574
chr16-45039652-INS-197
chr16-45043826-INS-3174
chr16-45061045-INS-211
chr16-45068350-INS-52
chr16-45173425-INS-3053
chr16-45173586-INS-96
chr16-45176308-INS-468
chr16-45194993-INS-134
chr16-45204044-INS-78
chr16-45246015-INS-311
chr16-45278858-INS-871
chr16-45306358-INS-434
chr16-45315133-INS-349
chr16-45331313-INS-6473
chr16-45363910-INS-112
chr16-45382370-INS-172
chr16-45409598-INS-114
chr16-45413250-INS-126
chr16-45427456-INS-233
chr16-45428393-INS-351
chr16-45444912-INS-74
chr16-45451863-INS-716
chr16-45475215-INS-230
chr16-45487378-INS-6808
chr16-45498845-INS-72
chr16-45516893-INS-51
chr16-45526914-INS-69
chr16-45604532-INS-111
chr16-45610252-INS-70
chr16-45615646-INS-74

chr16-45625116-INS-110
chr16-45703045-INS-6488
chr16-45715898-INS-6413
chr16-45718764-INS-577
chr16-45777782-INS-259
chr16-45824361-INS-99
chr16-45826607-INS-51
chr16-45835685-INS-242
chr16-45871698-INS-1073
chr16-45877184-INS-68
chr16-45899949-INS-1356
chr16-45917106-INS-51
chr16-45929003-INS-470
chr16-45970436-INS-401
chr16-45979977-INS-6186
chr16-45980487-INS-527
chr16-45984500-INS-6481
chr16-46001969-INS-61
chr16-46004048-INS-221
chr16-46010746-INS-1983
chr16-46013806-INS-6758
chr16-46020943-INS-96
chr16-46036226-INS-52
chr16-46044911-INS-180
chr16-46052297-INS-55
chr16-46056324-INS-1891
chr16-46067304-INS-55
chr16-46080168-INS-965
chr16-46089488-INS-68
chr16-46130521-INS-125
chr16-46166432-INS-149
chr16-46200792-INS-182
chr16-46218809-INS-131
chr16-46220492-INS-632
chr16-46226170-INS-81
chr16-46252002-INS-6689
chr16-46265048-INS-236
chr16-46285748-INS-543
chr16-46288861-INS-204
chr16-46323884-INS-6694
chr16-46338469-INS-667
chr16-46340919-INS-58
chr16-46350032-INS-207
chr16-46359961-INS-73
chr16-46364002-INS-368
chr16-46374252-INS-512
chr16-46376406-INS-145
chr16-46392883-INS-106
chr16-46393821-INS-205
chr16-46412907-INS-54
chr16-46426704-INS-136
chr16-46428675-INS-52
chr16-46442046-INS-78
chr16-46443671-INS-962
chr16-46477575-INS-7122
chr16-46477576-INS-377
chr16-46484059-INS-6233
chr16-46487687-INS-58
chr16-46491973-INS-63
chr16-46502962-INS-3228
chr16-46511366-INS-6164
chr16-46522433-INS-614
chr16-46525430-INS-83
chr16-46552789-INS-59
chr16-46561190-INS-52
chr16-46593644-INS-7539
chr16-46594633-INS-55
chr16-46604221-INS-72
chr16-46634327-INS-6781
chr16-46687355-INS-149
chr16-46701759-INS-122
chr16-46761375-INS-66
chr16-46783449-INS-582
chr16-46878604-INS-65
chr16-46891376-INS-53
chr16-46893113-INS-146
chr16-46895423-INS-64

chr16-46932838-INS-362
chr16-46950508-INS-467
chr16-47001853-INS-75
chr16-47090801-INS-137
chr16-47098269-INS-6101
chr16-47100265-INS-52
chr16-47103930-INS-168
chr16-47152393-INS-68
chr16-47169672-INS-95
chr16-47207931-INS-247
chr16-47218721-INS-408
chr16-47239713-INS-95
chr16-47272807-INS-159
chr16-47283355-INS-74
chr16-47295276-INS-171
chr16-47319384-INS-8985
chr16-47326954-INS-282
chr16-47335995-INS-990
chr16-47351388-INS-566
chr16-47354577-INS-359
chr16-47365742-INS-56
chr16-47385876-INS-88
chr16-47414920-INS-396
chr16-47422791-INS-52
chr16-47436546-INS-399
chr16-47446068-INS-169
chr16-47473240-INS-6833
chr16-47480748-INS-197
chr16-47513176-INS-64
chr16-47558659-INS-116
chr16-47579485-INS-88
chr16-47594315-INS-93
chr16-47614910-INS-54
chr16-47621279-INS-7256
chr16-47636479-INS-84
chr16-47656906-INS-289
chr16-47657148-INS-6065
chr16-47664230-INS-2570
chr16-47668297-INS-498
chr16-47694558-INS-7204
chr16-47721149-INS-10937
chr16-47724912-INS-309
chr16-47737687-INS-74
chr16-47753727-INS-77
chr16-47758719-INS-150
chr16-47795707-INS-152
chr16-47824151-INS-93
chr16-47824641-INS-248
chr16-47860025-INS-203
chr16-47861423-INS-5871
chr16-47874185-INS-62
chr16-47876312-INS-58
chr16-47884948-INS-6386
chr16-47911876-INS-111
chr16-47914297-INS-101
chr16-47915875-INS-933
chr16-47945488-INS-78
chr16-48017938-INS-211
chr16-48025844-INS-52
chr16-48033593-INS-600
chr16-48043661-INS-539
chr16-48055071-INS-62
chr16-48070109-INS-138
chr16-48099959-INS-52
chr16-48103042-INS-143
chr16-48121088-INS-183
chr16-48132160-INS-454
chr16-48151633-INS-432
chr16-48153844-INS-329
chr16-48155632-INS-191
chr16-48158566-INS-57
chr16-48161182-INS-109
chr16-48165157-INS-194
chr16-48171345-INS-55
chr16-48175741-INS-86
chr16-48176207-INS-102
chr16-48180884-INS-592

chr16-48182508-INS-352
chr16-48189959-INS-201
chr16-48193934-INS-101
chr16-48195713-INS-345
chr16-48199862-INS-301
chr16-48213686-INS-94
chr16-48228894-INS-58
chr16-48235407-INS-130
chr16-48243576-INS-254
chr16-48244952-INS-59
chr16-48251345-INS-52
chr16-48259297-INS-283
chr16-48265340-INS-161
chr16-48276441-INS-844
chr16-48278166-INS-201
chr16-48305658-INS-549
chr16-48333610-INS-167
chr16-48337372-INS-525
chr16-48355728-INS-86
chr16-48375305-INS-207
chr16-48401601-INS-146
chr16-48403565-INS-58
chr16-48411880-INS-215
chr16-48423263-INS-135
chr16-48425783-INS-384
chr16-48432411-INS-202
chr16-48460660-INS-5340
chr16-48470326-INS-172
chr16-48471794-INS-135
chr16-48486269-INS-74
chr16-48503090-INS-54
chr16-48508143-INS-763
chr16-48508964-INS-77
chr16-48517540-INS-66
chr16-48525912-INS-56
chr16-48562922-INS-55
chr16-48590104-INS-581
chr16-48609713-INS-6444
chr16-48615149-INS-70
chr16-48617678-INS-7149
chr16-48625557-INS-835
chr16-48627957-INS-365
chr16-48636227-INS-65
chr16-48648100-INS-52
chr16-48648542-INS-4138
chr16-48669423-INS-926
chr16-48692494-INS-107
chr16-48712921-INS-66
chr16-48723963-INS-11989
chr16-48732101-INS-566
chr16-48735021-INS-95
chr16-48766085-INS-111
chr16-48785602-INS-498
chr16-48801975-INS-683
chr16-48810070-INS-231
chr16-48814982-INS-131
chr16-48818107-INS-86
chr16-48823858-INS-400
chr16-48830224-INS-52
chr16-48831668-INS-78
chr16-48857349-INS-63
chr16-48859459-INS-926
chr16-48977964-INS-1141
chr16-48994078-INS-3478
chr16-49005182-INS-6253
chr16-49007159-INS-51
chr16-49012154-INS-72
chr16-49014055-INS-504
chr16-49034062-INS-211
chr16-49039528-INS-876
chr16-49041581-INS-52
chr16-49066867-INS-217
chr16-49086422-INS-71
chr16-49099396-INS-875
chr16-49101878-INS-219
chr16-49112048-INS-358
chr16-49132319-INS-68

chr16-49176609-INS-80
chr16-49217529-INS-66
chr16-49218866-INS-62
chr16-49247745-INS-6603
chr16-49268109-INS-207
chr16-49280156-INS-7005
chr16-49302292-INS-193
chr16-49303559-INS-6649
chr16-49304132-INS-2042
chr16-49318275-INS-135
chr16-49321952-INS-54
chr16-49337493-INS-6812
chr16-49342411-INS-58
chr16-49362564-INS-73
chr16-49370299-INS-102
chr16-49381282-INS-104
chr16-49479795-INS-6691
chr16-49487209-INS-76
chr16-49497247-INS-803
chr16-49526426-INS-51
chr16-49534084-INS-813
chr16-49543353-INS-88
chr16-49545216-INS-61
chr16-49556051-INS-340
chr16-49590749-INS-52
chr16-49603292-INS-218
chr16-49610608-INS-61
chr16-49650119-INS-326
chr16-49683510-INS-124
chr16-49726426-INS-58
chr16-49734156-INS-129
chr16-49758593-INS-68
chr16-49786926-INS-74
chr16-49810164-INS-148
chr16-49818978-INS-226
chr16-49834090-INS-410
chr16-49850835-INS-1313
chr16-49850925-INS-781
chr16-49864930-INS-132
chr16-49899208-INS-419
chr16-49939167-INS-125
chr16-49942811-INS-56
chr16-49946920-INS-58
chr16-49955396-INS-858
chr16-49969735-INS-54
chr16-49981385-INS-200
chr16-49989645-INS-58
chr16-49990415-INS-60
chr16-50077719-INS-1279
chr16-50103824-INS-108
chr16-50131487-INS-151
chr16-50144181-INS-2765
chr16-50199186-INS-199
chr16-50223750-INS-196
chr16-50242113-INS-216
chr16-50247993-INS-195
chr16-50264168-INS-55
chr16-50308590-INS-209
chr16-50315383-INS-111
chr16-50330457-INS-66
chr16-50352626-INS-90
chr16-50362895-INS-70
chr16-50364623-INS-60
chr16-50438658-INS-448
chr16-50458739-INS-197
chr16-50466475-INS-422
chr16-50468789-INS-64
chr16-50473107-INS-264
chr16-50483150-INS-172
chr16-50519049-INS-95
chr16-50578476-INS-322
chr16-50611378-INS-508
chr16-50623693-INS-216
chr16-50624912-INS-852
chr16-50639666-INS-340
chr16-50641693-INS-121
chr16-50650655-INS-358

chr16-50654615-INS-2968
chr16-50670243-INS-174
chr16-50679107-INS-454
chr16-50691292-INS-269
chr16-50698863-INS-65
chr16-50711001-INS-6622
chr16-50717687-INS-50
chr16-50721054-INS-2995
chr16-50724708-INS-6877
chr16-50726934-INS-185
chr16-50767161-INS-401
chr16-50783197-INS-1000
chr16-50788291-INS-121
chr16-50798217-INS-132
chr16-50818756-INS-65
chr16-50819781-INS-64
chr16-50898942-INS-6627
chr16-50924577-INS-6012
chr16-50926413-INS-88
chr16-50933680-INS-145
chr16-50937754-INS-74
chr16-50984116-INS-69
chr16-51005920-INS-204
chr16-51032669-INS-212
chr16-51050297-INS-6776
chr16-51052276-INS-407
chr16-51053716-INS-97
chr16-51055203-INS-124
chr16-51099141-INS-69
chr16-51117080-INS-1405
chr16-51122829-INS-127
chr16-51139545-INS-1041
chr16-51154187-INS-3511
chr16-51157784-INS-52
chr16-51164555-INS-414
chr16-51176117-INS-314
chr16-51178541-INS-72
chr16-51196066-INS-6756
chr16-51266987-INS-722
chr16-51286498-INS-654
chr16-51289714-INS-5352
chr16-51294020-INS-214
chr16-51294679-INS-6716
chr16-51301345-INS-65
chr16-51302437-INS-104
chr16-51303150-INS-1003
chr16-51303244-INS-7098
chr16-51314325-INS-675
chr16-51337888-INS-55
chr16-51346770-INS-145
chr16-51442441-INS-52
chr16-51465153-INS-60
chr16-51475098-INS-68
chr16-51480720-INS-914
chr16-51523908-INS-456
chr16-51525987-INS-149
chr16-51534173-INS-70
chr16-51538985-INS-84
chr16-51567443-INS-78
chr16-51600176-INS-185
chr16-51617423-INS-60
chr16-51635853-INS-68
chr16-51637698-INS-56
chr16-51660676-INS-644
chr16-51671689-INS-337
chr16-51704579-INS-6665
chr16-51710213-INS-281
chr16-51713157-INS-6148
chr16-51716714-INS-471
chr16-51747461-INS-98
chr16-51751364-INS-121
chr16-51784617-INS-3028
chr16-51804846-INS-56
chr16-51840085-INS-59
chr16-51858127-INS-55
chr16-51862211-INS-57
chr16-51864565-INS-98

chr16-51866894-INS-78
chr16-51887202-INS-198
chr16-51924366-INS-136
chr16-51962373-INS-136
chr16-51984466-INS-158
chr16-51996221-INS-64
chr16-52011553-INS-349
chr16-52040658-INS-191
chr16-52054405-INS-53
chr16-52085208-INS-75
chr16-52095960-INS-6694
chr16-52120327-INS-58
chr16-52124381-INS-3204
chr16-52141584-INS-6392
chr16-52162242-INS-63
chr16-52163941-INS-60
chr16-52168298-INS-379
chr16-52170166-INS-699
chr16-52185574-INS-2442
chr16-52199175-INS-57
chr16-52206136-INS-399
chr16-52224612-INS-81
chr16-52229892-INS-202
chr16-52235441-INS-200
chr16-52257570-INS-400
chr16-52286272-INS-6757
chr16-52293782-INS-408
chr16-52296507-INS-142
chr16-52369812-INS-5317
chr16-52401684-INS-171
chr16-52405687-INS-6330
chr16-52412713-INS-6208
chr16-52429932-INS-6452
chr16-52435049-INS-844
chr16-52478214-INS-858
chr16-52478891-INS-71
chr16-52567395-INS-394
chr16-52572353-INS-188
chr16-52583214-INS-6574
chr16-52593562-INS-60
chr16-52599868-INS-77
chr16-52603022-INS-161
chr16-52627967-INS-777
chr16-52630289-INS-209
chr16-52666370-INS-6368
chr16-52744864-INS-79
chr16-52770583-INS-187
chr16-52795459-INS-342
chr16-52828528-INS-339
chr16-52847426-INS-1607
chr16-52862386-INS-96
chr16-52864915-INS-432
chr16-52873332-INS-768
chr16-52886133-INS-372
chr16-52887110-INS-279
chr16-52887776-INS-758
chr16-52893553-INS-56
chr16-52912432-INS-62
chr16-52921309-INS-121
chr16-52922039-INS-6520
chr16-52924027-INS-6575
chr16-52927401-INS-270
chr16-52990156-INS-87
chr16-53025945-INS-63
chr16-53038516-INS-492
chr16-53060326-INS-484
chr16-53070829-INS-7122
chr16-53073967-INS-65
chr16-53082763-INS-62
chr16-53095169-INS-121
chr16-53102841-INS-292
chr16-53120342-INS-55
chr16-53124302-INS-56
chr16-53166082-INS-444
chr16-53168286-INS-208
chr16-53219670-INS-801
chr16-53220778-INS-528

chr16-53249453-INS-6396
chr16-53252893-INS-351
chr16-53253761-INS-59
chr16-53265949-INS-398
chr16-53296211-INS-6991
chr16-53334475-INS-61
chr16-53406027-INS-7206
chr16-53420052-INS-944
chr16-53480294-INS-787
chr16-53497779-INS-2514
chr16-53512534-INS-1191
chr16-53533234-INS-71
chr16-53537495-INS-5274
chr16-53580788-INS-135
chr16-53590062-INS-608
chr16-53603556-INS-847
chr16-53641204-INS-57
chr16-53688176-INS-156
chr16-53694796-INS-92
chr16-53704755-INS-194
chr16-53715508-INS-231
chr16-53747226-INS-138
chr16-53772201-INS-476
chr16-53775290-INS-112
chr16-53798733-INS-59
chr16-53847416-INS-171
chr16-53869799-INS-60
chr16-53898764-INS-77
chr16-53911315-INS-54
chr16-53922957-INS-69
chr16-53925795-INS-180
chr16-53931065-INS-6415
chr16-53938689-INS-68
chr16-53972329-INS-62
chr16-53984387-INS-702
chr16-53991155-INS-86
chr16-54007713-INS-52
chr16-54008911-INS-62
chr16-54034569-INS-171
chr16-54083899-INS-636
chr16-54086806-INS-251
chr16-54092960-INS-75
chr16-54132216-INS-5217
chr16-54146174-INS-68
chr16-54170952-INS-88
chr16-54177502-INS-6230
chr16-54199812-INS-6320
chr16-54203870-INS-248
chr16-54210662-INS-61
chr16-54224487-INS-4095
chr16-54225072-INS-527
chr16-54225807-INS-4328
chr16-54240010-INS-7173
chr16-54274425-INS-56
chr16-54276841-INS-117
chr16-54283600-INS-162
chr16-54337726-INS-1142
chr16-54350704-INS-871
chr16-54372612-INS-6216
chr16-54377525-INS-82
chr16-54421682-INS-227
chr16-54425410-INS-6540
chr16-54447258-INS-58
chr16-54516056-INS-187
chr16-54528418-INS-99
chr16-54544140-INS-53
chr16-54548753-INS-68
chr16-54616537-INS-181
chr16-54625621-INS-202
chr16-54651275-INS-7093
chr16-54660107-INS-573
chr16-54679710-INS-65
chr16-54680617-INS-1209
chr16-54687935-INS-212
chr16-54688358-INS-423
chr16-54692232-INS-6249
chr16-54694811-INS-52

chr16-54699297-INS-6506
chr16-54712515-INS-68
chr16-54719516-INS-608
chr16-54725087-INS-592
chr16-54738120-INS-95
chr16-54758709-INS-88
chr16-54802921-INS-6464
chr16-54820962-INS-105
chr16-54823400-INS-141
chr16-54852341-INS-60
chr16-54854939-INS-703
chr16-54857840-INS-4672
chr16-54858186-INS-842
chr16-54860698-INS-1987
chr16-54866901-INS-70
chr16-54890401-INS-543
chr16-54914461-INS-6747
chr16-54914933-INS-196
chr16-54923794-INS-278
chr16-54925839-INS-104
chr16-54933723-INS-6608
chr16-54947227-INS-496
chr16-54949908-INS-143
chr16-54957779-INS-53
chr16-54962807-INS-330
chr16-54997661-INS-56
chr16-55015015-INS-78
chr16-55057601-INS-229
chr16-55062340-INS-52
chr16-55068450-INS-252
chr16-55070591-INS-150
chr16-55096505-INS-76
chr16-55115630-INS-96
chr16-55126046-INS-133
chr16-55126911-INS-490
chr16-55137589-INS-104
chr16-55172168-INS-62
chr16-55180275-INS-132
chr16-55197680-INS-148
chr16-55204882-INS-68
chr16-55213544-INS-890
chr16-55215716-INS-84
chr16-55231979-INS-373
chr16-55235512-INS-498
chr16-55236597-INS-54
chr16-55237093-INS-6643
chr16-55238816-INS-449
chr16-55240310-INS-77
chr16-55242941-INS-1588
chr16-55251604-INS-51
chr16-55291630-INS-69
chr16-55297452-INS-269
chr16-55305339-INS-53
chr16-55307476-INS-6381
chr16-55312406-INS-8857
chr16-55315267-INS-141
chr16-55321622-INS-205
chr16-55329688-INS-454
chr16-55373911-INS-196
chr16-55381666-INS-441
chr16-55395204-INS-770
chr16-55406949-INS-55
chr16-55410567-INS-60
chr16-55422620-INS-6345
chr16-55452986-INS-156
chr16-55458438-INS-366
chr16-55489518-INS-222
chr16-55504334-INS-58
chr16-55508107-INS-3283
chr16-55514931-INS-455
chr16-55518136-INS-2822
chr16-55526294-INS-161
chr16-55530286-INS-141
chr16-55533703-INS-304
chr16-55534101-INS-304
chr16-55538585-INS-196
chr16-55549785-INS-386

chr16-55563541-INS-141
chr16-55581361-INS-51
chr16-55587342-INS-5690
chr16-55589547-INS-206
chr16-55594673-INS-127
chr16-55598176-INS-70
chr16-55614932-INS-303
chr16-55625897-INS-132
chr16-55639299-INS-942
chr16-55675406-INS-60
chr16-55689136-INS-194
chr16-55696326-INS-78
chr16-55708087-INS-392
chr16-55713429-INS-193
chr16-55718581-INS-177
chr16-55719983-INS-451
chr16-55729740-INS-63
chr16-55736952-INS-64
chr16-55744240-INS-74
chr16-55753970-INS-157
chr16-55766656-INS-52
chr16-55767033-INS-95
chr16-55768728-INS-366
chr16-55774929-INS-184
chr16-55785653-INS-184
chr16-55799390-INS-398
chr16-55806625-INS-51
chr16-55815782-INS-93
chr16-55876983-INS-770
chr16-55880107-INS-206
chr16-55886191-INS-209
chr16-55889671-INS-498
chr16-55901641-INS-199
chr16-55916945-INS-210
chr16-55918829-INS-57
chr16-55924827-INS-403
chr16-55955962-INS-95
chr16-55971995-INS-64
chr16-55973393-INS-665
chr16-55985211-INS-337
chr16-55994386-INS-188
chr16-56000000-INS-550
chr16-56029796-INS-261
chr16-56047522-INS-105
chr16-56051060-INS-72
chr16-56052641-INS-657
chr16-56055375-INS-93
chr16-56065532-INS-322
chr16-56085020-INS-391
chr16-56090718-INS-89
chr16-56101900-INS-55
chr16-56115278-INS-52
chr16-56115913-INS-93
chr16-56117336-INS-7523
chr16-56123237-INS-360
chr16-56133379-INS-283
chr16-56151979-INS-3174
chr16-56178400-INS-375
chr16-56181065-INS-60
chr16-56182466-INS-62
chr16-56188060-INS-1811
chr16-56195385-INS-287
chr16-56218961-INS-6047
chr16-56221288-INS-7099
chr16-56245976-INS-1589
chr16-56260956-INS-432
chr16-56264310-INS-6422
chr16-56277505-INS-180
chr16-56289319-INS-116
chr16-56302619-INS-400
chr16-56309103-INS-57
chr16-56319240-INS-745
chr16-56329494-INS-279
chr16-56347331-INS-152
chr16-56388665-INS-6202
chr16-56393828-INS-6809
chr16-56399552-INS-6899

chr16-56402534-INS-72
chr16-56405007-INS-3727
chr16-56408692-INS-6415
chr16-56425536-INS-57
chr16-56460729-INS-78
chr16-56464290-INS-252
chr16-56505185-INS-189
chr16-56520307-INS-4271
chr16-56536195-INS-5172
chr16-56538495-INS-1307
chr16-56540119-INS-432
chr16-56576010-INS-711
chr16-56582791-INS-168
chr16-56600740-INS-55
chr16-56617256-INS-558
chr16-56623854-INS-166
chr16-56628666-INS-66
chr16-56659835-INS-3912
chr16-56669296-INS-73
chr16-56679393-INS-6190
chr16-56680071-INS-60
chr16-56680542-INS-954
chr16-56685229-INS-63
chr16-56699035-INS-52
chr16-56704773-INS-6401
chr16-56740684-INS-7521
chr16-56743817-INS-88
chr16-56749355-INS-5148
chr16-56752495-INS-110
chr16-56753525-INS-372
chr16-56760957-INS-204
chr16-56763469-INS-356
chr16-56766502-INS-221
chr16-56769983-INS-1892
chr16-56787480-INS-254
chr16-56794081-INS-245
chr16-56794677-INS-196
chr16-56803618-INS-55
chr16-56808382-INS-140
chr16-56830094-INS-55
chr16-56830235-INS-1067
chr16-56833534-INS-56
chr16-56834548-INS-5545
chr16-56844521-INS-58
chr16-56858850-INS-996
chr16-56867107-INS-641
chr16-56873734-INS-507
chr16-56892199-INS-132
chr16-56893976-INS-287
chr16-56895890-INS-6445
chr16-56905268-INS-235
chr16-56917196-INS-414
chr16-56917706-INS-748
chr16-56930748-INS-122
chr16-56933631-INS-52
chr16-56941226-INS-996
chr16-56965936-INS-78
chr16-56988710-INS-498
chr16-56991384-INS-189
chr16-56992622-INS-200
chr16-56993840-INS-200
chr16-57013482-INS-244
chr16-57030977-INS-383
chr16-57055438-INS-274
chr16-57066313-INS-54
chr16-57077079-INS-51
chr16-57081676-INS-172
chr16-57083412-INS-7177
chr16-57087761-INS-74
chr16-57108101-INS-274
chr16-57112582-INS-178
chr16-57118025-INS-7082
chr16-57153319-INS-6686
chr16-57162703-INS-379
chr16-57172297-INS-273
chr16-57178497-INS-106
chr16-57188856-INS-370

chr16-57210450-INS-226
chr16-57228853-INS-66
chr16-57230236-INS-6200
chr16-57240706-INS-148
chr16-57253455-INS-2467
chr16-57262492-INS-54
chr16-57274884-INS-67
chr16-57319296-INS-53
chr16-57344093-INS-162
chr16-57346329-INS-259
chr16-57364112-INS-55
chr16-57415472-INS-85
chr16-57424858-INS-85
chr16-57430055-INS-109
chr16-57434076-INS-480
chr16-57436561-INS-94
chr16-57465816-INS-101
chr16-57493774-INS-72
chr16-57513797-INS-64
chr16-57542797-INS-323
chr16-57571201-INS-57
chr16-57592204-INS-372
chr16-57594752-INS-6589
chr16-57598529-INS-401
chr16-57605881-INS-103
chr16-57638329-INS-72
chr16-57652720-INS-2203
chr16-57653340-INS-465
chr16-57655874-INS-2258
chr16-57664261-INS-52
chr16-57666158-INS-70
chr16-57733710-INS-603
chr16-57747182-INS-179
chr16-57750514-INS-354
chr16-57759522-INS-75
chr16-57768012-INS-5267
chr16-57814164-INS-64
chr16-57861772-INS-6244
chr16-57903358-INS-992
chr16-57909371-INS-144
chr16-57921024-INS-53
chr16-57937309-INS-1336
chr16-57942790-INS-3416
chr16-57957060-INS-4157
chr16-57958802-INS-107
chr16-57976389-INS-232
chr16-57989512-INS-112
chr16-57995008-INS-159
chr16-58004224-INS-134
chr16-58027872-INS-83
chr16-58030324-INS-6769
chr16-58049254-INS-70
chr16-58052123-INS-71
chr16-58052689-INS-6929
chr16-58055667-INS-52
chr16-58056591-INS-6308
chr16-58066365-INS-159
chr16-58077713-INS-150
chr16-58089390-INS-235
chr16-58091854-INS-483
chr16-58095313-INS-2103
chr16-58096465-INS-392
chr16-58120856-INS-536
chr16-58148163-INS-82
chr16-58155583-INS-688
chr16-58176503-INS-230
chr16-58177952-INS-367
chr16-58181381-INS-143
chr16-58208314-INS-6624
chr16-58242845-INS-124
chr16-58257116-INS-92
chr16-58264875-INS-623
chr16-58305509-INS-130
chr16-58311507-INS-89
chr16-58328666-INS-146
chr16-58331521-INS-52
chr16-58356090-INS-894

chr16-58394384-INS-117
chr16-58401248-INS-873
chr16-58423352-INS-203
chr16-58446818-INS-68
chr16-58448824-INS-70
chr16-58451336-INS-204
chr16-58454225-INS-64
chr16-58454291-INS-630
chr16-58477847-INS-64
chr16-58496160-INS-56
chr16-58497139-INS-290
chr16-58498125-INS-1385
chr16-58504886-INS-66
chr16-58512546-INS-60
chr16-58526332-INS-253
chr16-58528290-INS-75
chr16-58543463-INS-1442
chr16-58557657-INS-55
chr16-58589326-INS-2924
chr16-58601368-INS-56
chr16-58604629-INS-6491
chr16-58607675-INS-6412
chr16-58619386-INS-70
chr16-58677807-INS-309
chr16-58702357-INS-554
chr16-58703718-INS-116
chr16-58709104-INS-201
chr16-58743785-INS-1909
chr16-58745840-INS-5301
chr16-58768375-INS-100
chr16-58795205-INS-62
chr16-58807750-INS-98
chr16-58811910-INS-59
chr16-58814100-INS-80
chr16-58817715-INS-122
chr16-58843885-INS-510
chr16-58862148-INS-226
chr16-58896480-INS-222
chr16-58930404-INS-6588
chr16-58935657-INS-418
chr16-58937114-INS-7025
chr16-58952901-INS-287
chr16-58962735-INS-211
chr16-58964032-INS-1582
chr16-58965895-INS-810
chr16-59013836-INS-100
chr16-59019245-INS-130
chr16-59023119-INS-831
chr16-59057733-INS-6754
chr16-59073275-INS-6381
chr16-59098968-INS-86
chr16-59105001-INS-10741
chr16-59129926-INS-312
chr16-59134137-INS-51
chr16-59224178-INS-7072
chr16-59252414-INS-52
chr16-59296459-INS-124
chr16-59317021-INS-891
chr16-59337200-INS-52
chr16-59344190-INS-405
chr16-59351699-INS-58
chr16-59360729-INS-163
chr16-59374157-INS-208
chr16-59396286-INS-84
chr16-59413028-INS-56
chr16-59431726-INS-57
chr16-59443820-INS-224
chr16-59445232-INS-238
chr16-59476365-INS-834
chr16-59479865-INS-4469
chr16-59496864-INS-770
chr16-59512762-INS-95
chr16-59514036-INS-208
chr16-59514464-INS-607
chr16-59516814-INS-3324
chr16-59529972-INS-70
chr16-59574890-INS-435

chr16-59580491-INS-1348
chr16-59591902-INS-6186
chr16-59630379-INS-7364
chr16-59656770-INS-448
chr16-59660143-INS-64
chr16-59700251-INS-7316
chr16-59700659-INS-65
chr16-59704142-INS-241
chr16-59706763-INS-963
chr16-59710610-INS-165
chr16-59733988-INS-992
chr16-59743118-INS-98
chr16-59744880-INS-655
chr16-59753690-INS-624
chr16-59772736-INS-275
chr16-59790659-INS-6481
chr16-59809475-INS-489
chr16-59813241-INS-7123
chr16-59814418-INS-57
chr16-59825281-INS-644
chr16-59856196-INS-469
chr16-59866459-INS-638
chr16-59869723-INS-497
chr16-59881677-INS-592
chr16-59896072-INS-203
chr16-59900333-INS-210
chr16-59906572-INS-132
chr16-59909052-INS-365
chr16-59920147-INS-57
chr16-59943306-INS-243
chr16-59955814-INS-390
chr16-59970757-INS-268
chr16-59975042-INS-6858
chr16-59982296-INS-177
chr16-59985463-INS-51
chr16-59993150-INS-66
chr16-59996376-INS-985
chr16-60024259-INS-883
chr16-60028202-INS-639
chr16-60033337-INS-475
chr16-60063280-INS-109
chr16-60069980-INS-138
chr16-60103026-INS-52
chr16-60118125-INS-174
chr16-60119303-INS-225
chr16-60121584-INS-81
chr16-60132345-INS-78
chr16-60136679-INS-76
chr16-60154665-INS-572
chr16-60187401-INS-57
chr16-60190545-INS-424
chr16-60191259-INS-499
chr16-60193814-INS-52
chr16-60208870-INS-428
chr16-60225990-INS-1054
chr16-60232123-INS-143
chr16-60233688-INS-52
chr16-60236997-INS-184
chr16-60310456-INS-184
chr16-60323556-INS-141
chr16-60368128-INS-272
chr16-60381019-INS-507
chr16-60398223-INS-53
chr16-60412858-INS-135
chr16-60419975-INS-174
chr16-60457596-INS-819
chr16-60467405-INS-781
chr16-60479242-INS-64
chr16-60525444-INS-285
chr16-60543140-INS-2062
chr16-60601004-INS-234
chr16-60623597-INS-189
chr16-60627457-INS-133
chr16-60632794-INS-341
chr16-60638230-INS-82
chr16-60660059-INS-83
chr16-60700112-INS-66

chr16-60725067-INS-270
chr16-60727076-INS-991
chr16-60737331-INS-94
chr16-60768781-INS-178
chr16-60815259-INS-417
chr16-60818471-INS-69
chr16-60819824-INS-104
chr16-60832033-INS-80
chr16-60837336-INS-187
chr16-60856701-INS-67
chr16-60866616-INS-96
chr16-60876685-INS-460
chr16-60900239-INS-64
chr16-60915404-INS-56
chr16-60929660-INS-928
chr16-60942653-INS-2869
chr16-60942679-INS-1861
chr16-60961246-INS-1955
chr16-61017192-INS-65
chr16-61018575-INS-416
chr16-61020558-INS-62
chr16-61040279-INS-74
chr16-61066594-INS-1358
chr16-61094873-INS-4535
chr16-61102265-INS-262
chr16-61138292-INS-98
chr16-61140293-INS-104
chr16-61143057-INS-445
chr16-61143831-INS-376
chr16-61177530-INS-374
chr16-61195591-INS-308
chr16-61207530-INS-418
chr16-61223584-INS-427
chr16-61233122-INS-6700
chr16-61242913-INS-3301
chr16-61250621-INS-1107
chr16-61264350-INS-109
chr16-61271125-INS-105
chr16-61280001-INS-319
chr16-61289302-INS-230
chr16-61314543-INS-52
chr16-61316424-INS-188
chr16-61330524-INS-190
chr16-61334789-INS-104
chr16-61339844-INS-66
chr16-61392149-INS-414
chr16-61395424-INS-123
chr16-61417074-INS-108
chr16-61432306-INS-400
chr16-61437485-INS-195
chr16-61458565-INS-891
chr16-61465315-INS-51
chr16-61468299-INS-74
chr16-61503958-INS-97
chr16-61527460-INS-629
chr16-61547848-INS-104
chr16-61583757-INS-353
chr16-61616627-INS-358
chr16-61621575-INS-337
chr16-61665245-INS-329
chr16-61668572-INS-6249
chr16-61701121-INS-141
chr16-61719118-INS-52
chr16-61720008-INS-6315
chr16-61820724-INS-281
chr16-61857523-INS-845
chr16-61859023-INS-54
chr16-61909023-INS-702
chr16-61928797-INS-6365
chr16-61944467-INS-6481
chr16-61945095-INS-156
chr16-61945705-INS-470
chr16-61951188-INS-7165
chr16-61981916-INS-54
chr16-61995908-INS-149
chr16-62044394-INS-542
chr16-62046895-INS-464

chr16-62051699-INS-120
chr16-62062497-INS-612
chr16-62074944-INS-776
chr16-62085188-INS-478
chr16-62147434-INS-87
chr16-62192772-INS-515
chr16-62206008-INS-84
chr16-62231799-INS-54
chr16-62261866-INS-1000
chr16-62265496-INS-72
chr16-62276901-INS-451
chr16-62283880-INS-2802
chr16-62288719-INS-978
chr16-62295163-INS-4378
chr16-62320186-INS-56
chr16-62322327-INS-431
chr16-62349022-INS-497
chr16-62363990-INS-54
chr16-62371562-INS-555
chr16-62390768-INS-212
chr16-62419117-INS-2137
chr16-62426985-INS-128
chr16-62466084-INS-302
chr16-62466916-INS-59
chr16-62468031-INS-52
chr16-62483210-INS-112
chr16-62504889-INS-697
chr16-62505673-INS-6596
chr16-62506268-INS-5271
chr16-62513358-INS-773
chr16-62520599-INS-6052
chr16-62535650-INS-188
chr16-62542900-INS-4148
chr16-62561411-INS-208
chr16-62578172-INS-51
chr16-62583628-INS-76
chr16-62583872-INS-7076
chr16-62588586-INS-6797
chr16-62593623-INS-150
chr16-62604936-INS-580
chr16-62683775-INS-442
chr16-62704793-INS-627
chr16-62717179-INS-204
chr16-62718293-INS-458
chr16-62720285-INS-86
chr16-62740293-INS-6746
chr16-62753402-INS-52
chr16-62783563-INS-162
chr16-62786205-INS-76
chr16-62789170-INS-550
chr16-62791096-INS-6673
chr16-62795607-INS-413
chr16-62798111-INS-6738
chr16-62800176-INS-269
chr16-62809196-INS-331
chr16-62812273-INS-6427
chr16-62827203-INS-2539
chr16-62834504-INS-162
chr16-62856880-INS-76
chr16-62866660-INS-62
chr16-62868781-INS-423
chr16-62875517-INS-7695
chr16-62877129-INS-489
chr16-62877788-INS-59
chr16-62880090-INS-337
chr16-62917515-INS-78
chr16-62923931-INS-65
chr16-62944618-INS-60
chr16-62965820-INS-243
chr16-62979255-INS-939
chr16-62982943-INS-60
chr16-62985125-INS-388
chr16-63027773-INS-61
chr16-63029497-INS-8988
chr16-63030483-INS-6181
chr16-63033127-INS-88
chr16-63065135-INS-7078

chr16-63067214-INS-212
chr16-63073885-INS-566
chr16-63096278-INS-6470
chr16-63098357-INS-120
chr16-63118733-INS-72
chr16-63130433-INS-271
chr16-63149571-INS-108
chr16-63158306-INS-81
chr16-63168987-INS-113
chr16-63176972-INS-356
chr16-63177926-INS-59
chr16-63185674-INS-279
chr16-63242757-INS-361
chr16-63252817-INS-549
chr16-63266532-INS-1860
chr16-63279054-INS-98
chr16-63282916-INS-555
chr16-63326005-INS-220
chr16-63369477-INS-95
chr16-63386210-INS-711
chr16-63446893-INS-861
chr16-63453726-INS-578
chr16-63468208-INS-176
chr16-63532873-INS-243
chr16-63537375-INS-86
chr16-63554010-INS-242
chr16-63589429-INS-78
chr16-63626435-INS-1785
chr16-63637011-INS-52
chr16-63638737-INS-288
chr16-63643319-INS-234
chr16-63654867-INS-86
chr16-63656041-INS-314
chr16-63658006-INS-80
chr16-63686334-INS-66
chr16-63705561-INS-54
chr16-63713867-INS-295
chr16-63734818-INS-208
chr16-63770024-INS-80
chr16-63782601-INS-52
chr16-63786177-INS-439
chr16-63863742-INS-108
chr16-63873500-INS-315
chr16-63881254-INS-100
chr16-63885242-INS-128
chr16-63903146-INS-285
chr16-63920554-INS-54
chr16-63927558-INS-254
chr16-63983331-INS-704
chr16-63986240-INS-269
chr16-64015090-INS-500
chr16-64028794-INS-507
chr16-64064943-INS-104
chr16-64068700-INS-7264
chr16-64069742-INS-997
chr16-64081282-INS-232
chr16-64088535-INS-144
chr16-64090059-INS-7118
chr16-64099299-INS-1227
chr16-64122858-INS-125
chr16-64154734-INS-95
chr16-64160641-INS-364
chr16-64196300-INS-148
chr16-64200643-INS-55
chr16-64214150-INS-303
chr16-64221116-INS-7115
chr16-64223402-INS-416
chr16-64226184-INS-99
chr16-64250996-INS-669
chr16-64261859-INS-54
chr16-64293054-INS-84
chr16-64314044-INS-87
chr16-64316410-INS-60
chr16-64329578-INS-72
chr16-64354279-INS-309
chr16-64357421-INS-253
chr16-64378785-INS-274

chr16-64380673-INS-71
chr16-64388826-INS-6746
chr16-64409075-INS-61
chr16-64429862-INS-136
chr16-64446853-INS-216
chr16-64489437-INS-56
chr16-64493712-INS-83
chr16-64494544-INS-55
chr16-64513877-INS-238
chr16-64521634-INS-69
chr16-64523471-INS-372
chr16-64535195-INS-6690
chr16-64543533-INS-102
chr16-64544594-INS-52
chr16-64548614-INS-86
chr16-64562139-INS-77
chr16-64573319-INS-317
chr16-64579552-INS-3010
chr16-64604561-INS-66
chr16-64665754-INS-427
chr16-64683293-INS-81
chr16-64697280-INS-498
chr16-64704070-INS-5590
chr16-64710393-INS-390
chr16-64719756-INS-58
chr16-64732929-INS-104
chr16-64736057-INS-197
chr16-64747852-INS-187
chr16-64749457-INS-56
chr16-64749762-INS-496
chr16-64754949-INS-948
chr16-64754990-INS-62
chr16-64771467-INS-208
chr16-64776626-INS-570
chr16-64813615-INS-207
chr16-64822258-INS-814
chr16-64822384-INS-2559
chr16-64828503-INS-544
chr16-64833795-INS-132
chr16-64841879-INS-9127
chr16-64848599-INS-54
chr16-64857251-INS-190
chr16-64859737-INS-104
chr16-64889503-INS-201
chr16-64903638-INS-66
chr16-64910294-INS-110
chr16-64922970-INS-232
chr16-64929367-INS-52
chr16-64933149-INS-62
chr16-64936162-INS-154
chr16-64970795-INS-82
chr16-64972347-INS-70
chr16-64994390-INS-56
chr16-65007817-INS-237
chr16-65009120-INS-58
chr16-65013637-INS-776
chr16-65105499-INS-54
chr16-65163961-INS-69
chr16-65165844-INS-74
chr16-65180428-INS-58
chr16-65181296-INS-347
chr16-65195886-INS-138
chr16-65196694-INS-721
chr16-65199699-INS-187
chr16-65203199-INS-409
chr16-65206252-INS-448
chr16-65207126-INS-101
chr16-65208134-INS-993
chr16-65230569-INS-218
chr16-65235424-INS-640
chr16-65247443-INS-340
chr16-65270704-INS-66
chr16-65276102-INS-485
chr16-65305911-INS-70
chr16-65379404-INS-141
chr16-65381511-INS-327
chr16-65410349-INS-781

chr16-65423201-INS-93
chr16-65432258-INS-107
chr16-65437340-INS-261
chr16-65438755-INS-52
chr16-65442629-INS-75
chr16-65446493-INS-5205
chr16-65448708-INS-402
chr16-65495856-INS-6999
chr16-65500862-INS-2284
chr16-65513154-INS-56
chr16-65516293-INS-3226
chr16-65533297-INS-407
chr16-65565262-INS-205
chr16-65574209-INS-286
chr16-65619295-INS-551
chr16-65620085-INS-166
chr16-65653498-INS-96
chr16-65707533-INS-116
chr16-65708932-INS-381
chr16-65737192-INS-7128
chr16-65741921-INS-54
chr16-65756758-INS-182
chr16-65758718-INS-58
chr16-65766681-INS-142
chr16-65774560-INS-489
chr16-65779992-INS-51
chr16-65802892-INS-74
chr16-65826824-INS-1886
chr16-65827597-INS-149
chr16-65828032-INS-5495
chr16-65831922-INS-497
chr16-65876449-INS-5289
chr16-65889134-INS-62
chr16-65893262-INS-51
chr16-65896180-INS-155
chr16-65921250-INS-146
chr16-66015033-INS-4308
chr16-66018731-INS-861
chr16-66051646-INS-6442
chr16-66054709-INS-151
chr16-66077155-INS-52
chr16-66101680-INS-479
chr16-66104026-INS-52
chr16-66106646-INS-6351
chr16-66117152-INS-5549
chr16-66148406-INS-549
chr16-66162429-INS-354
chr16-66170323-INS-76
chr16-66182567-INS-59
chr16-66214284-INS-94
chr16-66221663-INS-933
chr16-66235376-INS-399
chr16-66294681-INS-71
chr16-66311123-INS-110
chr16-66312342-INS-229
chr16-66323558-INS-6430
chr16-66324814-INS-55
chr16-66352571-INS-691
chr16-66358903-INS-180
chr16-66362536-INS-202
chr16-66374851-INS-105
chr16-66386841-INS-1627
chr16-66402049-INS-118
chr16-66495057-INS-587
chr16-66516385-INS-180
chr16-66518002-INS-60
chr16-66534581-INS-386
chr16-66560367-INS-53
chr16-66624791-INS-199
chr16-66637759-INS-259
chr16-66642735-INS-7352
chr16-66647108-INS-484
chr16-66656537-INS-201
chr16-66662109-INS-112
chr16-66667342-INS-101
chr16-66674464-INS-805
chr16-66686444-INS-405

chr16-66711203-INS-181
chr16-66734507-INS-401
chr16-66789892-INS-251
chr16-66791146-INS-291
chr16-66792018-INS-770
chr16-66810422-INS-92
chr16-66830288-INS-528
chr16-66855600-INS-91
chr16-66861787-INS-283
chr16-66861990-INS-487
chr16-66864251-INS-388
chr16-66865441-INS-62
chr16-66925968-INS-86
chr16-66937288-INS-523
chr16-66945404-INS-78
chr16-66982342-INS-459
chr16-66982953-INS-136
chr16-66993228-INS-506
chr16-66999756-INS-63
chr16-67002068-INS-51
chr16-67050400-INS-182
chr16-67139634-INS-75
chr16-67145083-INS-7123
chr16-67160575-INS-102
chr16-67175897-INS-450
chr16-67190279-INS-62
chr16-67195541-INS-720
chr16-67218272-INS-186
chr16-67237961-INS-500
chr16-67241976-INS-93
chr16-67244276-INS-165
chr16-67276553-INS-57
chr16-67309567-INS-506
chr16-67381536-INS-52
chr16-67392168-INS-593
chr16-67396112-INS-121
chr16-67414941-INS-62
chr16-67438779-INS-212
chr16-67449341-INS-87
chr16-67457179-INS-761
chr16-67459997-INS-183
chr16-67499507-INS-194
chr16-67508012-INS-598
chr16-67518634-INS-557
chr16-67519760-INS-141
chr16-67543201-INS-119
chr16-67544184-INS-56
chr16-67548637-INS-840
chr16-67549677-INS-990
chr16-67576148-INS-72
chr16-67580919-INS-51
chr16-67605394-INS-6785
chr16-67639506-INS-52
chr16-67678138-INS-56
chr16-67692843-INS-971
chr16-67694443-INS-53
chr16-67792382-INS-6427
chr16-67800940-INS-6725
chr16-67810233-INS-323
chr16-67823387-INS-74
chr16-67857159-INS-422
chr16-67862644-INS-1044
chr16-67890565-INS-533
chr16-67897463-INS-312
chr16-67905909-INS-256
chr16-67958145-INS-78
chr16-67967336-INS-100
chr16-68013542-INS-952
chr16-68029346-INS-312
chr16-68052703-INS-3147
chr16-68080260-INS-72
chr16-68104236-INS-104
chr16-68114473-INS-195
chr16-68152958-INS-129
chr16-68154163-INS-6830
chr16-68188207-INS-56
chr16-68202489-INS-96

chr16-68220277-INS-1563
chr16-68222654-INS-56
chr16-68247828-INS-176
chr16-68252848-INS-190
chr16-68266147-INS-75
chr16-68278328-INS-567
chr16-68299325-INS-51
chr16-68311024-INS-376
chr16-68320531-INS-52
chr16-68334958-INS-959
chr16-68351778-INS-799
chr16-68353640-INS-68
chr16-68373280-INS-54
chr16-68398291-INS-113
chr16-68412150-INS-251
chr16-68430563-INS-78
chr16-68442115-INS-178
chr16-68444446-INS-211
chr16-68447914-INS-400
chr16-68483385-INS-90
chr16-68535013-INS-1043
chr16-68542730-INS-54
chr16-68557994-INS-116
chr16-68598963-INS-639
chr16-68602315-INS-323
chr16-68616786-INS-339
chr16-68647993-INS-67
chr16-68661949-INS-604
chr16-68666726-INS-53
chr16-68670615-INS-99
chr16-68690781-INS-6813
chr16-68749642-INS-4022
chr16-68756298-INS-54
chr16-68806382-INS-57
chr16-68827259-INS-1661
chr16-69055219-INS-347
chr16-69074215-INS-58
chr16-69086414-INS-373
chr16-69115404-INS-70
chr16-69195946-INS-177
chr16-69197603-INS-63
chr16-69226764-INS-104
chr16-69235788-INS-176
chr16-69285602-INS-76
chr16-69308157-INS-2803
chr16-69329190-INS-755
chr16-69333291-INS-6994
chr16-69339757-INS-78
chr16-69341010-INS-387
chr16-69363084-INS-64
chr16-69375078-INS-96
chr16-69377378-INS-90
chr16-69393392-INS-133
chr16-69395734-INS-108
chr16-69401221-INS-466
chr16-69434723-INS-219
chr16-69451344-INS-975
chr16-69457771-INS-166
chr16-69474575-INS-81
chr16-69478900-INS-58
chr16-69488146-INS-629
chr16-69489693-INS-89
chr16-69493410-INS-432
chr16-69496849-INS-1375
chr16-69504824-INS-711
chr16-69507392-INS-212
chr16-69516682-INS-60
chr16-69525490-INS-60
chr16-69541832-INS-90
chr16-69556201-INS-98
chr16-69583126-INS-110
chr16-69605478-INS-72
chr16-69635514-INS-53
chr16-69649586-INS-104
chr16-69690167-INS-212
chr16-69700202-INS-62
chr16-69710823-INS-78

chr16-69725062-INS-6963
chr16-69732908-INS-60
chr16-69758615-INS-118
chr16-69772507-INS-8081
chr16-69774334-INS-328
chr16-69785256-INS-136
chr16-69785863-INS-188
chr16-69788314-INS-114
chr16-69791516-INS-62
chr16-69806137-INS-334
chr16-69807300-INS-76
chr16-69812434-INS-68
chr16-69816253-INS-120
chr16-69820588-INS-475
chr16-69845981-INS-449
chr16-69872372-INS-504
chr16-69879878-INS-106
chr16-69884074-INS-8419
chr16-69887350-INS-374
chr16-69889716-INS-175
chr16-69921063-INS-178
chr16-69943063-INS-336
chr16-69944192-INS-64
chr16-69962191-INS-95
chr16-69974928-INS-3094
chr16-69975432-INS-56
chr16-69994335-INS-296
chr16-69995672-INS-80
chr16-70017568-INS-2854
chr16-70022461-INS-11829
chr16-70028094-INS-169
chr16-70034612-INS-738
chr16-70038763-INS-329
chr16-70057131-INS-303
chr16-70065248-INS-56
chr16-70080149-INS-389
chr16-70095770-INS-163
chr16-70098449-INS-146
chr16-70098453-INS-57
chr16-70115740-INS-183
chr16-70125958-INS-736
chr16-70135191-INS-60
chr16-70139558-INS-6438
chr16-70155293-INS-56
chr16-70202450-INS-55
chr16-70218498-INS-56
chr16-70222120-INS-517
chr16-70277314-INS-93
chr16-70280465-INS-53
chr16-70297169-INS-943
chr16-70297533-INS-56
chr16-70312558-INS-415
chr16-70314778-INS-531
chr16-70323194-INS-445
chr16-70330794-INS-251
chr16-70355308-INS-724
chr16-70365109-INS-56
chr16-70368862-INS-626
chr16-70406909-INS-5398
chr16-70427988-INS-60
chr16-70457522-INS-227
chr16-70475878-INS-61
chr16-70513261-INS-1377
chr16-70554720-INS-722
chr16-70586384-INS-59
chr16-70589675-INS-715
chr16-70600693-INS-68
chr16-70614928-INS-201
chr16-70622647-INS-610
chr16-70648418-INS-198
chr16-70651514-INS-578
chr16-70656514-INS-941
chr16-70668655-INS-425
chr16-70669487-INS-129
chr16-70670057-INS-403
chr16-70672923-INS-371
chr16-70707282-INS-6713

chr16-70718903-INS-495
chr16-70751707-INS-229
chr16-70786122-INS-188
chr16-70815300-INS-135
chr16-70830445-INS-6293
chr16-70868054-INS-70
chr16-70878777-INS-798
chr16-70897250-INS-96
chr16-70899804-INS-906
chr16-70908707-INS-390
chr16-70920066-INS-102
chr16-70958635-INS-158
chr16-70996775-INS-289
chr16-71014643-INS-99
chr16-71022843-INS-104
chr16-71024542-INS-3679
chr16-71035768-INS-54
chr16-71038150-INS-493
chr16-71041907-INS-537
chr16-71065655-INS-83
chr16-71081628-INS-386
chr16-71084008-INS-63
chr16-71096247-INS-243
chr16-71103552-INS-7010
chr16-71120734-INS-196
chr16-71122774-INS-452
chr16-71129170-INS-84
chr16-71134560-INS-58
chr16-71153501-INS-611
chr16-71177159-INS-208
chr16-71183314-INS-444
chr16-71187049-INS-613
chr16-71200173-INS-147
chr16-71203972-INS-4705
chr16-71205071-INS-6832
chr16-71206252-INS-6284
chr16-71206512-INS-857
chr16-71229095-INS-311
chr16-71235714-INS-144
chr16-71239635-INS-132
chr16-71247139-INS-337
chr16-71276052-INS-6435
chr16-71301837-INS-365
chr16-71303395-INS-503
chr16-71305554-INS-198
chr16-71324895-INS-6441
chr16-71379208-INS-67
chr16-71410562-INS-111
chr16-71412025-INS-582
chr16-71443548-INS-52
chr16-71449274-INS-70
chr16-71504047-INS-482
chr16-71512956-INS-131
chr16-71524598-INS-182
chr16-71567339-INS-744
chr16-71567926-INS-82
chr16-71575163-INS-274
chr16-71596893-INS-97
chr16-71601121-INS-6609
chr16-71603106-INS-124
chr16-71605731-INS-156
chr16-71616624-INS-124
chr16-71624034-INS-347
chr16-71626211-INS-101
chr16-71633741-INS-237
chr16-71636784-INS-60
chr16-71641207-INS-1093
chr16-71663455-INS-76
chr16-71708594-INS-461
chr16-71735737-INS-174
chr16-71752203-INS-235
chr16-71755947-INS-545
chr16-71762410-INS-3330
chr16-71763947-INS-166
chr16-71768076-INS-6648
chr16-71786333-INS-109
chr16-71787308-INS-76

chr16-71800669-INS-886
chr16-71823176-INS-51
chr16-71839439-INS-86
chr16-71862311-INS-94
chr16-71895071-INS-540
chr16-71911527-INS-579
chr16-71953775-INS-380
chr16-71954511-INS-512
chr16-71986923-INS-208
chr16-71994282-INS-62
chr16-72025220-INS-61
chr16-72031658-INS-974
chr16-72042108-INS-56
chr16-72051060-INS-111
chr16-72053794-INS-404
chr16-72054425-INS-642
chr16-72054434-INS-52
chr16-72067328-INS-6437
chr16-72114976-INS-67
chr16-72123079-INS-54
chr16-72147475-INS-85
chr16-72155101-INS-52
chr16-72158051-INS-235
chr16-72174745-INS-187
chr16-72186664-INS-232
chr16-72189436-INS-129
chr16-72206651-INS-185
chr16-72219072-INS-58
chr16-72240237-INS-5301
chr16-72259641-INS-138
chr16-72261423-INS-67
chr16-72269060-INS-57
chr16-72274972-INS-108
chr16-72280663-INS-51
chr16-72290986-INS-98
chr16-72306081-INS-366
chr16-72311809-INS-84
chr16-72323358-INS-722
chr16-72333379-INS-3219
chr16-72335338-INS-117
chr16-72346196-INS-84
chr16-72353052-INS-52
chr16-72366797-INS-5127
chr16-72373365-INS-158
chr16-72377982-INS-58
chr16-72380793-INS-6940
chr16-72385194-INS-174
chr16-72431929-INS-132
chr16-72454216-INS-6649
chr16-72472741-INS-64
chr16-72491300-INS-138
chr16-72495610-INS-209
chr16-72499153-INS-96
chr16-72501797-INS-842
chr16-72507707-INS-59
chr16-72543251-INS-203
chr16-72562832-INS-810
chr16-72602422-INS-144
chr16-72612398-INS-65
chr16-72619290-INS-140
chr16-72634953-INS-201
chr16-72657448-INS-109
chr16-72664119-INS-469
chr16-72667799-INS-64
chr16-72678819-INS-189
chr16-72681449-INS-477
chr16-72705859-INS-1895
chr16-72750056-INS-709
chr16-72762559-INS-51
chr16-72777559-INS-54
chr16-72815974-INS-1901
chr16-72817193-INS-175
chr16-72818903-INS-105
chr16-72865236-INS-96
chr16-72868296-INS-132
chr16-72925989-INS-367
chr16-72927279-INS-185

chr16-72931813-INS-54
chr16-72958203-INS-533
chr16-72961981-INS-1781
chr16-72975347-INS-169
chr16-72984981-INS-62
chr16-73013440-INS-216
chr16-73017390-INS-6399
chr16-73024167-INS-903
chr16-73038451-INS-63
chr16-73069441-INS-84
chr16-73088011-INS-234
chr16-73094883-INS-533
chr16-73096712-INS-166
chr16-73104683-INS-137
chr16-73110671-INS-80
chr16-73120396-INS-507
chr16-73132653-INS-4344
chr16-73137049-INS-354
chr16-73176609-INS-123
chr16-73235054-INS-56
chr16-73241816-INS-66
chr16-73244065-INS-60
chr16-73246502-INS-77
chr16-73270195-INS-743
chr16-73277321-INS-97
chr16-73286147-INS-68
chr16-73305193-INS-112
chr16-73317948-INS-52
chr16-73320049-INS-158
chr16-73320592-INS-395
chr16-73342866-INS-6522
chr16-73355383-INS-173
chr16-73371681-INS-51
chr16-73373963-INS-530
chr16-73391410-INS-68
chr16-73394327-INS-292
chr16-73422764-INS-207
chr16-73444806-INS-107
chr16-73445438-INS-319
chr16-73446584-INS-8825
chr16-73446744-INS-97
chr16-73463267-INS-87
chr16-73469145-INS-70
chr16-73469353-INS-6207
chr16-73493727-INS-2826
chr16-73498310-INS-266
chr16-73510724-INS-212
chr16-73516989-INS-6590
chr16-73519322-INS-564
chr16-73533842-INS-212
chr16-73536057-INS-249
chr16-73539048-INS-158
chr16-73539990-INS-90
chr16-73576310-INS-468
chr16-73609592-INS-6692
chr16-73612449-INS-6875
chr16-73645249-INS-88
chr16-73662945-INS-84
chr16-73667992-INS-470
chr16-73723358-INS-78
chr16-73745411-INS-648
chr16-73812259-INS-372
chr16-73814385-INS-230
chr16-73819443-INS-68
chr16-73850042-INS-59
chr16-73880989-INS-79
chr16-73900746-INS-790
chr16-73904075-INS-123
chr16-73905295-INS-130
chr16-73922691-INS-519
chr16-73923703-INS-279
chr16-73925131-INS-550
chr16-73934199-INS-162
chr16-73944572-INS-1893
chr16-73967704-INS-56
chr16-73971291-INS-370
chr16-74030787-INS-55

chr16-74039646-INS-51
chr16-74074951-INS-57
chr16-74086916-INS-130
chr16-74128953-INS-623
chr16-74193014-INS-176
chr16-74218599-INS-5549
chr16-74227903-INS-124
chr16-74236313-INS-349
chr16-74245907-INS-58
chr16-74248559-INS-78
chr16-74250670-INS-183
chr16-74267762-INS-488
chr16-74286696-INS-261
chr16-74353563-INS-69
chr16-74381299-INS-54
chr16-74383132-INS-209
chr16-74395854-INS-104
chr16-74406552-INS-135
chr16-74417900-INS-144
chr16-74449453-INS-57
chr16-74478061-INS-95
chr16-74496650-INS-1887
chr16-74504524-INS-110
chr16-74522595-INS-446
chr16-74550324-INS-547
chr16-74564892-INS-5244
chr16-74580077-INS-100
chr16-74585395-INS-7498
chr16-74617735-INS-92
chr16-74621594-INS-195
chr16-74645638-INS-71
chr16-74653928-INS-121
chr16-74658571-INS-263
chr16-74735692-INS-157
chr16-74741892-INS-179
chr16-74758472-INS-1888
chr16-74782939-INS-7563
chr16-74788176-INS-159
chr16-74790245-INS-339
chr16-74824563-INS-52
chr16-74846080-INS-1929
chr16-74847446-INS-54
chr16-74855942-INS-92
chr16-74880423-INS-163
chr16-74905234-INS-1465
chr16-74907441-INS-52
chr16-74941859-INS-240
chr16-74972902-INS-134
chr16-74978038-INS-1339
chr16-74986314-INS-164
chr16-74997687-INS-184
chr16-75001667-INS-557
chr16-75023425-INS-175
chr16-75024403-INS-108
chr16-75075720-INS-196
chr16-75078576-INS-497
chr16-75085990-INS-72
chr16-75097268-INS-285
chr16-75102111-INS-114
chr16-75105782-INS-62
chr16-75148316-INS-186
chr16-75173093-INS-78
chr16-75190704-INS-216
chr16-75224434-INS-84
chr16-75256474-INS-125
chr16-75261451-INS-234
chr16-75282938-INS-60
chr16-75315237-INS-7747
chr16-75335258-INS-64
chr16-75346052-INS-72
chr16-75361847-INS-71
chr16-75372961-INS-140
chr16-75382343-INS-89
chr16-75390262-INS-282
chr16-75392500-INS-344
chr16-75410162-INS-6983
chr16-75430502-INS-612

chr16-75461512-INS-110
chr16-75470308-INS-183
chr16-75505241-INS-420
chr16-75507459-INS-230
chr16-75515200-INS-66
chr16-75531858-INS-485
chr16-75548475-INS-864
chr16-75550815-INS-146
chr16-75551867-INS-292
chr16-75578727-INS-90
chr16-75579869-INS-400
chr16-75602923-INS-222
chr16-75653914-INS-497
chr16-75668694-INS-60
chr16-75675162-INS-94
chr16-75679802-INS-1224
chr16-75688511-INS-1892
chr16-75699463-INS-168
chr16-75719848-INS-253
chr16-75740341-INS-137
chr16-75741800-INS-66
chr16-75756811-INS-6536
chr16-75769896-INS-6818
chr16-75800166-INS-689
chr16-75816510-INS-143
chr16-75832237-INS-71
chr16-75866260-INS-106
chr16-75870485-INS-400
chr16-75894796-INS-232
chr16-75921106-INS-474
chr16-75984433-INS-7086
chr16-75990978-INS-193
chr16-76013488-INS-100
chr16-76033794-INS-64
chr16-76033899-INS-488
chr16-76041765-INS-260
chr16-76054109-INS-231
chr16-76058135-INS-176
chr16-76073070-INS-81
chr16-76083890-INS-180
chr16-76122202-INS-76
chr16-76153641-INS-54
chr16-76174060-INS-884
chr16-76206161-INS-147
chr16-76237139-INS-52
chr16-76246528-INS-164
chr16-76253489-INS-144
chr16-76281491-INS-94
chr16-76309405-INS-108
chr16-76325954-INS-69
chr16-76328315-INS-77
chr16-76357834-INS-1353
chr16-76357988-INS-136
chr16-76367064-INS-70
chr16-76367587-INS-72
chr16-76369208-INS-60
chr16-76378377-INS-106
chr16-76401156-INS-414
chr16-76411993-INS-52
chr16-76432227-INS-634
chr16-76442153-INS-126
chr16-76448691-INS-3041
chr16-76455948-INS-873
chr16-76486791-INS-216
chr16-76504410-INS-512
chr16-76507110-INS-7483
chr16-76529616-INS-119
chr16-76563301-INS-254
chr16-76585935-INS-424
chr16-76596781-INS-152
chr16-76599460-INS-97
chr16-76603621-INS-201
chr16-76641930-INS-72
chr16-76656537-INS-67
chr16-76656621-INS-652
chr16-76662546-INS-66
chr16-76687571-INS-404

chr16-76704675-INS-56
chr16-76709712-INS-72
chr16-76727224-INS-327
chr16-76752112-INS-191
chr16-76770333-INS-76
chr16-76772245-INS-80
chr16-76776270-INS-154
chr16-76779966-INS-78
chr16-76793639-INS-62
chr16-76819788-INS-185
chr16-76823422-INS-263
chr16-76869542-INS-120
chr16-76877797-INS-348
chr16-76882737-INS-195
chr16-76891190-INS-102
chr16-76895989-INS-1886
chr16-76901871-INS-537
chr16-76919637-INS-6165
chr16-76920895-INS-52
chr16-76935185-INS-139
chr16-76936415-INS-581
chr16-76941860-INS-425
chr16-76978027-INS-448
chr16-77084033-INS-771
chr16-77084966-INS-191
chr16-77091926-INS-56
chr16-77121814-INS-140
chr16-77135303-INS-147
chr16-77140943-INS-62
chr16-77141656-INS-60
chr16-77212723-INS-190
chr16-77227588-INS-54
chr16-77229928-INS-52
chr16-77238611-INS-93
chr16-77263159-INS-68
chr16-77280275-INS-111
chr16-77281310-INS-101
chr16-77286152-INS-569
chr16-77288465-INS-7045
chr16-77307753-INS-179
chr16-77322587-INS-182
chr16-77337124-INS-59
chr16-77340038-INS-52
chr16-77362421-INS-1010
chr16-77364392-INS-505
chr16-77395489-INS-76
chr16-77410513-INS-162
chr16-77419882-INS-386
chr16-77434653-INS-178
chr16-77448296-INS-80
chr16-77467736-INS-621
chr16-77474835-INS-728
chr16-77488933-INS-77
chr16-77491877-INS-263
chr16-77509226-INS-126
chr16-77539246-INS-138
chr16-77544077-INS-6442
chr16-77561512-INS-56
chr16-77581842-INS-170
chr16-77599030-INS-552
chr16-77619466-INS-58
chr16-77627616-INS-737
chr16-77628458-INS-52
chr16-77653382-INS-348
chr16-77672520-INS-142
chr16-77702831-INS-387
chr16-77705561-INS-380
chr16-77706194-INS-752
chr16-77708200-INS-6694
chr16-77715058-INS-208
chr16-77722147-INS-62
chr16-77744892-INS-923
chr16-77745645-INS-106
chr16-77754409-INS-6438
chr16-77779365-INS-378
chr16-77788549-INS-124
chr16-77808571-INS-54

chr16-77816631-INS-439
chr16-77838282-INS-161
chr16-77840810-INS-69
chr16-77858879-INS-115
chr16-77875059-INS-63
chr16-77883128-INS-6589
chr16-77890427-INS-1613
chr16-77890974-INS-4157
chr16-77933381-INS-544
chr16-77935994-INS-468
chr16-77943225-INS-313
chr16-77955962-INS-5119
chr16-77971684-INS-61
chr16-77987512-INS-4553
chr16-77991784-INS-142
chr16-77997808-INS-128
chr16-78008957-INS-366
chr16-78033266-INS-230
chr16-78035050-INS-53
chr16-78067911-INS-60
chr16-78075046-INS-71
chr16-78081606-INS-256
chr16-78132474-INS-329
chr16-78148067-INS-214
chr16-78171045-INS-107
chr16-78177423-INS-987
chr16-78196197-INS-423
chr16-78199151-INS-206
chr16-78200991-INS-6902
chr16-78204442-INS-6847
chr16-78205387-INS-327
chr16-78214174-INS-106
chr16-78248015-INS-92
chr16-78253862-INS-506
chr16-78276281-INS-403
chr16-78286915-INS-54
chr16-78326771-INS-184
chr16-78329960-INS-132
chr16-78332296-INS-262
chr16-78345013-INS-211
chr16-78350636-INS-53
chr16-78361626-INS-250
chr16-78378454-INS-57
chr16-78381302-INS-84
chr16-78387138-INS-162
chr16-78399849-INS-100
chr16-78426695-INS-844
chr16-78437892-INS-53
chr16-78440062-INS-142
chr16-78448813-INS-54
chr16-78457662-INS-102
chr16-78477526-INS-183
chr16-78489430-INS-479
chr16-78495557-INS-200
chr16-78498270-INS-52
chr16-78506617-INS-572
chr16-78509730-INS-6520
chr16-78511240-INS-688
chr16-78517606-INS-449
chr16-78521737-INS-56
chr16-78535492-INS-75
chr16-78546546-INS-76
chr16-78569208-INS-72
chr16-78590252-INS-390
chr16-78622064-INS-648
chr16-78637260-INS-54
chr16-78638701-INS-76
chr16-78643266-INS-176
chr16-78647697-INS-70
chr16-78663419-INS-7965
chr16-78669616-INS-193
chr16-78681799-INS-188
chr16-78699677-INS-52
chr16-78702117-INS-56
chr16-78706449-INS-665
chr16-78759994-INS-181
chr16-78777811-INS-57

chr16-78783366-INS-222
chr16-78797843-INS-671
chr16-78862012-INS-218
chr16-78888792-INS-6415
chr16-78890259-INS-76
chr16-78895610-INS-67
chr16-78912944-INS-52
chr16-78926919-INS-165
chr16-78930471-INS-260
chr16-78936184-INS-386
chr16-78963428-INS-70
chr16-78984913-INS-149
chr16-78986329-INS-371
chr16-79007450-INS-711
chr16-79015164-INS-58
chr16-79034039-INS-64
chr16-79042878-INS-784
chr16-79045970-INS-4450
chr16-79050409-INS-405
chr16-79051876-INS-115
chr16-79052289-INS-509
chr16-79057737-INS-65
chr16-79127755-INS-5341
chr16-79142721-INS-740
chr16-79159545-INS-329
chr16-79232066-INS-101
chr16-79240064-INS-52
chr16-79266606-INS-66
chr16-79278468-INS-55
chr16-79346432-INS-57
chr16-79349612-INS-105
chr16-79351411-INS-5268
chr16-79368039-INS-959
chr16-79370023-INS-59
chr16-79395274-INS-110
chr16-79398829-INS-72
chr16-79448109-INS-61
chr16-79474154-INS-2782
chr16-79501057-INS-57
chr16-79505972-INS-206
chr16-79506850-INS-2844
chr16-79512932-INS-179
chr16-79520297-INS-52
chr16-79549725-INS-7084
chr16-79551188-INS-54
chr16-79593117-INS-106
chr16-79652730-INS-130
chr16-79656385-INS-58
chr16-79697070-INS-201
chr16-79708026-INS-384
chr16-79720302-INS-78
chr16-79732923-INS-58
chr16-79749818-INS-309
chr16-79756998-INS-69
chr16-79772643-INS-72
chr16-79891917-INS-6251
chr16-79910444-INS-503
chr16-79924735-INS-586
chr16-79950725-INS-279
chr16-79984381-INS-56
chr16-79997588-INS-392
chr16-80037428-INS-929
chr16-80050921-INS-56
chr16-80052674-INS-52
chr16-80060670-INS-140
chr16-80068420-INS-457
chr16-80075830-INS-63
chr16-80096926-INS-313
chr16-80098587-INS-6231
chr16-80116845-INS-4038
chr16-80122075-INS-1546
chr16-80130395-INS-191
chr16-80181035-INS-572
chr16-80182247-INS-409
chr16-80234037-INS-80
chr16-80236944-INS-60
chr16-80243035-INS-53

chr16-80251757-INS-114
chr16-80253553-INS-57
chr16-80267245-INS-56
chr16-80302743-INS-204
chr16-80313761-INS-111
chr16-80319211-INS-51
chr16-80323326-INS-133
chr16-80324336-INS-63
chr16-80326699-INS-67
chr16-80352676-INS-3046
chr16-80359398-INS-6355
chr16-80362323-INS-813
chr16-80392200-INS-51
chr16-80414372-INS-187
chr16-80433680-INS-756
chr16-80451971-INS-162
chr16-80492115-INS-60
chr16-80504021-INS-138
chr16-80530058-INS-414
chr16-80531444-INS-139
chr16-80535102-INS-199
chr16-80551147-INS-76
chr16-80564898-INS-113
chr16-80570038-INS-132
chr16-80626354-INS-218
chr16-80635207-INS-139
chr16-80645709-INS-210
chr16-80660118-INS-68
chr16-80698751-INS-401
chr16-80700179-INS-379
chr16-80709410-INS-133
chr16-80744369-INS-147
chr16-80746392-INS-51
chr16-80800379-INS-57
chr16-80808849-INS-72
chr16-80858351-INS-920
chr16-80882163-INS-476
chr16-80896455-INS-874
chr16-80907231-INS-870
chr16-80908398-INS-67
chr16-80937190-INS-683
chr16-80940713-INS-106
chr16-80961811-INS-65
chr16-80962678-INS-7003
chr16-80978496-INS-71
chr16-81010208-INS-455
chr16-81016225-INS-64
chr16-81046732-INS-108
chr16-81048106-INS-130
chr16-81049184-INS-210
chr16-81111752-INS-64
chr16-81114829-INS-364
chr16-81123222-INS-54
chr16-81127677-INS-671
chr16-81130015-INS-268
chr16-81145327-INS-556
chr16-81178048-INS-79
chr16-81179529-INS-336
chr16-81181251-INS-67
chr16-81200726-INS-484
chr16-81231312-INS-2659
chr16-81261076-INS-68
chr16-81266002-INS-884
chr16-81269055-INS-114
chr16-81287617-INS-253
chr16-81303910-INS-138
chr16-81310732-INS-228
chr16-81321116-INS-839
chr16-81335961-INS-6733
chr16-81360959-INS-504
chr16-81375273-INS-194
chr16-81383068-INS-71
chr16-81393413-INS-65
chr16-81408251-INS-172
chr16-81415554-INS-72
chr16-81501681-INS-345
chr16-81519185-INS-70

chr16-81522813-INS-317
chr16-81530710-INS-6528
chr16-81533351-INS-69
chr16-81554467-INS-50
chr16-81560107-INS-65
chr16-81561873-INS-479
chr16-81605953-INS-68
chr16-81627076-INS-71
chr16-81630081-INS-52
chr16-81632097-INS-104
chr16-81636027-INS-323
chr16-81656157-INS-212
chr16-81713125-INS-105
chr16-81722568-INS-224
chr16-81736274-INS-512
chr16-81750592-INS-991
chr16-81757173-INS-458
chr16-81761893-INS-97
chr16-81771417-INS-398
chr16-81780821-INS-59
chr16-81793069-INS-67
chr16-81815995-INS-78
chr16-81816948-INS-486
chr16-81825687-INS-5849
chr16-81852467-INS-143
chr16-81855787-INS-7761
chr16-81856217-INS-96
chr16-81897657-INS-345
chr16-81898787-INS-785
chr16-81935075-INS-6382
chr16-81944964-INS-52
chr16-81967262-INS-56
chr16-82027996-INS-507
chr16-82029893-INS-285
chr16-82040825-INS-820
chr16-82049055-INS-90
chr16-82051617-INS-86
chr16-82056116-INS-62
chr16-82058424-INS-99
chr16-82083450-INS-6017
chr16-82112122-INS-413
chr16-82172297-INS-962
chr16-82189851-INS-798
chr16-82192250-INS-181
chr16-82199115-INS-6263
chr16-82207012-INS-62
chr16-82272860-INS-402
chr16-82306764-INS-1864
chr16-82351287-INS-78
chr16-82354933-INS-54
chr16-82358529-INS-120
chr16-82389477-INS-248
chr16-82401195-INS-179
chr16-82401584-INS-4367
chr16-82479669-INS-358
chr16-82490796-INS-4335
chr16-82505184-INS-1281
chr16-82509214-INS-332
chr16-82534854-INS-4090
chr16-82549225-INS-7528
chr16-82555323-INS-100
chr16-82561912-INS-176
chr16-82562058-INS-6335
chr16-82565290-INS-7092
chr16-82581356-INS-6390
chr16-82606759-INS-6828
chr16-82608086-INS-650
chr16-82608599-INS-1799
chr16-82611280-INS-208
chr16-82621105-INS-640
chr16-82635334-INS-54
chr16-82656689-INS-792
chr16-82670409-INS-292
chr16-82732317-INS-56
chr16-82737854-INS-56
chr16-82752488-INS-204
chr16-82756593-INS-233

chr16-82770983-INS-76
chr16-82826915-INS-560
chr16-82876999-INS-198
chr16-82913379-INS-4936
chr16-82984995-INS-61
chr16-82992790-INS-195
chr16-83045838-INS-742
chr16-83074341-INS-467
chr16-83093102-INS-55
chr16-83114576-INS-208
chr16-83132070-INS-113
chr16-83165591-INS-399
chr16-83227045-INS-213
chr16-83288831-INS-2306
chr16-83289529-INS-241
chr16-83315167-INS-95
chr16-83345785-INS-400
chr16-83354818-INS-206
chr16-83372325-INS-186
chr16-83401703-INS-434
chr16-83424379-INS-56
chr16-83426573-INS-179
chr16-83431125-INS-404
chr16-83472444-INS-300
chr16-83474622-INS-842
chr16-83477467-INS-82
chr16-83510651-INS-92
chr16-83515129-INS-94
chr16-83523373-INS-56
chr16-83528222-INS-78
chr16-83601860-INS-84
chr16-83609282-INS-767
chr16-83610067-INS-590
chr16-83641066-INS-81
chr16-83647323-INS-190
chr16-83658024-INS-66
chr16-83676234-INS-63
chr16-83685986-INS-118
chr16-83716904-INS-138
chr16-83739115-INS-50
chr16-83769897-INS-7166
chr16-83771074-INS-134
chr16-83787575-INS-120
chr16-83791319-INS-470
chr16-83806916-INS-64
chr16-83822447-INS-482
chr16-83836302-INS-220
chr16-83840339-INS-337
chr16-83845244-INS-378
chr16-83846197-INS-131
chr16-83886344-INS-81
chr16-83887949-INS-265
chr16-83894956-INS-70
chr16-83903763-INS-217
chr16-83905455-INS-6157
chr16-83919780-INS-150
chr16-83955369-INS-74
chr16-83966868-INS-145
chr16-83983147-INS-890
chr16-83989301-INS-3219
chr16-84018663-INS-110
chr16-84028106-INS-6642
chr16-84034659-INS-58
chr16-84055815-INS-627
chr16-84087813-INS-167
chr16-84096235-INS-4172
chr16-84105554-INS-303
chr16-84110431-INS-6801
chr16-84113745-INS-6446
chr16-84161812-INS-50
chr16-84165196-INS-72
chr16-84167000-INS-445
chr16-84176872-INS-80
chr16-84203836-INS-225
chr16-84211234-INS-51
chr16-84215887-INS-504
chr16-84223604-INS-1535

chr16-84259894-INS-133
chr16-84296102-INS-585
chr16-84323064-INS-72
chr16-84339497-INS-614
chr16-84377787-INS-194
chr16-84379869-INS-636
chr16-84384305-INS-1003
chr16-84384879-INS-190
chr16-84394715-INS-705
chr16-84412670-INS-89
chr16-84414284-INS-52
chr16-84417726-INS-206
chr16-84439049-INS-177
chr16-84442241-INS-905
chr16-84450840-INS-103
chr16-84457897-INS-219
chr16-84485381-INS-407
chr16-84495805-INS-53
chr16-84507806-INS-100
chr16-84560109-INS-80
chr16-84587853-INS-85
chr16-84589526-INS-51
chr16-84593126-INS-54
chr16-84597947-INS-56
chr16-84602930-INS-87
chr16-84612289-INS-249
chr16-84619247-INS-98
chr16-84622328-INS-465
chr16-84628129-INS-220
chr16-84652305-INS-55
chr16-84653830-INS-207
chr16-84662793-INS-208
chr16-84670074-INS-70
chr16-84671322-INS-4419
chr16-84673850-INS-221
chr16-84686016-INS-5322
chr16-84693304-INS-6253
chr16-84711630-INS-236
chr16-84722382-INS-116
chr16-84723560-INS-64
chr16-84739175-INS-212
chr16-84755268-INS-486
chr16-84770698-INS-209
chr16-84825336-INS-141
chr16-84834070-INS-298
chr16-84857601-INS-1016
chr16-84869990-INS-197
chr16-84870741-INS-579
chr16-84896530-INS-7741
chr16-84915163-INS-1987
chr16-84926682-INS-6538
chr16-84944199-INS-398
chr16-84948431-INS-1040
chr16-84957954-INS-66
chr16-84961597-INS-85
chr16-84973296-INS-144
chr16-85002294-INS-51
chr16-85009316-INS-155
chr16-85030650-INS-107
chr16-85051601-INS-67
chr16-85084036-INS-587
chr16-85094040-INS-80
chr16-85106126-INS-108
chr16-85111761-INS-369
chr16-85128305-INS-978
chr16-85164338-INS-58
chr16-85194399-INS-172
chr16-85221395-INS-52
chr16-85223878-INS-329
chr16-85225172-INS-239
chr16-85275265-INS-437
chr16-85282008-INS-210
chr16-85319423-INS-58
chr16-85324386-INS-107
chr16-85326423-INS-206
chr16-85354328-INS-6566
chr16-85373708-INS-200

chr16-85382773-INS-1584
chr16-85394859-INS-2246
chr16-85398770-INS-430
chr16-85403687-INS-573
chr16-85424211-INS-92
chr16-85454129-INS-60
chr16-85464761-INS-60
chr16-85468956-INS-407
chr16-85476508-INS-69
chr16-85496143-INS-58
chr16-85497501-INS-60
chr16-85517510-INS-6726
chr16-85525472-INS-52
chr16-85534433-INS-82
chr16-85559872-INS-125
chr16-85572060-INS-323
chr16-85582548-INS-51
chr16-85618967-INS-98
chr16-85624811-INS-126
chr16-85661921-INS-519
chr16-85678418-INS-71
chr16-85681739-INS-143
chr16-85685792-INS-535
chr16-85701300-INS-66
chr16-85703458-INS-54
chr16-85718384-INS-236
chr16-85740212-INS-304
chr16-85785850-INS-371
chr16-85818709-INS-429
chr16-85818967-INS-6206
chr16-85826928-INS-120
chr16-85827181-INS-917
chr16-85859230-INS-246
chr16-85866155-INS-183
chr16-85871191-INS-172
chr16-85886862-INS-570
chr16-85903306-INS-59
chr16-85909691-INS-53
chr16-85914226-INS-4365
chr16-85954141-INS-81
chr16-85958647-INS-62
chr16-86010494-INS-844
chr16-86013235-INS-328
chr16-86047886-INS-110
chr16-86051260-INS-340
chr16-86065213-INS-67
chr16-86072037-INS-79
chr16-86074618-INS-62
chr16-86076603-INS-54
chr16-86100351-INS-400
chr16-86125931-INS-64
chr16-86146472-INS-106
chr16-86147544-INS-300
chr16-86156631-INS-119
chr16-86164855-INS-194
chr16-86180411-INS-75
chr16-86206868-INS-78
chr16-86219370-INS-72
chr16-86237348-INS-399
chr16-86252853-INS-139
chr16-86288845-INS-60
chr16-86308811-INS-991
chr16-86323701-INS-725
chr16-86358963-INS-220
chr16-86368525-INS-78
chr16-86371321-INS-60
chr16-86377985-INS-74
chr16-86397943-INS-199
chr16-86398896-INS-77
chr16-86403208-INS-401
chr16-86404713-INS-966
chr16-86438782-INS-179
chr16-86452682-INS-71
chr16-86462155-INS-121
chr16-86463966-INS-361
chr16-86489520-INS-143
chr16-86493239-INS-566

chr16-86504956-INS-219
chr16-86513231-INS-463
chr16-86547441-INS-370
chr16-86562405-INS-76
chr16-86567974-INS-299
chr16-86594860-INS-126
chr16-86617781-INS-3604
chr16-86626988-INS-82
chr16-86650366-INS-233
chr16-86668613-INS-6638
chr16-86674493-INS-438
chr16-86676939-INS-680
chr16-86692302-INS-80
chr16-86707217-INS-362
chr16-86735316-INS-135
chr16-86740905-INS-305
chr16-86744182-INS-6886
chr16-86764269-INS-105
chr16-86780553-INS-340
chr16-86781328-INS-70
chr16-86784058-INS-67
chr16-86799706-INS-163
chr16-86806745-INS-530
chr16-86825094-INS-117
chr16-86830155-INS-269
chr16-86837080-INS-66
chr16-86859203-INS-142
chr16-86861603-INS-7800
chr16-86885755-INS-451
chr16-86898328-INS-163
chr16-86904385-INS-3261
chr16-86907445-INS-76
chr16-86973700-INS-164
chr16-86984861-INS-93
chr16-86985443-INS-86
chr16-87003403-INS-618
chr16-87008720-INS-60
chr16-87011290-INS-799
chr16-87017884-INS-52
chr16-87038843-INS-60
chr16-87040313-INS-68
chr16-87048234-INS-189
chr16-87053309-INS-266
chr16-87059726-INS-52
chr16-87066178-INS-72
chr16-87071574-INS-881
chr16-87081629-INS-52
chr16-87111176-INS-255
chr16-87127736-INS-167
chr16-87139515-INS-60
chr16-87148906-INS-56
chr16-87154468-INS-52
chr16-87169304-INS-393
chr16-87183294-INS-278
chr16-87183321-INS-869
chr16-87207874-INS-326
chr16-87212232-INS-65
chr16-87219167-INS-164
chr16-87225812-INS-302
chr16-87228456-INS-807
chr16-87236053-INS-241
chr16-87239313-INS-85
chr16-87240879-INS-103
chr16-87245000-INS-325
chr16-87265210-INS-118
chr16-87278943-INS-323
chr16-87323172-INS-88
chr16-87323184-INS-156
chr16-87340779-INS-407
chr16-87347672-INS-175
chr16-87348655-INS-202
chr16-87357730-INS-61
chr16-87366217-INS-1276
chr16-87374559-INS-65
chr16-87389516-INS-72
chr16-87391566-INS-56
chr16-87399016-INS-144

chr16-87403076-INS-264
chr16-87405457-INS-52
chr16-87411128-INS-824
chr16-87413386-INS-377
chr16-87416666-INS-68
chr16-87448425-INS-8985
chr16-87478754-INS-69
chr16-87481992-INS-89
chr16-87483016-INS-145
chr16-87484412-INS-201
chr16-87487820-INS-492
chr16-87511348-INS-881
chr16-87526035-INS-213
chr16-87541238-INS-87
chr16-87559795-INS-407
chr16-87573371-INS-336
chr16-87575835-INS-120
chr16-87581036-INS-99
chr16-87582203-INS-74
chr16-87587845-INS-56
chr16-87591817-INS-737
chr16-87678593-INS-6676
chr16-87698680-INS-175
chr16-87711818-INS-76
chr16-87729537-INS-148
chr16-87765893-INS-276
chr16-87807388-INS-96
chr16-87809456-INS-913
chr16-87815965-INS-303
chr16-87857010-INS-69
chr16-87883582-INS-585
chr16-87886625-INS-304
chr16-87899164-INS-391
chr16-87900160-INS-108
chr16-87936831-INS-611
chr16-87942042-INS-4972
chr16-87946093-INS-410
chr16-87963302-INS-68
chr16-87972299-INS-97
chr16-87992743-INS-1356
chr16-87998914-INS-59
chr16-88003135-INS-271
chr16-88011145-INS-242
chr16-88040890-INS-549
chr16-88109283-INS-738
chr16-88122318-INS-74
chr16-88131325-INS-56
chr16-88135748-INS-55
chr16-88141975-INS-192
chr16-88156463-INS-336
chr16-88159629-INS-498
chr16-88162353-INS-1880
chr16-88163929-INS-6705
chr16-88180931-INS-57
chr16-88209269-INS-1331
chr16-88210552-INS-120
chr16-88214154-INS-198
chr16-88232008-INS-458
chr16-88240160-INS-1272
chr16-88241051-INS-1250
chr16-88265057-INS-294
chr16-88267322-INS-63
chr16-88268518-INS-504
chr16-88268778-INS-3073
chr16-88279483-INS-413
chr16-88286446-INS-349
chr16-88293539-INS-734
chr16-88300968-INS-125
chr16-88320969-INS-360
chr16-88324659-INS-212
chr16-88343719-INS-152
chr16-88347538-INS-145
chr16-88348348-INS-331
chr16-88351086-INS-58
chr16-88372261-INS-52
chr16-88401399-INS-52
chr16-88417785-INS-2906

chr16-88421944-INS-63
chr16-88436210-INS-7071
chr16-88437383-INS-74
chr16-88439163-INS-54
chr16-88442902-INS-59
chr16-88475026-INS-102
chr16-88506969-INS-66
chr16-88534327-INS-795
chr16-88535014-INS-102
chr16-88570451-INS-111
chr16-88603860-INS-170
chr16-88624041-INS-539
chr16-88653888-INS-63
chr16-88671936-INS-74
chr16-88713903-INS-1183
chr16-88750655-INS-280
chr16-88778063-INS-644
chr16-88794808-INS-474
chr16-88801528-INS-70
chr16-88845915-INS-4516
chr16-88857236-INS-54
chr16-88857533-INS-7848
chr16-88884037-INS-390
chr16-88932861-INS-828
chr16-88978382-INS-183
chr16-89001659-INS-6445
chr16-89002174-INS-208
chr16-89009058-INS-147
chr16-89021635-INS-6654
chr16-89023850-INS-51
chr16-89061700-INS-1345
chr16-89094098-INS-83
chr16-89113568-INS-204
chr16-89139639-INS-3211
chr16-89168511-INS-1084
chr16-89181847-INS-126
chr16-89189871-INS-69
chr16-89196106-INS-488
chr16-89209324-INS-92
chr16-89210884-INS-286
chr16-89248171-INS-149
chr16-89255809-INS-759
chr16-89262974-INS-339
chr16-89277819-INS-160
chr16-89279118-INS-57
chr16-89292354-INS-66
chr16-89294438-INS-60
chr16-89307282-INS-95
chr16-89313561-INS-153
chr16-89316422-INS-160
chr16-89316964-INS-71
chr16-89321524-INS-101
chr16-89325346-INS-330
chr16-89336532-INS-86
chr16-89340826-INS-191
chr16-89350267-INS-57
chr16-89354977-INS-214
chr16-89365628-INS-65
chr16-89374150-INS-836
chr16-89385323-INS-81
chr16-89417100-INS-127
chr16-89433207-INS-856
chr16-89456366-INS-554
chr16-89473186-INS-697
chr16-89493650-INS-129
chr16-89499064-INS-169
chr16-89505932-INS-342
chr16-89539181-INS-67
chr16-89550292-INS-210
chr16-89555026-INS-84
chr16-89572519-INS-89
chr16-89577189-INS-7385
chr16-89612885-INS-410
chr16-89662588-INS-62
chr16-89702625-INS-767
chr16-89729834-INS-357
chr16-89732650-INS-80

chr16-89746231-INS-170
chr16-89757874-INS-61
chr16-89761113-INS-51
chr16-89770461-INS-57
chr16-89840322-INS-734
chr16-89849948-INS-72
chr16-89887778-INS-101
chr16-89888617-INS-164
chr16-89897676-INS-290
chr16-89903256-INS-105
chr16-89903708-INS-814
chr16-89911173-INS-114
chr16-89918388-INS-80
chr16-89920630-INS-68
chr16-89921965-INS-471
chr16-89930338-INS-61
chr16-89937464-INS-66
chr16-89951395-INS-209
chr16-89953542-INS-216
chr16-89955683-INS-58
chr16-89958589-INS-137
chr16-89962190-INS-198
chr16-89965418-INS-106
chr16-89966296-INS-702
chr16-89966350-INS-72
chr16-89970678-INS-140
chr16-89975359-INS-209
chr16-89991348-INS-302
chr16-89999919-INS-56
chr16-90003939-INS-113
chr16-90008939-INS-84
chr16-90009985-INS-55
chr16-90014353-INS-199
chr16-90077917-INS-52
chr16-90083704-INS-93
chr16-90089681-INS-176
chr16-90101351-INS-138
chr16-90103583-INS-203
chr16-90122031-INS-201
chr16-90129438-INS-199
chr16-90137191-INS-52
chr16-90152594-INS-51
chr16-90156489-INS-101
chr16-90204871-INS-176
chr16-90212422-INS-128
chr16-90225065-INS-351
chr16-90251325-INS-60
chr16-90257873-INS-68
chr16-90262745-INS-221
chr16-90264555-INS-7145
chr16-90284115-INS-66
chr16-90287411-INS-60
chr16-90310488-INS-80
chr16-90393503-INS-445
chr16-90397126-INS-170
chr16-90399763-INS-1345
chr16-90410427-INS-218
chr16-90432684-INS-191
chr16-90438819-INS-123
chr16-90440037-INS-265
chr16-90454897-INS-58
chr16-90454897-INS-64
chr16-90457483-INS-277
chr16-90465055-INS-4539
chr16-90465827-INS-73
chr16-90469754-INS-70
chr16-90480718-INS-344
chr16-90486766-INS-60
chr16-90488551-INS-308
chr16-90498568-INS-60
chr16-90528416-INS-54
chr16-90531191-INS-232
chr16-90536515-INS-316
chr16-90540385-INS-90
chr16-90544988-INS-67
chr16-90555258-INS-62
chr16-90562220-INS-85

chr16-90567522-INS-70
chr16-90587819-INS-98
chr16-90600695-INS-498
chr16-90611720-INS-409
chr16-90619464-INS-5294
chr16-90662384-INS-51
chr16-90664042-INS-55
chr16-90674721-INS-54
chr16-90676448-INS-51
chr16-90682054-INS-321
chr16-90689620-INS-439
chr16-90691234-INS-289
chr16-90706836-INS-51
chr16-90710145-INS-68
chr16-90737627-INS-73
chr16-90737739-INS-73
chr16-90740709-INS-52
chr16-90771959-INS-184
chr16-90772248-INS-184
chr16-90782565-INS-350
chr16-90797533-INS-526
chr16-90834761-INS-166
chr16-90859243-INS-770
chr16-90864152-INS-80
chr16-90872435-INS-68
chr16-90889098-INS-541
chr16-90926839-INS-62
chr16-91028690-INS-59
chr16-91036870-INS-754
chr16-91037047-INS-104
chr16-91054323-INS-54
chr16-91078045-INS-60
chr16-91091934-INS-405
chr16-91103443-INS-66
chr16-91120461-INS-375
chr16-91122929-INS-129
chr16-91139704-INS-896
chr16-91140273-INS-192
chr16-91155189-INS-211
chr16-91155696-INS-89
chr16-91167716-INS-64
chr16-91177033-INS-52
chr16-91178442-INS-52
chr16-91189030-INS-93
chr16-91200293-INS-103
chr16-91214902-INS-385
chr16-91222297-INS-54
chr16-91229300-INS-328
chr16-91246273-INS-487
chr16-91251627-INS-130
chr16-91251812-INS-490
chr16-91260373-INS-72
chr16-91273504-INS-437
chr16-91286180-INS-244
chr16-91293238-INS-240
chr16-91300419-INS-84
chr16-91310002-INS-105
chr16-91314870-INS-64
chr16-91319727-INS-208
chr16-91321877-INS-528
chr16-91325178-INS-55
chr16-91329725-INS-71
chr16-91337642-INS-204
chr16-91346246-INS-66
chr16-91350910-INS-87
chr16-91364729-INS-274
chr16-91377346-INS-172
chr16-91377651-INS-408
chr16-91389006-INS-198
chr16-91400471-INS-96
chr16-91401546-INS-203
chr16-91428352-INS-58
chr16-91446713-INS-131
chr16-91449737-INS-212
chr16-91465632-INS-561
chr16-91493087-INS-580
chr16-91512858-INS-79

chr16-91518370-INS-184
chr16-91520878-INS-3416
chr16-91535084-INS-212
chr16-91547679-INS-772
chr16-91551512-INS-92
chr16-91571493-INS-942
chr16-91608069-INS-230
chr16-91612772-INS-213
chr16-91629334-INS-51
chr16-91652991-INS-220
chr16-91686207-INS-108
chr16-91693613-INS-53
chr16-91701692-INS-183
chr16-91710188-INS-54
chr16-91717782-INS-128
chr16-91726816-INS-204
chr16-91728719-INS-499
chr16-91730079-INS-417
chr16-91734480-INS-348
chr16-91737327-INS-1508
chr16-91737329-INS-876
chr16-91737334-INS-58
chr16-91740610-INS-62
chr16-91748037-INS-6401
chr16-91751796-INS-212
chr16-91769285-INS-78
chr16-91776304-INS-232
chr16-91784019-INS-203
chr16-91785511-INS-92
chr16-91786023-INS-859
chr16-91787389-INS-96
chr16-91791195-INS-62
chr16-91807182-INS-306
chr16-91819228-INS-54
chr16-91820805-INS-85
chr16-91821579-INS-6651
chr16-91825746-INS-55
chr16-91840263-INS-92
chr16-91842116-INS-85
chr16-91849923-INS-55
chr16-91915163-INS-388
chr16-91928560-INS-6042
chr16-91930468-INS-60
chr16-91934301-INS-154
chr16-91946012-INS-62
chr16-91985094-INS-175
chr16-91985839-INS-64
chr16-91987931-INS-56
chr16-92014733-INS-180
chr16-92017560-INS-750
chr16-92024767-INS-7108
chr16-92045634-INS-102
chr16-92048076-INS-56
chr16-92051981-INS-54
chr16-92052383-INS-77
chr16-92064219-INS-82
chr16-92078803-INS-330
chr16-92095631-INS-7076
chr16-92118638-INS-352
chr16-92133717-INS-79
chr16-92135477-INS-370
chr16-92139025-INS-122
chr16-92166288-INS-99
chr16-92172929-INS-172
chr16-92175927-INS-552
chr16-92182037-INS-60
chr16-92203933-INS-52
chr16-92229457-INS-131
chr16-92237169-INS-7079
chr16-92265911-INS-69
chr16-92271093-INS-123
chr16-92293126-INS-7158
chr16-92310732-INS-67
chr16-92316360-INS-75
chr16-92321431-INS-64
chr16-92326023-INS-72
chr16-92360693-INS-82

chr16-92371244-INS-180
chr16-92372391-INS-515
chr16-92374806-INS-178
chr16-92377906-INS-86
chr16-92382472-INS-354
chr16-92425468-INS-59
chr16-92457763-INS-261
chr16-92511239-INS-54
chr16-92547619-INS-108
chr16-92548805-INS-189
chr16-92558673-INS-73
chr16-92569102-INS-54
chr16-92569832-INS-52
chr16-92584554-INS-149
chr16-92586055-INS-69
chr16-92595784-INS-209
chr16-92597629-INS-64
chr16-92602266-INS-110
chr16-92603377-INS-83
chr16-92621075-INS-151
chr16-92634054-INS-498
chr16-92640060-INS-66
chr16-92644898-INS-83
chr16-92655706-INS-902
chr16-92657192-INS-273
chr16-92659598-INS-153
chr16-92662731-INS-68
chr16-92669981-INS-117
chr16-92685149-INS-87
chr16-92692365-INS-52
chr16-92699671-INS-86
chr16-92717143-INS-72
chr16-92730987-INS-277
chr16-92731159-INS-196
chr16-92741389-INS-265
chr16-92744422-INS-75
chr16-92744438-INS-999
chr16-92746930-INS-641
chr16-92752043-INS-58
chr16-92757649-INS-60
chr16-92761842-INS-418
chr16-92768591-INS-81
chr16-92777538-INS-64
chr16-92795218-INS-138
chr16-92801137-INS-51
chr16-92812964-INS-892
chr16-92814738-INS-168
chr16-92824951-INS-74
chr16-92831733-INS-263
chr16-92841396-INS-99
chr16-92857141-INS-84
chr16-92895608-INS-189
chr16-92897736-INS-200
chr16-92914862-INS-64
chr16-92924394-INS-114
chr16-92928263-INS-57
chr16-92931550-INS-92
chr16-92950804-INS-54
chr16-92961604-INS-273
chr16-92968136-INS-58
chr16-92972205-INS-53
chr16-92987835-INS-90
chr16-92992583-INS-108
chr16-93033835-INS-200
chr16-93045214-INS-66
chr16-93047273-INS-342
chr16-93056386-INS-288
chr16-93079913-INS-138
chr16-93086727-INS-136
chr16-93127716-INS-86
chr16-93135570-INS-54
chr16-93138877-INS-58
chr16-93145492-INS-541
chr16-93177393-INS-153
chr16-93199591-INS-116
chr16-93204203-INS-110
chr16-93205753-INS-288

chr16-93208467-INS-64
chr16-93242502-INS-76
chr16-93246759-INS-318
chr16-93249453-INS-55
chr16-93252676-INS-190
chr16-93259746-INS-88
chr16-93266172-INS-170
chr16-93280674-INS-54
chr16-93281742-INS-84
chr16-93284210-INS-142
chr16-93286708-INS-76
chr16-93302587-INS-230
chr16-93306254-INS-414
chr16-93318040-INS-222
chr16-93321257-INS-112
chr16-93333774-INS-83
chr16-93345243-INS-213
chr16-93364325-INS-464
chr16-93368007-INS-52
chr16-93369040-INS-100
chr16-93370694-INS-120
chr16-93398688-INS-84
chr16-93404247-INS-389
chr16-93443786-INS-66
chr16-93451635-INS-151
chr16-93455399-INS-60
chr16-93457282-INS-95
chr16-93468061-INS-118
chr16-93474045-INS-78
chr16-93474235-INS-462
chr16-93532435-INS-99
chr16-93533542-INS-60
chr16-93538498-INS-229
chr16-93552130-INS-69
chr16-93561661-INS-248
chr16-93569329-INS-160
chr16-93575469-INS-361
chr16-93575916-INS-197
chr16-93580518-INS-131
chr16-93583862-INS-318
chr16-93586173-INS-201
chr16-93595561-INS-440
chr16-93598926-INS-80
chr16-93600871-INS-168
chr16-93607699-INS-204
chr16-93608921-INS-397
chr16-93609893-INS-321
chr16-93623376-INS-65
chr16-93626044-INS-8681
chr16-93628239-INS-72
chr16-93631857-INS-199
chr16-93637706-INS-182
chr16-93660645-INS-117
chr16-93673094-INS-99
chr16-93682855-INS-209
chr16-93707499-INS-5270
chr16-93710155-INS-479
chr16-93712728-INS-102
chr16-93714361-INS-70
chr16-93718287-INS-192
chr16-93722422-INS-197
chr16-93724451-INS-170
chr16-93727966-INS-55
chr16-93734254-INS-254
chr16-93741696-INS-74
chr16-93803988-INS-359
chr16-93829080-INS-154
chr16-93836900-INS-65
chr16-93838674-INS-202
chr16-93844085-INS-56
chr16-93850780-INS-66
chr16-93858348-INS-196
chr16-93874761-INS-202
chr16-93883609-INS-53
chr16-93885553-INS-52
chr16-93899605-INS-242
chr16-93917756-INS-189

chr16-93937343-INS-5041
chr16-93960201-INS-160
chr16-93966457-INS-70
chr16-93984373-INS-810
chr16-93998032-INS-207
chr16-94003472-INS-138
chr16-94021913-INS-107
chr16-94025216-INS-5259
chr16-94032950-INS-102
chr16-94035910-INS-249
chr16-94038444-INS-78
chr16-94056951-INS-215
chr16-94057493-INS-201
chr16-94072558-INS-376
chr16-94085919-INS-204
chr16-94087837-INS-108
chr16-94096558-INS-196
chr16-94108787-INS-196
chr16-94112281-INS-989
chr16-94117314-INS-79
chr16-94145835-INS-198
chr16-94150134-INS-73
chr16-94159512-INS-124
chr16-94162103-INS-95
chr16-94163537-INS-295
chr16-94177926-INS-236
chr16-94264964-INS-355
chr16-94305413-INS-222
chr16-94340528-INS-51
chr16-94352818-INS-200
chr16-94376049-INS-64
chr16-94379685-INS-393
chr16-94381455-INS-209
chr16-94383134-INS-336
chr16-94405335-INS-202
chr16-94405807-INS-6054
chr16-94406766-INS-51
chr16-94415802-INS-350
chr16-94449913-INS-195
chr16-94455683-INS-60
chr16-94462750-INS-91
chr16-94462795-INS-6401
chr16-94476682-INS-360
chr16-94478138-INS-310
chr16-94495066-INS-57
chr16-94534109-INS-102
chr16-94537110-INS-228
chr16-94539603-INS-306
chr16-94539816-INS-2448
chr16-94577263-INS-95
chr16-94581278-INS-192
chr16-94586024-INS-70
chr16-94591803-INS-312
chr16-94599484-INS-111
chr16-94628529-INS-50
chr16-94632220-INS-351
chr16-94641179-INS-74
chr16-94703299-INS-128
chr16-94769342-INS-90
chr16-94785799-INS-71
chr16-94794178-INS-164
chr16-94801284-INS-59
chr16-94833436-INS-104
chr16-94860321-INS-98
chr16-94891015-INS-140
chr16-94897920-INS-61
chr16-94926981-INS-454
chr16-94937297-INS-558
chr16-94943566-INS-126
chr16-94945987-INS-63
chr16-94960405-INS-156
chr16-94961987-INS-190
chr16-94966745-INS-157
chr16-94973189-INS-58
chr16-94987245-INS-445
chr16-95010785-INS-653
chr16-95011477-INS-60

chr16-95017604-INS-82
chr16-95078835-INS-61
chr16-95091867-INS-106
chr16-95099854-INS-56
chr16-95109524-INS-211
chr16-95120951-INS-116
chr16-95127232-INS-64
chr16-95130166-INS-400
chr16-95145371-INS-270
chr16-95148104-INS-365
chr16-95156903-INS-80
chr16-95163317-INS-112
chr16-95184575-INS-118
chr16-95191228-INS-180
chr16-95199707-INS-79
chr16-95201251-INS-106
chr16-95230853-INS-206
chr16-95238364-INS-81
chr16-95244333-INS-52
chr16-95275799-INS-68
chr16-95282942-INS-200
chr16-95285558-INS-161
chr16-95286940-INS-119
chr16-95287810-INS-572
chr16-95302317-INS-339
chr16-95323371-INS-52
chr16-95337515-INS-51
chr16-95347584-INS-111
chr16-95350081-INS-175
chr16-95352873-INS-765
chr16-95360566-INS-537
chr16-95365849-INS-342
chr16-95374977-INS-135
chr16-95389612-INS-111
chr16-95396609-INS-68
chr16-95401149-INS-107
chr16-95404257-INS-74
chr16-95415446-INS-393
chr16-95475568-INS-67
chr16-95482496-INS-68
chr16-95483479-INS-196
chr16-95494312-INS-224
chr16-95533141-INS-55
chr16-95537494-INS-88
chr16-95539045-INS-52
chr16-95545681-INS-98
chr16-95591456-INS-96
chr16-95594048-INS-146
chr16-95605336-INS-114
chr16-95609055-INS-78
chr16-95614515-INS-119
chr16-95615717-INS-6440
chr16-95629494-INS-601
chr16-95630902-INS-76
chr16-95638319-INS-576
chr16-95669886-INS-228
chr16-95696575-INS-164
chr16-95697920-INS-68
chr16-95732073-INS-70
chr16-95735940-INS-60
chr16-95743659-INS-65
chr16-95760670-INS-96
chr16-95774937-INS-4186
chr16-95780841-INS-366
chr16-95780949-INS-1045
chr16-95786871-INS-74
chr16-95856780-INS-102
chr16-95872481-INS-56
chr16-95879174-INS-86
chr16-95897040-INS-175
chr16-95932516-INS-147
chr16-95933847-INS-203
chr16-95946482-INS-61
chr16-95952028-INS-164
chr16-95969091-INS-666
chr16-95994672-INS-420
chr16-96012970-INS-329

chr16-96014923-INS-82
chr16-96021197-INS-52
chr16-96037529-INS-108
chr16-96057179-INS-116
chr16-96068009-INS-953
chr16-96070001-INS-326
chr16-96078885-INS-295
chr16-96094978-INS-153
chr16-96106755-INS-144
chr16-96115082-INS-783
chr16-96138187-INS-956
chr16-96165440-INS-391
chr16-96167197-INS-401
chr16-96173471-INS-300
chr16-96197165-INS-351
chr16-96207580-INS-58
chr16-96243992-INS-76
chr16-96247865-INS-337
chr16-96263817-INS-371
chr16-96277029-INS-557
chr16-96277841-INS-97
chr16-96305552-INS-52
chr16-96311724-INS-126
chr16-96343065-INS-69
chr16-96371981-INS-64
chr16-96377669-INS-3106
chr16-96403944-INS-57
chr16-96462657-INS-52
chr16-96469804-INS-112
chr16-96501091-INS-210
chr16-96524806-INS-705
chr16-96541232-INS-211
chr16-96565535-INS-134
chr16-96576430-INS-66
chr16-96600816-INS-57
chr16-96615348-INS-327
chr16-96649552-INS-61
chr16-96676078-INS-186
chr16-96714513-INS-62
chr16-96732936-INS-290
chr16-96748049-INS-351
chr16-96761701-INS-1541
chr16-96762779-INS-81
chr16-96811640-INS-52
chr16-96814512-INS-52
chr16-96816533-INS-105
chr16-96817683-INS-69
chr16-96826860-INS-54
chr16-96838169-INS-400
chr16-96871116-INS-166
chr16-96874960-INS-141
chr16-96880027-INS-170
chr16-96882205-INS-52
chr16-96885560-INS-150
chr16-96914524-INS-489
chr16-96928221-INS-57
chr16-96964092-INS-52
chr16-96976024-INS-211
chr16-96992023-INS-108
chr16-97006863-INS-290
chr16-97015315-INS-53
chr16-97024794-INS-6655
chr16-97033055-INS-6487
chr16-97037886-INS-1941
chr16-97040685-INS-1869
chr16-97045748-INS-183
chr16-97050333-INS-343
chr16-97051245-INS-623
chr16-97056643-INS-290
chr16-97067111-INS-54
chr16-97068125-INS-438
chr16-97073937-INS-118
chr16-97091343-INS-52
chr16-97103431-INS-176
chr16-97111006-INS-620
chr16-97113159-INS-63
chr16-97122779-INS-2706

chr16-97122780-INS-415
chr16-97132139-INS-1889
chr16-97137615-INS-51
chr16-97138598-INS-88
chr16-97163288-INS-56
chr16-97181737-INS-81
chr16-97224171-INS-507
chr16-97244549-INS-9544
chr16-97253670-INS-3351
chr16-97261119-INS-5953
chr16-97279046-INS-156
chr16-97331143-INS-70
chr16-97362832-INS-2032
chr16-97379203-INS-52
chr16-97381123-INS-200
chr16-97391346-INS-53
chr16-97404369-INS-61
chr16-97442565-INS-180
chr16-97446843-INS-6584
chr16-97456061-INS-65
chr16-97461220-INS-745
chr16-97461494-INS-124
chr16-97485341-INS-72
chr16-97509090-INS-406
chr16-97553992-INS-229
chr16-97588377-INS-70
chr16-97596251-INS-274
chr16-97598333-INS-101
chr16-97647818-INS-1251
chr16-97659126-INS-136
chr16-97674234-INS-209
chr16-97698534-INS-69
chr16-97704640-INS-85
chr16-97733211-INS-359
chr16-97736051-INS-93
chr16-97787916-INS-55
chr16-97808339-INS-191
chr16-97812742-INS-588
chr16-97817267-INS-177
chr16-97820352-INS-54
chr16-97823614-INS-70
chr16-97834716-INS-314
chr16-97853565-INS-897
chr16-97873357-INS-1299
chr16-97873358-INS-1608
chr16-97873392-INS-213
chr17-3051869-INS-114
chr17-3053108-INS-134
chr17-3062188-INS-166
chr17-3075624-INS-294
chr17-3077085-INS-219
chr17-3123942-INS-109
chr17-3154766-INS-401
chr17-3160863-INS-204
chr17-3183180-INS-221
chr17-3204312-INS-283
chr17-3236231-INS-400
chr17-3268260-INS-916
chr17-3270002-INS-173
chr17-3288988-INS-100
chr17-3301796-INS-88
chr17-3303444-INS-117
chr17-3321425-INS-230
chr17-3324077-INS-219
chr17-3332994-INS-108
chr17-3338448-INS-183
chr17-3356246-INS-52
chr17-3369580-INS-221
chr17-3374570-INS-94
chr17-3392684-INS-540
chr17-3403207-INS-1062
chr17-3403339-INS-247
chr17-3403476-INS-956
chr17-3411598-INS-120
chr17-3433415-INS-314
chr17-3461366-INS-156
chr17-3466726-INS-162

chr17-3514171-INS-60
chr17-3525603-INS-489
chr17-3564358-INS-169
chr17-3581856-INS-213
chr17-3589330-INS-442
chr17-3619653-INS-99
chr17-3631353-INS-126
chr17-3640866-INS-399
chr17-3658462-INS-56
chr17-3704083-INS-60
chr17-3712663-INS-87
chr17-3733952-INS-196
chr17-3735463-INS-351
chr17-3748571-INS-353
chr17-3768378-INS-100
chr17-3803300-INS-87
chr17-3807407-INS-61
chr17-3833724-INS-101
chr17-3846007-INS-213
chr17-3882370-INS-266
chr17-3888140-INS-624
chr17-3890091-INS-155
chr17-3894068-INS-197
chr17-3921544-INS-160
chr17-3951288-INS-342
chr17-3954303-INS-887
chr17-4019538-INS-64
chr17-4061195-INS-13251
chr17-4066702-INS-103
chr17-4090169-INS-54
chr17-4122914-INS-156
chr17-4125198-INS-7117
chr17-4126175-INS-605
chr17-4157528-INS-133
chr17-4161134-INS-134
chr17-4186091-INS-676
chr17-4208464-INS-424
chr17-4238781-INS-230
chr17-4249298-INS-54
chr17-4252709-INS-238
chr17-4263775-INS-711
chr17-4276202-INS-52
chr17-4320331-INS-294
chr17-4348661-INS-111
chr17-4367134-INS-51
chr17-4367562-INS-125
chr17-4404684-INS-60
chr17-4427157-INS-173
chr17-4427808-INS-395
chr17-4447061-INS-79
chr17-4461396-INS-1928
chr17-4465903-INS-108
chr17-4468601-INS-628
chr17-4477620-INS-135
chr17-4479062-INS-60
chr17-4491087-INS-52
chr17-4494767-INS-56
chr17-4519411-INS-98
chr17-4549568-INS-98
chr17-4554414-INS-64
chr17-4557228-INS-88
chr17-4558760-INS-723
chr17-4568867-INS-52
chr17-4572784-INS-99
chr17-4587679-INS-4546
chr17-4592380-INS-125
chr17-4597774-INS-62
chr17-4604655-INS-60
chr17-4643696-INS-72
chr17-4652734-INS-159
chr17-4661634-INS-6516
chr17-4667614-INS-61
chr17-4670062-INS-110
chr17-4673436-INS-215
chr17-4676484-INS-305
chr17-4694975-INS-68
chr17-4710618-INS-1864

chr17-4710922-INS-60
chr17-4785491-INS-53
chr17-4790542-INS-51
chr17-4791696-INS-332
chr17-4798015-INS-432
chr17-4807917-INS-5263
chr17-4817580-INS-172
chr17-4830578-INS-157
chr17-4842558-INS-144
chr17-4843142-INS-399
chr17-4845940-INS-234
chr17-4847721-INS-59
chr17-4849440-INS-163
chr17-4855167-INS-116
chr17-4858751-INS-706
chr17-4861302-INS-142
chr17-4865493-INS-261
chr17-4897794-INS-54
chr17-4907079-INS-163
chr17-4907591-INS-131
chr17-4912005-INS-270
chr17-4914063-INS-236
chr17-4923080-INS-54
chr17-4924383-INS-54
chr17-4944235-INS-64
chr17-4961759-INS-230
chr17-4973932-INS-291
chr17-4975609-INS-607
chr17-4991663-INS-185
chr17-4993590-INS-218
chr17-5011869-INS-202
chr17-5020729-INS-102
chr17-5055904-INS-306
chr17-5080789-INS-200
chr17-5092752-INS-1890
chr17-5105116-INS-55
chr17-5106592-INS-108
chr17-5123519-INS-218
chr17-5132277-INS-261
chr17-5150171-INS-193
chr17-5153317-INS-143
chr17-5182975-INS-304
chr17-5190678-INS-139
chr17-5194858-INS-120
chr17-5197772-INS-60
chr17-5206940-INS-128
chr17-5208839-INS-506
chr17-5209283-INS-483
chr17-5217184-INS-70
chr17-5224661-INS-390
chr17-5228268-INS-130
chr17-5242651-INS-192
chr17-5247213-INS-541
chr17-5249604-INS-272
chr17-5291284-INS-3527
chr17-5291516-INS-132
chr17-5301879-INS-197
chr17-5304392-INS-612
chr17-5348838-INS-57
chr17-5352378-INS-206
chr17-5363832-INS-86
chr17-5363924-INS-778
chr17-5365246-INS-140
chr17-5383173-INS-52
chr17-5406511-INS-56
chr17-5410835-INS-318
chr17-5434829-INS-312
chr17-5442229-INS-233
chr17-5444002-INS-55
chr17-5459671-INS-5645
chr17-5464992-INS-497
chr17-5475970-INS-1038
chr17-5506584-INS-210
chr17-5510229-INS-52
chr17-5511135-INS-487
chr17-5512749-INS-68
chr17-5528813-INS-247

chr17-5548609-INS-71
chr17-5552290-INS-132
chr17-5579196-INS-114
chr17-5579851-INS-5659
chr17-5581966-INS-132
chr17-5607288-INS-76
chr17-5614269-INS-91
chr17-5617920-INS-52
chr17-5622017-INS-218
chr17-5631404-INS-324
chr17-5633101-INS-98
chr17-5641711-INS-6835
chr17-5645992-INS-734
chr17-5655719-INS-344
chr17-5673273-INS-538
chr17-5684133-INS-728
chr17-5691406-INS-80
chr17-5710339-INS-196
chr17-5717766-INS-192
chr17-5733770-INS-608
chr17-5767714-INS-52
chr17-5780670-INS-72
chr17-5791620-INS-144
chr17-5799452-INS-130
chr17-5807373-INS-72
chr17-5815498-INS-142
chr17-5827864-INS-57
chr17-5839154-INS-6433
chr17-5842755-INS-64
chr17-5847381-INS-141
chr17-5848561-INS-204
chr17-5859177-INS-86
chr17-5864530-INS-200
chr17-5865630-INS-997
chr17-5868135-INS-400
chr17-5883540-INS-204
chr17-5887795-INS-214
chr17-5894685-INS-275
chr17-5896903-INS-291
chr17-5933936-INS-163
chr17-5934343-INS-2402
chr17-5938620-INS-117
chr17-5943943-INS-273
chr17-5945173-INS-83
chr17-5951081-INS-225
chr17-5951632-INS-225
chr17-5967199-INS-59
chr17-5978708-INS-204
chr17-5978782-INS-736
chr17-5991532-INS-53
chr17-6015556-INS-169
chr17-6025361-INS-113
chr17-6064698-INS-86
chr17-6081156-INS-184
chr17-6084344-INS-55
chr17-6090026-INS-62
chr17-6103632-INS-152
chr17-6113236-INS-407
chr17-6115772-INS-213
chr17-6132350-INS-77
chr17-6154108-INS-214
chr17-6170638-INS-90
chr17-6180910-INS-106
chr17-6196753-INS-59
chr17-6232610-INS-320
chr17-6234946-INS-84
chr17-6246485-INS-634
chr17-6269895-INS-128
chr17-6286240-INS-65
chr17-6325823-INS-593
chr17-6375265-INS-3425
chr17-6394059-INS-332
chr17-6395269-INS-68
chr17-6441308-INS-196
chr17-6658335-INS-612
chr17-6693836-INS-51
chr17-6697755-INS-569

chr17-6731607-INS-963
chr17-6758191-INS-209
chr17-6771686-INS-64
chr17-6798681-INS-193
chr17-6847601-INS-5722
chr17-6849681-INS-205
chr17-6852341-INS-70
chr17-6854578-INS-196
chr17-6856930-INS-328
chr17-6946552-INS-434
chr17-6961196-INS-381
chr17-6971231-INS-749
chr17-6977025-INS-173
chr17-6984649-INS-171
chr17-6997747-INS-450
chr17-7016590-INS-90
chr17-7030712-INS-58
chr17-7048729-INS-110
chr17-7054834-INS-85
chr17-7058566-INS-65
chr17-7067210-INS-58
chr17-7078245-INS-90
chr17-7083026-INS-134
chr17-7101826-INS-5552
chr17-7106010-INS-5220
chr17-7109010-INS-52
chr17-7127478-INS-77
chr17-7260836-INS-110
chr17-7271267-INS-1866
chr17-7273800-INS-1890
chr17-7296786-INS-451
chr17-7357148-INS-147
chr17-7407947-INS-292
chr17-7409630-INS-65
chr17-7422547-INS-190
chr17-7423304-INS-278
chr17-7454489-INS-64
chr17-7467933-INS-160
chr17-7496368-INS-180
chr17-7505341-INS-100
chr17-7539728-INS-206
chr17-7555278-INS-203
chr17-7560329-INS-141
chr17-7590015-INS-118
chr17-7610385-INS-78
chr17-7616887-INS-1887
chr17-7623176-INS-6245
chr17-7633040-INS-68
chr17-7651166-INS-6468
chr17-7670495-INS-122
chr17-7681226-INS-5472
chr17-7706302-INS-66
chr17-7725792-INS-119
chr17-7745941-INS-189
chr17-7749188-INS-243
chr17-7759851-INS-134
chr17-7780235-INS-535
chr17-7795423-INS-80
chr17-7798394-INS-52
chr17-7807228-INS-63
chr17-7812771-INS-82
chr17-7824647-INS-95
chr17-7829094-INS-166
chr17-7829929-INS-267
chr17-7871533-INS-6941
chr17-7879792-INS-6400
chr17-7906176-INS-76
chr17-7925188-INS-218
chr17-7943138-INS-70
chr17-7949935-INS-2124
chr17-7962536-INS-94
chr17-7963690-INS-252
chr17-7970348-INS-74
chr17-7971104-INS-112
chr17-7978240-INS-200
chr17-7985969-INS-73
chr17-7997945-INS-63

chr17-7998587-INS-506
chr17-8052588-INS-351
chr17-8070466-INS-212
chr17-8080837-INS-461
chr17-8100442-INS-572
chr17-8105164-INS-52
chr17-8123367-INS-103
chr17-8135357-INS-875
chr17-8163979-INS-5219
chr17-8175444-INS-1450
chr17-8177834-INS-5301
chr17-8218617-INS-4232
chr17-8239531-INS-4663
chr17-8253487-INS-101
chr17-8254465-INS-7090
chr17-8260662-INS-2445
chr17-8273093-INS-72
chr17-8283500-INS-399
chr17-8290551-INS-7159
chr17-8291661-INS-118
chr17-8294031-INS-69
chr17-8316174-INS-451
chr17-8339803-INS-76
chr17-8349023-INS-80
chr17-8352197-INS-79
chr17-8365682-INS-55
chr17-8373656-INS-215
chr17-8394788-INS-984
chr17-8396940-INS-70
chr17-8412725-INS-57
chr17-8414745-INS-88
chr17-8419359-INS-87
chr17-8427059-INS-126
chr17-8446320-INS-129
chr17-8455829-INS-110
chr17-8460262-INS-249
chr17-8470386-INS-350
chr17-8477879-INS-54
chr17-8478923-INS-122
chr17-8494750-INS-1076
chr17-8500237-INS-172
chr17-8501448-INS-66
chr17-8526794-INS-57
chr17-8545475-INS-72
chr17-8548786-INS-154
chr17-8566117-INS-225
chr17-8613061-INS-728
chr17-8620719-INS-317
chr17-8640051-INS-180
chr17-8642071-INS-150
chr17-8644137-INS-118
chr17-8646417-INS-84
chr17-8649218-INS-122
chr17-8665178-INS-96
chr17-8667090-INS-206
chr17-8680927-INS-499
chr17-8683538-INS-5489
chr17-8686529-INS-6436
chr17-8689599-INS-92
chr17-8692970-INS-58
chr17-8719930-INS-77
chr17-8737995-INS-219
chr17-8760345-INS-254
chr17-8787924-INS-830
chr17-8801336-INS-147
chr17-8807875-INS-72
chr17-8823451-INS-355
chr17-8890112-INS-293
chr17-8894400-INS-162
chr17-8900453-INS-175
chr17-8916950-INS-52
chr17-8918017-INS-63
chr17-8920735-INS-201
chr17-8923179-INS-176
chr17-8934303-INS-164
chr17-8939187-INS-149
chr17-8959689-INS-62

chr17-8961994-INS-62
chr17-8973755-INS-259
chr17-8979881-INS-70
chr17-9044227-INS-121
chr17-9045265-INS-64
chr17-9046207-INS-60
chr17-9051767-INS-536
chr17-9071616-INS-144
chr17-9074681-INS-100
chr17-9096827-INS-730
chr17-9155051-INS-111
chr17-9155261-INS-500
chr17-9234167-INS-397
chr17-9285857-INS-100
chr17-9310690-INS-106
chr17-9319194-INS-1531
chr17-9322500-INS-74
chr17-9353968-INS-343
chr17-9360773-INS-247
chr17-9414382-INS-69
chr17-9416569-INS-130
chr17-9419877-INS-274
chr17-9420731-INS-66
chr17-9422668-INS-52
chr17-9424781-INS-7096
chr17-9434309-INS-966
chr17-9434910-INS-138
chr17-9437041-INS-69
chr17-9451171-INS-226
chr17-9457334-INS-68
chr17-9472884-INS-329
chr17-9496956-INS-50
chr17-9506732-INS-273
chr17-9508054-INS-244
chr17-9628172-INS-104
chr17-9632127-INS-56
chr17-9646149-INS-154
chr17-9650945-INS-579
chr17-9681118-INS-129
chr17-9736788-INS-64
chr17-9748244-INS-398
chr17-9761791-INS-559
chr17-9793560-INS-70
chr17-9800926-INS-58
chr17-9831211-INS-56
chr17-9849517-INS-72
chr17-9850514-INS-471
chr17-9895753-INS-60
chr17-9902447-INS-217
chr17-9924774-INS-966
chr17-9963687-INS-354
chr17-9980187-INS-92
chr17-9982591-INS-102
chr17-9986035-INS-64
chr17-9987166-INS-548
chr17-9989001-INS-64
chr17-9998083-INS-57
chr17-10033730-INS-117
chr17-10052423-INS-60
chr17-10057100-INS-99
chr17-10058046-INS-98
chr17-10076139-INS-488
chr17-10103291-INS-88
chr17-10128489-INS-61
chr17-10133661-INS-389
chr17-10147804-INS-114
chr17-10242423-INS-2151
chr17-10246977-INS-102
chr17-10256843-INS-89
chr17-10259098-INS-7584
chr17-10276177-INS-575
chr17-10337591-INS-54
chr17-10345500-INS-270
chr17-10382269-INS-6725
chr17-10453203-INS-275
chr17-10469696-INS-89
chr17-10473633-INS-93

chr17-10475808-INS-200
chr17-10499756-INS-59
chr17-10502193-INS-172
chr17-10520418-INS-200
chr17-10549866-INS-51
chr17-10586961-INS-399
chr17-10608826-INS-59
chr17-10617644-INS-64
chr17-10628408-INS-116
chr17-10636022-INS-5547
chr17-10646163-INS-158
chr17-10652494-INS-186
chr17-10663895-INS-198
chr17-10664002-INS-793
chr17-10665355-INS-62
chr17-10669908-INS-691
chr17-10683216-INS-161
chr17-10747429-INS-830
chr17-10809892-INS-351
chr17-10815161-INS-176
chr17-10826053-INS-67
chr17-10828235-INS-3111
chr17-10831318-INS-1746
chr17-10834513-INS-53
chr17-10841536-INS-92
chr17-10850715-INS-60
chr17-10910426-INS-67
chr17-10931927-INS-359
chr17-10948105-INS-75
chr17-10958270-INS-170
chr17-10976838-INS-75
chr17-10980885-INS-160
chr17-11020712-INS-883
chr17-11034666-INS-246
chr17-11047638-INS-56
chr17-11054242-INS-532
chr17-11087937-INS-372
chr17-11102540-INS-1685
chr17-11123922-INS-4358
chr17-11131742-INS-169
chr17-11149805-INS-200
chr17-11166095-INS-87
chr17-11179278-INS-53
chr17-11180123-INS-6738
chr17-11227650-INS-201
chr17-11237239-INS-60
chr17-11241208-INS-58
chr17-11243660-INS-64
chr17-11247757-INS-226
chr17-11262053-INS-70
chr17-11268828-INS-435
chr17-11279383-INS-410
chr17-11301744-INS-198
chr17-11326202-INS-252
chr17-11336913-INS-52
chr17-11366829-INS-5224
chr17-11398525-INS-157
chr17-11470080-INS-519
chr17-11471288-INS-2834
chr17-11493927-INS-60
chr17-11494472-INS-1331
chr17-11499922-INS-644
chr17-11502534-INS-52
chr17-11511727-INS-382
chr17-11533413-INS-1624
chr17-11538097-INS-6144
chr17-11565004-INS-84
chr17-11602156-INS-898
chr17-11603865-INS-98
chr17-11620146-INS-58
chr17-11620981-INS-126
chr17-11622213-INS-6485
chr17-11637318-INS-78
chr17-11647196-INS-449
chr17-11682587-INS-75
chr17-11728781-INS-7255
chr17-11737145-INS-139

chr17-11741217-INS-133
chr17-11743724-INS-163
chr17-11747031-INS-87
chr17-11763700-INS-72
chr17-11777914-INS-6552
chr17-11787794-INS-337
chr17-11796802-INS-52
chr17-11807835-INS-208
chr17-11839769-INS-554
chr17-11842960-INS-174
chr17-11850768-INS-51
chr17-11872523-INS-124
chr17-11874093-INS-66
chr17-11891326-INS-71
chr17-11905948-INS-240
chr17-11907330-INS-113
chr17-11934148-INS-54
chr17-11952039-INS-141
chr17-11966894-INS-80
chr17-11972840-INS-67
chr17-12033659-INS-70
chr17-12042709-INS-7234
chr17-12094271-INS-61
chr17-12114014-INS-74
chr17-12134317-INS-572
chr17-12137795-INS-244
chr17-12156826-INS-421
chr17-12158007-INS-328
chr17-12197868-INS-91
chr17-12269479-INS-3135
chr17-12282201-INS-5410
chr17-12292871-INS-7081
chr17-12298726-INS-6699
chr17-12307484-INS-51
chr17-12311013-INS-153
chr17-12323517-INS-92
chr17-12349369-INS-213
chr17-12392009-INS-333
chr17-12411375-INS-207
chr17-12471175-INS-99
chr17-12494472-INS-1764
chr17-12508533-INS-377
chr17-12533987-INS-61
chr17-12543781-INS-479
chr17-12572506-INS-76
chr17-12581645-INS-76
chr17-12584018-INS-132
chr17-12596229-INS-217
chr17-12598618-INS-78
chr17-12605299-INS-855
chr17-12612724-INS-59
chr17-12628902-INS-401
chr17-12630823-INS-120
chr17-12654295-INS-56
chr17-12655401-INS-98
chr17-12657769-INS-692
chr17-12666481-INS-133
chr17-12671176-INS-7330
chr17-12674467-INS-146
chr17-12676289-INS-143
chr17-12691451-INS-7115
chr17-12698561-INS-66
chr17-12705199-INS-253
chr17-12735403-INS-1862
chr17-12743436-INS-153
chr17-12744503-INS-339
chr17-12767242-INS-78
chr17-12798419-INS-59
chr17-12832783-INS-607
chr17-12835707-INS-118
chr17-12836766-INS-308
chr17-12851727-INS-71
chr17-12859675-INS-12115
chr17-12861043-INS-5304
chr17-12862817-INS-4845
chr17-12868253-INS-161
chr17-12932014-INS-191

chr17-12956800-INS-760
chr17-12972585-INS-1869
chr17-13024635-INS-149
chr17-13067990-INS-171
chr17-13079145-INS-179
chr17-13087858-INS-86
chr17-13090064-INS-130
chr17-13097509-INS-60
chr17-13106214-INS-55
chr17-13122714-INS-57
chr17-13131515-INS-203
chr17-13168555-INS-64
chr17-13169727-INS-223
chr17-13172236-INS-197
chr17-13172798-INS-734
chr17-13185565-INS-159
chr17-13197901-INS-214
chr17-13203569-INS-66
chr17-13220969-INS-7795
chr17-13225541-INS-421
chr17-13228177-INS-715
chr17-13288080-INS-52
chr17-13298495-INS-52
chr17-13391050-INS-71
chr17-13391342-INS-6585
chr17-13394038-INS-96
chr17-13403156-INS-66
chr17-13413143-INS-229
chr17-13418234-INS-72
chr17-13422986-INS-56
chr17-13426877-INS-54
chr17-13536028-INS-141
chr17-13556951-INS-6176
chr17-13598051-INS-6689
chr17-13623726-INS-81
chr17-13665854-INS-122
chr17-13673001-INS-7142
chr17-13678633-INS-82
chr17-13723866-INS-57
chr17-13736930-INS-6382
chr17-13772159-INS-90
chr17-13779185-INS-55
chr17-13786672-INS-54
chr17-13888098-INS-5565
chr17-13888100-INS-8985
chr17-13896373-INS-154
chr17-13901333-INS-90
chr17-13907417-INS-72
chr17-13914332-INS-908
chr17-13976012-INS-79
chr17-13994916-INS-84
chr17-13997679-INS-126
chr17-14010462-INS-98
chr17-14013075-INS-846
chr17-14022585-INS-394
chr17-14107183-INS-90
chr17-14121291-INS-60
chr17-14133614-INS-208
chr17-14137687-INS-60
chr17-14148072-INS-270
chr17-14181925-INS-542
chr17-14186402-INS-345
chr17-14203365-INS-184
chr17-14232224-INS-246
chr17-14261440-INS-345
chr17-14299789-INS-794
chr17-14345946-INS-58
chr17-14350936-INS-2064
chr17-14354032-INS-58
chr17-14441541-INS-159
chr17-14452208-INS-73
chr17-14472396-INS-61
chr17-14487000-INS-61
chr17-14490903-INS-199
chr17-14511424-INS-80
chr17-14511602-INS-272
chr17-14561387-INS-678

chr17-14582899-INS-186
chr17-14594307-INS-6679
chr17-14595067-INS-52
chr17-14601879-INS-53
chr17-14606282-INS-80
chr17-14620286-INS-5176
chr17-14647853-INS-51
chr17-14665004-INS-58
chr17-14665611-INS-792
chr17-14674382-INS-143
chr17-14694429-INS-469
chr17-14768689-INS-171
chr17-14771242-INS-80
chr17-14794348-INS-6720
chr17-14807876-INS-98
chr17-14862583-INS-59
chr17-14863617-INS-194
chr17-14865735-INS-53
chr17-14881864-INS-390
chr17-14907865-INS-6789
chr17-14950272-INS-51
chr17-14957735-INS-174
chr17-14962020-INS-148
chr17-14984053-INS-1696
chr17-14994820-INS-528
chr17-15012028-INS-73
chr17-15013515-INS-65
chr17-15015842-INS-296
chr17-15026943-INS-213
chr17-15044790-INS-308
chr17-15053755-INS-184
chr17-15060614-INS-979
chr17-15061842-INS-147
chr17-15070935-INS-64
chr17-15076884-INS-6858
chr17-15080183-INS-526
chr17-15082064-INS-273
chr17-15117869-INS-204
chr17-15136304-INS-66
chr17-15141806-INS-81
chr17-15153079-INS-318
chr17-15153973-INS-125
chr17-15158857-INS-522
chr17-15179829-INS-144
chr17-15185189-INS-195
chr17-15196880-INS-213
chr17-15223849-INS-212
chr17-15259621-INS-224
chr17-15270198-INS-75
chr17-15271549-INS-250
chr17-15274058-INS-369
chr17-15284548-INS-72
chr17-15310884-INS-98
chr17-15328399-INS-55
chr17-15349212-INS-158
chr17-15352689-INS-226
chr17-15361046-INS-221
chr17-15377847-INS-93
chr17-15378971-INS-59
chr17-15412201-INS-361
chr17-15425678-INS-66
chr17-15428917-INS-58
chr17-15432320-INS-401
chr17-15435883-INS-58
chr17-15439024-INS-52
chr17-15469090-INS-69
chr17-15481358-INS-243
chr17-15482315-INS-156
chr17-15485386-INS-6687
chr17-15512483-INS-101
chr17-15567651-INS-906
chr17-15580612-INS-202
chr17-15582595-INS-68
chr17-15610434-INS-385
chr17-15614656-INS-433
chr17-15632219-INS-208
chr17-15647326-INS-102

chr17-15659703-INS-199
chr17-15670617-INS-54
chr17-15700629-INS-60
chr17-15721481-INS-178
chr17-15753956-INS-273
chr17-15763105-INS-329
chr17-15767366-INS-325
chr17-15780679-INS-174
chr17-15803321-INS-56
chr17-15805510-INS-379
chr17-15809768-INS-506
chr17-15811401-INS-56
chr17-15814763-INS-178
chr17-15816610-INS-161
chr17-15822397-INS-789
chr17-15822400-INS-190
chr17-15825852-INS-5610
chr17-15832982-INS-143
chr17-15835821-INS-77
chr17-15838661-INS-213
chr17-15857584-INS-217
chr17-15862867-INS-54
chr17-15863957-INS-4544
chr17-15865873-INS-201
chr17-15885306-INS-6348
chr17-15903094-INS-54
chr17-15911472-INS-345
chr17-15914073-INS-64
chr17-15919155-INS-211
chr17-15940142-INS-212
chr17-15945757-INS-62
chr17-15994016-INS-200
chr17-16017245-INS-56
chr17-16023324-INS-90
chr17-16039026-INS-3840
chr17-16059254-INS-6440
chr17-16066203-INS-186
chr17-16077063-INS-390
chr17-16083804-INS-134
chr17-16105839-INS-64
chr17-16120029-INS-2555
chr17-16120161-INS-110
chr17-16122195-INS-250
chr17-16177206-INS-6462
chr17-16178438-INS-180
chr17-16189688-INS-151
chr17-16212307-INS-122
chr17-16226796-INS-147
chr17-16237395-INS-126
chr17-16241841-INS-87
chr17-16251058-INS-465
chr17-16273532-INS-216
chr17-16278721-INS-320
chr17-16312040-INS-6244
chr17-16312237-INS-281
chr17-16337728-INS-53
chr17-16351055-INS-64
chr17-16376957-INS-694
chr17-16402141-INS-663
chr17-16402811-INS-81
chr17-16443349-INS-251
chr17-16455045-INS-7045
chr17-16458289-INS-55
chr17-16467151-INS-443
chr17-16479124-INS-6939
chr17-16517267-INS-4453
chr17-16517588-INS-2864
chr17-16542165-INS-82
chr17-16544170-INS-104
chr17-16544706-INS-510
chr17-16558264-INS-150
chr17-16570700-INS-6865
chr17-16574282-INS-282
chr17-16600949-INS-2717
chr17-16632330-INS-783
chr17-16668708-INS-313
chr17-16696350-INS-56

chr17-16711989-INS-240
chr17-16713516-INS-52
chr17-16743886-INS-94
chr17-16746730-INS-114
chr17-16781483-INS-81
chr17-16834722-INS-448
chr17-16836140-INS-153
chr17-16846262-INS-70
chr17-16849606-INS-233
chr17-16854662-INS-67
chr17-16868022-INS-61
chr17-16896795-INS-318
chr17-16920319-INS-262
chr17-16928570-INS-310
chr17-16938404-INS-869
chr17-16940446-INS-203
chr17-16957651-INS-104
chr17-16966348-INS-6410
chr17-16995115-INS-70
chr17-16998255-INS-192
chr17-17009901-INS-106
chr17-17011080-INS-554
chr17-17033873-INS-6711
chr17-17039272-INS-60
chr17-17041442-INS-6436
chr17-17078382-INS-214
chr17-17082004-INS-130
chr17-17087033-INS-100
chr17-17111222-INS-125
chr17-17123130-INS-308
chr17-17124267-INS-586
chr17-17128824-INS-173
chr17-17144564-INS-52
chr17-17148218-INS-206
chr17-17164808-INS-54
chr17-17189900-INS-67
chr17-17191573-INS-51
chr17-17230643-INS-624
chr17-17238967-INS-192
chr17-17256074-INS-666
chr17-17298267-INS-7088
chr17-17344919-INS-2738
chr17-17390768-INS-55
chr17-17506460-INS-54
chr17-17512695-INS-502
chr17-17546623-INS-219
chr17-17561149-INS-98
chr17-17585572-INS-485
chr17-17634863-INS-92
chr17-17635651-INS-203
chr17-17639297-INS-96
chr17-17674880-INS-175
chr17-17679274-INS-71
chr17-17680961-INS-5220
chr17-17683208-INS-119
chr17-17684733-INS-193
chr17-17713019-INS-242
chr17-17721865-INS-61
chr17-17723733-INS-64
chr17-17746778-INS-54
chr17-17753604-INS-353
chr17-17788182-INS-219
chr17-17789451-INS-54
chr17-17797007-INS-80
chr17-17806094-INS-155
chr17-17807187-INS-56
chr17-17818493-INS-187
chr17-17829101-INS-75
chr17-17838480-INS-182
chr17-17840454-INS-194
chr17-17867123-INS-1709
chr17-17887770-INS-437
chr17-17891164-INS-7101
chr17-17891165-INS-227
chr17-17899743-INS-495
chr17-17905096-INS-4441
chr17-17922498-INS-599

chr17-17930894-INS-621
chr17-17935025-INS-329
chr17-17956621-INS-324
chr17-17977388-INS-812
chr17-17982365-INS-68
chr17-18009774-INS-160
chr17-18010446-INS-575
chr17-18031095-INS-338
chr17-18035188-INS-141
chr17-18038500-INS-52
chr17-18040779-INS-73
chr17-18079736-INS-66
chr17-18122366-INS-2020
chr17-18153876-INS-7399
chr17-18157261-INS-578
chr17-18160897-INS-514
chr17-18197469-INS-1894
chr17-18203276-INS-431
chr17-18213389-INS-6301
chr17-18239931-INS-56
chr17-18250468-INS-793
chr17-18260838-INS-6238
chr17-18273223-INS-698
chr17-18288935-INS-4083
chr17-18292398-INS-1224
chr17-18297924-INS-7216
chr17-18300758-INS-10060
chr17-18300859-INS-6670
chr17-18306216-INS-94
chr17-18307122-INS-2882
chr17-18362773-INS-961
chr17-18373071-INS-249
chr17-18377665-INS-379
chr17-18432330-INS-3949
chr17-18457169-INS-2225
chr17-18474344-INS-204
chr17-18483501-INS-56
chr17-18498481-INS-186
chr17-18505840-INS-66
chr17-18509941-INS-66
chr17-18581778-INS-85
chr17-18586350-INS-54
chr17-18593581-INS-132
chr17-18657781-INS-615
chr17-18658329-INS-196
chr17-18704416-INS-204
chr17-18747240-INS-8771
chr17-18800688-INS-3186
chr17-18837130-INS-59
chr17-18922327-INS-64
chr17-19051828-INS-99
chr17-19128795-INS-7326
chr17-19257736-INS-2794
chr17-19262864-INS-577
chr17-19294412-INS-154
chr17-19339036-INS-85
chr17-19355490-INS-614
chr17-19402944-INS-57
chr17-19409490-INS-380
chr17-19411208-INS-5475
chr17-19446168-INS-59
chr17-19458922-INS-355
chr17-19479153-INS-124
chr17-19501453-INS-6709
chr17-19619184-INS-494
chr17-19668958-INS-52
chr17-19774163-INS-6475
chr17-19806564-INS-195
chr17-19837624-INS-55
chr17-19851590-INS-53
chr17-19873363-INS-6424
chr17-19907218-INS-77
chr17-19922243-INS-499
chr17-19923711-INS-140
chr17-19945783-INS-52
chr17-19954443-INS-69
chr17-20054858-INS-257

chr17-20074939-INS-193
chr17-20146092-INS-6536
chr17-20154487-INS-127
chr17-20163578-INS-55
chr17-20194392-INS-935
chr17-20289911-INS-56
chr17-20304931-INS-52
chr17-20325044-INS-323
chr17-20345879-INS-130
chr17-20356203-INS-443
chr17-20391222-INS-57
chr17-20411792-INS-540
chr17-20414937-INS-58
chr17-20417409-INS-53
chr17-20421870-INS-68
chr17-20424319-INS-100
chr17-20434645-INS-52
chr17-20443089-INS-64
chr17-20455998-INS-55
chr17-20482792-INS-7752
chr17-20490592-INS-8326
chr17-20505381-INS-1318
chr17-20521869-INS-3099
chr17-20538771-INS-625
chr17-20542840-INS-6655
chr17-20543118-INS-3963
chr17-20543998-INS-6883
chr17-20555156-INS-108
chr17-20562421-INS-360
chr17-20588484-INS-7195
chr17-20594938-INS-224
chr17-20631481-INS-120
chr17-20644763-INS-110
chr17-20695749-INS-182
chr17-20792837-INS-584
chr17-20842409-INS-11492
chr17-20886705-INS-62
chr17-20903169-INS-1263
chr17-20919661-INS-62
chr17-20959621-INS-266
chr17-20965850-INS-129
chr17-20995804-INS-457
chr17-21001113-INS-660
chr17-21022247-INS-6482
chr17-21042152-INS-6839
chr17-21049561-INS-326
chr17-21072360-INS-501
chr17-21081221-INS-6501
chr17-21088075-INS-104
chr17-21197333-INS-7132
chr17-21218123-INS-937
chr17-21227017-INS-404
chr17-21235449-INS-3058
chr17-21242960-INS-82
chr17-21255916-INS-64
chr17-21255960-INS-2034
chr17-21256720-INS-2037
chr17-21284297-INS-749
chr17-21372773-INS-182
chr17-21406118-INS-922
chr17-21468971-INS-9036
chr17-21484703-INS-975
chr17-21536088-INS-908
chr17-21551501-INS-60
chr17-21554453-INS-190
chr17-21592280-INS-53
chr17-21629284-INS-197
chr17-21658121-INS-112
chr17-21682075-INS-181
chr17-21730514-INS-498
chr17-21732791-INS-244
chr17-21737487-INS-257
chr17-21761494-INS-73
chr17-21778296-INS-6465
chr17-21794527-INS-127
chr17-21806024-INS-135
chr17-21807353-INS-99

chr17-21817198-INS-304
chr17-21840930-INS-54
chr17-21874967-INS-401
chr17-21884791-INS-199
chr17-21910182-INS-84
chr17-21912985-INS-345
chr17-21914731-INS-6442
chr17-21914764-INS-3176
chr17-21920000-INS-293
chr17-21926178-INS-6504
chr17-21947834-INS-481
chr17-21963140-INS-84
chr17-21967126-INS-75
chr17-21985065-INS-54
chr17-21991903-INS-657
chr17-22005863-INS-120
chr17-22010277-INS-7372
chr17-22039372-INS-11687
chr17-22044924-INS-188
chr17-22058579-INS-64
chr17-22077652-INS-208
chr17-22088660-INS-830
chr17-22115522-INS-201
chr17-22117445-INS-6427
chr17-22129128-INS-51
chr17-22143110-INS-202
chr17-22170637-INS-303
chr17-22201870-INS-328
chr17-22205766-INS-6243
chr17-22219044-INS-92
chr17-22312743-INS-59
chr17-22393714-INS-5417
chr17-22394115-INS-411
chr17-22401564-INS-144
chr17-22410337-INS-154
chr17-22418930-INS-162
chr17-22459028-INS-407
chr17-22478015-INS-287
chr17-22497884-INS-175
chr17-22514393-INS-6440
chr17-22564324-INS-196
chr17-22582708-INS-250
chr17-22611918-INS-54
chr17-22651665-INS-341
chr17-22654046-INS-6615
chr17-22661233-INS-656
chr17-22665033-INS-54
chr17-22701309-INS-6240
chr17-22702364-INS-526
chr17-22705493-INS-5774
chr17-22723884-INS-572
chr17-22727447-INS-71
chr17-22735510-INS-66
chr17-22763130-INS-5497
chr17-22773766-INS-6801
chr17-22775469-INS-1163
chr17-22785707-INS-5161
chr17-22803605-INS-11267
chr17-22816356-INS-6215
chr17-22823296-INS-131
chr17-22909514-INS-84
chr17-22913254-INS-68
chr17-22930487-INS-6661
chr17-22948065-INS-63
chr17-22966550-INS-5311
chr17-22998649-INS-740
chr17-23006865-INS-6399
chr17-23025171-INS-787
chr17-23039055-INS-5150
chr17-23057044-INS-696
chr17-23060107-INS-342
chr17-23108509-INS-562
chr17-23140294-INS-497
chr17-23172632-INS-350
chr17-23183467-INS-115
chr17-23235394-INS-7200
chr17-23277639-INS-56

chr17-23332375-INS-135
chr17-23353706-INS-7483
chr17-23385236-INS-5029
chr17-23417017-INS-6452
chr17-23458919-INS-54
chr17-23494896-INS-1068
chr17-23666973-INS-208
chr17-23691896-INS-208
chr17-23730413-INS-74
chr17-23735838-INS-8018
chr17-23737529-INS-184
chr17-23744919-INS-218
chr17-23750205-INS-140
chr17-23756121-INS-133
chr17-23767940-INS-210
chr17-23776149-INS-112
chr17-23802385-INS-189
chr17-23809120-INS-99
chr17-23810683-INS-64
chr17-23840251-INS-90
chr17-23845137-INS-550
chr17-23871161-INS-108
chr17-23876716-INS-51
chr17-23877813-INS-53
chr17-23885179-INS-193
chr17-23899043-INS-165
chr17-23914389-INS-199
chr17-23922774-INS-54
chr17-23972984-INS-134
chr17-23974154-INS-210
chr17-23987620-INS-209
chr17-23989494-INS-152
chr17-23994479-INS-71
chr17-24007241-INS-124
chr17-24030185-INS-205
chr17-24059325-INS-205
chr17-24062134-INS-52
chr17-24064213-INS-404
chr17-24097506-INS-62
chr17-24146930-INS-188
chr17-24167167-INS-381
chr17-24183533-INS-269
chr17-24204985-INS-108
chr17-24214780-INS-82
chr17-24216106-INS-5215
chr17-24219848-INS-56
chr17-24242623-INS-212
chr17-24278316-INS-173
chr17-24303223-INS-129
chr17-24313041-INS-1012
chr17-24314807-INS-228
chr17-24326003-INS-135
chr17-24330787-INS-204
chr17-24336340-INS-60
chr17-24337380-INS-275
chr17-24352836-INS-6440
chr17-24366462-INS-869
chr17-24371681-INS-214
chr17-24404013-INS-266
chr17-24406762-INS-51
chr17-24409090-INS-241
chr17-24410013-INS-212
chr17-24446548-INS-56
chr17-24459490-INS-172
chr17-24466552-INS-160
chr17-24472123-INS-230
chr17-24478520-INS-69
chr17-24486359-INS-130
chr17-24489677-INS-193
chr17-24507260-INS-122
chr17-24520323-INS-61
chr17-24526333-INS-179
chr17-24528929-INS-89
chr17-24538943-INS-51
chr17-24561850-INS-154
chr17-24567618-INS-192
chr17-24575782-INS-146

chr17-24587825-INS-153
chr17-24592651-INS-124
chr17-24599178-INS-492
chr17-24601335-INS-198
chr17-24623629-INS-227
chr17-24651405-INS-73
chr17-24657556-INS-138
chr17-24672292-INS-213
chr17-24710226-INS-238
chr17-24711999-INS-164
chr17-24716770-INS-122
chr17-24740230-INS-207
chr17-24742074-INS-189
chr17-24749933-INS-89
chr17-24755448-INS-203
chr17-24760779-INS-248
chr17-24763950-INS-96
chr17-24766276-INS-283
chr17-24836221-INS-115
chr17-24836676-INS-156
chr17-24840758-INS-209
chr17-24844058-INS-826
chr17-24881548-INS-249
chr17-24882696-INS-203
chr17-24921957-INS-137
chr17-24923907-INS-338
chr17-24925959-INS-311
chr17-24929185-INS-209
chr17-24933276-INS-51
chr17-24935797-INS-157
chr17-24944490-INS-52
chr17-24944950-INS-175
chr17-24949232-INS-2393
chr17-24981409-INS-209
chr17-24984953-INS-78
chr17-24990228-INS-201
chr17-24993828-INS-96
chr17-24995672-INS-141
chr17-25005084-INS-721
chr17-25007541-INS-211
chr17-25011533-INS-203
chr17-25014329-INS-539
chr17-25017863-INS-52
chr17-25020362-INS-205
chr17-25025061-INS-68
chr17-25092351-INS-60
chr17-25092908-INS-461
chr17-25164672-INS-142
chr17-25190029-INS-197
chr17-25261593-INS-116
chr17-25267857-INS-171
chr17-25269815-INS-118
chr17-25290411-INS-654
chr17-25309133-INS-222
chr17-25310440-INS-136
chr17-25312497-INS-198
chr17-25323089-INS-77
chr17-25344408-INS-54
chr17-25348135-INS-89
chr17-25377920-INS-82
chr17-25414991-INS-58
chr17-25418354-INS-107
chr17-25429424-INS-173
chr17-25434373-INS-227
chr17-25457184-INS-66
chr17-25488860-INS-93
chr17-25495077-INS-230
chr17-25508863-INS-207
chr17-25522581-INS-76
chr17-25535267-INS-262
chr17-25540388-INS-195
chr17-25550896-INS-6413
chr17-25554641-INS-61
chr17-25565277-INS-439
chr17-25571664-INS-5669
chr17-25582970-INS-326
chr17-25621980-INS-212

chr17-25658901-INS-211
chr17-25689938-INS-70
chr17-25697404-INS-122
chr17-25705648-INS-364
chr17-25726228-INS-120
chr17-25744573-INS-106
chr17-25759136-INS-92
chr17-25763485-INS-7124
chr17-25769387-INS-66
chr17-25807601-INS-210
chr17-25824877-INS-1912
chr17-25837289-INS-80
chr17-25859747-INS-67
chr17-25861733-INS-90
chr17-25867425-INS-127
chr17-25877149-INS-141
chr17-25878420-INS-174
chr17-25879123-INS-54
chr17-25886209-INS-201
chr17-25889028-INS-1891
chr17-25896707-INS-55
chr17-25918980-INS-330
chr17-25923778-INS-379
chr17-25933874-INS-210
chr17-25952129-INS-167
chr17-25973851-INS-269
chr17-25978544-INS-72
chr17-25979690-INS-80
chr17-25981895-INS-178
chr17-25984103-INS-55
chr17-25988734-INS-99
chr17-26020300-INS-219
chr17-26033023-INS-66
chr17-26039831-INS-51
chr17-26074448-INS-5537
chr17-26076564-INS-275
chr17-26090088-INS-203
chr17-26092404-INS-227
chr17-26114322-INS-123
chr17-26121142-INS-210
chr17-26123801-INS-206
chr17-26142613-INS-83
chr17-26154993-INS-69
chr17-26189533-INS-248
chr17-26258689-INS-92
chr17-26279563-INS-212
chr17-26282013-INS-217
chr17-26291255-INS-168
chr17-26292032-INS-996
chr17-26296095-INS-211
chr17-26301460-INS-95
chr17-26301465-INS-56
chr17-26304683-INS-493
chr17-26305215-INS-181
chr17-26305879-INS-120
chr17-26308733-INS-55
chr17-26315019-INS-237
chr17-26319875-INS-3744
chr17-26322965-INS-105
chr17-26329633-INS-229
chr17-26331721-INS-171
chr17-26344769-INS-120
chr17-26386557-INS-463
chr17-26391171-INS-208
chr17-26393574-INS-189
chr17-26399005-INS-207
chr17-26442048-INS-192
chr17-26464969-INS-62
chr17-26484353-INS-177
chr17-26497090-INS-170
chr17-26535480-INS-251
chr17-26537114-INS-68
chr17-26539293-INS-98
chr17-26544849-INS-182
chr17-26547429-INS-313
chr17-26564525-INS-400
chr17-26568597-INS-214

chr17-26574313-INS-130
chr17-26605934-INS-1864
chr17-26606886-INS-187
chr17-26616177-INS-136
chr17-26617570-INS-92
chr17-26655854-INS-113
chr17-26657220-INS-182
chr17-26663457-INS-63
chr17-26691875-INS-763
chr17-26697063-INS-124
chr17-26730931-INS-250
chr17-26742290-INS-233
chr17-26757411-INS-1199
chr17-26771501-INS-228
chr17-26777269-INS-108
chr17-26797922-INS-132
chr17-26809889-INS-108
chr17-26811173-INS-363
chr17-26852131-INS-159
chr17-26867167-INS-216
chr17-26879550-INS-53
chr17-26889022-INS-529
chr17-26900646-INS-102
chr17-26914165-INS-286
chr17-26915178-INS-215
chr17-26920204-INS-195
chr17-26927468-INS-81
chr17-26942859-INS-161
chr17-26966393-INS-209
chr17-26983354-INS-276
chr17-26983544-INS-162
chr17-26983865-INS-3659
chr17-27013172-INS-222
chr17-27021289-INS-58
chr17-27031515-INS-191
chr17-27043170-INS-2707
chr17-27045492-INS-89
chr17-27050679-INS-4082
chr17-27050758-INS-1119
chr17-27055797-INS-209
chr17-27066248-INS-68
chr17-27070491-INS-300
chr17-27090716-INS-128
chr17-27102214-INS-620
chr17-27115171-INS-71
chr17-27200152-INS-143
chr17-27201047-INS-164
chr17-27203885-INS-90
chr17-27209319-INS-206
chr17-27214262-INS-129
chr17-27214747-INS-99
chr17-27220979-INS-217
chr17-27229484-INS-270
chr17-27233211-INS-210
chr17-27255495-INS-52
chr17-27292664-INS-192
chr17-27293768-INS-206
chr17-27310969-INS-120
chr17-27324675-INS-164
chr17-27351974-INS-119
chr17-27356753-INS-115
chr17-27365217-INS-218
chr17-27381063-INS-150
chr17-27416528-INS-115
chr17-27424182-INS-205
chr17-27430528-INS-209
chr17-27450502-INS-264
chr17-27453361-INS-497
chr17-27456221-INS-197
chr17-27478790-INS-505
chr17-27493338-INS-53
chr17-27509815-INS-187
chr17-27542670-INS-55
chr17-27561083-INS-56
chr17-27565700-INS-57
chr17-27610356-INS-65
chr17-27614619-INS-358

chr17-27655664-INS-82
chr17-27689349-INS-125
chr17-27694545-INS-137
chr17-27741884-INS-69
chr17-27742681-INS-126
chr17-27755143-INS-87
chr17-27759297-INS-186
chr17-27764349-INS-54
chr17-27769411-INS-188
chr17-27790690-INS-158
chr17-27800396-INS-2520
chr17-27809177-INS-135
chr17-27816473-INS-186
chr17-27818301-INS-439
chr17-27822015-INS-199
chr17-27827556-INS-209
chr17-27834882-INS-136
chr17-27836436-INS-72
chr17-27879605-INS-343
chr17-27886749-INS-166
chr17-27889472-INS-116
chr17-27890761-INS-349
chr17-27892154-INS-216
chr17-27897561-INS-62
chr17-27899580-INS-195
chr17-27934316-INS-62
chr17-27936924-INS-133
chr17-27941347-INS-169
chr17-27961062-INS-210
chr17-27963066-INS-83
chr17-27970648-INS-64
chr17-27977874-INS-134
chr17-27993302-INS-208
chr17-28009463-INS-208
chr17-28011505-INS-209
chr17-28026328-INS-737
chr17-28028303-INS-83
chr17-28041362-INS-108
chr17-28046294-INS-171
chr17-28069285-INS-330
chr17-28077026-INS-202
chr17-28080302-INS-200
chr17-28082596-INS-104
chr17-28092918-INS-192
chr17-28099499-INS-86
chr17-28107008-INS-72
chr17-28107836-INS-75
chr17-28125496-INS-218
chr17-28138019-INS-155
chr17-28146700-INS-700
chr17-28172507-INS-222
chr17-28240681-INS-353
chr17-28244667-INS-205
chr17-28245497-INS-290
chr17-28285105-INS-143
chr17-28291425-INS-157
chr17-28292851-INS-56
chr17-28295428-INS-52
chr17-28309286-INS-312
chr17-28315181-INS-64
chr17-28321445-INS-192
chr17-28400283-INS-217
chr17-28409509-INS-234
chr17-28441913-INS-87
chr17-28446322-INS-320
chr17-28448998-INS-218
chr17-28457715-INS-102
chr17-28468769-INS-82
chr17-28472628-INS-55
chr17-28481523-INS-206
chr17-28482413-INS-206
chr17-28487199-INS-98
chr17-28494902-INS-100
chr17-28504096-INS-292
chr17-28507778-INS-270
chr17-28557066-INS-217
chr17-28585091-INS-70

chr17-28599919-INS-204
chr17-28602496-INS-104
chr17-28608991-INS-51
chr17-28614060-INS-199
chr17-28615266-INS-174
chr17-28619187-INS-53
chr17-28622161-INS-996
chr17-28623508-INS-83
chr17-28628112-INS-187
chr17-28646638-INS-234
chr17-28656128-INS-72
chr17-28658071-INS-60
chr17-28673395-INS-109
chr17-28676398-INS-207
chr17-28698424-INS-203
chr17-28703832-INS-1010
chr17-28703854-INS-610
chr17-28708645-INS-54
chr17-28723989-INS-189
chr17-28748042-INS-52
chr17-28749218-INS-199
chr17-28773709-INS-180
chr17-28778977-INS-207
chr17-28782083-INS-85
chr17-28783356-INS-154
chr17-28788109-INS-72
chr17-28789189-INS-294
chr17-28790720-INS-868
chr17-28790871-INS-167
chr17-28796176-INS-126
chr17-28814233-INS-233
chr17-28815200-INS-217
chr17-28825918-INS-72
chr17-28828360-INS-319
chr17-28845078-INS-437
chr17-28849994-INS-116
chr17-28850814-INS-211
chr17-28853588-INS-143
chr17-28863355-INS-180
chr17-28864301-INS-261
chr17-28889995-INS-777
chr17-28895773-INS-146
chr17-28941948-INS-208
chr17-28951781-INS-60
chr17-28963840-INS-766
chr17-28967639-INS-372
chr17-28976017-INS-356
chr17-28979179-INS-142
chr17-28982963-INS-89
chr17-28983574-INS-204
chr17-28988806-INS-56
chr17-28994750-INS-64
chr17-28996041-INS-54
chr17-29002085-INS-749
chr17-29003710-INS-586
chr17-29004987-INS-558
chr17-29030603-INS-174
chr17-29034173-INS-256
chr17-29039889-INS-411
chr17-29046114-INS-192
chr17-29080276-INS-329
chr17-29090140-INS-197
chr17-29095542-INS-216
chr17-29106624-INS-467
chr17-29118838-INS-105
chr17-29120249-INS-203
chr17-29125862-INS-111
chr17-29137938-INS-286
chr17-29141021-INS-55
chr17-29151381-INS-505
chr17-29189688-INS-89
chr17-29195839-INS-76
chr17-29199818-INS-119
chr17-29204603-INS-215
chr17-29218410-INS-149
chr17-29222471-INS-85
chr17-29229593-INS-60

chr17-29232754-INS-291
chr17-29237326-INS-203
chr17-29239530-INS-251
chr17-29247443-INS-909
chr17-29248800-INS-80
chr17-29254111-INS-282
chr17-29262888-INS-241
chr17-29270716-INS-68
chr17-29282197-INS-62
chr17-29305116-INS-53
chr17-29321100-INS-116
chr17-29327157-INS-2094
chr17-29328318-INS-170
chr17-29337528-INS-54
chr17-29351476-INS-102
chr17-29365173-INS-76
chr17-29401611-INS-436
chr17-29403545-INS-179
chr17-29409415-INS-213
chr17-29410255-INS-88
chr17-29411713-INS-88
chr17-29414579-INS-208
chr17-29421110-INS-92
chr17-29423879-INS-211
chr17-29429249-INS-84
chr17-29431709-INS-227
chr17-29442782-INS-210
chr17-29474753-INS-74
chr17-29488280-INS-206
chr17-29495939-INS-79
chr17-29498810-INS-213
chr17-29503989-INS-142
chr17-29504927-INS-689
chr17-29505256-INS-690
chr17-29505828-INS-272
chr17-29508314-INS-64
chr17-29516790-INS-87
chr17-29533477-INS-499
chr17-29540966-INS-5741
chr17-29543071-INS-5466
chr17-29555032-INS-120
chr17-29563812-INS-186
chr17-29572665-INS-203
chr17-29578770-INS-212
chr17-29589828-INS-341
chr17-29622154-INS-182
chr17-29640163-INS-124
chr17-29644725-INS-200
chr17-29654031-INS-52
chr17-29660397-INS-234
chr17-29666892-INS-3041
chr17-29680808-INS-8418
chr17-29706083-INS-52
chr17-29717654-INS-164
chr17-29724267-INS-60
chr17-29725550-INS-978
chr17-29726549-INS-56
chr17-29753505-INS-60
chr17-29761396-INS-330
chr17-29764098-INS-327
chr17-29781258-INS-121
chr17-29813442-INS-270
chr17-29826057-INS-257
chr17-29853897-INS-210
chr17-29865199-INS-830
chr17-29875114-INS-229
chr17-29884202-INS-518
chr17-29888730-INS-209
chr17-29891047-INS-173
chr17-29923321-INS-51
chr17-29930914-INS-54
chr17-29938543-INS-159
chr17-29940961-INS-197
chr17-29974115-INS-171
chr17-29976737-INS-204
chr17-29983749-INS-132
chr17-29996415-INS-509

chr17-29997721-INS-244
chr17-29998988-INS-7237
chr17-30001685-INS-508
chr17-30003726-INS-51
chr17-30009924-INS-166
chr17-30025292-INS-422
chr17-30035803-INS-124
chr17-30044872-INS-56
chr17-30047664-INS-78
chr17-30051536-INS-405
chr17-30082025-INS-74
chr17-30091512-INS-94
chr17-30107903-INS-61
chr17-30133403-INS-150
chr17-30136057-INS-228
chr17-30139436-INS-68
chr17-30165250-INS-397
chr17-30171955-INS-226
chr17-30189719-INS-76
chr17-30197655-INS-181
chr17-30214836-INS-54
chr17-30235382-INS-68
chr17-30236566-INS-207
chr17-30237822-INS-494
chr17-30271790-INS-197
chr17-30302283-INS-145
chr17-30308206-INS-67
chr17-30311125-INS-97
chr17-30318422-INS-199
chr17-30325839-INS-64
chr17-30328729-INS-87
chr17-30338414-INS-230
chr17-30367613-INS-118
chr17-30384613-INS-65
chr17-30387866-INS-174
chr17-30405599-INS-65
chr17-30426175-INS-1882
chr17-30440551-INS-163
chr17-30452631-INS-6441
chr17-30490532-INS-746
chr17-30505376-INS-210
chr17-30569796-INS-306
chr17-30586193-INS-59
chr17-30665668-INS-2557
chr17-30702508-INS-1200
chr17-30702821-INS-375
chr17-30711691-INS-89
chr17-30718690-INS-204
chr17-30731052-INS-73
chr17-30737923-INS-58
chr17-30781655-INS-62
chr17-30789462-INS-5015
chr17-30793161-INS-129
chr17-30801560-INS-184
chr17-30802103-INS-1573
chr17-30806457-INS-5549
chr17-30835896-INS-4249
chr17-30846325-INS-62
chr17-30848437-INS-51
chr17-30848885-INS-186
chr17-30862438-INS-119
chr17-30868870-INS-60
chr17-30869375-INS-378
chr17-30879244-INS-200
chr17-30886224-INS-59
chr17-30887918-INS-372
chr17-30909807-INS-191
chr17-30917326-INS-177
chr17-30936864-INS-110
chr17-30956030-INS-51
chr17-30957881-INS-696
chr17-30962494-INS-76
chr17-30969121-INS-72
chr17-30975017-INS-111
chr17-30990996-INS-350
chr17-31000805-INS-185
chr17-31022747-INS-52

chr17-31040780-INS-54
chr17-31056851-INS-62
chr17-31059748-INS-73
chr17-31108311-INS-536
chr17-31110027-INS-58
chr17-31114520-INS-1417
chr17-31114520-INS-965
chr17-31114995-INS-260
chr17-31118398-INS-267
chr17-31128755-INS-58
chr17-31176797-INS-77
chr17-31182760-INS-75
chr17-31183714-INS-78
chr17-31193759-INS-588
chr17-31196485-INS-332
chr17-31220454-INS-60
chr17-31224294-INS-168
chr17-31342292-INS-554
chr17-31351472-INS-169
chr17-31353451-INS-362
chr17-31390924-INS-202
chr17-31431773-INS-358
chr17-31437906-INS-554
chr17-31449275-INS-66
chr17-31453427-INS-72
chr17-31464509-INS-59
chr17-31467298-INS-548
chr17-31499346-INS-211
chr17-31508640-INS-228
chr17-31510652-INS-66
chr17-31512254-INS-58
chr17-31518688-INS-5778
chr17-31537237-INS-170
chr17-31547900-INS-70
chr17-31550393-INS-352
chr17-31561007-INS-135
chr17-31580890-INS-254
chr17-31609334-INS-62
chr17-31614991-INS-130
chr17-31626461-INS-236
chr17-31644096-INS-75
chr17-31654325-INS-267
chr17-31659778-INS-63
chr17-31663219-INS-63
chr17-31665474-INS-51
chr17-31675075-INS-72
chr17-31700187-INS-198
chr17-31708894-INS-254
chr17-31716229-INS-114
chr17-31719899-INS-264
chr17-31745172-INS-1250
chr17-31750908-INS-212
chr17-31770938-INS-371
chr17-31792898-INS-191
chr17-31817461-INS-230
chr17-31830886-INS-189
chr17-31847799-INS-7083
chr17-31851426-INS-220
chr17-31858376-INS-113
chr17-31859432-INS-199
chr17-31885215-INS-59
chr17-31903710-INS-6442
chr17-31908381-INS-178
chr17-31930023-INS-72
chr17-31930553-INS-60
chr17-31961170-INS-66
chr17-31978390-INS-107
chr17-31984183-INS-60
chr17-31986660-INS-56
chr17-32019601-INS-350
chr17-32026993-INS-59
chr17-32030722-INS-93
chr17-32045239-INS-5283
chr17-32075427-INS-63
chr17-32113294-INS-70
chr17-32140052-INS-316
chr17-32140757-INS-115

chr17-32146439-INS-52
chr17-32160777-INS-118
chr17-32199658-INS-3130
chr17-32207035-INS-385
chr17-32213110-INS-60
chr17-32218080-INS-6476
chr17-32218495-INS-509
chr17-32233151-INS-212
chr17-32241253-INS-712
chr17-32246671-INS-247
chr17-32248807-INS-220
chr17-32261614-INS-95
chr17-32272648-INS-202
chr17-32276622-INS-163
chr17-32295106-INS-64
chr17-32301836-INS-128
chr17-32307839-INS-62
chr17-32317847-INS-163
chr17-32320189-INS-69
chr17-32342466-INS-56
chr17-32348168-INS-206
chr17-32349943-INS-771
chr17-32364908-INS-133
chr17-32392829-INS-58
chr17-32402286-INS-212
chr17-32411103-INS-6440
chr17-32413836-INS-204
chr17-32430517-INS-86
chr17-32430567-INS-1313
chr17-32434969-INS-132
chr17-32449834-INS-436
chr17-32453634-INS-370
chr17-32465882-INS-456
chr17-32467159-INS-153
chr17-32474734-INS-347
chr17-32483177-INS-213
chr17-32491068-INS-5309
chr17-32497058-INS-102
chr17-32510319-INS-162
chr17-32516010-INS-202
chr17-32521695-INS-200
chr17-32526113-INS-170
chr17-32546243-INS-214
chr17-32548706-INS-5548
chr17-32550639-INS-1311
chr17-32556371-INS-57
chr17-32560366-INS-201
chr17-32565766-INS-221
chr17-32619419-INS-108
chr17-32631150-INS-212
chr17-32635382-INS-102
chr17-32636306-INS-197
chr17-32641038-INS-72
chr17-32642053-INS-1090
chr17-32649073-INS-192
chr17-32658261-INS-160
chr17-32679330-INS-874
chr17-32704056-INS-278
chr17-32723676-INS-185
chr17-32733575-INS-893
chr17-32739842-INS-51
chr17-32744381-INS-106
chr17-32745554-INS-80
chr17-32754359-INS-989
chr17-32767108-INS-72
chr17-32792117-INS-84
chr17-32795858-INS-70
chr17-32796874-INS-566
chr17-32805577-INS-226
chr17-32821217-INS-388
chr17-32823879-INS-121
chr17-32825752-INS-58
chr17-32835065-INS-309
chr17-32855093-INS-203
chr17-32867795-INS-301
chr17-32872931-INS-7595
chr17-32872932-INS-976

chr17-32880865-INS-164
chr17-32931346-INS-771
chr17-32943301-INS-59
chr17-32945142-INS-205
chr17-32955161-INS-431
chr17-32973478-INS-111
chr17-32975428-INS-270
chr17-32975456-INS-912
chr17-32978864-INS-57
chr17-32995408-INS-104
chr17-33005112-INS-66
chr17-33010833-INS-326
chr17-33012249-INS-183
chr17-33015840-INS-60
chr17-33027713-INS-76
chr17-33034992-INS-105
chr17-33047686-INS-144
chr17-33055862-INS-203
chr17-33066830-INS-187
chr17-33087305-INS-343
chr17-33114536-INS-430
chr17-33117523-INS-972
chr17-33118962-INS-63
chr17-33137240-INS-116
chr17-33140812-INS-205
chr17-33145165-INS-101
chr17-33151184-INS-753
chr17-33179935-INS-65
chr17-33194044-INS-403
chr17-33195919-INS-208
chr17-33199626-INS-124
chr17-33214187-INS-104
chr17-33229725-INS-228
chr17-33239572-INS-164
chr17-33244421-INS-498
chr17-33254280-INS-6387
chr17-33277873-INS-440
chr17-33282946-INS-598
chr17-33300037-INS-198
chr17-33331281-INS-358
chr17-33337428-INS-4892
chr17-33338727-INS-364
chr17-33346673-INS-74
chr17-33347683-INS-4995
chr17-33372209-INS-3140
chr17-33373293-INS-7112
chr17-33411665-INS-52
chr17-33435470-INS-67
chr17-33529485-INS-60
chr17-33535059-INS-272
chr17-33552511-INS-98
chr17-33559544-INS-54
chr17-33568261-INS-198
chr17-33577351-INS-60
chr17-33581702-INS-3301
chr17-33622119-INS-7259
chr17-33622624-INS-388
chr17-33633318-INS-214
chr17-33657390-INS-364
chr17-33683932-INS-281
chr17-33686008-INS-72
chr17-33691073-INS-126
chr17-33693736-INS-8681
chr17-33703571-INS-335
chr17-33708250-INS-565
chr17-33711481-INS-144
chr17-33715519-INS-52
chr17-33738256-INS-1718
chr17-33738722-INS-77
chr17-33752043-INS-120
chr17-33765998-INS-283
chr17-33775678-INS-94
chr17-33776369-INS-66
chr17-33785685-INS-106
chr17-33791467-INS-845
chr17-33803658-INS-91
chr17-33806654-INS-100

chr17-33812854-INS-52
chr17-33815290-INS-138
chr17-33822727-INS-107
chr17-33854513-INS-242
chr17-33893266-INS-199
chr17-33920399-INS-199
chr17-33923293-INS-54
chr17-33941136-INS-72
chr17-33946711-INS-68
chr17-33950597-INS-238
chr17-33963305-INS-217
chr17-33980836-INS-168
chr17-33989777-INS-127
chr17-34007482-INS-168
chr17-34018451-INS-5180
chr17-34027933-INS-461
chr17-34045459-INS-192
chr17-34054464-INS-4821
chr17-34057442-INS-301
chr17-34058810-INS-99
chr17-34071717-INS-55
chr17-34097024-INS-135
chr17-34106425-INS-328
chr17-34115058-INS-226
chr17-34116841-INS-204
chr17-34140354-INS-222
chr17-34142211-INS-206
chr17-34146285-INS-248
chr17-34153149-INS-78
chr17-34192167-INS-148
chr17-34214290-INS-400
chr17-34239726-INS-179
chr17-34266423-INS-105
chr17-34283311-INS-218
chr17-34292061-INS-763
chr17-34293458-INS-333
chr17-34333301-INS-206
chr17-34362859-INS-458
chr17-34371179-INS-914
chr17-34376983-INS-90
chr17-34380799-INS-223
chr17-34407712-INS-55
chr17-34411620-INS-201
chr17-34468670-INS-158
chr17-34473194-INS-204
chr17-34484696-INS-301
chr17-34492473-INS-351
chr17-34547945-INS-198
chr17-34549590-INS-178
chr17-34572653-INS-84
chr17-34577329-INS-159
chr17-34591479-INS-72
chr17-34653982-INS-80
chr17-34669515-INS-128
chr17-34690635-INS-443
chr17-34697895-INS-90
chr17-34737193-INS-97
chr17-34738759-INS-61
chr17-34740255-INS-60
chr17-34752905-INS-105
chr17-34774761-INS-875
chr17-34834422-INS-145
chr17-34860784-INS-61
chr17-34917624-INS-80
chr17-34921071-INS-215
chr17-34930571-INS-2233
chr17-34942437-INS-548
chr17-34979408-INS-55
chr17-34998676-INS-4305
chr17-35046831-INS-324
chr17-35055670-INS-62
chr17-35088177-INS-1890
chr17-35091498-INS-208
chr17-35108214-INS-119
chr17-35120933-INS-1144
chr17-35133878-INS-77
chr17-35139741-INS-228

chr17-35158137-INS-207
chr17-35194368-INS-61
chr17-35213538-INS-225
chr17-35226109-INS-236
chr17-35294173-INS-82
chr17-35311165-INS-62
chr17-35348494-INS-182
chr17-35353637-INS-180
chr17-35356481-INS-190
chr17-35370048-INS-212
chr17-35412813-INS-120
chr17-35424745-INS-61
chr17-35425920-INS-60
chr17-35464616-INS-59
chr17-35475386-INS-4678
chr17-35476856-INS-200
chr17-35480602-INS-358
chr17-35490865-INS-184
chr17-35498672-INS-218
chr17-35500761-INS-493
chr17-35563235-INS-123
chr17-35572412-INS-444
chr17-35574268-INS-222
chr17-35600275-INS-77
chr17-35604138-INS-327
chr17-35605460-INS-543
chr17-35621705-INS-118
chr17-35623407-INS-937
chr17-35753854-INS-66
chr17-35760428-INS-90
chr17-35782778-INS-169
chr17-35798108-INS-94
chr17-35799149-INS-356
chr17-35806790-INS-195
chr17-35810324-INS-74
chr17-35822173-INS-233
chr17-35830259-INS-218
chr17-35842403-INS-81
chr17-35851471-INS-73
chr17-35852671-INS-208
chr17-35873050-INS-664
chr17-35875450-INS-211
chr17-35883415-INS-74
chr17-35898992-INS-111
chr17-35905472-INS-203
chr17-35913792-INS-188
chr17-35930978-INS-202
chr17-35931729-INS-135
chr17-35933865-INS-178
chr17-35951287-INS-139
chr17-35989459-INS-564
chr17-35990077-INS-226
chr17-36006052-INS-172
chr17-36013994-INS-58
chr17-36020644-INS-144
chr17-36024183-INS-542
chr17-36025636-INS-63
chr17-36027750-INS-218
chr17-36029621-INS-5498
chr17-36034456-INS-582
chr17-36042954-INS-414
chr17-36068241-INS-5217
chr17-36071535-INS-188
chr17-36089299-INS-225
chr17-36092009-INS-219
chr17-36099977-INS-216
chr17-36100126-INS-816
chr17-36107743-INS-192
chr17-36110365-INS-386
chr17-36111866-INS-844
chr17-36121820-INS-224
chr17-36137542-INS-191
chr17-36140127-INS-207
chr17-36155302-INS-158
chr17-36157717-INS-161
chr17-36173074-INS-73
chr17-36202944-INS-64

chr17-36207068-INS-181
chr17-36224530-INS-207
chr17-36245348-INS-7661
chr17-36246285-INS-142
chr17-36247898-INS-145
chr17-36251591-INS-65
chr17-36286137-INS-208
chr17-36316382-INS-511
chr17-36350632-INS-54
chr17-36358941-INS-228
chr17-36419030-INS-207
chr17-36433867-INS-193
chr17-36437493-INS-413
chr17-36530720-INS-211
chr17-36544850-INS-331
chr17-36548942-INS-58
chr17-36563892-INS-57
chr17-36601234-INS-187
chr17-36604239-INS-507
chr17-36604838-INS-180
chr17-36614646-INS-177
chr17-36627017-INS-54
chr17-36646590-INS-55
chr17-36663915-INS-457
chr17-36669721-INS-505
chr17-36716233-INS-74
chr17-36717493-INS-289
chr17-36733393-INS-6774
chr17-36734432-INS-1572
chr17-36735621-INS-7085
chr17-36765122-INS-8048
chr17-36774306-INS-6403
chr17-36812228-INS-52
chr17-36816520-INS-1700
chr17-36830088-INS-4557
chr17-36850065-INS-369
chr17-36856024-INS-9124
chr17-36863604-INS-59
chr17-36866587-INS-7148
chr17-36866819-INS-6546
chr17-36893601-INS-54
chr17-36901332-INS-74
chr17-36905772-INS-6440
chr17-36911479-INS-241
chr17-36916894-INS-996
chr17-36932212-INS-179
chr17-36933468-INS-8619
chr17-36943960-INS-6194
chr17-36955583-INS-6240
chr17-36960461-INS-279
chr17-36975271-INS-204
chr17-37000635-INS-867
chr17-37015889-INS-820
chr17-37025106-INS-97
chr17-37033380-INS-548
chr17-37048312-INS-79
chr17-37111002-INS-128
chr17-37184950-INS-221
chr17-37204757-INS-202
chr17-37228891-INS-218
chr17-37246543-INS-4458
chr17-37271776-INS-157
chr17-37279946-INS-540
chr17-37292406-INS-56
chr17-37294864-INS-216
chr17-37297235-INS-140
chr17-37300922-INS-84
chr17-37306991-INS-217
chr17-37311758-INS-245
chr17-37316636-INS-464
chr17-37324072-INS-53
chr17-37330094-INS-118
chr17-37348817-INS-502
chr17-37400370-INS-361
chr17-37428637-INS-201
chr17-37452776-INS-1763
chr17-37452926-INS-575

chr17-37461721-INS-172
chr17-37500406-INS-306
chr17-37514503-INS-185
chr17-37515076-INS-96
chr17-37520744-INS-3870
chr17-37525296-INS-915
chr17-37526681-INS-417
chr17-37529955-INS-201
chr17-37532163-INS-176
chr17-37542613-INS-6886
chr17-37597360-INS-498
chr17-37599213-INS-6464
chr17-37607697-INS-124
chr17-37673011-INS-6450
chr17-37683078-INS-7816
chr17-37683392-INS-7923
chr17-37684213-INS-54
chr17-37707984-INS-4378
chr17-37709832-INS-242
chr17-37712352-INS-544
chr17-37718666-INS-401
chr17-37722170-INS-6885
chr17-37725137-INS-66
chr17-37730710-INS-54
chr17-37745317-INS-51
chr17-37779511-INS-184
chr17-37779743-INS-7196
chr17-37783649-INS-204
chr17-37794717-INS-446
chr17-37804548-INS-52
chr17-37830890-INS-100
chr17-37846926-INS-7163
chr17-37855999-INS-119
chr17-37862325-INS-6385
chr17-37879893-INS-86
chr17-37902804-INS-6460
chr17-37908390-INS-582
chr17-37944221-INS-5461
chr17-37963836-INS-131
chr17-37967494-INS-51
chr17-37987729-INS-89
chr17-38012894-INS-110
chr17-38014096-INS-58
chr17-38029957-INS-7598
chr17-38060439-INS-117
chr17-38077207-INS-59
chr17-38079352-INS-75
chr17-38083865-INS-104
chr17-38090156-INS-6428
chr17-38095844-INS-7068
chr17-38109047-INS-257
chr17-38141180-INS-143
chr17-38153885-INS-76
chr17-38161370-INS-395
chr17-38164774-INS-624
chr17-38281603-INS-88
chr17-38322753-INS-209
chr17-38330164-INS-64
chr17-38334722-INS-118
chr17-38339274-INS-95
chr17-38340404-INS-250
chr17-38342041-INS-846
chr17-38348888-INS-944
chr17-38357494-INS-79
chr17-38363451-INS-626
chr17-38371345-INS-180
chr17-38405416-INS-62
chr17-38406199-INS-1640
chr17-38415012-INS-4063
chr17-38423901-INS-701
chr17-38452234-INS-212
chr17-38476749-INS-6496
chr17-38490169-INS-917
chr17-38494799-INS-6397
chr17-38508960-INS-6414
chr17-38520407-INS-268
chr17-38532384-INS-86

chr17-38556041-INS-2638
chr17-38625522-INS-968
chr17-38641668-INS-349
chr17-38651512-INS-8842
chr17-38651824-INS-67
chr17-38665534-INS-7098
chr17-38683544-INS-63
chr17-38709737-INS-3255
chr17-38722149-INS-56
chr17-38723769-INS-289
chr17-38740402-INS-99
chr17-38740856-INS-1893
chr17-38744050-INS-332
chr17-38750384-INS-496
chr17-38773783-INS-173
chr17-38783409-INS-378
chr17-38784413-INS-498
chr17-38860976-INS-4936
chr17-38861264-INS-7043
chr17-38883135-INS-7084
chr17-38982304-INS-118
chr17-38986553-INS-164
chr17-39037152-INS-77
chr17-39038747-INS-894
chr17-39055140-INS-151
chr17-39067547-INS-546
chr17-39114750-INS-87
chr17-39133512-INS-70
chr17-39164829-INS-5729
chr17-39242454-INS-6407
chr17-39255385-INS-110
chr17-39283980-INS-53
chr17-39298478-INS-56
chr17-39329149-INS-170
chr17-39358832-INS-1257
chr17-39411520-INS-51
chr17-39419498-INS-358
chr17-39578246-INS-82
chr17-39611046-INS-184
chr17-39614676-INS-378
chr17-39640572-INS-63
chr17-39738656-INS-1636
chr17-39767919-INS-1177
chr17-39804381-INS-3135
chr17-39883559-INS-7073
chr17-39976641-INS-143
chr17-40035806-INS-60
chr17-40053205-INS-6540
chr17-40104437-INS-6361
chr17-40148212-INS-204
chr17-40171758-INS-2933
chr17-40302947-INS-149
chr17-40321627-INS-440
chr17-40342955-INS-399
chr17-40357538-INS-193
chr17-40359420-INS-236
chr17-40363612-INS-58
chr17-40492096-INS-7194
chr17-40498953-INS-103
chr17-40510693-INS-185
chr17-40521985-INS-70
chr17-40536876-INS-75
chr17-40541402-INS-83
chr17-40543626-INS-84
chr17-40549277-INS-74
chr17-40571430-INS-188
chr17-40662974-INS-94
chr17-40669079-INS-176
chr17-40691878-INS-292
chr17-40718128-INS-245
chr17-40732848-INS-3693
chr17-40759823-INS-313
chr17-40784353-INS-133
chr17-40797179-INS-831
chr17-40807867-INS-147
chr17-40808850-INS-75
chr17-40815930-INS-66

chr17-40845313-INS-707
chr17-40861097-INS-323
chr17-40863177-INS-370
chr17-40874617-INS-58
chr17-40880823-INS-206
chr17-40886874-INS-533
chr17-40902953-INS-641
chr17-40907973-INS-7164
chr17-40958898-INS-2994
chr17-40985426-INS-6782
chr17-40991641-INS-111
chr17-41065656-INS-7111
chr17-41082466-INS-70
chr17-41087718-INS-51
chr17-41118998-INS-2984
chr17-41129715-INS-251
chr17-41137427-INS-69
chr17-41148518-INS-53
chr17-41149608-INS-245
chr17-41155781-INS-52
chr17-41158364-INS-439
chr17-41181949-INS-148
chr17-41195534-INS-92
chr17-41205488-INS-90
chr17-41226670-INS-96
chr17-41238233-INS-206
chr17-41311029-INS-57
chr17-41312682-INS-52
chr17-41325893-INS-6434
chr17-41351818-INS-445
chr17-41373502-INS-340
chr17-41377681-INS-63
chr17-41411192-INS-306
chr17-41417757-INS-69
chr17-41443094-INS-935
chr17-41443976-INS-56
chr17-41445641-INS-211
chr17-41452354-INS-976
chr17-41458763-INS-162
chr17-41469835-INS-118
chr17-41470456-INS-121
chr17-41473102-INS-74
chr17-41476103-INS-164
chr17-41491283-INS-157
chr17-41514899-INS-62
chr17-41524129-INS-389
chr17-41527997-INS-253
chr17-41532295-INS-316
chr17-41548954-INS-234
chr17-41568114-INS-6824
chr17-41628089-INS-58
chr17-41643875-INS-178
chr17-41645249-INS-6645
chr17-41646356-INS-209
chr17-41685732-INS-1860
chr17-41687478-INS-80
chr17-41718737-INS-108
chr17-41727681-INS-6529
chr17-41727776-INS-140
chr17-41745882-INS-425
chr17-41753834-INS-55
chr17-41761116-INS-57
chr17-41766625-INS-458
chr17-41773883-INS-177
chr17-41839745-INS-52
chr17-41842400-INS-826
chr17-41844868-INS-390
chr17-41849792-INS-58
chr17-41859856-INS-1014
chr17-41871545-INS-75
chr17-41884237-INS-107
chr17-41896675-INS-97
chr17-41901690-INS-121
chr17-41931596-INS-6457
chr17-41945569-INS-52
chr17-41950660-INS-675
chr17-41967779-INS-600

chr17-41992806-INS-3049
chr17-42004337-INS-52
chr17-42006844-INS-114
chr17-42020160-INS-70
chr17-42037958-INS-397
chr17-42047513-INS-72
chr17-42067238-INS-104
chr17-42129184-INS-77
chr17-42145521-INS-669
chr17-42169959-INS-2813
chr17-42179434-INS-990
chr17-42184456-INS-206
chr17-42191292-INS-184
chr17-42199425-INS-60
chr17-42205833-INS-337
chr17-42207702-INS-419
chr17-42235361-INS-52
chr17-42236515-INS-137
chr17-42279718-INS-76
chr17-42282426-INS-386
chr17-42283471-INS-64
chr17-42317752-INS-209
chr17-42323238-INS-1863
chr17-42342467-INS-6360
chr17-42349203-INS-136
chr17-42352209-INS-514
chr17-42365610-INS-53
chr17-42366498-INS-489
chr17-42375853-INS-392
chr17-42418261-INS-347
chr17-42453008-INS-70
chr17-42470374-INS-400
chr17-42478096-INS-501
chr17-42505593-INS-3207
chr17-42509744-INS-211
chr17-42510324-INS-555
chr17-42560836-INS-376
chr17-42567539-INS-399
chr17-42583370-INS-274
chr17-42585670-INS-72
chr17-42588724-INS-366
chr17-42610832-INS-7168
chr17-42617660-INS-6462
chr17-42662363-INS-186
chr17-42664905-INS-53
chr17-42687322-INS-197
chr17-42692647-INS-163
chr17-42713412-INS-288
chr17-42722849-INS-77
chr17-42732547-INS-84
chr17-42738590-INS-473
chr17-42745438-INS-6116
chr17-42768715-INS-71
chr17-42772005-INS-307
chr17-42778205-INS-53
chr17-42784232-INS-293
chr17-42793639-INS-76
chr17-42841708-INS-645
chr17-42880593-INS-65
chr17-42911060-INS-93
chr17-42915200-INS-164
chr17-42917409-INS-106
chr17-42981596-INS-340
chr17-42984549-INS-173
chr17-42989304-INS-525
chr17-43001178-INS-83
chr17-43005307-INS-55
chr17-43027442-INS-354
chr17-43038990-INS-52
chr17-43047236-INS-56
chr17-43054577-INS-1407
chr17-43055036-INS-360
chr17-43074857-INS-437
chr17-43080198-INS-1290
chr17-43153528-INS-60
chr17-43155547-INS-52
chr17-43207400-INS-189

chr17-43213612-INS-209
chr17-43216362-INS-175
chr17-43216646-INS-813
chr17-43237265-INS-80
chr17-43244104-INS-114
chr17-43244872-INS-100
chr17-43250101-INS-300
chr17-43254963-INS-75
chr17-43289575-INS-123
chr17-43299037-INS-159
chr17-43303607-INS-87
chr17-43305064-INS-233
chr17-43312346-INS-78
chr17-43362302-INS-72
chr17-43425859-INS-63
chr17-43501319-INS-435
chr17-43503956-INS-65
chr17-43522554-INS-189
chr17-43534347-INS-178
chr17-43563406-INS-224
chr17-43567071-INS-108
chr17-43569420-INS-78
chr17-43570948-INS-52
chr17-43594313-INS-91
chr17-43608443-INS-118
chr17-43626625-INS-60
chr17-43641029-INS-89
chr17-43650175-INS-62
chr17-43668868-INS-92
chr17-43680608-INS-181
chr17-43684189-INS-177
chr17-43704900-INS-52
chr17-43763500-INS-185
chr17-43764687-INS-215
chr17-43778049-INS-310
chr17-43779425-INS-53
chr17-43794092-INS-236
chr17-43811809-INS-187
chr17-43817543-INS-51
chr17-43840706-INS-148
chr17-43844622-INS-432
chr17-43846103-INS-668
chr17-43847441-INS-81
chr17-43863434-INS-390
chr17-43889293-INS-189
chr17-43896628-INS-404
chr17-43902715-INS-52
chr17-43904532-INS-64
chr17-43915689-INS-51
chr17-43959641-INS-51
chr17-43961822-INS-97
chr17-43973468-INS-70
chr17-43978322-INS-98
chr17-43983382-INS-57
chr17-44004323-INS-167
chr17-44008200-INS-340
chr17-44015605-INS-146
chr17-44024821-INS-80
chr17-44026520-INS-230
chr17-44034418-INS-6413
chr17-44038790-INS-51
chr17-44054768-INS-58
chr17-44056827-INS-80
chr17-44066527-INS-612
chr17-44066712-INS-2010
chr17-44083682-INS-109
chr17-44085545-INS-77
chr17-44089922-INS-392
chr17-44101465-INS-54
chr17-44106387-INS-56
chr17-44141831-INS-140
chr17-44167844-INS-62
chr17-44171679-INS-84
chr17-44182350-INS-5056
chr17-44185498-INS-528
chr17-44260119-INS-99
chr17-44261862-INS-357

chr17-44270257-INS-119
chr17-44272773-INS-84
chr17-44280246-INS-70
chr17-44296840-INS-52
chr17-44300892-INS-368
chr17-44322967-INS-105
chr17-44325614-INS-61
chr17-44330919-INS-52
chr17-44366927-INS-444
chr17-44400866-INS-171
chr17-44418362-INS-169
chr17-44441640-INS-69
chr17-44449573-INS-232
chr17-44470849-INS-67
chr17-44488949-INS-148
chr17-44499127-INS-53
chr17-44518532-INS-51
chr17-44560183-INS-1515
chr17-44563451-INS-113
chr17-44573034-INS-52
chr17-44592275-INS-68
chr17-44612274-INS-392
chr17-44616863-INS-125
chr17-44621471-INS-551
chr17-44631757-INS-523
chr17-44638157-INS-52
chr17-44642447-INS-54
chr17-44651745-INS-495
chr17-44687546-INS-912
chr17-44688943-INS-68
chr17-44706369-INS-74
chr17-44721546-INS-2588
chr17-44725806-INS-66
chr17-44743031-INS-200
chr17-44752942-INS-417
chr17-44756032-INS-52
chr17-44771597-INS-110
chr17-44778772-INS-54
chr17-44791909-INS-132
chr17-44810913-INS-249
chr17-44829843-INS-56
chr17-44887456-INS-140
chr17-44895164-INS-141
chr17-44913588-INS-155
chr17-44946366-INS-279
chr17-44946786-INS-450
chr17-44948759-INS-121
chr17-44957990-INS-53
chr17-44988073-INS-76
chr17-45004732-INS-59
chr17-45010512-INS-65
chr17-45019962-INS-180
chr17-45029598-INS-83
chr17-45037724-INS-116
chr17-45062296-INS-471
chr17-45067548-INS-226
chr17-45075580-INS-52
chr17-45079987-INS-180
chr17-45133039-INS-66
chr17-45180042-INS-57
chr17-45184012-INS-250
chr17-45229094-INS-436
chr17-45243883-INS-5335
chr17-45245019-INS-402
chr17-45248966-INS-77
chr17-45290567-INS-178
chr17-45316567-INS-74
chr17-45351684-INS-386
chr17-45384661-INS-6554
chr17-45409652-INS-86
chr17-45420692-INS-208
chr17-45529965-INS-88
chr17-45534712-INS-62
chr17-45554052-INS-90
chr17-45559762-INS-382
chr17-45568467-INS-335
chr17-45592431-INS-66

chr17-45595352-INS-58
chr17-45597865-INS-765
chr17-45600741-INS-70
chr17-45622542-INS-56
chr17-45646127-INS-132
chr17-45655084-INS-145
chr17-45675412-INS-335
chr17-45677271-INS-306
chr17-45690036-INS-348
chr17-45706185-INS-2807
chr17-45720222-INS-5974
chr17-45736420-INS-831
chr17-45740391-INS-180
chr17-45764310-INS-110
chr17-45770157-INS-175
chr17-45772725-INS-205
chr17-45791162-INS-94
chr17-45792318-INS-92
chr17-45794130-INS-125
chr17-45795461-INS-203
chr17-45797456-INS-184
chr17-45798820-INS-856
chr17-45799625-INS-93
chr17-45805990-INS-85
chr17-45845424-INS-78
chr17-45859750-INS-54
chr17-45865755-INS-209
chr17-45882637-INS-54
chr17-45917076-INS-5029
chr17-45917360-INS-132
chr17-45924059-INS-59
chr17-45927909-INS-203
chr17-45931990-INS-195
chr17-45935272-INS-210
chr17-45951271-INS-198
chr17-45954459-INS-133
chr17-45963642-INS-81
chr17-45966867-INS-203
chr17-45983110-INS-116
chr17-46002187-INS-609
chr17-46012262-INS-12386
chr17-46026097-INS-95
chr17-46079317-INS-60
chr17-46089239-INS-102
chr17-46092193-INS-394
chr17-46102243-INS-62
chr17-46104215-INS-6413
chr17-46105177-INS-2804
chr17-46110441-INS-104
chr17-46119892-INS-445
chr17-46150656-INS-250
chr17-46182624-INS-401
chr17-46187461-INS-53
chr17-46190111-INS-104
chr17-46194658-INS-922
chr17-46194711-INS-530
chr17-46195883-INS-60
chr17-46231811-INS-77
chr17-46249178-INS-144
chr17-46261273-INS-138
chr17-46288479-INS-54
chr17-46294420-INS-51
chr17-46300747-INS-149
chr17-46313776-INS-431
chr17-46316488-INS-208
chr17-46322090-INS-161
chr17-46322909-INS-368
chr17-46355452-INS-59
chr17-46390642-INS-71
chr17-46404257-INS-90
chr17-46410319-INS-229
chr17-46428522-INS-80
chr17-46433248-INS-6401
chr17-46444217-INS-84
chr17-46448133-INS-66
chr17-46452080-INS-94
chr17-46456567-INS-223

chr17-46465082-INS-72
chr17-46495084-INS-128
chr17-46507894-INS-270
chr17-46535143-INS-55
chr17-46545785-INS-61
chr17-46559448-INS-53
chr17-46590653-INS-397
chr17-46597762-INS-114
chr17-46611529-INS-121
chr17-46622952-INS-95
chr17-46640215-INS-209
chr17-46649094-INS-97
chr17-46670582-INS-56
chr17-46746628-INS-209
chr17-46750452-INS-181
chr17-46788099-INS-60
chr17-46801149-INS-457
chr17-46804319-INS-2866
chr17-46846102-INS-244
chr17-46853661-INS-64
chr17-46882752-INS-82
chr17-46886412-INS-72
chr17-46949832-INS-221
chr17-46968257-INS-64
chr17-46981164-INS-735
chr17-47001483-INS-934
chr17-47001598-INS-249
chr17-47002745-INS-296
chr17-47004956-INS-183
chr17-47024372-INS-91
chr17-47043095-INS-56
chr17-47044330-INS-178
chr17-47054420-INS-54
chr17-47054818-INS-309
chr17-47059104-INS-256
chr17-47070346-INS-57
chr17-47072704-INS-202
chr17-47077324-INS-66
chr17-47078223-INS-85
chr17-47081121-INS-56
chr17-47094654-INS-407
chr17-47096753-INS-56
chr17-47097422-INS-581
chr17-47099079-INS-70
chr17-47103618-INS-136
chr17-47116338-INS-831
chr17-47121044-INS-202
chr17-47122276-INS-214
chr17-47124425-INS-162
chr17-47139355-INS-127
chr17-47155919-INS-295
chr17-47160791-INS-56
chr17-47164437-INS-74
chr17-47166231-INS-195
chr17-47186613-INS-205
chr17-47187137-INS-197
chr17-47189966-INS-190
chr17-47191964-INS-57
chr17-47195722-INS-126
chr17-47199431-INS-86
chr17-47202914-INS-238
chr17-47210803-INS-158
chr17-47213043-INS-202
chr17-47216260-INS-210
chr17-47226142-INS-70
chr17-47247240-INS-87
chr17-47249444-INS-54
chr17-47272904-INS-63
chr17-47281213-INS-188
chr17-47351748-INS-509
chr17-47363828-INS-62
chr17-47372558-INS-66
chr17-47380209-INS-7089
chr17-47386309-INS-261
chr17-47400514-INS-64
chr17-47447850-INS-66
chr17-47448937-INS-124

chr17-47471227-INS-81
chr17-47484248-INS-62
chr17-47492372-INS-58
chr17-47508266-INS-72
chr17-47509750-INS-246
chr17-47513556-INS-88
chr17-47524547-INS-186
chr17-47526504-INS-85
chr17-47542099-INS-79
chr17-47543592-INS-134
chr17-47582787-INS-51
chr17-47583831-INS-57
chr17-47585722-INS-61
chr17-47595149-INS-354
chr17-47600767-INS-194
chr17-47602679-INS-465
chr17-47640473-INS-128
chr17-47653709-INS-463
chr17-47665466-INS-88
chr17-47681704-INS-89
chr17-47732069-INS-112
chr17-47736004-INS-390
chr17-47771789-INS-100
chr17-47792633-INS-242
chr17-47808588-INS-79
chr17-47820813-INS-86
chr17-47830006-INS-219
chr17-47858520-INS-155
chr17-47859586-INS-83
chr17-47865222-INS-52
chr17-47890424-INS-89
chr17-47891856-INS-52
chr17-47953296-INS-51
chr17-47967218-INS-132
chr17-47971168-INS-224
chr17-47981382-INS-82
chr17-48034086-INS-198
chr17-48085798-INS-197
chr17-48088786-INS-108
chr17-48109417-INS-51
chr17-48112840-INS-118
chr17-48159035-INS-51
chr17-48213111-INS-77
chr17-48216087-INS-223
chr17-48248091-INS-67
chr17-48248410-INS-388
chr17-48259062-INS-86
chr17-48296195-INS-212
chr17-48304186-INS-52
chr17-48307236-INS-2842
chr17-48314069-INS-52
chr17-48329297-INS-60
chr17-48336790-INS-220
chr17-48363784-INS-5548
chr17-48364302-INS-169
chr17-48372621-INS-141
chr17-48409457-INS-72
chr17-48411370-INS-91
chr17-48414472-INS-439
chr17-48435092-INS-60
chr17-48465881-INS-124
chr17-48478216-INS-1890
chr17-48493002-INS-72
chr17-48493303-INS-323
chr17-48494103-INS-217
chr17-48503647-INS-56
chr17-48505765-INS-75
chr17-48525992-INS-259
chr17-48527842-INS-75
chr17-48535612-INS-76
chr17-48540134-INS-112
chr17-48549560-INS-66
chr17-48556041-INS-380
chr17-48556642-INS-53
chr17-48559787-INS-123
chr17-48562097-INS-133
chr17-48597211-INS-58

chr17-48600338-INS-60
chr17-48604352-INS-617
chr17-48605173-INS-52
chr17-48608509-INS-60
chr17-48611398-INS-318
chr17-48614371-INS-6401
chr17-48618530-INS-125
chr17-48642345-INS-400
chr17-48664166-INS-86
chr17-48667687-INS-54
chr17-48679126-INS-783
chr17-48683664-INS-101
chr17-48690212-INS-66
chr17-48694945-INS-54
chr17-48709820-INS-58
chr17-48737951-INS-82
chr17-48752420-INS-318
chr17-48753379-INS-282
chr17-48759992-INS-85
chr17-48769376-INS-59
chr17-48772944-INS-134
chr17-48782551-INS-178
chr17-48792583-INS-901
chr17-48801934-INS-57
chr17-48822480-INS-81
chr17-48843651-INS-338
chr17-48847286-INS-173
chr17-48920942-INS-70
chr17-48927885-INS-158
chr17-48940593-INS-64
chr17-48949084-INS-329
chr17-48997102-INS-87
chr17-49011928-INS-140
chr17-49059255-INS-100
chr17-49085454-INS-56
chr17-49093984-INS-88
chr17-49116717-INS-56
chr17-49140608-INS-7109
chr17-49174841-INS-86
chr17-49176053-INS-152
chr17-49190958-INS-63
chr17-49197862-INS-118
chr17-49213691-INS-151
chr17-49267520-INS-84
chr17-49279149-INS-115
chr17-49344182-INS-118
chr17-49356648-INS-58
chr17-49383337-INS-104
chr17-49402812-INS-557
chr17-49411392-INS-174
chr17-49422455-INS-186
chr17-49432328-INS-60
chr17-49434559-INS-52
chr17-49482807-INS-51
chr17-49498426-INS-148
chr17-49507065-INS-56
chr17-49511654-INS-515
chr17-49525376-INS-62
chr17-49529244-INS-365
chr17-49565129-INS-61
chr17-49567152-INS-63
chr17-49575731-INS-157
chr17-49580530-INS-102
chr17-49584562-INS-431
chr17-49584993-INS-228
chr17-49588888-INS-471
chr17-49671157-INS-102
chr17-49690135-INS-207
chr17-49693826-INS-264
chr17-49700598-INS-452
chr17-49710957-INS-5380
chr17-49718692-INS-87
chr17-49768357-INS-2778
chr17-49823588-INS-151
chr17-49834148-INS-416
chr17-49837310-INS-85
chr17-49838413-INS-69

chr17-49842145-INS-52
chr17-49856091-INS-66
chr17-49858932-INS-113
chr17-49865842-INS-66
chr17-49866321-INS-219
chr17-49902566-INS-126
chr17-49909822-INS-927
chr17-49912192-INS-53
chr17-49924375-INS-74
chr17-49925365-INS-83
chr17-49930861-INS-121
chr17-49931975-INS-54
chr17-49938205-INS-199
chr17-49939702-INS-74
chr17-49983700-INS-143
chr17-49989198-INS-94
chr17-49998891-INS-114
chr17-50000239-INS-275
chr17-50019027-INS-440
chr17-50032786-INS-390
chr17-50036454-INS-51
chr17-50040515-INS-54
chr17-50057265-INS-577
chr17-50068094-INS-55
chr17-50089355-INS-52
chr17-50098742-INS-83
chr17-50122999-INS-434
chr17-50125703-INS-76
chr17-50139891-INS-53
chr17-50142123-INS-126
chr17-50161150-INS-339
chr17-50177008-INS-65
chr17-50185450-INS-94
chr17-50189684-INS-124
chr17-50199190-INS-76
chr17-50238200-INS-337
chr17-50243300-INS-102
chr17-50248003-INS-246
chr17-50278550-INS-426
chr17-50279665-INS-108
chr17-50283262-INS-228
chr17-50298402-INS-5549
chr17-50302882-INS-485
chr17-50323560-INS-752
chr17-50360238-INS-144
chr17-50362820-INS-104
chr17-50392416-INS-70
chr17-50403977-INS-83
chr17-50423207-INS-80
chr17-50427766-INS-88
chr17-50448563-INS-209
chr17-50451493-INS-524
chr17-50482976-INS-84
chr17-50514909-INS-189
chr17-50535912-INS-137
chr17-50541535-INS-111
chr17-50546342-INS-72
chr17-50552517-INS-202
chr17-50553771-INS-66
chr17-50564291-INS-67
chr17-50565717-INS-60
chr17-50567497-INS-85
chr17-50576543-INS-3064
chr17-50577115-INS-390
chr17-50608042-INS-439
chr17-50619839-INS-58
chr17-50628514-INS-220
chr17-50643630-INS-158
chr17-50651645-INS-129
chr17-50666839-INS-159
chr17-50667673-INS-575
chr17-50680084-INS-577
chr17-50685472-INS-205
chr17-50701916-INS-319
chr17-50705138-INS-318
chr17-50706343-INS-282
chr17-50738137-INS-77

chr17-50749418-INS-155
chr17-50751815-INS-94
chr17-50771869-INS-87
chr17-50781375-INS-214
chr17-50791502-INS-88
chr17-50800479-INS-72
chr17-50875491-INS-495
chr17-50890413-INS-853
chr17-50891214-INS-308
chr17-50903637-INS-114
chr17-50926429-INS-7109
chr17-50947805-INS-129
chr17-50960303-INS-122
chr17-50966838-INS-67
chr17-50983810-INS-62
chr17-51001902-INS-7125
chr17-51003793-INS-523
chr17-51034081-INS-56
chr17-51039824-INS-211
chr17-51077521-INS-183
chr17-51078600-INS-321
chr17-51096331-INS-126
chr17-51160193-INS-56
chr17-51231709-INS-52
chr17-51242994-INS-4745
chr17-51242995-INS-358
chr17-51252244-INS-6862
chr17-51257928-INS-4267
chr17-51266420-INS-145
chr17-51280608-INS-413
chr17-51297319-INS-146
chr17-51311494-INS-698
chr17-51344324-INS-479
chr17-51351387-INS-3871
chr17-51369355-INS-59
chr17-51373024-INS-6869
chr17-51375812-INS-99
chr17-51397404-INS-6795
chr17-51399856-INS-51
chr17-51406210-INS-214
chr17-51410246-INS-103
chr17-51421677-INS-74
chr17-51454834-INS-61
chr17-51504478-INS-86
chr17-51516175-INS-55
chr17-51635572-INS-58
chr17-51651026-INS-6519
chr17-51651980-INS-56
chr17-51657417-INS-7055
chr17-51660372-INS-87
chr17-51690250-INS-85
chr17-51696946-INS-126
chr17-51738010-INS-118
chr17-51759473-INS-549
chr17-51803886-INS-6582
chr17-51804787-INS-5269
chr17-51873069-INS-454
chr17-51887912-INS-88
chr17-51958602-INS-395
chr17-51966413-INS-56
chr17-51983298-INS-60
chr17-51990082-INS-200
chr17-52005315-INS-68
chr17-52015268-INS-514
chr17-52021625-INS-380
chr17-52035889-INS-716
chr17-52083998-INS-60
chr17-52102820-INS-115
chr17-52126346-INS-314
chr17-52145979-INS-212
chr17-52165623-INS-65
chr17-52179935-INS-176
chr17-52199023-INS-84
chr17-52225628-INS-75
chr17-52254061-INS-340
chr17-52262527-INS-76
chr17-52272171-INS-6603

chr17-52306063-INS-60
chr17-52322831-INS-310
chr17-52340977-INS-227
chr17-52346824-INS-293
chr17-52363270-INS-221
chr17-52374014-INS-99
chr17-52427548-INS-192
chr17-52430582-INS-148
chr17-52441795-INS-317
chr17-52448854-INS-132
chr17-52454197-INS-5101
chr17-52454749-INS-1491
chr17-52462639-INS-54
chr17-52463441-INS-571
chr17-52475753-INS-54
chr17-52481809-INS-52
chr17-52510151-INS-56
chr17-52518978-INS-108
chr17-52533708-INS-53
chr17-52537188-INS-199
chr17-52550170-INS-88
chr17-52578925-INS-894
chr17-52639647-INS-7897
chr17-52641840-INS-6378
chr17-52653343-INS-74
chr17-52678877-INS-7141
chr17-52691021-INS-60
chr17-52694791-INS-62
chr17-52708031-INS-671
chr17-52722001-INS-88
chr17-52726546-INS-322
chr17-52738087-INS-60
chr17-52801042-INS-722
chr17-52804844-INS-163
chr17-52819321-INS-208
chr17-52823923-INS-6478
chr17-52826373-INS-76
chr17-52874069-INS-270
chr17-52883167-INS-608
chr17-52893899-INS-492
chr17-52895757-INS-175
chr17-52911753-INS-200
chr17-52937327-INS-70
chr17-52952484-INS-82
chr17-52959042-INS-64
chr17-52962647-INS-52
chr17-52980913-INS-407
chr17-52990379-INS-100
chr17-53001533-INS-124
chr17-53012368-INS-94
chr17-53024326-INS-817
chr17-53038716-INS-78
chr17-53045246-INS-301
chr17-53085539-INS-448
chr17-53141398-INS-69
chr17-53152871-INS-403
chr17-53170543-INS-65
chr17-53173873-INS-210
chr17-53177298-INS-945
chr17-53181909-INS-2935
chr17-53184678-INS-158
chr17-53196975-INS-387
chr17-53202270-INS-572
chr17-53246143-INS-4004
chr17-53246456-INS-6445
chr17-53296364-INS-67
chr17-53358677-INS-220
chr17-53363470-INS-6269
chr17-53391530-INS-80
chr17-53400234-INS-70
chr17-53412194-INS-66
chr17-53441916-INS-242
chr17-53472246-INS-356
chr17-53502382-INS-131
chr17-53514000-INS-626
chr17-53541892-INS-7111
chr17-53545024-INS-55

chr17-53549711-INS-458
chr17-53554233-INS-742
chr17-53564719-INS-328
chr17-53577383-INS-730
chr17-53602293-INS-292
chr17-53605998-INS-74
chr17-53616764-INS-120
chr17-53631584-INS-131
chr17-53633404-INS-358
chr17-53635520-INS-666
chr17-53641430-INS-6887
chr17-53645946-INS-394
chr17-53646486-INS-76
chr17-53649530-INS-291
chr17-53651519-INS-6645
chr17-53654458-INS-51
chr17-53684044-INS-193
chr17-53689407-INS-674
chr17-53693259-INS-226
chr17-53694857-INS-52
chr17-53702716-INS-339
chr17-53713950-INS-204
chr17-53719663-INS-245
chr17-53748744-INS-92
chr17-53757043-INS-60
chr17-53764351-INS-78
chr17-53775184-INS-6524
chr17-53783078-INS-64
chr17-53812512-INS-356
chr17-53847626-INS-154
chr17-53865452-INS-138
chr17-53870237-INS-75
chr17-53871491-INS-267
chr17-53877822-INS-709
chr17-53878389-INS-70
chr17-53894249-INS-1888
chr17-53897822-INS-80
chr17-53900455-INS-199
chr17-53904936-INS-230
chr17-53928823-INS-84
chr17-53944400-INS-223
chr17-53956920-INS-86
chr17-53959279-INS-504
chr17-53964651-INS-473
chr17-54005924-INS-201
chr17-54034831-INS-3038
chr17-54044952-INS-80
chr17-54054314-INS-9907
chr17-54121061-INS-370
chr17-54142753-INS-7071
chr17-54144573-INS-77
chr17-54150022-INS-182
chr17-54154118-INS-142
chr17-54162967-INS-189
chr17-54171468-INS-86
chr17-54208182-INS-413
chr17-54233867-INS-119
chr17-54234132-INS-438
chr17-54243336-INS-210
chr17-54357016-INS-214
chr17-54364088-INS-87
chr17-54365088-INS-179
chr17-54370166-INS-397
chr17-54387106-INS-487
chr17-54390791-INS-56
chr17-54481394-INS-978
chr17-54496692-INS-135
chr17-54509922-INS-70
chr17-54512019-INS-80
chr17-54537994-INS-121
chr17-54548964-INS-54
chr17-54553108-INS-437
chr17-54554435-INS-236
chr17-54555223-INS-6442
chr17-54558589-INS-63
chr17-54579251-INS-325
chr17-54616251-INS-437

chr17-54643960-INS-52
chr17-54647657-INS-555
chr17-54648458-INS-87
chr17-54649244-INS-83
chr17-54676999-INS-4852
chr17-54741363-INS-837
chr17-54751954-INS-137
chr17-54766920-INS-531
chr17-54769497-INS-65
chr17-54782010-INS-201
chr17-54820388-INS-283
chr17-54866641-INS-100
chr17-54886930-INS-148
chr17-54891394-INS-340
chr17-54931077-INS-168
chr17-54981774-INS-64
chr17-54994183-INS-239
chr17-54996084-INS-110
chr17-55005905-INS-717
chr17-55014154-INS-9047
chr17-55018651-INS-78
chr17-55028364-INS-121
chr17-55032976-INS-98
chr17-55045492-INS-348
chr17-55046590-INS-7074
chr17-55052224-INS-107
chr17-55094138-INS-6641
chr17-55098661-INS-1159
chr17-55108977-INS-65
chr17-55135256-INS-78
chr17-55140780-INS-96
chr17-55146260-INS-166
chr17-55157391-INS-54
chr17-55167030-INS-906
chr17-55244593-INS-279
chr17-55274921-INS-328
chr17-55292372-INS-199
chr17-55325994-INS-90
chr17-55352085-INS-56
chr17-55355622-INS-1518
chr17-55361085-INS-6193
chr17-55371619-INS-6235
chr17-55375455-INS-65
chr17-55378105-INS-245
chr17-55380659-INS-6252
chr17-55388634-INS-65
chr17-55389923-INS-471
chr17-55415307-INS-979
chr17-55428586-INS-68
chr17-55473744-INS-59
chr17-55481049-INS-416
chr17-55528724-INS-64
chr17-55556924-INS-6357
chr17-55562163-INS-148
chr17-55563331-INS-363
chr17-55565303-INS-58
chr17-55576223-INS-91
chr17-55603225-INS-207
chr17-55704921-INS-75
chr17-55739375-INS-353
chr17-55783711-INS-292
chr17-55814662-INS-328
chr17-55823597-INS-86
chr17-55823753-INS-995
chr17-55871403-INS-6492
chr17-55875253-INS-420
chr17-55906929-INS-951
chr17-55909290-INS-166
chr17-55930365-INS-183
chr17-55949176-INS-123
chr17-55959962-INS-212
chr17-55993375-INS-127
chr17-55998535-INS-3489
chr17-55998931-INS-60
chr17-56014223-INS-389
chr17-56072554-INS-7179
chr17-56083112-INS-197

chr17-56103741-INS-102
chr17-56106834-INS-52
chr17-56128795-INS-1606
chr17-56131099-INS-962
chr17-56147496-INS-204
chr17-56169107-INS-3107
chr17-56189832-INS-60
chr17-56214418-INS-76
chr17-56239139-INS-111
chr17-56256202-INS-225
chr17-56294950-INS-168
chr17-56309264-INS-63
chr17-56311030-INS-101
chr17-56346744-INS-268
chr17-56348374-INS-2648
chr17-56348422-INS-303
chr17-56356728-INS-211
chr17-56401431-INS-198
chr17-56411826-INS-198
chr17-56473136-INS-54
chr17-56474118-INS-187
chr17-56522847-INS-71
chr17-56527576-INS-198
chr17-56531587-INS-195
chr17-56536827-INS-180
chr17-56545251-INS-100
chr17-56545853-INS-180
chr17-56549647-INS-68
chr17-56560640-INS-165
chr17-56588940-INS-208
chr17-56592083-INS-277
chr17-56606756-INS-219
chr17-56630968-INS-450
chr17-56710398-INS-128
chr17-56723081-INS-63
chr17-56729771-INS-72
chr17-56737323-INS-79
chr17-56744247-INS-204
chr17-56774192-INS-412
chr17-56774204-INS-1306
chr17-56790805-INS-556
chr17-56791503-INS-204
chr17-56804776-INS-335
chr17-56807635-INS-133
chr17-56815779-INS-56
chr17-56831243-INS-86
chr17-56844235-INS-66
chr17-56862040-INS-210
chr17-56863005-INS-85
chr17-56885034-INS-166
chr17-56963436-INS-152
chr17-56971897-INS-61
chr17-56976340-INS-85
chr17-56983849-INS-192
chr17-56985413-INS-196
chr17-57011291-INS-160
chr17-57026680-INS-225
chr17-57077118-INS-213
chr17-57081071-INS-258
chr17-57101833-INS-460
chr17-57103523-INS-200
chr17-57113946-INS-256
chr17-57114638-INS-109
chr17-57124677-INS-132
chr17-57141064-INS-210
chr17-57148713-INS-93
chr17-57158370-INS-438
chr17-57164404-INS-206
chr17-57168754-INS-328
chr17-57172108-INS-598
chr17-57204915-INS-52
chr17-57249001-INS-594
chr17-57267205-INS-86
chr17-57273818-INS-199
chr17-57276471-INS-179
chr17-57284602-INS-113
chr17-57296447-INS-82

chr17-57302428-INS-215
chr17-57310315-INS-133
chr17-57323479-INS-142
chr17-57355648-INS-82
chr17-57360179-INS-65
chr17-57392901-INS-400
chr17-57401581-INS-792
chr17-57452627-INS-339
chr17-57456416-INS-75
chr17-57463734-INS-125
chr17-57475669-INS-110
chr17-57476751-INS-57
chr17-57481236-INS-216
chr17-57491340-INS-170
chr17-57507498-INS-407
chr17-57511760-INS-980
chr17-57545651-INS-69
chr17-57553011-INS-135
chr17-57573975-INS-63
chr17-57589113-INS-72
chr17-57594271-INS-177
chr17-57607696-INS-550
chr17-57613774-INS-52
chr17-57623523-INS-130
chr17-57632361-INS-104
chr17-57633812-INS-52
chr17-57643835-INS-122
chr17-57654457-INS-472
chr17-57655628-INS-52
chr17-57667606-INS-88
chr17-57702242-INS-201
chr17-57713452-INS-1801
chr17-57726481-INS-372
chr17-57758774-INS-158
chr17-57763350-INS-98
chr17-57773713-INS-54
chr17-57784521-INS-107
chr17-57796770-INS-2872
chr17-57808319-INS-6135
chr17-57818644-INS-672
chr17-57819199-INS-6179
chr17-57823191-INS-72
chr17-57824681-INS-64
chr17-57890280-INS-6799
chr17-57909455-INS-190
chr17-57910058-INS-52
chr17-57949931-INS-199
chr17-57958453-INS-1766
chr17-58021478-INS-326
chr17-58036542-INS-518
chr17-58050382-INS-831
chr17-58051718-INS-126
chr17-58063108-INS-58
chr17-58075971-INS-72
chr17-58082831-INS-52
chr17-58086163-INS-57
chr17-58101250-INS-93
chr17-58109620-INS-73
chr17-58119746-INS-835
chr17-58139077-INS-96
chr17-58140053-INS-269
chr17-58183704-INS-119
chr17-58187146-INS-113
chr17-58192745-INS-151
chr17-58215349-INS-59
chr17-58217857-INS-588
chr17-58258449-INS-6843
chr17-58259334-INS-6703
chr17-58284243-INS-982
chr17-58295480-INS-162
chr17-58313409-INS-66
chr17-58333555-INS-83
chr17-58350348-INS-111
chr17-58355140-INS-65
chr17-58371550-INS-225
chr17-58384115-INS-390
chr17-58411386-INS-68

chr17-58471586-INS-112
chr17-58488782-INS-96
chr17-58490316-INS-108
chr17-58496402-INS-60
chr17-58509099-INS-435
chr17-58533969-INS-126
chr17-58545750-INS-98
chr17-58573322-INS-897
chr17-58578576-INS-209
chr17-58590339-INS-60
chr17-58603561-INS-52
chr17-58647166-INS-188
chr17-58650026-INS-52
chr17-58665155-INS-85
chr17-58677387-INS-938
chr17-58682650-INS-185
chr17-58720896-INS-93
chr17-58744921-INS-234
chr17-58768830-INS-54
chr17-58802498-INS-54
chr17-58810473-INS-52
chr17-58814250-INS-91
chr17-58833963-INS-180
chr17-58858758-INS-317
chr17-58906899-INS-160
chr17-58912931-INS-590
chr17-58914632-INS-52
chr17-58923687-INS-79
chr17-58962229-INS-789
chr17-58993659-INS-319
chr17-59082667-INS-956
chr17-59084690-INS-7716
chr17-59097843-INS-172
chr17-59116628-INS-79
chr17-59133681-INS-183
chr17-59139179-INS-128
chr17-59159830-INS-76
chr17-59164963-INS-75
chr17-59176445-INS-158
chr17-59177204-INS-515
chr17-59207283-INS-60
chr17-59208679-INS-1896
chr17-59217784-INS-153
chr17-59225243-INS-56
chr17-59233868-INS-345
chr17-59239098-INS-434
chr17-59243763-INS-197
chr17-59244464-INS-79
chr17-59254946-INS-273
chr17-59263442-INS-82
chr17-59270512-INS-107
chr17-59275445-INS-487
chr17-59279237-INS-196
chr17-59283290-INS-67
chr17-59301712-INS-830
chr17-59306227-INS-203
chr17-59312930-INS-329
chr17-59330906-INS-69
chr17-59333967-INS-669
chr17-59339122-INS-601
chr17-59348652-INS-202
chr17-59353052-INS-416
chr17-59364092-INS-64
chr17-59372035-INS-414
chr17-59380697-INS-127
chr17-59385694-INS-60
chr17-59394065-INS-864
chr17-59419839-INS-158
chr17-59426540-INS-6932
chr17-59427001-INS-913
chr17-59436023-INS-454
chr17-59489307-INS-753
chr17-59508690-INS-311
chr17-59539583-INS-342
chr17-59544633-INS-75
chr17-59567734-INS-217
chr17-59589203-INS-54

chr17-59600091-INS-5889
chr17-59606139-INS-63
chr17-59611076-INS-222
chr17-59618424-INS-56
chr17-59635948-INS-188
chr17-59639890-INS-61
chr17-59662561-INS-55
chr17-59664364-INS-333
chr17-59682656-INS-52
chr17-59727688-INS-160
chr17-59731978-INS-190
chr17-59741586-INS-167
chr17-59743530-INS-190
chr17-59746392-INS-127
chr17-59748998-INS-51
chr17-59799050-INS-954
chr17-59847791-INS-53
chr17-59859328-INS-142
chr17-59861338-INS-75
chr17-59866613-INS-139
chr17-59876774-INS-7147
chr17-59919252-INS-60
chr17-59922322-INS-54
chr17-59929845-INS-864
chr17-59945581-INS-79
chr17-59947133-INS-179
chr17-59957149-INS-62
chr17-59958820-INS-129
chr17-59973142-INS-54
chr17-59975011-INS-917
chr17-59985090-INS-130
chr17-59996228-INS-52
chr17-60020369-INS-393
chr17-60021499-INS-54
chr17-60071648-INS-7076
chr17-60076467-INS-101
chr17-60138637-INS-61
chr17-60140619-INS-93
chr17-60141695-INS-52
chr17-60181782-INS-54
chr17-60183771-INS-593
chr17-60209527-INS-158
chr17-60221455-INS-6806
chr17-60323540-INS-224
chr17-60345422-INS-78
chr17-60372650-INS-654
chr17-60375451-INS-101
chr17-60392993-INS-240
chr17-60434078-INS-112
chr17-60436277-INS-424
chr17-60477606-INS-80
chr17-60511152-INS-67
chr17-60514975-INS-3813
chr17-60523581-INS-75
chr17-60591735-INS-144
chr17-60604573-INS-5031
chr17-60626102-INS-7161
chr17-60639691-INS-215
chr17-60644303-INS-2095
chr17-60647354-INS-78
chr17-60827889-INS-51
chr17-60835123-INS-64
chr17-60850318-INS-299
chr17-60877548-INS-210
chr17-60878425-INS-62
chr17-60899501-INS-6422
chr17-60907718-INS-6515
chr17-60908993-INS-51
chr17-60917120-INS-7121
chr17-60922414-INS-232
chr17-60974413-INS-348
chr17-61028575-INS-60
chr17-61044647-INS-199
chr17-61051173-INS-350
chr17-61069617-INS-496
chr17-61110451-INS-473
chr17-61114183-INS-816

chr17-61129467-INS-289
chr17-61134988-INS-188
chr17-61152595-INS-94
chr17-61208394-INS-1084
chr17-61218846-INS-225
chr17-61223935-INS-134
chr17-61260646-INS-6872
chr17-61314662-INS-358
chr17-61326964-INS-106
chr17-61365902-INS-395
chr17-61379678-INS-114
chr17-61414036-INS-56
chr17-61415579-INS-105
chr17-61441155-INS-605
chr17-61443149-INS-6329
chr17-61457937-INS-1891
chr17-61458838-INS-113
chr17-61475979-INS-5337
chr17-61479434-INS-120
chr17-61534560-INS-150
chr17-61546048-INS-7717
chr17-61546194-INS-60
chr17-61575811-INS-52
chr17-61594594-INS-901
chr17-61609543-INS-117
chr17-61659325-INS-1159
chr17-61669997-INS-56
chr17-61718266-INS-51
chr17-61721769-INS-52
chr17-61753177-INS-105
chr17-61761150-INS-72
chr17-61767272-INS-108
chr17-61807463-INS-52
chr17-61811342-INS-372
chr17-61824264-INS-540
chr17-61867656-INS-557
chr17-61874529-INS-53
chr17-61880344-INS-272
chr17-61894508-INS-79
chr17-61915108-INS-52
chr17-61960125-INS-116
chr17-61961811-INS-52
chr17-61966497-INS-128
chr17-62041145-INS-89
chr17-62063242-INS-2728
chr17-62071518-INS-64
chr17-62076313-INS-55
chr17-62108156-INS-138
chr17-62133696-INS-487
chr17-62155246-INS-72
chr17-62168576-INS-147
chr17-62295033-INS-130
chr17-62301824-INS-698
chr17-62320012-INS-128
chr17-62366528-INS-183
chr17-62422426-INS-3520
chr17-62423476-INS-3711
chr17-62498313-INS-193
chr17-62519775-INS-100
chr17-62521310-INS-6242
chr17-62540318-INS-320
chr17-62553462-INS-239
chr17-62574465-INS-150
chr17-62577176-INS-66
chr17-62621451-INS-119
chr17-62624860-INS-59
chr17-62642471-INS-52
chr17-62654145-INS-116
chr17-62660509-INS-185
chr17-62686429-INS-70
chr17-62709537-INS-201
chr17-62713130-INS-66
chr17-62713830-INS-64
chr17-62722083-INS-72
chr17-62744290-INS-54
chr17-62764333-INS-276
chr17-62798805-INS-6440

chr17-62801209-INS-127
chr17-62830679-INS-359
chr17-62872425-INS-193
chr17-62889736-INS-59
chr17-62903948-INS-118
chr17-62957771-INS-61
chr17-62966296-INS-72
chr17-62971870-INS-198
chr17-62975367-INS-58
chr17-63019737-INS-124
chr17-63038910-INS-76
chr17-63097259-INS-100
chr17-63163018-INS-120
chr17-63223880-INS-108
chr17-63273813-INS-80
chr17-63330151-INS-544
chr17-63331223-INS-54
chr17-63343972-INS-170
chr17-63357907-INS-223
chr17-63379332-INS-64
chr17-63381333-INS-66
chr17-63390584-INS-6912
chr17-63404575-INS-706
chr17-63409428-INS-107
chr17-63415824-INS-224
chr17-63427557-INS-80
chr17-63448148-INS-368
chr17-63507073-INS-71
chr17-63519952-INS-382
chr17-63521654-INS-56
chr17-63543186-INS-68
chr17-63573685-INS-76
chr17-63587498-INS-70
chr17-63601529-INS-64
chr17-63643714-INS-4353
chr17-63647014-INS-76
chr17-63663140-INS-185
chr17-63666612-INS-224
chr17-63715916-INS-119
chr17-63718895-INS-218
chr17-63730523-INS-166
chr17-63761188-INS-88
chr17-63774035-INS-215
chr17-63789189-INS-189
chr17-63806660-INS-91
chr17-63814597-INS-77
chr17-63816989-INS-202
chr17-63820729-INS-129
chr17-63825130-INS-363
chr17-63840991-INS-163
chr17-63845149-INS-120
chr17-63870196-INS-68
chr17-63879760-INS-192
chr17-63907616-INS-518
chr17-63953952-INS-470
chr17-63957743-INS-575
chr17-63958924-INS-360
chr17-63962557-INS-97
chr17-63965674-INS-401
chr17-63981897-INS-539
chr17-63999900-INS-540
chr17-64010075-INS-58
chr17-64011371-INS-65
chr17-64019871-INS-358
chr17-64030708-INS-379
chr17-64033977-INS-184
chr17-64034638-INS-506
chr17-64042169-INS-253
chr17-64046981-INS-6193
chr17-64064254-INS-66
chr17-64079054-INS-52
chr17-64092355-INS-433
chr17-64116956-INS-54
chr17-64119989-INS-51
chr17-64160090-INS-91
chr17-64165521-INS-2270
chr17-64232542-INS-190

chr17-64238186-INS-1621
chr17-64250968-INS-1710
chr17-64256780-INS-79
chr17-64278300-INS-196
chr17-64320761-INS-110
chr17-64322000-INS-105
chr17-64349155-INS-55
chr17-64354376-INS-361
chr17-64355597-INS-69
chr17-64367324-INS-70
chr17-64390295-INS-287
chr17-64391580-INS-84
chr17-64402354-INS-54
chr17-64404589-INS-53
chr17-64406594-INS-111
chr17-64407993-INS-76
chr17-64411354-INS-488
chr17-64438731-INS-348
chr17-64458218-INS-1480
chr17-64468784-INS-372
chr17-64505231-INS-4799
chr17-64519417-INS-80
chr17-64522891-INS-828
chr17-64531139-INS-653
chr17-64565855-INS-440
chr17-64627878-INS-217
chr17-64636099-INS-52
chr17-64689826-INS-156
chr17-64706036-INS-542
chr17-64708458-INS-287
chr17-64745281-INS-162
chr17-64748324-INS-56
chr17-64756077-INS-615
chr17-64757500-INS-115
chr17-64759684-INS-69
chr17-64775105-INS-193
chr17-64784408-INS-98
chr17-64802086-INS-90
chr17-64807443-INS-80
chr17-64827095-INS-184
chr17-64828819-INS-451
chr17-64863285-INS-56
chr17-64875274-INS-60
chr17-64877052-INS-94
chr17-64877242-INS-454
chr17-64889256-INS-77
chr17-64895152-INS-561
chr17-64898252-INS-95
chr17-64918055-INS-199
chr17-64920358-INS-72
chr17-64923994-INS-1165
chr17-64923995-INS-120
chr17-64933462-INS-355
chr17-64938154-INS-104
chr17-64949812-INS-84
chr17-64979466-INS-306
chr17-64985674-INS-225
chr17-65006707-INS-401
chr17-65020386-INS-55
chr17-65024822-INS-6442
chr17-65044235-INS-68
chr17-65062890-INS-51
chr17-65089066-INS-358
chr17-65092833-INS-656
chr17-65110281-INS-549
chr17-65140665-INS-66
chr17-65147906-INS-309
chr17-65178298-INS-122
chr17-65251188-INS-192
chr17-65258779-INS-54
chr17-65263076-INS-445
chr17-65277451-INS-9327
chr17-65287706-INS-507
chr17-65304627-INS-64
chr17-65306066-INS-224
chr17-65309210-INS-51
chr17-65313565-INS-107

chr17-65319268-INS-6897
chr17-65333143-INS-166
chr17-65334157-INS-110
chr17-65351696-INS-60
chr17-65373039-INS-493
chr17-65374740-INS-52
chr17-65379067-INS-59
chr17-65384602-INS-76
chr17-65387342-INS-191
chr17-65429589-INS-486
chr17-65489137-INS-65
chr17-65493907-INS-66
chr17-65496951-INS-440
chr17-65499048-INS-52
chr17-65571245-INS-208
chr17-65588614-INS-136
chr17-65595483-INS-69
chr17-65612038-INS-300
chr17-65635104-INS-55
chr17-65653672-INS-396
chr17-65687425-INS-68
chr17-65687552-INS-668
chr17-65692515-INS-213
chr17-65725315-INS-210
chr17-65727056-INS-70
chr17-65731354-INS-73
chr17-65744148-INS-924
chr17-65751175-INS-55
chr17-65791165-INS-194
chr17-65793214-INS-72
chr17-65812084-INS-6401
chr17-65821261-INS-157
chr17-65836469-INS-143
chr17-65848989-INS-377
chr17-65849714-INS-98
chr17-65852604-INS-162
chr17-65868921-INS-138
chr17-65868921-INS-450
chr17-65875338-INS-51
chr17-65898355-INS-52
chr17-65980539-INS-85
chr17-66001968-INS-342
chr17-66014205-INS-428
chr17-66015514-INS-678
chr17-66042740-INS-265
chr17-66058133-INS-111
chr17-66063974-INS-196
chr17-66077574-INS-163
chr17-66110031-INS-399
chr17-66116872-INS-65
chr17-66121904-INS-191
chr17-66154403-INS-190
chr17-66154501-INS-682
chr17-66164897-INS-59
chr17-66202070-INS-195
chr17-66216034-INS-233
chr17-66245251-INS-314
chr17-66249154-INS-138
chr17-66387564-INS-136
chr17-66397600-INS-203
chr17-66405489-INS-202
chr17-66412588-INS-825
chr17-66417678-INS-78
chr17-66429416-INS-91
chr17-66449845-INS-666
chr17-66486404-INS-206
chr17-66494066-INS-234
chr17-66504057-INS-52
chr17-66517094-INS-92
chr17-66537643-INS-449
chr17-66571194-INS-123
chr17-66581576-INS-125
chr17-66587221-INS-74
chr17-66594850-INS-223
chr17-66595056-INS-6738
chr17-66606859-INS-94
chr17-66621428-INS-289

chr17-66631913-INS-480
chr17-66635425-INS-51
chr17-66680165-INS-72
chr17-66689673-INS-60
chr17-66721922-INS-198
chr17-66727415-INS-195
chr17-66748961-INS-55
chr17-66751872-INS-230
chr17-66777574-INS-142
chr17-66780036-INS-230
chr17-66783362-INS-75
chr17-66784570-INS-197
chr17-66792191-INS-435
chr17-66795903-INS-428
chr17-66797738-INS-166
chr17-66799691-INS-366
chr17-66805878-INS-840
chr17-66828054-INS-205
chr17-66831987-INS-151
chr17-66842381-INS-78
chr17-66847707-INS-324
chr17-66849810-INS-100
chr17-66853601-INS-146
chr17-66857969-INS-440
chr17-66867156-INS-98
chr17-66876686-INS-79
chr17-66902568-INS-268
chr17-66910738-INS-160
chr17-66931673-INS-105
chr17-66933482-INS-289
chr17-66935671-INS-67
chr17-66951841-INS-60
chr17-66980679-INS-140
chr17-67009919-INS-105
chr17-67021973-INS-69
chr17-67024851-INS-124
chr17-67028796-INS-318
chr17-67034037-INS-84
chr17-67129389-INS-212
chr17-67186180-INS-65
chr17-67207383-INS-66
chr17-67208813-INS-52
chr17-67260544-INS-122
chr17-67298970-INS-57
chr17-67304494-INS-578
chr17-67306147-INS-336
chr17-67332974-INS-85
chr17-67340518-INS-90
chr17-67345182-INS-136
chr17-67358068-INS-251
chr17-67366306-INS-63
chr17-67370347-INS-146
chr17-67387509-INS-236
chr17-67392384-INS-88
chr17-67395493-INS-272
chr17-67415317-INS-56
chr17-67419520-INS-191
chr17-67437185-INS-5938
chr17-67446551-INS-186
chr17-67475564-INS-67
chr17-67486588-INS-183
chr17-67494539-INS-262
chr17-67532505-INS-578
chr17-67540944-INS-497
chr17-67556060-INS-63
chr17-67638595-INS-329
chr17-67657115-INS-59
chr17-67685118-INS-102
chr17-67691436-INS-289
chr17-67717512-INS-130
chr17-67747317-INS-72
chr17-67782995-INS-214
chr17-67793543-INS-208
chr17-67800802-INS-233
chr17-67803891-INS-55
chr17-67819513-INS-80
chr17-67836355-INS-295

chr17-67850757-INS-126
chr17-67877389-INS-52
chr17-67884823-INS-198
chr17-67885543-INS-61
chr17-67894576-INS-714
chr17-67936690-INS-770
chr17-67948640-INS-61
chr17-67957072-INS-58
chr17-67964741-INS-291
chr17-67982099-INS-98
chr17-68022423-INS-64
chr17-68027489-INS-138
chr17-68031766-INS-93
chr17-68036684-INS-56
chr17-68048425-INS-141
chr17-68056419-INS-102
chr17-68066406-INS-440
chr17-68067555-INS-66
chr17-68070138-INS-55
chr17-68077367-INS-64
chr17-68085164-INS-55
chr17-68104017-INS-401
chr17-68130914-INS-228
chr17-68153179-INS-297
chr17-68161039-INS-112
chr17-68194372-INS-60
chr17-68210043-INS-226
chr17-68218710-INS-177
chr17-68303762-INS-51
chr17-68311927-INS-63
chr17-68328301-INS-52
chr17-68357990-INS-424
chr17-68363912-INS-263
chr17-68403922-INS-283
chr17-68415189-INS-56
chr17-68422096-INS-52
chr17-68435664-INS-53
chr17-68439643-INS-580
chr17-68444133-INS-96
chr17-68460245-INS-69
chr17-68468228-INS-214
chr17-68468668-INS-51
chr17-68469901-INS-6692
chr17-68499774-INS-364
chr17-68506836-INS-111
chr17-68520281-INS-66
chr17-68521567-INS-272
chr17-68543281-INS-130
chr17-68549210-INS-170
chr17-68571788-INS-74
chr17-68590511-INS-121
chr17-68598022-INS-58
chr17-68627142-INS-119
chr17-68627186-INS-60
chr17-68631488-INS-52
chr17-68633730-INS-533
chr17-68640060-INS-96
chr17-68670012-INS-99
chr17-68699011-INS-66
chr17-68705534-INS-64
chr17-68760684-INS-109
chr17-68763960-INS-54
chr17-68767286-INS-218
chr17-68789853-INS-977
chr17-68816653-INS-204
chr17-68844638-INS-3607
chr17-68847601-INS-417
chr17-68848654-INS-584
chr17-68894896-INS-54
chr17-68897874-INS-120
chr17-68920305-INS-159
chr17-68943938-INS-54
chr17-68981766-INS-62
chr17-69003940-INS-62
chr17-69012535-INS-58
chr17-69016283-INS-250
chr17-69018712-INS-74

chr17-69023056-INS-368
chr17-69025667-INS-140
chr17-69098997-INS-161
chr17-69141787-INS-517
chr17-69165546-INS-4456
chr17-69173156-INS-101
chr17-69175463-INS-312
chr17-69181740-INS-56
chr17-69201714-INS-639
chr17-69223949-INS-210
chr17-69309660-INS-749
chr17-69326486-INS-926
chr17-69335788-INS-66
chr17-69338958-INS-106
chr17-69365737-INS-130
chr17-69379197-INS-209
chr17-69387357-INS-89
chr17-69390207-INS-56
chr17-69392195-INS-124
chr17-69397536-INS-368
chr17-69402378-INS-69
chr17-69435721-INS-96
chr17-69450724-INS-106
chr17-69452498-INS-66
chr17-69522647-INS-54
chr17-69535492-INS-124
chr17-69542030-INS-193
chr17-69545937-INS-190
chr17-69546178-INS-872
chr17-69593430-INS-819
chr17-69594159-INS-207
chr17-69604048-INS-56
chr17-69610480-INS-235
chr17-69622882-INS-149
chr17-69630670-INS-212
chr17-69706702-INS-79
chr17-69717975-INS-68
chr17-69740887-INS-96
chr17-69752383-INS-190
chr17-69762750-INS-162
chr17-69797654-INS-3136
chr17-69807213-INS-75
chr17-69837136-INS-6450
chr17-69843642-INS-273
chr17-69850590-INS-318
chr17-69875534-INS-352
chr17-69876426-INS-134
chr17-69898278-INS-60
chr17-69910562-INS-116
chr17-69943303-INS-237
chr17-69969764-INS-716
chr17-69985947-INS-63
chr17-70019201-INS-161
chr17-70024143-INS-56
chr17-70042977-INS-61
chr17-70090063-INS-126
chr17-70115831-INS-2378
chr17-70147499-INS-429
chr17-70152037-INS-195
chr17-70161171-INS-287
chr17-70167521-INS-287
chr17-70182588-INS-209
chr17-70215005-INS-255
chr17-70233660-INS-993
chr17-70237580-INS-206
chr17-70241758-INS-252
chr17-70244097-INS-55
chr17-70299926-INS-1250
chr17-70315727-INS-645
chr17-70377262-INS-369
chr17-70412929-INS-1130
chr17-70428969-INS-207
chr17-70439062-INS-665
chr17-70450180-INS-99
chr17-70468792-INS-101
chr17-70476694-INS-81
chr17-70496038-INS-7928

chr17-70521235-INS-61
chr17-70529926-INS-57
chr17-70531774-INS-84
chr17-70578306-INS-139
chr17-70618437-INS-63
chr17-70624992-INS-6438
chr17-70649713-INS-110
chr17-70654157-INS-584
chr17-70677296-INS-216
chr17-70689326-INS-64
chr17-70698634-INS-185
chr17-70711387-INS-72
chr17-70720112-INS-119
chr17-70727496-INS-57
chr17-70728655-INS-55
chr17-70738536-INS-57
chr17-70752350-INS-229
chr17-70761549-INS-124
chr17-70773428-INS-310
chr17-70789667-INS-7085
chr17-70794415-INS-52
chr17-70807066-INS-90
chr17-70813961-INS-84
chr17-70815995-INS-87
chr17-70836357-INS-100
chr17-70843027-INS-78
chr17-70847223-INS-352
chr17-70864111-INS-52
chr17-70866360-INS-86
chr17-70881507-INS-190
chr17-70886627-INS-108
chr17-70898173-INS-72
chr17-70900568-INS-123
chr17-70932658-INS-55
chr17-70959552-INS-180
chr17-70968634-INS-161
chr17-71001951-INS-471
chr17-71012345-INS-192
chr17-71037547-INS-208
chr17-71050961-INS-232
chr17-71052716-INS-197
chr17-71061631-INS-726
chr17-71066979-INS-179
chr17-71078414-INS-66
chr17-71089952-INS-136
chr17-71110587-INS-158
chr17-71165624-INS-133
chr17-71166915-INS-68
chr17-71172924-INS-208
chr17-71175924-INS-74
chr17-71221721-INS-276
chr17-71235948-INS-326
chr17-71237291-INS-406
chr17-71241893-INS-81
chr17-71243149-INS-162
chr17-71245958-INS-168
chr17-71246258-INS-195
chr17-71248117-INS-201
chr17-71255142-INS-67
chr17-71262801-INS-193
chr17-71274219-INS-53
chr17-71305781-INS-5225
chr17-71315980-INS-204
chr17-71322555-INS-60
chr17-71335881-INS-313
chr17-71345068-INS-246
chr17-71355124-INS-121
chr17-71360219-INS-202
chr17-71361271-INS-74
chr17-71377828-INS-80
chr17-71385092-INS-232
chr17-71390231-INS-203
chr17-71421745-INS-328
chr17-71427546-INS-52
chr17-71441107-INS-327
chr17-71449718-INS-110
chr17-71458684-INS-71

chr17-71469506-INS-218
chr17-71480063-INS-62
chr17-71494376-INS-153
chr17-71516553-INS-54
chr17-71527501-INS-55
chr17-71552342-INS-52
chr17-71604717-INS-100
chr17-71607810-INS-110
chr17-71643436-INS-85
chr17-71646938-INS-59
chr17-71687490-INS-56
chr17-71699077-INS-60
chr17-71707026-INS-248
chr17-71744958-INS-351
chr17-71746227-INS-824
chr17-71765596-INS-149
chr17-71776421-INS-218
chr17-71780765-INS-171
chr17-71792194-INS-589
chr17-71796508-INS-405
chr17-71806192-INS-174
chr17-71807684-INS-199
chr17-71812489-INS-225
chr17-71829632-INS-55
chr17-71840907-INS-469
chr17-71843796-INS-531
chr17-71883043-INS-70
chr17-71896784-INS-216
chr17-71901242-INS-6452
chr17-71906525-INS-76
chr17-71929284-INS-90
chr17-71942441-INS-126
chr17-71954367-INS-88
chr17-71960697-INS-58
chr17-71970933-INS-215
chr17-71977725-INS-219
chr17-71989211-INS-161
chr17-72008388-INS-69
chr17-72011596-INS-54
chr17-72032648-INS-116
chr17-72037255-INS-340
chr17-72042384-INS-821
chr17-72076593-INS-6325
chr17-72078138-INS-54
chr17-72084374-INS-106
chr17-72094793-INS-464
chr17-72127538-INS-822
chr17-72132631-INS-390
chr17-72162256-INS-174
chr17-72168958-INS-445
chr17-72182595-INS-53
chr17-72200954-INS-69
chr17-72205360-INS-207
chr17-72206913-INS-120
chr17-72211009-INS-385
chr17-72211831-INS-164
chr17-72220818-INS-70
chr17-72242617-INS-108
chr17-72247349-INS-60
chr17-72290209-INS-380
chr17-72308903-INS-273
chr17-72315989-INS-82
chr17-72320318-INS-87
chr17-72339039-INS-115
chr17-72369213-INS-142
chr17-72376649-INS-115
chr17-72379547-INS-54
chr17-72421331-INS-277
chr17-72448290-INS-720
chr17-72461894-INS-51
chr17-72463113-INS-1823
chr17-72466968-INS-87
chr17-72482178-INS-131
chr17-72503717-INS-98
chr17-72583442-INS-6383
chr17-72598798-INS-79
chr17-72605783-INS-53

chr17-72660604-INS-473
chr17-72663510-INS-56
chr17-72672032-INS-57
chr17-72704693-INS-2992
chr17-72724174-INS-300
chr17-72732010-INS-108
chr17-72746815-INS-203
chr17-72765915-INS-581
chr17-72773463-INS-90
chr17-72803383-INS-159
chr17-72808113-INS-465
chr17-72834480-INS-66
chr17-72842554-INS-58
chr17-72846389-INS-75
chr17-72849300-INS-69
chr17-72857254-INS-108
chr17-72861590-INS-63
chr17-72877918-INS-65
chr17-72902971-INS-214
chr17-72909853-INS-60
chr17-72922151-INS-108
chr17-72932005-INS-64
chr17-72932805-INS-55
chr17-72974532-INS-505
chr17-72984324-INS-140
chr17-72991934-INS-52
chr17-73002210-INS-161
chr17-73013230-INS-118
chr17-73014441-INS-160
chr17-73041782-INS-78
chr17-73055550-INS-4270
chr17-73111900-INS-106
chr17-73126135-INS-64
chr17-73165216-INS-203
chr17-73173975-INS-666
chr17-73174110-INS-314
chr17-73194806-INS-196
chr17-73206887-INS-104
chr17-73231965-INS-193
chr17-73279355-INS-73
chr17-73292244-INS-5542
chr17-73305137-INS-1000
chr17-73308333-INS-223
chr17-73339631-INS-96
chr17-73341370-INS-169
chr17-73380873-INS-81
chr17-73397588-INS-93
chr17-73405444-INS-136
chr17-73424286-INS-1203
chr17-73447314-INS-649
chr17-73452106-INS-205
chr17-73473221-INS-190
chr17-73492011-INS-168
chr17-73524224-INS-78
chr17-73535607-INS-5254
chr17-73545061-INS-82
chr17-73546194-INS-76
chr17-73559404-INS-58
chr17-73586564-INS-73
chr17-73590885-INS-224
chr17-73652590-INS-115
chr17-73663455-INS-53
chr17-73670456-INS-127
chr17-73720218-INS-321
chr17-73764847-INS-116
chr17-73773271-INS-101
chr17-73788454-INS-120
chr17-73819794-INS-62
chr17-73840729-INS-101
chr17-73884678-INS-67
chr17-73930465-INS-718
chr17-73932692-INS-56
chr17-73933854-INS-132
chr17-73951215-INS-1920
chr17-73951863-INS-91
chr17-73958465-INS-366
chr17-73991690-INS-132

chr17-74076481-INS-6272
chr17-74078921-INS-93
chr17-74082153-INS-401
chr17-74136564-INS-55
chr17-74140777-INS-522
chr17-74146808-INS-85
chr17-74153290-INS-6602
chr17-74156959-INS-108
chr17-74159260-INS-176
chr17-74173764-INS-60
chr17-74174459-INS-62
chr17-74181850-INS-338
chr17-74215911-INS-86
chr17-74220223-INS-741
chr17-74238071-INS-129
chr17-74252810-INS-90
chr17-74271322-INS-514
chr17-74289933-INS-75
chr17-74340395-INS-102
chr17-74371866-INS-208
chr17-74377189-INS-96
chr17-74392007-INS-91
chr17-74392330-INS-743
chr17-74396864-INS-80
chr17-74398197-INS-185
chr17-74399156-INS-833
chr17-74405700-INS-55
chr17-74416774-INS-136
chr17-74433794-INS-589
chr17-74462050-INS-56
chr17-74479721-INS-721
chr17-74489166-INS-185
chr17-74494134-INS-90
chr17-74508947-INS-77
chr17-74513298-INS-129
chr17-74516820-INS-312
chr17-74557499-INS-78
chr17-74565181-INS-72
chr17-74573611-INS-209
chr17-74584318-INS-81
chr17-74589215-INS-202
chr17-74594292-INS-250
chr17-74596123-INS-118
chr17-74606889-INS-196
chr17-74609753-INS-114
chr17-74610823-INS-100
chr17-74614370-INS-57
chr17-74635427-INS-107
chr17-74655743-INS-155
chr17-74667750-INS-218
chr17-74672090-INS-94
chr17-74673460-INS-185
chr17-74691621-INS-213
chr17-74692214-INS-220
chr17-74693239-INS-59
chr17-74694404-INS-399
chr17-74715466-INS-254
chr17-74720467-INS-639
chr17-74735504-INS-116
chr17-74735504-INS-178
chr17-74757585-INS-60
chr17-74760706-INS-108
chr17-74771892-INS-55
chr17-74786736-INS-973
chr17-74813639-INS-56
chr17-74816747-INS-358
chr17-74819824-INS-213
chr17-74821508-INS-199
chr17-74827762-INS-327
chr17-74837862-INS-203
chr17-74848250-INS-498
chr17-74854889-INS-226
chr17-74856766-INS-104
chr17-74863592-INS-475
chr17-74878015-INS-297
chr17-74878402-INS-823
chr17-74888573-INS-95

chr17-74903423-INS-79
chr17-74965748-INS-213
chr17-75016718-INS-79
chr17-75029581-INS-856
chr17-75037068-INS-498
chr17-75048858-INS-219
chr17-75077159-INS-3431
chr17-75086031-INS-236
chr17-75148538-INS-129
chr17-75154366-INS-167
chr17-75167200-INS-565
chr17-75208701-INS-235
chr17-75209712-INS-274
chr17-75211636-INS-89
chr17-75241096-INS-179
chr17-75254944-INS-111
chr17-75257806-INS-67
chr17-75275899-INS-232
chr17-75306622-INS-276
chr17-75372014-INS-192
chr17-75372556-INS-972
chr17-75382012-INS-438
chr17-75390925-INS-78
chr17-75402817-INS-62
chr17-75411649-INS-102
chr17-75416366-INS-147
chr17-75440894-INS-56
chr17-75443387-INS-138
chr17-75454484-INS-8011
chr17-75462587-INS-188
chr17-75467569-INS-265
chr17-75483304-INS-112
chr17-75496920-INS-104
chr17-75511439-INS-723
chr17-75569517-INS-1874
chr17-75574427-INS-224
chr17-75588156-INS-540
chr17-75628613-INS-574
chr17-75661859-INS-78
chr17-75686825-INS-237
chr17-75699954-INS-304
chr17-75738828-INS-924
chr17-75750083-INS-214
chr17-75795243-INS-624
chr17-75817266-INS-204
chr17-75820563-INS-869
chr17-75827996-INS-110
chr17-75833115-INS-55
chr17-75839531-INS-302
chr17-75843626-INS-64
chr17-75849311-INS-110
chr17-75888549-INS-179
chr17-75908870-INS-316
chr17-75939647-INS-124
chr17-75952883-INS-487
chr17-75968188-INS-149
chr17-75998091-INS-214
chr17-76006984-INS-102
chr17-76028346-INS-6687
chr17-76030069-INS-746
chr17-76030391-INS-7936
chr17-76057569-INS-118
chr17-76058072-INS-7425
chr17-76073578-INS-95
chr17-76081780-INS-159
chr17-76084897-INS-94
chr17-76104179-INS-345
chr17-76109720-INS-211
chr17-76121138-INS-1436
chr17-76129102-INS-114
chr17-76137918-INS-954
chr17-76167777-INS-240
chr17-76178566-INS-65
chr17-76180221-INS-5293
chr17-76182772-INS-64
chr17-76221801-INS-90
chr17-76244320-INS-942

chr17-76280577-INS-360
chr17-76322288-INS-5699
chr17-76334980-INS-68
chr17-76368919-INS-399
chr17-76371754-INS-96
chr17-76393179-INS-399
chr17-76425847-INS-4273
chr17-76426603-INS-104
chr17-76430163-INS-12652
chr17-76464324-INS-6442
chr17-76466168-INS-75
chr17-76469470-INS-55
chr17-76481566-INS-922
chr17-76500556-INS-326
chr17-76510359-INS-731
chr17-76530688-INS-84
chr17-76537284-INS-98
chr17-76650296-INS-55
chr17-76685911-INS-426
chr17-76689809-INS-7101
chr17-76689810-INS-358
chr17-76695744-INS-56
chr17-76814951-INS-5081
chr17-76815923-INS-53
chr17-76843588-INS-3867
chr17-76846973-INS-52
chr17-76850571-INS-6219
chr17-76887360-INS-119
chr17-76928605-INS-54
chr17-76932797-INS-1282
chr17-76935993-INS-108
chr17-76947184-INS-1670
chr17-76954747-INS-5983
chr17-76959165-INS-285
chr17-76978733-INS-70
chr17-76986569-INS-464
chr17-76988124-INS-82
chr17-76997353-INS-779
chr17-77002806-INS-513
chr17-77008229-INS-157
chr17-77012526-INS-60
chr17-77022481-INS-86
chr17-77028359-INS-395
chr17-77031337-INS-329
chr17-77033047-INS-64
chr17-77034106-INS-70
chr17-77037915-INS-2151
chr17-77044049-INS-201
chr17-77047224-INS-6629
chr17-77057378-INS-75
chr17-77062251-INS-7094
chr17-77072707-INS-72
chr17-77073402-INS-4509
chr17-77084217-INS-64
chr17-77086001-INS-58
chr17-77090885-INS-408
chr17-77123324-INS-147
chr17-77140169-INS-88
chr17-77173655-INS-56
chr17-77178374-INS-115
chr17-77180282-INS-76
chr17-77185144-INS-445
chr17-77195044-INS-165
chr17-77214041-INS-77
chr17-77216282-INS-52
chr17-77259801-INS-191
chr17-77306040-INS-72
chr17-77353142-INS-516
chr17-77407329-INS-162
chr17-77426749-INS-169
chr17-77439603-INS-159
chr17-77442319-INS-542
chr17-77444434-INS-539
chr17-77502162-INS-276
chr17-77566574-INS-159
chr17-77578018-INS-6670
chr17-77578723-INS-154

chr17-77581714-INS-431
chr17-77584123-INS-850
chr17-77584814-INS-143
chr17-77620073-INS-5438
chr17-77628939-INS-5523
chr17-77635260-INS-601
chr17-77639279-INS-84
chr17-77658678-INS-80
chr17-77673190-INS-5094
chr17-77678123-INS-640
chr17-77683530-INS-267
chr17-77766001-INS-1887
chr17-77774057-INS-266
chr17-77778103-INS-111
chr17-77790147-INS-292
chr17-77792348-INS-107
chr17-77794538-INS-190
chr17-77824108-INS-92
chr17-77831058-INS-53
chr17-77837273-INS-123
chr17-77875975-INS-6649
chr17-77876857-INS-120
chr17-77885334-INS-126
chr17-77911045-INS-254
chr17-77921414-INS-256
chr17-77930364-INS-712
chr17-77943026-INS-375
chr17-77965757-INS-6633
chr17-77968131-INS-1611
chr17-77975590-INS-90
chr17-77979166-INS-84
chr17-77986163-INS-6589
chr17-78073874-INS-8227
chr17-78108893-INS-203
chr17-78112981-INS-56
chr17-78115535-INS-91
chr17-78119671-INS-116
chr17-78123869-INS-488
chr17-78145944-INS-76
chr17-78163690-INS-69
chr17-78177085-INS-89
chr17-78185426-INS-101
chr17-78205355-INS-192
chr17-78226239-INS-68
chr17-78235340-INS-61
chr17-78248367-INS-52
chr17-78250412-INS-369
chr17-78264072-INS-142
chr17-78273805-INS-4169
chr17-78301442-INS-973
chr17-78315050-INS-52
chr17-78329049-INS-93
chr17-78359834-INS-102
chr17-78364236-INS-95
chr17-78384580-INS-66
chr17-78397007-INS-210
chr17-78401797-INS-549
chr17-78405297-INS-1275
chr17-78426950-INS-1236
chr17-78437274-INS-78
chr17-78451829-INS-625
chr17-78454351-INS-60
chr17-78483746-INS-131
chr17-78490557-INS-189
chr17-78494230-INS-756
chr17-78559569-INS-60
chr17-78591461-INS-384
chr17-78631904-INS-96
chr17-78679603-INS-54
chr17-78721408-INS-72
chr17-78737459-INS-157
chr17-78740345-INS-1862
chr17-78755984-INS-194
chr17-78761217-INS-497
chr17-78766531-INS-152
chr17-78769414-INS-51
chr17-78785062-INS-6534

chr17-78806461-INS-118
chr17-78810664-INS-665
chr17-78814342-INS-166
chr17-78833089-INS-114
chr17-78839800-INS-96
chr17-78847904-INS-256
chr17-79021339-INS-114
chr17-79046523-INS-162
chr17-79058293-INS-486
chr17-79058293-INS-931
chr17-79071188-INS-182
chr17-79079429-INS-140
chr17-79079430-INS-157
chr17-79098044-INS-106
chr17-79139913-INS-78
chr17-79197253-INS-61
chr17-79202683-INS-69
chr17-79205732-INS-89
chr17-79211470-INS-114
chr17-79252879-INS-124
chr17-79265067-INS-295
chr17-79271258-INS-169
chr17-79281322-INS-268
chr17-79285058-INS-1586
chr17-79313115-INS-230
chr17-79316513-INS-64
chr17-79342433-INS-60
chr17-79366954-INS-106
chr17-79373602-INS-101
chr17-79396807-INS-463
chr17-79403510-INS-62
chr17-79431386-INS-139
chr17-79442314-INS-55
chr17-79466210-INS-232
chr17-79529840-INS-400
chr17-79562797-INS-474
chr17-79563598-INS-180
chr17-79570013-INS-721
chr17-79579008-INS-71
chr17-79590575-INS-82
chr17-79617855-INS-309
chr17-79623985-INS-70
chr17-79692520-INS-216
chr17-79696548-INS-64
chr17-79704316-INS-6419
chr17-79760215-INS-60
chr17-79761083-INS-54
chr17-79831573-INS-307
chr17-79839443-INS-322
chr17-79879922-INS-906
chr17-79879981-INS-1347
chr17-79912294-INS-70
chr17-79915345-INS-187
chr17-79946191-INS-54
chr17-79977267-INS-51
chr17-79987470-INS-944
chr17-80002359-INS-383
chr17-80004501-INS-56
chr17-80011634-INS-347
chr17-80025739-INS-64
chr17-80029590-INS-61
chr17-80049976-INS-73
chr17-80055960-INS-153
chr17-80061068-INS-81
chr17-80065600-INS-198
chr17-80101569-INS-740
chr17-80102203-INS-134
chr17-80108281-INS-63
chr17-80114123-INS-60
chr17-80124020-INS-201
chr17-80143925-INS-876
chr17-80153578-INS-290
chr17-80203900-INS-64
chr17-80209847-INS-198
chr17-80228280-INS-81
chr17-80235580-INS-506
chr17-80254451-INS-202

chr17-80263896-INS-530
chr17-80267018-INS-274
chr17-80267334-INS-163
chr17-80270096-INS-329
chr17-80271191-INS-90
chr17-80280182-INS-52
chr17-80282684-INS-114
chr17-80299643-INS-313
chr17-80303526-INS-193
chr17-80314517-INS-65
chr17-80321187-INS-223
chr17-80324561-INS-57
chr17-80341088-INS-59
chr17-80356739-INS-219
chr17-80359725-INS-181
chr17-80366769-INS-563
chr17-80373815-INS-419
chr17-80390703-INS-54
chr17-80391471-INS-60
chr17-80408912-INS-81
chr17-80412975-INS-73
chr17-80421569-INS-111
chr17-80428941-INS-62
chr17-80437280-INS-205
chr17-80449571-INS-552
chr17-80450780-INS-207
chr17-80455650-INS-59
chr17-80457043-INS-228
chr17-80458564-INS-150
chr17-80458790-INS-304
chr17-80460676-INS-55
chr17-80461201-INS-272
chr17-80469936-INS-178
chr17-80486241-INS-198
chr17-80488508-INS-250
chr17-80490539-INS-180
chr17-80495367-INS-204
chr17-80497662-INS-328
chr17-80499222-INS-111
chr17-80520268-INS-208
chr17-80526624-INS-214
chr17-80545019-INS-311
chr17-80548472-INS-209
chr17-80567241-INS-216
chr17-80569058-INS-188
chr17-80571612-INS-290
chr17-80605315-INS-207
chr17-80613212-INS-54
chr17-80655506-INS-358
chr17-80675409-INS-68
chr17-80723490-INS-218
chr17-80729189-INS-228
chr17-80733320-INS-54
chr17-80755396-INS-90
chr17-80794034-INS-197
chr17-80798746-INS-132
chr17-80803569-INS-246
chr17-80814189-INS-228
chr17-80816634-INS-220
chr17-80818529-INS-218
chr17-80829960-INS-669
chr17-80848355-INS-340
chr17-80857809-INS-64
chr17-80861792-INS-498
chr17-80906533-INS-262
chr17-80912100-INS-86
chr17-80920690-INS-54
chr17-80925487-INS-60
chr17-80928362-INS-56
chr17-80930347-INS-226
chr17-80946488-INS-182
chr17-80947562-INS-52
chr17-80948219-INS-431
chr17-80981559-INS-124
chr17-81010183-INS-55
chr17-81024235-INS-563
chr17-81036580-INS-150

chr17-81042830-INS-203
chr17-81064352-INS-98
chr17-81068666-INS-50
chr17-81075086-INS-128
chr17-81082229-INS-60
chr17-81098854-INS-72
chr17-81113653-INS-759
chr17-81118558-INS-66
chr17-81135833-INS-96
chr17-81157658-INS-63
chr17-81158355-INS-931
chr17-81160015-INS-62
chr17-81161010-INS-122
chr17-81172999-INS-50
chr17-81177328-INS-447
chr17-81201283-INS-313
chr17-81219448-INS-104
chr17-81261986-INS-374
chr17-81262694-INS-104
chr17-81283735-INS-136
chr17-81286787-INS-70
chr17-81292988-INS-126
chr17-81296139-INS-192
chr17-81306016-INS-184
chr17-81327152-INS-62
chr17-81329985-INS-116
chr17-81332085-INS-53
chr17-81341179-INS-54
chr17-81355870-INS-217
chr17-81382947-INS-204
chr17-81395770-INS-58
chr17-81399420-INS-223
chr17-81406990-INS-51
chr17-81433464-INS-129
chr17-81438629-INS-374
chr17-81457014-INS-189
chr17-81478287-INS-910
chr17-81478356-INS-336
chr17-81485158-INS-310
chr17-81491268-INS-336
chr17-81495608-INS-273
chr17-81502727-INS-292
chr17-81511029-INS-204
chr17-81522183-INS-400
chr17-81532397-INS-681
chr17-81571721-INS-225
chr17-81573705-INS-515
chr17-81578412-INS-63
chr17-81614693-INS-6511
chr17-81634400-INS-1639
chr17-81646103-INS-146
chr17-81646896-INS-897
chr17-81649997-INS-81
chr17-81702756-INS-52
chr17-81721651-INS-111
chr17-81723102-INS-512
chr17-81725768-INS-233
chr17-81738615-INS-111
chr17-81755574-INS-205
chr17-81758564-INS-6695
chr17-81764510-INS-255
chr17-81780699-INS-62
chr17-81795163-INS-62
chr17-81796834-INS-148
chr17-81806555-INS-237
chr17-81814880-INS-227
chr17-81819892-INS-104
chr17-81822977-INS-5240
chr17-81829505-INS-1890
chr17-81871318-INS-511
chr17-81877930-INS-246
chr17-81879315-INS-167
chr17-81884035-INS-91
chr17-81889989-INS-113
chr17-81915514-INS-380
chr17-81939544-INS-52
chr17-81991887-INS-212

chr17-81998311-INS-64
chr17-82010001-INS-58
chr17-82011506-INS-136
chr17-82019299-INS-639
chr17-82023974-INS-212
chr17-82033367-INS-51
chr17-82035479-INS-83
chr17-82038079-INS-111
chr17-82066202-INS-81
chr17-82069648-INS-80
chr17-82077082-INS-5544
chr17-82077917-INS-569
chr17-82089783-INS-328
chr17-82105408-INS-1897
chr17-82110945-INS-498
chr17-82144966-INS-85
chr17-82161389-INS-66
chr17-82169073-INS-386
chr17-82181236-INS-690
chr17-82183655-INS-860
chr17-82196528-INS-206
chr17-82247862-INS-148
chr17-82250254-INS-76
chr17-82258417-INS-6631
chr17-82260298-INS-106
chr17-82266342-INS-130
chr17-82293766-INS-1295
chr17-82312228-INS-663
chr17-82314806-INS-333
chr17-82332647-INS-991
chr17-82344944-INS-74
chr17-82346489-INS-632
chr17-82365592-INS-5299
chr17-82388593-INS-56
chr17-82423195-INS-52
chr17-82432059-INS-8119
chr17-82492350-INS-192
chr17-82542854-INS-282
chr17-82551487-INS-53
chr17-82561309-INS-6282
chr17-82561963-INS-61
chr17-82573671-INS-7220
chr17-82594603-INS-136
chr17-82605801-INS-650
chr17-82607340-INS-802
chr17-82619712-INS-6487
chr17-82623541-INS-96
chr17-82631398-INS-63
chr17-82658715-INS-51
chr17-82666080-INS-324
chr17-82667422-INS-6500
chr17-82668990-INS-438
chr17-82683004-INS-6399
chr17-82683005-INS-496
chr17-82690211-INS-355
chr17-82706106-INS-280
chr17-82755563-INS-84
chr17-82769527-INS-4561
chr17-82771382-INS-208
chr17-82817185-INS-149
chr17-82823292-INS-203
chr17-82826947-INS-518
chr17-82859867-INS-60
chr17-82896150-INS-335
chr17-82898733-INS-98
chr17-82914073-INS-56
chr17-82975742-INS-195
chr17-82988977-INS-268
chr17-82991667-INS-6941
chr17-83013624-INS-126
chr17-83017627-INS-72
chr17-83044380-INS-116
chr17-83060215-INS-5345
chr17-83060396-INS-4166
chr17-83069939-INS-124
chr17-83087171-INS-69
chr17-83087996-INS-164

chr17-83094622-INS-2599
chr17-83117370-INS-65
chr17-83141411-INS-191
chr17-83142745-INS-105
chr17-83148633-INS-418
chr17-83169614-INS-258
chr17-83182945-INS-107
chr17-83189619-INS-114
chr17-83241126-INS-376
chr17-83261831-INS-83
chr17-83284620-INS-1371
chr17-83289451-INS-214
chr17-83313572-INS-75
chr17-83314414-INS-100
chr17-83395922-INS-195
chr17-83412550-INS-62
chr17-83433263-INS-62
chr17-83462854-INS-7245
chr17-83465920-INS-108
chr17-83500464-INS-208
chr17-83519917-INS-90
chr17-83536081-INS-59
chr17-83536373-INS-1112
chr17-83539884-INS-78
chr17-83556353-INS-58
chr17-83557755-INS-74
chr17-83561733-INS-116
chr17-83572473-INS-75
chr17-83576143-INS-199
chr17-83580090-INS-111
chr17-83590968-INS-315
chr17-83594061-INS-665
chr17-83599905-INS-137
chr17-83601834-INS-90
chr17-83604294-INS-196
chr17-83605210-INS-60
chr17-83606013-INS-409
chr17-83607688-INS-300
chr17-83617647-INS-218
chr17-83645026-INS-165
chr17-83655740-INS-60
chr17-83658214-INS-2158
chr17-83664194-INS-564
chr17-83677758-INS-52
chr17-83683936-INS-648
chr17-83688727-INS-80
chr17-83701126-INS-205
chr17-83707373-INS-53
chr17-83728650-INS-408
chr17-83731987-INS-180
chr17-83738160-INS-57
chr17-83752179-INS-773
chr17-83757264-INS-222
chr17-83759476-INS-75
chr17-83797998-INS-353
chr17-83813728-INS-672
chr17-83829506-INS-489
chr17-83838349-INS-220
chr17-83850110-INS-91
chr17-83861733-INS-440
chr17-83864361-INS-172
chr17-83866158-INS-80
chr17-83876189-INS-168
chr17-83887205-INS-223
chr17-83887760-INS-158
chr17-83899418-INS-7124
chr17-83899435-INS-62
chr17-83926697-INS-369
chr17-83928684-INS-274
chr17-83945763-INS-78
chr17-83954639-INS-51
chr17-83958412-INS-205
chr17-83959239-INS-202
chr17-83976891-INS-264
chr17-83984522-INS-211
chr17-84001881-INS-128
chr17-84003069-INS-535

chr17-84032505-INS-52
chr17-84035881-INS-179
chr17-84039087-INS-300
chr17-84043861-INS-89
chr17-84045532-INS-209
chr17-84080062-INS-181
chr17-84085694-INS-175
chr17-84093431-INS-199
chr17-84131247-INS-84
chr17-84135040-INS-207
chr17-84150115-INS-86
chr17-84161263-INS-203
chr17-84167218-INS-64
chr17-84176302-INS-5267
chr17-84177757-INS-2894
chr17-84205820-INS-529
chr17-84230393-INS-55
chr17-84245556-INS-389
chr17-84254002-INS-71
chr17-84260562-INS-210
chr17-84284347-INS-210
chr17-84286141-INS-68
chr17-84324839-INS-164
chr17-84350036-INS-7119
chr17-84365294-INS-204
chr17-84377745-INS-63
chr17-84380766-INS-78
chr17-84398565-INS-406
chr17-84466371-INS-652
chr17-84473076-INS-230
chr17-84510640-INS-151
chr17-84564306-INS-906
chr17-84576168-INS-366
chr17-84576803-INS-71
chr17-84588032-INS-846
chr17-84601071-INS-78
chr17-84603532-INS-51
chr17-84627876-INS-137
chr17-84630716-INS-59
chr17-84635065-INS-78
chr17-84643382-INS-140
chr17-84652053-INS-138
chr17-84676446-INS-68
chr17-84712674-INS-210
chr17-84717569-INS-75
chr17-84726235-INS-120
chr17-84754759-INS-684
chr17-84767474-INS-192
chr17-84769799-INS-168
chr17-84778745-INS-195
chr17-84779720-INS-527
chr17-84789433-INS-325
chr17-84814168-INS-54
chr17-84818440-INS-60
chr17-84821757-INS-67
chr17-84823004-INS-109
chr17-84826415-INS-56
chr17-84846539-INS-87
chr17-84846694-INS-87
chr17-84885888-INS-119
chr17-84920245-INS-54
chr17-84921362-INS-166
chr17-84937484-INS-96
chr17-84941753-INS-56
chr17-84946196-INS-188
chr17-84961375-INS-52
chr17-84963624-INS-196
chr17-84973762-INS-52
chr17-84974419-INS-388
chr17-84986520-INS-166
chr17-84991414-INS-63
chr17-85077235-INS-102
chr17-85102889-INS-207
chr17-85111601-INS-733
chr17-85118199-INS-64
chr17-85128260-INS-143
chr17-85138547-INS-82

chr17-85147189-INS-215
chr17-85151187-INS-891
chr17-85165460-INS-76
chr17-85170310-INS-75
chr17-85179366-INS-131
chr17-85201660-INS-63
chr17-85209930-INS-174
chr17-85211321-INS-69
chr17-85212534-INS-293
chr17-85218049-INS-338
chr17-85238244-INS-359
chr17-85240600-INS-243
chr17-85254459-INS-296
chr17-85282773-INS-195
chr17-85291695-INS-57
chr17-85299020-INS-120
chr17-85308857-INS-90
chr17-85314178-INS-64
chr17-85329560-INS-437
chr17-85337003-INS-52
chr17-85339778-INS-244
chr17-85356729-INS-85
chr17-85358431-INS-104
chr17-85364398-INS-63
chr17-85365454-INS-198
chr17-85367773-INS-60
chr17-85407785-INS-82
chr17-85426683-INS-157
chr17-85427991-INS-568
chr17-85440021-INS-142
chr17-85441160-INS-124
chr17-85448285-INS-86
chr17-85457893-INS-70
chr17-85481768-INS-204
chr17-85492781-INS-176
chr17-85504781-INS-166
chr17-85527104-INS-84
chr17-85559317-INS-74
chr17-85575182-INS-119
chr17-85604581-INS-242
chr17-85619910-INS-264
chr17-85672543-INS-94
chr17-85677489-INS-221
chr17-85691309-INS-98
chr17-85692399-INS-76
chr17-85698357-INS-189
chr17-85717555-INS-758
chr17-85754586-INS-112
chr17-85761677-INS-52
chr17-85779462-INS-374
chr17-85784938-INS-381
chr17-85812102-INS-5409
chr17-85816627-INS-58
chr17-85818123-INS-183
chr17-85825827-INS-60
chr17-85859872-INS-324
chr17-85863361-INS-106
chr17-85876493-INS-228
chr17-85893699-INS-120
chr17-85896500-INS-86
chr17-85911461-INS-243
chr17-85945657-INS-912
chr17-85945841-INS-361
chr17-85946133-INS-1057
chr17-85948030-INS-187
chr17-85951345-INS-905
chr17-85961677-INS-126
chr17-85962976-INS-575
chr17-85966012-INS-224
chr17-85984507-INS-114
chr17-85984983-INS-232
chr17-86005542-INS-189
chr17-86009220-INS-148
chr17-86013350-INS-52
chr17-86061083-INS-63
chr17-86084451-INS-151
chr17-86086377-INS-2014

chr17-86092236-INS-151
chr17-86117277-INS-157
chr17-86141393-INS-106
chr17-86147061-INS-54
chr17-86148549-INS-217
chr17-86184752-INS-78
chr17-86186520-INS-318
chr17-86211419-INS-310
chr17-86212752-INS-540
chr17-86281234-INS-157
chr17-86288183-INS-179
chr17-86300761-INS-198
chr17-86315716-INS-129
chr17-86351952-INS-60
chr17-86359975-INS-4357
chr17-86362797-INS-6238
chr17-86421829-INS-208
chr17-86434079-INS-103
chr17-86439064-INS-84
chr17-86439178-INS-888
chr17-86455699-INS-200
chr17-86458588-INS-52
chr17-86463581-INS-674
chr17-86463590-INS-189
chr17-86467282-INS-73
chr17-86470714-INS-119
chr17-86472826-INS-268
chr17-86485034-INS-73
chr17-86491047-INS-84
chr17-86493828-INS-89
chr17-86498417-INS-329
chr17-86505487-INS-171
chr17-86508170-INS-80
chr17-86508947-INS-52
chr17-86524771-INS-213
chr17-86565212-INS-60
chr17-86567314-INS-58
chr17-86568332-INS-95
chr17-86576035-INS-54
chr17-86603675-INS-56
chr17-86605096-INS-62
chr17-86611870-INS-94
chr17-86619770-INS-616
chr17-86629983-INS-183
chr17-86631364-INS-89
chr17-86644514-INS-160
chr17-86651219-INS-882
chr17-86652095-INS-64
chr17-86667839-INS-171
chr17-86679834-INS-208
chr17-86700239-INS-409
chr17-86706162-INS-69
chr17-86723199-INS-64
chr17-86736395-INS-94
chr17-86758592-INS-174
chr17-86774038-INS-64
chr17-86775753-INS-207
chr17-86797957-INS-102
chr17-86800911-INS-52
chr17-86802255-INS-340
chr17-86814080-INS-208
chr17-86827280-INS-57
chr17-86833997-INS-342
chr17-86842545-INS-51
chr17-86846237-INS-54
chr17-86882191-INS-204
chr17-86885459-INS-170
chr17-86906338-INS-115
chr17-86913541-INS-156
chr17-86915306-INS-60
chr17-86921123-INS-72
chr17-86940409-INS-62
chr17-86950516-INS-190
chr17-86964409-INS-66
chr17-86970298-INS-137
chr17-86972081-INS-51
chr17-86980673-INS-68

chr17-86995964-INS-103
chr17-86998128-INS-96
chr17-87023139-INS-91
chr17-87031811-INS-108
chr17-87047051-INS-62
chr17-87069560-INS-358
chr17-87100234-INS-110
chr17-87124875-INS-129
chr17-87137907-INS-122
chr17-87142762-INS-69
chr17-87162588-INS-76
chr17-87167523-INS-184
chr17-87171262-INS-162
chr17-87191610-INS-146
chr17-87197890-INS-106
chr17-87204751-INS-175
chr17-87218854-INS-105
chr17-87249508-INS-72
chr17-87253634-INS-461
chr17-87257139-INS-431
chr17-87258793-INS-465
chr17-87278243-INS-72
chr17-87288468-INS-299
chr17-87301860-INS-98
chr17-87320816-INS-68
chr17-87321187-INS-1089
chr17-87321563-INS-198
chr17-87323318-INS-163
chr17-87329519-INS-2791
chr17-87334696-INS-405
chr17-87337678-INS-211
chr17-87339492-INS-197
chr17-87340948-INS-92
chr17-87343472-INS-58
chr17-87370328-INS-68
chr17-87379568-INS-99
chr17-87381012-INS-63
chr17-87396629-INS-583
chr17-87401596-INS-56
chr17-87425863-INS-200
chr17-87433002-INS-81
chr17-87460247-INS-62
chr17-87464398-INS-95
chr17-87468255-INS-127
chr17-87469566-INS-320
chr17-87491969-INS-52
chr17-87499831-INS-89
chr17-87503955-INS-168
chr17-87523564-INS-3575
chr17-87524337-INS-64
chr17-87540173-INS-55
chr17-87542264-INS-72
chr17-87575048-INS-198
chr17-87580122-INS-133
chr17-87584333-INS-229
chr17-87604529-INS-401
chr17-87609234-INS-224
chr17-87633588-INS-214
chr17-87660476-INS-3055
chr17-87664899-INS-221
chr17-87701560-INS-66
chr17-87702422-INS-158
chr17-87714134-INS-1245
chr17-87719491-INS-73
chr17-87726073-INS-94
chr17-87728876-INS-224
chr17-87735573-INS-174
chr17-87745561-INS-56
chr17-87758973-INS-163
chr17-87766252-INS-80
chr17-87771010-INS-275
chr17-87773808-INS-480
chr17-87776359-INS-66
chr17-87780216-INS-194
chr17-87783062-INS-78
chr17-87783469-INS-52
chr17-87784653-INS-96

chr17-87788832-INS-245
chr17-87805209-INS-7390
chr17-87805847-INS-688
chr17-87807899-INS-500
chr17-87816391-INS-165
chr17-87821097-INS-757
chr17-87822869-INS-71
chr17-87823019-INS-119
chr17-87831577-INS-96
chr17-87843199-INS-56
chr17-87848147-INS-79
chr17-87851213-INS-221
chr17-87856710-INS-456
chr17-87857589-INS-204
chr17-87859741-INS-213
chr17-87872956-INS-211
chr17-87875593-INS-66
chr17-87876468-INS-329
chr17-87882856-INS-66
chr17-87897801-INS-211
chr17-87899325-INS-220
chr17-87913611-INS-87
chr17-87914142-INS-87
chr17-87924836-INS-82
chr17-87925823-INS-215
chr17-87929581-INS-197
chr17-87934171-INS-83
chr17-87938249-INS-58
chr17-87945151-INS-169
chr17-87962436-INS-209
chr17-87984646-INS-57
chr17-87995755-INS-103
chr17-88002253-INS-150
chr17-88013684-INS-216
chr17-88015324-INS-504
chr17-88016401-INS-203
chr17-88029806-INS-215
chr17-88042729-INS-8732
chr17-88042739-INS-53
chr17-88062061-INS-167
chr17-88114882-INS-199
chr17-88115728-INS-231
chr17-88123794-INS-84
chr17-88136452-INS-58
chr17-88136831-INS-220
chr17-88144088-INS-154
chr17-88152916-INS-133
chr17-88159163-INS-154
chr17-88169511-INS-335
chr17-88173356-INS-124
chr17-88178990-INS-7126
chr17-88198210-INS-59
chr17-88200934-INS-92
chr17-88207755-INS-400
chr17-88211610-INS-56
chr17-88213598-INS-170
chr17-88218323-INS-192
chr17-88227751-INS-168
chr17-88234512-INS-57
chr17-88238299-INS-141
chr17-88244158-INS-63
chr17-88255099-INS-52
chr17-88260684-INS-59
chr17-88268381-INS-192
chr17-88284180-INS-123
chr17-88285527-INS-190
chr17-88308030-INS-52
chr17-88376108-INS-185
chr17-88376828-INS-162
chr17-88383819-INS-271
chr17-88394356-INS-960
chr17-88395019-INS-80
chr17-88398813-INS-62
chr17-88400153-INS-60
chr17-88402411-INS-506
chr17-88406342-INS-52
chr17-88407780-INS-106

chr17-88411263-INS-688
chr17-88413334-INS-190
chr17-88415099-INS-155
chr17-88423131-INS-94
chr17-88452545-INS-346
chr17-88473661-INS-279
chr17-88481626-INS-66
chr17-88484070-INS-131
chr17-88484212-INS-258
chr17-88488517-INS-92
chr17-88488682-INS-534
chr17-88495182-INS-65
chr17-88535794-INS-74
chr17-88535809-INS-545
chr17-88545490-INS-62
chr17-88546666-INS-388
chr17-88552157-INS-209
chr17-88561745-INS-247
chr17-88562355-INS-120
chr17-88565257-INS-58
chr17-88566441-INS-385
chr17-88567270-INS-52
chr17-88576548-INS-110
chr17-88587382-INS-126
chr17-88617622-INS-222
chr17-88619623-INS-60
chr17-88626831-INS-178
chr17-88630841-INS-1035
chr17-88630988-INS-87
chr17-88665640-INS-405
chr17-88673376-INS-871
chr17-88684212-INS-63
chr17-88691450-INS-311
chr17-88717379-INS-81
chr17-88724846-INS-195
chr17-88741308-INS-122
chr17-88772488-INS-102
chr17-88779905-INS-139
chr17-88815846-INS-54
chr17-88819167-INS-54
chr17-88840952-INS-54
chr17-88865806-INS-352
chr17-88866734-INS-163
chr17-88881028-INS-134
chr17-88889461-INS-369
chr17-88891872-INS-647
chr17-88901774-INS-170
chr17-88907589-INS-182
chr17-88910267-INS-466
chr17-88914895-INS-81
chr17-88944825-INS-149
chr17-88951073-INS-62
chr17-88951173-INS-740
chr17-88951465-INS-941
chr17-88953222-INS-206
chr17-88955161-INS-52
chr17-88961056-INS-4437
chr17-88964249-INS-93
chr17-88965917-INS-106
chr17-88971739-INS-55
chr17-88989031-INS-80
chr17-88995003-INS-262
chr17-89016032-INS-231
chr17-89021455-INS-797
chr17-89025091-INS-190
chr17-89042798-INS-64
chr17-89124444-INS-452
chr17-89146669-INS-179
chr17-89171300-INS-411
chr17-89198790-INS-60
chr17-89227239-INS-82
chr17-89230324-INS-104
chr17-89237055-INS-555
chr17-89240777-INS-1714
chr17-89255344-INS-55
chr17-89278971-INS-287
chr17-89305034-INS-2868

chr17-89308366-INS-93
chr17-89323983-INS-419
chr17-89330607-INS-5375
chr17-89360840-INS-57
chr17-89365683-INS-73
chr17-89405890-INS-363
chr17-89419211-INS-602
chr17-89420786-INS-112
chr17-89429519-INS-264
chr17-89475362-INS-172
chr17-89484075-INS-158
chr17-89504161-INS-121
chr17-89536399-INS-80
chr17-89595855-INS-206
chr17-89612380-INS-52
chr17-89627810-INS-54
chr17-89649768-INS-990
chr17-89684711-INS-54
chr17-89702069-INS-74
chr17-89721386-INS-69
chr17-89742547-INS-336
chr17-89758003-INS-123
chr17-89761708-INS-52
chr17-89778485-INS-125
chr17-89784263-INS-284
chr17-89831491-INS-316
chr17-89866626-INS-4573
chr17-89882911-INS-264
chr17-89895344-INS-319
chr17-89906306-INS-138
chr17-89907412-INS-505
chr17-89919443-INS-63
chr17-89931212-INS-347
chr17-89932598-INS-6375
chr17-89944529-INS-185
chr17-89946291-INS-183
chr17-89950893-INS-57
chr17-89959523-INS-100
chr17-89960822-INS-777
chr17-89970492-INS-60
chr17-89987744-INS-519
chr17-90008878-INS-751
chr17-90024365-INS-184
chr17-90028019-INS-346
chr17-90080700-INS-63
chr17-90092824-INS-54
chr17-90095720-INS-52
chr17-90107623-INS-404
chr17-90131476-INS-914
chr17-90138787-INS-52
chr17-90165428-INS-277
chr17-90169069-INS-138
chr17-90173664-INS-59
chr17-90211277-INS-202
chr17-90262433-INS-94
chr17-90264395-INS-492
chr17-90265762-INS-53
chr17-90271345-INS-380
chr17-90299414-INS-52
chr17-90307411-INS-751
chr17-90314300-INS-72
chr17-90337130-INS-58
chr17-90374479-INS-6731
chr17-90377599-INS-90
chr17-90383025-INS-127
chr17-90418896-INS-52
chr17-90514585-INS-64
chr17-90526543-INS-363
chr17-90537428-INS-136
chr17-90565059-INS-67
chr17-90565318-INS-55
chr17-90568430-INS-165
chr17-90569105-INS-575
chr17-90571203-INS-52
chr17-90586204-INS-1142
chr17-90586451-INS-343
chr17-90595546-INS-59

chr17-90598250-INS-322
chr17-90620708-INS-256
chr17-90647904-INS-321
chr17-90653259-INS-104
chr17-90656782-INS-5266
chr17-90657506-INS-350
chr17-90663983-INS-497
chr17-90680448-INS-58
chr17-90705862-INS-109
chr17-90711980-INS-240
chr17-90713844-INS-201
chr17-90739370-INS-65
chr17-90809174-INS-207
chr17-90811565-INS-127
chr17-90825802-INS-6398
chr17-90832988-INS-7087
chr17-90841199-INS-80
chr17-90845079-INS-482
chr17-90845873-INS-54
chr17-90900456-INS-241
chr17-90900771-INS-416
chr17-90938176-INS-52
chr17-90973035-INS-103
chr17-90977805-INS-148
chr17-91014562-INS-88
chr17-91027344-INS-509
chr17-91030349-INS-54
chr17-91079428-INS-104
chr17-91085891-INS-71
chr17-91132763-INS-478
chr17-91152204-INS-636
chr17-91183168-INS-181
chr17-91202683-INS-88
chr17-91203035-INS-7724
chr17-91204305-INS-92
chr17-91241970-INS-1453
chr17-91252501-INS-640
chr17-91262554-INS-56
chr17-91276556-INS-52
chr17-91286894-INS-514
chr17-91292450-INS-71
chr17-91302132-INS-68
chr17-91331753-INS-55
chr17-91334875-INS-469
chr17-91339985-INS-192
chr17-91356427-INS-507
chr17-91362863-INS-155
chr17-91370418-INS-119
chr17-91379319-INS-52
chr17-91388987-INS-402
chr17-91449397-INS-267
chr17-91475272-INS-114
chr17-91485190-INS-225
chr17-91494408-INS-869
chr17-91527464-INS-52
chr17-91554883-INS-6486
chr17-91566648-INS-505
chr17-91611956-INS-121
chr17-91631765-INS-73
chr17-91632656-INS-128
chr17-91635664-INS-729
chr17-91636963-INS-62
chr17-91637380-INS-125
chr17-91648748-INS-324
chr17-91662884-INS-4143
chr17-91663237-INS-133
chr17-91670738-INS-5602
chr17-91681673-INS-107
chr17-91697063-INS-67
chr17-91701412-INS-282
chr17-91753835-INS-741
chr17-91769772-INS-56
chr17-91777885-INS-818
chr17-91792948-INS-281
chr17-91823473-INS-278
chr17-91839450-INS-84
chr17-91842677-INS-258

chr17-91895128-INS-336
chr17-91907417-INS-120
chr17-91920930-INS-433
chr17-91943411-INS-6922
chr17-91943715-INS-384
chr17-91946409-INS-92
chr17-91969991-INS-409
chr17-91974005-INS-216
chr17-91981543-INS-68
chr17-91981579-INS-756
chr17-92009780-INS-737
chr17-92011805-INS-367
chr17-92042212-INS-424
chr17-92043283-INS-107
chr17-92049816-INS-195
chr17-92052161-INS-133
chr17-92064256-INS-98
chr17-92087025-INS-90
chr17-92100510-INS-59
chr17-92128231-INS-881
chr17-92136318-INS-65
chr17-92148108-INS-202
chr17-92153641-INS-5482
chr17-92159863-INS-724
chr17-92171654-INS-57
chr17-92178372-INS-504
chr17-92187625-INS-1198
chr17-92222236-INS-7086
chr17-92230164-INS-390
chr17-92242149-INS-70
chr17-92268631-INS-6396
chr17-92279643-INS-64
chr17-92303498-INS-68
chr17-92322567-INS-7120
chr17-92340468-INS-357
chr17-92343928-INS-84
chr17-92350607-INS-203
chr17-92355225-INS-378
chr17-92359148-INS-537
chr17-92361069-INS-660
chr17-92380602-INS-121
chr17-92388689-INS-132
chr17-92405056-INS-88
chr17-92412449-INS-64
chr17-92429570-INS-405
chr17-92446627-INS-196
chr17-92450336-INS-523
chr17-92488418-INS-2882
chr17-92500280-INS-141
chr17-92505415-INS-624
chr17-92508981-INS-78
chr17-92523260-INS-449
chr17-92576133-INS-79
chr17-92583517-INS-108
chr17-92584608-INS-545
chr17-92617903-INS-535
chr17-92636199-INS-626
chr17-92678095-INS-89
chr17-92712078-INS-6406
chr17-92721762-INS-52
chr17-92727160-INS-4452
chr17-92752266-INS-401
chr17-92755602-INS-109
chr17-92755615-INS-712
chr17-92769150-INS-125
chr17-92772179-INS-965
chr17-92784993-INS-64
chr17-92803465-INS-513
chr17-92818874-INS-231
chr17-92850923-INS-329
chr17-92891206-INS-6712
chr17-92892440-INS-53
chr17-92914679-INS-2260
chr17-92924565-INS-7083
chr17-92939605-INS-161
chr17-92942936-INS-656
chr17-92985947-INS-4742

chr17-93057047-INS-74
chr17-93069038-INS-292
chr17-93076099-INS-413
chr17-93095855-INS-55
chr17-93096501-INS-6413
chr17-93102774-INS-56
chr17-93126660-INS-69
chr17-93129565-INS-281
chr17-93131702-INS-58
chr17-93149821-INS-52
chr17-93157100-INS-225
chr17-93184542-INS-234
chr17-93205661-INS-491
chr17-93307679-INS-5948
chr17-93318708-INS-6930
chr17-93352316-INS-95
chr17-93405895-INS-88
chr17-93480137-INS-55
chr17-93491909-INS-417
chr17-93503255-INS-123
chr17-93577409-INS-137
chr17-93631351-INS-55
chr17-93680376-INS-180
chr17-93702059-INS-897
chr17-93707593-INS-871
chr17-93731385-INS-91
chr17-93733021-INS-348
chr17-93744240-INS-60
chr17-93759572-INS-210
chr17-93775206-INS-135
chr17-93778404-INS-95
chr17-93788935-INS-187
chr17-93836830-INS-5085
chr17-93838807-INS-227
chr17-93841152-INS-432
chr17-93845820-INS-203
chr17-93867142-INS-202
chr17-93893912-INS-126
chr17-93908041-INS-5509
chr17-93908281-INS-108
chr17-93920579-INS-460
chr17-93925885-INS-6450
chr17-93930490-INS-144
chr17-93945282-INS-6561
chr17-93947223-INS-6226
chr17-93962513-INS-422
chr17-93976146-INS-63
chr17-93999168-INS-108
chr17-94018694-INS-368
chr17-94021834-INS-7356
chr17-94048727-INS-2391
chr17-94071313-INS-60
chr17-94094132-INS-134
chr17-94095509-INS-626
chr17-94107172-INS-52
chr17-94110414-INS-7120
chr17-94123688-INS-242
chr17-94137876-INS-144
chr17-94161230-INS-66
chr17-94178540-INS-265
chr17-94182891-INS-52
chr17-94201706-INS-6679
chr17-94206530-INS-6495
chr17-94230500-INS-6214
chr17-94234834-INS-175
chr17-94263042-INS-232
chr17-94266841-INS-396
chr17-94303082-INS-240
chr17-94333491-INS-4327
chr17-94438186-INS-445
chr17-94464862-INS-217
chr17-94543643-INS-524
chr17-94578301-INS-521
chr17-94592611-INS-7080
chr17-94623460-INS-361
chr17-94645626-INS-60
chr17-94688160-INS-1004

chr17-94696818-INS-5279
chr17-94698560-INS-1083
chr17-94742832-INS-327
chr17-94760432-INS-7085
chr17-94775556-INS-167
chr17-94777509-INS-434
chr17-94803829-INS-6470
chr17-94826758-INS-51
chr17-94833847-INS-206
chr17-94876553-INS-247
chr17-94888689-INS-233
chr17-94930575-INS-55
chr17-94960937-INS-6173
chr17-94966686-INS-1833
chr17-94970164-INS-192
chr17-94984137-INS-214
chr17-94984797-INS-53
chr17-95025802-INS-492
chr17-95069989-INS-2927
chr17-95074356-INS-1830
chr17-95087741-INS-8085
chr17-95088065-INS-922
chr17-95131036-INS-681
chr17-95177450-INS-207
chr17-95189251-INS-530
chr17-95193172-INS-62
chr18-3118753-INS-4244
chr18-3169748-INS-223
chr18-3266988-INS-104
chr18-3275024-INS-69
chr18-3299928-INS-80
chr18-3315212-INS-230
chr18-3318482-INS-211
chr18-3324355-INS-240
chr18-3328877-INS-173
chr18-3354172-INS-5173
chr18-3368250-INS-389
chr18-3392608-INS-332
chr18-3416879-INS-215
chr18-3430820-INS-52
chr18-3441044-INS-442
chr18-3452343-INS-498
chr18-3490669-INS-202
chr18-3499878-INS-122
chr18-3501456-INS-62
chr18-3514498-INS-53
chr18-3549357-INS-57
chr18-3553181-INS-181
chr18-3557259-INS-81
chr18-3559429-INS-454
chr18-3561840-INS-110
chr18-3566333-INS-60
chr18-3571030-INS-207
chr18-3595119-INS-55
chr18-3598732-INS-7333
chr18-3601769-INS-52
chr18-3629225-INS-65
chr18-3637778-INS-444
chr18-3711724-INS-68
chr18-3725484-INS-60
chr18-3730851-INS-581
chr18-3742422-INS-80
chr18-3762528-INS-58
chr18-3770194-INS-54
chr18-3792019-INS-291
chr18-3799404-INS-82
chr18-3812489-INS-3096
chr18-3829736-INS-54
chr18-3863558-INS-148
chr18-3909216-INS-414
chr18-3911835-INS-78
chr18-3934656-INS-205
chr18-3938071-INS-378
chr18-3995916-INS-95
chr18-3996957-INS-59
chr18-4054367-INS-367
chr18-4060684-INS-545

chr18-4061633-INS-1323
chr18-4086968-INS-102
chr18-4089558-INS-471
chr18-4107369-INS-1131
chr18-4108939-INS-96
chr18-4120445-INS-205
chr18-4139116-INS-6916
chr18-4147294-INS-182
chr18-4160673-INS-56
chr18-4165568-INS-263
chr18-4173476-INS-93
chr18-4191907-INS-319
chr18-4213365-INS-559
chr18-4251225-INS-165
chr18-4290793-INS-147
chr18-4301470-INS-151
chr18-4316185-INS-100
chr18-4333690-INS-53
chr18-4340028-INS-125
chr18-4367563-INS-117
chr18-4376801-INS-66
chr18-4384086-INS-164
chr18-4503989-INS-97
chr18-4516217-INS-68
chr18-4525315-INS-53
chr18-4571124-INS-59
chr18-4580330-INS-69
chr18-4616389-INS-6181
chr18-4631100-INS-116
chr18-4676809-INS-60
chr18-4742168-INS-68
chr18-4745938-INS-69
chr18-4755726-INS-195
chr18-4757110-INS-86
chr18-4768684-INS-83
chr18-4815610-INS-56
chr18-4835130-INS-509
chr18-4841229-INS-74
chr18-4862120-INS-116
chr18-4916548-INS-2196
chr18-4916570-INS-1372
chr18-4937747-INS-258
chr18-4956105-INS-58
chr18-4963045-INS-413
chr18-4999054-INS-290
chr18-5000774-INS-413
chr18-5024243-INS-249
chr18-5027299-INS-341
chr18-5037519-INS-74
chr18-5057991-INS-102
chr18-5061372-INS-68
chr18-5062633-INS-222
chr18-5090589-INS-182
chr18-5102546-INS-78
chr18-5122802-INS-320
chr18-5134211-INS-52
chr18-5141872-INS-240
chr18-5158700-INS-146
chr18-5184323-INS-84
chr18-5218163-INS-52
chr18-5225064-INS-6895
chr18-5260297-INS-56
chr18-5267845-INS-85
chr18-5295619-INS-81
chr18-5298198-INS-6587
chr18-5303295-INS-695
chr18-5342196-INS-284
chr18-5345188-INS-76
chr18-5347795-INS-92
chr18-5377399-INS-171
chr18-5386579-INS-167
chr18-5402458-INS-83
chr18-5415972-INS-91
chr18-5440501-INS-441
chr18-5460044-INS-315
chr18-5465769-INS-760
chr18-5470078-INS-454

chr18-5498171-INS-80
chr18-5504718-INS-68
chr18-5509281-INS-310
chr18-5526161-INS-225
chr18-5527964-INS-198
chr18-5534629-INS-55
chr18-5584295-INS-55
chr18-5603539-INS-76
chr18-5648544-INS-96
chr18-5677788-INS-395
chr18-5681271-INS-247
chr18-5683952-INS-55
chr18-5694301-INS-75
chr18-5742848-INS-199
chr18-5748184-INS-64
chr18-5763849-INS-169
chr18-5796853-INS-720
chr18-5806278-INS-51
chr18-5813674-INS-498
chr18-5840767-INS-68
chr18-5845788-INS-4299
chr18-5858334-INS-65
chr18-5885577-INS-51
chr18-5900892-INS-79
chr18-5906187-INS-6485
chr18-5930892-INS-2693
chr18-5958507-INS-6245
chr18-5962359-INS-260
chr18-5979809-INS-238
chr18-5981787-INS-467
chr18-5986339-INS-78
chr18-5987778-INS-56
chr18-6023777-INS-342
chr18-6027089-INS-54
chr18-6033871-INS-315
chr18-6047795-INS-468
chr18-6060916-INS-986
chr18-6065584-INS-70
chr18-6067757-INS-746
chr18-6082348-INS-96
chr18-6090314-INS-56
chr18-6090891-INS-503
chr18-6098444-INS-158
chr18-6104462-INS-450
chr18-6124742-INS-87
chr18-6149017-INS-401
chr18-6154293-INS-506
chr18-6158946-INS-358
chr18-6163739-INS-73
chr18-6164139-INS-6565
chr18-6166143-INS-114
chr18-6169047-INS-88
chr18-6182703-INS-64
chr18-6210848-INS-132
chr18-6248724-INS-214
chr18-6265583-INS-192
chr18-6272971-INS-258
chr18-6283150-INS-4039
chr18-6287608-INS-126
chr18-6294765-INS-154
chr18-6299636-INS-204
chr18-6306368-INS-201
chr18-6314855-INS-62
chr18-6363991-INS-64
chr18-6365880-INS-84
chr18-6378006-INS-209
chr18-6382513-INS-507
chr18-6385085-INS-194
chr18-6410664-INS-1728
chr18-6411569-INS-1562
chr18-6412549-INS-52
chr18-6412964-INS-6290
chr18-6415903-INS-222
chr18-6432110-INS-7235
chr18-6439085-INS-399
chr18-6461979-INS-740
chr18-6493042-INS-1229

chr18-6495660-INS-496
chr18-6499444-INS-1889
chr18-6507236-INS-240
chr18-6509746-INS-68
chr18-6523784-INS-210
chr18-6540524-INS-397
chr18-6541415-INS-150
chr18-6555439-INS-55
chr18-6559100-INS-142
chr18-6564607-INS-351
chr18-6575105-INS-126
chr18-6625948-INS-58
chr18-6648510-INS-429
chr18-6652251-INS-99
chr18-6653652-INS-1893
chr18-6675770-INS-68
chr18-6691651-INS-304
chr18-6713881-INS-465
chr18-6720633-INS-237
chr18-6733113-INS-93
chr18-6736184-INS-112
chr18-6785199-INS-160
chr18-6787847-INS-91
chr18-6845610-INS-56
chr18-6869509-INS-374
chr18-6871927-INS-1886
chr18-6883725-INS-127
chr18-6894142-INS-6364
chr18-6915921-INS-201
chr18-6967328-INS-76
chr18-6998863-INS-68
chr18-7035088-INS-91
chr18-7046741-INS-778
chr18-7055615-INS-2729
chr18-7082003-INS-208
chr18-7091270-INS-55
chr18-7096738-INS-172
chr18-7146746-INS-84
chr18-7166588-INS-975
chr18-7211989-INS-75
chr18-7228701-INS-371
chr18-7235543-INS-206
chr18-7239269-INS-359
chr18-7240393-INS-210
chr18-7245537-INS-333
chr18-7255546-INS-310
chr18-7265310-INS-313
chr18-7303702-INS-5332
chr18-7384375-INS-116
chr18-7426080-INS-51
chr18-7452537-INS-134
chr18-7460517-INS-63
chr18-7490591-INS-1162
chr18-7493198-INS-186
chr18-7503094-INS-53
chr18-7504808-INS-190
chr18-7523072-INS-70
chr18-7546160-INS-1176
chr18-7569624-INS-387
chr18-7582228-INS-620
chr18-7608146-INS-185
chr18-7608948-INS-181
chr18-7609524-INS-305
chr18-7609783-INS-185
chr18-7612934-INS-54
chr18-7616721-INS-5218
chr18-7628697-INS-487
chr18-7637480-INS-85
chr18-7654444-INS-160
chr18-7662542-INS-540
chr18-7663433-INS-405
chr18-7673697-INS-161
chr18-7676924-INS-322
chr18-7698795-INS-52
chr18-7702871-INS-110
chr18-7735501-INS-325
chr18-7763422-INS-9658

chr18-7795618-INS-2614
chr18-7796131-INS-292
chr18-7805790-INS-100
chr18-7806571-INS-233
chr18-7812346-INS-1865
chr18-7835906-INS-418
chr18-7865451-INS-79
chr18-7873986-INS-63
chr18-7909705-INS-61
chr18-7915762-INS-58
chr18-7933701-INS-51
chr18-7952536-INS-610
chr18-7953798-INS-69
chr18-7955856-INS-639
chr18-7957299-INS-591
chr18-7957422-INS-4984
chr18-7974943-INS-6471
chr18-7975706-INS-52
chr18-7977615-INS-855
chr18-7987561-INS-323
chr18-7995695-INS-7099
chr18-8003509-INS-98
chr18-8011372-INS-57
chr18-8024547-INS-6450
chr18-8031833-INS-192
chr18-8042377-INS-70
chr18-8052336-INS-590
chr18-8105958-INS-70
chr18-8131324-INS-514
chr18-8134033-INS-564
chr18-8200572-INS-139
chr18-8272933-INS-1703
chr18-8283837-INS-52
chr18-8299967-INS-7105
chr18-8300153-INS-116
chr18-8344881-INS-84
chr18-8373598-INS-117
chr18-8377500-INS-6726
chr18-8431945-INS-938
chr18-8455410-INS-193
chr18-8482846-INS-176
chr18-8493715-INS-379
chr18-8498694-INS-5548
chr18-8533548-INS-184
chr18-8549261-INS-506
chr18-8575315-INS-51
chr18-8642841-INS-4458
chr18-8654077-INS-78
chr18-8657740-INS-6257
chr18-8674361-INS-6337
chr18-8678295-INS-98
chr18-8707347-INS-359
chr18-8724111-INS-55
chr18-8761813-INS-200
chr18-8770308-INS-331
chr18-8788123-INS-6492
chr18-8828306-INS-897
chr18-8834386-INS-88
chr18-8840078-INS-573
chr18-8854195-INS-82
chr18-8859389-INS-754
chr18-8864370-INS-68
chr18-8884600-INS-55
chr18-8906531-INS-1464
chr18-8911426-INS-51
chr18-8927695-INS-55
chr18-8976299-INS-170
chr18-8979701-INS-127
chr18-8989672-INS-68
chr18-8995805-INS-78
chr18-9064951-INS-889
chr18-9065667-INS-548
chr18-9071799-INS-390
chr18-9110096-INS-6927
chr18-9163408-INS-442
chr18-9178921-INS-56
chr18-9180998-INS-60

chr18-9201631-INS-238
chr18-9241215-INS-58
chr18-9255982-INS-5551
chr18-9296495-INS-115
chr18-9324946-INS-588
chr18-9334432-INS-70
chr18-9335470-INS-198
chr18-9350670-INS-504
chr18-9368226-INS-389
chr18-9398696-INS-167
chr18-9404183-INS-331
chr18-9433899-INS-70
chr18-9443090-INS-173
chr18-9451577-INS-174
chr18-9453453-INS-381
chr18-9482851-INS-328
chr18-9489499-INS-571
chr18-9504134-INS-64
chr18-9509838-INS-272
chr18-9530783-INS-165
chr18-9554370-INS-80
chr18-9559411-INS-80
chr18-9567480-INS-341
chr18-9602619-INS-289
chr18-9619567-INS-117
chr18-9626053-INS-324
chr18-9633217-INS-6695
chr18-9669591-INS-869
chr18-9671376-INS-195
chr18-9676178-INS-156
chr18-9700520-INS-380
chr18-9711578-INS-182
chr18-9720075-INS-515
chr18-9772307-INS-230
chr18-9774729-INS-338
chr18-9790073-INS-140
chr18-9803235-INS-6744
chr18-9812599-INS-4845
chr18-9815512-INS-7263
chr18-9827377-INS-56
chr18-9842024-INS-270
chr18-9861237-INS-966
chr18-9878344-INS-113
chr18-9897675-INS-4747
chr18-9899092-INS-4219
chr18-9901618-INS-195
chr18-9922936-INS-6210
chr18-9952991-INS-326
chr18-9965864-INS-70
chr18-9980661-INS-860
chr18-10037628-INS-130
chr18-10044471-INS-72
chr18-10053586-INS-94
chr18-10055951-INS-1503
chr18-10058008-INS-879
chr18-10205345-INS-75
chr18-10265066-INS-54
chr18-10265893-INS-467
chr18-10350090-INS-411
chr18-10353591-INS-249
chr18-10354496-INS-427
chr18-10412320-INS-212
chr18-10443643-INS-53
chr18-10505316-INS-339
chr18-10508463-INS-123
chr18-10530927-INS-63
chr18-10558203-INS-217
chr18-10634882-INS-117
chr18-10642679-INS-106
chr18-10653365-INS-288
chr18-10660941-INS-285
chr18-10665650-INS-109
chr18-10684203-INS-200
chr18-10703328-INS-223
chr18-10724042-INS-677
chr18-10744355-INS-210
chr18-10853416-INS-98

chr18-10856312-INS-199
chr18-10908122-INS-79
chr18-10918438-INS-102
chr18-10939065-INS-102
chr18-10949404-INS-57
chr18-10984863-INS-140
chr18-11031176-INS-66
chr18-11038060-INS-119
chr18-11043034-INS-128
chr18-11049041-INS-136
chr18-11061541-INS-118
chr18-11079111-INS-50
chr18-11087184-INS-63
chr18-11093335-INS-66
chr18-11113828-INS-91
chr18-11118677-INS-173
chr18-11134066-INS-132
chr18-11166904-INS-419
chr18-11198942-INS-218
chr18-11203611-INS-64
chr18-11206517-INS-134
chr18-11208302-INS-162
chr18-11236203-INS-144
chr18-11246995-INS-85
chr18-11287382-INS-105
chr18-11303136-INS-367
chr18-11312483-INS-166
chr18-11332775-INS-325
chr18-11347477-INS-70
chr18-11349765-INS-62
chr18-11361761-INS-145
chr18-11394854-INS-204
chr18-11402772-INS-187
chr18-11409237-INS-6700
chr18-11413906-INS-914
chr18-11464691-INS-111
chr18-11490975-INS-130
chr18-11517499-INS-132
chr18-11554127-INS-222
chr18-11557456-INS-200
chr18-11569103-INS-157
chr18-11578846-INS-515
chr18-11582775-INS-163
chr18-11624507-INS-61
chr18-11639135-INS-1108
chr18-11639136-INS-724
chr18-11651220-INS-54
chr18-11670010-INS-182
chr18-11680346-INS-58
chr18-11688255-INS-715
chr18-11702174-INS-817
chr18-11709011-INS-4725
chr18-11726543-INS-60
chr18-11729625-INS-5244
chr18-11751749-INS-52
chr18-11753039-INS-6231
chr18-11815967-INS-61
chr18-11822252-INS-384
chr18-11829583-INS-73
chr18-11863991-INS-360
chr18-11865848-INS-234
chr18-11866036-INS-2584
chr18-11876375-INS-6241
chr18-11881098-INS-1899
chr18-11885032-INS-3474
chr18-11948887-INS-76
chr18-11954160-INS-198
chr18-11978636-INS-88
chr18-12018854-INS-53
chr18-12033409-INS-119
chr18-12056880-INS-180
chr18-12079724-INS-622
chr18-12090562-INS-194
chr18-12120842-INS-328
chr18-12128042-INS-53
chr18-12139070-INS-75
chr18-12143602-INS-91

chr18-12145857-INS-263
chr18-12151024-INS-115
chr18-12171132-INS-197
chr18-12175591-INS-121
chr18-12187084-INS-105
chr18-12187522-INS-208
chr18-12256887-INS-519
chr18-12285359-INS-53
chr18-12292187-INS-5059
chr18-12292288-INS-5839
chr18-12294275-INS-94
chr18-12310698-INS-304
chr18-12311030-INS-108
chr18-12322594-INS-203
chr18-12330711-INS-59
chr18-12336596-INS-61
chr18-12352063-INS-218
chr18-12363687-INS-195
chr18-12391351-INS-54
chr18-12429753-INS-72
chr18-12462318-INS-78
chr18-12475020-INS-279
chr18-12507271-INS-101
chr18-12514259-INS-62
chr18-12520107-INS-85
chr18-12536996-INS-104
chr18-12547427-INS-971
chr18-12556981-INS-133
chr18-12591601-INS-244
chr18-12603504-INS-123
chr18-12604722-INS-64
chr18-12609670-INS-359
chr18-12619868-INS-73
chr18-12651630-INS-189
chr18-12655198-INS-107
chr18-12656405-INS-236
chr18-12659551-INS-58
chr18-12665806-INS-62
chr18-12668663-INS-1225
chr18-12668664-INS-762
chr18-12669351-INS-65
chr18-12670451-INS-63
chr18-12672426-INS-102
chr18-12679959-INS-497
chr18-12684041-INS-62
chr18-12694214-INS-199
chr18-12695364-INS-80
chr18-12735338-INS-72
chr18-12736316-INS-494
chr18-12819849-INS-198
chr18-12823674-INS-58
chr18-12834691-INS-92
chr18-12853555-INS-280
chr18-12856954-INS-106
chr18-12857908-INS-166
chr18-12890658-INS-190
chr18-12900902-INS-224
chr18-12901075-INS-860
chr18-12902706-INS-84
chr18-12925282-INS-110
chr18-12929472-INS-6186
chr18-12939683-INS-240
chr18-12947489-INS-72
chr18-12988820-INS-1276
chr18-12988853-INS-160
chr18-12993087-INS-110
chr18-13010861-INS-72
chr18-13013722-INS-80
chr18-13031066-INS-58
chr18-13048150-INS-319
chr18-13068589-INS-816
chr18-13070544-INS-68
chr18-13082726-INS-54
chr18-13086446-INS-78
chr18-13089570-INS-402
chr18-13106915-INS-60
chr18-13111527-INS-61

chr18-13136483-INS-754
chr18-13147442-INS-60
chr18-13157774-INS-198
chr18-13231854-INS-619
chr18-13239817-INS-850
chr18-13255771-INS-64
chr18-13257472-INS-101
chr18-13289400-INS-65
chr18-13291733-INS-211
chr18-13296955-INS-7295
chr18-13315633-INS-80
chr18-13324781-INS-566
chr18-13326897-INS-56
chr18-13334803-INS-6314
chr18-13339647-INS-1473
chr18-13368881-INS-452
chr18-13383037-INS-144
chr18-13418783-INS-609
chr18-13428186-INS-129
chr18-13430377-INS-300
chr18-13432545-INS-1349
chr18-13436723-INS-73
chr18-13450876-INS-96
chr18-13458802-INS-2519
chr18-13467171-INS-560
chr18-13495652-INS-116
chr18-13511599-INS-92
chr18-13524718-INS-208
chr18-13534064-INS-62
chr18-13544557-INS-98
chr18-13549237-INS-62
chr18-13561271-INS-72
chr18-13571169-INS-57
chr18-13575693-INS-127
chr18-13618951-INS-140
chr18-13676147-INS-99
chr18-13721268-INS-504
chr18-13741545-INS-52
chr18-13749400-INS-56
chr18-13766798-INS-138
chr18-13774711-INS-207
chr18-13779385-INS-479
chr18-13830094-INS-54
chr18-13833660-INS-87
chr18-13845994-INS-286
chr18-13866303-INS-87
chr18-13877648-INS-115
chr18-13883222-INS-81
chr18-13970518-INS-140
chr18-14014562-INS-74
chr18-14021099-INS-56
chr18-14090453-INS-116
chr18-14111136-INS-108
chr18-14150071-INS-72
chr18-14153211-INS-754
chr18-14166009-INS-58
chr18-14166807-INS-789
chr18-14196000-INS-83
chr18-14200166-INS-408
chr18-14201334-INS-70
chr18-14231887-INS-196
chr18-14235622-INS-75
chr18-14239244-INS-924
chr18-14265937-INS-161
chr18-14270668-INS-94
chr18-14295196-INS-362
chr18-14301424-INS-169
chr18-14318113-INS-524
chr18-14331961-INS-136
chr18-14358912-INS-477
chr18-14389875-INS-408
chr18-14420213-INS-78
chr18-14425326-INS-166
chr18-14433628-INS-251
chr18-14468138-INS-163
chr18-14480515-INS-256
chr18-14480601-INS-881

chr18-14497146-INS-542
chr18-14506496-INS-56
chr18-14511263-INS-96
chr18-14530427-INS-180
chr18-14532655-INS-6272
chr18-14551338-INS-436
chr18-14569882-INS-60
chr18-14587038-INS-128
chr18-14597543-INS-66
chr18-14604417-INS-784
chr18-14607617-INS-866
chr18-14609002-INS-246
chr18-14633894-INS-148
chr18-14644808-INS-137
chr18-14658028-INS-343
chr18-14698135-INS-150
chr18-14702332-INS-822
chr18-14707185-INS-2143
chr18-14720438-INS-217
chr18-14725756-INS-194
chr18-14742102-INS-86
chr18-14764689-INS-59
chr18-14778948-INS-351
chr18-14786604-INS-80
chr18-14791338-INS-216
chr18-14798388-INS-226
chr18-14825757-INS-212
chr18-14827407-INS-118
chr18-14832092-INS-54
chr18-14848558-INS-250
chr18-14888096-INS-788
chr18-14888731-INS-6442
chr18-14909385-INS-64
chr18-14914025-INS-8734
chr18-14957716-INS-139
chr18-14997036-INS-170
chr18-15028540-INS-241
chr18-15048599-INS-59
chr18-15056246-INS-171
chr18-15071318-INS-1891
chr18-15075118-INS-98
chr18-15127073-INS-84
chr18-15166105-INS-60
chr18-15172908-INS-306
chr18-15180669-INS-80
chr18-15187041-INS-137
chr18-15188604-INS-86
chr18-15191928-INS-198
chr18-15220254-INS-76
chr18-15235085-INS-54
chr18-15254581-INS-571
chr18-15259978-INS-84
chr18-15261288-INS-54
chr18-15262373-INS-843
chr18-15266889-INS-98
chr18-15269969-INS-541
chr18-15273217-INS-74
chr18-15280041-INS-162
chr18-15298735-INS-7089
chr18-15301931-INS-74
chr18-15316509-INS-549
chr18-15354000-INS-440
chr18-15372905-INS-153
chr18-15405070-INS-274
chr18-15418917-INS-52
chr18-15422948-INS-188
chr18-15426821-INS-264
chr18-15441461-INS-133
chr18-15454735-INS-616
chr18-15481895-INS-50
chr18-15491066-INS-232
chr18-15520792-INS-61
chr18-15567420-INS-635
chr18-15587717-INS-637
chr18-15595942-INS-67
chr18-15632403-INS-119
chr18-15636486-INS-82

chr18-15647788-INS-305
chr18-15653662-INS-72
chr18-15673634-INS-120
chr18-15677661-INS-218
chr18-15702377-INS-242
chr18-15713148-INS-225
chr18-15714668-INS-318
chr18-15740316-INS-867
chr18-15766065-INS-55
chr18-15768924-INS-110
chr18-15775373-INS-3326
chr18-15788753-INS-82
chr18-15811620-INS-396
chr18-15846335-INS-64
chr18-15931794-INS-120
chr18-15936018-INS-6696
chr18-15948311-INS-53
chr18-15989118-INS-429
chr18-15995390-INS-76
chr18-15998175-INS-139
chr18-16001822-INS-671
chr18-16011702-INS-60
chr18-16029595-INS-256
chr18-16041709-INS-172
chr18-16089216-INS-10737
chr18-16128462-INS-524
chr18-16136065-INS-207
chr18-16141007-INS-96
chr18-16157434-INS-220
chr18-16161616-INS-59
chr18-16185479-INS-58
chr18-16205925-INS-90
chr18-16225178-INS-106
chr18-16259471-INS-7302
chr18-16260763-INS-6680
chr18-16261927-INS-152
chr18-16265106-INS-60
chr18-16292819-INS-53
chr18-16307590-INS-137
chr18-16328407-INS-149
chr18-16383768-INS-1361
chr18-16411712-INS-6313
chr18-16452980-INS-76
chr18-16484924-INS-1865
chr18-16494126-INS-56
chr18-16496758-INS-10049
chr18-16510771-INS-80
chr18-16513417-INS-6987
chr18-16563398-INS-136
chr18-16604380-INS-159
chr18-16636164-INS-381
chr18-16644171-INS-604
chr18-16648138-INS-75
chr18-16676891-INS-4057
chr18-16699795-INS-83
chr18-16708100-INS-59
chr18-16787986-INS-66
chr18-16802802-INS-62
chr18-16847611-INS-115
chr18-16877363-INS-86
chr18-16888448-INS-233
chr18-16901642-INS-7144
chr18-16909412-INS-78
chr18-16922277-INS-104
chr18-16927489-INS-2607
chr18-16935544-INS-63
chr18-16968420-INS-8949
chr18-17016511-INS-321
chr18-17050360-INS-53
chr18-17063527-INS-308
chr18-17094595-INS-54
chr18-17095264-INS-211
chr18-17103236-INS-75
chr18-17128045-INS-214
chr18-17137166-INS-578
chr18-17139941-INS-70
chr18-17142502-INS-74

chr18-17150836-INS-5161
chr18-17167903-INS-324
chr18-17171073-INS-87
chr18-17177301-INS-334
chr18-17188219-INS-596
chr18-17206006-INS-6245
chr18-17217716-INS-567
chr18-17230572-INS-115
chr18-17255838-INS-6252
chr18-17267277-INS-485
chr18-17272049-INS-724
chr18-17285380-INS-3855
chr18-17296845-INS-88
chr18-17319551-INS-196
chr18-17351187-INS-531
chr18-17372039-INS-55
chr18-17427245-INS-499
chr18-17445293-INS-498
chr18-17466614-INS-738
chr18-17469058-INS-67
chr18-17472422-INS-6270
chr18-17490275-INS-56
chr18-17499229-INS-164
chr18-17527944-INS-94
chr18-17535700-INS-5306
chr18-17546311-INS-1009
chr18-17546955-INS-9749
chr18-17553681-INS-173
chr18-17578823-INS-127
chr18-17594429-INS-62
chr18-17606208-INS-59
chr18-17610929-INS-85
chr18-17641479-INS-150
chr18-17644330-INS-494
chr18-17652656-INS-56
chr18-17668148-INS-622
chr18-17683253-INS-76
chr18-17691471-INS-53
chr18-17693330-INS-6191
chr18-17698283-INS-826
chr18-17698405-INS-91
chr18-17702655-INS-71
chr18-17704880-INS-346
chr18-17711076-INS-58
chr18-17718686-INS-52
chr18-17738827-INS-64
chr18-17744751-INS-640
chr18-17761018-INS-514
chr18-17767889-INS-214
chr18-17768713-INS-2078
chr18-17775007-INS-73
chr18-17777833-INS-444
chr18-17786442-INS-6120
chr18-17792162-INS-1894
chr18-17822527-INS-2987
chr18-17823075-INS-6454
chr18-17835010-INS-6666
chr18-17836524-INS-201
chr18-17839254-INS-488
chr18-17844986-INS-5224
chr18-17853424-INS-6296
chr18-17856499-INS-6876
chr18-17860542-INS-160
chr18-17881012-INS-6280
chr18-17899350-INS-84
chr18-17907000-INS-59
chr18-17907995-INS-55
chr18-17930136-INS-6099
chr18-17944661-INS-52
chr18-17987568-INS-59
chr18-18037915-INS-7222
chr18-18038736-INS-421
chr18-18062391-INS-56
chr18-18063327-INS-498
chr18-18113889-INS-7124
chr18-18158863-INS-787
chr18-18196606-INS-6245

chr18-18235453-INS-829
chr18-18253987-INS-5549
chr18-18254847-INS-181
chr18-18261189-INS-65
chr18-18273840-INS-83
chr18-18326332-INS-92
chr18-18332044-INS-91
chr18-18334438-INS-61
chr18-18340833-INS-524
chr18-18370206-INS-96
chr18-18375977-INS-57
chr18-18384125-INS-345
chr18-18398614-INS-82
chr18-18400794-INS-102
chr18-18408149-INS-6779
chr18-18419854-INS-223
chr18-18430241-INS-141
chr18-18463457-INS-3174
chr18-18472127-INS-56
chr18-18495685-INS-67
chr18-18516612-INS-58
chr18-18548418-INS-433
chr18-18555857-INS-303
chr18-18597987-INS-404
chr18-18598912-INS-54
chr18-18606511-INS-74
chr18-18617639-INS-191
chr18-18625209-INS-5080
chr18-18639929-INS-528
chr18-18669842-INS-82
chr18-18695772-INS-4978
chr18-18718191-INS-1138
chr18-18736179-INS-449
chr18-18765925-INS-637
chr18-18770928-INS-100
chr18-18787107-INS-295
chr18-18793353-INS-56
chr18-18796262-INS-6369
chr18-18806583-INS-84
chr18-18819435-INS-6294
chr18-18820456-INS-576
chr18-18830816-INS-968
chr18-18848419-INS-2815
chr18-18850650-INS-56
chr18-18869299-INS-185
chr18-18885517-INS-6631
chr18-18911897-INS-331
chr18-18931739-INS-233
chr18-18936612-INS-6690
chr18-18946615-INS-4320
chr18-18948314-INS-1008
chr18-18963492-INS-497
chr18-18986859-INS-1510
chr18-19016466-INS-156
chr18-19061651-INS-6421
chr18-19101005-INS-5508
chr18-19105505-INS-6645
chr18-19134633-INS-175
chr18-19135867-INS-201
chr18-19148331-INS-579
chr18-19164772-INS-3881
chr18-19171302-INS-397
chr18-19178616-INS-457
chr18-19188651-INS-88
chr18-19200288-INS-206
chr18-19210658-INS-192
chr18-19215437-INS-212
chr18-19229593-INS-418
chr18-19238260-INS-91
chr18-19243451-INS-358
chr18-19249978-INS-225
chr18-19251154-INS-7147
chr18-19254850-INS-359
chr18-19255873-INS-6977
chr18-19278969-INS-3422
chr18-19302490-INS-160
chr18-19323824-INS-237

chr18-19348825-INS-2734
chr18-19359182-INS-599
chr18-19390589-INS-750
chr18-19393503-INS-62
chr18-19413359-INS-7500
chr18-19433496-INS-51
chr18-19439894-INS-5208
chr18-19458335-INS-183
chr18-19463049-INS-59
chr18-19512533-INS-345
chr18-19538518-INS-414
chr18-19555136-INS-91
chr18-19557913-INS-323
chr18-19560674-INS-104
chr18-19586363-INS-792
chr18-19607234-INS-193
chr18-19617878-INS-62
chr18-19660490-INS-771
chr18-19672594-INS-80
chr18-19677836-INS-72
chr18-19701856-INS-358
chr18-19712126-INS-1888
chr18-19728501-INS-542
chr18-19763015-INS-92
chr18-19775066-INS-484
chr18-19863412-INS-1684
chr18-19884987-INS-68
chr18-19922264-INS-287
chr18-19929982-INS-268
chr18-19954201-INS-398
chr18-19994909-INS-54
chr18-20011656-INS-70
chr18-20049084-INS-231
chr18-20052764-INS-62
chr18-20056461-INS-325
chr18-20063911-INS-61
chr18-20069996-INS-58
chr18-20075934-INS-72
chr18-20093696-INS-69
chr18-20129473-INS-167
chr18-20152175-INS-84
chr18-20178787-INS-127
chr18-20217837-INS-51
chr18-20258049-INS-6444
chr18-20263459-INS-74
chr18-20275887-INS-169
chr18-20280867-INS-90
chr18-20315736-INS-371
chr18-20318316-INS-450
chr18-20333431-INS-115
chr18-20333497-INS-1243
chr18-20349193-INS-62
chr18-20431306-INS-6344
chr18-20453699-INS-65
chr18-20471650-INS-528
chr18-20492502-INS-6707
chr18-20521660-INS-143
chr18-20541209-INS-299
chr18-20580590-INS-121
chr18-20587262-INS-69
chr18-20602147-INS-66
chr18-20611136-INS-328
chr18-20621402-INS-54
chr18-20638898-INS-3368
chr18-20659664-INS-144
chr18-20701667-INS-75
chr18-20707676-INS-925
chr18-20711772-INS-163
chr18-20723726-INS-95
chr18-20728421-INS-710
chr18-20735919-INS-70
chr18-20762461-INS-154
chr18-20770085-INS-547
chr18-20774593-INS-108
chr18-20801505-INS-72
chr18-20849249-INS-99
chr18-20852121-INS-62

chr18-20877435-INS-61
chr18-20896202-INS-114
chr18-20913955-INS-109
chr18-20916856-INS-3525
chr18-20918209-INS-113
chr18-20922180-INS-196
chr18-20923182-INS-203
chr18-20928302-INS-126
chr18-20929748-INS-481
chr18-20956010-INS-164
chr18-20961991-INS-219
chr18-20989884-INS-374
chr18-20991478-INS-196
chr18-21003942-INS-78
chr18-21006384-INS-203
chr18-21026361-INS-190
chr18-21031927-INS-973
chr18-21033634-INS-296
chr18-21083192-INS-100
chr18-21092198-INS-56
chr18-21112383-INS-214
chr18-21140604-INS-90
chr18-21188398-INS-81
chr18-21188814-INS-207
chr18-21195597-INS-5430
chr18-21242873-INS-228
chr18-21258229-INS-56
chr18-21287941-INS-181
chr18-21297028-INS-284
chr18-21312477-INS-54
chr18-21315372-INS-51
chr18-21334044-INS-190
chr18-21346280-INS-124
chr18-21382558-INS-160
chr18-21389557-INS-600
chr18-21407655-INS-6213
chr18-21454478-INS-132
chr18-21479711-INS-70
chr18-21498292-INS-178
chr18-21540117-INS-208
chr18-21541094-INS-512
chr18-21558833-INS-90
chr18-21563000-INS-150
chr18-21608757-INS-401
chr18-21610083-INS-102
chr18-21640427-INS-1286
chr18-21695538-INS-6303
chr18-21697240-INS-56
chr18-21711104-INS-264
chr18-21737154-INS-6403
chr18-21761109-INS-155
chr18-21771177-INS-199
chr18-21804907-INS-74
chr18-21822288-INS-549
chr18-21835783-INS-57
chr18-21841302-INS-400
chr18-21849645-INS-203
chr18-21858818-INS-55
chr18-21871355-INS-113
chr18-21907614-INS-256
chr18-21946648-INS-104
chr18-21952316-INS-397
chr18-21955286-INS-69
chr18-21976153-INS-76
chr18-21983939-INS-138
chr18-21989223-INS-6194
chr18-21993498-INS-107
chr18-21994610-INS-6292
chr18-22000105-INS-51
chr18-22025198-INS-249
chr18-22039597-INS-55
chr18-22049731-INS-6997
chr18-22081172-INS-51
chr18-22148898-INS-57
chr18-22183694-INS-52
chr18-22222409-INS-52
chr18-22234362-INS-6924

chr18-22257020-INS-3119
chr18-22271056-INS-51
chr18-22293701-INS-259
chr18-22298693-INS-393
chr18-22323919-INS-75
chr18-22328479-INS-202
chr18-22335568-INS-6433
chr18-22367394-INS-53
chr18-22384389-INS-855
chr18-22394302-INS-7078
chr18-22403849-INS-519
chr18-22426767-INS-56
chr18-22442675-INS-66
chr18-22450411-INS-6190
chr18-22463570-INS-52
chr18-22568325-INS-600
chr18-22579525-INS-52
chr18-22586092-INS-132
chr18-22593073-INS-52
chr18-22610116-INS-316
chr18-22614472-INS-92
chr18-22623259-INS-524
chr18-22697682-INS-491
chr18-22724248-INS-227
chr18-22735055-INS-81
chr18-22763136-INS-299
chr18-22764208-INS-128
chr18-22807429-INS-992
chr18-22808061-INS-1242
chr18-22818401-INS-183
chr18-22824787-INS-64
chr18-22831367-INS-197
chr18-22858019-INS-137
chr18-22875832-INS-882
chr18-22911153-INS-268
chr18-22916248-INS-6254
chr18-22916768-INS-400
chr18-22936043-INS-593
chr18-22946273-INS-122
chr18-22953708-INS-82
chr18-22999964-INS-62
chr18-23039409-INS-181
chr18-23044339-INS-211
chr18-23071533-INS-592
chr18-23087767-INS-56
chr18-23131173-INS-391
chr18-23142332-INS-104
chr18-23162395-INS-68
chr18-23193203-INS-291
chr18-23207708-INS-73
chr18-23230200-INS-506
chr18-23237432-INS-179
chr18-23239597-INS-385
chr18-23241434-INS-90
chr18-23245526-INS-178
chr18-23262133-INS-276
chr18-23262182-INS-6481
chr18-23264043-INS-56
chr18-23270317-INS-65
chr18-23285636-INS-150
chr18-23289373-INS-175
chr18-23298155-INS-100
chr18-23300509-INS-55
chr18-23330305-INS-6410
chr18-23393892-INS-292
chr18-23417442-INS-168
chr18-23445051-INS-1473
chr18-23447728-INS-194
chr18-23466820-INS-114
chr18-23476624-INS-290
chr18-23499222-INS-496
chr18-23506502-INS-97
chr18-23514084-INS-51
chr18-23536748-INS-60
chr18-23538089-INS-208
chr18-23607263-INS-50
chr18-23629260-INS-64

chr18-23649241-INS-83
chr18-23656578-INS-52
chr18-23658472-INS-146
chr18-23660465-INS-554
chr18-23664666-INS-72
chr18-23684480-INS-81
chr18-23726523-INS-82
chr18-23773992-INS-170
chr18-23793417-INS-449
chr18-23795247-INS-5232
chr18-23804308-INS-544
chr18-23805893-INS-162
chr18-23823449-INS-61
chr18-23872767-INS-4860
chr18-23899614-INS-56
chr18-23903131-INS-191
chr18-23912734-INS-93
chr18-23936164-INS-120
chr18-23945708-INS-72
chr18-23951796-INS-167
chr18-23977738-INS-64
chr18-23978510-INS-107
chr18-24049807-INS-169
chr18-24062264-INS-225
chr18-24065605-INS-634
chr18-24071475-INS-408
chr18-24086746-INS-573
chr18-24096681-INS-1143
chr18-24105015-INS-6458
chr18-24116469-INS-415
chr18-24132756-INS-1863
chr18-24132897-INS-106
chr18-24134176-INS-201
chr18-24148590-INS-301
chr18-24150254-INS-177
chr18-24203015-INS-201
chr18-24205705-INS-548
chr18-24210124-INS-62
chr18-24217076-INS-197
chr18-24220472-INS-234
chr18-24224869-INS-125
chr18-24239332-INS-209
chr18-24248797-INS-251
chr18-24255711-INS-101
chr18-24266012-INS-90
chr18-24278725-INS-217
chr18-24279838-INS-120
chr18-24280176-INS-1800
chr18-24280920-INS-135
chr18-24305635-INS-158
chr18-24312922-INS-114
chr18-24318310-INS-121
chr18-24328015-INS-395
chr18-24329142-INS-976
chr18-24367689-INS-100
chr18-24381196-INS-118
chr18-24392411-INS-110
chr18-24426686-INS-215
chr18-24434378-INS-222
chr18-24441856-INS-68
chr18-24445281-INS-112
chr18-24459089-INS-164
chr18-24462660-INS-60
chr18-24470938-INS-112
chr18-24519909-INS-220
chr18-24566581-INS-133
chr18-24598461-INS-86
chr18-24602085-INS-303
chr18-24614002-INS-87
chr18-24650902-INS-174
chr18-24656678-INS-179
chr18-24676465-INS-88
chr18-24678013-INS-678
chr18-24692232-INS-85
chr18-24694738-INS-557
chr18-24728152-INS-78
chr18-24755885-INS-91

chr18-24779171-INS-68
chr18-24788738-INS-132
chr18-24805108-INS-559
chr18-24806082-INS-207
chr18-24811355-INS-806
chr18-24813576-INS-1888
chr18-24814624-INS-701
chr18-24815440-INS-67
chr18-24823139-INS-164
chr18-24824853-INS-88
chr18-24845957-INS-194
chr18-24848991-INS-70
chr18-24859211-INS-336
chr18-24879906-INS-168
chr18-24916409-INS-687
chr18-24927603-INS-229
chr18-24934433-INS-233
chr18-24936806-INS-94
chr18-24938692-INS-52
chr18-24942865-INS-550
chr18-24953121-INS-340
chr18-24962054-INS-86
chr18-24968427-INS-68
chr18-24987474-INS-54
chr18-25019711-INS-243
chr18-25025648-INS-414
chr18-25034265-INS-200
chr18-25062077-INS-172
chr18-25086334-INS-84
chr18-25097119-INS-162
chr18-25119509-INS-113
chr18-25186325-INS-232
chr18-25186405-INS-6706
chr18-25188002-INS-70
chr18-25191503-INS-109
chr18-25225668-INS-107
chr18-25264713-INS-59
chr18-25270109-INS-7138
chr18-25306522-INS-566
chr18-25320455-INS-79
chr18-25384375-INS-315
chr18-25385100-INS-399
chr18-25385687-INS-1467
chr18-25391422-INS-5400
chr18-25405026-INS-68
chr18-25406958-INS-528
chr18-25413214-INS-173
chr18-25426587-INS-218
chr18-25434709-INS-249
chr18-25502372-INS-648
chr18-25520384-INS-62
chr18-25559740-INS-59
chr18-25579060-INS-474
chr18-25584000-INS-540
chr18-25585094-INS-109
chr18-25629431-INS-56
chr18-25633716-INS-63
chr18-25731403-INS-68
chr18-25735668-INS-58
chr18-25757708-INS-179
chr18-25765543-INS-66
chr18-25798148-INS-66
chr18-25843155-INS-272
chr18-25846608-INS-284
chr18-25877324-INS-116
chr18-25922696-INS-194
chr18-25990914-INS-80
chr18-25996127-INS-56
chr18-26002661-INS-84
chr18-26035552-INS-133
chr18-26077378-INS-6390
chr18-26103837-INS-94
chr18-26124319-INS-6396
chr18-26128243-INS-66
chr18-26151091-INS-353
chr18-26182979-INS-414
chr18-26208145-INS-62

chr18-26213095-INS-84
chr18-26216761-INS-397
chr18-26234960-INS-5633
chr18-26261827-INS-199
chr18-26282196-INS-1439
chr18-26302596-INS-60
chr18-26322268-INS-133
chr18-26329113-INS-296
chr18-26364273-INS-93
chr18-26364680-INS-78
chr18-26365087-INS-724
chr18-26370330-INS-58
chr18-26375900-INS-422
chr18-26387366-INS-68
chr18-26396797-INS-51
chr18-26424090-INS-198
chr18-26439797-INS-85
chr18-26449903-INS-330
chr18-26468440-INS-70
chr18-26482298-INS-64
chr18-26500088-INS-54
chr18-26522656-INS-6644
chr18-26524256-INS-323
chr18-26553371-INS-6175
chr18-26556317-INS-6492
chr18-26579785-INS-7273
chr18-26587191-INS-664
chr18-26598366-INS-67
chr18-26601126-INS-278
chr18-26628171-INS-447
chr18-26630192-INS-128
chr18-26631285-INS-442
chr18-26632358-INS-776
chr18-26636814-INS-3320
chr18-26647146-INS-176
chr18-26654202-INS-187
chr18-26678923-INS-635
chr18-26684112-INS-7544
chr18-26692195-INS-271
chr18-26704937-INS-76
chr18-26749193-INS-112
chr18-26758281-INS-6454
chr18-26766591-INS-204
chr18-26792669-INS-360
chr18-26804018-INS-510
chr18-26840057-INS-56
chr18-26842284-INS-103
chr18-26860536-INS-90
chr18-26863639-INS-58
chr18-26875334-INS-54
chr18-26886439-INS-54
chr18-26935010-INS-71
chr18-26952069-INS-66
chr18-26958888-INS-600
chr18-26961653-INS-585
chr18-26985599-INS-476
chr18-26997570-INS-80
chr18-26998391-INS-128
chr18-27002339-INS-74
chr18-27002771-INS-345
chr18-27011335-INS-533
chr18-27012213-INS-267
chr18-27069037-INS-136
chr18-27069058-INS-341
chr18-27069463-INS-108
chr18-27085125-INS-572
chr18-27086716-INS-486
chr18-27113601-INS-502
chr18-27114892-INS-84
chr18-27141628-INS-101
chr18-27162536-INS-56
chr18-27263691-INS-154
chr18-27281877-INS-157
chr18-27289130-INS-136
chr18-27302795-INS-97
chr18-27322655-INS-939
chr18-27341282-INS-114

chr18-27344194-INS-2826
chr18-27359078-INS-170
chr18-27386295-INS-650
chr18-27421896-INS-56
chr18-27423717-INS-104
chr18-27445580-INS-120
chr18-27464995-INS-142
chr18-27467231-INS-64
chr18-27499244-INS-275
chr18-27513585-INS-59
chr18-27528683-INS-100
chr18-27561377-INS-4832
chr18-27581472-INS-61
chr18-27584300-INS-5969
chr18-27608721-INS-666
chr18-27610942-INS-73
chr18-27633761-INS-55
chr18-27635379-INS-198
chr18-27641299-INS-159
chr18-27665017-INS-6405
chr18-27697151-INS-154
chr18-27698642-INS-289
chr18-27775731-INS-185
chr18-27808210-INS-264
chr18-27812978-INS-114
chr18-27829879-INS-401
chr18-27833851-INS-56
chr18-27837972-INS-70
chr18-27842323-INS-6295
chr18-27844570-INS-606
chr18-27849777-INS-72
chr18-27874616-INS-200
chr18-27883220-INS-104
chr18-27886905-INS-168
chr18-27890994-INS-73
chr18-27895878-INS-85
chr18-27924190-INS-70
chr18-27944662-INS-7119
chr18-27959015-INS-144
chr18-27960852-INS-209
chr18-27969219-INS-174
chr18-27985486-INS-209
chr18-27991999-INS-358
chr18-27998188-INS-62
chr18-28020629-INS-82
chr18-28024232-INS-52
chr18-28041252-INS-68
chr18-28066247-INS-52
chr18-28111584-INS-95
chr18-28133837-INS-6447
chr18-28173270-INS-279
chr18-28174164-INS-82
chr18-28199079-INS-2138
chr18-28212400-INS-469
chr18-28291506-INS-187
chr18-28294790-INS-75
chr18-28296889-INS-201
chr18-28304755-INS-614
chr18-28306918-INS-65
chr18-28311002-INS-3525
chr18-28314858-INS-350
chr18-28328089-INS-242
chr18-28333166-INS-52
chr18-28364975-INS-696
chr18-28379022-INS-208
chr18-28440879-INS-52
chr18-28453723-INS-5303
chr18-28475825-INS-59
chr18-28503038-INS-467
chr18-28505166-INS-68
chr18-28525782-INS-1520
chr18-28538292-INS-98
chr18-28561783-INS-54
chr18-28582886-INS-51
chr18-28629810-INS-86
chr18-28631179-INS-415
chr18-28653660-INS-84

chr18-28657753-INS-53
chr18-28668356-INS-62
chr18-28679960-INS-97
chr18-28698560-INS-52
chr18-28717358-INS-77
chr18-28772740-INS-54
chr18-28778582-INS-466
chr18-28782045-INS-561
chr18-28804171-INS-170
chr18-28810762-INS-80
chr18-28821700-INS-65
chr18-28856226-INS-71
chr18-28861141-INS-1002
chr18-28873223-INS-741
chr18-28896787-INS-70
chr18-28899449-INS-614
chr18-28907944-INS-57
chr18-28928937-INS-53
chr18-28959750-INS-531
chr18-28971569-INS-81
chr18-28982554-INS-52
chr18-29007047-INS-264
chr18-29053135-INS-241
chr18-29078664-INS-101
chr18-29113460-INS-54
chr18-29127685-INS-107
chr18-29135075-INS-108
chr18-29171303-INS-302
chr18-29193865-INS-72
chr18-29199013-INS-478
chr18-29209565-INS-991
chr18-29218509-INS-144
chr18-29244419-INS-55
chr18-29246035-INS-986
chr18-29247046-INS-136
chr18-29250625-INS-149
chr18-29254800-INS-68
chr18-29262740-INS-88
chr18-29271171-INS-127
chr18-29285719-INS-196
chr18-29313502-INS-85
chr18-29346491-INS-265
chr18-29379523-INS-2071
chr18-29413896-INS-52
chr18-29433157-INS-92
chr18-29472570-INS-62
chr18-29514956-INS-56
chr18-29532959-INS-80
chr18-29539067-INS-114
chr18-29559047-INS-148
chr18-29585085-INS-72
chr18-29600950-INS-898
chr18-29621155-INS-132
chr18-29624443-INS-78
chr18-29628037-INS-192
chr18-29631325-INS-270
chr18-29648996-INS-185
chr18-29657152-INS-464
chr18-29713395-INS-66
chr18-29747795-INS-202
chr18-29750945-INS-797
chr18-29754482-INS-97
chr18-29776763-INS-440
chr18-29787731-INS-136
chr18-29789123-INS-338
chr18-29799933-INS-846
chr18-29824646-INS-1397
chr18-29829960-INS-464
chr18-29835603-INS-1228
chr18-29845092-INS-1619
chr18-29868977-INS-116
chr18-29943258-INS-9874
chr18-29946297-INS-72
chr18-29976389-INS-425
chr18-29986747-INS-72
chr18-30002784-INS-413
chr18-30008549-INS-68

chr18-30015560-INS-68
chr18-30042961-INS-73
chr18-30047996-INS-260
chr18-30050652-INS-239
chr18-30072830-INS-761
chr18-30080669-INS-104
chr18-30081813-INS-62
chr18-30098063-INS-56
chr18-30098683-INS-7211
chr18-30114493-INS-183
chr18-30128150-INS-305
chr18-30129236-INS-72
chr18-30139980-INS-88
chr18-30144343-INS-206
chr18-30163910-INS-611
chr18-30195347-INS-232
chr18-30202525-INS-52
chr18-30208287-INS-828
chr18-30212660-INS-7311
chr18-30216292-INS-92
chr18-30294330-INS-6874
chr18-30295019-INS-281
chr18-30301070-INS-6496
chr18-30328300-INS-883
chr18-30359853-INS-327
chr18-30362515-INS-191
chr18-30366789-INS-108
chr18-30373617-INS-60
chr18-30377535-INS-59
chr18-30380936-INS-565
chr18-30409641-INS-168
chr18-30411303-INS-75
chr18-30419896-INS-886
chr18-30421556-INS-224
chr18-30458194-INS-171
chr18-30460006-INS-433
chr18-30491858-INS-52
chr18-30500938-INS-188
chr18-30603630-INS-82
chr18-30605879-INS-399
chr18-30608196-INS-119
chr18-30647235-INS-269
chr18-30651556-INS-624
chr18-30704523-INS-433
chr18-30717261-INS-76
chr18-30766500-INS-343
chr18-30768308-INS-62
chr18-30809009-INS-72
chr18-30822169-INS-84
chr18-30851112-INS-92
chr18-30871118-INS-146
chr18-30872822-INS-409
chr18-30891469-INS-59
chr18-30896626-INS-65
chr18-30930000-INS-55
chr18-30931049-INS-229
chr18-30933388-INS-105
chr18-30953500-INS-7227
chr18-30956885-INS-6964
chr18-30967541-INS-668
chr18-30968289-INS-60
chr18-30990108-INS-190
chr18-31000183-INS-5393
chr18-31001958-INS-8722
chr18-31041989-INS-75
chr18-31049496-INS-53
chr18-31050918-INS-161
chr18-31067261-INS-202
chr18-31079944-INS-126
chr18-31110601-INS-222
chr18-31111656-INS-749
chr18-31120420-INS-6545
chr18-31143501-INS-175
chr18-31170187-INS-284
chr18-31192377-INS-107
chr18-31194963-INS-354
chr18-31219960-INS-69

chr18-31242518-INS-163
chr18-31252209-INS-66
chr18-31269208-INS-211
chr18-31272797-INS-7102
chr18-31307180-INS-54
chr18-31309236-INS-51
chr18-31310104-INS-127
chr18-31328740-INS-269
chr18-31346279-INS-117
chr18-31378494-INS-5548
chr18-31386674-INS-945
chr18-31408214-INS-496
chr18-31426931-INS-214
chr18-31445507-INS-111
chr18-31448615-INS-62
chr18-31462403-INS-6697
chr18-31469082-INS-99
chr18-31485854-INS-344
chr18-31530349-INS-486
chr18-31544596-INS-246
chr18-31560702-INS-148
chr18-31564039-INS-6927
chr18-31564492-INS-853
chr18-31596416-INS-939
chr18-31599189-INS-108
chr18-31605039-INS-6735
chr18-31625309-INS-95
chr18-31668814-INS-118
chr18-31675742-INS-156
chr18-31680786-INS-362
chr18-31690743-INS-908
chr18-31730806-INS-393
chr18-31778126-INS-54
chr18-31785135-INS-144
chr18-31796366-INS-58
chr18-31822132-INS-95
chr18-31842856-INS-212
chr18-31847505-INS-182
chr18-31865012-INS-279
chr18-31874523-INS-4733
chr18-31881663-INS-5397
chr18-31884629-INS-7106
chr18-31888975-INS-5199
chr18-31923468-INS-204
chr18-31944658-INS-266
chr18-31945937-INS-195
chr18-31947611-INS-183
chr18-31949680-INS-535
chr18-31956729-INS-214
chr18-31979832-INS-60
chr18-31987896-INS-55
chr18-31994042-INS-214
chr18-31997679-INS-203
chr18-32047183-INS-184
chr18-32048798-INS-82
chr18-32052547-INS-573
chr18-32055017-INS-106
chr18-32061706-INS-241
chr18-32066184-INS-52
chr18-32093744-INS-88
chr18-32096324-INS-60
chr18-32144665-INS-77
chr18-32166866-INS-57
chr18-32188865-INS-209
chr18-32208594-INS-170
chr18-32214873-INS-60
chr18-32225918-INS-65
chr18-32285988-INS-89
chr18-32327310-INS-313
chr18-32332196-INS-58
chr18-32339440-INS-87
chr18-32347677-INS-342
chr18-32351291-INS-617
chr18-32375857-INS-148
chr18-32377769-INS-118
chr18-32391287-INS-312
chr18-32415217-INS-936

chr18-32419836-INS-61
chr18-32439925-INS-107
chr18-32455786-INS-54
chr18-32461681-INS-58
chr18-32466904-INS-343
chr18-32482881-INS-114
chr18-32497500-INS-4359
chr18-32501666-INS-51
chr18-32504735-INS-61
chr18-32514161-INS-70
chr18-32520106-INS-63
chr18-32521842-INS-128
chr18-32567167-INS-121
chr18-32583530-INS-373
chr18-32600345-INS-6245
chr18-32654031-INS-276
chr18-32655850-INS-337
chr18-32693927-INS-84
chr18-32695819-INS-130
chr18-32699595-INS-170
chr18-32705928-INS-55
chr18-32706436-INS-76
chr18-32718016-INS-644
chr18-32721773-INS-66
chr18-32730964-INS-523
chr18-32750924-INS-159
chr18-32758857-INS-647
chr18-32760552-INS-8994
chr18-32771566-INS-112
chr18-32809400-INS-6803
chr18-32812852-INS-1826
chr18-32833128-INS-188
chr18-32833316-INS-123
chr18-32833368-INS-154
chr18-32847762-INS-392
chr18-32849544-INS-987
chr18-32857542-INS-387
chr18-32858996-INS-90
chr18-32870371-INS-52
chr18-32928697-INS-186
chr18-32965324-INS-492
chr18-32983312-INS-130
chr18-32984618-INS-72
chr18-33030258-INS-137
chr18-33081117-INS-1261
chr18-33081118-INS-337
chr18-33136227-INS-68
chr18-33184464-INS-67
chr18-33225262-INS-130
chr18-33245918-INS-1038
chr18-33245919-INS-870
chr18-33266015-INS-298
chr18-33270413-INS-69
chr18-33272800-INS-661
chr18-33276522-INS-139
chr18-33287996-INS-196
chr18-33437529-INS-118
chr18-33446736-INS-54
chr18-33526493-INS-152
chr18-33555865-INS-337
chr18-33561422-INS-64
chr18-33584602-INS-57
chr18-33600166-INS-238
chr18-33602717-INS-52
chr18-33605299-INS-146
chr18-33641729-INS-194
chr18-33642946-INS-405
chr18-33654654-INS-147
chr18-33657862-INS-130
chr18-33661042-INS-595
chr18-33672460-INS-72
chr18-33675851-INS-126
chr18-33683133-INS-886
chr18-33697783-INS-5293
chr18-33707749-INS-3291
chr18-33718738-INS-914
chr18-33726626-INS-82

chr18-33728620-INS-1905
chr18-33728622-INS-75
chr18-33734106-INS-958
chr18-33747513-INS-668
chr18-33751826-INS-797
chr18-33804326-INS-125
chr18-33807433-INS-455
chr18-33817013-INS-6866
chr18-33843738-INS-7102
chr18-33860215-INS-64
chr18-33862077-INS-274
chr18-33869759-INS-57
chr18-33876648-INS-6716
chr18-33883733-INS-362
chr18-33884549-INS-64
chr18-33885714-INS-220
chr18-33896722-INS-111
chr18-33898091-INS-164
chr18-33902371-INS-93
chr18-33932305-INS-162
chr18-33943666-INS-70
chr18-33944500-INS-72
chr18-33954898-INS-71
chr18-33961029-INS-52
chr18-33974933-INS-61
chr18-33979572-INS-195
chr18-33985353-INS-188
chr18-34003504-INS-299
chr18-34020894-INS-206
chr18-34037843-INS-98
chr18-34058348-INS-64
chr18-34072396-INS-202
chr18-34085241-INS-90
chr18-34090417-INS-82
chr18-34141531-INS-102
chr18-34147926-INS-176
chr18-34152614-INS-140
chr18-34158473-INS-331
chr18-34161913-INS-58
chr18-34162880-INS-186
chr18-34163561-INS-435
chr18-34167845-INS-52
chr18-34170144-INS-434
chr18-34191313-INS-189
chr18-34205860-INS-408
chr18-34208055-INS-94
chr18-34226764-INS-479
chr18-34237207-INS-538
chr18-34238517-INS-326
chr18-34244339-INS-453
chr18-34255872-INS-445
chr18-34259715-INS-169
chr18-34271222-INS-195
chr18-34294932-INS-223
chr18-34318606-INS-464
chr18-34320190-INS-376
chr18-34338434-INS-108
chr18-34372337-INS-96
chr18-34373508-INS-458
chr18-34402987-INS-209
chr18-34428439-INS-58
chr18-34443390-INS-204
chr18-34457088-INS-222
chr18-34462286-INS-192
chr18-34464648-INS-56
chr18-34475115-INS-199
chr18-34476262-INS-328
chr18-34483372-INS-186
chr18-34495467-INS-121
chr18-34497005-INS-288
chr18-34516720-INS-52
chr18-34518932-INS-3278
chr18-34533683-INS-328
chr18-34548907-INS-555
chr18-34560737-INS-159
chr18-34563592-INS-84
chr18-34609615-INS-191

chr18-34654089-INS-164
chr18-34654814-INS-54
chr18-34662422-INS-67
chr18-34666763-INS-420
chr18-34674726-INS-133
chr18-34693722-INS-58
chr18-34699377-INS-100
chr18-34704307-INS-230
chr18-34708728-INS-66
chr18-34718294-INS-61
chr18-34719522-INS-199
chr18-34720970-INS-203
chr18-34721578-INS-203
chr18-34723488-INS-199
chr18-34725588-INS-202
chr18-34726574-INS-129
chr18-34757066-INS-164
chr18-34772258-INS-116
chr18-34777527-INS-124
chr18-34779853-INS-54
chr18-34783496-INS-374
chr18-34785335-INS-213
chr18-34786348-INS-84
chr18-34793437-INS-193
chr18-34803344-INS-170
chr18-34832766-INS-458
chr18-34838454-INS-337
chr18-34843864-INS-133
chr18-34862813-INS-170
chr18-34865088-INS-108
chr18-34867221-INS-204
chr18-34885960-INS-183
chr18-34896926-INS-70
chr18-34913520-INS-97
chr18-34917720-INS-59
chr18-34918960-INS-355
chr18-34919876-INS-200
chr18-34923234-INS-70
chr18-34953073-INS-62
chr18-34968545-INS-155
chr18-34976811-INS-209
chr18-34981276-INS-242
chr18-34982325-INS-156
chr18-35009142-INS-243
chr18-35011602-INS-214
chr18-35015489-INS-5554
chr18-35021587-INS-52
chr18-35023020-INS-74
chr18-35026347-INS-198
chr18-35039451-INS-205
chr18-35042413-INS-82
chr18-35060459-INS-234
chr18-35094342-INS-466
chr18-35109166-INS-597
chr18-35110140-INS-57
chr18-35114391-INS-420
chr18-35130002-INS-116
chr18-35136512-INS-54
chr18-35164006-INS-104
chr18-35170340-INS-125
chr18-35172614-INS-54
chr18-35174283-INS-63
chr18-35182318-INS-227
chr18-35186045-INS-65
chr18-35190467-INS-7251
chr18-35206506-INS-379
chr18-35239739-INS-623
chr18-35258380-INS-171
chr18-35259931-INS-69
chr18-35271733-INS-211
chr18-35306103-INS-175
chr18-35308656-INS-64
chr18-35329486-INS-264
chr18-35336096-INS-234
chr18-35364457-INS-92
chr18-35369553-INS-58
chr18-35395474-INS-64

chr18-35409706-INS-401
chr18-35416246-INS-72
chr18-35419806-INS-52
chr18-35430304-INS-99
chr18-35434152-INS-216
chr18-35435856-INS-54
chr18-35450812-INS-188
chr18-35462335-INS-66
chr18-35463659-INS-72
chr18-35488441-INS-174
chr18-35494919-INS-517
chr18-35540523-INS-381
chr18-35599649-INS-80
chr18-35645109-INS-186
chr18-35647943-INS-3583
chr18-35671484-INS-60
chr18-35697980-INS-185
chr18-35755217-INS-154
chr18-35765477-INS-52
chr18-35766544-INS-213
chr18-35767665-INS-71
chr18-35770907-INS-215
chr18-35773005-INS-212
chr18-35777274-INS-58
chr18-35795614-INS-582
chr18-35797779-INS-434
chr18-35813385-INS-5005
chr18-35817106-INS-336
chr18-35834792-INS-201
chr18-35838368-INS-732
chr18-35890187-INS-195
chr18-35891725-INS-431
chr18-35900742-INS-211
chr18-35925384-INS-169
chr18-35928692-INS-212
chr18-35931139-INS-197
chr18-35935550-INS-215
chr18-35936563-INS-198
chr18-35942008-INS-227
chr18-35950926-INS-252
chr18-35959839-INS-57
chr18-35984174-INS-75
chr18-36004687-INS-182
chr18-36029902-INS-216
chr18-36039639-INS-193
chr18-36044214-INS-458
chr18-36050304-INS-64
chr18-36064315-INS-54
chr18-36073618-INS-112
chr18-36087506-INS-186
chr18-36102609-INS-102
chr18-36117933-INS-205
chr18-36136376-INS-114
chr18-36142613-INS-56
chr18-36144999-INS-255
chr18-36183677-INS-209
chr18-36199248-INS-207
chr18-36270069-INS-98
chr18-36285783-INS-159
chr18-36301878-INS-60
chr18-36307454-INS-174
chr18-36324807-INS-68
chr18-36334548-INS-246
chr18-36338421-INS-228
chr18-36339966-INS-527
chr18-36340830-INS-508
chr18-36344222-INS-52
chr18-36349190-INS-87
chr18-36362110-INS-52
chr18-36376495-INS-173
chr18-36389678-INS-76
chr18-36445668-INS-124
chr18-36462702-INS-456
chr18-36470425-INS-89
chr18-36473079-INS-196
chr18-36481449-INS-57
chr18-36507558-INS-84

chr18-36510161-INS-179
chr18-36515750-INS-63
chr18-36539773-INS-164
chr18-36540655-INS-209
chr18-36548287-INS-459
chr18-36554020-INS-58
chr18-36566800-INS-160
chr18-36577228-INS-92
chr18-36581558-INS-7139
chr18-36584522-INS-133
chr18-36591166-INS-55
chr18-36642270-INS-181
chr18-36681252-INS-232
chr18-36684032-INS-216
chr18-36692158-INS-166
chr18-36767171-INS-75
chr18-36784836-INS-102
chr18-36805870-INS-128
chr18-36806296-INS-199
chr18-36806494-INS-118
chr18-36813807-INS-119
chr18-36840703-INS-204
chr18-36864372-INS-138
chr18-36885593-INS-98
chr18-36909059-INS-202
chr18-36910366-INS-149
chr18-36935697-INS-176
chr18-36942712-INS-241
chr18-36959313-INS-5468
chr18-37106360-INS-6428
chr18-37128570-INS-273
chr18-37152522-INS-163
chr18-37184388-INS-7067
chr18-37235778-INS-3466
chr18-37238728-INS-119
chr18-37257027-INS-1290
chr18-37333647-INS-217
chr18-37343533-INS-434
chr18-37349056-INS-60
chr18-37366728-INS-446
chr18-37411993-INS-55
chr18-37463591-INS-72
chr18-37511188-INS-73
chr18-37543615-INS-551
chr18-37593553-INS-59
chr18-37600096-INS-107
chr18-37609985-INS-74
chr18-37626589-INS-197
chr18-37674650-INS-679
chr18-37718209-INS-128
chr18-37730073-INS-7909
chr18-37732660-INS-192
chr18-37744156-INS-83
chr18-37809981-INS-829
chr18-37831670-INS-131
chr18-37858040-INS-80
chr18-37910175-INS-58
chr18-37941729-INS-5552
chr18-37945568-INS-195
chr18-37980500-INS-224
chr18-37983946-INS-202
chr18-38008085-INS-204
chr18-38012349-INS-6413
chr18-38084603-INS-194
chr18-38098231-INS-199
chr18-38113477-INS-71
chr18-38114829-INS-666
chr18-38116522-INS-56
chr18-38142986-INS-209
chr18-38144825-INS-143
chr18-38151554-INS-54
chr18-38198044-INS-84
chr18-38206884-INS-159
chr18-38240471-INS-51
chr18-38241113-INS-203
chr18-38243156-INS-92
chr18-38273918-INS-72

chr18-38364669-INS-52
chr18-38369527-INS-78
chr18-38378856-INS-151
chr18-38394045-INS-76
chr18-38398828-INS-64
chr18-38444111-INS-346
chr18-38446976-INS-658
chr18-38463316-INS-295
chr18-38464488-INS-202
chr18-38472621-INS-236
chr18-38482259-INS-226
chr18-38485117-INS-241
chr18-38489505-INS-206
chr18-38516421-INS-418
chr18-38518918-INS-448
chr18-38526140-INS-205
chr18-38530147-INS-55
chr18-38531165-INS-340
chr18-38538335-INS-202
chr18-38545369-INS-68
chr18-38550514-INS-7283
chr18-38556639-INS-165
chr18-38568747-INS-518
chr18-38572284-INS-87
chr18-38584885-INS-399
chr18-38609583-INS-192
chr18-38621846-INS-196
chr18-38627471-INS-175
chr18-38638990-INS-155
chr18-38646836-INS-990
chr18-38649005-INS-219
chr18-38659356-INS-52
chr18-38660077-INS-202
chr18-38686972-INS-277
chr18-38696131-INS-120
chr18-38729717-INS-52
chr18-38738491-INS-52
chr18-38740475-INS-100
chr18-38753210-INS-213
chr18-38759945-INS-89
chr18-38760848-INS-216
chr18-38780412-INS-402
chr18-38793631-INS-199
chr18-38801166-INS-196
chr18-38802484-INS-93
chr18-38838653-INS-52
chr18-38841891-INS-337
chr18-38845995-INS-93
chr18-38873512-INS-86
chr18-38892573-INS-137
chr18-38894209-INS-156
chr18-38920105-INS-290
chr18-38958102-INS-441
chr18-38967640-INS-211
chr18-39007021-INS-5549
chr18-39035855-INS-54
chr18-39049617-INS-104
chr18-39050274-INS-151
chr18-39053673-INS-167
chr18-39056193-INS-210
chr18-39067524-INS-1893
chr18-39069204-INS-265
chr18-39085688-INS-88
chr18-39091571-INS-344
chr18-39093738-INS-77
chr18-39113980-INS-110
chr18-39138815-INS-125
chr18-39155111-INS-216
chr18-39166279-INS-66
chr18-39203141-INS-87
chr18-39204154-INS-668
chr18-39209610-INS-709
chr18-39224366-INS-6477
chr18-39266149-INS-187
chr18-39308101-INS-213
chr18-39311817-INS-124
chr18-39314932-INS-206

chr18-39331329-INS-203
chr18-39340235-INS-920
chr18-39352698-INS-472
chr18-39375697-INS-4842
chr18-39383024-INS-7700
chr18-39404123-INS-2171
chr18-39441249-INS-287
chr18-39454810-INS-1466
chr18-39463055-INS-939
chr18-39484497-INS-588
chr18-39515170-INS-80
chr18-39583150-INS-210
chr18-39589500-INS-98
chr18-39683039-INS-198
chr18-39691061-INS-565
chr18-39707466-INS-400
chr18-39711480-INS-166
chr18-39751969-INS-84
chr18-39763702-INS-108
chr18-39802604-INS-104
chr18-39826037-INS-304
chr18-39855462-INS-162
chr18-39870845-INS-6307
chr18-39884327-INS-108
chr18-39909897-INS-120
chr18-39913204-INS-957
chr18-39940870-INS-169
chr18-39948870-INS-53
chr18-39949855-INS-79
chr18-39994920-INS-66
chr18-40018145-INS-146
chr18-40087537-INS-94
chr18-40107899-INS-599
chr18-40111550-INS-487
chr18-40123293-INS-634
chr18-40177687-INS-7191
chr18-40193787-INS-88
chr18-40200751-INS-178
chr18-40244994-INS-1188
chr18-40247670-INS-254
chr18-40247705-INS-2085
chr18-40264642-INS-6141
chr18-40290950-INS-69
chr18-40296812-INS-96
chr18-40305418-INS-6129
chr18-40325915-INS-656
chr18-40330056-INS-6422
chr18-40333642-INS-260
chr18-40347186-INS-212
chr18-40358902-INS-2743
chr18-40365584-INS-538
chr18-40367245-INS-183
chr18-40376109-INS-343
chr18-40388300-INS-57
chr18-40399055-INS-82
chr18-40416388-INS-7760
chr18-40432016-INS-74
chr18-40476930-INS-92
chr18-40496113-INS-304
chr18-40511295-INS-182
chr18-40522723-INS-785
chr18-40524267-INS-459
chr18-40529789-INS-55
chr18-40549691-INS-223
chr18-40555213-INS-323
chr18-40567689-INS-6832
chr18-40569723-INS-247
chr18-40608936-INS-374
chr18-40611217-INS-93
chr18-40612104-INS-6592
chr18-40616753-INS-55
chr18-40650390-INS-271
chr18-40691912-INS-327
chr18-40700787-INS-267
chr18-40707155-INS-6402
chr18-40713117-INS-2787
chr18-40723319-INS-6548

chr18-40734652-INS-93
chr18-40767894-INS-294
chr18-40791377-INS-56
chr18-40825225-INS-278
chr18-40839588-INS-6435
chr18-40907426-INS-133
chr18-40909624-INS-997
chr18-40932268-INS-138
chr18-40933502-INS-56
chr18-40948967-INS-768
chr18-40970351-INS-55
chr18-40982795-INS-513
chr18-41038875-INS-442
chr18-41045148-INS-99
chr18-41048914-INS-2131
chr18-41056654-INS-817
chr18-41087487-INS-169
chr18-41094303-INS-6252
chr18-41102537-INS-547
chr18-41106197-INS-2932
chr18-41121979-INS-1439
chr18-41126677-INS-6286
chr18-41127734-INS-76
chr18-41155407-INS-138
chr18-41171431-INS-182
chr18-41177115-INS-549
chr18-41193152-INS-209
chr18-41202280-INS-101
chr18-41210824-INS-5247
chr18-41213849-INS-233
chr18-41237165-INS-743
chr18-41243882-INS-50
chr18-41254131-INS-342
chr18-41265317-INS-51
chr18-41269360-INS-201
chr18-41309727-INS-52
chr18-41311826-INS-132
chr18-41314806-INS-763
chr18-41315457-INS-6413
chr18-41326089-INS-224
chr18-41355257-INS-780
chr18-41380168-INS-114
chr18-41401182-INS-66
chr18-41409698-INS-52
chr18-41445306-INS-92
chr18-41462506-INS-231
chr18-41484416-INS-486
chr18-41507495-INS-85
chr18-41541651-INS-281
chr18-41546290-INS-160
chr18-41559665-INS-102
chr18-41568779-INS-441
chr18-41608253-INS-51
chr18-41614630-INS-197
chr18-41617854-INS-200
chr18-41655218-INS-521
chr18-41662514-INS-6954
chr18-41677416-INS-63
chr18-41700126-INS-56
chr18-41720890-INS-892
chr18-41769381-INS-62
chr18-41775617-INS-230
chr18-41777500-INS-6583
chr18-41789123-INS-161
chr18-41805314-INS-227
chr18-41839432-INS-226
chr18-41846085-INS-353
chr18-41903923-INS-4589
chr18-41909350-INS-158
chr18-41914279-INS-6172
chr18-41918322-INS-907
chr18-41941066-INS-108
chr18-41944384-INS-56
chr18-41973653-INS-186
chr18-41976597-INS-373
chr18-42027503-INS-206
chr18-42037009-INS-195

chr18-42046718-INS-202
chr18-42050845-INS-617
chr18-42079023-INS-73
chr18-42090223-INS-171
chr18-42108528-INS-58
chr18-42126355-INS-175
chr18-42137649-INS-60
chr18-42139509-INS-91
chr18-42148585-INS-55
chr18-42149702-INS-132
chr18-42151796-INS-391
chr18-42193592-INS-54
chr18-42199168-INS-51
chr18-42207949-INS-1255
chr18-42225823-INS-106
chr18-42239269-INS-52
chr18-42242324-INS-52
chr18-42259946-INS-331
chr18-42270765-INS-288
chr18-42284416-INS-69
chr18-42302746-INS-55
chr18-42328247-INS-177
chr18-42331556-INS-174
chr18-42339299-INS-64
chr18-42347209-INS-134
chr18-42371900-INS-201
chr18-42373717-INS-123
chr18-42376617-INS-353
chr18-42393851-INS-102
chr18-42406481-INS-206
chr18-42445733-INS-207
chr18-42454314-INS-198
chr18-42500193-INS-329
chr18-42568278-INS-211
chr18-42591624-INS-108
chr18-42608579-INS-206
chr18-42635512-INS-375
chr18-42670855-INS-207
chr18-42691649-INS-226
chr18-42713660-INS-135
chr18-42714482-INS-410
chr18-42763421-INS-60
chr18-42774251-INS-367
chr18-42789642-INS-417
chr18-42801110-INS-331
chr18-42820438-INS-304
chr18-42845655-INS-51
chr18-42890525-INS-70
chr18-42907088-INS-93
chr18-42920480-INS-53
chr18-42937673-INS-75
chr18-42938762-INS-6640
chr18-42956330-INS-171
chr18-43010743-INS-134
chr18-43033299-INS-56
chr18-43039707-INS-86
chr18-43048155-INS-187
chr18-43055228-INS-74
chr18-43060962-INS-120
chr18-43068387-INS-57
chr18-43081958-INS-64
chr18-43108961-INS-56
chr18-43112875-INS-775
chr18-43154785-INS-66
chr18-43177517-INS-239
chr18-43217973-INS-54
chr18-43225093-INS-390
chr18-43233166-INS-561
chr18-43244613-INS-137
chr18-43249434-INS-142
chr18-43284768-INS-64
chr18-43291422-INS-86
chr18-43293000-INS-552
chr18-43309789-INS-69
chr18-43311397-INS-179
chr18-43326049-INS-164
chr18-43328632-INS-967

chr18-43337217-INS-552
chr18-43385072-INS-72
chr18-43399267-INS-81
chr18-43419912-INS-189
chr18-43440767-INS-123
chr18-43441860-INS-6403
chr18-43449949-INS-59
chr18-43461244-INS-66
chr18-43479011-INS-102
chr18-43485138-INS-318
chr18-43510383-INS-175
chr18-43534670-INS-204
chr18-43561796-INS-6263
chr18-43572443-INS-130
chr18-43591255-INS-319
chr18-43603622-INS-264
chr18-43615207-INS-85
chr18-43620002-INS-55
chr18-43623106-INS-697
chr18-43623775-INS-121
chr18-43633092-INS-178
chr18-43648192-INS-86
chr18-43649307-INS-6887
chr18-43651468-INS-326
chr18-43654624-INS-80
chr18-43672814-INS-61
chr18-43685446-INS-228
chr18-43728268-INS-185
chr18-43743888-INS-6456
chr18-43744915-INS-692
chr18-43747887-INS-7087
chr18-43765988-INS-76
chr18-43777187-INS-326
chr18-43780580-INS-555
chr18-43788196-INS-56
chr18-43789981-INS-60
chr18-43795871-INS-58
chr18-43831229-INS-324
chr18-43860027-INS-52
chr18-43886661-INS-52
chr18-43894037-INS-315
chr18-43931661-INS-804
chr18-43936727-INS-301
chr18-43945203-INS-60
chr18-43949105-INS-53
chr18-43980259-INS-401
chr18-43986927-INS-296
chr18-44007866-INS-752
chr18-44015079-INS-7487
chr18-44018948-INS-217
chr18-44021066-INS-255
chr18-44035275-INS-156
chr18-44073454-INS-96
chr18-44116182-INS-447
chr18-44126262-INS-581
chr18-44132236-INS-60
chr18-44242953-INS-56
chr18-44245010-INS-4413
chr18-44251659-INS-182
chr18-44257664-INS-186
chr18-44271397-INS-764
chr18-44289978-INS-56
chr18-44310074-INS-96
chr18-44327640-INS-93
chr18-44340790-INS-373
chr18-44353765-INS-6692
chr18-44357593-INS-465
chr18-44401152-INS-59
chr18-44419802-INS-299
chr18-44465248-INS-680
chr18-44537124-INS-338
chr18-44543985-INS-748
chr18-44544470-INS-312
chr18-44559399-INS-61
chr18-44589776-INS-55
chr18-44599331-INS-99
chr18-44614173-INS-376

chr18-44616455-INS-64
chr18-44649356-INS-52
chr18-44660637-INS-414
chr18-44716579-INS-6941
chr18-44721950-INS-188
chr18-44732134-INS-282
chr18-44760285-INS-52
chr18-44774277-INS-131
chr18-44800135-INS-223
chr18-44813775-INS-98
chr18-44820007-INS-52
chr18-44828644-INS-59
chr18-44837727-INS-85
chr18-44865135-INS-98
chr18-44893759-INS-6548
chr18-44900562-INS-203
chr18-44903594-INS-55
chr18-44911848-INS-73
chr18-44913018-INS-165
chr18-44914765-INS-82
chr18-44916945-INS-543
chr18-44931843-INS-66
chr18-44934877-INS-54
chr18-44936269-INS-56
chr18-44945597-INS-58
chr18-44960095-INS-55
chr18-44975742-INS-586
chr18-44997901-INS-176
chr18-45002225-INS-203
chr18-45013020-INS-8675
chr18-45051177-INS-210
chr18-45072947-INS-56
chr18-45085523-INS-92
chr18-45086188-INS-512
chr18-45101055-INS-124
chr18-45126760-INS-6174
chr18-45168008-INS-150
chr18-45172531-INS-60
chr18-45175011-INS-2882
chr18-45227608-INS-60
chr18-45231215-INS-239
chr18-45237585-INS-55
chr18-45244261-INS-638
chr18-45247682-INS-74
chr18-45257921-INS-279
chr18-45271914-INS-212
chr18-45275114-INS-136
chr18-45289134-INS-50
chr18-45311753-INS-79
chr18-45321302-INS-2461
chr18-45322188-INS-6178
chr18-45327625-INS-68
chr18-45337739-INS-130
chr18-45355907-INS-995
chr18-45363814-INS-122
chr18-45369528-INS-497
chr18-45377959-INS-72
chr18-45393923-INS-53
chr18-45424010-INS-369
chr18-45434449-INS-7000
chr18-45443019-INS-52
chr18-45456274-INS-64
chr18-45469128-INS-61
chr18-45480852-INS-498
chr18-45483505-INS-7585
chr18-45486229-INS-85
chr18-45490315-INS-88
chr18-45498961-INS-6586
chr18-45514665-INS-51
chr18-45538150-INS-64
chr18-45566059-INS-529
chr18-45569145-INS-6518
chr18-45573576-INS-63
chr18-45579223-INS-301
chr18-45580585-INS-54
chr18-45594828-INS-86
chr18-45606644-INS-128

chr18-45608894-INS-6199
chr18-45611501-INS-6475
chr18-45647694-INS-928
chr18-45681238-INS-89
chr18-45682419-INS-1394
chr18-45720539-INS-69
chr18-45729753-INS-73
chr18-45736081-INS-64
chr18-45736995-INS-6444
chr18-45756830-INS-5512
chr18-45793929-INS-772
chr18-45800754-INS-180
chr18-45818138-INS-197
chr18-45820085-INS-86
chr18-45835893-INS-3223
chr18-45847971-INS-283
chr18-45864545-INS-53
chr18-45865613-INS-429
chr18-45881437-INS-72
chr18-45889811-INS-72
chr18-45918949-INS-6627
chr18-45925682-INS-238
chr18-45944320-INS-56
chr18-45974451-INS-412
chr18-45978388-INS-65
chr18-45993158-INS-149
chr18-45995215-INS-124
chr18-46024848-INS-7087
chr18-46044582-INS-6895
chr18-46056347-INS-176
chr18-46082596-INS-56
chr18-46110020-INS-1888
chr18-46112678-INS-6950
chr18-46118301-INS-331
chr18-46132765-INS-64
chr18-46135217-INS-341
chr18-46173685-INS-56
chr18-46200049-INS-52
chr18-46212507-INS-575
chr18-46225194-INS-90
chr18-46226786-INS-912
chr18-46243346-INS-492
chr18-46244452-INS-51
chr18-46268124-INS-1110
chr18-46268125-INS-888
chr18-46285295-INS-341
chr18-46326222-INS-98
chr18-46340715-INS-64
chr18-46353783-INS-729
chr18-46362212-INS-108
chr18-46365622-INS-995
chr18-46389230-INS-68
chr18-46417211-INS-117
chr18-46417375-INS-170
chr18-46428354-INS-167
chr18-46448383-INS-525
chr18-46460489-INS-642
chr18-46461061-INS-84
chr18-46464606-INS-56
chr18-46466457-INS-60
chr18-46503544-INS-330
chr18-46519356-INS-80
chr18-46524145-INS-69
chr18-46532857-INS-61
chr18-46541110-INS-267
chr18-46545642-INS-5552
chr18-46559590-INS-54
chr18-46561799-INS-69
chr18-46578741-INS-60
chr18-46582388-INS-54
chr18-46584630-INS-66
chr18-46644487-INS-197
chr18-46652367-INS-262
chr18-46664972-INS-54
chr18-46685126-INS-487
chr18-46686312-INS-146
chr18-46694928-INS-897

chr18-46705984-INS-176
chr18-46706753-INS-7012
chr18-46716346-INS-74
chr18-46718587-INS-740
chr18-46728728-INS-221
chr18-46739862-INS-211
chr18-46742644-INS-54
chr18-46745243-INS-83
chr18-46795000-INS-527
chr18-46801411-INS-103
chr18-46825666-INS-212
chr18-46828769-INS-62
chr18-46835308-INS-467
chr18-46866041-INS-116
chr18-46869016-INS-234
chr18-46954689-INS-139
chr18-46991844-INS-172
chr18-47012148-INS-86
chr18-47037604-INS-341
chr18-47042616-INS-100
chr18-47049964-INS-515
chr18-47068183-INS-79
chr18-47076811-INS-2470
chr18-47106301-INS-134
chr18-47113747-INS-78
chr18-47131128-INS-825
chr18-47139757-INS-319
chr18-47145469-INS-210
chr18-47155598-INS-106
chr18-47162481-INS-79
chr18-47188072-INS-289
chr18-47216429-INS-489
chr18-47223317-INS-65
chr18-47251642-INS-1866
chr18-47253029-INS-4042
chr18-47260597-INS-198
chr18-47266232-INS-208
chr18-47284923-INS-75
chr18-47326565-INS-56
chr18-47360620-INS-62
chr18-47364688-INS-76
chr18-47400215-INS-53
chr18-47465146-INS-72
chr18-47467022-INS-64
chr18-47479845-INS-70
chr18-47484606-INS-63
chr18-47489957-INS-362
chr18-47518336-INS-103
chr18-47536830-INS-53
chr18-47557801-INS-238
chr18-47571916-INS-318
chr18-47577029-INS-445
chr18-47595516-INS-58
chr18-47613822-INS-3191
chr18-47615575-INS-305
chr18-47619734-INS-197
chr18-47627753-INS-384
chr18-47634948-INS-112
chr18-47641119-INS-235
chr18-47646399-INS-64
chr18-47653143-INS-56
chr18-47667538-INS-557
chr18-47675160-INS-530
chr18-47700056-INS-5317
chr18-47704534-INS-62
chr18-47705279-INS-7007
chr18-47720098-INS-100
chr18-47727979-INS-407
chr18-47734482-INS-56
chr18-47736178-INS-100
chr18-47751997-INS-613
chr18-47761175-INS-164
chr18-47763871-INS-831
chr18-47763882-INS-1149
chr18-47765307-INS-119
chr18-47793501-INS-145
chr18-47805498-INS-77

chr18-47806568-INS-82
chr18-47816816-INS-208
chr18-47828180-INS-204
chr18-47839825-INS-61
chr18-47840943-INS-60
chr18-47844392-INS-122
chr18-47857046-INS-501
chr18-47858930-INS-96
chr18-47862029-INS-482
chr18-47878971-INS-59
chr18-47899235-INS-182
chr18-47919757-INS-55
chr18-47926297-INS-220
chr18-47955005-INS-152
chr18-47958139-INS-188
chr18-47983763-INS-624
chr18-47993477-INS-6387
chr18-47994797-INS-6474
chr18-48017406-INS-121
chr18-48086174-INS-123
chr18-48107388-INS-231
chr18-48112539-INS-56
chr18-48141334-INS-57
chr18-48173085-INS-715
chr18-48177937-INS-311
chr18-48189970-INS-6772
chr18-48216897-INS-53
chr18-48225882-INS-446
chr18-48231577-INS-157
chr18-48249419-INS-297
chr18-48272924-INS-52
chr18-48273747-INS-56
chr18-48285587-INS-498
chr18-48286281-INS-62
chr18-48286874-INS-53
chr18-48298579-INS-592
chr18-48338402-INS-72
chr18-48356973-INS-111
chr18-48375209-INS-109
chr18-48389971-INS-56
chr18-48397905-INS-7235
chr18-48409208-INS-92
chr18-48415120-INS-326
chr18-48417193-INS-164
chr18-48419267-INS-311
chr18-48419633-INS-3132
chr18-48422758-INS-6442
chr18-48424094-INS-649
chr18-48434142-INS-655
chr18-48438898-INS-60
chr18-48439439-INS-774
chr18-48452626-INS-84
chr18-48460306-INS-325
chr18-48492056-INS-126
chr18-48508940-INS-429
chr18-48520954-INS-66
chr18-48527804-INS-55
chr18-48532029-INS-320
chr18-48542929-INS-68
chr18-48559725-INS-511
chr18-48561549-INS-256
chr18-48603836-INS-81
chr18-48715581-INS-422
chr18-48736909-INS-196
chr18-48747185-INS-66
chr18-48758439-INS-99
chr18-48766370-INS-394
chr18-48783172-INS-54
chr18-48793878-INS-209
chr18-48799425-INS-847
chr18-48817647-INS-216
chr18-48830761-INS-126
chr18-48837033-INS-202
chr18-48850702-INS-7253
chr18-48867592-INS-56
chr18-48877368-INS-1954
chr18-48880896-INS-3787

chr18-48901766-INS-6947
chr18-48904744-INS-682
chr18-48936305-INS-79
chr18-48939864-INS-741
chr18-48944703-INS-72
chr18-48949936-INS-97
chr18-48952900-INS-58
chr18-48955558-INS-125
chr18-48958033-INS-399
chr18-49004553-INS-6802
chr18-49008738-INS-90
chr18-49034184-INS-127
chr18-49043177-INS-974
chr18-49053536-INS-65
chr18-49079244-INS-60
chr18-49085718-INS-198
chr18-49089339-INS-479
chr18-49130186-INS-53
chr18-49172523-INS-240
chr18-49197519-INS-246
chr18-49225515-INS-157
chr18-49226930-INS-230
chr18-49249511-INS-390
chr18-49250235-INS-90
chr18-49256843-INS-8023
chr18-49290237-INS-73
chr18-49298428-INS-805
chr18-49313856-INS-850
chr18-49315082-INS-250
chr18-49319046-INS-992
chr18-49321903-INS-68
chr18-49336834-INS-364
chr18-49342795-INS-62
chr18-49351197-INS-54
chr18-49353676-INS-284
chr18-49357043-INS-215
chr18-49377445-INS-558
chr18-49387834-INS-127
chr18-49405920-INS-206
chr18-49462280-INS-64
chr18-49471108-INS-6257
chr18-49473048-INS-6186
chr18-49476139-INS-61
chr18-49546315-INS-789
chr18-49547925-INS-268
chr18-49551142-INS-93
chr18-49569220-INS-544
chr18-49585759-INS-77
chr18-49595405-INS-5205
chr18-49632937-INS-57
chr18-49704631-INS-264
chr18-49713229-INS-979
chr18-49726372-INS-498
chr18-49751147-INS-87
chr18-49758678-INS-223
chr18-49760091-INS-6907
chr18-49770188-INS-429
chr18-49772895-INS-180
chr18-49775780-INS-1124
chr18-49793091-INS-170
chr18-49794087-INS-92
chr18-49811260-INS-56
chr18-49820311-INS-77
chr18-49821702-INS-6868
chr18-49843118-INS-194
chr18-49867624-INS-252
chr18-49869947-INS-195
chr18-49880371-INS-435
chr18-49882608-INS-208
chr18-49908819-INS-386
chr18-49912855-INS-533
chr18-49923913-INS-87
chr18-49929903-INS-401
chr18-49949758-INS-252
chr18-49950936-INS-228
chr18-49971797-INS-206
chr18-49976179-INS-208

chr18-49981027-INS-164
chr18-49985773-INS-266
chr18-50068218-INS-445
chr18-50087103-INS-52
chr18-50107153-INS-95
chr18-50116632-INS-203
chr18-50142493-INS-78
chr18-50144416-INS-260
chr18-50154470-INS-58
chr18-50177455-INS-88
chr18-50209067-INS-177
chr18-50215594-INS-533
chr18-50232866-INS-202
chr18-50253328-INS-106
chr18-50269376-INS-551
chr18-50276014-INS-476
chr18-50280065-INS-217
chr18-50283301-INS-200
chr18-50291959-INS-50
chr18-50293705-INS-379
chr18-50307225-INS-672
chr18-50323521-INS-116
chr18-50325117-INS-347
chr18-50342311-INS-54
chr18-50354394-INS-64
chr18-50357468-INS-64
chr18-50377442-INS-989
chr18-50379111-INS-213
chr18-50409826-INS-208
chr18-50423605-INS-333
chr18-50426922-INS-304
chr18-50468154-INS-542
chr18-50484447-INS-65
chr18-50485544-INS-102
chr18-50497172-INS-82
chr18-50498952-INS-65
chr18-50525435-INS-479
chr18-50534046-INS-55
chr18-50544397-INS-55
chr18-50546765-INS-498
chr18-50567284-INS-6239
chr18-50569491-INS-632
chr18-50591268-INS-156
chr18-50609230-INS-6387
chr18-50613555-INS-288
chr18-50618642-INS-132
chr18-50629280-INS-401
chr18-50630366-INS-303
chr18-50637080-INS-1660
chr18-50641294-INS-6616
chr18-50661956-INS-108
chr18-50689209-INS-132
chr18-50694316-INS-472
chr18-50718580-INS-6157
chr18-50740713-INS-474
chr18-50799618-INS-851
chr18-50839941-INS-360
chr18-50869904-INS-228
chr18-50886404-INS-433
chr18-50903578-INS-64
chr18-50904939-INS-6404
chr18-50908940-INS-659
chr18-50932661-INS-393
chr18-50935720-INS-82
chr18-50941836-INS-81
chr18-50943299-INS-69
chr18-50952891-INS-6382
chr18-50953062-INS-74
chr18-50955122-INS-180
chr18-50974288-INS-82
chr18-50990271-INS-67
chr18-51000868-INS-7207
chr18-51009537-INS-149
chr18-51020291-INS-680
chr18-51035617-INS-309
chr18-51085037-INS-290
chr18-51091183-INS-6393

chr18-51097317-INS-55
chr18-51152298-INS-76
chr18-51152837-INS-82
chr18-51155567-INS-813
chr18-51166769-INS-93
chr18-51168391-INS-98
chr18-51191178-INS-88
chr18-51197327-INS-104
chr18-51211144-INS-53
chr18-51214234-INS-80
chr18-51235082-INS-72
chr18-51242021-INS-448
chr18-51246369-INS-328
chr18-51267872-INS-88
chr18-51283295-INS-57
chr18-51290530-INS-78
chr18-51299057-INS-63
chr18-51313753-INS-723
chr18-51348893-INS-656
chr18-51350152-INS-914
chr18-51360637-INS-3987
chr18-51361703-INS-164
chr18-51369016-INS-65
chr18-51369591-INS-142
chr18-51370108-INS-197
chr18-51389718-INS-79
chr18-51402219-INS-115
chr18-51403194-INS-216
chr18-51419549-INS-175
chr18-51434391-INS-162
chr18-51443965-INS-216
chr18-51451817-INS-145
chr18-51485929-INS-65
chr18-51487785-INS-539
chr18-51516933-INS-126
chr18-51522316-INS-154
chr18-51574153-INS-120
chr18-51576003-INS-1968
chr18-51583733-INS-6442
chr18-51584471-INS-77
chr18-51590205-INS-88
chr18-51606674-INS-81
chr18-51628768-INS-53
chr18-51648787-INS-507
chr18-51654282-INS-486
chr18-51677239-INS-256
chr18-51678005-INS-76
chr18-51697292-INS-151
chr18-51701787-INS-614
chr18-51714469-INS-65
chr18-51730152-INS-904
chr18-51734172-INS-219
chr18-51739902-INS-562
chr18-51778473-INS-478
chr18-51780701-INS-439
chr18-51800393-INS-51
chr18-51803423-INS-95
chr18-51808292-INS-189
chr18-51823261-INS-64
chr18-51841712-INS-69
chr18-51851959-INS-85
chr18-51897780-INS-64
chr18-51914349-INS-803
chr18-51920701-INS-64
chr18-51931738-INS-1888
chr18-51959933-INS-72
chr18-51962157-INS-121
chr18-51967466-INS-130
chr18-51976612-INS-185
chr18-52014275-INS-530
chr18-52060887-INS-153
chr18-52064410-INS-189
chr18-52066573-INS-238
chr18-52073499-INS-288
chr18-52112682-INS-90
chr18-52125927-INS-296
chr18-52127900-INS-54

chr18-52135157-INS-156
chr18-52192382-INS-174
chr18-52228962-INS-8437
chr18-52245139-INS-60
chr18-52250325-INS-157
chr18-52305395-INS-351
chr18-52317629-INS-177
chr18-52325808-INS-199
chr18-52339292-INS-293
chr18-52358280-INS-361
chr18-52361583-INS-122
chr18-52378000-INS-431
chr18-52400672-INS-2630
chr18-52424998-INS-84
chr18-52460811-INS-571
chr18-52477619-INS-92
chr18-52483397-INS-54
chr18-52488881-INS-92
chr18-52506863-INS-101
chr18-52530651-INS-178
chr18-52562548-INS-228
chr18-52607059-INS-62
chr18-52615326-INS-400
chr18-52639074-INS-72
chr18-52640324-INS-396
chr18-52645142-INS-59
chr18-52657668-INS-623
chr18-52665037-INS-461
chr18-52675531-INS-64
chr18-52698896-INS-100
chr18-52700219-INS-52
chr18-52732667-INS-56
chr18-52736596-INS-799
chr18-52799692-INS-225
chr18-52814385-INS-413
chr18-52837222-INS-500
chr18-52847194-INS-60
chr18-52852898-INS-74
chr18-52857876-INS-58
chr18-52869429-INS-63
chr18-52871526-INS-332
chr18-52894273-INS-196
chr18-52925972-INS-74
chr18-52987786-INS-331
chr18-52996565-INS-66
chr18-53006478-INS-998
chr18-53009442-INS-212
chr18-53017196-INS-60
chr18-53034069-INS-68
chr18-53064417-INS-258
chr18-53096809-INS-138
chr18-53114021-INS-80
chr18-53140180-INS-225
chr18-53145650-INS-133
chr18-53153391-INS-215
chr18-53175117-INS-1502
chr18-53175118-INS-217
chr18-53181253-INS-6417
chr18-53184193-INS-6615
chr18-53187969-INS-201
chr18-53198790-INS-296
chr18-53207115-INS-64
chr18-53223244-INS-155
chr18-53227520-INS-3237
chr18-53235146-INS-6782
chr18-53246056-INS-96
chr18-53265806-INS-192
chr18-53277778-INS-144
chr18-53283148-INS-208
chr18-53302435-INS-83
chr18-53315750-INS-133
chr18-53331487-INS-508
chr18-53353982-INS-295
chr18-53355739-INS-203
chr18-53370585-INS-51
chr18-53388215-INS-216
chr18-53393041-INS-208

chr18-53417753-INS-106
chr18-53423804-INS-93
chr18-53427483-INS-588
chr18-53444216-INS-100
chr18-53465659-INS-828
chr18-53475859-INS-61
chr18-53488253-INS-280
chr18-53502105-INS-53
chr18-53559042-INS-101
chr18-53565916-INS-145
chr18-53572382-INS-77
chr18-53600621-INS-53
chr18-53615898-INS-79
chr18-53651417-INS-73
chr18-53655726-INS-397
chr18-53683486-INS-190
chr18-53688763-INS-52
chr18-53715197-INS-128
chr18-53722527-INS-126
chr18-53750434-INS-66
chr18-53753946-INS-329
chr18-53756401-INS-102
chr18-53774447-INS-52
chr18-53778797-INS-64
chr18-53780893-INS-54
chr18-53788646-INS-845
chr18-53798249-INS-164
chr18-53799922-INS-60
chr18-53804818-INS-52
chr18-53808864-INS-362
chr18-53834554-INS-940
chr18-53846974-INS-881
chr18-53920148-INS-233
chr18-53934891-INS-567
chr18-53937819-INS-78
chr18-53951080-INS-330
chr18-53954044-INS-243
chr18-53976294-INS-8417
chr18-54026805-INS-77
chr18-54028229-INS-422
chr18-54112623-INS-76
chr18-54152615-INS-56
chr18-54156901-INS-201
chr18-54180617-INS-201
chr18-54196761-INS-119
chr18-54219393-INS-79
chr18-54230898-INS-100
chr18-54236622-INS-62
chr18-54247879-INS-577
chr18-54270461-INS-7296
chr18-54274155-INS-72
chr18-54321941-INS-61
chr18-54336195-INS-481
chr18-54351887-INS-125
chr18-54354792-INS-6519
chr18-54430700-INS-112
chr18-54431328-INS-183
chr18-54475133-INS-5269
chr18-54493777-INS-52
chr18-54499317-INS-86
chr18-54530251-INS-66
chr18-54565711-INS-578
chr18-54576960-INS-56
chr18-54583599-INS-68
chr18-54628212-INS-62
chr18-54646688-INS-629
chr18-54649414-INS-959
chr18-54649487-INS-131
chr18-54664971-INS-51
chr18-54667456-INS-252
chr18-54677709-INS-104
chr18-54714636-INS-60
chr18-54728204-INS-62
chr18-54749854-INS-617
chr18-54773333-INS-112
chr18-54776446-INS-84
chr18-54777643-INS-533

chr18-54807409-INS-82
chr18-54816931-INS-175
chr18-54829169-INS-68
chr18-54832944-INS-52
chr18-54843830-INS-67
chr18-54879008-INS-74
chr18-54894928-INS-79
chr18-54953324-INS-103
chr18-54969786-INS-57
chr18-54985240-INS-57
chr18-54986523-INS-9015
chr18-55002415-INS-58
chr18-55033009-INS-55
chr18-55047668-INS-205
chr18-55089364-INS-172
chr18-55097093-INS-186
chr18-55113851-INS-82
chr18-55118134-INS-56
chr18-55132933-INS-72
chr18-55164471-INS-684
chr18-55170102-INS-90
chr18-55177641-INS-51
chr18-55219934-INS-52
chr18-55225050-INS-198
chr18-55227181-INS-114
chr18-55232621-INS-62
chr18-55260821-INS-105
chr18-55270107-INS-51
chr18-55278999-INS-236
chr18-55292873-INS-64
chr18-55305481-INS-74
chr18-55322425-INS-60
chr18-55342000-INS-380
chr18-55379182-INS-64
chr18-55392955-INS-52
chr18-55400248-INS-175
chr18-55402217-INS-678
chr18-55403336-INS-152
chr18-55410672-INS-166
chr18-55458994-INS-164
chr18-55465957-INS-111
chr18-55466630-INS-6813
chr18-55504675-INS-633
chr18-55511324-INS-51
chr18-55598056-INS-366
chr18-55616399-INS-55
chr18-55656476-INS-70
chr18-55660275-INS-327
chr18-55660997-INS-56
chr18-55687921-INS-58
chr18-55704356-INS-176
chr18-55720027-INS-568
chr18-55784437-INS-62
chr18-55786839-INS-53
chr18-55808700-INS-148
chr18-55820288-INS-52
chr18-55821166-INS-330
chr18-55840459-INS-416
chr18-55869306-INS-71
chr18-55870776-INS-329
chr18-55878021-INS-76
chr18-55924495-INS-65
chr18-55954709-INS-7606
chr18-55967575-INS-120
chr18-56019491-INS-400
chr18-56030195-INS-7506
chr18-56039755-INS-64
chr18-56044048-INS-204
chr18-56067261-INS-315
chr18-56092510-INS-7362
chr18-56096017-INS-1285
chr18-56098969-INS-400
chr18-56124881-INS-770
chr18-56159539-INS-116
chr18-56174995-INS-158
chr18-56178657-INS-479
chr18-56187660-INS-492

chr18-56211622-INS-70
chr18-56270793-INS-93
chr18-56275940-INS-70
chr18-56287634-INS-58
chr18-56299374-INS-114
chr18-56306229-INS-280
chr18-56313488-INS-148
chr18-56319515-INS-364
chr18-56321745-INS-50
chr18-56330669-INS-86
chr18-56342668-INS-6449
chr18-56342877-INS-86
chr18-56375770-INS-150
chr18-56386204-INS-51
chr18-56396210-INS-57
chr18-56420047-INS-160
chr18-56430907-INS-60
chr18-56453971-INS-93
chr18-56486247-INS-85
chr18-56494360-INS-61
chr18-56513531-INS-87
chr18-56518032-INS-75
chr18-56520113-INS-96
chr18-56546550-INS-498
chr18-56576640-INS-200
chr18-56587934-INS-803
chr18-56605428-INS-92
chr18-56641841-INS-84
chr18-56648805-INS-152
chr18-56663866-INS-109
chr18-56685558-INS-167
chr18-56696820-INS-187
chr18-56716367-INS-203
chr18-56747429-INS-118
chr18-56749749-INS-81
chr18-56751043-INS-153
chr18-56753071-INS-153
chr18-56761175-INS-64
chr18-56761650-INS-396
chr18-56766771-INS-196
chr18-56771111-INS-81
chr18-56780301-INS-91
chr18-56788731-INS-399
chr18-56791957-INS-60
chr18-56798889-INS-206
chr18-56814696-INS-53
chr18-56819432-INS-69
chr18-56822484-INS-208
chr18-56824190-INS-220
chr18-56834778-INS-133
chr18-56836731-INS-375
chr18-56849834-INS-183
chr18-56855188-INS-220
chr18-56857391-INS-196
chr18-56866581-INS-198
chr18-56874360-INS-62
chr18-56882261-INS-249
chr18-56897330-INS-6667
chr18-56897331-INS-7305
chr18-56956788-INS-504
chr18-56991273-INS-156
chr18-57011317-INS-60
chr18-57027379-INS-51
chr18-57032495-INS-94
chr18-57038513-INS-56
chr18-57051924-INS-56
chr18-57059636-INS-82
chr18-57071445-INS-155
chr18-57074967-INS-94
chr18-57077429-INS-6981
chr18-57089769-INS-733
chr18-57114914-INS-53
chr18-57117103-INS-6644
chr18-57174643-INS-688
chr18-57181326-INS-172
chr18-57184046-INS-56
chr18-57192270-INS-64

chr18-57202006-INS-141
chr18-57209130-INS-53
chr18-57218600-INS-70
chr18-57223164-INS-52
chr18-57229282-INS-54
chr18-57233494-INS-174
chr18-57241788-INS-200
chr18-57273005-INS-70
chr18-57279128-INS-5221
chr18-57299555-INS-186
chr18-57359290-INS-85
chr18-57361215-INS-232
chr18-57377298-INS-192
chr18-57421693-INS-63
chr18-57452622-INS-76
chr18-57483111-INS-497
chr18-57491673-INS-71
chr18-57496873-INS-54
chr18-57497825-INS-120
chr18-57517645-INS-71
chr18-57519009-INS-94
chr18-57528102-INS-72
chr18-57529221-INS-81
chr18-57552690-INS-234
chr18-57565665-INS-66
chr18-57573343-INS-423
chr18-57576418-INS-306
chr18-57586570-INS-66
chr18-57607062-INS-51
chr18-57611527-INS-342
chr18-57618308-INS-64
chr18-57705223-INS-57
chr18-57759869-INS-71
chr18-57782791-INS-59
chr18-57806226-INS-87
chr18-57810656-INS-435
chr18-57811817-INS-685
chr18-57812666-INS-76
chr18-57817935-INS-262
chr18-57818451-INS-1178
chr18-57818608-INS-172
chr18-57842635-INS-55
chr18-57851979-INS-114
chr18-57857334-INS-400
chr18-57862550-INS-88
chr18-57863588-INS-111
chr18-57876575-INS-737
chr18-57878638-INS-186
chr18-57890371-INS-52
chr18-57954556-INS-60
chr18-57972699-INS-3183
chr18-57985682-INS-114
chr18-58001744-INS-140
chr18-58014825-INS-263
chr18-58029011-INS-817
chr18-58088806-INS-86
chr18-58091192-INS-307
chr18-58096437-INS-54
chr18-58104059-INS-80
chr18-58130997-INS-60
chr18-58157914-INS-85
chr18-58204041-INS-948
chr18-58217402-INS-84
chr18-58221954-INS-208
chr18-58241723-INS-160
chr18-58270998-INS-199
chr18-58323589-INS-52
chr18-58336618-INS-68
chr18-58340371-INS-132
chr18-58346083-INS-423
chr18-58359575-INS-171
chr18-58367587-INS-77
chr18-58371738-INS-405
chr18-58387407-INS-198
chr18-58415564-INS-171
chr18-58417170-INS-60
chr18-58422556-INS-564

chr18-58423942-INS-4418
chr18-58431919-INS-58
chr18-58442509-INS-171
chr18-58474054-INS-68
chr18-58486433-INS-74
chr18-58495349-INS-338
chr18-58498172-INS-400
chr18-58507628-INS-77
chr18-58510881-INS-474
chr18-58523186-INS-140
chr18-58534643-INS-103
chr18-58540333-INS-6443
chr18-58572741-INS-1092
chr18-58576685-INS-52
chr18-58592954-INS-52
chr18-58611864-INS-109
chr18-58619744-INS-286
chr18-58653229-INS-61
chr18-58682817-INS-134
chr18-58723349-INS-69
chr18-58732065-INS-55
chr18-58735132-INS-96
chr18-58789737-INS-56
chr18-58800090-INS-66
chr18-58814736-INS-101
chr18-58826308-INS-498
chr18-58828350-INS-63
chr18-58860191-INS-63
chr18-58871655-INS-190
chr18-58897603-INS-94
chr18-58915953-INS-162
chr18-58923618-INS-214
chr18-58940343-INS-454
chr18-58957960-INS-56
chr18-58995338-INS-203
chr18-59007430-INS-3710
chr18-59009773-INS-72
chr18-59037625-INS-122
chr18-59038892-INS-278
chr18-59043836-INS-187
chr18-59048426-INS-289
chr18-59090970-INS-59
chr18-59096226-INS-261
chr18-59119182-INS-194
chr18-59134362-INS-60
chr18-59136466-INS-54
chr18-59183112-INS-978
chr18-59200101-INS-84
chr18-59207087-INS-370
chr18-59215031-INS-806
chr18-59216184-INS-411
chr18-59242309-INS-82
chr18-59283191-INS-60
chr18-59289784-INS-336
chr18-59323669-INS-85
chr18-59337135-INS-56
chr18-59373905-INS-286
chr18-59386898-INS-52
chr18-59434362-INS-6486
chr18-59435166-INS-2408
chr18-59436559-INS-579
chr18-59436943-INS-5233
chr18-59458963-INS-1835
chr18-59494342-INS-648
chr18-59508925-INS-650
chr18-59509454-INS-120
chr18-59531500-INS-66
chr18-59532743-INS-400
chr18-59535487-INS-217
chr18-59539972-INS-774
chr18-59541121-INS-51
chr18-59558853-INS-101
chr18-59571044-INS-291
chr18-59586276-INS-8442
chr18-59602800-INS-376
chr18-59617338-INS-74
chr18-59637744-INS-201

chr18-59645739-INS-144
chr18-59655105-INS-179
chr18-59668816-INS-104
chr18-59672627-INS-6870
chr18-59697160-INS-240
chr18-59702578-INS-7023
chr18-59717413-INS-417
chr18-59735153-INS-797
chr18-59746454-INS-56
chr18-59755156-INS-55
chr18-59758584-INS-1802
chr18-59768124-INS-191
chr18-59777838-INS-101
chr18-59777838-INS-112
chr18-59779895-INS-51
chr18-59842880-INS-57
chr18-59876979-INS-238
chr18-59883572-INS-52
chr18-59924725-INS-59
chr18-59942075-INS-58
chr18-59980310-INS-96
chr18-60004935-INS-125
chr18-60007273-INS-708
chr18-60008908-INS-127
chr18-60032421-INS-410
chr18-60034405-INS-729
chr18-60034415-INS-1064
chr18-60048536-INS-54
chr18-60060821-INS-56
chr18-60090307-INS-207
chr18-60095925-INS-124
chr18-60129788-INS-593
chr18-60136725-INS-357
chr18-60154803-INS-70
chr18-60158203-INS-230
chr18-60160412-INS-648
chr18-60167431-INS-339
chr18-60219451-INS-166
chr18-60228992-INS-209
chr18-60230024-INS-358
chr18-60233239-INS-6471
chr18-60239827-INS-91
chr18-60272803-INS-6288
chr18-60298733-INS-400
chr18-60307138-INS-454
chr18-60351052-INS-125
chr18-60375910-INS-452
chr18-60383074-INS-194
chr18-60394940-INS-450
chr18-60418873-INS-162
chr18-60427930-INS-166
chr18-60453597-INS-331
chr18-60532260-INS-8324
chr18-60572639-INS-1890
chr18-60573186-INS-90
chr18-60574197-INS-106
chr18-60593531-INS-87
chr18-60623612-INS-235
chr18-60721684-INS-300
chr18-60758918-INS-52
chr18-60766451-INS-208
chr18-60778701-INS-51
chr18-60796613-INS-344
chr18-60867338-INS-253
chr18-60875402-INS-94
chr18-60877600-INS-189
chr18-60911805-INS-1851
chr18-60923200-INS-111
chr18-60923622-INS-637
chr18-60934381-INS-74
chr18-60940102-INS-1238
chr18-61001676-INS-2085
chr18-61069187-INS-120
chr18-61088106-INS-103
chr18-61095655-INS-168
chr18-61163796-INS-54
chr18-61165098-INS-82

chr18-61207226-INS-55
chr18-61262692-INS-293
chr18-61267272-INS-190
chr18-61275988-INS-62
chr18-61287801-INS-247
chr18-61315494-INS-220
chr18-61318378-INS-120
chr18-61337954-INS-367
chr18-61364214-INS-340
chr18-61411893-INS-433
chr18-61453571-INS-66
chr18-61463084-INS-54
chr18-61466916-INS-592
chr18-61489689-INS-96
chr18-61516059-INS-90
chr18-61548038-INS-284
chr18-61552061-INS-64
chr18-61562028-INS-72
chr18-61591400-INS-74
chr18-61595682-INS-84
chr18-61597409-INS-66
chr18-61602756-INS-59
chr18-61616241-INS-97
chr18-61652688-INS-203
chr18-61728885-INS-113
chr18-61732123-INS-60
chr18-61732956-INS-52
chr18-61740110-INS-232
chr18-61746176-INS-292
chr18-61752776-INS-455
chr18-61767406-INS-6473
chr18-61795971-INS-86
chr18-61801553-INS-60
chr18-61833824-INS-212
chr18-61842073-INS-59
chr18-61894042-INS-52
chr18-61897972-INS-64
chr18-61907042-INS-91
chr18-61907640-INS-192
chr18-61927891-INS-52
chr18-61947826-INS-1888
chr18-61953275-INS-53
chr18-61997334-INS-96
chr18-62005381-INS-775
chr18-62053412-INS-133
chr18-62059604-INS-146
chr18-62060725-INS-7192
chr18-62061903-INS-267
chr18-62077461-INS-102
chr18-62078960-INS-334
chr18-62081319-INS-1176
chr18-62081320-INS-798
chr18-62113742-INS-342
chr18-62122039-INS-77
chr18-62126126-INS-161
chr18-62153782-INS-1131
chr18-62173086-INS-54
chr18-62193722-INS-127
chr18-62219933-INS-79
chr18-62226189-INS-78
chr18-62284278-INS-669
chr18-62292852-INS-59
chr18-62294108-INS-55
chr18-62301454-INS-57
chr18-62361161-INS-82
chr18-62381444-INS-4134
chr18-62394247-INS-227
chr18-62406187-INS-203
chr18-62406363-INS-649
chr18-62440651-INS-233
chr18-62454297-INS-326
chr18-62470018-INS-179
chr18-62476699-INS-52
chr18-62525563-INS-91
chr18-62547452-INS-62
chr18-62559001-INS-3030
chr18-62565676-INS-548

chr18-62576292-INS-62
chr18-62601941-INS-728
chr18-62630115-INS-58
chr18-62631123-INS-75
chr18-62638214-INS-281
chr18-62647310-INS-102
chr18-62650412-INS-824
chr18-62653403-INS-94
chr18-62690140-INS-2841
chr18-62694637-INS-440
chr18-62705210-INS-104
chr18-62714057-INS-120
chr18-62730637-INS-262
chr18-62739275-INS-106
chr18-62769258-INS-65
chr18-62780878-INS-5549
chr18-62799917-INS-56
chr18-62800928-INS-672
chr18-62819242-INS-60
chr18-62845275-INS-72
chr18-62871425-INS-271
chr18-62872448-INS-89
chr18-62875367-INS-79
chr18-62887175-INS-448
chr18-62891743-INS-69
chr18-62895130-INS-73
chr18-62913150-INS-198
chr18-62914861-INS-59
chr18-62916716-INS-73
chr18-62926844-INS-325
chr18-62933647-INS-88
chr18-62978026-INS-60
chr18-63003631-INS-606
chr18-63008805-INS-52
chr18-63028132-INS-128
chr18-63030681-INS-240
chr18-63032175-INS-60
chr18-63099550-INS-82
chr18-63106119-INS-60
chr18-63107920-INS-65
chr18-63108902-INS-324
chr18-63117515-INS-87
chr18-63140529-INS-218
chr18-63158100-INS-150
chr18-63162720-INS-195
chr18-63176777-INS-56
chr18-63216583-INS-70
chr18-63240867-INS-272
chr18-63245021-INS-192
chr18-63249347-INS-128
chr18-63252636-INS-65
chr18-63262333-INS-58
chr18-63268209-INS-200
chr18-63270778-INS-70
chr18-63319471-INS-74
chr18-63321936-INS-77
chr18-63346950-INS-69
chr18-63353806-INS-286
chr18-63382517-INS-592
chr18-63388161-INS-126
chr18-63406393-INS-52
chr18-63410652-INS-198
chr18-63421179-INS-70
chr18-63449341-INS-122
chr18-63462602-INS-107
chr18-63505369-INS-591
chr18-63512527-INS-234
chr18-63526497-INS-72
chr18-63529067-INS-399
chr18-63537085-INS-163
chr18-63545132-INS-56
chr18-63553320-INS-139
chr18-63561598-INS-176
chr18-63595418-INS-58
chr18-63600605-INS-864
chr18-63674229-INS-415
chr18-63717136-INS-175

chr18-63743581-INS-400
chr18-63751341-INS-184
chr18-63766397-INS-313
chr18-63774437-INS-807
chr18-63777003-INS-84
chr18-63802129-INS-179
chr18-63808066-INS-750
chr18-63819975-INS-71
chr18-63831211-INS-231
chr18-63845600-INS-119
chr18-63854679-INS-268
chr18-63878894-INS-52
chr18-63933913-INS-4442
chr18-63937936-INS-72
chr18-63955253-INS-91
chr18-63958446-INS-88
chr18-63962273-INS-125
chr18-63999579-INS-194
chr18-64011699-INS-86
chr18-64029825-INS-207
chr18-64033695-INS-673
chr18-64039842-INS-53
chr18-64051488-INS-54
chr18-64096391-INS-228
chr18-64149097-INS-75
chr18-64156260-INS-54
chr18-64243455-INS-142
chr18-64253350-INS-118
chr18-64281317-INS-1175
chr18-64348497-INS-80
chr18-64350265-INS-509
chr18-64352020-INS-91
chr18-64362938-INS-60
chr18-64386539-INS-553
chr18-64393809-INS-293
chr18-64395446-INS-58
chr18-64447401-INS-68
chr18-64448131-INS-876
chr18-64454780-INS-80
chr18-64481466-INS-102
chr18-64524615-INS-78
chr18-64528851-INS-400
chr18-64544703-INS-146
chr18-64609319-INS-169
chr18-64617024-INS-94
chr18-64630274-INS-84
chr18-64636268-INS-203
chr18-64639244-INS-84
chr18-64650900-INS-231
chr18-64655377-INS-64
chr18-64669182-INS-84
chr18-64686515-INS-206
chr18-64687657-INS-382
chr18-64690348-INS-125
chr18-64703359-INS-745
chr18-64730288-INS-507
chr18-64750793-INS-86
chr18-64756936-INS-91
chr18-64760044-INS-68
chr18-64807862-INS-54
chr18-64810065-INS-262
chr18-64827265-INS-57
chr18-64830347-INS-52
chr18-64892015-INS-400
chr18-64893075-INS-203
chr18-64925251-INS-57
chr18-64934786-INS-104
chr18-64942156-INS-204
chr18-64947114-INS-1656
chr18-64949178-INS-199
chr18-64955962-INS-60
chr18-64986473-INS-398
chr18-64986646-INS-1875
chr18-64990012-INS-181
chr18-64995243-INS-221
chr18-65001421-INS-909
chr18-65032607-INS-79

chr18-65038022-INS-70
chr18-65040336-INS-207
chr18-65043195-INS-64
chr18-65048563-INS-108
chr18-65099490-INS-118
chr18-65116355-INS-181
chr18-65120888-INS-74
chr18-65131891-INS-116
chr18-65142939-INS-312
chr18-65170246-INS-69
chr18-65182992-INS-95
chr18-65201543-INS-97
chr18-65208415-INS-140
chr18-65233436-INS-269
chr18-65258981-INS-463
chr18-65272667-INS-179
chr18-65355472-INS-206
chr18-65380435-INS-461
chr18-65381496-INS-90
chr18-65383222-INS-53
chr18-65386058-INS-195
chr18-65406041-INS-259
chr18-65415304-INS-456
chr18-65470705-INS-76
chr18-65476643-INS-68
chr18-65544888-INS-308
chr18-65549304-INS-223
chr18-65551154-INS-286
chr18-65558393-INS-60
chr18-65628543-INS-199
chr18-65629589-INS-69
chr18-65647569-INS-255
chr18-65647951-INS-287
chr18-65668627-INS-331
chr18-65672292-INS-534
chr18-65677313-INS-63
chr18-65702599-INS-206
chr18-65719758-INS-207
chr18-65755304-INS-93
chr18-65755801-INS-188
chr18-65765576-INS-324
chr18-65799323-INS-174
chr18-65811906-INS-399
chr18-65827366-INS-5565
chr18-65856480-INS-235
chr18-65894659-INS-56
chr18-65906896-INS-75
chr18-65916126-INS-1099
chr18-65962619-INS-190
chr18-65986913-INS-88
chr18-65992478-INS-994
chr18-66002045-INS-814
chr18-66016063-INS-82
chr18-66055051-INS-53
chr18-66061685-INS-51
chr18-66119486-INS-150
chr18-66139049-INS-454
chr18-66144215-INS-75
chr18-66153803-INS-51
chr18-66168993-INS-72
chr18-66176969-INS-6592
chr18-66216077-INS-64
chr18-66224564-INS-121
chr18-66242786-INS-279
chr18-66274778-INS-66
chr18-66289469-INS-334
chr18-66351581-INS-61
chr18-66409539-INS-86
chr18-66429236-INS-64
chr18-66439911-INS-6704
chr18-66464397-INS-168
chr18-66475622-INS-56
chr18-66487772-INS-246
chr18-66496973-INS-84
chr18-66498674-INS-114
chr18-66501852-INS-281
chr18-66505598-INS-180

chr18-66518428-INS-207
chr18-66526681-INS-4397
chr18-66546688-INS-142
chr18-66586091-INS-68
chr18-66624655-INS-691
chr18-66644103-INS-203
chr18-66645246-INS-715
chr18-66646107-INS-52
chr18-66648489-INS-92
chr18-66695886-INS-116
chr18-66697305-INS-110
chr18-66702181-INS-330
chr18-66719950-INS-250
chr18-66726543-INS-251
chr18-66729065-INS-103
chr18-66747603-INS-118
chr18-66777044-INS-86
chr18-66805156-INS-419
chr18-66809727-INS-167
chr18-66815411-INS-6793
chr18-66822995-INS-395
chr18-66833769-INS-80
chr18-66842878-INS-76
chr18-66850916-INS-170
chr18-66853021-INS-540
chr18-66854760-INS-60
chr18-66856831-INS-542
chr18-66859601-INS-54
chr18-66860118-INS-6111
chr18-66892570-INS-108
chr18-66911752-INS-348
chr18-66912480-INS-69
chr18-66916954-INS-168
chr18-66918281-INS-465
chr18-66924847-INS-7278
chr18-66927810-INS-92
chr18-66968068-INS-358
chr18-67039109-INS-92
chr18-67045280-INS-58
chr18-67090709-INS-109
chr18-67100025-INS-331
chr18-67104575-INS-948
chr18-67107306-INS-60
chr18-67136016-INS-71
chr18-67137544-INS-153
chr18-67166829-INS-60
chr18-67181368-INS-163
chr18-67185643-INS-82
chr18-67207866-INS-236
chr18-67231074-INS-179
chr18-67261993-INS-150
chr18-67271698-INS-139
chr18-67280928-INS-169
chr18-67313949-INS-211
chr18-67317490-INS-60
chr18-67333068-INS-227
chr18-67355015-INS-74
chr18-67386622-INS-109
chr18-67390137-INS-66
chr18-67391287-INS-504
chr18-67404984-INS-216
chr18-67407921-INS-57
chr18-67411135-INS-207
chr18-67414754-INS-384
chr18-67438532-INS-168
chr18-67460172-INS-228
chr18-67468222-INS-109
chr18-67503631-INS-110
chr18-67517012-INS-66
chr18-67518478-INS-5476
chr18-67527027-INS-210
chr18-67531832-INS-187
chr18-67541434-INS-197
chr18-67554863-INS-74
chr18-67575038-INS-635
chr18-67577937-INS-88
chr18-67591901-INS-205

chr18-67592998-INS-380
chr18-67600124-INS-552
chr18-67614260-INS-193
chr18-67620965-INS-73
chr18-67629205-INS-381
chr18-67631719-INS-55
chr18-67633640-INS-57
chr18-67649199-INS-208
chr18-67652659-INS-106
chr18-67653572-INS-64
chr18-67666171-INS-502
chr18-67684552-INS-211
chr18-67693801-INS-225
chr18-67695085-INS-504
chr18-67696050-INS-71
chr18-67699098-INS-56
chr18-67701792-INS-242
chr18-67712995-INS-103
chr18-67714377-INS-6440
chr18-67720489-INS-145
chr18-67727089-INS-202
chr18-67730934-INS-229
chr18-67732980-INS-202
chr18-67746041-INS-212
chr18-67753688-INS-399
chr18-67756562-INS-231
chr18-67760566-INS-76
chr18-67767061-INS-655
chr18-67768175-INS-119
chr18-67779946-INS-126
chr18-67788296-INS-361
chr18-67793074-INS-201
chr18-67799577-INS-298
chr18-67817946-INS-1891
chr18-67838911-INS-200
chr18-67842897-INS-169
chr18-67844896-INS-272
chr18-67845462-INS-209
chr18-67846649-INS-198
chr18-67849545-INS-84
chr18-67853450-INS-772
chr18-67869812-INS-54
chr18-67870312-INS-192
chr18-67872926-INS-182
chr18-67878415-INS-52
chr18-67888126-INS-54
chr18-67901891-INS-170
chr18-67904769-INS-205
chr18-67922653-INS-175
chr18-67929574-INS-220
chr18-67946370-INS-330
chr18-67949163-INS-118
chr18-67951830-INS-64
chr18-67954186-INS-52
chr18-67959752-INS-187
chr18-67970159-INS-152
chr18-67988109-INS-52
chr18-68004253-INS-91
chr18-68022834-INS-206
chr18-68023588-INS-53
chr18-68033881-INS-557
chr18-68042183-INS-52
chr18-68053289-INS-378
chr18-68099450-INS-120
chr18-68193008-INS-348
chr18-68205175-INS-200
chr18-68248053-INS-484
chr18-68275170-INS-230
chr18-68299845-INS-288
chr18-68307215-INS-199
chr18-68325338-INS-58
chr18-68340345-INS-86
chr18-68365705-INS-51
chr18-68377145-INS-925
chr18-68378349-INS-208
chr18-68418299-INS-422
chr18-68448063-INS-300

chr18-68466840-INS-51
chr18-68472874-INS-531
chr18-68511541-INS-70
chr18-68512960-INS-404
chr18-68521853-INS-89
chr18-68554607-INS-486
chr18-68564752-INS-188
chr18-68566007-INS-121
chr18-68567060-INS-240
chr18-68579269-INS-208
chr18-68583320-INS-195
chr18-68626070-INS-65
chr18-68653733-INS-129
chr18-68665139-INS-74
chr18-68669300-INS-460
chr18-68677837-INS-69
chr18-68680424-INS-66
chr18-68687170-INS-82
chr18-68696290-INS-1818
chr18-68752798-INS-60
chr18-68753858-INS-96
chr18-68757064-INS-7097
chr18-68773031-INS-186
chr18-68800306-INS-78
chr18-68802616-INS-95
chr18-68817082-INS-851
chr18-68832596-INS-78
chr18-68854930-INS-8419
chr18-68862851-INS-81
chr18-68897103-INS-288
chr18-68941663-INS-65
chr18-68944004-INS-82
chr18-68953047-INS-51
chr18-68971418-INS-60
chr18-69018655-INS-52
chr18-69020008-INS-177
chr18-69027000-INS-222
chr18-69028931-INS-547
chr18-69030317-INS-120
chr18-69032941-INS-132
chr18-69043473-INS-86
chr18-69102256-INS-52
chr18-69112400-INS-66
chr18-69125032-INS-304
chr18-69152651-INS-52
chr18-69165881-INS-66
chr18-69175444-INS-462
chr18-69204244-INS-112
chr18-69269027-INS-80
chr18-69307787-INS-52
chr18-69324328-INS-127
chr18-69340618-INS-203
chr18-69354178-INS-93
chr18-69362012-INS-145
chr18-69363594-INS-166
chr18-69394406-INS-552
chr18-69414621-INS-146
chr18-69421261-INS-112
chr18-69438190-INS-544
chr18-69484240-INS-64
chr18-69485453-INS-67
chr18-69492829-INS-101
chr18-69516177-INS-202
chr18-69596829-INS-90
chr18-69602580-INS-117
chr18-69614133-INS-76
chr18-69617005-INS-51
chr18-69635925-INS-69
chr18-69644626-INS-264
chr18-69656717-INS-58
chr18-69830935-INS-953
chr18-69858671-INS-56
chr18-69877765-INS-181
chr18-69912275-INS-184
chr18-69935713-INS-52
chr18-69936540-INS-212
chr18-69948858-INS-7263

chr18-70012886-INS-6699
chr18-70020776-INS-7429
chr18-70027314-INS-101
chr18-70043219-INS-6442
chr18-70049586-INS-115
chr18-70076535-INS-56
chr18-70087197-INS-330
chr18-70103862-INS-200
chr18-70104352-INS-377
chr18-70106616-INS-429
chr18-70121085-INS-7318
chr18-70127404-INS-65
chr18-70134663-INS-200
chr18-70147590-INS-137
chr18-70149708-INS-332
chr18-70151186-INS-61
chr18-70190648-INS-152
chr18-70218101-INS-86
chr18-70332570-INS-90
chr18-70335587-INS-422
chr18-70348936-INS-78
chr18-70362315-INS-325
chr18-70363263-INS-366
chr18-70376042-INS-116
chr18-70389292-INS-142
chr18-70400654-INS-369
chr18-70436034-INS-907
chr18-70478113-INS-60
chr18-70484096-INS-52
chr18-70511249-INS-56
chr18-70517886-INS-5128
chr18-70518941-INS-107
chr18-70593168-INS-208
chr18-70604563-INS-665
chr18-70615038-INS-71
chr18-70641034-INS-919
chr18-70656579-INS-186
chr18-70663594-INS-54
chr18-70666722-INS-127
chr18-70671224-INS-55
chr18-70723844-INS-165
chr18-70740850-INS-116
chr18-70764320-INS-322
chr18-70777012-INS-130
chr18-70792694-INS-124
chr18-70795884-INS-88
chr18-70796709-INS-138
chr18-70890307-INS-559
chr18-70891304-INS-461
chr18-70893737-INS-526
chr18-70895954-INS-296
chr18-70905503-INS-443
chr18-70905963-INS-6168
chr18-70907616-INS-230
chr18-70918328-INS-418
chr18-70952348-INS-362
chr18-70965808-INS-714
chr18-71001336-INS-53
chr18-71025446-INS-56
chr18-71035915-INS-340
chr18-71038315-INS-126
chr18-71043855-INS-74
chr18-71053062-INS-6856
chr18-71062545-INS-167
chr18-71071777-INS-668
chr18-71072432-INS-88
chr18-71117884-INS-61
chr18-71126358-INS-541
chr18-71128495-INS-53
chr18-71135734-INS-257
chr18-71140659-INS-217
chr18-71172284-INS-139
chr18-71173742-INS-6992
chr18-71241826-INS-6601
chr18-71253658-INS-113
chr18-71286716-INS-2930
chr18-71295495-INS-52

chr18-71305825-INS-2868
chr18-71318099-INS-498
chr18-71348261-INS-60
chr18-71381657-INS-136
chr18-71411252-INS-6221
chr18-71425011-INS-176
chr18-71442462-INS-138
chr18-71444145-INS-269
chr18-71447598-INS-398
chr18-71484236-INS-122
chr18-71485756-INS-82
chr18-71513413-INS-182
chr18-71518867-INS-6599
chr18-71523692-INS-64
chr18-71525533-INS-702
chr18-71550567-INS-224
chr18-71572707-INS-6697
chr18-71591794-INS-51
chr18-71608993-INS-568
chr18-71614148-INS-382
chr18-71633552-INS-58
chr18-71647642-INS-125
chr18-71652738-INS-7116
chr18-71662829-INS-52
chr18-71678309-INS-6384
chr18-71691897-INS-140
chr18-71695166-INS-85
chr18-71707281-INS-75
chr18-71714176-INS-53
chr18-71737002-INS-62
chr18-71744918-INS-58
chr18-71789080-INS-54
chr18-71792561-INS-205
chr18-71798611-INS-540
chr18-71811891-INS-102
chr18-71829692-INS-95
chr18-71841389-INS-417
chr18-71881315-INS-839
chr18-71892209-INS-88
chr18-71898245-INS-70
chr18-71899593-INS-186
chr18-71917948-INS-166
chr18-71963672-INS-780
chr18-71977576-INS-183
chr18-71985108-INS-306
chr18-71997415-INS-6845
chr18-72006827-INS-112
chr18-72050650-INS-54
chr18-72071952-INS-53
chr18-72082591-INS-72
chr18-72101370-INS-122
chr18-72128368-INS-55
chr18-72130133-INS-66
chr18-72136447-INS-280
chr18-72189009-INS-61
chr18-72191969-INS-482
chr18-72197026-INS-52
chr18-72218321-INS-721
chr18-72260378-INS-56
chr18-72268187-INS-608
chr18-72287079-INS-58
chr18-72288058-INS-4232
chr18-72297118-INS-60
chr18-72305999-INS-54
chr18-72316736-INS-2072
chr18-72319056-INS-177
chr18-72324155-INS-53
chr18-72345054-INS-236
chr18-72357789-INS-51
chr18-72366586-INS-420
chr18-72379157-INS-1420
chr18-72428402-INS-71
chr18-72430483-INS-79
chr18-72442140-INS-5676
chr18-72459132-INS-1455
chr18-72466149-INS-53
chr18-72523817-INS-59

chr18-72530183-INS-229
chr18-72551910-INS-76
chr18-72576112-INS-205
chr18-72585206-INS-50
chr18-72604230-INS-74
chr18-72613951-INS-506
chr18-72617168-INS-689
chr18-72639090-INS-86
chr18-72680666-INS-215
chr18-72682222-INS-521
chr18-72684016-INS-82
chr18-72705748-INS-740
chr18-72718663-INS-72
chr18-72782674-INS-6871
chr18-72793850-INS-124
chr18-72822377-INS-67
chr18-72854980-INS-66
chr18-72859769-INS-4839
chr18-72875077-INS-157
chr18-72878719-INS-72
chr18-72897928-INS-346
chr18-72972213-INS-133
chr18-72979225-INS-167
chr18-73022641-INS-1394
chr18-73023026-INS-321
chr18-73028519-INS-58
chr18-73032079-INS-79
chr18-73037285-INS-114
chr18-73046530-INS-60
chr18-73052919-INS-137
chr18-73054286-INS-243
chr18-73082788-INS-5343
chr18-73085760-INS-4000
chr18-73086049-INS-278
chr18-73087220-INS-392
chr18-73120988-INS-297
chr18-73124392-INS-158
chr18-73144092-INS-52
chr18-73156334-INS-74
chr18-73168306-INS-420
chr18-73171123-INS-1443
chr18-73180034-INS-51
chr18-73180996-INS-61
chr18-73191735-INS-768
chr18-73193752-INS-51
chr18-73225780-INS-5975
chr18-73233949-INS-51
chr18-73239999-INS-7839
chr18-73240835-INS-6504
chr18-73242724-INS-62
chr18-73243567-INS-212
chr18-73251356-INS-57
chr18-73264386-INS-172
chr18-73272846-INS-188
chr18-73287163-INS-52
chr18-73308294-INS-362
chr18-73327844-INS-677
chr18-73331332-INS-55
chr18-73345055-INS-337
chr18-73355588-INS-573
chr18-73360073-INS-508
chr18-73361086-INS-4832
chr18-73370934-INS-56
chr18-73376135-INS-6222
chr18-73377767-INS-60
chr18-73390135-INS-775
chr18-73399701-INS-158
chr18-73410073-INS-203
chr18-73411803-INS-51
chr18-73422019-INS-1296
chr18-73422020-INS-190
chr18-73423826-INS-65
chr18-73424891-INS-191
chr18-73428963-INS-134
chr18-73459604-INS-68
chr18-73514088-INS-81
chr18-73525637-INS-433

chr18-73553144-INS-121
chr18-73562486-INS-197
chr18-73566847-INS-104
chr18-73580933-INS-414
chr18-73581126-INS-828
chr18-73581167-INS-1864
chr18-73581309-INS-1909
chr18-73582302-INS-155
chr18-73591486-INS-306
chr18-73596705-INS-57
chr18-73603692-INS-61
chr18-73621049-INS-243
chr18-73638472-INS-324
chr18-73642331-INS-202
chr18-73655442-INS-883
chr18-73683196-INS-208
chr18-73692960-INS-57
chr18-73719343-INS-212
chr18-73733206-INS-208
chr18-73734450-INS-1892
chr18-73739232-INS-241
chr18-73741729-INS-5294
chr18-73754042-INS-148
chr18-73767711-INS-202
chr18-73829277-INS-102
chr18-73847415-INS-164
chr18-73853406-INS-95
chr18-73859135-INS-54
chr18-73862952-INS-428
chr18-73895497-INS-114
chr18-73923799-INS-55
chr18-73944444-INS-144
chr18-73960322-INS-112
chr18-73960486-INS-1197
chr18-73964440-INS-767
chr18-73978550-INS-1127
chr18-73979357-INS-211
chr18-73980619-INS-54
chr18-73991799-INS-393
chr18-74009130-INS-79
chr18-74020715-INS-53
chr18-74036155-INS-583
chr18-74061156-INS-173
chr18-74070895-INS-558
chr18-74089406-INS-81
chr18-74119236-INS-4510
chr18-74121769-INS-85
chr18-74145045-INS-167
chr18-74168479-INS-169
chr18-74207552-INS-523
chr18-74208827-INS-4099
chr18-74230045-INS-198
chr18-74230875-INS-458
chr18-74232982-INS-81
chr18-74234247-INS-74
chr18-74236846-INS-90
chr18-74241321-INS-531
chr18-74254957-INS-54
chr18-74274865-INS-135
chr18-74276937-INS-178
chr18-74283215-INS-76
chr18-74288984-INS-59
chr18-74323074-INS-200
chr18-74338459-INS-58
chr18-74373182-INS-245
chr18-74381157-INS-4427
chr18-74387667-INS-6413
chr18-74428441-INS-453
chr18-74434039-INS-183
chr18-74450220-INS-341
chr18-74458500-INS-57
chr18-74504453-INS-66
chr18-74507461-INS-324
chr18-74524832-INS-3053
chr18-74541009-INS-158
chr18-74548046-INS-69
chr18-74555987-INS-245

chr18-74571444-INS-134
chr18-74584738-INS-54
chr18-74591474-INS-241
chr18-74593099-INS-242
chr18-74596984-INS-68
chr18-74617588-INS-272
chr18-74623331-INS-1888
chr18-74653719-INS-128
chr18-74663971-INS-1199
chr18-74676780-INS-83
chr18-74709359-INS-82
chr18-74714138-INS-66
chr18-74715116-INS-310
chr18-74721263-INS-248
chr18-74743935-INS-448
chr18-74757086-INS-249
chr18-74781965-INS-6511
chr18-74806491-INS-371
chr18-74819503-INS-150
chr18-74824588-INS-182
chr18-74835985-INS-79
chr18-74839233-INS-1283
chr18-74839622-INS-189
chr18-74896138-INS-155
chr18-74920666-INS-86
chr18-74924012-INS-51
chr18-74930203-INS-137
chr18-74933961-INS-579
chr18-74935829-INS-54
chr18-74943662-INS-397
chr18-74959887-INS-124
chr18-74969627-INS-150
chr18-74994771-INS-138
chr18-74996508-INS-270
chr18-75009110-INS-55
chr18-75013208-INS-55
chr18-75013829-INS-90
chr18-75023137-INS-64
chr18-75047374-INS-219
chr18-75049997-INS-148
chr18-75065058-INS-254
chr18-75065967-INS-197
chr18-75077416-INS-75
chr18-75108484-INS-82
chr18-75110406-INS-213
chr18-75128320-INS-56
chr18-75128352-INS-807
chr18-75139870-INS-324
chr18-75141252-INS-96
chr18-75156578-INS-106
chr18-75166489-INS-110
chr18-75174975-INS-598
chr18-75194531-INS-113
chr18-75203387-INS-206
chr18-75219500-INS-140
chr18-75229405-INS-966
chr18-75232690-INS-200
chr18-75244565-INS-255
chr18-75268879-INS-64
chr18-75276076-INS-588
chr18-75284727-INS-558
chr18-75294634-INS-112
chr18-75296805-INS-136
chr18-75312580-INS-77
chr18-75320571-INS-204
chr18-75323786-INS-62
chr18-75333273-INS-219
chr18-75362157-INS-60
chr18-75372744-INS-74
chr18-75383074-INS-52
chr18-75390754-INS-56
chr18-75397529-INS-211
chr18-75406254-INS-164
chr18-75415433-INS-194
chr18-75415459-INS-666
chr18-75429488-INS-132
chr18-75431961-INS-63

chr18-75446634-INS-78
chr18-75448357-INS-132
chr18-75491910-INS-205
chr18-75522105-INS-96
chr18-75551355-INS-70
chr18-75578429-INS-5406
chr18-75588312-INS-152
chr18-75589651-INS-529
chr18-75591312-INS-217
chr18-75594678-INS-58
chr18-75599497-INS-52
chr18-75601336-INS-56
chr18-75634431-INS-196
chr18-75663596-INS-118
chr18-75673603-INS-164
chr18-75675627-INS-344
chr18-75676888-INS-90
chr18-75700849-INS-57
chr18-75701640-INS-131
chr18-75709429-INS-261
chr18-75717818-INS-56
chr18-75723339-INS-76
chr18-75748782-INS-93
chr18-75762021-INS-85
chr18-75798508-INS-54
chr18-75834134-INS-813
chr18-75845994-INS-209
chr18-75850747-INS-394
chr18-75853344-INS-128
chr18-75855492-INS-481
chr18-75887068-INS-644
chr18-75889896-INS-102
chr18-75901438-INS-191
chr18-75908214-INS-86
chr18-75914485-INS-859
chr18-75918656-INS-95
chr18-75928965-INS-1048
chr18-75931957-INS-90
chr18-75937022-INS-64
chr18-75946953-INS-149
chr18-75959752-INS-543
chr18-75960328-INS-97
chr18-75987165-INS-103
chr18-76000793-INS-234
chr18-76070232-INS-69
chr18-76109004-INS-208
chr18-76118469-INS-62
chr18-76141789-INS-54
chr18-76154969-INS-178
chr18-76191560-INS-5404
chr18-76327523-INS-81
chr18-76368020-INS-378
chr18-76402028-INS-298
chr18-76480161-INS-163
chr18-76486766-INS-52
chr18-76504531-INS-54
chr18-76510029-INS-71
chr18-76515562-INS-54
chr18-76517414-INS-206
chr18-76526934-INS-160
chr18-76530825-INS-5294
chr18-76535565-INS-665
chr18-76549049-INS-345
chr18-76558531-INS-78
chr18-76559556-INS-206
chr18-76590116-INS-669
chr18-76614888-INS-60
chr18-76618621-INS-4838
chr18-76674122-INS-57
chr18-76679697-INS-656
chr18-76681150-INS-141
chr18-76681734-INS-84
chr18-76689835-INS-79
chr18-76708371-INS-430
chr18-76710778-INS-498
chr18-76752476-INS-364
chr18-76759972-INS-339

chr18-76762995-INS-97
chr18-76764563-INS-105
chr18-76765097-INS-488
chr18-76774008-INS-53
chr18-76781357-INS-380
chr18-76786937-INS-197
chr18-76794108-INS-78
chr18-76815373-INS-3266
chr18-76849905-INS-217
chr18-76852701-INS-67
chr18-76863692-INS-813
chr18-76865896-INS-387
chr18-76868715-INS-52
chr18-76935376-INS-64
chr18-76938402-INS-121
chr18-76956310-INS-52
chr18-76960518-INS-183
chr18-76971372-INS-92
chr18-76979077-INS-63
chr18-76995031-INS-62
chr18-77008291-INS-74
chr18-77009770-INS-67
chr18-77048092-INS-215
chr18-77054198-INS-454
chr18-77055863-INS-68
chr18-77056735-INS-211
chr18-77067899-INS-172
chr18-77072891-INS-54
chr18-77089401-INS-212
chr18-77090777-INS-182
chr18-77100798-INS-56
chr18-77106039-INS-189
chr18-77117480-INS-115
chr18-77119400-INS-7102
chr18-77128154-INS-54
chr18-77139379-INS-153
chr18-77147477-INS-51
chr18-77166543-INS-156
chr18-77173529-INS-64
chr18-77183931-INS-763
chr18-77194078-INS-56
chr18-77204459-INS-55
chr18-77270245-INS-215
chr18-77277529-INS-61
chr18-77300770-INS-53
chr18-77316071-INS-245
chr18-77317829-INS-68
chr18-77322298-INS-587
chr18-77348721-INS-548
chr18-77351519-INS-91
chr18-77360716-INS-69
chr18-77368263-INS-131
chr18-77401053-INS-150
chr18-77402380-INS-107
chr18-77424581-INS-796
chr18-77426922-INS-210
chr18-77445760-INS-110
chr18-77491579-INS-1349
chr18-77533415-INS-53
chr18-77556216-INS-90
chr18-77569125-INS-164
chr18-77585777-INS-56
chr18-77599088-INS-110
chr18-77606711-INS-104
chr18-77607730-INS-400
chr18-77673184-INS-214
chr18-77675263-INS-136
chr18-77675440-INS-197
chr18-77682291-INS-168
chr18-77702635-INS-415
chr18-77716289-INS-72
chr18-77745352-INS-208
chr18-77766588-INS-64
chr18-77770058-INS-6741
chr18-77771974-INS-52
chr18-77782515-INS-178
chr18-77784818-INS-56

chr18-77795556-INS-72
chr18-77806598-INS-148
chr18-77813384-INS-165
chr18-77815609-INS-86
chr18-77817013-INS-70
chr18-77821648-INS-231
chr18-77829300-INS-130
chr18-77837758-INS-984
chr18-77839213-INS-197
chr18-77853108-INS-52
chr18-77859192-INS-78
chr18-77862826-INS-207
chr18-77863971-INS-211
chr18-77864588-INS-214
chr18-77871118-INS-211
chr18-77880845-INS-55
chr18-77894821-INS-935
chr18-77895755-INS-297
chr18-77920202-INS-53
chr18-77927094-INS-385
chr18-77932149-INS-417
chr18-77935376-INS-201
chr18-77939507-INS-101
chr18-77945350-INS-300
chr18-77967173-INS-235
chr18-77967443-INS-761
chr18-77975312-INS-209
chr18-77993272-INS-206
chr18-77994198-INS-333
chr18-77995716-INS-209
chr18-78000332-INS-156
chr18-78002450-INS-57
chr18-78007489-INS-181
chr18-78014906-INS-206
chr18-78020137-INS-109
chr18-78020601-INS-269
chr18-78033928-INS-78
chr18-78044144-INS-207
chr18-78045334-INS-173
chr18-78055270-INS-239
chr18-78060284-INS-545
chr18-78080756-INS-1888
chr18-78080757-INS-399
chr18-78087924-INS-62
chr18-78095729-INS-67
chr18-78096941-INS-444
chr18-78117628-INS-167
chr18-78124144-INS-201
chr18-78170797-INS-289
chr18-78180699-INS-348
chr18-78182838-INS-646
chr18-78183857-INS-52
chr18-78185773-INS-199
chr18-78187031-INS-516
chr18-78209253-INS-72
chr18-78246858-INS-226
chr18-78248191-INS-259
chr18-78270292-INS-75
chr18-78272407-INS-391
chr18-78335247-INS-998
chr18-78360706-INS-945
chr18-78361969-INS-96
chr18-78370305-INS-737
chr18-78378510-INS-156
chr18-78398785-INS-273
chr18-78413681-INS-129
chr18-78417632-INS-212
chr18-78430143-INS-253
chr18-78456055-INS-2841
chr18-78479114-INS-72
chr18-78510662-INS-175
chr18-78524461-INS-181
chr18-78542011-INS-148
chr18-78542260-INS-148
chr18-78548098-INS-94
chr18-78553444-INS-60
chr18-78570079-INS-58

chr18-78591067-INS-64
chr18-78605130-INS-63
chr18-78606688-INS-52
chr18-78608922-INS-63
chr18-78629568-INS-807
chr18-78641180-INS-83
chr18-78648022-INS-1888
chr18-78652805-INS-205
chr18-78678137-INS-70
chr18-78681501-INS-59
chr18-78709060-INS-72
chr18-78736896-INS-52
chr18-78786905-INS-205
chr18-78802870-INS-91
chr18-78803984-INS-219
chr18-78807945-INS-997
chr18-78812659-INS-147
chr18-78814908-INS-158
chr18-78831756-INS-62
chr18-78848399-INS-59
chr18-78882230-INS-62
chr18-78889963-INS-58
chr18-78896459-INS-6443
chr18-78923465-INS-54
chr18-78935388-INS-52
chr18-78937557-INS-207
chr18-78939579-INS-61
chr18-78970953-INS-55
chr18-78984997-INS-180
chr18-79003658-INS-370
chr18-79006818-INS-6531
chr18-79007257-INS-184
chr18-79022519-INS-64
chr18-79070773-INS-63
chr18-79083637-INS-387
chr18-79088801-INS-78
chr18-79099408-INS-54
chr18-79105051-INS-315
chr18-79113434-INS-87
chr18-79149912-INS-64
chr18-79209390-INS-6447
chr18-79254679-INS-7282
chr18-79256777-INS-53
chr18-79268300-INS-104
chr18-79271607-INS-253
chr18-79272169-INS-115
chr18-79291212-INS-903
chr18-79293277-INS-376
chr18-79293942-INS-212
chr18-79298518-INS-53
chr18-79300397-INS-52
chr18-79302162-INS-59
chr18-79312613-INS-6928
chr18-79319557-INS-91
chr18-79320083-INS-804
chr18-79326800-INS-68
chr18-79368565-INS-1841
chr18-79409850-INS-52
chr18-79411270-INS-230
chr18-79457358-INS-137
chr18-79475198-INS-97
chr18-79500745-INS-483
chr18-79516730-INS-142
chr18-79527977-INS-174
chr18-79536923-INS-413
chr18-79550794-INS-211
chr18-79560869-INS-693
chr18-79564704-INS-94
chr18-79586006-INS-54
chr18-79588537-INS-221
chr18-79591364-INS-212
chr18-79607487-INS-402
chr18-79661434-INS-63
chr18-79688258-INS-203
chr18-79690811-INS-324
chr18-79702035-INS-223
chr18-79703871-INS-86

chr18-79736869-INS-121
chr18-79757684-INS-3674
chr18-79761314-INS-407
chr18-79775376-INS-338
chr18-79784994-INS-6621
chr18-79823284-INS-76
chr18-79824100-INS-376
chr18-79825026-INS-7075
chr18-79827502-INS-75
chr18-79829198-INS-7326
chr18-79868011-INS-240
chr18-79871630-INS-7068
chr18-79883317-INS-53
chr18-79890742-INS-63
chr18-79900933-INS-60
chr18-79902673-INS-162
chr18-79913937-INS-252
chr18-79928099-INS-202
chr18-79942267-INS-97
chr18-79973427-INS-53
chr18-79985262-INS-100
chr18-79989832-INS-101
chr18-79998605-INS-98
chr18-80008260-INS-318
chr18-80016316-INS-236
chr18-80035891-INS-389
chr18-80051900-INS-240
chr18-80052966-INS-521
chr18-80077642-INS-108
chr18-80078630-INS-514
chr18-80132955-INS-148
chr18-80148012-INS-99
chr18-80178228-INS-153
chr18-80184290-INS-135
chr18-80203085-INS-321
chr18-80204232-INS-83
chr18-80209675-INS-193
chr18-80214294-INS-65
chr18-80224079-INS-212
chr18-80239379-INS-238
chr18-80242454-INS-485
chr18-80259761-INS-135
chr18-80271750-INS-66
chr18-80285720-INS-198
chr18-80299133-INS-55
chr18-80303089-INS-168
chr18-80385718-INS-308
chr18-80391219-INS-52
chr18-80393942-INS-52
chr18-80396604-INS-69
chr18-80428451-INS-103
chr18-80453810-INS-63
chr18-80479729-INS-58
chr18-80497649-INS-155
chr18-80545231-INS-258
chr18-80549159-INS-212
chr18-80564410-INS-95
chr18-80568890-INS-53
chr18-80581110-INS-113
chr18-80633109-INS-241
chr18-80655814-INS-63
chr18-80701948-INS-97
chr18-80716718-INS-384
chr18-80793674-INS-52
chr18-80817546-INS-114
chr18-80837528-INS-145
chr18-80878424-INS-60
chr18-80886501-INS-1334
chr18-80886502-INS-735
chr18-80908527-INS-495
chr18-80936762-INS-84
chr18-80944088-INS-78
chr18-80961412-INS-52
chr18-80971502-INS-236
chr18-81006481-INS-115
chr18-81099179-INS-146
chr18-81102425-INS-717

chr18-81107661-INS-299
chr18-81109160-INS-56
chr18-81122258-INS-145
chr18-81136226-INS-92
chr18-81140764-INS-286
chr18-81147562-INS-142
chr18-81159613-INS-92
chr18-81207742-INS-62
chr18-81211485-INS-72
chr18-81213600-INS-69
chr18-81224916-INS-54
chr18-81228033-INS-594
chr18-81228041-INS-1950
chr18-81229008-INS-76
chr18-81257101-INS-464
chr18-81262719-INS-90
chr18-81265350-INS-110
chr18-81271970-INS-82
chr18-81288208-INS-108
chr18-81289571-INS-416
chr18-81311745-INS-138
chr18-81315954-INS-248
chr18-81317856-INS-68
chr18-81333876-INS-57
chr18-81337636-INS-903
chr18-81351208-INS-111
chr18-81364181-INS-68
chr18-81365037-INS-839
chr18-81389931-INS-977
chr18-81391501-INS-91
chr18-81393856-INS-161
chr18-81399582-INS-1676
chr18-81399593-INS-1830
chr18-81408751-INS-72
chr18-81418487-INS-368
chr18-81427127-INS-188
chr18-81444603-INS-135
chr18-81447195-INS-399
chr18-81480304-INS-151
chr18-81489459-INS-96
chr18-81502057-INS-209
chr18-81535539-INS-160
chr18-81557883-INS-118
chr18-81562476-INS-544
chr18-81600237-INS-80
chr18-81605442-INS-86
chr18-81608161-INS-118
chr18-81615607-INS-58
chr18-81647352-INS-63
chr18-81659024-INS-222
chr18-81665748-INS-52
chr18-81669337-INS-76
chr18-81679202-INS-176
chr18-81707887-INS-70
chr18-81712331-INS-215
chr18-81731801-INS-52
chr18-81734391-INS-53
chr18-81742506-INS-258
chr18-81744343-INS-56
chr18-81756417-INS-83
chr18-81758351-INS-56
chr18-81765259-INS-162
chr18-81778817-INS-186
chr18-81781852-INS-107
chr18-81800209-INS-109
chr18-81818625-INS-281
chr18-81821187-INS-69
chr18-81838324-INS-323
chr18-81846461-INS-75
chr18-81856341-INS-245
chr18-81864414-INS-102
chr18-81872832-INS-152
chr18-81876583-INS-69
chr18-81879052-INS-186
chr18-81893320-INS-62
chr18-81896932-INS-52
chr18-81899552-INS-377

chr18-81926200-INS-68
chr18-81967669-INS-252
chr18-81979144-INS-167
chr18-81984513-INS-168
chr18-82011441-INS-74
chr18-82023051-INS-161
chr18-82040312-INS-75
chr18-82043968-INS-211
chr18-82056075-INS-61
chr18-82057503-INS-131
chr18-82068149-INS-139
chr18-82085070-INS-64
chr18-82095395-INS-201
chr18-82105177-INS-363
chr18-82120359-INS-328
chr18-82124628-INS-88
chr18-82126231-INS-72
chr18-82129360-INS-7263
chr18-82130240-INS-82
chr18-82140565-INS-162
chr18-82153502-INS-67
chr18-82153764-INS-125
chr18-82156510-INS-213
chr18-82175981-INS-136
chr18-82178174-INS-149
chr18-82195030-INS-525
chr18-82207200-INS-121
chr18-82229951-INS-164
chr18-82230952-INS-142
chr18-82239782-INS-398
chr18-82259208-INS-58
chr18-82271361-INS-248
chr18-82281509-INS-80
chr18-82292165-INS-149
chr18-82297268-INS-7163
chr18-82300108-INS-125
chr18-82303679-INS-94
chr18-82304602-INS-2178
chr18-82318352-INS-51
chr18-82329875-INS-172
chr18-82333777-INS-221
chr18-82335463-INS-378
chr18-82349013-INS-202
chr18-82351132-INS-210
chr18-82354105-INS-152
chr18-82366660-INS-76
chr18-82394400-INS-272
chr18-82397953-INS-285
chr18-82398961-INS-106
chr18-82406148-INS-82
chr18-82418184-INS-783
chr18-82433410-INS-142
chr18-82453487-INS-172
chr18-82464999-INS-484
chr18-82474966-INS-96
chr18-82490190-INS-126
chr18-82499455-INS-403
chr18-82501520-INS-102
chr18-82514219-INS-85
chr18-82541677-INS-5344
chr18-82546569-INS-89
chr18-82547316-INS-604
chr18-82557108-INS-90
chr18-82562785-INS-171
chr18-82564571-INS-72
chr18-82566938-INS-75
chr18-82575080-INS-160
chr18-82604888-INS-197
chr18-82609500-INS-75
chr18-82618781-INS-58
chr18-82627776-INS-175
chr18-82633694-INS-246
chr18-82638159-INS-106
chr18-82650129-INS-312
chr18-82654607-INS-693
chr18-82673109-INS-99
chr18-82682256-INS-531

chr18-82690342-INS-368
chr18-82695753-INS-132
chr18-82702240-INS-410
chr18-82708023-INS-172
chr18-82747744-INS-185
chr18-82751419-INS-109
chr18-82766988-INS-157
chr18-82806398-INS-104
chr18-82826782-INS-244
chr18-82833053-INS-133
chr18-82840023-INS-70
chr18-82860363-INS-56
chr18-82870339-INS-211
chr18-82883477-INS-99
chr18-82894826-INS-143
chr18-82897071-INS-215
chr18-82921970-INS-58
chr18-82942191-INS-191
chr18-82952602-INS-102
chr18-82969805-INS-194
chr18-82989979-INS-255
chr18-82991868-INS-60
chr18-82994523-INS-54
chr18-83003796-INS-117
chr18-83008773-INS-98
chr18-83009495-INS-63
chr18-83018298-INS-58
chr18-83025944-INS-95
chr18-83032248-INS-57
chr18-83035331-INS-326
chr18-83036947-INS-83
chr18-83043110-INS-1887
chr18-83046644-INS-224
chr18-83070746-INS-399
chr18-83076031-INS-89
chr18-83080495-INS-435
chr18-83084164-INS-192
chr18-83089288-INS-52
chr18-83101343-INS-114
chr18-83120955-INS-257
chr18-83149539-INS-7011
chr18-83151697-INS-192
chr18-83164791-INS-72
chr18-83176938-INS-336
chr18-83207170-INS-96
chr18-83208179-INS-66
chr18-83214848-INS-127
chr18-83264456-INS-53
chr18-83269693-INS-110
chr18-83273511-INS-54
chr18-83284406-INS-211
chr18-83289324-INS-211
chr18-83297670-INS-431
chr18-83331930-INS-55
chr18-83334027-INS-179
chr18-83357317-INS-52
chr18-83376358-INS-80
chr18-83388319-INS-64
chr18-83389321-INS-82
chr18-83392897-INS-198
chr18-83394142-INS-194
chr18-83417248-INS-84
chr18-83449767-INS-116
chr18-83451812-INS-133
chr18-83463516-INS-118
chr18-83481760-INS-64
chr18-83491017-INS-62
chr18-83505952-INS-115
chr18-83521100-INS-204
chr18-83552779-INS-286
chr18-83559763-INS-176
chr18-83571150-INS-100
chr18-83572855-INS-88
chr18-83578128-INS-159
chr18-83582105-INS-68
chr18-83588706-INS-76
chr18-83598396-INS-348

chr18-83610646-INS-54
chr18-83679269-INS-60
chr18-83683360-INS-60
chr18-83699272-INS-57
chr18-83712965-INS-149
chr18-83715634-INS-70
chr18-83716097-INS-143
chr18-83736299-INS-144
chr18-83749799-INS-94
chr18-83757495-INS-325
chr18-83796728-INS-364
chr18-83821385-INS-91
chr18-83839876-INS-111
chr18-83844458-INS-60
chr18-83849550-INS-72
chr18-83860637-INS-210
chr18-83865527-INS-669
chr18-83866339-INS-52
chr18-83882806-INS-283
chr18-83892252-INS-56
chr18-83901040-INS-190
chr18-83913106-INS-99
chr18-83926554-INS-94
chr18-83942527-INS-75
chr18-83978708-INS-392
chr18-83986459-INS-204
chr18-83991776-INS-58
chr18-83998941-INS-58
chr18-84000700-INS-156
chr18-84048550-INS-75
chr18-84057937-INS-54
chr18-84068033-INS-93
chr18-84084019-INS-81
chr18-84097808-INS-50
chr18-84124255-INS-56
chr18-84142029-INS-82
chr18-84149488-INS-163
chr18-84154141-INS-216
chr18-84188827-INS-72
chr18-84211790-INS-55
chr18-84213761-INS-60
chr18-84238176-INS-179
chr18-84272068-INS-204
chr18-84294731-INS-194
chr18-84316121-INS-234
chr18-84326343-INS-112
chr18-84336273-INS-979
chr18-84340307-INS-53
chr18-84366170-INS-52
chr18-84383773-INS-206
chr18-84384791-INS-60
chr18-84402905-INS-5766
chr18-84406223-INS-800
chr18-84418017-INS-65
chr18-84442944-INS-80
chr18-84444543-INS-6832
chr18-84447142-INS-92
chr18-84503947-INS-494
chr18-84527340-INS-801
chr18-84536137-INS-72
chr18-84538970-INS-103
chr18-84553987-INS-56
chr18-84562844-INS-166
chr18-84582286-INS-230
chr18-84582922-INS-179
chr18-84582938-INS-1857
chr18-84588100-INS-274
chr18-84599452-INS-748
chr18-84620766-INS-199
chr18-84636572-INS-5173
chr18-84644436-INS-497
chr18-84651965-INS-103
chr18-84654119-INS-238
chr18-84656574-INS-52
chr18-84701496-INS-60
chr18-84712495-INS-52
chr18-84714866-INS-308

chr18-84730414-INS-324
chr18-84744784-INS-252
chr18-84751677-INS-535
chr18-84761194-INS-360
chr18-84786909-INS-343
chr18-84788022-INS-842
chr18-84794357-INS-119
chr18-84797491-INS-53
chr18-84797491-INS-689
chr18-84815706-INS-163
chr18-84860564-INS-498
chr18-84892933-INS-133
chr18-84903024-INS-147
chr18-84924167-INS-111
chr18-84997145-INS-112
chr18-85006833-INS-267
chr18-85034286-INS-341
chr18-85034571-INS-844
chr18-85041883-INS-656
chr18-85075129-INS-330
chr18-85083650-INS-769
chr18-85101874-INS-61
chr18-85109736-INS-138
chr18-85112423-INS-51
chr18-85115541-INS-6412
chr18-85125596-INS-2254
chr18-85145096-INS-212
chr18-85161749-INS-728
chr18-85197759-INS-94
chr18-85208829-INS-107
chr18-85211731-INS-80
chr18-85238448-INS-57
chr18-85256588-INS-108
chr18-85268678-INS-145
chr18-85272968-INS-282
chr18-85309656-INS-940
chr18-85319125-INS-356
chr18-85335099-INS-53
chr18-85343923-INS-181
chr18-85388392-INS-401
chr18-85416709-INS-684
chr18-85438471-INS-191
chr18-85439886-INS-102
chr18-85452282-INS-52
chr18-85470027-INS-113
chr18-85476615-INS-87
chr18-85547634-INS-55
chr18-85565517-INS-422
chr18-85572153-INS-51
chr18-85608473-INS-221
chr18-85615693-INS-64
chr18-85624399-INS-56
chr18-85639151-INS-54
chr18-85656065-INS-91
chr18-85665707-INS-548
chr18-85684203-INS-82
chr18-85714597-INS-5529
chr18-85733921-INS-79
chr18-85747671-INS-142
chr18-85761266-INS-216
chr18-85781602-INS-3169
chr18-85782287-INS-70
chr18-85786437-INS-116
chr18-85788121-INS-158
chr18-85791395-INS-261
chr18-85815790-INS-80
chr18-85850877-INS-63
chr18-85875905-INS-64
chr18-85887839-INS-588
chr18-85910818-INS-107
chr18-85952611-INS-645
chr18-85961273-INS-919
chr18-85964884-INS-313
chr18-86048859-INS-111
chr18-86049760-INS-116
chr18-86051867-INS-7183
chr18-86060637-INS-448

chr18-86100338-INS-122
chr18-86111209-INS-61
chr18-86132464-INS-58
chr18-86136057-INS-56
chr18-86194162-INS-125
chr18-86211096-INS-7142
chr18-86233619-INS-237
chr18-86239626-INS-418
chr18-86242526-INS-53
chr18-86273451-INS-133
chr18-86279643-INS-89
chr18-86287413-INS-59
chr18-86296564-INS-483
chr18-86320234-INS-120
chr18-86328542-INS-652
chr18-86365483-INS-54
chr18-86375403-INS-510
chr18-86395883-INS-963
chr18-86397157-INS-62
chr18-86399253-INS-61
chr18-86410108-INS-66
chr18-86459265-INS-55
chr18-86471480-INS-697
chr18-86499625-INS-56
chr18-86508480-INS-413
chr18-86509922-INS-53
chr18-86526969-INS-169
chr18-86528154-INS-144
chr18-86543394-INS-54
chr18-86568246-INS-148
chr18-86571659-INS-65
chr18-86578251-INS-102
chr18-86600756-INS-160
chr18-86608642-INS-52
chr18-86631066-INS-673
chr18-86749740-INS-978
chr18-86751104-INS-305
chr18-86755851-INS-277
chr18-86769399-INS-364
chr18-86771006-INS-92
chr18-86772219-INS-960
chr18-86800188-INS-451
chr18-86814373-INS-202
chr18-86855468-INS-81
chr18-86863965-INS-116
chr18-86867673-INS-60
chr18-86897369-INS-105
chr18-86904804-INS-110
chr18-86910027-INS-879
chr18-86911329-INS-346
chr18-86924826-INS-306
chr18-86925792-INS-6383
chr18-86953848-INS-601
chr18-86972230-INS-196
chr18-87019081-INS-377
chr18-87021508-INS-64
chr18-87035060-INS-6403
chr18-87040219-INS-223
chr18-87048501-INS-6198
chr18-87050129-INS-5324
chr18-87059767-INS-55
chr18-87078109-INS-194
chr18-87097674-INS-84
chr18-87113756-INS-52
chr18-87121052-INS-75
chr18-87126254-INS-599
chr18-87137325-INS-323
chr18-87146882-INS-540
chr18-87155560-INS-174
chr18-87188143-INS-83
chr18-87249293-INS-58
chr18-87254632-INS-76
chr18-87265971-INS-76
chr18-87283284-INS-54
chr18-87283826-INS-491
chr18-87335839-INS-387
chr18-87355240-INS-163

chr18-87361676-INS-222
chr18-87387248-INS-123
chr18-87397173-INS-56
chr18-87399964-INS-109
chr18-87406732-INS-6552
chr18-87415403-INS-64
chr18-87419560-INS-61
chr18-87469640-INS-666
chr18-87470552-INS-310
chr18-87503546-INS-1888
chr18-87535413-INS-431
chr18-87545048-INS-98
chr18-87547112-INS-64
chr18-87549359-INS-57
chr18-87557006-INS-114
chr18-87564466-INS-943
chr18-87576834-INS-5550
chr18-87592341-INS-301
chr18-87593376-INS-70
chr18-87608379-INS-54
chr18-87635862-INS-2694
chr18-87639943-INS-127
chr18-87677957-INS-505
chr18-87682840-INS-286
chr18-87694097-INS-6893
chr18-87696280-INS-199
chr18-87718029-INS-183
chr18-87718931-INS-60
chr18-87736272-INS-196
chr18-87743477-INS-546
chr18-87752053-INS-698
chr18-87758503-INS-641
chr18-87791713-INS-5548
chr18-87801195-INS-212
chr18-87801195-INS-848
chr18-87847832-INS-2565
chr18-87852061-INS-647
chr18-87853667-INS-66
chr18-87855944-INS-9581
chr18-87860511-INS-913
chr18-87885079-INS-2913
chr18-87923571-INS-143
chr18-87930246-INS-54
chr18-87960447-INS-105
chr18-87964398-INS-70
chr18-87998396-INS-411
chr18-88056900-INS-60
chr18-88060372-INS-241
chr18-88062390-INS-106
chr18-88067131-INS-959
chr18-88069387-INS-85
chr18-88074052-INS-485
chr18-88077195-INS-101
chr18-88080359-INS-54
chr18-88097175-INS-100
chr18-88115892-INS-974
chr18-88134597-INS-883
chr18-88156548-INS-695
chr18-88159001-INS-6105
chr18-88178482-INS-56
chr18-88180672-INS-219
chr18-88208409-INS-363
chr18-88261823-INS-58
chr18-88301039-INS-56
chr18-88303474-INS-84
chr18-88329257-INS-182
chr18-88337230-INS-306
chr18-88341194-INS-51
chr18-88342660-INS-102
chr18-88346866-INS-642
chr18-88348639-INS-170
chr18-88352480-INS-6387
chr18-88400489-INS-53
chr18-88401177-INS-356
chr18-88419389-INS-141
chr18-88450584-INS-58
chr18-88459540-INS-92

chr18-88511437-INS-248
chr18-88535285-INS-6383
chr18-88554984-INS-66
chr18-88573859-INS-152
chr18-88579594-INS-166
chr18-88582770-INS-117
chr18-88622334-INS-68
chr18-88625245-INS-81
chr18-88636414-INS-259
chr18-88658136-INS-616
chr18-88683323-INS-60
chr18-88715773-INS-259
chr18-88728417-INS-60
chr18-88788726-INS-226
chr18-88796328-INS-1008
chr18-88806771-INS-206
chr18-88812315-INS-8130
chr18-88830760-INS-6205
chr18-88853901-INS-5426
chr18-88861817-INS-356
chr18-88862877-INS-7068
chr18-88914812-INS-94
chr18-88924227-INS-390
chr18-88925149-INS-80
chr18-88927463-INS-114
chr18-88954812-INS-660
chr18-88968517-INS-6462
chr18-88969248-INS-4351
chr18-88976543-INS-425
chr18-88984969-INS-6280
chr18-88985288-INS-82
chr18-88989654-INS-91
chr18-89000606-INS-467
chr18-89001732-INS-2156
chr18-89004554-INS-559
chr18-89006937-INS-59
chr18-89023964-INS-79
chr18-89039946-INS-70
chr18-89050180-INS-201
chr18-89071570-INS-868
chr18-89081799-INS-6401
chr18-89083703-INS-210
chr18-89085738-INS-190
chr18-89090556-INS-466
chr18-89091603-INS-131
chr18-89132867-INS-141
chr18-89148497-INS-173
chr18-89151922-INS-79
chr18-89211224-INS-6480
chr18-89222997-INS-6205
chr18-89238546-INS-197
chr18-89244788-INS-83
chr18-89249227-INS-229
chr18-89356105-INS-64
chr18-89357819-INS-161
chr18-89369510-INS-230
chr18-89386955-INS-65
chr18-89418058-INS-6592
chr18-89443043-INS-6197
chr18-89474020-INS-332
chr18-89505165-INS-89
chr18-89512080-INS-363
chr18-89528378-INS-169
chr18-89531066-INS-58
chr18-89587263-INS-55
chr18-89643170-INS-190
chr18-89653834-INS-3461
chr18-89657015-INS-73
chr18-89659681-INS-4062
chr18-89663147-INS-2554
chr18-89678145-INS-518
chr18-89695188-INS-2938
chr18-89697349-INS-6452
chr18-89728400-INS-379
chr18-89737011-INS-67
chr18-89797730-INS-129
chr18-89799365-INS-1705

chr18-89819861-INS-6874
chr18-89899439-INS-3423
chr18-89900994-INS-57
chr18-89906250-INS-2658
chr18-89927717-INS-393
chr18-89964393-INS-7582
chr18-89989704-INS-195
chr18-90027208-INS-4860
chr18-90035591-INS-161
chr18-90040609-INS-290
chr18-90056205-INS-486
chr18-90092110-INS-6251
chr18-90121683-INS-6418
chr18-90153800-INS-88
chr18-90179356-INS-825
chr18-90180269-INS-76
chr18-90191467-INS-147
chr18-90193488-INS-4852
chr18-90198861-INS-7175
chr18-90229037-INS-6759
chr18-90259318-INS-550
chr18-90307084-INS-188
chr18-90309914-INS-137
chr18-90314323-INS-102
chr18-90330980-INS-412
chr18-90386763-INS-75
chr18-90387311-INS-436
chr18-90403205-INS-3228
chr18-90462860-INS-125
chr18-90464564-INS-399
chr18-90490986-INS-78
chr18-90495528-INS-4825
chr18-90496903-INS-54
chr18-90502741-INS-252
chr18-90517788-INS-329
chr18-90523518-INS-324
chr18-90577849-INS-205
chr18-90583681-INS-217
chr18-90586635-INS-1374
chr18-90587333-INS-76
chr18-90613580-INS-234
chr19-3094583-INS-212
chr19-3122576-INS-7179
chr19-3129906-INS-214
chr19-3168552-INS-725
chr19-3173958-INS-214
chr19-3174710-INS-79
chr19-3188394-INS-6401
chr19-3201760-INS-61
chr19-3236419-INS-74
chr19-3257199-INS-190
chr19-3316067-INS-125
chr19-3338191-INS-52
chr19-3357133-INS-203
chr19-3358668-INS-194
chr19-3368143-INS-978
chr19-3368155-INS-425
chr19-3387288-INS-194
chr19-3392391-INS-166
chr19-3403354-INS-987
chr19-3405039-INS-186
chr19-3409282-INS-53
chr19-3422464-INS-157
chr19-3424906-INS-57
chr19-3454318-INS-68
chr19-3485708-INS-5449
chr19-3491779-INS-90
chr19-3497588-INS-188
chr19-3502282-INS-125
chr19-3515258-INS-147
chr19-3517765-INS-196
chr19-3530214-INS-690
chr19-3547743-INS-103
chr19-3560265-INS-1947
chr19-3578547-INS-198
chr19-3579158-INS-438
chr19-3611767-INS-55

chr19-3631500-INS-341
chr19-3636023-INS-223
chr19-3638438-INS-62
chr19-3663475-INS-200
chr19-3677482-INS-63
chr19-3687985-INS-215
chr19-3693191-INS-244
chr19-3699243-INS-142
chr19-3711232-INS-319
chr19-3727338-INS-193
chr19-3749558-INS-72
chr19-3756068-INS-246
chr19-3770115-INS-180
chr19-3775577-INS-209
chr19-3776874-INS-139
chr19-3783362-INS-5929
chr19-3787116-INS-191
chr19-3787897-INS-193
chr19-3794499-INS-158
chr19-3803245-INS-169
chr19-3806907-INS-56
chr19-3813438-INS-179
chr19-3822886-INS-215
chr19-3825475-INS-216
chr19-3827450-INS-205
chr19-3833837-INS-219
chr19-3874403-INS-1112
chr19-3879717-INS-136
chr19-3896705-INS-194
chr19-3898367-INS-785
chr19-3911606-INS-55
chr19-3922719-INS-61
chr19-3939054-INS-72
chr19-3977472-INS-217
chr19-3984520-INS-903
chr19-3988410-INS-108
chr19-3996155-INS-201
chr19-3999671-INS-106
chr19-4001674-INS-135
chr19-4003263-INS-209
chr19-4007029-INS-82
chr19-4010934-INS-88
chr19-4012714-INS-355
chr19-4017640-INS-89
chr19-4022892-INS-81
chr19-4034230-INS-598
chr19-4082852-INS-196
chr19-4089574-INS-80
chr19-4124018-INS-100
chr19-4130228-INS-183
chr19-4168745-INS-163
chr19-4193537-INS-144
chr19-4211293-INS-203
chr19-4217849-INS-100
chr19-4229830-INS-195
chr19-4234544-INS-101
chr19-4249990-INS-213
chr19-4259713-INS-213
chr19-4265917-INS-176
chr19-4277977-INS-212
chr19-4283214-INS-161
chr19-4288684-INS-207
chr19-4292181-INS-221
chr19-4308368-INS-205
chr19-4311095-INS-133
chr19-4315796-INS-71
chr19-4317322-INS-119
chr19-4337543-INS-314
chr19-4360464-INS-142
chr19-4364216-INS-104
chr19-4417258-INS-192
chr19-4420039-INS-216
chr19-4478764-INS-83
chr19-4504176-INS-138
chr19-4510673-INS-207
chr19-4515635-INS-60
chr19-4520606-INS-52

chr19-4530467-INS-215
chr19-4536857-INS-407
chr19-4572563-INS-159
chr19-4596989-INS-232
chr19-4626672-INS-101
chr19-4637043-INS-245
chr19-4644597-INS-303
chr19-4681237-INS-197
chr19-4684883-INS-80
chr19-4685999-INS-93
chr19-4687037-INS-64
chr19-4705415-INS-57
chr19-4707590-INS-180
chr19-4716051-INS-289
chr19-4724020-INS-360
chr19-4744944-INS-434
chr19-4813330-INS-204
chr19-4833377-INS-232
chr19-4862561-INS-76
chr19-4864031-INS-102
chr19-4866290-INS-192
chr19-4868679-INS-231
chr19-4869581-INS-1167
chr19-4872399-INS-5113
chr19-4877572-INS-205
chr19-4886963-INS-78
chr19-4901820-INS-178
chr19-4905015-INS-183
chr19-4925357-INS-218
chr19-4931426-INS-441
chr19-4932259-INS-190
chr19-4938502-INS-160
chr19-4943240-INS-205
chr19-4961956-INS-272
chr19-4968748-INS-386
chr19-4984666-INS-54
chr19-4989533-INS-126
chr19-5001975-INS-224
chr19-5006531-INS-197
chr19-5007278-INS-4605
chr19-5007350-INS-2285
chr19-5007892-INS-248
chr19-5011171-INS-196
chr19-5012808-INS-260
chr19-5016755-INS-120
chr19-5019063-INS-7088
chr19-5024904-INS-51
chr19-5029635-INS-217
chr19-5030950-INS-208
chr19-5054305-INS-442
chr19-5077593-INS-59
chr19-5084911-INS-216
chr19-5095334-INS-65
chr19-5108774-INS-51
chr19-5115087-INS-202
chr19-5116940-INS-56
chr19-5123024-INS-100
chr19-5127270-INS-273
chr19-5199136-INS-480
chr19-5205558-INS-72
chr19-5210247-INS-134
chr19-5216938-INS-584
chr19-5240595-INS-52
chr19-5242351-INS-1888
chr19-5255344-INS-80
chr19-5258006-INS-196
chr19-5259970-INS-207
chr19-5266998-INS-603
chr19-5277901-INS-58
chr19-5295525-INS-217
chr19-5296897-INS-52
chr19-5306385-INS-115
chr19-5309687-INS-127
chr19-5310561-INS-591
chr19-5311674-INS-63
chr19-5330438-INS-184
chr19-5333032-INS-591

chr19-5346025-INS-200
chr19-5346620-INS-201
chr19-5358425-INS-60
chr19-5359230-INS-1739
chr19-5359246-INS-2170
chr19-5359254-INS-3240
chr19-5359255-INS-2879
chr19-5370873-INS-429
chr19-5371917-INS-262
chr19-5379276-INS-228
chr19-5387490-INS-88
chr19-5407324-INS-89
chr19-5426595-INS-503
chr19-5442634-INS-520
chr19-5451493-INS-434
chr19-5454589-INS-75
chr19-5468640-INS-923
chr19-5483130-INS-59
chr19-5491246-INS-88
chr19-5500721-INS-152
chr19-5519151-INS-119
chr19-5521150-INS-181
chr19-5553927-INS-4799
chr19-5555745-INS-226
chr19-5563827-INS-223
chr19-5566246-INS-211
chr19-5573345-INS-282
chr19-5591495-INS-64
chr19-5614306-INS-210
chr19-5632008-INS-58
chr19-5633666-INS-133
chr19-5639822-INS-529
chr19-5641034-INS-203
chr19-5647499-INS-111
chr19-5650485-INS-134
chr19-5656440-INS-50
chr19-5666870-INS-200
chr19-5727898-INS-82
chr19-5736911-INS-483
chr19-5736951-INS-924
chr19-5782701-INS-210
chr19-5825691-INS-53
chr19-5830450-INS-335
chr19-5835556-INS-165
chr19-5838325-INS-66
chr19-5857233-INS-198
chr19-5865580-INS-184
chr19-5880617-INS-176
chr19-5910288-INS-179
chr19-5934006-INS-51
chr19-5936153-INS-193
chr19-5937718-INS-214
chr19-5950197-INS-206
chr19-5955707-INS-533
chr19-5971338-INS-380
chr19-5983167-INS-57
chr19-5986567-INS-7451
chr19-5987104-INS-93
chr19-5997293-INS-203
chr19-6005387-INS-226
chr19-6007377-INS-168
chr19-6015873-INS-130
chr19-6017451-INS-129
chr19-6020038-INS-221
chr19-6027679-INS-72
chr19-6061559-INS-58
chr19-6067870-INS-199
chr19-6070152-INS-186
chr19-6071915-INS-54
chr19-6072607-INS-119
chr19-6080817-INS-217
chr19-6083636-INS-196
chr19-6109905-INS-178
chr19-6113093-INS-140
chr19-6131439-INS-206
chr19-6146091-INS-207
chr19-6152101-INS-964

chr19-6152819-INS-80
chr19-6157148-INS-655
chr19-6170960-INS-72
chr19-6171474-INS-99
chr19-6196362-INS-91
chr19-6198194-INS-204
chr19-6202313-INS-377
chr19-6204492-INS-112
chr19-6213897-INS-221
chr19-6222832-INS-414
chr19-6225294-INS-80
chr19-6226246-INS-207
chr19-6266521-INS-56
chr19-6267708-INS-184
chr19-6271403-INS-69
chr19-6278930-INS-55
chr19-6299121-INS-164
chr19-6337690-INS-130
chr19-6378406-INS-212
chr19-6379463-INS-425
chr19-6381718-INS-205
chr19-6382192-INS-203
chr19-6431060-INS-322
chr19-6444266-INS-219
chr19-6455784-INS-115
chr19-6482334-INS-56
chr19-6495347-INS-211
chr19-6510680-INS-166
chr19-6515931-INS-113
chr19-6530149-INS-62
chr19-6531973-INS-55
chr19-6558763-INS-132
chr19-6582687-INS-130
chr19-6628375-INS-66
chr19-6629432-INS-216
chr19-6640311-INS-53
chr19-6684289-INS-369
chr19-6693378-INS-226
chr19-6701708-INS-273
chr19-6706162-INS-55
chr19-6717723-INS-62
chr19-6721456-INS-162
chr19-6721800-INS-636
chr19-6726803-INS-170
chr19-6737367-INS-138
chr19-6821180-INS-232
chr19-6859357-INS-192
chr19-6871291-INS-215
chr19-6879757-INS-56
chr19-6880967-INS-5981
chr19-6914135-INS-155
chr19-6941428-INS-134
chr19-6961596-INS-106
chr19-6980953-INS-151
chr19-7007154-INS-174
chr19-7008542-INS-225
chr19-7010681-INS-135
chr19-7014205-INS-172
chr19-7023313-INS-209
chr19-7098809-INS-990
chr19-7101848-INS-52
chr19-7128118-INS-229
chr19-7177670-INS-204
chr19-7179492-INS-124
chr19-7180369-INS-207
chr19-7189805-INS-176
chr19-7197364-INS-193
chr19-7213988-INS-213
chr19-7216176-INS-92
chr19-7219841-INS-200
chr19-7222612-INS-188
chr19-7225320-INS-189
chr19-7226766-INS-78
chr19-7233211-INS-244
chr19-7274228-INS-170
chr19-7294712-INS-228
chr19-7304999-INS-194

chr19-7310747-INS-208
chr19-7312901-INS-198
chr19-7316360-INS-1098
chr19-7319999-INS-128
chr19-7324416-INS-513
chr19-7328066-INS-216
chr19-7333744-INS-296
chr19-7337216-INS-53
chr19-7340696-INS-51
chr19-7344110-INS-185
chr19-7354925-INS-60
chr19-7369529-INS-235
chr19-7374808-INS-577
chr19-7377234-INS-3691
chr19-7380374-INS-181
chr19-7416032-INS-209
chr19-7423272-INS-57
chr19-7452472-INS-662
chr19-7468232-INS-183
chr19-7475467-INS-393
chr19-7487638-INS-5549
chr19-7492170-INS-206
chr19-7507297-INS-216
chr19-7515247-INS-60
chr19-7527767-INS-8973
chr19-7542472-INS-240
chr19-7547990-INS-229
chr19-7594972-INS-219
chr19-7601219-INS-205
chr19-7607317-INS-253
chr19-7612351-INS-62
chr19-7615101-INS-628
chr19-7648024-INS-126
chr19-7658283-INS-402
chr19-7666082-INS-197
chr19-7673677-INS-145
chr19-7680211-INS-6202
chr19-7704513-INS-131
chr19-7776283-INS-6196
chr19-7811656-INS-7106
chr19-7827793-INS-174
chr19-7832253-INS-324
chr19-7835441-INS-284
chr19-7839310-INS-149
chr19-7842750-INS-212
chr19-7946782-INS-79
chr19-7974690-INS-7205
chr19-8032987-INS-314
chr19-8079614-INS-60
chr19-8099388-INS-1625
chr19-8119889-INS-6090
chr19-8162416-INS-1842
chr19-8162954-INS-1142
chr19-8183338-INS-56
chr19-8188700-INS-2895
chr19-8243150-INS-71
chr19-8255225-INS-6432
chr19-8319247-INS-78
chr19-8328261-INS-52
chr19-8376754-INS-684
chr19-8381673-INS-51
chr19-8411220-INS-5413
chr19-8426785-INS-53
chr19-8443942-INS-112
chr19-8455011-INS-400
chr19-8504615-INS-176
chr19-8524478-INS-6475
chr19-8527157-INS-147
chr19-8527346-INS-6441
chr19-8562662-INS-6686
chr19-8579738-INS-56
chr19-8593506-INS-312
chr19-8617793-INS-81
chr19-8621402-INS-80
chr19-8639096-INS-103
chr19-8647585-INS-530
chr19-8665572-INS-121

chr19-8677990-INS-200
chr19-8682182-INS-58
chr19-8689651-INS-55
chr19-8696878-INS-203
chr19-8708519-INS-185
chr19-8709121-INS-234
chr19-8712222-INS-192
chr19-8730110-INS-688
chr19-8731175-INS-197
chr19-8737601-INS-572
chr19-8738257-INS-133
chr19-8770253-INS-213
chr19-8793574-INS-794
chr19-8805065-INS-198
chr19-8814306-INS-202
chr19-8822342-INS-217
chr19-8824988-INS-88
chr19-8825631-INS-364
chr19-8826682-INS-165
chr19-8836129-INS-79
chr19-8838817-INS-238
chr19-8842060-INS-54
chr19-8852309-INS-189
chr19-8857115-INS-142
chr19-8882894-INS-435
chr19-8901939-INS-118
chr19-8917284-INS-145
chr19-8924236-INS-349
chr19-8947674-INS-164
chr19-8972966-INS-52
chr19-9034944-INS-64
chr19-9056158-INS-53
chr19-9064494-INS-168
chr19-9073758-INS-87
chr19-9077629-INS-160
chr19-9083894-INS-234
chr19-9097493-INS-210
chr19-9112445-INS-59
chr19-9116937-INS-6388
chr19-9119741-INS-7082
chr19-9153265-INS-189
chr19-9160235-INS-99
chr19-9166123-INS-206
chr19-9248010-INS-77
chr19-9253422-INS-282
chr19-9257730-INS-72
chr19-9274905-INS-302
chr19-9392293-INS-671
chr19-9402498-INS-83
chr19-9478252-INS-54
chr19-9499838-INS-687
chr19-9511583-INS-7182
chr19-9556115-INS-4830
chr19-9571100-INS-57
chr19-9602089-INS-3086
chr19-9611249-INS-78
chr19-9687216-INS-6063
chr19-9691422-INS-6380
chr19-9713054-INS-205
chr19-9787487-INS-465
chr19-9795354-INS-518
chr19-9826620-INS-78
chr19-9837139-INS-87
chr19-9843439-INS-6674
chr19-9845331-INS-243
chr19-9884630-INS-62
chr19-9911627-INS-79
chr19-9941192-INS-376
chr19-9965331-INS-204
chr19-9968466-INS-123
chr19-9971063-INS-418
chr19-9975992-INS-134
chr19-10025318-INS-52
chr19-10038446-INS-56
chr19-10054358-INS-182
chr19-10074999-INS-79
chr19-10081329-INS-6440

chr19-10095191-INS-120
chr19-10103951-INS-140
chr19-10107573-INS-54
chr19-10120801-INS-62
chr19-10123383-INS-92
chr19-10138977-INS-77
chr19-10142130-INS-184
chr19-10147258-INS-215
chr19-10159238-INS-52
chr19-10169752-INS-328
chr19-10206984-INS-67
chr19-10218774-INS-188
chr19-10231955-INS-88
chr19-10243199-INS-58
chr19-10252193-INS-487
chr19-10293824-INS-223
chr19-10295153-INS-297
chr19-10315469-INS-272
chr19-10332907-INS-60
chr19-10344247-INS-74
chr19-10356406-INS-286
chr19-10363627-INS-75
chr19-10385338-INS-73
chr19-10433404-INS-56
chr19-10442897-INS-114
chr19-10448750-INS-452
chr19-10450293-INS-110
chr19-10481025-INS-618
chr19-10486923-INS-81
chr19-10498455-INS-212
chr19-10527111-INS-229
chr19-10530786-INS-202
chr19-10535283-INS-63
chr19-10546199-INS-197
chr19-10550594-INS-202
chr19-10569921-INS-68
chr19-10572537-INS-300
chr19-10578394-INS-182
chr19-10587632-INS-134
chr19-10617025-INS-51
chr19-10637557-INS-297
chr19-10655460-INS-223
chr19-10659185-INS-59
chr19-10676796-INS-933
chr19-10699214-INS-211
chr19-10717776-INS-68
chr19-10723553-INS-51
chr19-10727222-INS-134
chr19-10730932-INS-54
chr19-10736595-INS-135
chr19-10744056-INS-66
chr19-10745712-INS-60
chr19-10764135-INS-254
chr19-10773519-INS-52
chr19-10798250-INS-101
chr19-10802104-INS-61
chr19-10824761-INS-200
chr19-10825671-INS-148
chr19-10856409-INS-149
chr19-10901872-INS-64
chr19-10903217-INS-199
chr19-10915580-INS-236
chr19-10935113-INS-217
chr19-10939992-INS-72
chr19-10952593-INS-7485
chr19-10961271-INS-168
chr19-10964826-INS-6475
chr19-10976555-INS-101
chr19-10979196-INS-66
chr19-10992284-INS-227
chr19-10998009-INS-73
chr19-11007451-INS-182
chr19-11050920-INS-60
chr19-11075338-INS-580
chr19-11084026-INS-57
chr19-11086328-INS-107
chr19-11108634-INS-72

chr19-11116248-INS-80
chr19-11154257-INS-56
chr19-11160491-INS-1890
chr19-11166276-INS-730
chr19-11186727-INS-8322
chr19-11189543-INS-3422
chr19-11192881-INS-122
chr19-11211666-INS-943
chr19-11212895-INS-205
chr19-11217655-INS-84
chr19-11222609-INS-163
chr19-11246063-INS-6661
chr19-11251811-INS-130
chr19-11251966-INS-1313
chr19-11299858-INS-61
chr19-11315895-INS-146
chr19-11320316-INS-54
chr19-11328303-INS-358
chr19-11329108-INS-79
chr19-11342266-INS-6198
chr19-11370285-INS-173
chr19-11393433-INS-286
chr19-11449303-INS-1093
chr19-11449466-INS-2653
chr19-11455406-INS-81
chr19-11459183-INS-85
chr19-11470886-INS-69
chr19-11500952-INS-7139
chr19-11546319-INS-62
chr19-11559074-INS-270
chr19-11563696-INS-302
chr19-11569669-INS-1512
chr19-11586833-INS-1114
chr19-11657764-INS-293
chr19-11658796-INS-307
chr19-11664056-INS-216
chr19-11672976-INS-58
chr19-11679263-INS-488
chr19-11695486-INS-492
chr19-11705370-INS-293
chr19-11718686-INS-83
chr19-11718799-INS-3347
chr19-11724687-INS-72
chr19-11744747-INS-380
chr19-11746946-INS-60
chr19-11751062-INS-510
chr19-11758649-INS-54
chr19-11761076-INS-264
chr19-11764846-INS-128
chr19-11765450-INS-175
chr19-11769449-INS-187
chr19-11771648-INS-97
chr19-11777565-INS-1210
chr19-11782961-INS-636
chr19-11792892-INS-337
chr19-11798723-INS-498
chr19-11817567-INS-172
chr19-11824158-INS-197
chr19-11829164-INS-328
chr19-11830868-INS-7190
chr19-11853650-INS-170
chr19-11855587-INS-126
chr19-11858448-INS-131
chr19-11867669-INS-128
chr19-11876716-INS-499
chr19-11893956-INS-60
chr19-11895757-INS-96
chr19-11901749-INS-88
chr19-11904393-INS-590
chr19-11908504-INS-322
chr19-11930240-INS-419
chr19-11935792-INS-53
chr19-11939948-INS-63
chr19-11980784-INS-132
chr19-11990489-INS-58
chr19-11991210-INS-74
chr19-11994880-INS-66

chr19-12005061-INS-54
chr19-12010130-INS-196
chr19-12010378-INS-118
chr19-12015600-INS-58
chr19-12054024-INS-110
chr19-12056548-INS-56
chr19-12058029-INS-67
chr19-12060357-INS-75
chr19-12062375-INS-54
chr19-12091937-INS-282
chr19-12104088-INS-199
chr19-12106937-INS-6481
chr19-12110798-INS-6213
chr19-12115374-INS-93
chr19-12148538-INS-421
chr19-12154832-INS-170
chr19-12165553-INS-69
chr19-12182622-INS-59
chr19-12182729-INS-6500
chr19-12199083-INS-345
chr19-12204459-INS-71
chr19-12252563-INS-140
chr19-12256776-INS-385
chr19-12260190-INS-74
chr19-12307200-INS-97
chr19-12318698-INS-55
chr19-12349375-INS-416
chr19-12354854-INS-993
chr19-12356646-INS-232
chr19-12397391-INS-529
chr19-12412997-INS-230
chr19-12418063-INS-126
chr19-12461944-INS-99
chr19-12471915-INS-102
chr19-12475608-INS-54
chr19-12499973-INS-105
chr19-12506784-INS-863
chr19-12587566-INS-240
chr19-12594441-INS-54
chr19-12596472-INS-6626
chr19-12596538-INS-5291
chr19-12602179-INS-51
chr19-12624412-INS-598
chr19-12650928-INS-366
chr19-12703118-INS-428
chr19-12703145-INS-6704
chr19-12708221-INS-313
chr19-12758982-INS-225
chr19-12789808-INS-170
chr19-12811556-INS-91
chr19-12829322-INS-55
chr19-12843630-INS-409
chr19-12855225-INS-136
chr19-12858812-INS-187
chr19-12894110-INS-58
chr19-12928114-INS-99
chr19-12935980-INS-172
chr19-12942851-INS-70
chr19-12980065-INS-392
chr19-12983073-INS-88
chr19-12994768-INS-57
chr19-13041142-INS-669
chr19-13049618-INS-1133
chr19-13081271-INS-200
chr19-13104255-INS-2890
chr19-13132250-INS-125
chr19-13175845-INS-6173
chr19-13214062-INS-130
chr19-13278786-INS-1885
chr19-13303159-INS-6677
chr19-13323586-INS-60
chr19-13338512-INS-601
chr19-13434462-INS-112
chr19-13530902-INS-84
chr19-13550539-INS-51
chr19-13605142-INS-5302
chr19-13650070-INS-389

chr19-13662020-INS-268
chr19-13678831-INS-160
chr19-13688229-INS-605
chr19-13723679-INS-70
chr19-13740980-INS-116
chr19-13771128-INS-78
chr19-13776832-INS-9486
chr19-13780183-INS-399
chr19-13795585-INS-61
chr19-13797794-INS-507
chr19-13822060-INS-96
chr19-13841305-INS-362
chr19-13849206-INS-2460
chr19-13859984-INS-7037
chr19-13936606-INS-51
chr19-13968446-INS-70
chr19-13991127-INS-720
chr19-14011714-INS-116
chr19-14020647-INS-654
chr19-14063296-INS-132
chr19-14098124-INS-56
chr19-14102125-INS-8419
chr19-14124190-INS-7024
chr19-14155067-INS-53
chr19-14215779-INS-122
chr19-14222177-INS-58
chr19-14235699-INS-284
chr19-14248204-INS-12830
chr19-14249523-INS-3129
chr19-14263979-INS-72
chr19-14264450-INS-64
chr19-14339022-INS-160
chr19-14341954-INS-76
chr19-14342688-INS-7240
chr19-14342959-INS-256
chr19-14343488-INS-102
chr19-14372914-INS-405
chr19-14374607-INS-2288
chr19-14445747-INS-317
chr19-14461282-INS-634
chr19-14470054-INS-401
chr19-14472012-INS-64
chr19-14472756-INS-894
chr19-14479525-INS-58
chr19-14523170-INS-439
chr19-14525941-INS-52
chr19-14535556-INS-302
chr19-14536475-INS-86
chr19-14552428-INS-249
chr19-14557050-INS-544
chr19-14587755-INS-241
chr19-14603403-INS-79
chr19-14611508-INS-195
chr19-14614151-INS-66
chr19-14640031-INS-198
chr19-14641150-INS-239
chr19-14655039-INS-781
chr19-14693554-INS-358
chr19-14697907-INS-1190
chr19-14699493-INS-4278
chr19-14701262-INS-163
chr19-14706944-INS-217
chr19-14720104-INS-59
chr19-14753410-INS-504
chr19-14844421-INS-52
chr19-14871035-INS-53
chr19-14871541-INS-60
chr19-14871878-INS-1897
chr19-14907429-INS-446
chr19-14909918-INS-60
chr19-14944971-INS-478
chr19-14946932-INS-1889
chr19-14962724-INS-358
chr19-14966113-INS-6440
chr19-14993690-INS-1489
chr19-15014005-INS-398
chr19-15019591-INS-7107

chr19-15078347-INS-2856
chr19-15094994-INS-390
chr19-15097437-INS-71
chr19-15114739-INS-6282
chr19-15119468-INS-546
chr19-15123743-INS-6605
chr19-15151550-INS-51
chr19-15158318-INS-265
chr19-15176630-INS-697
chr19-15179982-INS-94
chr19-15190016-INS-118
chr19-15195768-INS-419
chr19-15196671-INS-59
chr19-15219814-INS-66
chr19-15221661-INS-223
chr19-15225165-INS-56
chr19-15243604-INS-400
chr19-15255861-INS-60
chr19-15264713-INS-7185
chr19-15275329-INS-176
chr19-15278654-INS-356
chr19-15308859-INS-104
chr19-15367571-INS-6630
chr19-15394747-INS-932
chr19-15450681-INS-66
chr19-15451203-INS-396
chr19-15456957-INS-54
chr19-15479270-INS-51
chr19-15503894-INS-420
chr19-15513648-INS-6642
chr19-15524178-INS-51
chr19-15529169-INS-58
chr19-15530843-INS-80
chr19-15569250-INS-62
chr19-15586408-INS-254
chr19-15587810-INS-72
chr19-15591061-INS-62
chr19-15604856-INS-364
chr19-15641168-INS-107
chr19-15643188-INS-52
chr19-15707488-INS-449
chr19-15710419-INS-54
chr19-15721103-INS-60
chr19-15736846-INS-272
chr19-15763547-INS-93
chr19-15794604-INS-209
chr19-15796613-INS-164
chr19-15802114-INS-123
chr19-15840328-INS-73
chr19-15856292-INS-52
chr19-15866953-INS-217
chr19-15871573-INS-64
chr19-15879181-INS-880
chr19-15893099-INS-58
chr19-15900958-INS-217
chr19-15911126-INS-60
chr19-15914389-INS-61
chr19-15921028-INS-383
chr19-15938093-INS-227
chr19-15951614-INS-68
chr19-15966239-INS-114
chr19-15971178-INS-384
chr19-16006107-INS-54
chr19-16010735-INS-145
chr19-16027605-INS-505
chr19-16030537-INS-98
chr19-16032386-INS-7126
chr19-16050058-INS-79
chr19-16062077-INS-75
chr19-16085542-INS-497
chr19-16088001-INS-51
chr19-16095407-INS-78
chr19-16115558-INS-60
chr19-16131378-INS-81
chr19-16132539-INS-289
chr19-16175459-INS-200
chr19-16178820-INS-399

chr19-16180810-INS-159
chr19-16188238-INS-180
chr19-16190713-INS-238
chr19-16194281-INS-350
chr19-16195811-INS-133
chr19-16198669-INS-96
chr19-16207471-INS-60
chr19-16238674-INS-545
chr19-16251725-INS-93
chr19-16254594-INS-330
chr19-16258055-INS-55
chr19-16264631-INS-120
chr19-16294166-INS-5547
chr19-16307882-INS-7517
chr19-16311300-INS-171
chr19-16313166-INS-86
chr19-16383226-INS-5000
chr19-16383932-INS-212
chr19-16393073-INS-4984
chr19-16410999-INS-7282
chr19-16419124-INS-56
chr19-16423353-INS-122
chr19-16424746-INS-72
chr19-16441266-INS-166
chr19-16461122-INS-60
chr19-16465995-INS-52
chr19-16474548-INS-151
chr19-16519243-INS-480
chr19-16541949-INS-92
chr19-16545271-INS-345
chr19-16546481-INS-63
chr19-16574020-INS-308
chr19-16592189-INS-50
chr19-16599506-INS-57
chr19-16614215-INS-857
chr19-16621438-INS-142
chr19-16652469-INS-168
chr19-16658548-INS-7203
chr19-16677184-INS-686
chr19-16679659-INS-739
chr19-16694317-INS-123
chr19-16708268-INS-68
chr19-16730609-INS-201
chr19-16741056-INS-175
chr19-16747096-INS-63
chr19-16773336-INS-52
chr19-16791792-INS-72
chr19-16801166-INS-194
chr19-16808956-INS-391
chr19-16841774-INS-68
chr19-16863518-INS-62
chr19-16872594-INS-191
chr19-16874446-INS-90
chr19-16876606-INS-161
chr19-16878571-INS-5434
chr19-16889826-INS-68
chr19-16895674-INS-688
chr19-16946992-INS-108
chr19-16962702-INS-218
chr19-17040522-INS-220
chr19-17060331-INS-609
chr19-17069481-INS-541
chr19-17089995-INS-67
chr19-17111637-INS-54
chr19-17139007-INS-151
chr19-17150373-INS-400
chr19-17171026-INS-396
chr19-17175580-INS-626
chr19-17304347-INS-176
chr19-17319082-INS-118
chr19-17332762-INS-55
chr19-17343620-INS-68
chr19-17346594-INS-62
chr19-17357649-INS-322
chr19-17365067-INS-73
chr19-17377364-INS-76
chr19-17388945-INS-55

chr19-17410260-INS-84
chr19-17435623-INS-83
chr19-17465116-INS-615
chr19-17495174-INS-6383
chr19-17501800-INS-367
chr19-17517589-INS-72
chr19-17518938-INS-66
chr19-17523117-INS-52
chr19-17526607-INS-206
chr19-17541817-INS-88
chr19-17557088-INS-120
chr19-17571133-INS-78
chr19-17577201-INS-212
chr19-17578460-INS-52
chr19-17609891-INS-722
chr19-17695887-INS-117
chr19-17706577-INS-253
chr19-17734333-INS-108
chr19-17752251-INS-140
chr19-17769874-INS-390
chr19-17786901-INS-60
chr19-17795367-INS-123
chr19-17837626-INS-7532
chr19-17844251-INS-513
chr19-17862231-INS-394
chr19-17912780-INS-152
chr19-17932252-INS-53
chr19-17934995-INS-56
chr19-17939671-INS-471
chr19-17943939-INS-507
chr19-17988199-INS-422
chr19-17994477-INS-431
chr19-17997380-INS-79
chr19-18078725-INS-212
chr19-18112758-INS-315
chr19-18148879-INS-6574
chr19-18149864-INS-56
chr19-18178465-INS-140
chr19-18190253-INS-872
chr19-18212736-INS-60
chr19-18215951-INS-394
chr19-18227851-INS-64
chr19-18276212-INS-625
chr19-18277237-INS-72
chr19-18299443-INS-56
chr19-18306066-INS-498
chr19-18311279-INS-70
chr19-18313036-INS-93
chr19-18352756-INS-87
chr19-18411065-INS-57
chr19-18422704-INS-162
chr19-18436384-INS-52
chr19-18444029-INS-83
chr19-18469398-INS-179
chr19-18475461-INS-521
chr19-18476389-INS-55
chr19-18484230-INS-75
chr19-18488604-INS-853
chr19-18514215-INS-53
chr19-18533394-INS-577
chr19-18536582-INS-107
chr19-18537536-INS-51
chr19-18541911-INS-52
chr19-18575697-INS-73
chr19-18606945-INS-211
chr19-18620046-INS-431
chr19-18639700-INS-184
chr19-18645689-INS-6669
chr19-18646825-INS-87
chr19-18666144-INS-85
chr19-18689765-INS-493
chr19-18691498-INS-170
chr19-18701807-INS-54
chr19-18709066-INS-190
chr19-18719594-INS-2336
chr19-18722553-INS-96
chr19-18728564-INS-105

chr19-18732543-INS-240
chr19-18738008-INS-53
chr19-18752944-INS-52
chr19-18765368-INS-347
chr19-18770796-INS-134
chr19-18771211-INS-100
chr19-18774512-INS-627
chr19-18784236-INS-130
chr19-18785481-INS-958
chr19-18801919-INS-157
chr19-18802466-INS-60
chr19-18812110-INS-55
chr19-18837862-INS-217
chr19-18859803-INS-55
chr19-18862496-INS-411
chr19-18887945-INS-892
chr19-18989913-INS-704
chr19-18991793-INS-104
chr19-19034797-INS-654
chr19-19049686-INS-142
chr19-19067836-INS-171
chr19-19171346-INS-7083
chr19-19174166-INS-137
chr19-19176616-INS-193
chr19-19210838-INS-52
chr19-19212540-INS-144
chr19-19238043-INS-60
chr19-19244625-INS-391
chr19-19268944-INS-7088
chr19-19292102-INS-77
chr19-19363265-INS-88
chr19-19377166-INS-6004
chr19-19419891-INS-63
chr19-19439071-INS-219
chr19-19465212-INS-829
chr19-19467488-INS-250
chr19-19482764-INS-224
chr19-19503398-INS-120
chr19-19534397-INS-62
chr19-19543123-INS-56
chr19-19561987-INS-127
chr19-19566780-INS-87
chr19-19601308-INS-553
chr19-19623364-INS-277
chr19-19634949-INS-389
chr19-19647488-INS-54
chr19-19678988-INS-198
chr19-19686009-INS-5654
chr19-19713510-INS-622
chr19-19735555-INS-68
chr19-19766984-INS-97
chr19-19785479-INS-51
chr19-19793521-INS-56
chr19-19814312-INS-1982
chr19-19815305-INS-59
chr19-19828242-INS-2975
chr19-19831336-INS-62
chr19-19846836-INS-145
chr19-19884567-INS-129
chr19-19889351-INS-97
chr19-19938843-INS-165
chr19-19944116-INS-147
chr19-19945584-INS-680
chr19-19961407-INS-374
chr19-19968733-INS-191
chr19-19990035-INS-159
chr19-19995827-INS-104
chr19-20003912-INS-330
chr19-20024768-INS-356
chr19-20037216-INS-60
chr19-20043039-INS-5493
chr19-20056515-INS-106
chr19-20095112-INS-583
chr19-20107419-INS-520
chr19-20115064-INS-6790
chr19-20123885-INS-6415
chr19-20124401-INS-399

chr19-20130478-INS-65
chr19-20134515-INS-358
chr19-20141610-INS-219
chr19-20170040-INS-6936
chr19-20181318-INS-85
chr19-20206929-INS-164
chr19-20216781-INS-64
chr19-20243194-INS-401
chr19-20260527-INS-685
chr19-20288656-INS-120
chr19-20293354-INS-192
chr19-20302540-INS-157
chr19-20333662-INS-293
chr19-20335970-INS-130
chr19-20369623-INS-63
chr19-20370192-INS-62
chr19-20385986-INS-471
chr19-20400323-INS-66
chr19-20402780-INS-292
chr19-20415014-INS-218
chr19-20473422-INS-407
chr19-20476309-INS-65
chr19-20479341-INS-130
chr19-20495005-INS-585
chr19-20498255-INS-204
chr19-20531460-INS-96
chr19-20532022-INS-211
chr19-20537300-INS-200
chr19-20550854-INS-211
chr19-20590998-INS-68
chr19-20638383-INS-6440
chr19-20641942-INS-1563
chr19-20660409-INS-6604
chr19-20667509-INS-66
chr19-20671167-INS-1894
chr19-20733505-INS-232
chr19-20734053-INS-2845
chr19-20755518-INS-66
chr19-20767092-INS-517
chr19-20807674-INS-476
chr19-20822709-INS-56
chr19-20867558-INS-140
chr19-20886302-INS-113
chr19-20909498-INS-288
chr19-20927167-INS-88
chr19-20930581-INS-84
chr19-20964771-INS-54
chr19-20968380-INS-604
chr19-20994481-INS-158
chr19-20996773-INS-330
chr19-21001652-INS-52
chr19-21003331-INS-78
chr19-21019564-INS-210
chr19-21020928-INS-51
chr19-21067513-INS-94
chr19-21079359-INS-78
chr19-21090283-INS-142
chr19-21091252-INS-91
chr19-21098221-INS-133
chr19-21103096-INS-814
chr19-21106497-INS-162
chr19-21133442-INS-365
chr19-21155087-INS-775
chr19-21240247-INS-85
chr19-21265797-INS-284
chr19-21282491-INS-535
chr19-21296936-INS-942
chr19-21299162-INS-339
chr19-21344285-INS-174
chr19-21367184-INS-206
chr19-21370072-INS-163
chr19-21394550-INS-754
chr19-21405350-INS-54
chr19-21440432-INS-274
chr19-21455741-INS-3679
chr19-21481130-INS-178
chr19-21504475-INS-332

chr19-21526822-INS-321
chr19-21528040-INS-322
chr19-21568086-INS-60
chr19-21571888-INS-55
chr19-21573627-INS-730
chr19-21575570-INS-68
chr19-21580961-INS-153
chr19-21584618-INS-141
chr19-21613394-INS-74
chr19-21724084-INS-119
chr19-21728578-INS-505
chr19-21745767-INS-508
chr19-21748699-INS-53
chr19-21761098-INS-467
chr19-21781334-INS-87
chr19-21793725-INS-104
chr19-21837397-INS-91
chr19-21848537-INS-207
chr19-21854598-INS-52
chr19-21869057-INS-86
chr19-21885904-INS-7349
chr19-21886420-INS-56
chr19-21889301-INS-138
chr19-21923563-INS-72
chr19-21977770-INS-212
chr19-21989017-INS-967
chr19-21991491-INS-52
chr19-21997038-INS-428
chr19-22013057-INS-66
chr19-22097282-INS-56
chr19-22099021-INS-56
chr19-22101808-INS-401
chr19-22105355-INS-62
chr19-22130726-INS-58
chr19-22176311-INS-64
chr19-22205041-INS-193
chr19-22283068-INS-74
chr19-22304675-INS-1312
chr19-22316345-INS-57
chr19-22325121-INS-54
chr19-22326142-INS-6598
chr19-22330656-INS-108
chr19-22353042-INS-60
chr19-22371224-INS-126
chr19-22379957-INS-314
chr19-22390640-INS-173
chr19-22469764-INS-6189
chr19-22499536-INS-682
chr19-22501538-INS-82
chr19-22606154-INS-374
chr19-22617939-INS-189
chr19-22664715-INS-69
chr19-22688353-INS-97
chr19-22712929-INS-167
chr19-22756885-INS-344
chr19-22768528-INS-574
chr19-22782602-INS-7286
chr19-22789077-INS-340
chr19-22820073-INS-400
chr19-22827442-INS-86
chr19-22831372-INS-452
chr19-22849426-INS-629
chr19-22859457-INS-130
chr19-22863378-INS-102
chr19-22891020-INS-213
chr19-22893096-INS-120
chr19-22897592-INS-190
chr19-22914495-INS-56
chr19-22924836-INS-173
chr19-22931714-INS-376
chr19-22937529-INS-118
chr19-22963498-INS-61
chr19-22967785-INS-53
chr19-22968860-INS-60
chr19-22976230-INS-69
chr19-22988429-INS-6511
chr19-22989848-INS-108

chr19-22996740-INS-121
chr19-23008998-INS-86
chr19-23015513-INS-90
chr19-23028283-INS-264
chr19-23031432-INS-294
chr19-23036370-INS-65
chr19-23041330-INS-62
chr19-23044707-INS-58
chr19-23051256-INS-208
chr19-23055325-INS-112
chr19-23088178-INS-214
chr19-23138540-INS-444
chr19-23146203-INS-55
chr19-23181394-INS-148
chr19-23206114-INS-207
chr19-23212719-INS-76
chr19-23215874-INS-59
chr19-23217623-INS-341
chr19-23226993-INS-183
chr19-23243277-INS-214
chr19-23250385-INS-133
chr19-23262341-INS-59
chr19-23276309-INS-878
chr19-23314479-INS-951
chr19-23320590-INS-138
chr19-23347312-INS-187
chr19-23349591-INS-126
chr19-23353220-INS-196
chr19-23370532-INS-126
chr19-23396643-INS-248
chr19-23402610-INS-6468
chr19-23404961-INS-993
chr19-23406912-INS-148
chr19-23413212-INS-124
chr19-23423223-INS-74
chr19-23435817-INS-54
chr19-23443664-INS-78
chr19-23444258-INS-52
chr19-23473173-INS-236
chr19-23488748-INS-691
chr19-23495730-INS-85
chr19-23506143-INS-125
chr19-23531826-INS-295
chr19-23552977-INS-103
chr19-23595074-INS-59
chr19-23626836-INS-791
chr19-23635586-INS-228
chr19-23641814-INS-305
chr19-23650419-INS-56
chr19-23692719-INS-474
chr19-23696929-INS-64
chr19-23703166-INS-91
chr19-23713204-INS-87
chr19-23745673-INS-56
chr19-23795117-INS-455
chr19-23820732-INS-84
chr19-23825317-INS-70
chr19-23834755-INS-252
chr19-23857729-INS-74
chr19-23931085-INS-904
chr19-23939721-INS-103
chr19-23992585-INS-216
chr19-23995334-INS-87
chr19-24020357-INS-191
chr19-24040712-INS-52
chr19-24063958-INS-627
chr19-24079755-INS-79
chr19-24095249-INS-71
chr19-24132115-INS-186
chr19-24140165-INS-97
chr19-24163335-INS-443
chr19-24181945-INS-143
chr19-24186018-INS-60
chr19-24197318-INS-112
chr19-24219486-INS-205
chr19-24224911-INS-199
chr19-24237098-INS-177

chr19-24252275-INS-76
chr19-24256873-INS-200
chr19-24274218-INS-167
chr19-24286532-INS-229
chr19-24296793-INS-296
chr19-24308872-INS-54
chr19-24320021-INS-136
chr19-24323492-INS-51
chr19-24332686-INS-110
chr19-24374031-INS-155
chr19-24444815-INS-82
chr19-24449850-INS-146
chr19-24459795-INS-255
chr19-24465828-INS-66
chr19-24471621-INS-510
chr19-24506111-INS-98
chr19-24507093-INS-100
chr19-24523433-INS-202
chr19-24539806-INS-99
chr19-24541278-INS-197
chr19-24578069-INS-257
chr19-24583356-INS-6329
chr19-24608193-INS-102
chr19-24621244-INS-60
chr19-24636254-INS-205
chr19-24636470-INS-682
chr19-24661489-INS-82
chr19-24663701-INS-51
chr19-24673446-INS-66
chr19-24718535-INS-222
chr19-24721443-INS-369
chr19-24753368-INS-55
chr19-24800194-INS-54
chr19-24820719-INS-109
chr19-24831817-INS-88
chr19-24835718-INS-238
chr19-24870081-INS-6421
chr19-24875393-INS-65
chr19-24884419-INS-377
chr19-24933673-INS-52
chr19-24948046-INS-90
chr19-24960545-INS-56
chr19-24960761-INS-945
chr19-24963904-INS-357
chr19-24970280-INS-74
chr19-24987933-INS-60
chr19-24989030-INS-323
chr19-25004246-INS-63
chr19-25022356-INS-70
chr19-25038715-INS-148
chr19-25075187-INS-52
chr19-25091730-INS-174
chr19-25102211-INS-80
chr19-25194082-INS-67
chr19-25236985-INS-6449
chr19-25264839-INS-62
chr19-25274096-INS-93
chr19-25310724-INS-68
chr19-25322536-INS-68
chr19-25338741-INS-203
chr19-25349503-INS-340
chr19-25354041-INS-126
chr19-25422969-INS-130
chr19-25430065-INS-88
chr19-25436189-INS-6401
chr19-25438106-INS-252
chr19-25442273-INS-194
chr19-25449405-INS-153
chr19-25452047-INS-176
chr19-25457807-INS-197
chr19-25461840-INS-187
chr19-25500604-INS-448
chr19-25509107-INS-166
chr19-25509866-INS-129
chr19-25544984-INS-124
chr19-25547274-INS-213
chr19-25573549-INS-62

chr19-25596498-INS-64
chr19-25656696-INS-68
chr19-25738513-INS-198
chr19-25749039-INS-52
chr19-25753058-INS-6694
chr19-25754418-INS-118
chr19-25756855-INS-115
chr19-25763038-INS-559
chr19-25763054-INS-714
chr19-25792276-INS-420
chr19-25814697-INS-68
chr19-25835078-INS-69
chr19-25860905-INS-509
chr19-25869440-INS-56
chr19-25887426-INS-183
chr19-25929952-INS-175
chr19-25932423-INS-132
chr19-25944935-INS-69
chr19-25964976-INS-530
chr19-25986227-INS-58
chr19-26039584-INS-148
chr19-26048951-INS-91
chr19-26081498-INS-406
chr19-26094195-INS-376
chr19-26118671-INS-82
chr19-26126368-INS-326
chr19-26127539-INS-75
chr19-26134665-INS-86
chr19-26291215-INS-51
chr19-26304240-INS-96
chr19-26305738-INS-130
chr19-26314619-INS-512
chr19-26350873-INS-88
chr19-26409245-INS-60
chr19-26410245-INS-84
chr19-26420723-INS-658
chr19-26436976-INS-54
chr19-26446137-INS-157
chr19-26489648-INS-57
chr19-26514271-INS-78
chr19-26529636-INS-554
chr19-26593662-INS-86
chr19-26612398-INS-398
chr19-26615123-INS-85
chr19-26646058-INS-498
chr19-26648299-INS-68
chr19-26649175-INS-228
chr19-26658782-INS-51
chr19-26672933-INS-56
chr19-26687404-INS-80
chr19-26691986-INS-92
chr19-26707081-INS-110
chr19-26708576-INS-131
chr19-26725558-INS-59
chr19-26730734-INS-84
chr19-26738184-INS-78
chr19-26748066-INS-517
chr19-26789680-INS-128
chr19-26794838-INS-173
chr19-26796589-INS-130
chr19-26880938-INS-374
chr19-26900098-INS-143
chr19-26930451-INS-84
chr19-26938322-INS-3137
chr19-26939749-INS-204
chr19-26950727-INS-197
chr19-26954017-INS-57
chr19-26972528-INS-306
chr19-26988860-INS-95
chr19-26994632-INS-153
chr19-27006717-INS-53
chr19-27018101-INS-70
chr19-27021894-INS-68
chr19-27029695-INS-59
chr19-27051224-INS-137
chr19-27052866-INS-6585
chr19-27057211-INS-209

chr19-27058354-INS-260
chr19-27070309-INS-276
chr19-27086394-INS-477
chr19-27105551-INS-7276
chr19-27178238-INS-149
chr19-27184787-INS-428
chr19-27219358-INS-98
chr19-27220082-INS-399
chr19-27256766-INS-82
chr19-27258245-INS-210
chr19-27263764-INS-566
chr19-27274310-INS-86
chr19-27292423-INS-52
chr19-27299397-INS-191
chr19-27312233-INS-61
chr19-27328755-INS-4859
chr19-27334421-INS-62
chr19-27341612-INS-56
chr19-27387148-INS-193
chr19-27396966-INS-410
chr19-27411856-INS-663
chr19-27421122-INS-78
chr19-27426162-INS-132
chr19-27437194-INS-117
chr19-27446010-INS-150
chr19-27500116-INS-64
chr19-27507684-INS-72
chr19-27525330-INS-373
chr19-27542873-INS-61
chr19-27550186-INS-464
chr19-27550889-INS-54
chr19-27583555-INS-93
chr19-27588237-INS-51
chr19-27597285-INS-143
chr19-27603136-INS-233
chr19-27616695-INS-432
chr19-27622095-INS-342
chr19-27636826-INS-66
chr19-27642299-INS-148
chr19-27654405-INS-7194
chr19-27656296-INS-387
chr19-27659718-INS-56
chr19-27699544-INS-122
chr19-27747054-INS-204
chr19-27779035-INS-64
chr19-27816004-INS-134
chr19-27817191-INS-4076
chr19-27821178-INS-366
chr19-27831862-INS-90
chr19-27856223-INS-99
chr19-27861281-INS-57
chr19-27908472-INS-91
chr19-27913662-INS-215
chr19-27941377-INS-84
chr19-27950843-INS-427
chr19-27953439-INS-70
chr19-28025396-INS-58
chr19-28061027-INS-100
chr19-28084768-INS-185
chr19-28085643-INS-488
chr19-28097294-INS-52
chr19-28136585-INS-52
chr19-28171832-INS-65
chr19-28177579-INS-107
chr19-28178544-INS-222
chr19-28194163-INS-61
chr19-28255316-INS-141
chr19-28259135-INS-683
chr19-28270725-INS-105
chr19-28347995-INS-340
chr19-28349496-INS-52
chr19-28402487-INS-1803
chr19-28450155-INS-210
chr19-28465969-INS-94
chr19-28507115-INS-524
chr19-28521923-INS-71
chr19-28591286-INS-73

chr19-28606590-INS-68
chr19-28612738-INS-153
chr19-28614766-INS-55
chr19-28637947-INS-54
chr19-28658411-INS-136
chr19-28674153-INS-191
chr19-28677422-INS-95
chr19-28690453-INS-123
chr19-28715246-INS-260
chr19-28738480-INS-174
chr19-28751706-INS-421
chr19-28757340-INS-400
chr19-28762429-INS-110
chr19-28789180-INS-61
chr19-28839379-INS-96
chr19-28846217-INS-62
chr19-28859969-INS-60
chr19-28902008-INS-101
chr19-28910726-INS-103
chr19-28911534-INS-60
chr19-28931064-INS-316
chr19-28971949-INS-101
chr19-28980558-INS-135
chr19-29004271-INS-5558
chr19-29007899-INS-202
chr19-29012567-INS-54
chr19-29013709-INS-2857
chr19-29035185-INS-105
chr19-29075567-INS-224
chr19-29093553-INS-865
chr19-29094439-INS-140
chr19-29138804-INS-71
chr19-29140697-INS-79
chr19-29148106-INS-773
chr19-29155349-INS-385
chr19-29170938-INS-89
chr19-29174012-INS-295
chr19-29181077-INS-84
chr19-29206987-INS-226
chr19-29210431-INS-199
chr19-29237064-INS-191
chr19-29273313-INS-79
chr19-29305093-INS-60
chr19-29335142-INS-633
chr19-29335913-INS-120
chr19-29340350-INS-321
chr19-29384001-INS-65
chr19-29418159-INS-78
chr19-29427172-INS-99
chr19-29435473-INS-234
chr19-29439337-INS-53
chr19-29459047-INS-184
chr19-29474937-INS-135
chr19-29495825-INS-61
chr19-29508051-INS-495
chr19-29520386-INS-474
chr19-29548978-INS-194
chr19-29567016-INS-209
chr19-29600166-INS-76
chr19-29606577-INS-315
chr19-29612843-INS-62
chr19-29623822-INS-76
chr19-29629734-INS-182
chr19-29643076-INS-89
chr19-29655284-INS-110
chr19-29674297-INS-233
chr19-29679604-INS-79
chr19-29688625-INS-2869
chr19-29714703-INS-204
chr19-29753169-INS-73
chr19-29764810-INS-6126
chr19-29775886-INS-215
chr19-29778478-INS-99
chr19-29792673-INS-100
chr19-29799405-INS-358
chr19-29815322-INS-462
chr19-29837254-INS-196

chr19-29845819-INS-324
chr19-29853910-INS-310
chr19-29870570-INS-88
chr19-29872283-INS-342
chr19-29875804-INS-56
chr19-29930253-INS-449
chr19-29961200-INS-354
chr19-29980617-INS-280
chr19-29998334-INS-88
chr19-30003022-INS-699
chr19-30003951-INS-60
chr19-30007087-INS-59
chr19-30023030-INS-240
chr19-30051670-INS-113
chr19-30058259-INS-60
chr19-30060671-INS-207
chr19-30065274-INS-224
chr19-30067614-INS-76
chr19-30069974-INS-365
chr19-30088034-INS-66
chr19-30146449-INS-106
chr19-30147561-INS-56
chr19-30153878-INS-105
chr19-30161255-INS-205
chr19-30161627-INS-900
chr19-30182299-INS-267
chr19-30193233-INS-75
chr19-30194089-INS-588
chr19-30205146-INS-60
chr19-30213295-INS-135
chr19-30215693-INS-126
chr19-30229720-INS-108
chr19-30249735-INS-98
chr19-30251256-INS-321
chr19-30253277-INS-103
chr19-30254351-INS-79
chr19-30270558-INS-608
chr19-30274069-INS-140
chr19-30279313-INS-351
chr19-30283884-INS-62
chr19-30316302-INS-52
chr19-30323238-INS-301
chr19-30375578-INS-298
chr19-30397853-INS-51
chr19-30427950-INS-52
chr19-30440287-INS-116
chr19-30442942-INS-855
chr19-30445975-INS-342
chr19-30461493-INS-60
chr19-30476547-INS-150
chr19-30477151-INS-243
chr19-30480176-INS-51
chr19-30483561-INS-76
chr19-30488501-INS-194
chr19-30530327-INS-56
chr19-30535231-INS-538
chr19-30544041-INS-126
chr19-30573024-INS-101
chr19-30576747-INS-50
chr19-30598917-INS-131
chr19-30608270-INS-638
chr19-30622861-INS-124
chr19-30686029-INS-65
chr19-30740890-INS-211
chr19-30746993-INS-109
chr19-30751394-INS-846
chr19-30785678-INS-726
chr19-30793740-INS-76
chr19-30826151-INS-400
chr19-30841778-INS-75
chr19-30845254-INS-64
chr19-30848771-INS-83
chr19-30862801-INS-118
chr19-30870059-INS-90
chr19-30873959-INS-659
chr19-30888688-INS-163
chr19-30909097-INS-56

chr19-30936416-INS-423
chr19-30954891-INS-62
chr19-30961062-INS-667
chr19-30968835-INS-242
chr19-30983567-INS-185
chr19-31002253-INS-174
chr19-31031304-INS-54
chr19-31044204-INS-86
chr19-31089869-INS-64
chr19-31111414-INS-61
chr19-31116263-INS-424
chr19-31143480-INS-83
chr19-31154192-INS-7475
chr19-31160585-INS-52
chr19-31189918-INS-863
chr19-31209415-INS-869
chr19-31229040-INS-180
chr19-31238692-INS-103
chr19-31253959-INS-668
chr19-31260979-INS-98
chr19-31302736-INS-698
chr19-31307357-INS-185
chr19-31317334-INS-152
chr19-31325422-INS-190
chr19-31326822-INS-135
chr19-31338311-INS-5549
chr19-31340736-INS-160
chr19-31347909-INS-52
chr19-31396984-INS-86
chr19-31417553-INS-544
chr19-31434145-INS-723
chr19-31453984-INS-61
chr19-31458704-INS-52
chr19-31494777-INS-266
chr19-31505958-INS-57
chr19-31506946-INS-56
chr19-31520197-INS-6699
chr19-31546628-INS-200
chr19-31563394-INS-700
chr19-31573768-INS-410
chr19-31584430-INS-99
chr19-31591115-INS-245
chr19-31643269-INS-63
chr19-31676828-INS-61
chr19-31680030-INS-90
chr19-31700318-INS-156
chr19-31728249-INS-61
chr19-31771521-INS-302
chr19-31777596-INS-98
chr19-31778246-INS-452
chr19-31781373-INS-204
chr19-31792416-INS-910
chr19-31795996-INS-72
chr19-31821619-INS-5849
chr19-31851693-INS-111
chr19-31853310-INS-75
chr19-31875167-INS-97
chr19-31879075-INS-425
chr19-31879998-INS-203
chr19-31888426-INS-64
chr19-31897597-INS-128
chr19-31906708-INS-90
chr19-31921523-INS-51
chr19-31923705-INS-222
chr19-31924228-INS-84
chr19-31927478-INS-84
chr19-31956049-INS-104
chr19-31962363-INS-52
chr19-31977977-INS-973
chr19-32004759-INS-126
chr19-32018811-INS-101
chr19-32028767-INS-6720
chr19-32029487-INS-73
chr19-32030532-INS-74
chr19-32041358-INS-730
chr19-32056155-INS-197
chr19-32060255-INS-56

chr19-32071627-INS-705
chr19-32078991-INS-66
chr19-32084495-INS-546
chr19-32090852-INS-60
chr19-32118044-INS-201
chr19-32130988-INS-439
chr19-32151876-INS-239
chr19-32154967-INS-97
chr19-32157285-INS-75
chr19-32162247-INS-198
chr19-32175819-INS-58
chr19-32207587-INS-218
chr19-32210796-INS-55
chr19-32214586-INS-3414
chr19-32260851-INS-52
chr19-32278751-INS-53
chr19-32303125-INS-87
chr19-32311777-INS-56
chr19-32315801-INS-66
chr19-32317576-INS-59
chr19-32339650-INS-451
chr19-32341780-INS-101
chr19-32370243-INS-594
chr19-32371531-INS-208
chr19-32375768-INS-58
chr19-32378646-INS-69
chr19-32389275-INS-84
chr19-32390685-INS-52
chr19-32396055-INS-65
chr19-32429139-INS-117
chr19-32430988-INS-231
chr19-32434431-INS-503
chr19-32448487-INS-120
chr19-32448538-INS-590
chr19-32475179-INS-618
chr19-32480107-INS-448
chr19-32480569-INS-165
chr19-32512646-INS-202
chr19-32530126-INS-134
chr19-32544305-INS-100
chr19-32555758-INS-50
chr19-32616523-INS-64
chr19-32627638-INS-62
chr19-32635587-INS-390
chr19-32656247-INS-210
chr19-32664032-INS-260
chr19-32670050-INS-51
chr19-32677913-INS-61
chr19-32687465-INS-183
chr19-32698287-INS-247
chr19-32705143-INS-205
chr19-32705654-INS-56
chr19-32712021-INS-208
chr19-32713628-INS-93
chr19-32721618-INS-83
chr19-32728599-INS-271
chr19-32728599-INS-375
chr19-32732614-INS-221
chr19-32742695-INS-60
chr19-32768489-INS-54
chr19-32821377-INS-213
chr19-32836991-INS-146
chr19-32842935-INS-206
chr19-32848846-INS-209
chr19-32851201-INS-180
chr19-32852576-INS-82
chr19-32884239-INS-133
chr19-32902912-INS-64
chr19-32912696-INS-81
chr19-32945201-INS-51
chr19-32948426-INS-88
chr19-32972642-INS-59
chr19-32974639-INS-143
chr19-32977666-INS-187
chr19-32983079-INS-988
chr19-33027426-INS-76
chr19-33073113-INS-105

chr19-33073825-INS-456
chr19-33097934-INS-7104
chr19-33099774-INS-232
chr19-33126749-INS-606
chr19-33131906-INS-66
chr19-33136507-INS-60
chr19-33143820-INS-982
chr19-33148835-INS-55
chr19-33150132-INS-57
chr19-33150767-INS-878
chr19-33151179-INS-6190
chr19-33152003-INS-64
chr19-33161591-INS-125
chr19-33165404-INS-5547
chr19-33167357-INS-200
chr19-33179126-INS-56
chr19-33204074-INS-388
chr19-33208584-INS-53
chr19-33231837-INS-6118
chr19-33249737-INS-511
chr19-33252444-INS-68
chr19-33262160-INS-6870
chr19-33276512-INS-119
chr19-33281160-INS-594
chr19-33286637-INS-85
chr19-33319508-INS-536
chr19-33333592-INS-81
chr19-33451257-INS-6495
chr19-33463090-INS-7163
chr19-33473403-INS-5512
chr19-33496394-INS-79
chr19-33510558-INS-2038
chr19-33512577-INS-8851
chr19-33547107-INS-774
chr19-33635752-INS-79
chr19-33731581-INS-868
chr19-33761488-INS-346
chr19-33777123-INS-8668
chr19-33786587-INS-7149
chr19-33819691-INS-60
chr19-33827436-INS-2894
chr19-33847078-INS-2361
chr19-33880238-INS-5314
chr19-33887767-INS-2463
chr19-33944997-INS-354
chr19-33954981-INS-661
chr19-34028969-INS-195
chr19-34060701-INS-167
chr19-34067986-INS-837
chr19-34069160-INS-196
chr19-34071664-INS-899
chr19-34093379-INS-98
chr19-34097465-INS-61
chr19-34139878-INS-203
chr19-34154697-INS-420
chr19-34159086-INS-392
chr19-34160109-INS-207
chr19-34182975-INS-78
chr19-34209840-INS-151
chr19-34255227-INS-60
chr19-34277018-INS-152
chr19-34315524-INS-198
chr19-34331968-INS-52
chr19-34345213-INS-563
chr19-34377381-INS-189
chr19-34389511-INS-57
chr19-34393585-INS-70
chr19-34397223-INS-51
chr19-34426881-INS-52
chr19-34436002-INS-967
chr19-34440233-INS-55
chr19-34487100-INS-83
chr19-34531535-INS-200
chr19-34539402-INS-115
chr19-34547449-INS-95
chr19-34599189-INS-497
chr19-34616091-INS-70

chr19-34642325-INS-1239
chr19-34662358-INS-136
chr19-34666010-INS-600
chr19-34670463-INS-52
chr19-34708663-INS-334
chr19-34713107-INS-53
chr19-34727352-INS-72
chr19-34755434-INS-54
chr19-34768593-INS-65
chr19-34794433-INS-72
chr19-34800350-INS-328
chr19-34833164-INS-347
chr19-34837152-INS-51
chr19-34852806-INS-62
chr19-34865441-INS-89
chr19-34880071-INS-102
chr19-34882231-INS-103
chr19-34901580-INS-169
chr19-34923342-INS-186
chr19-34931770-INS-6452
chr19-34951523-INS-72
chr19-34970022-INS-449
chr19-34979148-INS-401
chr19-34985138-INS-52
chr19-34997658-INS-66
chr19-35001194-INS-55
chr19-35004987-INS-72
chr19-35016601-INS-357
chr19-35018106-INS-52
chr19-35022125-INS-895
chr19-35032201-INS-6395
chr19-35047621-INS-474
chr19-35067325-INS-371
chr19-35079776-INS-68
chr19-35096197-INS-397
chr19-35100132-INS-528
chr19-35107983-INS-926
chr19-35116270-INS-53
chr19-35120437-INS-58
chr19-35145564-INS-60
chr19-35174399-INS-401
chr19-35178183-INS-236
chr19-35186708-INS-6701
chr19-35194659-INS-6550
chr19-35212168-INS-214
chr19-35223161-INS-3734
chr19-35235947-INS-496
chr19-35239138-INS-255
chr19-35244273-INS-81
chr19-35262149-INS-301
chr19-35265727-INS-252
chr19-35284322-INS-6373
chr19-35286804-INS-86
chr19-35302569-INS-68
chr19-35307060-INS-6391
chr19-35317698-INS-190
chr19-35334939-INS-50
chr19-35357539-INS-411
chr19-35369733-INS-707
chr19-35391085-INS-189
chr19-35428222-INS-146
chr19-35430136-INS-6536
chr19-35438071-INS-2271
chr19-35453063-INS-2804
chr19-35458268-INS-6261
chr19-35474012-INS-3311
chr19-35479350-INS-185
chr19-35486777-INS-221
chr19-35499843-INS-99
chr19-35544026-INS-68
chr19-35571631-INS-75
chr19-35572673-INS-395
chr19-35576970-INS-90
chr19-35625752-INS-77
chr19-35634001-INS-207
chr19-35651161-INS-207
chr19-35652462-INS-122

chr19-35661938-INS-928
chr19-35666326-INS-64
chr19-35671756-INS-131
chr19-35713020-INS-69
chr19-35716421-INS-59
chr19-35717783-INS-640
chr19-35727047-INS-118
chr19-35746258-INS-8387
chr19-35757074-INS-6389
chr19-35758693-INS-2280
chr19-35775142-INS-667
chr19-35784558-INS-152
chr19-35788463-INS-54
chr19-35797474-INS-56
chr19-35812582-INS-76
chr19-35821579-INS-390
chr19-35825990-INS-6388
chr19-35849485-INS-804
chr19-35856045-INS-449
chr19-35858157-INS-53
chr19-35866658-INS-1403
chr19-35887317-INS-2071
chr19-35905072-INS-192
chr19-35918133-INS-436
chr19-35940981-INS-814
chr19-35956670-INS-64
chr19-35970786-INS-101
chr19-36025582-INS-78
chr19-36041014-INS-88
chr19-36047028-INS-392
chr19-36051730-INS-52
chr19-36054921-INS-612
chr19-36059896-INS-59
chr19-36079763-INS-55
chr19-36090503-INS-734
chr19-36130606-INS-221
chr19-36208711-INS-205
chr19-36240130-INS-1138
chr19-36252336-INS-339
chr19-36255204-INS-6477
chr19-36277542-INS-482
chr19-36285125-INS-7239
chr19-36306411-INS-57
chr19-36345519-INS-210
chr19-36370187-INS-7156
chr19-36382151-INS-56
chr19-36390556-INS-388
chr19-36401610-INS-498
chr19-36414211-INS-399
chr19-36424022-INS-61
chr19-36452480-INS-979
chr19-36471841-INS-371
chr19-36472246-INS-4833
chr19-36495310-INS-421
chr19-36540194-INS-1218
chr19-36546563-INS-6358
chr19-36549394-INS-52
chr19-36572266-INS-184
chr19-36583466-INS-793
chr19-36595138-INS-60
chr19-36616461-INS-104
chr19-36642393-INS-82
chr19-36648183-INS-54
chr19-36651968-INS-241
chr19-36674287-INS-54
chr19-36683269-INS-72
chr19-36683969-INS-62
chr19-36684902-INS-118
chr19-36696359-INS-135
chr19-36721663-INS-171
chr19-36725049-INS-622
chr19-36744778-INS-197
chr19-36749871-INS-54
chr19-36761156-INS-473
chr19-36772772-INS-7194
chr19-36804462-INS-198
chr19-36824951-INS-208

chr19-36827733-INS-241
chr19-36835904-INS-107
chr19-36843055-INS-54
chr19-36877978-INS-63
chr19-36882603-INS-226
chr19-36889640-INS-54
chr19-36898440-INS-203
chr19-36905158-INS-174
chr19-36913541-INS-701
chr19-36966047-INS-210
chr19-36969721-INS-140
chr19-36978883-INS-498
chr19-36979733-INS-55
chr19-36990899-INS-108
chr19-36996435-INS-77
chr19-37005437-INS-168
chr19-37015947-INS-54
chr19-37025628-INS-215
chr19-37070988-INS-510
chr19-37078214-INS-80
chr19-37087616-INS-54
chr19-37121323-INS-882
chr19-37129898-INS-159
chr19-37131888-INS-195
chr19-37149225-INS-339
chr19-37156476-INS-576
chr19-37174537-INS-431
chr19-37176637-INS-208
chr19-37189536-INS-108
chr19-37203625-INS-83
chr19-37205375-INS-201
chr19-37210524-INS-100
chr19-37212249-INS-184
chr19-37217944-INS-762
chr19-37218695-INS-115
chr19-37231022-INS-56
chr19-37238763-INS-72
chr19-37241905-INS-492
chr19-37250682-INS-65
chr19-37277786-INS-115
chr19-37278077-INS-57
chr19-37299171-INS-97
chr19-37300200-INS-229
chr19-37303492-INS-223
chr19-37305126-INS-65
chr19-37308543-INS-124
chr19-37322292-INS-84
chr19-37332842-INS-51
chr19-37349295-INS-188
chr19-37350531-INS-216
chr19-37365795-INS-353
chr19-37368128-INS-58
chr19-37370630-INS-244
chr19-37373382-INS-217
chr19-37374446-INS-58
chr19-37394800-INS-65
chr19-37403364-INS-182
chr19-37415947-INS-66
chr19-37443954-INS-52
chr19-37447295-INS-199
chr19-37459654-INS-185
chr19-37477043-INS-82
chr19-37489649-INS-175
chr19-37490977-INS-66
chr19-37504948-INS-193
chr19-37510574-INS-307
chr19-37550081-INS-234
chr19-37552120-INS-203
chr19-37553618-INS-629
chr19-37560109-INS-1420
chr19-37560110-INS-429
chr19-37565621-INS-61
chr19-37596233-INS-173
chr19-37631680-INS-122
chr19-37635826-INS-57
chr19-37659788-INS-457
chr19-37677545-INS-66

chr19-37711159-INS-135
chr19-37717039-INS-326
chr19-37730136-INS-62
chr19-37768902-INS-220
chr19-37776621-INS-385
chr19-37787279-INS-56
chr19-37790660-INS-52
chr19-37805547-INS-126
chr19-37821537-INS-624
chr19-37831858-INS-57
chr19-37852663-INS-88
chr19-37861668-INS-64
chr19-37865677-INS-199
chr19-37884274-INS-62
chr19-37913254-INS-56
chr19-37927812-INS-122
chr19-37936898-INS-6441
chr19-37954766-INS-55
chr19-37973142-INS-280
chr19-37991641-INS-80
chr19-37997487-INS-211
chr19-38012548-INS-103
chr19-38029285-INS-175
chr19-38041101-INS-130
chr19-38044817-INS-216
chr19-38047105-INS-386
chr19-38070932-INS-163
chr19-38081781-INS-56
chr19-38092962-INS-83
chr19-38094887-INS-74
chr19-38129683-INS-134
chr19-38148453-INS-335
chr19-38150934-INS-99
chr19-38154190-INS-58
chr19-38169834-INS-65
chr19-38184385-INS-77
chr19-38187819-INS-53
chr19-38210535-INS-51
chr19-38230310-INS-159
chr19-38246956-INS-105
chr19-38249371-INS-58
chr19-38267578-INS-73
chr19-38290365-INS-82
chr19-38306600-INS-56
chr19-38314628-INS-212
chr19-38314630-INS-810
chr19-38317507-INS-194
chr19-38322474-INS-297
chr19-38324326-INS-192
chr19-38340132-INS-97
chr19-38340744-INS-1111
chr19-38345875-INS-177
chr19-38356580-INS-185
chr19-38358963-INS-175
chr19-38375213-INS-5369
chr19-38383905-INS-199
chr19-38413002-INS-153
chr19-38418367-INS-208
chr19-38421322-INS-103
chr19-38502805-INS-51
chr19-38534668-INS-60
chr19-38538715-INS-115
chr19-38552292-INS-333
chr19-38555459-INS-63
chr19-38590970-INS-76
chr19-38593228-INS-82
chr19-38595285-INS-68
chr19-38600709-INS-341
chr19-38609701-INS-232
chr19-38652602-INS-51
chr19-38655883-INS-92
chr19-38669575-INS-987
chr19-38673728-INS-254
chr19-38690148-INS-52
chr19-38699140-INS-134
chr19-38724032-INS-230
chr19-38730389-INS-71

chr19-38748920-INS-60
chr19-38768813-INS-127
chr19-38770205-INS-194
chr19-38790546-INS-208
chr19-38791957-INS-207
chr19-38831814-INS-7099
chr19-38833422-INS-7112
chr19-38840890-INS-64
chr19-38848504-INS-104
chr19-38849969-INS-52
chr19-38850848-INS-64
chr19-38855610-INS-621
chr19-38870302-INS-59
chr19-38891844-INS-215
chr19-38892330-INS-2779
chr19-38903493-INS-56
chr19-38905324-INS-5314
chr19-38910370-INS-60
chr19-38924108-INS-201
chr19-38926234-INS-69
chr19-38934796-INS-208
chr19-38948363-INS-68
chr19-38973269-INS-188
chr19-38988275-INS-285
chr19-38992167-INS-594
chr19-38992179-INS-69
chr19-39011333-INS-210
chr19-39072078-INS-8621
chr19-39073363-INS-161
chr19-39076881-INS-79
chr19-39083055-INS-57
chr19-39084172-INS-419
chr19-39153939-INS-56
chr19-39170704-INS-970
chr19-39171881-INS-81
chr19-39176729-INS-61
chr19-39182986-INS-157
chr19-39227890-INS-463
chr19-39244150-INS-566
chr19-39282069-INS-120
chr19-39316585-INS-169
chr19-39325296-INS-218
chr19-39365837-INS-65
chr19-39386803-INS-178
chr19-39396339-INS-687
chr19-39437067-INS-74
chr19-39513769-INS-857
chr19-39526700-INS-534
chr19-39599662-INS-1917
chr19-39673983-INS-8743
chr19-39754393-INS-126
chr19-39795260-INS-168
chr19-39825066-INS-3773
chr19-39827216-INS-1524
chr19-39857683-INS-1185
chr19-39867680-INS-4918
chr19-39868849-INS-310
chr19-39886757-INS-58
chr19-39912974-INS-771
chr19-39914531-INS-498
chr19-39917401-INS-1976
chr19-39919571-INS-1104
chr19-39927864-INS-7170
chr19-39931705-INS-6688
chr19-39962599-INS-497
chr19-39970236-INS-581
chr19-39971006-INS-6443
chr19-39980320-INS-306
chr19-40018203-INS-76
chr19-40054767-INS-81
chr19-40077827-INS-72
chr19-40102968-INS-389
chr19-40107388-INS-750
chr19-40119667-INS-106
chr19-40170736-INS-51
chr19-40178998-INS-6403
chr19-40179143-INS-52

chr19-40217595-INS-486
chr19-40221704-INS-127
chr19-40244022-INS-174
chr19-40271859-INS-236
chr19-40304188-INS-400
chr19-40317603-INS-163
chr19-40326474-INS-188
chr19-40346840-INS-53
chr19-40402414-INS-347
chr19-40404875-INS-92
chr19-40420175-INS-92
chr19-40426006-INS-168
chr19-40444566-INS-178
chr19-40467511-INS-72
chr19-40497236-INS-308
chr19-40553594-INS-100
chr19-40562959-INS-138
chr19-40582997-INS-3091
chr19-40591523-INS-56
chr19-40605459-INS-123
chr19-40608932-INS-106
chr19-40614939-INS-6440
chr19-40615969-INS-307
chr19-40630751-INS-472
chr19-40658175-INS-111
chr19-40697574-INS-51
chr19-40702158-INS-118
chr19-40747723-INS-55
chr19-40748998-INS-83
chr19-40754756-INS-52
chr19-40776203-INS-130
chr19-40783816-INS-183
chr19-40787405-INS-129
chr19-40789524-INS-485
chr19-40805446-INS-166
chr19-40807695-INS-81
chr19-40812690-INS-99
chr19-40820792-INS-105
chr19-40827438-INS-399
chr19-40831011-INS-133
chr19-40835883-INS-365
chr19-40843349-INS-134
chr19-40853917-INS-276
chr19-40856655-INS-137
chr19-40858635-INS-75
chr19-40873120-INS-86
chr19-40890898-INS-53
chr19-40923439-INS-75
chr19-40929231-INS-215
chr19-40947367-INS-131
chr19-40952479-INS-419
chr19-40957520-INS-209
chr19-40977346-INS-98
chr19-41039431-INS-401
chr19-41047087-INS-184
chr19-41060091-INS-323
chr19-41078856-INS-52
chr19-41094742-INS-70
chr19-41096610-INS-205
chr19-41123191-INS-76
chr19-41124974-INS-226
chr19-41127910-INS-81
chr19-41152616-INS-84
chr19-41158129-INS-444
chr19-41175042-INS-71
chr19-41212188-INS-156
chr19-41236981-INS-115
chr19-41260044-INS-54
chr19-41276163-INS-72
chr19-41280718-INS-72
chr19-41344518-INS-62
chr19-41376331-INS-89
chr19-41377488-INS-143
chr19-41390448-INS-66
chr19-41406274-INS-70
chr19-41411367-INS-240
chr19-41426643-INS-206

chr19-41432248-INS-242
chr19-41436934-INS-163
chr19-41441610-INS-89
chr19-41442797-INS-170
chr19-41447380-INS-91
chr19-41485064-INS-451
chr19-41489402-INS-201
chr19-41493768-INS-3303
chr19-41508227-INS-100
chr19-41508275-INS-773
chr19-41516698-INS-432
chr19-41518758-INS-135
chr19-41538078-INS-63
chr19-41569707-INS-211
chr19-41571696-INS-126
chr19-41604267-INS-102
chr19-41608513-INS-65
chr19-41620609-INS-68
chr19-41621732-INS-55
chr19-41647400-INS-215
chr19-41653973-INS-54
chr19-41690862-INS-188
chr19-41702657-INS-162
chr19-41708724-INS-628
chr19-41711769-INS-125
chr19-41713726-INS-95
chr19-41741772-INS-51
chr19-41749977-INS-142
chr19-41758930-INS-149
chr19-41760198-INS-80
chr19-41765156-INS-92
chr19-41778744-INS-56
chr19-41794225-INS-604
chr19-41811046-INS-193
chr19-41817370-INS-313
chr19-41829750-INS-66
chr19-41832185-INS-253
chr19-41834254-INS-61
chr19-41840305-INS-240
chr19-41861538-INS-72
chr19-41872305-INS-308
chr19-41873641-INS-171
chr19-41892028-INS-216
chr19-41901485-INS-234
chr19-41909815-INS-201
chr19-41918078-INS-175
chr19-41943378-INS-209
chr19-41948477-INS-68
chr19-41974043-INS-52
chr19-42006168-INS-55
chr19-42019220-INS-400
chr19-42044378-INS-202
chr19-42069409-INS-168
chr19-42074045-INS-183
chr19-42087417-INS-59
chr19-42091943-INS-74
chr19-42093230-INS-96
chr19-42096968-INS-317
chr19-42121243-INS-114
chr19-42122026-INS-721
chr19-42143871-INS-72
chr19-42162762-INS-174
chr19-42165531-INS-64
chr19-42170670-INS-230
chr19-42171779-INS-174
chr19-42206364-INS-84
chr19-42243129-INS-238
chr19-42255830-INS-52
chr19-42263932-INS-111
chr19-42276122-INS-62
chr19-42309264-INS-380
chr19-42329878-INS-165
chr19-42389962-INS-141
chr19-42421993-INS-54
chr19-42460980-INS-202
chr19-42470117-INS-223
chr19-42488971-INS-56

chr19-42502020-INS-68
chr19-42509784-INS-52
chr19-42514958-INS-401
chr19-42519977-INS-63
chr19-42530641-INS-52
chr19-42537025-INS-638
chr19-42576120-INS-667
chr19-42603724-INS-73
chr19-42605712-INS-173
chr19-42610739-INS-64
chr19-42624485-INS-57
chr19-42624642-INS-1089
chr19-42658911-INS-52
chr19-42701127-INS-60
chr19-42704121-INS-210
chr19-42704954-INS-976
chr19-42729500-INS-52
chr19-42742438-INS-86
chr19-42762020-INS-129
chr19-42772582-INS-67
chr19-42789079-INS-71
chr19-42799432-INS-6440
chr19-42823150-INS-506
chr19-42827746-INS-191
chr19-42864120-INS-444
chr19-42867723-INS-530
chr19-42876077-INS-170
chr19-42878945-INS-66
chr19-42887218-INS-110
chr19-42889657-INS-849
chr19-42891993-INS-63
chr19-42897507-INS-164
chr19-42930354-INS-6439
chr19-42938371-INS-678
chr19-42939020-INS-116
chr19-42967798-INS-204
chr19-42983174-INS-7579
chr19-42994942-INS-498
chr19-43018216-INS-246
chr19-43023403-INS-354
chr19-43061490-INS-596
chr19-43096134-INS-62
chr19-43102319-INS-577
chr19-43131170-INS-262
chr19-43138029-INS-513
chr19-43175582-INS-2523
chr19-43184177-INS-70
chr19-43184727-INS-477
chr19-43210344-INS-99
chr19-43239051-INS-264
chr19-43246972-INS-60
chr19-43254428-INS-8762
chr19-43255992-INS-510
chr19-43267800-INS-569
chr19-43273592-INS-6724
chr19-43274342-INS-368
chr19-43357642-INS-98
chr19-43382202-INS-234
chr19-43386483-INS-525
chr19-43397733-INS-170
chr19-43414155-INS-308
chr19-43425022-INS-214
chr19-43436445-INS-183
chr19-43471862-INS-59
chr19-43475578-INS-52
chr19-43480088-INS-217
chr19-43486991-INS-5141
chr19-43499029-INS-140
chr19-43503922-INS-77
chr19-43509392-INS-84
chr19-43526445-INS-131
chr19-43570340-INS-339
chr19-43579382-INS-118
chr19-43585054-INS-107
chr19-43596164-INS-52
chr19-43619604-INS-110
chr19-43622708-INS-125

chr19-43639210-INS-221
chr19-43644886-INS-128
chr19-43671520-INS-60
chr19-43676088-INS-148
chr19-43680690-INS-60
chr19-43687892-INS-60
chr19-43695358-INS-58
chr19-43709572-INS-205
chr19-43723763-INS-56
chr19-43730899-INS-92
chr19-43750853-INS-197
chr19-43758632-INS-507
chr19-43765390-INS-71
chr19-43771562-INS-62
chr19-43794116-INS-58
chr19-43808924-INS-66
chr19-43818185-INS-81
chr19-43845447-INS-91
chr19-43879402-INS-53
chr19-43880987-INS-432
chr19-43883164-INS-368
chr19-43887341-INS-58
chr19-43892221-INS-229
chr19-43898809-INS-54
chr19-43910082-INS-166
chr19-43924220-INS-52
chr19-43932478-INS-207
chr19-43937933-INS-84
chr19-43953821-INS-60
chr19-43955813-INS-65
chr19-43956718-INS-222
chr19-43962617-INS-243
chr19-43970572-INS-192
chr19-44007764-INS-60
chr19-44038779-INS-146
chr19-44039550-INS-396
chr19-44065395-INS-506
chr19-44077805-INS-55
chr19-44086907-INS-184
chr19-44099743-INS-85
chr19-44104760-INS-341
chr19-44114554-INS-112
chr19-44119016-INS-214
chr19-44148759-INS-245
chr19-44153356-INS-214
chr19-44163807-INS-52
chr19-44174574-INS-129
chr19-44175116-INS-124
chr19-44193331-INS-4759
chr19-44206165-INS-53
chr19-44215291-INS-192
chr19-44216563-INS-337
chr19-44223677-INS-5348
chr19-44231699-INS-145
chr19-44239970-INS-53
chr19-44253714-INS-199
chr19-44257481-INS-242
chr19-44261271-INS-73
chr19-44264828-INS-98
chr19-44298278-INS-160
chr19-44307360-INS-88
chr19-44309340-INS-171
chr19-44319796-INS-127
chr19-44346443-INS-160
chr19-44362216-INS-957
chr19-44363234-INS-435
chr19-44363871-INS-253
chr19-44365387-INS-72
chr19-44367525-INS-460
chr19-44368300-INS-200
chr19-44369535-INS-260
chr19-44372233-INS-60
chr19-44375505-INS-166
chr19-44398282-INS-166
chr19-44401339-INS-64
chr19-44408632-INS-104
chr19-44422526-INS-189

chr19-44424970-INS-74
chr19-44428008-INS-61
chr19-44429465-INS-154
chr19-44431870-INS-205
chr19-44437364-INS-139
chr19-44442531-INS-322
chr19-44448708-INS-180
chr19-44451480-INS-138
chr19-44467692-INS-113
chr19-44472046-INS-293
chr19-44475451-INS-56
chr19-44486480-INS-305
chr19-44510190-INS-86
chr19-44517328-INS-129
chr19-44524279-INS-713
chr19-44598770-INS-158
chr19-44599872-INS-6442
chr19-44629348-INS-54
chr19-44635946-INS-146
chr19-44641487-INS-144
chr19-44642953-INS-108
chr19-44724300-INS-51
chr19-44733555-INS-61
chr19-44834540-INS-175
chr19-44837535-INS-56
chr19-44840765-INS-64
chr19-44841929-INS-70
chr19-44857134-INS-52
chr19-44866880-INS-227
chr19-44871748-INS-54
chr19-44883534-INS-114
chr19-44908722-INS-96
chr19-44922635-INS-176
chr19-44924413-INS-182
chr19-44948026-INS-238
chr19-44957948-INS-379
chr19-45006338-INS-55
chr19-45023521-INS-220
chr19-45060715-INS-174
chr19-45100923-INS-555
chr19-45107953-INS-64
chr19-45111866-INS-68
chr19-45126362-INS-172
chr19-45127418-INS-204
chr19-45140977-INS-72
chr19-45162866-INS-759
chr19-45166783-INS-58
chr19-45180981-INS-243
chr19-45187613-INS-149
chr19-45193497-INS-554
chr19-45198869-INS-5187
chr19-45207181-INS-92
chr19-45225522-INS-63
chr19-45249500-INS-112
chr19-45261875-INS-193
chr19-45274517-INS-1416
chr19-45277276-INS-156
chr19-45304811-INS-788
chr19-45308289-INS-207
chr19-45310020-INS-55
chr19-45320713-INS-102
chr19-45323293-INS-176
chr19-45338666-INS-51
chr19-45347927-INS-195
chr19-45363252-INS-81
chr19-45374270-INS-71
chr19-45380295-INS-126
chr19-45383693-INS-69
chr19-45388629-INS-75
chr19-45389978-INS-102
chr19-45390647-INS-180
chr19-45401806-INS-254
chr19-45404771-INS-209
chr19-45425463-INS-295
chr19-45440003-INS-198
chr19-45454084-INS-78
chr19-45509597-INS-268

chr19-45537016-INS-75
chr19-45546027-INS-209
chr19-45550534-INS-60
chr19-45553114-INS-224
chr19-45560552-INS-195
chr19-45600078-INS-68
chr19-45603248-INS-79
chr19-45610338-INS-201
chr19-45627232-INS-408
chr19-45627406-INS-874
chr19-45663433-INS-52
chr19-45671325-INS-52
chr19-45671847-INS-63
chr19-45676378-INS-75
chr19-45679304-INS-57
chr19-45696427-INS-134
chr19-45697142-INS-60
chr19-45698774-INS-214
chr19-45702683-INS-78
chr19-45705721-INS-120
chr19-45710523-INS-246
chr19-45718134-INS-153
chr19-45720933-INS-74
chr19-45755490-INS-96
chr19-45777412-INS-278
chr19-45779651-INS-302
chr19-45780551-INS-149
chr19-45789865-INS-208
chr19-45817913-INS-218
chr19-45824706-INS-206
chr19-45827958-INS-224
chr19-45838379-INS-65
chr19-45838683-INS-62
chr19-45839490-INS-206
chr19-45840455-INS-114
chr19-45842365-INS-263
chr19-45844594-INS-55
chr19-45845890-INS-60
chr19-45869584-INS-229
chr19-45871736-INS-202
chr19-45874699-INS-115
chr19-45877807-INS-220
chr19-45889915-INS-84
chr19-45932167-INS-51
chr19-45933522-INS-123
chr19-45939416-INS-133
chr19-45944311-INS-59
chr19-45953496-INS-70
chr19-45960996-INS-719
chr19-45964970-INS-162
chr19-45968596-INS-227
chr19-45973853-INS-132
chr19-45976658-INS-221
chr19-45980069-INS-146
chr19-45997829-INS-132
chr19-45999229-INS-61
chr19-46009445-INS-69
chr19-46015742-INS-128
chr19-46016247-INS-68
chr19-46019423-INS-130
chr19-46035568-INS-5548
chr19-46036608-INS-197
chr19-46038273-INS-817
chr19-46041933-INS-186
chr19-46043096-INS-5567
chr19-46044771-INS-8941
chr19-46050159-INS-220
chr19-46066094-INS-208
chr19-46077748-INS-505
chr19-46083802-INS-173
chr19-46094209-INS-59
chr19-46096868-INS-68
chr19-46103436-INS-63
chr19-46106289-INS-50
chr19-46110180-INS-722
chr19-46196852-INS-127
chr19-46227047-INS-265

chr19-46235692-INS-56
chr19-46278828-INS-198
chr19-46280441-INS-157
chr19-46284004-INS-169
chr19-46284972-INS-120
chr19-46290619-INS-126
chr19-46303759-INS-213
chr19-46351489-INS-243
chr19-46371194-INS-209
chr19-46381565-INS-207
chr19-46408299-INS-173
chr19-46485777-INS-214
chr19-46527565-INS-52
chr19-46532648-INS-134
chr19-46537327-INS-80
chr19-46540334-INS-124
chr19-46557552-INS-552
chr19-46569009-INS-572
chr19-46572619-INS-188
chr19-46572849-INS-632
chr19-46583742-INS-61
chr19-46599985-INS-53
chr19-46604252-INS-202
chr19-46622181-INS-52
chr19-46625652-INS-53
chr19-46632614-INS-57
chr19-46639496-INS-52
chr19-46642192-INS-998
chr19-46667307-INS-62
chr19-46672971-INS-203
chr19-46700424-INS-78
chr19-46703378-INS-1888
chr19-46706997-INS-248
chr19-46720488-INS-191
chr19-46742366-INS-815
chr19-46757293-INS-232
chr19-46759014-INS-224
chr19-46765721-INS-165
chr19-46775629-INS-524
chr19-46781918-INS-180
chr19-46795394-INS-76
chr19-46806624-INS-376
chr19-46838530-INS-181
chr19-46849161-INS-120
chr19-46860745-INS-56
chr19-46865972-INS-149
chr19-46874692-INS-491
chr19-46899194-INS-325
chr19-46906251-INS-199
chr19-46908542-INS-52
chr19-46910868-INS-184
chr19-46921049-INS-204
chr19-46926850-INS-51
chr19-46940185-INS-243
chr19-46955532-INS-207
chr19-46974555-INS-96
chr19-46981676-INS-201
chr19-46983885-INS-200
chr19-46993549-INS-62
chr19-46997767-INS-78
chr19-46999901-INS-254
chr19-47014569-INS-138
chr19-47021156-INS-363
chr19-47027822-INS-172
chr19-47049291-INS-52
chr19-47084115-INS-266
chr19-47090370-INS-64
chr19-47095234-INS-212
chr19-47138722-INS-58
chr19-47143598-INS-200
chr19-47175703-INS-208
chr19-47177914-INS-194
chr19-47194920-INS-218
chr19-47207115-INS-340
chr19-47223371-INS-102
chr19-47230093-INS-833
chr19-47244400-INS-211

chr19-47245454-INS-63
chr19-47264509-INS-162
chr19-47296761-INS-101
chr19-47298957-INS-52
chr19-47316870-INS-57
chr19-47395617-INS-71
chr19-47400433-INS-72
chr19-47413186-INS-60
chr19-47444396-INS-152
chr19-47454810-INS-51
chr19-47456766-INS-60
chr19-47458282-INS-164
chr19-47461981-INS-54
chr19-47466999-INS-218
chr19-47493905-INS-122
chr19-47514229-INS-619
chr19-47518447-INS-294
chr19-47560232-INS-310
chr19-47564975-INS-180
chr19-47569691-INS-210
chr19-47582442-INS-164
chr19-47624054-INS-479
chr19-47692367-INS-206
chr19-47704230-INS-56
chr19-47743086-INS-110
chr19-47756081-INS-51
chr19-47768898-INS-128
chr19-47771902-INS-314
chr19-47791494-INS-310
chr19-47816212-INS-52
chr19-47827600-INS-183
chr19-47845047-INS-445
chr19-47847276-INS-768
chr19-47879448-INS-1693
chr19-47901155-INS-97
chr19-47922208-INS-3759
chr19-47940499-INS-131
chr19-47958460-INS-61
chr19-47970727-INS-104
chr19-48082682-INS-1939
chr19-48096508-INS-413
chr19-48133260-INS-207
chr19-48146544-INS-68
chr19-48162347-INS-240
chr19-48178183-INS-358
chr19-48183900-INS-67
chr19-48212549-INS-56
chr19-48288039-INS-602
chr19-48307776-INS-244
chr19-48317165-INS-339
chr19-48352765-INS-234
chr19-48367535-INS-89
chr19-48369242-INS-429
chr19-48408945-INS-145
chr19-48420877-INS-144
chr19-48421384-INS-124
chr19-48440801-INS-254
chr19-48487371-INS-584
chr19-48515418-INS-430
chr19-48523624-INS-484
chr19-48568352-INS-462
chr19-48602328-INS-80
chr19-48635482-INS-80
chr19-48643230-INS-60
chr19-48681105-INS-150
chr19-48717004-INS-269
chr19-48761520-INS-214
chr19-48767304-INS-294
chr19-48802722-INS-147
chr19-48838701-INS-729
chr19-48859171-INS-68
chr19-48862778-INS-64
chr19-48875036-INS-354
chr19-48880198-INS-665
chr19-48898637-INS-466
chr19-48920999-INS-212
chr19-48956646-INS-2785

chr19-48965761-INS-1794
chr19-48965836-INS-197
chr19-48980577-INS-6747
chr19-49002656-INS-52
chr19-49010615-INS-55
chr19-49015149-INS-7141
chr19-49021617-INS-194
chr19-49051572-INS-173
chr19-49082821-INS-6876
chr19-49101440-INS-79
chr19-49108875-INS-144
chr19-49128784-INS-75
chr19-49132834-INS-372
chr19-49153837-INS-405
chr19-49179888-INS-739
chr19-49181788-INS-53
chr19-49185315-INS-157
chr19-49200051-INS-145
chr19-49254938-INS-241
chr19-49275018-INS-62
chr19-49283955-INS-238
chr19-49285557-INS-292
chr19-49290847-INS-69
chr19-49298623-INS-129
chr19-49306740-INS-56
chr19-49315086-INS-56
chr19-49367760-INS-74
chr19-49371213-INS-121
chr19-49380104-INS-53
chr19-49394757-INS-160
chr19-49410323-INS-385
chr19-49413432-INS-388
chr19-49413864-INS-456
chr19-49419028-INS-64
chr19-49434412-INS-93
chr19-49435064-INS-390
chr19-49445456-INS-306
chr19-49449625-INS-212
chr19-49467358-INS-78
chr19-49485894-INS-299
chr19-49490736-INS-120
chr19-49501600-INS-76
chr19-49502106-INS-379
chr19-49511778-INS-212
chr19-49521477-INS-52
chr19-49530888-INS-1549
chr19-49533354-INS-162
chr19-49549007-INS-70
chr19-49562019-INS-142
chr19-49566528-INS-212
chr19-49587411-INS-61
chr19-49606864-INS-62
chr19-49616499-INS-399
chr19-49625096-INS-59
chr19-49642077-INS-331
chr19-49646479-INS-198
chr19-49655315-INS-89
chr19-49697952-INS-192
chr19-49702694-INS-231
chr19-49710613-INS-101
chr19-49753642-INS-492
chr19-49769911-INS-492
chr19-49792914-INS-548
chr19-49811883-INS-323
chr19-49825226-INS-429
chr19-49832640-INS-121
chr19-49849172-INS-536
chr19-49907145-INS-365
chr19-49908263-INS-60
chr19-49919521-INS-313
chr19-49943096-INS-880
chr19-49945099-INS-104
chr19-49999933-INS-801
chr19-50003190-INS-392
chr19-50017106-INS-66
chr19-50023756-INS-234
chr19-50030878-INS-83

chr19-50044679-INS-113
chr19-50045454-INS-186
chr19-50047122-INS-162
chr19-50052115-INS-848
chr19-50060725-INS-551
chr19-50061998-INS-815
chr19-50070710-INS-197
chr19-50071978-INS-1861
chr19-50072344-INS-65
chr19-50080844-INS-174
chr19-50084768-INS-1893
chr19-50119825-INS-190
chr19-50129787-INS-77
chr19-50145665-INS-100
chr19-50150781-INS-51
chr19-50170157-INS-398
chr19-50172992-INS-54
chr19-50176424-INS-157
chr19-50230627-INS-382
chr19-50243315-INS-200
chr19-50259672-INS-2736
chr19-50260405-INS-190
chr19-50270972-INS-327
chr19-50279445-INS-63
chr19-50360920-INS-171
chr19-50372463-INS-177
chr19-50396221-INS-58
chr19-50406250-INS-316
chr19-50422392-INS-101
chr19-50445545-INS-234
chr19-50473962-INS-433
chr19-50506141-INS-248
chr19-50517435-INS-180
chr19-50525455-INS-58
chr19-50527216-INS-56
chr19-50558058-INS-71
chr19-50591652-INS-54
chr19-50678569-INS-162
chr19-50692948-INS-544
chr19-50696813-INS-357
chr19-50712753-INS-110
chr19-50725116-INS-123
chr19-50728987-INS-351
chr19-50730707-INS-6336
chr19-50757512-INS-64
chr19-50764204-INS-362
chr19-50831227-INS-62
chr19-50834141-INS-6279
chr19-50923363-INS-69
chr19-50954700-INS-76
chr19-50959054-INS-279
chr19-50969526-INS-124
chr19-50973539-INS-74
chr19-50976457-INS-68
chr19-50977647-INS-124
chr19-51041085-INS-66
chr19-51048509-INS-91
chr19-51103797-INS-135
chr19-51109290-INS-52
chr19-51140475-INS-56
chr19-51156901-INS-872
chr19-51168345-INS-5601
chr19-51174952-INS-260
chr19-51190198-INS-681
chr19-51274602-INS-306
chr19-51280241-INS-410
chr19-51289078-INS-144
chr19-51299074-INS-801
chr19-51309675-INS-69
chr19-51317670-INS-56
chr19-51326022-INS-74
chr19-51341128-INS-315
chr19-51413104-INS-120
chr19-51415312-INS-778
chr19-51434886-INS-6004
chr19-51476845-INS-7109
chr19-51497003-INS-84

chr19-51511511-INS-50
chr19-51576576-INS-538
chr19-51610116-INS-7638
chr19-51612076-INS-63
chr19-51669398-INS-398
chr19-51730212-INS-132
chr19-51740605-INS-112
chr19-51749939-INS-194
chr19-51750910-INS-52
chr19-51761130-INS-108
chr19-51776464-INS-60
chr19-51799952-INS-107
chr19-51830337-INS-52
chr19-51831971-INS-7114
chr19-51859913-INS-162
chr19-51867674-INS-80
chr19-51868049-INS-537
chr19-51870704-INS-140
chr19-51882962-INS-52
chr19-51895582-INS-194
chr19-51897490-INS-63
chr19-51911918-INS-456
chr19-51915026-INS-82
chr19-51916786-INS-502
chr19-51943830-INS-424
chr19-51945992-INS-62
chr19-51954410-INS-397
chr19-51969660-INS-54
chr19-52008314-INS-5296
chr19-52014118-INS-6282
chr19-52019990-INS-59
chr19-52055906-INS-64
chr19-52064604-INS-569
chr19-52130211-INS-708
chr19-52131375-INS-220
chr19-52147454-INS-232
chr19-52157050-INS-283
chr19-52159745-INS-64
chr19-52161935-INS-82
chr19-52181243-INS-92
chr19-52190153-INS-76
chr19-52209025-INS-224
chr19-52215095-INS-73
chr19-52223891-INS-6152
chr19-52226092-INS-56
chr19-52239533-INS-129
chr19-52260583-INS-150
chr19-52263828-INS-88
chr19-52328088-INS-86
chr19-52330733-INS-62
chr19-52358635-INS-1979
chr19-52385933-INS-62
chr19-52408495-INS-509
chr19-52419593-INS-331
chr19-52425075-INS-145
chr19-52433107-INS-419
chr19-52436556-INS-64
chr19-52476414-INS-306
chr19-52499572-INS-63
chr19-52503641-INS-753
chr19-52509793-INS-819
chr19-52545510-INS-158
chr19-52552310-INS-184
chr19-52590565-INS-399
chr19-52593035-INS-195
chr19-52595793-INS-110
chr19-52596529-INS-53
chr19-52603736-INS-76
chr19-52620938-INS-305
chr19-52623930-INS-402
chr19-52644743-INS-491
chr19-52651175-INS-169
chr19-52652701-INS-183
chr19-52661318-INS-806
chr19-52679641-INS-317
chr19-52707547-INS-380
chr19-52709023-INS-58

chr19-52711553-INS-146
chr19-52739406-INS-51
chr19-52754205-INS-124
chr19-52760521-INS-62
chr19-52769876-INS-2794
chr19-52780832-INS-76
chr19-52782342-INS-7469
chr19-52794700-INS-469
chr19-52802768-INS-100
chr19-52844632-INS-432
chr19-52861882-INS-162
chr19-52869187-INS-564
chr19-52880677-INS-104
chr19-52881119-INS-165
chr19-52900708-INS-154
chr19-52909920-INS-59
chr19-52940780-INS-359
chr19-52983761-INS-195
chr19-52994207-INS-160
chr19-52995313-INS-54
chr19-53009232-INS-379
chr19-53013024-INS-201
chr19-53028509-INS-114
chr19-53030568-INS-8940
chr19-53040850-INS-205
chr19-53051102-INS-334
chr19-53059749-INS-92
chr19-53061619-INS-436
chr19-53077221-INS-58
chr19-53080371-INS-458
chr19-53083264-INS-67
chr19-53136563-INS-501
chr19-53140616-INS-242
chr19-53141853-INS-248
chr19-53153020-INS-62
chr19-53169063-INS-171
chr19-53194074-INS-96
chr19-53203710-INS-347
chr19-53214222-INS-76
chr19-53229127-INS-58
chr19-53248879-INS-196
chr19-53275700-INS-117
chr19-53283266-INS-107
chr19-53291608-INS-229
chr19-53317363-INS-105
chr19-53352248-INS-200
chr19-53365233-INS-156
chr19-53367416-INS-186
chr19-53369435-INS-191
chr19-53371058-INS-88
chr19-53382489-INS-247
chr19-53409086-INS-80
chr19-53415229-INS-1860
chr19-53415576-INS-114
chr19-53430352-INS-86
chr19-53435181-INS-469
chr19-53440547-INS-52
chr19-53441587-INS-78
chr19-53481761-INS-104
chr19-53485125-INS-264
chr19-53488467-INS-57
chr19-53492141-INS-60
chr19-53506622-INS-70
chr19-53536677-INS-67
chr19-53538076-INS-52
chr19-53542981-INS-213
chr19-53549721-INS-207
chr19-53578965-INS-232
chr19-53610914-INS-198
chr19-53614909-INS-422
chr19-53618775-INS-200
chr19-53631768-INS-106
chr19-53634855-INS-58
chr19-53647134-INS-379
chr19-53650140-INS-103
chr19-53651368-INS-118
chr19-53662444-INS-105

chr19-53673589-INS-78
chr19-53689918-INS-57
chr19-53717032-INS-68
chr19-53730093-INS-152
chr19-53737253-INS-63
chr19-53738533-INS-260
chr19-53744379-INS-386
chr19-53750593-INS-94
chr19-53765664-INS-3927
chr19-53765727-INS-422
chr19-53767154-INS-561
chr19-53800060-INS-67
chr19-53804428-INS-52
chr19-53807463-INS-51
chr19-53823708-INS-375
chr19-53852253-INS-51
chr19-53855735-INS-64
chr19-53859424-INS-52
chr19-53868517-INS-52
chr19-53879514-INS-108
chr19-53880788-INS-60
chr19-53919158-INS-102
chr19-53928523-INS-264
chr19-53962890-INS-74
chr19-53988854-INS-214
chr19-53999980-INS-58
chr19-54053986-INS-341
chr19-54096662-INS-74
chr19-54138114-INS-194
chr19-54139504-INS-176
chr19-54144993-INS-394
chr19-54151187-INS-192
chr19-54170248-INS-289
chr19-54174205-INS-376
chr19-54178658-INS-411
chr19-54179788-INS-396
chr19-54181821-INS-452
chr19-54209367-INS-196
chr19-54251041-INS-233
chr19-54263004-INS-65
chr19-54280547-INS-157
chr19-54323727-INS-204
chr19-54326844-INS-62
chr19-54371175-INS-143
chr19-54397640-INS-52
chr19-54406095-INS-112
chr19-54408715-INS-157
chr19-54417542-INS-108
chr19-54422145-INS-84
chr19-54439646-INS-60
chr19-54461643-INS-154
chr19-54483420-INS-72
chr19-54489118-INS-129
chr19-54492986-INS-227
chr19-54503776-INS-118
chr19-54549417-INS-60
chr19-54559370-INS-138
chr19-54581768-INS-60
chr19-54621075-INS-51
chr19-54624308-INS-167
chr19-54637525-INS-124
chr19-54642762-INS-78
chr19-54651698-INS-51
chr19-54664697-INS-58
chr19-54686672-INS-146
chr19-54727279-INS-233
chr19-54737453-INS-72
chr19-54761711-INS-80
chr19-54777073-INS-69
chr19-54783241-INS-223
chr19-54788705-INS-96
chr19-54791226-INS-149
chr19-54806085-INS-52
chr19-54813195-INS-55
chr19-54839118-INS-94
chr19-54843489-INS-58
chr19-54857715-INS-146

chr19-54871196-INS-55
chr19-54876657-INS-62
chr19-54887482-INS-91
chr19-54889133-INS-103
chr19-54890061-INS-140
chr19-54913981-INS-108
chr19-54925195-INS-61
chr19-54935974-INS-64
chr19-54956416-INS-120
chr19-54965631-INS-59
chr19-54999079-INS-463
chr19-55010907-INS-200
chr19-55027367-INS-50
chr19-55037855-INS-57
chr19-55047105-INS-130
chr19-55050327-INS-798
chr19-55059922-INS-76
chr19-55083398-INS-159
chr19-55093601-INS-72
chr19-55100805-INS-155
chr19-55120855-INS-140
chr19-55135012-INS-63
chr19-55142615-INS-64
chr19-55151325-INS-51
chr19-55164561-INS-72
chr19-55186569-INS-425
chr19-55188334-INS-68
chr19-55198079-INS-197
chr19-55228769-INS-250
chr19-55235354-INS-190
chr19-55256218-INS-206
chr19-55268061-INS-199
chr19-55275485-INS-96
chr19-55294932-INS-199
chr19-55296587-INS-190
chr19-55306942-INS-405
chr19-55313878-INS-210
chr19-55317388-INS-58
chr19-55335537-INS-123
chr19-55357266-INS-112
chr19-55383654-INS-93
chr19-55397507-INS-59
chr19-55405344-INS-122
chr19-55421678-INS-71
chr19-55460633-INS-714
chr19-55516882-INS-2412
chr19-55517231-INS-386
chr19-55541630-INS-71
chr19-55568466-INS-66
chr19-55576854-INS-91
chr19-55582024-INS-55
chr19-55625035-INS-95
chr19-55641186-INS-347
chr19-55653821-INS-231
chr19-55666624-INS-156
chr19-55694792-INS-205
chr19-55701691-INS-88
chr19-55717561-INS-215
chr19-55724141-INS-108
chr19-55778953-INS-217
chr19-55802726-INS-82
chr19-55834415-INS-55
chr19-55835095-INS-60
chr19-55849221-INS-209
chr19-55857756-INS-104
chr19-55907036-INS-82
chr19-55941066-INS-126
chr19-55976650-INS-60
chr19-55993946-INS-612
chr19-56005627-INS-93
chr19-56019865-INS-52
chr19-56022060-INS-225
chr19-56025579-INS-236
chr19-56033793-INS-905
chr19-56052033-INS-80
chr19-56083680-INS-56
chr19-56090607-INS-142

chr19-56099689-INS-97
chr19-56112857-INS-139
chr19-56114708-INS-128
chr19-56151444-INS-78
chr19-56168528-INS-356
chr19-56196385-INS-54
chr19-56215354-INS-66
chr19-56233270-INS-77
chr19-56288389-INS-54
chr19-56291428-INS-168
chr19-56298776-INS-86
chr19-56304291-INS-183
chr19-56304302-INS-902
chr19-56311275-INS-55
chr19-56311533-INS-168
chr19-56333406-INS-82
chr19-56343859-INS-86
chr19-56390034-INS-67
chr19-56392119-INS-72
chr19-56405604-INS-91
chr19-56435082-INS-7235
chr19-56435108-INS-56
chr19-56452333-INS-171
chr19-56470180-INS-57
chr19-56471368-INS-573
chr19-56477473-INS-259
chr19-56478844-INS-503
chr19-56485309-INS-56
chr19-56491757-INS-58
chr19-56496275-INS-86
chr19-56501041-INS-72
chr19-56529410-INS-84
chr19-56542222-INS-338
chr19-56571499-INS-106
chr19-56605500-INS-51
chr19-56646293-INS-56
chr19-56665889-INS-58
chr19-56672182-INS-195
chr19-56684792-INS-133
chr19-56724505-INS-226
chr19-56767086-INS-72
chr19-56778374-INS-156
chr19-56786924-INS-129
chr19-56792432-INS-68
chr19-56835976-INS-1096
chr19-56835994-INS-754
chr19-56872213-INS-84
chr19-56876817-INS-51
chr19-56888074-INS-71
chr19-56922962-INS-60
chr19-56940078-INS-240
chr19-56941988-INS-76
chr19-56955992-INS-82
chr19-56965058-INS-159
chr19-56985055-INS-58
chr19-57046820-INS-123
chr19-57052877-INS-227
chr19-57065100-INS-142
chr19-57077816-INS-70
chr19-57090068-INS-7158
chr19-57090755-INS-401
chr19-57094653-INS-104
chr19-57100022-INS-54
chr19-57101840-INS-1148
chr19-57101840-INS-795
chr19-57101841-INS-60
chr19-57122507-INS-63
chr19-57128827-INS-91
chr19-57141551-INS-150
chr19-57148075-INS-56
chr19-57156658-INS-57
chr19-57165473-INS-134
chr19-57196056-INS-52
chr19-57215780-INS-73
chr19-57220551-INS-90
chr19-57236873-INS-84
chr19-57254175-INS-170

chr19-57283706-INS-205
chr19-57286696-INS-172
chr19-57293018-INS-196
chr19-57307617-INS-54
chr19-57323914-INS-203
chr19-57336004-INS-587
chr19-57336973-INS-162
chr19-57353483-INS-208
chr19-57362071-INS-53
chr19-57380786-INS-437
chr19-57383580-INS-132
chr19-57395661-INS-138
chr19-57404026-INS-148
chr19-57417048-INS-729
chr19-57427761-INS-142
chr19-57432168-INS-98
chr19-57442217-INS-569
chr19-57446991-INS-64
chr19-57466878-INS-225
chr19-57468113-INS-59
chr19-57485042-INS-72
chr19-57490487-INS-98
chr19-57496218-INS-54
chr19-57533813-INS-52
chr19-57537778-INS-161
chr19-57548035-INS-268
chr19-57551112-INS-61
chr19-57564018-INS-553
chr19-57586529-INS-233
chr19-57590109-INS-124
chr19-57596033-INS-194
chr19-57637151-INS-300
chr19-57638155-INS-208
chr19-57650420-INS-52
chr19-57654606-INS-76
chr19-57670113-INS-114
chr19-57691275-INS-53
chr19-57696294-INS-88
chr19-57719502-INS-185
chr19-57722293-INS-134
chr19-57724808-INS-238
chr19-57749845-INS-101
chr19-57783853-INS-156
chr19-57792872-INS-52
chr19-57798915-INS-54
chr19-57807847-INS-100
chr19-57813496-INS-134
chr19-57838027-INS-147
chr19-57915011-INS-2005
chr19-57915070-INS-647
chr19-57917013-INS-272
chr19-57917932-INS-58
chr19-57937437-INS-99
chr19-57955803-INS-998
chr19-57968626-INS-82
chr19-57977921-INS-497
chr19-57993933-INS-520
chr19-58005029-INS-6433
chr19-58026003-INS-143
chr19-58049361-INS-57
chr19-58067243-INS-175
chr19-58075658-INS-84
chr19-58105012-INS-197
chr19-58111063-INS-90
chr19-58126257-INS-53
chr19-58138023-INS-215
chr19-58142372-INS-420
chr19-58179561-INS-96
chr19-58215130-INS-60
chr19-58217953-INS-221
chr19-58241721-INS-827
chr19-58255060-INS-900
chr19-58280957-INS-102
chr19-58283378-INS-108
chr19-58291793-INS-175
chr19-58293473-INS-252
chr19-58312493-INS-134

chr19-58318629-INS-126
chr19-58321519-INS-148
chr19-58351144-INS-204
chr19-58406453-INS-128
chr19-58438690-INS-52
chr19-58461709-INS-272
chr19-58474534-INS-193
chr19-58478107-INS-407
chr19-58505629-INS-6773
chr19-58507030-INS-148
chr19-58509393-INS-2921
chr19-58514036-INS-415
chr19-58530105-INS-64
chr19-58541105-INS-70
chr19-58544202-INS-6820
chr19-58554910-INS-68
chr19-58557439-INS-193
chr19-58562557-INS-189
chr19-58590096-INS-466
chr19-58592488-INS-62
chr19-58601865-INS-112
chr19-58617233-INS-6691
chr19-58643925-INS-110
chr19-58648125-INS-114
chr19-58656074-INS-152
chr19-58744719-INS-60
chr19-58756776-INS-58
chr19-58758895-INS-74
chr19-58762963-INS-122
chr19-58801172-INS-124
chr19-58821116-INS-105
chr19-58844456-INS-62
chr19-58863402-INS-71
chr19-58876602-INS-120
chr19-58882291-INS-172
chr19-58889798-INS-387
chr19-58930451-INS-80
chr19-58976070-INS-142
chr19-58990061-INS-58
chr19-59019871-INS-66
chr19-59028259-INS-379
chr19-59032054-INS-200
chr19-59040535-INS-93
chr19-59064583-INS-56
chr19-59070594-INS-524
chr19-59099163-INS-54
chr19-59117431-INS-93
chr19-59118386-INS-7102
chr19-59124649-INS-133
chr19-59130487-INS-85
chr19-59142383-INS-7140
chr19-59153059-INS-52
chr19-59160723-INS-89
chr19-59163739-INS-58
chr19-59169945-INS-59
chr19-59183570-INS-77
chr19-59185159-INS-89
chr19-59200476-INS-815
chr19-59207535-INS-105
chr19-59214819-INS-177
chr19-59233168-INS-129
chr19-59243462-INS-54
chr19-59248659-INS-72
chr19-59342975-INS-71
chr19-59345699-INS-111
chr19-59347679-INS-160
chr19-59375672-INS-441
chr19-59391722-INS-85
chr19-59404657-INS-86
chr19-59410444-INS-82
chr19-59415626-INS-173
chr19-59416824-INS-68
chr19-59470753-INS-118
chr19-59488583-INS-110
chr19-59490103-INS-52
chr19-59491435-INS-57
chr19-59503754-INS-186

chr19-59516943-INS-639
chr19-59528427-INS-236
chr19-59538379-INS-54
chr19-59544472-INS-95
chr19-59551072-INS-56
chr19-59559693-INS-173
chr19-59575136-INS-54
chr19-59584760-INS-3427
chr19-59637030-INS-100
chr19-59680838-INS-217
chr19-59688604-INS-235
chr19-59724702-INS-158
chr19-59735916-INS-69
chr19-59745813-INS-66
chr19-59750540-INS-56
chr19-59769457-INS-119
chr19-59840400-INS-60
chr19-59845248-INS-277
chr19-59869104-INS-327
chr19-59940423-INS-64
chr19-59975999-INS-1080
chr19-59976161-INS-286
chr19-59977300-INS-72
chr19-59989471-INS-52
chr19-59995910-INS-517
chr19-60017623-INS-6922
chr19-60044733-INS-92
chr19-60062989-INS-139
chr19-60065772-INS-312
chr19-60082037-INS-54
chr19-60097188-INS-194
chr19-60127673-INS-6633
chr19-60128792-INS-51
chr19-60142991-INS-80
chr19-60165447-INS-54
chr19-60220833-INS-6207
chr19-60229836-INS-275
chr19-60283793-INS-60
chr19-60304946-INS-94
chr19-60308378-INS-160
chr19-60312620-INS-58
chr19-60333487-INS-54
chr19-60368889-INS-58
chr19-60380992-INS-56
chr19-60383253-INS-1368
chr19-60390816-INS-183
chr19-60408192-INS-70
chr19-60417456-INS-214
chr19-60443581-INS-118
chr19-60455108-INS-71
chr19-60468237-INS-124
chr19-60483512-INS-94
chr19-60502818-INS-5312
chr19-60557762-INS-71
chr19-60572412-INS-164
chr19-60585552-INS-4823
chr19-60605382-INS-90
chr19-60621806-INS-5667
chr19-60654625-INS-706
chr19-60655691-INS-246
chr19-60657162-INS-95
chr19-60664918-INS-130
chr19-60666149-INS-55
chr19-60708107-INS-158
chr19-60731261-INS-110
chr19-60736160-INS-213
chr19-60772368-INS-87
chr19-60789528-INS-175
chr19-60801050-INS-198
chr19-60809940-INS-55
chr19-60815466-INS-230
chr19-60836221-INS-402
chr19-60844693-INS-52
chr19-60869582-INS-364
chr19-60874436-INS-1889
chr19-60893603-INS-285
chr19-60913078-INS-100

chr19-60922885-INS-56
chr19-60935502-INS-162
chr19-60938325-INS-122
chr19-60958295-INS-99
chr19-60980573-INS-6900
chr19-60996303-INS-206
chr19-60998198-INS-116
chr19-61040023-INS-511
chr19-61047738-INS-222
chr19-61052888-INS-183
chr19-61113313-INS-127
chr19-61124345-INS-109
chr19-61166724-INS-9767
chr19-61224820-INS-202
chr19-61257342-INS-92
chr19-61285525-INS-168
chr2-3051051-INS-165
chr2-3095898-INS-333
chr2-3110279-INS-259
chr2-3139382-INS-87
chr2-3153665-INS-117
chr2-3174247-INS-159
chr2-3192557-INS-258
chr2-3199360-INS-72
chr2-3225402-INS-136
chr2-3233738-INS-492
chr2-3246965-INS-67
chr2-3265733-INS-98
chr2-3300702-INS-60
chr2-3356616-INS-58
chr2-3358615-INS-76
chr2-3386955-INS-88
chr2-3394963-INS-150
chr2-3406564-INS-669
chr2-3408413-INS-116
chr2-3434434-INS-131
chr2-3435006-INS-147
chr2-3440780-INS-359
chr2-3462645-INS-190
chr2-3538451-INS-528
chr2-3545481-INS-53
chr2-3568506-INS-116
chr2-3587073-INS-72
chr2-3602665-INS-138
chr2-3606610-INS-304
chr2-3614893-INS-79
chr2-3615698-INS-124
chr2-3630861-INS-80
chr2-3633177-INS-225
chr2-3654668-INS-323
chr2-3662725-INS-171
chr2-3672340-INS-350
chr2-3684103-INS-233
chr2-3688803-INS-400
chr2-3701906-INS-995
chr2-3704221-INS-245
chr2-3711471-INS-56
chr2-3738642-INS-490
chr2-3753970-INS-100
chr2-3779743-INS-616
chr2-3784902-INS-80
chr2-3786127-INS-167
chr2-3808501-INS-97
chr2-3814099-INS-54
chr2-3822016-INS-116
chr2-3834916-INS-294
chr2-3858569-INS-66
chr2-3860879-INS-645
chr2-3869532-INS-6416
chr2-3870554-INS-241
chr2-3872945-INS-82
chr2-3902910-INS-1862
chr2-3909831-INS-108
chr2-3921701-INS-228
chr2-3931029-INS-96
chr2-3958892-INS-190
chr2-4005515-INS-1036

chr2-4021118-INS-155
chr2-4040670-INS-93
chr2-4054269-INS-6494
chr2-4061281-INS-6037
chr2-4091434-INS-168
chr2-4102539-INS-68
chr2-4106945-INS-88
chr2-4155771-INS-524
chr2-4166551-INS-51
chr2-4173996-INS-198
chr2-4226570-INS-58
chr2-4236348-INS-389
chr2-4287195-INS-72
chr2-4306004-INS-52
chr2-4309975-INS-185
chr2-4314245-INS-60
chr2-4321481-INS-120
chr2-4324707-INS-204
chr2-4354726-INS-54
chr2-4387253-INS-174
chr2-4394132-INS-76
chr2-4398077-INS-104
chr2-4412861-INS-108
chr2-4421025-INS-252
chr2-4424180-INS-78
chr2-4425043-INS-549
chr2-4457994-INS-217
chr2-4458741-INS-414
chr2-4475984-INS-232
chr2-4493531-INS-167
chr2-4520904-INS-66
chr2-4523857-INS-70
chr2-4587126-INS-130
chr2-4598439-INS-132
chr2-4636390-INS-466
chr2-4671226-INS-56
chr2-4684157-INS-681
chr2-4704437-INS-400
chr2-4716668-INS-194
chr2-4734275-INS-374
chr2-4737656-INS-2835
chr2-4745306-INS-75
chr2-4759928-INS-68
chr2-4775828-INS-164
chr2-4796184-INS-126
chr2-4818238-INS-73
chr2-4850033-INS-269
chr2-4851177-INS-53
chr2-4853317-INS-251
chr2-4869122-INS-62
chr2-4875219-INS-373
chr2-4918173-INS-52
chr2-4958257-INS-118
chr2-4970182-INS-204
chr2-4971764-INS-400
chr2-4987647-INS-102
chr2-5001789-INS-411
chr2-5004166-INS-62
chr2-5014849-INS-359
chr2-5023841-INS-196
chr2-5036308-INS-214
chr2-5042866-INS-308
chr2-5053218-INS-2895
chr2-5060158-INS-78
chr2-5073686-INS-194
chr2-5082164-INS-247
chr2-5088337-INS-79
chr2-5104406-INS-227
chr2-5123222-INS-368
chr2-5124157-INS-6869
chr2-5149873-INS-343
chr2-5151857-INS-156
chr2-5159812-INS-52
chr2-5174637-INS-133
chr2-5178967-INS-137
chr2-5186561-INS-195
chr2-5231247-INS-78

chr2-5266473-INS-130
chr2-5316836-INS-140
chr2-5320380-INS-119
chr2-5323554-INS-666
chr2-5341421-INS-592
chr2-5344378-INS-52
chr2-5364040-INS-295
chr2-5411393-INS-89
chr2-5412804-INS-98
chr2-5417291-INS-186
chr2-5430048-INS-69
chr2-5432694-INS-538
chr2-5444498-INS-145
chr2-5456733-INS-799
chr2-5462106-INS-112
chr2-5467316-INS-55
chr2-5488162-INS-140
chr2-5496008-INS-130
chr2-5529926-INS-234
chr2-5543183-INS-257
chr2-5543868-INS-70
chr2-5587348-INS-387
chr2-5587350-INS-1075
chr2-5601784-INS-206
chr2-5606783-INS-426
chr2-5654353-INS-159
chr2-5672411-INS-156
chr2-5684488-INS-296
chr2-5685260-INS-64
chr2-5697088-INS-83
chr2-5697882-INS-6445
chr2-5702473-INS-181
chr2-5724482-INS-80
chr2-5728244-INS-51
chr2-5731424-INS-198
chr2-5758064-INS-5435
chr2-5790656-INS-965
chr2-5792612-INS-60
chr2-5799150-INS-109
chr2-5803295-INS-289
chr2-5805447-INS-55
chr2-5825914-INS-368
chr2-5831239-INS-111
chr2-5834577-INS-126
chr2-5844594-INS-257
chr2-5847185-INS-90
chr2-5884765-INS-62
chr2-5925643-INS-212
chr2-5927699-INS-66
chr2-5943583-INS-207
chr2-5944828-INS-82
chr2-5952706-INS-83
chr2-5983327-INS-232
chr2-6001705-INS-701
chr2-6003805-INS-125
chr2-6007371-INS-89
chr2-6074907-INS-72
chr2-6088972-INS-63
chr2-6106212-INS-465
chr2-6107452-INS-55
chr2-6108338-INS-66
chr2-6109447-INS-231
chr2-6110780-INS-91
chr2-6113903-INS-69
chr2-6157226-INS-200
chr2-6159668-INS-80
chr2-6201969-INS-88
chr2-6220885-INS-340
chr2-6243535-INS-55
chr2-6245575-INS-93
chr2-6251037-INS-54
chr2-6282690-INS-411
chr2-6285053-INS-60
chr2-6294098-INS-5273
chr2-6307932-INS-72
chr2-6332235-INS-301
chr2-6350074-INS-73

chr2-6360158-INS-60
chr2-6378508-INS-60
chr2-6390081-INS-397
chr2-6418598-INS-101
chr2-6474705-INS-52
chr2-6482180-INS-400
chr2-6492458-INS-108
chr2-6494141-INS-165
chr2-6513612-INS-342
chr2-6548271-INS-67
chr2-6582135-INS-117
chr2-6596426-INS-200
chr2-6633220-INS-84
chr2-6635219-INS-132
chr2-6637588-INS-114
chr2-6695533-INS-63
chr2-6718012-INS-757
chr2-6798219-INS-174
chr2-6831794-INS-569
chr2-6842345-INS-262
chr2-6862269-INS-547
chr2-6932772-INS-60
chr2-6937333-INS-401
chr2-6960151-INS-657
chr2-7006650-INS-105
chr2-7013973-INS-250
chr2-7055682-INS-362
chr2-7088002-INS-72
chr2-7100287-INS-588
chr2-7104073-INS-56
chr2-7121557-INS-78
chr2-7124712-INS-62
chr2-7133033-INS-68
chr2-7144706-INS-72
chr2-7146497-INS-484
chr2-7155640-INS-894
chr2-7168319-INS-95
chr2-7180001-INS-149
chr2-7191496-INS-154
chr2-7210797-INS-90
chr2-7239064-INS-6802
chr2-7267603-INS-55
chr2-7283572-INS-51
chr2-7309091-INS-106
chr2-7350771-INS-106
chr2-7419981-INS-242
chr2-7432461-INS-401
chr2-7441134-INS-58
chr2-7482959-INS-423
chr2-7497933-INS-573
chr2-7513530-INS-88
chr2-7547823-INS-97
chr2-7551885-INS-888
chr2-7591731-INS-404
chr2-7615487-INS-8270
chr2-7644426-INS-1316
chr2-7646421-INS-64
chr2-7677961-INS-368
chr2-7696201-INS-290
chr2-7699387-INS-195
chr2-7717331-INS-205
chr2-7725508-INS-67
chr2-7787491-INS-82
chr2-7791855-INS-849
chr2-7845924-INS-69
chr2-7851237-INS-627
chr2-7856268-INS-94
chr2-7859833-INS-293
chr2-7885601-INS-195
chr2-7888543-INS-56
chr2-7904750-INS-886
chr2-7911926-INS-6213
chr2-7912840-INS-220
chr2-7926875-INS-212
chr2-7931211-INS-131
chr2-7939972-INS-77
chr2-7942162-INS-87

chr2-7952120-INS-84
chr2-7962340-INS-4199
chr2-7980929-INS-906
chr2-8009066-INS-57
chr2-8057215-INS-58
chr2-8063082-INS-72
chr2-8067541-INS-182
chr2-8071873-INS-52
chr2-8083408-INS-56
chr2-8097620-INS-1781
chr2-8108490-INS-71
chr2-8114326-INS-607
chr2-8115451-INS-930
chr2-8115714-INS-61
chr2-8121888-INS-112
chr2-8139081-INS-53
chr2-8144091-INS-163
chr2-8177050-INS-648
chr2-8182195-INS-6465
chr2-8212598-INS-417
chr2-8238845-INS-106
chr2-8254404-INS-195
chr2-8261477-INS-941
chr2-8266382-INS-767
chr2-8292171-INS-122
chr2-8320190-INS-250
chr2-8370330-INS-61
chr2-8412613-INS-7320
chr2-8417774-INS-8694
chr2-8423108-INS-75
chr2-8443295-INS-68
chr2-8502462-INS-6390
chr2-8511271-INS-7310
chr2-8515768-INS-90
chr2-8526905-INS-781
chr2-8528947-INS-154
chr2-8573783-INS-443
chr2-8678580-INS-110
chr2-8700086-INS-69
chr2-8704164-INS-255
chr2-8706369-INS-400
chr2-8713657-INS-360
chr2-8740358-INS-206
chr2-8746673-INS-188
chr2-8748539-INS-58
chr2-8806005-INS-2499
chr2-8811932-INS-562
chr2-8813495-INS-74
chr2-8829536-INS-1871
chr2-8833662-INS-104
chr2-8855309-INS-72
chr2-8863836-INS-128
chr2-8880047-INS-239
chr2-8882519-INS-74
chr2-8894200-INS-8357
chr2-8918045-INS-62
chr2-8930667-INS-993
chr2-8939167-INS-84
chr2-8939533-INS-6360
chr2-8960822-INS-64
chr2-8977686-INS-176
chr2-8986483-INS-94
chr2-9016864-INS-65
chr2-9045380-INS-64
chr2-9078356-INS-159
chr2-9079429-INS-124
chr2-9094459-INS-2554
chr2-9118053-INS-52
chr2-9118073-INS-1419
chr2-9137474-INS-211
chr2-9139701-INS-332
chr2-9144697-INS-7130
chr2-9145221-INS-54
chr2-9166349-INS-644
chr2-9171122-INS-300
chr2-9201209-INS-76
chr2-9206382-INS-801

chr2-9228720-INS-75
chr2-9234010-INS-336
chr2-9250593-INS-54
chr2-9296595-INS-103
chr2-9301643-INS-200
chr2-9319950-INS-503
chr2-9320742-INS-53
chr2-9328400-INS-77
chr2-9353385-INS-60
chr2-9360962-INS-94
chr2-9404773-INS-157
chr2-9409873-INS-54
chr2-9458773-INS-60
chr2-9470960-INS-303
chr2-9498228-INS-76
chr2-9507419-INS-220
chr2-9532790-INS-56
chr2-9538369-INS-80
chr2-9553490-INS-60
chr2-9563202-INS-53
chr2-9577171-INS-98
chr2-9613689-INS-64
chr2-9618254-INS-104
chr2-9664227-INS-7485
chr2-9677599-INS-655
chr2-9679277-INS-6479
chr2-9705238-INS-60
chr2-9711371-INS-3695
chr2-9715822-INS-234
chr2-9741560-INS-55
chr2-9742553-INS-71
chr2-9780189-INS-62
chr2-9785361-INS-381
chr2-9788112-INS-108
chr2-9789705-INS-137
chr2-9798119-INS-476
chr2-9802712-INS-358
chr2-9804363-INS-255
chr2-9837958-INS-681
chr2-9844800-INS-217
chr2-9858761-INS-1888
chr2-9859623-INS-199
chr2-9893412-INS-74
chr2-9898640-INS-430
chr2-9909888-INS-122
chr2-9935530-INS-102
chr2-9938082-INS-64
chr2-9939235-INS-1593
chr2-9951323-INS-60
chr2-9970883-INS-370
chr2-9980682-INS-161
chr2-9991167-INS-192
chr2-10003401-INS-640
chr2-10019439-INS-54
chr2-10029815-INS-498
chr2-10043576-INS-194
chr2-10058477-INS-56
chr2-10061021-INS-177
chr2-10064187-INS-52
chr2-10067379-INS-207
chr2-10075982-INS-64
chr2-10086952-INS-108
chr2-10090800-INS-5036
chr2-10101115-INS-52
chr2-10111083-INS-67
chr2-10123594-INS-345
chr2-10127345-INS-76
chr2-10131481-INS-225
chr2-10137096-INS-351
chr2-10137536-INS-497
chr2-10148015-INS-66
chr2-10150938-INS-174
chr2-10151669-INS-707
chr2-10160047-INS-108
chr2-10162535-INS-79
chr2-10163088-INS-59
chr2-10172542-INS-195

chr2-10176691-INS-98
chr2-10190822-INS-87
chr2-10192566-INS-344
chr2-10203787-INS-66
chr2-10208216-INS-82
chr2-10210514-INS-196
chr2-10210887-INS-344
chr2-10218263-INS-59
chr2-10229542-INS-216
chr2-10258849-INS-212
chr2-10271470-INS-92
chr2-10282002-INS-257
chr2-10288450-INS-376
chr2-10295313-INS-189
chr2-10326625-INS-58
chr2-10330445-INS-724
chr2-10333502-INS-54
chr2-10336084-INS-70
chr2-10342682-INS-228
chr2-10348977-INS-127
chr2-10359137-INS-65
chr2-10363318-INS-187
chr2-10381007-INS-215
chr2-10405245-INS-130
chr2-10412632-INS-58
chr2-10423126-INS-56
chr2-10445967-INS-61
chr2-10454106-INS-910
chr2-10459999-INS-182
chr2-10462946-INS-1341
chr2-10472921-INS-114
chr2-10511595-INS-90
chr2-10532211-INS-76
chr2-10533383-INS-176
chr2-10539062-INS-64
chr2-10540196-INS-295
chr2-10570286-INS-87
chr2-10578974-INS-70
chr2-10601750-INS-426
chr2-10604744-INS-77
chr2-10607097-INS-234
chr2-10610450-INS-221
chr2-10613915-INS-993
chr2-10620926-INS-62
chr2-10625731-INS-753
chr2-10647901-INS-768
chr2-10667060-INS-83
chr2-10671166-INS-393
chr2-10704073-INS-419
chr2-10720261-INS-236
chr2-10732354-INS-421
chr2-10735683-INS-467
chr2-10755858-INS-164
chr2-10763658-INS-54
chr2-10776395-INS-52
chr2-10784266-INS-56
chr2-10786665-INS-351
chr2-10791668-INS-81
chr2-10807896-INS-60
chr2-10814828-INS-88
chr2-10816663-INS-58
chr2-10843262-INS-368
chr2-10855255-INS-76
chr2-10860648-INS-99
chr2-10869804-INS-162
chr2-10883511-INS-191
chr2-10884411-INS-58
chr2-10897771-INS-141
chr2-10905030-INS-59
chr2-10940692-INS-796
chr2-10973018-INS-813
chr2-10993018-INS-358
chr2-11019203-INS-131
chr2-11022689-INS-80
chr2-11024878-INS-56
chr2-11034834-INS-434
chr2-11037536-INS-59

chr2-11040967-INS-52
chr2-11065219-INS-300
chr2-11098001-INS-134
chr2-11099848-INS-440
chr2-11100888-INS-61
chr2-11116964-INS-60
chr2-11124142-INS-200
chr2-11180532-INS-724
chr2-11185911-INS-95
chr2-11187237-INS-60
chr2-11200862-INS-71
chr2-11206840-INS-205
chr2-11207692-INS-54
chr2-11214567-INS-193
chr2-11222679-INS-196
chr2-11232361-INS-140
chr2-11236389-INS-225
chr2-11242392-INS-87
chr2-11245942-INS-69
chr2-11256859-INS-234
chr2-11270235-INS-92
chr2-11284131-INS-160
chr2-11289040-INS-347
chr2-11303724-INS-57
chr2-11308257-INS-200
chr2-11322682-INS-88
chr2-11328251-INS-222
chr2-11330487-INS-506
chr2-11345339-INS-83
chr2-11348999-INS-68
chr2-11387070-INS-86
chr2-11399392-INS-1485
chr2-11407802-INS-64
chr2-11432747-INS-649
chr2-11440241-INS-253
chr2-11450353-INS-266
chr2-11460527-INS-124
chr2-11461969-INS-237
chr2-11466859-INS-177
chr2-11478428-INS-74
chr2-11493011-INS-304
chr2-11507239-INS-58
chr2-11512199-INS-52
chr2-11512724-INS-80
chr2-11531558-INS-164
chr2-11544130-INS-188
chr2-11548035-INS-207
chr2-11555749-INS-204
chr2-11562788-INS-872
chr2-11583690-INS-323
chr2-11599411-INS-168
chr2-11605962-INS-84
chr2-11625924-INS-145
chr2-11631685-INS-137
chr2-11650852-INS-200
chr2-11657454-INS-140
chr2-11659669-INS-51
chr2-11705901-INS-58
chr2-11750227-INS-440
chr2-11776255-INS-598
chr2-11777630-INS-132
chr2-11790160-INS-79
chr2-11816086-INS-196
chr2-11818870-INS-473
chr2-11900600-INS-58
chr2-11910840-INS-121
chr2-11916627-INS-705
chr2-11934614-INS-840
chr2-11941209-INS-6681
chr2-11948254-INS-200
chr2-11966206-INS-132
chr2-11969037-INS-6273
chr2-12026264-INS-251
chr2-12055944-INS-399
chr2-12060813-INS-70
chr2-12074128-INS-180
chr2-12084784-INS-102

chr2-12096232-INS-166
chr2-12120016-INS-338
chr2-12125097-INS-72
chr2-12147373-INS-589
chr2-12150178-INS-937
chr2-12160753-INS-56
chr2-12175495-INS-183
chr2-12176904-INS-147
chr2-12187224-INS-64
chr2-12193091-INS-110
chr2-12218174-INS-112
chr2-12230069-INS-80
chr2-12286144-INS-111
chr2-12317491-INS-51
chr2-12321885-INS-534
chr2-12323609-INS-121
chr2-12324754-INS-89
chr2-12334280-INS-73
chr2-12341839-INS-58
chr2-12348457-INS-141
chr2-12351871-INS-400
chr2-12364728-INS-208
chr2-12366133-INS-497
chr2-12380957-INS-454
chr2-12390262-INS-378
chr2-12401513-INS-54
chr2-12420831-INS-186
chr2-12431926-INS-54
chr2-12438071-INS-87
chr2-12453843-INS-100
chr2-12458489-INS-74
chr2-12466583-INS-3152
chr2-12469043-INS-123
chr2-12476460-INS-172
chr2-12484662-INS-425
chr2-12501437-INS-153
chr2-12518296-INS-57
chr2-12532022-INS-728
chr2-12553375-INS-143
chr2-12566789-INS-457
chr2-12575906-INS-78
chr2-12585231-INS-844
chr2-12586092-INS-92
chr2-12601110-INS-1598
chr2-12613759-INS-206
chr2-12617624-INS-87
chr2-12634283-INS-6204
chr2-12662099-INS-54
chr2-12663172-INS-126
chr2-12679799-INS-452
chr2-12681411-INS-95
chr2-12692983-INS-547
chr2-12694637-INS-136
chr2-12706824-INS-90
chr2-12722074-INS-524
chr2-12731294-INS-53
chr2-12732463-INS-401
chr2-12737527-INS-55
chr2-12763052-INS-53
chr2-12765362-INS-51
chr2-12772519-INS-436
chr2-12776132-INS-54
chr2-12778791-INS-190
chr2-12803180-INS-60
chr2-12807289-INS-7040
chr2-12807984-INS-74
chr2-12823896-INS-714
chr2-12841067-INS-6188
chr2-12841259-INS-139
chr2-12847013-INS-1094
chr2-12847130-INS-113
chr2-12881824-INS-8747
chr2-12890663-INS-195
chr2-12899072-INS-1113
chr2-12905361-INS-1630
chr2-12909459-INS-6264
chr2-12913369-INS-129

chr2-12920302-INS-359
chr2-12925128-INS-589
chr2-12962422-INS-6539
chr2-12984648-INS-72
chr2-12988509-INS-400
chr2-12999457-INS-66
chr2-13010113-INS-65
chr2-13013746-INS-52
chr2-13058523-INS-84
chr2-13082421-INS-58
chr2-13095638-INS-93
chr2-13115644-INS-217
chr2-13128725-INS-132
chr2-13131499-INS-84
chr2-13170064-INS-104
chr2-13176569-INS-150
chr2-13184897-INS-78
chr2-13218105-INS-5256
chr2-13218689-INS-988
chr2-13220276-INS-263
chr2-13224173-INS-243
chr2-13281985-INS-96
chr2-13306106-INS-1359
chr2-13319733-INS-3057
chr2-13342483-INS-64
chr2-13380902-INS-1894
chr2-13417387-INS-185
chr2-13442474-INS-613
chr2-13452359-INS-54
chr2-13457416-INS-153
chr2-13467545-INS-191
chr2-13476701-INS-235
chr2-13476701-INS-752
chr2-13492109-INS-76
chr2-13500396-INS-58
chr2-13516249-INS-68
chr2-13517756-INS-908
chr2-13538526-INS-148
chr2-13540713-INS-275
chr2-13601641-INS-111
chr2-13604798-INS-187
chr2-13612868-INS-95
chr2-13613857-INS-274
chr2-13643022-INS-105
chr2-13662623-INS-81
chr2-13674224-INS-747
chr2-13678173-INS-1436
chr2-13696322-INS-6176
chr2-13701929-INS-6677
chr2-13717755-INS-251
chr2-13747418-INS-412
chr2-13748767-INS-693
chr2-13756752-INS-53
chr2-13773259-INS-98
chr2-13802927-INS-66
chr2-13806280-INS-448
chr2-13822616-INS-985
chr2-13843636-INS-7890
chr2-13844985-INS-172
chr2-13856648-INS-58
chr2-13911719-INS-254
chr2-13920599-INS-6403
chr2-13922814-INS-73
chr2-13927834-INS-721
chr2-13935164-INS-52
chr2-13939984-INS-152
chr2-13961136-INS-440
chr2-13968504-INS-450
chr2-14046562-INS-80
chr2-14050660-INS-399
chr2-14069751-INS-1039
chr2-14072175-INS-229
chr2-14074048-INS-6972
chr2-14091665-INS-60
chr2-14095407-INS-943
chr2-14121559-INS-192
chr2-14155640-INS-88

chr2-14201465-INS-5810
chr2-14208962-INS-73
chr2-14215270-INS-89
chr2-14224520-INS-204
chr2-14229598-INS-393
chr2-14232318-INS-4712
chr2-14254702-INS-230
chr2-14272974-INS-132
chr2-14275626-INS-475
chr2-14344386-INS-427
chr2-14360955-INS-2993
chr2-14363481-INS-97
chr2-14369976-INS-227
chr2-14371630-INS-55
chr2-14373663-INS-249
chr2-14382839-INS-60
chr2-14408055-INS-6403
chr2-14427289-INS-154
chr2-14456596-INS-234
chr2-14459819-INS-192
chr2-14471522-INS-76
chr2-14475079-INS-153
chr2-14479321-INS-266
chr2-14504494-INS-136
chr2-14522460-INS-618
chr2-14555304-INS-452
chr2-14569632-INS-236
chr2-14572693-INS-97
chr2-14584180-INS-56
chr2-14665130-INS-654
chr2-14675791-INS-896
chr2-14676426-INS-295
chr2-14689881-INS-56
chr2-14705156-INS-52
chr2-14718299-INS-87
chr2-14762817-INS-456
chr2-14805203-INS-74
chr2-14806192-INS-714
chr2-14809569-INS-526
chr2-14810251-INS-54
chr2-14813058-INS-58
chr2-14834711-INS-70
chr2-14839432-INS-63
chr2-14880992-INS-55
chr2-14891380-INS-67
chr2-14899639-INS-100
chr2-14904249-INS-115
chr2-14940242-INS-304
chr2-14956576-INS-400
chr2-14974047-INS-53
chr2-15001949-INS-2935
chr2-15012312-INS-168
chr2-15019716-INS-96
chr2-15023751-INS-6456
chr2-15048604-INS-52
chr2-15107853-INS-58
chr2-15110043-INS-60
chr2-15117308-INS-57
chr2-15118143-INS-258
chr2-15126421-INS-418
chr2-15136540-INS-492
chr2-15141031-INS-58
chr2-15163978-INS-342
chr2-15170479-INS-70
chr2-15171638-INS-66
chr2-15182377-INS-2015
chr2-15213548-INS-60
chr2-15228357-INS-139
chr2-15243935-INS-310
chr2-15265368-INS-110
chr2-15275849-INS-473
chr2-15278152-INS-647
chr2-15282434-INS-3135
chr2-15292241-INS-6413
chr2-15315373-INS-201
chr2-15341977-INS-925
chr2-15345616-INS-6504

chr2-15356496-INS-356
chr2-15359528-INS-386
chr2-15360358-INS-570
chr2-15365095-INS-2433
chr2-15375528-INS-301
chr2-15375709-INS-95
chr2-15393122-INS-3349
chr2-15393457-INS-5281
chr2-15402931-INS-361
chr2-15422633-INS-4650
chr2-15449815-INS-1488
chr2-15454880-INS-55
chr2-15459823-INS-81
chr2-15465438-INS-91
chr2-15477447-INS-123
chr2-15479328-INS-59
chr2-15481404-INS-392
chr2-15492840-INS-363
chr2-15511434-INS-135
chr2-15545578-INS-722
chr2-15554346-INS-721
chr2-15558772-INS-167
chr2-15571538-INS-1550
chr2-15601721-INS-200
chr2-15611970-INS-86
chr2-15652941-INS-354
chr2-15677840-INS-6755
chr2-15684465-INS-7082
chr2-15719968-INS-675
chr2-15727847-INS-52
chr2-15730044-INS-6443
chr2-15745149-INS-781
chr2-15755617-INS-6259
chr2-15762043-INS-156
chr2-15778152-INS-466
chr2-15789825-INS-888
chr2-15834308-INS-521
chr2-15840945-INS-181
chr2-15848834-INS-152
chr2-15853839-INS-114
chr2-15879422-INS-2660
chr2-15905468-INS-446
chr2-15909464-INS-74
chr2-15927202-INS-934
chr2-15956153-INS-81
chr2-15981107-INS-52
chr2-15992707-INS-131
chr2-16005209-INS-78
chr2-16006892-INS-3293
chr2-16019120-INS-214
chr2-16020259-INS-7473
chr2-16020260-INS-325
chr2-16038257-INS-801
chr2-16041921-INS-1341
chr2-16048415-INS-939
chr2-16061253-INS-207
chr2-16088487-INS-57
chr2-16105423-INS-384
chr2-16118350-INS-555
chr2-16119411-INS-227
chr2-16129815-INS-6222
chr2-16131467-INS-7108
chr2-16140221-INS-498
chr2-16142642-INS-6918
chr2-16151546-INS-72
chr2-16153877-INS-543
chr2-16167698-INS-286
chr2-16168694-INS-66
chr2-16182603-INS-526
chr2-16193723-INS-82
chr2-16207110-INS-52
chr2-16250775-INS-261
chr2-16261093-INS-284
chr2-16277797-INS-1490
chr2-16286717-INS-6489
chr2-16301059-INS-56
chr2-16311796-INS-224

chr2-16316761-INS-236
chr2-16339229-INS-514
chr2-16344241-INS-2643
chr2-16349877-INS-2244
chr2-16355496-INS-959
chr2-16379405-INS-119
chr2-16421976-INS-251
chr2-16423976-INS-864
chr2-16465668-INS-194
chr2-16468270-INS-114
chr2-16489862-INS-6456
chr2-16491730-INS-6973
chr2-16504958-INS-4537
chr2-16524912-INS-54
chr2-16529824-INS-423
chr2-16538444-INS-185
chr2-16541945-INS-84
chr2-16546649-INS-6539
chr2-16547836-INS-1490
chr2-16556217-INS-257
chr2-16572022-INS-919
chr2-16573294-INS-418
chr2-16601614-INS-146
chr2-16602107-INS-6868
chr2-16603958-INS-199
chr2-16657874-INS-295
chr2-16680101-INS-310
chr2-16682354-INS-174
chr2-16693691-INS-211
chr2-16700881-INS-57
chr2-16724785-INS-271
chr2-16725880-INS-81
chr2-16728301-INS-127
chr2-16737796-INS-115
chr2-16737886-INS-745
chr2-16744320-INS-240
chr2-16752835-INS-618
chr2-16775118-INS-69
chr2-16819421-INS-72
chr2-16823767-INS-6329
chr2-16865003-INS-3172
chr2-16870146-INS-114
chr2-16880869-INS-444
chr2-16916965-INS-65
chr2-16924780-INS-62
chr2-16926868-INS-195
chr2-16928124-INS-97
chr2-16932004-INS-56
chr2-16967296-INS-1600
chr2-16977129-INS-252
chr2-16982971-INS-53
chr2-16998090-INS-2976
chr2-17013325-INS-9178
chr2-17013509-INS-441
chr2-17015347-INS-155
chr2-17019632-INS-1848
chr2-17021248-INS-101
chr2-17025108-INS-7596
chr2-17029039-INS-88
chr2-17029294-INS-2240
chr2-17035020-INS-87
chr2-17044208-INS-112
chr2-17055567-INS-58
chr2-17059285-INS-53
chr2-17108134-INS-56
chr2-17115358-INS-55
chr2-17127566-INS-6266
chr2-17138776-INS-6496
chr2-17140320-INS-202
chr2-17198873-INS-116
chr2-17201028-INS-670
chr2-17203863-INS-537
chr2-17214533-INS-6344
chr2-17219368-INS-78
chr2-17234334-INS-377
chr2-17238988-INS-1825
chr2-17248065-INS-7577

chr2-17257733-INS-77
chr2-17290946-INS-134
chr2-17293350-INS-196
chr2-17306822-INS-6684
chr2-17309008-INS-6797
chr2-17309501-INS-7350
chr2-17316196-INS-52
chr2-17321255-INS-174
chr2-17338606-INS-1281
chr2-17341458-INS-56
chr2-17386927-INS-185
chr2-17397009-INS-68
chr2-17408002-INS-251
chr2-17445760-INS-349
chr2-17446710-INS-100
chr2-17468157-INS-147
chr2-17481107-INS-257
chr2-17482890-INS-172
chr2-17498197-INS-109
chr2-17511736-INS-2294
chr2-17521579-INS-68
chr2-17528137-INS-72
chr2-17533596-INS-52
chr2-17547086-INS-394
chr2-17553334-INS-240
chr2-17564985-INS-206
chr2-17579089-INS-75
chr2-17593108-INS-2362
chr2-17601041-INS-616
chr2-17613652-INS-104
chr2-17614937-INS-54
chr2-17645281-INS-90
chr2-17653444-INS-58
chr2-17661309-INS-399
chr2-17696913-INS-102
chr2-17724316-INS-177
chr2-17746376-INS-5543
chr2-17750163-INS-53
chr2-17781991-INS-124
chr2-17802884-INS-223
chr2-17804726-INS-209
chr2-17812538-INS-935
chr2-17814832-INS-60
chr2-17820814-INS-99
chr2-17824846-INS-218
chr2-17826493-INS-105
chr2-17827841-INS-59
chr2-17829623-INS-52
chr2-17839196-INS-72
chr2-17861682-INS-149
chr2-17862471-INS-94
chr2-17864281-INS-75
chr2-17869228-INS-216
chr2-17874936-INS-94
chr2-17884286-INS-207
chr2-17898498-INS-227
chr2-17901678-INS-468
chr2-17902310-INS-7165
chr2-17905617-INS-189
chr2-17916788-INS-94
chr2-17919543-INS-52
chr2-17934846-INS-83
chr2-17940981-INS-264
chr2-17942866-INS-304
chr2-17949544-INS-104
chr2-17966625-INS-288
chr2-17969148-INS-94
chr2-17982757-INS-149
chr2-17988392-INS-165
chr2-18003620-INS-209
chr2-18008210-INS-513
chr2-18020951-INS-442
chr2-18024103-INS-202
chr2-18025790-INS-501
chr2-18027145-INS-56
chr2-18078759-INS-207
chr2-18084267-INS-170

chr2-18086656-INS-67
chr2-18088280-INS-171
chr2-18099245-INS-104
chr2-18100792-INS-231
chr2-18107443-INS-65
chr2-18110919-INS-81
chr2-18124374-INS-208
chr2-18137744-INS-153
chr2-18140697-INS-84
chr2-18159428-INS-360
chr2-18161929-INS-763
chr2-18183546-INS-492
chr2-18221937-INS-202
chr2-18229731-INS-219
chr2-18232081-INS-223
chr2-18246575-INS-62
chr2-18276028-INS-3333
chr2-18282658-INS-2732
chr2-18422888-INS-6941
chr2-18422897-INS-67
chr2-18436489-INS-72
chr2-18437241-INS-60
chr2-18479867-INS-715
chr2-18489983-INS-4995
chr2-18493282-INS-64
chr2-18500985-INS-268
chr2-18514230-INS-890
chr2-18518119-INS-110
chr2-18526253-INS-79
chr2-18542579-INS-66
chr2-18566829-INS-161
chr2-18570805-INS-51
chr2-18573052-INS-4298
chr2-18576593-INS-6415
chr2-18577066-INS-486
chr2-18584746-INS-55
chr2-18586782-INS-224
chr2-18589621-INS-54
chr2-18615255-INS-98
chr2-18635425-INS-76
chr2-18646653-INS-96
chr2-18651786-INS-458
chr2-18780216-INS-63
chr2-18793730-INS-98
chr2-18798934-INS-6966
chr2-18814503-INS-88
chr2-18824504-INS-89
chr2-18867138-INS-426
chr2-18898258-INS-642
chr2-18909061-INS-86
chr2-18979156-INS-108
chr2-19016991-INS-102
chr2-19024652-INS-65
chr2-19026894-INS-439
chr2-19030525-INS-419
chr2-19068883-INS-252
chr2-19080482-INS-5334
chr2-19082954-INS-825
chr2-19095955-INS-78
chr2-19113430-INS-381
chr2-19163890-INS-52
chr2-19194490-INS-64
chr2-19209528-INS-148
chr2-19274366-INS-131
chr2-19288834-INS-126
chr2-19310081-INS-76
chr2-19320803-INS-58
chr2-19326938-INS-407
chr2-19334831-INS-172
chr2-19367198-INS-172
chr2-19371752-INS-63
chr2-19375893-INS-802
chr2-19392648-INS-490
chr2-19423757-INS-139
chr2-19425097-INS-168
chr2-19426054-INS-284
chr2-19433326-INS-65

chr2-19448206-INS-62
chr2-19497610-INS-325
chr2-19513246-INS-68
chr2-19514174-INS-456
chr2-19519950-INS-62
chr2-19522035-INS-411
chr2-19532265-INS-61
chr2-19539525-INS-76
chr2-19566257-INS-93
chr2-19574689-INS-505
chr2-19577030-INS-198
chr2-19596708-INS-185
chr2-19629993-INS-205
chr2-19645008-INS-95
chr2-19692007-INS-114
chr2-19700287-INS-271
chr2-19702382-INS-150
chr2-19703435-INS-512
chr2-19732250-INS-342
chr2-19761178-INS-53
chr2-19766338-INS-286
chr2-19777212-INS-173
chr2-19781790-INS-114
chr2-19787090-INS-110
chr2-19804468-INS-61
chr2-19833950-INS-6295
chr2-19855149-INS-229
chr2-19857452-INS-56
chr2-19888765-INS-336
chr2-19900381-INS-525
chr2-19920433-INS-901
chr2-19994523-INS-6518
chr2-20007243-INS-73
chr2-20037849-INS-76
chr2-20063266-INS-201
chr2-20087186-INS-671
chr2-20100101-INS-6811
chr2-20112174-INS-113
chr2-20124533-INS-400
chr2-20135227-INS-134
chr2-20139617-INS-313
chr2-20141282-INS-868
chr2-20141371-INS-180
chr2-20159987-INS-80
chr2-20180971-INS-54
chr2-20181714-INS-326
chr2-20195200-INS-518
chr2-20217832-INS-580
chr2-20280412-INS-227
chr2-20283750-INS-86
chr2-20288206-INS-287
chr2-20302792-INS-6454
chr2-20312220-INS-6692
chr2-20336529-INS-100
chr2-20340296-INS-484
chr2-20347184-INS-61
chr2-20400208-INS-468
chr2-20449816-INS-200
chr2-20456081-INS-223
chr2-20456684-INS-359
chr2-20460240-INS-155
chr2-20492983-INS-85
chr2-20510811-INS-238
chr2-20515815-INS-162
chr2-20601182-INS-200
chr2-20611917-INS-154
chr2-20641582-INS-212
chr2-20694663-INS-471
chr2-20725448-INS-82
chr2-20747089-INS-77
chr2-20749272-INS-199
chr2-20754600-INS-597
chr2-20767337-INS-223
chr2-20773982-INS-68
chr2-20801239-INS-187
chr2-20803751-INS-291
chr2-20818623-INS-337

chr2-20823622-INS-5298
chr2-20867796-INS-267
chr2-20889589-INS-160
chr2-20900059-INS-81
chr2-20910557-INS-132
chr2-20913117-INS-282
chr2-20917959-INS-53
chr2-20918506-INS-429
chr2-20920776-INS-63
chr2-20942292-INS-197
chr2-20945815-INS-54
chr2-20964375-INS-211
chr2-20990761-INS-313
chr2-21012388-INS-82
chr2-21022287-INS-155
chr2-21030181-INS-247
chr2-21037142-INS-139
chr2-21071474-INS-118
chr2-21089811-INS-8723
chr2-21100623-INS-64
chr2-21110773-INS-4420
chr2-21113087-INS-72
chr2-21122150-INS-217
chr2-21124318-INS-2764
chr2-21126340-INS-85
chr2-21136794-INS-134
chr2-21143099-INS-184
chr2-21157251-INS-8128
chr2-21161222-INS-54
chr2-21224952-INS-202
chr2-21226264-INS-5542
chr2-21227977-INS-205
chr2-21258110-INS-230
chr2-21260493-INS-98
chr2-21276561-INS-4373
chr2-21312589-INS-402
chr2-21342682-INS-6624
chr2-21347004-INS-149
chr2-21348796-INS-6440
chr2-21352457-INS-60
chr2-21356225-INS-84
chr2-21419705-INS-188
chr2-21435134-INS-70
chr2-21442253-INS-52
chr2-21447511-INS-504
chr2-21453180-INS-382
chr2-21500344-INS-6428
chr2-21501984-INS-427
chr2-21505659-INS-201
chr2-21515074-INS-154
chr2-21525091-INS-2948
chr2-21535509-INS-977
chr2-21555221-INS-183
chr2-21600104-INS-752
chr2-21612741-INS-3749
chr2-21640960-INS-111
chr2-21657938-INS-1890
chr2-21674249-INS-67
chr2-21688240-INS-6381
chr2-21690542-INS-4896
chr2-21728402-INS-418
chr2-21747248-INS-536
chr2-21757451-INS-2810
chr2-21825733-INS-439
chr2-21826970-INS-677
chr2-21858399-INS-8149
chr2-21891966-INS-203
chr2-21946973-INS-368
chr2-21959960-INS-409
chr2-21979365-INS-690
chr2-21983100-INS-649
chr2-22058236-INS-3988
chr2-22074710-INS-126
chr2-22102875-INS-389
chr2-22104912-INS-61
chr2-22107162-INS-204
chr2-22111071-INS-183

chr2-22120339-INS-401
chr2-22140102-INS-331
chr2-22171798-INS-84
chr2-22178166-INS-73
chr2-22182475-INS-112
chr2-22185205-INS-2550
chr2-22235428-INS-8862
chr2-22245215-INS-56
chr2-22258390-INS-341
chr2-22260225-INS-488
chr2-22283288-INS-327
chr2-22376645-INS-385
chr2-22380435-INS-91
chr2-22437542-INS-230
chr2-22533543-INS-163
chr2-22552252-INS-272
chr2-22576065-INS-146
chr2-22585463-INS-705
chr2-22622193-INS-78
chr2-22630420-INS-216
chr2-22633247-INS-109
chr2-22646900-INS-161
chr2-22650463-INS-174
chr2-22658474-INS-89
chr2-22698795-INS-127
chr2-22725246-INS-338
chr2-22735517-INS-200
chr2-22754721-INS-102
chr2-22767805-INS-98
chr2-22778628-INS-113
chr2-22835723-INS-104
chr2-22839390-INS-52
chr2-22848952-INS-88
chr2-22856205-INS-75
chr2-22857499-INS-69
chr2-22875190-INS-101
chr2-22931129-INS-516
chr2-22946911-INS-66
chr2-22948280-INS-213
chr2-22955138-INS-361
chr2-22981375-INS-137
chr2-22987395-INS-477
chr2-22992630-INS-208
chr2-23017395-INS-200
chr2-23091415-INS-579
chr2-23117184-INS-186
chr2-23120296-INS-269
chr2-23123303-INS-204
chr2-23181257-INS-58
chr2-23206453-INS-386
chr2-23284213-INS-355
chr2-23302593-INS-55
chr2-23333029-INS-655
chr2-23361123-INS-88
chr2-23379255-INS-214
chr2-23389467-INS-128
chr2-23391685-INS-7097
chr2-23428077-INS-459
chr2-23429162-INS-235
chr2-23440477-INS-108
chr2-23444908-INS-904
chr2-23460213-INS-294
chr2-23476291-INS-100
chr2-23477727-INS-89
chr2-23479316-INS-588
chr2-23512889-INS-312
chr2-23534008-INS-207
chr2-23547749-INS-6295
chr2-23558089-INS-6797
chr2-23575228-INS-574
chr2-23576234-INS-292
chr2-23604291-INS-6423
chr2-23608345-INS-582
chr2-23613364-INS-390
chr2-23642445-INS-1893
chr2-23655485-INS-53
chr2-23666595-INS-147

chr2-23730737-INS-85
chr2-23751918-INS-67
chr2-23780276-INS-1003
chr2-23791160-INS-165
chr2-23805953-INS-6587
chr2-23817869-INS-199
chr2-23884222-INS-55
chr2-23953551-INS-6777
chr2-23961198-INS-1986
chr2-23963191-INS-1523
chr2-23999294-INS-7166
chr2-24003689-INS-6424
chr2-24016459-INS-6248
chr2-24044125-INS-280
chr2-24054013-INS-136
chr2-24082254-INS-52
chr2-24138874-INS-108
chr2-24152463-INS-628
chr2-24213380-INS-831
chr2-24228651-INS-142
chr2-24240782-INS-188
chr2-24257348-INS-118
chr2-24261336-INS-51
chr2-24349437-INS-102
chr2-24350915-INS-84
chr2-24356822-INS-67
chr2-24404778-INS-70
chr2-24411234-INS-81
chr2-24428012-INS-187
chr2-24435792-INS-62
chr2-24436520-INS-108
chr2-24437962-INS-104
chr2-24441046-INS-5333
chr2-24457635-INS-143
chr2-24476611-INS-256
chr2-24504892-INS-6787
chr2-24518617-INS-51
chr2-24553540-INS-112
chr2-24557025-INS-94
chr2-24575063-INS-110
chr2-24578082-INS-56
chr2-24581839-INS-97
chr2-24601952-INS-61
chr2-24605784-INS-3892
chr2-24608661-INS-199
chr2-24613602-INS-154
chr2-24614351-INS-128
chr2-24624253-INS-869
chr2-24624751-INS-188
chr2-24639072-INS-106
chr2-24639856-INS-679
chr2-24669495-INS-52
chr2-24702036-INS-61
chr2-24724877-INS-73
chr2-24769322-INS-171
chr2-24777976-INS-417
chr2-24802122-INS-115
chr2-24823026-INS-216
chr2-24826523-INS-210
chr2-24828489-INS-102
chr2-24831411-INS-161
chr2-24832863-INS-104
chr2-24834036-INS-101
chr2-24834756-INS-559
chr2-24844408-INS-341
chr2-24857173-INS-378
chr2-24863768-INS-59
chr2-24880866-INS-936
chr2-24881295-INS-341
chr2-24887819-INS-257
chr2-24894855-INS-258
chr2-24897734-INS-52
chr2-24903384-INS-203
chr2-24917065-INS-157
chr2-24917910-INS-174
chr2-24924667-INS-149
chr2-24925031-INS-203

chr2-24927008-INS-87
chr2-24930805-INS-163
chr2-24959686-INS-202
chr2-24963391-INS-568
chr2-24999332-INS-128
chr2-24999790-INS-334
chr2-25027859-INS-202
chr2-25045849-INS-140
chr2-25055882-INS-52
chr2-25078792-INS-76
chr2-25081819-INS-134
chr2-25098985-INS-217
chr2-25141408-INS-87
chr2-25142783-INS-126
chr2-25150113-INS-64
chr2-25171694-INS-445
chr2-25192681-INS-62
chr2-25196695-INS-120
chr2-25198708-INS-176
chr2-25218556-INS-192
chr2-25231172-INS-70
chr2-25232270-INS-173
chr2-25261397-INS-98
chr2-25310554-INS-294
chr2-25311596-INS-148
chr2-25334786-INS-74
chr2-25360978-INS-229
chr2-25379458-INS-72
chr2-25388591-INS-54
chr2-25426870-INS-193
chr2-25443506-INS-80
chr2-25462722-INS-390
chr2-25500025-INS-96
chr2-25549797-INS-52
chr2-25555550-INS-309
chr2-25555684-INS-3168
chr2-25558733-INS-944
chr2-25568093-INS-200
chr2-25588279-INS-58
chr2-25618884-INS-54
chr2-25632532-INS-105
chr2-25686702-INS-52
chr2-25687741-INS-64
chr2-25721912-INS-160
chr2-25747147-INS-52
chr2-25753715-INS-122
chr2-25822206-INS-2401
chr2-25877392-INS-180
chr2-25921261-INS-90
chr2-25969252-INS-71
chr2-26018469-INS-173
chr2-26037357-INS-77
chr2-26039945-INS-135
chr2-26049622-INS-51
chr2-26068711-INS-192
chr2-26089019-INS-78
chr2-26100029-INS-53
chr2-26101860-INS-203
chr2-26128922-INS-100
chr2-26159505-INS-86
chr2-26160604-INS-86
chr2-26173953-INS-91
chr2-26177059-INS-156
chr2-26188759-INS-54
chr2-26191320-INS-261
chr2-26192196-INS-56
chr2-26238323-INS-2864
chr2-26252258-INS-1895
chr2-26285365-INS-816
chr2-26294601-INS-52
chr2-26295524-INS-169
chr2-26307783-INS-801
chr2-26312952-INS-57
chr2-26351486-INS-528
chr2-26364622-INS-210
chr2-26371285-INS-119
chr2-26399081-INS-96

chr2-26404410-INS-193
chr2-26411785-INS-167
chr2-26439866-INS-99
chr2-26455195-INS-111
chr2-26464097-INS-164
chr2-26486819-INS-56
chr2-26488400-INS-210
chr2-26496068-INS-198
chr2-26513536-INS-86
chr2-26521695-INS-205
chr2-26537088-INS-136
chr2-26540200-INS-209
chr2-26548830-INS-59
chr2-26553052-INS-60
chr2-26558730-INS-197
chr2-26564257-INS-77
chr2-26598942-INS-938
chr2-26620668-INS-112
chr2-26638641-INS-184
chr2-26663925-INS-151
chr2-26678589-INS-506
chr2-26690896-INS-70
chr2-26694611-INS-187
chr2-26726396-INS-322
chr2-26738156-INS-146
chr2-26759982-INS-63
chr2-26772830-INS-195
chr2-26787351-INS-167
chr2-26799767-INS-311
chr2-26855413-INS-131
chr2-26870153-INS-561
chr2-26877802-INS-372
chr2-26887319-INS-56
chr2-26940266-INS-258
chr2-26954804-INS-312
chr2-27007726-INS-328
chr2-27017631-INS-148
chr2-27019927-INS-265
chr2-27024537-INS-57
chr2-27054789-INS-113
chr2-27058505-INS-56
chr2-27063835-INS-237
chr2-27083882-INS-55
chr2-27099022-INS-140
chr2-27104300-INS-56
chr2-27135806-INS-209
chr2-27148956-INS-200
chr2-27161125-INS-287
chr2-27165937-INS-483
chr2-27193515-INS-80
chr2-27210117-INS-67
chr2-27212408-INS-492
chr2-27214409-INS-58
chr2-27232783-INS-64
chr2-27252933-INS-196
chr2-27269640-INS-141
chr2-27325963-INS-67
chr2-27371080-INS-67
chr2-27381390-INS-485
chr2-27388088-INS-254
chr2-27391281-INS-5543
chr2-27396998-INS-289
chr2-27403536-INS-207
chr2-27407927-INS-3008
chr2-27419773-INS-195
chr2-27426839-INS-114
chr2-27441042-INS-147
chr2-27468137-INS-170
chr2-27476117-INS-106
chr2-27484393-INS-107
chr2-27499892-INS-60
chr2-27532514-INS-75
chr2-27538763-INS-461
chr2-27549405-INS-126
chr2-27658873-INS-144
chr2-27702567-INS-288
chr2-27719306-INS-52

chr2-27720837-INS-104
chr2-27823374-INS-215
chr2-27831450-INS-245
chr2-27930947-INS-52
chr2-27952864-INS-64
chr2-27953386-INS-68
chr2-27960964-INS-93
chr2-27993051-INS-77
chr2-27993969-INS-58
chr2-28039277-INS-56
chr2-28043178-INS-399
chr2-28061651-INS-56
chr2-28080377-INS-60
chr2-28118465-INS-100
chr2-28127323-INS-1874
chr2-28157406-INS-232
chr2-28186874-INS-212
chr2-28188606-INS-102
chr2-28196909-INS-53
chr2-28199036-INS-106
chr2-28210536-INS-458
chr2-28242775-INS-114
chr2-28250083-INS-278
chr2-28317085-INS-142
chr2-28321704-INS-59
chr2-28323879-INS-414
chr2-28331249-INS-199
chr2-28372422-INS-213
chr2-28384564-INS-426
chr2-28398231-INS-483
chr2-28401956-INS-358
chr2-28437967-INS-61
chr2-28443593-INS-206
chr2-28460968-INS-821
chr2-28469222-INS-740
chr2-28472370-INS-152
chr2-28476221-INS-191
chr2-28479526-INS-193
chr2-28480858-INS-52
chr2-28485679-INS-118
chr2-28520544-INS-70
chr2-28524778-INS-208
chr2-28533534-INS-245
chr2-28548588-INS-202
chr2-28549691-INS-120
chr2-28550912-INS-339
chr2-28565591-INS-279
chr2-28572663-INS-201
chr2-28586624-INS-136
chr2-28591907-INS-213
chr2-28606243-INS-853
chr2-28623236-INS-98
chr2-28638439-INS-59
chr2-28649516-INS-177
chr2-28656414-INS-235
chr2-28659913-INS-214
chr2-28665036-INS-200
chr2-28672594-INS-123
chr2-28680027-INS-158
chr2-28694665-INS-201
chr2-28707113-INS-428
chr2-28709097-INS-160
chr2-28768553-INS-197
chr2-28772471-INS-130
chr2-28811148-INS-72
chr2-28813207-INS-99
chr2-28818328-INS-79
chr2-28820887-INS-371
chr2-28842650-INS-111
chr2-28862437-INS-64
chr2-28866033-INS-266
chr2-28872607-INS-569
chr2-28875751-INS-360
chr2-28890723-INS-73
chr2-28905571-INS-102
chr2-28916305-INS-271
chr2-28933229-INS-233

chr2-28942287-INS-187
chr2-28966777-INS-136
chr2-28975877-INS-427
chr2-28993355-INS-51
chr2-28995966-INS-75
chr2-29009034-INS-219
chr2-29009939-INS-450
chr2-29017895-INS-91
chr2-29043139-INS-837
chr2-29065706-INS-110
chr2-29101092-INS-75
chr2-29105198-INS-201
chr2-29106109-INS-100
chr2-29111580-INS-80
chr2-29121655-INS-80
chr2-29123465-INS-59
chr2-29180405-INS-76
chr2-29206057-INS-187
chr2-29213539-INS-56
chr2-29235042-INS-53
chr2-29235861-INS-283
chr2-29247660-INS-68
chr2-29270323-INS-794
chr2-29316084-INS-152
chr2-29329394-INS-96
chr2-29333773-INS-480
chr2-29348025-INS-52
chr2-29349700-INS-212
chr2-29358045-INS-132
chr2-29377580-INS-76
chr2-29420030-INS-57
chr2-29430495-INS-216
chr2-29441365-INS-382
chr2-29446421-INS-74
chr2-29486115-INS-128
chr2-29502237-INS-68
chr2-29507080-INS-199
chr2-29535630-INS-222
chr2-29561040-INS-63
chr2-29580710-INS-214
chr2-29614550-INS-348
chr2-29627627-INS-57
chr2-29632958-INS-84
chr2-29645338-INS-234
chr2-29665496-INS-136
chr2-29679860-INS-310
chr2-29708618-INS-60
chr2-29713746-INS-379
chr2-29714556-INS-137
chr2-29742145-INS-84
chr2-29763457-INS-493
chr2-29767554-INS-305
chr2-29778738-INS-54
chr2-29784131-INS-292
chr2-29799458-INS-190
chr2-29834231-INS-101
chr2-29847226-INS-193
chr2-29885744-INS-80
chr2-29911001-INS-58
chr2-29913806-INS-204
chr2-29936116-INS-202
chr2-29936847-INS-239
chr2-29941235-INS-78
chr2-29943467-INS-194
chr2-29965015-INS-233
chr2-30015702-INS-210
chr2-30018967-INS-52
chr2-30035065-INS-192
chr2-30044299-INS-132
chr2-30049278-INS-367
chr2-30059744-INS-92
chr2-30071489-INS-53
chr2-30085120-INS-69
chr2-30088430-INS-310
chr2-30089735-INS-210
chr2-30105351-INS-54
chr2-30115982-INS-101

chr2-30137674-INS-94
chr2-30142473-INS-212
chr2-30178976-INS-484
chr2-30187778-INS-89
chr2-30214801-INS-182
chr2-30232029-INS-206
chr2-30246514-INS-57
chr2-30253385-INS-170
chr2-30259408-INS-206
chr2-30315726-INS-167
chr2-30324245-INS-149
chr2-30339794-INS-237
chr2-30342709-INS-725
chr2-30348870-INS-113
chr2-30353121-INS-225
chr2-30383212-INS-191
chr2-30384393-INS-89
chr2-30406725-INS-711
chr2-30407637-INS-56
chr2-30410352-INS-100
chr2-30430312-INS-64
chr2-30436345-INS-54
chr2-30447295-INS-586
chr2-30454305-INS-399
chr2-30470941-INS-191
chr2-30474049-INS-70
chr2-30475487-INS-284
chr2-30505464-INS-58
chr2-30528646-INS-144
chr2-30530199-INS-98
chr2-30534269-INS-222
chr2-30558250-INS-448
chr2-30562705-INS-64
chr2-30564680-INS-99
chr2-30578681-INS-103
chr2-30600634-INS-52
chr2-30603463-INS-201
chr2-30621217-INS-121
chr2-30629810-INS-418
chr2-30630608-INS-100
chr2-30638556-INS-260
chr2-30643712-INS-234
chr2-30650742-INS-100
chr2-30655511-INS-273
chr2-30656967-INS-178
chr2-30657309-INS-5521
chr2-30667344-INS-144
chr2-30669857-INS-68
chr2-30677299-INS-54
chr2-30679302-INS-70
chr2-30719321-INS-202
chr2-30736618-INS-63
chr2-30755783-INS-120
chr2-30759062-INS-54
chr2-30770125-INS-287
chr2-30776691-INS-165
chr2-30802752-INS-130
chr2-30802752-INS-640
chr2-30803698-INS-418
chr2-30809980-INS-250
chr2-30812897-INS-61
chr2-30814788-INS-52
chr2-30823158-INS-385
chr2-30862171-INS-51
chr2-30867279-INS-54
chr2-30891011-INS-200
chr2-30891746-INS-197
chr2-30899928-INS-209
chr2-30908037-INS-171
chr2-30933679-INS-55
chr2-30934716-INS-825
chr2-30941233-INS-56
chr2-30968832-INS-136
chr2-30971780-INS-5200
chr2-30973974-INS-219
chr2-30987486-INS-283
chr2-30993035-INS-77

chr2-31005891-INS-74
chr2-31007303-INS-72
chr2-31041420-INS-394
chr2-31044377-INS-214
chr2-31046571-INS-499
chr2-31062804-INS-243
chr2-31074544-INS-206
chr2-31110619-INS-1892
chr2-31118810-INS-59
chr2-31120293-INS-290
chr2-31131277-INS-164
chr2-31133343-INS-59
chr2-31143434-INS-54
chr2-31168887-INS-179
chr2-31181657-INS-69
chr2-31197295-INS-441
chr2-31208173-INS-60
chr2-31218737-INS-96
chr2-31245659-INS-62
chr2-31265529-INS-68
chr2-31299870-INS-97
chr2-31346910-INS-67
chr2-31352109-INS-105
chr2-31355568-INS-127
chr2-31368289-INS-1490
chr2-31393437-INS-190
chr2-31425109-INS-219
chr2-31440925-INS-52
chr2-31459565-INS-201
chr2-31494097-INS-55
chr2-31514525-INS-52
chr2-31516263-INS-52
chr2-31519723-INS-201
chr2-31554167-INS-181
chr2-31556836-INS-94
chr2-31563476-INS-53
chr2-31574608-INS-135
chr2-31584065-INS-157
chr2-31591538-INS-299
chr2-31592100-INS-96
chr2-31597199-INS-174
chr2-31612890-INS-170
chr2-31615028-INS-239
chr2-31648522-INS-129
chr2-31648874-INS-491
chr2-31657255-INS-135
chr2-31693592-INS-110
chr2-31720225-INS-53
chr2-31740827-INS-96
chr2-31742435-INS-51
chr2-31743586-INS-83
chr2-31823280-INS-450
chr2-31825495-INS-193
chr2-31837744-INS-159
chr2-31853806-INS-90
chr2-31856279-INS-57
chr2-31869416-INS-202
chr2-31881908-INS-202
chr2-31884252-INS-90
chr2-31931396-INS-213
chr2-31934787-INS-227
chr2-31967614-INS-66
chr2-31974797-INS-175
chr2-31990422-INS-184
chr2-31994872-INS-209
chr2-32001633-INS-88
chr2-32017643-INS-65
chr2-32023377-INS-210
chr2-32024999-INS-163
chr2-32026513-INS-52
chr2-32030384-INS-341
chr2-32032009-INS-171
chr2-32056196-INS-214
chr2-32066807-INS-211
chr2-32092579-INS-60
chr2-32100620-INS-283
chr2-32112702-INS-6684

chr2-32214899-INS-52
chr2-32220120-INS-193
chr2-32221551-INS-214
chr2-32234643-INS-176
chr2-32284761-INS-365
chr2-32302653-INS-108
chr2-32333583-INS-52
chr2-32349239-INS-155
chr2-32362199-INS-63
chr2-32370320-INS-219
chr2-32373188-INS-142
chr2-32408757-INS-74
chr2-32446351-INS-216
chr2-32487537-INS-1862
chr2-32493055-INS-929
chr2-32516163-INS-57
chr2-32528940-INS-191
chr2-32552983-INS-67
chr2-32585854-INS-141
chr2-32588918-INS-320
chr2-32606063-INS-272
chr2-32608530-INS-56
chr2-32613104-INS-58
chr2-32614741-INS-92
chr2-32616878-INS-143
chr2-32629669-INS-395
chr2-32642188-INS-91
chr2-32643642-INS-136
chr2-32646242-INS-200
chr2-32650865-INS-115
chr2-32675655-INS-85
chr2-32699827-INS-204
chr2-32704434-INS-205
chr2-32715176-INS-198
chr2-32719958-INS-62
chr2-32722683-INS-198
chr2-32727925-INS-410
chr2-32738781-INS-208
chr2-32739345-INS-831
chr2-32744314-INS-219
chr2-32745419-INS-96
chr2-32770998-INS-66
chr2-32779604-INS-200
chr2-32791345-INS-80
chr2-32820567-INS-203
chr2-32839780-INS-205
chr2-32841235-INS-185
chr2-32848606-INS-183
chr2-32860552-INS-56
chr2-32875242-INS-109
chr2-32886890-INS-330
chr2-32910095-INS-186
chr2-32933756-INS-219
chr2-32982896-INS-224
chr2-33029221-INS-201
chr2-33030191-INS-114
chr2-33032150-INS-96
chr2-33083144-INS-105
chr2-33088516-INS-212
chr2-33115597-INS-207
chr2-33146399-INS-215
chr2-33148063-INS-167
chr2-33184410-INS-137
chr2-33205103-INS-74
chr2-33207742-INS-76
chr2-33243915-INS-1234
chr2-33288329-INS-57
chr2-33307201-INS-143
chr2-33322249-INS-598
chr2-33323227-INS-190
chr2-33324470-INS-102
chr2-33346475-INS-74
chr2-33355088-INS-483
chr2-33374607-INS-562
chr2-33378621-INS-128
chr2-33394965-INS-185
chr2-33416371-INS-50

chr2-33424080-INS-433
chr2-33440951-INS-675
chr2-33464405-INS-63
chr2-33471909-INS-86
chr2-33498447-INS-74
chr2-33540865-INS-210
chr2-33573223-INS-80
chr2-33577235-INS-224
chr2-33594581-INS-117
chr2-33621717-INS-51
chr2-33670182-INS-227
chr2-33698306-INS-402
chr2-33710995-INS-113
chr2-33729078-INS-144
chr2-33769817-INS-400
chr2-33790669-INS-361
chr2-33804299-INS-99
chr2-33810903-INS-74
chr2-33813482-INS-56
chr2-33824244-INS-202
chr2-33844591-INS-745
chr2-33853912-INS-77
chr2-33867386-INS-222
chr2-33869620-INS-118
chr2-33874917-INS-565
chr2-33900768-INS-78
chr2-33960903-INS-400
chr2-33964346-INS-63
chr2-33976770-INS-728
chr2-33981273-INS-400
chr2-33993205-INS-278
chr2-34035831-INS-92
chr2-34082486-INS-74
chr2-34117448-INS-60
chr2-34185784-INS-69
chr2-34191132-INS-326
chr2-34278961-INS-5551
chr2-34288170-INS-54
chr2-34431672-INS-208
chr2-34443375-INS-215
chr2-34452020-INS-992
chr2-34508726-INS-1888
chr2-34535387-INS-187
chr2-34548877-INS-140
chr2-34558970-INS-76
chr2-34580738-INS-78
chr2-34584581-INS-149
chr2-34622589-INS-59
chr2-34624545-INS-411
chr2-34626707-INS-480
chr2-34639725-INS-134
chr2-34651798-INS-216
chr2-34657118-INS-101
chr2-34661311-INS-52
chr2-34671268-INS-325
chr2-34676874-INS-612
chr2-34721510-INS-71
chr2-34726760-INS-58
chr2-34730120-INS-429
chr2-34743423-INS-137
chr2-34745236-INS-52
chr2-34746380-INS-95
chr2-34761517-INS-103
chr2-34771817-INS-247
chr2-34778666-INS-218
chr2-34810746-INS-120
chr2-34830807-INS-63
chr2-34832954-INS-55
chr2-34851672-INS-71
chr2-34882564-INS-98
chr2-34890415-INS-200
chr2-34890966-INS-196
chr2-34913238-INS-201
chr2-34958096-INS-238
chr2-34965483-INS-6415
chr2-34965559-INS-103
chr2-35012300-INS-206

chr2-35016834-INS-448
chr2-35025028-INS-104
chr2-35095956-INS-98
chr2-35099273-INS-161
chr2-35101070-INS-172
chr2-35116931-INS-53
chr2-35134272-INS-196
chr2-35139182-INS-74
chr2-35203777-INS-334
chr2-35268396-INS-90
chr2-35269701-INS-61
chr2-35275245-INS-262
chr2-35280237-INS-522
chr2-35282526-INS-209
chr2-35285955-INS-217
chr2-35288327-INS-530
chr2-35303204-INS-51
chr2-35304372-INS-489
chr2-35318681-INS-67
chr2-35321845-INS-56
chr2-35360126-INS-4903
chr2-35388762-INS-348
chr2-35400426-INS-209
chr2-35425656-INS-831
chr2-35441037-INS-104
chr2-35454550-INS-119
chr2-35489398-INS-64
chr2-35526739-INS-495
chr2-35531610-INS-67
chr2-35544428-INS-204
chr2-35615136-INS-95
chr2-35639716-INS-172
chr2-35657325-INS-83
chr2-35664982-INS-108
chr2-35668755-INS-93
chr2-35692858-INS-71
chr2-35774704-INS-111
chr2-35822115-INS-189
chr2-35879153-INS-211
chr2-35892664-INS-97
chr2-35893710-INS-501
chr2-35910128-INS-96
chr2-35919476-INS-63
chr2-35936399-INS-423
chr2-35946284-INS-76
chr2-36014333-INS-6440
chr2-36029860-INS-192
chr2-36052105-INS-122
chr2-36082734-INS-188
chr2-36089179-INS-226
chr2-36102978-INS-112
chr2-36132024-INS-58
chr2-36164076-INS-198
chr2-36171537-INS-7137
chr2-36171540-INS-166
chr2-36181246-INS-63
chr2-36204846-INS-332
chr2-36208544-INS-3270
chr2-36210504-INS-96
chr2-36218763-INS-6663
chr2-36231119-INS-957
chr2-36237207-INS-5578
chr2-36256466-INS-6338
chr2-36263741-INS-64
chr2-36287076-INS-7520
chr2-36298121-INS-214
chr2-36306348-INS-331
chr2-36324159-INS-604
chr2-36325296-INS-325
chr2-36333737-INS-210
chr2-36335325-INS-1918
chr2-36349031-INS-178
chr2-36398521-INS-930
chr2-36411166-INS-105
chr2-36422586-INS-153
chr2-36451648-INS-64
chr2-36466038-INS-52

chr2-36472786-INS-60
chr2-36482238-INS-160
chr2-36485302-INS-572
chr2-36486839-INS-358
chr2-36489376-INS-54
chr2-36531302-INS-145
chr2-36537449-INS-1865
chr2-36538061-INS-64
chr2-36552878-INS-6725
chr2-36601382-INS-7295
chr2-36658125-INS-956
chr2-36678386-INS-14330
chr2-36683199-INS-5449
chr2-36705691-INS-642
chr2-36730491-INS-4561
chr2-36738082-INS-142
chr2-36741802-INS-219
chr2-36748377-INS-51
chr2-36794796-INS-7003
chr2-36828189-INS-9432
chr2-36854633-INS-203
chr2-36864473-INS-185
chr2-36865520-INS-6473
chr2-36891867-INS-57
chr2-36900214-INS-1987
chr2-36918694-INS-6600
chr2-36947029-INS-804
chr2-36962749-INS-1399
chr2-36984379-INS-320
chr2-36987112-INS-923
chr2-36990024-INS-1490
chr2-37019474-INS-74
chr2-37054104-INS-107
chr2-37068208-INS-557
chr2-37077518-INS-979
chr2-37106396-INS-56
chr2-37139484-INS-478
chr2-37202161-INS-2973
chr2-37204111-INS-404
chr2-37206965-INS-393
chr2-37207712-INS-856
chr2-37250313-INS-7978
chr2-37250822-INS-229
chr2-37256319-INS-388
chr2-37325068-INS-394
chr2-37358123-INS-73
chr2-37360029-INS-58
chr2-37372773-INS-748
chr2-37383604-INS-224
chr2-37385393-INS-118
chr2-37465079-INS-58
chr2-37481739-INS-63
chr2-37511822-INS-208
chr2-37550046-INS-83
chr2-37557067-INS-821
chr2-37617100-INS-200
chr2-37637226-INS-205
chr2-37650386-INS-86
chr2-37651991-INS-221
chr2-37667447-INS-90
chr2-37670990-INS-158
chr2-37703486-INS-155
chr2-37754216-INS-194
chr2-37804823-INS-360
chr2-37811676-INS-435
chr2-37842206-INS-107
chr2-37856079-INS-349
chr2-37965753-INS-734
chr2-37966287-INS-222
chr2-37977080-INS-70
chr2-37993654-INS-514
chr2-38076966-INS-60
chr2-38082662-INS-527
chr2-38088658-INS-356
chr2-38093225-INS-2128
chr2-38117845-INS-1430
chr2-38122430-INS-326

chr2-38125947-INS-527
chr2-38182830-INS-190
chr2-38184874-INS-772
chr2-38187176-INS-51
chr2-38192791-INS-124
chr2-38205101-INS-211
chr2-38214387-INS-95
chr2-38220580-INS-481
chr2-38247203-INS-60
chr2-38264344-INS-199
chr2-38283468-INS-217
chr2-38292157-INS-121
chr2-38306191-INS-89
chr2-38314776-INS-82
chr2-38336428-INS-215
chr2-38341345-INS-98
chr2-38354324-INS-62
chr2-38373725-INS-110
chr2-38378365-INS-7483
chr2-38386539-INS-205
chr2-38397259-INS-90
chr2-38414855-INS-202
chr2-38417254-INS-62
chr2-38457845-INS-88
chr2-38490862-INS-123
chr2-38518748-INS-748
chr2-38525379-INS-100
chr2-38545052-INS-78
chr2-38572176-INS-403
chr2-38612535-INS-72
chr2-38638825-INS-215
chr2-38645848-INS-169
chr2-38659977-INS-58
chr2-38675680-INS-91
chr2-38694397-INS-53
chr2-38711290-INS-120
chr2-38722337-INS-284
chr2-38729468-INS-562
chr2-38741836-INS-87
chr2-38769009-INS-205
chr2-38790558-INS-205
chr2-38803881-INS-91
chr2-38804396-INS-502
chr2-38824565-INS-68
chr2-38829614-INS-224
chr2-38836889-INS-60
chr2-38846531-INS-240
chr2-38858208-INS-112
chr2-38868819-INS-222
chr2-38893929-INS-69
chr2-38904773-INS-206
chr2-38917381-INS-348
chr2-38935336-INS-62
chr2-38938075-INS-285
chr2-38990259-INS-288
chr2-38998248-INS-53
chr2-39003037-INS-1328
chr2-39015544-INS-845
chr2-39097981-INS-96
chr2-39111229-INS-171
chr2-39153585-INS-194
chr2-39207029-INS-6454
chr2-39216240-INS-653
chr2-39237579-INS-126
chr2-39241322-INS-6257
chr2-39250339-INS-123
chr2-39259725-INS-52
chr2-39260993-INS-6378
chr2-39264521-INS-214
chr2-39322392-INS-61
chr2-39340303-INS-58
chr2-39378643-INS-86
chr2-39388830-INS-500
chr2-39389796-INS-982
chr2-39414503-INS-141
chr2-39436580-INS-790
chr2-39458660-INS-208

chr2-39505305-INS-334
chr2-39507766-INS-513
chr2-39524876-INS-166
chr2-39540018-INS-60
chr2-39547687-INS-129
chr2-39557645-INS-196
chr2-39618783-INS-180
chr2-39622272-INS-56
chr2-39641618-INS-254
chr2-39644343-INS-2196
chr2-39711732-INS-516
chr2-39712824-INS-337
chr2-39715699-INS-216
chr2-39773738-INS-7097
chr2-39781221-INS-534
chr2-39790426-INS-86
chr2-39791773-INS-60
chr2-39795407-INS-217
chr2-39800914-INS-369
chr2-39825538-INS-180
chr2-39829286-INS-6399
chr2-39829728-INS-409
chr2-39833780-INS-400
chr2-39838622-INS-68
chr2-39865889-INS-400
chr2-39867783-INS-2076
chr2-39868251-INS-462
chr2-39871049-INS-548
chr2-39876342-INS-53
chr2-39890237-INS-551
chr2-39899257-INS-930
chr2-39907073-INS-496
chr2-39916311-INS-57
chr2-39928433-INS-326
chr2-39942901-INS-549
chr2-39947952-INS-351
chr2-39951783-INS-4529
chr2-39976747-INS-55
chr2-39984950-INS-966
chr2-39992173-INS-318
chr2-39994555-INS-52
chr2-40017915-INS-2626
chr2-40075438-INS-3429
chr2-40076454-INS-665
chr2-40122856-INS-51
chr2-40143260-INS-494
chr2-40149909-INS-7621
chr2-40164998-INS-6488
chr2-40178157-INS-124
chr2-40192948-INS-149
chr2-40195583-INS-68
chr2-40225518-INS-75
chr2-40227851-INS-424
chr2-40230129-INS-63
chr2-40236054-INS-7018
chr2-40241006-INS-283
chr2-40265989-INS-6071
chr2-40271054-INS-147
chr2-40287445-INS-60
chr2-40297448-INS-6603
chr2-40311484-INS-347
chr2-40311672-INS-3705
chr2-40316617-INS-759
chr2-40320015-INS-200
chr2-40327721-INS-85
chr2-40339558-INS-216
chr2-40355080-INS-380
chr2-40356133-INS-57
chr2-40357318-INS-830
chr2-40387854-INS-3518
chr2-40387986-INS-323
chr2-40400279-INS-296
chr2-40428657-INS-1252
chr2-40449351-INS-100
chr2-40461708-INS-5790
chr2-40495165-INS-69
chr2-40498970-INS-344

chr2-40499696-INS-108
chr2-40511536-INS-283
chr2-40511691-INS-301
chr2-40512422-INS-59
chr2-40520618-INS-176
chr2-40528613-INS-421
chr2-40534527-INS-154
chr2-40541275-INS-67
chr2-40556740-INS-87
chr2-40557556-INS-298
chr2-40560429-INS-640
chr2-40588021-INS-57
chr2-40597777-INS-222
chr2-40611422-INS-161
chr2-40636783-INS-56
chr2-40637974-INS-52
chr2-40639043-INS-174
chr2-40650621-INS-321
chr2-40661101-INS-143
chr2-40664603-INS-246
chr2-40669411-INS-79
chr2-40687411-INS-54
chr2-40713117-INS-75
chr2-40727394-INS-952
chr2-40728515-INS-430
chr2-40730566-INS-67
chr2-40738192-INS-75
chr2-40744062-INS-66
chr2-40789273-INS-6675
chr2-40789579-INS-64
chr2-40807614-INS-117
chr2-40853815-INS-988
chr2-40865282-INS-508
chr2-40871966-INS-78
chr2-40898890-INS-522
chr2-40899515-INS-966
chr2-40911013-INS-180
chr2-40941370-INS-491
chr2-40981688-INS-282
chr2-40989758-INS-66
chr2-41014707-INS-720
chr2-41017082-INS-66
chr2-41019334-INS-1185
chr2-41025004-INS-7125
chr2-41046237-INS-811
chr2-41048868-INS-174
chr2-41066619-INS-440
chr2-41076513-INS-6473
chr2-41081565-INS-66
chr2-41084001-INS-56
chr2-41124497-INS-64
chr2-41135510-INS-115
chr2-41138682-INS-80
chr2-41146509-INS-90
chr2-41150098-INS-6178
chr2-41155444-INS-72
chr2-41169514-INS-69
chr2-41214837-INS-328
chr2-41227173-INS-199
chr2-41228639-INS-62
chr2-41244537-INS-1380
chr2-41274087-INS-128
chr2-41328014-INS-611
chr2-41330663-INS-62
chr2-41333118-INS-248
chr2-41351700-INS-58
chr2-41380995-INS-400
chr2-41386636-INS-6486
chr2-41389203-INS-274
chr2-41393295-INS-108
chr2-41394543-INS-1895
chr2-41398047-INS-52
chr2-41418236-INS-343
chr2-41422564-INS-978
chr2-41423478-INS-210
chr2-41450831-INS-52
chr2-41469799-INS-106

chr2-41494199-INS-517
chr2-41506326-INS-172
chr2-41508231-INS-51
chr2-41521191-INS-60
chr2-41527432-INS-212
chr2-41530268-INS-199
chr2-41556984-INS-54
chr2-41582300-INS-65
chr2-41621031-INS-228
chr2-41646016-INS-2664
chr2-41657218-INS-2188
chr2-41728886-INS-62
chr2-41762977-INS-130
chr2-41787005-INS-885
chr2-41808021-INS-62
chr2-41814700-INS-1075
chr2-41823344-INS-281
chr2-41871767-INS-363
chr2-41904487-INS-165
chr2-41918744-INS-734
chr2-41942391-INS-211
chr2-41943847-INS-60
chr2-41956320-INS-105
chr2-41983211-INS-496
chr2-41989691-INS-61
chr2-41998308-INS-62
chr2-42009959-INS-58
chr2-42012127-INS-231
chr2-42032952-INS-78
chr2-42035168-INS-182
chr2-42042283-INS-2715
chr2-42043312-INS-510
chr2-42049275-INS-202
chr2-42058456-INS-58
chr2-42064104-INS-154
chr2-42070847-INS-390
chr2-42136487-INS-219
chr2-42146713-INS-991
chr2-42165913-INS-373
chr2-42189004-INS-101
chr2-42198001-INS-6278
chr2-42201448-INS-96
chr2-42219120-INS-4782
chr2-42219353-INS-58
chr2-42221112-INS-136
chr2-42226046-INS-54
chr2-42246376-INS-78
chr2-42257986-INS-201
chr2-42263088-INS-57
chr2-42273209-INS-1496
chr2-42286331-INS-66
chr2-42305597-INS-56
chr2-42314168-INS-205
chr2-42318165-INS-329
chr2-42351979-INS-58
chr2-42380339-INS-62
chr2-42385469-INS-318
chr2-42422372-INS-52
chr2-42431995-INS-322
chr2-42433268-INS-462
chr2-42438091-INS-366
chr2-42446040-INS-825
chr2-42459609-INS-602
chr2-42477773-INS-74
chr2-42484694-INS-376
chr2-42508036-INS-52
chr2-42517703-INS-142
chr2-42533509-INS-74
chr2-42550778-INS-52
chr2-42566420-INS-7193
chr2-42574249-INS-159
chr2-42578478-INS-178
chr2-42583654-INS-95
chr2-42589654-INS-166
chr2-42612114-INS-613
chr2-42625638-INS-69
chr2-42630449-INS-111

chr2-42642692-INS-187
chr2-42662608-INS-192
chr2-42700614-INS-68
chr2-42704677-INS-148
chr2-42721040-INS-716
chr2-42726493-INS-82
chr2-42728390-INS-57
chr2-42742302-INS-532
chr2-42757530-INS-66
chr2-42758903-INS-948
chr2-42778013-INS-2273
chr2-42789660-INS-173
chr2-42790587-INS-8419
chr2-42792625-INS-94
chr2-42794387-INS-69
chr2-42802343-INS-401
chr2-42832074-INS-96
chr2-42863724-INS-497
chr2-42864930-INS-233
chr2-42882948-INS-7467
chr2-42883437-INS-73
chr2-42885517-INS-214
chr2-42924986-INS-80
chr2-42928717-INS-186
chr2-42945045-INS-6295
chr2-42965459-INS-54
chr2-42977792-INS-622
chr2-42992667-INS-1877
chr2-43032205-INS-410
chr2-43043945-INS-400
chr2-43044609-INS-1892
chr2-43065960-INS-81
chr2-43082069-INS-72
chr2-43083589-INS-151
chr2-43130436-INS-54
chr2-43133970-INS-283
chr2-43166317-INS-6788
chr2-43245765-INS-3486
chr2-43279366-INS-202
chr2-43297588-INS-6589
chr2-43306384-INS-84
chr2-43339103-INS-59
chr2-43354515-INS-144
chr2-43369560-INS-113
chr2-43377858-INS-71
chr2-43414892-INS-266
chr2-43441012-INS-185
chr2-43456228-INS-132
chr2-43465397-INS-4780
chr2-43468268-INS-149
chr2-43512905-INS-57
chr2-43542577-INS-78
chr2-43591720-INS-235
chr2-43596187-INS-100
chr2-43629882-INS-224
chr2-43636476-INS-104
chr2-43646587-INS-331
chr2-43649340-INS-1219
chr2-43655537-INS-111
chr2-43660166-INS-1280
chr2-43685981-INS-193
chr2-43687125-INS-98
chr2-43708245-INS-495
chr2-43715854-INS-81
chr2-43720096-INS-137
chr2-43727532-INS-342
chr2-43740422-INS-116
chr2-43744578-INS-79
chr2-43762548-INS-64
chr2-43813006-INS-53
chr2-43857852-INS-2547
chr2-43875138-INS-603
chr2-43898367-INS-404
chr2-43933955-INS-76
chr2-43967814-INS-207
chr2-43977125-INS-2332
chr2-43979223-INS-2799

chr2-43992121-INS-1890
chr2-43992905-INS-1822
chr2-44012750-INS-52
chr2-44016647-INS-7062
chr2-44018396-INS-6892
chr2-44026884-INS-104
chr2-44048894-INS-190
chr2-44064484-INS-215
chr2-44090545-INS-123
chr2-44120243-INS-220
chr2-44122174-INS-100
chr2-44126521-INS-117
chr2-44132510-INS-66
chr2-44144851-INS-82
chr2-44173124-INS-54
chr2-44210889-INS-90
chr2-44226197-INS-223
chr2-44248187-INS-192
chr2-44260408-INS-181
chr2-44291667-INS-68
chr2-44303494-INS-258
chr2-44325387-INS-5566
chr2-44336857-INS-61
chr2-44346471-INS-256
chr2-44366189-INS-6800
chr2-44367284-INS-8246
chr2-44377497-INS-6185
chr2-44379660-INS-341
chr2-44384344-INS-82
chr2-44385930-INS-410
chr2-44393701-INS-106
chr2-44398727-INS-115
chr2-44427016-INS-570
chr2-44441922-INS-929
chr2-44483809-INS-203
chr2-44484618-INS-207
chr2-44501593-INS-214
chr2-44527212-INS-52
chr2-44536019-INS-124
chr2-44637642-INS-890
chr2-44641809-INS-249
chr2-44662219-INS-366
chr2-44697541-INS-213
chr2-44706501-INS-3471
chr2-44711510-INS-98
chr2-44719453-INS-504
chr2-44738240-INS-877
chr2-44755405-INS-563
chr2-44778206-INS-56
chr2-44848579-INS-166
chr2-44852508-INS-96
chr2-44872891-INS-106
chr2-44876189-INS-115
chr2-44884899-INS-668
chr2-44894342-INS-244
chr2-44910859-INS-70
chr2-44997062-INS-61
chr2-45026695-INS-81
chr2-45055154-INS-93
chr2-45077214-INS-431
chr2-45081320-INS-641
chr2-45081333-INS-193
chr2-45091868-INS-64
chr2-45107961-INS-60
chr2-45130752-INS-486
chr2-45133354-INS-60
chr2-45159264-INS-1890
chr2-45164058-INS-669
chr2-45169300-INS-203
chr2-45256672-INS-936
chr2-45260799-INS-127
chr2-45264261-INS-53
chr2-45265552-INS-64
chr2-45310749-INS-54
chr2-45312267-INS-67
chr2-45314615-INS-509
chr2-45315888-INS-136

chr2-45368256-INS-444
chr2-45378163-INS-6680
chr2-45392438-INS-160
chr2-45407771-INS-117
chr2-45424210-INS-90
chr2-45429502-INS-56
chr2-45431817-INS-6492
chr2-45443086-INS-74
chr2-45460836-INS-84
chr2-45464287-INS-58
chr2-45487450-INS-225
chr2-45490163-INS-97
chr2-45506324-INS-294
chr2-45525953-INS-223
chr2-45541935-INS-6647
chr2-45545918-INS-52
chr2-45556723-INS-335
chr2-45557199-INS-6220
chr2-45565668-INS-54
chr2-45566779-INS-144
chr2-45576281-INS-68
chr2-45620108-INS-166
chr2-45624212-INS-6335
chr2-45635718-INS-100
chr2-45639829-INS-371
chr2-45657651-INS-6663
chr2-45664328-INS-84
chr2-45678769-INS-54
chr2-45748008-INS-119
chr2-45749117-INS-255
chr2-45751584-INS-56
chr2-45776101-INS-66
chr2-45784362-INS-72
chr2-45795482-INS-100
chr2-45817089-INS-111
chr2-45819765-INS-52
chr2-45832357-INS-705
chr2-45840429-INS-710
chr2-45865484-INS-837
chr2-45902611-INS-3515
chr2-45902759-INS-7443
chr2-45913792-INS-704
chr2-45956801-INS-159
chr2-45994620-INS-63
chr2-46035230-INS-445
chr2-46070594-INS-55
chr2-46087413-INS-148
chr2-46139768-INS-735
chr2-46149693-INS-278
chr2-46171831-INS-1020
chr2-46171889-INS-462
chr2-46210922-INS-819
chr2-46298817-INS-63
chr2-46352171-INS-199
chr2-46364213-INS-329
chr2-46371154-INS-276
chr2-46387083-INS-98
chr2-46389423-INS-62
chr2-46389783-INS-2416
chr2-46404014-INS-56
chr2-46417987-INS-81
chr2-46419717-INS-6231
chr2-46426954-INS-308
chr2-46460873-INS-7083
chr2-46523077-INS-70
chr2-46566795-INS-84
chr2-46568686-INS-3003
chr2-46635274-INS-155
chr2-46647492-INS-357
chr2-46652365-INS-52
chr2-46663746-INS-57
chr2-46667686-INS-186
chr2-46675682-INS-84
chr2-46683652-INS-245
chr2-46694715-INS-85
chr2-46704643-INS-110
chr2-46712606-INS-213

chr2-46716242-INS-927
chr2-46721486-INS-102
chr2-46758026-INS-64
chr2-46787207-INS-70
chr2-46802837-INS-56
chr2-46843612-INS-70
chr2-46897972-INS-343
chr2-46899319-INS-312
chr2-46900033-INS-208
chr2-46925217-INS-2966
chr2-46927922-INS-4540
chr2-46932105-INS-903
chr2-46934680-INS-56
chr2-46938686-INS-187
chr2-46950823-INS-2194
chr2-47032489-INS-56
chr2-47053920-INS-70
chr2-47061352-INS-102
chr2-47069368-INS-65
chr2-47070537-INS-444
chr2-47095970-INS-155
chr2-47111423-INS-52
chr2-47122219-INS-6529
chr2-47122977-INS-76
chr2-47145939-INS-508
chr2-47168887-INS-59
chr2-47200649-INS-70
chr2-47215489-INS-381
chr2-47230315-INS-141
chr2-47231182-INS-325
chr2-47256743-INS-152
chr2-47278692-INS-84
chr2-47289445-INS-104
chr2-47309352-INS-198
chr2-47310564-INS-6403
chr2-47313225-INS-798
chr2-47366377-INS-62
chr2-47377461-INS-255
chr2-47396627-INS-56
chr2-47429605-INS-113
chr2-47434760-INS-71
chr2-47475207-INS-57
chr2-47500287-INS-104
chr2-47509489-INS-66
chr2-47546108-INS-6372
chr2-47549174-INS-273
chr2-47624365-INS-494
chr2-47638178-INS-562
chr2-47665055-INS-668
chr2-47712185-INS-446
chr2-47720043-INS-84
chr2-47753185-INS-582
chr2-47759284-INS-72
chr2-47789907-INS-122
chr2-47793362-INS-555
chr2-47811439-INS-123
chr2-47823211-INS-96
chr2-47827807-INS-740
chr2-47844460-INS-443
chr2-47909106-INS-52
chr2-48026269-INS-113
chr2-48069846-INS-356
chr2-48073860-INS-456
chr2-48075610-INS-56
chr2-48108030-INS-54
chr2-48128392-INS-6640
chr2-48128629-INS-6232
chr2-48130040-INS-381
chr2-48137665-INS-550
chr2-48138621-INS-6643
chr2-48173425-INS-177
chr2-48182197-INS-7687
chr2-48198035-INS-7291
chr2-48235270-INS-185
chr2-48261898-INS-360
chr2-48276960-INS-64
chr2-48356680-INS-290

chr2-48396887-INS-146
chr2-48402549-INS-181
chr2-48414336-INS-152
chr2-48423688-INS-53
chr2-48438752-INS-433
chr2-48456019-INS-58
chr2-48459769-INS-6415
chr2-48501634-INS-288
chr2-48545611-INS-3106
chr2-48550001-INS-52
chr2-48561423-INS-6654
chr2-48580470-INS-60
chr2-48604476-INS-83
chr2-48611772-INS-239
chr2-48627699-INS-71
chr2-48632645-INS-130
chr2-48636493-INS-60
chr2-48688264-INS-72
chr2-48700490-INS-53
chr2-48734210-INS-368
chr2-48739323-INS-75
chr2-48787641-INS-702
chr2-48796600-INS-1216
chr2-48796600-INS-576
chr2-48796633-INS-187
chr2-48801830-INS-60
chr2-48805932-INS-52
chr2-48808304-INS-718
chr2-48813144-INS-7080
chr2-48852899-INS-858
chr2-48859038-INS-723
chr2-48872388-INS-83
chr2-48880715-INS-148
chr2-48916542-INS-124
chr2-48940198-INS-199
chr2-48959364-INS-4098
chr2-48984419-INS-122
chr2-49003808-INS-398
chr2-49012926-INS-78
chr2-49028941-INS-380
chr2-49038581-INS-116
chr2-49062983-INS-91
chr2-49102451-INS-494
chr2-49111407-INS-498
chr2-49124902-INS-53
chr2-49135213-INS-62
chr2-49168509-INS-102
chr2-49172177-INS-431
chr2-49194203-INS-672
chr2-49197554-INS-211
chr2-49201176-INS-117
chr2-49227149-INS-509
chr2-49238263-INS-81
chr2-49251055-INS-194
chr2-49268549-INS-295
chr2-49288839-INS-66
chr2-49296623-INS-319
chr2-49300184-INS-107
chr2-49357725-INS-140
chr2-49389714-INS-401
chr2-49403570-INS-94
chr2-49407826-INS-208
chr2-49436530-INS-6474
chr2-49503116-INS-183
chr2-49524940-INS-67
chr2-49536516-INS-83
chr2-49575660-INS-498
chr2-49582539-INS-514
chr2-49607857-INS-150
chr2-49669367-INS-61
chr2-49684271-INS-119
chr2-49686107-INS-484
chr2-49722916-INS-162
chr2-49723950-INS-167
chr2-49730349-INS-66
chr2-49749612-INS-52
chr2-49751251-INS-60

chr2-49761389-INS-143
chr2-49773977-INS-55
chr2-49776499-INS-52
chr2-49779533-INS-67
chr2-49809449-INS-60
chr2-49811708-INS-82
chr2-49817398-INS-115
chr2-49825699-INS-7293
chr2-49846624-INS-712
chr2-49866226-INS-60
chr2-49920079-INS-65
chr2-49942728-INS-93
chr2-49962526-INS-60
chr2-49964904-INS-266
chr2-49991032-INS-236
chr2-50009639-INS-112
chr2-50022504-INS-171
chr2-50053686-INS-242
chr2-50057280-INS-204
chr2-50085328-INS-5290
chr2-50107608-INS-206
chr2-50140841-INS-82
chr2-50155365-INS-94
chr2-50161482-INS-68
chr2-50177342-INS-276
chr2-50179089-INS-588
chr2-50190193-INS-60
chr2-50201397-INS-56
chr2-50203218-INS-646
chr2-50208913-INS-336
chr2-50227375-INS-53
chr2-50237209-INS-6442
chr2-50240181-INS-214
chr2-50272736-INS-538
chr2-50314833-INS-777
chr2-50319096-INS-66
chr2-50353412-INS-5705
chr2-50373367-INS-283
chr2-50381416-INS-761
chr2-50414955-INS-62
chr2-50423852-INS-100
chr2-50430385-INS-68
chr2-50436168-INS-307
chr2-50454195-INS-109
chr2-50471552-INS-6586
chr2-50484258-INS-59
chr2-50490098-INS-847
chr2-50508372-INS-92
chr2-50528341-INS-4207
chr2-50541244-INS-63
chr2-50543907-INS-478
chr2-50577758-INS-562
chr2-50608839-INS-7401
chr2-50622779-INS-52
chr2-50630916-INS-403
chr2-50631457-INS-173
chr2-50632724-INS-73
chr2-50688369-INS-109
chr2-50726057-INS-370
chr2-50765160-INS-440
chr2-50766934-INS-190
chr2-50802653-INS-64
chr2-50816242-INS-74
chr2-50823661-INS-108
chr2-50826410-INS-78
chr2-50829466-INS-935
chr2-50830686-INS-231
chr2-50887421-INS-118
chr2-50899160-INS-95
chr2-50901933-INS-332
chr2-50936267-INS-380
chr2-50965127-INS-73
chr2-50971153-INS-227
chr2-51000142-INS-54
chr2-51003640-INS-434
chr2-51042032-INS-110
chr2-51052306-INS-499

chr2-51061658-INS-54
chr2-51086522-INS-58
chr2-51111319-INS-66
chr2-51122644-INS-760
chr2-51187930-INS-354
chr2-51194611-INS-57
chr2-51212181-INS-57
chr2-51233144-INS-86
chr2-51263634-INS-54
chr2-51291490-INS-406
chr2-51329010-INS-2469
chr2-51351222-INS-437
chr2-51372712-INS-52
chr2-51402751-INS-56
chr2-51410898-INS-58
chr2-51418844-INS-60
chr2-51434635-INS-831
chr2-51437585-INS-60
chr2-51443213-INS-201
chr2-51450418-INS-128
chr2-51452496-INS-985
chr2-51469685-INS-82
chr2-51537654-INS-195
chr2-51539007-INS-52
chr2-51541068-INS-237
chr2-51543885-INS-101
chr2-51561841-INS-208
chr2-51566257-INS-64
chr2-51586989-INS-530
chr2-51655477-INS-84
chr2-51661401-INS-126
chr2-51661470-INS-621
chr2-51706957-INS-89
chr2-51709556-INS-134
chr2-51727396-INS-70
chr2-51752819-INS-112
chr2-51767623-INS-52
chr2-51779807-INS-1864
chr2-51783486-INS-125
chr2-51793949-INS-6342
chr2-51807050-INS-55
chr2-51811668-INS-5364
chr2-51836203-INS-153
chr2-51837777-INS-552
chr2-51854624-INS-593
chr2-51859105-INS-75
chr2-51873792-INS-132
chr2-51886790-INS-114
chr2-51895569-INS-78
chr2-51904891-INS-119
chr2-51913811-INS-709
chr2-51948002-INS-356
chr2-51955342-INS-55
chr2-51958284-INS-89
chr2-51960555-INS-158
chr2-51979160-INS-636
chr2-51987604-INS-206
chr2-52070162-INS-130
chr2-52081014-INS-271
chr2-52119329-INS-68
chr2-52129568-INS-5550
chr2-52178111-INS-56
chr2-52191837-INS-100
chr2-52208841-INS-236
chr2-52210212-INS-540
chr2-52231770-INS-1427
chr2-52244434-INS-211
chr2-52250495-INS-59
chr2-52259503-INS-56
chr2-52263684-INS-62
chr2-52268061-INS-71
chr2-52274697-INS-58
chr2-52276806-INS-63
chr2-52308850-INS-226
chr2-52328939-INS-945
chr2-52339581-INS-96
chr2-52388781-INS-54

chr2-52394430-INS-265
chr2-52398005-INS-56
chr2-52415248-INS-169
chr2-52438222-INS-56
chr2-52450722-INS-137
chr2-52462047-INS-866
chr2-52463221-INS-53
chr2-52470098-INS-492
chr2-52490591-INS-558
chr2-52503454-INS-153
chr2-52517347-INS-7402
chr2-52519554-INS-75
chr2-52567556-INS-112
chr2-52581930-INS-205
chr2-52586967-INS-4388
chr2-52587604-INS-4396
chr2-52587905-INS-4789
chr2-52588592-INS-3648
chr2-52599891-INS-643
chr2-52608224-INS-115
chr2-52618878-INS-116
chr2-52627350-INS-105
chr2-52634760-INS-219
chr2-52640034-INS-534
chr2-52645080-INS-873
chr2-52650321-INS-182
chr2-52655889-INS-150
chr2-52661723-INS-967
chr2-52666936-INS-471
chr2-52668388-INS-57
chr2-52689277-INS-390
chr2-52692176-INS-53
chr2-52697305-INS-791
chr2-52716210-INS-183
chr2-52747345-INS-121
chr2-52760540-INS-64
chr2-52791118-INS-76
chr2-52794710-INS-200
chr2-52808552-INS-220
chr2-52813436-INS-92
chr2-52845183-INS-63
chr2-52847175-INS-3055
chr2-52860086-INS-92
chr2-52861438-INS-197
chr2-52864930-INS-233
chr2-52878552-INS-209
chr2-52885325-INS-215
chr2-52906612-INS-177
chr2-52915656-INS-56
chr2-52916890-INS-190
chr2-52921221-INS-213
chr2-52967172-INS-64
chr2-53006182-INS-290
chr2-53006265-INS-444
chr2-53015462-INS-786
chr2-53066687-INS-124
chr2-53069611-INS-231
chr2-53074312-INS-211
chr2-53089666-INS-257
chr2-53125206-INS-56
chr2-53142250-INS-280
chr2-53157487-INS-152
chr2-53173589-INS-367
chr2-53186445-INS-1244
chr2-53191103-INS-6581
chr2-53199731-INS-128
chr2-53205218-INS-400
chr2-53223864-INS-2879
chr2-53235783-INS-286
chr2-53236362-INS-979
chr2-53239669-INS-5653
chr2-53244489-INS-64
chr2-53246984-INS-52
chr2-53260102-INS-115
chr2-53262061-INS-73
chr2-53283310-INS-2348
chr2-53345994-INS-207

chr2-53382124-INS-417
chr2-53384143-INS-53
chr2-53390043-INS-53
chr2-53399576-INS-202
chr2-53422387-INS-447
chr2-53434968-INS-352
chr2-53471225-INS-120
chr2-53474706-INS-5318
chr2-53479663-INS-143
chr2-53547018-INS-340
chr2-53559448-INS-53
chr2-53620057-INS-393
chr2-53662478-INS-291
chr2-53665509-INS-59
chr2-53695069-INS-6176
chr2-53704340-INS-101
chr2-53734537-INS-208
chr2-53740185-INS-264
chr2-53753947-INS-62
chr2-53758622-INS-631
chr2-53762555-INS-355
chr2-53779263-INS-75
chr2-53800508-INS-241
chr2-53817923-INS-86
chr2-53830238-INS-112
chr2-53830778-INS-6925
chr2-53856101-INS-7210
chr2-53860656-INS-7560
chr2-53863338-INS-6460
chr2-53865095-INS-60
chr2-53892729-INS-314
chr2-53910781-INS-78
chr2-53919230-INS-398
chr2-53931902-INS-1268
chr2-53935599-INS-77
chr2-53954934-INS-438
chr2-53982311-INS-2547
chr2-53986516-INS-6599
chr2-54005249-INS-56
chr2-54083799-INS-187
chr2-54085892-INS-106
chr2-54110742-INS-85
chr2-54112709-INS-99
chr2-54115958-INS-712
chr2-54131897-INS-332
chr2-54150233-INS-66
chr2-54159326-INS-58
chr2-54169390-INS-720
chr2-54209598-INS-69
chr2-54257370-INS-6513
chr2-54274335-INS-60
chr2-54293428-INS-119
chr2-54307647-INS-54
chr2-54323923-INS-7446
chr2-54333558-INS-112
chr2-54355897-INS-105
chr2-54360999-INS-197
chr2-54368713-INS-656
chr2-54388901-INS-56
chr2-54393481-INS-76
chr2-54413263-INS-7088
chr2-54416894-INS-203
chr2-54443375-INS-72
chr2-54545926-INS-1895
chr2-54549993-INS-297
chr2-54552903-INS-55
chr2-54556591-INS-103
chr2-54558583-INS-6702
chr2-54570999-INS-168
chr2-54580283-INS-8672
chr2-54621194-INS-72
chr2-54631315-INS-70
chr2-54662110-INS-84
chr2-54681264-INS-7087
chr2-54684618-INS-90
chr2-54687262-INS-8831
chr2-54695009-INS-902

chr2-54721797-INS-56
chr2-54789330-INS-537
chr2-54798569-INS-111
chr2-54804985-INS-327
chr2-54834935-INS-1240
chr2-54900095-INS-53
chr2-54907863-INS-64
chr2-54931402-INS-51
chr2-54931975-INS-7110
chr2-54951105-INS-83
chr2-54954006-INS-6720
chr2-54967674-INS-759
chr2-54971583-INS-6317
chr2-54980685-INS-154
chr2-55005401-INS-100
chr2-55028105-INS-289
chr2-55031439-INS-198
chr2-55066008-INS-84
chr2-55099810-INS-308
chr2-55115862-INS-200
chr2-55137873-INS-649
chr2-55174096-INS-376
chr2-55188280-INS-83
chr2-55191102-INS-321
chr2-55193559-INS-7393
chr2-55201550-INS-432
chr2-55205083-INS-104
chr2-55249267-INS-538
chr2-55281869-INS-7109
chr2-55294893-INS-5543
chr2-55300414-INS-60
chr2-55302631-INS-217
chr2-55313085-INS-135
chr2-55328980-INS-84
chr2-55360504-INS-54
chr2-55415515-INS-81
chr2-55428179-INS-109
chr2-55471919-INS-179
chr2-55503080-INS-75
chr2-55538561-INS-126
chr2-55559327-INS-480
chr2-55560306-INS-74
chr2-55575459-INS-73
chr2-55577941-INS-452
chr2-55598564-INS-126
chr2-55608806-INS-454
chr2-55661660-INS-6464
chr2-55666597-INS-95
chr2-55668660-INS-1491
chr2-55689790-INS-1563
chr2-55694113-INS-8792
chr2-55706445-INS-126
chr2-55753722-INS-6395
chr2-55769612-INS-354
chr2-55773260-INS-431
chr2-55814225-INS-656
chr2-55849634-INS-57
chr2-55858342-INS-221
chr2-55862423-INS-74
chr2-55872586-INS-6349
chr2-55874221-INS-62
chr2-55896655-INS-647
chr2-55903050-INS-7080
chr2-55903497-INS-6595
chr2-55905343-INS-74
chr2-55912466-INS-6367
chr2-55952777-INS-160
chr2-55974425-INS-379
chr2-55988063-INS-6433
chr2-56006843-INS-626
chr2-56010999-INS-339
chr2-56035123-INS-54
chr2-56059416-INS-175
chr2-56088423-INS-94
chr2-56095680-INS-120
chr2-56100131-INS-98
chr2-56114004-INS-320

chr2-56116346-INS-168
chr2-56128777-INS-359
chr2-56132068-INS-198
chr2-56136948-INS-104
chr2-56163415-INS-606
chr2-56175394-INS-6670
chr2-56177402-INS-261
chr2-56183171-INS-99
chr2-56204859-INS-5351
chr2-56218713-INS-56
chr2-56229275-INS-3412
chr2-56244593-INS-340
chr2-56273803-INS-4109
chr2-56277423-INS-440
chr2-56284901-INS-56
chr2-56322222-INS-51
chr2-56373845-INS-442
chr2-56402927-INS-180
chr2-56419297-INS-114
chr2-56420145-INS-452
chr2-56439772-INS-7099
chr2-56449443-INS-91
chr2-56461215-INS-347
chr2-56487677-INS-101
chr2-56489751-INS-95
chr2-56532595-INS-354
chr2-56548052-INS-482
chr2-56549927-INS-64
chr2-56552329-INS-64
chr2-56557303-INS-76
chr2-56574989-INS-170
chr2-56585606-INS-72
chr2-56606437-INS-597
chr2-56610254-INS-281
chr2-56615997-INS-141
chr2-56656015-INS-199
chr2-56663881-INS-6619
chr2-56665431-INS-197
chr2-56668074-INS-83
chr2-56673952-INS-208
chr2-56676379-INS-3691
chr2-56679613-INS-59
chr2-56704908-INS-410
chr2-56716003-INS-94
chr2-56730401-INS-6787
chr2-56731701-INS-1923
chr2-56773824-INS-61
chr2-56777551-INS-6623
chr2-56779527-INS-586
chr2-56793427-INS-200
chr2-56798375-INS-57
chr2-56813108-INS-410
chr2-56827922-INS-6524
chr2-56829205-INS-167
chr2-56840275-INS-76
chr2-56869674-INS-491
chr2-56871005-INS-358
chr2-56892911-INS-58
chr2-56909090-INS-198
chr2-56932399-INS-60
chr2-56946601-INS-1888
chr2-56946767-INS-623
chr2-56962353-INS-105
chr2-56978624-INS-444
chr2-57006304-INS-52
chr2-57020815-INS-3093
chr2-57025118-INS-117
chr2-57039062-INS-58
chr2-57042741-INS-72
chr2-57065199-INS-548
chr2-57080438-INS-51
chr2-57100731-INS-82
chr2-57116326-INS-81
chr2-57117702-INS-53
chr2-57133162-INS-105
chr2-57150783-INS-64
chr2-57153825-INS-94

chr2-57154479-INS-621
chr2-57162398-INS-238
chr2-57219040-INS-110
chr2-57220300-INS-251
chr2-57291723-INS-180
chr2-57301943-INS-641
chr2-57304447-INS-59
chr2-57342330-INS-64
chr2-57346979-INS-396
chr2-57351686-INS-107
chr2-57361659-INS-339
chr2-57419794-INS-242
chr2-57430163-INS-561
chr2-57461823-INS-130
chr2-57479743-INS-330
chr2-57485664-INS-6716
chr2-57497998-INS-388
chr2-57517594-INS-338
chr2-57533031-INS-395
chr2-57538488-INS-6696
chr2-57564721-INS-84
chr2-57566671-INS-92
chr2-57577273-INS-152
chr2-57577538-INS-6635
chr2-57631749-INS-819
chr2-57642327-INS-315
chr2-57667115-INS-529
chr2-57682971-INS-52
chr2-57691603-INS-65
chr2-57720959-INS-54
chr2-57723500-INS-6931
chr2-57726972-INS-391
chr2-57755516-INS-114
chr2-57760672-INS-129
chr2-57778471-INS-107
chr2-57786794-INS-128
chr2-57814178-INS-274
chr2-57821412-INS-353
chr2-57879093-INS-6510
chr2-57882131-INS-1553
chr2-57882259-INS-173
chr2-57913115-INS-105
chr2-57949959-INS-199
chr2-57987301-INS-85
chr2-58003761-INS-111
chr2-58041623-INS-352
chr2-58058647-INS-120
chr2-58089043-INS-79
chr2-58091791-INS-92
chr2-58138505-INS-54
chr2-58147012-INS-208
chr2-58153853-INS-204
chr2-58175994-INS-717
chr2-58203186-INS-240
chr2-58226878-INS-6601
chr2-58238179-INS-77
chr2-58249265-INS-640
chr2-58253004-INS-53
chr2-58261973-INS-52
chr2-58280213-INS-515
chr2-58289551-INS-154
chr2-58300013-INS-252
chr2-58330046-INS-226
chr2-58331841-INS-69
chr2-58343105-INS-161
chr2-58366019-INS-57
chr2-58382192-INS-86
chr2-58384585-INS-86
chr2-58423985-INS-446
chr2-58439637-INS-54
chr2-58468254-INS-54
chr2-58471474-INS-159
chr2-58506456-INS-217
chr2-58513801-INS-74
chr2-58516028-INS-488
chr2-58525391-INS-75
chr2-58525664-INS-132

chr2-58550309-INS-56
chr2-58609745-INS-72
chr2-58616284-INS-241
chr2-58619115-INS-197
chr2-58651887-INS-1526
chr2-58669355-INS-391
chr2-58672029-INS-72
chr2-58688495-INS-190
chr2-58690986-INS-91
chr2-58708600-INS-548
chr2-58714958-INS-6886
chr2-58728398-INS-6227
chr2-58814064-INS-80
chr2-58879824-INS-76
chr2-58883320-INS-52
chr2-58897449-INS-322
chr2-58912988-INS-2500
chr2-58915757-INS-51
chr2-58953132-INS-64
chr2-58984268-INS-74
chr2-59006835-INS-10319
chr2-59071006-INS-91
chr2-59083309-INS-379
chr2-59086128-INS-62
chr2-59104593-INS-90
chr2-59135482-INS-543
chr2-59173661-INS-208
chr2-59189808-INS-211
chr2-59220864-INS-400
chr2-59228799-INS-207
chr2-59241823-INS-413
chr2-59252219-INS-2825
chr2-59295160-INS-64
chr2-59316735-INS-66
chr2-59338696-INS-211
chr2-59354852-INS-78
chr2-59356361-INS-7488
chr2-59364352-INS-203
chr2-59373385-INS-131
chr2-59384271-INS-180
chr2-59385371-INS-201
chr2-59397819-INS-207
chr2-59410991-INS-90
chr2-59421767-INS-779
chr2-59427750-INS-215
chr2-59452600-INS-68
chr2-59535489-INS-403
chr2-59536840-INS-102
chr2-59545816-INS-120
chr2-59557883-INS-81
chr2-59580783-INS-119
chr2-59606981-INS-156
chr2-59616638-INS-64
chr2-59631156-INS-216
chr2-59631699-INS-261
chr2-59645484-INS-88
chr2-59649583-INS-172
chr2-59657768-INS-70
chr2-59658499-INS-459
chr2-59687761-INS-546
chr2-59695529-INS-399
chr2-59697570-INS-56
chr2-59698983-INS-399
chr2-59704232-INS-441
chr2-59717699-INS-444
chr2-59718854-INS-218
chr2-59719455-INS-180
chr2-59736390-INS-420
chr2-59745583-INS-56
chr2-59747515-INS-205
chr2-59749118-INS-369
chr2-59755447-INS-60
chr2-59761062-INS-6464
chr2-59803466-INS-329
chr2-59804618-INS-307
chr2-59824310-INS-117
chr2-59883150-INS-75

chr2-59903808-INS-212
chr2-59912064-INS-60
chr2-59939870-INS-406
chr2-59977157-INS-60
chr2-59983525-INS-57
chr2-59992593-INS-263
chr2-60008712-INS-129
chr2-60012375-INS-56
chr2-60017072-INS-431
chr2-60019806-INS-175
chr2-60022022-INS-90
chr2-60025102-INS-69
chr2-60038391-INS-75
chr2-60091704-INS-90
chr2-60100746-INS-1188
chr2-60100929-INS-698
chr2-60101668-INS-107
chr2-60119784-INS-56
chr2-60134085-INS-345
chr2-60136144-INS-440
chr2-60149160-INS-53
chr2-60150499-INS-226
chr2-60164551-INS-72
chr2-60174467-INS-80
chr2-60175874-INS-209
chr2-60193678-INS-136
chr2-60196588-INS-68
chr2-60231483-INS-132
chr2-60233123-INS-95
chr2-60236181-INS-305
chr2-60239265-INS-134
chr2-60253225-INS-94
chr2-60259504-INS-465
chr2-60266290-INS-180
chr2-60317917-INS-442
chr2-60327885-INS-969
chr2-60330449-INS-54
chr2-60356847-INS-199
chr2-60411019-INS-78
chr2-60430033-INS-372
chr2-60451042-INS-61
chr2-60468891-INS-2373
chr2-60497070-INS-182
chr2-60500563-INS-54
chr2-60565994-INS-388
chr2-60599275-INS-56
chr2-60623319-INS-63
chr2-60655691-INS-480
chr2-60717444-INS-71
chr2-60720732-INS-195
chr2-60756656-INS-51
chr2-60768784-INS-72
chr2-60773459-INS-61
chr2-60788027-INS-107
chr2-60790320-INS-345
chr2-60791468-INS-120
chr2-60805273-INS-75
chr2-60821920-INS-127
chr2-60824772-INS-5707
chr2-60826761-INS-117
chr2-60828093-INS-6804
chr2-60831465-INS-5377
chr2-60834099-INS-831
chr2-60843818-INS-175
chr2-60866698-INS-685
chr2-60894905-INS-118
chr2-60896359-INS-126
chr2-60920297-INS-169
chr2-60921597-INS-70
chr2-60926404-INS-63
chr2-60938364-INS-75
chr2-60942492-INS-120
chr2-60946110-INS-54
chr2-60952370-INS-346
chr2-60957358-INS-80
chr2-60973226-INS-186
chr2-60980244-INS-193

chr2-60983172-INS-291
chr2-60985519-INS-72
chr2-61014438-INS-68
chr2-61024723-INS-58
chr2-61040740-INS-321
chr2-61059865-INS-330
chr2-61064072-INS-75
chr2-61067424-INS-54
chr2-61084381-INS-486
chr2-61112741-INS-65
chr2-61122236-INS-66
chr2-61128458-INS-223
chr2-61162701-INS-64
chr2-61169676-INS-77
chr2-61219499-INS-96
chr2-61243175-INS-358
chr2-61267328-INS-2598
chr2-61299723-INS-290
chr2-61302861-INS-56
chr2-61306383-INS-60
chr2-61310003-INS-202
chr2-61320621-INS-140
chr2-61331464-INS-84
chr2-61342640-INS-167
chr2-61358642-INS-97
chr2-61359918-INS-163
chr2-61361322-INS-145
chr2-61375847-INS-114
chr2-61412823-INS-56
chr2-61424700-INS-195
chr2-61449470-INS-3160
chr2-61456236-INS-651
chr2-61457838-INS-55
chr2-61463491-INS-915
chr2-61505721-INS-440
chr2-61508292-INS-102
chr2-61526639-INS-67
chr2-61538412-INS-8985
chr2-61560540-INS-772
chr2-61571115-INS-60
chr2-61577965-INS-328
chr2-61626066-INS-107
chr2-61662114-INS-78
chr2-61668346-INS-1087
chr2-61668369-INS-840
chr2-61685238-INS-150
chr2-61689736-INS-6193
chr2-61715953-INS-6984
chr2-61720530-INS-6389
chr2-61721175-INS-5091
chr2-61729392-INS-2863
chr2-61738574-INS-6071
chr2-61779025-INS-163
chr2-61800316-INS-250
chr2-61837086-INS-116
chr2-61854076-INS-245
chr2-61859861-INS-6552
chr2-61862509-INS-121
chr2-61875618-INS-613
chr2-61893237-INS-105
chr2-61937237-INS-60
chr2-61941323-INS-470
chr2-61944643-INS-119
chr2-61948632-INS-3861
chr2-61951880-INS-68
chr2-61984635-INS-76
chr2-62035354-INS-6385
chr2-62048463-INS-2262
chr2-62052162-INS-72
chr2-62110980-INS-51
chr2-62120605-INS-354
chr2-62133050-INS-318
chr2-62144689-INS-125
chr2-62171854-INS-54
chr2-62186944-INS-477
chr2-62206765-INS-66
chr2-62219862-INS-206

chr2-62248876-INS-82
chr2-62258438-INS-95
chr2-62304505-INS-56
chr2-62321781-INS-52
chr2-62325859-INS-754
chr2-62329665-INS-5311
chr2-62331680-INS-239
chr2-62356075-INS-77
chr2-62374065-INS-128
chr2-62380793-INS-437
chr2-62395066-INS-131
chr2-62433030-INS-70
chr2-62478405-INS-184
chr2-62513789-INS-746
chr2-62541535-INS-440
chr2-62551038-INS-162
chr2-62562405-INS-9742
chr2-62567866-INS-476
chr2-62649070-INS-6268
chr2-62723304-INS-152
chr2-62730570-INS-654
chr2-62731088-INS-59
chr2-62738872-INS-351
chr2-62754069-INS-64
chr2-62769739-INS-83
chr2-62783611-INS-74
chr2-62787344-INS-220
chr2-62792259-INS-964
chr2-62804841-INS-55
chr2-62832635-INS-535
chr2-62855123-INS-276
chr2-62883887-INS-51
chr2-62887982-INS-256
chr2-62893961-INS-382
chr2-62910394-INS-104
chr2-62946822-INS-82
chr2-62962397-INS-693
chr2-63009778-INS-52
chr2-63030022-INS-6045
chr2-63039945-INS-56
chr2-63062683-INS-696
chr2-63067770-INS-167
chr2-63073954-INS-2845
chr2-63079796-INS-426
chr2-63089459-INS-109
chr2-63101594-INS-197
chr2-63114350-INS-916
chr2-63115613-INS-360
chr2-63119264-INS-3084
chr2-63144050-INS-83
chr2-63156017-INS-70
chr2-63183834-INS-70
chr2-63187251-INS-183
chr2-63195328-INS-8457
chr2-63275375-INS-107
chr2-63297478-INS-65
chr2-63332364-INS-5245
chr2-63353702-INS-7237
chr2-63356931-INS-6579
chr2-63374598-INS-131
chr2-63385650-INS-65
chr2-63386771-INS-524
chr2-63410428-INS-4806
chr2-63492557-INS-405
chr2-63499159-INS-194
chr2-63510845-INS-57
chr2-63518614-INS-68
chr2-63522684-INS-66
chr2-63530346-INS-222
chr2-63593278-INS-62
chr2-63629373-INS-147
chr2-63642960-INS-71
chr2-63647300-INS-156
chr2-63701116-INS-454
chr2-63719961-INS-110
chr2-63730021-INS-601
chr2-63737111-INS-108

chr2-63755453-INS-384
chr2-63764828-INS-208
chr2-63767469-INS-355
chr2-63782214-INS-6974
chr2-63811574-INS-106
chr2-63852795-INS-68
chr2-63882949-INS-64
chr2-63889334-INS-6466
chr2-63904051-INS-62
chr2-63955468-INS-146
chr2-63957540-INS-146
chr2-63963073-INS-312
chr2-63984686-INS-412
chr2-63990647-INS-58
chr2-64011751-INS-394
chr2-64022051-INS-6793
chr2-64032752-INS-554
chr2-64033777-INS-54
chr2-64040058-INS-7154
chr2-64084921-INS-162
chr2-64107203-INS-54
chr2-64116608-INS-94
chr2-64137833-INS-351
chr2-64171523-INS-86
chr2-64173490-INS-76
chr2-64174925-INS-245
chr2-64202402-INS-887
chr2-64206428-INS-144
chr2-64251967-INS-187
chr2-64301219-INS-631
chr2-64344524-INS-96
chr2-64347010-INS-326
chr2-64348118-INS-670
chr2-64374666-INS-7150
chr2-64393324-INS-506
chr2-64394202-INS-286
chr2-64406246-INS-252
chr2-64408334-INS-436
chr2-64472463-INS-506
chr2-64482161-INS-4293
chr2-64489582-INS-64
chr2-64499447-INS-6099
chr2-64519172-INS-2107
chr2-64524470-INS-209
chr2-64526908-INS-338
chr2-64573078-INS-2262
chr2-64665083-INS-279
chr2-64666295-INS-68
chr2-64675779-INS-566
chr2-64763006-INS-55
chr2-64789829-INS-78
chr2-64792369-INS-194
chr2-64823936-INS-427
chr2-64860641-INS-296
chr2-64861204-INS-197
chr2-64870237-INS-89
chr2-64876137-INS-136
chr2-64917077-INS-130
chr2-64947448-INS-564
chr2-64974273-INS-334
chr2-64995430-INS-991
chr2-65009200-INS-90
chr2-65010399-INS-4302
chr2-65014946-INS-105
chr2-65030674-INS-575
chr2-65031205-INS-62
chr2-65046898-INS-75
chr2-65058299-INS-462
chr2-65072972-INS-78
chr2-65080598-INS-62
chr2-65083019-INS-223
chr2-65086157-INS-60
chr2-65090414-INS-310
chr2-65094389-INS-58
chr2-65101832-INS-54
chr2-65110420-INS-431
chr2-65118493-INS-144

chr2-65121377-INS-194
chr2-65131963-INS-73
chr2-65140439-INS-98
chr2-65145919-INS-3138
chr2-65179322-INS-68
chr2-65206485-INS-169
chr2-65227456-INS-592
chr2-65238094-INS-140
chr2-65253156-INS-78
chr2-65260198-INS-474
chr2-65305435-INS-57
chr2-65334009-INS-191
chr2-65342226-INS-89
chr2-65365828-INS-369
chr2-65371465-INS-222
chr2-65437549-INS-126
chr2-65441291-INS-82
chr2-65458516-INS-258
chr2-65467848-INS-301
chr2-65482526-INS-7078
chr2-65490160-INS-7745
chr2-65493363-INS-62
chr2-65558397-INS-196
chr2-65575556-INS-92
chr2-65600776-INS-7642
chr2-65605445-INS-212
chr2-65626384-INS-81
chr2-65634772-INS-111
chr2-65658008-INS-56
chr2-65668304-INS-208
chr2-65671921-INS-56
chr2-65688605-INS-137
chr2-65692895-INS-883
chr2-65743082-INS-120
chr2-65744794-INS-60
chr2-65757050-INS-517
chr2-65794020-INS-6385
chr2-65794538-INS-69
chr2-65812218-INS-182
chr2-65828294-INS-7553
chr2-65843473-INS-785
chr2-65881567-INS-55
chr2-65886426-INS-201
chr2-65921938-INS-62
chr2-65946827-INS-71
chr2-65978293-INS-390
chr2-65982252-INS-98
chr2-66008214-INS-55
chr2-66011506-INS-525
chr2-66011856-INS-445
chr2-66025932-INS-242
chr2-66071092-INS-70
chr2-66089362-INS-574
chr2-66169804-INS-65
chr2-66170876-INS-94
chr2-66175996-INS-112
chr2-66177881-INS-72
chr2-66247685-INS-139
chr2-66291631-INS-400
chr2-66309914-INS-93
chr2-66315614-INS-542
chr2-66334930-INS-342
chr2-66372896-INS-208
chr2-66399525-INS-726
chr2-66408662-INS-526
chr2-66422632-INS-244
chr2-66423806-INS-4078
chr2-66430802-INS-6916
chr2-66441978-INS-260
chr2-66455029-INS-100
chr2-66457442-INS-536
chr2-66472031-INS-72
chr2-66485563-INS-62
chr2-66503293-INS-238
chr2-66516346-INS-72
chr2-66524512-INS-298
chr2-66581251-INS-90

chr2-66592848-INS-156
chr2-66593525-INS-431
chr2-66618382-INS-64
chr2-66654780-INS-58
chr2-66701045-INS-636
chr2-66715959-INS-218
chr2-66754587-INS-1194
chr2-66772962-INS-6180
chr2-66778336-INS-2023
chr2-66798640-INS-56
chr2-66806046-INS-298
chr2-66814139-INS-57
chr2-66816832-INS-64
chr2-66817973-INS-105
chr2-66824342-INS-326
chr2-66825700-INS-6133
chr2-66849708-INS-7191
chr2-66877575-INS-54
chr2-66910391-INS-222
chr2-66931712-INS-203
chr2-66943660-INS-101
chr2-66951228-INS-303
chr2-66975283-INS-184
chr2-66986139-INS-51
chr2-67009852-INS-630
chr2-67014611-INS-363
chr2-67030529-INS-73
chr2-67052534-INS-712
chr2-67065388-INS-360
chr2-67095411-INS-52
chr2-67123319-INS-130
chr2-67136313-INS-147
chr2-67143355-INS-182
chr2-67166253-INS-80
chr2-67197443-INS-449
chr2-67205105-INS-456
chr2-67261053-INS-83
chr2-67271165-INS-100
chr2-67314746-INS-366
chr2-67325701-INS-82
chr2-67332770-INS-478
chr2-67347980-INS-108
chr2-67391533-INS-62
chr2-67399441-INS-88
chr2-67417286-INS-62
chr2-67419385-INS-205
chr2-67424459-INS-75
chr2-67430073-INS-136
chr2-67439848-INS-108
chr2-67462037-INS-54
chr2-67476254-INS-52
chr2-67482184-INS-56
chr2-67490720-INS-56
chr2-67510579-INS-293
chr2-67518659-INS-59
chr2-67522359-INS-7028
chr2-67526751-INS-56
chr2-67544298-INS-64
chr2-67557359-INS-431
chr2-67563579-INS-919
chr2-67582711-INS-91
chr2-67594775-INS-310
chr2-67619227-INS-104
chr2-67622232-INS-874
chr2-67684929-INS-590
chr2-67688551-INS-61
chr2-67693787-INS-629
chr2-67699708-INS-170
chr2-67701566-INS-78
chr2-67718101-INS-126
chr2-67740931-INS-81
chr2-67782902-INS-110
chr2-67784791-INS-97
chr2-67797232-INS-192
chr2-67811382-INS-78
chr2-67814348-INS-158
chr2-67847543-INS-246

chr2-67857506-INS-79
chr2-67864956-INS-116
chr2-67869817-INS-109
chr2-67874427-INS-130
chr2-67891535-INS-327
chr2-67910576-INS-68
chr2-67917969-INS-784
chr2-67970622-INS-85
chr2-68010938-INS-307
chr2-68011150-INS-10569
chr2-68019503-INS-62
chr2-68152939-INS-70
chr2-68166971-INS-54
chr2-68185667-INS-145
chr2-68188166-INS-110
chr2-68202609-INS-70
chr2-68213941-INS-280
chr2-68227522-INS-160
chr2-68233174-INS-127
chr2-68242502-INS-148
chr2-68279444-INS-150
chr2-68280433-INS-195
chr2-68286471-INS-598
chr2-68290308-INS-224
chr2-68292898-INS-209
chr2-68298375-INS-115
chr2-68314207-INS-112
chr2-68373270-INS-120
chr2-68437584-INS-7829
chr2-68443313-INS-458
chr2-68447441-INS-333
chr2-68467186-INS-936
chr2-68470337-INS-701
chr2-68555882-INS-568
chr2-68564183-INS-5149
chr2-68584711-INS-336
chr2-68587133-INS-62
chr2-68600886-INS-212
chr2-68608046-INS-2183
chr2-68635447-INS-60
chr2-68646527-INS-56
chr2-68700932-INS-63
chr2-68711904-INS-117
chr2-68718226-INS-61
chr2-68740414-INS-56
chr2-68745082-INS-72
chr2-68780544-INS-212
chr2-68783850-INS-933
chr2-68799298-INS-58
chr2-68820323-INS-598
chr2-68843893-INS-6879
chr2-68859614-INS-61
chr2-68917580-INS-92
chr2-68923808-INS-495
chr2-68932149-INS-99
chr2-68985434-INS-103
chr2-68988512-INS-449
chr2-69004441-INS-70
chr2-69007460-INS-447
chr2-69011976-INS-57
chr2-69075667-INS-2801
chr2-69078648-INS-7087
chr2-69086164-INS-111
chr2-69088518-INS-7101
chr2-69094872-INS-64
chr2-69141536-INS-5404
chr2-69141614-INS-6903
chr2-69151589-INS-53
chr2-69155062-INS-174
chr2-69189745-INS-329
chr2-69198287-INS-123
chr2-69207158-INS-151
chr2-69209359-INS-581
chr2-69240832-INS-899
chr2-69272820-INS-137
chr2-69283001-INS-124
chr2-69301562-INS-869

chr2-69302601-INS-58
chr2-69331694-INS-84
chr2-69346115-INS-265
chr2-69351468-INS-209
chr2-69356993-INS-202
chr2-69381116-INS-367
chr2-69393137-INS-4883
chr2-69396019-INS-69
chr2-69415293-INS-52
chr2-69416882-INS-59
chr2-69426595-INS-143
chr2-69435213-INS-57
chr2-69440686-INS-106
chr2-69446481-INS-100
chr2-69449057-INS-55
chr2-69453271-INS-255
chr2-69457375-INS-96
chr2-69459353-INS-54
chr2-69472085-INS-57
chr2-69476927-INS-59
chr2-69482770-INS-84
chr2-69486726-INS-118
chr2-69490064-INS-139
chr2-69491322-INS-269
chr2-69492557-INS-97
chr2-69517877-INS-53
chr2-69527822-INS-465
chr2-69531211-INS-76
chr2-69538308-INS-206
chr2-69546423-INS-90
chr2-69550626-INS-77
chr2-69610538-INS-97
chr2-69615433-INS-67
chr2-69635415-INS-178
chr2-69639731-INS-179
chr2-69661853-INS-199
chr2-69669902-INS-118
chr2-69676099-INS-688
chr2-69679955-INS-6951
chr2-69718352-INS-205
chr2-69733741-INS-320
chr2-69735337-INS-208
chr2-69736449-INS-148
chr2-69742797-INS-172
chr2-69746520-INS-160
chr2-69760525-INS-218
chr2-69763658-INS-201
chr2-69765959-INS-91
chr2-69769838-INS-117
chr2-69785820-INS-205
chr2-69817342-INS-208
chr2-69818264-INS-60
chr2-69827912-INS-62
chr2-69833817-INS-196
chr2-69841956-INS-174
chr2-69876868-INS-870
chr2-69878744-INS-472
chr2-69886832-INS-84
chr2-69942034-INS-200
chr2-69942277-INS-344
chr2-69955031-INS-108
chr2-69963124-INS-397
chr2-69976496-INS-116
chr2-69995686-INS-54
chr2-69997369-INS-52
chr2-70014379-INS-124
chr2-70022268-INS-58
chr2-70024279-INS-62
chr2-70040706-INS-652
chr2-70056285-INS-66
chr2-70077231-INS-80
chr2-70089733-INS-168
chr2-70105837-INS-62
chr2-70106868-INS-111
chr2-70108495-INS-115
chr2-70113952-INS-120
chr2-70130606-INS-86

chr2-70155405-INS-117
chr2-70165796-INS-173
chr2-70201767-INS-6452
chr2-70216521-INS-717
chr2-70217804-INS-693
chr2-70227810-INS-116
chr2-70236220-INS-412
chr2-70240364-INS-169
chr2-70242111-INS-694
chr2-70262380-INS-305
chr2-70263011-INS-117
chr2-70268002-INS-555
chr2-70268894-INS-215
chr2-70283188-INS-88
chr2-70295631-INS-307
chr2-70319015-INS-116
chr2-70322138-INS-77
chr2-70351242-INS-66
chr2-70352866-INS-98
chr2-70357210-INS-60
chr2-70367580-INS-138
chr2-70368638-INS-200
chr2-70379701-INS-197
chr2-70380661-INS-62
chr2-70407522-INS-56
chr2-70419091-INS-155
chr2-70420802-INS-139
chr2-70422630-INS-52
chr2-70425702-INS-246
chr2-70433171-INS-59
chr2-70446768-INS-136
chr2-70479961-INS-760
chr2-70522483-INS-320
chr2-70524751-INS-78
chr2-70573994-INS-320
chr2-70634471-INS-144
chr2-70648244-INS-203
chr2-70668635-INS-52
chr2-70672181-INS-88
chr2-70689623-INS-60
chr2-70705291-INS-128
chr2-70716051-INS-133
chr2-70721665-INS-744
chr2-70725406-INS-192
chr2-70756797-INS-56
chr2-70761755-INS-67
chr2-70764835-INS-55
chr2-70787531-INS-88
chr2-70822284-INS-5548
chr2-70846118-INS-54
chr2-70932885-INS-153
chr2-70943850-INS-72
chr2-70981777-INS-532
chr2-70986379-INS-229
chr2-70989243-INS-149
chr2-71039048-INS-114
chr2-71040473-INS-204
chr2-71059195-INS-372
chr2-71072617-INS-209
chr2-71100998-INS-52
chr2-71118910-INS-206
chr2-71123018-INS-56
chr2-71136786-INS-195
chr2-71140212-INS-128
chr2-71149475-INS-184
chr2-71176465-INS-124
chr2-71180983-INS-136
chr2-71195190-INS-60
chr2-71200273-INS-196
chr2-71206223-INS-57
chr2-71255062-INS-162
chr2-71267664-INS-51
chr2-71273604-INS-201
chr2-71299721-INS-143
chr2-71309106-INS-198
chr2-71318679-INS-370
chr2-71328132-INS-1875

chr2-71406524-INS-208
chr2-71412362-INS-52
chr2-71420625-INS-103
chr2-71433339-INS-202
chr2-71434886-INS-53
chr2-71452646-INS-975
chr2-71491818-INS-173
chr2-71511166-INS-147
chr2-71534149-INS-917
chr2-71556980-INS-115
chr2-71583273-INS-223
chr2-71651272-INS-201
chr2-71678889-INS-352
chr2-71680013-INS-66
chr2-71696960-INS-392
chr2-71701895-INS-401
chr2-71728562-INS-76
chr2-71756567-INS-356
chr2-71764429-INS-56
chr2-71771195-INS-175
chr2-71823559-INS-456
chr2-71846950-INS-93
chr2-71850792-INS-581
chr2-71855408-INS-63
chr2-71857472-INS-542
chr2-71877694-INS-278
chr2-71879727-INS-268
chr2-71888281-INS-52
chr2-71889492-INS-60
chr2-71892999-INS-56
chr2-71904831-INS-60
chr2-71910415-INS-72
chr2-71940181-INS-145
chr2-71962259-INS-64
chr2-71968872-INS-83
chr2-72003734-INS-58
chr2-72057655-INS-67
chr2-72096828-INS-202
chr2-72119090-INS-1892
chr2-72121609-INS-366
chr2-72130568-INS-70
chr2-72140526-INS-52
chr2-72156193-INS-75
chr2-72175971-INS-144
chr2-72201436-INS-55
chr2-72226025-INS-332
chr2-72228850-INS-65
chr2-72244542-INS-524
chr2-72262729-INS-201
chr2-72302733-INS-5339
chr2-72318548-INS-898
chr2-72332095-INS-52
chr2-72363641-INS-1884
chr2-72389608-INS-96
chr2-72403371-INS-998
chr2-72405106-INS-142
chr2-72418956-INS-206
chr2-72424782-INS-206
chr2-72431607-INS-148
chr2-72451219-INS-324
chr2-72485466-INS-155
chr2-72519829-INS-251
chr2-72535306-INS-166
chr2-72539598-INS-59
chr2-72549948-INS-310
chr2-72556592-INS-368
chr2-72571386-INS-58
chr2-72578220-INS-51
chr2-72580292-INS-762
chr2-72585025-INS-84
chr2-72595428-INS-94
chr2-72623421-INS-56
chr2-72647940-INS-220
chr2-72651473-INS-52
chr2-72665198-INS-72
chr2-72673693-INS-102
chr2-72676043-INS-166

chr2-72685899-INS-57
chr2-72701956-INS-173
chr2-72710155-INS-363
chr2-72712864-INS-411
chr2-72755602-INS-210
chr2-72763744-INS-93
chr2-72781369-INS-575
chr2-72790379-INS-52
chr2-72795138-INS-196
chr2-72796499-INS-384
chr2-72824682-INS-70
chr2-72840900-INS-188
chr2-72854556-INS-397
chr2-72865318-INS-169
chr2-72869558-INS-56
chr2-72887967-INS-203
chr2-72908328-INS-52
chr2-72913183-INS-184
chr2-72934933-INS-152
chr2-72945762-INS-489
chr2-73013600-INS-56
chr2-73020912-INS-137
chr2-73034008-INS-7040
chr2-73037820-INS-202
chr2-73043128-INS-340
chr2-73055569-INS-58
chr2-73092772-INS-145
chr2-73150116-INS-154
chr2-73151901-INS-248
chr2-73153314-INS-100
chr2-73156738-INS-128
chr2-73182381-INS-399
chr2-73194496-INS-55
chr2-73225251-INS-344
chr2-73234090-INS-1864
chr2-73237886-INS-90
chr2-73257234-INS-62
chr2-73295370-INS-188
chr2-73355115-INS-60
chr2-73373805-INS-219
chr2-73376378-INS-1443
chr2-73376966-INS-81
chr2-73393585-INS-445
chr2-73413444-INS-6440
chr2-73416477-INS-56
chr2-73424977-INS-195
chr2-73435841-INS-240
chr2-73481056-INS-351
chr2-73483546-INS-64
chr2-73492371-INS-78
chr2-73507244-INS-135
chr2-73513291-INS-84
chr2-73517202-INS-96
chr2-73520149-INS-251
chr2-73528141-INS-6594
chr2-73539525-INS-123
chr2-73541745-INS-96
chr2-73553706-INS-793
chr2-73562112-INS-183
chr2-73592223-INS-187
chr2-73593840-INS-211
chr2-73595969-INS-506
chr2-73599150-INS-61
chr2-73614965-INS-96
chr2-73617465-INS-559
chr2-73626897-INS-59
chr2-73628409-INS-66
chr2-73684956-INS-149
chr2-73697692-INS-824
chr2-73737785-INS-916
chr2-73760247-INS-76
chr2-73770870-INS-65
chr2-73779189-INS-207
chr2-73830170-INS-166
chr2-73842967-INS-127
chr2-73843984-INS-54
chr2-73875204-INS-266

chr2-73920483-INS-1299
chr2-73935373-INS-157
chr2-73954886-INS-204
chr2-73964500-INS-6103
chr2-73973078-INS-440
chr2-73978829-INS-429
chr2-73985906-INS-915
chr2-73993893-INS-522
chr2-74027556-INS-216
chr2-74051313-INS-74
chr2-74081971-INS-690
chr2-74110504-INS-137
chr2-74115420-INS-409
chr2-74126948-INS-174
chr2-74182396-INS-82
chr2-74210468-INS-63
chr2-74211784-INS-51
chr2-74236566-INS-5386
chr2-74283949-INS-6178
chr2-74295247-INS-182
chr2-74329911-INS-431
chr2-74332966-INS-1537
chr2-74344809-INS-55
chr2-74397373-INS-82
chr2-74402057-INS-564
chr2-74430226-INS-256
chr2-74433585-INS-56
chr2-74573407-INS-164
chr2-74609435-INS-88
chr2-74616811-INS-55
chr2-74635197-INS-5538
chr2-74636979-INS-5060
chr2-74641684-INS-165
chr2-74661854-INS-291
chr2-74679818-INS-112
chr2-74682034-INS-56
chr2-74682238-INS-7003
chr2-74706164-INS-481
chr2-74728372-INS-7168
chr2-74731861-INS-132
chr2-74752127-INS-89
chr2-74786218-INS-78
chr2-74789560-INS-76
chr2-74791210-INS-111
chr2-74827518-INS-103
chr2-74829609-INS-145
chr2-74846503-INS-84
chr2-74876057-INS-6793
chr2-74894802-INS-51
chr2-74928835-INS-57
chr2-74940000-INS-64
chr2-74941189-INS-84
chr2-74970960-INS-5556
chr2-74986322-INS-512
chr2-74989395-INS-73
chr2-74995378-INS-86
chr2-75040913-INS-54
chr2-75049337-INS-399
chr2-75056619-INS-53
chr2-75077281-INS-452
chr2-75079212-INS-834
chr2-75109727-INS-227
chr2-75130279-INS-668
chr2-75130852-INS-52
chr2-75135192-INS-1881
chr2-75177315-INS-74
chr2-75188250-INS-400
chr2-75231447-INS-276
chr2-75235946-INS-472
chr2-75248122-INS-434
chr2-75249421-INS-53
chr2-75261789-INS-52
chr2-75275060-INS-348
chr2-75294661-INS-51
chr2-75308809-INS-644
chr2-75338158-INS-63
chr2-75360159-INS-60

chr2-75365602-INS-5518
chr2-75381739-INS-292
chr2-75401611-INS-1754
chr2-75442406-INS-207
chr2-75445301-INS-178
chr2-75448863-INS-274
chr2-75453157-INS-531
chr2-75465261-INS-5548
chr2-75478322-INS-124
chr2-75512210-INS-586
chr2-75520370-INS-179
chr2-75524427-INS-576
chr2-75539570-INS-197
chr2-75541428-INS-206
chr2-75547673-INS-196
chr2-75558192-INS-229
chr2-75563333-INS-108
chr2-75564494-INS-400
chr2-75573589-INS-144
chr2-75578406-INS-3177
chr2-75578831-INS-64
chr2-75578917-INS-1274
chr2-75600750-INS-128
chr2-75602111-INS-455
chr2-75604678-INS-116
chr2-75610285-INS-322
chr2-75642046-INS-58
chr2-75647070-INS-206
chr2-75655103-INS-192
chr2-75710145-INS-433
chr2-75771693-INS-498
chr2-75794042-INS-297
chr2-75826979-INS-52
chr2-75838435-INS-60
chr2-75852439-INS-56
chr2-75857990-INS-51
chr2-75912156-INS-55
chr2-75921438-INS-187
chr2-75932004-INS-196
chr2-75937013-INS-5219
chr2-75943298-INS-6176
chr2-75944311-INS-67
chr2-75958038-INS-204
chr2-75960273-INS-174
chr2-75985913-INS-125
chr2-75987740-INS-431
chr2-76019707-INS-56
chr2-76039836-INS-218
chr2-76059674-INS-1524
chr2-76132562-INS-80
chr2-76141586-INS-485
chr2-76184885-INS-127
chr2-76194104-INS-88
chr2-76207047-INS-63
chr2-76231025-INS-52
chr2-76233799-INS-118
chr2-76249201-INS-87
chr2-76271807-INS-105
chr2-76299293-INS-233
chr2-76317180-INS-160
chr2-76324256-INS-102
chr2-76326053-INS-138
chr2-76342712-INS-60
chr2-76349917-INS-156
chr2-76352839-INS-116
chr2-76419109-INS-147
chr2-76421574-INS-78
chr2-76435155-INS-831
chr2-76437508-INS-224
chr2-76475085-INS-222
chr2-76487485-INS-54
chr2-76512581-INS-149
chr2-76513965-INS-55
chr2-76535735-INS-205
chr2-76633310-INS-114
chr2-76635822-INS-92
chr2-76668206-INS-207

chr2-76714014-INS-60
chr2-76743705-INS-220
chr2-76768594-INS-126
chr2-76807253-INS-272
chr2-76832352-INS-185
chr2-76868632-INS-145
chr2-76901158-INS-229
chr2-76942209-INS-68
chr2-76970818-INS-120
chr2-76998517-INS-504
chr2-77030949-INS-125
chr2-77051138-INS-6739
chr2-77051901-INS-89
chr2-77087652-INS-128
chr2-77089721-INS-62
chr2-77105203-INS-52
chr2-77126060-INS-146
chr2-77139344-INS-101
chr2-77184721-INS-397
chr2-77193577-INS-54
chr2-77203709-INS-238
chr2-77238494-INS-54
chr2-77253733-INS-203
chr2-77285694-INS-136
chr2-77293265-INS-64
chr2-77295927-INS-81
chr2-77336008-INS-53
chr2-77348057-INS-114
chr2-77351257-INS-68
chr2-77355338-INS-72
chr2-77394683-INS-255
chr2-77414043-INS-242
chr2-77418999-INS-364
chr2-77453931-INS-142
chr2-77461569-INS-54
chr2-77466503-INS-465
chr2-77562069-INS-78
chr2-77571910-INS-51
chr2-77610898-INS-60
chr2-77639724-INS-54
chr2-77697705-INS-59
chr2-77705890-INS-56
chr2-77719507-INS-7137
chr2-77726734-INS-151
chr2-77750054-INS-6238
chr2-77766634-INS-149
chr2-77770929-INS-428
chr2-77777595-INS-53
chr2-77801510-INS-111
chr2-77801802-INS-4360
chr2-77802426-INS-179
chr2-77864223-INS-60
chr2-77866267-INS-236
chr2-77882974-INS-69
chr2-77906735-INS-658
chr2-77925152-INS-66
chr2-77963155-INS-160
chr2-77965353-INS-12013
chr2-77987848-INS-53
chr2-77989722-INS-98
chr2-78012635-INS-344
chr2-78031646-INS-1148
chr2-78033151-INS-777
chr2-78041746-INS-61
chr2-78045419-INS-80
chr2-78046592-INS-507
chr2-78051986-INS-2831
chr2-78061241-INS-166
chr2-78065547-INS-168
chr2-78078067-INS-6973
chr2-78087963-INS-54
chr2-78100606-INS-3856
chr2-78115925-INS-66
chr2-78134647-INS-322
chr2-78146297-INS-104
chr2-78178064-INS-1053
chr2-78186507-INS-6281

chr2-78187046-INS-52
chr2-78192305-INS-165
chr2-78213824-INS-182
chr2-78215971-INS-84
chr2-78245811-INS-51
chr2-78274233-INS-487
chr2-78312520-INS-2097
chr2-78319403-INS-98
chr2-78329940-INS-65
chr2-78352271-INS-61
chr2-78381752-INS-6609
chr2-78393665-INS-470
chr2-78408448-INS-6694
chr2-78425899-INS-100
chr2-78434156-INS-435
chr2-78463439-INS-111
chr2-78466382-INS-70
chr2-78524271-INS-166
chr2-78538322-INS-226
chr2-78588269-INS-62
chr2-78631834-INS-133
chr2-78632517-INS-102
chr2-78635456-INS-62
chr2-78637023-INS-52
chr2-78639438-INS-58
chr2-78663777-INS-955
chr2-78666863-INS-92
chr2-78679072-INS-6396
chr2-78680136-INS-174
chr2-78732131-INS-1868
chr2-78749494-INS-6473
chr2-78767048-INS-494
chr2-78787642-INS-306
chr2-78789242-INS-289
chr2-78813926-INS-177
chr2-78839799-INS-79
chr2-78850598-INS-56
chr2-78852457-INS-105
chr2-78868995-INS-774
chr2-78888339-INS-109
chr2-78950747-INS-110
chr2-78964378-INS-221
chr2-78972576-INS-529
chr2-78985764-INS-167
chr2-78987356-INS-1005
chr2-78987357-INS-60
chr2-78987357-INS-940
chr2-78988315-INS-121
chr2-79000687-INS-52
chr2-79000756-INS-803
chr2-79028550-INS-51
chr2-79047031-INS-52
chr2-79062515-INS-136
chr2-79082302-INS-58
chr2-79084596-INS-638
chr2-79102346-INS-59
chr2-79159351-INS-7369
chr2-79163316-INS-130
chr2-79168684-INS-66
chr2-79232590-INS-133
chr2-79273469-INS-262
chr2-79306043-INS-1019
chr2-79307834-INS-432
chr2-79311069-INS-210
chr2-79316788-INS-78
chr2-79349974-INS-6240
chr2-79365184-INS-62
chr2-79379723-INS-1543
chr2-79406195-INS-70
chr2-79408814-INS-183
chr2-79410056-INS-81
chr2-79436113-INS-109
chr2-79518930-INS-212
chr2-79529930-INS-843
chr2-79548805-INS-60
chr2-79555178-INS-330
chr2-79557880-INS-102

chr2-79569774-INS-271
chr2-79588890-INS-305
chr2-79599506-INS-52
chr2-79613591-INS-144
chr2-79627115-INS-155
chr2-79647471-INS-789
chr2-79649175-INS-601
chr2-79693186-INS-78
chr2-79704181-INS-78
chr2-79730978-INS-150
chr2-79737006-INS-413
chr2-79747568-INS-169
chr2-79764573-INS-6446
chr2-79846400-INS-124
chr2-79848549-INS-911
chr2-79848575-INS-273
chr2-79854350-INS-112
chr2-79860760-INS-304
chr2-79881090-INS-412
chr2-79894574-INS-95
chr2-79903875-INS-6721
chr2-79923180-INS-248
chr2-79929758-INS-4410
chr2-79930652-INS-294
chr2-79946550-INS-51
chr2-79948946-INS-118
chr2-79968414-INS-592
chr2-79971424-INS-81
chr2-79976957-INS-92
chr2-79987298-INS-51
chr2-80002891-INS-632
chr2-80049907-INS-6065
chr2-80055246-INS-6527
chr2-80087461-INS-1517
chr2-80127126-INS-825
chr2-80131916-INS-6457
chr2-80133538-INS-65
chr2-80169634-INS-120
chr2-80210084-INS-11960
chr2-80213829-INS-191
chr2-80221737-INS-615
chr2-80235533-INS-2684
chr2-80244474-INS-89
chr2-80251266-INS-58
chr2-80263596-INS-78
chr2-80280443-INS-134
chr2-80298283-INS-68
chr2-80316037-INS-80
chr2-80323420-INS-1345
chr2-80335678-INS-78
chr2-80337527-INS-6448
chr2-80352972-INS-204
chr2-80356840-INS-64
chr2-80369168-INS-173
chr2-80388631-INS-345
chr2-80406283-INS-62
chr2-80414693-INS-1893
chr2-80414786-INS-494
chr2-80416777-INS-356
chr2-80427270-INS-64
chr2-80429010-INS-328
chr2-80445091-INS-52
chr2-80458507-INS-57
chr2-80466029-INS-52
chr2-80489731-INS-68
chr2-80492005-INS-188
chr2-80500664-INS-219
chr2-80507278-INS-52
chr2-80508293-INS-114
chr2-80532529-INS-444
chr2-80587717-INS-6242
chr2-80592057-INS-87
chr2-80685929-INS-152
chr2-80700536-INS-51
chr2-80712795-INS-268
chr2-80764607-INS-58
chr2-80767432-INS-982

chr2-80768092-INS-1714
chr2-80794752-INS-92
chr2-80860553-INS-75
chr2-80862313-INS-57
chr2-80868195-INS-99
chr2-80874676-INS-332
chr2-80890217-INS-910
chr2-80927633-INS-937
chr2-80946237-INS-602
chr2-80950096-INS-4847
chr2-80960067-INS-771
chr2-80978125-INS-483
chr2-80990718-INS-435
chr2-81001118-INS-3061
chr2-81038786-INS-55
chr2-81064842-INS-54
chr2-81076751-INS-266
chr2-81077721-INS-765
chr2-81091798-INS-213
chr2-81118618-INS-60
chr2-81120687-INS-103
chr2-81125057-INS-72
chr2-81171729-INS-6845
chr2-81186041-INS-241
chr2-81186687-INS-74
chr2-81215654-INS-270
chr2-81231240-INS-172
chr2-81240709-INS-324
chr2-81272530-INS-117
chr2-81290045-INS-51
chr2-81298438-INS-306
chr2-81303661-INS-69
chr2-81305474-INS-445
chr2-81325447-INS-56
chr2-81345540-INS-207
chr2-81348322-INS-400
chr2-81356292-INS-128
chr2-81356733-INS-6452
chr2-81361733-INS-68
chr2-81363037-INS-56
chr2-81389798-INS-621
chr2-81400934-INS-74
chr2-81479934-INS-102
chr2-81482063-INS-6428
chr2-81490365-INS-603
chr2-81496269-INS-248
chr2-81522857-INS-6405
chr2-81565060-INS-586
chr2-81595666-INS-60
chr2-81601740-INS-125
chr2-81643187-INS-223
chr2-81646443-INS-521
chr2-81659687-INS-496
chr2-81659933-INS-835
chr2-81664361-INS-86
chr2-81666141-INS-56
chr2-81690607-INS-5309
chr2-81693283-INS-1000
chr2-81744563-INS-52
chr2-81754564-INS-76
chr2-81762084-INS-58
chr2-81766781-INS-80
chr2-81768010-INS-197
chr2-81779840-INS-273
chr2-81782887-INS-195
chr2-81817922-INS-731
chr2-81832553-INS-688
chr2-81849423-INS-56
chr2-81888754-INS-213
chr2-81951230-INS-207
chr2-81965299-INS-84
chr2-81975556-INS-173
chr2-81977270-INS-5473
chr2-82007619-INS-51
chr2-82053093-INS-101
chr2-82065143-INS-64
chr2-82070505-INS-69

chr2-82103821-INS-483
chr2-82104566-INS-229
chr2-82106230-INS-1871
chr2-82118402-INS-208
chr2-82134224-INS-55
chr2-82173794-INS-6586
chr2-82177645-INS-794
chr2-82188055-INS-99
chr2-82206351-INS-155
chr2-82208042-INS-288
chr2-82212071-INS-1515
chr2-82212943-INS-144
chr2-82224642-INS-53
chr2-82251200-INS-71
chr2-82251778-INS-1305
chr2-82310281-INS-3542
chr2-82338204-INS-172
chr2-82358208-INS-148
chr2-82376558-INS-6651
chr2-82390072-INS-55
chr2-82397630-INS-150
chr2-82401385-INS-219
chr2-82404007-INS-53
chr2-82409350-INS-244
chr2-82424321-INS-68
chr2-82429554-INS-7096
chr2-82432673-INS-7295
chr2-82449598-INS-1067
chr2-82449731-INS-142
chr2-82471978-INS-600
chr2-82474187-INS-573
chr2-82521800-INS-56
chr2-82526326-INS-52
chr2-82532967-INS-4093
chr2-82562014-INS-52
chr2-82574372-INS-895
chr2-82605388-INS-76
chr2-82609057-INS-100
chr2-82611560-INS-205
chr2-82624507-INS-106
chr2-82633235-INS-583
chr2-82641859-INS-56
chr2-82650432-INS-229
chr2-82661800-INS-356
chr2-82666918-INS-293
chr2-82676876-INS-798
chr2-82683326-INS-399
chr2-82691672-INS-325
chr2-82695862-INS-73
chr2-82726347-INS-79
chr2-82733055-INS-228
chr2-82738265-INS-195
chr2-82746005-INS-4390
chr2-82772226-INS-114
chr2-82790865-INS-597
chr2-82835986-INS-199
chr2-82839573-INS-67
chr2-82844259-INS-239
chr2-82887314-INS-54
chr2-82892228-INS-91
chr2-82914220-INS-137
chr2-82915705-INS-74
chr2-82918090-INS-63
chr2-82925042-INS-2215
chr2-82925855-INS-156
chr2-82931846-INS-999
chr2-82985244-INS-98
chr2-82994274-INS-71
chr2-83000660-INS-6256
chr2-83002790-INS-207
chr2-83003283-INS-986
chr2-83034458-INS-1478
chr2-83062182-INS-841
chr2-83073436-INS-186
chr2-83090506-INS-243
chr2-83110107-INS-198
chr2-83146685-INS-506

chr2-83168167-INS-52
chr2-83199991-INS-6660
chr2-83200107-INS-172
chr2-83201737-INS-52
chr2-83241800-INS-11912
chr2-83254472-INS-352
chr2-83269096-INS-338
chr2-83272151-INS-139
chr2-83337289-INS-74
chr2-83345459-INS-497
chr2-83346877-INS-401
chr2-83357596-INS-80
chr2-83378876-INS-6798
chr2-83381681-INS-6213
chr2-83384954-INS-834
chr2-83399617-INS-847
chr2-83420290-INS-12585
chr2-83423290-INS-153
chr2-83426841-INS-144
chr2-83427466-INS-72
chr2-83427995-INS-6201
chr2-83430785-INS-197
chr2-83439951-INS-499
chr2-83447529-INS-877
chr2-83463801-INS-258
chr2-83465044-INS-363
chr2-83470856-INS-58
chr2-83493719-INS-303
chr2-83532025-INS-362
chr2-83543490-INS-1875
chr2-83551653-INS-6440
chr2-83551693-INS-90
chr2-83554170-INS-67
chr2-83555699-INS-70
chr2-83560938-INS-171
chr2-83561920-INS-203
chr2-83608784-INS-64
chr2-83648128-INS-74
chr2-83753631-INS-52
chr2-83816023-INS-238
chr2-83826098-INS-56
chr2-83917450-INS-212
chr2-83951046-INS-450
chr2-83956007-INS-52
chr2-84031126-INS-602
chr2-84033361-INS-572
chr2-84058781-INS-6962
chr2-84078241-INS-910
chr2-84098571-INS-68
chr2-84137444-INS-6898
chr2-84137529-INS-1561
chr2-84144025-INS-182
chr2-84150022-INS-104
chr2-84158269-INS-593
chr2-84210661-INS-639
chr2-84224795-INS-81
chr2-84227482-INS-83
chr2-84229915-INS-578
chr2-84233760-INS-156
chr2-84249512-INS-56
chr2-84264791-INS-52
chr2-84285565-INS-241
chr2-84290888-INS-60
chr2-84300205-INS-120
chr2-84301261-INS-152
chr2-84310663-INS-207
chr2-84319436-INS-209
chr2-84325168-INS-206
chr2-84329591-INS-155
chr2-84347464-INS-50
chr2-84351685-INS-277
chr2-84362436-INS-110
chr2-84365778-INS-72
chr2-84367708-INS-74
chr2-84370731-INS-53
chr2-84372689-INS-96
chr2-84374084-INS-119

chr2-84380555-INS-64
chr2-84384289-INS-213
chr2-84412175-INS-175
chr2-84416777-INS-116
chr2-84419477-INS-57
chr2-84422756-INS-59
chr2-84425505-INS-6475
chr2-84429402-INS-72
chr2-84439767-INS-561
chr2-84443114-INS-102
chr2-84445237-INS-6399
chr2-84460176-INS-201
chr2-84485501-INS-210
chr2-84515158-INS-267
chr2-84529167-INS-60
chr2-84532298-INS-209
chr2-84547069-INS-204
chr2-84550779-INS-52
chr2-84560122-INS-156
chr2-84570169-INS-97
chr2-84604988-INS-192
chr2-84612652-INS-208
chr2-84624529-INS-5397
chr2-84637789-INS-143
chr2-84658402-INS-62
chr2-84663533-INS-219
chr2-84686444-INS-542
chr2-84695940-INS-86
chr2-84727626-INS-84
chr2-84728617-INS-185
chr2-84752285-INS-110
chr2-84762295-INS-84
chr2-84769555-INS-74
chr2-84789967-INS-5380
chr2-84792945-INS-183
chr2-84794309-INS-230
chr2-84799380-INS-214
chr2-84800994-INS-54
chr2-84804244-INS-75
chr2-84815949-INS-53
chr2-84827888-INS-116
chr2-84957325-INS-230
chr2-84964112-INS-94
chr2-84973483-INS-54
chr2-85001729-INS-90
chr2-85004538-INS-52
chr2-85010002-INS-6465
chr2-85014462-INS-83
chr2-85016662-INS-116
chr2-85034042-INS-441
chr2-85035670-INS-646
chr2-85042328-INS-64
chr2-85052250-INS-93
chr2-85069460-INS-129
chr2-85074301-INS-1527
chr2-85101025-INS-275
chr2-85130824-INS-206
chr2-85139993-INS-180
chr2-85147219-INS-924
chr2-85152797-INS-885
chr2-85155845-INS-56
chr2-85167070-INS-6853
chr2-85177512-INS-91
chr2-85215286-INS-6838
chr2-85223699-INS-84
chr2-85256022-INS-52
chr2-85271512-INS-997
chr2-85299654-INS-506
chr2-85304800-INS-1341
chr2-85332658-INS-225
chr2-85342407-INS-51
chr2-85385102-INS-581
chr2-85398169-INS-1897
chr2-85411396-INS-62
chr2-85431881-INS-66
chr2-85466540-INS-130
chr2-85498827-INS-60

chr2-85511225-INS-84
chr2-85514788-INS-7078
chr2-85545015-INS-407
chr2-85564430-INS-64
chr2-85580538-INS-209
chr2-85582100-INS-182
chr2-85599572-INS-244
chr2-85602160-INS-7157
chr2-85624546-INS-53
chr2-85628337-INS-64
chr2-85648565-INS-501
chr2-85651302-INS-100
chr2-85658305-INS-1354
chr2-85659091-INS-94
chr2-85664728-INS-213
chr2-85671181-INS-172
chr2-85689009-INS-133
chr2-85714394-INS-2104
chr2-85722688-INS-1287
chr2-85737363-INS-6557
chr2-85740381-INS-141
chr2-85747815-INS-80
chr2-85759006-INS-94
chr2-85760514-INS-52
chr2-85831466-INS-52
chr2-85866332-INS-202
chr2-85871402-INS-53
chr2-85887795-INS-68
chr2-85912855-INS-470
chr2-85948573-INS-58
chr2-85960175-INS-54
chr2-85970142-INS-7067
chr2-85994998-INS-92
chr2-86017617-INS-368
chr2-86018664-INS-207
chr2-86035863-INS-119
chr2-86049807-INS-340
chr2-86053896-INS-458
chr2-86055569-INS-53
chr2-86093369-INS-568
chr2-86097532-INS-60
chr2-86111144-INS-113
chr2-86136903-INS-670
chr2-86152686-INS-1892
chr2-86180436-INS-667
chr2-86220625-INS-920
chr2-86261563-INS-5283
chr2-86283548-INS-919
chr2-86317396-INS-85
chr2-86319500-INS-62
chr2-86335682-INS-60
chr2-86353185-INS-224
chr2-86359686-INS-677
chr2-86367992-INS-2949
chr2-86376956-INS-71
chr2-86405796-INS-58
chr2-86429722-INS-3415
chr2-86457668-INS-2679
chr2-86488959-INS-4585
chr2-86489986-INS-91
chr2-86508144-INS-90
chr2-86510696-INS-60
chr2-86512681-INS-4555
chr2-86513174-INS-152
chr2-86515013-INS-70
chr2-86552654-INS-6707
chr2-86564101-INS-54
chr2-86576637-INS-952
chr2-86577727-INS-88
chr2-86602491-INS-109
chr2-86633738-INS-398
chr2-86638427-INS-164
chr2-86643147-INS-117
chr2-86663679-INS-500
chr2-86686885-INS-6439
chr2-86710694-INS-652
chr2-86713826-INS-1266

chr2-86714353-INS-2517
chr2-86728311-INS-85
chr2-86760139-INS-83
chr2-86763256-INS-72
chr2-86763557-INS-6385
chr2-86807186-INS-1864
chr2-86842062-INS-6361
chr2-86859968-INS-140
chr2-86863115-INS-124
chr2-86907130-INS-397
chr2-86931109-INS-404
chr2-86936591-INS-773
chr2-87014898-INS-125
chr2-87021265-INS-130
chr2-87028987-INS-60
chr2-87034861-INS-542
chr2-87055464-INS-4501
chr2-87135423-INS-13281
chr2-87140239-INS-6239
chr2-87150201-INS-6205
chr2-87156525-INS-233
chr2-87175644-INS-358
chr2-87182287-INS-78
chr2-87189135-INS-131
chr2-87196781-INS-84
chr2-87222342-INS-203
chr2-87253553-INS-2674
chr2-87259477-INS-359
chr2-87290973-INS-201
chr2-87314781-INS-67
chr2-87323150-INS-62
chr2-87330868-INS-367
chr2-87356049-INS-205
chr2-87364138-INS-346
chr2-87369339-INS-92
chr2-87369776-INS-5228
chr2-87373463-INS-172
chr2-87374086-INS-232
chr2-87374783-INS-405
chr2-87376990-INS-134
chr2-87377979-INS-226
chr2-87379646-INS-6180
chr2-87383978-INS-336
chr2-87390091-INS-407
chr2-87390257-INS-2107
chr2-87399284-INS-352
chr2-87442119-INS-100
chr2-87452732-INS-6452
chr2-87453760-INS-53
chr2-87484252-INS-94
chr2-87484676-INS-6450
chr2-87499716-INS-226
chr2-87519496-INS-7188
chr2-87523092-INS-202
chr2-87531089-INS-117
chr2-87538932-INS-4660
chr2-87540998-INS-54
chr2-87544004-INS-56
chr2-87545345-INS-6822
chr2-87548559-INS-6389
chr2-87564512-INS-68
chr2-87569887-INS-6237
chr2-87595083-INS-234
chr2-87614057-INS-69
chr2-87633743-INS-306
chr2-87636482-INS-393
chr2-87658796-INS-691
chr2-87679778-INS-6535
chr2-87697059-INS-693
chr2-87729714-INS-294
chr2-87747527-INS-2168
chr2-87774739-INS-7283
chr2-87777311-INS-6398
chr2-87819322-INS-980
chr2-87821934-INS-2168
chr2-87841946-INS-300
chr2-87851110-INS-204

chr2-87861328-INS-109
chr2-87878498-INS-68
chr2-87884882-INS-329
chr2-87901689-INS-187
chr2-87904694-INS-310
chr2-87906585-INS-185
chr2-87933056-INS-557
chr2-87982840-INS-6679
chr2-87992114-INS-156
chr2-88028020-INS-2304
chr2-88039968-INS-6215
chr2-88042994-INS-201
chr2-88079884-INS-6233
chr2-88089397-INS-928
chr2-88094514-INS-7654
chr2-88124616-INS-76
chr2-88127049-INS-111
chr2-88208562-INS-844
chr2-88224627-INS-55
chr2-88230921-INS-117
chr2-88235173-INS-195
chr2-88283687-INS-399
chr2-88285185-INS-52
chr2-88302954-INS-249
chr2-88320980-INS-63
chr2-88361468-INS-802
chr2-88369027-INS-423
chr2-88376314-INS-389
chr2-88393804-INS-183
chr2-88404856-INS-323
chr2-88412133-INS-492
chr2-88432910-INS-167
chr2-88469334-INS-2986
chr2-88484359-INS-934
chr2-88491516-INS-6577
chr2-88512171-INS-54
chr2-88557701-INS-8223
chr2-88558710-INS-58
chr2-88560294-INS-6515
chr2-88591486-INS-5154
chr2-88613777-INS-4200
chr2-88616394-INS-60
chr2-88627334-INS-170
chr2-88631985-INS-66
chr2-88699449-INS-95
chr2-88707393-INS-76
chr2-88731530-INS-61
chr2-88755797-INS-96
chr2-88763070-INS-138
chr2-88764814-INS-80
chr2-88767371-INS-368
chr2-88809970-INS-481
chr2-88826807-INS-4600
chr2-88834851-INS-726
chr2-88847538-INS-416
chr2-88869124-INS-247
chr2-88879305-INS-152
chr2-88915835-INS-93
chr2-88921390-INS-52
chr2-88923271-INS-343
chr2-88941244-INS-90
chr2-88965687-INS-132
chr2-88974252-INS-185
chr2-88984155-INS-54
chr2-89011802-INS-109
chr2-89015124-INS-144
chr2-89016498-INS-430
chr2-89026363-INS-182
chr2-89032566-INS-1029
chr2-89033507-INS-243
chr2-89141389-INS-452
chr2-89153141-INS-513
chr2-89191671-INS-65
chr2-89215085-INS-218
chr2-89230902-INS-67
chr2-89231880-INS-57
chr2-89235377-INS-130

chr2-89238182-INS-90
chr2-89268659-INS-56
chr2-89284024-INS-59
chr2-89301990-INS-6440
chr2-89311221-INS-83
chr2-89335466-INS-354
chr2-89338520-INS-2598
chr2-89338755-INS-960
chr2-89338834-INS-105
chr2-89360142-INS-92
chr2-89394220-INS-415
chr2-89397620-INS-208
chr2-89430055-INS-7630
chr2-89435933-INS-457
chr2-89445656-INS-368
chr2-89447289-INS-7147
chr2-89468883-INS-82
chr2-89473694-INS-265
chr2-89486298-INS-6488
chr2-89490350-INS-772
chr2-89493499-INS-128
chr2-89503951-INS-174
chr2-89520431-INS-56
chr2-89529355-INS-290
chr2-89590892-INS-2384
chr2-89611077-INS-400
chr2-89631725-INS-104
chr2-89639363-INS-6790
chr2-89673927-INS-1444
chr2-89708485-INS-57
chr2-89746086-INS-6417
chr2-89780572-INS-70
chr2-89792625-INS-3805
chr2-89838307-INS-52
chr2-89844768-INS-69
chr2-89858931-INS-787
chr2-89883842-INS-64
chr2-89886979-INS-100
chr2-89888139-INS-204
chr2-89897060-INS-371
chr2-89944730-INS-192
chr2-89973078-INS-557
chr2-90101604-INS-492
chr2-90108111-INS-7811
chr2-90115209-INS-123
chr2-90125687-INS-65
chr2-90138474-INS-223
chr2-90145268-INS-872
chr2-90149520-INS-517
chr2-90151133-INS-121
chr2-90187469-INS-6226
chr2-90202755-INS-506
chr2-90211161-INS-129
chr2-90218303-INS-635
chr2-90221222-INS-874
chr2-90236162-INS-154
chr2-90239154-INS-955
chr2-90246814-INS-7112
chr2-90276041-INS-86
chr2-90277570-INS-52
chr2-90302495-INS-201
chr2-90309281-INS-689
chr2-90323601-INS-90
chr2-90347429-INS-52
chr2-90359497-INS-70
chr2-90393809-INS-82
chr2-90409297-INS-79
chr2-90432871-INS-64
chr2-90436653-INS-229
chr2-90438320-INS-7141
chr2-90441985-INS-432
chr2-90450607-INS-475
chr2-90508793-INS-64
chr2-90523629-INS-297
chr2-90532101-INS-179
chr2-90542449-INS-542
chr2-90544553-INS-98

chr2-90553433-INS-90
chr2-90561198-INS-314
chr2-90564981-INS-200
chr2-90569061-INS-392
chr2-90572764-INS-202
chr2-90578751-INS-51
chr2-90583751-INS-199
chr2-90595985-INS-141
chr2-90601825-INS-203
chr2-90619677-INS-51
chr2-90622932-INS-205
chr2-90625990-INS-66
chr2-90637892-INS-53
chr2-90641298-INS-60
chr2-90642964-INS-368
chr2-90648152-INS-119
chr2-90655383-INS-195
chr2-90660122-INS-157
chr2-90667889-INS-153
chr2-90676187-INS-201
chr2-90693095-INS-216
chr2-90703057-INS-204
chr2-90742763-INS-130
chr2-90763205-INS-201
chr2-90782461-INS-103
chr2-90793382-INS-69
chr2-90804014-INS-152
chr2-90824278-INS-208
chr2-90832029-INS-194
chr2-90851040-INS-4858
chr2-90852298-INS-163
chr2-90862684-INS-54
chr2-90865237-INS-145
chr2-90880890-INS-6415
chr2-90906219-INS-193
chr2-90925922-INS-104
chr2-90935656-INS-100
chr2-90935989-INS-51
chr2-90942394-INS-164
chr2-90947116-INS-117
chr2-90978495-INS-128
chr2-90979647-INS-424
chr2-91028762-INS-437
chr2-91050519-INS-112
chr2-91116506-INS-221
chr2-91123776-INS-303
chr2-91139733-INS-5197
chr2-91140885-INS-61
chr2-91154321-INS-142
chr2-91155977-INS-60
chr2-91182652-INS-215
chr2-91209275-INS-212
chr2-91212248-INS-217
chr2-91235487-INS-67
chr2-91274646-INS-156
chr2-91279010-INS-199
chr2-91323500-INS-160
chr2-91332545-INS-82
chr2-91334461-INS-255
chr2-91348013-INS-168
chr2-91374353-INS-175
chr2-91385299-INS-60
chr2-91414133-INS-193
chr2-91421601-INS-139
chr2-91455504-INS-289
chr2-91457701-INS-851
chr2-91513729-INS-99
chr2-91520504-INS-50
chr2-91524165-INS-88
chr2-91535439-INS-57
chr2-91545552-INS-219
chr2-91554739-INS-104
chr2-91559933-INS-52
chr2-91579945-INS-196
chr2-91584004-INS-209
chr2-91588903-INS-174
chr2-91590544-INS-293

chr2-91599717-INS-179
chr2-91617743-INS-62
chr2-91644621-INS-221
chr2-91692336-INS-264
chr2-91729291-INS-58
chr2-91743719-INS-53
chr2-91744729-INS-179
chr2-91772554-INS-194
chr2-91777754-INS-60
chr2-91781853-INS-846
chr2-91781873-INS-89
chr2-91786869-INS-135
chr2-91797725-INS-55
chr2-91807819-INS-65
chr2-91811957-INS-97
chr2-91829707-INS-715
chr2-91831059-INS-99
chr2-91855524-INS-70
chr2-91865936-INS-52
chr2-91905871-INS-169
chr2-91910162-INS-529
chr2-91911783-INS-75
chr2-91920210-INS-66
chr2-91924057-INS-60
chr2-91925337-INS-56
chr2-91933516-INS-56
chr2-91934171-INS-108
chr2-91936079-INS-434
chr2-91940120-INS-79
chr2-91956648-INS-255
chr2-91962895-INS-208
chr2-91971808-INS-244
chr2-91973930-INS-94
chr2-91982096-INS-252
chr2-92003507-INS-265
chr2-92038326-INS-85
chr2-92049378-INS-172
chr2-92074348-INS-52
chr2-92123110-INS-128
chr2-92143199-INS-574
chr2-92165152-INS-112
chr2-92170137-INS-90
chr2-92183101-INS-80
chr2-92233235-INS-52
chr2-92255957-INS-793
chr2-92304839-INS-281
chr2-92309817-INS-591
chr2-92312680-INS-60
chr2-92323527-INS-51
chr2-92342408-INS-126
chr2-92346543-INS-211
chr2-92349193-INS-341
chr2-92369471-INS-95
chr2-92383998-INS-246
chr2-92412914-INS-62
chr2-92479477-INS-56
chr2-92487463-INS-345
chr2-92491319-INS-293
chr2-92498525-INS-599
chr2-92500228-INS-2601
chr2-92541181-INS-187
chr2-92576401-INS-3377
chr2-92587407-INS-63
chr2-92610719-INS-103
chr2-92616807-INS-562
chr2-92621113-INS-102
chr2-92637604-INS-74
chr2-92660352-INS-88
chr2-92662756-INS-132
chr2-92665905-INS-100
chr2-92670912-INS-78
chr2-92672632-INS-64
chr2-92678039-INS-313
chr2-92694169-INS-306
chr2-92700563-INS-56
chr2-92702684-INS-55
chr2-92708223-INS-87

chr2-92710530-INS-207
chr2-92738442-INS-54
chr2-92749402-INS-237
chr2-92790718-INS-66
chr2-92794465-INS-171
chr2-92836895-INS-78
chr2-92864154-INS-52
chr2-92878920-INS-112
chr2-92887515-INS-62
chr2-92894072-INS-52
chr2-92916443-INS-397
chr2-92940504-INS-151
chr2-92959873-INS-150
chr2-92962097-INS-64
chr2-92970255-INS-84
chr2-92995284-INS-71
chr2-93014964-INS-139
chr2-93055901-INS-132
chr2-93073369-INS-96
chr2-93080409-INS-94
chr2-93109341-INS-89
chr2-93136714-INS-77
chr2-93183003-INS-83
chr2-93213794-INS-52
chr2-93225838-INS-188
chr2-93227323-INS-170
chr2-93274802-INS-205
chr2-93280484-INS-162
chr2-93299767-INS-515
chr2-93301866-INS-151
chr2-93311531-INS-189
chr2-93353648-INS-831
chr2-93364740-INS-56
chr2-93380826-INS-176
chr2-93389593-INS-2442
chr2-93411618-INS-213
chr2-93463876-INS-183
chr2-93496823-INS-87
chr2-93512308-INS-357
chr2-93553303-INS-80
chr2-93557664-INS-1151
chr2-93564615-INS-124
chr2-93572960-INS-563
chr2-93582343-INS-209
chr2-93604126-INS-246
chr2-93606691-INS-70
chr2-93634138-INS-174
chr2-93640044-INS-56
chr2-93645084-INS-118
chr2-93682504-INS-632
chr2-93682508-INS-505
chr2-93690623-INS-60
chr2-93707250-INS-113
chr2-93710508-INS-728
chr2-93713995-INS-144
chr2-93725681-INS-723
chr2-93750273-INS-1891
chr2-93801829-INS-240
chr2-93812107-INS-472
chr2-93820196-INS-70
chr2-93851831-INS-52
chr2-93947952-INS-5217
chr2-93969347-INS-5549
chr2-93969879-INS-52
chr2-93975682-INS-58
chr2-94035523-INS-83
chr2-94037186-INS-464
chr2-94049593-INS-5168
chr2-94051244-INS-196
chr2-94062544-INS-144
chr2-94116944-INS-5581
chr2-94122617-INS-398
chr2-94137359-INS-585
chr2-94141448-INS-240
chr2-94143697-INS-57
chr2-94175776-INS-238
chr2-94179327-INS-144

chr2-94220367-INS-270
chr2-94221670-INS-165
chr2-94230291-INS-183
chr2-94269952-INS-74
chr2-94278422-INS-219
chr2-94279662-INS-68
chr2-94323052-INS-215
chr2-94354852-INS-204
chr2-94373962-INS-476
chr2-94409759-INS-86
chr2-94443097-INS-6166
chr2-94443753-INS-146
chr2-94475471-INS-344
chr2-94477284-INS-64
chr2-94496857-INS-84
chr2-94503317-INS-7302
chr2-94504676-INS-55
chr2-94508477-INS-7297
chr2-94516174-INS-242
chr2-94518418-INS-6434
chr2-94518477-INS-182
chr2-94532583-INS-487
chr2-94544810-INS-8210
chr2-94555658-INS-54
chr2-94572078-INS-73
chr2-94582388-INS-83
chr2-94612896-INS-328
chr2-94643589-INS-68
chr2-94653045-INS-52
chr2-94659654-INS-590
chr2-94690246-INS-644
chr2-94714602-INS-653
chr2-94748729-INS-291
chr2-94787006-INS-60
chr2-94799825-INS-2732
chr2-94813093-INS-61
chr2-94813284-INS-5262
chr2-94830439-INS-6432
chr2-94840893-INS-149
chr2-94848586-INS-60
chr2-94883628-INS-616
chr2-94913109-INS-507
chr2-94935238-INS-263
chr2-94938224-INS-93
chr2-94950222-INS-770
chr2-95018743-INS-131
chr2-95021324-INS-260
chr2-95026341-INS-54
chr2-95041593-INS-2784
chr2-95068596-INS-358
chr2-95079870-INS-521
chr2-95090531-INS-4410
chr2-95132499-INS-5772
chr2-95158712-INS-4990
chr2-95161379-INS-250
chr2-95174500-INS-66
chr2-95183939-INS-387
chr2-95192087-INS-70
chr2-95208242-INS-188
chr2-95242886-INS-89
chr2-95244611-INS-1060
chr2-95253627-INS-7002
chr2-95256423-INS-60
chr2-95295113-INS-54
chr2-95302095-INS-208
chr2-95332103-INS-83
chr2-95363448-INS-584
chr2-95382379-INS-2255
chr2-95414305-INS-71
chr2-95433272-INS-74
chr2-95435549-INS-6190
chr2-95438549-INS-56
chr2-95490793-INS-74
chr2-95501005-INS-389
chr2-95511862-INS-102
chr2-95517738-INS-120
chr2-95524740-INS-64

chr2-95531451-INS-136
chr2-95539940-INS-224
chr2-95549123-INS-78
chr2-95572596-INS-445
chr2-95573566-INS-2976
chr2-95573567-INS-477
chr2-95580941-INS-82
chr2-95599700-INS-140
chr2-95607936-INS-58
chr2-95612920-INS-907
chr2-95621237-INS-330
chr2-95622943-INS-66
chr2-95688908-INS-237
chr2-95706771-INS-118
chr2-95732725-INS-92
chr2-95744859-INS-95
chr2-95747416-INS-70
chr2-95765567-INS-152
chr2-95776486-INS-126
chr2-95787733-INS-260
chr2-95814568-INS-1890
chr2-95820588-INS-51
chr2-95837455-INS-62
chr2-95847948-INS-52
chr2-95873137-INS-526
chr2-95888127-INS-70
chr2-95897013-INS-160
chr2-95912013-INS-528
chr2-95921579-INS-76
chr2-95939472-INS-51
chr2-95950727-INS-7324
chr2-95953564-INS-145
chr2-95967110-INS-385
chr2-95980723-INS-309
chr2-95997601-INS-72
chr2-96019288-INS-70
chr2-96031151-INS-623
chr2-96043967-INS-52
chr2-96044651-INS-498
chr2-96061986-INS-532
chr2-96068458-INS-71
chr2-96106802-INS-103
chr2-96126533-INS-458
chr2-96161468-INS-62
chr2-96179959-INS-74
chr2-96187312-INS-791
chr2-96191097-INS-88
chr2-96194943-INS-731
chr2-96200593-INS-54
chr2-96243325-INS-52
chr2-96253933-INS-60
chr2-96320132-INS-596
chr2-96325418-INS-70
chr2-96375362-INS-387
chr2-96383982-INS-384
chr2-96407430-INS-222
chr2-96451371-INS-105
chr2-96456420-INS-90
chr2-96487586-INS-64
chr2-96528452-INS-1593
chr2-96551761-INS-275
chr2-96565344-INS-209
chr2-96626936-INS-92
chr2-96652748-INS-6646
chr2-96673759-INS-341
chr2-96700239-INS-109
chr2-96709908-INS-147
chr2-96713699-INS-99
chr2-96716830-INS-358
chr2-96745094-INS-60
chr2-96757879-INS-52
chr2-96782239-INS-52
chr2-96783319-INS-97
chr2-96794228-INS-57
chr2-96795931-INS-90
chr2-96819148-INS-6672
chr2-96824265-INS-969

chr2-96827006-INS-59
chr2-96834024-INS-815
chr2-96841919-INS-52
chr2-96909918-INS-155
chr2-96930048-INS-203
chr2-96947104-INS-651
chr2-96979210-INS-64
chr2-97002454-INS-6312
chr2-97039721-INS-53
chr2-97050916-INS-354
chr2-97070219-INS-104
chr2-97078490-INS-758
chr2-97095186-INS-96
chr2-97105270-INS-51
chr2-97108896-INS-2207
chr2-97109834-INS-744
chr2-97115730-INS-627
chr2-97132235-INS-87
chr2-97170969-INS-818
chr2-97204898-INS-142
chr2-97212117-INS-122
chr2-97235003-INS-901
chr2-97261886-INS-983
chr2-97264537-INS-561
chr2-97279147-INS-872
chr2-97300226-INS-64
chr2-97302051-INS-84
chr2-97373311-INS-988
chr2-97381240-INS-94
chr2-97393394-INS-75
chr2-97414474-INS-76
chr2-97415808-INS-285
chr2-97417934-INS-55
chr2-97418690-INS-805
chr2-97419974-INS-6442
chr2-97452676-INS-65
chr2-97502179-INS-1132
chr2-97517425-INS-56
chr2-97521952-INS-120
chr2-97536702-INS-70
chr2-97547913-INS-236
chr2-97553771-INS-1233
chr2-97555355-INS-6401
chr2-97574059-INS-392
chr2-97588021-INS-56
chr2-97589549-INS-60
chr2-97602951-INS-2320
chr2-97606079-INS-342
chr2-97619832-INS-143
chr2-97647308-INS-298
chr2-97650642-INS-57
chr2-97661752-INS-502
chr2-97675009-INS-15515
chr2-97682299-INS-1421
chr2-97683247-INS-6842
chr2-97685488-INS-129
chr2-97688119-INS-6166
chr2-97692120-INS-4101
chr2-97701858-INS-434
chr2-97719354-INS-57
chr2-97783886-INS-155
chr2-97794995-INS-499
chr2-97830858-INS-305
chr2-97834671-INS-95
chr2-97891289-INS-71
chr2-97917200-INS-669
chr2-97936618-INS-640
chr2-97971120-INS-338
chr2-98042711-INS-72
chr2-98049516-INS-492
chr2-98079660-INS-776
chr2-98090908-INS-76
chr2-98125213-INS-365
chr2-98148469-INS-333
chr2-98266742-INS-155
chr2-98268993-INS-54
chr2-98316860-INS-93

chr2-98317515-INS-54
chr2-98358086-INS-54
chr2-98360460-INS-396
chr2-98389967-INS-525
chr2-98432353-INS-71
chr2-98448961-INS-51
chr2-98451211-INS-6407
chr2-98454615-INS-1115
chr2-98454649-INS-7581
chr2-98475192-INS-426
chr2-98485209-INS-1342
chr2-98510845-INS-219
chr2-98554552-INS-545
chr2-98576244-INS-596
chr2-98605637-INS-91
chr2-98619620-INS-561
chr2-98642127-INS-52
chr2-98669046-INS-84
chr2-98675973-INS-583
chr2-98735971-INS-677
chr2-98784544-INS-139
chr2-98791783-INS-210
chr2-98800404-INS-493
chr2-98812520-INS-58
chr2-98814121-INS-844
chr2-98828274-INS-600
chr2-98870397-INS-102
chr2-98895029-INS-509
chr2-98908071-INS-75
chr2-98913373-INS-137
chr2-98973522-INS-54
chr2-98975612-INS-714
chr2-99002351-INS-54
chr2-99037514-INS-231
chr2-99097113-INS-202
chr2-99102993-INS-100
chr2-99160756-INS-66
chr2-99178407-INS-58
chr2-99206338-INS-474
chr2-99208969-INS-178
chr2-99259002-INS-83
chr2-99299552-INS-66
chr2-99302688-INS-54
chr2-99307325-INS-121
chr2-99310683-INS-176
chr2-99331613-INS-471
chr2-99349972-INS-348
chr2-99356402-INS-117
chr2-99389629-INS-563
chr2-99455967-INS-6608
chr2-99462452-INS-6442
chr2-99490933-INS-850
chr2-99499318-INS-284
chr2-99503577-INS-867
chr2-99525357-INS-1386
chr2-99537394-INS-52
chr2-99547933-INS-297
chr2-99551668-INS-142
chr2-99554072-INS-50
chr2-99580896-INS-135
chr2-99585546-INS-168
chr2-99615989-INS-5549
chr2-99643804-INS-74
chr2-99647877-INS-195
chr2-99653989-INS-708
chr2-99690867-INS-92
chr2-99692377-INS-942
chr2-99694051-INS-118
chr2-99696116-INS-51
chr2-99698976-INS-52
chr2-99711267-INS-6331
chr2-99714942-INS-59
chr2-99720130-INS-102
chr2-99727372-INS-168
chr2-99746087-INS-6577
chr2-99749011-INS-804
chr2-99770542-INS-199

chr2-99773233-INS-431
chr2-99774322-INS-96
chr2-99785239-INS-227
chr2-99819114-INS-165
chr2-99826227-INS-62
chr2-99887497-INS-80
chr2-99915895-INS-112
chr2-99937324-INS-7110
chr2-99975670-INS-220
chr2-99977409-INS-130
chr2-100009025-INS-6615
chr2-100009523-INS-88
chr2-100009875-INS-6263
chr2-100017276-INS-223
chr2-100021685-INS-5157
chr2-100041600-INS-999
chr2-100077064-INS-187
chr2-100082205-INS-102
chr2-100089106-INS-80
chr2-100124345-INS-627
chr2-100129320-INS-6140
chr2-100154481-INS-358
chr2-100160483-INS-56
chr2-100166899-INS-408
chr2-100168376-INS-532
chr2-100175087-INS-796
chr2-100188876-INS-125
chr2-100197839-INS-312
chr2-100236846-INS-63
chr2-100269287-INS-54
chr2-100272403-INS-831
chr2-100286410-INS-1676
chr2-100319802-INS-1660
chr2-100382765-INS-100
chr2-100402280-INS-7293
chr2-100422256-INS-920
chr2-100446251-INS-155
chr2-100596640-INS-206
chr2-100637904-INS-737
chr2-100663838-INS-505
chr2-100679910-INS-65
chr2-100684118-INS-120
chr2-100686165-INS-66
chr2-100699439-INS-52
chr2-100707670-INS-998
chr2-100719850-INS-1483
chr2-100723020-INS-5092
chr2-100726983-INS-3288
chr2-100739752-INS-578
chr2-100777560-INS-63
chr2-100787957-INS-52
chr2-100800693-INS-206
chr2-100818669-INS-385
chr2-100826491-INS-243
chr2-100832163-INS-6384
chr2-100838838-INS-96
chr2-100858809-INS-56
chr2-100861786-INS-199
chr2-100877044-INS-68
chr2-100896888-INS-68
chr2-100914648-INS-299
chr2-100933230-INS-376
chr2-101008514-INS-92
chr2-101009199-INS-630
chr2-101081141-INS-75
chr2-101086397-INS-62
chr2-101091718-INS-72
chr2-101107272-INS-76
chr2-101109452-INS-377
chr2-101124049-INS-109
chr2-101128163-INS-70
chr2-101129840-INS-375
chr2-101131354-INS-7273
chr2-101132543-INS-6869
chr2-101150594-INS-6413
chr2-101172871-INS-645
chr2-101174126-INS-58

chr2-101201266-INS-87
chr2-101217089-INS-7129
chr2-101239769-INS-395
chr2-101243250-INS-75
chr2-101245051-INS-1983
chr2-101251533-INS-277
chr2-101258841-INS-949
chr2-101301599-INS-6683
chr2-101302087-INS-557
chr2-101307938-INS-155
chr2-101352686-INS-114
chr2-101357133-INS-187
chr2-101376925-INS-357
chr2-101399275-INS-126
chr2-101420737-INS-899
chr2-101426003-INS-669
chr2-101452201-INS-611
chr2-101488265-INS-122
chr2-101507777-INS-64
chr2-101509505-INS-60
chr2-101539401-INS-198
chr2-101541724-INS-53
chr2-101560628-INS-154
chr2-101574107-INS-193
chr2-101576425-INS-114
chr2-101591152-INS-233
chr2-101604074-INS-61
chr2-101619493-INS-74
chr2-101629119-INS-61
chr2-101639326-INS-53
chr2-101680428-INS-119
chr2-101699538-INS-121
chr2-101794554-INS-65
chr2-101826097-INS-2722
chr2-101828698-INS-6564
chr2-101841952-INS-52
chr2-101848559-INS-81
chr2-101857116-INS-51
chr2-101886327-INS-182
chr2-101907356-INS-108
chr2-101925851-INS-62
chr2-101927410-INS-165
chr2-101953216-INS-241
chr2-102007904-INS-53
chr2-102042986-INS-143
chr2-102089095-INS-275
chr2-102103219-INS-98
chr2-102114420-INS-391
chr2-102125860-INS-441
chr2-102155659-INS-106
chr2-102165634-INS-108
chr2-102196843-INS-6442
chr2-102199230-INS-255
chr2-102200958-INS-67
chr2-102245542-INS-5016
chr2-102250334-INS-93
chr2-102264052-INS-87
chr2-102266747-INS-166
chr2-102278102-INS-250
chr2-102308389-INS-112
chr2-102309756-INS-555
chr2-102313391-INS-128
chr2-102317911-INS-151
chr2-102330646-INS-97
chr2-102339676-INS-320
chr2-102347297-INS-80
chr2-102348514-INS-115
chr2-102352053-INS-339
chr2-102366529-INS-53
chr2-102371668-INS-122
chr2-102378811-INS-52
chr2-102397769-INS-311
chr2-102407053-INS-6520
chr2-102415073-INS-53
chr2-102450110-INS-222
chr2-102484039-INS-73
chr2-102492209-INS-54

chr2-102529045-INS-55
chr2-102545004-INS-82
chr2-102637912-INS-214
chr2-102640170-INS-302
chr2-102665511-INS-111
chr2-102688230-INS-6299
chr2-102710452-INS-124
chr2-102733845-INS-554
chr2-102742084-INS-75
chr2-102769310-INS-597
chr2-102782007-INS-57
chr2-102790371-INS-72
chr2-102795171-INS-927
chr2-102795885-INS-190
chr2-102796297-INS-6440
chr2-102804202-INS-159
chr2-102849858-INS-683
chr2-102925178-INS-57
chr2-102929143-INS-415
chr2-102936741-INS-119
chr2-102945860-INS-83
chr2-102966005-INS-59
chr2-102999835-INS-70
chr2-103031297-INS-118
chr2-103050668-INS-55
chr2-103054081-INS-218
chr2-103055522-INS-55
chr2-103071833-INS-65
chr2-103078101-INS-56
chr2-103102497-INS-51
chr2-103108486-INS-82
chr2-103126822-INS-283
chr2-103136858-INS-52
chr2-103144991-INS-267
chr2-103179885-INS-183
chr2-103182171-INS-58
chr2-103185958-INS-60
chr2-103212363-INS-61
chr2-103226050-INS-297
chr2-103235882-INS-59
chr2-103261065-INS-195
chr2-103301965-INS-84
chr2-103317098-INS-358
chr2-103327680-INS-501
chr2-103335073-INS-167
chr2-103343932-INS-122
chr2-103447509-INS-56
chr2-103460091-INS-171
chr2-103464375-INS-62
chr2-103491082-INS-1888
chr2-103510554-INS-273
chr2-103556343-INS-52
chr2-103570353-INS-75
chr2-103602501-INS-198
chr2-103607521-INS-119
chr2-103635707-INS-609
chr2-103650212-INS-94
chr2-103656166-INS-158
chr2-103664765-INS-223
chr2-103689508-INS-153
chr2-103699164-INS-192
chr2-103701754-INS-197
chr2-103727318-INS-99
chr2-103757073-INS-179
chr2-103790932-INS-532
chr2-103805052-INS-108
chr2-103848817-INS-314
chr2-103873656-INS-304
chr2-103877082-INS-51
chr2-103889267-INS-60
chr2-103900929-INS-62
chr2-103915009-INS-54
chr2-103941814-INS-51
chr2-103950583-INS-7482
chr2-103956717-INS-78
chr2-103957344-INS-532
chr2-103980047-INS-179

chr2-104017978-INS-235
chr2-104032658-INS-98
chr2-104036832-INS-623
chr2-104074664-INS-319
chr2-104095499-INS-67
chr2-104101820-INS-88
chr2-104126580-INS-136
chr2-104135039-INS-124
chr2-104136148-INS-249
chr2-104141941-INS-59
chr2-104153032-INS-108
chr2-104175288-INS-70
chr2-104205600-INS-126
chr2-104208432-INS-266
chr2-104211057-INS-82
chr2-104228086-INS-7140
chr2-104237651-INS-408
chr2-104246085-INS-532
chr2-104251962-INS-55
chr2-104252554-INS-210
chr2-104278559-INS-79
chr2-104283569-INS-74
chr2-104297980-INS-66
chr2-104299330-INS-172
chr2-104302878-INS-791
chr2-104330874-INS-86
chr2-104333102-INS-622
chr2-104342837-INS-106
chr2-104347796-INS-213
chr2-104348726-INS-68
chr2-104351607-INS-69
chr2-104352824-INS-890
chr2-104359958-INS-266
chr2-104372435-INS-562
chr2-104380793-INS-503
chr2-104393418-INS-58
chr2-104404954-INS-325
chr2-104416737-INS-95
chr2-104464677-INS-84
chr2-104471433-INS-266
chr2-104486795-INS-318
chr2-104489739-INS-63
chr2-104495673-INS-100
chr2-104502168-INS-212
chr2-104512674-INS-78
chr2-104530712-INS-567
chr2-104545895-INS-165
chr2-104548651-INS-172
chr2-104575953-INS-128
chr2-104608003-INS-72
chr2-104632769-INS-2757
chr2-104637544-INS-52
chr2-104642291-INS-184
chr2-104642875-INS-560
chr2-104655065-INS-51
chr2-104659880-INS-196
chr2-104669971-INS-212
chr2-104695425-INS-193
chr2-104705093-INS-92
chr2-104715115-INS-236
chr2-104721638-INS-164
chr2-104728932-INS-6358
chr2-104738465-INS-187
chr2-104758921-INS-95
chr2-104779811-INS-393
chr2-104815867-INS-76
chr2-104873401-INS-365
chr2-104890015-INS-52
chr2-104904249-INS-225
chr2-104915377-INS-121
chr2-104942844-INS-52
chr2-104955172-INS-67
chr2-104960174-INS-61
chr2-104982754-INS-54
chr2-104984344-INS-89
chr2-104990525-INS-53
chr2-105003264-INS-61

chr2-105017551-INS-60
chr2-105018353-INS-270
chr2-105042679-INS-70
chr2-105056091-INS-65
chr2-105065095-INS-222
chr2-105083299-INS-504
chr2-105105537-INS-54
chr2-105108485-INS-109
chr2-105113519-INS-70
chr2-105120124-INS-199
chr2-105139639-INS-549
chr2-105155096-INS-577
chr2-105156087-INS-192
chr2-105161626-INS-5299
chr2-105162867-INS-606
chr2-105168697-INS-93
chr2-105193380-INS-219
chr2-105199412-INS-697
chr2-105214910-INS-81
chr2-105225744-INS-64
chr2-105236646-INS-51
chr2-105239942-INS-143
chr2-105251074-INS-75
chr2-105257274-INS-264
chr2-105258693-INS-6442
chr2-105261951-INS-58
chr2-105292528-INS-520
chr2-105302027-INS-63
chr2-105306072-INS-136
chr2-105308235-INS-7151
chr2-105311625-INS-57
chr2-105325998-INS-52
chr2-105327251-INS-507
chr2-105334401-INS-233
chr2-105337381-INS-63
chr2-105338422-INS-1028
chr2-105366907-INS-230
chr2-105386433-INS-751
chr2-105394626-INS-6160
chr2-105449402-INS-62
chr2-105479132-INS-56
chr2-105503294-INS-54
chr2-105514019-INS-119
chr2-105526431-INS-87
chr2-105580544-INS-442
chr2-105597491-INS-139
chr2-105599265-INS-208
chr2-105674773-INS-6404
chr2-105699221-INS-6506
chr2-105713969-INS-6664
chr2-105741356-INS-109
chr2-105760044-INS-135
chr2-105781306-INS-399
chr2-105844329-INS-11759
chr2-105848547-INS-551
chr2-105870073-INS-58
chr2-105878578-INS-486
chr2-105879889-INS-274
chr2-105905913-INS-52
chr2-105971690-INS-222
chr2-105972984-INS-611
chr2-106001447-INS-52
chr2-106014613-INS-83
chr2-106075401-INS-1237
chr2-106080247-INS-200
chr2-106098505-INS-64
chr2-106109449-INS-295
chr2-106111238-INS-793
chr2-106120801-INS-94
chr2-106130880-INS-115
chr2-106146155-INS-54
chr2-106190096-INS-611
chr2-106243622-INS-66
chr2-106253937-INS-72
chr2-106324563-INS-117
chr2-106350517-INS-54
chr2-106365710-INS-55

chr2-106395130-INS-64
chr2-106434118-INS-68
chr2-106447037-INS-221
chr2-106452672-INS-730
chr2-106460894-INS-66
chr2-106470980-INS-4408
chr2-106484786-INS-7214
chr2-106487102-INS-7292
chr2-106511047-INS-81
chr2-106525792-INS-52
chr2-106556874-INS-62
chr2-106570333-INS-84
chr2-106581616-INS-98
chr2-106616644-INS-70
chr2-106636470-INS-213
chr2-106642570-INS-90
chr2-106645171-INS-67
chr2-106710126-INS-5133
chr2-106734205-INS-326
chr2-106741312-INS-84
chr2-106809408-INS-1533
chr2-106810263-INS-73
chr2-106813662-INS-51
chr2-106852230-INS-131
chr2-106877891-INS-180
chr2-106899279-INS-6895
chr2-106912602-INS-124
chr2-106914530-INS-219
chr2-106926691-INS-91
chr2-106940412-INS-7136
chr2-106946416-INS-56
chr2-106956331-INS-90
chr2-106992885-INS-7013
chr2-106995886-INS-500
chr2-107013422-INS-122
chr2-107021044-INS-326
chr2-107025972-INS-629
chr2-107034723-INS-65
chr2-107043663-INS-185
chr2-107058405-INS-94
chr2-107092016-INS-64
chr2-107099142-INS-196
chr2-107106256-INS-6216
chr2-107174300-INS-159
chr2-107242349-INS-68
chr2-107279252-INS-322
chr2-107317072-INS-162
chr2-107330893-INS-72
chr2-107393659-INS-70
chr2-107410729-INS-152
chr2-107423755-INS-64
chr2-107434823-INS-76
chr2-107459499-INS-62
chr2-107464158-INS-73
chr2-107470355-INS-618
chr2-107479878-INS-158
chr2-107491283-INS-91
chr2-107503772-INS-78
chr2-107512697-INS-50
chr2-107517216-INS-56
chr2-107546552-INS-336
chr2-107550668-INS-65
chr2-107557772-INS-6193
chr2-107612954-INS-400
chr2-107627480-INS-440
chr2-107638366-INS-54
chr2-107644510-INS-105
chr2-107658521-INS-315
chr2-107685677-INS-134
chr2-107705013-INS-52
chr2-107736863-INS-195
chr2-107772210-INS-7774
chr2-107805290-INS-837
chr2-107836357-INS-6240
chr2-107851547-INS-70
chr2-107859529-INS-218
chr2-107912187-INS-302

chr2-107920449-INS-51
chr2-107946341-INS-52
chr2-107961154-INS-383
chr2-107970344-INS-141
chr2-108000424-INS-7464
chr2-108000548-INS-176
chr2-108028135-INS-99
chr2-108032758-INS-431
chr2-108057689-INS-63
chr2-108059147-INS-650
chr2-108070336-INS-166
chr2-108082362-INS-228
chr2-108135672-INS-82
chr2-108151743-INS-2474
chr2-108196007-INS-76
chr2-108202259-INS-569
chr2-108205699-INS-426
chr2-108227040-INS-3091
chr2-108229034-INS-76
chr2-108291951-INS-66
chr2-108332813-INS-517
chr2-108337385-INS-207
chr2-108346858-INS-82
chr2-108351878-INS-359
chr2-108369177-INS-635
chr2-108394475-INS-149
chr2-108410305-INS-356
chr2-108429090-INS-11621
chr2-108429832-INS-1646
chr2-108459957-INS-6392
chr2-108474501-INS-68
chr2-108476408-INS-780
chr2-108485483-INS-432
chr2-108493054-INS-55
chr2-108494220-INS-317
chr2-108497599-INS-783
chr2-108533671-INS-490
chr2-108557517-INS-101
chr2-108583816-INS-76
chr2-108605251-INS-496
chr2-108618705-INS-58
chr2-108638168-INS-109
chr2-108653871-INS-656
chr2-108654120-INS-6391
chr2-108667182-INS-119
chr2-108683608-INS-555
chr2-108693953-INS-214
chr2-108700223-INS-271
chr2-108701084-INS-7326
chr2-108715977-INS-251
chr2-108734381-INS-159
chr2-108743380-INS-222
chr2-108744967-INS-67
chr2-108754266-INS-96
chr2-108763628-INS-7108
chr2-108770048-INS-227
chr2-108806651-INS-474
chr2-108806950-INS-691
chr2-108811196-INS-56
chr2-108813145-INS-525
chr2-108837910-INS-852
chr2-108862096-INS-241
chr2-108869394-INS-54
chr2-108885702-INS-76
chr2-108903950-INS-3168
chr2-108939741-INS-117
chr2-108948955-INS-227
chr2-108962562-INS-78
chr2-108983114-INS-168
chr2-108988374-INS-3760
chr2-108996987-INS-492
chr2-109064081-INS-490
chr2-109092298-INS-6548
chr2-109117714-INS-68
chr2-109141816-INS-100
chr2-109183174-INS-256
chr2-109192775-INS-54

chr2-109194980-INS-56
chr2-109203035-INS-295
chr2-109227452-INS-76
chr2-109232197-INS-106
chr2-109252866-INS-119
chr2-109258366-INS-88
chr2-109297372-INS-648
chr2-109317457-INS-70
chr2-109336809-INS-106
chr2-109400628-INS-72
chr2-109405946-INS-55
chr2-109410311-INS-137
chr2-109418484-INS-143
chr2-109425317-INS-62
chr2-109432305-INS-669
chr2-109487311-INS-98
chr2-109551603-INS-99
chr2-109561639-INS-108
chr2-109580121-INS-262
chr2-109586150-INS-266
chr2-109605726-INS-53
chr2-109620069-INS-215
chr2-109625227-INS-172
chr2-109637377-INS-59
chr2-109640172-INS-706
chr2-109645573-INS-6008
chr2-109648767-INS-103
chr2-109661481-INS-181
chr2-109670585-INS-64
chr2-109705235-INS-53
chr2-109710413-INS-7160
chr2-109715980-INS-147
chr2-109725582-INS-275
chr2-109740851-INS-267
chr2-109754713-INS-145
chr2-109780605-INS-57
chr2-109812036-INS-2654
chr2-109823043-INS-64
chr2-109869957-INS-628
chr2-109892318-INS-3989
chr2-109901282-INS-137
chr2-109908271-INS-52
chr2-109927682-INS-81
chr2-109941191-INS-245
chr2-109945870-INS-54
chr2-109946629-INS-186
chr2-109964693-INS-55
chr2-109981414-INS-157
chr2-109983916-INS-85
chr2-109996632-INS-91
chr2-109997687-INS-394
chr2-110035343-INS-997
chr2-110037560-INS-52
chr2-110050183-INS-53
chr2-110057867-INS-53
chr2-110058529-INS-456
chr2-110063501-INS-114
chr2-110065790-INS-400
chr2-110103210-INS-400
chr2-110111831-INS-293
chr2-110124006-INS-96
chr2-110136511-INS-480
chr2-110146519-INS-744
chr2-110177956-INS-5091
chr2-110214678-INS-1687
chr2-110216658-INS-374
chr2-110240887-INS-3561
chr2-110256261-INS-78
chr2-110269988-INS-561
chr2-110279195-INS-171
chr2-110282349-INS-51
chr2-110286115-INS-570
chr2-110296118-INS-1998
chr2-110308982-INS-6410
chr2-110312720-INS-63
chr2-110315659-INS-228
chr2-110331046-INS-7094

chr2-110339278-INS-313
chr2-110342742-INS-172
chr2-110348798-INS-6364
chr2-110353006-INS-739
chr2-110375379-INS-1281
chr2-110385308-INS-4873
chr2-110390460-INS-6942
chr2-110391975-INS-132
chr2-110393638-INS-182
chr2-110394976-INS-59
chr2-110396698-INS-175
chr2-110414518-INS-427
chr2-110455274-INS-221
chr2-110465590-INS-92
chr2-110480629-INS-249
chr2-110519377-INS-50
chr2-110522756-INS-52
chr2-110524921-INS-796
chr2-110530884-INS-157
chr2-110612917-INS-56
chr2-110620169-INS-78
chr2-110624978-INS-55
chr2-110648224-INS-511
chr2-110652362-INS-54
chr2-110663190-INS-65
chr2-110671277-INS-411
chr2-110672906-INS-686
chr2-110685463-INS-55
chr2-110714177-INS-169
chr2-110715521-INS-376
chr2-110716665-INS-113
chr2-110732184-INS-54
chr2-110786486-INS-653
chr2-110790830-INS-78
chr2-110798193-INS-4833
chr2-110810386-INS-280
chr2-110867074-INS-658
chr2-110935927-INS-83
chr2-110938280-INS-663
chr2-110955551-INS-6382
chr2-110967999-INS-239
chr2-110972137-INS-64
chr2-110992606-INS-261
chr2-110993344-INS-657
chr2-111012325-INS-83
chr2-111025620-INS-403
chr2-111072959-INS-368
chr2-111076353-INS-7068
chr2-111076354-INS-343
chr2-111076770-INS-7151
chr2-111091390-INS-194
chr2-111102549-INS-64
chr2-111150483-INS-65
chr2-111160723-INS-154
chr2-111248951-INS-68
chr2-111252282-INS-92
chr2-111275639-INS-11622
chr2-111298620-INS-483
chr2-111301225-INS-2380
chr2-111351224-INS-90
chr2-111363215-INS-52
chr2-111374317-INS-156
chr2-111405783-INS-354
chr2-111439090-INS-56
chr2-111444405-INS-394
chr2-111499625-INS-71
chr2-111500316-INS-963
chr2-111509756-INS-199
chr2-111526115-INS-115
chr2-111535922-INS-3509
chr2-111561365-INS-627
chr2-111577336-INS-6305
chr2-111594723-INS-6212
chr2-111597538-INS-60
chr2-111603913-INS-182
chr2-111605226-INS-68
chr2-111628736-INS-570

chr2-111654054-INS-70
chr2-111655668-INS-157
chr2-111700003-INS-61
chr2-111709234-INS-65
chr2-111731382-INS-5068
chr2-111776006-INS-235
chr2-111781374-INS-6686
chr2-111793934-INS-125
chr2-111809769-INS-74
chr2-111818753-INS-204
chr2-111837346-INS-60
chr2-111839834-INS-51
chr2-111857891-INS-236
chr2-111868509-INS-5623
chr2-111879466-INS-107
chr2-111883711-INS-56
chr2-111891278-INS-212
chr2-111907312-INS-2274
chr2-111923573-INS-75
chr2-111928184-INS-6021
chr2-111966863-INS-4559
chr2-111981022-INS-3419
chr2-112009875-INS-171
chr2-112045332-INS-104
chr2-112087989-INS-78
chr2-112093283-INS-980
chr2-112113555-INS-942
chr2-112118905-INS-77
chr2-112158393-INS-8734
chr2-112175467-INS-58
chr2-112183718-INS-76
chr2-112187624-INS-228
chr2-112230910-INS-154
chr2-112237297-INS-268
chr2-112255588-INS-438
chr2-112266552-INS-217
chr2-112314709-INS-242
chr2-112324577-INS-135
chr2-112346166-INS-201
chr2-112347726-INS-210
chr2-112354046-INS-120
chr2-112356034-INS-71
chr2-112401916-INS-2392
chr2-112409614-INS-5548
chr2-112421014-INS-367
chr2-112437919-INS-7167
chr2-112449625-INS-634
chr2-112464131-INS-197
chr2-112472788-INS-158
chr2-112512985-INS-120
chr2-112523514-INS-264
chr2-112530316-INS-6878
chr2-112542097-INS-105
chr2-112548303-INS-84
chr2-112626388-INS-390
chr2-112632740-INS-119
chr2-112636283-INS-589
chr2-112682985-INS-521
chr2-112693562-INS-235
chr2-112717051-INS-6628
chr2-112733909-INS-706
chr2-112758373-INS-56
chr2-112760177-INS-83
chr2-112768851-INS-185
chr2-112782519-INS-60
chr2-112793822-INS-102
chr2-112798470-INS-58
chr2-112847381-INS-486
chr2-112878755-INS-74
chr2-112889024-INS-752
chr2-112897390-INS-574
chr2-112904482-INS-387
chr2-112925429-INS-92
chr2-112977009-INS-6614
chr2-113022824-INS-57
chr2-113039188-INS-79
chr2-113050131-INS-403

chr2-113062716-INS-175
chr2-113110575-INS-5338
chr2-113119624-INS-66
chr2-113152618-INS-62
chr2-113153008-INS-306
chr2-113244901-INS-72
chr2-113254278-INS-83
chr2-113286152-INS-232
chr2-113320221-INS-66
chr2-113423719-INS-358
chr2-113479266-INS-140
chr2-113491817-INS-434
chr2-113501346-INS-315
chr2-113537371-INS-52
chr2-113548945-INS-156
chr2-113566624-INS-55
chr2-113593590-INS-168
chr2-113604088-INS-717
chr2-113607560-INS-2715
chr2-113613773-INS-52
chr2-113628693-INS-117
chr2-113634084-INS-92
chr2-113639728-INS-66
chr2-113674617-INS-306
chr2-113682493-INS-59
chr2-113689778-INS-6592
chr2-113689943-INS-6240
chr2-113712026-INS-56
chr2-113725101-INS-428
chr2-113732246-INS-86
chr2-113741649-INS-56
chr2-113812938-INS-107
chr2-113863414-INS-70
chr2-113891851-INS-560
chr2-113953264-INS-94
chr2-113985834-INS-450
chr2-114009265-INS-206
chr2-114058935-INS-199
chr2-114066296-INS-116
chr2-114071628-INS-63
chr2-114118138-INS-52
chr2-114139319-INS-646
chr2-114159320-INS-189
chr2-114162712-INS-126
chr2-114166939-INS-180
chr2-114182648-INS-52
chr2-114192810-INS-6830
chr2-114213660-INS-63
chr2-114218052-INS-412
chr2-114231962-INS-76
chr2-114236585-INS-394
chr2-114253815-INS-297
chr2-114257159-INS-124
chr2-114263050-INS-160
chr2-114283510-INS-73
chr2-114319856-INS-940
chr2-114406211-INS-141
chr2-114456454-INS-80
chr2-114486117-INS-202
chr2-114490778-INS-60
chr2-114504955-INS-66
chr2-114520833-INS-76
chr2-114523868-INS-52
chr2-114535816-INS-429
chr2-114545496-INS-263
chr2-114582325-INS-532
chr2-114596473-INS-115
chr2-114599235-INS-90
chr2-114611864-INS-816
chr2-114619676-INS-435
chr2-114649309-INS-94
chr2-114652948-INS-5091
chr2-114678366-INS-975
chr2-114686711-INS-10597
chr2-114695481-INS-177
chr2-114707912-INS-326
chr2-114722699-INS-253

chr2-114762364-INS-506
chr2-114786587-INS-6886
chr2-114811376-INS-1344
chr2-114811453-INS-129
chr2-114814743-INS-372
chr2-114823444-INS-2600
chr2-114824609-INS-7018
chr2-114830501-INS-232
chr2-114832711-INS-513
chr2-114832774-INS-7089
chr2-114851580-INS-132
chr2-114862175-INS-6181
chr2-114871568-INS-6506
chr2-114880371-INS-215
chr2-114883328-INS-6354
chr2-114903340-INS-205
chr2-114956766-INS-67
chr2-114971264-INS-6387
chr2-114977090-INS-78
chr2-114999057-INS-55
chr2-115016970-INS-924
chr2-115034688-INS-251
chr2-115037182-INS-75
chr2-115055897-INS-87
chr2-115062238-INS-777
chr2-115072255-INS-432
chr2-115105055-INS-303
chr2-115111462-INS-639
chr2-115122441-INS-59
chr2-115137031-INS-344
chr2-115137600-INS-54
chr2-115143274-INS-3346
chr2-115174483-INS-6811
chr2-115189749-INS-1864
chr2-115207752-INS-604
chr2-115254692-INS-81
chr2-115266269-INS-383
chr2-115271953-INS-190
chr2-115273681-INS-250
chr2-115279875-INS-808
chr2-115301372-INS-202
chr2-115306674-INS-208
chr2-115328987-INS-860
chr2-115354929-INS-114
chr2-115371931-INS-866
chr2-115376770-INS-82
chr2-115381968-INS-239
chr2-115384301-INS-6498
chr2-115391529-INS-376
chr2-115395529-INS-119
chr2-115402110-INS-1892
chr2-115419352-INS-826
chr2-115430379-INS-133
chr2-115486248-INS-796
chr2-115491540-INS-583
chr2-115527711-INS-179
chr2-115541027-INS-230
chr2-115569940-INS-62
chr2-115588203-INS-457
chr2-115615274-INS-300
chr2-115630639-INS-1853
chr2-115630640-INS-105
chr2-115630640-INS-800
chr2-115645392-INS-58
chr2-115734684-INS-70
chr2-115738736-INS-439
chr2-115740707-INS-76
chr2-115754698-INS-52
chr2-115801004-INS-478
chr2-115803933-INS-65
chr2-115804361-INS-180
chr2-115811330-INS-208
chr2-115830464-INS-85
chr2-115918573-INS-298
chr2-115924250-INS-129
chr2-115945997-INS-106
chr2-115953317-INS-550

chr2-115965783-INS-220
chr2-115972666-INS-56
chr2-115998578-INS-58
chr2-116030438-INS-65
chr2-116049585-INS-487
chr2-116052195-INS-220
chr2-116060651-INS-237
chr2-116082350-INS-152
chr2-116184692-INS-637
chr2-116186189-INS-88
chr2-116218280-INS-128
chr2-116222409-INS-89
chr2-116225697-INS-6786
chr2-116245407-INS-7966
chr2-116253340-INS-68
chr2-116254372-INS-55
chr2-116292764-INS-6172
chr2-116293309-INS-1019
chr2-116302781-INS-209
chr2-116306548-INS-173
chr2-116309871-INS-84
chr2-116344660-INS-13690
chr2-116344660-INS-83
chr2-116386088-INS-173
chr2-116403810-INS-554
chr2-116405536-INS-384
chr2-116413356-INS-277
chr2-116429412-INS-6271
chr2-116441744-INS-221
chr2-116442742-INS-3639
chr2-116450704-INS-342
chr2-116458305-INS-114
chr2-116466988-INS-181
chr2-116480179-INS-358
chr2-116490716-INS-95
chr2-116493417-INS-7239
chr2-116495441-INS-96
chr2-116498249-INS-309
chr2-116503647-INS-605
chr2-116527109-INS-110
chr2-116541816-INS-79
chr2-116544863-INS-192
chr2-116546427-INS-66
chr2-116552451-INS-6423
chr2-116615599-INS-5993
chr2-116618049-INS-85
chr2-116626998-INS-90
chr2-116633223-INS-176
chr2-116637192-INS-426
chr2-116658735-INS-105
chr2-116666111-INS-52
chr2-116673564-INS-222
chr2-116676930-INS-487
chr2-116683708-INS-53
chr2-116684074-INS-253
chr2-116686231-INS-52
chr2-116696890-INS-179
chr2-116721707-INS-532
chr2-116732936-INS-57
chr2-116734518-INS-614
chr2-116737577-INS-116
chr2-116756711-INS-62
chr2-116764764-INS-440
chr2-116765649-INS-62
chr2-116773862-INS-54
chr2-116775072-INS-933
chr2-116775245-INS-96
chr2-116788288-INS-52
chr2-116798150-INS-61
chr2-116808279-INS-56
chr2-116816983-INS-52
chr2-116837801-INS-106
chr2-116839303-INS-312
chr2-116865442-INS-51
chr2-116865516-INS-813
chr2-116885709-INS-113
chr2-116886795-INS-225

chr2-116891518-INS-95
chr2-116906006-INS-194
chr2-116906959-INS-63
chr2-116930459-INS-4695
chr2-116940577-INS-282
chr2-116945570-INS-363
chr2-116974749-INS-617
chr2-116975844-INS-71
chr2-117007492-INS-128
chr2-117034042-INS-328
chr2-117037391-INS-9163
chr2-117054837-INS-81
chr2-117079326-INS-51
chr2-117080067-INS-141
chr2-117086147-INS-64
chr2-117097132-INS-1793
chr2-117104287-INS-6217
chr2-117128380-INS-60
chr2-117134237-INS-60
chr2-117137836-INS-104
chr2-117188784-INS-90
chr2-117200721-INS-156
chr2-117218398-INS-143
chr2-117243422-INS-76
chr2-117262511-INS-71
chr2-117271173-INS-75
chr2-117274834-INS-561
chr2-117278833-INS-65
chr2-117288848-INS-339
chr2-117296031-INS-88
chr2-117311516-INS-150
chr2-117316170-INS-123
chr2-117330060-INS-82
chr2-117332481-INS-78
chr2-117333743-INS-126
chr2-117341945-INS-159
chr2-117353845-INS-200
chr2-117358633-INS-56
chr2-117363915-INS-663
chr2-117396071-INS-174
chr2-117399779-INS-138
chr2-117427389-INS-162
chr2-117430821-INS-6235
chr2-117444932-INS-54
chr2-117453898-INS-90
chr2-117475182-INS-864
chr2-117484577-INS-89
chr2-117496252-INS-70
chr2-117502537-INS-64
chr2-117518902-INS-77
chr2-117525416-INS-509
chr2-117527951-INS-55
chr2-117529143-INS-102
chr2-117565792-INS-648
chr2-117585472-INS-445
chr2-117607219-INS-579
chr2-117610265-INS-59
chr2-117616980-INS-400
chr2-117635086-INS-52
chr2-117668705-INS-62
chr2-117703681-INS-852
chr2-117730065-INS-65
chr2-117747660-INS-78
chr2-117782605-INS-72
chr2-117788997-INS-339
chr2-117791498-INS-78
chr2-117798994-INS-6442
chr2-117800240-INS-6379
chr2-117809068-INS-206
chr2-117853130-INS-82
chr2-117879783-INS-663
chr2-117888101-INS-115
chr2-117890111-INS-5990
chr2-117897670-INS-222
chr2-117909512-INS-97
chr2-117920944-INS-7261
chr2-117937405-INS-62

chr2-117976201-INS-140
chr2-118007710-INS-63
chr2-118029724-INS-292
chr2-118031660-INS-411
chr2-118038667-INS-58
chr2-118045094-INS-90
chr2-118052187-INS-61
chr2-118058176-INS-401
chr2-118072645-INS-165
chr2-118098907-INS-166
chr2-118137477-INS-52
chr2-118168021-INS-321
chr2-118182607-INS-83
chr2-118187205-INS-206
chr2-118191859-INS-141
chr2-118199827-INS-273
chr2-118210894-INS-125
chr2-118255010-INS-58
chr2-118267759-INS-478
chr2-118269381-INS-76
chr2-118275186-INS-866
chr2-118275273-INS-200
chr2-118275323-INS-250
chr2-118312818-INS-68
chr2-118326958-INS-312
chr2-118332910-INS-249
chr2-118333232-INS-624
chr2-118337064-INS-122
chr2-118347699-INS-67
chr2-118372072-INS-81
chr2-118395882-INS-208
chr2-118403992-INS-62
chr2-118405183-INS-66
chr2-118405824-INS-321
chr2-118424534-INS-157
chr2-118447415-INS-90
chr2-118465946-INS-62
chr2-118502571-INS-66
chr2-118512753-INS-66
chr2-118544125-INS-102
chr2-118570296-INS-196
chr2-118583494-INS-91
chr2-118601481-INS-198
chr2-118612532-INS-190
chr2-118637214-INS-244
chr2-118678773-INS-205
chr2-118683701-INS-361
chr2-118687809-INS-108
chr2-118693858-INS-311
chr2-118705000-INS-68
chr2-118707245-INS-196
chr2-118716425-INS-207
chr2-118767551-INS-50
chr2-118772767-INS-203
chr2-118774258-INS-68
chr2-118822424-INS-65
chr2-118830404-INS-364
chr2-118839164-INS-64
chr2-118852207-INS-216
chr2-118858087-INS-109
chr2-118858604-INS-224
chr2-118874613-INS-217
chr2-118893245-INS-174
chr2-118897819-INS-174
chr2-118908304-INS-321
chr2-118909875-INS-214
chr2-118944709-INS-198
chr2-118949595-INS-56
chr2-118961700-INS-88
chr2-118966994-INS-72
chr2-118968598-INS-225
chr2-118973312-INS-211
chr2-118979190-INS-76
chr2-118988523-INS-152
chr2-119008098-INS-189
chr2-119020668-INS-216
chr2-119025149-INS-388

chr2-119027724-INS-171
chr2-119038419-INS-211
chr2-119044966-INS-104
chr2-119048863-INS-226
chr2-119061267-INS-101
chr2-119079671-INS-1867
chr2-119083561-INS-123
chr2-119112105-INS-204
chr2-119116882-INS-198
chr2-119126041-INS-191
chr2-119149542-INS-86
chr2-119193397-INS-83
chr2-119209141-INS-144
chr2-119212252-INS-172
chr2-119219825-INS-273
chr2-119233815-INS-354
chr2-119236589-INS-203
chr2-119287888-INS-230
chr2-119290637-INS-71
chr2-119323951-INS-57
chr2-119328887-INS-193
chr2-119330798-INS-222
chr2-119336106-INS-57
chr2-119346738-INS-210
chr2-119358680-INS-197
chr2-119365281-INS-188
chr2-119389813-INS-204
chr2-119398093-INS-248
chr2-119402623-INS-62
chr2-119405820-INS-200
chr2-119411868-INS-201
chr2-119413469-INS-68
chr2-119419995-INS-154
chr2-119422078-INS-174
chr2-119432345-INS-86
chr2-119447103-INS-204
chr2-119479239-INS-261
chr2-119487483-INS-810
chr2-119487511-INS-206
chr2-119489672-INS-875
chr2-119516211-INS-203
chr2-119532966-INS-244
chr2-119542663-INS-199
chr2-119550243-INS-81
chr2-119558303-INS-191
chr2-119561767-INS-133
chr2-119611813-INS-85
chr2-119615548-INS-189
chr2-119618733-INS-361
chr2-119625644-INS-75
chr2-119670375-INS-219
chr2-119735977-INS-460
chr2-119763938-INS-178
chr2-119785692-INS-6401
chr2-119789206-INS-60
chr2-119791750-INS-225
chr2-119813481-INS-54
chr2-119830399-INS-112
chr2-119837517-INS-254
chr2-119906324-INS-147
chr2-119969351-INS-60
chr2-119976595-INS-419
chr2-119990628-INS-170
chr2-119997909-INS-283
chr2-120012687-INS-753
chr2-120038028-INS-65
chr2-120058253-INS-102
chr2-120063539-INS-196
chr2-120098382-INS-130
chr2-120106105-INS-68
chr2-120127765-INS-95
chr2-120131706-INS-199
chr2-120160566-INS-85
chr2-120173517-INS-204
chr2-120175844-INS-51
chr2-120185688-INS-215
chr2-120215848-INS-71

chr2-120242175-INS-60
chr2-120243983-INS-222
chr2-120263668-INS-202
chr2-120266665-INS-165
chr2-120317697-INS-215
chr2-120323718-INS-223
chr2-120325114-INS-6442
chr2-120372889-INS-176
chr2-120399025-INS-133
chr2-120427848-INS-148
chr2-120442414-INS-999
chr2-120449000-INS-832
chr2-120451360-INS-81
chr2-120481446-INS-302
chr2-120491403-INS-211
chr2-120498034-INS-259
chr2-120498363-INS-480
chr2-120500786-INS-252
chr2-120522293-INS-199
chr2-120558001-INS-95
chr2-120598114-INS-142
chr2-120609301-INS-174
chr2-120638992-INS-65
chr2-120664700-INS-99
chr2-120673570-INS-616
chr2-120675552-INS-60
chr2-120684514-INS-213
chr2-120687070-INS-90
chr2-120699747-INS-217
chr2-120753335-INS-683
chr2-120765436-INS-234
chr2-120770439-INS-67
chr2-120791270-INS-210
chr2-120817637-INS-399
chr2-120820506-INS-164
chr2-120827656-INS-95
chr2-120887312-INS-54
chr2-120899999-INS-102
chr2-120912525-INS-58
chr2-120918257-INS-104
chr2-120919592-INS-71
chr2-120920723-INS-129
chr2-120923122-INS-56
chr2-120979768-INS-249
chr2-120982085-INS-180
chr2-121015759-INS-169
chr2-121053108-INS-203
chr2-121060509-INS-103
chr2-121071191-INS-1000
chr2-121074303-INS-87
chr2-121096727-INS-224
chr2-121103692-INS-219
chr2-121106990-INS-166
chr2-121108914-INS-148
chr2-121129614-INS-197
chr2-121170729-INS-221
chr2-121174975-INS-53
chr2-121256625-INS-256
chr2-121259538-INS-63
chr2-121262329-INS-203
chr2-121266366-INS-68
chr2-121278106-INS-157
chr2-121282360-INS-5551
chr2-121285306-INS-56
chr2-121293258-INS-122
chr2-121299401-INS-210
chr2-121305202-INS-8940
chr2-121305386-INS-207
chr2-121308429-INS-213
chr2-121310470-INS-211
chr2-121315932-INS-230
chr2-121323404-INS-64
chr2-121331954-INS-210
chr2-121333996-INS-203
chr2-121343755-INS-100
chr2-121345319-INS-194
chr2-121365519-INS-311

chr2-121368847-INS-204
chr2-121405403-INS-60
chr2-121415280-INS-56
chr2-121429710-INS-56
chr2-121438299-INS-74
chr2-121442124-INS-159
chr2-121449985-INS-136
chr2-121456318-INS-78
chr2-121469762-INS-54
chr2-121477762-INS-493
chr2-121494542-INS-60
chr2-121537058-INS-113
chr2-121541819-INS-67
chr2-121566865-INS-94
chr2-121576529-INS-79
chr2-121589002-INS-765
chr2-121591667-INS-123
chr2-121611528-INS-204
chr2-121626126-INS-338
chr2-121629460-INS-171
chr2-121651008-INS-75
chr2-121664730-INS-52
chr2-121703333-INS-308
chr2-121707484-INS-836
chr2-121716932-INS-201
chr2-121726422-INS-62
chr2-121733791-INS-209
chr2-121739300-INS-161
chr2-121749678-INS-123
chr2-121764678-INS-136
chr2-121806957-INS-120
chr2-121817310-INS-181
chr2-121817710-INS-975
chr2-121840364-INS-205
chr2-121846503-INS-253
chr2-121851639-INS-199
chr2-121866370-INS-197
chr2-121881938-INS-60
chr2-121911204-INS-204
chr2-121928443-INS-488
chr2-121953755-INS-58
chr2-121972422-INS-108
chr2-121975184-INS-713
chr2-121976078-INS-51
chr2-122021959-INS-201
chr2-122023086-INS-323
chr2-122034633-INS-378
chr2-122064259-INS-245
chr2-122106624-INS-147
chr2-122118819-INS-54
chr2-122138626-INS-209
chr2-122172072-INS-79
chr2-122189117-INS-290
chr2-122222684-INS-64
chr2-122406694-INS-209
chr2-122441043-INS-325
chr2-122459917-INS-71
chr2-122470442-INS-63
chr2-122482733-INS-73
chr2-122495833-INS-70
chr2-122496126-INS-8985
chr2-122497452-INS-7096
chr2-122497988-INS-3246
chr2-122497989-INS-787
chr2-122518370-INS-183
chr2-122561154-INS-181
chr2-122572779-INS-704
chr2-122590418-INS-405
chr2-122592799-INS-137
chr2-122598544-INS-224
chr2-122620654-INS-89
chr2-122626607-INS-107
chr2-122629140-INS-209
chr2-122647270-INS-64
chr2-122653610-INS-463
chr2-122655888-INS-252
chr2-122712183-INS-124

chr2-122727423-INS-2138
chr2-122730875-INS-172
chr2-122751634-INS-76
chr2-122770003-INS-707
chr2-122830072-INS-601
chr2-122849398-INS-70
chr2-122858855-INS-319
chr2-122859681-INS-6459
chr2-122865114-INS-400
chr2-122879613-INS-6295
chr2-122908579-INS-128
chr2-122916730-INS-82
chr2-122919304-INS-68
chr2-122922009-INS-56
chr2-122970763-INS-52
chr2-122971357-INS-9344
chr2-122989195-INS-941
chr2-123007729-INS-228
chr2-123025914-INS-239
chr2-123031116-INS-117
chr2-123033159-INS-101
chr2-123037959-INS-749
chr2-123144663-INS-83
chr2-123149817-INS-83
chr2-123176314-INS-177
chr2-123196563-INS-529
chr2-123199920-INS-320
chr2-123205357-INS-575
chr2-123246868-INS-2585
chr2-123251413-INS-376
chr2-123253313-INS-275
chr2-123258212-INS-394
chr2-123291936-INS-2910
chr2-123294251-INS-76
chr2-123299926-INS-5149
chr2-123314508-INS-888
chr2-123352708-INS-7192
chr2-123374731-INS-591
chr2-123385637-INS-148
chr2-123388297-INS-76
chr2-123397553-INS-72
chr2-123467137-INS-694
chr2-123487362-INS-700
chr2-123503917-INS-92
chr2-123520044-INS-55
chr2-123555602-INS-238
chr2-123586864-INS-2943
chr2-123591068-INS-6535
chr2-123597638-INS-108
chr2-123598559-INS-79
chr2-123612383-INS-92
chr2-123671089-INS-241
chr2-123672323-INS-63
chr2-123673816-INS-98
chr2-123752472-INS-342
chr2-123765677-INS-395
chr2-123776367-INS-355
chr2-123779768-INS-6474
chr2-123783107-INS-751
chr2-123801016-INS-6404
chr2-123840118-INS-7135
chr2-123853623-INS-165
chr2-123862773-INS-54
chr2-123924086-INS-51
chr2-123936629-INS-60
chr2-123944594-INS-1371
chr2-123946744-INS-174
chr2-123947364-INS-893
chr2-124006685-INS-156
chr2-124037137-INS-67
chr2-124099543-INS-6106
chr2-124124139-INS-211
chr2-124155125-INS-2850
chr2-124163498-INS-64
chr2-124176991-INS-55
chr2-124217665-INS-994
chr2-124232956-INS-68

chr2-124266424-INS-87
chr2-124276054-INS-56
chr2-124293316-INS-394
chr2-124298749-INS-7280
chr2-124335914-INS-54
chr2-124411545-INS-78
chr2-124483822-INS-107
chr2-124516809-INS-170
chr2-124520762-INS-110
chr2-124521755-INS-4912
chr2-124523330-INS-52
chr2-124535334-INS-334
chr2-124536382-INS-78
chr2-124547678-INS-101
chr2-124554856-INS-100
chr2-124571290-INS-3161
chr2-124585695-INS-75
chr2-124587468-INS-86
chr2-124623319-INS-60
chr2-124651416-INS-6787
chr2-124659672-INS-293
chr2-124667308-INS-201
chr2-124693360-INS-403
chr2-124698004-INS-6400
chr2-124710470-INS-6538
chr2-124743474-INS-292
chr2-124752606-INS-308
chr2-124774088-INS-186
chr2-124787457-INS-112
chr2-124796225-INS-190
chr2-124828535-INS-68
chr2-124832702-INS-123
chr2-124855083-INS-7437
chr2-124876963-INS-225
chr2-124879615-INS-100
chr2-124881603-INS-58
chr2-124891038-INS-1207
chr2-124891256-INS-355
chr2-124900519-INS-469
chr2-124913875-INS-58
chr2-124921063-INS-54
chr2-124940238-INS-180
chr2-124963454-INS-184
chr2-124968952-INS-8940
chr2-125009530-INS-176
chr2-125021528-INS-102
chr2-125121174-INS-328
chr2-125200301-INS-72
chr2-125209370-INS-5862
chr2-125233228-INS-56
chr2-125239334-INS-796
chr2-125241272-INS-202
chr2-125262927-INS-64
chr2-125276078-INS-1628
chr2-125300986-INS-84
chr2-125336371-INS-69
chr2-125338085-INS-71
chr2-125343587-INS-148
chr2-125354960-INS-67
chr2-125435504-INS-65
chr2-125444065-INS-72
chr2-125449964-INS-65
chr2-125459601-INS-144
chr2-125479003-INS-53
chr2-125483236-INS-68
chr2-125488094-INS-54
chr2-125510711-INS-53
chr2-125521629-INS-221
chr2-125525893-INS-120
chr2-125535767-INS-141
chr2-125550344-INS-59
chr2-125552881-INS-52
chr2-125556290-INS-127
chr2-125560988-INS-55
chr2-125583640-INS-1531
chr2-125584601-INS-255
chr2-125586287-INS-108

chr2-125607818-INS-264
chr2-125608680-INS-128
chr2-125608774-INS-459
chr2-125645728-INS-120
chr2-125659509-INS-240
chr2-125659956-INS-120
chr2-125674327-INS-274
chr2-125689972-INS-206
chr2-125691211-INS-84
chr2-125725342-INS-767
chr2-125729912-INS-3350
chr2-125740907-INS-105
chr2-125770136-INS-13111
chr2-125782200-INS-98
chr2-125784329-INS-166
chr2-125789892-INS-6491
chr2-125792557-INS-5916
chr2-125846539-INS-68
chr2-125850479-INS-233
chr2-125863350-INS-59
chr2-125868700-INS-479
chr2-125881240-INS-280
chr2-125939689-INS-186
chr2-125993242-INS-479
chr2-126009820-INS-6227
chr2-126016557-INS-9737
chr2-126070471-INS-424
chr2-126120916-INS-838
chr2-126123650-INS-6581
chr2-126134314-INS-398
chr2-126136465-INS-597
chr2-126201261-INS-58
chr2-126207220-INS-111
chr2-126224967-INS-933
chr2-126239465-INS-96
chr2-126251933-INS-77
chr2-126263007-INS-134
chr2-126272875-INS-330
chr2-126290202-INS-202
chr2-126300092-INS-77
chr2-126312119-INS-428
chr2-126382933-INS-52
chr2-126383800-INS-247
chr2-126391338-INS-158
chr2-126444401-INS-885
chr2-126471182-INS-697
chr2-126490886-INS-217
chr2-126495181-INS-138
chr2-126497492-INS-211
chr2-126498973-INS-203
chr2-126509292-INS-1454
chr2-126540096-INS-53
chr2-126542182-INS-428
chr2-126545759-INS-55
chr2-126550356-INS-129
chr2-126558676-INS-203
chr2-126586506-INS-209
chr2-126587252-INS-215
chr2-126615213-INS-52
chr2-126638571-INS-208
chr2-126664234-INS-60
chr2-126680259-INS-54
chr2-126680714-INS-217
chr2-126692426-INS-91
chr2-126719989-INS-159
chr2-126721257-INS-809
chr2-126723716-INS-90
chr2-126730036-INS-105
chr2-126732155-INS-58
chr2-126741148-INS-133
chr2-126749525-INS-219
chr2-126751719-INS-59
chr2-126766919-INS-197
chr2-126795728-INS-169
chr2-126801677-INS-207
chr2-126855115-INS-891
chr2-126878270-INS-203

chr2-126882314-INS-52
chr2-126893011-INS-525
chr2-126896067-INS-208
chr2-126906096-INS-108
chr2-126919570-INS-243
chr2-126924108-INS-254
chr2-126928472-INS-203
chr2-126930655-INS-200
chr2-126931737-INS-206
chr2-126940802-INS-213
chr2-126942695-INS-502
chr2-126960254-INS-211
chr2-126965274-INS-54
chr2-126966993-INS-204
chr2-126977922-INS-112
chr2-126979330-INS-209
chr2-126994362-INS-146
chr2-126998860-INS-159
chr2-127021249-INS-82
chr2-127042117-INS-339
chr2-127048431-INS-451
chr2-127061452-INS-117
chr2-127065883-INS-94
chr2-127083040-INS-205
chr2-127093260-INS-506
chr2-127107957-INS-68
chr2-127119177-INS-208
chr2-127134938-INS-152
chr2-127145121-INS-155
chr2-127148425-INS-7482
chr2-127161420-INS-106
chr2-127197658-INS-116
chr2-127222367-INS-735
chr2-127250156-INS-1990
chr2-127260076-INS-56
chr2-127293584-INS-587
chr2-127297300-INS-179
chr2-127355984-INS-57
chr2-127368127-INS-72
chr2-127372911-INS-82
chr2-127391050-INS-7286
chr2-127392431-INS-81
chr2-127395673-INS-7188
chr2-127408248-INS-101
chr2-127409259-INS-194
chr2-127410752-INS-66
chr2-127415962-INS-53
chr2-127420623-INS-68
chr2-127421676-INS-68
chr2-127477016-INS-386
chr2-127478692-INS-76
chr2-127500527-INS-102
chr2-127503389-INS-141
chr2-127509089-INS-224
chr2-127512071-INS-108
chr2-127525933-INS-413
chr2-127552567-INS-728
chr2-127552568-INS-476
chr2-127561621-INS-165
chr2-127577266-INS-64
chr2-127579782-INS-180
chr2-127593581-INS-511
chr2-127600776-INS-68
chr2-127619114-INS-96
chr2-127624236-INS-52
chr2-127677599-INS-73
chr2-127684469-INS-265
chr2-127687368-INS-843
chr2-127697791-INS-72
chr2-127700768-INS-52
chr2-127737765-INS-92
chr2-127756940-INS-227
chr2-127771377-INS-130
chr2-127782434-INS-53
chr2-127796681-INS-60
chr2-127809930-INS-93
chr2-127848052-INS-69

chr2-127908035-INS-73
chr2-127916972-INS-69
chr2-127960158-INS-72
chr2-128030007-INS-244
chr2-128054752-INS-120
chr2-128074785-INS-178
chr2-128111634-INS-353
chr2-128115144-INS-228
chr2-128157385-INS-160
chr2-128185251-INS-400
chr2-128249321-INS-78
chr2-128251617-INS-168
chr2-128282595-INS-148
chr2-128284688-INS-289
chr2-128299199-INS-54
chr2-128344305-INS-73
chr2-128370698-INS-104
chr2-128398215-INS-51
chr2-128404383-INS-340
chr2-128416593-INS-353
chr2-128431651-INS-378
chr2-128437287-INS-274
chr2-128478570-INS-134
chr2-128523290-INS-209
chr2-128575142-INS-202
chr2-128583645-INS-174
chr2-128589989-INS-106
chr2-128595780-INS-52
chr2-128599724-INS-162
chr2-128631506-INS-62
chr2-128639434-INS-59
chr2-128652061-INS-121
chr2-128656782-INS-181
chr2-128666831-INS-165
chr2-128687757-INS-671
chr2-128690032-INS-188
chr2-128701874-INS-208
chr2-128702555-INS-148
chr2-128720196-INS-63
chr2-128728268-INS-71
chr2-128747207-INS-52
chr2-128754191-INS-349
chr2-128789723-INS-342
chr2-128795993-INS-365
chr2-128798983-INS-167
chr2-128822606-INS-65
chr2-128846577-INS-75
chr2-128883547-INS-128
chr2-128894108-INS-187
chr2-128902305-INS-150
chr2-128910651-INS-162
chr2-128914142-INS-60
chr2-128920071-INS-285
chr2-128926357-INS-77
chr2-128942666-INS-83
chr2-128975164-INS-400
chr2-128999987-INS-104
chr2-129015188-INS-367
chr2-129021239-INS-72
chr2-129037287-INS-391
chr2-129064708-INS-212
chr2-129079957-INS-163
chr2-129089686-INS-198
chr2-129090587-INS-55
chr2-129098980-INS-52
chr2-129103616-INS-281
chr2-129116849-INS-196
chr2-129123237-INS-163
chr2-129162635-INS-205
chr2-129168068-INS-52
chr2-129178693-INS-64
chr2-129183194-INS-7089
chr2-129184404-INS-3637
chr2-129212774-INS-71
chr2-129218684-INS-151
chr2-129260187-INS-112
chr2-129262117-INS-487

chr2-129278721-INS-1386
chr2-129280386-INS-6236
chr2-129284146-INS-1093
chr2-129284148-INS-617
chr2-129306130-INS-54
chr2-129388541-INS-64
chr2-129389993-INS-63
chr2-129423047-INS-68
chr2-129429239-INS-59
chr2-129445561-INS-704
chr2-129445564-INS-161
chr2-129482972-INS-60
chr2-129494801-INS-117
chr2-129568647-INS-159
chr2-129574893-INS-58
chr2-129611574-INS-60
chr2-129620186-INS-52
chr2-129625124-INS-61
chr2-129725514-INS-141
chr2-129779637-INS-3233
chr2-129782913-INS-120
chr2-129812350-INS-54
chr2-129821284-INS-189
chr2-129877459-INS-52
chr2-129901412-INS-62
chr2-129915289-INS-497
chr2-129940502-INS-6690
chr2-129951051-INS-68
chr2-129957500-INS-123
chr2-129972362-INS-112
chr2-129989266-INS-160
chr2-130002439-INS-64
chr2-130005126-INS-323
chr2-130027771-INS-76
chr2-130055892-INS-209
chr2-130078581-INS-168
chr2-130079407-INS-228
chr2-130081139-INS-200
chr2-130084879-INS-198
chr2-130108767-INS-334
chr2-130110897-INS-211
chr2-130115309-INS-90
chr2-130131940-INS-282
chr2-130144263-INS-56
chr2-130215285-INS-56
chr2-130242913-INS-135
chr2-130250788-INS-91
chr2-130296300-INS-102
chr2-130304604-INS-98
chr2-130305284-INS-1113
chr2-130321932-INS-207
chr2-130324973-INS-356
chr2-130330374-INS-440
chr2-130340329-INS-212
chr2-130351942-INS-75
chr2-130398651-INS-68
chr2-130409207-INS-271
chr2-130435811-INS-216
chr2-130436105-INS-1048
chr2-130439993-INS-63
chr2-130449263-INS-142
chr2-130454946-INS-203
chr2-130463302-INS-70
chr2-130463706-INS-187
chr2-130487946-INS-305
chr2-130489358-INS-180
chr2-130495201-INS-84
chr2-130497571-INS-410
chr2-130515671-INS-220
chr2-130544042-INS-114
chr2-130551324-INS-54
chr2-130588099-INS-193
chr2-130591973-INS-205
chr2-130600071-INS-54
chr2-130612993-INS-374
chr2-130624748-INS-53
chr2-130630738-INS-105

chr2-130638101-INS-151
chr2-130653325-INS-319
chr2-130672437-INS-683
chr2-130675991-INS-69
chr2-130680711-INS-78
chr2-130691008-INS-57
chr2-130715459-INS-347
chr2-130763217-INS-56
chr2-130781199-INS-388
chr2-130795276-INS-58
chr2-130801563-INS-166
chr2-130814882-INS-52
chr2-130825123-INS-113
chr2-130831742-INS-60
chr2-130847394-INS-131
chr2-130865956-INS-81
chr2-130902693-INS-54
chr2-130921445-INS-51
chr2-130937834-INS-87
chr2-130949678-INS-209
chr2-130962860-INS-196
chr2-130965503-INS-209
chr2-130972508-INS-238
chr2-130996676-INS-79
chr2-131042827-INS-136
chr2-131053659-INS-90
chr2-131056280-INS-99
chr2-131057240-INS-213
chr2-131060949-INS-72
chr2-131064769-INS-112
chr2-131072995-INS-55
chr2-131086037-INS-66
chr2-131115645-INS-125
chr2-131132591-INS-245
chr2-131136376-INS-104
chr2-131146927-INS-84
chr2-131153813-INS-53
chr2-131165340-INS-282
chr2-131210139-INS-81
chr2-131221174-INS-71
chr2-131227359-INS-62
chr2-131247509-INS-612
chr2-131256309-INS-391
chr2-131269946-INS-110
chr2-131293061-INS-166
chr2-131307507-INS-54
chr2-131311661-INS-180
chr2-131323476-INS-77
chr2-131350812-INS-63
chr2-131375757-INS-172
chr2-131377357-INS-333
chr2-131379577-INS-305
chr2-131406728-INS-195
chr2-131424667-INS-190
chr2-131465515-INS-100
chr2-131483244-INS-6385
chr2-131495754-INS-55
chr2-131504175-INS-54
chr2-131512172-INS-266
chr2-131514536-INS-138
chr2-131525512-INS-55
chr2-131528172-INS-395
chr2-131542135-INS-406
chr2-131543923-INS-468
chr2-131563678-INS-60
chr2-131584314-INS-126
chr2-131618643-INS-54
chr2-131644455-INS-6274
chr2-131652881-INS-118
chr2-131658720-INS-57
chr2-131669312-INS-141
chr2-131670260-INS-245
chr2-131685914-INS-228
chr2-131701592-INS-291
chr2-131709029-INS-140
chr2-131716691-INS-234
chr2-131781400-INS-703

chr2-131817708-INS-558
chr2-131849859-INS-96
chr2-131880354-INS-60
chr2-131904632-INS-135
chr2-131926758-INS-253
chr2-131930147-INS-64
chr2-131946688-INS-315
chr2-131958643-INS-54
chr2-131959671-INS-64
chr2-131966626-INS-279
chr2-131976510-INS-57
chr2-131979571-INS-139
chr2-131998285-INS-286
chr2-132000635-INS-94
chr2-132017262-INS-354
chr2-132025475-INS-62
chr2-132065454-INS-329
chr2-132075509-INS-206
chr2-132086319-INS-5348
chr2-132086784-INS-197
chr2-132109874-INS-225
chr2-132111750-INS-177
chr2-132121985-INS-6442
chr2-132123190-INS-236
chr2-132123244-INS-918
chr2-132134794-INS-104
chr2-132170966-INS-58
chr2-132180245-INS-861
chr2-132232332-INS-60
chr2-132237979-INS-217
chr2-132239028-INS-266
chr2-132243753-INS-56
chr2-132247628-INS-58
chr2-132285624-INS-275
chr2-132313029-INS-321
chr2-132317603-INS-51
chr2-132319211-INS-610
chr2-132323580-INS-56
chr2-132334067-INS-442
chr2-132341321-INS-62
chr2-132347967-INS-60
chr2-132350960-INS-258
chr2-132353636-INS-140
chr2-132369814-INS-229
chr2-132370163-INS-697
chr2-132380576-INS-91
chr2-132391582-INS-263
chr2-132396999-INS-52
chr2-132400625-INS-227
chr2-132440619-INS-162
chr2-132442363-INS-57
chr2-132446237-INS-451
chr2-132448273-INS-57
chr2-132450829-INS-54
chr2-132454761-INS-241
chr2-132469312-INS-169
chr2-132484725-INS-95
chr2-132493956-INS-191
chr2-132494454-INS-598
chr2-132501972-INS-51
chr2-132510920-INS-542
chr2-132521103-INS-165
chr2-132527489-INS-148
chr2-132534229-INS-52
chr2-132536721-INS-1770
chr2-132544547-INS-95
chr2-132545873-INS-206
chr2-132554188-INS-59
chr2-132560896-INS-6652
chr2-132569994-INS-70
chr2-132573840-INS-87
chr2-132587722-INS-212
chr2-132593234-INS-488
chr2-132607876-INS-64
chr2-132617347-INS-400
chr2-132618414-INS-2904
chr2-132621432-INS-52

chr2-132623000-INS-1008
chr2-132643161-INS-60
chr2-132674595-INS-294
chr2-132681275-INS-266
chr2-132681890-INS-52
chr2-132716715-INS-201
chr2-132747632-INS-132
chr2-132759079-INS-917
chr2-132761483-INS-95
chr2-132765254-INS-77
chr2-132799653-INS-139
chr2-132801781-INS-227
chr2-132823313-INS-80
chr2-132834039-INS-84
chr2-132843226-INS-61
chr2-132859803-INS-128
chr2-132883013-INS-1599
chr2-132886132-INS-53
chr2-132916287-INS-72
chr2-132954532-INS-70
chr2-132981723-INS-269
chr2-132989574-INS-64
chr2-132995504-INS-498
chr2-133000189-INS-828
chr2-133005722-INS-844
chr2-133011807-INS-69
chr2-133043693-INS-52
chr2-133072844-INS-935
chr2-133080194-INS-51
chr2-133087624-INS-7131
chr2-133093131-INS-89
chr2-133149719-INS-832
chr2-133153371-INS-1440
chr2-133154381-INS-741
chr2-133157756-INS-412
chr2-133174688-INS-64
chr2-133189828-INS-56
chr2-133195243-INS-123
chr2-133201610-INS-192
chr2-133205805-INS-75
chr2-133208050-INS-58
chr2-133228348-INS-196
chr2-133236538-INS-60
chr2-133242348-INS-132
chr2-133294288-INS-144
chr2-133296534-INS-60
chr2-133333449-INS-52
chr2-133388617-INS-86
chr2-133401266-INS-120
chr2-133428857-INS-323
chr2-133430870-INS-57
chr2-133444660-INS-172
chr2-133466711-INS-100
chr2-133472071-INS-60
chr2-133489227-INS-64
chr2-133497413-INS-217
chr2-133512953-INS-70
chr2-133529336-INS-445
chr2-133534551-INS-346
chr2-133542989-INS-433
chr2-133561484-INS-499
chr2-133562306-INS-753
chr2-133579535-INS-514
chr2-133659667-INS-51
chr2-133662736-INS-52
chr2-133700382-INS-393
chr2-133702811-INS-5186
chr2-133709862-INS-191
chr2-133732136-INS-835
chr2-133737993-INS-580
chr2-133748863-INS-302
chr2-133761578-INS-418
chr2-133783946-INS-131
chr2-133803353-INS-96
chr2-133815086-INS-152
chr2-133838368-INS-60
chr2-133840537-INS-174

chr2-133853794-INS-6615
chr2-133869590-INS-52
chr2-133892517-INS-76
chr2-133950720-INS-71
chr2-133966521-INS-7097
chr2-134004982-INS-68
chr2-134053576-INS-1838
chr2-134055833-INS-6277
chr2-134088181-INS-92
chr2-134096397-INS-2556
chr2-134097654-INS-197
chr2-134102356-INS-274
chr2-134113502-INS-891
chr2-134148098-INS-72
chr2-134163824-INS-417
chr2-134176554-INS-404
chr2-134181995-INS-669
chr2-134199236-INS-191
chr2-134205207-INS-191
chr2-134243140-INS-87
chr2-134255850-INS-199
chr2-134260975-INS-680
chr2-134262952-INS-207
chr2-134286162-INS-53
chr2-134292521-INS-6569
chr2-134312735-INS-52
chr2-134355897-INS-74
chr2-134356645-INS-259
chr2-134373434-INS-88
chr2-134435335-INS-466
chr2-134477499-INS-123
chr2-134479336-INS-70
chr2-134490577-INS-91
chr2-134498030-INS-54
chr2-134537828-INS-51
chr2-134563781-INS-372
chr2-134566904-INS-518
chr2-134603556-INS-213
chr2-134608124-INS-64
chr2-134633143-INS-647
chr2-134639405-INS-127
chr2-134659757-INS-431
chr2-134684362-INS-88
chr2-134710643-INS-388
chr2-134735015-INS-386
chr2-134736365-INS-58
chr2-134739329-INS-260
chr2-134791680-INS-50
chr2-134793331-INS-6811
chr2-134799385-INS-154
chr2-134801692-INS-118
chr2-134842071-INS-60
chr2-134843498-INS-366
chr2-134846844-INS-243
chr2-134853293-INS-342
chr2-134871262-INS-108
chr2-134873488-INS-179
chr2-134881643-INS-209
chr2-134883126-INS-271
chr2-134901507-INS-56
chr2-134960211-INS-136
chr2-134973441-INS-398
chr2-135009223-INS-66
chr2-135013798-INS-56
chr2-135020552-INS-145
chr2-135047107-INS-798
chr2-135051027-INS-5297
chr2-135060685-INS-285
chr2-135099879-INS-54
chr2-135101817-INS-199
chr2-135105483-INS-95
chr2-135108847-INS-6470
chr2-135119602-INS-72
chr2-135147500-INS-549
chr2-135152691-INS-527
chr2-135156272-INS-88
chr2-135174744-INS-185

chr2-135185970-INS-158
chr2-135198216-INS-6265
chr2-135252682-INS-6593
chr2-135259039-INS-112
chr2-135311868-INS-113
chr2-135312072-INS-3196
chr2-135312930-INS-181
chr2-135320097-INS-8492
chr2-135335409-INS-105
chr2-135348664-INS-56
chr2-135369138-INS-926
chr2-135382883-INS-84
chr2-135392893-INS-50
chr2-135400839-INS-328
chr2-135417786-INS-76
chr2-135434447-INS-208
chr2-135466742-INS-55
chr2-135479591-INS-434
chr2-135480735-INS-56
chr2-135488909-INS-174
chr2-135511188-INS-189
chr2-135520416-INS-210
chr2-135531004-INS-51
chr2-135533368-INS-53
chr2-135545655-INS-344
chr2-135552072-INS-72
chr2-135561773-INS-576
chr2-135577690-INS-102
chr2-135600874-INS-324
chr2-135620179-INS-267
chr2-135638458-INS-71
chr2-135649022-INS-172
chr2-135651068-INS-95
chr2-135658666-INS-91
chr2-135658878-INS-1630
chr2-135680798-INS-145
chr2-135693329-INS-83
chr2-135719747-INS-713
chr2-135722758-INS-60
chr2-135728406-INS-339
chr2-135741841-INS-488
chr2-135750666-INS-99
chr2-135795317-INS-510
chr2-135811639-INS-84
chr2-135824431-INS-159
chr2-135830461-INS-1124
chr2-135838084-INS-75
chr2-135842330-INS-84
chr2-135867668-INS-350
chr2-135869523-INS-81
chr2-135879173-INS-368
chr2-135881996-INS-69
chr2-135886395-INS-193
chr2-135894435-INS-196
chr2-135897478-INS-60
chr2-135936923-INS-59
chr2-135978524-INS-954
chr2-135988357-INS-58
chr2-136012059-INS-63
chr2-136034518-INS-587
chr2-136041562-INS-210
chr2-136046115-INS-6561
chr2-136069192-INS-176
chr2-136089671-INS-224
chr2-136091235-INS-75
chr2-136101166-INS-100
chr2-136108312-INS-5714
chr2-136114256-INS-56
chr2-136123529-INS-94
chr2-136124030-INS-97
chr2-136128885-INS-340
chr2-136144086-INS-56
chr2-136145447-INS-990
chr2-136154949-INS-92
chr2-136159734-INS-996
chr2-136182710-INS-304
chr2-136196780-INS-91

chr2-136258073-INS-1888
chr2-136261777-INS-336
chr2-136289601-INS-628
chr2-136306370-INS-7416
chr2-136320679-INS-51
chr2-136323392-INS-358
chr2-136325908-INS-187
chr2-136348683-INS-120
chr2-136371622-INS-547
chr2-136375190-INS-478
chr2-136415274-INS-490
chr2-136456501-INS-313
chr2-136493771-INS-239
chr2-136507223-INS-68
chr2-136522471-INS-514
chr2-136535643-INS-1891
chr2-136539692-INS-58
chr2-136554025-INS-82
chr2-136562131-INS-634
chr2-136573034-INS-619
chr2-136593687-INS-2333
chr2-136610894-INS-198
chr2-136617440-INS-58
chr2-136622494-INS-493
chr2-136635853-INS-146
chr2-136680417-INS-68
chr2-136694128-INS-7119
chr2-136731428-INS-238
chr2-136732628-INS-187
chr2-136752157-INS-5391
chr2-136753793-INS-6502
chr2-136777284-INS-6791
chr2-136782049-INS-53
chr2-136793935-INS-56
chr2-136825788-INS-938
chr2-136923947-INS-58
chr2-137001464-INS-448
chr2-137077813-INS-120
chr2-137105634-INS-204
chr2-137105845-INS-468
chr2-137125514-INS-116
chr2-137186900-INS-51
chr2-137195688-INS-6574
chr2-137214751-INS-6878
chr2-137218919-INS-787
chr2-137239348-INS-2550
chr2-137258380-INS-53
chr2-137260277-INS-595
chr2-137268336-INS-54
chr2-137279315-INS-482
chr2-137281656-INS-86
chr2-137289542-INS-53
chr2-137292451-INS-57
chr2-137299768-INS-68
chr2-137318322-INS-52
chr2-137345789-INS-80
chr2-137366866-INS-97
chr2-137399345-INS-242
chr2-137466218-INS-398
chr2-137470665-INS-254
chr2-137474920-INS-56
chr2-137520002-INS-6403
chr2-137567090-INS-195
chr2-137590734-INS-70
chr2-137593142-INS-352
chr2-137623627-INS-58
chr2-137651521-INS-904
chr2-137654084-INS-155
chr2-137657529-INS-57
chr2-137660748-INS-72
chr2-137700941-INS-74
chr2-137740804-INS-292
chr2-137788934-INS-651
chr2-137825466-INS-103
chr2-137827872-INS-6322
chr2-137899010-INS-60
chr2-137904776-INS-58

chr2-137918868-INS-81
chr2-137928354-INS-117
chr2-137957029-INS-525
chr2-137996720-INS-151
chr2-138000404-INS-369
chr2-138004158-INS-80
chr2-138026539-INS-53
chr2-138040005-INS-200
chr2-138060110-INS-157
chr2-138081538-INS-58
chr2-138105930-INS-61
chr2-138110517-INS-216
chr2-138117792-INS-62
chr2-138154096-INS-3291
chr2-138155893-INS-134
chr2-138165091-INS-67
chr2-138175746-INS-62
chr2-138186035-INS-6348
chr2-138187768-INS-446
chr2-138199229-INS-6587
chr2-138209681-INS-432
chr2-138260712-INS-2303
chr2-138266675-INS-469
chr2-138277596-INS-386
chr2-138333300-INS-60
chr2-138333767-INS-6547
chr2-138342175-INS-6632
chr2-138373713-INS-480
chr2-138392764-INS-6398
chr2-138399056-INS-57
chr2-138400439-INS-5320
chr2-138403203-INS-405
chr2-138449579-INS-364
chr2-138458915-INS-7081
chr2-138467889-INS-2804
chr2-138500104-INS-70
chr2-138501025-INS-327
chr2-138504524-INS-63
chr2-138515966-INS-782
chr2-138518271-INS-73
chr2-138519227-INS-195
chr2-138557300-INS-6843
chr2-138562555-INS-75
chr2-138574634-INS-527
chr2-138583910-INS-230
chr2-138587086-INS-53
chr2-138591275-INS-2699
chr2-138601009-INS-6670
chr2-138623873-INS-116
chr2-138626103-INS-93
chr2-138647223-INS-336
chr2-138668015-INS-153
chr2-138700748-INS-6390
chr2-138704837-INS-6525
chr2-138706085-INS-105
chr2-138710155-INS-68
chr2-138725683-INS-508
chr2-138732762-INS-905
chr2-138759944-INS-112
chr2-138763544-INS-196
chr2-138770701-INS-206
chr2-138771332-INS-201
chr2-138776824-INS-150
chr2-138784890-INS-122
chr2-138810184-INS-86
chr2-138816034-INS-4820
chr2-138824358-INS-115
chr2-138825107-INS-892
chr2-138826316-INS-109
chr2-138842566-INS-57
chr2-138849680-INS-401
chr2-138853075-INS-53
chr2-138859436-INS-508
chr2-138867615-INS-7483
chr2-138887321-INS-149
chr2-138892746-INS-755
chr2-138900627-INS-1962

chr2-138926123-INS-657
chr2-138946879-INS-166
chr2-138951744-INS-64
chr2-139023738-INS-187
chr2-139042850-INS-180
chr2-139046401-INS-331
chr2-139055278-INS-69
chr2-139076408-INS-604
chr2-139077247-INS-368
chr2-139084160-INS-122
chr2-139094385-INS-207
chr2-139127768-INS-6096
chr2-139129989-INS-1720
chr2-139171809-INS-88
chr2-139180208-INS-88
chr2-139185090-INS-100
chr2-139188521-INS-127
chr2-139206393-INS-97
chr2-139209517-INS-5424
chr2-139213542-INS-6590
chr2-139244557-INS-7141
chr2-139323687-INS-5139
chr2-139326345-INS-228
chr2-139335267-INS-263
chr2-139336855-INS-74
chr2-139342430-INS-1190
chr2-139393605-INS-205
chr2-139412875-INS-377
chr2-139430256-INS-211
chr2-139438412-INS-122
chr2-139464553-INS-106
chr2-139465408-INS-613
chr2-139486189-INS-8307
chr2-139498065-INS-190
chr2-139517090-INS-62
chr2-139526895-INS-261
chr2-139531047-INS-287
chr2-139536575-INS-183
chr2-139575043-INS-578
chr2-139601820-INS-108
chr2-139616510-INS-137
chr2-139703522-INS-85
chr2-139714777-INS-478
chr2-139734142-INS-141
chr2-139734292-INS-6712
chr2-139736486-INS-83
chr2-139768803-INS-517
chr2-139770723-INS-58
chr2-139773647-INS-73
chr2-139790951-INS-94
chr2-139860908-INS-178
chr2-139864032-INS-118
chr2-139874061-INS-127
chr2-139882317-INS-68
chr2-139921314-INS-541
chr2-139946806-INS-618
chr2-139953283-INS-127
chr2-139987551-INS-76
chr2-139995624-INS-56
chr2-140001976-INS-60
chr2-140012205-INS-95
chr2-140042118-INS-159
chr2-140049065-INS-155
chr2-140093842-INS-52
chr2-140126465-INS-340
chr2-140139656-INS-82
chr2-140179997-INS-456
chr2-140216822-INS-345
chr2-140224077-INS-415
chr2-140353027-INS-98
chr2-140419108-INS-228
chr2-140473174-INS-114
chr2-140489420-INS-401
chr2-140522368-INS-75
chr2-140541352-INS-220
chr2-140560070-INS-190
chr2-140572268-INS-51

chr2-140621302-INS-372
chr2-140646401-INS-53
chr2-140652178-INS-338
chr2-140660802-INS-81
chr2-140667853-INS-72
chr2-140689683-INS-1839
chr2-140706989-INS-400
chr2-140753630-INS-52
chr2-140754844-INS-84
chr2-140755350-INS-7178
chr2-140762833-INS-52
chr2-140779383-INS-975
chr2-140783923-INS-443
chr2-140789231-INS-54
chr2-140811241-INS-124
chr2-140819772-INS-175
chr2-140824150-INS-93
chr2-140838820-INS-172
chr2-140930466-INS-377
chr2-140937919-INS-1291
chr2-140963341-INS-92
chr2-140966081-INS-457
chr2-140991578-INS-200
chr2-140995722-INS-94
chr2-141001063-INS-400
chr2-141028330-INS-52
chr2-141081154-INS-365
chr2-141100891-INS-240
chr2-141119513-INS-117
chr2-141148320-INS-111
chr2-141177932-INS-6459
chr2-141210551-INS-198
chr2-141217860-INS-88
chr2-141222240-INS-54
chr2-141237782-INS-56
chr2-141239055-INS-60
chr2-141241366-INS-302
chr2-141271806-INS-56
chr2-141277671-INS-104
chr2-141288561-INS-76
chr2-141292682-INS-8792
chr2-141295719-INS-75
chr2-141328899-INS-108
chr2-141336763-INS-137
chr2-141364950-INS-76
chr2-141365998-INS-248
chr2-141368183-INS-469
chr2-141372566-INS-162
chr2-141408568-INS-87
chr2-141421205-INS-910
chr2-141460722-INS-102
chr2-141467842-INS-58
chr2-141471188-INS-94
chr2-141476494-INS-152
chr2-141477394-INS-849
chr2-141479302-INS-452
chr2-141487184-INS-176
chr2-141491786-INS-68
chr2-141516692-INS-72
chr2-141543680-INS-150
chr2-141568636-INS-66
chr2-141591115-INS-332
chr2-141614132-INS-602
chr2-141620275-INS-52
chr2-141624462-INS-142
chr2-141634736-INS-58
chr2-141655703-INS-390
chr2-141661328-INS-107
chr2-141667150-INS-2576
chr2-141668547-INS-6630
chr2-141669034-INS-68
chr2-141677865-INS-215
chr2-141740177-INS-56
chr2-141757254-INS-52
chr2-141760161-INS-3679
chr2-141771777-INS-7455
chr2-141806787-INS-64

chr2-141826402-INS-6401
chr2-141840727-INS-3528
chr2-141919363-INS-182
chr2-141923234-INS-59
chr2-141925944-INS-76
chr2-142054071-INS-510
chr2-142061323-INS-56
chr2-142069646-INS-98
chr2-142076822-INS-92
chr2-142078993-INS-52
chr2-142089275-INS-56
chr2-142103115-INS-857
chr2-142130507-INS-64
chr2-142131910-INS-664
chr2-142135789-INS-208
chr2-142160959-INS-57
chr2-142164484-INS-8458
chr2-142173832-INS-61
chr2-142181784-INS-64
chr2-142189190-INS-1348
chr2-142192851-INS-709
chr2-142208200-INS-631
chr2-142248828-INS-884
chr2-142261745-INS-1595
chr2-142290519-INS-466
chr2-142295596-INS-7148
chr2-142296861-INS-5553
chr2-142298725-INS-6003
chr2-142304422-INS-304
chr2-142339087-INS-571
chr2-142379457-INS-113
chr2-142416957-INS-145
chr2-142453816-INS-246
chr2-142459720-INS-98
chr2-142494381-INS-84
chr2-142512467-INS-810
chr2-142513710-INS-57
chr2-142524611-INS-52
chr2-142536513-INS-7132
chr2-142574209-INS-117
chr2-142581972-INS-135
chr2-142590428-INS-76
chr2-142620130-INS-183
chr2-142659236-INS-118
chr2-142687308-INS-100
chr2-142693779-INS-178
chr2-142703246-INS-205
chr2-142723099-INS-64
chr2-142729631-INS-200
chr2-142732557-INS-2915
chr2-142732659-INS-199
chr2-142735073-INS-182
chr2-142752762-INS-66
chr2-142760525-INS-59
chr2-142763798-INS-64
chr2-142766057-INS-217
chr2-142768001-INS-72
chr2-142780801-INS-92
chr2-142839141-INS-72
chr2-142868441-INS-530
chr2-142880512-INS-95
chr2-142896481-INS-103
chr2-142903325-INS-332
chr2-142906117-INS-187
chr2-142928547-INS-60
chr2-142944098-INS-192
chr2-142944998-INS-206
chr2-142946760-INS-93
chr2-142959625-INS-680
chr2-142967316-INS-190
chr2-142973858-INS-80
chr2-142986759-INS-204
chr2-142994550-INS-56
chr2-143008011-INS-6240
chr2-143013485-INS-55
chr2-143015585-INS-400
chr2-143055135-INS-719

chr2-143061085-INS-69
chr2-143111173-INS-51
chr2-143122941-INS-72
chr2-143129408-INS-101
chr2-143146346-INS-787
chr2-143160967-INS-67
chr2-143162491-INS-7373
chr2-143168036-INS-56
chr2-143172086-INS-65
chr2-143173236-INS-401
chr2-143179360-INS-54
chr2-143183559-INS-874
chr2-143187944-INS-85
chr2-143196875-INS-776
chr2-143201539-INS-466
chr2-143208965-INS-55
chr2-143224581-INS-647
chr2-143237629-INS-190
chr2-143241388-INS-498
chr2-143243170-INS-72
chr2-143266160-INS-377
chr2-143267361-INS-205
chr2-143282184-INS-6481
chr2-143307352-INS-6380
chr2-143308415-INS-199
chr2-143313191-INS-130
chr2-143360854-INS-54
chr2-143381171-INS-58
chr2-143400360-INS-60
chr2-143406323-INS-112
chr2-143435918-INS-132
chr2-143443728-INS-1025
chr2-143464437-INS-155
chr2-143475921-INS-54
chr2-143505192-INS-6470
chr2-143582884-INS-142
chr2-143596150-INS-276
chr2-143600550-INS-214
chr2-143606957-INS-241
chr2-143610758-INS-356
chr2-143662996-INS-2318
chr2-143671300-INS-57
chr2-143720504-INS-237
chr2-143835833-INS-1287
chr2-143843890-INS-56
chr2-143874247-INS-82
chr2-143880482-INS-71
chr2-143893294-INS-57
chr2-143899389-INS-244
chr2-143907990-INS-56
chr2-143908302-INS-50
chr2-143913889-INS-51
chr2-143923403-INS-62
chr2-143923766-INS-169
chr2-143933581-INS-147
chr2-143966984-INS-99
chr2-143986261-INS-261
chr2-144011126-INS-128
chr2-144013941-INS-90
chr2-144020234-INS-295
chr2-144044459-INS-74
chr2-144071525-INS-316
chr2-144081061-INS-154
chr2-144093789-INS-224
chr2-144118748-INS-173
chr2-144120238-INS-200
chr2-144124085-INS-200
chr2-144155312-INS-199
chr2-144178064-INS-142
chr2-144179065-INS-141
chr2-144182538-INS-220
chr2-144196378-INS-580
chr2-144237096-INS-76
chr2-144239472-INS-120
chr2-144284461-INS-54
chr2-144288023-INS-214
chr2-144288042-INS-6448

chr2-144293127-INS-507
chr2-144297610-INS-200
chr2-144339422-INS-70
chr2-144341914-INS-56
chr2-144349912-INS-258
chr2-144352662-INS-51
chr2-144362811-INS-77
chr2-144399400-INS-93
chr2-144421840-INS-693
chr2-144424299-INS-159
chr2-144437498-INS-219
chr2-144444502-INS-240
chr2-144450255-INS-5428
chr2-144454017-INS-52
chr2-144456321-INS-63
chr2-144515945-INS-54
chr2-144516303-INS-6595
chr2-144522594-INS-108
chr2-144534891-INS-110
chr2-144549230-INS-82
chr2-144555110-INS-395
chr2-144564347-INS-6648
chr2-144578060-INS-116
chr2-144622635-INS-374
chr2-144626569-INS-407
chr2-144637494-INS-90
chr2-144666534-INS-93
chr2-144679782-INS-91
chr2-144696823-INS-69
chr2-144755031-INS-123
chr2-144765499-INS-504
chr2-144786186-INS-102
chr2-144787586-INS-184
chr2-144793154-INS-494
chr2-144804974-INS-79
chr2-144825318-INS-80
chr2-144861950-INS-3148
chr2-144862936-INS-56
chr2-144865679-INS-90
chr2-144900826-INS-62
chr2-144909770-INS-973
chr2-144919287-INS-196
chr2-144929679-INS-66
chr2-144937859-INS-615
chr2-144940256-INS-281
chr2-144972095-INS-6495
chr2-144977334-INS-778
chr2-144981132-INS-57
chr2-145003568-INS-636
chr2-145017391-INS-6582
chr2-145038365-INS-84
chr2-145039579-INS-186
chr2-145062571-INS-194
chr2-145063091-INS-698
chr2-145071825-INS-60
chr2-145096044-INS-171
chr2-145103500-INS-399
chr2-145149772-INS-51
chr2-145176346-INS-188
chr2-145178057-INS-539
chr2-145192935-INS-359
chr2-145208150-INS-165
chr2-145234607-INS-226
chr2-145264235-INS-54
chr2-145324330-INS-78
chr2-145326391-INS-114
chr2-145327721-INS-237
chr2-145367964-INS-52
chr2-145382178-INS-108
chr2-145440923-INS-77
chr2-145444346-INS-63
chr2-145457057-INS-68
chr2-145460410-INS-52
chr2-145480010-INS-67
chr2-145490071-INS-179
chr2-145507843-INS-112
chr2-145523111-INS-833

chr2-145535331-INS-87
chr2-145537469-INS-4525
chr2-145541050-INS-203
chr2-145562685-INS-137
chr2-145614640-INS-5453
chr2-145615898-INS-7099
chr2-145670548-INS-189
chr2-145683216-INS-134
chr2-145723163-INS-6497
chr2-145733279-INS-155
chr2-145734991-INS-95
chr2-145738098-INS-119
chr2-145739212-INS-554
chr2-145763403-INS-398
chr2-145834039-INS-331
chr2-145839255-INS-74
chr2-145856041-INS-80
chr2-145874174-INS-507
chr2-145901120-INS-56
chr2-145928645-INS-1952
chr2-145928979-INS-6444
chr2-145936528-INS-143
chr2-145941842-INS-91
chr2-145944475-INS-137
chr2-145948676-INS-71
chr2-145964531-INS-436
chr2-145989567-INS-446
chr2-145996457-INS-7912
chr2-146015266-INS-82
chr2-146021061-INS-155
chr2-146027450-INS-148
chr2-146047031-INS-51
chr2-146048009-INS-378
chr2-146050327-INS-112
chr2-146057609-INS-60
chr2-146059782-INS-89
chr2-146085071-INS-64
chr2-146127433-INS-66
chr2-146212862-INS-211
chr2-146324365-INS-54
chr2-146338384-INS-608
chr2-146357568-INS-200
chr2-146366916-INS-63
chr2-146401791-INS-223
chr2-146414197-INS-147
chr2-146428816-INS-530
chr2-146446354-INS-250
chr2-146446367-INS-506
chr2-146456935-INS-186
chr2-146501454-INS-7195
chr2-146514130-INS-77
chr2-146517507-INS-56
chr2-146535767-INS-100
chr2-146589122-INS-197
chr2-146598124-INS-103
chr2-146612423-INS-90
chr2-146622849-INS-368
chr2-146632258-INS-181
chr2-146632381-INS-1843
chr2-146633108-INS-435
chr2-146646904-INS-550
chr2-146650174-INS-53
chr2-146663921-INS-62
chr2-146686153-INS-78
chr2-146687705-INS-112
chr2-146699315-INS-70
chr2-146756454-INS-233
chr2-146756734-INS-965
chr2-146771589-INS-104
chr2-146776686-INS-62
chr2-146797924-INS-62
chr2-146809701-INS-368
chr2-146815193-INS-328
chr2-146823268-INS-57
chr2-146828001-INS-1793
chr2-146850735-INS-204
chr2-146859614-INS-3412

chr2-146867255-INS-51
chr2-146912996-INS-237
chr2-146918173-INS-68
chr2-146934204-INS-130
chr2-146935786-INS-201
chr2-146951892-INS-288
chr2-146968837-INS-1864
chr2-146974296-INS-6197
chr2-146976566-INS-7168
chr2-147006227-INS-52
chr2-147050975-INS-71
chr2-147064807-INS-175
chr2-147099500-INS-63
chr2-147111110-INS-189
chr2-147144796-INS-98
chr2-147154365-INS-270
chr2-147164642-INS-220
chr2-147188722-INS-56
chr2-147214494-INS-94
chr2-147223177-INS-98
chr2-147231546-INS-139
chr2-147257835-INS-332
chr2-147272533-INS-81
chr2-147277019-INS-291
chr2-147315046-INS-194
chr2-147320897-INS-109
chr2-147351393-INS-289
chr2-147375590-INS-6446
chr2-147403584-INS-126
chr2-147405659-INS-212
chr2-147409945-INS-136
chr2-147466131-INS-654
chr2-147521109-INS-52
chr2-147524626-INS-219
chr2-147529188-INS-60
chr2-147534396-INS-54
chr2-147562053-INS-98
chr2-147616181-INS-180
chr2-147626187-INS-51
chr2-147639709-INS-105
chr2-147652006-INS-302
chr2-147685863-INS-7172
chr2-147695774-INS-71
chr2-147702122-INS-52
chr2-147705558-INS-140
chr2-147712765-INS-214
chr2-147732522-INS-152
chr2-147744736-INS-190
chr2-147745117-INS-394
chr2-147775883-INS-76
chr2-147825360-INS-8062
chr2-147848912-INS-51
chr2-147867020-INS-218
chr2-147946901-INS-196
chr2-147954951-INS-247
chr2-147965134-INS-84
chr2-147985314-INS-252
chr2-148005115-INS-6737
chr2-148030490-INS-1136
chr2-148036535-INS-5106
chr2-148048762-INS-6593
chr2-148053553-INS-3282
chr2-148062599-INS-223
chr2-148071950-INS-51
chr2-148077534-INS-178
chr2-148093553-INS-589
chr2-148105864-INS-51
chr2-148114443-INS-55
chr2-148128132-INS-270
chr2-148155941-INS-51
chr2-148171757-INS-106
chr2-148177018-INS-64
chr2-148263918-INS-7302
chr2-148264643-INS-228
chr2-148368622-INS-56
chr2-148377506-INS-406
chr2-148406224-INS-5387

chr2-148424572-INS-57
chr2-148447834-INS-72
chr2-148476141-INS-160
chr2-148487354-INS-293
chr2-148488894-INS-106
chr2-148491039-INS-359
chr2-148498399-INS-450
chr2-148499078-INS-55
chr2-148499962-INS-498
chr2-148550516-INS-194
chr2-148569072-INS-91
chr2-148581248-INS-949
chr2-148591781-INS-58
chr2-148607504-INS-87
chr2-148611640-INS-65
chr2-148629512-INS-168
chr2-148638221-INS-60
chr2-148679968-INS-503
chr2-148693678-INS-776
chr2-148708678-INS-696
chr2-148731689-INS-117
chr2-148737476-INS-210
chr2-148746210-INS-2462
chr2-148753418-INS-5402
chr2-148772481-INS-6094
chr2-148803504-INS-58
chr2-148822534-INS-3009
chr2-148838424-INS-332
chr2-148860249-INS-738
chr2-148888391-INS-504
chr2-148888456-INS-889
chr2-148928412-INS-135
chr2-148935888-INS-65
chr2-148978673-INS-214
chr2-149074982-INS-6671
chr2-149123228-INS-6345
chr2-149124231-INS-60
chr2-149174054-INS-599
chr2-149197886-INS-64
chr2-149233545-INS-6711
chr2-149255399-INS-143
chr2-149260209-INS-244
chr2-149260323-INS-977
chr2-149267162-INS-200
chr2-149274373-INS-7067
chr2-149276131-INS-534
chr2-149308949-INS-227
chr2-149312883-INS-70
chr2-149316737-INS-988
chr2-149333586-INS-1886
chr2-149336451-INS-142
chr2-149347997-INS-6398
chr2-149374436-INS-87
chr2-149394923-INS-144
chr2-149398501-INS-371
chr2-149426702-INS-84
chr2-149472234-INS-306
chr2-149495229-INS-204
chr2-149521513-INS-520
chr2-149532523-INS-4828
chr2-149545214-INS-262
chr2-149552853-INS-5463
chr2-149571799-INS-933
chr2-149578461-INS-56
chr2-149604655-INS-64
chr2-149650512-INS-722
chr2-149663852-INS-97
chr2-149679429-INS-6152
chr2-149709119-INS-572
chr2-149756723-INS-100
chr2-149774699-INS-400
chr2-149824286-INS-585
chr2-149824729-INS-6401
chr2-149830641-INS-274
chr2-149831417-INS-521
chr2-149861124-INS-475
chr2-149863457-INS-682

chr2-149883843-INS-610
chr2-149894141-INS-56
chr2-149904781-INS-493
chr2-149935850-INS-7385
chr2-149969608-INS-80
chr2-149990426-INS-486
chr2-150020413-INS-730
chr2-150110249-INS-168
chr2-150181949-INS-126
chr2-150188505-INS-321
chr2-150195735-INS-104
chr2-150248552-INS-71
chr2-150265526-INS-1541
chr2-150300021-INS-195
chr2-150301613-INS-5068
chr2-150319523-INS-525
chr2-150368397-INS-225
chr2-150381859-INS-52
chr2-150392346-INS-71
chr2-150405175-INS-153
chr2-150418835-INS-70
chr2-150445923-INS-71
chr2-150473071-INS-61
chr2-150474687-INS-167
chr2-150498809-INS-106
chr2-150519864-INS-138
chr2-150565703-INS-177
chr2-150567669-INS-413
chr2-150618588-INS-227
chr2-150620084-INS-238
chr2-150636183-INS-239
chr2-150641911-INS-212
chr2-150651229-INS-131
chr2-150663745-INS-220
chr2-150685232-INS-389
chr2-150723849-INS-199
chr2-150740534-INS-230
chr2-150754798-INS-185
chr2-150759045-INS-208
chr2-150763019-INS-205
chr2-150766885-INS-560
chr2-150774017-INS-189
chr2-150822843-INS-64
chr2-150836915-INS-70
chr2-150843731-INS-61
chr2-150875596-INS-832
chr2-150942066-INS-5468
chr2-150946022-INS-121
chr2-150949339-INS-101
chr2-151011338-INS-6332
chr2-151017662-INS-344
chr2-151032630-INS-3041
chr2-151043604-INS-358
chr2-151143441-INS-2465
chr2-151146464-INS-96
chr2-151165799-INS-1889
chr2-151184866-INS-201
chr2-151212207-INS-358
chr2-151215357-INS-6520
chr2-151223092-INS-89
chr2-151293374-INS-270
chr2-151312294-INS-160
chr2-151367420-INS-650
chr2-151423399-INS-642
chr2-151425207-INS-592
chr2-151479341-INS-205
chr2-151492910-INS-98
chr2-151535526-INS-878
chr2-151588705-INS-268
chr2-151592871-INS-499
chr2-151610383-INS-250
chr2-151623699-INS-158
chr2-151665839-INS-56
chr2-151685700-INS-91
chr2-151689470-INS-196
chr2-151695220-INS-6403
chr2-151695697-INS-525

chr2-151707276-INS-87
chr2-151734040-INS-315
chr2-151829472-INS-141
chr2-151863982-INS-163
chr2-151865369-INS-186
chr2-151873951-INS-206
chr2-151877040-INS-71
chr2-151882058-INS-98
chr2-151883478-INS-112
chr2-151925276-INS-149
chr2-151942916-INS-144
chr2-152013975-INS-111
chr2-152020841-INS-4518
chr2-152020842-INS-340
chr2-152022896-INS-123
chr2-152060274-INS-80
chr2-152062611-INS-98
chr2-152092840-INS-99
chr2-152100770-INS-178
chr2-152126884-INS-102
chr2-152148789-INS-388
chr2-152158843-INS-130
chr2-152172725-INS-200
chr2-152193541-INS-381
chr2-152208725-INS-65
chr2-152231292-INS-169
chr2-152232042-INS-351
chr2-152234001-INS-81
chr2-152235122-INS-105
chr2-152262614-INS-206
chr2-152264862-INS-54
chr2-152267552-INS-106
chr2-152272103-INS-80
chr2-152273146-INS-206
chr2-152279917-INS-74
chr2-152286922-INS-171
chr2-152289191-INS-104
chr2-152296747-INS-155
chr2-152298913-INS-151
chr2-152300111-INS-79
chr2-152302903-INS-535
chr2-152302936-INS-73
chr2-152309795-INS-715
chr2-152316176-INS-755
chr2-152335458-INS-192
chr2-152383934-INS-443
chr2-152410648-INS-169
chr2-152416143-INS-227
chr2-152437674-INS-56
chr2-152454277-INS-164
chr2-152455560-INS-258
chr2-152464864-INS-74
chr2-152487730-INS-6408
chr2-152490498-INS-104
chr2-152491535-INS-652
chr2-152493360-INS-329
chr2-152518428-INS-64
chr2-152590653-INS-198
chr2-152597796-INS-213
chr2-152610246-INS-56
chr2-152624986-INS-129
chr2-152628417-INS-51
chr2-152638408-INS-129
chr2-152643124-INS-195
chr2-152649242-INS-54
chr2-152661671-INS-216
chr2-152665238-INS-180
chr2-152703870-INS-213
chr2-152708334-INS-57
chr2-152744573-INS-260
chr2-152754930-INS-212
chr2-152763759-INS-64
chr2-152800109-INS-166
chr2-152803408-INS-122
chr2-152804745-INS-505
chr2-152823765-INS-201
chr2-152892177-INS-54

chr2-152911349-INS-68
chr2-152967714-INS-52
chr2-153014808-INS-103
chr2-153087553-INS-559
chr2-153102988-INS-52
chr2-153117885-INS-433
chr2-153124269-INS-294
chr2-153129085-INS-51
chr2-153136544-INS-177
chr2-153140476-INS-213
chr2-153143792-INS-524
chr2-153152998-INS-135
chr2-153168506-INS-54
chr2-153178747-INS-66
chr2-153182420-INS-103
chr2-153185778-INS-391
chr2-153215658-INS-646
chr2-153225381-INS-1551
chr2-153268722-INS-118
chr2-153281834-INS-87
chr2-153288770-INS-74
chr2-153300729-INS-203
chr2-153306772-INS-204
chr2-153346923-INS-52
chr2-153376466-INS-128
chr2-153381193-INS-58
chr2-153399261-INS-167
chr2-153404051-INS-82
chr2-153411245-INS-416
chr2-153413318-INS-198
chr2-153414973-INS-252
chr2-153419905-INS-54
chr2-153435929-INS-221
chr2-153440712-INS-76
chr2-153445452-INS-363
chr2-153452469-INS-8734
chr2-153456208-INS-143
chr2-153458329-INS-53
chr2-153466797-INS-357
chr2-153482683-INS-198
chr2-153486831-INS-202
chr2-153488396-INS-128
chr2-153515180-INS-683
chr2-153580108-INS-61
chr2-153606235-INS-881
chr2-153624360-INS-108
chr2-153647254-INS-103
chr2-153649444-INS-88
chr2-153650760-INS-52
chr2-153675286-INS-109
chr2-153677463-INS-77
chr2-153682141-INS-73
chr2-153699754-INS-1812
chr2-153708362-INS-68
chr2-153715981-INS-162
chr2-153738668-INS-187
chr2-153781122-INS-363
chr2-153793553-INS-91
chr2-153797035-INS-258
chr2-153821041-INS-129
chr2-153833191-INS-332
chr2-153865291-INS-175
chr2-153866562-INS-273
chr2-153891707-INS-252
chr2-153925705-INS-90
chr2-153955859-INS-1205
chr2-153994683-INS-375
chr2-153995610-INS-187
chr2-154028501-INS-113
chr2-154049684-INS-119
chr2-154078926-INS-227
chr2-154088515-INS-198
chr2-154089806-INS-2339
chr2-154111565-INS-88
chr2-154126487-INS-84
chr2-154135376-INS-76
chr2-154137129-INS-56

chr2-154147832-INS-94
chr2-154162684-INS-149
chr2-154166911-INS-7081
chr2-154169291-INS-57
chr2-154173340-INS-81
chr2-154174771-INS-445
chr2-154205081-INS-293
chr2-154205567-INS-127
chr2-154207190-INS-75
chr2-154212085-INS-214
chr2-154216766-INS-293
chr2-154230354-INS-124
chr2-154244165-INS-203
chr2-154251617-INS-169
chr2-154258553-INS-51
chr2-154265963-INS-211
chr2-154266963-INS-352
chr2-154274593-INS-149
chr2-154275990-INS-283
chr2-154283316-INS-583
chr2-154291135-INS-197
chr2-154299630-INS-53
chr2-154301925-INS-56
chr2-154311620-INS-459
chr2-154320261-INS-318
chr2-154321279-INS-212
chr2-154344300-INS-190
chr2-154348663-INS-243
chr2-154353319-INS-208
chr2-154359700-INS-417
chr2-154364481-INS-93
chr2-154414884-INS-121
chr2-154443915-INS-104
chr2-154458669-INS-258
chr2-154459824-INS-171
chr2-154472051-INS-121
chr2-154480584-INS-5112
chr2-154484175-INS-145
chr2-154508286-INS-914
chr2-154528233-INS-72
chr2-154540616-INS-106
chr2-154544276-INS-209
chr2-154548356-INS-88
chr2-154550511-INS-255
chr2-154555151-INS-63
chr2-154566635-INS-184
chr2-154571656-INS-128
chr2-154584237-INS-78
chr2-154593254-INS-108
chr2-154614701-INS-68
chr2-154625684-INS-75
chr2-154715979-INS-184
chr2-154719783-INS-174
chr2-154724395-INS-54
chr2-154732041-INS-175
chr2-154738085-INS-491
chr2-154740094-INS-8747
chr2-154753037-INS-60
chr2-154755328-INS-77
chr2-154761634-INS-258
chr2-154766922-INS-88
chr2-154769891-INS-112
chr2-154772832-INS-64
chr2-154780944-INS-375
chr2-154788373-INS-332
chr2-154796645-INS-7123
chr2-154809110-INS-197
chr2-154811003-INS-107
chr2-154818846-INS-349
chr2-154825279-INS-156
chr2-154844960-INS-218
chr2-154854596-INS-442
chr2-154881091-INS-275
chr2-154885650-INS-189
chr2-154903358-INS-341
chr2-154914731-INS-411
chr2-154927540-INS-202

chr2-154960130-INS-199
chr2-154979912-INS-204
chr2-155033496-INS-212
chr2-155037191-INS-205
chr2-155039676-INS-54
chr2-155049104-INS-204
chr2-155072045-INS-5974
chr2-155082183-INS-70
chr2-155083945-INS-66
chr2-155085122-INS-821
chr2-155088929-INS-100
chr2-155101482-INS-203
chr2-155111708-INS-98
chr2-155114626-INS-198
chr2-155115742-INS-164
chr2-155125095-INS-94
chr2-155127947-INS-74
chr2-155141865-INS-218
chr2-155152339-INS-192
chr2-155171425-INS-66
chr2-155187843-INS-157
chr2-155225434-INS-58
chr2-155269179-INS-973
chr2-155269801-INS-714
chr2-155274481-INS-52
chr2-155286347-INS-387
chr2-155290999-INS-155
chr2-155328822-INS-222
chr2-155387917-INS-126
chr2-155443335-INS-6442
chr2-155449768-INS-106
chr2-155463390-INS-52
chr2-155473911-INS-60
chr2-155475024-INS-440
chr2-155504814-INS-173
chr2-155506437-INS-421
chr2-155508660-INS-84
chr2-155512205-INS-1866
chr2-155527494-INS-189
chr2-155542135-INS-98
chr2-155550794-INS-117
chr2-155580099-INS-196
chr2-155583024-INS-57
chr2-155583466-INS-235
chr2-155594106-INS-143
chr2-155600092-INS-2764
chr2-155611525-INS-6442
chr2-155615291-INS-403
chr2-155635702-INS-119
chr2-155638448-INS-405
chr2-155685890-INS-52
chr2-155707646-INS-91
chr2-155709120-INS-581
chr2-155770743-INS-168
chr2-155776227-INS-211
chr2-155780221-INS-399
chr2-155797574-INS-221
chr2-155800651-INS-405
chr2-155804622-INS-162
chr2-155821876-INS-209
chr2-155846780-INS-194
chr2-155863189-INS-167
chr2-155876941-INS-280
chr2-155892592-INS-205
chr2-155892824-INS-549
chr2-155917028-INS-54
chr2-155963490-INS-154
chr2-155967404-INS-179
chr2-155982934-INS-90
chr2-156026308-INS-178
chr2-156031891-INS-233
chr2-156034920-INS-111
chr2-156036201-INS-216
chr2-156043461-INS-160
chr2-156046331-INS-111
chr2-156086514-INS-70
chr2-156103267-INS-104

chr2-156117207-INS-61
chr2-156121644-INS-65
chr2-156149190-INS-66
chr2-156165360-INS-572
chr2-156178639-INS-177
chr2-156196956-INS-57
chr2-156235004-INS-50
chr2-156236266-INS-110
chr2-156248469-INS-400
chr2-156252058-INS-240
chr2-156253188-INS-80
chr2-156254674-INS-204
chr2-156275829-INS-224
chr2-156281232-INS-65
chr2-156285705-INS-82
chr2-156287867-INS-54
chr2-156299753-INS-331
chr2-156321107-INS-207
chr2-156326237-INS-250
chr2-156351265-INS-108
chr2-156358690-INS-119
chr2-156386166-INS-79
chr2-156395598-INS-61
chr2-156397185-INS-72
chr2-156417203-INS-90
chr2-156444521-INS-117
chr2-156483491-INS-113
chr2-156533342-INS-292
chr2-156552172-INS-279
chr2-156592560-INS-208
chr2-156616292-INS-177
chr2-156626511-INS-230
chr2-156628818-INS-485
chr2-156679392-INS-57
chr2-156709130-INS-70
chr2-156712132-INS-54
chr2-156731275-INS-51
chr2-156735318-INS-223
chr2-156738235-INS-212
chr2-156740664-INS-100
chr2-156742565-INS-55
chr2-156744436-INS-54
chr2-156746774-INS-423
chr2-156749757-INS-7951
chr2-156753984-INS-66
chr2-156760483-INS-205
chr2-156785772-INS-209
chr2-156787604-INS-111
chr2-156806831-INS-209
chr2-156814242-INS-60
chr2-156827416-INS-203
chr2-156840177-INS-252
chr2-156844946-INS-199
chr2-156874375-INS-134
chr2-156932528-INS-162
chr2-156940944-INS-76
chr2-156950605-INS-270
chr2-156993462-INS-207
chr2-157007826-INS-72
chr2-157015039-INS-208
chr2-157028555-INS-260
chr2-157036285-INS-113
chr2-157050183-INS-203
chr2-157065669-INS-140
chr2-157066404-INS-918
chr2-157074011-INS-72
chr2-157078286-INS-327
chr2-157092395-INS-323
chr2-157097803-INS-97
chr2-157134646-INS-81
chr2-157140741-INS-194
chr2-157154017-INS-74
chr2-157168404-INS-52
chr2-157199486-INS-90
chr2-157221458-INS-943
chr2-157242709-INS-615
chr2-157250870-INS-153

chr2-157274759-INS-226
chr2-157310631-INS-388
chr2-157320895-INS-59
chr2-157323415-INS-180
chr2-157323415-INS-714
chr2-157330396-INS-59
chr2-157335921-INS-167
chr2-157341448-INS-117
chr2-157345710-INS-87
chr2-157354567-INS-205
chr2-157365844-INS-68
chr2-157376806-INS-386
chr2-157385661-INS-87
chr2-157386213-INS-162
chr2-157426633-INS-213
chr2-157435642-INS-625
chr2-157440741-INS-77
chr2-157444257-INS-246
chr2-157452753-INS-115
chr2-157469039-INS-116
chr2-157477523-INS-60
chr2-157483979-INS-153
chr2-157489250-INS-81
chr2-157491082-INS-541
chr2-157494010-INS-139
chr2-157505603-INS-154
chr2-157528839-INS-56
chr2-157531868-INS-343
chr2-157546554-INS-172
chr2-157549935-INS-60
chr2-157552014-INS-162
chr2-157571896-INS-227
chr2-157577075-INS-226
chr2-157607518-INS-204
chr2-157611081-INS-52
chr2-157622123-INS-76
chr2-157669697-INS-359
chr2-157674467-INS-3208
chr2-157684748-INS-598
chr2-157711073-INS-68
chr2-157721479-INS-222
chr2-157723024-INS-610
chr2-157729132-INS-160
chr2-157734475-INS-121
chr2-157749012-INS-401
chr2-157770030-INS-156
chr2-157775069-INS-61
chr2-157779996-INS-214
chr2-157800946-INS-90
chr2-157807746-INS-56
chr2-157818775-INS-119
chr2-157856113-INS-263
chr2-157875470-INS-68
chr2-157893286-INS-72
chr2-157927195-INS-431
chr2-157968310-INS-116
chr2-157997428-INS-139
chr2-158028341-INS-102
chr2-158092365-INS-58
chr2-158092940-INS-116
chr2-158112949-INS-68
chr2-158115608-INS-68
chr2-158116868-INS-247
chr2-158119600-INS-56
chr2-158120316-INS-53
chr2-158122065-INS-66
chr2-158164395-INS-5095
chr2-158172087-INS-114
chr2-158182498-INS-129
chr2-158194588-INS-162
chr2-158194604-INS-289
chr2-158219362-INS-55
chr2-158225523-INS-129
chr2-158225525-INS-202
chr2-158229643-INS-139
chr2-158235530-INS-231
chr2-158242851-INS-96

chr2-158245440-INS-4886
chr2-158259098-INS-70
chr2-158259181-INS-836
chr2-158310497-INS-762
chr2-158358726-INS-88
chr2-158371919-INS-151
chr2-158375985-INS-56
chr2-158377871-INS-232
chr2-158382805-INS-61
chr2-158398609-INS-82
chr2-158399106-INS-201
chr2-158404768-INS-80
chr2-158406802-INS-51
chr2-158409050-INS-173
chr2-158418547-INS-104
chr2-158419954-INS-69
chr2-158427688-INS-108
chr2-158429907-INS-75
chr2-158469287-INS-539
chr2-158470434-INS-288
chr2-158471754-INS-201
chr2-158473003-INS-498
chr2-158482664-INS-3642
chr2-158486167-INS-412
chr2-158501960-INS-63
chr2-158520939-INS-80
chr2-158524267-INS-68
chr2-158546157-INS-57
chr2-158552132-INS-6452
chr2-158566673-INS-55
chr2-158588777-INS-270
chr2-158612695-INS-287
chr2-158620231-INS-485
chr2-158644314-INS-327
chr2-158650682-INS-118
chr2-158682974-INS-204
chr2-158694390-INS-222
chr2-158707573-INS-107
chr2-158723413-INS-258
chr2-158725956-INS-88
chr2-158733764-INS-58
chr2-158738958-INS-365
chr2-158766527-INS-164
chr2-158777905-INS-672
chr2-158795044-INS-203
chr2-158807974-INS-58
chr2-158812794-INS-64
chr2-158827954-INS-498
chr2-158854008-INS-487
chr2-158869636-INS-74
chr2-158883805-INS-676
chr2-158889449-INS-54
chr2-158941810-INS-75
chr2-158972075-INS-68
chr2-158977769-INS-61
chr2-158982646-INS-87
chr2-159003357-INS-52
chr2-159026646-INS-52
chr2-159048403-INS-274
chr2-159061003-INS-6293
chr2-159064686-INS-72
chr2-159076519-INS-52
chr2-159086482-INS-82
chr2-159140326-INS-54
chr2-159149603-INS-65
chr2-159164718-INS-55
chr2-159172838-INS-2252
chr2-159207305-INS-84
chr2-159233994-INS-52
chr2-159238417-INS-408
chr2-159276984-INS-218
chr2-159313245-INS-6680
chr2-159318745-INS-151
chr2-159320288-INS-107
chr2-159331119-INS-66
chr2-159344295-INS-62
chr2-159360927-INS-106

chr2-159380663-INS-54
chr2-159392427-INS-622
chr2-159401181-INS-66
chr2-159450332-INS-58
chr2-159466945-INS-52
chr2-159506388-INS-134
chr2-159514945-INS-106
chr2-159546473-INS-61
chr2-159578014-INS-74
chr2-159582621-INS-60
chr2-159599574-INS-1533
chr2-159600910-INS-275
chr2-159613065-INS-754
chr2-159652949-INS-304
chr2-159662810-INS-152
chr2-159670083-INS-142
chr2-159682246-INS-125
chr2-159685370-INS-64
chr2-159696195-INS-241
chr2-159704145-INS-303
chr2-159723300-INS-160
chr2-159729330-INS-66
chr2-159772160-INS-58
chr2-159789492-INS-266
chr2-159790387-INS-400
chr2-159839216-INS-141
chr2-159866753-INS-67
chr2-159885141-INS-151
chr2-159917069-INS-410
chr2-159934065-INS-90
chr2-159945091-INS-144
chr2-159958647-INS-124
chr2-159981524-INS-64
chr2-160001848-INS-190
chr2-160010973-INS-7128
chr2-160034023-INS-102
chr2-160048764-INS-64
chr2-160063175-INS-109
chr2-160092870-INS-306
chr2-160094417-INS-407
chr2-160094949-INS-52
chr2-160101441-INS-90
chr2-160119500-INS-120
chr2-160123645-INS-59
chr2-160144759-INS-6538
chr2-160161131-INS-72
chr2-160165790-INS-54
chr2-160188576-INS-108
chr2-160193964-INS-60
chr2-160233839-INS-144
chr2-160246095-INS-92
chr2-160247043-INS-91
chr2-160280634-INS-171
chr2-160290215-INS-329
chr2-160294092-INS-7043
chr2-160300574-INS-58
chr2-160360610-INS-169
chr2-160365351-INS-65
chr2-160383230-INS-150
chr2-160389770-INS-200
chr2-160393209-INS-93
chr2-160408905-INS-293
chr2-160419456-INS-54
chr2-160430772-INS-120
chr2-160435670-INS-496
chr2-160444793-INS-378
chr2-160453194-INS-75
chr2-160459685-INS-180
chr2-160466452-INS-207
chr2-160467729-INS-1892
chr2-160467914-INS-53
chr2-160502235-INS-216
chr2-160521215-INS-281
chr2-160533570-INS-638
chr2-160538603-INS-200
chr2-160550295-INS-210
chr2-160556864-INS-68

chr2-160585235-INS-177
chr2-160615223-INS-60
chr2-160653554-INS-218
chr2-160668876-INS-116
chr2-160735549-INS-70
chr2-160756557-INS-130
chr2-160760425-INS-58
chr2-160763601-INS-65
chr2-160777677-INS-176
chr2-160804561-INS-446
chr2-160815713-INS-70
chr2-160834545-INS-213
chr2-160866354-INS-88
chr2-160879284-INS-75
chr2-160879788-INS-599
chr2-160891965-INS-110
chr2-160956082-INS-202
chr2-160968985-INS-242
chr2-160971646-INS-204
chr2-160986100-INS-6705
chr2-160992485-INS-232
chr2-160999540-INS-79
chr2-161012171-INS-252
chr2-161035882-INS-548
chr2-161039681-INS-7134
chr2-161047713-INS-289
chr2-161064570-INS-137
chr2-161081240-INS-76
chr2-161083081-INS-72
chr2-161092900-INS-73
chr2-161123666-INS-167
chr2-161126258-INS-65
chr2-161134736-INS-1874
chr2-161147492-INS-52
chr2-161160824-INS-490
chr2-161184981-INS-549
chr2-161193646-INS-62
chr2-161249902-INS-56
chr2-161263578-INS-64
chr2-161264310-INS-314
chr2-161267762-INS-646
chr2-161288574-INS-80
chr2-161340093-INS-107
chr2-161384360-INS-68
chr2-161391280-INS-82
chr2-161403903-INS-590
chr2-161404418-INS-779
chr2-161410007-INS-61
chr2-161414827-INS-51
chr2-161438974-INS-139
chr2-161479711-INS-6862
chr2-161513805-INS-52
chr2-161562821-INS-54
chr2-161565155-INS-6830
chr2-161613579-INS-200
chr2-161622242-INS-199
chr2-161627274-INS-78
chr2-161648978-INS-501
chr2-161663902-INS-55
chr2-161666628-INS-90
chr2-161691914-INS-1525
chr2-161708283-INS-3560
chr2-161718022-INS-67
chr2-161724943-INS-134
chr2-161761820-INS-122
chr2-161767564-INS-173
chr2-161806946-INS-54
chr2-161849412-INS-615
chr2-161871368-INS-58
chr2-161892083-INS-6233
chr2-161895188-INS-640
chr2-161909201-INS-226
chr2-161923881-INS-96
chr2-161931291-INS-211
chr2-161933689-INS-2389
chr2-161934511-INS-1204
chr2-161938364-INS-147

chr2-161946099-INS-97
chr2-161951025-INS-92
chr2-161964219-INS-70
chr2-161973843-INS-662
chr2-161999611-INS-2455
chr2-162071009-INS-372
chr2-162123817-INS-6677
chr2-162134475-INS-53
chr2-162138354-INS-64
chr2-162162704-INS-165
chr2-162163938-INS-338
chr2-162202957-INS-60
chr2-162213713-INS-69
chr2-162226679-INS-613
chr2-162238006-INS-362
chr2-162285712-INS-165
chr2-162286969-INS-57
chr2-162291520-INS-149
chr2-162297413-INS-1131
chr2-162319372-INS-347
chr2-162331910-INS-421
chr2-162346659-INS-56
chr2-162395685-INS-72
chr2-162453134-INS-132
chr2-162472603-INS-264
chr2-162474680-INS-401
chr2-162484218-INS-809
chr2-162498277-INS-86
chr2-162514890-INS-175
chr2-162524160-INS-64
chr2-162544457-INS-62
chr2-162546036-INS-59
chr2-162579879-INS-296
chr2-162592875-INS-181
chr2-162596312-INS-250
chr2-162613996-INS-54
chr2-162626511-INS-775
chr2-162638478-INS-199
chr2-162657669-INS-69
chr2-162661994-INS-262
chr2-162705858-INS-63
chr2-162720730-INS-68
chr2-162725483-INS-118
chr2-162736547-INS-88
chr2-162769631-INS-145
chr2-162772065-INS-298
chr2-162779735-INS-217
chr2-162804568-INS-193
chr2-162820354-INS-223
chr2-162832647-INS-58
chr2-162845659-INS-73
chr2-162853188-INS-185
chr2-162864059-INS-258
chr2-162879824-INS-165
chr2-162882147-INS-56
chr2-162883408-INS-586
chr2-162885696-INS-194
chr2-162895225-INS-198
chr2-162906502-INS-257
chr2-162915134-INS-200
chr2-162933566-INS-74
chr2-162942929-INS-194
chr2-162947844-INS-52
chr2-162958655-INS-780
chr2-162960212-INS-155
chr2-162966608-INS-211
chr2-162992906-INS-64
chr2-163027273-INS-280
chr2-163040027-INS-4170
chr2-163074474-INS-85
chr2-163102397-INS-177
chr2-163118415-INS-66
chr2-163154719-INS-98
chr2-163172474-INS-192
chr2-163182758-INS-68
chr2-163186897-INS-78
chr2-163198882-INS-111

chr2-163200887-INS-88
chr2-163211758-INS-112
chr2-163213481-INS-76
chr2-163251824-INS-829
chr2-163253158-INS-129
chr2-163265217-INS-183
chr2-163267323-INS-111
chr2-163308702-INS-221
chr2-163327882-INS-395
chr2-163333745-INS-65
chr2-163335921-INS-53
chr2-163346785-INS-58
chr2-163368308-INS-6378
chr2-163370156-INS-158
chr2-163373663-INS-188
chr2-163396591-INS-130
chr2-163417839-INS-58
chr2-163424451-INS-68
chr2-163428650-INS-274
chr2-163439521-INS-285
chr2-163439819-INS-302
chr2-163447733-INS-202
chr2-163469319-INS-182
chr2-163482051-INS-288
chr2-163489395-INS-387
chr2-163514155-INS-246
chr2-163544816-INS-205
chr2-163551638-INS-176
chr2-163577159-INS-213
chr2-163583568-INS-204
chr2-163599973-INS-356
chr2-163601813-INS-94
chr2-163604132-INS-64
chr2-163608670-INS-69
chr2-163615746-INS-56
chr2-163620602-INS-51
chr2-163625227-INS-190
chr2-163652174-INS-56
chr2-163698346-INS-105
chr2-163699331-INS-436
chr2-163729248-INS-52
chr2-163766964-INS-222
chr2-163773852-INS-150
chr2-163818009-INS-200
chr2-163824934-INS-60
chr2-163830913-INS-66
chr2-163855387-INS-96
chr2-163866114-INS-949
chr2-163870311-INS-151
chr2-163890553-INS-253
chr2-163893816-INS-229
chr2-163899848-INS-151
chr2-163905960-INS-54
chr2-163912238-INS-114
chr2-163917833-INS-85
chr2-163940175-INS-303
chr2-163943588-INS-340
chr2-163948256-INS-215
chr2-164014197-INS-5238
chr2-164039789-INS-511
chr2-164078221-INS-519
chr2-164087485-INS-52
chr2-164116003-INS-304
chr2-164119685-INS-66
chr2-164152936-INS-97
chr2-164154386-INS-52
chr2-164173641-INS-94
chr2-164185039-INS-559
chr2-164190663-INS-69
chr2-164206474-INS-429
chr2-164208939-INS-207
chr2-164216675-INS-169
chr2-164219226-INS-198
chr2-164240431-INS-262
chr2-164240433-INS-937
chr2-164264381-INS-201
chr2-164295772-INS-84

chr2-164316076-INS-6440
chr2-164341650-INS-94
chr2-164353998-INS-109
chr2-164384184-INS-210
chr2-164391807-INS-309
chr2-164400107-INS-101
chr2-164402325-INS-61
chr2-164413490-INS-51
chr2-164417218-INS-626
chr2-164424157-INS-66
chr2-164424956-INS-911
chr2-164431620-INS-184
chr2-164444023-INS-300
chr2-164455495-INS-301
chr2-164457672-INS-84
chr2-164485500-INS-75
chr2-164495973-INS-56
chr2-164500694-INS-70
chr2-164501368-INS-142
chr2-164508218-INS-88
chr2-164530344-INS-1482
chr2-164544787-INS-600
chr2-164547060-INS-1260
chr2-164547069-INS-94
chr2-164547218-INS-491
chr2-164568578-INS-224
chr2-164570443-INS-58
chr2-164583379-INS-88
chr2-164592010-INS-218
chr2-164601983-INS-208
chr2-164603276-INS-126
chr2-164639502-INS-63
chr2-164653656-INS-96
chr2-164657867-INS-59
chr2-164677635-INS-88
chr2-164702074-INS-253
chr2-164708308-INS-126
chr2-164715842-INS-213
chr2-164738995-INS-209
chr2-164742703-INS-156
chr2-164750632-INS-211
chr2-164786766-INS-201
chr2-164791082-INS-68
chr2-164803851-INS-75
chr2-164815949-INS-98
chr2-164819784-INS-60
chr2-164824130-INS-198
chr2-164825290-INS-86
chr2-164880987-INS-58
chr2-164892218-INS-56
chr2-164901496-INS-241
chr2-164908184-INS-73
chr2-164909254-INS-126
chr2-164923927-INS-95
chr2-164937564-INS-374
chr2-164977746-INS-51
chr2-164996161-INS-149
chr2-164996691-INS-172
chr2-165021943-INS-60
chr2-165050090-INS-90
chr2-165051054-INS-152
chr2-165091201-INS-55
chr2-165107644-INS-135
chr2-165120708-INS-98
chr2-165134615-INS-146
chr2-165142869-INS-7156
chr2-165176715-INS-130
chr2-165177909-INS-125
chr2-165181124-INS-1864
chr2-165188156-INS-312
chr2-165202874-INS-120
chr2-165228627-INS-54
chr2-165253552-INS-307
chr2-165293500-INS-56
chr2-165303948-INS-59
chr2-165307428-INS-182
chr2-165321622-INS-72

chr2-165356010-INS-162
chr2-165368863-INS-59
chr2-165372493-INS-207
chr2-165382799-INS-429
chr2-165397042-INS-55
chr2-165408529-INS-90
chr2-165453755-INS-63
chr2-165461689-INS-90
chr2-165476159-INS-59
chr2-165486987-INS-94
chr2-165543003-INS-238
chr2-165565374-INS-69
chr2-165577752-INS-139
chr2-165582601-INS-338
chr2-165585697-INS-70
chr2-165586181-INS-77
chr2-165590056-INS-52
chr2-165591253-INS-62
chr2-165604424-INS-443
chr2-165618973-INS-451
chr2-165630448-INS-74
chr2-165638226-INS-452
chr2-165644001-INS-213
chr2-165689449-INS-203
chr2-165737631-INS-1992
chr2-165744657-INS-169
chr2-165762272-INS-186
chr2-165776556-INS-180
chr2-165778870-INS-224
chr2-165780918-INS-218
chr2-165784492-INS-149
chr2-165786712-INS-185
chr2-165789659-INS-212
chr2-165790290-INS-208
chr2-165811370-INS-87
chr2-165814873-INS-173
chr2-165816816-INS-114
chr2-165818387-INS-5471
chr2-165818626-INS-215
chr2-165822614-INS-244
chr2-165823155-INS-72
chr2-165828980-INS-56
chr2-165832446-INS-221
chr2-165838644-INS-63
chr2-165846758-INS-96
chr2-165850098-INS-211
chr2-165858333-INS-481
chr2-165864797-INS-206
chr2-165868314-INS-99
chr2-165889057-INS-75
chr2-165938582-INS-99
chr2-165944277-INS-68
chr2-165951374-INS-218
chr2-165959993-INS-422
chr2-165969506-INS-110
chr2-166003038-INS-6467
chr2-166003754-INS-59
chr2-166011342-INS-6399
chr2-166013005-INS-166
chr2-166062770-INS-1888
chr2-166070360-INS-58
chr2-166076198-INS-60
chr2-166086340-INS-63
chr2-166104964-INS-60
chr2-166105464-INS-6442
chr2-166114460-INS-205
chr2-166136064-INS-69
chr2-166143586-INS-97
chr2-166149889-INS-7261
chr2-166193357-INS-574
chr2-166210172-INS-96
chr2-166244759-INS-233
chr2-166254929-INS-61
chr2-166255717-INS-889
chr2-166266234-INS-205
chr2-166270722-INS-82
chr2-166271793-INS-92

chr2-166277257-INS-54
chr2-166289974-INS-180
chr2-166309583-INS-64
chr2-166312596-INS-799
chr2-166331646-INS-210
chr2-166365696-INS-208
chr2-166371040-INS-115
chr2-166406390-INS-122
chr2-166421192-INS-64
chr2-166446207-INS-217
chr2-166456939-INS-73
chr2-166485698-INS-75
chr2-166501367-INS-428
chr2-166507241-INS-51
chr2-166509173-INS-611
chr2-166530971-INS-227
chr2-166544818-INS-186
chr2-166580855-INS-218
chr2-166612682-INS-181
chr2-166618844-INS-156
chr2-166621904-INS-194
chr2-166623258-INS-202
chr2-166627066-INS-86
chr2-166665545-INS-201
chr2-166671387-INS-90
chr2-166681352-INS-102
chr2-166690360-INS-140
chr2-166698966-INS-168
chr2-166702403-INS-187
chr2-166704183-INS-5289
chr2-166725471-INS-59
chr2-166727374-INS-64
chr2-166735774-INS-85
chr2-166743062-INS-120
chr2-166755152-INS-56
chr2-166758102-INS-223
chr2-166765342-INS-208
chr2-166797825-INS-120
chr2-166802063-INS-317
chr2-166810282-INS-66
chr2-166844769-INS-187
chr2-166845996-INS-484
chr2-166847308-INS-376
chr2-166872196-INS-201
chr2-166887739-INS-54
chr2-166911311-INS-212
chr2-166926848-INS-139
chr2-166932827-INS-63
chr2-166948710-INS-123
chr2-166966593-INS-212
chr2-166969596-INS-54
chr2-166979898-INS-101
chr2-166999250-INS-92
chr2-167002499-INS-204
chr2-167011256-INS-263
chr2-167012260-INS-53
chr2-167024588-INS-178
chr2-167027402-INS-63
chr2-167040050-INS-188
chr2-167055296-INS-213
chr2-167069011-INS-176
chr2-167075634-INS-279
chr2-167079566-INS-111
chr2-167080989-INS-206
chr2-167091142-INS-54
chr2-167096092-INS-58
chr2-167099335-INS-194
chr2-167111329-INS-1354
chr2-167111339-INS-429
chr2-167120548-INS-496
chr2-167123236-INS-197
chr2-167132782-INS-55
chr2-167164466-INS-331
chr2-167183854-INS-202
chr2-167195794-INS-60
chr2-167204198-INS-72
chr2-167207101-INS-228

chr2-167211076-INS-91
chr2-167220967-INS-354
chr2-167223326-INS-70
chr2-167226782-INS-399
chr2-167239755-INS-444
chr2-167240653-INS-367
chr2-167257382-INS-68
chr2-167260634-INS-315
chr2-167274094-INS-7214
chr2-167284304-INS-188
chr2-167297175-INS-102
chr2-167299955-INS-156
chr2-167308852-INS-106
chr2-167314303-INS-509
chr2-167347996-INS-56
chr2-167355173-INS-57
chr2-167369624-INS-167
chr2-167391401-INS-52
chr2-167411353-INS-66
chr2-167432046-INS-91
chr2-167435093-INS-259
chr2-167470741-INS-223
chr2-167478403-INS-61
chr2-167511303-INS-521
chr2-167518727-INS-57
chr2-167543709-INS-440
chr2-167549614-INS-286
chr2-167560399-INS-152
chr2-167581809-INS-368
chr2-167583224-INS-377
chr2-167626632-INS-72
chr2-167633283-INS-88
chr2-167646170-INS-113
chr2-167658773-INS-76
chr2-167660486-INS-54
chr2-167686207-INS-68
chr2-167688015-INS-80
chr2-167721637-INS-58
chr2-167744100-INS-194
chr2-167755617-INS-86
chr2-167829194-INS-64
chr2-167843334-INS-260
chr2-167846613-INS-722
chr2-167849595-INS-5319
chr2-167864533-INS-208
chr2-167869271-INS-5549
chr2-167869389-INS-101
chr2-167872276-INS-198
chr2-167888217-INS-127
chr2-167913755-INS-5340
chr2-167932762-INS-64
chr2-167954747-INS-104
chr2-167960787-INS-422
chr2-167968077-INS-106
chr2-167975833-INS-66
chr2-167995546-INS-56
chr2-168015673-INS-83
chr2-168017528-INS-198
chr2-168029123-INS-146
chr2-168044215-INS-198
chr2-168068521-INS-71
chr2-168084375-INS-199
chr2-168096032-INS-492
chr2-168106634-INS-52
chr2-168107692-INS-58
chr2-168129330-INS-83
chr2-168135667-INS-54
chr2-168138094-INS-61
chr2-168142910-INS-254
chr2-168151792-INS-107
chr2-168153732-INS-52
chr2-168165257-INS-89
chr2-168179086-INS-222
chr2-168196197-INS-60
chr2-168217410-INS-80
chr2-168220146-INS-915
chr2-168251555-INS-64

chr2-168265150-INS-92
chr2-168292507-INS-495
chr2-168321826-INS-118
chr2-168337888-INS-67
chr2-168341592-INS-112
chr2-168344868-INS-251
chr2-168349249-INS-73
chr2-168369801-INS-6404
chr2-168383121-INS-51
chr2-168387373-INS-91
chr2-168410557-INS-148
chr2-168434569-INS-369
chr2-168439174-INS-408
chr2-168461354-INS-676
chr2-168461453-INS-330
chr2-168480418-INS-177
chr2-168487370-INS-315
chr2-168498054-INS-67
chr2-168522091-INS-141
chr2-168565940-INS-87
chr2-168572883-INS-183
chr2-168574878-INS-221
chr2-168589266-INS-96
chr2-168590700-INS-128
chr2-168632632-INS-210
chr2-168634805-INS-64
chr2-168638603-INS-63
chr2-168640314-INS-117
chr2-168654616-INS-228
chr2-168659967-INS-56
chr2-168674669-INS-202
chr2-168677682-INS-93
chr2-168687179-INS-217
chr2-168690388-INS-556
chr2-168697000-INS-137
chr2-168699171-INS-60
chr2-168705404-INS-199
chr2-168722905-INS-56
chr2-168728742-INS-262
chr2-168731642-INS-302
chr2-168734361-INS-184
chr2-168738487-INS-60
chr2-168742186-INS-528
chr2-168747981-INS-52
chr2-168751225-INS-390
chr2-168764400-INS-181
chr2-168780433-INS-56
chr2-168792820-INS-459
chr2-168818574-INS-214
chr2-168822193-INS-176
chr2-168855701-INS-232
chr2-168861562-INS-69
chr2-168875683-INS-421
chr2-168916289-INS-7104
chr2-168944786-INS-172
chr2-168954813-INS-427
chr2-168956907-INS-77
chr2-168965513-INS-103
chr2-168986639-INS-121
chr2-168993107-INS-156
chr2-168995468-INS-58
chr2-169001316-INS-92
chr2-169004895-INS-430
chr2-169016109-INS-66
chr2-169025687-INS-190
chr2-169038841-INS-147
chr2-169041834-INS-1550
chr2-169052341-INS-100
chr2-169062913-INS-331
chr2-169079783-INS-55
chr2-169086933-INS-57
chr2-169091285-INS-290
chr2-169105424-INS-111
chr2-169108342-INS-86
chr2-169113526-INS-122
chr2-169127252-INS-80
chr2-169130134-INS-172

chr2-169141088-INS-312
chr2-169148340-INS-281
chr2-169177468-INS-111
chr2-169190118-INS-67
chr2-169218169-INS-503
chr2-169244109-INS-477
chr2-169249259-INS-137
chr2-169257067-INS-64
chr2-169269003-INS-72
chr2-169294034-INS-177
chr2-169297913-INS-54
chr2-169310597-INS-162
chr2-169322161-INS-98
chr2-169382421-INS-81
chr2-169404123-INS-508
chr2-169409527-INS-54
chr2-169411405-INS-233
chr2-169413407-INS-55
chr2-169436965-INS-94
chr2-169440777-INS-146
chr2-169450739-INS-822
chr2-169456888-INS-80
chr2-169460231-INS-88
chr2-169462312-INS-553
chr2-169472440-INS-176
chr2-169487055-INS-194
chr2-169510450-INS-59
chr2-169511497-INS-205
chr2-169573853-INS-130
chr2-169598695-INS-138
chr2-169623670-INS-303
chr2-169642510-INS-341
chr2-169646787-INS-90
chr2-169649020-INS-102
chr2-169659185-INS-116
chr2-169659768-INS-130
chr2-169691497-INS-752
chr2-169696731-INS-54
chr2-169699120-INS-152
chr2-169700914-INS-102
chr2-169715899-INS-55
chr2-169717137-INS-288
chr2-169746730-INS-157
chr2-169789598-INS-258
chr2-169806519-INS-84
chr2-169814031-INS-78
chr2-169832678-INS-538
chr2-169869153-INS-210
chr2-169880815-INS-69
chr2-169892844-INS-923
chr2-169918032-INS-57
chr2-169924304-INS-198
chr2-169926216-INS-52
chr2-169935445-INS-277
chr2-169997262-INS-279
chr2-170001571-INS-61
chr2-170005108-INS-56
chr2-170010592-INS-491
chr2-170012125-INS-183
chr2-170013913-INS-86
chr2-170022056-INS-129
chr2-170038901-INS-534
chr2-170045022-INS-209
chr2-170046531-INS-72
chr2-170050045-INS-106
chr2-170053646-INS-197
chr2-170068774-INS-478
chr2-170072603-INS-165
chr2-170073859-INS-104
chr2-170085775-INS-697
chr2-170099970-INS-102
chr2-170113717-INS-203
chr2-170116187-INS-121
chr2-170121502-INS-578
chr2-170140773-INS-52
chr2-170144849-INS-126
chr2-170159160-INS-54

chr2-170172235-INS-440
chr2-170173160-INS-198
chr2-170194013-INS-91
chr2-170199937-INS-194
chr2-170202435-INS-51
chr2-170205378-INS-94
chr2-170215864-INS-109
chr2-170261371-INS-151
chr2-170273730-INS-298
chr2-170276447-INS-722
chr2-170306156-INS-76
chr2-170320125-INS-112
chr2-170342064-INS-306
chr2-170346365-INS-295
chr2-170346961-INS-264
chr2-170354576-INS-101
chr2-170365834-INS-126
chr2-170400225-INS-321
chr2-170403428-INS-113
chr2-170407797-INS-75
chr2-170413491-INS-117
chr2-170414538-INS-122
chr2-170423234-INS-192
chr2-170426930-INS-96
chr2-170433171-INS-52
chr2-170475047-INS-281
chr2-170475827-INS-86
chr2-170476792-INS-68
chr2-170493074-INS-53
chr2-170519364-INS-5319
chr2-170536517-INS-96
chr2-170541698-INS-305
chr2-170556290-INS-149
chr2-170563213-INS-64
chr2-170566906-INS-189
chr2-170571345-INS-163
chr2-170575301-INS-123
chr2-170593635-INS-169
chr2-170601112-INS-137
chr2-170616388-INS-102
chr2-170651640-INS-425
chr2-170655426-INS-57
chr2-170659626-INS-60
chr2-170666164-INS-183
chr2-170667063-INS-154
chr2-170677857-INS-67
chr2-170684538-INS-86
chr2-170714323-INS-53
chr2-170728236-INS-83
chr2-170730151-INS-96
chr2-170732633-INS-451
chr2-170732653-INS-363
chr2-170734183-INS-153
chr2-170743933-INS-133
chr2-170745531-INS-62
chr2-170753277-INS-88
chr2-170765878-INS-374
chr2-170784439-INS-272
chr2-170795916-INS-55
chr2-170801804-INS-1152
chr2-170814090-INS-130
chr2-170846840-INS-60
chr2-170867422-INS-236
chr2-170876686-INS-195
chr2-170884235-INS-224
chr2-170932659-INS-142
chr2-170936442-INS-75
chr2-170966937-INS-68
chr2-171005534-INS-65
chr2-171046104-INS-830
chr2-171048278-INS-534
chr2-171056220-INS-79
chr2-171069939-INS-69
chr2-171074852-INS-60
chr2-171141290-INS-71
chr2-171147818-INS-125
chr2-171155309-INS-504

chr2-171182230-INS-155
chr2-171236325-INS-106
chr2-171239366-INS-499
chr2-171262255-INS-194
chr2-171272178-INS-7091
chr2-171276252-INS-99
chr2-171281020-INS-86
chr2-171286284-INS-112
chr2-171300287-INS-850
chr2-171339965-INS-94
chr2-171348105-INS-165
chr2-171410064-INS-280
chr2-171436841-INS-2636
chr2-171444292-INS-54
chr2-171463253-INS-374
chr2-171481018-INS-78
chr2-171506131-INS-55
chr2-171532422-INS-57
chr2-171538890-INS-52
chr2-171541842-INS-77
chr2-171545107-INS-195
chr2-171582026-INS-123
chr2-171599354-INS-176
chr2-171612901-INS-318
chr2-171640113-INS-201
chr2-171649213-INS-73
chr2-171657488-INS-218
chr2-171657566-INS-908
chr2-171657731-INS-1144
chr2-171658834-INS-76
chr2-171665669-INS-228
chr2-171674716-INS-105
chr2-171724611-INS-67
chr2-171725742-INS-186
chr2-171781843-INS-70
chr2-171804979-INS-468
chr2-171811281-INS-114
chr2-171814218-INS-190
chr2-171833237-INS-98
chr2-171847424-INS-126
chr2-171887399-INS-461
chr2-171891790-INS-6333
chr2-171900620-INS-340
chr2-171903073-INS-111
chr2-171905949-INS-2014
chr2-171907527-INS-1652
chr2-171908090-INS-1026
chr2-171908118-INS-80
chr2-171913537-INS-60
chr2-171921980-INS-713
chr2-171941262-INS-386
chr2-171950713-INS-344
chr2-171954888-INS-52
chr2-171999203-INS-163
chr2-172000265-INS-7080
chr2-172011044-INS-94
chr2-172016854-INS-7483
chr2-172017143-INS-200
chr2-172031469-INS-427
chr2-172046862-INS-408
chr2-172047096-INS-432
chr2-172048027-INS-463
chr2-172051832-INS-230
chr2-172066452-INS-72
chr2-172075619-INS-60
chr2-172081821-INS-6638
chr2-172099701-INS-84
chr2-172102388-INS-67
chr2-172109436-INS-901
chr2-172109573-INS-119
chr2-172118674-INS-51
chr2-172129505-INS-212
chr2-172138512-INS-765
chr2-172151517-INS-89
chr2-172172019-INS-264
chr2-172176238-INS-237
chr2-172189628-INS-56

chr2-172230214-INS-7159
chr2-172251459-INS-142
chr2-172266168-INS-169
chr2-172286278-INS-704
chr2-172306235-INS-231
chr2-172312850-INS-77
chr2-172323176-INS-116
chr2-172325100-INS-345
chr2-172325565-INS-386
chr2-172328331-INS-165
chr2-172343356-INS-474
chr2-172357042-INS-89
chr2-172361459-INS-71
chr2-172366040-INS-356
chr2-172374889-INS-230
chr2-172384491-INS-97
chr2-172418745-INS-64
chr2-172423257-INS-51
chr2-172429140-INS-213
chr2-172433689-INS-385
chr2-172440700-INS-177
chr2-172466576-INS-66
chr2-172468087-INS-5561
chr2-172469146-INS-193
chr2-172483885-INS-56
chr2-172499706-INS-120
chr2-172506601-INS-172
chr2-172510577-INS-54
chr2-172516026-INS-111
chr2-172557937-INS-186
chr2-172575172-INS-100
chr2-172581481-INS-140
chr2-172594099-INS-4300
chr2-172595820-INS-54
chr2-172604774-INS-214
chr2-172650645-INS-232
chr2-172664357-INS-60
chr2-172677079-INS-138
chr2-172683664-INS-159
chr2-172698484-INS-143
chr2-172701813-INS-192
chr2-172706434-INS-86
chr2-172755152-INS-51
chr2-172760152-INS-60
chr2-172762855-INS-144
chr2-172763186-INS-72
chr2-172766145-INS-430
chr2-172768567-INS-439
chr2-172794463-INS-209
chr2-172808803-INS-153
chr2-172817842-INS-53
chr2-172833500-INS-441
chr2-172857017-INS-66
chr2-172858382-INS-77
chr2-172893598-INS-87
chr2-172931223-INS-51
chr2-172939272-INS-95
chr2-172941581-INS-236
chr2-172959865-INS-56
chr2-172963374-INS-246
chr2-172969723-INS-214
chr2-172990225-INS-139
chr2-172992190-INS-64
chr2-172999636-INS-131
chr2-173006186-INS-286
chr2-173018275-INS-130
chr2-173024823-INS-456
chr2-173029647-INS-118
chr2-173046824-INS-75
chr2-173055102-INS-389
chr2-173061781-INS-294
chr2-173065314-INS-357
chr2-173112777-INS-205
chr2-173136400-INS-171
chr2-173156536-INS-58
chr2-173169019-INS-72
chr2-173171755-INS-363

chr2-173188159-INS-66
chr2-173190347-INS-137
chr2-173198030-INS-87
chr2-173219614-INS-52
chr2-173276570-INS-167
chr2-173315449-INS-150
chr2-173319911-INS-74
chr2-173327827-INS-140
chr2-173329904-INS-81
chr2-173365210-INS-343
chr2-173368222-INS-62
chr2-173376185-INS-453
chr2-173384028-INS-100
chr2-173384312-INS-298
chr2-173436972-INS-52
chr2-173451338-INS-444
chr2-173486656-INS-170
chr2-173497765-INS-575
chr2-173513758-INS-367
chr2-173529628-INS-197
chr2-173541870-INS-262
chr2-173558030-INS-119
chr2-173565483-INS-1063
chr2-173565495-INS-966
chr2-173568479-INS-97
chr2-173570858-INS-103
chr2-173577239-INS-52
chr2-173588480-INS-54
chr2-173598168-INS-99
chr2-173612057-INS-90
chr2-173628426-INS-55
chr2-173681468-INS-56
chr2-173682013-INS-124
chr2-173684445-INS-398
chr2-173703296-INS-133
chr2-173760657-INS-899
chr2-173786512-INS-73
chr2-173792897-INS-68
chr2-173803132-INS-6278
chr2-173803998-INS-346
chr2-173809695-INS-58
chr2-173839024-INS-95
chr2-173848578-INS-204
chr2-173906527-INS-1058
chr2-173911064-INS-102
chr2-173935059-INS-128
chr2-173972157-INS-60
chr2-173981056-INS-60
chr2-173984798-INS-203
chr2-173993869-INS-157
chr2-174003115-INS-61
chr2-174037135-INS-294
chr2-174060298-INS-315
chr2-174061492-INS-68
chr2-174081749-INS-96
chr2-174092907-INS-132
chr2-174131889-INS-62
chr2-174157489-INS-107
chr2-174189712-INS-208
chr2-174190685-INS-244
chr2-174194860-INS-92
chr2-174197630-INS-196
chr2-174241691-INS-109
chr2-174252972-INS-585
chr2-174260227-INS-387
chr2-174293590-INS-59
chr2-174303510-INS-520
chr2-174333486-INS-63
chr2-174343905-INS-52
chr2-174361758-INS-115
chr2-174365754-INS-76
chr2-174428587-INS-92
chr2-174437230-INS-93
chr2-174450861-INS-66
chr2-174473107-INS-114
chr2-174480116-INS-286
chr2-174503999-INS-510

chr2-174517106-INS-80
chr2-174525780-INS-60
chr2-174576491-INS-80
chr2-174598524-INS-56
chr2-174615509-INS-7167
chr2-174617260-INS-75
chr2-174627463-INS-411
chr2-174632859-INS-5568
chr2-174634123-INS-76
chr2-174643692-INS-166
chr2-174667139-INS-76
chr2-174669632-INS-165
chr2-174679587-INS-116
chr2-174750433-INS-67
chr2-174793427-INS-6017
chr2-174797400-INS-7823
chr2-174944131-INS-359
chr2-174975884-INS-195
chr2-175037635-INS-84
chr2-175655661-INS-486
chr2-175672079-INS-84
chr2-175770467-INS-507
chr2-175772093-INS-208
chr2-176393126-INS-500
chr2-176412779-INS-167
chr2-176631001-INS-358
chr2-176642158-INS-3556
chr2-176746622-INS-420
chr2-176749301-INS-362
chr2-176758927-INS-142
chr2-176817680-INS-420
chr2-176848859-INS-79
chr2-176883290-INS-209
chr2-176884916-INS-252
chr2-177006961-INS-2436
chr2-177022837-INS-228
chr2-177047830-INS-201
chr2-177093600-INS-7124
chr2-177112860-INS-250
chr2-177233667-INS-7189
chr2-177234389-INS-6047
chr2-177290379-INS-198
chr2-177479392-INS-1488
chr2-177486943-INS-5547
chr2-177527496-INS-7176
chr2-177542528-INS-1407
chr2-177607956-INS-7113
chr2-177622445-INS-168
chr2-177626022-INS-6440
chr2-177654559-INS-58
chr2-177704225-INS-347
chr2-177750854-INS-4646
chr2-177764262-INS-68
chr2-177770221-INS-367
chr2-177778828-INS-669
chr2-177782593-INS-216
chr2-177788763-INS-80
chr2-177793214-INS-57
chr2-177794242-INS-108
chr2-177802961-INS-99
chr2-177808566-INS-5564
chr2-177852774-INS-672
chr2-177864580-INS-623
chr2-177865088-INS-76
chr2-177871293-INS-54
chr2-177900340-INS-183
chr2-177914912-INS-58
chr2-177940446-INS-52
chr2-177952031-INS-56
chr2-177995932-INS-73
chr2-178039274-INS-963
chr2-178045689-INS-532
chr2-178064234-INS-54
chr2-178068091-INS-80
chr2-178104025-INS-147
chr2-178134456-INS-76
chr2-178150490-INS-195

chr2-178151484-INS-57
chr2-178159340-INS-67
chr2-178170680-INS-232
chr2-178173992-INS-363
chr2-178192127-INS-56
chr2-178194035-INS-3682
chr2-178206191-INS-66
chr2-178245671-INS-595
chr2-178247311-INS-63
chr2-178263256-INS-360
chr2-178280642-INS-125
chr2-178281234-INS-391
chr2-178295916-INS-62
chr2-178296878-INS-7113
chr2-178302513-INS-101
chr2-178304355-INS-70
chr2-178306778-INS-80
chr2-178319269-INS-81
chr2-178377267-INS-472
chr2-178378063-INS-125
chr2-178384998-INS-1050
chr2-178388280-INS-155
chr2-178396191-INS-122
chr2-178397505-INS-480
chr2-178405113-INS-142
chr2-178413559-INS-272
chr2-178428763-INS-679
chr2-178442410-INS-161
chr2-178494590-INS-2907
chr2-178504280-INS-58
chr2-178530191-INS-52
chr2-178536941-INS-168
chr2-178540026-INS-52
chr2-178552423-INS-138
chr2-178555041-INS-56
chr2-178614439-INS-119
chr2-178634007-INS-54
chr2-178644979-INS-418
chr2-178672618-INS-63
chr2-178690712-INS-54
chr2-178695131-INS-95
chr2-178697153-INS-150
chr2-178715952-INS-6401
chr2-178721982-INS-69
chr2-178739664-INS-59
chr2-178745202-INS-3140
chr2-178772512-INS-110
chr2-178779680-INS-208
chr2-178783467-INS-66
chr2-178785653-INS-66
chr2-178802767-INS-194
chr2-178831837-INS-59
chr2-178836562-INS-131
chr2-178843138-INS-112
chr2-178846233-INS-60
chr2-178858224-INS-189
chr2-178862890-INS-1520
chr2-178885679-INS-68
chr2-178916943-INS-95
chr2-178951749-INS-198
chr2-178951999-INS-429
chr2-178952567-INS-297
chr2-178954892-INS-59
chr2-178964269-INS-63
chr2-178971153-INS-57
chr2-178981810-INS-367
chr2-178985781-INS-78
chr2-178999409-INS-91
chr2-179021510-INS-240
chr2-179025838-INS-54
chr2-179045175-INS-69
chr2-179047583-INS-86
chr2-179081629-INS-74
chr2-179087565-INS-108
chr2-179134164-INS-72
chr2-179137819-INS-267
chr2-179143114-INS-86

chr2-179159134-INS-280
chr2-179163395-INS-64
chr2-179203542-INS-350
chr2-179214885-INS-157
chr2-179222585-INS-177
chr2-179230373-INS-119
chr2-179240109-INS-197
chr2-179249050-INS-1296
chr2-179262428-INS-63
chr2-179294281-INS-304
chr2-179320470-INS-731
chr2-179320995-INS-6428
chr2-179326486-INS-80
chr2-179332017-INS-84
chr2-179341808-INS-78
chr2-179345786-INS-137
chr2-179349972-INS-273
chr2-179361373-INS-508
chr2-179379658-INS-94
chr2-179406708-INS-62
chr2-179429938-INS-196
chr2-179436157-INS-590
chr2-179436985-INS-57
chr2-179447772-INS-64
chr2-179456496-INS-74
chr2-179472678-INS-67
chr2-179482775-INS-57
chr2-179609793-INS-201
chr2-179626070-INS-56
chr2-179635558-INS-194
chr2-179660757-INS-506
chr2-179692208-INS-206
chr2-179708644-INS-421
chr2-179714429-INS-73
chr2-179718794-INS-204
chr2-179773328-INS-208
chr2-179778187-INS-63
chr2-179786530-INS-230
chr2-179792425-INS-85
chr2-179802082-INS-211
chr2-179829922-INS-160
chr2-179839938-INS-208
chr2-179880525-INS-215
chr2-179889466-INS-127
chr2-179895836-INS-204
chr2-179897985-INS-229
chr2-179907984-INS-177
chr2-179916282-INS-129
chr2-179929596-INS-234
chr2-179937206-INS-170
chr2-179941296-INS-98
chr2-179955743-INS-62
chr2-179982605-INS-59
chr2-180014658-INS-320
chr2-180022371-INS-106
chr2-180033555-INS-99
chr2-180034871-INS-116
chr2-180065750-INS-102
chr2-180096436-INS-112
chr2-180116973-INS-243
chr2-180121583-INS-51
chr2-180164863-INS-53
chr2-180201989-INS-78
chr2-180215854-INS-58
chr2-180230166-INS-146
chr2-180254100-INS-196
chr2-180257917-INS-52
chr2-180296265-INS-354
chr2-180304725-INS-118
chr2-180333008-INS-60
chr2-180339089-INS-205
chr2-180373545-INS-218
chr2-180376055-INS-321
chr2-180389415-INS-193
chr2-180394008-INS-114
chr2-180395895-INS-195
chr2-180408057-INS-199

chr2-180408835-INS-205
chr2-180422338-INS-223
chr2-180429771-INS-64
chr2-180452786-INS-114
chr2-180464749-INS-57
chr2-180490729-INS-201
chr2-180491574-INS-58
chr2-180492925-INS-143
chr2-180494775-INS-101
chr2-180512849-INS-128
chr2-180555200-INS-203
chr2-180606145-INS-83
chr2-180644744-INS-62
chr2-180671981-INS-95
chr2-180692034-INS-84
chr2-180707018-INS-613
chr2-180711174-INS-4749
chr2-180748153-INS-135
chr2-180757026-INS-78
chr2-180768801-INS-243
chr2-180802175-INS-193
chr2-180823542-INS-5289
chr2-180833685-INS-52
chr2-180843072-INS-116
chr2-180857249-INS-474
chr2-180859224-INS-168
chr2-180860387-INS-80
chr2-180866862-INS-112
chr2-180883366-INS-114
chr2-180885407-INS-106
chr2-180898681-INS-180
chr2-180911342-INS-195
chr2-180918040-INS-58
chr2-180938767-INS-399
chr2-180966944-INS-399
chr2-181000451-INS-99
chr2-181002649-INS-65
chr2-181026811-INS-52
chr2-181056177-INS-198
chr2-181081603-INS-69
chr2-181159934-INS-275
chr2-181171395-INS-183
chr2-181177770-INS-55
chr2-181199957-INS-209
chr2-181236689-INS-195
chr2-181247179-INS-204
chr2-181249435-INS-208
chr2-181258034-INS-195
chr2-181264808-INS-451
chr2-181265667-INS-200
chr2-181268691-INS-188
chr2-181285376-INS-56
chr2-181306630-INS-56
chr2-181353442-INS-1900
chr2-181379017-INS-1251
chr2-181379550-INS-8822
chr2-181392712-INS-1102
chr2-181393263-INS-1086
chr2-181401796-INS-64
chr2-181424004-INS-103
chr2-181434969-INS-51
chr2-181450388-INS-508
chr2-181451502-INS-2450
chr2-181464338-INS-72
chr2-181491161-INS-159
chr2-181508649-INS-498
chr2-181523875-INS-7224
chr2-181530045-INS-225
chr2-181531447-INS-6795
chr2-181571042-INS-114
chr2-181602422-INS-201
chr2-181627730-INS-69
chr3-3041226-INS-5309
chr3-3043479-INS-606
chr3-3180902-INS-63
chr3-3196594-INS-141
chr3-3210732-INS-75

chr3-3251276-INS-99
chr3-3255451-INS-52
chr3-3281564-INS-323
chr3-3290096-INS-358
chr3-3312850-INS-87
chr3-3344948-INS-7926
chr3-3354950-INS-826
chr3-3367209-INS-4836
chr3-3369645-INS-2543
chr3-3373398-INS-10485
chr3-3406863-INS-1842
chr3-3415018-INS-4998
chr3-3435729-INS-773
chr3-3464110-INS-64
chr3-3521649-INS-52
chr3-3525143-INS-184
chr3-3537104-INS-494
chr3-3543141-INS-56
chr3-3549329-INS-633
chr3-3558247-INS-178
chr3-3560358-INS-115
chr3-3568429-INS-145
chr3-3598515-INS-162
chr3-3610775-INS-207
chr3-3628653-INS-140
chr3-3647922-INS-667
chr3-3647922-INS-72
chr3-3656761-INS-74
chr3-3663342-INS-91
chr3-3690719-INS-163
chr3-3699804-INS-119
chr3-3719269-INS-369
chr3-3737901-INS-329
chr3-3746796-INS-54
chr3-3775998-INS-339
chr3-3792320-INS-245
chr3-3793400-INS-2575
chr3-3797824-INS-295
chr3-3815909-INS-63
chr3-3844131-INS-6647
chr3-3860286-INS-54
chr3-3872269-INS-272
chr3-3882092-INS-268
chr3-3898609-INS-6542
chr3-3901824-INS-6503
chr3-3919848-INS-289
chr3-3921125-INS-85
chr3-3922115-INS-199
chr3-3954015-INS-62
chr3-3960289-INS-68
chr3-3970174-INS-195
chr3-3970899-INS-590
chr3-3981447-INS-328
chr3-4007733-INS-67
chr3-4010416-INS-457
chr3-4015384-INS-6761
chr3-4023144-INS-76
chr3-4101864-INS-6381
chr3-4120672-INS-60
chr3-4147626-INS-5941
chr3-4150307-INS-8564
chr3-4157351-INS-7210
chr3-4159445-INS-52
chr3-4176783-INS-7043
chr3-4185664-INS-66
chr3-4217407-INS-6547
chr3-4217518-INS-54
chr3-4222105-INS-7580
chr3-4275187-INS-373
chr3-4297073-INS-4228
chr3-4318507-INS-339
chr3-4325349-INS-6454
chr3-4332178-INS-56
chr3-4357652-INS-145
chr3-4359575-INS-76
chr3-4374925-INS-538
chr3-4394447-INS-7103

chr3-4399988-INS-150
chr3-4411830-INS-115
chr3-4415803-INS-7199
chr3-4415812-INS-6810
chr3-4420075-INS-54
chr3-4432976-INS-140
chr3-4445025-INS-6388
chr3-4456503-INS-7108
chr3-4480072-INS-492
chr3-4507975-INS-212
chr3-4517758-INS-59
chr3-4523575-INS-7440
chr3-4550333-INS-7122
chr3-4599187-INS-203
chr3-4612663-INS-426
chr3-4625368-INS-7064
chr3-4632948-INS-6763
chr3-4639412-INS-3134
chr3-4646661-INS-469
chr3-4671453-INS-6456
chr3-4691612-INS-457
chr3-4718351-INS-81
chr3-4719343-INS-204
chr3-4738964-INS-50
chr3-4744811-INS-225
chr3-4749971-INS-6787
chr3-4752825-INS-7405
chr3-4756485-INS-7099
chr3-4759908-INS-6485
chr3-4762764-INS-169
chr3-4783071-INS-2286
chr3-4811458-INS-252
chr3-4831709-INS-3151
chr3-4842523-INS-6471
chr3-4842714-INS-216
chr3-4850651-INS-82
chr3-4857573-INS-69
chr3-4864514-INS-7661
chr3-4867477-INS-189
chr3-4869421-INS-7018
chr3-4883183-INS-206
chr3-4906857-INS-6737
chr3-4913688-INS-179
chr3-4932700-INS-131
chr3-4942414-INS-53
chr3-4956942-INS-4953
chr3-4957356-INS-4370
chr3-4959879-INS-7125
chr3-4962251-INS-1518
chr3-4964637-INS-3055
chr3-4971301-INS-88
chr3-5005081-INS-2686
chr3-5010308-INS-2784
chr3-5015177-INS-104
chr3-5016699-INS-266
chr3-5019513-INS-414
chr3-5023305-INS-57
chr3-5043291-INS-292
chr3-5047004-INS-84
chr3-5059124-INS-1889
chr3-5081743-INS-885
chr3-5085199-INS-100
chr3-5163318-INS-161
chr3-5168461-INS-208
chr3-5202258-INS-88
chr3-5225669-INS-6452
chr3-5236763-INS-63
chr3-5250913-INS-486
chr3-5254209-INS-52
chr3-5267255-INS-70
chr3-5285337-INS-105
chr3-5340922-INS-286
chr3-5375448-INS-125
chr3-5439363-INS-52
chr3-5455087-INS-56
chr3-5459202-INS-104
chr3-5488923-INS-60

chr3-5499085-INS-6595
chr3-5519604-INS-627
chr3-5537331-INS-6242
chr3-5566948-INS-428
chr3-5580854-INS-55
chr3-5592684-INS-63
chr3-5609452-INS-160
chr3-5616340-INS-302
chr3-5702488-INS-51
chr3-5721134-INS-108
chr3-5739364-INS-309
chr3-5745226-INS-198
chr3-5765264-INS-79
chr3-5779020-INS-60
chr3-5794889-INS-72
chr3-5803706-INS-390
chr3-5809811-INS-214
chr3-5817789-INS-1848
chr3-5837410-INS-5366
chr3-5841033-INS-68
chr3-5844494-INS-59
chr3-5887746-INS-105
chr3-5898422-INS-5586
chr3-5916677-INS-84
chr3-5941753-INS-177
chr3-5963846-INS-7225
chr3-5973441-INS-502
chr3-5977079-INS-197
chr3-5981311-INS-452
chr3-6019174-INS-3428
chr3-6028608-INS-108
chr3-6033640-INS-114
chr3-6053117-INS-60
chr3-6090061-INS-401
chr3-6127071-INS-79
chr3-6146811-INS-153
chr3-6150418-INS-90
chr3-6157684-INS-183
chr3-6158781-INS-624
chr3-6186417-INS-159
chr3-6235825-INS-193
chr3-6245177-INS-522
chr3-6274182-INS-1821
chr3-6297432-INS-54
chr3-6299315-INS-2084
chr3-6299316-INS-392
chr3-6318058-INS-1313
chr3-6321511-INS-62
chr3-6327052-INS-6971
chr3-6329018-INS-78
chr3-6337562-INS-2125
chr3-6337938-INS-63
chr3-6344623-INS-7125
chr3-6408832-INS-4356
chr3-6416030-INS-6576
chr3-6424080-INS-60
chr3-6439046-INS-107
chr3-6453676-INS-51
chr3-6492876-INS-64
chr3-6501635-INS-58
chr3-6506629-INS-198
chr3-6531694-INS-62
chr3-6533248-INS-7490
chr3-6545619-INS-78
chr3-6558995-INS-427
chr3-6569344-INS-456
chr3-6586790-INS-202
chr3-6587965-INS-114
chr3-6631636-INS-70
chr3-6656747-INS-493
chr3-6718968-INS-108
chr3-6720310-INS-55
chr3-6724232-INS-2242
chr3-6736889-INS-168
chr3-6749639-INS-54
chr3-6785853-INS-850
chr3-6792546-INS-63

chr3-6815710-INS-77
chr3-6819018-INS-51
chr3-6857984-INS-65
chr3-6874887-INS-235
chr3-6879963-INS-62
chr3-6886226-INS-266
chr3-6928092-INS-387
chr3-6956129-INS-68
chr3-6959738-INS-224
chr3-6983034-INS-67
chr3-7015977-INS-71
chr3-7029133-INS-66
chr3-7030959-INS-111
chr3-7065887-INS-299
chr3-7077065-INS-62
chr3-7081837-INS-64
chr3-7083241-INS-950
chr3-7093545-INS-185
chr3-7095339-INS-833
chr3-7099380-INS-192
chr3-7111665-INS-416
chr3-7121949-INS-53
chr3-7125312-INS-218
chr3-7137084-INS-568
chr3-7156668-INS-214
chr3-7205572-INS-134
chr3-7209907-INS-62
chr3-7219049-INS-321
chr3-7269856-INS-73
chr3-7271300-INS-367
chr3-7282683-INS-212
chr3-7291299-INS-338
chr3-7327716-INS-590
chr3-7329303-INS-65
chr3-7332252-INS-52
chr3-7356948-INS-596
chr3-7385298-INS-7326
chr3-7415574-INS-105
chr3-7418509-INS-746
chr3-7420457-INS-75
chr3-7438162-INS-79
chr3-7466074-INS-4586
chr3-7469252-INS-66
chr3-7497481-INS-154
chr3-7501033-INS-198
chr3-7501509-INS-198
chr3-7501976-INS-824
chr3-7512202-INS-61
chr3-7514396-INS-146
chr3-7523807-INS-599
chr3-7526925-INS-57
chr3-7540605-INS-404
chr3-7551146-INS-54
chr3-7564416-INS-331
chr3-7625602-INS-516
chr3-7638112-INS-80
chr3-7654004-INS-54
chr3-7657099-INS-157
chr3-7675926-INS-240
chr3-7763472-INS-70
chr3-7797181-INS-130
chr3-7808184-INS-6964
chr3-7829925-INS-140
chr3-7863743-INS-4797
chr3-7869025-INS-56
chr3-7871872-INS-5727
chr3-7872755-INS-4058
chr3-7908601-INS-10637
chr3-7936578-INS-1840
chr3-7946080-INS-117
chr3-7950585-INS-197
chr3-7985803-INS-6448
chr3-8023211-INS-727
chr3-8026377-INS-549
chr3-8043677-INS-58
chr3-8067343-INS-2599
chr3-8107633-INS-194

chr3-8115769-INS-100
chr3-8126498-INS-554
chr3-8142710-INS-390
chr3-8155143-INS-70
chr3-8157472-INS-93
chr3-8164620-INS-140
chr3-8177953-INS-56
chr3-8210846-INS-399
chr3-8224385-INS-138
chr3-8228696-INS-113
chr3-8233185-INS-383
chr3-8233306-INS-6758
chr3-8235742-INS-258
chr3-8241078-INS-57
chr3-8256949-INS-1892
chr3-8258457-INS-84
chr3-8258885-INS-121
chr3-8267088-INS-6213
chr3-8315263-INS-173
chr3-8347071-INS-6675
chr3-8353345-INS-904
chr3-8356399-INS-607
chr3-8362800-INS-128
chr3-8363047-INS-986
chr3-8367728-INS-75
chr3-8372962-INS-184
chr3-8377292-INS-52
chr3-8424371-INS-5730
chr3-8468156-INS-123
chr3-8478584-INS-1192
chr3-8482827-INS-204
chr3-8490749-INS-73
chr3-8522221-INS-6411
chr3-8525860-INS-176
chr3-8559542-INS-74
chr3-8562229-INS-1893
chr3-8567708-INS-6225
chr3-8584002-INS-125
chr3-8585973-INS-58
chr3-8633458-INS-442
chr3-8634917-INS-443
chr3-8645242-INS-95
chr3-8672917-INS-573
chr3-8698808-INS-125
chr3-8716583-INS-206
chr3-8723973-INS-7086
chr3-8747993-INS-96
chr3-8749049-INS-52
chr3-8767026-INS-88
chr3-8771048-INS-473
chr3-8794302-INS-82
chr3-8795475-INS-277
chr3-8804108-INS-50
chr3-8824329-INS-322
chr3-8830772-INS-64
chr3-8834427-INS-1249
chr3-8841176-INS-78
chr3-8845619-INS-405
chr3-8853595-INS-107
chr3-8854193-INS-7224
chr3-8854980-INS-430
chr3-8858171-INS-70
chr3-8859320-INS-251
chr3-8866412-INS-325
chr3-8872568-INS-64
chr3-8873270-INS-172
chr3-8922945-INS-590
chr3-8934094-INS-204
chr3-8951984-INS-169
chr3-8953474-INS-52
chr3-8956299-INS-126
chr3-8961212-INS-147
chr3-8989264-INS-218
chr3-8991916-INS-96
chr3-8993538-INS-56
chr3-9010520-INS-6319
chr3-9013321-INS-86

chr3-9018268-INS-116
chr3-9043883-INS-130
chr3-9051354-INS-404
chr3-9065012-INS-202
chr3-9068048-INS-176
chr3-9072304-INS-175
chr3-9074732-INS-645
chr3-9075247-INS-71
chr3-9079372-INS-208
chr3-9085765-INS-53
chr3-9095595-INS-403
chr3-9112053-INS-369
chr3-9117778-INS-411
chr3-9118823-INS-203
chr3-9124451-INS-227
chr3-9126137-INS-169
chr3-9131749-INS-198
chr3-9140598-INS-214
chr3-9145914-INS-61
chr3-9149042-INS-75
chr3-9182178-INS-314
chr3-9193902-INS-101
chr3-9200217-INS-245
chr3-9237718-INS-1273
chr3-9237880-INS-138
chr3-9253459-INS-96
chr3-9294163-INS-140
chr3-9296060-INS-109
chr3-9296709-INS-87
chr3-9300918-INS-75
chr3-9307563-INS-349
chr3-9313725-INS-194
chr3-9320513-INS-133
chr3-9329382-INS-52
chr3-9336706-INS-440
chr3-9344778-INS-349
chr3-9363218-INS-358
chr3-9369726-INS-85
chr3-9406468-INS-369
chr3-9425115-INS-636
chr3-9429986-INS-52
chr3-9444209-INS-128
chr3-9462471-INS-220
chr3-9471885-INS-166
chr3-9485501-INS-4622
chr3-9507094-INS-196
chr3-9516128-INS-104
chr3-9519267-INS-69
chr3-9531363-INS-3500
chr3-9566847-INS-96
chr3-9572909-INS-133
chr3-9582975-INS-265
chr3-9593221-INS-150
chr3-9605765-INS-130
chr3-9610885-INS-90
chr3-9620400-INS-765
chr3-9625380-INS-231
chr3-9630735-INS-185
chr3-9639082-INS-67
chr3-9658705-INS-252
chr3-9666460-INS-100
chr3-9680138-INS-186
chr3-9696818-INS-226
chr3-9702092-INS-658
chr3-9714962-INS-312
chr3-9717661-INS-7209
chr3-9721840-INS-58
chr3-9724492-INS-92
chr3-9728463-INS-72
chr3-9734535-INS-337
chr3-9756624-INS-732
chr3-9789229-INS-141
chr3-9792532-INS-101
chr3-9794372-INS-501
chr3-9809584-INS-210
chr3-9827323-INS-422
chr3-9852159-INS-84

chr3-9853628-INS-64
chr3-9855475-INS-57
chr3-9865424-INS-56
chr3-9867853-INS-1158
chr3-9867853-INS-90
chr3-9867854-INS-968
chr3-9868580-INS-1314
chr3-9868656-INS-2444
chr3-9868662-INS-4248
chr3-9879396-INS-264
chr3-9909207-INS-273
chr3-9918828-INS-55
chr3-9963060-INS-74
chr3-9982927-INS-94
chr3-9983619-INS-81
chr3-9990716-INS-919
chr3-9990841-INS-52
chr3-9996984-INS-6440
chr3-10013439-INS-500
chr3-10021162-INS-100
chr3-10040571-INS-7180
chr3-10049203-INS-61
chr3-10103495-INS-183
chr3-10119050-INS-61
chr3-10170712-INS-191
chr3-10177986-INS-187
chr3-10185037-INS-122
chr3-10198755-INS-234
chr3-10213119-INS-214
chr3-10243438-INS-254
chr3-10249621-INS-54
chr3-10270005-INS-299
chr3-10275204-INS-152
chr3-10295773-INS-155
chr3-10308284-INS-260
chr3-10315596-INS-202
chr3-10322458-INS-55
chr3-10329323-INS-105
chr3-10367343-INS-206
chr3-10369062-INS-269
chr3-10370336-INS-131
chr3-10377772-INS-498
chr3-10393174-INS-257
chr3-10394292-INS-108
chr3-10404495-INS-434
chr3-10423101-INS-78
chr3-10423519-INS-199
chr3-10429836-INS-127
chr3-10442800-INS-166
chr3-10452887-INS-232
chr3-10469680-INS-518
chr3-10472018-INS-7412
chr3-10472496-INS-72
chr3-10479627-INS-358
chr3-10498650-INS-134
chr3-10520958-INS-5542
chr3-10586348-INS-6409
chr3-10603817-INS-7094
chr3-10606344-INS-447
chr3-10634812-INS-66
chr3-10639778-INS-129
chr3-10651286-INS-126
chr3-10679648-INS-64
chr3-10704497-INS-176
chr3-10729732-INS-136
chr3-10744256-INS-52
chr3-10745294-INS-306
chr3-10747193-INS-252
chr3-10779871-INS-277
chr3-10793067-INS-65
chr3-10795108-INS-115
chr3-10812942-INS-58
chr3-10816649-INS-510
chr3-10829125-INS-719
chr3-10846283-INS-6405
chr3-10864593-INS-58
chr3-10866103-INS-264

chr3-10878516-INS-115
chr3-10885807-INS-358
chr3-10903103-INS-532
chr3-10925378-INS-424
chr3-10971249-INS-215
chr3-10980522-INS-1655
chr3-10996560-INS-85
chr3-10997233-INS-70
chr3-11003229-INS-216
chr3-11005874-INS-264
chr3-11025188-INS-574
chr3-11060880-INS-157
chr3-11089976-INS-281
chr3-11097193-INS-467
chr3-11103317-INS-740
chr3-11108805-INS-857
chr3-11144239-INS-208
chr3-11160394-INS-544
chr3-11200521-INS-61
chr3-11215254-INS-77
chr3-11225204-INS-426
chr3-11228930-INS-125
chr3-11240323-INS-158
chr3-11258562-INS-55
chr3-11267496-INS-139
chr3-11276080-INS-70
chr3-11286092-INS-58
chr3-11287430-INS-419
chr3-11304295-INS-686
chr3-11318814-INS-561
chr3-11336285-INS-751
chr3-11379084-INS-54
chr3-11388371-INS-389
chr3-11390860-INS-162
chr3-11403455-INS-179
chr3-11453159-INS-174
chr3-11489545-INS-557
chr3-11492382-INS-68
chr3-11498316-INS-454
chr3-11499319-INS-385
chr3-11505965-INS-358
chr3-11510057-INS-105
chr3-11536902-INS-75
chr3-11543025-INS-686
chr3-11555812-INS-7036
chr3-11563601-INS-56
chr3-11577925-INS-1047
chr3-11583246-INS-3072
chr3-11594767-INS-382
chr3-11608902-INS-82
chr3-11670577-INS-288
chr3-11708369-INS-6573
chr3-11725474-INS-6388
chr3-11760205-INS-161
chr3-11796274-INS-365
chr3-11809005-INS-185
chr3-11825998-INS-107
chr3-11856222-INS-168
chr3-11911427-INS-52
chr3-11926233-INS-60
chr3-11956823-INS-494
chr3-11960511-INS-297
chr3-11962448-INS-90
chr3-11964444-INS-451
chr3-11970014-INS-6695
chr3-12017739-INS-6248
chr3-12081484-INS-635
chr3-12105927-INS-57
chr3-12108437-INS-548
chr3-12120687-INS-55
chr3-12226731-INS-7436
chr3-12284412-INS-228
chr3-12307020-INS-60
chr3-12312480-INS-819
chr3-12336554-INS-339
chr3-12346396-INS-55
chr3-12370618-INS-54

chr3-12393974-INS-118
chr3-12439165-INS-6672
chr3-12448905-INS-84
chr3-12457761-INS-2928
chr3-12565677-INS-84
chr3-12588023-INS-234
chr3-12592022-INS-5169
chr3-12636291-INS-580
chr3-12650870-INS-3948
chr3-12681197-INS-185
chr3-12732200-INS-400
chr3-12757984-INS-4612
chr3-12758065-INS-64
chr3-12762931-INS-225
chr3-12818794-INS-502
chr3-12826362-INS-220
chr3-12864263-INS-459
chr3-12877942-INS-152
chr3-12973834-INS-213
chr3-12982555-INS-85
chr3-12991113-INS-970
chr3-13026752-INS-499
chr3-13029417-INS-8738
chr3-13039164-INS-303
chr3-13043645-INS-63
chr3-13046151-INS-56
chr3-13056185-INS-498
chr3-13107186-INS-246
chr3-13108746-INS-117
chr3-13119858-INS-54
chr3-13147935-INS-986
chr3-13161003-INS-6791
chr3-13161618-INS-52
chr3-13180683-INS-1889
chr3-13190336-INS-6674
chr3-13190527-INS-2962
chr3-13202206-INS-74
chr3-13207319-INS-895
chr3-13218740-INS-2504
chr3-13239134-INS-1371
chr3-13259099-INS-130
chr3-13274511-INS-6418
chr3-13282627-INS-6266
chr3-13289341-INS-5191
chr3-13297753-INS-80
chr3-13339723-INS-7010
chr3-13352709-INS-393
chr3-13352762-INS-6403
chr3-13417701-INS-199
chr3-13455647-INS-136
chr3-13457646-INS-6677
chr3-13469322-INS-722
chr3-13481073-INS-567
chr3-13510974-INS-556
chr3-13527315-INS-547
chr3-13528151-INS-77
chr3-13553745-INS-184
chr3-13554641-INS-564
chr3-13601973-INS-52
chr3-13608022-INS-6892
chr3-13614324-INS-52
chr3-13640305-INS-435
chr3-13658436-INS-77
chr3-13710312-INS-74
chr3-13801715-INS-311
chr3-13817774-INS-428
chr3-13826592-INS-325
chr3-13837283-INS-146
chr3-13839712-INS-416
chr3-13844740-INS-178
chr3-13878831-INS-94
chr3-13884528-INS-80
chr3-13921471-INS-528
chr3-13923280-INS-62
chr3-13998652-INS-6421
chr3-14024479-INS-51
chr3-14082386-INS-65

chr3-14149236-INS-60
chr3-14153577-INS-128
chr3-14165878-INS-58
chr3-14177373-INS-598
chr3-14197539-INS-120
chr3-14200869-INS-401
chr3-14201410-INS-352
chr3-14218061-INS-95
chr3-14219004-INS-923
chr3-14289517-INS-68
chr3-14327928-INS-65
chr3-14330021-INS-559
chr3-14383617-INS-75
chr3-14487608-INS-67
chr3-14520059-INS-131
chr3-14604241-INS-247
chr3-14608871-INS-630
chr3-14666821-INS-239
chr3-14677693-INS-108
chr3-14722434-INS-2391
chr3-14723151-INS-187
chr3-14726019-INS-6745
chr3-14760072-INS-60
chr3-14849825-INS-82
chr3-14855304-INS-140
chr3-14882159-INS-372
chr3-14890811-INS-544
chr3-14907139-INS-204
chr3-14930309-INS-56
chr3-14946043-INS-66
chr3-14957953-INS-112
chr3-14964666-INS-100
chr3-15005429-INS-65
chr3-15032059-INS-315
chr3-15035740-INS-67
chr3-15041388-INS-110
chr3-15045056-INS-121
chr3-15065164-INS-4541
chr3-15114688-INS-76
chr3-15142504-INS-470
chr3-15182259-INS-164
chr3-15190297-INS-108
chr3-15203290-INS-2805
chr3-15205008-INS-327
chr3-15233273-INS-5627
chr3-15236235-INS-3576
chr3-15244382-INS-62
chr3-15295060-INS-96
chr3-15301516-INS-326
chr3-15302911-INS-667
chr3-15327052-INS-52
chr3-15329781-INS-800
chr3-15331794-INS-7098
chr3-15363517-INS-575
chr3-15389726-INS-76
chr3-15405568-INS-122
chr3-15470365-INS-474
chr3-15477095-INS-117
chr3-15539599-INS-567
chr3-15541561-INS-59
chr3-15544566-INS-534
chr3-15599457-INS-54
chr3-15601873-INS-388
chr3-15611293-INS-536
chr3-15614411-INS-110
chr3-15614411-INS-112
chr3-15633197-INS-82
chr3-15638466-INS-152
chr3-15650134-INS-175
chr3-15696474-INS-195
chr3-15700550-INS-225
chr3-15801252-INS-58
chr3-15887943-INS-150
chr3-15907264-INS-224
chr3-15985325-INS-7234
chr3-16003704-INS-506
chr3-16004557-INS-198

chr3-16059118-INS-62
chr3-16088319-INS-247
chr3-16089891-INS-61
chr3-16097323-INS-927
chr3-16106718-INS-6451
chr3-16122553-INS-62
chr3-16126775-INS-455
chr3-16129523-INS-53
chr3-16138687-INS-52
chr3-16142611-INS-208
chr3-16163441-INS-67
chr3-16174223-INS-94
chr3-16187081-INS-583
chr3-16203461-INS-475
chr3-16207996-INS-124
chr3-16210963-INS-409
chr3-16218671-INS-399
chr3-16263772-INS-76
chr3-16266786-INS-59
chr3-16279211-INS-360
chr3-16296817-INS-68
chr3-16300918-INS-5082
chr3-16323176-INS-66
chr3-16332773-INS-83
chr3-16385169-INS-171
chr3-16389557-INS-581
chr3-16390945-INS-665
chr3-16391592-INS-219
chr3-16418170-INS-194
chr3-16430557-INS-528
chr3-16433867-INS-94
chr3-16451149-INS-417
chr3-16462546-INS-130
chr3-16476357-INS-1748
chr3-16486352-INS-85
chr3-16506391-INS-131
chr3-16523766-INS-341
chr3-16560148-INS-1970
chr3-16595937-INS-982
chr3-16606317-INS-134
chr3-16613828-INS-6484
chr3-16618120-INS-740
chr3-16619495-INS-449
chr3-16631391-INS-6650
chr3-16641892-INS-309
chr3-16646543-INS-831
chr3-16664393-INS-7024
chr3-16684856-INS-9644
chr3-16708486-INS-552
chr3-16742597-INS-143
chr3-16780446-INS-72
chr3-16819025-INS-82
chr3-16842004-INS-106
chr3-16876486-INS-364
chr3-16895359-INS-4943
chr3-16910639-INS-55
chr3-16928305-INS-168
chr3-16930469-INS-67
chr3-16936552-INS-59
chr3-16937842-INS-78
chr3-16944301-INS-119
chr3-16972430-INS-68
chr3-16994781-INS-128
chr3-17035775-INS-84
chr3-17038459-INS-674
chr3-17097536-INS-132
chr3-17136260-INS-54
chr3-17143797-INS-150
chr3-17144410-INS-866
chr3-17172256-INS-5549
chr3-17229431-INS-157
chr3-17262060-INS-5962
chr3-17267119-INS-7147
chr3-17290442-INS-54
chr3-17301540-INS-54
chr3-17320909-INS-102
chr3-17329741-INS-622

chr3-17336917-INS-6635
chr3-17389668-INS-485
chr3-17392672-INS-232
chr3-17398462-INS-73
chr3-17402469-INS-839
chr3-17413236-INS-191
chr3-17415272-INS-453
chr3-17417166-INS-54
chr3-17427381-INS-4621
chr3-17433041-INS-90
chr3-17449266-INS-1927
chr3-17450102-INS-7326
chr3-17460175-INS-2167
chr3-17490270-INS-53
chr3-17492915-INS-183
chr3-17509766-INS-6154
chr3-17511400-INS-390
chr3-17540686-INS-126
chr3-17558396-INS-159
chr3-17571308-INS-1890
chr3-17581380-INS-60
chr3-17589053-INS-130
chr3-17598734-INS-739
chr3-17612851-INS-200
chr3-17623774-INS-6589
chr3-17626836-INS-81
chr3-17653813-INS-6592
chr3-17660579-INS-306
chr3-17667080-INS-124
chr3-17676597-INS-229
chr3-17679020-INS-226
chr3-17686460-INS-7107
chr3-17698658-INS-70
chr3-17700306-INS-6173
chr3-17748441-INS-229
chr3-17767903-INS-337
chr3-17797115-INS-83
chr3-17821876-INS-1887
chr3-17824447-INS-1102
chr3-17872857-INS-72
chr3-17875442-INS-58
chr3-17876470-INS-88
chr3-17883635-INS-164
chr3-17895390-INS-6524
chr3-17898457-INS-64
chr3-18000907-INS-70
chr3-18018054-INS-178
chr3-18051973-INS-127
chr3-18070403-INS-728
chr3-18112985-INS-92
chr3-18114032-INS-201
chr3-18119519-INS-226
chr3-18131769-INS-57
chr3-18167249-INS-195
chr3-18182171-INS-104
chr3-18189754-INS-102
chr3-18203882-INS-108
chr3-18205811-INS-514
chr3-18211085-INS-2769
chr3-18229743-INS-90
chr3-18257490-INS-910
chr3-18277677-INS-656
chr3-18289836-INS-54
chr3-18294879-INS-169
chr3-18300757-INS-220
chr3-18353947-INS-244
chr3-18387996-INS-54
chr3-18417068-INS-1725
chr3-18417779-INS-386
chr3-18477261-INS-5326
chr3-18478930-INS-230
chr3-18484862-INS-3637
chr3-18484862-INS-408
chr3-18500391-INS-357
chr3-18511196-INS-244
chr3-18544975-INS-168
chr3-18561088-INS-121

chr3-18589525-INS-6466
chr3-18590604-INS-5889
chr3-18593496-INS-6589
chr3-18598129-INS-154
chr3-18601297-INS-6182
chr3-18610815-INS-208
chr3-18622780-INS-5362
chr3-18624814-INS-101
chr3-18646233-INS-262
chr3-18654733-INS-7002
chr3-18659280-INS-260
chr3-18662403-INS-727
chr3-18703280-INS-56
chr3-18705420-INS-66
chr3-18749060-INS-92
chr3-18749872-INS-451
chr3-18795761-INS-175
chr3-18817340-INS-400
chr3-18819845-INS-178
chr3-18820370-INS-6393
chr3-18827756-INS-6593
chr3-18842837-INS-1899
chr3-18862406-INS-175
chr3-18867192-INS-3183
chr3-18872140-INS-678
chr3-18899714-INS-62
chr3-18941144-INS-6281
chr3-18964434-INS-2255
chr3-18978880-INS-1877
chr3-18981242-INS-771
chr3-19006360-INS-273
chr3-19017914-INS-84
chr3-19042912-INS-82
chr3-19047581-INS-464
chr3-19066311-INS-58
chr3-19089736-INS-483
chr3-19110924-INS-104
chr3-19126105-INS-6410
chr3-19130803-INS-5012
chr3-19134039-INS-194
chr3-19159738-INS-737
chr3-19218269-INS-97
chr3-19246277-INS-76
chr3-19261289-INS-52
chr3-19289464-INS-58
chr3-19307568-INS-81
chr3-19307630-INS-401
chr3-19321657-INS-734
chr3-19333883-INS-173
chr3-19341869-INS-56
chr3-19344127-INS-70
chr3-19349636-INS-51
chr3-19351869-INS-432
chr3-19379329-INS-64
chr3-19393345-INS-87
chr3-19407277-INS-367
chr3-19416119-INS-244
chr3-19430034-INS-81
chr3-19440422-INS-524
chr3-19461789-INS-398
chr3-19463725-INS-239
chr3-19468660-INS-67
chr3-19500118-INS-119
chr3-19509672-INS-639
chr3-19569117-INS-168
chr3-19582115-INS-6689
chr3-19595838-INS-6399
chr3-19597851-INS-6198
chr3-19602756-INS-5548
chr3-19618000-INS-123
chr3-19620587-INS-86
chr3-19658001-INS-192
chr3-19681287-INS-552
chr3-19700098-INS-415
chr3-19713685-INS-83
chr3-19724162-INS-72
chr3-19737171-INS-149

chr3-19743496-INS-800
chr3-19746585-INS-64
chr3-19753221-INS-54
chr3-19787491-INS-52
chr3-19789665-INS-619
chr3-19809681-INS-278
chr3-19813136-INS-59
chr3-19813889-INS-996
chr3-19824941-INS-54
chr3-19843795-INS-532
chr3-19853490-INS-329
chr3-19877647-INS-183
chr3-19878648-INS-607
chr3-19918952-INS-58
chr3-19921022-INS-177
chr3-19934913-INS-54
chr3-19938413-INS-80
chr3-19959839-INS-219
chr3-19986393-INS-670
chr3-19989011-INS-167
chr3-19992609-INS-6825
chr3-20001827-INS-147
chr3-20015830-INS-113
chr3-20017744-INS-1043
chr3-20052530-INS-70
chr3-20055377-INS-88
chr3-20062112-INS-413
chr3-20069212-INS-164
chr3-20077620-INS-565
chr3-20086705-INS-55
chr3-20091588-INS-185
chr3-20093359-INS-167
chr3-20110754-INS-67
chr3-20113823-INS-113
chr3-20131120-INS-83
chr3-20131439-INS-392
chr3-20140669-INS-519
chr3-20175854-INS-810
chr3-20181581-INS-195
chr3-20189010-INS-128
chr3-20197417-INS-142
chr3-20228529-INS-200
chr3-20233462-INS-2104
chr3-20241527-INS-145
chr3-20268436-INS-6440
chr3-20274602-INS-80
chr3-20292485-INS-116
chr3-20297567-INS-67
chr3-20300862-INS-90
chr3-20303251-INS-265
chr3-20323945-INS-375
chr3-20332291-INS-93
chr3-20347833-INS-58
chr3-20365484-INS-295
chr3-20379714-INS-360
chr3-20384649-INS-62
chr3-20400099-INS-90
chr3-20406316-INS-339
chr3-20414627-INS-457
chr3-20428312-INS-6647
chr3-20431174-INS-77
chr3-20436654-INS-842
chr3-20449818-INS-6466
chr3-20449837-INS-294
chr3-20455395-INS-428
chr3-20458739-INS-177
chr3-20487048-INS-408
chr3-20498327-INS-71
chr3-20508996-INS-558
chr3-20513837-INS-207
chr3-20516010-INS-498
chr3-20518306-INS-171
chr3-20552552-INS-390
chr3-20553026-INS-3010
chr3-20562726-INS-132
chr3-20563867-INS-80
chr3-20566107-INS-453

chr3-20570623-INS-241
chr3-20598535-INS-439
chr3-20599710-INS-1947
chr3-20611799-INS-54
chr3-20638823-INS-61
chr3-20642172-INS-111
chr3-20643644-INS-7096
chr3-20686949-INS-7339
chr3-20702829-INS-79
chr3-20711876-INS-164
chr3-20718643-INS-6730
chr3-20723858-INS-161
chr3-20750502-INS-445
chr3-20753242-INS-453
chr3-20759568-INS-593
chr3-20786387-INS-96
chr3-20786479-INS-900
chr3-20794821-INS-204
chr3-20806626-INS-7148
chr3-20810230-INS-52
chr3-20824083-INS-134
chr3-20835907-INS-65
chr3-20856758-INS-505
chr3-20860883-INS-71
chr3-20923115-INS-800
chr3-20930068-INS-65
chr3-20946143-INS-148
chr3-20982545-INS-105
chr3-21029176-INS-130
chr3-21031632-INS-73
chr3-21065982-INS-6455
chr3-21081176-INS-992
chr3-21129149-INS-179
chr3-21145118-INS-282
chr3-21149202-INS-58
chr3-21153414-INS-238
chr3-21162447-INS-6182
chr3-21165373-INS-923
chr3-21181061-INS-133
chr3-21240163-INS-89
chr3-21240306-INS-7301
chr3-21306191-INS-429
chr3-21307414-INS-201
chr3-21325443-INS-84
chr3-21328960-INS-184
chr3-21353245-INS-58
chr3-21379553-INS-60
chr3-21391004-INS-80
chr3-21413591-INS-361
chr3-21444423-INS-7451
chr3-21453294-INS-90
chr3-21463975-INS-357
chr3-21482414-INS-343
chr3-21494335-INS-10276
chr3-21500520-INS-64
chr3-21503298-INS-59
chr3-21517987-INS-464
chr3-21542282-INS-51
chr3-21544358-INS-509
chr3-21562645-INS-82
chr3-21568510-INS-1579
chr3-21584937-INS-6042
chr3-21635566-INS-80
chr3-21686450-INS-272
chr3-21689932-INS-76
chr3-21736544-INS-6835
chr3-21747154-INS-429
chr3-21755509-INS-6268
chr3-21788248-INS-123
chr3-21797878-INS-203
chr3-21822435-INS-63
chr3-21867504-INS-64
chr3-21874713-INS-68
chr3-21880634-INS-209
chr3-21887877-INS-84
chr3-21942138-INS-209
chr3-21948262-INS-153

chr3-21968732-INS-54
chr3-21971264-INS-122
chr3-21978580-INS-81
chr3-21983315-INS-472
chr3-21993411-INS-140
chr3-22011005-INS-864
chr3-22036409-INS-165
chr3-22043917-INS-227
chr3-22056036-INS-518
chr3-22061747-INS-278
chr3-22096997-INS-322
chr3-22098390-INS-7055
chr3-22101220-INS-185
chr3-22103188-INS-313
chr3-22275122-INS-736
chr3-22280632-INS-54
chr3-22289729-INS-596
chr3-22290828-INS-496
chr3-22292598-INS-68
chr3-22296186-INS-4541
chr3-22297539-INS-2802
chr3-22306367-INS-6452
chr3-22311379-INS-499
chr3-22371964-INS-54
chr3-22382960-INS-8846
chr3-22465883-INS-519
chr3-22496486-INS-160
chr3-22514305-INS-617
chr3-22523406-INS-58
chr3-22563027-INS-83
chr3-22575979-INS-80
chr3-22588580-INS-87
chr3-22590257-INS-68
chr3-22593446-INS-858
chr3-22634077-INS-63
chr3-22648842-INS-128
chr3-22651743-INS-324
chr3-22689293-INS-88
chr3-22742305-INS-126
chr3-22769580-INS-208
chr3-22770521-INS-6259
chr3-22901580-INS-67
chr3-22937746-INS-54
chr3-22950928-INS-2403
chr3-22954523-INS-675
chr3-22979455-INS-176
chr3-22985039-INS-7086
chr3-22990855-INS-1890
chr3-23002234-INS-82
chr3-23015045-INS-107
chr3-23064708-INS-2689
chr3-23081872-INS-362
chr3-23127942-INS-51
chr3-23138243-INS-166
chr3-23143868-INS-61
chr3-23147212-INS-115
chr3-23159011-INS-128
chr3-23182453-INS-55
chr3-23193760-INS-119
chr3-23216435-INS-6472
chr3-23222535-INS-144
chr3-23242299-INS-5458
chr3-23249319-INS-212
chr3-23255522-INS-193
chr3-23268328-INS-104
chr3-23295030-INS-168
chr3-23302084-INS-95
chr3-23340423-INS-106
chr3-23377503-INS-316
chr3-23389296-INS-80
chr3-23397772-INS-4834
chr3-23398836-INS-529
chr3-23406678-INS-828
chr3-23407872-INS-213
chr3-23464308-INS-1773
chr3-23480040-INS-56
chr3-23515201-INS-5213

chr3-23523512-INS-199
chr3-23535522-INS-598
chr3-23547248-INS-68
chr3-23550377-INS-1590
chr3-23580147-INS-6931
chr3-23619789-INS-7087
chr3-23650088-INS-58
chr3-23657624-INS-380
chr3-23670895-INS-880
chr3-23685236-INS-56
chr3-23718798-INS-84
chr3-23730244-INS-7302
chr3-23731638-INS-861
chr3-23736584-INS-498
chr3-23749594-INS-493
chr3-23758829-INS-277
chr3-23774492-INS-92
chr3-23788589-INS-11817
chr3-23817764-INS-164
chr3-23819025-INS-109
chr3-23828811-INS-2556
chr3-23877579-INS-227
chr3-23899876-INS-141
chr3-23902108-INS-961
chr3-23905229-INS-140
chr3-23932790-INS-850
chr3-24018150-INS-217
chr3-24023575-INS-521
chr3-24064638-INS-525
chr3-24131262-INS-2176
chr3-24137931-INS-6427
chr3-24155987-INS-1229
chr3-24158694-INS-60
chr3-24165511-INS-497
chr3-24173188-INS-56
chr3-24197595-INS-3189
chr3-24205729-INS-1196
chr3-24209005-INS-322
chr3-24233225-INS-358
chr3-24251610-INS-6166
chr3-24262274-INS-63
chr3-24293799-INS-207
chr3-24324828-INS-144
chr3-24372945-INS-558
chr3-24374305-INS-114
chr3-24379414-INS-443
chr3-24392628-INS-53
chr3-24411497-INS-84
chr3-24413393-INS-841
chr3-24419184-INS-130
chr3-24424881-INS-437
chr3-24430460-INS-56
chr3-24465255-INS-118
chr3-24469805-INS-190
chr3-24476105-INS-148
chr3-24503405-INS-399
chr3-24514025-INS-88
chr3-24532143-INS-99
chr3-24559224-INS-183
chr3-24568123-INS-56
chr3-24570705-INS-289
chr3-24571727-INS-665
chr3-24607335-INS-56
chr3-24609854-INS-144
chr3-24627883-INS-7127
chr3-24628020-INS-608
chr3-24636185-INS-538
chr3-24643510-INS-75
chr3-24648288-INS-190
chr3-24654382-INS-177
chr3-24672589-INS-203
chr3-24685014-INS-652
chr3-24705865-INS-54
chr3-24717827-INS-853
chr3-24724274-INS-218
chr3-24755426-INS-54
chr3-24771145-INS-66

chr3-24774177-INS-219
chr3-24806788-INS-58
chr3-24809235-INS-2296
chr3-24816859-INS-346
chr3-24836599-INS-271
chr3-24875205-INS-52
chr3-24880010-INS-802
chr3-24888516-INS-64
chr3-24896445-INS-489
chr3-24924692-INS-319
chr3-24940353-INS-154
chr3-24946977-INS-55
chr3-24949010-INS-76
chr3-24968580-INS-160
chr3-24983816-INS-55
chr3-24993589-INS-70
chr3-25048765-INS-7086
chr3-25050411-INS-64
chr3-25096852-INS-133
chr3-25118023-INS-399
chr3-25121736-INS-108
chr3-25131180-INS-90
chr3-25138556-INS-903
chr3-25172669-INS-1642
chr3-25172849-INS-230
chr3-25207191-INS-465
chr3-25241306-INS-68
chr3-25249428-INS-3131
chr3-25255097-INS-154
chr3-25288776-INS-62
chr3-25294967-INS-114
chr3-25304290-INS-214
chr3-25317645-INS-204
chr3-25411719-INS-62
chr3-25420347-INS-52
chr3-25428584-INS-759
chr3-25442150-INS-144
chr3-25446710-INS-173
chr3-25484846-INS-192
chr3-25499235-INS-348
chr3-25507285-INS-212
chr3-25538095-INS-124
chr3-25546667-INS-1796
chr3-25550011-INS-156
chr3-25571497-INS-832
chr3-25606748-INS-93
chr3-25622969-INS-51
chr3-25664862-INS-60
chr3-25673859-INS-56
chr3-25676449-INS-481
chr3-25678982-INS-217
chr3-25703170-INS-592
chr3-25710223-INS-153
chr3-25736425-INS-58
chr3-25745581-INS-78
chr3-25771252-INS-145
chr3-25775491-INS-522
chr3-25778804-INS-126
chr3-25830051-INS-91
chr3-25840804-INS-54
chr3-25844408-INS-155
chr3-25851088-INS-108
chr3-25864189-INS-77
chr3-25871151-INS-3980
chr3-25871748-INS-315
chr3-25892480-INS-129
chr3-25906983-INS-636
chr3-25955705-INS-279
chr3-25973114-INS-66
chr3-26030352-INS-53
chr3-26036534-INS-609
chr3-26056682-INS-61
chr3-26063731-INS-61
chr3-26100051-INS-152
chr3-26103962-INS-932
chr3-26109606-INS-115
chr3-26129718-INS-67

chr3-26167028-INS-422
chr3-26179994-INS-553
chr3-26226412-INS-7308
chr3-26255984-INS-6386
chr3-26280850-INS-365
chr3-26283468-INS-54
chr3-26294640-INS-115
chr3-26301041-INS-687
chr3-26306269-INS-140
chr3-26332519-INS-529
chr3-26337789-INS-116
chr3-26362066-INS-76
chr3-26381506-INS-264
chr3-26383653-INS-56
chr3-26415340-INS-182
chr3-26431394-INS-72
chr3-26439637-INS-126
chr3-26505683-INS-313
chr3-26517780-INS-6254
chr3-26522610-INS-96
chr3-26529245-INS-877
chr3-26560461-INS-161
chr3-26572881-INS-108
chr3-26578445-INS-94
chr3-26591869-INS-6936
chr3-26610227-INS-776
chr3-26649031-INS-117
chr3-26656206-INS-130
chr3-26677286-INS-69
chr3-26690371-INS-53
chr3-26699198-INS-321
chr3-26712807-INS-96
chr3-26742948-INS-320
chr3-26759262-INS-80
chr3-26767857-INS-56
chr3-26768864-INS-1155
chr3-26771305-INS-104
chr3-26774615-INS-163
chr3-26778268-INS-93
chr3-26779221-INS-113
chr3-26779605-INS-319
chr3-26883432-INS-432
chr3-26887842-INS-54
chr3-26936811-INS-60
chr3-26988715-INS-55
chr3-27021829-INS-172
chr3-27038517-INS-242
chr3-27042321-INS-77
chr3-27048084-INS-745
chr3-27051644-INS-250
chr3-27080724-INS-66
chr3-27087875-INS-58
chr3-27094417-INS-56
chr3-27100173-INS-56
chr3-27104566-INS-62
chr3-27129492-INS-379
chr3-27174684-INS-63
chr3-27181525-INS-69
chr3-27195719-INS-64
chr3-27201570-INS-99
chr3-27208336-INS-6527
chr3-27241456-INS-57
chr3-27269992-INS-170
chr3-27304244-INS-265
chr3-27333947-INS-71
chr3-27340294-INS-318
chr3-27340859-INS-7174
chr3-27355796-INS-203
chr3-27355973-INS-142
chr3-27400453-INS-140
chr3-27407881-INS-84
chr3-27408455-INS-176
chr3-27420068-INS-58
chr3-27435351-INS-197
chr3-27443549-INS-136
chr3-27445933-INS-327
chr3-27465736-INS-183

chr3-27474461-INS-95
chr3-27490663-INS-74
chr3-27495790-INS-207
chr3-27522134-INS-67
chr3-27540878-INS-160
chr3-27550211-INS-224
chr3-27562532-INS-55
chr3-27571492-INS-97
chr3-27582229-INS-206
chr3-27624676-INS-59
chr3-27644230-INS-72
chr3-27653960-INS-354
chr3-27655803-INS-68
chr3-27689253-INS-375
chr3-27696312-INS-172
chr3-27699916-INS-196
chr3-27701042-INS-56
chr3-27708254-INS-372
chr3-27727595-INS-220
chr3-27754453-INS-108
chr3-27775315-INS-58
chr3-27795218-INS-115
chr3-27811740-INS-114
chr3-27815522-INS-105
chr3-27835845-INS-53
chr3-27851059-INS-55
chr3-27857269-INS-519
chr3-27875094-INS-412
chr3-27876340-INS-168
chr3-27924591-INS-56
chr3-27932035-INS-317
chr3-27932648-INS-74
chr3-27937944-INS-230
chr3-27949850-INS-159
chr3-27954384-INS-70
chr3-27957046-INS-90
chr3-27981640-INS-129
chr3-28003275-INS-56
chr3-28020027-INS-144
chr3-28021581-INS-330
chr3-28080958-INS-74
chr3-28086322-INS-68
chr3-28090614-INS-435
chr3-28101123-INS-201
chr3-28152050-INS-338
chr3-28155509-INS-136
chr3-28174188-INS-7172
chr3-28226523-INS-112
chr3-28228785-INS-77
chr3-28263110-INS-79
chr3-28275693-INS-211
chr3-28277958-INS-198
chr3-28286840-INS-197
chr3-28290937-INS-57
chr3-28298921-INS-52
chr3-28299918-INS-623
chr3-28328669-INS-73
chr3-28332052-INS-76
chr3-28342649-INS-112
chr3-28352974-INS-76
chr3-28356771-INS-87
chr3-28360045-INS-66
chr3-28365046-INS-64
chr3-28403829-INS-56
chr3-28405762-INS-686
chr3-28418796-INS-154
chr3-28455404-INS-122
chr3-28457411-INS-546
chr3-28467840-INS-278
chr3-28487780-INS-108
chr3-28493099-INS-66
chr3-28513938-INS-78
chr3-28535586-INS-74
chr3-28539048-INS-120
chr3-28558773-INS-54
chr3-28584819-INS-396
chr3-28616407-INS-154

chr3-28618142-INS-126
chr3-28624414-INS-80
chr3-28649246-INS-56
chr3-28660700-INS-301
chr3-28687992-INS-1464
chr3-28695454-INS-59
chr3-28758318-INS-54
chr3-28787894-INS-54
chr3-28793301-INS-365
chr3-28819547-INS-1937
chr3-28857115-INS-138
chr3-28861032-INS-60
chr3-28866103-INS-58
chr3-28879488-INS-54
chr3-28895892-INS-537
chr3-28946425-INS-240
chr3-28962785-INS-184
chr3-29022795-INS-132
chr3-29060486-INS-51
chr3-29067557-INS-62
chr3-29068952-INS-208
chr3-29078124-INS-188
chr3-29079727-INS-118
chr3-29083304-INS-136
chr3-29102635-INS-6475
chr3-29104282-INS-133
chr3-29122049-INS-6465
chr3-29134710-INS-411
chr3-29168596-INS-228
chr3-29210808-INS-69
chr3-29217329-INS-608
chr3-29228851-INS-84
chr3-29254460-INS-69
chr3-29267456-INS-52
chr3-29279554-INS-194
chr3-29295440-INS-113
chr3-29320911-INS-194
chr3-29325844-INS-145
chr3-29329463-INS-144
chr3-29343479-INS-77
chr3-29356167-INS-88
chr3-29359347-INS-6593
chr3-29416001-INS-1175
chr3-29422453-INS-84
chr3-29450417-INS-218
chr3-29468334-INS-104
chr3-29495394-INS-54
chr3-29501456-INS-75
chr3-29506285-INS-57
chr3-29523978-INS-2563
chr3-29535204-INS-137
chr3-29555604-INS-60
chr3-29558868-INS-349
chr3-29571425-INS-54
chr3-29580412-INS-367
chr3-29608923-INS-421
chr3-29621011-INS-55
chr3-29640996-INS-56
chr3-29663943-INS-260
chr3-29672246-INS-54
chr3-29693043-INS-426
chr3-29697676-INS-1093
chr3-29697941-INS-187
chr3-29727270-INS-1706
chr3-29750667-INS-159
chr3-29771069-INS-4341
chr3-29772552-INS-454
chr3-29805280-INS-80
chr3-29816688-INS-65
chr3-29849299-INS-53
chr3-29856282-INS-180
chr3-29871613-INS-466
chr3-29896766-INS-54
chr3-29897976-INS-162
chr3-29898423-INS-180
chr3-29931274-INS-151
chr3-29933122-INS-192

chr3-29959505-INS-316
chr3-29982719-INS-128
chr3-30041588-INS-153
chr3-30043822-INS-160
chr3-30047990-INS-90
chr3-30084692-INS-205
chr3-30108519-INS-876
chr3-30119122-INS-217
chr3-30126884-INS-7259
chr3-30150754-INS-88
chr3-30184063-INS-60
chr3-30230644-INS-7483
chr3-30237216-INS-54
chr3-30238231-INS-115
chr3-30279714-INS-154
chr3-30302177-INS-74
chr3-30342386-INS-58
chr3-30382769-INS-201
chr3-30417795-INS-54
chr3-30418624-INS-290
chr3-30440403-INS-60
chr3-30444083-INS-58
chr3-30444585-INS-563
chr3-30448233-INS-313
chr3-30475742-INS-55
chr3-30480580-INS-6849
chr3-30500802-INS-123
chr3-30506491-INS-172
chr3-30514569-INS-170
chr3-30517789-INS-93
chr3-30524027-INS-290
chr3-30525268-INS-286
chr3-30525544-INS-108
chr3-30549923-INS-148
chr3-30557245-INS-315
chr3-30570979-INS-98
chr3-30586259-INS-6409
chr3-30587705-INS-154
chr3-30597523-INS-96
chr3-30626293-INS-111
chr3-30634390-INS-204
chr3-30675240-INS-114
chr3-30679392-INS-78
chr3-30707317-INS-60
chr3-30712807-INS-127
chr3-30728032-INS-327
chr3-30733627-INS-222
chr3-30751588-INS-210
chr3-30753460-INS-203
chr3-30755720-INS-63
chr3-30757353-INS-193
chr3-30777163-INS-6855
chr3-30779571-INS-214
chr3-30784501-INS-491
chr3-30786044-INS-799
chr3-30797699-INS-179
chr3-30799649-INS-365
chr3-30818279-INS-61
chr3-30834182-INS-385
chr3-30855472-INS-857
chr3-30856157-INS-6440
chr3-30882705-INS-144
chr3-30893562-INS-75
chr3-30898919-INS-196
chr3-30916575-INS-92
chr3-30925093-INS-51
chr3-30926310-INS-401
chr3-30938054-INS-292
chr3-30951105-INS-116
chr3-30969767-INS-114
chr3-30973118-INS-80
chr3-30978552-INS-209
chr3-30985524-INS-224
chr3-31020804-INS-199
chr3-31073091-INS-130
chr3-31091638-INS-206
chr3-31092579-INS-134

chr3-31095095-INS-206
chr3-31101011-INS-148
chr3-31124235-INS-1902
chr3-31124816-INS-2640
chr3-31126713-INS-74
chr3-31133348-INS-136
chr3-31139060-INS-200
chr3-31159720-INS-201
chr3-31192522-INS-431
chr3-31197594-INS-210
chr3-31203782-INS-119
chr3-31209442-INS-213
chr3-31224832-INS-95
chr3-31225756-INS-60
chr3-31234627-INS-909
chr3-31247585-INS-216
chr3-31247632-INS-2869
chr3-31248278-INS-261
chr3-31279247-INS-104
chr3-31312201-INS-522
chr3-31319845-INS-466
chr3-31324625-INS-544
chr3-31329814-INS-134
chr3-31336544-INS-123
chr3-31344397-INS-236
chr3-31346544-INS-340
chr3-31375779-INS-174
chr3-31383282-INS-149
chr3-31389260-INS-124
chr3-31431519-INS-315
chr3-31456466-INS-74
chr3-31482136-INS-5814
chr3-31486718-INS-579
chr3-31507990-INS-919
chr3-31520938-INS-6440
chr3-31521347-INS-62
chr3-31559505-INS-234
chr3-31608292-INS-51
chr3-31642413-INS-59
chr3-31644937-INS-122
chr3-31655521-INS-51
chr3-31701512-INS-67
chr3-31713664-INS-279
chr3-31721588-INS-60
chr3-31722521-INS-60
chr3-31740483-INS-56
chr3-31740803-INS-55
chr3-31766434-INS-6308
chr3-31769360-INS-1890
chr3-31769921-INS-6401
chr3-31773386-INS-160
chr3-31777550-INS-208
chr3-31785618-INS-604
chr3-31793087-INS-7211
chr3-31795885-INS-4133
chr3-31807254-INS-349
chr3-31815709-INS-90
chr3-31821216-INS-54
chr3-31821767-INS-428
chr3-31825809-INS-268
chr3-31865179-INS-80
chr3-31910313-INS-131
chr3-31916821-INS-216
chr3-31926833-INS-76
chr3-31937501-INS-400
chr3-31942680-INS-73
chr3-31961223-INS-5552
chr3-31967307-INS-120
chr3-31972412-INS-415
chr3-31978123-INS-58
chr3-31983245-INS-134
chr3-32009553-INS-410
chr3-32012664-INS-56
chr3-32042043-INS-322
chr3-32065845-INS-401
chr3-32068502-INS-264
chr3-32078565-INS-452

chr3-32085893-INS-256
chr3-32096821-INS-94
chr3-32103526-INS-189
chr3-32124438-INS-199
chr3-32181731-INS-624
chr3-32202212-INS-9047
chr3-32215302-INS-2297
chr3-32230078-INS-215
chr3-32237323-INS-1075
chr3-32244134-INS-4468
chr3-32268615-INS-490
chr3-32269202-INS-136
chr3-32280507-INS-227
chr3-32292562-INS-92
chr3-32305462-INS-131
chr3-32346608-INS-7879
chr3-32357537-INS-8062
chr3-32363676-INS-6849
chr3-32375318-INS-316
chr3-32377486-INS-188
chr3-32394572-INS-339
chr3-32397965-INS-185
chr3-32408895-INS-427
chr3-32424741-INS-60
chr3-32456753-INS-211
chr3-32500184-INS-96
chr3-32504921-INS-73
chr3-32533742-INS-399
chr3-32543783-INS-97
chr3-32577584-INS-199
chr3-32581353-INS-172
chr3-32584951-INS-63
chr3-32587962-INS-320
chr3-32603078-INS-185
chr3-32622125-INS-371
chr3-32643781-INS-753
chr3-32654066-INS-510
chr3-32680153-INS-207
chr3-32682540-INS-57
chr3-32694539-INS-252
chr3-32701096-INS-188
chr3-32703527-INS-629
chr3-32709234-INS-86
chr3-32715348-INS-184
chr3-32717117-INS-245
chr3-32744599-INS-342
chr3-32771461-INS-69
chr3-32781320-INS-322
chr3-32790245-INS-144
chr3-32802046-INS-202
chr3-32822266-INS-546
chr3-32837874-INS-60
chr3-32866861-INS-5075
chr3-32872442-INS-103
chr3-32875450-INS-6409
chr3-32882997-INS-82
chr3-32889074-INS-52
chr3-32900340-INS-70
chr3-32906099-INS-420
chr3-32919763-INS-68
chr3-32928590-INS-53
chr3-32945558-INS-151
chr3-32955542-INS-160
chr3-32976748-INS-364
chr3-32984127-INS-52
chr3-32993087-INS-612
chr3-33025018-INS-54
chr3-33027021-INS-52
chr3-33035427-INS-108
chr3-33038104-INS-71
chr3-33039908-INS-643
chr3-33057543-INS-63
chr3-33059822-INS-73
chr3-33079389-INS-280
chr3-33086730-INS-102
chr3-33092635-INS-120
chr3-33096436-INS-835

chr3-33103953-INS-109
chr3-33117437-INS-1857
chr3-33139684-INS-74
chr3-33159997-INS-256
chr3-33173483-INS-910
chr3-33211120-INS-261
chr3-33213814-INS-574
chr3-33223783-INS-616
chr3-33234235-INS-408
chr3-33240864-INS-5275
chr3-33241658-INS-433
chr3-33244220-INS-95
chr3-33250798-INS-147
chr3-33251206-INS-6481
chr3-33276462-INS-61
chr3-33280315-INS-4551
chr3-33284470-INS-61
chr3-33310288-INS-6489
chr3-33317339-INS-114
chr3-33320307-INS-813
chr3-33336325-INS-56
chr3-33372763-INS-60
chr3-33393124-INS-222
chr3-33405505-INS-72
chr3-33406208-INS-596
chr3-33409225-INS-337
chr3-33410060-INS-79
chr3-33453185-INS-106
chr3-33461019-INS-110
chr3-33498661-INS-6758
chr3-33510112-INS-430
chr3-33523247-INS-569
chr3-33527233-INS-3364
chr3-33527639-INS-52
chr3-33562232-INS-63
chr3-33571633-INS-108
chr3-33608332-INS-149
chr3-33636952-INS-89
chr3-33638947-INS-523
chr3-33644386-INS-6598
chr3-33671038-INS-135
chr3-33695075-INS-6738
chr3-33695812-INS-305
chr3-33704644-INS-94
chr3-33709864-INS-56
chr3-33711107-INS-161
chr3-33768911-INS-470
chr3-33792788-INS-52
chr3-33808376-INS-6883
chr3-33810284-INS-1877
chr3-33810506-INS-358
chr3-33811885-INS-55
chr3-33812983-INS-1286
chr3-33817228-INS-6409
chr3-33830192-INS-70
chr3-33894485-INS-449
chr3-33914359-INS-448
chr3-33916993-INS-591
chr3-33922416-INS-199
chr3-33935379-INS-214
chr3-33945372-INS-577
chr3-33982247-INS-198
chr3-33993619-INS-78
chr3-33993755-INS-96
chr3-34011934-INS-97
chr3-34016005-INS-87
chr3-34018535-INS-1862
chr3-34024131-INS-103
chr3-34024227-INS-641
chr3-34031361-INS-206
chr3-34050304-INS-599
chr3-34061712-INS-637
chr3-34116793-INS-204
chr3-34125830-INS-76
chr3-34142926-INS-191
chr3-34151241-INS-416
chr3-34163260-INS-105

chr3-34164638-INS-61
chr3-34172865-INS-483
chr3-34181138-INS-212
chr3-34194269-INS-584
chr3-34203828-INS-126
chr3-34205559-INS-59
chr3-34223671-INS-211
chr3-34241313-INS-110
chr3-34262470-INS-188
chr3-34277802-INS-70
chr3-34290150-INS-60
chr3-34293245-INS-355
chr3-34297672-INS-7248
chr3-34309857-INS-1374
chr3-34359042-INS-145
chr3-34372132-INS-7025
chr3-34372707-INS-256
chr3-34399171-INS-6442
chr3-34404490-INS-422
chr3-34432079-INS-56
chr3-34445431-INS-6494
chr3-34448492-INS-58
chr3-34463341-INS-3188
chr3-34485652-INS-66
chr3-34493179-INS-120
chr3-34494507-INS-69
chr3-34508384-INS-70
chr3-34548032-INS-316
chr3-34559320-INS-177
chr3-34578785-INS-76
chr3-34582112-INS-354
chr3-34623494-INS-60
chr3-34625319-INS-70
chr3-34646267-INS-93
chr3-34659128-INS-79
chr3-34661473-INS-61
chr3-34666727-INS-107
chr3-34687556-INS-52
chr3-34728687-INS-126
chr3-34762688-INS-204
chr3-34768444-INS-182
chr3-34777223-INS-54
chr3-34796415-INS-87
chr3-34826748-INS-61
chr3-34831406-INS-51
chr3-34842468-INS-154
chr3-34844258-INS-201
chr3-34849315-INS-88
chr3-34855617-INS-93
chr3-34860053-INS-248
chr3-34869634-INS-535
chr3-34882064-INS-54
chr3-34908105-INS-56
chr3-34911130-INS-70
chr3-34921066-INS-63
chr3-34940172-INS-54
chr3-34942507-INS-364
chr3-34946403-INS-79
chr3-34969468-INS-89
chr3-34996863-INS-781
chr3-34999183-INS-63
chr3-35003786-INS-60
chr3-35015246-INS-174
chr3-35035079-INS-69
chr3-35040150-INS-60
chr3-35056730-INS-385
chr3-35061694-INS-262
chr3-35065649-INS-169
chr3-35078902-INS-64
chr3-35083902-INS-6403
chr3-35091274-INS-62
chr3-35099526-INS-4911
chr3-35126611-INS-76
chr3-35135075-INS-191
chr3-35161777-INS-52
chr3-35224367-INS-499
chr3-35245521-INS-327

chr3-35247800-INS-59
chr3-35258845-INS-96
chr3-35264114-INS-54
chr3-35264391-INS-5259
chr3-35268819-INS-6034
chr3-35279181-INS-800
chr3-35288941-INS-184
chr3-35347142-INS-272
chr3-35350762-INS-57
chr3-35354227-INS-143
chr3-35355314-INS-355
chr3-35362573-INS-115
chr3-35379597-INS-67
chr3-35385726-INS-201
chr3-35412710-INS-324
chr3-35428355-INS-52
chr3-35443027-INS-65
chr3-35473902-INS-187
chr3-35474611-INS-6489
chr3-35506967-INS-336
chr3-35524244-INS-99
chr3-35525546-INS-60
chr3-35539498-INS-60
chr3-35548379-INS-85
chr3-35554143-INS-297
chr3-35560627-INS-171
chr3-35575358-INS-419
chr3-35608537-INS-53
chr3-35621505-INS-574
chr3-35629842-INS-61
chr3-35677415-INS-408
chr3-35695406-INS-435
chr3-35704054-INS-52
chr3-35706638-INS-7328
chr3-35727526-INS-104
chr3-35729805-INS-200
chr3-35731995-INS-60
chr3-35734754-INS-72
chr3-35736517-INS-141
chr3-35769636-INS-213
chr3-35770685-INS-220
chr3-35775573-INS-94
chr3-35783092-INS-360
chr3-35785461-INS-64
chr3-35813759-INS-132
chr3-35816738-INS-227
chr3-35833907-INS-288
chr3-35880060-INS-78
chr3-35893744-INS-488
chr3-35912544-INS-984
chr3-35930302-INS-162
chr3-35931878-INS-105
chr3-35950831-INS-59
chr3-35957093-INS-124
chr3-35958563-INS-52
chr3-35961374-INS-6403
chr3-35963887-INS-98
chr3-35966222-INS-678
chr3-36010008-INS-105
chr3-36029681-INS-546
chr3-36035395-INS-54
chr3-36052123-INS-4604
chr3-36060575-INS-6442
chr3-36066227-INS-140
chr3-36068426-INS-218
chr3-36077154-INS-282
chr3-36082630-INS-3863
chr3-36082638-INS-483
chr3-36085224-INS-348
chr3-36116653-INS-173
chr3-36118561-INS-56
chr3-36124777-INS-208
chr3-36133466-INS-400
chr3-36137808-INS-117
chr3-36145884-INS-316
chr3-36150723-INS-5645
chr3-36170560-INS-62

chr3-36171559-INS-1890
chr3-36210630-INS-422
chr3-36217141-INS-183
chr3-36231621-INS-90
chr3-36259617-INS-94
chr3-36266934-INS-138
chr3-36270822-INS-326
chr3-36278362-INS-65
chr3-36305231-INS-54
chr3-36308505-INS-364
chr3-36312786-INS-54
chr3-36320396-INS-86
chr3-36326682-INS-5236
chr3-36333877-INS-454
chr3-36350492-INS-96
chr3-36361639-INS-119
chr3-36373086-INS-340
chr3-36397921-INS-79
chr3-36401200-INS-2064
chr3-36403150-INS-188
chr3-36416339-INS-7091
chr3-36441867-INS-4913
chr3-36454287-INS-108
chr3-36455494-INS-472
chr3-36486096-INS-51
chr3-36499793-INS-188
chr3-36501132-INS-934
chr3-36522689-INS-64
chr3-36539534-INS-482
chr3-36542556-INS-212
chr3-36588735-INS-249
chr3-36589530-INS-331
chr3-36601514-INS-64
chr3-36634458-INS-388
chr3-36656283-INS-252
chr3-36664081-INS-70
chr3-36666403-INS-231
chr3-36692820-INS-315
chr3-36740724-INS-59
chr3-36771973-INS-206
chr3-36774335-INS-131
chr3-36792912-INS-78
chr3-36804265-INS-345
chr3-36807602-INS-54
chr3-36816260-INS-222
chr3-36844176-INS-52
chr3-36914119-INS-126
chr3-36959528-INS-193
chr3-36972907-INS-8239
chr3-36978891-INS-153
chr3-37079685-INS-237
chr3-37108808-INS-5915
chr3-37119939-INS-6692
chr3-37131156-INS-347
chr3-37158509-INS-86
chr3-37162396-INS-4599
chr3-37162414-INS-6893
chr3-37200443-INS-265
chr3-37204867-INS-70
chr3-37234693-INS-63
chr3-37247454-INS-399
chr3-37298110-INS-166
chr3-37318077-INS-104
chr3-37322895-INS-66
chr3-37323234-INS-374
chr3-37324718-INS-64
chr3-37340418-INS-525
chr3-37365477-INS-2073
chr3-37413889-INS-53
chr3-37416118-INS-90
chr3-37417459-INS-105
chr3-37420004-INS-294
chr3-37440975-INS-82
chr3-37455593-INS-186
chr3-37470680-INS-4125
chr3-37491526-INS-498
chr3-37504057-INS-70

chr3-37506993-INS-119
chr3-37636150-INS-419
chr3-37658792-INS-76
chr3-37664660-INS-203
chr3-37666013-INS-952
chr3-37688894-INS-90
chr3-37699918-INS-54
chr3-37700456-INS-212
chr3-37709480-INS-51
chr3-37731773-INS-296
chr3-37735286-INS-338
chr3-37740682-INS-349
chr3-37768153-INS-82
chr3-37826080-INS-115
chr3-37853879-INS-154
chr3-37861246-INS-282
chr3-37903857-INS-106
chr3-37913866-INS-143
chr3-37924953-INS-94
chr3-37944664-INS-221
chr3-37947976-INS-374
chr3-37972571-INS-922
chr3-37974555-INS-110
chr3-38020511-INS-216
chr3-38043315-INS-274
chr3-38068727-INS-161
chr3-38085835-INS-124
chr3-38110698-INS-346
chr3-38153891-INS-3982
chr3-38155621-INS-3212
chr3-38188115-INS-132
chr3-38210390-INS-191
chr3-38227107-INS-513
chr3-38246354-INS-208
chr3-38250937-INS-126
chr3-38301706-INS-64
chr3-38324039-INS-61
chr3-38352078-INS-76
chr3-38356835-INS-104
chr3-38396436-INS-287
chr3-38419168-INS-70
chr3-38428836-INS-6440
chr3-38429728-INS-79
chr3-38442563-INS-82
chr3-38450731-INS-205
chr3-38467178-INS-97
chr3-38480334-INS-783
chr3-38485146-INS-110
chr3-38493135-INS-107
chr3-38496553-INS-86
chr3-38552137-INS-340
chr3-38554657-INS-578
chr3-38577329-INS-614
chr3-38586652-INS-398
chr3-38617621-INS-434
chr3-38623689-INS-75
chr3-38629699-INS-332
chr3-38704770-INS-498
chr3-38705524-INS-53
chr3-38711232-INS-70
chr3-38719175-INS-278
chr3-38726168-INS-114
chr3-38733654-INS-404
chr3-38760752-INS-6445
chr3-38766629-INS-82
chr3-38792398-INS-920
chr3-38833406-INS-55
chr3-38861567-INS-6665
chr3-38863708-INS-77
chr3-38876612-INS-87
chr3-38878418-INS-77
chr3-38898434-INS-58
chr3-38900578-INS-141
chr3-38907378-INS-53
chr3-39002579-INS-52
chr3-39015424-INS-88
chr3-39017953-INS-389

chr3-39018906-INS-147
chr3-39033496-INS-400
chr3-39052763-INS-376
chr3-39057312-INS-51
chr3-39062898-INS-54
chr3-39069585-INS-826
chr3-39073454-INS-105
chr3-39111246-INS-153
chr3-39117398-INS-194
chr3-39161274-INS-110
chr3-39178085-INS-6431
chr3-39197675-INS-54
chr3-39198451-INS-158
chr3-39207851-INS-608
chr3-39233909-INS-2459
chr3-39244516-INS-70
chr3-39271060-INS-53
chr3-39286090-INS-92
chr3-39315205-INS-4571
chr3-39389260-INS-56
chr3-39461512-INS-52
chr3-39469652-INS-80
chr3-39479171-INS-55
chr3-39487563-INS-5530
chr3-39498519-INS-343
chr3-39511400-INS-88
chr3-39530839-INS-56
chr3-39544542-INS-64
chr3-39622063-INS-168
chr3-39622350-INS-615
chr3-39627030-INS-202
chr3-39629810-INS-320
chr3-39643867-INS-134
chr3-39654227-INS-6261
chr3-39668920-INS-203
chr3-39676564-INS-646
chr3-39695578-INS-51
chr3-39711062-INS-2153
chr3-39714588-INS-426
chr3-39720309-INS-56
chr3-39722781-INS-55
chr3-39726203-INS-76
chr3-39763633-INS-387
chr3-39769559-INS-65
chr3-39783310-INS-181
chr3-39788098-INS-767
chr3-39817692-INS-7268
chr3-39838822-INS-354
chr3-39910564-INS-3166
chr3-39924854-INS-1038
chr3-39965976-INS-246
chr3-39981386-INS-414
chr3-39988654-INS-398
chr3-39998287-INS-127
chr3-40000667-INS-63
chr3-40004259-INS-131
chr3-40047682-INS-7110
chr3-40048709-INS-124
chr3-40051644-INS-161
chr3-40061441-INS-72
chr3-40062939-INS-79
chr3-40065641-INS-54
chr3-40077570-INS-64
chr3-40108346-INS-8142
chr3-40155250-INS-79
chr3-40159042-INS-423
chr3-40203669-INS-654
chr3-40248013-INS-267
chr3-40256128-INS-342
chr3-40271761-INS-408
chr3-40284983-INS-80
chr3-40291893-INS-78
chr3-40293437-INS-66
chr3-40295336-INS-7092
chr3-40338753-INS-152
chr3-40342312-INS-63
chr3-40390709-INS-215

chr3-40436465-INS-573
chr3-40479990-INS-5199
chr3-40488248-INS-496
chr3-40489841-INS-6538
chr3-40497374-INS-52
chr3-40520260-INS-238
chr3-40526046-INS-117
chr3-40529674-INS-134
chr3-40598120-INS-59
chr3-40604738-INS-516
chr3-40623208-INS-122
chr3-40636286-INS-6459
chr3-40655125-INS-104
chr3-40662246-INS-800
chr3-40682576-INS-3843
chr3-40710846-INS-80
chr3-40713866-INS-102
chr3-40729088-INS-686
chr3-40735176-INS-92
chr3-40767642-INS-54
chr3-40774700-INS-443
chr3-40780714-INS-225
chr3-40783682-INS-55
chr3-40790525-INS-446
chr3-40803534-INS-52
chr3-40806785-INS-860
chr3-40878112-INS-316
chr3-40895303-INS-132
chr3-40942827-INS-58
chr3-40949092-INS-831
chr3-40952313-INS-170
chr3-40958191-INS-557
chr3-40973094-INS-340
chr3-40974109-INS-190
chr3-40991683-INS-176
chr3-41000450-INS-51
chr3-41042685-INS-64
chr3-41063694-INS-801
chr3-41067076-INS-182
chr3-41080390-INS-56
chr3-41098940-INS-139
chr3-41130108-INS-232
chr3-41142079-INS-176
chr3-41148032-INS-240
chr3-41170757-INS-254
chr3-41173491-INS-211
chr3-41223628-INS-54
chr3-41273516-INS-74
chr3-41282099-INS-3389
chr3-41289796-INS-60
chr3-41315811-INS-329
chr3-41322460-INS-62
chr3-41355597-INS-70
chr3-41357613-INS-390
chr3-41376952-INS-238
chr3-41436901-INS-200
chr3-41502750-INS-2784
chr3-41504642-INS-246
chr3-41508789-INS-76
chr3-41549692-INS-146
chr3-41599295-INS-407
chr3-41610023-INS-311
chr3-41616966-INS-507
chr3-41622051-INS-212
chr3-41637455-INS-193
chr3-41670269-INS-3373
chr3-41675674-INS-1032
chr3-41710593-INS-6691
chr3-41724596-INS-201
chr3-41733958-INS-884
chr3-41773875-INS-832
chr3-41784780-INS-6352
chr3-41792439-INS-423
chr3-41795988-INS-84
chr3-41796513-INS-3195
chr3-41809705-INS-402
chr3-41822923-INS-191

chr3-41855846-INS-6203
chr3-41872054-INS-167
chr3-41877796-INS-78
chr3-41928466-INS-6392
chr3-41934078-INS-6180
chr3-41963047-INS-399
chr3-41975486-INS-70
chr3-41988427-INS-108
chr3-41990211-INS-6809
chr3-41998850-INS-677
chr3-42003063-INS-6425
chr3-42005759-INS-7518
chr3-42015103-INS-148
chr3-42026399-INS-379
chr3-42070475-INS-214
chr3-42081717-INS-63
chr3-42095268-INS-69
chr3-42096463-INS-6377
chr3-42103650-INS-270
chr3-42124421-INS-779
chr3-42125648-INS-68
chr3-42128153-INS-7155
chr3-42180877-INS-151
chr3-42190996-INS-6733
chr3-42206326-INS-6419
chr3-42246834-INS-56
chr3-42254970-INS-3077
chr3-42267253-INS-94
chr3-42302602-INS-83
chr3-42303571-INS-179
chr3-42316838-INS-6801
chr3-42320569-INS-723
chr3-42330041-INS-58
chr3-42343759-INS-145
chr3-42368234-INS-741
chr3-42370525-INS-877
chr3-42383321-INS-2849
chr3-42390126-INS-52
chr3-42428713-INS-205
chr3-42448833-INS-599
chr3-42450375-INS-137
chr3-42462445-INS-417
chr3-42496100-INS-217
chr3-42500443-INS-786
chr3-42522098-INS-129
chr3-42540279-INS-342
chr3-42545667-INS-320
chr3-42557705-INS-7034
chr3-42595370-INS-220
chr3-42613838-INS-7690
chr3-42617354-INS-56
chr3-42626885-INS-91
chr3-42634107-INS-437
chr3-42635372-INS-142
chr3-42647016-INS-60
chr3-42693872-INS-76
chr3-42716471-INS-575
chr3-42734145-INS-339
chr3-42739067-INS-341
chr3-42785973-INS-156
chr3-42808988-INS-55
chr3-42816327-INS-221
chr3-42846211-INS-6279
chr3-42853014-INS-658
chr3-42897771-INS-134
chr3-42915304-INS-5567
chr3-42951500-INS-582
chr3-42952666-INS-79
chr3-42956949-INS-52
chr3-42958826-INS-7230
chr3-42973433-INS-88
chr3-43014506-INS-199
chr3-43025987-INS-185
chr3-43069269-INS-52
chr3-43075794-INS-783
chr3-43154648-INS-907
chr3-43180835-INS-6609

chr3-43215526-INS-56
chr3-43222128-INS-55
chr3-43225352-INS-208
chr3-43253246-INS-1891
chr3-43286944-INS-220
chr3-43311754-INS-54
chr3-43337070-INS-53
chr3-43365204-INS-52
chr3-43368665-INS-4541
chr3-43407240-INS-68
chr3-43436693-INS-2650
chr3-43458193-INS-80
chr3-43493090-INS-84
chr3-43511898-INS-424
chr3-43524984-INS-68
chr3-43526057-INS-7095
chr3-43531936-INS-549
chr3-43555495-INS-1213
chr3-43594361-INS-60
chr3-43629814-INS-162
chr3-43631003-INS-867
chr3-43639279-INS-181
chr3-43663628-INS-56
chr3-43675620-INS-687
chr3-43708534-INS-66
chr3-43720006-INS-151
chr3-43731782-INS-7927
chr3-43756634-INS-56
chr3-43763885-INS-457
chr3-43777535-INS-5328
chr3-43793889-INS-57
chr3-43806540-INS-116
chr3-43843597-INS-147
chr3-43848227-INS-274
chr3-43855058-INS-835
chr3-43874316-INS-52
chr3-43885171-INS-113
chr3-43932626-INS-56
chr3-43956211-INS-336
chr3-43959051-INS-51
chr3-43968839-INS-437
chr3-43970199-INS-105
chr3-43992422-INS-54
chr3-43992599-INS-646
chr3-44008243-INS-119
chr3-44021809-INS-54
chr3-44048207-INS-53
chr3-44053192-INS-529
chr3-44065434-INS-67
chr3-44079150-INS-5232
chr3-44110907-INS-6235
chr3-44125259-INS-74
chr3-44131232-INS-96
chr3-44161485-INS-72
chr3-44205425-INS-6436
chr3-44225855-INS-64
chr3-44228047-INS-58
chr3-44240684-INS-138
chr3-44266083-INS-197
chr3-44295650-INS-934
chr3-44305263-INS-170
chr3-44330911-INS-6614
chr3-44333336-INS-261
chr3-44403717-INS-337
chr3-44413355-INS-6368
chr3-44421166-INS-781
chr3-44545458-INS-90
chr3-44549387-INS-6388
chr3-44601687-INS-292
chr3-44637994-INS-83
chr3-44664863-INS-340
chr3-44694443-INS-110
chr3-44697592-INS-6798
chr3-44701204-INS-6518
chr3-44707727-INS-5163
chr3-44727589-INS-74
chr3-44730029-INS-155

chr3-44737471-INS-6662
chr3-44739050-INS-81
chr3-44744914-INS-4638
chr3-44751846-INS-99
chr3-44753843-INS-498
chr3-44780987-INS-60
chr3-44782213-INS-405
chr3-44791431-INS-228
chr3-44823287-INS-184
chr3-44871243-INS-80
chr3-44901549-INS-56
chr3-44926229-INS-6609
chr3-44935503-INS-506
chr3-44940389-INS-68
chr3-44943546-INS-148
chr3-44960313-INS-403
chr3-44973385-INS-200
chr3-44983814-INS-54
chr3-44994499-INS-6174
chr3-45016414-INS-255
chr3-45016955-INS-6905
chr3-45044007-INS-112
chr3-45064146-INS-467
chr3-45074787-INS-3066
chr3-45083763-INS-356
chr3-45092782-INS-596
chr3-45102477-INS-74
chr3-45140245-INS-289
chr3-45143070-INS-54
chr3-45146582-INS-54
chr3-45165214-INS-336
chr3-45167287-INS-50
chr3-45186560-INS-7163
chr3-45204090-INS-423
chr3-45206124-INS-8568
chr3-45207662-INS-360
chr3-45231589-INS-118
chr3-45233670-INS-886
chr3-45248346-INS-6582
chr3-45248956-INS-6632
chr3-45252458-INS-5144
chr3-45261123-INS-71
chr3-45272018-INS-87
chr3-45279973-INS-532
chr3-45287190-INS-71
chr3-45300582-INS-60
chr3-45314298-INS-7055
chr3-45365018-INS-1996
chr3-45392518-INS-80
chr3-45408055-INS-1112
chr3-45411829-INS-6152
chr3-45424646-INS-7130
chr3-45428756-INS-262
chr3-45430151-INS-130
chr3-45431156-INS-58
chr3-45440003-INS-7185
chr3-45444076-INS-6044
chr3-45445665-INS-66
chr3-45463535-INS-437
chr3-45465238-INS-4472
chr3-45468235-INS-347
chr3-45473507-INS-112
chr3-45479913-INS-96
chr3-45484496-INS-56
chr3-45488597-INS-485
chr3-45492782-INS-109
chr3-45498594-INS-3898
chr3-45503002-INS-1021
chr3-45511656-INS-67
chr3-45525244-INS-830
chr3-45527597-INS-8777
chr3-45548243-INS-5864
chr3-45552730-INS-72
chr3-45570842-INS-994
chr3-45578958-INS-54
chr3-45580104-INS-439
chr3-45592175-INS-993

chr3-45593693-INS-2930
chr3-45597743-INS-280
chr3-45624404-INS-239
chr3-45637199-INS-54
chr3-45661792-INS-124
chr3-45689058-INS-492
chr3-45689743-INS-98
chr3-45709693-INS-202
chr3-45719442-INS-60
chr3-45724234-INS-56
chr3-45743514-INS-7160
chr3-45763443-INS-448
chr3-45767278-INS-250
chr3-45778425-INS-278
chr3-45799190-INS-165
chr3-45800917-INS-412
chr3-45846013-INS-7107
chr3-45877579-INS-7445
chr3-45886592-INS-6253
chr3-45893949-INS-6404
chr3-45901020-INS-132
chr3-45918266-INS-243
chr3-45946455-INS-364
chr3-45949636-INS-1302
chr3-45950322-INS-1038
chr3-46001033-INS-2812
chr3-46006505-INS-51
chr3-46014930-INS-911
chr3-46016387-INS-65
chr3-46025413-INS-64
chr3-46033382-INS-2215
chr3-46040565-INS-773
chr3-46041168-INS-55
chr3-46057105-INS-108
chr3-46165975-INS-161
chr3-46170357-INS-331
chr3-46188582-INS-8390
chr3-46193871-INS-291
chr3-46204413-INS-7249
chr3-46217274-INS-74
chr3-46226338-INS-157
chr3-46245249-INS-400
chr3-46251638-INS-75
chr3-46259683-INS-6900
chr3-46280954-INS-85
chr3-46296058-INS-51
chr3-46308315-INS-51
chr3-46315860-INS-256
chr3-46323923-INS-63
chr3-46349608-INS-307
chr3-46369124-INS-197
chr3-46371431-INS-53
chr3-46371843-INS-63
chr3-46375298-INS-8660
chr3-46393394-INS-58
chr3-46405134-INS-74
chr3-46440360-INS-452
chr3-46462136-INS-6932
chr3-46501042-INS-104
chr3-46543967-INS-52
chr3-46550123-INS-296
chr3-46555074-INS-7211
chr3-46560929-INS-57
chr3-46561569-INS-477
chr3-46576417-INS-56
chr3-46587768-INS-63
chr3-46615814-INS-162
chr3-46628074-INS-171
chr3-46630295-INS-81
chr3-46660138-INS-6145
chr3-46665526-INS-195
chr3-46673760-INS-56
chr3-46674986-INS-319
chr3-46702996-INS-63
chr3-46744085-INS-70
chr3-46770151-INS-170
chr3-46795396-INS-66

chr3-46799108-INS-97
chr3-46804989-INS-409
chr3-46847157-INS-6909
chr3-46847986-INS-313
chr3-46854717-INS-55
chr3-46859176-INS-614
chr3-46860396-INS-870
chr3-46861219-INS-58
chr3-46872867-INS-56
chr3-46872994-INS-6615
chr3-46926642-INS-565
chr3-46930348-INS-6288
chr3-46945964-INS-317
chr3-46946573-INS-76
chr3-46953531-INS-520
chr3-46954402-INS-53
chr3-46961390-INS-6415
chr3-46967714-INS-8325
chr3-47014211-INS-161
chr3-47017332-INS-223
chr3-47046762-INS-208
chr3-47062896-INS-245
chr3-47063661-INS-150
chr3-47133176-INS-129
chr3-47152262-INS-170
chr3-47170337-INS-2782
chr3-47182783-INS-69
chr3-47183965-INS-108
chr3-47198393-INS-113
chr3-47200329-INS-130
chr3-47221112-INS-470
chr3-47230989-INS-166
chr3-47237216-INS-326
chr3-47289886-INS-472
chr3-47301135-INS-333
chr3-47314397-INS-6070
chr3-47323855-INS-77
chr3-47334717-INS-218
chr3-47351358-INS-976
chr3-47351542-INS-231
chr3-47354445-INS-170
chr3-47363056-INS-412
chr3-47370137-INS-495
chr3-47381801-INS-64
chr3-47383694-INS-56
chr3-47396256-INS-7162
chr3-47442953-INS-120
chr3-47452325-INS-59
chr3-47461329-INS-56
chr3-47473745-INS-999
chr3-47489489-INS-157
chr3-47567688-INS-68
chr3-47574514-INS-6355
chr3-47585027-INS-220
chr3-47608711-INS-53
chr3-47614934-INS-954
chr3-47616139-INS-879
chr3-47620192-INS-56
chr3-47632608-INS-67
chr3-47639360-INS-463
chr3-47640443-INS-2365
chr3-47682937-INS-104
chr3-47686760-INS-54
chr3-47688550-INS-52
chr3-47690621-INS-271
chr3-47692009-INS-129
chr3-47712664-INS-113
chr3-47734651-INS-52
chr3-47756414-INS-150
chr3-47763373-INS-962
chr3-47766857-INS-119
chr3-47804022-INS-227
chr3-47805992-INS-655
chr3-47839826-INS-87
chr3-47936701-INS-108
chr3-47937913-INS-384
chr3-47971724-INS-195

chr3-47989958-INS-196
chr3-47994508-INS-978
chr3-48020979-INS-140
chr3-48039320-INS-7087
chr3-48040167-INS-390
chr3-48054095-INS-657
chr3-48154652-INS-351
chr3-48163920-INS-114
chr3-48188364-INS-81
chr3-48195597-INS-868
chr3-48197349-INS-6547
chr3-48201353-INS-458
chr3-48203033-INS-434
chr3-48207821-INS-485
chr3-48214085-INS-841
chr3-48214998-INS-107
chr3-48216460-INS-84
chr3-48225289-INS-480
chr3-48227282-INS-327
chr3-48236471-INS-500
chr3-48269532-INS-98
chr3-48279372-INS-110
chr3-48283155-INS-283
chr3-48294091-INS-139
chr3-48325304-INS-69
chr3-48329886-INS-997
chr3-48343166-INS-200
chr3-48354054-INS-537
chr3-48397109-INS-2576
chr3-48411429-INS-57
chr3-48412349-INS-981
chr3-48440511-INS-76
chr3-48458002-INS-334
chr3-48472904-INS-70
chr3-48518022-INS-3400
chr3-48550321-INS-5998
chr3-48580598-INS-238
chr3-48593298-INS-6728
chr3-48601173-INS-67
chr3-48608355-INS-52
chr3-48640224-INS-251
chr3-48668998-INS-2882
chr3-48669268-INS-352
chr3-48683169-INS-96
chr3-48685562-INS-87
chr3-48704116-INS-416
chr3-48727312-INS-232
chr3-48737777-INS-208
chr3-48746874-INS-447
chr3-48754858-INS-343
chr3-48763283-INS-369
chr3-48765719-INS-53
chr3-48777218-INS-54
chr3-48778573-INS-71
chr3-48790071-INS-748
chr3-48799440-INS-995
chr3-48816377-INS-234
chr3-48856461-INS-284
chr3-48860826-INS-108
chr3-48874814-INS-692
chr3-48876428-INS-259
chr3-48889934-INS-52
chr3-48919639-INS-400
chr3-48940512-INS-52
chr3-48954689-INS-114
chr3-48967936-INS-562
chr3-49013345-INS-55
chr3-49044774-INS-134
chr3-49049576-INS-237
chr3-49099928-INS-499
chr3-49122065-INS-154
chr3-49145747-INS-54
chr3-49201133-INS-6385
chr3-49238405-INS-159
chr3-49272116-INS-93
chr3-49275933-INS-181
chr3-49453721-INS-229

chr3-49459716-INS-78
chr3-49473783-INS-78
chr3-49484336-INS-486
chr3-49515636-INS-76
chr3-49529422-INS-56
chr3-49535805-INS-52
chr3-49539283-INS-5291
chr3-49565246-INS-415
chr3-49613679-INS-7079
chr3-49614216-INS-70
chr3-49634258-INS-59
chr3-49650488-INS-221
chr3-49659040-INS-110
chr3-49669284-INS-2003
chr3-49681296-INS-1358
chr3-49693957-INS-94
chr3-49729088-INS-62
chr3-49738499-INS-104
chr3-49741220-INS-270
chr3-49761818-INS-177
chr3-49793900-INS-104
chr3-49802645-INS-129
chr3-49810382-INS-728
chr3-49838044-INS-231
chr3-49874486-INS-327
chr3-49905491-INS-186
chr3-49921337-INS-52
chr3-49924031-INS-86
chr3-49953177-INS-88
chr3-49957466-INS-91
chr3-49973175-INS-356
chr3-49980435-INS-123
chr3-49987873-INS-343
chr3-50004688-INS-692
chr3-50040024-INS-75
chr3-50048488-INS-578
chr3-50057136-INS-362
chr3-50059046-INS-6727
chr3-50060960-INS-1834
chr3-50082329-INS-225
chr3-50091799-INS-114
chr3-50108362-INS-789
chr3-50109502-INS-968
chr3-50110723-INS-53
chr3-50114255-INS-84
chr3-50114759-INS-56
chr3-50116906-INS-226
chr3-50137199-INS-53
chr3-50210807-INS-191
chr3-50235819-INS-240
chr3-50242481-INS-313
chr3-50246673-INS-279
chr3-50259597-INS-66
chr3-50265268-INS-87
chr3-50282865-INS-52
chr3-50297769-INS-325
chr3-50340412-INS-70
chr3-50370331-INS-62
chr3-50399584-INS-75
chr3-50402587-INS-92
chr3-50416033-INS-59
chr3-50435635-INS-152
chr3-50482423-INS-400
chr3-50506657-INS-108
chr3-50517570-INS-54
chr3-50527005-INS-290
chr3-50538746-INS-92
chr3-50594636-INS-527
chr3-50597660-INS-6260
chr3-50597983-INS-68
chr3-50602446-INS-216
chr3-50630740-INS-496
chr3-50631825-INS-314
chr3-50640670-INS-6323
chr3-50666804-INS-108
chr3-50687233-INS-57
chr3-50725860-INS-6223

chr3-50737898-INS-638
chr3-50748998-INS-104
chr3-50776119-INS-3141
chr3-50779798-INS-76
chr3-50818448-INS-556
chr3-50825021-INS-150
chr3-50859256-INS-208
chr3-50893155-INS-127
chr3-50896369-INS-127
chr3-50907870-INS-73
chr3-50911329-INS-96
chr3-50913438-INS-345
chr3-50916295-INS-6368
chr3-50917671-INS-97
chr3-50980327-INS-64
chr3-50982177-INS-161
chr3-50991080-INS-54
chr3-50998646-INS-798
chr3-51015309-INS-406
chr3-51017444-INS-56
chr3-51030799-INS-357
chr3-51054512-INS-68
chr3-51055749-INS-296
chr3-51060321-INS-194
chr3-51100640-INS-84
chr3-51107184-INS-53
chr3-51111832-INS-249
chr3-51113539-INS-497
chr3-51176573-INS-150
chr3-51180285-INS-119
chr3-51182074-INS-267
chr3-51201203-INS-144
chr3-51203019-INS-243
chr3-51209366-INS-142
chr3-51212188-INS-512
chr3-51226640-INS-200
chr3-51229071-INS-201
chr3-51240175-INS-216
chr3-51243566-INS-873
chr3-51244310-INS-204
chr3-51253142-INS-303
chr3-51257538-INS-236
chr3-51263854-INS-115
chr3-51274331-INS-6729
chr3-51276020-INS-564
chr3-51282267-INS-220
chr3-51326110-INS-56
chr3-51333804-INS-326
chr3-51341167-INS-201
chr3-51351020-INS-945
chr3-51352240-INS-216
chr3-51373947-INS-77
chr3-51375540-INS-376
chr3-51387357-INS-736
chr3-51394560-INS-137
chr3-51412615-INS-577
chr3-51414148-INS-744
chr3-51414571-INS-832
chr3-51416949-INS-58
chr3-51418239-INS-91
chr3-51421636-INS-68
chr3-51443263-INS-111
chr3-51454200-INS-487
chr3-51460591-INS-204
chr3-51464244-INS-54
chr3-51488477-INS-196
chr3-51507403-INS-90
chr3-51508999-INS-60
chr3-51518264-INS-82
chr3-51532976-INS-126
chr3-51535729-INS-56
chr3-51540934-INS-903
chr3-51552866-INS-167
chr3-51554678-INS-120
chr3-51563561-INS-192
chr3-51578564-INS-197
chr3-51580819-INS-210

chr3-51603903-INS-333
chr3-51606024-INS-196
chr3-51621746-INS-81
chr3-51649705-INS-296
chr3-51673580-INS-1222
chr3-51673961-INS-1616
chr3-51673990-INS-2656
chr3-51674234-INS-2916
chr3-51676158-INS-54
chr3-51684536-INS-174
chr3-51691385-INS-263
chr3-51692508-INS-144
chr3-51712512-INS-138
chr3-51731067-INS-62
chr3-51741799-INS-117
chr3-51757935-INS-223
chr3-51765165-INS-180
chr3-51794843-INS-78
chr3-51861829-INS-229
chr3-51864419-INS-103
chr3-51872097-INS-389
chr3-51888769-INS-64
chr3-51921133-INS-62
chr3-51949790-INS-76
chr3-52013505-INS-60
chr3-52016837-INS-54
chr3-52035682-INS-65
chr3-52055550-INS-691
chr3-52057337-INS-58
chr3-52077995-INS-137
chr3-52085855-INS-79
chr3-52088833-INS-117
chr3-52096325-INS-84
chr3-52102312-INS-400
chr3-52111653-INS-51
chr3-52115922-INS-101
chr3-52147294-INS-5005
chr3-52164184-INS-156
chr3-52184262-INS-213
chr3-52202135-INS-58
chr3-52208267-INS-87
chr3-52220654-INS-423
chr3-52224251-INS-84
chr3-52235807-INS-474
chr3-52247025-INS-473
chr3-52254089-INS-52
chr3-52261399-INS-141
chr3-52262409-INS-399
chr3-52264027-INS-201
chr3-52285388-INS-498
chr3-52287608-INS-60
chr3-52309823-INS-95
chr3-52331126-INS-342
chr3-52356916-INS-80
chr3-52387887-INS-78
chr3-52399717-INS-61
chr3-52412775-INS-240
chr3-52425264-INS-122
chr3-52427562-INS-54
chr3-52428832-INS-201
chr3-52431248-INS-148
chr3-52471066-INS-202
chr3-52477083-INS-171
chr3-52488690-INS-65
chr3-52492835-INS-74
chr3-52504327-INS-54
chr3-52509894-INS-338
chr3-52516576-INS-55
chr3-52566242-INS-895
chr3-52581213-INS-6147
chr3-52584535-INS-214
chr3-52596743-INS-53
chr3-52607096-INS-126
chr3-52608055-INS-180
chr3-52611771-INS-68
chr3-52634541-INS-216
chr3-52642584-INS-273

chr3-52643227-INS-250
chr3-52646695-INS-135
chr3-52652298-INS-59
chr3-52662854-INS-83
chr3-52665141-INS-135
chr3-52666197-INS-756
chr3-52675627-INS-7221
chr3-52679042-INS-7184
chr3-52749301-INS-222
chr3-52762002-INS-400
chr3-52762645-INS-862
chr3-52788098-INS-62
chr3-52807947-INS-371
chr3-52809037-INS-79
chr3-52813958-INS-74
chr3-52857710-INS-233
chr3-52881971-INS-254
chr3-52885300-INS-79
chr3-52887617-INS-92
chr3-52893079-INS-90
chr3-52918798-INS-8376
chr3-52936559-INS-6595
chr3-52969997-INS-166
chr3-52988078-INS-278
chr3-52997782-INS-84
chr3-53012378-INS-401
chr3-53035983-INS-571
chr3-53049106-INS-127
chr3-53053863-INS-66
chr3-53067771-INS-1007
chr3-53081900-INS-81
chr3-53082842-INS-55
chr3-53087201-INS-52
chr3-53103720-INS-5218
chr3-53119410-INS-76
chr3-53121843-INS-238
chr3-53134679-INS-116
chr3-53173094-INS-449
chr3-53191990-INS-91
chr3-53222285-INS-385
chr3-53231012-INS-88
chr3-53268718-INS-75
chr3-53280163-INS-94
chr3-53310127-INS-141
chr3-53321736-INS-108
chr3-53362417-INS-60
chr3-53363599-INS-72
chr3-53391377-INS-58
chr3-53392676-INS-198
chr3-53409213-INS-142
chr3-53419090-INS-59
chr3-53427537-INS-96
chr3-53428309-INS-138
chr3-53436755-INS-82
chr3-53439185-INS-350
chr3-53442312-INS-60
chr3-53470631-INS-74
chr3-53481162-INS-6251
chr3-53513016-INS-210
chr3-53517695-INS-94
chr3-53520954-INS-51
chr3-53533703-INS-142
chr3-53544905-INS-326
chr3-53558966-INS-324
chr3-53587101-INS-883
chr3-53588453-INS-3618
chr3-53590906-INS-223
chr3-53607134-INS-92
chr3-53617832-INS-64
chr3-53619316-INS-69
chr3-53632903-INS-86
chr3-53641808-INS-3123
chr3-53676548-INS-106
chr3-53683284-INS-455
chr3-53703178-INS-1862
chr3-53720455-INS-55
chr3-53742321-INS-132

chr3-53743468-INS-180
chr3-53784479-INS-446
chr3-53810101-INS-6508
chr3-53814396-INS-74
chr3-53823086-INS-323
chr3-53831150-INS-7163
chr3-53873995-INS-193
chr3-53881784-INS-84
chr3-53912864-INS-174
chr3-53948903-INS-73
chr3-53991590-INS-58
chr3-54011997-INS-132
chr3-54038614-INS-920
chr3-54048271-INS-504
chr3-54054616-INS-141
chr3-54067124-INS-133
chr3-54069312-INS-6426
chr3-54074585-INS-497
chr3-54095224-INS-67
chr3-54102595-INS-310
chr3-54150133-INS-144
chr3-54172543-INS-54
chr3-54183933-INS-1412
chr3-54201522-INS-57
chr3-54208745-INS-393
chr3-54244537-INS-60
chr3-54255455-INS-305
chr3-54257478-INS-427
chr3-54310324-INS-76
chr3-54319315-INS-175
chr3-54322668-INS-92
chr3-54326951-INS-139
chr3-54334186-INS-66
chr3-54346507-INS-271
chr3-54380531-INS-160
chr3-54382464-INS-76
chr3-54401701-INS-67
chr3-54417735-INS-151
chr3-54435842-INS-985
chr3-54437899-INS-82
chr3-54439658-INS-51
chr3-54441013-INS-200
chr3-54450301-INS-72
chr3-54461124-INS-180
chr3-54499901-INS-744
chr3-54502914-INS-54
chr3-54503876-INS-69
chr3-54592024-INS-115
chr3-54615094-INS-146
chr3-54635070-INS-54
chr3-54667173-INS-58
chr3-54736169-INS-178
chr3-54740998-INS-1067
chr3-54775896-INS-390
chr3-54792746-INS-58
chr3-54798755-INS-121
chr3-54811722-INS-260
chr3-54824845-INS-400
chr3-54829899-INS-58
chr3-54905693-INS-240
chr3-54915755-INS-410
chr3-54931073-INS-148
chr3-54957716-INS-95
chr3-54975466-INS-472
chr3-54999602-INS-946
chr3-55005071-INS-53
chr3-55007066-INS-230
chr3-55010062-INS-56
chr3-55017645-INS-106
chr3-55039503-INS-86
chr3-55087496-INS-153
chr3-55100191-INS-56
chr3-55100970-INS-70
chr3-55146235-INS-162
chr3-55172260-INS-60
chr3-55195097-INS-201
chr3-55235830-INS-57

chr3-55262310-INS-208
chr3-55271019-INS-66
chr3-55281062-INS-177
chr3-55292509-INS-72
chr3-55321527-INS-128
chr3-55323741-INS-187
chr3-55341143-INS-201
chr3-55343718-INS-459
chr3-55357640-INS-333
chr3-55381520-INS-102
chr3-55435450-INS-132
chr3-55460917-INS-278
chr3-55496306-INS-56
chr3-55502120-INS-59
chr3-55505374-INS-182
chr3-55506330-INS-195
chr3-55542015-INS-297
chr3-55568734-INS-52
chr3-55631717-INS-1270
chr3-55640612-INS-216
chr3-55644941-INS-153
chr3-55659778-INS-3482
chr3-55686517-INS-93
chr3-55705208-INS-929
chr3-55710560-INS-382
chr3-55710609-INS-56
chr3-55729577-INS-83
chr3-55743847-INS-350
chr3-55757180-INS-252
chr3-55759180-INS-509
chr3-55794491-INS-284
chr3-55836870-INS-144
chr3-55902027-INS-122
chr3-55917934-INS-198
chr3-55920373-INS-6442
chr3-55960844-INS-667
chr3-56003405-INS-6529
chr3-56007021-INS-532
chr3-56033501-INS-397
chr3-56037791-INS-52
chr3-56071311-INS-385
chr3-56097757-INS-56
chr3-56119536-INS-1158
chr3-56126094-INS-67
chr3-56139192-INS-725
chr3-56202978-INS-67
chr3-56221472-INS-6566
chr3-56222132-INS-57
chr3-56249066-INS-440
chr3-56253364-INS-548
chr3-56254254-INS-1960
chr3-56255795-INS-487
chr3-56257893-INS-4522
chr3-56261911-INS-6508
chr3-56266089-INS-58
chr3-56289566-INS-7083
chr3-56291419-INS-108
chr3-56301132-INS-586
chr3-56308223-INS-582
chr3-56320277-INS-74
chr3-56323712-INS-100
chr3-56324428-INS-167
chr3-56345904-INS-5338
chr3-56350646-INS-210
chr3-56362240-INS-159
chr3-56376121-INS-150
chr3-56408341-INS-462
chr3-56445771-INS-546
chr3-56482993-INS-59
chr3-56486859-INS-57
chr3-56498753-INS-6611
chr3-56500548-INS-105
chr3-56529516-INS-449
chr3-56533050-INS-58
chr3-56575566-INS-332
chr3-56604288-INS-65
chr3-56649310-INS-5436

chr3-56672354-INS-439
chr3-56690016-INS-544
chr3-56703781-INS-3781
chr3-56739133-INS-88
chr3-56744698-INS-111
chr3-56782732-INS-5256
chr3-56795185-INS-572
chr3-56825512-INS-358
chr3-56839135-INS-358
chr3-56847262-INS-102
chr3-56852268-INS-96
chr3-56896598-INS-196
chr3-56939639-INS-169
chr3-56942242-INS-184
chr3-56943372-INS-52
chr3-56962881-INS-142
chr3-56996678-INS-158
chr3-57004802-INS-60
chr3-57019579-INS-744
chr3-57024216-INS-1423
chr3-57073374-INS-276
chr3-57079117-INS-66
chr3-57109534-INS-66
chr3-57126143-INS-51
chr3-57157297-INS-6494
chr3-57170327-INS-673
chr3-57184319-INS-832
chr3-57185652-INS-6758
chr3-57223476-INS-550
chr3-57242398-INS-7106
chr3-57242917-INS-979
chr3-57253827-INS-60
chr3-57270477-INS-7214
chr3-57299048-INS-80
chr3-57311131-INS-100
chr3-57315631-INS-95
chr3-57317166-INS-100
chr3-57319615-INS-140
chr3-57331390-INS-240
chr3-57336524-INS-59
chr3-57345505-INS-83
chr3-57350545-INS-54
chr3-57376417-INS-980
chr3-57405579-INS-132
chr3-57492915-INS-51
chr3-57498862-INS-62
chr3-57520766-INS-248
chr3-57523398-INS-130
chr3-57532208-INS-156
chr3-57534274-INS-114
chr3-57550951-INS-5313
chr3-57561496-INS-516
chr3-57581351-INS-51
chr3-57601299-INS-90
chr3-57643334-INS-847
chr3-57678997-INS-52
chr3-57682190-INS-197
chr3-57688943-INS-62
chr3-57691147-INS-562
chr3-57692616-INS-492
chr3-57696340-INS-179
chr3-57700893-INS-764
chr3-57703184-INS-54
chr3-57707673-INS-1890
chr3-57720166-INS-457
chr3-57744541-INS-779
chr3-57762643-INS-865
chr3-57777282-INS-204
chr3-57783287-INS-478
chr3-57799046-INS-683
chr3-57806371-INS-203
chr3-57811317-INS-400
chr3-57817292-INS-340
chr3-57825231-INS-59
chr3-57839802-INS-330
chr3-57846696-INS-133
chr3-57870228-INS-84

chr3-57884757-INS-7081
chr3-57893208-INS-60
chr3-57897563-INS-146
chr3-57909778-INS-52
chr3-57920242-INS-301
chr3-57939591-INS-115
chr3-57955959-INS-184
chr3-57976565-INS-389
chr3-57977533-INS-79
chr3-57986900-INS-182
chr3-58008575-INS-64
chr3-58023042-INS-54
chr3-58028146-INS-338
chr3-58033276-INS-355
chr3-58045973-INS-70
chr3-58046927-INS-3082
chr3-58053118-INS-130
chr3-58058786-INS-228
chr3-58060478-INS-181
chr3-58067343-INS-276
chr3-58072769-INS-665
chr3-58078936-INS-68
chr3-58090300-INS-178
chr3-58091651-INS-789
chr3-58115300-INS-663
chr3-58115321-INS-117
chr3-58117458-INS-81
chr3-58122397-INS-241
chr3-58128555-INS-133
chr3-58158525-INS-608
chr3-58159508-INS-290
chr3-58160528-INS-54
chr3-58188462-INS-81
chr3-58190283-INS-64
chr3-58210154-INS-67
chr3-58225476-INS-213
chr3-58252011-INS-7147
chr3-58261069-INS-435
chr3-58268417-INS-386
chr3-58281917-INS-95
chr3-58309728-INS-51
chr3-58360454-INS-74
chr3-58390170-INS-723
chr3-58396278-INS-200
chr3-58409311-INS-79
chr3-58410616-INS-686
chr3-58439337-INS-206
chr3-58442806-INS-183
chr3-58444801-INS-280
chr3-58457447-INS-152
chr3-58470351-INS-468
chr3-58487581-INS-61
chr3-58514436-INS-221
chr3-58522107-INS-179
chr3-58526729-INS-97
chr3-58530722-INS-5557
chr3-58535293-INS-898
chr3-58562344-INS-166
chr3-58576384-INS-86
chr3-58598658-INS-71
chr3-58602539-INS-73
chr3-58608847-INS-690
chr3-58635687-INS-6401
chr3-58640722-INS-294
chr3-58642915-INS-1890
chr3-58644650-INS-64
chr3-58645731-INS-267
chr3-58661082-INS-57
chr3-58666141-INS-236
chr3-58667823-INS-52
chr3-58688158-INS-6399
chr3-58689707-INS-7073
chr3-58711551-INS-202
chr3-58719186-INS-638
chr3-58742250-INS-74
chr3-58752558-INS-76
chr3-58767722-INS-118

chr3-58805165-INS-6704
chr3-58810364-INS-219
chr3-58812932-INS-52
chr3-58842171-INS-68
chr3-58848731-INS-8439
chr3-58853648-INS-1372
chr3-58870267-INS-367
chr3-58872724-INS-91
chr3-58876930-INS-400
chr3-58890230-INS-56
chr3-58900205-INS-484
chr3-58903723-INS-377
chr3-58921341-INS-230
chr3-58925053-INS-181
chr3-58935072-INS-74
chr3-58940123-INS-150
chr3-58962823-INS-2296
chr3-58985593-INS-82
chr3-58987354-INS-55
chr3-58988151-INS-604
chr3-58990004-INS-81
chr3-59017637-INS-52
chr3-59019081-INS-6460
chr3-59053106-INS-276
chr3-59062402-INS-148
chr3-59066876-INS-724
chr3-59082112-INS-153
chr3-59098620-INS-84
chr3-59102870-INS-397
chr3-59112212-INS-60
chr3-59134319-INS-189
chr3-59175159-INS-163
chr3-59177635-INS-640
chr3-59216134-INS-1493
chr3-59234205-INS-82
chr3-59298163-INS-89
chr3-59299605-INS-6977
chr3-59321983-INS-86
chr3-59352535-INS-60
chr3-59359710-INS-542
chr3-59367301-INS-113
chr3-59376747-INS-148
chr3-59378344-INS-540
chr3-59471436-INS-3278
chr3-59600691-INS-52
chr3-59602487-INS-7098
chr3-59611874-INS-555
chr3-59640442-INS-602
chr3-59646606-INS-117
chr3-59657376-INS-413
chr3-59668446-INS-7051
chr3-59703873-INS-204
chr3-59724375-INS-64
chr3-59737247-INS-206
chr3-59740592-INS-104
chr3-59755030-INS-132
chr3-59775244-INS-264
chr3-59782449-INS-76
chr3-59785281-INS-408
chr3-59810481-INS-274
chr3-59822308-INS-989
chr3-59824618-INS-71
chr3-59853369-INS-296
chr3-59911286-INS-6634
chr3-59949438-INS-673
chr3-59953381-INS-137
chr3-59960100-INS-62
chr3-59972985-INS-354
chr3-59992821-INS-60
chr3-59996688-INS-943
chr3-60006370-INS-6816
chr3-60010288-INS-102
chr3-60015251-INS-201
chr3-60023484-INS-589
chr3-60024302-INS-406
chr3-60025375-INS-88
chr3-60044547-INS-61

chr3-60057936-INS-52
chr3-60064085-INS-639
chr3-60067405-INS-91
chr3-60073795-INS-77
chr3-60090010-INS-770
chr3-60103065-INS-78
chr3-60111618-INS-272
chr3-60125780-INS-111
chr3-60126976-INS-6278
chr3-60135766-INS-61
chr3-60156142-INS-2854
chr3-60158531-INS-1829
chr3-60180468-INS-400
chr3-60186852-INS-6401
chr3-60210627-INS-392
chr3-60215307-INS-95
chr3-60236799-INS-6399
chr3-60260571-INS-90
chr3-60293865-INS-226
chr3-60298114-INS-54
chr3-60314503-INS-7359
chr3-60315749-INS-2527
chr3-60404018-INS-563
chr3-60421068-INS-56
chr3-60432612-INS-124
chr3-60439529-INS-11917
chr3-60479238-INS-52
chr3-60489011-INS-54
chr3-60549652-INS-372
chr3-60559732-INS-74
chr3-60582283-INS-722
chr3-60615829-INS-112
chr3-60625571-INS-147
chr3-60633538-INS-556
chr3-60660938-INS-117
chr3-60677859-INS-116
chr3-60713594-INS-1528
chr3-60731524-INS-498
chr3-60763604-INS-52
chr3-60788880-INS-653
chr3-60805972-INS-79
chr3-60810832-INS-6431
chr3-60820183-INS-66
chr3-60825503-INS-139
chr3-60831022-INS-6635
chr3-60850764-INS-304
chr3-60856220-INS-1891
chr3-60868254-INS-52
chr3-60868676-INS-331
chr3-60916839-INS-95
chr3-60921006-INS-65
chr3-60932906-INS-3039
chr3-60937660-INS-2596
chr3-60944914-INS-382
chr3-60960557-INS-4274
chr3-60999879-INS-122
chr3-61002389-INS-810
chr3-61024382-INS-1410
chr3-61045730-INS-6356
chr3-61046501-INS-6592
chr3-61048889-INS-120
chr3-61074369-INS-62
chr3-61094139-INS-153
chr3-61097144-INS-86
chr3-61101692-INS-266
chr3-61121396-INS-1342
chr3-61139766-INS-385
chr3-61163885-INS-86
chr3-61189780-INS-158
chr3-61195268-INS-92
chr3-61219697-INS-4844
chr3-61250968-INS-184
chr3-61260894-INS-2698
chr3-61282444-INS-361
chr3-61286637-INS-55
chr3-61289407-INS-6194
chr3-61294360-INS-54

chr3-61299768-INS-69
chr3-61312034-INS-330
chr3-61331725-INS-140
chr3-61397634-INS-463
chr3-61445622-INS-51
chr3-61464425-INS-2251
chr3-61468471-INS-6671
chr3-61496487-INS-400
chr3-61504893-INS-6195
chr3-61509045-INS-6390
chr3-61543167-INS-452
chr3-61570926-INS-92
chr3-61586687-INS-6320
chr3-61590239-INS-183
chr3-61605381-INS-1843
chr3-61636832-INS-56
chr3-61641560-INS-7358
chr3-61661332-INS-6372
chr3-61672261-INS-135
chr3-61702848-INS-61
chr3-61715094-INS-736
chr3-61716400-INS-276
chr3-61721735-INS-112
chr3-61727833-INS-62
chr3-61730012-INS-931
chr3-61810259-INS-208
chr3-61814455-INS-6122
chr3-61822716-INS-695
chr3-61831110-INS-6222
chr3-61839988-INS-646
chr3-61868575-INS-190
chr3-61878861-INS-74
chr3-61906429-INS-127
chr3-61924309-INS-933
chr3-61931979-INS-55
chr3-61952588-INS-756
chr3-61957260-INS-833
chr3-61966045-INS-52
chr3-61974180-INS-6910
chr3-61975666-INS-72
chr3-61997810-INS-74
chr3-62005430-INS-284
chr3-62007159-INS-81
chr3-62037237-INS-1928
chr3-62045600-INS-69
chr3-62048555-INS-6810
chr3-62082883-INS-53
chr3-62091183-INS-205
chr3-62108828-INS-3845
chr3-62123136-INS-173
chr3-62130389-INS-497
chr3-62132690-INS-127
chr3-62145493-INS-124
chr3-62178455-INS-270
chr3-62199786-INS-540
chr3-62202222-INS-87
chr3-62206850-INS-60
chr3-62217623-INS-57
chr3-62218672-INS-124
chr3-62243055-INS-64
chr3-62253846-INS-244
chr3-62259413-INS-6434
chr3-62288694-INS-51
chr3-62304474-INS-65
chr3-62314849-INS-1890
chr3-62316819-INS-321
chr3-62340392-INS-130
chr3-62357401-INS-125
chr3-62358684-INS-82
chr3-62378406-INS-399
chr3-62415925-INS-92
chr3-62418702-INS-52
chr3-62504700-INS-66
chr3-62506628-INS-59
chr3-62508513-INS-59
chr3-62512450-INS-57
chr3-62537101-INS-146

chr3-62575717-INS-63
chr3-62598740-INS-521
chr3-62617854-INS-7709
chr3-62619069-INS-933
chr3-62631522-INS-63
chr3-62639481-INS-415
chr3-62652932-INS-146
chr3-62664723-INS-134
chr3-62669807-INS-308
chr3-62696821-INS-847
chr3-62699932-INS-431
chr3-62701090-INS-4044
chr3-62709902-INS-204
chr3-62725820-INS-54
chr3-62748074-INS-6577
chr3-62758336-INS-858
chr3-62761762-INS-71
chr3-62762704-INS-1892
chr3-62764047-INS-204
chr3-62778132-INS-1912
chr3-62798330-INS-384
chr3-62799695-INS-6396
chr3-62817670-INS-156
chr3-62834314-INS-178
chr3-62856243-INS-7042
chr3-62902275-INS-6727
chr3-62931703-INS-241
chr3-62970807-INS-104
chr3-62994775-INS-86
chr3-63012562-INS-357
chr3-63021837-INS-871
chr3-63035094-INS-184
chr3-63041245-INS-110
chr3-63057183-INS-231
chr3-63061456-INS-6402
chr3-63123004-INS-6497
chr3-63130651-INS-351
chr3-63140229-INS-6233
chr3-63159768-INS-528
chr3-63189095-INS-410
chr3-63189336-INS-6851
chr3-63194962-INS-52
chr3-63195530-INS-5142
chr3-63201675-INS-52
chr3-63222399-INS-6678
chr3-63232863-INS-54
chr3-63236005-INS-538
chr3-63258804-INS-5332
chr3-63263518-INS-58
chr3-63307941-INS-137
chr3-63342723-INS-56
chr3-63387507-INS-6189
chr3-63389684-INS-740
chr3-63395756-INS-80
chr3-63396397-INS-7336
chr3-63399860-INS-62
chr3-63437454-INS-201
chr3-63469865-INS-6907
chr3-63476758-INS-62
chr3-63492098-INS-1450
chr3-63522864-INS-458
chr3-63559351-INS-66
chr3-63562212-INS-342
chr3-63573929-INS-102
chr3-63583216-INS-212
chr3-63588446-INS-99
chr3-63602521-INS-472
chr3-63619198-INS-97
chr3-63643519-INS-84
chr3-63645692-INS-944
chr3-63650848-INS-7079
chr3-63673283-INS-6792
chr3-63673787-INS-817
chr3-63675819-INS-54
chr3-63684213-INS-65
chr3-63698147-INS-72
chr3-63708289-INS-580

chr3-63721456-INS-58
chr3-63744479-INS-6691
chr3-63757019-INS-553
chr3-63769472-INS-83
chr3-63828074-INS-130
chr3-63829558-INS-186
chr3-63848003-INS-69
chr3-63865996-INS-817
chr3-63894052-INS-63
chr3-63917161-INS-105
chr3-63936804-INS-230
chr3-63944976-INS-5385
chr3-63957314-INS-7183
chr3-63988341-INS-57
chr3-64005892-INS-58
chr3-64017160-INS-7504
chr3-64048477-INS-840
chr3-64059997-INS-61
chr3-64078946-INS-194
chr3-64128428-INS-147
chr3-64161424-INS-7121
chr3-64177690-INS-415
chr3-64204496-INS-52
chr3-64205239-INS-841
chr3-64225807-INS-66
chr3-64230730-INS-52
chr3-64258420-INS-75
chr3-64265003-INS-6396
chr3-64333121-INS-278
chr3-64390437-INS-7078
chr3-64418598-INS-1432
chr3-64448919-INS-116
chr3-64506451-INS-56
chr3-64509972-INS-218
chr3-64526500-INS-4462
chr3-64534757-INS-979
chr3-64576078-INS-55
chr3-64648045-INS-60
chr3-64650265-INS-6403
chr3-64695443-INS-107
chr3-64700802-INS-458
chr3-64774317-INS-113
chr3-64776875-INS-84
chr3-64778172-INS-454
chr3-64787166-INS-422
chr3-64815239-INS-296
chr3-64858025-INS-69
chr3-64869418-INS-6288
chr3-64910246-INS-88
chr3-64948102-INS-196
chr3-64962047-INS-6919
chr3-64967817-INS-176
chr3-64977930-INS-1556
chr3-64988271-INS-63
chr3-64988936-INS-499
chr3-64990443-INS-540
chr3-64998838-INS-236
chr3-65000279-INS-52
chr3-65002375-INS-107
chr3-65026049-INS-6912
chr3-65034704-INS-413
chr3-65073375-INS-68
chr3-65088577-INS-6812
chr3-65128025-INS-125
chr3-65135300-INS-98
chr3-65136941-INS-433
chr3-65142285-INS-76
chr3-65152573-INS-104
chr3-65159147-INS-417
chr3-65177153-INS-54
chr3-65182015-INS-279
chr3-65192841-INS-6226
chr3-65197846-INS-165
chr3-65210812-INS-59
chr3-65213503-INS-56
chr3-65219598-INS-757
chr3-65220781-INS-81

chr3-65224541-INS-66
chr3-65227581-INS-7111
chr3-65233807-INS-785
chr3-65247144-INS-227
chr3-65249858-INS-400
chr3-65254954-INS-85
chr3-65258718-INS-94
chr3-65272814-INS-234
chr3-65300618-INS-57
chr3-65316808-INS-635
chr3-65328963-INS-210
chr3-65346577-INS-197
chr3-65350258-INS-89
chr3-65352351-INS-70
chr3-65374214-INS-115
chr3-65385709-INS-212
chr3-65392168-INS-52
chr3-65406293-INS-7327
chr3-65407427-INS-642
chr3-65437318-INS-194
chr3-65476269-INS-448
chr3-65492151-INS-97
chr3-65503575-INS-60
chr3-65514509-INS-133
chr3-65548893-INS-476
chr3-65552542-INS-125
chr3-65589848-INS-409
chr3-65603748-INS-119
chr3-65649506-INS-3280
chr3-65655028-INS-83
chr3-65661738-INS-407
chr3-65675995-INS-184
chr3-65697666-INS-90
chr3-65715137-INS-57
chr3-65792713-INS-7076
chr3-65812608-INS-52
chr3-65814415-INS-185
chr3-65823395-INS-126
chr3-65827788-INS-197
chr3-65839014-INS-450
chr3-65843867-INS-217
chr3-65848014-INS-187
chr3-65849980-INS-79
chr3-65936697-INS-53
chr3-65949353-INS-55
chr3-65950448-INS-6481
chr3-65972674-INS-102
chr3-65975867-INS-1833
chr3-65993538-INS-218
chr3-66010460-INS-7175
chr3-66023232-INS-197
chr3-66047474-INS-214
chr3-66077868-INS-85
chr3-66112750-INS-52
chr3-66152707-INS-110
chr3-66176298-INS-135
chr3-66188120-INS-468
chr3-66208167-INS-115
chr3-66232122-INS-2095
chr3-66246178-INS-1324
chr3-66265288-INS-6198
chr3-66269206-INS-282
chr3-66285860-INS-56
chr3-66288331-INS-51
chr3-66291526-INS-75
chr3-66351177-INS-224
chr3-66400351-INS-486
chr3-66426396-INS-93
chr3-66462954-INS-112
chr3-66470416-INS-1838
chr3-66491109-INS-56
chr3-66520598-INS-58
chr3-66559744-INS-62
chr3-66594016-INS-86
chr3-66599539-INS-1519
chr3-66601599-INS-141
chr3-66605883-INS-94

chr3-66630335-INS-137
chr3-66638792-INS-59
chr3-66670278-INS-290
chr3-66677104-INS-56
chr3-66686403-INS-55
chr3-66688682-INS-567
chr3-66710976-INS-3839
chr3-66712393-INS-6631
chr3-66716827-INS-99
chr3-66758593-INS-53
chr3-66761191-INS-7641
chr3-66789558-INS-118
chr3-66796838-INS-7892
chr3-66803532-INS-509
chr3-66818754-INS-131
chr3-66849475-INS-7128
chr3-66853510-INS-134
chr3-66863955-INS-382
chr3-66892431-INS-140
chr3-66894685-INS-195
chr3-66898573-INS-886
chr3-66907438-INS-2177
chr3-66948495-INS-2081
chr3-66970681-INS-52
chr3-66980981-INS-1138
chr3-66981045-INS-394
chr3-66995449-INS-90
chr3-67014687-INS-1059
chr3-67029117-INS-9212
chr3-67038703-INS-274
chr3-67041874-INS-550
chr3-67090232-INS-6180
chr3-67101375-INS-392
chr3-67105134-INS-353
chr3-67116820-INS-358
chr3-67192925-INS-295
chr3-67194653-INS-398
chr3-67195197-INS-5039
chr3-67209909-INS-7405
chr3-67233741-INS-51
chr3-67238754-INS-830
chr3-67252184-INS-288
chr3-67253231-INS-56
chr3-67278232-INS-4438
chr3-67279857-INS-6380
chr3-67297513-INS-53
chr3-67343505-INS-7716
chr3-67385018-INS-207
chr3-67388411-INS-60
chr3-67400935-INS-101
chr3-67419518-INS-99
chr3-67441543-INS-768
chr3-67478925-INS-6442
chr3-67481110-INS-285
chr3-67481731-INS-352
chr3-67503423-INS-7140
chr3-67508964-INS-52
chr3-67511678-INS-1087
chr3-67521468-INS-338
chr3-67525871-INS-205
chr3-67529780-INS-152
chr3-67538042-INS-73
chr3-67544831-INS-93
chr3-67549468-INS-1797
chr3-67559025-INS-56
chr3-67562876-INS-240
chr3-67563212-INS-6690
chr3-67571496-INS-70
chr3-67604158-INS-63
chr3-67605626-INS-225
chr3-67614734-INS-399
chr3-67624798-INS-148
chr3-67648306-INS-52
chr3-67668914-INS-142
chr3-67674708-INS-97
chr3-67679718-INS-417
chr3-67690773-INS-222

chr3-67717931-INS-7266
chr3-67726856-INS-536
chr3-67757364-INS-649
chr3-67763680-INS-300
chr3-67764941-INS-269
chr3-67795381-INS-651
chr3-67804856-INS-700
chr3-67819203-INS-57
chr3-67831912-INS-52
chr3-67849104-INS-1440
chr3-67883591-INS-640
chr3-67884583-INS-79
chr3-67928654-INS-108
chr3-67948355-INS-1202
chr3-67953618-INS-66
chr3-68005802-INS-895
chr3-68032721-INS-62
chr3-68033254-INS-75
chr3-68034919-INS-7329
chr3-68058362-INS-6471
chr3-68060854-INS-62
chr3-68061610-INS-399
chr3-68073856-INS-58
chr3-68084560-INS-76
chr3-68119211-INS-440
chr3-68156051-INS-83
chr3-68203636-INS-57
chr3-68241529-INS-295
chr3-68268727-INS-90
chr3-68274207-INS-73
chr3-68304822-INS-151
chr3-68338385-INS-58
chr3-68347319-INS-71
chr3-68362182-INS-199
chr3-68364098-INS-8455
chr3-68411027-INS-207
chr3-68420668-INS-144
chr3-68422108-INS-70
chr3-68422898-INS-66
chr3-68436134-INS-6702
chr3-68440399-INS-216
chr3-68457293-INS-412
chr3-68475258-INS-123
chr3-68485145-INS-155
chr3-68500848-INS-154
chr3-68501668-INS-87
chr3-68512773-INS-65
chr3-68521045-INS-53
chr3-68547506-INS-282
chr3-68563736-INS-58
chr3-68566862-INS-149
chr3-68571232-INS-68
chr3-68611781-INS-181
chr3-68618755-INS-151
chr3-68621060-INS-583
chr3-68625609-INS-191
chr3-68647289-INS-87
chr3-68664434-INS-893
chr3-68691573-INS-1202
chr3-68716240-INS-4069
chr3-68725484-INS-6683
chr3-68755229-INS-133
chr3-68765517-INS-3827
chr3-68767012-INS-147
chr3-68814525-INS-62
chr3-68833637-INS-468
chr3-68858102-INS-215
chr3-68858489-INS-8986
chr3-68861264-INS-63
chr3-68881187-INS-6048
chr3-68888878-INS-67
chr3-68896205-INS-293
chr3-68901744-INS-124
chr3-68908626-INS-180
chr3-68909731-INS-204
chr3-68932591-INS-194
chr3-68956256-INS-374

chr3-68964484-INS-140
chr3-68995125-INS-209
chr3-69000601-INS-199
chr3-69003275-INS-488
chr3-69045589-INS-243
chr3-69059546-INS-97
chr3-69061589-INS-176
chr3-69069037-INS-107
chr3-69083634-INS-62
chr3-69096791-INS-80
chr3-69109385-INS-128
chr3-69134455-INS-204
chr3-69142729-INS-6166
chr3-69162805-INS-205
chr3-69187864-INS-52
chr3-69201131-INS-192
chr3-69216809-INS-74
chr3-69220775-INS-89
chr3-69236575-INS-461
chr3-69243355-INS-2335
chr3-69279012-INS-202
chr3-69284178-INS-207
chr3-69288718-INS-60
chr3-69319359-INS-94
chr3-69319989-INS-76
chr3-69331479-INS-71
chr3-69332310-INS-303
chr3-69333916-INS-86
chr3-69357851-INS-769
chr3-69373110-INS-216
chr3-69380389-INS-176
chr3-69404491-INS-100
chr3-69419689-INS-111
chr3-69433155-INS-66
chr3-69437810-INS-203
chr3-69454859-INS-742
chr3-69477640-INS-251
chr3-69489189-INS-168
chr3-69513885-INS-139
chr3-69522367-INS-60
chr3-69542940-INS-72
chr3-69551873-INS-91
chr3-69595065-INS-700
chr3-69623287-INS-5386
chr3-69633489-INS-51
chr3-69683046-INS-565
chr3-69684816-INS-6415
chr3-69686695-INS-667
chr3-69693031-INS-367
chr3-69703184-INS-64
chr3-69709200-INS-782
chr3-69711464-INS-767
chr3-69717413-INS-64
chr3-69732738-INS-92
chr3-69740481-INS-170
chr3-69751980-INS-52
chr3-69776686-INS-105
chr3-69787909-INS-300
chr3-69791935-INS-132
chr3-69792704-INS-616
chr3-69808613-INS-60
chr3-69816054-INS-228
chr3-69862385-INS-691
chr3-69880062-INS-5804
chr3-69900067-INS-340
chr3-69903829-INS-64
chr3-69905958-INS-196
chr3-69933307-INS-303
chr3-69962414-INS-334
chr3-69964469-INS-306
chr3-69970860-INS-54
chr3-69996511-INS-114
chr3-70021702-INS-118
chr3-70033406-INS-179
chr3-70046696-INS-68
chr3-70050274-INS-6705
chr3-70069796-INS-70

chr3-70083168-INS-75
chr3-70096504-INS-56
chr3-70100523-INS-6377
chr3-70106031-INS-2527
chr3-70108574-INS-73
chr3-70108846-INS-955
chr3-70112132-INS-173
chr3-70117510-INS-102
chr3-70122018-INS-917
chr3-70123037-INS-52
chr3-70130029-INS-230
chr3-70131798-INS-563
chr3-70136693-INS-551
chr3-70138265-INS-311
chr3-70139253-INS-1088
chr3-70143824-INS-86
chr3-70158612-INS-679
chr3-70221192-INS-65
chr3-70242169-INS-235
chr3-70250240-INS-195
chr3-70252542-INS-302
chr3-70269856-INS-521
chr3-70314866-INS-55
chr3-70333719-INS-6372
chr3-70354249-INS-1492
chr3-70355479-INS-96
chr3-70358718-INS-221
chr3-70362604-INS-52
chr3-70376513-INS-110
chr3-70382612-INS-456
chr3-70396927-INS-93
chr3-70398688-INS-70
chr3-70411129-INS-326
chr3-70425900-INS-644
chr3-70430263-INS-152
chr3-70449098-INS-501
chr3-70453772-INS-301
chr3-70456737-INS-70
chr3-70471069-INS-329
chr3-70484900-INS-214
chr3-70492186-INS-288
chr3-70551739-INS-80
chr3-70567481-INS-136
chr3-70571444-INS-459
chr3-70589947-INS-801
chr3-70602948-INS-6495
chr3-70608643-INS-79
chr3-70641047-INS-442
chr3-70670669-INS-111
chr3-70673518-INS-562
chr3-70691445-INS-498
chr3-70695452-INS-385
chr3-70715920-INS-262
chr3-70751331-INS-101
chr3-70766131-INS-115
chr3-70774681-INS-542
chr3-70787540-INS-4047
chr3-70787999-INS-153
chr3-70788424-INS-507
chr3-70800052-INS-354
chr3-70801041-INS-68
chr3-70826949-INS-224
chr3-70828468-INS-143
chr3-70830106-INS-164
chr3-70832024-INS-655
chr3-70835534-INS-62
chr3-70837427-INS-192
chr3-70848270-INS-88
chr3-70864537-INS-181
chr3-70864537-INS-3294
chr3-70891718-INS-690
chr3-70909105-INS-138
chr3-70916266-INS-517
chr3-70974026-INS-445
chr3-70985700-INS-328
chr3-71000082-INS-65
chr3-71000672-INS-1864

chr3-71007413-INS-498
chr3-71034859-INS-71
chr3-71077649-INS-354
chr3-71081844-INS-256
chr3-71142670-INS-158
chr3-71143453-INS-209
chr3-71145330-INS-177
chr3-71190372-INS-78
chr3-71191069-INS-5081
chr3-71197547-INS-208
chr3-71206190-INS-54
chr3-71227966-INS-778
chr3-71240167-INS-146
chr3-71250040-INS-359
chr3-71256170-INS-382
chr3-71263267-INS-341
chr3-71289875-INS-55
chr3-71313533-INS-493
chr3-71326325-INS-6800
chr3-71329842-INS-6398
chr3-71331432-INS-53
chr3-71355750-INS-121
chr3-71375061-INS-199
chr3-71379957-INS-84
chr3-71384514-INS-62
chr3-71389941-INS-917
chr3-71402795-INS-390
chr3-71402826-INS-1025
chr3-71402851-INS-661
chr3-71402938-INS-1390
chr3-71413289-INS-6365
chr3-71434378-INS-832
chr3-71455941-INS-6386
chr3-71565534-INS-466
chr3-71572069-INS-52
chr3-71602010-INS-203
chr3-71630783-INS-92
chr3-71652127-INS-4569
chr3-71664349-INS-8029
chr3-71670456-INS-454
chr3-71675251-INS-581
chr3-71688122-INS-214
chr3-71702284-INS-502
chr3-71728777-INS-68
chr3-71734470-INS-73
chr3-71741021-INS-78
chr3-71776162-INS-292
chr3-71816186-INS-390
chr3-71838899-INS-59
chr3-71855079-INS-75
chr3-71905290-INS-214
chr3-71906527-INS-83
chr3-71908472-INS-167
chr3-71928797-INS-68
chr3-71964990-INS-6190
chr3-71982725-INS-95
chr3-72006326-INS-53
chr3-72031191-INS-58
chr3-72044387-INS-104
chr3-72053598-INS-6290
chr3-72073460-INS-60
chr3-72082351-INS-56
chr3-72107787-INS-1455
chr3-72132894-INS-55
chr3-72150990-INS-645
chr3-72153154-INS-6180
chr3-72154516-INS-62
chr3-72161152-INS-2038
chr3-72162240-INS-347
chr3-72171332-INS-6330
chr3-72183585-INS-5868
chr3-72187021-INS-199
chr3-72188096-INS-136
chr3-72192085-INS-4328
chr3-72213541-INS-852
chr3-72250950-INS-3918
chr3-72277684-INS-880

chr3-72284582-INS-6324
chr3-72291759-INS-61
chr3-72293884-INS-97
chr3-72353288-INS-446
chr3-72364467-INS-822
chr3-72370789-INS-7784
chr3-72466043-INS-207
chr3-72467288-INS-399
chr3-72481313-INS-52
chr3-72512030-INS-72
chr3-72514650-INS-118
chr3-72521597-INS-5549
chr3-72548143-INS-339
chr3-72582619-INS-92
chr3-72599215-INS-265
chr3-72651731-INS-99
chr3-72660402-INS-6472
chr3-72704417-INS-159
chr3-72716453-INS-84
chr3-72726853-INS-120
chr3-72734383-INS-82
chr3-72747473-INS-673
chr3-72773811-INS-1887
chr3-72775247-INS-82
chr3-72785949-INS-52
chr3-72812537-INS-102
chr3-72833368-INS-210
chr3-72849360-INS-51
chr3-72854119-INS-749
chr3-72871219-INS-334
chr3-72882644-INS-7513
chr3-72936620-INS-98
chr3-72939344-INS-116
chr3-72942653-INS-100
chr3-72945935-INS-55
chr3-73003511-INS-536
chr3-73033802-INS-299
chr3-73046254-INS-77
chr3-73056394-INS-196
chr3-73071282-INS-653
chr3-73076793-INS-83
chr3-73079411-INS-399
chr3-73084809-INS-58
chr3-73087818-INS-54
chr3-73111594-INS-898
chr3-73118495-INS-92
chr3-73124062-INS-3075
chr3-73124727-INS-6376
chr3-73139585-INS-191
chr3-73142040-INS-62
chr3-73168346-INS-334
chr3-73180692-INS-6394
chr3-73224635-INS-51
chr3-73289020-INS-1891
chr3-73327207-INS-438
chr3-73341592-INS-7483
chr3-73387868-INS-6442
chr3-73453242-INS-219
chr3-73456255-INS-411
chr3-73458973-INS-66
chr3-73465437-INS-83
chr3-73495150-INS-230
chr3-73510041-INS-7076
chr3-73512447-INS-1344
chr3-73513845-INS-2799
chr3-73515908-INS-4604
chr3-73518608-INS-165
chr3-73521045-INS-545
chr3-73529934-INS-5355
chr3-73559836-INS-263
chr3-73579907-INS-6736
chr3-73593995-INS-580
chr3-73603723-INS-258
chr3-73607246-INS-181
chr3-73609972-INS-88
chr3-73631261-INS-319
chr3-73661339-INS-80

chr3-73662432-INS-474
chr3-73686129-INS-53
chr3-73691332-INS-494
chr3-73725657-INS-1359
chr3-73736734-INS-440
chr3-73745237-INS-82
chr3-73750518-INS-6512
chr3-73755820-INS-80
chr3-73757837-INS-382
chr3-73769628-INS-530
chr3-73786178-INS-201
chr3-73851913-INS-60
chr3-73860763-INS-591
chr3-73864101-INS-358
chr3-73871561-INS-166
chr3-73883842-INS-579
chr3-73892928-INS-90
chr3-73947142-INS-138
chr3-73967414-INS-6185
chr3-73979207-INS-63
chr3-73988891-INS-119
chr3-73996824-INS-274
chr3-74003128-INS-86
chr3-74009705-INS-567
chr3-74021419-INS-314
chr3-74023914-INS-236
chr3-74044478-INS-512
chr3-74063602-INS-81
chr3-74068124-INS-193
chr3-74068508-INS-6335
chr3-74077516-INS-56
chr3-74112023-INS-741
chr3-74114041-INS-935
chr3-74134736-INS-110
chr3-74215525-INS-170
chr3-74318967-INS-344
chr3-74321592-INS-568
chr3-74339647-INS-69
chr3-74360477-INS-1493
chr3-74376411-INS-73
chr3-74391623-INS-341
chr3-74413762-INS-122
chr3-74416826-INS-189
chr3-74441746-INS-51
chr3-74452444-INS-55
chr3-74488072-INS-262
chr3-74497801-INS-139
chr3-74512011-INS-444
chr3-74514160-INS-6384
chr3-74515068-INS-694
chr3-74518496-INS-4423
chr3-74576738-INS-7103
chr3-74621893-INS-6110
chr3-74630220-INS-460
chr3-74645655-INS-340
chr3-74676963-INS-129
chr3-74680402-INS-502
chr3-74684480-INS-366
chr3-74686515-INS-301
chr3-74713171-INS-61
chr3-74732255-INS-671
chr3-74751910-INS-80
chr3-74792545-INS-8328
chr3-74882907-INS-172
chr3-74888773-INS-113
chr3-74898073-INS-70
chr3-74925023-INS-58
chr3-74966773-INS-1413
chr3-74974583-INS-7715
chr3-75023982-INS-128
chr3-75040678-INS-1033
chr3-75042437-INS-52
chr3-75044295-INS-56
chr3-75069970-INS-6415
chr3-75073388-INS-70
chr3-75081644-INS-2442
chr3-75092746-INS-642

chr3-75133552-INS-102
chr3-75154340-INS-186
chr3-75158112-INS-749
chr3-75161007-INS-57
chr3-75188709-INS-306
chr3-75200156-INS-51
chr3-75207414-INS-836
chr3-75208142-INS-85
chr3-75240208-INS-422
chr3-75276669-INS-158
chr3-75305633-INS-462
chr3-75306812-INS-6490
chr3-75306982-INS-142
chr3-75347626-INS-383
chr3-75379604-INS-69
chr3-75383284-INS-65
chr3-75411220-INS-60
chr3-75418604-INS-58
chr3-75425386-INS-234
chr3-75459796-INS-67
chr3-75476442-INS-654
chr3-75488206-INS-231
chr3-75495290-INS-328
chr3-75499273-INS-52
chr3-75504171-INS-171
chr3-75530822-INS-115
chr3-75533291-INS-401
chr3-75549913-INS-64
chr3-75551841-INS-90
chr3-75579526-INS-69
chr3-75598070-INS-88
chr3-75613826-INS-2166
chr3-75620182-INS-378
chr3-75621686-INS-56
chr3-75625830-INS-202
chr3-75630323-INS-635
chr3-75661386-INS-60
chr3-75668633-INS-318
chr3-75672895-INS-56
chr3-75680097-INS-162
chr3-75681511-INS-64
chr3-75683893-INS-6428
chr3-75684656-INS-6198
chr3-75707834-INS-6382
chr3-75727921-INS-176
chr3-75734250-INS-348
chr3-75788905-INS-72
chr3-75790689-INS-54
chr3-75815671-INS-76
chr3-75820678-INS-121
chr3-75824750-INS-232
chr3-75836956-INS-83
chr3-75940458-INS-974
chr3-75948024-INS-753
chr3-75959136-INS-486
chr3-75967845-INS-2812
chr3-75979728-INS-72
chr3-76003981-INS-496
chr3-76011064-INS-92
chr3-76015406-INS-63
chr3-76030347-INS-980
chr3-76044335-INS-276
chr3-76079466-INS-1786
chr3-76089522-INS-701
chr3-76108307-INS-675
chr3-76148637-INS-407
chr3-76167376-INS-54
chr3-76189612-INS-82
chr3-76196645-INS-190
chr3-76233392-INS-6360
chr3-76240382-INS-120
chr3-76248438-INS-101
chr3-76296750-INS-54
chr3-76302236-INS-139
chr3-76311448-INS-197
chr3-76340202-INS-4167
chr3-76341721-INS-119

chr3-76352882-INS-320
chr3-76392867-INS-60
chr3-76395687-INS-62
chr3-76399276-INS-2792
chr3-76404349-INS-5695
chr3-76407126-INS-213
chr3-76456531-INS-2337
chr3-76489818-INS-228
chr3-76496046-INS-66
chr3-76498362-INS-82
chr3-76541165-INS-603
chr3-76557578-INS-56
chr3-76609977-INS-78
chr3-76650087-INS-120
chr3-76681827-INS-56
chr3-76683196-INS-166
chr3-76753822-INS-252
chr3-76791421-INS-291
chr3-76809525-INS-7217
chr3-76811312-INS-129
chr3-76837540-INS-286
chr3-76850010-INS-52
chr3-76878876-INS-500
chr3-76885596-INS-473
chr3-76889755-INS-6394
chr3-76903688-INS-59
chr3-76906904-INS-66
chr3-76908955-INS-116
chr3-76960916-INS-357
chr3-76977996-INS-70
chr3-76988135-INS-52
chr3-77017975-INS-171
chr3-77022830-INS-253
chr3-77047912-INS-323
chr3-77049207-INS-347
chr3-77056597-INS-517
chr3-77060902-INS-202
chr3-77087668-INS-1488
chr3-77088356-INS-872
chr3-77090588-INS-74
chr3-77097539-INS-214
chr3-77103204-INS-56
chr3-77120317-INS-109
chr3-77188365-INS-52
chr3-77198164-INS-403
chr3-77201343-INS-688
chr3-77205884-INS-177
chr3-77233130-INS-145
chr3-77251721-INS-323
chr3-77262016-INS-7506
chr3-77273594-INS-608
chr3-77301405-INS-76
chr3-77325659-INS-634
chr3-77362368-INS-286
chr3-77368453-INS-86
chr3-77396826-INS-546
chr3-77400318-INS-315
chr3-77402552-INS-6311
chr3-77486786-INS-211
chr3-77500509-INS-403
chr3-77513142-INS-4358
chr3-77633075-INS-64
chr3-77643738-INS-88
chr3-77652944-INS-56
chr3-77655487-INS-106
chr3-77668400-INS-269
chr3-77699897-INS-276
chr3-77704903-INS-136
chr3-77707392-INS-6480
chr3-77760075-INS-134
chr3-77774236-INS-623
chr3-77847387-INS-76
chr3-77852521-INS-3048
chr3-77862986-INS-215
chr3-77869170-INS-473
chr3-77903582-INS-7575
chr3-77917816-INS-52

chr3-77994433-INS-679
chr3-77996375-INS-153
chr3-77996741-INS-394
chr3-77997550-INS-6871
chr3-78000110-INS-440
chr3-78004737-INS-61
chr3-78014379-INS-76
chr3-78043358-INS-100
chr3-78043403-INS-700
chr3-78045082-INS-61
chr3-78063005-INS-122
chr3-78065271-INS-576
chr3-78069865-INS-2276
chr3-78071144-INS-68
chr3-78088402-INS-260
chr3-78091247-INS-65
chr3-78123476-INS-60
chr3-78143764-INS-70
chr3-78174581-INS-396
chr3-78179265-INS-4059
chr3-78222973-INS-54
chr3-78223426-INS-948
chr3-78235758-INS-258
chr3-78238407-INS-6488
chr3-78246626-INS-1218
chr3-78247639-INS-1864
chr3-78249386-INS-150
chr3-78256179-INS-7581
chr3-78261172-INS-50
chr3-78280252-INS-497
chr3-78293045-INS-552
chr3-78366274-INS-81
chr3-78373783-INS-6552
chr3-78378413-INS-6640
chr3-78383888-INS-64
chr3-78422767-INS-54
chr3-78429245-INS-800
chr3-78442435-INS-66
chr3-78443805-INS-399
chr3-78468082-INS-64
chr3-78469240-INS-76
chr3-78510487-INS-158
chr3-78523568-INS-96
chr3-78524947-INS-68
chr3-78526495-INS-376
chr3-78531500-INS-57
chr3-78534112-INS-92
chr3-78537831-INS-5249
chr3-78541018-INS-92
chr3-78554650-INS-175
chr3-78556860-INS-102
chr3-78573616-INS-57
chr3-78574070-INS-518
chr3-78576608-INS-273
chr3-78635983-INS-56
chr3-78692548-INS-54
chr3-78696995-INS-358
chr3-78700924-INS-70
chr3-78743421-INS-99
chr3-78763882-INS-226
chr3-78782293-INS-52
chr3-78791946-INS-76
chr3-78795447-INS-189
chr3-78821374-INS-59
chr3-78826730-INS-135
chr3-78845370-INS-160
chr3-78848043-INS-57
chr3-78857013-INS-328
chr3-78886856-INS-566
chr3-78906153-INS-57
chr3-78917927-INS-321
chr3-78949115-INS-6401
chr3-78983764-INS-238
chr3-79025273-INS-160
chr3-79049017-INS-72
chr3-79091348-INS-177
chr3-79094783-INS-76

chr3-79104672-INS-577
chr3-79107043-INS-498
chr3-79122715-INS-7115
chr3-79125111-INS-6440
chr3-79129352-INS-402
chr3-79133049-INS-498
chr3-79169927-INS-250
chr3-79228659-INS-64
chr3-79264023-INS-737
chr3-79271736-INS-74
chr3-79273161-INS-464
chr3-79285147-INS-173
chr3-79306062-INS-137
chr3-79307828-INS-507
chr3-79347944-INS-180
chr3-79360526-INS-253
chr3-79373963-INS-78
chr3-79381326-INS-351
chr3-79405637-INS-74
chr3-79421623-INS-556
chr3-79454021-INS-72
chr3-79462150-INS-52
chr3-79482202-INS-91
chr3-79492944-INS-292
chr3-79501149-INS-305
chr3-79502369-INS-836
chr3-79531922-INS-197
chr3-79540153-INS-193
chr3-79554498-INS-64
chr3-79558792-INS-206
chr3-79585436-INS-62
chr3-79586827-INS-1888
chr3-79598194-INS-99
chr3-79610201-INS-673
chr3-79612129-INS-200
chr3-79615597-INS-166
chr3-79661083-INS-73
chr3-79663144-INS-54
chr3-79727686-INS-434
chr3-79729690-INS-116
chr3-79751219-INS-239
chr3-79764119-INS-621
chr3-79766174-INS-879
chr3-79775603-INS-95
chr3-79776438-INS-360
chr3-79794909-INS-142
chr3-79799439-INS-199
chr3-79818975-INS-52
chr3-79828576-INS-150
chr3-79835117-INS-340
chr3-79839759-INS-5549
chr3-79881301-INS-6344
chr3-79881818-INS-466
chr3-79885890-INS-62
chr3-79888427-INS-667
chr3-79898940-INS-79
chr3-79909722-INS-6509
chr3-79926681-INS-201
chr3-79932416-INS-192
chr3-79933553-INS-859
chr3-79942953-INS-218
chr3-80005667-INS-6952
chr3-80014949-INS-892
chr3-80015707-INS-66
chr3-80065846-INS-433
chr3-80066954-INS-92
chr3-80096986-INS-967
chr3-80101465-INS-54
chr3-80125956-INS-6350
chr3-80170962-INS-60
chr3-80188537-INS-59
chr3-80209772-INS-151
chr3-80222989-INS-165
chr3-80243815-INS-150
chr3-80267762-INS-6456
chr3-80302028-INS-91
chr3-80329420-INS-86

chr3-80330730-INS-58
chr3-80333745-INS-534
chr3-80334689-INS-225
chr3-80387510-INS-102
chr3-80389483-INS-481
chr3-80394077-INS-350
chr3-80409024-INS-56
chr3-80411679-INS-87
chr3-80417240-INS-327
chr3-80434360-INS-209
chr3-80495142-INS-66
chr3-80519000-INS-9081
chr3-80520353-INS-866
chr3-80525978-INS-110
chr3-80563568-INS-118
chr3-80588223-INS-359
chr3-80598322-INS-63
chr3-80612640-INS-259
chr3-80638212-INS-70
chr3-80641639-INS-444
chr3-80674629-INS-800
chr3-80681248-INS-853
chr3-80689656-INS-6502
chr3-80691138-INS-299
chr3-80695812-INS-82
chr3-80727294-INS-104
chr3-80734359-INS-89
chr3-80741140-INS-62
chr3-80770958-INS-395
chr3-80774500-INS-200
chr3-80779132-INS-7910
chr3-80781212-INS-76
chr3-80783821-INS-6377
chr3-80799455-INS-130
chr3-80810188-INS-8894
chr3-80836686-INS-128
chr3-80838105-INS-236
chr3-80859586-INS-7145
chr3-80862007-INS-7207
chr3-80903217-INS-112
chr3-80936742-INS-104
chr3-80939030-INS-184
chr3-80941271-INS-136
chr3-80951691-INS-407
chr3-80957011-INS-283
chr3-80963609-INS-64
chr3-81028960-INS-52
chr3-81075766-INS-113
chr3-81076530-INS-68
chr3-81095920-INS-120
chr3-81126147-INS-5724
chr3-81127233-INS-522
chr3-81127980-INS-85
chr3-81153898-INS-490
chr3-81178340-INS-6563
chr3-81187267-INS-6427
chr3-81211395-INS-142
chr3-81235968-INS-72
chr3-81243317-INS-146
chr3-81248973-INS-78
chr3-81260484-INS-522
chr3-81281426-INS-338
chr3-81298461-INS-6535
chr3-81315936-INS-128
chr3-81324833-INS-165
chr3-81329564-INS-80
chr3-81341593-INS-189
chr3-81408201-INS-196
chr3-81415570-INS-51
chr3-81462087-INS-83
chr3-81495153-INS-110
chr3-81513848-INS-358
chr3-81535877-INS-265
chr3-81538650-INS-59
chr3-81542459-INS-6257
chr3-81554952-INS-62
chr3-81605848-INS-197

chr3-81626171-INS-416
chr3-81627393-INS-2314
chr3-81640876-INS-59
chr3-81646477-INS-934
chr3-81656125-INS-54
chr3-81661276-INS-206
chr3-81662469-INS-3367
chr3-81673659-INS-6649
chr3-81681489-INS-250
chr3-81689309-INS-154
chr3-81708782-INS-55
chr3-81709571-INS-7568
chr3-81711809-INS-6383
chr3-81715111-INS-323
chr3-81725771-INS-3310
chr3-81730508-INS-210
chr3-81748169-INS-52
chr3-81772342-INS-4495
chr3-81777984-INS-299
chr3-81790870-INS-498
chr3-81804945-INS-54
chr3-81846074-INS-98
chr3-81850782-INS-92
chr3-81888164-INS-920
chr3-81928769-INS-1195
chr3-81932544-INS-104
chr3-81960317-INS-198
chr3-81971946-INS-315
chr3-81983798-INS-93
chr3-82022602-INS-225
chr3-82032120-INS-66
chr3-82039196-INS-60
chr3-82042187-INS-250
chr3-82046413-INS-69
chr3-82057181-INS-98
chr3-82062608-INS-74
chr3-82075358-INS-58
chr3-82080160-INS-106
chr3-82116715-INS-52
chr3-82127266-INS-78
chr3-82136338-INS-212
chr3-82147267-INS-80
chr3-82168644-INS-1243
chr3-82189301-INS-78
chr3-82200786-INS-56
chr3-82208236-INS-6296
chr3-82213357-INS-89
chr3-82215834-INS-665
chr3-82217541-INS-2889
chr3-82217560-INS-325
chr3-82228126-INS-6176
chr3-82231765-INS-6415
chr3-82252787-INS-169
chr3-82254208-INS-111
chr3-82259472-INS-68
chr3-82287903-INS-201
chr3-82313126-INS-51
chr3-82336345-INS-216
chr3-82347897-INS-584
chr3-82355742-INS-56
chr3-82364570-INS-157
chr3-82380721-INS-6454
chr3-82407176-INS-792
chr3-82417629-INS-184
chr3-82427139-INS-89
chr3-82429020-INS-197
chr3-82442282-INS-593
chr3-82449344-INS-52
chr3-82458282-INS-556
chr3-82490721-INS-64
chr3-82496402-INS-100
chr3-82530251-INS-372
chr3-82537188-INS-498
chr3-82539205-INS-94
chr3-82571754-INS-399
chr3-82590278-INS-723
chr3-82624910-INS-74

chr3-82648702-INS-482
chr3-82659375-INS-119
chr3-82666031-INS-60
chr3-82667306-INS-75
chr3-82680755-INS-113
chr3-82687136-INS-96
chr3-82692944-INS-996
chr3-82711460-INS-100
chr3-82714787-INS-479
chr3-82728206-INS-922
chr3-82732104-INS-516
chr3-82748737-INS-732
chr3-82777422-INS-439
chr3-82806753-INS-262
chr3-82831043-INS-631
chr3-82847656-INS-74
chr3-82867700-INS-53
chr3-82888502-INS-67
chr3-82894663-INS-7108
chr3-82895051-INS-148
chr3-82915343-INS-85
chr3-82924112-INS-54
chr3-82931163-INS-57
chr3-82936031-INS-52
chr3-82957816-INS-769
chr3-82968167-INS-427
chr3-82971730-INS-292
chr3-82972447-INS-681
chr3-82978317-INS-57
chr3-82988944-INS-214
chr3-83008884-INS-195
chr3-83010573-INS-57
chr3-83031411-INS-831
chr3-83054099-INS-81
chr3-83056432-INS-86
chr3-83060633-INS-109
chr3-83101226-INS-769
chr3-83109241-INS-114
chr3-83137684-INS-54
chr3-83164699-INS-593
chr3-83165928-INS-74
chr3-83185198-INS-233
chr3-83188695-INS-6394
chr3-83198044-INS-66
chr3-83224456-INS-87
chr3-83225027-INS-8284
chr3-83231037-INS-224
chr3-83335465-INS-52
chr3-83335890-INS-52
chr3-83347125-INS-427
chr3-83389909-INS-116
chr3-83392155-INS-268
chr3-83406252-INS-6459
chr3-83413814-INS-195
chr3-83419869-INS-54
chr3-83422990-INS-213
chr3-83468481-INS-68
chr3-83478799-INS-219
chr3-83478833-INS-392
chr3-83479447-INS-53
chr3-83506730-INS-212
chr3-83511831-INS-86
chr3-83541623-INS-68
chr3-83556875-INS-440
chr3-83561986-INS-6427
chr3-83566239-INS-5904
chr3-83568523-INS-55
chr3-83598326-INS-3290
chr3-83608996-INS-191
chr3-83637925-INS-277
chr3-83649695-INS-169
chr3-83659014-INS-115
chr3-83668693-INS-607
chr3-83711092-INS-54
chr3-83713818-INS-233
chr3-83738734-INS-215
chr3-83754038-INS-82

chr3-83761635-INS-882
chr3-83768489-INS-193
chr3-83791347-INS-4457
chr3-83823159-INS-107
chr3-83827441-INS-64
chr3-83830243-INS-77
chr3-83861550-INS-65
chr3-83871316-INS-3222
chr3-83872945-INS-215
chr3-83879017-INS-215
chr3-83905736-INS-58
chr3-83913325-INS-167
chr3-83916384-INS-60
chr3-83937110-INS-148
chr3-83951383-INS-134
chr3-83955827-INS-52
chr3-83957520-INS-201
chr3-83966382-INS-88
chr3-83979652-INS-74
chr3-84005608-INS-93
chr3-84022105-INS-176
chr3-84057805-INS-55
chr3-84061892-INS-72
chr3-84077433-INS-63
chr3-84081950-INS-198
chr3-84083446-INS-52
chr3-84107188-INS-132
chr3-84118725-INS-124
chr3-84129058-INS-528
chr3-84130330-INS-64
chr3-84135414-INS-136
chr3-84194980-INS-60
chr3-84205198-INS-75
chr3-84228820-INS-236
chr3-84245832-INS-193
chr3-84263262-INS-71
chr3-84280015-INS-343
chr3-84286142-INS-408
chr3-84308685-INS-274
chr3-84318546-INS-266
chr3-84325352-INS-55
chr3-84372242-INS-84
chr3-84391403-INS-322
chr3-84406119-INS-918
chr3-84424034-INS-252
chr3-84439389-INS-749
chr3-84489156-INS-66
chr3-84492835-INS-75
chr3-84496201-INS-138
chr3-84499578-INS-150
chr3-84522456-INS-162
chr3-84527374-INS-351
chr3-84528265-INS-64
chr3-84531533-INS-211
chr3-84532496-INS-146
chr3-84533908-INS-192
chr3-84537046-INS-66
chr3-84538587-INS-224
chr3-84540487-INS-70
chr3-84541662-INS-201
chr3-84544671-INS-60
chr3-84551475-INS-221
chr3-84574390-INS-754
chr3-84604880-INS-197
chr3-84616881-INS-151
chr3-84623873-INS-62
chr3-84625749-INS-916
chr3-84626593-INS-90
chr3-84639064-INS-56
chr3-84646848-INS-60
chr3-84678939-INS-54
chr3-84727187-INS-214
chr3-84750100-INS-52
chr3-84806252-INS-229
chr3-84819921-INS-59
chr3-84893274-INS-190
chr3-84901906-INS-70

chr3-84937070-INS-399
chr3-84942195-INS-5316
chr3-84948067-INS-53
chr3-84978419-INS-105
chr3-85002304-INS-60
chr3-85016067-INS-86
chr3-85037398-INS-221
chr3-85047326-INS-56
chr3-85083207-INS-333
chr3-85097755-INS-92
chr3-85126813-INS-681
chr3-85128501-INS-163
chr3-85173794-INS-270
chr3-85185238-INS-62
chr3-85188990-INS-104
chr3-85194284-INS-62
chr3-85209278-INS-82
chr3-85250451-INS-118
chr3-85308644-INS-56
chr3-85312321-INS-66
chr3-85327110-INS-68
chr3-85344953-INS-104
chr3-85347275-INS-56
chr3-85354951-INS-62
chr3-85409215-INS-96
chr3-85422228-INS-263
chr3-85449771-INS-386
chr3-85488258-INS-227
chr3-85495201-INS-104
chr3-85514841-INS-114
chr3-85540965-INS-113
chr3-85579064-INS-76
chr3-85588408-INS-184
chr3-85589186-INS-135
chr3-85613082-INS-303
chr3-85669609-INS-6403
chr3-85683181-INS-54
chr3-85714964-INS-57
chr3-85759645-INS-240
chr3-85763953-INS-141
chr3-85789845-INS-124
chr3-85807691-INS-135
chr3-85848651-INS-118
chr3-85850295-INS-204
chr3-85857396-INS-56
chr3-85864421-INS-203
chr3-85887350-INS-195
chr3-85892054-INS-192
chr3-85906011-INS-64
chr3-85910016-INS-199
chr3-85919900-INS-180
chr3-85940150-INS-2821
chr3-85978386-INS-6440
chr3-86001576-INS-467
chr3-86009182-INS-136
chr3-86052023-INS-60
chr3-86082233-INS-453
chr3-86112616-INS-1426
chr3-86117442-INS-147
chr3-86143117-INS-158
chr3-86143202-INS-282
chr3-86153962-INS-129
chr3-86157349-INS-1640
chr3-86161403-INS-192
chr3-86176733-INS-222
chr3-86187226-INS-148
chr3-86194111-INS-140
chr3-86206998-INS-7146
chr3-86216435-INS-259
chr3-86257270-INS-201
chr3-86263388-INS-689
chr3-86325765-INS-207
chr3-86328576-INS-6231
chr3-86328634-INS-154
chr3-86333085-INS-59
chr3-86359050-INS-184
chr3-86374153-INS-60

chr3-86391508-INS-114
chr3-86397619-INS-73
chr3-86398988-INS-131
chr3-86410477-INS-110
chr3-86434191-INS-68
chr3-86475391-INS-244
chr3-86493114-INS-231
chr3-86522496-INS-239
chr3-86528544-INS-7430
chr3-86544530-INS-527
chr3-86563493-INS-480
chr3-86591492-INS-261
chr3-86634643-INS-7088
chr3-86645595-INS-116
chr3-86671745-INS-123
chr3-86681552-INS-215
chr3-86690376-INS-240
chr3-86691387-INS-399
chr3-86700178-INS-192
chr3-86706096-INS-64
chr3-86716872-INS-325
chr3-86764238-INS-114
chr3-86771145-INS-90
chr3-86772698-INS-106
chr3-86785436-INS-54
chr3-86797221-INS-141
chr3-86803713-INS-337
chr3-86815251-INS-266
chr3-86817670-INS-376
chr3-86817961-INS-8470
chr3-86837793-INS-466
chr3-86844265-INS-405
chr3-86847339-INS-96
chr3-86888969-INS-55
chr3-86900459-INS-135
chr3-86904241-INS-206
chr3-86933811-INS-482
chr3-86947731-INS-53
chr3-87039170-INS-73
chr3-87064537-INS-142
chr3-87069325-INS-185
chr3-87125551-INS-362
chr3-87134592-INS-205
chr3-87153556-INS-330
chr3-87161021-INS-219
chr3-87202766-INS-66
chr3-87211742-INS-263
chr3-87215974-INS-56
chr3-87229792-INS-6320
chr3-87234161-INS-60
chr3-87249442-INS-3694
chr3-87253665-INS-186
chr3-87257712-INS-199
chr3-87264493-INS-767
chr3-87325395-INS-574
chr3-87482689-INS-433
chr3-87495015-INS-612
chr3-87575267-INS-71
chr3-87589269-INS-531
chr3-87662855-INS-81
chr3-87664701-INS-90
chr3-87665995-INS-108
chr3-87682392-INS-72
chr3-87699156-INS-210
chr3-87711578-INS-54
chr3-87733163-INS-476
chr3-87748359-INS-124
chr3-87749495-INS-238
chr3-87763113-INS-208
chr3-87770275-INS-61
chr3-87773765-INS-179
chr3-87779463-INS-88
chr3-87783972-INS-211
chr3-87796108-INS-254
chr3-87800066-INS-104
chr3-87803071-INS-68
chr3-87807378-INS-177

chr3-87828855-INS-120
chr3-87830052-INS-444
chr3-87841292-INS-742
chr3-87861972-INS-54
chr3-87871080-INS-57
chr3-87882379-INS-215
chr3-87912403-INS-879
chr3-87920745-INS-68
chr3-87928063-INS-102
chr3-87956423-INS-80
chr3-87964454-INS-98
chr3-87985087-INS-889
chr3-87991542-INS-81
chr3-88008402-INS-107
chr3-88036095-INS-200
chr3-88037793-INS-400
chr3-88039210-INS-104
chr3-88064845-INS-104
chr3-88091730-INS-211
chr3-88096633-INS-114
chr3-88098536-INS-399
chr3-88102499-INS-91
chr3-88134937-INS-52
chr3-88147286-INS-79
chr3-88166064-INS-190
chr3-88170022-INS-201
chr3-88176715-INS-202
chr3-88177674-INS-118
chr3-88178401-INS-2150
chr3-88186230-INS-5671
chr3-88229206-INS-235
chr3-88273913-INS-59
chr3-88293403-INS-279
chr3-88294912-INS-129
chr3-88296749-INS-209
chr3-88309524-INS-160
chr3-88323961-INS-203
chr3-88340659-INS-562
chr3-88341819-INS-161
chr3-88347752-INS-217
chr3-88351554-INS-139
chr3-88376645-INS-395
chr3-88385733-INS-133
chr3-88391679-INS-65
chr3-88399354-INS-84
chr3-88402014-INS-69
chr3-88413787-INS-470
chr3-88481486-INS-170
chr3-88483016-INS-234
chr3-88493829-INS-230
chr3-88499437-INS-167
chr3-88503100-INS-60
chr3-88503993-INS-279
chr3-88506800-INS-190
chr3-88512073-INS-243
chr3-88513077-INS-205
chr3-88517191-INS-99
chr3-88519629-INS-138
chr3-88527365-INS-68
chr3-88544856-INS-191
chr3-88549652-INS-66
chr3-88551844-INS-219
chr3-88558348-INS-1894
chr3-88561147-INS-66
chr3-88563285-INS-224
chr3-88609959-INS-221
chr3-88632893-INS-214
chr3-88686098-INS-272
chr3-88687474-INS-110
chr3-88688868-INS-215
chr3-88700226-INS-134
chr3-88718470-INS-84
chr3-88736563-INS-95
chr3-88749588-INS-110
chr3-88750983-INS-214
chr3-88754248-INS-2842
chr3-88756487-INS-54

chr3-88759750-INS-6442
chr3-88770970-INS-436
chr3-88771519-INS-84
chr3-88773647-INS-397
chr3-88791262-INS-1279
chr3-88833081-INS-147
chr3-88839220-INS-204
chr3-88839971-INS-69
chr3-88875268-INS-97
chr3-88877205-INS-758
chr3-88881300-INS-1888
chr3-88902456-INS-221
chr3-88921669-INS-74
chr3-88926719-INS-195
chr3-88930094-INS-200
chr3-88945344-INS-52
chr3-88948423-INS-74
chr3-88981843-INS-508
chr3-89014892-INS-170
chr3-89020957-INS-170
chr3-89025275-INS-62
chr3-89047738-INS-83
chr3-89050855-INS-63
chr3-89059533-INS-316
chr3-89064353-INS-205
chr3-89068564-INS-165
chr3-89082198-INS-65
chr3-89086080-INS-219
chr3-89098659-INS-397
chr3-89105396-INS-209
chr3-89107845-INS-107
chr3-89133179-INS-70
chr3-89135169-INS-58
chr3-89148494-INS-57
chr3-89162066-INS-2502
chr3-89162915-INS-198
chr3-89167472-INS-197
chr3-89172001-INS-210
chr3-89190732-INS-191
chr3-89197688-INS-791
chr3-89199230-INS-52
chr3-89208594-INS-104
chr3-89209688-INS-100
chr3-89224619-INS-147
chr3-89232741-INS-192
chr3-89233563-INS-217
chr3-89237132-INS-65
chr3-89241886-INS-209
chr3-89252321-INS-106
chr3-89253908-INS-170
chr3-89258692-INS-59
chr3-89264429-INS-229
chr3-89305419-INS-64
chr3-89332489-INS-222
chr3-89368798-INS-190
chr3-89377754-INS-166
chr3-89387274-INS-51
chr3-89388827-INS-86
chr3-89404287-INS-91
chr3-89413244-INS-249
chr3-89434565-INS-85
chr3-89462259-INS-64
chr3-89504197-INS-1365
chr3-89537544-INS-363
chr3-89546204-INS-513
chr3-89548746-INS-183
chr3-89552341-INS-174
chr3-89554485-INS-230
chr3-89568952-INS-62
chr3-89589101-INS-2967
chr3-89596333-INS-56
chr3-89600447-INS-502
chr3-89605369-INS-552
chr3-89608993-INS-515
chr3-89656852-INS-95
chr3-89673433-INS-135
chr3-89713298-INS-56

chr3-89733777-INS-103
chr3-89751361-INS-208
chr3-89770116-INS-68
chr3-89799296-INS-52
chr3-89802123-INS-198
chr3-89809808-INS-132
chr3-89810544-INS-378
chr3-89825083-INS-207
chr3-89831042-INS-54
chr3-89832349-INS-56
chr3-89849981-INS-265
chr3-89873944-INS-148
chr3-89877738-INS-157
chr3-89881974-INS-271
chr3-89883540-INS-334
chr3-89894050-INS-74
chr3-89937083-INS-191
chr3-89950539-INS-202
chr3-89967836-INS-208
chr3-89980763-INS-108
chr3-90010583-INS-196
chr3-90014055-INS-201
chr3-90017463-INS-216
chr3-90019346-INS-122
chr3-90024777-INS-52
chr3-90044593-INS-195
chr3-90047557-INS-5518
chr3-90047977-INS-147
chr3-90052545-INS-79
chr3-90056906-INS-343
chr3-90094241-INS-369
chr3-90104056-INS-358
chr3-90111648-INS-214
chr3-90151334-INS-170
chr3-90153429-INS-81
chr3-90154100-INS-5548
chr3-90154850-INS-1236
chr3-90174216-INS-64
chr3-90179987-INS-200
chr3-90193639-INS-417
chr3-90195063-INS-194
chr3-90197781-INS-396
chr3-90219297-INS-204
chr3-90221978-INS-171
chr3-90222919-INS-213
chr3-90222931-INS-5542
chr3-90225634-INS-228
chr3-90231555-INS-87
chr3-90242565-INS-111
chr3-90249924-INS-460
chr3-90271951-INS-8985
chr3-90272715-INS-52
chr3-90275408-INS-204
chr3-90281003-INS-482
chr3-90287281-INS-178
chr3-90287548-INS-196
chr3-90312497-INS-68
chr3-90332741-INS-208
chr3-90334205-INS-206
chr3-90337670-INS-78
chr3-90385937-INS-51
chr3-90403790-INS-439
chr3-90429025-INS-893
chr3-90447363-INS-122
chr3-90457346-INS-3526
chr3-90458802-INS-55
chr3-90468032-INS-205
chr3-90474000-INS-221
chr3-90475502-INS-115
chr3-90481316-INS-204
chr3-90495623-INS-741
chr3-90505604-INS-140
chr3-90539508-INS-870
chr3-90549500-INS-334
chr3-90552598-INS-156
chr3-90604568-INS-67
chr3-90618134-INS-7973

chr3-90620638-INS-7170
chr3-90629090-INS-7080
chr3-90679212-INS-537
chr3-90686737-INS-401
chr3-90700686-INS-75
chr3-90720062-INS-6991
chr3-90825791-INS-401
chr3-90876951-INS-212
chr3-90898187-INS-76
chr3-90906533-INS-243
chr3-90936299-INS-7128
chr3-90964050-INS-280
chr3-91003224-INS-209
chr3-91042623-INS-180
chr3-91112135-INS-56
chr3-91163188-INS-8534
chr3-91172404-INS-7132
chr3-91244455-INS-1003
chr3-91247487-INS-68
chr3-91271404-INS-78
chr3-91272893-INS-60
chr3-91313806-INS-68
chr3-91322606-INS-324
chr3-91323702-INS-207
chr3-91339527-INS-277
chr3-91370363-INS-451
chr3-91373860-INS-1508
chr3-91415262-INS-505
chr3-91452199-INS-86
chr3-91454352-INS-114
chr3-91459953-INS-92
chr3-91460891-INS-324
chr3-91497249-INS-60
chr3-91503212-INS-6477
chr3-91529748-INS-10582
chr3-91534320-INS-59
chr3-91535612-INS-1471
chr3-91621304-INS-1020
chr3-91631360-INS-585
chr3-91723081-INS-3253
chr3-91757803-INS-1776
chr3-91760486-INS-64
chr3-91844446-INS-6196
chr3-91872021-INS-868
chr3-91878233-INS-144
chr3-91884473-INS-58
chr3-91888171-INS-347
chr3-91918957-INS-162
chr3-91932399-INS-6912
chr3-91949855-INS-62
chr3-91972954-INS-528
chr3-91978601-INS-90
chr3-91993251-INS-6352
chr3-92035495-INS-292
chr3-92047578-INS-396
chr3-92052892-INS-64
chr3-92066423-INS-86
chr3-92080389-INS-110
chr3-92084121-INS-54
chr3-92088355-INS-116
chr3-92104061-INS-54
chr3-92108903-INS-80
chr3-92114542-INS-109
chr3-92121429-INS-62
chr3-92175867-INS-654
chr3-92178047-INS-53
chr3-92214671-INS-52
chr3-92224556-INS-465
chr3-92227266-INS-601
chr3-92233643-INS-196
chr3-92237921-INS-981
chr3-92247333-INS-79
chr3-92250880-INS-6391
chr3-92251810-INS-673
chr3-92264200-INS-56
chr3-92273313-INS-189
chr3-92274105-INS-593

chr3-92275511-INS-176
chr3-92289295-INS-889
chr3-92294845-INS-194
chr3-92299997-INS-68
chr3-92301060-INS-220
chr3-92303973-INS-333
chr3-92307247-INS-459
chr3-92313636-INS-158
chr3-92324453-INS-70
chr3-92324627-INS-1000
chr3-92347171-INS-87
chr3-92380429-INS-829
chr3-92402529-INS-486
chr3-92414065-INS-145
chr3-92436993-INS-193
chr3-92467461-INS-53
chr3-92479165-INS-432
chr3-92481872-INS-76
chr3-92482903-INS-177
chr3-92485082-INS-70
chr3-92489235-INS-93
chr3-92503269-INS-6731
chr3-92517659-INS-58
chr3-92537366-INS-56
chr3-92565893-INS-255
chr3-92591826-INS-202
chr3-92604312-INS-119
chr3-92619607-INS-204
chr3-92622392-INS-67
chr3-92622776-INS-932
chr3-92624445-INS-68
chr3-92628739-INS-1672
chr3-92638071-INS-6609
chr3-92657469-INS-198
chr3-92665799-INS-376
chr3-92692785-INS-83
chr3-92694295-INS-221
chr3-92715649-INS-555
chr3-92731645-INS-84
chr3-92770588-INS-7757
chr3-92774619-INS-61
chr3-92800234-INS-142
chr3-92807106-INS-60
chr3-92811808-INS-99
chr3-92813360-INS-148
chr3-92826989-INS-6685
chr3-92859418-INS-6488
chr3-92859653-INS-72
chr3-92865688-INS-935
chr3-92885830-INS-52
chr3-92893001-INS-384
chr3-92903512-INS-6676
chr3-92904800-INS-6705
chr3-92916802-INS-839
chr3-92942374-INS-6403
chr3-92946278-INS-108
chr3-92951390-INS-348
chr3-93002098-INS-128
chr3-93003668-INS-6371
chr3-93013183-INS-8774
chr3-93019278-INS-197
chr3-93028277-INS-1479
chr3-93059885-INS-6516
chr3-93154029-INS-124
chr3-93205304-INS-6443
chr3-93209707-INS-66
chr3-93241702-INS-278
chr3-93243576-INS-51
chr3-93251231-INS-168
chr3-93253211-INS-70
chr3-93257797-INS-96
chr3-93258194-INS-369
chr3-93280849-INS-394
chr3-93294209-INS-66
chr3-93307702-INS-147
chr3-93310034-INS-106
chr3-93312166-INS-64

chr3-93318302-INS-132
chr3-93368994-INS-430
chr3-93379611-INS-71
chr3-93386433-INS-158
chr3-93387569-INS-206
chr3-93393668-INS-397
chr3-93398456-INS-118
chr3-93402528-INS-105
chr3-93408583-INS-6622
chr3-93432608-INS-79
chr3-93434831-INS-332
chr3-93466823-INS-399
chr3-93495313-INS-60
chr3-93502912-INS-66
chr3-93535696-INS-384
chr3-93544625-INS-6387
chr3-93609470-INS-70
chr3-93623156-INS-6580
chr3-93656470-INS-204
chr3-93787161-INS-112
chr3-93861901-INS-8706
chr3-93883154-INS-6442
chr3-93930652-INS-217
chr3-93946924-INS-188
chr3-93959361-INS-81
chr3-93973233-INS-7660
chr3-93981841-INS-70
chr3-94084603-INS-220
chr3-94086168-INS-3307
chr3-94087652-INS-6707
chr3-94099602-INS-110
chr3-94161193-INS-486
chr3-94206156-INS-72
chr3-94228512-INS-196
chr3-94234705-INS-145
chr3-94265749-INS-282
chr3-94266297-INS-74
chr3-94293337-INS-92
chr3-94311307-INS-60
chr3-94329153-INS-199
chr3-94337103-INS-53
chr3-94340312-INS-109
chr3-94359805-INS-96
chr3-94368223-INS-715
chr3-94374305-INS-148
chr3-94375278-INS-821
chr3-94376048-INS-56
chr3-94401402-INS-72
chr3-94428110-INS-313
chr3-94443422-INS-197
chr3-94453268-INS-545
chr3-94464500-INS-162
chr3-94497018-INS-95
chr3-94509444-INS-82
chr3-94510713-INS-201
chr3-94513067-INS-223
chr3-94516291-INS-209
chr3-94535791-INS-51
chr3-94550948-INS-711
chr3-94552658-INS-52
chr3-94560075-INS-229
chr3-94574121-INS-333
chr3-94593770-INS-170
chr3-94596578-INS-63
chr3-94608174-INS-107
chr3-94644314-INS-3887
chr3-94666562-INS-95
chr3-94697128-INS-178
chr3-94705881-INS-104
chr3-94708938-INS-94
chr3-94722924-INS-165
chr3-94726039-INS-202
chr3-94733438-INS-5215
chr3-94765345-INS-113
chr3-94783670-INS-429
chr3-94817644-INS-128
chr3-94841384-INS-83

chr3-94855148-INS-78
chr3-94897829-INS-193
chr3-94904293-INS-54
chr3-94927137-INS-205
chr3-94930743-INS-1384
chr3-94933990-INS-445
chr3-94950635-INS-185
chr3-94954320-INS-2865
chr3-94959574-INS-371
chr3-94970859-INS-673
chr3-94978594-INS-210
chr3-94980242-INS-61
chr3-94996950-INS-441
chr3-94999224-INS-202
chr3-95014648-INS-195
chr3-95019299-INS-207
chr3-95057650-INS-52
chr3-95058446-INS-71
chr3-95058906-INS-212
chr3-95085204-INS-60
chr3-95085702-INS-716
chr3-95088178-INS-104
chr3-95109336-INS-213
chr3-95110706-INS-99
chr3-95113958-INS-111
chr3-95115180-INS-61
chr3-95121213-INS-205
chr3-95128497-INS-52
chr3-95149398-INS-183
chr3-95150657-INS-338
chr3-95157009-INS-285
chr3-95158654-INS-54
chr3-95183546-INS-54
chr3-95211107-INS-135
chr3-95236631-INS-172
chr3-95266471-INS-132
chr3-95287122-INS-296
chr3-95298416-INS-216
chr3-95317603-INS-67
chr3-95350092-INS-90
chr3-95352622-INS-675
chr3-95354856-INS-193
chr3-95379467-INS-226
chr3-95416808-INS-72
chr3-95425110-INS-62
chr3-95431996-INS-70
chr3-95432793-INS-55
chr3-95435395-INS-160
chr3-95449781-INS-172
chr3-95454419-INS-401
chr3-95462304-INS-205
chr3-95464028-INS-447
chr3-95477749-INS-106
chr3-95487385-INS-103
chr3-95489015-INS-129
chr3-95490043-INS-1298
chr3-95511050-INS-209
chr3-95517752-INS-55
chr3-95521034-INS-273
chr3-95527165-INS-200
chr3-95548153-INS-205
chr3-95550497-INS-204
chr3-95554459-INS-175
chr3-95564469-INS-64
chr3-95597293-INS-179
chr3-95612730-INS-104
chr3-95636015-INS-174
chr3-95670833-INS-84
chr3-95699658-INS-195
chr3-95703006-INS-60
chr3-95710293-INS-218
chr3-95733001-INS-106
chr3-95734621-INS-927
chr3-95734636-INS-1150
chr3-95745189-INS-326
chr3-95769544-INS-52
chr3-95772987-INS-284

chr3-95773826-INS-56
chr3-95794915-INS-92
chr3-95795877-INS-212
chr3-95820574-INS-77
chr3-95843491-INS-57
chr3-95861035-INS-211
chr3-95900613-INS-56
chr3-95920977-INS-171
chr3-95923945-INS-60
chr3-95933534-INS-195
chr3-95936092-INS-627
chr3-95944607-INS-910
chr3-95951819-INS-470
chr3-95952271-INS-8158
chr3-95952339-INS-3606
chr3-95967324-INS-201
chr3-95970272-INS-198
chr3-95993077-INS-52
chr3-95996221-INS-75
chr3-96038361-INS-173
chr3-96050296-INS-58
chr3-96051915-INS-199
chr3-96059074-INS-1886
chr3-96093447-INS-63
chr3-96136570-INS-755
chr3-96164070-INS-82
chr3-96198440-INS-188
chr3-96268078-INS-204
chr3-96269514-INS-434
chr3-96325998-INS-100
chr3-96338721-INS-72
chr3-96345603-INS-187
chr3-96351772-INS-59
chr3-96381992-INS-321
chr3-96394543-INS-194
chr3-96405765-INS-7484
chr3-96413655-INS-187
chr3-96429087-INS-212
chr3-96443703-INS-83
chr3-96451383-INS-195
chr3-96480065-INS-256
chr3-96482784-INS-221
chr3-96496287-INS-132
chr3-96511119-INS-182
chr3-96514659-INS-56
chr3-96526121-INS-137
chr3-96545551-INS-708
chr3-96548552-INS-57
chr3-96549521-INS-199
chr3-96568130-INS-60
chr3-96576065-INS-306
chr3-96579891-INS-54
chr3-96610243-INS-205
chr3-96625762-INS-211
chr3-96641378-INS-69
chr3-96647675-INS-110
chr3-96650315-INS-52
chr3-96652248-INS-93
chr3-96658449-INS-99
chr3-96658976-INS-115
chr3-96660980-INS-212
chr3-96666195-INS-210
chr3-96677412-INS-75
chr3-96679763-INS-826
chr3-96702150-INS-268
chr3-96714794-INS-220
chr3-96717148-INS-223
chr3-96730135-INS-216
chr3-96747875-INS-58
chr3-96751961-INS-1456
chr3-96753963-INS-51
chr3-96756124-INS-264
chr3-96763477-INS-66
chr3-96785224-INS-74
chr3-96791401-INS-208
chr3-96809176-INS-58
chr3-96818971-INS-58

chr3-96820591-INS-855
chr3-96869839-INS-299
chr3-96873098-INS-131
chr3-96876898-INS-104
chr3-96878293-INS-81
chr3-96879548-INS-186
chr3-96893444-INS-150
chr3-96899741-INS-81
chr3-96918861-INS-3331
chr3-96951474-INS-58
chr3-96956048-INS-61
chr3-96967767-INS-70
chr3-96976914-INS-105
chr3-96986921-INS-219
chr3-97003974-INS-112
chr3-97005635-INS-508
chr3-97032404-INS-535
chr3-97039003-INS-507
chr3-97041571-INS-7443
chr3-97050170-INS-7226
chr3-97052901-INS-498
chr3-97068838-INS-264
chr3-97069925-INS-101
chr3-97078093-INS-310
chr3-97088453-INS-179
chr3-97092996-INS-141
chr3-97106277-INS-149
chr3-97162781-INS-51
chr3-97168198-INS-96
chr3-97168290-INS-351
chr3-97179371-INS-400
chr3-97197916-INS-198
chr3-97215107-INS-98
chr3-97222419-INS-154
chr3-97228567-INS-146
chr3-97235022-INS-435
chr3-97236376-INS-147
chr3-97286454-INS-321
chr3-97326414-INS-51
chr3-97367116-INS-522
chr3-97369582-INS-202
chr3-97370824-INS-392
chr3-97402917-INS-103
chr3-97435653-INS-6218
chr3-97439479-INS-142
chr3-97442035-INS-121
chr3-97442287-INS-310
chr3-97443057-INS-68
chr3-97452727-INS-244
chr3-97454909-INS-110
chr3-97457068-INS-61
chr3-97458269-INS-186
chr3-97460210-INS-444
chr3-97476019-INS-68
chr3-97497217-INS-444
chr3-97512178-INS-250
chr3-97529552-INS-52
chr3-97560524-INS-156
chr3-97584082-INS-73
chr3-97595862-INS-141
chr3-97633452-INS-68
chr3-97651674-INS-268
chr3-97680546-INS-1166
chr3-97680715-INS-81
chr3-97696629-INS-101
chr3-97714541-INS-147
chr3-97732655-INS-78
chr3-97736738-INS-62
chr3-97739516-INS-746
chr3-97740078-INS-53
chr3-97772816-INS-360
chr3-97774534-INS-81
chr3-97785372-INS-997
chr3-97793834-INS-162
chr3-97797406-INS-204
chr3-97837974-INS-60
chr3-97860878-INS-111

chr3-97863542-INS-150
chr3-97924095-INS-1010
chr3-97924096-INS-467
chr3-97932245-INS-104
chr3-97941718-INS-71
chr3-97963395-INS-67
chr3-97965927-INS-67
chr3-97976186-INS-142
chr3-98003028-INS-60
chr3-98004569-INS-125
chr3-98021785-INS-409
chr3-98039304-INS-190
chr3-98071382-INS-53
chr3-98100835-INS-68
chr3-98107474-INS-155
chr3-98133893-INS-61
chr3-98134811-INS-406
chr3-98159417-INS-177
chr3-98168377-INS-231
chr3-98171013-INS-56
chr3-98172413-INS-398
chr3-98175571-INS-72
chr3-98205042-INS-70
chr3-98210881-INS-55
chr3-98220295-INS-936
chr3-98235537-INS-100
chr3-98251665-INS-358
chr3-98273969-INS-55
chr3-98339810-INS-81
chr3-98341630-INS-95
chr3-98343783-INS-78
chr3-98430450-INS-96
chr3-98453124-INS-308
chr3-98455640-INS-185
chr3-98482227-INS-189
chr3-98487340-INS-343
chr3-98519381-INS-81
chr3-98522886-INS-64
chr3-98532190-INS-309
chr3-98550531-INS-413
chr3-98566001-INS-6298
chr3-98567812-INS-204
chr3-98569821-INS-159
chr3-98571251-INS-6714
chr3-98573952-INS-6792
chr3-98579633-INS-7079
chr3-98645312-INS-6452
chr3-98691965-INS-392
chr3-98715685-INS-70
chr3-98732967-INS-101
chr3-98735597-INS-166
chr3-98801797-INS-340
chr3-98819464-INS-204
chr3-98822765-INS-66
chr3-98824869-INS-549
chr3-98856747-INS-966
chr3-98902035-INS-52
chr3-98905056-INS-664
chr3-98910065-INS-111
chr3-98932031-INS-5313
chr3-98957311-INS-251
chr3-98976391-INS-2782
chr3-98991554-INS-198
chr3-98997007-INS-292
chr3-99001383-INS-86
chr3-99041089-INS-1545
chr3-99044010-INS-198
chr3-99051944-INS-204
chr3-99070643-INS-276
chr3-99090074-INS-156
chr3-99098308-INS-72
chr3-99113535-INS-89
chr3-99166812-INS-80
chr3-99169757-INS-140
chr3-99192734-INS-168
chr3-99200542-INS-104
chr3-99242232-INS-7555

chr3-99296940-INS-270
chr3-99333251-INS-350
chr3-99333901-INS-821
chr3-99336793-INS-6349
chr3-99336795-INS-165
chr3-99349835-INS-3397
chr3-99370585-INS-8787
chr3-99375403-INS-52
chr3-99402907-INS-538
chr3-99428515-INS-136
chr3-99444503-INS-100
chr3-99445639-INS-335
chr3-99473585-INS-54
chr3-99480612-INS-58
chr3-99501997-INS-1030
chr3-99501998-INS-635
chr3-99503287-INS-6733
chr3-99510595-INS-97
chr3-99512981-INS-72
chr3-99543144-INS-908
chr3-99580850-INS-6997
chr3-99649268-INS-80
chr3-99689123-INS-91
chr3-99692673-INS-169
chr3-99734901-INS-6652
chr3-99737697-INS-212
chr3-99745017-INS-354
chr3-99746522-INS-1929
chr3-99752214-INS-555
chr3-99780762-INS-97
chr3-99797589-INS-57
chr3-99821721-INS-69
chr3-99823685-INS-82
chr3-99843805-INS-377
chr3-99845070-INS-210
chr3-99874726-INS-54
chr3-99882346-INS-319
chr3-99887488-INS-414
chr3-99892393-INS-140
chr3-99905620-INS-138
chr3-99917952-INS-630
chr3-99938871-INS-309
chr3-99944632-INS-164
chr3-99950502-INS-74
chr3-99956271-INS-115
chr3-99959626-INS-637
chr3-99982145-INS-221
chr3-99994028-INS-350
chr3-100015951-INS-52
chr3-100021433-INS-7113
chr3-100021592-INS-373
chr3-100043841-INS-6316
chr3-100061061-INS-56
chr3-100067436-INS-195
chr3-100082121-INS-76
chr3-100084837-INS-336
chr3-100092201-INS-121
chr3-100110135-INS-54
chr3-100129395-INS-686
chr3-100131085-INS-519
chr3-100134326-INS-155
chr3-100189782-INS-73
chr3-100194104-INS-89
chr3-100198579-INS-656
chr3-100200556-INS-605
chr3-100204264-INS-98
chr3-100212248-INS-335
chr3-100215321-INS-200
chr3-100227989-INS-96
chr3-100238520-INS-66
chr3-100246134-INS-123
chr3-100253068-INS-6928
chr3-100263202-INS-317
chr3-100280454-INS-304
chr3-100294890-INS-545
chr3-100295495-INS-1950
chr3-100322819-INS-338

chr3-100360810-INS-810
chr3-100381331-INS-80
chr3-100400033-INS-56
chr3-100450505-INS-152
chr3-100450917-INS-133
chr3-100453254-INS-153
chr3-100458240-INS-240
chr3-100536641-INS-295
chr3-100614989-INS-90
chr3-100619848-INS-378
chr3-100666101-INS-112
chr3-100669113-INS-87
chr3-100670938-INS-59
chr3-100679715-INS-52
chr3-100686315-INS-158
chr3-100719413-INS-70
chr3-100760513-INS-139
chr3-100790579-INS-204
chr3-100794222-INS-63
chr3-100801033-INS-52
chr3-100810085-INS-530
chr3-100811428-INS-496
chr3-100819698-INS-54
chr3-100849509-INS-69
chr3-100856081-INS-68
chr3-100863313-INS-106
chr3-100864329-INS-153
chr3-100877753-INS-233
chr3-100887356-INS-115
chr3-100901484-INS-72
chr3-100917666-INS-58
chr3-100942899-INS-60
chr3-100961741-INS-176
chr3-101031582-INS-51
chr3-101039795-INS-235
chr3-101056867-INS-87
chr3-101080404-INS-174
chr3-101082764-INS-176
chr3-101095871-INS-1888
chr3-101135056-INS-168
chr3-101166990-INS-90
chr3-101197286-INS-351
chr3-101201020-INS-58
chr3-101226801-INS-318
chr3-101276165-INS-84
chr3-101308026-INS-92
chr3-101321573-INS-213
chr3-101379778-INS-51
chr3-101400579-INS-104
chr3-101416866-INS-68
chr3-101422677-INS-244
chr3-101449568-INS-865
chr3-101489839-INS-55
chr3-101520827-INS-50
chr3-101554591-INS-89
chr3-101557782-INS-286
chr3-101565999-INS-70
chr3-101570631-INS-420
chr3-101606807-INS-440
chr3-101621248-INS-507
chr3-101622815-INS-132
chr3-101635598-INS-73
chr3-101637067-INS-53
chr3-101645891-INS-76
chr3-101648876-INS-83
chr3-101652319-INS-232
chr3-101656970-INS-80
chr3-101669454-INS-52
chr3-101716996-INS-203
chr3-101719324-INS-3382
chr3-101728575-INS-117
chr3-101806731-INS-80
chr3-101878232-INS-218
chr3-101897155-INS-58
chr3-101901975-INS-61
chr3-101938918-INS-60
chr3-101951688-INS-69

chr3-101968684-INS-80
chr3-101973601-INS-57
chr3-101976006-INS-56
chr3-101987446-INS-56
chr3-101996679-INS-6460
chr3-102000775-INS-105
chr3-102030706-INS-79
chr3-102040457-INS-53
chr3-102044533-INS-180
chr3-102077306-INS-126
chr3-102097995-INS-506
chr3-102125804-INS-74
chr3-102197866-INS-52
chr3-102200324-INS-54
chr3-102208625-INS-112
chr3-102220884-INS-111
chr3-102222540-INS-2449
chr3-102226488-INS-367
chr3-102235853-INS-102
chr3-102253049-INS-382
chr3-102269764-INS-54
chr3-102274523-INS-70
chr3-102309355-INS-115
chr3-102324438-INS-579
chr3-102350554-INS-485
chr3-102356212-INS-440
chr3-102359384-INS-96
chr3-102399385-INS-56
chr3-102399683-INS-108
chr3-102432034-INS-52
chr3-102453428-INS-218
chr3-102455213-INS-51
chr3-102465975-INS-198
chr3-102474605-INS-1567
chr3-102477299-INS-2325
chr3-102482779-INS-754
chr3-102485850-INS-85
chr3-102488134-INS-263
chr3-102514920-INS-147
chr3-102553500-INS-126
chr3-102556858-INS-167
chr3-102614517-INS-444
chr3-102629041-INS-272
chr3-102631511-INS-56
chr3-102636411-INS-133
chr3-102644641-INS-81
chr3-102665553-INS-329
chr3-102684606-INS-356
chr3-102693682-INS-198
chr3-102700980-INS-631
chr3-102702681-INS-272
chr3-102710827-INS-673
chr3-102717535-INS-2953
chr3-102757669-INS-995
chr3-102782543-INS-118
chr3-102828151-INS-386
chr3-102835575-INS-839
chr3-102851283-INS-202
chr3-102861508-INS-128
chr3-102866653-INS-5523
chr3-102867109-INS-108
chr3-102899196-INS-118
chr3-102916140-INS-458
chr3-102917244-INS-166
chr3-102919858-INS-55
chr3-102923673-INS-76
chr3-102925276-INS-80
chr3-102985656-INS-73
chr3-102999291-INS-307
chr3-103001024-INS-75
chr3-103013951-INS-254
chr3-103032445-INS-192
chr3-103039828-INS-63
chr3-103041188-INS-2890
chr3-103042134-INS-95
chr3-103045502-INS-73
chr3-103047981-INS-540

chr3-103047989-INS-56
chr3-103050908-INS-68
chr3-103065210-INS-165
chr3-103072985-INS-170
chr3-103079594-INS-88
chr3-103083217-INS-288
chr3-103092184-INS-57
chr3-103095440-INS-77
chr3-103102847-INS-55
chr3-103109547-INS-574
chr3-103113894-INS-176
chr3-103122759-INS-514
chr3-103123278-INS-198
chr3-103124475-INS-215
chr3-103129331-INS-210
chr3-103130835-INS-174
chr3-103142104-INS-75
chr3-103144349-INS-301
chr3-103161946-INS-210
chr3-103168611-INS-76
chr3-103182395-INS-159
chr3-103197936-INS-61
chr3-103205270-INS-139
chr3-103207130-INS-193
chr3-103210120-INS-82
chr3-103229875-INS-227
chr3-103235719-INS-56
chr3-103248526-INS-56
chr3-103288093-INS-251
chr3-103295842-INS-300
chr3-103310984-INS-109
chr3-103317672-INS-203
chr3-103324012-INS-66
chr3-103326058-INS-216
chr3-103339129-INS-60
chr3-103351891-INS-901
chr3-103375759-INS-610
chr3-103446333-INS-212
chr3-103450437-INS-56
chr3-103465262-INS-372
chr3-103504601-INS-151
chr3-103534969-INS-170
chr3-103581005-INS-583
chr3-103583038-INS-380
chr3-103598613-INS-156
chr3-103613105-INS-63
chr3-103640211-INS-56
chr3-103664773-INS-134
chr3-103668790-INS-350
chr3-103701825-INS-5226
chr3-103702569-INS-64
chr3-103725265-INS-244
chr3-103737340-INS-124
chr3-103737341-INS-140
chr3-103745189-INS-204
chr3-103762073-INS-116
chr3-103766692-INS-193
chr3-103783078-INS-54
chr3-103789977-INS-115
chr3-103789977-INS-117
chr3-103798052-INS-103
chr3-103808891-INS-76
chr3-103826205-INS-54
chr3-103846833-INS-204
chr3-103850875-INS-63
chr3-103859833-INS-105
chr3-103865316-INS-52
chr3-103868387-INS-322
chr3-103901700-INS-52
chr3-103916156-INS-664
chr3-103918586-INS-622
chr3-103935741-INS-66
chr3-103949161-INS-625
chr3-103972559-INS-619
chr3-103984242-INS-63
chr3-104013630-INS-837
chr3-104042189-INS-195

chr3-104064429-INS-216
chr3-104082180-INS-216
chr3-104092079-INS-113
chr3-104109715-INS-459
chr3-104112635-INS-1322
chr3-104132290-INS-56
chr3-104137197-INS-94
chr3-104138584-INS-75
chr3-104151401-INS-1891
chr3-104159069-INS-118
chr3-104182683-INS-6412
chr3-104186055-INS-51
chr3-104215448-INS-108
chr3-104219700-INS-318
chr3-104221598-INS-83
chr3-104254492-INS-77
chr3-104258818-INS-84
chr3-104269026-INS-331
chr3-104279623-INS-64
chr3-104299518-INS-523
chr3-104311970-INS-68
chr3-104318679-INS-58
chr3-104326302-INS-51
chr3-104334947-INS-98
chr3-104368371-INS-137
chr3-104374823-INS-81
chr3-104391596-INS-61
chr3-104400041-INS-62
chr3-104421267-INS-357
chr3-104441625-INS-96
chr3-104443496-INS-124
chr3-104446510-INS-94
chr3-104446722-INS-391
chr3-104457672-INS-74
chr3-104464975-INS-56
chr3-104465966-INS-190
chr3-104467372-INS-87
chr3-104469936-INS-102
chr3-104477103-INS-906
chr3-104477203-INS-170
chr3-104513101-INS-220
chr3-104530916-INS-60
chr3-104575405-INS-207
chr3-104593365-INS-78
chr3-104597525-INS-60
chr3-104604605-INS-52
chr3-104611204-INS-150
chr3-104635582-INS-320
chr3-104639071-INS-144
chr3-104660431-INS-98
chr3-104663217-INS-92
chr3-104672642-INS-203
chr3-104699371-INS-212
chr3-104705971-INS-174
chr3-104745458-INS-6460
chr3-104745597-INS-60
chr3-104807578-INS-1227
chr3-104820535-INS-76
chr3-104824660-INS-52
chr3-104862264-INS-202
chr3-104866823-INS-136
chr3-104894428-INS-70
chr3-104894769-INS-120
chr3-104929338-INS-767
chr3-104935766-INS-112
chr3-104935799-INS-812
chr3-104936501-INS-340
chr3-104936912-INS-177
chr3-104941759-INS-356
chr3-104972671-INS-235
chr3-104978977-INS-215
chr3-104981149-INS-5357
chr3-104991461-INS-206
chr3-104996947-INS-183
chr3-104999573-INS-164
chr3-105001866-INS-52
chr3-105011031-INS-74

chr3-105018866-INS-193
chr3-105028630-INS-62
chr3-105057163-INS-6690
chr3-105063471-INS-6429
chr3-105064671-INS-165
chr3-105066468-INS-698
chr3-105067753-INS-6227
chr3-105071211-INS-116
chr3-105128758-INS-202
chr3-105139444-INS-723
chr3-105146747-INS-72
chr3-105169052-INS-242
chr3-105191252-INS-78
chr3-105201265-INS-692
chr3-105204103-INS-59
chr3-105208078-INS-6941
chr3-105234011-INS-360
chr3-105239040-INS-90
chr3-105254979-INS-131
chr3-105266899-INS-7120
chr3-105300114-INS-75
chr3-105301659-INS-8190
chr3-105302098-INS-63
chr3-105316645-INS-427
chr3-105355768-INS-78
chr3-105390254-INS-71
chr3-105398993-INS-110
chr3-105427546-INS-1617
chr3-105438580-INS-94
chr3-105463243-INS-90
chr3-105470813-INS-192
chr3-105486138-INS-110
chr3-105521662-INS-6838
chr3-105525754-INS-106
chr3-105526227-INS-84
chr3-105542036-INS-401
chr3-105563894-INS-54
chr3-105596834-INS-368
chr3-105598555-INS-5249
chr3-105608047-INS-86
chr3-105623261-INS-94
chr3-105626786-INS-302
chr3-105652439-INS-242
chr3-105667026-INS-67
chr3-105693515-INS-201
chr3-105701801-INS-86
chr3-105704938-INS-145
chr3-105723810-INS-718
chr3-105731488-INS-108
chr3-105732172-INS-62
chr3-105737111-INS-102
chr3-105748081-INS-128
chr3-105756626-INS-73
chr3-105788383-INS-76
chr3-105811008-INS-124
chr3-105811208-INS-113
chr3-105813659-INS-60
chr3-105830956-INS-58
chr3-105886354-INS-655
chr3-105893021-INS-55
chr3-105893744-INS-147
chr3-105952174-INS-186
chr3-105961533-INS-429
chr3-105962320-INS-6442
chr3-105968989-INS-180
chr3-105988600-INS-442
chr3-105998557-INS-684
chr3-106009440-INS-65
chr3-106062555-INS-2362
chr3-106150033-INS-6064
chr3-106153544-INS-515
chr3-106173877-INS-208
chr3-106217393-INS-478
chr3-106222252-INS-527
chr3-106225696-INS-58
chr3-106238581-INS-108
chr3-106243043-INS-651

chr3-106290141-INS-52
chr3-106313076-INS-112
chr3-106314464-INS-457
chr3-106348099-INS-6853
chr3-106377179-INS-52
chr3-106387299-INS-60
chr3-106402706-INS-99
chr3-106404936-INS-211
chr3-106422228-INS-66
chr3-106439657-INS-184
chr3-106491272-INS-397
chr3-106506501-INS-64
chr3-106539746-INS-57
chr3-106595611-INS-6919
chr3-106608500-INS-691
chr3-106644661-INS-204
chr3-106654727-INS-139
chr3-106675419-INS-74
chr3-106685975-INS-901
chr3-106698072-INS-99
chr3-106698728-INS-2371
chr3-106722325-INS-143
chr3-106737054-INS-507
chr3-106738954-INS-186
chr3-106740167-INS-74
chr3-106747921-INS-75
chr3-106755093-INS-266
chr3-106759921-INS-4958
chr3-106782095-INS-6621
chr3-106815443-INS-319
chr3-106816186-INS-96
chr3-106840399-INS-74
chr3-106864120-INS-122
chr3-106870766-INS-109
chr3-106904771-INS-738
chr3-106907964-INS-602
chr3-106911641-INS-95
chr3-106920181-INS-52
chr3-106921440-INS-82
chr3-106958811-INS-176
chr3-106966938-INS-3120
chr3-107015182-INS-74
chr3-107027663-INS-190
chr3-107041729-INS-1138
chr3-107078896-INS-56
chr3-107105130-INS-221
chr3-107114451-INS-73
chr3-107133843-INS-170
chr3-107159489-INS-108
chr3-107180764-INS-64
chr3-107181497-INS-201
chr3-107196036-INS-80
chr3-107198851-INS-52
chr3-107212715-INS-74
chr3-107219765-INS-74
chr3-107264925-INS-82
chr3-107298241-INS-145
chr3-107317845-INS-86
chr3-107342238-INS-68
chr3-107356166-INS-82
chr3-107386027-INS-406
chr3-107387087-INS-420
chr3-107387088-INS-168
chr3-107389118-INS-148
chr3-107414553-INS-66
chr3-107430576-INS-1645
chr3-107447349-INS-141
chr3-107562707-INS-56
chr3-107629014-INS-84
chr3-107732040-INS-64
chr3-107734674-INS-90
chr3-107764924-INS-187
chr3-107769584-INS-424
chr3-107785783-INS-92
chr3-107797212-INS-121
chr3-107842577-INS-54
chr3-107848733-INS-357

chr3-107887790-INS-99
chr3-107922659-INS-220
chr3-107940559-INS-200
chr3-107942567-INS-152
chr3-107945289-INS-60
chr3-107959693-INS-398
chr3-107966301-INS-138
chr3-107966905-INS-682
chr3-107970558-INS-189
chr3-108000083-INS-424
chr3-108017173-INS-75
chr3-108055337-INS-54
chr3-108077339-INS-202
chr3-108097271-INS-60
chr3-108102509-INS-52
chr3-108125893-INS-120
chr3-108136962-INS-1215
chr3-108154631-INS-87
chr3-108156096-INS-198
chr3-108157969-INS-170
chr3-108167367-INS-74
chr3-108182737-INS-120
chr3-108197155-INS-200
chr3-108198182-INS-447
chr3-108207026-INS-121
chr3-108223210-INS-61
chr3-108238809-INS-206
chr3-108245910-INS-207
chr3-108251060-INS-198
chr3-108276191-INS-5252
chr3-108277884-INS-53
chr3-108288010-INS-70
chr3-108292063-INS-260
chr3-108297882-INS-331
chr3-108298240-INS-121
chr3-108344184-INS-64
chr3-108367660-INS-52
chr3-108439450-INS-59
chr3-108448847-INS-107
chr3-108451664-INS-211
chr3-108453177-INS-362
chr3-108454422-INS-81
chr3-108493286-INS-536
chr3-108522074-INS-193
chr3-108530869-INS-202
chr3-108533344-INS-434
chr3-108543935-INS-66
chr3-108557049-INS-534
chr3-108596281-INS-123
chr3-108605496-INS-299
chr3-108610769-INS-96
chr3-108612376-INS-200
chr3-108625043-INS-2571
chr3-108660427-INS-61
chr3-108712096-INS-162
chr3-108724940-INS-671
chr3-108730598-INS-172
chr3-108737611-INS-428
chr3-108771098-INS-69
chr3-108825164-INS-401
chr3-108827533-INS-300
chr3-108887654-INS-56
chr3-108898247-INS-58
chr3-108906799-INS-82
chr3-108906866-INS-522
chr3-108906991-INS-1680
chr3-108923418-INS-64
chr3-108927091-INS-52
chr3-108933243-INS-60
chr3-108944449-INS-400
chr3-108951496-INS-521
chr3-108956678-INS-120
chr3-108964399-INS-113
chr3-108965832-INS-56
chr3-108995413-INS-242
chr3-109000931-INS-203
chr3-109025855-INS-6640

chr3-109044837-INS-69
chr3-109045473-INS-129
chr3-109061557-INS-205
chr3-109065167-INS-70
chr3-109065992-INS-305
chr3-109095600-INS-68
chr3-109096560-INS-230
chr3-109101902-INS-215
chr3-109122906-INS-6715
chr3-109159219-INS-2256
chr3-109176002-INS-376
chr3-109195572-INS-341
chr3-109202047-INS-83
chr3-109215877-INS-139
chr3-109218086-INS-371
chr3-109235123-INS-68
chr3-109239474-INS-313
chr3-109245430-INS-329
chr3-109249499-INS-56
chr3-109266146-INS-150
chr3-109275569-INS-242
chr3-109276653-INS-68
chr3-109288317-INS-379
chr3-109294393-INS-199
chr3-109300919-INS-90
chr3-109310912-INS-6824
chr3-109312877-INS-60
chr3-109333256-INS-183
chr3-109339098-INS-56
chr3-109349309-INS-52
chr3-109369991-INS-64
chr3-109390210-INS-166
chr3-109403228-INS-95
chr3-109404911-INS-229
chr3-109427525-INS-134
chr3-109461728-INS-574
chr3-109480196-INS-77
chr3-109496143-INS-4892
chr3-109529529-INS-76
chr3-109532634-INS-70
chr3-109537177-INS-58
chr3-109584478-INS-76
chr3-109601217-INS-624
chr3-109606546-INS-407
chr3-109611122-INS-4746
chr3-109625953-INS-81
chr3-109662453-INS-90
chr3-109698543-INS-543
chr3-109703513-INS-55
chr3-109707136-INS-116
chr3-109720182-INS-120
chr3-109798983-INS-576
chr3-109809989-INS-101
chr3-109814625-INS-828
chr3-109829258-INS-210
chr3-109862471-INS-64
chr3-109871389-INS-7136
chr3-109880620-INS-210
chr3-109882400-INS-108
chr3-109915860-INS-7195
chr3-109926312-INS-184
chr3-109933849-INS-261
chr3-109957230-INS-442
chr3-109974626-INS-6918
chr3-109987549-INS-190
chr3-109988654-INS-80
chr3-109993833-INS-556
chr3-110011098-INS-64
chr3-110022936-INS-536
chr3-110053992-INS-51
chr3-110078509-INS-87
chr3-110085729-INS-238
chr3-110090599-INS-52
chr3-110126029-INS-302
chr3-110146150-INS-62
chr3-110156115-INS-969
chr3-110161589-INS-484

chr3-110194686-INS-269
chr3-110210046-INS-971
chr3-110218135-INS-62
chr3-110218294-INS-1291
chr3-110226777-INS-56
chr3-110256559-INS-52
chr3-110287872-INS-78
chr3-110316468-INS-677
chr3-110318476-INS-115
chr3-110331368-INS-60
chr3-110343008-INS-197
chr3-110402125-INS-6363
chr3-110403578-INS-52
chr3-110410504-INS-56
chr3-110452332-INS-234
chr3-110463297-INS-487
chr3-110468265-INS-56
chr3-110495555-INS-5250
chr3-110513207-INS-344
chr3-110526932-INS-55
chr3-110549729-INS-80
chr3-110577586-INS-398
chr3-110590596-INS-58
chr3-110620392-INS-7256
chr3-110663782-INS-434
chr3-110676558-INS-598
chr3-110693665-INS-7196
chr3-110694137-INS-122
chr3-110709182-INS-389
chr3-110748639-INS-52
chr3-110770482-INS-173
chr3-110775608-INS-58
chr3-110777327-INS-56
chr3-110780121-INS-163
chr3-110798477-INS-64
chr3-110804318-INS-57
chr3-110813638-INS-270
chr3-110837893-INS-55
chr3-110853054-INS-87
chr3-110864461-INS-6954
chr3-110870333-INS-518
chr3-110885098-INS-59
chr3-110891512-INS-392
chr3-110903101-INS-100
chr3-110907099-INS-167
chr3-110916204-INS-291
chr3-110930190-INS-96
chr3-110980961-INS-104
chr3-111030989-INS-118
chr3-111065715-INS-73
chr3-111067958-INS-112
chr3-111069233-INS-191
chr3-111104069-INS-96
chr3-111117133-INS-173
chr3-111144921-INS-199
chr3-111169417-INS-199
chr3-111187794-INS-66
chr3-111203925-INS-282
chr3-111206664-INS-77
chr3-111212667-INS-6392
chr3-111242115-INS-4507
chr3-111255119-INS-80
chr3-111260552-INS-57
chr3-111274121-INS-80
chr3-111293102-INS-110
chr3-111298626-INS-94
chr3-111310031-INS-52
chr3-111322001-INS-131
chr3-111328928-INS-67
chr3-111382905-INS-933
chr3-111395328-INS-699
chr3-111404846-INS-59
chr3-111405584-INS-447
chr3-111407587-INS-86
chr3-111444653-INS-4595
chr3-111456106-INS-63
chr3-111496604-INS-67

chr3-111501024-INS-148
chr3-111527079-INS-105
chr3-111544335-INS-144
chr3-111560794-INS-66
chr3-111568265-INS-457
chr3-111594268-INS-6371
chr3-111600013-INS-206
chr3-111613286-INS-6274
chr3-111619953-INS-160
chr3-111628897-INS-200
chr3-111662396-INS-64
chr3-111679467-INS-121
chr3-111690735-INS-114
chr3-111696156-INS-489
chr3-111697496-INS-4925
chr3-111698243-INS-7653
chr3-111699585-INS-60
chr3-111699971-INS-6806
chr3-111713326-INS-88
chr3-111737066-INS-592
chr3-111751213-INS-4017
chr3-111752650-INS-351
chr3-111754012-INS-491
chr3-111756997-INS-1840
chr3-111765716-INS-398
chr3-111781423-INS-54
chr3-111791930-INS-77
chr3-111801932-INS-524
chr3-111839611-INS-922
chr3-111850130-INS-6392
chr3-111860987-INS-686
chr3-111866660-INS-203
chr3-111923226-INS-4364
chr3-111932215-INS-567
chr3-111940878-INS-6326
chr3-111950914-INS-1025
chr3-111964555-INS-200
chr3-112004498-INS-537
chr3-112013733-INS-52
chr3-112047630-INS-349
chr3-112050487-INS-158
chr3-112098566-INS-3996
chr3-112099772-INS-213
chr3-112166680-INS-8402
chr3-112179476-INS-9947
chr3-112252633-INS-98
chr3-112292487-INS-81
chr3-112321792-INS-6509
chr3-112323669-INS-562
chr3-112329846-INS-6246
chr3-112329971-INS-469
chr3-112335066-INS-52
chr3-112337015-INS-6934
chr3-112340726-INS-6718
chr3-112355570-INS-55
chr3-112370190-INS-159
chr3-112399471-INS-315
chr3-112441481-INS-6478
chr3-112465203-INS-500
chr3-112513062-INS-5544
chr3-112522955-INS-137
chr3-112533837-INS-357
chr3-112557060-INS-53
chr3-112561359-INS-88
chr3-112574435-INS-58
chr3-112609276-INS-107
chr3-112611904-INS-484
chr3-112613365-INS-66
chr3-112647299-INS-6485
chr3-112700459-INS-52
chr3-112712658-INS-600
chr3-112726592-INS-70
chr3-112732189-INS-84
chr3-112778693-INS-61
chr3-112797764-INS-1212
chr3-112805094-INS-58
chr3-112820336-INS-60

chr3-112856126-INS-109
chr3-112856559-INS-116
chr3-112879841-INS-72
chr3-112896307-INS-52
chr3-112904652-INS-153
chr3-112970601-INS-6948
chr3-112971925-INS-322
chr3-112988310-INS-731
chr3-113016223-INS-86
chr3-113038384-INS-727
chr3-113075485-INS-1010
chr3-113076362-INS-7931
chr3-113082830-INS-377
chr3-113089945-INS-472
chr3-113155555-INS-390
chr3-113317395-INS-212
chr3-113321191-INS-641
chr3-113330748-INS-234
chr3-113333382-INS-498
chr3-113336616-INS-761
chr3-113340957-INS-407
chr3-113345113-INS-176
chr3-113349010-INS-57
chr3-113372584-INS-914
chr3-113376860-INS-82
chr3-113396438-INS-6246
chr3-113402835-INS-56
chr3-113428377-INS-1822
chr3-113443605-INS-493
chr3-113453715-INS-165
chr3-113458484-INS-64
chr3-113460966-INS-69
chr3-113482323-INS-358
chr3-113500947-INS-806
chr3-113547340-INS-57
chr3-113563057-INS-94
chr3-113573044-INS-809
chr3-113586457-INS-336
chr3-113604448-INS-1272
chr3-113607872-INS-1731
chr3-113617359-INS-11048
chr3-113621622-INS-56
chr3-113635379-INS-188
chr3-113655506-INS-5995
chr3-113668091-INS-78
chr3-113692683-INS-6180
chr3-113694331-INS-6591
chr3-113716483-INS-56
chr3-113717780-INS-281
chr3-113723793-INS-80
chr3-113741766-INS-6375
chr3-113742553-INS-6625
chr3-113745828-INS-60
chr3-113748492-INS-343
chr3-113766554-INS-149
chr3-113771079-INS-257
chr3-113781431-INS-89
chr3-113783071-INS-52
chr3-113815466-INS-147
chr3-113851031-INS-52
chr3-113864127-INS-6456
chr3-113897301-INS-176
chr3-113898451-INS-75
chr3-113902435-INS-332
chr3-113917564-INS-698
chr3-113941691-INS-78
chr3-113949485-INS-298
chr3-113960557-INS-292
chr3-113961436-INS-144
chr3-113963091-INS-971
chr3-113969887-INS-179
chr3-114034699-INS-495
chr3-114042675-INS-118
chr3-114047781-INS-598
chr3-114051570-INS-112
chr3-114077889-INS-6489
chr3-114097959-INS-5003

chr3-114109630-INS-53
chr3-114133695-INS-65
chr3-114149375-INS-444
chr3-114180842-INS-766
chr3-114195830-INS-54
chr3-114197822-INS-737
chr3-114227848-INS-452
chr3-114240078-INS-2666
chr3-114256346-INS-79
chr3-114259787-INS-88
chr3-114278533-INS-61
chr3-114278853-INS-6717
chr3-114281287-INS-112
chr3-114285239-INS-58
chr3-114308375-INS-65
chr3-114310458-INS-86
chr3-114346959-INS-1032
chr3-114361818-INS-118
chr3-114364661-INS-58
chr3-114404205-INS-1864
chr3-114459479-INS-72
chr3-114552771-INS-3224
chr3-114564230-INS-101
chr3-114569957-INS-350
chr3-114579005-INS-338
chr3-114585133-INS-479
chr3-114613350-INS-60
chr3-114638524-INS-393
chr3-114653383-INS-231
chr3-114670155-INS-233
chr3-114686090-INS-53
chr3-114737651-INS-437
chr3-114752345-INS-95
chr3-114785227-INS-885
chr3-114808516-INS-57
chr3-114830126-INS-404
chr3-114839826-INS-506
chr3-114851961-INS-5339
chr3-114901226-INS-360
chr3-114905745-INS-6894
chr3-114912283-INS-64
chr3-114920519-INS-107
chr3-114958346-INS-57
chr3-114976901-INS-5720
chr3-115002252-INS-250
chr3-115008981-INS-2490
chr3-115018404-INS-568
chr3-115024650-INS-114
chr3-115039879-INS-58
chr3-115052233-INS-59
chr3-115058973-INS-7240
chr3-115085783-INS-197
chr3-115096972-INS-60
chr3-115102619-INS-405
chr3-115112952-INS-205
chr3-115122843-INS-164
chr3-115138472-INS-133
chr3-115184227-INS-108
chr3-115186131-INS-387
chr3-115206036-INS-209
chr3-115210216-INS-7206
chr3-115212069-INS-1685
chr3-115220338-INS-203
chr3-115227212-INS-651
chr3-115228688-INS-67
chr3-115239121-INS-67
chr3-115248471-INS-58
chr3-115260877-INS-5838
chr3-115264108-INS-680
chr3-115271026-INS-54
chr3-115282643-INS-217
chr3-115300611-INS-4651
chr3-115305015-INS-1052
chr3-115342911-INS-97
chr3-115345489-INS-7115
chr3-115359620-INS-526
chr3-115364124-INS-84

chr3-115372845-INS-110
chr3-115373198-INS-60
chr3-115407029-INS-6632
chr3-115418581-INS-54
chr3-115421055-INS-207
chr3-115434150-INS-78
chr3-115446815-INS-486
chr3-115452746-INS-52
chr3-115460659-INS-157
chr3-115465147-INS-285
chr3-115475872-INS-809
chr3-115475964-INS-196
chr3-115480323-INS-399
chr3-115501064-INS-311
chr3-115514312-INS-358
chr3-115517366-INS-206
chr3-115528001-INS-203
chr3-115538503-INS-72
chr3-115545969-INS-164
chr3-115560430-INS-66
chr3-115585889-INS-104
chr3-115593686-INS-66
chr3-115604453-INS-60
chr3-115614996-INS-318
chr3-115624976-INS-131
chr3-115632904-INS-203
chr3-115645924-INS-7111
chr3-115653541-INS-136
chr3-115661923-INS-753
chr3-115676126-INS-77
chr3-115684462-INS-200
chr3-115700028-INS-897
chr3-115700267-INS-3334
chr3-115700520-INS-1661
chr3-115700718-INS-2174
chr3-115700893-INS-197
chr3-115722018-INS-58
chr3-115726241-INS-6240
chr3-115732931-INS-273
chr3-115744334-INS-96
chr3-115762108-INS-6401
chr3-115783601-INS-76
chr3-115794639-INS-109
chr3-115803318-INS-64
chr3-115823038-INS-61
chr3-115831177-INS-6863
chr3-115837430-INS-63
chr3-115846437-INS-851
chr3-115852411-INS-51
chr3-115853208-INS-66
chr3-115857331-INS-77
chr3-115866093-INS-138
chr3-115934199-INS-391
chr3-116000332-INS-890
chr3-116029454-INS-267
chr3-116041778-INS-117
chr3-116048190-INS-51
chr3-116079672-INS-61
chr3-116085201-INS-64
chr3-116107408-INS-55
chr3-116116670-INS-140
chr3-116153085-INS-65
chr3-116162443-INS-163
chr3-116166646-INS-61
chr3-116189455-INS-58
chr3-116207444-INS-55
chr3-116211461-INS-154
chr3-116263580-INS-123
chr3-116266884-INS-310
chr3-116280982-INS-201
chr3-116283874-INS-789
chr3-116286518-INS-172
chr3-116287468-INS-69
chr3-116290817-INS-425
chr3-116312167-INS-106
chr3-116313182-INS-386
chr3-116325403-INS-60

chr3-116343070-INS-60
chr3-116362321-INS-194
chr3-116368507-INS-105
chr3-116374492-INS-105
chr3-116385930-INS-212
chr3-116442957-INS-60
chr3-116460003-INS-212
chr3-116485083-INS-220
chr3-116486370-INS-62
chr3-116510558-INS-428
chr3-116512229-INS-51
chr3-116512899-INS-112
chr3-116522696-INS-151
chr3-116527247-INS-498
chr3-116530950-INS-1796
chr3-116536621-INS-51
chr3-116544720-INS-172
chr3-116553083-INS-144
chr3-116586859-INS-179
chr3-116604602-INS-79
chr3-116606179-INS-728
chr3-116635371-INS-165
chr3-116636047-INS-68
chr3-116642283-INS-90
chr3-116655963-INS-230
chr3-116656710-INS-185
chr3-116659535-INS-138
chr3-116662642-INS-110
chr3-116670481-INS-492
chr3-116692051-INS-204
chr3-116720382-INS-263
chr3-116726685-INS-147
chr3-116743566-INS-153
chr3-116752524-INS-168
chr3-116756842-INS-195
chr3-116757361-INS-85
chr3-116768353-INS-453
chr3-116769868-INS-6417
chr3-116771018-INS-208
chr3-116774380-INS-293
chr3-116777958-INS-322
chr3-116794873-INS-54
chr3-116822683-INS-52
chr3-116844191-INS-173
chr3-116844383-INS-195
chr3-116878445-INS-2914
chr3-116878551-INS-6532
chr3-116881253-INS-568
chr3-116885303-INS-78
chr3-116887834-INS-72
chr3-116888667-INS-490
chr3-116900576-INS-52
chr3-116912554-INS-98
chr3-116916909-INS-75
chr3-116919543-INS-366
chr3-116924781-INS-640
chr3-116929583-INS-150
chr3-116943914-INS-7295
chr3-116955244-INS-223
chr3-116978357-INS-52
chr3-116980522-INS-214
chr3-116990056-INS-6255
chr3-117025859-INS-369
chr3-117034398-INS-93
chr3-117045879-INS-118
chr3-117076020-INS-603
chr3-117089466-INS-315
chr3-117090894-INS-64
chr3-117102180-INS-61
chr3-117125914-INS-895
chr3-117149661-INS-63
chr3-117170086-INS-67
chr3-117186680-INS-399
chr3-117208383-INS-107
chr3-117218047-INS-98
chr3-117221920-INS-534
chr3-117237261-INS-200

chr3-117248702-INS-143
chr3-117255851-INS-54
chr3-117269795-INS-58
chr3-117281259-INS-510
chr3-117297101-INS-267
chr3-117298464-INS-6470
chr3-117298497-INS-6725
chr3-117301325-INS-6567
chr3-117314933-INS-414
chr3-117331886-INS-497
chr3-117362887-INS-229
chr3-117364216-INS-106
chr3-117374446-INS-553
chr3-117388037-INS-60
chr3-117390758-INS-175
chr3-117400865-INS-212
chr3-117412022-INS-72
chr3-117413402-INS-91
chr3-117423374-INS-427
chr3-117438308-INS-150
chr3-117450263-INS-197
chr3-117458493-INS-874
chr3-117471546-INS-330
chr3-117506997-INS-156
chr3-117543413-INS-104
chr3-117556943-INS-178
chr3-117565029-INS-56
chr3-117566205-INS-360
chr3-117604016-INS-342
chr3-117606861-INS-60
chr3-117631097-INS-749
chr3-117663399-INS-205
chr3-117669506-INS-576
chr3-117676903-INS-63
chr3-117678257-INS-155
chr3-117681866-INS-817
chr3-117696589-INS-232
chr3-117699408-INS-519
chr3-117753504-INS-339
chr3-117771399-INS-714
chr3-117781989-INS-75
chr3-117788188-INS-237
chr3-117816339-INS-53
chr3-117823904-INS-6916
chr3-117824561-INS-148
chr3-117827468-INS-4652
chr3-117831729-INS-65
chr3-117855405-INS-64
chr3-117884509-INS-92
chr3-117887238-INS-137
chr3-117895903-INS-62
chr3-117918438-INS-1460
chr3-117953381-INS-78
chr3-117955041-INS-172
chr3-117961209-INS-68
chr3-117979030-INS-201
chr3-118003697-INS-6348
chr3-118004144-INS-62
chr3-118016358-INS-197
chr3-118017271-INS-3088
chr3-118034853-INS-125
chr3-118045510-INS-63
chr3-118047272-INS-299
chr3-118071906-INS-78
chr3-118104997-INS-60
chr3-118114596-INS-412
chr3-118145949-INS-6652
chr3-118205111-INS-831
chr3-118209201-INS-301
chr3-118238312-INS-109
chr3-118241636-INS-53
chr3-118268077-INS-52
chr3-118315043-INS-88
chr3-118331765-INS-60
chr3-118334411-INS-974
chr3-118335001-INS-66
chr3-118340455-INS-848

chr3-118343172-INS-90
chr3-118355340-INS-227
chr3-118403388-INS-6224
chr3-118410223-INS-576
chr3-118414804-INS-78
chr3-118420875-INS-559
chr3-118460005-INS-204
chr3-118463681-INS-52
chr3-118478744-INS-90
chr3-118481386-INS-299
chr3-118493725-INS-126
chr3-118501452-INS-131
chr3-118539402-INS-53
chr3-118548174-INS-87
chr3-118558420-INS-220
chr3-118572689-INS-93
chr3-118577222-INS-174
chr3-118587275-INS-95
chr3-118588919-INS-60
chr3-118598290-INS-359
chr3-118626525-INS-544
chr3-118633643-INS-153
chr3-118654028-INS-6930
chr3-118703679-INS-64
chr3-118711263-INS-320
chr3-118776205-INS-56
chr3-118790881-INS-241
chr3-118820618-INS-64
chr3-118824791-INS-176
chr3-118843094-INS-351
chr3-118853861-INS-68
chr3-118863296-INS-539
chr3-118888367-INS-65
chr3-118896802-INS-54
chr3-118914219-INS-5327
chr3-118931365-INS-114
chr3-118960960-INS-586
chr3-118978259-INS-66
chr3-118984612-INS-119
chr3-119007142-INS-273
chr3-119016055-INS-991
chr3-119021501-INS-55
chr3-119057248-INS-201
chr3-119060448-INS-58
chr3-119065453-INS-92
chr3-119093759-INS-227
chr3-119111144-INS-52
chr3-119116061-INS-55
chr3-119132514-INS-114
chr3-119147864-INS-914
chr3-119168979-INS-88
chr3-119226996-INS-177
chr3-119230539-INS-52
chr3-119263449-INS-499
chr3-119272256-INS-55
chr3-119275161-INS-52
chr3-119276213-INS-303
chr3-119276810-INS-59
chr3-119288565-INS-56
chr3-119301236-INS-267
chr3-119313032-INS-222
chr3-119319551-INS-96
chr3-119329984-INS-760
chr3-119413752-INS-249
chr3-119424711-INS-215
chr3-119438702-INS-70
chr3-119445661-INS-341
chr3-119446708-INS-65
chr3-119464037-INS-69
chr3-119477174-INS-74
chr3-119488202-INS-1069
chr3-119495565-INS-63
chr3-119506129-INS-52
chr3-119570873-INS-313
chr3-119607721-INS-169
chr3-119609713-INS-105
chr3-119641956-INS-72

chr3-119675528-INS-402
chr3-119703415-INS-270
chr3-119719509-INS-193
chr3-119729962-INS-68
chr3-119746118-INS-178
chr3-119754765-INS-52
chr3-119761541-INS-84
chr3-119767836-INS-6581
chr3-119794071-INS-516
chr3-119816714-INS-181
chr3-119820750-INS-84
chr3-119821231-INS-1259
chr3-119825255-INS-86
chr3-119852497-INS-104
chr3-119862573-INS-78
chr3-119873167-INS-81
chr3-119881777-INS-564
chr3-119894389-INS-6452
chr3-119898105-INS-118
chr3-119945952-INS-60
chr3-119954130-INS-123
chr3-119955416-INS-207
chr3-119967725-INS-353
chr3-119977075-INS-65
chr3-119988567-INS-93
chr3-119991659-INS-323
chr3-120003613-INS-191
chr3-120020296-INS-90
chr3-120023285-INS-176
chr3-120026750-INS-153
chr3-120045185-INS-214
chr3-120051660-INS-6757
chr3-120087596-INS-254
chr3-120130337-INS-226
chr3-120133250-INS-254
chr3-120170739-INS-4393
chr3-120171332-INS-87
chr3-120192039-INS-671
chr3-120217624-INS-960
chr3-120220136-INS-2695
chr3-120222852-INS-160
chr3-120255201-INS-62
chr3-120277827-INS-774
chr3-120284650-INS-54
chr3-120293715-INS-51
chr3-120296081-INS-106
chr3-120322026-INS-102
chr3-120324484-INS-181
chr3-120331274-INS-187
chr3-120372044-INS-191
chr3-120379069-INS-146
chr3-120387368-INS-73
chr3-120391411-INS-337
chr3-120394986-INS-398
chr3-120416933-INS-76
chr3-120427039-INS-54
chr3-120433934-INS-347
chr3-120508170-INS-260
chr3-120535433-INS-424
chr3-120539744-INS-819
chr3-120548371-INS-795
chr3-120592255-INS-104
chr3-120602050-INS-196
chr3-120605538-INS-103
chr3-120645874-INS-184
chr3-120649223-INS-95
chr3-120661678-INS-98
chr3-120684455-INS-80
chr3-120697017-INS-667
chr3-120721847-INS-208
chr3-120745482-INS-66
chr3-120752132-INS-454
chr3-120756645-INS-6687
chr3-120758674-INS-813
chr3-120775248-INS-73
chr3-120784354-INS-192
chr3-120785807-INS-64

chr3-120786646-INS-85
chr3-120805656-INS-54
chr3-120833738-INS-70
chr3-120838208-INS-6657
chr3-120860718-INS-5499
chr3-120870272-INS-96
chr3-120872456-INS-143
chr3-120935182-INS-792
chr3-120945865-INS-995
chr3-120946712-INS-60
chr3-120988590-INS-222
chr3-121016706-INS-234
chr3-121018634-INS-149
chr3-121023855-INS-136
chr3-121043190-INS-598
chr3-121066878-INS-59
chr3-121073382-INS-53
chr3-121079117-INS-96
chr3-121093360-INS-206
chr3-121129267-INS-433
chr3-121140071-INS-105
chr3-121156371-INS-130
chr3-121172157-INS-105
chr3-121175331-INS-82
chr3-121178491-INS-345
chr3-121213635-INS-62
chr3-121238574-INS-56
chr3-121270558-INS-67
chr3-121291085-INS-75
chr3-121314100-INS-189
chr3-121321059-INS-72
chr3-121336075-INS-184
chr3-121358819-INS-207
chr3-121374685-INS-90
chr3-121386113-INS-223
chr3-121388962-INS-197
chr3-121393852-INS-173
chr3-121410386-INS-103
chr3-121418875-INS-206
chr3-121433032-INS-51
chr3-121487972-INS-70
chr3-121523557-INS-486
chr3-121595030-INS-225
chr3-121617864-INS-215
chr3-121619083-INS-99
chr3-121630125-INS-284
chr3-121660702-INS-209
chr3-121663261-INS-335
chr3-121706230-INS-66
chr3-121709969-INS-178
chr3-121715087-INS-104
chr3-121725618-INS-225
chr3-121756115-INS-57
chr3-121819132-INS-104
chr3-121830943-INS-64
chr3-121840837-INS-204
chr3-121851198-INS-123
chr3-121856156-INS-231
chr3-121870947-INS-72
chr3-121876625-INS-52
chr3-121876986-INS-987
chr3-121889418-INS-52
chr3-121916750-INS-340
chr3-121923615-INS-72
chr3-121929132-INS-430
chr3-121979185-INS-219
chr3-121979964-INS-51
chr3-121992023-INS-212
chr3-122005392-INS-874
chr3-122022343-INS-81
chr3-122073906-INS-61
chr3-122094780-INS-488
chr3-122096157-INS-90
chr3-122101861-INS-885
chr3-122102051-INS-2524
chr3-122109151-INS-329
chr3-122120914-INS-216

chr3-122122718-INS-120
chr3-122133648-INS-67
chr3-122143895-INS-6440
chr3-122157908-INS-72
chr3-122167031-INS-1963
chr3-122193942-INS-177
chr3-122207341-INS-201
chr3-122228703-INS-323
chr3-122237612-INS-60
chr3-122241756-INS-80
chr3-122290428-INS-51
chr3-122295171-INS-235
chr3-122307720-INS-68
chr3-122318301-INS-120
chr3-122361300-INS-234
chr3-122371016-INS-180
chr3-122379977-INS-204
chr3-122387895-INS-195
chr3-122388693-INS-209
chr3-122392143-INS-52
chr3-122398396-INS-52
chr3-122404557-INS-1701
chr3-122412567-INS-52
chr3-122418853-INS-297
chr3-122426031-INS-588
chr3-122465263-INS-224
chr3-122469625-INS-180
chr3-122519295-INS-68
chr3-122571213-INS-358
chr3-122628585-INS-172
chr3-122642513-INS-400
chr3-122646040-INS-226
chr3-122655460-INS-55
chr3-122681194-INS-72
chr3-122684209-INS-50
chr3-122712400-INS-151
chr3-122721757-INS-177
chr3-122735056-INS-302
chr3-122764562-INS-149
chr3-122787090-INS-52
chr3-122825253-INS-111
chr3-122857259-INS-165
chr3-122878213-INS-107
chr3-122939256-INS-115
chr3-122978734-INS-52
chr3-123004646-INS-7084
chr3-123005919-INS-52
chr3-123007362-INS-200
chr3-123020928-INS-103
chr3-123024036-INS-176
chr3-123053397-INS-399
chr3-123059592-INS-193
chr3-123095244-INS-336
chr3-123098328-INS-328
chr3-123101178-INS-208
chr3-123111443-INS-101
chr3-123124797-INS-64
chr3-123156821-INS-175
chr3-123167166-INS-1018
chr3-123189713-INS-86
chr3-123192599-INS-90
chr3-123193615-INS-192
chr3-123209585-INS-405
chr3-123215219-INS-5000
chr3-123227050-INS-80
chr3-123246114-INS-87
chr3-123247812-INS-76
chr3-123253651-INS-57
chr3-123259814-INS-493
chr3-123271196-INS-554
chr3-123273095-INS-54
chr3-123277595-INS-65
chr3-123281307-INS-209
chr3-123290967-INS-561
chr3-123305132-INS-523
chr3-123317156-INS-54
chr3-123326974-INS-78

chr3-123328701-INS-66
chr3-123333194-INS-6465
chr3-123334426-INS-65
chr3-123334910-INS-6525
chr3-123369433-INS-290
chr3-123371990-INS-325
chr3-123377782-INS-164
chr3-123389561-INS-56
chr3-123414492-INS-69
chr3-123422011-INS-128
chr3-123425367-INS-146
chr3-123435366-INS-5158
chr3-123466637-INS-963
chr3-123473969-INS-845
chr3-123485617-INS-126
chr3-123501752-INS-400
chr3-123537412-INS-155
chr3-123540855-INS-60
chr3-123545017-INS-259
chr3-123547174-INS-54
chr3-123552773-INS-207
chr3-123581891-INS-52
chr3-123593312-INS-227
chr3-123648284-INS-176
chr3-123665978-INS-547
chr3-123669544-INS-51
chr3-123691871-INS-205
chr3-123696054-INS-56
chr3-123709222-INS-398
chr3-123785412-INS-170
chr3-123789687-INS-5219
chr3-123804719-INS-52
chr3-123842284-INS-327
chr3-123843104-INS-68
chr3-123846520-INS-59
chr3-123880580-INS-328
chr3-123894138-INS-88
chr3-123896482-INS-66
chr3-123897867-INS-58
chr3-123902777-INS-66
chr3-123980821-INS-469
chr3-124009378-INS-53
chr3-124027467-INS-7228
chr3-124042094-INS-399
chr3-124058807-INS-597
chr3-124084890-INS-878
chr3-124103855-INS-69
chr3-124120338-INS-146
chr3-124138187-INS-869
chr3-124152202-INS-739
chr3-124158769-INS-875
chr3-124166345-INS-203
chr3-124169098-INS-106
chr3-124171651-INS-115
chr3-124171919-INS-5468
chr3-124204061-INS-77
chr3-124286450-INS-68
chr3-124346989-INS-601
chr3-124347420-INS-828
chr3-124359135-INS-141
chr3-124363629-INS-293
chr3-124366006-INS-54
chr3-124381932-INS-224
chr3-124388801-INS-236
chr3-124393694-INS-82
chr3-124394818-INS-354
chr3-124414564-INS-6401
chr3-124434270-INS-76
chr3-124447928-INS-59
chr3-124452235-INS-132
chr3-124462466-INS-194
chr3-124470380-INS-112
chr3-124476673-INS-420
chr3-124484818-INS-144
chr3-124518144-INS-66
chr3-124586194-INS-447
chr3-124620725-INS-166

chr3-124655579-INS-51
chr3-124715533-INS-55
chr3-124753718-INS-91
chr3-124765682-INS-5356
chr3-124780606-INS-222
chr3-124782867-INS-98
chr3-124795480-INS-6583
chr3-124839079-INS-54
chr3-124842690-INS-279
chr3-124899536-INS-5375
chr3-124906222-INS-121
chr3-124932489-INS-52
chr3-124960100-INS-6938
chr3-124993870-INS-64
chr3-125000041-INS-77
chr3-125039082-INS-206
chr3-125047675-INS-572
chr3-125068305-INS-110
chr3-125080435-INS-65
chr3-125087826-INS-892
chr3-125088568-INS-52
chr3-125089798-INS-59
chr3-125100039-INS-135
chr3-125111420-INS-95
chr3-125116333-INS-63
chr3-125122413-INS-435
chr3-125143563-INS-225
chr3-125156070-INS-100
chr3-125178358-INS-401
chr3-125180107-INS-53
chr3-125198811-INS-66
chr3-125207724-INS-3989
chr3-125215606-INS-664
chr3-125222542-INS-54
chr3-125244265-INS-556
chr3-125263366-INS-63
chr3-125282315-INS-7166
chr3-125305370-INS-763
chr3-125318997-INS-732
chr3-125341752-INS-52
chr3-125347849-INS-183
chr3-125383046-INS-55
chr3-125439352-INS-187
chr3-125452215-INS-52
chr3-125504711-INS-453
chr3-125532043-INS-52
chr3-125560507-INS-441
chr3-125562829-INS-76
chr3-125622536-INS-347
chr3-125632541-INS-58
chr3-125639058-INS-58
chr3-125640154-INS-587
chr3-125653333-INS-493
chr3-125654608-INS-53
chr3-125677508-INS-615
chr3-125687904-INS-94
chr3-125710942-INS-927
chr3-125726141-INS-193
chr3-125759396-INS-208
chr3-125804976-INS-60
chr3-125807502-INS-55
chr3-125809933-INS-795
chr3-125813649-INS-84
chr3-125871787-INS-380
chr3-125875577-INS-923
chr3-125885942-INS-65
chr3-125917143-INS-6229
chr3-125985277-INS-422
chr3-126022781-INS-60
chr3-126032673-INS-457
chr3-126083145-INS-56
chr3-126085040-INS-4351
chr3-126105895-INS-65
chr3-126128427-INS-229
chr3-126130363-INS-76
chr3-126154840-INS-72
chr3-126162626-INS-633

chr3-126184654-INS-8151
chr3-126214712-INS-73
chr3-126226838-INS-56
chr3-126231824-INS-5261
chr3-126241919-INS-61
chr3-126244592-INS-57
chr3-126274780-INS-7049
chr3-126291631-INS-64
chr3-126303004-INS-832
chr3-126307178-INS-112
chr3-126313494-INS-408
chr3-126324199-INS-154
chr3-126331445-INS-174
chr3-126368073-INS-88
chr3-126439764-INS-250
chr3-126480087-INS-248
chr3-126514542-INS-144
chr3-126538306-INS-336
chr3-126542800-INS-85
chr3-126573952-INS-109
chr3-126581584-INS-8067
chr3-126618350-INS-101
chr3-126631071-INS-54
chr3-126650172-INS-3847
chr3-126650477-INS-1941
chr3-126664308-INS-76
chr3-126684231-INS-103
chr3-126692169-INS-52
chr3-126702570-INS-58
chr3-126820928-INS-124
chr3-126831849-INS-112
chr3-126875936-INS-64
chr3-126876783-INS-80
chr3-126878185-INS-727
chr3-126890289-INS-63
chr3-126899822-INS-75
chr3-126910375-INS-198
chr3-126947897-INS-60
chr3-126960265-INS-59
chr3-127025355-INS-431
chr3-127057904-INS-94
chr3-127081746-INS-324
chr3-127089811-INS-57
chr3-127095908-INS-197
chr3-127098404-INS-70
chr3-127121469-INS-69
chr3-127129027-INS-190
chr3-127157999-INS-53
chr3-127161914-INS-395
chr3-127164329-INS-160
chr3-127179360-INS-351
chr3-127188619-INS-208
chr3-127197862-INS-121
chr3-127220837-INS-85
chr3-127227439-INS-264
chr3-127230744-INS-58
chr3-127234291-INS-214
chr3-127237536-INS-79
chr3-127255533-INS-309
chr3-127283604-INS-194
chr3-127291673-INS-225
chr3-127298512-INS-827
chr3-127307012-INS-125
chr3-127308946-INS-104
chr3-127314599-INS-56
chr3-127318766-INS-400
chr3-127320104-INS-52
chr3-127320682-INS-212
chr3-127324851-INS-121
chr3-127329453-INS-211
chr3-127333703-INS-483
chr3-127343641-INS-569
chr3-127348347-INS-214
chr3-127358356-INS-217
chr3-127368802-INS-158
chr3-127374715-INS-104
chr3-127378290-INS-126

chr3-127384637-INS-202
chr3-127385660-INS-198
chr3-127401694-INS-904
chr3-127406000-INS-214
chr3-127409795-INS-102
chr3-127423809-INS-198
chr3-127444581-INS-278
chr3-127448087-INS-381
chr3-127461384-INS-63
chr3-127469102-INS-52
chr3-127469880-INS-120
chr3-127471866-INS-108
chr3-127477515-INS-535
chr3-127481532-INS-193
chr3-127484671-INS-877
chr3-127491863-INS-127
chr3-127517674-INS-57
chr3-127526123-INS-85
chr3-127559571-INS-570
chr3-127560310-INS-52
chr3-127565673-INS-208
chr3-127580359-INS-196
chr3-127601624-INS-58
chr3-127602799-INS-200
chr3-127608216-INS-105
chr3-127608373-INS-305
chr3-127640328-INS-72
chr3-127642356-INS-185
chr3-127654595-INS-203
chr3-127661293-INS-325
chr3-127699938-INS-70
chr3-127701566-INS-548
chr3-127712321-INS-53
chr3-127724822-INS-458
chr3-127749170-INS-227
chr3-127764130-INS-300
chr3-127784404-INS-54
chr3-127793666-INS-54
chr3-127801310-INS-51
chr3-127803853-INS-7073
chr3-127813731-INS-189
chr3-127814517-INS-130
chr3-127822919-INS-351
chr3-127826205-INS-159
chr3-127826241-INS-1044
chr3-127827712-INS-64
chr3-127860236-INS-152
chr3-127878036-INS-73
chr3-127887832-INS-5207
chr3-127919798-INS-659
chr3-127921367-INS-680
chr3-127921767-INS-80
chr3-127956926-INS-112
chr3-127972221-INS-52
chr3-128001579-INS-67
chr3-128001937-INS-751
chr3-128018536-INS-267
chr3-128064013-INS-120
chr3-128126735-INS-56
chr3-128133311-INS-80
chr3-128136235-INS-57
chr3-128159249-INS-244
chr3-128167620-INS-342
chr3-128186714-INS-78
chr3-128203175-INS-482
chr3-128290726-INS-198
chr3-128306396-INS-274
chr3-128316117-INS-68
chr3-128322921-INS-2855
chr3-128327028-INS-146
chr3-128347860-INS-492
chr3-128361795-INS-122
chr3-128363003-INS-441
chr3-128375680-INS-1614
chr3-128380696-INS-440
chr3-128403082-INS-386
chr3-128412336-INS-540

chr3-128413165-INS-67
chr3-128432559-INS-112
chr3-128435450-INS-764
chr3-128436211-INS-144
chr3-128438569-INS-58
chr3-128545634-INS-61
chr3-128589479-INS-1888
chr3-128594963-INS-456
chr3-128635936-INS-7129
chr3-128640303-INS-1831
chr3-128660569-INS-107
chr3-128671047-INS-5319
chr3-128694594-INS-742
chr3-128729802-INS-184
chr3-128736262-INS-57
chr3-128791941-INS-640
chr3-128810412-INS-209
chr3-128839061-INS-151
chr3-128840211-INS-64
chr3-128866529-INS-6268
chr3-128892871-INS-162
chr3-128905833-INS-63
chr3-128933288-INS-179
chr3-128970381-INS-118
chr3-128971680-INS-86
chr3-129004074-INS-56
chr3-129005607-INS-76
chr3-129038190-INS-174
chr3-129046769-INS-58
chr3-129053452-INS-409
chr3-129054772-INS-7235
chr3-129073836-INS-105
chr3-129084836-INS-107
chr3-129086117-INS-52
chr3-129113247-INS-205
chr3-129118995-INS-176
chr3-129149814-INS-208
chr3-129153151-INS-204
chr3-129171993-INS-178
chr3-129176475-INS-158
chr3-129176482-INS-762
chr3-129181769-INS-124
chr3-129202847-INS-60
chr3-129206208-INS-399
chr3-129214843-INS-52
chr3-129259181-INS-52
chr3-129260800-INS-51
chr3-129280764-INS-132
chr3-129285304-INS-76
chr3-129308487-INS-148
chr3-129312731-INS-273
chr3-129323611-INS-204
chr3-129347223-INS-198
chr3-129354363-INS-393
chr3-129378931-INS-68
chr3-129397780-INS-60
chr3-129402631-INS-63
chr3-129409876-INS-363
chr3-129410004-INS-584
chr3-129438445-INS-210
chr3-129440500-INS-93
chr3-129441466-INS-514
chr3-129452709-INS-63
chr3-129453872-INS-208
chr3-129454750-INS-442
chr3-129457885-INS-59
chr3-129474722-INS-181
chr3-129477616-INS-630
chr3-129514326-INS-5195
chr3-129539657-INS-87
chr3-129542677-INS-186
chr3-129543554-INS-152
chr3-129570581-INS-178
chr3-129575309-INS-115
chr3-129577490-INS-67
chr3-129582455-INS-63
chr3-129592688-INS-72

chr3-129599305-INS-407
chr3-129608257-INS-52
chr3-129609878-INS-493
chr3-129623921-INS-286
chr3-129627050-INS-358
chr3-129645168-INS-50
chr3-129651144-INS-57
chr3-129654023-INS-845
chr3-129664715-INS-194
chr3-129669689-INS-57
chr3-129701315-INS-78
chr3-129716418-INS-149
chr3-129735232-INS-196
chr3-129744756-INS-423
chr3-129759330-INS-115
chr3-129773297-INS-110
chr3-129783243-INS-81
chr3-129785942-INS-64
chr3-129867650-INS-85
chr3-129869896-INS-165
chr3-129884936-INS-168
chr3-129899846-INS-340
chr3-129902830-INS-245
chr3-129906501-INS-91
chr3-129943030-INS-60
chr3-129949613-INS-312
chr3-129967542-INS-154
chr3-129993238-INS-62
chr3-130014212-INS-66
chr3-130020566-INS-54
chr3-130030550-INS-742
chr3-130031922-INS-6709
chr3-130051803-INS-55
chr3-130061641-INS-231
chr3-130068619-INS-88
chr3-130073874-INS-54
chr3-130077836-INS-701
chr3-130119275-INS-99
chr3-130120697-INS-62
chr3-130137828-INS-400
chr3-130153347-INS-364
chr3-130163658-INS-110
chr3-130170315-INS-58
chr3-130178738-INS-104
chr3-130223143-INS-141
chr3-130237330-INS-127
chr3-130275977-INS-205
chr3-130282542-INS-7446
chr3-130338688-INS-155
chr3-130367821-INS-63
chr3-130398154-INS-466
chr3-130402786-INS-186
chr3-130407332-INS-118
chr3-130417992-INS-80
chr3-130419518-INS-268
chr3-130444455-INS-237
chr3-130469681-INS-205
chr3-130487232-INS-193
chr3-130495475-INS-57
chr3-130536317-INS-177
chr3-130537390-INS-152
chr3-130577735-INS-216
chr3-130594867-INS-398
chr3-130606100-INS-152
chr3-130617273-INS-881
chr3-130646021-INS-212
chr3-130700276-INS-62
chr3-130719638-INS-75
chr3-130728631-INS-400
chr3-130743183-INS-549
chr3-130775757-INS-113
chr3-130778176-INS-72
chr3-130808946-INS-91
chr3-130837660-INS-842
chr3-130855998-INS-69
chr3-130858498-INS-183
chr3-130859488-INS-124

chr3-130861798-INS-66
chr3-130864978-INS-278
chr3-130877612-INS-133
chr3-130902463-INS-88
chr3-130934270-INS-116
chr3-130956375-INS-83
chr3-130994107-INS-54
chr3-131025784-INS-184
chr3-131049033-INS-156
chr3-131050785-INS-56
chr3-131063575-INS-52
chr3-131071432-INS-137
chr3-131077306-INS-61
chr3-131080058-INS-61
chr3-131086557-INS-60
chr3-131173367-INS-60
chr3-131175352-INS-79
chr3-131202609-INS-398
chr3-131240738-INS-157
chr3-131245295-INS-51
chr3-131265314-INS-181
chr3-131277461-INS-231
chr3-131297454-INS-55
chr3-131315112-INS-52
chr3-131318665-INS-350
chr3-131321236-INS-779
chr3-131328773-INS-140
chr3-131338118-INS-149
chr3-131381047-INS-56
chr3-131383663-INS-64
chr3-131387354-INS-788
chr3-131463418-INS-303
chr3-131488959-INS-518
chr3-131515428-INS-251
chr3-131516881-INS-66
chr3-131518338-INS-76
chr3-131531534-INS-243
chr3-131534990-INS-50
chr3-131569309-INS-80
chr3-131573046-INS-52
chr3-131577380-INS-79
chr3-131591788-INS-357
chr3-131608523-INS-53
chr3-131632491-INS-141
chr3-131641730-INS-94
chr3-131650467-INS-473
chr3-131657814-INS-142
chr3-131667723-INS-172
chr3-131681361-INS-323
chr3-131705016-INS-306
chr3-131711725-INS-214
chr3-131744627-INS-108
chr3-131746439-INS-126
chr3-131775430-INS-594
chr3-131839759-INS-52
chr3-131873045-INS-52
chr3-131875181-INS-314
chr3-131895082-INS-71
chr3-131898363-INS-96
chr3-131971702-INS-54
chr3-131994145-INS-198
chr3-132055381-INS-56
chr3-132058506-INS-58
chr3-132064657-INS-86
chr3-132079783-INS-7004
chr3-132082955-INS-914
chr3-132110130-INS-107
chr3-132118091-INS-411
chr3-132118255-INS-411
chr3-132127946-INS-6631
chr3-132132852-INS-239
chr3-132135575-INS-352
chr3-132160404-INS-127
chr3-132170417-INS-55
chr3-132171908-INS-60
chr3-132207136-INS-180
chr3-132211819-INS-56

chr3-132214579-INS-53
chr3-132237577-INS-90
chr3-132258899-INS-132
chr3-132265621-INS-58
chr3-132278565-INS-932
chr3-132289176-INS-409
chr3-132327758-INS-359
chr3-132328366-INS-689
chr3-132334832-INS-86
chr3-132422450-INS-70
chr3-132505414-INS-273
chr3-132523066-INS-95
chr3-132525506-INS-6318
chr3-132559181-INS-59
chr3-132585770-INS-64
chr3-132625799-INS-141
chr3-132630733-INS-159
chr3-132648733-INS-122
chr3-132654912-INS-64
chr3-132666449-INS-298
chr3-132674165-INS-583
chr3-132683733-INS-74
chr3-132691043-INS-68
chr3-132723417-INS-454
chr3-132733654-INS-65
chr3-132746845-INS-74
chr3-132754454-INS-294
chr3-132768696-INS-851
chr3-132779119-INS-65
chr3-132841629-INS-146
chr3-132853202-INS-54
chr3-132855702-INS-3840
chr3-132864666-INS-67
chr3-132907819-INS-6997
chr3-132947134-INS-764
chr3-132970060-INS-195
chr3-132976503-INS-6405
chr3-132989352-INS-51
chr3-133012844-INS-188
chr3-133021761-INS-185
chr3-133043782-INS-174
chr3-133058542-INS-266
chr3-133079023-INS-6913
chr3-133081085-INS-185
chr3-133093971-INS-211
chr3-133139688-INS-60
chr3-133147235-INS-210
chr3-133164023-INS-135
chr3-133174566-INS-67
chr3-133186489-INS-60
chr3-133210565-INS-196
chr3-133222661-INS-56
chr3-133241383-INS-655
chr3-133245517-INS-118
chr3-133250197-INS-129
chr3-133258383-INS-503
chr3-133262228-INS-105
chr3-133262782-INS-114
chr3-133267942-INS-292
chr3-133279380-INS-52
chr3-133315803-INS-76
chr3-133323390-INS-189
chr3-133341536-INS-204
chr3-133355182-INS-160
chr3-133373223-INS-57
chr3-133383543-INS-174
chr3-133439358-INS-55
chr3-133452202-INS-367
chr3-133470712-INS-334
chr3-133480872-INS-56
chr3-133495413-INS-209
chr3-133513426-INS-102
chr3-133518530-INS-62
chr3-133522076-INS-117
chr3-133546831-INS-7185
chr3-133576879-INS-676
chr3-133586394-INS-317

chr3-133608627-INS-66
chr3-133610951-INS-462
chr3-133619221-INS-53
chr3-133692698-INS-52
chr3-133697907-INS-155
chr3-133703081-INS-52
chr3-133747790-INS-164
chr3-133749125-INS-71
chr3-133763670-INS-132
chr3-133772209-INS-380
chr3-133787016-INS-102
chr3-133815884-INS-786
chr3-133819749-INS-6273
chr3-133829361-INS-191
chr3-133837438-INS-96
chr3-133854988-INS-835
chr3-133877220-INS-172
chr3-133882479-INS-586
chr3-133885743-INS-535
chr3-133901681-INS-466
chr3-133913716-INS-575
chr3-133934052-INS-410
chr3-133983156-INS-71
chr3-133986350-INS-51
chr3-134005874-INS-52
chr3-134013959-INS-173
chr3-134034833-INS-250
chr3-134036412-INS-180
chr3-134041998-INS-577
chr3-134045071-INS-204
chr3-134047723-INS-131
chr3-134052310-INS-317
chr3-134082730-INS-827
chr3-134105448-INS-68
chr3-134107920-INS-164
chr3-134124585-INS-64
chr3-134126085-INS-494
chr3-134130763-INS-152
chr3-134140607-INS-537
chr3-134183657-INS-70
chr3-134186427-INS-380
chr3-134187925-INS-915
chr3-134189243-INS-71
chr3-134196479-INS-939
chr3-134200451-INS-411
chr3-134200824-INS-6534
chr3-134204779-INS-76
chr3-134223297-INS-72
chr3-134251827-INS-416
chr3-134284457-INS-196
chr3-134296481-INS-128
chr3-134315482-INS-53
chr3-134316358-INS-146
chr3-134330303-INS-53
chr3-134357479-INS-2592
chr3-134367067-INS-2656
chr3-134375649-INS-330
chr3-134430091-INS-3159
chr3-134450140-INS-100
chr3-134451919-INS-962
chr3-134460973-INS-895
chr3-134483155-INS-824
chr3-134483895-INS-169
chr3-134516036-INS-7689
chr3-134545404-INS-103
chr3-134553563-INS-230
chr3-134603176-INS-7359
chr3-134606112-INS-6454
chr3-134623453-INS-354
chr3-134627875-INS-130
chr3-134638689-INS-152
chr3-134643833-INS-521
chr3-134683676-INS-3201
chr3-134691761-INS-1402
chr3-134697566-INS-6901
chr3-134721261-INS-377
chr3-134724139-INS-762

chr3-134729599-INS-52
chr3-134752848-INS-81
chr3-134754313-INS-68
chr3-134775009-INS-725
chr3-134777990-INS-691
chr3-134786284-INS-675
chr3-134816642-INS-210
chr3-134820407-INS-60
chr3-134821960-INS-6960
chr3-134827085-INS-65
chr3-134858843-INS-56
chr3-134878773-INS-8459
chr3-134900098-INS-138
chr3-134909126-INS-6452
chr3-134910632-INS-5219
chr3-134940154-INS-119
chr3-134966204-INS-111
chr3-134996158-INS-106
chr3-135006271-INS-52
chr3-135015031-INS-153
chr3-135019542-INS-165
chr3-135042582-INS-76
chr3-135049220-INS-5302
chr3-135053142-INS-881
chr3-135091558-INS-54
chr3-135120554-INS-68
chr3-135125954-INS-198
chr3-135158771-INS-192
chr3-135180402-INS-230
chr3-135189508-INS-162
chr3-135194138-INS-51
chr3-135201695-INS-72
chr3-135213281-INS-67
chr3-135213722-INS-58
chr3-135217939-INS-6473
chr3-135222951-INS-63
chr3-135227600-INS-107
chr3-135236235-INS-146
chr3-135237219-INS-5304
chr3-135245066-INS-68
chr3-135256152-INS-377
chr3-135266234-INS-356
chr3-135267419-INS-367
chr3-135279305-INS-7123
chr3-135279798-INS-499
chr3-135282972-INS-76
chr3-135297661-INS-187
chr3-135325400-INS-793
chr3-135328082-INS-93
chr3-135329047-INS-56
chr3-135345916-INS-1891
chr3-135369607-INS-56
chr3-135374027-INS-521
chr3-135381603-INS-60
chr3-135386260-INS-836
chr3-135390163-INS-324
chr3-135419598-INS-116
chr3-135424683-INS-391
chr3-135447543-INS-82
chr3-135448087-INS-164
chr3-135452396-INS-57
chr3-135454646-INS-252
chr3-135455720-INS-54
chr3-135460609-INS-158
chr3-135472122-INS-106
chr3-135485593-INS-511
chr3-135489116-INS-285
chr3-135494459-INS-6188
chr3-135509912-INS-183
chr3-135513253-INS-284
chr3-135514030-INS-2817
chr3-135537462-INS-198
chr3-135543014-INS-402
chr3-135557358-INS-63
chr3-135590272-INS-56
chr3-135603478-INS-135
chr3-135609215-INS-61

chr3-135617732-INS-244
chr3-135618788-INS-64
chr3-135643244-INS-68
chr3-135660074-INS-62
chr3-135664481-INS-54
chr3-135681960-INS-133
chr3-135740217-INS-105
chr3-135763285-INS-228
chr3-135767332-INS-332
chr3-135806952-INS-644
chr3-135808994-INS-397
chr3-135819558-INS-156
chr3-135859380-INS-277
chr3-135861338-INS-190
chr3-135862995-INS-90
chr3-135901790-INS-1175
chr3-135905477-INS-152
chr3-135921044-INS-212
chr3-135928326-INS-1035
chr3-135941794-INS-79
chr3-135954782-INS-115
chr3-135978405-INS-117
chr3-135988890-INS-76
chr3-135999724-INS-123
chr3-136023256-INS-5910
chr3-136046890-INS-72
chr3-136126307-INS-344
chr3-136138790-INS-6721
chr3-136145119-INS-194
chr3-136150143-INS-94
chr3-136154608-INS-148
chr3-136166864-INS-6679
chr3-136175031-INS-6666
chr3-136177776-INS-83
chr3-136226891-INS-194
chr3-136241900-INS-5265
chr3-136261177-INS-59
chr3-136263000-INS-61
chr3-136273198-INS-169
chr3-136293433-INS-474
chr3-136332579-INS-123
chr3-136334192-INS-354
chr3-136340320-INS-152
chr3-136363309-INS-69
chr3-136365191-INS-234
chr3-136367933-INS-60
chr3-136369248-INS-154
chr3-136383127-INS-141
chr3-136389419-INS-57
chr3-136393835-INS-52
chr3-136395992-INS-308
chr3-136398047-INS-772
chr3-136449248-INS-106
chr3-136473236-INS-54
chr3-136521318-INS-52
chr3-136544508-INS-1885
chr3-136551020-INS-145
chr3-136556486-INS-118
chr3-136561441-INS-204
chr3-136563482-INS-141
chr3-136581613-INS-57
chr3-136592852-INS-110
chr3-136595777-INS-657
chr3-136648085-INS-1104
chr3-136689792-INS-697
chr3-136694445-INS-600
chr3-136702862-INS-570
chr3-136768424-INS-144
chr3-136770381-INS-7077
chr3-136788845-INS-567
chr3-136823116-INS-6376
chr3-136842236-INS-66
chr3-136857243-INS-223
chr3-136881187-INS-185
chr3-136883124-INS-64
chr3-136914765-INS-111
chr3-136940128-INS-54

chr3-137005958-INS-115
chr3-137015755-INS-54
chr3-137016858-INS-54
chr3-137022696-INS-63
chr3-137026134-INS-273
chr3-137030190-INS-92
chr3-137034677-INS-73
chr3-137039448-INS-270
chr3-137052760-INS-64
chr3-137081662-INS-121
chr3-137127738-INS-72
chr3-137139122-INS-326
chr3-137162020-INS-706
chr3-137177975-INS-81
chr3-137187647-INS-65
chr3-137196192-INS-239
chr3-137201017-INS-56
chr3-137238899-INS-5567
chr3-137277743-INS-150
chr3-137298060-INS-54
chr3-137298636-INS-54
chr3-137316067-INS-86
chr3-137319837-INS-50
chr3-137335189-INS-353
chr3-137356654-INS-189
chr3-137392852-INS-197
chr3-137411633-INS-570
chr3-137413723-INS-115
chr3-137418900-INS-73
chr3-137480281-INS-60
chr3-137483280-INS-118
chr3-137495813-INS-132
chr3-137499891-INS-352
chr3-137511212-INS-68
chr3-137514509-INS-177
chr3-137518507-INS-77
chr3-137538737-INS-66
chr3-137541255-INS-63
chr3-137547565-INS-209
chr3-137549326-INS-108
chr3-137550125-INS-376
chr3-137558929-INS-101
chr3-137575146-INS-58
chr3-137577491-INS-76
chr3-137592508-INS-52
chr3-137605656-INS-66
chr3-137615461-INS-52
chr3-137625214-INS-101
chr3-137646349-INS-174
chr3-137671248-INS-53
chr3-137672467-INS-122
chr3-137679142-INS-54
chr3-137689075-INS-52
chr3-137812097-INS-202
chr3-137813493-INS-417
chr3-137814449-INS-2815
chr3-137820415-INS-96
chr3-137834009-INS-260
chr3-137838126-INS-52
chr3-137864328-INS-98
chr3-137872473-INS-197
chr3-137878583-INS-66
chr3-137890947-INS-6475
chr3-137901912-INS-199
chr3-137934811-INS-63
chr3-137935787-INS-198
chr3-137955336-INS-390
chr3-137963722-INS-492
chr3-137985298-INS-98
chr3-137999590-INS-134
chr3-138001961-INS-99
chr3-138007309-INS-634
chr3-138025859-INS-54
chr3-138042352-INS-441
chr3-138046330-INS-84
chr3-138067642-INS-62
chr3-138075943-INS-72

chr3-138080170-INS-56
chr3-138091696-INS-147
chr3-138107854-INS-212
chr3-138122239-INS-330
chr3-138155060-INS-126
chr3-138186254-INS-186
chr3-138191103-INS-916
chr3-138202455-INS-161
chr3-138204777-INS-80
chr3-138218783-INS-739
chr3-138240693-INS-163
chr3-138260116-INS-157
chr3-138272938-INS-189
chr3-138275956-INS-351
chr3-138291447-INS-369
chr3-138294806-INS-51
chr3-138305328-INS-80
chr3-138306683-INS-575
chr3-138329514-INS-58
chr3-138330368-INS-57
chr3-138345337-INS-84
chr3-138348289-INS-2393
chr3-138384518-INS-6501
chr3-138384625-INS-142
chr3-138387777-INS-153
chr3-138398135-INS-55
chr3-138441903-INS-140
chr3-138459824-INS-100
chr3-138464315-INS-52
chr3-138475075-INS-145
chr3-138478391-INS-244
chr3-138491373-INS-131
chr3-138529059-INS-77
chr3-138549311-INS-7118
chr3-138555256-INS-60
chr3-138571837-INS-643
chr3-138588070-INS-227
chr3-138603214-INS-6110
chr3-138610731-INS-216
chr3-138625008-INS-531
chr3-138649623-INS-137
chr3-138756545-INS-4063
chr3-138781867-INS-57
chr3-138796146-INS-166
chr3-138803810-INS-564
chr3-138817293-INS-116
chr3-138822213-INS-77
chr3-138870186-INS-64
chr3-138904005-INS-104
chr3-138922030-INS-67
chr3-138928252-INS-400
chr3-138934565-INS-194
chr3-138955705-INS-87
chr3-138968908-INS-53
chr3-139014255-INS-55
chr3-139026221-INS-51
chr3-139067136-INS-99
chr3-139071296-INS-183
chr3-139081436-INS-433
chr3-139087139-INS-6363
chr3-139088345-INS-56
chr3-139090737-INS-625
chr3-139102570-INS-138
chr3-139126853-INS-74
chr3-139142437-INS-162
chr3-139146225-INS-1481
chr3-139195458-INS-518
chr3-139205690-INS-51
chr3-139209284-INS-72
chr3-139225878-INS-3055
chr3-139229322-INS-964
chr3-139231927-INS-298
chr3-139271491-INS-212
chr3-139288204-INS-57
chr3-139322383-INS-227
chr3-139347514-INS-4448
chr3-139371755-INS-511

chr3-139382527-INS-845
chr3-139390586-INS-6434
chr3-139424596-INS-52
chr3-139448936-INS-147
chr3-139454330-INS-1075
chr3-139454872-INS-6787
chr3-139474185-INS-639
chr3-139490673-INS-355
chr3-139496472-INS-102
chr3-139502067-INS-190
chr3-139509599-INS-430
chr3-139545423-INS-135
chr3-139565384-INS-782
chr3-139574874-INS-58
chr3-139584314-INS-353
chr3-139605410-INS-152
chr3-139616623-INS-986
chr3-139629460-INS-83
chr3-139692724-INS-539
chr3-139715255-INS-80
chr3-139736713-INS-72
chr3-139768410-INS-213
chr3-139772208-INS-487
chr3-139781281-INS-76
chr3-139785648-INS-1091
chr3-139800846-INS-232
chr3-139818193-INS-109
chr3-139833804-INS-308
chr3-139880639-INS-77
chr3-139896949-INS-85
chr3-139957140-INS-88
chr3-139994844-INS-175
chr3-140042365-INS-231
chr3-140053150-INS-362
chr3-140067913-INS-437
chr3-140068657-INS-913
chr3-140073651-INS-531
chr3-140080079-INS-175
chr3-140129525-INS-642
chr3-140136979-INS-76
chr3-140151244-INS-406
chr3-140157738-INS-6796
chr3-140171284-INS-7134
chr3-140181570-INS-759
chr3-140307841-INS-786
chr3-140320660-INS-591
chr3-140337569-INS-972
chr3-140356170-INS-665
chr3-140400509-INS-74
chr3-140402086-INS-138
chr3-140405461-INS-697
chr3-140478931-INS-3120
chr3-140478936-INS-2704
chr3-140496053-INS-53
chr3-140511014-INS-1136
chr3-140515481-INS-54
chr3-140527560-INS-557
chr3-140598332-INS-84
chr3-140606042-INS-60
chr3-140609523-INS-78
chr3-140621107-INS-6648
chr3-140638769-INS-56
chr3-140677434-INS-52
chr3-140705892-INS-831
chr3-140730678-INS-78
chr3-140750752-INS-116
chr3-140806648-INS-53
chr3-140819147-INS-833
chr3-140824384-INS-62
chr3-140826391-INS-659
chr3-140839525-INS-662
chr3-140841200-INS-200
chr3-140865046-INS-680
chr3-140885891-INS-874
chr3-140914911-INS-81
chr3-140952422-INS-92
chr3-141018675-INS-56

chr3-141084441-INS-449
chr3-141098036-INS-137
chr3-141138543-INS-197
chr3-141155810-INS-642
chr3-141210333-INS-241
chr3-141211720-INS-7109
chr3-141244807-INS-120
chr3-141247421-INS-68
chr3-141259848-INS-72
chr3-141275172-INS-418
chr3-141287640-INS-8027
chr3-141289035-INS-74
chr3-141297095-INS-210
chr3-141298497-INS-126
chr3-141340232-INS-139
chr3-141350017-INS-267
chr3-141380601-INS-373
chr3-141391759-INS-160
chr3-141395799-INS-249
chr3-141443270-INS-510
chr3-141475748-INS-67
chr3-141479617-INS-70
chr3-141498027-INS-72
chr3-141517363-INS-1527
chr3-141533228-INS-1892
chr3-141551578-INS-93
chr3-141552641-INS-60
chr3-141575323-INS-63
chr3-141588907-INS-881
chr3-141614592-INS-74
chr3-141626565-INS-122
chr3-141660415-INS-78
chr3-141664401-INS-118
chr3-141681872-INS-711
chr3-141684230-INS-202
chr3-141718173-INS-52
chr3-141732934-INS-170
chr3-141738725-INS-97
chr3-141753663-INS-52
chr3-141769035-INS-92
chr3-141783467-INS-66
chr3-141787175-INS-249
chr3-141825917-INS-56
chr3-141836776-INS-866
chr3-141855625-INS-120
chr3-141880168-INS-76
chr3-141881307-INS-98
chr3-141897286-INS-349
chr3-141897427-INS-70
chr3-141919216-INS-68
chr3-141923505-INS-56
chr3-141930650-INS-140
chr3-141969623-INS-6854
chr3-141981999-INS-185
chr3-141985850-INS-199
chr3-141990970-INS-294
chr3-141992362-INS-642
chr3-141995458-INS-205
chr3-141995740-INS-6185
chr3-142000672-INS-769
chr3-142003824-INS-52
chr3-142004453-INS-4360
chr3-142035019-INS-854
chr3-142085370-INS-1351
chr3-142088696-INS-51
chr3-142106918-INS-396
chr3-142115542-INS-230
chr3-142116536-INS-111
chr3-142118412-INS-392
chr3-142126817-INS-84
chr3-142138428-INS-64
chr3-142190298-INS-85
chr3-142198216-INS-459
chr3-142201989-INS-338
chr3-142203729-INS-57
chr3-142228031-INS-258
chr3-142232118-INS-100

chr3-142233045-INS-879
chr3-142241835-INS-603
chr3-142262893-INS-427
chr3-142268128-INS-54
chr3-142331538-INS-505
chr3-142354982-INS-323
chr3-142369556-INS-243
chr3-142383109-INS-558
chr3-142387589-INS-58
chr3-142388066-INS-197
chr3-142429793-INS-843
chr3-142432776-INS-66
chr3-142473120-INS-77
chr3-142475294-INS-309
chr3-142495600-INS-3104
chr3-142542502-INS-204
chr3-142619642-INS-392
chr3-142629272-INS-62
chr3-142637204-INS-61
chr3-142643043-INS-825
chr3-142675299-INS-66
chr3-142688676-INS-857
chr3-142701116-INS-365
chr3-142761867-INS-6523
chr3-142817560-INS-2977
chr3-142820683-INS-1657
chr3-142841226-INS-3648
chr3-142857138-INS-80
chr3-142863299-INS-333
chr3-142894536-INS-54
chr3-142894585-INS-1216
chr3-142895076-INS-119
chr3-142953888-INS-307
chr3-142957749-INS-73
chr3-143006371-INS-157
chr3-143015301-INS-100
chr3-143016707-INS-165
chr3-143019001-INS-80
chr3-143029661-INS-159
chr3-143059882-INS-7173
chr3-143067545-INS-381
chr3-143071349-INS-81
chr3-143099230-INS-52
chr3-143138096-INS-159
chr3-143144839-INS-426
chr3-143152176-INS-407
chr3-143198399-INS-516
chr3-143240011-INS-54
chr3-143243154-INS-57
chr3-143254196-INS-109
chr3-143274092-INS-78
chr3-143277315-INS-4100
chr3-143286944-INS-4356
chr3-143292501-INS-98
chr3-143367370-INS-132
chr3-143384199-INS-5187
chr3-143413196-INS-882
chr3-143420852-INS-90
chr3-143474770-INS-57
chr3-143490991-INS-64
chr3-143497592-INS-62
chr3-143507688-INS-401
chr3-143577164-INS-120
chr3-143580260-INS-104
chr3-143607336-INS-7967
chr3-143612237-INS-96
chr3-143626230-INS-60
chr3-143646071-INS-126
chr3-143656421-INS-105
chr3-143678454-INS-117
chr3-143698257-INS-7129
chr3-143708052-INS-531
chr3-143728447-INS-344
chr3-143732750-INS-170
chr3-143743274-INS-320
chr3-143746374-INS-51
chr3-143754250-INS-56

chr3-143760784-INS-400
chr3-143762116-INS-875
chr3-143763753-INS-65
chr3-143807414-INS-205
chr3-143849212-INS-535
chr3-143862062-INS-112
chr3-143869866-INS-56
chr3-143871669-INS-438
chr3-143897602-INS-160
chr3-143919338-INS-408
chr3-143959652-INS-126
chr3-143985872-INS-307
chr3-144025969-INS-54
chr3-144047811-INS-58
chr3-144061553-INS-543
chr3-144091356-INS-124
chr3-144121972-INS-72
chr3-144135490-INS-187
chr3-144161030-INS-61
chr3-144172600-INS-851
chr3-144175335-INS-206
chr3-144189052-INS-65
chr3-144195873-INS-728
chr3-144195878-INS-195
chr3-144196654-INS-110
chr3-144213840-INS-88
chr3-144245802-INS-58
chr3-144252865-INS-477
chr3-144282032-INS-119
chr3-144283023-INS-84
chr3-144304904-INS-437
chr3-144309101-INS-434
chr3-144349791-INS-952
chr3-144366677-INS-51
chr3-144388224-INS-65
chr3-144415869-INS-65
chr3-144451475-INS-231
chr3-144494353-INS-105
chr3-144510081-INS-66
chr3-144546379-INS-113
chr3-144554787-INS-216
chr3-144570449-INS-89
chr3-144615239-INS-69
chr3-144633259-INS-65
chr3-144657812-INS-56
chr3-144659252-INS-54
chr3-144689006-INS-82
chr3-144693449-INS-483
chr3-144717403-INS-145
chr3-144733484-INS-60
chr3-144741168-INS-63
chr3-144743103-INS-3168
chr3-144752102-INS-214
chr3-144757138-INS-52
chr3-144760972-INS-6879
chr3-144843433-INS-287
chr3-144854583-INS-153
chr3-144864990-INS-81
chr3-144868849-INS-158
chr3-144898201-INS-370
chr3-144907292-INS-813
chr3-144923417-INS-89
chr3-144973079-INS-190
chr3-144974749-INS-7014
chr3-144988582-INS-80
chr3-144991306-INS-294
chr3-144992088-INS-58
chr3-145009513-INS-701
chr3-145019276-INS-54
chr3-145033086-INS-58
chr3-145050968-INS-202
chr3-145092589-INS-142
chr3-145127106-INS-182
chr3-145131476-INS-589
chr3-145146626-INS-55
chr3-145152504-INS-6688
chr3-145153701-INS-120

chr3-145154812-INS-116
chr3-145161453-INS-194
chr3-145162513-INS-6603
chr3-145166808-INS-79
chr3-145183300-INS-186
chr3-145192812-INS-636
chr3-145199620-INS-6476
chr3-145201920-INS-123
chr3-145220440-INS-352
chr3-145233557-INS-141
chr3-145240747-INS-404
chr3-145249816-INS-525
chr3-145255010-INS-140
chr3-145259024-INS-90
chr3-145268591-INS-840
chr3-145277392-INS-70
chr3-145280658-INS-140
chr3-145296479-INS-52
chr3-145307947-INS-61
chr3-145325705-INS-171
chr3-145343574-INS-479
chr3-145368262-INS-86
chr3-145376283-INS-76
chr3-145378344-INS-4492
chr3-145398854-INS-391
chr3-145401818-INS-60
chr3-145405186-INS-356
chr3-145409906-INS-227
chr3-145418146-INS-94
chr3-145450309-INS-94
chr3-145452515-INS-202
chr3-145453889-INS-165
chr3-145455518-INS-53
chr3-145497980-INS-949
chr3-145505164-INS-314
chr3-145525008-INS-1892
chr3-145527250-INS-95
chr3-145534334-INS-120
chr3-145537267-INS-1816
chr3-145549561-INS-60
chr3-145551676-INS-256
chr3-145556472-INS-386
chr3-145559901-INS-131
chr3-145579395-INS-464
chr3-145587052-INS-80
chr3-145604294-INS-78
chr3-145604443-INS-555
chr3-145606617-INS-52
chr3-145609752-INS-54
chr3-145622889-INS-1892
chr3-145623707-INS-413
chr3-145628569-INS-5282
chr3-145633288-INS-3012
chr3-145637715-INS-93
chr3-145646011-INS-265
chr3-145651048-INS-703
chr3-145652902-INS-285
chr3-145660270-INS-136
chr3-145661940-INS-181
chr3-145667733-INS-204
chr3-145668471-INS-164
chr3-145683799-INS-458
chr3-145697149-INS-833
chr3-145699379-INS-170
chr3-145714198-INS-964
chr3-145714830-INS-194
chr3-145731364-INS-190
chr3-145735666-INS-338
chr3-145773590-INS-58
chr3-145806923-INS-420
chr3-145806973-INS-1721
chr3-145807128-INS-659
chr3-145834758-INS-53
chr3-145844698-INS-132
chr3-145845738-INS-80
chr3-145852294-INS-508
chr3-145868056-INS-410

chr3-145886251-INS-60
chr3-145887679-INS-171
chr3-145899663-INS-282
chr3-145901881-INS-301
chr3-145902922-INS-637
chr3-145912834-INS-338
chr3-145928325-INS-231
chr3-145941217-INS-56
chr3-145949698-INS-122
chr3-145950955-INS-130
chr3-145976779-INS-63
chr3-145983493-INS-386
chr3-145987725-INS-693
chr3-145999518-INS-214
chr3-146013877-INS-605
chr3-146039473-INS-114
chr3-146040835-INS-298
chr3-146044169-INS-162
chr3-146060284-INS-363
chr3-146074791-INS-234
chr3-146079444-INS-515
chr3-146081904-INS-201
chr3-146088359-INS-139
chr3-146100793-INS-378
chr3-146104081-INS-92
chr3-146131208-INS-91
chr3-146147540-INS-164
chr3-146159244-INS-140
chr3-146170415-INS-63
chr3-146176710-INS-710
chr3-146183585-INS-2467
chr3-146183822-INS-237
chr3-146184047-INS-295
chr3-146244667-INS-192
chr3-146246364-INS-68
chr3-146252233-INS-82
chr3-146265981-INS-386
chr3-146290772-INS-78
chr3-146300508-INS-223
chr3-146307429-INS-228
chr3-146318007-INS-131
chr3-146329541-INS-154
chr3-146342445-INS-53
chr3-146351619-INS-52
chr3-146353802-INS-53
chr3-146377489-INS-185
chr3-146434829-INS-84
chr3-146459960-INS-64
chr3-146465609-INS-76
chr3-146526451-INS-91
chr3-146537295-INS-391
chr3-146565703-INS-92
chr3-146569155-INS-766
chr3-146574686-INS-224
chr3-146602942-INS-76
chr3-146614060-INS-300
chr3-146616659-INS-89
chr3-146657366-INS-85
chr3-146671116-INS-389
chr3-146675581-INS-53
chr3-146690514-INS-230
chr3-146695574-INS-179
chr3-146698409-INS-157
chr3-146702150-INS-140
chr3-146707905-INS-831
chr3-146728567-INS-105
chr3-146751562-INS-82
chr3-146754495-INS-264
chr3-146762709-INS-5321
chr3-146781128-INS-80
chr3-146784299-INS-222
chr3-146821088-INS-357
chr3-146852216-INS-56
chr3-146853713-INS-100
chr3-146864616-INS-210
chr3-146893259-INS-329
chr3-146930850-INS-212

chr3-146980731-INS-60
chr3-147017339-INS-524
chr3-147023922-INS-120
chr3-147050077-INS-173
chr3-147055044-INS-3092
chr3-147072432-INS-131
chr3-147075645-INS-6721
chr3-147090137-INS-313
chr3-147111722-INS-93
chr3-147120205-INS-165
chr3-147185917-INS-429
chr3-147225827-INS-55
chr3-147233857-INS-221
chr3-147254580-INS-68
chr3-147273167-INS-135
chr3-147276849-INS-417
chr3-147279910-INS-6425
chr3-147288018-INS-59
chr3-147288590-INS-81
chr3-147301125-INS-200
chr3-147306685-INS-128
chr3-147308638-INS-218
chr3-147319624-INS-142
chr3-147325063-INS-98
chr3-147333556-INS-280
chr3-147428325-INS-114
chr3-147430951-INS-106
chr3-147437324-INS-296
chr3-147443052-INS-578
chr3-147461914-INS-365
chr3-147515457-INS-148
chr3-147539242-INS-75
chr3-147558343-INS-52
chr3-147561812-INS-174
chr3-147574145-INS-219
chr3-147600050-INS-86
chr3-147604272-INS-113
chr3-147697432-INS-64
chr3-147700267-INS-988
chr3-147701671-INS-440
chr3-147721617-INS-57
chr3-147727552-INS-173
chr3-147735395-INS-55
chr3-147743624-INS-56
chr3-147744648-INS-55
chr3-147745212-INS-123
chr3-147774021-INS-14064
chr3-147799588-INS-386
chr3-147800513-INS-80
chr3-147808333-INS-707
chr3-147827193-INS-86
chr3-147840640-INS-65
chr3-147912845-INS-119
chr3-147935254-INS-116
chr3-147942703-INS-63
chr3-147948950-INS-512
chr3-147959083-INS-1432
chr3-147967011-INS-53
chr3-148034323-INS-76
chr3-148052866-INS-88
chr3-148103400-INS-166
chr3-148107683-INS-265
chr3-148108808-INS-498
chr3-148122532-INS-60
chr3-148126340-INS-72
chr3-148129631-INS-76
chr3-148146160-INS-72
chr3-148167865-INS-96
chr3-148176369-INS-213
chr3-148190827-INS-120
chr3-148192202-INS-97
chr3-148194385-INS-165
chr3-148239493-INS-52
chr3-148264195-INS-144
chr3-148281164-INS-62
chr3-148283952-INS-60
chr3-148299741-INS-64

chr3-148313574-INS-58
chr3-148319968-INS-196
chr3-148328732-INS-189
chr3-148345551-INS-185
chr3-148345764-INS-159
chr3-148357910-INS-514
chr3-148368531-INS-573
chr3-148381587-INS-61
chr3-148391050-INS-162
chr3-148391409-INS-231
chr3-148392958-INS-762
chr3-148426858-INS-371
chr3-148464753-INS-114
chr3-148484683-INS-61
chr3-148498703-INS-58
chr3-148507992-INS-78
chr3-148556186-INS-80
chr3-148584760-INS-84
chr3-148608555-INS-226
chr3-148737345-INS-198
chr3-148738987-INS-84
chr3-148738987-INS-989
chr3-148739591-INS-62
chr3-148745804-INS-148
chr3-148751454-INS-236
chr3-148755579-INS-98
chr3-148763640-INS-201
chr3-148792855-INS-522
chr3-148833610-INS-96
chr3-148841560-INS-195
chr3-148843610-INS-64
chr3-148843610-INS-739
chr3-148851678-INS-369
chr3-148873139-INS-807
chr3-148878990-INS-70
chr3-148881397-INS-55
chr3-148892510-INS-116
chr3-148909610-INS-58
chr3-148912900-INS-88
chr3-148915248-INS-90
chr3-148934466-INS-162
chr3-148938067-INS-60
chr3-148973381-INS-56
chr3-148985927-INS-61
chr3-149003699-INS-193
chr3-149066644-INS-208
chr3-149068293-INS-135
chr3-149078273-INS-171
chr3-149080662-INS-5541
chr3-149089949-INS-509
chr3-149098066-INS-275
chr3-149110271-INS-60
chr3-149135522-INS-67
chr3-149157647-INS-60
chr3-149166050-INS-115
chr3-149183935-INS-1187
chr3-149188495-INS-401
chr3-149227288-INS-147
chr3-149278238-INS-51
chr3-149280615-INS-71
chr3-149293022-INS-63
chr3-149308485-INS-132
chr3-149311160-INS-72
chr3-149320033-INS-488
chr3-149348962-INS-954
chr3-149420281-INS-96
chr3-149425110-INS-3530
chr3-149450924-INS-132
chr3-149452163-INS-129
chr3-149457737-INS-315
chr3-149485999-INS-54
chr3-149488160-INS-54
chr3-149491378-INS-6305
chr3-149505685-INS-122
chr3-149514524-INS-102
chr3-149525911-INS-51
chr3-149530056-INS-58

chr3-149531402-INS-269
chr3-149532922-INS-56
chr3-149557437-INS-137
chr3-149558139-INS-127
chr3-149589156-INS-79
chr3-149612120-INS-94
chr3-149627581-INS-5121
chr3-149660771-INS-622
chr3-149662969-INS-479
chr3-149688857-INS-6383
chr3-149726378-INS-118
chr3-149731779-INS-58
chr3-149734690-INS-78
chr3-149748680-INS-362
chr3-149764424-INS-200
chr3-149773159-INS-95
chr3-149779397-INS-73
chr3-149781890-INS-113
chr3-149795752-INS-65
chr3-149814802-INS-54
chr3-149837276-INS-519
chr3-149844149-INS-879
chr3-149857634-INS-107
chr3-149907322-INS-927
chr3-149909640-INS-370
chr3-149921632-INS-65
chr3-149991190-INS-53
chr3-150004066-INS-53
chr3-150013239-INS-108
chr3-150020126-INS-746
chr3-150094225-INS-60
chr3-150095647-INS-226
chr3-150119425-INS-5626
chr3-150158971-INS-125
chr3-150206181-INS-187
chr3-150212998-INS-324
chr3-150294295-INS-503
chr3-150300836-INS-110
chr3-150304414-INS-66
chr3-150317047-INS-7528
chr3-150336503-INS-75
chr3-150354543-INS-6668
chr3-150361344-INS-723
chr3-150364129-INS-348
chr3-150366114-INS-927
chr3-150375259-INS-488
chr3-150386288-INS-275
chr3-150417867-INS-381
chr3-150437190-INS-78
chr3-150437539-INS-892
chr3-150439197-INS-274
chr3-150457815-INS-55
chr3-150464795-INS-72
chr3-150469699-INS-52
chr3-150479302-INS-423
chr3-150485807-INS-1501
chr3-150503103-INS-70
chr3-150509983-INS-6730
chr3-150570947-INS-165
chr3-150583443-INS-232
chr3-150586286-INS-52
chr3-150594543-INS-152
chr3-150647840-INS-54
chr3-150650291-INS-339
chr3-150668923-INS-423
chr3-150682207-INS-61
chr3-150683780-INS-216
chr3-150772367-INS-5235
chr3-150778564-INS-123
chr3-150795519-INS-53
chr3-150824616-INS-76
chr3-150835441-INS-54
chr3-150852984-INS-579
chr3-150905915-INS-52
chr3-150927232-INS-280
chr3-150934938-INS-240
chr3-150957256-INS-56

chr3-150960071-INS-165
chr3-150974825-INS-392
chr3-150985164-INS-80
chr3-151066756-INS-53
chr3-151072985-INS-95
chr3-151080160-INS-150
chr3-151111914-INS-118
chr3-151135673-INS-94
chr3-151143964-INS-968
chr3-151152766-INS-60
chr3-151170362-INS-57
chr3-151190594-INS-77
chr3-151227774-INS-242
chr3-151239244-INS-72
chr3-151242687-INS-149
chr3-151257634-INS-590
chr3-151265394-INS-428
chr3-151290567-INS-60
chr3-151292227-INS-139
chr3-151298286-INS-400
chr3-151318147-INS-95
chr3-151320527-INS-85
chr3-151369420-INS-7526
chr3-151401889-INS-230
chr3-151407046-INS-120
chr3-151438872-INS-80
chr3-151469376-INS-141
chr3-151491884-INS-63
chr3-151525941-INS-55
chr3-151538681-INS-780
chr3-151567352-INS-68
chr3-151619137-INS-53
chr3-151625917-INS-835
chr3-151644966-INS-315
chr3-151656215-INS-470
chr3-151673337-INS-126
chr3-151674212-INS-177
chr3-151682516-INS-317
chr3-151693865-INS-6625
chr3-151694900-INS-57
chr3-151793620-INS-162
chr3-151802836-INS-540
chr3-151817819-INS-65
chr3-151821138-INS-78
chr3-151824962-INS-225
chr3-151830578-INS-75
chr3-151830940-INS-119
chr3-151837236-INS-65
chr3-151843421-INS-56
chr3-151849877-INS-59
chr3-151851817-INS-287
chr3-151858861-INS-102
chr3-151863333-INS-191
chr3-151879337-INS-77
chr3-151904707-INS-588
chr3-151909667-INS-95
chr3-151941632-INS-76
chr3-151956365-INS-174
chr3-151965358-INS-79
chr3-151986291-INS-63
chr3-151991123-INS-342
chr3-152005378-INS-249
chr3-152020271-INS-1229
chr3-152020356-INS-164
chr3-152047274-INS-78
chr3-152049830-INS-62
chr3-152060914-INS-228
chr3-152080655-INS-130
chr3-152118453-INS-90
chr3-152120867-INS-134
chr3-152123338-INS-60
chr3-152187548-INS-337
chr3-152192187-INS-119
chr3-152199206-INS-82
chr3-152213137-INS-844
chr3-152263450-INS-536
chr3-152271154-INS-598

chr3-152301860-INS-619
chr3-152304671-INS-170
chr3-152313536-INS-198
chr3-152319826-INS-175
chr3-152387621-INS-202
chr3-152392988-INS-641
chr3-152400060-INS-670
chr3-152432210-INS-65
chr3-152445575-INS-185
chr3-152454098-INS-238
chr3-152461108-INS-157
chr3-152464669-INS-812
chr3-152477160-INS-3091
chr3-152495993-INS-396
chr3-152532147-INS-91
chr3-152537416-INS-783
chr3-152546662-INS-52
chr3-152550862-INS-86
chr3-152557023-INS-51
chr3-152584767-INS-75
chr3-152593167-INS-261
chr3-152601252-INS-102
chr3-152647477-INS-150
chr3-152659039-INS-94
chr3-152666896-INS-93
chr3-152699906-INS-97
chr3-152701171-INS-82
chr3-152704274-INS-348
chr3-152705911-INS-561
chr3-152709675-INS-68
chr3-152712186-INS-95
chr3-152715516-INS-51
chr3-152722751-INS-74
chr3-152735354-INS-187
chr3-152736348-INS-810
chr3-152739855-INS-90
chr3-152760657-INS-431
chr3-152776473-INS-113
chr3-152777373-INS-514
chr3-152784538-INS-106
chr3-152813963-INS-258
chr3-152830916-INS-58
chr3-152832806-INS-212
chr3-152853896-INS-6442
chr3-152854426-INS-204
chr3-152861116-INS-53
chr3-152866525-INS-136
chr3-152879324-INS-199
chr3-152885246-INS-65
chr3-152919065-INS-324
chr3-152931019-INS-437
chr3-152934367-INS-120
chr3-152948120-INS-51
chr3-152963644-INS-67
chr3-152972719-INS-64
chr3-153020734-INS-143
chr3-153048729-INS-80
chr3-153062530-INS-99
chr3-153112344-INS-195
chr3-153123302-INS-78
chr3-153126175-INS-65
chr3-153134332-INS-96
chr3-153141250-INS-72
chr3-153156552-INS-225
chr3-153168870-INS-105
chr3-153199160-INS-62
chr3-153229145-INS-202
chr3-153278911-INS-192
chr3-153302533-INS-177
chr3-153312860-INS-341
chr3-153332426-INS-132
chr3-153338823-INS-53
chr3-153356635-INS-870
chr3-153380900-INS-64
chr3-153385873-INS-59
chr3-153403186-INS-191
chr3-153415973-INS-72

chr3-153424973-INS-63
chr3-153433143-INS-179
chr3-153434934-INS-316
chr3-153443573-INS-798
chr3-153450950-INS-138
chr3-153475941-INS-62
chr3-153490611-INS-363
chr3-153498469-INS-154
chr3-153500178-INS-52
chr3-153504307-INS-377
chr3-153517159-INS-57
chr3-153520737-INS-401
chr3-153524691-INS-136
chr3-153534960-INS-221
chr3-153543764-INS-179
chr3-153555800-INS-56
chr3-153557593-INS-164
chr3-153584137-INS-214
chr3-153585863-INS-217
chr3-153590020-INS-188
chr3-153604027-INS-198
chr3-153605448-INS-205
chr3-153625395-INS-169
chr3-153627684-INS-226
chr3-153641136-INS-440
chr3-153645054-INS-5552
chr3-153656831-INS-223
chr3-153664955-INS-60
chr3-153685717-INS-104
chr3-153709009-INS-203
chr3-153717867-INS-55
chr3-153721773-INS-92
chr3-153740624-INS-195
chr3-153765076-INS-107
chr3-153767481-INS-539
chr3-153769719-INS-257
chr3-153780082-INS-60
chr3-153822146-INS-365
chr3-153823098-INS-52
chr3-153827863-INS-112
chr3-153840374-INS-55
chr3-153845959-INS-253
chr3-153881169-INS-221
chr3-153884962-INS-135
chr3-153901527-INS-140
chr3-153926723-INS-74
chr3-153929495-INS-54
chr3-153934071-INS-66
chr3-154013927-INS-67
chr3-154024567-INS-68
chr3-154030836-INS-54
chr3-154044084-INS-108
chr3-154050090-INS-95
chr3-154075499-INS-54
chr3-154078391-INS-458
chr3-154084587-INS-381
chr3-154090396-INS-308
chr3-154095913-INS-516
chr3-154101986-INS-83
chr3-154124484-INS-65
chr3-154134979-INS-323
chr3-154160832-INS-98
chr3-154178939-INS-112
chr3-154179439-INS-192
chr3-154192298-INS-362
chr3-154194521-INS-152
chr3-154242864-INS-190
chr3-154246817-INS-134
chr3-154249766-INS-782
chr3-154251730-INS-673
chr3-154259834-INS-437
chr3-154269652-INS-68
chr3-154283843-INS-592
chr3-154284099-INS-740
chr3-154286311-INS-54
chr3-154292585-INS-60
chr3-154292885-INS-260

chr3-154314844-INS-284
chr3-154322541-INS-53
chr3-154323590-INS-86
chr3-154325496-INS-76
chr3-154334535-INS-297
chr3-154344365-INS-206
chr3-154356194-INS-72
chr3-154372717-INS-99
chr3-154394663-INS-273
chr3-154429366-INS-96
chr3-154447526-INS-6161
chr3-154459114-INS-57
chr3-154465114-INS-4546
chr3-154467089-INS-5493
chr3-154468181-INS-66
chr3-154484502-INS-385
chr3-154509088-INS-169
chr3-154510484-INS-5543
chr3-154518097-INS-87
chr3-154551537-INS-80
chr3-154552161-INS-241
chr3-154584186-INS-60
chr3-154613697-INS-471
chr3-154615985-INS-90
chr3-154618242-INS-555
chr3-154629060-INS-432
chr3-154630736-INS-196
chr3-154642145-INS-262
chr3-154663039-INS-411
chr3-154713538-INS-347
chr3-154734949-INS-78
chr3-154739132-INS-119
chr3-154779493-INS-291
chr3-154809112-INS-65
chr3-154834445-INS-61
chr3-154858454-INS-6997
chr3-154868725-INS-72
chr3-154869832-INS-168
chr3-154872782-INS-5750
chr3-154909335-INS-128
chr3-154936428-INS-79
chr3-154964391-INS-72
chr3-154967759-INS-399
chr3-154999356-INS-86
chr3-155004515-INS-273
chr3-155024357-INS-237
chr3-155025525-INS-196
chr3-155051859-INS-483
chr3-155074731-INS-204
chr3-155092916-INS-57
chr3-155094742-INS-137
chr3-155117665-INS-6280
chr3-155118178-INS-96
chr3-155124433-INS-58
chr3-155156615-INS-261
chr3-155178463-INS-336
chr3-155229469-INS-81
chr3-155232686-INS-248
chr3-155234938-INS-54
chr3-155236961-INS-76
chr3-155256176-INS-167
chr3-155260642-INS-393
chr3-155273618-INS-52
chr3-155327394-INS-64
chr3-155329996-INS-66
chr3-155332560-INS-440
chr3-155364166-INS-2722
chr3-155379644-INS-56
chr3-155421847-INS-87
chr3-155423585-INS-6769
chr3-155452341-INS-167
chr3-155498701-INS-338
chr3-155511574-INS-129
chr3-155522700-INS-80
chr3-155550043-INS-97
chr3-155556058-INS-63
chr3-155560262-INS-66

chr3-155576945-INS-160
chr3-155596666-INS-6333
chr3-155618608-INS-801
chr3-155663406-INS-68
chr3-155689711-INS-6527
chr3-155692912-INS-1992
chr3-155703266-INS-133
chr3-155704330-INS-170
chr3-155730514-INS-99
chr3-155731029-INS-1511
chr3-155750303-INS-218
chr3-155752531-INS-579
chr3-155773107-INS-81
chr3-155789352-INS-1661
chr3-155789365-INS-3919
chr3-155802594-INS-87
chr3-155810858-INS-56
chr3-155825921-INS-3355
chr3-155827902-INS-72
chr3-155830193-INS-86
chr3-155836310-INS-52
chr3-155862332-INS-8734
chr3-155865819-INS-54
chr3-155892384-INS-88
chr3-155925461-INS-206
chr3-155941864-INS-273
chr3-155952994-INS-6675
chr3-155955059-INS-4057
chr3-155955897-INS-51
chr3-155959266-INS-175
chr3-155987063-INS-66
chr3-156004255-INS-103
chr3-156043874-INS-66
chr3-156087783-INS-7769
chr3-156092573-INS-78
chr3-156111253-INS-2786
chr3-156111810-INS-83
chr3-156115708-INS-67
chr3-156118308-INS-123
chr3-156121434-INS-67
chr3-156125092-INS-1550
chr3-156139695-INS-1578
chr3-156156650-INS-64
chr3-156218208-INS-6567
chr3-156245668-INS-477
chr3-156271656-INS-197
chr3-156306421-INS-87
chr3-156319004-INS-195
chr3-156320901-INS-2682
chr3-156325077-INS-589
chr3-156360378-INS-399
chr3-156387066-INS-58
chr3-156391050-INS-61
chr3-156442303-INS-72
chr3-156447854-INS-6183
chr3-156472699-INS-95
chr3-156503657-INS-353
chr3-156517420-INS-62
chr3-156564003-INS-127
chr3-156599671-INS-106
chr3-156621233-INS-7211
chr3-156623766-INS-1042
chr3-156659137-INS-54
chr3-156697612-INS-845
chr3-156706713-INS-569
chr3-156727056-INS-70
chr3-156745112-INS-145
chr3-156750878-INS-109
chr3-156758637-INS-1485
chr3-156771227-INS-6590
chr3-156788535-INS-89
chr3-156818251-INS-76
chr3-156834571-INS-275
chr3-156889419-INS-1355
chr3-156900622-INS-6872
chr3-156939487-INS-219
chr3-156947900-INS-76

chr3-156956640-INS-578
chr3-156980184-INS-80
chr3-156987981-INS-54
chr3-156994784-INS-63
chr3-156998891-INS-70
chr3-157002541-INS-61
chr3-157032392-INS-58
chr3-157034374-INS-146
chr3-157066975-INS-5403
chr3-157074419-INS-180
chr3-157075541-INS-358
chr3-157079015-INS-389
chr3-157085388-INS-358
chr3-157087213-INS-113
chr3-157094284-INS-127
chr3-157112660-INS-6901
chr3-157130774-INS-1890
chr3-157137999-INS-6054
chr3-157140809-INS-745
chr3-157142204-INS-7290
chr3-157147759-INS-302
chr3-157166179-INS-63
chr3-157181996-INS-57
chr3-157190941-INS-76
chr3-157192131-INS-309
chr3-157220673-INS-339
chr3-157226825-INS-258
chr3-157231002-INS-7143
chr3-157232368-INS-55
chr3-157281788-INS-313
chr3-157304844-INS-701
chr3-157309879-INS-52
chr3-157397480-INS-722
chr3-157427707-INS-6953
chr3-157438903-INS-655
chr3-157455209-INS-187
chr3-157472437-INS-310
chr3-157478352-INS-278
chr3-157496886-INS-831
chr3-157501269-INS-5547
chr3-157502014-INS-259
chr3-157502897-INS-442
chr3-157517859-INS-92
chr3-157540552-INS-4891
chr3-157549493-INS-167
chr3-157574765-INS-430
chr3-157575772-INS-1194
chr3-157578789-INS-208
chr3-157582640-INS-6632
chr3-157591860-INS-173
chr3-157614231-INS-243
chr3-157618215-INS-229
chr3-157649862-INS-167
chr3-157662404-INS-56
chr3-157703839-INS-156
chr3-157746493-INS-163
chr3-157755366-INS-75
chr3-157813984-INS-1751
chr3-157845349-INS-51
chr3-157890433-INS-652
chr3-157899405-INS-55
chr3-157906075-INS-522
chr3-157916046-INS-769
chr3-157922104-INS-599
chr3-157924851-INS-74
chr3-157930997-INS-58
chr3-157943976-INS-476
chr3-157949751-INS-438
chr3-157952654-INS-330
chr3-157956587-INS-467
chr3-157969693-INS-86
chr3-158035067-INS-201
chr3-158040712-INS-206
chr3-158045532-INS-1139
chr3-158068808-INS-58
chr3-158069801-INS-569
chr3-158084687-INS-449

chr3-158095364-INS-497
chr3-158158233-INS-81
chr3-158179955-INS-88
chr3-158237417-INS-6518
chr3-158295402-INS-687
chr3-158314851-INS-951
chr3-158315629-INS-178
chr3-158346540-INS-94
chr3-158368386-INS-280
chr3-158377798-INS-190
chr3-158383058-INS-816
chr3-158395836-INS-142
chr3-158418233-INS-627
chr3-158441929-INS-208
chr3-158519554-INS-133
chr3-158540732-INS-486
chr3-158588130-INS-187
chr3-158625758-INS-60
chr3-158655382-INS-185
chr3-158662204-INS-306
chr3-158680212-INS-54
chr3-158687663-INS-6507
chr3-158702751-INS-400
chr3-158711511-INS-319
chr3-158764182-INS-625
chr3-158774135-INS-151
chr3-158784535-INS-76
chr3-158788274-INS-138
chr3-158813829-INS-628
chr3-158825187-INS-508
chr3-158838682-INS-64
chr3-158859636-INS-218
chr3-158917760-INS-310
chr3-158922983-INS-380
chr3-158959774-INS-136
chr3-158971832-INS-6143
chr3-158978643-INS-56
chr3-158991458-INS-1076
chr3-159005427-INS-396
chr3-159016263-INS-5377
chr3-159016681-INS-977
chr3-159030941-INS-284
chr3-159071749-INS-1890
chr3-159077968-INS-497
chr3-159090015-INS-257
chr3-159100780-INS-265
chr3-159104486-INS-69
chr3-159113482-INS-74
chr3-159121085-INS-440
chr3-159163085-INS-68
chr3-159165798-INS-118
chr3-159178522-INS-6479
chr3-159217293-INS-69
chr3-159307925-INS-56
chr3-159321620-INS-242
chr3-159351180-INS-374
chr3-159356165-INS-56
chr3-159369453-INS-297
chr3-159390952-INS-7167
chr3-159416453-INS-4739
chr3-159418472-INS-6214
chr3-159421898-INS-377
chr3-159442203-INS-462
chr3-159496570-INS-59
chr3-159497711-INS-267
chr3-159577009-INS-96
chr3-159578031-INS-359
chr3-159579900-INS-170
chr3-159582590-INS-550
chr3-159613797-INS-68
chr3-159637049-INS-591
chr4-3069282-INS-169
chr4-3082199-INS-63
chr4-3090555-INS-52
chr4-3216359-INS-5557
chr4-3229150-INS-7244
chr4-3231739-INS-150

chr4-3250304-INS-5252
chr4-3271135-INS-72
chr4-3271621-INS-187
chr4-3275600-INS-6228
chr4-3275800-INS-52
chr4-3296731-INS-5188
chr4-3302596-INS-107
chr4-3320314-INS-81
chr4-3354759-INS-445
chr4-3367067-INS-412
chr4-3387233-INS-5081
chr4-3412091-INS-72
chr4-3416662-INS-197
chr4-3418540-INS-361
chr4-3430291-INS-415
chr4-3444107-INS-7047
chr4-3453751-INS-194
chr4-3471156-INS-99
chr4-3483499-INS-373
chr4-3502144-INS-266
chr4-3511352-INS-874
chr4-3528056-INS-122
chr4-3538942-INS-6315
chr4-3542895-INS-1075
chr4-3548371-INS-82
chr4-3593673-INS-78
chr4-3593918-INS-5446
chr4-3606925-INS-61
chr4-3613707-INS-141
chr4-3643495-INS-437
chr4-3651261-INS-726
chr4-3669528-INS-68
chr4-3685271-INS-60
chr4-3704264-INS-823
chr4-3717467-INS-59
chr4-3726736-INS-70
chr4-3764493-INS-385
chr4-3771522-INS-6487
chr4-3780390-INS-119
chr4-3785183-INS-58
chr4-3817216-INS-8401
chr4-3842464-INS-66
chr4-3861555-INS-342
chr4-3879741-INS-7265
chr4-3885507-INS-8161
chr4-3894307-INS-844
chr4-3927009-INS-75
chr4-3945098-INS-572
chr4-3948020-INS-51
chr4-3952109-INS-68
chr4-3978691-INS-731
chr4-4046193-INS-6697
chr4-4061507-INS-52
chr4-4072985-INS-205
chr4-4080442-INS-86
chr4-4090535-INS-82
chr4-4098052-INS-736
chr4-4108646-INS-109
chr4-4147244-INS-100
chr4-4156240-INS-60
chr4-4158727-INS-144
chr4-4176689-INS-165
chr4-4185019-INS-190
chr4-4190685-INS-181
chr4-4201035-INS-7093
chr4-4209673-INS-59
chr4-4257962-INS-62
chr4-4283183-INS-7847
chr4-4291307-INS-56
chr4-4340087-INS-56
chr4-4349532-INS-6678
chr4-4387566-INS-65
chr4-4403110-INS-3193
chr4-4417728-INS-74
chr4-4423087-INS-99
chr4-4434515-INS-6447
chr4-4447178-INS-81

chr4-4448356-INS-103
chr4-4467373-INS-623
chr4-4483013-INS-3870
chr4-4502332-INS-5841
chr4-4524583-INS-437
chr4-4527673-INS-1297
chr4-4535143-INS-212
chr4-4545346-INS-120
chr4-4584679-INS-128
chr4-4593340-INS-169
chr4-4599002-INS-962
chr4-4620944-INS-209
chr4-4625116-INS-5545
chr4-4632653-INS-220
chr4-4637652-INS-105
chr4-4709354-INS-6507
chr4-4710036-INS-902
chr4-4715099-INS-62
chr4-4719398-INS-216
chr4-4723265-INS-69
chr4-4724767-INS-191
chr4-4789644-INS-197
chr4-4798640-INS-266
chr4-4832260-INS-498
chr4-4839706-INS-51
chr4-4906910-INS-86
chr4-4907588-INS-691
chr4-4930039-INS-75
chr4-4952559-INS-422
chr4-4961155-INS-398
chr4-5038157-INS-79
chr4-5038372-INS-758
chr4-5074641-INS-168
chr4-5081740-INS-68
chr4-5089590-INS-390
chr4-5135881-INS-73
chr4-5168872-INS-111
chr4-5173858-INS-6839
chr4-5183122-INS-6217
chr4-5199638-INS-458
chr4-5205515-INS-60
chr4-5211454-INS-78
chr4-5286456-INS-197
chr4-5336991-INS-76
chr4-5358706-INS-161
chr4-5361406-INS-401
chr4-5385990-INS-185
chr4-5399299-INS-81
chr4-5422946-INS-151
chr4-5427706-INS-430
chr4-5433143-INS-401
chr4-5473159-INS-6070
chr4-5498028-INS-485
chr4-5532999-INS-126
chr4-5539386-INS-147
chr4-5543432-INS-194
chr4-5571487-INS-96
chr4-5583768-INS-354
chr4-5604304-INS-878
chr4-5611678-INS-83
chr4-5616533-INS-52
chr4-5626529-INS-607
chr4-5630403-INS-765
chr4-5638771-INS-94
chr4-5654217-INS-675
chr4-5670341-INS-6836
chr4-5673512-INS-184
chr4-5678371-INS-238
chr4-5685925-INS-58
chr4-5697395-INS-102
chr4-5703261-INS-340
chr4-5705572-INS-108
chr4-5716420-INS-54
chr4-5727382-INS-206
chr4-5748485-INS-280
chr4-5750286-INS-238
chr4-5784735-INS-54

chr4-5790565-INS-150
chr4-5817865-INS-126
chr4-5847838-INS-357
chr4-5860760-INS-285
chr4-5918487-INS-80
chr4-5921824-INS-64
chr4-5924234-INS-539
chr4-5966877-INS-192
chr4-5978546-INS-100
chr4-5981744-INS-855
chr4-6082487-INS-64
chr4-6083323-INS-476
chr4-6100801-INS-6503
chr4-6101937-INS-347
chr4-6102477-INS-2120
chr4-6113699-INS-191
chr4-6115644-INS-129
chr4-6143051-INS-330
chr4-6163182-INS-132
chr4-6164709-INS-71
chr4-6204864-INS-162
chr4-6206238-INS-62
chr4-6233612-INS-1888
chr4-6245859-INS-143
chr4-6259049-INS-176
chr4-6274063-INS-56
chr4-6279289-INS-51
chr4-6323563-INS-77
chr4-6377463-INS-80
chr4-6383358-INS-988
chr4-6387242-INS-201
chr4-6390305-INS-530
chr4-6421610-INS-924
chr4-6432315-INS-365
chr4-6439632-INS-105
chr4-6440524-INS-144
chr4-6444077-INS-184
chr4-6478323-INS-373
chr4-6490632-INS-427
chr4-6505623-INS-62
chr4-6507637-INS-83
chr4-6510464-INS-106
chr4-6515483-INS-383
chr4-6520315-INS-7559
chr4-6533482-INS-211
chr4-6538594-INS-65
chr4-6540836-INS-55
chr4-6556504-INS-97
chr4-6559620-INS-345
chr4-6560891-INS-343
chr4-6566213-INS-128
chr4-6628070-INS-7575
chr4-6648246-INS-85
chr4-6663550-INS-535
chr4-6670911-INS-72
chr4-6673684-INS-164
chr4-6683896-INS-69
chr4-6685410-INS-54
chr4-6770022-INS-81
chr4-6798926-INS-60
chr4-6800552-INS-430
chr4-6829387-INS-78
chr4-6837726-INS-213
chr4-6846702-INS-174
chr4-6853922-INS-165
chr4-6879729-INS-257
chr4-6912372-INS-64
chr4-6964785-INS-69
chr4-7010098-INS-1125
chr4-7013283-INS-87
chr4-7024692-INS-120
chr4-7026123-INS-56
chr4-7030585-INS-64
chr4-7033378-INS-80
chr4-7076054-INS-105
chr4-7076844-INS-464
chr4-7094110-INS-58

chr4-7109450-INS-4805
chr4-7110943-INS-6158
chr4-7113927-INS-54
chr4-7127779-INS-113
chr4-7129858-INS-142
chr4-7134424-INS-430
chr4-7145720-INS-52
chr4-7154750-INS-290
chr4-7183913-INS-76
chr4-7194605-INS-182
chr4-7232147-INS-58
chr4-7234856-INS-59
chr4-7259152-INS-6401
chr4-7282991-INS-67
chr4-7293092-INS-214
chr4-7304097-INS-52
chr4-7325306-INS-81
chr4-7335342-INS-736
chr4-7344387-INS-56
chr4-7345973-INS-103
chr4-7383014-INS-609
chr4-7403686-INS-56
chr4-7418457-INS-274
chr4-7428294-INS-65
chr4-7467153-INS-956
chr4-7468915-INS-273
chr4-7511812-INS-962
chr4-7553854-INS-102
chr4-7577361-INS-257
chr4-7588368-INS-78
chr4-7590807-INS-56
chr4-7592663-INS-52
chr4-7628742-INS-407
chr4-7646012-INS-646
chr4-7666490-INS-116
chr4-7697280-INS-353
chr4-7748642-INS-8677
chr4-7792216-INS-552
chr4-7798152-INS-109
chr4-7799796-INS-94
chr4-7842457-INS-60
chr4-7862711-INS-59
chr4-7878383-INS-872
chr4-7879452-INS-134
chr4-7922372-INS-6543
chr4-7923556-INS-2004
chr4-7933668-INS-363
chr4-7941824-INS-141
chr4-7957098-INS-152
chr4-7976902-INS-59
chr4-7991198-INS-144
chr4-7998954-INS-326
chr4-8018973-INS-70
chr4-8036582-INS-413
chr4-8038258-INS-60
chr4-8071027-INS-514
chr4-8107480-INS-190
chr4-8109433-INS-407
chr4-8144991-INS-53
chr4-8152066-INS-73
chr4-8212872-INS-331
chr4-8223103-INS-400
chr4-8230097-INS-152
chr4-8235905-INS-65
chr4-8265993-INS-1890
chr4-8273863-INS-104
chr4-8279321-INS-213
chr4-8280711-INS-74
chr4-8283607-INS-419
chr4-8306524-INS-61
chr4-8307799-INS-148
chr4-8334275-INS-80
chr4-8359906-INS-128
chr4-8392336-INS-304
chr4-8416302-INS-90
chr4-8421136-INS-472
chr4-8423520-INS-145

chr4-8431762-INS-598
chr4-8443097-INS-214
chr4-8447730-INS-275
chr4-8456136-INS-270
chr4-8466086-INS-243
chr4-8487335-INS-138
chr4-8497941-INS-123
chr4-8509573-INS-339
chr4-8529759-INS-245
chr4-8566896-INS-107
chr4-8604636-INS-72
chr4-8629970-INS-106
chr4-8630209-INS-703
chr4-8654252-INS-116
chr4-8686268-INS-400
chr4-8821620-INS-444
chr4-8824546-INS-118
chr4-8842630-INS-141
chr4-8883302-INS-545
chr4-8902631-INS-60
chr4-8978463-INS-56
chr4-8982142-INS-89
chr4-9002409-INS-66
chr4-9029389-INS-491
chr4-9032995-INS-87
chr4-9061359-INS-72
chr4-9070913-INS-382
chr4-9071867-INS-123
chr4-9072895-INS-359
chr4-9086646-INS-146
chr4-9091018-INS-197
chr4-9117525-INS-149
chr4-9142568-INS-379
chr4-9146950-INS-154
chr4-9150162-INS-400
chr4-9157885-INS-57
chr4-9215512-INS-456
chr4-9231222-INS-52
chr4-9242692-INS-165
chr4-9284097-INS-56
chr4-9299415-INS-167
chr4-9320593-INS-1890
chr4-9325914-INS-1450
chr4-9349963-INS-155
chr4-9364145-INS-56
chr4-9378946-INS-80
chr4-9385885-INS-231
chr4-9388774-INS-6729
chr4-9394945-INS-529
chr4-9397666-INS-508
chr4-9407594-INS-154
chr4-9431252-INS-66
chr4-9442196-INS-56
chr4-9476620-INS-135
chr4-9488623-INS-59
chr4-9493857-INS-214
chr4-9526229-INS-235
chr4-9540659-INS-105
chr4-9552774-INS-52
chr4-9555306-INS-294
chr4-9584330-INS-155
chr4-9605496-INS-120
chr4-9613733-INS-60
chr4-9654427-INS-256
chr4-9665150-INS-56
chr4-9681332-INS-431
chr4-9689801-INS-777
chr4-9703733-INS-206
chr4-9715577-INS-624
chr4-9784210-INS-87
chr4-9790253-INS-615
chr4-9791238-INS-68
chr4-9827597-INS-140
chr4-9838039-INS-338
chr4-9891349-INS-56
chr4-9954354-INS-366
chr4-9964968-INS-56

chr4-9985353-INS-84
chr4-9989919-INS-95
chr4-9992020-INS-366
chr4-10001723-INS-284
chr4-10028772-INS-292
chr4-10030746-INS-61
chr4-10042050-INS-165
chr4-10085330-INS-4686
chr4-10090917-INS-2581
chr4-10091385-INS-6595
chr4-10099505-INS-140
chr4-10121152-INS-2316
chr4-10122790-INS-694
chr4-10126397-INS-440
chr4-10137522-INS-1030
chr4-10157756-INS-61
chr4-10167753-INS-327
chr4-10170811-INS-397
chr4-10174967-INS-138
chr4-10181743-INS-259
chr4-10208104-INS-169
chr4-10225114-INS-56
chr4-10246906-INS-552
chr4-10273216-INS-134
chr4-10278277-INS-788
chr4-10284978-INS-90
chr4-10298536-INS-178
chr4-10306528-INS-6922
chr4-10311377-INS-6677
chr4-10355058-INS-195
chr4-10361268-INS-69
chr4-10386331-INS-165
chr4-10404055-INS-92
chr4-10426978-INS-912
chr4-10438000-INS-225
chr4-10463738-INS-51
chr4-10509842-INS-73
chr4-10517225-INS-83
chr4-10518306-INS-64
chr4-10522172-INS-238
chr4-10525375-INS-852
chr4-10534863-INS-340
chr4-10551717-INS-1752
chr4-10577205-INS-241
chr4-10580517-INS-146
chr4-10650681-INS-51
chr4-10656401-INS-82
chr4-10664548-INS-4478
chr4-10694272-INS-93
chr4-10743855-INS-68
chr4-10748098-INS-8552
chr4-10752929-INS-387
chr4-10771832-INS-649
chr4-10802513-INS-2115
chr4-10806495-INS-556
chr4-10823728-INS-150
chr4-10845574-INS-564
chr4-10872803-INS-734
chr4-10903921-INS-80
chr4-10935792-INS-180
chr4-10939587-INS-63
chr4-10945409-INS-196
chr4-10951328-INS-325
chr4-10978135-INS-581
chr4-10987059-INS-69
chr4-10996572-INS-140
chr4-11000675-INS-210
chr4-11008663-INS-155
chr4-11010629-INS-376
chr4-11017548-INS-268
chr4-11031743-INS-519
chr4-11055282-INS-89
chr4-11076964-INS-200
chr4-11078724-INS-70
chr4-11083560-INS-209
chr4-11086669-INS-212
chr4-11106316-INS-407

chr4-11107973-INS-162
chr4-11113778-INS-155
chr4-11139668-INS-76
chr4-11162657-INS-422
chr4-11197892-INS-194
chr4-11215910-INS-62
chr4-11222736-INS-274
chr4-11231895-INS-72
chr4-11238721-INS-90
chr4-11256394-INS-507
chr4-11259780-INS-124
chr4-11266429-INS-566
chr4-11328283-INS-64
chr4-11347309-INS-285
chr4-11364424-INS-401
chr4-11426727-INS-90
chr4-11453629-INS-153
chr4-11532215-INS-155
chr4-11536152-INS-193
chr4-11553400-INS-112
chr4-11567400-INS-938
chr4-11571207-INS-114
chr4-11584491-INS-66
chr4-11605443-INS-561
chr4-11608897-INS-82
chr4-11609510-INS-112
chr4-11634502-INS-244
chr4-11679116-INS-161
chr4-11695004-INS-1587
chr4-11695068-INS-297
chr4-11697339-INS-53
chr4-11717035-INS-286
chr4-11757650-INS-80
chr4-11759017-INS-715
chr4-11786344-INS-5542
chr4-11793394-INS-70
chr4-11805422-INS-197
chr4-11810464-INS-70
chr4-11831451-INS-328
chr4-11843116-INS-541
chr4-11875591-INS-121
chr4-11884238-INS-553
chr4-11889379-INS-492
chr4-11898293-INS-60
chr4-11901757-INS-431
chr4-11903831-INS-217
chr4-11906937-INS-104
chr4-11913635-INS-58
chr4-11938879-INS-364
chr4-11947138-INS-207
chr4-11970090-INS-267
chr4-11981773-INS-765
chr4-12004790-INS-72
chr4-12037472-INS-1235
chr4-12037556-INS-58
chr4-12037635-INS-1514
chr4-12041912-INS-141
chr4-12047974-INS-266
chr4-12058059-INS-68
chr4-12097246-INS-72
chr4-12139498-INS-191
chr4-12190540-INS-493
chr4-12197940-INS-108
chr4-12224432-INS-426
chr4-12259005-INS-63
chr4-12265448-INS-123
chr4-12267826-INS-222
chr4-12280801-INS-6240
chr4-12290840-INS-210
chr4-12315968-INS-185
chr4-12317486-INS-6796
chr4-12369348-INS-5078
chr4-12370418-INS-59
chr4-12379821-INS-7332
chr4-12387228-INS-113
chr4-12414357-INS-6474
chr4-12423343-INS-66

chr4-12429907-INS-354
chr4-12458978-INS-283
chr4-12483023-INS-115
chr4-12499652-INS-394
chr4-12514286-INS-369
chr4-12529239-INS-103
chr4-12557136-INS-52
chr4-12558324-INS-372
chr4-12592715-INS-82
chr4-12633579-INS-53
chr4-12638645-INS-1876
chr4-12679919-INS-280
chr4-12684434-INS-2969
chr4-12699728-INS-659
chr4-12705385-INS-78
chr4-12706333-INS-163
chr4-12721915-INS-129
chr4-12738318-INS-56
chr4-12740582-INS-119
chr4-12746544-INS-55
chr4-12764475-INS-291
chr4-12785693-INS-66
chr4-12789146-INS-6738
chr4-12791565-INS-732
chr4-12854174-INS-195
chr4-12891486-INS-119
chr4-12894333-INS-322
chr4-12894507-INS-8850
chr4-12900950-INS-480
chr4-12909720-INS-57
chr4-12911884-INS-120
chr4-12922883-INS-61
chr4-12962819-INS-305
chr4-12977965-INS-6253
chr4-13003427-INS-167
chr4-13029977-INS-79
chr4-13073531-INS-328
chr4-13074552-INS-52
chr4-13087672-INS-98
chr4-13097813-INS-64
chr4-13101022-INS-480
chr4-13104458-INS-60
chr4-13113590-INS-52
chr4-13134629-INS-329
chr4-13146389-INS-63
chr4-13194233-INS-60
chr4-13220486-INS-98
chr4-13245025-INS-893
chr4-13251284-INS-7079
chr4-13265744-INS-77
chr4-13267044-INS-60
chr4-13364170-INS-164
chr4-13401801-INS-133
chr4-13416372-INS-84
chr4-13458472-INS-1451
chr4-13464160-INS-177
chr4-13478076-INS-75
chr4-13489151-INS-6644
chr4-13518036-INS-234
chr4-13564575-INS-114
chr4-13568458-INS-109
chr4-13584372-INS-130
chr4-13643599-INS-126
chr4-13656957-INS-101
chr4-13664521-INS-179
chr4-13701123-INS-115
chr4-13712324-INS-101
chr4-13772846-INS-201
chr4-13808515-INS-228
chr4-13824382-INS-83
chr4-13838042-INS-60
chr4-13866961-INS-210
chr4-13946872-INS-2801
chr4-13952804-INS-6409
chr4-13965020-INS-78
chr4-13983230-INS-2992
chr4-13986165-INS-4245

chr4-13986560-INS-6236
chr4-13997188-INS-63
chr4-14005204-INS-128
chr4-14006428-INS-360
chr4-14015932-INS-551
chr4-14034532-INS-6241
chr4-14068658-INS-6413
chr4-14078439-INS-5393
chr4-14081704-INS-328
chr4-14098556-INS-73
chr4-14146315-INS-66
chr4-14186977-INS-350
chr4-14189671-INS-129
chr4-14198750-INS-398
chr4-14233908-INS-985
chr4-14265574-INS-125
chr4-14299493-INS-188
chr4-14361491-INS-6797
chr4-14369477-INS-6272
chr4-14390906-INS-54
chr4-14421841-INS-51
chr4-14447728-INS-160
chr4-14489104-INS-122
chr4-14517605-INS-124
chr4-14527062-INS-897
chr4-14528704-INS-8419
chr4-14541397-INS-162
chr4-14592407-INS-510
chr4-14610723-INS-186
chr4-14613540-INS-172
chr4-14648648-INS-234
chr4-14651780-INS-7549
chr4-14657108-INS-6673
chr4-14668234-INS-6662
chr4-14695174-INS-345
chr4-14711601-INS-545
chr4-14714765-INS-1892
chr4-14744815-INS-79
chr4-14758997-INS-212
chr4-14782710-INS-95
chr4-14820669-INS-140
chr4-14884822-INS-65
chr4-14887912-INS-85
chr4-14898164-INS-612
chr4-14905863-INS-117
chr4-14908845-INS-80
chr4-14913043-INS-206
chr4-14924219-INS-94
chr4-14949244-INS-200
chr4-14994151-INS-249
chr4-14999522-INS-64
chr4-15010128-INS-136
chr4-15031568-INS-409
chr4-15036044-INS-138
chr4-15038303-INS-6004
chr4-15045012-INS-86
chr4-15047365-INS-1074
chr4-15061744-INS-235
chr4-15083618-INS-454
chr4-15100581-INS-340
chr4-15124671-INS-70
chr4-15130060-INS-132
chr4-15148092-INS-54
chr4-15151555-INS-193
chr4-15153940-INS-76
chr4-15169131-INS-108
chr4-15172415-INS-407
chr4-15176569-INS-55
chr4-15176703-INS-552
chr4-15201151-INS-623
chr4-15212767-INS-938
chr4-15227365-INS-68
chr4-15232979-INS-224
chr4-15261451-INS-66
chr4-15299489-INS-79
chr4-15315441-INS-72
chr4-15320216-INS-56

chr4-15330118-INS-91
chr4-15359121-INS-78
chr4-15365468-INS-139
chr4-15367392-INS-76
chr4-15386411-INS-71
chr4-15404033-INS-75
chr4-15425204-INS-952
chr4-15428416-INS-790
chr4-15439417-INS-114
chr4-15471069-INS-332
chr4-15481472-INS-7095
chr4-15500962-INS-130
chr4-15504993-INS-963
chr4-15507950-INS-80
chr4-15518875-INS-314
chr4-15530771-INS-329
chr4-15556930-INS-75
chr4-15567807-INS-95
chr4-15574892-INS-60
chr4-15595583-INS-151
chr4-15654116-INS-62
chr4-15678414-INS-57
chr4-15697065-INS-106
chr4-15704462-INS-836
chr4-15733937-INS-481
chr4-15744483-INS-6402
chr4-15757930-INS-96
chr4-15817607-INS-962
chr4-15819886-INS-106
chr4-15821489-INS-82
chr4-15831300-INS-177
chr4-15875383-INS-100
chr4-15903660-INS-344
chr4-15911478-INS-65
chr4-15928235-INS-140
chr4-15961704-INS-58
chr4-15986010-INS-406
chr4-15991902-INS-332
chr4-16040413-INS-319
chr4-16107313-INS-72
chr4-16120634-INS-5836
chr4-16122290-INS-53
chr4-16125807-INS-105
chr4-16203788-INS-546
chr4-16258049-INS-840
chr4-16266734-INS-203
chr4-16309938-INS-207
chr4-16313573-INS-6604
chr4-16314190-INS-537
chr4-16402356-INS-6220
chr4-16425344-INS-204
chr4-16426781-INS-405
chr4-16445023-INS-6897
chr4-16462690-INS-790
chr4-16467698-INS-121
chr4-16490820-INS-73
chr4-16500977-INS-2659
chr4-16509514-INS-531
chr4-16524975-INS-150
chr4-16542037-INS-451
chr4-16561758-INS-358
chr4-16570867-INS-453
chr4-16593667-INS-149
chr4-16613711-INS-351
chr4-16624069-INS-54
chr4-16628962-INS-114
chr4-16667608-INS-92
chr4-16681655-INS-52
chr4-16693923-INS-241
chr4-16712901-INS-486
chr4-16713688-INS-84
chr4-16729927-INS-61
chr4-16734319-INS-181
chr4-16734319-INS-984
chr4-16748088-INS-583
chr4-16751749-INS-348
chr4-16764630-INS-52

chr4-16767010-INS-115
chr4-16768338-INS-52
chr4-16824977-INS-84
chr4-16845488-INS-62
chr4-16858291-INS-493
chr4-16870938-INS-216
chr4-16894330-INS-56
chr4-16935209-INS-51
chr4-16948927-INS-330
chr4-16971218-INS-551
chr4-16977404-INS-66
chr4-17045162-INS-55
chr4-17053712-INS-62
chr4-17062959-INS-61
chr4-17078230-INS-760
chr4-17083849-INS-54
chr4-17105433-INS-354
chr4-17108614-INS-223
chr4-17117220-INS-588
chr4-17142150-INS-520
chr4-17188133-INS-90
chr4-17210199-INS-384
chr4-17215698-INS-322
chr4-17227186-INS-6844
chr4-17234618-INS-636
chr4-17251125-INS-204
chr4-17258639-INS-504
chr4-17279641-INS-54
chr4-17309307-INS-213
chr4-17366299-INS-56
chr4-17379763-INS-107
chr4-17382389-INS-152
chr4-17423672-INS-399
chr4-17436665-INS-1895
chr4-17450742-INS-520
chr4-17467863-INS-777
chr4-17499420-INS-136
chr4-17512346-INS-198
chr4-17516957-INS-554
chr4-17533495-INS-260
chr4-17562557-INS-397
chr4-17583890-INS-207
chr4-17602268-INS-62
chr4-17633230-INS-244
chr4-17652800-INS-79
chr4-17659056-INS-128
chr4-17685226-INS-118
chr4-17701525-INS-143
chr4-17701589-INS-3380
chr4-17720471-INS-181
chr4-17739574-INS-6708
chr4-17741864-INS-132
chr4-17774203-INS-78
chr4-17800604-INS-867
chr4-17806132-INS-7233
chr4-17860307-INS-52
chr4-17884877-INS-52
chr4-17898166-INS-492
chr4-17900625-INS-494
chr4-17922274-INS-613
chr4-17953289-INS-61
chr4-17964162-INS-242
chr4-17965157-INS-204
chr4-17974550-INS-892
chr4-17993207-INS-64
chr4-17999708-INS-224
chr4-18058670-INS-990
chr4-18063126-INS-453
chr4-18111079-INS-370
chr4-18135127-INS-402
chr4-18137732-INS-842
chr4-18163115-INS-711
chr4-18165863-INS-4373
chr4-18202010-INS-60
chr4-18208751-INS-66
chr4-18217014-INS-58
chr4-18253219-INS-857

chr4-18291485-INS-60
chr4-18322107-INS-6458
chr4-18322926-INS-362
chr4-18329530-INS-616
chr4-18340069-INS-135
chr4-18355427-INS-924
chr4-18362025-INS-139
chr4-18415976-INS-745
chr4-18458737-INS-936
chr4-18460671-INS-902
chr4-18599348-INS-121
chr4-18659020-INS-106
chr4-18699812-INS-6403
chr4-18755666-INS-51
chr4-18796754-INS-89
chr4-18808117-INS-201
chr4-18825892-INS-125
chr4-18828844-INS-68
chr4-18828859-INS-186
chr4-18830276-INS-52
chr4-18832145-INS-236
chr4-18841006-INS-123
chr4-18852641-INS-60
chr4-18864135-INS-82
chr4-18871694-INS-5549
chr4-18893473-INS-974
chr4-18895654-INS-69
chr4-18913950-INS-64
chr4-18917010-INS-71
chr4-18919598-INS-53
chr4-18924055-INS-630
chr4-18929901-INS-106
chr4-18968753-INS-7047
chr4-18969382-INS-6571
chr4-18973298-INS-51
chr4-19002495-INS-76
chr4-19018953-INS-3056
chr4-19058598-INS-86
chr4-19072476-INS-60
chr4-19096116-INS-976
chr4-19140105-INS-89
chr4-19181447-INS-431
chr4-19183424-INS-1872
chr4-19252484-INS-59
chr4-19276186-INS-70
chr4-19278534-INS-203
chr4-19296562-INS-1890
chr4-19299496-INS-153
chr4-19302094-INS-69
chr4-19317503-INS-886
chr4-19353353-INS-659
chr4-19358452-INS-58
chr4-19386746-INS-214
chr4-19444984-INS-6640
chr4-19445781-INS-190
chr4-19457470-INS-267
chr4-19460680-INS-132
chr4-19538601-INS-366
chr4-19544404-INS-443
chr4-19555714-INS-72
chr4-19590292-INS-346
chr4-19592099-INS-217
chr4-19602622-INS-135
chr4-19630193-INS-66
chr4-19680093-INS-92
chr4-19682156-INS-197
chr4-19682225-INS-5213
chr4-19740946-INS-54
chr4-19785941-INS-162
chr4-19807538-INS-1266
chr4-19821542-INS-156
chr4-19828205-INS-54
chr4-19833785-INS-6474
chr4-19837227-INS-209
chr4-19846847-INS-64
chr4-19850407-INS-72
chr4-19852976-INS-4526

chr4-19859290-INS-646
chr4-19888552-INS-129
chr4-19890138-INS-65
chr4-19913672-INS-695
chr4-19938988-INS-289
chr4-19951653-INS-58
chr4-19952219-INS-749
chr4-19959308-INS-605
chr4-19967591-INS-68
chr4-19982428-INS-77
chr4-19991363-INS-203
chr4-19999615-INS-915
chr4-20006337-INS-404
chr4-20024719-INS-276
chr4-20050789-INS-184
chr4-20054695-INS-117
chr4-20060544-INS-199
chr4-20062515-INS-112
chr4-20070895-INS-91
chr4-20137267-INS-302
chr4-20154880-INS-90
chr4-20175102-INS-207
chr4-20187542-INS-93
chr4-20195454-INS-53
chr4-20238769-INS-6804
chr4-20272101-INS-64
chr4-20281741-INS-52
chr4-20330728-INS-72
chr4-20333184-INS-258
chr4-20370441-INS-94
chr4-20393542-INS-119
chr4-20394728-INS-206
chr4-20400891-INS-137
chr4-20404699-INS-55
chr4-20406791-INS-58
chr4-20442476-INS-387
chr4-20449523-INS-256
chr4-20485262-INS-65
chr4-20504691-INS-236
chr4-20528697-INS-3830
chr4-20548182-INS-251
chr4-20550902-INS-217
chr4-20563622-INS-3228
chr4-20591908-INS-4137
chr4-20595060-INS-227
chr4-20615477-INS-80
chr4-20640431-INS-388
chr4-20648199-INS-6390
chr4-20651616-INS-63
chr4-20656443-INS-845
chr4-20663647-INS-86
chr4-20707481-INS-297
chr4-20735246-INS-52
chr4-20740675-INS-7311
chr4-20741762-INS-61
chr4-20748370-INS-226
chr4-20748617-INS-6401
chr4-20753956-INS-268
chr4-20770512-INS-66
chr4-20812073-INS-66
chr4-20814441-INS-89
chr4-20892809-INS-206
chr4-20902705-INS-926
chr4-20913107-INS-534
chr4-20913625-INS-2804
chr4-20922136-INS-318
chr4-20955094-INS-238
chr4-20957466-INS-55
chr4-20958028-INS-930
chr4-20964240-INS-90
chr4-20965365-INS-7209
chr4-20975236-INS-197
chr4-20977142-INS-114
chr4-20982518-INS-146
chr4-21002983-INS-56
chr4-21003933-INS-398
chr4-21006141-INS-91

chr4-21056643-INS-329
chr4-21064807-INS-6230
chr4-21068075-INS-54
chr4-21100615-INS-56
chr4-21195666-INS-55
chr4-21200633-INS-63
chr4-21206303-INS-201
chr4-21238846-INS-148
chr4-21248157-INS-328
chr4-21262690-INS-54
chr4-21274242-INS-377
chr4-21280244-INS-58
chr4-21328190-INS-732
chr4-21330869-INS-66
chr4-21331666-INS-63
chr4-21345046-INS-473
chr4-21347152-INS-66
chr4-21360779-INS-70
chr4-21393326-INS-221
chr4-21408498-INS-6189
chr4-21448251-INS-211
chr4-21494058-INS-53
chr4-21498820-INS-1410
chr4-21502695-INS-201
chr4-21534187-INS-230
chr4-21576769-INS-6815
chr4-21579134-INS-402
chr4-21609161-INS-135
chr4-21619431-INS-56
chr4-21629523-INS-400
chr4-21638479-INS-55
chr4-21638915-INS-129
chr4-21649384-INS-52
chr4-21669400-INS-6464
chr4-21692744-INS-2132
chr4-21706250-INS-3166
chr4-21725245-INS-87
chr4-21745047-INS-3938
chr4-21758725-INS-98
chr4-21760468-INS-128
chr4-21810462-INS-228
chr4-21840177-INS-54
chr4-21843536-INS-660
chr4-21857904-INS-212
chr4-21876824-INS-195
chr4-21885749-INS-85
chr4-21888667-INS-92
chr4-21892553-INS-56
chr4-21926572-INS-440
chr4-21939832-INS-350
chr4-21944955-INS-417
chr4-21962297-INS-91
chr4-21978829-INS-432
chr4-21985389-INS-102
chr4-22002181-INS-126
chr4-22009577-INS-66
chr4-22011198-INS-58
chr4-22013290-INS-400
chr4-22021074-INS-242
chr4-22043625-INS-119
chr4-22050917-INS-2967
chr4-22058136-INS-179
chr4-22075091-INS-669
chr4-22078358-INS-175
chr4-22097653-INS-4415
chr4-22105009-INS-320
chr4-22123312-INS-66
chr4-22131513-INS-6712
chr4-22135236-INS-173
chr4-22157089-INS-84
chr4-22162209-INS-6384
chr4-22182678-INS-252
chr4-22213669-INS-361
chr4-22244402-INS-354
chr4-22246735-INS-211
chr4-22253079-INS-6599
chr4-22273473-INS-387

chr4-22280104-INS-102
chr4-22284013-INS-180
chr4-22290198-INS-118
chr4-22297046-INS-66
chr4-22333135-INS-399
chr4-22364384-INS-217
chr4-22382870-INS-70
chr4-22431623-INS-175
chr4-22443751-INS-90
chr4-22450671-INS-63
chr4-22509769-INS-450
chr4-22514060-INS-175
chr4-22523046-INS-863
chr4-22523191-INS-1636
chr4-22529972-INS-6482
chr4-22549524-INS-224
chr4-22554377-INS-7223
chr4-22561856-INS-72
chr4-22562427-INS-3332
chr4-22577208-INS-103
chr4-22654670-INS-78
chr4-22661607-INS-62
chr4-22664396-INS-82
chr4-22673754-INS-6894
chr4-22707429-INS-52
chr4-22715338-INS-804
chr4-22752416-INS-1868
chr4-22761581-INS-52
chr4-22764404-INS-491
chr4-22790195-INS-897
chr4-22794985-INS-422
chr4-22814756-INS-93
chr4-22842607-INS-7180
chr4-22868765-INS-3832
chr4-22886101-INS-513
chr4-22909715-INS-297
chr4-22920882-INS-233
chr4-22933569-INS-567
chr4-23002264-INS-150
chr4-23007863-INS-3177
chr4-23009426-INS-211
chr4-23010522-INS-753
chr4-23016284-INS-506
chr4-23017775-INS-6396
chr4-23064351-INS-2127
chr4-23105210-INS-492
chr4-23113938-INS-208
chr4-23132478-INS-50
chr4-23146792-INS-128
chr4-23170616-INS-205
chr4-23187896-INS-6884
chr4-23212751-INS-121
chr4-23239507-INS-429
chr4-23254619-INS-51
chr4-23257068-INS-65
chr4-23298903-INS-6668
chr4-23300565-INS-56
chr4-23301687-INS-1753
chr4-23302281-INS-52
chr4-23308181-INS-863
chr4-23317283-INS-74
chr4-23325582-INS-52
chr4-23332175-INS-146
chr4-23348053-INS-235
chr4-23350666-INS-213
chr4-23354920-INS-401
chr4-23368550-INS-706
chr4-23390979-INS-66
chr4-23407637-INS-358
chr4-23408719-INS-106
chr4-23456145-INS-522
chr4-23459930-INS-584
chr4-23510242-INS-100
chr4-23517496-INS-86
chr4-23552641-INS-140
chr4-23562227-INS-100
chr4-23566077-INS-616

chr4-23577169-INS-600
chr4-23578221-INS-89
chr4-23578969-INS-6253
chr4-23621003-INS-479
chr4-23641956-INS-673
chr4-23644776-INS-7122
chr4-23663049-INS-6887
chr4-23664736-INS-134
chr4-23664911-INS-3947
chr4-23675508-INS-6952
chr4-23683555-INS-690
chr4-23687946-INS-82
chr4-23736286-INS-887
chr4-23749602-INS-85
chr4-23759063-INS-60
chr4-23768436-INS-439
chr4-23775685-INS-180
chr4-23787433-INS-548
chr4-23793023-INS-77
chr4-23794100-INS-7125
chr4-23798911-INS-6963
chr4-23803607-INS-593
chr4-23816658-INS-220
chr4-23842554-INS-1060
chr4-23857684-INS-416
chr4-23877247-INS-68
chr4-23886172-INS-266
chr4-23890705-INS-207
chr4-23899433-INS-6247
chr4-23915054-INS-218
chr4-23916568-INS-85
chr4-23919519-INS-61
chr4-23925042-INS-309
chr4-23930662-INS-56
chr4-23953185-INS-92
chr4-23955267-INS-474
chr4-24013081-INS-277
chr4-24113130-INS-294
chr4-24122833-INS-80
chr4-24133362-INS-5151
chr4-24150896-INS-210
chr4-24163367-INS-188
chr4-24177185-INS-256
chr4-24194031-INS-212
chr4-24244362-INS-154
chr4-24279584-INS-557
chr4-24288116-INS-4076
chr4-24369866-INS-54
chr4-24398463-INS-56
chr4-24402011-INS-80
chr4-24422138-INS-1726
chr4-24439557-INS-100
chr4-24443692-INS-128
chr4-24486832-INS-173
chr4-24497287-INS-207
chr4-24500720-INS-184
chr4-24517974-INS-130
chr4-24545911-INS-202
chr4-24576444-INS-104
chr4-24579709-INS-301
chr4-24588430-INS-182
chr4-24603843-INS-521
chr4-24624963-INS-310
chr4-24638852-INS-969
chr4-24668787-INS-80
chr4-24680983-INS-100
chr4-24685578-INS-530
chr4-24689262-INS-286
chr4-24704221-INS-4573
chr4-24758384-INS-358
chr4-24760560-INS-670
chr4-24768792-INS-118
chr4-24788589-INS-891
chr4-24788803-INS-75
chr4-24800475-INS-132
chr4-24801172-INS-779
chr4-24810664-INS-686

chr4-24812319-INS-378
chr4-24814820-INS-694
chr4-24823422-INS-593
chr4-24846655-INS-228
chr4-24854637-INS-500
chr4-24875843-INS-623
chr4-24881950-INS-7091
chr4-24918502-INS-158
chr4-24922539-INS-2034
chr4-24938292-INS-78
chr4-24961439-INS-634
chr4-24964754-INS-394
chr4-24973500-INS-8125
chr4-25026099-INS-506
chr4-25035012-INS-84
chr4-25049235-INS-139
chr4-25081891-INS-1500
chr4-25097283-INS-409
chr4-25120351-INS-58
chr4-25199137-INS-204
chr4-25232610-INS-178
chr4-25245761-INS-78
chr4-25275664-INS-152
chr4-25305789-INS-809
chr4-25309139-INS-318
chr4-25319077-INS-74
chr4-25342663-INS-407
chr4-25363997-INS-926
chr4-25372574-INS-59
chr4-25393216-INS-214
chr4-25398274-INS-261
chr4-25428371-INS-136
chr4-25461657-INS-156
chr4-25471909-INS-139
chr4-25475716-INS-175
chr4-25487845-INS-380
chr4-25595461-INS-6435
chr4-25597132-INS-7428
chr4-25618531-INS-55
chr4-25630368-INS-108
chr4-25658284-INS-262
chr4-25659037-INS-596
chr4-25667231-INS-864
chr4-25676686-INS-150
chr4-25689042-INS-614
chr4-25692542-INS-129
chr4-25717364-INS-333
chr4-25724409-INS-76
chr4-25725823-INS-123
chr4-25728234-INS-209
chr4-25770726-INS-147
chr4-25777788-INS-358
chr4-25793170-INS-750
chr4-25800817-INS-116
chr4-25824391-INS-206
chr4-25891549-INS-142
chr4-25899996-INS-6566
chr4-25906198-INS-121
chr4-25940476-INS-80
chr4-25941995-INS-790
chr4-25944838-INS-130
chr4-25949031-INS-55
chr4-25974227-INS-107
chr4-25988841-INS-6845
chr4-25996922-INS-6739
chr4-26007510-INS-133
chr4-26011039-INS-357
chr4-26031731-INS-124
chr4-26048095-INS-281
chr4-26069737-INS-4999
chr4-26086290-INS-7136
chr4-26086518-INS-388
chr4-26127727-INS-51
chr4-26133747-INS-84
chr4-26182352-INS-201
chr4-26210025-INS-526
chr4-26212604-INS-216

chr4-26222907-INS-494
chr4-26224890-INS-401
chr4-26248564-INS-76
chr4-26267085-INS-414
chr4-26268448-INS-292
chr4-26280990-INS-232
chr4-26289861-INS-56
chr4-26323222-INS-124
chr4-26337266-INS-83
chr4-26341733-INS-853
chr4-26376354-INS-72
chr4-26381754-INS-129
chr4-26456417-INS-151
chr4-26473986-INS-450
chr4-26475245-INS-130
chr4-26513053-INS-59
chr4-26513818-INS-558
chr4-26517783-INS-410
chr4-26574805-INS-167
chr4-26592673-INS-573
chr4-26621043-INS-202
chr4-26636791-INS-76
chr4-26687411-INS-71
chr4-26705761-INS-248
chr4-26712381-INS-160
chr4-26754797-INS-243
chr4-26772820-INS-75
chr4-26791350-INS-56
chr4-26815570-INS-469
chr4-26835518-INS-484
chr4-26839095-INS-106
chr4-26855795-INS-102
chr4-26863386-INS-189
chr4-26867695-INS-7071
chr4-26879621-INS-6944
chr4-26881493-INS-68
chr4-26883113-INS-6682
chr4-26886530-INS-6951
chr4-26897172-INS-127
chr4-26926096-INS-461
chr4-26950309-INS-55
chr4-26974438-INS-108
chr4-26983759-INS-58
chr4-26988431-INS-100
chr4-27122873-INS-164
chr4-27125080-INS-363
chr4-27131876-INS-55
chr4-27178750-INS-199
chr4-27205711-INS-421
chr4-27250908-INS-68
chr4-27253225-INS-52
chr4-27279964-INS-64
chr4-27315251-INS-87
chr4-27322461-INS-1811
chr4-27325898-INS-759
chr4-27365927-INS-56
chr4-27422249-INS-119
chr4-27449170-INS-88
chr4-27488212-INS-186
chr4-27500612-INS-52
chr4-27502838-INS-125
chr4-27510885-INS-248
chr4-27527096-INS-55
chr4-27534156-INS-880
chr4-27573006-INS-277
chr4-27594012-INS-199
chr4-27656088-INS-165
chr4-27661896-INS-54
chr4-27672829-INS-121
chr4-27721845-INS-466
chr4-27725658-INS-68
chr4-27824069-INS-89
chr4-27832332-INS-146
chr4-27838520-INS-60
chr4-27841640-INS-63
chr4-27849332-INS-56
chr4-27883943-INS-2156

chr4-27912894-INS-353
chr4-27926661-INS-378
chr4-27928917-INS-210
chr4-27946261-INS-245
chr4-27959297-INS-242
chr4-27972003-INS-88
chr4-28003519-INS-3376
chr4-28030035-INS-417
chr4-28124805-INS-52
chr4-28127508-INS-359
chr4-28129222-INS-356
chr4-28162263-INS-6800
chr4-28163779-INS-52
chr4-28172081-INS-84
chr4-28178261-INS-339
chr4-28202541-INS-5240
chr4-28214692-INS-703
chr4-28224365-INS-2010
chr4-28249427-INS-55
chr4-28259542-INS-109
chr4-28283451-INS-234
chr4-28293502-INS-351
chr4-28295445-INS-6505
chr4-28307787-INS-64
chr4-28332412-INS-175
chr4-28333716-INS-598
chr4-28341985-INS-78
chr4-28382248-INS-172
chr4-28393378-INS-705
chr4-28400992-INS-133
chr4-28427992-INS-431
chr4-28429275-INS-366
chr4-28434144-INS-412
chr4-28439914-INS-234
chr4-28455339-INS-241
chr4-28549374-INS-67
chr4-28550702-INS-72
chr4-28572832-INS-83
chr4-28573528-INS-591
chr4-28628442-INS-52
chr4-28634302-INS-2522
chr4-28672847-INS-699
chr4-28702433-INS-131
chr4-28710231-INS-60
chr4-28715036-INS-58
chr4-28719465-INS-2373
chr4-28761202-INS-488
chr4-28765339-INS-196
chr4-28776634-INS-111
chr4-28799571-INS-54
chr4-28827666-INS-127
chr4-28855514-INS-743
chr4-28858855-INS-369
chr4-28902686-INS-129
chr4-28912140-INS-756
chr4-28916209-INS-64
chr4-28921684-INS-250
chr4-28928328-INS-691
chr4-28929408-INS-56
chr4-28942991-INS-400
chr4-28972562-INS-57
chr4-28975402-INS-60
chr4-28990200-INS-863
chr4-29002563-INS-189
chr4-29029042-INS-70
chr4-29043399-INS-52
chr4-29045865-INS-6976
chr4-29085339-INS-66
chr4-29123436-INS-293
chr4-29134988-INS-84
chr4-29140959-INS-344
chr4-29160924-INS-80
chr4-29181215-INS-80
chr4-29260864-INS-172
chr4-29289322-INS-282
chr4-29309234-INS-668
chr4-29320529-INS-72

chr4-29332900-INS-306
chr4-29372001-INS-7095
chr4-29374283-INS-54
chr4-29383493-INS-55
chr4-29414138-INS-5210
chr4-29460296-INS-124
chr4-29496999-INS-74
chr4-29521527-INS-525
chr4-29545221-INS-88
chr4-29551105-INS-60
chr4-29582080-INS-55
chr4-29585021-INS-200
chr4-29620870-INS-6378
chr4-29632009-INS-56
chr4-29636431-INS-60
chr4-29643456-INS-228
chr4-29644678-INS-74
chr4-29653257-INS-6407
chr4-29679568-INS-2142
chr4-29758842-INS-50
chr4-29760462-INS-450
chr4-29764517-INS-72
chr4-29774416-INS-448
chr4-29783851-INS-80
chr4-29785296-INS-6932
chr4-29815585-INS-210
chr4-29817875-INS-92
chr4-29820815-INS-176
chr4-29825129-INS-55
chr4-29846047-INS-97
chr4-29856998-INS-5325
chr4-29880786-INS-438
chr4-29891304-INS-72
chr4-29906481-INS-144
chr4-29933490-INS-582
chr4-29934949-INS-94
chr4-29939737-INS-2223
chr4-29959349-INS-52
chr4-29963722-INS-209
chr4-29967855-INS-4479
chr4-29985579-INS-7337
chr4-30000183-INS-480
chr4-30015113-INS-113
chr4-30021955-INS-106
chr4-30030179-INS-73
chr4-30048964-INS-1229
chr4-30082194-INS-4447
chr4-30093787-INS-431
chr4-30097123-INS-60
chr4-30111721-INS-134
chr4-30114266-INS-164
chr4-30122447-INS-116
chr4-30140603-INS-307
chr4-30157762-INS-75
chr4-30162262-INS-111
chr4-30167638-INS-298
chr4-30207575-INS-55
chr4-30238224-INS-80
chr4-30272989-INS-6466
chr4-30299648-INS-51
chr4-30305334-INS-88
chr4-30395818-INS-340
chr4-30419723-INS-360
chr4-30424579-INS-112
chr4-30456153-INS-84
chr4-30460082-INS-61
chr4-30469080-INS-61
chr4-30475385-INS-78
chr4-30478691-INS-53
chr4-30486721-INS-103
chr4-30487379-INS-706
chr4-30492580-INS-180
chr4-30496494-INS-74
chr4-30507163-INS-879
chr4-30507755-INS-86
chr4-30509086-INS-447
chr4-30541824-INS-212

chr4-30545295-INS-67
chr4-30556589-INS-212
chr4-30570925-INS-6864
chr4-30573463-INS-111
chr4-30588982-INS-6209
chr4-30612086-INS-1311
chr4-30621213-INS-275
chr4-30630402-INS-382
chr4-30657250-INS-65
chr4-30676561-INS-103
chr4-30705199-INS-735
chr4-30749967-INS-418
chr4-30763264-INS-897
chr4-30764283-INS-52
chr4-30772602-INS-55
chr4-30796267-INS-192
chr4-30821090-INS-57
chr4-30849194-INS-69
chr4-30850414-INS-619
chr4-30876155-INS-76
chr4-30885997-INS-214
chr4-30910417-INS-66
chr4-30930568-INS-187
chr4-30963025-INS-444
chr4-30973888-INS-169
chr4-30999441-INS-83
chr4-31004841-INS-61
chr4-31008094-INS-525
chr4-31021400-INS-231
chr4-31045191-INS-177
chr4-31048145-INS-63
chr4-31050335-INS-437
chr4-31051653-INS-61
chr4-31074749-INS-231
chr4-31087511-INS-563
chr4-31110637-INS-63
chr4-31143829-INS-145
chr4-31150072-INS-175
chr4-31166261-INS-70
chr4-31178330-INS-262
chr4-31187724-INS-400
chr4-31191767-INS-268
chr4-31194640-INS-254
chr4-31219150-INS-207
chr4-31233969-INS-52
chr4-31237471-INS-323
chr4-31253237-INS-75
chr4-31268117-INS-54
chr4-31277172-INS-163
chr4-31284124-INS-780
chr4-31303630-INS-1311
chr4-31309771-INS-6401
chr4-31322219-INS-140
chr4-31340624-INS-140
chr4-31341494-INS-349
chr4-31348687-INS-103
chr4-31355265-INS-1720
chr4-31380430-INS-397
chr4-31388828-INS-703
chr4-31399613-INS-129
chr4-31416050-INS-79
chr4-31432948-INS-146
chr4-31462909-INS-506
chr4-31470998-INS-146
chr4-31473376-INS-58
chr4-31492695-INS-6332
chr4-31499951-INS-463
chr4-31521196-INS-6411
chr4-31526738-INS-604
chr4-31538302-INS-83
chr4-31539039-INS-480
chr4-31547757-INS-230
chr4-31552983-INS-322
chr4-31574751-INS-87
chr4-31579877-INS-609
chr4-31654753-INS-58
chr4-31695948-INS-84

chr4-31719413-INS-645
chr4-31732040-INS-163
chr4-31775516-INS-128
chr4-31777698-INS-624
chr4-31801727-INS-923
chr4-31807158-INS-68
chr4-31814926-INS-50
chr4-31828421-INS-177
chr4-31855236-INS-62
chr4-31884195-INS-366
chr4-31918614-INS-332
chr4-31987102-INS-136
chr4-31987614-INS-270
chr4-32026958-INS-392
chr4-32044305-INS-360
chr4-32048160-INS-51
chr4-32049661-INS-80
chr4-32078991-INS-66
chr4-32096400-INS-178
chr4-32106119-INS-113
chr4-32108985-INS-251
chr4-32119966-INS-134
chr4-32140950-INS-76
chr4-32152926-INS-65
chr4-32215794-INS-56
chr4-32246782-INS-112
chr4-32276155-INS-60
chr4-32353595-INS-110
chr4-32356763-INS-3413
chr4-32362449-INS-6936
chr4-32378317-INS-57
chr4-32412295-INS-54
chr4-32483982-INS-84
chr4-32515502-INS-891
chr4-32540014-INS-331
chr4-32566979-INS-72
chr4-32569711-INS-112
chr4-32574659-INS-204
chr4-32578630-INS-410
chr4-32592921-INS-208
chr4-32594049-INS-588
chr4-32605001-INS-59
chr4-32629293-INS-98
chr4-32665478-INS-213
chr4-32673639-INS-56
chr4-32721896-INS-59
chr4-32780700-INS-61
chr4-32782287-INS-236
chr4-32850066-INS-208
chr4-32852065-INS-69
chr4-32858157-INS-74
chr4-32860175-INS-67
chr4-32865247-INS-96
chr4-32868783-INS-101
chr4-32871130-INS-165
chr4-32897391-INS-125
chr4-32899117-INS-70
chr4-32940074-INS-108
chr4-32958405-INS-60
chr4-32966691-INS-280
chr4-32969948-INS-113
chr4-32988876-INS-441
chr4-33007045-INS-204
chr4-33010402-INS-84
chr4-33011779-INS-428
chr4-33016533-INS-336
chr4-33039034-INS-80
chr4-33052467-INS-218
chr4-33054920-INS-68
chr4-33057712-INS-378
chr4-33078224-INS-442
chr4-33080055-INS-225
chr4-33083046-INS-1205
chr4-33084758-INS-204
chr4-33089150-INS-400
chr4-33102133-INS-80
chr4-33112137-INS-391

chr4-33153878-INS-138
chr4-33166279-INS-218
chr4-33173788-INS-92
chr4-33177535-INS-983
chr4-33203867-INS-138
chr4-33207815-INS-209
chr4-33229519-INS-72
chr4-33232541-INS-62
chr4-33254373-INS-200
chr4-33259925-INS-203
chr4-33273327-INS-192
chr4-33282745-INS-60
chr4-33296276-INS-876
chr4-33299702-INS-264
chr4-33309731-INS-148
chr4-33342135-INS-55
chr4-33344785-INS-155
chr4-33346347-INS-289
chr4-33358539-INS-286
chr4-33392131-INS-135
chr4-33409981-INS-76
chr4-33422510-INS-55
chr4-33492133-INS-397
chr4-33536187-INS-3116
chr4-33540464-INS-118
chr4-33550331-INS-58
chr4-33579716-INS-775
chr4-33586013-INS-646
chr4-33602221-INS-66
chr4-33609539-INS-285
chr4-33618029-INS-141
chr4-33624025-INS-555
chr4-33629977-INS-84
chr4-33647471-INS-78
chr4-33653473-INS-62
chr4-33665730-INS-100
chr4-33671935-INS-239
chr4-33675027-INS-150
chr4-33678668-INS-233
chr4-33681512-INS-421
chr4-33690019-INS-60
chr4-33691553-INS-109
chr4-33701543-INS-114
chr4-33710722-INS-89
chr4-33720149-INS-142
chr4-33729976-INS-313
chr4-33777334-INS-66
chr4-33783674-INS-117
chr4-33787924-INS-338
chr4-33805442-INS-216
chr4-33812896-INS-66
chr4-33828739-INS-60
chr4-33867774-INS-191
chr4-33874123-INS-164
chr4-33918407-INS-192
chr4-33987922-INS-70
chr4-34005807-INS-3267
chr4-34037083-INS-449
chr4-34058846-INS-880
chr4-34060227-INS-64
chr4-34065395-INS-120
chr4-34084960-INS-165
chr4-34091299-INS-58
chr4-34107876-INS-456
chr4-34176682-INS-177
chr4-34181092-INS-192
chr4-34182266-INS-471
chr4-34189038-INS-227
chr4-34197306-INS-72
chr4-34205682-INS-54
chr4-34218753-INS-151
chr4-34220477-INS-160
chr4-34242575-INS-248
chr4-34249156-INS-589
chr4-34298673-INS-86
chr4-34302651-INS-527
chr4-34324072-INS-310

chr4-34329997-INS-383
chr4-34340818-INS-56
chr4-34377297-INS-84
chr4-34388584-INS-657
chr4-34429379-INS-480
chr4-34457363-INS-651
chr4-34461327-INS-440
chr4-34497469-INS-63
chr4-34511532-INS-199
chr4-34682403-INS-930
chr4-34687829-INS-5121
chr4-34689179-INS-265
chr4-34691261-INS-148
chr4-34709784-INS-282
chr4-34718134-INS-57
chr4-34741450-INS-98
chr4-34783797-INS-56
chr4-34786344-INS-152
chr4-34915296-INS-150
chr4-34921945-INS-195
chr4-34924517-INS-169
chr4-34928531-INS-627
chr4-34934762-INS-281
chr4-34966730-INS-76
chr4-34968518-INS-192
chr4-34975910-INS-116
chr4-34992765-INS-103
chr4-34994689-INS-159
chr4-35011483-INS-301
chr4-35028586-INS-105
chr4-35041696-INS-494
chr4-35052017-INS-187
chr4-35057324-INS-72
chr4-35093637-INS-146
chr4-35097658-INS-98
chr4-35101727-INS-137
chr4-35138499-INS-54
chr4-35178133-INS-172
chr4-35259806-INS-203
chr4-35285596-INS-60
chr4-35302849-INS-54
chr4-35310874-INS-99
chr4-35332381-INS-369
chr4-35335346-INS-3160
chr4-35345327-INS-125
chr4-35356639-INS-7100
chr4-35387245-INS-84
chr4-35388314-INS-332
chr4-35411344-INS-512
chr4-35412470-INS-639
chr4-35422903-INS-78
chr4-35437298-INS-449
chr4-35444217-INS-168
chr4-35474869-INS-52
chr4-35477155-INS-57
chr4-35478472-INS-56
chr4-35512455-INS-2991
chr4-35523899-INS-60
chr4-35534155-INS-781
chr4-35535275-INS-6219
chr4-35546760-INS-98
chr4-35564186-INS-173
chr4-35581957-INS-1978
chr4-35582715-INS-6117
chr4-35591817-INS-6788
chr4-35596720-INS-53
chr4-35607579-INS-178
chr4-35609663-INS-6236
chr4-35647160-INS-317
chr4-35678329-INS-497
chr4-35706150-INS-62
chr4-35720304-INS-102
chr4-35736117-INS-399
chr4-35774402-INS-53
chr4-35806514-INS-109
chr4-35819861-INS-68
chr4-35875451-INS-54

chr4-35878701-INS-102
chr4-35904075-INS-100
chr4-35926493-INS-81
chr4-35934261-INS-112
chr4-35941352-INS-80
chr4-35955675-INS-92
chr4-35988522-INS-443
chr4-35992343-INS-53
chr4-35998218-INS-68
chr4-36089846-INS-68
chr4-36090955-INS-76
chr4-36105892-INS-92
chr4-36132946-INS-55
chr4-36149825-INS-489
chr4-36154706-INS-66
chr4-36166486-INS-73
chr4-36168255-INS-191
chr4-36176108-INS-303
chr4-36177124-INS-245
chr4-36205176-INS-388
chr4-36268530-INS-137
chr4-36285184-INS-80
chr4-36297658-INS-256
chr4-36351132-INS-436
chr4-36372338-INS-582
chr4-36376189-INS-6890
chr4-36392485-INS-158
chr4-36407365-INS-60
chr4-36411779-INS-989
chr4-36429053-INS-780
chr4-36471176-INS-234
chr4-36518309-INS-174
chr4-36546430-INS-52
chr4-36564939-INS-218
chr4-36568011-INS-54
chr4-36602189-INS-253
chr4-36626378-INS-752
chr4-36627343-INS-213
chr4-36672508-INS-367
chr4-36679085-INS-435
chr4-36681627-INS-381
chr4-36690192-INS-428
chr4-36736925-INS-645
chr4-36739205-INS-106
chr4-36757401-INS-193
chr4-36768488-INS-624
chr4-36780261-INS-59
chr4-36787349-INS-328
chr4-36844654-INS-66
chr4-36853127-INS-459
chr4-36867428-INS-98
chr4-36878196-INS-310
chr4-36888345-INS-51
chr4-36970885-INS-7061
chr4-37064958-INS-317
chr4-37093691-INS-3424
chr4-37099982-INS-6894
chr4-37149310-INS-60
chr4-37176666-INS-212
chr4-37179280-INS-6506
chr4-37190160-INS-268
chr4-37191225-INS-337
chr4-37192704-INS-289
chr4-37227717-INS-54
chr4-37270214-INS-72
chr4-37271121-INS-5240
chr4-37305487-INS-52
chr4-37308448-INS-212
chr4-37309513-INS-723
chr4-37315637-INS-332
chr4-37324811-INS-303
chr4-37341826-INS-6483
chr4-37390916-INS-328
chr4-37396411-INS-508
chr4-37407754-INS-92
chr4-37412861-INS-56
chr4-37428948-INS-6460

chr4-37463906-INS-267
chr4-37473272-INS-557
chr4-37490700-INS-138
chr4-37541660-INS-102
chr4-37601586-INS-141
chr4-37623826-INS-188
chr4-37642871-INS-68
chr4-37706820-INS-1888
chr4-37712199-INS-6581
chr4-37732782-INS-144
chr4-37743699-INS-53
chr4-37755185-INS-102
chr4-37769411-INS-400
chr4-37777628-INS-322
chr4-37852094-INS-192
chr4-37854575-INS-98
chr4-37858526-INS-7471
chr4-37861999-INS-94
chr4-37863267-INS-55
chr4-37882444-INS-104
chr4-37887998-INS-158
chr4-37925245-INS-72
chr4-37926909-INS-239
chr4-37947618-INS-7135
chr4-37960152-INS-5282
chr4-37962649-INS-1133
chr4-37963835-INS-921
chr4-37972420-INS-383
chr4-37979074-INS-2140
chr4-37983417-INS-6165
chr4-37996085-INS-4952
chr4-37996176-INS-517
chr4-38008524-INS-192
chr4-38026145-INS-1494
chr4-38072124-INS-907
chr4-38074279-INS-85
chr4-38076930-INS-52
chr4-38101540-INS-404
chr4-38131039-INS-819
chr4-38139158-INS-2747
chr4-38141727-INS-54
chr4-38164347-INS-6226
chr4-38173197-INS-67
chr4-38176994-INS-65
chr4-38196262-INS-96
chr4-38199693-INS-6602
chr4-38248979-INS-100
chr4-38257539-INS-961
chr4-38281918-INS-746
chr4-38296906-INS-58
chr4-38301675-INS-248
chr4-38310466-INS-478
chr4-38341177-INS-9134
chr4-38353515-INS-96
chr4-38382275-INS-477
chr4-38393429-INS-432
chr4-38448835-INS-401
chr4-38485959-INS-167
chr4-38513283-INS-5129
chr4-38574724-INS-500
chr4-38620544-INS-78
chr4-38650107-INS-481
chr4-38658289-INS-391
chr4-38664201-INS-6646
chr4-38747403-INS-52
chr4-38757211-INS-7187
chr4-38827072-INS-80
chr4-38830978-INS-340
chr4-38835314-INS-57
chr4-38843221-INS-845
chr4-38844325-INS-331
chr4-38882142-INS-2577
chr4-38894255-INS-203
chr4-38901979-INS-6835
chr4-38943637-INS-80
chr4-38949387-INS-3050
chr4-38968316-INS-54

chr4-38994681-INS-52
chr4-39002944-INS-189
chr4-39030751-INS-181
chr4-39037310-INS-794
chr4-39048654-INS-190
chr4-39055392-INS-4831
chr4-39067009-INS-140
chr4-39069999-INS-66
chr4-39080635-INS-281
chr4-39116788-INS-80
chr4-39133210-INS-6451
chr4-39151394-INS-75
chr4-39222974-INS-777
chr4-39243302-INS-444
chr4-39245705-INS-64
chr4-39248040-INS-397
chr4-39307849-INS-222
chr4-39332294-INS-120
chr4-39353945-INS-191
chr4-39363015-INS-134
chr4-39392594-INS-193
chr4-39440111-INS-492
chr4-39444233-INS-52
chr4-39522255-INS-192
chr4-39548775-INS-52
chr4-39554432-INS-176
chr4-39590933-INS-339
chr4-39598986-INS-264
chr4-39605216-INS-146
chr4-39666569-INS-422
chr4-39674950-INS-200
chr4-39692243-INS-725
chr4-39718706-INS-245
chr4-39731675-INS-108
chr4-39750321-INS-6983
chr4-39800084-INS-163
chr4-39833616-INS-983
chr4-39843851-INS-390
chr4-39863942-INS-54
chr4-39875167-INS-64
chr4-39878710-INS-66
chr4-39893747-INS-59
chr4-39924390-INS-195
chr4-39930095-INS-142
chr4-39959517-INS-68
chr4-39969598-INS-399
chr4-39975788-INS-209
chr4-39981699-INS-272
chr4-40020911-INS-345
chr4-40040071-INS-79
chr4-40044551-INS-1197
chr4-40050761-INS-5620
chr4-40064803-INS-6483
chr4-40091608-INS-80
chr4-40106753-INS-120
chr4-40116135-INS-6935
chr4-40122192-INS-400
chr4-40138508-INS-2473
chr4-40140621-INS-439
chr4-40154673-INS-62
chr4-40156705-INS-667
chr4-40162276-INS-445
chr4-40171709-INS-92
chr4-40176643-INS-68
chr4-40178477-INS-205
chr4-40207631-INS-170
chr4-40216736-INS-198
chr4-40230982-INS-173
chr4-40249157-INS-133
chr4-40250577-INS-360
chr4-40251587-INS-206
chr4-40256645-INS-200
chr4-40263424-INS-57
chr4-40281427-INS-212
chr4-40291885-INS-78
chr4-40323132-INS-107
chr4-40325253-INS-432

chr4-40358384-INS-64
chr4-40380459-INS-52
chr4-40389726-INS-86
chr4-40422546-INS-273
chr4-40489105-INS-93
chr4-40505626-INS-51
chr4-40516289-INS-93
chr4-40522493-INS-96
chr4-40542412-INS-215
chr4-40552950-INS-509
chr4-40574276-INS-898
chr4-40615967-INS-783
chr4-40623250-INS-597
chr4-40638137-INS-108
chr4-40662840-INS-495
chr4-40672600-INS-341
chr4-40677764-INS-476
chr4-40710880-INS-63
chr4-40726558-INS-108
chr4-40752262-INS-55
chr4-40754986-INS-78
chr4-40756933-INS-80
chr4-40764882-INS-214
chr4-40773141-INS-54
chr4-40778498-INS-496
chr4-40799586-INS-208
chr4-40836798-INS-72
chr4-40839943-INS-215
chr4-40850408-INS-132
chr4-40878864-INS-82
chr4-40880981-INS-205
chr4-40889764-INS-80
chr4-40907122-INS-658
chr4-40914389-INS-196
chr4-40919217-INS-336
chr4-40942179-INS-54
chr4-41015903-INS-80
chr4-41030555-INS-115
chr4-41042579-INS-91
chr4-41057758-INS-354
chr4-41059216-INS-75
chr4-41074879-INS-189
chr4-41079432-INS-298
chr4-41100294-INS-56
chr4-41104910-INS-238
chr4-41105894-INS-156
chr4-41110269-INS-219
chr4-41125230-INS-214
chr4-41128764-INS-224
chr4-41172670-INS-6475
chr4-41174791-INS-134
chr4-41175827-INS-6523
chr4-41176180-INS-65
chr4-41201197-INS-67
chr4-41208487-INS-60
chr4-41227810-INS-101
chr4-41234762-INS-906
chr4-41238184-INS-302
chr4-41249330-INS-78
chr4-41276775-INS-203
chr4-41277840-INS-150
chr4-41281514-INS-92
chr4-41300794-INS-224
chr4-41303931-INS-59
chr4-41334953-INS-51
chr4-41338121-INS-70
chr4-41342867-INS-188
chr4-41346661-INS-166
chr4-41367506-INS-188
chr4-41393116-INS-138
chr4-41412966-INS-52
chr4-41438430-INS-265
chr4-41516506-INS-61
chr4-41533358-INS-176
chr4-41534637-INS-102
chr4-41540814-INS-80
chr4-41575020-INS-136

chr4-41589606-INS-401
chr4-41590684-INS-269
chr4-41633372-INS-203
chr4-41645024-INS-70
chr4-41666140-INS-60
chr4-41678866-INS-209
chr4-41690777-INS-83
chr4-41783040-INS-357
chr4-41857146-INS-520
chr4-42205819-INS-131
chr4-42368324-INS-62
chr4-42454639-INS-498
chr4-42560496-INS-2776
chr4-42633842-INS-5238
chr4-42720025-INS-498
chr4-42739835-INS-879
chr4-42742055-INS-467
chr4-42743838-INS-257
chr4-42745834-INS-99
chr4-42775195-INS-156
chr4-42776215-INS-71
chr4-42778476-INS-202
chr4-42785069-INS-52
chr4-42801089-INS-76
chr4-42810265-INS-6669
chr4-42818352-INS-558
chr4-42862058-INS-234
chr4-42964579-INS-328
chr4-42966860-INS-192
chr4-43019015-INS-139
chr4-43079379-INS-312
chr4-43108893-INS-541
chr4-43291892-INS-354
chr4-43296551-INS-114
chr4-43319825-INS-5549
chr4-43324047-INS-54
chr4-43359052-INS-3621
chr4-43411250-INS-64
chr4-43429546-INS-232
chr4-43463527-INS-880
chr4-43471477-INS-348
chr4-43477510-INS-56
chr4-43481542-INS-57
chr4-43487725-INS-90
chr4-43529623-INS-234
chr4-43541124-INS-82
chr4-43596526-INS-78
chr4-43681342-INS-265
chr4-43683769-INS-52
chr4-43702517-INS-83
chr4-43713467-INS-52
chr4-43717350-INS-75
chr4-43719780-INS-81
chr4-43758108-INS-264
chr4-43763938-INS-120
chr4-43779518-INS-82
chr4-43803133-INS-150
chr4-43827372-INS-517
chr4-43830110-INS-90
chr4-43844212-INS-343
chr4-43846571-INS-767
chr4-43871240-INS-80
chr4-43872369-INS-60
chr4-43882456-INS-60
chr4-43917730-INS-345
chr4-43945298-INS-59
chr4-43950952-INS-660
chr4-43951835-INS-54
chr4-43967193-INS-78
chr4-43969084-INS-296
chr4-43990482-INS-422
chr4-44011556-INS-129
chr4-44012120-INS-626
chr4-44035984-INS-116
chr4-44038064-INS-148
chr4-44045255-INS-489
chr4-44057025-INS-425

chr4-44071450-INS-51
chr4-44078229-INS-53
chr4-44099525-INS-63
chr4-44111826-INS-207
chr4-44120795-INS-284
chr4-44138156-INS-66
chr4-44172925-INS-199
chr4-44193550-INS-121
chr4-44201778-INS-617
chr4-44202320-INS-52
chr4-44203837-INS-147
chr4-44220468-INS-281
chr4-44226734-INS-55
chr4-44241433-INS-64
chr4-44242752-INS-57
chr4-44244660-INS-68
chr4-44247003-INS-76
chr4-44255220-INS-126
chr4-44258426-INS-64
chr4-44268088-INS-450
chr4-44268847-INS-52
chr4-44274724-INS-92
chr4-44295717-INS-204
chr4-44303217-INS-370
chr4-44313999-INS-474
chr4-44318417-INS-419
chr4-44326434-INS-199
chr4-44330815-INS-206
chr4-44337152-INS-51
chr4-44346292-INS-62
chr4-44350227-INS-128
chr4-44366488-INS-557
chr4-44368580-INS-5955
chr4-44376977-INS-111
chr4-44389737-INS-70
chr4-44399291-INS-227
chr4-44409613-INS-214
chr4-44437684-INS-76
chr4-44439578-INS-184
chr4-44451862-INS-126
chr4-44492448-INS-101
chr4-44499332-INS-240
chr4-44505075-INS-65
chr4-44507692-INS-104
chr4-44517826-INS-375
chr4-44568192-INS-70
chr4-44574417-INS-78
chr4-44600609-INS-339
chr4-44627683-INS-115
chr4-44637609-INS-169
chr4-44644268-INS-72
chr4-44667726-INS-208
chr4-44683048-INS-226
chr4-44701314-INS-75
chr4-44721754-INS-652
chr4-44725351-INS-85
chr4-44744937-INS-104
chr4-44764542-INS-94
chr4-44799452-INS-52
chr4-44879078-INS-6464
chr4-44882714-INS-658
chr4-44898346-INS-915
chr4-44966709-INS-213
chr4-44972686-INS-104
chr4-44975391-INS-208
chr4-44979272-INS-308
chr4-44986396-INS-71
chr4-44999210-INS-6427
chr4-45013705-INS-196
chr4-45014134-INS-6468
chr4-45015736-INS-580
chr4-45073403-INS-200
chr4-45103966-INS-67
chr4-45110101-INS-53
chr4-45130870-INS-60
chr4-45132461-INS-174
chr4-45151292-INS-198

chr4-45153451-INS-226
chr4-45155130-INS-78
chr4-45165554-INS-128
chr4-45178852-INS-1231
chr4-45189543-INS-499
chr4-45190785-INS-180
chr4-45195345-INS-108
chr4-45232047-INS-127
chr4-45240967-INS-91
chr4-45257681-INS-185
chr4-45276673-INS-318
chr4-45276772-INS-662
chr4-45286651-INS-115
chr4-45303925-INS-506
chr4-45321012-INS-225
chr4-45339119-INS-76
chr4-45384699-INS-204
chr4-45386540-INS-374
chr4-45393618-INS-67
chr4-45402127-INS-213
chr4-45404652-INS-115
chr4-45404652-INS-150
chr4-45416323-INS-204
chr4-45441061-INS-178
chr4-45447788-INS-103
chr4-45472076-INS-82
chr4-45475800-INS-509
chr4-45483602-INS-216
chr4-45527284-INS-52
chr4-45553734-INS-206
chr4-45559521-INS-5305
chr4-45575278-INS-211
chr4-45579768-INS-146
chr4-45613704-INS-294
chr4-45619812-INS-1035
chr4-45637722-INS-243
chr4-45641123-INS-196
chr4-45644974-INS-193
chr4-45659852-INS-74
chr4-45664188-INS-74
chr4-45677547-INS-572
chr4-45686840-INS-374
chr4-45714602-INS-114
chr4-45728450-INS-190
chr4-45740148-INS-139
chr4-45743866-INS-51
chr4-45757979-INS-55
chr4-45774319-INS-203
chr4-45776278-INS-77
chr4-45799685-INS-92
chr4-45805000-INS-218
chr4-45847453-INS-81
chr4-45864430-INS-76
chr4-45883799-INS-155
chr4-45890140-INS-198
chr4-45891866-INS-51
chr4-45899395-INS-93
chr4-45911870-INS-235
chr4-45922333-INS-76
chr4-45926978-INS-123
chr4-45939683-INS-86
chr4-45942241-INS-81
chr4-46046490-INS-80
chr4-46052943-INS-736
chr4-46058763-INS-80
chr4-46075173-INS-98
chr4-46088474-INS-207
chr4-46091268-INS-58
chr4-46104318-INS-100
chr4-46112351-INS-51
chr4-46125199-INS-105
chr4-46128216-INS-66
chr4-46129528-INS-88
chr4-46144406-INS-304
chr4-46145029-INS-121
chr4-46148381-INS-544
chr4-46177285-INS-74

chr4-46178937-INS-134
chr4-46185032-INS-190
chr4-46187292-INS-53
chr4-46188713-INS-601
chr4-46194059-INS-85
chr4-46217935-INS-54
chr4-46223977-INS-90
chr4-46232765-INS-163
chr4-46235609-INS-341
chr4-46252829-INS-127
chr4-46259349-INS-186
chr4-46274423-INS-65
chr4-46285504-INS-171
chr4-46289491-INS-181
chr4-46300620-INS-241
chr4-46314076-INS-216
chr4-46317695-INS-280
chr4-46342803-INS-60
chr4-46354596-INS-71
chr4-46357391-INS-76
chr4-46372120-INS-82
chr4-46373095-INS-230
chr4-46382038-INS-51
chr4-46408331-INS-108
chr4-46426573-INS-55
chr4-46432332-INS-52
chr4-46446584-INS-402
chr4-46447500-INS-85
chr4-46455491-INS-177
chr4-46461282-INS-201
chr4-46466945-INS-51
chr4-46482122-INS-84
chr4-46493456-INS-393
chr4-46494199-INS-62
chr4-46507810-INS-65
chr4-46521183-INS-138
chr4-46525868-INS-78
chr4-46539823-INS-54
chr4-46629305-INS-160
chr4-46634976-INS-103
chr4-46659372-INS-194
chr4-46666791-INS-58
chr4-46673391-INS-52
chr4-46685074-INS-64
chr4-46702195-INS-75
chr4-46705051-INS-135
chr4-46716041-INS-243
chr4-46746040-INS-88
chr4-46757138-INS-511
chr4-46768053-INS-218
chr4-46769696-INS-751
chr4-46769740-INS-160
chr4-46777741-INS-178
chr4-46864404-INS-68
chr4-46872570-INS-400
chr4-46893456-INS-89
chr4-46922426-INS-185
chr4-46939464-INS-131
chr4-46940936-INS-175
chr4-46994875-INS-84
chr4-47054450-INS-186
chr4-47085236-INS-592
chr4-47096377-INS-269
chr4-47125282-INS-117
chr4-47134899-INS-52
chr4-47185509-INS-98
chr4-47189546-INS-93
chr4-47192101-INS-151
chr4-47194119-INS-594
chr4-47194119-INS-660
chr4-47215006-INS-1931
chr4-47216470-INS-58
chr4-47241605-INS-237
chr4-47270589-INS-64
chr4-47275628-INS-52
chr4-47332179-INS-70
chr4-47335363-INS-108

chr4-47343951-INS-221
chr4-47346657-INS-64
chr4-47349674-INS-55
chr4-47397649-INS-135
chr4-47425565-INS-65
chr4-47429439-INS-52
chr4-47442051-INS-58
chr4-47478531-INS-61
chr4-47488389-INS-102
chr4-47513924-INS-335
chr4-47534663-INS-90
chr4-47547740-INS-2760
chr4-47558901-INS-185
chr4-47569023-INS-2721
chr4-47569559-INS-257
chr4-47582063-INS-170
chr4-47623999-INS-60
chr4-47687849-INS-85
chr4-47707940-INS-57
chr4-47715335-INS-6898
chr4-47722542-INS-593
chr4-47723612-INS-490
chr4-47766415-INS-170
chr4-47797434-INS-663
chr4-47818746-INS-66
chr4-47828706-INS-101
chr4-47838481-INS-375
chr4-47844813-INS-52
chr4-47852372-INS-319
chr4-47861595-INS-78
chr4-47883666-INS-130
chr4-47887096-INS-228
chr4-47895613-INS-54
chr4-47897243-INS-617
chr4-47898300-INS-84
chr4-47900951-INS-578
chr4-47910331-INS-52
chr4-47921956-INS-186
chr4-47931981-INS-497
chr4-47992497-INS-51
chr4-48005711-INS-69
chr4-48030063-INS-948
chr4-48123654-INS-69
chr4-48146970-INS-62
chr4-48187604-INS-196
chr4-48203831-INS-1859
chr4-48221305-INS-235
chr4-48223285-INS-6971
chr4-48239405-INS-499
chr4-48254785-INS-732
chr4-48259803-INS-814
chr4-48261176-INS-581
chr4-48274923-INS-437
chr4-48300695-INS-134
chr4-48330731-INS-207
chr4-48340576-INS-64
chr4-48342474-INS-53
chr4-48349934-INS-614
chr4-48380323-INS-506
chr4-48398977-INS-164
chr4-48405642-INS-78
chr4-48409547-INS-224
chr4-48409561-INS-920
chr4-48412532-INS-176
chr4-48420015-INS-248
chr4-48434618-INS-189
chr4-48450887-INS-492
chr4-48451648-INS-188
chr4-48482380-INS-230
chr4-48486977-INS-194
chr4-48489831-INS-175
chr4-48493796-INS-60
chr4-48495384-INS-58
chr4-48497799-INS-196
chr4-48520787-INS-398
chr4-48528482-INS-53
chr4-48532879-INS-187

chr4-48566294-INS-173
chr4-48572297-INS-198
chr4-48575060-INS-141
chr4-48591600-INS-58
chr4-48593808-INS-54
chr4-48621717-INS-279
chr4-48636273-INS-900
chr4-48646582-INS-218
chr4-48685759-INS-132
chr4-48717724-INS-6886
chr4-48722922-INS-198
chr4-48734360-INS-211
chr4-48736472-INS-64
chr4-48746836-INS-381
chr4-48760273-INS-400
chr4-48827421-INS-466
chr4-48830394-INS-165
chr4-48839936-INS-71
chr4-48870343-INS-189
chr4-48879268-INS-87
chr4-48884002-INS-135
chr4-48915197-INS-820
chr4-48916633-INS-147
chr4-48917571-INS-503
chr4-48936093-INS-6173
chr4-48964449-INS-75
chr4-48973948-INS-64
chr4-49057641-INS-358
chr4-49074737-INS-281
chr4-49077517-INS-70
chr4-49089569-INS-82
chr4-49109298-INS-52
chr4-49116760-INS-62
chr4-49150803-INS-928
chr4-49210466-INS-397
chr4-49211606-INS-734
chr4-49227208-INS-650
chr4-49233462-INS-91
chr4-49240421-INS-55
chr4-49240826-INS-564
chr4-49243102-INS-56
chr4-49247584-INS-5101
chr4-49252619-INS-6140
chr4-49294141-INS-492
chr4-49297619-INS-248
chr4-49300841-INS-565
chr4-49323263-INS-71
chr4-49323738-INS-1435
chr4-49348363-INS-14098
chr4-49350793-INS-6725
chr4-49356471-INS-328
chr4-49373543-INS-163
chr4-49451086-INS-1893
chr4-49458533-INS-82
chr4-49479997-INS-548
chr4-49505762-INS-130
chr4-49518095-INS-54
chr4-49527406-INS-463
chr4-49532214-INS-210
chr4-49533433-INS-73
chr4-49571418-INS-308
chr4-49614841-INS-99
chr4-49620369-INS-51
chr4-49637471-INS-172
chr4-49648753-INS-63
chr4-49675230-INS-146
chr4-49700962-INS-391
chr4-49712638-INS-150
chr4-49738476-INS-112
chr4-49747126-INS-609
chr4-49781109-INS-354
chr4-49813180-INS-57
chr4-49826646-INS-149
chr4-49832383-INS-322
chr4-49835159-INS-67
chr4-49849979-INS-265
chr4-49883194-INS-323

chr4-49903852-INS-7151
chr4-49917406-INS-79
chr4-49923213-INS-430
chr4-49924900-INS-79
chr4-49959190-INS-378
chr4-49979144-INS-824
chr4-49992535-INS-52
chr4-50018414-INS-58
chr4-50027590-INS-166
chr4-50048388-INS-103
chr4-50054863-INS-317
chr4-50065828-INS-129
chr4-50066661-INS-340
chr4-50068964-INS-140
chr4-50092753-INS-5339
chr4-50137616-INS-156
chr4-50138039-INS-7023
chr4-50145137-INS-52
chr4-50184215-INS-110
chr4-50188260-INS-464
chr4-50204893-INS-670
chr4-50221977-INS-301
chr4-50264847-INS-67
chr4-50280542-INS-635
chr4-50285721-INS-54
chr4-50302483-INS-978
chr4-50306360-INS-6154
chr4-50357980-INS-142
chr4-50371070-INS-204
chr4-50372395-INS-511
chr4-50391150-INS-7209
chr4-50416417-INS-714
chr4-50473395-INS-6424
chr4-50487065-INS-68
chr4-50491778-INS-132
chr4-50498237-INS-76
chr4-50525074-INS-201
chr4-50573018-INS-56
chr4-50575777-INS-64
chr4-50595477-INS-247
chr4-50662456-INS-76
chr4-50668108-INS-1268
chr4-50678784-INS-6709
chr4-50748064-INS-362
chr4-50772212-INS-597
chr4-50792885-INS-959
chr4-50804702-INS-64
chr4-50815735-INS-60
chr4-50854646-INS-60
chr4-50910303-INS-150
chr4-50910479-INS-1115
chr4-50935957-INS-239
chr4-50951475-INS-336
chr4-50958298-INS-2926
chr4-50971276-INS-60
chr4-50978776-INS-54
chr4-50984196-INS-113
chr4-51008082-INS-157
chr4-51020076-INS-472
chr4-51027074-INS-70
chr4-51028138-INS-57
chr4-51035592-INS-355
chr4-51047440-INS-145
chr4-51053182-INS-183
chr4-51082265-INS-51
chr4-51087496-INS-95
chr4-51093868-INS-6440
chr4-51098098-INS-985
chr4-51110919-INS-1248
chr4-51116472-INS-61
chr4-51118208-INS-1674
chr4-51118844-INS-444
chr4-51126311-INS-95
chr4-51141117-INS-52
chr4-51150163-INS-87
chr4-51167650-INS-145
chr4-51185099-INS-65

chr4-51186915-INS-612
chr4-51215551-INS-94
chr4-51220146-INS-259
chr4-51230737-INS-454
chr4-51267165-INS-4370
chr4-51268448-INS-142
chr4-51280178-INS-100
chr4-51316357-INS-247
chr4-51345639-INS-2594
chr4-51359739-INS-6398
chr4-51371461-INS-106
chr4-51392065-INS-320
chr4-51412645-INS-2712
chr4-51432181-INS-145
chr4-51451629-INS-400
chr4-51473435-INS-56
chr4-51480133-INS-6689
chr4-51486247-INS-418
chr4-51488774-INS-226
chr4-51520487-INS-445
chr4-51527948-INS-60
chr4-51534994-INS-795
chr4-51545176-INS-956
chr4-51557286-INS-407
chr4-51570783-INS-65
chr4-51581638-INS-97
chr4-51585091-INS-74
chr4-51591453-INS-715
chr4-51625849-INS-57
chr4-51639969-INS-6386
chr4-51676135-INS-6224
chr4-51685498-INS-6156
chr4-51704620-INS-2626
chr4-51731587-INS-87
chr4-51758533-INS-865
chr4-51771588-INS-59
chr4-51817868-INS-530
chr4-51832727-INS-162
chr4-51843574-INS-6441
chr4-51888120-INS-443
chr4-51908610-INS-54
chr4-51964428-INS-81
chr4-51995738-INS-71
chr4-51997151-INS-56
chr4-52061275-INS-564
chr4-52103608-INS-6215
chr4-52111057-INS-90
chr4-52139961-INS-7272
chr4-52151545-INS-327
chr4-52154058-INS-406
chr4-52158323-INS-755
chr4-52177992-INS-6224
chr4-52242278-INS-56
chr4-52249610-INS-6387
chr4-52274204-INS-11982
chr4-52275760-INS-108
chr4-52318435-INS-82
chr4-52354499-INS-209
chr4-52367643-INS-496
chr4-52403872-INS-242
chr4-52406795-INS-121
chr4-52429909-INS-211
chr4-52453522-INS-51
chr4-52494022-INS-192
chr4-52533145-INS-76
chr4-52551506-INS-667
chr4-52616693-INS-282
chr4-52635875-INS-52
chr4-52640149-INS-52
chr4-52748918-INS-211
chr4-52752512-INS-3412
chr4-52781325-INS-65
chr4-52802728-INS-875
chr4-52813582-INS-83
chr4-52823490-INS-79
chr4-52829470-INS-169
chr4-52849587-INS-401

chr4-52886929-INS-145
chr4-52942599-INS-6802
chr4-52944032-INS-6490
chr4-52960332-INS-216
chr4-52986585-INS-64
chr4-53111673-INS-114
chr4-53112905-INS-284
chr4-53126716-INS-117
chr4-53130326-INS-225
chr4-53210773-INS-99
chr4-53226769-INS-257
chr4-53263835-INS-549
chr4-53275591-INS-62
chr4-53277195-INS-103
chr4-53293097-INS-54
chr4-53309289-INS-751
chr4-53312121-INS-383
chr4-53326384-INS-210
chr4-53342859-INS-253
chr4-53346529-INS-118
chr4-53364120-INS-50
chr4-53375226-INS-136
chr4-53412909-INS-184
chr4-53417736-INS-105
chr4-53424272-INS-54
chr4-53432819-INS-472
chr4-53446877-INS-316
chr4-53448873-INS-61
chr4-53469929-INS-96
chr4-53480053-INS-51
chr4-53489230-INS-144
chr4-53489937-INS-434
chr4-53492708-INS-167
chr4-53531284-INS-214
chr4-53543202-INS-74
chr4-53546302-INS-70
chr4-53579771-INS-736
chr4-53617780-INS-66
chr4-53623747-INS-62
chr4-53631052-INS-182
chr4-53708796-INS-916
chr4-53712302-INS-6501
chr4-53729109-INS-220
chr4-53752702-INS-128
chr4-53757863-INS-993
chr4-53773232-INS-224
chr4-53775216-INS-67
chr4-53791781-INS-55
chr4-53801033-INS-60
chr4-53838575-INS-195
chr4-53849912-INS-109
chr4-53863556-INS-340
chr4-53871365-INS-66
chr4-53875656-INS-128
chr4-53900315-INS-60
chr4-53914846-INS-407
chr4-53929193-INS-55
chr4-53936952-INS-178
chr4-53948671-INS-526
chr4-53959075-INS-212
chr4-53986810-INS-151
chr4-53992809-INS-92
chr4-53996036-INS-290
chr4-54006348-INS-74
chr4-54042940-INS-127
chr4-54047897-INS-330
chr4-54114527-INS-58
chr4-54138192-INS-63
chr4-54148359-INS-337
chr4-54166637-INS-101
chr4-54207508-INS-952
chr4-54224520-INS-73
chr4-54252363-INS-343
chr4-54315347-INS-311
chr4-54326118-INS-70
chr4-54346474-INS-152
chr4-54393983-INS-64

chr4-54402863-INS-6483
chr4-54478218-INS-185
chr4-54605826-INS-232
chr4-54664244-INS-6475
chr4-54689963-INS-76
chr4-54706578-INS-1024
chr4-54706799-INS-349
chr4-54776630-INS-82
chr4-54778612-INS-186
chr4-54842011-INS-90
chr4-54877904-INS-139
chr4-54887532-INS-52
chr4-54891669-INS-98
chr4-54912601-INS-114
chr4-54922699-INS-52
chr4-54930064-INS-50
chr4-54931091-INS-169
chr4-54940371-INS-482
chr4-54984367-INS-73
chr4-54996342-INS-200
chr4-55037804-INS-62
chr4-55086471-INS-59
chr4-55104127-INS-61
chr4-55125159-INS-140
chr4-55127640-INS-5547
chr4-55156400-INS-65
chr4-55207167-INS-116
chr4-55224286-INS-293
chr4-55226218-INS-99
chr4-55283815-INS-266
chr4-55295605-INS-80
chr4-55305667-INS-178
chr4-55356571-INS-52
chr4-55384557-INS-52
chr4-55386420-INS-63
chr4-55414380-INS-78
chr4-55414983-INS-6544
chr4-55427642-INS-138
chr4-55446297-INS-79
chr4-55463376-INS-204
chr4-55475218-INS-73
chr4-55512941-INS-203
chr4-55521516-INS-78
chr4-55555279-INS-194
chr4-55579913-INS-61
chr4-55583060-INS-114
chr4-55584714-INS-126
chr4-55592028-INS-87
chr4-55608032-INS-75
chr4-55631558-INS-70
chr4-55663080-INS-92
chr4-55717875-INS-171
chr4-55730937-INS-122
chr4-55742557-INS-59
chr4-55756540-INS-53
chr4-55761089-INS-54
chr4-55765241-INS-287
chr4-55776759-INS-52
chr4-55789075-INS-68
chr4-55789803-INS-64
chr4-55823438-INS-835
chr4-55858636-INS-190
chr4-55868477-INS-239
chr4-55874120-INS-2709
chr4-55880364-INS-204
chr4-55929792-INS-54
chr4-55936122-INS-6481
chr4-55940157-INS-6379
chr4-55979296-INS-52
chr4-56012327-INS-204
chr4-56015066-INS-195
chr4-56020790-INS-75
chr4-56056640-INS-2616
chr4-56100497-INS-399
chr4-56116635-INS-536
chr4-56143049-INS-104
chr4-56156166-INS-938

chr4-56171586-INS-458
chr4-56182172-INS-585
chr4-56184087-INS-500
chr4-56194498-INS-60
chr4-56211241-INS-54
chr4-56227657-INS-168
chr4-56240041-INS-52
chr4-56262337-INS-130
chr4-56265406-INS-51
chr4-56267414-INS-100
chr4-56283730-INS-177
chr4-56383977-INS-247
chr4-56391074-INS-69
chr4-56402414-INS-70
chr4-56431394-INS-212
chr4-56435626-INS-600
chr4-56444290-INS-238
chr4-56461190-INS-144
chr4-56484732-INS-72
chr4-56488605-INS-671
chr4-56501399-INS-703
chr4-56503294-INS-61
chr4-56542892-INS-55
chr4-56565160-INS-927
chr4-56625288-INS-253
chr4-56641890-INS-66
chr4-56643406-INS-385
chr4-56652113-INS-432
chr4-56675195-INS-438
chr4-56680472-INS-168
chr4-56693279-INS-380
chr4-56693536-INS-400
chr4-56695313-INS-55
chr4-56698042-INS-201
chr4-56698119-INS-808
chr4-56705481-INS-206
chr4-56727247-INS-144
chr4-56728068-INS-267
chr4-56776942-INS-228
chr4-56792640-INS-95
chr4-56821505-INS-266
chr4-56826720-INS-189
chr4-56828555-INS-77
chr4-56842914-INS-138
chr4-56858539-INS-231
chr4-56879115-INS-51
chr4-56893649-INS-123
chr4-56927608-INS-218
chr4-56933899-INS-121
chr4-56956025-INS-231
chr4-56975143-INS-884
chr4-56978140-INS-167
chr4-56997040-INS-62
chr4-56998383-INS-217
chr4-57005531-INS-134
chr4-57011861-INS-202
chr4-57020194-INS-57
chr4-57023237-INS-462
chr4-57034065-INS-126
chr4-57052478-INS-635
chr4-57091276-INS-614
chr4-57120473-INS-51
chr4-57172871-INS-68
chr4-57188100-INS-128
chr4-57219823-INS-175
chr4-57253671-INS-95
chr4-57260796-INS-60
chr4-57313491-INS-66
chr4-57313663-INS-579
chr4-57332587-INS-67
chr4-57340361-INS-619
chr4-57352183-INS-51
chr4-57357087-INS-218
chr4-57364635-INS-70
chr4-57376085-INS-125
chr4-57384890-INS-78
chr4-57412206-INS-80

chr4-57463729-INS-88
chr4-57483231-INS-401
chr4-57490590-INS-398
chr4-57497608-INS-651
chr4-57523506-INS-401
chr4-57537217-INS-54
chr4-57555789-INS-58
chr4-57590757-INS-138
chr4-57597308-INS-96
chr4-57615302-INS-211
chr4-57620097-INS-119
chr4-57625729-INS-56
chr4-57659520-INS-299
chr4-57695802-INS-137
chr4-57723968-INS-83
chr4-57725333-INS-508
chr4-57736976-INS-52
chr4-57753944-INS-239
chr4-57761620-INS-70
chr4-57772112-INS-97
chr4-57774028-INS-127
chr4-57783814-INS-61
chr4-57786748-INS-195
chr4-57803771-INS-58
chr4-57807309-INS-240
chr4-57811602-INS-222
chr4-57814575-INS-224
chr4-57817800-INS-193
chr4-57839518-INS-53
chr4-57850464-INS-51
chr4-57864736-INS-82
chr4-57868806-INS-55
chr4-57872849-INS-76
chr4-57878418-INS-58
chr4-57927470-INS-60
chr4-57932385-INS-507
chr4-57936103-INS-63
chr4-57938314-INS-72
chr4-57962794-INS-58
chr4-58007595-INS-72
chr4-58061498-INS-720
chr4-58082341-INS-52
chr4-58083513-INS-6450
chr4-58092471-INS-606
chr4-58094281-INS-83
chr4-58097214-INS-162
chr4-58098441-INS-113
chr4-58115252-INS-154
chr4-58122742-INS-54
chr4-58148593-INS-1933
chr4-58161521-INS-60
chr4-58177949-INS-409
chr4-58179385-INS-114
chr4-58198693-INS-367
chr4-58219892-INS-210
chr4-58223645-INS-6594
chr4-58236601-INS-65
chr4-58251774-INS-127
chr4-58253805-INS-476
chr4-58270549-INS-298
chr4-58272424-INS-401
chr4-58273958-INS-201
chr4-58302339-INS-138
chr4-58312102-INS-199
chr4-58321366-INS-155
chr4-58322844-INS-150
chr4-58326706-INS-56
chr4-58361119-INS-237
chr4-58385195-INS-75
chr4-58404761-INS-96
chr4-58422439-INS-240
chr4-58447004-INS-58
chr4-58484769-INS-220
chr4-58489946-INS-218
chr4-58505001-INS-330
chr4-58561349-INS-293
chr4-58592318-INS-64

chr4-58606397-INS-52
chr4-58623765-INS-166
chr4-58639592-INS-64
chr4-58660263-INS-196
chr4-58670021-INS-262
chr4-58681680-INS-106
chr4-58716360-INS-354
chr4-58718440-INS-66
chr4-58725099-INS-333
chr4-58730410-INS-325
chr4-58760150-INS-432
chr4-58767696-INS-209
chr4-58776912-INS-467
chr4-58780285-INS-70
chr4-58783517-INS-985
chr4-58788551-INS-6179
chr4-58800650-INS-847
chr4-58877332-INS-712
chr4-58898290-INS-214
chr4-58903876-INS-399
chr4-58916213-INS-116
chr4-58930314-INS-6442
chr4-59020906-INS-280
chr4-59029867-INS-238
chr4-59045809-INS-77
chr4-59057570-INS-52
chr4-59064081-INS-218
chr4-59067684-INS-104
chr4-59077710-INS-58
chr4-59086753-INS-660
chr4-59095997-INS-198
chr4-59107405-INS-506
chr4-59110923-INS-503
chr4-59111971-INS-927
chr4-59127402-INS-60
chr4-59133849-INS-247
chr4-59139411-INS-52
chr4-59166353-INS-84
chr4-59171053-INS-514
chr4-59172948-INS-871
chr4-59185489-INS-416
chr4-59185992-INS-54
chr4-59216585-INS-181
chr4-59232574-INS-240
chr4-59242936-INS-6445
chr4-59245270-INS-101
chr4-59250238-INS-60
chr4-59267131-INS-343
chr4-59270230-INS-1415
chr4-59275550-INS-68
chr4-59277147-INS-280
chr4-59277814-INS-590
chr4-59279891-INS-78
chr4-59281932-INS-51
chr4-59288183-INS-343
chr4-59288364-INS-4756
chr4-59291679-INS-104
chr4-59293011-INS-197
chr4-59302008-INS-93
chr4-59305737-INS-209
chr4-59317099-INS-197
chr4-59373414-INS-138
chr4-59383000-INS-204
chr4-59392632-INS-192
chr4-59397626-INS-118
chr4-59414391-INS-68
chr4-59418494-INS-212
chr4-59432805-INS-131
chr4-59445454-INS-92
chr4-59455048-INS-56
chr4-59483832-INS-6427
chr4-59554012-INS-158
chr4-59556822-INS-188
chr4-59560516-INS-81
chr4-59573665-INS-96
chr4-59577201-INS-202
chr4-59578549-INS-114

chr4-59586035-INS-715
chr4-59586282-INS-231
chr4-59604688-INS-6744
chr4-59606157-INS-212
chr4-59608379-INS-1237
chr4-59616021-INS-129
chr4-59664802-INS-60
chr4-59667927-INS-52
chr4-59668745-INS-258
chr4-59679730-INS-462
chr4-59687822-INS-148
chr4-59701163-INS-3231
chr4-59740839-INS-56
chr4-59757794-INS-108
chr4-59764346-INS-5569
chr4-59776107-INS-204
chr4-59788294-INS-1449
chr4-59789693-INS-193
chr4-59790716-INS-217
chr4-59791437-INS-199
chr4-59815915-INS-213
chr4-59822522-INS-196
chr4-59827701-INS-60
chr4-59832512-INS-108
chr4-59837143-INS-212
chr4-59839272-INS-223
chr4-59859987-INS-312
chr4-59864663-INS-7162
chr4-59870787-INS-197
chr4-59873281-INS-80
chr4-59875666-INS-210
chr4-59882184-INS-59
chr4-59890969-INS-180
chr4-59921619-INS-121
chr4-59923817-INS-64
chr4-59944085-INS-112
chr4-59946737-INS-116
chr4-59997402-INS-55
chr4-60017233-INS-92
chr4-60018742-INS-84
chr4-60018811-INS-1334
chr4-60023839-INS-1523
chr4-60051446-INS-346
chr4-60069416-INS-327
chr4-60075597-INS-109
chr4-60081881-INS-74
chr4-60085785-INS-55
chr4-60102927-INS-3601
chr4-60161951-INS-395
chr4-60174096-INS-1304
chr4-60229985-INS-785
chr4-60249326-INS-51
chr4-60270344-INS-54
chr4-60414091-INS-424
chr4-60469287-INS-60
chr4-61108956-INS-136
chr4-61114985-INS-182
chr4-61171199-INS-5054
chr4-61181912-INS-3356
chr4-61191771-INS-223
chr4-61364443-INS-2938
chr4-61408583-INS-1344
chr4-61479322-INS-1890
chr4-61501044-INS-5052
chr4-61579258-INS-5052
chr4-61594434-INS-6401
chr4-61645453-INS-144
chr4-61717231-INS-52
chr4-61741552-INS-140
chr4-61743197-INS-94
chr4-61817223-INS-4608
chr4-61835650-INS-7633
chr4-61847739-INS-326
chr4-61865810-INS-5555
chr4-61884844-INS-6473
chr4-61916829-INS-122
chr4-61918430-INS-7191

chr4-61963793-INS-135
chr4-62008955-INS-181
chr4-62014234-INS-68
chr4-62019572-INS-6413
chr4-62091378-INS-134
chr4-62095022-INS-76
chr4-62113814-INS-98
chr4-62136756-INS-80
chr4-62171723-INS-54
chr4-62195707-INS-86
chr4-62219708-INS-3562
chr4-62229357-INS-52
chr4-62235075-INS-464
chr4-62274371-INS-64
chr4-62284205-INS-52
chr4-62329477-INS-129
chr4-62384073-INS-100
chr4-62384985-INS-231
chr4-62396921-INS-183
chr4-62401469-INS-52
chr4-62404622-INS-201
chr4-62407794-INS-218
chr4-62418673-INS-459
chr4-62434665-INS-587
chr4-62453526-INS-81
chr4-62462906-INS-87
chr4-62464877-INS-66
chr4-62470001-INS-239
chr4-62476207-INS-159
chr4-62498954-INS-65
chr4-62505086-INS-72
chr4-62511050-INS-62
chr4-62515800-INS-73
chr4-62530761-INS-193
chr4-62532647-INS-216
chr4-62552949-INS-54
chr4-62555874-INS-65
chr4-62562617-INS-116
chr4-62628519-INS-103
chr4-62636731-INS-208
chr4-62644814-INS-60
chr4-62684165-INS-6442
chr4-62686812-INS-64
chr4-62691474-INS-79
chr4-62703575-INS-164
chr4-62705228-INS-7180
chr4-62729955-INS-188
chr4-62735955-INS-828
chr4-62736274-INS-6452
chr4-62744389-INS-190
chr4-62764070-INS-105
chr4-62801136-INS-100
chr4-62805124-INS-151
chr4-62806285-INS-889
chr4-62816943-INS-57
chr4-62816947-INS-903
chr4-62821459-INS-112
chr4-62836012-INS-102
chr4-62862077-INS-55
chr4-62876294-INS-176
chr4-62877810-INS-112
chr4-62895307-INS-52
chr4-62899730-INS-532
chr4-62906023-INS-57
chr4-62933865-INS-124
chr4-62959717-INS-212
chr4-62968249-INS-60
chr4-62970220-INS-1862
chr4-63074688-INS-51
chr4-63076157-INS-68
chr4-63111050-INS-74
chr4-63128123-INS-52
chr4-63154470-INS-266
chr4-63158179-INS-196
chr4-63171140-INS-139
chr4-63180946-INS-67
chr4-63194964-INS-61

chr4-63304413-INS-251
chr4-63310189-INS-55
chr4-63325262-INS-99
chr4-63337486-INS-212
chr4-63353858-INS-55
chr4-63359161-INS-58
chr4-63384579-INS-158
chr4-63404587-INS-52
chr4-63431613-INS-60
chr4-63449056-INS-223
chr4-63465461-INS-127
chr4-63505099-INS-52
chr4-63520321-INS-151
chr4-63527216-INS-6871
chr4-63545730-INS-6680
chr4-63573075-INS-67
chr4-63573696-INS-789
chr4-63577765-INS-197
chr4-63577865-INS-3739
chr4-63577868-INS-416
chr4-63582259-INS-57
chr4-63612839-INS-216
chr4-63664607-INS-60
chr4-63708522-INS-52
chr4-63746892-INS-59
chr4-63775451-INS-591
chr4-63896721-INS-52
chr4-63928053-INS-552
chr4-63948025-INS-526
chr4-63958948-INS-312
chr4-64014336-INS-108
chr4-64021252-INS-531
chr4-64037476-INS-799
chr4-64038236-INS-6169
chr4-64040469-INS-62
chr4-64049918-INS-490
chr4-64055229-INS-6639
chr4-64055230-INS-202
chr4-64070459-INS-6261
chr4-64133783-INS-205
chr4-64251738-INS-6174
chr4-64261750-INS-369
chr4-64268021-INS-61
chr4-64299591-INS-90
chr4-64302629-INS-76
chr4-64312832-INS-65
chr4-64335059-INS-81
chr4-64346606-INS-113
chr4-64347799-INS-402
chr4-64348781-INS-68
chr4-64367485-INS-54
chr4-64405910-INS-81
chr4-64427769-INS-254
chr4-64432051-INS-89
chr4-64435843-INS-64
chr4-64440271-INS-383
chr4-64468577-INS-197
chr4-64480757-INS-99
chr4-64483207-INS-65
chr4-64509926-INS-399
chr4-64527378-INS-153
chr4-64593471-INS-6205
chr4-64634456-INS-396
chr4-64639281-INS-716
chr4-64649616-INS-1900
chr4-64663567-INS-6386
chr4-64666611-INS-6155
chr4-64703846-INS-148
chr4-64714356-INS-271
chr4-64716025-INS-56
chr4-64717881-INS-111
chr4-64721070-INS-75
chr4-64724350-INS-705
chr4-64749784-INS-474
chr4-64778511-INS-102
chr4-64785679-INS-154
chr4-64786844-INS-745

chr4-64822703-INS-184
chr4-64832830-INS-6419
chr4-64844940-INS-56
chr4-64872841-INS-68
chr4-64875177-INS-80
chr4-64886959-INS-94
chr4-64896167-INS-693
chr4-64920265-INS-122
chr4-64937188-INS-6428
chr4-64944276-INS-200
chr4-64962362-INS-70
chr4-64969821-INS-87
chr4-64981278-INS-271
chr4-64981817-INS-60
chr4-64986708-INS-6765
chr4-64991957-INS-52
chr4-65002483-INS-593
chr4-65013352-INS-131
chr4-65026543-INS-348
chr4-65035616-INS-77
chr4-65038225-INS-92
chr4-65044421-INS-80
chr4-65065841-INS-54
chr4-65091031-INS-667
chr4-65096352-INS-113
chr4-65131793-INS-238
chr4-65155747-INS-117
chr4-65184747-INS-65
chr4-65190432-INS-55
chr4-65198777-INS-89
chr4-65211921-INS-196
chr4-65227420-INS-815
chr4-65232595-INS-6839
chr4-65287694-INS-5328
chr4-65298757-INS-6923
chr4-65312187-INS-384
chr4-65338396-INS-52
chr4-65363119-INS-124
chr4-65363800-INS-481
chr4-65375205-INS-200
chr4-65381612-INS-58
chr4-65388864-INS-400
chr4-65405308-INS-108
chr4-65417289-INS-6401
chr4-65420205-INS-6688
chr4-65421791-INS-123
chr4-65429465-INS-52
chr4-65431896-INS-62
chr4-65468010-INS-92
chr4-65508375-INS-56
chr4-65530034-INS-91
chr4-65547262-INS-72
chr4-65550281-INS-199
chr4-65550893-INS-6776
chr4-65560414-INS-93
chr4-65573403-INS-158
chr4-65606023-INS-52
chr4-65608678-INS-513
chr4-65628809-INS-347
chr4-65629847-INS-54
chr4-65635584-INS-61
chr4-65640766-INS-84
chr4-65655935-INS-364
chr4-65664784-INS-126
chr4-65690004-INS-85
chr4-65692986-INS-217
chr4-65705499-INS-72
chr4-65709654-INS-56
chr4-65710120-INS-2399
chr4-65771737-INS-1253
chr4-65774499-INS-336
chr4-65800938-INS-6326
chr4-65848287-INS-114
chr4-65865322-INS-192
chr4-65870239-INS-57
chr4-65874064-INS-6830
chr4-65876965-INS-651

chr4-65882985-INS-7315
chr4-65886750-INS-55
chr4-65914107-INS-6591
chr4-65923808-INS-99
chr4-65931489-INS-7163
chr4-65939409-INS-54
chr4-65940525-INS-6448
chr4-65961052-INS-139
chr4-65981135-INS-91
chr4-65995146-INS-140
chr4-66004687-INS-90
chr4-66005426-INS-503
chr4-66012267-INS-58
chr4-66016990-INS-187
chr4-66018415-INS-71
chr4-66030741-INS-58
chr4-66066602-INS-339
chr4-66077882-INS-78
chr4-66084705-INS-230
chr4-66102974-INS-6496
chr4-66109049-INS-85
chr4-66123199-INS-399
chr4-66131199-INS-180
chr4-66159302-INS-120
chr4-66162373-INS-56
chr4-66168266-INS-80
chr4-66215336-INS-168
chr4-66218189-INS-6884
chr4-66219768-INS-58
chr4-66222687-INS-6176
chr4-66229454-INS-921
chr4-66230084-INS-6971
chr4-66243952-INS-506
chr4-66260385-INS-53
chr4-66302672-INS-168
chr4-66305773-INS-195
chr4-66329380-INS-1893
chr4-66331038-INS-59
chr4-66334070-INS-115
chr4-66335483-INS-7049
chr4-66345872-INS-52
chr4-66369924-INS-57
chr4-66425689-INS-192
chr4-66458031-INS-5831
chr4-66465435-INS-1888
chr4-66479711-INS-159
chr4-66502536-INS-240
chr4-66525724-INS-326
chr4-66528366-INS-329
chr4-66530003-INS-203
chr4-66546685-INS-967
chr4-66560451-INS-677
chr4-66567954-INS-6723
chr4-66602950-INS-6889
chr4-66604957-INS-3010
chr4-66657536-INS-6112
chr4-66662714-INS-54
chr4-66673219-INS-831
chr4-66685820-INS-210
chr4-66723461-INS-109
chr4-66738846-INS-8451
chr4-66742617-INS-6498
chr4-66770667-INS-221
chr4-66772601-INS-719
chr4-66788943-INS-157
chr4-66802251-INS-819
chr4-66813621-INS-342
chr4-66814941-INS-51
chr4-66816418-INS-133
chr4-66841964-INS-96
chr4-66861190-INS-62
chr4-66878206-INS-69
chr4-66891522-INS-1482
chr4-66898155-INS-72
chr4-66922600-INS-403
chr4-66927477-INS-271
chr4-66932801-INS-7367

chr4-66956198-INS-4766
chr4-66973814-INS-62
chr4-67010761-INS-67
chr4-67019471-INS-405
chr4-67037725-INS-6960
chr4-67041787-INS-248
chr4-67046053-INS-89
chr4-67051455-INS-326
chr4-67077584-INS-344
chr4-67093709-INS-60
chr4-67121137-INS-83
chr4-67129553-INS-62
chr4-67130136-INS-181
chr4-67136527-INS-6441
chr4-67144382-INS-58
chr4-67150436-INS-74
chr4-67159239-INS-294
chr4-67161636-INS-65
chr4-67192456-INS-4325
chr4-67199379-INS-748
chr4-67225027-INS-944
chr4-67265941-INS-775
chr4-67271500-INS-6699
chr4-67315762-INS-88
chr4-67317971-INS-6753
chr4-67332555-INS-72
chr4-67361126-INS-176
chr4-67363226-INS-6862
chr4-67392645-INS-213
chr4-67405461-INS-52
chr4-67409795-INS-67
chr4-67424555-INS-400
chr4-67427394-INS-258
chr4-67428199-INS-215
chr4-67448450-INS-71
chr4-67456174-INS-3387
chr4-67478366-INS-164
chr4-67479445-INS-134
chr4-67489779-INS-6176
chr4-67505883-INS-504
chr4-67509404-INS-130
chr4-67514545-INS-162
chr4-67529752-INS-54
chr4-67531108-INS-2774
chr4-67547783-INS-114
chr4-67555186-INS-251
chr4-67576489-INS-901
chr4-67593705-INS-940
chr4-67599617-INS-91
chr4-67620678-INS-62
chr4-67653647-INS-7232
chr4-67656773-INS-449
chr4-67660139-INS-111
chr4-67731669-INS-70
chr4-67753332-INS-498
chr4-67754261-INS-868
chr4-67785335-INS-85
chr4-67823179-INS-640
chr4-67823713-INS-1644
chr4-67869291-INS-700
chr4-67883559-INS-84
chr4-67949047-INS-208
chr4-67966335-INS-106
chr4-67976742-INS-52
chr4-67987209-INS-212
chr4-68024253-INS-61
chr4-68041999-INS-6366
chr4-68042744-INS-52
chr4-68044391-INS-240
chr4-68046968-INS-116
chr4-68061161-INS-6611
chr4-68122652-INS-4198
chr4-68133783-INS-401
chr4-68136458-INS-696
chr4-68161645-INS-52
chr4-68222620-INS-313
chr4-68248906-INS-103

chr4-68270034-INS-129
chr4-68284620-INS-1063
chr4-68297929-INS-72
chr4-68307051-INS-103
chr4-68313077-INS-544
chr4-68339918-INS-57
chr4-68384391-INS-152
chr4-68406216-INS-203
chr4-68412579-INS-84
chr4-68424899-INS-496
chr4-68429730-INS-5185
chr4-68438547-INS-838
chr4-68445792-INS-58
chr4-68450817-INS-77
chr4-68486104-INS-70
chr4-68490297-INS-100
chr4-68491417-INS-74
chr4-68498577-INS-2898
chr4-68558710-INS-6454
chr4-68614625-INS-111
chr4-68660234-INS-110
chr4-68670583-INS-66
chr4-68675776-INS-118
chr4-68685055-INS-76
chr4-68688374-INS-62
chr4-68691430-INS-177
chr4-68744378-INS-52
chr4-68745256-INS-309
chr4-68749733-INS-206
chr4-68763141-INS-156
chr4-68775039-INS-109
chr4-68777478-INS-77
chr4-68782300-INS-62
chr4-68786967-INS-247
chr4-68797366-INS-351
chr4-68800044-INS-60
chr4-68803656-INS-105
chr4-68812806-INS-476
chr4-68834818-INS-1632
chr4-68841586-INS-6485
chr4-68876123-INS-54
chr4-68893641-INS-782
chr4-68927725-INS-1178
chr4-68950053-INS-166
chr4-68952341-INS-67
chr4-68963313-INS-590
chr4-68975848-INS-614
chr4-68999861-INS-63
chr4-69009592-INS-51
chr4-69017839-INS-60
chr4-69048060-INS-110
chr4-69059109-INS-376
chr4-69113628-INS-105
chr4-69119567-INS-285
chr4-69142449-INS-413
chr4-69210190-INS-299
chr4-69211688-INS-60
chr4-69262145-INS-455
chr4-69274783-INS-4194
chr4-69277825-INS-188
chr4-69293735-INS-209
chr4-69318867-INS-51
chr4-69384638-INS-7341
chr4-69385787-INS-132
chr4-69400012-INS-1231
chr4-69407977-INS-6517
chr4-69448851-INS-72
chr4-69451688-INS-106
chr4-69460802-INS-261
chr4-69461942-INS-281
chr4-69466216-INS-6284
chr4-69470454-INS-150
chr4-69476414-INS-99
chr4-69490161-INS-1084
chr4-69522002-INS-72
chr4-69551591-INS-9875
chr4-69558541-INS-8028

chr4-69569502-INS-344
chr4-69576671-INS-79
chr4-69585629-INS-462
chr4-69596450-INS-76
chr4-69596883-INS-3267
chr4-69597918-INS-682
chr4-69599417-INS-6224
chr4-69612772-INS-911
chr4-69644063-INS-6819
chr4-69644589-INS-502
chr4-69652695-INS-98
chr4-69683267-INS-328
chr4-69729239-INS-496
chr4-69765502-INS-532
chr4-69783605-INS-296
chr4-69802947-INS-380
chr4-69827094-INS-201
chr4-69847738-INS-180
chr4-69862960-INS-170
chr4-69877085-INS-60
chr4-69879099-INS-56
chr4-69881525-INS-156
chr4-69905935-INS-116
chr4-69927166-INS-832
chr4-69927668-INS-95
chr4-69934756-INS-1527
chr4-69971463-INS-85
chr4-69975049-INS-63
chr4-69990230-INS-1892
chr4-70040934-INS-2715
chr4-70053172-INS-4361
chr4-70060186-INS-76
chr4-70096261-INS-510
chr4-70106987-INS-72
chr4-70130881-INS-768
chr4-70134562-INS-53
chr4-70253500-INS-564
chr4-70255354-INS-197
chr4-70265006-INS-1894
chr4-70293111-INS-224
chr4-70303736-INS-208
chr4-70332382-INS-55
chr4-70334745-INS-126
chr4-70338480-INS-107
chr4-70341347-INS-55
chr4-70383535-INS-125
chr4-70399174-INS-342
chr4-70420593-INS-65
chr4-70422824-INS-197
chr4-70432112-INS-133
chr4-70441112-INS-444
chr4-70452081-INS-138
chr4-70467314-INS-6459
chr4-70471387-INS-103
chr4-70482227-INS-51
chr4-70511826-INS-2024
chr4-70531395-INS-230
chr4-70533023-INS-71
chr4-70541839-INS-52
chr4-70542149-INS-6386
chr4-70557014-INS-51
chr4-70558705-INS-114
chr4-70564087-INS-400
chr4-70573175-INS-324
chr4-70575104-INS-133
chr4-70601373-INS-4451
chr4-70625548-INS-71
chr4-70656971-INS-370
chr4-70657489-INS-104
chr4-70660881-INS-54
chr4-70666990-INS-75
chr4-70673623-INS-526
chr4-70700684-INS-60
chr4-70725053-INS-6371
chr4-70739092-INS-5182
chr4-70747104-INS-7004
chr4-70758367-INS-117

chr4-70774989-INS-497
chr4-70780288-INS-60
chr4-70802569-INS-53
chr4-70804745-INS-7129
chr4-70804887-INS-52
chr4-70811378-INS-487
chr4-70833345-INS-6594
chr4-70841582-INS-587
chr4-70872782-INS-673
chr4-70877291-INS-195
chr4-70888675-INS-103
chr4-70934768-INS-1022
chr4-71003656-INS-206
chr4-71026146-INS-7099
chr4-71035987-INS-509
chr4-71038630-INS-102
chr4-71081833-INS-270
chr4-71106741-INS-127
chr4-71114503-INS-69
chr4-71124509-INS-56
chr4-71169242-INS-277
chr4-71217619-INS-764
chr4-71221372-INS-76
chr4-71259887-INS-58
chr4-71279531-INS-6376
chr4-71291712-INS-214
chr4-71293236-INS-947
chr4-71297031-INS-69
chr4-71297681-INS-1864
chr4-71318328-INS-75
chr4-71320215-INS-469
chr4-71340147-INS-6493
chr4-71342953-INS-305
chr4-71374434-INS-6736
chr4-71383042-INS-80
chr4-71409982-INS-69
chr4-71434200-INS-1988
chr4-71447792-INS-279
chr4-71452723-INS-411
chr4-71469642-INS-116
chr4-71519693-INS-58
chr4-71575589-INS-195
chr4-71580236-INS-76
chr4-71597962-INS-108
chr4-71646427-INS-680
chr4-71653488-INS-66
chr4-71671955-INS-788
chr4-71723696-INS-576
chr4-71762015-INS-493
chr4-71791800-INS-6615
chr4-71818533-INS-7031
chr4-71829319-INS-72
chr4-71831385-INS-51
chr4-71842932-INS-78
chr4-71844674-INS-63
chr4-71845141-INS-6151
chr4-71853756-INS-493
chr4-71874372-INS-6923
chr4-71891096-INS-6389
chr4-71906106-INS-64
chr4-71925342-INS-4613
chr4-71927848-INS-980
chr4-71938383-INS-237
chr4-71965227-INS-639
chr4-71983513-INS-1175
chr4-72049418-INS-73
chr4-72106020-INS-60
chr4-72108385-INS-201
chr4-72158944-INS-1861
chr4-72162102-INS-219
chr4-72166270-INS-53
chr4-72172289-INS-206
chr4-72205068-INS-164
chr4-72216894-INS-80
chr4-72279811-INS-9355
chr4-72285038-INS-688
chr4-72288552-INS-4980

chr4-72348337-INS-544
chr4-72375938-INS-6447
chr4-72432703-INS-54
chr4-72436801-INS-92
chr4-72506436-INS-4784
chr4-72575600-INS-467
chr4-72586626-INS-137
chr4-72612295-INS-315
chr4-72613122-INS-56
chr4-72617525-INS-634
chr4-72622034-INS-179
chr4-72635089-INS-602
chr4-72653250-INS-6737
chr4-72671744-INS-128
chr4-72673004-INS-148
chr4-72677672-INS-139
chr4-72693817-INS-61
chr4-72704390-INS-6481
chr4-72722540-INS-6382
chr4-72731799-INS-249
chr4-72748194-INS-272
chr4-72764056-INS-123
chr4-72776928-INS-138
chr4-72779681-INS-77
chr4-72801466-INS-373
chr4-72813351-INS-88
chr4-72815365-INS-772
chr4-72828253-INS-163
chr4-72837619-INS-400
chr4-72838471-INS-110
chr4-72840866-INS-183
chr4-72862099-INS-946
chr4-72880064-INS-211
chr4-72902937-INS-60
chr4-72915230-INS-795
chr4-72927870-INS-180
chr4-72953806-INS-359
chr4-72997002-INS-54
chr4-73014621-INS-6842
chr4-73020055-INS-3227
chr4-73021805-INS-294
chr4-73029369-INS-6454
chr4-73048651-INS-856
chr4-73052690-INS-414
chr4-73073927-INS-559
chr4-73092156-INS-5284
chr4-73092949-INS-145
chr4-73118282-INS-275
chr4-73147319-INS-464
chr4-73154992-INS-3554
chr4-73174498-INS-74
chr4-73200690-INS-235
chr4-73268473-INS-2132
chr4-73270434-INS-407
chr4-73274851-INS-327
chr4-73295868-INS-56
chr4-73296988-INS-868
chr4-73299030-INS-64
chr4-73301548-INS-194
chr4-73352780-INS-2145
chr4-73359760-INS-6350
chr4-73361429-INS-103
chr4-73363589-INS-61
chr4-73365084-INS-57
chr4-73371447-INS-72
chr4-73375415-INS-58
chr4-73409086-INS-6431
chr4-73419072-INS-441
chr4-73422418-INS-88
chr4-73425243-INS-127
chr4-73500651-INS-111
chr4-73512439-INS-63
chr4-73536773-INS-99
chr4-73621988-INS-268
chr4-73635246-INS-178
chr4-73661838-INS-962
chr4-73667964-INS-63

chr4-73685924-INS-81
chr4-73701418-INS-54
chr4-73740706-INS-477
chr4-73858634-INS-163
chr4-73864953-INS-88
chr4-73882957-INS-889
chr4-73897185-INS-400
chr4-73900936-INS-6379
chr4-73920427-INS-2665
chr4-73930484-INS-150
chr4-73944121-INS-388
chr4-73965058-INS-117
chr4-73982952-INS-2078
chr4-73988436-INS-6160
chr4-73998723-INS-57
chr4-74012241-INS-155
chr4-74055319-INS-122
chr4-74065947-INS-111
chr4-74092905-INS-225
chr4-74094308-INS-6637
chr4-74097532-INS-6549
chr4-74116512-INS-68
chr4-74117919-INS-413
chr4-74127792-INS-68
chr4-74130463-INS-88
chr4-74136994-INS-780
chr4-74148441-INS-58
chr4-74149113-INS-506
chr4-74169058-INS-192
chr4-74179046-INS-89
chr4-74184718-INS-214
chr4-74190553-INS-198
chr4-74191141-INS-276
chr4-74228086-INS-57
chr4-74255517-INS-202
chr4-74256275-INS-203
chr4-74261342-INS-213
chr4-74273011-INS-9412
chr4-74279388-INS-4392
chr4-74292599-INS-445
chr4-74301330-INS-60
chr4-74302994-INS-211
chr4-74310312-INS-112
chr4-74387799-INS-172
chr4-74389484-INS-64
chr4-74402354-INS-53
chr4-74403389-INS-56
chr4-74405233-INS-329
chr4-74431479-INS-60
chr4-74441736-INS-268
chr4-74493938-INS-7007
chr4-74499399-INS-68
chr4-74507784-INS-66
chr4-74508896-INS-354
chr4-74548060-INS-255
chr4-74549301-INS-880
chr4-74551907-INS-4949
chr4-74626544-INS-62
chr4-74695113-INS-783
chr4-74699069-INS-52
chr4-74734476-INS-870
chr4-74748571-INS-929
chr4-74765128-INS-53
chr4-74771194-INS-115
chr4-74773636-INS-111
chr4-74779138-INS-8472
chr4-74797871-INS-363
chr4-74815982-INS-70
chr4-74823354-INS-56
chr4-74843904-INS-452
chr4-74940026-INS-136
chr4-74942935-INS-7090
chr4-75007297-INS-316
chr4-75015556-INS-52
chr4-75039900-INS-1205
chr4-75059369-INS-6328
chr4-75075658-INS-765

chr4-75102407-INS-6424
chr4-75104152-INS-6319
chr4-75108580-INS-92
chr4-75111113-INS-497
chr4-75112518-INS-286
chr4-75133862-INS-354
chr4-75186723-INS-256
chr4-75218793-INS-972
chr4-75232802-INS-64
chr4-75244524-INS-859
chr4-75255541-INS-52
chr4-75261736-INS-6384
chr4-75262448-INS-6366
chr4-75263192-INS-344
chr4-75273376-INS-201
chr4-75301038-INS-679
chr4-75314215-INS-487
chr4-75334874-INS-86
chr4-75347962-INS-6440
chr4-75390318-INS-78
chr4-75399846-INS-977
chr4-75429818-INS-250
chr4-75434527-INS-56
chr4-75455828-INS-176
chr4-75467595-INS-130
chr4-75477763-INS-73
chr4-75493681-INS-409
chr4-75505292-INS-179
chr4-75507022-INS-1893
chr4-75512845-INS-5474
chr4-75544745-INS-197
chr4-75553981-INS-789
chr4-75556754-INS-54
chr4-75564129-INS-62
chr4-75567275-INS-144
chr4-75571799-INS-57
chr4-75573719-INS-1494
chr4-75586568-INS-165
chr4-75602972-INS-51
chr4-75624245-INS-65
chr4-75661069-INS-6605
chr4-75676130-INS-6249
chr4-75678658-INS-54
chr4-75688775-INS-233
chr4-75693973-INS-51
chr4-75736876-INS-64
chr4-75762883-INS-54
chr4-75780928-INS-993
chr4-75805564-INS-6639
chr4-75816143-INS-54
chr4-75831888-INS-90
chr4-75840126-INS-89
chr4-75931411-INS-186
chr4-75945435-INS-245
chr4-75953205-INS-735
chr4-75956805-INS-76
chr4-75981753-INS-1191
chr4-75999250-INS-76
chr4-76001937-INS-64
chr4-76023183-INS-97
chr4-76049547-INS-86
chr4-76069164-INS-265
chr4-76099535-INS-652
chr4-76134485-INS-563
chr4-76137338-INS-64
chr4-76138120-INS-2216
chr4-76175546-INS-60
chr4-76184214-INS-400
chr4-76199375-INS-6450
chr4-76199624-INS-56
chr4-76202175-INS-6702
chr4-76209955-INS-6501
chr4-76218052-INS-6309
chr4-76253479-INS-64
chr4-76283325-INS-8816
chr4-76290642-INS-130
chr4-76401203-INS-75

chr4-76452446-INS-60
chr4-76487591-INS-60
chr4-76503453-INS-52
chr4-76567478-INS-65
chr4-76575941-INS-142
chr4-76585639-INS-686
chr4-76585642-INS-1090
chr4-76588856-INS-2909
chr4-76595114-INS-358
chr4-76598907-INS-579
chr4-76617781-INS-115
chr4-76619647-INS-161
chr4-76620258-INS-85
chr4-76632023-INS-672
chr4-76641780-INS-52
chr4-76671984-INS-1078
chr4-76679948-INS-6241
chr4-76727411-INS-872
chr4-76728635-INS-64
chr4-76742465-INS-68
chr4-76754103-INS-1852
chr4-76762476-INS-227
chr4-76771579-INS-61
chr4-76779244-INS-314
chr4-76797187-INS-260
chr4-76811869-INS-960
chr4-76813577-INS-59
chr4-76818898-INS-73
chr4-76882269-INS-631
chr4-76887296-INS-503
chr4-76900621-INS-60
chr4-76907560-INS-311
chr4-76913449-INS-218
chr4-76921467-INS-628
chr4-76948331-INS-72
chr4-76961968-INS-70
chr4-76972006-INS-154
chr4-76975842-INS-53
chr4-76998375-INS-63
chr4-76998985-INS-499
chr4-77003417-INS-585
chr4-77027166-INS-51
chr4-77056596-INS-243
chr4-77074807-INS-1558
chr4-77080247-INS-81
chr4-77146167-INS-120
chr4-77177583-INS-56
chr4-77185407-INS-90
chr4-77187724-INS-6905
chr4-77196048-INS-6467
chr4-77210982-INS-56
chr4-77263552-INS-377
chr4-77271490-INS-62
chr4-77282268-INS-510
chr4-77329359-INS-88
chr4-77330770-INS-7102
chr4-77341379-INS-58
chr4-77359239-INS-201
chr4-77364348-INS-60
chr4-77366091-INS-773
chr4-77372432-INS-840
chr4-77445167-INS-855
chr4-77450005-INS-83
chr4-77463896-INS-6219
chr4-77479412-INS-147
chr4-77518810-INS-300
chr4-77523779-INS-6379
chr4-77539839-INS-4073
chr4-77549854-INS-86
chr4-77557104-INS-63
chr4-77558829-INS-955
chr4-77560661-INS-60
chr4-77581203-INS-6500
chr4-77589897-INS-653
chr4-77593631-INS-147
chr4-77596183-INS-362
chr4-77614533-INS-60

chr4-77617141-INS-114
chr4-77647058-INS-290
chr4-77673186-INS-232
chr4-77681507-INS-831
chr4-77684285-INS-820
chr4-77685318-INS-6417
chr4-77685832-INS-971
chr4-77727474-INS-102
chr4-77736382-INS-478
chr4-77748305-INS-400
chr4-77758135-INS-815
chr4-77777083-INS-2130
chr4-77794508-INS-70
chr4-77805587-INS-737
chr4-77812099-INS-64
chr4-77826985-INS-66
chr4-77829562-INS-82
chr4-77838993-INS-112
chr4-77846624-INS-293
chr4-77855002-INS-208
chr4-77857313-INS-949
chr4-77885620-INS-52
chr4-77914064-INS-212
chr4-77916460-INS-7573
chr4-77950299-INS-66
chr4-77955255-INS-6554
chr4-77957961-INS-64
chr4-77975572-INS-96
chr4-77977948-INS-63
chr4-78000541-INS-210
chr4-78014354-INS-60
chr4-78022098-INS-443
chr4-78023142-INS-220
chr4-78027306-INS-246
chr4-78063322-INS-130
chr4-78070353-INS-60
chr4-78088517-INS-4162
chr4-78101132-INS-400
chr4-78105613-INS-160
chr4-78122397-INS-96
chr4-78146450-INS-76
chr4-78161004-INS-138
chr4-78184495-INS-96
chr4-78240621-INS-192
chr4-78283592-INS-60
chr4-78328426-INS-65
chr4-78332462-INS-395
chr4-78406572-INS-64
chr4-78420303-INS-258
chr4-78421804-INS-216
chr4-78439291-INS-109
chr4-78450481-INS-6816
chr4-78481443-INS-5299
chr4-78553641-INS-73
chr4-78582181-INS-57
chr4-78588915-INS-2764
chr4-78606890-INS-75
chr4-78620590-INS-2990
chr4-78698274-INS-283
chr4-78729649-INS-65
chr4-78749571-INS-829
chr4-78810315-INS-7235
chr4-78824188-INS-53
chr4-78840657-INS-6792
chr4-78920098-INS-1420
chr4-78936989-INS-279
chr4-78939511-INS-56
chr4-78942798-INS-360
chr4-78958225-INS-62
chr4-78959470-INS-5547
chr4-79021888-INS-519
chr4-79038985-INS-784
chr4-79046193-INS-6462
chr4-79057669-INS-74
chr4-79077220-INS-423
chr4-79093242-INS-70
chr4-79139114-INS-6674

chr4-79208007-INS-54
chr4-79237077-INS-52
chr4-79247687-INS-369
chr4-79321254-INS-317
chr4-79338645-INS-58
chr4-79353368-INS-58
chr4-79355178-INS-64
chr4-79375838-INS-52
chr4-79382195-INS-92
chr4-79411018-INS-83
chr4-79422707-INS-374
chr4-79451136-INS-120
chr4-79453077-INS-495
chr4-79468643-INS-6921
chr4-79472952-INS-416
chr4-79509650-INS-1890
chr4-79518008-INS-86
chr4-79521275-INS-72
chr4-79555558-INS-138
chr4-79566088-INS-6706
chr4-79577399-INS-345
chr4-79598513-INS-88
chr4-79617741-INS-58
chr4-79651135-INS-70
chr4-79654771-INS-57
chr4-79662421-INS-380
chr4-79677736-INS-96
chr4-79721228-INS-80
chr4-79741502-INS-1608
chr4-79769362-INS-1034
chr4-79794307-INS-202
chr4-79817410-INS-54
chr4-79823235-INS-387
chr4-79848040-INS-55
chr4-79849873-INS-53
chr4-79905559-INS-87
chr4-79919892-INS-454
chr4-79925091-INS-4743
chr4-79939208-INS-363
chr4-79941167-INS-3620
chr4-79951197-INS-116
chr4-79951771-INS-704
chr4-79976154-INS-60
chr4-79980898-INS-230
chr4-80040833-INS-160
chr4-80043851-INS-5246
chr4-80056079-INS-64
chr4-80069341-INS-5300
chr4-80075143-INS-782
chr4-80081959-INS-60
chr4-80120785-INS-362
chr4-80143196-INS-392
chr4-80174225-INS-67
chr4-80222002-INS-165
chr4-80263851-INS-68
chr4-80302237-INS-66
chr4-80312382-INS-168
chr4-80314414-INS-421
chr4-80316528-INS-57
chr4-80339319-INS-189
chr4-80363295-INS-634
chr4-80373938-INS-460
chr4-80377572-INS-442
chr4-80404033-INS-855
chr4-80405229-INS-256
chr4-80418517-INS-132
chr4-80436729-INS-6773
chr4-80451062-INS-6643
chr4-80462481-INS-95
chr4-80470035-INS-400
chr4-80470141-INS-1264
chr4-80472456-INS-999
chr4-80474648-INS-1294
chr4-80478844-INS-639
chr4-80505689-INS-7133
chr4-80506419-INS-54
chr4-80525622-INS-61

chr4-80527813-INS-343
chr4-80533819-INS-6367
chr4-80537620-INS-90
chr4-80545648-INS-479
chr4-80594876-INS-88
chr4-80600939-INS-787
chr4-80603271-INS-244
chr4-80604941-INS-52
chr4-80608169-INS-8947
chr4-80613452-INS-100
chr4-80623846-INS-6429
chr4-80643827-INS-943
chr4-80666051-INS-410
chr4-80667976-INS-294
chr4-80672331-INS-548
chr4-80712806-INS-6820
chr4-80737669-INS-69
chr4-80739797-INS-84
chr4-80770855-INS-178
chr4-80780438-INS-70
chr4-80782614-INS-55
chr4-80798084-INS-89
chr4-80828411-INS-69
chr4-80849715-INS-125
chr4-80869021-INS-408
chr4-80879437-INS-464
chr4-80886766-INS-57
chr4-80890540-INS-499
chr4-80892053-INS-254
chr4-80894274-INS-1022
chr4-80900948-INS-365
chr4-80903795-INS-137
chr4-80915262-INS-571
chr4-80925333-INS-248
chr4-80962717-INS-84
chr4-81011928-INS-5996
chr4-81018693-INS-1619
chr4-81031194-INS-904
chr4-81055460-INS-116
chr4-81112626-INS-194
chr4-81165780-INS-185
chr4-81181538-INS-304
chr4-81205526-INS-6516
chr4-81210288-INS-204
chr4-81220773-INS-57
chr4-81252528-INS-75
chr4-81270324-INS-133
chr4-81276046-INS-257
chr4-81307896-INS-203
chr4-81318458-INS-117
chr4-81370913-INS-79
chr4-81387912-INS-303
chr4-81389684-INS-148
chr4-81395739-INS-730
chr4-81401382-INS-874
chr4-81419307-INS-286
chr4-81424371-INS-108
chr4-81442182-INS-365
chr4-81445234-INS-180
chr4-81460830-INS-146
chr4-81477098-INS-57
chr4-81477944-INS-229
chr4-81521936-INS-86
chr4-81522271-INS-915
chr4-81529172-INS-390
chr4-81541951-INS-70
chr4-81550963-INS-6247
chr4-81554062-INS-65
chr4-81567164-INS-89
chr4-81596933-INS-191
chr4-81598050-INS-109
chr4-81632418-INS-202
chr4-81650578-INS-738
chr4-81659253-INS-231
chr4-81684177-INS-6635
chr4-81693681-INS-6692
chr4-81713392-INS-70

chr4-81743419-INS-54
chr4-81751015-INS-137
chr4-81770746-INS-54
chr4-81806750-INS-1273
chr4-81815009-INS-115
chr4-81816266-INS-246
chr4-81826941-INS-636
chr4-81836430-INS-54
chr4-81864051-INS-1727
chr4-81868020-INS-401
chr4-81892671-INS-530
chr4-81904450-INS-73
chr4-81914099-INS-341
chr4-81918694-INS-64
chr4-81946482-INS-72
chr4-81950665-INS-6543
chr4-81956696-INS-7415
chr4-81970468-INS-51
chr4-81986693-INS-86
chr4-81996099-INS-6600
chr4-82000780-INS-92
chr4-82001692-INS-6986
chr4-82020125-INS-1276
chr4-82029193-INS-99
chr4-82030276-INS-6839
chr4-82033450-INS-527
chr4-82042526-INS-106
chr4-82045051-INS-104
chr4-82045534-INS-3621
chr4-82050993-INS-4369
chr4-82056875-INS-67
chr4-82063644-INS-91
chr4-82088945-INS-79
chr4-82163597-INS-433
chr4-82165271-INS-666
chr4-82171021-INS-296
chr4-82261688-INS-239
chr4-82272778-INS-134
chr4-82273794-INS-76
chr4-82280025-INS-75
chr4-82295593-INS-723
chr4-82300467-INS-86
chr4-82313614-INS-62
chr4-82354691-INS-718
chr4-82399696-INS-424
chr4-82436634-INS-674
chr4-82482897-INS-73
chr4-82501964-INS-92
chr4-82529883-INS-448
chr4-82545901-INS-52
chr4-82551537-INS-56
chr4-82556255-INS-348
chr4-82564307-INS-75
chr4-82587048-INS-76
chr4-82590541-INS-214
chr4-82593622-INS-507
chr4-82600653-INS-229
chr4-82619289-INS-7111
chr4-82627147-INS-151
chr4-82629509-INS-203
chr4-82630253-INS-194
chr4-82635897-INS-62
chr4-82668477-INS-389
chr4-82675203-INS-6821
chr4-82676986-INS-521
chr4-82687317-INS-604
chr4-82752796-INS-87
chr4-82806946-INS-6255
chr4-82811255-INS-193
chr4-82829471-INS-127
chr4-82867350-INS-630
chr4-82893106-INS-482
chr4-82897808-INS-98
chr4-82919486-INS-106
chr4-82931921-INS-92
chr4-82951756-INS-106
chr4-82970711-INS-64

chr4-82981228-INS-560
chr4-83005602-INS-134
chr4-83010474-INS-490
chr4-83018362-INS-103
chr4-83023534-INS-61
chr4-83030433-INS-1802
chr4-83039053-INS-6581
chr4-83065913-INS-578
chr4-83067874-INS-661
chr4-83071816-INS-479
chr4-83091183-INS-88
chr4-83095023-INS-742
chr4-83105900-INS-70
chr4-83119274-INS-341
chr4-83129714-INS-148
chr4-83139996-INS-54
chr4-83141892-INS-155
chr4-83157592-INS-88
chr4-83161749-INS-231
chr4-83196971-INS-236
chr4-83199616-INS-157
chr4-83201328-INS-81
chr4-83207977-INS-942
chr4-83208139-INS-179
chr4-83221985-INS-210
chr4-83228926-INS-220
chr4-83243996-INS-209
chr4-83249457-INS-76
chr4-83253237-INS-206
chr4-83272037-INS-199
chr4-83277318-INS-219
chr4-83293177-INS-6638
chr4-83318925-INS-198
chr4-83321661-INS-420
chr4-83328785-INS-420
chr4-83329385-INS-108
chr4-83330476-INS-736
chr4-83333984-INS-197
chr4-83349717-INS-70
chr4-83384570-INS-200
chr4-83386924-INS-63
chr4-83387688-INS-180
chr4-83390109-INS-213
chr4-83393071-INS-202
chr4-83393341-INS-838
chr4-83413584-INS-426
chr4-83415678-INS-379
chr4-83426329-INS-213
chr4-83437939-INS-81
chr4-83439378-INS-237
chr4-83451546-INS-192
chr4-83462043-INS-136
chr4-83466287-INS-166
chr4-83473990-INS-112
chr4-83494262-INS-5548
chr4-83496306-INS-771
chr4-83497975-INS-662
chr4-83500566-INS-80
chr4-83506155-INS-196
chr4-83511982-INS-6517
chr4-83588584-INS-64
chr4-83593665-INS-86
chr4-83603454-INS-816
chr4-83605740-INS-257
chr4-83669682-INS-123
chr4-83692566-INS-537
chr4-83699687-INS-335
chr4-83719286-INS-572
chr4-83723655-INS-1892
chr4-83738931-INS-168
chr4-83742981-INS-199
chr4-83745869-INS-396
chr4-83760356-INS-152
chr4-83761057-INS-139
chr4-83762567-INS-6254
chr4-83765907-INS-76
chr4-83811682-INS-69

chr4-83813184-INS-78
chr4-83840048-INS-7346
chr4-83840990-INS-71
chr4-83853282-INS-7138
chr4-83854677-INS-677
chr4-83885881-INS-7069
chr4-84031689-INS-5547
chr4-84036747-INS-108
chr4-84041997-INS-141
chr4-84062196-INS-114
chr4-84062876-INS-157
chr4-84080122-INS-80
chr4-84116249-INS-98
chr4-84158497-INS-80
chr4-84184394-INS-220
chr4-84248722-INS-69
chr4-84251857-INS-540
chr4-84255463-INS-589
chr4-84270441-INS-56
chr4-84271552-INS-73
chr4-84360639-INS-6679
chr4-84382040-INS-117
chr4-84391607-INS-617
chr4-84397315-INS-1754
chr4-84398109-INS-77
chr4-84403804-INS-5551
chr4-84505491-INS-109
chr4-84507654-INS-57
chr4-84526488-INS-102
chr4-84575820-INS-820
chr4-84589363-INS-181
chr4-84594702-INS-82
chr4-84601572-INS-101
chr4-84607414-INS-6994
chr4-84610376-INS-832
chr4-84614024-INS-60
chr4-84614572-INS-140
chr4-84617392-INS-52
chr4-84622406-INS-86
chr4-84635286-INS-71
chr4-84656512-INS-67
chr4-84659078-INS-590
chr4-84690126-INS-191
chr4-84698607-INS-152
chr4-84729454-INS-208
chr4-84735517-INS-1958
chr4-84735615-INS-4734
chr4-84742885-INS-2881
chr4-84744381-INS-497
chr4-84748831-INS-83
chr4-84767837-INS-171
chr4-84777544-INS-52
chr4-84789546-INS-332
chr4-84790773-INS-616
chr4-84801210-INS-1890
chr4-84831574-INS-234
chr4-84880396-INS-65
chr4-84892803-INS-978
chr4-84895732-INS-118
chr4-84898341-INS-183
chr4-84937754-INS-62
chr4-84940593-INS-51
chr4-84966287-INS-97
chr4-84971963-INS-337
chr4-85007960-INS-191
chr4-85017684-INS-79
chr4-85019550-INS-840
chr4-85023698-INS-7437
chr4-85115392-INS-268
chr4-85125561-INS-2496
chr4-85136247-INS-117
chr4-85140215-INS-76
chr4-85142021-INS-56
chr4-85148786-INS-112
chr4-85151794-INS-98
chr4-85168313-INS-6927
chr4-85177492-INS-125

chr4-85197802-INS-476
chr4-85248856-INS-425
chr4-85257581-INS-59
chr4-85302280-INS-432
chr4-85306687-INS-54
chr4-85311984-INS-144
chr4-85325012-INS-150
chr4-85346321-INS-220
chr4-85350368-INS-70
chr4-85355588-INS-62
chr4-85365504-INS-56
chr4-85372167-INS-54
chr4-85373283-INS-51
chr4-85411050-INS-52
chr4-85451531-INS-139
chr4-85458738-INS-226
chr4-85467572-INS-996
chr4-85468762-INS-74
chr4-85547438-INS-485
chr4-85560316-INS-77
chr4-85586154-INS-7075
chr4-85589909-INS-99
chr4-85625313-INS-52
chr4-85641149-INS-63
chr4-85661210-INS-428
chr4-85665805-INS-76
chr4-85668186-INS-56
chr4-85687312-INS-368
chr4-85747779-INS-171
chr4-85750904-INS-174
chr4-85753265-INS-68
chr4-85760901-INS-509
chr4-85774639-INS-198
chr4-85776638-INS-399
chr4-85777526-INS-130
chr4-85831060-INS-52
chr4-85862227-INS-293
chr4-85893054-INS-108
chr4-85902058-INS-107
chr4-85927384-INS-117
chr4-85928132-INS-398
chr4-85935350-INS-893
chr4-85936800-INS-106
chr4-85939313-INS-52
chr4-85966368-INS-151
chr4-85970106-INS-52
chr4-85982643-INS-56
chr4-86000978-INS-141
chr4-86027935-INS-6181
chr4-86029627-INS-5284
chr4-86046951-INS-55
chr4-86078705-INS-75
chr4-86089048-INS-64
chr4-86102586-INS-6470
chr4-86148544-INS-66
chr4-86168494-INS-3114
chr4-86200811-INS-60
chr4-86212417-INS-391
chr4-86219716-INS-60
chr4-86265361-INS-200
chr4-86280249-INS-52
chr4-86356260-INS-113
chr4-86402985-INS-52
chr4-86409393-INS-58
chr4-86411127-INS-4527
chr4-86417514-INS-62
chr4-86425343-INS-330
chr4-86446914-INS-52
chr4-86456108-INS-227
chr4-86479182-INS-759
chr4-86481328-INS-158
chr4-86520118-INS-56
chr4-86524180-INS-404
chr4-86527796-INS-55
chr4-86539720-INS-4092
chr4-86554490-INS-211
chr4-86569486-INS-277

chr4-86590076-INS-178
chr4-86594119-INS-70
chr4-86596830-INS-84
chr4-86601583-INS-125
chr4-86607277-INS-208
chr4-86610856-INS-53
chr4-86626143-INS-493
chr4-86630907-INS-51
chr4-86639526-INS-99
chr4-86645071-INS-205
chr4-86648845-INS-317
chr4-86651886-INS-194
chr4-86659396-INS-74
chr4-86664760-INS-217
chr4-86681626-INS-70
chr4-86717286-INS-68
chr4-86722468-INS-60
chr4-86727713-INS-6401
chr4-86727759-INS-226
chr4-86728351-INS-51
chr4-86738873-INS-415
chr4-86764802-INS-88
chr4-86773695-INS-80
chr4-86789378-INS-56
chr4-86799994-INS-108
chr4-86808456-INS-186
chr4-86816610-INS-55
chr4-86828835-INS-75
chr4-86856879-INS-223
chr4-86860800-INS-214
chr4-86861699-INS-420
chr4-86863904-INS-85
chr4-86870305-INS-226
chr4-86874718-INS-173
chr4-86876073-INS-484
chr4-86879274-INS-84
chr4-86891441-INS-214
chr4-86899979-INS-118
chr4-86902280-INS-65
chr4-86911135-INS-109
chr4-86915737-INS-100
chr4-86919114-INS-74
chr4-86928427-INS-96
chr4-86944870-INS-372
chr4-86959208-INS-52
chr4-86987760-INS-70
chr4-86994296-INS-4636
chr4-87004361-INS-359
chr4-87035160-INS-7560
chr4-87042460-INS-127
chr4-87064839-INS-5453
chr4-87110329-INS-51
chr4-87113064-INS-70
chr4-87139724-INS-6624
chr4-87140305-INS-51
chr4-87143246-INS-199
chr4-87151892-INS-6268
chr4-87178750-INS-6316
chr4-87188622-INS-63
chr4-87192932-INS-7766
chr4-87211639-INS-554
chr4-87213764-INS-7708
chr4-87218680-INS-6285
chr4-87261377-INS-273
chr4-87294701-INS-399
chr4-87301149-INS-1200
chr4-87304174-INS-448
chr4-87306421-INS-245
chr4-87321555-INS-112
chr4-87347610-INS-55
chr4-87382417-INS-51
chr4-87392599-INS-64
chr4-87430121-INS-116
chr4-87450764-INS-141
chr4-87465475-INS-58
chr4-87482237-INS-492
chr4-87516794-INS-423

chr4-87520233-INS-463
chr4-87524722-INS-6649
chr4-87527799-INS-205
chr4-87542871-INS-255
chr4-87544964-INS-81
chr4-87550774-INS-6182
chr4-87552447-INS-6303
chr4-87561840-INS-159
chr4-87646460-INS-237
chr4-87648160-INS-90
chr4-87659299-INS-326
chr4-87693100-INS-6436
chr4-87705362-INS-56
chr4-87732663-INS-7250
chr4-87735043-INS-125
chr4-87767428-INS-64
chr4-87779466-INS-199
chr4-87781776-INS-151
chr4-87807085-INS-740
chr4-87832080-INS-72
chr4-87840408-INS-175
chr4-87861535-INS-192
chr4-87889842-INS-160
chr4-87900392-INS-60
chr4-87908016-INS-52
chr4-87916085-INS-601
chr4-87920310-INS-675
chr4-87922598-INS-194
chr4-87941394-INS-61
chr4-87948355-INS-132
chr4-87952784-INS-303
chr4-87960866-INS-65
chr4-87966720-INS-53
chr4-87968706-INS-90
chr4-87970024-INS-130
chr4-87970906-INS-2895
chr4-87973431-INS-82
chr4-87977887-INS-1060
chr4-87980785-INS-72
chr4-87984356-INS-87
chr4-87997130-INS-148
chr4-88025531-INS-64
chr4-88044299-INS-199
chr4-88049790-INS-124
chr4-88066524-INS-1403
chr4-88068246-INS-241
chr4-88068747-INS-240
chr4-88081172-INS-92
chr4-88091777-INS-59
chr4-88135731-INS-175
chr4-88138248-INS-77
chr4-88149986-INS-611
chr4-88166052-INS-99
chr4-88178961-INS-127
chr4-88181225-INS-831
chr4-88187885-INS-6358
chr4-88189722-INS-61
chr4-88209088-INS-472
chr4-88221883-INS-6593
chr4-88222022-INS-209
chr4-88237675-INS-72
chr4-88249494-INS-61
chr4-88250332-INS-6457
chr4-88251889-INS-59
chr4-88257878-INS-7774
chr4-88273048-INS-180
chr4-88292762-INS-6700
chr4-88293919-INS-524
chr4-88308652-INS-206
chr4-88310549-INS-213
chr4-88378900-INS-193
chr4-88383135-INS-1255
chr4-88399694-INS-199
chr4-88429544-INS-90
chr4-88450177-INS-857
chr4-88468912-INS-124
chr4-88499828-INS-6590

chr4-88501199-INS-344
chr4-88511425-INS-58
chr4-88515704-INS-139
chr4-88524241-INS-55
chr4-88544205-INS-881
chr4-88549718-INS-54
chr4-88559708-INS-268
chr4-88560550-INS-57
chr4-88566094-INS-82
chr4-88576113-INS-210
chr4-88595838-INS-923
chr4-88605137-INS-431
chr4-88625721-INS-168
chr4-88653663-INS-465
chr4-88662897-INS-269
chr4-88665171-INS-1878
chr4-88669714-INS-7077
chr4-88679502-INS-7345
chr4-88745704-INS-60
chr4-88761494-INS-427
chr4-88766929-INS-208
chr4-88779739-INS-89
chr4-88788286-INS-88
chr4-88791060-INS-64
chr4-88800819-INS-51
chr4-88831715-INS-94
chr4-88890225-INS-402
chr4-88911632-INS-138
chr4-88928136-INS-54
chr4-88931500-INS-66
chr4-88981831-INS-440
chr4-88985038-INS-162
chr4-88987923-INS-461
chr4-88991580-INS-53
chr4-89014988-INS-90
chr4-89024271-INS-186
chr4-89032849-INS-58
chr4-89041137-INS-54
chr4-89065490-INS-54
chr4-89074409-INS-52
chr4-89091669-INS-58
chr4-89101795-INS-1691
chr4-89124655-INS-1546
chr4-89127319-INS-200
chr4-89167069-INS-220
chr4-89175590-INS-68
chr4-89182936-INS-3830
chr4-89183141-INS-1241
chr4-89183143-INS-1891
chr4-89183144-INS-908
chr4-89192816-INS-56
chr4-89201678-INS-66
chr4-89203313-INS-973
chr4-89203541-INS-523
chr4-89205866-INS-58
chr4-89230452-INS-163
chr4-89235035-INS-3298
chr4-89236868-INS-203
chr4-89253604-INS-58
chr4-89269167-INS-95
chr4-89275988-INS-308
chr4-89289211-INS-202
chr4-89291018-INS-439
chr4-89297569-INS-96
chr4-89311377-INS-1458
chr4-89320444-INS-6638
chr4-89320648-INS-196
chr4-89328480-INS-82
chr4-89359016-INS-490
chr4-89365739-INS-263
chr4-89376528-INS-98
chr4-89408564-INS-115
chr4-89425385-INS-185
chr4-89443440-INS-81
chr4-89452926-INS-292
chr4-89461020-INS-315
chr4-89472111-INS-65

chr4-89477218-INS-67
chr4-89537557-INS-76
chr4-89538377-INS-844
chr4-89603402-INS-162
chr4-89618911-INS-58
chr4-89628385-INS-86
chr4-89645501-INS-882
chr4-89648490-INS-70
chr4-89649831-INS-358
chr4-89662225-INS-7111
chr4-89670560-INS-6155
chr4-89677981-INS-1890
chr4-89691344-INS-313
chr4-89789564-INS-6430
chr4-89814196-INS-1126
chr4-89818181-INS-510
chr4-89821038-INS-571
chr4-89825857-INS-6210
chr4-89833554-INS-4447
chr4-89841838-INS-348
chr4-89847599-INS-209
chr4-89862480-INS-107
chr4-89866218-INS-294
chr4-89899543-INS-293
chr4-89905684-INS-6323
chr4-89918460-INS-1127
chr4-89944709-INS-60
chr4-89954192-INS-65
chr4-89957906-INS-144
chr4-89960749-INS-6264
chr4-89962570-INS-63
chr4-89968267-INS-6873
chr4-89979079-INS-6395
chr4-89981570-INS-539
chr4-89984822-INS-60
chr4-89994917-INS-7206
chr4-90012274-INS-6422
chr4-90015716-INS-6359
chr4-90025907-INS-6602
chr4-90026230-INS-498
chr4-90035221-INS-67
chr4-90038378-INS-6425
chr4-90061152-INS-783
chr4-90064973-INS-401
chr4-90077260-INS-200
chr4-90081114-INS-60
chr4-90084731-INS-401
chr4-90089295-INS-117
chr4-90118962-INS-393
chr4-90137876-INS-115
chr4-90139190-INS-6208
chr4-90141347-INS-54
chr4-90151109-INS-217
chr4-90154849-INS-64
chr4-90186912-INS-597
chr4-90188710-INS-461
chr4-90197015-INS-379
chr4-90252028-INS-454
chr4-90258980-INS-56
chr4-90279818-INS-4747
chr4-90319215-INS-99
chr4-90346321-INS-102
chr4-90376822-INS-52
chr4-90378543-INS-109
chr4-90384389-INS-79
chr4-90400802-INS-295
chr4-90410410-INS-503
chr4-90433005-INS-52
chr4-90461704-INS-54
chr4-90468290-INS-586
chr4-90473364-INS-332
chr4-90478260-INS-157
chr4-90480553-INS-60
chr4-90488313-INS-51
chr4-90571662-INS-196
chr4-90581160-INS-205
chr4-90588641-INS-88

chr4-90593198-INS-62
chr4-90601157-INS-265
chr4-90648940-INS-111
chr4-90657911-INS-307
chr4-90690118-INS-56
chr4-90714568-INS-87
chr4-90714835-INS-6321
chr4-90738313-INS-187
chr4-90744567-INS-2253
chr4-90776707-INS-58
chr4-90778523-INS-58
chr4-90801063-INS-6860
chr4-90809728-INS-58
chr4-90811766-INS-54
chr4-90835528-INS-51
chr4-90838609-INS-238
chr4-90848685-INS-350
chr4-90867957-INS-338
chr4-90874982-INS-382
chr4-90897845-INS-6423
chr4-90908442-INS-70
chr4-90912909-INS-6450
chr4-90938973-INS-6989
chr4-90975833-INS-54
chr4-91004204-INS-74
chr4-91017014-INS-6679
chr4-91052162-INS-79
chr4-91054869-INS-2042
chr4-91070137-INS-149
chr4-91071181-INS-4240
chr4-91096707-INS-400
chr4-91137146-INS-152
chr4-91150594-INS-319
chr4-91167619-INS-116
chr4-91181103-INS-142
chr4-91278421-INS-76
chr4-91280965-INS-153
chr4-91298773-INS-342
chr4-91301472-INS-202
chr4-91308788-INS-197
chr4-91315614-INS-56
chr4-91336116-INS-419
chr4-91338440-INS-141
chr4-91340398-INS-88
chr4-91345400-INS-204
chr4-91354227-INS-6440
chr4-91369587-INS-1513
chr4-91388403-INS-6590
chr4-91396529-INS-68
chr4-91402095-INS-643
chr4-91424672-INS-162
chr4-91427323-INS-497
chr4-91450728-INS-59
chr4-91454223-INS-3501
chr4-91455400-INS-4754
chr4-91475633-INS-98
chr4-91482960-INS-190
chr4-91509720-INS-528
chr4-91551766-INS-52
chr4-91565924-INS-6909
chr4-91589947-INS-965
chr4-91593936-INS-82
chr4-91599813-INS-118
chr4-91609477-INS-52
chr4-91610356-INS-2843
chr4-91615316-INS-6439
chr4-91659561-INS-352
chr4-91669046-INS-127
chr4-91669255-INS-6616
chr4-91701491-INS-6473
chr4-91710345-INS-92
chr4-91711484-INS-571
chr4-91734329-INS-163
chr4-91753009-INS-58
chr4-91812473-INS-177
chr4-91816155-INS-142
chr4-91825269-INS-68

chr4-91835787-INS-518
chr4-91836484-INS-59
chr4-91884873-INS-565
chr4-91890716-INS-51
chr4-91923473-INS-441
chr4-91928478-INS-309
chr4-91932392-INS-60
chr4-91949899-INS-6394
chr4-91957488-INS-68
chr4-91977571-INS-57
chr4-91987087-INS-551
chr4-91995107-INS-970
chr4-92003169-INS-69
chr4-92010048-INS-6583
chr4-92030074-INS-252
chr4-92031189-INS-1420
chr4-92047748-INS-352
chr4-92074403-INS-399
chr4-92078815-INS-319
chr4-92087750-INS-6473
chr4-92094158-INS-6449
chr4-92106039-INS-605
chr4-92145416-INS-151
chr4-92151277-INS-323
chr4-92168941-INS-7160
chr4-92226877-INS-3405
chr4-92241794-INS-57
chr4-92251022-INS-106
chr4-92267977-INS-602
chr4-92268006-INS-1193
chr4-92281025-INS-68
chr4-92306124-INS-66
chr4-92334040-INS-68
chr4-92339135-INS-253
chr4-92355743-INS-72
chr4-92365020-INS-56
chr4-92366290-INS-90
chr4-92403668-INS-509
chr4-92419598-INS-994
chr4-92440192-INS-66
chr4-92442614-INS-5101
chr4-92452881-INS-448
chr4-92454992-INS-239
chr4-92473355-INS-666
chr4-92476682-INS-562
chr4-92494286-INS-70
chr4-92496946-INS-343
chr4-92538628-INS-7075
chr4-92541090-INS-731
chr4-92545496-INS-77
chr4-92574542-INS-127
chr4-92574542-INS-938
chr4-92589460-INS-765
chr4-92600759-INS-185
chr4-92605186-INS-6180
chr4-92620618-INS-58
chr4-92628148-INS-6826
chr4-92674393-INS-344
chr4-92698798-INS-317
chr4-92742504-INS-617
chr4-92754684-INS-6648
chr4-92760019-INS-1895
chr4-92760664-INS-1064
chr4-92768487-INS-128
chr4-92781882-INS-509
chr4-92797479-INS-593
chr4-92801675-INS-70
chr4-92802962-INS-6186
chr4-92844901-INS-4819
chr4-92847423-INS-538
chr4-92848914-INS-72
chr4-92852362-INS-68
chr4-92885057-INS-52
chr4-92895692-INS-72
chr4-92898101-INS-173
chr4-92902252-INS-6415
chr4-92918274-INS-57

chr4-92919440-INS-716
chr4-92927039-INS-7228
chr4-92938101-INS-477
chr4-92964144-INS-56
chr4-92969915-INS-87
chr4-93002855-INS-6474
chr4-93061719-INS-76
chr4-93078512-INS-690
chr4-93112364-INS-105
chr4-93113035-INS-58
chr4-93114544-INS-394
chr4-93118819-INS-3177
chr4-93122781-INS-81
chr4-93127028-INS-196
chr4-93150397-INS-190
chr4-93151832-INS-2910
chr4-93199348-INS-120
chr4-93204580-INS-3855
chr4-93247433-INS-395
chr4-93249991-INS-420
chr4-93278012-INS-136
chr4-93300900-INS-7136
chr4-93313540-INS-6069
chr4-93371142-INS-66
chr4-93390780-INS-652
chr4-93394441-INS-101
chr4-93397548-INS-57
chr4-93410826-INS-206
chr4-93414971-INS-6335
chr4-93417307-INS-5337
chr4-93425623-INS-234
chr4-93440723-INS-2816
chr4-93446498-INS-57
chr4-93448757-INS-97
chr4-93468991-INS-270
chr4-93469656-INS-76
chr4-93484522-INS-73
chr4-93490743-INS-115
chr4-93527580-INS-7439
chr4-93530941-INS-62
chr4-93558584-INS-93
chr4-93559448-INS-554
chr4-93568568-INS-54
chr4-93581844-INS-214
chr4-93638992-INS-458
chr4-93646077-INS-2740
chr4-93650796-INS-101
chr4-93662706-INS-200
chr4-93663147-INS-80
chr4-93685146-INS-200
chr4-93720316-INS-6669
chr4-93723245-INS-285
chr4-93726249-INS-1894
chr4-93737721-INS-356
chr4-93746982-INS-949
chr4-93753044-INS-56
chr4-93759920-INS-697
chr4-93772210-INS-52
chr4-93800485-INS-54
chr4-93839562-INS-138
chr4-93863502-INS-6603
chr4-93886194-INS-202
chr4-93898879-INS-62
chr4-93919894-INS-182
chr4-93921721-INS-3744
chr4-93928346-INS-7769
chr4-93929024-INS-5393
chr4-93932593-INS-95
chr4-93950201-INS-6666
chr4-93956417-INS-219
chr4-93976121-INS-3202
chr4-94026658-INS-6400
chr4-94026709-INS-67
chr4-94034671-INS-6436
chr4-94040739-INS-616
chr4-94043081-INS-941
chr4-94060606-INS-1000

chr4-94091769-INS-768
chr4-94101605-INS-62
chr4-94102585-INS-104
chr4-94104798-INS-332
chr4-94124667-INS-60
chr4-94134269-INS-207
chr4-94139203-INS-1137
chr4-94139740-INS-398
chr4-94145323-INS-6376
chr4-94161774-INS-190
chr4-94181113-INS-6582
chr4-94203070-INS-1375
chr4-94276792-INS-123
chr4-94287700-INS-152
chr4-94293405-INS-52
chr4-94297652-INS-62
chr4-94338559-INS-117
chr4-94339118-INS-54
chr4-94355877-INS-54
chr4-94378379-INS-90
chr4-94404516-INS-181
chr4-94464277-INS-135
chr4-94501882-INS-197
chr4-94508920-INS-187
chr4-94519561-INS-123
chr4-94525409-INS-183
chr4-94540576-INS-86
chr4-94563467-INS-208
chr4-94573690-INS-118
chr4-94593565-INS-212
chr4-94595834-INS-5991
chr4-94597378-INS-1208
chr4-94600563-INS-1889
chr4-94602150-INS-194
chr4-94604068-INS-522
chr4-94608239-INS-6445
chr4-94613059-INS-54
chr4-94634352-INS-286
chr4-94637270-INS-54
chr4-94650422-INS-401
chr4-94657711-INS-6431
chr4-94676734-INS-6948
chr4-94677230-INS-6472
chr4-94689277-INS-464
chr4-94697978-INS-72
chr4-94698787-INS-237
chr4-94736707-INS-111
chr4-94760793-INS-6452
chr4-94794201-INS-7228
chr4-94821418-INS-101
chr4-94822671-INS-246
chr4-94855295-INS-616
chr4-94856437-INS-51
chr4-94874499-INS-349
chr4-94920806-INS-3068
chr4-94927283-INS-256
chr4-94933262-INS-208
chr4-94948932-INS-160
chr4-94959318-INS-358
chr4-95008284-INS-232
chr4-95018232-INS-7491
chr4-95028403-INS-92
chr4-95037757-INS-100
chr4-95065380-INS-85
chr4-95087145-INS-125
chr4-95107574-INS-492
chr4-95111849-INS-248
chr4-95115513-INS-3045
chr4-95127938-INS-549
chr4-95129459-INS-72
chr4-95158171-INS-6440
chr4-95171072-INS-292
chr4-95199578-INS-54
chr4-95204771-INS-221
chr4-95241985-INS-61
chr4-95289868-INS-102
chr4-95314691-INS-302

chr4-95326097-INS-482
chr4-95361327-INS-53
chr4-95368937-INS-76
chr4-95373611-INS-785
chr4-95381542-INS-143
chr4-95383274-INS-82
chr4-95392805-INS-6627
chr4-95395874-INS-6923
chr4-95396654-INS-204
chr4-95407752-INS-486
chr4-95434980-INS-229
chr4-95447108-INS-57
chr4-95458858-INS-1158
chr4-95483441-INS-76
chr4-95488857-INS-54
chr4-95491489-INS-6831
chr4-95505041-INS-231
chr4-95509059-INS-341
chr4-95518595-INS-6379
chr4-95536206-INS-92
chr4-95542125-INS-84
chr4-95548938-INS-63
chr4-95551671-INS-796
chr4-95551675-INS-160
chr4-95561955-INS-222
chr4-95567895-INS-163
chr4-95580961-INS-749
chr4-95613657-INS-6682
chr4-95614966-INS-54
chr4-95650413-INS-603
chr4-95668137-INS-248
chr4-95690652-INS-723
chr4-95711873-INS-106
chr4-95717635-INS-200
chr4-95767865-INS-7211
chr4-95769627-INS-63
chr4-95773658-INS-277
chr4-95777528-INS-354
chr4-95833306-INS-420
chr4-95837233-INS-7344
chr4-95853001-INS-166
chr4-95858667-INS-175
chr4-95930370-INS-6500
chr4-95937315-INS-296
chr4-95941858-INS-51
chr4-95945480-INS-69
chr4-95947369-INS-1862
chr4-95947370-INS-385
chr4-95960391-INS-172
chr4-95989645-INS-325
chr4-95990687-INS-1437
chr4-96035742-INS-6371
chr4-96058703-INS-6295
chr4-96069603-INS-63
chr4-96072044-INS-809
chr4-96114767-INS-331
chr4-96116431-INS-213
chr4-96120081-INS-6991
chr4-96130635-INS-54
chr4-96133211-INS-135
chr4-96135645-INS-147
chr4-96184014-INS-189
chr4-96188989-INS-284
chr4-96226010-INS-177
chr4-96227097-INS-60
chr4-96227609-INS-97
chr4-96235335-INS-1398
chr4-96260372-INS-140
chr4-96313374-INS-53
chr4-96327101-INS-60
chr4-96339499-INS-669
chr4-96352448-INS-7418
chr4-96354352-INS-122
chr4-96359389-INS-65
chr4-96372413-INS-364
chr4-96388860-INS-57
chr4-96407639-INS-434

chr4-96449105-INS-72
chr4-96477572-INS-295
chr4-96484927-INS-58
chr4-96497953-INS-430
chr4-96509389-INS-53
chr4-96578101-INS-257
chr4-96581647-INS-60
chr4-96595379-INS-51
chr4-96600896-INS-176
chr4-96605451-INS-659
chr4-96642173-INS-66
chr4-96645016-INS-581
chr4-96703041-INS-401
chr4-96723594-INS-262
chr4-96769151-INS-112
chr4-96810076-INS-456
chr4-96813910-INS-67
chr4-96823468-INS-247
chr4-96832204-INS-73
chr4-96869325-INS-143
chr4-96871673-INS-164
chr4-96873648-INS-175
chr4-96904148-INS-443
chr4-96905693-INS-118
chr4-96908812-INS-439
chr4-96919075-INS-75
chr4-96924008-INS-53
chr4-96953927-INS-55
chr4-96960135-INS-80
chr4-97022314-INS-912
chr4-97041477-INS-148
chr4-97080684-INS-432
chr4-97082826-INS-106
chr4-97100657-INS-64
chr4-97105983-INS-461
chr4-97132828-INS-6737
chr4-97133681-INS-75
chr4-97152104-INS-98
chr4-97156662-INS-90
chr4-97191970-INS-55
chr4-97196158-INS-7089
chr4-97197205-INS-6168
chr4-97234934-INS-62
chr4-97253585-INS-6701
chr4-97273198-INS-252
chr4-97320480-INS-6729
chr4-97320812-INS-139
chr4-97325427-INS-322
chr4-97347469-INS-5243
chr4-97358072-INS-52
chr4-97359345-INS-7210
chr4-97375732-INS-156
chr4-97386703-INS-384
chr4-97389086-INS-66
chr4-97407788-INS-1864
chr4-97413555-INS-492
chr4-97429120-INS-400
chr4-97434635-INS-6995
chr4-97437242-INS-543
chr4-97455102-INS-6207
chr4-97477542-INS-62
chr4-97507673-INS-98
chr4-97518361-INS-135
chr4-97537432-INS-52
chr4-97566816-INS-183
chr4-97570694-INS-6452
chr4-97605566-INS-3769
chr4-97645416-INS-68
chr4-97660855-INS-53
chr4-97695289-INS-123
chr4-97716328-INS-144
chr4-97735337-INS-102
chr4-97736330-INS-74
chr4-97774230-INS-1451
chr4-97800968-INS-130
chr4-97802399-INS-67
chr4-97821310-INS-105

chr4-97859838-INS-55
chr4-97932712-INS-204
chr4-97936043-INS-52
chr4-97981344-INS-120
chr4-98041708-INS-77
chr4-98052303-INS-719
chr4-98059818-INS-1272
chr4-98085931-INS-129
chr4-98088950-INS-72
chr4-98096002-INS-1586
chr4-98113533-INS-565
chr4-98125164-INS-63
chr4-98135550-INS-800
chr4-98223219-INS-220
chr4-98223838-INS-94
chr4-98240749-INS-377
chr4-98251028-INS-510
chr4-98268192-INS-116
chr4-98271622-INS-60
chr4-98292109-INS-54
chr4-98304114-INS-211
chr4-98307094-INS-7399
chr4-98310693-INS-205
chr4-98325252-INS-56
chr4-98327723-INS-247
chr4-98346674-INS-492
chr4-98366817-INS-73
chr4-98371126-INS-7914
chr4-98421888-INS-194
chr4-98432998-INS-156
chr4-98477385-INS-102
chr4-98492687-INS-634
chr4-98492764-INS-55
chr4-98502115-INS-52
chr4-98505054-INS-260
chr4-98506459-INS-141
chr4-98517403-INS-260
chr4-98538867-INS-54
chr4-98546825-INS-52
chr4-98555082-INS-71
chr4-98557807-INS-65
chr4-98584091-INS-208
chr4-98595289-INS-172
chr4-98609623-INS-72
chr4-98628231-INS-77
chr4-98635955-INS-745
chr4-98642465-INS-170
chr4-98651466-INS-722
chr4-98674049-INS-556
chr4-98681029-INS-114
chr4-98714739-INS-131
chr4-98716033-INS-64
chr4-98718741-INS-203
chr4-98720237-INS-85
chr4-98733847-INS-495
chr4-98744847-INS-186
chr4-98746256-INS-1562
chr4-98749870-INS-108
chr4-98754486-INS-205
chr4-98756614-INS-219
chr4-98772621-INS-194
chr4-98787331-INS-209
chr4-98792571-INS-196
chr4-98793438-INS-186
chr4-98800961-INS-521
chr4-98802217-INS-75
chr4-98804370-INS-128
chr4-98830781-INS-7315
chr4-98882271-INS-203
chr4-98889922-INS-127
chr4-98935891-INS-175
chr4-98948602-INS-365
chr4-98964023-INS-399
chr4-98971528-INS-76
chr4-98975656-INS-84
chr4-98979869-INS-196
chr4-98986698-INS-1092

chr4-99025508-INS-554
chr4-99057182-INS-184
chr4-99059302-INS-111
chr4-99067700-INS-54
chr4-99071634-INS-4042
chr4-99077180-INS-207
chr4-99077813-INS-390
chr4-99084759-INS-76
chr4-99089406-INS-78
chr4-99090242-INS-234
chr4-99099095-INS-226
chr4-99108729-INS-483
chr4-99136056-INS-73
chr4-99136733-INS-70
chr4-99137013-INS-699
chr4-99150302-INS-51
chr4-99153937-INS-6184
chr4-99181944-INS-84
chr4-99191046-INS-60
chr4-99232162-INS-104
chr4-99235165-INS-230
chr4-99240538-INS-108
chr4-99271356-INS-59
chr4-99281782-INS-398
chr4-99294366-INS-52
chr4-99298638-INS-68
chr4-99302046-INS-496
chr4-99303007-INS-240
chr4-99307448-INS-181
chr4-99314923-INS-107
chr4-99323130-INS-5571
chr4-99345621-INS-52
chr4-99365399-INS-397
chr4-99394423-INS-78
chr4-99420662-INS-101
chr4-99420662-INS-737
chr4-99421069-INS-293
chr4-99427509-INS-53
chr4-99470766-INS-913
chr4-99470839-INS-297
chr4-99481554-INS-104
chr4-99485133-INS-190
chr4-99488487-INS-90
chr4-99505635-INS-65
chr4-99507823-INS-78
chr4-99547693-INS-67
chr4-99559200-INS-408
chr4-99573582-INS-55
chr4-99574998-INS-197
chr4-99591156-INS-71
chr4-99594134-INS-52
chr4-99596136-INS-53
chr4-99596664-INS-52
chr4-99629292-INS-66
chr4-99633866-INS-56
chr4-99695508-INS-52
chr4-99698341-INS-52
chr4-99713247-INS-597
chr4-99716388-INS-559
chr4-99748193-INS-4457
chr4-99749693-INS-54
chr4-99751831-INS-181
chr4-99762073-INS-4896
chr4-99794957-INS-105
chr4-99819603-INS-52
chr4-99830005-INS-287
chr4-99848313-INS-158
chr4-99884750-INS-343
chr4-99892766-INS-164
chr4-99922296-INS-429
chr4-99934921-INS-6723
chr4-99940717-INS-401
chr4-99959939-INS-7803
chr4-99970423-INS-117
chr4-99972347-INS-52
chr4-99980637-INS-57
chr4-99993346-INS-95

chr4-100023153-INS-54
chr4-100044452-INS-81
chr4-100059576-INS-204
chr4-100065308-INS-198
chr4-100077248-INS-54
chr4-100086170-INS-7639
chr4-100106917-INS-51
chr4-100114063-INS-463
chr4-100114254-INS-865
chr4-100167513-INS-139
chr4-100190061-INS-148
chr4-100213264-INS-7443
chr4-100224646-INS-6707
chr4-100275246-INS-63
chr4-100279206-INS-73
chr4-100291365-INS-6997
chr4-100307557-INS-208
chr4-100317101-INS-5354
chr4-100324482-INS-401
chr4-100328506-INS-88
chr4-100338457-INS-108
chr4-100345815-INS-109
chr4-100346803-INS-509
chr4-100354604-INS-234
chr4-100407484-INS-764
chr4-100434119-INS-122
chr4-100449234-INS-7248
chr4-100460149-INS-288
chr4-100485646-INS-272
chr4-100500814-INS-3420
chr4-100519918-INS-170
chr4-100524647-INS-76
chr4-100532672-INS-65
chr4-100554690-INS-80
chr4-100558153-INS-80
chr4-100566698-INS-203
chr4-100569444-INS-501
chr4-100589588-INS-212
chr4-100595006-INS-232
chr4-100605773-INS-59
chr4-100619122-INS-132
chr4-100623472-INS-191
chr4-100629759-INS-68
chr4-100649171-INS-194
chr4-100662422-INS-1864
chr4-100667682-INS-161
chr4-100695245-INS-122
chr4-100699394-INS-81
chr4-100717005-INS-6560
chr4-100718288-INS-83
chr4-100731481-INS-120
chr4-100739217-INS-84
chr4-100759526-INS-622
chr4-100800170-INS-67
chr4-100808342-INS-57
chr4-100824620-INS-293
chr4-100830269-INS-56
chr4-100859760-INS-130
chr4-100860939-INS-63
chr4-100882209-INS-69
chr4-100911985-INS-201
chr4-100921123-INS-6854
chr4-100923648-INS-53
chr4-100949288-INS-324
chr4-100963541-INS-130
chr4-100980907-INS-156
chr4-100998236-INS-80
chr4-101000127-INS-54
chr4-101010497-INS-68
chr4-101018914-INS-155
chr4-101033877-INS-90
chr4-101038773-INS-62
chr4-101046105-INS-857
chr4-101052121-INS-199
chr4-101062305-INS-102
chr4-101082864-INS-5219
chr4-101122792-INS-187

chr4-101140596-INS-87
chr4-101148188-INS-163
chr4-101151920-INS-239
chr4-101170308-INS-212
chr4-101170389-INS-2174
chr4-101170850-INS-191
chr4-101171138-INS-107
chr4-101220548-INS-98
chr4-101226987-INS-60
chr4-101234501-INS-200
chr4-101236229-INS-221
chr4-101243925-INS-224
chr4-101284354-INS-199
chr4-101285072-INS-192
chr4-101305599-INS-161
chr4-101354428-INS-53
chr4-101397320-INS-62
chr4-101406681-INS-166
chr4-101424558-INS-138
chr4-101431173-INS-144
chr4-101466461-INS-150
chr4-101477379-INS-278
chr4-101481062-INS-54
chr4-101484970-INS-66
chr4-101496717-INS-179
chr4-101511811-INS-200
chr4-101529002-INS-400
chr4-101570394-INS-105
chr4-101586966-INS-160
chr4-101623007-INS-83
chr4-101626522-INS-1967
chr4-101626642-INS-222
chr4-101627476-INS-1248
chr4-101634941-INS-60
chr4-101663285-INS-195
chr4-101688507-INS-133
chr4-101690275-INS-56
chr4-101697385-INS-64
chr4-101699200-INS-272
chr4-101710308-INS-620
chr4-101719919-INS-158
chr4-101733394-INS-192
chr4-101735436-INS-5292
chr4-101741489-INS-206
chr4-101745972-INS-101
chr4-101756666-INS-190
chr4-101758889-INS-134
chr4-101764448-INS-199
chr4-101771623-INS-208
chr4-101788039-INS-147
chr4-101820391-INS-232
chr4-101830968-INS-93
chr4-101848769-INS-216
chr4-101850911-INS-519
chr4-101851881-INS-483
chr4-101860976-INS-5155
chr4-101861233-INS-321
chr4-101876327-INS-1892
chr4-101876328-INS-400
chr4-101886884-INS-199
chr4-101896553-INS-229
chr4-101924223-INS-62
chr4-101975634-INS-344
chr4-102025822-INS-420
chr4-102045117-INS-64
chr4-102046385-INS-422
chr4-102054186-INS-84
chr4-102063256-INS-131
chr4-102072078-INS-449
chr4-102081665-INS-400
chr4-102092361-INS-6524
chr4-102135636-INS-85
chr4-102151688-INS-351
chr4-102174839-INS-99
chr4-102192358-INS-6685
chr4-102194886-INS-64
chr4-102225948-INS-6649

chr4-102252095-INS-61
chr4-102253611-INS-54
chr4-102256908-INS-74
chr4-102257697-INS-721
chr4-102268760-INS-6667
chr4-102276663-INS-57
chr4-102294558-INS-3304
chr4-102306314-INS-142
chr4-102318124-INS-108
chr4-102351331-INS-1378
chr4-102351349-INS-613
chr4-102351886-INS-1885
chr4-102351982-INS-129
chr4-102406917-INS-909
chr4-102419836-INS-7255
chr4-102424599-INS-1075
chr4-102464777-INS-51
chr4-102466429-INS-144
chr4-102472294-INS-102
chr4-102490117-INS-384
chr4-102496125-INS-655
chr4-102522063-INS-252
chr4-102563130-INS-201
chr4-102586266-INS-200
chr4-102626020-INS-157
chr4-102643558-INS-74
chr4-102654288-INS-82
chr4-102655022-INS-91
chr4-102659219-INS-359
chr4-102672071-INS-66
chr4-102676627-INS-58
chr4-102722507-INS-212
chr4-102763798-INS-100
chr4-102793797-INS-1016
chr4-102804656-INS-141
chr4-102808556-INS-6625
chr4-102818031-INS-314
chr4-102826701-INS-258
chr4-102838042-INS-279
chr4-102874695-INS-184
chr4-102897654-INS-425
chr4-102904501-INS-607
chr4-102907923-INS-123
chr4-102913908-INS-90
chr4-102915778-INS-66
chr4-102927190-INS-154
chr4-102942623-INS-564
chr4-102950922-INS-68
chr4-102954795-INS-1880
chr4-102957303-INS-929
chr4-102995153-INS-148
chr4-103030463-INS-52
chr4-103066597-INS-215
chr4-103082337-INS-5546
chr4-103104307-INS-58
chr4-103121039-INS-6440
chr4-103123214-INS-94
chr4-103126860-INS-1580
chr4-103149148-INS-56
chr4-103151789-INS-6489
chr4-103157386-INS-93
chr4-103159689-INS-702
chr4-103164446-INS-497
chr4-103169869-INS-176
chr4-103197225-INS-54
chr4-103199434-INS-371
chr4-103241522-INS-58
chr4-103295626-INS-53
chr4-103303518-INS-84
chr4-103352091-INS-6675
chr4-103360158-INS-52
chr4-103378564-INS-2910
chr4-103382643-INS-1225
chr4-103386227-INS-606
chr4-103402235-INS-192
chr4-103422766-INS-75
chr4-103482308-INS-57

chr4-103511481-INS-54
chr4-103539148-INS-52
chr4-103541585-INS-172
chr4-103554214-INS-110
chr4-103577048-INS-6922
chr4-103606750-INS-295
chr4-103628124-INS-172
chr4-103636934-INS-570
chr4-103665894-INS-56
chr4-103673649-INS-63
chr4-103682645-INS-6459
chr4-103702408-INS-184
chr4-103718519-INS-149
chr4-103740995-INS-72
chr4-103743107-INS-419
chr4-103751073-INS-2648
chr4-103761663-INS-54
chr4-103793366-INS-104
chr4-103797259-INS-282
chr4-103802956-INS-177
chr4-103816227-INS-385
chr4-103831032-INS-1678
chr4-103832564-INS-1764
chr4-103859532-INS-100
chr4-103860402-INS-64
chr4-103885911-INS-101
chr4-103888708-INS-105
chr4-103905573-INS-2606
chr4-103934981-INS-129
chr4-103941315-INS-6599
chr4-103944409-INS-973
chr4-103959655-INS-247
chr4-103959950-INS-5277
chr4-103979479-INS-7338
chr4-104032532-INS-130
chr4-104040531-INS-6489
chr4-104061072-INS-607
chr4-104080941-INS-520
chr4-104085804-INS-7578
chr4-104103376-INS-6669
chr4-104128222-INS-7543
chr4-104190986-INS-67
chr4-104193052-INS-172
chr4-104207251-INS-104
chr4-104264764-INS-57
chr4-104280847-INS-80
chr4-104301002-INS-130
chr4-104317536-INS-56
chr4-104344134-INS-358
chr4-104351339-INS-346
chr4-104363900-INS-70
chr4-104365757-INS-55
chr4-104372553-INS-466
chr4-104380201-INS-70
chr4-104390181-INS-596
chr4-104400824-INS-54
chr4-104402339-INS-57
chr4-104418503-INS-64
chr4-104420575-INS-51
chr4-104428921-INS-58
chr4-104431137-INS-68
chr4-104461083-INS-6440
chr4-104469523-INS-57
chr4-104484048-INS-94
chr4-104506069-INS-128
chr4-104512142-INS-53
chr4-104523012-INS-61
chr4-104556839-INS-70
chr4-104568586-INS-858
chr4-104619381-INS-60
chr4-104622733-INS-239
chr4-104624375-INS-348
chr4-104634942-INS-68
chr4-104721141-INS-349
chr4-104737965-INS-90
chr4-104755167-INS-59
chr4-104755792-INS-53

chr4-104766107-INS-2927
chr4-104775658-INS-875
chr4-104800813-INS-56
chr4-104804213-INS-222
chr4-104822496-INS-63
chr4-104823457-INS-58
chr4-104828662-INS-92
chr4-104840919-INS-173
chr4-104852524-INS-471
chr4-104863561-INS-171
chr4-104884113-INS-157
chr4-104900366-INS-117
chr4-104904755-INS-6444
chr4-104927813-INS-109
chr4-104939249-INS-58
chr4-104962601-INS-60
chr4-104969413-INS-192
chr4-104970113-INS-63
chr4-104973173-INS-56
chr4-104993775-INS-59
chr4-104999064-INS-52
chr4-105023954-INS-60
chr4-105051094-INS-6965
chr4-105071264-INS-109
chr4-105091681-INS-217
chr4-105100348-INS-60
chr4-105127362-INS-166
chr4-105161942-INS-74
chr4-105187542-INS-1327
chr4-105201028-INS-88
chr4-105204753-INS-773
chr4-105208286-INS-75
chr4-105212913-INS-489
chr4-105221252-INS-400
chr4-105231027-INS-75
chr4-105246618-INS-63
chr4-105254149-INS-66
chr4-105259615-INS-288
chr4-105268615-INS-144
chr4-105382133-INS-52
chr4-105383042-INS-469
chr4-105405293-INS-260
chr4-105411885-INS-316
chr4-105433663-INS-5497
chr4-105461996-INS-694
chr4-105467862-INS-66
chr4-105468638-INS-6360
chr4-105507241-INS-54
chr4-105538957-INS-6603
chr4-105544895-INS-70
chr4-105550051-INS-176
chr4-105568232-INS-994
chr4-105569032-INS-6440
chr4-105575344-INS-6692
chr4-105607992-INS-429
chr4-105641100-INS-52
chr4-105645220-INS-2061
chr4-105667492-INS-52
chr4-105718972-INS-86
chr4-105729921-INS-148
chr4-105822471-INS-57
chr4-105842354-INS-53
chr4-105854084-INS-120
chr4-105887320-INS-447
chr4-105898146-INS-58
chr4-105909451-INS-58
chr4-105936355-INS-226
chr4-105974691-INS-70
chr4-105981421-INS-61
chr4-105986777-INS-6914
chr4-106012760-INS-110
chr4-106030889-INS-4083
chr4-106045984-INS-6395
chr4-106059097-INS-280
chr4-106060371-INS-52
chr4-106063759-INS-8457
chr4-106071674-INS-64

chr4-106074774-INS-242
chr4-106085609-INS-52
chr4-106091761-INS-180
chr4-106103815-INS-140
chr4-106108246-INS-132
chr4-106143591-INS-670
chr4-106152166-INS-336
chr4-106160911-INS-64
chr4-106162885-INS-496
chr4-106172806-INS-80
chr4-106181676-INS-535
chr4-106183469-INS-217
chr4-106185514-INS-59
chr4-106200148-INS-699
chr4-106207912-INS-205
chr4-106224574-INS-170
chr4-106231888-INS-94
chr4-106235824-INS-238
chr4-106236756-INS-91
chr4-106241618-INS-195
chr4-106245915-INS-58
chr4-106254218-INS-90
chr4-106277314-INS-151
chr4-106316349-INS-112
chr4-106317495-INS-442
chr4-106327077-INS-102
chr4-106379769-INS-230
chr4-106437413-INS-76
chr4-106452814-INS-619
chr4-106456608-INS-206
chr4-106470944-INS-83
chr4-106483647-INS-121
chr4-106517853-INS-95
chr4-106540758-INS-76
chr4-106575573-INS-171
chr4-106583776-INS-203
chr4-106598837-INS-196
chr4-106607467-INS-192
chr4-106618148-INS-194
chr4-106631631-INS-156
chr4-106680006-INS-106
chr4-106683423-INS-161
chr4-106684519-INS-57
chr4-106688827-INS-63
chr4-106715713-INS-203
chr4-106731143-INS-5389
chr4-106797482-INS-143
chr4-106815036-INS-68
chr4-106817667-INS-492
chr4-106825712-INS-66
chr4-106828662-INS-229
chr4-106832070-INS-214
chr4-106834212-INS-206
chr4-106853876-INS-55
chr4-106951454-INS-75
chr4-106968539-INS-61
chr4-106986999-INS-107
chr4-107027876-INS-812
chr4-107028904-INS-541
chr4-107033212-INS-1891
chr4-107052220-INS-85
chr4-107064746-INS-217
chr4-107067961-INS-182
chr4-107077918-INS-56
chr4-107084574-INS-201
chr4-107085973-INS-163
chr4-107087943-INS-139
chr4-107115328-INS-54
chr4-107139390-INS-148
chr4-107140434-INS-401
chr4-107153138-INS-57
chr4-107165484-INS-54
chr4-107177674-INS-479
chr4-107181409-INS-212
chr4-107188219-INS-507
chr4-107202791-INS-91
chr4-107220167-INS-209

chr4-107227291-INS-219
chr4-107234702-INS-503
chr4-107253356-INS-131
chr4-107256151-INS-160
chr4-107277152-INS-338
chr4-107288433-INS-254
chr4-107289672-INS-71
chr4-107408810-INS-172
chr4-107432473-INS-127
chr4-107448932-INS-128
chr4-107514473-INS-315
chr4-107518308-INS-51
chr4-107526920-INS-458
chr4-107573119-INS-57
chr4-107591124-INS-73
chr4-107603804-INS-60
chr4-107623019-INS-150
chr4-107648574-INS-62
chr4-107683592-INS-54
chr4-107716670-INS-205
chr4-107735791-INS-229
chr4-107738814-INS-242
chr4-107743943-INS-60
chr4-107746775-INS-60
chr4-107749982-INS-153
chr4-107752922-INS-169
chr4-107760916-INS-168
chr4-107774892-INS-62
chr4-107791708-INS-64
chr4-107793433-INS-82
chr4-107796092-INS-354
chr4-107830534-INS-97
chr4-107835024-INS-524
chr4-107858323-INS-198
chr4-107862792-INS-80
chr4-107883080-INS-106
chr4-107902144-INS-51
chr4-107906711-INS-169
chr4-107935790-INS-80
chr4-107937712-INS-196
chr4-107940034-INS-616
chr4-107949738-INS-54
chr4-107972851-INS-121
chr4-107993326-INS-284
chr4-107994923-INS-339
chr4-108001266-INS-8985
chr4-108020246-INS-129
chr4-108021091-INS-633
chr4-108026370-INS-234
chr4-108031942-INS-123
chr4-108037101-INS-54
chr4-108045874-INS-185
chr4-108054324-INS-167
chr4-108071319-INS-86
chr4-108077577-INS-62
chr4-108080800-INS-209
chr4-108114056-INS-72
chr4-108115281-INS-212
chr4-108119255-INS-199
chr4-108120593-INS-96
chr4-108127851-INS-55
chr4-108129662-INS-433
chr4-108131303-INS-187
chr4-108137685-INS-466
chr4-108139782-INS-431
chr4-108155905-INS-121
chr4-108176762-INS-199
chr4-108210199-INS-51
chr4-108210820-INS-58
chr4-108212030-INS-209
chr4-108220617-INS-360
chr4-108242393-INS-225
chr4-108254725-INS-158
chr4-108257366-INS-143
chr4-108262755-INS-178
chr4-108275130-INS-1859
chr4-108288698-INS-235

chr4-108291100-INS-60
chr4-108293833-INS-78
chr4-108299909-INS-55
chr4-108303031-INS-239
chr4-108312635-INS-218
chr4-108322908-INS-409
chr4-108387241-INS-211
chr4-108390495-INS-212
chr4-108429498-INS-73
chr4-108442017-INS-497
chr4-108469044-INS-225
chr4-108478306-INS-62
chr4-108502894-INS-197
chr4-108518143-INS-204
chr4-108518999-INS-191
chr4-108542832-INS-84
chr4-108543416-INS-153
chr4-108547841-INS-54
chr4-108569268-INS-82
chr4-108578440-INS-221
chr4-108598882-INS-561
chr4-108610669-INS-2242
chr4-108616525-INS-651
chr4-108621954-INS-180
chr4-108630111-INS-249
chr4-108634605-INS-209
chr4-108639712-INS-6793
chr4-108644444-INS-179
chr4-108650076-INS-203
chr4-108669314-INS-206
chr4-108677288-INS-121
chr4-108696989-INS-206
chr4-108713228-INS-64
chr4-108758957-INS-70
chr4-108761788-INS-8986
chr4-108773964-INS-54
chr4-108776463-INS-64
chr4-108777693-INS-60
chr4-108780436-INS-169
chr4-108785417-INS-810
chr4-108787957-INS-63
chr4-108796565-INS-812
chr4-108803284-INS-62
chr4-108807003-INS-196
chr4-108823063-INS-140
chr4-108824446-INS-66
chr4-108835315-INS-404
chr4-108844170-INS-54
chr4-108847850-INS-481
chr4-108864961-INS-132
chr4-108866720-INS-5549
chr4-108887053-INS-140
chr4-108905219-INS-72
chr4-108908987-INS-74
chr4-108935257-INS-59
chr4-108979499-INS-62
chr4-108984500-INS-176
chr4-108995132-INS-51
chr4-109000236-INS-112
chr4-109010848-INS-85
chr4-109045920-INS-52
chr4-109052159-INS-978
chr4-109069034-INS-383
chr4-109076724-INS-130
chr4-109081702-INS-208
chr4-109082201-INS-1888
chr4-109084660-INS-380
chr4-109087864-INS-355
chr4-109097718-INS-68
chr4-109115264-INS-83
chr4-109119547-INS-351
chr4-109120277-INS-54
chr4-109144538-INS-320
chr4-109175723-INS-373
chr4-109176236-INS-873
chr4-109192583-INS-403
chr4-109246106-INS-113

chr4-109254915-INS-274
chr4-109256762-INS-68
chr4-109280760-INS-52
chr4-109286355-INS-295
chr4-109290170-INS-108
chr4-109323967-INS-432
chr4-109351497-INS-915
chr4-109351912-INS-83
chr4-109368210-INS-194
chr4-109379742-INS-102
chr4-109382928-INS-227
chr4-109395531-INS-210
chr4-109404939-INS-129
chr4-109419319-INS-112
chr4-109436606-INS-99
chr4-109439838-INS-132
chr4-109444166-INS-6325
chr4-109448014-INS-135
chr4-109453481-INS-92
chr4-109455889-INS-125
chr4-109459682-INS-200
chr4-109464270-INS-4971
chr4-109473208-INS-98
chr4-109474499-INS-66
chr4-109481383-INS-7909
chr4-109482980-INS-404
chr4-109529937-INS-58
chr4-109533255-INS-1267
chr4-109545455-INS-1697
chr4-109557974-INS-4604
chr4-109589833-INS-66
chr4-109591960-INS-213
chr4-109607327-INS-52
chr4-109608115-INS-7158
chr4-109638071-INS-64
chr4-109656240-INS-334
chr4-109667135-INS-1305
chr4-109675260-INS-68
chr4-109753801-INS-148
chr4-109776552-INS-52
chr4-109847200-INS-204
chr4-109848520-INS-52
chr4-109873923-INS-181
chr4-109895614-INS-57
chr4-109901977-INS-486
chr4-109952382-INS-130
chr4-109970580-INS-7135
chr4-109975390-INS-2883
chr4-109987062-INS-56
chr4-109989657-INS-54
chr4-109993591-INS-6782
chr4-109994680-INS-115
chr4-110030209-INS-243
chr4-110038681-INS-78
chr4-110041435-INS-3956
chr4-110054468-INS-8046
chr4-110068874-INS-200
chr4-110104410-INS-306
chr4-110124585-INS-796
chr4-110127342-INS-566
chr4-110155313-INS-60
chr4-110156068-INS-146
chr4-110194330-INS-52
chr4-110214719-INS-255
chr4-110243892-INS-966
chr4-110259093-INS-630
chr4-110263129-INS-90
chr4-110271672-INS-7421
chr4-110291044-INS-81
chr4-110293292-INS-6084
chr4-110310375-INS-448
chr4-110313192-INS-1145
chr4-110320163-INS-325
chr4-110337033-INS-517
chr4-110338585-INS-393
chr4-110377695-INS-2426
chr4-110378018-INS-928

chr4-110390213-INS-265
chr4-110404251-INS-355
chr4-110405320-INS-53
chr4-110407974-INS-216
chr4-110409268-INS-65
chr4-110418976-INS-544
chr4-110421951-INS-3582
chr4-110467095-INS-532
chr4-110493728-INS-449
chr4-110521051-INS-51
chr4-110521250-INS-6897
chr4-110532991-INS-446
chr4-110539526-INS-142
chr4-110539738-INS-268
chr4-110581112-INS-285
chr4-110611245-INS-6468
chr4-110611911-INS-159
chr4-110613233-INS-495
chr4-110624643-INS-236
chr4-110630491-INS-3462
chr4-110640603-INS-958
chr4-110659446-INS-674
chr4-110672442-INS-92
chr4-110695213-INS-64
chr4-110702575-INS-437
chr4-110731687-INS-72
chr4-110750545-INS-589
chr4-110764548-INS-249
chr4-110764734-INS-864
chr4-110784537-INS-476
chr4-110813422-INS-6595
chr4-110825632-INS-373
chr4-110826212-INS-6230
chr4-110827450-INS-5407
chr4-110827710-INS-799
chr4-110853211-INS-7350
chr4-110863060-INS-2224
chr4-110880966-INS-6422
chr4-110931074-INS-66
chr4-110931725-INS-53
chr4-110932655-INS-6572
chr4-110965900-INS-6487
chr4-110973230-INS-400
chr4-110988753-INS-148
chr4-110995927-INS-84
chr4-111000030-INS-987
chr4-111012737-INS-6670
chr4-111026717-INS-1293
chr4-111034465-INS-3465
chr4-111050633-INS-175
chr4-111062054-INS-158
chr4-111069887-INS-76
chr4-111082798-INS-930
chr4-111084253-INS-4844
chr4-111093972-INS-63
chr4-111161700-INS-6975
chr4-111172355-INS-219
chr4-111204876-INS-56
chr4-111233492-INS-6320
chr4-111250372-INS-217
chr4-111323239-INS-322
chr4-111336715-INS-462
chr4-111355022-INS-135
chr4-111421681-INS-829
chr4-111427833-INS-84
chr4-111436981-INS-66
chr4-111459766-INS-54
chr4-111467612-INS-7513
chr4-111470463-INS-51
chr4-111490380-INS-111
chr4-111496321-INS-104
chr4-111558161-INS-77
chr4-111567055-INS-64
chr4-111572821-INS-58
chr4-111591913-INS-6477
chr4-111603037-INS-8662
chr4-111606406-INS-121

chr4-111609435-INS-223
chr4-111648080-INS-86
chr4-111649113-INS-458
chr4-111650967-INS-969
chr4-111653300-INS-90
chr4-111658306-INS-400
chr4-111683546-INS-4355
chr4-111721716-INS-6192
chr4-111728407-INS-69
chr4-111731809-INS-95
chr4-111738682-INS-53
chr4-111762090-INS-153
chr4-111777819-INS-78
chr4-111777940-INS-622
chr4-111786711-INS-66
chr4-111840356-INS-776
chr4-111996352-INS-139
chr4-112000638-INS-53
chr4-112114125-INS-65
chr4-112142363-INS-919
chr4-112146228-INS-830
chr4-112151763-INS-444
chr4-112153694-INS-221
chr4-112172580-INS-497
chr4-112174410-INS-71
chr4-112314360-INS-207
chr4-112318817-INS-63
chr4-112327989-INS-143
chr4-112350730-INS-638
chr4-112406043-INS-199
chr4-112416324-INS-688
chr4-112474596-INS-702
chr4-112479481-INS-198
chr4-112500411-INS-503
chr4-112510005-INS-448
chr4-112536403-INS-6531
chr4-112536668-INS-8257
chr4-112590381-INS-7082
chr4-112590975-INS-120
chr4-112606231-INS-104
chr4-112628368-INS-66
chr4-112641519-INS-191
chr4-113090476-INS-1802
chr4-113983958-INS-340
chr4-114100672-INS-308
chr4-114163380-INS-252
chr4-114166011-INS-6391
chr4-114197516-INS-419
chr4-114222048-INS-66
chr4-114223501-INS-117
chr4-114243013-INS-148
chr4-114250091-INS-152
chr4-114254653-INS-63
chr4-114285610-INS-78
chr4-114299592-INS-2955
chr4-114352447-INS-430
chr4-114368728-INS-222
chr4-114417847-INS-68
chr4-114454701-INS-646
chr4-114497757-INS-65
chr4-114518824-INS-60
chr4-114525424-INS-93
chr4-114527378-INS-99
chr4-114564697-INS-99
chr4-114582699-INS-508
chr4-114612131-INS-213
chr4-114664823-INS-148
chr4-114693942-INS-1289
chr4-114697146-INS-88
chr4-114713753-INS-7073
chr4-114726332-INS-56
chr4-114734392-INS-62
chr4-114743223-INS-876
chr4-114793870-INS-67
chr4-114802004-INS-210
chr4-114803183-INS-67
chr4-114809229-INS-72

chr4-114815260-INS-96
chr4-114872671-INS-192
chr4-114897321-INS-94
chr4-114904542-INS-52
chr4-114910999-INS-91
chr4-114940864-INS-558
chr4-114957171-INS-297
chr4-114959294-INS-525
chr4-114973609-INS-120
chr4-115007017-INS-446
chr4-115017076-INS-316
chr4-115017995-INS-197
chr4-115020506-INS-72
chr4-115030838-INS-51
chr4-115031853-INS-54
chr4-115051770-INS-630
chr4-115073222-INS-128
chr4-115079092-INS-7096
chr4-115085615-INS-62
chr4-115132306-INS-98
chr4-115145519-INS-988
chr4-115152609-INS-84
chr4-115159479-INS-51
chr4-115181837-INS-117
chr4-115185152-INS-635
chr4-115272123-INS-65
chr4-115309895-INS-6704
chr4-115341433-INS-76
chr4-115375262-INS-61
chr4-115409986-INS-7382
chr4-115441036-INS-340
chr4-115478583-INS-2762
chr4-115486218-INS-5377
chr4-115492142-INS-204
chr4-115495252-INS-7757
chr4-115499687-INS-457
chr4-115510833-INS-111
chr4-115520134-INS-198
chr4-115529796-INS-6396
chr4-115558271-INS-270
chr4-115570618-INS-58
chr4-115572825-INS-209
chr4-115576683-INS-214
chr4-115586077-INS-221
chr4-115588774-INS-210
chr4-115620494-INS-57
chr4-115625332-INS-52
chr4-115626615-INS-64
chr4-115640046-INS-217
chr4-115646005-INS-51
chr4-115672832-INS-208
chr4-115709627-INS-358
chr4-115724555-INS-268
chr4-115774095-INS-118
chr4-115774672-INS-54
chr4-115776687-INS-70
chr4-115795284-INS-91
chr4-115797860-INS-52
chr4-115804022-INS-229
chr4-115805274-INS-401
chr4-115806371-INS-172
chr4-115813063-INS-51
chr4-115845398-INS-359
chr4-115847097-INS-572
chr4-115876667-INS-218
chr4-115882560-INS-103
chr4-115885514-INS-117
chr4-115886456-INS-216
chr4-115893900-INS-196
chr4-115900331-INS-260
chr4-115922888-INS-627
chr4-115931099-INS-204
chr4-115934584-INS-219
chr4-115941464-INS-205
chr4-115973138-INS-51
chr4-115990157-INS-204
chr4-116040875-INS-193

chr4-116050989-INS-205
chr4-116053316-INS-418
chr4-116055049-INS-214
chr4-116058039-INS-1892
chr4-116059746-INS-3648
chr4-116062722-INS-56
chr4-116064487-INS-132
chr4-116068392-INS-129
chr4-116076778-INS-133
chr4-116086286-INS-167
chr4-116092876-INS-213
chr4-116095064-INS-162
chr4-116097996-INS-243
chr4-116119960-INS-222
chr4-116129445-INS-68
chr4-116133612-INS-208
chr4-116136902-INS-560
chr4-116154689-INS-220
chr4-116155892-INS-96
chr4-116176626-INS-153
chr4-116202320-INS-147
chr4-116209256-INS-92
chr4-116219056-INS-158
chr4-116226607-INS-612
chr4-116247075-INS-113
chr4-116274462-INS-56
chr4-116275804-INS-8656
chr4-116296540-INS-231
chr4-116317714-INS-128
chr4-116327410-INS-64
chr4-116343898-INS-80
chr4-116374673-INS-68
chr4-116375389-INS-267
chr4-116387118-INS-2295
chr4-116387998-INS-207
chr4-116392917-INS-197
chr4-116447051-INS-215
chr4-116448618-INS-184
chr4-116453816-INS-230
chr4-116491806-INS-185
chr4-116496769-INS-205
chr4-116500524-INS-200
chr4-116507751-INS-226
chr4-116553266-INS-224
chr4-116567800-INS-155
chr4-116580416-INS-203
chr4-116590867-INS-156
chr4-116592935-INS-211
chr4-116598347-INS-125
chr4-116601431-INS-386
chr4-116604869-INS-52
chr4-116604913-INS-92
chr4-116625626-INS-211
chr4-116628209-INS-204
chr4-116631309-INS-213
chr4-116632594-INS-200
chr4-116636417-INS-123
chr4-116670177-INS-67
chr4-116672255-INS-90
chr4-116683199-INS-163
chr4-116687971-INS-52
chr4-116698053-INS-100
chr4-116700409-INS-185
chr4-116706186-INS-100
chr4-116709145-INS-320
chr4-116712518-INS-152
chr4-116715045-INS-111
chr4-116719621-INS-400
chr4-116727335-INS-53
chr4-116727746-INS-129
chr4-116729887-INS-181
chr4-116729969-INS-608
chr4-116731694-INS-5549
chr4-116735480-INS-69
chr4-116738019-INS-74
chr4-116743854-INS-223
chr4-116751722-INS-102

chr4-116765969-INS-225
chr4-116795036-INS-217
chr4-116798077-INS-68
chr4-116798705-INS-525
chr4-116804475-INS-164
chr4-116811084-INS-262
chr4-116826648-INS-160
chr4-116834852-INS-185
chr4-116850398-INS-228
chr4-116870705-INS-205
chr4-116888982-INS-83
chr4-116894147-INS-301
chr4-116897065-INS-112
chr4-116903505-INS-103
chr4-116909497-INS-54
chr4-116944638-INS-59
chr4-116948204-INS-186
chr4-116978946-INS-230
chr4-116997192-INS-186
chr4-117005607-INS-80
chr4-117007733-INS-200
chr4-117026850-INS-441
chr4-117042789-INS-197
chr4-117045554-INS-52
chr4-117059870-INS-200
chr4-117063808-INS-218
chr4-117064819-INS-204
chr4-117071094-INS-139
chr4-117073793-INS-188
chr4-117076612-INS-132
chr4-117082663-INS-166
chr4-117098504-INS-574
chr4-117101926-INS-118
chr4-117102279-INS-182
chr4-117118367-INS-120
chr4-117147502-INS-138
chr4-117184550-INS-194
chr4-117189284-INS-57
chr4-117196591-INS-169
chr4-117207686-INS-155
chr4-117253334-INS-52
chr4-117257570-INS-70
chr4-117271830-INS-55
chr4-117287757-INS-140
chr4-117292505-INS-60
chr4-117304142-INS-6442
chr4-117313858-INS-198
chr4-117317913-INS-79
chr4-117324707-INS-213
chr4-117326430-INS-262
chr4-117328743-INS-54
chr4-117333858-INS-122
chr4-117364612-INS-130
chr4-117366240-INS-60
chr4-117369916-INS-110
chr4-117372888-INS-56
chr4-117374436-INS-120
chr4-117379341-INS-63
chr4-117383689-INS-58
chr4-117405937-INS-197
chr4-117410973-INS-317
chr4-117434572-INS-56
chr4-117451749-INS-207
chr4-117454115-INS-111
chr4-117473399-INS-430
chr4-117489950-INS-100
chr4-117541060-INS-68
chr4-117543998-INS-66
chr4-117546401-INS-219
chr4-117600461-INS-280
chr4-117627832-INS-208
chr4-117629474-INS-96
chr4-117639727-INS-186
chr4-117645559-INS-293
chr4-117652194-INS-130
chr4-117674651-INS-52
chr4-117677593-INS-200

chr4-117739100-INS-186
chr4-117788476-INS-199
chr4-117822327-INS-116
chr4-117830207-INS-242
chr4-117856212-INS-128
chr4-117869762-INS-271
chr4-117875303-INS-89
chr4-117918024-INS-56
chr4-117920032-INS-62
chr4-117923349-INS-60
chr4-117945588-INS-6401
chr4-117959721-INS-118
chr4-117975799-INS-90
chr4-118001067-INS-327
chr4-118051620-INS-59
chr4-118056431-INS-216
chr4-118155939-INS-54
chr4-118169346-INS-51
chr4-118224086-INS-55
chr4-118297384-INS-90
chr4-118330442-INS-102
chr4-118345688-INS-389
chr4-118361967-INS-228
chr4-118365884-INS-53
chr4-118368189-INS-178
chr4-118380557-INS-173
chr4-118387677-INS-64
chr4-118405047-INS-1624
chr4-118405532-INS-2988
chr4-118405887-INS-8833
chr4-118405924-INS-2111
chr4-118409228-INS-93
chr4-118414022-INS-76
chr4-118468043-INS-82
chr4-118481874-INS-63
chr4-118499098-INS-228
chr4-118499159-INS-444
chr4-118519332-INS-216
chr4-118525950-INS-293
chr4-118551212-INS-53
chr4-118595371-INS-68
chr4-118629833-INS-343
chr4-118631739-INS-349
chr4-118646230-INS-100
chr4-118648022-INS-391
chr4-118656390-INS-155
chr4-118659326-INS-6463
chr4-118694505-INS-62
chr4-118735533-INS-135
chr4-118740465-INS-6608
chr4-118764072-INS-324
chr4-118780136-INS-120
chr4-118797459-INS-112
chr4-118810738-INS-96
chr4-118811274-INS-128
chr4-118827941-INS-312
chr4-118832155-INS-108
chr4-118832913-INS-129
chr4-118835729-INS-157
chr4-118838163-INS-8347
chr4-118838164-INS-828
chr4-118846633-INS-1074
chr4-118849030-INS-667
chr4-118857741-INS-110
chr4-118873522-INS-138
chr4-118879989-INS-110
chr4-118885140-INS-185
chr4-118900052-INS-85
chr4-118906238-INS-127
chr4-118913402-INS-334
chr4-118917957-INS-207
chr4-118920074-INS-267
chr4-118941793-INS-56
chr4-118943516-INS-59
chr4-118988332-INS-240
chr4-119002946-INS-202
chr4-119021609-INS-74

chr4-119042630-INS-5549
chr4-119060046-INS-507
chr4-119068359-INS-285
chr4-119121194-INS-64
chr4-119123455-INS-214
chr4-119124865-INS-206
chr4-119141152-INS-215
chr4-119157170-INS-236
chr4-119173831-INS-164
chr4-119198124-INS-195
chr4-119203931-INS-57
chr4-119213778-INS-120
chr4-119214345-INS-79
chr4-119250231-INS-104
chr4-119273142-INS-186
chr4-119308411-INS-86
chr4-119311874-INS-56
chr4-119314184-INS-186
chr4-119369114-INS-417
chr4-119371309-INS-418
chr4-119385894-INS-123
chr4-119404504-INS-200
chr4-119469838-INS-55
chr4-119473352-INS-200
chr4-119476668-INS-189
chr4-119483007-INS-213
chr4-119489293-INS-58
chr4-119524078-INS-98
chr4-119527836-INS-804
chr4-119532695-INS-62
chr4-119571543-INS-179
chr4-119587426-INS-158
chr4-119620104-INS-54
chr4-119633060-INS-64
chr4-119651507-INS-94
chr4-119697328-INS-195
chr4-119758418-INS-80
chr4-119777485-INS-108
chr4-119812501-INS-756
chr4-119833985-INS-79
chr4-119837191-INS-52
chr4-119858111-INS-62
chr4-119861072-INS-175
chr4-119877277-INS-7167
chr4-119895492-INS-67
chr4-119931945-INS-203
chr4-119936642-INS-95
chr4-119943946-INS-203
chr4-119950450-INS-88
chr4-119970569-INS-72
chr4-120017692-INS-84
chr4-120074907-INS-57
chr4-120081466-INS-400
chr4-120108243-INS-78
chr4-120135559-INS-260
chr4-120150545-INS-57
chr4-120165246-INS-106
chr4-120176568-INS-125
chr4-120203068-INS-87
chr4-120206243-INS-221
chr4-120229559-INS-130
chr4-120270174-INS-210
chr4-120316361-INS-292
chr4-120327340-INS-84
chr4-120361057-INS-3618
chr4-120369724-INS-198
chr4-120380448-INS-116
chr4-120385102-INS-386
chr4-120404953-INS-400
chr4-120426868-INS-153
chr4-120430489-INS-198
chr4-120444645-INS-850
chr4-120454699-INS-52
chr4-120465018-INS-95
chr4-120494760-INS-8697
chr4-120502570-INS-321
chr4-120508925-INS-84

chr4-120545321-INS-209
chr4-120548018-INS-160
chr4-120591737-INS-52
chr4-120596645-INS-53
chr4-120607899-INS-69
chr4-120611799-INS-68
chr4-120620458-INS-68
chr4-120623764-INS-117
chr4-120666083-INS-87
chr4-120677016-INS-177
chr4-120690583-INS-192
chr4-120692752-INS-8734
chr4-120707381-INS-204
chr4-120722264-INS-69
chr4-120754825-INS-201
chr4-120758507-INS-1080
chr4-120766071-INS-111
chr4-120780886-INS-88
chr4-120788849-INS-211
chr4-120789972-INS-893
chr4-120804582-INS-195
chr4-120805326-INS-406
chr4-120807816-INS-205
chr4-120813487-INS-60
chr4-120820917-INS-56
chr4-120834775-INS-124
chr4-120841783-INS-204
chr4-120850562-INS-198
chr4-120861974-INS-6755
chr4-120865610-INS-114
chr4-120883610-INS-60
chr4-120890561-INS-81
chr4-120899302-INS-92
chr4-120906729-INS-356
chr4-120946447-INS-60
chr4-120967325-INS-175
chr4-120980754-INS-6705
chr4-120987310-INS-6502
chr4-120991232-INS-182
chr4-120993756-INS-211
chr4-121026035-INS-77
chr4-121030465-INS-752
chr4-121033728-INS-2215
chr4-121035187-INS-59
chr4-121056209-INS-53
chr4-121081076-INS-568
chr4-121089637-INS-156
chr4-121117714-INS-236
chr4-121138992-INS-242
chr4-121152383-INS-168
chr4-121181950-INS-2871
chr4-121203796-INS-114
chr4-121228580-INS-119
chr4-121243190-INS-92
chr4-121245809-INS-60
chr4-121247896-INS-557
chr4-121251238-INS-56
chr4-121253509-INS-3799
chr4-121258031-INS-135
chr4-121258056-INS-1888
chr4-121259050-INS-68
chr4-121327755-INS-4198
chr4-121334468-INS-6441
chr4-121499307-INS-99
chr4-121509845-INS-100
chr4-121536183-INS-6403
chr4-121548932-INS-1625
chr4-121571017-INS-7133
chr4-121625853-INS-7285
chr4-121630908-INS-172
chr4-121756032-INS-5266
chr4-121905554-INS-5171
chr4-122298495-INS-5171
chr4-122385746-INS-523
chr4-122407267-INS-6432
chr4-122412437-INS-608
chr4-122505735-INS-1888

chr4-122528156-INS-64
chr4-122569943-INS-780
chr4-122606349-INS-424
chr4-122623419-INS-81
chr4-122637232-INS-160
chr4-122662998-INS-52
chr4-122691267-INS-878
chr4-122709517-INS-59
chr4-122730447-INS-65
chr4-122734376-INS-378
chr4-122739002-INS-73
chr4-122748484-INS-84
chr4-122755253-INS-56
chr4-122786222-INS-772
chr4-122788303-INS-545
chr4-122793394-INS-236
chr4-122795696-INS-75
chr4-122807723-INS-224
chr4-122814340-INS-405
chr4-122835414-INS-329
chr4-122845118-INS-60
chr4-122858197-INS-564
chr4-122865844-INS-66
chr4-122867918-INS-84
chr4-122908580-INS-96
chr4-122914062-INS-211
chr4-122932116-INS-208
chr4-122941704-INS-67
chr4-122949789-INS-207
chr4-122954281-INS-82
chr4-122961339-INS-52
chr4-122965831-INS-68
chr4-122972287-INS-225
chr4-122973730-INS-231
chr4-122979593-INS-84
chr4-122980994-INS-126
chr4-122983174-INS-118
chr4-123040036-INS-134
chr4-123046389-INS-56
chr4-123050685-INS-200
chr4-123059191-INS-61
chr4-123061058-INS-109
chr4-123098308-INS-201
chr4-123111395-INS-184
chr4-123117567-INS-100
chr4-123147035-INS-79
chr4-123153096-INS-117
chr4-123159258-INS-491
chr4-123166812-INS-75
chr4-123172643-INS-200
chr4-123200243-INS-231
chr4-123211853-INS-57
chr4-123214738-INS-158
chr4-123238267-INS-256
chr4-123261334-INS-180
chr4-123267822-INS-136
chr4-123277062-INS-90
chr4-123285362-INS-192
chr4-123314717-INS-54
chr4-123317281-INS-146
chr4-123318603-INS-183
chr4-123320461-INS-95
chr4-123332669-INS-447
chr4-123336344-INS-69
chr4-123347251-INS-80
chr4-123362555-INS-90
chr4-123392765-INS-71
chr4-123413886-INS-104
chr4-123419348-INS-63
chr4-123424262-INS-63
chr4-123432197-INS-448
chr4-123444921-INS-56
chr4-123469809-INS-201
chr4-123493766-INS-60
chr4-123497218-INS-216
chr4-123529689-INS-184
chr4-123540781-INS-209

chr4-123542226-INS-217
chr4-123543245-INS-171
chr4-123548230-INS-51
chr4-123554097-INS-198
chr4-123560261-INS-52
chr4-123566044-INS-271
chr4-123572163-INS-78
chr4-123573801-INS-180
chr4-123588073-INS-197
chr4-123595714-INS-434
chr4-123596481-INS-93
chr4-123597647-INS-537
chr4-123599587-INS-556
chr4-123605069-INS-83
chr4-123615486-INS-395
chr4-123647280-INS-167
chr4-123663026-INS-363
chr4-123685818-INS-180
chr4-123693508-INS-398
chr4-123694406-INS-233
chr4-123696937-INS-184
chr4-123711117-INS-245
chr4-123711851-INS-68
chr4-123712901-INS-166
chr4-123718454-INS-72
chr4-123719511-INS-212
chr4-123720946-INS-207
chr4-123721748-INS-72
chr4-123753204-INS-800
chr4-123770731-INS-513
chr4-123793339-INS-7184
chr4-123796399-INS-56
chr4-123800175-INS-110
chr4-123822178-INS-180
chr4-123824436-INS-168
chr4-123834047-INS-133
chr4-123847854-INS-60
chr4-123868617-INS-60
chr4-123873501-INS-981
chr4-123877128-INS-261
chr4-123894997-INS-53
chr4-123918000-INS-280
chr4-123932876-INS-266
chr4-123934831-INS-81
chr4-123939107-INS-76
chr4-123940585-INS-110
chr4-123968850-INS-446
chr4-123972666-INS-431
chr4-123984306-INS-102
chr4-123998664-INS-484
chr4-124008519-INS-171
chr4-124013890-INS-377
chr4-124039415-INS-185
chr4-124049340-INS-152
chr4-124053446-INS-102
chr4-124082279-INS-89
chr4-124088180-INS-269
chr4-124106193-INS-80
chr4-124128758-INS-68
chr4-124143101-INS-758
chr4-124183033-INS-90
chr4-124189720-INS-113
chr4-124196103-INS-100
chr4-124239677-INS-305
chr4-124243540-INS-69
chr4-124258109-INS-53
chr4-124272509-INS-166
chr4-124290431-INS-209
chr4-124304530-INS-72
chr4-124315734-INS-196
chr4-124351561-INS-187
chr4-124361949-INS-428
chr4-124368668-INS-337
chr4-124378861-INS-67
chr4-124386882-INS-66
chr4-124391134-INS-53
chr4-124425886-INS-60

chr4-124435520-INS-256
chr4-124447324-INS-262
chr4-124461602-INS-417
chr4-124470986-INS-63
chr4-124476067-INS-70
chr4-124480021-INS-52
chr4-124498617-INS-357
chr4-124507884-INS-118
chr4-124525460-INS-6440
chr4-124547640-INS-6808
chr4-124600519-INS-225
chr4-124611267-INS-411
chr4-124617786-INS-224
chr4-124621462-INS-227
chr4-124633919-INS-187
chr4-124639615-INS-215
chr4-124651771-INS-220
chr4-124663752-INS-962
chr4-124665219-INS-54
chr4-124683414-INS-2222
chr4-124689700-INS-184
chr4-124712737-INS-393
chr4-124716314-INS-139
chr4-124718697-INS-165
chr4-124758128-INS-206
chr4-124764103-INS-120
chr4-124781004-INS-56
chr4-124784233-INS-233
chr4-124791232-INS-52
chr4-124797043-INS-141
chr4-124802659-INS-84
chr4-124819432-INS-170
chr4-124832211-INS-207
chr4-124834592-INS-250
chr4-124840097-INS-171
chr4-124854779-INS-56
chr4-124857708-INS-247
chr4-124878938-INS-408
chr4-124887073-INS-54
chr4-124903142-INS-58
chr4-124905613-INS-70
chr4-124923360-INS-108
chr4-124929878-INS-178
chr4-124976548-INS-194
chr4-125031165-INS-51
chr4-125033909-INS-55
chr4-125035692-INS-177
chr4-125046729-INS-412
chr4-125055735-INS-67
chr4-125059835-INS-273
chr4-125062039-INS-86
chr4-125065611-INS-53
chr4-125071417-INS-184
chr4-125073009-INS-318
chr4-125075930-INS-212
chr4-125077378-INS-314
chr4-125108998-INS-308
chr4-125145160-INS-84
chr4-125166571-INS-110
chr4-125197634-INS-58
chr4-125206218-INS-7078
chr4-125227159-INS-5553
chr4-125228551-INS-172
chr4-125245088-INS-73
chr4-125271157-INS-60
chr4-125275824-INS-66
chr4-125282340-INS-96
chr4-125294406-INS-319
chr4-125296025-INS-410
chr4-125333351-INS-133
chr4-125337372-INS-804
chr4-125342511-INS-171
chr4-125369141-INS-378
chr4-125378855-INS-133
chr4-125456533-INS-195
chr4-125462486-INS-208
chr4-125490846-INS-60

chr4-125504496-INS-75
chr4-125510462-INS-69
chr4-125511341-INS-58
chr4-125515413-INS-76
chr4-125520947-INS-323
chr4-125526794-INS-57
chr4-125563813-INS-54
chr4-125597074-INS-72
chr4-125614405-INS-520
chr4-125615600-INS-5280
chr4-125619081-INS-62
chr4-125670127-INS-81
chr4-125699560-INS-202
chr4-125712344-INS-65
chr4-125715040-INS-54
chr4-125722730-INS-94
chr4-125761990-INS-301
chr4-125775501-INS-71
chr4-125788763-INS-60
chr4-125809712-INS-446
chr4-125828484-INS-635
chr4-125845584-INS-284
chr4-125857670-INS-171
chr4-125870220-INS-90
chr4-125903575-INS-192
chr4-125906893-INS-228
chr4-125926530-INS-69
chr4-125946242-INS-176
chr4-125953083-INS-153
chr4-125967047-INS-105
chr4-125970898-INS-223
chr4-125973210-INS-90
chr4-125975656-INS-160
chr4-125983928-INS-209
chr4-126037021-INS-2924
chr4-126049408-INS-179
chr4-126053475-INS-73
chr4-126082463-INS-377
chr4-126112995-INS-232
chr4-126113487-INS-254
chr4-126114495-INS-430
chr4-126115883-INS-206
chr4-126123469-INS-190
chr4-126124876-INS-644
chr4-126136797-INS-209
chr4-126163821-INS-205
chr4-126164559-INS-202
chr4-126165832-INS-178
chr4-126168537-INS-436
chr4-126192660-INS-200
chr4-126197712-INS-60
chr4-126201666-INS-52
chr4-126224151-INS-199
chr4-126225864-INS-149
chr4-126229814-INS-92
chr4-126231620-INS-392
chr4-126246112-INS-209
chr4-126256110-INS-8731
chr4-126257840-INS-146
chr4-126264259-INS-114
chr4-126285258-INS-57
chr4-126308287-INS-52
chr4-126321022-INS-56
chr4-126345360-INS-128
chr4-126346483-INS-56
chr4-126366459-INS-104
chr4-126379328-INS-62
chr4-126382764-INS-581
chr4-126391998-INS-208
chr4-126394104-INS-52
chr4-126395446-INS-64
chr4-126399130-INS-78
chr4-126401593-INS-205
chr4-126409903-INS-569
chr4-126414433-INS-281
chr4-126428983-INS-195
chr4-126434890-INS-52

chr4-126439151-INS-525
chr4-126445507-INS-175
chr4-126459366-INS-207
chr4-126482788-INS-403
chr4-126483495-INS-541
chr4-126485201-INS-94
chr4-126489575-INS-55
chr4-126516142-INS-1895
chr4-126517110-INS-70
chr4-126523184-INS-223
chr4-126525339-INS-52
chr4-126525801-INS-254
chr4-126546487-INS-55
chr4-126593137-INS-225
chr4-126604334-INS-1107
chr4-126618151-INS-221
chr4-126648569-INS-143
chr4-126672503-INS-398
chr4-126680396-INS-93
chr4-126698717-INS-152
chr4-126740652-INS-71
chr4-126750358-INS-169
chr4-126753216-INS-451
chr4-126767621-INS-121
chr4-126792990-INS-203
chr4-126794592-INS-210
chr4-126825957-INS-52
chr4-126845327-INS-199
chr4-126849242-INS-227
chr4-126850744-INS-88
chr4-126871977-INS-1893
chr4-126872303-INS-1257
chr4-126872304-INS-624
chr4-126877586-INS-1295
chr4-126879770-INS-198
chr4-126884108-INS-214
chr4-126893372-INS-585
chr4-126893696-INS-6845
chr4-126897618-INS-54
chr4-126901422-INS-227
chr4-126912080-INS-216
chr4-126931276-INS-190
chr4-126956013-INS-68
chr4-126957191-INS-82
chr4-126962255-INS-86
chr4-126963173-INS-245
chr4-126967447-INS-212
chr4-126991706-INS-95
chr4-126997431-INS-113
chr4-127008451-INS-181
chr4-127039297-INS-202
chr4-127042390-INS-5538
chr4-127050483-INS-240
chr4-127052230-INS-96
chr4-127054091-INS-677
chr4-127105098-INS-78
chr4-127115115-INS-56
chr4-127123118-INS-62
chr4-127150253-INS-119
chr4-127170858-INS-93
chr4-127174934-INS-253
chr4-127176994-INS-53
chr4-127188608-INS-58
chr4-127215492-INS-679
chr4-127229317-INS-53
chr4-127229784-INS-628
chr4-127247898-INS-357
chr4-127254921-INS-209
chr4-127256394-INS-52
chr4-127268510-INS-57
chr4-127270656-INS-52
chr4-127290762-INS-75
chr4-127295115-INS-80
chr4-127296492-INS-249
chr4-127298892-INS-5280
chr4-127313800-INS-62
chr4-127330846-INS-52

chr4-127332724-INS-59
chr4-127337702-INS-2824
chr4-127363885-INS-671
chr4-127379081-INS-72
chr4-127383337-INS-53
chr4-127393459-INS-58
chr4-127393897-INS-69
chr4-127418512-INS-79
chr4-127438126-INS-156
chr4-127451229-INS-195
chr4-127455831-INS-235
chr4-127496053-INS-7108
chr4-127499245-INS-152
chr4-127512561-INS-395
chr4-127517370-INS-791
chr4-127526026-INS-52
chr4-127526144-INS-418
chr4-127529141-INS-811
chr4-127544047-INS-175
chr4-127557535-INS-4781
chr4-127563394-INS-63
chr4-127563843-INS-426
chr4-127565821-INS-785
chr4-127568987-INS-345
chr4-127597622-INS-122
chr4-127621252-INS-90
chr4-127626059-INS-222
chr4-127629895-INS-96
chr4-127638517-INS-76
chr4-127674608-INS-5509
chr4-127680896-INS-70
chr4-127682728-INS-91
chr4-127696372-INS-111
chr4-127699835-INS-141
chr4-127725833-INS-532
chr4-127773612-INS-6764
chr4-127777455-INS-91
chr4-127778419-INS-400
chr4-127784541-INS-74
chr4-127797654-INS-298
chr4-127800528-INS-763
chr4-127808336-INS-422
chr4-127818233-INS-206
chr4-127827961-INS-86
chr4-127842138-INS-58
chr4-127871941-INS-248
chr4-127893857-INS-59
chr4-127918042-INS-108
chr4-127918902-INS-110
chr4-127929990-INS-490
chr4-127950032-INS-57
chr4-127957619-INS-181
chr4-127963188-INS-72
chr4-127989838-INS-66
chr4-127999867-INS-1888
chr4-128003858-INS-83
chr4-128031682-INS-52
chr4-128040791-INS-57
chr4-128061201-INS-56
chr4-128083729-INS-103
chr4-128098836-INS-134
chr4-128115378-INS-1007
chr4-128115396-INS-2367
chr4-128115416-INS-130
chr4-128124225-INS-448
chr4-128134333-INS-51
chr4-128153832-INS-70
chr4-128178200-INS-523
chr4-128195464-INS-81
chr4-128217625-INS-131
chr4-128238405-INS-70
chr4-128255474-INS-63
chr4-128257395-INS-60
chr4-128295076-INS-103
chr4-128296056-INS-64
chr4-128317508-INS-151
chr4-128357507-INS-943

chr4-128414750-INS-56
chr4-128441622-INS-206
chr4-128463602-INS-69
chr4-128467435-INS-157
chr4-128475762-INS-78
chr4-128481622-INS-201
chr4-128496651-INS-68
chr4-128520709-INS-265
chr4-128586374-INS-77
chr4-128594256-INS-77
chr4-128604687-INS-105
chr4-128631504-INS-54
chr4-128647022-INS-96
chr4-128721203-INS-182
chr4-128721894-INS-58
chr4-128729574-INS-279
chr4-128731401-INS-56
chr4-128742239-INS-210
chr4-128752089-INS-144
chr4-128754455-INS-1238
chr4-128758636-INS-52
chr4-128766626-INS-53
chr4-128797569-INS-89
chr4-128809372-INS-112
chr4-128816176-INS-839
chr4-128837531-INS-87
chr4-128846077-INS-238
chr4-128850353-INS-106
chr4-128850353-INS-632
chr4-128860089-INS-129
chr4-128865924-INS-129
chr4-128901812-INS-403
chr4-128908753-INS-90
chr4-128933250-INS-86
chr4-128935727-INS-216
chr4-128947842-INS-206
chr4-128950184-INS-208
chr4-128954626-INS-207
chr4-128958991-INS-201
chr4-128971065-INS-56
chr4-128987200-INS-175
chr4-129010455-INS-63
chr4-129021026-INS-172
chr4-129029627-INS-5554
chr4-129039798-INS-73
chr4-129059676-INS-81
chr4-129073065-INS-215
chr4-129079229-INS-203
chr4-129093027-INS-101
chr4-129095913-INS-56
chr4-129127699-INS-192
chr4-129130639-INS-207
chr4-129137460-INS-207
chr4-129145350-INS-173
chr4-129148481-INS-520
chr4-129163231-INS-614
chr4-129169094-INS-108
chr4-129171330-INS-113
chr4-129172378-INS-69
chr4-129179240-INS-273
chr4-129191140-INS-213
chr4-129220203-INS-60
chr4-129231863-INS-169
chr4-129252087-INS-309
chr4-129265097-INS-248
chr4-129266987-INS-68
chr4-129275638-INS-377
chr4-129283050-INS-224
chr4-129306618-INS-204
chr4-129307988-INS-124
chr4-129317406-INS-142
chr4-129319180-INS-746
chr4-129332959-INS-98
chr4-129333460-INS-84
chr4-129349503-INS-60
chr4-129365109-INS-93
chr4-129367005-INS-259

chr4-129371088-INS-170
chr4-129372701-INS-59
chr4-129373166-INS-202
chr4-129377172-INS-72
chr4-129386160-INS-116
chr4-129405178-INS-64
chr4-129426396-INS-217
chr4-129442985-INS-363
chr4-129445467-INS-77
chr4-129446756-INS-97
chr4-129485010-INS-66
chr4-129498429-INS-201
chr4-129500458-INS-107
chr4-129501477-INS-217
chr4-129529425-INS-193
chr4-129545881-INS-225
chr4-129565257-INS-241
chr4-129567930-INS-127
chr4-129570737-INS-125
chr4-129574248-INS-267
chr4-129582491-INS-195
chr4-129613704-INS-170
chr4-129614844-INS-199
chr4-129617641-INS-199
chr4-129626399-INS-204
chr4-129633187-INS-234
chr4-129638901-INS-71
chr4-129639774-INS-244
chr4-129647535-INS-195
chr4-129659497-INS-288
chr4-129668147-INS-174
chr4-129681726-INS-207
chr4-129683910-INS-182
chr4-129704033-INS-191
chr4-129709144-INS-196
chr4-129727665-INS-212
chr4-129757865-INS-62
chr4-129777554-INS-233
chr4-129788271-INS-176
chr4-129792436-INS-57
chr4-129797135-INS-196
chr4-129798690-INS-836
chr4-129808940-INS-313
chr4-129826449-INS-702
chr4-129890682-INS-388
chr4-129926626-INS-53
chr4-129968351-INS-96
chr4-130006888-INS-82
chr4-130013149-INS-110
chr4-130039688-INS-216
chr4-130046510-INS-163
chr4-130046679-INS-1146
chr4-130046753-INS-710
chr4-130072054-INS-65
chr4-130087442-INS-60
chr4-130090316-INS-246
chr4-130098221-INS-255
chr4-130129825-INS-56
chr4-130134810-INS-54
chr4-130135447-INS-127
chr4-130137459-INS-55
chr4-130139346-INS-115
chr4-130144243-INS-195
chr4-130150055-INS-333
chr4-130151237-INS-104
chr4-130164317-INS-99
chr4-130170266-INS-54
chr4-130185021-INS-190
chr4-130187362-INS-163
chr4-130215783-INS-56
chr4-130220216-INS-200
chr4-130249982-INS-164
chr4-130288319-INS-198
chr4-130291501-INS-169
chr4-130301441-INS-101
chr4-130303860-INS-53
chr4-130328885-INS-58

chr4-130341469-INS-181
chr4-130342692-INS-192
chr4-130344411-INS-156
chr4-130347441-INS-200
chr4-130355004-INS-78
chr4-130357162-INS-210
chr4-130361767-INS-201
chr4-130370181-INS-62
chr4-130372396-INS-213
chr4-130377156-INS-202
chr4-130379417-INS-173
chr4-130384060-INS-87
chr4-130394076-INS-168
chr4-130397717-INS-202
chr4-130441293-INS-64
chr4-130441912-INS-456
chr4-130466306-INS-208
chr4-130481012-INS-1886
chr4-130500287-INS-56
chr4-130504423-INS-52
chr4-130516582-INS-115
chr4-130521181-INS-69
chr4-130564309-INS-53
chr4-130566779-INS-51
chr4-130569963-INS-143
chr4-130575539-INS-168
chr4-130599208-INS-130
chr4-130608555-INS-72
chr4-130613924-INS-122
chr4-130657892-INS-130
chr4-130660030-INS-1127
chr4-130660382-INS-97
chr4-130662744-INS-108
chr4-130663308-INS-70
chr4-130667742-INS-106
chr4-130669458-INS-57
chr4-130678952-INS-52
chr4-130697707-INS-128
chr4-130708995-INS-90
chr4-130732991-INS-80
chr4-130736548-INS-68
chr4-130745447-INS-123
chr4-130754217-INS-262
chr4-130782193-INS-59
chr4-130783969-INS-114
chr4-130794255-INS-196
chr4-130806484-INS-548
chr4-130808260-INS-79
chr4-130809753-INS-60
chr4-130826506-INS-90
chr4-130827318-INS-149
chr4-130830984-INS-54
chr4-130836634-INS-280
chr4-130859468-INS-53
chr4-130873234-INS-81
chr4-130890061-INS-61
chr4-130897006-INS-56
chr4-130911961-INS-82
chr4-131000226-INS-135
chr4-131024859-INS-59
chr4-131041938-INS-66
chr4-131048997-INS-55
chr4-131075194-INS-251
chr4-131077256-INS-205
chr4-131079271-INS-384
chr4-131153048-INS-180
chr4-131180121-INS-129
chr4-131190301-INS-66
chr4-131191971-INS-56
chr4-131197545-INS-98
chr4-131236401-INS-180
chr4-131236581-INS-1076
chr4-131237643-INS-211
chr4-131250408-INS-331
chr4-131288808-INS-400
chr4-131294475-INS-114
chr4-131311553-INS-72

chr4-131318708-INS-51
chr4-131320195-INS-207
chr4-131336306-INS-90
chr4-131339855-INS-107
chr4-131355209-INS-312
chr4-131357079-INS-92
chr4-131389000-INS-61
chr4-131402687-INS-76
chr4-131420524-INS-61
chr4-131431933-INS-52
chr4-131448621-INS-57
chr4-131455018-INS-91
chr4-131473137-INS-147
chr4-131479787-INS-74
chr4-131532065-INS-213
chr4-131532907-INS-363
chr4-131536370-INS-251
chr4-131541905-INS-59
chr4-131550271-INS-207
chr4-131554573-INS-58
chr4-131556135-INS-210
chr4-131567115-INS-66
chr4-131596219-INS-243
chr4-131631637-INS-254
chr4-131660478-INS-66
chr4-131668027-INS-212
chr4-131696297-INS-108
chr4-131700380-INS-1889
chr4-131713964-INS-294
chr4-131730075-INS-52
chr4-131745155-INS-204
chr4-131752507-INS-211
chr4-131752961-INS-5289
chr4-131754753-INS-204
chr4-131758487-INS-83
chr4-131770692-INS-51
chr4-131781866-INS-206
chr4-131786050-INS-93
chr4-131799921-INS-209
chr4-131801245-INS-108
chr4-131817333-INS-213
chr4-131829694-INS-52
chr4-131831429-INS-739
chr4-131835263-INS-562
chr4-131852637-INS-1120
chr4-131861607-INS-209
chr4-131863301-INS-212
chr4-131877008-INS-8195
chr4-131878524-INS-130
chr4-131883199-INS-200
chr4-131888408-INS-151
chr4-131889142-INS-73
chr4-131890562-INS-193
chr4-131898870-INS-51
chr4-131906153-INS-56
chr4-131921029-INS-205
chr4-131926119-INS-224
chr4-131949085-INS-153
chr4-131950649-INS-56
chr4-131951379-INS-145
chr4-131968057-INS-67
chr4-131975517-INS-94
chr4-131992371-INS-136
chr4-131999517-INS-126
chr4-132012951-INS-530
chr4-132016394-INS-481
chr4-132018070-INS-83
chr4-132024467-INS-199
chr4-132038756-INS-183
chr4-132042492-INS-248
chr4-132049520-INS-242
chr4-132057885-INS-198
chr4-132064382-INS-134
chr4-132076020-INS-189
chr4-132082226-INS-207
chr4-132083758-INS-96
chr4-132089483-INS-195

chr4-132090574-INS-390
chr4-132091392-INS-290
chr4-132096798-INS-224
chr4-132111659-INS-52
chr4-132135941-INS-58
chr4-132141712-INS-226
chr4-132142977-INS-144
chr4-132144122-INS-57
chr4-132151113-INS-212
chr4-132162350-INS-383
chr4-132164861-INS-57
chr4-132166225-INS-183
chr4-132174682-INS-71
chr4-132175133-INS-241
chr4-132177546-INS-103
chr4-132183786-INS-94
chr4-132194411-INS-60
chr4-132195989-INS-54
chr4-132197015-INS-205
chr4-132205467-INS-208
chr4-132230268-INS-5289
chr4-132240287-INS-54
chr4-132243379-INS-215
chr4-132262772-INS-159
chr4-132275649-INS-57
chr4-132282610-INS-225
chr4-132283914-INS-126
chr4-132288671-INS-78
chr4-132293775-INS-116
chr4-132301892-INS-68
chr4-132304224-INS-54
chr4-132305796-INS-187
chr4-132314182-INS-54
chr4-132317699-INS-113
chr4-132332580-INS-191
chr4-132334772-INS-571
chr4-132337527-INS-51
chr4-132345871-INS-62
chr4-132349108-INS-95
chr4-132352327-INS-238
chr4-132356288-INS-183
chr4-132357987-INS-518
chr4-132360370-INS-205
chr4-132373866-INS-200
chr4-132383234-INS-212
chr4-132386994-INS-170
chr4-132396447-INS-195
chr4-132399106-INS-54
chr4-132402454-INS-201
chr4-132416661-INS-209
chr4-132422697-INS-186
chr4-132423395-INS-254
chr4-132429787-INS-223
chr4-132438076-INS-87
chr4-132442430-INS-111
chr4-132446060-INS-179
chr4-132457709-INS-52
chr4-132470742-INS-223
chr4-132472627-INS-98
chr4-132479154-INS-669
chr4-132481975-INS-115
chr4-132508763-INS-900
chr4-132512078-INS-60
chr4-132513946-INS-85
chr4-132528547-INS-224
chr4-132538143-INS-518
chr4-132541591-INS-204
chr4-132560476-INS-186
chr4-132589371-INS-70
chr4-132597369-INS-205
chr4-132604343-INS-205
chr4-132617081-INS-57
chr4-132618495-INS-251
chr4-132620086-INS-180
chr4-132621203-INS-205
chr4-132628863-INS-214
chr4-132634128-INS-204

chr4-132638506-INS-106
chr4-132655170-INS-128
chr4-132656283-INS-206
chr4-132658740-INS-55
chr4-132659702-INS-132
chr4-132671863-INS-75
chr4-132679889-INS-532
chr4-132683056-INS-57
chr4-132693758-INS-233
chr4-132697364-INS-54
chr4-132708044-INS-52
chr4-132708731-INS-63
chr4-132725927-INS-80
chr4-132744290-INS-142
chr4-132757339-INS-53
chr4-132829746-INS-199
chr4-132848036-INS-220
chr4-132864222-INS-221
chr4-132880076-INS-206
chr4-132881864-INS-60
chr4-132885757-INS-166
chr4-132890099-INS-245
chr4-132890419-INS-120
chr4-132913122-INS-84
chr4-132942391-INS-214
chr4-132956690-INS-138
chr4-132962702-INS-52
chr4-132964931-INS-111
chr4-132965663-INS-197
chr4-132993384-INS-6462
chr4-132998697-INS-232
chr4-133001268-INS-163
chr4-133011291-INS-89
chr4-133014963-INS-403
chr4-133015883-INS-107
chr4-133018752-INS-210
chr4-133048586-INS-152
chr4-133072582-INS-248
chr4-133074001-INS-214
chr4-133075445-INS-110
chr4-133085905-INS-6335
chr4-133095933-INS-207
chr4-133120770-INS-120
chr4-133128494-INS-209
chr4-133144801-INS-197
chr4-133163691-INS-208
chr4-133180460-INS-408
chr4-133190483-INS-175
chr4-133227223-INS-112
chr4-133230350-INS-206
chr4-133241398-INS-117
chr4-133245812-INS-205
chr4-133256682-INS-75
chr4-133275374-INS-196
chr4-133282187-INS-149
chr4-133282757-INS-215
chr4-133285189-INS-66
chr4-133288115-INS-55
chr4-133291931-INS-323
chr4-133304407-INS-200
chr4-133313007-INS-396
chr4-133321447-INS-78
chr4-133323783-INS-992
chr4-133346658-INS-68
chr4-133368064-INS-71
chr4-133372604-INS-223
chr4-133374025-INS-146
chr4-133375080-INS-147
chr4-133380630-INS-178
chr4-133384572-INS-100
chr4-133389205-INS-204
chr4-133395249-INS-250
chr4-133424489-INS-235
chr4-133445611-INS-148
chr4-133454911-INS-209
chr4-133475477-INS-207
chr4-133510114-INS-83

chr4-133522139-INS-132
chr4-133527775-INS-164
chr4-133549878-INS-118
chr4-133557590-INS-192
chr4-133559251-INS-135
chr4-133566078-INS-323
chr4-133574347-INS-54
chr4-133580181-INS-216
chr4-133609745-INS-70
chr4-133628364-INS-175
chr4-133631082-INS-1515
chr4-133631082-INS-632
chr4-133632312-INS-72
chr4-133651790-INS-185
chr4-133657916-INS-67
chr4-133661505-INS-207
chr4-133699769-INS-233
chr4-133758393-INS-168
chr4-133760678-INS-215
chr4-133790355-INS-94
chr4-133797792-INS-72
chr4-133805217-INS-202
chr4-133810053-INS-726
chr4-133825102-INS-5062
chr4-133828761-INS-201
chr4-133869964-INS-119
chr4-133893684-INS-208
chr4-133916221-INS-91
chr4-133931320-INS-57
chr4-133947529-INS-693
chr4-133953758-INS-68
chr4-133958598-INS-195
chr4-133986324-INS-219
chr4-133990942-INS-124
chr4-133991724-INS-88
chr4-134059209-INS-78
chr4-134064103-INS-161
chr4-134071557-INS-52
chr4-134073744-INS-221
chr4-134080419-INS-204
chr4-134105004-INS-114
chr4-134108097-INS-322
chr4-134108384-INS-340
chr4-134108785-INS-75
chr4-134116132-INS-219
chr4-134120201-INS-175
chr4-134128201-INS-99
chr4-134138358-INS-228
chr4-134146320-INS-197
chr4-134159067-INS-80
chr4-134185464-INS-401
chr4-134191688-INS-74
chr4-134239518-INS-203
chr4-134246210-INS-357
chr4-134256779-INS-62
chr4-134279791-INS-186
chr4-134290526-INS-544
chr4-134300932-INS-196
chr4-134301371-INS-106
chr4-134312495-INS-98
chr4-134335590-INS-58
chr4-134339362-INS-225
chr4-134340766-INS-100
chr4-134349514-INS-240
chr4-134394003-INS-287
chr4-134400317-INS-231
chr4-134408966-INS-74
chr4-134413373-INS-75
chr4-134419832-INS-156
chr4-134422237-INS-115
chr4-134457894-INS-185
chr4-134505134-INS-55
chr4-134505914-INS-219
chr4-134508580-INS-211
chr4-134515666-INS-193
chr4-134518209-INS-78
chr4-134546627-INS-212

chr4-134573383-INS-57
chr4-134574927-INS-219
chr4-134577334-INS-79
chr4-134593965-INS-73
chr4-134594074-INS-126
chr4-134598706-INS-66
chr4-134600564-INS-217
chr4-134621783-INS-62
chr4-134631560-INS-191
chr4-134633179-INS-133
chr4-134649665-INS-390
chr4-134679180-INS-110
chr4-134693630-INS-225
chr4-134735483-INS-110
chr4-134745746-INS-201
chr4-134747648-INS-59
chr4-134769740-INS-364
chr4-134770378-INS-72
chr4-134862560-INS-58
chr4-134916685-INS-126
chr4-134924434-INS-58
chr4-134929983-INS-614
chr4-134956503-INS-226
chr4-134971265-INS-82
chr4-135023076-INS-249
chr4-135026877-INS-4652
chr4-135035428-INS-150
chr4-135044884-INS-497
chr4-135053007-INS-59
chr4-135053619-INS-779
chr4-135057342-INS-214
chr4-135077110-INS-175
chr4-135089637-INS-240
chr4-135091906-INS-73
chr4-135094708-INS-2237
chr4-135096602-INS-242
chr4-135102317-INS-55
chr4-135134427-INS-188
chr4-135136811-INS-132
chr4-135152256-INS-175
chr4-135166340-INS-67
chr4-135171299-INS-201
chr4-135180864-INS-380
chr4-135200249-INS-418
chr4-135202645-INS-78
chr4-135211315-INS-124
chr4-135214994-INS-161
chr4-135217767-INS-77
chr4-135228224-INS-548
chr4-135228827-INS-198
chr4-135240839-INS-63
chr4-135263408-INS-177
chr4-135308263-INS-61
chr4-135326252-INS-104
chr4-135329667-INS-212
chr4-135370442-INS-70
chr4-135376859-INS-85
chr4-135379490-INS-138
chr4-135405895-INS-546
chr4-135460129-INS-62
chr4-135485986-INS-331
chr4-135513734-INS-123
chr4-135527683-INS-72
chr4-135533899-INS-293
chr4-135557151-INS-199
chr4-135575821-INS-80
chr4-135580353-INS-78
chr4-135582278-INS-502
chr4-135605969-INS-202
chr4-135606458-INS-427
chr4-135612757-INS-505
chr4-135633360-INS-203
chr4-135654937-INS-183
chr4-135670753-INS-124
chr4-135702168-INS-231
chr4-135703925-INS-205
chr4-135713120-INS-351

chr4-135720870-INS-504
chr4-135722405-INS-207
chr4-135724641-INS-204
chr4-135729246-INS-231
chr4-135735582-INS-114
chr4-135737388-INS-187
chr4-135745303-INS-51
chr4-135751017-INS-64
chr4-135766675-INS-194
chr4-135785143-INS-297
chr4-135786284-INS-469
chr4-135796005-INS-190
chr4-135801211-INS-4630
chr4-135810899-INS-171
chr4-135815339-INS-172
chr4-135821487-INS-209
chr4-135831545-INS-168
chr4-135836314-INS-87
chr4-135838999-INS-548
chr4-135840751-INS-254
chr4-135843743-INS-66
chr4-135845713-INS-51
chr4-135849457-INS-193
chr4-135851682-INS-272
chr4-135883965-INS-96
chr4-135885691-INS-167
chr4-135894557-INS-200
chr4-135898493-INS-173
chr4-135910413-INS-207
chr4-135933260-INS-132
chr4-135939977-INS-208
chr4-135940520-INS-203
chr4-135957344-INS-6795
chr4-135958461-INS-215
chr4-135990060-INS-71
chr4-136031521-INS-626
chr4-136036941-INS-56
chr4-136047731-INS-52
chr4-136049950-INS-919
chr4-136076419-INS-54
chr4-136097423-INS-257
chr4-136136772-INS-215
chr4-136139490-INS-52
chr4-136140565-INS-52
chr4-136156421-INS-214
chr4-136158938-INS-70
chr4-136180186-INS-178
chr4-136181629-INS-203
chr4-136183308-INS-66
chr4-136239554-INS-324
chr4-136249873-INS-101
chr4-136258398-INS-62
chr4-136284886-INS-52
chr4-136338641-INS-123
chr4-136342244-INS-124
chr4-136343417-INS-164
chr4-136351928-INS-432
chr4-136381170-INS-96
chr4-136383842-INS-68
chr4-136422593-INS-104
chr4-136427248-INS-215
chr4-136431004-INS-53
chr4-136433323-INS-208
chr4-136443188-INS-80
chr4-136462319-INS-76
chr4-136496152-INS-70
chr4-136542120-INS-204
chr4-136568128-INS-70
chr4-136581637-INS-52
chr4-136605107-INS-104
chr4-136610368-INS-56
chr4-136688658-INS-59
chr4-136734145-INS-354
chr4-136742678-INS-52
chr4-136753076-INS-164
chr4-136779383-INS-58
chr4-136785897-INS-83

chr4-136801254-INS-150
chr4-136807517-INS-206
chr4-136873182-INS-52
chr4-136912935-INS-399
chr4-136918724-INS-80
chr4-136943451-INS-329
chr4-136961742-INS-72
chr4-136962244-INS-138
chr4-136969993-INS-224
chr4-136988700-INS-169
chr4-136993552-INS-70
chr4-137000963-INS-78
chr4-137053694-INS-202
chr4-137067217-INS-76
chr4-137074807-INS-105
chr4-137083762-INS-192
chr4-137085182-INS-186
chr4-137107821-INS-143
chr4-137118863-INS-373
chr4-137125073-INS-150
chr4-137161027-INS-326
chr4-137174589-INS-57
chr4-137174651-INS-75
chr4-137176693-INS-163
chr4-137187610-INS-230
chr4-137217987-INS-265
chr4-137223712-INS-88
chr4-137227053-INS-56
chr4-137282098-INS-215
chr4-137295237-INS-625
chr4-137306083-INS-147
chr4-137306369-INS-230
chr4-137314202-INS-584
chr4-137314251-INS-706
chr4-137325453-INS-204
chr4-137329938-INS-84
chr4-137335262-INS-226
chr4-137362770-INS-209
chr4-137379136-INS-79
chr4-137381748-INS-64
chr4-137382629-INS-577
chr4-137389898-INS-87
chr4-137436173-INS-219
chr4-137442521-INS-59
chr4-137453675-INS-203
chr4-137461441-INS-214
chr4-137464291-INS-158
chr4-137467870-INS-206
chr4-137486750-INS-60
chr4-137489980-INS-56
chr4-137543143-INS-57
chr4-137548923-INS-88
chr4-137559287-INS-127
chr4-137559287-INS-131
chr4-137568224-INS-343
chr4-137574891-INS-222
chr4-137576649-INS-678
chr4-137595561-INS-204
chr4-137625814-INS-198
chr4-137632972-INS-143
chr4-137638054-INS-198
chr4-137639478-INS-193
chr4-137643970-INS-128
chr4-137648838-INS-272
chr4-137651957-INS-610
chr4-137688582-INS-60
chr4-137694838-INS-69
chr4-137710393-INS-132
chr4-137753925-INS-182
chr4-137758686-INS-160
chr4-137759698-INS-51
chr4-137766108-INS-74
chr4-137768761-INS-130
chr4-137769343-INS-139
chr4-137777044-INS-247
chr4-137790112-INS-176
chr4-137804134-INS-121

chr4-137861176-INS-87
chr4-137885191-INS-202
chr4-137889259-INS-168
chr4-137890997-INS-189
chr4-137897465-INS-482
chr4-137921257-INS-156
chr4-137924818-INS-78
chr4-137929008-INS-410
chr4-137932263-INS-71
chr4-137973707-INS-176
chr4-137975246-INS-209
chr4-137976533-INS-77
chr4-137993833-INS-3409
chr4-138014485-INS-91
chr4-138021891-INS-161
chr4-138025075-INS-66
chr4-138031464-INS-531
chr4-138054625-INS-159
chr4-138064422-INS-56
chr4-138097566-INS-63
chr4-138107606-INS-85
chr4-138119794-INS-72
chr4-138125777-INS-343
chr4-138137088-INS-350
chr4-138143861-INS-300
chr4-138193721-INS-140
chr4-138206901-INS-230
chr4-138242103-INS-172
chr4-138248961-INS-65
chr4-138258370-INS-232
chr4-138268878-INS-68
chr4-138275067-INS-123
chr4-138291354-INS-593
chr4-138304325-INS-53
chr4-138309535-INS-156
chr4-138318902-INS-60
chr4-138322868-INS-224
chr4-138342917-INS-105
chr4-138366771-INS-162
chr4-138388582-INS-59
chr4-138390116-INS-88
chr4-138399402-INS-70
chr4-138400204-INS-302
chr4-138409565-INS-376
chr4-138418139-INS-52
chr4-138464011-INS-107
chr4-138483136-INS-56
chr4-138486902-INS-112
chr4-138489324-INS-81
chr4-138501458-INS-63
chr4-138502730-INS-80
chr4-138516624-INS-598
chr4-138551127-INS-270
chr4-138572280-INS-160
chr4-138582349-INS-182
chr4-138599993-INS-167
chr4-138614629-INS-87
chr4-138624658-INS-140
chr4-138642994-INS-208
chr4-138662405-INS-86
chr4-138675805-INS-70
chr4-138683560-INS-66
chr4-138715216-INS-70
chr4-138716606-INS-170
chr4-138739853-INS-62
chr4-138771996-INS-426
chr4-138779173-INS-213
chr4-138804600-INS-67
chr4-138805187-INS-106
chr4-138811951-INS-51
chr4-138813577-INS-63
chr4-138833378-INS-4362
chr4-138833379-INS-359
chr4-138871821-INS-107
chr4-138872761-INS-64
chr4-138876636-INS-129
chr4-138879239-INS-174

chr4-138889556-INS-336
chr4-138896425-INS-540
chr4-138905662-INS-234
chr4-138919129-INS-201
chr4-138932379-INS-90
chr4-138934194-INS-680
chr4-138940107-INS-109
chr4-138947731-INS-243
chr4-138951989-INS-5245
chr4-138952982-INS-782
chr4-138968954-INS-206
chr4-139006831-INS-57
chr4-139042471-INS-197
chr4-139051038-INS-427
chr4-139052345-INS-424
chr4-139053720-INS-57
chr4-139056255-INS-422
chr4-139059288-INS-204
chr4-139066605-INS-104
chr4-139068586-INS-161
chr4-139082928-INS-137
chr4-139090048-INS-86
chr4-139094914-INS-217
chr4-139111953-INS-213
chr4-139115906-INS-98
chr4-139137269-INS-336
chr4-139188291-INS-54
chr4-139230795-INS-428
chr4-139231792-INS-63
chr4-139243611-INS-69
chr4-139251903-INS-52
chr4-139260693-INS-64
chr4-139263142-INS-124
chr4-139272167-INS-150
chr4-139286903-INS-99
chr4-139309815-INS-157
chr4-139316463-INS-83
chr4-139316499-INS-959
chr4-139357244-INS-60
chr4-139361971-INS-119
chr4-139414990-INS-693
chr4-139420095-INS-60
chr4-139444619-INS-66
chr4-139445357-INS-70
chr4-139446138-INS-65
chr4-139457455-INS-469
chr4-139468611-INS-176
chr4-139492303-INS-210
chr4-139509207-INS-74
chr4-139542334-INS-82
chr4-139556498-INS-62
chr4-139584457-INS-200
chr4-139593019-INS-950
chr4-139623585-INS-53
chr4-139707864-INS-179
chr4-139710404-INS-114
chr4-139724194-INS-377
chr4-139737949-INS-400
chr4-139747898-INS-74
chr4-139757738-INS-56
chr4-139770865-INS-418
chr4-139774838-INS-97
chr4-139824974-INS-198
chr4-139856451-INS-357
chr4-139874066-INS-62
chr4-139897528-INS-76
chr4-139898849-INS-98
chr4-139901624-INS-166
chr4-139909952-INS-93
chr4-139944920-INS-84
chr4-139969401-INS-53
chr4-140004022-INS-178
chr4-140016363-INS-62
chr4-140019984-INS-199
chr4-140031156-INS-68
chr4-140034571-INS-116
chr4-140038866-INS-242

chr4-140042236-INS-265
chr4-140070679-INS-112
chr4-140102287-INS-55
chr4-140116658-INS-118
chr4-140121309-INS-102
chr4-140133210-INS-81
chr4-140140855-INS-96
chr4-140150290-INS-141
chr4-140155274-INS-82
chr4-140158993-INS-85
chr4-140164637-INS-62
chr4-140175350-INS-87
chr4-140178316-INS-58
chr4-140214665-INS-62
chr4-140220140-INS-94
chr4-140236712-INS-146
chr4-140238166-INS-75
chr4-140244362-INS-830
chr4-140284729-INS-111
chr4-140298640-INS-261
chr4-140316498-INS-52
chr4-140368109-INS-114
chr4-140379244-INS-84
chr4-140410010-INS-199
chr4-140423277-INS-5649
chr4-140439058-INS-401
chr4-140442630-INS-203
chr4-140446792-INS-198
chr4-140450560-INS-706
chr4-140456961-INS-57
chr4-140461952-INS-206
chr4-140465916-INS-200
chr4-140476611-INS-64
chr4-140494360-INS-98
chr4-140509880-INS-200
chr4-140535356-INS-74
chr4-140578788-INS-80
chr4-140581271-INS-64
chr4-140593007-INS-248
chr4-140646952-INS-56
chr4-140735165-INS-279
chr4-140786764-INS-80
chr4-140806890-INS-224
chr4-140808802-INS-84
chr4-140811883-INS-255
chr4-140831456-INS-72
chr4-140834274-INS-209
chr4-140856103-INS-243
chr4-140895663-INS-798
chr4-140896200-INS-107
chr4-140897584-INS-62
chr4-140909717-INS-64
chr4-140933460-INS-73
chr4-140943496-INS-337
chr4-140968901-INS-205
chr4-141001392-INS-118
chr4-141023656-INS-129
chr4-141024947-INS-206
chr4-141029871-INS-228
chr4-141031927-INS-169
chr4-141046924-INS-201
chr4-141086684-INS-166
chr4-141103137-INS-193
chr4-141114178-INS-209
chr4-141116517-INS-180
chr4-141121568-INS-96
chr4-141139757-INS-193
chr4-141157108-INS-54
chr4-141158179-INS-166
chr4-141159586-INS-389
chr4-141160812-INS-54
chr4-141164811-INS-55
chr4-141168623-INS-97
chr4-141169822-INS-220
chr4-141208583-INS-444
chr4-141212462-INS-70
chr4-141224593-INS-204

chr4-141242395-INS-208
chr4-141271545-INS-191
chr4-141278424-INS-207
chr4-141288454-INS-211
chr4-141289482-INS-153
chr4-141294214-INS-72
chr4-141312026-INS-67
chr4-141332265-INS-223
chr4-141334391-INS-610
chr4-141334571-INS-53
chr4-141340447-INS-59
chr4-141364124-INS-164
chr4-141401048-INS-70
chr4-141402500-INS-206
chr4-141451820-INS-54
chr4-141452814-INS-197
chr4-141454119-INS-248
chr4-141455988-INS-52
chr4-141464462-INS-77
chr4-141478124-INS-247
chr4-141482421-INS-64
chr4-141485302-INS-212
chr4-141500744-INS-97
chr4-141519236-INS-227
chr4-141561124-INS-78
chr4-141569637-INS-203
chr4-141572738-INS-55
chr4-141613154-INS-5285
chr4-141616084-INS-124
chr4-141625961-INS-60
chr4-141632199-INS-54
chr4-141634654-INS-129
chr4-141676138-INS-123
chr4-141679984-INS-58
chr4-141692223-INS-653
chr4-141715646-INS-232
chr4-141721407-INS-79
chr4-141731434-INS-55
chr4-141732729-INS-694
chr4-141736067-INS-59
chr4-141757435-INS-70
chr4-141762204-INS-284
chr4-141783828-INS-399
chr4-141821360-INS-53
chr4-141826931-INS-76
chr4-141827073-INS-584
chr4-141862847-INS-5412
chr4-141882297-INS-198
chr4-141909716-INS-217
chr4-141927302-INS-54
chr4-141932475-INS-144
chr4-141935180-INS-73
chr4-141968024-INS-80
chr4-141972900-INS-138
chr4-141995443-INS-246
chr4-141997449-INS-50
chr4-141999886-INS-1022
chr4-141999887-INS-157
chr4-142000505-INS-957
chr4-142021585-INS-51
chr4-142030543-INS-51
chr4-142032527-INS-62
chr4-142036479-INS-111
chr4-142044867-INS-153
chr4-142065630-INS-56
chr4-142072991-INS-93
chr4-142073742-INS-467
chr4-142105643-INS-268
chr4-142110645-INS-734
chr4-142122866-INS-400
chr4-142151014-INS-246
chr4-142181160-INS-376
chr4-142194582-INS-135
chr4-142213848-INS-78
chr4-142222759-INS-105
chr4-142226115-INS-95
chr4-142257693-INS-85

chr4-142258841-INS-318
chr4-142267480-INS-63
chr4-142269528-INS-72
chr4-142278070-INS-58
chr4-142292814-INS-60
chr4-142296724-INS-160
chr4-142317369-INS-238
chr4-142318219-INS-410
chr4-142319022-INS-312
chr4-142321111-INS-92
chr4-142325270-INS-68
chr4-142332363-INS-135
chr4-142341074-INS-504
chr4-142347837-INS-59
chr4-142362645-INS-486
chr4-142383744-INS-2136
chr4-142398181-INS-61
chr4-142400610-INS-75
chr4-142409108-INS-53
chr4-142447450-INS-306
chr4-142458946-INS-71
chr4-142482653-INS-114
chr4-142505898-INS-85
chr4-142508348-INS-60
chr4-142527612-INS-538
chr4-142544006-INS-52
chr4-142550057-INS-373
chr4-142559935-INS-282
chr4-142567931-INS-233
chr4-142616743-INS-343
chr4-142629974-INS-87
chr4-142635159-INS-171
chr4-142653338-INS-330
chr4-142674807-INS-94
chr4-142678727-INS-360
chr4-142702858-INS-239
chr4-142716348-INS-60
chr4-142719043-INS-132
chr4-142755984-INS-410
chr4-142766368-INS-7522
chr4-142808591-INS-655
chr4-142817679-INS-186
chr4-142832061-INS-76
chr4-142850438-INS-440
chr4-142850698-INS-843
chr4-142852283-INS-83
chr4-142910319-INS-106
chr4-142911662-INS-362
chr4-142925868-INS-51
chr4-142930275-INS-312
chr4-142942689-INS-112
chr4-142945297-INS-74
chr4-142947019-INS-185
chr4-142949949-INS-213
chr4-142954581-INS-508
chr4-142958475-INS-95
chr4-142967797-INS-196
chr4-142973444-INS-70
chr4-142974689-INS-169
chr4-142978645-INS-210
chr4-142985087-INS-124
chr4-142986997-INS-241
chr4-142997032-INS-214
chr4-143002084-INS-66
chr4-143007644-INS-78
chr4-143009735-INS-54
chr4-143014454-INS-61
chr4-143034339-INS-106
chr4-143036109-INS-276
chr4-143037700-INS-51
chr4-143044107-INS-76
chr4-143054059-INS-547
chr4-143091242-INS-291
chr4-143110470-INS-185
chr4-143111307-INS-385
chr4-143118482-INS-85
chr4-143128775-INS-889

chr4-143130103-INS-113
chr4-143132871-INS-52
chr4-143184573-INS-347
chr4-143218602-INS-614
chr4-143219846-INS-5604
chr4-143221094-INS-52
chr4-143291011-INS-204
chr4-143386788-INS-76
chr4-143395836-INS-602
chr4-143452104-INS-6386
chr4-143471486-INS-586
chr4-143503601-INS-5158
chr4-143510505-INS-201
chr4-143574754-INS-497
chr4-143631117-INS-59
chr4-143698071-INS-6220
chr4-143714726-INS-75
chr4-143742826-INS-64
chr4-143775755-INS-189
chr4-143788840-INS-78
chr4-143806583-INS-187
chr4-143842502-INS-167
chr4-143868379-INS-8219
chr4-143875370-INS-201
chr4-143891578-INS-62
chr4-143913037-INS-212
chr4-143918154-INS-5609
chr4-143942479-INS-5345
chr4-143966343-INS-871
chr4-143972665-INS-57
chr4-143974073-INS-82
chr4-143981681-INS-102
chr4-143989579-INS-680
chr4-143995617-INS-201
chr4-144012055-INS-87
chr4-144017332-INS-224
chr4-144028287-INS-60
chr4-144035095-INS-195
chr4-144036145-INS-208
chr4-144049292-INS-209
chr4-144080531-INS-343
chr4-144091659-INS-445
chr4-144150699-INS-515
chr4-144156178-INS-151
chr4-144158407-INS-220
chr4-144194505-INS-74
chr4-144206191-INS-144
chr4-144227718-INS-486
chr4-144288839-INS-6982
chr4-144302703-INS-6244
chr4-144381723-INS-179
chr4-144424371-INS-477
chr4-144487618-INS-7026
chr4-144562802-INS-182
chr4-144584796-INS-94
chr4-144587569-INS-202
chr4-144613591-INS-225
chr4-144622293-INS-146
chr4-144662353-INS-80
chr4-144663049-INS-849
chr4-144671897-INS-84
chr4-144696234-INS-53
chr4-144697032-INS-88
chr4-144772727-INS-80
chr4-144799620-INS-79
chr4-144836938-INS-973
chr4-144893540-INS-114
chr4-144897485-INS-69
chr4-144930513-INS-51
chr4-144938110-INS-5347
chr4-144956177-INS-327
chr4-144986257-INS-90
chr4-144999056-INS-58
chr4-145000228-INS-53
chr4-145014528-INS-190
chr4-145038431-INS-64
chr4-145046451-INS-1455

chr4-145046452-INS-866
chr4-145047532-INS-328
chr4-145049649-INS-100
chr4-145073540-INS-757
chr4-145078570-INS-193
chr4-145088361-INS-321
chr4-145128729-INS-5413
chr4-145305812-INS-5233
chr4-145316193-INS-301
chr4-145334132-INS-348
chr4-145386029-INS-7116
chr4-145428989-INS-209
chr4-145438199-INS-62
chr4-145445365-INS-171
chr4-145481980-INS-4336
chr4-145538961-INS-7113
chr4-146050959-INS-291
chr4-146051998-INS-86
chr4-146069228-INS-77
chr4-146076030-INS-72
chr4-146153383-INS-1227
chr4-146153384-INS-120
chr4-146154293-INS-55
chr4-146158005-INS-361
chr4-146186289-INS-4336
chr4-146187455-INS-7075
chr4-146191960-INS-136
chr4-146205757-INS-76
chr4-146228796-INS-7113
chr4-146258684-INS-53
chr4-146266755-INS-348
chr4-146318570-INS-421
chr4-146355037-INS-92
chr4-146358078-INS-221
chr4-146376292-INS-6817
chr4-146376390-INS-2083
chr4-146381603-INS-276
chr4-146389018-INS-218
chr4-146412564-INS-345
chr4-146415545-INS-110
chr4-146425705-INS-189
chr4-146512394-INS-7094
chr4-146548276-INS-7123
chr4-146566284-INS-5179
chr4-146574276-INS-222
chr4-146577814-INS-198
chr4-146620038-INS-218
chr4-146622061-INS-336
chr4-146641215-INS-140
chr4-146646308-INS-211
chr4-146799339-INS-202
chr4-146813475-INS-75
chr4-146821920-INS-485
chr4-146831312-INS-68
chr4-146893620-INS-79
chr4-146896984-INS-65
chr4-146913219-INS-84
chr4-146995966-INS-755
chr4-147088905-INS-52
chr4-147127660-INS-56
chr4-147143173-INS-252
chr4-147194325-INS-387
chr4-147201961-INS-137
chr4-147254851-INS-3990
chr4-147272480-INS-181
chr4-147278473-INS-221
chr4-147315454-INS-8578
chr4-147317941-INS-51
chr4-147327157-INS-582
chr4-147328190-INS-348
chr4-147337208-INS-7100
chr4-147401371-INS-2083
chr4-147451055-INS-5621
chr4-147467488-INS-55
chr4-147511368-INS-99
chr4-147520148-INS-225
chr4-147529370-INS-970

chr4-147577260-INS-513
chr4-147661137-INS-3290
chr4-147719897-INS-3649
chr4-147758912-INS-84
chr4-147759389-INS-167
chr4-147769004-INS-150
chr4-147823710-INS-620
chr4-147839024-INS-126
chr4-147855333-INS-199
chr4-147855877-INS-3208
chr4-147878952-INS-7040
chr4-147904447-INS-172
chr4-147928784-INS-148
chr4-147937668-INS-661
chr4-147944771-INS-215
chr4-147948940-INS-176
chr4-147954886-INS-90
chr4-147978762-INS-164
chr4-148011691-INS-168
chr4-148012746-INS-92
chr4-148014741-INS-186
chr4-148019743-INS-97
chr4-148020771-INS-8454
chr4-148042417-INS-224
chr4-148056639-INS-58
chr4-148061118-INS-244
chr4-148076289-INS-197
chr4-148088040-INS-203
chr4-148105075-INS-76
chr4-148129845-INS-56
chr4-148131021-INS-166
chr4-148131667-INS-242
chr4-148150318-INS-249
chr4-148160405-INS-190
chr4-148168506-INS-801
chr4-148179817-INS-912
chr4-148193840-INS-205
chr4-148195806-INS-54
chr4-148204339-INS-210
chr4-148205759-INS-90
chr4-148232298-INS-174
chr4-148282873-INS-75
chr4-148285553-INS-326
chr4-148298779-INS-392
chr4-148301450-INS-463
chr4-148310172-INS-114
chr4-148325647-INS-67
chr4-148331241-INS-124
chr4-148349744-INS-441
chr4-148361163-INS-86
chr4-148392865-INS-84
chr4-148408242-INS-55
chr4-148415969-INS-56
chr4-148423463-INS-95
chr4-148455652-INS-151
chr4-148471862-INS-85
chr4-148505752-INS-113
chr4-148507266-INS-188
chr4-148509153-INS-401
chr4-148546286-INS-212
chr4-148601695-INS-79
chr4-148607036-INS-221
chr4-148611336-INS-91
chr4-148625733-INS-197
chr4-148633448-INS-201
chr4-148652511-INS-237
chr4-148674664-INS-174
chr4-148680126-INS-253
chr4-148691393-INS-224
chr4-148712236-INS-235
chr4-148714333-INS-202
chr4-148746374-INS-203
chr4-148765540-INS-168
chr4-148767723-INS-83
chr4-148773211-INS-51
chr4-148774191-INS-420
chr4-148781998-INS-53

chr4-148788758-INS-94
chr4-148795214-INS-63
chr4-148798943-INS-52
chr4-148805596-INS-186
chr4-148836562-INS-115
chr4-148842757-INS-212
chr4-148849906-INS-443
chr4-148854346-INS-1893
chr4-148897632-INS-58
chr4-149012334-INS-130
chr4-149038110-INS-54
chr4-149067229-INS-237
chr4-149080577-INS-308
chr4-149099447-INS-70
chr4-149103734-INS-54
chr4-149127011-INS-213
chr4-149165485-INS-148
chr4-149173002-INS-266
chr4-149177983-INS-135
chr4-149187952-INS-54
chr4-149190480-INS-219
chr4-149211807-INS-87
chr4-149224828-INS-880
chr4-149231875-INS-53
chr4-149236909-INS-239
chr4-149237084-INS-568
chr4-149245994-INS-217
chr4-149253368-INS-195
chr4-149254302-INS-60
chr4-149259162-INS-174
chr4-149267821-INS-176
chr4-149273761-INS-216
chr4-149276549-INS-186
chr4-149281868-INS-965
chr4-149283537-INS-463
chr4-149286267-INS-170
chr4-149292200-INS-202
chr4-149294021-INS-61
chr4-149296832-INS-208
chr4-149298985-INS-216
chr4-149314751-INS-175
chr4-149317990-INS-187
chr4-149328347-INS-184
chr4-149360151-INS-158
chr4-149369735-INS-61
chr4-149371366-INS-76
chr4-149394429-INS-203
chr4-149396241-INS-56
chr4-149411415-INS-213
chr4-149416563-INS-79
chr4-149432926-INS-110
chr4-149439340-INS-356
chr4-149465101-INS-97
chr4-149479393-INS-489
chr4-149480551-INS-97
chr4-149481038-INS-579
chr4-149485944-INS-196
chr4-149494647-INS-353
chr4-149495796-INS-216
chr4-149496394-INS-192
chr4-149496912-INS-465
chr4-149498111-INS-136
chr4-149512730-INS-6525
chr4-149515016-INS-222
chr4-149515942-INS-5245
chr4-149516626-INS-198
chr4-149526282-INS-202
chr4-149536094-INS-199
chr4-149537502-INS-163
chr4-149538237-INS-198
chr4-149543927-INS-441
chr4-149551352-INS-117
chr4-149554742-INS-197
chr4-149555788-INS-208
chr4-149557829-INS-181
chr4-149558569-INS-56
chr4-149562738-INS-141

chr4-149567644-INS-174
chr4-149581633-INS-93
chr4-149583143-INS-153
chr4-149586423-INS-54
chr4-149591778-INS-64
chr4-149593628-INS-193
chr4-149594320-INS-202
chr4-149599238-INS-216
chr4-149619734-INS-83
chr4-149628251-INS-167
chr4-149665921-INS-204
chr4-149674244-INS-75
chr4-149677216-INS-129
chr4-149702198-INS-210
chr4-149707560-INS-289
chr4-149720491-INS-172
chr4-149721982-INS-201
chr4-149728717-INS-218
chr4-149754099-INS-227
chr4-149768810-INS-190
chr4-149770315-INS-212
chr4-149771571-INS-161
chr4-149779896-INS-232
chr4-149788573-INS-165
chr4-149791075-INS-71
chr4-149793081-INS-211
chr4-149797794-INS-54
chr4-149814615-INS-171
chr4-149834862-INS-212
chr4-149841826-INS-118
chr4-149844317-INS-242
chr4-149850759-INS-114
chr4-149867447-INS-142
chr4-149873531-INS-192
chr4-149885958-INS-55
chr4-149895212-INS-1224
chr4-149895269-INS-571
chr4-149903607-INS-154
chr4-149909101-INS-64
chr4-149913366-INS-738
chr4-149920646-INS-307
chr4-149927275-INS-60
chr4-149951112-INS-116
chr4-149956974-INS-136
chr4-149976266-INS-80
chr4-149984429-INS-105
chr4-150058334-INS-244
chr4-150076247-INS-119
chr4-150105681-INS-576
chr4-150111787-INS-233
chr4-150131462-INS-63
chr4-150143978-INS-129
chr4-150148386-INS-201
chr4-150159068-INS-108
chr4-150180259-INS-168
chr4-150184310-INS-215
chr4-150190828-INS-90
chr4-150211392-INS-145
chr4-150236196-INS-240
chr4-150254370-INS-891
chr4-150254584-INS-52
chr4-150255096-INS-266
chr4-150264591-INS-225
chr4-150267970-INS-174
chr4-150274648-INS-62
chr4-150279042-INS-208
chr4-150287779-INS-210
chr4-150300196-INS-58
chr4-150303634-INS-150
chr4-150304694-INS-198
chr4-150309082-INS-211
chr4-150310032-INS-183
chr4-150337053-INS-62
chr4-150352443-INS-204
chr4-150353969-INS-307
chr4-150361189-INS-178
chr4-150362064-INS-182

chr4-150364412-INS-214
chr4-150377039-INS-132
chr4-150388117-INS-221
chr4-150405227-INS-216
chr4-150409221-INS-123
chr4-150412226-INS-127
chr4-150426088-INS-59
chr4-150429848-INS-278
chr4-150435395-INS-219
chr4-150441720-INS-92
chr4-150445505-INS-204
chr4-150464946-INS-54
chr4-150472641-INS-464
chr4-150482478-INS-371
chr4-150486982-INS-192
chr4-150506172-INS-498
chr4-150517701-INS-94
chr4-150525236-INS-111
chr4-150529742-INS-56
chr4-150550645-INS-286
chr4-150556343-INS-570
chr4-150562811-INS-123
chr4-150570351-INS-209
chr4-150582870-INS-174
chr4-150588494-INS-177
chr4-150589005-INS-93
chr4-150590183-INS-58
chr4-150598522-INS-526
chr4-150608925-INS-1499
chr4-150615050-INS-205
chr4-150624567-INS-724
chr4-150643910-INS-193
chr4-150692189-INS-215
chr4-150721519-INS-52
chr4-150723561-INS-296
chr4-150738382-INS-67
chr4-150752063-INS-212
chr4-150753800-INS-201
chr4-150758499-INS-473
chr4-150770421-INS-59
chr4-150776818-INS-128
chr4-150778838-INS-176
chr4-150786546-INS-392
chr4-150799985-INS-80
chr4-150811319-INS-868
chr4-150824002-INS-193
chr4-150829105-INS-209
chr4-150831293-INS-196
chr4-150838288-INS-116
chr4-150846576-INS-52
chr4-150859523-INS-51
chr4-150861922-INS-72
chr4-150886800-INS-534
chr4-150953708-INS-76
chr4-150966658-INS-153
chr4-150976048-INS-436
chr4-150978140-INS-113
chr4-151000656-INS-204
chr4-151004522-INS-76
chr4-151014767-INS-103
chr4-151022303-INS-8326
chr4-151032881-INS-203
chr4-151039794-INS-411
chr4-151058729-INS-84
chr4-151072772-INS-259
chr4-151165694-INS-55
chr4-151169818-INS-148
chr4-151172038-INS-76
chr4-151208260-INS-170
chr4-151238525-INS-186
chr4-151263956-INS-380
chr4-151274672-INS-118
chr4-151317928-INS-109
chr4-151323910-INS-70
chr4-151331342-INS-467
chr4-151361090-INS-135
chr4-151393634-INS-106

chr4-151402655-INS-55
chr4-151415586-INS-70
chr4-151434818-INS-76
chr4-151459651-INS-67
chr4-151490964-INS-66
chr4-151516609-INS-390
chr4-151554828-INS-114
chr4-151569548-INS-263
chr4-151571973-INS-72
chr4-151584394-INS-118
chr4-151616690-INS-132
chr4-151617129-INS-6074
chr4-151656281-INS-174
chr4-151683612-INS-54
chr4-151706350-INS-81
chr4-151708739-INS-74
chr4-151726711-INS-104
chr4-151733961-INS-148
chr4-151737196-INS-55
chr4-151742240-INS-84
chr4-151752333-INS-151
chr4-151769497-INS-80
chr4-151784032-INS-179
chr4-151790294-INS-126
chr4-151822725-INS-542
chr4-151829552-INS-174
chr4-151880752-INS-69
chr4-151885578-INS-231
chr4-151887233-INS-168
chr4-151895780-INS-51
chr4-151926554-INS-213
chr4-151929946-INS-267
chr4-151931088-INS-103
chr4-151950489-INS-78
chr4-151957184-INS-126
chr4-151973763-INS-232
chr4-151975650-INS-580
chr4-151979374-INS-268
chr4-151982704-INS-58
chr4-151998600-INS-79
chr4-152006940-INS-131
chr4-152008959-INS-58
chr4-152018003-INS-69
chr4-152025225-INS-81
chr4-152026806-INS-99
chr4-152042041-INS-167
chr4-152043006-INS-196
chr4-152059296-INS-168
chr4-152066928-INS-208
chr4-152072660-INS-84
chr4-152085639-INS-624
chr4-152085648-INS-931
chr4-152088821-INS-6319
chr4-152101404-INS-54
chr4-152120671-INS-197
chr4-152139558-INS-396
chr4-152144208-INS-260
chr4-152165189-INS-52
chr4-152166185-INS-229
chr4-152167170-INS-89
chr4-152179827-INS-276
chr4-152180850-INS-62
chr4-152191969-INS-223
chr4-152196448-INS-151
chr4-152240504-INS-188
chr4-152241928-INS-113
chr4-152244209-INS-173
chr4-152247189-INS-180
chr4-152256337-INS-197
chr4-152256716-INS-456
chr4-152269723-INS-201
chr4-152271716-INS-72
chr4-152282794-INS-162
chr4-152323548-INS-141
chr4-152328606-INS-62
chr4-152330195-INS-127
chr4-152331792-INS-72

chr4-152346134-INS-208
chr4-152373624-INS-152
chr4-152373687-INS-220
chr4-152380822-INS-60
chr4-152395566-INS-216
chr4-152401859-INS-115
chr4-152406681-INS-56
chr4-152412937-INS-212
chr4-152426420-INS-213
chr4-152448295-INS-169
chr4-152452062-INS-113
chr4-152470744-INS-168
chr4-152528557-INS-59
chr4-152553268-INS-530
chr4-152556470-INS-170
chr4-152559935-INS-83
chr4-152612233-INS-73
chr4-152612966-INS-670
chr4-152640216-INS-102
chr4-152640377-INS-188
chr4-152651706-INS-97
chr4-152654813-INS-264
chr4-152669164-INS-76
chr4-152675591-INS-668
chr4-152678042-INS-333
chr4-152686391-INS-63
chr4-152693854-INS-90
chr4-152703178-INS-195
chr4-152743777-INS-327
chr4-152752806-INS-122
chr4-152781889-INS-60
chr4-152787134-INS-315
chr4-152790765-INS-79
chr4-152841880-INS-344
chr4-152847050-INS-322
chr4-152873564-INS-437
chr4-152898137-INS-311
chr4-152899762-INS-54
chr4-152900682-INS-77
chr4-152956606-INS-57
chr4-152965234-INS-127
chr4-152982104-INS-285
chr4-152994487-INS-54
chr4-152999059-INS-107
chr4-153012133-INS-75
chr4-153016367-INS-184
chr4-153024142-INS-54
chr4-153040922-INS-52
chr4-153043563-INS-72
chr4-153093011-INS-1020
chr4-153131194-INS-532
chr4-153141878-INS-57
chr4-153143523-INS-74
chr4-153143895-INS-401
chr4-153150298-INS-69
chr4-153151395-INS-73
chr4-153153307-INS-54
chr4-153172186-INS-149
chr4-153179864-INS-86
chr4-153205993-INS-56
chr4-153210949-INS-80
chr4-153215295-INS-186
chr4-153218609-INS-151
chr4-153246993-INS-128
chr4-153258387-INS-234
chr4-153263412-INS-214
chr4-153268773-INS-53
chr4-153284762-INS-323
chr4-153298865-INS-78
chr4-153303878-INS-61
chr4-153318172-INS-115
chr4-153321208-INS-197
chr4-153324953-INS-256
chr4-153337849-INS-111
chr4-153349377-INS-290
chr4-153351838-INS-72
chr4-153352859-INS-102

chr4-153357500-INS-134
chr4-153368047-INS-120
chr4-153377884-INS-72
chr4-153395444-INS-118
chr4-153402189-INS-94
chr4-153411896-INS-62
chr4-153422924-INS-109
chr4-153433470-INS-119
chr4-153434295-INS-86
chr4-153437155-INS-149
chr4-153453692-INS-67
chr4-153467502-INS-230
chr4-153481982-INS-174
chr4-153488643-INS-212
chr4-153502741-INS-223
chr4-153526043-INS-134
chr4-153529207-INS-81
chr4-153570327-INS-61
chr4-153572530-INS-97
chr4-153573753-INS-65
chr4-153581499-INS-59
chr4-153593408-INS-138
chr4-153595214-INS-56
chr4-153601230-INS-108
chr4-153639747-INS-66
chr4-153651800-INS-99
chr4-153652515-INS-100
chr4-153658859-INS-343
chr4-153664138-INS-55
chr4-153665205-INS-191
chr4-153669723-INS-54
chr4-153671809-INS-52
chr4-153679103-INS-440
chr4-153691000-INS-1790
chr4-153702419-INS-120
chr4-153703640-INS-85
chr4-153708084-INS-51
chr4-153741753-INS-64
chr4-153748787-INS-54
chr4-153753440-INS-82
chr4-153754762-INS-62
chr4-153756058-INS-65
chr4-153758212-INS-103
chr4-153760011-INS-172
chr4-153767238-INS-54
chr4-153784468-INS-497
chr4-153803575-INS-52
chr4-153804706-INS-289
chr4-153817129-INS-77
chr4-153846287-INS-294
chr4-153854863-INS-404
chr4-153863665-INS-59
chr4-153867665-INS-150
chr4-153876589-INS-116
chr4-153884759-INS-77
chr4-153890104-INS-236
chr4-153894772-INS-259
chr4-153903009-INS-73
chr4-153905520-INS-89
chr4-153908420-INS-430
chr4-153912593-INS-197
chr4-153927942-INS-60
chr4-153981589-INS-64
chr4-153983691-INS-341
chr4-153987219-INS-389
chr4-154004137-INS-177
chr4-154005868-INS-145
chr4-154011465-INS-114
chr4-154013178-INS-91
chr4-154016967-INS-68
chr4-154030273-INS-205
chr4-154033492-INS-57
chr4-154052591-INS-250
chr4-154054808-INS-910
chr4-154066505-INS-970
chr4-154068196-INS-66
chr4-154073277-INS-133

chr4-154097109-INS-56
chr4-154100079-INS-126
chr4-154116410-INS-60
chr4-154118855-INS-76
chr4-154122398-INS-120
chr4-154124379-INS-165
chr4-154135145-INS-149
chr4-154157534-INS-56
chr4-154158454-INS-775
chr4-154158454-INS-776
chr4-154177368-INS-72
chr4-154178079-INS-831
chr4-154192848-INS-233
chr4-154209309-INS-56
chr4-154218220-INS-82
chr4-154219784-INS-5544
chr4-154221584-INS-70
chr4-154232642-INS-229
chr4-154233750-INS-232
chr4-154281743-INS-77
chr4-154292770-INS-182
chr4-154338459-INS-102
chr4-154359259-INS-129
chr4-154373195-INS-167
chr4-154373779-INS-139
chr4-154396741-INS-119
chr4-154398691-INS-118
chr4-154462647-INS-92
chr4-154516149-INS-513
chr4-154560548-INS-146
chr4-154589685-INS-204
chr4-154610695-INS-129
chr4-154622504-INS-248
chr4-154647166-INS-66
chr4-154707551-INS-56
chr4-154757301-INS-64
chr4-154764885-INS-187
chr4-154767509-INS-108
chr4-154776207-INS-57
chr4-154779729-INS-86
chr4-154818480-INS-52
chr4-154828654-INS-5318
chr4-154898560-INS-99
chr4-154903915-INS-140
chr4-154907047-INS-54
chr4-154924166-INS-88
chr4-154957121-INS-227
chr4-154960206-INS-452
chr4-154963983-INS-340
chr4-154969741-INS-705
chr4-154996692-INS-229
chr4-155022016-INS-69
chr4-155023484-INS-236
chr4-155026641-INS-1008
chr4-155026642-INS-793
chr4-155029344-INS-83
chr4-155029923-INS-60
chr4-155073331-INS-78
chr4-155085820-INS-5434
chr4-155088160-INS-204
chr4-155101158-INS-52
chr4-155111022-INS-257
chr4-155112470-INS-98
chr4-155119795-INS-211
chr4-155121631-INS-219
chr4-155123643-INS-67
chr4-155195765-INS-86
chr4-155201063-INS-52
chr4-155221256-INS-146
chr4-155253383-INS-129
chr4-155265882-INS-224
chr4-155273976-INS-93
chr4-155281052-INS-199
chr4-155301174-INS-444
chr4-155313433-INS-270
chr4-155325124-INS-55
chr4-155343931-INS-218

chr4-155358300-INS-1220
chr4-155358387-INS-319
chr4-155364187-INS-288
chr4-155381848-INS-177
chr4-155390756-INS-58
chr4-155405889-INS-98
chr4-155411701-INS-64
chr4-155419158-INS-229
chr4-155423135-INS-203
chr4-155434985-INS-1445
chr4-155439930-INS-400
chr4-155455193-INS-222
chr4-155458183-INS-348
chr4-155480621-INS-192
chr4-155487456-INS-141
chr4-155488399-INS-182
chr4-155489751-INS-76
chr4-155494467-INS-244
chr4-155510991-INS-78
chr4-155526456-INS-103
chr4-155527864-INS-51
chr4-155535178-INS-182
chr4-155537688-INS-102
chr4-155543613-INS-76
chr4-155548934-INS-233
chr4-155551829-INS-105
chr4-155559987-INS-204
chr4-155564104-INS-70
chr4-155599463-INS-122
chr4-155624121-INS-60
chr4-155635076-INS-53
chr4-155635600-INS-215
chr4-155651492-INS-131
chr4-155656719-INS-71
chr4-155664297-INS-84
chr4-155678537-INS-324
chr4-155682080-INS-347
chr4-155689253-INS-537
chr4-155694090-INS-107
chr4-155698919-INS-143
chr4-155719369-INS-193
chr4-155729802-INS-557
chr4-155747690-INS-1268
chr4-155756314-INS-213
chr4-155764360-INS-1860
chr4-155772701-INS-74
chr4-155775002-INS-201
chr4-155784840-INS-631
chr4-155806178-INS-75
chr4-155842731-INS-215
chr4-155865124-INS-361
chr4-155872807-INS-203
chr4-155900511-INS-52
chr4-155915081-INS-184
chr4-155922125-INS-55
chr4-155932895-INS-214
chr4-155944152-INS-563
chr4-155954542-INS-223
chr4-155958979-INS-209
chr4-155960206-INS-202
chr4-155961486-INS-189
chr4-155966557-INS-209
chr4-155982401-INS-191
chr4-155991742-INS-55
chr4-155993975-INS-198
chr4-156043815-INS-455
chr4-156061341-INS-207
chr4-156077211-INS-196
chr4-156086244-INS-468
chr4-156095351-INS-90
chr4-156126132-INS-140
chr4-156135549-INS-54
chr4-156171590-INS-314
chr4-156180460-INS-74
chr4-156194837-INS-201
chr4-156202254-INS-200
chr4-156222225-INS-99

chr4-156245653-INS-122
chr4-156296569-INS-246
chr4-156302625-INS-396
chr4-156321228-INS-396
chr4-156322765-INS-383
chr4-156337941-INS-147
chr4-156387808-INS-205
chr4-156398665-INS-146
chr4-156404212-INS-55
chr4-156452096-INS-356
chr4-156485857-INS-220
chr4-156505945-INS-68
chr4-156593130-INS-141
chr4-156603744-INS-197
chr4-156702169-INS-220
chr5-3066995-INS-90
chr5-3086606-INS-66
chr5-3197426-INS-58
chr5-3203998-INS-65
chr5-3207217-INS-95
chr5-3217160-INS-70
chr5-3236617-INS-402
chr5-3267369-INS-59
chr5-3269616-INS-3454
chr5-3271604-INS-542
chr5-3291805-INS-58
chr5-3297174-INS-7178
chr5-3316497-INS-178
chr5-3317998-INS-698
chr5-3333819-INS-71
chr5-3361960-INS-406
chr5-3419693-INS-162
chr5-3429053-INS-82
chr5-3430870-INS-75
chr5-3435296-INS-202
chr5-3440336-INS-170
chr5-3449471-INS-54
chr5-3453243-INS-354
chr5-3459052-INS-267
chr5-3475445-INS-683
chr5-3476961-INS-85
chr5-3484612-INS-250
chr5-3512624-INS-118
chr5-3528569-INS-144
chr5-3539182-INS-848
chr5-3551770-INS-56
chr5-3565891-INS-160
chr5-3569868-INS-4108
chr5-3585073-INS-600
chr5-3585472-INS-73
chr5-3610456-INS-137
chr5-3652846-INS-153
chr5-3653919-INS-258
chr5-3657001-INS-197
chr5-3659756-INS-86
chr5-3663194-INS-260
chr5-3663559-INS-95
chr5-3695071-INS-56
chr5-3696775-INS-106
chr5-3705847-INS-115
chr5-3709980-INS-105
chr5-3721881-INS-213
chr5-3732934-INS-151
chr5-3737530-INS-171
chr5-3766869-INS-65
chr5-3771698-INS-210
chr5-3787805-INS-60
chr5-3798826-INS-6354
chr5-3924738-INS-7056
chr5-3930685-INS-196
chr5-3960361-INS-109
chr5-3983819-INS-505
chr5-3999191-INS-72
chr5-4012835-INS-416
chr5-4015398-INS-100
chr5-4027773-INS-442
chr5-4037546-INS-848

chr5-4071119-INS-82
chr5-4080242-INS-504
chr5-4102389-INS-3926
chr5-4102612-INS-106
chr5-4102613-INS-311
chr5-4102777-INS-4698
chr5-4103418-INS-2715
chr5-4104177-INS-379
chr5-4107863-INS-123
chr5-4108175-INS-692
chr5-4109857-INS-400
chr5-4117311-INS-945
chr5-4117323-INS-53
chr5-4119641-INS-124
chr5-4138362-INS-186
chr5-4141333-INS-59
chr5-4145516-INS-74
chr5-4174899-INS-204
chr5-4219510-INS-324
chr5-4234571-INS-665
chr5-4243931-INS-401
chr5-4277232-INS-182
chr5-4282246-INS-289
chr5-4283760-INS-64
chr5-4286804-INS-87
chr5-4288424-INS-68
chr5-4309809-INS-100
chr5-4328451-INS-481
chr5-4351579-INS-86
chr5-4356377-INS-58
chr5-4388707-INS-231
chr5-4413641-INS-185
chr5-4417091-INS-2792
chr5-4429830-INS-54
chr5-4447110-INS-7173
chr5-4475581-INS-52
chr5-4518880-INS-324
chr5-4526074-INS-85
chr5-4539110-INS-52
chr5-4550307-INS-55
chr5-4588363-INS-50
chr5-4588418-INS-64
chr5-4597956-INS-151
chr5-4607215-INS-60
chr5-4613484-INS-361
chr5-4616223-INS-487
chr5-4711498-INS-380
chr5-4713022-INS-665
chr5-4733142-INS-65
chr5-4773767-INS-336
chr5-4818239-INS-401
chr5-4843489-INS-70
chr5-4863240-INS-153
chr5-4916650-INS-115
chr5-4953944-INS-407
chr5-4962676-INS-197
chr5-5005480-INS-57
chr5-5014481-INS-795
chr5-5026168-INS-100
chr5-5037744-INS-243
chr5-5078144-INS-176
chr5-5096036-INS-94
chr5-5133400-INS-64
chr5-5136131-INS-105
chr5-5193193-INS-912
chr5-5198829-INS-88
chr5-5204357-INS-61
chr5-5215394-INS-55
chr5-5237150-INS-2802
chr5-5258918-INS-88
chr5-5269996-INS-7924
chr5-5279107-INS-228
chr5-5325812-INS-175
chr5-5332239-INS-53
chr5-5355929-INS-59
chr5-5361623-INS-88
chr5-5367133-INS-54

chr5-5371474-INS-739
chr5-5378671-INS-159
chr5-5382341-INS-95
chr5-5393211-INS-207
chr5-5405978-INS-328
chr5-5406965-INS-69
chr5-5432153-INS-127
chr5-5441279-INS-68
chr5-5443285-INS-107
chr5-5445679-INS-326
chr5-5448320-INS-107
chr5-5458159-INS-427
chr5-5461374-INS-52
chr5-5463939-INS-78
chr5-5548089-INS-70
chr5-5549992-INS-1491
chr5-5578462-INS-94
chr5-5582284-INS-660
chr5-5592224-INS-259
chr5-5610564-INS-288
chr5-5628523-INS-516
chr5-5667624-INS-169
chr5-5672370-INS-7347
chr5-5676695-INS-73
chr5-5677424-INS-406
chr5-5711314-INS-234
chr5-5735668-INS-403
chr5-5802546-INS-470
chr5-5834193-INS-87
chr5-5835768-INS-159
chr5-5861656-INS-60
chr5-5867589-INS-59
chr5-5885975-INS-67
chr5-5888648-INS-506
chr5-5909228-INS-58
chr5-5910080-INS-537
chr5-5920322-INS-252
chr5-5930146-INS-124
chr5-5966367-INS-82
chr5-5968039-INS-81
chr5-5968282-INS-626
chr5-5989269-INS-483
chr5-6005146-INS-6390
chr5-6010648-INS-52
chr5-6010984-INS-55
chr5-6012233-INS-125
chr5-6030257-INS-63
chr5-6030874-INS-343
chr5-6042312-INS-2622
chr5-6044002-INS-67
chr5-6048257-INS-6361
chr5-6052983-INS-60
chr5-6066266-INS-65
chr5-6066371-INS-6378
chr5-6071882-INS-553
chr5-6081607-INS-80
chr5-6089578-INS-103
chr5-6103849-INS-377
chr5-6118548-INS-80
chr5-6120779-INS-120
chr5-6197099-INS-188
chr5-6211791-INS-832
chr5-6226044-INS-84
chr5-6228836-INS-6261
chr5-6245251-INS-58
chr5-6257990-INS-646
chr5-6268907-INS-573
chr5-6269631-INS-874
chr5-6270562-INS-142
chr5-6288164-INS-52
chr5-6294666-INS-8350
chr5-6304306-INS-60
chr5-6329525-INS-62
chr5-6331856-INS-405
chr5-6337994-INS-219
chr5-6343306-INS-8000
chr5-6348957-INS-519

chr5-6369940-INS-509
chr5-6390196-INS-5513
chr5-6394524-INS-10891
chr5-6455468-INS-6383
chr5-6490903-INS-3412
chr5-6501317-INS-505
chr5-6515013-INS-119
chr5-6521950-INS-198
chr5-6524523-INS-1460
chr5-6530350-INS-250
chr5-6543541-INS-173
chr5-6580642-INS-178
chr5-6583423-INS-6440
chr5-6595860-INS-5164
chr5-6600674-INS-374
chr5-6618412-INS-357
chr5-6623097-INS-539
chr5-6624067-INS-7109
chr5-6636616-INS-6209
chr5-6650130-INS-6463
chr5-6670441-INS-165
chr5-6678305-INS-184
chr5-6697639-INS-503
chr5-6719778-INS-3280
chr5-6765521-INS-400
chr5-6779195-INS-528
chr5-6803037-INS-401
chr5-6863431-INS-51
chr5-6883689-INS-82
chr5-6900282-INS-320
chr5-6928515-INS-447
chr5-6933774-INS-243
chr5-6935882-INS-295
chr5-6980815-INS-355
chr5-6992418-INS-62
chr5-7010626-INS-188
chr5-7032906-INS-52
chr5-7036142-INS-52
chr5-7042919-INS-51
chr5-7053991-INS-506
chr5-7060879-INS-7258
chr5-7062293-INS-108
chr5-7074815-INS-2870
chr5-7077982-INS-56
chr5-7093999-INS-1412
chr5-7101424-INS-95
chr5-7103917-INS-78
chr5-7108083-INS-1779
chr5-7122767-INS-6238
chr5-7123389-INS-74
chr5-7128672-INS-926
chr5-7178378-INS-60
chr5-7265580-INS-100
chr5-7432062-INS-60
chr5-7463936-INS-200
chr5-7494349-INS-5233
chr5-7503661-INS-157
chr5-7556680-INS-7102
chr5-7571511-INS-181
chr5-7571587-INS-6477
chr5-7588471-INS-60
chr5-7626131-INS-4523
chr5-7635043-INS-105
chr5-7675612-INS-464
chr5-7726959-INS-789
chr5-7761409-INS-7102
chr5-7764569-INS-116
chr5-7767528-INS-52
chr5-7789122-INS-419
chr5-7795282-INS-980
chr5-7796781-INS-2825
chr5-7824304-INS-952
chr5-7869418-INS-64
chr5-7881625-INS-189
chr5-7899852-INS-185
chr5-7925732-INS-548
chr5-7940529-INS-140

chr5-7977397-INS-6608
chr5-8003398-INS-573
chr5-8051781-INS-215
chr5-8105610-INS-54
chr5-8133576-INS-98
chr5-8147017-INS-455
chr5-8178899-INS-448
chr5-8192507-INS-101
chr5-8200675-INS-95
chr5-8210474-INS-83
chr5-8232842-INS-64
chr5-8235366-INS-225
chr5-8239448-INS-131
chr5-8243338-INS-8265
chr5-8250688-INS-1943
chr5-8253801-INS-81
chr5-8259233-INS-430
chr5-8283079-INS-337
chr5-8296284-INS-100
chr5-8304815-INS-3491
chr5-8307869-INS-225
chr5-8316602-INS-423
chr5-8324557-INS-6520
chr5-8338289-INS-913
chr5-8346960-INS-62
chr5-8376747-INS-198
chr5-8391817-INS-490
chr5-8412278-INS-55
chr5-8429942-INS-212
chr5-8484387-INS-66
chr5-8530660-INS-70
chr5-8553617-INS-910
chr5-8564476-INS-52
chr5-8571663-INS-238
chr5-8576914-INS-96
chr5-8614257-INS-2696
chr5-8614686-INS-1252
chr5-8631790-INS-351
chr5-8632756-INS-52
chr5-8634458-INS-106
chr5-8645544-INS-1551
chr5-8650450-INS-7465
chr5-8680837-INS-60
chr5-8692277-INS-97
chr5-8699308-INS-56
chr5-8718394-INS-119
chr5-8721267-INS-60
chr5-8738831-INS-498
chr5-8739787-INS-5169
chr5-8753559-INS-107
chr5-8761989-INS-212
chr5-8762341-INS-82
chr5-8764156-INS-213
chr5-8778183-INS-72
chr5-8871809-INS-515
chr5-8872663-INS-156
chr5-8882668-INS-315
chr5-8888779-INS-877
chr5-8901813-INS-976
chr5-8926412-INS-66
chr5-8964472-INS-460
chr5-8976237-INS-291
chr5-8989288-INS-260
chr5-9002298-INS-946
chr5-9006587-INS-205
chr5-9060151-INS-56
chr5-9062652-INS-169
chr5-9065274-INS-67
chr5-9087031-INS-199
chr5-9089174-INS-60
chr5-9095592-INS-84
chr5-9096611-INS-207
chr5-9099445-INS-212
chr5-9119295-INS-200
chr5-9131491-INS-3808
chr5-9132649-INS-326
chr5-9143423-INS-194

chr5-9156022-INS-611
chr5-9163914-INS-310
chr5-9198092-INS-54
chr5-9218691-INS-512
chr5-9223285-INS-83
chr5-9226881-INS-845
chr5-9230289-INS-235
chr5-9253926-INS-855
chr5-9257767-INS-2121
chr5-9260852-INS-60
chr5-9291850-INS-75
chr5-9303525-INS-145
chr5-9307237-INS-742
chr5-9312902-INS-320
chr5-9325705-INS-1337
chr5-9348803-INS-6487
chr5-9352470-INS-1980
chr5-9352471-INS-504
chr5-9390138-INS-76
chr5-9391145-INS-6429
chr5-9437840-INS-164
chr5-9439153-INS-339
chr5-9476772-INS-53
chr5-9501727-INS-58
chr5-9502651-INS-220
chr5-9523664-INS-400
chr5-9534290-INS-217
chr5-9540647-INS-270
chr5-9551843-INS-432
chr5-9580031-INS-152
chr5-9590209-INS-81
chr5-9610302-INS-58
chr5-9621876-INS-86
chr5-9692076-INS-590
chr5-9694518-INS-6463
chr5-9714563-INS-518
chr5-9723487-INS-986
chr5-9773524-INS-294
chr5-9781154-INS-98
chr5-9795944-INS-75
chr5-9803429-INS-438
chr5-9809060-INS-100
chr5-9815828-INS-6356
chr5-9818416-INS-6292
chr5-9823014-INS-2071
chr5-9835077-INS-6801
chr5-9836346-INS-8359
chr5-9846846-INS-83
chr5-9917928-INS-128
chr5-9924286-INS-62
chr5-9925993-INS-59
chr5-9937817-INS-980
chr5-9944889-INS-306
chr5-9948132-INS-69
chr5-9957595-INS-53
chr5-9965754-INS-96
chr5-9990467-INS-60
chr5-10001424-INS-64
chr5-10011869-INS-2904
chr5-10075314-INS-113
chr5-10090774-INS-981
chr5-10128092-INS-69
chr5-10139139-INS-226
chr5-10172360-INS-494
chr5-10179167-INS-70
chr5-10193279-INS-683
chr5-10200228-INS-401
chr5-10285556-INS-1821
chr5-10288466-INS-92
chr5-10321831-INS-6266
chr5-10336761-INS-229
chr5-10348995-INS-272
chr5-10367423-INS-349
chr5-10385191-INS-725
chr5-10392728-INS-5999
chr5-10424635-INS-63
chr5-10435713-INS-52

chr5-10472741-INS-66
chr5-10523392-INS-51
chr5-10611390-INS-140
chr5-10628147-INS-56
chr5-10631638-INS-473
chr5-10632780-INS-52
chr5-10637927-INS-57
chr5-10643154-INS-62
chr5-10649268-INS-164
chr5-10663383-INS-105
chr5-10697793-INS-398
chr5-10708385-INS-56
chr5-10712193-INS-296
chr5-10744148-INS-357
chr5-10775711-INS-135
chr5-10785262-INS-79
chr5-10807833-INS-172
chr5-10824899-INS-3727
chr5-10830597-INS-54
chr5-10849776-INS-462
chr5-10861541-INS-153
chr5-10879476-INS-6806
chr5-10882892-INS-842
chr5-10887648-INS-6933
chr5-10915468-INS-1166
chr5-10970633-INS-493
chr5-11199737-INS-331
chr5-11236726-INS-5367
chr5-11241237-INS-112
chr5-11257116-INS-480
chr5-11279854-INS-606
chr5-11297016-INS-347
chr5-11455826-INS-69
chr5-11750199-INS-60
chr5-11771444-INS-344
chr5-11803989-INS-867
chr5-11821405-INS-70
chr5-11930319-INS-3609
chr5-11946347-INS-6600
chr5-11951612-INS-212
chr5-11961121-INS-173
chr5-12024622-INS-213
chr5-12091550-INS-314
chr5-12106951-INS-75
chr5-12113576-INS-447
chr5-12153447-INS-451
chr5-12190298-INS-8419
chr5-12227855-INS-98
chr5-12231596-INS-144
chr5-12232982-INS-2034
chr5-12259818-INS-521
chr5-12262429-INS-7249
chr5-12267989-INS-58
chr5-12278194-INS-52
chr5-12282068-INS-196
chr5-12286939-INS-62
chr5-12289497-INS-6190
chr5-12309748-INS-437
chr5-12313671-INS-70
chr5-12319227-INS-6254
chr5-12324046-INS-56
chr5-12325166-INS-68
chr5-12377015-INS-102
chr5-12424148-INS-164
chr5-12432727-INS-56
chr5-12436779-INS-358
chr5-12469818-INS-205
chr5-12474087-INS-164
chr5-12478236-INS-62
chr5-12481678-INS-144
chr5-12517972-INS-498
chr5-12530474-INS-54
chr5-12551364-INS-236
chr5-12561776-INS-176
chr5-12572411-INS-66
chr5-12581877-INS-62
chr5-12604108-INS-102

chr5-12606224-INS-51
chr5-12606799-INS-902
chr5-12609443-INS-61
chr5-12617244-INS-346
chr5-12649822-INS-67
chr5-12668097-INS-69
chr5-12668966-INS-52
chr5-12686576-INS-84
chr5-12687434-INS-830
chr5-12695300-INS-352
chr5-12699864-INS-192
chr5-12703993-INS-448
chr5-12737255-INS-60
chr5-12741427-INS-6204
chr5-12777516-INS-264
chr5-12785021-INS-54
chr5-12803034-INS-208
chr5-12805431-INS-52
chr5-12863247-INS-61
chr5-12877076-INS-66
chr5-12884544-INS-52
chr5-12910198-INS-834
chr5-12912075-INS-7410
chr5-12913423-INS-167
chr5-12914968-INS-520
chr5-12916424-INS-344
chr5-12938163-INS-64
chr5-12943024-INS-75
chr5-12954855-INS-187
chr5-12972826-INS-53
chr5-12975655-INS-73
chr5-12984711-INS-231
chr5-12997601-INS-96
chr5-13005575-INS-3851
chr5-13005870-INS-365
chr5-13054073-INS-234
chr5-13064013-INS-168
chr5-13073095-INS-82
chr5-13115201-INS-275
chr5-13124001-INS-466
chr5-13131727-INS-391
chr5-13143087-INS-357
chr5-13162146-INS-67
chr5-13169243-INS-242
chr5-13169954-INS-1344
chr5-13183290-INS-426
chr5-13197386-INS-52
chr5-13211587-INS-56
chr5-13215431-INS-56
chr5-13215591-INS-1277
chr5-13219363-INS-166
chr5-13233828-INS-6226
chr5-13238567-INS-6208
chr5-13249065-INS-107
chr5-13249355-INS-522
chr5-13256949-INS-540
chr5-13338160-INS-114
chr5-13377519-INS-63
chr5-13462817-INS-90
chr5-13468492-INS-201
chr5-13475930-INS-53
chr5-13490956-INS-108
chr5-13509601-INS-295
chr5-13513163-INS-1070
chr5-13518272-INS-104
chr5-13547297-INS-818
chr5-13574384-INS-62
chr5-13575269-INS-64
chr5-13602764-INS-118
chr5-13604076-INS-400
chr5-13663154-INS-105
chr5-13696058-INS-6582
chr5-13704121-INS-227
chr5-13714264-INS-1159
chr5-13722072-INS-188
chr5-13733756-INS-58
chr5-13744319-INS-494

chr5-13787648-INS-685
chr5-13813175-INS-56
chr5-13825541-INS-52
chr5-13827193-INS-442
chr5-13880228-INS-5373
chr5-13914933-INS-174
chr5-13963714-INS-7050
chr5-13964669-INS-126
chr5-13984767-INS-405
chr5-13987084-INS-6442
chr5-14013752-INS-131
chr5-14017642-INS-6484
chr5-14033444-INS-77
chr5-14048130-INS-58
chr5-14055187-INS-111
chr5-14056781-INS-534
chr5-14059642-INS-400
chr5-14070131-INS-176
chr5-14090568-INS-573
chr5-14096551-INS-439
chr5-14110907-INS-400
chr5-14147280-INS-824
chr5-14179513-INS-52
chr5-14185509-INS-230
chr5-14187295-INS-128
chr5-14190730-INS-163
chr5-14204004-INS-51
chr5-14217994-INS-825
chr5-14230103-INS-212
chr5-14231505-INS-330
chr5-14238452-INS-57
chr5-14242336-INS-606
chr5-14249832-INS-51
chr5-14275116-INS-6574
chr5-14286495-INS-173
chr5-14287541-INS-180
chr5-14305239-INS-70
chr5-14308640-INS-66
chr5-14313882-INS-55
chr5-14320633-INS-159
chr5-14328166-INS-246
chr5-14357120-INS-56
chr5-14371136-INS-469
chr5-14378471-INS-143
chr5-14420561-INS-6869
chr5-14425670-INS-155
chr5-14435682-INS-679
chr5-14437429-INS-144
chr5-14443671-INS-54
chr5-14457852-INS-557
chr5-14466019-INS-254
chr5-14521622-INS-64
chr5-14522811-INS-144
chr5-14541261-INS-51
chr5-14544140-INS-66
chr5-14552565-INS-290
chr5-14619416-INS-664
chr5-14639963-INS-255
chr5-14651756-INS-51
chr5-14653457-INS-695
chr5-14654483-INS-68
chr5-14670508-INS-176
chr5-14697402-INS-7858
chr5-14697760-INS-452
chr5-14708123-INS-55
chr5-14708842-INS-132
chr5-14721865-INS-429
chr5-14747585-INS-7687
chr5-14787468-INS-177
chr5-14790203-INS-192
chr5-14814707-INS-336
chr5-14830049-INS-2350
chr5-14843538-INS-305
chr5-14855320-INS-5285
chr5-14858184-INS-157
chr5-14859241-INS-170
chr5-14901533-INS-416

chr5-14906419-INS-538
chr5-15000723-INS-433
chr5-15037459-INS-6469
chr5-15081217-INS-722
chr5-15091572-INS-6468
chr5-15701528-INS-77
chr5-15704185-INS-394
chr5-15751629-INS-77
chr5-15960782-INS-521
chr5-15976206-INS-150
chr5-15982517-INS-7231
chr5-15992827-INS-256
chr5-15997305-INS-6442
chr5-16003230-INS-343
chr5-16004319-INS-880
chr5-16031588-INS-88
chr5-16064838-INS-51
chr5-16067441-INS-702
chr5-16074816-INS-498
chr5-16094067-INS-75
chr5-16102299-INS-85
chr5-16120833-INS-124
chr5-16160315-INS-179
chr5-16162346-INS-92
chr5-16193182-INS-54
chr5-16198852-INS-60
chr5-16208298-INS-56
chr5-16209025-INS-364
chr5-16210891-INS-472
chr5-16242267-INS-123
chr5-16270102-INS-101
chr5-16294751-INS-74
chr5-16321433-INS-132
chr5-16341931-INS-239
chr5-16361503-INS-108
chr5-16367116-INS-7604
chr5-16441408-INS-388
chr5-16446357-INS-126
chr5-16450872-INS-656
chr5-16489162-INS-783
chr5-16501603-INS-135
chr5-16501759-INS-175
chr5-16514708-INS-979
chr5-16530941-INS-169
chr5-16600856-INS-64
chr5-16625831-INS-76
chr5-16634973-INS-150
chr5-16635500-INS-88
chr5-16669328-INS-534
chr5-16669844-INS-62
chr5-16687937-INS-73
chr5-16699601-INS-70
chr5-16715720-INS-68
chr5-16745697-INS-175
chr5-16771145-INS-52
chr5-16808416-INS-280
chr5-16827998-INS-529
chr5-16842142-INS-78
chr5-16855242-INS-868
chr5-16888689-INS-59
chr5-16906524-INS-7319
chr5-16909484-INS-6863
chr5-16918021-INS-681
chr5-16919765-INS-64
chr5-16938450-INS-60
chr5-16953298-INS-6536
chr5-16954004-INS-616
chr5-16972086-INS-6219
chr5-16980526-INS-53
chr5-16989757-INS-414
chr5-17008601-INS-769
chr5-17014770-INS-232
chr5-17055670-INS-6553
chr5-17070499-INS-67
chr5-17096782-INS-504
chr5-17106896-INS-439
chr5-17107805-INS-328

chr5-17133275-INS-54
chr5-17136411-INS-94
chr5-17207619-INS-89
chr5-17208573-INS-287
chr5-17220193-INS-85
chr5-17246907-INS-70
chr5-17256179-INS-6970
chr5-17264804-INS-6264
chr5-17273714-INS-6378
chr5-17277816-INS-944
chr5-17287950-INS-1000
chr5-17300070-INS-70
chr5-17322212-INS-146
chr5-17341637-INS-411
chr5-17346536-INS-411
chr5-17383273-INS-205
chr5-17390578-INS-505
chr5-17411582-INS-52
chr5-17429735-INS-6951
chr5-17430755-INS-98
chr5-17433609-INS-193
chr5-17463582-INS-52
chr5-17482698-INS-54
chr5-17508228-INS-68
chr5-17509438-INS-7114
chr5-17512748-INS-737
chr5-17536829-INS-214
chr5-17548437-INS-794
chr5-17588306-INS-810
chr5-17601076-INS-138
chr5-17603006-INS-115
chr5-17610880-INS-5787
chr5-17617953-INS-1240
chr5-17654039-INS-369
chr5-17672768-INS-59
chr5-17698965-INS-361
chr5-17716133-INS-175
chr5-17805524-INS-60
chr5-17831180-INS-54
chr5-17857731-INS-128
chr5-17879349-INS-60
chr5-17906402-INS-369
chr5-17907575-INS-204
chr5-17952093-INS-56
chr5-17984547-INS-171
chr5-17991566-INS-84
chr5-18001739-INS-56
chr5-18015556-INS-414
chr5-18034914-INS-205
chr5-18055025-INS-349
chr5-18058094-INS-113
chr5-18066152-INS-120
chr5-18103255-INS-275
chr5-18134835-INS-6272
chr5-18155099-INS-135
chr5-18179476-INS-112
chr5-18198136-INS-360
chr5-18200458-INS-7171
chr5-18201216-INS-207
chr5-18202126-INS-116
chr5-18222728-INS-54
chr5-18238554-INS-406
chr5-18247920-INS-51
chr5-18256349-INS-6612
chr5-18260853-INS-5607
chr5-18264770-INS-56
chr5-18290311-INS-824
chr5-18294967-INS-6421
chr5-18312738-INS-88
chr5-18322083-INS-53
chr5-18324320-INS-169
chr5-18344865-INS-287
chr5-18347443-INS-128
chr5-18366775-INS-79
chr5-18372992-INS-202
chr5-18393797-INS-1019
chr5-18409380-INS-215

chr5-18415173-INS-384
chr5-18431691-INS-497
chr5-18443247-INS-52
chr5-18444442-INS-61
chr5-18465328-INS-418
chr5-18478133-INS-1044
chr5-18511026-INS-462
chr5-18534046-INS-62
chr5-18544791-INS-160
chr5-18547700-INS-70
chr5-18552237-INS-51
chr5-18593277-INS-144
chr5-18610303-INS-6573
chr5-18625238-INS-295
chr5-18629982-INS-138
chr5-18634204-INS-178
chr5-18654564-INS-72
chr5-18706298-INS-458
chr5-18716782-INS-684
chr5-18753467-INS-931
chr5-18757119-INS-334
chr5-18767879-INS-8941
chr5-18769131-INS-507
chr5-18777036-INS-132
chr5-18783499-INS-208
chr5-18794794-INS-359
chr5-18809864-INS-389
chr5-18816750-INS-152
chr5-18818138-INS-546
chr5-18833585-INS-94
chr5-18834211-INS-8656
chr5-18846057-INS-6636
chr5-18899484-INS-188
chr5-18917789-INS-230
chr5-18976715-INS-64
chr5-19001903-INS-923
chr5-19065050-INS-54
chr5-19095406-INS-68
chr5-19115042-INS-103
chr5-19129040-INS-72
chr5-19150613-INS-329
chr5-19160629-INS-313
chr5-19167682-INS-6413
chr5-19174901-INS-67
chr5-19216392-INS-6825
chr5-19218566-INS-270
chr5-19219603-INS-55
chr5-19223808-INS-293
chr5-19272289-INS-504
chr5-19279346-INS-386
chr5-19282662-INS-57
chr5-19284078-INS-476
chr5-19328939-INS-950
chr5-19343399-INS-62
chr5-19347951-INS-228
chr5-19349171-INS-525
chr5-19384406-INS-132
chr5-19413868-INS-638
chr5-19425600-INS-398
chr5-19436687-INS-429
chr5-19441154-INS-53
chr5-19444396-INS-742
chr5-19462270-INS-52
chr5-19498887-INS-163
chr5-19510331-INS-2733
chr5-19530307-INS-255
chr5-19572482-INS-502
chr5-19594648-INS-553
chr5-19602848-INS-6289
chr5-19603680-INS-87
chr5-19607519-INS-119
chr5-19650707-INS-178
chr5-19677505-INS-55
chr5-19677808-INS-673
chr5-19681230-INS-52
chr5-19681606-INS-214
chr5-19696766-INS-7375

chr5-19704226-INS-51
chr5-19707906-INS-332
chr5-19710017-INS-183
chr5-19715648-INS-191
chr5-19727358-INS-225
chr5-19767539-INS-73
chr5-19772134-INS-581
chr5-19815370-INS-6935
chr5-19825888-INS-214
chr5-19839683-INS-90
chr5-19852187-INS-52
chr5-19866332-INS-164
chr5-19871403-INS-7170
chr5-19876444-INS-463
chr5-19880992-INS-60
chr5-19883456-INS-2576
chr5-19887353-INS-152
chr5-19899744-INS-68
chr5-19956406-INS-94
chr5-19967648-INS-439
chr5-19981544-INS-71
chr5-19995585-INS-51
chr5-20019915-INS-139
chr5-20078728-INS-2794
chr5-20080851-INS-407
chr5-20095654-INS-92
chr5-20102552-INS-69
chr5-20159771-INS-169
chr5-20192622-INS-226
chr5-20193069-INS-84
chr5-20196599-INS-87
chr5-20199156-INS-328
chr5-20214082-INS-2118
chr5-20226613-INS-927
chr5-20236279-INS-1882
chr5-20240876-INS-5541
chr5-20244291-INS-640
chr5-20246032-INS-52
chr5-20274470-INS-406
chr5-20282233-INS-54
chr5-20316495-INS-104
chr5-20334810-INS-74
chr5-20341052-INS-208
chr5-20407026-INS-136
chr5-20410741-INS-213
chr5-20442116-INS-895
chr5-20451435-INS-148
chr5-20463194-INS-68
chr5-20473170-INS-117
chr5-20546757-INS-131
chr5-20561218-INS-6481
chr5-20576629-INS-74
chr5-20604784-INS-64
chr5-20619729-INS-146
chr5-20643089-INS-530
chr5-20655332-INS-721
chr5-20656179-INS-75
chr5-20668590-INS-64
chr5-20675331-INS-69
chr5-20705694-INS-76
chr5-20719688-INS-215
chr5-20798869-INS-422
chr5-20810735-INS-66
chr5-20813181-INS-242
chr5-20814585-INS-120
chr5-20843561-INS-158
chr5-20843561-INS-56
chr5-20849685-INS-80
chr5-20850746-INS-112
chr5-20877890-INS-494
chr5-20878003-INS-188
chr5-20886447-INS-202
chr5-20887797-INS-202
chr5-20891706-INS-60
chr5-20895394-INS-173
chr5-20898912-INS-204
chr5-20917226-INS-167

chr5-20918426-INS-514
chr5-20922405-INS-58
chr5-20922808-INS-164
chr5-20929140-INS-226
chr5-20935037-INS-397
chr5-20944300-INS-208
chr5-20971662-INS-379
chr5-20988703-INS-215
chr5-20991819-INS-315
chr5-20999255-INS-4204
chr5-21034331-INS-1649
chr5-21039976-INS-302
chr5-21044709-INS-53
chr5-21061478-INS-64
chr5-21061787-INS-64
chr5-21063109-INS-95
chr5-21075183-INS-61
chr5-21078664-INS-186
chr5-21084340-INS-54
chr5-21105115-INS-110
chr5-21120504-INS-76
chr5-21143301-INS-154
chr5-21151681-INS-80
chr5-21152730-INS-175
chr5-21159637-INS-72
chr5-21164821-INS-271
chr5-21174753-INS-148
chr5-21179146-INS-117
chr5-21179480-INS-152
chr5-21204532-INS-76
chr5-21206799-INS-112
chr5-21219653-INS-238
chr5-21230677-INS-52
chr5-21233775-INS-66
chr5-21246765-INS-280
chr5-21279447-INS-203
chr5-21285454-INS-59
chr5-21303985-INS-464
chr5-21320329-INS-196
chr5-21326427-INS-558
chr5-21330643-INS-72
chr5-21342317-INS-211
chr5-21352044-INS-213
chr5-21392363-INS-2330
chr5-21404594-INS-60
chr5-21415605-INS-50
chr5-21418294-INS-52
chr5-21423600-INS-60
chr5-21429034-INS-82
chr5-21465558-INS-164
chr5-21470013-INS-110
chr5-21471404-INS-283
chr5-21471552-INS-1007
chr5-21479757-INS-144
chr5-21495487-INS-433
chr5-21529682-INS-96
chr5-21533473-INS-96
chr5-21551998-INS-430
chr5-21563716-INS-712
chr5-21569673-INS-88
chr5-21575983-INS-255
chr5-21599546-INS-101
chr5-21614155-INS-67
chr5-21661974-INS-330
chr5-21670639-INS-507
chr5-21673988-INS-88
chr5-21693496-INS-210
chr5-21700823-INS-65
chr5-21706300-INS-60
chr5-21720520-INS-57
chr5-21722803-INS-56
chr5-21741263-INS-189
chr5-21746689-INS-251
chr5-21750403-INS-257
chr5-21777580-INS-83
chr5-21781282-INS-285
chr5-21792068-INS-217

chr5-21844122-INS-64
chr5-21847394-INS-424
chr5-21866964-INS-5219
chr5-21881564-INS-154
chr5-21890021-INS-209
chr5-21898356-INS-53
chr5-21900712-INS-171
chr5-21916544-INS-132
chr5-21921224-INS-572
chr5-21928413-INS-204
chr5-21931792-INS-201
chr5-21935808-INS-260
chr5-21950005-INS-166
chr5-21957213-INS-73
chr5-21972911-INS-80
chr5-21985804-INS-210
chr5-21992015-INS-68
chr5-22000590-INS-418
chr5-22009850-INS-127
chr5-22019158-INS-112
chr5-22026962-INS-132
chr5-22040855-INS-504
chr5-22059647-INS-64
chr5-22060532-INS-104
chr5-22061591-INS-209
chr5-22088988-INS-516
chr5-22100269-INS-66
chr5-22114191-INS-127
chr5-22120265-INS-54
chr5-22121466-INS-55
chr5-22126194-INS-409
chr5-22129786-INS-524
chr5-22136988-INS-588
chr5-22139675-INS-5302
chr5-22195407-INS-629
chr5-22225351-INS-102
chr5-22251236-INS-322
chr5-22262137-INS-440
chr5-22287754-INS-51
chr5-22355733-INS-60
chr5-22363790-INS-112
chr5-22394981-INS-906
chr5-22430685-INS-102
chr5-22441178-INS-476
chr5-22444282-INS-215
chr5-22450981-INS-208
chr5-22468107-INS-105
chr5-22474822-INS-2697
chr5-22496113-INS-6484
chr5-22520649-INS-170
chr5-22533425-INS-120
chr5-22584850-INS-6641
chr5-22591116-INS-895
chr5-22604252-INS-331
chr5-22617182-INS-60
chr5-22632874-INS-106
chr5-22646136-INS-56
chr5-22648980-INS-722
chr5-22654242-INS-320
chr5-22673885-INS-298
chr5-22690982-INS-52
chr5-22709914-INS-116
chr5-22815681-INS-84
chr5-22830986-INS-58
chr5-22833534-INS-404
chr5-22840230-INS-503
chr5-22849745-INS-515
chr5-22863979-INS-5452
chr5-22874798-INS-157
chr5-22890692-INS-150
chr5-22908546-INS-76
chr5-22910228-INS-70
chr5-22956591-INS-53
chr5-22987270-INS-634
chr5-23009992-INS-51
chr5-23019214-INS-87
chr5-23020367-INS-321

chr5-23030903-INS-120
chr5-23067371-INS-151
chr5-23097622-INS-141
chr5-23138884-INS-96
chr5-23140794-INS-170
chr5-23227686-INS-64
chr5-23242722-INS-144
chr5-23256340-INS-500
chr5-23261975-INS-161
chr5-23263589-INS-58
chr5-23269437-INS-53
chr5-23304025-INS-58
chr5-23317331-INS-455
chr5-23328230-INS-485
chr5-23331778-INS-777
chr5-23364709-INS-182
chr5-23383035-INS-210
chr5-23406346-INS-80
chr5-23424899-INS-178
chr5-23432512-INS-91
chr5-23450424-INS-400
chr5-23456496-INS-52
chr5-23462074-INS-747
chr5-23518356-INS-437
chr5-23519349-INS-91
chr5-23548227-INS-184
chr5-23553823-INS-251
chr5-23556109-INS-640
chr5-23556120-INS-241
chr5-23556134-INS-1084
chr5-23577599-INS-431
chr5-23587649-INS-153
chr5-23589725-INS-4156
chr5-23607945-INS-120
chr5-23609493-INS-76
chr5-23627609-INS-63
chr5-23644598-INS-158
chr5-23663668-INS-68
chr5-23675778-INS-53
chr5-23728665-INS-52
chr5-23754245-INS-76
chr5-23757415-INS-182
chr5-23810765-INS-170
chr5-23831839-INS-111
chr5-23833870-INS-53
chr5-23839311-INS-240
chr5-23842167-INS-55
chr5-23844413-INS-252
chr5-23877164-INS-644
chr5-23882888-INS-846
chr5-23885901-INS-224
chr5-23891075-INS-197
chr5-23895378-INS-585
chr5-23903217-INS-525
chr5-23918094-INS-382
chr5-23920459-INS-524
chr5-23929491-INS-209
chr5-23961847-INS-182
chr5-23966400-INS-253
chr5-24003877-INS-417
chr5-24020097-INS-198
chr5-24025077-INS-217
chr5-24026367-INS-418
chr5-24032518-INS-322
chr5-24052410-INS-214
chr5-24078704-INS-506
chr5-24080065-INS-157
chr5-24094498-INS-196
chr5-24107170-INS-67
chr5-24110333-INS-80
chr5-24113845-INS-372
chr5-24170815-INS-77
chr5-24200726-INS-206
chr5-24236867-INS-170
chr5-24243594-INS-255
chr5-24269368-INS-206
chr5-24283721-INS-55

chr5-24288258-INS-203
chr5-24320216-INS-56
chr5-24340662-INS-441
chr5-24345717-INS-644
chr5-24356045-INS-793
chr5-24365907-INS-199
chr5-24394129-INS-8420
chr5-24409930-INS-172
chr5-24410918-INS-60
chr5-24428495-INS-658
chr5-24430624-INS-52
chr5-24438642-INS-62
chr5-24450100-INS-60
chr5-24466060-INS-75
chr5-24510024-INS-146
chr5-24568441-INS-215
chr5-24575482-INS-136
chr5-24580651-INS-92
chr5-24608569-INS-196
chr5-24712039-INS-100
chr5-24716517-INS-133
chr5-24717179-INS-1891
chr5-24752055-INS-283
chr5-24762271-INS-203
chr5-24764279-INS-171
chr5-24768704-INS-62
chr5-24781758-INS-5671
chr5-24788379-INS-105
chr5-24793512-INS-84
chr5-24818473-INS-85
chr5-24819394-INS-228
chr5-24863475-INS-52
chr5-24867407-INS-615
chr5-24887306-INS-288
chr5-24912096-INS-178
chr5-24912456-INS-428
chr5-24919502-INS-57
chr5-24927243-INS-171
chr5-24928146-INS-102
chr5-24932619-INS-53
chr5-24950674-INS-239
chr5-24957187-INS-117
chr5-24982317-INS-152
chr5-25006143-INS-116
chr5-25029926-INS-207
chr5-25056641-INS-590
chr5-25058673-INS-927
chr5-25073633-INS-134
chr5-25098263-INS-100
chr5-25129600-INS-79
chr5-25169585-INS-211
chr5-25192471-INS-409
chr5-25194219-INS-156
chr5-25196627-INS-90
chr5-25208407-INS-60
chr5-25219065-INS-51
chr5-25220549-INS-271
chr5-25236605-INS-210
chr5-25245337-INS-7061
chr5-25246709-INS-460
chr5-25288311-INS-2225
chr5-25296539-INS-6483
chr5-25301016-INS-52
chr5-25307437-INS-134
chr5-25312918-INS-443
chr5-25314573-INS-206
chr5-25369824-INS-178
chr5-25379960-INS-9587
chr5-25385452-INS-57
chr5-25395815-INS-7367
chr5-25401491-INS-423
chr5-25408611-INS-76
chr5-25420540-INS-445
chr5-25422875-INS-533
chr5-25423733-INS-186
chr5-25432929-INS-276
chr5-25538359-INS-304

chr5-25540568-INS-62
chr5-25551428-INS-249
chr5-25564050-INS-451
chr5-25567281-INS-609
chr5-25598117-INS-577
chr5-25605989-INS-70
chr5-25631356-INS-322
chr5-25636997-INS-183
chr5-25642222-INS-131
chr5-25654872-INS-184
chr5-25660489-INS-133
chr5-25689246-INS-411
chr5-25700033-INS-351
chr5-25724949-INS-98
chr5-25755991-INS-728
chr5-25759705-INS-493
chr5-25769016-INS-138
chr5-25806721-INS-121
chr5-25823363-INS-54
chr5-25824349-INS-311
chr5-25831354-INS-134
chr5-25856413-INS-246
chr5-25916879-INS-79
chr5-25925049-INS-52
chr5-25939276-INS-64
chr5-25970994-INS-72
chr5-25985533-INS-156
chr5-26015440-INS-984
chr5-26017748-INS-122
chr5-26018412-INS-144
chr5-26020238-INS-281
chr5-26023766-INS-136
chr5-26026535-INS-920
chr5-26030609-INS-158
chr5-26038108-INS-109
chr5-26050173-INS-98
chr5-26063348-INS-89
chr5-26079024-INS-722
chr5-26081937-INS-86
chr5-26114632-INS-156
chr5-26191736-INS-92
chr5-26229481-INS-164
chr5-26282972-INS-336
chr5-26386262-INS-501
chr5-26421052-INS-6663
chr5-26421359-INS-914
chr5-26433734-INS-58
chr5-26442881-INS-6203
chr5-26482583-INS-902
chr5-26487437-INS-523
chr5-26500013-INS-234
chr5-26504248-INS-498
chr5-26537390-INS-226
chr5-26614423-INS-87
chr5-26679018-INS-72
chr5-26690958-INS-57
chr5-26715009-INS-53
chr5-26730954-INS-453
chr5-26736597-INS-91
chr5-26783327-INS-54
chr5-26788509-INS-102
chr5-26827128-INS-601
chr5-26846565-INS-56
chr5-26877294-INS-78
chr5-26903206-INS-136
chr5-26920606-INS-365
chr5-26943046-INS-250
chr5-26946325-INS-152
chr5-26955258-INS-62
chr5-26956695-INS-400
chr5-26987462-INS-800
chr5-27012405-INS-138
chr5-27036790-INS-341
chr5-27052071-INS-99
chr5-27057289-INS-418
chr5-27062029-INS-109
chr5-27084001-INS-340

chr5-27154959-INS-186
chr5-27158804-INS-73
chr5-27167283-INS-60
chr5-27211003-INS-6336
chr5-27222242-INS-190
chr5-27234004-INS-6441
chr5-27264151-INS-68
chr5-27275822-INS-270
chr5-27278006-INS-66
chr5-27303976-INS-80
chr5-27339051-INS-66
chr5-27357140-INS-398
chr5-27386135-INS-204
chr5-27390466-INS-70
chr5-27402274-INS-82
chr5-27495799-INS-56
chr5-27498931-INS-130
chr5-27499993-INS-78
chr5-27501381-INS-212
chr5-27520948-INS-52
chr5-27524451-INS-158
chr5-27538474-INS-54
chr5-27576743-INS-196
chr5-27598466-INS-261
chr5-27602118-INS-67
chr5-27617818-INS-203
chr5-27679251-INS-621
chr5-27743868-INS-6588
chr5-27748041-INS-111
chr5-27751379-INS-148
chr5-27763136-INS-919
chr5-27765294-INS-775
chr5-27778859-INS-154
chr5-27809229-INS-4592
chr5-27810733-INS-400
chr5-27860502-INS-514
chr5-27867591-INS-679
chr5-27871927-INS-70
chr5-27873312-INS-56
chr5-27881939-INS-90
chr5-27887081-INS-135
chr5-27900486-INS-400
chr5-27921972-INS-62
chr5-27927835-INS-56
chr5-27930059-INS-69
chr5-27950091-INS-103
chr5-27966874-INS-60
chr5-27980279-INS-56
chr5-27992737-INS-213
chr5-27997944-INS-127
chr5-28025778-INS-950
chr5-28043044-INS-5324
chr5-28051760-INS-61
chr5-28054122-INS-52
chr5-28075755-INS-110
chr5-28077818-INS-151
chr5-28080724-INS-231
chr5-28094279-INS-474
chr5-28114733-INS-208
chr5-28162864-INS-130
chr5-28164816-INS-399
chr5-28198463-INS-59
chr5-28217178-INS-216
chr5-28220710-INS-116
chr5-28241327-INS-74
chr5-28252717-INS-672
chr5-28273898-INS-123
chr5-28309277-INS-54
chr5-28317723-INS-123
chr5-28323938-INS-56
chr5-28324712-INS-98
chr5-28435992-INS-140
chr5-28438165-INS-143
chr5-28442921-INS-3382
chr5-28445327-INS-67
chr5-28462353-INS-251
chr5-28476730-INS-213

chr5-28493014-INS-111
chr5-28496867-INS-84
chr5-28498629-INS-64
chr5-28501474-INS-328
chr5-28507047-INS-234
chr5-28512999-INS-180
chr5-28531337-INS-62
chr5-28549534-INS-203
chr5-28582788-INS-60
chr5-28597481-INS-66
chr5-28615148-INS-946
chr5-28629721-INS-82
chr5-28633121-INS-130
chr5-28684713-INS-185
chr5-28709559-INS-394
chr5-28720198-INS-2182
chr5-28739075-INS-6883
chr5-28779991-INS-7117
chr5-28806186-INS-381
chr5-28817966-INS-101
chr5-28841857-INS-6703
chr5-28868310-INS-157
chr5-28875654-INS-596
chr5-28904635-INS-120
chr5-28912116-INS-265
chr5-28921941-INS-64
chr5-28934509-INS-54
chr5-28951609-INS-61
chr5-28984817-INS-126
chr5-28985791-INS-174
chr5-29000802-INS-619
chr5-29028072-INS-399
chr5-29055401-INS-284
chr5-29059640-INS-55
chr5-29075618-INS-100
chr5-29099849-INS-62
chr5-29129178-INS-972
chr5-29133538-INS-245
chr5-29198459-INS-135
chr5-29220354-INS-76
chr5-29245419-INS-6664
chr5-29303845-INS-51
chr5-29310135-INS-597
chr5-29311999-INS-951
chr5-29340302-INS-430
chr5-29357936-INS-60
chr5-29361492-INS-499
chr5-29363590-INS-105
chr5-29371021-INS-64
chr5-29427799-INS-704
chr5-29435337-INS-377
chr5-29436756-INS-178
chr5-29448730-INS-2629
chr5-29466971-INS-228
chr5-29490389-INS-60
chr5-29515173-INS-224
chr5-29530139-INS-226
chr5-29544499-INS-53
chr5-29578319-INS-249
chr5-29580555-INS-72
chr5-29593017-INS-197
chr5-29634163-INS-286
chr5-29660025-INS-869
chr5-29668922-INS-100
chr5-29672568-INS-232
chr5-29685357-INS-231
chr5-29731606-INS-549
chr5-29739777-INS-197
chr5-29766398-INS-178
chr5-29766892-INS-538
chr5-29772666-INS-62
chr5-29800412-INS-87
chr5-29810235-INS-58
chr5-29854381-INS-207
chr5-29883857-INS-278
chr5-29890037-INS-90
chr5-29903145-INS-189

chr5-29910419-INS-6432
chr5-29919676-INS-6468
chr5-29922361-INS-849
chr5-29924182-INS-210
chr5-29928150-INS-174
chr5-29930880-INS-251
chr5-29963729-INS-196
chr5-29992519-INS-203
chr5-29994221-INS-157
chr5-29999245-INS-401
chr5-30016771-INS-207
chr5-30027395-INS-133
chr5-30027867-INS-215
chr5-30028720-INS-573
chr5-30035838-INS-87
chr5-30040835-INS-90
chr5-30050452-INS-54
chr5-30068824-INS-190
chr5-30075913-INS-975
chr5-30107779-INS-55
chr5-30116649-INS-66
chr5-30121622-INS-823
chr5-30124627-INS-188
chr5-30125536-INS-78
chr5-30152639-INS-277
chr5-30163988-INS-90
chr5-30169157-INS-407
chr5-30169767-INS-168
chr5-30171538-INS-298
chr5-30248204-INS-227
chr5-30263959-INS-491
chr5-30269386-INS-214
chr5-30281478-INS-394
chr5-30283157-INS-240
chr5-30285042-INS-223
chr5-30322625-INS-59
chr5-30327903-INS-171
chr5-30330622-INS-236
chr5-30399642-INS-107
chr5-30410070-INS-60
chr5-30419984-INS-200
chr5-30423903-INS-117
chr5-30426679-INS-269
chr5-30456858-INS-622
chr5-30460432-INS-123
chr5-30462160-INS-202
chr5-30467684-INS-267
chr5-30478943-INS-205
chr5-30509717-INS-54
chr5-30511642-INS-60
chr5-30519127-INS-462
chr5-30568053-INS-112
chr5-30573019-INS-52
chr5-30574697-INS-60
chr5-30604028-INS-272
chr5-30621246-INS-166
chr5-30649698-INS-346
chr5-30651770-INS-124
chr5-30653277-INS-422
chr5-30669161-INS-61
chr5-30671794-INS-79
chr5-30696193-INS-111
chr5-30704738-INS-52
chr5-30723806-INS-51
chr5-30727336-INS-103
chr5-30737263-INS-52
chr5-30820806-INS-492
chr5-30823144-INS-52
chr5-30827406-INS-79
chr5-30856159-INS-171
chr5-30876314-INS-54
chr5-30906963-INS-401
chr5-30908149-INS-202
chr5-30921603-INS-143
chr5-30946563-INS-80
chr5-30958988-INS-200
chr5-30997763-INS-51

chr5-30998712-INS-231
chr5-31058077-INS-204
chr5-31091449-INS-204
chr5-31119038-INS-214
chr5-31137599-INS-190
chr5-31148881-INS-189
chr5-31155005-INS-120
chr5-31168105-INS-217
chr5-31170471-INS-289
chr5-31184755-INS-383
chr5-31185558-INS-84
chr5-31257578-INS-219
chr5-31263721-INS-4458
chr5-31273429-INS-140
chr5-31284259-INS-209
chr5-31288893-INS-213
chr5-31299929-INS-84
chr5-31306744-INS-95
chr5-31322894-INS-77
chr5-31345710-INS-205
chr5-31354924-INS-121
chr5-31383970-INS-120
chr5-31388871-INS-181
chr5-31389412-INS-65
chr5-31400638-INS-450
chr5-31418243-INS-90
chr5-31429936-INS-81
chr5-31479642-INS-229
chr5-31487384-INS-6738
chr5-31504181-INS-215
chr5-31524626-INS-58
chr5-31539064-INS-6345
chr5-31545545-INS-358
chr5-31560625-INS-522
chr5-31592457-INS-70
chr5-31636279-INS-111
chr5-31642303-INS-57
chr5-31649046-INS-187
chr5-31651315-INS-403
chr5-31661837-INS-73
chr5-31665705-INS-376
chr5-31666374-INS-814
chr5-31691565-INS-168
chr5-31715010-INS-139
chr5-31726827-INS-55
chr5-31729983-INS-173
chr5-31730648-INS-60
chr5-31758236-INS-138
chr5-31780025-INS-195
chr5-31844697-INS-235
chr5-31852034-INS-299
chr5-31859632-INS-516
chr5-31899850-INS-81
chr5-31928163-INS-55
chr5-31966043-INS-281
chr5-31994970-INS-973
chr5-32030722-INS-387
chr5-32036961-INS-60
chr5-32053829-INS-64
chr5-32056830-INS-72
chr5-32059306-INS-500
chr5-32066706-INS-578
chr5-32111599-INS-202
chr5-32127934-INS-74
chr5-32167326-INS-692
chr5-32169490-INS-76
chr5-32195706-INS-54
chr5-32202883-INS-92
chr5-32236752-INS-190
chr5-32265095-INS-58
chr5-32272044-INS-139
chr5-32285373-INS-69
chr5-32289607-INS-883
chr5-32297150-INS-66
chr5-32302086-INS-76
chr5-32317560-INS-111
chr5-32327019-INS-90

chr5-32340206-INS-120
chr5-32344357-INS-1870
chr5-32346467-INS-86
chr5-32361538-INS-94
chr5-32367472-INS-61
chr5-32381849-INS-358
chr5-32388279-INS-135
chr5-32399951-INS-62
chr5-32433197-INS-229
chr5-32467663-INS-64
chr5-32471627-INS-94
chr5-32499030-INS-136
chr5-32573460-INS-72
chr5-32574896-INS-52
chr5-32618771-INS-270
chr5-32642759-INS-148
chr5-32676608-INS-534
chr5-32707657-INS-567
chr5-32710578-INS-60
chr5-32717197-INS-101
chr5-32728570-INS-472
chr5-32776612-INS-68
chr5-32829909-INS-207
chr5-32840810-INS-386
chr5-32847212-INS-84
chr5-32868515-INS-338
chr5-32874424-INS-2202
chr5-32901057-INS-577
chr5-32922699-INS-80
chr5-32930710-INS-166
chr5-32945321-INS-658
chr5-32957567-INS-51
chr5-32961023-INS-52
chr5-32993454-INS-205
chr5-33003093-INS-698
chr5-33006474-INS-312
chr5-33018916-INS-197
chr5-33034027-INS-174
chr5-33043448-INS-172
chr5-33046237-INS-175
chr5-33060365-INS-250
chr5-33063526-INS-203
chr5-33076411-INS-394
chr5-33089990-INS-201
chr5-33093457-INS-199
chr5-33108088-INS-828
chr5-33127880-INS-60
chr5-33161751-INS-570
chr5-33187620-INS-212
chr5-33188836-INS-408
chr5-33198354-INS-109
chr5-33199792-INS-5861
chr5-33201853-INS-120
chr5-33211639-INS-283
chr5-33228015-INS-7474
chr5-33251242-INS-366
chr5-33259798-INS-8994
chr5-33284513-INS-395
chr5-33305953-INS-64
chr5-33326289-INS-6235
chr5-33364064-INS-154
chr5-33379035-INS-84
chr5-33414937-INS-394
chr5-33423317-INS-152
chr5-33438681-INS-66
chr5-33441297-INS-178
chr5-33443296-INS-7068
chr5-33446101-INS-86
chr5-33450229-INS-104
chr5-33512160-INS-147
chr5-33528429-INS-57
chr5-33552509-INS-144
chr5-33565699-INS-333
chr5-33576487-INS-527
chr5-33585805-INS-189
chr5-33593719-INS-194
chr5-33644232-INS-54

chr5-33675786-INS-93
chr5-33688098-INS-156
chr5-33695607-INS-69
chr5-33769008-INS-128
chr5-33821797-INS-180
chr5-33827154-INS-217
chr5-33869646-INS-137
chr5-33911116-INS-58
chr5-33914622-INS-96
chr5-33915953-INS-167
chr5-33934128-INS-86
chr5-33945565-INS-127
chr5-33952371-INS-211
chr5-33953509-INS-196
chr5-33960150-INS-191
chr5-33962946-INS-72
chr5-33981759-INS-60
chr5-33985379-INS-857
chr5-33989372-INS-158
chr5-33992544-INS-70
chr5-33992744-INS-485
chr5-34014321-INS-217
chr5-34058853-INS-95
chr5-34076624-INS-203
chr5-34167964-INS-7172
chr5-34170620-INS-3033
chr5-34183665-INS-594
chr5-34196002-INS-633
chr5-34214680-INS-8007
chr5-34251259-INS-250
chr5-34265058-INS-96
chr5-34279937-INS-671
chr5-34290043-INS-666
chr5-34302054-INS-66
chr5-34317324-INS-77
chr5-34317558-INS-947
chr5-34320254-INS-222
chr5-34322062-INS-174
chr5-34324557-INS-112
chr5-34350036-INS-183
chr5-34363141-INS-443
chr5-34368413-INS-204
chr5-34501690-INS-178
chr5-34535795-INS-386
chr5-34547163-INS-205
chr5-34548344-INS-206
chr5-34562708-INS-193
chr5-34598470-INS-162
chr5-34600803-INS-99
chr5-34629330-INS-168
chr5-34650248-INS-207
chr5-34657296-INS-345
chr5-34661627-INS-111
chr5-34662861-INS-66
chr5-34666724-INS-123
chr5-34675943-INS-123
chr5-34724268-INS-192
chr5-34738446-INS-270
chr5-34753061-INS-301
chr5-34763426-INS-111
chr5-34812900-INS-200
chr5-34846844-INS-340
chr5-34849472-INS-481
chr5-34860759-INS-131
chr5-34902731-INS-214
chr5-34907531-INS-131
chr5-34910660-INS-94
chr5-34915158-INS-149
chr5-34925078-INS-207
chr5-34925178-INS-728
chr5-34936491-INS-264
chr5-34951915-INS-178
chr5-34975622-INS-69
chr5-34990744-INS-663
chr5-35001658-INS-175
chr5-35143520-INS-4993
chr5-35161471-INS-170

chr5-35180861-INS-51
chr5-35216195-INS-152
chr5-35230396-INS-111
chr5-35233334-INS-123
chr5-35234916-INS-52
chr5-35285395-INS-164
chr5-35293051-INS-667
chr5-35301446-INS-434
chr5-35337276-INS-183
chr5-35351157-INS-58
chr5-35368946-INS-84
chr5-35376594-INS-66
chr5-35382191-INS-74
chr5-35392290-INS-51
chr5-35394774-INS-120
chr5-35431598-INS-54
chr5-35473460-INS-147
chr5-35487693-INS-390
chr5-35492536-INS-386
chr5-35499196-INS-364
chr5-35511489-INS-154
chr5-35513109-INS-127
chr5-35532620-INS-55
chr5-35533608-INS-244
chr5-35538914-INS-152
chr5-35617239-INS-385
chr5-35633867-INS-206
chr5-35641797-INS-143
chr5-35665974-INS-186
chr5-35675727-INS-6458
chr5-35690677-INS-440
chr5-35696532-INS-242
chr5-35697540-INS-67
chr5-35712220-INS-326
chr5-35728291-INS-2974
chr5-35746298-INS-202
chr5-35758448-INS-84
chr5-35760676-INS-130
chr5-35796691-INS-87
chr5-35909610-INS-2676
chr5-35948936-INS-91
chr5-35951052-INS-549
chr5-35974352-INS-75
chr5-35981568-INS-54
chr5-35988841-INS-106
chr5-35991161-INS-176
chr5-36001487-INS-155
chr5-36012379-INS-196
chr5-36018191-INS-95
chr5-36091767-INS-179
chr5-36120310-INS-333
chr5-36132254-INS-56
chr5-36150091-INS-227
chr5-36155567-INS-64
chr5-36226562-INS-101
chr5-36252558-INS-88
chr5-36262274-INS-380
chr5-36265371-INS-201
chr5-36296299-INS-77
chr5-36298544-INS-52
chr5-36376928-INS-927
chr5-36396750-INS-1059
chr5-36396948-INS-212
chr5-36420936-INS-400
chr5-36450346-INS-53
chr5-36455927-INS-67
chr5-36461274-INS-51
chr5-36508191-INS-64
chr5-36522525-INS-93
chr5-36555661-INS-64
chr5-36559480-INS-67
chr5-36589702-INS-400
chr5-36604813-INS-2529
chr5-36624984-INS-250
chr5-36634373-INS-234
chr5-36636898-INS-53
chr5-36638015-INS-761

chr5-36646412-INS-269
chr5-36672550-INS-1192
chr5-36692786-INS-73
chr5-36700165-INS-234
chr5-36711057-INS-199
chr5-36736300-INS-62
chr5-36746380-INS-56
chr5-36784211-INS-54
chr5-36818920-INS-330
chr5-36834996-INS-128
chr5-36859630-INS-77
chr5-36892210-INS-563
chr5-36898710-INS-69
chr5-36936458-INS-120
chr5-36946200-INS-206
chr5-36954466-INS-124
chr5-36961884-INS-96
chr5-36963020-INS-64
chr5-36968876-INS-86
chr5-36983679-INS-198
chr5-36983739-INS-1330
chr5-36994235-INS-138
chr5-37056300-INS-63
chr5-37058707-INS-108
chr5-37113120-INS-237
chr5-37140236-INS-67
chr5-37156882-INS-52
chr5-37177900-INS-286
chr5-37203800-INS-132
chr5-37215694-INS-171
chr5-37246796-INS-513
chr5-37251766-INS-162
chr5-37260286-INS-88
chr5-37267989-INS-70
chr5-37292659-INS-77
chr5-37312745-INS-116
chr5-37345212-INS-91
chr5-37359704-INS-320
chr5-37384051-INS-66
chr5-37477367-INS-55
chr5-37504134-INS-71
chr5-37518743-INS-193
chr5-37536986-INS-295
chr5-37538843-INS-81
chr5-37551413-INS-66
chr5-37553450-INS-58
chr5-37554750-INS-495
chr5-37564671-INS-61
chr5-37568860-INS-177
chr5-37636792-INS-242
chr5-37648393-INS-60
chr5-37669641-INS-89
chr5-37690358-INS-496
chr5-37704489-INS-108
chr5-37716240-INS-58
chr5-37753625-INS-1264
chr5-37753658-INS-624
chr5-37873000-INS-109
chr5-37884152-INS-52
chr5-37906036-INS-431
chr5-37939698-INS-480
chr5-37951591-INS-243
chr5-37968527-INS-116
chr5-37968834-INS-87
chr5-38023981-INS-792
chr5-38028746-INS-7113
chr5-38043882-INS-144
chr5-38069780-INS-211
chr5-38092664-INS-115
chr5-38161074-INS-57
chr5-38179563-INS-87
chr5-38193751-INS-71
chr5-38272613-INS-342
chr5-38304365-INS-70
chr5-38329813-INS-60
chr5-38342736-INS-400
chr5-38467927-INS-443

chr5-38624988-INS-162
chr5-38636065-INS-68
chr5-38643631-INS-53
chr5-38669597-INS-331
chr5-38674807-INS-181
chr5-38675906-INS-581
chr5-38682135-INS-618
chr5-38694596-INS-97
chr5-38705998-INS-282
chr5-38708578-INS-106
chr5-38749330-INS-399
chr5-38755673-INS-158
chr5-38758487-INS-61
chr5-38759707-INS-73
chr5-38781825-INS-79
chr5-38789580-INS-61
chr5-38799288-INS-390
chr5-38804011-INS-166
chr5-38812002-INS-82
chr5-38814031-INS-171
chr5-38835704-INS-211
chr5-38852508-INS-209
chr5-38857794-INS-99
chr5-38889251-INS-176
chr5-38900680-INS-100
chr5-38987412-INS-272
chr5-38987767-INS-543
chr5-38988376-INS-68
chr5-39004272-INS-80
chr5-39080750-INS-52
chr5-39084797-INS-144
chr5-39089732-INS-304
chr5-39096784-INS-346
chr5-39111975-INS-774
chr5-39150547-INS-268
chr5-39153520-INS-54
chr5-39172566-INS-92
chr5-39206787-INS-89
chr5-39224569-INS-189
chr5-39232873-INS-86
chr5-39237641-INS-210
chr5-39247673-INS-484
chr5-39261864-INS-291
chr5-39267619-INS-541
chr5-39271365-INS-53
chr5-39281537-INS-6932
chr5-39298030-INS-701
chr5-39347687-INS-244
chr5-39359275-INS-201
chr5-39421866-INS-63
chr5-39425682-INS-113
chr5-39427884-INS-304
chr5-39430364-INS-69
chr5-39435833-INS-52
chr5-39438831-INS-144
chr5-39440276-INS-685
chr5-39441136-INS-827
chr5-39447934-INS-373
chr5-39485930-INS-209
chr5-39487469-INS-60
chr5-39498739-INS-6183
chr5-39529884-INS-81
chr5-39536882-INS-56
chr5-39543567-INS-677
chr5-39552007-INS-57
chr5-39608241-INS-198
chr5-39641177-INS-167
chr5-39650132-INS-246
chr5-39652433-INS-70
chr5-39652928-INS-171
chr5-39707264-INS-649
chr5-39746326-INS-707
chr5-39749076-INS-82
chr5-39759670-INS-622
chr5-39760236-INS-64
chr5-39764879-INS-607
chr5-39785327-INS-58

chr5-39816902-INS-5548
chr5-39828852-INS-80
chr5-39838584-INS-118
chr5-39840603-INS-868
chr5-39890263-INS-55
chr5-39890837-INS-5074
chr5-39893114-INS-276
chr5-39926952-INS-56
chr5-39976909-INS-286
chr5-39987996-INS-246
chr5-39998238-INS-214
chr5-40010222-INS-3131
chr5-40045564-INS-148
chr5-40070004-INS-52
chr5-40079850-INS-467
chr5-40092646-INS-436
chr5-40115179-INS-219
chr5-40129628-INS-361
chr5-40132839-INS-84
chr5-40143457-INS-276
chr5-40159805-INS-150
chr5-40190258-INS-7170
chr5-40262215-INS-52
chr5-40295517-INS-359
chr5-40307179-INS-77
chr5-40341766-INS-631
chr5-40351178-INS-128
chr5-40353477-INS-143
chr5-40354417-INS-400
chr5-40381820-INS-246
chr5-40407175-INS-171
chr5-40413716-INS-450
chr5-40418129-INS-63
chr5-40421938-INS-60
chr5-40441007-INS-932
chr5-40446199-INS-168
chr5-40455463-INS-72
chr5-40473713-INS-337
chr5-40537795-INS-124
chr5-40538880-INS-740
chr5-40599604-INS-135
chr5-40609619-INS-153
chr5-40614816-INS-63
chr5-40659060-INS-112
chr5-40669269-INS-54
chr5-40686749-INS-841
chr5-40708496-INS-6596
chr5-40715446-INS-928
chr5-40716806-INS-969
chr5-40734189-INS-199
chr5-40744770-INS-72
chr5-40748130-INS-916
chr5-40768726-INS-73
chr5-40769875-INS-234
chr5-40770566-INS-859
chr5-40771068-INS-50
chr5-40780965-INS-266
chr5-40831798-INS-54
chr5-40836006-INS-51
chr5-40847579-INS-94
chr5-40885823-INS-698
chr5-40920762-INS-116
chr5-40950462-INS-3419
chr5-40972454-INS-569
chr5-41029169-INS-518
chr5-41030982-INS-54
chr5-41043921-INS-217
chr5-41047555-INS-104
chr5-41048320-INS-559
chr5-41067436-INS-84
chr5-41086844-INS-64
chr5-41087960-INS-347
chr5-41115609-INS-192
chr5-41116323-INS-6165
chr5-41116975-INS-788
chr5-41119519-INS-372
chr5-41146721-INS-66

chr5-41150665-INS-6490
chr5-41164332-INS-79
chr5-41166947-INS-112
chr5-41183368-INS-509
chr5-41194375-INS-851
chr5-41200525-INS-253
chr5-41203577-INS-68
chr5-41237230-INS-414
chr5-41246230-INS-58
chr5-41278389-INS-110
chr5-41302109-INS-132
chr5-41346383-INS-86
chr5-41387759-INS-838
chr5-41413807-INS-89
chr5-41420970-INS-202
chr5-41422128-INS-666
chr5-41431007-INS-672
chr5-41486154-INS-52
chr5-41509565-INS-556
chr5-41525880-INS-760
chr5-41534458-INS-69
chr5-41542281-INS-106
chr5-41548982-INS-63
chr5-41562894-INS-866
chr5-41565249-INS-58
chr5-41570292-INS-126
chr5-41573403-INS-7211
chr5-41590367-INS-158
chr5-41597854-INS-6714
chr5-41641658-INS-72
chr5-41644053-INS-72
chr5-41664517-INS-6282
chr5-41707057-INS-68
chr5-41718938-INS-317
chr5-41721844-INS-327
chr5-41737725-INS-497
chr5-41769898-INS-386
chr5-41792746-INS-161
chr5-41808271-INS-6927
chr5-41813258-INS-913
chr5-41824172-INS-56
chr5-41827236-INS-632
chr5-41833993-INS-199
chr5-41871602-INS-296
chr5-41872890-INS-347
chr5-41872987-INS-827
chr5-41886235-INS-82
chr5-41888128-INS-51
chr5-41891181-INS-69
chr5-41903968-INS-214
chr5-41910147-INS-545
chr5-41914116-INS-451
chr5-41917729-INS-55
chr5-41924406-INS-60
chr5-41930819-INS-81
chr5-41935438-INS-62
chr5-41952933-INS-654
chr5-41966274-INS-6268
chr5-41981133-INS-54
chr5-41981779-INS-417
chr5-41991968-INS-432
chr5-42031683-INS-66
chr5-42038739-INS-125
chr5-42045200-INS-899
chr5-42052231-INS-959
chr5-42061808-INS-171
chr5-42070075-INS-155
chr5-42088502-INS-680
chr5-42090385-INS-281
chr5-42113193-INS-1370
chr5-42117819-INS-127
chr5-42119934-INS-156
chr5-42123356-INS-1590
chr5-42125282-INS-245
chr5-42129632-INS-52
chr5-42147925-INS-6024
chr5-42202185-INS-112

chr5-42238190-INS-748
chr5-42257930-INS-3180
chr5-42259059-INS-53
chr5-42305715-INS-75
chr5-42313569-INS-293
chr5-42315359-INS-65
chr5-42324100-INS-62
chr5-42368924-INS-154
chr5-42370424-INS-831
chr5-42373771-INS-345
chr5-42384857-INS-1014
chr5-42387038-INS-135
chr5-42397435-INS-937
chr5-42401435-INS-321
chr5-42408425-INS-55
chr5-42437360-INS-678
chr5-42462823-INS-222
chr5-42467536-INS-58
chr5-42488367-INS-407
chr5-42489539-INS-1707
chr5-42495610-INS-72
chr5-42498677-INS-741
chr5-42525823-INS-6173
chr5-42591585-INS-394
chr5-42600038-INS-721
chr5-42610601-INS-180
chr5-42615557-INS-662
chr5-42655339-INS-109
chr5-42673988-INS-54
chr5-42708166-INS-942
chr5-42718738-INS-56
chr5-42734243-INS-153
chr5-42756712-INS-51
chr5-42773939-INS-94
chr5-42774715-INS-68
chr5-42805485-INS-95
chr5-42825981-INS-245
chr5-42836093-INS-66
chr5-42883665-INS-81
chr5-42898683-INS-498
chr5-42925101-INS-64
chr5-42956763-INS-488
chr5-42995190-INS-593
chr5-42997570-INS-199
chr5-43006032-INS-266
chr5-43012130-INS-68
chr5-43017941-INS-144
chr5-43142409-INS-56
chr5-43194451-INS-180
chr5-43212506-INS-111
chr5-43219420-INS-83
chr5-43235822-INS-421
chr5-43277804-INS-235
chr5-43284593-INS-70
chr5-43293648-INS-959
chr5-43321012-INS-700
chr5-43330702-INS-184
chr5-43334099-INS-892
chr5-43340325-INS-8734
chr5-43421025-INS-635
chr5-43472186-INS-420
chr5-43491320-INS-56
chr5-43501773-INS-101
chr5-43511665-INS-141
chr5-43513312-INS-114
chr5-43527944-INS-187
chr5-43561056-INS-173
chr5-43592118-INS-54
chr5-43593363-INS-7009
chr5-43597585-INS-6270
chr5-43599391-INS-62
chr5-43603585-INS-292
chr5-43611571-INS-332
chr5-43642916-INS-7009
chr5-43643817-INS-3365
chr5-43657642-INS-157
chr5-43670701-INS-76

chr5-43690874-INS-475
chr5-43711647-INS-6568
chr5-43712689-INS-57
chr5-43732948-INS-227
chr5-43756723-INS-179
chr5-43775222-INS-56
chr5-43777732-INS-54
chr5-43787469-INS-6563
chr5-43794111-INS-6808
chr5-43832347-INS-62
chr5-43836973-INS-58
chr5-43851072-INS-382
chr5-43854048-INS-58
chr5-43866620-INS-70
chr5-43867598-INS-126
chr5-43883927-INS-121
chr5-43907409-INS-52
chr5-43935579-INS-62
chr5-43942473-INS-73
chr5-43960791-INS-210
chr5-43964828-INS-114
chr5-43966104-INS-72
chr5-43976041-INS-179
chr5-44004412-INS-71
chr5-44014760-INS-78
chr5-44016539-INS-66
chr5-44022210-INS-468
chr5-44024479-INS-215
chr5-44028591-INS-228
chr5-44068370-INS-66
chr5-44075590-INS-57
chr5-44086277-INS-588
chr5-44086811-INS-54
chr5-44142006-INS-444
chr5-44175810-INS-1089
chr5-44184464-INS-314
chr5-44212945-INS-70
chr5-44228526-INS-122
chr5-44229497-INS-125
chr5-44260976-INS-267
chr5-44267594-INS-140
chr5-44273398-INS-197
chr5-44277964-INS-383
chr5-44329577-INS-152
chr5-44332202-INS-1888
chr5-44366341-INS-191
chr5-44368064-INS-160
chr5-44404485-INS-216
chr5-44421629-INS-243
chr5-44426785-INS-88
chr5-44438081-INS-73
chr5-44449109-INS-547
chr5-44476110-INS-6781
chr5-44477825-INS-325
chr5-44527145-INS-52
chr5-44547741-INS-400
chr5-44548820-INS-7124
chr5-44549720-INS-69
chr5-44571230-INS-6422
chr5-44591940-INS-132
chr5-44593374-INS-6708
chr5-44600905-INS-12965
chr5-44601078-INS-6406
chr5-44604029-INS-550
chr5-44604325-INS-6668
chr5-44614745-INS-7381
chr5-44637953-INS-56
chr5-44655986-INS-55
chr5-44675747-INS-495
chr5-44684042-INS-52
chr5-44706413-INS-191
chr5-44709700-INS-773
chr5-44725898-INS-556
chr5-44730252-INS-419
chr5-44752266-INS-342
chr5-44759570-INS-56
chr5-44779723-INS-138

chr5-44795283-INS-106
chr5-44818679-INS-6503
chr5-44860352-INS-56
chr5-44861672-INS-55
chr5-44882836-INS-60
chr5-44916239-INS-182
chr5-44918046-INS-2918
chr5-44972107-INS-60
chr5-44992361-INS-86
chr5-44995007-INS-64
chr5-45019559-INS-393
chr5-45038464-INS-445
chr5-45066894-INS-282
chr5-45074335-INS-249
chr5-45113097-INS-814
chr5-45114940-INS-498
chr5-45132104-INS-207
chr5-45138480-INS-147
chr5-45152740-INS-51
chr5-45195239-INS-51
chr5-45196728-INS-110
chr5-45227426-INS-828
chr5-45236722-INS-80
chr5-45270584-INS-52
chr5-45273222-INS-101
chr5-45286767-INS-223
chr5-45296408-INS-1110
chr5-45323185-INS-7355
chr5-45356503-INS-176
chr5-45361465-INS-116
chr5-45372416-INS-169
chr5-45385017-INS-274
chr5-45422144-INS-63
chr5-45437987-INS-134
chr5-45450636-INS-201
chr5-45561704-INS-626
chr5-45562268-INS-65
chr5-45589891-INS-541
chr5-45609913-INS-77
chr5-45612772-INS-58
chr5-45658611-INS-699
chr5-45686716-INS-194
chr5-45702844-INS-102
chr5-45733583-INS-3075
chr5-45784067-INS-1880
chr5-45803134-INS-1027
chr5-45812931-INS-793
chr5-45831517-INS-192
chr5-45900268-INS-80
chr5-45911914-INS-491
chr5-45988336-INS-1002
chr5-46044719-INS-66
chr5-46062358-INS-59
chr5-46116359-INS-160
chr5-46117010-INS-6727
chr5-46121649-INS-6440
chr5-46124651-INS-6426
chr5-46134075-INS-161
chr5-46212449-INS-89
chr5-46265738-INS-508
chr5-46283692-INS-620
chr5-46290260-INS-520
chr5-46341288-INS-56
chr5-46350763-INS-65
chr5-46355912-INS-441
chr5-46356381-INS-714
chr5-46373891-INS-899
chr5-46388553-INS-51
chr5-46408016-INS-6278
chr5-46416895-INS-52
chr5-46427995-INS-451
chr5-46435074-INS-680
chr5-46463079-INS-6692
chr5-46487530-INS-3581
chr5-46507509-INS-81
chr5-46512381-INS-85
chr5-46545773-INS-54

chr5-46558312-INS-212
chr5-46562999-INS-6267
chr5-46571708-INS-1888
chr5-46582688-INS-399
chr5-46607468-INS-830
chr5-46611701-INS-507
chr5-46625449-INS-369
chr5-46671103-INS-78
chr5-46702073-INS-76
chr5-46743899-INS-90
chr5-46762163-INS-204
chr5-46766703-INS-64
chr5-46812449-INS-310
chr5-46817697-INS-98
chr5-46850601-INS-629
chr5-46902216-INS-567
chr5-46905161-INS-72
chr5-46916420-INS-93
chr5-46963265-INS-695
chr5-46971707-INS-4796
chr5-46978171-INS-64
chr5-46984358-INS-75
chr5-46996644-INS-80
chr5-47022921-INS-186
chr5-47025681-INS-114
chr5-47026195-INS-67
chr5-47033870-INS-71
chr5-47068996-INS-6393
chr5-47077826-INS-6401
chr5-47089290-INS-163
chr5-47093019-INS-7114
chr5-47124809-INS-130
chr5-47127661-INS-220
chr5-47134851-INS-757
chr5-47141231-INS-6647
chr5-47166455-INS-5325
chr5-47247331-INS-319
chr5-47274215-INS-5057
chr5-47289749-INS-6590
chr5-47290807-INS-54
chr5-47296191-INS-7085
chr5-47297439-INS-61
chr5-47306309-INS-6654
chr5-47323524-INS-53
chr5-47326983-INS-6461
chr5-47397321-INS-204
chr5-47402705-INS-271
chr5-47444655-INS-424
chr5-47450506-INS-202
chr5-47454828-INS-148
chr5-47469293-INS-6903
chr5-47484114-INS-52
chr5-47543322-INS-348
chr5-47580982-INS-152
chr5-47594832-INS-126
chr5-47606775-INS-249
chr5-47678521-INS-68
chr5-47717893-INS-478
chr5-47732640-INS-528
chr5-47741529-INS-257
chr5-47751927-INS-54
chr5-47755667-INS-252
chr5-47767948-INS-207
chr5-47784999-INS-511
chr5-47787068-INS-326
chr5-47792725-INS-60
chr5-47843814-INS-93
chr5-47859991-INS-6237
chr5-47941602-INS-461
chr5-47945157-INS-6299
chr5-47977778-INS-330
chr5-48031791-INS-61
chr5-48040368-INS-6449
chr5-48049889-INS-400
chr5-48071352-INS-124
chr5-48074472-INS-66
chr5-48128061-INS-75

chr5-48134383-INS-57
chr5-48149260-INS-449
chr5-48152519-INS-112
chr5-48170270-INS-58
chr5-48183902-INS-66
chr5-48194105-INS-4083
chr5-48223976-INS-4679
chr5-48248399-INS-180
chr5-48261701-INS-115
chr5-48306547-INS-700
chr5-48320222-INS-426
chr5-48330373-INS-121
chr5-48345990-INS-1893
chr5-48348367-INS-52
chr5-48376224-INS-179
chr5-48408043-INS-7308
chr5-48414058-INS-289
chr5-48416373-INS-68
chr5-48438375-INS-60
chr5-48464366-INS-1170
chr5-48479797-INS-6678
chr5-48485189-INS-559
chr5-48487991-INS-94
chr5-48505103-INS-86
chr5-48505876-INS-88
chr5-48510387-INS-407
chr5-48519032-INS-7307
chr5-48531942-INS-5550
chr5-48558102-INS-74
chr5-48575181-INS-140
chr5-48597915-INS-276
chr5-48602591-INS-435
chr5-48611419-INS-144
chr5-48611903-INS-768
chr5-48618206-INS-323
chr5-48642821-INS-322
chr5-48651022-INS-54
chr5-48660816-INS-414
chr5-48664787-INS-687
chr5-48672168-INS-94
chr5-48696945-INS-164
chr5-48699077-INS-765
chr5-48727763-INS-318
chr5-48753159-INS-202
chr5-48770023-INS-135
chr5-48785600-INS-69
chr5-48851691-INS-117
chr5-48926473-INS-191
chr5-48949024-INS-514
chr5-48954565-INS-62
chr5-48961897-INS-204
chr5-48983276-INS-143
chr5-48991742-INS-6441
chr5-49000522-INS-604
chr5-49017836-INS-774
chr5-49058095-INS-54
chr5-49079517-INS-110
chr5-49101561-INS-835
chr5-49120832-INS-356
chr5-49125319-INS-139
chr5-49127603-INS-609
chr5-49156071-INS-59
chr5-49176362-INS-309
chr5-49194439-INS-68
chr5-49244117-INS-279
chr5-49268611-INS-221
chr5-49269935-INS-406
chr5-49330662-INS-64
chr5-49338194-INS-283
chr5-49340454-INS-505
chr5-49344964-INS-822
chr5-49398370-INS-8450
chr5-49412912-INS-558
chr5-49462920-INS-6442
chr5-49493130-INS-162
chr5-49500023-INS-838
chr5-49507395-INS-52

chr5-49557776-INS-67
chr5-49590432-INS-183
chr5-49605652-INS-72
chr5-49611580-INS-57
chr5-49714773-INS-160
chr5-49716847-INS-118
chr5-49723331-INS-352
chr5-49726492-INS-85
chr5-49742221-INS-215
chr5-49756019-INS-271
chr5-49774161-INS-580
chr5-49782862-INS-56
chr5-49785540-INS-77
chr5-49800202-INS-336
chr5-49809175-INS-171
chr5-49834271-INS-6447
chr5-49835656-INS-5478
chr5-49844888-INS-56
chr5-49863649-INS-182
chr5-49889264-INS-832
chr5-49931696-INS-664
chr5-49938815-INS-550
chr5-49948518-INS-165
chr5-49977706-INS-425
chr5-49977779-INS-424
chr5-49989547-INS-555
chr5-50034469-INS-582
chr5-50045987-INS-544
chr5-50046824-INS-319
chr5-50054687-INS-231
chr5-50070251-INS-238
chr5-50137092-INS-7386
chr5-50189977-INS-130
chr5-50210811-INS-122
chr5-50250300-INS-701
chr5-50299736-INS-54
chr5-50303023-INS-434
chr5-50316570-INS-1819
chr5-50329056-INS-98
chr5-50344941-INS-149
chr5-50345639-INS-60
chr5-50384065-INS-694
chr5-50388068-INS-78
chr5-50448940-INS-51
chr5-50450120-INS-112
chr5-50462492-INS-6402
chr5-50466968-INS-55
chr5-50500287-INS-70
chr5-50502520-INS-68
chr5-50528090-INS-659
chr5-50529662-INS-100
chr5-50541633-INS-90
chr5-50549289-INS-52
chr5-50552846-INS-54
chr5-50558494-INS-112
chr5-50581158-INS-236
chr5-50583012-INS-64
chr5-50586884-INS-702
chr5-50609659-INS-192
chr5-50643348-INS-72
chr5-50668315-INS-135
chr5-50672505-INS-58
chr5-50674471-INS-462
chr5-50716703-INS-70
chr5-50767193-INS-248
chr5-50778146-INS-72
chr5-50783460-INS-156
chr5-50800868-INS-100
chr5-50808017-INS-100
chr5-50823255-INS-320
chr5-50825526-INS-128
chr5-50835476-INS-135
chr5-50859644-INS-472
chr5-50872083-INS-258
chr5-50876278-INS-90
chr5-50957536-INS-802
chr5-50974920-INS-659

chr5-50981733-INS-115
chr5-51000409-INS-6330
chr5-51033816-INS-456
chr5-51061976-INS-155
chr5-51082638-INS-66
chr5-51090126-INS-87
chr5-51103649-INS-359
chr5-51140656-INS-386
chr5-51149854-INS-6447
chr5-51177772-INS-204
chr5-51205153-INS-317
chr5-51217115-INS-52
chr5-51220935-INS-189
chr5-51225629-INS-6889
chr5-51241989-INS-568
chr5-51251602-INS-85
chr5-51256460-INS-228
chr5-51264295-INS-64
chr5-51288046-INS-275
chr5-51291556-INS-57
chr5-51310886-INS-60
chr5-51316934-INS-111
chr5-51323676-INS-79
chr5-51331846-INS-508
chr5-51343941-INS-70
chr5-51358843-INS-487
chr5-51359555-INS-426
chr5-51411637-INS-151
chr5-51435734-INS-53
chr5-51443016-INS-197
chr5-51475169-INS-248
chr5-51505275-INS-68
chr5-51577565-INS-358
chr5-51580178-INS-54
chr5-51614796-INS-114
chr5-51664228-INS-276
chr5-51707216-INS-182
chr5-51713106-INS-58
chr5-51786263-INS-133
chr5-51793449-INS-80
chr5-51826076-INS-7056
chr5-51847168-INS-134
chr5-51886087-INS-58
chr5-51910084-INS-68
chr5-51912370-INS-96
chr5-51933240-INS-214
chr5-51956342-INS-74
chr5-51968248-INS-53
chr5-51997826-INS-93
chr5-52009802-INS-62
chr5-52023229-INS-53
chr5-52051228-INS-725
chr5-52064613-INS-350
chr5-52080378-INS-943
chr5-52080529-INS-74
chr5-52143408-INS-70
chr5-52167460-INS-73
chr5-52170513-INS-52
chr5-52176324-INS-400
chr5-52206877-INS-198
chr5-52217443-INS-65
chr5-52228758-INS-116
chr5-52231732-INS-445
chr5-52256752-INS-62
chr5-52288594-INS-86
chr5-52305135-INS-62
chr5-52362597-INS-117
chr5-52366970-INS-83
chr5-52428271-INS-62
chr5-52433401-INS-571
chr5-52446595-INS-99
chr5-52450304-INS-112
chr5-52460422-INS-7082
chr5-52465522-INS-136
chr5-52488741-INS-439
chr5-52495468-INS-82
chr5-52508741-INS-56

chr5-52545273-INS-177
chr5-52562255-INS-70
chr5-52580033-INS-334
chr5-52590260-INS-90
chr5-52594057-INS-91
chr5-52621586-INS-118
chr5-52633035-INS-195
chr5-52637658-INS-388
chr5-52645969-INS-68
chr5-52660082-INS-77
chr5-52661794-INS-198
chr5-52663236-INS-413
chr5-52679137-INS-83
chr5-52730025-INS-6452
chr5-52735804-INS-102
chr5-52749984-INS-54
chr5-52750420-INS-523
chr5-52753436-INS-60
chr5-52770189-INS-56
chr5-52782779-INS-66
chr5-52784422-INS-57
chr5-52834601-INS-57
chr5-52848804-INS-130
chr5-52854334-INS-56
chr5-52866095-INS-353
chr5-52885424-INS-486
chr5-52897705-INS-223
chr5-52983146-INS-52
chr5-53009021-INS-62
chr5-53048554-INS-82
chr5-53090168-INS-166
chr5-53096709-INS-251
chr5-53105085-INS-160
chr5-53106316-INS-66
chr5-53115925-INS-104
chr5-53123051-INS-51
chr5-53127783-INS-91
chr5-53140753-INS-253
chr5-53141965-INS-1000
chr5-53162101-INS-132
chr5-53164307-INS-307
chr5-53185553-INS-444
chr5-53191976-INS-239
chr5-53193866-INS-125
chr5-53195781-INS-186
chr5-53208951-INS-171
chr5-53270455-INS-173
chr5-53307661-INS-81
chr5-53308667-INS-54
chr5-53333159-INS-113
chr5-53342774-INS-75
chr5-53349252-INS-102
chr5-53387179-INS-61
chr5-53392583-INS-56
chr5-53402358-INS-156
chr5-53404057-INS-58
chr5-53454559-INS-5270
chr5-53457490-INS-238
chr5-53463571-INS-84
chr5-53494536-INS-66
chr5-53496011-INS-60
chr5-53496362-INS-58
chr5-53499328-INS-198
chr5-53505897-INS-182
chr5-53510830-INS-312
chr5-53520184-INS-562
chr5-53562045-INS-508
chr5-53563179-INS-76
chr5-53571836-INS-91
chr5-53598016-INS-60
chr5-53599645-INS-52
chr5-53616726-INS-84
chr5-53617358-INS-180
chr5-53627782-INS-79
chr5-53650122-INS-248
chr5-53673547-INS-83
chr5-53679692-INS-60

chr5-53681075-INS-51
chr5-53686134-INS-178
chr5-53692112-INS-195
chr5-53719009-INS-78
chr5-53729307-INS-153
chr5-53731707-INS-133
chr5-53736519-INS-351
chr5-53738712-INS-72
chr5-53740016-INS-62
chr5-53756902-INS-207
chr5-53764143-INS-8770
chr5-53778792-INS-6705
chr5-53783915-INS-341
chr5-53808843-INS-106
chr5-53821149-INS-4964
chr5-53833583-INS-337
chr5-53839593-INS-6524
chr5-53841277-INS-203
chr5-53845113-INS-82
chr5-53853229-INS-68
chr5-53875058-INS-117
chr5-53882140-INS-330
chr5-53891122-INS-59
chr5-53902513-INS-92
chr5-53916662-INS-54
chr5-53931041-INS-288
chr5-53939297-INS-327
chr5-53954741-INS-7343
chr5-53969558-INS-53
chr5-54003518-INS-65
chr5-54016522-INS-62
chr5-54025819-INS-384
chr5-54045832-INS-136
chr5-54054992-INS-181
chr5-54087786-INS-448
chr5-54117120-INS-141
chr5-54120619-INS-91
chr5-54135502-INS-81
chr5-54138032-INS-2186
chr5-54221574-INS-175
chr5-54223283-INS-102
chr5-54284605-INS-77
chr5-54307680-INS-54
chr5-54325423-INS-7153
chr5-54327913-INS-5868
chr5-54347443-INS-80
chr5-54367866-INS-6475
chr5-54372835-INS-60
chr5-54389095-INS-98
chr5-54404079-INS-52
chr5-54431895-INS-4577
chr5-54440272-INS-6596
chr5-54458657-INS-1480
chr5-54466916-INS-132
chr5-54498668-INS-73
chr5-54520127-INS-2276
chr5-54522353-INS-1890
chr5-54538385-INS-56
chr5-54542272-INS-124
chr5-54545070-INS-6195
chr5-54575007-INS-63
chr5-54582214-INS-6460
chr5-54595809-INS-208
chr5-54599591-INS-52
chr5-54632036-INS-271
chr5-54637937-INS-193
chr5-54654063-INS-80
chr5-54669871-INS-421
chr5-54681963-INS-64
chr5-54685993-INS-777
chr5-54694538-INS-6314
chr5-54711606-INS-2780
chr5-54742301-INS-300
chr5-54760646-INS-60
chr5-54767531-INS-65
chr5-54804453-INS-61
chr5-54805863-INS-253

chr5-54821921-INS-483
chr5-54826524-INS-55
chr5-54836186-INS-144
chr5-54841802-INS-179
chr5-54864948-INS-154
chr5-54877937-INS-6276
chr5-54882928-INS-76
chr5-54888087-INS-6314
chr5-54889416-INS-194
chr5-54890018-INS-69
chr5-54925618-INS-77
chr5-54958736-INS-51
chr5-54959762-INS-175
chr5-54965537-INS-441
chr5-54986867-INS-92
chr5-54988143-INS-79
chr5-54991907-INS-95
chr5-55004263-INS-73
chr5-55020870-INS-52
chr5-55046943-INS-956
chr5-55054039-INS-211
chr5-55067232-INS-362
chr5-55069101-INS-94
chr5-55091526-INS-115
chr5-55145158-INS-78
chr5-55152385-INS-381
chr5-55162437-INS-194
chr5-55180957-INS-197
chr5-55183444-INS-56
chr5-55195261-INS-60
chr5-55216179-INS-938
chr5-55257159-INS-52
chr5-55268422-INS-56
chr5-55312490-INS-52
chr5-55348606-INS-54
chr5-55367093-INS-229
chr5-55388672-INS-104
chr5-55405837-INS-717
chr5-55419702-INS-78
chr5-55437258-INS-537
chr5-55452039-INS-52
chr5-55463670-INS-52
chr5-55471125-INS-70
chr5-55499562-INS-201
chr5-55506058-INS-55
chr5-55508633-INS-71
chr5-55531703-INS-88
chr5-55534287-INS-6438
chr5-55536434-INS-76
chr5-55572285-INS-6390
chr5-55591014-INS-141
chr5-55595940-INS-68
chr5-55681539-INS-6743
chr5-55690335-INS-691
chr5-55696853-INS-69
chr5-55724371-INS-318
chr5-55748214-INS-294
chr5-55752675-INS-113
chr5-55760398-INS-135
chr5-55764207-INS-298
chr5-55768088-INS-155
chr5-55770078-INS-322
chr5-55786118-INS-198
chr5-55843375-INS-119
chr5-55865258-INS-100
chr5-55912141-INS-52
chr5-55927059-INS-5266
chr5-55943889-INS-6724
chr5-55971966-INS-63
chr5-56007248-INS-749
chr5-56010718-INS-166
chr5-56040163-INS-150
chr5-56042901-INS-113
chr5-56064381-INS-258
chr5-56066151-INS-92
chr5-56085810-INS-80
chr5-56090523-INS-5543

chr5-56128029-INS-377
chr5-56143829-INS-369
chr5-56144393-INS-54
chr5-56145660-INS-52
chr5-56156327-INS-10324
chr5-56174915-INS-74
chr5-56191254-INS-2892
chr5-56194650-INS-66
chr5-56196602-INS-64
chr5-56199205-INS-196
chr5-56286630-INS-214
chr5-56322294-INS-855
chr5-56330286-INS-152
chr5-56331335-INS-7918
chr5-56333678-INS-494
chr5-56418144-INS-59
chr5-56435448-INS-55
chr5-56437224-INS-522
chr5-56443156-INS-5082
chr5-56509004-INS-299
chr5-56564869-INS-6783
chr5-56571207-INS-67
chr5-56585998-INS-75
chr5-56594718-INS-181
chr5-56595819-INS-167
chr5-56602200-INS-60
chr5-56620813-INS-520
chr5-56626350-INS-54
chr5-56646064-INS-53
chr5-56673185-INS-338
chr5-56694643-INS-72
chr5-56731905-INS-63
chr5-56742084-INS-52
chr5-56760409-INS-118
chr5-56771401-INS-693
chr5-56773275-INS-65
chr5-56828131-INS-396
chr5-56834949-INS-313
chr5-56850084-INS-68
chr5-56876531-INS-765
chr5-56877553-INS-380
chr5-56890493-INS-665
chr5-56905808-INS-69
chr5-56934897-INS-77
chr5-56944424-INS-56
chr5-56944785-INS-808
chr5-56949349-INS-201
chr5-56965423-INS-62
chr5-57014737-INS-88
chr5-57027149-INS-7355
chr5-57038995-INS-59
chr5-57062382-INS-75
chr5-57068134-INS-247
chr5-57068945-INS-420
chr5-57069596-INS-54
chr5-57076865-INS-662
chr5-57206684-INS-792
chr5-57223644-INS-665
chr5-57229639-INS-230
chr5-57240571-INS-71
chr5-57254754-INS-487
chr5-57264876-INS-104
chr5-57269416-INS-80
chr5-57281350-INS-64
chr5-57282808-INS-54
chr5-57293352-INS-88
chr5-57302399-INS-62
chr5-57319677-INS-6721
chr5-57320264-INS-663
chr5-57324862-INS-133
chr5-57327093-INS-666
chr5-57356253-INS-361
chr5-57373409-INS-56
chr5-57375095-INS-128
chr5-57378205-INS-73
chr5-57381046-INS-792
chr5-57382541-INS-946

chr5-57384342-INS-8426
chr5-57408844-INS-383
chr5-57442305-INS-390
chr5-57453248-INS-171
chr5-57455408-INS-322
chr5-57492329-INS-130
chr5-57503723-INS-60
chr5-57507661-INS-467
chr5-57512349-INS-71
chr5-57519981-INS-69
chr5-57541817-INS-63
chr5-57627631-INS-393
chr5-57628983-INS-400
chr5-57653177-INS-55
chr5-57654477-INS-389
chr5-57663799-INS-113
chr5-57668964-INS-57
chr5-57672054-INS-65
chr5-57688919-INS-733
chr5-57695726-INS-6210
chr5-57702675-INS-73
chr5-57720887-INS-72
chr5-57798903-INS-799
chr5-57841993-INS-6675
chr5-57861186-INS-496
chr5-57867815-INS-199
chr5-57906541-INS-208
chr5-57952681-INS-58
chr5-57957946-INS-51
chr5-57982981-INS-152
chr5-57987924-INS-211
chr5-58034543-INS-185
chr5-58044842-INS-256
chr5-58048149-INS-178
chr5-58096258-INS-112
chr5-58097028-INS-411
chr5-58105438-INS-57
chr5-58110138-INS-65
chr5-58138589-INS-64
chr5-58190587-INS-118
chr5-58204532-INS-52
chr5-58208495-INS-311
chr5-58209476-INS-791
chr5-58210454-INS-610
chr5-58213501-INS-58
chr5-58247557-INS-238
chr5-58251864-INS-78
chr5-58281686-INS-179
chr5-58291733-INS-544
chr5-58297975-INS-923
chr5-58302412-INS-60
chr5-58316643-INS-54
chr5-58364777-INS-316
chr5-58398101-INS-358
chr5-58418445-INS-58
chr5-58507606-INS-6429
chr5-58531488-INS-170
chr5-58542301-INS-53
chr5-58545580-INS-472
chr5-58555388-INS-141
chr5-58580556-INS-69
chr5-58619903-INS-90
chr5-58623308-INS-55
chr5-58658879-INS-342
chr5-58671008-INS-56
chr5-58678825-INS-81
chr5-58689430-INS-716
chr5-58699455-INS-506
chr5-58752475-INS-127
chr5-58753923-INS-87
chr5-58772290-INS-412
chr5-58777643-INS-462
chr5-58795091-INS-56
chr5-58857394-INS-54
chr5-58860083-INS-109
chr5-58869412-INS-6334
chr5-58878246-INS-65

chr5-58881890-INS-60
chr5-58892255-INS-72
chr5-58901450-INS-216
chr5-58917477-INS-373
chr5-58972648-INS-78
chr5-58983626-INS-63
chr5-59018729-INS-614
chr5-59029024-INS-162
chr5-59039484-INS-289
chr5-59054770-INS-401
chr5-59073686-INS-3170
chr5-59152388-INS-968
chr5-59154012-INS-6433
chr5-59168607-INS-7047
chr5-59171918-INS-54
chr5-59208670-INS-56
chr5-59222242-INS-64
chr5-59253109-INS-64
chr5-59258496-INS-113
chr5-59267394-INS-66
chr5-59334212-INS-279
chr5-59338631-INS-61
chr5-59396500-INS-6607
chr5-59427425-INS-60
chr5-59463093-INS-15549
chr5-59490313-INS-4549
chr5-59510123-INS-6859
chr5-59539187-INS-58
chr5-59549990-INS-792
chr5-59550902-INS-86
chr5-59583874-INS-1894
chr5-59588744-INS-53
chr5-59645250-INS-284
chr5-59653214-INS-486
chr5-59667620-INS-166
chr5-59691468-INS-67
chr5-59705042-INS-89
chr5-59707786-INS-206
chr5-59714872-INS-434
chr5-59721748-INS-202
chr5-59745089-INS-88
chr5-59792412-INS-2192
chr5-59797604-INS-1706
chr5-59845655-INS-209
chr5-59859559-INS-56
chr5-59870345-INS-358
chr5-59952862-INS-98
chr5-59956651-INS-57
chr5-59958629-INS-288
chr5-59959277-INS-68
chr5-59960751-INS-61
chr5-59972708-INS-6665
chr5-60007100-INS-94
chr5-60015739-INS-158
chr5-60020532-INS-406
chr5-60024724-INS-650
chr5-60038304-INS-88
chr5-60053197-INS-2295
chr5-60064122-INS-565
chr5-60073137-INS-68
chr5-60084944-INS-214
chr5-60094491-INS-7301
chr5-60099151-INS-61
chr5-60120783-INS-86
chr5-60218156-INS-51
chr5-60222391-INS-117
chr5-60261778-INS-54
chr5-60301124-INS-91
chr5-60354295-INS-191
chr5-60365703-INS-465
chr5-60391471-INS-55
chr5-60395721-INS-296
chr5-60407138-INS-84
chr5-60425091-INS-52
chr5-60441394-INS-54
chr5-60459278-INS-1949
chr5-60460634-INS-260

chr5-60473939-INS-588
chr5-60533088-INS-67
chr5-60605548-INS-66
chr5-60618338-INS-986
chr5-60638537-INS-369
chr5-60659200-INS-53
chr5-60663790-INS-53
chr5-60670538-INS-136
chr5-60672242-INS-245
chr5-60737413-INS-198
chr5-60739855-INS-108
chr5-60771127-INS-100
chr5-60776860-INS-248
chr5-60809240-INS-208
chr5-60846959-INS-83
chr5-60861898-INS-676
chr5-60869743-INS-169
chr5-60871100-INS-153
chr5-60881006-INS-5275
chr5-60902177-INS-7146
chr5-60905396-INS-905
chr5-60908522-INS-122
chr5-60922032-INS-102
chr5-60923072-INS-108
chr5-60930149-INS-246
chr5-60940113-INS-52
chr5-60980147-INS-157
chr5-60987631-INS-123
chr5-60991625-INS-874
chr5-61015408-INS-63
chr5-61015576-INS-7105
chr5-61063348-INS-162
chr5-61089306-INS-62
chr5-61090929-INS-128
chr5-61105211-INS-439
chr5-61113312-INS-411
chr5-61155416-INS-116
chr5-61156421-INS-7076
chr5-61167772-INS-6392
chr5-61168666-INS-6348
chr5-61182168-INS-486
chr5-61240749-INS-82
chr5-61253630-INS-375
chr5-61296128-INS-678
chr5-61330960-INS-120
chr5-61337939-INS-6641
chr5-61349717-INS-446
chr5-61350599-INS-575
chr5-61377158-INS-62
chr5-61388501-INS-114
chr5-61402714-INS-182
chr5-61408008-INS-459
chr5-61437310-INS-195
chr5-61438015-INS-234
chr5-61446843-INS-6826
chr5-61518780-INS-411
chr5-61525265-INS-947
chr5-61577437-INS-589
chr5-61598440-INS-82
chr5-61609156-INS-310
chr5-61639252-INS-155
chr5-61642607-INS-572
chr5-61653022-INS-72
chr5-61684227-INS-506
chr5-61717991-INS-346
chr5-61738940-INS-409
chr5-61759682-INS-186
chr5-61767606-INS-58
chr5-61777751-INS-2029
chr5-61787610-INS-123
chr5-61790320-INS-7242
chr5-61805190-INS-76
chr5-61809917-INS-176
chr5-61820354-INS-6002
chr5-61821802-INS-82
chr5-61827451-INS-70
chr5-61857184-INS-756

chr5-61913224-INS-227
chr5-61931789-INS-441
chr5-61973002-INS-123
chr5-61991889-INS-501
chr5-62024045-INS-342
chr5-62030517-INS-719
chr5-62031430-INS-100
chr5-62034851-INS-229
chr5-62051956-INS-53
chr5-62071600-INS-624
chr5-62086116-INS-876
chr5-62087688-INS-5379
chr5-62111378-INS-108
chr5-62144683-INS-892
chr5-62146022-INS-51
chr5-62149583-INS-167
chr5-62212612-INS-92
chr5-62219065-INS-146
chr5-62230288-INS-713
chr5-62232208-INS-86
chr5-62244851-INS-55
chr5-62266702-INS-286
chr5-62267990-INS-3014
chr5-62274094-INS-163
chr5-62286612-INS-437
chr5-62303332-INS-113
chr5-62307977-INS-52
chr5-62316926-INS-52
chr5-62327428-INS-60
chr5-62337505-INS-764
chr5-62338715-INS-295
chr5-62344925-INS-508
chr5-62346883-INS-400
chr5-62362739-INS-7487
chr5-62364263-INS-2217
chr5-62364790-INS-1963
chr5-62383352-INS-214
chr5-62424407-INS-239
chr5-62489645-INS-407
chr5-62492241-INS-79
chr5-62502237-INS-518
chr5-62543054-INS-68
chr5-62557317-INS-74
chr5-62595282-INS-177
chr5-62597904-INS-688
chr5-62620485-INS-438
chr5-62639976-INS-145
chr5-62643964-INS-628
chr5-62649617-INS-189
chr5-62649773-INS-290
chr5-62654541-INS-180
chr5-62709704-INS-70
chr5-62756132-INS-188
chr5-62766805-INS-437
chr5-62789452-INS-62
chr5-62794727-INS-120
chr5-62832713-INS-110
chr5-62838974-INS-123
chr5-62839463-INS-395
chr5-62844401-INS-57
chr5-62852470-INS-146
chr5-62866580-INS-143
chr5-62893833-INS-129
chr5-62895903-INS-497
chr5-62898939-INS-150
chr5-62911701-INS-361
chr5-62917349-INS-116
chr5-62950215-INS-6758
chr5-62986900-INS-72
chr5-62995312-INS-6475
chr5-62997597-INS-770
chr5-63001053-INS-287
chr5-63009539-INS-195
chr5-63026101-INS-408
chr5-63026542-INS-332
chr5-63042546-INS-52
chr5-63044165-INS-132

chr5-63049007-INS-253
chr5-63072269-INS-129
chr5-63085538-INS-403
chr5-63096441-INS-54
chr5-63111429-INS-56
chr5-63119431-INS-321
chr5-63154707-INS-72
chr5-63166305-INS-6574
chr5-63185319-INS-6377
chr5-63214784-INS-72
chr5-63227343-INS-88
chr5-63227816-INS-221
chr5-63266396-INS-6465
chr5-63283421-INS-118
chr5-63307448-INS-201
chr5-63327077-INS-5551
chr5-63328762-INS-6986
chr5-63334902-INS-505
chr5-63339138-INS-6552
chr5-63355984-INS-293
chr5-63371429-INS-275
chr5-63398087-INS-164
chr5-63399698-INS-269
chr5-63418118-INS-101
chr5-63419150-INS-1862
chr5-63419254-INS-461
chr5-63430088-INS-54
chr5-63439824-INS-171
chr5-63444350-INS-904
chr5-63474061-INS-302
chr5-63489804-INS-286
chr5-63505935-INS-56
chr5-63524600-INS-71
chr5-63559060-INS-667
chr5-63563838-INS-153
chr5-63603216-INS-439
chr5-63605486-INS-195
chr5-63618246-INS-100
chr5-63620789-INS-93
chr5-63638483-INS-75
chr5-63646594-INS-2513
chr5-63664598-INS-673
chr5-63665509-INS-262
chr5-63692004-INS-85
chr5-63700059-INS-52
chr5-63708712-INS-5251
chr5-63720164-INS-68
chr5-63742866-INS-167
chr5-63746017-INS-7225
chr5-63760600-INS-5543
chr5-63766505-INS-1996
chr5-63767674-INS-6472
chr5-63770958-INS-6504
chr5-63784714-INS-50
chr5-63792546-INS-110
chr5-63822291-INS-72
chr5-63840480-INS-236
chr5-63856813-INS-60
chr5-63860272-INS-98
chr5-63863947-INS-6470
chr5-63904866-INS-325
chr5-63906132-INS-6474
chr5-63916779-INS-175
chr5-63926322-INS-92
chr5-63929575-INS-53
chr5-63931433-INS-167
chr5-63933693-INS-497
chr5-63934774-INS-538
chr5-63956346-INS-62
chr5-63972857-INS-66
chr5-63976820-INS-451
chr5-63995136-INS-674
chr5-64004746-INS-125
chr5-64021444-INS-96
chr5-64058957-INS-6731
chr5-64059433-INS-56
chr5-64064307-INS-293

chr5-64084986-INS-177
chr5-64096154-INS-102
chr5-64099449-INS-191
chr5-64122453-INS-196
chr5-64129819-INS-151
chr5-64142840-INS-212
chr5-64172954-INS-5136
chr5-64184962-INS-6307
chr5-64226380-INS-182
chr5-64231313-INS-598
chr5-64240326-INS-204
chr5-64268052-INS-639
chr5-64273965-INS-172
chr5-64283000-INS-64
chr5-64286673-INS-204
chr5-64295824-INS-376
chr5-64297194-INS-417
chr5-64322184-INS-882
chr5-64333140-INS-54
chr5-64336221-INS-211
chr5-64354669-INS-370
chr5-64357841-INS-179
chr5-64360681-INS-82
chr5-64402827-INS-491
chr5-64405189-INS-111
chr5-64408122-INS-92
chr5-64416489-INS-192
chr5-64419783-INS-111
chr5-64448041-INS-350
chr5-64448961-INS-115
chr5-64480008-INS-213
chr5-64501517-INS-233
chr5-64519656-INS-384
chr5-64522462-INS-66
chr5-64542049-INS-65
chr5-64551374-INS-73
chr5-64558737-INS-204
chr5-64593765-INS-54
chr5-64610031-INS-54
chr5-64611193-INS-99
chr5-64613598-INS-193
chr5-64614816-INS-104
chr5-64631885-INS-300
chr5-64633308-INS-195
chr5-64662035-INS-78
chr5-64700163-INS-1890
chr5-64709205-INS-102
chr5-64711017-INS-424
chr5-64712560-INS-185
chr5-64726046-INS-56
chr5-64746132-INS-56
chr5-64754308-INS-729
chr5-64792096-INS-219
chr5-64793783-INS-75
chr5-64798348-INS-54
chr5-64848689-INS-77
chr5-64850027-INS-56
chr5-64897733-INS-92
chr5-64908267-INS-66
chr5-64916232-INS-53
chr5-64938967-INS-62
chr5-64971087-INS-377
chr5-64993560-INS-5494
chr5-65001147-INS-1472
chr5-65036683-INS-168
chr5-65037056-INS-237
chr5-65057338-INS-60
chr5-65059328-INS-352
chr5-65201944-INS-278
chr5-65201953-INS-212
chr5-65202041-INS-524
chr5-65218094-INS-54
chr5-65226637-INS-60
chr5-65234146-INS-203
chr5-65244311-INS-168
chr5-65250623-INS-5167
chr5-65285545-INS-278

chr5-65287247-INS-55
chr5-65305637-INS-434
chr5-65306515-INS-52
chr5-65334264-INS-57
chr5-65352777-INS-358
chr5-65366670-INS-434
chr5-65372646-INS-164
chr5-65377381-INS-366
chr5-65378551-INS-64
chr5-65389690-INS-326
chr5-65402201-INS-75
chr5-65406805-INS-227
chr5-65428336-INS-335
chr5-65459740-INS-62
chr5-65462408-INS-828
chr5-65503894-INS-208
chr5-65560729-INS-165
chr5-65586720-INS-62
chr5-65603757-INS-209
chr5-65613405-INS-55
chr5-65630553-INS-395
chr5-65635632-INS-395
chr5-65640937-INS-149
chr5-65663550-INS-81
chr5-65665386-INS-223
chr5-65675826-INS-5059
chr5-65678323-INS-225
chr5-65690314-INS-401
chr5-65697864-INS-261
chr5-65728488-INS-561
chr5-65730320-INS-56
chr5-65739280-INS-52
chr5-65747365-INS-149
chr5-65750165-INS-56
chr5-65786444-INS-335
chr5-65793769-INS-602
chr5-65808090-INS-776
chr5-65809640-INS-362
chr5-65820741-INS-144
chr5-65845515-INS-205
chr5-65849920-INS-447
chr5-65859582-INS-66
chr5-65881464-INS-326
chr5-65891566-INS-252
chr5-65893386-INS-278
chr5-65900671-INS-54
chr5-65901585-INS-4615
chr5-65903853-INS-65
chr5-65927048-INS-204
chr5-65946232-INS-101
chr5-65953733-INS-368
chr5-65968058-INS-174
chr5-65989048-INS-117
chr5-65991382-INS-105
chr5-65994064-INS-74
chr5-66000109-INS-138
chr5-66005199-INS-71
chr5-66025723-INS-52
chr5-66052883-INS-93
chr5-66064682-INS-395
chr5-66065886-INS-449
chr5-66090739-INS-56
chr5-66099006-INS-162
chr5-66107033-INS-80
chr5-66121737-INS-6473
chr5-66134714-INS-1886
chr5-66138051-INS-139
chr5-66160865-INS-90
chr5-66170571-INS-209
chr5-66202836-INS-132
chr5-66204951-INS-313
chr5-66215315-INS-72
chr5-66249063-INS-127
chr5-66258767-INS-299
chr5-66265440-INS-52
chr5-66266738-INS-482
chr5-66276573-INS-385

chr5-66284166-INS-165
chr5-66291316-INS-97
chr5-66318463-INS-99
chr5-66329001-INS-197
chr5-66333852-INS-62
chr5-66340943-INS-238
chr5-66354009-INS-195
chr5-66357911-INS-168
chr5-66361431-INS-165
chr5-66372528-INS-360
chr5-66383161-INS-80
chr5-66391415-INS-54
chr5-66391731-INS-249
chr5-66398213-INS-82
chr5-66407500-INS-75
chr5-66407519-INS-305
chr5-66413159-INS-173
chr5-66419774-INS-53
chr5-66426139-INS-102
chr5-66439963-INS-712
chr5-66440674-INS-117
chr5-66446372-INS-1295
chr5-66447689-INS-138
chr5-66535432-INS-68
chr5-66540654-INS-251
chr5-66571191-INS-55
chr5-66577791-INS-70
chr5-66586342-INS-290
chr5-66592335-INS-89
chr5-66611195-INS-386
chr5-66667577-INS-96
chr5-66688016-INS-57
chr5-66702338-INS-69
chr5-66714341-INS-172
chr5-66754915-INS-55
chr5-66757722-INS-83
chr5-66773594-INS-211
chr5-66782805-INS-120
chr5-66804457-INS-329
chr5-66813645-INS-179
chr5-66818706-INS-140
chr5-66824259-INS-120
chr5-66854141-INS-114
chr5-66860909-INS-308
chr5-66870331-INS-96
chr5-66870912-INS-99
chr5-66915629-INS-103
chr5-66922282-INS-63
chr5-66949248-INS-72
chr5-66961743-INS-261
chr5-66963582-INS-70
chr5-66974031-INS-393
chr5-66986226-INS-140
chr5-66999039-INS-78
chr5-67012684-INS-91
chr5-67059431-INS-55
chr5-67070698-INS-740
chr5-67076646-INS-847
chr5-67120104-INS-52
chr5-67135747-INS-291
chr5-67147547-INS-141
chr5-67155984-INS-54
chr5-67174303-INS-518
chr5-67177306-INS-227
chr5-67178758-INS-87
chr5-67207218-INS-115
chr5-67241003-INS-434
chr5-67267406-INS-230
chr5-67268644-INS-52
chr5-67326673-INS-205
chr5-67337302-INS-377
chr5-67344219-INS-7900
chr5-67355799-INS-415
chr5-67374893-INS-102
chr5-67382973-INS-67
chr5-67425052-INS-71
chr5-67452123-INS-168

chr5-67463798-INS-80
chr5-67481394-INS-342
chr5-67487129-INS-397
chr5-67495551-INS-209
chr5-67514499-INS-497
chr5-67517354-INS-222
chr5-67565634-INS-449
chr5-67576550-INS-217
chr5-67595489-INS-54
chr5-67606101-INS-393
chr5-67618552-INS-67
chr5-67627743-INS-61
chr5-67649330-INS-52
chr5-67662923-INS-52
chr5-67681274-INS-400
chr5-67686197-INS-291
chr5-67696745-INS-921
chr5-67749472-INS-382
chr5-67761587-INS-71
chr5-67806229-INS-1916
chr5-67818670-INS-105
chr5-67821259-INS-60
chr5-67888973-INS-61
chr5-67923825-INS-181
chr5-67949382-INS-384
chr5-67985247-INS-85
chr5-67994944-INS-115
chr5-68012886-INS-601
chr5-68024083-INS-162
chr5-68152859-INS-234
chr5-68186638-INS-160
chr5-68207600-INS-65
chr5-68210060-INS-675
chr5-68219893-INS-52
chr5-68225606-INS-90
chr5-68231725-INS-314
chr5-68235739-INS-188
chr5-68241441-INS-2109
chr5-68248630-INS-150
chr5-68321989-INS-64
chr5-68347018-INS-727
chr5-68352605-INS-196
chr5-68396060-INS-444
chr5-68527851-INS-201
chr5-68554671-INS-91
chr5-68599655-INS-7224
chr5-68600567-INS-6487
chr5-68604825-INS-94
chr5-68661942-INS-166
chr5-68746174-INS-438
chr5-68783893-INS-726
chr5-68793831-INS-498
chr5-68808003-INS-811
chr5-68847984-INS-407
chr5-68852495-INS-60
chr5-68860767-INS-157
chr5-68872176-INS-58
chr5-68890046-INS-76
chr5-68909245-INS-652
chr5-68911274-INS-65
chr5-68915094-INS-283
chr5-68918805-INS-119
chr5-68932244-INS-336
chr5-68939281-INS-123
chr5-69001325-INS-439
chr5-69002443-INS-60
chr5-69004702-INS-54
chr5-69016254-INS-64
chr5-69030464-INS-435
chr5-69032272-INS-106
chr5-69073235-INS-55
chr5-69111797-INS-723
chr5-69151135-INS-67
chr5-69173356-INS-65
chr5-69186353-INS-783
chr5-69227468-INS-54
chr5-69230663-INS-634

chr5-69262361-INS-74
chr5-69269763-INS-690
chr5-69300323-INS-189
chr5-69320000-INS-118
chr5-69328196-INS-62
chr5-69360386-INS-63
chr5-69373573-INS-57
chr5-69397576-INS-185
chr5-69421290-INS-66
chr5-69433620-INS-100
chr5-69439442-INS-164
chr5-69461700-INS-201
chr5-69470425-INS-6475
chr5-69495696-INS-88
chr5-69558346-INS-56
chr5-69562668-INS-52
chr5-69568300-INS-491
chr5-69593884-INS-64
chr5-69602801-INS-118
chr5-69620860-INS-78
chr5-69625564-INS-630
chr5-69640874-INS-184
chr5-69668873-INS-60
chr5-69672515-INS-218
chr5-69689204-INS-214
chr5-69703959-INS-92
chr5-69711158-INS-64
chr5-69713996-INS-116
chr5-69754000-INS-55
chr5-69777216-INS-409
chr5-69787524-INS-651
chr5-69826159-INS-364
chr5-69830728-INS-65
chr5-69832642-INS-408
chr5-69849373-INS-148
chr5-69867938-INS-52
chr5-69870599-INS-61
chr5-69894912-INS-98
chr5-69900184-INS-104
chr5-69902181-INS-64
chr5-69922305-INS-466
chr5-69945439-INS-491
chr5-69962004-INS-676
chr5-70004049-INS-401
chr5-70006045-INS-259
chr5-70017730-INS-525
chr5-70047946-INS-899
chr5-70099492-INS-304
chr5-70102624-INS-277
chr5-70113943-INS-247
chr5-70136567-INS-748
chr5-70139441-INS-55
chr5-70169190-INS-61
chr5-70170314-INS-52
chr5-70188418-INS-243
chr5-70198909-INS-455
chr5-70203050-INS-346
chr5-70331608-INS-391
chr5-70374216-INS-88
chr5-70375901-INS-6785
chr5-70404154-INS-56
chr5-70411720-INS-64
chr5-70417417-INS-60
chr5-70434541-INS-61
chr5-70460751-INS-88
chr5-70466203-INS-56
chr5-70477183-INS-300
chr5-70631663-INS-747
chr5-70643390-INS-82
chr5-70645397-INS-5551
chr5-70646624-INS-137
chr5-70669369-INS-52
chr5-70689665-INS-196
chr5-70690948-INS-823
chr5-70716956-INS-8260
chr5-70803353-INS-53
chr5-70861536-INS-218

chr5-70868443-INS-543
chr5-70886452-INS-830
chr5-70918841-INS-618
chr5-70919417-INS-618
chr5-70947421-INS-270
chr5-70950970-INS-120
chr5-70985342-INS-61
chr5-70989346-INS-300
chr5-70996380-INS-591
chr5-71003449-INS-829
chr5-71007004-INS-137
chr5-71026788-INS-80
chr5-71044043-INS-155
chr5-71056822-INS-136
chr5-71066388-INS-54
chr5-71118148-INS-221
chr5-71125851-INS-60
chr5-71129991-INS-222
chr5-71153460-INS-52
chr5-71190499-INS-443
chr5-71200832-INS-208
chr5-71202138-INS-339
chr5-71222338-INS-106
chr5-71231609-INS-576
chr5-71239535-INS-144
chr5-71263921-INS-380
chr5-71292361-INS-97
chr5-71308049-INS-71
chr5-71352875-INS-52
chr5-71409678-INS-883
chr5-71421247-INS-170
chr5-71458845-INS-119
chr5-71465434-INS-75
chr5-71496290-INS-173
chr5-71514850-INS-509
chr5-71526731-INS-122
chr5-71578252-INS-735
chr5-71582751-INS-104
chr5-71602860-INS-80
chr5-71613981-INS-1166
chr5-71625779-INS-67
chr5-71634521-INS-571
chr5-71644492-INS-63
chr5-71670005-INS-4974
chr5-71709712-INS-6096
chr5-71723962-INS-57
chr5-71729859-INS-164
chr5-71758557-INS-134
chr5-71763867-INS-163
chr5-71787994-INS-73
chr5-71822958-INS-86
chr5-71831301-INS-314
chr5-71843847-INS-719
chr5-71852601-INS-133
chr5-71853264-INS-537
chr5-71865903-INS-123
chr5-71876587-INS-143
chr5-71889977-INS-556
chr5-71926147-INS-66
chr5-71927176-INS-1864
chr5-71931591-INS-214
chr5-71954621-INS-626
chr5-71977127-INS-92
chr5-71987728-INS-6229
chr5-71998288-INS-517
chr5-72006061-INS-80
chr5-72013267-INS-56
chr5-72013803-INS-6487
chr5-72017334-INS-6242
chr5-72040294-INS-138
chr5-72040661-INS-207
chr5-72054042-INS-126
chr5-72056156-INS-495
chr5-72064140-INS-64
chr5-72068904-INS-110
chr5-72128514-INS-365
chr5-72133363-INS-175

chr5-72152394-INS-54
chr5-72160555-INS-66
chr5-72164178-INS-309
chr5-72182775-INS-242
chr5-72195315-INS-54
chr5-72221873-INS-112
chr5-72281590-INS-378
chr5-72308186-INS-102
chr5-72314913-INS-76
chr5-72321102-INS-203
chr5-72346864-INS-969
chr5-72356693-INS-68
chr5-72374386-INS-56
chr5-72402717-INS-849
chr5-72427098-INS-181
chr5-72429905-INS-58
chr5-72465739-INS-967
chr5-72467513-INS-6906
chr5-72472799-INS-61
chr5-72477661-INS-7446
chr5-72526277-INS-58
chr5-72540217-INS-70
chr5-72545517-INS-56
chr5-72549073-INS-148
chr5-72570502-INS-144
chr5-72574857-INS-52
chr5-72580896-INS-794
chr5-72589584-INS-446
chr5-72597774-INS-225
chr5-72608889-INS-315
chr5-72634519-INS-539
chr5-72640987-INS-357
chr5-72721089-INS-129
chr5-72741549-INS-74
chr5-72771832-INS-173
chr5-72866821-INS-55
chr5-72885344-INS-397
chr5-72890318-INS-109
chr5-72890626-INS-186
chr5-72892776-INS-56
chr5-72939128-INS-207
chr5-72965220-INS-72
chr5-72991238-INS-74
chr5-72998592-INS-129
chr5-73004844-INS-76
chr5-73015452-INS-208
chr5-73026869-INS-77
chr5-73030234-INS-280
chr5-73034505-INS-51
chr5-73039449-INS-258
chr5-73058657-INS-134
chr5-73099455-INS-187
chr5-73109735-INS-749
chr5-73112375-INS-193
chr5-73146773-INS-100
chr5-73150868-INS-134
chr5-73181086-INS-64
chr5-73215816-INS-278
chr5-73216255-INS-212
chr5-73230497-INS-52
chr5-73276622-INS-58
chr5-73309974-INS-52
chr5-73317645-INS-116
chr5-73322827-INS-69
chr5-73340911-INS-328
chr5-73350953-INS-53
chr5-73373388-INS-127
chr5-73378847-INS-940
chr5-73384623-INS-101
chr5-73385525-INS-1888
chr5-73391025-INS-53
chr5-73397712-INS-82
chr5-73405037-INS-72
chr5-73406950-INS-129
chr5-73420193-INS-66
chr5-73487200-INS-361
chr5-73488837-INS-200

chr5-73505265-INS-116
chr5-73513778-INS-74
chr5-73547293-INS-181
chr5-73551993-INS-414
chr5-73556577-INS-1044
chr5-73556591-INS-981
chr5-73556611-INS-163
chr5-73566719-INS-270
chr5-73580183-INS-174
chr5-73600352-INS-223
chr5-73610248-INS-98
chr5-73613058-INS-181
chr5-73653021-INS-172
chr5-73691092-INS-122
chr5-73705939-INS-207
chr5-73709227-INS-116
chr5-73725655-INS-66
chr5-73732842-INS-79
chr5-73754951-INS-4569
chr5-73774494-INS-209
chr5-73775993-INS-282
chr5-73857903-INS-121
chr5-73860715-INS-52
chr5-73877638-INS-72
chr5-73883043-INS-1567
chr5-73883062-INS-197
chr5-73918633-INS-240
chr5-73925242-INS-70
chr5-73970645-INS-56
chr5-73989147-INS-246
chr5-74022108-INS-128
chr5-74027029-INS-65
chr5-74035703-INS-147
chr5-74059499-INS-548
chr5-74071983-INS-226
chr5-74079965-INS-485
chr5-74116290-INS-105
chr5-74122546-INS-71
chr5-74138725-INS-614
chr5-74156216-INS-91
chr5-74181633-INS-208
chr5-74189472-INS-90
chr5-74202233-INS-180
chr5-74204182-INS-308
chr5-74214483-INS-209
chr5-74218388-INS-152
chr5-74233695-INS-115
chr5-74241768-INS-215
chr5-74246783-INS-72
chr5-74250719-INS-293
chr5-74260241-INS-199
chr5-74264267-INS-225
chr5-74265985-INS-57
chr5-74278677-INS-82
chr5-74284945-INS-395
chr5-74325945-INS-106
chr5-74332043-INS-58
chr5-74365618-INS-58
chr5-74375193-INS-60
chr5-74435491-INS-958
chr5-74462028-INS-6020
chr5-74462114-INS-70
chr5-74468411-INS-107
chr5-74470718-INS-304
chr5-74510069-INS-76
chr5-74510847-INS-56
chr5-74517795-INS-267
chr5-74545584-INS-135
chr5-74553332-INS-166
chr5-74553600-INS-2671
chr5-74554728-INS-4812
chr5-74567559-INS-56
chr5-74581744-INS-928
chr5-74583854-INS-61
chr5-74585390-INS-174
chr5-74603124-INS-173
chr5-74607351-INS-164

chr5-74683497-INS-490
chr5-74687834-INS-197
chr5-74694452-INS-216
chr5-74697335-INS-327
chr5-74700806-INS-54
chr5-74707724-INS-222
chr5-74775586-INS-8101
chr5-74778282-INS-63
chr5-74799056-INS-54
chr5-74817808-INS-67
chr5-74848528-INS-4549
chr5-74852237-INS-394
chr5-74922003-INS-312
chr5-74938343-INS-52
chr5-74973572-INS-52
chr5-74974954-INS-72
chr5-74990526-INS-58
chr5-75017745-INS-77
chr5-75028771-INS-524
chr5-75040994-INS-224
chr5-75048951-INS-206
chr5-75052134-INS-288
chr5-75071524-INS-209
chr5-75104215-INS-91
chr5-75116546-INS-66
chr5-75127253-INS-60
chr5-75137168-INS-413
chr5-75197450-INS-60
chr5-75209615-INS-107
chr5-75239758-INS-50
chr5-75249646-INS-151
chr5-75250743-INS-60
chr5-75261549-INS-52
chr5-75273086-INS-219
chr5-75281926-INS-1638
chr5-75308343-INS-63
chr5-75322091-INS-62
chr5-75327905-INS-223
chr5-75372091-INS-52
chr5-75421838-INS-6635
chr5-75434126-INS-621
chr5-75436741-INS-310
chr5-75444000-INS-266
chr5-75465296-INS-3059
chr5-75495669-INS-100
chr5-75514810-INS-58
chr5-75599515-INS-71
chr5-75601067-INS-59
chr5-75606365-INS-66
chr5-75613534-INS-127
chr5-75622808-INS-154
chr5-75625865-INS-71
chr5-75642837-INS-65
chr5-75647402-INS-296
chr5-75656901-INS-347
chr5-75658079-INS-121
chr5-75674378-INS-70
chr5-75689048-INS-103
chr5-75696458-INS-608
chr5-75699573-INS-56
chr5-75702268-INS-111
chr5-75706552-INS-72
chr5-75759674-INS-64
chr5-75783767-INS-69
chr5-75828482-INS-106
chr5-75829821-INS-186
chr5-75833216-INS-97
chr5-75840321-INS-74
chr5-75888160-INS-210
chr5-75893100-INS-193
chr5-75921356-INS-1621
chr5-75959846-INS-96
chr5-75980407-INS-115
chr5-75981953-INS-52
chr5-75994582-INS-197
chr5-76009919-INS-121
chr5-76050032-INS-477

chr5-76068705-INS-6615
chr5-76074463-INS-56
chr5-76074837-INS-6524
chr5-76077144-INS-4374
chr5-76088466-INS-87
chr5-76111646-INS-52
chr5-76137984-INS-120
chr5-76141592-INS-77
chr5-76143978-INS-213
chr5-76149959-INS-115
chr5-76170602-INS-6594
chr5-76181216-INS-128
chr5-76204620-INS-249
chr5-76277821-INS-111
chr5-76300864-INS-56
chr5-76342333-INS-118
chr5-76358799-INS-53
chr5-76371724-INS-204
chr5-76468844-INS-83
chr5-76479496-INS-52
chr5-76491052-INS-57
chr5-76512588-INS-93
chr5-76527653-INS-57
chr5-76546839-INS-82
chr5-76569057-INS-95
chr5-76580160-INS-6943
chr5-76636205-INS-184
chr5-76642858-INS-58
chr5-76680536-INS-212
chr5-76690948-INS-51
chr5-76718588-INS-205
chr5-76722776-INS-528
chr5-76734479-INS-52
chr5-76740166-INS-181
chr5-76759781-INS-134
chr5-76810978-INS-59
chr5-76823873-INS-58
chr5-76826624-INS-433
chr5-76826784-INS-516
chr5-76830281-INS-62
chr5-76831676-INS-106
chr5-76838871-INS-60
chr5-76861999-INS-198
chr5-76893506-INS-68
chr5-76920509-INS-88
chr5-76933620-INS-66
chr5-76945934-INS-218
chr5-76958667-INS-60
chr5-76968674-INS-70
chr5-76970143-INS-76
chr5-76986101-INS-7130
chr5-77037076-INS-208
chr5-77038742-INS-553
chr5-77041235-INS-229
chr5-77042549-INS-190
chr5-77049678-INS-92
chr5-77050776-INS-108
chr5-77091794-INS-208
chr5-77094317-INS-125
chr5-77104651-INS-196
chr5-77110052-INS-198
chr5-77126013-INS-551
chr5-77160899-INS-210
chr5-77182382-INS-327
chr5-77197982-INS-325
chr5-77235071-INS-60
chr5-77246126-INS-73
chr5-77292045-INS-703
chr5-77298135-INS-788
chr5-77345497-INS-238
chr5-77356006-INS-1296
chr5-77391315-INS-282
chr5-77396977-INS-367
chr5-77403824-INS-201
chr5-77408518-INS-170
chr5-77420482-INS-212
chr5-77437439-INS-211

chr5-77447685-INS-187
chr5-77452024-INS-216
chr5-77490018-INS-88
chr5-77521047-INS-126
chr5-77526509-INS-55
chr5-77549975-INS-87
chr5-77587284-INS-880
chr5-77588249-INS-265
chr5-77602856-INS-960
chr5-77617583-INS-176
chr5-77650482-INS-56
chr5-77657298-INS-306
chr5-77659857-INS-211
chr5-77667810-INS-62
chr5-77671421-INS-129
chr5-77683204-INS-57
chr5-77692046-INS-133
chr5-77694112-INS-83
chr5-77734480-INS-78
chr5-77800431-INS-322
chr5-77807584-INS-7226
chr5-77808917-INS-108
chr5-77810743-INS-168
chr5-77814161-INS-3018
chr5-77841090-INS-800
chr5-77843275-INS-200
chr5-77859072-INS-286
chr5-77864322-INS-6418
chr5-77868282-INS-68
chr5-77871416-INS-81
chr5-77872667-INS-133
chr5-77881621-INS-7534
chr5-77926079-INS-137
chr5-77939485-INS-142
chr5-77948826-INS-128
chr5-77955741-INS-458
chr5-78009432-INS-207
chr5-78009569-INS-5750
chr5-78010822-INS-142
chr5-78035085-INS-133
chr5-78045003-INS-112
chr5-78062756-INS-347
chr5-78105523-INS-364
chr5-78132802-INS-136
chr5-78156259-INS-662
chr5-78156900-INS-172
chr5-78171728-INS-343
chr5-78175472-INS-6655
chr5-78196888-INS-124
chr5-78211155-INS-67
chr5-78253170-INS-361
chr5-78288255-INS-653
chr5-78309309-INS-470
chr5-78309534-INS-208
chr5-78313061-INS-63
chr5-78383202-INS-166
chr5-78454237-INS-170
chr5-78486743-INS-372
chr5-78490574-INS-892
chr5-78502138-INS-244
chr5-78503807-INS-447
chr5-78506059-INS-5251
chr5-78509712-INS-6439
chr5-78521965-INS-6913
chr5-78565433-INS-375
chr5-78569908-INS-67
chr5-78583284-INS-239
chr5-78588158-INS-398
chr5-78594123-INS-77
chr5-78596440-INS-653
chr5-78610968-INS-66
chr5-78678987-INS-97
chr5-78689682-INS-340
chr5-78734369-INS-69
chr5-78741278-INS-785
chr5-78743411-INS-66
chr5-78782525-INS-171

chr5-78783970-INS-113
chr5-78787853-INS-56
chr5-78788431-INS-541
chr5-78820312-INS-339
chr5-78889315-INS-56
chr5-78891092-INS-68
chr5-78891818-INS-6664
chr5-78912830-INS-101
chr5-78921248-INS-79
chr5-78946150-INS-100
chr5-78974996-INS-572
chr5-78982502-INS-155
chr5-78986002-INS-55
chr5-79015948-INS-348
chr5-79017128-INS-827
chr5-79028409-INS-11580
chr5-79051228-INS-74
chr5-79054323-INS-52
chr5-79058676-INS-51
chr5-79065241-INS-284
chr5-79070478-INS-174
chr5-79096700-INS-521
chr5-79103879-INS-358
chr5-79113953-INS-426
chr5-79115891-INS-99
chr5-79131702-INS-243
chr5-79134662-INS-114
chr5-79163513-INS-348
chr5-79180865-INS-212
chr5-79198424-INS-56
chr5-79198928-INS-964
chr5-79200751-INS-6942
chr5-79285689-INS-311
chr5-79293580-INS-202
chr5-79302869-INS-67
chr5-79326648-INS-212
chr5-79347630-INS-304
chr5-79362505-INS-134
chr5-79404653-INS-79
chr5-79416368-INS-1029
chr5-79452588-INS-5312
chr5-79506259-INS-390
chr5-79538775-INS-5678
chr5-79569573-INS-245
chr5-79584397-INS-76
chr5-79587613-INS-285
chr5-79607837-INS-114
chr5-79615046-INS-819
chr5-79620883-INS-796
chr5-79627180-INS-99
chr5-79633465-INS-55
chr5-79654812-INS-445
chr5-79655936-INS-448
chr5-79657444-INS-186
chr5-79664103-INS-215
chr5-79701132-INS-213
chr5-79703568-INS-148
chr5-79721348-INS-128
chr5-79724245-INS-1864
chr5-79742811-INS-993
chr5-79763074-INS-60
chr5-79769426-INS-70
chr5-79788748-INS-376
chr5-79809587-INS-648
chr5-79828220-INS-6088
chr5-79828649-INS-267
chr5-79836510-INS-451
chr5-79844530-INS-516
chr5-79899248-INS-351
chr5-79899986-INS-138
chr5-79901378-INS-163
chr5-79926053-INS-78
chr5-79992749-INS-830
chr5-80012358-INS-699
chr5-80041875-INS-6297
chr5-80066399-INS-313
chr5-80066509-INS-7281

chr5-80073447-INS-89
chr5-80098852-INS-111
chr5-80151426-INS-86
chr5-80182576-INS-507
chr5-80193327-INS-924
chr5-80212648-INS-1632
chr5-80214872-INS-52
chr5-80236673-INS-2086
chr5-80236852-INS-52
chr5-80238724-INS-931
chr5-80256177-INS-4362
chr5-80260104-INS-1489
chr5-80302236-INS-412
chr5-80307394-INS-6567
chr5-80324481-INS-170
chr5-80327847-INS-1864
chr5-80348117-INS-56
chr5-80360493-INS-740
chr5-80374877-INS-702
chr5-80377614-INS-94
chr5-80417246-INS-57
chr5-80451559-INS-350
chr5-80460593-INS-1551
chr5-80473531-INS-113
chr5-80476574-INS-1073
chr5-80494546-INS-72
chr5-80512924-INS-6290
chr5-80515596-INS-504
chr5-80519290-INS-699
chr5-80549938-INS-84
chr5-80580332-INS-75
chr5-80612825-INS-54
chr5-80625641-INS-212
chr5-80643847-INS-331
chr5-80650167-INS-121
chr5-80651073-INS-281
chr5-80652797-INS-132
chr5-80669345-INS-2997
chr5-80672426-INS-629
chr5-80675351-INS-82
chr5-80680728-INS-7124
chr5-80699277-INS-704
chr5-80719607-INS-1892
chr5-80737886-INS-3277
chr5-80740473-INS-173
chr5-80767966-INS-72
chr5-80776336-INS-117
chr5-80783278-INS-94
chr5-80793001-INS-6639
chr5-80801080-INS-69
chr5-80814177-INS-238
chr5-80817909-INS-427
chr5-80829217-INS-82
chr5-80831082-INS-312
chr5-80851057-INS-110
chr5-80885637-INS-113
chr5-80896861-INS-6355
chr5-80900783-INS-350
chr5-80914697-INS-681
chr5-80927716-INS-972
chr5-80930909-INS-58
chr5-80944983-INS-118
chr5-80955893-INS-133
chr5-80958483-INS-548
chr5-80962389-INS-751
chr5-80971213-INS-360
chr5-80974135-INS-7796
chr5-80983918-INS-6420
chr5-81033561-INS-58
chr5-81034251-INS-284
chr5-81072446-INS-1892
chr5-81075980-INS-7143
chr5-81081114-INS-1362
chr5-81090652-INS-1096
chr5-81107604-INS-372
chr5-81111460-INS-694
chr5-81115228-INS-455

chr5-81119518-INS-167
chr5-81120002-INS-524
chr5-81124162-INS-64
chr5-81150334-INS-174
chr5-81155314-INS-344
chr5-81160589-INS-88
chr5-81174790-INS-5414
chr5-81176515-INS-52
chr5-81204911-INS-728
chr5-81218214-INS-115
chr5-81239563-INS-56
chr5-81269504-INS-628
chr5-81286321-INS-6444
chr5-81286367-INS-1866
chr5-81293830-INS-54
chr5-81303411-INS-6480
chr5-81309010-INS-634
chr5-81332738-INS-64
chr5-81385805-INS-95
chr5-81394279-INS-66
chr5-81401925-INS-109
chr5-81427526-INS-68
chr5-81491461-INS-54
chr5-81508703-INS-73
chr5-81524247-INS-183
chr5-81537279-INS-431
chr5-81554152-INS-198
chr5-81565487-INS-451
chr5-81577681-INS-56
chr5-81603819-INS-72
chr5-81628272-INS-524
chr5-81632082-INS-1891
chr5-81634987-INS-67
chr5-81640989-INS-241
chr5-81642746-INS-203
chr5-81647206-INS-2308
chr5-81677252-INS-879
chr5-81702062-INS-284
chr5-81710934-INS-202
chr5-81739212-INS-193
chr5-81752314-INS-65
chr5-81759130-INS-58
chr5-81771285-INS-6473
chr5-81784020-INS-561
chr5-81809234-INS-282
chr5-81828383-INS-64
chr5-81847928-INS-7321
chr5-81866909-INS-391
chr5-81867850-INS-65
chr5-81895403-INS-360
chr5-81907949-INS-65
chr5-81918538-INS-62
chr5-81941520-INS-3694
chr5-81948679-INS-203
chr5-81960827-INS-55
chr5-81961833-INS-147
chr5-81988324-INS-171
chr5-81990422-INS-66
chr5-82033740-INS-498
chr5-82062742-INS-124
chr5-82070461-INS-399
chr5-82095859-INS-422
chr5-82104954-INS-372
chr5-82118718-INS-7640
chr5-82129960-INS-174
chr5-82141170-INS-83
chr5-82202376-INS-743
chr5-82208973-INS-591
chr5-82217556-INS-198
chr5-82227506-INS-225
chr5-82240847-INS-217
chr5-82244286-INS-99
chr5-82250927-INS-69
chr5-82254364-INS-225
chr5-82268065-INS-536
chr5-82276164-INS-316
chr5-82294767-INS-589

chr5-82300233-INS-372
chr5-82309127-INS-94
chr5-82346762-INS-8113
chr5-82365020-INS-590
chr5-82383932-INS-283
chr5-82401133-INS-339
chr5-82402369-INS-6327
chr5-82436808-INS-6214
chr5-82498171-INS-733
chr5-82503119-INS-51
chr5-82537883-INS-80
chr5-82572771-INS-52
chr5-82572938-INS-891
chr5-82586196-INS-678
chr5-82590173-INS-360
chr5-82594508-INS-521
chr5-82601427-INS-78
chr5-82644510-INS-84
chr5-82670339-INS-296
chr5-82690469-INS-107
chr5-82694888-INS-6723
chr5-82857745-INS-542
chr5-82864616-INS-89
chr5-82887673-INS-7236
chr5-82913620-INS-97
chr5-82938561-INS-259
chr5-82945922-INS-597
chr5-82965437-INS-642
chr5-82968124-INS-288
chr5-82987383-INS-405
chr5-82989095-INS-199
chr5-82989657-INS-6880
chr5-82990457-INS-5320
chr5-82998413-INS-492
chr5-83077746-INS-610
chr5-83082237-INS-384
chr5-83106243-INS-476
chr5-83151074-INS-343
chr5-83152514-INS-132
chr5-83182560-INS-91
chr5-83270717-INS-6625
chr5-83276822-INS-5292
chr5-83328959-INS-99
chr5-83329169-INS-861
chr5-83347248-INS-7125
chr5-83470176-INS-7141
chr5-83484661-INS-392
chr5-83519419-INS-369
chr5-83539783-INS-54
chr5-83545342-INS-116
chr5-83558372-INS-2224
chr5-83570534-INS-70
chr5-83571776-INS-114
chr5-83662831-INS-400
chr5-83678565-INS-425
chr5-83686878-INS-235
chr5-83690463-INS-73
chr5-83731287-INS-68
chr5-83742916-INS-1338
chr5-83776247-INS-231
chr5-83788520-INS-181
chr5-83827724-INS-156
chr5-83835926-INS-204
chr5-83844514-INS-77
chr5-83850553-INS-63
chr5-83872415-INS-331
chr5-83928595-INS-124
chr5-83947680-INS-280
chr5-83959475-INS-229
chr5-83988597-INS-111
chr5-83994790-INS-205
chr5-84000467-INS-56
chr5-84057980-INS-489
chr5-84153607-INS-190
chr5-84160563-INS-679
chr5-84162977-INS-7276
chr5-84166697-INS-457

chr5-84167696-INS-390
chr5-84186937-INS-6675
chr5-84192636-INS-103
chr5-84224899-INS-926
chr5-84233031-INS-74
chr5-84246147-INS-280
chr5-84275687-INS-112
chr5-84305366-INS-68
chr5-84314781-INS-288
chr5-84320924-INS-334
chr5-84391785-INS-213
chr5-84394623-INS-6934
chr5-84408248-INS-108
chr5-84419624-INS-143
chr5-84453494-INS-7198
chr5-84456880-INS-199
chr5-84496932-INS-58
chr5-84509931-INS-116
chr5-84518819-INS-550
chr5-84552126-INS-243
chr5-84558767-INS-105
chr5-84559670-INS-350
chr5-84574426-INS-208
chr5-84578946-INS-441
chr5-84607441-INS-52
chr5-84631672-INS-116
chr5-84633526-INS-939
chr5-84666723-INS-72
chr5-84681577-INS-539
chr5-84682642-INS-86
chr5-84686272-INS-7397
chr5-84721906-INS-6473
chr5-84726240-INS-285
chr5-84727130-INS-100
chr5-84731232-INS-343
chr5-84734617-INS-212
chr5-84741913-INS-52
chr5-84743913-INS-212
chr5-84746313-INS-63
chr5-84748144-INS-395
chr5-84749213-INS-197
chr5-84751395-INS-65
chr5-84760283-INS-324
chr5-84776048-INS-211
chr5-84833479-INS-3624
chr5-84856240-INS-301
chr5-84989677-INS-252
chr5-85040638-INS-664
chr5-85056336-INS-6826
chr5-85058559-INS-6441
chr5-85080518-INS-133
chr5-85099481-INS-62
chr5-85149713-INS-134
chr5-85156756-INS-384
chr5-85164837-INS-407
chr5-85219343-INS-653
chr5-85233877-INS-60
chr5-85247069-INS-52
chr5-85347914-INS-269
chr5-85350442-INS-59
chr5-85351495-INS-564
chr5-85357989-INS-54
chr5-85369365-INS-186
chr5-85381576-INS-742
chr5-85439324-INS-301
chr5-85457717-INS-329
chr5-85499835-INS-701
chr5-85512749-INS-52
chr5-85514774-INS-7085
chr5-85560516-INS-74
chr5-85588359-INS-74
chr5-85632849-INS-341
chr5-85637731-INS-92
chr5-85668872-INS-1862
chr5-85684635-INS-56
chr5-85688249-INS-423
chr5-85696098-INS-52

chr5-85698670-INS-494
chr5-85732568-INS-171
chr5-85759145-INS-928
chr5-85777730-INS-7042
chr5-85800437-INS-487
chr5-85810672-INS-214
chr5-85836001-INS-108
chr5-85861213-INS-129
chr5-85878379-INS-110
chr5-85890012-INS-126
chr5-85910902-INS-205
chr5-85914514-INS-551
chr5-85922505-INS-358
chr5-85923858-INS-193
chr5-85957567-INS-489
chr5-86046767-INS-138
chr5-86049013-INS-328
chr5-86050873-INS-246
chr5-86083854-INS-373
chr5-86084873-INS-346
chr5-86109506-INS-90
chr5-86113768-INS-281
chr5-86122816-INS-183
chr5-86130456-INS-222
chr5-86142306-INS-55
chr5-86161662-INS-102
chr5-86190174-INS-980
chr5-86223133-INS-361
chr5-86233230-INS-65
chr5-86245778-INS-59
chr5-86255494-INS-1231
chr5-86265226-INS-52
chr5-86297165-INS-234
chr5-86321460-INS-514
chr5-86347098-INS-118
chr5-86382342-INS-171
chr5-86395170-INS-63
chr5-86401432-INS-103
chr5-86408539-INS-72
chr5-86460184-INS-63
chr5-86477344-INS-51
chr5-86492393-INS-66
chr5-86494240-INS-76
chr5-86500012-INS-334
chr5-86547204-INS-108
chr5-86552886-INS-466
chr5-86556937-INS-672
chr5-86564610-INS-202
chr5-86593914-INS-410
chr5-86607333-INS-202
chr5-86629318-INS-6909
chr5-86654840-INS-470
chr5-86659327-INS-53
chr5-86679479-INS-162
chr5-86684350-INS-6384
chr5-86691805-INS-64
chr5-86711075-INS-659
chr5-86757772-INS-400
chr5-86761943-INS-391
chr5-86811769-INS-258
chr5-86812685-INS-721
chr5-86869425-INS-894
chr5-86871588-INS-206
chr5-86880289-INS-75
chr5-86916840-INS-66
chr5-86924597-INS-136
chr5-86930336-INS-140
chr5-86936043-INS-187
chr5-86953478-INS-58
chr5-86959820-INS-614
chr5-87003633-INS-128
chr5-87051217-INS-69
chr5-87088677-INS-298
chr5-87125100-INS-13805
chr5-87134353-INS-461
chr5-87177160-INS-54
chr5-87177991-INS-494

chr5-87212333-INS-357
chr5-87217617-INS-52
chr5-87219435-INS-83
chr5-87232118-INS-79
chr5-87232540-INS-719
chr5-87245807-INS-1891
chr5-87248421-INS-58
chr5-87340826-INS-474
chr5-87343628-INS-531
chr5-87392201-INS-152
chr5-87424872-INS-6511
chr5-87452159-INS-332
chr5-87527972-INS-199
chr5-87565330-INS-368
chr5-87586641-INS-9745
chr5-87601685-INS-6868
chr5-87610431-INS-7186
chr5-87648179-INS-176
chr5-87666054-INS-324
chr5-87667721-INS-330
chr5-87684877-INS-106
chr5-87702114-INS-9247
chr5-87718473-INS-737
chr5-87733505-INS-131
chr5-87742112-INS-104
chr5-87761558-INS-56
chr5-87765811-INS-66
chr5-87803712-INS-127
chr5-87823815-INS-258
chr5-87849853-INS-53
chr5-87858073-INS-138
chr5-87867120-INS-55
chr5-87920967-INS-349
chr5-87948207-INS-63
chr5-87950277-INS-344
chr5-87952098-INS-217
chr5-87952461-INS-154
chr5-87958900-INS-209
chr5-87965877-INS-54
chr5-87975146-INS-92
chr5-87980075-INS-87
chr5-87999457-INS-826
chr5-88006253-INS-77
chr5-88038583-INS-66
chr5-88078810-INS-290
chr5-88088904-INS-667
chr5-88094563-INS-60
chr5-88150004-INS-1095
chr5-88158558-INS-90
chr5-88181348-INS-2919
chr5-88182266-INS-7036
chr5-88189758-INS-1152
chr5-88203655-INS-326
chr5-88209522-INS-150
chr5-88248647-INS-429
chr5-88362261-INS-687
chr5-88421499-INS-514
chr5-88422444-INS-6710
chr5-88434861-INS-102
chr5-88470686-INS-142
chr5-88527833-INS-206
chr5-88546026-INS-384
chr5-88567184-INS-489
chr5-88570753-INS-52
chr5-88601634-INS-131
chr5-88614362-INS-550
chr5-88637186-INS-385
chr5-88643626-INS-536
chr5-88661074-INS-626
chr5-88676421-INS-77
chr5-88678348-INS-139
chr5-88684162-INS-169
chr5-88690554-INS-465
chr5-88719075-INS-51
chr5-88726264-INS-103
chr5-88742700-INS-51
chr5-88777548-INS-247

chr5-88782697-INS-322
chr5-88819276-INS-176
chr5-88824869-INS-171
chr5-88829229-INS-54
chr5-88834604-INS-225
chr5-88842848-INS-366
chr5-88846936-INS-195
chr5-88854633-INS-99
chr5-88855926-INS-801
chr5-88858554-INS-95
chr5-88869328-INS-64
chr5-88873386-INS-242
chr5-88876129-INS-212
chr5-88878058-INS-816
chr5-88895313-INS-872
chr5-88926618-INS-51
chr5-88944305-INS-699
chr5-88960627-INS-88
chr5-88970575-INS-6396
chr5-88985776-INS-363
chr5-88994809-INS-7124
chr5-88999837-INS-60
chr5-89009401-INS-164
chr5-89028456-INS-6629
chr5-89097214-INS-524
chr5-89139781-INS-52
chr5-89142302-INS-166
chr5-89156439-INS-82
chr5-89164853-INS-6828
chr5-89189933-INS-80
chr5-89211950-INS-186
chr5-89214157-INS-1893
chr5-89215720-INS-68
chr5-89224134-INS-51
chr5-89233889-INS-186
chr5-89237121-INS-52
chr5-89250207-INS-110
chr5-89280977-INS-100
chr5-89288287-INS-170
chr5-89328846-INS-52
chr5-89360565-INS-7290
chr5-89389872-INS-5369
chr5-89392951-INS-55
chr5-89396047-INS-6978
chr5-89415573-INS-6297
chr5-89466994-INS-164
chr5-89467510-INS-638
chr5-89468880-INS-6679
chr5-89470756-INS-4199
chr5-89473381-INS-365
chr5-89497370-INS-329
chr5-89530532-INS-126
chr5-89531245-INS-1329
chr5-89539343-INS-99
chr5-89555267-INS-474
chr5-89588807-INS-492
chr5-89594159-INS-6481
chr5-89614447-INS-129
chr5-89636330-INS-354
chr5-89638969-INS-144
chr5-89644376-INS-201
chr5-89648545-INS-8723
chr5-89670186-INS-190
chr5-89672132-INS-213
chr5-89677445-INS-583
chr5-89715251-INS-6437
chr5-89739978-INS-157
chr5-89754219-INS-324
chr5-89768808-INS-252
chr5-89821953-INS-6618
chr5-89828947-INS-535
chr5-89840642-INS-296
chr5-89847089-INS-70
chr5-89891239-INS-327
chr5-89898943-INS-124
chr5-89924915-INS-432
chr5-89939145-INS-85

chr5-89964939-INS-121
chr5-89974521-INS-136
chr5-89974570-INS-200
chr5-89997907-INS-84
chr5-90000260-INS-128
chr5-90005898-INS-917
chr5-90057569-INS-72
chr5-90065455-INS-324
chr5-90084584-INS-9547
chr5-90095813-INS-505
chr5-90101007-INS-204
chr5-90111734-INS-315
chr5-90143990-INS-112
chr5-90147688-INS-141
chr5-90163056-INS-1988
chr5-90165375-INS-729
chr5-90170032-INS-70
chr5-90181877-INS-86
chr5-90186906-INS-2495
chr5-90189432-INS-171
chr5-90213287-INS-58
chr5-90232072-INS-121
chr5-90237973-INS-155
chr5-90263411-INS-64
chr5-90273709-INS-130
chr5-90294709-INS-52
chr5-90303313-INS-7095
chr5-90305321-INS-214
chr5-90308530-INS-121
chr5-90326501-INS-508
chr5-90328436-INS-231
chr5-90335750-INS-264
chr5-90341988-INS-243
chr5-90352375-INS-65
chr5-90354270-INS-66
chr5-90358928-INS-754
chr5-90368827-INS-194
chr5-90372329-INS-104
chr5-90373303-INS-206
chr5-90373966-INS-77
chr5-90389094-INS-328
chr5-90402098-INS-503
chr5-90464568-INS-80
chr5-90475208-INS-72
chr5-90481461-INS-53
chr5-90525547-INS-6582
chr5-90531957-INS-577
chr5-90558884-INS-159
chr5-90561288-INS-172
chr5-90582149-INS-84
chr5-90585210-INS-59
chr5-90589133-INS-55
chr5-90606473-INS-79
chr5-90664959-INS-100
chr5-90668587-INS-151
chr5-90741374-INS-55
chr5-90758608-INS-132
chr5-90766094-INS-114
chr5-90778400-INS-54
chr5-90834193-INS-129
chr5-90845991-INS-424
chr5-90858058-INS-55
chr5-90860481-INS-73
chr5-90873974-INS-646
chr5-90900877-INS-421
chr5-90902821-INS-80
chr5-90906492-INS-52
chr5-90949070-INS-58
chr5-90969524-INS-91
chr5-90989082-INS-165
chr5-91025674-INS-209
chr5-91030024-INS-87
chr5-91034041-INS-382
chr5-91036244-INS-178
chr5-91054523-INS-386
chr5-91067634-INS-7252
chr5-91099488-INS-145

chr5-91100871-INS-93
chr5-91111124-INS-560
chr5-91121087-INS-78
chr5-91130206-INS-110
chr5-91146548-INS-113
chr5-91151358-INS-132
chr5-91173665-INS-52
chr5-91191444-INS-304
chr5-91197126-INS-507
chr5-91198672-INS-52
chr5-91203029-INS-173
chr5-91228415-INS-638
chr5-91257777-INS-350
chr5-91289651-INS-69
chr5-91306311-INS-104
chr5-91320282-INS-115
chr5-91323161-INS-581
chr5-91333948-INS-68
chr5-91371153-INS-84
chr5-91397700-INS-4558
chr5-91410336-INS-392
chr5-91438278-INS-115
chr5-91440502-INS-76
chr5-91444109-INS-145
chr5-91483702-INS-149
chr5-91492298-INS-390
chr5-91543265-INS-152
chr5-91547254-INS-99
chr5-91564483-INS-551
chr5-91574940-INS-128
chr5-91587451-INS-493
chr5-91592051-INS-134
chr5-91598900-INS-166
chr5-91601972-INS-117
chr5-91611110-INS-7220
chr5-91631281-INS-166
chr5-91635124-INS-55
chr5-91641009-INS-6405
chr5-91642860-INS-6441
chr5-91688844-INS-70
chr5-91693332-INS-584
chr5-91701301-INS-432
chr5-91752448-INS-70
chr5-91817839-INS-456
chr5-91824498-INS-205
chr5-91855664-INS-187
chr5-91881900-INS-6363
chr5-91891404-INS-59
chr5-91897707-INS-122
chr5-91920358-INS-380
chr5-91927551-INS-114
chr5-91932189-INS-6652
chr5-91942285-INS-453
chr5-92005571-INS-64
chr5-92037508-INS-176
chr5-92042147-INS-154
chr5-92044039-INS-594
chr5-92058074-INS-6601
chr5-92097131-INS-74
chr5-92122443-INS-359
chr5-92151231-INS-309
chr5-92179899-INS-339
chr5-92183947-INS-203
chr5-92197245-INS-204
chr5-92224482-INS-193
chr5-92237804-INS-54
chr5-92240824-INS-74
chr5-92248158-INS-230
chr5-92254015-INS-197
chr5-92271495-INS-5364
chr5-92274082-INS-209
chr5-92293515-INS-55
chr5-92295484-INS-224
chr5-92303676-INS-172
chr5-92308157-INS-203
chr5-92309020-INS-207
chr5-92311734-INS-324

chr5-92370993-INS-129
chr5-92377939-INS-1880
chr5-92421991-INS-57
chr5-92426771-INS-534
chr5-92428391-INS-105
chr5-92440816-INS-143
chr5-92449229-INS-93
chr5-92481704-INS-60
chr5-92498848-INS-72
chr5-92518146-INS-103
chr5-92531620-INS-211
chr5-92560643-INS-113
chr5-92568214-INS-72
chr5-92577880-INS-219
chr5-92588394-INS-495
chr5-92610501-INS-328
chr5-92645741-INS-91
chr5-92654907-INS-173
chr5-92675372-INS-232
chr5-92683060-INS-402
chr5-92693332-INS-6401
chr5-92704732-INS-532
chr5-92709353-INS-68
chr5-92711720-INS-100
chr5-92739489-INS-888
chr5-92741084-INS-180
chr5-92754687-INS-117
chr5-92775946-INS-127
chr5-92791923-INS-96
chr5-92792472-INS-167
chr5-92794406-INS-283
chr5-92832187-INS-63
chr5-92832634-INS-2601
chr5-92864623-INS-107
chr5-92882382-INS-495
chr5-92883800-INS-3105
chr5-92885353-INS-88
chr5-92895743-INS-462
chr5-92896515-INS-70
chr5-92900197-INS-195
chr5-92910160-INS-62
chr5-92920194-INS-53
chr5-92927214-INS-263
chr5-92931867-INS-54
chr5-92933563-INS-171
chr5-92937978-INS-125
chr5-92941380-INS-64
chr5-92945742-INS-64
chr5-92960548-INS-71
chr5-92965674-INS-162
chr5-92967410-INS-68
chr5-92968795-INS-203
chr5-92975616-INS-71
chr5-92978821-INS-604
chr5-92995689-INS-199
chr5-93005490-INS-1601
chr5-93037858-INS-205
chr5-93060066-INS-206
chr5-93070437-INS-106
chr5-93075518-INS-75
chr5-93091486-INS-198
chr5-93127537-INS-232
chr5-93133328-INS-358
chr5-93169888-INS-68
chr5-93186464-INS-238
chr5-93205817-INS-60
chr5-93207573-INS-218
chr5-93211108-INS-2523
chr5-93220057-INS-58
chr5-93221195-INS-82
chr5-93222172-INS-239
chr5-93234548-INS-173
chr5-93259242-INS-128
chr5-93280832-INS-68
chr5-93308165-INS-131
chr5-93325918-INS-116
chr5-93350846-INS-102

chr5-93391622-INS-65
chr5-93392446-INS-2968
chr5-93408784-INS-66
chr5-93457638-INS-201
chr5-93469264-INS-7722
chr5-93475159-INS-6885
chr5-93478036-INS-1891
chr5-93483027-INS-7379
chr5-93612679-INS-539
chr5-93621438-INS-6413
chr5-93674344-INS-7086
chr5-93736521-INS-4513
chr5-93787993-INS-232
chr5-93801852-INS-7129
chr5-93885648-INS-6671
chr5-93928519-INS-296
chr5-93929684-INS-170
chr5-93939229-INS-321
chr5-93943347-INS-50
chr5-94008494-INS-56
chr5-94511934-INS-58
chr5-94801712-INS-6539
chr5-95595221-INS-2972
chr5-95626804-INS-238
chr5-95643942-INS-6249
chr5-95654780-INS-150
chr5-95657130-INS-113
chr5-95742331-INS-69
chr5-95924656-INS-7105
chr5-95964657-INS-169
chr5-95966087-INS-3523
chr5-95967225-INS-7094
chr5-95975821-INS-1845
chr5-95988637-INS-4972
chr5-96022814-INS-82
chr5-96036306-INS-2099
chr5-96059366-INS-6792
chr5-96066573-INS-98
chr5-96067158-INS-1483
chr5-96070979-INS-94
chr5-96092439-INS-76
chr5-96129375-INS-6565
chr5-96138455-INS-357
chr5-96146133-INS-498
chr5-96153789-INS-149
chr5-96161167-INS-3909
chr5-96172566-INS-147
chr5-96178577-INS-126
chr5-96189056-INS-357
chr5-96191541-INS-6199
chr5-96191835-INS-574
chr5-96200886-INS-2118
chr5-96212204-INS-64
chr5-96321574-INS-99
chr5-96332215-INS-84
chr5-96343281-INS-60
chr5-96344760-INS-697
chr5-96350128-INS-7165
chr5-96352078-INS-75
chr5-96382240-INS-59
chr5-96395957-INS-162
chr5-96418171-INS-871
chr5-96504297-INS-362
chr5-96508176-INS-143
chr5-96517727-INS-365
chr5-96541586-INS-108
chr5-96550000-INS-54
chr5-96578583-INS-86
chr5-96584088-INS-192
chr5-96594901-INS-87
chr5-96658584-INS-647
chr5-96698982-INS-87
chr5-96708916-INS-204
chr5-96717792-INS-67
chr5-96721922-INS-248
chr5-96735757-INS-171
chr5-96738381-INS-76

chr5-96809945-INS-105
chr5-96858248-INS-248
chr5-96875325-INS-78
chr5-96878413-INS-433
chr5-96899433-INS-74
chr5-96913706-INS-102
chr5-96939326-INS-52
chr5-96948887-INS-63
chr5-96961300-INS-249
chr5-96962836-INS-2088
chr5-96963650-INS-329
chr5-96994896-INS-501
chr5-97011881-INS-56
chr5-97031363-INS-82
chr5-97034552-INS-152
chr5-97051418-INS-68
chr5-97056601-INS-250
chr5-97058111-INS-56
chr5-97060105-INS-353
chr5-97083051-INS-51
chr5-97100159-INS-538
chr5-97105303-INS-60
chr5-97167146-INS-336
chr5-97169408-INS-325
chr5-97175057-INS-467
chr5-97257387-INS-668
chr5-97318872-INS-461
chr5-97320385-INS-92
chr5-97322913-INS-190
chr5-97339395-INS-219
chr5-97344626-INS-3274
chr5-97363109-INS-68
chr5-97364714-INS-58
chr5-97378319-INS-54
chr5-97445441-INS-202
chr5-97451335-INS-200
chr5-97462680-INS-54
chr5-97473180-INS-294
chr5-97487344-INS-54
chr5-97514851-INS-80
chr5-97517471-INS-182
chr5-97551040-INS-549
chr5-97565100-INS-380
chr5-97565862-INS-7471
chr5-97611533-INS-522
chr5-97627729-INS-161
chr5-97635635-INS-76
chr5-97642919-INS-212
chr5-97653649-INS-54
chr5-97673980-INS-424
chr5-97676385-INS-53
chr5-97676530-INS-200
chr5-97695606-INS-120
chr5-97715431-INS-72
chr5-97720279-INS-53
chr5-97729751-INS-64
chr5-97738345-INS-464
chr5-97740123-INS-7321
chr5-97754091-INS-275
chr5-97799236-INS-116
chr5-97829112-INS-114
chr5-97853592-INS-62
chr5-97861694-INS-7907
chr5-97865344-INS-170
chr5-97866765-INS-212
chr5-97896373-INS-128
chr5-97925119-INS-219
chr5-97929741-INS-117
chr5-97958063-INS-7002
chr5-97962446-INS-134
chr5-97964694-INS-6204
chr5-97967936-INS-527
chr5-97996671-INS-407
chr5-98014523-INS-97
chr5-98040998-INS-52
chr5-98069330-INS-6418
chr5-98081006-INS-110

chr5-98099426-INS-897
chr5-98109750-INS-268
chr5-98129224-INS-6543
chr5-98162904-INS-9117
chr5-98165578-INS-179
chr5-98172224-INS-144
chr5-98208291-INS-371
chr5-98255434-INS-64
chr5-98261487-INS-53
chr5-98287893-INS-922
chr5-98299305-INS-247
chr5-98300927-INS-78
chr5-98301459-INS-830
chr5-98308478-INS-228
chr5-98310897-INS-177
chr5-98322838-INS-231
chr5-98337025-INS-7111
chr5-98350936-INS-251
chr5-98364289-INS-168
chr5-98384348-INS-203
chr5-98396269-INS-72
chr5-98428620-INS-731
chr5-98435467-INS-227
chr5-98444780-INS-424
chr5-98456095-INS-87
chr5-98488251-INS-56
chr5-98495076-INS-54
chr5-98518648-INS-303
chr5-98538197-INS-639
chr5-98560420-INS-225
chr5-98565204-INS-2923
chr5-98585255-INS-174
chr5-98609394-INS-54
chr5-98610579-INS-333
chr5-98646737-INS-964
chr5-98661714-INS-1316
chr5-98669356-INS-111
chr5-98671034-INS-182
chr5-98675974-INS-248
chr5-98688157-INS-63
chr5-98692403-INS-61
chr5-98697737-INS-232
chr5-98747935-INS-52
chr5-98843742-INS-321
chr5-98857297-INS-72
chr5-98857948-INS-189
chr5-98864572-INS-372
chr5-98902282-INS-503
chr5-98905286-INS-84
chr5-98942241-INS-269
chr5-98945963-INS-66
chr5-98949998-INS-103
chr5-98972871-INS-56
chr5-98974719-INS-64
chr5-98978645-INS-60
chr5-98988899-INS-56
chr5-99003986-INS-55
chr5-99022273-INS-108
chr5-99032786-INS-66
chr5-99042556-INS-83
chr5-99050852-INS-417
chr5-99063312-INS-212
chr5-99082962-INS-148
chr5-99086857-INS-86
chr5-99107798-INS-87
chr5-99150475-INS-117
chr5-99158879-INS-7025
chr5-99171700-INS-50
chr5-99173556-INS-187
chr5-99176129-INS-826
chr5-99227663-INS-68
chr5-99233841-INS-76
chr5-99254804-INS-76
chr5-99287113-INS-393
chr5-99299062-INS-6592
chr5-99329035-INS-265
chr5-99408854-INS-61

chr5-99411481-INS-89
chr5-99441332-INS-124
chr5-99478248-INS-78
chr5-99494915-INS-352
chr5-99510542-INS-70
chr5-99516508-INS-61
chr5-99549493-INS-211
chr5-99560829-INS-70
chr5-99563412-INS-331
chr5-99566912-INS-102
chr5-99573250-INS-252
chr5-99582757-INS-65
chr5-99603237-INS-89
chr5-99612969-INS-99
chr5-99618095-INS-109
chr5-99636056-INS-195
chr5-99644181-INS-57
chr5-99694501-INS-76
chr5-99699529-INS-88
chr5-99718160-INS-57
chr5-99718205-INS-556
chr5-99720336-INS-54
chr5-99720948-INS-189
chr5-99765367-INS-78
chr5-99778889-INS-280
chr5-99815574-INS-271
chr5-99844440-INS-164
chr5-99879144-INS-100
chr5-99881468-INS-52
chr5-99883119-INS-117
chr5-99903844-INS-56
chr5-99925090-INS-119
chr5-99941699-INS-314
chr5-99964224-INS-315
chr5-99971317-INS-79
chr5-99987129-INS-125
chr5-99995568-INS-550
chr5-100062860-INS-295
chr5-100091751-INS-686
chr5-100094518-INS-230
chr5-100098240-INS-126
chr5-100130513-INS-153
chr5-100147196-INS-131
chr5-100150644-INS-279
chr5-100161107-INS-198
chr5-100168431-INS-117
chr5-100175867-INS-125
chr5-100191004-INS-54
chr5-100193010-INS-87
chr5-100222727-INS-221
chr5-100236524-INS-101
chr5-100244345-INS-581
chr5-100247316-INS-62
chr5-100271732-INS-192
chr5-100309256-INS-67
chr5-100311955-INS-176
chr5-100314374-INS-86
chr5-100316317-INS-60
chr5-100330606-INS-434
chr5-100350000-INS-60
chr5-100354935-INS-56
chr5-100364152-INS-751
chr5-100370132-INS-357
chr5-100373826-INS-719
chr5-100396346-INS-144
chr5-100398669-INS-126
chr5-100400705-INS-213
chr5-100440485-INS-164
chr5-100448047-INS-498
chr5-100450742-INS-55
chr5-100454800-INS-7148
chr5-100459429-INS-189
chr5-100461498-INS-587
chr5-100472820-INS-94
chr5-100498750-INS-137
chr5-100522779-INS-142
chr5-100545191-INS-216

chr5-100560846-INS-52
chr5-100571546-INS-204
chr5-100573881-INS-497
chr5-100638354-INS-138
chr5-100659699-INS-52
chr5-100676664-INS-61
chr5-100686691-INS-179
chr5-100690196-INS-451
chr5-100697535-INS-449
chr5-100718437-INS-498
chr5-100721081-INS-128
chr5-100731448-INS-60
chr5-100737709-INS-98
chr5-100743409-INS-402
chr5-100806983-INS-134
chr5-100817687-INS-126
chr5-100824601-INS-58
chr5-100844908-INS-222
chr5-100854975-INS-178
chr5-100909300-INS-58
chr5-100927447-INS-120
chr5-100927981-INS-796
chr5-100933809-INS-95
chr5-100934597-INS-63
chr5-100935358-INS-6801
chr5-100949927-INS-67
chr5-100956115-INS-4522
chr5-101017460-INS-52
chr5-101026315-INS-127
chr5-101036077-INS-109
chr5-101071642-INS-58
chr5-101083982-INS-372
chr5-101101983-INS-116
chr5-101104176-INS-74
chr5-101105107-INS-157
chr5-101106507-INS-148
chr5-101111713-INS-52
chr5-101132900-INS-754
chr5-101186166-INS-407
chr5-101214744-INS-760
chr5-101223303-INS-61
chr5-101227203-INS-66
chr5-101233318-INS-99
chr5-101234627-INS-120
chr5-101244574-INS-225
chr5-101246637-INS-379
chr5-101249321-INS-321
chr5-101282125-INS-130
chr5-101297119-INS-114
chr5-101300056-INS-112
chr5-101301097-INS-503
chr5-101308347-INS-118
chr5-101313176-INS-84
chr5-101317611-INS-1058
chr5-101317612-INS-897
chr5-101324640-INS-146
chr5-101357294-INS-215
chr5-101358756-INS-73
chr5-101363623-INS-52
chr5-101369612-INS-919
chr5-101370881-INS-82
chr5-101383760-INS-90
chr5-101420493-INS-126
chr5-101427279-INS-79
chr5-101479565-INS-177
chr5-101503308-INS-106
chr5-101529562-INS-457
chr5-101554230-INS-58
chr5-101567684-INS-94
chr5-101572178-INS-88
chr5-101581296-INS-240
chr5-101647321-INS-124
chr5-101677136-INS-153
chr5-101684884-INS-407
chr5-101731839-INS-68
chr5-101746966-INS-56
chr5-101750364-INS-202

chr5-101761610-INS-3501
chr5-101785563-INS-712
chr5-101794144-INS-76
chr5-101845047-INS-62
chr5-101845849-INS-245
chr5-101847863-INS-80
chr5-101897942-INS-213
chr5-101924131-INS-320
chr5-101929651-INS-982
chr5-101932626-INS-155
chr5-101933669-INS-391
chr5-101953267-INS-75
chr5-101968265-INS-52
chr5-101970994-INS-140
chr5-101974089-INS-174
chr5-101989025-INS-203
chr5-102027849-INS-265
chr5-102039698-INS-92
chr5-102046643-INS-52
chr5-102054825-INS-66
chr5-102057078-INS-167
chr5-102087427-INS-309
chr5-102090474-INS-367
chr5-102115066-INS-177
chr5-102122354-INS-52
chr5-102184031-INS-104
chr5-102185929-INS-221
chr5-102187850-INS-142
chr5-102199033-INS-282
chr5-102222285-INS-142
chr5-102238968-INS-333
chr5-102240592-INS-95
chr5-102245565-INS-6511
chr5-102264861-INS-225
chr5-102278113-INS-293
chr5-102295222-INS-6334
chr5-102300124-INS-1044
chr5-102306848-INS-111
chr5-102316904-INS-227
chr5-102361267-INS-128
chr5-102362557-INS-60
chr5-102423849-INS-75
chr5-102455565-INS-175
chr5-102457270-INS-106
chr5-102466810-INS-212
chr5-102473289-INS-354
chr5-102483126-INS-129
chr5-102511060-INS-443
chr5-102525701-INS-184
chr5-102544611-INS-53
chr5-102616292-INS-95
chr5-102623256-INS-233
chr5-102623901-INS-204
chr5-102627569-INS-96
chr5-102639172-INS-255
chr5-102680203-INS-63
chr5-102683159-INS-67
chr5-102701240-INS-56
chr5-102737423-INS-6681
chr5-102743322-INS-60
chr5-102754169-INS-71
chr5-102782866-INS-67
chr5-102828127-INS-73
chr5-102855791-INS-72
chr5-102870365-INS-138
chr5-102891811-INS-60
chr5-102907465-INS-75
chr5-102911653-INS-183
chr5-102926817-INS-261
chr5-102936736-INS-108
chr5-102945690-INS-72
chr5-103008783-INS-489
chr5-103021134-INS-64
chr5-103023240-INS-803
chr5-103030234-INS-198
chr5-103076043-INS-60
chr5-103112454-INS-117

chr5-103113020-INS-1864
chr5-103159964-INS-54
chr5-103207291-INS-775
chr5-103215771-INS-7239
chr5-103222623-INS-156
chr5-103231745-INS-349
chr5-103242141-INS-62
chr5-103264769-INS-175
chr5-103276856-INS-416
chr5-103286647-INS-169
chr5-103290676-INS-97
chr5-103323894-INS-462
chr5-103328381-INS-90
chr5-103346603-INS-107
chr5-103363299-INS-114
chr5-103389670-INS-6416
chr5-103396655-INS-496
chr5-103404010-INS-339
chr5-103413255-INS-524
chr5-103431589-INS-160
chr5-103453885-INS-213
chr5-103457092-INS-148
chr5-103457943-INS-711
chr5-103459372-INS-356
chr5-103490320-INS-100
chr5-103504941-INS-64
chr5-103506442-INS-122
chr5-103508862-INS-59
chr5-103518246-INS-55
chr5-103540598-INS-71
chr5-103551324-INS-116
chr5-103561886-INS-56
chr5-103578322-INS-220
chr5-103593193-INS-110
chr5-103603407-INS-203
chr5-103618684-INS-76
chr5-103625189-INS-552
chr5-103631485-INS-88
chr5-103638330-INS-56
chr5-103656213-INS-130
chr5-103686727-INS-76
chr5-103714682-INS-97
chr5-103718291-INS-177
chr5-103731753-INS-89
chr5-103751682-INS-210
chr5-103766531-INS-93
chr5-103787787-INS-221
chr5-103790450-INS-85
chr5-103791763-INS-51
chr5-103822405-INS-210
chr5-103834550-INS-184
chr5-103844839-INS-108
chr5-103848671-INS-73
chr5-103851502-INS-198
chr5-103856378-INS-226
chr5-103899759-INS-749
chr5-103915499-INS-58
chr5-103921517-INS-327
chr5-103927484-INS-200
chr5-103930699-INS-56
chr5-103938500-INS-614
chr5-103946338-INS-60
chr5-103948524-INS-54
chr5-103968663-INS-97
chr5-103974510-INS-54
chr5-103988335-INS-228
chr5-104006989-INS-159
chr5-104016943-INS-189
chr5-104040192-INS-435
chr5-104045389-INS-54
chr5-104055330-INS-71
chr5-104072848-INS-164
chr5-104087781-INS-68
chr5-104091816-INS-204
chr5-104104295-INS-7487
chr5-104106811-INS-64
chr5-104109465-INS-197

chr5-104110289-INS-51
chr5-104115856-INS-54
chr5-104144940-INS-172
chr5-104146314-INS-56
chr5-104162073-INS-205
chr5-104162618-INS-112
chr5-104176726-INS-339
chr5-104180095-INS-206
chr5-104201352-INS-202
chr5-104204161-INS-227
chr5-104250808-INS-125
chr5-104251165-INS-84
chr5-104278071-INS-62
chr5-104287174-INS-305
chr5-104300824-INS-180
chr5-104310890-INS-118
chr5-104325526-INS-270
chr5-104413416-INS-55
chr5-104479557-INS-188
chr5-104480092-INS-5320
chr5-104496709-INS-275
chr5-104520266-INS-952
chr5-104520321-INS-244
chr5-104521298-INS-265
chr5-104536918-INS-328
chr5-104542188-INS-210
chr5-104567051-INS-9385
chr5-104570882-INS-117
chr5-104656154-INS-56
chr5-104716041-INS-62
chr5-104723358-INS-171
chr5-104991341-INS-52
chr5-105032773-INS-84
chr5-105067398-INS-3706
chr5-105106756-INS-52
chr5-105125244-INS-469
chr5-105128409-INS-127
chr5-105137331-INS-62
chr5-105152212-INS-6827
chr5-105161563-INS-70
chr5-105191232-INS-205
chr5-105191257-INS-207
chr5-105200255-INS-9396
chr5-105252695-INS-168
chr5-105283760-INS-133
chr5-105346255-INS-122
chr5-105417654-INS-336
chr5-105451840-INS-171
chr5-105574520-INS-150
chr5-105606447-INS-307
chr5-105617583-INS-922
chr5-105655789-INS-679
chr5-105668159-INS-92
chr5-105687124-INS-168
chr5-105696633-INS-65
chr5-105710342-INS-70
chr5-105737093-INS-65
chr5-105742335-INS-54
chr5-105744124-INS-76
chr5-105764398-INS-66
chr5-105780718-INS-94
chr5-105786907-INS-6401
chr5-105808803-INS-224
chr5-105816915-INS-203
chr5-105836941-INS-77
chr5-105847402-INS-232
chr5-105854759-INS-193
chr5-105858213-INS-63
chr5-105864477-INS-54
chr5-105883500-INS-320
chr5-105884638-INS-204
chr5-105903165-INS-212
chr5-105929684-INS-124
chr5-105938348-INS-225
chr5-105949054-INS-378
chr5-105977704-INS-7065
chr5-105993451-INS-413

chr5-105999292-INS-102
chr5-106058643-INS-514
chr5-106077453-INS-327
chr5-106093348-INS-75
chr5-106116708-INS-79
chr5-106135431-INS-104
chr5-106138301-INS-65
chr5-106179209-INS-57
chr5-106191513-INS-280
chr5-106210995-INS-83
chr5-106213666-INS-68
chr5-106233062-INS-58
chr5-106241688-INS-244
chr5-106243110-INS-204
chr5-106247344-INS-239
chr5-106262501-INS-89
chr5-106275770-INS-127
chr5-106276450-INS-396
chr5-106294635-INS-127
chr5-106303629-INS-214
chr5-106303902-INS-538
chr5-106305506-INS-94
chr5-106316771-INS-193
chr5-106379071-INS-52
chr5-106382633-INS-521
chr5-106395457-INS-6252
chr5-106396062-INS-108
chr5-106417198-INS-70
chr5-106436426-INS-93
chr5-106493576-INS-423
chr5-106503902-INS-69
chr5-106516549-INS-906
chr5-106532617-INS-69
chr5-106539816-INS-341
chr5-106555745-INS-140
chr5-106563436-INS-441
chr5-106576148-INS-59
chr5-106578477-INS-249
chr5-106588860-INS-488
chr5-106589648-INS-51
chr5-106627419-INS-63
chr5-106633454-INS-425
chr5-106668961-INS-64
chr5-106717329-INS-75
chr5-106742315-INS-72
chr5-106744128-INS-200
chr5-106773598-INS-52
chr5-106812618-INS-217
chr5-106836173-INS-182
chr5-106852018-INS-52
chr5-106855327-INS-66
chr5-106878921-INS-345
chr5-106880435-INS-142
chr5-106898014-INS-327
chr5-106934723-INS-228
chr5-106938651-INS-390
chr5-106956038-INS-170
chr5-106956241-INS-548
chr5-106960008-INS-285
chr5-106969130-INS-62
chr5-106976961-INS-88
chr5-106991875-INS-357
chr5-107007800-INS-95
chr5-107019629-INS-328
chr5-107032364-INS-68
chr5-107042458-INS-219
chr5-107091109-INS-359
chr5-107101521-INS-109
chr5-107105890-INS-198
chr5-107107336-INS-283
chr5-107109800-INS-4997
chr5-107115819-INS-189
chr5-107118171-INS-62
chr5-107128228-INS-206
chr5-107143358-INS-199
chr5-107145388-INS-54
chr5-107148801-INS-77

chr5-107149697-INS-125
chr5-107171550-INS-86
chr5-107185510-INS-224
chr5-107200949-INS-110
chr5-107239564-INS-145
chr5-107251968-INS-445
chr5-107260141-INS-54
chr5-107269354-INS-166
chr5-107306330-INS-52
chr5-107328299-INS-235
chr5-107339913-INS-78
chr5-107355033-INS-210
chr5-107360261-INS-500
chr5-107362136-INS-93
chr5-107375833-INS-502
chr5-107390565-INS-96
chr5-107413464-INS-62
chr5-107415026-INS-152
chr5-107442148-INS-124
chr5-107483573-INS-2189
chr5-107497194-INS-111
chr5-107501737-INS-52
chr5-107560171-INS-56
chr5-107573287-INS-200
chr5-107583016-INS-165
chr5-107637829-INS-86
chr5-107668662-INS-341
chr5-107697640-INS-51
chr5-107705387-INS-463
chr5-107721080-INS-61
chr5-107750379-INS-697
chr5-107758037-INS-97
chr5-107839510-INS-149
chr5-107849295-INS-138
chr5-107850264-INS-129
chr5-107855023-INS-63
chr5-107898014-INS-196
chr5-107908292-INS-125
chr5-108027401-INS-161
chr5-108042391-INS-292
chr5-108054327-INS-200
chr5-108061622-INS-100
chr5-108095016-INS-121
chr5-108115906-INS-199
chr5-108145522-INS-54
chr5-108166012-INS-201
chr5-108182518-INS-141
chr5-108184233-INS-198
chr5-108229575-INS-54
chr5-108233045-INS-202
chr5-108257677-INS-60
chr5-108264361-INS-188
chr5-108274998-INS-92
chr5-108276195-INS-164
chr5-108318133-INS-303
chr5-108326530-INS-185
chr5-108364744-INS-57
chr5-108382326-INS-1894
chr5-108404389-INS-234
chr5-108413808-INS-66
chr5-108433645-INS-6536
chr5-108437564-INS-855
chr5-108449026-INS-62
chr5-108482656-INS-205
chr5-108502777-INS-74
chr5-108505071-INS-114
chr5-108544449-INS-60
chr5-108555390-INS-54
chr5-108561694-INS-73
chr5-108563472-INS-147
chr5-108592109-INS-66
chr5-108601139-INS-122
chr5-108602514-INS-56
chr5-108606543-INS-400
chr5-108628755-INS-67
chr5-108665976-INS-254
chr5-108680627-INS-171

chr5-108712239-INS-194
chr5-108731719-INS-54
chr5-108742450-INS-688
chr5-108830318-INS-94
chr5-108874590-INS-66
chr5-108897941-INS-1886
chr5-108946667-INS-6349
chr5-108987050-INS-6396
chr5-109032869-INS-17856
chr5-109072753-INS-77
chr5-109145915-INS-54
chr5-109168642-INS-60
chr5-109206535-INS-61
chr5-109251484-INS-66
chr5-109259385-INS-96
chr5-109284709-INS-170
chr5-109339204-INS-78
chr5-109355470-INS-794
chr5-109365913-INS-55
chr5-109383908-INS-6599
chr5-109467593-INS-108
chr5-109514834-INS-853
chr5-109526918-INS-7000
chr5-109571764-INS-343
chr5-109586093-INS-62
chr5-109668868-INS-624
chr5-109670518-INS-418
chr5-109742663-INS-74
chr5-109757008-INS-88
chr5-109761093-INS-6214
chr5-109762466-INS-1256
chr5-109786329-INS-54
chr5-109841545-INS-886
chr5-109847554-INS-542
chr5-109853436-INS-161
chr5-109857812-INS-303
chr5-109861391-INS-6175
chr5-109881512-INS-71
chr5-109883372-INS-51
chr5-109925365-INS-292
chr5-109965877-INS-173
chr5-109974110-INS-185
chr5-109975141-INS-380
chr5-110005372-INS-224
chr5-110043400-INS-212
chr5-110057672-INS-7069
chr5-110073090-INS-232
chr5-110173022-INS-200
chr5-110193050-INS-179
chr5-110211612-INS-5337
chr5-110221483-INS-119
chr5-110225458-INS-128
chr5-110235323-INS-403
chr5-110238858-INS-240
chr5-110240557-INS-198
chr5-110280850-INS-4528
chr5-110286956-INS-203
chr5-110288697-INS-219
chr5-110314793-INS-132
chr5-110329284-INS-207
chr5-110338194-INS-220
chr5-110341316-INS-1256
chr5-110394542-INS-367
chr5-110395411-INS-204
chr5-110403780-INS-220
chr5-110419716-INS-198
chr5-110425681-INS-147
chr5-110432154-INS-170
chr5-110444279-INS-204
chr5-110449582-INS-131
chr5-110456201-INS-88
chr5-110478535-INS-210
chr5-110480834-INS-181
chr5-110493196-INS-216
chr5-110496253-INS-78
chr5-110507336-INS-428
chr5-110599645-INS-3228

chr5-110625705-INS-185
chr5-110640548-INS-62
chr5-110680676-INS-69
chr5-110684939-INS-205
chr5-110700705-INS-84
chr5-110705146-INS-217
chr5-110714042-INS-177
chr5-110715186-INS-60
chr5-110743127-INS-65
chr5-110747960-INS-51
chr5-110789065-INS-441
chr5-110792045-INS-174
chr5-110800594-INS-54
chr5-110855965-INS-399
chr5-110858493-INS-69
chr5-110881166-INS-242
chr5-110882304-INS-222
chr5-110898808-INS-118
chr5-110906188-INS-170
chr5-110916294-INS-212
chr5-110924962-INS-64
chr5-110952884-INS-128
chr5-110960911-INS-294
chr5-110991064-INS-55
chr5-111006604-INS-201
chr5-111012716-INS-226
chr5-111015197-INS-210
chr5-111041157-INS-252
chr5-111042495-INS-64
chr5-111052917-INS-203
chr5-111078572-INS-831
chr5-111084393-INS-256
chr5-111111744-INS-189
chr5-111122944-INS-51
chr5-111137699-INS-264
chr5-111143512-INS-90
chr5-111193796-INS-231
chr5-111195803-INS-72
chr5-111227581-INS-80
chr5-111231124-INS-93
chr5-111266583-INS-127
chr5-111278535-INS-96
chr5-111293025-INS-419
chr5-111294368-INS-99
chr5-111297470-INS-102
chr5-111313483-INS-564
chr5-111318895-INS-175
chr5-111335594-INS-60
chr5-111340756-INS-170
chr5-111348687-INS-64
chr5-111401373-INS-185
chr5-111407769-INS-122
chr5-111447506-INS-422
chr5-111464833-INS-106
chr5-111515624-INS-407
chr5-111532625-INS-5259
chr5-111552819-INS-220
chr5-111555224-INS-106
chr5-111592931-INS-85
chr5-111598378-INS-78
chr5-111620737-INS-92
chr5-111641045-INS-191
chr5-111642882-INS-306
chr5-111672278-INS-331
chr5-111674346-INS-59
chr5-111709765-INS-94
chr5-111731785-INS-381
chr5-111740725-INS-439
chr5-111743411-INS-99
chr5-111764670-INS-63
chr5-111770110-INS-59
chr5-111777444-INS-657
chr5-111802392-INS-114
chr5-111810593-INS-82
chr5-111812428-INS-73
chr5-111867187-INS-65
chr5-111871293-INS-61

chr5-111874312-INS-58
chr5-111900960-INS-275
chr5-111915836-INS-204
chr5-111945914-INS-100
chr5-111966246-INS-63
chr5-111972267-INS-78
chr5-111973723-INS-75
chr5-111979278-INS-323
chr5-111991166-INS-102
chr5-112011826-INS-110
chr5-112015270-INS-174
chr5-112015578-INS-102
chr5-112020004-INS-66
chr5-112024172-INS-310
chr5-112087317-INS-251
chr5-112088947-INS-92
chr5-112151581-INS-160
chr5-112162700-INS-441
chr5-112174755-INS-251
chr5-112191282-INS-197
chr5-112203786-INS-150
chr5-112226020-INS-206
chr5-112325418-INS-56
chr5-112351657-INS-449
chr5-112379046-INS-208
chr5-112387597-INS-186
chr5-112389900-INS-64
chr5-112391919-INS-51
chr5-112405581-INS-189
chr5-112410002-INS-76
chr5-112473671-INS-153
chr5-112515286-INS-56
chr5-112612184-INS-160
chr5-112617024-INS-160
chr5-112629299-INS-86
chr5-112630694-INS-62
chr5-112677966-INS-75
chr5-112689326-INS-62
chr5-112691252-INS-75
chr5-112747425-INS-66
chr5-112748940-INS-108
chr5-112770201-INS-100
chr5-112791276-INS-65
chr5-112794957-INS-60
chr5-112796644-INS-168
chr5-112802857-INS-69
chr5-112814315-INS-136
chr5-112824233-INS-251
chr5-112863978-INS-60
chr5-112870949-INS-62
chr5-112877185-INS-62
chr5-112901065-INS-231
chr5-112933686-INS-134
chr5-112949681-INS-533
chr5-112976213-INS-83
chr5-113038241-INS-302
chr5-113061325-INS-57
chr5-113085920-INS-98
chr5-113108412-INS-167
chr5-113111034-INS-97
chr5-113135500-INS-67
chr5-113149149-INS-136
chr5-113178529-INS-972
chr5-113196062-INS-53
chr5-113205828-INS-51
chr5-113219806-INS-123
chr5-113225350-INS-53
chr5-113252788-INS-60
chr5-113256012-INS-64
chr5-113258568-INS-300
chr5-113300537-INS-157
chr5-113304813-INS-60
chr5-113348414-INS-70
chr5-113358701-INS-213
chr5-113407765-INS-460
chr5-113420570-INS-118
chr5-113426829-INS-81

chr5-113429598-INS-148
chr5-113476043-INS-197
chr5-113515783-INS-60
chr5-113518017-INS-264
chr5-113524177-INS-359
chr5-113525674-INS-113
chr5-113548398-INS-176
chr5-113582330-INS-1267
chr5-113586323-INS-116
chr5-113598625-INS-84
chr5-113631171-INS-68
chr5-113643868-INS-156
chr5-113648010-INS-395
chr5-113713536-INS-335
chr5-113736788-INS-281
chr5-113738700-INS-140
chr5-113742414-INS-152
chr5-113742556-INS-208
chr5-113757814-INS-72
chr5-113784717-INS-179
chr5-113787169-INS-539
chr5-113807462-INS-52
chr5-113814545-INS-262
chr5-113824380-INS-951
chr5-113830968-INS-625
chr5-113842980-INS-66
chr5-113857003-INS-52
chr5-113863096-INS-169
chr5-113869976-INS-278
chr5-113881831-INS-97
chr5-113908629-INS-145
chr5-113910594-INS-58
chr5-113917420-INS-158
chr5-113926565-INS-6440
chr5-113946865-INS-128
chr5-113959683-INS-196
chr5-113969721-INS-1891
chr5-113980465-INS-434
chr5-113983215-INS-69
chr5-113997919-INS-197
chr5-114010450-INS-986
chr5-114021926-INS-116
chr5-114026516-INS-209
chr5-114103792-INS-148
chr5-114111582-INS-81
chr5-114141377-INS-70
chr5-114144097-INS-88
chr5-114150377-INS-230
chr5-114151292-INS-108
chr5-114172403-INS-64
chr5-114178900-INS-110
chr5-114182204-INS-320
chr5-114209235-INS-499
chr5-114232018-INS-354
chr5-114234723-INS-879
chr5-114240399-INS-231
chr5-114249556-INS-201
chr5-114254645-INS-226
chr5-114258692-INS-56
chr5-114264659-INS-247
chr5-114281189-INS-5528
chr5-114284981-INS-85
chr5-114298339-INS-192
chr5-114304635-INS-222
chr5-114306452-INS-204
chr5-114386115-INS-224
chr5-114390871-INS-331
chr5-114394387-INS-66
chr5-114426391-INS-176
chr5-114441038-INS-354
chr5-114459482-INS-64
chr5-114462183-INS-134
chr5-114471574-INS-52
chr5-114479399-INS-229
chr5-114491280-INS-303
chr5-114515300-INS-68
chr5-114524747-INS-61

chr5-114552606-INS-209
chr5-114555782-INS-355
chr5-114569745-INS-207
chr5-114593764-INS-95
chr5-114602835-INS-69
chr5-114606217-INS-207
chr5-114617613-INS-277
chr5-114650249-INS-87
chr5-114682391-INS-170
chr5-114693318-INS-214
chr5-114695021-INS-140
chr5-114702531-INS-177
chr5-114715845-INS-90
chr5-114716788-INS-207
chr5-114721803-INS-6440
chr5-114725984-INS-198
chr5-114735237-INS-209
chr5-114736785-INS-254
chr5-114746231-INS-54
chr5-114800522-INS-189
chr5-114809468-INS-207
chr5-114810802-INS-1018
chr5-114838803-INS-84
chr5-114891891-INS-79
chr5-114898885-INS-826
chr5-114911739-INS-174
chr5-114930546-INS-205
chr5-114939078-INS-176
chr5-114950311-INS-165
chr5-114971283-INS-60
chr5-114983524-INS-207
chr5-114997384-INS-55
chr5-115006090-INS-180
chr5-115023701-INS-54
chr5-115025892-INS-8734
chr5-115049680-INS-69
chr5-115052380-INS-312
chr5-115067141-INS-224
chr5-115121358-INS-173
chr5-115128749-INS-188
chr5-115132157-INS-185
chr5-115153099-INS-380
chr5-115155942-INS-238
chr5-115178189-INS-130
chr5-115181068-INS-57
chr5-115182734-INS-160
chr5-115204762-INS-208
chr5-115209293-INS-59
chr5-115210321-INS-54
chr5-115217216-INS-54
chr5-115225734-INS-227
chr5-115238903-INS-53
chr5-115247406-INS-349
chr5-115252337-INS-151
chr5-115253880-INS-78
chr5-115257519-INS-176
chr5-115265014-INS-96
chr5-115276231-INS-176
chr5-115289799-INS-58
chr5-115309298-INS-112
chr5-115335420-INS-180
chr5-115337906-INS-283
chr5-115348899-INS-82
chr5-115359881-INS-130
chr5-115364114-INS-52
chr5-115365216-INS-200
chr5-115365250-INS-674
chr5-115374149-INS-210
chr5-115401841-INS-136
chr5-115412360-INS-171
chr5-115419742-INS-209
chr5-115431940-INS-210
chr5-115432515-INS-211
chr5-115453971-INS-972
chr5-115479673-INS-201
chr5-115481744-INS-210
chr5-115491231-INS-157

chr5-115495879-INS-67
chr5-115496919-INS-211
chr5-115526074-INS-130
chr5-115533659-INS-52
chr5-115539576-INS-116
chr5-115563761-INS-87
chr5-115572779-INS-52
chr5-115582218-INS-273
chr5-115608483-INS-216
chr5-115614473-INS-150
chr5-115627087-INS-181
chr5-115628634-INS-665
chr5-115637664-INS-221
chr5-115674484-INS-5546
chr5-115711078-INS-2785
chr5-115733721-INS-212
chr5-115807690-INS-207
chr5-115814773-INS-166
chr5-115816267-INS-145
chr5-115828951-INS-193
chr5-115832506-INS-66
chr5-115865024-INS-254
chr5-115888887-INS-175
chr5-115895062-INS-203
chr5-115898591-INS-493
chr5-115912114-INS-77
chr5-115920833-INS-4363
chr5-115922376-INS-999
chr5-115947501-INS-246
chr5-115947823-INS-176
chr5-115948794-INS-82
chr5-115954674-INS-135
chr5-115959326-INS-55
chr5-115965275-INS-61
chr5-116005117-INS-212
chr5-116028869-INS-182
chr5-116087980-INS-231
chr5-116128634-INS-175
chr5-116130962-INS-79
chr5-116134038-INS-52
chr5-116137020-INS-107
chr5-116175469-INS-105
chr5-116182969-INS-205
chr5-116187429-INS-78
chr5-116191787-INS-138
chr5-116215472-INS-92
chr5-116225826-INS-58
chr5-116230527-INS-178
chr5-116243694-INS-114
chr5-116251334-INS-54
chr5-116258012-INS-311
chr5-116268613-INS-6481
chr5-116279840-INS-596
chr5-116302913-INS-218
chr5-116305558-INS-131
chr5-116319691-INS-399
chr5-116337745-INS-55
chr5-116343198-INS-125
chr5-116364219-INS-512
chr5-116418762-INS-60
chr5-116436126-INS-148
chr5-116458555-INS-192
chr5-116469378-INS-76
chr5-116471123-INS-106
chr5-116474858-INS-73
chr5-116482107-INS-59
chr5-116498592-INS-244
chr5-116501957-INS-54
chr5-116525059-INS-66
chr5-116559847-INS-89
chr5-116566061-INS-64
chr5-116567321-INS-78
chr5-116586808-INS-58
chr5-116590798-INS-394
chr5-116592402-INS-79
chr5-116625403-INS-63
chr5-116634477-INS-655

chr5-116643050-INS-5545
chr5-116652813-INS-136
chr5-116655152-INS-157
chr5-116664335-INS-84
chr5-116669696-INS-1146
chr5-116671742-INS-64
chr5-116674280-INS-263
chr5-116713162-INS-168
chr5-116732193-INS-545
chr5-116735573-INS-57
chr5-116760499-INS-86
chr5-116765492-INS-179
chr5-116781869-INS-106
chr5-116832499-INS-74
chr5-116838340-INS-99
chr5-116842933-INS-76
chr5-116848390-INS-144
chr5-116891163-INS-892
chr5-116912230-INS-51
chr5-116922048-INS-116
chr5-116941492-INS-115
chr5-116945323-INS-74
chr5-116950165-INS-63
chr5-116983335-INS-65
chr5-116996510-INS-212
chr5-117015243-INS-403
chr5-117022374-INS-56
chr5-117025215-INS-387
chr5-117034139-INS-120
chr5-117049844-INS-288
chr5-117066273-INS-740
chr5-117074179-INS-125
chr5-117079115-INS-67
chr5-117085094-INS-52
chr5-117135043-INS-52
chr5-117154115-INS-507
chr5-117160613-INS-165
chr5-117171297-INS-582
chr5-117175978-INS-53
chr5-117200913-INS-65
chr5-117201975-INS-119
chr5-117209533-INS-156
chr5-117211936-INS-159
chr5-117220241-INS-498
chr5-117262081-INS-140
chr5-117267768-INS-104
chr5-117272796-INS-51
chr5-117317485-INS-170
chr5-117319748-INS-64
chr5-117321234-INS-172
chr5-117340421-INS-285
chr5-117343742-INS-260
chr5-117357721-INS-197
chr5-117360789-INS-52
chr5-117369998-INS-207
chr5-117402817-INS-83
chr5-117416622-INS-293
chr5-117431665-INS-872
chr5-117444670-INS-158
chr5-117454391-INS-607
chr5-117528114-INS-169
chr5-117534075-INS-89
chr5-117536053-INS-114
chr5-117555360-INS-57
chr5-117586247-INS-68
chr5-117590425-INS-56
chr5-117611563-INS-105
chr5-117618653-INS-343
chr5-117620477-INS-549
chr5-117622871-INS-76
chr5-117630908-INS-54
chr5-117645916-INS-507
chr5-117722513-INS-80
chr5-117735837-INS-467
chr5-117751504-INS-283
chr5-117763206-INS-52
chr5-117772445-INS-551

chr5-117790759-INS-52
chr5-117804909-INS-57
chr5-117810221-INS-56
chr5-117819001-INS-52
chr5-117835397-INS-528
chr5-117864886-INS-168
chr5-117916715-INS-84
chr5-117918009-INS-132
chr5-117922378-INS-383
chr5-117925703-INS-51
chr5-117934730-INS-857
chr5-117941444-INS-56
chr5-117973071-INS-334
chr5-117996263-INS-458
chr5-118023717-INS-70
chr5-118029771-INS-313
chr5-118038045-INS-2258
chr5-118038137-INS-1230
chr5-118038138-INS-517
chr5-118100069-INS-102
chr5-118109108-INS-183
chr5-118110494-INS-922
chr5-118113459-INS-191
chr5-118116084-INS-164
chr5-118151088-INS-190
chr5-118155842-INS-52
chr5-118161083-INS-7111
chr5-118170604-INS-64
chr5-118173568-INS-537
chr5-118208740-INS-52
chr5-118224413-INS-96
chr5-118264537-INS-191
chr5-118280315-INS-1845
chr5-118280650-INS-57
chr5-118290904-INS-96
chr5-118303574-INS-52
chr5-118320460-INS-170
chr5-118321481-INS-96
chr5-118337566-INS-572
chr5-118340334-INS-100
chr5-118356605-INS-67
chr5-118380429-INS-208
chr5-118385316-INS-62
chr5-118388494-INS-51
chr5-118394244-INS-109
chr5-118401413-INS-458
chr5-118408637-INS-66
chr5-118414038-INS-130
chr5-118417398-INS-67
chr5-118434338-INS-339
chr5-118443504-INS-63
chr5-118452715-INS-306
chr5-118462112-INS-178
chr5-118475448-INS-126
chr5-118477681-INS-64
chr5-118484388-INS-60
chr5-118489757-INS-65
chr5-118491678-INS-74
chr5-118530532-INS-98
chr5-118561309-INS-115
chr5-118572284-INS-200
chr5-118577089-INS-62
chr5-118580485-INS-395
chr5-118587630-INS-58
chr5-118591480-INS-52
chr5-118596739-INS-54
chr5-118613114-INS-103
chr5-118633844-INS-124
chr5-118649922-INS-230
chr5-118659814-INS-113
chr5-118660839-INS-7135
chr5-118676093-INS-62
chr5-118706530-INS-87
chr5-118721901-INS-165
chr5-118726564-INS-54
chr5-118764382-INS-61
chr5-118801393-INS-158

chr5-118815101-INS-87
chr5-118852051-INS-76
chr5-118853290-INS-62
chr5-118875634-INS-468
chr5-118886306-INS-87
chr5-118912929-INS-114
chr5-118914176-INS-60
chr5-118918471-INS-205
chr5-118926608-INS-184
chr5-118937946-INS-97
chr5-118955152-INS-203
chr5-118965306-INS-466
chr5-118969063-INS-88
chr5-118974342-INS-72
chr5-118982903-INS-138
chr5-119005329-INS-55
chr5-119007851-INS-52
chr5-119011450-INS-345
chr5-119023684-INS-56
chr5-119032409-INS-120
chr5-119066873-INS-181
chr5-119085923-INS-158
chr5-119101048-INS-57
chr5-119124734-INS-238
chr5-119126535-INS-111
chr5-119130642-INS-171
chr5-119131812-INS-76
chr5-119146803-INS-391
chr5-119155649-INS-505
chr5-119159111-INS-102
chr5-119159876-INS-114
chr5-119166757-INS-90
chr5-119207771-INS-81
chr5-119215867-INS-121
chr5-119221636-INS-54
chr5-119231581-INS-54
chr5-119247578-INS-105
chr5-119265526-INS-122
chr5-119282148-INS-96
chr5-119285697-INS-88
chr5-119302512-INS-254
chr5-119303776-INS-326
chr5-119323370-INS-186
chr5-119336306-INS-67
chr5-119337258-INS-50
chr5-119342504-INS-70
chr5-119346255-INS-76
chr5-119370246-INS-82
chr5-119373190-INS-121
chr5-119394931-INS-70
chr5-119449455-INS-51
chr5-119474463-INS-60
chr5-119604061-INS-125
chr5-119615679-INS-123
chr5-119635157-INS-151
chr5-119636717-INS-197
chr5-119650474-INS-280
chr5-119661515-INS-156
chr5-119677479-INS-92
chr5-119684555-INS-162
chr5-119693182-INS-64
chr5-119700363-INS-59
chr5-119703226-INS-63
chr5-119709777-INS-226
chr5-119712345-INS-198
chr5-119716341-INS-68
chr5-119724293-INS-140
chr5-119729199-INS-266
chr5-119742474-INS-195
chr5-119794226-INS-96
chr5-119829453-INS-118
chr5-119848693-INS-83
chr5-119860213-INS-319
chr5-119866360-INS-106
chr5-119872999-INS-195
chr5-119892504-INS-58
chr5-119894908-INS-61

chr5-119899979-INS-58
chr5-119904012-INS-54
chr5-119921353-INS-186
chr5-119922591-INS-178
chr5-119923652-INS-214
chr5-119928576-INS-664
chr5-119992933-INS-103
chr5-120007266-INS-60
chr5-120014820-INS-148
chr5-120016848-INS-72
chr5-120032918-INS-179
chr5-120048903-INS-101
chr5-120063316-INS-140
chr5-120074765-INS-117
chr5-120078558-INS-611
chr5-120079354-INS-63
chr5-120089195-INS-110
chr5-120102212-INS-108
chr5-120108003-INS-84
chr5-120118639-INS-113
chr5-120121668-INS-6581
chr5-120130101-INS-76
chr5-120141365-INS-68
chr5-120145882-INS-4850
chr5-120163050-INS-8980
chr5-120175775-INS-6257
chr5-120185931-INS-54
chr5-120206910-INS-54
chr5-120219998-INS-65
chr5-120221393-INS-66
chr5-120225007-INS-184
chr5-120233355-INS-498
chr5-120268525-INS-111
chr5-120306939-INS-144
chr5-120341538-INS-621
chr5-120365595-INS-790
chr5-120388950-INS-120
chr5-120413796-INS-121
chr5-120445017-INS-398
chr5-120465311-INS-198
chr5-120477117-INS-106
chr5-120480516-INS-423
chr5-120481027-INS-522
chr5-120483081-INS-58
chr5-120487289-INS-56
chr5-120501322-INS-87
chr5-120513330-INS-84
chr5-120533814-INS-64
chr5-120546256-INS-503
chr5-120564198-INS-55
chr5-120587938-INS-76
chr5-120603061-INS-66
chr5-120606131-INS-165
chr5-120608182-INS-207
chr5-120611212-INS-63
chr5-120618054-INS-106
chr5-120656618-INS-75
chr5-120690550-INS-56
chr5-120695280-INS-120
chr5-120724104-INS-102
chr5-120731960-INS-68
chr5-120755450-INS-207
chr5-120807001-INS-56
chr5-120835173-INS-140
chr5-120853585-INS-54
chr5-120857565-INS-212
chr5-120863722-INS-122
chr5-120883403-INS-132
chr5-120892814-INS-451
chr5-120896315-INS-54
chr5-120897655-INS-60
chr5-120918659-INS-125
chr5-120920108-INS-51
chr5-120932117-INS-134
chr5-120971908-INS-423
chr5-121012837-INS-92
chr5-121020902-INS-52

chr5-121024042-INS-133
chr5-121053682-INS-703
chr5-121103974-INS-127
chr5-121107223-INS-122
chr5-121119023-INS-478
chr5-121136597-INS-307
chr5-121144367-INS-72
chr5-121157125-INS-58
chr5-121159856-INS-60
chr5-121161394-INS-54
chr5-121162064-INS-935
chr5-121190034-INS-108
chr5-121193214-INS-58
chr5-121199278-INS-79
chr5-121218430-INS-451
chr5-121225692-INS-58
chr5-121251007-INS-5318
chr5-121251112-INS-597
chr5-121256202-INS-68
chr5-121262151-INS-128
chr5-121303510-INS-201
chr5-121310251-INS-63
chr5-121324453-INS-735
chr5-121335748-INS-236
chr5-121340797-INS-139
chr5-121342324-INS-215
chr5-121352651-INS-455
chr5-121365611-INS-68
chr5-121374048-INS-312
chr5-121376087-INS-198
chr5-121378567-INS-198
chr5-121385208-INS-64
chr5-121399289-INS-1138
chr5-121421530-INS-2628
chr5-121465549-INS-536
chr5-121470661-INS-125
chr5-121485525-INS-204
chr5-121511756-INS-208
chr5-121534505-INS-191
chr5-121542578-INS-202
chr5-121547344-INS-138
chr5-121559937-INS-224
chr5-121580004-INS-66
chr5-121580741-INS-469
chr5-121593520-INS-8789
chr5-121602363-INS-161
chr5-121621142-INS-322
chr5-121653410-INS-199
chr5-121683726-INS-51
chr5-121703059-INS-155
chr5-121709992-INS-186
chr5-121771749-INS-52
chr5-121787225-INS-96
chr5-121826648-INS-50
chr5-121829890-INS-127
chr5-121833286-INS-686
chr5-121840118-INS-62
chr5-121845320-INS-210
chr5-121858367-INS-57
chr5-121859985-INS-222
chr5-121871322-INS-459
chr5-121877245-INS-188
chr5-121906080-INS-199
chr5-121925580-INS-206
chr5-121931208-INS-88
chr5-121933405-INS-352
chr5-121971353-INS-190
chr5-121985198-INS-224
chr5-122019056-INS-181
chr5-122071877-INS-212
chr5-122102511-INS-448
chr5-122146471-INS-64
chr5-122151856-INS-162
chr5-122184083-INS-63
chr5-122194626-INS-72
chr5-122204059-INS-901
chr5-122205017-INS-470

chr5-122244099-INS-121
chr5-122249979-INS-70
chr5-122261183-INS-245
chr5-122280934-INS-112
chr5-122283315-INS-76
chr5-122307819-INS-167
chr5-122314256-INS-223
chr5-122316116-INS-73
chr5-122343845-INS-76
chr5-122365288-INS-56
chr5-122377814-INS-117
chr5-122398757-INS-377
chr5-122416500-INS-212
chr5-122417363-INS-135
chr5-122419639-INS-244
chr5-122434337-INS-224
chr5-122438982-INS-59
chr5-122443172-INS-136
chr5-122447545-INS-203
chr5-122486261-INS-154
chr5-122488576-INS-130
chr5-122502342-INS-196
chr5-122504354-INS-62
chr5-122511266-INS-173
chr5-122513742-INS-205
chr5-122517565-INS-66
chr5-122533682-INS-56
chr5-122537653-INS-204
chr5-122546248-INS-227
chr5-122550415-INS-144
chr5-122556500-INS-1667
chr5-122576330-INS-201
chr5-122608637-INS-154
chr5-122614295-INS-63
chr5-122626479-INS-197
chr5-122643321-INS-114
chr5-122649665-INS-71
chr5-122656370-INS-724
chr5-122658247-INS-201
chr5-122697317-INS-210
chr5-122700400-INS-82
chr5-122700892-INS-156
chr5-122769473-INS-56
chr5-122778985-INS-220
chr5-122811011-INS-52
chr5-122815271-INS-54
chr5-122823157-INS-91
chr5-122827928-INS-144
chr5-122867592-INS-655
chr5-122907274-INS-201
chr5-122914322-INS-82
chr5-122921364-INS-441
chr5-122943980-INS-68
chr5-122945119-INS-1890
chr5-122947277-INS-60
chr5-122962501-INS-76
chr5-122967889-INS-52
chr5-122968664-INS-199
chr5-122976974-INS-116
chr5-122979112-INS-134
chr5-122980705-INS-127
chr5-122986129-INS-5326
chr5-122996447-INS-135
chr5-123005949-INS-460
chr5-123010925-INS-102
chr5-123022621-INS-58
chr5-123049616-INS-66
chr5-123050715-INS-701
chr5-123092242-INS-185
chr5-123104602-INS-166
chr5-123107775-INS-216
chr5-123109033-INS-211
chr5-123119578-INS-129
chr5-123140062-INS-84
chr5-123145337-INS-384
chr5-123158413-INS-90
chr5-123176828-INS-198

chr5-123179716-INS-211
chr5-123194052-INS-94
chr5-123208414-INS-60
chr5-123210983-INS-202
chr5-123225793-INS-108
chr5-123243589-INS-195
chr5-123260334-INS-210
chr5-123262537-INS-205
chr5-123283161-INS-1890
chr5-123312952-INS-208
chr5-123318664-INS-62
chr5-123322080-INS-71
chr5-123330948-INS-203
chr5-123336983-INS-101
chr5-123358812-INS-78
chr5-123376636-INS-217
chr5-123391078-INS-104
chr5-123415040-INS-52
chr5-123418213-INS-153
chr5-123430607-INS-206
chr5-123444997-INS-103
chr5-123446373-INS-137
chr5-123465542-INS-211
chr5-123499714-INS-266
chr5-123501961-INS-277
chr5-123518179-INS-51
chr5-123530406-INS-430
chr5-123538671-INS-66
chr5-123549612-INS-122
chr5-123553662-INS-140
chr5-123561728-INS-104
chr5-123568950-INS-213
chr5-123580096-INS-333
chr5-123597850-INS-205
chr5-123613591-INS-207
chr5-123623911-INS-204
chr5-123626604-INS-212
chr5-123628980-INS-143
chr5-123643887-INS-54
chr5-123645927-INS-192
chr5-123659671-INS-60
chr5-123666410-INS-103
chr5-123698179-INS-208
chr5-123703318-INS-201
chr5-123705793-INS-206
chr5-123729730-INS-98
chr5-123730752-INS-135
chr5-123734267-INS-128
chr5-123736466-INS-85
chr5-123756961-INS-163
chr5-123758546-INS-309
chr5-123771146-INS-115
chr5-123781520-INS-58
chr5-123789914-INS-277
chr5-123796081-INS-75
chr5-123806920-INS-68
chr5-123816545-INS-125
chr5-123818593-INS-360
chr5-123825485-INS-198
chr5-123829524-INS-108
chr5-123836758-INS-138
chr5-123846472-INS-325
chr5-123848679-INS-197
chr5-123850880-INS-65
chr5-123860550-INS-54
chr5-123861311-INS-506
chr5-123863906-INS-112
chr5-123911194-INS-401
chr5-123912151-INS-202
chr5-123955507-INS-132
chr5-123981336-INS-52
chr5-123993810-INS-200
chr5-124007378-INS-56
chr5-124019181-INS-125
chr5-124025611-INS-100
chr5-124044045-INS-110
chr5-124047251-INS-289

chr5-124066631-INS-171
chr5-124072909-INS-197
chr5-124075234-INS-164
chr5-124090930-INS-112
chr5-124167990-INS-221
chr5-124178943-INS-261
chr5-124190638-INS-86
chr5-124204006-INS-187
chr5-124207387-INS-62
chr5-124224113-INS-5512
chr5-124237928-INS-205
chr5-124241032-INS-171
chr5-124242272-INS-209
chr5-124306412-INS-1204
chr5-124306935-INS-87
chr5-124315727-INS-302
chr5-124372927-INS-61
chr5-124401423-INS-54
chr5-124437909-INS-56
chr5-124439912-INS-5365
chr5-124440784-INS-87
chr5-124464084-INS-67
chr5-124473725-INS-197
chr5-124479112-INS-148
chr5-124482427-INS-310
chr5-124517738-INS-61
chr5-124552294-INS-317
chr5-124559867-INS-242
chr5-124572630-INS-207
chr5-124606130-INS-159
chr5-124613259-INS-170
chr5-124614257-INS-64
chr5-124619146-INS-182
chr5-124651471-INS-242
chr5-124655667-INS-192
chr5-124670150-INS-219
chr5-124675894-INS-249
chr5-124697714-INS-227
chr5-124702410-INS-117
chr5-124703477-INS-78
chr5-124737877-INS-232
chr5-124750427-INS-92
chr5-124752671-INS-439
chr5-124762108-INS-199
chr5-124764593-INS-178
chr5-124765979-INS-66
chr5-124771220-INS-99
chr5-124774181-INS-62
chr5-124786975-INS-125
chr5-124788181-INS-87
chr5-124804387-INS-605
chr5-124813593-INS-57
chr5-124814612-INS-140
chr5-124820256-INS-194
chr5-124824029-INS-377
chr5-124829834-INS-54
chr5-124832341-INS-888
chr5-124836533-INS-108
chr5-124840738-INS-945
chr5-124851280-INS-54
chr5-124852703-INS-215
chr5-124856490-INS-1018
chr5-124856518-INS-628
chr5-124876400-INS-205
chr5-124890843-INS-90
chr5-124923030-INS-146
chr5-124943462-INS-221
chr5-124945152-INS-212
chr5-124968291-INS-56
chr5-124992121-INS-252
chr5-125003268-INS-4117
chr5-125051874-INS-204
chr5-125056029-INS-125
chr5-125077662-INS-68
chr5-125098729-INS-64
chr5-125100821-INS-54
chr5-125139571-INS-74

chr5-125150220-INS-72
chr5-125160835-INS-62
chr5-125182875-INS-115
chr5-125243532-INS-335
chr5-125253405-INS-186
chr5-125265193-INS-53
chr5-125279121-INS-555
chr5-125288052-INS-123
chr5-125303104-INS-198
chr5-125304290-INS-61
chr5-125307255-INS-4161
chr5-125308130-INS-213
chr5-125309393-INS-54
chr5-125313108-INS-54
chr5-125317108-INS-116
chr5-125318140-INS-112
chr5-125320273-INS-125
chr5-125320394-INS-899
chr5-125347751-INS-215
chr5-125349291-INS-355
chr5-125359543-INS-201
chr5-125369556-INS-97
chr5-125372139-INS-64
chr5-125372811-INS-394
chr5-125374876-INS-50
chr5-125394304-INS-205
chr5-125412701-INS-164
chr5-125424755-INS-133
chr5-125432363-INS-390
chr5-125439211-INS-112
chr5-125451239-INS-5346
chr5-125464955-INS-228
chr5-125471612-INS-54
chr5-125473294-INS-204
chr5-125474728-INS-199
chr5-125475317-INS-74
chr5-125477910-INS-169
chr5-125513903-INS-102
chr5-125533912-INS-202
chr5-125547701-INS-144
chr5-125559136-INS-60
chr5-125574673-INS-578
chr5-125601642-INS-1892
chr5-125607881-INS-57
chr5-125614204-INS-51
chr5-125630303-INS-96
chr5-125672937-INS-104
chr5-125678213-INS-54
chr5-125685722-INS-172
chr5-125731439-INS-58
chr5-125733637-INS-55
chr5-125738190-INS-58
chr5-125746970-INS-54
chr5-125753858-INS-91
chr5-125780096-INS-964
chr5-125803028-INS-69
chr5-125824889-INS-81
chr5-125845071-INS-55
chr5-125853777-INS-87
chr5-125872374-INS-102
chr5-125907377-INS-58
chr5-125923919-INS-1000
chr5-125928212-INS-207
chr5-125982133-INS-57
chr5-125987257-INS-58
chr5-126009253-INS-171
chr5-126015914-INS-97
chr5-126022941-INS-213
chr5-126079261-INS-5318
chr5-126095420-INS-120
chr5-126101435-INS-185
chr5-126116158-INS-153
chr5-126157719-INS-878
chr5-126179973-INS-612
chr5-126194388-INS-465
chr5-126235699-INS-203
chr5-126260700-INS-379

chr5-126270940-INS-91
chr5-126308416-INS-69
chr5-126315739-INS-210
chr5-126323040-INS-442
chr5-126330697-INS-155
chr5-126342769-INS-525
chr5-126343313-INS-58
chr5-126388745-INS-872
chr5-126409191-INS-54
chr5-126411147-INS-85
chr5-126414516-INS-487
chr5-126426063-INS-60
chr5-126449511-INS-928
chr5-126473757-INS-107
chr5-126486853-INS-892
chr5-126534758-INS-389
chr5-126540481-INS-166
chr5-126543080-INS-363
chr5-126575021-INS-132
chr5-126606825-INS-52
chr5-126610289-INS-135
chr5-126610291-INS-205
chr5-126613290-INS-120
chr5-126616728-INS-133
chr5-126629835-INS-55
chr5-126632141-INS-204
chr5-126636568-INS-140
chr5-126662812-INS-223
chr5-126688708-INS-58
chr5-126693351-INS-362
chr5-126720859-INS-51
chr5-126730287-INS-68
chr5-126733113-INS-68
chr5-126765739-INS-54
chr5-126775685-INS-828
chr5-126794447-INS-51
chr5-126794901-INS-52
chr5-126807414-INS-7589
chr5-126814609-INS-555
chr5-126828318-INS-62
chr5-126832156-INS-370
chr5-126840175-INS-55
chr5-126851947-INS-54
chr5-126866511-INS-162
chr5-126871865-INS-54
chr5-126875097-INS-67
chr5-126883280-INS-103
chr5-126889046-INS-135
chr5-126904669-INS-4533
chr5-126912021-INS-108
chr5-126925016-INS-129
chr5-126930862-INS-63
chr5-126943385-INS-175
chr5-126943914-INS-126
chr5-126947241-INS-269
chr5-126996483-INS-58
chr5-126997498-INS-106
chr5-127000147-INS-63
chr5-127004465-INS-276
chr5-127017963-INS-56
chr5-127019010-INS-51
chr5-127032386-INS-62
chr5-127094999-INS-104
chr5-127097945-INS-57
chr5-127131560-INS-285
chr5-127160813-INS-85
chr5-127187819-INS-51
chr5-127204291-INS-72
chr5-127204435-INS-314
chr5-127207248-INS-441
chr5-127262731-INS-279
chr5-127274817-INS-267
chr5-127286538-INS-60
chr5-127287579-INS-142
chr5-127334190-INS-6452
chr5-127371830-INS-210
chr5-127383442-INS-51

chr5-127387273-INS-247
chr5-127408562-INS-179
chr5-127413603-INS-67
chr5-127430113-INS-92
chr5-127457219-INS-96
chr5-127465342-INS-141
chr5-127475450-INS-59
chr5-127547200-INS-91
chr5-127571155-INS-52
chr5-127577324-INS-174
chr5-127589418-INS-77
chr5-127595460-INS-156
chr5-127606492-INS-137
chr5-127612752-INS-202
chr5-127636374-INS-297
chr5-127649953-INS-489
chr5-127660424-INS-197
chr5-127675003-INS-178
chr5-127700342-INS-267
chr5-127703880-INS-340
chr5-127705780-INS-248
chr5-127708073-INS-190
chr5-127714336-INS-7087
chr5-127719140-INS-198
chr5-127723194-INS-131
chr5-127730840-INS-230
chr5-127735769-INS-194
chr5-127749245-INS-56
chr5-127758819-INS-200
chr5-127780198-INS-193
chr5-127782069-INS-351
chr5-127782916-INS-202
chr5-127791303-INS-54
chr5-127797799-INS-472
chr5-127809376-INS-116
chr5-127819111-INS-173
chr5-127831103-INS-149
chr5-127836717-INS-4394
chr5-127846264-INS-120
chr5-127849933-INS-130
chr5-127850852-INS-55
chr5-127867779-INS-227
chr5-127878280-INS-52
chr5-127879303-INS-60
chr5-127893718-INS-132
chr5-127899580-INS-603
chr5-127935455-INS-105
chr5-127945039-INS-613
chr5-127978397-INS-175
chr5-127990526-INS-62
chr5-127992181-INS-55
chr5-128009627-INS-546
chr5-128012556-INS-363
chr5-128038289-INS-108
chr5-128039924-INS-489
chr5-128055412-INS-130
chr5-128064082-INS-70
chr5-128070159-INS-51
chr5-128096758-INS-358
chr5-128101062-INS-64
chr5-128130807-INS-72
chr5-128142469-INS-62
chr5-128146679-INS-941
chr5-128151813-INS-56
chr5-128154802-INS-1841
chr5-128194877-INS-74
chr5-128199435-INS-220
chr5-128250615-INS-122
chr5-128262986-INS-56
chr5-128270954-INS-71
chr5-128296768-INS-68
chr5-128304082-INS-99
chr5-128312143-INS-69
chr5-128313068-INS-56
chr5-128342241-INS-72
chr5-128377765-INS-56
chr5-128419521-INS-433

chr5-128477689-INS-1595
chr5-128480848-INS-197
chr5-128483614-INS-70
chr5-128497942-INS-135
chr5-128531845-INS-322
chr5-128545212-INS-66
chr5-128549176-INS-449
chr5-128551073-INS-207
chr5-128556703-INS-209
chr5-128571368-INS-226
chr5-128588973-INS-57
chr5-128594927-INS-54
chr5-128598346-INS-88
chr5-128599291-INS-301
chr5-128614616-INS-56
chr5-128643078-INS-165
chr5-128645846-INS-198
chr5-128647553-INS-443
chr5-128650540-INS-72
chr5-128665368-INS-81
chr5-128692526-INS-56
chr5-128704879-INS-64
chr5-128708345-INS-82
chr5-128710948-INS-66
chr5-128711659-INS-64
chr5-128743549-INS-122
chr5-128759209-INS-66
chr5-128776826-INS-50
chr5-128805135-INS-300
chr5-128811769-INS-86
chr5-128831697-INS-73
chr5-128852734-INS-92
chr5-128862415-INS-92
chr5-128867151-INS-57
chr5-128903506-INS-474
chr5-128905513-INS-145
chr5-128935234-INS-58
chr5-128954794-INS-251
chr5-128961813-INS-60
chr5-128970243-INS-88
chr5-128985562-INS-63
chr5-128995211-INS-60
chr5-128997509-INS-52
chr5-129000559-INS-84
chr5-129024375-INS-95
chr5-129034438-INS-52
chr5-129039576-INS-125
chr5-129050206-INS-51
chr5-129054715-INS-133
chr5-129054737-INS-366
chr5-129056496-INS-73
chr5-129073200-INS-60
chr5-129093635-INS-363
chr5-129103973-INS-207
chr5-129114752-INS-53
chr5-129121745-INS-89
chr5-129126767-INS-172
chr5-129132922-INS-60
chr5-129157174-INS-57
chr5-129181550-INS-151
chr5-129182490-INS-6327
chr5-129202737-INS-660
chr5-129212728-INS-90
chr5-129240419-INS-67
chr5-129263797-INS-63
chr5-129287360-INS-203
chr5-129317125-INS-70
chr5-129324073-INS-84
chr5-129331075-INS-1966
chr5-129331273-INS-142
chr5-129331775-INS-195
chr5-129350203-INS-1034
chr5-129350222-INS-988
chr5-129350762-INS-165
chr5-129376960-INS-590
chr5-129380043-INS-216
chr5-129382484-INS-103

chr5-129433835-INS-203
chr5-129438081-INS-354
chr5-129443399-INS-369
chr5-129460620-INS-110
chr5-129471372-INS-58
chr5-129489087-INS-158
chr5-129490128-INS-56
chr5-129492694-INS-148
chr5-129494164-INS-84
chr5-129522326-INS-5772
chr5-129527688-INS-165
chr5-129541836-INS-76
chr5-129545891-INS-65
chr5-129548570-INS-77
chr5-129551168-INS-283
chr5-129575258-INS-492
chr5-129587019-INS-480
chr5-129633515-INS-130
chr5-129652245-INS-112
chr5-129669797-INS-110
chr5-129678359-INS-387
chr5-129679536-INS-154
chr5-129680585-INS-53
chr5-129688773-INS-173
chr5-129695026-INS-270
chr5-129707418-INS-719
chr5-129710367-INS-517
chr5-129729885-INS-59
chr5-129766491-INS-78
chr5-129775248-INS-188
chr5-129780194-INS-172
chr5-129785916-INS-302
chr5-129792122-INS-70
chr5-129799936-INS-191
chr5-129849659-INS-50
chr5-129898938-INS-178
chr5-129905645-INS-180
chr5-129911162-INS-161
chr5-129918472-INS-60
chr5-129922773-INS-978
chr5-129939360-INS-171
chr5-129951759-INS-164
chr5-129972164-INS-193
chr5-129985329-INS-227
chr5-129990053-INS-119
chr5-130020482-INS-158
chr5-130035296-INS-214
chr5-130043331-INS-202
chr5-130062085-INS-359
chr5-130064766-INS-111
chr5-130071568-INS-211
chr5-130072070-INS-280
chr5-130089746-INS-70
chr5-130094857-INS-239
chr5-130095833-INS-116
chr5-130100024-INS-80
chr5-130102648-INS-52
chr5-130124041-INS-202
chr5-130131396-INS-51
chr5-130157030-INS-160
chr5-130162507-INS-63
chr5-130166244-INS-173
chr5-130179339-INS-112
chr5-130180149-INS-970
chr5-130187480-INS-124
chr5-130195355-INS-576
chr5-130199126-INS-87
chr5-130201993-INS-204
chr5-130209091-INS-182
chr5-130226209-INS-671
chr5-130232710-INS-164
chr5-130243317-INS-493
chr5-130245033-INS-206
chr5-130255121-INS-329
chr5-130263711-INS-221
chr5-130284774-INS-204
chr5-130288324-INS-110

chr5-130295587-INS-59
chr5-130296113-INS-64
chr5-130297441-INS-51
chr5-130325291-INS-86
chr5-130331372-INS-280
chr5-130351612-INS-110
chr5-130356535-INS-110
chr5-130371837-INS-78
chr5-130374442-INS-159
chr5-130384793-INS-84
chr5-130401804-INS-51
chr5-130406074-INS-204
chr5-130406886-INS-501
chr5-130410563-INS-84
chr5-130425792-INS-343
chr5-130431324-INS-108
chr5-130441290-INS-62
chr5-130444532-INS-583
chr5-130482966-INS-401
chr5-130485113-INS-62
chr5-130498893-INS-72
chr5-130522229-INS-70
chr5-130524330-INS-52
chr5-130529827-INS-66
chr5-130533490-INS-184
chr5-130545814-INS-241
chr5-130554390-INS-54
chr5-130560569-INS-62
chr5-130574119-INS-115
chr5-130600915-INS-116
chr5-130607817-INS-75
chr5-130614573-INS-63
chr5-130622186-INS-52
chr5-130626804-INS-114
chr5-130628023-INS-164
chr5-130635326-INS-79
chr5-130692109-INS-63
chr5-130728196-INS-59
chr5-130742371-INS-62
chr5-130751796-INS-3222
chr5-130775482-INS-54
chr5-130793596-INS-72
chr5-130812676-INS-63
chr5-130828051-INS-160
chr5-130830024-INS-83
chr5-130838310-INS-130
chr5-130849246-INS-210
chr5-130850767-INS-64
chr5-130887083-INS-55
chr5-130893759-INS-777
chr5-130900485-INS-108
chr5-130914498-INS-335
chr5-130919784-INS-56
chr5-130925962-INS-106
chr5-130934599-INS-265
chr5-130942127-INS-2940
chr5-130949199-INS-109
chr5-130950480-INS-211
chr5-130961538-INS-135
chr5-130963931-INS-51
chr5-130971479-INS-54
chr5-130987112-INS-218
chr5-130995350-INS-83
chr5-131008884-INS-324
chr5-131034275-INS-201
chr5-131074356-INS-192
chr5-131076660-INS-104
chr5-131111542-INS-5556
chr5-131119845-INS-6927
chr5-131122571-INS-232
chr5-131127324-INS-132
chr5-131128743-INS-52
chr5-131167203-INS-264
chr5-131182317-INS-62
chr5-131209421-INS-81
chr5-131218814-INS-58
chr5-131223823-INS-56

chr5-131242234-INS-3069
chr5-131248608-INS-159
chr5-131287951-INS-127
chr5-131290319-INS-52
chr5-131344848-INS-401
chr5-131351070-INS-86
chr5-131379713-INS-565
chr5-131385616-INS-96
chr5-131396241-INS-56
chr5-131410123-INS-192
chr5-131416781-INS-72
chr5-131417601-INS-6853
chr5-131422866-INS-212
chr5-131458116-INS-145
chr5-131460289-INS-130
chr5-131522630-INS-369
chr5-131525629-INS-231
chr5-131577230-INS-190
chr5-131581197-INS-54
chr5-131610867-INS-54
chr5-131634402-INS-141
chr5-131648687-INS-63
chr5-131652254-INS-343
chr5-131669168-INS-276
chr5-131670835-INS-55
chr5-131673113-INS-52
chr5-131675898-INS-155
chr5-131698770-INS-67
chr5-131701639-INS-53
chr5-131720767-INS-425
chr5-131721830-INS-965
chr5-131722077-INS-55
chr5-131734844-INS-6401
chr5-131752578-INS-1069
chr5-131754712-INS-504
chr5-131780032-INS-74
chr5-131784431-INS-51
chr5-131803223-INS-67
chr5-131826889-INS-120
chr5-131856146-INS-56
chr5-131856854-INS-159
chr5-131876580-INS-7146
chr5-131877792-INS-374
chr5-131902550-INS-79
chr5-131939060-INS-248
chr5-131971377-INS-122
chr5-131998213-INS-99
chr5-132025691-INS-116
chr5-132032936-INS-59
chr5-132036182-INS-298
chr5-132056976-INS-125
chr5-132077997-INS-57
chr5-132093283-INS-6504
chr5-132107432-INS-159
chr5-132143077-INS-53
chr5-132174629-INS-132
chr5-132192265-INS-186
chr5-132257914-INS-160
chr5-132264727-INS-100
chr5-132281709-INS-80
chr5-132303933-INS-83
chr5-132313123-INS-313
chr5-132330831-INS-128
chr5-132344905-INS-74
chr5-132354544-INS-64
chr5-132358458-INS-55
chr5-132380988-INS-63
chr5-132391643-INS-84
chr5-132419264-INS-957
chr5-132419327-INS-479
chr5-132424325-INS-327
chr5-132427582-INS-129
chr5-132440413-INS-70
chr5-132441891-INS-91
chr5-132443037-INS-62
chr5-132454005-INS-90
chr5-132470559-INS-111

chr5-132478168-INS-212
chr5-132490752-INS-63
chr5-132497606-INS-132
chr5-132525979-INS-628
chr5-132549688-INS-709
chr5-132575566-INS-54
chr5-132582747-INS-63
chr5-132587701-INS-53
chr5-132603729-INS-129
chr5-132611018-INS-334
chr5-132624673-INS-159
chr5-132630214-INS-100
chr5-132635039-INS-197
chr5-132640834-INS-84
chr5-132644958-INS-506
chr5-132655955-INS-833
chr5-132655957-INS-74
chr5-132664246-INS-194
chr5-132664352-INS-677
chr5-132670442-INS-52
chr5-132690424-INS-63
chr5-132698319-INS-54
chr5-132721105-INS-72
chr5-132725643-INS-54
chr5-132728130-INS-53
chr5-132735232-INS-132
chr5-132737739-INS-88
chr5-132753088-INS-142
chr5-132757538-INS-62
chr5-132777595-INS-55
chr5-132779019-INS-449
chr5-132783344-INS-99
chr5-132785291-INS-52
chr5-132800796-INS-60
chr5-132851055-INS-122
chr5-132861244-INS-76
chr5-132863879-INS-54
chr5-132865812-INS-206
chr5-132866601-INS-119
chr5-132891784-INS-59
chr5-132905150-INS-178
chr5-132906662-INS-159
chr5-132912504-INS-99
chr5-132921201-INS-51
chr5-132929748-INS-79
chr5-132933852-INS-53
chr5-132948616-INS-7134
chr5-132972840-INS-94
chr5-132977037-INS-159
chr5-132992887-INS-61
chr5-132994629-INS-70
chr5-132996533-INS-78
chr5-132997547-INS-91
chr5-133001303-INS-7390
chr5-133003758-INS-54
chr5-133013729-INS-55
chr5-133013914-INS-106
chr5-133017471-INS-939
chr5-133022622-INS-239
chr5-133029937-INS-78
chr5-133052690-INS-618
chr5-133060283-INS-52
chr5-133074175-INS-146
chr5-133086897-INS-51
chr5-133126252-INS-106
chr5-133152036-INS-109
chr5-133156676-INS-444
chr5-133158748-INS-106
chr5-133168265-INS-80
chr5-133172592-INS-216
chr5-133174095-INS-52
chr5-133196886-INS-613
chr5-133227579-INS-6183
chr5-133231948-INS-6469
chr5-133242235-INS-69
chr5-133246241-INS-93
chr5-133252529-INS-197

chr5-133283601-INS-129
chr5-133342245-INS-80
chr5-133368121-INS-6661
chr5-133369103-INS-594
chr5-133389451-INS-194
chr5-133411997-INS-180
chr5-133423892-INS-341
chr5-133429035-INS-60
chr5-133456402-INS-65
chr5-133478763-INS-70
chr5-133519153-INS-55
chr5-133521683-INS-69
chr5-133531980-INS-84
chr5-133535918-INS-397
chr5-133538161-INS-346
chr5-133541796-INS-60
chr5-133553008-INS-82
chr5-133599272-INS-58
chr5-133621272-INS-56
chr5-133652414-INS-369
chr5-133664953-INS-126
chr5-133686697-INS-355
chr5-133686710-INS-418
chr5-133692625-INS-101
chr5-133693625-INS-158
chr5-133702336-INS-63
chr5-133703925-INS-57
chr5-133709916-INS-78
chr5-133720052-INS-107
chr5-133722778-INS-54
chr5-133738726-INS-60
chr5-133752495-INS-599
chr5-133764220-INS-80
chr5-133799338-INS-92
chr5-133823935-INS-62
chr5-133829594-INS-691
chr5-133849578-INS-160
chr5-133854414-INS-81
chr5-133857904-INS-68
chr5-133862875-INS-73
chr5-133886189-INS-159
chr5-133891044-INS-281
chr5-133893998-INS-96
chr5-133895779-INS-68
chr5-133913435-INS-164
chr5-133929154-INS-110
chr5-133937188-INS-930
chr5-133940709-INS-90
chr5-133957348-INS-356
chr5-133961062-INS-58
chr5-133972380-INS-67
chr5-133973976-INS-169
chr5-133976374-INS-298
chr5-134017986-INS-334
chr5-134022895-INS-285
chr5-134036283-INS-161
chr5-134036773-INS-112
chr5-134039135-INS-53
chr5-134045130-INS-74
chr5-134047790-INS-543
chr5-134050800-INS-112
chr5-134055113-INS-72
chr5-134072484-INS-52
chr5-134076541-INS-75
chr5-134079373-INS-142
chr5-134080778-INS-558
chr5-134090430-INS-106
chr5-134094796-INS-80
chr5-134096410-INS-365
chr5-134101099-INS-53
chr5-134111267-INS-203
chr5-134113469-INS-200
chr5-134118852-INS-107
chr5-134131189-INS-100
chr5-134136258-INS-56
chr5-134146227-INS-422
chr5-134147221-INS-104

chr5-134150644-INS-214
chr5-134155414-INS-54
chr5-134180674-INS-409
chr5-134181806-INS-97
chr5-134197525-INS-281
chr5-134203919-INS-208
chr5-134208197-INS-208
chr5-134209386-INS-226
chr5-134210836-INS-52
chr5-134225642-INS-215
chr5-134234914-INS-57
chr5-134241353-INS-104
chr5-134247288-INS-396
chr5-134252562-INS-98
chr5-134285129-INS-289
chr5-134285610-INS-75
chr5-134287380-INS-102
chr5-134306980-INS-80
chr5-134314629-INS-258
chr5-134316678-INS-148
chr5-134375312-INS-127
chr5-134382704-INS-210
chr5-134383580-INS-354
chr5-134388624-INS-205
chr5-134394445-INS-92
chr5-134395401-INS-81
chr5-134396730-INS-210
chr5-134404002-INS-53
chr5-134406959-INS-223
chr5-134409341-INS-107
chr5-134415390-INS-203
chr5-134417721-INS-52
chr5-134422235-INS-217
chr5-134424002-INS-53
chr5-134427727-INS-64
chr5-134448052-INS-179
chr5-134448889-INS-7364
chr5-134451236-INS-67
chr5-134468511-INS-212
chr5-134470181-INS-52
chr5-134494691-INS-133
chr5-134511919-INS-52
chr5-134513975-INS-64
chr5-134515764-INS-208
chr5-134540629-INS-223
chr5-134547377-INS-106
chr5-134593583-INS-140
chr5-134612276-INS-364
chr5-134614099-INS-175
chr5-134624103-INS-85
chr5-134634089-INS-199
chr5-134639618-INS-53
chr5-134640206-INS-176
chr5-134641940-INS-52
chr5-134646477-INS-175
chr5-134651888-INS-76
chr5-134658981-INS-328
chr5-134672013-INS-64
chr5-134678213-INS-234
chr5-134687055-INS-436
chr5-134689931-INS-169
chr5-134718604-INS-240
chr5-134733719-INS-265
chr5-134735710-INS-106
chr5-134743430-INS-133
chr5-134755655-INS-53
chr5-134761626-INS-52
chr5-134764431-INS-371
chr5-134786767-INS-111
chr5-134799867-INS-84
chr5-134802358-INS-107
chr5-134808861-INS-153
chr5-134825385-INS-87
chr5-134836683-INS-151
chr5-134836732-INS-1549
chr5-134839718-INS-377
chr5-134840770-INS-609

chr5-134843631-INS-106
chr5-134846123-INS-83
chr5-134848618-INS-52
chr5-134852497-INS-53
chr5-134863906-INS-8455
chr5-134870157-INS-499
chr5-134873244-INS-52
chr5-134899542-INS-81
chr5-134915540-INS-106
chr5-134934327-INS-174
chr5-134938723-INS-82
chr5-134949618-INS-969
chr5-134950391-INS-73
chr5-134954676-INS-60
chr5-134956048-INS-205
chr5-134960881-INS-256
chr5-134967663-INS-228
chr5-134973144-INS-178
chr5-134991387-INS-55
chr5-135005511-INS-106
chr5-135012543-INS-84
chr5-135022777-INS-54
chr5-135025126-INS-7174
chr5-135030481-INS-211
chr5-135037240-INS-207
chr5-135067472-INS-121
chr5-135096830-INS-179
chr5-135115455-INS-61
chr5-135128236-INS-203
chr5-135131839-INS-166
chr5-135146502-INS-670
chr5-135152208-INS-295
chr5-135160660-INS-64
chr5-135177549-INS-210
chr5-135213505-INS-208
chr5-135227844-INS-350
chr5-135247430-INS-56
chr5-135256549-INS-97
chr5-135296145-INS-544
chr5-135296238-INS-1781
chr5-135301621-INS-168
chr5-135316663-INS-355
chr5-135326459-INS-231
chr5-135329071-INS-7484
chr5-135332050-INS-925
chr5-135335101-INS-7146
chr5-135353572-INS-207
chr5-135357404-INS-192
chr5-135374340-INS-87
chr5-135384128-INS-6452
chr5-135398159-INS-53
chr5-135402607-INS-214
chr5-135417357-INS-122
chr5-135426252-INS-78
chr5-135427492-INS-56
chr5-135446098-INS-137
chr5-135447715-INS-130
chr5-135460991-INS-203
chr5-135467538-INS-260
chr5-135468333-INS-105
chr5-135470860-INS-85
chr5-135475273-INS-212
chr5-135520023-INS-56
chr5-135523164-INS-53
chr5-135527430-INS-74
chr5-135529587-INS-105
chr5-135532972-INS-68
chr5-135534806-INS-6282
chr5-135540992-INS-280
chr5-135543199-INS-191
chr5-135556309-INS-110
chr5-135558348-INS-216
chr5-135563250-INS-265
chr5-135567041-INS-145
chr5-135570454-INS-61
chr5-135581282-INS-56
chr5-135583917-INS-860

chr5-135620187-INS-60
chr5-135644555-INS-140
chr5-135645955-INS-4827
chr5-135646348-INS-59
chr5-135648613-INS-329
chr5-135653500-INS-76
chr5-135667601-INS-109
chr5-135678614-INS-217
chr5-135679507-INS-84
chr5-135681591-INS-117
chr5-135688578-INS-64
chr5-135690626-INS-207
chr5-135691745-INS-76
chr5-135694311-INS-327
chr5-135695233-INS-55
chr5-135701552-INS-53
chr5-135702974-INS-77
chr5-135707974-INS-94
chr5-135722693-INS-181
chr5-135734345-INS-201
chr5-135745664-INS-267
chr5-135750305-INS-162
chr5-135778516-INS-73
chr5-135778963-INS-6475
chr5-135796143-INS-163
chr5-135801258-INS-2194
chr5-135815548-INS-228
chr5-135822080-INS-199
chr5-135827916-INS-102
chr5-135828895-INS-135
chr5-135838736-INS-314
chr5-135859491-INS-957
chr5-135869759-INS-52
chr5-135877425-INS-76
chr5-135882430-INS-74
chr5-135890149-INS-63
chr5-135906361-INS-81
chr5-135913396-INS-211
chr5-135920696-INS-377
chr5-135923320-INS-185
chr5-135926143-INS-79
chr5-135939421-INS-61
chr5-135944727-INS-98
chr5-135954136-INS-68
chr5-135967070-INS-199
chr5-135969009-INS-140
chr5-135971241-INS-109
chr5-135971555-INS-94
chr5-135976062-INS-116
chr5-135983085-INS-170
chr5-135995166-INS-164
chr5-135998847-INS-5313
chr5-136015739-INS-203
chr5-136020847-INS-153
chr5-136029404-INS-4752
chr5-136033097-INS-241
chr5-136050518-INS-118
chr5-136058376-INS-53
chr5-136060574-INS-111
chr5-136077881-INS-215
chr5-136083267-INS-208
chr5-136084353-INS-140
chr5-136096581-INS-120
chr5-136109314-INS-106
chr5-136110632-INS-209
chr5-136136941-INS-59
chr5-136142421-INS-53
chr5-136172625-INS-11833
chr5-136177363-INS-107
chr5-136189912-INS-56
chr5-136193513-INS-161
chr5-136219570-INS-62
chr5-136229686-INS-106
chr5-136231454-INS-89
chr5-136241122-INS-476
chr5-136254724-INS-187
chr5-136278744-INS-265

chr5-136283321-INS-209
chr5-136288177-INS-204
chr5-136374544-INS-56
chr5-136377371-INS-72
chr5-136381178-INS-90
chr5-136406879-INS-53
chr5-136415021-INS-196
chr5-136486282-INS-156
chr5-136496261-INS-109
chr5-136513468-INS-1864
chr5-136523797-INS-229
chr5-136526469-INS-64
chr5-136589411-INS-197
chr5-136616101-INS-72
chr5-136618649-INS-210
chr5-136621997-INS-149
chr5-136626004-INS-58
chr5-136628164-INS-55
chr5-136641350-INS-76
chr5-136647064-INS-55
chr5-136653730-INS-51
chr5-136655233-INS-185
chr5-136662000-INS-106
chr5-136665755-INS-265
chr5-136674373-INS-52
chr5-136675328-INS-53
chr5-136679932-INS-530
chr5-136686331-INS-134
chr5-136692133-INS-71
chr5-136698600-INS-69
chr5-136718631-INS-200
chr5-136740804-INS-110
chr5-136741808-INS-64
chr5-136745157-INS-106
chr5-136746750-INS-376
chr5-136765409-INS-53
chr5-136788225-INS-64
chr5-136795066-INS-72
chr5-136795998-INS-53
chr5-136799303-INS-208
chr5-136808265-INS-179
chr5-136814570-INS-187
chr5-136820292-INS-81
chr5-136824155-INS-211
chr5-136838923-INS-87
chr5-136846790-INS-218
chr5-136848084-INS-74
chr5-136849895-INS-192
chr5-136851515-INS-221
chr5-136856258-INS-158
chr5-136859140-INS-582
chr5-136861228-INS-122
chr5-136870755-INS-248
chr5-136876584-INS-76
chr5-136879251-INS-53
chr5-136881731-INS-90
chr5-136889633-INS-105
chr5-136900866-INS-226
chr5-136904424-INS-252
chr5-136909320-INS-54
chr5-136927061-INS-196
chr5-136933039-INS-217
chr5-136943024-INS-219
chr5-136970486-INS-542
chr5-136971250-INS-274
chr5-136973102-INS-393
chr5-136974652-INS-91
chr5-136976793-INS-83
chr5-136981516-INS-227
chr5-136982639-INS-223
chr5-136988986-INS-208
chr5-137002350-INS-78
chr5-137039908-INS-50
chr5-137042946-INS-226
chr5-137045590-INS-222
chr5-137047474-INS-59
chr5-137048739-INS-215

chr5-137051258-INS-67
chr5-137071135-INS-205
chr5-137081648-INS-197
chr5-137097614-INS-59
chr5-137109363-INS-87
chr5-137122394-INS-171
chr5-137123824-INS-234
chr5-137130773-INS-123
chr5-137153391-INS-317
chr5-137155280-INS-80
chr5-137169103-INS-56
chr5-137171927-INS-109
chr5-137173469-INS-223
chr5-137180234-INS-54
chr5-137195188-INS-55
chr5-137199421-INS-121
chr5-137205386-INS-54
chr5-137221152-INS-65
chr5-137224349-INS-53
chr5-137226010-INS-66
chr5-137233584-INS-70
chr5-137240135-INS-122
chr5-137242928-INS-406
chr5-137257933-INS-642
chr5-137259416-INS-180
chr5-137268142-INS-63
chr5-137274251-INS-186
chr5-137275376-INS-64
chr5-137287150-INS-54
chr5-137328252-INS-82
chr5-137344778-INS-60
chr5-137351987-INS-52
chr5-137353861-INS-240
chr5-137361129-INS-53
chr5-137366157-INS-54
chr5-137366488-INS-584
chr5-137368300-INS-130
chr5-137379311-INS-206
chr5-137393120-INS-80
chr5-137407355-INS-210
chr5-137413873-INS-68
chr5-137421311-INS-54
chr5-137430780-INS-64
chr5-137435388-INS-212
chr5-137440565-INS-59
chr5-137442503-INS-150
chr5-137444692-INS-141
chr5-137449816-INS-134
chr5-137457878-INS-174
chr5-137459619-INS-106
chr5-137461033-INS-91
chr5-137484975-INS-262
chr5-137487253-INS-201
chr5-137488765-INS-59
chr5-137493651-INS-116
chr5-137499290-INS-2965
chr5-137504934-INS-192
chr5-137536618-INS-107
chr5-137539637-INS-56
chr5-137547682-INS-307
chr5-137551188-INS-54
chr5-137559080-INS-231
chr5-137586533-INS-72
chr5-137592009-INS-255
chr5-137593317-INS-51
chr5-137599787-INS-121
chr5-137600802-INS-198
chr5-137615417-INS-106
chr5-137618807-INS-301
chr5-137629324-INS-115
chr5-137641877-INS-213
chr5-137656762-INS-401
chr5-137657822-INS-104
chr5-137663119-INS-316
chr5-137666702-INS-148
chr5-137696924-INS-121
chr5-137697916-INS-92

chr5-137709321-INS-82
chr5-137713631-INS-212
chr5-137724106-INS-178
chr5-137731621-INS-52
chr5-137754938-INS-121
chr5-137769663-INS-439
chr5-137806024-INS-384
chr5-137809372-INS-165
chr5-137830667-INS-7439
chr5-137854088-INS-72
chr5-137866516-INS-96
chr5-137882604-INS-214
chr5-137894951-INS-111
chr5-137898477-INS-64
chr5-137920180-INS-89
chr5-137934017-INS-620
chr5-137935786-INS-210
chr5-137937186-INS-137
chr5-137958387-INS-51
chr5-137959921-INS-87
chr5-137964718-INS-233
chr5-137968225-INS-283
chr5-137981326-INS-197
chr5-137981469-INS-215
chr5-137982694-INS-69
chr5-137986304-INS-53
chr5-137994496-INS-431
chr5-138013938-INS-72
chr5-138038317-INS-149
chr5-138046325-INS-225
chr5-138065951-INS-53
chr5-138071875-INS-53
chr5-138089191-INS-175
chr5-138119330-INS-212
chr5-138121921-INS-196
chr5-138146659-INS-216
chr5-138154528-INS-159
chr5-138155822-INS-268
chr5-138156518-INS-721
chr5-138173964-INS-183
chr5-138186892-INS-215
chr5-138217143-INS-57
chr5-138233770-INS-391
chr5-138256590-INS-158
chr5-138271888-INS-53
chr5-138300452-INS-65
chr5-138334971-INS-114
chr5-138360612-INS-250
chr5-138364459-INS-6279
chr5-138365046-INS-453
chr5-138379675-INS-6183
chr5-138385378-INS-6178
chr5-138388869-INS-136
chr5-138391093-INS-72
chr5-138394145-INS-74
chr5-138397426-INS-6426
chr5-138401160-INS-5559
chr5-138432690-INS-340
chr5-138433369-INS-203
chr5-138442111-INS-138
chr5-138443989-INS-146
chr5-138461882-INS-439
chr5-138468032-INS-71
chr5-138509768-INS-375
chr5-138525808-INS-5299
chr5-138544216-INS-70
chr5-138613699-INS-249
chr5-138646471-INS-76
chr5-138674307-INS-158
chr5-138678233-INS-52
chr5-138689806-INS-385
chr5-138707129-INS-4890
chr5-138722044-INS-200
chr5-138746877-INS-54
chr5-138748492-INS-81
chr5-138753171-INS-800
chr5-138766891-INS-99

chr5-138767587-INS-54
chr5-138824590-INS-70
chr5-138850368-INS-92
chr5-138890151-INS-664
chr5-138895758-INS-151
chr5-138929548-INS-103
chr5-138932496-INS-222
chr5-138950680-INS-266
chr5-138989741-INS-157
chr5-138992297-INS-546
chr5-139002203-INS-267
chr5-139007238-INS-191
chr5-139026407-INS-75
chr5-139038392-INS-54
chr5-139051271-INS-105
chr5-139058037-INS-52
chr5-139060819-INS-267
chr5-139069067-INS-54
chr5-139073693-INS-739
chr5-139079003-INS-126
chr5-139080920-INS-354
chr5-139084514-INS-67
chr5-139086319-INS-52
chr5-139090479-INS-96
chr5-139094573-INS-472
chr5-139112022-INS-56
chr5-139119538-INS-105
chr5-139132728-INS-73
chr5-139149757-INS-69
chr5-139159649-INS-209
chr5-139167703-INS-62
chr5-139178995-INS-207
chr5-139181786-INS-117
chr5-139182219-INS-87
chr5-139193192-INS-55
chr5-139236211-INS-199
chr5-139242396-INS-70
chr5-139247763-INS-6263
chr5-139250475-INS-54
chr5-139264627-INS-63
chr5-139264932-INS-112
chr5-139277394-INS-211
chr5-139285806-INS-279
chr5-139292092-INS-135
chr5-139293122-INS-108
chr5-139294069-INS-52
chr5-139301858-INS-120
chr5-139312723-INS-212
chr5-139316505-INS-60
chr5-139317402-INS-213
chr5-139321778-INS-256
chr5-139323258-INS-174
chr5-139326731-INS-102
chr5-139331893-INS-87
chr5-139333659-INS-262
chr5-139345696-INS-151
chr5-139360936-INS-166
chr5-139364158-INS-577
chr5-139392535-INS-65
chr5-139420703-INS-166
chr5-139436364-INS-112
chr5-139457720-INS-54
chr5-139459361-INS-6401
chr5-139475304-INS-208
chr5-139475490-INS-1299
chr5-139479925-INS-122
chr5-139486362-INS-6440
chr5-139496516-INS-86
chr5-139505987-INS-52
chr5-139509975-INS-222
chr5-139512235-INS-105
chr5-139513168-INS-59
chr5-139520697-INS-61
chr5-139548085-INS-73
chr5-139550563-INS-51
chr5-139553333-INS-399
chr5-139556851-INS-162

chr5-139561497-INS-189
chr5-139576361-INS-52
chr5-139593192-INS-102
chr5-139594425-INS-53
chr5-139597483-INS-52
chr5-139622659-INS-70
chr5-139626082-INS-212
chr5-139636522-INS-143
chr5-139638066-INS-219
chr5-139639938-INS-59
chr5-139641801-INS-1865
chr5-139656563-INS-54
chr5-139666456-INS-52
chr5-139685527-INS-52
chr5-139689389-INS-320
chr5-139700668-INS-313
chr5-139705476-INS-54
chr5-139719081-INS-56
chr5-139727551-INS-268
chr5-139739512-INS-51
chr5-139774890-INS-62
chr5-139776317-INS-101
chr5-139778212-INS-201
chr5-139801174-INS-52
chr5-139814470-INS-106
chr5-139824467-INS-105
chr5-139829261-INS-497
chr5-139858089-INS-265
chr5-139863010-INS-105
chr5-139875972-INS-162
chr5-139879066-INS-6403
chr5-139892612-INS-195
chr5-139906136-INS-242
chr5-139913688-INS-200
chr5-139935043-INS-74
chr5-139952901-INS-135
chr5-139962506-INS-176
chr5-139967897-INS-990
chr5-139968139-INS-111
chr5-139968910-INS-78
chr5-139973177-INS-65
chr5-139993044-INS-202
chr5-140001024-INS-105
chr5-140044218-INS-52
chr5-140081401-INS-156
chr5-140085150-INS-54
chr5-140093150-INS-189
chr5-140098020-INS-467
chr5-140107342-INS-104
chr5-140121449-INS-96
chr5-140133555-INS-209
chr5-140135340-INS-205
chr5-140138182-INS-183
chr5-140163982-INS-112
chr5-140181481-INS-64
chr5-140193988-INS-73
chr5-140207498-INS-54
chr5-140209501-INS-58
chr5-140214759-INS-99
chr5-140216835-INS-435
chr5-140221368-INS-313
chr5-140229551-INS-157
chr5-140259301-INS-309
chr5-140263495-INS-61
chr5-140267369-INS-53
chr5-140276524-INS-91
chr5-140279318-INS-254
chr5-140282855-INS-171
chr5-140291830-INS-438
chr5-140292635-INS-67
chr5-140313100-INS-92
chr5-140328920-INS-708
chr5-140346303-INS-110
chr5-140357994-INS-54
chr5-140362144-INS-69
chr5-140366306-INS-198
chr5-140371395-INS-222

chr5-140377229-INS-199
chr5-140379529-INS-123
chr5-140379811-INS-5253
chr5-140383101-INS-408
chr5-140384282-INS-106
chr5-140395059-INS-5564
chr5-140396562-INS-198
chr5-140398376-INS-96
chr5-140412927-INS-91
chr5-140432697-INS-86
chr5-140438955-INS-142
chr5-140439733-INS-61
chr5-140442288-INS-165
chr5-140444640-INS-198
chr5-140449939-INS-52
chr5-140460316-INS-198
chr5-140467032-INS-87
chr5-140471875-INS-308
chr5-140473195-INS-144
chr5-140473905-INS-942
chr5-140474047-INS-102
chr5-140486430-INS-203
chr5-140498251-INS-370
chr5-140499178-INS-210
chr5-140501338-INS-104
chr5-140504437-INS-222
chr5-140506414-INS-116
chr5-140508842-INS-132
chr5-140514471-INS-206
chr5-140533990-INS-189
chr5-140538536-INS-54
chr5-140554760-INS-109
chr5-140562085-INS-55
chr5-140563763-INS-114
chr5-140568538-INS-197
chr5-140575681-INS-660
chr5-140639936-INS-57
chr5-140654001-INS-76
chr5-140658613-INS-157
chr5-140673214-INS-80
chr5-140676584-INS-104
chr5-140692903-INS-212
chr5-140693905-INS-207
chr5-140696163-INS-398
chr5-140716222-INS-108
chr5-140736947-INS-421
chr5-140742362-INS-449
chr5-140783112-INS-202
chr5-140797665-INS-319
chr5-140807774-INS-303
chr5-140844983-INS-53
chr5-140862829-INS-291
chr5-140884458-INS-738
chr5-140898390-INS-51
chr5-140905002-INS-106
chr5-140912010-INS-499
chr5-140916981-INS-240
chr5-140925650-INS-221
chr5-140934722-INS-184
chr5-140949644-INS-106
chr5-140961193-INS-110
chr5-140977806-INS-275
chr5-140981380-INS-66
chr5-140984454-INS-791
chr5-140984473-INS-209
chr5-140987789-INS-51
chr5-141000736-INS-197
chr5-141004552-INS-189
chr5-141016303-INS-53
chr5-141034552-INS-158
chr5-141038820-INS-58
chr5-141041546-INS-106
chr5-141049277-INS-156
chr5-141057737-INS-75
chr5-141060929-INS-106
chr5-141062704-INS-265
chr5-141101059-INS-51

chr5-141121711-INS-66
chr5-141142266-INS-69
chr5-141152115-INS-122
chr5-141172702-INS-144
chr5-141197890-INS-54
chr5-141208252-INS-72
chr5-141223844-INS-61
chr5-141236813-INS-5568
chr5-141244069-INS-6034
chr5-141255400-INS-74
chr5-141278053-INS-647
chr5-141289302-INS-117
chr5-141320215-INS-207
chr5-141334927-INS-457
chr5-141352956-INS-72
chr5-141362024-INS-58
chr5-141379530-INS-460
chr5-141403893-INS-472
chr5-141404725-INS-474
chr5-141418706-INS-554
chr5-141477550-INS-94
chr5-141485776-INS-56
chr5-141504670-INS-355
chr5-141527044-INS-89
chr5-141528027-INS-104
chr5-141532018-INS-51
chr5-141561196-INS-305
chr5-141565237-INS-103
chr5-141571669-INS-108
chr5-141575968-INS-97
chr5-141579002-INS-173
chr5-141606871-INS-6822
chr5-141608280-INS-497
chr5-141609739-INS-256
chr5-141669999-INS-57
chr5-141685214-INS-224
chr5-141689453-INS-230
chr5-141697265-INS-66
chr5-141728918-INS-54
chr5-141741533-INS-103
chr5-141758089-INS-54
chr5-141768559-INS-106
chr5-141789098-INS-128
chr5-141797388-INS-76
chr5-141813942-INS-56
chr5-141825517-INS-265
chr5-141826527-INS-55
chr5-141840945-INS-370
chr5-141847059-INS-244
chr5-141857441-INS-104
chr5-141871010-INS-503
chr5-141873767-INS-373
chr5-141894651-INS-58
chr5-141906109-INS-55
chr5-141907448-INS-238
chr5-141949118-INS-132
chr5-141966104-INS-76
chr5-141968102-INS-459
chr5-142001316-INS-66
chr5-142018917-INS-73
chr5-142019774-INS-52
chr5-142023502-INS-213
chr5-142033793-INS-167
chr5-142043209-INS-105
chr5-142049111-INS-204
chr5-142051230-INS-52
chr5-142062091-INS-126
chr5-142103980-INS-275
chr5-142106532-INS-55
chr5-142110672-INS-140
chr5-142112805-INS-123
chr5-142116210-INS-105
chr5-142119272-INS-199
chr5-142130326-INS-222
chr5-142138274-INS-329
chr5-142140488-INS-54
chr5-142143900-INS-7135

chr5-142154706-INS-696
chr5-142158408-INS-112
chr5-142164792-INS-52
chr5-142166904-INS-240
chr5-142179894-INS-180
chr5-142192550-INS-492
chr5-142205123-INS-107
chr5-142209506-INS-227
chr5-142217179-INS-5546
chr5-142229846-INS-53
chr5-142246713-INS-119
chr5-142253594-INS-53
chr5-142256263-INS-649
chr5-142266094-INS-52
chr5-142267273-INS-63
chr5-142270673-INS-87
chr5-142298767-INS-68
chr5-142302235-INS-102
chr5-142304606-INS-64
chr5-142322899-INS-124
chr5-142331398-INS-116
chr5-142335807-INS-113
chr5-142346610-INS-156
chr5-142368205-INS-58
chr5-142379576-INS-106
chr5-142380911-INS-197
chr5-142408829-INS-258
chr5-142415872-INS-105
chr5-142429043-INS-76
chr5-142437660-INS-123
chr5-142441951-INS-243
chr5-142456450-INS-53
chr5-142461658-INS-196
chr5-142466651-INS-51
chr5-142505722-INS-160
chr5-142525823-INS-52
chr5-142541089-INS-57
chr5-142543165-INS-220
chr5-142544663-INS-98
chr5-142557243-INS-163
chr5-142561884-INS-62
chr5-142596853-INS-262
chr5-142601902-INS-156
chr5-142607578-INS-125
chr5-142610022-INS-225
chr5-142648563-INS-167
chr5-142654018-INS-52
chr5-142660142-INS-152
chr5-142663062-INS-637
chr5-142665123-INS-89
chr5-142669514-INS-415
chr5-142670810-INS-107
chr5-142685662-INS-52
chr5-142690906-INS-156
chr5-142697391-INS-77
chr5-142699200-INS-92
chr5-142700230-INS-358
chr5-142716086-INS-1893
chr5-142724056-INS-106
chr5-142730314-INS-156
chr5-142744844-INS-137
chr5-142756527-INS-528
chr5-142763653-INS-54
chr5-142768853-INS-65
chr5-142791484-INS-66
chr5-142813140-INS-182
chr5-142850116-INS-96
chr5-142865155-INS-177
chr5-142873253-INS-146
chr5-142875046-INS-128
chr5-142899852-INS-416
chr5-142918478-INS-209
chr5-142922839-INS-91
chr5-142931612-INS-65
chr5-142939720-INS-52
chr5-142960196-INS-197
chr5-142967839-INS-215

chr5-142968353-INS-519
chr5-142971457-INS-106
chr5-143046785-INS-89
chr5-143066353-INS-86
chr5-143089356-INS-60
chr5-143101405-INS-57
chr5-143103607-INS-611
chr5-143109211-INS-8985
chr5-143113431-INS-343
chr5-143114428-INS-135
chr5-143125758-INS-463
chr5-143132056-INS-168
chr5-143134878-INS-86
chr5-143138102-INS-69
chr5-143148367-INS-3391
chr5-143162910-INS-321
chr5-143168737-INS-209
chr5-143202023-INS-56
chr5-143216495-INS-407
chr5-143224808-INS-157
chr5-143228682-INS-78
chr5-143233789-INS-83
chr5-143245801-INS-287
chr5-143245873-INS-738
chr5-143246739-INS-356
chr5-143257165-INS-364
chr5-143269162-INS-288
chr5-143270605-INS-56
chr5-143283062-INS-329
chr5-143311231-INS-104
chr5-143332695-INS-158
chr5-143344318-INS-175
chr5-143346975-INS-264
chr5-143352715-INS-101
chr5-143353224-INS-1890
chr5-143363557-INS-132
chr5-143364443-INS-107
chr5-143376298-INS-184
chr5-143391998-INS-218
chr5-143394173-INS-63
chr5-143399719-INS-54
chr5-143404617-INS-92
chr5-143418587-INS-57
chr5-143440788-INS-60
chr5-143452546-INS-204
chr5-143457116-INS-378
chr5-143464954-INS-202
chr5-143480953-INS-480
chr5-143497053-INS-172
chr5-143519250-INS-96
chr5-143522175-INS-264
chr5-143544916-INS-122
chr5-143561838-INS-169
chr5-143614010-INS-108
chr5-143624780-INS-74
chr5-143626512-INS-76
chr5-143639750-INS-820
chr5-143639766-INS-127
chr5-143658190-INS-198
chr5-143692202-INS-87
chr5-143747097-INS-820
chr5-143791203-INS-72
chr5-143792582-INS-572
chr5-143809333-INS-102
chr5-143820299-INS-211
chr5-143820299-INS-372
chr5-143864428-INS-185
chr5-143867603-INS-1885
chr5-143904783-INS-300
chr5-143909836-INS-279
chr5-143919964-INS-53
chr5-143981628-INS-247
chr5-143999034-INS-628
chr5-144004296-INS-126
chr5-144007665-INS-266
chr5-144031385-INS-404
chr5-144032713-INS-219

chr5-144050335-INS-196
chr5-144060220-INS-145
chr5-144099881-INS-323
chr5-144104709-INS-95
chr5-144114532-INS-53
chr5-144117307-INS-120
chr5-144120518-INS-53
chr5-144129236-INS-66
chr5-144144062-INS-168
chr5-144161243-INS-57
chr5-144166211-INS-426
chr5-144174299-INS-217
chr5-144175281-INS-56
chr5-144178610-INS-206
chr5-144228781-INS-57
chr5-144233818-INS-201
chr5-144237764-INS-276
chr5-144244701-INS-60
chr5-144264034-INS-54
chr5-144279893-INS-509
chr5-144296007-INS-52
chr5-144333547-INS-157
chr5-144340975-INS-676
chr5-144350451-INS-186
chr5-144351226-INS-204
chr5-144362345-INS-82
chr5-144365978-INS-53
chr5-144367773-INS-287
chr5-144377142-INS-142
chr5-144382911-INS-106
chr5-144383005-INS-210
chr5-144384506-INS-198
chr5-144388950-INS-1864
chr5-144395141-INS-58
chr5-144408855-INS-173
chr5-144415211-INS-58
chr5-144431726-INS-146
chr5-144432863-INS-56
chr5-144434360-INS-182
chr5-144436296-INS-80
chr5-144441872-INS-210
chr5-144451510-INS-52
chr5-144459812-INS-62
chr5-144469819-INS-396
chr5-144473293-INS-400
chr5-144478968-INS-62
chr5-144493796-INS-62
chr5-144494819-INS-114
chr5-144497142-INS-56
chr5-144530958-INS-235
chr5-144542910-INS-316
chr5-144555187-INS-59
chr5-144561937-INS-52
chr5-144572508-INS-213
chr5-144574895-INS-332
chr5-144581333-INS-264
chr5-144591272-INS-54
chr5-144597283-INS-205
chr5-144619841-INS-105
chr5-144623733-INS-144
chr5-144631990-INS-128
chr5-144640571-INS-235
chr5-144643656-INS-455
chr5-144645105-INS-53
chr5-144658099-INS-69
chr5-144659794-INS-368
chr5-144693622-INS-147
chr5-144708710-INS-153
chr5-144727154-INS-379
chr5-144732707-INS-480
chr5-144733452-INS-262
chr5-144742857-INS-149
chr5-144783274-INS-77
chr5-144785645-INS-173
chr5-144797873-INS-123
chr5-144803205-INS-64
chr5-144818797-INS-129

chr5-144826892-INS-370
chr5-144832155-INS-322
chr5-144843554-INS-8293
chr5-144852596-INS-90
chr5-144854443-INS-126
chr5-144869465-INS-197
chr5-144877760-INS-68
chr5-144907098-INS-66
chr5-144909376-INS-204
chr5-144924015-INS-66
chr5-144937406-INS-111
chr5-144963412-INS-80
chr5-144972978-INS-706
chr5-144981018-INS-78
chr5-144995629-INS-165
chr5-145019076-INS-226
chr5-145019755-INS-79
chr5-145030591-INS-386
chr5-145039448-INS-80
chr5-145040645-INS-64
chr5-145093941-INS-227
chr5-145113997-INS-5257
chr5-145130017-INS-89
chr5-145131458-INS-84
chr5-145138907-INS-56
chr5-145143896-INS-439
chr5-145151121-INS-508
chr5-145164485-INS-204
chr5-145196226-INS-207
chr5-145223856-INS-69
chr5-145229550-INS-6784
chr5-145253598-INS-7093
chr5-145283739-INS-66
chr5-145284925-INS-283
chr5-145330244-INS-6819
chr5-145380248-INS-380
chr5-145442822-INS-201
chr5-145460804-INS-52
chr5-145474147-INS-6379
chr5-145476020-INS-2750
chr5-145535418-INS-562
chr5-145547833-INS-292
chr5-145576703-INS-6687
chr5-145607360-INS-7966
chr5-145628499-INS-558
chr5-145760046-INS-64
chr5-145801385-INS-82
chr5-145842312-INS-73
chr5-145879170-INS-417
chr5-145949727-INS-263
chr5-145968501-INS-198
chr5-146019431-INS-229
chr5-146048101-INS-498
chr5-146069741-INS-170
chr5-146199595-INS-6401
chr5-146206290-INS-170
chr5-146215598-INS-54
chr5-146217614-INS-844
chr5-146218576-INS-183
chr5-146226587-INS-278
chr5-146235046-INS-286
chr5-146243191-INS-105
chr5-146272674-INS-119
chr5-146305196-INS-136
chr5-146340394-INS-96
chr5-146343441-INS-165
chr5-146351317-INS-337
chr5-146371165-INS-162
chr5-146391014-INS-318
chr5-146486430-INS-92
chr5-146499748-INS-327
chr5-146503984-INS-213
chr5-146589735-INS-191
chr5-146603063-INS-124
chr5-146642697-INS-67
chr5-146644321-INS-56
chr5-146651499-INS-142

chr5-146653672-INS-54
chr5-146659711-INS-151
chr5-146664129-INS-210
chr5-146667602-INS-64
chr5-146675400-INS-195
chr5-146676830-INS-305
chr5-146689765-INS-58
chr5-146696941-INS-4390
chr5-146706367-INS-60
chr5-146716234-INS-91
chr5-146722513-INS-241
chr5-146734744-INS-114
chr5-146757953-INS-369
chr5-146792204-INS-129
chr5-146802805-INS-54
chr5-146806756-INS-3819
chr5-146812396-INS-557
chr5-146825115-INS-180
chr5-146844248-INS-5293
chr5-146846271-INS-142
chr5-146860515-INS-180
chr5-146883565-INS-440
chr5-146894874-INS-232
chr5-146900850-INS-201
chr5-146902590-INS-229
chr5-146904742-INS-212
chr5-146912994-INS-178
chr5-146915573-INS-75
chr5-146917492-INS-180
chr5-146920450-INS-415
chr5-146923413-INS-102
chr5-146931588-INS-60
chr5-146948888-INS-123
chr5-146985009-INS-209
chr5-146987427-INS-230
chr5-146989677-INS-440
chr5-146995435-INS-179
chr5-146998675-INS-223
chr5-147002738-INS-157
chr5-147005122-INS-157
chr5-147029382-INS-161
chr5-147030871-INS-7228
chr5-147036136-INS-202
chr5-147044759-INS-202
chr5-147053153-INS-51
chr5-147107758-INS-62
chr5-147126502-INS-60
chr5-147139561-INS-84
chr5-147144409-INS-341
chr5-147157586-INS-64
chr5-147167246-INS-60
chr5-147185007-INS-116
chr5-147193782-INS-260
chr5-147194750-INS-304
chr5-147200336-INS-66
chr5-147255499-INS-120
chr5-147257567-INS-8989
chr5-147266073-INS-60
chr5-147272936-INS-60
chr5-147276862-INS-54
chr5-147278504-INS-305
chr5-147279776-INS-68
chr5-147283740-INS-51
chr5-147302457-INS-134
chr5-147307963-INS-72
chr5-147316089-INS-457
chr5-147321590-INS-203
chr5-147322186-INS-205
chr5-147334287-INS-300
chr5-147335377-INS-251
chr5-147342330-INS-70
chr5-147343517-INS-344
chr5-147357931-INS-319
chr5-147359526-INS-232
chr5-147368265-INS-82
chr5-147373637-INS-206
chr5-147397761-INS-214

chr5-147414828-INS-60
chr5-147418946-INS-284
chr5-147433207-INS-51
chr5-147433952-INS-7205
chr5-147443863-INS-64
chr5-147453343-INS-53
chr5-147481202-INS-58
chr5-147488113-INS-102
chr5-147511958-INS-108
chr5-147514597-INS-107
chr5-147523790-INS-51
chr5-147534507-INS-271
chr5-147548260-INS-264
chr5-147553041-INS-407
chr5-147560569-INS-97
chr5-147576826-INS-159
chr5-147578696-INS-56
chr5-147591675-INS-92
chr5-147605122-INS-81
chr5-147617171-INS-55
chr5-147648546-INS-68
chr5-147653132-INS-6485
chr5-147674363-INS-100
chr5-147677965-INS-66
chr5-147682083-INS-108
chr5-147683999-INS-54
chr5-147688994-INS-233
chr5-147694617-INS-278
chr5-147699354-INS-112
chr5-147701317-INS-687
chr5-147721611-INS-65
chr5-147729284-INS-60
chr5-147732429-INS-246
chr5-147752441-INS-179
chr5-147760417-INS-60
chr5-147778260-INS-587
chr5-147788963-INS-211
chr5-147815752-INS-107
chr5-147833015-INS-132
chr5-147838103-INS-225
chr5-147842169-INS-102
chr5-147850615-INS-188
chr5-147864218-INS-88
chr5-147871669-INS-192
chr5-147875567-INS-202
chr5-147888986-INS-506
chr5-147910902-INS-83
chr5-147929255-INS-211
chr5-147939265-INS-6295
chr5-147942580-INS-111
chr5-147960987-INS-59
chr5-147967525-INS-102
chr5-147977879-INS-64
chr5-147979738-INS-54
chr5-147983671-INS-145
chr5-147990933-INS-56
chr5-148003516-INS-158
chr5-148012416-INS-54
chr5-148017507-INS-162
chr5-148030243-INS-128
chr5-148054940-INS-210
chr5-148065563-INS-685
chr5-148076523-INS-372
chr5-148080367-INS-63
chr5-148098205-INS-68
chr5-148100632-INS-64
chr5-148136366-INS-255
chr5-148154939-INS-681
chr5-148157662-INS-75
chr5-148205219-INS-52
chr5-148221491-INS-3654
chr5-148222820-INS-7331
chr5-148222821-INS-425
chr5-148227871-INS-1888
chr5-148249586-INS-62
chr5-148270598-INS-365
chr5-148271730-INS-76

chr5-148288007-INS-401
chr5-148292088-INS-66
chr5-148294081-INS-188
chr5-148297085-INS-100
chr5-148305758-INS-213
chr5-148307682-INS-139
chr5-148345025-INS-54
chr5-148350084-INS-103
chr5-148351971-INS-80
chr5-148359088-INS-84
chr5-148367985-INS-104
chr5-148369605-INS-246
chr5-148411860-INS-117
chr5-148419395-INS-118
chr5-148425084-INS-221
chr5-148434201-INS-63
chr5-148456866-INS-71
chr5-148499415-INS-54
chr5-148503152-INS-51
chr5-148503734-INS-84
chr5-148547114-INS-137
chr5-148576370-INS-54
chr5-148663455-INS-66
chr5-148671409-INS-124
chr5-148673132-INS-400
chr5-148688053-INS-122
chr5-148688738-INS-58
chr5-148713682-INS-76
chr5-148720805-INS-322
chr5-148732275-INS-1888
chr5-148761679-INS-645
chr5-148770588-INS-291
chr5-148819614-INS-428
chr5-148830173-INS-113
chr5-148852875-INS-111
chr5-148857032-INS-720
chr5-148857202-INS-260
chr5-148898147-INS-67
chr5-148914351-INS-304
chr5-148924518-INS-57
chr5-148938389-INS-485
chr5-148941700-INS-211
chr5-148946610-INS-88
chr5-148955795-INS-135
chr5-148960009-INS-90
chr5-148964660-INS-107
chr5-148976253-INS-58
chr5-149000938-INS-481
chr5-149017845-INS-197
chr5-149020250-INS-750
chr5-149022151-INS-75
chr5-149035825-INS-71
chr5-149051400-INS-366
chr5-149055698-INS-62
chr5-149096682-INS-55
chr5-149108892-INS-220
chr5-149195432-INS-140
chr5-149200882-INS-719
chr5-149209141-INS-54
chr5-149210904-INS-270
chr5-149215520-INS-245
chr5-149229885-INS-92
chr5-149244932-INS-68
chr5-149257726-INS-80
chr5-149263027-INS-104
chr5-149274025-INS-144
chr5-149281047-INS-136
chr5-149281715-INS-507
chr5-149290408-INS-91
chr5-149297154-INS-144
chr5-149336959-INS-52
chr5-149351274-INS-124
chr5-149374460-INS-209
chr5-149395809-INS-222
chr5-149397871-INS-94
chr5-149400953-INS-62
chr5-149403353-INS-54

chr5-149423920-INS-54
chr5-149425905-INS-241
chr5-149434549-INS-675
chr5-149435683-INS-126
chr5-149451092-INS-70
chr5-149472220-INS-68
chr5-149482132-INS-573
chr5-149483535-INS-74
chr5-149489447-INS-208
chr5-149498650-INS-270
chr5-149502597-INS-76
chr5-149516702-INS-133
chr5-149524984-INS-287
chr5-149526731-INS-6965
chr5-149527643-INS-229
chr5-149531503-INS-121
chr5-149548798-INS-215
chr5-149560522-INS-225
chr5-149567766-INS-217
chr5-149581379-INS-186
chr5-149609945-INS-390
chr5-149613743-INS-138
chr5-149625172-INS-127
chr5-149641195-INS-59
chr5-149649482-INS-217
chr5-149659748-INS-127
chr5-149701843-INS-92
chr5-149708697-INS-62
chr5-149716710-INS-433
chr5-149735121-INS-177
chr5-149755621-INS-212
chr5-149794016-INS-331
chr5-149799766-INS-86
chr5-149818314-INS-200
chr5-149835357-INS-53
chr5-149841678-INS-833
chr5-149901664-INS-441
chr5-149903925-INS-7260
chr5-149913220-INS-57
chr5-149930227-INS-53
chr5-149931278-INS-192
chr5-149950696-INS-5112
chr5-149951272-INS-76
chr5-149973176-INS-58
chr5-149985911-INS-185
chr5-149994554-INS-83
chr5-150000813-INS-198
chr5-150008568-INS-51
chr5-150012782-INS-169
chr5-150036633-INS-54
chr5-150071580-INS-377
chr5-150080643-INS-1862
chr5-150101601-INS-61
chr5-150132744-INS-70
chr5-150132837-INS-84
chr5-150161689-INS-147
chr5-150165798-INS-191
chr5-150189334-INS-201
chr5-150191656-INS-107
chr5-150243260-INS-54
chr5-150335190-INS-110
chr5-150363628-INS-403
chr5-150391215-INS-57
chr5-150398620-INS-6415
chr5-150402469-INS-78
chr5-150406854-INS-117
chr5-150426736-INS-205
chr5-150431833-INS-93
chr5-150445632-INS-90
chr5-150479746-INS-216
chr5-150504940-INS-203
chr5-150512230-INS-5229
chr5-150554325-INS-61
chr5-150565406-INS-173
chr5-150574335-INS-234
chr5-150577281-INS-309
chr5-150581970-INS-56

chr5-150605055-INS-76
chr5-150615415-INS-157
chr5-150617403-INS-327
chr5-150699390-INS-55
chr5-150749990-INS-6425
chr5-150751069-INS-53
chr5-150777983-INS-181
chr5-150808568-INS-158
chr5-150825021-INS-63
chr5-150837772-INS-57
chr5-150843773-INS-1481
chr5-150865353-INS-364
chr5-150872220-INS-7918
chr5-150885891-INS-502
chr5-150920964-INS-358
chr5-150925058-INS-394
chr5-150945178-INS-302
chr5-150947339-INS-215
chr5-150953912-INS-131
chr5-150958513-INS-195
chr5-150994071-INS-74
chr5-151016801-INS-76
chr5-151029187-INS-402
chr5-151076391-INS-1064
chr5-151160571-INS-167
chr5-151196760-INS-112
chr5-151198549-INS-119
chr5-151234182-INS-176
chr5-151263666-INS-7093
chr5-151285466-INS-182
chr5-151298990-INS-80
chr5-151326802-INS-112
chr5-151365732-INS-6256
chr5-151417842-INS-986
chr5-151468901-INS-150
chr5-151490740-INS-326
chr5-151524839-INS-7270
chr5-151527319-INS-5407
chr5-151539066-INS-9752
chr5-151547331-INS-671
chr5-151560559-INS-6188
chr5-151575202-INS-6252
chr5-151578870-INS-214
chr5-151593522-INS-7036
chr5-151594115-INS-93
chr5-151596685-INS-3709
chr5-151614415-INS-363
chr5-151621043-INS-406
chr5-151630855-INS-235
chr5-151653717-INS-327
chr5-151656073-INS-108
chr6-3121450-INS-153
chr6-3140031-INS-77
chr6-3141132-INS-680
chr6-3193828-INS-91
chr6-3206703-INS-7160
chr6-3214442-INS-75
chr6-3216768-INS-1512
chr6-3220724-INS-5526
chr6-3231553-INS-6473
chr6-3236168-INS-74
chr6-3247423-INS-104
chr6-3275972-INS-6527
chr6-3286612-INS-159
chr6-3290662-INS-927
chr6-3291363-INS-294
chr6-3291792-INS-249
chr6-3302746-INS-104
chr6-3306362-INS-539
chr6-3328502-INS-72
chr6-3344777-INS-112
chr6-3384021-INS-83
chr6-3439043-INS-235
chr6-3461394-INS-52
chr6-3469486-INS-462
chr6-3501229-INS-124
chr6-3501229-INS-187

chr6-3539767-INS-73
chr6-3558022-INS-148
chr6-3568591-INS-54
chr6-3572823-INS-77
chr6-3588928-INS-58
chr6-3620157-INS-576
chr6-3631396-INS-333
chr6-3637749-INS-390
chr6-3683109-INS-630
chr6-3688416-INS-1333
chr6-3722705-INS-3362
chr6-3747000-INS-64
chr6-3792988-INS-6071
chr6-3796906-INS-713
chr6-3801686-INS-80
chr6-3810869-INS-74
chr6-3827926-INS-55
chr6-3879570-INS-427
chr6-3887254-INS-763
chr6-3894632-INS-80
chr6-3904812-INS-1021
chr6-3934831-INS-7120
chr6-3935342-INS-597
chr6-3938592-INS-338
chr6-3956299-INS-190
chr6-3981265-INS-5108
chr6-3999192-INS-4933
chr6-4003223-INS-587
chr6-4003894-INS-88
chr6-4019923-INS-122
chr6-4024484-INS-6844
chr6-4025088-INS-4408
chr6-4034083-INS-57
chr6-4039751-INS-280
chr6-4049960-INS-6245
chr6-4052699-INS-7147
chr6-4062811-INS-7556
chr6-4065658-INS-54
chr6-4083837-INS-128
chr6-4089089-INS-203
chr6-4091230-INS-4359
chr6-4111328-INS-195
chr6-4114706-INS-538
chr6-4145078-INS-89
chr6-4146137-INS-372
chr6-4156302-INS-237
chr6-4156670-INS-400
chr6-4171455-INS-55
chr6-4185750-INS-498
chr6-4215742-INS-51
chr6-4216201-INS-646
chr6-4223181-INS-439
chr6-4235506-INS-347
chr6-4241731-INS-185
chr6-4269076-INS-6529
chr6-4292812-INS-112
chr6-4296514-INS-61
chr6-4312536-INS-64
chr6-4315350-INS-865
chr6-4346169-INS-6199
chr6-4350701-INS-72
chr6-4370236-INS-68
chr6-4374850-INS-53
chr6-4389813-INS-497
chr6-4391425-INS-225
chr6-4391836-INS-961
chr6-4394514-INS-53
chr6-4408596-INS-329
chr6-4417803-INS-104
chr6-4461909-INS-125
chr6-4464958-INS-643
chr6-4465628-INS-57
chr6-4467475-INS-274
chr6-4469301-INS-440
chr6-4503848-INS-69
chr6-4515442-INS-51
chr6-4517669-INS-68

chr6-4547241-INS-5207
chr6-4579121-INS-109
chr6-4583382-INS-130
chr6-4585257-INS-168
chr6-4586473-INS-145
chr6-4589714-INS-551
chr6-4593652-INS-54
chr6-4598629-INS-154
chr6-4606528-INS-92
chr6-4611552-INS-819
chr6-4613799-INS-53
chr6-4648003-INS-113
chr6-4658508-INS-415
chr6-4666890-INS-204
chr6-4686131-INS-169
chr6-4712528-INS-204
chr6-4737385-INS-242
chr6-4777742-INS-888
chr6-4778415-INS-204
chr6-4790506-INS-60
chr6-4814381-INS-136
chr6-4843079-INS-146
chr6-4845756-INS-78
chr6-4885700-INS-381
chr6-4911514-INS-480
chr6-4917544-INS-69
chr6-4940294-INS-367
chr6-4959292-INS-76
chr6-4961488-INS-96
chr6-4962062-INS-334
chr6-4965665-INS-208
chr6-4987566-INS-134
chr6-4992906-INS-70
chr6-4995631-INS-5549
chr6-5002136-INS-106
chr6-5003798-INS-70
chr6-5008455-INS-155
chr6-5022658-INS-211
chr6-5032875-INS-88
chr6-5050052-INS-96
chr6-5051270-INS-52
chr6-5063173-INS-5547
chr6-5075275-INS-57
chr6-5150926-INS-197
chr6-5164175-INS-78
chr6-5188116-INS-230
chr6-5193338-INS-52
chr6-5208079-INS-412
chr6-5232619-INS-118
chr6-5239703-INS-65
chr6-5260642-INS-60
chr6-5263731-INS-315
chr6-5273775-INS-4576
chr6-5307332-INS-6954
chr6-5325246-INS-618
chr6-5332915-INS-7130
chr6-5334973-INS-55
chr6-5337770-INS-128
chr6-5345115-INS-473
chr6-5355009-INS-60
chr6-5363811-INS-385
chr6-5366932-INS-562
chr6-5378606-INS-101
chr6-5393960-INS-211
chr6-5401122-INS-65
chr6-5412451-INS-276
chr6-5413054-INS-207
chr6-5415827-INS-476
chr6-5436988-INS-79
chr6-5445668-INS-582
chr6-5457165-INS-108
chr6-5460863-INS-221
chr6-5465693-INS-507
chr6-5495742-INS-108
chr6-5498112-INS-57
chr6-5501046-INS-67
chr6-5527275-INS-76

chr6-5550818-INS-125
chr6-5578827-INS-398
chr6-5586828-INS-269
chr6-5599154-INS-56
chr6-5638739-INS-142
chr6-5639898-INS-71
chr6-5674832-INS-92
chr6-5693281-INS-177
chr6-5713005-INS-238
chr6-5720297-INS-202
chr6-5739116-INS-820
chr6-5744481-INS-94
chr6-5779498-INS-120
chr6-5781654-INS-52
chr6-5805640-INS-550
chr6-5805640-INS-78
chr6-5814579-INS-349
chr6-5820855-INS-137
chr6-5823696-INS-315
chr6-5833345-INS-74
chr6-5835874-INS-164
chr6-5862680-INS-249
chr6-5866527-INS-61
chr6-5893813-INS-161
chr6-5897710-INS-85
chr6-5903563-INS-254
chr6-5911560-INS-88
chr6-5915051-INS-312
chr6-5930154-INS-90
chr6-5955078-INS-61
chr6-5966070-INS-126
chr6-6003904-INS-179
chr6-6012088-INS-82
chr6-6032708-INS-184
chr6-6032731-INS-1037
chr6-6090392-INS-68
chr6-6115892-INS-89
chr6-6123450-INS-148
chr6-6129938-INS-175
chr6-6133036-INS-349
chr6-6136538-INS-546
chr6-6145571-INS-261
chr6-6160616-INS-113
chr6-6179478-INS-111
chr6-6215282-INS-315
chr6-6231925-INS-62
chr6-6237017-INS-328
chr6-6242914-INS-198
chr6-6244542-INS-312
chr6-6274951-INS-536
chr6-6323498-INS-7245
chr6-6334241-INS-2857
chr6-6341687-INS-400
chr6-6347862-INS-904
chr6-6391696-INS-412
chr6-6439388-INS-53
chr6-6446750-INS-104
chr6-6459377-INS-183
chr6-6462407-INS-291
chr6-6469309-INS-328
chr6-6497748-INS-74
chr6-6506517-INS-847
chr6-6509011-INS-75
chr6-6524541-INS-383
chr6-6540156-INS-98
chr6-6579626-INS-248
chr6-6640654-INS-80
chr6-6683249-INS-4721
chr6-6687392-INS-6702
chr6-6688460-INS-102
chr6-6722446-INS-947
chr6-6731093-INS-54
chr6-6760082-INS-142
chr6-6771823-INS-64
chr6-6779929-INS-54
chr6-6790065-INS-94
chr6-6798668-INS-63

chr6-6806040-INS-101
chr6-6816168-INS-162
chr6-6825734-INS-53
chr6-6867017-INS-83
chr6-6901861-INS-72
chr6-6925468-INS-57
chr6-6934873-INS-6592
chr6-6962126-INS-1139
chr6-6962153-INS-719
chr6-6963271-INS-151
chr6-6972286-INS-678
chr6-6995070-INS-7341
chr6-6995633-INS-73
chr6-7063948-INS-374
chr6-7067334-INS-70
chr6-7117094-INS-271
chr6-7162802-INS-781
chr6-7186360-INS-572
chr6-7202815-INS-246
chr6-7246958-INS-201
chr6-7248872-INS-199
chr6-7256460-INS-96
chr6-7262101-INS-7089
chr6-7326742-INS-10509
chr6-7375052-INS-140
chr6-7379092-INS-485
chr6-7480537-INS-113
chr6-7491580-INS-2987
chr6-7495358-INS-142
chr6-7535245-INS-148
chr6-7536090-INS-7077
chr6-7537566-INS-56
chr6-7567292-INS-76
chr6-7568017-INS-113
chr6-7576420-INS-640
chr6-7577449-INS-92
chr6-7579826-INS-279
chr6-7600378-INS-123
chr6-7615108-INS-66
chr6-7649075-INS-584
chr6-7659309-INS-66
chr6-7662195-INS-60
chr6-7667609-INS-187
chr6-7684407-INS-554
chr6-7704682-INS-5601
chr6-7715216-INS-90
chr6-7750614-INS-86
chr6-7752277-INS-579
chr6-7757861-INS-289
chr6-7766992-INS-562
chr6-7783620-INS-66
chr6-7804303-INS-358
chr6-7829538-INS-6632
chr6-7876895-INS-927
chr6-7886555-INS-6762
chr6-7888308-INS-53
chr6-7904791-INS-68
chr6-7958333-INS-906
chr6-7960612-INS-90
chr6-7961905-INS-853
chr6-7968895-INS-136
chr6-7975621-INS-505
chr6-7995150-INS-84
chr6-8010985-INS-888
chr6-8018460-INS-51
chr6-8027619-INS-6267
chr6-8057307-INS-1250
chr6-8074945-INS-6685
chr6-8074957-INS-594
chr6-8077460-INS-54
chr6-8079237-INS-2205
chr6-8090877-INS-6260
chr6-8101382-INS-6338
chr6-8140238-INS-388
chr6-8165550-INS-130
chr6-8171723-INS-294
chr6-8184637-INS-880

chr6-8186028-INS-190
chr6-8191167-INS-112
chr6-8203097-INS-56
chr6-8207849-INS-70
chr6-8210885-INS-343
chr6-8225215-INS-181
chr6-8240091-INS-309
chr6-8269692-INS-806
chr6-8284537-INS-377
chr6-8355814-INS-3563
chr6-8409235-INS-64
chr6-8419189-INS-104
chr6-8420916-INS-57
chr6-8432313-INS-85
chr6-8459385-INS-131
chr6-8464018-INS-338
chr6-8483888-INS-70
chr6-8502311-INS-369
chr6-8526817-INS-156
chr6-8600056-INS-168
chr6-8631270-INS-153
chr6-8640292-INS-146
chr6-8681032-INS-78
chr6-8681882-INS-136
chr6-8684330-INS-216
chr6-8685990-INS-390
chr6-8733400-INS-98
chr6-8738337-INS-277
chr6-8746052-INS-148
chr6-8763603-INS-103
chr6-8766711-INS-492
chr6-8773461-INS-70
chr6-8774623-INS-146
chr6-8809509-INS-52
chr6-8818488-INS-7105
chr6-8819600-INS-967
chr6-8862945-INS-76
chr6-8885745-INS-76
chr6-8902250-INS-232
chr6-8936294-INS-91
chr6-8971074-INS-143
chr6-9017629-INS-120
chr6-9035675-INS-51
chr6-9053278-INS-138
chr6-9055138-INS-427
chr6-9069323-INS-64
chr6-9073902-INS-164
chr6-9096325-INS-705
chr6-9096507-INS-2688
chr6-9106438-INS-1702
chr6-9134498-INS-86
chr6-9136044-INS-6457
chr6-9145238-INS-212
chr6-9153487-INS-68
chr6-9157034-INS-65
chr6-9162093-INS-6193
chr6-9166966-INS-64
chr6-9173036-INS-119
chr6-9178157-INS-85
chr6-9200526-INS-1720
chr6-9225203-INS-384
chr6-9242201-INS-6212
chr6-9242403-INS-399
chr6-9316516-INS-52
chr6-9325176-INS-56
chr6-9357199-INS-356
chr6-9359971-INS-275
chr6-9367502-INS-110
chr6-9390052-INS-145
chr6-9414908-INS-6627
chr6-9439739-INS-370
chr6-9445791-INS-289
chr6-9487297-INS-53
chr6-9489581-INS-498
chr6-9492417-INS-106
chr6-9494314-INS-217
chr6-9497304-INS-98

chr6-9537232-INS-129
chr6-9592309-INS-52
chr6-9639820-INS-69
chr6-9657189-INS-651
chr6-9694344-INS-55
chr6-9702724-INS-52
chr6-9706035-INS-64
chr6-9719492-INS-79
chr6-9742773-INS-451
chr6-9761143-INS-171
chr6-9771958-INS-372
chr6-9788448-INS-286
chr6-9789351-INS-454
chr6-9812116-INS-146
chr6-9834392-INS-62
chr6-9841714-INS-191
chr6-9844479-INS-57
chr6-9870438-INS-423
chr6-9873362-INS-396
chr6-9887331-INS-116
chr6-9895721-INS-213
chr6-9896799-INS-68
chr6-9907457-INS-120
chr6-9918711-INS-212
chr6-9919212-INS-63
chr6-9953322-INS-228
chr6-9955027-INS-100
chr6-9959705-INS-405
chr6-9997948-INS-615
chr6-10044249-INS-7187
chr6-10105001-INS-156
chr6-10106360-INS-303
chr6-10125568-INS-51
chr6-10132682-INS-78
chr6-10183576-INS-4802
chr6-10198676-INS-6942
chr6-10214435-INS-214
chr6-10223741-INS-1992
chr6-10230009-INS-3856
chr6-10230835-INS-6634
chr6-10239236-INS-7123
chr6-10251829-INS-62
chr6-10252174-INS-66
chr6-10258677-INS-655
chr6-10278225-INS-113
chr6-10285730-INS-8234
chr6-10347488-INS-51
chr6-10350999-INS-413
chr6-10351433-INS-6664
chr6-10375909-INS-1882
chr6-10382670-INS-6096
chr6-10386496-INS-7071
chr6-10413953-INS-264
chr6-10418962-INS-203
chr6-10442848-INS-4722
chr6-10447124-INS-2688
chr6-10453730-INS-266
chr6-10471359-INS-107
chr6-10477812-INS-327
chr6-10480739-INS-190
chr6-10491328-INS-54
chr6-10495956-INS-4271
chr6-10510665-INS-1107
chr6-10530777-INS-401
chr6-10536279-INS-77
chr6-10600435-INS-357
chr6-10607191-INS-185
chr6-10610545-INS-57
chr6-10611193-INS-133
chr6-10613326-INS-787
chr6-10615484-INS-411
chr6-10641294-INS-93
chr6-10647517-INS-3227
chr6-10654923-INS-52
chr6-10667042-INS-168
chr6-10684903-INS-940
chr6-10698296-INS-53

chr6-10707376-INS-93
chr6-10724840-INS-58
chr6-10727612-INS-82
chr6-10749741-INS-627
chr6-10757047-INS-156
chr6-10757059-INS-674
chr6-10794846-INS-95
chr6-10812849-INS-4709
chr6-10830426-INS-875
chr6-10850322-INS-164
chr6-10871896-INS-106
chr6-10891378-INS-192
chr6-10894553-INS-575
chr6-10899527-INS-6641
chr6-10941097-INS-64
chr6-10981845-INS-406
chr6-10989453-INS-128
chr6-10991792-INS-189
chr6-11014296-INS-2908
chr6-11016261-INS-1467
chr6-11067503-INS-192
chr6-11074871-INS-7087
chr6-11081617-INS-6643
chr6-11113042-INS-7318
chr6-11136867-INS-331
chr6-11143970-INS-270
chr6-11146811-INS-81
chr6-11150976-INS-129
chr6-11155591-INS-246
chr6-11179633-INS-78
chr6-11191802-INS-57
chr6-11234157-INS-60
chr6-11238577-INS-183
chr6-11281788-INS-120
chr6-11296660-INS-919
chr6-11326157-INS-262
chr6-11344465-INS-66
chr6-11354805-INS-498
chr6-11358935-INS-54
chr6-11388107-INS-498
chr6-11397274-INS-6482
chr6-11397449-INS-137
chr6-11413529-INS-57
chr6-11420649-INS-942
chr6-11428618-INS-199
chr6-11458643-INS-70
chr6-11472359-INS-74
chr6-11492052-INS-252
chr6-11511868-INS-840
chr6-11520426-INS-54
chr6-11523192-INS-60
chr6-11538011-INS-193
chr6-11552532-INS-6724
chr6-11566988-INS-7131
chr6-11587252-INS-6401
chr6-11591745-INS-184
chr6-11599258-INS-63
chr6-11609894-INS-708
chr6-11611101-INS-51
chr6-11663207-INS-127
chr6-11666377-INS-510
chr6-11675676-INS-553
chr6-11689604-INS-156
chr6-11692648-INS-715
chr6-11730632-INS-3319
chr6-11736918-INS-52
chr6-11766513-INS-7082
chr6-11770193-INS-280
chr6-11773577-INS-6598
chr6-11774130-INS-60
chr6-11779124-INS-6656
chr6-11786929-INS-400
chr6-11790315-INS-748
chr6-11797547-INS-70
chr6-11799162-INS-148
chr6-11803799-INS-358
chr6-11845639-INS-214

chr6-11883906-INS-979
chr6-11899428-INS-55
chr6-11916377-INS-354
chr6-11922942-INS-144
chr6-11937228-INS-125
chr6-11948969-INS-431
chr6-11971947-INS-359
chr6-11979568-INS-449
chr6-12013728-INS-59
chr6-12059447-INS-291
chr6-12076815-INS-56
chr6-12082055-INS-872
chr6-12097735-INS-92
chr6-12102488-INS-185
chr6-12105921-INS-68
chr6-12107693-INS-432
chr6-12116850-INS-142
chr6-12125978-INS-431
chr6-12139593-INS-58
chr6-12152244-INS-596
chr6-12159695-INS-57
chr6-12162404-INS-144
chr6-12178126-INS-208
chr6-12182406-INS-66
chr6-12203307-INS-190
chr6-12204628-INS-339
chr6-12215467-INS-7241
chr6-12227915-INS-60
chr6-12244741-INS-4555
chr6-12261947-INS-172
chr6-12293520-INS-947
chr6-12307844-INS-362
chr6-12329561-INS-55
chr6-12334443-INS-59
chr6-12346419-INS-211
chr6-12359125-INS-128
chr6-12370878-INS-221
chr6-12382244-INS-201
chr6-12385629-INS-496
chr6-12402344-INS-182
chr6-12404282-INS-724
chr6-12425416-INS-809
chr6-12432129-INS-204
chr6-12433456-INS-160
chr6-12448424-INS-642
chr6-12457437-INS-459
chr6-12482603-INS-92
chr6-12484180-INS-114
chr6-12514120-INS-487
chr6-12531239-INS-528
chr6-12563651-INS-320
chr6-12569182-INS-62
chr6-12571839-INS-359
chr6-12579674-INS-385
chr6-12590328-INS-53
chr6-12599259-INS-150
chr6-12604609-INS-87
chr6-12611547-INS-83
chr6-12634392-INS-98
chr6-12637380-INS-6927
chr6-12639914-INS-101
chr6-12649462-INS-295
chr6-12650664-INS-175
chr6-12658464-INS-370
chr6-12685943-INS-72
chr6-12703847-INS-116
chr6-12714954-INS-158
chr6-12725836-INS-80
chr6-12752134-INS-62
chr6-12759274-INS-76
chr6-12800224-INS-155
chr6-12804965-INS-65
chr6-12817031-INS-499
chr6-12818794-INS-53
chr6-12846210-INS-307
chr6-12864443-INS-71
chr6-12880991-INS-7086

chr6-12900878-INS-123
chr6-12909366-INS-8393
chr6-12940332-INS-90
chr6-12944114-INS-64
chr6-12963316-INS-329
chr6-12964547-INS-92
chr6-13001944-INS-104
chr6-13012229-INS-538
chr6-13025015-INS-280
chr6-13062695-INS-6116
chr6-13153908-INS-420
chr6-13154273-INS-265
chr6-13219619-INS-315
chr6-13220250-INS-200
chr6-13223581-INS-368
chr6-13226611-INS-87
chr6-13232319-INS-219
chr6-13235380-INS-6945
chr6-13290281-INS-342
chr6-13293623-INS-62
chr6-13300826-INS-70
chr6-13303821-INS-928
chr6-13311661-INS-6672
chr6-13367590-INS-178
chr6-13380158-INS-54
chr6-13402751-INS-434
chr6-13404622-INS-340
chr6-13417620-INS-56
chr6-13475642-INS-569
chr6-13505064-INS-85
chr6-13507818-INS-6315
chr6-13517923-INS-336
chr6-13518845-INS-6129
chr6-13572181-INS-204
chr6-13575276-INS-55
chr6-13576287-INS-227
chr6-13589712-INS-312
chr6-13594663-INS-52
chr6-13600467-INS-1327
chr6-13605402-INS-66
chr6-13621538-INS-237
chr6-13623825-INS-75
chr6-13642453-INS-266
chr6-13646888-INS-165
chr6-13663174-INS-308
chr6-13672540-INS-358
chr6-13682172-INS-206
chr6-13695556-INS-154
chr6-13699442-INS-64
chr6-13699730-INS-778
chr6-13748606-INS-88
chr6-13755663-INS-68
chr6-13761014-INS-6803
chr6-13776785-INS-163
chr6-13802419-INS-675
chr6-13808815-INS-52
chr6-13821185-INS-336
chr6-13822646-INS-78
chr6-13846068-INS-76
chr6-13881106-INS-53
chr6-13898154-INS-84
chr6-13901520-INS-376
chr6-13969180-INS-64
chr6-13977547-INS-60
chr6-13982182-INS-91
chr6-13993984-INS-74
chr6-14006506-INS-177
chr6-14022012-INS-339
chr6-14066169-INS-310
chr6-14084384-INS-105
chr6-14091989-INS-65
chr6-14149720-INS-350
chr6-14151360-INS-399
chr6-14155590-INS-2660
chr6-14156516-INS-194
chr6-14173909-INS-273
chr6-14190142-INS-746

chr6-14215007-INS-4409
chr6-14219808-INS-79
chr6-14220731-INS-135
chr6-14230826-INS-169
chr6-14246095-INS-1181
chr6-14248573-INS-3865
chr6-14261153-INS-569
chr6-14282113-INS-996
chr6-14368995-INS-880
chr6-14372373-INS-500
chr6-14404443-INS-331
chr6-14437014-INS-396
chr6-14447794-INS-757
chr6-14458118-INS-7017
chr6-14474955-INS-131
chr6-14478509-INS-6415
chr6-14485907-INS-73
chr6-14498573-INS-64
chr6-14501746-INS-435
chr6-14511415-INS-1541
chr6-14517798-INS-54
chr6-14575222-INS-56
chr6-14605458-INS-77
chr6-14644035-INS-54
chr6-14683200-INS-80
chr6-14706209-INS-192
chr6-14723362-INS-59
chr6-14738300-INS-225
chr6-14745511-INS-148
chr6-14748158-INS-76
chr6-14752777-INS-62
chr6-14775402-INS-132
chr6-14791815-INS-984
chr6-14886765-INS-100
chr6-14896488-INS-153
chr6-14902810-INS-104
chr6-14913588-INS-108
chr6-14915497-INS-60
chr6-14939294-INS-6540
chr6-14948178-INS-85
chr6-14948766-INS-565
chr6-14973443-INS-320
chr6-15016449-INS-72
chr6-15021743-INS-924
chr6-15023422-INS-117
chr6-15086031-INS-309
chr6-15103578-INS-57
chr6-15146595-INS-158
chr6-15224030-INS-68
chr6-15260025-INS-502
chr6-15263441-INS-249
chr6-15265214-INS-86
chr6-15272705-INS-65
chr6-15294711-INS-3097
chr6-15297432-INS-108
chr6-15312862-INS-1499
chr6-15327136-INS-56
chr6-15359333-INS-61
chr6-15364796-INS-78
chr6-15370935-INS-161
chr6-15385368-INS-73
chr6-15392713-INS-240
chr6-15407893-INS-215
chr6-15496569-INS-4528
chr6-15498441-INS-1067
chr6-15502168-INS-59
chr6-15518561-INS-401
chr6-15560896-INS-108
chr6-15582872-INS-61
chr6-15596561-INS-231
chr6-15615160-INS-5453
chr6-15616741-INS-200
chr6-15618342-INS-130
chr6-15633238-INS-6915
chr6-15676471-INS-202
chr6-15712991-INS-7739
chr6-15755175-INS-154

chr6-15760329-INS-390
chr6-15788373-INS-70
chr6-15843640-INS-424
chr6-15877522-INS-6308
chr6-15879732-INS-3259
chr6-15892040-INS-292
chr6-15910971-INS-184
chr6-15924278-INS-61
chr6-15950118-INS-520
chr6-15956646-INS-134
chr6-15988583-INS-6903
chr6-15994099-INS-218
chr6-16014112-INS-404
chr6-16058347-INS-335
chr6-16083148-INS-115
chr6-16098301-INS-436
chr6-16108117-INS-171
chr6-16138240-INS-224
chr6-16148740-INS-66
chr6-16173079-INS-400
chr6-16179589-INS-384
chr6-16192739-INS-198
chr6-16202477-INS-602
chr6-16232563-INS-195
chr6-16236262-INS-73
chr6-16241660-INS-6476
chr6-16251961-INS-6438
chr6-16264200-INS-289
chr6-16265199-INS-508
chr6-16269602-INS-6217
chr6-16279184-INS-390
chr6-16279900-INS-1289
chr6-16282112-INS-7510
chr6-16285281-INS-74
chr6-16286900-INS-104
chr6-16291140-INS-3264
chr6-16296126-INS-156
chr6-16340770-INS-54
chr6-16351607-INS-666
chr6-16403201-INS-115
chr6-16426350-INS-211
chr6-16439791-INS-401
chr6-16447791-INS-202
chr6-16476225-INS-83
chr6-16512553-INS-60
chr6-16570429-INS-1932
chr6-16589121-INS-61
chr6-16594338-INS-54
chr6-16612097-INS-195
chr6-16629841-INS-55
chr6-16649198-INS-112
chr6-16658726-INS-102
chr6-16676594-INS-76
chr6-16686907-INS-54
chr6-16695394-INS-60
chr6-16722505-INS-838
chr6-16759851-INS-7792
chr6-16771175-INS-157
chr6-16775929-INS-553
chr6-16782680-INS-160
chr6-16801931-INS-564
chr6-16815007-INS-75
chr6-16830323-INS-156
chr6-16836592-INS-107
chr6-16841233-INS-65
chr6-16866932-INS-98
chr6-16892315-INS-294
chr6-16905254-INS-88
chr6-16920254-INS-1809
chr6-16928926-INS-286
chr6-16961533-INS-220
chr6-16976222-INS-154
chr6-16989393-INS-168
chr6-17023860-INS-345
chr6-17051914-INS-7070
chr6-17058340-INS-201
chr6-17067258-INS-6544

chr6-17093569-INS-74
chr6-17113104-INS-274
chr6-17125514-INS-359
chr6-17140668-INS-272
chr6-17145873-INS-65
chr6-17185959-INS-140
chr6-17192838-INS-53
chr6-17214480-INS-278
chr6-17233451-INS-167
chr6-17243826-INS-6714
chr6-17274006-INS-393
chr6-17301446-INS-66
chr6-17302969-INS-497
chr6-17311771-INS-333
chr6-17312725-INS-613
chr6-17318416-INS-56
chr6-17322890-INS-199
chr6-17326403-INS-159
chr6-17334808-INS-54
chr6-17355572-INS-4546
chr6-17376260-INS-85
chr6-17389894-INS-250
chr6-17391109-INS-198
chr6-17427877-INS-76
chr6-17487283-INS-7222
chr6-17493021-INS-1114
chr6-17493085-INS-1402
chr6-17493332-INS-357
chr6-17501186-INS-144
chr6-17502320-INS-328
chr6-17570333-INS-946
chr6-17578979-INS-80
chr6-17590228-INS-78
chr6-17591214-INS-54
chr6-17665665-INS-95
chr6-17685017-INS-156
chr6-17693514-INS-199
chr6-17696679-INS-140
chr6-17711484-INS-234
chr6-17727728-INS-409
chr6-17734901-INS-502
chr6-17738432-INS-56
chr6-17753606-INS-139
chr6-17766172-INS-207
chr6-17774783-INS-397
chr6-17794392-INS-140
chr6-17860000-INS-170
chr6-17878187-INS-166
chr6-17921093-INS-146
chr6-17935874-INS-297
chr6-17936893-INS-175
chr6-17977006-INS-100
chr6-18020574-INS-229
chr6-18046907-INS-203
chr6-18136706-INS-7014
chr6-18176328-INS-57
chr6-18178196-INS-111
chr6-18194847-INS-152
chr6-18221176-INS-67
chr6-18240182-INS-60
chr6-18244252-INS-79
chr6-18247661-INS-750
chr6-18262963-INS-89
chr6-18311646-INS-148
chr6-18312778-INS-64
chr6-18315776-INS-423
chr6-18374680-INS-84
chr6-18383231-INS-198
chr6-18397099-INS-704
chr6-18517540-INS-68
chr6-18536423-INS-51
chr6-18552995-INS-501
chr6-18558339-INS-53
chr6-18574788-INS-317
chr6-18604522-INS-80
chr6-18626151-INS-464
chr6-18627297-INS-108

chr6-18647493-INS-628
chr6-18652133-INS-748
chr6-18669560-INS-399
chr6-18734558-INS-99
chr6-18745664-INS-64
chr6-18761148-INS-84
chr6-18828661-INS-498
chr6-18831997-INS-492
chr6-18837041-INS-3381
chr6-18840184-INS-6392
chr6-18896676-INS-6451
chr6-18908941-INS-204
chr6-18936287-INS-64
chr6-18943964-INS-321
chr6-18944892-INS-71
chr6-18945361-INS-5283
chr6-18946737-INS-184
chr6-18949181-INS-52
chr6-18966693-INS-671
chr6-19009671-INS-633
chr6-19022696-INS-72
chr6-19026144-INS-219
chr6-19046243-INS-73
chr6-19057922-INS-52
chr6-19060440-INS-62
chr6-19062577-INS-70
chr6-19066240-INS-67
chr6-19076867-INS-6812
chr6-19163231-INS-454
chr6-19171183-INS-140
chr6-19202345-INS-60
chr6-19204466-INS-84
chr6-19238508-INS-62
chr6-19257035-INS-6845
chr6-19268017-INS-54
chr6-19277728-INS-84
chr6-19303222-INS-4362
chr6-19329970-INS-1265
chr6-19332502-INS-88
chr6-19409489-INS-6909
chr6-19420434-INS-399
chr6-19426063-INS-380
chr6-19427305-INS-60
chr6-19448303-INS-55
chr6-19479278-INS-180
chr6-19530462-INS-467
chr6-19532814-INS-56
chr6-19546357-INS-100
chr6-19547462-INS-206
chr6-19626192-INS-77
chr6-19628562-INS-154
chr6-19638553-INS-296
chr6-19698603-INS-416
chr6-19703983-INS-123
chr6-19767692-INS-354
chr6-19797362-INS-701
chr6-19813578-INS-205
chr6-19819044-INS-70
chr6-19826136-INS-817
chr6-19827036-INS-2958
chr6-19864101-INS-379
chr6-19901419-INS-84
chr6-19920298-INS-54
chr6-19960992-INS-713
chr6-20002455-INS-58
chr6-20005402-INS-6239
chr6-20010508-INS-494
chr6-20010909-INS-504
chr6-20023807-INS-424
chr6-20029391-INS-54
chr6-20031152-INS-183
chr6-20050681-INS-54
chr6-20069600-INS-392
chr6-20138303-INS-83
chr6-20188196-INS-201
chr6-20207471-INS-627
chr6-20242302-INS-72

chr6-20289226-INS-962
chr6-20292920-INS-242
chr6-20306050-INS-794
chr6-20357407-INS-82
chr6-20395248-INS-6382
chr6-20404703-INS-7291
chr6-20423773-INS-361
chr6-20478376-INS-54
chr6-20479949-INS-176
chr6-20503943-INS-52
chr6-20519357-INS-65
chr6-20560731-INS-6373
chr6-20603315-INS-82
chr6-20626604-INS-57
chr6-20632935-INS-310
chr6-20639348-INS-490
chr6-20639582-INS-6253
chr6-20651161-INS-4880
chr6-20654962-INS-6484
chr6-20661791-INS-5693
chr6-20663982-INS-64
chr6-20668426-INS-68
chr6-20686462-INS-436
chr6-20707180-INS-99
chr6-20739469-INS-216
chr6-20744818-INS-6245
chr6-20792783-INS-3450
chr6-20806351-INS-396
chr6-20826775-INS-52
chr6-20844539-INS-587
chr6-20901498-INS-119
chr6-20913719-INS-162
chr6-20951900-INS-143
chr6-20952054-INS-249
chr6-20959390-INS-130
chr6-20967353-INS-82
chr6-20983983-INS-54
chr6-21007065-INS-337
chr6-21009095-INS-244
chr6-21024079-INS-55
chr6-21028224-INS-52
chr6-21029159-INS-273
chr6-21054049-INS-642
chr6-21110982-INS-6600
chr6-21111941-INS-150
chr6-21132348-INS-6692
chr6-21167055-INS-863
chr6-21187616-INS-55
chr6-21224460-INS-52
chr6-21317646-INS-1891
chr6-21323204-INS-65
chr6-21373673-INS-344
chr6-21466912-INS-143
chr6-21539280-INS-60
chr6-21552650-INS-60
chr6-21562892-INS-147
chr6-21578049-INS-506
chr6-21584822-INS-6870
chr6-21595382-INS-3727
chr6-21602116-INS-515
chr6-21609221-INS-8575
chr6-21648338-INS-208
chr6-21662359-INS-60
chr6-21667904-INS-207
chr6-21685615-INS-78
chr6-21690382-INS-267
chr6-21710627-INS-58
chr6-21729850-INS-252
chr6-21778159-INS-73
chr6-21806066-INS-154
chr6-21821146-INS-52
chr6-21825027-INS-69
chr6-21867802-INS-232
chr6-21869287-INS-160
chr6-21876926-INS-134
chr6-21880377-INS-120
chr6-21888241-INS-102

chr6-21901297-INS-51
chr6-21955449-INS-190
chr6-21971435-INS-126
chr6-21983175-INS-78
chr6-21985431-INS-1150
chr6-21987403-INS-121
chr6-22001945-INS-1688
chr6-22027363-INS-96
chr6-22045844-INS-54
chr6-22046967-INS-69
chr6-22055953-INS-56
chr6-22059351-INS-101
chr6-22065723-INS-78
chr6-22084265-INS-274
chr6-22095786-INS-100
chr6-22098627-INS-122
chr6-22107033-INS-520
chr6-22110422-INS-344
chr6-22112765-INS-154
chr6-22118730-INS-66
chr6-22153598-INS-897
chr6-22172622-INS-231
chr6-22177608-INS-56
chr6-22185333-INS-136
chr6-22190842-INS-2755
chr6-22199038-INS-232
chr6-22216223-INS-702
chr6-22219170-INS-52
chr6-22219963-INS-92
chr6-22264482-INS-494
chr6-22273870-INS-69
chr6-22275831-INS-132
chr6-22285646-INS-52
chr6-22298732-INS-149
chr6-22305242-INS-190
chr6-22352645-INS-65
chr6-22359729-INS-58
chr6-22360491-INS-53
chr6-22360993-INS-89
chr6-22365925-INS-208
chr6-22377346-INS-69
chr6-22379705-INS-70
chr6-22390354-INS-369
chr6-22391232-INS-635
chr6-22397481-INS-212
chr6-22401644-INS-128
chr6-22402899-INS-60
chr6-22405033-INS-330
chr6-22421862-INS-181
chr6-22423365-INS-52
chr6-22430115-INS-1891
chr6-22441176-INS-251
chr6-22449628-INS-180
chr6-22460831-INS-84
chr6-22470067-INS-464
chr6-22490138-INS-68
chr6-22506307-INS-213
chr6-22516294-INS-123
chr6-22536033-INS-562
chr6-22537417-INS-385
chr6-22540737-INS-212
chr6-22547681-INS-78
chr6-22551644-INS-149
chr6-22558086-INS-475
chr6-22558854-INS-7158
chr6-22569025-INS-60
chr6-22575430-INS-839
chr6-22583083-INS-6290
chr6-22596625-INS-8341
chr6-22621585-INS-62
chr6-22628899-INS-55
chr6-22631954-INS-7993
chr6-22634493-INS-6893
chr6-22638478-INS-427
chr6-22644782-INS-4945
chr6-22651467-INS-3167
chr6-22655037-INS-89

chr6-22670368-INS-167
chr6-22680015-INS-198
chr6-22708633-INS-114
chr6-22719203-INS-6411
chr6-22728745-INS-4411
chr6-22729818-INS-6743
chr6-22835926-INS-254
chr6-22847161-INS-92
chr6-22853891-INS-180
chr6-22864283-INS-125
chr6-22867599-INS-65
chr6-22877857-INS-53
chr6-22902514-INS-67
chr6-22917881-INS-397
chr6-22947174-INS-288
chr6-22951511-INS-51
chr6-22969665-INS-591
chr6-22987975-INS-123
chr6-22995889-INS-73
chr6-23005161-INS-116
chr6-23027172-INS-813
chr6-23029546-INS-253
chr6-23032677-INS-54
chr6-23042103-INS-156
chr6-23045170-INS-84
chr6-23066514-INS-691
chr6-23067732-INS-178
chr6-23097157-INS-113
chr6-23121904-INS-162
chr6-23130334-INS-55
chr6-23141815-INS-60
chr6-23155601-INS-54
chr6-23171145-INS-400
chr6-23178413-INS-148
chr6-23180771-INS-77
chr6-23192478-INS-952
chr6-23206403-INS-383
chr6-23223559-INS-64
chr6-23227251-INS-792
chr6-23231689-INS-189
chr6-23246307-INS-122
chr6-23266514-INS-84
chr6-23284378-INS-134
chr6-23320406-INS-310
chr6-23333631-INS-78
chr6-23336404-INS-132
chr6-23349802-INS-176
chr6-23395997-INS-94
chr6-23399995-INS-330
chr6-23405223-INS-194
chr6-23430151-INS-455
chr6-23434881-INS-254
chr6-23458831-INS-108
chr6-23464980-INS-134
chr6-23482619-INS-279
chr6-23534182-INS-162
chr6-23570903-INS-117
chr6-23580372-INS-419
chr6-23582201-INS-70
chr6-23587913-INS-676
chr6-23589568-INS-322
chr6-23597989-INS-609
chr6-23615290-INS-348
chr6-23641812-INS-167
chr6-23642763-INS-64
chr6-23643481-INS-592
chr6-23646705-INS-237
chr6-23666944-INS-305
chr6-23679345-INS-9095
chr6-23699867-INS-80
chr6-23740447-INS-59
chr6-23759387-INS-637
chr6-23775412-INS-56
chr6-23788618-INS-605
chr6-23805922-INS-681
chr6-23819254-INS-70
chr6-23867846-INS-51

chr6-23887513-INS-586
chr6-23904414-INS-191
chr6-23908224-INS-5090
chr6-23918977-INS-132
chr6-23924367-INS-65
chr6-23980970-INS-6543
chr6-24020712-INS-82
chr6-24043936-INS-118
chr6-24048219-INS-54
chr6-24056407-INS-53
chr6-24066563-INS-5302
chr6-24098054-INS-122
chr6-24110516-INS-106
chr6-24128675-INS-153
chr6-24129622-INS-84
chr6-24189626-INS-235
chr6-24202264-INS-108
chr6-24203882-INS-805
chr6-24215151-INS-401
chr6-24218028-INS-124
chr6-24228681-INS-5464
chr6-24233132-INS-107
chr6-24254282-INS-120
chr6-24307831-INS-139
chr6-24323046-INS-110
chr6-24330517-INS-501
chr6-24344496-INS-152
chr6-24361703-INS-52
chr6-24367754-INS-62
chr6-24369596-INS-60
chr6-24381788-INS-351
chr6-24450188-INS-351
chr6-24454568-INS-71
chr6-24455594-INS-1890
chr6-24455804-INS-68
chr6-24472586-INS-108
chr6-24476290-INS-2136
chr6-24510194-INS-2534
chr6-24533183-INS-6497
chr6-24535017-INS-84
chr6-24541503-INS-54
chr6-24547519-INS-120
chr6-24563902-INS-386
chr6-24565443-INS-482
chr6-24566660-INS-145
chr6-24579694-INS-58
chr6-24583941-INS-2982
chr6-24630554-INS-52
chr6-24634740-INS-89
chr6-24683894-INS-1028
chr6-24684929-INS-660
chr6-24686972-INS-381
chr6-24732441-INS-64
chr6-24754581-INS-51
chr6-24760260-INS-983
chr6-24765420-INS-134
chr6-24773714-INS-657
chr6-24779905-INS-104
chr6-24799062-INS-214
chr6-24802340-INS-54
chr6-24822702-INS-463
chr6-24834476-INS-60
chr6-24853641-INS-214
chr6-24864240-INS-87
chr6-24889328-INS-99
chr6-24894793-INS-122
chr6-24911226-INS-82
chr6-24925263-INS-2890
chr6-24941158-INS-4542
chr6-24952618-INS-95
chr6-24976819-INS-53
chr6-24977648-INS-53
chr6-24980616-INS-77
chr6-24980945-INS-849
chr6-24982552-INS-194
chr6-24986138-INS-81
chr6-25001265-INS-265

chr6-25016017-INS-72
chr6-25027198-INS-171
chr6-25032532-INS-57
chr6-25039964-INS-105
chr6-25058845-INS-54
chr6-25066881-INS-112
chr6-25069404-INS-792
chr6-25168156-INS-245
chr6-25170137-INS-108
chr6-25206151-INS-694
chr6-25217618-INS-6254
chr6-25247134-INS-67
chr6-25257080-INS-722
chr6-25265079-INS-401
chr6-25276519-INS-386
chr6-25286213-INS-72
chr6-25307270-INS-64
chr6-25317500-INS-6424
chr6-25323359-INS-1417
chr6-25377478-INS-94
chr6-25379134-INS-107
chr6-25386390-INS-61
chr6-25387203-INS-92
chr6-25388550-INS-79
chr6-25401380-INS-102
chr6-25427929-INS-56
chr6-25444603-INS-282
chr6-25459047-INS-205
chr6-25475131-INS-219
chr6-25487960-INS-7207
chr6-25491255-INS-158
chr6-25498119-INS-448
chr6-25640268-INS-5204
chr6-25644532-INS-101
chr6-25648354-INS-951
chr6-25657442-INS-55
chr6-25714406-INS-5308
chr6-25718203-INS-1372
chr6-25730387-INS-6716
chr6-25739676-INS-204
chr6-25748115-INS-6647
chr6-25775113-INS-509
chr6-25779393-INS-171
chr6-25801330-INS-275
chr6-25826813-INS-937
chr6-25837173-INS-115
chr6-25862290-INS-588
chr6-25868568-INS-323
chr6-25917704-INS-159
chr6-25918446-INS-555
chr6-25926812-INS-62
chr6-25929092-INS-66
chr6-25931616-INS-119
chr6-25932350-INS-6724
chr6-25936581-INS-385
chr6-25950163-INS-307
chr6-25967709-INS-296
chr6-25970828-INS-64
chr6-25981376-INS-275
chr6-26022574-INS-96
chr6-26027734-INS-82
chr6-26033908-INS-80
chr6-26106956-INS-57
chr6-26182343-INS-138
chr6-26197253-INS-90
chr6-26207695-INS-352
chr6-26218849-INS-749
chr6-26233832-INS-203
chr6-26242138-INS-68
chr6-26243780-INS-164
chr6-26268019-INS-199
chr6-26288876-INS-131
chr6-26309166-INS-83
chr6-26324977-INS-638
chr6-26358557-INS-804
chr6-26364532-INS-335
chr6-26380428-INS-159

chr6-26381051-INS-7346
chr6-26388627-INS-2604
chr6-26391263-INS-154
chr6-26422360-INS-375
chr6-26425878-INS-139
chr6-26456790-INS-6899
chr6-26499077-INS-69
chr6-26515318-INS-65
chr6-26564211-INS-151
chr6-26571918-INS-90
chr6-26577475-INS-633
chr6-26616031-INS-549
chr6-26646157-INS-56
chr6-26648297-INS-52
chr6-26672812-INS-378
chr6-26690647-INS-185
chr6-26704711-INS-200
chr6-26721652-INS-116
chr6-26731736-INS-536
chr6-26741592-INS-5595
chr6-26742305-INS-416
chr6-26746845-INS-352
chr6-26767711-INS-66
chr6-26777557-INS-6253
chr6-26781316-INS-109
chr6-26795748-INS-175
chr6-26801156-INS-66
chr6-26828857-INS-55
chr6-26856420-INS-68
chr6-26858658-INS-387
chr6-26887518-INS-558
chr6-26899403-INS-77
chr6-26932978-INS-470
chr6-27026543-INS-446
chr6-27058572-INS-72
chr6-27137246-INS-142
chr6-27155863-INS-190
chr6-27159484-INS-680
chr6-27167415-INS-56
chr6-27191807-INS-62
chr6-27216773-INS-815
chr6-27266272-INS-498
chr6-27280238-INS-61
chr6-27312108-INS-229
chr6-27316792-INS-60
chr6-27325208-INS-170
chr6-27336626-INS-6320
chr6-27339732-INS-58
chr6-27367943-INS-1911
chr6-27372336-INS-159
chr6-27374430-INS-54
chr6-27375331-INS-367
chr6-27377730-INS-112
chr6-27380110-INS-56
chr6-27388990-INS-135
chr6-27397393-INS-78
chr6-27421648-INS-6427
chr6-27437465-INS-6209
chr6-27457250-INS-304
chr6-27483530-INS-90
chr6-27494378-INS-201
chr6-27497782-INS-7218
chr6-27500322-INS-774
chr6-27503670-INS-100
chr6-27513279-INS-358
chr6-27515069-INS-54
chr6-27517072-INS-67
chr6-27521095-INS-6934
chr6-27545252-INS-7017
chr6-27576972-INS-52
chr6-27582313-INS-6535
chr6-27584985-INS-52
chr6-27592606-INS-76
chr6-27610959-INS-268
chr6-27628977-INS-77
chr6-27630113-INS-809
chr6-27634356-INS-87

chr6-27665680-INS-166
chr6-27667727-INS-144
chr6-27672082-INS-54
chr6-27674261-INS-6633
chr6-27675676-INS-100
chr6-27683215-INS-199
chr6-27687678-INS-179
chr6-27719703-INS-210
chr6-27758908-INS-108
chr6-27771604-INS-108
chr6-27801965-INS-771
chr6-27816064-INS-177
chr6-27836074-INS-6722
chr6-27843929-INS-113
chr6-27847546-INS-497
chr6-27857594-INS-130
chr6-27867739-INS-340
chr6-27876632-INS-1076
chr6-27917913-INS-550
chr6-27918920-INS-61
chr6-27921254-INS-98
chr6-28000305-INS-88
chr6-28027757-INS-79
chr6-28036069-INS-492
chr6-28046202-INS-118
chr6-28047261-INS-130
chr6-28047431-INS-302
chr6-28055491-INS-168
chr6-28074436-INS-124
chr6-28090565-INS-55
chr6-28090794-INS-2013
chr6-28094006-INS-75
chr6-28101116-INS-100
chr6-28109381-INS-202
chr6-28126770-INS-1891
chr6-28127112-INS-589
chr6-28175944-INS-6328
chr6-28189959-INS-198
chr6-28192259-INS-53
chr6-28232949-INS-92
chr6-28274532-INS-546
chr6-28278967-INS-313
chr6-28295657-INS-140
chr6-28306260-INS-404
chr6-28325706-INS-194
chr6-28332034-INS-497
chr6-28340193-INS-684
chr6-28377416-INS-59
chr6-28384697-INS-213
chr6-28385141-INS-401
chr6-28391955-INS-365
chr6-28412083-INS-215
chr6-28428557-INS-70
chr6-28431444-INS-66
chr6-28458284-INS-51
chr6-28462314-INS-259
chr6-28462403-INS-871
chr6-28475894-INS-105
chr6-28486176-INS-70
chr6-28502381-INS-152
chr6-28531955-INS-64
chr6-28582068-INS-473
chr6-28590647-INS-58
chr6-28594887-INS-1619
chr6-28735820-INS-146
chr6-28738515-INS-203
chr6-28755074-INS-137
chr6-28772724-INS-162
chr6-28792705-INS-82
chr6-28825713-INS-208
chr6-28840160-INS-204
chr6-28841245-INS-218
chr6-28889623-INS-60
chr6-28891462-INS-58
chr6-28911180-INS-401
chr6-28934267-INS-146
chr6-28937873-INS-332

chr6-28946139-INS-500
chr6-28954050-INS-76
chr6-28985100-INS-119
chr6-29006875-INS-80
chr6-29028419-INS-482
chr6-29082636-INS-96
chr6-29091573-INS-81
chr6-29148727-INS-215
chr6-29151995-INS-679
chr6-29152637-INS-195
chr6-29153912-INS-203
chr6-29162802-INS-346
chr6-29175071-INS-55
chr6-29219548-INS-205
chr6-29221487-INS-61
chr6-29236162-INS-64
chr6-29240307-INS-242
chr6-29249243-INS-307
chr6-29250111-INS-84
chr6-29258135-INS-66
chr6-29271963-INS-159
chr6-29278321-INS-1869
chr6-29288481-INS-206
chr6-29289938-INS-54
chr6-29293043-INS-201
chr6-29300776-INS-7062
chr6-29302011-INS-6399
chr6-29317256-INS-409
chr6-29327058-INS-56
chr6-29383023-INS-76
chr6-29394343-INS-192
chr6-29408940-INS-96
chr6-29466570-INS-223
chr6-29476193-INS-573
chr6-29508512-INS-76
chr6-29508638-INS-1887
chr6-29525298-INS-249
chr6-29551401-INS-301
chr6-29597591-INS-414
chr6-29610219-INS-93
chr6-29635966-INS-176
chr6-29640231-INS-179
chr6-29649729-INS-58
chr6-29660445-INS-64
chr6-29671748-INS-232
chr6-29678394-INS-222
chr6-29682962-INS-81
chr6-29686566-INS-68
chr6-29698942-INS-109
chr6-29708962-INS-196
chr6-29724371-INS-265
chr6-29731221-INS-105
chr6-29731397-INS-410
chr6-29733277-INS-148
chr6-29747953-INS-3038
chr6-29747970-INS-963
chr6-29748655-INS-2523
chr6-29748977-INS-101
chr6-29774292-INS-202
chr6-29789923-INS-219
chr6-29816678-INS-208
chr6-29827038-INS-68
chr6-29830677-INS-187
chr6-29848449-INS-633
chr6-29900249-INS-205
chr6-29912777-INS-233
chr6-29916078-INS-173
chr6-29925310-INS-87
chr6-29968186-INS-207
chr6-29979957-INS-210
chr6-29991059-INS-64
chr6-29991593-INS-194
chr6-29992859-INS-59
chr6-29996890-INS-80
chr6-30006234-INS-154
chr6-30023783-INS-127
chr6-30024574-INS-262

chr6-30026243-INS-243
chr6-30044376-INS-124
chr6-30058016-INS-771
chr6-30060538-INS-6440
chr6-30063541-INS-68
chr6-30068772-INS-148
chr6-30099364-INS-630
chr6-30119184-INS-88
chr6-30126941-INS-172
chr6-30133762-INS-100
chr6-30185095-INS-262
chr6-30193180-INS-57
chr6-30215243-INS-130
chr6-30224759-INS-205
chr6-30230616-INS-134
chr6-30246246-INS-287
chr6-30255343-INS-450
chr6-30269855-INS-253
chr6-30273645-INS-131
chr6-30277661-INS-476
chr6-30287355-INS-60
chr6-30296469-INS-293
chr6-30318319-INS-56
chr6-30331878-INS-209
chr6-30335149-INS-382
chr6-30337174-INS-78
chr6-30339413-INS-339
chr6-30341748-INS-98
chr6-30354424-INS-60
chr6-30367777-INS-235
chr6-30371678-INS-214
chr6-30378612-INS-181
chr6-30394730-INS-124
chr6-30398374-INS-136
chr6-30398741-INS-6440
chr6-30413878-INS-515
chr6-30420633-INS-176
chr6-30429978-INS-90
chr6-30433685-INS-201
chr6-30443497-INS-225
chr6-30445620-INS-422
chr6-30459442-INS-235
chr6-30462860-INS-89
chr6-30467798-INS-7170
chr6-30471092-INS-112
chr6-30483484-INS-171
chr6-30505082-INS-206
chr6-30527326-INS-54
chr6-30566905-INS-73
chr6-30586801-INS-260
chr6-30588505-INS-150
chr6-30601024-INS-135
chr6-30614992-INS-52
chr6-30629921-INS-136
chr6-30707358-INS-75
chr6-30755058-INS-110
chr6-30762054-INS-83
chr6-30808572-INS-581
chr6-30845506-INS-775
chr6-30862898-INS-297
chr6-30874692-INS-131
chr6-30885112-INS-2873
chr6-30895872-INS-259
chr6-30921885-INS-125
chr6-30922915-INS-121
chr6-30953559-INS-79
chr6-30956680-INS-306
chr6-30969427-INS-112
chr6-30985809-INS-220
chr6-31012875-INS-545
chr6-31017801-INS-1648
chr6-31053073-INS-57
chr6-31056937-INS-83
chr6-31060013-INS-72
chr6-31100428-INS-52
chr6-31132190-INS-101
chr6-31145140-INS-267

chr6-31149685-INS-52
chr6-31151301-INS-58
chr6-31165643-INS-202
chr6-31205104-INS-54
chr6-31208982-INS-52
chr6-31211675-INS-125
chr6-31234107-INS-123
chr6-31306372-INS-401
chr6-31326320-INS-303
chr6-31331553-INS-761
chr6-31351443-INS-5327
chr6-31363677-INS-90
chr6-31485167-INS-568
chr6-31562699-INS-76
chr6-31572967-INS-188
chr6-31574287-INS-51
chr6-31575924-INS-231
chr6-31581618-INS-52
chr6-31587314-INS-151
chr6-31602480-INS-120
chr6-31608725-INS-52
chr6-31645758-INS-6213
chr6-31650499-INS-58
chr6-31654958-INS-84
chr6-31665039-INS-54
chr6-31703584-INS-133
chr6-31731801-INS-54
chr6-31757414-INS-86
chr6-31767059-INS-345
chr6-31790069-INS-579
chr6-31811141-INS-58
chr6-31833493-INS-621
chr6-31836308-INS-100
chr6-31841609-INS-377
chr6-31883035-INS-75
chr6-31893915-INS-52
chr6-31924307-INS-228
chr6-31937893-INS-184
chr6-31939416-INS-54
chr6-31946800-INS-132
chr6-31973863-INS-62
chr6-32004821-INS-1477
chr6-32057637-INS-833
chr6-32070925-INS-1060
chr6-32101851-INS-410
chr6-32155902-INS-58
chr6-32170416-INS-410
chr6-32179433-INS-216
chr6-32194539-INS-74
chr6-32210847-INS-185
chr6-32229574-INS-56
chr6-32279013-INS-100
chr6-32281668-INS-52
chr6-32373903-INS-76
chr6-32375590-INS-230
chr6-32379202-INS-215
chr6-32395946-INS-137
chr6-32446219-INS-57
chr6-32463216-INS-1605
chr6-32498826-INS-6870
chr6-32509150-INS-6553
chr6-32522795-INS-168
chr6-32601835-INS-733
chr6-32616725-INS-289
chr6-32694524-INS-267
chr6-32724293-INS-220
chr6-32733602-INS-359
chr6-32757392-INS-160
chr6-32778125-INS-421
chr6-32784295-INS-87
chr6-32841533-INS-312
chr6-32858555-INS-56
chr6-32948003-INS-623
chr6-32957441-INS-96
chr6-32970415-INS-200
chr6-32975466-INS-531
chr6-32982739-INS-6777

chr6-32982887-INS-7064
chr6-32992579-INS-353
chr6-33003351-INS-1890
chr6-33082556-INS-643
chr6-33084904-INS-400
chr6-33091845-INS-58
chr6-33094831-INS-66
chr6-33096470-INS-236
chr6-33103365-INS-318
chr6-33112362-INS-208
chr6-33168483-INS-92
chr6-33186620-INS-199
chr6-33205944-INS-90
chr6-33212670-INS-1732
chr6-33215652-INS-216
chr6-33224575-INS-223
chr6-33226727-INS-203
chr6-33245630-INS-179
chr6-33271088-INS-110
chr6-33295909-INS-69
chr6-33302794-INS-1438
chr6-33332204-INS-873
chr6-33352480-INS-195
chr6-33369192-INS-185
chr6-33377733-INS-325
chr6-33405589-INS-59
chr6-33411237-INS-316
chr6-33418691-INS-52
chr6-33419752-INS-60
chr6-33447280-INS-174
chr6-33456920-INS-78
chr6-33476543-INS-51
chr6-33486669-INS-74
chr6-33490849-INS-244
chr6-33519664-INS-92
chr6-33535378-INS-199
chr6-33595726-INS-1227
chr6-33607595-INS-2604
chr6-33615335-INS-52
chr6-33653485-INS-54
chr6-33673640-INS-55
chr6-33687824-INS-72
chr6-33697063-INS-2938
chr6-33700653-INS-2464
chr6-33708399-INS-117
chr6-33728362-INS-562
chr6-33761361-INS-133
chr6-33797315-INS-120
chr6-33803074-INS-552
chr6-33813639-INS-172
chr6-33820044-INS-82
chr6-33827670-INS-82
chr6-33845200-INS-316
chr6-33889402-INS-2204
chr6-33895586-INS-217
chr6-33898129-INS-1447
chr6-33914730-INS-645
chr6-33943982-INS-138
chr6-33977934-INS-428
chr6-33995063-INS-103
chr6-34003544-INS-110
chr6-34020761-INS-60
chr6-34078018-INS-77
chr6-34097375-INS-1492
chr6-34111478-INS-53
chr6-34148845-INS-5543
chr6-34151162-INS-204
chr6-34179664-INS-72
chr6-34181284-INS-73
chr6-34181372-INS-532
chr6-34203835-INS-332
chr6-34204038-INS-7077
chr6-34218621-INS-200
chr6-34237107-INS-235
chr6-34239946-INS-81
chr6-34240394-INS-841
chr6-34250607-INS-547

chr6-34268689-INS-52
chr6-34276179-INS-96
chr6-34278201-INS-503
chr6-34296442-INS-168
chr6-34300513-INS-97
chr6-34316274-INS-187
chr6-34335330-INS-72
chr6-34337088-INS-168
chr6-34372782-INS-104
chr6-34374115-INS-105
chr6-34393835-INS-7171
chr6-34400279-INS-70
chr6-34412087-INS-400
chr6-34434391-INS-5348
chr6-34444541-INS-51
chr6-34461617-INS-56
chr6-34476237-INS-241
chr6-34485771-INS-55
chr6-34501923-INS-66
chr6-34505748-INS-94
chr6-34507620-INS-52
chr6-34521861-INS-198
chr6-34524131-INS-98
chr6-34536108-INS-85
chr6-34542540-INS-203
chr6-34555298-INS-364
chr6-34585367-INS-55
chr6-34607486-INS-104
chr6-34625894-INS-89
chr6-34645606-INS-64
chr6-34646085-INS-522
chr6-34657060-INS-206
chr6-34667111-INS-136
chr6-34678630-INS-99
chr6-34706051-INS-71
chr6-34715613-INS-92
chr6-34741048-INS-140
chr6-34758787-INS-204
chr6-34766359-INS-62
chr6-34802557-INS-167
chr6-34817740-INS-66
chr6-34821273-INS-64
chr6-34825915-INS-712
chr6-34826910-INS-559
chr6-34832978-INS-87
chr6-34838248-INS-520
chr6-34841187-INS-98
chr6-34847796-INS-498
chr6-34849404-INS-492
chr6-34858414-INS-64
chr6-34860977-INS-169
chr6-34882380-INS-117
chr6-34888269-INS-66
chr6-34890478-INS-121
chr6-34897093-INS-218
chr6-34912850-INS-288
chr6-34918233-INS-744
chr6-34923919-INS-460
chr6-34928947-INS-87
chr6-34935347-INS-128
chr6-34944377-INS-96
chr6-34947139-INS-80
chr6-34961299-INS-108
chr6-34962301-INS-78
chr6-34988761-INS-132
chr6-34990406-INS-74
chr6-35031710-INS-196
chr6-35052445-INS-166
chr6-35069337-INS-84
chr6-35076441-INS-64
chr6-35116114-INS-54
chr6-35117569-INS-78
chr6-35125945-INS-211
chr6-35131937-INS-179
chr6-35139231-INS-194
chr6-35150667-INS-87
chr6-35179302-INS-175

chr6-35182828-INS-236
chr6-35227705-INS-604
chr6-35232373-INS-114
chr6-35238807-INS-835
chr6-35238889-INS-215
chr6-35268717-INS-60
chr6-35285851-INS-235
chr6-35326629-INS-258
chr6-35332581-INS-524
chr6-35344614-INS-72
chr6-35352701-INS-54
chr6-35365078-INS-255
chr6-35374117-INS-90
chr6-35377357-INS-2438
chr6-35389645-INS-80
chr6-35391501-INS-2596
chr6-35406334-INS-152
chr6-35417885-INS-90
chr6-35438421-INS-211
chr6-35454606-INS-68
chr6-35461110-INS-238
chr6-35491316-INS-52
chr6-35507716-INS-174
chr6-35539707-INS-75
chr6-35541139-INS-172
chr6-35545514-INS-72
chr6-35568363-INS-84
chr6-35576220-INS-53
chr6-35583763-INS-58
chr6-35585938-INS-54
chr6-35601797-INS-513
chr6-35606081-INS-56
chr6-35645331-INS-433
chr6-35659386-INS-84
chr6-35667614-INS-52
chr6-35694133-INS-551
chr6-35715516-INS-179
chr6-35720390-INS-56
chr6-35743621-INS-308
chr6-35778283-INS-193
chr6-35788575-INS-60
chr6-35792264-INS-102
chr6-35808460-INS-55
chr6-35818798-INS-56
chr6-35824496-INS-164
chr6-35831591-INS-126
chr6-35847236-INS-129
chr6-35854124-INS-220
chr6-35855249-INS-96
chr6-35863630-INS-709
chr6-35882926-INS-86
chr6-35923261-INS-84
chr6-35943016-INS-80
chr6-35955537-INS-52
chr6-35974655-INS-155
chr6-35990031-INS-148
chr6-36001236-INS-96
chr6-36002116-INS-170
chr6-36004283-INS-291
chr6-36016387-INS-114
chr6-36037397-INS-340
chr6-36051874-INS-7032
chr6-36078290-INS-74
chr6-36079595-INS-74
chr6-36105144-INS-250
chr6-36105584-INS-385
chr6-36116537-INS-480
chr6-36138078-INS-428
chr6-36138825-INS-68
chr6-36143453-INS-89
chr6-36190233-INS-56
chr6-36200867-INS-98
chr6-36208713-INS-159
chr6-36216764-INS-52
chr6-36220804-INS-741
chr6-36231890-INS-51
chr6-36252711-INS-401

chr6-36284722-INS-624
chr6-36289187-INS-170
chr6-36319453-INS-52
chr6-36323918-INS-231
chr6-36340924-INS-1269
chr6-36395024-INS-779
chr6-36404016-INS-68
chr6-36406229-INS-54
chr6-36424357-INS-176
chr6-36431154-INS-189
chr6-36456121-INS-522
chr6-36502833-INS-66
chr6-36523210-INS-52
chr6-36545464-INS-84
chr6-36556229-INS-199
chr6-36566058-INS-74
chr6-36580269-INS-53
chr6-36596178-INS-68
chr6-36610843-INS-543
chr6-36615167-INS-433
chr6-36656611-INS-132
chr6-36717738-INS-93
chr6-36722429-INS-56
chr6-36731032-INS-56
chr6-36734079-INS-200
chr6-36737284-INS-60
chr6-36742954-INS-546
chr6-36745875-INS-398
chr6-36769083-INS-224
chr6-36812148-INS-66
chr6-36842484-INS-86
chr6-36844001-INS-70
chr6-36866130-INS-60
chr6-36876152-INS-310
chr6-36878155-INS-6323
chr6-36884854-INS-125
chr6-36909972-INS-86
chr6-36950084-INS-613
chr6-36974382-INS-68
chr6-36979656-INS-206
chr6-36986639-INS-69
chr6-36991658-INS-440
chr6-37010788-INS-535
chr6-37022549-INS-176
chr6-37052222-INS-1177
chr6-37059923-INS-115
chr6-37080751-INS-118
chr6-37085627-INS-113
chr6-37092276-INS-89
chr6-37105703-INS-403
chr6-37159869-INS-488
chr6-37182008-INS-478
chr6-37190526-INS-78
chr6-37197020-INS-501
chr6-37201320-INS-117
chr6-37247745-INS-56
chr6-37280024-INS-105
chr6-37295552-INS-57
chr6-37330420-INS-79
chr6-37332998-INS-53
chr6-37342190-INS-146
chr6-37343428-INS-7127
chr6-37356195-INS-388
chr6-37363618-INS-199
chr6-37389251-INS-296
chr6-37403429-INS-235
chr6-37410726-INS-328
chr6-37438520-INS-265
chr6-37512583-INS-200
chr6-37513807-INS-72
chr6-37521302-INS-107
chr6-37531961-INS-184
chr6-37554217-INS-220
chr6-37557315-INS-391
chr6-37565059-INS-56
chr6-37572316-INS-192
chr6-37582331-INS-228

chr6-37608298-INS-119
chr6-37624347-INS-79
chr6-37629518-INS-482
chr6-37668021-INS-179
chr6-37677296-INS-130
chr6-37681894-INS-138
chr6-37688256-INS-60
chr6-37690541-INS-503
chr6-37707069-INS-435
chr6-37708716-INS-83
chr6-37715413-INS-513
chr6-37721154-INS-186
chr6-37732541-INS-400
chr6-37742037-INS-335
chr6-37752612-INS-140
chr6-37753573-INS-5556
chr6-37795021-INS-260
chr6-37810460-INS-443
chr6-37835430-INS-61
chr6-37836588-INS-160
chr6-37844724-INS-567
chr6-37851609-INS-122
chr6-37870023-INS-142
chr6-37883516-INS-75
chr6-37898039-INS-105
chr6-37921007-INS-116
chr6-37931086-INS-77
chr6-37941352-INS-66
chr6-37975517-INS-62
chr6-37979424-INS-176
chr6-37984474-INS-52
chr6-37985021-INS-75
chr6-37991064-INS-143
chr6-38075767-INS-94
chr6-38097359-INS-209
chr6-38106184-INS-89
chr6-38110336-INS-106
chr6-38130687-INS-217
chr6-38133079-INS-151
chr6-38137807-INS-783
chr6-38157352-INS-56
chr6-38177263-INS-109
chr6-38189557-INS-71
chr6-38278412-INS-114
chr6-38294421-INS-201
chr6-38297292-INS-68
chr6-38300100-INS-170
chr6-38302001-INS-510
chr6-38320813-INS-165
chr6-38330021-INS-141
chr6-38334272-INS-118
chr6-38337832-INS-52
chr6-38342277-INS-191
chr6-38344798-INS-466
chr6-38386066-INS-279
chr6-38387597-INS-65
chr6-38420829-INS-209
chr6-38423547-INS-63
chr6-38424709-INS-56
chr6-38427490-INS-161
chr6-38436045-INS-748
chr6-38456730-INS-182
chr6-38460492-INS-107
chr6-38474948-INS-68
chr6-38476282-INS-53
chr6-38493848-INS-56
chr6-38517948-INS-354
chr6-38518759-INS-180
chr6-38523189-INS-96
chr6-38552194-INS-145
chr6-38591057-INS-90
chr6-38599521-INS-206
chr6-38606514-INS-54
chr6-38616770-INS-410
chr6-38630824-INS-515
chr6-38632409-INS-405
chr6-38637174-INS-176

chr6-38659791-INS-310
chr6-38662894-INS-211
chr6-38711382-INS-206
chr6-38721103-INS-55
chr6-38751255-INS-72
chr6-38787699-INS-66
chr6-38800040-INS-52
chr6-38804846-INS-206
chr6-38808874-INS-325
chr6-38871938-INS-414
chr6-38888693-INS-232
chr6-38900199-INS-925
chr6-38916759-INS-86
chr6-38937391-INS-68
chr6-38961074-INS-107
chr6-38968678-INS-62
chr6-38971533-INS-144
chr6-39009739-INS-613
chr6-39036040-INS-156
chr6-39039530-INS-73
chr6-39048116-INS-231
chr6-39056138-INS-212
chr6-39057129-INS-140
chr6-39073353-INS-113
chr6-39084379-INS-68
chr6-39113576-INS-238
chr6-39146862-INS-200
chr6-39157828-INS-206
chr6-39164143-INS-203
chr6-39168735-INS-133
chr6-39176927-INS-174
chr6-39177417-INS-202
chr6-39179149-INS-197
chr6-39197664-INS-202
chr6-39206806-INS-207
chr6-39208014-INS-83
chr6-39212330-INS-297
chr6-39216246-INS-199
chr6-39225648-INS-119
chr6-39235409-INS-129
chr6-39242137-INS-81
chr6-39243217-INS-52
chr6-39252751-INS-216
chr6-39277302-INS-114
chr6-39292692-INS-222
chr6-39304417-INS-110
chr6-39328198-INS-241
chr6-39331317-INS-188
chr6-39336325-INS-65
chr6-39340453-INS-100
chr6-39345160-INS-336
chr6-39345662-INS-80
chr6-39358696-INS-57
chr6-39362209-INS-73
chr6-39363306-INS-65
chr6-39367706-INS-184
chr6-39370171-INS-94
chr6-39374676-INS-214
chr6-39385510-INS-94
chr6-39386805-INS-210
chr6-39393612-INS-344
chr6-39405326-INS-82
chr6-39407866-INS-179
chr6-39413702-INS-1860
chr6-39421249-INS-60
chr6-39438629-INS-343
chr6-39442726-INS-177
chr6-39444784-INS-99
chr6-39466504-INS-172
chr6-39476443-INS-180
chr6-39481094-INS-62
chr6-39487789-INS-554
chr6-39490409-INS-187
chr6-39503779-INS-206
chr6-39508337-INS-209
chr6-39528315-INS-78
chr6-39530199-INS-400

chr6-39540708-INS-215
chr6-39568656-INS-218
chr6-39578527-INS-53
chr6-39609121-INS-216
chr6-39618821-INS-51
chr6-39630242-INS-245
chr6-39644717-INS-387
chr6-39653149-INS-136
chr6-39674085-INS-80
chr6-39706202-INS-7141
chr6-39708090-INS-374
chr6-39708211-INS-620
chr6-39711459-INS-278
chr6-39735375-INS-67
chr6-39745403-INS-194
chr6-39747500-INS-187
chr6-39760292-INS-90
chr6-39762617-INS-195
chr6-39765852-INS-123
chr6-39767215-INS-177
chr6-39774720-INS-58
chr6-39804359-INS-366
chr6-39833442-INS-67
chr6-39834367-INS-839
chr6-39886596-INS-6522
chr6-39894984-INS-405
chr6-39941581-INS-58
chr6-39946180-INS-58
chr6-39957814-INS-1932
chr6-39960229-INS-1344
chr6-39960361-INS-242
chr6-39975408-INS-113
chr6-39979662-INS-91
chr6-39985690-INS-122
chr6-39999230-INS-53
chr6-40034835-INS-1823
chr6-40036224-INS-399
chr6-40046251-INS-128
chr6-40058493-INS-328
chr6-40099463-INS-60
chr6-40104345-INS-52
chr6-40105602-INS-56
chr6-40127669-INS-86
chr6-40153770-INS-61
chr6-40197164-INS-146
chr6-40201428-INS-6245
chr6-40219382-INS-1794
chr6-40220628-INS-289
chr6-40328193-INS-72
chr6-40351414-INS-192
chr6-40425601-INS-206
chr6-40444615-INS-70
chr6-40447479-INS-161
chr6-40465188-INS-66
chr6-40479952-INS-354
chr6-40502883-INS-199
chr6-40505236-INS-6440
chr6-40508736-INS-56
chr6-40510580-INS-753
chr6-40519625-INS-59
chr6-40531423-INS-134
chr6-40559519-INS-800
chr6-40586411-INS-63
chr6-40593488-INS-57
chr6-40594984-INS-61
chr6-40625187-INS-66
chr6-40667625-INS-184
chr6-40690111-INS-556
chr6-40717985-INS-56
chr6-40729669-INS-101
chr6-40731280-INS-145
chr6-40738739-INS-181
chr6-40743364-INS-52
chr6-40744842-INS-84
chr6-40754245-INS-126
chr6-40768227-INS-56
chr6-40770648-INS-89

chr6-40778342-INS-63
chr6-40786505-INS-62
chr6-40799546-INS-161
chr6-40812764-INS-58
chr6-40828843-INS-186
chr6-40829807-INS-132
chr6-40841217-INS-385
chr6-40844545-INS-120
chr6-40844849-INS-357
chr6-40856815-INS-64
chr6-40857602-INS-441
chr6-40870922-INS-202
chr6-40879757-INS-288
chr6-40881483-INS-451
chr6-40881815-INS-6613
chr6-40894134-INS-7003
chr6-40944566-INS-68
chr6-40945955-INS-2830
chr6-40973556-INS-70
chr6-40981637-INS-347
chr6-41022831-INS-104
chr6-41032224-INS-64
chr6-41035020-INS-182
chr6-41043389-INS-271
chr6-41071112-INS-91
chr6-41083857-INS-89
chr6-41093795-INS-98
chr6-41111151-INS-147
chr6-41123452-INS-110
chr6-41140988-INS-2623
chr6-41142558-INS-1773
chr6-41143973-INS-139
chr6-41148138-INS-276
chr6-41152557-INS-569
chr6-41153857-INS-71
chr6-41154874-INS-89
chr6-41168609-INS-172
chr6-41172720-INS-194
chr6-41199040-INS-363
chr6-41209002-INS-1616
chr6-41216593-INS-7964
chr6-41216951-INS-292
chr6-41219615-INS-70
chr6-41225312-INS-6928
chr6-41244107-INS-330
chr6-41290543-INS-84
chr6-41316423-INS-91
chr6-41326580-INS-55
chr6-41329961-INS-6452
chr6-41336825-INS-56
chr6-41360100-INS-439
chr6-41385564-INS-64
chr6-41450103-INS-279
chr6-41463172-INS-963
chr6-41480858-INS-679
chr6-41483154-INS-232
chr6-41500620-INS-85
chr6-41504198-INS-205
chr6-41513720-INS-226
chr6-41617385-INS-54
chr6-41640759-INS-132
chr6-41652137-INS-86
chr6-41668640-INS-374
chr6-41669569-INS-128
chr6-41687943-INS-99
chr6-41690676-INS-354
chr6-41725823-INS-125
chr6-41736534-INS-198
chr6-41741670-INS-492
chr6-41751424-INS-351
chr6-41758274-INS-126
chr6-41764766-INS-518
chr6-41769204-INS-375
chr6-41772967-INS-78
chr6-41773400-INS-686
chr6-41793844-INS-54
chr6-41804085-INS-74

chr6-41876502-INS-330
chr6-41895448-INS-128
chr6-41900525-INS-57
chr6-41929734-INS-734
chr6-41937569-INS-469
chr6-41941374-INS-874
chr6-42001552-INS-61
chr6-42003482-INS-88
chr6-42006624-INS-6442
chr6-42008970-INS-90
chr6-42021175-INS-6882
chr6-42079012-INS-6413
chr6-42092856-INS-63
chr6-42100049-INS-207
chr6-42114475-INS-86
chr6-42158063-INS-76
chr6-42168107-INS-7203
chr6-42190393-INS-84
chr6-42209797-INS-224
chr6-42211950-INS-236
chr6-42224246-INS-87
chr6-42368314-INS-538
chr6-42378624-INS-156
chr6-42383599-INS-151
chr6-42397488-INS-58
chr6-42399029-INS-52
chr6-42411698-INS-8256
chr6-42414096-INS-57
chr6-42422486-INS-7176
chr6-42424641-INS-138
chr6-42451362-INS-52
chr6-42457017-INS-6690
chr6-42478831-INS-5968
chr6-42498837-INS-103
chr6-42501303-INS-6318
chr6-42510660-INS-62
chr6-42511210-INS-7176
chr6-42528164-INS-997
chr6-42535619-INS-84
chr6-42556253-INS-359
chr6-42572411-INS-60
chr6-42598013-INS-8115
chr6-42604549-INS-142
chr6-42617917-INS-90
chr6-42633368-INS-148
chr6-42636970-INS-514
chr6-42658908-INS-274
chr6-42694735-INS-212
chr6-42732492-INS-64
chr6-42734148-INS-6805
chr6-42734929-INS-6924
chr6-42738882-INS-592
chr6-42738957-INS-6699
chr6-42757624-INS-4426
chr6-42776462-INS-112
chr6-42781613-INS-134
chr6-42787596-INS-63
chr6-42797692-INS-97
chr6-42826286-INS-90
chr6-42830264-INS-1893
chr6-42843488-INS-69
chr6-42880568-INS-327
chr6-42890356-INS-60
chr6-42906828-INS-261
chr6-42910572-INS-135
chr6-42972145-INS-118
chr6-42975922-INS-257
chr6-42976507-INS-1312
chr6-42977660-INS-291
chr6-43005569-INS-464
chr6-43035543-INS-220
chr6-43040967-INS-71
chr6-43051693-INS-54
chr6-43065604-INS-62
chr6-43075191-INS-549
chr6-43079501-INS-117
chr6-43112400-INS-98

chr6-43125095-INS-79
chr6-43134899-INS-326
chr6-43142297-INS-58
chr6-43148255-INS-329
chr6-43188132-INS-6336
chr6-43213539-INS-362
chr6-43245169-INS-112
chr6-43292060-INS-639
chr6-43293738-INS-60
chr6-43294476-INS-6245
chr6-43310982-INS-546
chr6-43337645-INS-6879
chr6-43340813-INS-71
chr6-43342730-INS-2843
chr6-43342852-INS-198
chr6-43351781-INS-6356
chr6-43409582-INS-58
chr6-43420850-INS-6260
chr6-43443907-INS-2557
chr6-43450178-INS-8348
chr6-43463177-INS-1773
chr6-43473653-INS-3684
chr6-43485896-INS-199
chr6-43509842-INS-578
chr6-43534702-INS-61
chr6-43569054-INS-7179
chr6-43616815-INS-92
chr6-43624511-INS-6460
chr6-43626185-INS-92
chr6-43646071-INS-866
chr6-43650319-INS-1117
chr6-43662263-INS-116
chr6-43679606-INS-52
chr6-43682228-INS-59
chr6-43689350-INS-2467
chr6-43704397-INS-53
chr6-43705852-INS-140
chr6-43729102-INS-172
chr6-43742095-INS-6625
chr6-43743274-INS-191
chr6-43747129-INS-6269
chr6-43756815-INS-299
chr6-43773132-INS-86
chr6-43802705-INS-6615
chr6-43804936-INS-341
chr6-43845541-INS-228
chr6-43898852-INS-377
chr6-43944908-INS-54
chr6-43989043-INS-218
chr6-43996952-INS-100
chr6-44010920-INS-6967
chr6-44017541-INS-122
chr6-44048730-INS-55
chr6-44121865-INS-174
chr6-44158448-INS-52
chr6-44175313-INS-92
chr6-44191789-INS-74
chr6-44291157-INS-664
chr6-44299347-INS-59
chr6-44306313-INS-564
chr6-44339717-INS-64
chr6-44340972-INS-60
chr6-44342644-INS-220
chr6-44347025-INS-172
chr6-44353807-INS-6396
chr6-44362809-INS-169
chr6-44366974-INS-164
chr6-44414521-INS-154
chr6-44454659-INS-106
chr6-44456472-INS-89
chr6-44479703-INS-88
chr6-44483643-INS-233
chr6-44495273-INS-272
chr6-44519051-INS-51
chr6-44635746-INS-69
chr6-44644745-INS-55
chr6-44658554-INS-321

chr6-44664306-INS-54
chr6-44665679-INS-528
chr6-44666915-INS-104
chr6-44676716-INS-177
chr6-44679915-INS-5960
chr6-44680260-INS-126
chr6-44692421-INS-82
chr6-44720032-INS-3128
chr6-44724865-INS-84
chr6-44733963-INS-51
chr6-44789829-INS-346
chr6-44793642-INS-160
chr6-44819883-INS-56
chr6-44837543-INS-72
chr6-44851534-INS-926
chr6-44852745-INS-201
chr6-44874482-INS-185
chr6-44884761-INS-70
chr6-44901589-INS-52
chr6-44912300-INS-52
chr6-44921239-INS-228
chr6-44924214-INS-7112
chr6-44926767-INS-376
chr6-44944529-INS-976
chr6-44970961-INS-77
chr6-44994980-INS-126
chr6-45018304-INS-90
chr6-45028670-INS-446
chr6-45041575-INS-68
chr6-45045704-INS-64
chr6-45050194-INS-68
chr6-45118584-INS-70
chr6-45124492-INS-150
chr6-45156981-INS-81
chr6-45167109-INS-95
chr6-45202084-INS-343
chr6-45205233-INS-60
chr6-45209297-INS-59
chr6-45233341-INS-866
chr6-45244451-INS-77
chr6-45256024-INS-54
chr6-45281283-INS-262
chr6-45285491-INS-120
chr6-45300561-INS-295
chr6-45337530-INS-52
chr6-45342777-INS-76
chr6-45360573-INS-762
chr6-45364065-INS-127
chr6-45378795-INS-117
chr6-45392594-INS-304
chr6-45417528-INS-606
chr6-45481626-INS-88
chr6-45485347-INS-644
chr6-45499559-INS-105
chr6-45515876-INS-736
chr6-45517620-INS-58
chr6-45535467-INS-54
chr6-45541580-INS-208
chr6-45581618-INS-482
chr6-45589987-INS-551
chr6-45644483-INS-123
chr6-45658482-INS-60
chr6-45664504-INS-587
chr6-45665889-INS-497
chr6-45673207-INS-124
chr6-45676902-INS-87
chr6-45677804-INS-199
chr6-45720109-INS-51
chr6-45761967-INS-275
chr6-45765627-INS-54
chr6-45771501-INS-537
chr6-45776788-INS-52
chr6-45779864-INS-146
chr6-45780618-INS-74
chr6-45785885-INS-88
chr6-45792669-INS-148
chr6-45857372-INS-102

chr6-45865456-INS-638
chr6-45866885-INS-80
chr6-45920194-INS-92
chr6-45927675-INS-6579
chr6-45992553-INS-52
chr6-46004287-INS-247
chr6-46024799-INS-226
chr6-46026040-INS-68
chr6-46072713-INS-908
chr6-46096383-INS-192
chr6-46117201-INS-8065
chr6-46121959-INS-52
chr6-46125695-INS-403
chr6-46140548-INS-120
chr6-46183443-INS-64
chr6-46197692-INS-420
chr6-46220860-INS-269
chr6-46263312-INS-202
chr6-46264456-INS-214
chr6-46297275-INS-110
chr6-46314357-INS-6487
chr6-46321786-INS-499
chr6-46332782-INS-6831
chr6-46399697-INS-288
chr6-46512002-INS-62
chr6-46519925-INS-363
chr6-46546616-INS-122
chr6-46550748-INS-6162
chr6-46594224-INS-581
chr6-46602279-INS-182
chr6-46642077-INS-6487
chr6-46664180-INS-6247
chr6-46720526-INS-72
chr6-46727433-INS-2802
chr6-46759158-INS-561
chr6-46760477-INS-52
chr6-46770766-INS-388
chr6-46773102-INS-370
chr6-46775921-INS-972
chr6-46788223-INS-203
chr6-46794613-INS-4524
chr6-46803657-INS-118
chr6-46817284-INS-127
chr6-46819707-INS-100
chr6-46825097-INS-54
chr6-46842966-INS-201
chr6-46880390-INS-283
chr6-46895314-INS-3981
chr6-46896688-INS-183
chr6-46902586-INS-86
chr6-46906014-INS-326
chr6-46924265-INS-99
chr6-46927338-INS-163
chr6-46929796-INS-6442
chr6-46934817-INS-215
chr6-46959026-INS-6807
chr6-46989752-INS-690
chr6-46999679-INS-8408
chr6-47004739-INS-62
chr6-47013859-INS-60
chr6-47018197-INS-60
chr6-47023558-INS-258
chr6-47048091-INS-6672
chr6-47048212-INS-960
chr6-47055497-INS-69
chr6-47075809-INS-6276
chr6-47081814-INS-1237
chr6-47098905-INS-257
chr6-47099101-INS-916
chr6-47116721-INS-58
chr6-47165501-INS-200
chr6-47208967-INS-61
chr6-47214645-INS-6432
chr6-47239647-INS-74
chr6-47250610-INS-51
chr6-47257993-INS-72
chr6-47260203-INS-199

chr6-47263571-INS-113
chr6-47302410-INS-68
chr6-47305728-INS-822
chr6-47306703-INS-910
chr6-47308254-INS-94
chr6-47312593-INS-163
chr6-47328696-INS-71
chr6-47335977-INS-429
chr6-47369206-INS-186
chr6-47371102-INS-1866
chr6-47383856-INS-513
chr6-47391864-INS-70
chr6-47415436-INS-52
chr6-47417925-INS-70
chr6-47447575-INS-880
chr6-47503308-INS-164
chr6-47560459-INS-206
chr6-47565582-INS-657
chr6-47590123-INS-193
chr6-47591148-INS-179
chr6-47593273-INS-62
chr6-47593694-INS-152
chr6-47600189-INS-56
chr6-47612719-INS-6405
chr6-47631642-INS-51
chr6-47635924-INS-102
chr6-47717327-INS-54
chr6-47738496-INS-3054
chr6-47739828-INS-56
chr6-47740737-INS-93
chr6-47744175-INS-60
chr6-47745073-INS-63
chr6-47748526-INS-57
chr6-47749479-INS-82
chr6-47754810-INS-90
chr6-47768329-INS-414
chr6-47791043-INS-135
chr6-47808206-INS-70
chr6-47824804-INS-212
chr6-47838550-INS-6442
chr6-47838863-INS-86
chr6-47865103-INS-134
chr6-47901600-INS-162
chr6-47912533-INS-91
chr6-47912976-INS-218
chr6-47925157-INS-401
chr6-47929196-INS-104
chr6-47931812-INS-57
chr6-47954286-INS-506
chr6-47990782-INS-104
chr6-48031537-INS-57
chr6-48070185-INS-6428
chr6-48074738-INS-60
chr6-48075983-INS-796
chr6-48091390-INS-223
chr6-48092050-INS-238
chr6-48096223-INS-70
chr6-48118262-INS-83
chr6-48141231-INS-52
chr6-48155245-INS-64
chr6-48158291-INS-277
chr6-48159606-INS-80
chr6-48176541-INS-217
chr6-48190027-INS-101
chr6-48200743-INS-68
chr6-48203656-INS-204
chr6-48205157-INS-78
chr6-48221169-INS-7156
chr6-48239709-INS-209
chr6-48245391-INS-1515
chr6-48246531-INS-78
chr6-48290276-INS-796
chr6-48296048-INS-4636
chr6-48346250-INS-86
chr6-48353060-INS-58
chr6-48361267-INS-206
chr6-48398591-INS-51

chr6-48483785-INS-200
chr6-48490225-INS-963
chr6-48494944-INS-106
chr6-48512814-INS-204
chr6-48520158-INS-53
chr6-48530706-INS-58
chr6-48539093-INS-204
chr6-48541362-INS-52
chr6-48552838-INS-91
chr6-48556980-INS-193
chr6-48559029-INS-780
chr6-48570361-INS-462
chr6-48615685-INS-193
chr6-48620290-INS-106
chr6-48646471-INS-390
chr6-48660950-INS-119
chr6-48703388-INS-235
chr6-48706754-INS-484
chr6-48708036-INS-76
chr6-48715579-INS-162
chr6-48716644-INS-68
chr6-48719267-INS-163
chr6-48744361-INS-108
chr6-48752313-INS-323
chr6-48758460-INS-171
chr6-48760761-INS-221
chr6-48769618-INS-138
chr6-48770355-INS-54
chr6-48775050-INS-431
chr6-48797014-INS-64
chr6-48823614-INS-203
chr6-48828406-INS-82
chr6-48836160-INS-8397
chr6-48846616-INS-74
chr6-48897560-INS-109
chr6-48932537-INS-322
chr6-48934136-INS-444
chr6-48935642-INS-67
chr6-48944533-INS-62
chr6-48970421-INS-88
chr6-48977015-INS-53
chr6-48979389-INS-302
chr6-48984646-INS-972
chr6-48995483-INS-148
chr6-48997504-INS-56
chr6-49004282-INS-91
chr6-49036425-INS-717
chr6-49038017-INS-326
chr6-49045670-INS-97
chr6-49124396-INS-196
chr6-49126512-INS-366
chr6-49132604-INS-266
chr6-49137370-INS-204
chr6-49138295-INS-51
chr6-49158667-INS-95
chr6-49168964-INS-105
chr6-49178323-INS-212
chr6-49199504-INS-222
chr6-49202898-INS-109
chr6-49209492-INS-706
chr6-49215625-INS-283
chr6-49225331-INS-53
chr6-49249053-INS-533
chr6-49281980-INS-400
chr6-49285323-INS-207
chr6-49298879-INS-204
chr6-49301422-INS-143
chr6-49315026-INS-225
chr6-49327484-INS-185
chr6-49329073-INS-103
chr6-49332214-INS-203
chr6-49370522-INS-365
chr6-49383892-INS-558
chr6-49393034-INS-116
chr6-49448339-INS-81
chr6-49465183-INS-52
chr6-49470367-INS-58

chr6-49480573-INS-427
chr6-49482719-INS-372
chr6-49489011-INS-52
chr6-49493832-INS-1897
chr6-49535774-INS-212
chr6-49544030-INS-87
chr6-49555803-INS-117
chr6-49556602-INS-51
chr6-49561561-INS-179
chr6-49573756-INS-69
chr6-49600548-INS-57
chr6-49625205-INS-92
chr6-49640935-INS-292
chr6-49645521-INS-120
chr6-49676221-INS-528
chr6-49684959-INS-51
chr6-49691227-INS-64
chr6-49711036-INS-613
chr6-49712771-INS-60
chr6-49737865-INS-54
chr6-49755103-INS-1766
chr6-49764980-INS-194
chr6-49770863-INS-189
chr6-49855767-INS-96
chr6-49859278-INS-361
chr6-49873315-INS-58
chr6-49878491-INS-108
chr6-49896772-INS-140
chr6-49942254-INS-88
chr6-49948611-INS-127
chr6-49949688-INS-81
chr6-49961801-INS-70
chr6-49978798-INS-66
chr6-49991011-INS-299
chr6-49992389-INS-153
chr6-50014555-INS-86
chr6-50029036-INS-156
chr6-50036006-INS-71
chr6-50042853-INS-62
chr6-50081576-INS-162
chr6-50121277-INS-117
chr6-50131063-INS-184
chr6-50140731-INS-252
chr6-50146182-INS-116
chr6-50174058-INS-277
chr6-50174213-INS-624
chr6-50219555-INS-184
chr6-50259387-INS-165
chr6-50314792-INS-60
chr6-50319066-INS-109
chr6-50406156-INS-58
chr6-50424634-INS-75
chr6-50444082-INS-84
chr6-50444844-INS-577
chr6-50458879-INS-174
chr6-50462766-INS-62
chr6-50463955-INS-308
chr6-50469062-INS-253
chr6-50473734-INS-174
chr6-50482369-INS-976
chr6-50485039-INS-266
chr6-50487223-INS-95
chr6-50499301-INS-56
chr6-50531545-INS-66
chr6-50535496-INS-199
chr6-50577292-INS-385
chr6-50579681-INS-60
chr6-50613167-INS-168
chr6-50613512-INS-524
chr6-50614590-INS-4351
chr6-50615983-INS-51
chr6-50657776-INS-65
chr6-50678052-INS-54
chr6-50689747-INS-58
chr6-50733031-INS-58
chr6-50781420-INS-84
chr6-50832724-INS-107

chr6-50856627-INS-147
chr6-50862011-INS-88
chr6-50869029-INS-5286
chr6-50870091-INS-70
chr6-50877676-INS-105
chr6-50882232-INS-55
chr6-50896996-INS-98
chr6-50911818-INS-181
chr6-50916542-INS-974
chr6-50930717-INS-54
chr6-50965989-INS-333
chr6-50975393-INS-192
chr6-50978910-INS-113
chr6-50982308-INS-174
chr6-50985923-INS-412
chr6-50995524-INS-54
chr6-51005008-INS-134
chr6-51026501-INS-248
chr6-51036797-INS-123
chr6-51049421-INS-222
chr6-51120816-INS-85
chr6-51159228-INS-51
chr6-51160322-INS-54
chr6-51163080-INS-166
chr6-51163869-INS-56
chr6-51191334-INS-632
chr6-51194221-INS-74
chr6-51215641-INS-60
chr6-51237108-INS-194
chr6-51237891-INS-6419
chr6-51263824-INS-374
chr6-51272970-INS-138
chr6-51295972-INS-82
chr6-51297380-INS-134
chr6-51304050-INS-52
chr6-51307533-INS-452
chr6-51335540-INS-311
chr6-51341191-INS-69
chr6-51358038-INS-101
chr6-51384201-INS-54
chr6-51385141-INS-358
chr6-51393896-INS-921
chr6-51395301-INS-190
chr6-51399362-INS-96
chr6-51401625-INS-223
chr6-51402913-INS-5512
chr6-51418607-INS-143
chr6-51424048-INS-81
chr6-51432718-INS-536
chr6-51455984-INS-220
chr6-51459201-INS-106
chr6-51462300-INS-190
chr6-51473725-INS-174
chr6-51487588-INS-220
chr6-51497123-INS-208
chr6-51498218-INS-255
chr6-51501689-INS-119
chr6-51514261-INS-63
chr6-51560184-INS-196
chr6-51579200-INS-69
chr6-51627016-INS-5261
chr6-51627167-INS-66
chr6-51629783-INS-103
chr6-51636725-INS-73
chr6-51638072-INS-76
chr6-51642689-INS-71
chr6-51651507-INS-153
chr6-51665820-INS-54
chr6-51678947-INS-87
chr6-51683327-INS-234
chr6-51690648-INS-452
chr6-51692366-INS-68
chr6-51701521-INS-183
chr6-51702178-INS-59
chr6-51716129-INS-69
chr6-51724370-INS-204
chr6-51740246-INS-129

chr6-51741309-INS-52
chr6-51761329-INS-212
chr6-51762296-INS-162
chr6-51765797-INS-53
chr6-51787317-INS-54
chr6-51804082-INS-212
chr6-51847663-INS-61
chr6-51919153-INS-400
chr6-51949846-INS-84
chr6-51972861-INS-168
chr6-51990947-INS-70
chr6-52005574-INS-51
chr6-52011177-INS-1051
chr6-52026066-INS-68
chr6-52046124-INS-140
chr6-52046997-INS-112
chr6-52086913-INS-192
chr6-52089638-INS-142
chr6-52096729-INS-64
chr6-52098111-INS-54
chr6-52106060-INS-95
chr6-52109727-INS-51
chr6-52111505-INS-52
chr6-52121095-INS-93
chr6-52143398-INS-54
chr6-52184710-INS-90
chr6-52187045-INS-118
chr6-52246329-INS-117
chr6-52254547-INS-170
chr6-52261378-INS-55
chr6-52264447-INS-74
chr6-52266881-INS-51
chr6-52281679-INS-212
chr6-52304537-INS-55
chr6-52319149-INS-204
chr6-52335202-INS-463
chr6-52377740-INS-54
chr6-52386538-INS-430
chr6-52393376-INS-219
chr6-52396894-INS-200
chr6-52408890-INS-400
chr6-52430781-INS-58
chr6-52432505-INS-64
chr6-52444578-INS-56
chr6-52446105-INS-447
chr6-52450033-INS-92
chr6-52457057-INS-116
chr6-52468777-INS-68
chr6-52493354-INS-78
chr6-52497149-INS-88
chr6-52499672-INS-57
chr6-52508941-INS-61
chr6-52510066-INS-154
chr6-52518433-INS-51
chr6-52524471-INS-500
chr6-52525682-INS-56
chr6-52527755-INS-195
chr6-52557182-INS-952
chr6-52559880-INS-89
chr6-52565540-INS-385
chr6-52576122-INS-166
chr6-52602050-INS-117
chr6-52604317-INS-179
chr6-52655983-INS-260
chr6-52663723-INS-405
chr6-52676159-INS-459
chr6-52686533-INS-370
chr6-52688287-INS-83
chr6-52688933-INS-831
chr6-52740186-INS-58
chr6-52770247-INS-80
chr6-52801266-INS-270
chr6-52813823-INS-425
chr6-52825146-INS-437
chr6-52865547-INS-92
chr6-52868422-INS-76
chr6-52870978-INS-80

chr6-52882816-INS-82
chr6-52907971-INS-110
chr6-52908990-INS-74
chr6-52980360-INS-286
chr6-53006088-INS-756
chr6-53024478-INS-182
chr6-53061309-INS-155
chr6-53076618-INS-55
chr6-53102784-INS-438
chr6-53105171-INS-87
chr6-53110349-INS-220
chr6-53125556-INS-58
chr6-53133988-INS-230
chr6-53136684-INS-1866
chr6-53144101-INS-78
chr6-53145437-INS-54
chr6-53147180-INS-567
chr6-53226162-INS-302
chr6-53228323-INS-246
chr6-53244049-INS-252
chr6-53258143-INS-121
chr6-53293329-INS-52
chr6-53303260-INS-400
chr6-53305824-INS-90
chr6-53331162-INS-193
chr6-53332458-INS-97
chr6-53333810-INS-88
chr6-53341783-INS-103
chr6-53401197-INS-907
chr6-53402412-INS-254
chr6-53443568-INS-125
chr6-53448787-INS-130
chr6-53473063-INS-82
chr6-53492321-INS-210
chr6-53512122-INS-268
chr6-53524698-INS-52
chr6-53536362-INS-55
chr6-53584736-INS-78
chr6-53642541-INS-57
chr6-53647981-INS-116
chr6-53684715-INS-56
chr6-53685720-INS-489
chr6-53708623-INS-60
chr6-53723496-INS-275
chr6-53728037-INS-94
chr6-53734916-INS-68
chr6-53738989-INS-363
chr6-53742592-INS-52
chr6-53774480-INS-255
chr6-53775982-INS-638
chr6-53786059-INS-469
chr6-53788386-INS-96
chr6-53803320-INS-60
chr6-53828198-INS-769
chr6-53841237-INS-52
chr6-53843199-INS-212
chr6-53863046-INS-51
chr6-53868888-INS-94
chr6-53891015-INS-170
chr6-53914605-INS-395
chr6-53942445-INS-146
chr6-53982379-INS-426
chr6-53993875-INS-114
chr6-54006426-INS-131
chr6-54023424-INS-568
chr6-54056322-INS-254
chr6-54059951-INS-74
chr6-54063239-INS-96
chr6-54108818-INS-76
chr6-54133053-INS-60
chr6-54138732-INS-328
chr6-54143438-INS-255
chr6-54145681-INS-134
chr6-54166160-INS-61
chr6-54166938-INS-72
chr6-54167863-INS-262
chr6-54183094-INS-152

chr6-54189885-INS-64
chr6-54191717-INS-62
chr6-54194893-INS-64
chr6-54206792-INS-1533
chr6-54233414-INS-131
chr6-54243695-INS-73
chr6-54244719-INS-343
chr6-54268199-INS-158
chr6-54321790-INS-9394
chr6-54360142-INS-78
chr6-54360422-INS-1487
chr6-54366958-INS-109
chr6-54373797-INS-215
chr6-54381385-INS-69
chr6-54424742-INS-64
chr6-54426798-INS-60
chr6-54445905-INS-271
chr6-54454674-INS-92
chr6-54464108-INS-540
chr6-54517662-INS-306
chr6-54539861-INS-61
chr6-54556997-INS-83
chr6-54565611-INS-714
chr6-54571603-INS-203
chr6-54581604-INS-143
chr6-54610171-INS-175
chr6-54619376-INS-210
chr6-54626946-INS-92
chr6-54631486-INS-2203
chr6-54643599-INS-70
chr6-54674451-INS-80
chr6-54684351-INS-104
chr6-54749872-INS-56
chr6-54771841-INS-965
chr6-54791722-INS-98
chr6-54887922-INS-114
chr6-54906146-INS-71
chr6-54909953-INS-142
chr6-54931707-INS-8419
chr6-54936970-INS-288
chr6-55008520-INS-116
chr6-55024763-INS-56
chr6-55039703-INS-208
chr6-55108904-INS-429
chr6-55112870-INS-58
chr6-55118591-INS-400
chr6-55122157-INS-172
chr6-55135700-INS-214
chr6-55144902-INS-498
chr6-55169380-INS-285
chr6-55189771-INS-66
chr6-55193842-INS-358
chr6-55196958-INS-252
chr6-55219999-INS-711
chr6-55235304-INS-142
chr6-55243921-INS-6638
chr6-55260207-INS-235
chr6-55267651-INS-433
chr6-55328031-INS-58
chr6-55348128-INS-66
chr6-55370486-INS-253
chr6-55432564-INS-132
chr6-55460333-INS-56
chr6-55509550-INS-1009
chr6-55512398-INS-56
chr6-55523026-INS-56
chr6-55561359-INS-5178
chr6-55568260-INS-100
chr6-55599507-INS-144
chr6-55607622-INS-133
chr6-55613435-INS-53
chr6-55616787-INS-101
chr6-55625257-INS-60
chr6-55677426-INS-52
chr6-55688044-INS-87
chr6-55696717-INS-250
chr6-55711878-INS-428

chr6-55770321-INS-81
chr6-55779269-INS-111
chr6-55793895-INS-66
chr6-55806952-INS-75
chr6-55838270-INS-66
chr6-55839793-INS-61
chr6-55845281-INS-64
chr6-55858571-INS-67
chr6-55910229-INS-610
chr6-55922064-INS-378
chr6-55930675-INS-952
chr6-55988499-INS-60
chr6-55990409-INS-489
chr6-55998684-INS-150
chr6-56024313-INS-83
chr6-56043795-INS-91
chr6-56065273-INS-160
chr6-56103684-INS-696
chr6-56111909-INS-155
chr6-56143493-INS-64
chr6-56170704-INS-126
chr6-56197639-INS-577
chr6-56198661-INS-311
chr6-56209336-INS-469
chr6-56225067-INS-206
chr6-56229452-INS-523
chr6-56233155-INS-692
chr6-56235118-INS-83
chr6-56237397-INS-507
chr6-56242530-INS-389
chr6-56261082-INS-403
chr6-56276759-INS-212
chr6-56312849-INS-201
chr6-56322639-INS-118
chr6-56351951-INS-114
chr6-56355524-INS-400
chr6-56380674-INS-6681
chr6-56386929-INS-55
chr6-56389521-INS-2212
chr6-56412289-INS-5390
chr6-56418753-INS-492
chr6-56428709-INS-6768
chr6-56437038-INS-58
chr6-56456969-INS-55
chr6-56478962-INS-522
chr6-56479925-INS-85
chr6-56527268-INS-143
chr6-56551683-INS-57
chr6-56555461-INS-759
chr6-56569732-INS-468
chr6-56574395-INS-126
chr6-56586744-INS-51
chr6-56600389-INS-216
chr6-56605176-INS-88
chr6-56612591-INS-108
chr6-56654925-INS-249
chr6-56658031-INS-123
chr6-56671661-INS-2337
chr6-56685729-INS-532
chr6-56690813-INS-56
chr6-56694944-INS-140
chr6-56702627-INS-200
chr6-56746619-INS-180
chr6-56754081-INS-325
chr6-56781356-INS-296
chr6-56787711-INS-596
chr6-56797367-INS-811
chr6-56824015-INS-52
chr6-56839213-INS-1261
chr6-56912002-INS-151
chr6-56929270-INS-286
chr6-56994876-INS-6381
chr6-57014492-INS-6410
chr6-57014640-INS-137
chr6-57022065-INS-120
chr6-57024961-INS-6467
chr6-57027250-INS-1887

chr6-57057028-INS-136
chr6-57072276-INS-6202
chr6-57077326-INS-55
chr6-57105599-INS-332
chr6-57106937-INS-188
chr6-57109229-INS-524
chr6-57138718-INS-528
chr6-57156699-INS-6298
chr6-57159184-INS-6590
chr6-57174618-INS-186
chr6-57216206-INS-6333
chr6-57226705-INS-52
chr6-57231171-INS-3804
chr6-57233163-INS-191
chr6-57254181-INS-6418
chr6-57281365-INS-214
chr6-57391179-INS-2794
chr6-57410872-INS-355
chr6-57476839-INS-57
chr6-57508290-INS-166
chr6-57530346-INS-96
chr6-57583633-INS-54
chr6-57621417-INS-179
chr6-57664257-INS-4548
chr6-57710914-INS-99
chr6-57726601-INS-54
chr6-57738303-INS-196
chr6-57751371-INS-72
chr6-57775729-INS-596
chr6-57800269-INS-80
chr6-57828373-INS-124
chr6-57840478-INS-52
chr6-57885576-INS-7398
chr6-57896682-INS-214
chr6-57914216-INS-7462
chr6-57917801-INS-1426
chr6-57948050-INS-6454
chr6-57958469-INS-177
chr6-57996149-INS-638
chr6-58034557-INS-6511
chr6-58041084-INS-590
chr6-58044791-INS-118
chr6-58069651-INS-2266
chr6-58076765-INS-309
chr6-58080016-INS-1260
chr6-58087720-INS-577
chr6-58098659-INS-492
chr6-58110856-INS-3277
chr6-58132757-INS-353
chr6-58147739-INS-333
chr6-58191583-INS-52
chr6-58231559-INS-60
chr6-58240422-INS-77
chr6-58253336-INS-67
chr6-58305678-INS-54
chr6-58315935-INS-67
chr6-58318913-INS-9142
chr6-58365574-INS-291
chr6-58370500-INS-68
chr6-58372216-INS-379
chr6-58394357-INS-6991
chr6-58397564-INS-6236
chr6-58423430-INS-216
chr6-58434102-INS-56
chr6-58457955-INS-401
chr6-58472470-INS-1035
chr6-58494266-INS-67
chr6-58502972-INS-112
chr6-58518310-INS-280
chr6-58531438-INS-114
chr6-58532542-INS-93
chr6-58570811-INS-70
chr6-58575437-INS-70
chr6-58597852-INS-6177
chr6-58672239-INS-358
chr6-58676230-INS-1286
chr6-58699531-INS-69

chr6-58701244-INS-141
chr6-58706729-INS-132
chr6-58709711-INS-507
chr6-58716257-INS-54
chr6-58724086-INS-500
chr6-58727780-INS-2513
chr6-58733054-INS-2917
chr6-58776042-INS-90
chr6-58783994-INS-186
chr6-58790748-INS-67
chr6-58809346-INS-69
chr6-58812349-INS-70
chr6-58821539-INS-365
chr6-58829339-INS-222
chr6-58832314-INS-151
chr6-58839060-INS-965
chr6-58842377-INS-446
chr6-58855915-INS-329
chr6-58862593-INS-57
chr6-58869242-INS-75
chr6-58881621-INS-60
chr6-58909037-INS-52
chr6-58929900-INS-75
chr6-58962241-INS-54
chr6-58979862-INS-70
chr6-59005468-INS-5543
chr6-59019168-INS-6199
chr6-59058318-INS-6569
chr6-59065491-INS-298
chr6-59082880-INS-89
chr6-59133834-INS-1024
chr6-59154832-INS-191
chr6-59161142-INS-177
chr6-59185500-INS-205
chr6-59200164-INS-751
chr6-59228936-INS-180
chr6-59254306-INS-7034
chr6-59289442-INS-193
chr6-59293531-INS-119
chr6-59304218-INS-1353
chr6-59315922-INS-1210
chr6-59345258-INS-296
chr6-59350179-INS-492
chr6-59371854-INS-64
chr6-59378609-INS-70
chr6-59407607-INS-164
chr6-59432125-INS-50
chr6-59445129-INS-102
chr6-59458884-INS-4856
chr6-59474709-INS-147
chr6-59479365-INS-109
chr6-59546414-INS-65
chr6-59579134-INS-898
chr6-59591160-INS-119
chr6-59635710-INS-92
chr6-59688759-INS-89
chr6-59716134-INS-333
chr6-59733330-INS-110
chr6-60286956-INS-138
chr6-60287330-INS-727
chr6-60300192-INS-6914
chr6-60306653-INS-1477
chr6-60311101-INS-58
chr6-60320114-INS-69
chr6-60324370-INS-530
chr6-60399448-INS-2858
chr6-60446732-INS-329
chr6-60458614-INS-5234
chr6-60481887-INS-76
chr6-60544372-INS-202
chr6-60565431-INS-60
chr6-60579490-INS-204
chr6-60593731-INS-944
chr6-60594678-INS-61
chr6-60603317-INS-69
chr6-60626812-INS-115
chr6-60637884-INS-53

chr6-60647220-INS-1891
chr6-60655855-INS-50
chr6-60675574-INS-529
chr6-60676446-INS-206
chr6-60677826-INS-640
chr6-60694637-INS-60
chr6-60699868-INS-98
chr6-60706522-INS-162
chr6-60756066-INS-62
chr6-60811951-INS-60
chr6-60843804-INS-52
chr6-60871376-INS-152
chr6-60876546-INS-874
chr6-60894337-INS-911
chr6-60900006-INS-177
chr6-60923974-INS-92
chr6-60942091-INS-69
chr6-60962416-INS-266
chr6-60971116-INS-314
chr6-60978619-INS-398
chr6-60991567-INS-7164
chr6-60994396-INS-6384
chr6-61007350-INS-2953
chr6-61008100-INS-5547
chr6-61018448-INS-197
chr6-61022405-INS-298
chr6-61035500-INS-80
chr6-61044729-INS-81
chr6-61055170-INS-181
chr6-61063476-INS-73
chr6-61079832-INS-155
chr6-61090888-INS-64
chr6-61092140-INS-64
chr6-61115114-INS-57
chr6-61123988-INS-905
chr6-61148961-INS-147
chr6-61152768-INS-109
chr6-61166462-INS-59
chr6-61167811-INS-68
chr6-61172146-INS-295
chr6-61176111-INS-76
chr6-61179388-INS-89
chr6-61206227-INS-728
chr6-61220214-INS-54
chr6-61227502-INS-60
chr6-61238851-INS-70
chr6-61257778-INS-781
chr6-61259512-INS-757
chr6-61287538-INS-405
chr6-61295132-INS-405
chr6-61314022-INS-94
chr6-61325643-INS-115
chr6-61347537-INS-7528
chr6-61392792-INS-80
chr6-61399255-INS-97
chr6-61477898-INS-211
chr6-61481225-INS-830
chr6-61496576-INS-158
chr6-61514835-INS-905
chr6-61521601-INS-56
chr6-61541109-INS-99
chr6-61622229-INS-109
chr6-61626566-INS-58
chr6-61635422-INS-70
chr6-61666509-INS-107
chr6-61668026-INS-204
chr6-61672858-INS-318
chr6-61678265-INS-436
chr6-61724133-INS-57
chr6-61747223-INS-252
chr6-61750232-INS-760
chr6-61754265-INS-58
chr6-61765662-INS-153
chr6-61779296-INS-75
chr6-61791449-INS-137
chr6-61805848-INS-79
chr6-61807151-INS-663

chr6-61809476-INS-70
chr6-61834920-INS-645
chr6-61851608-INS-157
chr6-61867396-INS-109
chr6-61878256-INS-106
chr6-61969001-INS-441
chr6-62005320-INS-265
chr6-62021914-INS-321
chr6-62027944-INS-60
chr6-62126157-INS-1887
chr6-62171739-INS-113
chr6-62234309-INS-66
chr6-62252401-INS-176
chr6-62254883-INS-76
chr6-62256821-INS-69
chr6-62281779-INS-148
chr6-62292188-INS-58
chr6-62317901-INS-263
chr6-62355381-INS-136
chr6-62398945-INS-209
chr6-62456567-INS-226
chr6-62480597-INS-56
chr6-62497529-INS-53
chr6-62520221-INS-73
chr6-62528677-INS-204
chr6-62531416-INS-56
chr6-62551442-INS-83
chr6-62552820-INS-221
chr6-62555416-INS-920
chr6-62633429-INS-847
chr6-62663622-INS-86
chr6-62692741-INS-7120
chr6-62725155-INS-90
chr6-62752891-INS-498
chr6-62762920-INS-819
chr6-62767516-INS-3235
chr6-62767801-INS-1254
chr6-62783225-INS-103
chr6-62785982-INS-7270
chr6-62811430-INS-392
chr6-62815007-INS-64
chr6-62837416-INS-213
chr6-62839255-INS-152
chr6-62850329-INS-857
chr6-62854224-INS-129
chr6-62861932-INS-51
chr6-62873226-INS-96
chr6-62902372-INS-722
chr6-62911190-INS-137
chr6-62926406-INS-619
chr6-62947797-INS-80
chr6-62952207-INS-58
chr6-62978121-INS-6695
chr6-63014446-INS-675
chr6-63040090-INS-5591
chr6-63080554-INS-404
chr6-63087034-INS-142
chr6-63089955-INS-54
chr6-63107231-INS-98
chr6-63111246-INS-5246
chr6-63118117-INS-224
chr6-63124548-INS-6796
chr6-63169441-INS-64
chr6-63192149-INS-5323
chr6-63209008-INS-257
chr6-63254779-INS-60
chr6-63260698-INS-54
chr6-63271816-INS-81
chr6-63289718-INS-152
chr6-63304744-INS-282
chr6-63308563-INS-1070
chr6-63344947-INS-160
chr6-63349767-INS-94
chr6-63370178-INS-221
chr6-63373352-INS-583
chr6-63388570-INS-687
chr6-63406249-INS-214

chr6-63411558-INS-100
chr6-63420313-INS-91
chr6-63435013-INS-562
chr6-63437540-INS-57
chr6-63439970-INS-996
chr6-63458472-INS-438
chr6-63476582-INS-180
chr6-63569260-INS-1587
chr6-63574664-INS-73
chr6-63583718-INS-2198
chr6-63596107-INS-100
chr6-63600041-INS-6490
chr6-63638243-INS-90
chr6-63668120-INS-213
chr6-63692002-INS-53
chr6-63697311-INS-276
chr6-63704384-INS-472
chr6-63790998-INS-204
chr6-63802196-INS-229
chr6-63813182-INS-52
chr6-63824849-INS-429
chr6-63834395-INS-463
chr6-63838757-INS-6688
chr6-63843923-INS-381
chr6-63856827-INS-72
chr6-63886951-INS-95
chr6-63887075-INS-471
chr6-63917266-INS-422
chr6-63925817-INS-142
chr6-63943225-INS-77
chr6-63955539-INS-201
chr6-63972271-INS-67
chr6-64003622-INS-1609
chr6-64003711-INS-55
chr6-64052122-INS-170
chr6-64059293-INS-399
chr6-64059920-INS-218
chr6-64065930-INS-72
chr6-64077958-INS-449
chr6-64104210-INS-66
chr6-64110089-INS-453
chr6-64118855-INS-69
chr6-64142381-INS-102
chr6-64167237-INS-204
chr6-64197970-INS-869
chr6-64228502-INS-96
chr6-64244558-INS-69
chr6-64259213-INS-639
chr6-64265450-INS-7200
chr6-64269241-INS-218
chr6-64272744-INS-356
chr6-64282974-INS-55
chr6-64302394-INS-58
chr6-64315726-INS-68
chr6-64318439-INS-326
chr6-64334816-INS-190
chr6-64349695-INS-224
chr6-64377290-INS-95
chr6-64378491-INS-102
chr6-64385003-INS-643
chr6-64427357-INS-52
chr6-64466659-INS-56
chr6-64497088-INS-238
chr6-64503798-INS-118
chr6-64529605-INS-140
chr6-64546748-INS-501
chr6-64564364-INS-146
chr6-64582671-INS-1888
chr6-64590244-INS-67
chr6-64628286-INS-157
chr6-64650612-INS-6406
chr6-64657653-INS-80
chr6-64667563-INS-181
chr6-64679948-INS-293
chr6-64684912-INS-124
chr6-64690071-INS-72
chr6-64700428-INS-7309

chr6-64702161-INS-2697
chr6-64717470-INS-104
chr6-64734179-INS-74
chr6-64773448-INS-197
chr6-64790241-INS-98
chr6-64817159-INS-557
chr6-64817980-INS-70
chr6-64819545-INS-66
chr6-64820485-INS-307
chr6-64849463-INS-276
chr6-64856672-INS-88
chr6-64882593-INS-790
chr6-64913388-INS-203
chr6-64923638-INS-518
chr6-64925580-INS-122
chr6-64944992-INS-76
chr6-64945884-INS-120
chr6-64951019-INS-78
chr6-64981148-INS-698
chr6-64982130-INS-166
chr6-64987573-INS-123
chr6-64988515-INS-121
chr6-64989778-INS-52
chr6-65002285-INS-199
chr6-65007511-INS-63
chr6-65009947-INS-155
chr6-65040745-INS-216
chr6-65041415-INS-801
chr6-65043864-INS-100
chr6-65087629-INS-201
chr6-65094723-INS-401
chr6-65104069-INS-96
chr6-65109796-INS-206
chr6-65114367-INS-6688
chr6-65126058-INS-915
chr6-65131611-INS-1070
chr6-65132948-INS-299
chr6-65196675-INS-7351
chr6-65203733-INS-432
chr6-65211666-INS-441
chr6-65223370-INS-206
chr6-65225917-INS-88
chr6-65230595-INS-528
chr6-65230990-INS-4142
chr6-65231453-INS-59
chr6-65287238-INS-760
chr6-65303618-INS-160
chr6-65342157-INS-57
chr6-65343492-INS-105
chr6-65350850-INS-373
chr6-65353022-INS-63
chr6-65373540-INS-6506
chr6-65385999-INS-6385
chr6-65386861-INS-476
chr6-65390104-INS-1441
chr6-65408282-INS-89
chr6-65409269-INS-371
chr6-65425295-INS-83
chr6-65435228-INS-66
chr6-65451756-INS-473
chr6-65464750-INS-134
chr6-65494760-INS-80
chr6-65505001-INS-62
chr6-65506623-INS-272
chr6-65515928-INS-62
chr6-65553273-INS-85
chr6-65578820-INS-515
chr6-65580701-INS-175
chr6-65584771-INS-239
chr6-65589727-INS-82
chr6-65651430-INS-105
chr6-65695783-INS-163
chr6-65703949-INS-60
chr6-65723461-INS-110
chr6-65727077-INS-517
chr6-65768779-INS-59
chr6-65786338-INS-163

chr6-65813222-INS-419
chr6-65856984-INS-56
chr6-65882256-INS-2796
chr6-65888410-INS-442
chr6-65901930-INS-184
chr6-65972412-INS-81
chr6-65992380-INS-91
chr6-66033864-INS-966
chr6-66087997-INS-76
chr6-66094077-INS-86
chr6-66103818-INS-2827
chr6-66140963-INS-130
chr6-66181307-INS-57
chr6-66186722-INS-600
chr6-66191070-INS-520
chr6-66194292-INS-185
chr6-66197132-INS-1032
chr6-66282814-INS-6200
chr6-66289220-INS-197
chr6-66308577-INS-115
chr6-66316801-INS-9636
chr6-66337421-INS-85
chr6-66369379-INS-125
chr6-66371131-INS-277
chr6-66374825-INS-6786
chr6-66392820-INS-94
chr6-66442350-INS-76
chr6-66455033-INS-126
chr6-66459092-INS-6245
chr6-66462819-INS-65
chr6-66475209-INS-52
chr6-66478461-INS-203
chr6-66483161-INS-72
chr6-66519241-INS-683
chr6-66520391-INS-92
chr6-66536406-INS-499
chr6-66550640-INS-2435
chr6-66598717-INS-4021
chr6-66601204-INS-58
chr6-66662012-INS-7195
chr6-66683816-INS-70
chr6-66689913-INS-378
chr6-66724478-INS-730
chr6-66725750-INS-60
chr6-66736119-INS-54
chr6-66812647-INS-63
chr6-66815868-INS-938
chr6-66819653-INS-73
chr6-66865639-INS-181
chr6-66894251-INS-99
chr6-66904175-INS-99
chr6-66947169-INS-228
chr6-66998907-INS-52
chr6-67000734-INS-2755
chr6-67005718-INS-85
chr6-67016288-INS-165
chr6-67028707-INS-198
chr6-67043405-INS-148
chr6-67049292-INS-499
chr6-67066511-INS-200
chr6-67077030-INS-86
chr6-67080971-INS-52
chr6-67093243-INS-260
chr6-67095146-INS-66
chr6-67102438-INS-52
chr6-67137622-INS-5313
chr6-67151918-INS-233
chr6-67174961-INS-98
chr6-67178264-INS-106
chr6-67180757-INS-313
chr6-67206550-INS-172
chr6-67210240-INS-434
chr6-67229239-INS-211
chr6-67234499-INS-52
chr6-67247743-INS-186
chr6-67254661-INS-181
chr6-67259300-INS-217

chr6-67259849-INS-107
chr6-67263640-INS-96
chr6-67278659-INS-209
chr6-67281259-INS-79
chr6-67284541-INS-161
chr6-67284541-INS-273
chr6-67304123-INS-363
chr6-67307411-INS-205
chr6-67310215-INS-207
chr6-67339624-INS-123
chr6-67343921-INS-287
chr6-67345796-INS-150
chr6-67376313-INS-103
chr6-67377313-INS-56
chr6-67379885-INS-64
chr6-67382950-INS-190
chr6-67385000-INS-55
chr6-67393712-INS-465
chr6-67403681-INS-95
chr6-67440302-INS-375
chr6-67452787-INS-209
chr6-67485584-INS-212
chr6-67506731-INS-5544
chr6-67517853-INS-201
chr6-67524181-INS-318
chr6-67527190-INS-205
chr6-67530662-INS-5586
chr6-67536083-INS-54
chr6-67536328-INS-6335
chr6-67539156-INS-7061
chr6-67546678-INS-856
chr6-67572205-INS-197
chr6-67578157-INS-6902
chr6-67603146-INS-212
chr6-67607933-INS-2381
chr6-67635035-INS-6401
chr6-67692922-INS-6428
chr6-67697746-INS-6471
chr6-67735687-INS-4687
chr6-67789509-INS-514
chr6-67820506-INS-3868
chr6-67846838-INS-2297
chr6-67883701-INS-248
chr6-67908289-INS-181
chr6-67928335-INS-6384
chr6-67933183-INS-135
chr6-67947426-INS-309
chr6-67957622-INS-55
chr6-67960929-INS-624
chr6-67988659-INS-8419
chr6-68005184-INS-6274
chr6-68070095-INS-6436
chr6-68076105-INS-7283
chr6-68077553-INS-286
chr6-68096474-INS-321
chr6-68145306-INS-1288
chr6-68177384-INS-52
chr6-68193771-INS-6659
chr6-68356761-INS-52
chr6-68367985-INS-81
chr6-68419885-INS-325
chr6-68444545-INS-84
chr6-68612304-INS-89
chr6-68628833-INS-276
chr6-68644117-INS-69
chr6-68653109-INS-159
chr6-68704282-INS-66
chr6-68706219-INS-180
chr6-68795873-INS-69
chr6-68803467-INS-540
chr6-68818104-INS-56
chr6-68818716-INS-2262
chr6-68832977-INS-6201
chr6-68959022-INS-881
chr6-69013097-INS-213
chr6-69148676-INS-6548
chr6-69153644-INS-376

chr6-69173886-INS-64
chr6-69267847-INS-199
chr6-69292279-INS-134
chr6-69568834-INS-5549
chr6-69671580-INS-364
chr6-69673794-INS-453
chr6-69699185-INS-114
chr6-69789889-INS-6185
chr6-69793512-INS-6186
chr6-69805907-INS-5828
chr6-69806338-INS-57
chr6-69817571-INS-6924
chr6-69896646-INS-6231
chr6-70023887-INS-168
chr6-70064633-INS-107
chr6-70089765-INS-51
chr6-70096160-INS-51
chr6-70173462-INS-808
chr6-70187052-INS-61
chr6-70198985-INS-80
chr6-70209649-INS-253
chr6-70264810-INS-56
chr6-70266293-INS-78
chr6-70334048-INS-189
chr6-70336064-INS-808
chr6-70367770-INS-199
chr6-70389812-INS-723
chr6-70473509-INS-427
chr6-70489328-INS-79
chr6-70680343-INS-593
chr6-70692600-INS-305
chr6-70708475-INS-186
chr6-70726176-INS-210
chr6-70728043-INS-63
chr6-70746700-INS-6453
chr6-70750392-INS-484
chr6-70767557-INS-150
chr6-70786687-INS-204
chr6-70788342-INS-53
chr6-70790256-INS-188
chr6-70796665-INS-329
chr6-70798332-INS-57
chr6-70839543-INS-95
chr6-70844401-INS-64
chr6-70858718-INS-112
chr6-70867645-INS-150
chr6-70870560-INS-97
chr6-70873731-INS-200
chr6-70892348-INS-56
chr6-70913976-INS-63
chr6-70921475-INS-181
chr6-70922552-INS-99
chr6-70926011-INS-52
chr6-70929111-INS-60
chr6-70932083-INS-194
chr6-70938207-INS-77
chr6-70943240-INS-211
chr6-70952345-INS-81
chr6-70964853-INS-53
chr6-70971009-INS-199
chr6-70986980-INS-61
chr6-70994270-INS-399
chr6-71007228-INS-51
chr6-71015432-INS-394
chr6-71032243-INS-613
chr6-71033488-INS-78
chr6-71051009-INS-84
chr6-71055106-INS-64
chr6-71089347-INS-180
chr6-71095019-INS-182
chr6-71100656-INS-57
chr6-71161024-INS-5548
chr6-71167316-INS-54
chr6-71167971-INS-599
chr6-71175914-INS-6442
chr6-71179693-INS-52
chr6-71181656-INS-73

chr6-71205149-INS-68
chr6-71211119-INS-192
chr6-71222404-INS-135
chr6-71242074-INS-52
chr6-71285256-INS-167
chr6-71347980-INS-276
chr6-71359667-INS-177
chr6-71371264-INS-182
chr6-71399459-INS-203
chr6-71401551-INS-66
chr6-71429042-INS-6069
chr6-71434179-INS-67
chr6-71441067-INS-146
chr6-71454529-INS-58
chr6-71458730-INS-72
chr6-71460481-INS-506
chr6-71462574-INS-192
chr6-71505687-INS-172
chr6-71508492-INS-402
chr6-71520463-INS-51
chr6-71568168-INS-214
chr6-71583024-INS-445
chr6-71585457-INS-137
chr6-71622608-INS-327
chr6-71640237-INS-177
chr6-71649689-INS-208
chr6-71660046-INS-57
chr6-71667221-INS-362
chr6-71680353-INS-53
chr6-71695963-INS-75
chr6-71706538-INS-75
chr6-71723920-INS-155
chr6-71746214-INS-52
chr6-71750928-INS-98
chr6-71762323-INS-125
chr6-71774959-INS-75
chr6-71776010-INS-202
chr6-71802313-INS-200
chr6-71818188-INS-160
chr6-71838894-INS-52
chr6-71873524-INS-199
chr6-71882604-INS-105
chr6-71909099-INS-222
chr6-71916630-INS-70
chr6-71951860-INS-90
chr6-71977858-INS-57
chr6-71987808-INS-6659
chr6-71996896-INS-51
chr6-72020103-INS-635
chr6-72036224-INS-64
chr6-72067848-INS-88
chr6-72085788-INS-107
chr6-72093763-INS-250
chr6-72096090-INS-137
chr6-72098878-INS-230
chr6-72117988-INS-57
chr6-72133818-INS-80
chr6-72140466-INS-58
chr6-72146671-INS-440
chr6-72169182-INS-67
chr6-72177356-INS-90
chr6-72179399-INS-58
chr6-72223725-INS-149
chr6-72226766-INS-108
chr6-72234631-INS-212
chr6-72248479-INS-118
chr6-72256210-INS-212
chr6-72260229-INS-65
chr6-72271081-INS-92
chr6-72288188-INS-60
chr6-72309278-INS-222
chr6-72329092-INS-61
chr6-72358061-INS-71
chr6-72392835-INS-87
chr6-72394124-INS-62
chr6-72441009-INS-128
chr6-72447556-INS-60

chr6-72457615-INS-310
chr6-72459733-INS-54
chr6-72471987-INS-199
chr6-72472386-INS-351
chr6-72502057-INS-400
chr6-72531287-INS-283
chr6-72557190-INS-225
chr6-72563145-INS-51
chr6-72602973-INS-274
chr6-72680514-INS-183
chr6-72684260-INS-126
chr6-72699347-INS-138
chr6-72703079-INS-82
chr6-72703079-INS-969
chr6-72703850-INS-71
chr6-72711442-INS-182
chr6-72728058-INS-234
chr6-72735079-INS-120
chr6-72736400-INS-212
chr6-72758783-INS-51
chr6-72767750-INS-209
chr6-72774168-INS-102
chr6-72779855-INS-54
chr6-72781704-INS-216
chr6-72782557-INS-382
chr6-72784190-INS-264
chr6-72785931-INS-235
chr6-72804682-INS-276
chr6-72840870-INS-373
chr6-72851127-INS-234
chr6-72870780-INS-156
chr6-72878925-INS-835
chr6-72895078-INS-97
chr6-72911819-INS-197
chr6-72918500-INS-56
chr6-72920302-INS-54
chr6-72962524-INS-56
chr6-72985247-INS-147
chr6-72989210-INS-123
chr6-73014719-INS-66
chr6-73019099-INS-52
chr6-73027209-INS-102
chr6-73041284-INS-61
chr6-73055511-INS-420
chr6-73060687-INS-66
chr6-73068632-INS-55
chr6-73076805-INS-78
chr6-73095926-INS-86
chr6-73099654-INS-86
chr6-73110173-INS-202
chr6-73141601-INS-445
chr6-73166896-INS-52
chr6-73178416-INS-981
chr6-73207382-INS-209
chr6-73214604-INS-129
chr6-73219365-INS-226
chr6-73240502-INS-60
chr6-73242577-INS-75
chr6-73260918-INS-222
chr6-73281044-INS-455
chr6-73294694-INS-531
chr6-73304514-INS-66
chr6-73316464-INS-76
chr6-73321376-INS-338
chr6-73332204-INS-508
chr6-73337238-INS-293
chr6-73407511-INS-275
chr6-73413133-INS-115
chr6-73434595-INS-54
chr6-73437074-INS-190
chr6-73453754-INS-219
chr6-73464619-INS-63
chr6-73468676-INS-197
chr6-73487277-INS-129
chr6-73497155-INS-68
chr6-73538394-INS-320
chr6-73540929-INS-336

chr6-73557150-INS-636
chr6-73595363-INS-169
chr6-73600190-INS-609
chr6-73618912-INS-563
chr6-73621114-INS-488
chr6-73625423-INS-85
chr6-73628359-INS-76
chr6-73633679-INS-854
chr6-73640403-INS-66
chr6-73645334-INS-57
chr6-73660102-INS-325
chr6-73676034-INS-70
chr6-73689302-INS-66
chr6-73693603-INS-6198
chr6-73700087-INS-6668
chr6-73706404-INS-1719
chr6-73722970-INS-7338
chr6-73729191-INS-524
chr6-73730384-INS-150
chr6-73750694-INS-354
chr6-73765914-INS-79
chr6-73774823-INS-56
chr6-73795269-INS-508
chr6-73814204-INS-56
chr6-73819950-INS-377
chr6-73849362-INS-378
chr6-73865796-INS-755
chr6-73874176-INS-6590
chr6-73883585-INS-56
chr6-73902309-INS-88
chr6-73911429-INS-7340
chr6-73929405-INS-996
chr6-73933837-INS-276
chr6-73971238-INS-621
chr6-73975635-INS-190
chr6-73994283-INS-187
chr6-73999364-INS-78
chr6-74008210-INS-284
chr6-74013267-INS-60
chr6-74028223-INS-61
chr6-74040630-INS-373
chr6-74042211-INS-6528
chr6-74050349-INS-177
chr6-74062000-INS-308
chr6-74066351-INS-64
chr6-74177281-INS-54
chr6-74187662-INS-269
chr6-74193987-INS-6489
chr6-74195982-INS-174
chr6-74214278-INS-274
chr6-74234522-INS-62
chr6-74238458-INS-176
chr6-74296681-INS-693
chr6-74311918-INS-326
chr6-74315956-INS-181
chr6-74328043-INS-88
chr6-74366320-INS-570
chr6-74367856-INS-6405
chr6-74368061-INS-159
chr6-74369919-INS-1358
chr6-74393535-INS-57
chr6-74401639-INS-55
chr6-74466560-INS-422
chr6-74478378-INS-223
chr6-74492995-INS-123
chr6-74528366-INS-339
chr6-74535048-INS-108
chr6-74539175-INS-188
chr6-74546326-INS-66
chr6-74552642-INS-64
chr6-74566968-INS-667
chr6-74601228-INS-57
chr6-74610502-INS-166
chr6-74639236-INS-153
chr6-74645167-INS-420
chr6-74672192-INS-386
chr6-74676332-INS-659

chr6-74696523-INS-981
chr6-74709850-INS-170
chr6-74727045-INS-362
chr6-74743695-INS-63
chr6-74784118-INS-56
chr6-74789795-INS-133
chr6-74812499-INS-54
chr6-74923731-INS-721
chr6-74942652-INS-226
chr6-74962745-INS-4906
chr6-75046733-INS-68
chr6-75074681-INS-63
chr6-75090077-INS-54
chr6-75096896-INS-6398
chr6-75118447-INS-62
chr6-75124349-INS-58
chr6-75147372-INS-664
chr6-75150317-INS-471
chr6-75157496-INS-283
chr6-75173779-INS-6454
chr6-75202865-INS-78
chr6-75208995-INS-2853
chr6-75219182-INS-70
chr6-75243234-INS-184
chr6-75253643-INS-335
chr6-75259453-INS-254
chr6-75263123-INS-5667
chr6-75315793-INS-65
chr6-75366321-INS-58
chr6-75400967-INS-181
chr6-75451170-INS-6455
chr6-75459102-INS-525
chr6-75462161-INS-608
chr6-75469172-INS-352
chr6-75485395-INS-377
chr6-75503426-INS-573
chr6-75504106-INS-102
chr6-75512457-INS-75
chr6-75558081-INS-115
chr6-75581263-INS-77
chr6-75585437-INS-174
chr6-75599743-INS-2452
chr6-75615890-INS-762
chr6-75636967-INS-91
chr6-75666825-INS-1892
chr6-75680424-INS-561
chr6-75703548-INS-6178
chr6-75733715-INS-60
chr6-75744987-INS-4448
chr6-75781476-INS-53
chr6-75809277-INS-2431
chr6-75819107-INS-50
chr6-75821600-INS-79
chr6-75882188-INS-843
chr6-75883659-INS-267
chr6-75887668-INS-98
chr6-75898365-INS-77
chr6-75930465-INS-6538
chr6-75950985-INS-1593
chr6-76012506-INS-54
chr6-76029461-INS-726
chr6-76034777-INS-579
chr6-76040635-INS-136
chr6-76070361-INS-582
chr6-76085985-INS-52
chr6-76095949-INS-58
chr6-76110235-INS-163
chr6-76136916-INS-58
chr6-76143339-INS-202
chr6-76144844-INS-257
chr6-76184152-INS-60
chr6-76216650-INS-6427
chr6-76219264-INS-819
chr6-76249996-INS-501
chr6-76296464-INS-5502
chr6-76298819-INS-7073
chr6-76313638-INS-162

chr6-76369002-INS-54
chr6-76432506-INS-465
chr6-76455396-INS-64
chr6-76463940-INS-479
chr6-76484706-INS-5276
chr6-76496130-INS-96
chr6-76504478-INS-369
chr6-76508842-INS-792
chr6-76546836-INS-104
chr6-76549297-INS-662
chr6-76550448-INS-520
chr6-76563816-INS-480
chr6-76580943-INS-105
chr6-76585107-INS-52
chr6-76592726-INS-6648
chr6-76632555-INS-144
chr6-76646207-INS-60
chr6-76676286-INS-58
chr6-76713561-INS-6401
chr6-76721738-INS-260
chr6-76742052-INS-128
chr6-76785904-INS-5541
chr6-76795346-INS-643
chr6-76806274-INS-6249
chr6-76813814-INS-6415
chr6-76814463-INS-187
chr6-76816447-INS-6444
chr6-76820577-INS-206
chr6-76823096-INS-74
chr6-76844105-INS-205
chr6-76851313-INS-165
chr6-76886500-INS-813
chr6-76905076-INS-317
chr6-76909051-INS-947
chr6-76926259-INS-55
chr6-76941901-INS-52
chr6-77015037-INS-55
chr6-77053244-INS-303
chr6-77084659-INS-429
chr6-77089999-INS-126
chr6-77104752-INS-62
chr6-77126719-INS-673
chr6-77159200-INS-118
chr6-77167212-INS-105
chr6-77170000-INS-54
chr6-77183344-INS-210
chr6-77187011-INS-55
chr6-77190620-INS-54
chr6-77206827-INS-279
chr6-77219121-INS-188
chr6-77239820-INS-193
chr6-77250639-INS-126
chr6-77293408-INS-595
chr6-77294482-INS-142
chr6-77318722-INS-2612
chr6-77323960-INS-480
chr6-77334188-INS-81
chr6-77352690-INS-309
chr6-77379558-INS-60
chr6-77385918-INS-7084
chr6-77405041-INS-60
chr6-77432381-INS-879
chr6-77465152-INS-301
chr6-77475607-INS-200
chr6-77479491-INS-6537
chr6-77480471-INS-1558
chr6-77511511-INS-2580
chr6-77557570-INS-715
chr6-77612583-INS-147
chr6-77628282-INS-723
chr6-77669168-INS-404
chr6-77689520-INS-368
chr6-77694658-INS-330
chr6-77701380-INS-54
chr6-77717731-INS-174
chr6-77747417-INS-119
chr6-77798571-INS-808

chr6-77802189-INS-6834
chr6-77803284-INS-894
chr6-77803305-INS-1410
chr6-77811460-INS-373
chr6-77832173-INS-56
chr6-77843289-INS-333
chr6-77887711-INS-107
chr6-77900692-INS-991
chr6-77914566-INS-114
chr6-77917568-INS-1164
chr6-77933986-INS-104
chr6-77939001-INS-56
chr6-77962342-INS-207
chr6-77962752-INS-801
chr6-77974144-INS-6400
chr6-77977727-INS-313
chr6-77983163-INS-326
chr6-77995063-INS-79
chr6-78011231-INS-52
chr6-78020612-INS-191
chr6-78021991-INS-7104
chr6-78045276-INS-6593
chr6-78046354-INS-2678
chr6-78065335-INS-58
chr6-78086672-INS-111
chr6-78091569-INS-122
chr6-78100191-INS-69
chr6-78161236-INS-128
chr6-78172130-INS-107
chr6-78181772-INS-469
chr6-78224306-INS-59
chr6-78268487-INS-455
chr6-78269754-INS-110
chr6-78284096-INS-291
chr6-78295378-INS-110
chr6-78301157-INS-6146
chr6-78317014-INS-3445
chr6-78383879-INS-1890
chr6-78392640-INS-8830
chr6-78410661-INS-226
chr6-78426519-INS-363
chr6-78429581-INS-52
chr6-78432948-INS-123
chr6-78457786-INS-191
chr6-78489028-INS-428
chr6-78513325-INS-56
chr6-78536299-INS-554
chr6-78590966-INS-62
chr6-78593034-INS-75
chr6-78599619-INS-67
chr6-78617277-INS-1888
chr6-78640304-INS-337
chr6-78650520-INS-6204
chr6-78699502-INS-69
chr6-78711543-INS-1164
chr6-78737499-INS-58
chr6-78738055-INS-96
chr6-78743817-INS-1674
chr6-78744954-INS-56
chr6-78751333-INS-691
chr6-78784302-INS-652
chr6-78787976-INS-76
chr6-78824216-INS-89
chr6-78830449-INS-1086
chr6-78847306-INS-204
chr6-78870655-INS-187
chr6-78873369-INS-7080
chr6-78888400-INS-82
chr6-78906233-INS-116
chr6-78929617-INS-328
chr6-78962196-INS-6492
chr6-78985722-INS-510
chr6-79019222-INS-61
chr6-79029372-INS-675
chr6-79030553-INS-433
chr6-79039057-INS-63
chr6-79045631-INS-68

chr6-79057455-INS-54
chr6-79058108-INS-834
chr6-79065967-INS-826
chr6-79076997-INS-88
chr6-79089400-INS-1866
chr6-79093735-INS-62
chr6-79099226-INS-1375
chr6-79101520-INS-287
chr6-79121475-INS-234
chr6-79177969-INS-571
chr6-79207677-INS-193
chr6-79216321-INS-483
chr6-79239846-INS-56
chr6-79246428-INS-1630
chr6-79250547-INS-194
chr6-79268573-INS-129
chr6-79329119-INS-217
chr6-79341186-INS-72
chr6-79343596-INS-79
chr6-79364520-INS-549
chr6-79427978-INS-235
chr6-79437678-INS-6442
chr6-79447191-INS-273
chr6-79452875-INS-2302
chr6-79456699-INS-264
chr6-79470010-INS-962
chr6-79474789-INS-251
chr6-79477220-INS-58
chr6-79529353-INS-7231
chr6-79566388-INS-464
chr6-79575273-INS-204
chr6-79595773-INS-6486
chr6-79616882-INS-448
chr6-79617460-INS-951
chr6-79623411-INS-160
chr6-79629872-INS-813
chr6-79631778-INS-175
chr6-79634334-INS-6378
chr6-79659994-INS-59
chr6-79669611-INS-449
chr6-79680110-INS-83
chr6-79697449-INS-432
chr6-79709515-INS-162
chr6-79756686-INS-70
chr6-79760322-INS-112
chr6-79779335-INS-6742
chr6-79822234-INS-54
chr6-79853007-INS-270
chr6-79857697-INS-6208
chr6-79870330-INS-358
chr6-79872920-INS-4807
chr6-79888589-INS-401
chr6-79891550-INS-84
chr6-79909138-INS-58
chr6-79948189-INS-354
chr6-79969299-INS-144
chr6-79986500-INS-208
chr6-80010078-INS-51
chr6-80021851-INS-56
chr6-80027014-INS-370
chr6-80036482-INS-76
chr6-80050820-INS-56
chr6-80053482-INS-58
chr6-80067021-INS-103
chr6-80102980-INS-179
chr6-80143929-INS-236
chr6-80151252-INS-56
chr6-80168483-INS-3659
chr6-80172565-INS-81
chr6-80174863-INS-54
chr6-80188986-INS-151
chr6-80212734-INS-53
chr6-80234832-INS-521
chr6-80247959-INS-759
chr6-80255975-INS-86
chr6-80273324-INS-84
chr6-80275312-INS-52

chr6-80310645-INS-56
chr6-80312115-INS-204
chr6-80319434-INS-522
chr6-80350951-INS-108
chr6-80363547-INS-555
chr6-80366663-INS-210
chr6-80369317-INS-80
chr6-80380611-INS-276
chr6-80381753-INS-883
chr6-80394639-INS-488
chr6-80416071-INS-150
chr6-80440875-INS-488
chr6-80444885-INS-141
chr6-80446660-INS-57
chr6-80461624-INS-2661
chr6-80469505-INS-447
chr6-80494390-INS-683
chr6-80499677-INS-174
chr6-80501687-INS-237
chr6-80507601-INS-52
chr6-80508716-INS-129
chr6-80515633-INS-73
chr6-80537085-INS-4993
chr6-80538846-INS-70
chr6-80541684-INS-60
chr6-80552006-INS-530
chr6-80567355-INS-98
chr6-80567542-INS-6186
chr6-80572393-INS-410
chr6-80598512-INS-800
chr6-80626918-INS-687
chr6-80642429-INS-5195
chr6-80667571-INS-319
chr6-80681057-INS-2582
chr6-80682959-INS-53
chr6-80697728-INS-60
chr6-80736095-INS-93
chr6-80775475-INS-104
chr6-80787959-INS-843
chr6-80791074-INS-119
chr6-80818166-INS-54
chr6-80825644-INS-877
chr6-80869534-INS-6740
chr6-80875991-INS-71
chr6-80880725-INS-8648
chr6-80884345-INS-91
chr6-80920234-INS-6487
chr6-80929479-INS-582
chr6-80934974-INS-53
chr6-80954921-INS-100
chr6-80958035-INS-531
chr6-80962059-INS-258
chr6-80969814-INS-153
chr6-80973040-INS-132
chr6-80975944-INS-94
chr6-80976983-INS-76
chr6-81003606-INS-916
chr6-81010912-INS-66
chr6-81013330-INS-214
chr6-81028451-INS-70
chr6-81036316-INS-152
chr6-81041617-INS-53
chr6-81044054-INS-92
chr6-81061924-INS-579
chr6-81071961-INS-845
chr6-81073670-INS-125
chr6-81081741-INS-116
chr6-81083969-INS-90
chr6-81110137-INS-64
chr6-81136535-INS-521
chr6-81160624-INS-148
chr6-81170780-INS-6198
chr6-81171280-INS-58
chr6-81177200-INS-61
chr6-81177783-INS-767
chr6-81177841-INS-66
chr6-81180303-INS-829

chr6-81188226-INS-175
chr6-81201719-INS-596
chr6-81208142-INS-52
chr6-81229127-INS-59
chr6-81234306-INS-69
chr6-81244392-INS-159
chr6-81247934-INS-563
chr6-81249581-INS-700
chr6-81271498-INS-60
chr6-81277287-INS-60
chr6-81295512-INS-51
chr6-81309406-INS-79
chr6-81318257-INS-6706
chr6-81323900-INS-570
chr6-81327387-INS-6261
chr6-81333124-INS-275
chr6-81347240-INS-54
chr6-81355635-INS-118
chr6-81359227-INS-69
chr6-81360521-INS-66
chr6-81371550-INS-54
chr6-81390166-INS-102
chr6-81391207-INS-56
chr6-81392612-INS-494
chr6-81430313-INS-113
chr6-81464414-INS-359
chr6-81467293-INS-399
chr6-81479899-INS-57
chr6-81488982-INS-93
chr6-81509187-INS-78
chr6-81516124-INS-68
chr6-81549419-INS-2804
chr6-81563769-INS-61
chr6-81564746-INS-214
chr6-81586343-INS-54
chr6-81595483-INS-54
chr6-81606568-INS-756
chr6-81608045-INS-76
chr6-81614926-INS-51
chr6-81622189-INS-207
chr6-81626233-INS-125
chr6-81633590-INS-7115
chr6-81646868-INS-63
chr6-81663071-INS-469
chr6-81664455-INS-108
chr6-81666742-INS-58
chr6-81675464-INS-58
chr6-81679798-INS-80
chr6-81709305-INS-209
chr6-81720165-INS-86
chr6-81755663-INS-65
chr6-81795692-INS-127
chr6-81816228-INS-414
chr6-81824206-INS-98
chr6-81889326-INS-162
chr6-81896833-INS-733
chr6-81906435-INS-394
chr6-81917559-INS-197
chr6-81932444-INS-6818
chr6-81945279-INS-505
chr6-81990890-INS-740
chr6-81993568-INS-67
chr6-81996530-INS-203
chr6-81997803-INS-859
chr6-82001406-INS-51
chr6-82006452-INS-78
chr6-82042880-INS-106
chr6-82090265-INS-56
chr6-82097471-INS-54
chr6-82111605-INS-125
chr6-82116498-INS-63
chr6-82132726-INS-65
chr6-82137918-INS-1057
chr6-82170797-INS-7321
chr6-82180937-INS-557
chr6-82185221-INS-6499
chr6-82206796-INS-154

chr6-82208681-INS-57
chr6-82221344-INS-84
chr6-82245139-INS-120
chr6-82275003-INS-54
chr6-82278993-INS-957
chr6-82312000-INS-144
chr6-82344038-INS-51
chr6-82355358-INS-58
chr6-82362372-INS-949
chr6-82367631-INS-5286
chr6-82376327-INS-62
chr6-82387216-INS-60
chr6-82399709-INS-297
chr6-82421724-INS-317
chr6-82421813-INS-748
chr6-82475463-INS-115
chr6-82483315-INS-6654
chr6-82519003-INS-128
chr6-82547757-INS-897
chr6-82586555-INS-141
chr6-82587833-INS-67
chr6-82596273-INS-5542
chr6-82625969-INS-95
chr6-82657416-INS-70
chr6-82658724-INS-101
chr6-82663588-INS-173
chr6-82665266-INS-370
chr6-82676659-INS-99
chr6-82682098-INS-578
chr6-82684422-INS-548
chr6-82706873-INS-382
chr6-82718444-INS-161
chr6-82726600-INS-84
chr6-82754871-INS-279
chr6-82756822-INS-141
chr6-82765115-INS-88
chr6-82780171-INS-54
chr6-82790663-INS-82
chr6-82793514-INS-340
chr6-82799400-INS-7296
chr6-82803186-INS-60
chr6-82805868-INS-117
chr6-82808208-INS-121
chr6-82811337-INS-104
chr6-82823100-INS-214
chr6-82848398-INS-84
chr6-82859214-INS-122
chr6-82866416-INS-93
chr6-82871660-INS-89
chr6-82891514-INS-112
chr6-82893990-INS-66
chr6-82901428-INS-108
chr6-82902667-INS-57
chr6-82930750-INS-578
chr6-82950002-INS-135
chr6-82953417-INS-178
chr6-82957086-INS-208
chr6-82964761-INS-105
chr6-82999963-INS-96
chr6-83066030-INS-104
chr6-83076537-INS-68
chr6-83107024-INS-207
chr6-83107621-INS-429
chr6-83116594-INS-404
chr6-83118044-INS-198
chr6-83123145-INS-76
chr6-83132916-INS-54
chr6-83141462-INS-215
chr6-83151048-INS-204
chr6-83158276-INS-90
chr6-83178931-INS-231
chr6-83193684-INS-128
chr6-83208370-INS-204
chr6-83246814-INS-290
chr6-83257805-INS-218
chr6-83259710-INS-374
chr6-83271669-INS-64

chr6-83275453-INS-52
chr6-83285520-INS-115
chr6-83292998-INS-201
chr6-83324919-INS-98
chr6-83361049-INS-126
chr6-83362362-INS-95
chr6-83378447-INS-168
chr6-83413511-INS-179
chr6-83425038-INS-58
chr6-83484156-INS-508
chr6-83485685-INS-167
chr6-83492675-INS-88
chr6-83497921-INS-136
chr6-83508285-INS-98
chr6-83531898-INS-92
chr6-83553483-INS-50
chr6-83582448-INS-313
chr6-83602320-INS-58
chr6-83629306-INS-56
chr6-83644025-INS-439
chr6-83670157-INS-58
chr6-83677669-INS-60
chr6-83678309-INS-75
chr6-83686746-INS-62
chr6-83694759-INS-94
chr6-83698428-INS-56
chr6-83718894-INS-56
chr6-83729474-INS-73
chr6-83741145-INS-421
chr6-83755356-INS-444
chr6-83811098-INS-347
chr6-83837690-INS-54
chr6-83852867-INS-51
chr6-83854138-INS-147
chr6-83856552-INS-51
chr6-83868440-INS-242
chr6-83915913-INS-242
chr6-83922532-INS-192
chr6-83925307-INS-205
chr6-83933075-INS-52
chr6-83945839-INS-260
chr6-83965422-INS-55
chr6-83967491-INS-1628
chr6-83971554-INS-222
chr6-83974907-INS-66
chr6-84004012-INS-210
chr6-84013687-INS-801
chr6-84019465-INS-227
chr6-84020859-INS-234
chr6-84022555-INS-56
chr6-84098753-INS-212
chr6-84102650-INS-361
chr6-84105276-INS-117
chr6-84107753-INS-182
chr6-84140314-INS-68
chr6-84166468-INS-105
chr6-84173863-INS-52
chr6-84178613-INS-197
chr6-84191111-INS-56
chr6-84191810-INS-88
chr6-84210175-INS-154
chr6-84222138-INS-54
chr6-84264205-INS-138
chr6-84275380-INS-552
chr6-84295347-INS-133
chr6-84297726-INS-177
chr6-84303233-INS-100
chr6-84319549-INS-97
chr6-84323516-INS-283
chr6-84345266-INS-165
chr6-84400600-INS-87
chr6-84401177-INS-52
chr6-84415281-INS-115
chr6-84429929-INS-74
chr6-84433754-INS-135
chr6-84445966-INS-80
chr6-84455848-INS-109

chr6-84491610-INS-362
chr6-84517150-INS-57
chr6-84628609-INS-106
chr6-84710902-INS-106
chr6-84726217-INS-6686
chr6-84755932-INS-58
chr6-84757359-INS-5275
chr6-84778546-INS-6201
chr6-84791683-INS-118
chr6-84801177-INS-1344
chr6-84801452-INS-1488
chr6-84813705-INS-208
chr6-84846092-INS-68
chr6-84877195-INS-228
chr6-84921149-INS-137
chr6-84933637-INS-402
chr6-84943055-INS-56
chr6-84950479-INS-96
chr6-84956856-INS-77
chr6-85034163-INS-228
chr6-85048978-INS-61
chr6-85051090-INS-102
chr6-85122116-INS-959
chr6-85150100-INS-202
chr6-85157520-INS-332
chr6-85159907-INS-52
chr6-85169819-INS-115
chr6-85186097-INS-104
chr6-85224967-INS-60
chr6-85287058-INS-163
chr6-85294402-INS-204
chr6-85329941-INS-68
chr6-85332474-INS-72
chr6-85336903-INS-165
chr6-85367169-INS-127
chr6-85368791-INS-90
chr6-85373221-INS-245
chr6-85374583-INS-81
chr6-85381028-INS-68
chr6-85393176-INS-104
chr6-85401735-INS-207
chr6-85466280-INS-131
chr6-85472915-INS-66
chr6-85503972-INS-76
chr6-85532858-INS-592
chr6-85537918-INS-55
chr6-85574157-INS-191
chr6-85578927-INS-134
chr6-85579276-INS-237
chr6-85592641-INS-148
chr6-85807449-INS-198
chr6-85844672-INS-7720
chr6-85860078-INS-5699
chr6-85889131-INS-508
chr6-85925399-INS-187
chr6-85933021-INS-99
chr6-85943440-INS-92
chr6-85944747-INS-53
chr6-85949255-INS-5478
chr6-85955238-INS-251
chr6-85967325-INS-212
chr6-85989265-INS-54
chr6-86054050-INS-379
chr6-86065889-INS-66
chr6-86091353-INS-216
chr6-86111462-INS-92
chr6-86117000-INS-72
chr6-86135899-INS-151
chr6-86140090-INS-212
chr6-86140943-INS-74
chr6-86160371-INS-445
chr6-86206898-INS-100
chr6-86230135-INS-56
chr6-86256032-INS-89
chr6-86258760-INS-60
chr6-86264428-INS-114
chr6-86267778-INS-563

chr6-86268628-INS-126
chr6-86283728-INS-648
chr6-86301988-INS-59
chr6-86308742-INS-4230
chr6-86308922-INS-1761
chr6-86329642-INS-102
chr6-86339787-INS-65
chr6-86352509-INS-226
chr6-86369655-INS-230
chr6-86388180-INS-110
chr6-86407050-INS-169
chr6-86422262-INS-255
chr6-86429199-INS-187
chr6-86430270-INS-220
chr6-86444941-INS-178
chr6-86447773-INS-321
chr6-86458683-INS-221
chr6-86462951-INS-274
chr6-86476312-INS-74
chr6-86487433-INS-171
chr6-86489676-INS-243
chr6-86508169-INS-74
chr6-86516347-INS-161
chr6-86522124-INS-64
chr6-86525656-INS-134
chr6-86534432-INS-185
chr6-86547411-INS-59
chr6-86564973-INS-74
chr6-86569754-INS-59
chr6-86593013-INS-252
chr6-86620166-INS-78
chr6-86632445-INS-56
chr6-86648214-INS-206
chr6-86651067-INS-194
chr6-86657244-INS-400
chr6-86667835-INS-200
chr6-86676245-INS-134
chr6-86685167-INS-938
chr6-86695791-INS-77
chr6-86723894-INS-220
chr6-86734101-INS-191
chr6-86762780-INS-70
chr6-86768650-INS-94
chr6-86774979-INS-323
chr6-86779795-INS-215
chr6-86792656-INS-159
chr6-86795572-INS-85
chr6-86823615-INS-70
chr6-86830074-INS-277
chr6-86858367-INS-519
chr6-86860435-INS-54
chr6-86876933-INS-51
chr6-86889081-INS-105
chr6-86904363-INS-52
chr6-86932519-INS-210
chr6-86956279-INS-121
chr6-86992589-INS-131
chr6-86999220-INS-183
chr6-87003635-INS-486
chr6-87015503-INS-57
chr6-87017623-INS-192
chr6-87025583-INS-104
chr6-87037129-INS-171
chr6-87052250-INS-96
chr6-87059669-INS-65
chr6-87061537-INS-67
chr6-87063075-INS-67
chr6-87067470-INS-64
chr6-87073918-INS-58
chr6-87080667-INS-418
chr6-87096478-INS-111
chr6-87127505-INS-60
chr6-87203273-INS-181
chr6-87211547-INS-149
chr6-87216285-INS-57
chr6-87218384-INS-146
chr6-87220747-INS-62

chr6-87222189-INS-54
chr6-87228940-INS-59
chr6-87288707-INS-128
chr6-87326845-INS-99
chr6-87331033-INS-90
chr6-87348470-INS-544
chr6-87356031-INS-54
chr6-87377525-INS-106
chr6-87434202-INS-197
chr6-87434821-INS-124
chr6-87449808-INS-81
chr6-87451061-INS-129
chr6-87457601-INS-89
chr6-87482784-INS-184
chr6-87503248-INS-229
chr6-87513733-INS-72
chr6-87547006-INS-64
chr6-87599417-INS-420
chr6-87612636-INS-505
chr6-87643825-INS-104
chr6-87664732-INS-51
chr6-87674323-INS-266
chr6-87685643-INS-52
chr6-87718952-INS-52
chr6-87732966-INS-83
chr6-87744969-INS-311
chr6-87746394-INS-70
chr6-87752202-INS-2484
chr6-87752680-INS-2296
chr6-87752729-INS-1467
chr6-87752730-INS-869
chr6-87754792-INS-78
chr6-87762753-INS-198
chr6-87774287-INS-110
chr6-87780037-INS-76
chr6-87781460-INS-64
chr6-87791450-INS-198
chr6-87808586-INS-219
chr6-87810562-INS-103
chr6-87813710-INS-191
chr6-87832095-INS-101
chr6-87847802-INS-52
chr6-87862720-INS-995
chr6-87909684-INS-144
chr6-87919205-INS-5263
chr6-87938443-INS-65
chr6-87944424-INS-443
chr6-87947632-INS-203
chr6-87991494-INS-146
chr6-88000294-INS-305
chr6-88010947-INS-52
chr6-88027097-INS-191
chr6-88031735-INS-203
chr6-88050589-INS-158
chr6-88058685-INS-153
chr6-88095123-INS-349
chr6-88112167-INS-132
chr6-88134875-INS-55
chr6-88155618-INS-180
chr6-88187995-INS-540
chr6-88205081-INS-51
chr6-88253118-INS-661
chr6-88266143-INS-192
chr6-88286846-INS-51
chr6-88310723-INS-261
chr6-88314604-INS-6645
chr6-88337255-INS-984
chr6-88338081-INS-106
chr6-88345636-INS-207
chr6-88407773-INS-463
chr6-88420647-INS-87
chr6-88420932-INS-1301
chr6-88423748-INS-56
chr6-88424001-INS-834
chr6-88451359-INS-579
chr6-88471632-INS-163
chr6-88471773-INS-792

chr6-88482537-INS-225
chr6-88483301-INS-70
chr6-88489056-INS-151
chr6-88498478-INS-174
chr6-88500016-INS-93
chr6-88507240-INS-66
chr6-88510468-INS-1158
chr6-88510693-INS-939
chr6-88518925-INS-598
chr6-88564880-INS-668
chr6-88637201-INS-587
chr6-88664974-INS-62
chr6-88797689-INS-76
chr6-88846015-INS-78
chr6-88881479-INS-52
chr6-88891129-INS-78
chr6-88922669-INS-228
chr6-88969640-INS-333
chr6-88972738-INS-101
chr6-88981374-INS-172
chr6-89000375-INS-214
chr6-89016009-INS-96
chr6-89029988-INS-6878
chr6-89055632-INS-53
chr6-89058440-INS-76
chr6-89069454-INS-65
chr6-89073719-INS-150
chr6-89104225-INS-61
chr6-89142069-INS-107
chr6-89164289-INS-681
chr6-89185247-INS-492
chr6-89185952-INS-132
chr6-89232917-INS-54
chr6-89352964-INS-56
chr6-89369458-INS-187
chr6-89465294-INS-352
chr6-89539945-INS-102
chr6-89556267-INS-164
chr6-89567623-INS-188
chr6-89599719-INS-64
chr6-89618318-INS-64
chr6-89630184-INS-55
chr6-89653377-INS-7204
chr6-89688320-INS-64
chr6-89696422-INS-54
chr6-89721543-INS-52
chr6-89722891-INS-54
chr6-89729781-INS-57
chr6-89747394-INS-105
chr6-89750318-INS-305
chr6-89766924-INS-68
chr6-89771925-INS-7158
chr6-89776351-INS-112
chr6-89802416-INS-562
chr6-89854156-INS-234
chr6-89856029-INS-192
chr6-89864426-INS-81
chr6-89887922-INS-624
chr6-89913428-INS-662
chr6-89944147-INS-52
chr6-89986287-INS-60
chr6-90011228-INS-424
chr6-90020028-INS-6272
chr6-90023245-INS-56
chr6-90043175-INS-52
chr6-90047218-INS-121
chr6-90140495-INS-6875
chr6-90143467-INS-7235
chr6-90146782-INS-2342
chr6-90172232-INS-555
chr6-90186630-INS-247
chr6-90199227-INS-5159
chr6-90199998-INS-89
chr6-90238843-INS-171
chr6-90244689-INS-58
chr6-90276579-INS-124
chr6-90298004-INS-59

chr6-90334857-INS-54
chr6-90391077-INS-295
chr6-90413992-INS-100
chr6-90414159-INS-5433
chr6-90436284-INS-444
chr6-90441894-INS-210
chr6-90470221-INS-229
chr6-90486715-INS-768
chr6-90495652-INS-2400
chr6-90521435-INS-120
chr6-90522595-INS-405
chr6-90534494-INS-401
chr6-90551069-INS-74
chr6-90553896-INS-297
chr6-90572185-INS-78
chr6-90597654-INS-611
chr6-90604747-INS-56
chr6-90633165-INS-196
chr6-90635724-INS-196
chr6-90651965-INS-52
chr6-90752442-INS-191
chr6-90771455-INS-60
chr6-90816865-INS-7159
chr6-90824198-INS-364
chr6-90829934-INS-2227
chr6-90836008-INS-114
chr6-90841261-INS-90
chr6-90878795-INS-52
chr6-90892504-INS-79
chr6-90904341-INS-204
chr6-90911172-INS-70
chr6-90934458-INS-330
chr6-90958250-INS-264
chr6-90995151-INS-64
chr6-91010999-INS-59
chr6-91012468-INS-131
chr6-91027519-INS-153
chr6-91035273-INS-54
chr6-91041109-INS-104
chr6-91067654-INS-726
chr6-91073271-INS-231
chr6-91103449-INS-581
chr6-91104691-INS-64
chr6-91113044-INS-205
chr6-91121852-INS-120
chr6-91154303-INS-81
chr6-91172252-INS-60
chr6-91189429-INS-172
chr6-91214544-INS-159
chr6-91224855-INS-54
chr6-91250887-INS-513
chr6-91282177-INS-58
chr6-91294693-INS-852
chr6-91310356-INS-77
chr6-91340023-INS-227
chr6-91348064-INS-106
chr6-91349366-INS-143
chr6-91351317-INS-94
chr6-91363359-INS-110
chr6-91396101-INS-2414
chr6-91402454-INS-198
chr6-91417044-INS-704
chr6-91417746-INS-5595
chr6-91425603-INS-218
chr6-91436582-INS-60
chr6-91447943-INS-200
chr6-91477658-INS-209
chr6-91508077-INS-215
chr6-91513953-INS-144
chr6-91522566-INS-2649
chr6-91532486-INS-1888
chr6-91586383-INS-152
chr6-91591683-INS-289
chr6-91645110-INS-835
chr6-91653346-INS-212
chr6-91657878-INS-62
chr6-91675710-INS-84

chr6-91694819-INS-206
chr6-91698127-INS-53
chr6-91723634-INS-98
chr6-91754804-INS-136
chr6-91758632-INS-102
chr6-91812416-INS-419
chr6-91852395-INS-1864
chr6-91856143-INS-57
chr6-91872078-INS-65
chr6-91879575-INS-526
chr6-91885071-INS-257
chr6-91886912-INS-59
chr6-91893904-INS-211
chr6-91916721-INS-63
chr6-91947685-INS-101
chr6-91974354-INS-57
chr6-91982803-INS-374
chr6-91983543-INS-58
chr6-92002867-INS-453
chr6-92006928-INS-70
chr6-92026325-INS-72
chr6-92045819-INS-1864
chr6-92075270-INS-60
chr6-92101997-INS-198
chr6-92151624-INS-200
chr6-92157695-INS-93
chr6-92175922-INS-163
chr6-92177715-INS-781
chr6-92187191-INS-66
chr6-92203816-INS-203
chr6-92206647-INS-68
chr6-92214459-INS-70
chr6-92223857-INS-201
chr6-92227506-INS-217
chr6-92231706-INS-507
chr6-92265850-INS-115
chr6-92266683-INS-173
chr6-92274275-INS-228
chr6-92292641-INS-56
chr6-92297167-INS-191
chr6-92312590-INS-1048
chr6-92332003-INS-6564
chr6-92339027-INS-6201
chr6-92346271-INS-112
chr6-92409821-INS-51
chr6-92447440-INS-6771
chr6-92475220-INS-175
chr6-92480745-INS-77
chr6-92484659-INS-255
chr6-92489249-INS-400
chr6-92506898-INS-92
chr6-92618430-INS-6364
chr6-92672499-INS-63
chr6-92673930-INS-202
chr6-92683707-INS-52
chr6-92695511-INS-167
chr6-92699964-INS-214
chr6-92722659-INS-617
chr6-92759145-INS-266
chr6-92811967-INS-54
chr6-92862728-INS-64
chr6-92875082-INS-874
chr6-92877003-INS-80
chr6-92882026-INS-185
chr6-92884774-INS-53
chr6-92926268-INS-364
chr6-92934456-INS-83
chr6-92937523-INS-649
chr6-92941035-INS-277
chr6-92960297-INS-137
chr6-93014325-INS-102
chr6-93019283-INS-386
chr6-93086414-INS-64
chr6-93100011-INS-83
chr6-93100429-INS-189
chr6-93101926-INS-102
chr6-93113814-INS-104

chr6-93126519-INS-61
chr6-93138495-INS-145
chr6-93150367-INS-51
chr6-93158093-INS-102
chr6-93166956-INS-148
chr6-93171608-INS-107
chr6-93204578-INS-203
chr6-93232587-INS-967
chr6-93244993-INS-200
chr6-93275967-INS-82
chr6-93294191-INS-406
chr6-93313178-INS-52
chr6-93336299-INS-55
chr6-93337037-INS-2143
chr6-93341129-INS-200
chr6-93344484-INS-569
chr6-93349702-INS-6877
chr6-93363088-INS-112
chr6-93404330-INS-7475
chr6-93406489-INS-903
chr6-93407207-INS-87
chr6-93408912-INS-52
chr6-93486544-INS-7398
chr6-93489789-INS-62
chr6-93499386-INS-267
chr6-93510005-INS-572
chr6-93545282-INS-251
chr6-93547188-INS-56
chr6-93551643-INS-398
chr6-93588050-INS-66
chr6-93598037-INS-65
chr6-93600248-INS-104
chr6-93628532-INS-142
chr6-93646983-INS-612
chr6-93648126-INS-56
chr6-93684591-INS-375
chr6-93717393-INS-75
chr6-93718857-INS-81
chr6-93724663-INS-108
chr6-93760170-INS-432
chr6-93789230-INS-52
chr6-93789845-INS-262
chr6-93804118-INS-275
chr6-93860657-INS-549
chr6-93860924-INS-196
chr6-93864997-INS-74
chr6-93869270-INS-66
chr6-93877334-INS-58
chr6-93888070-INS-56
chr6-93911305-INS-98
chr6-93916307-INS-77
chr6-93917940-INS-2972
chr6-93941621-INS-184
chr6-93954196-INS-192
chr6-93972495-INS-86
chr6-93993373-INS-114
chr6-94000528-INS-64
chr6-94016353-INS-193
chr6-94018771-INS-145
chr6-94024849-INS-257
chr6-94031803-INS-170
chr6-94041562-INS-208
chr6-94106306-INS-198
chr6-94150338-INS-212
chr6-94152576-INS-91
chr6-94169021-INS-54
chr6-94194576-INS-127
chr6-94212349-INS-169
chr6-94253135-INS-216
chr6-94297928-INS-227
chr6-94305190-INS-4827
chr6-94320135-INS-400
chr6-94320248-INS-6583
chr6-94320249-INS-6152
chr6-94323174-INS-6653
chr6-94326122-INS-60
chr6-94382320-INS-171

chr6-94432454-INS-640
chr6-94443849-INS-84
chr6-94471842-INS-348
chr6-94485848-INS-385
chr6-94491208-INS-214
chr6-94534447-INS-68
chr6-94539294-INS-6596
chr6-94557449-INS-96
chr6-94561360-INS-99
chr6-94568011-INS-51
chr6-94607293-INS-58
chr6-94616876-INS-56
chr6-94667573-INS-113
chr6-94685716-INS-111
chr6-94694000-INS-128
chr6-94719203-INS-206
chr6-94728745-INS-85
chr6-94742533-INS-80
chr6-94742955-INS-91
chr6-94767938-INS-7103
chr6-94772995-INS-197
chr6-94775343-INS-76
chr6-94817181-INS-137
chr6-94840248-INS-207
chr6-94846094-INS-70
chr6-94851546-INS-55
chr6-94871266-INS-51
chr6-94904609-INS-68
chr6-94906480-INS-75
chr6-94913276-INS-92
chr6-94926382-INS-54
chr6-94932156-INS-470
chr6-94937161-INS-128
chr6-94957888-INS-216
chr6-95006360-INS-114
chr6-95014194-INS-213
chr6-95024570-INS-64
chr6-95032989-INS-136
chr6-95068823-INS-692
chr6-95083817-INS-237
chr6-95129128-INS-64
chr6-95171524-INS-58
chr6-95178645-INS-398
chr6-95182821-INS-587
chr6-95185779-INS-420
chr6-95195305-INS-930
chr6-95216737-INS-58
chr6-95332475-INS-68
chr6-95375917-INS-8061
chr6-95393679-INS-120
chr6-95415348-INS-112
chr6-95429479-INS-96
chr6-95449005-INS-7127
chr6-95462116-INS-85
chr6-95470786-INS-336
chr6-95514071-INS-6689
chr6-95565560-INS-52
chr6-95565565-INS-666
chr6-95578029-INS-513
chr6-95585617-INS-82
chr6-95591654-INS-64
chr6-95599968-INS-94
chr6-95603344-INS-318
chr6-95607011-INS-64
chr6-95616315-INS-983
chr6-95616996-INS-142
chr6-95628171-INS-136
chr6-95661123-INS-178
chr6-95670567-INS-507
chr6-95675818-INS-454
chr6-95677613-INS-54
chr6-95708604-INS-73
chr6-95724258-INS-504
chr6-95729097-INS-54
chr6-95736209-INS-70
chr6-95756702-INS-54
chr6-95757135-INS-4616

chr6-95761450-INS-693
chr6-95763944-INS-84
chr6-95781247-INS-739
chr6-95799746-INS-6880
chr6-95809419-INS-621
chr6-95834649-INS-68
chr6-95836180-INS-72
chr6-95858578-INS-56
chr6-95859042-INS-589
chr6-95863995-INS-56
chr6-95921253-INS-195
chr6-95952034-INS-706
chr6-96006523-INS-1888
chr6-96030473-INS-7057
chr6-96044853-INS-233
chr6-96047225-INS-80
chr6-96072279-INS-58
chr6-96078204-INS-62
chr6-96080546-INS-76
chr6-96107545-INS-87
chr6-96169777-INS-125
chr6-96182466-INS-66
chr6-96184039-INS-304
chr6-96193884-INS-268
chr6-96201430-INS-81
chr6-96206181-INS-54
chr6-96210727-INS-52
chr6-96225862-INS-288
chr6-96232822-INS-251
chr6-96253644-INS-163
chr6-96259581-INS-7008
chr6-96273676-INS-212
chr6-96273935-INS-1060
chr6-96281079-INS-641
chr6-96284467-INS-180
chr6-96289822-INS-5545
chr6-96315343-INS-289
chr6-96364240-INS-179
chr6-96377125-INS-69
chr6-96393398-INS-145
chr6-96400014-INS-68
chr6-96431621-INS-416
chr6-96436891-INS-593
chr6-96461171-INS-93
chr6-96514345-INS-79
chr6-96537193-INS-56
chr6-96549706-INS-58
chr6-96550407-INS-388
chr6-96553753-INS-66
chr6-96562389-INS-9744
chr6-96591083-INS-419
chr6-96592711-INS-159
chr6-96652323-INS-60
chr6-96679829-INS-197
chr6-96715980-INS-352
chr6-96720491-INS-51
chr6-96723918-INS-144
chr6-96747764-INS-342
chr6-96749024-INS-897
chr6-96760226-INS-1486
chr6-96779922-INS-76
chr6-96812801-INS-92
chr6-96824246-INS-474
chr6-96861242-INS-124
chr6-96898418-INS-1167
chr6-96914341-INS-371
chr6-96923846-INS-191
chr6-96925236-INS-61
chr6-96938281-INS-2751
chr6-96946743-INS-214
chr6-96955179-INS-80
chr6-96960125-INS-60
chr6-96962228-INS-188
chr6-96967851-INS-56
chr6-96968929-INS-624
chr6-96975342-INS-394
chr6-96988216-INS-483

chr6-96995573-INS-62
chr6-97011402-INS-112
chr6-97013201-INS-54
chr6-97033871-INS-1179
chr6-97046849-INS-112
chr6-97053745-INS-136
chr6-97072717-INS-116
chr6-97085162-INS-710
chr6-97085530-INS-5563
chr6-97133650-INS-222
chr6-97136093-INS-84
chr6-97139241-INS-259
chr6-97151627-INS-84
chr6-97171366-INS-123
chr6-97184078-INS-58
chr6-97194666-INS-75
chr6-97201246-INS-52
chr6-97205791-INS-4194
chr6-97222972-INS-6409
chr6-97226293-INS-114
chr6-97249189-INS-95
chr6-97257274-INS-294
chr6-97260117-INS-56
chr6-97275312-INS-7184
chr6-97301914-INS-88
chr6-97313483-INS-399
chr6-97314764-INS-224
chr6-97343499-INS-53
chr6-97345148-INS-258
chr6-97346325-INS-68
chr6-97365904-INS-54
chr6-97387716-INS-111
chr6-97397916-INS-62
chr6-97426519-INS-132
chr6-97445605-INS-400
chr6-97448717-INS-296
chr6-97455852-INS-54
chr6-97466827-INS-68
chr6-97470282-INS-69
chr6-97480778-INS-143
chr6-97481666-INS-6423
chr6-97488615-INS-207
chr6-97520453-INS-416
chr6-97525192-INS-146
chr6-97527831-INS-106
chr6-97540614-INS-74
chr6-97545501-INS-176
chr6-97562337-INS-463
chr6-97611873-INS-400
chr6-97613057-INS-637
chr6-97615619-INS-62
chr6-97635866-INS-957
chr6-97649496-INS-58
chr6-97661825-INS-191
chr6-97665744-INS-71
chr6-97686927-INS-125
chr6-97693398-INS-197
chr6-97698579-INS-98
chr6-97699812-INS-66
chr6-97703759-INS-173
chr6-97711298-INS-370
chr6-97732878-INS-346
chr6-97742392-INS-428
chr6-97743039-INS-449
chr6-97743362-INS-101
chr6-97751494-INS-242
chr6-97758643-INS-253
chr6-97764931-INS-375
chr6-97769774-INS-62
chr6-97781612-INS-108
chr6-97787872-INS-59
chr6-97800105-INS-58
chr6-97802805-INS-90
chr6-97830972-INS-52
chr6-97840177-INS-6442
chr6-97851826-INS-54
chr6-97852289-INS-8106

chr6-97857846-INS-313
chr6-97859829-INS-6169
chr6-97866719-INS-86
chr6-97897059-INS-7331
chr6-97902394-INS-5255
chr6-97909054-INS-7034
chr6-97930446-INS-219
chr6-97948004-INS-222
chr6-97952347-INS-7239
chr6-97953679-INS-120
chr6-97966710-INS-368
chr6-97977109-INS-60
chr6-98011351-INS-6992
chr6-98020155-INS-516
chr6-98041731-INS-738
chr6-98059567-INS-72
chr6-98063051-INS-345
chr6-98092471-INS-198
chr6-98105598-INS-6520
chr6-98113212-INS-192
chr6-98119029-INS-260
chr6-98139619-INS-350
chr6-98173794-INS-78
chr6-98193192-INS-134
chr6-98205218-INS-60
chr6-98222883-INS-123
chr6-98227418-INS-480
chr6-98258636-INS-3180
chr6-98260495-INS-75
chr6-98286781-INS-382
chr6-98293122-INS-158
chr6-98294053-INS-387
chr6-98301106-INS-419
chr6-98311928-INS-178
chr6-98336958-INS-6578
chr6-98345236-INS-108
chr6-98358922-INS-5219
chr6-98383335-INS-6099
chr6-98392116-INS-342
chr6-98402281-INS-1481
chr6-98409169-INS-68
chr6-98434645-INS-61
chr6-98435038-INS-6442
chr6-98437412-INS-308
chr6-98439004-INS-604
chr6-98442964-INS-127
chr6-98472371-INS-229
chr6-98495434-INS-64
chr6-98527958-INS-52
chr6-98543628-INS-52
chr6-98552565-INS-64
chr6-98572496-INS-87
chr6-98579731-INS-8467
chr6-98582370-INS-72
chr6-98591876-INS-341
chr6-98599640-INS-100
chr6-98619800-INS-56
chr6-98644277-INS-8419
chr6-98672727-INS-200
chr6-98712064-INS-360
chr6-98775532-INS-528
chr6-98779565-INS-172
chr6-98784831-INS-149
chr6-98792527-INS-147
chr6-98810112-INS-195
chr6-98819171-INS-54
chr6-98824674-INS-6587
chr6-98825489-INS-56
chr6-98835541-INS-1886
chr6-98861305-INS-3219
chr6-98869768-INS-271
chr6-98900667-INS-90
chr6-98910711-INS-171
chr6-98921422-INS-57
chr6-98963401-INS-60
chr6-98970223-INS-53
chr6-99015983-INS-233

chr6-99027922-INS-54
chr6-99029521-INS-85
chr6-99054900-INS-281
chr6-99075601-INS-224
chr6-99110693-INS-51
chr6-99128132-INS-56
chr6-99133188-INS-8216
chr6-99192940-INS-170
chr6-99259852-INS-62
chr6-99275349-INS-208
chr6-99314000-INS-100
chr6-99331278-INS-65
chr6-99335309-INS-131
chr6-99392960-INS-302
chr6-99423047-INS-110
chr6-99470507-INS-331
chr6-99488194-INS-7159
chr6-99524394-INS-400
chr6-99574297-INS-150
chr6-99611799-INS-122
chr6-99615795-INS-116
chr6-99685269-INS-202
chr6-99715413-INS-60
chr6-99746267-INS-64
chr6-99763280-INS-87
chr6-99768067-INS-118
chr6-99777774-INS-134
chr6-99783268-INS-57
chr6-99787389-INS-77
chr6-99790705-INS-429
chr6-99810762-INS-52
chr6-99816898-INS-980
chr6-99825116-INS-374
chr6-99837912-INS-69
chr6-99843598-INS-67
chr6-99872177-INS-184
chr6-99876172-INS-72
chr6-99892603-INS-103
chr6-99942395-INS-66
chr6-99942610-INS-261
chr6-99974080-INS-63
chr6-99979416-INS-66
chr6-99984571-INS-88
chr6-99994989-INS-234
chr6-100009590-INS-145
chr6-100032518-INS-54
chr6-100034809-INS-53
chr6-100038444-INS-60
chr6-100043774-INS-76
chr6-100065419-INS-53
chr6-100066561-INS-80
chr6-100091539-INS-133
chr6-100098768-INS-56
chr6-100131502-INS-72
chr6-100139708-INS-204
chr6-100145182-INS-209
chr6-100168947-INS-182
chr6-100176600-INS-296
chr6-100181840-INS-262
chr6-100252948-INS-198
chr6-100276717-INS-62
chr6-100297736-INS-90
chr6-100300506-INS-221
chr6-100305680-INS-188
chr6-100306524-INS-8734
chr6-100314355-INS-52
chr6-100316732-INS-178
chr6-100320873-INS-480
chr6-100321622-INS-130
chr6-100330692-INS-54
chr6-100360414-INS-6442
chr6-100396188-INS-69
chr6-100398231-INS-255
chr6-100400815-INS-108
chr6-100411846-INS-229
chr6-100413271-INS-87
chr6-100437816-INS-72

chr6-100440562-INS-55
chr6-100444707-INS-201
chr6-100458130-INS-88
chr6-100486175-INS-496
chr6-100489387-INS-219
chr6-100495288-INS-63
chr6-100498007-INS-67
chr6-100511392-INS-124
chr6-100516226-INS-90
chr6-100521398-INS-341
chr6-100522699-INS-68
chr6-100538465-INS-206
chr6-100556302-INS-52
chr6-100587774-INS-441
chr6-100590978-INS-56
chr6-100600976-INS-78
chr6-100627429-INS-400
chr6-100630788-INS-397
chr6-100642798-INS-489
chr6-100668247-INS-190
chr6-100669869-INS-5372
chr6-100674219-INS-328
chr6-100676996-INS-83
chr6-100695789-INS-52
chr6-100725694-INS-6423
chr6-100753040-INS-55
chr6-100754674-INS-84
chr6-100772275-INS-280
chr6-100780525-INS-52
chr6-100802189-INS-166
chr6-100806476-INS-214
chr6-100810047-INS-613
chr6-100816022-INS-78
chr6-100830518-INS-53
chr6-100835519-INS-608
chr6-100863587-INS-54
chr6-100873666-INS-6327
chr6-100879713-INS-54
chr6-100880197-INS-100
chr6-100889817-INS-64
chr6-100897235-INS-120
chr6-100914654-INS-144
chr6-100926660-INS-70
chr6-100928513-INS-411
chr6-100932563-INS-66
chr6-100940082-INS-141
chr6-100956907-INS-72
chr6-100962072-INS-853
chr6-100977234-INS-6676
chr6-101000261-INS-91
chr6-101014866-INS-524
chr6-101046270-INS-104
chr6-101051167-INS-109
chr6-101055165-INS-54
chr6-101061003-INS-53
chr6-101078210-INS-395
chr6-101099897-INS-207
chr6-101102078-INS-70
chr6-101140484-INS-78
chr6-101156056-INS-102
chr6-101170584-INS-639
chr6-101179631-INS-72
chr6-101188713-INS-228
chr6-101191193-INS-116
chr6-101223437-INS-428
chr6-101224749-INS-138
chr6-101243708-INS-58
chr6-101259382-INS-80
chr6-101293487-INS-59
chr6-101336136-INS-106
chr6-101341341-INS-212
chr6-101357454-INS-60
chr6-101368703-INS-117
chr6-101383013-INS-51
chr6-101437298-INS-77
chr6-101448565-INS-419
chr6-101454081-INS-52

chr6-101459133-INS-101
chr6-101472878-INS-68
chr6-101486119-INS-54
chr6-101497243-INS-57
chr6-101506550-INS-7435
chr6-101511628-INS-569
chr6-101521048-INS-204
chr6-101530864-INS-87
chr6-101547064-INS-6388
chr6-101548099-INS-971
chr6-101550227-INS-2220
chr6-101556824-INS-6520
chr6-101557346-INS-708
chr6-101559258-INS-92
chr6-101573509-INS-65
chr6-101591431-INS-154
chr6-101598264-INS-244
chr6-101613390-INS-366
chr6-101616181-INS-248
chr6-101631430-INS-7201
chr6-101670295-INS-165
chr6-101705622-INS-461
chr6-101715934-INS-59
chr6-101716954-INS-214
chr6-101727301-INS-72
chr6-101727494-INS-454
chr6-101754791-INS-76
chr6-101757688-INS-62
chr6-101759055-INS-8008
chr6-101784312-INS-510
chr6-101785559-INS-257
chr6-101793618-INS-927
chr6-101802198-INS-402
chr6-101802566-INS-159
chr6-101831862-INS-99
chr6-101839201-INS-5412
chr6-101846876-INS-6594
chr6-101854596-INS-60
chr6-101857086-INS-60
chr6-101863207-INS-95
chr6-101886460-INS-6401
chr6-101895144-INS-137
chr6-101921909-INS-6085
chr6-101966612-INS-58
chr6-101969747-INS-71
chr6-102010418-INS-52
chr6-102024968-INS-52
chr6-102037723-INS-498
chr6-102070410-INS-668
chr6-102084470-INS-178
chr6-102123096-INS-7796
chr6-102147989-INS-51
chr6-102188108-INS-399
chr6-102202043-INS-1556
chr6-102219384-INS-60
chr6-102239303-INS-273
chr6-102240964-INS-59
chr6-102256040-INS-50
chr6-102257306-INS-78
chr6-102260871-INS-94
chr6-102262222-INS-362
chr6-102264066-INS-6166
chr6-102269616-INS-420
chr6-102284822-INS-93
chr6-102295384-INS-80
chr6-102310704-INS-696
chr6-102315648-INS-968
chr6-102334439-INS-189
chr6-102337211-INS-377
chr6-102343718-INS-123
chr6-102374204-INS-5549
chr6-102376618-INS-58
chr6-102377282-INS-1892
chr6-102401288-INS-60
chr6-102414151-INS-72
chr6-102417007-INS-71
chr6-102424502-INS-139

chr6-102430292-INS-6771
chr6-102434071-INS-56
chr6-102438284-INS-63
chr6-102438285-INS-604
chr6-102476362-INS-56
chr6-102484710-INS-6670
chr6-102485761-INS-491
chr6-102489602-INS-55
chr6-102491206-INS-882
chr6-102501544-INS-318
chr6-102505169-INS-195
chr6-102508629-INS-96
chr6-102515597-INS-133
chr6-102545045-INS-62
chr6-102571491-INS-7166
chr6-102581066-INS-180
chr6-102595337-INS-212
chr6-102600886-INS-75
chr6-102629208-INS-342
chr6-102635242-INS-56
chr6-102650871-INS-198
chr6-102671017-INS-102
chr6-102673303-INS-145
chr6-102684447-INS-166
chr6-102686554-INS-96
chr6-102738657-INS-229
chr6-102753116-INS-103
chr6-102776619-INS-4162
chr6-102785031-INS-66
chr6-102797365-INS-1862
chr6-102848553-INS-261
chr6-102888960-INS-2850
chr6-102892370-INS-185
chr6-102910234-INS-190
chr6-102917503-INS-93
chr6-102927563-INS-7143
chr6-102934674-INS-52
chr6-102939120-INS-201
chr6-102952619-INS-2765
chr6-102974747-INS-52
chr6-103003622-INS-91
chr6-103045429-INS-212
chr6-103049173-INS-120
chr6-103120117-INS-55
chr6-103131962-INS-56
chr6-103154064-INS-66
chr6-103162345-INS-75
chr6-103187567-INS-220
chr6-103207572-INS-67
chr6-103234821-INS-446
chr6-103247874-INS-55
chr6-103256261-INS-259
chr6-103280246-INS-74
chr6-103286290-INS-217
chr6-103287244-INS-52
chr6-103308826-INS-204
chr6-103313513-INS-180
chr6-103381803-INS-523
chr6-103473314-INS-57
chr6-103476291-INS-60
chr6-103479245-INS-268
chr6-103508167-INS-51
chr6-103513262-INS-75
chr6-103528266-INS-324
chr6-103530406-INS-405
chr6-103543441-INS-223
chr6-103570138-INS-6577
chr6-103595932-INS-74
chr6-103598438-INS-84
chr6-103612977-INS-446
chr6-103630210-INS-72
chr6-103632105-INS-107
chr6-103635645-INS-396
chr6-103640355-INS-914
chr6-103643715-INS-76
chr6-103650456-INS-229
chr6-103652870-INS-65

chr6-103660507-INS-60
chr6-103723948-INS-64
chr6-103731341-INS-55
chr6-103733235-INS-74
chr6-103742606-INS-788
chr6-103790811-INS-147
chr6-103805861-INS-144
chr6-103819225-INS-6685
chr6-103831300-INS-84
chr6-103856054-INS-6465
chr6-103869614-INS-252
chr6-103885322-INS-602
chr6-103902361-INS-5582
chr6-103905897-INS-80
chr6-103909105-INS-127
chr6-103920138-INS-105
chr6-103923274-INS-69
chr6-103925977-INS-535
chr6-103932082-INS-152
chr6-103945944-INS-255
chr6-103987739-INS-535
chr6-103998669-INS-61
chr6-104021776-INS-64
chr6-104057234-INS-851
chr6-104072913-INS-320
chr6-104100901-INS-100
chr6-104109049-INS-79
chr6-104116828-INS-82
chr6-104147371-INS-83
chr6-104154390-INS-54
chr6-104157370-INS-60
chr6-104168023-INS-410
chr6-104210753-INS-377
chr6-104213443-INS-1888
chr6-104228639-INS-70
chr6-104232710-INS-104
chr6-104276678-INS-196
chr6-104314728-INS-98
chr6-104328012-INS-159
chr6-104331734-INS-6485
chr6-104344926-INS-106
chr6-104353278-INS-711
chr6-104368997-INS-67
chr6-104396305-INS-286
chr6-104403906-INS-415
chr6-104404924-INS-329
chr6-104408312-INS-6295
chr6-104408908-INS-84
chr6-104414535-INS-3327
chr6-104437394-INS-56
chr6-104440154-INS-135
chr6-104548976-INS-559
chr6-104564691-INS-217
chr6-104591477-INS-117
chr6-104603438-INS-72
chr6-104609633-INS-233
chr6-104622687-INS-77
chr6-104625507-INS-395
chr6-104627884-INS-80
chr6-104647505-INS-179
chr6-104649419-INS-83
chr6-104658423-INS-338
chr6-104689673-INS-120
chr6-104695599-INS-160
chr6-104699220-INS-60
chr6-104712822-INS-995
chr6-104714155-INS-293
chr6-104728512-INS-153
chr6-104740399-INS-951
chr6-104750747-INS-375
chr6-104756122-INS-76
chr6-104760138-INS-92
chr6-104786751-INS-314
chr6-104817963-INS-255
chr6-104819139-INS-111
chr6-104824478-INS-449
chr6-104828153-INS-225

chr6-104854810-INS-99
chr6-104858077-INS-1708
chr6-104864093-INS-60
chr6-104874473-INS-5277
chr6-104942453-INS-501
chr6-104945846-INS-91
chr6-104955408-INS-441
chr6-104964825-INS-114
chr6-104969608-INS-387
chr6-104979383-INS-240
chr6-104981129-INS-62
chr6-104985802-INS-643
chr6-105000802-INS-63
chr6-105000818-INS-1755
chr6-105001845-INS-3723
chr6-105013565-INS-342
chr6-105015836-INS-65
chr6-105017976-INS-459
chr6-105044019-INS-1792
chr6-105055496-INS-208
chr6-105060220-INS-265
chr6-105084464-INS-114
chr6-105096060-INS-342
chr6-105101844-INS-76
chr6-105103033-INS-1844
chr6-105109705-INS-1762
chr6-105125031-INS-62
chr6-105147433-INS-76
chr6-105167151-INS-6667
chr6-105179212-INS-261
chr6-105185796-INS-2152
chr6-105192871-INS-60
chr6-105213576-INS-201
chr6-105255126-INS-423
chr6-105282427-INS-386
chr6-105322352-INS-584
chr6-105323819-INS-6332
chr6-105325508-INS-98
chr6-105338949-INS-102
chr6-105346279-INS-140
chr6-105364077-INS-587
chr6-105373373-INS-260
chr6-105376347-INS-128
chr6-105378049-INS-420
chr6-105388139-INS-389
chr6-105404309-INS-207
chr6-105450051-INS-525
chr6-105463871-INS-222
chr6-105473893-INS-89
chr6-105506098-INS-136
chr6-105511999-INS-75
chr6-105520614-INS-193
chr6-105531784-INS-939
chr6-105585042-INS-72
chr6-105586588-INS-235
chr6-105595447-INS-428
chr6-105616637-INS-65
chr6-105629619-INS-90
chr6-105658445-INS-7633
chr6-105660736-INS-634
chr6-105666336-INS-114
chr6-105668111-INS-398
chr6-105669445-INS-52
chr6-105690419-INS-341
chr6-105702535-INS-72
chr6-105703874-INS-153
chr6-105710733-INS-2492
chr6-105747865-INS-153
chr6-105757473-INS-326
chr6-105782658-INS-84
chr6-105789229-INS-6298
chr6-105794024-INS-135
chr6-105802004-INS-52
chr6-105825485-INS-380
chr6-105828144-INS-62
chr6-105837661-INS-55
chr6-105857065-INS-122

chr6-105861922-INS-6912
chr6-105872261-INS-58
chr6-105882690-INS-71
chr6-105885225-INS-451
chr6-105910131-INS-52
chr6-105920386-INS-54
chr6-105921590-INS-58
chr6-105974793-INS-98
chr6-105982465-INS-110
chr6-106017827-INS-124
chr6-106020657-INS-615
chr6-106037758-INS-170
chr6-106046160-INS-269
chr6-106048654-INS-452
chr6-106052799-INS-841
chr6-106057835-INS-56
chr6-106089452-INS-343
chr6-106100960-INS-584
chr6-106120051-INS-55
chr6-106129128-INS-5998
chr6-106137277-INS-57
chr6-106143032-INS-132
chr6-106146282-INS-7073
chr6-106147621-INS-7329
chr6-106152090-INS-6460
chr6-106172056-INS-86
chr6-106184645-INS-528
chr6-106188208-INS-324
chr6-106206832-INS-112
chr6-106210111-INS-104
chr6-106240189-INS-76
chr6-106250402-INS-66
chr6-106252659-INS-68
chr6-106257809-INS-64
chr6-106283087-INS-238
chr6-106320381-INS-64
chr6-106323302-INS-548
chr6-106323326-INS-4646
chr6-106325790-INS-317
chr6-106342287-INS-434
chr6-106364383-INS-652
chr6-106385056-INS-307
chr6-106389508-INS-56
chr6-106419429-INS-198
chr6-106422446-INS-270
chr6-106438872-INS-75
chr6-106447252-INS-440
chr6-106451201-INS-60
chr6-106454084-INS-767
chr6-106464268-INS-56
chr6-106488674-INS-90
chr6-106509524-INS-101
chr6-106533168-INS-6686
chr6-106549648-INS-177
chr6-106569728-INS-55
chr6-106600246-INS-76
chr6-106613113-INS-400
chr6-106629885-INS-52
chr6-106634255-INS-487
chr6-106641012-INS-753
chr6-106669605-INS-563
chr6-106690449-INS-280
chr6-106702321-INS-58
chr6-106714430-INS-64
chr6-106777988-INS-5292
chr6-106800798-INS-866
chr6-106803206-INS-112
chr6-106816958-INS-52
chr6-106836889-INS-458
chr6-106848412-INS-57
chr6-106850919-INS-563
chr6-106907416-INS-253
chr6-106911637-INS-7086
chr6-106914658-INS-6677
chr6-106926242-INS-55
chr6-106927054-INS-82
chr6-106932857-INS-67

chr6-106935349-INS-6486
chr6-106949148-INS-60
chr6-106960579-INS-358
chr6-106965085-INS-68
chr6-106969594-INS-216
chr6-106986827-INS-6461
chr6-107009764-INS-6396
chr6-107025329-INS-93
chr6-107026518-INS-54
chr6-107058448-INS-175
chr6-107086225-INS-3964
chr6-107109077-INS-56
chr6-107116019-INS-1883
chr6-107194338-INS-745
chr6-107197140-INS-633
chr6-107216760-INS-85
chr6-107221841-INS-309
chr6-107234605-INS-62
chr6-107237218-INS-74
chr6-107244743-INS-509
chr6-107245944-INS-520
chr6-107250593-INS-98
chr6-107257683-INS-160
chr6-107284442-INS-126
chr6-107288306-INS-75
chr6-107296590-INS-4505
chr6-107298986-INS-51
chr6-107304922-INS-65
chr6-107322459-INS-54
chr6-107377098-INS-268
chr6-107387113-INS-430
chr6-107389374-INS-58
chr6-107406590-INS-728
chr6-107412592-INS-96
chr6-107438068-INS-208
chr6-107456534-INS-75
chr6-107457936-INS-102
chr6-107487576-INS-73
chr6-107490306-INS-83
chr6-107510939-INS-74
chr6-107514489-INS-64
chr6-107516144-INS-156
chr6-107524654-INS-71
chr6-107532894-INS-6664
chr6-107589009-INS-54
chr6-107625257-INS-283
chr6-107641504-INS-1599
chr6-107666998-INS-444
chr6-107672609-INS-454
chr6-107674039-INS-53
chr6-107680163-INS-60
chr6-107693706-INS-982
chr6-107697462-INS-68
chr6-107704571-INS-170
chr6-107721260-INS-51
chr6-107743424-INS-86
chr6-107777135-INS-56
chr6-107843672-INS-784
chr6-107850272-INS-302
chr6-107866690-INS-520
chr6-107889274-INS-424
chr6-107912529-INS-6219
chr6-107930208-INS-6237
chr6-107936880-INS-139
chr6-107955761-INS-764
chr6-107995251-INS-7142
chr6-107999883-INS-763
chr6-108005501-INS-399
chr6-108104671-INS-343
chr6-108128834-INS-60
chr6-108136753-INS-673
chr6-108157285-INS-144
chr6-108163495-INS-54
chr6-108173252-INS-524
chr6-108201369-INS-197
chr6-108217576-INS-134
chr6-108220721-INS-199

chr6-108274138-INS-50
chr6-108280700-INS-125
chr6-108321127-INS-185
chr6-108323874-INS-255
chr6-108331780-INS-56
chr6-108337912-INS-219
chr6-108339672-INS-63
chr6-108374956-INS-83
chr6-108381474-INS-78
chr6-108384559-INS-60
chr6-108413845-INS-52
chr6-108473412-INS-397
chr6-108498180-INS-398
chr6-108511159-INS-54
chr6-108521105-INS-72
chr6-108529452-INS-234
chr6-108543457-INS-92
chr6-108571054-INS-110
chr6-108581441-INS-121
chr6-108593173-INS-61
chr6-108603563-INS-125
chr6-108645320-INS-54
chr6-108647091-INS-764
chr6-108648073-INS-76
chr6-108650091-INS-223
chr6-108653480-INS-155
chr6-108670556-INS-62
chr6-108695116-INS-84
chr6-108701861-INS-230
chr6-108704077-INS-237
chr6-108720029-INS-798
chr6-108734705-INS-151
chr6-108738871-INS-55
chr6-108742064-INS-124
chr6-108751065-INS-69
chr6-108755991-INS-192
chr6-108780133-INS-68
chr6-108817053-INS-345
chr6-108826562-INS-57
chr6-108826965-INS-60
chr6-108840528-INS-75
chr6-108846799-INS-52
chr6-108850140-INS-137
chr6-108855039-INS-148
chr6-108881837-INS-178
chr6-108889371-INS-76
chr6-108893135-INS-124
chr6-108893266-INS-1008
chr6-108893292-INS-953
chr6-108906007-INS-266
chr6-108926921-INS-184
chr6-108950243-INS-6594
chr6-108959517-INS-2380
chr6-108972466-INS-443
chr6-108976568-INS-167
chr6-108978655-INS-302
chr6-109015003-INS-308
chr6-109058659-INS-203
chr6-109062364-INS-1253
chr6-109064380-INS-6109
chr6-109076112-INS-128
chr6-109082928-INS-6837
chr6-109101083-INS-725
chr6-109132792-INS-707
chr6-109135482-INS-617
chr6-109156824-INS-204
chr6-109183794-INS-126
chr6-109214690-INS-106
chr6-109223488-INS-658
chr6-109243650-INS-230
chr6-109284649-INS-407
chr6-109287660-INS-244
chr6-109328364-INS-8116
chr6-109329520-INS-64
chr6-109333041-INS-2268
chr6-109352079-INS-751
chr6-109355219-INS-1866

chr6-109363021-INS-6213
chr6-109393498-INS-1489
chr6-109418063-INS-559
chr6-109422600-INS-54
chr6-109428542-INS-6362
chr6-109431565-INS-5591
chr6-109453430-INS-102
chr6-109488962-INS-119
chr6-109496047-INS-53
chr6-109511336-INS-53
chr6-109512000-INS-626
chr6-109515600-INS-98
chr6-109568504-INS-214
chr6-109579689-INS-99
chr6-109632680-INS-501
chr6-109641512-INS-56
chr6-109642826-INS-76
chr6-109670212-INS-489
chr6-109693201-INS-4584
chr6-109698232-INS-80
chr6-109709528-INS-115
chr6-109719404-INS-373
chr6-109723204-INS-220
chr6-109751840-INS-157
chr6-109770852-INS-64
chr6-109778847-INS-54
chr6-109787042-INS-64
chr6-109792636-INS-60
chr6-109857390-INS-76
chr6-109874361-INS-65
chr6-109893872-INS-946
chr6-109910838-INS-53
chr6-109919266-INS-6424
chr6-109937922-INS-106
chr6-110006140-INS-253
chr6-110014768-INS-65
chr6-110018171-INS-112
chr6-110033993-INS-52
chr6-110050859-INS-130
chr6-110111489-INS-442
chr6-110121607-INS-53
chr6-110134539-INS-52
chr6-110169061-INS-96
chr6-110173173-INS-98
chr6-110272504-INS-60
chr6-110284560-INS-52
chr6-110317960-INS-126
chr6-110333193-INS-75
chr6-110372944-INS-68
chr6-110441676-INS-72
chr6-110456985-INS-60
chr6-110487329-INS-180
chr6-110489877-INS-195
chr6-110535878-INS-121
chr6-110538176-INS-93
chr6-110543170-INS-197
chr6-110546539-INS-75
chr6-110573643-INS-129
chr6-110582952-INS-633
chr6-110585772-INS-820
chr6-110637172-INS-148
chr6-110668576-INS-497
chr6-110682906-INS-195
chr6-110685991-INS-780
chr6-110693273-INS-153
chr6-110704760-INS-456
chr6-110706217-INS-66
chr6-110731457-INS-68
chr6-110763292-INS-262
chr6-110764593-INS-6461
chr6-110767843-INS-393
chr6-110774395-INS-159
chr6-110776025-INS-176
chr6-110779374-INS-59
chr6-110796616-INS-52
chr6-110821947-INS-498
chr6-110824533-INS-2331

chr6-110837088-INS-964
chr6-110840077-INS-79
chr6-110844855-INS-158
chr6-110854975-INS-1426
chr6-110862474-INS-141
chr6-110875970-INS-51
chr6-110958328-INS-986
chr6-110965322-INS-52
chr6-110968360-INS-403
chr6-110976546-INS-126
chr6-110987172-INS-429
chr6-110990504-INS-1272
chr6-111003721-INS-58
chr6-111009215-INS-80
chr6-111015928-INS-68
chr6-111019177-INS-627
chr6-111054142-INS-72
chr6-111080393-INS-6660
chr6-111084531-INS-7105
chr6-111087480-INS-102
chr6-111093002-INS-102
chr6-111109789-INS-52
chr6-111112875-INS-467
chr6-111148699-INS-575
chr6-111150694-INS-248
chr6-111158734-INS-164
chr6-111175528-INS-51
chr6-111197524-INS-196
chr6-111199942-INS-3341
chr6-111216403-INS-52
chr6-111236113-INS-54
chr6-111238124-INS-108
chr6-111239273-INS-385
chr6-111250362-INS-60
chr6-111266311-INS-174
chr6-111270237-INS-223
chr6-111275722-INS-323
chr6-111282742-INS-866
chr6-111285465-INS-115
chr6-111303379-INS-825
chr6-111326842-INS-52
chr6-111327847-INS-94
chr6-111338096-INS-62
chr6-111341085-INS-50
chr6-111366139-INS-6503
chr6-111379804-INS-108
chr6-111382763-INS-365
chr6-111384978-INS-99
chr6-111433322-INS-276
chr6-111484229-INS-74
chr6-111492799-INS-86
chr6-111499506-INS-60
chr6-111516279-INS-72
chr6-111517984-INS-52
chr6-111545831-INS-6646
chr6-111586597-INS-334
chr6-111629610-INS-343
chr6-111662563-INS-56
chr6-111672146-INS-61
chr6-111682441-INS-515
chr6-111744334-INS-457
chr6-111767297-INS-875
chr6-111780715-INS-327
chr6-111784321-INS-239
chr6-111796413-INS-2788
chr6-111803735-INS-1864
chr6-111803849-INS-430
chr6-111806848-INS-66
chr6-111818646-INS-7137
chr6-111837698-INS-55
chr6-111855046-INS-64
chr6-111860457-INS-104
chr6-111865009-INS-55
chr6-111898445-INS-64
chr6-111919368-INS-92
chr6-111992529-INS-142
chr6-112034597-INS-871

chr6-112058518-INS-52
chr6-112062593-INS-86
chr6-112076663-INS-52
chr6-112088646-INS-54
chr6-112089860-INS-64
chr6-112105532-INS-61
chr6-112111696-INS-80
chr6-112114287-INS-104
chr6-112116526-INS-254
chr6-112119483-INS-72
chr6-112164481-INS-6205
chr6-112182059-INS-341
chr6-112220323-INS-238
chr6-112237139-INS-285
chr6-112244461-INS-68
chr6-112299059-INS-66
chr6-112302621-INS-333
chr6-112317155-INS-77
chr6-112326740-INS-329
chr6-112345062-INS-139
chr6-112346567-INS-7090
chr6-112368843-INS-490
chr6-112375579-INS-123
chr6-112385883-INS-58
chr6-112391271-INS-280
chr6-112403974-INS-2219
chr6-112420112-INS-3029
chr6-112420267-INS-198
chr6-112428843-INS-174
chr6-112452787-INS-95
chr6-112457430-INS-169
chr6-112459238-INS-120
chr6-112472166-INS-125
chr6-112474202-INS-77
chr6-112477772-INS-65
chr6-112482604-INS-51
chr6-112493026-INS-579
chr6-112503305-INS-277
chr6-112505020-INS-116
chr6-112507880-INS-359
chr6-112508652-INS-158
chr6-112522203-INS-411
chr6-112536083-INS-114
chr6-112549885-INS-162
chr6-112592035-INS-125
chr6-112618490-INS-186
chr6-112647914-INS-400
chr6-112663465-INS-248
chr6-112672970-INS-77
chr6-112676722-INS-67
chr6-112719954-INS-820
chr6-112724566-INS-52
chr6-112748604-INS-193
chr6-112820699-INS-325
chr6-112837087-INS-164
chr6-112880886-INS-59
chr6-112882316-INS-54
chr6-112887387-INS-202
chr6-112891486-INS-324
chr6-112892478-INS-890
chr6-112904527-INS-5311
chr6-112911478-INS-58
chr6-112938594-INS-465
chr6-112942614-INS-200
chr6-112954964-INS-201
chr6-112956996-INS-178
chr6-112959391-INS-77
chr6-112971662-INS-69
chr6-112991891-INS-439
chr6-112998771-INS-70
chr6-113007253-INS-182
chr6-113016488-INS-144
chr6-113016558-INS-888
chr6-113041198-INS-221
chr6-113067764-INS-1888
chr6-113130544-INS-498
chr6-113190459-INS-59

chr6-113196207-INS-52
chr6-113200403-INS-445
chr6-113203656-INS-214
chr6-113204365-INS-70
chr6-113207163-INS-155
chr6-113226611-INS-81
chr6-113229908-INS-56
chr6-113272979-INS-207
chr6-113307704-INS-173
chr6-113314301-INS-211
chr6-113319275-INS-70
chr6-113323346-INS-212
chr6-113329369-INS-126
chr6-113332605-INS-201
chr6-113366881-INS-84
chr6-113374483-INS-7191
chr6-113379866-INS-202
chr6-113399378-INS-57
chr6-113403695-INS-104
chr6-113406384-INS-122
chr6-113414511-INS-170
chr6-113433283-INS-58
chr6-113445219-INS-52
chr6-113454824-INS-320
chr6-113501330-INS-212
chr6-113513548-INS-190
chr6-113515426-INS-178
chr6-113563136-INS-108
chr6-113596712-INS-184
chr6-113597570-INS-182
chr6-113598726-INS-440
chr6-113605782-INS-55
chr6-113610413-INS-193
chr6-113629107-INS-203
chr6-113634388-INS-92
chr6-113643099-INS-153
chr6-113645077-INS-68
chr6-113647801-INS-267
chr6-113649653-INS-139
chr6-113654700-INS-197
chr6-113658829-INS-175
chr6-113660971-INS-55
chr6-113676457-INS-52
chr6-113693601-INS-140
chr6-113696224-INS-62
chr6-113702197-INS-196
chr6-113741572-INS-136
chr6-113795599-INS-135
chr6-113796728-INS-1903
chr6-113797017-INS-491
chr6-113823390-INS-55
chr6-113825879-INS-6657
chr6-113826792-INS-84
chr6-113843831-INS-52
chr6-113878453-INS-75
chr6-113888676-INS-317
chr6-113918681-INS-60
chr6-113987806-INS-139
chr6-113988257-INS-78
chr6-114006841-INS-80
chr6-114010293-INS-120
chr6-114038204-INS-259
chr6-114046188-INS-84
chr6-114060865-INS-55
chr6-114097729-INS-402
chr6-114115609-INS-84
chr6-114117712-INS-277
chr6-114145216-INS-59
chr6-114145771-INS-849
chr6-114218911-INS-443
chr6-114232855-INS-200
chr6-114273299-INS-100
chr6-114310049-INS-66
chr6-114339210-INS-60
chr6-114372639-INS-76
chr6-114430410-INS-268
chr6-114438111-INS-328

chr6-114440543-INS-187
chr6-114443478-INS-75
chr6-114452529-INS-5129
chr6-114466451-INS-181
chr6-114482906-INS-62
chr6-114485277-INS-64
chr6-114558195-INS-58
chr6-114560542-INS-62
chr6-114590044-INS-55
chr6-114622418-INS-234
chr6-114626953-INS-214
chr6-114632262-INS-211
chr6-114649629-INS-60
chr6-114651775-INS-56
chr6-114656399-INS-179
chr6-114687526-INS-210
chr6-114692825-INS-84
chr6-114699001-INS-63
chr6-114732299-INS-64
chr6-114734004-INS-6894
chr6-114750649-INS-73
chr6-114767664-INS-274
chr6-114788807-INS-205
chr6-114795958-INS-5186
chr6-114804506-INS-52
chr6-114813245-INS-265
chr6-114820689-INS-892
chr6-114853492-INS-54
chr6-114854395-INS-225
chr6-114875626-INS-260
chr6-114887778-INS-183
chr6-114902738-INS-204
chr6-114916144-INS-66
chr6-114933300-INS-1802
chr6-114952781-INS-227
chr6-114955173-INS-124
chr6-114982068-INS-199
chr6-114990553-INS-58
chr6-115005875-INS-109
chr6-115027345-INS-6906
chr6-115035747-INS-72
chr6-115052562-INS-124
chr6-115072291-INS-84
chr6-115087195-INS-7882
chr6-115097540-INS-61
chr6-115106326-INS-68
chr6-115143969-INS-2634
chr6-115154942-INS-6731
chr6-115181307-INS-311
chr6-115199484-INS-60
chr6-115204815-INS-168
chr6-115211278-INS-56
chr6-115217519-INS-123
chr6-115217601-INS-645
chr6-115230780-INS-201
chr6-115233336-INS-72
chr6-115233690-INS-91
chr6-115250142-INS-188
chr6-115262673-INS-1886
chr6-115289577-INS-63
chr6-115290363-INS-60
chr6-115321074-INS-211
chr6-115323571-INS-604
chr6-115331921-INS-60
chr6-115347840-INS-124
chr6-115364892-INS-969
chr6-115378937-INS-400
chr6-115408522-INS-65
chr6-115425101-INS-680
chr6-115434093-INS-82
chr6-115510698-INS-51
chr6-115512635-INS-64
chr6-115517444-INS-218
chr6-115522150-INS-57
chr6-115527301-INS-203
chr6-115532482-INS-93
chr6-115539145-INS-6607

chr6-115539836-INS-103
chr6-115561523-INS-100
chr6-115563164-INS-66
chr6-115565324-INS-73
chr6-115582294-INS-179
chr6-115584187-INS-91
chr6-115643453-INS-54
chr6-115651382-INS-52
chr6-115683606-INS-7103
chr6-115684605-INS-51
chr6-115692942-INS-102
chr6-115695954-INS-1542
chr6-115696078-INS-69
chr6-115697027-INS-1890
chr6-115704400-INS-192
chr6-115712827-INS-88
chr6-115734409-INS-113
chr6-115737389-INS-5249
chr6-115738712-INS-415
chr6-115743381-INS-498
chr6-115748413-INS-164
chr6-115751334-INS-60
chr6-115751378-INS-58
chr6-115753283-INS-126
chr6-115779430-INS-217
chr6-115788470-INS-209
chr6-115809104-INS-105
chr6-115810799-INS-84
chr6-115819180-INS-210
chr6-115827068-INS-171
chr6-115861907-INS-102
chr6-115862396-INS-1084
chr6-115862798-INS-133
chr6-115867319-INS-129
chr6-115875067-INS-54
chr6-115898029-INS-264
chr6-115898619-INS-68
chr6-115899347-INS-67
chr6-115972251-INS-137
chr6-115974865-INS-68
chr6-115977520-INS-66
chr6-115992938-INS-150
chr6-115995448-INS-192
chr6-116005737-INS-517
chr6-116061097-INS-202
chr6-116065594-INS-195
chr6-116066908-INS-6432
chr6-116068267-INS-446
chr6-116106714-INS-184
chr6-116131776-INS-896
chr6-116146879-INS-5402
chr6-116153297-INS-539
chr6-116184147-INS-206
chr6-116188052-INS-224
chr6-116193548-INS-166
chr6-116201171-INS-223
chr6-116203467-INS-59
chr6-116207978-INS-379
chr6-116216181-INS-236
chr6-116219065-INS-891
chr6-116227423-INS-204
chr6-116228590-INS-155
chr6-116230456-INS-178
chr6-116231248-INS-183
chr6-116238139-INS-200
chr6-116247971-INS-103
chr6-116258805-INS-335
chr6-116259675-INS-51
chr6-116268233-INS-256
chr6-116299660-INS-2537
chr6-116305467-INS-291
chr6-116320082-INS-64
chr6-116323450-INS-78
chr6-116333740-INS-2621
chr6-116372186-INS-161
chr6-116374338-INS-58
chr6-116432726-INS-56

chr6-116448212-INS-7135
chr6-116456339-INS-5406
chr6-116457743-INS-211
chr6-116463998-INS-2226
chr6-116475983-INS-796
chr6-116492166-INS-296
chr6-116494745-INS-11288
chr6-116557792-INS-320
chr6-116600836-INS-382
chr6-116623614-INS-90
chr6-116643864-INS-199
chr6-116669456-INS-104
chr6-116673319-INS-115
chr6-116712092-INS-74
chr6-116717044-INS-79
chr6-116758622-INS-868
chr6-116786272-INS-132
chr6-116815894-INS-167
chr6-116816631-INS-366
chr6-116865943-INS-119
chr6-116903963-INS-425
chr6-116914131-INS-5326
chr6-116961522-INS-51
chr6-116965377-INS-313
chr6-116972071-INS-697
chr6-116972984-INS-456
chr6-116976328-INS-285
chr6-116976919-INS-66
chr6-116977920-INS-55
chr6-117024193-INS-66
chr6-117031105-INS-68
chr6-117036493-INS-107
chr6-117096663-INS-361
chr6-117105886-INS-126
chr6-117154173-INS-62
chr6-117171651-INS-99
chr6-117173465-INS-63
chr6-117175444-INS-302
chr6-117181460-INS-94
chr6-117190488-INS-109
chr6-117193311-INS-214
chr6-117210519-INS-203
chr6-117250240-INS-401
chr6-117265972-INS-82
chr6-117272289-INS-96
chr6-117280308-INS-435
chr6-117280752-INS-6238
chr6-117298876-INS-87
chr6-117327411-INS-651
chr6-117347139-INS-56
chr6-117353667-INS-52
chr6-117378827-INS-135
chr6-117418980-INS-6660
chr6-117419893-INS-400
chr6-117421188-INS-114
chr6-117429839-INS-743
chr6-117438726-INS-66
chr6-117454583-INS-110
chr6-117463558-INS-59
chr6-117489381-INS-6375
chr6-117530768-INS-400
chr6-117536067-INS-4891
chr6-117538324-INS-132
chr6-117541802-INS-168
chr6-117542044-INS-6149
chr6-117581556-INS-10436
chr6-117610090-INS-158
chr6-117610501-INS-1878
chr6-117611178-INS-76
chr6-117632241-INS-71
chr6-117635397-INS-134
chr6-117639740-INS-92
chr6-117653086-INS-64
chr6-117660488-INS-51
chr6-117662078-INS-89
chr6-117680799-INS-54
chr6-117684876-INS-146

chr6-117719641-INS-194
chr6-117728526-INS-70
chr6-117749332-INS-811
chr6-117754740-INS-413
chr6-117759041-INS-101
chr6-117776576-INS-154
chr6-117781538-INS-211
chr6-117793506-INS-561
chr6-117795858-INS-56
chr6-117810533-INS-52
chr6-117821717-INS-122
chr6-117824996-INS-149
chr6-117826621-INS-56
chr6-117837732-INS-174
chr6-117856026-INS-487
chr6-117861385-INS-5696
chr6-117874660-INS-113
chr6-117886804-INS-214
chr6-117918250-INS-57
chr6-117932519-INS-145
chr6-117944481-INS-351
chr6-117951894-INS-562
chr6-117958261-INS-116
chr6-117962342-INS-100
chr6-117987063-INS-186
chr6-118007284-INS-188
chr6-118019170-INS-218
chr6-118022891-INS-5452
chr6-118027808-INS-194
chr6-118029012-INS-55
chr6-118055555-INS-156
chr6-118068577-INS-55
chr6-118110028-INS-63
chr6-118110778-INS-346
chr6-118127093-INS-208
chr6-118135292-INS-7601
chr6-118150048-INS-190
chr6-118175826-INS-74
chr6-118195853-INS-462
chr6-118233187-INS-86
chr6-118281125-INS-51
chr6-118307463-INS-6194
chr6-118315925-INS-7239
chr6-118330175-INS-182
chr6-118363187-INS-124
chr6-118372597-INS-6349
chr6-118378332-INS-60
chr6-118387859-INS-498
chr6-118423231-INS-58
chr6-118436977-INS-874
chr6-118443524-INS-129
chr6-118457875-INS-148
chr6-118473093-INS-174
chr6-118474645-INS-197
chr6-118490200-INS-55
chr6-118491204-INS-88
chr6-118494726-INS-156
chr6-118543472-INS-317
chr6-118562673-INS-80
chr6-118640108-INS-52
chr6-118670925-INS-67
chr6-118689682-INS-60
chr6-118692379-INS-208
chr6-118762523-INS-97
chr6-118769710-INS-878
chr6-118770968-INS-868
chr6-118773167-INS-74
chr6-118787000-INS-62
chr6-118835063-INS-58
chr6-118841274-INS-100
chr6-118894112-INS-58
chr6-118934568-INS-515
chr6-118944751-INS-97
chr6-118949523-INS-140
chr6-118953520-INS-71
chr6-118968033-INS-79
chr6-118980450-INS-66

chr6-118999221-INS-68
chr6-119014785-INS-80
chr6-119017086-INS-60
chr6-119025558-INS-812
chr6-119035920-INS-110
chr6-119062011-INS-68
chr6-119069073-INS-375
chr6-119100907-INS-79
chr6-119104296-INS-52
chr6-119136002-INS-52
chr6-119140384-INS-246
chr6-119165076-INS-498
chr6-119168656-INS-997
chr6-119207274-INS-140
chr6-119220621-INS-130
chr6-119225643-INS-258
chr6-119229952-INS-434
chr6-119230725-INS-138
chr6-119231957-INS-56
chr6-119234753-INS-65
chr6-119238061-INS-180
chr6-119241690-INS-57
chr6-119247602-INS-92
chr6-119257559-INS-240
chr6-119271082-INS-353
chr6-119290920-INS-81
chr6-119380559-INS-211
chr6-119381888-INS-272
chr6-119386054-INS-135
chr6-119394804-INS-106
chr6-119400492-INS-58
chr6-119426140-INS-57
chr6-119439478-INS-63
chr6-119477484-INS-200
chr6-119489170-INS-112
chr6-119497820-INS-8674
chr6-119523135-INS-278
chr6-119530370-INS-6440
chr6-119532103-INS-1889
chr6-119534364-INS-80
chr6-119568738-INS-553
chr6-119592081-INS-60
chr6-119592254-INS-755
chr6-119598913-INS-890
chr6-119606441-INS-176
chr6-119606680-INS-6413
chr6-119613820-INS-2408
chr6-119642514-INS-65
chr6-119645995-INS-80
chr6-119655858-INS-7118
chr6-119698672-INS-81
chr6-119704780-INS-85
chr6-119709461-INS-6431
chr6-119763095-INS-196
chr6-119772623-INS-87
chr6-119797329-INS-474
chr6-119826602-INS-183
chr6-119827645-INS-63
chr6-119839602-INS-366
chr6-119859745-INS-203
chr6-119872661-INS-54
chr6-119888199-INS-142
chr6-119912127-INS-527
chr6-119913623-INS-101
chr6-119933532-INS-53
chr6-119950632-INS-62
chr6-119963705-INS-186
chr6-119965352-INS-70
chr6-119966935-INS-164
chr6-120041490-INS-198
chr6-120094904-INS-72
chr6-120097562-INS-215
chr6-120113418-INS-92
chr6-120114766-INS-167
chr6-120128457-INS-184
chr6-120131804-INS-170
chr6-120132373-INS-72

chr6-120134302-INS-75
chr6-120135590-INS-99
chr6-120155794-INS-209
chr6-120185373-INS-313
chr6-120186776-INS-57
chr6-120191077-INS-66
chr6-120206847-INS-170
chr6-120212576-INS-212
chr6-120219097-INS-203
chr6-120228972-INS-244
chr6-120250684-INS-217
chr6-120267698-INS-77
chr6-120270445-INS-225
chr6-120280809-INS-57
chr6-120309278-INS-210
chr6-120312672-INS-220
chr6-120318444-INS-200
chr6-120325517-INS-225
chr6-120334797-INS-50
chr6-120336839-INS-216
chr6-120344550-INS-168
chr6-120356205-INS-120
chr6-120395968-INS-229
chr6-120405932-INS-217
chr6-120429013-INS-149
chr6-120437216-INS-182
chr6-120451037-INS-72
chr6-120479845-INS-216
chr6-120491132-INS-53
chr6-120497426-INS-141
chr6-120517698-INS-57
chr6-120528288-INS-102
chr6-120532401-INS-204
chr6-120535193-INS-399
chr6-120538796-INS-212
chr6-120544246-INS-540
chr6-120546637-INS-181
chr6-120556220-INS-70
chr6-120563610-INS-61
chr6-120570705-INS-204
chr6-120574772-INS-400
chr6-120582767-INS-309
chr6-120603384-INS-199
chr6-120604497-INS-214
chr6-120606262-INS-226
chr6-120608917-INS-180
chr6-120612001-INS-344
chr6-120626267-INS-67
chr6-120629489-INS-234
chr6-120636964-INS-784
chr6-120641825-INS-204
chr6-120650630-INS-71
chr6-120662296-INS-202
chr6-120668037-INS-59
chr6-120672354-INS-53
chr6-120724238-INS-174
chr6-120755067-INS-211
chr6-120759770-INS-207
chr6-120764158-INS-125
chr6-120766685-INS-138
chr6-120771963-INS-343
chr6-120778267-INS-418
chr6-120780099-INS-158
chr6-120788458-INS-6440
chr6-120789050-INS-202
chr6-120802270-INS-206
chr6-120805975-INS-193
chr6-120807528-INS-195
chr6-120815288-INS-67
chr6-120822864-INS-77
chr6-120824724-INS-74
chr6-120829697-INS-183
chr6-120852078-INS-214
chr6-120854548-INS-65
chr6-120878532-INS-280
chr6-120895276-INS-196
chr6-120901157-INS-78

chr6-120902890-INS-98
chr6-120904434-INS-71
chr6-120916480-INS-202
chr6-120939153-INS-59
chr6-120945632-INS-77
chr6-120962356-INS-145
chr6-121000043-INS-136
chr6-121012847-INS-54
chr6-121025293-INS-120
chr6-121036268-INS-54
chr6-121051423-INS-154
chr6-121065015-INS-782
chr6-121083839-INS-266
chr6-121090743-INS-178
chr6-121099516-INS-138
chr6-121115715-INS-6475
chr6-121127662-INS-192
chr6-121134154-INS-582
chr6-121138553-INS-199
chr6-121140171-INS-215
chr6-121142060-INS-341
chr6-121165964-INS-406
chr6-121175411-INS-67
chr6-121176865-INS-145
chr6-121189842-INS-94
chr6-121219594-INS-269
chr6-121229992-INS-67
chr6-121246602-INS-259
chr6-121250808-INS-484
chr6-121255449-INS-231
chr6-121262581-INS-7001
chr6-121265334-INS-66
chr6-121268078-INS-68
chr6-121288666-INS-62
chr6-121335837-INS-79
chr6-121337091-INS-130
chr6-121343747-INS-433
chr6-121377195-INS-270
chr6-121397989-INS-5400
chr6-121402673-INS-2676
chr6-121418622-INS-104
chr6-121439491-INS-58
chr6-121453910-INS-202
chr6-121456034-INS-290
chr6-121496837-INS-75
chr6-121504965-INS-123
chr6-121509914-INS-56
chr6-121517514-INS-82
chr6-121519727-INS-193
chr6-121524697-INS-106
chr6-121579031-INS-54
chr6-121579779-INS-173
chr6-121589235-INS-106
chr6-121605925-INS-52
chr6-121608898-INS-103
chr6-121610446-INS-86
chr6-121633956-INS-63
chr6-121659240-INS-203
chr6-121700146-INS-185
chr6-121722154-INS-6695
chr6-121754552-INS-6594
chr6-121789327-INS-79
chr6-121887214-INS-180
chr6-121930905-INS-726
chr6-121980929-INS-100
chr6-121983024-INS-274
chr6-121994748-INS-4515
chr6-122000657-INS-1625
chr6-122032947-INS-6832
chr6-122035047-INS-269
chr6-122062873-INS-1862
chr6-122086291-INS-4440
chr6-122105350-INS-686
chr6-122226785-INS-572
chr6-122232847-INS-429
chr6-122235616-INS-5291
chr6-122238292-INS-70

chr6-122246788-INS-1886
chr6-122249556-INS-287
chr6-122251177-INS-76
chr6-122276358-INS-148
chr6-122307723-INS-113
chr6-122310514-INS-59
chr6-122326552-INS-202
chr6-122332463-INS-220
chr6-122336041-INS-949
chr6-122336493-INS-5287
chr6-122340834-INS-125
chr6-122345156-INS-56
chr6-122356853-INS-319
chr6-122358834-INS-66
chr6-122365940-INS-448
chr6-122373557-INS-58
chr6-122374945-INS-5566
chr6-122375759-INS-975
chr6-122398504-INS-55
chr6-122421963-INS-345
chr6-122437406-INS-961
chr6-122453071-INS-365
chr6-122468051-INS-159
chr6-122478435-INS-238
chr6-122480318-INS-64
chr6-122493690-INS-62
chr6-122499238-INS-218
chr6-122508596-INS-181
chr6-122521770-INS-94
chr6-122524971-INS-416
chr6-122537218-INS-99
chr6-122564047-INS-60
chr6-122566651-INS-66
chr6-122569737-INS-62
chr6-122570800-INS-509
chr6-122573580-INS-173
chr6-122580739-INS-195
chr6-122584270-INS-214
chr6-122587700-INS-51
chr6-122587784-INS-950
chr6-122592794-INS-3864
chr6-122605423-INS-171
chr6-122619711-INS-499
chr6-122621113-INS-201
chr6-122624629-INS-200
chr6-122635689-INS-201
chr6-122641345-INS-57
chr6-122642266-INS-754
chr6-122643613-INS-129
chr6-122654938-INS-198
chr6-122668181-INS-203
chr6-122676942-INS-189
chr6-122685886-INS-57
chr6-122690059-INS-244
chr6-122694352-INS-193
chr6-122703508-INS-405
chr6-122715141-INS-198
chr6-122743054-INS-79
chr6-122746238-INS-80
chr6-122748528-INS-327
chr6-122751557-INS-169
chr6-122759911-INS-331
chr6-122785341-INS-217
chr6-122808714-INS-437
chr6-122816068-INS-134
chr6-122817794-INS-199
chr6-122830235-INS-226
chr6-122833844-INS-76
chr6-122848265-INS-366
chr6-122854423-INS-226
chr6-122861856-INS-208
chr6-122866769-INS-68
chr6-122876545-INS-5452
chr6-122883643-INS-62
chr6-122909028-INS-64
chr6-122927374-INS-98
chr6-122979349-INS-514

chr6-122983233-INS-134
chr6-123003525-INS-316
chr6-123013067-INS-179
chr6-123025071-INS-197
chr6-123044732-INS-52
chr6-123050129-INS-7253
chr6-123050349-INS-8644
chr6-123074786-INS-484
chr6-123075846-INS-60
chr6-123076986-INS-424
chr6-123096491-INS-240
chr6-123099537-INS-68
chr6-123106706-INS-5188
chr6-123110239-INS-1340
chr6-123134293-INS-128
chr6-123153694-INS-6863
chr6-123167553-INS-498
chr6-123179722-INS-6185
chr6-123187368-INS-8155
chr6-123201855-INS-4380
chr6-123209268-INS-129
chr6-123315024-INS-714
chr6-123330110-INS-173
chr6-123331175-INS-105
chr6-123352787-INS-56
chr6-123368079-INS-7541
chr6-123380026-INS-169
chr6-123589497-INS-263
chr6-123607385-INS-54
chr6-123638357-INS-60
chr6-123639486-INS-847
chr6-123645712-INS-6594
chr6-123648891-INS-194
chr6-123710826-INS-74
chr6-123786278-INS-205
chr6-123794634-INS-60
chr6-123800922-INS-443
chr6-123813188-INS-232
chr6-123861924-INS-105
chr6-123881193-INS-513
chr6-123910454-INS-5482
chr6-123938706-INS-6453
chr6-123951111-INS-358
chr6-123965883-INS-868
chr6-123998536-INS-583
chr6-124000602-INS-352
chr6-124123283-INS-51
chr6-124143606-INS-429
chr6-124147907-INS-831
chr6-124175709-INS-110
chr6-124180948-INS-203
chr6-124183177-INS-6282
chr6-124196211-INS-6854
chr6-124215181-INS-214
chr6-124219620-INS-504
chr6-124255251-INS-86
chr6-124304487-INS-604
chr6-124320208-INS-92
chr6-124326309-INS-7286
chr6-124334706-INS-61
chr6-124338737-INS-63
chr6-124350821-INS-101
chr6-124365611-INS-56
chr6-124369208-INS-894
chr6-124411474-INS-859
chr6-124419415-INS-240
chr6-124462956-INS-67
chr6-124468675-INS-358
chr6-124495194-INS-65
chr6-124497008-INS-243
chr6-124501033-INS-52
chr6-124503711-INS-72
chr6-124506756-INS-157
chr6-124520780-INS-60
chr6-124531048-INS-210
chr6-124539412-INS-736
chr6-124547863-INS-98

chr6-124549821-INS-130
chr6-124558501-INS-138
chr6-124579384-INS-5496
chr6-124599323-INS-71
chr6-124667937-INS-64
chr6-124674216-INS-166
chr6-124682225-INS-854
chr6-124725876-INS-248
chr6-124750224-INS-159
chr6-124754239-INS-122
chr6-124763203-INS-729
chr6-124781203-INS-416
chr6-124785110-INS-198
chr6-124812054-INS-108
chr6-124821124-INS-197
chr6-124831103-INS-138
chr6-124833370-INS-170
chr6-124861659-INS-112
chr6-124874861-INS-58
chr6-124882588-INS-59
chr6-124918655-INS-88
chr6-124923321-INS-186
chr6-124925687-INS-136
chr6-124930998-INS-205
chr6-124947808-INS-167
chr6-124958554-INS-185
chr6-124960264-INS-190
chr6-124964737-INS-143
chr6-124966906-INS-94
chr6-124970253-INS-434
chr6-124991706-INS-514
chr6-124995677-INS-204
chr6-125029204-INS-193
chr6-125036952-INS-222
chr6-125045962-INS-209
chr6-125049096-INS-51
chr6-125056955-INS-175
chr6-125064019-INS-148
chr6-125069548-INS-209
chr6-125103489-INS-212
chr6-125145148-INS-57
chr6-125151825-INS-269
chr6-125159032-INS-132
chr6-125173700-INS-164
chr6-125176727-INS-203
chr6-125179911-INS-204
chr6-125184031-INS-229
chr6-125185430-INS-326
chr6-125190298-INS-215
chr6-125212431-INS-64
chr6-125217444-INS-303
chr6-125219268-INS-67
chr6-125222509-INS-72
chr6-125231312-INS-196
chr6-125235533-INS-60
chr6-125239750-INS-497
chr6-125246742-INS-1889
chr6-125269030-INS-203
chr6-125285570-INS-164
chr6-125288049-INS-182
chr6-125306522-INS-56
chr6-125317048-INS-303
chr6-125318400-INS-69
chr6-125329100-INS-333
chr6-125333030-INS-213
chr6-125368540-INS-181
chr6-125379378-INS-93
chr6-125386566-INS-178
chr6-125390916-INS-201
chr6-125392821-INS-1127
chr6-125411233-INS-263
chr6-125425042-INS-65
chr6-125432393-INS-221
chr6-125437141-INS-166
chr6-125438449-INS-175
chr6-125442386-INS-251
chr6-125456274-INS-70

chr6-125481719-INS-605
chr6-125485608-INS-196
chr6-125489285-INS-508
chr6-125492432-INS-54
chr6-125501394-INS-345
chr6-125513180-INS-64
chr6-125516876-INS-273
chr6-125520698-INS-72
chr6-125523986-INS-182
chr6-125538552-INS-183
chr6-125544590-INS-168
chr6-125553083-INS-901
chr6-125558844-INS-227
chr6-125568358-INS-951
chr6-125572168-INS-101
chr6-125586796-INS-62
chr6-125602975-INS-75
chr6-125621851-INS-91
chr6-125641790-INS-58
chr6-125683627-INS-52
chr6-125697987-INS-158
chr6-125725241-INS-164
chr6-125726131-INS-199
chr6-125740749-INS-360
chr6-125768442-INS-51
chr6-125776875-INS-242
chr6-125791083-INS-569
chr6-125812582-INS-54
chr6-125839032-INS-199
chr6-125866889-INS-794
chr6-125889126-INS-127
chr6-125953883-INS-116
chr6-125967160-INS-70
chr6-125969759-INS-156
chr6-125973044-INS-112
chr6-125976314-INS-70
chr6-125984087-INS-193
chr6-126012535-INS-66
chr6-126066268-INS-187
chr6-126127267-INS-52
chr6-126129896-INS-85
chr6-126172504-INS-279
chr6-126241610-INS-359
chr6-126242805-INS-450
chr6-126269451-INS-6469
chr6-126310390-INS-104
chr6-126319637-INS-285
chr6-126350509-INS-339
chr6-126408954-INS-236
chr6-126411805-INS-258
chr6-126467027-INS-498
chr6-126469024-INS-146
chr6-126478270-INS-76
chr6-126487461-INS-463
chr6-126490634-INS-60
chr6-126495495-INS-105
chr6-126514817-INS-53
chr6-126532728-INS-112
chr6-126558304-INS-73
chr6-126565604-INS-92
chr6-126585388-INS-414
chr6-126602208-INS-158
chr6-126607625-INS-449
chr6-126611534-INS-92
chr6-126659553-INS-730
chr6-126675809-INS-54
chr6-126694301-INS-64
chr6-126707233-INS-178
chr6-126734884-INS-165
chr6-126738940-INS-892
chr6-126742905-INS-262
chr6-126752861-INS-99
chr6-126794467-INS-210
chr6-126808787-INS-140
chr6-126811904-INS-53
chr6-126820323-INS-205
chr6-126823649-INS-106

chr6-126838609-INS-200
chr6-126843830-INS-88
chr6-126845844-INS-109
chr6-126857008-INS-66
chr6-126873609-INS-189
chr6-126888785-INS-52
chr6-126927371-INS-240
chr6-126963922-INS-65
chr6-126965348-INS-236
chr6-126966493-INS-176
chr6-126971722-INS-219
chr6-126973714-INS-54
chr6-126976110-INS-51
chr6-126979904-INS-191
chr6-126986259-INS-66
chr6-126999549-INS-211
chr6-127005741-INS-87
chr6-127009086-INS-172
chr6-127012753-INS-118
chr6-127026420-INS-112
chr6-127040138-INS-125
chr6-127060598-INS-57
chr6-127087107-INS-95
chr6-127110571-INS-556
chr6-127138719-INS-72
chr6-127138835-INS-288
chr6-127158068-INS-5520
chr6-127160681-INS-825
chr6-127203726-INS-70
chr6-127234601-INS-59
chr6-127283760-INS-96
chr6-127300036-INS-936
chr6-127304935-INS-91
chr6-127317207-INS-54
chr6-127324585-INS-171
chr6-127325227-INS-151
chr6-127335749-INS-93
chr6-127337178-INS-316
chr6-127344233-INS-60
chr6-127357274-INS-102
chr6-127370437-INS-56
chr6-127375184-INS-60
chr6-127376255-INS-263
chr6-127398268-INS-148
chr6-127404451-INS-88
chr6-127418197-INS-264
chr6-127419361-INS-164
chr6-127434932-INS-54
chr6-127445695-INS-80
chr6-127459329-INS-65
chr6-127467809-INS-436
chr6-127482221-INS-148
chr6-127507819-INS-106
chr6-127511249-INS-53
chr6-127524749-INS-98
chr6-127528803-INS-220
chr6-127547911-INS-76
chr6-127562103-INS-964
chr6-127567189-INS-94
chr6-127576134-INS-76
chr6-127585143-INS-173
chr6-127596572-INS-284
chr6-127602022-INS-90
chr6-127609164-INS-179
chr6-127616965-INS-54
chr6-127633422-INS-133
chr6-127651480-INS-196
chr6-127658634-INS-75
chr6-127665710-INS-401
chr6-127719979-INS-156
chr6-127720755-INS-57
chr6-127743528-INS-77
chr6-127761260-INS-77
chr6-127802552-INS-114
chr6-127828048-INS-331
chr6-127836851-INS-295
chr6-127850982-INS-60

chr6-127857535-INS-62
chr6-127875765-INS-53
chr6-127879367-INS-64
chr6-127897062-INS-148
chr6-127908712-INS-92
chr6-127910602-INS-66
chr6-127917599-INS-58
chr6-127931781-INS-52
chr6-127974841-INS-56
chr6-127979660-INS-59
chr6-127982365-INS-219
chr6-127985873-INS-96
chr6-128017715-INS-95
chr6-128064745-INS-114
chr6-128070039-INS-75
chr6-128089975-INS-122
chr6-128093108-INS-58
chr6-128096333-INS-126
chr6-128109162-INS-116
chr6-128143832-INS-207
chr6-128158907-INS-86
chr6-128175413-INS-215
chr6-128180893-INS-86
chr6-128219604-INS-332
chr6-128235341-INS-6937
chr6-128250123-INS-56
chr6-128251443-INS-194
chr6-128255677-INS-52
chr6-128268251-INS-199
chr6-128270266-INS-88
chr6-128271759-INS-466
chr6-128279900-INS-72
chr6-128291418-INS-221
chr6-128305506-INS-254
chr6-128306266-INS-218
chr6-128320866-INS-247
chr6-128321550-INS-2396
chr6-128323795-INS-231
chr6-128326024-INS-363
chr6-128363358-INS-78
chr6-128365345-INS-157
chr6-128366725-INS-215
chr6-128369817-INS-97
chr6-128376071-INS-59
chr6-128400395-INS-227
chr6-128404584-INS-55
chr6-128431211-INS-161
chr6-128438416-INS-141
chr6-128448950-INS-309
chr6-128467361-INS-52
chr6-128468337-INS-397
chr6-128470190-INS-102
chr6-128473857-INS-237
chr6-128476583-INS-60
chr6-128483145-INS-65
chr6-128486299-INS-77
chr6-128509170-INS-61
chr6-128512823-INS-487
chr6-128514958-INS-156
chr6-128516754-INS-444
chr6-128518832-INS-147
chr6-128560123-INS-53
chr6-128576002-INS-70
chr6-128582586-INS-136
chr6-128584362-INS-74
chr6-128611640-INS-190
chr6-128628198-INS-567
chr6-128666919-INS-81
chr6-128672445-INS-6475
chr6-128705829-INS-91
chr6-128718229-INS-74
chr6-128740580-INS-129
chr6-128748717-INS-135
chr6-128751434-INS-564
chr6-128806640-INS-325
chr6-128834454-INS-90
chr6-128868325-INS-621

chr6-128897175-INS-377
chr6-128907514-INS-119
chr6-128928217-INS-5507
chr6-128948169-INS-52
chr6-128949368-INS-580
chr6-128955904-INS-192
chr6-128963418-INS-326
chr6-128974185-INS-213
chr6-128975857-INS-54
chr6-128982845-INS-209
chr6-129023463-INS-277
chr6-129062335-INS-624
chr6-129086594-INS-141
chr6-129100641-INS-69
chr6-129113376-INS-2207
chr6-129117285-INS-155
chr6-129126115-INS-1720
chr6-129127841-INS-325
chr6-129134569-INS-439
chr6-129147691-INS-204
chr6-129155187-INS-107
chr6-129158771-INS-140
chr6-129163185-INS-110
chr6-129180284-INS-993
chr6-129187972-INS-72
chr6-129189154-INS-530
chr6-129195260-INS-194
chr6-129215672-INS-485
chr6-129243222-INS-62
chr6-129248024-INS-72
chr6-129255012-INS-58
chr6-129257137-INS-114
chr6-129271011-INS-471
chr6-129290012-INS-84
chr6-129293062-INS-405
chr6-129348898-INS-100
chr6-129388190-INS-114
chr6-129394773-INS-97
chr6-129420654-INS-174
chr6-129429776-INS-315
chr6-129460562-INS-400
chr6-129469409-INS-190
chr6-129513656-INS-92
chr6-129517826-INS-63
chr6-129526669-INS-52
chr6-129528863-INS-74
chr6-129531924-INS-66
chr6-129541252-INS-72
chr6-129553131-INS-124
chr6-129584211-INS-143
chr6-129589514-INS-402
chr6-129592639-INS-54
chr6-129614105-INS-473
chr6-129626877-INS-60
chr6-129628097-INS-89
chr6-129630345-INS-978
chr6-129638274-INS-684
chr6-129642507-INS-52
chr6-129671823-INS-958
chr6-129700177-INS-126
chr6-129795113-INS-6203
chr6-129804267-INS-4794
chr6-129805785-INS-7702
chr6-129834140-INS-212
chr6-129962700-INS-65
chr6-129993351-INS-55
chr6-130006533-INS-68
chr6-130020131-INS-94
chr6-130059343-INS-672
chr6-130076492-INS-892
chr6-130201645-INS-127
chr6-130294273-INS-163
chr6-130296297-INS-900
chr6-130309902-INS-318
chr6-130390015-INS-7440
chr6-130411390-INS-65
chr6-130415688-INS-103

chr6-130422808-INS-5207
chr6-130431954-INS-776
chr6-130448641-INS-67
chr6-130460341-INS-84
chr6-130467381-INS-734
chr6-130515900-INS-4891
chr6-130532887-INS-12124
chr6-130539833-INS-3264
chr6-130543867-INS-415
chr6-130544871-INS-1927
chr6-130552719-INS-757
chr6-130555930-INS-84
chr6-130606588-INS-168
chr6-130695241-INS-63
chr6-130697439-INS-148
chr6-130699864-INS-290
chr6-130725047-INS-6842
chr6-130763675-INS-168
chr6-130789572-INS-519
chr6-130794485-INS-77
chr6-130821710-INS-1291
chr6-130840474-INS-2595
chr6-130897882-INS-257
chr6-130915397-INS-168
chr6-130925760-INS-2817
chr6-131043605-INS-3061
chr6-131045989-INS-8571
chr6-131049947-INS-53
chr6-131059373-INS-56
chr6-131143415-INS-588
chr6-131171893-INS-508
chr6-131185287-INS-7291
chr6-131196177-INS-1210
chr6-131196779-INS-6598
chr6-131223463-INS-51
chr6-131267647-INS-865
chr6-131269922-INS-122
chr6-131276974-INS-65
chr6-131281939-INS-123
chr6-131285844-INS-244
chr6-131294795-INS-166
chr6-131297276-INS-90
chr6-131304120-INS-58
chr6-131318293-INS-6679
chr6-131319794-INS-2127
chr6-131324759-INS-68
chr6-131324763-INS-682
chr6-131326074-INS-631
chr6-131327417-INS-82
chr6-131333320-INS-209
chr6-131335198-INS-72
chr6-131336203-INS-84
chr6-131380565-INS-64
chr6-131419723-INS-6920
chr6-131440196-INS-55
chr6-131445408-INS-5102
chr6-131455913-INS-195
chr6-131460370-INS-86
chr6-131463501-INS-6474
chr6-131480403-INS-5096
chr6-131490370-INS-167
chr6-131517880-INS-136
chr6-131521288-INS-7097
chr6-131567418-INS-5325
chr6-131584859-INS-54
chr6-131596955-INS-6794
chr6-131603035-INS-6298
chr6-131620072-INS-52
chr6-131625983-INS-55
chr6-131644051-INS-85
chr6-131660528-INS-54
chr6-131725195-INS-91
chr6-131753567-INS-59
chr6-131783530-INS-283
chr6-131888534-INS-3850
chr6-131908497-INS-311
chr6-131919023-INS-51

chr6-132010847-INS-139
chr6-132036991-INS-326
chr6-132167575-INS-147
chr6-132214351-INS-838
chr6-132232221-INS-6393
chr6-132266260-INS-418
chr6-132272687-INS-86
chr6-132289152-INS-152
chr6-132366876-INS-165
chr6-132403095-INS-5082
chr6-132415581-INS-54
chr6-132420352-INS-83
chr6-132440682-INS-60
chr6-132452358-INS-105
chr6-132485493-INS-53
chr6-132606246-INS-407
chr6-132607400-INS-162
chr6-132624108-INS-207
chr6-132634599-INS-5275
chr6-132670911-INS-680
chr6-132703261-INS-65
chr6-132721659-INS-115
chr6-132756119-INS-517
chr6-132766135-INS-69
chr6-132783447-INS-434
chr6-132847422-INS-131
chr6-132865921-INS-94
chr6-132905155-INS-95
chr6-132951861-INS-51
chr6-132958582-INS-69
chr6-132962463-INS-52
chr6-132998987-INS-330
chr6-133005510-INS-492
chr6-133029545-INS-157
chr6-133044547-INS-7089
chr6-133074733-INS-206
chr6-133086792-INS-64
chr6-133099332-INS-62
chr6-133192853-INS-2859
chr6-133226786-INS-112
chr6-133230006-INS-2868
chr6-133241256-INS-55
chr6-133241457-INS-6214
chr6-133252767-INS-893
chr6-133253610-INS-7435
chr6-133298493-INS-4446
chr6-133325419-INS-1535
chr6-133338795-INS-6320
chr6-133346650-INS-105
chr6-133349343-INS-7031
chr6-133352301-INS-65
chr6-133361150-INS-141
chr6-133386584-INS-64
chr6-133389729-INS-52
chr6-133397438-INS-68
chr6-133397977-INS-317
chr6-133398953-INS-486
chr6-133410841-INS-203
chr6-133474281-INS-7507
chr6-133485608-INS-6685
chr6-133502904-INS-2497
chr6-133516635-INS-386
chr6-133548102-INS-87
chr6-133559031-INS-52
chr6-133565208-INS-745
chr6-133600467-INS-614
chr6-133699056-INS-59
chr6-133710963-INS-4697
chr6-133778036-INS-1897
chr6-133781021-INS-6632
chr6-133782820-INS-205
chr6-133803547-INS-57
chr6-133806529-INS-119
chr6-133815727-INS-195
chr6-133817669-INS-6292
chr6-133829964-INS-68
chr6-133836112-INS-6575

chr6-133845679-INS-6587
chr6-133878685-INS-525
chr6-133889138-INS-355
chr6-133891512-INS-86
chr6-133928044-INS-522
chr6-133929493-INS-228
chr6-134015692-INS-52
chr6-134017284-INS-110
chr6-134036682-INS-249
chr6-134043518-INS-59
chr6-134059106-INS-157
chr6-134061977-INS-84
chr6-134063183-INS-671
chr6-134101977-INS-60
chr6-134103585-INS-131
chr6-134112737-INS-128
chr6-134171618-INS-905
chr6-134198700-INS-7478
chr6-134203340-INS-138
chr6-134222639-INS-553
chr6-134228108-INS-104
chr6-134231634-INS-915
chr6-134247360-INS-57
chr6-134256346-INS-67
chr6-134265868-INS-264
chr6-134276180-INS-65
chr6-134280609-INS-59
chr6-134292917-INS-618
chr6-134293934-INS-3183
chr6-134305657-INS-197
chr6-134333356-INS-72
chr6-134337155-INS-72
chr6-134345748-INS-91
chr6-134358197-INS-120
chr6-134368845-INS-114
chr6-134371968-INS-76
chr6-134381354-INS-63
chr6-134383183-INS-84
chr6-134393996-INS-262
chr6-134403261-INS-56
chr6-134420040-INS-55
chr6-134442665-INS-230
chr6-134449177-INS-65
chr6-134465782-INS-181
chr6-134493628-INS-180
chr6-134499131-INS-105
chr6-134539567-INS-255
chr6-134547282-INS-288
chr6-134554343-INS-78
chr6-134563663-INS-67
chr6-134567921-INS-89
chr6-134568332-INS-52
chr6-134573391-INS-56
chr6-134574378-INS-644
chr6-134580016-INS-65
chr6-134585711-INS-95
chr6-134612001-INS-72
chr6-134642406-INS-86
chr6-134644224-INS-330
chr6-134681940-INS-6454
chr6-134720085-INS-169
chr6-134723024-INS-146
chr6-134748916-INS-78
chr6-134764201-INS-121
chr6-134774923-INS-287
chr6-134777341-INS-54
chr6-134788909-INS-339
chr6-134820821-INS-491
chr6-134842791-INS-93
chr6-134852932-INS-270
chr6-134871216-INS-203
chr6-134880253-INS-51
chr6-134887939-INS-177
chr6-134912175-INS-200
chr6-134916897-INS-194
chr6-134924235-INS-226
chr6-134927218-INS-143

chr6-134933547-INS-52
chr6-134973212-INS-250
chr6-134974392-INS-92
chr6-134976188-INS-83
chr6-134999207-INS-205
chr6-134999500-INS-529
chr6-135018301-INS-467
chr6-135024638-INS-480
chr6-135035941-INS-74
chr6-135039192-INS-264
chr6-135058176-INS-319
chr6-135063760-INS-56
chr6-135082580-INS-499
chr6-135090838-INS-51
chr6-135106007-INS-89
chr6-135117118-INS-72
chr6-135128628-INS-100
chr6-135133320-INS-106
chr6-135148765-INS-54
chr6-135150196-INS-56
chr6-135152720-INS-634
chr6-135159528-INS-65
chr6-135197853-INS-76
chr6-135203307-INS-204
chr6-135222117-INS-92
chr6-135278099-INS-60
chr6-135280563-INS-92
chr6-135286151-INS-193
chr6-135291343-INS-498
chr6-135356869-INS-203
chr6-135383593-INS-1274
chr6-135383667-INS-235
chr6-135393349-INS-79
chr6-135405865-INS-52
chr6-135429154-INS-287
chr6-135455280-INS-56
chr6-135460452-INS-77
chr6-135474931-INS-61
chr6-135475622-INS-5400
chr6-135476003-INS-56
chr6-135506597-INS-69
chr6-135515272-INS-63
chr6-135550824-INS-190
chr6-135571653-INS-623
chr6-135598247-INS-55
chr6-135603248-INS-107
chr6-135615797-INS-132
chr6-135650000-INS-329
chr6-135654540-INS-71
chr6-135662558-INS-467
chr6-135671600-INS-457
chr6-135677648-INS-503
chr6-135714464-INS-126
chr6-135730840-INS-1896
chr6-135779576-INS-115
chr6-135783572-INS-83
chr6-135835504-INS-169
chr6-135837312-INS-210
chr6-135845198-INS-189
chr6-135855564-INS-244
chr6-135866072-INS-70
chr6-135872435-INS-206
chr6-135896691-INS-225
chr6-135904057-INS-116
chr6-135937812-INS-86
chr6-135944831-INS-96
chr6-135953494-INS-461
chr6-135955586-INS-108
chr6-135961720-INS-474
chr6-135972271-INS-72
chr6-136015925-INS-56
chr6-136040616-INS-60
chr6-136060278-INS-700
chr6-136095951-INS-420
chr6-136125333-INS-1746
chr6-136132115-INS-107
chr6-136141037-INS-242

chr6-136172224-INS-1594
chr6-136190002-INS-180
chr6-136199465-INS-157
chr6-136226044-INS-53
chr6-136229493-INS-136
chr6-136232767-INS-51
chr6-136243714-INS-54
chr6-136302443-INS-217
chr6-136316407-INS-500
chr6-136329909-INS-54
chr6-136333574-INS-379
chr6-136337953-INS-872
chr6-136372794-INS-351
chr6-136377038-INS-202
chr6-136382499-INS-530
chr6-136382523-INS-1199
chr6-136390216-INS-70
chr6-136404441-INS-126
chr6-136410816-INS-131
chr6-136430529-INS-184
chr6-136432338-INS-890
chr6-136436203-INS-424
chr6-136438876-INS-198
chr6-136440361-INS-215
chr6-136481661-INS-207
chr6-136491559-INS-60
chr6-136494260-INS-127
chr6-136501485-INS-212
chr6-136504571-INS-129
chr6-136535934-INS-336
chr6-136539280-INS-294
chr6-136565902-INS-60
chr6-136579874-INS-83
chr6-136613977-INS-232
chr6-136643034-INS-52
chr6-136648424-INS-51
chr6-136654660-INS-188
chr6-136659307-INS-469
chr6-136660144-INS-182
chr6-136679041-INS-375
chr6-136687071-INS-136
chr6-136695152-INS-217
chr6-136695645-INS-164
chr6-136706064-INS-151
chr6-136741051-INS-991
chr6-136755464-INS-1892
chr6-136777415-INS-324
chr6-136793741-INS-651
chr6-136828289-INS-242
chr6-136828754-INS-670
chr6-136829984-INS-128
chr6-136856225-INS-92
chr6-136867076-INS-80
chr6-136871037-INS-125
chr6-136886451-INS-793
chr6-136894969-INS-101
chr6-136909496-INS-81
chr6-136922782-INS-98
chr6-136928703-INS-51
chr6-136934625-INS-316
chr6-136947657-INS-165
chr6-136977512-INS-117
chr6-136979743-INS-211
chr6-137004910-INS-107
chr6-137005996-INS-399
chr6-137100440-INS-140
chr6-137105760-INS-111
chr6-137142154-INS-220
chr6-137155564-INS-76
chr6-137165100-INS-80
chr6-137173381-INS-351
chr6-137194390-INS-60
chr6-137251092-INS-398
chr6-137279499-INS-247
chr6-137288799-INS-53
chr6-137297766-INS-84
chr6-137309677-INS-55

chr6-137313922-INS-219
chr6-137317488-INS-186
chr6-137339834-INS-358
chr6-137355166-INS-215
chr6-137383577-INS-86
chr6-137405506-INS-105
chr6-137411283-INS-498
chr6-137430708-INS-59
chr6-137435692-INS-140
chr6-137453974-INS-250
chr6-137525122-INS-5313
chr6-137573145-INS-72
chr6-137580100-INS-63
chr6-137598444-INS-192
chr6-137624543-INS-136
chr6-137641141-INS-454
chr6-137648602-INS-168
chr6-137655570-INS-196
chr6-137673035-INS-87
chr6-137685714-INS-78
chr6-137688412-INS-4587
chr6-137718606-INS-162
chr6-137726859-INS-198
chr6-137741330-INS-205
chr6-137752088-INS-102
chr6-137767521-INS-82
chr6-137770180-INS-471
chr6-137772309-INS-174
chr6-137788856-INS-296
chr6-137799800-INS-65
chr6-137804321-INS-54
chr6-137823240-INS-110
chr6-137875793-INS-1890
chr6-137875933-INS-188
chr6-137891051-INS-144
chr6-137900137-INS-91
chr6-137907539-INS-150
chr6-137919531-INS-55
chr6-137930204-INS-85
chr6-137934364-INS-60
chr6-137936878-INS-74
chr6-137948577-INS-60
chr6-137997842-INS-6342
chr6-138025206-INS-94
chr6-138025811-INS-8555
chr6-138039002-INS-95
chr6-138062927-INS-273
chr6-138071516-INS-99
chr6-138092611-INS-200
chr6-138113866-INS-62
chr6-138125447-INS-512
chr6-138128531-INS-57
chr6-138191370-INS-565
chr6-138219295-INS-350
chr6-138264110-INS-384
chr6-138283638-INS-56
chr6-138298971-INS-96
chr6-138301895-INS-65
chr6-138306295-INS-210
chr6-138374156-INS-89
chr6-138375238-INS-216
chr6-138385030-INS-134
chr6-138400660-INS-105
chr6-138408967-INS-238
chr6-138424353-INS-498
chr6-138432342-INS-56
chr6-138454373-INS-100
chr6-138511569-INS-70
chr6-138513588-INS-944
chr6-138540974-INS-70
chr6-138553801-INS-56
chr6-138561975-INS-272
chr6-138568742-INS-60
chr6-138577130-INS-504
chr6-138601694-INS-559
chr6-138601799-INS-930
chr6-138606732-INS-68

chr6-138609207-INS-54
chr6-138628675-INS-476
chr6-138640833-INS-207
chr6-138667901-INS-484
chr6-138680101-INS-58
chr6-138739858-INS-633
chr6-138750600-INS-82
chr6-138793050-INS-136
chr6-138807242-INS-162
chr6-138809181-INS-52
chr6-138826358-INS-136
chr6-138827329-INS-70
chr6-138835546-INS-106
chr6-138840364-INS-783
chr6-138842682-INS-203
chr6-138844113-INS-115
chr6-138853836-INS-6963
chr6-138862258-INS-645
chr6-138870428-INS-127
chr6-138872866-INS-66
chr6-138875676-INS-612
chr6-138883635-INS-51
chr6-138885614-INS-6898
chr6-138899261-INS-88
chr6-138934073-INS-270
chr6-138935190-INS-146
chr6-138946130-INS-290
chr6-138961534-INS-182
chr6-139005857-INS-164
chr6-139014680-INS-220
chr6-139014918-INS-391
chr6-139065680-INS-848
chr6-139075196-INS-138
chr6-139077329-INS-52
chr6-139097981-INS-140
chr6-139129587-INS-6542
chr6-139134613-INS-52
chr6-139189323-INS-60
chr6-139213943-INS-300
chr6-139233087-INS-52
chr6-139239848-INS-56
chr6-139245450-INS-71
chr6-139274204-INS-76
chr6-139290353-INS-70
chr6-139293011-INS-358
chr6-139298124-INS-192
chr6-139301102-INS-53
chr6-139319242-INS-109
chr6-139332367-INS-142
chr6-139361275-INS-6840
chr6-139392598-INS-179
chr6-139436700-INS-51
chr6-139443758-INS-403
chr6-139480794-INS-78
chr6-139491783-INS-297
chr6-139504477-INS-56
chr6-139512535-INS-72
chr6-139534202-INS-52
chr6-139539410-INS-72
chr6-139550562-INS-187
chr6-139558262-INS-60
chr6-139560442-INS-113
chr6-139581625-INS-80
chr6-139585335-INS-55
chr6-139608819-INS-70
chr6-139624691-INS-575
chr6-139633631-INS-83
chr6-139648593-INS-52
chr6-139662377-INS-388
chr6-139667464-INS-235
chr6-139667879-INS-615
chr6-139713960-INS-72
chr6-139730385-INS-74
chr6-139738058-INS-463
chr6-139778291-INS-84
chr6-139813738-INS-84
chr6-139851914-INS-164

chr6-139855766-INS-608
chr6-139860590-INS-407
chr6-139873526-INS-52
chr6-139897707-INS-86
chr6-139929705-INS-52
chr6-139931021-INS-88
chr6-139942100-INS-230
chr6-139962582-INS-72
chr6-139976173-INS-100
chr6-140016313-INS-300
chr6-140031073-INS-64
chr6-140054250-INS-124
chr6-140060169-INS-225
chr6-140102025-INS-66
chr6-140105535-INS-400
chr6-140124506-INS-53
chr6-140127242-INS-74
chr6-140143577-INS-132
chr6-140145950-INS-54
chr6-140170643-INS-4168
chr6-140180503-INS-148
chr6-140183084-INS-119
chr6-140213663-INS-76
chr6-140233609-INS-121
chr6-140235275-INS-56
chr6-140236337-INS-591
chr6-140249457-INS-528
chr6-140286403-INS-285
chr6-140307772-INS-404
chr6-140313359-INS-54
chr6-140323607-INS-709
chr6-140330180-INS-53
chr6-140357185-INS-233
chr6-140359824-INS-66
chr6-140360718-INS-60
chr6-140389805-INS-110
chr6-140419058-INS-148
chr6-140419100-INS-878
chr6-140439414-INS-164
chr6-140446254-INS-131
chr6-140450366-INS-7201
chr6-140462984-INS-183
chr6-140472934-INS-60
chr6-140478121-INS-118
chr6-140484480-INS-132
chr6-140486761-INS-60
chr6-140490584-INS-204
chr6-140507140-INS-51
chr6-140528853-INS-384
chr6-140530025-INS-169
chr6-140531081-INS-178
chr6-140534193-INS-120
chr6-140537000-INS-230
chr6-140563040-INS-809
chr6-140579085-INS-94
chr6-140584438-INS-53
chr6-140587648-INS-78
chr6-140592232-INS-419
chr6-140623101-INS-211
chr6-140623945-INS-226
chr6-140626136-INS-54
chr6-140631599-INS-71
chr6-140634975-INS-118
chr6-140639737-INS-203
chr6-140661737-INS-173
chr6-140662813-INS-58
chr6-140669251-INS-79
chr6-140670208-INS-207
chr6-140687664-INS-65
chr6-140732657-INS-201
chr6-140803572-INS-85
chr6-140820322-INS-54
chr6-140828527-INS-203
chr6-140830053-INS-67
chr6-140843773-INS-221
chr6-140864381-INS-859
chr6-140864411-INS-867

chr6-140905638-INS-344
chr6-140979809-INS-277
chr6-140994260-INS-156
chr6-141033829-INS-460
chr6-141034997-INS-82
chr6-141036147-INS-72
chr6-141045112-INS-143
chr6-141058360-INS-67
chr6-141082335-INS-53
chr6-141140894-INS-106
chr6-141164947-INS-54
chr6-141184676-INS-66
chr6-141187929-INS-134
chr6-141260333-INS-57
chr6-141263938-INS-144
chr6-141280284-INS-545
chr6-141331759-INS-233
chr6-141348155-INS-7423
chr6-141353741-INS-546
chr6-141367832-INS-220
chr6-141368547-INS-1888
chr6-141396786-INS-168
chr6-141403087-INS-229
chr6-141407408-INS-97
chr6-141424582-INS-172
chr6-141432546-INS-843
chr6-141466733-INS-70
chr6-141470609-INS-136
chr6-141475758-INS-326
chr6-141510138-INS-143
chr6-141510481-INS-808
chr6-141523566-INS-238
chr6-141533387-INS-676
chr6-141559853-INS-2539
chr6-141572119-INS-608
chr6-141590668-INS-526
chr6-141626149-INS-974
chr6-141626993-INS-60
chr6-141651729-INS-56
chr6-141655213-INS-148
chr6-141685014-INS-204
chr6-141704936-INS-485
chr6-141737130-INS-540
chr6-141748666-INS-52
chr6-141772734-INS-336
chr6-141802635-INS-201
chr6-141853442-INS-55
chr6-141865229-INS-426
chr6-141869580-INS-74
chr6-141874027-INS-619
chr6-141929753-INS-7195
chr6-141933133-INS-427
chr6-141934600-INS-5290
chr6-141936229-INS-5888
chr6-141946208-INS-87
chr6-141951805-INS-393
chr6-141969083-INS-626
chr6-141970071-INS-1649
chr6-141975143-INS-620
chr6-141987466-INS-6213
chr6-142058144-INS-175
chr6-142069032-INS-338
chr6-142071349-INS-725
chr6-142105090-INS-543
chr6-142131844-INS-559
chr6-142134275-INS-6257
chr6-142140433-INS-72
chr6-142188013-INS-913
chr6-142195054-INS-116
chr6-142205277-INS-62
chr6-142209288-INS-84
chr6-142215307-INS-54
chr6-142254400-INS-60
chr6-142266268-INS-181
chr6-142299860-INS-54
chr6-142302042-INS-60
chr6-142308005-INS-292

chr6-142319524-INS-96
chr6-142322561-INS-122
chr6-142326531-INS-135
chr6-142337988-INS-93
chr6-142345141-INS-204
chr6-142354286-INS-124
chr6-142365203-INS-210
chr6-142375142-INS-169
chr6-142376780-INS-385
chr6-142410219-INS-147
chr6-142414680-INS-51
chr6-142439338-INS-76
chr6-142443475-INS-55
chr6-142462831-INS-59
chr6-142464766-INS-887
chr6-142465299-INS-53
chr6-142471528-INS-64
chr6-142478155-INS-549
chr6-142506619-INS-305
chr6-142510838-INS-74
chr6-142518702-INS-58
chr6-142524548-INS-160
chr6-142531621-INS-672
chr6-142545635-INS-68
chr6-142602725-INS-410
chr6-142613496-INS-178
chr6-142657476-INS-83
chr6-142669470-INS-236
chr6-142679326-INS-148
chr6-142687854-INS-320
chr6-142696345-INS-158
chr6-142698285-INS-262
chr6-142713364-INS-259
chr6-142716584-INS-2937
chr6-142716603-INS-1584
chr6-142716615-INS-311
chr6-142723014-INS-184
chr6-142724015-INS-75
chr6-142730907-INS-1141
chr6-142755519-INS-73
chr6-142759218-INS-67
chr6-142763329-INS-52
chr6-142777319-INS-80
chr6-142784184-INS-85
chr6-142827477-INS-52
chr6-142835468-INS-6208
chr6-142836262-INS-89
chr6-142857406-INS-6661
chr6-142858313-INS-200
chr6-142872707-INS-576
chr6-142889807-INS-90
chr6-142903048-INS-71
chr6-142914165-INS-4137
chr6-142914690-INS-401
chr6-142935961-INS-129
chr6-142937595-INS-56
chr6-142939054-INS-742
chr6-142942918-INS-205
chr6-142946500-INS-411
chr6-142954378-INS-78
chr6-142955945-INS-716
chr6-142968506-INS-211
chr6-143001605-INS-52
chr6-143003192-INS-362
chr6-143035094-INS-80
chr6-143038747-INS-88
chr6-143044406-INS-93
chr6-143056390-INS-211
chr6-143079626-INS-155
chr6-143085285-INS-219
chr6-143091794-INS-218
chr6-143099142-INS-62
chr6-143120584-INS-73
chr6-143126753-INS-56
chr6-143131606-INS-62
chr6-143145869-INS-126
chr6-143166035-INS-227

chr6-143171499-INS-1156
chr6-143174127-INS-200
chr6-143210791-INS-171
chr6-143212447-INS-1150
chr6-143237374-INS-92
chr6-143253669-INS-124
chr6-143263510-INS-105
chr6-143263949-INS-439
chr6-143271429-INS-52
chr6-143302528-INS-276
chr6-143316464-INS-64
chr6-143333891-INS-280
chr6-143336951-INS-144
chr6-143339500-INS-498
chr6-143356899-INS-223
chr6-143374402-INS-306
chr6-143381598-INS-138
chr6-143401526-INS-57
chr6-143408784-INS-58
chr6-143410457-INS-103
chr6-143410486-INS-1152
chr6-143426297-INS-334
chr6-143441179-INS-97
chr6-143489971-INS-56
chr6-143494662-INS-881
chr6-143501466-INS-62
chr6-143509931-INS-384
chr6-143583528-INS-67
chr6-143631293-INS-86
chr6-143670024-INS-827
chr6-143704187-INS-300
chr6-143760124-INS-50
chr6-143788457-INS-89
chr6-143790882-INS-445
chr6-143827205-INS-54
chr6-143855092-INS-58
chr6-143860650-INS-1876
chr6-143866987-INS-6929
chr6-143868601-INS-121
chr6-143872145-INS-64
chr6-143872174-INS-504
chr6-143874968-INS-78
chr6-143900344-INS-153
chr6-143930714-INS-66
chr6-143933767-INS-198
chr6-143941526-INS-70
chr6-143955406-INS-62
chr6-143963335-INS-97
chr6-143997120-INS-272
chr6-144009672-INS-156
chr6-144014448-INS-62
chr6-144036502-INS-151
chr6-144054981-INS-52
chr6-144056071-INS-6842
chr6-144086159-INS-129
chr6-144110359-INS-404
chr6-144123971-INS-53
chr6-144129743-INS-115
chr6-144141637-INS-193
chr6-144160648-INS-62
chr6-144185660-INS-177
chr6-144193669-INS-54
chr6-144207754-INS-120
chr6-144224786-INS-96
chr6-144227851-INS-64
chr6-144232390-INS-59
chr6-144240092-INS-5347
chr6-144337626-INS-64
chr6-144366179-INS-162
chr6-144367746-INS-60
chr6-144432338-INS-373
chr6-144437499-INS-55
chr6-144455486-INS-156
chr6-144460217-INS-384
chr6-144465191-INS-215
chr6-144495279-INS-54
chr6-144507485-INS-385

chr6-144512148-INS-220
chr6-144518665-INS-58
chr6-144538795-INS-64
chr6-144540817-INS-76
chr6-144545370-INS-132
chr6-144547173-INS-579
chr6-144558317-INS-176
chr6-144561043-INS-256
chr6-144592101-INS-54
chr6-144596660-INS-56
chr6-144603269-INS-132
chr6-144605293-INS-374
chr6-144614225-INS-128
chr6-144615069-INS-64
chr6-144617111-INS-152
chr6-144620340-INS-334
chr6-144626250-INS-409
chr6-144633029-INS-422
chr6-144640933-INS-112
chr6-144647486-INS-70
chr6-144700146-INS-58
chr6-144711989-INS-82
chr6-144721647-INS-606
chr6-144731778-INS-110
chr6-144743077-INS-97
chr6-144770470-INS-219
chr6-144798982-INS-51
chr6-144821452-INS-293
chr6-144844257-INS-915
chr6-144856990-INS-54
chr6-144864849-INS-54
chr6-144871847-INS-65
chr6-144906477-INS-207
chr6-144915099-INS-209
chr6-144931043-INS-724
chr6-144950268-INS-199
chr6-144975609-INS-166
chr6-144979196-INS-191
chr6-144979694-INS-326
chr6-144982972-INS-452
chr6-144989022-INS-164
chr6-145001912-INS-201
chr6-145004297-INS-152
chr6-145012665-INS-170
chr6-145016308-INS-714
chr6-145019769-INS-74
chr6-145021166-INS-89
chr6-145035245-INS-200
chr6-145044817-INS-161
chr6-145061280-INS-77
chr6-145075845-INS-85
chr6-145076744-INS-234
chr6-145080067-INS-96
chr6-145089975-INS-68
chr6-145096370-INS-133
chr6-145104847-INS-53
chr6-145114672-INS-298
chr6-145144118-INS-98
chr6-145152348-INS-156
chr6-145180497-INS-268
chr6-145188928-INS-234
chr6-145197950-INS-64
chr6-145201409-INS-124
chr6-145210082-INS-90
chr6-145226354-INS-81
chr6-145227147-INS-379
chr6-145234608-INS-226
chr6-145244568-INS-60
chr6-145278220-INS-103
chr6-145279999-INS-71
chr6-145283541-INS-836
chr6-145283542-INS-539
chr6-145304138-INS-142
chr6-145319345-INS-401
chr6-145322646-INS-105
chr6-145325025-INS-210
chr6-145328014-INS-360

chr6-145347856-INS-160
chr6-145371691-INS-54
chr6-145385907-INS-275
chr6-145398625-INS-59
chr6-145400156-INS-1111
chr6-145400344-INS-142
chr6-145439603-INS-116
chr6-145444565-INS-99
chr6-145475392-INS-294
chr6-145484385-INS-88
chr6-145488664-INS-64
chr6-145508315-INS-69
chr6-145516282-INS-87
chr6-145523281-INS-139
chr6-145568954-INS-55
chr6-145581369-INS-354
chr6-145582874-INS-204
chr6-145595753-INS-81
chr6-145608545-INS-55
chr6-145614213-INS-357
chr6-145624870-INS-150
chr6-145629779-INS-109
chr6-145657699-INS-171
chr6-145684538-INS-81
chr6-145715839-INS-55
chr6-145743334-INS-63
chr6-145744294-INS-528
chr6-145762149-INS-81
chr6-145790911-INS-62
chr6-145794978-INS-106
chr6-145816769-INS-72
chr6-145818853-INS-58
chr6-145825556-INS-60
chr6-145849499-INS-62
chr6-145851656-INS-52
chr6-145855407-INS-97
chr6-145886862-INS-112
chr6-145896108-INS-57
chr6-145897154-INS-54
chr6-145929988-INS-5520
chr6-145966337-INS-176
chr6-145983670-INS-60
chr6-146005446-INS-252
chr6-146006389-INS-5199
chr6-146014872-INS-53
chr6-146133754-INS-108
chr6-146154018-INS-793
chr6-146182413-INS-618
chr6-146230008-INS-80
chr6-146235949-INS-401
chr6-146247158-INS-165
chr6-146252869-INS-396
chr6-146258348-INS-92
chr6-146280403-INS-310
chr6-146304130-INS-494
chr6-146307460-INS-64
chr6-146313119-INS-78
chr6-146331602-INS-365
chr6-146335523-INS-54
chr6-146337959-INS-68
chr6-146341570-INS-112
chr6-146413732-INS-192
chr6-146432843-INS-59
chr6-146453401-INS-171
chr6-146468237-INS-3366
chr6-146485549-INS-211
chr6-146491925-INS-183
chr6-146493070-INS-200
chr6-146500721-INS-1031
chr6-146533151-INS-77
chr6-146539754-INS-196
chr6-146540323-INS-224
chr6-146558639-INS-937
chr6-146582448-INS-146
chr6-146583463-INS-209
chr6-146615476-INS-83
chr6-146619721-INS-388

chr6-146635593-INS-992
chr6-146651326-INS-350
chr6-146657937-INS-225
chr6-146703683-INS-200
chr6-146710966-INS-224
chr6-146715038-INS-315
chr6-146739291-INS-356
chr6-146756862-INS-154
chr6-146763298-INS-434
chr6-146773435-INS-70
chr6-146785291-INS-66
chr6-146838661-INS-54
chr6-146877623-INS-399
chr6-146915478-INS-198
chr6-146938482-INS-112
chr6-146965936-INS-51
chr6-146975278-INS-85
chr6-146977950-INS-113
chr6-147007475-INS-53
chr6-147020655-INS-478
chr6-147020705-INS-481
chr6-147032046-INS-76
chr6-147058846-INS-582
chr6-147128132-INS-571
chr6-147182797-INS-212
chr6-147286479-INS-6405
chr6-147333691-INS-356
chr6-147358733-INS-6227
chr6-147426534-INS-7080
chr6-147426854-INS-148
chr6-147428247-INS-235
chr6-147468148-INS-71
chr6-147483061-INS-6376
chr6-147484270-INS-77
chr6-147501467-INS-68
chr6-147514931-INS-493
chr6-147522242-INS-6629
chr6-147530675-INS-333
chr6-147549307-INS-183
chr6-147564965-INS-123
chr6-147578892-INS-440
chr6-147586847-INS-132
chr6-147592764-INS-556
chr6-147593953-INS-116
chr6-147599966-INS-7092
chr6-147620498-INS-132
chr6-147625973-INS-69
chr6-147633221-INS-56
chr6-147666454-INS-264
chr6-147669872-INS-325
chr6-147673531-INS-52
chr6-147708738-INS-667
chr6-147716911-INS-58
chr6-147736213-INS-70
chr6-147737772-INS-605
chr6-147758864-INS-296
chr6-147769202-INS-435
chr6-147770415-INS-4077
chr6-147770562-INS-199
chr6-147772088-INS-107
chr6-147785584-INS-107
chr6-147786236-INS-97
chr6-147787215-INS-224
chr6-147820438-INS-54
chr6-147825103-INS-92
chr6-147840365-INS-60
chr6-147853244-INS-76
chr6-147856080-INS-93
chr6-147875338-INS-7388
chr6-147875627-INS-91
chr6-147889919-INS-122
chr6-147895794-INS-52
chr6-147898204-INS-109
chr6-147899908-INS-1151
chr6-147916414-INS-64
chr6-147921480-INS-207
chr6-147998338-INS-69

chr6-148045182-INS-152
chr6-148050377-INS-100
chr6-148054270-INS-51
chr6-148096315-INS-204
chr6-148100387-INS-88
chr6-148122368-INS-209
chr6-148166316-INS-69
chr6-148191941-INS-6372
chr6-148193277-INS-160
chr6-148195989-INS-231
chr6-148198474-INS-7249
chr6-148214743-INS-377
chr6-148229396-INS-52
chr6-148277873-INS-125
chr6-148281478-INS-58
chr6-148316442-INS-597
chr6-148359534-INS-1426
chr6-148361490-INS-7146
chr6-148364642-INS-97
chr6-148382798-INS-53
chr6-148391705-INS-443
chr6-148395197-INS-90
chr6-148408700-INS-558
chr6-148418310-INS-276
chr6-148418947-INS-138
chr6-148421137-INS-1633
chr6-148434666-INS-98
chr6-148453593-INS-346
chr6-148465394-INS-172
chr6-148475823-INS-715
chr6-148481036-INS-65
chr6-148487490-INS-71
chr6-148527547-INS-300
chr6-148531541-INS-794
chr6-148535162-INS-240
chr6-148536056-INS-194
chr6-148546561-INS-5514
chr6-148557461-INS-52
chr6-148558856-INS-52
chr6-148569828-INS-4875
chr6-148597817-INS-500
chr6-148600430-INS-541
chr6-148607831-INS-64
chr6-148634901-INS-338
chr6-148645412-INS-1814
chr6-148647467-INS-222
chr6-148670714-INS-123
chr6-148670987-INS-65
chr6-148682303-INS-818
chr6-148704114-INS-992
chr6-148712517-INS-89
chr6-148721066-INS-122
chr6-148729708-INS-251
chr6-148733421-INS-100
chr6-148736986-INS-1213
chr6-148741361-INS-206
chr6-148745408-INS-91
chr6-148764228-INS-55
chr6-148772469-INS-198
chr6-148780287-INS-208
chr6-148784811-INS-187
chr6-148786470-INS-212
chr6-148787806-INS-202
chr6-148793288-INS-384
chr6-148817745-INS-239
chr6-148828630-INS-198
chr6-148832934-INS-182
chr6-148840395-INS-237
chr6-148855391-INS-227
chr6-148866373-INS-204
chr6-148871124-INS-387
chr6-148886567-INS-7170
chr6-148886884-INS-63
chr6-148888196-INS-113
chr6-148903924-INS-210
chr6-148928463-INS-343
chr6-148935828-INS-209

chr6-148941318-INS-70
chr6-148943799-INS-7084
chr6-148957570-INS-51
chr6-148966764-INS-171
chr6-148972058-INS-60
chr6-148982873-INS-215
chr6-148985418-INS-157
chr6-149018225-INS-187
chr6-149022362-INS-198
chr6-149026015-INS-154
chr6-149028938-INS-126
chr6-149042276-INS-56
chr6-149047335-INS-419
chr6-149057886-INS-4567
chr6-149092527-INS-88
chr6-149096805-INS-5304
chr6-149097509-INS-745
chr6-149101724-INS-204
chr6-149105032-INS-342
chr6-149105456-INS-1324
chr6-149108349-INS-74
chr6-149124036-INS-190
chr6-149134913-INS-69
chr6-149153375-INS-394
chr6-149157658-INS-101
chr6-149165717-INS-390
chr6-149177043-INS-3242
chr6-149178976-INS-506
chr6-149189448-INS-5397
chr6-149249639-INS-6951
chr6-149249835-INS-4825
chr6-149250110-INS-226
chr6-149254348-INS-60
chr6-149257268-INS-459
chr6-149271841-INS-245
chr6-149275124-INS-6870
chr6-149278980-INS-106
chr6-149298732-INS-190
chr6-149298829-INS-1890
chr6-149302028-INS-275
chr6-149313344-INS-209
chr6-149325826-INS-91
chr6-149330411-INS-124
chr6-149340655-INS-62
chr6-149342657-INS-175
chr6-149343785-INS-257
chr6-149355728-INS-150
chr6-149378780-INS-52
chr6-149386161-INS-52
chr6-149475178-INS-816
chr7-3060546-INS-52
chr7-3148886-INS-189
chr7-3150574-INS-202
chr7-3158308-INS-92
chr7-3188916-INS-506
chr7-3197456-INS-164
chr7-3201310-INS-160
chr7-3206837-INS-60
chr7-3217062-INS-194
chr7-3219343-INS-211
chr7-3221417-INS-200
chr7-3237382-INS-214
chr7-3238039-INS-213
chr7-3240993-INS-227
chr7-3244884-INS-3228
chr7-3247998-INS-51
chr7-3253241-INS-218
chr7-3254725-INS-103
chr7-3256054-INS-76
chr7-3264744-INS-96
chr7-3275081-INS-463
chr7-3275616-INS-7478
chr7-3290417-INS-383
chr7-3300182-INS-284
chr7-3305385-INS-52
chr7-3307598-INS-396
chr7-3310685-INS-130

chr7-3374426-INS-61
chr7-3381057-INS-57
chr7-3388885-INS-87
chr7-3393616-INS-70
chr7-3399446-INS-203
chr7-3400931-INS-122
chr7-3410255-INS-169
chr7-3420959-INS-71
chr7-3433094-INS-128
chr7-3504172-INS-60
chr7-3509727-INS-88
chr7-3515318-INS-62
chr7-3549691-INS-52
chr7-3551321-INS-164
chr7-3558198-INS-813
chr7-3565279-INS-186
chr7-3567850-INS-80
chr7-3576085-INS-138
chr7-3584416-INS-98
chr7-3593281-INS-208
chr7-3604748-INS-218
chr7-3605603-INS-120
chr7-3606658-INS-170
chr7-3610837-INS-520
chr7-3679374-INS-59
chr7-3684405-INS-77
chr7-3689270-INS-164
chr7-3717344-INS-52
chr7-3725227-INS-329
chr7-3726069-INS-187
chr7-3726279-INS-572
chr7-3777790-INS-129
chr7-3824543-INS-220
chr7-3826899-INS-146
chr7-3831451-INS-382
chr7-3831492-INS-519
chr7-3837005-INS-282
chr7-3851528-INS-75
chr7-3875971-INS-62
chr7-3932258-INS-182
chr7-3936307-INS-6662
chr7-3947300-INS-6474
chr7-3950155-INS-1093
chr7-3983725-INS-52
chr7-3989684-INS-70
chr7-3999003-INS-184
chr7-4007749-INS-6250
chr7-4065858-INS-124
chr7-4067052-INS-70
chr7-4083460-INS-492
chr7-4086907-INS-105
chr7-4113202-INS-403
chr7-4124239-INS-60
chr7-4171473-INS-54
chr7-4177093-INS-184
chr7-4178508-INS-201
chr7-4205263-INS-684
chr7-4213325-INS-1087
chr7-4217349-INS-5717
chr7-4219681-INS-4671
chr7-4240319-INS-565
chr7-4275071-INS-200
chr7-4290253-INS-87
chr7-4301109-INS-542
chr7-4317208-INS-112
chr7-4319558-INS-444
chr7-4345049-INS-90
chr7-4361633-INS-54
chr7-4375055-INS-131
chr7-4397849-INS-143
chr7-4409086-INS-7662
chr7-4421689-INS-976
chr7-4424380-INS-194
chr7-4425646-INS-295
chr7-4427661-INS-404
chr7-4442362-INS-148
chr7-4448881-INS-56

chr7-4450126-INS-238
chr7-4456630-INS-354
chr7-4458092-INS-418
chr7-4464409-INS-70
chr7-4482739-INS-275
chr7-4490134-INS-114
chr7-4490857-INS-86
chr7-4492227-INS-217
chr7-4503511-INS-57
chr7-4509152-INS-488
chr7-4511699-INS-67
chr7-4519827-INS-79
chr7-4522307-INS-54
chr7-4531361-INS-268
chr7-4544753-INS-240
chr7-4556070-INS-70
chr7-4626883-INS-206
chr7-4630291-INS-128
chr7-4652927-INS-155
chr7-4654246-INS-337
chr7-4683508-INS-216
chr7-4698107-INS-66
chr7-4716501-INS-58
chr7-4722562-INS-129
chr7-4744673-INS-225
chr7-4763546-INS-86
chr7-4777438-INS-215
chr7-4778668-INS-296
chr7-4807170-INS-86
chr7-4817093-INS-56
chr7-4819610-INS-115
chr7-4823502-INS-76
chr7-4855490-INS-3790
chr7-4855665-INS-118
chr7-4856636-INS-197
chr7-4877060-INS-99
chr7-4887903-INS-2813
chr7-4887950-INS-394
chr7-4888128-INS-1302
chr7-4889749-INS-59
chr7-4891152-INS-282
chr7-4892664-INS-162
chr7-4898570-INS-54
chr7-4900046-INS-61
chr7-4911058-INS-102
chr7-4915892-INS-64
chr7-4941798-INS-92
chr7-4953152-INS-401
chr7-4971903-INS-62
chr7-5008804-INS-178
chr7-5028017-INS-572
chr7-5046884-INS-545
chr7-5085863-INS-108
chr7-5090439-INS-58
chr7-5097344-INS-292
chr7-5106676-INS-108
chr7-5112785-INS-386
chr7-5123051-INS-101
chr7-5129558-INS-54
chr7-5145610-INS-7077
chr7-5172655-INS-441
chr7-5177296-INS-344
chr7-5179395-INS-1292
chr7-5218768-INS-686
chr7-5242276-INS-3331
chr7-5282402-INS-328
chr7-5286288-INS-52
chr7-5307293-INS-605
chr7-5308417-INS-132
chr7-5313997-INS-227
chr7-5324352-INS-356
chr7-5326730-INS-179
chr7-5341352-INS-63
chr7-5342163-INS-966
chr7-5357458-INS-114
chr7-5368548-INS-202
chr7-5391755-INS-182

chr7-5401770-INS-3034
chr7-5414501-INS-84
chr7-5450925-INS-304
chr7-5458032-INS-84
chr7-5464015-INS-52
chr7-5473532-INS-432
chr7-5501878-INS-777
chr7-5509016-INS-6721
chr7-5523510-INS-2643
chr7-5528111-INS-199
chr7-5534044-INS-5854
chr7-5543437-INS-235
chr7-5562351-INS-208
chr7-5599114-INS-7155
chr7-5612449-INS-60
chr7-5673275-INS-6473
chr7-5769920-INS-142
chr7-5771819-INS-573
chr7-5772797-INS-200
chr7-5810792-INS-6917
chr7-5900215-INS-504
chr7-5952466-INS-2849
chr7-5963825-INS-64
chr7-5977285-INS-326
chr7-5978023-INS-3001
chr7-5985490-INS-1894
chr7-5991557-INS-6214
chr7-6046272-INS-171
chr7-6046865-INS-104
chr7-6059032-INS-212
chr7-6081487-INS-3980
chr7-6084412-INS-214
chr7-6107329-INS-205
chr7-6118166-INS-7166
chr7-6134005-INS-52
chr7-6164708-INS-82
chr7-6198614-INS-143
chr7-6202297-INS-51
chr7-6216722-INS-114
chr7-6217637-INS-120
chr7-6222951-INS-217
chr7-6228661-INS-209
chr7-6251013-INS-75
chr7-6259991-INS-80
chr7-6263025-INS-93
chr7-6264223-INS-598
chr7-6270289-INS-135
chr7-6271553-INS-180
chr7-6275372-INS-616
chr7-6283395-INS-53
chr7-6299104-INS-96
chr7-6302067-INS-88
chr7-6304190-INS-72
chr7-6312457-INS-231
chr7-6315418-INS-200
chr7-6321666-INS-275
chr7-6322933-INS-55
chr7-6342115-INS-7097
chr7-6354869-INS-78
chr7-6361880-INS-55
chr7-6368846-INS-72
chr7-6373643-INS-186
chr7-6376840-INS-275
chr7-6377141-INS-7139
chr7-6405404-INS-681
chr7-6411727-INS-58
chr7-6414434-INS-51
chr7-6416070-INS-52
chr7-6427951-INS-431
chr7-6453333-INS-550
chr7-6457219-INS-116
chr7-6475563-INS-1870
chr7-6476115-INS-104
chr7-6501505-INS-82
chr7-6510103-INS-769
chr7-6514065-INS-202
chr7-6522455-INS-403

chr7-6527471-INS-846
chr7-6532735-INS-107
chr7-6561779-INS-51
chr7-6564035-INS-123
chr7-6585987-INS-144
chr7-6591170-INS-63
chr7-6630418-INS-82
chr7-6641966-INS-134
chr7-6684356-INS-60
chr7-6705523-INS-110
chr7-6725281-INS-66
chr7-6749794-INS-70
chr7-6750686-INS-551
chr7-6752811-INS-334
chr7-6763656-INS-322
chr7-6776724-INS-80
chr7-6790742-INS-505
chr7-6791958-INS-6442
chr7-6839653-INS-286
chr7-6843675-INS-61
chr7-6851360-INS-246
chr7-6864799-INS-158
chr7-6895674-INS-5217
chr7-6897064-INS-57
chr7-6899447-INS-51
chr7-6905694-INS-277
chr7-6907666-INS-55
chr7-6910062-INS-118
chr7-6938504-INS-80
chr7-6939574-INS-228
chr7-6945238-INS-4753
chr7-6971256-INS-129
chr7-6985946-INS-222
chr7-7042471-INS-221
chr7-7056657-INS-53
chr7-7063247-INS-67
chr7-7128254-INS-162
chr7-7133663-INS-781
chr7-7133836-INS-3592
chr7-7160696-INS-78
chr7-7176470-INS-89
chr7-7181312-INS-156
chr7-7192189-INS-269
chr7-7214252-INS-54
chr7-7251005-INS-663
chr7-7259078-INS-302
chr7-7262906-INS-6421
chr7-7266243-INS-251
chr7-7274274-INS-64
chr7-7282132-INS-6224
chr7-7287578-INS-70
chr7-7288638-INS-116
chr7-7295611-INS-517
chr7-7303447-INS-95
chr7-7378939-INS-86
chr7-7483935-INS-569
chr7-7545887-INS-88
chr7-7582634-INS-6819
chr7-7600456-INS-82
chr7-7619191-INS-54
chr7-7627169-INS-68
chr7-7648263-INS-2851
chr7-7702504-INS-3117
chr7-7706404-INS-2771
chr7-7707480-INS-6560
chr7-7753850-INS-84
chr7-7770072-INS-209
chr7-7804613-INS-110
chr7-7809198-INS-111
chr7-7917779-INS-1883
chr7-8191162-INS-1061
chr7-8215842-INS-886
chr7-8244978-INS-7007
chr7-8285025-INS-6194
chr7-8542181-INS-72
chr7-8715863-INS-73
chr7-8728284-INS-6440

chr7-8776164-INS-7074
chr7-8877197-INS-3993
chr7-9010280-INS-290
chr7-9455676-INS-60
chr7-9603830-INS-1802
chr7-9724860-INS-146
chr7-9751231-INS-6501
chr7-9772768-INS-6414
chr7-9865857-INS-163
chr7-9867229-INS-5093
chr7-9913381-INS-52
chr7-9918530-INS-72
chr7-9934034-INS-454
chr7-9963315-INS-6591
chr7-9984050-INS-56
chr7-10038894-INS-56
chr7-10067944-INS-4321
chr7-10067945-INS-474
chr7-10092987-INS-56
chr7-10101821-INS-277
chr7-10114890-INS-4403
chr7-10132182-INS-7086
chr7-10152861-INS-7312
chr7-10156756-INS-137
chr7-10158607-INS-925
chr7-10192146-INS-6735
chr7-10193297-INS-344
chr7-10218786-INS-51
chr7-10220469-INS-501
chr7-10230438-INS-198
chr7-10248162-INS-145
chr7-10297376-INS-154
chr7-10341734-INS-52
chr7-10384000-INS-5391
chr7-10397008-INS-860
chr7-10425692-INS-63
chr7-10446423-INS-106
chr7-10450881-INS-191
chr7-10462901-INS-72
chr7-10470416-INS-6444
chr7-10488932-INS-550
chr7-10620610-INS-186
chr7-10640705-INS-258
chr7-10664255-INS-215
chr7-10728610-INS-498
chr7-10858945-INS-173
chr7-10882409-INS-485
chr7-10892883-INS-445
chr7-10965358-INS-99
chr7-11046334-INS-256
chr7-11051225-INS-828
chr7-11123564-INS-5261
chr7-11208916-INS-52
chr7-11357432-INS-7252
chr7-11368159-INS-8545
chr7-11376510-INS-385
chr7-11409206-INS-52
chr7-11433224-INS-163
chr7-11475061-INS-485
chr7-11495664-INS-63
chr7-11556847-INS-59
chr7-11571534-INS-84
chr7-11583843-INS-329
chr7-11586296-INS-197
chr7-11632432-INS-221
chr7-11632442-INS-6364
chr7-11639432-INS-6215
chr7-11643962-INS-190
chr7-11644361-INS-4660
chr7-11678325-INS-498
chr7-11695969-INS-605
chr7-11716347-INS-296
chr7-11723455-INS-655
chr7-11723471-INS-87
chr7-11744353-INS-10350
chr7-11751411-INS-7096
chr7-11760571-INS-7531

chr7-11799464-INS-7492
chr7-11802042-INS-627
chr7-11810710-INS-77
chr7-11858628-INS-78
chr7-11867355-INS-7367
chr7-11872940-INS-365
chr7-11912839-INS-328
chr7-11932668-INS-67
chr7-11985137-INS-6216
chr7-12013548-INS-6661
chr7-12042016-INS-197
chr7-12081542-INS-351
chr7-12093793-INS-58
chr7-12098006-INS-743
chr7-12148100-INS-186
chr7-12171658-INS-328
chr7-12180274-INS-175
chr7-12196578-INS-74
chr7-12198534-INS-73
chr7-12201652-INS-74
chr7-12211038-INS-401
chr7-12219263-INS-62
chr7-12234065-INS-81
chr7-12257102-INS-5405
chr7-12260511-INS-214
chr7-12270350-INS-400
chr7-12333762-INS-110
chr7-12339235-INS-166
chr7-12352582-INS-67
chr7-12355077-INS-6712
chr7-12363238-INS-8215
chr7-12389032-INS-73
chr7-12394123-INS-60
chr7-12407038-INS-2358
chr7-12438906-INS-3294
chr7-12447382-INS-61
chr7-12452135-INS-199
chr7-12473711-INS-65
chr7-12486502-INS-165
chr7-12497301-INS-142
chr7-12501390-INS-453
chr7-12520239-INS-195
chr7-12547196-INS-473
chr7-12549926-INS-175
chr7-12558940-INS-1143
chr7-12562820-INS-498
chr7-12586628-INS-227
chr7-12613752-INS-175
chr7-12627971-INS-101
chr7-12629100-INS-209
chr7-12644689-INS-167
chr7-12645318-INS-578
chr7-12648941-INS-51
chr7-12654961-INS-82
chr7-12658457-INS-271
chr7-12665988-INS-79
chr7-12670099-INS-516
chr7-12685785-INS-6961
chr7-12696902-INS-120
chr7-12706566-INS-224
chr7-12744880-INS-513
chr7-12753918-INS-78
chr7-12781669-INS-54
chr7-12783391-INS-224
chr7-12790960-INS-253
chr7-12794084-INS-264
chr7-12797161-INS-7247
chr7-12798726-INS-219
chr7-12821540-INS-329
chr7-12822179-INS-59
chr7-12884036-INS-2115
chr7-12887554-INS-577
chr7-12888164-INS-7078
chr7-12897111-INS-91
chr7-12899490-INS-548
chr7-12900283-INS-7426
chr7-12900284-INS-3391

chr7-12937622-INS-5030
chr7-12949801-INS-377
chr7-12958710-INS-133
chr7-13044183-INS-261
chr7-13050465-INS-6718
chr7-13053977-INS-63
chr7-13055488-INS-154
chr7-13076852-INS-56
chr7-13085618-INS-480
chr7-13113604-INS-54
chr7-13132926-INS-63
chr7-13142498-INS-86
chr7-13169130-INS-122
chr7-13255736-INS-5653
chr7-13268575-INS-62
chr7-13269227-INS-200
chr7-13279759-INS-56
chr7-13294670-INS-93
chr7-13341273-INS-2667
chr7-13356775-INS-6594
chr7-13370774-INS-140
chr7-13373157-INS-284
chr7-13377572-INS-5239
chr7-13405288-INS-52
chr7-13420700-INS-89
chr7-13438438-INS-1685
chr7-13456788-INS-109
chr7-13459914-INS-823
chr7-13479548-INS-6271
chr7-13521522-INS-4594
chr7-13642537-INS-469
chr7-13776963-INS-56
chr7-13817770-INS-77
chr7-13857779-INS-492
chr7-13905426-INS-8454
chr7-13925698-INS-154
chr7-13960870-INS-551
chr7-13970492-INS-164
chr7-14008347-INS-212
chr7-14013710-INS-5278
chr7-14020870-INS-87
chr7-14072491-INS-203
chr7-14137614-INS-201
chr7-14149272-INS-88
chr7-14204912-INS-214
chr7-14210452-INS-98
chr7-14222795-INS-1528
chr7-14240826-INS-348
chr7-14261418-INS-258
chr7-14303969-INS-167
chr7-14316411-INS-266
chr7-14329403-INS-338
chr7-14358154-INS-1956
chr7-14413260-INS-213
chr7-14464202-INS-1893
chr7-14474725-INS-7156
chr7-14496820-INS-470
chr7-14503036-INS-6853
chr7-14538828-INS-52
chr7-14584548-INS-2266
chr7-14600697-INS-1964
chr7-14611436-INS-6440
chr7-14612186-INS-6440
chr7-14629880-INS-5390
chr7-14766295-INS-6638
chr7-14782371-INS-6015
chr7-14785762-INS-1505
chr7-14802268-INS-5403
chr7-14836740-INS-1434
chr7-14859994-INS-6424
chr7-14892599-INS-98
chr7-14901639-INS-259
chr7-15116625-INS-4337
chr7-15210865-INS-147
chr7-15316595-INS-1347
chr7-15367218-INS-89
chr7-15368025-INS-66

chr7-15382251-INS-52
chr7-15393638-INS-5182
chr7-15407093-INS-307
chr7-15465577-INS-609
chr7-15552024-INS-7096
chr7-15553360-INS-68
chr7-15559910-INS-52
chr7-15560638-INS-62
chr7-15561736-INS-197
chr7-15565210-INS-64
chr7-15587319-INS-222
chr7-15644388-INS-65
chr7-15648774-INS-83
chr7-15655586-INS-65
chr7-15660052-INS-193
chr7-15665056-INS-94
chr7-15666548-INS-111
chr7-15666875-INS-89
chr7-15679112-INS-52
chr7-15680263-INS-169
chr7-15704946-INS-58
chr7-15712441-INS-8734
chr7-15715784-INS-79
chr7-15726589-INS-95
chr7-15739653-INS-92
chr7-15742832-INS-77
chr7-15748250-INS-53
chr7-15771270-INS-240
chr7-15785559-INS-108
chr7-15787168-INS-387
chr7-15799515-INS-171
chr7-15801972-INS-126
chr7-15852862-INS-217
chr7-15870909-INS-202
chr7-15907748-INS-250
chr7-15910712-INS-266
chr7-15927883-INS-7226
chr7-15940550-INS-98
chr7-15946497-INS-54
chr7-15960974-INS-249
chr7-15973242-INS-59
chr7-15974610-INS-108
chr7-15978386-INS-86
chr7-15988058-INS-64
chr7-15988691-INS-224
chr7-15994209-INS-422
chr7-16000183-INS-180
chr7-16021774-INS-58
chr7-16027789-INS-2045
chr7-16035976-INS-106
chr7-16036345-INS-98
chr7-16041309-INS-70
chr7-16048979-INS-222
chr7-16054663-INS-68
chr7-16055036-INS-51
chr7-16076678-INS-132
chr7-16087908-INS-424
chr7-16120384-INS-217
chr7-16166050-INS-212
chr7-16172637-INS-184
chr7-16176271-INS-176
chr7-16180716-INS-310
chr7-16207534-INS-74
chr7-16214199-INS-162
chr7-16217045-INS-541
chr7-16270528-INS-76
chr7-16291095-INS-252
chr7-16310348-INS-109
chr7-16314060-INS-52
chr7-16317132-INS-283
chr7-16337029-INS-72
chr7-16339904-INS-55
chr7-16363353-INS-552
chr7-16370329-INS-52
chr7-16395507-INS-212
chr7-16432026-INS-571
chr7-16438773-INS-383

chr7-16440786-INS-193
chr7-16446133-INS-75
chr7-16454729-INS-416
chr7-16455752-INS-121
chr7-16459558-INS-70
chr7-16461449-INS-930
chr7-16482410-INS-52
chr7-16484430-INS-525
chr7-16505007-INS-207
chr7-16528729-INS-216
chr7-16533205-INS-211
chr7-16536566-INS-320
chr7-16537256-INS-125
chr7-16546497-INS-150
chr7-16588812-INS-53
chr7-16615702-INS-213
chr7-16631994-INS-94
chr7-16645976-INS-175
chr7-16653134-INS-60
chr7-16668770-INS-367
chr7-16674684-INS-438
chr7-16682904-INS-209
chr7-16698904-INS-105
chr7-16729484-INS-96
chr7-16774896-INS-157
chr7-16784295-INS-65
chr7-16785510-INS-59
chr7-16798397-INS-212
chr7-16802375-INS-3102
chr7-16822801-INS-52
chr7-16857666-INS-1767
chr7-16864413-INS-341
chr7-16904112-INS-187
chr7-16906425-INS-83
chr7-16912565-INS-400
chr7-16922437-INS-201
chr7-16952469-INS-331
chr7-16954491-INS-90
chr7-16974854-INS-453
chr7-16982416-INS-622
chr7-16994451-INS-479
chr7-17036199-INS-569
chr7-17215811-INS-74
chr7-17223717-INS-6734
chr7-17274595-INS-99
chr7-17313175-INS-149
chr7-17334532-INS-304
chr7-17370063-INS-7585
chr7-17388463-INS-166
chr7-17437881-INS-204
chr7-17452654-INS-1036
chr7-17492803-INS-65
chr7-17498393-INS-52
chr7-17552045-INS-152
chr7-17553393-INS-66
chr7-17572485-INS-6254
chr7-17574778-INS-9096
chr7-17608702-INS-65
chr7-17614480-INS-193
chr7-17661645-INS-592
chr7-17665857-INS-419
chr7-17672024-INS-193
chr7-17705413-INS-164
chr7-17717088-INS-2222
chr7-17783375-INS-6331
chr7-17808198-INS-7324
chr7-17817573-INS-1329
chr7-17836928-INS-272
chr7-17838577-INS-6535
chr7-17840594-INS-52
chr7-17895747-INS-401
chr7-17922065-INS-211
chr7-17922070-INS-411
chr7-17943113-INS-1341
chr7-17955982-INS-208
chr7-17971735-INS-322
chr7-17976083-INS-88

chr7-17977649-INS-823
chr7-17988976-INS-89
chr7-17994831-INS-101
chr7-18004899-INS-83
chr7-18010826-INS-102
chr7-18031846-INS-69
chr7-18033752-INS-58
chr7-18043618-INS-6443
chr7-18056580-INS-56
chr7-18095648-INS-410
chr7-18098228-INS-4666
chr7-18104075-INS-138
chr7-18137618-INS-54
chr7-18142066-INS-135
chr7-18181916-INS-96
chr7-18183465-INS-70
chr7-18270292-INS-186
chr7-18349459-INS-131
chr7-18368591-INS-204
chr7-18389525-INS-51
chr7-18414034-INS-506
chr7-18486237-INS-434
chr7-18489657-INS-537
chr7-18538236-INS-56
chr7-18540970-INS-124
chr7-18554693-INS-60
chr7-18558093-INS-105
chr7-18583332-INS-224
chr7-18599884-INS-400
chr7-18607533-INS-142
chr7-18612279-INS-238
chr7-18616793-INS-83
chr7-18635490-INS-132
chr7-18637178-INS-292
chr7-18646933-INS-215
chr7-18669875-INS-52
chr7-18675109-INS-106
chr7-18678326-INS-123
chr7-18679795-INS-53
chr7-18682994-INS-53
chr7-18703414-INS-51
chr7-18713595-INS-64
chr7-18715028-INS-72
chr7-18731182-INS-54
chr7-18752925-INS-101
chr7-18766331-INS-223
chr7-18802960-INS-51
chr7-18857723-INS-153
chr7-18859766-INS-65
chr7-18888253-INS-116
chr7-18901984-INS-159
chr7-18923318-INS-51
chr7-18937540-INS-53
chr7-18944735-INS-51
chr7-18954095-INS-194
chr7-18964109-INS-148
chr7-18969732-INS-209
chr7-18971429-INS-166
chr7-18976034-INS-433
chr7-18997072-INS-52
chr7-19008987-INS-168
chr7-19012724-INS-54
chr7-19020081-INS-51
chr7-19021999-INS-269
chr7-19022276-INS-491
chr7-19029213-INS-55
chr7-19034186-INS-52
chr7-19053228-INS-92
chr7-19057386-INS-60
chr7-19070287-INS-200
chr7-19098550-INS-196
chr7-19099118-INS-432
chr7-19104802-INS-104
chr7-19106588-INS-56
chr7-19123479-INS-218
chr7-19176420-INS-105
chr7-19179912-INS-95

chr7-19200351-INS-181
chr7-19210592-INS-122
chr7-19219809-INS-106
chr7-19222743-INS-189
chr7-19226273-INS-94
chr7-19228459-INS-159
chr7-19236493-INS-189
chr7-19244440-INS-106
chr7-19245653-INS-219
chr7-19279103-INS-88
chr7-19342445-INS-56
chr7-19344820-INS-203
chr7-19350010-INS-209
chr7-19352082-INS-233
chr7-19359344-INS-207
chr7-19374169-INS-318
chr7-19387174-INS-92
chr7-19402252-INS-165
chr7-19405143-INS-53
chr7-19407888-INS-52
chr7-19411194-INS-56
chr7-19425763-INS-62
chr7-19460746-INS-143
chr7-19462965-INS-105
chr7-19464357-INS-658
chr7-19468789-INS-314
chr7-19469731-INS-130
chr7-19494595-INS-106
chr7-19519908-INS-202
chr7-19522332-INS-265
chr7-19532248-INS-350
chr7-19535469-INS-325
chr7-19536850-INS-98
chr7-19538648-INS-208
chr7-19555058-INS-80
chr7-19560473-INS-147
chr7-19561819-INS-102
chr7-19574143-INS-408
chr7-19583777-INS-130
chr7-19593109-INS-138
chr7-19608575-INS-69
chr7-19614928-INS-224
chr7-19617166-INS-65
chr7-19618736-INS-107
chr7-19626388-INS-320
chr7-19627927-INS-497
chr7-19642884-INS-316
chr7-19657819-INS-51
chr7-19677110-INS-307
chr7-19712344-INS-587
chr7-19721290-INS-435
chr7-19732236-INS-60
chr7-19736771-INS-169
chr7-19743899-INS-70
chr7-19751030-INS-205
chr7-19755022-INS-216
chr7-19756335-INS-860
chr7-19756339-INS-51
chr7-19778504-INS-152
chr7-19781274-INS-86
chr7-19807409-INS-68
chr7-19852891-INS-3159
chr7-20280519-INS-212
chr7-20531312-INS-6819
chr7-20559730-INS-5372
chr7-20561100-INS-1083
chr7-20581439-INS-793
chr7-20628057-INS-7028
chr7-20668112-INS-7108
chr7-20704412-INS-213
chr7-20710639-INS-82
chr7-20812969-INS-64
chr7-20846925-INS-682
chr7-20906171-INS-358
chr7-20962036-INS-85
chr7-20977552-INS-117
chr7-21154977-INS-68

chr7-21290157-INS-497
chr7-21398831-INS-358
chr7-21631885-INS-506
chr7-21738465-INS-498
chr7-22116778-INS-107
chr7-22464401-INS-812
chr7-22556081-INS-3396
chr7-22815448-INS-64
chr7-22852951-INS-57
chr7-22855860-INS-77
chr7-22861891-INS-255
chr7-22870250-INS-921
chr7-22883841-INS-104
chr7-22987649-INS-358
chr7-22998358-INS-296
chr7-23019460-INS-8323
chr7-23037551-INS-66
chr7-23052272-INS-76
chr7-23091521-INS-627
chr7-23130738-INS-237
chr7-23133662-INS-53
chr7-23139239-INS-171
chr7-23148962-INS-886
chr7-23151382-INS-5544
chr7-23162701-INS-114
chr7-23170736-INS-5448
chr7-23172710-INS-5305
chr7-23184909-INS-6299
chr7-23233065-INS-60
chr7-23234225-INS-7081
chr7-23237572-INS-51
chr7-23263652-INS-6952
chr7-23369486-INS-7060
chr7-23461753-INS-209
chr7-23484906-INS-7142
chr7-23534278-INS-7607
chr7-23583354-INS-119
chr7-23597796-INS-63
chr7-23599930-INS-924
chr7-23665443-INS-571
chr7-23667366-INS-94
chr7-23674443-INS-396
chr7-23688612-INS-914
chr7-23691234-INS-315
chr7-23700468-INS-6467
chr7-23718943-INS-580
chr7-23724062-INS-458
chr7-23774810-INS-70
chr7-23785541-INS-481
chr7-23786275-INS-5165
chr7-23796444-INS-64
chr7-23797935-INS-100
chr7-23799638-INS-53
chr7-23803246-INS-66
chr7-23831963-INS-342
chr7-23844086-INS-617
chr7-23847876-INS-56
chr7-23922697-INS-66
chr7-23924277-INS-858
chr7-23929919-INS-322
chr7-23931010-INS-393
chr7-23935206-INS-62
chr7-23958430-INS-60
chr7-23979147-INS-335
chr7-23988875-INS-395
chr7-24007206-INS-100
chr7-24019937-INS-343
chr7-24033857-INS-356
chr7-24040781-INS-88
chr7-24048861-INS-205
chr7-24049115-INS-346
chr7-24061421-INS-144
chr7-24062268-INS-415
chr7-24065619-INS-171
chr7-24071791-INS-288
chr7-24091113-INS-90
chr7-24100314-INS-208

chr7-24138993-INS-120
chr7-24145566-INS-211
chr7-24147389-INS-213
chr7-24160714-INS-106
chr7-24168953-INS-157
chr7-24204190-INS-199
chr7-24205832-INS-253
chr7-24217136-INS-55
chr7-24229268-INS-207
chr7-24256866-INS-60
chr7-24261104-INS-219
chr7-24267113-INS-223
chr7-24276822-INS-88
chr7-24295464-INS-437
chr7-24296790-INS-318
chr7-24299648-INS-127
chr7-24302110-INS-156
chr7-24312832-INS-103
chr7-24315557-INS-104
chr7-24322146-INS-66
chr7-24332872-INS-157
chr7-24334880-INS-212
chr7-24352720-INS-159
chr7-24368417-INS-441
chr7-24382828-INS-89
chr7-24388628-INS-51
chr7-24400950-INS-2110
chr7-24402982-INS-640
chr7-24444010-INS-68
chr7-24483724-INS-76
chr7-24488539-INS-66
chr7-24500683-INS-90
chr7-24510383-INS-155
chr7-24512517-INS-235
chr7-24543951-INS-338
chr7-24553941-INS-88
chr7-24563064-INS-961
chr7-24563157-INS-198
chr7-24565850-INS-78
chr7-24574076-INS-92
chr7-24588613-INS-169
chr7-24589762-INS-125
chr7-24591389-INS-168
chr7-24594354-INS-58
chr7-24620888-INS-118
chr7-24653263-INS-70
chr7-24659685-INS-2910
chr7-24659922-INS-1120
chr7-24659963-INS-150
chr7-24661587-INS-136
chr7-24663206-INS-346
chr7-24664705-INS-86
chr7-24667135-INS-63
chr7-24688164-INS-160
chr7-24691551-INS-100
chr7-24695041-INS-89
chr7-24720004-INS-334
chr7-24726487-INS-211
chr7-24735906-INS-162
chr7-24736818-INS-474
chr7-24738848-INS-71
chr7-24761064-INS-82
chr7-24804686-INS-719
chr7-24808359-INS-183
chr7-24812958-INS-210
chr7-24817349-INS-116
chr7-24823343-INS-54
chr7-24824747-INS-104
chr7-24831489-INS-201
chr7-24840666-INS-222
chr7-24843226-INS-910
chr7-24843294-INS-172
chr7-24884717-INS-59
chr7-24888568-INS-87
chr7-24910162-INS-339
chr7-24912628-INS-211
chr7-24956162-INS-197

chr7-24961646-INS-62
chr7-25024762-INS-155
chr7-25030737-INS-215
chr7-25038269-INS-205
chr7-25047107-INS-51
chr7-25052533-INS-59
chr7-25061891-INS-345
chr7-25087117-INS-206
chr7-25096631-INS-67
chr7-25107874-INS-201
chr7-25125157-INS-83
chr7-25136680-INS-59
chr7-25155592-INS-52
chr7-25178547-INS-84
chr7-25180588-INS-202
chr7-25181674-INS-242
chr7-25185983-INS-100
chr7-25210671-INS-218
chr7-25228412-INS-57
chr7-25230185-INS-171
chr7-25232914-INS-126
chr7-25257191-INS-225
chr7-25259118-INS-229
chr7-25259670-INS-109
chr7-25279683-INS-765
chr7-25282364-INS-101
chr7-25294652-INS-188
chr7-25295519-INS-58
chr7-25304628-INS-163
chr7-25310398-INS-735
chr7-25368052-INS-220
chr7-25378768-INS-81
chr7-25392525-INS-316
chr7-25399414-INS-82
chr7-25399875-INS-120
chr7-25406199-INS-57
chr7-25447241-INS-62
chr7-25464496-INS-423
chr7-25473194-INS-268
chr7-25474533-INS-212
chr7-25479507-INS-60
chr7-25487314-INS-52
chr7-25503781-INS-438
chr7-25506818-INS-306
chr7-25512068-INS-118
chr7-25521353-INS-59
chr7-25527545-INS-127
chr7-25533600-INS-50
chr7-25536989-INS-572
chr7-25549895-INS-797
chr7-25569217-INS-182
chr7-25572879-INS-476
chr7-25581264-INS-178
chr7-25585048-INS-112
chr7-25594563-INS-250
chr7-25599136-INS-52
chr7-25600526-INS-299
chr7-25601853-INS-114
chr7-25603052-INS-123
chr7-25604624-INS-52
chr7-25620439-INS-869
chr7-25635463-INS-300
chr7-25638215-INS-161
chr7-25640762-INS-354
chr7-25649313-INS-621
chr7-25665144-INS-93
chr7-25739628-INS-216
chr7-25832806-INS-114
chr7-25858910-INS-745
chr7-25860072-INS-133
chr7-25900807-INS-209
chr7-25901629-INS-6877
chr7-25910627-INS-166
chr7-25953081-INS-267
chr7-25955629-INS-57
chr7-25967932-INS-6531
chr7-25973119-INS-844

chr7-25981440-INS-6590
chr7-26000262-INS-60
chr7-26021279-INS-6750
chr7-26060855-INS-1890
chr7-26063204-INS-874
chr7-26136809-INS-196
chr7-26138636-INS-65
chr7-26151861-INS-66
chr7-26160093-INS-63
chr7-26170770-INS-59
chr7-26177157-INS-484
chr7-26180644-INS-7478
chr7-26181013-INS-517
chr7-26226679-INS-7734
chr7-26242734-INS-156
chr7-26255649-INS-54
chr7-26272449-INS-406
chr7-26278374-INS-71
chr7-26284332-INS-325
chr7-26313620-INS-232
chr7-26322806-INS-208
chr7-26326178-INS-7095
chr7-26328758-INS-6166
chr7-26342139-INS-52
chr7-26359443-INS-124
chr7-26362003-INS-570
chr7-26362838-INS-71
chr7-26367315-INS-77
chr7-26374084-INS-120
chr7-26384093-INS-6983
chr7-26406342-INS-291
chr7-26419731-INS-80
chr7-26425571-INS-1349
chr7-26430154-INS-6740
chr7-26435919-INS-529
chr7-26448583-INS-605
chr7-26476826-INS-70
chr7-26478336-INS-60
chr7-26485436-INS-325
chr7-26505365-INS-192
chr7-26512162-INS-438
chr7-26514722-INS-62
chr7-26528233-INS-1194
chr7-26528234-INS-426
chr7-26528236-INS-120
chr7-26547476-INS-4985
chr7-26550023-INS-56
chr7-26555820-INS-937
chr7-26629410-INS-85
chr7-26647016-INS-55
chr7-26656049-INS-74
chr7-26662345-INS-74
chr7-26672032-INS-871
chr7-26680175-INS-6418
chr7-26680176-INS-509
chr7-26692530-INS-73
chr7-26738565-INS-57
chr7-26767044-INS-982
chr7-26777491-INS-332
chr7-26782074-INS-158
chr7-26789127-INS-64
chr7-26790482-INS-390
chr7-26803198-INS-72
chr7-26809284-INS-263
chr7-26810414-INS-281
chr7-26816957-INS-54
chr7-26823304-INS-98
chr7-26825507-INS-212
chr7-26836640-INS-265
chr7-26856407-INS-56
chr7-26881233-INS-207
chr7-26906579-INS-210
chr7-26909539-INS-195
chr7-26918530-INS-161
chr7-26936109-INS-380
chr7-26937007-INS-56
chr7-26965347-INS-318

chr7-26969086-INS-62
chr7-26978432-INS-5554
chr7-26996518-INS-212
chr7-27000410-INS-175
chr7-27011848-INS-266
chr7-27049450-INS-124
chr7-27057927-INS-204
chr7-27062486-INS-57
chr7-27073034-INS-52
chr7-27076985-INS-124
chr7-27078999-INS-208
chr7-27080033-INS-52
chr7-27081336-INS-87
chr7-27084634-INS-93
chr7-27086012-INS-149
chr7-27088129-INS-60
chr7-27110502-INS-93
chr7-27118914-INS-62
chr7-27122630-INS-105
chr7-27126168-INS-446
chr7-27128138-INS-53
chr7-27130565-INS-194
chr7-27135287-INS-199
chr7-27138166-INS-160
chr7-27153802-INS-193
chr7-27160711-INS-179
chr7-27187326-INS-53
chr7-27210215-INS-123
chr7-27220517-INS-69
chr7-27229770-INS-53
chr7-27234258-INS-197
chr7-27246960-INS-56
chr7-27275264-INS-519
chr7-27284555-INS-78
chr7-27285568-INS-582
chr7-27287854-INS-206
chr7-27311245-INS-194
chr7-27312052-INS-393
chr7-27351910-INS-50
chr7-27375276-INS-213
chr7-27377580-INS-736
chr7-27383515-INS-61
chr7-27392500-INS-208
chr7-27397293-INS-148
chr7-27413770-INS-208
chr7-27413998-INS-4224
chr7-27427537-INS-938
chr7-27428529-INS-370
chr7-27430492-INS-101
chr7-27439350-INS-187
chr7-27450569-INS-213
chr7-27455369-INS-176
chr7-27457232-INS-5044
chr7-27461310-INS-338
chr7-27489969-INS-152
chr7-27491219-INS-175
chr7-27508630-INS-96
chr7-27519740-INS-5349
chr7-27544517-INS-190
chr7-27572422-INS-143
chr7-27599162-INS-272
chr7-27606774-INS-442
chr7-27622865-INS-70
chr7-27623667-INS-52
chr7-27629117-INS-217
chr7-27630910-INS-89
chr7-27631107-INS-136
chr7-27644392-INS-432
chr7-27665007-INS-125
chr7-27672868-INS-66
chr7-27681998-INS-209
chr7-27690310-INS-134
chr7-27712273-INS-522
chr7-27716185-INS-209
chr7-27731899-INS-194
chr7-27744126-INS-260
chr7-27751524-INS-583

chr7-27756484-INS-91
chr7-27759841-INS-60
chr7-27760323-INS-122
chr7-27762700-INS-58
chr7-27788137-INS-291
chr7-27794475-INS-207
chr7-27795830-INS-220
chr7-27796219-INS-316
chr7-27826078-INS-78
chr7-27827326-INS-232
chr7-27849879-INS-275
chr7-27850282-INS-204
chr7-27904040-INS-264
chr7-27905370-INS-87
chr7-27923129-INS-145
chr7-27936556-INS-80
chr7-27945237-INS-99
chr7-27946510-INS-264
chr7-27949545-INS-138
chr7-27952414-INS-240
chr7-27962746-INS-63
chr7-27973049-INS-315
chr7-27975000-INS-81
chr7-27985504-INS-215
chr7-27987500-INS-52
chr7-27994710-INS-243
chr7-27996214-INS-52
chr7-28005602-INS-162
chr7-28012612-INS-287
chr7-28024361-INS-205
chr7-28040435-INS-96
chr7-28041479-INS-209
chr7-28042056-INS-88
chr7-28046064-INS-221
chr7-28060864-INS-158
chr7-28065238-INS-72
chr7-28081803-INS-64
chr7-28115381-INS-201
chr7-28129611-INS-207
chr7-28144816-INS-93
chr7-28155226-INS-106
chr7-28159990-INS-215
chr7-28168147-INS-209
chr7-28172484-INS-83
chr7-28185266-INS-128
chr7-28185795-INS-58
chr7-28220205-INS-100
chr7-28226638-INS-144
chr7-28236159-INS-213
chr7-28242022-INS-53
chr7-28251553-INS-208
chr7-28301976-INS-265
chr7-28305839-INS-97
chr7-28308101-INS-102
chr7-28310913-INS-89
chr7-28312239-INS-94
chr7-28315412-INS-52
chr7-28319312-INS-183
chr7-28320975-INS-192
chr7-28332966-INS-118
chr7-28345752-INS-54
chr7-28349013-INS-440
chr7-28354830-INS-121
chr7-28367883-INS-53
chr7-28369131-INS-204
chr7-28380359-INS-70
chr7-28402457-INS-196
chr7-28404190-INS-257
chr7-28423525-INS-130
chr7-28450681-INS-66
chr7-28452595-INS-106
chr7-28453939-INS-971
chr7-28454532-INS-194
chr7-28471186-INS-70
chr7-28475490-INS-127
chr7-28478451-INS-63
chr7-28479975-INS-837

chr7-28481265-INS-223
chr7-28504577-INS-334
chr7-28523694-INS-360
chr7-28540406-INS-64
chr7-28541329-INS-212
chr7-28559461-INS-64
chr7-28562187-INS-253
chr7-28572678-INS-221
chr7-28574546-INS-201
chr7-28576921-INS-216
chr7-28581639-INS-78
chr7-28583561-INS-53
chr7-28604114-INS-142
chr7-28614371-INS-208
chr7-28642879-INS-56
chr7-28647213-INS-200
chr7-28648531-INS-237
chr7-28651990-INS-206
chr7-28654196-INS-195
chr7-28675649-INS-123
chr7-28684138-INS-206
chr7-28688618-INS-56
chr7-28689898-INS-91
chr7-28714268-INS-220
chr7-28720341-INS-496
chr7-28721051-INS-51
chr7-28723300-INS-102
chr7-28733733-INS-136
chr7-28734357-INS-106
chr7-28756021-INS-69
chr7-28760412-INS-159
chr7-28809807-INS-88
chr7-28910677-INS-72
chr7-28912202-INS-360
chr7-28925769-INS-212
chr7-28934342-INS-68
chr7-28949129-INS-74
chr7-28953685-INS-52
chr7-28967050-INS-62
chr7-28998748-INS-233
chr7-29001063-INS-51
chr7-29024627-INS-89
chr7-29025950-INS-787
chr7-29027054-INS-112
chr7-29029798-INS-83
chr7-29058960-INS-66
chr7-29061859-INS-115
chr7-29063783-INS-66
chr7-29075453-INS-476
chr7-29085026-INS-504
chr7-29089136-INS-151
chr7-29090310-INS-64
chr7-29097542-INS-181
chr7-29104199-INS-191
chr7-29105317-INS-252
chr7-29117923-INS-61
chr7-29134025-INS-119
chr7-29140366-INS-363
chr7-29143096-INS-102
chr7-29147349-INS-212
chr7-29171196-INS-273
chr7-29240015-INS-5340
chr7-29241274-INS-207
chr7-29245665-INS-211
chr7-29246234-INS-3147
chr7-29249580-INS-6454
chr7-29258809-INS-603
chr7-29265541-INS-385
chr7-29268121-INS-145
chr7-29275709-INS-63
chr7-29279312-INS-144
chr7-29308162-INS-721
chr7-29311741-INS-59
chr7-29328934-INS-5180
chr7-29349490-INS-218
chr7-29359576-INS-395
chr7-29378895-INS-419

chr7-29380316-INS-6475
chr7-29398837-INS-72
chr7-29407721-INS-5991
chr7-29459506-INS-1685
chr7-29470332-INS-59
chr7-29494037-INS-184
chr7-29500758-INS-577
chr7-29516595-INS-88
chr7-29524696-INS-56
chr7-29526903-INS-132
chr7-29547091-INS-184
chr7-29549045-INS-68
chr7-29551885-INS-57
chr7-29564281-INS-466
chr7-29572672-INS-196
chr7-29579093-INS-199
chr7-29613012-INS-55
chr7-29642144-INS-143
chr7-29647575-INS-226
chr7-29657771-INS-68
chr7-29663147-INS-150
chr7-29712173-INS-148
chr7-29729033-INS-157
chr7-29733874-INS-220
chr7-29759642-INS-175
chr7-29763230-INS-219
chr7-29771048-INS-186
chr7-29796023-INS-176
chr7-29802737-INS-104
chr7-29820471-INS-236
chr7-29826024-INS-490
chr7-29829243-INS-196
chr7-29844954-INS-206
chr7-29851605-INS-214
chr7-29852862-INS-396
chr7-29856921-INS-135
chr7-29865540-INS-209
chr7-29870299-INS-201
chr7-29890242-INS-59
chr7-29901448-INS-223
chr7-29902674-INS-210
chr7-29921018-INS-203
chr7-29921601-INS-397
chr7-29942127-INS-172
chr7-29955805-INS-222
chr7-29966742-INS-167
chr7-29969010-INS-211
chr7-29995766-INS-100
chr7-30009631-INS-3410
chr7-30016672-INS-52
chr7-30029828-INS-200
chr7-30064889-INS-157
chr7-30095571-INS-370
chr7-30096442-INS-259
chr7-30097333-INS-232
chr7-30103292-INS-204
chr7-30115217-INS-170
chr7-30128665-INS-106
chr7-30136768-INS-97
chr7-30150599-INS-202
chr7-30170956-INS-58
chr7-30187254-INS-104
chr7-30190839-INS-119
chr7-30195141-INS-77
chr7-30207219-INS-53
chr7-30243640-INS-96
chr7-30246797-INS-199
chr7-30252295-INS-247
chr7-30277180-INS-68
chr7-30300624-INS-200
chr7-30311375-INS-53
chr7-30314603-INS-211
chr7-30321082-INS-65
chr7-30324742-INS-674
chr7-30343342-INS-132
chr7-30344694-INS-154
chr7-30357218-INS-72

chr7-30366316-INS-162
chr7-30367362-INS-84
chr7-30376678-INS-217
chr7-30386613-INS-2826
chr7-30434757-INS-137
chr7-30451812-INS-93
chr7-30469579-INS-106
chr7-30483258-INS-67
chr7-30492521-INS-203
chr7-30496188-INS-389
chr7-30501945-INS-52
chr7-30508265-INS-67
chr7-30518388-INS-77
chr7-30526221-INS-226
chr7-30531209-INS-831
chr7-30531324-INS-315
chr7-30544835-INS-189
chr7-30557530-INS-54
chr7-30568496-INS-60
chr7-30587885-INS-128
chr7-30621795-INS-572
chr7-30624804-INS-201
chr7-30670435-INS-53
chr7-30689650-INS-62
chr7-30691572-INS-84
chr7-30702867-INS-74
chr7-30723584-INS-64
chr7-30734773-INS-216
chr7-30761671-INS-139
chr7-30800733-INS-202
chr7-30806602-INS-264
chr7-30821240-INS-51
chr7-30861448-INS-53
chr7-30873200-INS-81
chr7-30875027-INS-54
chr7-30882782-INS-52
chr7-30900267-INS-1864
chr7-30901589-INS-908
chr7-30914776-INS-6767
chr7-31016914-INS-2290
chr7-31035605-INS-52
chr7-31043452-INS-464
chr7-31055780-INS-115
chr7-31077475-INS-52
chr7-31170152-INS-234
chr7-31173551-INS-131
chr7-31258723-INS-340
chr7-31261810-INS-68
chr7-31285638-INS-7705
chr7-31335725-INS-3203
chr7-31352376-INS-2343
chr7-31384138-INS-4724
chr7-31429883-INS-4841
chr7-31461734-INS-68
chr7-31541161-INS-230
chr7-31544347-INS-51
chr7-31722644-INS-226
chr7-31802532-INS-148
chr7-31813698-INS-864
chr7-31815464-INS-307
chr7-31849087-INS-54
chr7-31862178-INS-866
chr7-31904716-INS-2337
chr7-31907101-INS-2387
chr7-32462952-INS-2873
chr7-32867436-INS-2337
chr7-32941272-INS-2840
chr7-32971974-INS-206
chr7-33078997-INS-7020
chr7-33121515-INS-577
chr7-33155998-INS-255
chr7-33159944-INS-204
chr7-33184033-INS-55
chr7-33235618-INS-178
chr7-33256875-INS-112
chr7-33273783-INS-91
chr7-33333713-INS-52

chr7-33432308-INS-6841
chr7-33438367-INS-354
chr7-33440176-INS-6161
chr7-33445247-INS-66
chr7-33453990-INS-6452
chr7-33490349-INS-58
chr7-33493662-INS-82
chr7-33499563-INS-7417
chr7-33506495-INS-6415
chr7-33509405-INS-70
chr7-33545356-INS-669
chr7-33576671-INS-84
chr7-33579035-INS-178
chr7-33590826-INS-430
chr7-33597443-INS-345
chr7-33604815-INS-6679
chr7-33609852-INS-78
chr7-33614066-INS-60
chr7-33636016-INS-121
chr7-33660294-INS-62
chr7-33664662-INS-114
chr7-33666236-INS-351
chr7-33691736-INS-7520
chr7-33714456-INS-967
chr7-33723056-INS-119
chr7-33785237-INS-204
chr7-33808447-INS-87
chr7-33816264-INS-6306
chr7-33824779-INS-62
chr7-33827035-INS-365
chr7-33835738-INS-60
chr7-33839097-INS-57
chr7-33866584-INS-197
chr7-33870774-INS-598
chr7-33877877-INS-3565
chr7-33890538-INS-78
chr7-33902964-INS-64
chr7-33904539-INS-72
chr7-33906651-INS-212
chr7-33909961-INS-74
chr7-33951666-INS-214
chr7-33959295-INS-212
chr7-34024650-INS-87
chr7-34026518-INS-113
chr7-34027217-INS-1156
chr7-34029545-INS-204
chr7-34035585-INS-60
chr7-34036392-INS-395
chr7-34046721-INS-283
chr7-34069248-INS-174
chr7-34091235-INS-872
chr7-34110294-INS-76
chr7-34114477-INS-733
chr7-34115815-INS-172
chr7-34117056-INS-9046
chr7-34122472-INS-127
chr7-34122577-INS-90
chr7-34123602-INS-118
chr7-34124563-INS-222
chr7-34130740-INS-161
chr7-34147832-INS-56
chr7-34152887-INS-181
chr7-34171577-INS-175
chr7-34174557-INS-527
chr7-34183128-INS-82
chr7-34190680-INS-149
chr7-34194314-INS-113
chr7-34195766-INS-88
chr7-34207362-INS-919
chr7-34238735-INS-107
chr7-34268826-INS-51
chr7-34272375-INS-853
chr7-34297095-INS-374
chr7-34312705-INS-154
chr7-34336117-INS-185
chr7-34337987-INS-256
chr7-34351361-INS-188

chr7-34361776-INS-65
chr7-34381592-INS-106
chr7-34388917-INS-84
chr7-34397153-INS-57
chr7-34413323-INS-53
chr7-34429342-INS-52
chr7-34431241-INS-137
chr7-34432557-INS-158
chr7-34439369-INS-59
chr7-34448853-INS-65
chr7-34460822-INS-223
chr7-34481681-INS-7091
chr7-34486186-INS-205
chr7-34488867-INS-210
chr7-34490999-INS-113
chr7-34502200-INS-398
chr7-34506828-INS-319
chr7-34515484-INS-79
chr7-34605648-INS-51
chr7-34622249-INS-527
chr7-34668138-INS-304
chr7-34677951-INS-82
chr7-34680481-INS-409
chr7-34680621-INS-7149
chr7-34681937-INS-96
chr7-34693051-INS-56
chr7-34707358-INS-549
chr7-34711827-INS-68
chr7-34735266-INS-93
chr7-34752557-INS-209
chr7-34769828-INS-464
chr7-34778102-INS-215
chr7-34794446-INS-294
chr7-34811176-INS-105
chr7-34814383-INS-78
chr7-34824821-INS-132
chr7-34833008-INS-118
chr7-34834089-INS-106
chr7-34859214-INS-1266
chr7-34859215-INS-902
chr7-34869359-INS-202
chr7-34889891-INS-90
chr7-34904714-INS-169
chr7-34921393-INS-62
chr7-34922633-INS-989
chr7-34929206-INS-53
chr7-34937481-INS-93
chr7-34950873-INS-98
chr7-34968216-INS-265
chr7-34977337-INS-3009
chr7-34978033-INS-768
chr7-34978120-INS-1132
chr7-34981520-INS-308
chr7-34983893-INS-199
chr7-34987750-INS-1897
chr7-34989097-INS-87
chr7-34995461-INS-72
chr7-34997213-INS-236
chr7-35003734-INS-53
chr7-35019024-INS-581
chr7-35051680-INS-207
chr7-35059996-INS-64
chr7-35064518-INS-70
chr7-35075275-INS-94
chr7-35077944-INS-432
chr7-35082525-INS-53
chr7-35094310-INS-103
chr7-35121148-INS-362
chr7-35124411-INS-168
chr7-35129908-INS-213
chr7-35136535-INS-282
chr7-35138145-INS-58
chr7-35141667-INS-384
chr7-35163622-INS-218
chr7-35180801-INS-242
chr7-35200193-INS-200
chr7-35204963-INS-323

chr7-35208694-INS-187
chr7-35212255-INS-51
chr7-35225682-INS-102
chr7-35227807-INS-166
chr7-35239800-INS-322
chr7-35244417-INS-95
chr7-35250680-INS-185
chr7-35263827-INS-58
chr7-35288465-INS-201
chr7-35307392-INS-210
chr7-35310602-INS-105
chr7-35313720-INS-960
chr7-35320709-INS-308
chr7-35331829-INS-215
chr7-35335307-INS-283
chr7-35341838-INS-217
chr7-35356430-INS-66
chr7-35358634-INS-56
chr7-35378598-INS-87
chr7-35384071-INS-153
chr7-35387258-INS-138
chr7-35404531-INS-203
chr7-35414261-INS-6442
chr7-35414793-INS-56
chr7-35416485-INS-142
chr7-35440065-INS-80
chr7-35458747-INS-210
chr7-35492709-INS-56
chr7-35496832-INS-141
chr7-35539507-INS-7077
chr7-35542136-INS-400
chr7-35548609-INS-56
chr7-35574354-INS-87
chr7-35580481-INS-53
chr7-35589755-INS-108
chr7-35605712-INS-113
chr7-35607516-INS-102
chr7-35612397-INS-294
chr7-35639722-INS-105
chr7-35653427-INS-178
chr7-35655812-INS-158
chr7-35661180-INS-100
chr7-35679481-INS-84
chr7-35699393-INS-184
chr7-35700507-INS-64
chr7-35707094-INS-132
chr7-35718778-INS-84
chr7-35724417-INS-53
chr7-35750654-INS-416
chr7-35764303-INS-57
chr7-35808286-INS-62
chr7-35821322-INS-144
chr7-35846914-INS-92
chr7-35848336-INS-228
chr7-35870457-INS-83
chr7-35872484-INS-76
chr7-35889478-INS-83
chr7-35923711-INS-52
chr7-35932255-INS-135
chr7-35957185-INS-277
chr7-35995554-INS-64
chr7-36003906-INS-440
chr7-36059647-INS-430
chr7-36078644-INS-363
chr7-36109200-INS-76
chr7-36114914-INS-621
chr7-36130546-INS-57
chr7-36140632-INS-92
chr7-36164451-INS-140
chr7-36199136-INS-568
chr7-36216350-INS-183
chr7-36217758-INS-53
chr7-36249085-INS-468
chr7-36250821-INS-129
chr7-36259717-INS-105
chr7-36263804-INS-63
chr7-36264835-INS-72

chr7-36296065-INS-80
chr7-36300663-INS-218
chr7-36309198-INS-60
chr7-36340301-INS-325
chr7-36344487-INS-55
chr7-36365053-INS-152
chr7-36366304-INS-820
chr7-36372886-INS-56
chr7-36410902-INS-57
chr7-36447911-INS-53
chr7-36477581-INS-624
chr7-36481688-INS-712
chr7-36492785-INS-249
chr7-36541600-INS-142
chr7-36550255-INS-234
chr7-36562570-INS-88
chr7-36572039-INS-832
chr7-36591575-INS-170
chr7-36623108-INS-56
chr7-36636627-INS-68
chr7-36682783-INS-52
chr7-36697411-INS-366
chr7-36698647-INS-111
chr7-36732649-INS-146
chr7-36733200-INS-79
chr7-36767079-INS-92
chr7-36771919-INS-60
chr7-36816942-INS-165
chr7-36825639-INS-56
chr7-36835439-INS-64
chr7-36839516-INS-62
chr7-36854573-INS-106
chr7-36866850-INS-243
chr7-36876715-INS-119
chr7-36883623-INS-76
chr7-36908301-INS-92
chr7-36923394-INS-110
chr7-36982506-INS-51
chr7-37008895-INS-323
chr7-37010126-INS-53
chr7-37052932-INS-55
chr7-37096441-INS-123
chr7-37104344-INS-57
chr7-37213340-INS-891
chr7-37226855-INS-401
chr7-37241319-INS-58
chr7-37256952-INS-53
chr7-37332374-INS-122
chr7-37335348-INS-330
chr7-37413586-INS-76
chr7-37436553-INS-84
chr7-37493325-INS-67
chr7-37512279-INS-51
chr7-37526323-INS-91
chr7-37562701-INS-65
chr7-37577387-INS-681
chr7-37600004-INS-56
chr7-37612156-INS-218
chr7-37614030-INS-585
chr7-37627527-INS-126
chr7-37647308-INS-321
chr7-37661326-INS-243
chr7-37681543-INS-196
chr7-37687221-INS-137
chr7-37690163-INS-176
chr7-37717847-INS-201
chr7-37726422-INS-193
chr7-37734703-INS-183
chr7-37737613-INS-67
chr7-37741338-INS-259
chr7-37745791-INS-100
chr7-37747576-INS-268
chr7-37762374-INS-190
chr7-37765559-INS-4555
chr7-37769905-INS-5081
chr7-37770914-INS-68
chr7-37779659-INS-52

chr7-37819794-INS-261
chr7-37822333-INS-128
chr7-37860889-INS-203
chr7-37862762-INS-190
chr7-37864238-INS-85
chr7-37891649-INS-170
chr7-37893152-INS-137
chr7-37945438-INS-557
chr7-37950943-INS-180
chr7-37953638-INS-329
chr7-37957935-INS-466
chr7-37959468-INS-93
chr7-37971823-INS-86
chr7-37990726-INS-166
chr7-38036665-INS-2356
chr7-38067580-INS-7803
chr7-38089698-INS-6473
chr7-38096476-INS-8834
chr7-38208839-INS-179
chr7-38316220-INS-71
chr7-38318724-INS-525
chr7-38325442-INS-397
chr7-38390503-INS-71
chr7-38391777-INS-116
chr7-38396919-INS-142
chr7-38412584-INS-55
chr7-38417470-INS-605
chr7-38451009-INS-630
chr7-38802066-INS-671
chr7-38849503-INS-212
chr7-38866546-INS-208
chr7-39062309-INS-6751
chr7-39151244-INS-62
chr7-39200448-INS-136
chr7-39212861-INS-127
chr7-39244936-INS-597
chr7-39257892-INS-402
chr7-39286060-INS-73
chr7-39300028-INS-459
chr7-39313953-INS-706
chr7-39326457-INS-74
chr7-39339377-INS-207
chr7-39360873-INS-3074
chr7-39360948-INS-694
chr7-39382768-INS-87
chr7-39388201-INS-61
chr7-39389201-INS-255
chr7-39390151-INS-775
chr7-39394358-INS-275
chr7-39398083-INS-125
chr7-39409072-INS-52
chr7-39417265-INS-388
chr7-39419065-INS-80
chr7-39420532-INS-81
chr7-39421302-INS-70
chr7-39426560-INS-5643
chr7-39435666-INS-13534
chr7-39463720-INS-63
chr7-39474183-INS-51
chr7-39492330-INS-197
chr7-39495879-INS-422
chr7-39508101-INS-90
chr7-39514715-INS-401
chr7-39521420-INS-52
chr7-39528492-INS-724
chr7-39549079-INS-6687
chr7-39571855-INS-95
chr7-39588983-INS-1890
chr7-39595874-INS-64
chr7-39652276-INS-214
chr7-39672926-INS-77
chr7-39683729-INS-135
chr7-39714197-INS-128
chr7-39721494-INS-132
chr7-39747936-INS-109
chr7-39757959-INS-400
chr7-39781182-INS-66

chr7-39808774-INS-487
chr7-39829491-INS-92
chr7-39936983-INS-636
chr7-39943493-INS-159
chr7-39956696-INS-144
chr7-39965222-INS-609
chr7-39980401-INS-194
chr7-40005271-INS-5549
chr7-40039162-INS-52
chr7-40079709-INS-56
chr7-40096466-INS-666
chr7-40159149-INS-73
chr7-40170660-INS-200
chr7-40180668-INS-280
chr7-40181882-INS-61
chr7-40210369-INS-236
chr7-40224937-INS-115
chr7-40227004-INS-1081
chr7-40231011-INS-460
chr7-40241377-INS-1349
chr7-40277000-INS-115
chr7-40298327-INS-69
chr7-40322501-INS-847
chr7-40339537-INS-56
chr7-40349870-INS-126
chr7-40364312-INS-74
chr7-40410173-INS-116
chr7-40419924-INS-235
chr7-40432091-INS-163
chr7-40463557-INS-354
chr7-40468581-INS-399
chr7-40479217-INS-51
chr7-40494446-INS-74
chr7-40553611-INS-60
chr7-40571841-INS-104
chr7-40583791-INS-75
chr7-40624835-INS-324
chr7-40663120-INS-6460
chr7-40679317-INS-970
chr7-40690988-INS-63
chr7-40698657-INS-416
chr7-40711019-INS-504
chr7-40737911-INS-50
chr7-40800846-INS-5407
chr7-40829642-INS-197
chr7-40877649-INS-83
chr7-41015643-INS-103
chr7-41027541-INS-6383
chr7-41035249-INS-60
chr7-41043523-INS-245
chr7-41057134-INS-230
chr7-41089140-INS-430
chr7-41104402-INS-991
chr7-41116645-INS-58
chr7-41135052-INS-89
chr7-41137125-INS-81
chr7-41145659-INS-246
chr7-41170102-INS-5002
chr7-41186711-INS-148
chr7-41206924-INS-63
chr7-41230866-INS-4395
chr7-41233092-INS-129
chr7-41238494-INS-6415
chr7-41263009-INS-130
chr7-41266492-INS-91
chr7-41270243-INS-324
chr7-41288984-INS-216
chr7-41320450-INS-831
chr7-41323444-INS-6477
chr7-41330279-INS-56
chr7-41338605-INS-52
chr7-41340204-INS-6454
chr7-41342972-INS-934
chr7-41346320-INS-5251
chr7-41346862-INS-7484
chr7-41365295-INS-55
chr7-41365793-INS-2929

chr7-41424562-INS-100
chr7-41491193-INS-1234
chr7-41544145-INS-4515
chr7-41584051-INS-989
chr7-41604169-INS-56
chr7-41666475-INS-153
chr7-41698718-INS-421
chr7-41721053-INS-57
chr7-41734144-INS-652
chr7-41738063-INS-155
chr7-41806326-INS-2684
chr7-41956746-INS-66
chr7-42000038-INS-108
chr7-42055709-INS-56
chr7-42088874-INS-2275
chr7-42117365-INS-906
chr7-42122069-INS-678
chr7-42134253-INS-273
chr7-42153833-INS-210
chr7-42194541-INS-198
chr7-42394966-INS-74
chr7-42397996-INS-546
chr7-42559739-INS-120
chr7-42709861-INS-175
chr7-42722541-INS-93
chr7-42781129-INS-282
chr7-42786578-INS-4460
chr7-42807932-INS-102
chr7-42808983-INS-93
chr7-42820740-INS-383
chr7-42841210-INS-91
chr7-42849229-INS-328
chr7-42856715-INS-189
chr7-42918079-INS-81
chr7-42926250-INS-641
chr7-42926308-INS-70
chr7-42927754-INS-66
chr7-42932213-INS-167
chr7-42984052-INS-809
chr7-42988408-INS-552
chr7-42991372-INS-101
chr7-42994878-INS-429
chr7-43009347-INS-173
chr7-43018865-INS-287
chr7-43028715-INS-216
chr7-43035859-INS-530
chr7-43070345-INS-82
chr7-43074573-INS-180
chr7-43077244-INS-215
chr7-43099883-INS-147
chr7-43142044-INS-200
chr7-43146620-INS-106
chr7-43154117-INS-52
chr7-43158038-INS-127
chr7-43160318-INS-60
chr7-43167056-INS-7112
chr7-43175234-INS-213
chr7-43199589-INS-548
chr7-43201863-INS-196
chr7-43215693-INS-51
chr7-43216155-INS-57
chr7-43218843-INS-111
chr7-43264592-INS-62
chr7-43273848-INS-90
chr7-43276799-INS-52
chr7-43279677-INS-60
chr7-43290327-INS-259
chr7-43294157-INS-80
chr7-43295705-INS-207
chr7-43304939-INS-162
chr7-43326426-INS-114
chr7-43348462-INS-322
chr7-43354859-INS-76
chr7-43383003-INS-405
chr7-43394907-INS-72
chr7-43401184-INS-54
chr7-43414200-INS-61

chr7-43451124-INS-74
chr7-43454065-INS-167
chr7-43471835-INS-160
chr7-43502157-INS-144
chr7-43503194-INS-67
chr7-43511010-INS-60
chr7-43538847-INS-61
chr7-43544034-INS-7156
chr7-43549155-INS-59
chr7-43554975-INS-241
chr7-43561695-INS-94
chr7-43592732-INS-258
chr7-43603025-INS-80
chr7-43609862-INS-53
chr7-43632740-INS-62
chr7-43652163-INS-326
chr7-43671188-INS-53
chr7-43724196-INS-112
chr7-43731985-INS-664
chr7-43783995-INS-64
chr7-43829793-INS-8755
chr7-43850505-INS-150
chr7-43858601-INS-204
chr7-43882986-INS-54
chr7-43889861-INS-737
chr7-43920770-INS-143
chr7-43922767-INS-224
chr7-43927062-INS-186
chr7-43937471-INS-72
chr7-43944111-INS-58
chr7-43948027-INS-206
chr7-43951380-INS-130
chr7-43980249-INS-72
chr7-43996574-INS-122
chr7-44038756-INS-68
chr7-44039185-INS-170
chr7-44064933-INS-63
chr7-44074465-INS-109
chr7-44102074-INS-96
chr7-44104567-INS-52
chr7-44123108-INS-54
chr7-44133622-INS-141
chr7-44141126-INS-61
chr7-44157035-INS-102
chr7-44163762-INS-176
chr7-44165527-INS-131
chr7-44173455-INS-72
chr7-44198752-INS-182
chr7-44227552-INS-208
chr7-44268159-INS-193
chr7-44270299-INS-1288
chr7-44272242-INS-56
chr7-44276266-INS-416
chr7-44276990-INS-255
chr7-44280444-INS-194
chr7-44350972-INS-53
chr7-44354263-INS-50
chr7-44354725-INS-53
chr7-44362488-INS-168
chr7-44364582-INS-114
chr7-44366067-INS-117
chr7-44368738-INS-92
chr7-44372905-INS-214
chr7-44385782-INS-371
chr7-44387256-INS-69
chr7-44388251-INS-77
chr7-44417690-INS-62
chr7-44421683-INS-126
chr7-44427366-INS-52
chr7-44430632-INS-678
chr7-44441942-INS-52
chr7-44454640-INS-51
chr7-44457680-INS-89
chr7-44482960-INS-176
chr7-44497486-INS-205
chr7-44505352-INS-219
chr7-44524923-INS-52

chr7-44555415-INS-55
chr7-44570973-INS-51
chr7-44575575-INS-65
chr7-44584006-INS-271
chr7-44597845-INS-60
chr7-44622209-INS-167
chr7-44659429-INS-80
chr7-44694597-INS-197
chr7-44714680-INS-54
chr7-44729251-INS-103
chr7-44746420-INS-62
chr7-44762406-INS-51
chr7-44788916-INS-112
chr7-44810311-INS-547
chr7-44823163-INS-135
chr7-44854690-INS-71
chr7-44860399-INS-51
chr7-44873896-INS-71
chr7-44887112-INS-218
chr7-44894781-INS-114
chr7-44896761-INS-552
chr7-44897702-INS-140
chr7-44906960-INS-65
chr7-44919178-INS-79
chr7-44919561-INS-54
chr7-44943918-INS-204
chr7-44956044-INS-261
chr7-44963726-INS-214
chr7-44968913-INS-209
chr7-44970756-INS-84
chr7-44999090-INS-80
chr7-45002141-INS-154
chr7-45013228-INS-74
chr7-45024646-INS-53
chr7-45028519-INS-204
chr7-45029297-INS-204
chr7-45038147-INS-884
chr7-45040600-INS-100
chr7-45050498-INS-57
chr7-45090042-INS-140
chr7-45098537-INS-351
chr7-45099518-INS-215
chr7-45103084-INS-201
chr7-45127183-INS-83
chr7-45135788-INS-5523
chr7-45146457-INS-207
chr7-45156103-INS-214
chr7-45173863-INS-471
chr7-45178221-INS-52
chr7-45183148-INS-796
chr7-45185772-INS-122
chr7-45192385-INS-172
chr7-45194699-INS-206
chr7-45218489-INS-210
chr7-45221792-INS-54
chr7-45232950-INS-189
chr7-45243914-INS-63
chr7-45256421-INS-181
chr7-45258535-INS-58
chr7-45261087-INS-880
chr7-45265425-INS-94
chr7-45281255-INS-102
chr7-45331810-INS-100
chr7-45386110-INS-78
chr7-45389475-INS-166
chr7-45411879-INS-71
chr7-45416647-INS-5542
chr7-45419883-INS-1893
chr7-45428219-INS-184
chr7-45430167-INS-113
chr7-45449042-INS-338
chr7-45456389-INS-72
chr7-45528518-INS-206
chr7-45548649-INS-159
chr7-45552699-INS-166
chr7-45553170-INS-521
chr7-45559456-INS-82

chr7-45581832-INS-206
chr7-45588738-INS-218
chr7-45589543-INS-67
chr7-45593708-INS-200
chr7-45605676-INS-312
chr7-45638726-INS-144
chr7-45640263-INS-54
chr7-45655776-INS-209
chr7-45660741-INS-236
chr7-45665369-INS-78
chr7-45685873-INS-178
chr7-45698639-INS-223
chr7-45710646-INS-923
chr7-45716720-INS-54
chr7-45741121-INS-53
chr7-45751395-INS-201
chr7-45764982-INS-162
chr7-45780012-INS-73
chr7-45791380-INS-60
chr7-45800611-INS-80
chr7-45801567-INS-168
chr7-45807144-INS-232
chr7-45817448-INS-72
chr7-45830444-INS-197
chr7-45849292-INS-55
chr7-45863038-INS-64
chr7-45873279-INS-215
chr7-45927063-INS-81
chr7-45940720-INS-65
chr7-45943423-INS-206
chr7-45979516-INS-87
chr7-45981788-INS-115
chr7-45985705-INS-118
chr7-46005051-INS-193
chr7-46009985-INS-290
chr7-46079660-INS-80
chr7-46098050-INS-156
chr7-46102861-INS-459
chr7-46133362-INS-365
chr7-46136565-INS-875
chr7-46138591-INS-58
chr7-46138631-INS-1060
chr7-46143197-INS-57
chr7-46147882-INS-118
chr7-46161721-INS-2320
chr7-46181449-INS-61
chr7-46181759-INS-4392
chr7-46198772-INS-54
chr7-46204434-INS-207
chr7-46205877-INS-68
chr7-46211131-INS-68
chr7-46213921-INS-56
chr7-46226068-INS-196
chr7-46247550-INS-127
chr7-46264642-INS-98
chr7-46268972-INS-247
chr7-46273462-INS-640
chr7-46273465-INS-123
chr7-46293164-INS-235
chr7-46301930-INS-114
chr7-46305104-INS-786
chr7-46341080-INS-207
chr7-46350585-INS-207
chr7-46360280-INS-54
chr7-46376700-INS-80
chr7-46380069-INS-251
chr7-46389521-INS-503
chr7-46395514-INS-12915
chr7-46400371-INS-201
chr7-46418958-INS-216
chr7-46420273-INS-512
chr7-46422864-INS-221
chr7-46469180-INS-52
chr7-46478211-INS-173
chr7-46481286-INS-125
chr7-46483640-INS-202
chr7-46507641-INS-441

chr7-46517249-INS-358
chr7-46521066-INS-214
chr7-46530692-INS-916
chr7-46539624-INS-204
chr7-46546871-INS-206
chr7-46548788-INS-208
chr7-46575252-INS-73
chr7-46585773-INS-86
chr7-46594690-INS-476
chr7-46596178-INS-602
chr7-46597835-INS-72
chr7-46609982-INS-219
chr7-46619153-INS-374
chr7-46644440-INS-54
chr7-46652833-INS-205
chr7-46664734-INS-69
chr7-46668734-INS-110
chr7-46671196-INS-211
chr7-46676838-INS-154
chr7-46682682-INS-61
chr7-46709637-INS-66
chr7-46715141-INS-256
chr7-46720718-INS-54
chr7-46724116-INS-400
chr7-46742389-INS-176
chr7-46753983-INS-60
chr7-46774598-INS-57
chr7-46788340-INS-179
chr7-46797892-INS-418
chr7-46821719-INS-87
chr7-46826741-INS-2181
chr7-46839251-INS-70
chr7-46844728-INS-76
chr7-46865262-INS-555
chr7-46878516-INS-382
chr7-46892439-INS-8459
chr7-46893857-INS-392
chr7-46896303-INS-54
chr7-46897926-INS-9005
chr7-46968764-INS-168
chr7-46991699-INS-935
chr7-47004874-INS-229
chr7-47011147-INS-400
chr7-47050286-INS-297
chr7-47060625-INS-124
chr7-47070488-INS-646
chr7-47106044-INS-200
chr7-47146661-INS-557
chr7-47150037-INS-215
chr7-47180660-INS-61
chr7-47200287-INS-56
chr7-47252890-INS-54
chr7-47274296-INS-6180
chr7-47278416-INS-1529
chr7-47284871-INS-60
chr7-47302634-INS-68
chr7-47352335-INS-6399
chr7-47370911-INS-82
chr7-47396089-INS-256
chr7-47455471-INS-347
chr7-47462620-INS-194
chr7-47485063-INS-92
chr7-47504069-INS-8005
chr7-47583234-INS-427
chr7-47624401-INS-129
chr7-47638336-INS-209
chr7-47639953-INS-123
chr7-47799136-INS-1757
chr7-47812022-INS-96
chr7-47871931-INS-213
chr7-47892825-INS-7119
chr7-47946033-INS-106
chr7-47949650-INS-2274
chr7-47961437-INS-5277
chr7-47970237-INS-7089
chr7-48000754-INS-6256
chr7-48015834-INS-75

chr7-48035675-INS-485
chr7-48047168-INS-1588
chr7-48051218-INS-695
chr7-48076952-INS-6446
chr7-48132987-INS-57
chr7-48160178-INS-64
chr7-48164938-INS-115
chr7-48170076-INS-211
chr7-48269757-INS-6139
chr7-48270838-INS-6669
chr7-48299796-INS-1391
chr7-48299863-INS-2228
chr7-48313346-INS-69
chr7-48322156-INS-128
chr7-48324314-INS-752
chr7-48325808-INS-3855
chr7-48339062-INS-54
chr7-48353207-INS-1682
chr7-48355469-INS-96
chr7-48357740-INS-109
chr7-48378056-INS-271
chr7-48386566-INS-400
chr7-48406655-INS-450
chr7-48413532-INS-395
chr7-48450187-INS-126
chr7-48458742-INS-234
chr7-48468608-INS-6494
chr7-48493251-INS-254
chr7-48540314-INS-186
chr7-48549568-INS-56
chr7-48554087-INS-182
chr7-48582418-INS-220
chr7-48595864-INS-380
chr7-48635000-INS-52
chr7-48643587-INS-165
chr7-48646014-INS-338
chr7-48648601-INS-114
chr7-48657622-INS-137
chr7-48712379-INS-373
chr7-48729721-INS-82
chr7-48737510-INS-211
chr7-48777863-INS-56
chr7-48791889-INS-55
chr7-48795202-INS-175
chr7-48815464-INS-3964
chr7-48822301-INS-84
chr7-48841962-INS-707
chr7-48862727-INS-166
chr7-48881482-INS-343
chr7-48922067-INS-86
chr7-48939726-INS-56
chr7-48960570-INS-56
chr7-48979451-INS-58
chr7-48980405-INS-106
chr7-48983691-INS-152
chr7-48985409-INS-54
chr7-48989816-INS-5215
chr7-49000095-INS-69
chr7-49009495-INS-409
chr7-49016500-INS-184
chr7-49030246-INS-231
chr7-49045786-INS-131
chr7-49077165-INS-248
chr7-49098281-INS-199
chr7-49103793-INS-126
chr7-49117215-INS-68
chr7-49138279-INS-52
chr7-49140248-INS-400
chr7-49142816-INS-246
chr7-49164608-INS-96
chr7-49174193-INS-64
chr7-49187144-INS-53
chr7-49210724-INS-65
chr7-49218967-INS-135
chr7-49220567-INS-203
chr7-49238012-INS-55
chr7-49244463-INS-210

chr7-49271372-INS-74
chr7-49285534-INS-70
chr7-49287486-INS-61
chr7-49302237-INS-86
chr7-49309202-INS-64
chr7-49310830-INS-204
chr7-49334504-INS-204
chr7-49345071-INS-377
chr7-49353183-INS-88
chr7-49362700-INS-60
chr7-49398505-INS-54
chr7-49407902-INS-70
chr7-49410885-INS-66
chr7-49451768-INS-109
chr7-49511293-INS-170
chr7-49514363-INS-147
chr7-49531138-INS-2932
chr7-49551666-INS-60
chr7-49552820-INS-52
chr7-49568424-INS-60
chr7-49578764-INS-57
chr7-49597066-INS-84
chr7-49609827-INS-56
chr7-49615570-INS-263
chr7-49639999-INS-88
chr7-49652233-INS-54
chr7-49666027-INS-115
chr7-49687072-INS-399
chr7-49688917-INS-249
chr7-49722239-INS-950
chr7-49725610-INS-62
chr7-49742037-INS-204
chr7-49745350-INS-354
chr7-49746426-INS-545
chr7-49747854-INS-83
chr7-49769499-INS-343
chr7-49780903-INS-2172
chr7-49805853-INS-80
chr7-49808127-INS-6668
chr7-49811024-INS-6923
chr7-49860115-INS-366
chr7-49881628-INS-126
chr7-49893939-INS-120
chr7-49925971-INS-393
chr7-49926932-INS-5128
chr7-49942707-INS-7156
chr7-49955878-INS-130
chr7-49956124-INS-442
chr7-49956647-INS-6367
chr7-49966292-INS-351
chr7-49989162-INS-215
chr7-49993611-INS-58
chr7-49998939-INS-549
chr7-50008829-INS-76
chr7-50013669-INS-520
chr7-50032070-INS-10459
chr7-50036598-INS-79
chr7-50064757-INS-96
chr7-50073202-INS-116
chr7-50093671-INS-285
chr7-50099884-INS-50
chr7-50118236-INS-108
chr7-50123312-INS-661
chr7-50125475-INS-6687
chr7-50125561-INS-229
chr7-50149763-INS-54
chr7-50153417-INS-185
chr7-50159353-INS-597
chr7-50159984-INS-6273
chr7-50162009-INS-116
chr7-50163648-INS-97
chr7-50182061-INS-6420
chr7-50206604-INS-152
chr7-50224716-INS-66
chr7-50226669-INS-421
chr7-50236845-INS-92
chr7-50241455-INS-88

chr7-50249178-INS-325
chr7-50250699-INS-364
chr7-50285496-INS-52
chr7-50299796-INS-177
chr7-50304201-INS-340
chr7-50310471-INS-102
chr7-50329049-INS-56
chr7-50332499-INS-179
chr7-50333737-INS-82
chr7-50348885-INS-141
chr7-50359824-INS-727
chr7-50378034-INS-7291
chr7-50388341-INS-167
chr7-50389508-INS-590
chr7-50392379-INS-6703
chr7-50414519-INS-168
chr7-50419287-INS-136
chr7-50420339-INS-126
chr7-50428453-INS-163
chr7-50436322-INS-75
chr7-50442363-INS-86
chr7-50449530-INS-116
chr7-50518105-INS-94
chr7-50522583-INS-374
chr7-50530668-INS-166
chr7-50556397-INS-87
chr7-50580798-INS-79
chr7-50582802-INS-115
chr7-50596680-INS-76
chr7-50605827-INS-58
chr7-50622941-INS-74
chr7-50631694-INS-70
chr7-50649193-INS-110
chr7-50664327-INS-144
chr7-50667576-INS-417
chr7-50708412-INS-108
chr7-50709949-INS-94
chr7-50715269-INS-6443
chr7-50719114-INS-51
chr7-50721243-INS-89
chr7-50734259-INS-59
chr7-50761412-INS-54
chr7-50765389-INS-7278
chr7-50784849-INS-158
chr7-50836687-INS-160
chr7-50961379-INS-53
chr7-50968893-INS-76
chr7-50988662-INS-100
chr7-50992160-INS-272
chr7-51004364-INS-454
chr7-51010805-INS-55
chr7-51013841-INS-92
chr7-51017522-INS-57
chr7-51027488-INS-433
chr7-51034910-INS-66
chr7-51053056-INS-78
chr7-51054038-INS-181
chr7-51113571-INS-63
chr7-51125848-INS-547
chr7-51149408-INS-106
chr7-51151075-INS-101
chr7-51153381-INS-54
chr7-51158164-INS-174
chr7-51163579-INS-80
chr7-51173348-INS-53
chr7-51175002-INS-505
chr7-51251093-INS-60
chr7-51257790-INS-164
chr7-51263908-INS-54
chr7-51303823-INS-566
chr7-51309342-INS-530
chr7-51313308-INS-102
chr7-51336700-INS-158
chr7-51353038-INS-52
chr7-51362559-INS-529
chr7-51370736-INS-105
chr7-51377276-INS-3783

chr7-51379245-INS-622
chr7-51390786-INS-104
chr7-51391723-INS-4948
chr7-51405515-INS-61
chr7-51408159-INS-51
chr7-51414193-INS-488
chr7-51441838-INS-65
chr7-51442649-INS-6641
chr7-51463693-INS-197
chr7-51474656-INS-56
chr7-51477962-INS-294
chr7-51484728-INS-135
chr7-51486127-INS-93
chr7-51518918-INS-152
chr7-51547588-INS-175
chr7-51605242-INS-1316
chr7-51605432-INS-77
chr7-51616545-INS-448
chr7-51622406-INS-221
chr7-51628764-INS-201
chr7-51633638-INS-7130
chr7-51636594-INS-194
chr7-51639181-INS-6454
chr7-51666316-INS-118
chr7-51684745-INS-6594
chr7-51707921-INS-52
chr7-51764851-INS-6402
chr7-51765726-INS-712
chr7-51768887-INS-685
chr7-51769603-INS-6874
chr7-51782323-INS-303
chr7-51804504-INS-83
chr7-51831273-INS-223
chr7-51833509-INS-64
chr7-51836236-INS-530
chr7-51839086-INS-103
chr7-51851031-INS-88
chr7-51852042-INS-372
chr7-51859728-INS-81
chr7-51884054-INS-228
chr7-51902998-INS-59
chr7-51910294-INS-503
chr7-51938994-INS-201
chr7-51942061-INS-384
chr7-51944402-INS-499
chr7-51954351-INS-352
chr7-51991964-INS-52
chr7-52013367-INS-63
chr7-52017353-INS-899
chr7-52029038-INS-58
chr7-52086534-INS-66
chr7-52117800-INS-343
chr7-52144929-INS-231
chr7-52145892-INS-53
chr7-52197390-INS-125
chr7-52198663-INS-522
chr7-52257211-INS-497
chr7-52283194-INS-140
chr7-52313861-INS-6493
chr7-52320691-INS-10045
chr7-52345862-INS-824
chr7-52347846-INS-6154
chr7-52348400-INS-138
chr7-52353884-INS-3103
chr7-52367525-INS-54
chr7-52390884-INS-58
chr7-52395722-INS-130
chr7-52398720-INS-79
chr7-52430730-INS-55
chr7-52430730-INS-796
chr7-52463853-INS-610
chr7-52474731-INS-70
chr7-52475826-INS-55
chr7-52485535-INS-214
chr7-52491756-INS-6214
chr7-52493993-INS-311
chr7-52494419-INS-6449

chr7-52509553-INS-193
chr7-52519631-INS-67
chr7-52530263-INS-78
chr7-52572627-INS-109
chr7-52583938-INS-435
chr7-52588042-INS-2187
chr7-52591759-INS-153
chr7-52717761-INS-322
chr7-52726402-INS-119
chr7-52738298-INS-142
chr7-52772405-INS-528
chr7-52797912-INS-6204
chr7-52812433-INS-189
chr7-52820851-INS-446
chr7-52856438-INS-396
chr7-52879692-INS-218
chr7-52890085-INS-532
chr7-52896085-INS-6478
chr7-52910699-INS-900
chr7-52911652-INS-210
chr7-52913283-INS-58
chr7-52920617-INS-6602
chr7-52932936-INS-199
chr7-52977648-INS-1892
chr7-52984008-INS-61
chr7-52998994-INS-6238
chr7-53011700-INS-58
chr7-53016373-INS-320
chr7-53023197-INS-132
chr7-53025348-INS-52
chr7-53027655-INS-144
chr7-53044873-INS-720
chr7-53079760-INS-60
chr7-53121831-INS-74
chr7-53174428-INS-66
chr7-53193068-INS-79
chr7-53195633-INS-75
chr7-53207652-INS-424
chr7-53271859-INS-204
chr7-53296171-INS-1359
chr7-53305516-INS-735
chr7-53312503-INS-219
chr7-53354044-INS-52
chr7-53354641-INS-87
chr7-53364232-INS-485
chr7-53374012-INS-56
chr7-53381206-INS-456
chr7-53407415-INS-397
chr7-53421605-INS-74
chr7-53433909-INS-99
chr7-53470486-INS-2396
chr7-53490710-INS-51
chr7-53505308-INS-1040
chr7-53522030-INS-52
chr7-53582778-INS-608
chr7-53584688-INS-334
chr7-53606620-INS-56
chr7-53622253-INS-288
chr7-53624760-INS-355
chr7-53636046-INS-141
chr7-53680363-INS-219
chr7-53682267-INS-351
chr7-53684224-INS-6573
chr7-53693111-INS-436
chr7-53696844-INS-80
chr7-53715319-INS-152
chr7-53722988-INS-7012
chr7-53723559-INS-7317
chr7-53750796-INS-535
chr7-53774039-INS-230
chr7-53776040-INS-219
chr7-53816212-INS-9857
chr7-53862540-INS-216
chr7-53874037-INS-481
chr7-53890814-INS-518
chr7-53900760-INS-927
chr7-53915918-INS-227

chr7-53932620-INS-60
chr7-53960623-INS-371
chr7-53965899-INS-827
chr7-53977389-INS-57
chr7-54013210-INS-692
chr7-54074466-INS-54
chr7-54121955-INS-161
chr7-54131160-INS-496
chr7-54135471-INS-55
chr7-54148811-INS-72
chr7-54159419-INS-786
chr7-54160463-INS-498
chr7-54172332-INS-722
chr7-54183828-INS-59
chr7-54188002-INS-6980
chr7-54189301-INS-54
chr7-54202980-INS-149
chr7-54208153-INS-387
chr7-54218053-INS-127
chr7-54223970-INS-694
chr7-54247069-INS-54
chr7-54268481-INS-6609
chr7-54270722-INS-88
chr7-54281274-INS-200
chr7-54292958-INS-231
chr7-54294642-INS-58
chr7-54299728-INS-935
chr7-54313462-INS-101
chr7-54351730-INS-500
chr7-54362136-INS-291
chr7-54366846-INS-90
chr7-54380092-INS-64
chr7-54382465-INS-72
chr7-54386396-INS-447
chr7-54450316-INS-82
chr7-54458691-INS-315
chr7-54507965-INS-942
chr7-54508723-INS-65
chr7-54515042-INS-712
chr7-54536129-INS-66
chr7-54552031-INS-533
chr7-54553611-INS-72
chr7-54568637-INS-308
chr7-54571212-INS-60
chr7-54580194-INS-52
chr7-54584765-INS-63
chr7-54588208-INS-623
chr7-54636194-INS-68
chr7-54688285-INS-156
chr7-54692976-INS-144
chr7-54703098-INS-6386
chr7-54727647-INS-1456
chr7-54760259-INS-176
chr7-54791926-INS-422
chr7-54834405-INS-240
chr7-54852330-INS-675
chr7-54852353-INS-107
chr7-54865674-INS-612
chr7-54876635-INS-52
chr7-54879502-INS-82
chr7-54884653-INS-6345
chr7-54979012-INS-78
chr7-54985209-INS-193
chr7-55026606-INS-657
chr7-55033821-INS-190
chr7-55071400-INS-57
chr7-55077415-INS-66
chr7-55096063-INS-521
chr7-55098518-INS-654
chr7-55111834-INS-6432
chr7-55113446-INS-6952
chr7-55114112-INS-7509
chr7-55122826-INS-92
chr7-55138540-INS-6170
chr7-55144899-INS-6621
chr7-55186609-INS-59
chr7-55189284-INS-876

chr7-55206117-INS-58
chr7-55226108-INS-192
chr7-55239180-INS-72
chr7-55261157-INS-57
chr7-55284540-INS-671
chr7-55295369-INS-161
chr7-55301855-INS-204
chr7-55316771-INS-201
chr7-55328512-INS-155
chr7-55351599-INS-625
chr7-55365490-INS-153
chr7-55407070-INS-285
chr7-55438406-INS-57
chr7-55441802-INS-188
chr7-55444157-INS-183
chr7-55482216-INS-223
chr7-55507160-INS-52
chr7-55510246-INS-74
chr7-55516458-INS-216
chr7-55541270-INS-51
chr7-55542827-INS-165
chr7-55543958-INS-579
chr7-55564249-INS-187
chr7-55584416-INS-80
chr7-55590757-INS-177
chr7-55604600-INS-441
chr7-55620134-INS-202
chr7-55633997-INS-51
chr7-55641311-INS-399
chr7-55662238-INS-56
chr7-55673848-INS-261
chr7-55687492-INS-203
chr7-55691882-INS-72
chr7-55720057-INS-693
chr7-55735045-INS-2793
chr7-55745339-INS-62
chr7-55753093-INS-108
chr7-55765989-INS-429
chr7-55780855-INS-164
chr7-55797478-INS-115
chr7-55801906-INS-317
chr7-55808286-INS-118
chr7-55812437-INS-1216
chr7-55856731-INS-234
chr7-55899826-INS-823
chr7-55903668-INS-52
chr7-55909765-INS-86
chr7-55922693-INS-117
chr7-55935336-INS-98
chr7-55940789-INS-62
chr7-55946041-INS-91
chr7-55956920-INS-5290
chr7-55961741-INS-106
chr7-55965429-INS-116
chr7-55971876-INS-221
chr7-55977269-INS-304
chr7-55982894-INS-6579
chr7-56015179-INS-6381
chr7-56057082-INS-85
chr7-56076296-INS-496
chr7-56083467-INS-172
chr7-56088267-INS-337
chr7-56091256-INS-158
chr7-56093103-INS-6401
chr7-56113345-INS-64
chr7-56177485-INS-55
chr7-56186363-INS-201
chr7-56215000-INS-358
chr7-56215644-INS-6266
chr7-56216862-INS-6347
chr7-56298272-INS-1377
chr7-56304053-INS-121
chr7-56323375-INS-6859
chr7-56327681-INS-1613
chr7-56358885-INS-104
chr7-56361691-INS-109
chr7-56365202-INS-83

chr7-56385293-INS-394
chr7-56401936-INS-318
chr7-56412485-INS-5057
chr7-56467365-INS-503
chr7-56509898-INS-102
chr7-56514129-INS-53
chr7-56601013-INS-867
chr7-56643991-INS-6705
chr7-56652928-INS-316
chr7-56674403-INS-95
chr7-56687316-INS-177
chr7-56713403-INS-120
chr7-56719391-INS-232
chr7-56724346-INS-6392
chr7-56733740-INS-400
chr7-56742396-INS-65
chr7-56807603-INS-71
chr7-56812075-INS-509
chr7-56839375-INS-445
chr7-56865599-INS-316
chr7-56882637-INS-70
chr7-56885273-INS-974
chr7-56893417-INS-116
chr7-56926392-INS-295
chr7-56945635-INS-466
chr7-56953775-INS-2298
chr7-56953776-INS-71
chr7-56960717-INS-57
chr7-56988545-INS-60
chr7-57004199-INS-60
chr7-57027703-INS-52
chr7-57045521-INS-4432
chr7-57045622-INS-410
chr7-57050185-INS-207
chr7-57061755-INS-6603
chr7-57064504-INS-7429
chr7-57065442-INS-53
chr7-57074177-INS-121
chr7-57084469-INS-72
chr7-57086003-INS-6430
chr7-57096448-INS-334
chr7-57107647-INS-326
chr7-57110948-INS-401
chr7-57178352-INS-106
chr7-57182877-INS-61
chr7-57196128-INS-764
chr7-57215563-INS-3874
chr7-57244283-INS-86
chr7-57294672-INS-55
chr7-57300204-INS-639
chr7-57317111-INS-2888
chr7-57322036-INS-359
chr7-57341603-INS-6401
chr7-57348084-INS-2874
chr7-57349720-INS-573
chr7-57365423-INS-119
chr7-57369231-INS-269
chr7-57387578-INS-296
chr7-57389895-INS-812
chr7-57401364-INS-194
chr7-57417825-INS-196
chr7-57420103-INS-51
chr7-57430552-INS-628
chr7-57449609-INS-52
chr7-57485597-INS-115
chr7-57493620-INS-7741
chr7-57494442-INS-824
chr7-57499789-INS-6241
chr7-57608355-INS-435
chr7-57613461-INS-134
chr7-57613749-INS-513
chr7-57625388-INS-201
chr7-57628007-INS-84
chr7-57630083-INS-102
chr7-57649962-INS-112
chr7-57660205-INS-604
chr7-57663718-INS-100

chr7-57665376-INS-316
chr7-57683636-INS-118
chr7-57697654-INS-6220
chr7-57718950-INS-620
chr7-57735640-INS-54
chr7-57738797-INS-6441
chr7-57738904-INS-155
chr7-57760402-INS-70
chr7-57764241-INS-593
chr7-57776836-INS-2678
chr7-57800787-INS-822
chr7-57805112-INS-64
chr7-57824457-INS-12160
chr7-57841635-INS-6230
chr7-57849169-INS-144
chr7-57853378-INS-161
chr7-57859107-INS-189
chr7-57863797-INS-6477
chr7-57869830-INS-217
chr7-57870569-INS-568
chr7-57876648-INS-6496
chr7-57882816-INS-211
chr7-57882937-INS-5241
chr7-57884410-INS-3345
chr7-57904736-INS-6060
chr7-57908872-INS-4354
chr7-57914855-INS-6203
chr7-57922706-INS-5113
chr7-57963844-INS-60
chr7-57964766-INS-96
chr7-58000430-INS-6249
chr7-58006384-INS-253
chr7-58006537-INS-1288
chr7-58009488-INS-648
chr7-58036398-INS-6345
chr7-58078949-INS-54
chr7-58083527-INS-311
chr7-58085848-INS-60
chr7-58096611-INS-56
chr7-58104324-INS-355
chr7-58111100-INS-106
chr7-58120549-INS-260
chr7-58130741-INS-938
chr7-58139992-INS-380
chr7-58159718-INS-6446
chr7-58164036-INS-586
chr7-58195250-INS-7806
chr7-58204840-INS-77
chr7-58211150-INS-84
chr7-58216616-INS-94
chr7-58234384-INS-184
chr7-58237211-INS-1976
chr7-58258301-INS-811
chr7-58260255-INS-6564
chr7-58276100-INS-683
chr7-58286808-INS-3147
chr7-58293545-INS-68
chr7-58318182-INS-100
chr7-58332559-INS-74
chr7-58373930-INS-74
chr7-58433347-INS-147
chr7-58444818-INS-191
chr7-58461989-INS-86
chr7-58482960-INS-64
chr7-58484865-INS-58
chr7-58518542-INS-3319
chr7-58551995-INS-155
chr7-58608487-INS-260
chr7-58635896-INS-7183
chr7-58662139-INS-107
chr7-58662891-INS-327
chr7-58670489-INS-413
chr7-58673542-INS-611
chr7-58676498-INS-563
chr7-58700543-INS-6672
chr7-58726031-INS-213
chr7-58811849-INS-62

chr7-58843112-INS-92
chr7-58849697-INS-84
chr7-58884713-INS-66
chr7-58889876-INS-865
chr7-58910763-INS-55
chr7-58929050-INS-75
chr7-58937965-INS-153
chr7-58950104-INS-113
chr7-58964736-INS-146
chr7-59003431-INS-1872
chr7-59063791-INS-2451
chr7-59267933-INS-188
chr7-59271996-INS-77
chr7-59295940-INS-236
chr7-59324386-INS-5093
chr7-59327326-INS-2517
chr7-59450372-INS-70
chr7-59555710-INS-237
chr7-59605496-INS-60
chr7-59615255-INS-427
chr7-59647327-INS-180
chr7-59678690-INS-488
chr7-59707037-INS-126
chr7-59715099-INS-466
chr7-59729232-INS-218
chr7-59732095-INS-317
chr7-59769488-INS-7032
chr7-59770923-INS-1359
chr7-59775127-INS-67
chr7-59775490-INS-939
chr7-59777649-INS-6713
chr7-59939069-INS-469
chr7-60001290-INS-2114
chr7-60005096-INS-421
chr7-60006343-INS-6573
chr7-60054729-INS-6711
chr7-60142052-INS-143
chr7-60158733-INS-152
chr7-60231733-INS-6400
chr7-60235396-INS-1960
chr7-60252412-INS-7107
chr7-60258770-INS-1894
chr7-60316200-INS-1161
chr7-60380693-INS-441
chr7-60388617-INS-124
chr7-60458387-INS-457
chr7-60492637-INS-846
chr7-60497215-INS-68
chr7-60595381-INS-6586
chr7-60604257-INS-6191
chr7-60622538-INS-526
chr7-60653665-INS-805
chr7-60656469-INS-584
chr7-60657754-INS-233
chr7-60703922-INS-55
chr7-60718365-INS-7015
chr7-60732669-INS-713
chr7-60775244-INS-877
chr7-60779977-INS-168
chr7-60820361-INS-703
chr7-60842317-INS-6251
chr7-60856398-INS-6850
chr7-60866015-INS-7090
chr7-60883737-INS-3078
chr7-60914827-INS-54
chr7-60921997-INS-213
chr7-60928435-INS-54
chr7-60963598-INS-84
chr7-60969092-INS-1000
chr7-60989187-INS-8018
chr7-61028122-INS-136
chr7-61030796-INS-57
chr7-61042031-INS-62
chr7-61049593-INS-151
chr7-61052818-INS-801
chr7-61137766-INS-72
chr7-61178017-INS-62

chr7-61192933-INS-113
chr7-61228952-INS-70
chr7-61272835-INS-150
chr7-61336744-INS-644
chr7-61344613-INS-2936
chr7-61362276-INS-53
chr7-61387345-INS-8036
chr7-61404874-INS-76
chr7-61423544-INS-313
chr7-61462225-INS-891
chr7-61465889-INS-52
chr7-61530942-INS-73
chr7-61557550-INS-385
chr7-61567547-INS-214
chr7-61571114-INS-881
chr7-61576287-INS-119
chr7-61581675-INS-59
chr7-61617098-INS-6309
chr7-61622587-INS-6393
chr7-61633528-INS-190
chr7-61675012-INS-174
chr7-61729170-INS-151
chr7-61797139-INS-87
chr7-61857074-INS-100
chr7-61907209-INS-88
chr7-61923944-INS-58
chr7-61958269-INS-59
chr7-61961175-INS-51
chr7-61993788-INS-592
chr7-62053345-INS-183
chr7-62074185-INS-94
chr7-62134371-INS-81
chr7-62143432-INS-11598
chr7-62152789-INS-6862
chr7-62174473-INS-7073
chr7-62181943-INS-742
chr7-62282209-INS-55
chr7-62292162-INS-1982
chr7-62295324-INS-382
chr7-62320017-INS-200
chr7-62323454-INS-7224
chr7-62420862-INS-180
chr7-62454918-INS-525
chr7-62456497-INS-103
chr7-62466265-INS-59
chr7-62547720-INS-98
chr7-62591141-INS-113
chr7-62605464-INS-61
chr7-62640616-INS-1891
chr7-62642385-INS-696
chr7-62650295-INS-760
chr7-62653253-INS-532
chr7-62671705-INS-211
chr7-62672288-INS-7111
chr7-62673719-INS-416
chr7-62681432-INS-60
chr7-62682839-INS-128
chr7-62693041-INS-63
chr7-62711786-INS-354
chr7-62716815-INS-79
chr7-62727745-INS-117
chr7-62738995-INS-3323
chr7-62743175-INS-112
chr7-62748424-INS-72
chr7-62751764-INS-6855
chr7-62763089-INS-89
chr7-62774849-INS-351
chr7-62821581-INS-2993
chr7-62826259-INS-2704
chr7-62864618-INS-3280
chr7-62875146-INS-1994
chr7-62882591-INS-63
chr7-62883674-INS-331
chr7-62903776-INS-69
chr7-62922228-INS-428
chr7-62984642-INS-60
chr7-62987932-INS-111

chr7-62999511-INS-1135
chr7-63010949-INS-11036
chr7-63069742-INS-269
chr7-63069997-INS-6910
chr7-63074316-INS-156
chr7-63079008-INS-6297
chr7-63110451-INS-86
chr7-63120859-INS-387
chr7-63123219-INS-219
chr7-63129998-INS-60
chr7-63136929-INS-6098
chr7-63137659-INS-7110
chr7-63152197-INS-79
chr7-63205951-INS-816
chr7-63211610-INS-126
chr7-63224361-INS-497
chr7-63249622-INS-608
chr7-63268145-INS-7399
chr7-63289366-INS-63
chr7-63289701-INS-1975
chr7-63290373-INS-6316
chr7-63303273-INS-57
chr7-63381251-INS-74
chr7-63398999-INS-69
chr7-63425031-INS-1139
chr7-63429922-INS-212
chr7-63446791-INS-433
chr7-63462014-INS-386
chr7-63496984-INS-131
chr7-63501095-INS-167
chr7-63504895-INS-295
chr7-63527027-INS-181
chr7-63544104-INS-242
chr7-63545597-INS-55
chr7-63549118-INS-159
chr7-63593356-INS-146
chr7-63609067-INS-112
chr7-63625079-INS-176
chr7-63650454-INS-96
chr7-63658338-INS-983
chr7-63683953-INS-70
chr7-63694420-INS-263
chr7-63710106-INS-133
chr7-63718128-INS-63
chr7-63732189-INS-134
chr7-63746226-INS-200
chr7-63747166-INS-74
chr7-63748791-INS-134
chr7-63752970-INS-174
chr7-63785714-INS-514
chr7-63792095-INS-150
chr7-63865265-INS-66
chr7-63866217-INS-74
chr7-63866462-INS-1893
chr7-63883102-INS-52
chr7-63916408-INS-6613
chr7-63920663-INS-176
chr7-63937932-INS-6730
chr7-63960979-INS-205
chr7-63992157-INS-57
chr7-64053585-INS-205
chr7-64055229-INS-132
chr7-64067352-INS-55
chr7-64074260-INS-112
chr7-64094863-INS-153
chr7-64150062-INS-120
chr7-64177234-INS-474
chr7-64184963-INS-419
chr7-64201636-INS-503
chr7-64257706-INS-70
chr7-64278607-INS-292
chr7-64302331-INS-327
chr7-64305772-INS-52
chr7-64340555-INS-66
chr7-64348468-INS-55
chr7-64389405-INS-180
chr7-64400664-INS-82

chr7-64425734-INS-82
chr7-64426927-INS-399
chr7-64461498-INS-110
chr7-64484729-INS-61
chr7-64494189-INS-1866
chr7-64500111-INS-403
chr7-64505154-INS-113
chr7-64507386-INS-126
chr7-64516789-INS-201
chr7-64531301-INS-609
chr7-64551333-INS-69
chr7-64596976-INS-327
chr7-64609743-INS-284
chr7-64613318-INS-93
chr7-64627933-INS-784
chr7-64628738-INS-319
chr7-64648355-INS-380
chr7-64651399-INS-54
chr7-64712704-INS-583
chr7-64800037-INS-85
chr7-64811542-INS-162
chr7-64814304-INS-327
chr7-64861763-INS-52
chr7-64869867-INS-6892
chr7-64874834-INS-281
chr7-64934562-INS-74
chr7-64966923-INS-56
chr7-64970987-INS-209
chr7-64975058-INS-115
chr7-65005494-INS-55
chr7-65008443-INS-111
chr7-65033320-INS-8940
chr7-65035886-INS-178
chr7-65053675-INS-51
chr7-65074872-INS-167
chr7-65087504-INS-66
chr7-65098409-INS-217
chr7-65121315-INS-60
chr7-65150119-INS-58
chr7-65170975-INS-64
chr7-65178767-INS-220
chr7-65196866-INS-100
chr7-65206291-INS-56
chr7-65221135-INS-61
chr7-65234675-INS-88
chr7-65268094-INS-66
chr7-65270323-INS-98
chr7-65273016-INS-262
chr7-65283731-INS-85
chr7-65308964-INS-247
chr7-65323587-INS-54
chr7-65339601-INS-358
chr7-65355922-INS-7127
chr7-65364005-INS-54
chr7-65381085-INS-341
chr7-65385529-INS-72
chr7-65388929-INS-94
chr7-65400630-INS-60
chr7-65402860-INS-51
chr7-65425815-INS-70
chr7-65431728-INS-566
chr7-65441218-INS-175
chr7-65444098-INS-382
chr7-65471167-INS-486
chr7-65485065-INS-122
chr7-65486592-INS-669
chr7-65487707-INS-200
chr7-65495274-INS-377
chr7-65517793-INS-120
chr7-65523667-INS-63
chr7-65524812-INS-183
chr7-65576773-INS-94
chr7-65682720-INS-63
chr7-65700347-INS-655
chr7-65723843-INS-64
chr7-65727748-INS-53
chr7-65737635-INS-172

chr7-65774141-INS-94
chr7-65778218-INS-96
chr7-65789463-INS-866
chr7-65795591-INS-236
chr7-65796687-INS-1892
chr7-65837977-INS-51
chr7-65838804-INS-175
chr7-65844994-INS-56
chr7-65853426-INS-52
chr7-65873236-INS-400
chr7-65913456-INS-180
chr7-65976514-INS-67
chr7-65984761-INS-56
chr7-66000363-INS-57
chr7-66001685-INS-52
chr7-66011649-INS-124
chr7-66012325-INS-868
chr7-66014678-INS-60
chr7-66022138-INS-77
chr7-66029801-INS-324
chr7-66035893-INS-676
chr7-66080238-INS-282
chr7-66145861-INS-80
chr7-66160824-INS-124
chr7-66160899-INS-271
chr7-66210255-INS-259
chr7-66215442-INS-617
chr7-66263211-INS-54
chr7-66273407-INS-5180
chr7-66277683-INS-60
chr7-66293095-INS-404
chr7-66297327-INS-56
chr7-66302872-INS-79
chr7-66348928-INS-300
chr7-66362493-INS-327
chr7-66376975-INS-497
chr7-66378265-INS-80
chr7-66389682-INS-160
chr7-66409170-INS-55
chr7-66443208-INS-80
chr7-66445336-INS-69
chr7-66450516-INS-69
chr7-66453643-INS-100
chr7-66483380-INS-54
chr7-66496401-INS-142
chr7-66551752-INS-80
chr7-66558707-INS-362
chr7-66578547-INS-52
chr7-66623703-INS-254
chr7-66687788-INS-402
chr7-66691855-INS-54
chr7-66691913-INS-153
chr7-66705249-INS-364
chr7-66713108-INS-260
chr7-66719299-INS-294
chr7-66747355-INS-56
chr7-66769214-INS-615
chr7-66774961-INS-64
chr7-66795312-INS-70
chr7-66830166-INS-213
chr7-66837638-INS-152
chr7-66847126-INS-81
chr7-66920066-INS-86
chr7-66930477-INS-158
chr7-66957664-INS-1893
chr7-66992572-INS-1399
chr7-66993957-INS-899
chr7-66997532-INS-74
chr7-67004992-INS-6622
chr7-67023799-INS-1551
chr7-67028751-INS-96
chr7-67063273-INS-202
chr7-67077765-INS-186
chr7-67109090-INS-74
chr7-67123884-INS-71
chr7-67151045-INS-648
chr7-67160090-INS-179

chr7-67178126-INS-153
chr7-67212184-INS-70
chr7-67237803-INS-85
chr7-67248485-INS-80
chr7-67256179-INS-214
chr7-67276354-INS-498
chr7-67299096-INS-3210
chr7-67312362-INS-66
chr7-67324627-INS-251
chr7-67328286-INS-173
chr7-67346735-INS-72
chr7-67348161-INS-59
chr7-67350137-INS-52
chr7-67405065-INS-75
chr7-67420306-INS-310
chr7-67423200-INS-54
chr7-67427306-INS-73
chr7-67438821-INS-217
chr7-67441022-INS-624
chr7-67456468-INS-71
chr7-67459204-INS-184
chr7-67466531-INS-137
chr7-67477130-INS-477
chr7-67512572-INS-4830
chr7-67531203-INS-94
chr7-67534602-INS-217
chr7-67539458-INS-86
chr7-67571364-INS-5638
chr7-67660390-INS-52
chr7-67683031-INS-66
chr7-67685486-INS-183
chr7-67690510-INS-53
chr7-67704362-INS-355
chr7-67707209-INS-3013
chr7-67745630-INS-485
chr7-67769347-INS-53
chr7-67793217-INS-212
chr7-67801715-INS-166
chr7-67809094-INS-60
chr7-67820130-INS-236
chr7-67821580-INS-158
chr7-67828610-INS-119
chr7-67855350-INS-202
chr7-67900525-INS-56
chr7-67909920-INS-211
chr7-67916158-INS-201
chr7-67919404-INS-196
chr7-67922451-INS-226
chr7-67959503-INS-60
chr7-67986257-INS-56
chr7-67988412-INS-157
chr7-67993296-INS-457
chr7-67998692-INS-245
chr7-68002086-INS-7091
chr7-68024962-INS-61
chr7-68038395-INS-6247
chr7-68045065-INS-174
chr7-68048245-INS-54
chr7-68051911-INS-74
chr7-68066046-INS-153
chr7-68070023-INS-6221
chr7-68070938-INS-1979
chr7-68077221-INS-6165
chr7-68083937-INS-1384
chr7-68121521-INS-60
chr7-68142599-INS-177
chr7-68146475-INS-414
chr7-68148409-INS-80
chr7-68156918-INS-55
chr7-68166855-INS-236
chr7-68174303-INS-6738
chr7-68179405-INS-6654
chr7-68179942-INS-108
chr7-68223748-INS-99
chr7-68231633-INS-98
chr7-68237848-INS-283
chr7-68263159-INS-68

chr7-68294757-INS-5351
chr7-68302879-INS-226
chr7-68303636-INS-65
chr7-68308167-INS-76
chr7-68315400-INS-59
chr7-68323740-INS-87
chr7-68324937-INS-434
chr7-68334289-INS-1748
chr7-68341256-INS-59
chr7-68342981-INS-92
chr7-68352450-INS-71
chr7-68353618-INS-91
chr7-68354037-INS-174
chr7-68362126-INS-54
chr7-68364128-INS-1663
chr7-68364180-INS-2234
chr7-68364545-INS-364
chr7-68425933-INS-7265
chr7-68428973-INS-217
chr7-68436402-INS-171
chr7-68470941-INS-88
chr7-68474862-INS-78
chr7-68476909-INS-318
chr7-68497675-INS-60
chr7-68558741-INS-493
chr7-68581381-INS-6738
chr7-68601776-INS-63
chr7-68604335-INS-71
chr7-68607775-INS-1414
chr7-68611378-INS-59
chr7-68620558-INS-186
chr7-68626165-INS-7176
chr7-68688528-INS-58
chr7-68698320-INS-504
chr7-68725741-INS-78
chr7-68734783-INS-357
chr7-68760016-INS-6575
chr7-68777595-INS-1394
chr7-68784089-INS-1034
chr7-68784429-INS-89
chr7-68786826-INS-7560
chr7-68796787-INS-7987
chr7-68809740-INS-52
chr7-68815984-INS-75
chr7-68816525-INS-353
chr7-68836490-INS-194
chr7-68852703-INS-257
chr7-68858356-INS-188
chr7-68866992-INS-76
chr7-68889906-INS-70
chr7-68897090-INS-978
chr7-68920761-INS-52
chr7-68936097-INS-155
chr7-68942086-INS-82
chr7-68983515-INS-252
chr7-69027626-INS-67
chr7-69030595-INS-512
chr7-69033586-INS-481
chr7-69053295-INS-1260
chr7-69092178-INS-401
chr7-69111824-INS-318
chr7-69118368-INS-8838
chr7-69119323-INS-121
chr7-69134034-INS-160
chr7-69135978-INS-533
chr7-69162731-INS-61
chr7-69187090-INS-489
chr7-69208014-INS-619
chr7-69213852-INS-66
chr7-69219753-INS-399
chr7-69237287-INS-266
chr7-69251312-INS-6273
chr7-69255394-INS-154
chr7-69270700-INS-144
chr7-69274182-INS-304
chr7-69286411-INS-239
chr7-69289775-INS-6467

chr7-69309380-INS-64
chr7-69326808-INS-284
chr7-69333290-INS-155
chr7-69334496-INS-3732
chr7-69381702-INS-276
chr7-69396482-INS-6440
chr7-69397607-INS-201
chr7-69410290-INS-840
chr7-69411337-INS-96
chr7-69434248-INS-6836
chr7-69435483-INS-499
chr7-69493568-INS-60
chr7-69504171-INS-65
chr7-69514848-INS-73
chr7-69516516-INS-489
chr7-69519847-INS-52
chr7-69530015-INS-157
chr7-69574271-INS-209
chr7-69622378-INS-758
chr7-69636714-INS-727
chr7-69647204-INS-54
chr7-69654675-INS-59
chr7-69658005-INS-459
chr7-69661988-INS-4080
chr7-69662442-INS-6979
chr7-69664350-INS-366
chr7-69664907-INS-54
chr7-69667973-INS-1246
chr7-69672419-INS-58
chr7-69673206-INS-146
chr7-69676636-INS-126
chr7-69677251-INS-859
chr7-69683256-INS-95
chr7-69696615-INS-4542
chr7-69708488-INS-330
chr7-69724598-INS-67
chr7-69733272-INS-996
chr7-69733331-INS-78
chr7-69742158-INS-55
chr7-69764313-INS-60
chr7-69774157-INS-133
chr7-69776455-INS-66
chr7-69792485-INS-138
chr7-69798869-INS-1820
chr7-69806053-INS-75
chr7-69811374-INS-78
chr7-69813975-INS-117
chr7-69850659-INS-108
chr7-69852088-INS-113
chr7-69865038-INS-94
chr7-69892461-INS-104
chr7-69909901-INS-100
chr7-69937361-INS-190
chr7-69950166-INS-56
chr7-69951872-INS-57
chr7-69959271-INS-2204
chr7-69968744-INS-70
chr7-69980385-INS-245
chr7-70030728-INS-51
chr7-70032831-INS-76
chr7-70045320-INS-57
chr7-70053100-INS-96
chr7-70056346-INS-205
chr7-70069487-INS-196
chr7-70079595-INS-68
chr7-70127462-INS-84
chr7-70133027-INS-240
chr7-70150861-INS-64
chr7-70168545-INS-100
chr7-70175833-INS-54
chr7-70179557-INS-144
chr7-70208735-INS-4329
chr7-70218069-INS-7154
chr7-70229184-INS-58
chr7-70256156-INS-525
chr7-70267029-INS-1284
chr7-70276491-INS-143

chr7-70277622-INS-66
chr7-70327476-INS-1426
chr7-70335364-INS-272
chr7-70368913-INS-198
chr7-70370346-INS-52
chr7-70404639-INS-11886
chr7-70427310-INS-210
chr7-70484250-INS-61
chr7-70505004-INS-6465
chr7-70526881-INS-6440
chr7-70530144-INS-6812
chr7-70536018-INS-178
chr7-70567137-INS-52
chr7-70627088-INS-888
chr7-70637535-INS-7734
chr7-70637536-INS-226
chr7-70650886-INS-128
chr7-70655870-INS-394
chr7-70678657-INS-102
chr7-70721548-INS-664
chr7-70807364-INS-106
chr7-70852573-INS-52
chr7-70904352-INS-94
chr7-70923123-INS-624
chr7-70925216-INS-51
chr7-70932645-INS-7139
chr7-70936514-INS-247
chr7-70945681-INS-64
chr7-70960656-INS-664
chr7-70966039-INS-2780
chr7-71008269-INS-186
chr7-71052933-INS-189
chr7-71055477-INS-246
chr7-71060138-INS-54
chr7-71068482-INS-65
chr7-71074622-INS-927
chr7-71116274-INS-477
chr7-71134207-INS-56
chr7-71152300-INS-80
chr7-71160566-INS-88
chr7-71212350-INS-56
chr7-71244506-INS-67
chr7-71245075-INS-222
chr7-71259083-INS-52
chr7-71264206-INS-72
chr7-71296526-INS-137
chr7-71309653-INS-562
chr7-71336830-INS-450
chr7-71373105-INS-735
chr7-71406010-INS-423
chr7-71439578-INS-82
chr7-71455115-INS-7505
chr7-71460064-INS-6871
chr7-71462131-INS-545
chr7-71538340-INS-4520
chr7-71568302-INS-56
chr7-71574941-INS-6215
chr7-71605603-INS-64
chr7-71615437-INS-56
chr7-71697463-INS-52
chr7-71704352-INS-6710
chr7-71723499-INS-6996
chr7-71754064-INS-596
chr7-71765767-INS-56
chr7-71781161-INS-7105
chr7-71792110-INS-63
chr7-71797903-INS-57
chr7-71837396-INS-110
chr7-71854665-INS-62
chr7-71860613-INS-475
chr7-71876655-INS-762
chr7-71880932-INS-78
chr7-71890932-INS-58
chr7-71893119-INS-204
chr7-71893438-INS-7483
chr7-71919799-INS-2836
chr7-71924426-INS-52

chr7-71928686-INS-138
chr7-71937644-INS-663
chr7-71963403-INS-6569
chr7-71978098-INS-72
chr7-71979312-INS-8414
chr7-71982207-INS-1684
chr7-72044977-INS-90
chr7-72049474-INS-536
chr7-72058108-INS-203
chr7-72061277-INS-57
chr7-72086369-INS-8570
chr7-72092717-INS-3205
chr7-72098253-INS-6431
chr7-72123413-INS-6744
chr7-72147863-INS-664
chr7-72166001-INS-70
chr7-72178302-INS-72
chr7-72200233-INS-202
chr7-72206172-INS-79
chr7-72250921-INS-52
chr7-72316277-INS-289
chr7-72321011-INS-6683
chr7-72354912-INS-154
chr7-72359272-INS-106
chr7-72386144-INS-67
chr7-72390557-INS-259
chr7-72397029-INS-57
chr7-72416921-INS-70
chr7-72428195-INS-259
chr7-72450179-INS-238
chr7-72459877-INS-52
chr7-72461826-INS-60
chr7-72463346-INS-121
chr7-72470384-INS-140
chr7-72477161-INS-1887
chr7-72482849-INS-60
chr7-72505835-INS-136
chr7-72505941-INS-1551
chr7-72507032-INS-398
chr7-72508356-INS-60
chr7-72542029-INS-397
chr7-72550150-INS-686
chr7-72574990-INS-111
chr7-72607107-INS-89
chr7-72621535-INS-1240
chr7-72631767-INS-136
chr7-72636138-INS-65
chr7-72642605-INS-88
chr7-72648543-INS-1047
chr7-72650140-INS-5984
chr7-72652872-INS-2775
chr7-72665170-INS-974
chr7-72670464-INS-52
chr7-72677888-INS-6615
chr7-72694824-INS-242
chr7-72696693-INS-198
chr7-72696893-INS-6705
chr7-72698539-INS-203
chr7-72714379-INS-788
chr7-72721165-INS-1214
chr7-72725848-INS-112
chr7-72732007-INS-201
chr7-72777688-INS-74
chr7-72796940-INS-69
chr7-72810330-INS-53
chr7-72855163-INS-266
chr7-72869614-INS-192
chr7-72883418-INS-56
chr7-72892832-INS-57
chr7-72913306-INS-51
chr7-72929550-INS-126
chr7-72947611-INS-51
chr7-72973833-INS-198
chr7-72974457-INS-2326
chr7-72974465-INS-61
chr7-72974471-INS-4368
chr7-72981181-INS-155

chr7-72989396-INS-85
chr7-72995804-INS-141
chr7-73003151-INS-53
chr7-73031487-INS-261
chr7-73034537-INS-227
chr7-73051850-INS-157
chr7-73060678-INS-57
chr7-73061191-INS-202
chr7-73062166-INS-63
chr7-73072302-INS-121
chr7-73081468-INS-70
chr7-73120124-INS-288
chr7-73133592-INS-63
chr7-73135787-INS-2265
chr7-73161232-INS-198
chr7-73162580-INS-227
chr7-73170189-INS-214
chr7-73176040-INS-92
chr7-73184877-INS-203
chr7-73192333-INS-104
chr7-73194559-INS-233
chr7-73198952-INS-217
chr7-73215253-INS-84
chr7-73218850-INS-161
chr7-73248906-INS-260
chr7-73276994-INS-231
chr7-73279591-INS-236
chr7-73292345-INS-466
chr7-73295324-INS-185
chr7-73319259-INS-6906
chr7-73330745-INS-199
chr7-73334376-INS-159
chr7-73336393-INS-60
chr7-73337413-INS-538
chr7-73356151-INS-340
chr7-73358462-INS-685
chr7-73363174-INS-396
chr7-73372514-INS-53
chr7-73399540-INS-618
chr7-73405518-INS-627
chr7-73411709-INS-74
chr7-73412636-INS-331
chr7-73412791-INS-1547
chr7-73489386-INS-137
chr7-73497967-INS-574
chr7-73500219-INS-52
chr7-73521760-INS-497
chr7-73601410-INS-51
chr7-73660999-INS-54
chr7-73664658-INS-101
chr7-73665972-INS-149
chr7-73692022-INS-297
chr7-73699473-INS-56
chr7-73716997-INS-116
chr7-73751844-INS-54
chr7-73756571-INS-63
chr7-73772988-INS-141
chr7-73782353-INS-83
chr7-73783921-INS-323
chr7-73786302-INS-1895
chr7-73796728-INS-66
chr7-73822287-INS-782
chr7-73843231-INS-507
chr7-73863593-INS-266
chr7-73869783-INS-60
chr7-73911940-INS-220
chr7-73917240-INS-7506
chr7-73974610-INS-136
chr7-73984357-INS-1491
chr7-73994054-INS-536
chr7-74007117-INS-6434
chr7-74011233-INS-230
chr7-74019131-INS-91
chr7-74048715-INS-86
chr7-74054373-INS-60
chr7-74062683-INS-254
chr7-74075722-INS-123

chr7-74097041-INS-554
chr7-74139959-INS-62
chr7-74186598-INS-134
chr7-74213674-INS-129
chr7-74227746-INS-6203
chr7-74227865-INS-186
chr7-74241622-INS-2565
chr7-74302330-INS-540
chr7-74326190-INS-54
chr7-74331925-INS-84
chr7-74334834-INS-84
chr7-74344911-INS-6599
chr7-74359565-INS-274
chr7-74362557-INS-97
chr7-74365281-INS-6640
chr7-74367687-INS-59
chr7-74405546-INS-70
chr7-74442197-INS-528
chr7-74443746-INS-60
chr7-74475889-INS-57
chr7-74486282-INS-52
chr7-74523901-INS-78
chr7-74539808-INS-448
chr7-74542749-INS-197
chr7-74552074-INS-60
chr7-74556076-INS-7089
chr7-74558207-INS-146
chr7-74559221-INS-58
chr7-74572347-INS-77
chr7-74582839-INS-90
chr7-74587866-INS-400
chr7-74591656-INS-62
chr7-74621591-INS-96
chr7-74637922-INS-57
chr7-74660589-INS-135
chr7-74661599-INS-6883
chr7-74661853-INS-2426
chr7-74684695-INS-57
chr7-74713077-INS-65
chr7-74721626-INS-220
chr7-74729317-INS-84
chr7-74730371-INS-7681
chr7-74736434-INS-214
chr7-74753389-INS-66
chr7-74755106-INS-429
chr7-74757477-INS-148
chr7-74778935-INS-6515
chr7-74803046-INS-54
chr7-74820762-INS-59
chr7-74846671-INS-620
chr7-74850255-INS-8790
chr7-74866156-INS-516
chr7-74874264-INS-364
chr7-74882758-INS-564
chr7-74896949-INS-128
chr7-74919314-INS-220
chr7-74944335-INS-97
chr7-74960926-INS-52
chr7-74968982-INS-135
chr7-74971861-INS-206
chr7-74995011-INS-176
chr7-74996818-INS-75
chr7-74997783-INS-176
chr7-75012930-INS-554
chr7-75023823-INS-169
chr7-75028490-INS-106
chr7-75039080-INS-578
chr7-75041306-INS-202
chr7-75046195-INS-60
chr7-75059016-INS-61
chr7-75064893-INS-217
chr7-75088767-INS-81
chr7-75101899-INS-150
chr7-75115579-INS-351
chr7-75125394-INS-364
chr7-75128518-INS-56
chr7-75150539-INS-867

chr7-75157516-INS-6689
chr7-75182386-INS-209
chr7-75184404-INS-252
chr7-75202648-INS-386
chr7-75273989-INS-66
chr7-75281635-INS-121
chr7-75323573-INS-1183
chr7-75323693-INS-96
chr7-75326299-INS-276
chr7-75326829-INS-62
chr7-75358304-INS-75
chr7-75368709-INS-211
chr7-75376590-INS-65
chr7-75381053-INS-210
chr7-75391011-INS-275
chr7-75415100-INS-194
chr7-75433433-INS-51
chr7-75434108-INS-172
chr7-75443321-INS-122
chr7-75446933-INS-533
chr7-75457609-INS-103
chr7-75470934-INS-125
chr7-75478370-INS-195
chr7-75480743-INS-203
chr7-75490961-INS-56
chr7-75534746-INS-152
chr7-75536651-INS-88
chr7-75542738-INS-75
chr7-75575777-INS-6903
chr7-75619305-INS-114
chr7-75636759-INS-64
chr7-75646994-INS-60
chr7-75686791-INS-6899
chr7-75694938-INS-69
chr7-75704819-INS-672
chr7-75715918-INS-207
chr7-75739879-INS-393
chr7-75741111-INS-194
chr7-75767685-INS-107
chr7-75788637-INS-131
chr7-75798623-INS-1103
chr7-75799361-INS-1461
chr7-75817569-INS-67
chr7-75852482-INS-54
chr7-75864992-INS-2852
chr7-75867713-INS-190
chr7-75890478-INS-7089
chr7-75921863-INS-56
chr7-75947272-INS-191
chr7-75956480-INS-139
chr7-75957534-INS-56
chr7-75963029-INS-77
chr7-75966397-INS-571
chr7-75984895-INS-739
chr7-75993747-INS-136
chr7-76004557-INS-1015
chr7-76031441-INS-932
chr7-76041453-INS-870
chr7-76058306-INS-77
chr7-76067606-INS-3909
chr7-76071292-INS-169
chr7-76085408-INS-539
chr7-76103345-INS-110
chr7-76125624-INS-433
chr7-76130682-INS-72
chr7-76135114-INS-60
chr7-76154950-INS-274
chr7-76164413-INS-458
chr7-76179553-INS-501
chr7-76187517-INS-6340
chr7-76191710-INS-85
chr7-76192522-INS-6385
chr7-76199298-INS-1195
chr7-76232119-INS-56
chr7-76233240-INS-4868
chr7-76240699-INS-64
chr7-76256162-INS-57

chr7-76271050-INS-146
chr7-76276744-INS-182
chr7-76277832-INS-78
chr7-76293507-INS-53
chr7-76295338-INS-86
chr7-76296942-INS-7074
chr7-76308214-INS-93
chr7-76312861-INS-74
chr7-76315195-INS-6392
chr7-76326971-INS-259
chr7-76327913-INS-54
chr7-76328875-INS-409
chr7-76338592-INS-84
chr7-76373296-INS-629
chr7-76375690-INS-199
chr7-76378338-INS-233
chr7-76384091-INS-688
chr7-76406092-INS-82
chr7-76412728-INS-51
chr7-76421799-INS-876
chr7-76425228-INS-64
chr7-76445381-INS-7256
chr7-76446015-INS-6208
chr7-76455476-INS-6188
chr7-76509087-INS-2228
chr7-76512843-INS-474
chr7-76593679-INS-108
chr7-76597417-INS-1913
chr7-76598325-INS-56
chr7-76625059-INS-127
chr7-76631824-INS-399
chr7-76634723-INS-197
chr7-76641969-INS-6517
chr7-76657713-INS-86
chr7-76670025-INS-659
chr7-76677490-INS-274
chr7-76690665-INS-285
chr7-76690735-INS-5198
chr7-76698201-INS-59
chr7-76703176-INS-983
chr7-76711016-INS-92
chr7-76724325-INS-137
chr7-76725492-INS-91
chr7-76738404-INS-126
chr7-76780011-INS-602
chr7-76786734-INS-148
chr7-76791968-INS-160
chr7-76794842-INS-106
chr7-76800341-INS-69
chr7-76807142-INS-535
chr7-76809542-INS-164
chr7-76819250-INS-172
chr7-76825655-INS-230
chr7-76832029-INS-85
chr7-76837776-INS-241
chr7-76840373-INS-869
chr7-76844851-INS-92
chr7-76856161-INS-167
chr7-76874387-INS-203
chr7-76884664-INS-106
chr7-76940000-INS-52
chr7-76961192-INS-131
chr7-76977726-INS-162
chr7-76987925-INS-6324
chr7-76990198-INS-324
chr7-76997785-INS-75
chr7-77006250-INS-6245
chr7-77019802-INS-84
chr7-77021928-INS-860
chr7-77024174-INS-414
chr7-77031396-INS-690
chr7-77045470-INS-64
chr7-77078711-INS-6432
chr7-77088580-INS-6287
chr7-77116051-INS-462
chr7-77118930-INS-134
chr7-77166746-INS-63

chr7-77212058-INS-109
chr7-77212700-INS-827
chr7-77270287-INS-58
chr7-77279788-INS-128
chr7-77304564-INS-213
chr7-77312620-INS-728
chr7-77313816-INS-62
chr7-77329147-INS-123
chr7-77358434-INS-803
chr7-77359543-INS-466
chr7-77364842-INS-618
chr7-77367063-INS-55
chr7-77373054-INS-220
chr7-77392468-INS-52
chr7-77409708-INS-431
chr7-77426795-INS-113
chr7-77454311-INS-310
chr7-77460137-INS-68
chr7-77479425-INS-63
chr7-77490166-INS-58
chr7-77504010-INS-150
chr7-77507101-INS-72
chr7-77579899-INS-98
chr7-77588862-INS-60
chr7-77598296-INS-222
chr7-77600854-INS-331
chr7-77617752-INS-249
chr7-77626526-INS-63
chr7-77638766-INS-153
chr7-77679682-INS-266
chr7-77703423-INS-424
chr7-77706532-INS-56
chr7-77741253-INS-357
chr7-77746262-INS-90
chr7-77749161-INS-132
chr7-77750244-INS-155
chr7-77777856-INS-56
chr7-77790313-INS-170
chr7-77809706-INS-438
chr7-77830704-INS-202
chr7-77853132-INS-1720
chr7-77856028-INS-660
chr7-77860786-INS-73
chr7-77900226-INS-173
chr7-77947490-INS-168
chr7-77957210-INS-80
chr7-77962362-INS-95
chr7-77973533-INS-80
chr7-78008907-INS-520
chr7-78011755-INS-52
chr7-78036251-INS-54
chr7-78048663-INS-2481
chr7-78074222-INS-6520
chr7-78079940-INS-340
chr7-78089912-INS-54
chr7-78151048-INS-60
chr7-78151962-INS-124
chr7-78164088-INS-51
chr7-78170337-INS-68
chr7-78187612-INS-717
chr7-78241345-INS-71
chr7-78242924-INS-65
chr7-78245152-INS-219
chr7-78264714-INS-78
chr7-78283109-INS-830
chr7-78304809-INS-80
chr7-78338911-INS-147
chr7-78394706-INS-143
chr7-78407772-INS-54
chr7-78429727-INS-232
chr7-78452742-INS-94
chr7-78504507-INS-66
chr7-78518092-INS-188
chr7-78541472-INS-282
chr7-78593775-INS-102
chr7-78601367-INS-72
chr7-78606085-INS-414

chr7-78607780-INS-173
chr7-78607953-INS-155
chr7-78609818-INS-350
chr7-78618362-INS-68
chr7-78625721-INS-109
chr7-78628470-INS-107
chr7-78633319-INS-135
chr7-78649869-INS-61
chr7-78676039-INS-189
chr7-78679319-INS-415
chr7-78687676-INS-58
chr7-78694223-INS-214
chr7-78696732-INS-60
chr7-78713708-INS-320
chr7-78715686-INS-128
chr7-78718955-INS-88
chr7-78728658-INS-192
chr7-78730890-INS-54
chr7-78774356-INS-140
chr7-78799893-INS-126
chr7-78806977-INS-101
chr7-78817532-INS-182
chr7-78820628-INS-60
chr7-78840923-INS-256
chr7-78845662-INS-991
chr7-78856982-INS-426
chr7-78865597-INS-103
chr7-78871669-INS-315
chr7-78873434-INS-5704
chr7-78915868-INS-56
chr7-78915877-INS-7590
chr7-78916467-INS-213
chr7-78917546-INS-846
chr7-78942124-INS-481
chr7-78945548-INS-209
chr7-78959559-INS-59
chr7-78962574-INS-98
chr7-78968822-INS-176
chr7-78996846-INS-134
chr7-79003297-INS-180
chr7-79022007-INS-201
chr7-79035701-INS-295
chr7-79039851-INS-118
chr7-79048413-INS-82
chr7-79055785-INS-195
chr7-79059137-INS-217
chr7-79060825-INS-173
chr7-79067508-INS-183
chr7-79089094-INS-256
chr7-79091471-INS-62
chr7-79096138-INS-210
chr7-79127764-INS-5548
chr7-79142616-INS-136
chr7-79187949-INS-206
chr7-79190565-INS-189
chr7-79192237-INS-201
chr7-79232435-INS-112
chr7-79261441-INS-200
chr7-79266782-INS-182
chr7-79282367-INS-1564
chr7-79300273-INS-96
chr7-79301680-INS-53
chr7-79309569-INS-108
chr7-79315694-INS-171
chr7-79326057-INS-119
chr7-79327552-INS-157
chr7-79331164-INS-141
chr7-79333695-INS-85
chr7-79336399-INS-60
chr7-79337731-INS-231
chr7-79362530-INS-264
chr7-79364571-INS-248
chr7-79380228-INS-324
chr7-79383271-INS-296
chr7-79405985-INS-203
chr7-79416213-INS-213
chr7-79418835-INS-138

chr7-79430197-INS-12178
chr7-79441138-INS-204
chr7-79452855-INS-60
chr7-79456727-INS-202
chr7-79466016-INS-572
chr7-79478664-INS-1296
chr7-79478664-INS-864
chr7-79479404-INS-1350
chr7-79487848-INS-217
chr7-79498357-INS-109
chr7-79505760-INS-201
chr7-79513984-INS-212
chr7-79514742-INS-201
chr7-79515140-INS-449
chr7-79515814-INS-198
chr7-79527986-INS-79
chr7-79552520-INS-266
chr7-79555598-INS-69
chr7-79558672-INS-250
chr7-79575868-INS-139
chr7-79585003-INS-54
chr7-79603557-INS-189
chr7-79622668-INS-104
chr7-79626031-INS-140
chr7-79632688-INS-81
chr7-79634138-INS-60
chr7-79637034-INS-200
chr7-79645468-INS-92
chr7-79660904-INS-175
chr7-79665781-INS-68
chr7-79669365-INS-254
chr7-79690832-INS-106
chr7-79703324-INS-167
chr7-79710837-INS-1101
chr7-79727172-INS-72
chr7-79729970-INS-404
chr7-79751896-INS-50
chr7-79769028-INS-6524
chr7-79775707-INS-170
chr7-79789023-INS-202
chr7-79803939-INS-54
chr7-79807453-INS-60
chr7-79830880-INS-223
chr7-79832833-INS-161
chr7-79846542-INS-122
chr7-79860241-INS-206
chr7-79871830-INS-939
chr7-79872873-INS-649
chr7-79887685-INS-218
chr7-79905672-INS-194
chr7-79917854-INS-3279
chr7-79921805-INS-111
chr7-79928850-INS-52
chr7-79931591-INS-216
chr7-79932742-INS-223
chr7-79939849-INS-202
chr7-79946403-INS-212
chr7-79950501-INS-60
chr7-79952556-INS-176
chr7-79956785-INS-209
chr7-80004972-INS-57
chr7-80024707-INS-1002
chr7-80071663-INS-1011
chr7-80073168-INS-168
chr7-80077990-INS-199
chr7-80095377-INS-6452
chr7-80117827-INS-304
chr7-80135945-INS-208
chr7-80137583-INS-731
chr7-80168929-INS-196
chr7-80170837-INS-55
chr7-80181603-INS-142
chr7-80194513-INS-174
chr7-80200060-INS-63
chr7-80205144-INS-60
chr7-80218186-INS-4345
chr7-80224061-INS-272

chr7-80260310-INS-51
chr7-80280282-INS-347
chr7-80290362-INS-64
chr7-80296141-INS-168
chr7-80303161-INS-347
chr7-80330642-INS-60
chr7-80358223-INS-260
chr7-80359029-INS-520
chr7-80359486-INS-78
chr7-80364529-INS-229
chr7-80374287-INS-60
chr7-80390416-INS-54
chr7-80396216-INS-63
chr7-80400716-INS-382
chr7-80419674-INS-788
chr7-80419912-INS-167
chr7-80431204-INS-226
chr7-80444195-INS-101
chr7-80449964-INS-56
chr7-80450289-INS-233
chr7-80458975-INS-60
chr7-80463940-INS-137
chr7-80465098-INS-280
chr7-80470213-INS-54
chr7-80473619-INS-64
chr7-80478843-INS-221
chr7-80508364-INS-218
chr7-80517974-INS-233
chr7-80526750-INS-332
chr7-80527602-INS-163
chr7-80531066-INS-123
chr7-80533314-INS-1123
chr7-80544281-INS-119
chr7-80555347-INS-198
chr7-80569787-INS-208
chr7-80580069-INS-123
chr7-80583564-INS-190
chr7-80593619-INS-298
chr7-80602398-INS-209
chr7-80605776-INS-52
chr7-80612414-INS-98
chr7-80613560-INS-218
chr7-80615061-INS-242
chr7-80615375-INS-286
chr7-80623495-INS-205
chr7-80645493-INS-196
chr7-80656571-INS-261
chr7-80678571-INS-202
chr7-80681750-INS-1892
chr7-80686089-INS-119
chr7-80689375-INS-64
chr7-80705243-INS-86
chr7-80706411-INS-66
chr7-80746622-INS-125
chr7-80780043-INS-463
chr7-80783084-INS-83
chr7-80790618-INS-212
chr7-80808238-INS-190
chr7-80810004-INS-142
chr7-80815246-INS-168
chr7-80818534-INS-55
chr7-80820718-INS-59
chr7-80823096-INS-191
chr7-80828639-INS-96
chr7-80843589-INS-58
chr7-80857674-INS-140
chr7-80880413-INS-78
chr7-80884532-INS-556
chr7-80885942-INS-247
chr7-80891999-INS-913
chr7-80924785-INS-66
chr7-80973073-INS-115
chr7-80987745-INS-425
chr7-80988505-INS-121
chr7-81009337-INS-132
chr7-81012856-INS-289
chr7-81017439-INS-62

chr7-81030658-INS-251
chr7-81038484-INS-132
chr7-81056294-INS-208
chr7-81084345-INS-209
chr7-81085402-INS-6234
chr7-81111936-INS-501
chr7-81129884-INS-200
chr7-81139095-INS-85
chr7-81146308-INS-131
chr7-81157640-INS-242
chr7-81167654-INS-582
chr7-81187084-INS-150
chr7-81196296-INS-387
chr7-81205661-INS-547
chr7-81206861-INS-216
chr7-81224615-INS-199
chr7-81228694-INS-84
chr7-81242437-INS-222
chr7-81273230-INS-58
chr7-81314108-INS-466
chr7-81320405-INS-90
chr7-81352361-INS-188
chr7-81361597-INS-256
chr7-81408990-INS-199
chr7-81431499-INS-566
chr7-81440195-INS-56
chr7-81483283-INS-573
chr7-81489107-INS-9045
chr7-81491963-INS-418
chr7-81494302-INS-196
chr7-81494445-INS-4532
chr7-81503869-INS-596
chr7-81505086-INS-72
chr7-81528344-INS-128
chr7-81559016-INS-239
chr7-81578736-INS-158
chr7-81626051-INS-115
chr7-81631605-INS-2281
chr7-81653850-INS-65
chr7-81657784-INS-5548
chr7-81669648-INS-75
chr7-81705301-INS-85
chr7-81722344-INS-54
chr7-81724153-INS-7156
chr7-81745070-INS-173
chr7-81756031-INS-187
chr7-81781124-INS-188
chr7-81813529-INS-189
chr7-81832027-INS-213
chr7-81835297-INS-52
chr7-81839051-INS-147
chr7-81841844-INS-984
chr7-81855744-INS-490
chr7-81863742-INS-212
chr7-81883057-INS-51
chr7-81883550-INS-97
chr7-81890707-INS-149
chr7-81901274-INS-514
chr7-81902850-INS-68
chr7-81917444-INS-150
chr7-81922462-INS-63
chr7-81928091-INS-52
chr7-81933404-INS-183
chr7-81947727-INS-2808
chr7-81951923-INS-86
chr7-81966122-INS-144
chr7-81971433-INS-120
chr7-81980425-INS-91
chr7-81993136-INS-69
chr7-81998366-INS-156
chr7-82001625-INS-461
chr7-82047238-INS-222
chr7-82058777-INS-4330
chr7-82062060-INS-61
chr7-82076071-INS-72
chr7-82107998-INS-102
chr7-82117295-INS-212

chr7-82125278-INS-83
chr7-82143892-INS-382
chr7-82155684-INS-82
chr7-82156099-INS-205
chr7-82157775-INS-433
chr7-82159670-INS-54
chr7-82171781-INS-60
chr7-82176830-INS-349
chr7-82193793-INS-7543
chr7-82203997-INS-52
chr7-82207904-INS-610
chr7-82208089-INS-2000
chr7-82211349-INS-138
chr7-82215258-INS-659
chr7-82224632-INS-186
chr7-82229635-INS-593
chr7-82233643-INS-166
chr7-82235238-INS-169
chr7-82238597-INS-188
chr7-82242322-INS-747
chr7-82247228-INS-6466
chr7-82267637-INS-105
chr7-82272129-INS-380
chr7-82301870-INS-101
chr7-82321154-INS-847
chr7-82326239-INS-4835
chr7-82346597-INS-464
chr7-82349263-INS-209
chr7-82353383-INS-130
chr7-82358884-INS-147
chr7-82373322-INS-70
chr7-82432672-INS-5546
chr7-82459847-INS-5556
chr7-82489214-INS-74
chr7-82523305-INS-213
chr7-82525321-INS-177
chr7-82544258-INS-3169
chr7-82569083-INS-6709
chr7-82572657-INS-51
chr7-82589056-INS-6357
chr7-82590360-INS-202
chr7-82636208-INS-131
chr7-82657685-INS-603
chr7-82663701-INS-262
chr7-82679092-INS-402
chr7-82695378-INS-54
chr7-82698497-INS-53
chr7-82713881-INS-56
chr7-82715437-INS-70
chr7-82730630-INS-63
chr7-82732775-INS-153
chr7-82747803-INS-182
chr7-82792587-INS-123
chr7-82803773-INS-330
chr7-82810098-INS-2245
chr7-82814329-INS-61
chr7-82821581-INS-406
chr7-82852152-INS-5544
chr7-82856590-INS-51
chr7-82883097-INS-62
chr7-82960029-INS-441
chr7-82966970-INS-2830
chr7-82968483-INS-53
chr7-82975175-INS-58
chr7-83047225-INS-1843
chr7-83065403-INS-267
chr7-83084255-INS-129
chr7-83090642-INS-88
chr7-83104375-INS-52
chr7-83148603-INS-66
chr7-83154844-INS-51
chr7-83162174-INS-439
chr7-83167013-INS-131
chr7-83170437-INS-322
chr7-83190660-INS-53
chr7-83204328-INS-60
chr7-83215084-INS-55

chr7-83224008-INS-75
chr7-83231607-INS-469
chr7-83237772-INS-92
chr7-83291059-INS-120
chr7-83308346-INS-65
chr7-83357001-INS-84
chr7-83409292-INS-89
chr7-83416279-INS-100
chr7-83427541-INS-139
chr7-83432646-INS-214
chr7-83470946-INS-399
chr7-83475275-INS-195
chr7-83475620-INS-557
chr7-83479886-INS-122
chr7-83480876-INS-60
chr7-83487552-INS-187
chr7-83498088-INS-98
chr7-83502670-INS-56
chr7-83517932-INS-76
chr7-83520666-INS-67
chr7-83528915-INS-58
chr7-83538910-INS-188
chr7-83545661-INS-215
chr7-83565696-INS-412
chr7-83565918-INS-6189
chr7-83577874-INS-204
chr7-83600050-INS-195
chr7-83620959-INS-54
chr7-83636018-INS-222
chr7-83639787-INS-55
chr7-83654177-INS-60
chr7-83655623-INS-70
chr7-83666794-INS-307
chr7-83673464-INS-82
chr7-83679161-INS-82
chr7-83680523-INS-445
chr7-83684278-INS-297
chr7-83691142-INS-3193
chr7-83720670-INS-56
chr7-83722885-INS-65
chr7-83732537-INS-146
chr7-83742343-INS-54
chr7-83751310-INS-54
chr7-83751843-INS-94
chr7-83755164-INS-765
chr7-83767935-INS-68
chr7-83772720-INS-707
chr7-83780424-INS-194
chr7-83790495-INS-222
chr7-83823443-INS-399
chr7-83833443-INS-203
chr7-83852779-INS-825
chr7-83865407-INS-285
chr7-83878477-INS-60
chr7-83903219-INS-107
chr7-83905179-INS-183
chr7-83915499-INS-207
chr7-83917083-INS-176
chr7-83925886-INS-60
chr7-83938707-INS-424
chr7-83951942-INS-52
chr7-83955699-INS-78
chr7-83987495-INS-215
chr7-84004160-INS-223
chr7-84006714-INS-78
chr7-84062485-INS-66
chr7-84106208-INS-6093
chr7-84109676-INS-6320
chr7-84114059-INS-6293
chr7-84123147-INS-416
chr7-84138089-INS-337
chr7-84141852-INS-175
chr7-84154319-INS-470
chr7-84161896-INS-787
chr7-84169809-INS-56
chr7-84229229-INS-128
chr7-84234036-INS-86

chr7-84307263-INS-471
chr7-84323702-INS-100
chr7-84335190-INS-79
chr7-84399861-INS-217
chr7-84436942-INS-6460
chr7-84451631-INS-54
chr7-84482340-INS-358
chr7-84491420-INS-5538
chr7-84495416-INS-812
chr7-84546943-INS-118
chr7-84556756-INS-75
chr7-84558325-INS-919
chr7-84587416-INS-995
chr7-84595263-INS-399
chr7-84612346-INS-66
chr7-84682860-INS-332
chr7-84753029-INS-831
chr7-84896298-INS-53
chr7-84928028-INS-549
chr7-85099682-INS-4866
chr7-85227907-INS-285
chr7-85228675-INS-5355
chr7-85266817-INS-921
chr7-85271673-INS-126
chr7-85332002-INS-58
chr7-85448247-INS-1957
chr7-85554947-INS-52
chr7-85561674-INS-1892
chr7-85574574-INS-6780
chr7-85607950-INS-6235
chr7-85615074-INS-6368
chr7-85618983-INS-202
chr7-85647066-INS-58
chr7-85668750-INS-347
chr7-85671733-INS-6465
chr7-85675722-INS-4844
chr7-85692784-INS-58
chr7-85694272-INS-6270
chr7-85796144-INS-888
chr7-85824489-INS-56
chr7-85874046-INS-60
chr7-85877685-INS-99
chr7-85892121-INS-625
chr7-85936500-INS-331
chr7-85947722-INS-123
chr7-85964592-INS-956
chr7-85972304-INS-232
chr7-85998881-INS-95
chr7-86003122-INS-8226
chr7-86010456-INS-629
chr7-86016832-INS-6222
chr7-86031923-INS-66
chr7-86077798-INS-130
chr7-86094200-INS-199
chr7-86096538-INS-530
chr7-86108182-INS-860
chr7-86119265-INS-116
chr7-86131079-INS-217
chr7-86165461-INS-983
chr7-86172104-INS-54
chr7-86173079-INS-116
chr7-86180469-INS-138
chr7-86228757-INS-6394
chr7-86244892-INS-55
chr7-86333904-INS-544
chr7-86365458-INS-58
chr7-86367480-INS-513
chr7-86367731-INS-6342
chr7-86387493-INS-86
chr7-86390250-INS-79
chr7-86401372-INS-97
chr7-86443094-INS-304
chr7-86486480-INS-5549
chr7-86566330-INS-6270
chr7-86623229-INS-54
chr7-86655159-INS-52
chr7-86712275-INS-580

chr7-86719144-INS-329
chr7-86786942-INS-13077
chr7-86843749-INS-5579
chr7-86843886-INS-215
chr7-86849124-INS-284
chr7-86862207-INS-202
chr7-86872933-INS-57
chr7-86894343-INS-185
chr7-86897148-INS-64
chr7-86903291-INS-116
chr7-86950523-INS-503
chr7-86962620-INS-70
chr7-86965359-INS-421
chr7-87004142-INS-689
chr7-87012313-INS-1891
chr7-87043620-INS-70
chr7-87052977-INS-6487
chr7-87088882-INS-85
chr7-87093379-INS-64
chr7-87098452-INS-6483
chr7-87099250-INS-7483
chr7-87112336-INS-104
chr7-87122625-INS-155
chr7-87130133-INS-155
chr7-87132798-INS-65
chr7-87160369-INS-6871
chr7-87189703-INS-6416
chr7-87196566-INS-630
chr7-87201825-INS-1971
chr7-87208063-INS-142
chr7-87208552-INS-856
chr7-87220386-INS-78
chr7-87240453-INS-54
chr7-87302641-INS-318
chr7-87329399-INS-177
chr7-87338750-INS-69
chr7-87373925-INS-6781
chr7-87406028-INS-401
chr7-87428943-INS-6285
chr7-87433683-INS-54
chr7-87475766-INS-64
chr7-87496778-INS-317
chr7-87501002-INS-387
chr7-87513195-INS-64
chr7-87515263-INS-518
chr7-87522099-INS-52
chr7-87523786-INS-92
chr7-87531786-INS-289
chr7-87577948-INS-178
chr7-87582787-INS-108
chr7-87585227-INS-66
chr7-87594924-INS-211
chr7-87656660-INS-451
chr7-87668590-INS-56
chr7-87690033-INS-77
chr7-87708255-INS-94
chr7-87720349-INS-372
chr7-87730599-INS-64
chr7-87837479-INS-8732
chr7-87838092-INS-131
chr7-87840908-INS-1607
chr7-87844697-INS-637
chr7-87870140-INS-7322
chr7-87883748-INS-121
chr7-87887537-INS-110
chr7-87916123-INS-475
chr7-87917292-INS-72
chr7-87918426-INS-1531
chr7-87919465-INS-172
chr7-87947924-INS-130
chr7-87965342-INS-52
chr7-87972414-INS-3844
chr7-87973528-INS-118
chr7-87974361-INS-338
chr7-87979060-INS-327
chr7-87990766-INS-209
chr7-88020274-INS-199

chr7-88040277-INS-60
chr7-88072765-INS-182
chr7-88106273-INS-359
chr7-88118304-INS-1814
chr7-88127881-INS-6275
chr7-88139300-INS-55
chr7-88153904-INS-92
chr7-88160293-INS-62
chr7-88166829-INS-65
chr7-88167916-INS-190
chr7-88170444-INS-183
chr7-88204962-INS-56
chr7-88211875-INS-60
chr7-88214628-INS-54
chr7-88240614-INS-148
chr7-88247342-INS-364
chr7-88266487-INS-247
chr7-88274167-INS-78
chr7-88288251-INS-3318
chr7-88302456-INS-59
chr7-88305819-INS-220
chr7-88309245-INS-6482
chr7-88339795-INS-240
chr7-88341556-INS-599
chr7-88375363-INS-551
chr7-88405109-INS-307
chr7-88407304-INS-58
chr7-88410112-INS-130
chr7-88411979-INS-83
chr7-88413401-INS-6226
chr7-88477948-INS-192
chr7-88477955-INS-2846
chr7-88494215-INS-7480
chr7-88503017-INS-437
chr7-88522261-INS-400
chr7-88557079-INS-476
chr7-88561060-INS-510
chr7-88580893-INS-163
chr7-88583336-INS-64
chr7-88590035-INS-506
chr7-88656432-INS-2151
chr7-88662031-INS-98
chr7-88673008-INS-286
chr7-88679568-INS-83
chr7-88688299-INS-202
chr7-88706537-INS-343
chr7-88764173-INS-586
chr7-88778669-INS-77
chr7-88886824-INS-232
chr7-88899935-INS-5152
chr7-88912364-INS-225
chr7-88913460-INS-482
chr7-88917105-INS-500
chr7-88919673-INS-72
chr7-88927542-INS-55
chr7-88929053-INS-237
chr7-88934271-INS-113
chr7-88967115-INS-68
chr7-88969115-INS-82
chr7-88992882-INS-211
chr7-89004140-INS-110
chr7-89044405-INS-56
chr7-89049134-INS-306
chr7-89109902-INS-424
chr7-89121751-INS-158
chr7-89124204-INS-54
chr7-89176234-INS-82
chr7-89222739-INS-61
chr7-89224015-INS-1725
chr7-89224423-INS-287
chr7-89239350-INS-333
chr7-89252676-INS-823
chr7-89267342-INS-60
chr7-89293936-INS-420
chr7-89317184-INS-803
chr7-89363237-INS-687
chr7-89364961-INS-300

chr7-89421611-INS-302
chr7-89462683-INS-249
chr7-89466863-INS-223
chr7-89482415-INS-330
chr7-89492224-INS-345
chr7-89495739-INS-65
chr7-89502170-INS-241
chr7-89503727-INS-324
chr7-89505357-INS-416
chr7-89527578-INS-152
chr7-89574403-INS-196
chr7-89582649-INS-178
chr7-89590682-INS-292
chr7-89592160-INS-154
chr7-89593152-INS-53
chr7-89595627-INS-51
chr7-89605159-INS-205
chr7-89615156-INS-691
chr7-89637711-INS-122
chr7-89648492-INS-206
chr7-89653021-INS-166
chr7-89654175-INS-402
chr7-89662197-INS-272
chr7-89664700-INS-87
chr7-89691363-INS-104
chr7-89698841-INS-4767
chr7-89729059-INS-219
chr7-89729463-INS-605
chr7-89732642-INS-164
chr7-89744187-INS-55
chr7-89746211-INS-176
chr7-89760134-INS-122
chr7-89775904-INS-307
chr7-89780376-INS-64
chr7-89812385-INS-6614
chr7-89822179-INS-4409
chr7-89833716-INS-52
chr7-89834857-INS-52
chr7-89846765-INS-126
chr7-89901518-INS-546
chr7-89907655-INS-444
chr7-89955019-INS-250
chr7-89970354-INS-244
chr7-89974284-INS-350
chr7-90051244-INS-200
chr7-90119587-INS-117
chr7-90130659-INS-272
chr7-90150305-INS-1267
chr7-90151141-INS-1910
chr7-90167271-INS-344
chr7-90190400-INS-549
chr7-90237403-INS-70
chr7-90245199-INS-64
chr7-90258260-INS-55
chr7-90286043-INS-208
chr7-90299083-INS-1461
chr7-90318969-INS-77
chr7-90322890-INS-204
chr7-90336939-INS-86
chr7-90353726-INS-132
chr7-90374060-INS-386
chr7-90384347-INS-72
chr7-90388632-INS-76
chr7-90394159-INS-640
chr7-90397534-INS-55
chr7-90414843-INS-162
chr7-90441520-INS-5678
chr7-90451774-INS-6351
chr7-90453906-INS-68
chr7-90457882-INS-6450
chr7-90476139-INS-166
chr7-90500516-INS-131
chr7-90526935-INS-105
chr7-90561446-INS-58
chr7-90564062-INS-107
chr7-90614754-INS-51
chr7-90615260-INS-6768

chr7-90638271-INS-81
chr7-90644521-INS-511
chr7-90646819-INS-933
chr7-90680403-INS-659
chr7-90690741-INS-554
chr7-90695514-INS-311
chr7-90697961-INS-114
chr7-90732800-INS-6334
chr7-90737073-INS-5988
chr7-90756776-INS-7486
chr7-90777885-INS-58
chr7-90778998-INS-131
chr7-90796580-INS-578
chr7-90810193-INS-391
chr7-90812479-INS-248
chr7-90819834-INS-207
chr7-90820869-INS-930
chr7-90827578-INS-735
chr7-90842978-INS-166
chr7-90853397-INS-90
chr7-90874844-INS-69
chr7-90943011-INS-96
chr7-90979103-INS-284
chr7-90981920-INS-5181
chr7-90982394-INS-364
chr7-91002313-INS-1064
chr7-91005662-INS-216
chr7-91015257-INS-279
chr7-91025270-INS-627
chr7-91042123-INS-585
chr7-91054821-INS-298
chr7-91064933-INS-114
chr7-91067393-INS-160
chr7-91111734-INS-336
chr7-91138694-INS-114
chr7-91167394-INS-7129
chr7-91197629-INS-107
chr7-91209879-INS-58
chr7-91213538-INS-179
chr7-91219924-INS-388
chr7-91229231-INS-173
chr7-91244255-INS-52
chr7-91257891-INS-346
chr7-91259947-INS-830
chr7-91284401-INS-437
chr7-91290555-INS-57
chr7-91292320-INS-54
chr7-91296118-INS-5249
chr7-91301201-INS-530
chr7-91305481-INS-58
chr7-91310857-INS-659
chr7-91327801-INS-414
chr7-91336262-INS-614
chr7-91338755-INS-576
chr7-91354305-INS-68
chr7-91365834-INS-54
chr7-91368303-INS-5289
chr7-91406291-INS-101
chr7-91418494-INS-494
chr7-91438634-INS-207
chr7-91450070-INS-58
chr7-91485219-INS-299
chr7-91530241-INS-139
chr7-91533504-INS-3098
chr7-91556894-INS-460
chr7-91589849-INS-218
chr7-91590980-INS-67
chr7-91603882-INS-139
chr7-91610775-INS-229
chr7-91618873-INS-89
chr7-91652387-INS-134
chr7-91663387-INS-364
chr7-91666084-INS-108
chr7-91693790-INS-54
chr7-91697924-INS-53
chr7-91707626-INS-201
chr7-91710576-INS-6455

chr7-91714292-INS-5332
chr7-91728311-INS-217
chr7-91732419-INS-127
chr7-91736721-INS-57
chr7-91783361-INS-54
chr7-91790673-INS-6390
chr7-91792622-INS-848
chr7-91816546-INS-94
chr7-91843917-INS-56
chr7-91863739-INS-59
chr7-91916282-INS-54
chr7-91967740-INS-52
chr7-91984500-INS-4032
chr7-92019060-INS-7030
chr7-92042292-INS-406
chr7-92047738-INS-227
chr7-92083440-INS-167
chr7-92098631-INS-67
chr7-92099928-INS-110
chr7-92113396-INS-99
chr7-92127162-INS-4308
chr7-92136403-INS-187
chr7-92139566-INS-386
chr7-92153877-INS-1370
chr7-92194044-INS-361
chr7-92217343-INS-199
chr7-92236618-INS-53
chr7-92281822-INS-203
chr7-92283436-INS-96
chr7-92297724-INS-290
chr7-92330460-INS-82
chr7-92337804-INS-62
chr7-92352080-INS-212
chr7-92358175-INS-122
chr7-92364389-INS-63
chr7-92366947-INS-56
chr7-92368845-INS-80
chr7-92376941-INS-223
chr7-92398626-INS-196
chr7-92423376-INS-188
chr7-92426868-INS-54
chr7-92431226-INS-58
chr7-92433800-INS-446
chr7-92445910-INS-297
chr7-92446919-INS-296
chr7-92451234-INS-6589
chr7-92495304-INS-85
chr7-92498208-INS-75
chr7-92538539-INS-402
chr7-92549987-INS-220
chr7-92554630-INS-76
chr7-92566202-INS-214
chr7-92566766-INS-117
chr7-92567963-INS-212
chr7-92593904-INS-61
chr7-92615247-INS-147
chr7-92641268-INS-531
chr7-92685985-INS-1606
chr7-92686720-INS-103
chr7-92688026-INS-121
chr7-92697732-INS-53
chr7-92707758-INS-691
chr7-92707810-INS-1079
chr7-92717885-INS-162
chr7-92744637-INS-486
chr7-92747514-INS-80
chr7-92773675-INS-6093
chr7-92781312-INS-6741
chr7-92784212-INS-60
chr7-92796379-INS-507
chr7-92800428-INS-203
chr7-92816844-INS-629
chr7-92830165-INS-54
chr7-92843877-INS-208
chr7-92901656-INS-88
chr7-92908833-INS-4600
chr7-92909982-INS-1867

chr7-92915294-INS-1864
chr7-92927200-INS-398
chr7-92929876-INS-80
chr7-92958689-INS-6367
chr7-92965894-INS-320
chr7-92969323-INS-175
chr7-92978719-INS-129
chr7-92980196-INS-65
chr7-92989553-INS-708
chr7-92995490-INS-4645
chr7-93014402-INS-148
chr7-93048329-INS-63
chr7-93058827-INS-6795
chr7-93071657-INS-107
chr7-93072517-INS-709
chr7-93074913-INS-244
chr7-93079048-INS-650
chr7-93109631-INS-54
chr7-93112426-INS-267
chr7-93126425-INS-374
chr7-93136573-INS-57
chr7-93143754-INS-284
chr7-93157829-INS-52
chr7-93167461-INS-509
chr7-93168902-INS-193
chr7-93193472-INS-52
chr7-93196591-INS-78
chr7-93205866-INS-627
chr7-93252457-INS-68
chr7-93286014-INS-208
chr7-93299683-INS-374
chr7-93312544-INS-386
chr7-93317648-INS-92
chr7-93344546-INS-116
chr7-93372559-INS-200
chr7-93373499-INS-51
chr7-93401585-INS-164
chr7-93417390-INS-704
chr7-93423614-INS-71
chr7-93432938-INS-340
chr7-93485980-INS-70
chr7-93490074-INS-4791
chr7-93494652-INS-668
chr7-93500303-INS-95
chr7-93510472-INS-184
chr7-93513546-INS-97
chr7-93516212-INS-277
chr7-93522094-INS-434
chr7-93601425-INS-52
chr7-93605684-INS-178
chr7-93612825-INS-6883
chr7-93634866-INS-430
chr7-93657266-INS-86
chr7-93667279-INS-69
chr7-93674058-INS-84
chr7-93677744-INS-296
chr7-93682275-INS-399
chr7-93696120-INS-243
chr7-93705701-INS-203
chr7-93711990-INS-57
chr7-93714505-INS-144
chr7-93734509-INS-102
chr7-93761538-INS-64
chr7-93798041-INS-118
chr7-93857134-INS-127
chr7-93863704-INS-57
chr7-93886211-INS-114
chr7-93899888-INS-2309
chr7-93959501-INS-738
chr7-94051084-INS-726
chr7-94052472-INS-219
chr7-94069022-INS-6749
chr7-94075732-INS-524
chr7-94085238-INS-54
chr7-94094882-INS-61
chr7-94106558-INS-172
chr7-94135258-INS-4207

chr7-94135907-INS-676
chr7-94167731-INS-53
chr7-94201466-INS-192
chr7-94229048-INS-6896
chr7-94254928-INS-6955
chr7-94274401-INS-121
chr7-94286258-INS-714
chr7-94328021-INS-6799
chr7-94377243-INS-6289
chr7-94412837-INS-6596
chr7-94429633-INS-60
chr7-94452205-INS-2704
chr7-94479376-INS-421
chr7-94484008-INS-90
chr7-94526824-INS-401
chr7-94533821-INS-89
chr7-94549279-INS-314
chr7-94556414-INS-649
chr7-94557889-INS-51
chr7-94570441-INS-2285
chr7-94574892-INS-507
chr7-94575306-INS-1701
chr7-94580879-INS-145
chr7-94588076-INS-6217
chr7-94598715-INS-356
chr7-94606392-INS-7622
chr7-94614584-INS-417
chr7-94617570-INS-80
chr7-94620012-INS-6900
chr7-94621436-INS-443
chr7-94690317-INS-52
chr7-94699574-INS-186
chr7-94727636-INS-208
chr7-94755621-INS-866
chr7-94794318-INS-59
chr7-94797491-INS-160
chr7-94799630-INS-115
chr7-94845516-INS-195
chr7-94850084-INS-67
chr7-94879956-INS-666
chr7-94900563-INS-63
chr7-94902684-INS-198
chr7-94949912-INS-149
chr7-94969644-INS-58
chr7-94986155-INS-6897
chr7-94997575-INS-76
chr7-95004423-INS-274
chr7-95011160-INS-1210
chr7-95016122-INS-269
chr7-95020360-INS-1376
chr7-95048474-INS-54
chr7-95070417-INS-72
chr7-95085948-INS-908
chr7-95101912-INS-76
chr7-95138577-INS-155
chr7-95150447-INS-376
chr7-95164972-INS-61
chr7-95189622-INS-67
chr7-95191295-INS-69
chr7-95211861-INS-122
chr7-95252884-INS-923
chr7-95266459-INS-54
chr7-95281177-INS-60
chr7-95284102-INS-61
chr7-95284718-INS-301
chr7-95300477-INS-60
chr7-95304698-INS-604
chr7-95309508-INS-102
chr7-95336970-INS-105
chr7-95373056-INS-191
chr7-95376917-INS-1107
chr7-95383093-INS-60
chr7-95392696-INS-195
chr7-95430863-INS-237
chr7-95442873-INS-55
chr7-95450371-INS-434
chr7-95475139-INS-60

chr7-95481893-INS-156
chr7-95494067-INS-61
chr7-95495906-INS-55
chr7-95496329-INS-236
chr7-95526600-INS-70
chr7-95537498-INS-743
chr7-95560135-INS-187
chr7-95581104-INS-201
chr7-95582711-INS-62
chr7-95596413-INS-215
chr7-95606382-INS-563
chr7-95641645-INS-56
chr7-95649176-INS-57
chr7-95669450-INS-357
chr7-95693300-INS-116
chr7-95724402-INS-197
chr7-95742647-INS-66
chr7-95746931-INS-922
chr7-95781409-INS-115
chr7-95807030-INS-204
chr7-95855592-INS-56
chr7-95904378-INS-662
chr7-95913882-INS-51
chr7-95928652-INS-54
chr7-95957827-INS-268
chr7-96022423-INS-7004
chr7-96058733-INS-509
chr7-96094166-INS-196
chr7-96096099-INS-68
chr7-96110895-INS-1504
chr7-96129171-INS-69
chr7-96146706-INS-993
chr7-96189507-INS-187
chr7-96228314-INS-112
chr7-96232400-INS-56
chr7-96237673-INS-66
chr7-96258651-INS-125
chr7-96286903-INS-112
chr7-96316378-INS-102
chr7-96341422-INS-598
chr7-96348384-INS-122
chr7-96393785-INS-127
chr7-96469127-INS-287
chr7-96475255-INS-52
chr7-96524818-INS-166
chr7-96538579-INS-283
chr7-96540892-INS-931
chr7-96594312-INS-65
chr7-96613494-INS-377
chr7-96640964-INS-508
chr7-96690022-INS-53
chr7-96719327-INS-196
chr7-96773066-INS-74
chr7-96778571-INS-99
chr7-96778648-INS-682
chr7-96843579-INS-752
chr7-96844107-INS-52
chr7-96917939-INS-397
chr7-96941416-INS-6440
chr7-97000777-INS-90
chr7-97008078-INS-201
chr7-97018223-INS-204
chr7-97036922-INS-66
chr7-97041963-INS-111
chr7-97082380-INS-52
chr7-97103587-INS-53
chr7-97156610-INS-186
chr7-97169055-INS-392
chr7-97171174-INS-408
chr7-97256652-INS-60
chr7-97271940-INS-64
chr7-97299104-INS-93
chr7-97301562-INS-59
chr7-97330797-INS-193
chr7-97344731-INS-156
chr7-97363460-INS-203
chr7-97369282-INS-138

chr7-97381322-INS-60
chr7-97385279-INS-214
chr7-97397720-INS-52
chr7-97408284-INS-203
chr7-97424375-INS-271
chr7-97445781-INS-526
chr7-97477183-INS-174
chr7-97482391-INS-88
chr7-97501312-INS-5543
chr7-97506480-INS-69
chr7-97519892-INS-57
chr7-97524151-INS-161
chr7-97536642-INS-58
chr7-97552035-INS-192
chr7-97562019-INS-115
chr7-97574413-INS-52
chr7-97591168-INS-182
chr7-97591923-INS-5550
chr7-97625536-INS-200
chr7-97650966-INS-220
chr7-97664849-INS-95
chr7-97685191-INS-62
chr7-97693003-INS-340
chr7-97749050-INS-214
chr7-97798260-INS-82
chr7-97804135-INS-300
chr7-97817569-INS-265
chr7-97845254-INS-202
chr7-97846656-INS-90
chr7-97856120-INS-527
chr7-97930840-INS-612
chr7-97989078-INS-162
chr7-98006230-INS-732
chr7-98010921-INS-82
chr7-98059887-INS-54
chr7-98107196-INS-178
chr7-98109051-INS-194
chr7-98131530-INS-81
chr7-98155220-INS-59
chr7-98163130-INS-66
chr7-98169669-INS-111
chr7-98177011-INS-224
chr7-98210011-INS-202
chr7-98230598-INS-2761
chr7-98262035-INS-85
chr7-98263249-INS-160
chr7-98274675-INS-178
chr7-98278212-INS-198
chr7-98315500-INS-938
chr7-98381817-INS-203
chr7-98401195-INS-109
chr7-98412758-INS-84
chr7-98444532-INS-122
chr7-98448792-INS-134
chr7-98480802-INS-210
chr7-98503380-INS-5921
chr7-98525853-INS-89
chr7-98534443-INS-152
chr7-98539012-INS-52
chr7-98564150-INS-100
chr7-98595862-INS-208
chr7-98601679-INS-74
chr7-98608088-INS-432
chr7-98617443-INS-61
chr7-98624628-INS-56
chr7-98646948-INS-390
chr7-98667165-INS-274
chr7-98680078-INS-115
chr7-98718122-INS-215
chr7-98722114-INS-66
chr7-98736484-INS-5259
chr7-98738390-INS-76
chr7-98741724-INS-216
chr7-98743680-INS-204
chr7-98759248-INS-112
chr7-98768831-INS-358
chr7-98787843-INS-64

chr7-98800483-INS-118
chr7-98858598-INS-960
chr7-98858671-INS-150
chr7-98858838-INS-234
chr7-98885486-INS-143
chr7-98905997-INS-246
chr7-98907324-INS-203
chr7-98907557-INS-7088
chr7-98918049-INS-138
chr7-98929861-INS-51
chr7-98934194-INS-60
chr7-98943134-INS-622
chr7-98958316-INS-70
chr7-98959075-INS-501
chr7-98961651-INS-68
chr7-98989056-INS-104
chr7-98992497-INS-67
chr7-99019327-INS-60
chr7-99023624-INS-66
chr7-99045969-INS-220
chr7-99054948-INS-116
chr7-99058370-INS-57
chr7-99061149-INS-2094
chr7-99064720-INS-62
chr7-99081778-INS-424
chr7-99133440-INS-314
chr7-99137895-INS-153
chr7-99140756-INS-339
chr7-99142629-INS-56
chr7-99154999-INS-164
chr7-99172105-INS-56
chr7-99200048-INS-416
chr7-99204259-INS-98
chr7-99214368-INS-400
chr7-99225562-INS-419
chr7-99244040-INS-129
chr7-99257308-INS-280
chr7-99291753-INS-151
chr7-99295631-INS-56
chr7-99363580-INS-50
chr7-99371580-INS-93
chr7-99378197-INS-251
chr7-99385967-INS-225
chr7-99394270-INS-55
chr7-99405830-INS-223
chr7-99413655-INS-141
chr7-99447091-INS-63
chr7-99450085-INS-6442
chr7-99491447-INS-83
chr7-99499066-INS-63
chr7-99526013-INS-375
chr7-99528547-INS-89
chr7-99535795-INS-208
chr7-99591773-INS-1264
chr7-99591924-INS-284
chr7-99609657-INS-59
chr7-99612476-INS-107
chr7-99624561-INS-6415
chr7-99637199-INS-96
chr7-99670746-INS-76
chr7-99687666-INS-421
chr7-99707776-INS-192
chr7-99718865-INS-143
chr7-99719422-INS-70
chr7-99748181-INS-203
chr7-99806665-INS-136
chr7-99812884-INS-165
chr7-99819949-INS-374
chr7-99826024-INS-88
chr7-99826704-INS-7102
chr7-99831135-INS-193
chr7-99835458-INS-139
chr7-99851179-INS-114
chr7-99869462-INS-1428
chr7-99890755-INS-97
chr7-99901156-INS-1892
chr7-99916625-INS-322

chr7-99944453-INS-226
chr7-99962856-INS-201
chr7-99965383-INS-176
chr7-99980402-INS-52
chr7-99990633-INS-105
chr7-100028035-INS-247
chr7-100064324-INS-167
chr7-100080869-INS-747
chr7-100085615-INS-51
chr7-100088593-INS-119
chr7-100090828-INS-950
chr7-100103431-INS-100
chr7-100133205-INS-389
chr7-100139260-INS-213
chr7-100159494-INS-311
chr7-100166608-INS-72
chr7-100204032-INS-382
chr7-100220883-INS-187
chr7-100230113-INS-95
chr7-100239593-INS-229
chr7-100271903-INS-162
chr7-100281751-INS-203
chr7-100301998-INS-187
chr7-100319447-INS-599
chr7-100320424-INS-56
chr7-100334711-INS-53
chr7-100338501-INS-62
chr7-100342184-INS-71
chr7-100345136-INS-142
chr7-100348642-INS-53
chr7-100351571-INS-69
chr7-100351836-INS-909
chr7-100364920-INS-268
chr7-100397127-INS-238
chr7-100413192-INS-200
chr7-100413864-INS-220
chr7-100421134-INS-88
chr7-100423935-INS-178
chr7-100428191-INS-163
chr7-100454960-INS-322
chr7-100478777-INS-62
chr7-100540596-INS-192
chr7-100541690-INS-83
chr7-100546912-INS-196
chr7-100551493-INS-247
chr7-100554692-INS-57
chr7-100600338-INS-81
chr7-100600536-INS-836
chr7-100605539-INS-275
chr7-100620264-INS-141
chr7-100624195-INS-96
chr7-100631650-INS-604
chr7-100668405-INS-51
chr7-100680377-INS-108
chr7-100685158-INS-599
chr7-100702978-INS-187
chr7-100708228-INS-176
chr7-100710008-INS-163
chr7-100729945-INS-70
chr7-100731165-INS-159
chr7-100732680-INS-233
chr7-100777532-INS-53
chr7-100787842-INS-60
chr7-100792554-INS-65
chr7-100858297-INS-153
chr7-100883946-INS-215
chr7-100885340-INS-537
chr7-100887684-INS-139
chr7-100911914-INS-162
chr7-100941952-INS-92
chr7-100973872-INS-72
chr7-101000356-INS-177
chr7-101008546-INS-195
chr7-101020760-INS-132
chr7-101036773-INS-84
chr7-101041149-INS-66
chr7-101059053-INS-185

chr7-101059053-INS-280
chr7-101069806-INS-168
chr7-101088906-INS-60
chr7-101097357-INS-107
chr7-101098479-INS-133
chr7-101169085-INS-52
chr7-101232111-INS-393
chr7-101248302-INS-56
chr7-101287201-INS-59
chr7-101299575-INS-5444
chr7-101302707-INS-52
chr7-101340748-INS-200
chr7-101381730-INS-94
chr7-101383629-INS-300
chr7-101430450-INS-176
chr7-101451605-INS-304
chr7-101479034-INS-252
chr7-101488994-INS-188
chr7-101505965-INS-339
chr7-101524098-INS-78
chr7-101537962-INS-199
chr7-101563474-INS-74
chr7-101579486-INS-247
chr7-101587393-INS-367
chr7-101625672-INS-202
chr7-101627453-INS-108
chr7-101627844-INS-147
chr7-101678081-INS-60
chr7-101683966-INS-74
chr7-101707010-INS-127
chr7-101722732-INS-59
chr7-101727401-INS-306
chr7-101729334-INS-203
chr7-101758832-INS-200
chr7-101774037-INS-216
chr7-101791851-INS-686
chr7-101805474-INS-204
chr7-101810975-INS-161
chr7-101827548-INS-221
chr7-101842644-INS-301
chr7-101844851-INS-893
chr7-101848781-INS-306
chr7-101850364-INS-209
chr7-101852432-INS-182
chr7-101878245-INS-233
chr7-101896055-INS-127
chr7-101911063-INS-126
chr7-101960406-INS-84
chr7-102018801-INS-64
chr7-102034390-INS-99
chr7-102037734-INS-121
chr7-102047254-INS-125
chr7-102053965-INS-186
chr7-102067346-INS-56
chr7-102072689-INS-225
chr7-102140325-INS-226
chr7-102169627-INS-106
chr7-102231365-INS-11663
chr7-102253067-INS-128
chr7-102269876-INS-117
chr7-102315137-INS-76
chr7-102320663-INS-53
chr7-102381686-INS-768
chr7-102384169-INS-5474
chr7-102390313-INS-517
chr7-102431926-INS-875
chr7-102439587-INS-7241
chr7-102453941-INS-160
chr7-102461988-INS-187
chr7-102473519-INS-93
chr7-102504826-INS-59
chr7-102531309-INS-5298
chr7-102531651-INS-3682
chr7-102557424-INS-803
chr7-102587725-INS-6716
chr7-102610352-INS-120
chr7-102610755-INS-181

chr7-102641627-INS-56
chr7-102659896-INS-378
chr7-102688227-INS-92
chr7-102719351-INS-110
chr7-102725451-INS-400
chr7-102750452-INS-421
chr7-102788679-INS-212
chr7-102796672-INS-405
chr7-102817115-INS-6363
chr7-102878926-INS-61
chr7-102908183-INS-518
chr7-103005753-INS-193
chr7-103022757-INS-52
chr7-103031205-INS-263
chr7-103065134-INS-12521
chr7-103123280-INS-5399
chr7-103188125-INS-6463
chr7-103193437-INS-4986
chr7-103238820-INS-51
chr7-103255273-INS-60
chr7-103353050-INS-1606
chr7-103367005-INS-221
chr7-103388440-INS-58
chr7-103464638-INS-88
chr7-103509440-INS-145
chr7-103535760-INS-72
chr7-103545833-INS-581
chr7-103570600-INS-206
chr7-103598816-INS-126
chr7-103622163-INS-1063
chr7-103644113-INS-368
chr7-103647397-INS-619
chr7-103648362-INS-85
chr7-103712757-INS-981
chr7-103737651-INS-6656
chr7-103764590-INS-349
chr7-103775943-INS-84
chr7-103781221-INS-210
chr7-103851365-INS-781
chr7-103869251-INS-200
chr7-103887465-INS-170
chr7-103911127-INS-204
chr7-103926985-INS-52
chr7-103935865-INS-112
chr7-103945808-INS-324
chr7-103950073-INS-91
chr7-103967776-INS-497
chr7-103983491-INS-178
chr7-103992715-INS-1346
chr7-104001012-INS-88
chr7-104022531-INS-64
chr7-104033332-INS-84
chr7-104038148-INS-500
chr7-104044714-INS-126
chr7-104054579-INS-548
chr7-104060223-INS-104
chr7-104086345-INS-500
chr7-104087534-INS-357
chr7-104114403-INS-664
chr7-104119027-INS-8209
chr7-104125933-INS-1285
chr7-104146080-INS-247
chr7-104228368-INS-108
chr7-104231490-INS-70
chr7-104248410-INS-164
chr7-104257405-INS-7785
chr7-104260412-INS-6439
chr7-104271323-INS-6630
chr7-104300411-INS-146
chr7-104339766-INS-6676
chr7-104343820-INS-7238
chr7-104372025-INS-72
chr7-104393493-INS-399
chr7-104404918-INS-141
chr7-104419897-INS-4591
chr7-104419912-INS-2321
chr7-104457426-INS-4337

chr7-104491124-INS-470
chr7-104505726-INS-56
chr7-104516283-INS-574
chr7-104519972-INS-372
chr7-104558645-INS-896
chr7-104582437-INS-5951
chr7-104589732-INS-87
chr7-104591695-INS-130
chr7-104593383-INS-2768
chr7-104654873-INS-6631
chr7-104671777-INS-56
chr7-104707263-INS-6756
chr7-104720805-INS-52
chr7-104725769-INS-62
chr7-104730606-INS-135
chr7-104757219-INS-114
chr7-104766575-INS-63
chr7-104768215-INS-52
chr7-104777198-INS-134
chr7-104828283-INS-8126
chr7-104835727-INS-220
chr7-104860951-INS-107
chr7-104867509-INS-5362
chr7-104873130-INS-57
chr7-104882779-INS-108
chr7-104888798-INS-204
chr7-104894781-INS-354
chr7-104906196-INS-296
chr7-104910880-INS-437
chr7-104912727-INS-201
chr7-104919020-INS-71
chr7-104924346-INS-607
chr7-104932477-INS-83
chr7-104934246-INS-228
chr7-104942844-INS-116
chr7-104951408-INS-341
chr7-104959809-INS-1827
chr7-104967496-INS-98
chr7-104971711-INS-393
chr7-104977950-INS-7337
chr7-104993325-INS-198
chr7-105011230-INS-449
chr7-105042707-INS-366
chr7-105050818-INS-56
chr7-105052434-INS-144
chr7-105082622-INS-107
chr7-105122117-INS-843
chr7-105127847-INS-158
chr7-105140881-INS-605
chr7-105152003-INS-199
chr7-105161855-INS-358
chr7-105179296-INS-150
chr7-105180438-INS-366
chr7-105182325-INS-142
chr7-105192401-INS-72
chr7-105201618-INS-485
chr7-105247114-INS-402
chr7-105279536-INS-147
chr7-105311227-INS-307
chr7-105320187-INS-582
chr7-105338010-INS-201
chr7-105349529-INS-57
chr7-105377301-INS-271
chr7-105431849-INS-63
chr7-105466948-INS-8734
chr7-105468780-INS-528
chr7-105470928-INS-265
chr7-105799660-INS-6402
chr7-105810709-INS-6267
chr7-105813475-INS-7991
chr7-105821437-INS-5554
chr7-105855527-INS-5022
chr7-105886747-INS-60
chr7-105920208-INS-171
chr7-105977095-INS-7130
chr7-105987189-INS-1335
chr7-105994491-INS-411

chr7-106005341-INS-97
chr7-106052953-INS-2155
chr7-106110254-INS-5384
chr7-106112760-INS-6644
chr7-106130938-INS-493
chr7-106206758-INS-568
chr7-106211826-INS-252
chr7-106225050-INS-2768
chr7-106227540-INS-66
chr7-106241056-INS-9136
chr7-106242960-INS-481
chr7-106253719-INS-5283
chr7-106439093-INS-769
chr7-106493752-INS-93
chr7-106518896-INS-78
chr7-106525049-INS-56
chr7-106539189-INS-6402
chr7-106564238-INS-55
chr7-106586363-INS-6572
chr7-106589762-INS-81
chr7-106595985-INS-9402
chr7-106599825-INS-73
chr7-106645697-INS-683
chr7-106677863-INS-6690
chr7-106678493-INS-199
chr7-106712389-INS-171
chr7-106775756-INS-315
chr7-106778723-INS-189
chr7-106778904-INS-1379
chr7-106794163-INS-357
chr7-106853845-INS-58
chr7-106886141-INS-84
chr7-106888613-INS-60
chr7-106891180-INS-194
chr7-106899889-INS-103
chr7-106912813-INS-187
chr7-106952058-INS-5278
chr7-106966654-INS-4852
chr7-106987470-INS-990
chr7-107025906-INS-70
chr7-107038756-INS-765
chr7-107068127-INS-51
chr7-107075185-INS-1888
chr7-107104051-INS-101
chr7-107111345-INS-292
chr7-107117614-INS-92
chr7-107132739-INS-87
chr7-107151199-INS-209
chr7-107161591-INS-72
chr7-107181125-INS-208
chr7-107183679-INS-54
chr7-107245977-INS-565
chr7-107299128-INS-54
chr7-107324233-INS-120
chr7-107354881-INS-260
chr7-107373512-INS-94
chr7-107374832-INS-6820
chr7-107382586-INS-486
chr7-107387130-INS-491
chr7-107408167-INS-504
chr7-107408930-INS-52
chr7-107415567-INS-102
chr7-107417745-INS-164
chr7-107427668-INS-543
chr7-107427683-INS-6797
chr7-107442483-INS-4864
chr7-107443583-INS-9234
chr7-107477577-INS-203
chr7-107500300-INS-6473
chr7-107535325-INS-146
chr7-107553642-INS-135
chr7-107619666-INS-254
chr7-107639059-INS-5680
chr7-107667640-INS-84
chr7-107684792-INS-66
chr7-107729519-INS-54
chr7-107729931-INS-167

chr7-107752415-INS-826
chr7-107762576-INS-89
chr7-107827356-INS-845
chr7-107849306-INS-118
chr7-107901077-INS-192
chr7-107905559-INS-105
chr7-107919663-INS-250
chr7-107931467-INS-503
chr7-107956363-INS-172
chr7-108006063-INS-1627
chr7-108033562-INS-782
chr7-108057834-INS-95
chr7-108078853-INS-98
chr7-108093973-INS-416
chr7-108209725-INS-580
chr7-108226174-INS-78
chr7-108238529-INS-201
chr7-108249950-INS-423
chr7-108254351-INS-60
chr7-108259645-INS-6704
chr7-108266207-INS-7405
chr7-108290725-INS-6576
chr7-108312040-INS-90
chr7-108317386-INS-171
chr7-108328209-INS-910
chr7-108389786-INS-164
chr7-108393575-INS-82
chr7-108397343-INS-202
chr7-108400917-INS-130
chr7-108412805-INS-7109
chr7-108414091-INS-56
chr7-108425662-INS-93
chr7-108445533-INS-6378
chr7-108471214-INS-119
chr7-108515969-INS-60
chr7-108531569-INS-66
chr7-108554764-INS-150
chr7-108556294-INS-218
chr7-108569727-INS-61
chr7-108606698-INS-898
chr7-108620985-INS-52
chr7-108645500-INS-6314
chr7-108702750-INS-60
chr7-108728687-INS-74
chr7-108734042-INS-62
chr7-108752687-INS-76
chr7-108780693-INS-58
chr7-108797512-INS-6155
chr7-108799368-INS-52
chr7-108817777-INS-56
chr7-108828207-INS-61
chr7-108829748-INS-7164
chr7-108862442-INS-132
chr7-108892471-INS-900
chr7-108901862-INS-74
chr7-108902778-INS-8205
chr7-108933996-INS-61
chr7-108944545-INS-7119
chr7-108956875-INS-193
chr7-108973794-INS-106
chr7-108980749-INS-196
chr7-108981703-INS-364
chr7-108988081-INS-1561
chr7-108996684-INS-1490
chr7-109033259-INS-1929
chr7-109041048-INS-533
chr7-109042358-INS-87
chr7-109047391-INS-75
chr7-109065115-INS-162
chr7-109071991-INS-55
chr7-109098536-INS-174
chr7-109099415-INS-222
chr7-109102856-INS-52
chr7-109127202-INS-52
chr7-109131087-INS-223
chr7-109143095-INS-76
chr7-109158373-INS-367

chr7-109202425-INS-81
chr7-109212703-INS-315
chr7-109223524-INS-221
chr7-109225784-INS-54
chr7-109294527-INS-175
chr7-109312521-INS-155
chr7-109327651-INS-67
chr7-109329532-INS-228
chr7-109332266-INS-400
chr7-109353616-INS-197
chr7-109358293-INS-101
chr7-109366559-INS-340
chr7-109387920-INS-156
chr7-109436420-INS-172
chr7-109440715-INS-236
chr7-109444419-INS-147
chr7-109445083-INS-474
chr7-109445333-INS-98
chr7-109446902-INS-88
chr7-109469298-INS-86
chr7-109472916-INS-132
chr7-109479241-INS-96
chr7-109502545-INS-206
chr7-109509178-INS-200
chr7-109510885-INS-980
chr7-109511861-INS-205
chr7-109513349-INS-214
chr7-109538596-INS-56
chr7-109552053-INS-147
chr7-109553555-INS-116
chr7-109555036-INS-203
chr7-109560809-INS-208
chr7-109562216-INS-209
chr7-109577624-INS-153
chr7-109594244-INS-175
chr7-109597221-INS-174
chr7-109601740-INS-835
chr7-109605651-INS-428
chr7-109607403-INS-198
chr7-109625689-INS-208
chr7-109628847-INS-170
chr7-109634379-INS-207
chr7-109636465-INS-95
chr7-109658054-INS-163
chr7-109666576-INS-497
chr7-109699375-INS-306
chr7-109704791-INS-229
chr7-109708220-INS-68
chr7-109710058-INS-177
chr7-109717537-INS-164
chr7-109732203-INS-230
chr7-109751053-INS-211
chr7-109755746-INS-57
chr7-109759896-INS-195
chr7-109763107-INS-194
chr7-109764850-INS-114
chr7-109777720-INS-112
chr7-109786947-INS-159
chr7-109789014-INS-205
chr7-109796520-INS-221
chr7-109804408-INS-98
chr7-109804485-INS-113
chr7-109806949-INS-775
chr7-109813428-INS-234
chr7-109816209-INS-231
chr7-109817544-INS-264
chr7-109827083-INS-288
chr7-109840742-INS-118
chr7-109864610-INS-226
chr7-109890844-INS-91
chr7-109892910-INS-368
chr7-109894340-INS-181
chr7-109909775-INS-5150
chr7-109941755-INS-270
chr7-109965608-INS-70
chr7-109998399-INS-78
chr7-110031399-INS-381

chr7-110069302-INS-125
chr7-110133267-INS-82
chr7-110145771-INS-389
chr7-110149572-INS-87
chr7-110195977-INS-284
chr7-110205276-INS-120
chr7-110224709-INS-51
chr7-110250202-INS-124
chr7-110251761-INS-55
chr7-110311139-INS-403
chr7-110351442-INS-143
chr7-110358725-INS-358
chr7-110415273-INS-78
chr7-110422710-INS-1246
chr7-110437454-INS-6686
chr7-110463725-INS-166
chr7-110466811-INS-424
chr7-110484663-INS-221
chr7-110494447-INS-5319
chr7-110533638-INS-6248
chr7-110552196-INS-251
chr7-110571114-INS-195
chr7-110598872-INS-260
chr7-110626111-INS-7341
chr7-110638647-INS-217
chr7-110658683-INS-535
chr7-110660679-INS-169
chr7-110700388-INS-55
chr7-110728853-INS-122
chr7-110732517-INS-141
chr7-110753063-INS-65
chr7-110761471-INS-59
chr7-110820786-INS-70
chr7-110828629-INS-103
chr7-110835927-INS-7628
chr7-110838181-INS-215
chr7-110855067-INS-7277
chr7-110857130-INS-117
chr7-110865255-INS-58
chr7-110880930-INS-276
chr7-110887036-INS-1059
chr7-110938591-INS-6228
chr7-110944660-INS-54
chr7-110957685-INS-609
chr7-110990873-INS-54
chr7-111012924-INS-138
chr7-111014433-INS-7782
chr7-111016428-INS-51
chr7-111024226-INS-120
chr7-111037514-INS-66
chr7-111057398-INS-614
chr7-111065058-INS-69
chr7-111126054-INS-110
chr7-111183824-INS-890
chr7-111199277-INS-357
chr7-111225803-INS-429
chr7-111254697-INS-58
chr7-111260041-INS-52
chr7-111292612-INS-383
chr7-111304625-INS-102
chr7-111344507-INS-80
chr7-111380011-INS-55
chr7-111393766-INS-401
chr7-111395321-INS-170
chr7-111411649-INS-315
chr7-111418819-INS-94
chr7-111424086-INS-124
chr7-111458864-INS-186
chr7-111479075-INS-72
chr7-111492035-INS-467
chr7-111498428-INS-663
chr7-111499460-INS-94
chr7-111507161-INS-6694
chr7-111508289-INS-291
chr7-111526839-INS-223
chr7-111544649-INS-224
chr7-111548894-INS-194

chr7-111552935-INS-55
chr7-111557008-INS-89
chr7-111583329-INS-6343
chr7-111597318-INS-69
chr7-111608286-INS-163
chr7-111684817-INS-61
chr7-111739513-INS-220
chr7-111759049-INS-55
chr7-111762249-INS-721
chr7-111762956-INS-76
chr7-111781415-INS-84
chr7-111841700-INS-149
chr7-111864759-INS-55
chr7-111880484-INS-56
chr7-111891783-INS-164
chr7-111894241-INS-234
chr7-111926671-INS-52
chr7-111949955-INS-152
chr7-111966037-INS-121
chr7-111972176-INS-458
chr7-111981564-INS-66
chr7-111982530-INS-72
chr7-111997832-INS-690
chr7-112005212-INS-104
chr7-112009979-INS-1895
chr7-112016649-INS-5407
chr7-112036187-INS-52
chr7-112043340-INS-256
chr7-112051889-INS-89
chr7-112056434-INS-80
chr7-112117745-INS-58
chr7-112131475-INS-275
chr7-112176800-INS-87
chr7-112180606-INS-60
chr7-112214871-INS-486
chr7-112216623-INS-199
chr7-112233168-INS-72
chr7-112236915-INS-300
chr7-112237436-INS-1290
chr7-112237641-INS-210
chr7-112253725-INS-128
chr7-112264554-INS-63
chr7-112288988-INS-173
chr7-112301134-INS-266
chr7-112303879-INS-54
chr7-112329041-INS-86
chr7-112382837-INS-1798
chr7-112388203-INS-216
chr7-112406133-INS-334
chr7-112409004-INS-61
chr7-112410778-INS-201
chr7-112420828-INS-199
chr7-112447353-INS-213
chr7-112460167-INS-177
chr7-112467454-INS-174
chr7-112494465-INS-138
chr7-112539424-INS-933
chr7-112541513-INS-83
chr7-112557698-INS-195
chr7-112565927-INS-129
chr7-112582304-INS-113
chr7-112582740-INS-5571
chr7-112602099-INS-136
chr7-112632799-INS-1028
chr7-112662007-INS-57
chr7-112669980-INS-52
chr7-112692531-INS-201
chr7-112719060-INS-56
chr7-112727719-INS-99
chr7-112729316-INS-77
chr7-112733789-INS-104
chr7-112743459-INS-90
chr7-112751764-INS-53
chr7-112770065-INS-51
chr7-112774453-INS-272
chr7-112779454-INS-203
chr7-112802042-INS-59

chr7-112830827-INS-103
chr7-112854123-INS-53
chr7-112874863-INS-90
chr7-112925560-INS-124
chr7-112928190-INS-82
chr7-112943643-INS-371
chr7-112961312-INS-228
chr7-113021619-INS-108
chr7-113031356-INS-86
chr7-113041212-INS-148
chr7-113055191-INS-76
chr7-113098576-INS-267
chr7-113101238-INS-374
chr7-113102193-INS-164
chr7-113135139-INS-168
chr7-113164959-INS-56
chr7-113201281-INS-7248
chr7-113229696-INS-368
chr7-113268771-INS-54
chr7-113276848-INS-421
chr7-113337432-INS-78
chr7-113363295-INS-112
chr7-113368172-INS-54
chr7-113376051-INS-98
chr7-113384654-INS-165
chr7-113440451-INS-410
chr7-113495824-INS-105
chr7-113505239-INS-63
chr7-113509433-INS-195
chr7-113517985-INS-158
chr7-113539427-INS-201
chr7-113575230-INS-316
chr7-113626510-INS-106
chr7-113644999-INS-140
chr7-113659993-INS-201
chr7-113660616-INS-164
chr7-113661113-INS-328
chr7-113729594-INS-59
chr7-113741220-INS-68
chr7-113751835-INS-120
chr7-113752472-INS-56
chr7-113764058-INS-214
chr7-113806484-INS-213
chr7-113814283-INS-608
chr7-113815959-INS-212
chr7-113827700-INS-54
chr7-113850666-INS-271
chr7-113878934-INS-399
chr7-113883077-INS-240
chr7-113897188-INS-63
chr7-113920974-INS-433
chr7-113924750-INS-289
chr7-113932403-INS-400
chr7-113942867-INS-199
chr7-113943671-INS-651
chr7-113944659-INS-67
chr7-113947128-INS-171
chr7-113952119-INS-608
chr7-113970215-INS-500
chr7-113974131-INS-855
chr7-113989811-INS-6499
chr7-114034817-INS-92
chr7-114053692-INS-702
chr7-114061983-INS-214
chr7-114086577-INS-156
chr7-114114210-INS-53
chr7-114123683-INS-124
chr7-114155808-INS-204
chr7-114198433-INS-130
chr7-114201689-INS-70
chr7-114241748-INS-431
chr7-114251854-INS-360
chr7-114252715-INS-6216
chr7-114267366-INS-64
chr7-114273747-INS-1324
chr7-114338793-INS-400
chr7-114340530-INS-106

chr7-114381025-INS-512
chr7-114384454-INS-322
chr7-114388774-INS-223
chr7-114396129-INS-435
chr7-114405571-INS-54
chr7-114454370-INS-402
chr7-114462940-INS-663
chr7-114530439-INS-2037
chr7-114530627-INS-68
chr7-114536301-INS-114
chr7-114585160-INS-425
chr7-114592801-INS-304
chr7-114612941-INS-717
chr7-114620952-INS-239
chr7-114656216-INS-64
chr7-114665102-INS-103
chr7-114709963-INS-52
chr7-114722825-INS-99
chr7-114732021-INS-666
chr7-114768811-INS-56
chr7-114804065-INS-72
chr7-114818959-INS-4848
chr7-114852843-INS-155
chr7-114855409-INS-60
chr7-114862029-INS-402
chr7-114869720-INS-60
chr7-114879769-INS-6440
chr7-114889799-INS-68
chr7-114906442-INS-528
chr7-114921824-INS-128
chr7-114933675-INS-54
chr7-114943708-INS-7534
chr7-114971093-INS-54
chr7-114979537-INS-74
chr7-114985158-INS-882
chr7-115003108-INS-57
chr7-115008935-INS-369
chr7-115017119-INS-194
chr7-115030248-INS-225
chr7-115056204-INS-66
chr7-115057502-INS-92
chr7-115102706-INS-60
chr7-115102977-INS-202
chr7-115103642-INS-700
chr7-115118613-INS-116
chr7-115166269-INS-72
chr7-115169303-INS-159
chr7-115188582-INS-204
chr7-115231204-INS-199
chr7-115243874-INS-1769
chr7-115244257-INS-53
chr7-115261133-INS-59
chr7-115295798-INS-70
chr7-115317062-INS-217
chr7-115339823-INS-377
chr7-115366531-INS-201
chr7-115480042-INS-82
chr7-115487457-INS-97
chr7-115530750-INS-56
chr7-115548730-INS-104
chr7-115552967-INS-275
chr7-115648683-INS-84
chr7-115652453-INS-341
chr7-115658095-INS-336
chr7-115667347-INS-76
chr7-115708310-INS-54
chr7-115715606-INS-281
chr7-115771273-INS-52
chr7-115791891-INS-230
chr7-115794921-INS-70
chr7-115801033-INS-195
chr7-115816773-INS-202
chr7-115831241-INS-106
chr7-115844423-INS-77
chr7-115851408-INS-218
chr7-115852163-INS-52
chr7-115858202-INS-68

chr7-115883580-INS-249
chr7-115889895-INS-225
chr7-115892584-INS-86
chr7-115896078-INS-128
chr7-115912356-INS-198
chr7-115924477-INS-165
chr7-115924477-INS-166
chr7-115925468-INS-192
chr7-115927599-INS-385
chr7-115939791-INS-212
chr7-115951659-INS-371
chr7-115952244-INS-202
chr7-115960016-INS-55
chr7-115977406-INS-202
chr7-116015065-INS-348
chr7-116049434-INS-69
chr7-116053851-INS-399
chr7-116082695-INS-119
chr7-116085082-INS-83
chr7-116095796-INS-60
chr7-116107379-INS-160
chr7-116143526-INS-88
chr7-116161661-INS-593
chr7-116195153-INS-6246
chr7-116201904-INS-6843
chr7-116222298-INS-76
chr7-116245478-INS-295
chr7-116271994-INS-187
chr7-116277092-INS-128
chr7-116278126-INS-191
chr7-116290001-INS-161
chr7-116309361-INS-58
chr7-116317688-INS-222
chr7-116325422-INS-273
chr7-116338016-INS-389
chr7-116366340-INS-1326
chr7-116366678-INS-860
chr7-116368689-INS-6292
chr7-116393402-INS-54
chr7-116405994-INS-56
chr7-116418547-INS-957
chr7-116432730-INS-103
chr7-116436713-INS-70
chr7-116453552-INS-174
chr7-116454436-INS-54
chr7-116483765-INS-58
chr7-116515678-INS-154
chr7-116536551-INS-114
chr7-116587887-INS-51
chr7-116602162-INS-69
chr7-116606818-INS-110
chr7-116617422-INS-665
chr7-116623166-INS-60
chr7-116634321-INS-470
chr7-116642505-INS-7086
chr7-116652024-INS-54
chr7-116671447-INS-52
chr7-116680596-INS-167
chr7-116698096-INS-118
chr7-116699027-INS-889
chr7-116720488-INS-373
chr7-116737690-INS-949
chr7-116739549-INS-80
chr7-116743720-INS-172
chr7-116747708-INS-62
chr7-116776037-INS-103
chr7-116834286-INS-460
chr7-116839204-INS-62
chr7-116864755-INS-895
chr7-116924437-INS-55
chr7-116968050-INS-84
chr7-116969718-INS-93
chr7-116972814-INS-84
chr7-116991909-INS-122
chr7-117012255-INS-92
chr7-117040943-INS-522
chr7-117052346-INS-98

chr7-117073371-INS-138
chr7-117077786-INS-85
chr7-117119196-INS-60
chr7-117159147-INS-5543
chr7-117181609-INS-71
chr7-117195163-INS-6989
chr7-117195895-INS-56
chr7-117209241-INS-53
chr7-117252840-INS-364
chr7-117276233-INS-52
chr7-117368447-INS-68
chr7-117426098-INS-215
chr7-117451486-INS-189
chr7-117463768-INS-1888
chr7-117471628-INS-1892
chr7-117480945-INS-57
chr7-117491957-INS-85
chr7-117501693-INS-60
chr7-117506705-INS-318
chr7-117508520-INS-69
chr7-117518224-INS-947
chr7-117522090-INS-78
chr7-117523842-INS-103
chr7-117526699-INS-106
chr7-117535756-INS-53
chr7-117541297-INS-93
chr7-117552128-INS-88
chr7-117555051-INS-76
chr7-117558652-INS-212
chr7-117560275-INS-60
chr7-117616814-INS-323
chr7-117618657-INS-54
chr7-117647957-INS-84
chr7-117652796-INS-90
chr7-117655880-INS-70
chr7-117683103-INS-213
chr7-117688632-INS-3077
chr7-117693146-INS-102
chr7-117727623-INS-92
chr7-117737529-INS-171
chr7-117758565-INS-112
chr7-117782743-INS-452
chr7-117786251-INS-721
chr7-117786763-INS-91
chr7-117798715-INS-594
chr7-117807742-INS-264
chr7-117808137-INS-406
chr7-117828003-INS-209
chr7-117851168-INS-140
chr7-117858072-INS-220
chr7-117859731-INS-77
chr7-117875645-INS-80
chr7-117897172-INS-143
chr7-117900943-INS-126
chr7-117916836-INS-170
chr7-117929319-INS-938
chr7-117931413-INS-184
chr7-117952944-INS-64
chr7-117962710-INS-5221
chr7-117979183-INS-62
chr7-118002682-INS-225
chr7-118011821-INS-391
chr7-118028697-INS-54
chr7-118054572-INS-153
chr7-118073958-INS-94
chr7-118084245-INS-68
chr7-118130999-INS-76
chr7-118141617-INS-89
chr7-118157620-INS-84
chr7-118166388-INS-91
chr7-118187630-INS-331
chr7-118202070-INS-180
chr7-118203301-INS-70
chr7-118206323-INS-59
chr7-118215849-INS-179
chr7-118218332-INS-439
chr7-118223147-INS-201

chr7-118242042-INS-52
chr7-118287253-INS-58
chr7-118309092-INS-208
chr7-118339516-INS-74
chr7-118350144-INS-187
chr7-118382895-INS-68
chr7-118396759-INS-165
chr7-118397194-INS-233
chr7-118398689-INS-511
chr7-118411529-INS-137
chr7-118413941-INS-136
chr7-118420619-INS-6413
chr7-118453822-INS-942
chr7-118480422-INS-122
chr7-118485070-INS-1043
chr7-118495618-INS-967
chr7-118498076-INS-69
chr7-118499531-INS-52
chr7-118504432-INS-220
chr7-118532605-INS-332
chr7-118534344-INS-215
chr7-118540095-INS-66
chr7-118550587-INS-145
chr7-118552488-INS-280
chr7-118612370-INS-84
chr7-118618583-INS-215
chr7-118624633-INS-546
chr7-118635451-INS-221
chr7-118641108-INS-80
chr7-118666776-INS-57
chr7-118685107-INS-134
chr7-118704424-INS-75
chr7-118746033-INS-72
chr7-118842956-INS-7114
chr7-118845407-INS-451
chr7-118850847-INS-96
chr7-118873385-INS-1015
chr7-118873386-INS-454
chr7-118874262-INS-570
chr7-118876765-INS-108
chr7-118895887-INS-78
chr7-118899108-INS-676
chr7-118931105-INS-114
chr7-118957860-INS-107
chr7-118959776-INS-64
chr7-118963929-INS-348
chr7-119010501-INS-140
chr7-119011609-INS-52
chr7-119066260-INS-150
chr7-119094958-INS-88
chr7-119096375-INS-90
chr7-119111934-INS-202
chr7-119130777-INS-170
chr7-119151643-INS-220
chr7-119183942-INS-146
chr7-119187029-INS-898
chr7-119204945-INS-124
chr7-119223306-INS-789
chr7-119228206-INS-270
chr7-119242225-INS-137
chr7-119265351-INS-334
chr7-119273312-INS-541
chr7-119309733-INS-78
chr7-119318886-INS-66
chr7-119321462-INS-84
chr7-119330722-INS-58
chr7-119338527-INS-483
chr7-119348449-INS-6233
chr7-119356892-INS-149
chr7-119358780-INS-120
chr7-119379312-INS-61
chr7-119387843-INS-1730
chr7-119402617-INS-59
chr7-119480872-INS-60
chr7-119514334-INS-92
chr7-119520390-INS-407
chr7-119533223-INS-108

chr7-119535948-INS-229
chr7-119541217-INS-270
chr7-119542332-INS-163
chr7-119552375-INS-56
chr7-119556504-INS-56
chr7-119569338-INS-461
chr7-119575798-INS-60
chr7-119577106-INS-118
chr7-119586682-INS-151
chr7-119587555-INS-7173
chr7-119590152-INS-316
chr7-119594378-INS-166
chr7-119606369-INS-622
chr7-119626112-INS-73
chr7-119651385-INS-243
chr7-119676368-INS-62
chr7-119727426-INS-1892
chr7-119728009-INS-51
chr7-119733897-INS-166
chr7-119740058-INS-53
chr7-119784151-INS-96
chr7-119792620-INS-58
chr7-119811133-INS-59
chr7-119813344-INS-85
chr7-119818645-INS-5549
chr7-119825419-INS-512
chr7-119835977-INS-78
chr7-119841614-INS-174
chr7-119850669-INS-229
chr7-119852459-INS-68
chr7-119854942-INS-117
chr7-119870918-INS-428
chr7-119874072-INS-99
chr7-119899748-INS-215
chr7-119910678-INS-211
chr7-119924543-INS-367
chr7-119958747-INS-250
chr7-119975758-INS-85
chr7-119981096-INS-88
chr7-119991228-INS-403
chr7-119992015-INS-382
chr7-120001228-INS-248
chr7-120003557-INS-1589
chr7-120015222-INS-1890
chr7-120019160-INS-205
chr7-120041080-INS-7131
chr7-120043788-INS-919
chr7-120050346-INS-88
chr7-120074034-INS-132
chr7-120099092-INS-168
chr7-120123798-INS-387
chr7-120129927-INS-84
chr7-120150490-INS-187
chr7-120154442-INS-594
chr7-120166290-INS-62
chr7-120168495-INS-392
chr7-120180258-INS-185
chr7-120204724-INS-202
chr7-120211202-INS-60
chr7-120218649-INS-128
chr7-120219874-INS-328
chr7-120220452-INS-187
chr7-120223862-INS-153
chr7-120226740-INS-158
chr7-120229132-INS-171
chr7-120249251-INS-200
chr7-120249264-INS-759
chr7-120254533-INS-6567
chr7-120260599-INS-127
chr7-120264832-INS-224
chr7-120274272-INS-698
chr7-120282371-INS-184
chr7-120306758-INS-230
chr7-120334994-INS-169
chr7-120337423-INS-55
chr7-120337614-INS-3125
chr7-120369191-INS-336

chr7-120377180-INS-81
chr7-120380527-INS-51
chr7-120399932-INS-409
chr7-120419500-INS-166
chr7-120436645-INS-60
chr7-120443853-INS-74
chr7-120446411-INS-55
chr7-120449211-INS-122
chr7-120449652-INS-5345
chr7-120453783-INS-68
chr7-120475575-INS-240
chr7-120476476-INS-51
chr7-120489510-INS-222
chr7-120491731-INS-55
chr7-120495947-INS-7099
chr7-120500239-INS-207
chr7-120512629-INS-5314
chr7-120514264-INS-102
chr7-120528923-INS-301
chr7-120554476-INS-167
chr7-120560174-INS-215
chr7-120569183-INS-54
chr7-120571303-INS-177
chr7-120572658-INS-56
chr7-120597298-INS-6285
chr7-120648145-INS-180
chr7-120669116-INS-299
chr7-120679003-INS-597
chr7-120679683-INS-67
chr7-120686235-INS-201
chr7-120695818-INS-196
chr7-120706647-INS-966
chr7-120723720-INS-164
chr7-120726014-INS-145
chr7-120728214-INS-156
chr7-120754758-INS-83
chr7-120769489-INS-760
chr7-120773763-INS-287
chr7-120773836-INS-920
chr7-120785327-INS-52
chr7-120799472-INS-504
chr7-120808271-INS-59
chr7-120812601-INS-74
chr7-120829900-INS-148
chr7-120835966-INS-101
chr7-120845406-INS-75
chr7-120853078-INS-235
chr7-120861350-INS-55
chr7-120917397-INS-72
chr7-120922971-INS-519
chr7-120924330-INS-975
chr7-120975847-INS-522
chr7-120996884-INS-52
chr7-121031207-INS-462
chr7-121041991-INS-208
chr7-121042389-INS-5883
chr7-121071878-INS-97
chr7-121086357-INS-352
chr7-121088295-INS-88
chr7-121104622-INS-858
chr7-121109095-INS-150
chr7-121128927-INS-220
chr7-121167139-INS-93
chr7-121174130-INS-60
chr7-121193055-INS-3174
chr7-121201304-INS-54
chr7-121215837-INS-6527
chr7-121223731-INS-148
chr7-121229241-INS-698
chr7-121255081-INS-120
chr7-121256270-INS-56
chr7-121267888-INS-199
chr7-121271616-INS-51
chr7-121280311-INS-67
chr7-121301108-INS-91
chr7-121313073-INS-85
chr7-121317089-INS-52

chr7-121333144-INS-196
chr7-121340444-INS-56
chr7-121343132-INS-106
chr7-121396325-INS-125
chr7-121398718-INS-153
chr7-121402343-INS-1886
chr7-121404880-INS-92
chr7-121443802-INS-493
chr7-121475949-INS-87
chr7-121500574-INS-226
chr7-121544563-INS-1212
chr7-121546707-INS-94
chr7-121547872-INS-136
chr7-121564803-INS-118
chr7-121566668-INS-52
chr7-121596771-INS-66
chr7-121598793-INS-209
chr7-121600249-INS-210
chr7-121608532-INS-222
chr7-121624279-INS-124
chr7-121629386-INS-394
chr7-121630861-INS-51
chr7-121640237-INS-205
chr7-121651379-INS-156
chr7-121651928-INS-62
chr7-121664252-INS-171
chr7-121716226-INS-56
chr7-121730751-INS-202
chr7-121736862-INS-854
chr7-121767918-INS-74
chr7-121773997-INS-95
chr7-121810506-INS-60
chr7-121819183-INS-195
chr7-121870148-INS-88
chr7-121871070-INS-418
chr7-121876876-INS-343
chr7-121885618-INS-123
chr7-121900514-INS-313
chr7-121918079-INS-143
chr7-121929433-INS-60
chr7-121947075-INS-110
chr7-121953747-INS-130
chr7-121961200-INS-286
chr7-121967021-INS-122
chr7-122000915-INS-73
chr7-122009849-INS-70
chr7-122019588-INS-83
chr7-122021137-INS-171
chr7-122026369-INS-254
chr7-122033554-INS-108
chr7-122038175-INS-80
chr7-122039406-INS-237
chr7-122047499-INS-279
chr7-122054450-INS-71
chr7-122087801-INS-76
chr7-122098723-INS-88
chr7-122113663-INS-129
chr7-122118289-INS-59
chr7-122122254-INS-74
chr7-122153604-INS-162
chr7-122202122-INS-64
chr7-122247926-INS-396
chr7-122282163-INS-171
chr7-122290879-INS-124
chr7-122293144-INS-525
chr7-122299217-INS-115
chr7-122325190-INS-65
chr7-122354891-INS-52
chr7-122356876-INS-51
chr7-122366684-INS-180
chr7-122371768-INS-117
chr7-122373645-INS-236
chr7-122377507-INS-227
chr7-122383808-INS-192
chr7-122397816-INS-716
chr7-122408273-INS-60
chr7-122415385-INS-1106

chr7-122441001-INS-72
chr7-122441679-INS-226
chr7-122450414-INS-201
chr7-122459316-INS-66
chr7-122468849-INS-88
chr7-122481821-INS-62
chr7-122482264-INS-179
chr7-122483101-INS-182
chr7-122487356-INS-104
chr7-122502083-INS-202
chr7-122514844-INS-1831
chr7-122531210-INS-58
chr7-122541927-INS-105
chr7-122555716-INS-631
chr7-122563228-INS-92
chr7-122585752-INS-58
chr7-122617239-INS-201
chr7-122622091-INS-120
chr7-122625146-INS-81
chr7-122628081-INS-102
chr7-122636232-INS-104
chr7-122640974-INS-203
chr7-122652503-INS-159
chr7-122653587-INS-225
chr7-122659072-INS-82
chr7-122669935-INS-52
chr7-122676900-INS-207
chr7-122702676-INS-291
chr7-122703285-INS-156
chr7-122713720-INS-62
chr7-122739826-INS-384
chr7-122758520-INS-98
chr7-122775170-INS-188
chr7-122786129-INS-97
chr7-122802353-INS-322
chr7-122821365-INS-182
chr7-122825297-INS-60
chr7-122838314-INS-126
chr7-122848710-INS-83
chr7-122855755-INS-238
chr7-122861045-INS-130
chr7-122884453-INS-282
chr7-122886750-INS-176
chr7-122893026-INS-128
chr7-122901016-INS-58
chr7-122902170-INS-201
chr7-122912470-INS-205
chr7-122918624-INS-130
chr7-122928316-INS-62
chr7-122957711-INS-261
chr7-122979209-INS-212
chr7-122998361-INS-400
chr7-123032352-INS-60
chr7-123037541-INS-72
chr7-123050110-INS-52
chr7-123073130-INS-207
chr7-123076797-INS-358
chr7-123104371-INS-60
chr7-123111388-INS-431
chr7-123127630-INS-63
chr7-123131571-INS-54
chr7-123134676-INS-89
chr7-123135704-INS-56
chr7-123146704-INS-466
chr7-123182944-INS-6464
chr7-123193713-INS-297
chr7-123195461-INS-129
chr7-123196741-INS-239
chr7-123248428-INS-199
chr7-123249458-INS-70
chr7-123252790-INS-61
chr7-123264759-INS-52
chr7-123266149-INS-221
chr7-123290872-INS-9840
chr7-123304495-INS-67
chr7-123310656-INS-218
chr7-123328448-INS-610

chr7-123336536-INS-324
chr7-123373376-INS-68
chr7-123386678-INS-68
chr7-123400260-INS-158
chr7-123481756-INS-95
chr7-123486071-INS-8459
chr7-123497513-INS-613
chr7-123504427-INS-2100
chr7-123509960-INS-68
chr7-123516667-INS-455
chr7-123528958-INS-174
chr7-123532586-INS-65
chr7-123533347-INS-79
chr7-123551112-INS-2600
chr7-123557801-INS-266
chr7-123572617-INS-151
chr7-123632388-INS-55
chr7-123636054-INS-52
chr7-123646213-INS-122
chr7-123655853-INS-61
chr7-123671892-INS-87
chr7-123694516-INS-222
chr7-123695095-INS-178
chr7-123698113-INS-172
chr7-123700798-INS-192
chr7-123737466-INS-461
chr7-123743989-INS-457
chr7-123753741-INS-155
chr7-123763791-INS-68
chr7-123764842-INS-112
chr7-123780084-INS-138
chr7-123790853-INS-6618
chr7-123791601-INS-155
chr7-123798873-INS-646
chr7-123802384-INS-3373
chr7-123844978-INS-52
chr7-123845040-INS-647
chr7-123851785-INS-326
chr7-123871250-INS-203
chr7-123877550-INS-437
chr7-123884524-INS-74
chr7-123905508-INS-125
chr7-123908726-INS-332
chr7-123954132-INS-53
chr7-123966792-INS-608
chr7-123986644-INS-96
chr7-123988558-INS-336
chr7-124003588-INS-213
chr7-124022363-INS-238
chr7-124039981-INS-58
chr7-124093842-INS-148
chr7-124111676-INS-113
chr7-124113229-INS-65
chr7-124138100-INS-294
chr7-124162449-INS-938
chr7-124176262-INS-51
chr7-124239387-INS-1843
chr7-124246326-INS-398
chr7-124248179-INS-7252
chr7-124249432-INS-123
chr7-124249610-INS-1856
chr7-124253283-INS-50
chr7-124302650-INS-312
chr7-124352055-INS-465
chr7-124386741-INS-77
chr7-124402062-INS-56
chr7-124420839-INS-95
chr7-124450716-INS-58
chr7-124472061-INS-96
chr7-124475676-INS-213
chr7-124477763-INS-117
chr7-124479242-INS-81
chr7-124493520-INS-367
chr7-124516433-INS-56
chr7-124527343-INS-130
chr7-124542772-INS-60
chr7-124544905-INS-952

chr7-124594342-INS-135
chr7-124607874-INS-7158
chr7-124625246-INS-212
chr7-124630783-INS-127
chr7-124665160-INS-93
chr7-124717240-INS-78
chr7-124719789-INS-179
chr7-124728553-INS-578
chr7-124728946-INS-6285
chr7-124732389-INS-385
chr7-124736483-INS-60
chr7-124742529-INS-1890
chr7-124761177-INS-64
chr7-124772031-INS-152
chr7-124774887-INS-130
chr7-124813137-INS-104
chr7-124830537-INS-81
chr7-124848030-INS-382
chr7-124867254-INS-76
chr7-124906991-INS-89
chr7-124907461-INS-196
chr7-124909142-INS-54
chr7-124910352-INS-1951
chr7-124981750-INS-249
chr7-124985531-INS-200
chr7-124986631-INS-52
chr7-125001062-INS-623
chr7-125003807-INS-64
chr7-125013926-INS-54
chr7-125039123-INS-366
chr7-125057610-INS-169
chr7-125073296-INS-124
chr7-125106784-INS-492
chr7-125111832-INS-169
chr7-125128153-INS-7082
chr7-125128458-INS-51
chr7-125167413-INS-82
chr7-125185943-INS-62
chr7-125194910-INS-111
chr7-125231305-INS-66
chr7-125233334-INS-52
chr7-125284774-INS-238
chr7-125288539-INS-199
chr7-125356478-INS-96
chr7-125360785-INS-79
chr7-125377821-INS-444
chr7-125378782-INS-57
chr7-125391320-INS-60
chr7-125402080-INS-108
chr7-125440268-INS-107
chr7-125462513-INS-54
chr7-125480938-INS-596
chr7-125482052-INS-54
chr7-125508887-INS-192
chr7-125516368-INS-200
chr7-125525489-INS-73
chr7-125531063-INS-74
chr7-125552342-INS-153
chr7-125575022-INS-182
chr7-125590094-INS-59
chr7-125603410-INS-256
chr7-125627124-INS-71
chr7-125670309-INS-88
chr7-125679674-INS-78
chr7-125691768-INS-256
chr7-125709712-INS-132
chr7-125739346-INS-55
chr7-125753497-INS-74
chr7-125757901-INS-189
chr7-125758771-INS-224
chr7-125766185-INS-200
chr7-125769431-INS-274
chr7-125775894-INS-831
chr7-125778332-INS-201
chr7-125815064-INS-177
chr7-125820600-INS-202
chr7-125822625-INS-139

chr7-125825734-INS-58
chr7-125831184-INS-221
chr7-125832964-INS-964
chr7-125832967-INS-72
chr7-125834672-INS-555
chr7-125835811-INS-221
chr7-125844045-INS-206
chr7-125862711-INS-54
chr7-125895237-INS-177
chr7-125912797-INS-244
chr7-125913967-INS-150
chr7-125914999-INS-347
chr7-125918224-INS-183
chr7-125920183-INS-126
chr7-125926413-INS-869
chr7-125937814-INS-67
chr7-125943362-INS-151
chr7-125956554-INS-82
chr7-125964753-INS-218
chr7-126004174-INS-7296
chr7-126020696-INS-183
chr7-126039801-INS-220
chr7-126050496-INS-56
chr7-126080361-INS-141
chr7-126081270-INS-914
chr7-126083584-INS-407
chr7-126117486-INS-206
chr7-126122858-INS-2989
chr7-126123974-INS-328
chr7-126135475-INS-102
chr7-126141501-INS-503
chr7-126144449-INS-188
chr7-126170690-INS-414
chr7-126177871-INS-234
chr7-126191167-INS-128
chr7-126203246-INS-5547
chr7-126212935-INS-97
chr7-126233990-INS-581
chr7-126237214-INS-54
chr7-126250994-INS-270
chr7-126257014-INS-202
chr7-126268075-INS-174
chr7-126280436-INS-2703
chr7-126280811-INS-167
chr7-126284287-INS-75
chr7-126318589-INS-177
chr7-126322438-INS-236
chr7-126325798-INS-3836
chr7-126325901-INS-430
chr7-126328682-INS-224
chr7-126329900-INS-119
chr7-126332248-INS-199
chr7-126337437-INS-131
chr7-126343067-INS-68
chr7-126413841-INS-132
chr7-126416966-INS-56
chr7-126435259-INS-484
chr7-126437525-INS-113
chr7-126475660-INS-247
chr7-126498858-INS-206
chr7-126503423-INS-238
chr7-126539947-INS-123
chr7-126602736-INS-430
chr7-126609278-INS-146
chr7-126642222-INS-204
chr7-126664244-INS-111
chr7-126730030-INS-181
chr7-126750391-INS-211
chr7-126759432-INS-244
chr7-126770009-INS-97
chr7-126772782-INS-126
chr7-126778094-INS-138
chr7-126785475-INS-52
chr7-126823160-INS-81
chr7-126825937-INS-77
chr7-126838802-INS-51
chr7-126863164-INS-919

chr7-126883962-INS-113
chr7-126902217-INS-116
chr7-126907280-INS-76
chr7-126925834-INS-65
chr7-126955786-INS-373
chr7-126967973-INS-237
chr7-126977274-INS-476
chr7-126985293-INS-213
chr7-126989120-INS-180
chr7-127001811-INS-331
chr7-127006620-INS-203
chr7-127027880-INS-206
chr7-127032036-INS-243
chr7-127038503-INS-57
chr7-127066334-INS-505
chr7-127069132-INS-142
chr7-127081260-INS-208
chr7-127092266-INS-196
chr7-127098018-INS-860
chr7-127115077-INS-292
chr7-127132490-INS-210
chr7-127180713-INS-219
chr7-127193283-INS-225
chr7-127226423-INS-96
chr7-127227457-INS-209
chr7-127233691-INS-147
chr7-127238138-INS-98
chr7-127239118-INS-214
chr7-127249378-INS-192
chr7-127258902-INS-191
chr7-127268606-INS-202
chr7-127278959-INS-250
chr7-127282658-INS-128
chr7-127285058-INS-170
chr7-127286659-INS-87
chr7-127294687-INS-207
chr7-127306861-INS-145
chr7-127309125-INS-220
chr7-127313650-INS-223
chr7-127329372-INS-425
chr7-127339524-INS-5220
chr7-127339663-INS-59
chr7-127342165-INS-173
chr7-127360949-INS-182
chr7-127364473-INS-54
chr7-127392435-INS-55
chr7-127405792-INS-117
chr7-127419485-INS-105
chr7-127430260-INS-226
chr7-127434963-INS-254
chr7-127451346-INS-230
chr7-127453233-INS-55
chr7-127458023-INS-205
chr7-127463663-INS-155
chr7-127473239-INS-204
chr7-127487133-INS-200
chr7-127494297-INS-206
chr7-127495227-INS-56
chr7-127500321-INS-208
chr7-127501562-INS-208
chr7-127518442-INS-63
chr7-127524638-INS-227
chr7-127538463-INS-76
chr7-127550676-INS-200
chr7-127556676-INS-197
chr7-127586267-INS-84
chr7-127618716-INS-60
chr7-127624361-INS-204
chr7-127625574-INS-102
chr7-127627523-INS-6400
chr7-127630226-INS-79
chr7-127634217-INS-60
chr7-127654423-INS-197
chr7-127655612-INS-196
chr7-127664086-INS-144
chr7-127738154-INS-175
chr7-127748693-INS-110

chr7-127760500-INS-79
chr7-127764776-INS-59
chr7-127784475-INS-242
chr7-127786677-INS-792
chr7-127786678-INS-167
chr7-127787181-INS-1466
chr7-127787289-INS-312
chr7-127794463-INS-979
chr7-127795173-INS-225
chr7-127809654-INS-92
chr7-127827934-INS-72
chr7-127832416-INS-184
chr7-127842988-INS-231
chr7-127844551-INS-148
chr7-127860769-INS-58
chr7-127864177-INS-59
chr7-127870808-INS-112
chr7-127882691-INS-101
chr7-127909120-INS-476
chr7-127910218-INS-741
chr7-127912742-INS-174
chr7-127939705-INS-268
chr7-127948565-INS-75
chr7-127954068-INS-70
chr7-127955572-INS-166
chr7-127956829-INS-420
chr7-127967229-INS-73
chr7-127981900-INS-190
chr7-127987045-INS-203
chr7-127995081-INS-70
chr7-128007837-INS-172
chr7-128024931-INS-341
chr7-128082969-INS-192
chr7-128091889-INS-316
chr7-128103485-INS-57
chr7-128111744-INS-204
chr7-128132281-INS-113
chr7-128151442-INS-271
chr7-128152392-INS-52
chr7-128158619-INS-54
chr7-128175220-INS-92
chr7-128177123-INS-66
chr7-128181887-INS-5503
chr7-128187290-INS-172
chr7-128193028-INS-164
chr7-128206132-INS-113
chr7-128234026-INS-201
chr7-128250556-INS-60
chr7-128272557-INS-145
chr7-128297801-INS-53
chr7-128323668-INS-121
chr7-128328235-INS-52
chr7-128337370-INS-223
chr7-128359037-INS-209
chr7-128360109-INS-243
chr7-128364747-INS-63
chr7-128382566-INS-51
chr7-128387953-INS-6471
chr7-128391849-INS-1673
chr7-128418792-INS-118
chr7-128421868-INS-54
chr7-128426425-INS-106
chr7-128439425-INS-286
chr7-128454558-INS-64
chr7-128474254-INS-181
chr7-128474791-INS-339
chr7-128502189-INS-399
chr7-128505451-INS-448
chr7-128507326-INS-406
chr7-128536397-INS-184
chr7-128536840-INS-6358
chr7-128543688-INS-176
chr7-128545279-INS-7080
chr7-128554943-INS-341
chr7-128560888-INS-63
chr7-128565623-INS-52
chr7-128578113-INS-7152

chr7-128597400-INS-346
chr7-128603265-INS-116
chr7-128609864-INS-1008
chr7-128611284-INS-90
chr7-128657100-INS-139
chr7-128676079-INS-412
chr7-128678773-INS-6010
chr7-128685784-INS-80
chr7-128703199-INS-74
chr7-128709151-INS-93
chr7-128711962-INS-59
chr7-128784672-INS-180
chr7-128786228-INS-105
chr7-128799532-INS-157
chr7-128828809-INS-626
chr7-128844556-INS-54
chr7-128856448-INS-120
chr7-128880497-INS-57
chr7-128897879-INS-799
chr7-128905789-INS-157
chr7-128907001-INS-110
chr7-128924341-INS-52
chr7-128928266-INS-6217
chr7-128951350-INS-55
chr7-129002697-INS-243
chr7-129006196-INS-93
chr7-129009701-INS-110
chr7-129023749-INS-81
chr7-129036414-INS-60
chr7-129038014-INS-78
chr7-129044796-INS-55
chr7-129058966-INS-652
chr7-129105047-INS-88
chr7-129121281-INS-4605
chr7-129129230-INS-80
chr7-129132568-INS-107
chr7-129164057-INS-5590
chr7-129174333-INS-124
chr7-129183700-INS-497
chr7-129185522-INS-137
chr7-129201051-INS-63
chr7-129209196-INS-2544
chr7-129212707-INS-1356
chr7-129248705-INS-195
chr7-129266754-INS-86
chr7-129277131-INS-81
chr7-129319326-INS-61
chr7-129319980-INS-80
chr7-129374235-INS-128
chr7-129383986-INS-96
chr7-129385985-INS-718
chr7-129398633-INS-603
chr7-129417972-INS-75
chr7-129419201-INS-53
chr7-129443551-INS-797
chr7-129456351-INS-107
chr7-129463670-INS-891
chr7-129463711-INS-328
chr7-129465818-INS-314
chr7-129473393-INS-458
chr7-129473418-INS-840
chr7-129488596-INS-92
chr7-129501847-INS-225
chr7-129505785-INS-89
chr7-129556301-INS-59
chr7-129560525-INS-186
chr7-129571564-INS-55
chr7-129628109-INS-400
chr7-129653229-INS-7072
chr7-129669739-INS-63
chr7-129676960-INS-67
chr7-129697369-INS-142
chr7-129728622-INS-399
chr7-129734708-INS-52
chr7-129739155-INS-51
chr7-129741537-INS-59
chr7-129783829-INS-106

chr7-129790306-INS-203
chr7-129873358-INS-51
chr7-129882165-INS-68
chr7-129886187-INS-186
chr7-129898977-INS-108
chr7-129937915-INS-120
chr7-129943647-INS-84
chr7-129972268-INS-86
chr7-129977397-INS-215
chr7-129987592-INS-178
chr7-130010718-INS-467
chr7-130015639-INS-228
chr7-130023345-INS-481
chr7-130059457-INS-747
chr7-130133556-INS-99
chr7-130148003-INS-128
chr7-130156542-INS-51
chr7-130161040-INS-222
chr7-130183333-INS-298
chr7-130193315-INS-54
chr7-130198187-INS-6307
chr7-130200693-INS-399
chr7-130214563-INS-7483
chr7-130241600-INS-164
chr7-130244813-INS-280
chr7-130312772-INS-575
chr7-130373055-INS-51
chr7-130377109-INS-62
chr7-130377778-INS-170
chr7-130391220-INS-81
chr7-130404786-INS-152
chr7-130417797-INS-793
chr7-130429181-INS-510
chr7-130433552-INS-136
chr7-130441262-INS-391
chr7-130453365-INS-80
chr7-130455901-INS-203
chr7-130490320-INS-206
chr7-130500019-INS-67
chr7-130519909-INS-62
chr7-130530304-INS-81
chr7-130543537-INS-83
chr7-130552123-INS-370
chr7-130557414-INS-91
chr7-130580134-INS-233
chr7-130592523-INS-68
chr7-130615627-INS-216
chr7-130616451-INS-876
chr7-130620708-INS-52
chr7-130626916-INS-96
chr7-130672632-INS-657
chr7-130694511-INS-52
chr7-130709119-INS-62
chr7-130711836-INS-55
chr7-130733825-INS-208
chr7-130774035-INS-114
chr7-130792595-INS-301
chr7-130800294-INS-474
chr7-130822094-INS-280
chr7-130847259-INS-139
chr7-130848589-INS-582
chr7-130872544-INS-91
chr7-130932173-INS-60
chr7-130936973-INS-108
chr7-130947654-INS-114
chr7-130961528-INS-203
chr7-130967733-INS-498
chr7-130973574-INS-54
chr7-130974984-INS-1858
chr7-131015117-INS-402
chr7-131024706-INS-164
chr7-131028631-INS-251
chr7-131049260-INS-116
chr7-131065749-INS-76
chr7-131069183-INS-60
chr7-131078288-INS-149
chr7-131111998-INS-66

chr7-131114598-INS-209
chr7-131127550-INS-520
chr7-131160776-INS-52
chr7-131167316-INS-64
chr7-131169367-INS-197
chr7-131184451-INS-64
chr7-131185115-INS-56
chr7-131198523-INS-157
chr7-131199060-INS-65
chr7-131238489-INS-68
chr7-131248573-INS-227
chr7-131267984-INS-115
chr7-131278899-INS-154
chr7-131297375-INS-196
chr7-131299388-INS-55
chr7-131304243-INS-80
chr7-131309446-INS-69
chr7-131322018-INS-86
chr7-131362107-INS-127
chr7-131385203-INS-80
chr7-131413518-INS-123
chr7-131422269-INS-122
chr7-131422734-INS-139
chr7-131450388-INS-52
chr7-131461422-INS-126
chr7-131473506-INS-52
chr7-131488102-INS-59
chr7-131497323-INS-256
chr7-131545686-INS-66
chr7-131554410-INS-54
chr7-131564665-INS-304
chr7-131603676-INS-145
chr7-131606989-INS-119
chr7-131616081-INS-572
chr7-131618425-INS-172
chr7-131622313-INS-60
chr7-131648489-INS-290
chr7-131659865-INS-174
chr7-131673038-INS-114
chr7-131678521-INS-60
chr7-131687922-INS-51
chr7-131694453-INS-68
chr7-131706069-INS-6405
chr7-131728554-INS-5465
chr7-131734747-INS-6472
chr7-131740563-INS-100
chr7-131742935-INS-121
chr7-131797764-INS-68
chr7-131811849-INS-103
chr7-131820257-INS-106
chr7-131852911-INS-97
chr7-131864257-INS-260
chr7-131873899-INS-52
chr7-131875891-INS-380
chr7-131883102-INS-231
chr7-131899961-INS-128
chr7-131901849-INS-53
chr7-131913291-INS-54
chr7-131913594-INS-96
chr7-131924706-INS-200
chr7-131932293-INS-62
chr7-131979277-INS-73
chr7-131995801-INS-144
chr7-132017680-INS-736
chr7-132024861-INS-51
chr7-132048111-INS-64
chr7-132048980-INS-595
chr7-132052931-INS-130
chr7-132074281-INS-54
chr7-132075854-INS-77
chr7-132084133-INS-63
chr7-132126169-INS-155
chr7-132142091-INS-64
chr7-132154591-INS-306
chr7-132160692-INS-73
chr7-132163477-INS-53
chr7-132164975-INS-54

chr7-132169525-INS-65
chr7-132187127-INS-177
chr7-132214298-INS-53
chr7-132215827-INS-197
chr7-132254489-INS-206
chr7-132264389-INS-68
chr7-132275144-INS-68
chr7-132278010-INS-52
chr7-132288177-INS-79
chr7-132304868-INS-63
chr7-132320005-INS-217
chr7-132327633-INS-149
chr7-132367801-INS-315
chr7-132372092-INS-53
chr7-132423687-INS-185
chr7-132428002-INS-243
chr7-132434084-INS-154
chr7-132435356-INS-5272
chr7-132442347-INS-53
chr7-132443440-INS-182
chr7-132447929-INS-298
chr7-132449550-INS-64
chr7-132481520-INS-399
chr7-132482682-INS-246
chr7-132546841-INS-306
chr7-132737885-INS-5542
chr7-132749972-INS-386
chr7-132760307-INS-200
chr7-132765127-INS-131
chr7-132770167-INS-94
chr7-132778057-INS-131
chr7-132806943-INS-265
chr7-132808127-INS-50
chr7-132813334-INS-117
chr7-132817249-INS-287
chr7-132845881-INS-75
chr7-132880462-INS-98
chr7-132909449-INS-212
chr7-132918560-INS-5549
chr7-132926115-INS-418
chr7-132932020-INS-104
chr7-132938259-INS-82
chr7-132970883-INS-90
chr7-132976301-INS-878
chr7-132978891-INS-57
chr7-132984754-INS-154
chr7-133006571-INS-181
chr7-133053898-INS-56
chr7-133090816-INS-98
chr7-133091076-INS-381
chr7-133114540-INS-174
chr7-133129156-INS-72
chr7-133143233-INS-82
chr7-133165664-INS-79
chr7-133193827-INS-134
chr7-133215824-INS-252
chr7-133226556-INS-118
chr7-133250530-INS-945
chr7-133269655-INS-91
chr7-133271038-INS-154
chr7-133277898-INS-207
chr7-133285011-INS-85
chr7-133306240-INS-152
chr7-133359774-INS-58
chr7-133360902-INS-68
chr7-133380562-INS-98
chr7-133407130-INS-72
chr7-133415892-INS-60
chr7-133417157-INS-72
chr7-133424496-INS-52
chr7-133447080-INS-160
chr7-133448282-INS-140
chr7-133456901-INS-122
chr7-133467803-INS-80
chr7-133469964-INS-68
chr7-133474543-INS-314
chr7-133493869-INS-6415

chr7-133495669-INS-871
chr7-133506933-INS-6605
chr7-133529781-INS-78
chr7-133541900-INS-54
chr7-133547818-INS-611
chr7-133575872-INS-94
chr7-133581794-INS-104
chr7-133612911-INS-91
chr7-133616163-INS-90
chr7-133633523-INS-117
chr7-133634647-INS-206
chr7-133681463-INS-81
chr7-133686811-INS-1086
chr7-133697480-INS-633
chr7-133722251-INS-6452
chr7-133723337-INS-62
chr7-133736687-INS-117
chr7-133756047-INS-58
chr7-133761255-INS-58
chr7-133762916-INS-321
chr7-133806427-INS-57
chr7-133807922-INS-6379
chr7-133818992-INS-150
chr7-133862541-INS-219
chr7-133877982-INS-65
chr7-133894358-INS-150
chr7-133906034-INS-50
chr7-133954621-INS-72
chr7-133974451-INS-53
chr7-133992579-INS-52
chr7-134017720-INS-240
chr7-134038812-INS-92
chr7-134042797-INS-2815
chr7-134045147-INS-516
chr7-134072905-INS-548
chr7-134079097-INS-66
chr7-134085972-INS-198
chr7-134098236-INS-2175
chr7-134122097-INS-405
chr7-134128475-INS-143
chr7-134129540-INS-64
chr7-134186189-INS-971
chr7-134196006-INS-6441
chr7-134207281-INS-442
chr7-134211287-INS-52
chr7-134218120-INS-6162
chr7-134234581-INS-146
chr7-134242708-INS-134
chr7-134268218-INS-87
chr7-134278274-INS-127
chr7-134288495-INS-90
chr7-134294712-INS-68
chr7-134313393-INS-543
chr7-134330713-INS-371
chr7-134341611-INS-116
chr7-134357689-INS-7344
chr7-134382345-INS-64
chr7-134397961-INS-75
chr7-134415026-INS-150
chr7-134446140-INS-240
chr7-134446909-INS-120
chr7-134452263-INS-484
chr7-134455737-INS-138
chr7-134461144-INS-147
chr7-134461358-INS-1273
chr7-134488903-INS-212
chr7-134499962-INS-136
chr7-134512911-INS-77
chr7-134528061-INS-58
chr7-134529479-INS-62
chr7-134548056-INS-58
chr7-134578643-INS-68
chr7-134581576-INS-152
chr7-134620898-INS-52
chr7-134681812-INS-164
chr7-134711917-INS-54
chr7-134782441-INS-125

chr7-134790826-INS-57
chr7-134795130-INS-91
chr7-134825580-INS-140
chr7-134891270-INS-52
chr7-134897804-INS-142
chr7-134902200-INS-133
chr7-134931469-INS-207
chr7-134936202-INS-171
chr7-134957824-INS-363
chr7-134966775-INS-112
chr7-134969912-INS-91
chr7-134976948-INS-104
chr7-135007576-INS-72
chr7-135017197-INS-53
chr7-135021604-INS-393
chr7-135043849-INS-208
chr7-135045818-INS-63
chr7-135051344-INS-104
chr7-135063432-INS-204
chr7-135064752-INS-125
chr7-135065808-INS-60
chr7-135066888-INS-234
chr7-135075238-INS-6156
chr7-135087904-INS-1892
chr7-135094344-INS-99
chr7-135132178-INS-98
chr7-135147353-INS-59
chr7-135168794-INS-165
chr7-135189150-INS-56
chr7-135191134-INS-146
chr7-135207274-INS-81
chr7-135266463-INS-118
chr7-135275739-INS-69
chr7-135284909-INS-177
chr7-135319455-INS-58
chr7-135366532-INS-79
chr7-135373548-INS-199
chr7-135375345-INS-64
chr7-135387745-INS-866
chr7-135387786-INS-593
chr7-135392092-INS-69
chr7-135401962-INS-174
chr7-135413245-INS-100
chr7-135417057-INS-88
chr7-135429937-INS-2830
chr7-135466212-INS-131
chr7-135487253-INS-60
chr7-135510028-INS-4484
chr7-135512924-INS-60
chr7-135514390-INS-67
chr7-135515628-INS-149
chr7-135526033-INS-52
chr7-135527877-INS-67
chr7-135552109-INS-6384
chr7-135566139-INS-88
chr7-135579353-INS-307
chr7-135591578-INS-110
chr7-135592906-INS-218
chr7-135613946-INS-76
chr7-135623876-INS-339
chr7-135641831-INS-68
chr7-135681086-INS-218
chr7-135701571-INS-54
chr7-135713224-INS-200
chr7-135744848-INS-53
chr7-135780923-INS-59
chr7-135805433-INS-1164
chr7-135812492-INS-56
chr7-135823067-INS-205
chr7-135874768-INS-90
chr7-135955714-INS-54
chr7-135969682-INS-6401
chr7-135978614-INS-228
chr7-135978723-INS-674
chr7-135986906-INS-111
chr7-135996552-INS-136
chr7-136026761-INS-168

chr7-136028684-INS-196
chr7-136061053-INS-66
chr7-136088423-INS-63
chr7-136175329-INS-559
chr7-136193424-INS-3298
chr7-136193705-INS-120
chr7-136214806-INS-56
chr7-136229054-INS-74
chr7-136255928-INS-354
chr7-136263962-INS-73
chr7-136286215-INS-52
chr7-136300476-INS-135
chr7-136315484-INS-51
chr7-136320114-INS-80
chr7-136327084-INS-108
chr7-136331399-INS-57
chr7-136347378-INS-222
chr7-136372057-INS-313
chr7-136375739-INS-456
chr7-136378029-INS-80
chr7-136388340-INS-66
chr7-136389035-INS-117
chr7-136404942-INS-108
chr7-136409434-INS-296
chr7-136431492-INS-62
chr7-136477468-INS-6606
chr7-136483544-INS-72
chr7-136498440-INS-114
chr7-136512504-INS-244
chr7-136535441-INS-58
chr7-136551536-INS-136
chr7-136620129-INS-120
chr7-136623938-INS-148
chr7-136658893-INS-94
chr7-136697560-INS-70
chr7-136699944-INS-79
chr7-136716604-INS-420
chr7-136737471-INS-111
chr7-136740790-INS-274
chr7-136777009-INS-65
chr7-136787561-INS-826
chr7-136791222-INS-227
chr7-136884138-INS-72
chr7-136893553-INS-117
chr7-136899411-INS-79
chr7-136943962-INS-57
chr7-136996512-INS-363
chr7-136997602-INS-209
chr7-137011497-INS-6401
chr7-137015710-INS-83
chr7-137023048-INS-402
chr7-137038156-INS-188
chr7-137041511-INS-202
chr7-137043363-INS-305
chr7-137052625-INS-221
chr7-137055685-INS-56
chr7-137057945-INS-176
chr7-137060719-INS-98
chr7-137070995-INS-64
chr7-137077774-INS-196
chr7-137134793-INS-189
chr7-137140730-INS-127
chr7-137145156-INS-522
chr7-137167614-INS-51
chr7-137191591-INS-72
chr7-137196608-INS-61
chr7-137204752-INS-361
chr7-137236623-INS-76
chr7-137254357-INS-621
chr7-137260820-INS-102
chr7-137295593-INS-444
chr7-137304356-INS-283
chr7-137321288-INS-349
chr7-137358028-INS-51
chr7-137378229-INS-193
chr7-137387348-INS-572
chr7-137388908-INS-6577

chr7-137435335-INS-2224
chr7-137444860-INS-334
chr7-137458006-INS-108
chr7-137488904-INS-51
chr7-137510738-INS-103
chr7-137520662-INS-61
chr7-137521329-INS-154
chr7-137540034-INS-400
chr7-137607576-INS-125
chr7-137618299-INS-173
chr7-137653767-INS-184
chr7-137656858-INS-98
chr7-137664250-INS-59
chr7-137675851-INS-62
chr7-137706369-INS-188
chr7-137709958-INS-162
chr7-137712344-INS-132
chr7-137716427-INS-56
chr7-137733297-INS-56
chr7-137745167-INS-114
chr7-137746929-INS-60
chr7-137770989-INS-60
chr7-137784311-INS-81
chr7-137831991-INS-160
chr7-137838502-INS-80
chr7-137888713-INS-82
chr7-137961967-INS-231
chr7-137980611-INS-170
chr7-137997432-INS-76
chr7-138012142-INS-168
chr7-138018783-INS-80
chr7-138034061-INS-53
chr7-138091777-INS-135
chr7-138102576-INS-6876
chr7-138147091-INS-55
chr7-138154890-INS-116
chr7-138173271-INS-56
chr7-138174083-INS-64
chr7-138191627-INS-147
chr7-138207364-INS-68
chr7-138228673-INS-54
chr7-138232859-INS-80
chr7-138263823-INS-117
chr7-138278356-INS-111
chr7-138283424-INS-828
chr7-138299779-INS-135
chr7-138326856-INS-909
chr7-138337862-INS-666
chr7-138343097-INS-177
chr7-138346189-INS-132
chr7-138364623-INS-148
chr7-138374611-INS-84
chr7-138377825-INS-512
chr7-138386869-INS-180
chr7-138392723-INS-51
chr7-138400324-INS-85
chr7-138403076-INS-61
chr7-138431064-INS-140
chr7-138431843-INS-153
chr7-138452757-INS-163
chr7-138472540-INS-933
chr7-138485882-INS-521
chr7-138495018-INS-114
chr7-138506214-INS-271
chr7-138514597-INS-126
chr7-138527527-INS-400
chr7-138539055-INS-84
chr7-138553246-INS-61
chr7-138555152-INS-89
chr7-138557137-INS-100
chr7-138573989-INS-70
chr7-138575816-INS-150
chr7-138619775-INS-59
chr7-138653198-INS-88
chr7-138762480-INS-326
chr7-138778726-INS-110
chr7-138783936-INS-271

chr7-138785861-INS-348
chr7-138812845-INS-625
chr7-138857506-INS-142
chr7-138901661-INS-172
chr7-138906326-INS-846
chr7-138927553-INS-6360
chr7-138937878-INS-93
chr7-138983393-INS-196
chr7-139003677-INS-633
chr7-139025255-INS-54
chr7-139027877-INS-347
chr7-139038112-INS-66
chr7-139041223-INS-680
chr7-139042853-INS-201
chr7-139058137-INS-639
chr7-139085418-INS-75
chr7-139105164-INS-503
chr7-139132216-INS-52
chr7-139133190-INS-67
chr7-139139461-INS-56
chr7-139169213-INS-129
chr7-139171869-INS-7156
chr7-139180089-INS-335
chr7-139188474-INS-329
chr7-139232798-INS-60
chr7-139237608-INS-152
chr7-139268360-INS-51
chr7-139288501-INS-56
chr7-139290015-INS-774
chr7-139290560-INS-175
chr7-139296638-INS-117
chr7-139333668-INS-61
chr7-139372447-INS-52
chr7-139382561-INS-180
chr7-139388571-INS-334
chr7-139394835-INS-52
chr7-139401815-INS-423
chr7-139425749-INS-61
chr7-139473704-INS-165
chr7-139483533-INS-99
chr7-139487105-INS-95
chr7-139505217-INS-168
chr7-139527585-INS-200
chr7-139570502-INS-120
chr7-139579261-INS-286
chr7-139582369-INS-454
chr7-139623766-INS-357
chr7-139647559-INS-566
chr7-139650914-INS-502
chr7-139656006-INS-212
chr7-139669041-INS-199
chr7-139723792-INS-55
chr7-139737659-INS-159
chr7-139740901-INS-889
chr7-139758947-INS-5061
chr7-139759065-INS-56
chr7-139762832-INS-60
chr7-139790098-INS-60
chr7-139801162-INS-694
chr7-139806109-INS-458
chr7-139828257-INS-59
chr7-139840623-INS-180
chr7-139844878-INS-149
chr7-139850418-INS-183
chr7-139855007-INS-67
chr7-139860813-INS-199
chr7-139890835-INS-3527
chr7-139891045-INS-3910
chr7-139917699-INS-124
chr7-139920453-INS-89
chr7-139922745-INS-58
chr7-139924440-INS-213
chr7-139943790-INS-6819
chr7-139955607-INS-73
chr7-139968173-INS-72
chr7-139989760-INS-6442
chr7-140027210-INS-6390

chr7-140028821-INS-3968
chr7-140037112-INS-51
chr7-140043790-INS-68
chr7-140055437-INS-1893
chr7-140056782-INS-6226
chr7-140060475-INS-3149
chr7-140125729-INS-6311
chr7-140136430-INS-775
chr7-140151023-INS-6393
chr7-140172914-INS-1103
chr7-140204674-INS-6908
chr7-140221252-INS-4586
chr7-140252348-INS-6853
chr7-140253094-INS-64
chr7-140259807-INS-1887
chr7-140285750-INS-99
chr7-140306541-INS-57
chr7-140313922-INS-958
chr7-140355284-INS-140
chr7-140393489-INS-92
chr7-140399365-INS-324
chr7-140444064-INS-580
chr7-140446513-INS-240
chr7-140451373-INS-192
chr7-140453615-INS-203
chr7-140455634-INS-83
chr7-140473444-INS-52
chr7-140505951-INS-186
chr7-140510208-INS-176
chr7-140533567-INS-169
chr7-140533567-INS-844
chr7-140537801-INS-52
chr7-140538977-INS-535
chr7-140544398-INS-407
chr7-140546365-INS-359
chr7-140548410-INS-149
chr7-140552387-INS-173
chr7-140558237-INS-391
chr7-140559739-INS-112
chr7-140565460-INS-163
chr7-140569408-INS-189
chr7-140575337-INS-259
chr7-140576849-INS-112
chr7-140580521-INS-179
chr7-140587713-INS-149
chr7-140588177-INS-200
chr7-140594954-INS-337
chr7-140602128-INS-93
chr7-140616578-INS-204
chr7-140666286-INS-52
chr7-140686132-INS-370
chr7-140692341-INS-206
chr7-140706213-INS-78
chr7-140709192-INS-73
chr7-140717560-INS-230
chr7-140737754-INS-193
chr7-140742106-INS-70
chr7-140750917-INS-72
chr7-140751669-INS-226
chr7-140760238-INS-247
chr7-140767665-INS-163
chr7-140780698-INS-68
chr7-140783472-INS-498
chr7-140831853-INS-52
chr7-140878179-INS-221
chr7-140879113-INS-214
chr7-140892219-INS-580
chr7-140899274-INS-147
chr7-140911870-INS-194
chr7-140939023-INS-70
chr7-140939417-INS-205
chr7-140950906-INS-278
chr7-140958306-INS-98
chr7-140959455-INS-219
chr7-140960394-INS-201
chr7-141005031-INS-198
chr7-141051443-INS-260

chr7-141086813-INS-158
chr7-141093004-INS-212
chr7-141107171-INS-188
chr7-141111659-INS-119
chr7-141131611-INS-68
chr7-141133936-INS-199
chr7-141134660-INS-203
chr7-141151951-INS-190
chr7-141163973-INS-303
chr7-141179179-INS-227
chr7-141217946-INS-1326
chr7-141219084-INS-251
chr7-141219847-INS-1845
chr7-141221227-INS-300
chr7-141225447-INS-801
chr7-141247228-INS-98
chr7-141253506-INS-483
chr7-141278553-INS-222
chr7-141285251-INS-138
chr7-141292037-INS-85
chr7-141297147-INS-288
chr7-141301449-INS-74
chr7-141304031-INS-60
chr7-141310645-INS-128
chr7-141336891-INS-60
chr7-141357320-INS-72
chr7-141362384-INS-143
chr7-141379133-INS-58
chr7-141397140-INS-124
chr7-141412631-INS-525
chr7-141450505-INS-150
chr7-141454888-INS-5088
chr7-141469844-INS-56
chr7-141511908-INS-58
chr7-141564759-INS-92
chr7-141584416-INS-68
chr7-141598644-INS-203
chr7-141625417-INS-184
chr7-141641613-INS-76
chr7-141657970-INS-331
chr7-141662309-INS-54
chr7-141665822-INS-63
chr7-141689104-INS-60
chr7-141701153-INS-313
chr7-141707478-INS-6485
chr7-141710654-INS-60
chr7-141715486-INS-56
chr7-141718768-INS-84
chr7-141736221-INS-208
chr7-141743604-INS-203
chr7-141755630-INS-77
chr7-141759150-INS-476
chr7-141760591-INS-476
chr7-141783349-INS-183
chr7-141786511-INS-130
chr7-141794512-INS-63
chr7-141805174-INS-192
chr7-141809215-INS-143
chr7-141850268-INS-54
chr7-141861649-INS-124
chr7-141960486-INS-55
chr7-141963015-INS-236
chr7-141968504-INS-70
chr7-142001003-INS-213
chr7-142020070-INS-127
chr7-142073273-INS-180
chr7-142114658-INS-349
chr7-142139980-INS-267
chr7-142168590-INS-346
chr7-142170505-INS-424
chr7-142174436-INS-178
chr7-142198518-INS-110
chr7-142225960-INS-130
chr7-142268537-INS-80
chr7-142290622-INS-288
chr7-142292169-INS-205
chr7-142344986-INS-508

chr7-142389755-INS-345
chr7-142404302-INS-932
chr7-142418385-INS-52
chr7-142429890-INS-6340
chr7-142440678-INS-324
chr7-142498196-INS-87
chr7-142502731-INS-52
chr7-142509388-INS-60
chr7-142530091-INS-1409
chr7-142543612-INS-75
chr7-142578843-INS-54
chr7-142597398-INS-114
chr7-142604386-INS-191
chr7-142658862-INS-72
chr7-142679338-INS-372
chr7-142691498-INS-56
chr7-142700560-INS-74
chr7-142783778-INS-65
chr7-142799361-INS-206
chr7-142868792-INS-56
chr7-142890328-INS-57
chr7-142909290-INS-81
chr7-142945145-INS-63
chr7-142958836-INS-76
chr7-143064038-INS-125
chr7-143082494-INS-202
chr7-143090412-INS-222
chr7-143111877-INS-199
chr7-143127659-INS-226
chr7-143134236-INS-65
chr7-143158787-INS-64
chr7-143174920-INS-55
chr7-143178417-INS-401
chr7-143181193-INS-124
chr7-143190695-INS-211
chr7-143198897-INS-51
chr7-143216400-INS-244
chr7-143236606-INS-173
chr7-143251761-INS-60
chr7-143302841-INS-80
chr7-143325378-INS-865
chr7-143327088-INS-63
chr7-143341009-INS-102
chr7-143342071-INS-53
chr7-143355749-INS-72
chr7-143363995-INS-6412
chr7-143420860-INS-60
chr7-143431731-INS-93
chr7-143439462-INS-61
chr7-143446638-INS-104
chr7-143452603-INS-3583
chr7-143475740-INS-441
chr7-143497911-INS-241
chr7-143498743-INS-6969
chr7-143512616-INS-890
chr7-143523744-INS-345
chr7-143524634-INS-4827
chr7-143577887-INS-56
chr7-143611081-INS-59
chr7-143614385-INS-54
chr7-143630696-INS-54
chr7-143658437-INS-61
chr7-143663195-INS-148
chr7-143675995-INS-213
chr7-143687886-INS-74
chr7-143712178-INS-130
chr7-143753170-INS-51
chr7-143759154-INS-82
chr7-143809112-INS-67
chr7-143818296-INS-702
chr7-143864191-INS-126
chr7-143874474-INS-72
chr7-143901691-INS-62
chr7-143937549-INS-93
chr7-143983349-INS-52
chr7-144022336-INS-209
chr7-144076992-INS-91

chr7-144084306-INS-63
chr7-144106988-INS-5897
chr7-144109741-INS-171
chr7-144113432-INS-202
chr7-144113532-INS-428
chr7-144157569-INS-76
chr7-144179514-INS-55
chr7-144194430-INS-52
chr7-144249758-INS-103
chr7-144320036-INS-531
chr7-144321878-INS-5627
chr7-144322685-INS-128
chr7-144339278-INS-88
chr7-144346186-INS-270
chr7-144363161-INS-53
chr7-144370356-INS-205
chr7-144385405-INS-80
chr7-144404266-INS-203
chr7-144406418-INS-97
chr7-144420261-INS-69
chr7-144425788-INS-498
chr7-144430072-INS-171
chr7-144448193-INS-62
chr7-144463180-INS-160
chr7-144464473-INS-282
chr7-144480030-INS-66
chr7-144502118-INS-5191
chr7-144515118-INS-201
chr7-144521531-INS-121
chr7-144527714-INS-2037
chr7-144550623-INS-1890
chr7-144577261-INS-198
chr7-144622986-INS-101
chr7-144627574-INS-85
chr7-144689357-INS-59
chr7-144694295-INS-1894
chr7-144757807-INS-138
chr7-144777338-INS-80
chr7-144800819-INS-6444
chr7-144859148-INS-7069
chr7-144873162-INS-200
chr7-144874243-INS-58
chr7-144882862-INS-193
chr7-144884239-INS-61
chr8-3055260-INS-52
chr8-3071401-INS-5412
chr8-3105762-INS-4952
chr8-3125650-INS-100
chr8-3156080-INS-59
chr8-3163895-INS-5553
chr8-3168125-INS-53
chr8-3175974-INS-296
chr8-3191042-INS-87
chr8-3205325-INS-124
chr8-3230455-INS-52
chr8-3267277-INS-499
chr8-3270671-INS-77
chr8-3307495-INS-68
chr8-3331336-INS-178
chr8-3342343-INS-110
chr8-3350668-INS-519
chr8-3381123-INS-132
chr8-3397654-INS-120
chr8-3414158-INS-56
chr8-3471057-INS-235
chr8-3481631-INS-149
chr8-3483744-INS-67
chr8-3490714-INS-616
chr8-3492472-INS-95
chr8-3522211-INS-212
chr8-3536186-INS-105
chr8-3580674-INS-496
chr8-3596576-INS-272
chr8-3604251-INS-568
chr8-3606288-INS-564
chr8-3611122-INS-168
chr8-3628667-INS-197

chr8-3679806-INS-184
chr8-3700060-INS-400
chr8-3709046-INS-205
chr8-3716327-INS-53
chr8-3742011-INS-212
chr8-3746556-INS-69
chr8-3755863-INS-336
chr8-3800135-INS-6413
chr8-3820009-INS-117
chr8-3867510-INS-400
chr8-3901839-INS-493
chr8-3919114-INS-6459
chr8-3986152-INS-484
chr8-3992785-INS-199
chr8-4056596-INS-241
chr8-4069166-INS-66
chr8-4134232-INS-267
chr8-4136091-INS-338
chr8-4138913-INS-82
chr8-4144674-INS-238
chr8-4145866-INS-390
chr8-4161368-INS-384
chr8-4190515-INS-52
chr8-4203503-INS-332
chr8-4323695-INS-138
chr8-4341166-INS-207
chr8-4368513-INS-214
chr8-4414170-INS-122
chr8-4417136-INS-139
chr8-4434120-INS-209
chr8-4439681-INS-5247
chr8-4468490-INS-2853
chr8-4479121-INS-523
chr8-4518579-INS-227
chr8-4544978-INS-63
chr8-4563379-INS-454
chr8-4577865-INS-291
chr8-4582068-INS-315
chr8-4589897-INS-200
chr8-4619574-INS-52
chr8-4664003-INS-549
chr8-4676394-INS-280
chr8-4703752-INS-133
chr8-4707289-INS-64
chr8-4711208-INS-221
chr8-4720384-INS-84
chr8-4724514-INS-565
chr8-4736717-INS-1864
chr8-4744848-INS-101
chr8-4756418-INS-332
chr8-4780107-INS-228
chr8-4781027-INS-64
chr8-4781826-INS-488
chr8-4843048-INS-97
chr8-4863245-INS-108
chr8-4873393-INS-1890
chr8-4910551-INS-51
chr8-4912075-INS-5903
chr8-4925387-INS-465
chr8-4928189-INS-6531
chr8-4930792-INS-6672
chr8-4930894-INS-7096
chr8-5054492-INS-60
chr8-5084456-INS-75
chr8-5087825-INS-196
chr8-5092094-INS-355
chr8-5105685-INS-152
chr8-5120605-INS-512
chr8-5124228-INS-197
chr8-5126109-INS-2029
chr8-5129815-INS-266
chr8-5134129-INS-104
chr8-5139927-INS-56
chr8-5196074-INS-344
chr8-5285475-INS-53
chr8-5287877-INS-122
chr8-5334447-INS-198

chr8-5373161-INS-111
chr8-5453118-INS-482
chr8-5473397-INS-262
chr8-5524946-INS-117
chr8-5527965-INS-349
chr8-5576911-INS-161
chr8-5593009-INS-123
chr8-5618202-INS-98
chr8-5630492-INS-74
chr8-5642923-INS-204
chr8-5647800-INS-493
chr8-5649326-INS-3174
chr8-5656195-INS-6389
chr8-5675362-INS-119
chr8-5730862-INS-286
chr8-5791426-INS-167
chr8-5799481-INS-102
chr8-5805866-INS-332
chr8-5867919-INS-83
chr8-5891317-INS-194
chr8-5899127-INS-190
chr8-5924378-INS-52
chr8-5941767-INS-180
chr8-5946966-INS-258
chr8-5947098-INS-7112
chr8-5948350-INS-72
chr8-5999269-INS-58
chr8-6030873-INS-84
chr8-6062088-INS-90
chr8-6098860-INS-86
chr8-6118657-INS-125
chr8-6123358-INS-92
chr8-6156669-INS-109
chr8-6158771-INS-3346
chr8-6160702-INS-535
chr8-6178867-INS-65
chr8-6179820-INS-250
chr8-6218499-INS-728
chr8-6231729-INS-154
chr8-6233155-INS-56
chr8-6234248-INS-92
chr8-6252045-INS-66
chr8-6266204-INS-5548
chr8-6287841-INS-330
chr8-6304460-INS-549
chr8-6333309-INS-120
chr8-6334406-INS-358
chr8-6345949-INS-73
chr8-6400802-INS-178
chr8-6401222-INS-115
chr8-6403324-INS-56
chr8-6420678-INS-109
chr8-6430253-INS-71
chr8-6461022-INS-507
chr8-6503272-INS-51
chr8-6505438-INS-67
chr8-6520751-INS-505
chr8-6543335-INS-93
chr8-6556306-INS-6642
chr8-6602146-INS-156
chr8-6633471-INS-134
chr8-6637191-INS-6687
chr8-6637457-INS-96
chr8-6659479-INS-565
chr8-6673964-INS-7064
chr8-6718784-INS-8277
chr8-6726960-INS-52
chr8-6738988-INS-186
chr8-6746150-INS-407
chr8-6765933-INS-6196
chr8-6835179-INS-102
chr8-6896042-INS-54
chr8-6924964-INS-120
chr8-6930571-INS-499
chr8-6961608-INS-51
chr8-7009615-INS-197
chr8-7054082-INS-754

chr8-7058345-INS-221
chr8-7061108-INS-268
chr8-7078517-INS-307
chr8-7095783-INS-197
chr8-7103458-INS-194
chr8-7127235-INS-295
chr8-7204071-INS-491
chr8-7215090-INS-208
chr8-7216528-INS-96
chr8-7217475-INS-321
chr8-7225525-INS-68
chr8-7249166-INS-371
chr8-7257057-INS-809
chr8-7261124-INS-183
chr8-7272248-INS-72
chr8-7295584-INS-413
chr8-7309457-INS-6341
chr8-7331270-INS-132
chr8-7332734-INS-59
chr8-7358007-INS-176
chr8-7384205-INS-6310
chr8-7414049-INS-7016
chr8-7428294-INS-142
chr8-7473669-INS-226
chr8-7479967-INS-95
chr8-7482327-INS-201
chr8-7521299-INS-112
chr8-7524531-INS-99
chr8-7533824-INS-184
chr8-7554712-INS-103
chr8-7565805-INS-56
chr8-7601615-INS-334
chr8-7604489-INS-254
chr8-7642962-INS-654
chr8-7649620-INS-111
chr8-7654851-INS-270
chr8-7677507-INS-56
chr8-7698259-INS-959
chr8-7753757-INS-72
chr8-7773822-INS-64
chr8-7848563-INS-6189
chr8-7857679-INS-6640
chr8-7860642-INS-6259
chr8-7877219-INS-6443
chr8-7893263-INS-156
chr8-7920838-INS-52
chr8-7934130-INS-145
chr8-7955666-INS-285
chr8-7990587-INS-138
chr8-7992781-INS-95
chr8-7999735-INS-254
chr8-8103913-INS-78
chr8-8104171-INS-7139
chr8-8109208-INS-6927
chr8-8109591-INS-532
chr8-8160488-INS-64
chr8-8172304-INS-417
chr8-8266514-INS-6921
chr8-8266747-INS-213
chr8-8274968-INS-57
chr8-8293911-INS-70
chr8-8307241-INS-60
chr8-8313245-INS-785
chr8-8349159-INS-219
chr8-8351647-INS-78
chr8-8353851-INS-80
chr8-8401757-INS-236
chr8-8462462-INS-132
chr8-8482787-INS-124
chr8-8488159-INS-397
chr8-8496900-INS-247
chr8-8504675-INS-391
chr8-8509288-INS-52
chr8-8519868-INS-392
chr8-8545212-INS-185
chr8-8554581-INS-104
chr8-8555984-INS-123

chr8-8575305-INS-66
chr8-8592279-INS-70
chr8-8610830-INS-66
chr8-8630798-INS-87
chr8-8656800-INS-76
chr8-8734056-INS-213
chr8-8797822-INS-53
chr8-8829240-INS-895
chr8-8859149-INS-60
chr8-8878568-INS-764
chr8-8901326-INS-140
chr8-8927943-INS-3666
chr8-8938947-INS-489
chr8-8966801-INS-96
chr8-8977423-INS-75
chr8-8983860-INS-251
chr8-8985234-INS-139
chr8-8994636-INS-57
chr8-8996547-INS-186
chr8-8998937-INS-62
chr8-9007201-INS-56
chr8-9016840-INS-227
chr8-9030518-INS-721
chr8-9046532-INS-52
chr8-9059699-INS-423
chr8-9082046-INS-193
chr8-9099266-INS-506
chr8-9107227-INS-74
chr8-9126452-INS-156
chr8-9129682-INS-244
chr8-9157879-INS-105
chr8-9160046-INS-506
chr8-9205475-INS-99
chr8-9213432-INS-52
chr8-9235917-INS-67
chr8-9235930-INS-7474
chr8-9249740-INS-86
chr8-9252456-INS-66
chr8-9270453-INS-72
chr8-9286641-INS-152
chr8-9317323-INS-358
chr8-9319162-INS-158
chr8-9347389-INS-161
chr8-9347577-INS-1350
chr8-9351068-INS-57
chr8-9357583-INS-86
chr8-9362848-INS-195
chr8-9368456-INS-90
chr8-9375935-INS-828
chr8-9412382-INS-416
chr8-9419090-INS-56
chr8-9420831-INS-51
chr8-9452421-INS-554
chr8-9458329-INS-156
chr8-9466377-INS-145
chr8-9477189-INS-348
chr8-9519517-INS-524
chr8-9542258-INS-63
chr8-9565181-INS-145
chr8-9572557-INS-516
chr8-9610506-INS-64
chr8-9614833-INS-141
chr8-9620093-INS-62
chr8-9644267-INS-342
chr8-9666111-INS-67
chr8-9693603-INS-75
chr8-9702284-INS-64
chr8-9719869-INS-89
chr8-9765115-INS-340
chr8-9779289-INS-305
chr8-9786508-INS-199
chr8-9806580-INS-108
chr8-9812325-INS-73
chr8-9824686-INS-58
chr8-9840464-INS-237
chr8-9876090-INS-115
chr8-9876903-INS-358

chr8-9917385-INS-565
chr8-9924092-INS-405
chr8-9926698-INS-294
chr8-9956920-INS-455
chr8-9967245-INS-7078
chr8-9973401-INS-133
chr8-9975556-INS-79
chr8-9979457-INS-120
chr8-9986367-INS-6903
chr8-9989137-INS-52
chr8-10003247-INS-64
chr8-10027595-INS-268
chr8-10045044-INS-52
chr8-10053290-INS-350
chr8-10089726-INS-76
chr8-10125088-INS-52
chr8-10155379-INS-269
chr8-10158144-INS-202
chr8-10161704-INS-1160
chr8-10161718-INS-120
chr8-10168526-INS-3973
chr8-10177126-INS-389
chr8-10177984-INS-222
chr8-10178245-INS-1295
chr8-10182411-INS-6595
chr8-10185764-INS-184
chr8-10203335-INS-68
chr8-10218404-INS-65
chr8-10234135-INS-66
chr8-10235402-INS-359
chr8-10235562-INS-623
chr8-10238664-INS-121
chr8-10242348-INS-75
chr8-10258688-INS-181
chr8-10315572-INS-101
chr8-10325426-INS-65
chr8-10334675-INS-1033
chr8-10336589-INS-979
chr8-10360913-INS-68
chr8-10374155-INS-144
chr8-10392985-INS-51
chr8-10398579-INS-93
chr8-10402972-INS-239
chr8-10410696-INS-60
chr8-10415060-INS-53
chr8-10418937-INS-122
chr8-10428618-INS-119
chr8-10436349-INS-69
chr8-10455005-INS-141
chr8-10473079-INS-53
chr8-10539190-INS-3839
chr8-10544479-INS-6586
chr8-10548690-INS-110
chr8-10560820-INS-194
chr8-10625297-INS-129
chr8-10634510-INS-618
chr8-10649949-INS-925
chr8-10662441-INS-54
chr8-10671805-INS-93
chr8-10682228-INS-160
chr8-10689380-INS-104
chr8-10718323-INS-259
chr8-10723517-INS-120
chr8-10746596-INS-60
chr8-10766709-INS-331
chr8-10775667-INS-147
chr8-10789812-INS-54
chr8-10802160-INS-51
chr8-10812736-INS-56
chr8-10830792-INS-64
chr8-10834993-INS-64
chr8-10836404-INS-124
chr8-10841491-INS-63
chr8-10847190-INS-78
chr8-10853702-INS-54
chr8-10856900-INS-440
chr8-10858904-INS-687

chr8-10892150-INS-72
chr8-10900402-INS-73
chr8-10903664-INS-734
chr8-10905416-INS-56
chr8-10910523-INS-77
chr8-10918906-INS-78
chr8-10925584-INS-53
chr8-10931596-INS-94
chr8-10932132-INS-2495
chr8-10932185-INS-403
chr8-10948886-INS-54
chr8-10950699-INS-53
chr8-10956135-INS-145
chr8-10962810-INS-490
chr8-10980174-INS-60
chr8-10981786-INS-64
chr8-10984038-INS-64
chr8-10989232-INS-148
chr8-10990575-INS-175
chr8-10991313-INS-401
chr8-10996983-INS-90
chr8-11006236-INS-147
chr8-11013298-INS-198
chr8-11014098-INS-66
chr8-11018426-INS-399
chr8-11022565-INS-137
chr8-11035636-INS-60
chr8-11045401-INS-260
chr8-11047645-INS-329
chr8-11048760-INS-180
chr8-11065206-INS-92
chr8-11081189-INS-245
chr8-11100149-INS-133
chr8-11103591-INS-75
chr8-11111413-INS-112
chr8-11114615-INS-108
chr8-11120973-INS-96
chr8-11123278-INS-54
chr8-11134141-INS-139
chr8-11137398-INS-55
chr8-11219424-INS-52
chr8-11237447-INS-164
chr8-11264350-INS-54
chr8-11290152-INS-261
chr8-11307992-INS-80
chr8-11357390-INS-96
chr8-11368066-INS-113
chr8-11384534-INS-60
chr8-11405584-INS-52
chr8-11415683-INS-83
chr8-11422856-INS-66
chr8-11440947-INS-75
chr8-11457831-INS-203
chr8-11459270-INS-104
chr8-11474419-INS-54
chr8-11486127-INS-98
chr8-11487239-INS-524
chr8-11497443-INS-198
chr8-11509531-INS-193
chr8-11521274-INS-118
chr8-11524760-INS-168
chr8-11541997-INS-72
chr8-11544830-INS-439
chr8-11553972-INS-62
chr8-11564965-INS-103
chr8-11575718-INS-147
chr8-11592894-INS-200
chr8-11595852-INS-252
chr8-11617573-INS-105
chr8-11636216-INS-196
chr8-11671988-INS-205
chr8-11680443-INS-178
chr8-11694709-INS-508
chr8-11696292-INS-243
chr8-11698351-INS-6439
chr8-11717141-INS-222
chr8-11721521-INS-401

chr8-11738948-INS-169
chr8-11756892-INS-183
chr8-11760768-INS-127
chr8-11762777-INS-177
chr8-11764388-INS-116
chr8-11793988-INS-324
chr8-11816190-INS-73
chr8-11816622-INS-206
chr8-11835307-INS-52
chr8-11855896-INS-133
chr8-11862071-INS-171
chr8-11887365-INS-68
chr8-11900624-INS-237
chr8-11939965-INS-110
chr8-11958176-INS-153
chr8-11977704-INS-65
chr8-12005119-INS-155
chr8-12012303-INS-391
chr8-12017427-INS-65
chr8-12029905-INS-176
chr8-12045694-INS-206
chr8-12049766-INS-127
chr8-12059292-INS-138
chr8-12069704-INS-2315
chr8-12071781-INS-179
chr8-12102856-INS-57
chr8-12105066-INS-54
chr8-12158775-INS-260
chr8-12160493-INS-65
chr8-12191585-INS-133
chr8-12218672-INS-449
chr8-12245788-INS-54
chr8-12250522-INS-58
chr8-12292882-INS-97
chr8-12295691-INS-66
chr8-12298746-INS-110
chr8-12319292-INS-106
chr8-12337321-INS-377
chr8-12386190-INS-3298
chr8-12389319-INS-75
chr8-12393110-INS-180
chr8-12409407-INS-880
chr8-12416616-INS-58
chr8-12426547-INS-806
chr8-12428270-INS-165
chr8-12451907-INS-200
chr8-12454575-INS-115
chr8-12459770-INS-90
chr8-12464965-INS-54
chr8-12467571-INS-229
chr8-12469524-INS-209
chr8-12556628-INS-80
chr8-12561187-INS-52
chr8-12571637-INS-165
chr8-12582852-INS-249
chr8-12616567-INS-204
chr8-12624113-INS-94
chr8-12631804-INS-206
chr8-12637537-INS-200
chr8-12640085-INS-61
chr8-12656041-INS-193
chr8-12666528-INS-60
chr8-12679997-INS-65
chr8-12693270-INS-75
chr8-12707334-INS-76
chr8-12716305-INS-263
chr8-12729197-INS-330
chr8-12737298-INS-435
chr8-12760201-INS-201
chr8-12771886-INS-112
chr8-12772990-INS-92
chr8-12779992-INS-123
chr8-12794099-INS-509
chr8-12819747-INS-68
chr8-12845814-INS-190
chr8-12907508-INS-286
chr8-12954500-INS-5543

chr8-12969137-INS-140
chr8-13021595-INS-50
chr8-13028590-INS-78
chr8-13073194-INS-75
chr8-13104918-INS-208
chr8-13111308-INS-179
chr8-13121070-INS-72
chr8-13125751-INS-120
chr8-13133573-INS-296
chr8-13134672-INS-807
chr8-13138916-INS-57
chr8-13150996-INS-426
chr8-13168674-INS-55
chr8-13207386-INS-117
chr8-13236355-INS-517
chr8-13257317-INS-412
chr8-13314800-INS-202
chr8-13319730-INS-60
chr8-13362971-INS-578
chr8-13366938-INS-316
chr8-13374205-INS-56
chr8-13377835-INS-63
chr8-13397865-INS-213
chr8-13433747-INS-697
chr8-13500766-INS-572
chr8-13530665-INS-71
chr8-13535102-INS-103
chr8-13553364-INS-190
chr8-13589735-INS-125
chr8-13596809-INS-270
chr8-13616095-INS-204
chr8-13621646-INS-103
chr8-13658757-INS-101
chr8-13662361-INS-158
chr8-13666721-INS-89
chr8-13666871-INS-6387
chr8-13668931-INS-348
chr8-13678037-INS-94
chr8-13691834-INS-984
chr8-13742167-INS-52
chr8-13751754-INS-165
chr8-13767684-INS-55
chr8-13780543-INS-76
chr8-13806435-INS-99
chr8-13834631-INS-232
chr8-13844289-INS-74
chr8-13851279-INS-373
chr8-13876912-INS-52
chr8-13893679-INS-280
chr8-13903438-INS-223
chr8-13907924-INS-5314
chr8-13941039-INS-66
chr8-13942106-INS-52
chr8-13945641-INS-96
chr8-13961841-INS-212
chr8-13966248-INS-85
chr8-13979565-INS-325
chr8-13990800-INS-419
chr8-13993546-INS-155
chr8-13994155-INS-268
chr8-13996717-INS-76
chr8-14017178-INS-269
chr8-14019267-INS-57
chr8-14037876-INS-178
chr8-14043317-INS-70
chr8-14044083-INS-208
chr8-14058319-INS-93
chr8-14084017-INS-60
chr8-14088553-INS-108
chr8-14104564-INS-174
chr8-14168840-INS-149
chr8-14169867-INS-95
chr8-14176546-INS-471
chr8-14210649-INS-52
chr8-14215534-INS-92
chr8-14259437-INS-201
chr8-14262751-INS-588

chr8-14266054-INS-60
chr8-14268773-INS-238
chr8-14298219-INS-98
chr8-14307436-INS-6050
chr8-14333809-INS-365
chr8-14343437-INS-176
chr8-14356834-INS-1953
chr8-14356886-INS-217
chr8-14357165-INS-1519
chr8-14419637-INS-185
chr8-14428395-INS-68
chr8-14502188-INS-52
chr8-14510943-INS-280
chr8-14515001-INS-68
chr8-14527817-INS-74
chr8-14603123-INS-761
chr8-14609436-INS-133
chr8-14612816-INS-51
chr8-14624755-INS-79
chr8-14654832-INS-94
chr8-14677796-INS-918
chr8-14685455-INS-131
chr8-14695858-INS-58
chr8-14726301-INS-83
chr8-14727929-INS-285
chr8-14735609-INS-56
chr8-14771433-INS-1048
chr8-14779370-INS-183
chr8-14789034-INS-74
chr8-14799345-INS-6966
chr8-14826123-INS-241
chr8-14844261-INS-471
chr8-14859550-INS-131
chr8-14926594-INS-121
chr8-14962108-INS-632
chr8-14986940-INS-382
chr8-14996486-INS-996
chr8-14998470-INS-128
chr8-15013372-INS-132
chr8-15026551-INS-68
chr8-15044915-INS-85
chr8-15056359-INS-72
chr8-15076836-INS-58
chr8-15087750-INS-62
chr8-15107233-INS-71
chr8-15128113-INS-199
chr8-15165681-INS-1875
chr8-15188104-INS-655
chr8-15192205-INS-1171
chr8-15192276-INS-586
chr8-15194065-INS-621
chr8-15240659-INS-583
chr8-15252922-INS-156
chr8-15261860-INS-239
chr8-15268601-INS-159
chr8-15273011-INS-159
chr8-15278621-INS-210
chr8-15281939-INS-156
chr8-15291672-INS-358
chr8-15309674-INS-101
chr8-15323067-INS-168
chr8-15371122-INS-54
chr8-15384300-INS-399
chr8-15420442-INS-51
chr8-15429523-INS-147
chr8-15429973-INS-484
chr8-15441503-INS-401
chr8-15443955-INS-77
chr8-15520463-INS-356
chr8-15526972-INS-337
chr8-15531998-INS-6395
chr8-15571812-INS-840
chr8-15681808-INS-51
chr8-15725871-INS-72
chr8-15729551-INS-114
chr8-15801311-INS-198
chr8-15834010-INS-72

chr8-15836579-INS-7477
chr8-15888012-INS-166
chr8-15892510-INS-440
chr8-15900475-INS-54
chr8-15901832-INS-212
chr8-15904245-INS-52
chr8-15929279-INS-458
chr8-15983351-INS-121
chr8-16008197-INS-84
chr8-16010975-INS-6586
chr8-16031471-INS-66
chr8-16109530-INS-211
chr8-16111698-INS-57
chr8-16131539-INS-51
chr8-16135955-INS-268
chr8-16141448-INS-53
chr8-16151971-INS-54
chr8-16163101-INS-60
chr8-16187008-INS-55
chr8-16202128-INS-242
chr8-16203977-INS-526
chr8-16238147-INS-94
chr8-16261563-INS-660
chr8-16300387-INS-412
chr8-16307246-INS-78
chr8-16336015-INS-317
chr8-16356950-INS-399
chr8-16383869-INS-53
chr8-16389963-INS-108
chr8-16408925-INS-314
chr8-16418373-INS-6881
chr8-16528785-INS-680
chr8-16564773-INS-116
chr8-16600613-INS-168
chr8-16633213-INS-351
chr8-16635571-INS-359
chr8-16648682-INS-53
chr8-16680953-INS-64
chr8-16709512-INS-60
chr8-16718918-INS-375
chr8-16822062-INS-211
chr8-16838388-INS-64
chr8-16869637-INS-385
chr8-16878911-INS-400
chr8-16885307-INS-583
chr8-16894907-INS-153
chr8-16895373-INS-71
chr8-16911369-INS-58
chr8-16916477-INS-60
chr8-16924270-INS-70
chr8-16928828-INS-200
chr8-16988633-INS-52
chr8-16992755-INS-191
chr8-17005008-INS-58
chr8-17062942-INS-421
chr8-17091233-INS-236
chr8-17093119-INS-124
chr8-17096567-INS-124
chr8-17172281-INS-338
chr8-17179485-INS-62
chr8-17199639-INS-236
chr8-17203422-INS-65
chr8-17225661-INS-158
chr8-17235345-INS-527
chr8-17255525-INS-70
chr8-17258794-INS-218
chr8-17264677-INS-141
chr8-17271165-INS-809
chr8-17282905-INS-54
chr8-17288140-INS-772
chr8-17290923-INS-80
chr8-17300924-INS-54
chr8-17315967-INS-230
chr8-17319897-INS-189
chr8-17324589-INS-103
chr8-17326085-INS-128
chr8-17326690-INS-236

chr8-17334120-INS-122
chr8-17399099-INS-336
chr8-17418431-INS-126
chr8-17430788-INS-60
chr8-17440832-INS-62
chr8-17456078-INS-184
chr8-17465129-INS-62
chr8-17474375-INS-72
chr8-17584004-INS-121
chr8-17605457-INS-400
chr8-17613173-INS-148
chr8-17627896-INS-72
chr8-17642530-INS-61
chr8-17646561-INS-216
chr8-17648095-INS-60
chr8-17746728-INS-717
chr8-17753290-INS-341
chr8-17775503-INS-307
chr8-17783342-INS-723
chr8-17821841-INS-385
chr8-17842727-INS-604
chr8-17843727-INS-102
chr8-17874985-INS-6815
chr8-17985154-INS-60
chr8-18024491-INS-81
chr8-18037104-INS-5990
chr8-18062768-INS-66
chr8-18068745-INS-282
chr8-18103905-INS-60
chr8-18106772-INS-286
chr8-18136098-INS-237
chr8-18146000-INS-100
chr8-18233922-INS-116
chr8-18248450-INS-197
chr8-18263632-INS-160
chr8-18265514-INS-365
chr8-18270485-INS-76
chr8-18378120-INS-76
chr8-18414916-INS-530
chr8-18427067-INS-52
chr8-18442892-INS-325
chr8-18450369-INS-394
chr8-18465217-INS-785
chr8-18480658-INS-7918
chr8-18496704-INS-61
chr8-18508700-INS-82
chr8-18524020-INS-114
chr8-18538717-INS-469
chr8-18543930-INS-53
chr8-18563288-INS-102
chr8-18569579-INS-59
chr8-18575207-INS-745
chr8-18575248-INS-73
chr8-18589448-INS-198
chr8-18594271-INS-59
chr8-18617157-INS-126
chr8-18667385-INS-213
chr8-18669114-INS-51
chr8-18674378-INS-54
chr8-18675425-INS-401
chr8-18701038-INS-117
chr8-18720639-INS-59
chr8-18723476-INS-93
chr8-18725368-INS-5546
chr8-18730144-INS-1289
chr8-18772367-INS-150
chr8-18777786-INS-60
chr8-18853311-INS-202
chr8-18865449-INS-180
chr8-18868738-INS-991
chr8-18900147-INS-68
chr8-18912370-INS-50
chr8-18925033-INS-604
chr8-18951264-INS-201
chr8-18952855-INS-400
chr8-18954179-INS-60
chr8-18977318-INS-109

chr8-19015582-INS-58
chr8-19027744-INS-516
chr8-19032088-INS-829
chr8-19050559-INS-198
chr8-19059064-INS-60
chr8-19071233-INS-1244
chr8-19072049-INS-6273
chr8-19080835-INS-468
chr8-19081735-INS-6599
chr8-19083531-INS-208
chr8-19084738-INS-60
chr8-19086310-INS-962
chr8-19114194-INS-108
chr8-19130305-INS-82
chr8-19186339-INS-6988
chr8-19194203-INS-98
chr8-19195725-INS-53
chr8-19198565-INS-54
chr8-19241501-INS-649
chr8-19265864-INS-493
chr8-19327389-INS-106
chr8-19340699-INS-124
chr8-19362959-INS-379
chr8-19363932-INS-137
chr8-19366936-INS-60
chr8-19402445-INS-88
chr8-19417963-INS-149
chr8-19462416-INS-66
chr8-19480534-INS-358
chr8-19514403-INS-120
chr8-19526857-INS-96
chr8-19623770-INS-690
chr8-19676527-INS-316
chr8-19676900-INS-549
chr8-19716562-INS-53
chr8-19728207-INS-53
chr8-19744868-INS-6457
chr8-19777997-INS-3573
chr8-19810243-INS-240
chr8-20486747-INS-94
chr8-20507947-INS-1875
chr8-20508081-INS-104
chr8-20540411-INS-1891
chr8-20732023-INS-167
chr8-21126969-INS-67
chr8-21360971-INS-8515
chr8-21377625-INS-426
chr8-21394880-INS-58
chr8-21397262-INS-56
chr8-21416369-INS-80
chr8-21417376-INS-71
chr8-21438999-INS-2800
chr8-21447705-INS-208
chr8-21459184-INS-91
chr8-21470208-INS-55
chr8-21499771-INS-98
chr8-21563074-INS-58
chr8-21585888-INS-213
chr8-21615840-INS-617
chr8-21762488-INS-155
chr8-22087757-INS-159
chr8-22089405-INS-6516
chr8-22116756-INS-51
chr8-22130639-INS-115
chr8-22136977-INS-10722
chr8-22165153-INS-83
chr8-22169216-INS-606
chr8-22173910-INS-270
chr8-22183507-INS-6688
chr8-22196883-INS-159
chr8-22227215-INS-199
chr8-22294519-INS-80
chr8-22301454-INS-71
chr8-22301996-INS-110
chr8-22302305-INS-181
chr8-22311557-INS-386
chr8-22339508-INS-517

chr8-22349408-INS-231
chr8-22410187-INS-131
chr8-22414352-INS-118
chr8-22416279-INS-174
chr8-22424022-INS-143
chr8-22432669-INS-5974
chr8-22444234-INS-498
chr8-22491247-INS-52
chr8-22493447-INS-54
chr8-22522712-INS-640
chr8-22577464-INS-211
chr8-22580137-INS-544
chr8-22580757-INS-134
chr8-22595988-INS-72
chr8-22599548-INS-89
chr8-22650829-INS-71
chr8-22686284-INS-205
chr8-22688278-INS-190
chr8-22692811-INS-124
chr8-22715596-INS-486
chr8-22737604-INS-56
chr8-22746869-INS-58
chr8-22755853-INS-77
chr8-22765513-INS-136
chr8-22771353-INS-325
chr8-22775116-INS-53
chr8-22787401-INS-92
chr8-22842471-INS-106
chr8-22845285-INS-95
chr8-22871055-INS-336
chr8-22876269-INS-96
chr8-22891964-INS-59
chr8-22900438-INS-207
chr8-22902830-INS-170
chr8-22909005-INS-328
chr8-22916434-INS-56
chr8-22951838-INS-85
chr8-22955959-INS-52
chr8-22971627-INS-72
chr8-22980840-INS-147
chr8-22986552-INS-76
chr8-23007322-INS-76
chr8-23012124-INS-62
chr8-23018015-INS-206
chr8-23056252-INS-5314
chr8-23056569-INS-99
chr8-23088103-INS-60
chr8-23095921-INS-341
chr8-23130175-INS-199
chr8-23137203-INS-207
chr8-23165264-INS-62
chr8-23191784-INS-79
chr8-23201084-INS-54
chr8-23225860-INS-124
chr8-23239321-INS-70
chr8-23249847-INS-80
chr8-23252879-INS-82
chr8-23269348-INS-106
chr8-23283373-INS-76
chr8-23285335-INS-54
chr8-23308622-INS-263
chr8-23326313-INS-147
chr8-23355209-INS-266
chr8-23355396-INS-671
chr8-23362331-INS-405
chr8-23377708-INS-239
chr8-23397434-INS-51
chr8-23436947-INS-399
chr8-23441987-INS-50
chr8-23452429-INS-94
chr8-23456668-INS-100
chr8-23494820-INS-113
chr8-23542084-INS-54
chr8-23546148-INS-216
chr8-23561245-INS-56
chr8-23603747-INS-81
chr8-23605598-INS-70

chr8-23617904-INS-138
chr8-23633352-INS-64
chr8-23678130-INS-76
chr8-23685049-INS-102
chr8-23693163-INS-204
chr8-23704392-INS-239
chr8-23726485-INS-53
chr8-23730156-INS-172
chr8-23743487-INS-145
chr8-23755228-INS-64
chr8-23768013-INS-196
chr8-23769376-INS-247
chr8-23784978-INS-79
chr8-23787036-INS-217
chr8-23792977-INS-80
chr8-23816605-INS-234
chr8-23840121-INS-137
chr8-23858854-INS-204
chr8-23864240-INS-4787
chr8-23872601-INS-57
chr8-23877422-INS-691
chr8-23903951-INS-97
chr8-23905648-INS-95
chr8-23906174-INS-151
chr8-23931765-INS-51
chr8-23943319-INS-86
chr8-23956944-INS-481
chr8-23964269-INS-3901
chr8-23971164-INS-6432
chr8-23975732-INS-140
chr8-23981324-INS-211
chr8-23982521-INS-1840
chr8-23984205-INS-239
chr8-24001089-INS-102
chr8-24010509-INS-81
chr8-24014120-INS-92
chr8-24043494-INS-287
chr8-24046247-INS-222
chr8-24063331-INS-63
chr8-24064826-INS-401
chr8-24065539-INS-82
chr8-24094613-INS-208
chr8-24101091-INS-381
chr8-24114295-INS-199
chr8-24132661-INS-324
chr8-24203645-INS-54
chr8-24205933-INS-354
chr8-24211535-INS-56
chr8-24223766-INS-110
chr8-24230061-INS-89
chr8-24242209-INS-489
chr8-24249990-INS-454
chr8-24257276-INS-145
chr8-24258524-INS-243
chr8-24269499-INS-168
chr8-24285433-INS-307
chr8-24311370-INS-61
chr8-24334305-INS-118
chr8-24381689-INS-190
chr8-24397664-INS-304
chr8-24409311-INS-153
chr8-24411736-INS-58
chr8-24416169-INS-231
chr8-24424741-INS-76
chr8-24434273-INS-118
chr8-24457974-INS-64
chr8-24478194-INS-114
chr8-24484701-INS-186
chr8-24494749-INS-98
chr8-24505258-INS-578
chr8-24512739-INS-91
chr8-24517521-INS-490
chr8-24518162-INS-138
chr8-24525705-INS-876
chr8-24541450-INS-349
chr8-24561978-INS-248
chr8-24581877-INS-84

chr8-24589524-INS-134
chr8-24596642-INS-1850
chr8-24615049-INS-120
chr8-24621661-INS-171
chr8-24623398-INS-81
chr8-24623919-INS-122
chr8-24705271-INS-66
chr8-24735180-INS-65
chr8-24752607-INS-51
chr8-24784566-INS-106
chr8-24787929-INS-105
chr8-24799526-INS-300
chr8-24818485-INS-53
chr8-24840911-INS-69
chr8-24846523-INS-92
chr8-24862666-INS-199
chr8-24870572-INS-264
chr8-24871853-INS-532
chr8-24882757-INS-52
chr8-24891341-INS-495
chr8-24896231-INS-58
chr8-24900799-INS-136
chr8-24916449-INS-108
chr8-24935132-INS-96
chr8-24959780-INS-605
chr8-24980359-INS-757
chr8-24983423-INS-387
chr8-25026393-INS-498
chr8-25045554-INS-90
chr8-25093019-INS-81
chr8-25123082-INS-68
chr8-25126964-INS-275
chr8-25129684-INS-859
chr8-25131980-INS-52
chr8-25133283-INS-74
chr8-25145104-INS-210
chr8-25171513-INS-60
chr8-25187745-INS-76
chr8-25194376-INS-79
chr8-25213109-INS-379
chr8-25213218-INS-1111
chr8-25238722-INS-148
chr8-25265395-INS-84
chr8-25272821-INS-64
chr8-25298643-INS-249
chr8-25300492-INS-210
chr8-25331936-INS-5551
chr8-25344124-INS-193
chr8-25353990-INS-408
chr8-25361481-INS-269
chr8-25385304-INS-99
chr8-25392217-INS-54
chr8-25393382-INS-536
chr8-25402087-INS-125
chr8-25442197-INS-88
chr8-25472786-INS-163
chr8-25487186-INS-445
chr8-25492234-INS-111
chr8-25492541-INS-218
chr8-25494445-INS-230
chr8-25514271-INS-52
chr8-25531894-INS-101
chr8-25538627-INS-51
chr8-25556695-INS-85
chr8-25561430-INS-58
chr8-25571182-INS-172
chr8-25580130-INS-109
chr8-25584027-INS-5385
chr8-25602257-INS-348
chr8-25628762-INS-142
chr8-25631483-INS-58
chr8-25633560-INS-52
chr8-25640972-INS-72
chr8-25666172-INS-57
chr8-25712743-INS-67
chr8-25717787-INS-772
chr8-25732272-INS-204

chr8-25744491-INS-56
chr8-25745190-INS-70
chr8-25751431-INS-672
chr8-25759079-INS-56
chr8-25808605-INS-399
chr8-25825435-INS-477
chr8-25827459-INS-178
chr8-25845128-INS-61
chr8-25850950-INS-204
chr8-25854165-INS-84
chr8-25889642-INS-63
chr8-25899052-INS-59
chr8-25903622-INS-241
chr8-25910503-INS-52
chr8-25918295-INS-99
chr8-25929070-INS-236
chr8-25935754-INS-186
chr8-25954560-INS-210
chr8-25961765-INS-100
chr8-25964875-INS-68
chr8-25980038-INS-94
chr8-25999407-INS-96
chr8-26000746-INS-150
chr8-26028914-INS-186
chr8-26051212-INS-103
chr8-26066999-INS-83
chr8-26072348-INS-56
chr8-26097091-INS-57
chr8-26117706-INS-70
chr8-26132827-INS-5548
chr8-26143461-INS-890
chr8-26147488-INS-650
chr8-26148806-INS-132
chr8-26170510-INS-116
chr8-26184126-INS-104
chr8-26192456-INS-53
chr8-26198863-INS-91
chr8-26223829-INS-147
chr8-26234532-INS-443
chr8-26260604-INS-373
chr8-26262067-INS-184
chr8-26263239-INS-52
chr8-26267823-INS-208
chr8-26295570-INS-218
chr8-26313779-INS-193
chr8-26325199-INS-181
chr8-26335822-INS-54
chr8-26359603-INS-210
chr8-26361315-INS-507
chr8-26398076-INS-214
chr8-26402334-INS-205
chr8-26405386-INS-564
chr8-26408657-INS-203
chr8-26412993-INS-712
chr8-26415080-INS-209
chr8-26417055-INS-70
chr8-26418218-INS-67
chr8-26420315-INS-222
chr8-26423403-INS-86
chr8-26425620-INS-107
chr8-26426193-INS-333
chr8-26428822-INS-385
chr8-26438832-INS-84
chr8-26449261-INS-395
chr8-26459568-INS-53
chr8-26471208-INS-129
chr8-26481532-INS-273
chr8-26493669-INS-297
chr8-26494868-INS-230
chr8-26515715-INS-172
chr8-26517819-INS-86
chr8-26524686-INS-76
chr8-26531484-INS-537
chr8-26553442-INS-214
chr8-26556236-INS-120
chr8-26563235-INS-76
chr8-26581089-INS-826

chr8-26583353-INS-448
chr8-26586377-INS-184
chr8-26617286-INS-56
chr8-26624403-INS-205
chr8-26635365-INS-71
chr8-26660648-INS-323
chr8-26664591-INS-57
chr8-26674669-INS-76
chr8-26683614-INS-68
chr8-26726601-INS-275
chr8-26726989-INS-257
chr8-26731630-INS-109
chr8-26735555-INS-168
chr8-26735721-INS-947
chr8-26760067-INS-398
chr8-26773772-INS-63
chr8-26775878-INS-90
chr8-26776042-INS-925
chr8-26778841-INS-592
chr8-26791795-INS-67
chr8-26796464-INS-56
chr8-26801429-INS-51
chr8-26824959-INS-115
chr8-26855346-INS-444
chr8-26864975-INS-52
chr8-26869504-INS-605
chr8-26873179-INS-147
chr8-26906571-INS-536
chr8-26914054-INS-534
chr8-26916028-INS-352
chr8-26918139-INS-442
chr8-26941524-INS-88
chr8-27008428-INS-6168
chr8-27043161-INS-246
chr8-27065893-INS-178
chr8-27066653-INS-344
chr8-27097192-INS-480
chr8-27125537-INS-61
chr8-27135933-INS-201
chr8-27192901-INS-93
chr8-27198945-INS-168
chr8-27217388-INS-63
chr8-27248363-INS-68
chr8-27270344-INS-157
chr8-27290582-INS-58
chr8-27299262-INS-72
chr8-27329059-INS-61
chr8-27336593-INS-177
chr8-27363622-INS-59
chr8-27372401-INS-67
chr8-27426089-INS-120
chr8-27462390-INS-58
chr8-27467202-INS-57
chr8-27480243-INS-51
chr8-27495611-INS-154
chr8-27501823-INS-206
chr8-27517402-INS-75
chr8-27553114-INS-91
chr8-27563898-INS-438
chr8-27568965-INS-218
chr8-27585504-INS-60
chr8-27588466-INS-68
chr8-27596158-INS-71
chr8-27605370-INS-53
chr8-27622660-INS-3312
chr8-27623124-INS-325
chr8-27634828-INS-542
chr8-27637612-INS-206
chr8-27639467-INS-251
chr8-27649605-INS-180
chr8-27662753-INS-174
chr8-27673930-INS-197
chr8-27676477-INS-284
chr8-27677544-INS-783
chr8-27701152-INS-223
chr8-27726408-INS-179
chr8-27735890-INS-173

chr8-27740644-INS-581
chr8-27744313-INS-205
chr8-27752807-INS-52
chr8-27757132-INS-63
chr8-27810031-INS-214
chr8-27811573-INS-75
chr8-27830209-INS-51
chr8-27842023-INS-468
chr8-27847194-INS-6467
chr8-27850336-INS-702
chr8-27855619-INS-395
chr8-27863784-INS-218
chr8-27866618-INS-945
chr8-27869302-INS-62
chr8-27878824-INS-82
chr8-27880762-INS-76
chr8-27890181-INS-56
chr8-27891199-INS-57
chr8-27902908-INS-58
chr8-27928629-INS-340
chr8-27966149-INS-6442
chr8-27978062-INS-482
chr8-27993312-INS-189
chr8-28009967-INS-256
chr8-28032056-INS-59
chr8-28032965-INS-6496
chr8-28038393-INS-5281
chr8-28042478-INS-6553
chr8-28044619-INS-6354
chr8-28050298-INS-73
chr8-28063509-INS-378
chr8-28094010-INS-128
chr8-28168865-INS-629
chr8-28177678-INS-110
chr8-28181668-INS-6169
chr8-28185690-INS-310
chr8-28198642-INS-77
chr8-28202613-INS-6377
chr8-28202847-INS-632
chr8-28207483-INS-108
chr8-28243465-INS-217
chr8-28273291-INS-56
chr8-28288983-INS-118
chr8-28316744-INS-4660
chr8-28324920-INS-58
chr8-28327067-INS-1380
chr8-28370158-INS-60
chr8-28380681-INS-82
chr8-28382102-INS-937
chr8-28385909-INS-79
chr8-28392655-INS-6463
chr8-28401590-INS-80
chr8-28414253-INS-488
chr8-28435161-INS-74
chr8-28456860-INS-816
chr8-28475776-INS-52
chr8-28483993-INS-252
chr8-28525716-INS-446
chr8-28538961-INS-91
chr8-28545203-INS-238
chr8-28560082-INS-400
chr8-28582288-INS-204
chr8-28588748-INS-144
chr8-28612477-INS-833
chr8-28613748-INS-52
chr8-28626635-INS-72
chr8-28631632-INS-641
chr8-28649522-INS-141
chr8-28654231-INS-220
chr8-28673348-INS-76
chr8-28697589-INS-57
chr8-28726574-INS-106
chr8-28743787-INS-469
chr8-28750740-INS-639
chr8-28752154-INS-527
chr8-28763993-INS-54
chr8-28793485-INS-545

chr8-28804930-INS-70
chr8-28839112-INS-838
chr8-28858146-INS-84
chr8-28872695-INS-72
chr8-28908622-INS-6927
chr8-28932657-INS-96
chr8-28937591-INS-59
chr8-29004795-INS-62
chr8-29016013-INS-96
chr8-29017644-INS-99
chr8-29057586-INS-475
chr8-29083305-INS-68
chr8-29130399-INS-91
chr8-29136194-INS-633
chr8-29155377-INS-69
chr8-29172179-INS-272
chr8-29175702-INS-253
chr8-29237760-INS-52
chr8-29238426-INS-102
chr8-29252384-INS-68
chr8-29259625-INS-60
chr8-29275660-INS-266
chr8-29308559-INS-68
chr8-29328465-INS-622
chr8-29344669-INS-54
chr8-29385603-INS-206
chr8-29390269-INS-62
chr8-29420082-INS-67
chr8-29428277-INS-2236
chr8-29428301-INS-1608
chr8-29428309-INS-1900
chr8-29428329-INS-1140
chr8-29462881-INS-456
chr8-29499016-INS-301
chr8-29507425-INS-447
chr8-29508913-INS-61
chr8-29511332-INS-388
chr8-29514491-INS-77
chr8-29523622-INS-68
chr8-29560116-INS-69
chr8-29576236-INS-240
chr8-29577857-INS-2904
chr8-29598888-INS-64
chr8-29614473-INS-90
chr8-29620873-INS-51
chr8-29647384-INS-508
chr8-29657226-INS-58
chr8-29658177-INS-6560
chr8-29703538-INS-58
chr8-29720716-INS-60
chr8-29724589-INS-52
chr8-29732331-INS-57
chr8-29743639-INS-53
chr8-29793805-INS-98
chr8-29795504-INS-200
chr8-29842579-INS-459
chr8-29854310-INS-52
chr8-29855736-INS-220
chr8-29877870-INS-120
chr8-29981407-INS-63
chr8-30019547-INS-53
chr8-30029700-INS-62
chr8-30031303-INS-80
chr8-30098592-INS-6894
chr8-30128791-INS-140
chr8-30132089-INS-71
chr8-30162027-INS-100
chr8-30175769-INS-100
chr8-30177136-INS-66
chr8-30194727-INS-52
chr8-30236176-INS-68
chr8-30246059-INS-906
chr8-30247210-INS-117
chr8-30256607-INS-742
chr8-30260898-INS-84
chr8-30274976-INS-6526
chr8-30280900-INS-100

chr8-30312502-INS-215
chr8-30324690-INS-64
chr8-30343752-INS-191
chr8-30388847-INS-178
chr8-30394813-INS-973
chr8-30409762-INS-200
chr8-30443073-INS-52
chr8-30447141-INS-54
chr8-30461102-INS-604
chr8-30519081-INS-6572
chr8-30519330-INS-660
chr8-30522018-INS-203
chr8-30536842-INS-68
chr8-30576305-INS-893
chr8-30583266-INS-52
chr8-30594197-INS-191
chr8-30608274-INS-539
chr8-30668856-INS-59
chr8-30676696-INS-608
chr8-30686751-INS-338
chr8-30755465-INS-69
chr8-30756959-INS-60
chr8-30757165-INS-4866
chr8-30843832-INS-69
chr8-30850251-INS-2240
chr8-30861947-INS-6201
chr8-30876556-INS-1317
chr8-30899465-INS-6887
chr8-30903893-INS-229
chr8-30904596-INS-645
chr8-30907646-INS-134
chr8-30963124-INS-191
chr8-30965045-INS-154
chr8-30993902-INS-180
chr8-30998344-INS-202
chr8-31003997-INS-60
chr8-31020214-INS-448
chr8-31021843-INS-55
chr8-31077337-INS-487
chr8-31090130-INS-901
chr8-31099114-INS-63
chr8-31116153-INS-6431
chr8-31121126-INS-378
chr8-31139598-INS-51
chr8-31156088-INS-116
chr8-31162890-INS-64
chr8-31194957-INS-53
chr8-31262625-INS-699
chr8-31267018-INS-81
chr8-31270147-INS-397
chr8-31299660-INS-566
chr8-31311112-INS-54
chr8-31319072-INS-55
chr8-31382297-INS-408
chr8-31386951-INS-726
chr8-31415373-INS-355
chr8-31416228-INS-194
chr8-31450455-INS-418
chr8-31475026-INS-210
chr8-31484929-INS-184
chr8-31499376-INS-65
chr8-31517125-INS-76
chr8-31534030-INS-63
chr8-31553004-INS-121
chr8-31563962-INS-163
chr8-31590131-INS-262
chr8-31608667-INS-5139
chr8-31625050-INS-197
chr8-31626583-INS-311
chr8-31633113-INS-441
chr8-31644037-INS-197
chr8-31649302-INS-52
chr8-31672854-INS-58
chr8-31674969-INS-149
chr8-31686712-INS-124
chr8-31688799-INS-368
chr8-31711478-INS-138

chr8-31720239-INS-57
chr8-31731839-INS-60
chr8-31753680-INS-87
chr8-31777261-INS-114
chr8-31808326-INS-52
chr8-31815527-INS-75
chr8-31829926-INS-610
chr8-31831804-INS-73
chr8-31873151-INS-95
chr8-31890630-INS-206
chr8-31904617-INS-316
chr8-31968751-INS-6737
chr8-31973900-INS-333
chr8-31984435-INS-66
chr8-31994784-INS-6232
chr8-32010937-INS-62
chr8-32017531-INS-6592
chr8-32067995-INS-6730
chr8-32072456-INS-152
chr8-32110484-INS-6414
chr8-32115027-INS-169
chr8-32119612-INS-292
chr8-32128157-INS-91
chr8-32175913-INS-6592
chr8-32177377-INS-435
chr8-32179896-INS-6627
chr8-32192148-INS-218
chr8-32197709-INS-60
chr8-32230679-INS-635
chr8-32249576-INS-66
chr8-32265505-INS-743
chr8-32268660-INS-400
chr8-32281988-INS-96
chr8-32286904-INS-528
chr8-32355956-INS-447
chr8-32400261-INS-74
chr8-32509066-INS-109
chr8-32536014-INS-76
chr8-32560933-INS-78
chr8-32586030-INS-149
chr8-32593561-INS-906
chr8-32597139-INS-64
chr8-32675914-INS-184
chr8-32732803-INS-850
chr8-32733831-INS-1211
chr8-32769880-INS-189
chr8-32790778-INS-404
chr8-32799776-INS-720
chr8-32805217-INS-98
chr8-32833025-INS-370
chr8-32872676-INS-81
chr8-32887019-INS-159
chr8-32889915-INS-99
chr8-32944697-INS-242
chr8-32946192-INS-828
chr8-33054219-INS-54
chr8-33085478-INS-285
chr8-33163343-INS-1070
chr8-33167604-INS-100
chr8-33175754-INS-104
chr8-33193663-INS-375
chr8-33200031-INS-229
chr8-33252140-INS-66
chr8-33254920-INS-56
chr8-33270868-INS-174
chr8-33274788-INS-140
chr8-33300681-INS-67
chr8-33302323-INS-133
chr8-33312368-INS-274
chr8-33314158-INS-91
chr8-33320111-INS-54
chr8-33331540-INS-296
chr8-33341221-INS-99
chr8-33364613-INS-80
chr8-33365375-INS-442
chr8-33405189-INS-256
chr8-33408356-INS-65

chr8-33417115-INS-216
chr8-33513548-INS-165
chr8-33519876-INS-365
chr8-33521046-INS-390
chr8-33523849-INS-53
chr8-33526025-INS-6232
chr8-33533613-INS-64
chr8-33547428-INS-780
chr8-33549665-INS-52
chr8-33559870-INS-78
chr8-33564704-INS-204
chr8-33574893-INS-428
chr8-33579339-INS-144
chr8-33608017-INS-616
chr8-33615672-INS-199
chr8-33625601-INS-7084
chr8-33626975-INS-56
chr8-33686760-INS-72
chr8-33691201-INS-660
chr8-33699456-INS-648
chr8-33705356-INS-60
chr8-33707946-INS-6440
chr8-33708256-INS-217
chr8-33721198-INS-7107
chr8-33723568-INS-223
chr8-33736897-INS-146
chr8-33739460-INS-107
chr8-33745670-INS-62
chr8-33749023-INS-109
chr8-33750287-INS-188
chr8-33755178-INS-125
chr8-33781729-INS-168
chr8-33789179-INS-80
chr8-33812038-INS-1058
chr8-33817820-INS-58
chr8-33851515-INS-76
chr8-33852563-INS-324
chr8-33865854-INS-101
chr8-33873181-INS-61
chr8-33999595-INS-59
chr8-34059196-INS-426
chr8-34060491-INS-5536
chr8-34075847-INS-98
chr8-34096488-INS-130
chr8-34124346-INS-149
chr8-34125205-INS-190
chr8-34126362-INS-53
chr8-34149837-INS-219
chr8-34162203-INS-205
chr8-34168107-INS-199
chr8-34171199-INS-207
chr8-34194259-INS-7460
chr8-34197041-INS-51
chr8-34198961-INS-235
chr8-34200012-INS-94
chr8-34217066-INS-184
chr8-34244956-INS-869
chr8-34246579-INS-206
chr8-34251936-INS-78
chr8-34265619-INS-2626
chr8-34270449-INS-114
chr8-34290612-INS-52
chr8-34305822-INS-55
chr8-34307372-INS-206
chr8-34362318-INS-112
chr8-34371174-INS-212
chr8-34377719-INS-227
chr8-34433129-INS-1095
chr8-34437185-INS-56
chr8-34440507-INS-184
chr8-34444973-INS-172
chr8-34464105-INS-814
chr8-34466895-INS-77
chr8-34487481-INS-237
chr8-34489209-INS-138
chr8-34494671-INS-118
chr8-34498048-INS-210

chr8-34498403-INS-342
chr8-34509159-INS-934
chr8-34533263-INS-177
chr8-34546591-INS-201
chr8-34562462-INS-109
chr8-34570115-INS-88
chr8-34574634-INS-106
chr8-34594077-INS-513
chr8-34615803-INS-87
chr8-34704728-INS-89
chr8-34707545-INS-63
chr8-34718370-INS-58
chr8-34754657-INS-70
chr8-34757517-INS-108
chr8-34764481-INS-688
chr8-34778254-INS-69
chr8-34844845-INS-56
chr8-34910680-INS-60
chr8-34943747-INS-504
chr8-34978865-INS-374
chr8-34979464-INS-144
chr8-34979567-INS-308
chr8-34981184-INS-167
chr8-35001341-INS-207
chr8-35013118-INS-60
chr8-35039004-INS-144
chr8-35083004-INS-51
chr8-35102156-INS-673
chr8-35104388-INS-165
chr8-35134892-INS-77
chr8-35140314-INS-62
chr8-35157031-INS-407
chr8-35189893-INS-116
chr8-35195753-INS-52
chr8-35211186-INS-365
chr8-35231508-INS-67
chr8-35257852-INS-133
chr8-35258018-INS-1890
chr8-35280999-INS-200
chr8-35292846-INS-507
chr8-35352029-INS-444
chr8-35437068-INS-77
chr8-35451031-INS-55
chr8-35456885-INS-74
chr8-35459328-INS-153
chr8-35471275-INS-400
chr8-35472200-INS-108
chr8-35475419-INS-406
chr8-35486968-INS-75
chr8-35531562-INS-106
chr8-35539470-INS-435
chr8-35571955-INS-80
chr8-35578138-INS-212
chr8-35599977-INS-203
chr8-35603743-INS-208
chr8-35611442-INS-540
chr8-35613270-INS-125
chr8-35622699-INS-90
chr8-35623306-INS-129
chr8-35638318-INS-173
chr8-35658036-INS-412
chr8-35662730-INS-254
chr8-35670600-INS-54
chr8-35673179-INS-52
chr8-35686252-INS-56
chr8-35719060-INS-144
chr8-35728986-INS-94
chr8-35755497-INS-116
chr8-35770945-INS-99
chr8-35771763-INS-223
chr8-35775157-INS-67
chr8-35783014-INS-97
chr8-35789683-INS-60
chr8-35798709-INS-274
chr8-35805811-INS-356
chr8-35815114-INS-280
chr8-35817845-INS-52

chr8-35833346-INS-2897
chr8-35843474-INS-5574
chr8-35848634-INS-57
chr8-35860442-INS-222
chr8-35994949-INS-55
chr8-36011576-INS-835
chr8-36025495-INS-55
chr8-36053947-INS-813
chr8-36102904-INS-213
chr8-36108060-INS-75
chr8-36123328-INS-125
chr8-36135584-INS-273
chr8-36150460-INS-580
chr8-36154291-INS-60
chr8-36160689-INS-77
chr8-36167500-INS-56
chr8-36210933-INS-77
chr8-36229782-INS-420
chr8-36241281-INS-54
chr8-36249326-INS-379
chr8-36313877-INS-164
chr8-36317997-INS-186
chr8-36358698-INS-85
chr8-36361031-INS-62
chr8-36368446-INS-221
chr8-36407159-INS-268
chr8-36419955-INS-180
chr8-36420902-INS-1830
chr8-36471428-INS-86
chr8-36532320-INS-68
chr8-36542496-INS-66
chr8-36546813-INS-172
chr8-36549502-INS-52
chr8-36554417-INS-240
chr8-36594789-INS-392
chr8-36625570-INS-108
chr8-36647744-INS-92
chr8-36717711-INS-54
chr8-36723006-INS-190
chr8-36786130-INS-56
chr8-36801419-INS-3086
chr8-36881034-INS-644
chr8-36900156-INS-67
chr8-36900945-INS-61
chr8-36930381-INS-56
chr8-36934831-INS-156
chr8-36973309-INS-197
chr8-36973560-INS-998
chr8-36991535-INS-323
chr8-36993399-INS-197
chr8-37003908-INS-169
chr8-37005281-INS-130
chr8-37007445-INS-195
chr8-37031996-INS-68
chr8-37062632-INS-131
chr8-37083657-INS-235
chr8-37087820-INS-59
chr8-37160063-INS-71
chr8-37168757-INS-85
chr8-37186542-INS-359
chr8-37189289-INS-441
chr8-37202423-INS-314
chr8-37209729-INS-78
chr8-37265425-INS-52
chr8-37281748-INS-76
chr8-37299878-INS-56
chr8-37314775-INS-96
chr8-37316761-INS-1138
chr8-37362707-INS-920
chr8-37377803-INS-202
chr8-37385718-INS-134
chr8-37394936-INS-67
chr8-37400785-INS-371
chr8-37403080-INS-58
chr8-37408344-INS-984
chr8-37409218-INS-494
chr8-37430265-INS-61

chr8-37451642-INS-96
chr8-37454021-INS-329
chr8-37457651-INS-151
chr8-37475303-INS-377
chr8-37477766-INS-135
chr8-37482007-INS-62
chr8-37494951-INS-63
chr8-37508983-INS-68
chr8-37511732-INS-133
chr8-37593751-INS-69
chr8-37600531-INS-88
chr8-37609179-INS-60
chr8-37620277-INS-224
chr8-37637382-INS-80
chr8-37666673-INS-5657
chr8-37684258-INS-65
chr8-37693344-INS-180
chr8-37746511-INS-69
chr8-37772120-INS-52
chr8-37795753-INS-75
chr8-37801890-INS-381
chr8-37807200-INS-65
chr8-37820747-INS-566
chr8-37838834-INS-555
chr8-37840763-INS-60
chr8-37972361-INS-4524
chr8-37995317-INS-1892
chr8-38029890-INS-56
chr8-38032535-INS-256
chr8-38036256-INS-62
chr8-38078687-INS-6753
chr8-38079578-INS-285
chr8-38085872-INS-1017
chr8-38087770-INS-338
chr8-38094190-INS-498
chr8-38102509-INS-479
chr8-38104417-INS-6566
chr8-38123969-INS-52
chr8-38164997-INS-59
chr8-38173816-INS-388
chr8-38199797-INS-282
chr8-38201927-INS-56
chr8-38227916-INS-72
chr8-38241499-INS-935
chr8-38322160-INS-72
chr8-38335969-INS-251
chr8-38482257-INS-165
chr8-38503895-INS-772
chr8-38532952-INS-239
chr8-38542922-INS-377
chr8-38547453-INS-110
chr8-38549579-INS-52
chr8-38550355-INS-74
chr8-38562126-INS-2776
chr8-38630139-INS-54
chr8-38664002-INS-55
chr8-38666504-INS-156
chr8-38695158-INS-459
chr8-38707172-INS-498
chr8-38716723-INS-1926
chr8-38752417-INS-56
chr8-38768066-INS-282
chr8-38772936-INS-976
chr8-38792797-INS-225
chr8-38812449-INS-228
chr8-38829717-INS-196
chr8-38892654-INS-209
chr8-38913459-INS-516
chr8-38958989-INS-904
chr8-39002539-INS-57
chr8-39007634-INS-100
chr8-39011998-INS-58
chr8-39013508-INS-107
chr8-39023807-INS-60
chr8-39062533-INS-106
chr8-39085231-INS-85
chr8-39092685-INS-86

chr8-39194642-INS-7178
chr8-39224480-INS-160
chr8-39236538-INS-400
chr8-39240306-INS-66
chr8-39240669-INS-926
chr8-39244644-INS-256
chr8-39265621-INS-124
chr8-39278759-INS-6519
chr8-39282122-INS-121
chr8-39295659-INS-6816
chr8-39316990-INS-56
chr8-39322613-INS-486
chr8-39333629-INS-65
chr8-39360419-INS-474
chr8-39361430-INS-1885
chr8-39368268-INS-7291
chr8-39392046-INS-6545
chr8-39394650-INS-2682
chr8-39443196-INS-7329
chr8-39446749-INS-180
chr8-39525749-INS-635
chr8-39536365-INS-231
chr8-39570730-INS-56
chr8-39590244-INS-259
chr8-39618525-INS-131
chr8-39623201-INS-52
chr8-39636788-INS-462
chr8-39639058-INS-883
chr8-39673330-INS-60
chr8-39681581-INS-59
chr8-39712316-INS-66
chr8-39717860-INS-860
chr8-39750917-INS-113
chr8-39798754-INS-441
chr8-39815549-INS-147
chr8-39863712-INS-186
chr8-39865346-INS-111
chr8-39892511-INS-192
chr8-39897891-INS-389
chr8-39931646-INS-626
chr8-39947088-INS-240
chr8-39959772-INS-684
chr8-39994557-INS-118
chr8-40048772-INS-266
chr8-40051988-INS-527
chr8-40080032-INS-671
chr8-40084817-INS-136
chr8-40189611-INS-256
chr8-40191541-INS-64
chr8-40206486-INS-679
chr8-40208420-INS-667
chr8-40214991-INS-665
chr8-40271215-INS-6451
chr8-40292561-INS-476
chr8-40293827-INS-248
chr8-40297973-INS-89
chr8-40320817-INS-395
chr8-40337093-INS-1860
chr8-40394622-INS-92
chr8-40400875-INS-380
chr8-40420242-INS-64
chr8-40455014-INS-296
chr8-40471029-INS-602
chr8-40500386-INS-163
chr8-40507377-INS-79
chr8-40528632-INS-149
chr8-40533627-INS-280
chr8-40603321-INS-6178
chr8-40606045-INS-104
chr8-40613072-INS-400
chr8-40636482-INS-400
chr8-40641973-INS-399
chr8-40675884-INS-100
chr8-40679859-INS-919
chr8-40695630-INS-360
chr8-40698485-INS-58
chr8-40714165-INS-112

chr8-40729315-INS-184
chr8-40743856-INS-340
chr8-40758004-INS-380
chr8-40773839-INS-6994
chr8-40813982-INS-6286
chr8-40866625-INS-116
chr8-40867649-INS-513
chr8-40884612-INS-55
chr8-40961674-INS-74
chr8-40963716-INS-213
chr8-40968625-INS-56
chr8-40985410-INS-131
chr8-40991633-INS-225
chr8-41018488-INS-5247
chr8-41018901-INS-74
chr8-41033871-INS-163
chr8-41040899-INS-376
chr8-41043959-INS-107
chr8-41099976-INS-120
chr8-41113426-INS-51
chr8-41137307-INS-128
chr8-41140936-INS-463
chr8-41146920-INS-6230
chr8-41148142-INS-243
chr8-41167685-INS-183
chr8-41182668-INS-329
chr8-41219122-INS-6664
chr8-41221267-INS-7078
chr8-41257259-INS-416
chr8-41259798-INS-77
chr8-41300919-INS-55
chr8-41310101-INS-847
chr8-41314651-INS-67
chr8-41319850-INS-145
chr8-41387292-INS-604
chr8-41393294-INS-82
chr8-41395577-INS-65
chr8-41413113-INS-112
chr8-41414790-INS-51
chr8-41420443-INS-211
chr8-41425276-INS-353
chr8-41426723-INS-6401
chr8-41472961-INS-153
chr8-41502526-INS-587
chr8-41513265-INS-64
chr8-41513926-INS-332
chr8-41523269-INS-73
chr8-41598137-INS-197
chr8-41616688-INS-90
chr8-41642467-INS-66
chr8-41656768-INS-181
chr8-41677538-INS-578
chr8-41686926-INS-51
chr8-41691169-INS-173
chr8-41742954-INS-452
chr8-41857702-INS-53
chr8-41863803-INS-219
chr8-41897263-INS-212
chr8-41983037-INS-771
chr8-41991051-INS-52
chr8-41994087-INS-280
chr8-41999669-INS-574
chr8-42009422-INS-53
chr8-42017306-INS-171
chr8-42029340-INS-199
chr8-42038496-INS-78
chr8-42052033-INS-54
chr8-42054546-INS-139
chr8-42076704-INS-141
chr8-42084788-INS-56
chr8-42088871-INS-738
chr8-42094432-INS-665
chr8-42107509-INS-7240
chr8-42137689-INS-911
chr8-42191788-INS-82
chr8-42195275-INS-178
chr8-42197138-INS-661

chr8-42243163-INS-2288
chr8-42248218-INS-5990
chr8-42268216-INS-134
chr8-42272056-INS-112
chr8-42281675-INS-7103
chr8-42282950-INS-95
chr8-42323479-INS-54
chr8-42420253-INS-6510
chr8-42424820-INS-72
chr8-42460046-INS-359
chr8-42473629-INS-60
chr8-42476442-INS-6298
chr8-42492084-INS-88
chr8-42499357-INS-183
chr8-42504513-INS-427
chr8-42522216-INS-6900
chr8-42571928-INS-3690
chr8-42575472-INS-6343
chr8-42628211-INS-101
chr8-42670466-INS-584
chr8-42692267-INS-103
chr8-42696254-INS-548
chr8-42707895-INS-132
chr8-42774630-INS-180
chr8-42786101-INS-82
chr8-42842771-INS-265
chr8-42849043-INS-170
chr8-42880162-INS-126
chr8-42911030-INS-74
chr8-42918656-INS-264
chr8-42927270-INS-58
chr8-42929826-INS-120
chr8-42949387-INS-52
chr8-42951044-INS-491
chr8-42959332-INS-60
chr8-42966201-INS-842
chr8-42973384-INS-737
chr8-42982050-INS-102
chr8-42987962-INS-752
chr8-43000145-INS-6381
chr8-43005662-INS-491
chr8-43021317-INS-1575
chr8-43031822-INS-169
chr8-43053723-INS-351
chr8-43070310-INS-228
chr8-43087831-INS-184
chr8-43095721-INS-377
chr8-43124632-INS-296
chr8-43143218-INS-74
chr8-43204422-INS-110
chr8-43256239-INS-4411
chr8-43282751-INS-6616
chr8-43288865-INS-664
chr8-43325554-INS-123
chr8-43340842-INS-69
chr8-43348566-INS-384
chr8-43355930-INS-6223
chr8-43359992-INS-86
chr8-43366568-INS-243
chr8-43413542-INS-389
chr8-43448850-INS-619
chr8-43452181-INS-6186
chr8-43469144-INS-1390
chr8-43575531-INS-937
chr8-43616981-INS-992
chr8-43626764-INS-468
chr8-43627474-INS-52
chr8-43688030-INS-573
chr8-43691368-INS-768
chr8-43691368-INS-960
chr8-43691401-INS-127
chr8-43712325-INS-62
chr8-43722270-INS-243
chr8-43786985-INS-65
chr8-43789790-INS-52
chr8-43791199-INS-55
chr8-43823480-INS-63

chr8-43842039-INS-56
chr8-43846013-INS-386
chr8-43875401-INS-60
chr8-43885924-INS-58
chr8-43894323-INS-54
chr8-43911323-INS-136
chr8-43927545-INS-132
chr8-43930795-INS-908
chr8-43940815-INS-512
chr8-43967938-INS-186
chr8-43970372-INS-69
chr8-43992966-INS-3238
chr8-44024925-INS-510
chr8-44051180-INS-6843
chr8-44060935-INS-655
chr8-44062462-INS-226
chr8-44065989-INS-150
chr8-44091092-INS-6501
chr8-44096177-INS-3997
chr8-44102669-INS-125
chr8-44119970-INS-3972
chr8-44148956-INS-585
chr8-44163772-INS-60
chr8-44174034-INS-66
chr8-44224807-INS-55
chr8-44254071-INS-82
chr8-44283240-INS-419
chr8-44284693-INS-106
chr8-44352945-INS-6387
chr8-44356357-INS-102
chr8-44362327-INS-72
chr8-44402112-INS-683
chr8-44404704-INS-100
chr8-44407306-INS-6460
chr8-44433479-INS-80
chr8-44466932-INS-865
chr8-44479861-INS-183
chr8-44482098-INS-54
chr8-44486471-INS-112
chr8-44517198-INS-257
chr8-44537942-INS-79
chr8-44578449-INS-352
chr8-44580886-INS-74
chr8-44593261-INS-96
chr8-44597170-INS-144
chr8-44633857-INS-60
chr8-44642854-INS-114
chr8-44645727-INS-423
chr8-44649181-INS-98
chr8-44652434-INS-216
chr8-44668245-INS-190
chr8-44703135-INS-118
chr8-44709352-INS-74
chr8-44713887-INS-224
chr8-44723402-INS-129
chr8-44727912-INS-168
chr8-44775808-INS-151
chr8-44783014-INS-80
chr8-44787973-INS-477
chr8-44845189-INS-88
chr8-44854414-INS-908
chr8-44874844-INS-74
chr8-44875207-INS-167
chr8-44892729-INS-154
chr8-44906772-INS-53
chr8-44909183-INS-280
chr8-44929098-INS-174
chr8-44981217-INS-498
chr8-44987676-INS-224
chr8-45006017-INS-54
chr8-45009431-INS-541
chr8-45017910-INS-117
chr8-45028803-INS-61
chr8-45054667-INS-59
chr8-45060594-INS-89
chr8-45067408-INS-81
chr8-45086829-INS-116

chr8-45100287-INS-170
chr8-45129385-INS-75
chr8-45160123-INS-134
chr8-45163180-INS-204
chr8-45184182-INS-124
chr8-45269495-INS-271
chr8-45291335-INS-52
chr8-45320092-INS-97
chr8-45347118-INS-412
chr8-45356905-INS-193
chr8-45359177-INS-67
chr8-45373010-INS-503
chr8-45381193-INS-271
chr8-45384152-INS-53
chr8-45400681-INS-76
chr8-45442466-INS-218
chr8-45443746-INS-83
chr8-45464152-INS-65
chr8-45497637-INS-5569
chr8-45511818-INS-431
chr8-45535880-INS-65
chr8-45537650-INS-60
chr8-45553037-INS-116
chr8-45571615-INS-81
chr8-45645060-INS-63
chr8-45663590-INS-108
chr8-45668543-INS-513
chr8-45673755-INS-249
chr8-45682360-INS-327
chr8-45686350-INS-51
chr8-45687345-INS-88
chr8-45707641-INS-56
chr8-45762083-INS-143
chr8-45802638-INS-210
chr8-45803808-INS-51
chr8-45831035-INS-58
chr8-45835673-INS-172
chr8-45848438-INS-99
chr8-45858152-INS-93
chr8-45863113-INS-60
chr8-45892124-INS-128
chr8-45910755-INS-175
chr8-45915254-INS-673
chr8-45915793-INS-123
chr8-45918414-INS-54
chr8-45918940-INS-92
chr8-45946020-INS-372
chr8-45956498-INS-382
chr8-45965598-INS-412
chr8-46001907-INS-347
chr8-46032811-INS-58
chr8-46046333-INS-78
chr8-46047989-INS-456
chr8-46069289-INS-56
chr8-46097255-INS-60
chr8-46101835-INS-65
chr8-46106960-INS-60
chr8-46124609-INS-714
chr8-46131742-INS-257
chr8-46174847-INS-70
chr8-46181674-INS-241
chr8-46191751-INS-52
chr8-46216712-INS-73
chr8-46221842-INS-322
chr8-46222405-INS-52
chr8-46240668-INS-126
chr8-46245043-INS-94
chr8-46261814-INS-108
chr8-46269366-INS-60
chr8-46321356-INS-202
chr8-46336569-INS-82
chr8-46346183-INS-390
chr8-46353965-INS-55
chr8-46384432-INS-127
chr8-46416443-INS-479
chr8-46427220-INS-69
chr8-46442030-INS-196

chr8-46447451-INS-60
chr8-46459735-INS-54
chr8-46460304-INS-188
chr8-46465205-INS-156
chr8-46481334-INS-163
chr8-46488486-INS-52
chr8-46498688-INS-307
chr8-46515203-INS-131
chr8-46534392-INS-52
chr8-46551412-INS-7358
chr8-46566956-INS-128
chr8-46584409-INS-187
chr8-46673493-INS-204
chr8-46690942-INS-253
chr8-46713174-INS-419
chr8-46714871-INS-188
chr8-46734463-INS-62
chr8-46736144-INS-401
chr8-46742101-INS-122
chr8-46755528-INS-98
chr8-46758091-INS-361
chr8-46761241-INS-56
chr8-46775706-INS-178
chr8-46776540-INS-192
chr8-46778753-INS-368
chr8-46784893-INS-317
chr8-46805821-INS-51
chr8-46806300-INS-225
chr8-46808721-INS-669
chr8-46810258-INS-7091
chr8-46814347-INS-95
chr8-46816622-INS-775
chr8-46855350-INS-836
chr8-46858311-INS-103
chr8-46874049-INS-70
chr8-46880586-INS-75
chr8-46895541-INS-99
chr8-46903747-INS-131
chr8-46929934-INS-59
chr8-46953866-INS-6320
chr8-46968528-INS-222
chr8-46998543-INS-55
chr8-47049559-INS-56
chr8-47058469-INS-451
chr8-47067684-INS-280
chr8-47068702-INS-165
chr8-47074539-INS-139
chr8-47079129-INS-52
chr8-47104212-INS-406
chr8-47125515-INS-108
chr8-47135537-INS-6840
chr8-47143976-INS-78
chr8-47145184-INS-55
chr8-47167332-INS-65
chr8-47220453-INS-53
chr8-47228414-INS-195
chr8-47247294-INS-62
chr8-47340956-INS-248
chr8-47353195-INS-67
chr8-47362389-INS-99
chr8-47367309-INS-227
chr8-47432170-INS-66
chr8-47452463-INS-87
chr8-47457338-INS-123
chr8-47457417-INS-129
chr8-47459694-INS-190
chr8-47473342-INS-124
chr8-47481292-INS-252
chr8-47494209-INS-60
chr8-47499983-INS-105
chr8-47500071-INS-130
chr8-47516138-INS-69
chr8-47554973-INS-69
chr8-47558419-INS-137
chr8-47575129-INS-125
chr8-47579738-INS-211
chr8-47689404-INS-98

chr8-47697908-INS-80
chr8-47700502-INS-176
chr8-47706682-INS-57
chr8-47711374-INS-90
chr8-47715599-INS-233
chr8-47717590-INS-59
chr8-47726160-INS-72
chr8-47731781-INS-196
chr8-47734755-INS-100
chr8-47744865-INS-225
chr8-47772238-INS-54
chr8-47782378-INS-157
chr8-47808832-INS-72
chr8-47814607-INS-70
chr8-47831474-INS-73
chr8-47857513-INS-7080
chr8-47860807-INS-54
chr8-47867503-INS-98
chr8-47925503-INS-66
chr8-47932178-INS-66
chr8-47933545-INS-55
chr8-47939627-INS-60
chr8-47941598-INS-178
chr8-47951954-INS-143
chr8-47961951-INS-362
chr8-47978144-INS-180
chr8-47996133-INS-63
chr8-47997551-INS-55
chr8-48010757-INS-427
chr8-48025894-INS-979
chr8-48033390-INS-132
chr8-48053245-INS-224
chr8-48063576-INS-60
chr8-48065139-INS-53
chr8-48094768-INS-930
chr8-48101214-INS-142
chr8-48106044-INS-498
chr8-48107025-INS-304
chr8-48111237-INS-121
chr8-48116987-INS-56
chr8-48123215-INS-124
chr8-48132145-INS-208
chr8-48135694-INS-229
chr8-48155056-INS-60
chr8-48173014-INS-100
chr8-48179202-INS-109
chr8-48181954-INS-192
chr8-48182827-INS-51
chr8-48185780-INS-355
chr8-48202299-INS-272
chr8-48208348-INS-200
chr8-48212232-INS-224
chr8-48229134-INS-173
chr8-48233743-INS-77
chr8-48245098-INS-188
chr8-48251537-INS-151
chr8-48260951-INS-85
chr8-48269176-INS-61
chr8-48270890-INS-2695
chr8-48272861-INS-77
chr8-48289230-INS-122
chr8-48304061-INS-550
chr8-48310014-INS-60
chr8-48346461-INS-406
chr8-48367541-INS-100
chr8-48375879-INS-495
chr8-48394526-INS-120
chr8-48398329-INS-64
chr8-48405755-INS-148
chr8-48434737-INS-60
chr8-48440845-INS-147
chr8-48463520-INS-76
chr8-48484106-INS-167
chr8-48485054-INS-220
chr8-48490647-INS-217
chr8-48511354-INS-400
chr8-48536176-INS-76

chr8-48538851-INS-74
chr8-48544153-INS-193
chr8-48549924-INS-236
chr8-48574699-INS-540
chr8-48614631-INS-251
chr8-48633596-INS-54
chr8-48651907-INS-93
chr8-48664273-INS-180
chr8-48699569-INS-81
chr8-48717031-INS-104
chr8-48722655-INS-214
chr8-48746229-INS-75
chr8-48778313-INS-506
chr8-48814696-INS-206
chr8-48849440-INS-56
chr8-48853654-INS-55
chr8-48857435-INS-68
chr8-48895117-INS-223
chr8-48924588-INS-6705
chr8-48936009-INS-62
chr8-48956307-INS-524
chr8-48975798-INS-92
chr8-48992596-INS-237
chr8-49032284-INS-197
chr8-49036372-INS-60
chr8-49040388-INS-63
chr8-49061559-INS-57
chr8-49071953-INS-399
chr8-49104354-INS-77
chr8-49107191-INS-80
chr8-49112429-INS-272
chr8-49113166-INS-120
chr8-49116007-INS-72
chr8-49121945-INS-54
chr8-49124928-INS-123
chr8-49135251-INS-213
chr8-49171471-INS-220
chr8-49221383-INS-7089
chr8-49228252-INS-122
chr8-49328920-INS-293
chr8-49363378-INS-83
chr8-49365335-INS-68
chr8-49367384-INS-56
chr8-49373896-INS-240
chr8-49401533-INS-702
chr8-49410608-INS-223
chr8-49416903-INS-6252
chr8-49421613-INS-52
chr8-49424417-INS-6699
chr8-49433505-INS-206
chr8-49445407-INS-63
chr8-49448208-INS-57
chr8-49456943-INS-76
chr8-49474292-INS-392
chr8-49492175-INS-94
chr8-49499971-INS-57
chr8-49548504-INS-52
chr8-49590778-INS-62
chr8-49633591-INS-928
chr8-49643569-INS-51
chr8-49700881-INS-92
chr8-49719446-INS-52
chr8-49722689-INS-275
chr8-49728928-INS-62
chr8-49750409-INS-56
chr8-49763776-INS-292
chr8-49778209-INS-6603
chr8-49784092-INS-175
chr8-49803422-INS-167
chr8-49805377-INS-2034
chr8-49805769-INS-341
chr8-49812606-INS-65
chr8-49835863-INS-91
chr8-49844030-INS-62
chr8-49845264-INS-57
chr8-49855111-INS-983
chr8-49892553-INS-289

chr8-49904970-INS-141
chr8-49906644-INS-240
chr8-49921618-INS-56
chr8-49925262-INS-113
chr8-49926662-INS-120
chr8-49985016-INS-654
chr8-50007197-INS-52
chr8-50025474-INS-54
chr8-50040129-INS-484
chr8-50042621-INS-153
chr8-50064930-INS-100
chr8-50065825-INS-7009
chr8-50067606-INS-61
chr8-50070583-INS-127
chr8-50084186-INS-129
chr8-50092783-INS-270
chr8-50124592-INS-124
chr8-50126607-INS-500
chr8-50192284-INS-239
chr8-50208211-INS-511
chr8-50228583-INS-56
chr8-50246712-INS-954
chr8-50276430-INS-54
chr8-50300999-INS-64
chr8-50302939-INS-75
chr8-50344577-INS-70
chr8-50346581-INS-516
chr8-50357473-INS-90
chr8-50367364-INS-53
chr8-50368291-INS-575
chr8-50393994-INS-68
chr8-50420797-INS-53
chr8-50477468-INS-60
chr8-50491498-INS-70
chr8-50507567-INS-85
chr8-50513506-INS-52
chr8-50533469-INS-208
chr8-50544086-INS-224
chr8-50547963-INS-66
chr8-50558676-INS-6178
chr8-50592787-INS-315
chr8-50598119-INS-2135
chr8-50640262-INS-110
chr8-50659768-INS-70
chr8-50660947-INS-76
chr8-50667105-INS-178
chr8-50708413-INS-1892
chr8-50712003-INS-164
chr8-50723133-INS-61
chr8-50754832-INS-2832
chr8-50765080-INS-58
chr8-50770440-INS-111
chr8-50771929-INS-555
chr8-50791597-INS-73
chr8-50812068-INS-88
chr8-50820872-INS-59
chr8-50833791-INS-905
chr8-50833811-INS-5313
chr8-50838917-INS-132
chr8-50842269-INS-602
chr8-50845525-INS-151
chr8-50859693-INS-180
chr8-50865935-INS-173
chr8-50870942-INS-6452
chr8-50880274-INS-66
chr8-50881328-INS-862
chr8-50887861-INS-334
chr8-50903948-INS-188
chr8-50934782-INS-343
chr8-50939818-INS-126
chr8-50952800-INS-789
chr8-50977694-INS-72
chr8-51005108-INS-621
chr8-51013535-INS-118
chr8-51026510-INS-106
chr8-51041480-INS-73
chr8-51044102-INS-72

chr8-51044924-INS-475
chr8-51058537-INS-170
chr8-51066854-INS-123
chr8-51091784-INS-108
chr8-51098868-INS-600
chr8-51115844-INS-80
chr8-51135558-INS-763
chr8-51145089-INS-54
chr8-51146475-INS-497
chr8-51148646-INS-173
chr8-51172121-INS-498
chr8-51182249-INS-56
chr8-51187915-INS-308
chr8-51231468-INS-422
chr8-51235789-INS-122
chr8-51292590-INS-57
chr8-51295163-INS-58
chr8-51302996-INS-131
chr8-51325928-INS-212
chr8-51360218-INS-131
chr8-51384738-INS-253
chr8-51390004-INS-3605
chr8-51398510-INS-52
chr8-51400531-INS-363
chr8-51402403-INS-54
chr8-51417165-INS-665
chr8-51444776-INS-77
chr8-51477160-INS-84
chr8-51481108-INS-90
chr8-51514235-INS-94
chr8-51519238-INS-58
chr8-51546593-INS-52
chr8-51554011-INS-56
chr8-51559994-INS-628
chr8-51569766-INS-52
chr8-51610458-INS-7587
chr8-51610458-INS-876
chr8-51642354-INS-506
chr8-51644286-INS-60
chr8-51656582-INS-61
chr8-51686066-INS-68
chr8-51692905-INS-104
chr8-51695141-INS-95
chr8-51700667-INS-835
chr8-51716325-INS-102
chr8-51732176-INS-59
chr8-51737571-INS-666
chr8-51738852-INS-6972
chr8-51746948-INS-676
chr8-51788414-INS-6920
chr8-51790362-INS-76
chr8-51808562-INS-81
chr8-51809964-INS-92
chr8-51813379-INS-604
chr8-51817106-INS-54
chr8-51819688-INS-108
chr8-51824355-INS-76
chr8-51826298-INS-86
chr8-51858669-INS-101
chr8-51863531-INS-452
chr8-51867291-INS-306
chr8-51929677-INS-52
chr8-52015916-INS-959
chr8-52023519-INS-76
chr8-52030202-INS-390
chr8-52043584-INS-61
chr8-52073545-INS-4174
chr8-52084975-INS-280
chr8-52090569-INS-236
chr8-52122042-INS-52
chr8-52124520-INS-162
chr8-52142561-INS-216
chr8-52151349-INS-200
chr8-52172985-INS-58
chr8-52191974-INS-54
chr8-52198270-INS-6537
chr8-52222993-INS-1726

chr8-52272904-INS-507
chr8-52308534-INS-89
chr8-52311066-INS-402
chr8-52331090-INS-999
chr8-52353186-INS-72
chr8-52363255-INS-126
chr8-52366226-INS-683
chr8-52421850-INS-6399
chr8-52438939-INS-198
chr8-52471686-INS-7439
chr8-52507906-INS-7407
chr8-52533969-INS-874
chr8-52582809-INS-61
chr8-52588660-INS-80
chr8-52648296-INS-55
chr8-52691713-INS-485
chr8-52696652-INS-54
chr8-52773141-INS-6414
chr8-52774336-INS-65
chr8-52788970-INS-51
chr8-52820899-INS-72
chr8-52835392-INS-67
chr8-52861297-INS-497
chr8-52863101-INS-414
chr8-52874440-INS-6882
chr8-52906431-INS-141
chr8-52916065-INS-58
chr8-52926796-INS-296
chr8-52935917-INS-138
chr8-52949100-INS-76
chr8-52983498-INS-400
chr8-53000380-INS-991
chr8-53003662-INS-149
chr8-53026622-INS-390
chr8-53028508-INS-474
chr8-53054109-INS-415
chr8-53071434-INS-6185
chr8-53073806-INS-4211
chr8-53091592-INS-407
chr8-53113305-INS-2841
chr8-53118055-INS-54
chr8-53154452-INS-566
chr8-53164266-INS-510
chr8-53178289-INS-51
chr8-53196583-INS-123
chr8-53212768-INS-54
chr8-53235390-INS-5258
chr8-53243400-INS-188
chr8-53271087-INS-6452
chr8-53286951-INS-201
chr8-53308044-INS-2782
chr8-53321067-INS-5733
chr8-53321944-INS-379
chr8-53332522-INS-224
chr8-53340419-INS-6431
chr8-53343808-INS-2910
chr8-53346934-INS-131
chr8-53356682-INS-53
chr8-53364409-INS-340
chr8-53385960-INS-357
chr8-53403099-INS-2537
chr8-53457633-INS-81
chr8-53473220-INS-352
chr8-53482050-INS-6193
chr8-53495702-INS-1194
chr8-53498919-INS-663
chr8-53503143-INS-1429
chr8-53506561-INS-8325
chr8-53516836-INS-124
chr8-53524486-INS-132
chr8-53543284-INS-977
chr8-53554064-INS-64
chr8-53563385-INS-63
chr8-53583860-INS-229
chr8-53588965-INS-408
chr8-53593590-INS-357
chr8-53596207-INS-5243

chr8-53607767-INS-120
chr8-53636293-INS-103
chr8-53639024-INS-6420
chr8-53639671-INS-128
chr8-53640880-INS-224
chr8-53645518-INS-198
chr8-53660572-INS-363
chr8-53672345-INS-124
chr8-53698796-INS-359
chr8-53772057-INS-209
chr8-53794655-INS-68
chr8-53840895-INS-893
chr8-53843639-INS-7086
chr8-53846569-INS-156
chr8-53851292-INS-52
chr8-53873172-INS-848
chr8-53882847-INS-56
chr8-53917469-INS-75
chr8-53923524-INS-628
chr8-53948225-INS-64
chr8-53950959-INS-578
chr8-53962876-INS-54
chr8-53972037-INS-92
chr8-53989780-INS-126
chr8-53992013-INS-70
chr8-54003717-INS-194
chr8-54029401-INS-198
chr8-54029492-INS-6443
chr8-54030282-INS-59
chr8-54032803-INS-209
chr8-54043683-INS-172
chr8-54045776-INS-58
chr8-54046284-INS-92
chr8-54048756-INS-79
chr8-54053745-INS-60
chr8-54056795-INS-87
chr8-54061476-INS-625
chr8-54063437-INS-7143
chr8-54073231-INS-473
chr8-54080278-INS-6257
chr8-54095050-INS-400
chr8-54100087-INS-82
chr8-54128876-INS-178
chr8-54131623-INS-546
chr8-54136994-INS-351
chr8-54140659-INS-6390
chr8-54155006-INS-67
chr8-54175719-INS-6385
chr8-54183394-INS-60
chr8-54211569-INS-333
chr8-54223367-INS-830
chr8-54231347-INS-246
chr8-54236127-INS-799
chr8-54239443-INS-166
chr8-54250799-INS-7146
chr8-54276242-INS-125
chr8-54295638-INS-224
chr8-54297967-INS-59
chr8-54298489-INS-52
chr8-54314118-INS-108
chr8-54316016-INS-7112
chr8-54319102-INS-62
chr8-54336940-INS-54
chr8-54348443-INS-6900
chr8-54364731-INS-547
chr8-54374123-INS-111
chr8-54382195-INS-94
chr8-54396934-INS-6877
chr8-54399127-INS-154
chr8-54437274-INS-78
chr8-54444751-INS-364
chr8-54449191-INS-5468
chr8-54455521-INS-74
chr8-54462659-INS-133
chr8-54473362-INS-5335
chr8-54474626-INS-5379
chr8-54479627-INS-646

chr8-54493626-INS-72
chr8-54502308-INS-105
chr8-54502342-INS-5549
chr8-54524605-INS-94
chr8-54526624-INS-357
chr8-54539168-INS-127
chr8-54563461-INS-70
chr8-54608374-INS-62
chr8-54629757-INS-54
chr8-54640328-INS-139
chr8-54644093-INS-471
chr8-54662019-INS-60
chr8-54672293-INS-102
chr8-54708555-INS-276
chr8-54709479-INS-72
chr8-54718259-INS-7957
chr8-54742611-INS-6691
chr8-54760530-INS-426
chr8-54761643-INS-202
chr8-54780358-INS-399
chr8-54787676-INS-313
chr8-54807970-INS-352
chr8-54821499-INS-76
chr8-54829491-INS-959
chr8-54835979-INS-56
chr8-54838166-INS-5259
chr8-54844973-INS-326
chr8-54855581-INS-426
chr8-54890182-INS-701
chr8-54905483-INS-58
chr8-54906473-INS-284
chr8-54929209-INS-147
chr8-54931609-INS-293
chr8-54938915-INS-72
chr8-54951027-INS-830
chr8-54956019-INS-57
chr8-54965274-INS-7192
chr8-54969998-INS-475
chr8-55028493-INS-165
chr8-55055911-INS-224
chr8-55056624-INS-786
chr8-55058389-INS-120
chr8-55097516-INS-69
chr8-55124610-INS-60
chr8-55139957-INS-103
chr8-55147506-INS-74
chr8-55152472-INS-391
chr8-55172033-INS-656
chr8-55193021-INS-573
chr8-55194541-INS-2561
chr8-55202288-INS-204
chr8-55205633-INS-157
chr8-55217536-INS-56
chr8-55237808-INS-247
chr8-55250879-INS-951
chr8-55261992-INS-143
chr8-55285463-INS-56
chr8-55336864-INS-86
chr8-55353800-INS-569
chr8-55362028-INS-71
chr8-55372988-INS-55
chr8-55377379-INS-56
chr8-55406165-INS-68
chr8-55418771-INS-805
chr8-55443596-INS-59
chr8-55459221-INS-467
chr8-55477894-INS-590
chr8-55483787-INS-130
chr8-55486445-INS-68
chr8-55496282-INS-190
chr8-55503399-INS-118
chr8-55505846-INS-55
chr8-55510991-INS-707
chr8-55862240-INS-81
chr8-56007541-INS-3136
chr8-56011933-INS-7612
chr8-56046475-INS-6536

chr8-56052060-INS-6657
chr8-56069367-INS-401
chr8-56073430-INS-127
chr8-56097583-INS-144
chr8-56099550-INS-919
chr8-56114191-INS-150
chr8-56115279-INS-874
chr8-56116872-INS-6208
chr8-56196170-INS-201
chr8-56231435-INS-61
chr8-56254338-INS-599
chr8-56259018-INS-214
chr8-56267521-INS-307
chr8-56281957-INS-7813
chr8-56327085-INS-223
chr8-56358763-INS-317
chr8-56370672-INS-928
chr8-56405477-INS-314
chr8-56407706-INS-62
chr8-56410933-INS-76
chr8-56420598-INS-379
chr8-56494271-INS-64
chr8-56511235-INS-61
chr8-56521722-INS-367
chr8-56522688-INS-102
chr8-56528181-INS-103
chr8-56534047-INS-74
chr8-56543703-INS-6797
chr8-56551266-INS-6795
chr8-56562414-INS-82
chr8-56569112-INS-58
chr8-56571948-INS-296
chr8-56600968-INS-319
chr8-56607467-INS-433
chr8-56642166-INS-492
chr8-56660860-INS-343
chr8-56707878-INS-196
chr8-56711692-INS-255
chr8-56716665-INS-112
chr8-56772568-INS-94
chr8-56779750-INS-1720
chr8-56801417-INS-359
chr8-56833414-INS-94
chr8-56834808-INS-2469
chr8-56872113-INS-933
chr8-56883263-INS-493
chr8-56901527-INS-324
chr8-56905113-INS-353
chr8-56915161-INS-156
chr8-56919910-INS-1862
chr8-56923544-INS-414
chr8-56928720-INS-54
chr8-56937894-INS-214
chr8-56940014-INS-79
chr8-56959896-INS-80
chr8-56968712-INS-6584
chr8-56977359-INS-7267
chr8-56981309-INS-62
chr8-56987184-INS-52
chr8-56990286-INS-68
chr8-57002427-INS-55
chr8-57013435-INS-4520
chr8-57016448-INS-1211
chr8-57018461-INS-68
chr8-57034121-INS-210
chr8-57044053-INS-143
chr8-57052500-INS-450
chr8-57088924-INS-134
chr8-57095956-INS-588
chr8-57101910-INS-76
chr8-57103623-INS-363
chr8-57115291-INS-105
chr8-57122975-INS-440
chr8-57140188-INS-52
chr8-57143230-INS-51
chr8-57149557-INS-71
chr8-57151141-INS-96

chr8-57155881-INS-56
chr8-57177904-INS-108
chr8-57196257-INS-132
chr8-57224508-INS-531
chr8-57227450-INS-709
chr8-57265962-INS-289
chr8-57267507-INS-79
chr8-57269598-INS-126
chr8-57282106-INS-5983
chr8-57340913-INS-68
chr8-57357171-INS-71
chr8-57371887-INS-66
chr8-57391199-INS-6248
chr8-57410153-INS-109
chr8-57412445-INS-1405
chr8-57414240-INS-232
chr8-57426050-INS-58
chr8-57426620-INS-697
chr8-57430127-INS-125
chr8-57449514-INS-189
chr8-57452516-INS-712
chr8-57452532-INS-69
chr8-57464529-INS-2256
chr8-57479231-INS-115
chr8-57482167-INS-202
chr8-57507915-INS-156
chr8-57513765-INS-1927
chr8-57529135-INS-604
chr8-57535038-INS-60
chr8-57546445-INS-739
chr8-57559382-INS-52
chr8-57589495-INS-71
chr8-57604696-INS-6539
chr8-57606742-INS-186
chr8-57609735-INS-69
chr8-57616959-INS-272
chr8-57619071-INS-6355
chr8-57654897-INS-72
chr8-57656162-INS-82
chr8-57665029-INS-51
chr8-57677698-INS-168
chr8-57690683-INS-6482
chr8-57698701-INS-94
chr8-57702878-INS-73
chr8-57706585-INS-100
chr8-57719931-INS-132
chr8-57744208-INS-332
chr8-57784364-INS-54
chr8-57819856-INS-61
chr8-57869504-INS-66
chr8-57882893-INS-101
chr8-57893044-INS-352
chr8-57927657-INS-53
chr8-57928598-INS-281
chr8-57931927-INS-76
chr8-57956742-INS-881
chr8-57957600-INS-60
chr8-57969783-INS-247
chr8-57976643-INS-207
chr8-57981540-INS-1887
chr8-57998077-INS-6442
chr8-58020081-INS-291
chr8-58024629-INS-505
chr8-58027363-INS-201
chr8-58029916-INS-177
chr8-58033632-INS-232
chr8-58049784-INS-185
chr8-58069919-INS-61
chr8-58081958-INS-54
chr8-58083013-INS-529
chr8-58121552-INS-52
chr8-58182853-INS-77
chr8-58191441-INS-463
chr8-58193245-INS-55
chr8-58209766-INS-115
chr8-58216675-INS-57
chr8-58217914-INS-804

chr8-58221198-INS-52
chr8-58262620-INS-472
chr8-58310093-INS-70
chr8-58314955-INS-176
chr8-58325429-INS-76
chr8-58334198-INS-56
chr8-58387073-INS-186
chr8-58396054-INS-53
chr8-58501720-INS-6129
chr8-58503161-INS-620
chr8-58532763-INS-205
chr8-58534556-INS-386
chr8-58546042-INS-119
chr8-58558565-INS-131
chr8-58567605-INS-560
chr8-58579927-INS-53
chr8-58591901-INS-69
chr8-58595930-INS-65
chr8-58615029-INS-9969
chr8-58615030-INS-284
chr8-58616050-INS-122
chr8-58645297-INS-65
chr8-58647864-INS-60
chr8-58656934-INS-228
chr8-58662255-INS-66
chr8-58663885-INS-200
chr8-58668512-INS-243
chr8-58671590-INS-51
chr8-58688739-INS-226
chr8-58705068-INS-484
chr8-58712251-INS-2742
chr8-58730653-INS-932
chr8-58739890-INS-470
chr8-58753186-INS-58
chr8-58763069-INS-959
chr8-58768827-INS-2929
chr8-58820258-INS-212
chr8-58851702-INS-261
chr8-58884489-INS-962
chr8-58931274-INS-124
chr8-58973524-INS-210
chr8-58999808-INS-6185
chr8-59002731-INS-676
chr8-59012661-INS-1042
chr8-59031965-INS-409
chr8-59038777-INS-490
chr8-59053645-INS-135
chr8-59053910-INS-329
chr8-59057515-INS-768
chr8-59102289-INS-201
chr8-59119601-INS-60
chr8-59144653-INS-58
chr8-59146078-INS-525
chr8-59157046-INS-289
chr8-59216615-INS-66
chr8-59222871-INS-1282
chr8-59224800-INS-552
chr8-59226003-INS-765
chr8-59230690-INS-95
chr8-59232879-INS-67
chr8-59240672-INS-404
chr8-59246764-INS-132
chr8-59286951-INS-51
chr8-59308742-INS-338
chr8-59331571-INS-925
chr8-59333571-INS-56
chr8-59359419-INS-53
chr8-59372320-INS-114
chr8-59409465-INS-209
chr8-59416054-INS-186
chr8-59421021-INS-54
chr8-59435556-INS-90
chr8-59456646-INS-52
chr8-59458744-INS-6884
chr8-59465734-INS-56
chr8-59527597-INS-316
chr8-59552817-INS-56

chr8-59583536-INS-92
chr8-59598724-INS-6186
chr8-59624586-INS-255
chr8-59629749-INS-54
chr8-59638869-INS-181
chr8-59655506-INS-6383
chr8-59655507-INS-356
chr8-59674998-INS-6277
chr8-59787013-INS-85
chr8-59833625-INS-88
chr8-59839674-INS-959
chr8-59853047-INS-56
chr8-59854732-INS-154
chr8-59890304-INS-275
chr8-59895638-INS-147
chr8-59917267-INS-80
chr8-59945743-INS-78
chr8-60011069-INS-160
chr8-60061283-INS-56
chr8-60094162-INS-2893
chr8-60103067-INS-212
chr8-60125168-INS-197
chr8-60183784-INS-186
chr8-60200880-INS-5522
chr8-60217274-INS-55
chr8-60271663-INS-60
chr8-60276004-INS-161
chr8-60284445-INS-200
chr8-60286472-INS-70
chr8-60299628-INS-6219
chr8-60301421-INS-223
chr8-60347158-INS-232
chr8-60413917-INS-58
chr8-60434698-INS-189
chr8-60437192-INS-928
chr8-60442118-INS-896
chr8-60495229-INS-77
chr8-60537478-INS-254
chr8-60537829-INS-6257
chr8-60545161-INS-354
chr8-60547313-INS-195
chr8-60555817-INS-616
chr8-60608372-INS-509
chr8-60627887-INS-61
chr8-60646924-INS-146
chr8-60649203-INS-112
chr8-60665764-INS-1342
chr8-60669395-INS-319
chr8-60677206-INS-213
chr8-60707261-INS-963
chr8-60718128-INS-206
chr8-60721244-INS-120
chr8-60729547-INS-445
chr8-60746285-INS-137
chr8-60753117-INS-96
chr8-60767602-INS-67
chr8-60770870-INS-6379
chr8-60809194-INS-5950
chr8-60834993-INS-56
chr8-60917042-INS-269
chr8-60943891-INS-636
chr8-60953339-INS-58
chr8-60974612-INS-328
chr8-60978301-INS-228
chr8-60994789-INS-90
chr8-60997681-INS-62
chr8-61003896-INS-261
chr8-61015712-INS-1376
chr8-61094264-INS-189
chr8-61134094-INS-154
chr8-61168663-INS-56
chr8-61173193-INS-56
chr8-61187705-INS-130
chr8-61200124-INS-55
chr8-61202361-INS-59
chr8-61221954-INS-402
chr8-61252904-INS-473

chr8-61255082-INS-132
chr8-61295097-INS-56
chr8-61325612-INS-129
chr8-61339640-INS-193
chr8-61354433-INS-423
chr8-61363295-INS-502
chr8-61383166-INS-1092
chr8-61409455-INS-69
chr8-61417574-INS-51
chr8-61418098-INS-6475
chr8-61461051-INS-7210
chr8-61474468-INS-60
chr8-61477327-INS-1470
chr8-61478063-INS-54
chr8-61491242-INS-59
chr8-61541862-INS-211
chr8-61585555-INS-169
chr8-61596599-INS-96
chr8-61632868-INS-57
chr8-61641999-INS-197
chr8-61648209-INS-499
chr8-61658616-INS-105
chr8-61663845-INS-62
chr8-61690928-INS-155
chr8-61694157-INS-57
chr8-61715478-INS-62
chr8-61719132-INS-390
chr8-61731149-INS-54
chr8-61762412-INS-51
chr8-61773630-INS-63
chr8-61779015-INS-169
chr8-61779799-INS-137
chr8-61781438-INS-337
chr8-61820185-INS-59
chr8-61824338-INS-59
chr8-61829598-INS-140
chr8-61843531-INS-5218
chr8-61853259-INS-93
chr8-61879087-INS-92
chr8-61899543-INS-6599
chr8-61906790-INS-98
chr8-61925527-INS-788
chr8-61926201-INS-285
chr8-61951303-INS-404
chr8-61957362-INS-67
chr8-61959421-INS-75
chr8-61966967-INS-75
chr8-61985617-INS-56
chr8-62035123-INS-84
chr8-62060274-INS-136
chr8-62075065-INS-55
chr8-62103326-INS-54
chr8-62112411-INS-134
chr8-62116384-INS-78
chr8-62133150-INS-159
chr8-62136415-INS-64
chr8-62168258-INS-84
chr8-62182015-INS-204
chr8-62197165-INS-83
chr8-62220376-INS-62
chr8-62245335-INS-358
chr8-62264446-INS-58
chr8-62267323-INS-180
chr8-62305860-INS-286
chr8-62358613-INS-6605
chr8-62379732-INS-448
chr8-62391112-INS-242
chr8-62398199-INS-454
chr8-62443874-INS-68
chr8-62445492-INS-153
chr8-62446976-INS-53
chr8-62479847-INS-798
chr8-62480392-INS-59
chr8-62507220-INS-1150
chr8-62518454-INS-78
chr8-62524911-INS-264
chr8-62532787-INS-560

chr8-62546779-INS-7107
chr8-62548239-INS-99
chr8-62549471-INS-7030
chr8-62564978-INS-51
chr8-62565351-INS-7140
chr8-62585310-INS-114
chr8-62604862-INS-6734
chr8-62625897-INS-90
chr8-62640627-INS-5769
chr8-62642399-INS-55
chr8-62672634-INS-57
chr8-62686475-INS-93
chr8-62692130-INS-173
chr8-62697036-INS-67
chr8-62735083-INS-144
chr8-62742243-INS-61
chr8-62763593-INS-6286
chr8-62764272-INS-53
chr8-62768319-INS-295
chr8-62787882-INS-200
chr8-62788465-INS-154
chr8-62812185-INS-3788
chr8-62820236-INS-859
chr8-62848915-INS-127
chr8-62863644-INS-399
chr8-62875939-INS-2694
chr8-62880247-INS-1328
chr8-62880316-INS-117
chr8-62884517-INS-72
chr8-62886306-INS-834
chr8-62888109-INS-351
chr8-62906513-INS-470
chr8-62908518-INS-270
chr8-62911466-INS-105
chr8-62918812-INS-6999
chr8-62919261-INS-53
chr8-62920326-INS-2875
chr8-62959373-INS-57
chr8-62977281-INS-825
chr8-62994516-INS-103
chr8-63065964-INS-727
chr8-63092077-INS-71
chr8-63111224-INS-152
chr8-63113486-INS-174
chr8-63144459-INS-164
chr8-63146882-INS-285
chr8-63150138-INS-55
chr8-63176588-INS-442
chr8-63192446-INS-80
chr8-63197795-INS-389
chr8-63210700-INS-642
chr8-63211217-INS-543
chr8-63236629-INS-557
chr8-63288106-INS-4667
chr8-63324510-INS-6885
chr8-63333234-INS-71
chr8-63346095-INS-6572
chr8-63377548-INS-395
chr8-63380486-INS-4751
chr8-63416127-INS-231
chr8-63443726-INS-82
chr8-63451139-INS-1280
chr8-63453469-INS-153
chr8-63458899-INS-805
chr8-63460237-INS-621
chr8-63467788-INS-53
chr8-63477476-INS-376
chr8-63478764-INS-160
chr8-63489462-INS-218
chr8-63494370-INS-413
chr8-63508673-INS-172
chr8-63529895-INS-80
chr8-63539221-INS-241
chr8-63544001-INS-53
chr8-63554238-INS-538
chr8-63569278-INS-73
chr8-63591407-INS-51

chr8-63592591-INS-5616
chr8-63616892-INS-94
chr8-63623765-INS-70
chr8-63631400-INS-6770
chr8-63638170-INS-1708
chr8-63638351-INS-56
chr8-63664318-INS-178
chr8-63666675-INS-113
chr8-63672608-INS-633
chr8-63673631-INS-141
chr8-63687992-INS-498
chr8-63740827-INS-113
chr8-63759512-INS-200
chr8-63774054-INS-6517
chr8-63782444-INS-387
chr8-63795829-INS-74
chr8-63816305-INS-51
chr8-63846842-INS-56
chr8-63849321-INS-791
chr8-63851861-INS-350
chr8-63860008-INS-7552
chr8-63881172-INS-60
chr8-63892054-INS-696
chr8-63905338-INS-363
chr8-63920838-INS-6440
chr8-63985703-INS-67
chr8-63985891-INS-610
chr8-64002665-INS-63
chr8-64009218-INS-468
chr8-64049139-INS-144
chr8-64054973-INS-77
chr8-64059387-INS-600
chr8-64061667-INS-413
chr8-64063942-INS-120
chr8-64109499-INS-619
chr8-64145964-INS-6434
chr8-64150867-INS-181
chr8-64153674-INS-78
chr8-64176709-INS-116
chr8-64179197-INS-204
chr8-64181648-INS-126
chr8-64221167-INS-504
chr8-64253384-INS-561
chr8-64282981-INS-430
chr8-64296591-INS-66
chr8-64317146-INS-485
chr8-64321229-INS-329
chr8-64322464-INS-54
chr8-64325480-INS-157
chr8-64393243-INS-195
chr8-64403181-INS-85
chr8-64410242-INS-440
chr8-64424022-INS-94
chr8-64434684-INS-67
chr8-64469997-INS-251
chr8-64487746-INS-487
chr8-64490213-INS-79
chr8-64492765-INS-403
chr8-64515053-INS-523
chr8-64525628-INS-66
chr8-64539589-INS-887
chr8-64565869-INS-719
chr8-64570604-INS-86
chr8-64580503-INS-83
chr8-64585733-INS-52
chr8-64590975-INS-141
chr8-64601951-INS-154
chr8-64621233-INS-530
chr8-64626326-INS-398
chr8-64632541-INS-72
chr8-64642510-INS-72
chr8-64670594-INS-93
chr8-64685713-INS-648
chr8-64709385-INS-443
chr8-64748410-INS-62
chr8-64749636-INS-52
chr8-64753206-INS-253

chr8-64776814-INS-90
chr8-64783347-INS-5156
chr8-64809662-INS-139
chr8-64816078-INS-242
chr8-64847083-INS-56
chr8-64856405-INS-449
chr8-64862534-INS-59
chr8-64864755-INS-115
chr8-64871974-INS-514
chr8-64873768-INS-80
chr8-64899365-INS-423
chr8-64909946-INS-115
chr8-64922460-INS-787
chr8-64931470-INS-6648
chr8-64935042-INS-1440
chr8-64936778-INS-72
chr8-64938395-INS-5512
chr8-64956289-INS-577
chr8-64966165-INS-386
chr8-64968448-INS-59
chr8-64974389-INS-227
chr8-64977302-INS-75
chr8-64977530-INS-690
chr8-64981626-INS-281
chr8-64991859-INS-60
chr8-64998623-INS-239
chr8-65006767-INS-58
chr8-65006836-INS-210
chr8-65017166-INS-72
chr8-65058124-INS-99
chr8-65060933-INS-81
chr8-65070014-INS-241
chr8-65086623-INS-55
chr8-65117618-INS-53
chr8-65142499-INS-58
chr8-65216511-INS-479
chr8-65219791-INS-60
chr8-65249565-INS-176
chr8-65253729-INS-189
chr8-65257467-INS-60
chr8-65259410-INS-508
chr8-65260934-INS-224
chr8-65263646-INS-52
chr8-65266000-INS-60
chr8-65320768-INS-141
chr8-65342200-INS-58
chr8-65342628-INS-183
chr8-65352852-INS-6296
chr8-65382512-INS-149
chr8-65384206-INS-84
chr8-65397554-INS-62
chr8-65426715-INS-6403
chr8-65432964-INS-400
chr8-65434887-INS-130
chr8-65456327-INS-160
chr8-65469622-INS-124
chr8-65476474-INS-70
chr8-65485652-INS-400
chr8-65533566-INS-276
chr8-65541040-INS-345
chr8-65553983-INS-153
chr8-65568752-INS-58
chr8-65572541-INS-104
chr8-65608617-INS-62
chr8-65610272-INS-81
chr8-65623838-INS-218
chr8-65635591-INS-482
chr8-65637411-INS-90
chr8-65648889-INS-82
chr8-65650366-INS-115
chr8-65656090-INS-65
chr8-65659233-INS-67
chr8-65661739-INS-117
chr8-65665420-INS-92
chr8-65669715-INS-974
chr8-65679116-INS-192
chr8-65694601-INS-1890

chr8-65723249-INS-114
chr8-65726139-INS-492
chr8-65764124-INS-156
chr8-65764840-INS-64
chr8-65768250-INS-5091
chr8-65773354-INS-378
chr8-65776609-INS-6360
chr8-65776684-INS-54
chr8-65793947-INS-6297
chr8-65808152-INS-635
chr8-65811086-INS-99
chr8-65830513-INS-578
chr8-65832039-INS-70
chr8-65837821-INS-867
chr8-65841397-INS-245
chr8-65841698-INS-416
chr8-65842532-INS-52
chr8-65846330-INS-199
chr8-65847478-INS-1984
chr8-65850497-INS-56
chr8-65852359-INS-408
chr8-65888120-INS-214
chr8-65897227-INS-140
chr8-65898108-INS-56
chr8-65904652-INS-173
chr8-65918319-INS-6873
chr8-65925500-INS-579
chr8-65945632-INS-785
chr8-65955223-INS-4776
chr8-65963182-INS-498
chr8-66013654-INS-232
chr8-66075326-INS-7050
chr8-66079243-INS-6439
chr8-66081960-INS-774
chr8-66089899-INS-203
chr8-66130138-INS-6401
chr8-66134360-INS-6198
chr8-66179587-INS-75
chr8-66191832-INS-103
chr8-66196194-INS-162
chr8-66215469-INS-58
chr8-66230629-INS-60
chr8-66254363-INS-213
chr8-66260254-INS-119
chr8-66264137-INS-136
chr8-66265290-INS-215
chr8-66292715-INS-977
chr8-66294095-INS-160
chr8-66297026-INS-118
chr8-66298018-INS-87
chr8-66302475-INS-51
chr8-66334686-INS-380
chr8-66338407-INS-60
chr8-66339142-INS-525
chr8-66343780-INS-203
chr8-66349375-INS-149
chr8-66402497-INS-400
chr8-66410989-INS-512
chr8-66435907-INS-6699
chr8-66485182-INS-4582
chr8-66487675-INS-56
chr8-66501757-INS-60
chr8-66507902-INS-380
chr8-66507949-INS-1174
chr8-66507950-INS-868
chr8-66535778-INS-72
chr8-66541184-INS-189
chr8-66544514-INS-831
chr8-66571506-INS-329
chr8-66572527-INS-258
chr8-66573487-INS-56
chr8-66585508-INS-6600
chr8-66602786-INS-539
chr8-66614927-INS-123
chr8-66650613-INS-154
chr8-66668557-INS-1000
chr8-66689270-INS-60

chr8-66694605-INS-535
chr8-66705491-INS-826
chr8-66710002-INS-76
chr8-66731780-INS-4634
chr8-66733570-INS-54
chr8-66734101-INS-102
chr8-66744932-INS-1864
chr8-66757984-INS-9630
chr8-66780790-INS-72
chr8-66786291-INS-60
chr8-66823131-INS-4290
chr8-66848942-INS-140
chr8-66901284-INS-83
chr8-66909515-INS-112
chr8-66909920-INS-71
chr8-66911455-INS-141
chr8-66942624-INS-316
chr8-66948600-INS-92
chr8-66982420-INS-6792
chr8-67008150-INS-105
chr8-67036294-INS-400
chr8-67073372-INS-401
chr8-67081674-INS-230
chr8-67091470-INS-225
chr8-67102833-INS-78
chr8-67123569-INS-61
chr8-67132201-INS-7124
chr8-67209798-INS-400
chr8-67213483-INS-6198
chr8-67219432-INS-185
chr8-67229368-INS-80
chr8-67229707-INS-7359
chr8-67255107-INS-83
chr8-67274521-INS-56
chr8-67301322-INS-400
chr8-67302420-INS-329
chr8-67304527-INS-448
chr8-67341447-INS-53
chr8-67344351-INS-52
chr8-67345655-INS-76
chr8-67380709-INS-67
chr8-67391800-INS-449
chr8-67398279-INS-830
chr8-67415645-INS-61
chr8-67416854-INS-55
chr8-67454649-INS-283
chr8-67477337-INS-56
chr8-67498427-INS-390
chr8-67502248-INS-265
chr8-67509790-INS-55
chr8-67537406-INS-5189
chr8-67553829-INS-90
chr8-67592176-INS-276
chr8-67664165-INS-6501
chr8-67716008-INS-102
chr8-67721967-INS-55
chr8-67725997-INS-339
chr8-67740439-INS-6611
chr8-67755802-INS-99
chr8-67810231-INS-153
chr8-67810799-INS-152
chr8-67835060-INS-81
chr8-67851951-INS-612
chr8-67855033-INS-530
chr8-67892264-INS-436
chr8-67912589-INS-275
chr8-67912596-INS-56
chr8-67929049-INS-59
chr8-67941738-INS-6309
chr8-67942409-INS-58
chr8-67952412-INS-1682
chr8-67979560-INS-821
chr8-67985813-INS-524
chr8-67997147-INS-154
chr8-68026494-INS-73
chr8-68027749-INS-77
chr8-68029794-INS-497

chr8-68030537-INS-54
chr8-68059488-INS-6837
chr8-68074931-INS-7068
chr8-68077491-INS-3296
chr8-68173269-INS-53
chr8-68174500-INS-56
chr8-68201802-INS-139
chr8-68225736-INS-184
chr8-68239300-INS-6960
chr8-68266242-INS-756
chr8-68271819-INS-420
chr8-68325101-INS-82
chr8-68371659-INS-76
chr8-68374271-INS-144
chr8-68378025-INS-494
chr8-68432179-INS-6520
chr8-68433535-INS-96
chr8-68449222-INS-1098
chr8-68449223-INS-840
chr8-68469241-INS-712
chr8-68484144-INS-54
chr8-68502230-INS-179
chr8-68523621-INS-169
chr8-68577814-INS-84
chr8-68583463-INS-128
chr8-68592107-INS-84
chr8-68605034-INS-166
chr8-68681080-INS-5388
chr8-68686393-INS-55
chr8-68711218-INS-51
chr8-68734227-INS-322
chr8-68743115-INS-119
chr8-68781222-INS-356
chr8-68789833-INS-421
chr8-68793840-INS-224
chr8-68831300-INS-7344
chr8-68840596-INS-3627
chr8-68849382-INS-1471
chr8-68851523-INS-458
chr8-68866044-INS-57
chr8-68870449-INS-162
chr8-68876588-INS-67
chr8-68882339-INS-76
chr8-68935952-INS-438
chr8-68948176-INS-507
chr8-68960581-INS-437
chr8-68968234-INS-60
chr8-68971923-INS-484
chr8-68992771-INS-201
chr8-69011462-INS-254
chr8-69022094-INS-654
chr8-69033255-INS-77
chr8-69039469-INS-180
chr8-69075527-INS-345
chr8-69088147-INS-347
chr8-69106206-INS-81
chr8-69113125-INS-6428
chr8-69136077-INS-1040
chr8-69157891-INS-6420
chr8-69161217-INS-91
chr8-69172594-INS-86
chr8-69194305-INS-1603
chr8-69222020-INS-50
chr8-69270800-INS-91
chr8-69281315-INS-166
chr8-69320277-INS-60
chr8-69328068-INS-99
chr8-69373520-INS-131
chr8-69373967-INS-307
chr8-69397546-INS-53
chr8-69413563-INS-64
chr8-69451039-INS-71
chr8-69461161-INS-216
chr8-69482178-INS-384
chr8-69487396-INS-113
chr8-69503063-INS-127
chr8-69521805-INS-175

chr8-69525138-INS-210
chr8-69549241-INS-158
chr8-69570053-INS-225
chr8-69584611-INS-182
chr8-69589718-INS-204
chr8-69592157-INS-206
chr8-69597966-INS-175
chr8-69605575-INS-6389
chr8-69619175-INS-84
chr8-69689179-INS-206
chr8-69746965-INS-3075
chr8-69749820-INS-134
chr8-69755842-INS-571
chr8-69770338-INS-4805
chr8-69778969-INS-98
chr8-69806262-INS-6643
chr8-69810831-INS-6892
chr8-69814274-INS-212
chr8-69818131-INS-7770
chr8-69832103-INS-108
chr8-69842480-INS-7107
chr8-69858361-INS-530
chr8-69874564-INS-104
chr8-69885408-INS-9559
chr8-69959408-INS-68
chr8-70003887-INS-647
chr8-70007426-INS-128
chr8-70033132-INS-4881
chr8-70079312-INS-598
chr8-70117125-INS-400
chr8-70150720-INS-690
chr8-70154499-INS-7114
chr8-70157767-INS-204
chr8-70165287-INS-233
chr8-70174014-INS-221
chr8-70187082-INS-74
chr8-70202043-INS-509
chr8-70219848-INS-182
chr8-70230490-INS-156
chr8-70232582-INS-487
chr8-70240928-INS-3479
chr8-70265345-INS-53
chr8-70281367-INS-384
chr8-70291326-INS-204
chr8-70308250-INS-213
chr8-70309992-INS-68
chr8-70310502-INS-480
chr8-70314492-INS-174
chr8-70316848-INS-146
chr8-70319917-INS-129
chr8-70325462-INS-116
chr8-70353549-INS-271
chr8-70383021-INS-85
chr8-70406440-INS-611
chr8-70421511-INS-54
chr8-70435534-INS-203
chr8-70436846-INS-193
chr8-70447676-INS-193
chr8-70456594-INS-624
chr8-70466779-INS-205
chr8-70484471-INS-51
chr8-70507024-INS-114
chr8-70535025-INS-65
chr8-70541798-INS-94
chr8-70566628-INS-53
chr8-70577041-INS-60
chr8-70577605-INS-124
chr8-70586647-INS-175
chr8-70604116-INS-274
chr8-70604863-INS-60
chr8-70641733-INS-211
chr8-70668860-INS-213
chr8-70670695-INS-1095
chr8-70671010-INS-996
chr8-70730050-INS-155
chr8-70732390-INS-347
chr8-70745517-INS-233

chr8-70774027-INS-203
chr8-70778271-INS-252
chr8-70818695-INS-134
chr8-70883113-INS-73
chr8-70885867-INS-105
chr8-70986339-INS-180
chr8-70989038-INS-53
chr8-70999171-INS-227
chr8-71007484-INS-246
chr8-71009327-INS-5808
chr8-71089885-INS-59
chr8-71106600-INS-220
chr8-71109766-INS-83
chr8-71116883-INS-215
chr8-71118797-INS-115
chr8-71119695-INS-208
chr8-71122726-INS-260
chr8-71174294-INS-106
chr8-71188899-INS-175
chr8-71210399-INS-170
chr8-71255675-INS-55
chr8-71270646-INS-370
chr8-71277339-INS-51
chr8-71279010-INS-202
chr8-71284891-INS-197
chr8-71295045-INS-208
chr8-71319196-INS-399
chr8-71324164-INS-173
chr8-71331326-INS-56
chr8-71355442-INS-565
chr8-71375377-INS-56
chr8-71453231-INS-73
chr8-71483068-INS-239
chr8-71555825-INS-6839
chr8-71560620-INS-5158
chr8-71577887-INS-102
chr8-71581030-INS-95
chr8-71642079-INS-69
chr8-71677455-INS-570
chr8-71698328-INS-217
chr8-71732789-INS-7099
chr8-71776392-INS-137
chr8-71805656-INS-207
chr8-71832815-INS-172
chr8-71840963-INS-80
chr8-71843445-INS-53
chr8-71863105-INS-195
chr8-71867340-INS-84
chr8-71873487-INS-574
chr8-71880160-INS-432
chr8-71892558-INS-106
chr8-71893621-INS-212
chr8-71899361-INS-187
chr8-71908187-INS-53
chr8-71912575-INS-205
chr8-71953677-INS-210
chr8-71968189-INS-183
chr8-71977373-INS-52
chr8-71984507-INS-64
chr8-72001199-INS-219
chr8-72005017-INS-602
chr8-72006270-INS-182
chr8-72006858-INS-74
chr8-72028031-INS-282
chr8-72036758-INS-176
chr8-72040663-INS-284
chr8-72056767-INS-179
chr8-72086288-INS-56
chr8-72090732-INS-244
chr8-72093500-INS-302
chr8-72144752-INS-205
chr8-72158407-INS-75
chr8-72209462-INS-149
chr8-72209941-INS-221
chr8-72235386-INS-2579
chr8-72572607-INS-457
chr8-72634830-INS-392

chr8-72636642-INS-70
chr8-72685552-INS-60
chr8-72689531-INS-199
chr8-72698002-INS-79
chr8-72702818-INS-6448
chr8-72712714-INS-6488
chr8-72725898-INS-162
chr8-72733123-INS-120
chr8-72759818-INS-491
chr8-72768745-INS-75
chr8-72797546-INS-68
chr8-72811377-INS-204
chr8-72814529-INS-83
chr8-72817601-INS-114
chr8-72827569-INS-461
chr8-72833111-INS-72
chr8-72834941-INS-294
chr8-72835870-INS-1877
chr8-72841437-INS-73
chr8-72844171-INS-75
chr8-72848188-INS-200
chr8-72867919-INS-94
chr8-72888076-INS-52
chr8-72922182-INS-68
chr8-72934873-INS-118
chr8-72956147-INS-249
chr8-72966632-INS-5368
chr8-72981540-INS-79
chr8-72986545-INS-59
chr8-72987926-INS-58
chr8-72991330-INS-59
chr8-73002421-INS-199
chr8-73017245-INS-206
chr8-73018897-INS-118
chr8-73030361-INS-56
chr8-73040250-INS-185
chr8-73045452-INS-82
chr8-73052347-INS-199
chr8-73069517-INS-142
chr8-73083659-INS-255
chr8-73135362-INS-418
chr8-73157321-INS-220
chr8-73161608-INS-400
chr8-73165813-INS-196
chr8-73169949-INS-56
chr8-73181501-INS-87
chr8-73195140-INS-201
chr8-73199314-INS-228
chr8-73202671-INS-54
chr8-73211456-INS-309
chr8-73211941-INS-359
chr8-73230548-INS-71
chr8-73235894-INS-4829
chr8-73258972-INS-66
chr8-73305794-INS-173
chr8-73333067-INS-163
chr8-73343164-INS-154
chr8-73344395-INS-7118
chr8-73344625-INS-117
chr8-73346403-INS-120
chr8-73352619-INS-120
chr8-73357280-INS-397
chr8-73362696-INS-60
chr8-73365044-INS-210
chr8-73366003-INS-688
chr8-73366801-INS-60
chr8-73367543-INS-715
chr8-73369007-INS-60
chr8-73398702-INS-86
chr8-73415455-INS-52
chr8-73417176-INS-63
chr8-73442020-INS-55
chr8-73459621-INS-180
chr8-73465217-INS-208
chr8-73481166-INS-158
chr8-73523189-INS-351
chr8-73530890-INS-498

chr8-73603246-INS-51
chr8-73608522-INS-6760
chr8-73642090-INS-7673
chr8-73658878-INS-68
chr8-73690865-INS-210
chr8-73718902-INS-182
chr8-73721517-INS-358
chr8-73732636-INS-576
chr8-73757053-INS-51
chr8-73759335-INS-131
chr8-73783723-INS-6654
chr8-73798513-INS-126
chr8-73831190-INS-226
chr8-73844720-INS-6182
chr8-73848228-INS-76
chr8-73862341-INS-155
chr8-73866982-INS-128
chr8-73871849-INS-238
chr8-73877505-INS-82
chr8-73917983-INS-903
chr8-73924329-INS-398
chr8-73930662-INS-144
chr8-74092047-INS-109
chr8-74127736-INS-195
chr8-74158812-INS-349
chr8-74168530-INS-135
chr8-74169157-INS-6676
chr8-74175855-INS-7251
chr8-74248571-INS-166
chr8-74296129-INS-5247
chr8-74298504-INS-52
chr8-74337737-INS-56
chr8-74343915-INS-72
chr8-74347533-INS-1746
chr8-74357357-INS-193
chr8-74363931-INS-70
chr8-74374065-INS-54
chr8-74403707-INS-201
chr8-74439176-INS-70
chr8-74449082-INS-6735
chr8-74469934-INS-821
chr8-74472789-INS-107
chr8-74479215-INS-133
chr8-74527583-INS-568
chr8-74536620-INS-5348
chr8-74553123-INS-203
chr8-74555125-INS-89
chr8-74564861-INS-6657
chr8-74569171-INS-60
chr8-74606539-INS-115
chr8-74631505-INS-82
chr8-74643429-INS-683
chr8-74665118-INS-538
chr8-74668094-INS-118
chr8-74672237-INS-4154
chr8-74693861-INS-3504
chr8-74703313-INS-6264
chr8-74713494-INS-6493
chr8-74715972-INS-492
chr8-74784616-INS-64
chr8-74797641-INS-603
chr8-74806461-INS-129
chr8-74830218-INS-297
chr8-74832374-INS-229
chr8-74855794-INS-112
chr8-74862751-INS-731
chr8-74872887-INS-66
chr8-74881212-INS-89
chr8-74936398-INS-66
chr8-74944848-INS-269
chr8-74953367-INS-6192
chr8-74990747-INS-324
chr8-75006185-INS-495
chr8-75011476-INS-117
chr8-75025599-INS-124
chr8-75032771-INS-794
chr8-75042520-INS-66

chr8-75046318-INS-342
chr8-75048904-INS-260
chr8-75067176-INS-168
chr8-75069052-INS-57
chr8-75071513-INS-100
chr8-75074134-INS-125
chr8-75091756-INS-2787
chr8-75102205-INS-7278
chr8-75115770-INS-6374
chr8-75119962-INS-66
chr8-75131887-INS-198
chr8-75146760-INS-112
chr8-75152346-INS-63
chr8-75182410-INS-83
chr8-75187503-INS-73
chr8-75287132-INS-518
chr8-75288350-INS-60
chr8-75310052-INS-145
chr8-75339947-INS-996
chr8-75346225-INS-395
chr8-75348873-INS-60
chr8-75366613-INS-6956
chr8-75369825-INS-93
chr8-75383026-INS-4853
chr8-75400725-INS-6458
chr8-75401628-INS-344
chr8-75405603-INS-7319
chr8-75409186-INS-151
chr8-75481416-INS-207
chr8-75492930-INS-58
chr8-75521362-INS-6442
chr8-75527071-INS-507
chr8-75538512-INS-241
chr8-75556465-INS-7230
chr8-75584997-INS-2488
chr8-75587604-INS-100
chr8-75610385-INS-197
chr8-75615429-INS-100
chr8-75616480-INS-215
chr8-75646388-INS-106
chr8-75657075-INS-892
chr8-75657836-INS-170
chr8-75661030-INS-195
chr8-75664747-INS-2066
chr8-75665170-INS-86
chr8-75675207-INS-185
chr8-75687586-INS-437
chr8-75691954-INS-194
chr8-75697175-INS-203
chr8-75701303-INS-201
chr8-75704888-INS-60
chr8-75732847-INS-196
chr8-75738697-INS-98
chr8-75741186-INS-219
chr8-75747861-INS-197
chr8-75756255-INS-68
chr8-75771938-INS-143
chr8-75799087-INS-226
chr8-75806014-INS-206
chr8-75807050-INS-398
chr8-75807551-INS-269
chr8-75810400-INS-7495
chr8-75813697-INS-75
chr8-75824864-INS-243
chr8-75834068-INS-187
chr8-75839747-INS-212
chr8-75857761-INS-57
chr8-75863208-INS-7078
chr8-75873060-INS-200
chr8-75882179-INS-282
chr8-75888228-INS-355
chr8-75892498-INS-7120
chr8-75900360-INS-236
chr8-75909740-INS-198
chr8-75946240-INS-138
chr8-75960966-INS-851
chr8-75962168-INS-161

chr8-75966666-INS-52
chr8-75974329-INS-57
chr8-75975667-INS-342
chr8-75993048-INS-135
chr8-76007710-INS-392
chr8-76013507-INS-209
chr8-76016821-INS-438
chr8-76017756-INS-119
chr8-76020953-INS-221
chr8-76024215-INS-209
chr8-76036171-INS-175
chr8-76039567-INS-208
chr8-76042961-INS-6847
chr8-76058462-INS-5044
chr8-76076319-INS-809
chr8-76077030-INS-506
chr8-76101117-INS-55
chr8-76115559-INS-6350
chr8-76124690-INS-419
chr8-76142027-INS-10006
chr8-76152431-INS-115
chr8-76160072-INS-7260
chr8-76162834-INS-127
chr8-76164347-INS-131
chr8-76171373-INS-639
chr8-76176299-INS-210
chr8-76193381-INS-209
chr8-76197036-INS-176
chr8-76218787-INS-787
chr8-76222000-INS-111
chr8-76223490-INS-937
chr8-76241523-INS-94
chr8-76245856-INS-88
chr8-76247454-INS-55
chr8-76269999-INS-269
chr8-76316202-INS-52
chr8-76337028-INS-53
chr8-76342003-INS-72
chr8-76361297-INS-58
chr8-76365897-INS-3400
chr8-76377872-INS-410
chr8-76388001-INS-62
chr8-76391793-INS-63
chr8-76401652-INS-80
chr8-76426916-INS-98
chr8-76449020-INS-556
chr8-76455945-INS-56
chr8-76482024-INS-435
chr8-76483676-INS-490
chr8-76490373-INS-6492
chr8-76518939-INS-5621
chr8-76519653-INS-54
chr8-76523868-INS-64
chr8-76536313-INS-80
chr8-76539257-INS-5406
chr8-76558787-INS-164
chr8-76564981-INS-108
chr8-76566590-INS-60
chr8-76589300-INS-4172
chr8-76598140-INS-1887
chr8-76602293-INS-66
chr8-76604364-INS-400
chr8-76606535-INS-7088
chr8-76614633-INS-74
chr8-76621550-INS-484
chr8-76624401-INS-54
chr8-76653904-INS-499
chr8-76679575-INS-191
chr8-76705208-INS-619
chr8-76717998-INS-1989
chr8-76735442-INS-981
chr8-76755086-INS-70
chr8-76762800-INS-54
chr8-76768850-INS-65
chr8-76774586-INS-696
chr8-76788596-INS-693
chr8-76807128-INS-187

chr8-76812532-INS-6582
chr8-76885591-INS-178
chr8-76925885-INS-1676
chr8-77006156-INS-92
chr8-77016159-INS-93
chr8-77023189-INS-6415
chr8-77035700-INS-412
chr8-77060178-INS-796
chr8-77086832-INS-306
chr8-77135063-INS-68
chr8-77138418-INS-99
chr8-77165998-INS-348
chr8-77166996-INS-185
chr8-77207957-INS-52
chr8-77213700-INS-114
chr8-77241696-INS-73
chr8-77243474-INS-77
chr8-77252862-INS-7131
chr8-77267665-INS-234
chr8-77272284-INS-502
chr8-77288265-INS-545
chr8-77292668-INS-56
chr8-77298154-INS-79
chr8-77308457-INS-6860
chr8-77317462-INS-1264
chr8-77321341-INS-60
chr8-77330741-INS-202
chr8-77368953-INS-1085
chr8-77385586-INS-837
chr8-77391195-INS-539
chr8-77423836-INS-132
chr8-77426056-INS-185
chr8-77434166-INS-66
chr8-77437149-INS-187
chr8-77477416-INS-98
chr8-77479886-INS-60
chr8-77502552-INS-66
chr8-77509245-INS-698
chr8-77539227-INS-961
chr8-77544930-INS-95
chr8-77587858-INS-75
chr8-77614832-INS-1114
chr8-77619860-INS-7250
chr8-77624916-INS-52
chr8-77630157-INS-61
chr8-77723442-INS-66
chr8-77791101-INS-72
chr8-77824251-INS-180
chr8-77837816-INS-52
chr8-77854301-INS-98
chr8-77854494-INS-149
chr8-77875405-INS-1362
chr8-77885687-INS-465
chr8-77890720-INS-52
chr8-77901707-INS-121
chr8-77956659-INS-53
chr8-77971602-INS-2828
chr8-77980723-INS-59
chr8-78022874-INS-94
chr8-78026686-INS-62
chr8-78047722-INS-400
chr8-78055565-INS-260
chr8-78062657-INS-51
chr8-78134031-INS-63
chr8-78142368-INS-207
chr8-78147665-INS-66
chr8-78149083-INS-72
chr8-78149457-INS-688
chr8-78150480-INS-68
chr8-78152098-INS-191
chr8-78158206-INS-74
chr8-78172045-INS-60
chr8-78175708-INS-117
chr8-78218466-INS-75
chr8-78219826-INS-299
chr8-78246592-INS-139
chr8-78248627-INS-207

chr8-78257067-INS-9045
chr8-78271425-INS-181
chr8-78290115-INS-50
chr8-78293683-INS-58
chr8-78298401-INS-500
chr8-78311108-INS-581
chr8-78322614-INS-219
chr8-78348299-INS-60
chr8-78355567-INS-246
chr8-78358099-INS-148
chr8-78379019-INS-76
chr8-78380428-INS-6649
chr8-78397481-INS-86
chr8-78428133-INS-60
chr8-78440971-INS-89
chr8-78445659-INS-410
chr8-78447613-INS-77
chr8-78454380-INS-158
chr8-78462081-INS-56
chr8-78471117-INS-143
chr8-78477611-INS-154
chr8-78485036-INS-282
chr8-78517302-INS-88
chr8-78534876-INS-1178
chr8-78547485-INS-157
chr8-78548817-INS-92
chr8-78559182-INS-76
chr8-78565313-INS-118
chr8-78572924-INS-121
chr8-78607673-INS-349
chr8-78619375-INS-206
chr8-78624588-INS-181
chr8-78641601-INS-263
chr8-78646181-INS-323
chr8-78647364-INS-312
chr8-78652044-INS-58
chr8-78654638-INS-442
chr8-78655376-INS-5108
chr8-78677264-INS-152
chr8-78738640-INS-200
chr8-78759148-INS-8640
chr8-78775389-INS-226
chr8-78779173-INS-51
chr8-78830704-INS-51
chr8-78853258-INS-99
chr8-78857304-INS-53
chr8-78860567-INS-71
chr8-78882558-INS-65
chr8-78884763-INS-131
chr8-78948473-INS-175
chr8-78982685-INS-80
chr8-78995381-INS-75
chr8-79015678-INS-100
chr8-79022293-INS-6365
chr8-79023430-INS-98
chr8-79026488-INS-62
chr8-79039403-INS-74
chr8-79051737-INS-58
chr8-79061036-INS-65
chr8-79062323-INS-138
chr8-79109653-INS-200
chr8-79110893-INS-320
chr8-79134637-INS-334
chr8-79136500-INS-154
chr8-79138402-INS-352
chr8-79140366-INS-346
chr8-79150287-INS-212
chr8-79168419-INS-88
chr8-79176859-INS-172
chr8-79182077-INS-240
chr8-79190966-INS-367
chr8-79200730-INS-76
chr8-79216191-INS-358
chr8-79227701-INS-130
chr8-79240132-INS-210
chr8-79247931-INS-207
chr8-79250400-INS-70

chr8-79259133-INS-322
chr8-79263857-INS-70
chr8-79267500-INS-228
chr8-79270016-INS-236
chr8-79284445-INS-76
chr8-79292185-INS-90
chr8-79294074-INS-73
chr8-79361625-INS-384
chr8-79365546-INS-71
chr8-79396322-INS-480
chr8-79399126-INS-59
chr8-79429285-INS-204
chr8-79433063-INS-224
chr8-79451499-INS-76
chr8-79461708-INS-634
chr8-79468044-INS-212
chr8-79494552-INS-415
chr8-79506509-INS-54
chr8-79528911-INS-321
chr8-79532283-INS-56
chr8-79535033-INS-87
chr8-79548303-INS-8011
chr8-79551825-INS-327
chr8-79553730-INS-128
chr8-79603826-INS-145
chr8-79650480-INS-58
chr8-79661391-INS-122
chr8-79671861-INS-352
chr8-79699651-INS-186
chr8-79702835-INS-59
chr8-79733133-INS-2495
chr8-79742293-INS-151
chr8-79744569-INS-97
chr8-79765256-INS-461
chr8-79791959-INS-260
chr8-79795607-INS-79
chr8-79824968-INS-69
chr8-79826228-INS-667
chr8-79858449-INS-344
chr8-79860843-INS-111
chr8-79870946-INS-70
chr8-79902150-INS-108
chr8-79905148-INS-55
chr8-79952134-INS-147
chr8-79987551-INS-99
chr8-79998456-INS-76
chr8-80006918-INS-192
chr8-80048920-INS-63
chr8-80057255-INS-668
chr8-80084613-INS-52
chr8-80109315-INS-197
chr8-80121271-INS-76
chr8-80197883-INS-893
chr8-80209581-INS-60
chr8-80216315-INS-60
chr8-80218144-INS-65
chr8-80225853-INS-71
chr8-80252162-INS-102
chr8-80263008-INS-99
chr8-80285253-INS-152
chr8-80288639-INS-326
chr8-80294891-INS-86
chr8-80322909-INS-51
chr8-80329878-INS-282
chr8-80335805-INS-52
chr8-80337675-INS-120
chr8-80340008-INS-54
chr8-80349497-INS-70
chr8-80356313-INS-186
chr8-80357976-INS-73
chr8-80405806-INS-988
chr8-80421454-INS-189
chr8-80431673-INS-202
chr8-80439609-INS-262
chr8-80445271-INS-182
chr8-80449805-INS-6708
chr8-80450798-INS-193

chr8-80454564-INS-267
chr8-80469158-INS-195
chr8-80530903-INS-197
chr8-80552085-INS-120
chr8-80568803-INS-782
chr8-80583373-INS-245
chr8-80601371-INS-649
chr8-80632579-INS-365
chr8-80646648-INS-1290
chr8-80669326-INS-78
chr8-80673969-INS-461
chr8-80678632-INS-204
chr8-80688648-INS-139
chr8-80703224-INS-1037
chr8-80712773-INS-52
chr8-80726506-INS-6187
chr8-80731748-INS-1228
chr8-80740048-INS-63
chr8-80742543-INS-56
chr8-80746957-INS-85
chr8-80763772-INS-106
chr8-80781154-INS-54
chr8-80804923-INS-6460
chr8-80816622-INS-498
chr8-80830526-INS-796
chr8-80835768-INS-52
chr8-80838914-INS-98
chr8-80849182-INS-124
chr8-80864445-INS-218
chr8-80880807-INS-92
chr8-80902655-INS-69
chr8-80912527-INS-66
chr8-80914313-INS-7080
chr8-80929593-INS-170
chr8-80939416-INS-55
chr8-80961183-INS-75
chr8-80986789-INS-53
chr8-81014615-INS-141
chr8-81031043-INS-85
chr8-81051424-INS-7139
chr8-81087259-INS-2387
chr8-81097936-INS-285
chr8-81108804-INS-159
chr8-81118303-INS-403
chr8-81123127-INS-55
chr8-81127867-INS-848
chr8-81148342-INS-757
chr8-81166832-INS-6587
chr8-81188450-INS-527
chr8-81200100-INS-3417
chr8-81200737-INS-448
chr8-81209394-INS-57
chr8-81213812-INS-52
chr8-81304757-INS-167
chr8-81320604-INS-121
chr8-81322748-INS-6368
chr8-81323429-INS-6771
chr8-81326423-INS-315
chr8-81345158-INS-105
chr8-81349851-INS-422
chr8-81360089-INS-206
chr8-81377959-INS-532
chr8-81403869-INS-6309
chr8-81414689-INS-115
chr8-81449279-INS-649
chr8-81450652-INS-133
chr8-81453918-INS-204
chr8-81476600-INS-498
chr8-81483760-INS-110
chr8-81488111-INS-73
chr8-81488748-INS-162
chr8-81500574-INS-505
chr8-81544902-INS-229
chr8-81546947-INS-198
chr8-81562887-INS-366
chr8-81564219-INS-54
chr8-81579366-INS-81

chr8-81602580-INS-303
chr8-81624988-INS-7080
chr8-81722139-INS-173
chr8-81750153-INS-355
chr8-81756415-INS-498
chr8-81758274-INS-358
chr8-81769828-INS-146
chr8-81770454-INS-84
chr8-81774109-INS-52
chr8-81789050-INS-7410
chr8-81817620-INS-402
chr8-81881854-INS-198
chr8-81889730-INS-215
chr8-81907271-INS-85
chr8-81933697-INS-87
chr8-81955360-INS-479
chr8-81976235-INS-57
chr8-81998294-INS-261
chr8-82022435-INS-65
chr8-82029356-INS-795
chr8-82058636-INS-278
chr8-82073706-INS-92
chr8-82076959-INS-204
chr8-82091070-INS-147
chr8-82177868-INS-238
chr8-82180428-INS-300
chr8-82182448-INS-251
chr8-82191303-INS-115
chr8-82207663-INS-6589
chr8-82216995-INS-801
chr8-82220430-INS-99
chr8-82224240-INS-310
chr8-82233674-INS-7556
chr8-82234662-INS-250
chr8-82298001-INS-57
chr8-82340671-INS-217
chr8-82347462-INS-127
chr8-82356913-INS-68
chr8-82380704-INS-6518
chr8-82402048-INS-60
chr8-82417789-INS-6145
chr8-82421367-INS-533
chr8-82433077-INS-292
chr8-82455442-INS-75
chr8-82472820-INS-58
chr8-82475932-INS-77
chr8-82498987-INS-210
chr8-82528353-INS-393
chr8-82569052-INS-522
chr8-82585054-INS-824
chr8-82626319-INS-72
chr8-82652712-INS-618
chr8-82688226-INS-52
chr8-82689996-INS-752
chr8-82704499-INS-54
chr8-82748522-INS-133
chr8-82757600-INS-167
chr8-82758283-INS-64
chr8-82775000-INS-312
chr8-82780912-INS-340
chr8-82789294-INS-53
chr8-82805025-INS-334
chr8-82824771-INS-132
chr8-82840515-INS-400
chr8-82841929-INS-6614
chr8-82881231-INS-265
chr8-82892530-INS-112
chr8-82914828-INS-6155
chr8-82915165-INS-546
chr8-82917522-INS-52
chr8-82979376-INS-95
chr8-82980536-INS-206
chr8-82983598-INS-6410
chr8-82993790-INS-117
chr8-83052309-INS-660
chr8-83065598-INS-70
chr8-83067168-INS-1494

chr8-83077435-INS-6406
chr8-83081689-INS-52
chr8-83130770-INS-106
chr8-83132693-INS-5768
chr8-83187884-INS-66
chr8-83205370-INS-72
chr8-83216916-INS-392
chr8-83245819-INS-1611
chr8-83269783-INS-53
chr8-83294370-INS-74
chr8-83319402-INS-375
chr8-83322412-INS-417
chr8-83329727-INS-1239
chr8-83335773-INS-253
chr8-83337911-INS-979
chr8-83339379-INS-7004
chr8-83391322-INS-94
chr8-83398347-INS-63
chr8-83410678-INS-60
chr8-83429868-INS-248
chr8-83432424-INS-73
chr8-83448187-INS-108
chr8-83470224-INS-189
chr8-83471609-INS-6776
chr8-83501835-INS-77
chr8-83511359-INS-84
chr8-83544604-INS-66
chr8-83564520-INS-66
chr8-83568029-INS-68
chr8-83576589-INS-188
chr8-83578360-INS-306
chr8-83593898-INS-60
chr8-83607006-INS-2276
chr8-83609601-INS-102
chr8-83613279-INS-51
chr8-83644654-INS-1425
chr8-83648792-INS-90
chr8-83664440-INS-124
chr8-83675056-INS-307
chr8-83676705-INS-69
chr8-83687456-INS-54
chr8-83725033-INS-80
chr8-83776638-INS-498
chr8-83786951-INS-55
chr8-83803408-INS-73
chr8-83811828-INS-56
chr8-83824739-INS-7101
chr8-83839406-INS-77
chr8-83841959-INS-223
chr8-83843228-INS-76
chr8-83843436-INS-439
chr8-83864649-INS-199
chr8-83870701-INS-52
chr8-83873657-INS-62
chr8-83885894-INS-210
chr8-83899032-INS-52
chr8-83902436-INS-740
chr8-83930742-INS-83
chr8-83933239-INS-74
chr8-83939922-INS-52
chr8-83944929-INS-80
chr8-83949460-INS-56
chr8-83955520-INS-245
chr8-83956083-INS-782
chr8-83957930-INS-136
chr8-83969947-INS-114
chr8-83976308-INS-585
chr8-83986365-INS-159
chr8-83993019-INS-97
chr8-84002834-INS-52
chr8-84005745-INS-77
chr8-84008014-INS-60
chr8-84031320-INS-233
chr8-84038132-INS-163
chr8-84051345-INS-453
chr8-84053021-INS-639
chr8-84062331-INS-7627

chr8-84080021-INS-112
chr8-84085759-INS-76
chr8-84093636-INS-194
chr8-84109767-INS-61
chr8-84110416-INS-7482
chr8-84113305-INS-197
chr8-84114902-INS-425
chr8-84128266-INS-413
chr8-84152123-INS-553
chr8-84157386-INS-130
chr8-84184316-INS-52
chr8-84244251-INS-436
chr8-84286234-INS-211
chr8-84287318-INS-267
chr8-84288170-INS-123
chr8-84296313-INS-68
chr8-84312715-INS-216
chr8-84322226-INS-960
chr8-84328116-INS-75
chr8-84329681-INS-116
chr8-84330214-INS-387
chr8-84346012-INS-163
chr8-84351265-INS-174
chr8-84353879-INS-204
chr8-84361673-INS-176
chr8-84372878-INS-193
chr8-84381027-INS-52
chr8-84422569-INS-222
chr8-84434678-INS-191
chr8-84468523-INS-211
chr8-84472152-INS-362
chr8-84517435-INS-329
chr8-84521760-INS-67
chr8-84523642-INS-257
chr8-84536259-INS-192
chr8-84552641-INS-181
chr8-84555216-INS-538
chr8-84557796-INS-72
chr8-84561355-INS-94
chr8-84571675-INS-218
chr8-84577425-INS-54
chr8-84579830-INS-54
chr8-84585869-INS-373
chr8-84586586-INS-5900
chr8-84672661-INS-210
chr8-84674899-INS-217
chr8-84680226-INS-62
chr8-84685418-INS-114
chr8-84693405-INS-218
chr8-84714052-INS-188
chr8-84720871-INS-65
chr8-84733243-INS-161
chr8-84757888-INS-80
chr8-84764250-INS-152
chr8-84777281-INS-439
chr8-84813886-INS-68
chr8-84820123-INS-52
chr8-84830225-INS-130
chr8-84835726-INS-84
chr8-84839928-INS-225
chr8-84841942-INS-173
chr8-84847133-INS-60
chr8-84864989-INS-208
chr8-84917808-INS-300
chr8-84993074-INS-63
chr8-85000290-INS-215
chr8-85033567-INS-60
chr8-85039056-INS-68
chr8-85043055-INS-89
chr8-85059751-INS-235
chr8-85075645-INS-289
chr8-85084081-INS-205
chr8-85102553-INS-102
chr8-85107247-INS-100
chr8-85111708-INS-57
chr8-85125870-INS-488
chr8-85134608-INS-107

chr8-85149415-INS-152
chr8-85157127-INS-126
chr8-85166202-INS-201
chr8-85184967-INS-58
chr8-85214215-INS-237
chr8-85229914-INS-57
chr8-85243155-INS-52
chr8-85302269-INS-53
chr8-85339841-INS-373
chr8-85340853-INS-184
chr8-85350606-INS-201
chr8-85381200-INS-67
chr8-85381665-INS-214
chr8-85390284-INS-224
chr8-85453586-INS-209
chr8-85472735-INS-298
chr8-85475437-INS-300
chr8-85484001-INS-213
chr8-85538763-INS-62
chr8-85553887-INS-197
chr8-85584773-INS-180
chr8-85586692-INS-209
chr8-85592340-INS-210
chr8-85596290-INS-202
chr8-85601305-INS-1366
chr8-85601389-INS-210
chr8-85604916-INS-236
chr8-85609796-INS-134
chr8-85621040-INS-223
chr8-85649998-INS-64
chr8-85651131-INS-53
chr8-85654900-INS-60
chr8-85656670-INS-211
chr8-85658114-INS-230
chr8-85658995-INS-121
chr8-85659887-INS-222
chr8-85676274-INS-319
chr8-85682159-INS-200
chr8-85691308-INS-249
chr8-85752835-INS-238
chr8-85787186-INS-129
chr8-85832658-INS-548
chr8-85840431-INS-96
chr8-85842776-INS-56
chr8-85848706-INS-371
chr8-85859919-INS-577
chr8-85860593-INS-203
chr8-85862193-INS-506
chr8-85871658-INS-898
chr8-85883447-INS-62
chr8-85884005-INS-8905
chr8-85901739-INS-647
chr8-85913050-INS-272
chr8-85919421-INS-327
chr8-85933603-INS-827
chr8-85948657-INS-222
chr8-86037569-INS-5467
chr8-86068109-INS-54
chr8-86076220-INS-208
chr8-86094900-INS-329
chr8-86100823-INS-179
chr8-86141108-INS-60
chr8-86143651-INS-727
chr8-86146856-INS-200
chr8-86148047-INS-260
chr8-86172445-INS-56
chr8-86179086-INS-55
chr8-86186738-INS-93
chr8-86198099-INS-56
chr8-86208192-INS-268
chr8-86210603-INS-66
chr8-86217990-INS-64
chr8-86220958-INS-92
chr8-86236558-INS-269
chr8-86237109-INS-108
chr8-86267473-INS-724
chr8-86287552-INS-230

chr8-86289545-INS-7143
chr8-86297013-INS-200
chr8-86419505-INS-94
chr8-86422683-INS-332
chr8-86437911-INS-6931
chr8-86437961-INS-198
chr8-86443100-INS-70
chr8-86446349-INS-216
chr8-86446889-INS-72
chr8-86472114-INS-173
chr8-86481487-INS-67
chr8-86504228-INS-54
chr8-86525828-INS-92
chr8-86533821-INS-486
chr8-86576627-INS-498
chr8-86584376-INS-105
chr8-86595577-INS-929
chr8-86613466-INS-2565
chr8-86617643-INS-204
chr8-86630284-INS-142
chr8-86656079-INS-116
chr8-86660404-INS-112
chr8-86666283-INS-88
chr8-86672090-INS-532
chr8-86675187-INS-428
chr8-86710552-INS-56
chr8-86721461-INS-54
chr8-86724578-INS-261
chr8-86733071-INS-56
chr8-86745436-INS-154
chr8-86753765-INS-191
chr8-86774714-INS-95
chr8-86775929-INS-129
chr8-86788962-INS-6595
chr8-86804282-INS-130
chr8-86809816-INS-2620
chr8-86813400-INS-468
chr8-86851259-INS-7248
chr8-86853486-INS-187
chr8-86858770-INS-513
chr8-86886558-INS-7137
chr8-86895272-INS-182
chr8-86924384-INS-51
chr8-86930016-INS-63
chr8-86950578-INS-200
chr8-86954559-INS-6676
chr8-86955026-INS-54
chr8-86955991-INS-165
chr8-86959721-INS-423
chr8-86963894-INS-84
chr8-86964464-INS-507
chr8-86966795-INS-6795
chr8-87008252-INS-469
chr8-87012727-INS-329
chr8-87018806-INS-65
chr8-87031812-INS-429
chr8-87040740-INS-56
chr8-87053521-INS-58
chr8-87086931-INS-160
chr8-87094415-INS-189
chr8-87115868-INS-109
chr8-87119471-INS-110
chr8-87135797-INS-6441
chr8-87137217-INS-164
chr8-87153525-INS-69
chr8-87154688-INS-1377
chr8-87160397-INS-60
chr8-87161233-INS-381
chr8-87234125-INS-416
chr8-87242688-INS-112
chr8-87334439-INS-59
chr8-87345938-INS-353
chr8-87351394-INS-69
chr8-87396693-INS-52
chr8-87416951-INS-230
chr8-87421176-INS-169
chr8-87458975-INS-215

chr8-87466903-INS-197
chr8-87468763-INS-3145
chr8-87490085-INS-306
chr8-87498744-INS-89
chr8-87504491-INS-68
chr8-87511341-INS-56
chr8-87542183-INS-191
chr8-87545433-INS-124
chr8-87568772-INS-58
chr8-87627080-INS-184
chr8-87637028-INS-153
chr8-87639922-INS-155
chr8-87644049-INS-184
chr8-87653252-INS-104
chr8-87663625-INS-387
chr8-87674341-INS-69
chr8-87686072-INS-212
chr8-87691333-INS-158
chr8-87707184-INS-120
chr8-87747550-INS-96
chr8-87751428-INS-6595
chr8-87753859-INS-259
chr8-87768214-INS-324
chr8-87779841-INS-177
chr8-87846715-INS-55
chr8-87852175-INS-400
chr8-87902003-INS-508
chr8-87903540-INS-237
chr8-87935292-INS-312
chr8-87974240-INS-328
chr8-87980241-INS-103
chr8-87990174-INS-60
chr8-87997893-INS-100
chr8-88047171-INS-286
chr8-88054602-INS-223
chr8-88083101-INS-84
chr8-88178090-INS-6296
chr8-88183038-INS-52
chr8-88189258-INS-88
chr8-88216710-INS-151
chr8-88223137-INS-87
chr8-88226554-INS-122
chr8-88245611-INS-144
chr8-88250498-INS-75
chr8-88258673-INS-60
chr8-88265200-INS-237
chr8-88271605-INS-124
chr8-88293833-INS-115
chr8-88299144-INS-1450
chr8-88359018-INS-323
chr8-88361182-INS-88
chr8-88368212-INS-302
chr8-88377438-INS-93
chr8-88381360-INS-694
chr8-88447056-INS-65
chr8-88475166-INS-76
chr8-88501054-INS-62
chr8-88522195-INS-128
chr8-88543851-INS-165
chr8-88551181-INS-60
chr8-88552485-INS-228
chr8-88556632-INS-204
chr8-88578993-INS-60
chr8-88608580-INS-188
chr8-88647861-INS-79
chr8-88651602-INS-261
chr8-88654790-INS-99
chr8-88698790-INS-185
chr8-88700381-INS-124
chr8-88710482-INS-52
chr8-88712716-INS-409
chr8-88714762-INS-179
chr8-88716656-INS-201
chr8-88719620-INS-222
chr8-88741642-INS-164
chr8-88744360-INS-87
chr8-88774248-INS-375

chr8-88800524-INS-170
chr8-88805499-INS-693
chr8-88809158-INS-74
chr8-88880248-INS-219
chr8-88899621-INS-53
chr8-88903158-INS-52
chr8-88907786-INS-135
chr8-88923117-INS-745
chr8-88961755-INS-69
chr8-88963109-INS-201
chr8-88976413-INS-279
chr8-88995600-INS-109
chr8-89005216-INS-56
chr8-89019462-INS-52
chr8-89032141-INS-630
chr8-89041069-INS-57
chr8-89091220-INS-71
chr8-89098333-INS-140
chr8-89104629-INS-83
chr8-89151316-INS-51
chr8-89154415-INS-57
chr8-89163154-INS-81
chr8-89183255-INS-187
chr8-89219521-INS-223
chr8-89232626-INS-84
chr8-89298824-INS-75
chr8-89329733-INS-6662
chr8-89333232-INS-169
chr8-89335216-INS-106
chr8-89339470-INS-59
chr8-89364080-INS-99
chr8-89371883-INS-58
chr8-89393662-INS-360
chr8-89492751-INS-416
chr8-89525993-INS-89
chr8-89529797-INS-89
chr8-89530779-INS-230
chr8-89545018-INS-61
chr8-89547587-INS-101
chr8-89573815-INS-76
chr8-89574189-INS-61
chr8-89576914-INS-272
chr8-89593592-INS-235
chr8-89626455-INS-215
chr8-89656679-INS-371
chr8-89676154-INS-475
chr8-89685414-INS-57
chr8-89697716-INS-176
chr8-89707535-INS-552
chr8-89711957-INS-53
chr8-89742968-INS-136
chr8-89813167-INS-209
chr8-89814735-INS-200
chr8-89815645-INS-106
chr8-89822103-INS-200
chr8-89853633-INS-8457
chr8-89854801-INS-52
chr8-89899060-INS-410
chr8-89940639-INS-436
chr8-89949974-INS-10258
chr8-89966543-INS-110
chr8-89979231-INS-210
chr8-89993071-INS-248
chr8-90007113-INS-57
chr8-90047909-INS-66
chr8-90050796-INS-62
chr8-90102134-INS-65
chr8-90106760-INS-60
chr8-90119117-INS-76
chr8-90127885-INS-195
chr8-90147699-INS-787
chr8-90163455-INS-51
chr8-90174195-INS-207
chr8-90175946-INS-64
chr8-90192681-INS-334
chr8-90201840-INS-68
chr8-90208414-INS-124

chr8-90211053-INS-204
chr8-90230178-INS-60
chr8-90265496-INS-124
chr8-90322334-INS-181
chr8-90354347-INS-939
chr8-90371973-INS-108
chr8-90387638-INS-131
chr8-90402769-INS-315
chr8-90409138-INS-539
chr8-90426243-INS-464
chr8-90445820-INS-218
chr8-90503208-INS-54
chr8-90517128-INS-68
chr8-90518401-INS-69
chr8-90521968-INS-169
chr8-90525300-INS-54
chr8-90561848-INS-1887
chr8-90566424-INS-146
chr8-90580783-INS-262
chr8-90584826-INS-201
chr8-90593957-INS-53
chr8-90610940-INS-422
chr8-90615276-INS-7195
chr8-90624674-INS-7325
chr8-90636896-INS-6223
chr8-90643634-INS-387
chr8-90672407-INS-207
chr8-90676472-INS-401
chr8-90678009-INS-72
chr8-90695794-INS-183
chr8-90717063-INS-202
chr8-90784967-INS-56
chr8-90789601-INS-105
chr8-90792460-INS-56
chr8-90794271-INS-94
chr8-90811808-INS-605
chr8-90840641-INS-360
chr8-90925063-INS-10017
chr8-90956921-INS-6401
chr8-90964459-INS-6628
chr8-90966010-INS-605
chr8-91005072-INS-52
chr8-91038577-INS-51
chr8-91069226-INS-223
chr8-91132740-INS-259
chr8-91164468-INS-1373
chr8-91184118-INS-100
chr8-91209979-INS-354
chr8-91210775-INS-100
chr8-91291462-INS-715
chr8-91311951-INS-1660
chr8-91352792-INS-92
chr8-91358447-INS-188
chr8-91370874-INS-77
chr8-91372338-INS-136
chr8-91387572-INS-54
chr8-91392375-INS-200
chr8-91396605-INS-52
chr8-91432947-INS-181
chr8-91438024-INS-196
chr8-91442134-INS-460
chr8-91455278-INS-207
chr8-91486942-INS-206
chr8-91504633-INS-71
chr8-91517929-INS-52
chr8-91520048-INS-205
chr8-91521788-INS-256
chr8-91534170-INS-98
chr8-91537543-INS-290
chr8-91540301-INS-229
chr8-91558308-INS-202
chr8-91559093-INS-245
chr8-91587536-INS-181
chr8-91592774-INS-183
chr8-91597548-INS-869
chr8-91605438-INS-121
chr8-91629941-INS-51

chr8-91671098-INS-211
chr8-91674217-INS-56
chr8-91679229-INS-66
chr8-91680818-INS-54
chr8-91695936-INS-210
chr8-91724032-INS-252
chr8-91733912-INS-280
chr8-91746519-INS-103
chr8-91765618-INS-118
chr8-91789372-INS-55
chr8-91790311-INS-855
chr8-91791180-INS-456
chr8-91814427-INS-73
chr8-91827335-INS-54
chr8-91831322-INS-57
chr8-91857550-INS-205
chr8-91863547-INS-133
chr8-91865434-INS-481
chr8-91914823-INS-159
chr8-91931875-INS-223
chr8-91941249-INS-1890
chr8-91962261-INS-138
chr8-91974854-INS-3124
chr8-91975055-INS-1576
chr8-91978286-INS-474
chr8-92003298-INS-399
chr8-92006069-INS-388
chr8-92007195-INS-62
chr8-92026333-INS-202
chr8-92027863-INS-194
chr8-92046223-INS-61
chr8-92052503-INS-81
chr8-92060547-INS-147
chr8-92062474-INS-60
chr8-92155828-INS-96
chr8-92165435-INS-148
chr8-92167506-INS-52
chr8-92174387-INS-56
chr8-92203324-INS-132
chr8-92210514-INS-58
chr8-92226392-INS-261
chr8-92255307-INS-88
chr8-92276311-INS-206
chr8-92305493-INS-1894
chr8-92306058-INS-83
chr8-92307773-INS-75
chr8-92313922-INS-71
chr8-92318367-INS-77
chr8-92407852-INS-85
chr8-92408626-INS-240
chr8-92442035-INS-55
chr8-92455628-INS-569
chr8-92465488-INS-79
chr8-92483965-INS-147
chr8-92491439-INS-177
chr8-92507827-INS-62
chr8-92516456-INS-54
chr8-92517331-INS-141
chr8-92521785-INS-98
chr8-92566455-INS-80
chr8-92607448-INS-386
chr8-92622752-INS-55
chr8-92645279-INS-6448
chr8-92645608-INS-65
chr8-92683694-INS-379
chr8-92732867-INS-58
chr8-92800933-INS-792
chr8-92805433-INS-205
chr8-92830955-INS-211
chr8-92868971-INS-162
chr8-92869934-INS-52
chr8-92884040-INS-502
chr8-92904030-INS-539
chr8-92912162-INS-210
chr8-92915264-INS-172
chr8-92938066-INS-72
chr8-92950447-INS-318

chr8-92982499-INS-86
chr8-92987885-INS-60
chr8-92996062-INS-78
chr8-92997084-INS-81
chr8-92998237-INS-88
chr8-93005788-INS-52
chr8-93008428-INS-633
chr8-93009581-INS-70
chr8-93013311-INS-60
chr8-93026303-INS-98
chr8-93057349-INS-212
chr8-93078887-INS-150
chr8-93108377-INS-401
chr8-93120777-INS-160
chr8-93157013-INS-823
chr8-93167328-INS-968
chr8-93189703-INS-99
chr8-93197690-INS-180
chr8-93270753-INS-200
chr8-93276808-INS-724
chr8-93292411-INS-302
chr8-93306084-INS-63
chr8-93319809-INS-63
chr8-93323579-INS-218
chr8-93370126-INS-155
chr8-93373955-INS-200
chr8-93432595-INS-54
chr8-93441796-INS-70
chr8-93453181-INS-358
chr8-93456350-INS-99
chr8-93459713-INS-88
chr8-93471334-INS-52
chr8-93485428-INS-97
chr8-93520433-INS-65
chr8-93541263-INS-374
chr8-93589173-INS-114
chr8-93595172-INS-492
chr8-93604368-INS-58
chr8-93634224-INS-2693
chr8-93646618-INS-61
chr8-93654900-INS-120
chr8-93657912-INS-66
chr8-93664079-INS-107
chr8-93665208-INS-659
chr8-93683455-INS-56
chr8-93715035-INS-258
chr8-93730829-INS-767
chr8-93735901-INS-67
chr8-93767249-INS-195
chr8-93767571-INS-266
chr8-93792155-INS-5026
chr8-93795963-INS-1354
chr8-93831276-INS-439
chr8-93853989-INS-53
chr8-93889806-INS-704
chr8-93891069-INS-400
chr8-93895721-INS-59
chr8-93953552-INS-703
chr8-93982683-INS-6274
chr8-94034699-INS-483
chr8-94035921-INS-292
chr8-94036378-INS-703
chr8-94048009-INS-956
chr8-94049525-INS-220
chr8-94052768-INS-495
chr8-94077896-INS-791
chr8-94090692-INS-191
chr8-94098758-INS-52
chr8-94110319-INS-295
chr8-94155432-INS-62
chr8-94157994-INS-376
chr8-94161069-INS-1892
chr8-94172180-INS-7434
chr8-94217921-INS-270
chr8-94219357-INS-260
chr8-94222352-INS-646
chr8-94224264-INS-358

chr8-94226948-INS-154
chr8-94241023-INS-52
chr8-94253738-INS-246
chr8-94257116-INS-103
chr8-94260532-INS-54
chr8-94286890-INS-271
chr8-94292203-INS-475
chr8-94323423-INS-72
chr8-94329937-INS-6970
chr8-94368915-INS-144
chr8-94402193-INS-6485
chr8-94413866-INS-310
chr8-94430132-INS-185
chr8-94436104-INS-58
chr8-94441899-INS-414
chr8-94447949-INS-57
chr8-94460483-INS-1480
chr8-94469125-INS-322
chr8-94501968-INS-124
chr8-94503937-INS-148
chr8-94523285-INS-105
chr8-94534869-INS-54
chr8-94546433-INS-6399
chr8-94563119-INS-52
chr8-94582702-INS-68
chr8-94584286-INS-485
chr8-94633237-INS-486
chr8-94641020-INS-221
chr8-94653215-INS-296
chr8-94685722-INS-58
chr8-94690246-INS-71
chr8-94780203-INS-130
chr8-94795679-INS-341
chr8-94832664-INS-191
chr8-94841761-INS-106
chr8-94843665-INS-102
chr8-94857380-INS-155
chr8-94872084-INS-160
chr8-94880503-INS-56
chr8-94884840-INS-92
chr8-94886497-INS-820
chr8-94891092-INS-255
chr8-94908962-INS-210
chr8-94926721-INS-725
chr8-94931592-INS-557
chr8-94932889-INS-51
chr8-94933845-INS-792
chr8-94955707-INS-179
chr8-94984638-INS-5263
chr8-94986629-INS-374
chr8-95009717-INS-226
chr8-95018607-INS-126
chr8-95044109-INS-158
chr8-95074394-INS-92
chr8-95093012-INS-118
chr8-95122905-INS-136
chr8-95129522-INS-6727
chr8-95135056-INS-978
chr8-95137228-INS-185
chr8-95141160-INS-61
chr8-95156457-INS-103
chr8-95171971-INS-194
chr8-95200282-INS-211
chr8-95205801-INS-109
chr8-95243156-INS-110
chr8-95244025-INS-548
chr8-95257706-INS-60
chr8-95320515-INS-118
chr8-95356992-INS-314
chr8-95380450-INS-231
chr8-95389318-INS-52
chr8-95414593-INS-169
chr8-95439406-INS-192
chr8-95459267-INS-237
chr8-95485768-INS-6422
chr8-95496654-INS-122
chr8-95515729-INS-138

chr8-95527907-INS-54
chr8-95543084-INS-773
chr8-95545768-INS-56
chr8-95552155-INS-54
chr8-95570576-INS-187
chr8-95603985-INS-88
chr8-95613135-INS-55
chr8-95656024-INS-254
chr8-95656943-INS-52
chr8-95662041-INS-970
chr8-95665702-INS-80
chr8-95666831-INS-135
chr8-95680173-INS-342
chr8-95681290-INS-400
chr8-95682889-INS-101
chr8-95686077-INS-67
chr8-95693066-INS-69
chr8-95709831-INS-126
chr8-95710449-INS-909
chr8-95713210-INS-68
chr8-95754650-INS-73
chr8-95757681-INS-220
chr8-95778877-INS-52
chr8-95790834-INS-61
chr8-95795688-INS-71
chr8-95811606-INS-113
chr8-95849792-INS-205
chr8-95874646-INS-203
chr8-95894183-INS-431
chr8-95896816-INS-88
chr8-95928602-INS-161
chr8-95939495-INS-548
chr8-95945473-INS-56
chr8-95960849-INS-202
chr8-95981516-INS-295
chr8-95983236-INS-207
chr8-95992955-INS-210
chr8-95995853-INS-117
chr8-96000162-INS-63
chr8-96002661-INS-304
chr8-96012007-INS-346
chr8-96042716-INS-91
chr8-96097119-INS-314
chr8-96105479-INS-138
chr8-96129390-INS-218
chr8-96148203-INS-54
chr8-96157597-INS-393
chr8-96205960-INS-207
chr8-96221427-INS-88
chr8-96228060-INS-64
chr8-96258083-INS-57
chr8-96259412-INS-275
chr8-96283272-INS-204
chr8-96309202-INS-5369
chr8-96324265-INS-306
chr8-96326116-INS-57
chr8-96332687-INS-195
chr8-96343475-INS-326
chr8-96356831-INS-198
chr8-96406075-INS-56
chr8-96421829-INS-119
chr8-96452905-INS-213
chr8-96456984-INS-171
chr8-96476938-INS-696
chr8-96505447-INS-205
chr8-96525146-INS-59
chr8-96539966-INS-267
chr8-96542974-INS-78
chr8-96545490-INS-221
chr8-96549234-INS-165
chr8-96558459-INS-173
chr8-96564646-INS-139
chr8-96580648-INS-146
chr8-96582388-INS-194
chr8-96592441-INS-186
chr8-96603856-INS-200
chr8-96633775-INS-234

chr8-96643905-INS-61
chr8-96657258-INS-741
chr8-96704543-INS-120
chr8-96728739-INS-294
chr8-96768506-INS-55
chr8-96777228-INS-6269
chr8-96777577-INS-55
chr8-96811802-INS-236
chr8-96823592-INS-6671
chr8-96824625-INS-92
chr8-96828181-INS-158
chr8-96832960-INS-6688
chr8-96875563-INS-54
chr8-96876506-INS-875
chr8-96887079-INS-71
chr8-96896728-INS-112
chr8-96950371-INS-438
chr8-96964395-INS-106
chr8-96991686-INS-203
chr8-97009754-INS-254
chr8-97013767-INS-228
chr8-97030037-INS-536
chr8-97038096-INS-353
chr8-97046007-INS-294
chr8-97048817-INS-666
chr8-97059263-INS-6454
chr8-97066529-INS-69
chr8-97091904-INS-383
chr8-97095464-INS-118
chr8-97153540-INS-132
chr8-97158553-INS-634
chr8-97167490-INS-288
chr8-97192256-INS-52
chr8-97196545-INS-119
chr8-97238975-INS-2659
chr8-97259790-INS-881
chr8-97262220-INS-60
chr8-97277529-INS-54
chr8-97285265-INS-496
chr8-97293979-INS-73
chr8-97301585-INS-158
chr8-97305471-INS-56
chr8-97322175-INS-639
chr8-97361088-INS-4380
chr8-97375760-INS-60
chr8-97385953-INS-87
chr8-97395661-INS-231
chr8-97397226-INS-189
chr8-97439534-INS-68
chr8-97453770-INS-54
chr8-97456123-INS-55
chr8-97473265-INS-52
chr8-97486185-INS-102
chr8-97510672-INS-742
chr8-97512507-INS-344
chr8-97531906-INS-100
chr8-97553468-INS-170
chr8-97582600-INS-61
chr8-97590309-INS-163
chr8-97622077-INS-92
chr8-97625786-INS-633
chr8-97645972-INS-711
chr8-97658279-INS-270
chr8-97668592-INS-54
chr8-97675172-INS-145
chr8-97688324-INS-68
chr8-97699382-INS-175
chr8-97710421-INS-260
chr8-97713920-INS-410
chr8-97757314-INS-228
chr8-97781217-INS-6655
chr8-97781948-INS-6379
chr8-97790357-INS-4478
chr8-97810233-INS-513
chr8-97814002-INS-59
chr8-97815598-INS-135
chr8-97871589-INS-66

chr8-97873079-INS-295
chr8-97880418-INS-717
chr8-97946820-INS-225
chr8-97976422-INS-189
chr8-97977465-INS-6291
chr8-98008333-INS-193
chr8-98009638-INS-206
chr8-98022411-INS-613
chr8-98038833-INS-244
chr8-98044373-INS-63
chr8-98070245-INS-202
chr8-98075071-INS-92
chr8-98079635-INS-53
chr8-98082302-INS-163
chr8-98106310-INS-120
chr8-98125050-INS-62
chr8-98142893-INS-70
chr8-98157254-INS-6208
chr8-98184164-INS-61
chr8-98200195-INS-942
chr8-98202304-INS-55
chr8-98207807-INS-460
chr8-98249384-INS-197
chr8-98256816-INS-2451
chr8-98262593-INS-56
chr8-98266111-INS-6846
chr8-98285643-INS-1202
chr8-98288106-INS-638
chr8-98321903-INS-541
chr8-98323249-INS-932
chr8-98345398-INS-57
chr8-98387561-INS-344
chr8-98394346-INS-7486
chr8-98402379-INS-315
chr8-98414519-INS-79
chr8-98425366-INS-61
chr8-98432428-INS-848
chr8-98434359-INS-154
chr8-98463846-INS-6384
chr8-98465423-INS-58
chr8-98470420-INS-79
chr8-98486623-INS-136
chr8-98499556-INS-88
chr8-98513601-INS-142
chr8-98514766-INS-57
chr8-98563448-INS-204
chr8-98570439-INS-100
chr8-98573970-INS-319
chr8-98584750-INS-524
chr8-98601776-INS-94
chr8-98628425-INS-493
chr8-98631463-INS-7044
chr8-98653933-INS-175
chr8-98657424-INS-112
chr8-98664583-INS-70
chr8-98672167-INS-56
chr8-98689177-INS-613
chr8-98699865-INS-66
chr8-98716463-INS-310
chr8-98742256-INS-96
chr8-98749059-INS-82
chr8-98758821-INS-286
chr8-98825641-INS-136
chr8-98826956-INS-287
chr8-98844927-INS-343
chr8-98890121-INS-108
chr8-98924348-INS-99
chr8-98960867-INS-66
chr8-98971302-INS-65
chr8-98974674-INS-978
chr8-98988662-INS-564
chr8-98997764-INS-301
chr8-98998716-INS-235
chr8-99021001-INS-420
chr8-99110358-INS-54
chr8-99144124-INS-184
chr8-99166727-INS-68

chr8-99179038-INS-52
chr8-99187701-INS-6612
chr8-99193892-INS-241
chr8-99208929-INS-88
chr8-99260964-INS-133
chr8-99287373-INS-66
chr8-99301166-INS-230
chr8-99317970-INS-767
chr8-99351308-INS-54
chr8-99352875-INS-52
chr8-99371524-INS-72
chr8-99401331-INS-502
chr8-99451085-INS-117
chr8-99457950-INS-197
chr8-99485719-INS-61
chr8-99497120-INS-73
chr8-99513499-INS-103
chr8-99542616-INS-216
chr8-99559837-INS-52
chr8-99571683-INS-832
chr8-99571907-INS-1456
chr8-99600056-INS-66
chr8-99621519-INS-189
chr8-99622889-INS-5294
chr8-99638360-INS-57
chr8-99645419-INS-240
chr8-99646147-INS-6411
chr8-99656448-INS-636
chr8-99727563-INS-78
chr8-99752068-INS-56
chr8-99780895-INS-52
chr8-99810313-INS-628
chr8-99841112-INS-65
chr8-99848328-INS-138
chr8-99851212-INS-54
chr8-99866464-INS-182
chr8-99909904-INS-354
chr8-99924212-INS-57
chr8-99938780-INS-64
chr8-99949225-INS-550
chr8-99981170-INS-63
chr8-99984811-INS-97
chr8-100004321-INS-54
chr8-100026126-INS-143
chr8-100029771-INS-458
chr8-100046556-INS-86
chr8-100051216-INS-259
chr8-100074927-INS-52
chr8-100097367-INS-61
chr8-100100302-INS-54
chr8-100117709-INS-86
chr8-100126191-INS-268
chr8-100128516-INS-65
chr8-100137501-INS-602
chr8-100147681-INS-128
chr8-100150654-INS-791
chr8-100190855-INS-66
chr8-100200867-INS-515
chr8-100207446-INS-318
chr8-100213851-INS-68
chr8-100232650-INS-547
chr8-100235359-INS-7310
chr8-100255072-INS-332
chr8-100259152-INS-75
chr8-100261662-INS-358
chr8-100263364-INS-761
chr8-100289858-INS-213
chr8-100330268-INS-200
chr8-100345773-INS-97
chr8-100350155-INS-642
chr8-100386836-INS-59
chr8-100404828-INS-367
chr8-100417022-INS-54
chr8-100422900-INS-9938
chr8-100426613-INS-71
chr8-100467133-INS-52
chr8-100540219-INS-78

chr8-100561516-INS-64
chr8-100583259-INS-409
chr8-100593291-INS-412
chr8-100607862-INS-2002
chr8-100617333-INS-74
chr8-100648335-INS-59
chr8-100677362-INS-74
chr8-100683399-INS-74
chr8-100683848-INS-241
chr8-100685993-INS-56
chr8-100708125-INS-132
chr8-100709983-INS-660
chr8-100716770-INS-60
chr8-100752332-INS-552
chr8-100800171-INS-249
chr8-100807185-INS-171
chr8-100813385-INS-59
chr8-100817424-INS-591
chr8-100830340-INS-737
chr8-100876037-INS-170
chr8-100941165-INS-52
chr8-100945162-INS-53
chr8-101033687-INS-56
chr8-101051196-INS-199
chr8-101065953-INS-245
chr8-101108956-INS-793
chr8-101120492-INS-115
chr8-101192532-INS-4671
chr8-101193622-INS-6395
chr8-101202011-INS-93
chr8-101216147-INS-7125
chr8-101223809-INS-699
chr8-101235795-INS-76
chr8-101280679-INS-228
chr8-101285486-INS-404
chr8-101290348-INS-80
chr8-101341027-INS-67
chr8-101350514-INS-200
chr8-101391091-INS-382
chr8-101437356-INS-475
chr8-101466932-INS-69
chr8-101480022-INS-52
chr8-101495439-INS-74
chr8-101503167-INS-52
chr8-101506706-INS-105
chr8-101529321-INS-98
chr8-101532767-INS-196
chr8-101536121-INS-120
chr8-101555795-INS-90
chr8-101581816-INS-449
chr8-101604295-INS-181
chr8-101611296-INS-67
chr8-101633198-INS-225
chr8-101657908-INS-66
chr8-101702850-INS-641
chr8-101723757-INS-195
chr8-101735432-INS-5008
chr8-101743474-INS-339
chr8-101755601-INS-475
chr8-101800977-INS-733
chr8-101820945-INS-548
chr8-101851697-INS-351
chr8-101910821-INS-68
chr8-101934981-INS-208
chr8-101966339-INS-96
chr8-101986839-INS-66
chr8-101997871-INS-276
chr8-102030620-INS-54
chr8-102040168-INS-53
chr8-102075246-INS-311
chr8-102120891-INS-58
chr8-102187794-INS-105
chr8-102244878-INS-6380
chr8-102248035-INS-66
chr8-102254415-INS-6770
chr8-102266061-INS-269
chr8-102285307-INS-391

chr8-102291315-INS-439
chr8-102297708-INS-51
chr8-102303401-INS-2168
chr8-102354128-INS-863
chr8-102360118-INS-246
chr8-102378566-INS-111
chr8-102474590-INS-53
chr8-102475473-INS-291
chr8-102493926-INS-706
chr8-102499073-INS-65
chr8-102516748-INS-2782
chr8-102548641-INS-67
chr8-102566338-INS-65
chr8-102610887-INS-214
chr8-102624286-INS-4379
chr8-102632076-INS-693
chr8-102632783-INS-70
chr8-102634035-INS-492
chr8-102653370-INS-80
chr8-102655323-INS-182
chr8-102686321-INS-104
chr8-102699932-INS-126
chr8-102718944-INS-217
chr8-102765461-INS-5747
chr8-102782020-INS-145
chr8-102847672-INS-58
chr8-102888936-INS-135
chr8-102890118-INS-328
chr8-102898952-INS-220
chr8-102903767-INS-85
chr8-102905928-INS-549
chr8-102923459-INS-76
chr8-102930311-INS-7180
chr8-102942222-INS-99
chr8-102955885-INS-245
chr8-103003645-INS-238
chr8-103012153-INS-2964
chr8-103021811-INS-60
chr8-103034097-INS-543
chr8-103038271-INS-332
chr8-103063228-INS-206
chr8-103076963-INS-52
chr8-103081582-INS-273
chr8-103089508-INS-56
chr8-103091316-INS-187
chr8-103151380-INS-233
chr8-103162862-INS-351
chr8-103175332-INS-58
chr8-103179227-INS-71
chr8-103181230-INS-117
chr8-103184365-INS-68
chr8-103191751-INS-7067
chr8-103201266-INS-6401
chr8-103211113-INS-69
chr8-103219339-INS-74
chr8-103222514-INS-261
chr8-103254853-INS-108
chr8-103263677-INS-57
chr8-103268191-INS-75
chr8-103273458-INS-236
chr8-103282115-INS-66
chr8-103300442-INS-136
chr8-103305746-INS-403
chr8-103338412-INS-160
chr8-103340152-INS-505
chr8-103341386-INS-7091
chr8-103372136-INS-103
chr8-103471510-INS-111
chr8-103479861-INS-606
chr8-103496141-INS-198
chr8-103531492-INS-88
chr8-103532438-INS-200
chr8-103591265-INS-5520
chr8-103611881-INS-60
chr8-103620707-INS-622
chr8-103628988-INS-105
chr8-103662852-INS-114

chr8-103695264-INS-147
chr8-103700682-INS-108
chr8-103708124-INS-129
chr8-103722994-INS-110
chr8-103727437-INS-6401
chr8-103761498-INS-3126
chr8-103785559-INS-144
chr8-103810121-INS-214
chr8-103819664-INS-148
chr8-103824149-INS-50
chr8-103832128-INS-201
chr8-103837329-INS-84
chr8-103839456-INS-70
chr8-103839963-INS-53
chr8-103847181-INS-114
chr8-103867170-INS-60
chr8-103868186-INS-56
chr8-103869771-INS-4102
chr8-103913193-INS-59
chr8-103923851-INS-299
chr8-103944238-INS-54
chr8-103954477-INS-165
chr8-103970470-INS-136
chr8-103992657-INS-67
chr8-104001939-INS-521
chr8-104010943-INS-523
chr8-104012085-INS-106
chr8-104023020-INS-290
chr8-104039454-INS-4394
chr8-104087308-INS-84
chr8-104093592-INS-63
chr8-104096590-INS-54
chr8-104112447-INS-515
chr8-104141876-INS-52
chr8-104165150-INS-158
chr8-104174395-INS-114
chr8-104238510-INS-206
chr8-104247762-INS-7102
chr8-104267668-INS-75
chr8-104276861-INS-61
chr8-104308140-INS-83
chr8-104325055-INS-52
chr8-104334318-INS-75
chr8-104339103-INS-373
chr8-104342693-INS-56
chr8-104354886-INS-147
chr8-104389232-INS-467
chr8-104390078-INS-52
chr8-104392713-INS-55
chr8-104436365-INS-97
chr8-104453178-INS-1045
chr8-104463069-INS-1467
chr8-104464883-INS-64
chr8-104479492-INS-266
chr8-104507649-INS-59
chr8-104532064-INS-87
chr8-104561805-INS-79
chr8-104569222-INS-95
chr8-104570715-INS-70
chr8-104593645-INS-240
chr8-104622883-INS-80
chr8-104654691-INS-144
chr8-104657132-INS-54
chr8-104668569-INS-184
chr8-104676936-INS-76
chr8-104681759-INS-53
chr8-104684940-INS-155
chr8-104690995-INS-67
chr8-104719420-INS-62
chr8-104726643-INS-106
chr8-104815278-INS-60
chr8-104819860-INS-213
chr8-104822296-INS-358
chr8-104843031-INS-248
chr8-104874513-INS-137
chr8-104878120-INS-335
chr8-104886298-INS-68

chr8-104889477-INS-169
chr8-104953070-INS-184
chr8-104961737-INS-60
chr8-104966200-INS-72
chr8-104991331-INS-99
chr8-104995727-INS-971
chr8-105007507-INS-70
chr8-105034922-INS-159
chr8-105042353-INS-290
chr8-105061951-INS-118
chr8-105087369-INS-55
chr8-105098004-INS-6436
chr8-105100290-INS-76
chr8-105105234-INS-213
chr8-105115984-INS-326
chr8-105117274-INS-104
chr8-105129262-INS-58
chr8-105156742-INS-215
chr8-105161047-INS-75
chr8-105184979-INS-527
chr8-105203670-INS-53
chr8-105223665-INS-57
chr8-105229622-INS-1888
chr8-105240111-INS-54
chr8-105243224-INS-196
chr8-105245133-INS-132
chr8-105248122-INS-142
chr8-105257925-INS-174
chr8-105267274-INS-1086
chr8-105326120-INS-90
chr8-105345373-INS-64
chr8-105358220-INS-74
chr8-105392703-INS-425
chr8-105424442-INS-102
chr8-105427063-INS-506
chr8-105444386-INS-292
chr8-105445020-INS-381
chr8-105457688-INS-201
chr8-105463195-INS-58
chr8-105475187-INS-4236
chr8-105497072-INS-69
chr8-105514880-INS-1493
chr8-105517510-INS-60
chr8-105518063-INS-58
chr8-105545116-INS-6440
chr8-105563638-INS-6236
chr8-105568142-INS-129
chr8-105606194-INS-207
chr8-105614864-INS-322
chr8-105634233-INS-189
chr8-105639448-INS-477
chr8-105639560-INS-6458
chr8-105644898-INS-1071
chr8-105684520-INS-397
chr8-105686348-INS-7452
chr8-105687940-INS-144
chr8-105702171-INS-73
chr8-105703467-INS-137
chr8-105714512-INS-67
chr8-105746463-INS-172
chr8-105763755-INS-426
chr8-105794042-INS-505
chr8-105836555-INS-92
chr8-105843426-INS-154
chr8-105846096-INS-140
chr8-105853090-INS-57
chr8-105867973-INS-52
chr8-105887113-INS-67
chr8-105912501-INS-192
chr8-105920552-INS-122
chr8-105924358-INS-198
chr8-105930327-INS-193
chr8-105935917-INS-210
chr8-105936930-INS-68
chr8-105949349-INS-5312
chr8-106001910-INS-394
chr8-106010996-INS-132

chr8-106018731-INS-189
chr8-106047479-INS-1215
chr8-106047486-INS-90
chr8-106123681-INS-972
chr8-106126661-INS-120
chr8-106136612-INS-122
chr8-106137776-INS-137
chr8-106147981-INS-204
chr8-106152436-INS-99
chr8-106155154-INS-199
chr8-106162277-INS-167
chr8-106171391-INS-311
chr8-106173092-INS-188
chr8-106224709-INS-56
chr8-106230743-INS-101
chr8-106242291-INS-167
chr8-106267952-INS-150
chr8-106276112-INS-212
chr8-106296102-INS-6438
chr8-106303843-INS-8735
chr8-106329307-INS-66
chr8-106347322-INS-98
chr8-106351511-INS-68
chr8-106356624-INS-156
chr8-106360486-INS-200
chr8-106392213-INS-213
chr8-106403782-INS-356
chr8-106420278-INS-232
chr8-106426147-INS-56
chr8-106441069-INS-202
chr8-106445531-INS-136
chr8-106449841-INS-994
chr8-106453807-INS-83
chr8-106457220-INS-216
chr8-106459296-INS-209
chr8-106463696-INS-99
chr8-106480968-INS-271
chr8-106482909-INS-164
chr8-106487542-INS-167
chr8-106503609-INS-218
chr8-106514334-INS-199
chr8-106525649-INS-55
chr8-106527891-INS-75
chr8-106533527-INS-55
chr8-106536489-INS-247
chr8-106540611-INS-900
chr8-106543385-INS-88
chr8-106544028-INS-144
chr8-106548470-INS-116
chr8-106550178-INS-166
chr8-106551826-INS-68
chr8-106557094-INS-193
chr8-106561176-INS-206
chr8-106577846-INS-59
chr8-106579475-INS-203
chr8-106592368-INS-317
chr8-106629564-INS-407
chr8-106637330-INS-290
chr8-106654481-INS-213
chr8-106663262-INS-583
chr8-106665663-INS-129
chr8-106679794-INS-214
chr8-106694519-INS-309
chr8-106700653-INS-115
chr8-106743369-INS-71
chr8-106748787-INS-76
chr8-106756072-INS-176
chr8-106759800-INS-125
chr8-106767921-INS-83
chr8-106771690-INS-442
chr8-106809180-INS-266
chr8-106814995-INS-175
chr8-106817476-INS-197
chr8-106843973-INS-176
chr8-106848410-INS-206
chr8-106850656-INS-201
chr8-106865927-INS-433

chr8-106876385-INS-134
chr8-106880777-INS-204
chr8-106882019-INS-164
chr8-106882535-INS-52
chr8-106884705-INS-131
chr8-106904888-INS-75
chr8-106913769-INS-134
chr8-106926846-INS-177
chr8-106959288-INS-504
chr8-106960180-INS-213
chr8-106963805-INS-825
chr8-106966176-INS-396
chr8-107022988-INS-98
chr8-107073525-INS-347
chr8-107088634-INS-502
chr8-107123915-INS-400
chr8-107135994-INS-75
chr8-107137699-INS-149
chr8-107153079-INS-143
chr8-107159617-INS-258
chr8-107168875-INS-139
chr8-107177288-INS-54
chr8-107189001-INS-3141
chr8-107197358-INS-68
chr8-107199993-INS-4097
chr8-107200295-INS-135
chr8-107251156-INS-197
chr8-107261056-INS-7139
chr8-107286429-INS-272
chr8-107294154-INS-277
chr8-107296776-INS-197
chr8-107302169-INS-157
chr8-107304516-INS-203
chr8-107307432-INS-136
chr8-107310988-INS-4376
chr8-107318058-INS-52
chr8-107320751-INS-166
chr8-107328589-INS-206
chr8-107331553-INS-202
chr8-107355472-INS-496
chr8-107367207-INS-307
chr8-107411777-INS-184
chr8-107417583-INS-79
chr8-107423165-INS-214
chr8-107423550-INS-3128
chr8-107425099-INS-1141
chr8-107425261-INS-158
chr8-107426578-INS-107
chr8-107434523-INS-197
chr8-107435718-INS-343
chr8-107470206-INS-51
chr8-107475178-INS-224
chr8-107492937-INS-68
chr8-107498533-INS-209
chr8-107501828-INS-551
chr8-107503447-INS-55
chr8-107513172-INS-1278
chr8-107519642-INS-382
chr8-107520848-INS-179
chr8-107527635-INS-155
chr8-107536648-INS-924
chr8-107541140-INS-183
chr8-107564524-INS-185
chr8-107570210-INS-156
chr8-107572824-INS-399
chr8-107578812-INS-212
chr8-107587927-INS-7128
chr8-107590257-INS-392
chr8-107593039-INS-357
chr8-107634350-INS-210
chr8-107636226-INS-236
chr8-107640909-INS-420
chr8-107644140-INS-64
chr8-107648570-INS-238
chr8-107691754-INS-205
chr8-107692866-INS-228
chr8-107693754-INS-60

chr8-107696333-INS-202
chr8-107702312-INS-4008
chr8-107721085-INS-193
chr8-107722902-INS-101
chr8-107736933-INS-217
chr8-107741831-INS-188
chr8-107743364-INS-206
chr8-107748506-INS-60
chr8-107752293-INS-5508
chr8-107757208-INS-53
chr8-107760011-INS-198
chr8-107768346-INS-205
chr8-107770636-INS-391
chr8-107773249-INS-188
chr8-107790883-INS-103
chr8-107792419-INS-384
chr8-107794062-INS-66
chr8-107824158-INS-548
chr8-107826996-INS-7099
chr8-107828131-INS-392
chr8-107833988-INS-202
chr8-107837145-INS-215
chr8-107838694-INS-75
chr8-107851773-INS-432
chr8-107855431-INS-63
chr8-107861230-INS-72
chr8-107863828-INS-212
chr8-107878131-INS-78
chr8-107887108-INS-59
chr8-107889825-INS-155
chr8-107917632-INS-170
chr8-107948155-INS-52
chr8-107953502-INS-168
chr8-107962280-INS-54
chr8-107968719-INS-196
chr8-107982932-INS-211
chr8-108008406-INS-125
chr8-108029330-INS-58
chr8-108033994-INS-56
chr8-108040043-INS-202
chr8-108042948-INS-211
chr8-108081254-INS-213
chr8-108087206-INS-210
chr8-108092143-INS-56
chr8-108103104-INS-151
chr8-108103767-INS-212
chr8-108110350-INS-217
chr8-108118462-INS-570
chr8-108123660-INS-206
chr8-108124616-INS-280
chr8-108144073-INS-109
chr8-108150802-INS-75
chr8-108154076-INS-51
chr8-108158508-INS-384
chr8-108159323-INS-176
chr8-108161054-INS-131
chr8-108200220-INS-174
chr8-108203333-INS-692
chr8-108204558-INS-196
chr8-108215225-INS-116
chr8-108229958-INS-73
chr8-108236324-INS-402
chr8-108258747-INS-116
chr8-108259507-INS-84
chr8-108261781-INS-646
chr8-108268847-INS-78
chr8-108269368-INS-54
chr8-108273716-INS-138
chr8-108289541-INS-208
chr8-108309948-INS-106
chr8-108315245-INS-108
chr8-108317769-INS-77
chr8-108321093-INS-56
chr8-108322448-INS-103
chr8-108327934-INS-123
chr8-108345752-INS-220
chr8-108346975-INS-166

chr8-108355957-INS-57
chr8-108360270-INS-330
chr8-108360319-INS-358
chr8-108404440-INS-60
chr8-108416172-INS-76
chr8-108422570-INS-267
chr8-108441825-INS-304
chr8-108447235-INS-325
chr8-108452148-INS-401
chr8-108456178-INS-137
chr8-108484193-INS-453
chr8-108495420-INS-200
chr8-108505237-INS-235
chr8-108536238-INS-52
chr8-108550436-INS-68
chr8-108561909-INS-106
chr8-108577325-INS-184
chr8-108580877-INS-219
chr8-108592270-INS-63
chr8-108606314-INS-52
chr8-108607533-INS-75
chr8-108610078-INS-57
chr8-108614631-INS-52
chr8-108620236-INS-76
chr8-108657911-INS-246
chr8-108663118-INS-64
chr8-108670571-INS-56
chr8-108715805-INS-59
chr8-108743016-INS-61
chr8-108774951-INS-223
chr8-108781164-INS-1419
chr8-108835137-INS-3003
chr8-108890774-INS-70
chr8-108891534-INS-366
chr8-108929445-INS-96
chr8-108957215-INS-103
chr8-108964750-INS-80
chr8-109009691-INS-261
chr8-109017076-INS-54
chr8-109024544-INS-64
chr8-109032490-INS-52
chr8-109148371-INS-359
chr8-109201723-INS-68
chr8-109202804-INS-114
chr8-109258423-INS-269
chr8-109270684-INS-213
chr8-109285581-INS-221
chr8-109320870-INS-72
chr8-109331813-INS-64
chr8-109337590-INS-137
chr8-109361510-INS-72
chr8-109367768-INS-62
chr8-109381243-INS-66
chr8-109408501-INS-396
chr8-109478286-INS-124
chr8-109498937-INS-64
chr8-109516699-INS-59
chr8-109532861-INS-180
chr8-109534472-INS-160
chr8-109614958-INS-60
chr8-109692791-INS-67
chr8-109704069-INS-154
chr8-109759023-INS-156
chr8-109775090-INS-756
chr8-109824411-INS-54
chr8-109826032-INS-52
chr8-109838833-INS-73
chr8-109923316-INS-95
chr8-109930227-INS-1492
chr8-109934748-INS-66
chr8-109952852-INS-80
chr8-109961201-INS-459
chr8-109984262-INS-66
chr8-110004392-INS-343
chr8-110006795-INS-189
chr8-110024313-INS-58
chr8-110039329-INS-63

chr8-110081241-INS-54
chr8-110113772-INS-172
chr8-110138753-INS-146
chr8-110150349-INS-52
chr8-110157426-INS-515
chr8-110176356-INS-100
chr8-110178974-INS-126
chr8-110217980-INS-58
chr8-110238749-INS-100
chr8-110242151-INS-397
chr8-110289919-INS-136
chr8-110304740-INS-57
chr8-110318911-INS-64
chr8-110327068-INS-396
chr8-110350673-INS-234
chr8-110355937-INS-64
chr8-110363481-INS-160
chr8-110389663-INS-178
chr8-110422367-INS-603
chr8-110427204-INS-60
chr8-110440259-INS-245
chr8-110494742-INS-127
chr8-110499722-INS-176
chr8-110503321-INS-256
chr8-110513359-INS-110
chr8-110519235-INS-90
chr8-110581144-INS-94
chr8-110597577-INS-276
chr8-110599269-INS-75
chr8-110611031-INS-649
chr8-110612856-INS-96
chr8-110653065-INS-298
chr8-110668614-INS-63
chr8-110677273-INS-52
chr8-110711149-INS-52
chr8-110715783-INS-196
chr8-110725750-INS-97
chr8-110732794-INS-312
chr8-110739831-INS-165
chr8-110783425-INS-56
chr8-110792657-INS-362
chr8-110853200-INS-64
chr8-110871659-INS-76
chr8-110877208-INS-53
chr8-110891493-INS-52
chr8-110916101-INS-54
chr8-110920007-INS-76
chr8-110943211-INS-1680
chr8-110945660-INS-142
chr8-110961939-INS-430
chr8-110971781-INS-563
chr8-110979052-INS-654
chr8-110988794-INS-59
chr8-110997132-INS-136
chr8-111011950-INS-177
chr8-111018176-INS-100
chr8-111020386-INS-6365
chr8-111025488-INS-640
chr8-111027934-INS-167
chr8-111033417-INS-463
chr8-111038473-INS-66
chr8-111042149-INS-109
chr8-111056105-INS-64
chr8-111061717-INS-86
chr8-111063947-INS-201
chr8-111088528-INS-200
chr8-111093577-INS-73
chr8-111096877-INS-52
chr8-111111404-INS-97
chr8-111112755-INS-69
chr8-111114022-INS-349
chr8-111118212-INS-57
chr8-111128561-INS-90
chr8-111136207-INS-1136
chr8-111155156-INS-4183
chr8-111156100-INS-64
chr8-111167633-INS-514

chr8-111172695-INS-69
chr8-111178472-INS-301
chr8-111197072-INS-57
chr8-111210753-INS-83
chr8-111214595-INS-82
chr8-111232252-INS-102
chr8-111284941-INS-60
chr8-111288408-INS-75
chr8-111308319-INS-401
chr8-111330663-INS-101
chr8-111335179-INS-65
chr8-111365877-INS-280
chr8-111419963-INS-1680
chr8-111424272-INS-165
chr8-111426970-INS-208
chr8-111444643-INS-60
chr8-111448577-INS-81
chr8-111472252-INS-217
chr8-111480515-INS-184
chr8-111488558-INS-227
chr8-111495269-INS-62
chr8-111497790-INS-177
chr8-111501868-INS-270
chr8-111528511-INS-186
chr8-111531458-INS-246
chr8-111582928-INS-131
chr8-111594798-INS-262
chr8-111599865-INS-59
chr8-111604962-INS-212
chr8-111611351-INS-191
chr8-111619523-INS-199
chr8-111631824-INS-100
chr8-111670719-INS-116
chr8-111675849-INS-325
chr8-111690750-INS-537
chr8-111701252-INS-192
chr8-111708448-INS-455
chr8-111717616-INS-62
chr8-111718071-INS-123
chr8-111720765-INS-203
chr8-111726119-INS-108
chr8-111727455-INS-209
chr8-111741244-INS-225
chr8-111744969-INS-252
chr8-111763889-INS-209
chr8-111825942-INS-201
chr8-111859407-INS-203
chr8-111885568-INS-210
chr8-111887771-INS-330
chr8-111897339-INS-95
chr8-111899601-INS-85
chr8-111927074-INS-183
chr8-111958128-INS-322
chr8-111970120-INS-70
chr8-111978864-INS-59
chr8-111980245-INS-161
chr8-111982695-INS-333
chr8-111983285-INS-54
chr8-111986219-INS-69
chr8-111992294-INS-181
chr8-111993375-INS-242
chr8-111995158-INS-185
chr8-112000296-INS-118
chr8-112005680-INS-60
chr8-112023124-INS-202
chr8-112029122-INS-147
chr8-112030166-INS-212
chr8-112042387-INS-54
chr8-112082035-INS-251
chr8-112094607-INS-392
chr8-112096316-INS-200
chr8-112097941-INS-332
chr8-112111941-INS-155
chr8-112114374-INS-307
chr8-112124222-INS-297
chr8-112151619-INS-214
chr8-112178689-INS-391

chr8-112180809-INS-99
chr8-112210232-INS-100
chr8-112221589-INS-62
chr8-112224948-INS-490
chr8-112250890-INS-82
chr8-112252823-INS-181
chr8-112289501-INS-282
chr8-112308172-INS-68
chr8-112329920-INS-207
chr8-112335006-INS-91
chr8-112341338-INS-112
chr8-112342402-INS-138
chr8-112343335-INS-210
chr8-112347448-INS-105
chr8-112358736-INS-152
chr8-112367864-INS-176
chr8-112388374-INS-459
chr8-112406635-INS-194
chr8-112414845-INS-414
chr8-112452128-INS-71
chr8-112474084-INS-137
chr8-112477578-INS-195
chr8-112483520-INS-556
chr8-112491854-INS-141
chr8-112497098-INS-56
chr8-112510905-INS-194
chr8-112522791-INS-73
chr8-112546135-INS-291
chr8-112570518-INS-207
chr8-112573769-INS-82
chr8-112578427-INS-212
chr8-112582238-INS-208
chr8-112583143-INS-51
chr8-112595113-INS-200
chr8-112604527-INS-123
chr8-112607490-INS-669
chr8-112614131-INS-143
chr8-112626744-INS-170
chr8-112633340-INS-221
chr8-112648409-INS-216
chr8-112663903-INS-55
chr8-112744024-INS-217
chr8-112745970-INS-191
chr8-112765413-INS-467
chr8-112772516-INS-136
chr8-112781542-INS-184
chr8-112784727-INS-161
chr8-112788802-INS-54
chr8-112802429-INS-63
chr8-112814935-INS-582
chr8-112818347-INS-52
chr8-112829476-INS-120
chr8-112843197-INS-89
chr8-112849251-INS-196
chr8-112855244-INS-474
chr8-112882574-INS-120
chr8-112884270-INS-151
chr8-112892521-INS-290
chr8-112921993-INS-75
chr8-112925949-INS-207
chr8-112933527-INS-188
chr8-112955743-INS-275
chr8-112959292-INS-121
chr8-112960436-INS-80
chr8-112974604-INS-4413
chr8-112988435-INS-180
chr8-112996571-INS-132
chr8-113009832-INS-120
chr8-113012670-INS-6485
chr8-113030517-INS-4647
chr8-113030769-INS-99
chr8-113051226-INS-409
chr8-113058481-INS-639
chr8-113063628-INS-525
chr8-113083784-INS-177
chr8-113105151-INS-746
chr8-113146967-INS-491

chr8-113185475-INS-697
chr8-113227316-INS-184
chr8-113234718-INS-623
chr8-113258951-INS-72
chr8-113262923-INS-54
chr8-113319254-INS-101
chr8-113329784-INS-972
chr8-113339871-INS-639
chr8-113357787-INS-56
chr8-113432209-INS-52
chr8-113444133-INS-72
chr8-113461230-INS-157
chr8-113461961-INS-57
chr8-113468723-INS-8714
chr8-113481114-INS-72
chr8-113544499-INS-57
chr8-113564061-INS-535
chr8-113599208-INS-471
chr8-113615398-INS-1264
chr8-113635111-INS-401
chr8-113656282-INS-53
chr8-113677298-INS-1891
chr8-113680377-INS-653
chr8-113682436-INS-54
chr8-113686976-INS-283
chr8-113702326-INS-6508
chr8-113716833-INS-186
chr8-113725876-INS-403
chr8-113749062-INS-95
chr8-113749549-INS-892
chr8-113774193-INS-60
chr8-113785990-INS-329
chr8-113820385-INS-108
chr8-113830903-INS-231
chr8-113834513-INS-98
chr8-113844954-INS-56
chr8-113887792-INS-54
chr8-113911832-INS-76
chr8-113914397-INS-420
chr8-113924141-INS-151
chr8-113937476-INS-55
chr8-113963578-INS-714
chr8-114035594-INS-56
chr8-114038931-INS-6473
chr8-114042427-INS-54
chr8-114046714-INS-64
chr8-114049598-INS-6395
chr8-114049926-INS-179
chr8-114080758-INS-90
chr8-114084758-INS-71
chr8-114095151-INS-150
chr8-114110127-INS-67
chr8-114112770-INS-51
chr8-114133475-INS-725
chr8-114172688-INS-609
chr8-114177457-INS-200
chr8-114186838-INS-96
chr8-114189759-INS-75
chr8-114196860-INS-164
chr8-114212858-INS-85
chr8-114216719-INS-172
chr8-114237903-INS-126
chr8-114258938-INS-99
chr8-114262818-INS-7064
chr8-114265878-INS-104
chr8-114266730-INS-5430
chr8-114279989-INS-60
chr8-114312679-INS-56
chr8-114314380-INS-435
chr8-114320768-INS-94
chr8-114322335-INS-62
chr8-114352547-INS-58
chr8-114374846-INS-94
chr8-114416854-INS-472
chr8-114453067-INS-137
chr8-114461843-INS-60
chr8-114475746-INS-55

chr8-114505312-INS-72
chr8-114512669-INS-112
chr8-114520641-INS-69
chr8-114551221-INS-544
chr8-114563132-INS-68
chr8-114566560-INS-239
chr8-114599282-INS-110
chr8-114610341-INS-56
chr8-114612781-INS-385
chr8-114616043-INS-56
chr8-114626400-INS-547
chr8-114630984-INS-88
chr8-114658685-INS-55
chr8-114666197-INS-80
chr8-114674148-INS-60
chr8-114681908-INS-159
chr8-114686552-INS-64
chr8-114689555-INS-497
chr8-114700181-INS-78
chr8-114711680-INS-82
chr8-114731606-INS-64
chr8-114736918-INS-806
chr8-114756801-INS-246
chr8-114761025-INS-549
chr8-114762958-INS-89
chr8-114772507-INS-68
chr8-114814654-INS-59
chr8-114818970-INS-136
chr8-114837480-INS-182
chr8-114850048-INS-62
chr8-114859846-INS-63
chr8-114870328-INS-67
chr8-114896945-INS-92
chr8-114908087-INS-51
chr8-114918647-INS-186
chr8-114924507-INS-94
chr8-114928659-INS-532
chr8-114933090-INS-233
chr8-114937432-INS-691
chr8-114943274-INS-209
chr8-114948337-INS-529
chr8-114949618-INS-133
chr8-114962869-INS-353
chr8-114970851-INS-75
chr8-114986398-INS-100
chr8-114988630-INS-878
chr8-115000735-INS-103
chr8-115027770-INS-84
chr8-115029408-INS-53
chr8-115031352-INS-178
chr8-115041943-INS-96
chr8-115069907-INS-97
chr8-115073232-INS-135
chr8-115086893-INS-104
chr8-115126620-INS-293
chr8-115144125-INS-58
chr8-115150971-INS-620
chr8-115183936-INS-410
chr8-115192625-INS-61
chr8-115194439-INS-416
chr8-115209704-INS-270
chr8-115227540-INS-56
chr8-115243487-INS-81
chr8-115251852-INS-58
chr8-115299783-INS-54
chr8-115308510-INS-457
chr8-115320883-INS-68
chr8-115323609-INS-103
chr8-115330120-INS-56
chr8-115338880-INS-78
chr8-115355581-INS-68
chr8-115389267-INS-400
chr8-115390282-INS-58
chr8-115414648-INS-72
chr8-115423359-INS-6440
chr8-115432187-INS-142
chr8-115449355-INS-282

chr8-115454597-INS-130
chr8-115500609-INS-225
chr8-115501870-INS-432
chr8-115534249-INS-82
chr8-115546585-INS-593
chr8-115568810-INS-99
chr8-115574869-INS-78
chr8-115584980-INS-386
chr8-115590643-INS-143
chr8-115596112-INS-54
chr8-115608870-INS-139
chr8-115641659-INS-53
chr8-115647166-INS-51
chr8-115658062-INS-506
chr8-115670929-INS-85
chr8-115681512-INS-6594
chr8-115689276-INS-187
chr8-115690641-INS-182
chr8-115708102-INS-66
chr8-115709083-INS-2379
chr8-115716248-INS-196
chr8-115730816-INS-143
chr8-115761228-INS-78
chr8-115774159-INS-481
chr8-115775540-INS-67
chr8-115820898-INS-606
chr8-115833348-INS-145
chr8-115853343-INS-55
chr8-115858391-INS-53
chr8-115872025-INS-152
chr8-115875961-INS-57
chr8-115879911-INS-252
chr8-115903889-INS-210
chr8-115966725-INS-73
chr8-115971986-INS-354
chr8-115978182-INS-56
chr8-115983843-INS-95
chr8-115991554-INS-422
chr8-116018697-INS-80
chr8-116028014-INS-86
chr8-116038966-INS-52
chr8-116050313-INS-270
chr8-116108410-INS-703
chr8-116111108-INS-53
chr8-116127476-INS-179
chr8-116136150-INS-291
chr8-116243601-INS-61
chr8-116274461-INS-73
chr8-116280329-INS-54
chr8-116283282-INS-53
chr8-116292140-INS-94
chr8-116301469-INS-565
chr8-116320943-INS-410
chr8-116329423-INS-222
chr8-116340261-INS-174
chr8-116342242-INS-539
chr8-116345533-INS-57
chr8-116371743-INS-89
chr8-116395663-INS-74
chr8-116402464-INS-60
chr8-116413914-INS-137
chr8-116476571-INS-51
chr8-116500926-INS-95
chr8-116503443-INS-374
chr8-116542210-INS-361
chr8-116549518-INS-61
chr8-116576953-INS-58
chr8-116599564-INS-180
chr8-116621448-INS-149
chr8-116647822-INS-175
chr8-116653111-INS-73
chr8-116674395-INS-90
chr8-116689245-INS-138
chr8-116694801-INS-191
chr8-116699571-INS-106
chr8-116717940-INS-180
chr8-116734101-INS-246

chr8-116773662-INS-115
chr8-116784000-INS-113
chr8-116807639-INS-80
chr8-116824725-INS-400
chr8-116828602-INS-54
chr8-116843819-INS-133
chr8-116850441-INS-56
chr8-116878230-INS-313
chr8-116888771-INS-117
chr8-116892209-INS-111
chr8-116893875-INS-52
chr8-116899526-INS-123
chr8-116926609-INS-132
chr8-116935733-INS-73
chr8-116944747-INS-159
chr8-116952080-INS-177
chr8-116956127-INS-72
chr8-117004509-INS-58
chr8-117025169-INS-537
chr8-117028286-INS-88
chr8-117034464-INS-386
chr8-117036501-INS-225
chr8-117055426-INS-76
chr8-117060318-INS-1864
chr8-117074380-INS-54
chr8-117083257-INS-5477
chr8-117093075-INS-51
chr8-117116119-INS-344
chr8-117117144-INS-180
chr8-117125690-INS-120
chr8-117153380-INS-95
chr8-117171579-INS-231
chr8-117177319-INS-7112
chr8-117183489-INS-367
chr8-117186304-INS-7078
chr8-117190160-INS-130
chr8-117198722-INS-8732
chr8-117204594-INS-66
chr8-117208154-INS-107
chr8-117252544-INS-52
chr8-117253939-INS-62
chr8-117257307-INS-114
chr8-117261731-INS-297
chr8-117263155-INS-118
chr8-117264073-INS-135
chr8-117268535-INS-393
chr8-117290933-INS-161
chr8-117302416-INS-146
chr8-117303176-INS-305
chr8-117327623-INS-116
chr8-117339454-INS-162
chr8-117343827-INS-60
chr8-117358470-INS-68
chr8-117379866-INS-104
chr8-117389705-INS-66
chr8-117392690-INS-811
chr8-117393396-INS-92
chr8-117410411-INS-65
chr8-117416016-INS-484
chr8-117419024-INS-477
chr8-117421417-INS-50
chr8-117430920-INS-223
chr8-117431444-INS-262
chr8-117432024-INS-306
chr8-117432678-INS-93
chr8-117466554-INS-56
chr8-117486469-INS-366
chr8-117493636-INS-5340
chr8-117496488-INS-156
chr8-117497094-INS-7293
chr8-117502439-INS-201
chr8-117505052-INS-270
chr8-117519460-INS-52
chr8-117539582-INS-75
chr8-117541914-INS-112
chr8-117551359-INS-328
chr8-117553093-INS-77

chr8-117559604-INS-85
chr8-117568030-INS-175
chr8-117587941-INS-70
chr8-117604044-INS-654
chr8-117609355-INS-61
chr8-117616884-INS-277
chr8-117618744-INS-98
chr8-117620760-INS-62
chr8-117622394-INS-136
chr8-117635149-INS-699
chr8-117647173-INS-56
chr8-117710980-INS-244
chr8-117727268-INS-146
chr8-117738792-INS-321
chr8-117746484-INS-9045
chr8-117748131-INS-450
chr8-117762222-INS-234
chr8-117766722-INS-60
chr8-117777350-INS-68
chr8-117779050-INS-60
chr8-117783725-INS-130
chr8-117796430-INS-199
chr8-117805960-INS-534
chr8-117821291-INS-200
chr8-117832605-INS-237
chr8-117833845-INS-87
chr8-117836226-INS-53
chr8-117857307-INS-343
chr8-117872185-INS-182
chr8-117892824-INS-74
chr8-117897548-INS-56
chr8-117900400-INS-194
chr8-117922973-INS-180
chr8-117946113-INS-205
chr8-117963422-INS-171
chr8-117972754-INS-79
chr8-117977028-INS-72
chr8-117996348-INS-228
chr8-118003800-INS-80
chr8-118011213-INS-204
chr8-118038944-INS-103
chr8-118040737-INS-104
chr8-118053831-INS-56
chr8-118077013-INS-423
chr8-118094148-INS-51
chr8-118119383-INS-56
chr8-118131708-INS-52
chr8-118136846-INS-169
chr8-118166904-INS-116
chr8-118209634-INS-97
chr8-118216151-INS-52
chr8-118235929-INS-114
chr8-118237875-INS-56
chr8-118240343-INS-141
chr8-118260718-INS-103
chr8-118272316-INS-52
chr8-118276087-INS-215
chr8-118287564-INS-69
chr8-118297639-INS-79
chr8-118305404-INS-170
chr8-118317902-INS-75
chr8-118321325-INS-68
chr8-118346652-INS-223
chr8-118349217-INS-112
chr8-118371015-INS-195
chr8-118383964-INS-56
chr8-118429897-INS-54
chr8-118433918-INS-54
chr8-118438822-INS-114
chr8-118445957-INS-52
chr8-118453851-INS-61
chr8-118460826-INS-60
chr8-118462584-INS-66
chr8-118465948-INS-54
chr8-118469972-INS-133
chr8-118471180-INS-80
chr8-118473174-INS-51

chr8-118475213-INS-107
chr8-118503764-INS-71
chr8-118516709-INS-224
chr8-118520352-INS-180
chr8-118561242-INS-189
chr8-118593276-INS-62
chr8-118600966-INS-129
chr8-118611591-INS-71
chr8-118623658-INS-133
chr8-118633124-INS-66
chr8-118636874-INS-79
chr8-118665112-INS-300
chr8-118686979-INS-209
chr8-118694428-INS-161
chr8-118700365-INS-84
chr8-118704976-INS-214
chr8-118712770-INS-120
chr8-118727059-INS-182
chr8-118740690-INS-1837
chr8-118799780-INS-110
chr8-118813166-INS-66
chr8-118819737-INS-54
chr8-118830497-INS-375
chr8-118849378-INS-86
chr8-118886125-INS-52
chr8-118906745-INS-84
chr8-118919697-INS-106
chr8-118932851-INS-532
chr8-118949833-INS-82
chr8-118961117-INS-401
chr8-118973024-INS-90
chr8-118975325-INS-164
chr8-118979761-INS-906
chr8-118986031-INS-552
chr8-119035504-INS-101
chr8-119037934-INS-308
chr8-119049681-INS-123
chr8-119082115-INS-354
chr8-119102465-INS-98
chr8-119108300-INS-132
chr8-119116904-INS-60
chr8-119118603-INS-55
chr8-119120279-INS-88
chr8-119155243-INS-70
chr8-119196738-INS-887
chr8-119197694-INS-87
chr8-119219308-INS-54
chr8-119241666-INS-168
chr8-119263276-INS-429
chr8-119274716-INS-91
chr8-119276785-INS-932
chr8-119303793-INS-65
chr8-119307851-INS-194
chr8-119340667-INS-600
chr8-119353439-INS-63
chr8-119356566-INS-151
chr8-119377158-INS-65
chr8-119404657-INS-70
chr8-119424345-INS-76
chr8-119434763-INS-216
chr8-119454146-INS-510
chr8-119472560-INS-69
chr8-119482943-INS-76
chr8-119510350-INS-52
chr8-119517545-INS-761
chr8-119572099-INS-150
chr8-119574769-INS-210
chr8-119578370-INS-58
chr8-119581024-INS-236
chr8-119608708-INS-235
chr8-119613811-INS-56
chr8-119615417-INS-58
chr8-119625838-INS-54
chr8-119630429-INS-183
chr8-119633554-INS-60
chr8-119649586-INS-76
chr8-119651610-INS-121

chr8-119682019-INS-77
chr8-119685391-INS-98
chr8-119687195-INS-60
chr8-119697368-INS-52
chr8-119711401-INS-220
chr8-119723339-INS-137
chr8-119733085-INS-214
chr8-119740106-INS-181
chr8-119759112-INS-65
chr8-119779602-INS-89
chr8-119784248-INS-5316
chr8-119797376-INS-323
chr8-119803659-INS-60
chr8-119806257-INS-126
chr8-119837235-INS-86
chr8-119847066-INS-85
chr8-119879363-INS-56
chr8-119879847-INS-85
chr8-119882908-INS-287
chr8-119897091-INS-58
chr8-119909440-INS-96
chr8-119920418-INS-67
chr8-119980633-INS-366
chr8-119985463-INS-60
chr8-119988099-INS-62
chr8-120009199-INS-64
chr8-120010296-INS-164
chr8-120020339-INS-86
chr8-120031860-INS-254
chr8-120039614-INS-60
chr8-120067581-INS-84
chr8-120068694-INS-57
chr8-120069167-INS-155
chr8-120093780-INS-196
chr8-120102638-INS-230
chr8-120103856-INS-84
chr8-120111957-INS-331
chr8-120142437-INS-72
chr8-120143069-INS-119
chr8-120143820-INS-249
chr8-120153888-INS-172
chr8-120185269-INS-71
chr8-120187949-INS-291
chr8-120204882-INS-200
chr8-120242963-INS-138
chr8-120243949-INS-75
chr8-120245114-INS-123
chr8-120251930-INS-57
chr8-120275924-INS-105
chr8-120281004-INS-506
chr8-120303254-INS-58
chr8-120308723-INS-59
chr8-120318804-INS-140
chr8-120338375-INS-229
chr8-120346048-INS-291
chr8-120347533-INS-171
chr8-120355884-INS-105
chr8-120367370-INS-190
chr8-120374550-INS-506
chr8-120378049-INS-384
chr8-120400570-INS-294
chr8-120411624-INS-152
chr8-120433993-INS-94
chr8-120440359-INS-96
chr8-120458603-INS-283
chr8-120467130-INS-150
chr8-120485727-INS-193
chr8-120509138-INS-182
chr8-120519719-INS-171
chr8-120528360-INS-367
chr8-120532990-INS-195
chr8-120568717-INS-157
chr8-120578243-INS-64
chr8-120585568-INS-177
chr8-120609720-INS-52
chr8-120613376-INS-94
chr8-120624409-INS-140

chr8-120645190-INS-251
chr8-120657711-INS-94
chr8-120661447-INS-81
chr8-120672837-INS-77
chr8-120688916-INS-202
chr8-120708916-INS-7082
chr8-120726847-INS-84
chr8-120759233-INS-77
chr8-120766916-INS-182
chr8-120775748-INS-88
chr8-120796074-INS-100
chr8-120815214-INS-289
chr8-120816743-INS-100
chr8-120822067-INS-568
chr8-120847444-INS-132
chr8-120852434-INS-111
chr8-120909939-INS-167
chr8-120915997-INS-123
chr8-120928170-INS-261
chr8-120930066-INS-56
chr8-120947390-INS-527
chr8-120966679-INS-64
chr8-120969215-INS-127
chr8-120972602-INS-53
chr8-120973003-INS-342
chr8-120974458-INS-210
chr8-120976140-INS-214
chr8-120982552-INS-137
chr8-120984566-INS-106
chr8-120986962-INS-258
chr8-120989803-INS-102
chr8-121012179-INS-376
chr8-121032315-INS-939
chr8-121032338-INS-58
chr8-121048458-INS-194
chr8-121060508-INS-76
chr8-121063579-INS-342
chr8-121094794-INS-148
chr8-121099321-INS-51
chr8-121147626-INS-97
chr8-121170933-INS-197
chr8-121174083-INS-121
chr8-121175560-INS-112
chr8-121202009-INS-196
chr8-121208740-INS-211
chr8-121211164-INS-233
chr8-121223046-INS-114
chr8-121228925-INS-208
chr8-121230539-INS-194
chr8-121253000-INS-52
chr8-121256394-INS-83
chr8-121257779-INS-229
chr8-121352249-INS-142
chr8-121357129-INS-380
chr8-121361789-INS-65
chr8-121403594-INS-124
chr8-121407525-INS-237
chr8-121426703-INS-354
chr8-121441085-INS-206
chr8-121457711-INS-158
chr8-121498765-INS-105
chr8-121511651-INS-57
chr8-121515347-INS-52
chr8-121555909-INS-200
chr8-121562846-INS-142
chr8-121567201-INS-168
chr8-121579621-INS-99
chr8-121581853-INS-172
chr8-121591582-INS-68
chr8-121597371-INS-97
chr8-121605406-INS-160
chr8-121617399-INS-68
chr8-121633234-INS-325
chr8-121671580-INS-580
chr8-121675120-INS-146
chr8-121680409-INS-60
chr8-121688016-INS-62

chr8-121691783-INS-154
chr8-121692611-INS-61
chr8-121708222-INS-72
chr8-121711392-INS-72
chr8-121713396-INS-158
chr8-121751459-INS-156
chr8-121752747-INS-56
chr8-121766573-INS-349
chr8-121773980-INS-184
chr8-121786063-INS-85
chr8-121795656-INS-117
chr8-121804535-INS-72
chr8-121821645-INS-108
chr8-121837082-INS-76
chr8-121850788-INS-58
chr8-121895804-INS-74
chr8-121908890-INS-807
chr8-121916816-INS-507
chr8-121927325-INS-105
chr8-121933305-INS-112
chr8-121947751-INS-160
chr8-121967591-INS-51
chr8-121972575-INS-84
chr8-121980966-INS-99
chr8-121984040-INS-509
chr8-121998541-INS-60
chr8-122008110-INS-51
chr8-122035267-INS-68
chr8-122066199-INS-386
chr8-122079178-INS-224
chr8-122086875-INS-152
chr8-122112651-INS-52
chr8-122156843-INS-62
chr8-122171200-INS-368
chr8-122176758-INS-236
chr8-122177813-INS-116
chr8-122190311-INS-118
chr8-122191373-INS-204
chr8-122199827-INS-150
chr8-122220797-INS-485
chr8-122222886-INS-198
chr8-122230769-INS-138
chr8-122237829-INS-51
chr8-122241659-INS-190
chr8-122251485-INS-287
chr8-122257019-INS-213
chr8-122262728-INS-84
chr8-122265176-INS-165
chr8-122265465-INS-609
chr8-122270531-INS-58
chr8-122273583-INS-142
chr8-122274599-INS-178
chr8-122282669-INS-155
chr8-122286363-INS-94
chr8-122308727-INS-204
chr8-122329903-INS-58
chr8-122361916-INS-108
chr8-122382726-INS-195
chr8-122405413-INS-126
chr8-122405829-INS-149
chr8-122427417-INS-53
chr8-122432706-INS-108
chr8-122438615-INS-113
chr8-122465945-INS-132
chr8-122475534-INS-62
chr8-122491182-INS-56
chr8-122572522-INS-55
chr8-122579185-INS-5340
chr8-122591188-INS-139
chr8-122621606-INS-134
chr8-122640089-INS-199
chr8-122643026-INS-51
chr8-122650729-INS-54
chr8-122652470-INS-5306
chr8-122675950-INS-3412
chr8-122684134-INS-214
chr8-122687356-INS-169

chr8-122690037-INS-102
chr8-122706889-INS-100
chr8-122711886-INS-210
chr8-122729704-INS-100
chr8-122730767-INS-72
chr8-122750186-INS-204
chr8-122751669-INS-214
chr8-122757100-INS-52
chr8-122780825-INS-444
chr8-122787713-INS-159
chr8-122789317-INS-57
chr8-122790472-INS-212
chr8-122809698-INS-199
chr8-122811036-INS-256
chr8-122827765-INS-394
chr8-122847078-INS-129
chr8-122855220-INS-202
chr8-122878317-INS-52
chr8-122883695-INS-54
chr8-122893293-INS-322
chr8-122901894-INS-170
chr8-122909824-INS-65
chr8-122955342-INS-199
chr8-122962109-INS-204
chr8-122968674-INS-241
chr8-122972221-INS-505
chr8-122983579-INS-120
chr8-123022545-INS-325
chr8-123058948-INS-68
chr8-123059601-INS-412
chr8-123095338-INS-487
chr8-123118522-INS-129
chr8-123124208-INS-217
chr8-123161281-INS-181
chr8-123223950-INS-294
chr8-123231781-INS-129
chr8-123236576-INS-154
chr8-123267777-INS-202
chr8-123280105-INS-214
chr8-123287520-INS-268
chr8-123328330-INS-333
chr8-123374231-INS-135
chr8-123379656-INS-56
chr8-123381360-INS-59
chr8-123383704-INS-158
chr8-123388895-INS-240
chr8-123416876-INS-665
chr8-123430894-INS-176
chr8-123434334-INS-213
chr8-123444374-INS-71
chr8-123448338-INS-82
chr8-123458522-INS-257
chr8-123481425-INS-130
chr8-123495157-INS-67
chr8-123526890-INS-58
chr8-123536526-INS-943
chr8-123555627-INS-164
chr8-123570633-INS-205
chr8-123573314-INS-234
chr8-123575298-INS-5294
chr8-123600914-INS-173
chr8-123602602-INS-206
chr8-123609764-INS-71
chr8-123650046-INS-702
chr8-123686390-INS-238
chr8-123687272-INS-370
chr8-123707896-INS-210
chr8-123710385-INS-190
chr8-123721728-INS-480
chr8-123725699-INS-4870
chr8-123728633-INS-70
chr8-123729860-INS-446
chr8-123741371-INS-631
chr8-123760762-INS-62
chr8-123760795-INS-1054
chr8-123760796-INS-310
chr8-123781313-INS-129

chr8-123813089-INS-129
chr8-123824838-INS-62
chr8-123827668-INS-79
chr8-123838058-INS-490
chr8-123851624-INS-176
chr8-123860105-INS-150
chr8-123868239-INS-54
chr8-123908807-INS-399
chr8-123944587-INS-232
chr8-123947012-INS-255
chr8-123990154-INS-204
chr8-124004188-INS-164
chr8-124011146-INS-70
chr8-124023301-INS-123
chr8-124026513-INS-236
chr8-124049832-INS-63
chr8-124078408-INS-239
chr8-124091480-INS-214
chr8-124093909-INS-257
chr8-124111028-INS-176
chr8-124121473-INS-54
chr8-124125792-INS-53
chr8-124127351-INS-82
chr8-124145124-INS-163
chr8-124163579-INS-138
chr8-124177014-INS-153
chr8-124179433-INS-202
chr8-124184505-INS-61
chr8-124194542-INS-216
chr8-124196561-INS-72
chr8-124198305-INS-139
chr8-124199374-INS-174
chr8-124201642-INS-270
chr8-124208717-INS-54
chr8-124213320-INS-207
chr8-124225114-INS-748
chr8-124238686-INS-56
chr8-124248060-INS-206
chr8-124250670-INS-87
chr8-124265213-INS-1721
chr8-124354875-INS-178
chr8-124359375-INS-112
chr8-124385706-INS-164
chr8-124402773-INS-266
chr8-124450312-INS-478
chr8-124470183-INS-76
chr8-124485592-INS-143
chr8-124492374-INS-83
chr8-124508975-INS-220
chr8-124529022-INS-60
chr8-124562332-INS-126
chr8-124564897-INS-79
chr8-124577668-INS-115
chr8-124588401-INS-166
chr8-124594267-INS-70
chr8-124626919-INS-204
chr8-124642334-INS-197
chr8-124664127-INS-189
chr8-124668327-INS-220
chr8-124686726-INS-123
chr8-124692775-INS-56
chr8-124713744-INS-178
chr8-124728677-INS-87
chr8-124753350-INS-494
chr8-124762939-INS-147
chr8-124799646-INS-217
chr8-124825698-INS-196
chr8-124833134-INS-56
chr8-124872113-INS-62
chr8-124885967-INS-132
chr8-124885986-INS-925
chr8-124888023-INS-137
chr8-124894987-INS-52
chr8-124897905-INS-63
chr8-124904254-INS-63
chr8-124914653-INS-115
chr8-124937199-INS-318

chr8-124995472-INS-213
chr8-125007529-INS-303
chr8-125041790-INS-945
chr8-125042753-INS-80
chr8-125082360-INS-694
chr8-125110149-INS-138
chr8-125137038-INS-80
chr8-125144812-INS-58
chr8-125149987-INS-150
chr8-125177971-INS-299
chr8-125191128-INS-76
chr8-125198431-INS-347
chr8-125200133-INS-3102
chr8-125202051-INS-156
chr8-125209223-INS-59
chr8-125215444-INS-366
chr8-125233920-INS-214
chr8-125239743-INS-433
chr8-125250909-INS-73
chr8-125262977-INS-615
chr8-125283287-INS-510
chr8-125299590-INS-359
chr8-125301719-INS-222
chr8-125304149-INS-167
chr8-125329923-INS-53
chr8-125341817-INS-532
chr8-125353746-INS-58
chr8-125377519-INS-213
chr8-125384117-INS-795
chr8-125384503-INS-141
chr8-125386102-INS-61
chr8-125399757-INS-68
chr8-125425266-INS-174
chr8-125430704-INS-218
chr8-125442266-INS-54
chr8-125459247-INS-54
chr8-125462735-INS-64
chr8-125482989-INS-217
chr8-125490903-INS-156
chr8-125493351-INS-244
chr8-125494279-INS-748
chr8-125518039-INS-53
chr8-125528352-INS-210
chr8-125532707-INS-212
chr8-125535200-INS-121
chr8-125553096-INS-170
chr8-125558153-INS-92
chr8-125561853-INS-97
chr8-125563670-INS-205
chr8-125585450-INS-808
chr8-125591190-INS-62
chr8-125610653-INS-69
chr8-125693605-INS-138
chr8-125710354-INS-61
chr8-125711246-INS-82
chr8-125711643-INS-70
chr8-125714755-INS-54
chr8-125732948-INS-167
chr8-125737634-INS-2932
chr8-125738242-INS-57
chr8-125746641-INS-64
chr8-125756560-INS-140
chr8-125758309-INS-792
chr8-125805051-INS-182
chr8-125819771-INS-121
chr8-125821481-INS-132
chr8-125824392-INS-260
chr8-125863270-INS-952
chr8-125888288-INS-54
chr8-125888989-INS-81
chr8-125898204-INS-2660
chr8-125908183-INS-74
chr8-125916161-INS-147
chr8-125918162-INS-249
chr8-125918627-INS-59
chr8-125919723-INS-127
chr8-125931117-INS-62

chr8-125942509-INS-82
chr8-125974688-INS-64
chr8-125988311-INS-90
chr8-125999849-INS-51
chr8-126027920-INS-73
chr8-126031849-INS-485
chr8-126067978-INS-128
chr8-126069185-INS-273
chr8-126098635-INS-148
chr8-126128140-INS-570
chr8-126137385-INS-383
chr8-126148238-INS-207
chr8-126153717-INS-200
chr8-126163488-INS-132
chr8-126182476-INS-6475
chr8-126183198-INS-72
chr8-126186279-INS-54
chr8-126192645-INS-94
chr8-126198562-INS-208
chr8-126204794-INS-156
chr8-126210079-INS-161
chr8-126214260-INS-122
chr8-126223051-INS-201
chr8-126236279-INS-84
chr8-126264586-INS-196
chr8-126281732-INS-78
chr8-126326453-INS-172
chr8-126340893-INS-66
chr8-126364824-INS-208
chr8-126384590-INS-173
chr8-126399158-INS-7483
chr8-126415331-INS-54
chr8-126424064-INS-437
chr8-126430046-INS-468
chr8-126433279-INS-411
chr8-126454460-INS-341
chr8-126501477-INS-102
chr8-126505550-INS-61
chr8-126510637-INS-113
chr8-126532564-INS-96
chr8-126535206-INS-83
chr8-126536184-INS-94
chr8-126564532-INS-104
chr8-126579895-INS-102
chr8-126592063-INS-184
chr8-126609918-INS-126
chr8-126630157-INS-468
chr8-126669605-INS-468
chr8-126695224-INS-70
chr8-126701583-INS-365
chr8-126747464-INS-242
chr8-126757252-INS-114
chr8-126765727-INS-63
chr8-126777896-INS-59
chr8-126804161-INS-237
chr8-126805845-INS-54
chr8-126817018-INS-196
chr8-126853688-INS-126
chr8-126859709-INS-172
chr8-126899738-INS-118
chr8-126913403-INS-128
chr8-126923714-INS-2668
chr8-126924893-INS-146
chr8-126926330-INS-2083
chr8-126930249-INS-100
chr8-126937966-INS-52
chr8-126964078-INS-130
chr8-126975590-INS-560
chr8-126982582-INS-270
chr8-127007966-INS-132
chr8-127010230-INS-101
chr8-127015456-INS-137
chr8-127015987-INS-160
chr8-127025960-INS-72
chr8-127042880-INS-673
chr8-127050653-INS-86
chr8-127065671-INS-160

chr8-127069615-INS-415
chr8-127094153-INS-411
chr8-127111950-INS-211
chr8-127166720-INS-180
chr8-127183460-INS-338
chr8-127194550-INS-55
chr8-127209043-INS-214
chr8-127217785-INS-218
chr8-127219093-INS-198
chr8-127238976-INS-239
chr8-127242885-INS-60
chr8-127245674-INS-712
chr8-127259072-INS-819
chr8-127261010-INS-66
chr8-127287677-INS-92
chr8-127288990-INS-72
chr8-127329519-INS-57
chr8-127335869-INS-145
chr8-127356181-INS-177
chr8-127370053-INS-84
chr8-127389479-INS-56
chr8-127395740-INS-160
chr8-127402394-INS-64
chr8-127476075-INS-148
chr8-127517469-INS-162
chr8-127525897-INS-53
chr8-127528424-INS-94
chr8-127544595-INS-80
chr8-127550643-INS-261
chr8-127567304-INS-63
chr8-127583156-INS-448
chr8-127587274-INS-160
chr8-127593341-INS-141
chr8-127600811-INS-64
chr8-127605592-INS-64
chr8-127632598-INS-72
chr8-127633970-INS-58
chr8-127638777-INS-325
chr8-127642433-INS-127
chr8-127648178-INS-366
chr8-127676544-INS-7195
chr8-127680646-INS-246
chr8-127681925-INS-63
chr8-127686105-INS-200
chr8-127699460-INS-511
chr8-127709343-INS-51
chr8-127711139-INS-193
chr8-127743970-INS-54
chr8-127752577-INS-517
chr8-127766981-INS-209
chr8-127773864-INS-345
chr8-127779733-INS-297
chr8-127804961-INS-969
chr8-127827615-INS-100
chr8-127846981-INS-271
chr8-127849672-INS-176
chr8-127865497-INS-400
chr8-127869134-INS-92
chr8-127879661-INS-206
chr8-127897198-INS-54
chr8-127907186-INS-428
chr8-127914846-INS-72
chr8-127927435-INS-72
chr8-127950957-INS-87
chr8-127953735-INS-72
chr8-127960173-INS-70
chr8-127988183-INS-74
chr8-127989424-INS-876
chr8-128006154-INS-181
chr8-128055285-INS-70
chr8-128104174-INS-238
chr8-128113721-INS-62
chr8-128198317-INS-56
chr8-128224952-INS-190
chr8-128239908-INS-249
chr8-128250387-INS-58
chr8-128263739-INS-69

chr8-128273979-INS-226
chr8-128281668-INS-54
chr8-128292888-INS-191
chr8-128306101-INS-265
chr8-128309912-INS-72
chr8-128315876-INS-79
chr8-128325416-INS-85
chr8-128334240-INS-650
chr8-128348068-INS-393
chr8-128378532-INS-937
chr8-128404275-INS-63
chr8-128404986-INS-2501
chr8-128405408-INS-60
chr8-128456907-INS-57
chr8-128461824-INS-7084
chr8-128473295-INS-401
chr8-128482013-INS-64
chr8-128493584-INS-411
chr8-128494142-INS-7095
chr8-128525910-INS-62
chr8-128544745-INS-54
chr8-128557661-INS-186
chr8-128574573-INS-153
chr8-128581447-INS-824
chr8-128588562-INS-150
chr8-128601902-INS-71
chr8-128605697-INS-354
chr8-128659263-INS-218
chr8-128661532-INS-400
chr8-128674175-INS-64
chr8-128689284-INS-75
chr8-128701425-INS-160
chr8-128708381-INS-85
chr8-128735157-INS-70
chr8-128782286-INS-147
chr8-128800809-INS-645
chr8-128808836-INS-84
chr8-128813296-INS-323
chr8-128813762-INS-757
chr8-128815147-INS-90
chr8-128818642-INS-7324
chr8-128821995-INS-56
chr8-128825031-INS-215
chr8-128838278-INS-70
chr8-128841013-INS-211
chr8-128848198-INS-204
chr8-128903126-INS-471
chr8-128910566-INS-55
chr8-128937218-INS-58
chr8-128946621-INS-189
chr8-129001598-INS-51
chr8-129003378-INS-100
chr8-129004671-INS-57
chr8-129020490-INS-56
chr8-129040280-INS-369
chr8-129057523-INS-65
chr8-129075647-INS-6214
chr8-129077835-INS-7654
chr8-129082250-INS-107
chr8-129158095-INS-1072
chr8-129179316-INS-65
chr8-129233807-INS-102
chr8-129239129-INS-6889
chr8-129260263-INS-56
chr8-129297380-INS-52
chr8-129299041-INS-6393
chr8-129307588-INS-566
chr8-129317976-INS-6442
chr8-129332020-INS-55
chr8-129335723-INS-400
chr8-129343980-INS-54
chr8-129369765-INS-879
chr8-129395381-INS-74
chr8-129397658-INS-161
chr8-129404042-INS-107
chr8-129407657-INS-1204
chr8-129430670-INS-917

chr8-129430765-INS-209
chr8-129445120-INS-130
chr8-129447787-INS-84
chr8-129481274-INS-506
chr8-129519494-INS-348
chr8-129522567-INS-221
chr8-129565962-INS-440
chr8-129580485-INS-68
chr8-129594184-INS-226
chr8-129596809-INS-482
chr8-129608594-INS-1202
chr8-129624326-INS-512
chr8-129626469-INS-1442
chr8-129631559-INS-98
chr8-129658132-INS-1006
chr8-129659968-INS-450
chr8-129690643-INS-1493
chr8-129701305-INS-109
chr8-129716504-INS-123
chr8-129751630-INS-248
chr8-129863182-INS-6280
chr8-129873845-INS-116
chr8-129885315-INS-1888
chr8-129921914-INS-79
chr8-129922893-INS-7305
chr8-129924034-INS-63
chr8-129934729-INS-2915
chr8-129963496-INS-179
chr8-130019639-INS-454
chr9-3038015-INS-1873
chr9-3062058-INS-72
chr9-3077077-INS-182
chr9-3100672-INS-392
chr9-3104966-INS-160
chr9-3111870-INS-72
chr9-3116781-INS-117
chr9-3196110-INS-99
chr9-3209694-INS-198
chr9-3218653-INS-611
chr9-3238028-INS-198
chr9-3253575-INS-180
chr9-3281216-INS-494
chr9-3296249-INS-2379
chr9-3308273-INS-6475
chr9-3336941-INS-126
chr9-3358377-INS-358
chr9-3386972-INS-7355
chr9-3412788-INS-6484
chr9-3441262-INS-52
chr9-3460923-INS-133
chr9-3497345-INS-72
chr9-3508966-INS-831
chr9-3511178-INS-354
chr9-3553089-INS-137
chr9-3570405-INS-187
chr9-3575847-INS-894
chr9-3603816-INS-64
chr9-3614790-INS-69
chr9-3629711-INS-342
chr9-3650995-INS-114
chr9-3663642-INS-1643
chr9-3665611-INS-6517
chr9-3672282-INS-379
chr9-3682745-INS-6674
chr9-3692924-INS-6770
chr9-3715287-INS-738
chr9-3716168-INS-68
chr9-3743185-INS-150
chr9-3770533-INS-357
chr9-3772303-INS-966
chr9-3823379-INS-195
chr9-3856865-INS-205
chr9-3864698-INS-56
chr9-3866971-INS-187
chr9-3873932-INS-5510
chr9-3879364-INS-203
chr9-3933356-INS-6364

chr9-3965790-INS-572
chr9-3986756-INS-468
chr9-4011100-INS-60
chr9-4034703-INS-2818
chr9-4060380-INS-70
chr9-4087278-INS-96
chr9-4089299-INS-131
chr9-4332077-INS-204
chr9-4378439-INS-6473
chr9-4394219-INS-52
chr9-4408470-INS-144
chr9-4446194-INS-494
chr9-4457239-INS-196
chr9-4465229-INS-171
chr9-4471412-INS-82
chr9-4500563-INS-527
chr9-4507351-INS-461
chr9-4522546-INS-52
chr9-4532448-INS-164
chr9-4534982-INS-60
chr9-4539753-INS-110
chr9-4540840-INS-56
chr9-4541841-INS-209
chr9-4546492-INS-52
chr9-4572361-INS-968
chr9-4575360-INS-554
chr9-4577577-INS-53
chr9-4599626-INS-52
chr9-4616690-INS-2449
chr9-4646038-INS-167
chr9-4660185-INS-65
chr9-4669610-INS-556
chr9-4673804-INS-6592
chr9-4683466-INS-7232
chr9-4683984-INS-68
chr9-4686204-INS-187
chr9-4702757-INS-204
chr9-4703753-INS-5091
chr9-4705018-INS-70
chr9-4708215-INS-324
chr9-4737360-INS-400
chr9-4771911-INS-212
chr9-4776577-INS-6348
chr9-4807645-INS-132
chr9-4814739-INS-58
chr9-4871067-INS-78
chr9-4873638-INS-200
chr9-4878923-INS-477
chr9-4888721-INS-54
chr9-4889699-INS-195
chr9-4895908-INS-52
chr9-4909977-INS-3127
chr9-4910788-INS-148
chr9-4920146-INS-6462
chr9-4933205-INS-286
chr9-4941801-INS-6218
chr9-4954790-INS-1989
chr9-4979777-INS-6402
chr9-4997926-INS-6151
chr9-4999087-INS-771
chr9-5006297-INS-344
chr9-5023417-INS-105
chr9-5023417-INS-815
chr9-5032202-INS-829
chr9-5045955-INS-260
chr9-5048891-INS-681
chr9-5052346-INS-5522
chr9-5058896-INS-408
chr9-5083312-INS-523
chr9-5090444-INS-122
chr9-5094167-INS-92
chr9-5116380-INS-53
chr9-5123383-INS-6912
chr9-5131355-INS-180
chr9-5143903-INS-891
chr9-5177917-INS-52
chr9-5197704-INS-112

chr9-5215230-INS-6728
chr9-5219117-INS-249
chr9-5229090-INS-486
chr9-5230754-INS-61
chr9-5247631-INS-86
chr9-5256395-INS-503
chr9-5292761-INS-56
chr9-5299147-INS-52
chr9-5305355-INS-632
chr9-5309255-INS-336
chr9-5312262-INS-710
chr9-5324022-INS-88
chr9-5343324-INS-489
chr9-5363365-INS-439
chr9-5390289-INS-344
chr9-5432476-INS-56
chr9-5440868-INS-71
chr9-5450667-INS-6233
chr9-5452365-INS-1888
chr9-5470634-INS-58
chr9-5488856-INS-67
chr9-5490211-INS-604
chr9-5490942-INS-237
chr9-5521232-INS-56
chr9-5540579-INS-6458
chr9-5546145-INS-6601
chr9-5558940-INS-5777
chr9-5561952-INS-145
chr9-5562788-INS-421
chr9-5584389-INS-63
chr9-5586751-INS-328
chr9-5605107-INS-78
chr9-5608706-INS-2838
chr9-5610428-INS-325
chr9-5630437-INS-71
chr9-5681963-INS-359
chr9-5700963-INS-110
chr9-5746240-INS-396
chr9-5769166-INS-7194
chr9-5773403-INS-6090
chr9-5842309-INS-5384
chr9-5853981-INS-7171
chr9-5873161-INS-265
chr9-5873288-INS-157
chr9-5900030-INS-7248
chr9-5951854-INS-330
chr9-5977481-INS-117
chr9-5980440-INS-116
chr9-5985008-INS-6190
chr9-5991919-INS-5002
chr9-5992278-INS-267
chr9-6000231-INS-52
chr9-6013151-INS-175
chr9-6035876-INS-264
chr9-6050565-INS-75
chr9-6085342-INS-59
chr9-6087735-INS-6230
chr9-6096122-INS-6274
chr9-6147080-INS-64
chr9-6216939-INS-1891
chr9-6250559-INS-400
chr9-6252264-INS-2602
chr9-6252809-INS-6599
chr9-6258389-INS-65
chr9-6263498-INS-84
chr9-6284224-INS-157
chr9-6295157-INS-198
chr9-6324066-INS-76
chr9-6350085-INS-6573
chr9-6352998-INS-468
chr9-6379175-INS-328
chr9-6391472-INS-245
chr9-6400995-INS-1367
chr9-6404485-INS-147
chr9-6409238-INS-156
chr9-6423897-INS-6479
chr9-6440410-INS-146

chr9-6454980-INS-120
chr9-6479756-INS-72
chr9-6509123-INS-419
chr9-6524141-INS-5606
chr9-6537691-INS-399
chr9-6551170-INS-132
chr9-6555700-INS-198
chr9-6607870-INS-290
chr9-6618610-INS-5541
chr9-6639328-INS-125
chr9-6665685-INS-120
chr9-6712625-INS-520
chr9-6717471-INS-344
chr9-6718567-INS-142
chr9-6732774-INS-464
chr9-6734251-INS-66
chr9-6737204-INS-539
chr9-6747008-INS-7564
chr9-6821937-INS-58
chr9-6827581-INS-199
chr9-6849265-INS-542
chr9-6853465-INS-297
chr9-6860145-INS-276
chr9-6863746-INS-55
chr9-6890848-INS-322
chr9-6902328-INS-405
chr9-6938738-INS-176
chr9-6941000-INS-186
chr9-6943470-INS-241
chr9-6984549-INS-2138
chr9-6991021-INS-5962
chr9-7024084-INS-326
chr9-7063623-INS-69
chr9-7072438-INS-181
chr9-7074469-INS-76
chr9-7157280-INS-272
chr9-7163020-INS-55
chr9-7171643-INS-506
chr9-7229920-INS-60
chr9-7235261-INS-537
chr9-7236963-INS-994
chr9-7304273-INS-348
chr9-7332970-INS-339
chr9-7353787-INS-505
chr9-7387291-INS-185
chr9-7403448-INS-197
chr9-7444312-INS-5332
chr9-7467422-INS-90
chr9-7476708-INS-183
chr9-7477711-INS-333
chr9-7512167-INS-527
chr9-7519751-INS-64
chr9-7527298-INS-246
chr9-7528664-INS-2113
chr9-7542609-INS-207
chr9-7562010-INS-517
chr9-7577258-INS-750
chr9-7581700-INS-205
chr9-7589794-INS-734
chr9-7591961-INS-56
chr9-7600763-INS-393
chr9-7607262-INS-738
chr9-7626765-INS-181
chr9-7627706-INS-64
chr9-7652396-INS-77
chr9-7655961-INS-432
chr9-7662271-INS-125
chr9-7685398-INS-51
chr9-7716797-INS-72
chr9-7734160-INS-51
chr9-7755267-INS-201
chr9-7761627-INS-66
chr9-7773876-INS-498
chr9-7834925-INS-114
chr9-7840371-INS-94
chr9-7850204-INS-81
chr9-7854011-INS-73

chr9-7875631-INS-5539
chr9-7876932-INS-437
chr9-7879232-INS-139
chr9-7911259-INS-204
chr9-7913466-INS-6451
chr9-7917046-INS-7333
chr9-7934901-INS-108
chr9-7934917-INS-432
chr9-7945016-INS-120
chr9-7977221-INS-54
chr9-7977810-INS-54
chr9-7997415-INS-66
chr9-8005224-INS-77
chr9-8009028-INS-149
chr9-8011927-INS-221
chr9-8028849-INS-172
chr9-8036205-INS-651
chr9-8040169-INS-144
chr9-8070934-INS-229
chr9-8079937-INS-146
chr9-8099081-INS-245
chr9-8109457-INS-200
chr9-8138264-INS-154
chr9-8182493-INS-3525
chr9-8226184-INS-7444
chr9-8269899-INS-311
chr9-8295967-INS-89
chr9-8303580-INS-531
chr9-8311170-INS-72
chr9-8347137-INS-6490
chr9-8351933-INS-100
chr9-8353958-INS-498
chr9-8367126-INS-6765
chr9-8400563-INS-186
chr9-8411782-INS-446
chr9-8428623-INS-99
chr9-8430094-INS-161
chr9-8440192-INS-95
chr9-8455791-INS-180
chr9-8489488-INS-91
chr9-8504732-INS-64
chr9-8556295-INS-204
chr9-8570192-INS-220
chr9-8575298-INS-78
chr9-8598184-INS-66
chr9-8620555-INS-6486
chr9-8636326-INS-423
chr9-8652734-INS-99
chr9-8658462-INS-62
chr9-8702027-INS-86
chr9-8714157-INS-627
chr9-8737904-INS-81
chr9-8747790-INS-56
chr9-8800288-INS-6182
chr9-8805735-INS-295
chr9-8811603-INS-217
chr9-8813271-INS-140
chr9-8830731-INS-399
chr9-8854043-INS-179
chr9-8858047-INS-5039
chr9-8864466-INS-6925
chr9-8905521-INS-906
chr9-8910402-INS-181
chr9-8946093-INS-149
chr9-8954435-INS-64
chr9-8960644-INS-131
chr9-8976735-INS-338
chr9-8986217-INS-62
chr9-9004328-INS-68
chr9-9035684-INS-188
chr9-9043064-INS-212
chr9-9107638-INS-105
chr9-9109465-INS-589
chr9-9133013-INS-731
chr9-9135806-INS-104
chr9-9143000-INS-126
chr9-9152352-INS-68

chr9-9154972-INS-120
chr9-9163166-INS-66
chr9-9174111-INS-362
chr9-9178643-INS-64
chr9-9183968-INS-181
chr9-9192650-INS-982
chr9-9193443-INS-176
chr9-9202786-INS-51
chr9-9205106-INS-84
chr9-9226443-INS-64
chr9-9228694-INS-136
chr9-9231427-INS-65
chr9-9258885-INS-200
chr9-9268659-INS-85
chr9-9283014-INS-224
chr9-9292960-INS-66
chr9-9295498-INS-55
chr9-9337965-INS-110
chr9-9365204-INS-56
chr9-9385987-INS-199
chr9-9430668-INS-70
chr9-9436049-INS-3157
chr9-9455735-INS-6224
chr9-9457289-INS-63
chr9-9468294-INS-214
chr9-9512315-INS-347
chr9-9534238-INS-52
chr9-9566005-INS-124
chr9-9572363-INS-6798
chr9-9701958-INS-206
chr9-9725893-INS-165
chr9-9729363-INS-5388
chr9-9732246-INS-4941
chr9-9766586-INS-74
chr9-9770262-INS-7722
chr9-9770387-INS-52
chr9-9776863-INS-285
chr9-9779557-INS-207
chr9-9784293-INS-92
chr9-9808778-INS-286
chr9-9822660-INS-113
chr9-9852982-INS-184
chr9-9944826-INS-8063
chr9-9964537-INS-252
chr9-9976970-INS-64
chr9-9991465-INS-52
chr9-10008053-INS-486
chr9-10040097-INS-307
chr9-10042972-INS-501
chr9-10069832-INS-106
chr9-10080524-INS-92
chr9-10091268-INS-186
chr9-10097444-INS-57
chr9-10182643-INS-6804
chr9-10202047-INS-113
chr9-10280344-INS-175
chr9-10288670-INS-55
chr9-10351094-INS-68
chr9-10362336-INS-1888
chr9-10364776-INS-201
chr9-10397204-INS-357
chr9-10402312-INS-891
chr9-10409286-INS-465
chr9-10484818-INS-6370
chr9-10509070-INS-61
chr9-10534872-INS-4001
chr9-10537338-INS-842
chr9-10539627-INS-104
chr9-10569361-INS-86
chr9-10604412-INS-81
chr9-10619405-INS-80
chr9-10624218-INS-86
chr9-10639704-INS-259
chr9-10643017-INS-96
chr9-10649781-INS-7358
chr9-10649841-INS-815
chr9-10660726-INS-54

chr9-10674030-INS-5025
chr9-10690822-INS-336
chr9-10699511-INS-52
chr9-10716320-INS-6173
chr9-10726039-INS-959
chr9-10763869-INS-70
chr9-10766659-INS-989
chr9-10779040-INS-218
chr9-10800724-INS-973
chr9-10807806-INS-721
chr9-10819878-INS-276
chr9-10823093-INS-358
chr9-10836937-INS-281
chr9-10881757-INS-55
chr9-10953364-INS-890
chr9-10956416-INS-302
chr9-10960571-INS-227
chr9-10964052-INS-53
chr9-10976545-INS-70
chr9-11006026-INS-150
chr9-11025181-INS-1512
chr9-11036258-INS-6550
chr9-11046932-INS-527
chr9-11061539-INS-76
chr9-11107416-INS-3897
chr9-11121861-INS-105
chr9-11125286-INS-209
chr9-11142927-INS-90
chr9-11146930-INS-82
chr9-11155240-INS-63
chr9-11170189-INS-103
chr9-11184171-INS-352
chr9-11186015-INS-54
chr9-11194652-INS-55
chr9-11234252-INS-209
chr9-11266351-INS-76
chr9-11268506-INS-61
chr9-11283979-INS-58
chr9-11297507-INS-155
chr9-11324848-INS-418
chr9-11329583-INS-62
chr9-11350556-INS-74
chr9-11369538-INS-111
chr9-11385477-INS-464
chr9-11390717-INS-184
chr9-11408518-INS-56
chr9-11428159-INS-90
chr9-11450706-INS-74
chr9-11464570-INS-988
chr9-11498768-INS-416
chr9-11529377-INS-7260
chr9-11533421-INS-446
chr9-11557430-INS-3278
chr9-11559311-INS-2186
chr9-11649355-INS-550
chr9-11665687-INS-731
chr9-11701950-INS-79
chr9-11777924-INS-64
chr9-11789501-INS-6783
chr9-11808174-INS-64
chr9-11815647-INS-1683
chr9-11838092-INS-617
chr9-11848332-INS-186
chr9-11870316-INS-53
chr9-11889357-INS-386
chr9-11896518-INS-6177
chr9-11897697-INS-86
chr9-11937722-INS-1893
chr9-11941272-INS-80
chr9-11982783-INS-103
chr9-11985932-INS-1455
chr9-11990826-INS-123
chr9-12001530-INS-73
chr9-12020776-INS-172
chr9-12037475-INS-204
chr9-12059488-INS-134
chr9-12096666-INS-327

chr9-12115331-INS-56
chr9-12206085-INS-62
chr9-12244639-INS-134
chr9-12264215-INS-147
chr9-12266199-INS-59
chr9-12336961-INS-1227
chr9-12344250-INS-1031
chr9-12357880-INS-182
chr9-12362366-INS-117
chr9-12381662-INS-76
chr9-12392970-INS-56
chr9-12405172-INS-201
chr9-12424453-INS-81
chr9-12428131-INS-52
chr9-12435367-INS-191
chr9-12473867-INS-62
chr9-12475092-INS-94
chr9-12497823-INS-359
chr9-12541411-INS-8533
chr9-12548216-INS-400
chr9-12583758-INS-67
chr9-12587402-INS-671
chr9-12605799-INS-6530
chr9-12610363-INS-77
chr9-12621501-INS-190
chr9-12642687-INS-87
chr9-12674607-INS-1662
chr9-12681063-INS-476
chr9-12693673-INS-204
chr9-12695821-INS-100
chr9-12715368-INS-511
chr9-12725947-INS-58
chr9-12730425-INS-4536
chr9-12755688-INS-69
chr9-12763215-INS-725
chr9-12789932-INS-202
chr9-12797848-INS-295
chr9-12814852-INS-200
chr9-12825223-INS-568
chr9-12830210-INS-53
chr9-12846036-INS-1942
chr9-12863491-INS-422
chr9-12878361-INS-565
chr9-12897985-INS-5249
chr9-12926421-INS-86
chr9-12927871-INS-6163
chr9-12932183-INS-101
chr9-12945880-INS-54
chr9-12946961-INS-54
chr9-12963103-INS-6609
chr9-12973553-INS-6318
chr9-12991245-INS-176
chr9-12995999-INS-115
chr9-13013742-INS-66
chr9-13015527-INS-237
chr9-13019302-INS-469
chr9-13066535-INS-66
chr9-13079548-INS-354
chr9-13085808-INS-57
chr9-13101948-INS-54
chr9-13104347-INS-62
chr9-13108190-INS-263
chr9-13158177-INS-58
chr9-13168663-INS-132
chr9-13173520-INS-83
chr9-13174027-INS-1485
chr9-13184603-INS-1900
chr9-13195136-INS-93
chr9-13207138-INS-133
chr9-13246449-INS-160
chr9-13293167-INS-88
chr9-13349078-INS-86
chr9-13354571-INS-198
chr9-13359696-INS-85
chr9-13374670-INS-56
chr9-13377627-INS-52
chr9-13389520-INS-678

chr9-13397832-INS-56
chr9-13409794-INS-207
chr9-13416754-INS-96
chr9-13421748-INS-333
chr9-13425731-INS-51
chr9-13441067-INS-126
chr9-13446014-INS-360
chr9-13464573-INS-52
chr9-13473861-INS-211
chr9-13478089-INS-352
chr9-13490329-INS-107
chr9-13496871-INS-78
chr9-13517744-INS-186
chr9-13522037-INS-172
chr9-13526253-INS-55
chr9-13527100-INS-230
chr9-13538370-INS-58
chr9-13561245-INS-73
chr9-13568186-INS-60
chr9-13599105-INS-221
chr9-13600748-INS-67
chr9-13623524-INS-326
chr9-13625208-INS-358
chr9-13650899-INS-787
chr9-13658324-INS-60
chr9-13689161-INS-162
chr9-13735515-INS-125
chr9-13746536-INS-413
chr9-13796241-INS-61
chr9-13821241-INS-78
chr9-13829387-INS-90
chr9-13861346-INS-121
chr9-13864679-INS-146
chr9-13867037-INS-985
chr9-13884505-INS-872
chr9-13885193-INS-130
chr9-13932459-INS-341
chr9-13933472-INS-80
chr9-13936657-INS-56
chr9-13943020-INS-135
chr9-13946419-INS-497
chr9-13955771-INS-62
chr9-13967910-INS-61
chr9-13969436-INS-483
chr9-13981792-INS-62
chr9-13986331-INS-197
chr9-13992007-INS-64
chr9-13998696-INS-68
chr9-14021497-INS-494
chr9-14040722-INS-560
chr9-14055980-INS-297
chr9-14075901-INS-399
chr9-14086665-INS-52
chr9-14093876-INS-62
chr9-14101391-INS-584
chr9-14103220-INS-155
chr9-14105018-INS-56
chr9-14116654-INS-2810
chr9-14120009-INS-1256
chr9-14134149-INS-91
chr9-14191019-INS-260
chr9-14208571-INS-643
chr9-14225157-INS-67
chr9-14226829-INS-191
chr9-14233079-INS-125
chr9-14245562-INS-201
chr9-14249145-INS-391
chr9-14256188-INS-442
chr9-14301339-INS-64
chr9-14304100-INS-51
chr9-14321230-INS-7079
chr9-14387345-INS-64
chr9-14408867-INS-191
chr9-14415791-INS-204
chr9-14463774-INS-76
chr9-14466191-INS-268
chr9-14483820-INS-236

chr9-14497407-INS-79
chr9-14507410-INS-74
chr9-14511757-INS-232
chr9-14521192-INS-106
chr9-14523922-INS-80
chr9-14539956-INS-136
chr9-14543561-INS-315
chr9-14556183-INS-475
chr9-14591691-INS-527
chr9-14597002-INS-400
chr9-14598913-INS-403
chr9-14607719-INS-53
chr9-14619744-INS-143
chr9-14629819-INS-62
chr9-14635625-INS-56
chr9-14657987-INS-70
chr9-14676872-INS-278
chr9-14679381-INS-6509
chr9-14697792-INS-205
chr9-14738506-INS-90
chr9-14755147-INS-244
chr9-14758473-INS-479
chr9-14796232-INS-126
chr9-14807609-INS-720
chr9-14809515-INS-62
chr9-14852101-INS-131
chr9-14865644-INS-66
chr9-14875018-INS-57
chr9-14879632-INS-132
chr9-14885581-INS-66
chr9-14889114-INS-264
chr9-14892467-INS-341
chr9-14906526-INS-201
chr9-14915606-INS-69
chr9-14925284-INS-56
chr9-14932011-INS-935
chr9-14959184-INS-82
chr9-14967348-INS-440
chr9-14974810-INS-221
chr9-15010975-INS-207
chr9-15021855-INS-334
chr9-15026387-INS-198
chr9-15045283-INS-1890
chr9-15060148-INS-362
chr9-15065613-INS-187
chr9-15089269-INS-56
chr9-15098315-INS-83
chr9-15112470-INS-56
chr9-15113910-INS-272
chr9-15125296-INS-825
chr9-15133451-INS-589
chr9-15145857-INS-60
chr9-15147389-INS-175
chr9-15213187-INS-301
chr9-15223329-INS-164
chr9-15227379-INS-165
chr9-15273774-INS-75
chr9-15280755-INS-200
chr9-15290825-INS-528
chr9-15294084-INS-429
chr9-15306503-INS-60
chr9-15306832-INS-54
chr9-15346266-INS-52
chr9-15359156-INS-329
chr9-15361641-INS-382
chr9-15366582-INS-100
chr9-15367430-INS-281
chr9-15373329-INS-441
chr9-15401727-INS-110
chr9-15408778-INS-207
chr9-15422610-INS-104
chr9-15449501-INS-58
chr9-15468707-INS-62
chr9-15475983-INS-225
chr9-15498343-INS-114
chr9-15502233-INS-62
chr9-15502956-INS-194

chr9-15519508-INS-476
chr9-15521110-INS-162
chr9-15531337-INS-235
chr9-15551843-INS-114
chr9-15561058-INS-118
chr9-15561337-INS-74
chr9-15574777-INS-58
chr9-15592626-INS-54
chr9-15604612-INS-3064
chr9-15610818-INS-226
chr9-15618401-INS-60
chr9-15628034-INS-88
chr9-15637683-INS-672
chr9-15641142-INS-524
chr9-15643979-INS-738
chr9-15647447-INS-111
chr9-15649039-INS-71
chr9-15677018-INS-89
chr9-15755872-INS-3187
chr9-15769605-INS-519
chr9-15782194-INS-54
chr9-15788742-INS-193
chr9-15806521-INS-144
chr9-15843806-INS-55
chr9-15889431-INS-100
chr9-15914332-INS-56
chr9-15923652-INS-372
chr9-15923652-INS-608
chr9-15968051-INS-60
chr9-15971833-INS-58
chr9-15997705-INS-120
chr9-15999426-INS-54
chr9-16000426-INS-70
chr9-16034161-INS-52
chr9-16045929-INS-91
chr9-16056793-INS-56
chr9-16060586-INS-171
chr9-16071124-INS-53
chr9-16088754-INS-5575
chr9-16213068-INS-66
chr9-16235783-INS-207
chr9-16259644-INS-70
chr9-16262213-INS-52
chr9-16296620-INS-207
chr9-16320437-INS-70
chr9-16329448-INS-78
chr9-16331623-INS-104
chr9-16336295-INS-90
chr9-16341613-INS-195
chr9-16344345-INS-71
chr9-16355859-INS-308
chr9-16360719-INS-6488
chr9-16370670-INS-108
chr9-16377674-INS-152
chr9-16380674-INS-179
chr9-16382700-INS-64
chr9-16387822-INS-4209
chr9-16419600-INS-206
chr9-16423195-INS-688
chr9-16429607-INS-365
chr9-16435884-INS-172
chr9-16480067-INS-400
chr9-16495212-INS-2707
chr9-16499826-INS-236
chr9-16511064-INS-6684
chr9-16552095-INS-6427
chr9-16559842-INS-542
chr9-16667915-INS-214
chr9-16711957-INS-150
chr9-16721414-INS-2821
chr9-16775898-INS-491
chr9-16784536-INS-56
chr9-16812144-INS-202
chr9-16828957-INS-176
chr9-16831852-INS-325
chr9-16908265-INS-810
chr9-16909805-INS-74

chr9-16926375-INS-52
chr9-16958361-INS-108
chr9-16963222-INS-54
chr9-17009220-INS-58
chr9-17014369-INS-102
chr9-17022880-INS-337
chr9-17040185-INS-146
chr9-17053743-INS-138
chr9-17076967-INS-138
chr9-17080698-INS-60
chr9-17101233-INS-55
chr9-17102866-INS-56
chr9-17135431-INS-64
chr9-17153165-INS-64
chr9-17160186-INS-66
chr9-17164727-INS-81
chr9-17175767-INS-62
chr9-17189732-INS-194
chr9-17193219-INS-66
chr9-17201173-INS-53
chr9-17238189-INS-287
chr9-17245852-INS-183
chr9-17280408-INS-890
chr9-17293389-INS-3871
chr9-17336890-INS-1864
chr9-17342940-INS-472
chr9-17379593-INS-226
chr9-17398099-INS-67
chr9-17406229-INS-682
chr9-17438035-INS-220
chr9-17449464-INS-473
chr9-17473974-INS-6478
chr9-17479789-INS-62
chr9-17483874-INS-51
chr9-17501873-INS-519
chr9-17532001-INS-51
chr9-17555399-INS-415
chr9-17578115-INS-57
chr9-17588779-INS-6725
chr9-17597989-INS-65
chr9-17605593-INS-638
chr9-17607908-INS-214
chr9-17647170-INS-951
chr9-17714806-INS-56
chr9-17721624-INS-96
chr9-17740449-INS-93
chr9-17750088-INS-983
chr9-17756448-INS-166
chr9-17760496-INS-398
chr9-17790343-INS-765
chr9-17796839-INS-56
chr9-17813147-INS-60
chr9-17814993-INS-375
chr9-17820587-INS-139
chr9-17852464-INS-132
chr9-17861260-INS-204
chr9-17862628-INS-276
chr9-17874291-INS-149
chr9-17896205-INS-228
chr9-17899404-INS-1573
chr9-17926506-INS-5281
chr9-17936039-INS-107
chr9-17945526-INS-181
chr9-17969403-INS-328
chr9-17984046-INS-188
chr9-18025961-INS-249
chr9-18061636-INS-7758
chr9-18083678-INS-193
chr9-18162537-INS-2733
chr9-18186542-INS-790
chr9-18212576-INS-384
chr9-18233630-INS-703
chr9-18249802-INS-86
chr9-18319139-INS-321
chr9-18327727-INS-56
chr9-18382242-INS-358
chr9-18383588-INS-55

chr9-18391106-INS-78
chr9-18424238-INS-7506
chr9-18468529-INS-77
chr9-18474911-INS-62
chr9-18495197-INS-70
chr9-18511940-INS-60
chr9-18581183-INS-960
chr9-18584716-INS-52
chr9-18588892-INS-184
chr9-18608443-INS-60
chr9-18611746-INS-78
chr9-18614057-INS-547
chr9-18616901-INS-54
chr9-18630977-INS-496
chr9-18637864-INS-696
chr9-18654695-INS-96
chr9-18657014-INS-88
chr9-18657987-INS-380
chr9-18686061-INS-52
chr9-18708483-INS-231
chr9-18709655-INS-86
chr9-18729748-INS-55
chr9-18732776-INS-5983
chr9-18734740-INS-7203
chr9-18796111-INS-141
chr9-18817748-INS-325
chr9-18871536-INS-7034
chr9-18887626-INS-236
chr9-19057806-INS-398
chr9-19070287-INS-4150
chr9-19113043-INS-62
chr9-19129974-INS-6385
chr9-19190856-INS-241
chr9-19240149-INS-208
chr9-19268742-INS-55
chr9-19306579-INS-831
chr9-19326554-INS-6500
chr9-19339752-INS-102
chr9-19340647-INS-104
chr9-19380133-INS-117
chr9-19388248-INS-8704
chr9-19390166-INS-172
chr9-19410579-INS-192
chr9-19434587-INS-86
chr9-19470369-INS-7085
chr9-19529771-INS-143
chr9-19540203-INS-173
chr9-19584831-INS-56
chr9-19596584-INS-150
chr9-19603484-INS-998
chr9-19623172-INS-211
chr9-19700301-INS-498
chr9-19729726-INS-332
chr9-19756287-INS-1367
chr9-19758008-INS-64
chr9-19762173-INS-106
chr9-19804201-INS-549
chr9-19821418-INS-203
chr9-19825607-INS-157
chr9-19841964-INS-708
chr9-19905140-INS-713
chr9-19907188-INS-495
chr9-19909869-INS-72
chr9-19914065-INS-56
chr9-19921827-INS-52
chr9-19987955-INS-264
chr9-20027808-INS-5665
chr9-20029195-INS-156
chr9-20043990-INS-95
chr9-20057822-INS-86
chr9-20075887-INS-165
chr9-20077350-INS-93
chr9-20088852-INS-740
chr9-20110710-INS-76
chr9-20138679-INS-425
chr9-20141704-INS-332
chr9-20178651-INS-63

chr9-20183616-INS-2676
chr9-20189252-INS-700
chr9-20191150-INS-2461
chr9-20203402-INS-4621
chr9-20214027-INS-8674
chr9-20221661-INS-2923
chr9-20233091-INS-208
chr9-20243046-INS-230
chr9-20255218-INS-63
chr9-20289402-INS-256
chr9-20326747-INS-203
chr9-20336170-INS-53
chr9-20373303-INS-71
chr9-20384113-INS-80
chr9-20386575-INS-54
chr9-20388237-INS-359
chr9-20399221-INS-326
chr9-20402800-INS-106
chr9-20405039-INS-200
chr9-20436686-INS-571
chr9-20438844-INS-53
chr9-20469663-INS-64
chr9-20493583-INS-51
chr9-20503064-INS-8735
chr9-20504069-INS-56
chr9-20511258-INS-407
chr9-20520070-INS-100
chr9-20585943-INS-68
chr9-20590445-INS-340
chr9-20605580-INS-398
chr9-20610429-INS-70
chr9-20618268-INS-104
chr9-20641482-INS-6452
chr9-20646576-INS-59
chr9-20665147-INS-532
chr9-20669498-INS-71
chr9-20677299-INS-551
chr9-20679145-INS-207
chr9-20729143-INS-147
chr9-20730477-INS-95
chr9-20732439-INS-84
chr9-20739758-INS-68
chr9-20743591-INS-62
chr9-20788810-INS-197
chr9-20813162-INS-264
chr9-20818198-INS-213
chr9-20832229-INS-132
chr9-20849912-INS-236
chr9-20865342-INS-174
chr9-20866345-INS-106
chr9-20895189-INS-127
chr9-20896519-INS-844
chr9-20903031-INS-754
chr9-20935080-INS-190
chr9-20938020-INS-104
chr9-20952603-INS-144
chr9-20959879-INS-152
chr9-20971513-INS-124
chr9-21004371-INS-128
chr9-21012551-INS-54
chr9-21015108-INS-52
chr9-21017896-INS-70
chr9-21055676-INS-56
chr9-21068121-INS-55
chr9-21085455-INS-206
chr9-21098768-INS-202
chr9-21111348-INS-178
chr9-21113894-INS-384
chr9-21119520-INS-52
chr9-21121700-INS-206
chr9-21125144-INS-211
chr9-21129544-INS-263
chr9-21132451-INS-409
chr9-21133167-INS-85
chr9-21157799-INS-52
chr9-21166232-INS-91
chr9-21168271-INS-176

chr9-21179292-INS-200
chr9-21214151-INS-140
chr9-21222448-INS-108
chr9-21225868-INS-63
chr9-21252402-INS-241
chr9-21260429-INS-103
chr9-21271127-INS-198
chr9-21275640-INS-51
chr9-21295006-INS-426
chr9-21307656-INS-64
chr9-21332532-INS-173
chr9-21332900-INS-482
chr9-21333506-INS-221
chr9-21345263-INS-76
chr9-21395770-INS-690
chr9-21400402-INS-74
chr9-21412114-INS-180
chr9-21414700-INS-190
chr9-21449487-INS-137
chr9-21450726-INS-213
chr9-21453277-INS-200
chr9-21455249-INS-62
chr9-21469479-INS-223
chr9-21474797-INS-382
chr9-21481391-INS-91
chr9-21509550-INS-153
chr9-21512307-INS-63
chr9-21519063-INS-82
chr9-21519869-INS-5276
chr9-21528932-INS-179
chr9-21530292-INS-212
chr9-21537658-INS-297
chr9-21550350-INS-85
chr9-21581374-INS-214
chr9-21599721-INS-200
chr9-21601007-INS-210
chr9-21615062-INS-210
chr9-21617752-INS-165
chr9-21624221-INS-222
chr9-21637460-INS-55
chr9-21644195-INS-301
chr9-21656705-INS-213
chr9-21658706-INS-66
chr9-21661997-INS-124
chr9-21684421-INS-207
chr9-21686515-INS-174
chr9-21689786-INS-155
chr9-21710649-INS-97
chr9-21720452-INS-198
chr9-21751522-INS-195
chr9-21769134-INS-201
chr9-21774656-INS-197
chr9-21784884-INS-182
chr9-21786922-INS-197
chr9-21816428-INS-355
chr9-21841467-INS-206
chr9-21849428-INS-315
chr9-21863549-INS-304
chr9-21864331-INS-56
chr9-21875840-INS-207
chr9-21895733-INS-196
chr9-21896778-INS-54
chr9-21951010-INS-51
chr9-21955022-INS-108
chr9-21959891-INS-200
chr9-22000747-INS-52
chr9-22005537-INS-156
chr9-22007940-INS-87
chr9-22053997-INS-125
chr9-22066568-INS-52
chr9-22080538-INS-296
chr9-22101214-INS-314
chr9-22103062-INS-93
chr9-22104313-INS-55
chr9-22107185-INS-132
chr9-22110992-INS-84
chr9-22113882-INS-702

chr9-22132577-INS-143
chr9-22136105-INS-134
chr9-22160092-INS-64
chr9-22169775-INS-172
chr9-22183363-INS-230
chr9-22228262-INS-369
chr9-22233743-INS-921
chr9-22238709-INS-326
chr9-22240793-INS-121
chr9-22249368-INS-469
chr9-22251483-INS-195
chr9-22283931-INS-53
chr9-22314119-INS-155
chr9-22356468-INS-175
chr9-22372653-INS-5373
chr9-22411517-INS-104
chr9-22426179-INS-66
chr9-22426844-INS-96
chr9-22438842-INS-229
chr9-22464694-INS-50
chr9-22477723-INS-153
chr9-22481697-INS-76
chr9-22486844-INS-280
chr9-22510215-INS-567
chr9-22511323-INS-88
chr9-22532144-INS-87
chr9-22542934-INS-69
chr9-22555228-INS-386
chr9-22570873-INS-60
chr9-22586815-INS-329
chr9-22588980-INS-65
chr9-22591981-INS-807
chr9-22596797-INS-102
chr9-22597853-INS-74
chr9-22600988-INS-58
chr9-22654917-INS-529
chr9-22664927-INS-120
chr9-22670342-INS-7152
chr9-22670475-INS-61
chr9-22676562-INS-94
chr9-22683131-INS-299
chr9-22684611-INS-330
chr9-22693503-INS-421
chr9-22728055-INS-269
chr9-22769964-INS-383
chr9-22775321-INS-79
chr9-22783363-INS-72
chr9-22788389-INS-126
chr9-22791410-INS-58
chr9-22814668-INS-416
chr9-22855721-INS-479
chr9-22857753-INS-394
chr9-22870062-INS-6473
chr9-22903502-INS-68
chr9-22913333-INS-401
chr9-22925629-INS-274
chr9-22933606-INS-573
chr9-22964643-INS-87
chr9-22998538-INS-83
chr9-23007038-INS-403
chr9-23014497-INS-60
chr9-23031585-INS-300
chr9-23066705-INS-67
chr9-23073484-INS-143
chr9-23104991-INS-211
chr9-23143259-INS-52
chr9-23151667-INS-58
chr9-23169274-INS-68
chr9-23211564-INS-60
chr9-23222419-INS-95
chr9-23230204-INS-466
chr9-23233010-INS-108
chr9-23266352-INS-96
chr9-23306726-INS-54
chr9-23316085-INS-168
chr9-23321054-INS-56
chr9-23354901-INS-52

chr9-23398174-INS-56
chr9-23412102-INS-548
chr9-23426436-INS-5313
chr9-23428824-INS-502
chr9-23450512-INS-90
chr9-23466327-INS-3013
chr9-23468860-INS-60
chr9-23476485-INS-362
chr9-23492029-INS-288
chr9-23498881-INS-386
chr9-23507257-INS-253
chr9-23513589-INS-1136
chr9-23537701-INS-153
chr9-23582561-INS-150
chr9-23631172-INS-537
chr9-23637533-INS-165
chr9-23645662-INS-107
chr9-23651457-INS-51
chr9-23663510-INS-382
chr9-23674744-INS-96
chr9-23711871-INS-107
chr9-23713007-INS-59
chr9-23714223-INS-664
chr9-23761955-INS-8174
chr9-23792774-INS-205
chr9-23794790-INS-68
chr9-23804211-INS-72
chr9-23831318-INS-64
chr9-23842636-INS-67
chr9-23925100-INS-60
chr9-23942491-INS-6395
chr9-23948347-INS-117
chr9-23951313-INS-76
chr9-23962065-INS-233
chr9-23967406-INS-72
chr9-24041102-INS-430
chr9-24067585-INS-90
chr9-24110749-INS-57
chr9-24174695-INS-464
chr9-24178318-INS-206
chr9-24229118-INS-80
chr9-24230958-INS-127
chr9-24268154-INS-344
chr9-24279637-INS-147
chr9-24280024-INS-683
chr9-24286572-INS-66
chr9-24307215-INS-399
chr9-24327347-INS-64
chr9-24370219-INS-160
chr9-24375177-INS-215
chr9-24379919-INS-54
chr9-24381628-INS-206
chr9-24392916-INS-286
chr9-24416750-INS-248
chr9-24433650-INS-103
chr9-24447339-INS-6458
chr9-24465817-INS-191
chr9-24524649-INS-781
chr9-24528400-INS-498
chr9-24548192-INS-1026
chr9-24559968-INS-64
chr9-24573986-INS-507
chr9-24579435-INS-75
chr9-24686877-INS-1150
chr9-24693582-INS-153
chr9-24719452-INS-52
chr9-24745877-INS-110
chr9-24749555-INS-120
chr9-24760075-INS-187
chr9-24774431-INS-116
chr9-24793488-INS-945
chr9-24806662-INS-908
chr9-24814078-INS-72
chr9-24822410-INS-290
chr9-24842262-INS-445
chr9-24852728-INS-655
chr9-24877962-INS-401

chr9-24895652-INS-52
chr9-24899458-INS-63
chr9-24912785-INS-123
chr9-24917883-INS-203
chr9-24931463-INS-60
chr9-24935414-INS-399
chr9-24950550-INS-195
chr9-24973459-INS-437
chr9-24992807-INS-844
chr9-25003462-INS-111
chr9-25085954-INS-88
chr9-25099467-INS-151
chr9-25102799-INS-1323
chr9-25116764-INS-198
chr9-25129000-INS-213
chr9-25138115-INS-1039
chr9-25196288-INS-659
chr9-25217924-INS-990
chr9-25222720-INS-76
chr9-25229206-INS-54
chr9-25238504-INS-6649
chr9-25251944-INS-61
chr9-25286323-INS-53
chr9-25291538-INS-220
chr9-25305242-INS-188
chr9-25342805-INS-429
chr9-25355283-INS-238
chr9-25376261-INS-232
chr9-25384072-INS-214
chr9-25387391-INS-66
chr9-25406001-INS-63
chr9-25425061-INS-457
chr9-25440858-INS-78
chr9-25470483-INS-225
chr9-25482228-INS-52
chr9-25494796-INS-127
chr9-25496434-INS-245
chr9-25498472-INS-639
chr9-25504838-INS-236
chr9-25523715-INS-2092
chr9-25559566-INS-557
chr9-25565443-INS-325
chr9-25571021-INS-6420
chr9-25579714-INS-6761
chr9-25595985-INS-520
chr9-25603225-INS-501
chr9-25628730-INS-56
chr9-25630843-INS-68
chr9-25641073-INS-6630
chr9-25702105-INS-475
chr9-25721149-INS-359
chr9-25749072-INS-859
chr9-25775724-INS-5944
chr9-25810329-INS-53
chr9-25848385-INS-60
chr9-25854273-INS-194
chr9-25863805-INS-53
chr9-25929138-INS-250
chr9-25947620-INS-52
chr9-26003283-INS-448
chr9-26023531-INS-6354
chr9-26044603-INS-99
chr9-26068254-INS-894
chr9-26078265-INS-324
chr9-26103671-INS-69
chr9-26140520-INS-6694
chr9-26167775-INS-124
chr9-26168728-INS-828
chr9-26198666-INS-86
chr9-26233379-INS-464
chr9-26247197-INS-81
chr9-26267362-INS-51
chr9-26274112-INS-284
chr9-26280534-INS-71
chr9-26284271-INS-913
chr9-26284333-INS-1476
chr9-26321402-INS-72

chr9-26322735-INS-51
chr9-26347712-INS-516
chr9-26361096-INS-382
chr9-26371756-INS-136
chr9-26402661-INS-52
chr9-26440717-INS-152
chr9-26461767-INS-240
chr9-26476849-INS-54
chr9-26492231-INS-269
chr9-26509581-INS-638
chr9-26510256-INS-55
chr9-26559794-INS-54
chr9-26570507-INS-2884
chr9-26599574-INS-199
chr9-26640630-INS-96
chr9-26683804-INS-165
chr9-26689153-INS-89
chr9-26699163-INS-185
chr9-26704577-INS-60
chr9-26709037-INS-66
chr9-26714943-INS-77
chr9-26727357-INS-53
chr9-26743800-INS-492
chr9-26748314-INS-79
chr9-26755390-INS-283
chr9-26757848-INS-533
chr9-26779542-INS-339
chr9-26789264-INS-106
chr9-26794142-INS-106
chr9-26835771-INS-7085
chr9-26861293-INS-122
chr9-26863690-INS-78
chr9-26883581-INS-104
chr9-26884977-INS-56
chr9-26898214-INS-144
chr9-26899417-INS-742
chr9-26907782-INS-89
chr9-26928560-INS-52
chr9-26930323-INS-240
chr9-26942692-INS-201
chr9-26945299-INS-6431
chr9-26967362-INS-161
chr9-26972723-INS-206
chr9-26984518-INS-84
chr9-26991970-INS-107
chr9-27023386-INS-985
chr9-27026720-INS-65
chr9-27043769-INS-2841
chr9-27047123-INS-85
chr9-27051136-INS-56
chr9-27059212-INS-201
chr9-27065033-INS-78
chr9-27069887-INS-193
chr9-27072913-INS-181
chr9-27094465-INS-93
chr9-27098204-INS-55
chr9-27101168-INS-329
chr9-27107818-INS-66
chr9-27135375-INS-7040
chr9-27138981-INS-6817
chr9-27156394-INS-235
chr9-27184969-INS-98
chr9-27191989-INS-104
chr9-27221728-INS-57
chr9-27271599-INS-119
chr9-27272242-INS-131
chr9-27275675-INS-103
chr9-27296031-INS-58
chr9-27307152-INS-58
chr9-27310482-INS-729
chr9-27315680-INS-108
chr9-27382602-INS-104
chr9-27385201-INS-745
chr9-27394055-INS-902
chr9-27409716-INS-344
chr9-27431812-INS-179
chr9-27452529-INS-52

chr9-27477872-INS-73
chr9-27492516-INS-68
chr9-27502221-INS-249
chr9-27506186-INS-52
chr9-27513934-INS-167
chr9-27523740-INS-67
chr9-27545091-INS-119
chr9-27557023-INS-191
chr9-27617179-INS-343
chr9-27617921-INS-449
chr9-27637393-INS-128
chr9-27646158-INS-72
chr9-27675313-INS-64
chr9-27710968-INS-146
chr9-27724237-INS-131
chr9-27727893-INS-330
chr9-27756808-INS-221
chr9-27760541-INS-784
chr9-27770572-INS-192
chr9-27785550-INS-58
chr9-27786603-INS-100
chr9-27793878-INS-324
chr9-27841264-INS-87
chr9-27843986-INS-79
chr9-27845279-INS-643
chr9-27863853-INS-83
chr9-27901020-INS-6233
chr9-27906245-INS-204
chr9-27909050-INS-169
chr9-27928054-INS-269
chr9-27968234-INS-397
chr9-28047390-INS-156
chr9-28116632-INS-1564
chr9-28117849-INS-272
chr9-28139138-INS-1166
chr9-28173371-INS-301
chr9-28184660-INS-56
chr9-28201778-INS-64
chr9-28224599-INS-2075
chr9-28242862-INS-394
chr9-28246349-INS-54
chr9-28246584-INS-59
chr9-28255816-INS-6199
chr9-28274275-INS-84
chr9-28295102-INS-821
chr9-28305974-INS-95
chr9-28306212-INS-53
chr9-28312736-INS-519
chr9-28317339-INS-82
chr9-28321361-INS-69
chr9-28331401-INS-6777
chr9-28337185-INS-70
chr9-28350683-INS-62
chr9-28362323-INS-3219
chr9-28362521-INS-89
chr9-28379113-INS-120
chr9-28410362-INS-64
chr9-28429286-INS-141
chr9-28492102-INS-147
chr9-28510494-INS-124
chr9-28514571-INS-170
chr9-28516752-INS-190
chr9-28522624-INS-78
chr9-28526529-INS-421
chr9-28527463-INS-56
chr9-28533785-INS-7075
chr9-28556481-INS-1496
chr9-28570389-INS-65
chr9-28574892-INS-6272
chr9-28616791-INS-1441
chr9-28663211-INS-280
chr9-28668566-INS-372
chr9-28675671-INS-446
chr9-28699740-INS-457
chr9-28709508-INS-66
chr9-28730495-INS-3024
chr9-28737999-INS-523

chr9-28741913-INS-141
chr9-28753230-INS-63
chr9-28780671-INS-7394
chr9-28791802-INS-432
chr9-28799047-INS-120
chr9-28802606-INS-196
chr9-28807872-INS-154
chr9-28810207-INS-134
chr9-28824095-INS-67
chr9-28883519-INS-68
chr9-28896287-INS-197
chr9-28922752-INS-7246
chr9-28929027-INS-132
chr9-28945707-INS-162
chr9-29007863-INS-323
chr9-29011781-INS-237
chr9-29015243-INS-56
chr9-29043150-INS-167
chr9-29069439-INS-70
chr9-29085889-INS-128
chr9-29115435-INS-74
chr9-29124898-INS-81
chr9-29137954-INS-532
chr9-29153414-INS-137
chr9-29160572-INS-66
chr9-29184096-INS-439
chr9-29210569-INS-5188
chr9-29219394-INS-2997
chr9-29256392-INS-331
chr9-29257615-INS-80
chr9-29262946-INS-54
chr9-29266002-INS-1894
chr9-29279303-INS-77
chr9-29291582-INS-120
chr9-29296591-INS-7131
chr9-29304877-INS-284
chr9-29339598-INS-51
chr9-29360574-INS-202
chr9-29371423-INS-55
chr9-29375719-INS-300
chr9-29386520-INS-774
chr9-29388060-INS-52
chr9-29416770-INS-74
chr9-29433858-INS-54
chr9-29474846-INS-192
chr9-29499455-INS-103
chr9-29541940-INS-52
chr9-29610992-INS-53
chr9-29659728-INS-2157
chr9-29679521-INS-163
chr9-29689906-INS-53
chr9-29695224-INS-62
chr9-29708433-INS-80
chr9-29739602-INS-61
chr9-29815134-INS-58
chr9-29853347-INS-81
chr9-29880027-INS-192
chr9-29921423-INS-177
chr9-29930009-INS-138
chr9-29966615-INS-204
chr9-29999098-INS-105
chr9-30008561-INS-74
chr9-30019871-INS-3503
chr9-30022874-INS-62
chr9-30027137-INS-56
chr9-30033286-INS-160
chr9-30074593-INS-1095
chr9-30074596-INS-1589
chr9-30074599-INS-1703
chr9-30074635-INS-591
chr9-30079490-INS-935
chr9-30093131-INS-6006
chr9-30107413-INS-232
chr9-30110203-INS-54
chr9-30147056-INS-7084
chr9-30161926-INS-433
chr9-30165998-INS-396

chr9-30180223-INS-64
chr9-30216711-INS-967
chr9-30266908-INS-1836
chr9-30269524-INS-90
chr9-30314251-INS-136
chr9-30315161-INS-280
chr9-30319830-INS-56
chr9-30345249-INS-78
chr9-30368489-INS-594
chr9-30379663-INS-210
chr9-30396552-INS-52
chr9-30425042-INS-73
chr9-30430262-INS-54
chr9-30432709-INS-68
chr9-30453715-INS-91
chr9-30454065-INS-140
chr9-30462468-INS-60
chr9-30471801-INS-105
chr9-30474484-INS-281
chr9-30482141-INS-241
chr9-30500825-INS-122
chr9-30567277-INS-90
chr9-30640628-INS-58
chr9-30651954-INS-66
chr9-30664244-INS-88
chr9-30689759-INS-362
chr9-30705408-INS-6399
chr9-30736443-INS-54
chr9-30744360-INS-285
chr9-30767562-INS-587
chr9-30771009-INS-507
chr9-30788921-INS-218
chr9-30794600-INS-123
chr9-30796054-INS-261
chr9-30805635-INS-86
chr9-30817747-INS-60
chr9-30824479-INS-239
chr9-30825077-INS-124
chr9-30838765-INS-52
chr9-30847713-INS-339
chr9-30849698-INS-92
chr9-30850801-INS-52
chr9-30877744-INS-54
chr9-30887439-INS-60
chr9-30895001-INS-67
chr9-30936339-INS-195
chr9-30966868-INS-123
chr9-30972598-INS-204
chr9-30992581-INS-84
chr9-31018287-INS-68
chr9-31024574-INS-157
chr9-31030210-INS-147
chr9-31057213-INS-68
chr9-31064328-INS-84
chr9-31066806-INS-206
chr9-31136215-INS-54
chr9-31138972-INS-362
chr9-31148263-INS-289
chr9-31161946-INS-53
chr9-31167231-INS-165
chr9-31173133-INS-110
chr9-31182026-INS-417
chr9-31189186-INS-197
chr9-31202351-INS-173
chr9-31215556-INS-56
chr9-31218356-INS-140
chr9-31257650-INS-100
chr9-31262898-INS-153
chr9-31296150-INS-62
chr9-31297061-INS-2031
chr9-31303520-INS-82
chr9-31305898-INS-542
chr9-31335075-INS-175
chr9-31336542-INS-72
chr9-31346510-INS-147
chr9-31379181-INS-76
chr9-31395087-INS-326

chr9-31418771-INS-83
chr9-31420949-INS-208
chr9-31426467-INS-67
chr9-31428938-INS-66
chr9-31429079-INS-606
chr9-31436007-INS-726
chr9-31456267-INS-179
chr9-31466162-INS-106
chr9-31467204-INS-554
chr9-31474873-INS-336
chr9-31487035-INS-98
chr9-31499051-INS-196
chr9-31504046-INS-66
chr9-31506016-INS-154
chr9-31522817-INS-468
chr9-31571261-INS-199
chr9-31595406-INS-242
chr9-31606014-INS-55
chr9-31615738-INS-8379
chr9-31626887-INS-6697
chr9-31630452-INS-53
chr9-31670792-INS-56
chr9-31679258-INS-66
chr9-31709714-INS-297
chr9-31716945-INS-122
chr9-31720808-INS-85
chr9-31774592-INS-121
chr9-31841118-INS-261
chr9-31865439-INS-204
chr9-31868204-INS-61
chr9-31869310-INS-104
chr9-31886892-INS-123
chr9-31887643-INS-76
chr9-31894659-INS-207
chr9-31910973-INS-174
chr9-31915847-INS-382
chr9-31939901-INS-127
chr9-31941877-INS-73
chr9-31961548-INS-86
chr9-31966799-INS-7129
chr9-31984227-INS-85
chr9-32023049-INS-157
chr9-32025251-INS-140
chr9-32026527-INS-55
chr9-32028745-INS-56
chr9-32042910-INS-185
chr9-32058388-INS-1890
chr9-32075371-INS-192
chr9-32099721-INS-189
chr9-32109889-INS-208
chr9-32156126-INS-72
chr9-32157911-INS-226
chr9-32177263-INS-1534
chr9-32195645-INS-289
chr9-32198308-INS-283
chr9-32199183-INS-84
chr9-32209192-INS-499
chr9-32266675-INS-164
chr9-32279351-INS-173
chr9-32291098-INS-449
chr9-32296571-INS-260
chr9-32300032-INS-68
chr9-32306501-INS-120
chr9-32312606-INS-191
chr9-32343930-INS-130
chr9-32354213-INS-261
chr9-32377956-INS-100
chr9-32390964-INS-58
chr9-32393199-INS-150
chr9-32427104-INS-63
chr9-32436472-INS-150
chr9-32451172-INS-54
chr9-32465142-INS-82
chr9-32481446-INS-121
chr9-32518899-INS-60
chr9-32532422-INS-87
chr9-32536001-INS-922

chr9-32552155-INS-52
chr9-32555957-INS-569
chr9-32566335-INS-117
chr9-32573894-INS-100
chr9-32590260-INS-52
chr9-32595008-INS-51
chr9-32599542-INS-148
chr9-32601155-INS-724
chr9-32640161-INS-262
chr9-32649120-INS-70
chr9-32677331-INS-381
chr9-32681172-INS-64
chr9-32684906-INS-156
chr9-32695216-INS-56
chr9-32702105-INS-77
chr9-32762673-INS-399
chr9-32792223-INS-371
chr9-32848493-INS-52
chr9-32871095-INS-193
chr9-32910233-INS-1892
chr9-32934237-INS-1814
chr9-32968078-INS-6528
chr9-32989087-INS-199
chr9-33010529-INS-64
chr9-33018120-INS-82
chr9-33036527-INS-6242
chr9-33050190-INS-383
chr9-33076590-INS-6242
chr9-33092234-INS-569
chr9-33165528-INS-1896
chr9-33167507-INS-6798
chr9-33176346-INS-292
chr9-33183587-INS-6385
chr9-33190241-INS-144
chr9-33221727-INS-176
chr9-33252923-INS-159
chr9-33258148-INS-126
chr9-33262850-INS-978
chr9-33284708-INS-6442
chr9-33304720-INS-214
chr9-33338265-INS-440
chr9-33339870-INS-859
chr9-33350780-INS-60
chr9-33376201-INS-53
chr9-33386716-INS-123
chr9-33393127-INS-152
chr9-33437238-INS-2991
chr9-33443466-INS-57
chr9-33448884-INS-2185
chr9-33462747-INS-325
chr9-33469305-INS-4800
chr9-33469387-INS-99
chr9-33482971-INS-219
chr9-33490709-INS-64
chr9-33492155-INS-110
chr9-33504495-INS-162
chr9-33518564-INS-503
chr9-33521412-INS-54
chr9-33550964-INS-59
chr9-33561818-INS-96
chr9-33565495-INS-110
chr9-33566745-INS-90
chr9-33592717-INS-56
chr9-33607894-INS-3521
chr9-33614967-INS-100
chr9-33621276-INS-6602
chr9-33634969-INS-76
chr9-33669069-INS-1892
chr9-33681126-INS-65
chr9-33695411-INS-126
chr9-33698118-INS-171
chr9-33706217-INS-651
chr9-33727935-INS-67
chr9-33730741-INS-821
chr9-33743164-INS-53
chr9-33744439-INS-218
chr9-33788848-INS-199

chr9-33844920-INS-317
chr9-33849971-INS-66
chr9-33889438-INS-401
chr9-33907930-INS-104
chr9-33922435-INS-65
chr9-33943631-INS-190
chr9-33970987-INS-87
chr9-33998562-INS-60
chr9-34003453-INS-60
chr9-34008245-INS-629
chr9-34016999-INS-52
chr9-34030545-INS-218
chr9-34033489-INS-653
chr9-34042339-INS-196
chr9-34050862-INS-62
chr9-34062618-INS-64
chr9-34088697-INS-161
chr9-34091619-INS-2260
chr9-34107616-INS-6378
chr9-34139388-INS-55
chr9-34207596-INS-68
chr9-34212015-INS-6281
chr9-34216510-INS-56
chr9-34239137-INS-259
chr9-34273262-INS-81
chr9-34275295-INS-90
chr9-34299499-INS-54
chr9-34318112-INS-96
chr9-34321006-INS-141
chr9-34381980-INS-172
chr9-34401802-INS-85
chr9-34438644-INS-52
chr9-34468249-INS-57
chr9-34477045-INS-153
chr9-34477951-INS-458
chr9-34492126-INS-1862
chr9-34506281-INS-283
chr9-34566030-INS-517
chr9-34573945-INS-126
chr9-34631191-INS-303
chr9-34633982-INS-59
chr9-34644444-INS-1014
chr9-34699852-INS-58
chr9-34700770-INS-130
chr9-34717173-INS-6413
chr9-34726602-INS-56
chr9-34742273-INS-55
chr9-34746400-INS-414
chr9-34750062-INS-6908
chr9-34752018-INS-55
chr9-34797133-INS-71
chr9-34808077-INS-1756
chr9-34815244-INS-122
chr9-34841119-INS-172
chr9-34901953-INS-872
chr9-34915130-INS-117
chr9-34934094-INS-6441
chr9-34934159-INS-62
chr9-34961102-INS-160
chr9-34964790-INS-111
chr9-34980157-INS-654
chr9-35117099-INS-266
chr9-35147947-INS-260
chr9-35172636-INS-68
chr9-35173484-INS-198
chr9-35201725-INS-369
chr9-35241654-INS-672
chr9-35250024-INS-60
chr9-35254431-INS-106
chr9-35262015-INS-517
chr9-35281943-INS-145
chr9-35285604-INS-90
chr9-35298966-INS-52
chr9-35330150-INS-311
chr9-35375598-INS-692
chr9-35490831-INS-324
chr9-35503968-INS-116

chr9-35516866-INS-4375
chr9-35546361-INS-9282
chr9-35549938-INS-182
chr9-35555025-INS-68
chr9-35564581-INS-215
chr9-35594527-INS-60
chr9-35618783-INS-98
chr9-35632159-INS-416
chr9-35658852-INS-225
chr9-35708547-INS-821
chr9-35719435-INS-981
chr9-35768902-INS-446
chr9-35770434-INS-86
chr9-35775158-INS-68
chr9-35800149-INS-157
chr9-35801780-INS-965
chr9-35802675-INS-6626
chr9-35824623-INS-6627
chr9-35833113-INS-5364
chr9-35840380-INS-202
chr9-35843916-INS-7544
chr9-35857831-INS-202
chr9-35863687-INS-139
chr9-35874394-INS-104
chr9-35874853-INS-55
chr9-35911749-INS-4017
chr9-35931117-INS-208
chr9-35964115-INS-398
chr9-35989978-INS-63
chr9-36009837-INS-812
chr9-36013550-INS-91
chr9-36015023-INS-6155
chr9-36024055-INS-3049
chr9-36027769-INS-7099
chr9-36108345-INS-371
chr9-36130654-INS-145
chr9-36156160-INS-10496
chr9-36164313-INS-256
chr9-36180480-INS-72
chr9-36210212-INS-60
chr9-36224130-INS-88
chr9-36234674-INS-204
chr9-36253539-INS-105
chr9-36320929-INS-6236
chr9-36329862-INS-7955
chr9-36340547-INS-119
chr9-36342835-INS-818
chr9-36344193-INS-1867
chr9-36345763-INS-6492
chr9-36353561-INS-1076
chr9-36361231-INS-1061
chr9-36363631-INS-335
chr9-36378922-INS-191
chr9-36381626-INS-210
chr9-36411355-INS-971
chr9-36426051-INS-120
chr9-36434799-INS-58
chr9-36450487-INS-122
chr9-36462786-INS-136
chr9-36470117-INS-4009
chr9-36470485-INS-6227
chr9-36492987-INS-7097
chr9-36495185-INS-91
chr9-36508998-INS-65
chr9-36560054-INS-6469
chr9-36577990-INS-985
chr9-36580450-INS-350
chr9-36606356-INS-56
chr9-36611897-INS-406
chr9-36623223-INS-225
chr9-36650819-INS-100
chr9-36653449-INS-248
chr9-36667466-INS-122
chr9-36668293-INS-139
chr9-36692316-INS-4779
chr9-36697153-INS-118
chr9-36700548-INS-174

chr9-36724414-INS-346
chr9-36732054-INS-71
chr9-36749102-INS-72
chr9-36758720-INS-96
chr9-36769418-INS-802
chr9-36775586-INS-104
chr9-36790343-INS-530
chr9-36833038-INS-92
chr9-36860098-INS-94
chr9-36871112-INS-189
chr9-36876848-INS-110
chr9-36906958-INS-98
chr9-36932801-INS-104
chr9-37016362-INS-55
chr9-37017198-INS-165
chr9-37017448-INS-6454
chr9-37092388-INS-158
chr9-37103901-INS-103
chr9-37105052-INS-95
chr9-37107006-INS-76
chr9-37138912-INS-102
chr9-37162511-INS-627
chr9-37178241-INS-85
chr9-37193728-INS-52
chr9-37208249-INS-219
chr9-37221542-INS-138
chr9-37264701-INS-183
chr9-37283188-INS-102
chr9-37290100-INS-103
chr9-37292537-INS-428
chr9-37352344-INS-322
chr9-37370462-INS-176
chr9-37378697-INS-154
chr9-37393633-INS-249
chr9-37421042-INS-422
chr9-37432202-INS-171
chr9-37435925-INS-198
chr9-37444876-INS-219
chr9-37462522-INS-66
chr9-37465766-INS-225
chr9-37474641-INS-122
chr9-37491860-INS-287
chr9-37508466-INS-88
chr9-37526034-INS-755
chr9-37528840-INS-287
chr9-37539373-INS-80
chr9-37539390-INS-225
chr9-37553900-INS-119
chr9-37555241-INS-157
chr9-37569631-INS-6473
chr9-37615527-INS-86
chr9-37626994-INS-55
chr9-37632880-INS-150
chr9-37634856-INS-289
chr9-37638503-INS-175
chr9-37651175-INS-107
chr9-37659517-INS-443
chr9-37669072-INS-76
chr9-37672208-INS-99
chr9-37680189-INS-112
chr9-37680322-INS-5272
chr9-37706568-INS-7153
chr9-37730928-INS-90
chr9-37733649-INS-174
chr9-37739579-INS-5473
chr9-37768016-INS-466
chr9-37774068-INS-182
chr9-37782221-INS-1426
chr9-37801504-INS-5544
chr9-37804217-INS-6816
chr9-37820194-INS-6414
chr9-37828978-INS-6560
chr9-37833692-INS-6646
chr9-37837428-INS-134
chr9-37841689-INS-1245
chr9-37841690-INS-846
chr9-37841789-INS-679

chr9-37845213-INS-344
chr9-37855027-INS-53
chr9-37863436-INS-346
chr9-37868048-INS-106
chr9-37876440-INS-120
chr9-37876741-INS-954
chr9-37883218-INS-6436
chr9-37885551-INS-6732
chr9-37887287-INS-341
chr9-37891522-INS-153
chr9-37900450-INS-6222
chr9-37902578-INS-403
chr9-37904690-INS-395
chr9-37909292-INS-77
chr9-37911628-INS-60
chr9-37914135-INS-503
chr9-37924852-INS-108
chr9-37926303-INS-7104
chr9-37936002-INS-52
chr9-37946788-INS-79
chr9-37964130-INS-6686
chr9-37975662-INS-52
chr9-37986386-INS-5544
chr9-37988901-INS-5207
chr9-37997577-INS-74
chr9-38009468-INS-92
chr9-38029653-INS-6719
chr9-38031902-INS-83
chr9-38039897-INS-117
chr9-38097440-INS-110
chr9-38110043-INS-182
chr9-38134195-INS-6117
chr9-38138947-INS-327
chr9-38140954-INS-6843
chr9-38141955-INS-153
chr9-38148062-INS-115
chr9-38161026-INS-70
chr9-38165866-INS-115
chr9-38167795-INS-6384
chr9-38183589-INS-1283
chr9-38198735-INS-133
chr9-38237246-INS-540
chr9-38244247-INS-857
chr9-38305377-INS-57
chr9-38327355-INS-6884
chr9-38358144-INS-6759
chr9-38359643-INS-100
chr9-38385024-INS-724
chr9-38390485-INS-73
chr9-38395884-INS-140
chr9-38419493-INS-6599
chr9-38419712-INS-4350
chr9-38448104-INS-213
chr9-38452156-INS-3938
chr9-38457647-INS-1115
chr9-38470525-INS-334
chr9-38488994-INS-399
chr9-38491926-INS-125
chr9-38500628-INS-6609
chr9-38502268-INS-102
chr9-38505030-INS-54
chr9-38509905-INS-956
chr9-38573177-INS-148
chr9-38597190-INS-486
chr9-38598888-INS-7318
chr9-38601022-INS-191
chr9-38612373-INS-254
chr9-38644542-INS-207
chr9-38669262-INS-6425
chr9-38678423-INS-6311
chr9-38681800-INS-2386
chr9-38682622-INS-289
chr9-38692538-INS-132
chr9-38695589-INS-70
chr9-38729469-INS-64
chr9-38733591-INS-416
chr9-38735470-INS-92

chr9-38744605-INS-367
chr9-38761418-INS-389
chr9-38774734-INS-89
chr9-38823470-INS-217
chr9-38845990-INS-205
chr9-38903004-INS-94
chr9-38922281-INS-3151
chr9-38938265-INS-64
chr9-38939253-INS-6325
chr9-38976369-INS-6688
chr9-38990859-INS-6584
chr9-38994557-INS-1674
chr9-38995973-INS-142
chr9-39008954-INS-6384
chr9-39018130-INS-57
chr9-39036165-INS-54
chr9-39036716-INS-7212
chr9-39042586-INS-6588
chr9-39067915-INS-7458
chr9-39076195-INS-519
chr9-39163875-INS-51
chr9-39187924-INS-63
chr9-39189815-INS-76
chr9-39208010-INS-231
chr9-39221839-INS-296
chr9-39224713-INS-932
chr9-39232134-INS-72
chr9-39260014-INS-68
chr9-39285373-INS-91
chr9-39323191-INS-442
chr9-39336226-INS-7427
chr9-39354659-INS-424
chr9-39362083-INS-60
chr9-39377864-INS-673
chr9-39387796-INS-6561
chr9-39404166-INS-6896
chr9-39444739-INS-83
chr9-39463315-INS-56
chr9-39471608-INS-124
chr9-39482517-INS-102
chr9-39495559-INS-60
chr9-39517143-INS-52
chr9-39521662-INS-300
chr9-39548921-INS-337
chr9-39551660-INS-224
chr9-39608650-INS-696
chr9-39610275-INS-400
chr9-39612859-INS-6836
chr9-39635870-INS-6795
chr9-39637094-INS-123
chr9-39655830-INS-76
chr9-39667129-INS-257
chr9-39668098-INS-344
chr9-39668831-INS-915
chr9-39705506-INS-168
chr9-39711968-INS-74
chr9-39719478-INS-63
chr9-39743095-INS-915
chr9-39762817-INS-513
chr9-39766752-INS-108
chr9-39776913-INS-82
chr9-39786315-INS-338
chr9-39809529-INS-154
chr9-39812768-INS-753
chr9-39813425-INS-57
chr9-39815587-INS-100
chr9-39842823-INS-72
chr9-39864825-INS-141
chr9-39878625-INS-59
chr9-39883319-INS-431
chr9-39884779-INS-51
chr9-39903138-INS-6456
chr9-39908912-INS-240
chr9-39909489-INS-3171
chr9-39926409-INS-86
chr9-39934749-INS-166
chr9-39943201-INS-226

chr9-39948780-INS-84
chr9-39973068-INS-544
chr9-39989289-INS-52
chr9-40029772-INS-651
chr9-40036223-INS-93
chr9-40043189-INS-7062
chr9-40058980-INS-415
chr9-40065179-INS-1904
chr9-40075679-INS-258
chr9-40131724-INS-58
chr9-40134062-INS-84
chr9-40137032-INS-327
chr9-40139251-INS-64
chr9-40159088-INS-3433
chr9-40164337-INS-57
chr9-40173009-INS-7217
chr9-40202577-INS-52
chr9-40222244-INS-211
chr9-40262553-INS-498
chr9-40263496-INS-78
chr9-40269543-INS-58
chr9-40271406-INS-77
chr9-40314140-INS-175
chr9-40321785-INS-218
chr9-40345286-INS-222
chr9-40359454-INS-237
chr9-40363325-INS-401
chr9-40382807-INS-380
chr9-40388034-INS-88
chr9-40401988-INS-156
chr9-40413271-INS-74
chr9-40417116-INS-91
chr9-40455311-INS-229
chr9-40471089-INS-1866
chr9-40479029-INS-51
chr9-40489324-INS-4681
chr9-40491536-INS-401
chr9-40493568-INS-216
chr9-40501105-INS-276
chr9-40507747-INS-229
chr9-40513955-INS-91
chr9-40520726-INS-285
chr9-40531726-INS-60
chr9-40534353-INS-50
chr9-40552558-INS-435
chr9-40558052-INS-128
chr9-40563502-INS-425
chr9-40565114-INS-56
chr9-40566633-INS-54
chr9-40569949-INS-195
chr9-40578858-INS-66
chr9-40580088-INS-76
chr9-40592387-INS-210
chr9-40601189-INS-99
chr9-40648420-INS-379
chr9-40652766-INS-165
chr9-40658904-INS-77
chr9-40661220-INS-209
chr9-40664705-INS-58
chr9-40669871-INS-241
chr9-40674265-INS-225
chr9-40675285-INS-214
chr9-40676149-INS-400
chr9-40676263-INS-1690
chr9-40681596-INS-145
chr9-40684241-INS-111
chr9-40686523-INS-58
chr9-40703160-INS-56
chr9-40708657-INS-66
chr9-40742770-INS-1102
chr9-40746992-INS-57
chr9-40758713-INS-136
chr9-40769282-INS-68
chr9-40826175-INS-218
chr9-40847566-INS-116
chr9-40893084-INS-217
chr9-40966344-INS-247

chr9-40971512-INS-165
chr9-40985721-INS-52
chr9-41003710-INS-248
chr9-41034724-INS-745
chr9-41065079-INS-141
chr9-41078734-INS-976
chr9-41083547-INS-67
chr9-41090116-INS-117
chr9-41113536-INS-112
chr9-41144643-INS-244
chr9-41146317-INS-473
chr9-41159086-INS-69
chr9-41174595-INS-161
chr9-41201495-INS-51
chr9-41216197-INS-54
chr9-41254826-INS-89
chr9-41268080-INS-493
chr9-41310196-INS-112
chr9-41316731-INS-56
chr9-41322810-INS-259
chr9-41373881-INS-6487
chr9-41376358-INS-62
chr9-41383306-INS-52
chr9-41428025-INS-58
chr9-41477593-INS-1265
chr9-41479694-INS-100
chr9-41487958-INS-58
chr9-41526964-INS-992
chr9-41531900-INS-88
chr9-41532373-INS-6886
chr9-41534123-INS-174
chr9-41545508-INS-90
chr9-41558596-INS-83
chr9-41587522-INS-176
chr9-41656613-INS-113
chr9-41712325-INS-72
chr9-41714662-INS-56
chr9-41739835-INS-54
chr9-41769487-INS-75
chr9-41781960-INS-51
chr9-41795132-INS-185
chr9-41799198-INS-54
chr9-41808754-INS-315
chr9-41826391-INS-159
chr9-41829533-INS-162
chr9-41855021-INS-88
chr9-41901896-INS-703
chr9-41907369-INS-122
chr9-41912618-INS-181
chr9-41919790-INS-54
chr9-41921723-INS-127
chr9-41932701-INS-155
chr9-41977485-INS-139
chr9-41979651-INS-108
chr9-42003502-INS-123
chr9-42008594-INS-189
chr9-42017156-INS-297
chr9-42020362-INS-119
chr9-42028800-INS-66
chr9-42063713-INS-1042
chr9-42066018-INS-115
chr9-42080458-INS-140
chr9-42088671-INS-60
chr9-42090828-INS-113
chr9-42110425-INS-83
chr9-42111087-INS-129
chr9-42131039-INS-2102
chr9-42137157-INS-6487
chr9-42153722-INS-6165
chr9-42181064-INS-88
chr9-42183902-INS-64
chr9-42190914-INS-72
chr9-42310249-INS-87
chr9-42313630-INS-52
chr9-42319505-INS-96
chr9-42329448-INS-305
chr9-42376309-INS-93

chr9-42387479-INS-180
chr9-42393544-INS-355
chr9-42422807-INS-400
chr9-42444035-INS-71
chr9-42451094-INS-326
chr9-42475433-INS-110
chr9-42521967-INS-153
chr9-42530782-INS-80
chr9-42545818-INS-78
chr9-42555519-INS-218
chr9-42561077-INS-75
chr9-42572165-INS-821
chr9-42603403-INS-7151
chr9-42608114-INS-112
chr9-42630628-INS-254
chr9-42636689-INS-57
chr9-42662021-INS-212
chr9-42722374-INS-58
chr9-42727644-INS-72
chr9-42739847-INS-66
chr9-42757207-INS-210
chr9-42775373-INS-104
chr9-42785854-INS-7240
chr9-42792141-INS-131
chr9-42811017-INS-54
chr9-42929994-INS-57
chr9-42930320-INS-299
chr9-42930887-INS-767
chr9-42931665-INS-190
chr9-42938315-INS-196
chr9-42959287-INS-82
chr9-42965511-INS-6416
chr9-42965936-INS-216
chr9-42969065-INS-2226
chr9-42972425-INS-150
chr9-42978759-INS-266
chr9-43002617-INS-65
chr9-43021323-INS-76
chr9-43023270-INS-197
chr9-43037241-INS-353
chr9-43075569-INS-158
chr9-43091286-INS-67
chr9-43101290-INS-167
chr9-43110358-INS-126
chr9-43128362-INS-203
chr9-43138622-INS-338
chr9-43149464-INS-65
chr9-43156518-INS-64
chr9-43159527-INS-165
chr9-43165302-INS-50
chr9-43169406-INS-201
chr9-43179569-INS-60
chr9-43186386-INS-164
chr9-43219801-INS-117
chr9-43256547-INS-51
chr9-43283776-INS-84
chr9-43301395-INS-256
chr9-43318955-INS-63
chr9-43341487-INS-343
chr9-43359671-INS-67
chr9-43364157-INS-353
chr9-43369863-INS-228
chr9-43383253-INS-204
chr9-43405339-INS-56
chr9-43426865-INS-98
chr9-43464167-INS-71
chr9-43486518-INS-180
chr9-43488377-INS-133
chr9-43492796-INS-62
chr9-43498976-INS-3302
chr9-43561361-INS-66
chr9-43568598-INS-57
chr9-43573782-INS-60
chr9-43588291-INS-66
chr9-43595735-INS-195
chr9-43619460-INS-63
chr9-43667946-INS-216

chr9-43695707-INS-56
chr9-43700792-INS-65
chr9-43731935-INS-74
chr9-43747026-INS-184
chr9-43765102-INS-55
chr9-43766299-INS-58
chr9-43767989-INS-62
chr9-43784595-INS-102
chr9-43797991-INS-60
chr9-43805205-INS-408
chr9-43813949-INS-75
chr9-43842061-INS-61
chr9-43849337-INS-412
chr9-43850254-INS-120
chr9-43861766-INS-125
chr9-43868236-INS-140
chr9-43937293-INS-55
chr9-43956803-INS-94
chr9-43965938-INS-52
chr9-43969087-INS-213
chr9-43969797-INS-1860
chr9-43973937-INS-207
chr9-44012151-INS-209
chr9-44057485-INS-90
chr9-44065704-INS-156
chr9-44068501-INS-295
chr9-44072111-INS-713
chr9-44074069-INS-198
chr9-44117690-INS-136
chr9-44122671-INS-248
chr9-44141086-INS-137
chr9-44147704-INS-188
chr9-44162464-INS-178
chr9-44191537-INS-180
chr9-44193898-INS-118
chr9-44205786-INS-207
chr9-44213441-INS-180
chr9-44223728-INS-155
chr9-44259981-INS-70
chr9-44284711-INS-202
chr9-44285564-INS-221
chr9-44301702-INS-270
chr9-44306179-INS-142
chr9-44315199-INS-84
chr9-44335957-INS-100
chr9-44339186-INS-205
chr9-44355627-INS-51
chr9-44359457-INS-67
chr9-44367929-INS-80
chr9-44372487-INS-57
chr9-44387056-INS-81
chr9-44425641-INS-51
chr9-44441025-INS-281
chr9-44441459-INS-451
chr9-44466238-INS-68
chr9-44473420-INS-173
chr9-44481035-INS-200
chr9-44491216-INS-56
chr9-44495205-INS-168
chr9-44501193-INS-63
chr9-44508355-INS-208
chr9-44527305-INS-109
chr9-44540225-INS-456
chr9-44577036-INS-121
chr9-44577672-INS-55
chr9-44659425-INS-169
chr9-44680095-INS-207
chr9-44694177-INS-181
chr9-44708477-INS-58
chr9-44739254-INS-77
chr9-44751198-INS-74
chr9-44801218-INS-212
chr9-44806735-INS-176
chr9-44814168-INS-191
chr9-44863108-INS-207
chr9-44873351-INS-204
chr9-44875316-INS-200

chr9-44883626-INS-54
chr9-44888357-INS-480
chr9-44889126-INS-75
chr9-44898148-INS-106
chr9-44929413-INS-5061
chr9-44930947-INS-80
chr9-44942384-INS-83
chr9-44945823-INS-82
chr9-44990287-INS-237
chr9-45010559-INS-3025
chr9-45017962-INS-53
chr9-45023480-INS-1365
chr9-45023481-INS-290
chr9-45023672-INS-1752
chr9-45023673-INS-582
chr9-45024673-INS-60
chr9-45055088-INS-147
chr9-45064224-INS-94
chr9-45066136-INS-142
chr9-45076922-INS-58
chr9-45088044-INS-202
chr9-45100970-INS-529
chr9-45167961-INS-100
chr9-45175232-INS-239
chr9-45203137-INS-70
chr9-45215061-INS-51
chr9-45239816-INS-81
chr9-45245129-INS-82
chr9-45245626-INS-197
chr9-45249269-INS-56
chr9-45275763-INS-212
chr9-45284174-INS-138
chr9-45304378-INS-369
chr9-45306226-INS-507
chr9-45316226-INS-136
chr9-45318470-INS-58
chr9-45327997-INS-433
chr9-45330640-INS-59
chr9-45339419-INS-60
chr9-45339873-INS-102
chr9-45377772-INS-127
chr9-45384293-INS-52
chr9-45391538-INS-68
chr9-45412857-INS-3008
chr9-45436535-INS-181
chr9-45467134-INS-1889
chr9-45475220-INS-214
chr9-45536562-INS-112
chr9-45570117-INS-52
chr9-45607221-INS-90
chr9-45610428-INS-58
chr9-45615243-INS-128
chr9-45629644-INS-124
chr9-45637764-INS-57
chr9-45641416-INS-112
chr9-45652859-INS-98
chr9-45668466-INS-52
chr9-45677020-INS-92
chr9-45705901-INS-81
chr9-45707637-INS-92
chr9-45720140-INS-72
chr9-45722227-INS-164
chr9-45725873-INS-64
chr9-45735410-INS-909
chr9-45735446-INS-54
chr9-45738583-INS-81
chr9-45741801-INS-266
chr9-45789281-INS-116
chr9-45871083-INS-61
chr9-45892094-INS-453
chr9-45893416-INS-72
chr9-45913415-INS-192
chr9-45934432-INS-232
chr9-45938047-INS-194
chr9-45945200-INS-198
chr9-45947640-INS-5522
chr9-45964907-INS-189

chr9-45967892-INS-354
chr9-45968038-INS-1220
chr9-45968349-INS-240
chr9-45983592-INS-63
chr9-45989112-INS-216
chr9-46077494-INS-53
chr9-46078183-INS-119
chr9-46079075-INS-72
chr9-46083239-INS-169
chr9-46097277-INS-95
chr9-46100891-INS-86
chr9-46110989-INS-104
chr9-46161533-INS-264
chr9-46164380-INS-276
chr9-46194852-INS-619
chr9-46204305-INS-438
chr9-46214146-INS-75
chr9-46219202-INS-103
chr9-46221720-INS-52
chr9-46228795-INS-221
chr9-46233764-INS-57
chr9-46238093-INS-401
chr9-46241914-INS-208
chr9-46245269-INS-121
chr9-46247306-INS-68
chr9-46311466-INS-130
chr9-46313851-INS-60
chr9-46321464-INS-70
chr9-46336138-INS-90
chr9-46336988-INS-76
chr9-46342163-INS-210
chr9-46370379-INS-70
chr9-46384659-INS-102
chr9-46388512-INS-561
chr9-46397303-INS-100
chr9-46400045-INS-73
chr9-46415152-INS-55
chr9-46458710-INS-498
chr9-46512286-INS-2985
chr9-46515297-INS-102
chr9-46524401-INS-72
chr9-46532327-INS-52
chr9-46569868-INS-145
chr9-46603045-INS-59
chr9-46605027-INS-58
chr9-46612567-INS-3904
chr9-46638978-INS-714
chr9-46670849-INS-518
chr9-46703519-INS-139
chr9-46720886-INS-119
chr9-46727482-INS-56
chr9-46737951-INS-116
chr9-46752680-INS-66
chr9-46763458-INS-235
chr9-46765720-INS-64
chr9-46773542-INS-492
chr9-46784185-INS-144
chr9-46795172-INS-81
chr9-46822800-INS-56
chr9-46823849-INS-331
chr9-46828973-INS-6206
chr9-46839120-INS-226
chr9-46864743-INS-7191
chr9-46868363-INS-437
chr9-46870984-INS-210
chr9-46894192-INS-306
chr9-46921107-INS-120
chr9-46927394-INS-107
chr9-46930457-INS-94
chr9-46962995-INS-5521
chr9-46966625-INS-59
chr9-46994667-INS-81
chr9-47008532-INS-75
chr9-47017667-INS-270
chr9-47019120-INS-742
chr9-47024301-INS-458
chr9-47029892-INS-138

chr9-47049377-INS-54
chr9-47088600-INS-128
chr9-47100492-INS-60
chr9-47103271-INS-227
chr9-47131496-INS-171
chr9-47145408-INS-3116
chr9-47156902-INS-76
chr9-47160280-INS-234
chr9-47171195-INS-150
chr9-47187613-INS-52
chr9-47194531-INS-51
chr9-47200505-INS-390
chr9-47210283-INS-148
chr9-47221529-INS-1890
chr9-47244746-INS-60
chr9-47338722-INS-330
chr9-47365822-INS-111
chr9-47376301-INS-52
chr9-47509081-INS-130
chr9-47550689-INS-420
chr9-47567375-INS-56
chr9-47666986-INS-54
chr9-47670249-INS-160
chr9-47764810-INS-72
chr9-47780502-INS-54
chr9-47808611-INS-57
chr9-47857118-INS-172
chr9-47859131-INS-67
chr9-47867243-INS-2171
chr9-47883850-INS-135
chr9-47886590-INS-64
chr9-47948432-INS-52
chr9-47950757-INS-64
chr9-47957891-INS-62
chr9-47967576-INS-284
chr9-47995189-INS-53
chr9-48036585-INS-51
chr9-48046886-INS-1591
chr9-48083586-INS-98
chr9-48118982-INS-196
chr9-48123272-INS-86
chr9-48188490-INS-2216
chr9-48226550-INS-175
chr9-48227339-INS-6210
chr9-48245089-INS-52
chr9-48279692-INS-55
chr9-48302131-INS-86
chr9-48325329-INS-64
chr9-48333094-INS-65
chr9-48352023-INS-54
chr9-48354422-INS-53
chr9-48366259-INS-498
chr9-48391389-INS-68
chr9-48411804-INS-52
chr9-48443645-INS-6721
chr9-48466940-INS-94
chr9-48486221-INS-146
chr9-48495335-INS-85
chr9-48509754-INS-88
chr9-48510625-INS-51
chr9-48513322-INS-53
chr9-48518442-INS-330
chr9-48530463-INS-55
chr9-48555550-INS-108
chr9-48577290-INS-341
chr9-48608083-INS-156
chr9-48633543-INS-58
chr9-48694347-INS-63
chr9-48696144-INS-4488
chr9-48718749-INS-58
chr9-48722862-INS-880
chr9-48750751-INS-76
chr9-48814551-INS-64
chr9-48842415-INS-51
chr9-48863513-INS-121
chr9-48865547-INS-55
chr9-48869121-INS-399

chr9-48884166-INS-76
chr9-48887281-INS-203
chr9-48905330-INS-212
chr9-48907054-INS-110
chr9-48908208-INS-154
chr9-48909365-INS-241
chr9-48913821-INS-338
chr9-48926843-INS-228
chr9-48929815-INS-206
chr9-48937641-INS-173
chr9-48938288-INS-83
chr9-48973083-INS-352
chr9-48975026-INS-196
chr9-49000244-INS-215
chr9-49001796-INS-169
chr9-49003672-INS-290
chr9-49008967-INS-129
chr9-49010619-INS-245
chr9-49015037-INS-117
chr9-49015566-INS-74
chr9-49039374-INS-582
chr9-49041733-INS-218
chr9-49056915-INS-321
chr9-49087967-INS-199
chr9-49131932-INS-142
chr9-49166492-INS-649
chr9-49199089-INS-713
chr9-49211148-INS-966
chr9-49216826-INS-427
chr9-49219821-INS-428
chr9-49232243-INS-6480
chr9-49246885-INS-284
chr9-49250198-INS-51
chr9-49311298-INS-52
chr9-49328879-INS-50
chr9-49336938-INS-84
chr9-49347536-INS-87
chr9-49350769-INS-5253
chr9-49352506-INS-54
chr9-49379595-INS-185
chr9-49379643-INS-558
chr9-49386033-INS-951
chr9-49394107-INS-215
chr9-49407817-INS-54
chr9-49446706-INS-73
chr9-49465473-INS-214
chr9-49468987-INS-65
chr9-49472191-INS-87
chr9-49500675-INS-622
chr9-49520109-INS-529
chr9-49533001-INS-1203
chr9-49554079-INS-62
chr9-49572617-INS-56
chr9-49604016-INS-66
chr9-49614099-INS-157
chr9-49617037-INS-756
chr9-49622313-INS-4143
chr9-49632475-INS-73
chr9-49655226-INS-892
chr9-49671116-INS-223
chr9-49681198-INS-80
chr9-49697476-INS-1894
chr9-49728975-INS-112
chr9-49731717-INS-314
chr9-49732179-INS-306
chr9-49746986-INS-65
chr9-49778441-INS-758
chr9-49838356-INS-498
chr9-49856410-INS-133
chr9-49914428-INS-57
chr9-49916612-INS-59
chr9-49932285-INS-6502
chr9-49933870-INS-503
chr9-49936185-INS-164
chr9-49938101-INS-184
chr9-49990406-INS-63
chr9-50006285-INS-908

chr9-50007895-INS-79
chr9-50016923-INS-72
chr9-50031824-INS-52
chr9-50042073-INS-194
chr9-50055739-INS-54
chr9-50064988-INS-142
chr9-50092657-INS-567
chr9-50099838-INS-56
chr9-50116046-INS-127
chr9-50128271-INS-99
chr9-50206407-INS-116
chr9-50251793-INS-293
chr9-50281128-INS-186
chr9-50284854-INS-202
chr9-50309196-INS-64
chr9-50327515-INS-273
chr9-50338366-INS-52
chr9-50356154-INS-63
chr9-50361345-INS-227
chr9-50377388-INS-80
chr9-50395790-INS-100
chr9-50401354-INS-213
chr9-50448495-INS-128
chr9-50453886-INS-343
chr9-50458549-INS-203
chr9-50462074-INS-131
chr9-50487064-INS-164
chr9-50488757-INS-283
chr9-50553200-INS-120
chr9-50566396-INS-221
chr9-50589871-INS-6400
chr9-50623593-INS-235
chr9-50665150-INS-55
chr9-50672078-INS-184
chr9-50676640-INS-233
chr9-50691482-INS-398
chr9-50733123-INS-145
chr9-50744276-INS-72
chr9-50769621-INS-202
chr9-50781051-INS-64
chr9-50787892-INS-58
chr9-50825410-INS-56
chr9-50852430-INS-768
chr9-50857283-INS-6399
chr9-50860161-INS-6230
chr9-50886633-INS-60
chr9-51011831-INS-192
chr9-51019957-INS-69
chr9-51032790-INS-106
chr9-51039488-INS-240
chr9-51045895-INS-94
chr9-51109864-INS-53
chr9-51117802-INS-68
chr9-51121591-INS-5981
chr9-51156522-INS-145
chr9-51167554-INS-82
chr9-51209825-INS-1886
chr9-51227838-INS-103
chr9-51232019-INS-557
chr9-51232825-INS-66
chr9-51252678-INS-78
chr9-51270142-INS-80
chr9-51271343-INS-158
chr9-51291911-INS-56
chr9-51299106-INS-548
chr9-51303510-INS-71
chr9-51311943-INS-118
chr9-51314557-INS-1758
chr9-51329479-INS-61
chr9-51369560-INS-109
chr9-51381518-INS-4731
chr9-51408326-INS-498
chr9-51457901-INS-56
chr9-51464328-INS-770
chr9-51484218-INS-106
chr9-51515082-INS-303
chr9-51548336-INS-51

chr9-51548522-INS-3039
chr9-51562806-INS-255
chr9-51579904-INS-177
chr9-51605544-INS-6201
chr9-51609966-INS-92
chr9-51626689-INS-535
chr9-51634433-INS-53
chr9-51654430-INS-140
chr9-51666123-INS-226
chr9-51672788-INS-54
chr9-51681273-INS-108
chr9-51697214-INS-74
chr9-51723751-INS-141
chr9-51808321-INS-52
chr9-51823401-INS-527
chr9-51838374-INS-112
chr9-51846458-INS-225
chr9-51857752-INS-229
chr9-51889913-INS-528
chr9-51906097-INS-82
chr9-51914716-INS-250
chr9-51946387-INS-268
chr9-51947040-INS-82
chr9-51949710-INS-199
chr9-51953927-INS-64
chr9-51967600-INS-61
chr9-51997431-INS-74
chr9-52034619-INS-504
chr9-52049063-INS-6546
chr9-52068780-INS-58
chr9-52090612-INS-76
chr9-52145376-INS-356
chr9-52152516-INS-7312
chr9-52168109-INS-191
chr9-52176264-INS-236
chr9-52178728-INS-134
chr9-52186899-INS-1885
chr9-52216346-INS-55
chr9-52221137-INS-51
chr9-52230518-INS-6396
chr9-52261646-INS-73
chr9-52264154-INS-66
chr9-52323304-INS-1502
chr9-52324762-INS-78
chr9-52354144-INS-5233
chr9-52360183-INS-54
chr9-52361512-INS-5997
chr9-52371532-INS-79
chr9-52372777-INS-457
chr9-52386235-INS-52
chr9-52406456-INS-66
chr9-52413006-INS-142
chr9-52419086-INS-636
chr9-52423791-INS-64
chr9-52430556-INS-8002
chr9-52436700-INS-72
chr9-52444513-INS-2429
chr9-52455842-INS-52
chr9-52457360-INS-4562
chr9-52486881-INS-838
chr9-52497242-INS-95
chr9-52497304-INS-130
chr9-52521813-INS-274
chr9-52531562-INS-669
chr9-52543128-INS-70
chr9-52577054-INS-68
chr9-52613939-INS-73
chr9-52646491-INS-537
chr9-52660277-INS-615
chr9-52661243-INS-212
chr9-52675712-INS-82
chr9-52680303-INS-6714
chr9-52687681-INS-113
chr9-52701212-INS-272
chr9-52710710-INS-100
chr9-52721918-INS-858
chr9-52729475-INS-462

chr9-52827656-INS-97
chr9-52833215-INS-342
chr9-52839734-INS-434
chr9-52841222-INS-240
chr9-52843530-INS-154
chr9-52870572-INS-56
chr9-52874589-INS-82
chr9-52889540-INS-390
chr9-52914244-INS-53
chr9-52916879-INS-52
chr9-52921428-INS-7115
chr9-52930933-INS-306
chr9-52942878-INS-496
chr9-52965463-INS-569
chr9-53005637-INS-2008
chr9-53011832-INS-329
chr9-53039908-INS-185
chr9-53046770-INS-245
chr9-53051586-INS-424
chr9-53065749-INS-2152
chr9-53081735-INS-94
chr9-53093489-INS-804
chr9-53094622-INS-206
chr9-53117462-INS-56
chr9-53118884-INS-99
chr9-53137049-INS-203
chr9-53178168-INS-82
chr9-53191779-INS-389
chr9-53199194-INS-132
chr9-53226008-INS-77
chr9-53240578-INS-72
chr9-53258175-INS-180
chr9-53268994-INS-87
chr9-53319886-INS-54
chr9-53328022-INS-235
chr9-53339762-INS-53
chr9-53345304-INS-132
chr9-53347674-INS-192
chr9-53357018-INS-194
chr9-53359555-INS-56
chr9-53363916-INS-124
chr9-53372439-INS-180
chr9-53380046-INS-364
chr9-53384559-INS-377
chr9-53388043-INS-95
chr9-53412836-INS-720
chr9-53414397-INS-209
chr9-53418116-INS-131
chr9-53426934-INS-100
chr9-53476244-INS-233
chr9-53477375-INS-493
chr9-53480808-INS-205
chr9-53486718-INS-79
chr9-53511342-INS-247
chr9-53512668-INS-60
chr9-53550530-INS-405
chr9-53567187-INS-620
chr9-53577150-INS-141
chr9-53584241-INS-83
chr9-53590449-INS-105
chr9-53591193-INS-84
chr9-53592123-INS-464
chr9-53594937-INS-66
chr9-53606277-INS-70
chr9-53615732-INS-187
chr9-53622513-INS-64
chr9-53626485-INS-58
chr9-53629276-INS-208
chr9-53634239-INS-66
chr9-53635138-INS-186
chr9-53662879-INS-177
chr9-53677535-INS-137
chr9-53689437-INS-202
chr9-53701321-INS-225
chr9-53705634-INS-158
chr9-53714480-INS-128
chr9-53732487-INS-94

chr9-53735167-INS-55
chr9-53736721-INS-59
chr9-53741756-INS-88
chr9-53743968-INS-753
chr9-53770975-INS-464
chr9-53778830-INS-58
chr9-53815610-INS-414
chr9-53820681-INS-74
chr9-53824244-INS-68
chr9-53888600-INS-61
chr9-53892264-INS-433
chr9-53897369-INS-131
chr9-53899633-INS-544
chr9-53911753-INS-157
chr9-53918668-INS-152
chr9-53919539-INS-3910
chr9-53919540-INS-498
chr9-53952518-INS-131
chr9-53966581-INS-100
chr9-53990403-INS-174
chr9-53999664-INS-60
chr9-54022257-INS-623
chr9-54039718-INS-2998
chr9-54061926-INS-54
chr9-54064194-INS-498
chr9-54113507-INS-96
chr9-54144214-INS-348
chr9-54161750-INS-7122
chr9-54188140-INS-235
chr9-54232419-INS-830
chr9-54235191-INS-97
chr9-54249898-INS-1889
chr9-54250268-INS-5528
chr9-54372248-INS-6365
chr9-54376040-INS-174
chr9-54390002-INS-923
chr9-54391558-INS-1361
chr9-54407101-INS-53
chr9-54415853-INS-6687
chr9-54424508-INS-759
chr9-54445170-INS-692
chr9-54464797-INS-62
chr9-54475265-INS-79
chr9-54543241-INS-74
chr9-54555719-INS-242
chr9-54571557-INS-6271
chr9-54574208-INS-77
chr9-54578791-INS-290
chr9-54596483-INS-92
chr9-54604837-INS-83
chr9-54629951-INS-177
chr9-54637566-INS-62
chr9-54644020-INS-55
chr9-54661451-INS-51
chr9-54688074-INS-55
chr9-54698556-INS-72
chr9-54705488-INS-97
chr9-54722012-INS-178
chr9-54729972-INS-214
chr9-54733866-INS-191
chr9-54746548-INS-6402
chr9-54754849-INS-354
chr9-54756244-INS-176
chr9-54769096-INS-70
chr9-54777743-INS-108
chr9-54780252-INS-747
chr9-54784901-INS-208
chr9-54846702-INS-52
chr9-54868805-INS-118
chr9-54870665-INS-149
chr9-54873823-INS-194
chr9-54884518-INS-224
chr9-54902527-INS-516
chr9-54910202-INS-83
chr9-55050906-INS-108
chr9-55073352-INS-178
chr9-55105904-INS-203

chr9-55109453-INS-110
chr9-55124021-INS-66
chr9-55128825-INS-200
chr9-55132193-INS-204
chr9-55134800-INS-394
chr9-55145196-INS-94
chr9-55151397-INS-427
chr9-55154665-INS-403
chr9-55157287-INS-88
chr9-55170230-INS-279
chr9-55180263-INS-56
chr9-55182383-INS-78
chr9-55190656-INS-70
chr9-55206704-INS-58
chr9-55214112-INS-90
chr9-55232657-INS-216
chr9-55261187-INS-430
chr9-55266015-INS-73
chr9-55267498-INS-72
chr9-55273453-INS-72
chr9-55293498-INS-72
chr9-55301090-INS-560
chr9-55310843-INS-55
chr9-55315113-INS-60
chr9-55325597-INS-73
chr9-55354523-INS-6919
chr9-55360266-INS-323
chr9-55364621-INS-60
chr9-55408336-INS-6065
chr9-55409950-INS-207
chr9-55420507-INS-213
chr9-55435460-INS-6362
chr9-55440009-INS-177
chr9-55480883-INS-53
chr9-55486332-INS-1236
chr9-55499280-INS-113
chr9-55523578-INS-6425
chr9-55545367-INS-210
chr9-55570610-INS-6761
chr9-55582921-INS-517
chr9-55585194-INS-52
chr9-55604318-INS-722
chr9-55609165-INS-7155
chr9-55705377-INS-400
chr9-55717584-INS-77
chr9-55727795-INS-694
chr9-55754148-INS-6783
chr9-55772689-INS-278
chr9-55780939-INS-1779
chr9-55815072-INS-56
chr9-55826007-INS-567
chr9-55829226-INS-6669
chr9-55834104-INS-68
chr9-55863944-INS-8581
chr9-55884455-INS-55
chr9-55901580-INS-1340
chr9-55910872-INS-101
chr9-55932786-INS-88
chr9-55943016-INS-95
chr9-55955075-INS-112
chr9-55958827-INS-193
chr9-55977338-INS-147
chr9-55991355-INS-207
chr9-55991436-INS-7099
chr9-56004697-INS-53
chr9-56010840-INS-130
chr9-56018486-INS-52
chr9-56026841-INS-77
chr9-56057594-INS-146
chr9-56060400-INS-127
chr9-56067281-INS-189
chr9-56075793-INS-60
chr9-56095281-INS-73
chr9-56133731-INS-188
chr9-56181380-INS-89
chr9-56182218-INS-222
chr9-56189566-INS-3244

chr9-56198981-INS-91
chr9-56238438-INS-66
chr9-56263994-INS-161
chr9-56332847-INS-240
chr9-56389161-INS-64
chr9-56396585-INS-60
chr9-56397792-INS-160
chr9-56452329-INS-61
chr9-56495561-INS-267
chr9-56511503-INS-144
chr9-56512649-INS-77
chr9-56570436-INS-142
chr9-56628494-INS-56
chr9-56666903-INS-51
chr9-56697447-INS-219
chr9-56716390-INS-103
chr9-56730318-INS-60
chr9-56739480-INS-200
chr9-56783630-INS-194
chr9-56797969-INS-215
chr9-56810114-INS-63
chr9-56818870-INS-197
chr9-56829919-INS-400
chr9-56841238-INS-97
chr9-56852385-INS-334
chr9-56867550-INS-93
chr9-56871005-INS-209
chr9-56886078-INS-83
chr9-56909137-INS-204
chr9-56917101-INS-503
chr9-56931785-INS-124
chr9-56937384-INS-222
chr9-56939674-INS-67
chr9-56947089-INS-147
chr9-56948423-INS-351
chr9-56949986-INS-211
chr9-56975170-INS-5541
chr9-56976628-INS-206
chr9-57001220-INS-57
chr9-57071187-INS-229
chr9-57071188-INS-204
chr9-57073284-INS-61
chr9-57078024-INS-216
chr9-57082180-INS-406
chr9-57084330-INS-233
chr9-57120885-INS-70
chr9-57145071-INS-339
chr9-57172433-INS-168
chr9-57181116-INS-86
chr9-57183217-INS-53
chr9-57184042-INS-237
chr9-57220612-INS-59
chr9-57222628-INS-206
chr9-57232647-INS-189
chr9-57235016-INS-681
chr9-57237843-INS-1887
chr9-57243879-INS-99
chr9-57247633-INS-112
chr9-57273599-INS-326
chr9-57281860-INS-211
chr9-57287645-INS-464
chr9-57300734-INS-198
chr9-57304931-INS-405
chr9-57317408-INS-576
chr9-57322548-INS-197
chr9-57330292-INS-6513
chr9-57334128-INS-209
chr9-57347561-INS-70
chr9-57361440-INS-596
chr9-57365564-INS-7107
chr9-57365963-INS-55
chr9-57367944-INS-206
chr9-57377858-INS-208
chr9-57389945-INS-435
chr9-57401289-INS-213
chr9-57408370-INS-58
chr9-57416457-INS-207

chr9-57419713-INS-52
chr9-57420972-INS-205
chr9-57463681-INS-188
chr9-57472614-INS-168
chr9-57483683-INS-59
chr9-57508067-INS-204
chr9-57510938-INS-5580
chr9-57568726-INS-197
chr9-57570526-INS-52
chr9-57571116-INS-208
chr9-57581257-INS-88
chr9-57607103-INS-235
chr9-57608488-INS-116
chr9-57611651-INS-60
chr9-57616160-INS-231
chr9-57619240-INS-216
chr9-57635975-INS-857
chr9-57636041-INS-322
chr9-57652391-INS-62
chr9-57677926-INS-89
chr9-57680205-INS-79
chr9-57685298-INS-239
chr9-57686729-INS-121
chr9-57722480-INS-162
chr9-57728152-INS-55
chr9-57730874-INS-150
chr9-57751807-INS-65
chr9-57754764-INS-108
chr9-57756123-INS-196
chr9-57757912-INS-462
chr9-57772430-INS-115
chr9-57782444-INS-156
chr9-57790703-INS-172
chr9-57793900-INS-339
chr9-57806623-INS-136
chr9-57808332-INS-204
chr9-57808863-INS-231
chr9-57836179-INS-179
chr9-57899800-INS-382
chr9-57914744-INS-221
chr9-57917559-INS-108
chr9-57924654-INS-70
chr9-57935128-INS-168
chr9-57972279-INS-108
chr9-58029166-INS-51
chr9-58035206-INS-118
chr9-58068648-INS-164
chr9-58079062-INS-205
chr9-58091418-INS-204
chr9-58093532-INS-446
chr9-58103192-INS-64
chr9-58116201-INS-64
chr9-58158929-INS-211
chr9-58161683-INS-54
chr9-58162441-INS-205
chr9-58164897-INS-147
chr9-58172879-INS-186
chr9-58178927-INS-862
chr9-58184725-INS-113
chr9-58191156-INS-221
chr9-58198781-INS-185
chr9-58202970-INS-171
chr9-58226676-INS-128
chr9-58246175-INS-78
chr9-58284536-INS-104
chr9-58286302-INS-65
chr9-58305092-INS-6543
chr9-58312390-INS-140
chr9-58323138-INS-57
chr9-58336862-INS-86
chr9-58394864-INS-74
chr9-58449182-INS-497
chr9-58470646-INS-53
chr9-58472022-INS-209
chr9-58480580-INS-80
chr9-58483230-INS-60
chr9-58487671-INS-156

chr9-58493217-INS-206
chr9-58515344-INS-278
chr9-58518146-INS-183
chr9-58609267-INS-998
chr9-58670468-INS-68
chr9-58699844-INS-235
chr9-58706143-INS-147
chr9-58764387-INS-54
chr9-58836934-INS-53
chr9-58861695-INS-84
chr9-58869141-INS-108
chr9-58899456-INS-660
chr9-58929063-INS-140
chr9-58961661-INS-148
chr9-58985529-INS-120
chr9-59007633-INS-89
chr9-59013744-INS-158
chr9-59052552-INS-54
chr9-59057577-INS-497
chr9-59059095-INS-84
chr9-59067902-INS-134
chr9-59069541-INS-5634
chr9-59086605-INS-348
chr9-59087715-INS-147
chr9-59128264-INS-60
chr9-59143104-INS-864
chr9-59153395-INS-130
chr9-59159541-INS-57
chr9-59160902-INS-78
chr9-59169254-INS-506
chr9-59182190-INS-119
chr9-59189581-INS-89
chr9-59197760-INS-60
chr9-59203633-INS-210
chr9-59212447-INS-456
chr9-59212952-INS-73
chr9-59226267-INS-188
chr9-59282369-INS-210
chr9-59286897-INS-212
chr9-59306596-INS-208
chr9-59312421-INS-138
chr9-59322610-INS-71
chr9-59348072-INS-66
chr9-59362666-INS-202
chr9-59397237-INS-958
chr9-59399460-INS-116
chr9-59403384-INS-506
chr9-59420581-INS-168
chr9-59439402-INS-51
chr9-59451947-INS-101
chr9-59482024-INS-211
chr9-59492684-INS-290
chr9-59507260-INS-99
chr9-59533738-INS-60
chr9-59546529-INS-207
chr9-59569900-INS-62
chr9-59581149-INS-64
chr9-59584446-INS-54
chr9-59590127-INS-88
chr9-59592366-INS-75
chr9-59601101-INS-428
chr9-59608105-INS-246
chr9-59609241-INS-55
chr9-59612793-INS-143
chr9-59625790-INS-1472
chr9-59651371-INS-214
chr9-59675618-INS-76
chr9-59678923-INS-59
chr9-59690704-INS-219
chr9-59700888-INS-279
chr9-59708649-INS-3326
chr9-59724714-INS-52
chr9-59758803-INS-197
chr9-59809291-INS-620
chr9-59839909-INS-181
chr9-59845666-INS-163
chr9-59873720-INS-404

chr9-59884052-INS-260
chr9-59897501-INS-794
chr9-59909086-INS-165
chr9-59911335-INS-72
chr9-59920375-INS-232
chr9-59936004-INS-4506
chr9-59962919-INS-106
chr9-59973460-INS-6475
chr9-59982223-INS-6252
chr9-60002410-INS-58
chr9-60046520-INS-83
chr9-60075017-INS-52
chr9-60088608-INS-88
chr9-60089591-INS-52
chr9-60119401-INS-51
chr9-60122643-INS-193
chr9-60166386-INS-245
chr9-60177050-INS-6721
chr9-60202076-INS-147
chr9-60211130-INS-292
chr9-60240482-INS-54
chr9-60267493-INS-52
chr9-60272919-INS-54
chr9-60293419-INS-53
chr9-60307417-INS-209
chr9-60315296-INS-72
chr9-60331215-INS-96
chr9-60336837-INS-94
chr9-60338562-INS-88
chr9-60348396-INS-150
chr9-60413913-INS-133
chr9-60446878-INS-166
chr9-60448076-INS-77
chr9-60466654-INS-58
chr9-60491261-INS-258
chr9-60508330-INS-56
chr9-60510777-INS-62
chr9-60536488-INS-62
chr9-60560584-INS-78
chr9-60564266-INS-112
chr9-60566442-INS-88
chr9-60580477-INS-504
chr9-60605658-INS-325
chr9-60606445-INS-196
chr9-60631790-INS-64
chr9-60654794-INS-523
chr9-60697195-INS-64
chr9-60704356-INS-64
chr9-60716106-INS-60
chr9-60720836-INS-58
chr9-60750652-INS-61
chr9-60755156-INS-130
chr9-60769997-INS-240
chr9-60785086-INS-76
chr9-60793039-INS-185
chr9-60805312-INS-66
chr9-60810900-INS-101
chr9-60816019-INS-389
chr9-60817345-INS-56
chr9-60846807-INS-56
chr9-60848939-INS-1891
chr9-60853207-INS-67
chr9-60899198-INS-56
chr9-60904178-INS-223
chr9-60909232-INS-72
chr9-60913708-INS-388
chr9-60935404-INS-51
chr9-60945168-INS-213
chr9-60947413-INS-138
chr9-60968634-INS-68
chr9-60993311-INS-108
chr9-61006214-INS-60
chr9-61035661-INS-250
chr9-61046978-INS-64
chr9-61060435-INS-64
chr9-61074485-INS-52
chr9-61078459-INS-614

chr9-61150299-INS-60
chr9-61161760-INS-124
chr9-61169936-INS-361
chr9-61171192-INS-67
chr9-61201345-INS-6486
chr9-61209946-INS-97
chr9-61228384-INS-79
chr9-61236499-INS-197
chr9-61270378-INS-223
chr9-61335081-INS-427
chr9-61361840-INS-77
chr9-61372208-INS-77
chr9-61379523-INS-203
chr9-61380885-INS-345
chr9-61385496-INS-71
chr9-61393003-INS-56
chr9-61399664-INS-68
chr9-61407812-INS-52
chr9-61418149-INS-57
chr9-61430950-INS-96
chr9-61448558-INS-523
chr9-61452754-INS-53
chr9-61459571-INS-70
chr9-61468188-INS-400
chr9-61469520-INS-68
chr9-61474719-INS-532
chr9-61478108-INS-124
chr9-61493016-INS-142
chr9-61532254-INS-60
chr9-61550724-INS-232
chr9-61566651-INS-96
chr9-61571259-INS-172
chr9-61575400-INS-156
chr9-61577447-INS-130
chr9-61589238-INS-286
chr9-61596335-INS-70
chr9-61610077-INS-1864
chr9-61617432-INS-70
chr9-61633168-INS-58
chr9-61652012-INS-74
chr9-61665948-INS-1860
chr9-61675364-INS-1040
chr9-61718048-INS-231
chr9-61733042-INS-636
chr9-61744256-INS-68
chr9-61749265-INS-66
chr9-61801010-INS-76
chr9-61802729-INS-372
chr9-61808221-INS-53
chr9-61852904-INS-201
chr9-61879397-INS-108
chr9-61887510-INS-60
chr9-61930691-INS-113
chr9-61933345-INS-95
chr9-61943060-INS-451
chr9-61955959-INS-105
chr9-61987429-INS-60
chr9-62004302-INS-511
chr9-62015205-INS-720
chr9-62027957-INS-61
chr9-62034069-INS-511
chr9-62039266-INS-68
chr9-62095480-INS-234
chr9-62103185-INS-2766
chr9-62106345-INS-290
chr9-62125133-INS-293
chr9-62145366-INS-612
chr9-62154494-INS-164
chr9-62187587-INS-241
chr9-62196719-INS-82
chr9-62208101-INS-51
chr9-62217916-INS-54
chr9-62222598-INS-177
chr9-62227500-INS-364
chr9-62229983-INS-59
chr9-62265618-INS-380
chr9-62296676-INS-198

chr9-62306026-INS-215
chr9-62315614-INS-6248
chr9-62318855-INS-132
chr9-62344005-INS-196
chr9-62375545-INS-128
chr9-62379900-INS-117
chr9-62385324-INS-53
chr9-62389358-INS-99
chr9-62395659-INS-261
chr9-62395725-INS-754
chr9-62397418-INS-439
chr9-62401320-INS-134
chr9-62417233-INS-134
chr9-62419857-INS-213
chr9-62466498-INS-218
chr9-62497332-INS-440
chr9-62516153-INS-65
chr9-62529850-INS-247
chr9-62531714-INS-139
chr9-62537923-INS-106
chr9-62544073-INS-173
chr9-62554790-INS-148
chr9-62561309-INS-160
chr9-62564056-INS-82
chr9-62575995-INS-323
chr9-62580361-INS-72
chr9-62597653-INS-126
chr9-62598198-INS-231
chr9-62612662-INS-310
chr9-62638943-INS-69
chr9-62640254-INS-120
chr9-62644877-INS-97
chr9-62653847-INS-185
chr9-62672030-INS-356
chr9-62676254-INS-52
chr9-62692295-INS-239
chr9-62693145-INS-79
chr9-62714646-INS-201
chr9-62721054-INS-239
chr9-62727811-INS-178
chr9-62732132-INS-222
chr9-62735341-INS-196
chr9-62756743-INS-76
chr9-62776768-INS-54
chr9-62782075-INS-60
chr9-62798415-INS-118
chr9-62800069-INS-203
chr9-62837153-INS-104
chr9-62855181-INS-487
chr9-62867265-INS-131
chr9-62874765-INS-84
chr9-62890615-INS-136
chr9-62893863-INS-106
chr9-62898266-INS-277
chr9-62926720-INS-197
chr9-62928418-INS-53
chr9-62932608-INS-329
chr9-62937486-INS-57
chr9-62940768-INS-75
chr9-62948122-INS-312
chr9-62954949-INS-140
chr9-62980869-INS-54
chr9-63001052-INS-101
chr9-63005012-INS-338
chr9-63017895-INS-120
chr9-63027153-INS-96
chr9-63063687-INS-222
chr9-63078588-INS-66
chr9-63144263-INS-51
chr9-63147140-INS-204
chr9-63160571-INS-74
chr9-63186373-INS-76
chr9-63215482-INS-84
chr9-63232497-INS-5068
chr9-63238548-INS-106
chr9-63238900-INS-268
chr9-63243056-INS-93

chr9-63265584-INS-469
chr9-63289177-INS-52
chr9-63298905-INS-140
chr9-63301579-INS-177
chr9-63386608-INS-829
chr9-63388570-INS-359
chr9-63423122-INS-3434
chr9-63435666-INS-56
chr9-63458141-INS-129
chr9-63463020-INS-150
chr9-63473019-INS-277
chr9-63492346-INS-73
chr9-63496925-INS-98
chr9-63522524-INS-255
chr9-63523063-INS-170
chr9-63547299-INS-1885
chr9-63567001-INS-163
chr9-63582946-INS-76
chr9-63597365-INS-201
chr9-63601993-INS-146
chr9-63607217-INS-58
chr9-63607850-INS-160
chr9-63631292-INS-70
chr9-63642190-INS-279
chr9-63655964-INS-224
chr9-63692293-INS-66
chr9-63704268-INS-927
chr9-63719628-INS-3068
chr9-63721327-INS-65
chr9-63731325-INS-402
chr9-63773037-INS-126
chr9-63774335-INS-53
chr9-63790774-INS-96
chr9-63804396-INS-114
chr9-63871866-INS-79
chr9-63922122-INS-143
chr9-63944414-INS-197
chr9-63947300-INS-63
chr9-63948741-INS-232
chr9-63959269-INS-59
chr9-63971008-INS-60
chr9-63972366-INS-248
chr9-63973577-INS-211
chr9-63983265-INS-138
chr9-64012451-INS-200
chr9-64019316-INS-183
chr9-64023026-INS-187
chr9-64033160-INS-190
chr9-64042358-INS-209
chr9-64060452-INS-108
chr9-64069403-INS-52
chr9-64077585-INS-84
chr9-64127296-INS-129
chr9-64128695-INS-63
chr9-64140799-INS-68
chr9-64167507-INS-172
chr9-64183654-INS-197
chr9-64188208-INS-156
chr9-64191363-INS-212
chr9-64197266-INS-204
chr9-64199672-INS-51
chr9-64238908-INS-51
chr9-64256716-INS-105
chr9-64280185-INS-118
chr9-64338791-INS-195
chr9-64354194-INS-1988
chr9-64374505-INS-84
chr9-64388767-INS-67
chr9-64397651-INS-206
chr9-64403968-INS-72
chr9-64404742-INS-312
chr9-64414362-INS-52
chr9-64422053-INS-101
chr9-64424925-INS-90
chr9-64447425-INS-137
chr9-64454900-INS-164
chr9-64456789-INS-178

chr9-64464234-INS-98
chr9-64479359-INS-66
chr9-64484992-INS-128
chr9-64494716-INS-63
chr9-64496584-INS-212
chr9-64510846-INS-90
chr9-64520828-INS-60
chr9-64529534-INS-100
chr9-64547529-INS-124
chr9-64568456-INS-184
chr9-64594076-INS-61
chr9-64624601-INS-57
chr9-64658693-INS-80
chr9-64667109-INS-128
chr9-64688052-INS-765
chr9-64706287-INS-492
chr9-64728877-INS-179
chr9-64751298-INS-166
chr9-64768108-INS-60
chr9-64776493-INS-66
chr9-64780649-INS-74
chr9-64785980-INS-173
chr9-64793074-INS-96
chr9-64806246-INS-55
chr9-64846881-INS-52
chr9-64850673-INS-236
chr9-64863618-INS-97
chr9-64874501-INS-987
chr9-64890908-INS-200
chr9-64910970-INS-62
chr9-64939353-INS-85
chr9-64945390-INS-83
chr9-64959708-INS-429
chr9-64978662-INS-51
chr9-64983805-INS-207
chr9-65024343-INS-52
chr9-65053288-INS-282
chr9-65058752-INS-205
chr9-65066530-INS-108
chr9-65068186-INS-199
chr9-65077863-INS-169
chr9-65082175-INS-741
chr9-65108226-INS-234
chr9-65108978-INS-165
chr9-65109800-INS-68
chr9-65130862-INS-199
chr9-65132810-INS-195
chr9-65139426-INS-78
chr9-65155985-INS-52
chr9-65198019-INS-186
chr9-65208165-INS-57
chr9-65215952-INS-75
chr9-65239116-INS-327
chr9-65240622-INS-376
chr9-65242015-INS-132
chr9-65251371-INS-137
chr9-65255631-INS-187
chr9-65273473-INS-172
chr9-65276003-INS-308
chr9-65286712-INS-198
chr9-65290630-INS-120
chr9-65304864-INS-56
chr9-65310251-INS-130
chr9-65328577-INS-75
chr9-65331143-INS-64
chr9-65340714-INS-6649
chr9-65344577-INS-144
chr9-65346097-INS-179
chr9-65371576-INS-1892
chr9-65391168-INS-186
chr9-65395554-INS-274
chr9-65409837-INS-159
chr9-65416148-INS-4989
chr9-65420992-INS-84
chr9-65429141-INS-208
chr9-65430613-INS-120
chr9-65452346-INS-104

chr9-65481799-INS-223
chr9-65492494-INS-197
chr9-65499969-INS-66
chr9-65502748-INS-260
chr9-65505634-INS-198
chr9-65510155-INS-208
chr9-65517053-INS-155
chr9-65519220-INS-200
chr9-65524194-INS-65
chr9-65526140-INS-208
chr9-65528049-INS-65
chr9-65531117-INS-52
chr9-65535826-INS-150
chr9-65554426-INS-208
chr9-65560867-INS-68
chr9-65573408-INS-8842
chr9-65593316-INS-54
chr9-65619658-INS-63
chr9-65624179-INS-725
chr9-65626626-INS-101
chr9-65630687-INS-202
chr9-65655654-INS-6473
chr9-65706461-INS-52
chr9-65709868-INS-210
chr9-65728397-INS-156
chr9-65744775-INS-209
chr9-65763354-INS-214
chr9-65769120-INS-91
chr9-65784050-INS-177
chr9-65799935-INS-157
chr9-65829954-INS-52
chr9-65869266-INS-181
chr9-65877827-INS-588
chr9-65883441-INS-170
chr9-65926373-INS-198
chr9-65935529-INS-64
chr9-65938580-INS-186
chr9-65940796-INS-190
chr9-65960389-INS-231
chr9-65980722-INS-234
chr9-66012279-INS-101
chr9-66019710-INS-101
chr9-66036838-INS-202
chr9-66041462-INS-269
chr9-66042654-INS-240
chr9-66049828-INS-77
chr9-66054141-INS-651
chr9-66093629-INS-158
chr9-66102366-INS-87
chr9-66111371-INS-68
chr9-66118043-INS-191
chr9-66124148-INS-804
chr9-66130671-INS-288
chr9-66140241-INS-83
chr9-66170762-INS-142
chr9-66180604-INS-55
chr9-66182764-INS-5307
chr9-66215369-INS-220
chr9-66228292-INS-58
chr9-66235188-INS-55
chr9-66253026-INS-140
chr9-66297828-INS-738
chr9-66297836-INS-58
chr9-66302137-INS-83
chr9-66317082-INS-102
chr9-66365089-INS-102
chr9-66371584-INS-82
chr9-66402138-INS-56
chr9-66410489-INS-178
chr9-66458665-INS-69
chr9-66460955-INS-6170
chr9-66475072-INS-55
chr9-66493126-INS-359
chr9-66504396-INS-54
chr9-66521317-INS-59
chr9-66531574-INS-1740
chr9-66551374-INS-299

chr9-66561800-INS-100
chr9-66570966-INS-298
chr9-66582289-INS-59
chr9-66600773-INS-51
chr9-66612545-INS-235
chr9-66616075-INS-211
chr9-66631164-INS-96
chr9-66645026-INS-62
chr9-66674152-INS-7105
chr9-66710596-INS-5138
chr9-66734913-INS-173
chr9-66757940-INS-226
chr9-66791230-INS-150
chr9-66811773-INS-132
chr9-66830846-INS-215
chr9-66850593-INS-117
chr9-66856814-INS-66
chr9-66872964-INS-87
chr9-66885160-INS-146
chr9-66910424-INS-498
chr9-66925949-INS-121
chr9-66931471-INS-52
chr9-66941794-INS-107
chr9-66961814-INS-234
chr9-66968989-INS-195
chr9-66978407-INS-209
chr9-66979753-INS-56
chr9-66983172-INS-1886
chr9-66995842-INS-96
chr9-67034464-INS-190
chr9-67035711-INS-7740
chr9-67051155-INS-7174
chr9-67058320-INS-52
chr9-67064583-INS-217
chr9-67074136-INS-55
chr9-67080306-INS-2982
chr9-67088889-INS-58
chr9-67090284-INS-75
chr9-67104897-INS-104
chr9-67167395-INS-52
chr9-67169018-INS-82
chr9-67180913-INS-541
chr9-67194665-INS-56
chr9-67207573-INS-166
chr9-67209184-INS-4434
chr9-67224736-INS-257
chr9-67227432-INS-188
chr9-67250497-INS-216
chr9-67270793-INS-199
chr9-67280809-INS-200
chr9-67329188-INS-767
chr9-67332380-INS-134
chr9-67344509-INS-54
chr9-67347831-INS-81
chr9-67347899-INS-560
chr9-67348637-INS-81
chr9-67357089-INS-112
chr9-67391864-INS-188
chr9-67405644-INS-106
chr9-67407633-INS-195
chr9-67417483-INS-73
chr9-67470660-INS-154
chr9-67471792-INS-221
chr9-67527541-INS-92
chr9-67535461-INS-216
chr9-67581415-INS-122
chr9-67582695-INS-525
chr9-67587515-INS-62
chr9-67601235-INS-55
chr9-67614100-INS-74
chr9-67632242-INS-328
chr9-67640236-INS-182
chr9-67665638-INS-58
chr9-67678312-INS-164
chr9-67694144-INS-288
chr9-67700210-INS-107
chr9-67702933-INS-158

chr9-67713199-INS-222
chr9-67714063-INS-135
chr9-67714991-INS-238
chr9-67718350-INS-183
chr9-67741330-INS-86
chr9-67745540-INS-51
chr9-67747485-INS-56
chr9-67749771-INS-63
chr9-67763687-INS-119
chr9-67772753-INS-72
chr9-67781524-INS-55
chr9-67785341-INS-93
chr9-67796435-INS-61
chr9-67798727-INS-208
chr9-67800972-INS-6440
chr9-67838059-INS-55
chr9-67876108-INS-129
chr9-67898892-INS-96
chr9-67920979-INS-146
chr9-67956525-INS-2734
chr9-67959305-INS-6245
chr9-67976088-INS-229
chr9-68019152-INS-60
chr9-68025594-INS-70
chr9-68047452-INS-3064
chr9-68062356-INS-6503
chr9-68073268-INS-96
chr9-68109754-INS-203
chr9-68132967-INS-2451
chr9-68133310-INS-52
chr9-68137220-INS-617
chr9-68141609-INS-60
chr9-68142858-INS-278
chr9-68156115-INS-107
chr9-68156679-INS-199
chr9-68159283-INS-142
chr9-68215651-INS-132
chr9-68235408-INS-110
chr9-68258523-INS-6711
chr9-68263218-INS-9726
chr9-68266158-INS-1342
chr9-68295789-INS-707
chr9-68300196-INS-77
chr9-68332827-INS-54
chr9-68337354-INS-777
chr9-68339348-INS-602
chr9-68353353-INS-52
chr9-68377785-INS-151
chr9-68390621-INS-94
chr9-68396677-INS-2604
chr9-68400069-INS-113
chr9-68429199-INS-1893
chr9-68442577-INS-57
chr9-68446818-INS-101
chr9-68471631-INS-235
chr9-68497320-INS-158
chr9-68499417-INS-100
chr9-68506135-INS-563
chr9-68547202-INS-251
chr9-68554744-INS-95
chr9-68564822-INS-70
chr9-68578883-INS-119
chr9-68587930-INS-95
chr9-68601629-INS-116
chr9-68623991-INS-125
chr9-68633423-INS-225
chr9-68635869-INS-214
chr9-68660591-INS-66
chr9-68664975-INS-198
chr9-68686944-INS-536
chr9-68768377-INS-61
chr9-68797035-INS-77
chr9-68801200-INS-52
chr9-68816663-INS-112
chr9-68819586-INS-104
chr9-68857010-INS-89
chr9-68865881-INS-1816

chr9-68893436-INS-52
chr9-68912694-INS-123
chr9-68917480-INS-159
chr9-68923638-INS-168
chr9-68967803-INS-58
chr9-69001336-INS-108
chr9-69048468-INS-170
chr9-69064768-INS-64
chr9-69067548-INS-113
chr9-69090976-INS-139
chr9-69115814-INS-66
chr9-69150643-INS-152
chr9-69157750-INS-157
chr9-69189571-INS-429
chr9-69209864-INS-94
chr9-69237765-INS-67
chr9-69243142-INS-189
chr9-69249622-INS-112
chr9-69262377-INS-292
chr9-69277716-INS-60
chr9-69295792-INS-50
chr9-69328487-INS-270
chr9-69339043-INS-129
chr9-69344332-INS-566
chr9-69345344-INS-5223
chr9-69350508-INS-58
chr9-69368685-INS-139
chr9-69381979-INS-79
chr9-69386646-INS-215
chr9-69388225-INS-53
chr9-69393631-INS-431
chr9-69420419-INS-103
chr9-69449839-INS-104
chr9-69493164-INS-132
chr9-69497838-INS-135
chr9-69526956-INS-52
chr9-69572487-INS-117
chr9-69584798-INS-139
chr9-69625145-INS-97
chr9-69627802-INS-57
chr9-69644559-INS-127
chr9-69691185-INS-151
chr9-69698374-INS-680
chr9-69708462-INS-108
chr9-69758848-INS-60
chr9-69760813-INS-6479
chr9-69767717-INS-81
chr9-69784372-INS-55
chr9-69816669-INS-55
chr9-69833613-INS-80
chr9-69846505-INS-67
chr9-69857527-INS-543
chr9-69870048-INS-74
chr9-69900524-INS-68
chr9-69908712-INS-52
chr9-69935404-INS-74
chr9-69963684-INS-114
chr9-69968383-INS-52
chr9-69972066-INS-384
chr9-69983948-INS-64
chr9-70019993-INS-68
chr9-70074614-INS-259
chr9-70165890-INS-196
chr9-70221614-INS-212
chr9-70252518-INS-115
chr9-70257283-INS-550
chr9-70265376-INS-53
chr9-70288443-INS-94
chr9-70300686-INS-110
chr9-70324410-INS-175
chr9-70366580-INS-177
chr9-70382953-INS-195
chr9-70391565-INS-277
chr9-70396394-INS-103
chr9-70424581-INS-394
chr9-70475517-INS-182
chr9-70527031-INS-84

chr9-70547983-INS-179
chr9-70553624-INS-93
chr9-70566082-INS-132
chr9-70595367-INS-100
chr9-70706239-INS-60
chr9-70716212-INS-64
chr9-70718077-INS-166
chr9-70721183-INS-72
chr9-70740406-INS-524
chr9-70768340-INS-846
chr9-70884495-INS-114
chr9-70898372-INS-54
chr9-70921845-INS-82
chr9-70936928-INS-52
chr9-70938354-INS-305
chr9-70975735-INS-924
chr9-70977770-INS-54
chr9-71009771-INS-56
chr9-71069020-INS-86
chr9-71071310-INS-86
chr9-71095254-INS-422
chr9-71105881-INS-131
chr9-71110632-INS-197
chr9-71142997-INS-70
chr9-71165789-INS-365
chr9-71184077-INS-53
chr9-71194911-INS-72
chr9-71231263-INS-59
chr9-71234416-INS-70
chr9-71242441-INS-69
chr9-71250481-INS-170
chr9-71252185-INS-56
chr9-71273558-INS-173
chr9-71278270-INS-119
chr9-71279510-INS-161
chr9-71291796-INS-62
chr9-71315976-INS-94
chr9-71356102-INS-207
chr9-71374562-INS-97
chr9-71383354-INS-209
chr9-71389123-INS-169
chr9-71391198-INS-207
chr9-71405066-INS-64
chr9-71421557-INS-202
chr9-71423704-INS-112
chr9-71447303-INS-54
chr9-71448756-INS-62
chr9-71460625-INS-537
chr9-71465935-INS-122
chr9-71474245-INS-59
chr9-71477161-INS-211
chr9-71528342-INS-505
chr9-71658286-INS-52
chr9-71675028-INS-221
chr9-71695124-INS-77
chr9-71707929-INS-236
chr9-71721147-INS-138
chr9-71722619-INS-1493
chr9-71736865-INS-90
chr9-71748044-INS-267
chr9-71811919-INS-132
chr9-71915451-INS-352
chr9-72001325-INS-129
chr9-72007681-INS-793
chr9-72012129-INS-85
chr9-72024320-INS-198
chr9-72066178-INS-154
chr9-72126930-INS-52
chr9-72129269-INS-55
chr9-72139242-INS-63
chr9-72166966-INS-246
chr9-72177304-INS-200
chr9-72218894-INS-201
chr9-72272337-INS-212
chr9-72308060-INS-56
chr9-72309060-INS-57
chr9-72320296-INS-242

chr9-72327706-INS-154
chr9-72332030-INS-90
chr9-72350298-INS-102
chr9-72358344-INS-163
chr9-72361434-INS-198
chr9-72361786-INS-357
chr9-72420388-INS-463
chr9-72422008-INS-208
chr9-72435435-INS-5540
chr9-72508344-INS-174
chr9-72513054-INS-197
chr9-72538868-INS-182
chr9-72546002-INS-59
chr9-72553868-INS-395
chr9-72557555-INS-73
chr9-72561185-INS-262
chr9-72580428-INS-181
chr9-72582519-INS-104
chr9-72585061-INS-327
chr9-72590043-INS-167
chr9-72633867-INS-85
chr9-72635847-INS-53
chr9-72658767-INS-313
chr9-72678895-INS-99
chr9-72680447-INS-206
chr9-72684237-INS-212
chr9-72704378-INS-208
chr9-72708572-INS-212
chr9-72731115-INS-160
chr9-72754325-INS-52
chr9-72757280-INS-197
chr9-72771142-INS-485
chr9-72774152-INS-62
chr9-72793034-INS-218
chr9-72800095-INS-66
chr9-72822673-INS-97
chr9-72843068-INS-56
chr9-72898729-INS-324
chr9-72902450-INS-53
chr9-72910577-INS-214
chr9-72913943-INS-60
chr9-72933796-INS-108
chr9-72940668-INS-76
chr9-72945820-INS-197
chr9-72960702-INS-81
chr9-72970470-INS-244
chr9-72977922-INS-192
chr9-73010723-INS-78
chr9-73021706-INS-614
chr9-73042888-INS-191
chr9-73051199-INS-201
chr9-73059110-INS-68
chr9-73067081-INS-80
chr9-73080523-INS-62
chr9-73091767-INS-62
chr9-73095258-INS-341
chr9-73096858-INS-52
chr9-73105279-INS-52
chr9-73112666-INS-207
chr9-73128939-INS-54
chr9-73151350-INS-180
chr9-73152792-INS-104
chr9-73157206-INS-146
chr9-73164040-INS-1888
chr9-73173438-INS-6904
chr9-73180380-INS-55
chr9-73191290-INS-405
chr9-73252747-INS-104
chr9-73283093-INS-206
chr9-73286831-INS-167
chr9-73327812-INS-133
chr9-73359897-INS-71
chr9-73381329-INS-72
chr9-73408449-INS-60
chr9-73418237-INS-79
chr9-73448915-INS-67
chr9-73451278-INS-53

chr9-73451398-INS-815
chr9-73468917-INS-124
chr9-73474979-INS-7005
chr9-73475370-INS-52
chr9-73504677-INS-237
chr9-73519216-INS-64
chr9-73549949-INS-493
chr9-73555840-INS-159
chr9-73555900-INS-983
chr9-73567084-INS-144
chr9-73578413-INS-437
chr9-73580360-INS-74
chr9-73581847-INS-160
chr9-73582470-INS-6186
chr9-73586205-INS-6437
chr9-73615300-INS-52
chr9-73623884-INS-50
chr9-73660022-INS-66
chr9-73662589-INS-78
chr9-73675226-INS-803
chr9-73677806-INS-142
chr9-73695271-INS-122
chr9-73738566-INS-57
chr9-73747973-INS-119
chr9-73753848-INS-147
chr9-73766290-INS-75
chr9-73775875-INS-208
chr9-73780135-INS-96
chr9-73822583-INS-6694
chr9-73857282-INS-711
chr9-73861632-INS-702
chr9-73865670-INS-207
chr9-73870580-INS-387
chr9-73889977-INS-77
chr9-73891659-INS-234
chr9-73923459-INS-395
chr9-73963592-INS-188
chr9-73984261-INS-52
chr9-74000425-INS-75
chr9-74001888-INS-64
chr9-74005707-INS-406
chr9-74014254-INS-1873
chr9-74016804-INS-238
chr9-74018851-INS-135
chr9-74031012-INS-61
chr9-74050283-INS-82
chr9-74096228-INS-381
chr9-74099146-INS-57
chr9-74131820-INS-310
chr9-74163408-INS-65
chr9-74192828-INS-227
chr9-74224936-INS-64
chr9-74228389-INS-51
chr9-74230852-INS-64
chr9-74240710-INS-55
chr9-74248072-INS-197
chr9-74264661-INS-102
chr9-74314617-INS-193
chr9-74317013-INS-52
chr9-74405050-INS-548
chr9-74435812-INS-140
chr9-74464179-INS-56
chr9-74480329-INS-106
chr9-74491749-INS-3523
chr9-74517244-INS-484
chr9-74534936-INS-51
chr9-74552432-INS-793
chr9-74553766-INS-6217
chr9-74566834-INS-128
chr9-74607530-INS-304
chr9-74614747-INS-60
chr9-74636380-INS-5382
chr9-74653674-INS-203
chr9-74687631-INS-2645
chr9-74687937-INS-840
chr9-74709346-INS-6699
chr9-74713865-INS-60

chr9-74732338-INS-421
chr9-74753878-INS-74
chr9-74904999-INS-448
chr9-74909572-INS-108
chr9-74912754-INS-459
chr9-74925698-INS-430
chr9-74926832-INS-61
chr9-74942382-INS-58
chr9-74948110-INS-57
chr9-74984096-INS-207
chr9-74989460-INS-1764
chr9-74990468-INS-1905
chr9-74999321-INS-54
chr9-75033583-INS-219
chr9-75058596-INS-598
chr9-75072607-INS-78
chr9-75123083-INS-379
chr9-75148938-INS-95
chr9-75206134-INS-294
chr9-75215976-INS-227
chr9-75220166-INS-74
chr9-75225672-INS-61
chr9-75227130-INS-62
chr9-75234050-INS-98
chr9-75281193-INS-179
chr9-75284137-INS-97
chr9-75288578-INS-211
chr9-75291211-INS-608
chr9-75332799-INS-211
chr9-75345159-INS-190
chr9-75345526-INS-288
chr9-75350789-INS-215
chr9-75356829-INS-164
chr9-75372895-INS-72
chr9-75374971-INS-204
chr9-75375887-INS-92
chr9-75381272-INS-176
chr9-75382010-INS-68
chr9-75384071-INS-58
chr9-75384859-INS-466
chr9-75386036-INS-165
chr9-75424513-INS-70
chr9-75434096-INS-171
chr9-75436309-INS-255
chr9-75447985-INS-154
chr9-75455253-INS-56
chr9-75515698-INS-137
chr9-75527343-INS-58
chr9-75544203-INS-834
chr9-75551599-INS-531
chr9-75561325-INS-60
chr9-75563799-INS-128
chr9-75566880-INS-53
chr9-75571741-INS-52
chr9-75578170-INS-258
chr9-75579559-INS-257
chr9-75580476-INS-1392
chr9-75599407-INS-241
chr9-75603343-INS-462
chr9-75615328-INS-979
chr9-75645477-INS-6586
chr9-75653712-INS-155
chr9-75664973-INS-188
chr9-75666973-INS-427
chr9-75670731-INS-64
chr9-75715051-INS-52
chr9-75722028-INS-84
chr9-75767201-INS-611
chr9-75783563-INS-160
chr9-75815523-INS-6390
chr9-75820646-INS-399
chr9-75852769-INS-177
chr9-75863901-INS-60
chr9-75900468-INS-361
chr9-75912048-INS-161
chr9-75928718-INS-197
chr9-75943157-INS-204

chr9-75944136-INS-110
chr9-75953157-INS-2044
chr9-75963621-INS-6254
chr9-75977357-INS-52
chr9-75983857-INS-82
chr9-76009275-INS-52
chr9-76043652-INS-2881
chr9-76052443-INS-77
chr9-76066893-INS-426
chr9-76070673-INS-51
chr9-76106634-INS-68
chr9-76117635-INS-1089
chr9-76132038-INS-126
chr9-76156101-INS-67
chr9-76167497-INS-54
chr9-76169089-INS-78
chr9-76182031-INS-7084
chr9-76183203-INS-51
chr9-76183523-INS-6605
chr9-76183863-INS-377
chr9-76186420-INS-489
chr9-76196239-INS-121
chr9-76205425-INS-590
chr9-76221984-INS-57
chr9-76239370-INS-6416
chr9-76264020-INS-6599
chr9-76264136-INS-5552
chr9-76271831-INS-148
chr9-76274604-INS-68
chr9-76276929-INS-7086
chr9-76288935-INS-69
chr9-76311856-INS-222
chr9-76323014-INS-187
chr9-76324034-INS-76
chr9-76353516-INS-322
chr9-76369680-INS-189
chr9-76384509-INS-108
chr9-76387327-INS-58
chr9-76418069-INS-119
chr9-76420464-INS-214
chr9-76423947-INS-64
chr9-76431655-INS-68
chr9-76471602-INS-212
chr9-76497281-INS-66
chr9-76510844-INS-68
chr9-76533566-INS-166
chr9-76557448-INS-78
chr9-76567647-INS-70
chr9-76573383-INS-124
chr9-76588057-INS-132
chr9-76590777-INS-69
chr9-76608671-INS-61
chr9-76630706-INS-132
chr9-76633300-INS-1281
chr9-76657362-INS-483
chr9-76690498-INS-101
chr9-76694692-INS-645
chr9-76729042-INS-5770
chr9-76730484-INS-100
chr9-76733510-INS-112
chr9-76758025-INS-804
chr9-76772056-INS-231
chr9-76776105-INS-7044
chr9-76783187-INS-68
chr9-76798287-INS-487
chr9-76817735-INS-75
chr9-76847987-INS-7756
chr9-76847991-INS-616
chr9-76852693-INS-79
chr9-76855534-INS-272
chr9-76864344-INS-54
chr9-76866824-INS-198
chr9-76896546-INS-288
chr9-76917604-INS-97
chr9-76943376-INS-56
chr9-76945665-INS-59
chr9-77013854-INS-168

chr9-77022611-INS-1881
chr9-77041715-INS-103
chr9-77115980-INS-1452
chr9-77143086-INS-352
chr9-77173996-INS-586
chr9-77178934-INS-55
chr9-77192392-INS-106
chr9-77200304-INS-65
chr9-77208303-INS-182
chr9-77209283-INS-218
chr9-77237432-INS-212
chr9-77271881-INS-61
chr9-77272527-INS-666
chr9-77277555-INS-260
chr9-77287711-INS-341
chr9-77289272-INS-264
chr9-77302924-INS-60
chr9-77315182-INS-51
chr9-77353817-INS-150
chr9-77371814-INS-67
chr9-77372822-INS-76
chr9-77388347-INS-6882
chr9-77397116-INS-100
chr9-77427724-INS-70
chr9-77430743-INS-144
chr9-77438459-INS-228
chr9-77440153-INS-120
chr9-77463242-INS-134
chr9-77493171-INS-95
chr9-77494961-INS-7116
chr9-77501222-INS-74
chr9-77514422-INS-70
chr9-77517070-INS-222
chr9-77518350-INS-713
chr9-77521917-INS-311
chr9-77560140-INS-1043
chr9-77575321-INS-126
chr9-77592073-INS-127
chr9-77599234-INS-2948
chr9-77617178-INS-55
chr9-77630578-INS-831
chr9-77643358-INS-104
chr9-77644312-INS-54
chr9-77661573-INS-166
chr9-77692071-INS-165
chr9-77710927-INS-114
chr9-77713651-INS-85
chr9-77718152-INS-263
chr9-77718496-INS-65
chr9-77724811-INS-68
chr9-77727479-INS-100
chr9-77734181-INS-144
chr9-77795081-INS-120
chr9-77803393-INS-62
chr9-77807875-INS-112
chr9-77815223-INS-298
chr9-77816640-INS-636
chr9-77840542-INS-95
chr9-77840814-INS-201
chr9-77842941-INS-78
chr9-77850258-INS-60
chr9-77920099-INS-125
chr9-77926223-INS-88
chr9-77940701-INS-68
chr9-77947731-INS-221
chr9-77967431-INS-261
chr9-77978742-INS-176
chr9-78003607-INS-4636
chr9-78012110-INS-136
chr9-78014593-INS-185
chr9-78029289-INS-379
chr9-78035575-INS-249
chr9-78042036-INS-101
chr9-78068246-INS-181
chr9-78072998-INS-64
chr9-78079543-INS-472
chr9-78082077-INS-89

chr9-78092006-INS-233
chr9-78122076-INS-333
chr9-78123698-INS-212
chr9-78128468-INS-220
chr9-78146201-INS-222
chr9-78173054-INS-203
chr9-78174428-INS-62
chr9-78177333-INS-54
chr9-78180415-INS-174
chr9-78199006-INS-249
chr9-78201569-INS-79
chr9-78214718-INS-54
chr9-78217953-INS-92
chr9-78226847-INS-74
chr9-78238051-INS-68
chr9-78249767-INS-171
chr9-78252196-INS-168
chr9-78271620-INS-103
chr9-78279685-INS-109
chr9-78283945-INS-202
chr9-78287249-INS-172
chr9-78288047-INS-87
chr9-78312409-INS-353
chr9-78323884-INS-325
chr9-78324439-INS-161
chr9-78345917-INS-209
chr9-78352590-INS-375
chr9-78362285-INS-190
chr9-78367695-INS-203
chr9-78370875-INS-182
chr9-78375106-INS-196
chr9-78376416-INS-191
chr9-78379452-INS-198
chr9-78384455-INS-539
chr9-78393006-INS-218
chr9-78402756-INS-127
chr9-78405412-INS-213
chr9-78433326-INS-193
chr9-78433897-INS-191
chr9-78449375-INS-192
chr9-78456778-INS-94
chr9-78460033-INS-69
chr9-78479198-INS-51
chr9-78502596-INS-441
chr9-78516181-INS-195
chr9-78525694-INS-108
chr9-78531290-INS-1472
chr9-78531389-INS-84
chr9-78549523-INS-7138
chr9-78579597-INS-52
chr9-78581312-INS-189
chr9-78599569-INS-90
chr9-78599991-INS-522
chr9-78607933-INS-106
chr9-78647585-INS-251
chr9-78672406-INS-84
chr9-78723236-INS-308
chr9-78738923-INS-8294
chr9-78741471-INS-6469
chr9-78741487-INS-714
chr9-78756152-INS-2109
chr9-78786134-INS-276
chr9-78800726-INS-302
chr9-78801369-INS-239
chr9-78803159-INS-72
chr9-78806140-INS-214
chr9-78818285-INS-116
chr9-78828787-INS-82
chr9-78854549-INS-300
chr9-78855801-INS-54
chr9-78862313-INS-132
chr9-78870845-INS-60
chr9-78889168-INS-52
chr9-78892917-INS-311
chr9-78894365-INS-54
chr9-78905309-INS-309
chr9-78932326-INS-1456

chr9-78934435-INS-6960
chr9-78935767-INS-5448
chr9-78947316-INS-104
chr9-78962089-INS-54
chr9-78966312-INS-83
chr9-78976344-INS-68
chr9-78980679-INS-55
chr9-78982577-INS-448
chr9-78983194-INS-51
chr9-78999783-INS-52
chr9-79013951-INS-288
chr9-79015683-INS-56
chr9-79027196-INS-64
chr9-79033602-INS-167
chr9-79055714-INS-331
chr9-79077712-INS-498
chr9-79090473-INS-558
chr9-79093161-INS-121
chr9-79111280-INS-57
chr9-79129001-INS-81
chr9-79137673-INS-104
chr9-79206931-INS-6681
chr9-79245846-INS-61
chr9-79260248-INS-226
chr9-79270560-INS-72
chr9-79291317-INS-70
chr9-79295335-INS-144
chr9-79305196-INS-461
chr9-79309541-INS-117
chr9-79320245-INS-68
chr9-79342213-INS-58
chr9-79346538-INS-57
chr9-79351256-INS-67
chr9-79368342-INS-61
chr9-79398844-INS-54
chr9-79409161-INS-187
chr9-79421194-INS-312
chr9-79440978-INS-4893
chr9-79462872-INS-306
chr9-79464455-INS-58
chr9-79466011-INS-64
chr9-79468623-INS-68
chr9-79493191-INS-104
chr9-79510146-INS-51
chr9-79512136-INS-89
chr9-79519548-INS-662
chr9-79525975-INS-98
chr9-79541708-INS-68
chr9-79545012-INS-108
chr9-79560346-INS-400
chr9-79575536-INS-91
chr9-79601982-INS-66
chr9-79640103-INS-278
chr9-79643610-INS-268
chr9-79647213-INS-102
chr9-79680437-INS-759
chr9-79696305-INS-168
chr9-79703090-INS-197
chr9-79731128-INS-161
chr9-79760203-INS-82
chr9-79763122-INS-123
chr9-79778592-INS-935
chr9-79779337-INS-382
chr9-79785725-INS-68
chr9-79794928-INS-175
chr9-79839779-INS-460
chr9-79843514-INS-196
chr9-79854530-INS-66
chr9-79855774-INS-54
chr9-79869713-INS-129
chr9-79870153-INS-514
chr9-79893227-INS-211
chr9-79973393-INS-158
chr9-79984688-INS-70
chr9-79992445-INS-5203
chr9-80019165-INS-287
chr9-80026646-INS-230

chr9-80027263-INS-644
chr9-80056874-INS-58
chr9-80059259-INS-54
chr9-80061665-INS-295
chr9-80065963-INS-116
chr9-80074400-INS-76
chr9-80080160-INS-84
chr9-80101132-INS-52
chr9-80103024-INS-56
chr9-80125048-INS-70
chr9-80147371-INS-199
chr9-80187812-INS-165
chr9-80239066-INS-196
chr9-80251470-INS-177
chr9-80259806-INS-218
chr9-80265672-INS-3286
chr9-80268393-INS-60
chr9-80278710-INS-542
chr9-80297829-INS-282
chr9-80349256-INS-232
chr9-80360647-INS-204
chr9-80362646-INS-563
chr9-80368973-INS-52
chr9-80398597-INS-219
chr9-80399088-INS-369
chr9-80423650-INS-439
chr9-80438803-INS-112
chr9-80447939-INS-107
chr9-80457576-INS-220
chr9-80496689-INS-81
chr9-80556542-INS-67
chr9-80586668-INS-186
chr9-80597900-INS-99
chr9-80627137-INS-375
chr9-80659925-INS-76
chr9-80689378-INS-94
chr9-80704163-INS-54
chr9-80705592-INS-70
chr9-80718945-INS-64
chr9-80720341-INS-4597
chr9-80726207-INS-59
chr9-80740248-INS-673
chr9-80763321-INS-196
chr9-80763321-INS-967
chr9-80778752-INS-57
chr9-80789845-INS-69
chr9-80792907-INS-371
chr9-80801324-INS-531
chr9-80812322-INS-131
chr9-80851206-INS-6298
chr9-80980170-INS-728
chr9-81005516-INS-185
chr9-81056066-INS-112
chr9-81073280-INS-668
chr9-81083892-INS-6568
chr9-81148705-INS-74
chr9-81157158-INS-85
chr9-81204670-INS-203
chr9-81286476-INS-257
chr9-81300902-INS-1945
chr9-81337014-INS-108
chr9-81341944-INS-439
chr9-81343689-INS-402
chr9-81355426-INS-66
chr9-81363598-INS-219
chr9-81380685-INS-54
chr9-81391856-INS-10424
chr9-81397421-INS-1761
chr9-81412053-INS-69
chr9-81452580-INS-106
chr9-81493040-INS-456
chr9-81550816-INS-60
chr9-81552247-INS-96
chr9-81555355-INS-229
chr9-81605623-INS-58
chr9-81607385-INS-264
chr9-81624351-INS-6561

chr9-81629377-INS-6574
chr9-81629511-INS-110
chr9-81630935-INS-4555
chr9-81643500-INS-93
chr9-81654310-INS-55
chr9-81659847-INS-118
chr9-81669033-INS-170
chr9-81674057-INS-180
chr9-81698641-INS-54
chr9-81704701-INS-63
chr9-81713521-INS-228
chr9-81732542-INS-59
chr9-81736306-INS-214
chr9-81785964-INS-968
chr9-81882483-INS-125
chr9-81887261-INS-1461
chr9-81888154-INS-7017
chr9-81898673-INS-227
chr9-81902316-INS-259
chr9-81933284-INS-857
chr9-81946853-INS-531
chr9-81958630-INS-128
chr9-81965855-INS-382
chr9-81972860-INS-59
chr9-81979198-INS-207
chr9-81987655-INS-359
chr9-82018230-INS-64
chr9-82028010-INS-216
chr9-82040376-INS-54
chr9-82090146-INS-272
chr9-82105350-INS-79
chr9-82107363-INS-192
chr9-82119434-INS-6798
chr9-82139548-INS-865
chr9-82146779-INS-141
chr9-82190912-INS-66
chr9-82203038-INS-62
chr9-82215648-INS-222
chr9-82225032-INS-464
chr9-82233023-INS-109
chr9-82248637-INS-52
chr9-82296743-INS-64
chr9-82299728-INS-85
chr9-82310382-INS-980
chr9-82350513-INS-531
chr9-82387061-INS-498
chr9-82398685-INS-55
chr9-82406878-INS-181
chr9-82411263-INS-74
chr9-82412923-INS-75
chr9-82415643-INS-131
chr9-82431768-INS-52
chr9-82437355-INS-84
chr9-82467693-INS-52
chr9-82487013-INS-53
chr9-82493518-INS-140
chr9-82498131-INS-125
chr9-82527914-INS-605
chr9-82531855-INS-459
chr9-82548259-INS-70
chr9-82553357-INS-55
chr9-82557113-INS-54
chr9-82562923-INS-62
chr9-82570545-INS-55
chr9-82601356-INS-54
chr9-82626436-INS-749
chr9-82631614-INS-2764
chr9-82645552-INS-998
chr9-82646869-INS-54
chr9-82708241-INS-155
chr9-82712677-INS-132
chr9-82732412-INS-6759
chr9-82735975-INS-212
chr9-82779106-INS-57
chr9-82816018-INS-145
chr9-82819485-INS-100
chr9-82820931-INS-139

chr9-82842321-INS-56
chr9-82852210-INS-56
chr9-82862114-INS-147
chr9-82878840-INS-145
chr9-82902701-INS-7252
chr9-82912820-INS-442
chr9-82937559-INS-151
chr9-82944226-INS-6252
chr9-82961762-INS-6787
chr9-82962598-INS-64
chr9-82970604-INS-717
chr9-83012624-INS-76
chr9-83037762-INS-64
chr9-83107761-INS-371
chr9-83136391-INS-52
chr9-83136688-INS-877
chr9-83137837-INS-558
chr9-83141136-INS-441
chr9-83144541-INS-54
chr9-83144886-INS-794
chr9-83155203-INS-68
chr9-83175249-INS-73
chr9-83200918-INS-5138
chr9-83212673-INS-140
chr9-83221341-INS-60
chr9-83226977-INS-309
chr9-83233826-INS-73
chr9-83240752-INS-120
chr9-83270966-INS-138
chr9-83299288-INS-54
chr9-83334367-INS-419
chr9-83350372-INS-56
chr9-83364488-INS-205
chr9-83373234-INS-103
chr9-83383669-INS-100
chr9-83396546-INS-59
chr9-83414211-INS-83
chr9-83417169-INS-486
chr9-83421908-INS-370
chr9-83441991-INS-158
chr9-83521558-INS-52
chr9-83537079-INS-409
chr9-83542627-INS-441
chr9-83555968-INS-199
chr9-83570497-INS-285
chr9-83608170-INS-72
chr9-83647132-INS-166
chr9-83657209-INS-768
chr9-83679416-INS-60
chr9-83684193-INS-92
chr9-83697329-INS-115
chr9-83742850-INS-54
chr9-83788529-INS-122
chr9-83794389-INS-504
chr9-83803212-INS-96
chr9-83809184-INS-52
chr9-83821468-INS-89
chr9-83833138-INS-52
chr9-83871129-INS-916
chr9-83920521-INS-58
chr9-83937148-INS-61
chr9-84000696-INS-72
chr9-84008215-INS-64
chr9-84011823-INS-54
chr9-84020303-INS-139
chr9-84033080-INS-440
chr9-84072906-INS-5554
chr9-84083668-INS-51
chr9-84095920-INS-355
chr9-84151183-INS-157
chr9-84203122-INS-192
chr9-84220362-INS-88
chr9-84237977-INS-196
chr9-84292808-INS-52
chr9-84294496-INS-56
chr9-84299297-INS-1894
chr9-84345574-INS-86

chr9-84381678-INS-768
chr9-84382672-INS-244
chr9-84389035-INS-5253
chr9-84414930-INS-119
chr9-84416957-INS-112
chr9-84419635-INS-56
chr9-84425214-INS-5054
chr9-84483842-INS-3187
chr9-84490641-INS-110
chr9-84536790-INS-871
chr9-84554298-INS-60
chr9-84563147-INS-4592
chr9-84567698-INS-67
chr9-84571853-INS-715
chr9-84588819-INS-3544
chr9-84594977-INS-5178
chr9-84603321-INS-52
chr9-84609823-INS-206
chr9-84634407-INS-166
chr9-84669601-INS-98
chr9-84700361-INS-52
chr9-84711735-INS-3315
chr9-84724043-INS-702
chr9-84740276-INS-438
chr9-84751462-INS-61
chr9-84755212-INS-168
chr9-84762556-INS-5875
chr9-84773533-INS-79
chr9-84776837-INS-6591
chr9-84796151-INS-245
chr9-84802033-INS-102
chr9-84804734-INS-7376
chr9-84812509-INS-80
chr9-84822821-INS-91
chr9-84840481-INS-489
chr9-84845765-INS-336
chr9-84852898-INS-88
chr9-84860177-INS-2823
chr9-84867200-INS-613
chr9-84880937-INS-6185
chr9-84884028-INS-5632
chr9-84892167-INS-259
chr9-84943227-INS-3810
chr9-84943904-INS-1603
chr9-84997109-INS-128
chr9-84998468-INS-120
chr9-85004732-INS-919
chr9-85012498-INS-58
chr9-85016048-INS-60
chr9-85019900-INS-194
chr9-85020895-INS-53
chr9-85041897-INS-66
chr9-85053517-INS-1166
chr9-85053804-INS-307
chr9-85063888-INS-74
chr9-85066469-INS-54
chr9-85080771-INS-720
chr9-85114987-INS-124
chr9-85136782-INS-203
chr9-85142859-INS-134
chr9-85160361-INS-298
chr9-85193503-INS-393
chr9-85195463-INS-183
chr9-85253794-INS-73
chr9-85255057-INS-53
chr9-85262091-INS-72
chr9-85265414-INS-52
chr9-85267814-INS-54
chr9-85285584-INS-90
chr9-85297498-INS-6436
chr9-85306932-INS-5358
chr9-85324077-INS-55
chr9-85327314-INS-58
chr9-85336261-INS-81
chr9-85340357-INS-455
chr9-85353054-INS-120
chr9-85355990-INS-692

chr9-85356829-INS-3906
chr9-85363932-INS-6375
chr9-85389588-INS-338
chr9-85399782-INS-56
chr9-85406034-INS-297
chr9-85416174-INS-660
chr9-85422868-INS-6492
chr9-85438008-INS-7698
chr9-85460569-INS-51
chr9-85483413-INS-67
chr9-85532031-INS-62
chr9-85545236-INS-66
chr9-85546370-INS-67
chr9-85566358-INS-372
chr9-85572441-INS-2532
chr9-85583799-INS-148
chr9-85608347-INS-812
chr9-85622434-INS-352
chr9-85633537-INS-385
chr9-85636018-INS-399
chr9-85638021-INS-110
chr9-85641817-INS-60
chr9-85649957-INS-831
chr9-85666525-INS-560
chr9-85670141-INS-246
chr9-85733738-INS-167
chr9-85736148-INS-104
chr9-85772897-INS-77
chr9-85778366-INS-88
chr9-85781239-INS-212
chr9-85790067-INS-202
chr9-85813752-INS-59
chr9-85877992-INS-146
chr9-85882206-INS-276
chr9-85961420-INS-87
chr9-85999782-INS-6956
chr9-86010508-INS-6893
chr9-86091808-INS-66
chr9-86103234-INS-1880
chr9-86109605-INS-293
chr9-86137048-INS-469
chr9-86139507-INS-438
chr9-86144533-INS-426
chr9-86159062-INS-232
chr9-86164598-INS-56
chr9-86202288-INS-82
chr9-86216208-INS-152
chr9-86219823-INS-76
chr9-86240245-INS-445
chr9-86253152-INS-72
chr9-86266773-INS-6592
chr9-86278404-INS-84
chr9-86279515-INS-198
chr9-86290328-INS-555
chr9-86303862-INS-216
chr9-86313277-INS-109
chr9-86316630-INS-179
chr9-86326374-INS-82
chr9-86331921-INS-62
chr9-86337154-INS-62
chr9-86348774-INS-297
chr9-86447898-INS-108
chr9-86470856-INS-55
chr9-86496099-INS-290
chr9-86519380-INS-462
chr9-86542951-INS-331
chr9-86543549-INS-131
chr9-86545484-INS-223
chr9-86547143-INS-108
chr9-86553121-INS-822
chr9-86556466-INS-185
chr9-86563572-INS-200
chr9-86572204-INS-59
chr9-86597165-INS-52
chr9-86617156-INS-104
chr9-86621122-INS-59
chr9-86623086-INS-714

chr9-86642871-INS-445
chr9-86653548-INS-254
chr9-86726154-INS-378
chr9-86787371-INS-60
chr9-86794254-INS-492
chr9-86807228-INS-361
chr9-86807394-INS-6686
chr9-86816369-INS-238
chr9-86822245-INS-1899
chr9-86841953-INS-62
chr9-86846553-INS-62
chr9-86906331-INS-266
chr9-86957731-INS-100
chr9-86977922-INS-232
chr9-87025499-INS-99
chr9-87033401-INS-99
chr9-87035168-INS-77
chr9-87058538-INS-6711
chr9-87083991-INS-366
chr9-87087719-INS-203
chr9-87098664-INS-6690
chr9-87101420-INS-1773
chr9-87148792-INS-1332
chr9-87150405-INS-52
chr9-87154363-INS-254
chr9-87173110-INS-65
chr9-87186461-INS-215
chr9-87212530-INS-219
chr9-87218120-INS-239
chr9-87231790-INS-328
chr9-87275348-INS-92
chr9-87281181-INS-865
chr9-87292489-INS-7436
chr9-87294721-INS-60
chr9-87305906-INS-224
chr9-87321924-INS-56
chr9-87356722-INS-52
chr9-87385779-INS-7256
chr9-87390751-INS-398
chr9-87395291-INS-394
chr9-87417847-INS-126
chr9-87442797-INS-244
chr9-87447386-INS-156
chr9-87467230-INS-100
chr9-87491041-INS-96
chr9-87491592-INS-224
chr9-87498391-INS-234
chr9-87529963-INS-6544
chr9-87541529-INS-1892
chr9-87568119-INS-498
chr9-87632870-INS-230
chr9-87634473-INS-67
chr9-87691849-INS-64
chr9-87708119-INS-222
chr9-87708904-INS-77
chr9-87725780-INS-189
chr9-87760460-INS-6222
chr9-87768738-INS-364
chr9-87769598-INS-4435
chr9-87789141-INS-670
chr9-87792315-INS-682
chr9-87792972-INS-312
chr9-87800775-INS-127
chr9-87820099-INS-105
chr9-87824412-INS-201
chr9-87845699-INS-58
chr9-87849292-INS-70
chr9-87855853-INS-651
chr9-87856861-INS-60
chr9-87863709-INS-121
chr9-87879168-INS-6492
chr9-87884656-INS-104
chr9-87899610-INS-5378
chr9-87901658-INS-147
chr9-87916132-INS-3128
chr9-87917542-INS-93
chr9-87933150-INS-145

chr9-87972532-INS-124
chr9-87976448-INS-559
chr9-87978453-INS-534
chr9-88006737-INS-68
chr9-88053693-INS-54
chr9-88066307-INS-344
chr9-88091779-INS-596
chr9-88091799-INS-52
chr9-88092338-INS-7080
chr9-88105493-INS-8081
chr9-88105769-INS-88
chr9-88115393-INS-396
chr9-88139932-INS-233
chr9-88152676-INS-449
chr9-88155143-INS-124
chr9-88162479-INS-70
chr9-88165934-INS-113
chr9-88173240-INS-54
chr9-88177926-INS-58
chr9-88194151-INS-153
chr9-88218536-INS-434
chr9-88240077-INS-76
chr9-88254347-INS-94
chr9-88285039-INS-298
chr9-88297249-INS-187
chr9-88306775-INS-223
chr9-88315310-INS-141
chr9-88352811-INS-213
chr9-88370349-INS-362
chr9-88392350-INS-67
chr9-88397160-INS-802
chr9-88398981-INS-66
chr9-88402652-INS-546
chr9-88420809-INS-194
chr9-88427175-INS-198
chr9-88437293-INS-340
chr9-88442848-INS-776
chr9-88444748-INS-52
chr9-88503804-INS-53
chr9-88578553-INS-6420
chr9-88621066-INS-861
chr9-88637507-INS-67
chr9-88814606-INS-572
chr9-88818629-INS-7107
chr9-88910566-INS-248
chr9-88945990-INS-342
chr9-88972764-INS-5371
chr9-88980555-INS-61
chr9-88996383-INS-5559
chr9-89044548-INS-65
chr9-89091539-INS-186
chr9-89160200-INS-127
chr9-89168331-INS-60
chr9-89174904-INS-429
chr9-89220719-INS-111
chr9-89228867-INS-506
chr9-89230175-INS-115
chr9-89251552-INS-749
chr9-89261992-INS-622
chr9-89262021-INS-63
chr9-89272344-INS-127
chr9-89279851-INS-105
chr9-89296155-INS-135
chr9-89309088-INS-727
chr9-89323180-INS-6876
chr9-89331109-INS-6276
chr9-89338182-INS-84
chr9-89365200-INS-220
chr9-89387272-INS-52
chr9-89388108-INS-279
chr9-89429843-INS-293
chr9-89470342-INS-224
chr9-89512919-INS-170
chr9-89527969-INS-67
chr9-89555153-INS-103
chr9-89576060-INS-245
chr9-89601663-INS-6543

chr9-89629705-INS-433
chr9-89632439-INS-7089
chr9-89637203-INS-60
chr9-89657765-INS-54
chr9-89660704-INS-76
chr9-89711011-INS-146
chr9-89721254-INS-51
chr9-89738897-INS-59
chr9-89752058-INS-52
chr9-89762023-INS-92
chr9-89762502-INS-272
chr9-89763444-INS-143
chr9-89769321-INS-52
chr9-89781613-INS-94
chr9-89785933-INS-196
chr9-89795860-INS-59
chr9-89811256-INS-68
chr9-89842014-INS-134
chr9-89864789-INS-6416
chr9-89871424-INS-206
chr9-89929947-INS-99
chr9-89932376-INS-177
chr9-89942228-INS-5271
chr9-89953512-INS-8734
chr9-89954907-INS-70
chr9-89965052-INS-197
chr9-89987968-INS-198
chr9-89989714-INS-205
chr9-89994845-INS-640
chr9-90003798-INS-1737
chr9-90012038-INS-208
chr9-90046395-INS-168
chr9-90093298-INS-63
chr9-90125595-INS-55
chr9-90136800-INS-400
chr9-90138389-INS-118
chr9-90183947-INS-370
chr9-90228594-INS-3371
chr9-90255775-INS-72
chr9-90273259-INS-7120
chr9-90282257-INS-91
chr9-90300656-INS-358
chr9-90323535-INS-61
chr9-90327579-INS-6667
chr9-90328949-INS-174
chr9-90349407-INS-570
chr9-90372023-INS-630
chr9-90382715-INS-7061
chr9-90408387-INS-207
chr9-90425934-INS-234
chr9-90426596-INS-2416
chr9-90427356-INS-190
chr9-90427592-INS-6236
chr9-90450257-INS-68
chr9-90450897-INS-439
chr9-90451448-INS-13648
chr9-90451721-INS-543
chr9-90454124-INS-227
chr9-90455939-INS-241
chr9-90458607-INS-88
chr9-90459950-INS-82
chr9-90481876-INS-236
chr9-90495198-INS-93
chr9-90536612-INS-678
chr9-90555369-INS-140
chr9-90561519-INS-371
chr9-90568366-INS-7095
chr9-90605721-INS-60
chr9-90609232-INS-877
chr9-90618606-INS-88
chr9-90628368-INS-106
chr9-90637456-INS-163
chr9-90649617-INS-105
chr9-90656901-INS-252
chr9-90660151-INS-2430
chr9-90676197-INS-204
chr9-90688127-INS-144

chr9-90702870-INS-343
chr9-90719104-INS-51
chr9-90734563-INS-168
chr9-90765737-INS-84
chr9-90766901-INS-5342
chr9-90770967-INS-82
chr9-90808609-INS-441
chr9-90821382-INS-76
chr9-90830010-INS-230
chr9-90848579-INS-88
chr9-90851714-INS-139
chr9-90875093-INS-58
chr9-90899236-INS-1570
chr9-90918990-INS-7115
chr9-90923859-INS-104
chr9-90931038-INS-69
chr9-90937031-INS-78
chr9-90955256-INS-53
chr9-90962068-INS-51
chr9-90974858-INS-360
chr9-90993755-INS-7813
chr9-91002899-INS-55
chr9-91004342-INS-7263
chr9-91007560-INS-276
chr9-91018051-INS-6671
chr9-91026160-INS-6456
chr9-91042637-INS-182
chr9-91081434-INS-6906
chr9-91083911-INS-9762
chr9-91125454-INS-206
chr9-91128203-INS-6472
chr9-91135389-INS-6211
chr9-91136414-INS-6141
chr9-91136924-INS-4521
chr9-91139083-INS-52
chr9-91151919-INS-7144
chr9-91180879-INS-614
chr9-91195395-INS-1017
chr9-91203329-INS-1212
chr9-91207475-INS-6412
chr9-91212068-INS-120
chr9-91222233-INS-7575
chr9-91228139-INS-80
chr9-91263141-INS-56
chr9-91306833-INS-366
chr9-91323121-INS-6475
chr9-91334478-INS-1185
chr9-91341948-INS-52
chr9-91372419-INS-261
chr9-91379129-INS-58
chr9-91385140-INS-4175
chr9-91400199-INS-6821
chr9-91404794-INS-82
chr9-91421472-INS-132
chr9-91431315-INS-57
chr9-91436983-INS-7637
chr9-91437702-INS-60
chr9-91482287-INS-120
chr9-91485215-INS-207
chr9-91494194-INS-8162
chr9-91530003-INS-63
chr9-91530924-INS-432
chr9-91547314-INS-6331
chr9-91558826-INS-80
chr9-91562201-INS-116
chr9-91567087-INS-108
chr9-91567624-INS-4953
chr9-91584587-INS-71
chr9-91589810-INS-63
chr9-91599692-INS-1491
chr9-91638986-INS-295
chr9-91642531-INS-97
chr9-91657039-INS-52
chr9-91684407-INS-1308
chr9-91715556-INS-62
chr9-91727641-INS-325
chr9-91734209-INS-70

chr9-91752887-INS-6182
chr9-91757460-INS-267
chr9-91764244-INS-471
chr9-91812670-INS-52
chr9-91817111-INS-536
chr9-91820162-INS-405
chr9-91851854-INS-9720
chr9-91853394-INS-1211
chr9-91858222-INS-337
chr9-91873750-INS-484
chr9-91890424-INS-143
chr9-91922291-INS-53
chr9-91932283-INS-6689
chr9-91944130-INS-54
chr9-91951400-INS-351
chr9-92011632-INS-827
chr9-92031411-INS-1334
chr9-92034969-INS-439
chr9-92040606-INS-158
chr9-92050413-INS-780
chr9-92073334-INS-217
chr9-92116596-INS-240
chr9-92116784-INS-389
chr9-92146796-INS-72
chr9-92161193-INS-120
chr9-92167731-INS-87
chr9-92198513-INS-312
chr9-92206488-INS-922
chr9-92220840-INS-6297
chr9-92225122-INS-163
chr9-92228633-INS-740
chr9-92232692-INS-56
chr9-92236372-INS-190
chr9-92251368-INS-6543
chr9-92257577-INS-558
chr9-92261202-INS-5241
chr9-92295462-INS-331
chr9-92334945-INS-424
chr9-92360467-INS-77
chr9-92368949-INS-529
chr9-92371569-INS-378
chr9-92383368-INS-50
chr9-92405482-INS-406
chr9-92415682-INS-6401
chr9-92418934-INS-57
chr9-92482301-INS-63
chr9-92491445-INS-66
chr9-92492659-INS-2959
chr9-92522740-INS-211
chr9-92524856-INS-73
chr9-92539821-INS-130
chr9-92543181-INS-525
chr9-92552348-INS-150
chr9-92568569-INS-3513
chr9-92574027-INS-227
chr9-92606883-INS-502
chr9-92611939-INS-87
chr9-92635694-INS-433
chr9-92644704-INS-64
chr9-92646246-INS-273
chr9-92649289-INS-66
chr9-92663062-INS-59
chr9-92710511-INS-190
chr9-92712868-INS-7111
chr9-92713956-INS-411
chr9-92755811-INS-362
chr9-92757938-INS-822
chr9-92809441-INS-54
chr9-92814729-INS-438
chr9-92851776-INS-85
chr9-92864679-INS-2606
chr9-92883800-INS-4908
chr9-92906952-INS-83
chr9-92908043-INS-225
chr9-92908839-INS-56
chr9-92926453-INS-727
chr9-92930081-INS-6176

chr9-92947943-INS-54
chr9-92984385-INS-12482
chr9-92984869-INS-988
chr9-92990826-INS-481
chr9-92993724-INS-54
chr9-93009440-INS-555
chr9-93017114-INS-4606
chr9-93029335-INS-52
chr9-93032914-INS-549
chr9-93058400-INS-76
chr9-93077585-INS-254
chr9-93089535-INS-56
chr9-93093991-INS-68
chr9-93097423-INS-64
chr9-93107066-INS-109
chr9-93113337-INS-60
chr9-93118918-INS-6788
chr9-93126742-INS-58
chr9-93148998-INS-64
chr9-93150664-INS-196
chr9-93183590-INS-540
chr9-93217946-INS-6734
chr9-93257852-INS-2301
chr9-93271650-INS-388
chr9-93307181-INS-6645
chr9-93347571-INS-138
chr9-93424663-INS-92
chr9-93447198-INS-114
chr9-93449890-INS-96
chr9-93458742-INS-71
chr9-93465008-INS-391
chr9-93480440-INS-559
chr9-93515782-INS-65
chr9-93534557-INS-54
chr9-93542164-INS-78
chr9-93555236-INS-187
chr9-93570993-INS-6893
chr9-93574291-INS-3402
chr9-93602470-INS-524
chr9-93634962-INS-143
chr9-93638926-INS-5264
chr9-93650986-INS-537
chr9-93652449-INS-524
chr9-93665069-INS-5357
chr9-93680143-INS-548
chr9-93682112-INS-6517
chr9-93724551-INS-298
chr9-93744468-INS-690
chr9-93750336-INS-58
chr9-93754237-INS-6594
chr9-93755077-INS-678
chr9-93756875-INS-229
chr9-93764638-INS-6612
chr9-93765203-INS-269
chr9-93769487-INS-6227
chr9-93772490-INS-6428
chr9-93783007-INS-1547
chr9-93789737-INS-2960
chr9-93790323-INS-383
chr9-93817409-INS-408
chr9-93833900-INS-123
chr9-93838950-INS-799
chr9-93855230-INS-193
chr9-93864735-INS-1131
chr9-93875218-INS-2945
chr9-93880787-INS-4352
chr9-93884009-INS-643
chr9-93885484-INS-59
chr9-93888091-INS-77
chr9-93897380-INS-137
chr9-93956787-INS-6407
chr9-93965560-INS-549
chr9-93975257-INS-56
chr9-93989792-INS-186
chr9-94015488-INS-5543
chr9-94023449-INS-954
chr9-94030978-INS-399

chr9-94039846-INS-107
chr9-94045631-INS-53
chr9-94063243-INS-94
chr9-94093438-INS-94
chr9-94103496-INS-56
chr9-94133731-INS-562
chr9-94141778-INS-82
chr9-94185823-INS-84
chr9-94267881-INS-5211
chr9-94304884-INS-594
chr9-94337329-INS-53
chr9-94387166-INS-332
chr9-94423565-INS-55
chr9-94463429-INS-52
chr9-94485243-INS-62
chr9-94486715-INS-360
chr9-94505942-INS-201
chr9-94536532-INS-947
chr9-94541620-INS-124
chr9-94580906-INS-6239
chr9-94586510-INS-4890
chr9-94592286-INS-74
chr9-94610392-INS-63
chr9-94616271-INS-54
chr9-94631954-INS-218
chr9-94643733-INS-91
chr9-94681345-INS-6732
chr9-94702073-INS-109
chr9-94725328-INS-537
chr9-94741784-INS-78
chr9-94769287-INS-136
chr9-94771549-INS-65
chr9-94774105-INS-114
chr9-94794970-INS-132
chr9-94812685-INS-117
chr9-94875715-INS-113
chr9-94876359-INS-7658
chr9-94878349-INS-357
chr9-94887651-INS-5294
chr9-94939294-INS-6401
chr9-94943019-INS-293
chr9-94951762-INS-6769
chr9-94953762-INS-68
chr9-94964106-INS-180
chr9-94989469-INS-99
chr9-94991769-INS-76
chr9-94992493-INS-1890
chr9-95030705-INS-6258
chr9-95036205-INS-454
chr9-95046582-INS-665
chr9-95049386-INS-91
chr9-95052038-INS-7018
chr9-95054381-INS-4053
chr9-95083768-INS-6211
chr9-95098037-INS-1615
chr9-95101429-INS-60
chr9-95110689-INS-112
chr9-95127236-INS-141
chr9-95139042-INS-7332
chr9-95140915-INS-6474
chr9-95147692-INS-144
chr9-95147987-INS-7037
chr9-95181892-INS-168
chr9-95204090-INS-66
chr9-95212627-INS-52
chr9-95224578-INS-100
chr9-95227023-INS-324
chr9-95229721-INS-105
chr9-95246273-INS-53
chr9-95246520-INS-367
chr9-95290495-INS-199
chr9-95296962-INS-6374
chr9-95304758-INS-60
chr9-95312364-INS-204
chr9-95348994-INS-262
chr9-95367999-INS-68
chr9-95386952-INS-269

chr9-95410659-INS-198
chr9-95412319-INS-203
chr9-95415515-INS-166
chr9-95420290-INS-321
chr9-95425825-INS-84
chr9-95426806-INS-241
chr9-95432821-INS-188
chr9-95464126-INS-139
chr9-95470797-INS-278
chr9-95485563-INS-147
chr9-95486809-INS-5385
chr9-95488982-INS-153
chr9-95490302-INS-184
chr9-95496621-INS-251
chr9-95497714-INS-70
chr9-95535272-INS-75
chr9-95556822-INS-2941
chr9-95568429-INS-358
chr9-95581060-INS-82
chr9-95617732-INS-71
chr9-95648156-INS-216
chr9-95658420-INS-52
chr9-95659485-INS-105
chr9-95672062-INS-314
chr9-95676743-INS-58
chr9-95681322-INS-498
chr9-95687497-INS-161
chr9-95694604-INS-58
chr9-95713714-INS-51
chr9-95718585-INS-196
chr9-95720876-INS-172
chr9-95729392-INS-209
chr9-95733353-INS-204
chr9-95784352-INS-76
chr9-95794788-INS-54
chr9-95852018-INS-380
chr9-95871597-INS-660
chr9-95872708-INS-75
chr9-95882916-INS-230
chr9-95887533-INS-58
chr9-95892823-INS-52
chr9-95896794-INS-115
chr9-95898073-INS-516
chr9-95904005-INS-210
chr9-95950842-INS-636
chr9-95954289-INS-410
chr9-95959728-INS-199
chr9-95962203-INS-411
chr9-95968177-INS-200
chr9-95969414-INS-84
chr9-95969660-INS-654
chr9-95986719-INS-584
chr9-96002694-INS-54
chr9-96008352-INS-213
chr9-96008907-INS-80
chr9-96012909-INS-135
chr9-96022978-INS-52
chr9-96027962-INS-129
chr9-96031335-INS-1831
chr9-96077338-INS-202
chr9-96085183-INS-176
chr9-96094600-INS-5099
chr9-96129276-INS-228
chr9-96133273-INS-717
chr9-96134963-INS-189
chr9-96158405-INS-492
chr9-96165718-INS-64
chr9-96174257-INS-167
chr9-96180479-INS-68
chr9-96195666-INS-206
chr9-96258695-INS-60
chr9-96277919-INS-194
chr9-96289533-INS-505
chr9-96300382-INS-657
chr9-96304838-INS-207
chr9-96315494-INS-55
chr9-96317247-INS-52

chr9-96345156-INS-266
chr9-96355485-INS-228
chr9-96364129-INS-210
chr9-96372709-INS-76
chr9-96387127-INS-167
chr9-96391953-INS-116
chr9-96401321-INS-56
chr9-96410113-INS-55
chr9-96417249-INS-283
chr9-96452172-INS-96
chr9-96461131-INS-101
chr9-96524853-INS-210
chr9-96559492-INS-73
chr9-96605075-INS-56
chr9-96608508-INS-57
chr9-96639783-INS-161
chr9-96641780-INS-320
chr9-96647997-INS-52
chr9-96674999-INS-51
chr9-96675777-INS-54
chr9-96676985-INS-55
chr9-96681927-INS-72
chr9-96689951-INS-7166
chr9-96695447-INS-57
chr9-96702913-INS-190
chr9-96713786-INS-202
chr9-96734494-INS-199
chr9-96735116-INS-195
chr9-96744488-INS-108
chr9-96745978-INS-1888
chr9-96748962-INS-98
chr9-96751182-INS-82
chr9-96760413-INS-6476
chr9-96784539-INS-86
chr9-96828878-INS-342
chr9-96904798-INS-122
chr9-96906907-INS-304
chr9-96931435-INS-181
chr9-96944545-INS-140
chr9-96962198-INS-82
chr9-96965805-INS-65
chr9-96965836-INS-921
chr9-96992514-INS-55
chr9-97000290-INS-250
chr9-97004078-INS-755
chr9-97020539-INS-241
chr9-97033980-INS-156
chr9-97056591-INS-95
chr9-97076880-INS-6458
chr9-97085197-INS-550
chr9-97086494-INS-445
chr9-97093619-INS-482
chr9-97099213-INS-79
chr9-97147503-INS-52
chr9-97158022-INS-72
chr9-97172405-INS-820
chr9-97226422-INS-5729
chr9-97233511-INS-54
chr9-97259635-INS-113
chr9-97300333-INS-80
chr9-97358915-INS-145
chr9-97381666-INS-344
chr9-97395880-INS-145
chr9-97412021-INS-400
chr9-97464112-INS-75
chr9-97491834-INS-88
chr9-97502230-INS-76
chr9-97520089-INS-69
chr9-97526524-INS-343
chr9-97530060-INS-177
chr9-97601343-INS-56
chr9-97683219-INS-1772
chr9-97685767-INS-248
chr9-97760469-INS-5630
chr9-97763448-INS-265
chr9-97898895-INS-850
chr9-97900336-INS-942

chr9-97934104-INS-7129
chr9-97939066-INS-86
chr9-97948466-INS-336
chr9-97966615-INS-226
chr9-97976389-INS-6380
chr9-97989842-INS-104
chr9-98001176-INS-56
chr9-98009081-INS-1864
chr9-98028685-INS-56
chr9-98038262-INS-87
chr9-98064281-INS-60
chr9-98078691-INS-267
chr9-98081579-INS-118
chr9-98089087-INS-373
chr9-98103198-INS-181
chr9-98114977-INS-398
chr9-98120271-INS-54
chr9-98122944-INS-212
chr9-98146295-INS-124
chr9-98158897-INS-63
chr9-98204940-INS-66
chr9-98205490-INS-9130
chr9-98209320-INS-197
chr9-98217835-INS-7146
chr9-98296325-INS-977
chr9-98296365-INS-123
chr9-98299830-INS-100
chr9-98313753-INS-94
chr9-98385300-INS-72
chr9-98403381-INS-56
chr9-98405253-INS-104
chr9-98412916-INS-422
chr9-98428455-INS-155
chr9-98439023-INS-60
chr9-98442193-INS-601
chr9-98459594-INS-52
chr9-98485266-INS-140
chr9-98491376-INS-1893
chr9-98491827-INS-288
chr9-98495938-INS-86
chr9-98543643-INS-453
chr9-98575101-INS-67
chr9-98584757-INS-53
chr9-98592997-INS-58
chr9-98614525-INS-59
chr9-98615066-INS-166
chr9-98680868-INS-150
chr9-98688452-INS-1864
chr9-98706108-INS-417
chr9-98745251-INS-2063
chr9-98772489-INS-7596
chr9-98799915-INS-108
chr9-98831623-INS-198
chr9-98835758-INS-58
chr9-98843482-INS-52
chr9-98853271-INS-227
chr9-98887868-INS-74
chr9-98899522-INS-66
chr9-98910540-INS-214
chr9-98925848-INS-79
chr9-98967231-INS-55
chr9-98973824-INS-1893
chr9-98975868-INS-257
chr9-98991068-INS-165
chr9-99005653-INS-176
chr9-99008681-INS-202
chr9-99015192-INS-209
chr9-99029912-INS-85
chr9-99047482-INS-54
chr9-99055701-INS-215
chr9-99056970-INS-154
chr9-99099497-INS-67
chr9-99105351-INS-214
chr9-99116130-INS-171
chr9-99118484-INS-103
chr9-99157135-INS-164
chr9-99158494-INS-180

chr9-99161399-INS-176
chr9-99176829-INS-218
chr9-99208835-INS-386
chr9-99221207-INS-246
chr9-99246473-INS-91
chr9-99249684-INS-173
chr9-99259503-INS-192
chr9-99320254-INS-64
chr9-99322797-INS-351
chr9-99324870-INS-172
chr9-99353112-INS-88
chr9-99359541-INS-79
chr9-99359978-INS-81
chr9-99368985-INS-554
chr9-99376538-INS-222
chr9-99377180-INS-6655
chr9-99427843-INS-170
chr9-99441565-INS-98
chr9-99444734-INS-313
chr9-99445286-INS-127
chr9-99446136-INS-216
chr9-99473691-INS-7349
chr9-99530804-INS-82
chr9-99540227-INS-84
chr9-99554080-INS-162
chr9-99557740-INS-204
chr9-99561352-INS-165
chr9-99596756-INS-108
chr9-99651496-INS-63
chr9-99677558-INS-7066
chr9-99679111-INS-54
chr9-99690390-INS-428
chr9-99693817-INS-64
chr9-99697321-INS-231
chr9-99703016-INS-122
chr9-99705244-INS-86
chr9-99714359-INS-62
chr9-99731149-INS-123
chr9-99763059-INS-68
chr9-99770281-INS-58
chr9-99778320-INS-6366
chr9-99811625-INS-3636
chr9-99813996-INS-5974
chr9-99846244-INS-56
chr9-99860082-INS-200
chr9-99882228-INS-55
chr9-99884938-INS-573
chr9-99897271-INS-276
chr9-99901194-INS-119
chr9-99922833-INS-56
chr9-99932074-INS-400
chr9-99937842-INS-199
chr9-99939904-INS-136
chr9-99941710-INS-66
chr9-99945040-INS-54
chr9-99946845-INS-59
chr9-99949680-INS-83
chr9-100004672-INS-72
chr9-100025231-INS-365
chr9-100059203-INS-459
chr9-100090791-INS-105
chr9-100106500-INS-64
chr9-100159001-INS-214
chr9-100165868-INS-76
chr9-100171679-INS-54
chr9-100211951-INS-60
chr9-100225544-INS-264
chr9-100233071-INS-194
chr9-100242872-INS-95
chr9-100247936-INS-171
chr9-100252905-INS-180
chr9-100255035-INS-830
chr9-100268275-INS-159
chr9-100274731-INS-611
chr9-100297061-INS-117
chr9-100307900-INS-524
chr9-100329354-INS-85

chr9-100338265-INS-53
chr9-100346940-INS-76
chr9-100392440-INS-58
chr9-100405592-INS-177
chr9-100415834-INS-70
chr9-100418947-INS-82
chr9-100439084-INS-58
chr9-100441136-INS-72
chr9-100459899-INS-108
chr9-100497382-INS-189
chr9-100499770-INS-879
chr9-100510787-INS-150
chr9-100517031-INS-7119
chr9-100583044-INS-182
chr9-100585735-INS-58
chr9-100609228-INS-204
chr9-100627852-INS-454
chr9-100643975-INS-202
chr9-100647535-INS-5357
chr9-100662380-INS-87
chr9-100688546-INS-118
chr9-100710984-INS-298
chr9-100741854-INS-218
chr9-100748952-INS-2999
chr9-100760157-INS-270
chr9-100775641-INS-88
chr9-100802095-INS-52
chr9-100827007-INS-550
chr9-100842738-INS-211
chr9-100860493-INS-93
chr9-100879079-INS-160
chr9-100926607-INS-56
chr9-100934327-INS-102
chr9-100937573-INS-204
chr9-100939975-INS-140
chr9-100941307-INS-56
chr9-100968411-INS-89
chr9-100974914-INS-420
chr9-100974938-INS-90
chr9-100986603-INS-113
chr9-101023467-INS-541
chr9-101025949-INS-82
chr9-101038289-INS-71
chr9-101051242-INS-101
chr9-101065861-INS-54
chr9-101066928-INS-125
chr9-101118969-INS-113
chr9-101130356-INS-204
chr9-101153344-INS-137
chr9-101153565-INS-6411
chr9-101154804-INS-177
chr9-101189996-INS-51
chr9-101221834-INS-215
chr9-101240603-INS-125
chr9-101241238-INS-6484
chr9-101243928-INS-96
chr9-101245590-INS-54
chr9-101278184-INS-53
chr9-101281339-INS-2841
chr9-101292977-INS-300
chr9-101337748-INS-63
chr9-101353042-INS-53
chr9-101357707-INS-400
chr9-101381077-INS-227
chr9-101416278-INS-797
chr9-101429317-INS-75
chr9-101431761-INS-55
chr9-101433118-INS-82
chr9-101465013-INS-3516
chr9-101474010-INS-6639
chr9-101482353-INS-79
chr9-101485346-INS-2578
chr9-101490042-INS-54
chr9-101492999-INS-300
chr9-101514729-INS-119
chr9-101521326-INS-5423
chr9-101550365-INS-72

chr9-101558263-INS-112
chr9-101574619-INS-132
chr9-101590317-INS-60
chr9-101599751-INS-474
chr9-101602081-INS-52
chr9-101606563-INS-55
chr9-101610060-INS-67
chr9-101619576-INS-68
chr9-101621761-INS-7233
chr9-101642422-INS-6403
chr9-101648191-INS-122
chr9-101666752-INS-54
chr9-101680563-INS-52
chr9-101682609-INS-121
chr9-101687901-INS-66
chr9-101741121-INS-114
chr9-101756251-INS-52
chr9-101803313-INS-60
chr9-101835783-INS-70
chr9-101847400-INS-321
chr9-101871054-INS-66
chr9-101900428-INS-498
chr9-101902300-INS-102
chr9-101929147-INS-202
chr9-101947260-INS-51
chr9-101951864-INS-52
chr9-101954783-INS-57
chr9-101960075-INS-51
chr9-101965616-INS-187
chr9-101980870-INS-65
chr9-101983657-INS-92
chr9-101994628-INS-63
chr9-102015739-INS-105
chr9-102048256-INS-67
chr9-102101732-INS-54
chr9-102116615-INS-109
chr9-102142344-INS-189
chr9-102181975-INS-108
chr9-102215151-INS-57
chr9-102264556-INS-143
chr9-102277299-INS-326
chr9-102293520-INS-60
chr9-102304773-INS-51
chr9-102315340-INS-221
chr9-102329739-INS-1888
chr9-102343973-INS-504
chr9-102367457-INS-291
chr9-102369519-INS-56
chr9-102397891-INS-52
chr9-102418551-INS-406
chr9-102437427-INS-70
chr9-102456207-INS-4796
chr9-102472188-INS-205
chr9-102556749-INS-73
chr9-102558301-INS-167
chr9-102559759-INS-54
chr9-102566262-INS-84
chr9-102568609-INS-68
chr9-102575354-INS-94
chr9-102580448-INS-192
chr9-102592620-INS-59
chr9-102628301-INS-70
chr9-102630369-INS-215
chr9-102641776-INS-57
chr9-102651130-INS-146
chr9-102671989-INS-52
chr9-102676768-INS-208
chr9-102678857-INS-408
chr9-102687261-INS-276
chr9-102690950-INS-459
chr9-102693385-INS-146
chr9-102694128-INS-214
chr9-102697637-INS-65
chr9-102732856-INS-130
chr9-102748701-INS-127
chr9-102768548-INS-556
chr9-102778594-INS-71

chr9-102787768-INS-88
chr9-102791107-INS-7047
chr9-102795452-INS-102
chr9-102797467-INS-58
chr9-102815979-INS-135
chr9-102826355-INS-7202
chr9-102826473-INS-178
chr9-102830240-INS-85
chr9-102830842-INS-460
chr9-102837819-INS-7136
chr9-102839949-INS-129
chr9-102849152-INS-113
chr9-102866850-INS-52
chr9-102875672-INS-146
chr9-102893257-INS-235
chr9-102908302-INS-158
chr9-102926944-INS-61
chr9-102965388-INS-84
chr9-102966548-INS-86
chr9-102972254-INS-108
chr9-102981922-INS-553
chr9-103077341-INS-128
chr9-103092659-INS-201
chr9-103098632-INS-212
chr9-103101489-INS-626
chr9-103124334-INS-66
chr9-103127146-INS-186
chr9-103132230-INS-263
chr9-103135078-INS-72
chr9-103154141-INS-78
chr9-103161342-INS-80
chr9-103190688-INS-96
chr9-103201146-INS-241
chr9-103208824-INS-267
chr9-103216480-INS-110
chr9-103218543-INS-196
chr9-103226121-INS-66
chr9-103228774-INS-76
chr9-103231359-INS-58
chr9-103236766-INS-55
chr9-103262742-INS-144
chr9-103278144-INS-202
chr9-103302060-INS-61
chr9-103312488-INS-75
chr9-103324034-INS-403
chr9-103329989-INS-307
chr9-103335895-INS-160
chr9-103339605-INS-66
chr9-103351437-INS-60
chr9-103362262-INS-74
chr9-103375945-INS-556
chr9-103378460-INS-182
chr9-103388281-INS-74
chr9-103401873-INS-79
chr9-103406124-INS-400
chr9-103423577-INS-191
chr9-103425184-INS-213
chr9-103425932-INS-1888
chr9-103489135-INS-76
chr9-103491765-INS-265
chr9-103503870-INS-54
chr9-103510384-INS-958
chr9-103526369-INS-218
chr9-103536103-INS-63
chr9-103626420-INS-309
chr9-103631729-INS-71
chr9-103634117-INS-60
chr9-103635850-INS-79
chr9-103640299-INS-108
chr9-103646578-INS-192
chr9-103669010-INS-98
chr9-103689326-INS-109
chr9-103694750-INS-439
chr9-103704210-INS-152
chr9-103717297-INS-216
chr9-103722106-INS-399
chr9-103727417-INS-153

chr9-103812701-INS-971
chr9-103821045-INS-289
chr9-103845321-INS-206
chr9-103847754-INS-300
chr9-103851109-INS-623
chr9-103859307-INS-143
chr9-103863497-INS-102
chr9-103877113-INS-128
chr9-103883466-INS-146
chr9-103884418-INS-68
chr9-103891221-INS-445
chr9-103903865-INS-52
chr9-103911115-INS-70
chr9-103934333-INS-135
chr9-103942789-INS-191
chr9-103957728-INS-54
chr9-103964056-INS-250
chr9-104022589-INS-66
chr9-104025206-INS-55
chr9-104041894-INS-54
chr9-104043495-INS-62
chr9-104047894-INS-55
chr9-104050774-INS-65
chr9-104054893-INS-516
chr9-104058467-INS-202
chr9-104078913-INS-69
chr9-104146995-INS-395
chr9-104167499-INS-61
chr9-104180608-INS-300
chr9-104187333-INS-150
chr9-104194231-INS-83
chr9-104196287-INS-198
chr9-104209641-INS-52
chr9-104226495-INS-80
chr9-104234796-INS-112
chr9-104257666-INS-91
chr9-104263757-INS-195
chr9-104272063-INS-174
chr9-104283696-INS-52
chr9-104297316-INS-77
chr9-104298590-INS-108
chr9-104301882-INS-60
chr9-104319771-INS-174
chr9-104321083-INS-196
chr9-104324818-INS-183
chr9-104329297-INS-206
chr9-104335303-INS-112
chr9-104349641-INS-57
chr9-104366262-INS-54
chr9-104382350-INS-66
chr9-104397578-INS-60
chr9-104399888-INS-60
chr9-104406161-INS-282
chr9-104407375-INS-52
chr9-104425794-INS-372
chr9-104425805-INS-966
chr9-104441419-INS-199
chr9-104449737-INS-68
chr9-104453740-INS-198
chr9-104456699-INS-166
chr9-104461262-INS-101
chr9-104480394-INS-63
chr9-104492623-INS-572
chr9-104497450-INS-122
chr9-104497887-INS-136
chr9-104504693-INS-98
chr9-104524938-INS-554
chr9-104533449-INS-104
chr9-104576546-INS-115
chr9-104585427-INS-76
chr9-104588687-INS-54
chr9-104589252-INS-71
chr9-104599034-INS-67
chr9-104653931-INS-93
chr9-104661374-INS-575
chr9-104677829-INS-7633
chr9-104695179-INS-53

chr9-104710522-INS-64
chr9-104725343-INS-201
chr9-104730718-INS-634
chr9-104745949-INS-135
chr9-104763487-INS-243
chr9-104769299-INS-185
chr9-104846879-INS-135
chr9-104851761-INS-100
chr9-104860285-INS-207
chr9-104865508-INS-130
chr9-104880359-INS-391
chr9-104892955-INS-66
chr9-104902004-INS-410
chr9-104950425-INS-162
chr9-104979156-INS-58
chr9-105038804-INS-54
chr9-105071292-INS-73
chr9-105075727-INS-51
chr9-105080877-INS-73
chr9-105080921-INS-636
chr9-105092268-INS-169
chr9-105097497-INS-216
chr9-105125911-INS-223
chr9-105173815-INS-252
chr9-105174761-INS-87
chr9-105177949-INS-401
chr9-105184552-INS-312
chr9-105192104-INS-74
chr9-105207489-INS-419
chr9-105222423-INS-6492
chr9-105236105-INS-234
chr9-105237147-INS-348
chr9-105252407-INS-67
chr9-105305231-INS-71
chr9-105308476-INS-200
chr9-105315080-INS-128
chr9-105319136-INS-238
chr9-105326396-INS-286
chr9-105332877-INS-162
chr9-105336246-INS-51
chr9-105344192-INS-124
chr9-105362189-INS-386
chr9-105393304-INS-211
chr9-105408024-INS-68
chr9-105413596-INS-434
chr9-105429654-INS-184
chr9-105449024-INS-286
chr9-105471234-INS-93
chr9-105474193-INS-148
chr9-105479735-INS-58
chr9-105481574-INS-101
chr9-105494919-INS-155
chr9-105511954-INS-400
chr9-105537795-INS-474
chr9-105544217-INS-169
chr9-105546408-INS-180
chr9-105552650-INS-112
chr9-105584085-INS-141
chr9-105607407-INS-160
chr9-105627205-INS-664
chr9-105668356-INS-392
chr9-105670362-INS-421
chr9-105702162-INS-88
chr9-105706304-INS-68
chr9-105706887-INS-572
chr9-105708956-INS-66
chr9-105717149-INS-8740
chr9-105719932-INS-492
chr9-105754229-INS-133
chr9-105763097-INS-198
chr9-105777185-INS-468
chr9-105778648-INS-208
chr9-105820762-INS-78
chr9-105833282-INS-91
chr9-105839234-INS-81
chr9-105843545-INS-164
chr9-105846554-INS-72

chr9-105860612-INS-130
chr9-105879052-INS-52
chr9-105903437-INS-816
chr9-105907777-INS-120
chr9-105928073-INS-56
chr9-105929740-INS-56
chr9-105963277-INS-212
chr9-105963467-INS-678
chr9-106008109-INS-852
chr9-106013921-INS-216
chr9-106015599-INS-268
chr9-106024042-INS-64
chr9-106071573-INS-103
chr9-106078720-INS-52
chr9-106107112-INS-298
chr9-106129417-INS-56
chr9-106132914-INS-83
chr9-106163320-INS-53
chr9-106171479-INS-61
chr9-106176700-INS-557
chr9-106198466-INS-56
chr9-106230675-INS-93
chr9-106235988-INS-60
chr9-106254990-INS-182
chr9-106266597-INS-400
chr9-106281294-INS-157
chr9-106301707-INS-698
chr9-106304330-INS-201
chr9-106318191-INS-427
chr9-106319971-INS-211
chr9-106356270-INS-199
chr9-106366146-INS-325
chr9-106367945-INS-197
chr9-106378438-INS-162
chr9-106383132-INS-57
chr9-106414809-INS-258
chr9-106427569-INS-63
chr9-106446212-INS-120
chr9-106451157-INS-134
chr9-106519133-INS-434
chr9-106559732-INS-84
chr9-106574026-INS-72
chr9-106587124-INS-53
chr9-106614127-INS-69
chr9-106637948-INS-118
chr9-106651988-INS-521
chr9-106662206-INS-54
chr9-106667252-INS-56
chr9-106687835-INS-222
chr9-106728824-INS-443
chr9-106731881-INS-301
chr9-106734747-INS-206
chr9-106750524-INS-72
chr9-106814261-INS-56
chr9-106817155-INS-236
chr9-106823218-INS-89
chr9-106830015-INS-109
chr9-106834530-INS-54
chr9-106839444-INS-390
chr9-106840098-INS-6498
chr9-106919307-INS-65
chr9-106928045-INS-53
chr9-106929601-INS-420
chr9-106939801-INS-51
chr9-106949557-INS-546
chr9-106965292-INS-4088
chr9-106965963-INS-2045
chr9-106994861-INS-54
chr9-106995786-INS-6248
chr9-107014170-INS-51
chr9-107014811-INS-233
chr9-107016062-INS-139
chr9-107023711-INS-6657
chr9-107042689-INS-149
chr9-107058287-INS-224
chr9-107089515-INS-203
chr9-107100969-INS-198

chr9-107117274-INS-135
chr9-107122182-INS-61
chr9-107123910-INS-168
chr9-107155098-INS-1959
chr9-107183853-INS-64
chr9-107239903-INS-130
chr9-107291449-INS-60
chr9-107296129-INS-97
chr9-107335950-INS-262
chr9-107351076-INS-6413
chr9-107351077-INS-503
chr9-107364363-INS-494
chr9-107405060-INS-1870
chr9-107407042-INS-324
chr9-107407049-INS-1312
chr9-107474804-INS-215
chr9-107526143-INS-204
chr9-107538736-INS-214
chr9-107564245-INS-94
chr9-107571517-INS-180
chr9-107596540-INS-164
chr9-107603804-INS-201
chr9-107631556-INS-200
chr9-107639523-INS-96
chr9-107643666-INS-183
chr9-107644709-INS-236
chr9-107670338-INS-221
chr9-107674743-INS-189
chr9-107677393-INS-189
chr9-107688460-INS-202
chr9-107726550-INS-189
chr9-107734787-INS-60
chr9-107739310-INS-216
chr9-107741296-INS-387
chr9-107741631-INS-226
chr9-107752293-INS-109
chr9-107755164-INS-201
chr9-107762674-INS-166
chr9-107764492-INS-63
chr9-107796981-INS-161
chr9-107799123-INS-52
chr9-107812392-INS-223
chr9-107821346-INS-223
chr9-107845435-INS-221
chr9-107890799-INS-106
chr9-107893993-INS-202
chr9-107910574-INS-125
chr9-107917475-INS-245
chr9-107966433-INS-203
chr9-107977713-INS-126
chr9-107997121-INS-74
chr9-108009364-INS-197
chr9-108011168-INS-58
chr9-108021763-INS-108
chr9-108026808-INS-227
chr9-108044833-INS-319
chr9-108074450-INS-172
chr9-108087944-INS-256
chr9-108108181-INS-72
chr9-108147471-INS-202
chr9-108149002-INS-112
chr9-108169655-INS-312
chr9-108169842-INS-85
chr9-108175633-INS-209
chr9-108181926-INS-295
chr9-108186550-INS-205
chr9-108212977-INS-163
chr9-108228010-INS-268
chr9-108231477-INS-194
chr9-108234564-INS-72
chr9-108236832-INS-276
chr9-108252728-INS-137
chr9-108254919-INS-1864
chr9-108261054-INS-63
chr9-108265154-INS-52
chr9-108266558-INS-196
chr9-108292261-INS-124

chr9-108308597-INS-111
chr9-108309650-INS-494
chr9-108310857-INS-151
chr9-108311274-INS-282
chr9-108318407-INS-182
chr9-108350667-INS-70
chr9-108399127-INS-207
chr9-108463205-INS-52
chr9-108475707-INS-348
chr9-108480037-INS-56
chr9-108488356-INS-213
chr9-108499095-INS-765
chr9-108507567-INS-425
chr9-108530925-INS-54
chr9-108534231-INS-5292
chr9-108536189-INS-324
chr9-108544033-INS-114
chr9-108567667-INS-237
chr9-108572885-INS-124
chr9-108597679-INS-56
chr9-108599151-INS-230
chr9-108620367-INS-71
chr9-108628121-INS-173
chr9-108638885-INS-60
chr9-108647230-INS-320
chr9-108658778-INS-200
chr9-108679316-INS-210
chr9-108681512-INS-659
chr9-108723802-INS-73
chr9-108748694-INS-88
chr9-108766587-INS-202
chr9-108768988-INS-59
chr9-108788971-INS-198
chr9-108805892-INS-210
chr9-108814871-INS-176
chr9-108830877-INS-174
chr9-108848527-INS-188
chr9-108849133-INS-130
chr9-108852700-INS-94
chr9-108878041-INS-740
chr9-108879080-INS-270
chr9-108881101-INS-61
chr9-108881515-INS-215
chr9-108895206-INS-64
chr9-108913506-INS-441
chr9-108914046-INS-118
chr9-108914727-INS-54
chr9-108950608-INS-204
chr9-108957518-INS-175
chr9-108961884-INS-866
chr9-108965603-INS-81
chr9-108968200-INS-641
chr9-108976105-INS-511
chr9-108985883-INS-92
chr9-108994409-INS-498
chr9-109004138-INS-89
chr9-109018228-INS-310
chr9-109023335-INS-58
chr9-109069284-INS-56
chr9-109166866-INS-53
chr9-109203521-INS-238
chr9-109206250-INS-1891
chr9-109239798-INS-5979
chr9-109256402-INS-331
chr9-109288285-INS-4171
chr9-109362277-INS-52
chr9-109363626-INS-138
chr9-109370241-INS-6241
chr9-109370503-INS-3544
chr9-109372256-INS-59
chr9-109378881-INS-497
chr9-109439996-INS-614
chr9-109491827-INS-64
chr9-109519210-INS-90
chr9-109521685-INS-200
chr9-109531393-INS-55
chr9-109564180-INS-578

chr9-109566015-INS-6642
chr9-109566595-INS-400
chr9-109603005-INS-1887
chr9-109603892-INS-526
chr9-109607015-INS-466
chr9-109633132-INS-530
chr9-109643859-INS-67
chr9-109644590-INS-127
chr9-109661088-INS-68
chr9-109669373-INS-212
chr9-109673927-INS-205
chr9-109686501-INS-54
chr9-109693596-INS-5369
chr9-109693601-INS-372
chr9-109697690-INS-51
chr9-109728055-INS-54
chr9-109741725-INS-149
chr9-109744703-INS-136
chr9-109748714-INS-102
chr9-109752410-INS-246
chr9-109755241-INS-58
chr9-109765467-INS-214
chr9-109777401-INS-211
chr9-109778616-INS-296
chr9-109782748-INS-111
chr9-109796875-INS-71
chr9-109811138-INS-158
chr9-109819399-INS-204
chr9-109822729-INS-213
chr9-109827228-INS-212
chr9-109836326-INS-260
chr9-109837139-INS-6452
chr9-109851101-INS-4837
chr9-109851813-INS-124
chr9-109852167-INS-142
chr9-109903096-INS-101
chr9-109922844-INS-123
chr9-109924302-INS-203
chr9-109927517-INS-211
chr9-109942770-INS-200
chr9-109948338-INS-199
chr9-109949514-INS-196
chr9-109953915-INS-58
chr9-109958927-INS-215
chr9-109960066-INS-210
chr9-109967398-INS-193
chr9-109972457-INS-54
chr9-109973602-INS-165
chr9-109975946-INS-59
chr9-109980130-INS-147
chr9-109982667-INS-209
chr9-109990114-INS-96
chr9-110007538-INS-203
chr9-110016329-INS-106
chr9-110018779-INS-254
chr9-110025795-INS-218
chr9-110037861-INS-215
chr9-110039463-INS-181
chr9-110041514-INS-246
chr9-110054627-INS-172
chr9-110059244-INS-60
chr9-110090077-INS-480
chr9-110105936-INS-69
chr9-110114973-INS-219
chr9-110119531-INS-96
chr9-110121087-INS-125
chr9-110129022-INS-234
chr9-110131313-INS-197
chr9-110147464-INS-161
chr9-110152328-INS-211
chr9-110168190-INS-114
chr9-110182873-INS-218
chr9-110185750-INS-206
chr9-110188323-INS-195
chr9-110197834-INS-161
chr9-110229617-INS-215
chr9-110239835-INS-64

chr9-110244235-INS-132
chr9-110252623-INS-131
chr9-110271544-INS-486
chr9-110273258-INS-188
chr9-110274957-INS-114
chr9-110279344-INS-197
chr9-110280900-INS-584
chr9-110290839-INS-123
chr9-110294800-INS-102
chr9-110311077-INS-333
chr9-110342380-INS-221
chr9-110347574-INS-84
chr9-110349335-INS-52
chr9-110405382-INS-359
chr9-110429534-INS-165
chr9-110492199-INS-372
chr9-110493474-INS-70
chr9-110494864-INS-211
chr9-110506094-INS-193
chr9-110517360-INS-52
chr9-110519048-INS-136
chr9-110547207-INS-200
chr9-110552348-INS-90
chr9-110566813-INS-64
chr9-110567592-INS-194
chr9-110584943-INS-55
chr9-110588818-INS-203
chr9-110594066-INS-53
chr9-110605936-INS-152
chr9-110618383-INS-124
chr9-110622914-INS-194
chr9-110633289-INS-201
chr9-110637567-INS-780
chr9-110641548-INS-123
chr9-110661866-INS-308
chr9-110662874-INS-5324
chr9-110683818-INS-179
chr9-110699217-INS-211
chr9-110733168-INS-220
chr9-110769543-INS-180
chr9-110775489-INS-213
chr9-110794459-INS-64
chr9-110806150-INS-206
chr9-110808441-INS-116
chr9-110814665-INS-193
chr9-110815636-INS-188
chr9-110825485-INS-57
chr9-110834466-INS-150
chr9-110840252-INS-54
chr9-110845663-INS-210
chr9-110846247-INS-114
chr9-110849003-INS-90
chr9-110858300-INS-1686
chr9-110861307-INS-165
chr9-110866698-INS-198
chr9-110871947-INS-67
chr9-110872816-INS-1892
chr9-110905260-INS-406
chr9-110912403-INS-51
chr9-110923138-INS-633
chr9-110959895-INS-79
chr9-111005465-INS-335
chr9-111038046-INS-109
chr9-111050926-INS-202
chr9-111075094-INS-281
chr9-111097999-INS-58
chr9-111114553-INS-199
chr9-111120785-INS-82
chr9-111143540-INS-185
chr9-111156661-INS-392
chr9-111164386-INS-125
chr9-111180735-INS-148
chr9-111186795-INS-166
chr9-111187898-INS-84
chr9-111189345-INS-519
chr9-111236491-INS-131
chr9-111278526-INS-72

chr9-111283315-INS-520
chr9-111283880-INS-65
chr9-111300354-INS-167
chr9-111374136-INS-52
chr9-111388245-INS-88
chr9-111396851-INS-52
chr9-111445966-INS-96
chr9-111446424-INS-512
chr9-111451288-INS-65
chr9-111456253-INS-52
chr9-111464813-INS-86
chr9-111481649-INS-54
chr9-111488999-INS-51
chr9-111515478-INS-232
chr9-111528610-INS-120
chr9-111585867-INS-224
chr9-111607797-INS-81
chr9-111645712-INS-631
chr9-111654429-INS-651
chr9-111699395-INS-102
chr9-111705887-INS-167
chr9-111707441-INS-339
chr9-111729760-INS-52
chr9-111744029-INS-442
chr9-111754816-INS-473
chr9-111756506-INS-95
chr9-111762247-INS-102
chr9-111769795-INS-89
chr9-111777585-INS-87
chr9-111786358-INS-52
chr9-111786869-INS-1430
chr9-111790447-INS-387
chr9-111807031-INS-83
chr9-111814908-INS-2981
chr9-111837443-INS-222
chr9-111850077-INS-184
chr9-111863032-INS-67
chr9-111869043-INS-123
chr9-111878522-INS-214
chr9-111918427-INS-86
chr9-111921046-INS-951
chr9-111926838-INS-783
chr9-111928180-INS-110
chr9-111977939-INS-137
chr9-111993209-INS-54
chr9-112014539-INS-59
chr9-112058913-INS-54
chr9-112064119-INS-69
chr9-112068845-INS-96
chr9-112083646-INS-180
chr9-112099241-INS-106
chr9-112120460-INS-67
chr9-112148960-INS-259
chr9-112150888-INS-278
chr9-112151360-INS-76
chr9-112152111-INS-614
chr9-112178754-INS-243
chr9-112191337-INS-278
chr9-112192164-INS-4650
chr9-112225873-INS-62
chr9-112231579-INS-58
chr9-112237595-INS-398
chr9-112257490-INS-152
chr9-112264281-INS-52
chr9-112267556-INS-59
chr9-112268436-INS-135
chr9-112274717-INS-60
chr9-112283560-INS-70
chr9-112344322-INS-442
chr9-112347286-INS-84
chr9-112358894-INS-75
chr9-112373695-INS-56
chr9-112387751-INS-103
chr9-112390075-INS-379
chr9-112400612-INS-693
chr9-112409318-INS-143
chr9-112425278-INS-52

chr9-112430048-INS-68
chr9-112446127-INS-2359
chr9-112446414-INS-321
chr9-112465921-INS-436
chr9-112467621-INS-397
chr9-112501038-INS-112
chr9-112509285-INS-98
chr9-112535594-INS-84
chr9-112558739-INS-791
chr9-112564941-INS-633
chr9-112572975-INS-72
chr9-112588759-INS-62
chr9-112608133-INS-151
chr9-112611532-INS-77
chr9-112628597-INS-72
chr9-112637660-INS-51
chr9-112660986-INS-264
chr9-112667886-INS-66
chr9-112699292-INS-204
chr9-112701884-INS-242
chr9-112706433-INS-146
chr9-112709277-INS-169
chr9-112711618-INS-134
chr9-112714747-INS-62
chr9-112732033-INS-133
chr9-112753440-INS-365
chr9-112757815-INS-68
chr9-112781915-INS-53
chr9-112798048-INS-167
chr9-112820993-INS-879
chr9-112846567-INS-487
chr9-112854911-INS-86
chr9-112872749-INS-6480
chr9-112875187-INS-68
chr9-112937799-INS-208
chr9-112944560-INS-63
chr9-112952465-INS-100
chr9-112953260-INS-89
chr9-112964779-INS-875
chr9-112981456-INS-55
chr9-113022808-INS-69
chr9-113024628-INS-167
chr9-113041409-INS-94
chr9-113048398-INS-59
chr9-113052714-INS-90
chr9-113090549-INS-102
chr9-113111305-INS-379
chr9-113148806-INS-4883
chr9-113155547-INS-4920
chr9-113162640-INS-53
chr9-113210287-INS-283
chr9-113219479-INS-52
chr9-113227951-INS-497
chr9-113232121-INS-376
chr9-113239939-INS-882
chr9-113258098-INS-384
chr9-113260063-INS-106
chr9-113316501-INS-96
chr9-113322590-INS-214
chr9-113325379-INS-205
chr9-113331058-INS-198
chr9-113342144-INS-148
chr9-113356348-INS-59
chr9-113365656-INS-203
chr9-113398939-INS-283
chr9-113420711-INS-2719
chr9-113447201-INS-166
chr9-113451840-INS-157
chr9-113453128-INS-95
chr9-113459274-INS-219
chr9-113478759-INS-737
chr9-113505378-INS-195
chr9-113527314-INS-250
chr9-113547521-INS-81
chr9-113548735-INS-402
chr9-113553158-INS-706
chr9-113557498-INS-60

chr9-113596242-INS-78
chr9-113607319-INS-180
chr9-113652525-INS-68
chr9-113698782-INS-54
chr9-113723645-INS-198
chr9-113741315-INS-259
chr9-113830524-INS-79
chr9-113845862-INS-209
chr9-113878113-INS-168
chr9-113880063-INS-131
chr9-113882084-INS-516
chr9-113892276-INS-128
chr9-113894902-INS-107
chr9-113899959-INS-354
chr9-113901191-INS-198
chr9-113938864-INS-159
chr9-113981707-INS-1363
chr9-114041656-INS-89
chr9-114089273-INS-6401
chr9-114213328-INS-210
chr9-114245264-INS-742
chr9-114282715-INS-218
chr9-114316500-INS-226
chr9-114320110-INS-179
chr9-114326212-INS-180
chr9-114335215-INS-819
chr9-114384681-INS-215
chr9-114393682-INS-70
chr9-114398005-INS-205
chr9-114403762-INS-212
chr9-114404409-INS-201
chr9-114406301-INS-528
chr9-114407179-INS-172
chr9-114409156-INS-54
chr9-114412876-INS-386
chr9-114417040-INS-209
chr9-114423057-INS-94
chr9-114432178-INS-211
chr9-114438466-INS-218
chr9-114445750-INS-200
chr9-114447835-INS-67
chr9-114470828-INS-141
chr9-114471884-INS-84
chr9-114473532-INS-64
chr9-114487949-INS-368
chr9-114495291-INS-50
chr9-114502967-INS-119
chr9-114506011-INS-476
chr9-114507073-INS-3023
chr9-114524280-INS-203
chr9-114539368-INS-213
chr9-114546884-INS-58
chr9-114555013-INS-117
chr9-114555688-INS-204
chr9-114568334-INS-199
chr9-114570584-INS-409
chr9-114572674-INS-1026
chr9-114575309-INS-163
chr9-114594732-INS-58
chr9-114606183-INS-217
chr9-114611341-INS-70
chr9-114614027-INS-58
chr9-114638224-INS-319
chr9-114642828-INS-157
chr9-114662425-INS-140
chr9-114671075-INS-446
chr9-114674232-INS-196
chr9-114685822-INS-110
chr9-114692631-INS-60
chr9-114694924-INS-174
chr9-114702486-INS-204
chr9-114711818-INS-701
chr9-114716388-INS-220
chr9-114719971-INS-400
chr9-114728811-INS-129
chr9-114732701-INS-80
chr9-114746510-INS-203

chr9-114750166-INS-119
chr9-114751361-INS-152
chr9-114753335-INS-281
chr9-114756632-INS-77
chr9-114767548-INS-57
chr9-114774135-INS-8985
chr9-114784989-INS-291
chr9-114794701-INS-284
chr9-114795762-INS-60
chr9-114812929-INS-216
chr9-114820183-INS-122
chr9-114822773-INS-178
chr9-114849455-INS-56
chr9-114852027-INS-324
chr9-114868732-INS-87
chr9-114894631-INS-78
chr9-114934513-INS-113
chr9-114935880-INS-408
chr9-114957505-INS-108
chr9-114965636-INS-492
chr9-114970576-INS-66
chr9-115043098-INS-204
chr9-115044339-INS-434
chr9-115054158-INS-207
chr9-115092044-INS-132
chr9-115097813-INS-121
chr9-115109634-INS-7024
chr9-115112301-INS-142
chr9-115128684-INS-142
chr9-115141294-INS-191
chr9-115145946-INS-71
chr9-115154761-INS-62
chr9-115187103-INS-440
chr9-115206658-INS-136
chr9-115219100-INS-64
chr9-115225640-INS-168
chr9-115279437-INS-291
chr9-115306323-INS-56
chr9-115312581-INS-50
chr9-115313109-INS-51
chr9-115337100-INS-54
chr9-115389249-INS-131
chr9-115402555-INS-52
chr9-115403984-INS-60
chr9-115405662-INS-57
chr9-115406387-INS-96
chr9-115466068-INS-72
chr9-115474011-INS-90
chr9-115477791-INS-440
chr9-115536016-INS-175
chr9-115543198-INS-3325
chr9-115563488-INS-110
chr9-115570596-INS-54
chr9-115575215-INS-574
chr9-115587418-INS-204
chr9-115594402-INS-200
chr9-115596599-INS-62
chr9-115638530-INS-61
chr9-115642886-INS-327
chr9-115646353-INS-71
chr9-115659582-INS-71
chr9-115694392-INS-62
chr9-115724462-INS-70
chr9-115754137-INS-53
chr9-115760145-INS-54
chr9-115764149-INS-174
chr9-115765476-INS-428
chr9-115774460-INS-87
chr9-115775037-INS-264
chr9-115792578-INS-400
chr9-115803609-INS-205
chr9-115812697-INS-62
chr9-115835256-INS-427
chr9-115842426-INS-238
chr9-115849735-INS-230
chr9-115855162-INS-899
chr9-115863909-INS-324

chr9-115869121-INS-154
chr9-115871712-INS-372
chr9-115901016-INS-60
chr9-115908981-INS-85
chr9-115915157-INS-52
chr9-115923550-INS-66
chr9-115944744-INS-167
chr9-115989097-INS-88
chr9-115997818-INS-78
chr9-116011568-INS-65
chr9-116029005-INS-51
chr9-116038499-INS-222
chr9-116048316-INS-55
chr9-116055840-INS-329
chr9-116070284-INS-252
chr9-116079196-INS-83
chr9-116080602-INS-191
chr9-116087712-INS-58
chr9-116091854-INS-288
chr9-116098393-INS-60
chr9-116103523-INS-172
chr9-116179027-INS-313
chr9-116184480-INS-108
chr9-116191824-INS-55
chr9-116227136-INS-242
chr9-116244915-INS-371
chr9-116262885-INS-178
chr9-116263741-INS-51
chr9-116264764-INS-4759
chr9-116285323-INS-114
chr9-116354011-INS-150
chr9-116380907-INS-135
chr9-116401478-INS-81
chr9-116412744-INS-56
chr9-116424188-INS-90
chr9-116436188-INS-920
chr9-116439088-INS-98
chr9-116445360-INS-220
chr9-116471356-INS-205
chr9-116502112-INS-151
chr9-116549028-INS-56
chr9-116582930-INS-60
chr9-116588742-INS-51
chr9-116610743-INS-80
chr9-116619236-INS-130
chr9-116634179-INS-88
chr9-116648210-INS-120
chr9-116653962-INS-84
chr9-116674282-INS-60
chr9-116682807-INS-412
chr9-116694999-INS-57
chr9-116707676-INS-171
chr9-116719125-INS-450
chr9-116745359-INS-60
chr9-116757590-INS-297
chr9-116766690-INS-50
chr9-116778564-INS-54
chr9-116798357-INS-80
chr9-116831510-INS-132
chr9-116834581-INS-68
chr9-116841006-INS-58
chr9-116868070-INS-433
chr9-116868848-INS-59
chr9-116883960-INS-127
chr9-116903342-INS-665
chr9-116904025-INS-52
chr9-116913006-INS-54
chr9-116914930-INS-193
chr9-116917217-INS-62
chr9-116926996-INS-52
chr9-116928152-INS-56
chr9-116937966-INS-238
chr9-116943311-INS-56
chr9-116950031-INS-52
chr9-116954455-INS-71
chr9-116958278-INS-230
chr9-116965134-INS-137

chr9-117010679-INS-74
chr9-117032895-INS-145
chr9-117041623-INS-58
chr9-117051242-INS-60
chr9-117059888-INS-348
chr9-117061107-INS-1165
chr9-117065230-INS-53
chr9-117087126-INS-68
chr9-117088233-INS-64
chr9-117090928-INS-64
chr9-117095873-INS-206
chr9-117116643-INS-148
chr9-117123789-INS-52
chr9-117138737-INS-324
chr9-117151509-INS-968
chr9-117157785-INS-381
chr9-117162407-INS-210
chr9-117173396-INS-122
chr9-117183319-INS-63
chr9-117188855-INS-5233
chr9-117190382-INS-188
chr9-117218657-INS-56
chr9-117233182-INS-99
chr9-117242263-INS-78
chr9-117242967-INS-75
chr9-117283792-INS-67
chr9-117328823-INS-193
chr9-117331438-INS-87
chr9-117352679-INS-159
chr9-117358276-INS-54
chr9-117371392-INS-62
chr9-117372571-INS-66
chr9-117386789-INS-71
chr9-117398663-INS-96
chr9-117428909-INS-73
chr9-117431929-INS-171
chr9-117455576-INS-77
chr9-117457482-INS-52
chr9-117471587-INS-220
chr9-117499145-INS-60
chr9-117520950-INS-264
chr9-117537106-INS-86
chr9-117541935-INS-56
chr9-117545372-INS-52
chr9-117546071-INS-434
chr9-117547537-INS-124
chr9-117566778-INS-400
chr9-117589978-INS-56
chr9-117619241-INS-154
chr9-117623229-INS-124
chr9-117636734-INS-67
chr9-117651216-INS-184
chr9-117667929-INS-186
chr9-117678900-INS-360
chr9-117689996-INS-76
chr9-117693343-INS-412
chr9-117695100-INS-134
chr9-117698213-INS-6372
chr9-117743340-INS-66
chr9-117746603-INS-794
chr9-117759664-INS-56
chr9-117764048-INS-144
chr9-117798086-INS-4051
chr9-117800537-INS-114
chr9-117801945-INS-307
chr9-117806555-INS-3624
chr9-117818662-INS-83
chr9-117823336-INS-57
chr9-117824575-INS-440
chr9-117832488-INS-766
chr9-117839982-INS-415
chr9-117854886-INS-545
chr9-117879762-INS-301
chr9-117907925-INS-122
chr9-117923566-INS-57
chr9-117933797-INS-98
chr9-117934595-INS-844

chr9-117949321-INS-62
chr9-117953399-INS-160
chr9-117996823-INS-615
chr9-118009686-INS-112
chr9-118015223-INS-190
chr9-118019472-INS-63
chr9-118021712-INS-77
chr9-118033367-INS-351
chr9-118044596-INS-54
chr9-118045800-INS-965
chr9-118049956-INS-169
chr9-118052332-INS-90
chr9-118060160-INS-205
chr9-118072301-INS-142
chr9-118085736-INS-56
chr9-118100238-INS-206
chr9-118114187-INS-103
chr9-118119815-INS-72
chr9-118144466-INS-70
chr9-118151928-INS-182
chr9-118181959-INS-237
chr9-118183998-INS-51
chr9-118191607-INS-108
chr9-118205271-INS-4403
chr9-118239140-INS-405
chr9-118244248-INS-77
chr9-118251219-INS-128
chr9-118269225-INS-65
chr9-118286923-INS-67
chr9-118289805-INS-104
chr9-118291409-INS-228
chr9-118292592-INS-74
chr9-118329836-INS-55
chr9-118338861-INS-173
chr9-118354543-INS-5292
chr9-118391323-INS-66
chr9-118398577-INS-109
chr9-118417498-INS-117
chr9-118428952-INS-74
chr9-118438830-INS-59
chr9-118447260-INS-315
chr9-118485323-INS-401
chr9-118493142-INS-303
chr9-118504107-INS-359
chr9-118516673-INS-169
chr9-118531755-INS-178
chr9-118533038-INS-112
chr9-118536348-INS-54
chr9-118559228-INS-88
chr9-118572693-INS-52
chr9-118579852-INS-98
chr9-118587301-INS-238
chr9-118600638-INS-245
chr9-118622618-INS-99
chr9-118627261-INS-155
chr9-118632156-INS-81
chr9-118641956-INS-184
chr9-118652040-INS-272
chr9-118676480-INS-60
chr9-118678492-INS-148
chr9-118687573-INS-54
chr9-118696593-INS-164
chr9-118703543-INS-8184
chr9-118719450-INS-2040
chr9-118734472-INS-63
chr9-118750281-INS-140
chr9-118784693-INS-73
chr9-118806217-INS-865
chr9-118815337-INS-507
chr9-118826249-INS-67
chr9-118831928-INS-84
chr9-118832160-INS-680
chr9-118842770-INS-166
chr9-118847622-INS-53
chr9-118867290-INS-112
chr9-118900045-INS-152
chr9-118918070-INS-149

chr9-118946416-INS-125
chr9-118948609-INS-203
chr9-118951449-INS-58
chr9-118986387-INS-2202
chr9-118991304-INS-100
chr9-118999199-INS-151
chr9-119011800-INS-483
chr9-119016798-INS-295
chr9-119023149-INS-125
chr9-119043265-INS-204
chr9-119053770-INS-186
chr9-119071658-INS-215
chr9-119100421-INS-102
chr9-119104632-INS-575
chr9-119161279-INS-205
chr9-119182173-INS-237
chr9-119187724-INS-64
chr9-119190624-INS-63
chr9-119195054-INS-167
chr9-119206919-INS-60
chr9-119215570-INS-136
chr9-119305784-INS-93
chr9-119333856-INS-168
chr9-119339190-INS-117
chr9-119339764-INS-58
chr9-119349673-INS-288
chr9-119357085-INS-250
chr9-119357087-INS-903
chr9-119368220-INS-177
chr9-119424868-INS-223
chr9-119425771-INS-122
chr9-119433019-INS-82
chr9-119437057-INS-66
chr9-119462359-INS-636
chr9-119470674-INS-96
chr9-119477786-INS-66
chr9-119483542-INS-77
chr9-119497451-INS-3300
chr9-119509835-INS-51
chr9-119540903-INS-51
chr9-119557788-INS-110
chr9-119559971-INS-174
chr9-119580960-INS-268
chr9-119597792-INS-766
chr9-119604049-INS-91
chr9-119621382-INS-201
chr9-119623877-INS-82
chr9-119639700-INS-276
chr9-119679306-INS-106
chr9-119685571-INS-379
chr9-119709724-INS-62
chr9-119715504-INS-467
chr9-119749896-INS-205
chr9-119768968-INS-185
chr9-119770167-INS-59
chr9-119776971-INS-337
chr9-119798805-INS-180
chr9-119822499-INS-429
chr9-119832574-INS-188
chr9-119855412-INS-197
chr9-119862805-INS-196
chr9-119885796-INS-128
chr9-119899381-INS-239
chr9-119906607-INS-573
chr9-119914978-INS-209
chr9-119938928-INS-224
chr9-119971596-INS-248
chr9-119979842-INS-122
chr9-119988393-INS-57
chr9-119990228-INS-193
chr9-120001998-INS-74
chr9-120009174-INS-426
chr9-120013429-INS-1888
chr9-120025015-INS-187
chr9-120033842-INS-139
chr9-120038394-INS-55
chr9-120041322-INS-135

chr9-120044476-INS-115
chr9-120049240-INS-2881
chr9-120065268-INS-475
chr9-120069238-INS-100
chr9-120072975-INS-60
chr9-120114951-INS-52
chr9-120116313-INS-956
chr9-120152703-INS-69
chr9-120176943-INS-96
chr9-120184895-INS-58
chr9-120186931-INS-102
chr9-120188072-INS-141
chr9-120193956-INS-66
chr9-120196467-INS-170
chr9-120216750-INS-56
chr9-120220163-INS-228
chr9-120236520-INS-124
chr9-120241792-INS-84
chr9-120262164-INS-81
chr9-120305339-INS-122
chr9-120310750-INS-114
chr9-120315999-INS-589
chr9-120329962-INS-137
chr9-120340688-INS-83
chr9-120345632-INS-61
chr9-120355932-INS-97
chr9-120370261-INS-255
chr9-120378104-INS-160
chr9-120383527-INS-55
chr9-120389175-INS-126
chr9-120392230-INS-92
chr9-120414776-INS-252
chr9-120416746-INS-474
chr9-120417779-INS-337
chr9-120422635-INS-55
chr9-120427681-INS-205
chr9-120436085-INS-217
chr9-120443023-INS-138
chr9-120449763-INS-58
chr9-120452216-INS-120
chr9-120457334-INS-151
chr9-120487636-INS-90
chr9-120494122-INS-90
chr9-120516495-INS-80
chr9-120517666-INS-61
chr9-120522478-INS-64
chr9-120549444-INS-380
chr9-120554125-INS-166
chr9-120560456-INS-65
chr9-120570536-INS-67
chr9-120572130-INS-62
chr9-120573631-INS-67
chr9-120583747-INS-192
chr9-120596275-INS-206
chr9-120596877-INS-183
chr9-120606463-INS-4374
chr9-120606571-INS-1479
chr9-120612427-INS-202
chr9-120629506-INS-105
chr9-120633252-INS-190
chr9-120657751-INS-109
chr9-120694055-INS-52
chr9-120708784-INS-58
chr9-120709091-INS-508
chr9-120726099-INS-200
chr9-120732829-INS-68
chr9-120735681-INS-53
chr9-120771665-INS-95
chr9-120796592-INS-169
chr9-120813438-INS-66
chr9-120814516-INS-84
chr9-120861088-INS-140
chr9-120870757-INS-168
chr9-120871077-INS-2352
chr9-120871081-INS-84
chr9-120884008-INS-5654
chr9-120887786-INS-699

chr9-120903324-INS-210
chr9-120904350-INS-228
chr9-120918735-INS-75
chr9-120924527-INS-1015
chr9-120933586-INS-63
chr9-120938087-INS-62
chr9-120949901-INS-215
chr9-120960917-INS-120
chr9-120971499-INS-53
chr9-120975626-INS-108
chr9-120976650-INS-114
chr9-120987526-INS-516
chr9-121014241-INS-90
chr9-121017576-INS-52
chr9-121019548-INS-60
chr9-121026699-INS-398
chr9-121044317-INS-212
chr9-121069568-INS-210
chr9-121079624-INS-348
chr9-121093947-INS-65
chr9-121103865-INS-207
chr9-121108630-INS-55
chr9-121119355-INS-200
chr9-121124910-INS-74
chr9-121160215-INS-221
chr9-121162347-INS-155
chr9-121167669-INS-143
chr9-121168709-INS-52
chr9-121188119-INS-91
chr9-121192126-INS-204
chr9-121194304-INS-298
chr9-121204731-INS-104
chr9-121215457-INS-221
chr9-121219607-INS-73
chr9-121228878-INS-1036
chr9-121246999-INS-117
chr9-121260026-INS-585
chr9-121283041-INS-492
chr9-121305777-INS-369
chr9-121323976-INS-52
chr9-121333448-INS-601
chr9-121342118-INS-128
chr9-121344436-INS-987
chr9-121351674-INS-90
chr9-121363930-INS-74
chr9-121380096-INS-62
chr9-121383038-INS-717
chr9-121388684-INS-170
chr9-121390717-INS-842
chr9-121407658-INS-161
chr9-121417674-INS-85
chr9-121423600-INS-4555
chr9-121429048-INS-158
chr9-121449522-INS-51
chr9-121461641-INS-180
chr9-121464200-INS-198
chr9-121466384-INS-78
chr9-121506063-INS-69
chr9-121514069-INS-52
chr9-121523534-INS-72
chr9-121527310-INS-176
chr9-121528916-INS-62
chr9-121534804-INS-175
chr9-121602377-INS-222
chr9-121616418-INS-60
chr9-121624926-INS-53
chr9-121635441-INS-90
chr9-121638686-INS-66
chr9-121670972-INS-730
chr9-121672363-INS-345
chr9-121674097-INS-404
chr9-121678187-INS-59
chr9-121682689-INS-60
chr9-121701738-INS-127
chr9-121706891-INS-217
chr9-121715711-INS-162
chr9-121728344-INS-272

chr9-121732570-INS-205
chr9-121758539-INS-1552
chr9-121758617-INS-191
chr9-121761009-INS-1034
chr9-121775177-INS-653
chr9-121783162-INS-132
chr9-121788267-INS-202
chr9-121805529-INS-112
chr9-121831867-INS-90
chr9-121834562-INS-7081
chr9-121836215-INS-652
chr9-121845917-INS-491
chr9-121854236-INS-69
chr9-121897619-INS-60
chr9-121915556-INS-326
chr9-121952720-INS-184
chr9-121984745-INS-258
chr9-121988910-INS-343
chr9-121991183-INS-1610
chr9-122027855-INS-97
chr9-122054055-INS-337
chr9-122063049-INS-58
chr9-122125968-INS-63
chr9-122126642-INS-201
chr9-122142258-INS-386
chr9-122151288-INS-72
chr9-122169778-INS-383
chr9-122176059-INS-84
chr9-122190989-INS-75
chr9-122197516-INS-197
chr9-122217762-INS-153
chr9-122219673-INS-85
chr9-122220600-INS-75
chr9-122233727-INS-5542
chr9-122242709-INS-130
chr9-122273936-INS-327
chr9-122284859-INS-4749
chr9-122286663-INS-2361
chr9-122297596-INS-52
chr9-122315084-INS-259
chr9-122335239-INS-216
chr9-122347696-INS-307
chr9-122351692-INS-116
chr9-122354043-INS-72
chr9-122359634-INS-511
chr9-122360337-INS-89
chr9-122392829-INS-61
chr9-122435806-INS-74
chr9-122437099-INS-261
chr9-122444105-INS-72
chr9-122445792-INS-85
chr9-122497864-INS-633
chr9-122515061-INS-195
chr9-122519629-INS-691
chr9-122552987-INS-88
chr9-122559517-INS-402
chr9-122561425-INS-117
chr9-122583532-INS-96
chr9-122590797-INS-92
chr9-122601049-INS-68
chr9-122606169-INS-54
chr9-122606725-INS-741
chr9-122609375-INS-97
chr9-122614113-INS-96
chr9-122628335-INS-201
chr9-122641988-INS-292
chr9-122657316-INS-93
chr9-122663509-INS-541
chr9-122664677-INS-209
chr9-122677452-INS-426
chr9-122694457-INS-60
chr9-122702165-INS-377
chr9-122703371-INS-557
chr9-122723588-INS-208
chr9-122727128-INS-190
chr9-122735578-INS-91
chr9-122749428-INS-124

chr9-122750163-INS-1930
chr9-122761722-INS-3861
chr9-122764392-INS-144
chr9-122772032-INS-6464
chr9-122779054-INS-202
chr9-122784369-INS-166
chr9-122789501-INS-506
chr9-122802399-INS-239
chr9-122809821-INS-80
chr9-122823355-INS-875
chr9-122828180-INS-206
chr9-122872323-INS-106
chr9-122880052-INS-145
chr9-122943602-INS-172
chr9-122944963-INS-214
chr9-122957220-INS-206
chr9-122982279-INS-94
chr9-123004689-INS-192
chr9-123018262-INS-221
chr9-123032562-INS-142
chr9-123037571-INS-108
chr9-123070638-INS-116
chr9-123077568-INS-150
chr9-123080155-INS-324
chr9-123114305-INS-213
chr9-123141993-INS-56
chr9-123153739-INS-64
chr9-123159213-INS-2703
chr9-123169984-INS-64
chr9-123178639-INS-117
chr9-123180045-INS-289
chr9-123181802-INS-55
chr9-123182994-INS-144
chr9-123202952-INS-64
chr9-123251392-INS-118
chr9-123281096-INS-78
chr9-123327751-INS-51
chr9-123336164-INS-212
chr9-123338748-INS-56
chr9-123353658-INS-53
chr9-123397262-INS-204
chr9-123404201-INS-96
chr9-123421992-INS-492
chr9-123448343-INS-109
chr9-123453767-INS-167
chr9-123469142-INS-145
chr9-123476118-INS-57
chr9-123501940-INS-80
chr9-123545352-INS-180
chr9-123546517-INS-56
chr9-123559918-INS-229
chr9-123593575-INS-72
chr9-123604438-INS-634
chr9-123606430-INS-627
chr9-123609993-INS-88
chr9-123614857-INS-693
chr9-123622291-INS-58
chr9-123627196-INS-77
chr9-123628539-INS-708
chr9-123653871-INS-52
chr9-123680961-INS-120
chr9-123696454-INS-54
chr9-123721983-INS-127
chr9-123723504-INS-142
chr9-123724882-INS-57
chr9-123744627-INS-54
chr9-123760044-INS-181
chr9-123761110-INS-53
chr9-123783808-INS-6450
chr9-123792139-INS-54
chr9-123807703-INS-282
chr9-123816060-INS-149
chr9-123817233-INS-59
chr9-123878944-INS-254
chr9-123886473-INS-332
chr9-123891290-INS-52
chr9-123940285-INS-108

chr9-123955778-INS-2261
chr9-124021363-INS-124
chr9-124027385-INS-76
chr9-124058295-INS-417
chr9-124069114-INS-744
chr9-124118977-INS-5550
chr9-124129202-INS-109
chr9-124194488-INS-150
chr9-124211027-INS-72
chr9-124218267-INS-209
chrX-3058199-INS-616
chrX-3164235-INS-616
chrX-3290525-INS-6413
chrX-3684763-INS-65
chrX-3850477-INS-146
chrX-3983301-INS-1004
chrX-4366858-INS-118
chrX-4371393-INS-82
chrX-4378380-INS-449
chrX-4445447-INS-7300
chrX-4521759-INS-111
chrX-4540266-INS-57
chrX-5317934-INS-6064
chrX-5318714-INS-1385
chrX-5322785-INS-72
chrX-5332703-INS-6851
chrX-5382642-INS-1312
chrX-5416746-INS-64
chrX-5442890-INS-233
chrX-5490199-INS-7092
chrX-5493859-INS-9232
chrX-5512641-INS-60
chrX-5518992-INS-329
chrX-5533266-INS-90
chrX-5569427-INS-2247
chrX-5644268-INS-400
chrX-5652323-INS-234
chrX-5661814-INS-913
chrX-5672606-INS-69
chrX-5678600-INS-76
chrX-5681612-INS-122
chrX-5695669-INS-408
chrX-5761895-INS-254
chrX-5766682-INS-689
chrX-5824314-INS-2864
chrX-5824352-INS-1447
chrX-5846764-INS-1119
chrX-5850436-INS-416
chrX-5915819-INS-52
chrX-5929713-INS-113
chrX-5958677-INS-106
chrX-5989128-INS-1538
chrX-5990856-INS-101
chrX-5994379-INS-78
chrX-6017887-INS-76
chrX-6032348-INS-495
chrX-6034946-INS-386
chrX-6037789-INS-728
chrX-6086111-INS-65
chrX-6094660-INS-413
chrX-6121857-INS-295
chrX-6179256-INS-495
chrX-6206171-INS-195
chrX-6218746-INS-314
chrX-6255480-INS-6185
chrX-6266366-INS-55
chrX-6272915-INS-106
chrX-6275838-INS-70
chrX-6304762-INS-60
chrX-6316053-INS-153
chrX-6338548-INS-120
chrX-6364896-INS-554
chrX-6389845-INS-6637
chrX-6390871-INS-567
chrX-6394725-INS-72
chrX-6486978-INS-69
chrX-6501449-INS-74

chrX-6521781-INS-608
chrX-6529546-INS-200
chrX-6532830-INS-185
chrX-6548630-INS-56
chrX-6568422-INS-7363
chrX-6571711-INS-5991
chrX-6587371-INS-6178
chrX-6589416-INS-365
chrX-6616498-INS-77
chrX-6630196-INS-191
chrX-6647901-INS-6256
chrX-6655348-INS-6625
chrX-6673813-INS-93
chrX-6697130-INS-230
chrX-6715325-INS-94
chrX-6736395-INS-54
chrX-6755672-INS-83
chrX-6764513-INS-68
chrX-6775433-INS-102
chrX-6800407-INS-371
chrX-6805558-INS-402
chrX-6806630-INS-615
chrX-6809655-INS-532
chrX-6815308-INS-204
chrX-6817905-INS-354
chrX-6832760-INS-454
chrX-6834084-INS-76
chrX-6880704-INS-66
chrX-6886977-INS-390
chrX-6935455-INS-133
chrX-6937736-INS-120
chrX-6956115-INS-6228
chrX-7011124-INS-123
chrX-7040631-INS-62
chrX-7062212-INS-108
chrX-7070149-INS-221
chrX-7073353-INS-109
chrX-7115285-INS-6440
chrX-7151013-INS-218
chrX-7170997-INS-171
chrX-7183862-INS-599
chrX-7189401-INS-289
chrX-7190664-INS-81
chrX-7197317-INS-374
chrX-7211257-INS-253
chrX-7229726-INS-54
chrX-7282281-INS-52
chrX-7291483-INS-61
chrX-7300061-INS-74
chrX-7304970-INS-203
chrX-7307788-INS-6363
chrX-7331399-INS-607
chrX-7342487-INS-516
chrX-7356250-INS-412
chrX-7358124-INS-66
chrX-7373144-INS-165
chrX-7396601-INS-52
chrX-7402726-INS-90
chrX-7408588-INS-73
chrX-7418633-INS-179
chrX-7463229-INS-427
chrX-7469034-INS-54
chrX-7523246-INS-108
chrX-7548138-INS-52
chrX-7552666-INS-210
chrX-7554575-INS-197
chrX-7569513-INS-176
chrX-7578916-INS-166
chrX-7587974-INS-308
chrX-7590879-INS-207
chrX-7615167-INS-147
chrX-7619351-INS-180
chrX-7619949-INS-172
chrX-7622281-INS-207
chrX-7625955-INS-207
chrX-7651037-INS-74
chrX-7664749-INS-197

chrX-7675698-INS-182
chrX-7705007-INS-55
chrX-7760832-INS-163
chrX-7770740-INS-3434
chrX-7773420-INS-215
chrX-7791837-INS-201
chrX-7844847-INS-175
chrX-7854752-INS-88
chrX-7855935-INS-757
chrX-7857220-INS-146
chrX-7861373-INS-147
chrX-7870816-INS-196
chrX-7877090-INS-70
chrX-7896456-INS-513
chrX-7899065-INS-195
chrX-7903673-INS-440
chrX-7914317-INS-207
chrX-7926439-INS-68
chrX-7959365-INS-226
chrX-7961974-INS-88
chrX-7965097-INS-537
chrX-7965162-INS-930
chrX-7966726-INS-56
chrX-7980462-INS-79
chrX-8018491-INS-183
chrX-8024443-INS-81
chrX-8075438-INS-206
chrX-8082721-INS-52
chrX-8097452-INS-205
chrX-8133391-INS-965
chrX-8139591-INS-61
chrX-8175761-INS-1886
chrX-8342340-INS-201
chrX-8393108-INS-93
chrX-8394880-INS-1431
chrX-8435125-INS-1922
chrX-8446237-INS-1166
chrX-8458974-INS-315
chrX-8503384-INS-2919
chrX-8552947-INS-86
chrX-8649493-INS-7699
chrX-8703174-INS-361
chrX-8731511-INS-234
chrX-8756612-INS-69
chrX-8787141-INS-86
chrX-8791548-INS-81
chrX-8814619-INS-706
chrX-8843196-INS-252
chrX-8875605-INS-402
chrX-8895873-INS-163
chrX-8899252-INS-566
chrX-8945621-INS-71
chrX-8975083-INS-389
chrX-8989462-INS-56
chrX-9004548-INS-324
chrX-9007749-INS-107
chrX-9010144-INS-62
chrX-9029514-INS-319
chrX-9063129-INS-480
chrX-9159873-INS-64
chrX-9169359-INS-768
chrX-9188135-INS-5061
chrX-9199588-INS-504
chrX-9202532-INS-176
chrX-9203919-INS-6271
chrX-9234797-INS-6983
chrX-9242970-INS-108
chrX-9246067-INS-171
chrX-9266658-INS-229
chrX-9270137-INS-647
chrX-9273127-INS-916
chrX-9275896-INS-106
chrX-9287160-INS-320
chrX-9309744-INS-3428
chrX-9320496-INS-89
chrX-9328737-INS-154
chrX-9329531-INS-237

chrX-9339197-INS-248
chrX-9350472-INS-953
chrX-9356905-INS-52
chrX-9394751-INS-7438
chrX-9411427-INS-55
chrX-9450875-INS-52
chrX-9452077-INS-54
chrX-9454169-INS-215
chrX-9487148-INS-53
chrX-9524649-INS-124
chrX-9553309-INS-487
chrX-9559135-INS-1088
chrX-9565283-INS-752
chrX-9576766-INS-4867
chrX-9598432-INS-878
chrX-9598980-INS-57
chrX-9620653-INS-226
chrX-9654756-INS-96
chrX-9669531-INS-3083
chrX-9714411-INS-6304
chrX-9723377-INS-128
chrX-9750556-INS-245
chrX-9777973-INS-229
chrX-9791385-INS-214
chrX-9814153-INS-213
chrX-9840963-INS-155
chrX-9850999-INS-383
chrX-9872109-INS-54
chrX-9877497-INS-614
chrX-9878102-INS-744
chrX-9881010-INS-72
chrX-9904090-INS-60
chrX-9910001-INS-60
chrX-9913181-INS-462
chrX-9936903-INS-117
chrX-9945042-INS-712
chrX-9952692-INS-92
chrX-9954265-INS-55
chrX-9974649-INS-95
chrX-10001836-INS-168
chrX-10023040-INS-117
chrX-10025680-INS-68
chrX-10072865-INS-239
chrX-10077105-INS-210
chrX-10083832-INS-6215
chrX-10113506-INS-100
chrX-10126162-INS-57
chrX-10145889-INS-125
chrX-10183683-INS-307
chrX-10204960-INS-6382
chrX-10219568-INS-235
chrX-10236149-INS-344
chrX-10261361-INS-274
chrX-10264175-INS-91
chrX-10278536-INS-146
chrX-10282976-INS-120
chrX-10297595-INS-1068
chrX-10307814-INS-53
chrX-10334445-INS-192
chrX-10350298-INS-187
chrX-10360968-INS-65
chrX-10366925-INS-60
chrX-10398645-INS-54
chrX-10401767-INS-90
chrX-10427942-INS-467
chrX-10456869-INS-58
chrX-10458978-INS-67
chrX-10497123-INS-72
chrX-10505586-INS-141
chrX-10512323-INS-104
chrX-10538848-INS-2435
chrX-10549297-INS-196
chrX-10551680-INS-399
chrX-10565564-INS-106
chrX-10570065-INS-90
chrX-10588488-INS-216
chrX-10612188-INS-329

chrX-10616975-INS-53
chrX-10639377-INS-465
chrX-10651504-INS-251
chrX-10788272-INS-125
chrX-10789761-INS-76
chrX-10792560-INS-269
chrX-10828200-INS-56
chrX-10853955-INS-53
chrX-10868856-INS-655
chrX-10871962-INS-58
chrX-10889676-INS-126
chrX-10969104-INS-138
chrX-10973985-INS-79
chrX-10979417-INS-100
chrX-11021789-INS-6665
chrX-11028485-INS-66
chrX-11046866-INS-223
chrX-11053685-INS-4064
chrX-11055584-INS-349
chrX-11071089-INS-3603
chrX-11078148-INS-168
chrX-11091619-INS-62
chrX-11119816-INS-83
chrX-11194933-INS-56
chrX-11209439-INS-57
chrX-11242730-INS-3157
chrX-11247893-INS-561
chrX-11254526-INS-562
chrX-11257554-INS-6352
chrX-11280499-INS-5542
chrX-11292212-INS-421
chrX-11355445-INS-52
chrX-11396124-INS-82
chrX-11405087-INS-53
chrX-11436194-INS-51
chrX-11500353-INS-229
chrX-11502844-INS-160
chrX-11571073-INS-184
chrX-11577470-INS-292
chrX-11615279-INS-87
chrX-11658428-INS-158
chrX-11708334-INS-752
chrX-11754449-INS-61
chrX-11800976-INS-101
chrX-11825526-INS-349
chrX-11833844-INS-62
chrX-11847018-INS-51
chrX-11863387-INS-55
chrX-11876782-INS-57
chrX-11880602-INS-103
chrX-11890202-INS-215
chrX-11897969-INS-55
chrX-11899047-INS-93
chrX-11960335-INS-53
chrX-11962017-INS-886
chrX-11974890-INS-52
chrX-11979169-INS-151
chrX-11984868-INS-171
chrX-11986442-INS-206
chrX-12000483-INS-56
chrX-12033051-INS-54
chrX-12035545-INS-64
chrX-12038428-INS-144
chrX-12039449-INS-204
chrX-12045055-INS-159
chrX-12077154-INS-196
chrX-12088133-INS-211
chrX-12092355-INS-102
chrX-12128066-INS-198
chrX-12169616-INS-64
chrX-12172358-INS-572
chrX-12184937-INS-117
chrX-12194858-INS-201
chrX-12200088-INS-137
chrX-12252630-INS-7246
chrX-12271475-INS-607
chrX-12301727-INS-56

chrX-12311646-INS-194
chrX-12322605-INS-73
chrX-12327903-INS-470
chrX-12330892-INS-522
chrX-12340799-INS-180
chrX-12369011-INS-664
chrX-12392831-INS-52
chrX-12408608-INS-144
chrX-12417054-INS-201
chrX-12423983-INS-82
chrX-12461271-INS-213
chrX-12470983-INS-209
chrX-12479234-INS-200
chrX-12490451-INS-61
chrX-12492266-INS-200
chrX-12499778-INS-5165
chrX-12517281-INS-350
chrX-12558107-INS-84
chrX-12564878-INS-199
chrX-12575948-INS-92
chrX-12577029-INS-200
chrX-12631864-INS-213
chrX-12641679-INS-163
chrX-12643786-INS-202
chrX-12656217-INS-150
chrX-12658894-INS-197
chrX-12667706-INS-54
chrX-12694479-INS-235
chrX-12699353-INS-205
chrX-12705255-INS-82
chrX-12715249-INS-701
chrX-12721149-INS-209
chrX-12722720-INS-208
chrX-12738725-INS-91
chrX-12761063-INS-57
chrX-12775064-INS-189
chrX-12819806-INS-201
chrX-12820774-INS-151
chrX-12825062-INS-122
chrX-12830288-INS-183
chrX-12840346-INS-82
chrX-12869408-INS-198
chrX-12881266-INS-150
chrX-12895349-INS-74
chrX-12896816-INS-401
chrX-12916778-INS-202
chrX-12929904-INS-232
chrX-12952870-INS-52
chrX-12957967-INS-65
chrX-12961456-INS-200
chrX-12971153-INS-164
chrX-12981290-INS-208
chrX-12982030-INS-229
chrX-12989796-INS-51
chrX-13005456-INS-209
chrX-13016460-INS-210
chrX-13025614-INS-221
chrX-13028680-INS-215
chrX-13080911-INS-126
chrX-13083883-INS-111
chrX-13087421-INS-165
chrX-13089092-INS-2098
chrX-13089632-INS-65
chrX-13090398-INS-417
chrX-13091820-INS-239
chrX-13110324-INS-58
chrX-13150745-INS-202
chrX-13166847-INS-120
chrX-13169827-INS-53
chrX-13175964-INS-291
chrX-13198968-INS-149
chrX-13225298-INS-58
chrX-13270939-INS-96
chrX-13273253-INS-70
chrX-13311941-INS-354
chrX-13321586-INS-111
chrX-13323748-INS-6582

chrX-13347964-INS-61
chrX-13348838-INS-225
chrX-13372702-INS-52
chrX-13456858-INS-55
chrX-13489409-INS-64
chrX-13524172-INS-147
chrX-13596369-INS-58
chrX-13599344-INS-6156
chrX-13644353-INS-605
chrX-13666265-INS-3621
chrX-13676227-INS-7112
chrX-13687000-INS-86
chrX-13739842-INS-51
chrX-13748638-INS-54
chrX-13792189-INS-569
chrX-13800490-INS-144
chrX-13834504-INS-98
chrX-13857148-INS-507
chrX-13921578-INS-84
chrX-13936489-INS-89
chrX-13938688-INS-274
chrX-13980337-INS-124
chrX-13989589-INS-82
chrX-13992977-INS-148
chrX-13999600-INS-516
chrX-14029236-INS-6217
chrX-14037049-INS-498
chrX-14048331-INS-6334
chrX-14050991-INS-6777
chrX-14055331-INS-6279
chrX-14055410-INS-6703
chrX-14078806-INS-6793
chrX-14090419-INS-211
chrX-14105738-INS-327
chrX-14114348-INS-1899
chrX-14116299-INS-132
chrX-14165493-INS-64
chrX-14188869-INS-54
chrX-14198631-INS-420
chrX-14235780-INS-125
chrX-14257212-INS-118
chrX-14287202-INS-139
chrX-14289969-INS-182
chrX-14328950-INS-428
chrX-14362654-INS-428
chrX-14470747-INS-71
chrX-14478122-INS-65
chrX-14487322-INS-844
chrX-14576405-INS-1288
chrX-14579825-INS-505
chrX-14580838-INS-571
chrX-14587944-INS-741
chrX-14602066-INS-66
chrX-14607901-INS-718
chrX-14624446-INS-5387
chrX-14635804-INS-358
chrX-14639729-INS-2305
chrX-14658380-INS-6800
chrX-14737792-INS-128
chrX-14752280-INS-161
chrX-14758473-INS-6368
chrX-14770321-INS-248
chrX-14771660-INS-6244
chrX-14795695-INS-86
chrX-14802027-INS-251
chrX-14856007-INS-174
chrX-14857115-INS-144
chrX-14882456-INS-2776
chrX-14892452-INS-4372
chrX-14908499-INS-1357
chrX-14922886-INS-263
chrX-14959236-INS-7979
chrX-14971196-INS-877
chrX-14976713-INS-139
chrX-15006081-INS-385
chrX-15025314-INS-64
chrX-15057660-INS-58

chrX-15058934-INS-129
chrX-15062386-INS-136
chrX-15080185-INS-2319
chrX-15100229-INS-79
chrX-15124854-INS-95
chrX-15128086-INS-4375
chrX-15138195-INS-139
chrX-15189065-INS-121
chrX-15209496-INS-210
chrX-15246552-INS-305
chrX-15274255-INS-69
chrX-15280648-INS-80
chrX-15317017-INS-216
chrX-15319288-INS-788
chrX-15331895-INS-519
chrX-15338639-INS-443
chrX-15361756-INS-2144
chrX-15380797-INS-72
chrX-15385106-INS-60
chrX-15445840-INS-76
chrX-15473033-INS-53
chrX-15507065-INS-154
chrX-15511058-INS-60
chrX-15514473-INS-546
chrX-15543605-INS-56
chrX-15555133-INS-176
chrX-15568859-INS-594
chrX-15581641-INS-486
chrX-15590172-INS-5794
chrX-15594547-INS-131
chrX-15662286-INS-167
chrX-15669101-INS-244
chrX-15709495-INS-214
chrX-15717237-INS-60
chrX-15754470-INS-104
chrX-15788625-INS-56
chrX-15789296-INS-226
chrX-15797572-INS-6635
chrX-15852457-INS-210
chrX-15886793-INS-77
chrX-15909618-INS-7109
chrX-15917743-INS-157
chrX-15928191-INS-168
chrX-15933459-INS-1148
chrX-15933844-INS-6306
chrX-15936830-INS-2281
chrX-15957758-INS-206
chrX-15988985-INS-87
chrX-16016567-INS-335
chrX-16028099-INS-232
chrX-16029982-INS-204
chrX-16066638-INS-107
chrX-16067726-INS-166
chrX-16073479-INS-69
chrX-16102132-INS-529
chrX-16112037-INS-53
chrX-16165463-INS-400
chrX-16195472-INS-51
chrX-16201321-INS-119
chrX-16219952-INS-823
chrX-16223427-INS-428
chrX-16227288-INS-6397
chrX-16228805-INS-60
chrX-16259282-INS-1903
chrX-16269086-INS-209
chrX-16303971-INS-458
chrX-16344792-INS-275
chrX-16351714-INS-163
chrX-16378349-INS-78
chrX-16427283-INS-90
chrX-16451574-INS-62
chrX-16465019-INS-80
chrX-16466164-INS-266
chrX-16473316-INS-86
chrX-16475083-INS-62
chrX-16479831-INS-178
chrX-16501721-INS-98

chrX-16512131-INS-59
chrX-16519082-INS-1019
chrX-16588960-INS-56
chrX-16598665-INS-170
chrX-16614114-INS-6483
chrX-16649266-INS-505
chrX-16650589-INS-1663
chrX-16710324-INS-70
chrX-16717487-INS-53
chrX-16749134-INS-91
chrX-16756140-INS-768
chrX-16783074-INS-395
chrX-16803416-INS-175
chrX-16818112-INS-285
chrX-16832222-INS-200
chrX-16844639-INS-4098
chrX-16858172-INS-493
chrX-16918471-INS-52
chrX-16943081-INS-62
chrX-16984415-INS-236
chrX-16984764-INS-924
chrX-17017760-INS-108
chrX-17076444-INS-339
chrX-17109245-INS-69
chrX-17112902-INS-409
chrX-17118517-INS-54
chrX-17179146-INS-400
chrX-17183100-INS-85
chrX-17189655-INS-71
chrX-17206523-INS-6365
chrX-17227357-INS-466
chrX-17242512-INS-60
chrX-17242721-INS-331
chrX-17260744-INS-134
chrX-17261212-INS-63
chrX-17273505-INS-192
chrX-17274019-INS-829
chrX-17322290-INS-4349
chrX-17346132-INS-127
chrX-17379579-INS-137
chrX-17391732-INS-136
chrX-17456457-INS-521
chrX-17487802-INS-70
chrX-17495789-INS-96
chrX-17505785-INS-66
chrX-17540215-INS-124
chrX-17554535-INS-996
chrX-17598506-INS-398
chrX-17600960-INS-5508
chrX-17624652-INS-180
chrX-17646598-INS-540
chrX-17667566-INS-59
chrX-17683961-INS-492
chrX-17684766-INS-6473
chrX-17689079-INS-3867
chrX-17697099-INS-153
chrX-17716251-INS-537
chrX-17717765-INS-69
chrX-17719115-INS-1053
chrX-17733156-INS-914
chrX-17748158-INS-207
chrX-17759466-INS-221
chrX-17780752-INS-420
chrX-17784117-INS-238
chrX-17831598-INS-175
chrX-17897077-INS-204
chrX-17923541-INS-248
chrX-17924900-INS-7140
chrX-17973536-INS-64
chrX-17985340-INS-695
chrX-18000890-INS-54
chrX-18018645-INS-60
chrX-18020445-INS-207
chrX-18098187-INS-64
chrX-18180287-INS-153
chrX-18187819-INS-82
chrX-18194497-INS-886

chrX-18238779-INS-70
chrX-18240213-INS-7127
chrX-18245890-INS-133
chrX-18268383-INS-58
chrX-18297218-INS-1424
chrX-18303049-INS-65
chrX-18306538-INS-280
chrX-18323171-INS-52
chrX-18326748-INS-164
chrX-18346585-INS-393
chrX-18355180-INS-186
chrX-18356525-INS-527
chrX-18397606-INS-74
chrX-18417017-INS-400
chrX-18459441-INS-430
chrX-18464927-INS-6330
chrX-18484909-INS-7323
chrX-18489012-INS-3129
chrX-18504959-INS-311
chrX-18542818-INS-68
chrX-18552874-INS-6370
chrX-18573822-INS-2042
chrX-18581088-INS-3526
chrX-18593656-INS-1890
chrX-18656058-INS-73
chrX-18658118-INS-400
chrX-18690471-INS-108
chrX-18711158-INS-191
chrX-18720971-INS-341
chrX-18748292-INS-206
chrX-18753394-INS-6313
chrX-18757248-INS-86
chrX-18785950-INS-60
chrX-18825836-INS-162
chrX-18846478-INS-378
chrX-18874774-INS-176
chrX-18879285-INS-6902
chrX-18883677-INS-284
chrX-18894982-INS-59
chrX-18904478-INS-3040
chrX-18922116-INS-856
chrX-18932509-INS-54
chrX-18947549-INS-152
chrX-18999730-INS-276
chrX-19025825-INS-335
chrX-19064477-INS-63
chrX-19079662-INS-302
chrX-19104705-INS-63
chrX-19125999-INS-423
chrX-19207372-INS-6573
chrX-19215290-INS-66
chrX-19224135-INS-182
chrX-19244420-INS-108
chrX-19249749-INS-94
chrX-19261916-INS-4353
chrX-19318818-INS-7107
chrX-19320831-INS-115
chrX-19322389-INS-416
chrX-19351819-INS-673
chrX-19375108-INS-6392
chrX-19383668-INS-54
chrX-19440336-INS-51
chrX-19506316-INS-4407
chrX-19571678-INS-87
chrX-19574042-INS-403
chrX-19602307-INS-7358
chrX-19612554-INS-133
chrX-19685042-INS-76
chrX-19705481-INS-402
chrX-19707908-INS-6586
chrX-19721902-INS-100
chrX-19776204-INS-6679
chrX-19776876-INS-112
chrX-19778106-INS-143
chrX-19804479-INS-346
chrX-19823260-INS-63
chrX-19852665-INS-902

chrX-19891369-INS-994
chrX-19907620-INS-58
chrX-19925084-INS-52
chrX-19935671-INS-289
chrX-19936916-INS-190
chrX-19983658-INS-120
chrX-19996885-INS-931
chrX-20016743-INS-51
chrX-20037831-INS-6614
chrX-20086282-INS-577
chrX-20122103-INS-214
chrX-20136345-INS-551
chrX-20139864-INS-204
chrX-20164600-INS-77
chrX-20174647-INS-232
chrX-20183613-INS-204
chrX-20189694-INS-214
chrX-20195658-INS-53
chrX-20196475-INS-400
chrX-20198367-INS-130
chrX-20250098-INS-201
chrX-20255144-INS-225
chrX-20261887-INS-70
chrX-20262987-INS-151
chrX-20269300-INS-152
chrX-20327969-INS-330
chrX-20370874-INS-76
chrX-20372867-INS-235
chrX-20382248-INS-145
chrX-20451230-INS-112
chrX-20519976-INS-202
chrX-20526626-INS-59
chrX-20594840-INS-177
chrX-20599800-INS-61
chrX-20611734-INS-174
chrX-20614584-INS-217
chrX-20618965-INS-55
chrX-20619975-INS-136
chrX-20621343-INS-211
chrX-20636729-INS-199
chrX-20638865-INS-5672
chrX-20659746-INS-195
chrX-20670953-INS-191
chrX-20680817-INS-532
chrX-20684961-INS-75
chrX-20690690-INS-172
chrX-20692564-INS-211
chrX-20702377-INS-110
chrX-20722145-INS-53
chrX-20741895-INS-387
chrX-20746432-INS-52
chrX-20760438-INS-114
chrX-20764592-INS-202
chrX-20767767-INS-95
chrX-20771903-INS-197
chrX-20773702-INS-870
chrX-20815656-INS-207
chrX-20823335-INS-256
chrX-20839472-INS-281
chrX-20845758-INS-623
chrX-20848599-INS-198
chrX-20864122-INS-6694
chrX-20927490-INS-136
chrX-20961948-INS-109
chrX-20965570-INS-74
chrX-21006298-INS-93
chrX-21049439-INS-280
chrX-21114407-INS-104
chrX-21136087-INS-498
chrX-21140493-INS-6299
chrX-21163617-INS-478
chrX-21208413-INS-211
chrX-21213226-INS-408
chrX-21236894-INS-100
chrX-21268859-INS-136
chrX-21281123-INS-52
chrX-21283359-INS-363

chrX-21345344-INS-555
chrX-21385104-INS-228
chrX-21386618-INS-143
chrX-21411029-INS-1192
chrX-21416988-INS-569
chrX-21437981-INS-1990
chrX-21468617-INS-283
chrX-21482295-INS-52
chrX-21529483-INS-491
chrX-21541914-INS-593
chrX-21543009-INS-53
chrX-21550028-INS-56
chrX-21553053-INS-71
chrX-21557437-INS-624
chrX-21561167-INS-114
chrX-21584547-INS-56
chrX-21602426-INS-118
chrX-21619045-INS-213
chrX-21639457-INS-256
chrX-21655086-INS-53
chrX-21661449-INS-336
chrX-21668998-INS-58
chrX-21679302-INS-313
chrX-21685635-INS-560
chrX-21780475-INS-3707
chrX-21806295-INS-150
chrX-21815704-INS-146
chrX-21839914-INS-245
chrX-21884459-INS-97
chrX-21943586-INS-6353
chrX-22033200-INS-326
chrX-22060543-INS-388
chrX-22063953-INS-55
chrX-22078469-INS-52
chrX-22100940-INS-6600
chrX-22127280-INS-131
chrX-22128303-INS-117
chrX-22144045-INS-141
chrX-22188975-INS-595
chrX-22229493-INS-3143
chrX-22235119-INS-183
chrX-22282301-INS-256
chrX-22288031-INS-66
chrX-22318245-INS-120
chrX-22323514-INS-294
chrX-22344138-INS-85
chrX-22351858-INS-6729
chrX-22355583-INS-334
chrX-22388556-INS-203
chrX-22389476-INS-58
chrX-22413784-INS-181
chrX-22425140-INS-125
chrX-22453614-INS-120
chrX-22478445-INS-94
chrX-22481140-INS-759
chrX-22507378-INS-471
chrX-22519976-INS-329
chrX-22521471-INS-485
chrX-22610385-INS-64
chrX-22655756-INS-58
chrX-22661513-INS-6946
chrX-22702187-INS-182
chrX-22774461-INS-59
chrX-22776862-INS-745
chrX-22824802-INS-72
chrX-22829479-INS-126
chrX-22875394-INS-958
chrX-22901869-INS-6386
chrX-22955154-INS-200
chrX-22966524-INS-4965
chrX-22999760-INS-6883
chrX-23000569-INS-482
chrX-23044597-INS-134
chrX-23070679-INS-519
chrX-23083692-INS-101
chrX-23186298-INS-701
chrX-23208946-INS-379

chrX-23217680-INS-59
chrX-23223500-INS-52
chrX-23226235-INS-72
chrX-23241069-INS-949
chrX-23249394-INS-68
chrX-23257442-INS-69
chrX-23276672-INS-354
chrX-23281064-INS-175
chrX-23325259-INS-446
chrX-23374612-INS-184
chrX-23386015-INS-2888
chrX-23387331-INS-443
chrX-23410854-INS-4366
chrX-23427221-INS-54
chrX-23432558-INS-86
chrX-23439839-INS-6202
chrX-23471172-INS-55
chrX-23601967-INS-51
chrX-23604437-INS-139
chrX-23610935-INS-641
chrX-23655850-INS-58
chrX-23677780-INS-480
chrX-23682110-INS-134
chrX-23716801-INS-6508
chrX-23725705-INS-272
chrX-23745296-INS-68
chrX-23758750-INS-52
chrX-23759675-INS-403
chrX-23776800-INS-196
chrX-23818707-INS-52
chrX-23867571-INS-1363
chrX-23868466-INS-213
chrX-23878790-INS-62
chrX-23900395-INS-1492
chrX-23915820-INS-125
chrX-24033330-INS-194
chrX-24201325-INS-400
chrX-24208541-INS-947
chrX-24259757-INS-280
chrX-24356154-INS-6730
chrX-24387777-INS-212
chrX-24702258-INS-4783
chrX-25339405-INS-1891
chrX-25391236-INS-354
chrX-26006577-INS-162
chrX-26010932-INS-6403
chrX-26141164-INS-1875
chrX-26210262-INS-66
chrX-26388038-INS-1890
chrX-26504625-INS-310
chrX-26505544-INS-59
chrX-26543151-INS-98
chrX-26560014-INS-53
chrX-26561314-INS-305
chrX-26686430-INS-6757
chrX-26714568-INS-53
chrX-26780075-INS-71
chrX-26830861-INS-400
chrX-26839482-INS-66
chrX-29942560-INS-921
chrX-30845972-INS-2246
chrX-30865242-INS-3124
chrX-30880775-INS-3749
chrX-30971011-INS-66
chrX-31216647-INS-81
chrX-31370032-INS-6226
chrX-31638581-INS-751
chrX-32054500-INS-81
chrX-32114399-INS-7097
chrX-32125539-INS-1016
chrX-32247584-INS-56
chrX-32253828-INS-51
chrX-32383880-INS-8780
chrX-32529844-INS-2603
chrX-32542632-INS-53
chrX-32559368-INS-212
chrX-32568984-INS-127

chrX-32651505-INS-72
chrX-32956409-INS-66
chrX-32980714-INS-398
chrX-33090682-INS-6399
chrX-33265083-INS-12380
chrX-33281586-INS-111
chrX-33408465-INS-66
chrX-33489261-INS-283
chrX-33565323-INS-2514
chrX-33740414-INS-52
chrX-34593864-INS-1895
chrX-34660022-INS-313
chrX-34909497-INS-6765
chrX-34941246-INS-180
chrX-35021385-INS-833
chrX-35043917-INS-2387
chrX-35054924-INS-813
chrX-35058034-INS-485
chrX-35061078-INS-548
chrX-35075171-INS-117
chrX-35078462-INS-267
chrX-35096402-INS-51
chrX-35103068-INS-7080
chrX-35143037-INS-943
chrX-35176253-INS-1413
chrX-35192217-INS-938
chrX-35213342-INS-52
chrX-35217654-INS-101
chrX-35280429-INS-59
chrX-35300773-INS-92
chrX-35316614-INS-59
chrX-35349742-INS-113
chrX-35367865-INS-208
chrX-35378026-INS-162
chrX-35397341-INS-70
chrX-35401016-INS-5541
chrX-35411936-INS-235
chrX-35427448-INS-70
chrX-35433928-INS-69
chrX-35442637-INS-68
chrX-35449277-INS-196
chrX-35455918-INS-400
chrX-35463324-INS-608
chrX-35465654-INS-68
chrX-35478531-INS-150
chrX-35483559-INS-88
chrX-35490918-INS-83
chrX-35497996-INS-200
chrX-35509515-INS-141
chrX-35515236-INS-52
chrX-35530269-INS-55
chrX-35543064-INS-105
chrX-35563284-INS-182
chrX-35581190-INS-610
chrX-35598617-INS-586
chrX-35600343-INS-63
chrX-35617267-INS-213
chrX-35638983-INS-433
chrX-35643795-INS-351
chrX-35690667-INS-177
chrX-35718694-INS-962
chrX-35721485-INS-107
chrX-35735934-INS-843
chrX-35761250-INS-86
chrX-35764385-INS-211
chrX-35777205-INS-54
chrX-35797357-INS-266
chrX-35811030-INS-120
chrX-35835792-INS-59
chrX-35846506-INS-449
chrX-35963470-INS-451
chrX-35964572-INS-184
chrX-36010166-INS-133
chrX-36023866-INS-54
chrX-36044547-INS-177
chrX-36045888-INS-207
chrX-36047943-INS-177

chrX-36055203-INS-64
chrX-36067493-INS-61
chrX-36103548-INS-168
chrX-36106231-INS-82
chrX-36110208-INS-61
chrX-36119403-INS-143
chrX-36121323-INS-182
chrX-36125735-INS-147
chrX-36129585-INS-176
chrX-36141626-INS-174
chrX-36143332-INS-84
chrX-36158376-INS-223
chrX-36168483-INS-655
chrX-36169082-INS-205
chrX-36170954-INS-381
chrX-36198589-INS-608
chrX-36201978-INS-582
chrX-36203386-INS-60
chrX-36223648-INS-96
chrX-36228320-INS-250
chrX-36284867-INS-93
chrX-36299270-INS-344
chrX-36319544-INS-54
chrX-36325384-INS-104
chrX-36330002-INS-105
chrX-36351375-INS-194
chrX-36397514-INS-199
chrX-36430560-INS-86
chrX-36432218-INS-68
chrX-36434823-INS-72
chrX-36456124-INS-119
chrX-36459320-INS-56
chrX-36465332-INS-53
chrX-36492619-INS-69
chrX-36501149-INS-399
chrX-36509473-INS-60
chrX-36586219-INS-399
chrX-36627871-INS-399
chrX-36629520-INS-1321
chrX-36747228-INS-1321
chrX-36759670-INS-3484
chrX-36778945-INS-169
chrX-36861318-INS-90
chrX-36862085-INS-361
chrX-36865610-INS-122
chrX-36875836-INS-95
chrX-36879739-INS-124
chrX-36889883-INS-90
chrX-36935073-INS-159
chrX-36937137-INS-95
chrX-36950351-INS-68
chrX-36967277-INS-81
chrX-36987012-INS-56
chrX-37017811-INS-1276
chrX-37028197-INS-179
chrX-37170523-INS-54
chrX-37177152-INS-100
chrX-37191300-INS-221
chrX-37201214-INS-666
chrX-37211414-INS-58
chrX-37217327-INS-237
chrX-37231064-INS-120
chrX-37238427-INS-142
chrX-37242206-INS-57
chrX-37294859-INS-54
chrX-37309040-INS-204
chrX-37334134-INS-104
chrX-37338276-INS-930
chrX-37362683-INS-66
chrX-37397517-INS-6456
chrX-37414423-INS-7965
chrX-37429369-INS-211
chrX-37434899-INS-261
chrX-37452641-INS-6334
chrX-37473669-INS-6881
chrX-37480244-INS-400
chrX-37483373-INS-889

chrX-37494995-INS-1358
chrX-37495194-INS-98
chrX-37532931-INS-84
chrX-37544470-INS-90
chrX-37572385-INS-66
chrX-37581814-INS-102
chrX-37590181-INS-77
chrX-37613755-INS-714
chrX-37641764-INS-320
chrX-37664818-INS-177
chrX-37670409-INS-66
chrX-37706053-INS-123
chrX-37712266-INS-157
chrX-37766911-INS-456
chrX-37785966-INS-399
chrX-37789099-INS-163
chrX-37795908-INS-6639
chrX-37829030-INS-60
chrX-37840034-INS-92
chrX-37852355-INS-198
chrX-37895354-INS-6657
chrX-37909451-INS-563
chrX-37963044-INS-7529
chrX-37987557-INS-51
chrX-38000209-INS-5826
chrX-38003304-INS-7273
chrX-38006559-INS-60
chrX-38007318-INS-6905
chrX-38012103-INS-55
chrX-38024012-INS-1364
chrX-38078174-INS-77
chrX-38095497-INS-97
chrX-38098016-INS-188
chrX-38122941-INS-285
chrX-38138546-INS-5559
chrX-38146979-INS-6875
chrX-38182526-INS-326
chrX-38186191-INS-52
chrX-38204889-INS-6235
chrX-38207736-INS-6465
chrX-38280415-INS-106
chrX-38314738-INS-828
chrX-38336750-INS-65
chrX-38348455-INS-6594
chrX-38357338-INS-6749
chrX-38397559-INS-166
chrX-38444821-INS-53
chrX-38513089-INS-102
chrX-38518977-INS-5561
chrX-38611675-INS-58
chrX-38627610-INS-1888
chrX-38637919-INS-1852
chrX-38676446-INS-122
chrX-38711411-INS-881
chrX-38767507-INS-322
chrX-38785774-INS-156
chrX-38816882-INS-345
chrX-38821360-INS-6592
chrX-38822725-INS-53
chrX-38828193-INS-859
chrX-38829375-INS-326
chrX-38843425-INS-69
chrX-38847791-INS-280
chrX-38860228-INS-400
chrX-38867275-INS-116
chrX-38888579-INS-167
chrX-38988683-INS-994
chrX-39029081-INS-417
chrX-39041861-INS-68
chrX-39053184-INS-83
chrX-39054296-INS-73
chrX-39058196-INS-236
chrX-39068519-INS-498
chrX-39080902-INS-2742
chrX-39117399-INS-540
chrX-39125006-INS-61
chrX-39134957-INS-52

chrX-39137395-INS-690
chrX-39138926-INS-68
chrX-39139474-INS-3673
chrX-39158773-INS-271
chrX-39163771-INS-52
chrX-39193550-INS-56
chrX-39194121-INS-539
chrX-39195977-INS-6612
chrX-39225800-INS-286
chrX-39238178-INS-131
chrX-39249710-INS-686
chrX-39258133-INS-620
chrX-39267976-INS-1682
chrX-39269555-INS-62
chrX-39303383-INS-82
chrX-39310563-INS-90
chrX-39371363-INS-2305
chrX-39371800-INS-209
chrX-39376375-INS-496
chrX-39379953-INS-287
chrX-39383082-INS-873
chrX-39432421-INS-62
chrX-39442398-INS-2900
chrX-39451000-INS-1549
chrX-39475306-INS-2506
chrX-39475473-INS-51
chrX-39489284-INS-3120
chrX-39499392-INS-422
chrX-39502166-INS-86
chrX-39511988-INS-68
chrX-39512932-INS-144
chrX-39524002-INS-2888
chrX-39532386-INS-181
chrX-39547420-INS-5300
chrX-39552729-INS-6371
chrX-39603757-INS-96
chrX-39611139-INS-6802
chrX-39670044-INS-125
chrX-39674293-INS-731
chrX-39699335-INS-235
chrX-39714105-INS-6227
chrX-39740195-INS-6391
chrX-39759668-INS-71
chrX-39831595-INS-296
chrX-39832895-INS-1884
chrX-39835445-INS-112
chrX-39848911-INS-431
chrX-39869950-INS-195
chrX-39876410-INS-72
chrX-39883105-INS-6686
chrX-39915456-INS-116
chrX-39917498-INS-126
chrX-39952972-INS-670
chrX-39953834-INS-198
chrX-40032233-INS-179
chrX-40045560-INS-90
chrX-40062396-INS-65
chrX-40066793-INS-130
chrX-40067445-INS-788
chrX-40070346-INS-737
chrX-40095700-INS-132
chrX-40132248-INS-130
chrX-40164819-INS-1139
chrX-40167111-INS-275
chrX-40199542-INS-441
chrX-40212249-INS-257
chrX-40228761-INS-187
chrX-40233106-INS-579
chrX-40234075-INS-6537
chrX-40246282-INS-70
chrX-40259828-INS-72
chrX-40277176-INS-195
chrX-40278214-INS-338
chrX-40281821-INS-360
chrX-40303714-INS-100
chrX-40310784-INS-507
chrX-40319383-INS-56

chrX-40329050-INS-6158
chrX-40339044-INS-2095
chrX-40369238-INS-255
chrX-40380308-INS-53
chrX-40406863-INS-345
chrX-40408519-INS-6259
chrX-40409818-INS-123
chrX-40415499-INS-58
chrX-40422783-INS-88
chrX-40429658-INS-6381
chrX-40429669-INS-55
chrX-40468631-INS-105
chrX-40510100-INS-68
chrX-40512684-INS-349
chrX-40516623-INS-125
chrX-40519123-INS-822
chrX-40547649-INS-68
chrX-40548732-INS-84
chrX-40562061-INS-2521
chrX-40567746-INS-347
chrX-40580736-INS-64
chrX-40588463-INS-732
chrX-40624767-INS-368
chrX-40636721-INS-275
chrX-40638724-INS-84
chrX-40649287-INS-108
chrX-40656055-INS-256
chrX-40659029-INS-171
chrX-40664757-INS-269
chrX-40670938-INS-7876
chrX-40672810-INS-1672
chrX-40685443-INS-568
chrX-40707103-INS-1099
chrX-40738403-INS-58
chrX-40768290-INS-802
chrX-40804602-INS-5783
chrX-40831864-INS-90
chrX-40843177-INS-322
chrX-40870902-INS-194
chrX-40899164-INS-62
chrX-40905248-INS-151
chrX-40949385-INS-225
chrX-40958150-INS-71
chrX-40961341-INS-513
chrX-41019840-INS-620
chrX-41027323-INS-127
chrX-41054202-INS-220
chrX-41057802-INS-66
chrX-41059358-INS-7190
chrX-41060984-INS-202
chrX-41070183-INS-65
chrX-41089180-INS-63
chrX-41099984-INS-179
chrX-41104444-INS-160
chrX-41123706-INS-2190
chrX-41137461-INS-107
chrX-41141347-INS-68
chrX-41153753-INS-84
chrX-41170029-INS-165
chrX-41179931-INS-395
chrX-41192709-INS-1860
chrX-41208879-INS-89
chrX-41216671-INS-5353
chrX-41218927-INS-83
chrX-41224655-INS-2742
chrX-41230390-INS-238
chrX-41262976-INS-58
chrX-41265528-INS-652
chrX-41282722-INS-54
chrX-41295752-INS-98
chrX-41304021-INS-60
chrX-41319964-INS-776
chrX-41330375-INS-88
chrX-41374787-INS-73
chrX-41403609-INS-399
chrX-41410813-INS-165
chrX-41417878-INS-52

chrX-41434567-INS-112
chrX-41439926-INS-136
chrX-41457132-INS-378
chrX-41464659-INS-333
chrX-41466750-INS-112
chrX-41479266-INS-57
chrX-41481699-INS-68
chrX-41497030-INS-6454
chrX-41515042-INS-6201
chrX-41539728-INS-122
chrX-41560411-INS-87
chrX-41575474-INS-62
chrX-41673454-INS-296
chrX-41676518-INS-142
chrX-41700231-INS-385
chrX-41730850-INS-412
chrX-41732496-INS-212
chrX-41752555-INS-151
chrX-41768852-INS-806
chrX-41788124-INS-544
chrX-41810623-INS-81
chrX-41831555-INS-481
chrX-41840790-INS-52
chrX-41865097-INS-463
chrX-41872738-INS-103
chrX-41872970-INS-6499
chrX-41880099-INS-52
chrX-41882377-INS-199
chrX-41884608-INS-64
chrX-41895180-INS-872
chrX-41902097-INS-206
chrX-41927482-INS-68
chrX-41958512-INS-268
chrX-41960467-INS-909
chrX-41962858-INS-713
chrX-41965844-INS-1852
chrX-41970487-INS-6703
chrX-41989802-INS-99
chrX-42005590-INS-87
chrX-42037193-INS-841
chrX-42071112-INS-320
chrX-42073576-INS-6689
chrX-42088932-INS-64
chrX-42100654-INS-52
chrX-42136554-INS-59
chrX-42137449-INS-182
chrX-42157950-INS-582
chrX-42170154-INS-143
chrX-42206395-INS-2006
chrX-42210339-INS-473
chrX-42259298-INS-336
chrX-42269010-INS-6190
chrX-42288687-INS-66
chrX-42309367-INS-211
chrX-42346585-INS-6492
chrX-42353161-INS-721
chrX-42361355-INS-700
chrX-42424047-INS-202
chrX-42429324-INS-7142
chrX-42452507-INS-57
chrX-42460162-INS-66
chrX-42473576-INS-265
chrX-42517608-INS-70
chrX-42550761-INS-222
chrX-42558388-INS-142
chrX-42560638-INS-851
chrX-42561285-INS-64
chrX-42582127-INS-294
chrX-42673658-INS-6499
chrX-42682618-INS-6432
chrX-42686522-INS-911
chrX-42701305-INS-58
chrX-42709936-INS-416
chrX-42712540-INS-521
chrX-42712721-INS-3582
chrX-42721202-INS-6926
chrX-42728700-INS-78

chrX-42738565-INS-6114
chrX-42775189-INS-594
chrX-42792751-INS-52
chrX-42822825-INS-250
chrX-42863000-INS-106
chrX-42892129-INS-406
chrX-42894559-INS-270
chrX-42900721-INS-583
chrX-42918420-INS-77
chrX-42964544-INS-7539
chrX-43007725-INS-143
chrX-43013683-INS-643
chrX-43016596-INS-172
chrX-43082514-INS-204
chrX-43093603-INS-214
chrX-43113602-INS-106
chrX-43141885-INS-133
chrX-43143539-INS-102
chrX-43149497-INS-535
chrX-43173521-INS-59
chrX-43224911-INS-1015
chrX-43234883-INS-288
chrX-43235448-INS-740
chrX-43240815-INS-60
chrX-43244470-INS-52
chrX-43248762-INS-71
chrX-43271631-INS-184
chrX-43284784-INS-65
chrX-43296146-INS-72
chrX-43324607-INS-198
chrX-43335179-INS-72
chrX-43337243-INS-6401
chrX-43346866-INS-256
chrX-43349259-INS-55
chrX-43350371-INS-818
chrX-43358006-INS-624
chrX-43410418-INS-285
chrX-43425327-INS-6426
chrX-43449490-INS-304
chrX-43485707-INS-76
chrX-43488420-INS-449
chrX-43493796-INS-2875
chrX-43494115-INS-166
chrX-43558107-INS-581
chrX-43618464-INS-136
chrX-43622209-INS-106
chrX-43650155-INS-98
chrX-43671299-INS-67
chrX-43727848-INS-119
chrX-43737205-INS-328
chrX-43738403-INS-897
chrX-43752943-INS-6881
chrX-43789021-INS-103
chrX-43829967-INS-1889
chrX-43862761-INS-4089
chrX-43868042-INS-415
chrX-43912538-INS-57
chrX-43935993-INS-278
chrX-43948084-INS-152
chrX-43968083-INS-67
chrX-43985478-INS-68
chrX-43994475-INS-60
chrX-44003676-INS-404
chrX-44014965-INS-885
chrX-44017678-INS-831
chrX-44034437-INS-55
chrX-44036920-INS-800
chrX-44050897-INS-121
chrX-44064399-INS-676
chrX-44084526-INS-494
chrX-44115825-INS-95
chrX-44120684-INS-117
chrX-44133526-INS-976
chrX-44158357-INS-201
chrX-44172132-INS-499
chrX-44229118-INS-57
chrX-44236408-INS-400

chrX-44271978-INS-204
chrX-44277671-INS-1181
chrX-44297383-INS-397
chrX-44298801-INS-60
chrX-44341405-INS-1078
chrX-44355491-INS-127
chrX-44360328-INS-177
chrX-44408493-INS-192
chrX-44413188-INS-6679
chrX-44441904-INS-698
chrX-44524767-INS-100
chrX-44529542-INS-75
chrX-44565827-INS-7073
chrX-44575847-INS-60
chrX-44576952-INS-198
chrX-44586027-INS-950
chrX-44617651-INS-239
chrX-44635360-INS-551
chrX-44657840-INS-56
chrX-44722755-INS-90
chrX-44776469-INS-227
chrX-44802405-INS-56
chrX-44812114-INS-94
chrX-44828913-INS-666
chrX-44831177-INS-471
chrX-44844013-INS-466
chrX-44848254-INS-60
chrX-44853666-INS-714
chrX-44855271-INS-62
chrX-44871651-INS-956
chrX-44912606-INS-129
chrX-44946723-INS-182
chrX-44952764-INS-70
chrX-44968227-INS-230
chrX-45016449-INS-2192
chrX-45017526-INS-6527
chrX-45028890-INS-1155
chrX-45033208-INS-151
chrX-45043339-INS-6592
chrX-45066151-INS-8532
chrX-45069077-INS-113
chrX-45080946-INS-926
chrX-45142162-INS-85
chrX-45156926-INS-435
chrX-45175974-INS-55
chrX-45222354-INS-6358
chrX-45229905-INS-1366
chrX-45230510-INS-1539
chrX-45245459-INS-539
chrX-45258601-INS-51
chrX-45376724-INS-70
chrX-45414107-INS-452
chrX-45447015-INS-59
chrX-45454342-INS-56
chrX-45478910-INS-164
chrX-45484144-INS-402
chrX-45501779-INS-54
chrX-45521877-INS-424
chrX-45544744-INS-359
chrX-45587321-INS-6445
chrX-45596409-INS-115
chrX-45600505-INS-575
chrX-45626365-INS-898
chrX-45699338-INS-499
chrX-45723849-INS-288
chrX-45753590-INS-7212
chrX-45773607-INS-66
chrX-45793252-INS-60
chrX-45814169-INS-603
chrX-45818909-INS-590
chrX-45823036-INS-398
chrX-45839057-INS-360
chrX-45860951-INS-55
chrX-45868877-INS-515
chrX-45881181-INS-176
chrX-45897751-INS-92
chrX-45937263-INS-73

chrX-45945356-INS-222
chrX-45958080-INS-6919
chrX-45963357-INS-393
chrX-45970094-INS-107
chrX-46057501-INS-9695
chrX-46073883-INS-121
chrX-46095156-INS-56
chrX-46113634-INS-400
chrX-46127249-INS-472
chrX-46173371-INS-331
chrX-46178245-INS-131
chrX-46185927-INS-79
chrX-46240658-INS-73
chrX-46342769-INS-57
chrX-46359789-INS-97
chrX-46383529-INS-72
chrX-46389423-INS-4890
chrX-46401973-INS-89
chrX-46467521-INS-6028
chrX-46487414-INS-206
chrX-46488330-INS-1532
chrX-46491232-INS-1788
chrX-46518091-INS-1923
chrX-46552798-INS-203
chrX-46564388-INS-172
chrX-46591428-INS-92
chrX-46606561-INS-54
chrX-46629011-INS-52
chrX-46636243-INS-803
chrX-46638959-INS-155
chrX-46651569-INS-418
chrX-46657158-INS-56
chrX-46688003-INS-400
chrX-46696459-INS-1045
chrX-46718661-INS-127
chrX-46732583-INS-456
chrX-46738718-INS-148
chrX-46740550-INS-1644
chrX-46787089-INS-346
chrX-46832988-INS-505
chrX-46837002-INS-519
chrX-46848560-INS-340
chrX-46862099-INS-56
chrX-46878678-INS-201
chrX-46892204-INS-182
chrX-46895702-INS-7241
chrX-46910030-INS-557
chrX-46924408-INS-822
chrX-46937551-INS-141
chrX-46945937-INS-731
chrX-46953818-INS-59
chrX-46964017-INS-159
chrX-46965664-INS-52
chrX-46985329-INS-1169
chrX-47018193-INS-1278
chrX-47034380-INS-284
chrX-47049785-INS-171
chrX-47050820-INS-124
chrX-47085540-INS-138
chrX-47091586-INS-436
chrX-47141628-INS-98
chrX-47142628-INS-57
chrX-47152486-INS-54
chrX-47165626-INS-204
chrX-47178135-INS-60
chrX-47203099-INS-59
chrX-47238770-INS-63
chrX-47241987-INS-108
chrX-47259828-INS-53
chrX-47280422-INS-90
chrX-47304956-INS-60
chrX-47328651-INS-173
chrX-47342068-INS-7862
chrX-47377343-INS-241
chrX-47392162-INS-214
chrX-47414805-INS-161
chrX-47419454-INS-389

chrX-47421225-INS-230
chrX-47423753-INS-191
chrX-47535664-INS-260
chrX-47542019-INS-64
chrX-47544583-INS-198
chrX-47557862-INS-520
chrX-47580104-INS-203
chrX-47589353-INS-270
chrX-47603102-INS-426
chrX-47620420-INS-58
chrX-47626971-INS-55
chrX-47645645-INS-213
chrX-47662578-INS-704
chrX-47684595-INS-148
chrX-47728536-INS-65
chrX-47733685-INS-201
chrX-47736274-INS-188
chrX-47785973-INS-208
chrX-47796929-INS-56
chrX-47834228-INS-211
chrX-47844242-INS-235
chrX-47861303-INS-6458
chrX-47869118-INS-239
chrX-47904617-INS-275
chrX-47919084-INS-177
chrX-47925026-INS-138
chrX-47931204-INS-52
chrX-47958568-INS-400
chrX-47963731-INS-207
chrX-47985847-INS-1039
chrX-47993865-INS-320
chrX-48015122-INS-86
chrX-48028786-INS-400
chrX-48031901-INS-60
chrX-48083191-INS-377
chrX-48105098-INS-84
chrX-48160939-INS-6252
chrX-48162969-INS-6261
chrX-48173125-INS-884
chrX-48191350-INS-3372
chrX-48211910-INS-1090
chrX-48214437-INS-1679
chrX-48268994-INS-888
chrX-48275305-INS-344
chrX-48307835-INS-54
chrX-48376203-INS-124
chrX-48470074-INS-55
chrX-48472995-INS-66
chrX-48478177-INS-57
chrX-48509675-INS-6552
chrX-48545563-INS-61
chrX-48560846-INS-80
chrX-48564927-INS-129
chrX-48576367-INS-72
chrX-48592244-INS-6948
chrX-48610171-INS-156
chrX-48624689-INS-136
chrX-48629129-INS-267
chrX-48630963-INS-459
chrX-48776415-INS-8410
chrX-48778399-INS-58
chrX-48784460-INS-4420
chrX-48825595-INS-1145
chrX-48856283-INS-111
chrX-48892467-INS-116
chrX-48897160-INS-335
chrX-48902952-INS-5749
chrX-48936628-INS-393
chrX-48988084-INS-57
chrX-48990566-INS-132
chrX-48995421-INS-73
chrX-49002256-INS-155
chrX-49036469-INS-64
chrX-49070155-INS-551
chrX-49074364-INS-6665
chrX-49111704-INS-252
chrX-49116907-INS-301

chrX-49119608-INS-55
chrX-49121665-INS-84
chrX-49172406-INS-411
chrX-49189357-INS-2224
chrX-49194734-INS-239
chrX-49220965-INS-586
chrX-49229981-INS-6336
chrX-49257741-INS-557
chrX-49265664-INS-6614
chrX-49299404-INS-7636
chrX-49347170-INS-331
chrX-49370153-INS-470
chrX-49385705-INS-152
chrX-49385809-INS-7092
chrX-49386840-INS-1827
chrX-49390524-INS-486
chrX-49395979-INS-6273
chrX-49403547-INS-6855
chrX-49421671-INS-7331
chrX-49460226-INS-133
chrX-49463265-INS-152
chrX-49477911-INS-129
chrX-49492777-INS-59
chrX-49523664-INS-403
chrX-49543420-INS-147
chrX-49552232-INS-7635
chrX-49564968-INS-326
chrX-49569019-INS-110
chrX-49572564-INS-254
chrX-49574359-INS-52
chrX-49582354-INS-530
chrX-49589825-INS-609
chrX-49605042-INS-996
chrX-49606079-INS-660
chrX-49672567-INS-200
chrX-49712825-INS-68
chrX-49787446-INS-7619
chrX-49825451-INS-136
chrX-49833132-INS-58
chrX-49850576-INS-115
chrX-49911274-INS-53
chrX-49948785-INS-62
chrX-49956254-INS-63
chrX-49962922-INS-146
chrX-49974986-INS-7129
chrX-49996907-INS-78
chrX-50058213-INS-56
chrX-50063235-INS-222
chrX-50084642-INS-120
chrX-50097918-INS-409
chrX-50125839-INS-466
chrX-50130773-INS-198
chrX-50151893-INS-59
chrX-50160741-INS-77
chrX-50162646-INS-74
chrX-50167111-INS-56
chrX-50177434-INS-60
chrX-50235728-INS-89
chrX-50245869-INS-112
chrX-50263884-INS-441
chrX-50281214-INS-98
chrX-50336478-INS-189
chrX-50382053-INS-131
chrX-50403994-INS-668
chrX-50448675-INS-1604
chrX-50456112-INS-2131
chrX-50473546-INS-147
chrX-50488872-INS-542
chrX-50551309-INS-966
chrX-50577723-INS-53
chrX-50624055-INS-52
chrX-50638045-INS-118
chrX-50642688-INS-128
chrX-50646351-INS-72
chrX-50669358-INS-69
chrX-50684578-INS-561
chrX-50691661-INS-520

chrX-50708840-INS-123
chrX-50712149-INS-235
chrX-50714933-INS-1858
chrX-50719411-INS-243
chrX-50758403-INS-56
chrX-50791266-INS-53
chrX-50796179-INS-62
chrX-50830210-INS-104
chrX-50890932-INS-270
chrX-50899197-INS-54
chrX-50939662-INS-52
chrX-50950674-INS-1726
chrX-50954121-INS-543
chrX-50970627-INS-82
chrX-51025813-INS-71
chrX-51050375-INS-358
chrX-51055388-INS-58
chrX-51093775-INS-424
chrX-51098577-INS-190
chrX-51103533-INS-481
chrX-51108313-INS-73
chrX-51109151-INS-53
chrX-51112512-INS-230
chrX-51122752-INS-273
chrX-51155859-INS-139
chrX-51156162-INS-6477
chrX-51169105-INS-60
chrX-51219312-INS-182
chrX-51220733-INS-116
chrX-51234423-INS-51
chrX-51247745-INS-553
chrX-51267424-INS-5160
chrX-51272122-INS-470
chrX-51292497-INS-7079
chrX-51293222-INS-53
chrX-51295066-INS-4972
chrX-51300722-INS-56
chrX-51312751-INS-76
chrX-51322236-INS-68
chrX-51358840-INS-4467
chrX-51379902-INS-139
chrX-51389164-INS-52
chrX-51406113-INS-7066
chrX-51443620-INS-196
chrX-51457346-INS-175
chrX-51466977-INS-60
chrX-51475459-INS-90
chrX-51489415-INS-6446
chrX-51490016-INS-242
chrX-51525937-INS-1127
chrX-51551777-INS-185
chrX-51599271-INS-225
chrX-51611608-INS-1958
chrX-51631063-INS-1028
chrX-51632805-INS-62
chrX-51711772-INS-53
chrX-51755158-INS-100
chrX-51761022-INS-54
chrX-51783006-INS-60
chrX-51787394-INS-6422
chrX-51788682-INS-629
chrX-51796597-INS-3244
chrX-51797642-INS-204
chrX-51802418-INS-6527
chrX-51812634-INS-318
chrX-51817212-INS-211
chrX-51824021-INS-8443
chrX-51824040-INS-8746
chrX-51857858-INS-79
chrX-51860521-INS-80
chrX-51875258-INS-190
chrX-51896601-INS-6146
chrX-51897934-INS-105
chrX-51901312-INS-1499
chrX-51909430-INS-219
chrX-51949856-INS-65
chrX-51956300-INS-201

chrX-51965686-INS-5269
chrX-51973652-INS-1323
chrX-51973698-INS-610
chrX-52033098-INS-116
chrX-52052481-INS-106
chrX-52068615-INS-198
chrX-52075640-INS-204
chrX-52078326-INS-198
chrX-52092729-INS-219
chrX-52093718-INS-175
chrX-52098121-INS-636
chrX-52123228-INS-56
chrX-52186233-INS-503
chrX-52201549-INS-227
chrX-52208320-INS-498
chrX-52278989-INS-56
chrX-52294407-INS-54
chrX-52331211-INS-165
chrX-52402246-INS-134
chrX-52408438-INS-78
chrX-52419363-INS-1986
chrX-52452900-INS-109
chrX-52471099-INS-80
chrX-52485797-INS-156
chrX-52499534-INS-65
chrX-52510011-INS-7327
chrX-52514519-INS-2235
chrX-52530898-INS-199
chrX-52547213-INS-62
chrX-52552093-INS-62
chrX-52556858-INS-144
chrX-52566009-INS-54
chrX-52579981-INS-128
chrX-52589954-INS-94
chrX-52591394-INS-716
chrX-52595708-INS-150
chrX-52601561-INS-128
chrX-52638417-INS-290
chrX-52654899-INS-128
chrX-52679726-INS-298
chrX-52695799-INS-215
chrX-52723392-INS-60
chrX-52729060-INS-100
chrX-52734943-INS-311
chrX-52791843-INS-6350
chrX-52793919-INS-847
chrX-52800692-INS-949
chrX-52812693-INS-6445
chrX-52869507-INS-585
chrX-52870781-INS-1890
chrX-52879486-INS-191
chrX-52897543-INS-76
chrX-52907518-INS-5549
chrX-52911054-INS-112
chrX-52952196-INS-4258
chrX-52960910-INS-190
chrX-52961605-INS-2702
chrX-52962353-INS-6520
chrX-52969439-INS-1651
chrX-52970835-INS-6828
chrX-52974827-INS-65
chrX-52990801-INS-6443
chrX-53002924-INS-7031
chrX-53032399-INS-65
chrX-53060794-INS-476
chrX-53074193-INS-5331
chrX-53078391-INS-5549
chrX-53085864-INS-5006
chrX-53330255-INS-1942
chrX-53361552-INS-5417
chrX-53463007-INS-1888
chrX-53470122-INS-1378
chrX-53805587-INS-4537
chrX-53893804-INS-239
chrX-53904556-INS-498
chrX-54032371-INS-1084
chrX-54335080-INS-102

chrX-54340627-INS-76
chrX-54398234-INS-228
chrX-54431526-INS-2756
chrX-54436449-INS-172
chrX-54482278-INS-2672
chrX-54725036-INS-374
chrX-54733258-INS-1891
chrX-54752134-INS-70
chrX-54769927-INS-3705
chrX-54820991-INS-88
chrX-54852334-INS-1991
chrX-54878687-INS-105
chrX-54937672-INS-156
chrX-54942861-INS-67
chrX-54947058-INS-78
chrX-55063001-INS-6362
chrX-55160976-INS-199
chrX-55180393-INS-183
chrX-55304459-INS-526
chrX-55307410-INS-258
chrX-55315213-INS-81
chrX-55317004-INS-287
chrX-55345046-INS-196
chrX-55351918-INS-55
chrX-55379330-INS-81
chrX-55380208-INS-969
chrX-55398405-INS-170
chrX-55413457-INS-220
chrX-55438317-INS-544
chrX-55439330-INS-221
chrX-55449362-INS-93
chrX-55478750-INS-6982
chrX-55502920-INS-154
chrX-55519039-INS-523
chrX-55522226-INS-2877
chrX-55564513-INS-6683
chrX-55565013-INS-105
chrX-55566863-INS-57
chrX-55574901-INS-220
chrX-55587197-INS-53
chrX-55612743-INS-713
chrX-55624775-INS-103
chrX-55626076-INS-63
chrX-55645010-INS-57
chrX-55733822-INS-76
chrX-55801383-INS-112
chrX-55852805-INS-52
chrX-55865025-INS-201
chrX-55865303-INS-5542
chrX-55874797-INS-56
chrX-55879945-INS-215
chrX-55916097-INS-56
chrX-55921035-INS-309
chrX-55936237-INS-122
chrX-55966118-INS-389
chrX-55969993-INS-140
chrX-55971848-INS-2777
chrX-55980284-INS-200
chrX-55999354-INS-118
chrX-56019279-INS-571
chrX-56060191-INS-322
chrX-56060926-INS-63
chrX-56064599-INS-6215
chrX-56102128-INS-150
chrX-56113783-INS-70
chrX-56154578-INS-56
chrX-56188361-INS-98
chrX-56223272-INS-156
chrX-56239739-INS-112
chrX-56244241-INS-3175
chrX-56313865-INS-60
chrX-56316608-INS-60
chrX-56319498-INS-72
chrX-56321349-INS-56
chrX-56360470-INS-1892
chrX-56362509-INS-183
chrX-56368064-INS-174

chrX-56376648-INS-77
chrX-56389322-INS-110
chrX-56390786-INS-528
chrX-56393634-INS-68
chrX-56409113-INS-88
chrX-56439449-INS-425
chrX-56450064-INS-100
chrX-56453391-INS-179
chrX-56461383-INS-66
chrX-56483018-INS-6210
chrX-56490598-INS-473
chrX-56501523-INS-7421
chrX-56531762-INS-56
chrX-56532650-INS-226
chrX-56540040-INS-538
chrX-56554752-INS-853
chrX-56579503-INS-1017
chrX-56580527-INS-363
chrX-56612167-INS-67
chrX-56627005-INS-70
chrX-56631275-INS-877
chrX-56632371-INS-2800
chrX-56634239-INS-560
chrX-56635556-INS-775
chrX-56712488-INS-213
chrX-56714176-INS-104
chrX-56716926-INS-321
chrX-56730138-INS-1893
chrX-56731003-INS-366
chrX-56746079-INS-84
chrX-56768117-INS-198
chrX-56790527-INS-339
chrX-56797133-INS-95
chrX-56855754-INS-54
chrX-56900249-INS-1389
chrX-56928228-INS-50
chrX-56957521-INS-2597
chrX-57025146-INS-7064
chrX-57039072-INS-63
chrX-57047101-INS-102
chrX-57062814-INS-73
chrX-57067120-INS-209
chrX-57069666-INS-105
chrX-57104777-INS-135
chrX-57115604-INS-78
chrX-57121628-INS-117
chrX-57126063-INS-414
chrX-57136004-INS-58
chrX-57150072-INS-76
chrX-57174823-INS-216
chrX-57177238-INS-133
chrX-57183052-INS-71
chrX-57217272-INS-149
chrX-57221796-INS-224
chrX-57229190-INS-54
chrX-57230418-INS-199
chrX-57243458-INS-123
chrX-57267028-INS-3166
chrX-57271147-INS-110
chrX-57300941-INS-74
chrX-57313879-INS-6682
chrX-57360441-INS-555
chrX-57435501-INS-310
chrX-57455944-INS-153
chrX-57478885-INS-6493
chrX-57488058-INS-644
chrX-57558471-INS-6454
chrX-57567768-INS-499
chrX-57584598-INS-74
chrX-57589388-INS-6860
chrX-57603055-INS-2910
chrX-57610763-INS-231
chrX-57619224-INS-312
chrX-57621207-INS-765
chrX-57643090-INS-301
chrX-57644495-INS-197
chrX-57653091-INS-501

chrX-57661696-INS-3555
chrX-57708557-INS-1075
chrX-57750321-INS-56
chrX-57750961-INS-3098
chrX-57766919-INS-6048
chrX-57774862-INS-3916
chrX-57776304-INS-433
chrX-57778597-INS-6432
chrX-57803204-INS-3165
chrX-57803205-INS-60
chrX-57824145-INS-164
chrX-57825293-INS-6373
chrX-57879469-INS-998
chrX-57904370-INS-56
chrX-57932123-INS-80
chrX-57980671-INS-828
chrX-57990516-INS-331
chrX-58030191-INS-426
chrX-58033698-INS-479
chrX-58035334-INS-358
chrX-58050729-INS-6424
chrX-58051009-INS-578
chrX-58055519-INS-1542
chrX-58055985-INS-337
chrX-58058672-INS-225
chrX-58069178-INS-4975
chrX-58073384-INS-215
chrX-58082216-INS-174
chrX-58090747-INS-868
chrX-58125914-INS-112
chrX-58144282-INS-6476
chrX-58144835-INS-72
chrX-58153287-INS-809
chrX-58185509-INS-413
chrX-58219247-INS-106
chrX-58239328-INS-6202
chrX-58242641-INS-6249
chrX-58284221-INS-56
chrX-58296111-INS-56
chrX-58312699-INS-519
chrX-58346323-INS-58
chrX-58355378-INS-6591
chrX-58376253-INS-603
chrX-58388465-INS-81
chrX-58409543-INS-401
chrX-58411345-INS-451
chrX-58426698-INS-6256
chrX-58428537-INS-778
chrX-58434371-INS-380
chrX-58435436-INS-52
chrX-58439417-INS-6171
chrX-58446741-INS-6651
chrX-58452005-INS-76
chrX-58461501-INS-509
chrX-58479910-INS-966
chrX-58501264-INS-153
chrX-58518773-INS-147
chrX-58521903-INS-106
chrX-58531148-INS-56
chrX-58586937-INS-66
chrX-58598546-INS-80
chrX-58637470-INS-67
chrX-58638230-INS-444
chrX-58641556-INS-358
chrX-58646545-INS-151
chrX-58659688-INS-70
chrX-58662352-INS-58
chrX-58663393-INS-673
chrX-58676698-INS-442
chrX-58682153-INS-69
chrX-58709936-INS-842
chrX-58728211-INS-113
chrX-58732617-INS-93
chrX-58733723-INS-87
chrX-58748409-INS-492
chrX-58778002-INS-6469
chrX-58825885-INS-6757

chrX-58826731-INS-115
chrX-58852024-INS-6434
chrX-58862359-INS-6249
chrX-58874757-INS-1298
chrX-58915158-INS-56
chrX-58923479-INS-115
chrX-58937929-INS-6881
chrX-58941497-INS-3933
chrX-58944279-INS-412
chrX-58949180-INS-7172
chrX-58952390-INS-55
chrX-58956835-INS-66
chrX-58988530-INS-6844
chrX-59003242-INS-139
chrX-59008528-INS-82
chrX-59020566-INS-640
chrX-59025626-INS-1949
chrX-59030156-INS-209
chrX-59061673-INS-6479
chrX-59063820-INS-120
chrX-59103495-INS-114
chrX-59123861-INS-378
chrX-59132842-INS-687
chrX-59135985-INS-6430
chrX-59152697-INS-120
chrX-59178591-INS-214
chrX-59178768-INS-339
chrX-59186590-INS-149
chrX-59205169-INS-184
chrX-59206387-INS-203
chrX-59209580-INS-67
chrX-59212727-INS-272
chrX-59234080-INS-279
chrX-59255008-INS-107
chrX-59260945-INS-137
chrX-59304987-INS-52
chrX-59317284-INS-5980
chrX-59327249-INS-778
chrX-59329438-INS-179
chrX-59330809-INS-64
chrX-59336208-INS-199
chrX-59384861-INS-51
chrX-59426176-INS-358
chrX-59429115-INS-55
chrX-59432158-INS-106
chrX-59463639-INS-332
chrX-59479263-INS-3657
chrX-59479333-INS-3035
chrX-59494518-INS-131
chrX-59502086-INS-6442
chrX-59504652-INS-3190
chrX-59509630-INS-59
chrX-59519157-INS-56
chrX-59536857-INS-274
chrX-59551533-INS-55
chrX-59579403-INS-393
chrX-59610061-INS-161
chrX-59630190-INS-492
chrX-59671231-INS-6455
chrX-59676412-INS-62
chrX-59681181-INS-342
chrX-59682307-INS-316
chrX-59683411-INS-128
chrX-59746544-INS-383
chrX-59747922-INS-8420
chrX-59749459-INS-6246
chrX-59780874-INS-481
chrX-59799578-INS-155
chrX-59800602-INS-52
chrX-59819583-INS-202
chrX-59833006-INS-68
chrX-59847537-INS-968
chrX-59864304-INS-52
chrX-59867598-INS-55
chrX-59883904-INS-111
chrX-59899093-INS-178
chrX-59906999-INS-105

chrX-59970122-INS-52
chrX-60005891-INS-6482
chrX-60010337-INS-6736
chrX-60017078-INS-943
chrX-60017286-INS-6410
chrX-60026378-INS-242
chrX-60032496-INS-162
chrX-60065279-INS-172
chrX-60067527-INS-488
chrX-60071524-INS-6738
chrX-60083122-INS-95
chrX-60087802-INS-62
chrX-60094749-INS-118
chrX-60097273-INS-63
chrX-60102898-INS-121
chrX-60117060-INS-6606
chrX-60117063-INS-519
chrX-60126344-INS-766
chrX-60130889-INS-1159
chrX-60141387-INS-953
chrX-60149234-INS-444
chrX-60173999-INS-93
chrX-60228458-INS-54
chrX-60228659-INS-71
chrX-60230518-INS-258
chrX-60237396-INS-62
chrX-60250857-INS-318
chrX-60290531-INS-66
chrX-60311758-INS-58
chrX-60318160-INS-701
chrX-60330379-INS-128
chrX-60346087-INS-52
chrX-60389358-INS-424
chrX-60428600-INS-144
chrX-60437226-INS-211
chrX-60480519-INS-6261
chrX-60508306-INS-526
chrX-60534204-INS-345
chrX-60578203-INS-56
chrX-60579603-INS-73
chrX-60583253-INS-6354
chrX-60602950-INS-199
chrX-60605467-INS-160
chrX-60605597-INS-6983
chrX-60646034-INS-824
chrX-60651140-INS-207
chrX-60668702-INS-1180
chrX-60684115-INS-2789
chrX-60691832-INS-7760
chrX-60704324-INS-531
chrX-60705926-INS-6395
chrX-60803979-INS-133
chrX-60810124-INS-3266
chrX-60819573-INS-54
chrX-60822554-INS-124
chrX-60824326-INS-393
chrX-60831522-INS-7642
chrX-60844781-INS-289
chrX-60848217-INS-115
chrX-60869429-INS-68
chrX-60874362-INS-6593
chrX-60877213-INS-55
chrX-60891499-INS-72
chrX-60922167-INS-72
chrX-60973329-INS-204
chrX-60997092-INS-6404
chrX-61057226-INS-424
chrX-61062609-INS-6615
chrX-61066538-INS-87
chrX-61087028-INS-82
chrX-61090217-INS-59
chrX-61101369-INS-195
chrX-61103089-INS-61
chrX-61103437-INS-57
chrX-61135720-INS-98
chrX-61146194-INS-560
chrX-61163674-INS-92

chrX-61175722-INS-56
chrX-61200231-INS-158
chrX-61205949-INS-674
chrX-61221277-INS-60
chrX-61253121-INS-110
chrX-61276123-INS-57
chrX-61279496-INS-443
chrX-61315365-INS-60
chrX-61320673-INS-57
chrX-61412443-INS-68
chrX-61453037-INS-329
chrX-61497351-INS-70
chrX-61533713-INS-352
chrX-61542416-INS-983
chrX-61560342-INS-178
chrX-61580009-INS-53
chrX-61585299-INS-74
chrX-61610698-INS-586
chrX-61617638-INS-150
chrX-61694605-INS-7070
chrX-61696434-INS-431
chrX-61700557-INS-62
chrX-61714207-INS-57
chrX-61748822-INS-5544
chrX-61763383-INS-2906
chrX-61774020-INS-108
chrX-61778861-INS-455
chrX-61779458-INS-1280
chrX-61811286-INS-393
chrX-61811806-INS-3382
chrX-61821284-INS-52
chrX-61823740-INS-60
chrX-61824038-INS-213
chrX-61853631-INS-98
chrX-61863933-INS-121
chrX-61867993-INS-6414
chrX-61871367-INS-528
chrX-61878434-INS-64
chrX-61905885-INS-446
chrX-61922594-INS-345
chrX-61922644-INS-6463
chrX-61927481-INS-330
chrX-61943700-INS-4749
chrX-61945336-INS-2467
chrX-61947891-INS-201
chrX-61958701-INS-636
chrX-61963334-INS-563
chrX-61971812-INS-3544
chrX-61983754-INS-122
chrX-62003069-INS-114
chrX-62010096-INS-6623
chrX-62044344-INS-7091
chrX-62072619-INS-152
chrX-62090109-INS-64
chrX-62117895-INS-215
chrX-62195227-INS-182
chrX-62199289-INS-56
chrX-62213204-INS-219
chrX-62216207-INS-108
chrX-62229407-INS-395
chrX-62237357-INS-314
chrX-62254548-INS-385
chrX-62255471-INS-95
chrX-62267897-INS-93
chrX-62280642-INS-85
chrX-62309849-INS-108
chrX-62329061-INS-109
chrX-62371063-INS-218
chrX-62390055-INS-400
chrX-62390678-INS-831
chrX-62398655-INS-989
chrX-62470679-INS-208
chrX-62485064-INS-346
chrX-62489113-INS-86
chrX-62495197-INS-1329
chrX-62504752-INS-88
chrX-62542742-INS-7147

chrX-62546813-INS-3123
chrX-62549449-INS-842
chrX-62552101-INS-421
chrX-62563677-INS-86
chrX-62585561-INS-247
chrX-62601486-INS-200
chrX-62615610-INS-962
chrX-62630728-INS-528
chrX-62647771-INS-1891
chrX-62651068-INS-633
chrX-62656214-INS-134
chrX-62678478-INS-214
chrX-62693367-INS-102
chrX-62701910-INS-920
chrX-62702958-INS-3100
chrX-62709817-INS-3412
chrX-62722771-INS-5804
chrX-62725689-INS-7114
chrX-62729153-INS-149
chrX-62743248-INS-187
chrX-62745110-INS-875
chrX-62769797-INS-60
chrX-62798679-INS-86
chrX-62823504-INS-688
chrX-62827324-INS-2977
chrX-62840874-INS-508
chrX-62873360-INS-101
chrX-62891955-INS-76
chrX-62909149-INS-6610
chrX-62909528-INS-177
chrX-62996925-INS-6825
chrX-63014056-INS-200
chrX-63059091-INS-266
chrX-63069952-INS-549
chrX-63090410-INS-69
chrX-63130884-INS-459
chrX-63142935-INS-60
chrX-63167557-INS-120
chrX-63198168-INS-110
chrX-63209053-INS-1949
chrX-63210455-INS-91
chrX-63225880-INS-51
chrX-63247996-INS-60
chrX-63256797-INS-463
chrX-63261878-INS-894
chrX-63280680-INS-399
chrX-63281366-INS-56
chrX-63322395-INS-256
chrX-63346454-INS-76
chrX-63354796-INS-204
chrX-63362724-INS-51
chrX-63365792-INS-230
chrX-63381252-INS-64
chrX-63394599-INS-6662
chrX-63436063-INS-399
chrX-63518118-INS-401
chrX-63523206-INS-109
chrX-63576971-INS-51
chrX-63621132-INS-1862
chrX-63648537-INS-77
chrX-63708502-INS-196
chrX-63743085-INS-6412
chrX-63745192-INS-112
chrX-63779282-INS-115
chrX-63783395-INS-171
chrX-63794552-INS-132
chrX-63797248-INS-51
chrX-63858090-INS-60
chrX-63880404-INS-401
chrX-63888087-INS-348
chrX-63949537-INS-6401
chrX-63957732-INS-338
chrX-63967942-INS-6775
chrX-63991582-INS-417
chrX-63992624-INS-62
chrX-64021753-INS-7440
chrX-64052039-INS-6295

chrX-64054169-INS-134
chrX-64070192-INS-7145
chrX-64081895-INS-52
chrX-64112116-INS-7513
chrX-64126530-INS-6494
chrX-64130392-INS-957
chrX-64140624-INS-185
chrX-64171738-INS-434
chrX-64172932-INS-1330
chrX-64173633-INS-997
chrX-64193800-INS-413
chrX-64206350-INS-490
chrX-64229890-INS-69
chrX-64249725-INS-111
chrX-64270127-INS-6180
chrX-64273376-INS-5115
chrX-64283358-INS-173
chrX-64287677-INS-1519
chrX-64297259-INS-4243
chrX-64310094-INS-1989
chrX-64319109-INS-6209
chrX-64330404-INS-124
chrX-64331617-INS-135
chrX-64368968-INS-64
chrX-64384421-INS-54
chrX-64397224-INS-1024
chrX-64400026-INS-288
chrX-64419395-INS-467
chrX-64464021-INS-6444
chrX-64476409-INS-467
chrX-64479204-INS-62
chrX-64495499-INS-135
chrX-64504765-INS-90
chrX-64599655-INS-3020
chrX-64628638-INS-854
chrX-64632872-INS-75
chrX-64637522-INS-6622
chrX-64657006-INS-382
chrX-64675136-INS-52
chrX-64768771-INS-579
chrX-64769580-INS-63
chrX-64769789-INS-2792
chrX-64776408-INS-905
chrX-64787761-INS-57
chrX-64801886-INS-6883
chrX-64812667-INS-84
chrX-64816943-INS-68
chrX-64857706-INS-64
chrX-64893320-INS-131
chrX-64903172-INS-498
chrX-64913127-INS-597
chrX-64915046-INS-385
chrX-64937095-INS-68
chrX-64939341-INS-438
chrX-64951687-INS-80
chrX-64955421-INS-3039
chrX-64965612-INS-1319
chrX-64972785-INS-222
chrX-65001564-INS-911
chrX-65058245-INS-482
chrX-65063519-INS-198
chrX-65106830-INS-128
chrX-65111817-INS-54
chrX-65114466-INS-77
chrX-65136842-INS-6705
chrX-65154372-INS-178
chrX-65173271-INS-275
chrX-65180833-INS-95
chrX-65186147-INS-453
chrX-65203402-INS-6151
chrX-65203477-INS-715
chrX-65211456-INS-66
chrX-65220886-INS-3572
chrX-65227977-INS-6694
chrX-65230356-INS-136
chrX-65232238-INS-274
chrX-65247724-INS-7504

chrX-65282561-INS-708
chrX-65303926-INS-54
chrX-65344215-INS-6244
chrX-65439404-INS-204
chrX-65449461-INS-669
chrX-65456486-INS-150
chrX-65475878-INS-400
chrX-65485280-INS-54
chrX-65502465-INS-100
chrX-65530046-INS-509
chrX-65567730-INS-76
chrX-65580032-INS-871
chrX-65604993-INS-776
chrX-65619665-INS-401
chrX-65633541-INS-52
chrX-65662734-INS-411
chrX-65671804-INS-60
chrX-65675749-INS-186
chrX-65682511-INS-2906
chrX-65686055-INS-80
chrX-65710357-INS-5547
chrX-65729663-INS-231
chrX-65747649-INS-5503
chrX-65749981-INS-96
chrX-65755258-INS-66
chrX-65779677-INS-373
chrX-65784955-INS-4880
chrX-65809689-INS-189
chrX-65851865-INS-244
chrX-65862153-INS-168
chrX-65863848-INS-334
chrX-65916220-INS-199
chrX-65928883-INS-340
chrX-65975602-INS-217
chrX-66009802-INS-68
chrX-66010850-INS-714
chrX-66012153-INS-307
chrX-66015903-INS-59
chrX-66044105-INS-427
chrX-66070134-INS-2193
chrX-66108438-INS-315
chrX-66119985-INS-2134
chrX-66122448-INS-166
chrX-66123692-INS-63
chrX-66152871-INS-779
chrX-66172630-INS-199
chrX-66177498-INS-89
chrX-66184235-INS-82
chrX-66185061-INS-5722
chrX-66203928-INS-64
chrX-66220826-INS-78
chrX-66243572-INS-64
chrX-66247116-INS-6171
chrX-66256348-INS-56
chrX-66261698-INS-6280
chrX-66314394-INS-333
chrX-66320348-INS-113
chrX-66365798-INS-128
chrX-66373883-INS-114
chrX-66429647-INS-507
chrX-66441639-INS-176
chrX-66473664-INS-70
chrX-66500773-INS-52
chrX-66503569-INS-65
chrX-66514710-INS-214
chrX-66521636-INS-5256
chrX-66530389-INS-145
chrX-66575328-INS-200
chrX-66617921-INS-282
chrX-66620553-INS-364
chrX-66685195-INS-88
chrX-66722489-INS-184
chrX-66783346-INS-5432
chrX-66787408-INS-90
chrX-66811671-INS-514
chrX-66826497-INS-443
chrX-66839106-INS-324

chrX-66844213-INS-7027
chrX-66845624-INS-282
chrX-66876189-INS-338
chrX-66902813-INS-200
chrX-66905958-INS-64
chrX-66919273-INS-120
chrX-66923762-INS-61
chrX-66928772-INS-219
chrX-66931342-INS-313
chrX-66936560-INS-60
chrX-66942887-INS-323
chrX-67040736-INS-56
chrX-67064083-INS-93
chrX-67073394-INS-358
chrX-67083350-INS-55
chrX-67093159-INS-275
chrX-67108876-INS-7125
chrX-67116655-INS-148
chrX-67136999-INS-1154
chrX-67159595-INS-642
chrX-67231693-INS-346
chrX-67253727-INS-52
chrX-67291615-INS-6399
chrX-67314630-INS-53
chrX-67316486-INS-495
chrX-67331163-INS-5464
chrX-67332712-INS-5749
chrX-67335013-INS-420
chrX-67341076-INS-285
chrX-67375128-INS-158
chrX-67395243-INS-75
chrX-67407942-INS-6590
chrX-67421323-INS-132
chrX-67455181-INS-54
chrX-67455261-INS-6225
chrX-67463207-INS-4316
chrX-67535943-INS-126
chrX-67537183-INS-7437
chrX-67537422-INS-58
chrX-67540514-INS-6260
chrX-67559755-INS-3336
chrX-67576178-INS-158
chrX-67629920-INS-84
chrX-67631967-INS-209
chrX-67643575-INS-162
chrX-67650491-INS-219
chrX-67664255-INS-492
chrX-67671568-INS-1888
chrX-67684380-INS-410
chrX-67770927-INS-5227
chrX-67774954-INS-873
chrX-67841397-INS-1089
chrX-67903783-INS-1647
chrX-67932027-INS-3269
chrX-67937032-INS-78
chrX-67958554-INS-206
chrX-67971614-INS-6321
chrX-68007667-INS-1008
chrX-68008555-INS-62
chrX-68015630-INS-62
chrX-68026491-INS-56
chrX-68079157-INS-56
chrX-68140897-INS-550
chrX-68169720-INS-98
chrX-68175993-INS-6213
chrX-68178741-INS-6371
chrX-68188609-INS-147
chrX-68194506-INS-70
chrX-68225789-INS-4080
chrX-68233275-INS-75
chrX-68246317-INS-416
chrX-68262559-INS-106
chrX-68270968-INS-6594
chrX-68274464-INS-511
chrX-68282804-INS-104
chrX-68289694-INS-1543
chrX-68302628-INS-1477

chrX-68307668-INS-6484
chrX-68325718-INS-1141
chrX-68330152-INS-96
chrX-68363525-INS-1937
chrX-68367658-INS-1928
chrX-68369892-INS-74
chrX-68373450-INS-80
chrX-68433907-INS-140
chrX-68467386-INS-694
chrX-68565768-INS-68
chrX-68644325-INS-274
chrX-68654510-INS-263
chrX-68657419-INS-501
chrX-68665085-INS-888
chrX-68670596-INS-6557
chrX-68684428-INS-55
chrX-68715073-INS-412
chrX-68718579-INS-91
chrX-68749046-INS-56
chrX-68760989-INS-54
chrX-68764671-INS-531
chrX-68825672-INS-1886
chrX-68866904-INS-380
chrX-68874150-INS-1449
chrX-68886025-INS-204
chrX-68891921-INS-1839
chrX-68893253-INS-76
chrX-68924036-INS-260
chrX-68924610-INS-131
chrX-68924641-INS-881
chrX-68930734-INS-318
chrX-68943064-INS-421
chrX-68960515-INS-112
chrX-68961504-INS-362
chrX-68966634-INS-75
chrX-68988509-INS-194
chrX-68989877-INS-107
chrX-68991077-INS-1225
chrX-68991275-INS-7303
chrX-68996096-INS-6278
chrX-69003160-INS-1190
chrX-69024926-INS-228
chrX-69029222-INS-56
chrX-69040494-INS-262
chrX-69078530-INS-463
chrX-69079909-INS-67
chrX-69149005-INS-395
chrX-69172376-INS-68
chrX-69177009-INS-62
chrX-69210823-INS-239
chrX-69224766-INS-813
chrX-69235299-INS-199
chrX-69237159-INS-329
chrX-69238996-INS-6831
chrX-69239694-INS-122
chrX-69241184-INS-52
chrX-69247060-INS-1307
chrX-69249927-INS-112
chrX-69255951-INS-320
chrX-69286459-INS-689
chrX-69289055-INS-5425
chrX-69292947-INS-2972
chrX-69356054-INS-528
chrX-69359441-INS-760
chrX-69398424-INS-162
chrX-69510391-INS-540
chrX-69618557-INS-1193
chrX-69637004-INS-476
chrX-69651259-INS-196
chrX-69683668-INS-85
chrX-69731808-INS-113
chrX-69739301-INS-687
chrX-69744842-INS-141
chrX-69757911-INS-370
chrX-69772397-INS-148
chrX-69839773-INS-70
chrX-69855819-INS-1210

chrX-69883639-INS-78
chrX-69918302-INS-60
chrX-69958736-INS-187
chrX-70019531-INS-90
chrX-70059265-INS-71
chrX-70070450-INS-72
chrX-70079154-INS-120
chrX-70101994-INS-653
chrX-70125973-INS-90
chrX-70135791-INS-108
chrX-70159625-INS-1891
chrX-70177515-INS-401
chrX-70201490-INS-695
chrX-70220121-INS-2360
chrX-70241396-INS-375
chrX-70284093-INS-196
chrX-70287291-INS-401
chrX-70311969-INS-90
chrX-70332608-INS-407
chrX-70335866-INS-113
chrX-70352092-INS-547
chrX-70385836-INS-572
chrX-70388618-INS-58
chrX-70401842-INS-152
chrX-70405579-INS-80
chrX-70408939-INS-127
chrX-70418485-INS-570
chrX-70436473-INS-368
chrX-70456755-INS-63
chrX-70483579-INS-832
chrX-70507823-INS-162
chrX-70513122-INS-777
chrX-70525436-INS-194
chrX-70539270-INS-115
chrX-70637477-INS-68
chrX-70688004-INS-76
chrX-70732681-INS-6843
chrX-70735731-INS-150
chrX-70750011-INS-529
chrX-70762534-INS-400
chrX-70772784-INS-498
chrX-70774480-INS-64
chrX-70812465-INS-54
chrX-70846087-INS-401
chrX-70874565-INS-199
chrX-70875276-INS-125
chrX-70877345-INS-67
chrX-70908943-INS-492
chrX-70917297-INS-60
chrX-70960160-INS-128
chrX-70961835-INS-360
chrX-70969392-INS-133
chrX-70980166-INS-499
chrX-70990209-INS-92
chrX-71018806-INS-119
chrX-71026144-INS-61
chrX-71058685-INS-1127
chrX-71075892-INS-126
chrX-71090128-INS-175
chrX-71110159-INS-280
chrX-71113509-INS-499
chrX-71121368-INS-426
chrX-71158985-INS-1864
chrX-71186094-INS-114
chrX-71187600-INS-214
chrX-71198427-INS-500
chrX-71211666-INS-280
chrX-71215518-INS-280
chrX-71235326-INS-1890
chrX-71242721-INS-274
chrX-71248060-INS-84
chrX-71274060-INS-96
chrX-71292235-INS-119
chrX-71300593-INS-52
chrX-71320183-INS-187
chrX-71357075-INS-67
chrX-71377173-INS-230

chrX-71382636-INS-690
chrX-71444646-INS-344
chrX-71454656-INS-6214
chrX-71458233-INS-1891
chrX-71475194-INS-587
chrX-71486844-INS-103
chrX-71540634-INS-214
chrX-71556744-INS-308
chrX-71563049-INS-62
chrX-71587248-INS-56
chrX-71593216-INS-1056
chrX-71593259-INS-451
chrX-71612895-INS-8883
chrX-71616811-INS-426
chrX-71670791-INS-6251
chrX-71677508-INS-54
chrX-71703177-INS-385
chrX-71716103-INS-392
chrX-71806616-INS-144
chrX-71815566-INS-227
chrX-71842916-INS-230
chrX-71865535-INS-78
chrX-71876617-INS-759
chrX-71890643-INS-165
chrX-71891636-INS-60
chrX-71916521-INS-120
chrX-71938241-INS-891
chrX-71948496-INS-202
chrX-71974183-INS-1458
chrX-71985028-INS-66
chrX-72015901-INS-1203
chrX-72023668-INS-76
chrX-72038049-INS-62
chrX-72058546-INS-483
chrX-72068176-INS-197
chrX-72069773-INS-63
chrX-72077033-INS-211
chrX-72080335-INS-6401
chrX-72098422-INS-65
chrX-72103902-INS-59
chrX-72161433-INS-137
chrX-72163352-INS-109
chrX-72169307-INS-139
chrX-72184384-INS-69
chrX-72192063-INS-137
chrX-72220578-INS-116
chrX-72257127-INS-103
chrX-72318219-INS-1696
chrX-72326272-INS-3137
chrX-72326721-INS-944
chrX-72338213-INS-134
chrX-72354450-INS-428
chrX-72363964-INS-190
chrX-72366351-INS-228
chrX-72381502-INS-154
chrX-72391219-INS-205
chrX-72421522-INS-66
chrX-72495562-INS-52
chrX-72499994-INS-74
chrX-72534284-INS-61
chrX-72541691-INS-884
chrX-72562002-INS-51
chrX-72574439-INS-112
chrX-72579384-INS-401
chrX-72583002-INS-59
chrX-72613334-INS-123
chrX-72633981-INS-122
chrX-72635680-INS-1884
chrX-72636721-INS-67
chrX-72676598-INS-235
chrX-72690801-INS-976
chrX-72695754-INS-194
chrX-72739929-INS-188
chrX-72740529-INS-79
chrX-72771791-INS-211
chrX-72786276-INS-637
chrX-72789298-INS-86

chrX-72854388-INS-155
chrX-72869811-INS-197
chrX-72873414-INS-52
chrX-72907368-INS-151
chrX-72929477-INS-163
chrX-72951419-INS-325
chrX-72961848-INS-224
chrX-72976418-INS-215
chrX-72980367-INS-197
chrX-72983148-INS-196
chrX-73030786-INS-204
chrX-73047393-INS-104
chrX-73053016-INS-201
chrX-73061392-INS-192
chrX-73140219-INS-2640
chrX-73156048-INS-58
chrX-73179013-INS-376
chrX-73194601-INS-374
chrX-73200006-INS-195
chrX-73202453-INS-3330
chrX-73212696-INS-70
chrX-73224663-INS-53
chrX-73228266-INS-260
chrX-73266490-INS-401
chrX-73296315-INS-179
chrX-73318312-INS-101
chrX-73341934-INS-86
chrX-73367829-INS-99
chrX-73369507-INS-739
chrX-73393492-INS-58
chrX-73397960-INS-282
chrX-73399870-INS-174
chrX-73408634-INS-198
chrX-73418724-INS-218
chrX-73433228-INS-484
chrX-73442685-INS-303
chrX-73497246-INS-166
chrX-73516035-INS-1377
chrX-73522120-INS-6663
chrX-73522965-INS-123
chrX-73532373-INS-958
chrX-73561676-INS-158
chrX-73596370-INS-161
chrX-73606175-INS-459
chrX-73636423-INS-127
chrX-73673017-INS-436
chrX-73712358-INS-66
chrX-73731754-INS-61
chrX-73740559-INS-63
chrX-73760012-INS-125
chrX-73830287-INS-149
chrX-73847525-INS-58
chrX-73888249-INS-274
chrX-73895491-INS-4802
chrX-73956896-INS-61
chrX-73966716-INS-3223
chrX-73991137-INS-3104
chrX-74015337-INS-409
chrX-74019535-INS-7013
chrX-74028393-INS-390
chrX-74042736-INS-6647
chrX-74058843-INS-6818
chrX-74073285-INS-52
chrX-74093731-INS-135
chrX-74111918-INS-66
chrX-74136085-INS-54
chrX-74181023-INS-220
chrX-74234542-INS-174
chrX-74250946-INS-93
chrX-74273530-INS-436
chrX-74283946-INS-87
chrX-74335118-INS-6192
chrX-74338098-INS-105
chrX-74339007-INS-6470
chrX-74355151-INS-70
chrX-74370111-INS-145
chrX-74374387-INS-6670

chrX-74380455-INS-421
chrX-74381430-INS-674
chrX-74384097-INS-7162
chrX-74385432-INS-964
chrX-74392277-INS-6786
chrX-74500205-INS-2788
chrX-74503414-INS-66
chrX-74544156-INS-217
chrX-74570640-INS-68
chrX-74646494-INS-595
chrX-74663363-INS-2388
chrX-74679665-INS-52
chrX-74688022-INS-60
chrX-74689371-INS-294
chrX-74699051-INS-328
chrX-74722666-INS-401
chrX-74751843-INS-6785
chrX-74779970-INS-6440
chrX-74812859-INS-57
chrX-74833192-INS-144
chrX-74834196-INS-134
chrX-74858084-INS-1893
chrX-74876599-INS-7115
chrX-74888367-INS-196
chrX-74891215-INS-524
chrX-74931079-INS-482
chrX-74961299-INS-321
chrX-74969579-INS-56
chrX-74979776-INS-498
chrX-74989431-INS-56
chrX-75015753-INS-505
chrX-75035011-INS-83
chrX-75074937-INS-489
chrX-75096603-INS-66
chrX-75115796-INS-1702
chrX-75119146-INS-6439
chrX-75123476-INS-117
chrX-75183749-INS-6530
chrX-75184093-INS-193
chrX-75188412-INS-498
chrX-75193267-INS-215
chrX-75215362-INS-240
chrX-75219609-INS-2926
chrX-75229509-INS-1204
chrX-75232483-INS-371
chrX-75233754-INS-646
chrX-75296255-INS-258
chrX-75332065-INS-8369
chrX-75338227-INS-7094
chrX-75349250-INS-2411
chrX-75378885-INS-117
chrX-75395874-INS-7115
chrX-75400674-INS-157
chrX-75414592-INS-7268
chrX-75416506-INS-3057
chrX-75484173-INS-60
chrX-75526326-INS-6475
chrX-75555336-INS-76
chrX-75583545-INS-454
chrX-75605390-INS-95
chrX-75608257-INS-530
chrX-75613026-INS-6428
chrX-75628382-INS-7118
chrX-75631878-INS-341
chrX-75684781-INS-623
chrX-75685919-INS-52
chrX-75725637-INS-52
chrX-75763447-INS-371
chrX-75794371-INS-3156
chrX-75817055-INS-969
chrX-75823276-INS-6147
chrX-75876517-INS-358
chrX-75923587-INS-70
chrX-75951749-INS-191
chrX-76016913-INS-2815
chrX-76017941-INS-385
chrX-76037456-INS-124

chrX-76059975-INS-511
chrX-76077378-INS-104
chrX-76080248-INS-955
chrX-76084927-INS-509
chrX-76087413-INS-68
chrX-76102766-INS-313
chrX-76112130-INS-207
chrX-76113865-INS-82
chrX-76116761-INS-1890
chrX-76135303-INS-62
chrX-76143460-INS-6248
chrX-76145627-INS-5613
chrX-76165884-INS-7091
chrX-76170452-INS-108
chrX-76234047-INS-134
chrX-76278330-INS-113
chrX-76281141-INS-52
chrX-76286696-INS-6693
chrX-76343732-INS-7247
chrX-76349501-INS-150
chrX-76387369-INS-208
chrX-76390581-INS-625
chrX-76408831-INS-554
chrX-76412263-INS-177
chrX-76420534-INS-235
chrX-76430791-INS-66
chrX-76434292-INS-88
chrX-76441702-INS-429
chrX-76464611-INS-54
chrX-76510160-INS-188
chrX-76520145-INS-160
chrX-76558523-INS-102
chrX-76565303-INS-52
chrX-76576383-INS-96
chrX-76581364-INS-177
chrX-76647982-INS-446
chrX-76656283-INS-135
chrX-76668089-INS-56
chrX-76685150-INS-53
chrX-76721249-INS-1443
chrX-76721250-INS-1947
chrX-76722026-INS-2018
chrX-76730840-INS-867
chrX-76740607-INS-200
chrX-76774999-INS-6457
chrX-76782429-INS-605
chrX-76812714-INS-2474
chrX-76821850-INS-64
chrX-76829470-INS-7111
chrX-76830741-INS-326
chrX-76845981-INS-328
chrX-76848311-INS-52
chrX-76860974-INS-106
chrX-76879874-INS-54
chrX-76911797-INS-55
chrX-76934707-INS-495
chrX-76965195-INS-88
chrX-77000454-INS-808
chrX-77035353-INS-87
chrX-77054180-INS-524
chrX-77063041-INS-1482
chrX-77066855-INS-238
chrX-77089947-INS-6345
chrX-77115188-INS-202
chrX-77121894-INS-1923
chrX-77146366-INS-218
chrX-77155393-INS-4931
chrX-77163602-INS-7768
chrX-77168338-INS-610
chrX-77246924-INS-84
chrX-77261095-INS-539
chrX-77263528-INS-279
chrX-77266060-INS-6934
chrX-77302642-INS-280
chrX-77329583-INS-193
chrX-77399477-INS-5356
chrX-77462541-INS-6422

chrX-77477908-INS-86
chrX-77484726-INS-128
chrX-77488966-INS-362
chrX-77542201-INS-6621
chrX-77549100-INS-439
chrX-77549202-INS-7146
chrX-77561908-INS-611
chrX-77564753-INS-7526
chrX-77584749-INS-990
chrX-77591425-INS-116
chrX-77626413-INS-78
chrX-77640866-INS-293
chrX-77656327-INS-150
chrX-77724899-INS-332
chrX-77734068-INS-109
chrX-77773724-INS-90
chrX-77776953-INS-417
chrX-77823460-INS-106
chrX-77854618-INS-1980
chrX-77865817-INS-653
chrX-77868789-INS-1891
chrX-77879896-INS-375
chrX-77890757-INS-6266
chrX-77897064-INS-622
chrX-77906889-INS-51
chrX-77912962-INS-240
chrX-77925267-INS-72
chrX-77976733-INS-70
chrX-78008062-INS-92
chrX-78037076-INS-111
chrX-78100881-INS-337
chrX-78134729-INS-6790
chrX-78163743-INS-66
chrX-78251718-INS-79
chrX-78255960-INS-83
chrX-78285745-INS-56
chrX-78342350-INS-1009
chrX-78379035-INS-7304
chrX-78409594-INS-6282
chrX-78416888-INS-587
chrX-78421822-INS-2860
chrX-78451295-INS-197
chrX-78460800-INS-403
chrX-78464159-INS-2846
chrX-78480479-INS-1191
chrX-78494743-INS-112
chrX-78499748-INS-6464
chrX-78538416-INS-537
chrX-78548978-INS-96
chrX-78580320-INS-1276
chrX-78598440-INS-108
chrX-78625050-INS-5744
chrX-78647401-INS-12537
chrX-78684534-INS-158
chrX-78690700-INS-1648
chrX-78691212-INS-55
chrX-78697748-INS-75
chrX-78699046-INS-6413
chrX-78702396-INS-671
chrX-78710079-INS-839
chrX-78716124-INS-156
chrX-78721389-INS-1890
chrX-78730002-INS-6602
chrX-78746002-INS-579
chrX-78764569-INS-6718
chrX-78767906-INS-357
chrX-78791502-INS-85
chrX-78801409-INS-6639
chrX-78803371-INS-64
chrX-78823734-INS-126
chrX-78824074-INS-758
chrX-78826652-INS-7959
chrX-78829191-INS-820
chrX-78878116-INS-68
chrX-78881335-INS-6399
chrX-78894854-INS-491
chrX-78904457-INS-191

chrX-78915868-INS-1900
chrX-78915896-INS-436
chrX-78919596-INS-505
chrX-78926155-INS-73
chrX-78951687-INS-400
chrX-78964217-INS-58
chrX-78965311-INS-201
chrX-78979842-INS-7602
chrX-78999992-INS-65
chrX-79006099-INS-110
chrX-79008526-INS-375
chrX-79017341-INS-838
chrX-79018827-INS-7431
chrX-79024250-INS-210
chrX-79046494-INS-145
chrX-79048517-INS-2081
chrX-79050365-INS-8465
chrX-79070022-INS-52
chrX-79075651-INS-6269
chrX-79079001-INS-3222
chrX-79091076-INS-401
chrX-79094703-INS-3635
chrX-79105665-INS-366
chrX-79126787-INS-669
chrX-79160786-INS-119
chrX-79165494-INS-65
chrX-79174311-INS-129
chrX-79180553-INS-641
chrX-79195890-INS-82
chrX-79221463-INS-57
chrX-79249029-INS-59
chrX-79265201-INS-69
chrX-79274021-INS-141
chrX-79284459-INS-52
chrX-79285535-INS-2837
chrX-79305548-INS-6503
chrX-79307727-INS-11841
chrX-79344898-INS-6473
chrX-79355532-INS-360
chrX-79376483-INS-142
chrX-79399710-INS-9339
chrX-79476877-INS-6796
chrX-79484214-INS-6684
chrX-79485769-INS-231
chrX-79489560-INS-109
chrX-79499104-INS-155
chrX-79504425-INS-116
chrX-79536728-INS-4084
chrX-79601009-INS-362
chrX-79610114-INS-3463
chrX-79628858-INS-208
chrX-79661829-INS-550
chrX-79666357-INS-73
chrX-79765767-INS-2754
chrX-79779932-INS-60
chrX-79802110-INS-65
chrX-79816765-INS-60
chrX-79818477-INS-75
chrX-79835607-INS-1891
chrX-79838006-INS-56
chrX-79851639-INS-179
chrX-79859412-INS-416
chrX-79904913-INS-4945
chrX-79914959-INS-958
chrX-79930818-INS-998
chrX-79955900-INS-6358
chrX-79963918-INS-7083
chrX-79966617-INS-6839
chrX-79972573-INS-66
chrX-79984842-INS-909
chrX-79993213-INS-6823
chrX-80040997-INS-333
chrX-80057623-INS-255
chrX-80069207-INS-493
chrX-80070001-INS-55
chrX-80081149-INS-831
chrX-80111441-INS-213

chrX-80121953-INS-69
chrX-80131879-INS-52
chrX-80143697-INS-195
chrX-80159117-INS-53
chrX-80175656-INS-1488
chrX-80177081-INS-153
chrX-80187030-INS-347
chrX-80201661-INS-4513
chrX-80210184-INS-4535
chrX-80212478-INS-141
chrX-80215652-INS-7258
chrX-80264031-INS-55
chrX-80266766-INS-6204
chrX-80285278-INS-7076
chrX-80298549-INS-5486
chrX-80314765-INS-115
chrX-80382554-INS-70
chrX-80398411-INS-6599
chrX-80456866-INS-208
chrX-80460981-INS-6501
chrX-80473205-INS-1864
chrX-80507048-INS-3318
chrX-80547335-INS-433
chrX-80576135-INS-97
chrX-80585354-INS-70
chrX-80605915-INS-741
chrX-80618795-INS-1487
chrX-80638788-INS-409
chrX-80779291-INS-221
chrX-80798383-INS-121
chrX-80822665-INS-79
chrX-80862238-INS-7135
chrX-80936923-INS-4469
chrX-80950534-INS-210
chrX-80951802-INS-144
chrX-80976549-INS-476
chrX-80987487-INS-221
chrX-81002330-INS-735
chrX-81016720-INS-58
chrX-81038774-INS-7107
chrX-81061398-INS-171
chrX-81068217-INS-218
chrX-81073574-INS-376
chrX-81074447-INS-54
chrX-81114941-INS-190
chrX-81120662-INS-1352
chrX-81141388-INS-8322
chrX-81166852-INS-1176
chrX-81181187-INS-57
chrX-81182457-INS-6340
chrX-81183487-INS-71
chrX-81209020-INS-122
chrX-81220629-INS-209
chrX-81241691-INS-202
chrX-81249597-INS-4759
chrX-81270799-INS-68
chrX-81271278-INS-1053
chrX-81303852-INS-150
chrX-81312690-INS-1503
chrX-81335482-INS-200
chrX-81337085-INS-6439
chrX-81370838-INS-52
chrX-81375693-INS-830
chrX-81376296-INS-76
chrX-81378170-INS-70
chrX-81396766-INS-1891
chrX-81399761-INS-117
chrX-81401031-INS-5284
chrX-81443140-INS-194
chrX-81457595-INS-58
chrX-81465878-INS-59
chrX-81491856-INS-60
chrX-81498786-INS-52
chrX-81527808-INS-2713
chrX-81560859-INS-282
chrX-81570991-INS-342
chrX-81602691-INS-93

chrX-81605131-INS-64
chrX-81653411-INS-60
chrX-81654448-INS-203
chrX-81688604-INS-69
chrX-81709249-INS-6600
chrX-81709401-INS-60
chrX-81714855-INS-6209
chrX-81724701-INS-6924
chrX-81726386-INS-6305
chrX-81739085-INS-52
chrX-81746078-INS-244
chrX-81769513-INS-6075
chrX-81779935-INS-99
chrX-81784007-INS-100
chrX-81811474-INS-840
chrX-81822273-INS-5354
chrX-81839805-INS-526
chrX-81873989-INS-410
chrX-81911999-INS-75
chrX-81919278-INS-3981
chrX-81965451-INS-374
chrX-82010566-INS-1363
chrX-82020720-INS-190
chrX-82024743-INS-7240
chrX-82030775-INS-192
chrX-82034144-INS-400
chrX-82038632-INS-122
chrX-82039096-INS-505
chrX-82121693-INS-52
chrX-82125335-INS-6892
chrX-82126354-INS-60
chrX-82149336-INS-578
chrX-82150846-INS-100
chrX-82170584-INS-354
chrX-82174230-INS-1348
chrX-82190394-INS-384
chrX-82208543-INS-1231
chrX-82209591-INS-1255
chrX-82228044-INS-584
chrX-82230805-INS-76
chrX-82231433-INS-51
chrX-82231624-INS-6876
chrX-82233428-INS-58
chrX-82251230-INS-722
chrX-82265487-INS-540
chrX-82265721-INS-6850
chrX-82270229-INS-89
chrX-82273658-INS-1274
chrX-82286512-INS-781
chrX-82299012-INS-2058
chrX-82300359-INS-1096
chrX-82314768-INS-3284
chrX-82319064-INS-4644
chrX-82356808-INS-6452
chrX-82361509-INS-491
chrX-82399123-INS-136
chrX-82422039-INS-5525
chrX-82425492-INS-6715
chrX-82445216-INS-68
chrX-82463853-INS-262
chrX-82478438-INS-179
chrX-82482217-INS-299
chrX-82491240-INS-383
chrX-82506995-INS-83
chrX-82530664-INS-117
chrX-82537020-INS-65
chrX-82615545-INS-818
chrX-82645500-INS-109
chrX-82647475-INS-120
chrX-82665747-INS-1099
chrX-82696442-INS-116
chrX-82697759-INS-73
chrX-82707322-INS-229
chrX-82709298-INS-52
chrX-82722138-INS-171
chrX-82750092-INS-437
chrX-82767002-INS-62

chrX-82786931-INS-782
chrX-82791985-INS-63
chrX-82804787-INS-56
chrX-82823579-INS-343
chrX-82838251-INS-220
chrX-82842788-INS-2529
chrX-82866161-INS-82
chrX-82880092-INS-491
chrX-82891699-INS-61
chrX-82918179-INS-6503
chrX-82945530-INS-72
chrX-82967723-INS-90
chrX-82975621-INS-52
chrX-82985583-INS-86
chrX-83019617-INS-6865
chrX-83023950-INS-339
chrX-83033406-INS-72
chrX-83043972-INS-627
chrX-83052537-INS-62
chrX-83054150-INS-492
chrX-83081323-INS-6385
chrX-83098382-INS-711
chrX-83123343-INS-410
chrX-83127431-INS-171
chrX-83146155-INS-56
chrX-83172246-INS-52
chrX-83173771-INS-817
chrX-83175060-INS-301
chrX-83189191-INS-68
chrX-83195440-INS-281
chrX-83215604-INS-605
chrX-83254396-INS-302
chrX-83261904-INS-170
chrX-83266958-INS-619
chrX-83279473-INS-95
chrX-83287187-INS-6928
chrX-83288132-INS-5217
chrX-83301129-INS-235
chrX-83314073-INS-164
chrX-83319124-INS-306
chrX-83328359-INS-609
chrX-83331878-INS-111
chrX-83333716-INS-443
chrX-83342135-INS-851
chrX-83344454-INS-10902
chrX-83348484-INS-6737
chrX-83364381-INS-765
chrX-83392012-INS-8001
chrX-83411962-INS-52
chrX-83424942-INS-406
chrX-83444701-INS-68
chrX-83461601-INS-76
chrX-83481339-INS-54
chrX-83482910-INS-242
chrX-83495700-INS-68
chrX-83498131-INS-51
chrX-83524212-INS-199
chrX-83561670-INS-79
chrX-83565425-INS-68
chrX-83578004-INS-112
chrX-83589864-INS-65
chrX-83591108-INS-199
chrX-83592422-INS-6537
chrX-83593712-INS-65
chrX-83595143-INS-57
chrX-83601201-INS-451
chrX-83601579-INS-6851
chrX-83605403-INS-1795
chrX-83623853-INS-186
chrX-83635693-INS-4353
chrX-83643906-INS-6459
chrX-83651346-INS-416
chrX-83682437-INS-431
chrX-83702455-INS-261
chrX-83712923-INS-80
chrX-83720816-INS-682
chrX-83726553-INS-86

chrX-83743409-INS-6440
chrX-83750463-INS-1143
chrX-83753383-INS-1525
chrX-83810714-INS-1607
chrX-83833550-INS-6460
chrX-83838560-INS-6922
chrX-83842453-INS-134
chrX-83850475-INS-497
chrX-83858489-INS-57
chrX-83860188-INS-186
chrX-83866884-INS-1314
chrX-83869901-INS-7113
chrX-83870123-INS-6652
chrX-83870127-INS-220
chrX-83875808-INS-142
chrX-83881175-INS-147
chrX-83906638-INS-126
chrX-83928692-INS-183
chrX-83937038-INS-5389
chrX-83940259-INS-75
chrX-83951955-INS-58
chrX-83954092-INS-96
chrX-83993319-INS-211
chrX-83999120-INS-482
chrX-84015052-INS-702
chrX-84075665-INS-176
chrX-84076518-INS-541
chrX-84079080-INS-267
chrX-84102235-INS-660
chrX-84102235-INS-67
chrX-84185543-INS-56
chrX-84186830-INS-109
chrX-84198219-INS-60
chrX-84224840-INS-914
chrX-84229917-INS-6524
chrX-84273546-INS-68
chrX-84277987-INS-6008
chrX-84295824-INS-382
chrX-84311683-INS-60
chrX-84338116-INS-52
chrX-84356470-INS-819
chrX-84375582-INS-52
chrX-84377941-INS-238
chrX-84394185-INS-385
chrX-84412901-INS-506
chrX-84448296-INS-112
chrX-84477187-INS-1726
chrX-84489416-INS-1366
chrX-84510306-INS-1870
chrX-84577140-INS-528
chrX-84599225-INS-101
chrX-84605460-INS-65
chrX-84679613-INS-876
chrX-84699058-INS-5352
chrX-84704660-INS-113
chrX-84711301-INS-216
chrX-84728676-INS-2820
chrX-84730379-INS-518
chrX-84772972-INS-77
chrX-84780362-INS-156
chrX-84791967-INS-92
chrX-84805615-INS-67
chrX-84808952-INS-6487
chrX-84810464-INS-54
chrX-84811440-INS-462
chrX-84821572-INS-143
chrX-84844017-INS-135
chrX-84844770-INS-1788
chrX-84873650-INS-89
chrX-84885177-INS-95
chrX-84895298-INS-551
chrX-84899467-INS-249
chrX-84900013-INS-179
chrX-84902570-INS-133
chrX-84933109-INS-497
chrX-85042039-INS-60
chrX-85047002-INS-490

chrX-85047824-INS-59
chrX-85051589-INS-6570
chrX-85061534-INS-683
chrX-85079532-INS-779
chrX-85080429-INS-472
chrX-85103225-INS-219
chrX-85121409-INS-1718
chrX-85123949-INS-2704
chrX-85124765-INS-6187
chrX-85131210-INS-2884
chrX-85161225-INS-99
chrX-85170672-INS-547
chrX-85177960-INS-124
chrX-85187140-INS-174
chrX-85197451-INS-7159
chrX-85215007-INS-379
chrX-85218376-INS-112
chrX-85233290-INS-69
chrX-85258310-INS-194
chrX-85343695-INS-264
chrX-85402034-INS-55
chrX-85412854-INS-104
chrX-85420733-INS-802
chrX-85434525-INS-317
chrX-85438269-INS-668
chrX-85443397-INS-408
chrX-85487890-INS-52
chrX-85509371-INS-60
chrX-85517500-INS-110
chrX-85548085-INS-453
chrX-85558501-INS-6891
chrX-85603630-INS-200
chrX-85609377-INS-494
chrX-85648516-INS-76
chrX-85686170-INS-247
chrX-85716607-INS-6768
chrX-85718948-INS-56
chrX-85724565-INS-175
chrX-85732437-INS-948
chrX-85759698-INS-70
chrX-85762336-INS-165
chrX-85764259-INS-367
chrX-85767546-INS-201
chrX-85782155-INS-277
chrX-85802245-INS-6471
chrX-85802378-INS-96
chrX-85817607-INS-58
chrX-85836406-INS-6677
chrX-85843922-INS-175
chrX-85849298-INS-469
chrX-85859002-INS-6387
chrX-85935373-INS-766
chrX-85974900-INS-52
chrX-85988186-INS-6254
chrX-86000303-INS-56
chrX-86058483-INS-6312
chrX-86080595-INS-200
chrX-86085414-INS-611
chrX-86087720-INS-63
chrX-86087861-INS-6610
chrX-86102324-INS-101
chrX-86130845-INS-83
chrX-86139346-INS-8340
chrX-86146854-INS-6440
chrX-86152821-INS-374
chrX-86157837-INS-199
chrX-86161355-INS-6375
chrX-86164057-INS-67
chrX-86170010-INS-410
chrX-86173945-INS-108
chrX-86182771-INS-561
chrX-86195778-INS-100
chrX-86212246-INS-96
chrX-86229528-INS-234
chrX-86248199-INS-195
chrX-86268031-INS-6245
chrX-86272678-INS-54

chrX-86283103-INS-829
chrX-86286092-INS-358
chrX-86298778-INS-201
chrX-86316555-INS-842
chrX-86320813-INS-420
chrX-86331663-INS-3861
chrX-86346790-INS-62
chrX-86353773-INS-1644
chrX-86356643-INS-54
chrX-86377552-INS-58
chrX-86413987-INS-58
chrX-86416752-INS-400
chrX-86425905-INS-455
chrX-86438904-INS-956
chrX-86441343-INS-530
chrX-86453966-INS-152
chrX-86488128-INS-3137
chrX-86495626-INS-54
chrX-86513144-INS-122
chrX-86518184-INS-79
chrX-86518924-INS-6379
chrX-86534056-INS-188
chrX-86558793-INS-73
chrX-86567570-INS-60
chrX-86588247-INS-51
chrX-86612184-INS-74
chrX-86632484-INS-320
chrX-86656507-INS-62
chrX-86693801-INS-235
chrX-86710035-INS-86
chrX-86726141-INS-56
chrX-86732557-INS-173
chrX-86743157-INS-77
chrX-86747897-INS-284
chrX-86752249-INS-62
chrX-86760858-INS-157
chrX-86777645-INS-5509
chrX-86783594-INS-179
chrX-86785541-INS-976
chrX-86815394-INS-81
chrX-86837168-INS-63
chrX-86865600-INS-182
chrX-86871008-INS-156
chrX-86896650-INS-992
chrX-86909208-INS-107
chrX-86946007-INS-74
chrX-86949140-INS-168
chrX-86986553-INS-6174
chrX-87009877-INS-66
chrX-87059117-INS-425
chrX-87060056-INS-145
chrX-87064672-INS-612
chrX-87066244-INS-54
chrX-87106173-INS-67
chrX-87141856-INS-657
chrX-87176489-INS-56
chrX-87176883-INS-969
chrX-87188203-INS-95
chrX-87208809-INS-57
chrX-87210611-INS-291
chrX-87251916-INS-2137
chrX-87261292-INS-52
chrX-87287684-INS-214
chrX-87296836-INS-66
chrX-87320314-INS-73
chrX-87340964-INS-6033
chrX-87347907-INS-268
chrX-87370225-INS-6780
chrX-87377218-INS-120
chrX-87388183-INS-114
chrX-87389640-INS-235
chrX-87446511-INS-6167
chrX-87454542-INS-51
chrX-87472005-INS-95
chrX-87537540-INS-99
chrX-87585718-INS-6833
chrX-87595436-INS-368

chrX-87629071-INS-116
chrX-87630613-INS-54
chrX-87684558-INS-197
chrX-87690594-INS-54
chrX-87731981-INS-356
chrX-87740096-INS-6180
chrX-87762213-INS-197
chrX-87817434-INS-109
chrX-87825752-INS-438
chrX-87842218-INS-55
chrX-87900528-INS-204
chrX-87907016-INS-429
chrX-87908005-INS-71
chrX-87917386-INS-518
chrX-87927877-INS-66
chrX-87936339-INS-411
chrX-87941370-INS-343
chrX-87944300-INS-169
chrX-87964255-INS-59
chrX-88076327-INS-108
chrX-88103089-INS-770
chrX-88127540-INS-1879
chrX-88133807-INS-3165
chrX-88160467-INS-55
chrX-88188962-INS-178
chrX-88199670-INS-187
chrX-88209490-INS-401
chrX-88213804-INS-6543
chrX-88230047-INS-55
chrX-88302089-INS-60
chrX-88312691-INS-1532
chrX-88313062-INS-106
chrX-88323218-INS-107
chrX-88334376-INS-64
chrX-88382705-INS-624
chrX-88385864-INS-384
chrX-88388012-INS-85
chrX-88392050-INS-650
chrX-88409746-INS-52
chrX-88440594-INS-439
chrX-88450669-INS-261
chrX-88516938-INS-108
chrX-88547964-INS-74
chrX-88549789-INS-163
chrX-88553723-INS-75
chrX-88581207-INS-68
chrX-88587433-INS-65
chrX-88598600-INS-227
chrX-88610723-INS-248
chrX-88612556-INS-160
chrX-88627250-INS-74
chrX-88640458-INS-743
chrX-88652880-INS-390
chrX-88704161-INS-453
chrX-88715089-INS-134
chrX-88730200-INS-339
chrX-88735460-INS-912
chrX-88752181-INS-201
chrX-88764979-INS-68
chrX-88862166-INS-133
chrX-88874206-INS-507
chrX-88876284-INS-71
chrX-88904274-INS-226
chrX-88944973-INS-100
chrX-88983800-INS-230
chrX-89032204-INS-91
chrX-89038884-INS-236
chrX-89095347-INS-6402
chrX-89097978-INS-5580
chrX-89104154-INS-91
chrX-89107828-INS-112
chrX-89145034-INS-5339
chrX-89149642-INS-1888
chrX-89159530-INS-676
chrX-89179478-INS-107
chrX-89180841-INS-153
chrX-89186872-INS-1853

chrX-89202295-INS-7250
chrX-89223850-INS-465
chrX-89226197-INS-7841
chrX-89232169-INS-773
chrX-89241692-INS-91
chrX-89256632-INS-6472
chrX-89284877-INS-92
chrX-89312698-INS-442
chrX-89323871-INS-830
chrX-89328466-INS-68
chrX-89329046-INS-6486
chrX-89365994-INS-6360
chrX-89369906-INS-1483
chrX-89429777-INS-200
chrX-89432838-INS-196
chrX-89439378-INS-395
chrX-89502435-INS-5153
chrX-89504319-INS-163
chrX-89505338-INS-692
chrX-89584422-INS-386
chrX-89656713-INS-83
chrX-89663279-INS-159
chrX-89673926-INS-56
chrX-89689484-INS-631
chrX-89765319-INS-78
chrX-89776049-INS-61
chrX-89814242-INS-2719
chrX-89832653-INS-68
chrX-89859206-INS-640
chrX-89881958-INS-1988
chrX-89894346-INS-987
chrX-89910089-INS-877
chrX-89953959-INS-1493
chrX-89978773-INS-7159
chrX-90016802-INS-93
chrX-90036835-INS-5462
chrX-90068535-INS-273
chrX-90082963-INS-181
chrX-90085058-INS-231
chrX-90091553-INS-7206
chrX-90128337-INS-4339
chrX-90145127-INS-242
chrX-90146890-INS-197
chrX-90215565-INS-54
chrX-90295303-INS-52
chrX-90328396-INS-399
chrX-90335012-INS-59
chrX-90349921-INS-164
chrX-90364052-INS-1252
chrX-90399702-INS-874
chrX-90555572-INS-55
chrX-90583227-INS-645
chrX-90708609-INS-1802
chrX-90738345-INS-251
chrX-90766232-INS-76
chrX-90802247-INS-5971
chrX-90871317-INS-6467
chrX-90874607-INS-215
chrX-90911253-INS-64
chrX-91069796-INS-88
chrX-91090891-INS-141
chrX-91093285-INS-439
chrX-91128610-INS-463
chrX-91135446-INS-821
chrX-91140410-INS-90
chrX-91167878-INS-198
chrX-91190679-INS-6483
chrX-91203396-INS-6699
chrX-91229164-INS-1391
chrX-91247724-INS-5973
chrX-91271740-INS-196
chrX-91287985-INS-5221
chrX-91303967-INS-377
chrX-91362456-INS-120
chrX-91376262-INS-101
chrX-91380377-INS-71
chrX-91389568-INS-160

chrX-91390655-INS-168
chrX-91405229-INS-344
chrX-91438775-INS-608
chrX-91442978-INS-5542
chrX-91444208-INS-104
chrX-91444859-INS-618
chrX-91465805-INS-6565
chrX-91484265-INS-75
chrX-91535926-INS-104
chrX-91552867-INS-128
chrX-91578282-INS-668
chrX-91601725-INS-182
chrX-91629253-INS-83
chrX-91643943-INS-7113
chrX-91684145-INS-367
chrX-91712350-INS-311
chrX-91755265-INS-214
chrX-91775940-INS-481
chrX-91788730-INS-159
chrX-91796983-INS-52
chrX-91801894-INS-3075
chrX-91808884-INS-173
chrX-91865341-INS-167
chrX-91912456-INS-490
chrX-91918318-INS-558
chrX-91919893-INS-64
chrX-91929013-INS-275
chrX-91930058-INS-7032
chrX-91936925-INS-6680
chrX-91945053-INS-333
chrX-91946718-INS-6705
chrX-91948831-INS-191
chrX-91971979-INS-641
chrX-91995139-INS-6376
chrX-92026601-INS-353
chrX-92032911-INS-1737
chrX-92057235-INS-2717
chrX-92063702-INS-139
chrX-92072845-INS-92
chrX-92075311-INS-1725
chrX-92098047-INS-320
chrX-92099335-INS-177
chrX-92103594-INS-51
chrX-92109722-INS-702
chrX-92112999-INS-63
chrX-92127561-INS-435
chrX-92132841-INS-1131
chrX-92137238-INS-177
chrX-92139839-INS-286
chrX-92159123-INS-1873
chrX-92171473-INS-4483
chrX-92174808-INS-1096
chrX-92189426-INS-60
chrX-92191198-INS-1179
chrX-92192661-INS-505
chrX-92198283-INS-230
chrX-92216121-INS-234
chrX-92245423-INS-58
chrX-92256424-INS-384
chrX-92268690-INS-52
chrX-92269521-INS-52
chrX-92305679-INS-543
chrX-92312755-INS-117
chrX-92340230-INS-64
chrX-92358215-INS-1864
chrX-92383662-INS-68
chrX-92436717-INS-143
chrX-92456488-INS-756
chrX-92467265-INS-551
chrX-92560558-INS-134
chrX-92578645-INS-197
chrX-92580900-INS-273
chrX-92582369-INS-228
chrX-92616035-INS-52
chrX-92653372-INS-6464
chrX-92655337-INS-85
chrX-92660029-INS-400

chrX-92666959-INS-446
chrX-92691181-INS-351
chrX-92695209-INS-238
chrX-92706450-INS-133
chrX-92714422-INS-186
chrX-92726025-INS-137
chrX-92738646-INS-375
chrX-92764192-INS-375
chrX-92778838-INS-310
chrX-92782265-INS-196
chrX-92797992-INS-204
chrX-92804092-INS-7329
chrX-92882675-INS-52
chrX-92892617-INS-123
chrX-92896655-INS-155
chrX-92915658-INS-52
chrX-92926781-INS-128
chrX-92959953-INS-164
chrX-92963279-INS-195
chrX-92966105-INS-52
chrX-92981911-INS-179
chrX-92984615-INS-176
chrX-93000001-INS-140
chrX-93016187-INS-62
chrX-93040894-INS-63
chrX-93043690-INS-258
chrX-93068718-INS-160
chrX-93081835-INS-107
chrX-93083108-INS-102
chrX-93097068-INS-221
chrX-93105731-INS-244
chrX-93140187-INS-92
chrX-93140630-INS-251
chrX-93153568-INS-65
chrX-93182233-INS-105
chrX-93200045-INS-177
chrX-93202111-INS-150
chrX-93205177-INS-201
chrX-93232416-INS-200
chrX-93236282-INS-322
chrX-93238548-INS-171
chrX-93250088-INS-208
chrX-93261586-INS-176
chrX-93269106-INS-6438
chrX-93284186-INS-216
chrX-93294004-INS-184
chrX-93302985-INS-550
chrX-93359600-INS-115
chrX-93376402-INS-94
chrX-93391381-INS-104
chrX-93409601-INS-53
chrX-93436276-INS-116
chrX-93437459-INS-52
chrX-93450102-INS-329
chrX-93452629-INS-67
chrX-93472837-INS-62
chrX-93487714-INS-68
chrX-93505213-INS-86
chrX-93605865-INS-55
chrX-93607449-INS-519
chrX-93628959-INS-178
chrX-93657354-INS-202
chrX-93660895-INS-461
chrX-93664514-INS-538
chrX-93677645-INS-311
chrX-93687151-INS-249
chrX-93692079-INS-93
chrX-93786535-INS-229
chrX-93802306-INS-680
chrX-93809086-INS-413
chrX-93834891-INS-204
chrX-93838302-INS-69
chrX-93886339-INS-62
chrX-93953815-INS-2072
chrX-93958677-INS-2069
chrX-94038553-INS-347
chrX-94064880-INS-92

chrX-94079253-INS-310
chrX-94085355-INS-443
chrX-94102322-INS-7593
chrX-94146267-INS-102
chrX-94156590-INS-738
chrX-94166949-INS-4389
chrX-94240866-INS-57
chrX-94246042-INS-383
chrX-94250314-INS-2640
chrX-94260009-INS-115
chrX-94282049-INS-191
chrX-94296261-INS-81
chrX-94304663-INS-433
chrX-94308379-INS-120
chrX-94317631-INS-96
chrX-94319345-INS-385
chrX-94336369-INS-233
chrX-94339921-INS-71
chrX-94361689-INS-60
chrX-94391793-INS-123
chrX-94413568-INS-1225
chrX-94423950-INS-5204
chrX-94425359-INS-84
chrX-94431956-INS-3735
chrX-94436365-INS-7859
chrX-94453192-INS-5649
chrX-94493215-INS-52
chrX-94506765-INS-5715
chrX-94531395-INS-69
chrX-94537676-INS-52
chrX-94545575-INS-579
chrX-94554557-INS-156
chrX-94572293-INS-167
chrX-94578218-INS-6388
chrX-94591149-INS-613
chrX-94610694-INS-171
chrX-94639953-INS-3043
chrX-94652514-INS-52
chrX-94676849-INS-629
chrX-94679516-INS-6484
chrX-94684689-INS-122
chrX-94686275-INS-88
chrX-94687998-INS-369
chrX-94698784-INS-176
chrX-94723066-INS-52
chrX-94775207-INS-461
chrX-94795450-INS-469
chrX-94827906-INS-426
chrX-94829766-INS-574
chrX-94832283-INS-391
chrX-94851101-INS-400
chrX-94876946-INS-1891
chrX-94884341-INS-288
chrX-94922154-INS-400
chrX-94958456-INS-90
chrX-95050224-INS-602
chrX-95052302-INS-106
chrX-95081881-INS-397
chrX-95122255-INS-623
chrX-95123724-INS-70
chrX-95126098-INS-380
chrX-95131328-INS-115
chrX-95147123-INS-681
chrX-95199436-INS-167
chrX-95208387-INS-149
chrX-95232930-INS-60
chrX-95236818-INS-290
chrX-95245942-INS-6260
chrX-95246185-INS-68
chrX-95258445-INS-194
chrX-95281495-INS-919
chrX-95286107-INS-217
chrX-95299093-INS-6341
chrX-95303958-INS-105
chrX-95358399-INS-7297
chrX-95366470-INS-194
chrX-95367932-INS-64

chrX-95370134-INS-52
chrX-95389483-INS-567
chrX-95402524-INS-160
chrX-95404746-INS-60
chrX-95413024-INS-6659
chrX-95464194-INS-73
chrX-95475794-INS-82
chrX-95478935-INS-82
chrX-95495903-INS-74
chrX-95507689-INS-57
chrX-95509936-INS-80
chrX-95512338-INS-67
chrX-95517435-INS-74
chrX-95536795-INS-7336
chrX-95551346-INS-769
chrX-95557662-INS-94
chrX-95565803-INS-104
chrX-95568425-INS-64
chrX-95577021-INS-116
chrX-95613192-INS-6226
chrX-95615462-INS-259
chrX-95621300-INS-6889
chrX-95623082-INS-57
chrX-95650774-INS-405
chrX-95662920-INS-721
chrX-95678764-INS-770
chrX-95694483-INS-76
chrX-95749070-INS-1892
chrX-95770258-INS-6711
chrX-95780684-INS-203
chrX-95798162-INS-123
chrX-95805041-INS-492
chrX-95884349-INS-6425
chrX-95902198-INS-70
chrX-95913187-INS-225
chrX-95921785-INS-6528
chrX-95970111-INS-2930
chrX-95984082-INS-7076
chrX-95987012-INS-745
chrX-96037577-INS-168
chrX-96041135-INS-72
chrX-96057664-INS-276
chrX-96069914-INS-97
chrX-96074312-INS-801
chrX-96101557-INS-6190
chrX-96114331-INS-284
chrX-96143604-INS-1080
chrX-96156469-INS-700
chrX-96160477-INS-80
chrX-96177540-INS-86
chrX-96186528-INS-298
chrX-96188482-INS-362
chrX-96208818-INS-216
chrX-96242457-INS-203
chrX-96274336-INS-6739
chrX-96286177-INS-57
chrX-96312683-INS-56
chrX-96315243-INS-240
chrX-96317719-INS-526
chrX-96319237-INS-68
chrX-96327174-INS-100
chrX-96337955-INS-55
chrX-96356000-INS-6187
chrX-96431422-INS-396
chrX-96435790-INS-281
chrX-96446574-INS-93
chrX-96461447-INS-59
chrX-96471213-INS-3112
chrX-96480292-INS-3155
chrX-96483123-INS-7322
chrX-96492452-INS-521
chrX-96514747-INS-1061
chrX-96535309-INS-199
chrX-96616538-INS-1244
chrX-96620324-INS-142
chrX-96623682-INS-358
chrX-96635163-INS-612

chrX-96673004-INS-1912
chrX-96716918-INS-1875
chrX-96745507-INS-84
chrX-96755612-INS-6198
chrX-96764078-INS-1363
chrX-96795921-INS-199
chrX-96831954-INS-697
chrX-96836028-INS-322
chrX-96854139-INS-112
chrX-96856893-INS-3874
chrX-96869485-INS-54
chrX-96943075-INS-10304
chrX-96943531-INS-110
chrX-96964357-INS-93
chrX-96984587-INS-201
chrX-96992365-INS-8480
chrX-96996918-INS-788
chrX-97023437-INS-885
chrX-97035017-INS-6477
chrX-97041853-INS-12907
chrX-97103707-INS-54
chrX-97116080-INS-354
chrX-97121062-INS-95
chrX-97139988-INS-209
chrX-97147312-INS-212
chrX-97149408-INS-115
chrX-97164982-INS-594
chrX-97172397-INS-87
chrX-97189974-INS-54
chrX-97226103-INS-764
chrX-97255366-INS-52
chrX-97258183-INS-68
chrX-97278091-INS-5550
chrX-97317543-INS-87
chrX-97326623-INS-68
chrX-97339843-INS-3262
chrX-97344946-INS-1867
chrX-97372647-INS-54
chrX-97372888-INS-671
chrX-97374594-INS-6808
chrX-97376668-INS-53
chrX-97410526-INS-331
chrX-97411799-INS-4801
chrX-97428467-INS-51
chrX-97433308-INS-667
chrX-97438321-INS-6864
chrX-97463680-INS-834
chrX-97497659-INS-429
chrX-97500081-INS-60
chrX-97533458-INS-353
chrX-97543827-INS-6399
chrX-97582268-INS-138
chrX-97583182-INS-6595
chrX-97591214-INS-6934
chrX-97592434-INS-469
chrX-97605317-INS-605
chrX-97634851-INS-66
chrX-97649239-INS-1101
chrX-97657842-INS-206
chrX-97661507-INS-8983
chrX-97669563-INS-456
chrX-97676498-INS-6184
chrX-97701598-INS-6204
chrX-97712843-INS-56
chrX-97714178-INS-7270
chrX-97714543-INS-84
chrX-97716891-INS-4733
chrX-97747244-INS-401
chrX-97754826-INS-400
chrX-97798542-INS-932
chrX-97816536-INS-198
chrX-97871009-INS-112
chrX-97880583-INS-100
chrX-97888195-INS-7486
chrX-97893890-INS-53
chrX-97894840-INS-56
chrX-97906843-INS-156

chrX-97913920-INS-124
chrX-97930838-INS-334
chrX-97965056-INS-88
chrX-97970424-INS-155
chrX-97984682-INS-206
chrX-97992403-INS-570
chrX-98021453-INS-169
chrX-98065867-INS-328
chrX-98137862-INS-6482
chrX-98146320-INS-110
chrX-98172306-INS-112
chrX-98217776-INS-216
chrX-98254101-INS-143
chrX-98261987-INS-5162
chrX-98266534-INS-98
chrX-98281940-INS-151
chrX-98287923-INS-162
chrX-98311589-INS-75
chrX-98369604-INS-74
chrX-98385673-INS-55
chrX-98395615-INS-51
chrX-98415315-INS-137
chrX-98429932-INS-6639
chrX-98493497-INS-107
chrX-98540562-INS-127
chrX-98562930-INS-113
chrX-98599009-INS-62
chrX-98669610-INS-98
chrX-98711948-INS-52
chrX-98744509-INS-136
chrX-98777945-INS-268
chrX-98793595-INS-335
chrX-98799564-INS-6220
chrX-98821315-INS-112
chrX-98856416-INS-1349
chrX-98895681-INS-7235
chrX-98899583-INS-161
chrX-98912479-INS-56
chrX-98937343-INS-6381
chrX-98956897-INS-63
chrX-98957362-INS-6393
chrX-98963549-INS-51
chrX-98980057-INS-326
chrX-98986680-INS-119
chrX-98994691-INS-401
chrX-99014739-INS-132
chrX-99030948-INS-106
chrX-99036598-INS-6339
chrX-99071084-INS-54
chrX-99090729-INS-77
chrX-99113401-INS-556
chrX-99128063-INS-319
chrX-99191433-INS-444
chrX-99261785-INS-508
chrX-99266866-INS-6217
chrX-99294600-INS-571
chrX-99323057-INS-330
chrX-99327026-INS-878
chrX-99350364-INS-52
chrX-99393567-INS-2541
chrX-99419911-INS-7031
chrX-99426237-INS-6477
chrX-99429745-INS-1792
chrX-99463244-INS-7298
chrX-99467527-INS-6843
chrX-99494647-INS-54
chrX-99495612-INS-93
chrX-99508821-INS-183
chrX-99511644-INS-60
chrX-99536408-INS-195
chrX-99588765-INS-183
chrX-99594039-INS-685
chrX-99653572-INS-205
chrX-99665905-INS-226
chrX-99679440-INS-104
chrX-99699474-INS-111
chrX-99702793-INS-516

chrX-99706809-INS-116
chrX-99715585-INS-202
chrX-99724024-INS-73
chrX-99774434-INS-88
chrX-99781711-INS-120
chrX-99784062-INS-60
chrX-99787478-INS-426
chrX-99801694-INS-53
chrX-99819217-INS-206
chrX-99842157-INS-95
chrX-99874532-INS-118
chrX-99885235-INS-209
chrX-99901685-INS-60
chrX-99918630-INS-212
chrX-99925978-INS-1097
chrX-99947438-INS-228
chrX-99962098-INS-62
chrX-99972202-INS-6505
chrX-99977227-INS-81
chrX-99984930-INS-5994
chrX-99995150-INS-690
chrX-99997183-INS-61
chrX-100052545-INS-205
chrX-100070361-INS-53
chrX-100076835-INS-430
chrX-100079613-INS-75
chrX-100097868-INS-62
chrX-100106186-INS-54
chrX-100108586-INS-213
chrX-100138970-INS-213
chrX-100142127-INS-273
chrX-100149782-INS-412
chrX-100153360-INS-128
chrX-100160791-INS-91
chrX-100163930-INS-365
chrX-100184433-INS-191
chrX-100243497-INS-263
chrX-100250075-INS-78
chrX-100261451-INS-229
chrX-100283887-INS-396
chrX-100356959-INS-532
chrX-100370959-INS-196
chrX-100400749-INS-168
chrX-100431121-INS-203
chrX-100444430-INS-109
chrX-100452575-INS-223
chrX-100480708-INS-855
chrX-100483361-INS-205
chrX-100484530-INS-199
chrX-100508908-INS-190
chrX-100509439-INS-1052
chrX-100519009-INS-164
chrX-100547940-INS-55
chrX-100551347-INS-626
chrX-100552640-INS-189
chrX-100564950-INS-167
chrX-100589308-INS-204
chrX-100609603-INS-218
chrX-100622989-INS-122
chrX-100658317-INS-132
chrX-100659528-INS-211
chrX-100667978-INS-1978
chrX-100674382-INS-212
chrX-100678635-INS-145
chrX-100709660-INS-205
chrX-100723387-INS-88
chrX-100723872-INS-139
chrX-100737344-INS-107
chrX-100739759-INS-163
chrX-100741117-INS-105
chrX-100763556-INS-52
chrX-100796764-INS-67
chrX-100876566-INS-76
chrX-100882638-INS-7168
chrX-100887492-INS-81
chrX-100921946-INS-528
chrX-100969464-INS-80

chrX-100982434-INS-327
chrX-101025091-INS-1713
chrX-101048598-INS-51
chrX-101055510-INS-60
chrX-101070083-INS-55
chrX-101076729-INS-217
chrX-101082776-INS-72
chrX-101125174-INS-1842
chrX-101162243-INS-104
chrX-101191501-INS-54
chrX-101194468-INS-238
chrX-101200038-INS-194
chrX-101203367-INS-140
chrX-101221549-INS-197
chrX-101226884-INS-231
chrX-101229869-INS-384
chrX-101237340-INS-199
chrX-101239823-INS-72
chrX-101255790-INS-75
chrX-101258470-INS-72
chrX-101259442-INS-109
chrX-101261733-INS-64
chrX-101262423-INS-97
chrX-101265912-INS-252
chrX-101283071-INS-183
chrX-101315051-INS-216
chrX-101323658-INS-191
chrX-101329935-INS-174
chrX-101339018-INS-56
chrX-101370968-INS-761
chrX-101374506-INS-106
chrX-101400416-INS-61
chrX-101409554-INS-90
chrX-101423839-INS-1936
chrX-101434283-INS-54
chrX-101439897-INS-186
chrX-101453299-INS-5173
chrX-101456439-INS-52
chrX-101500183-INS-439
chrX-101513386-INS-112
chrX-101527933-INS-6247
chrX-101534158-INS-158
chrX-101544040-INS-60
chrX-101550184-INS-58
chrX-101555851-INS-1140
chrX-101555920-INS-929
chrX-101556464-INS-2340
chrX-101556558-INS-990
chrX-101556612-INS-195
chrX-101556652-INS-1260
chrX-101556828-INS-2310
chrX-101589376-INS-107
chrX-101604861-INS-1040
chrX-101671177-INS-117
chrX-101679068-INS-187
chrX-101743399-INS-184
chrX-101780846-INS-284
chrX-101799712-INS-100
chrX-101837894-INS-140
chrX-101947470-INS-52
chrX-101980375-INS-108
chrX-101984820-INS-393
chrX-101995950-INS-71
chrX-102005485-INS-6343
chrX-102016096-INS-1278
chrX-102022996-INS-125
chrX-102026734-INS-169
chrX-102030963-INS-53
chrX-102035218-INS-61
chrX-102036132-INS-491
chrX-102080933-INS-202
chrX-102106171-INS-230
chrX-102153490-INS-97
chrX-102163642-INS-153
chrX-102203189-INS-62
chrX-102223121-INS-245
chrX-102236584-INS-94

chrX-102250846-INS-6200
chrX-102266940-INS-231
chrX-102287641-INS-57
chrX-102295409-INS-107
chrX-102300549-INS-205
chrX-102327688-INS-139
chrX-102328592-INS-851
chrX-102329129-INS-1206
chrX-102342619-INS-5154
chrX-102346684-INS-215
chrX-102364295-INS-85
chrX-102377540-INS-361
chrX-102379675-INS-220
chrX-102387538-INS-105
chrX-102390086-INS-539
chrX-102396080-INS-328
chrX-102425882-INS-152
chrX-102445207-INS-872
chrX-102450727-INS-209
chrX-102467678-INS-192
chrX-102469929-INS-129
chrX-102487812-INS-174
chrX-102499044-INS-68
chrX-102500125-INS-340
chrX-102500468-INS-544
chrX-102502099-INS-56
chrX-102535810-INS-52
chrX-102542399-INS-91
chrX-102547154-INS-386
chrX-102555842-INS-75
chrX-102558942-INS-124
chrX-102563759-INS-225
chrX-102600717-INS-556
chrX-102611059-INS-206
chrX-102612096-INS-177
chrX-102628564-INS-200
chrX-102640682-INS-101
chrX-102642154-INS-207
chrX-102645157-INS-206
chrX-102669236-INS-54
chrX-102680271-INS-154
chrX-102683030-INS-231
chrX-102683849-INS-202
chrX-102689194-INS-269
chrX-102719365-INS-132
chrX-102724296-INS-7130
chrX-102772156-INS-66
chrX-102779257-INS-72
chrX-102793679-INS-191
chrX-102836266-INS-66
chrX-102842497-INS-66
chrX-102855521-INS-158
chrX-102876186-INS-76
chrX-102909137-INS-147
chrX-102919796-INS-343
chrX-102955664-INS-1410
chrX-102977680-INS-115
chrX-102994740-INS-692
chrX-103008565-INS-199
chrX-103039335-INS-132
chrX-103076465-INS-51
chrX-103079332-INS-55
chrX-103105008-INS-1521
chrX-103105204-INS-58
chrX-103140331-INS-64
chrX-103140848-INS-58
chrX-103146242-INS-406
chrX-103164801-INS-61
chrX-103171595-INS-706
chrX-103178369-INS-51
chrX-103181594-INS-288
chrX-103194257-INS-315
chrX-103233045-INS-3714
chrX-103237080-INS-209
chrX-103238280-INS-205
chrX-103241330-INS-120
chrX-103286649-INS-86

chrX-103300713-INS-6704
chrX-103306482-INS-6338
chrX-103306810-INS-199
chrX-103323086-INS-6401
chrX-103329010-INS-102
chrX-103330411-INS-361
chrX-103338895-INS-98
chrX-103393806-INS-226
chrX-103406890-INS-358
chrX-103413134-INS-115
chrX-103422517-INS-59
chrX-103431059-INS-614
chrX-103438131-INS-66
chrX-103452029-INS-7146
chrX-103452801-INS-175
chrX-103463789-INS-69
chrX-103480500-INS-691
chrX-103497006-INS-101
chrX-103500722-INS-9071
chrX-103513169-INS-1085
chrX-103519315-INS-58
chrX-103559184-INS-124
chrX-103569243-INS-7084
chrX-103582491-INS-6613
chrX-103587601-INS-1884
chrX-103596067-INS-8068
chrX-103598660-INS-60
chrX-103628589-INS-134
chrX-103633651-INS-66
chrX-103656223-INS-1747
chrX-103704380-INS-766
chrX-103811121-INS-6792
chrX-103859492-INS-7006
chrX-103862180-INS-63
chrX-103866319-INS-65
chrX-103869419-INS-204
chrX-103878971-INS-52
chrX-103896324-INS-397
chrX-103936402-INS-6456
chrX-103968681-INS-428
chrX-103983394-INS-233
chrX-103990278-INS-1516
chrX-103990532-INS-629
chrX-103998396-INS-890
chrX-104003407-INS-213
chrX-104014092-INS-100
chrX-104030290-INS-159
chrX-104035492-INS-381
chrX-104038024-INS-60
chrX-104044424-INS-113
chrX-104052165-INS-800
chrX-104058328-INS-75
chrX-104063704-INS-658
chrX-104071727-INS-363
chrX-104073459-INS-59
chrX-104076953-INS-324
chrX-104088701-INS-210
chrX-104093888-INS-86
chrX-104112718-INS-851
chrX-104130544-INS-219
chrX-104141919-INS-398
chrX-104187376-INS-143
chrX-104187448-INS-1888
chrX-104188433-INS-11608
chrX-104212553-INS-481
chrX-104225586-INS-60
chrX-104226476-INS-110
chrX-104228073-INS-123
chrX-104274329-INS-52
chrX-104291519-INS-80
chrX-104316127-INS-295
chrX-104325003-INS-4064
chrX-104327756-INS-52
chrX-104346430-INS-735
chrX-104346993-INS-5943
chrX-104350301-INS-6461
chrX-104354254-INS-6028

chrX-104355704-INS-55
chrX-104364640-INS-6237
chrX-104369836-INS-6415
chrX-104416455-INS-68
chrX-104443517-INS-156
chrX-104474564-INS-145
chrX-104481642-INS-728
chrX-104502847-INS-9393
chrX-104527426-INS-1770
chrX-104553842-INS-78
chrX-104577035-INS-76
chrX-104579125-INS-54
chrX-104595635-INS-331
chrX-104601186-INS-101
chrX-104615231-INS-232
chrX-104639634-INS-198
chrX-104669903-INS-222
chrX-104671862-INS-59
chrX-104715959-INS-57
chrX-104809606-INS-64
chrX-104830805-INS-74
chrX-104845728-INS-71
chrX-104857679-INS-71
chrX-104860074-INS-128
chrX-104921729-INS-73
chrX-104932529-INS-67
chrX-104936848-INS-542
chrX-104964309-INS-306
chrX-104965373-INS-411
chrX-104967741-INS-113
chrX-104990773-INS-220
chrX-104992325-INS-195
chrX-105007670-INS-214
chrX-105043321-INS-224
chrX-105047388-INS-202
chrX-105051819-INS-197
chrX-105053001-INS-85
chrX-105075290-INS-404
chrX-105083838-INS-174
chrX-105089599-INS-133
chrX-105125771-INS-393
chrX-105139025-INS-420
chrX-105160526-INS-280
chrX-105163674-INS-195
chrX-105177805-INS-758
chrX-105194740-INS-96
chrX-105210901-INS-175
chrX-105275211-INS-944
chrX-105281235-INS-66
chrX-105290038-INS-182
chrX-105355930-INS-70
chrX-105384950-INS-644
chrX-105412837-INS-72
chrX-105459146-INS-212
chrX-105460736-INS-397
chrX-105464812-INS-151
chrX-105484384-INS-790
chrX-105503446-INS-642
chrX-105532877-INS-2183
chrX-105533569-INS-291
chrX-105536650-INS-371
chrX-105542860-INS-506
chrX-105552103-INS-103
chrX-105567270-INS-755
chrX-105634704-INS-235
chrX-105635933-INS-343
chrX-105647713-INS-52
chrX-105661005-INS-78
chrX-105684109-INS-61
chrX-105689432-INS-345
chrX-105690974-INS-64
chrX-105701882-INS-6170
chrX-105708451-INS-53
chrX-105745083-INS-166
chrX-105759058-INS-54
chrX-105816062-INS-56
chrX-105817510-INS-173

chrX-105831641-INS-210
chrX-105833117-INS-4324
chrX-105863067-INS-55
chrX-105870086-INS-120
chrX-105898612-INS-7107
chrX-105935584-INS-6795
chrX-105935964-INS-72
chrX-105958892-INS-212
chrX-105987856-INS-373
chrX-106028329-INS-6062
chrX-106035197-INS-230
chrX-106036374-INS-111
chrX-106036560-INS-4386
chrX-106065685-INS-315
chrX-106068835-INS-1961
chrX-106069691-INS-528
chrX-106070824-INS-6397
chrX-106076178-INS-83
chrX-106081443-INS-281
chrX-106088987-INS-6288
chrX-106158380-INS-184
chrX-106166055-INS-57
chrX-106177618-INS-3155
chrX-106189018-INS-104
chrX-106196689-INS-1422
chrX-106198604-INS-56
chrX-106223377-INS-95
chrX-106270045-INS-195
chrX-106270658-INS-6292
chrX-106286034-INS-282
chrX-106316271-INS-56
chrX-106341233-INS-499
chrX-106366768-INS-5217
chrX-106391695-INS-120
chrX-106405653-INS-404
chrX-106453121-INS-433
chrX-106461974-INS-54
chrX-106521150-INS-71
chrX-106542537-INS-195
chrX-106576672-INS-75
chrX-106581914-INS-5822
chrX-106593087-INS-60
chrX-106603380-INS-69
chrX-106604507-INS-262
chrX-106645796-INS-209
chrX-106661452-INS-5288
chrX-106705300-INS-177
chrX-106724223-INS-383
chrX-106736786-INS-4697
chrX-106744089-INS-1649
chrX-106756999-INS-196
chrX-106762270-INS-824
chrX-106790377-INS-278
chrX-106794595-INS-93
chrX-106832426-INS-769
chrX-106840983-INS-1150
chrX-106844678-INS-655
chrX-106858738-INS-89
chrX-106882522-INS-6677
chrX-106945477-INS-2011
chrX-106950944-INS-779
chrX-106952334-INS-116
chrX-106984166-INS-1706
chrX-106986370-INS-2613
chrX-106987864-INS-428
chrX-106988626-INS-7158
chrX-107015537-INS-7539
chrX-107042822-INS-143
chrX-107047858-INS-95
chrX-107058592-INS-197
chrX-107096777-INS-55
chrX-107114945-INS-201
chrX-107135639-INS-113
chrX-107217022-INS-440
chrX-107224297-INS-71
chrX-107264270-INS-828
chrX-107333471-INS-6509

chrX-107383355-INS-196
chrX-107432004-INS-310
chrX-107433237-INS-86
chrX-107471050-INS-60
chrX-107548282-INS-863
chrX-107562633-INS-7130
chrX-107563861-INS-155
chrX-107576481-INS-104
chrX-107590839-INS-55
chrX-107623284-INS-63
chrX-107639053-INS-63
chrX-107722711-INS-55
chrX-107746264-INS-6589
chrX-107770790-INS-839
chrX-107853204-INS-209
chrX-107880605-INS-56
chrX-107897145-INS-6171
chrX-107899141-INS-701
chrX-107908215-INS-477
chrX-107928938-INS-74
chrX-107933595-INS-326
chrX-107942078-INS-112
chrX-107950232-INS-138
chrX-107951773-INS-251
chrX-107988800-INS-625
chrX-107999679-INS-6434
chrX-108071683-INS-6143
chrX-108077319-INS-103
chrX-108081291-INS-203
chrX-108084358-INS-2954
chrX-108105336-INS-3875
chrX-108125581-INS-70
chrX-108147689-INS-90
chrX-108153347-INS-487
chrX-108176905-INS-124
chrX-108177661-INS-579
chrX-108202300-INS-510
chrX-108212190-INS-6326
chrX-108250386-INS-182
chrX-108251825-INS-74
chrX-108252743-INS-183
chrX-108274190-INS-70
chrX-108279699-INS-1482
chrX-108290454-INS-57
chrX-108296363-INS-597
chrX-108317136-INS-56
chrX-108334453-INS-379
chrX-108370844-INS-6622
chrX-108372210-INS-333
chrX-108374795-INS-1886
chrX-108385309-INS-331
chrX-108387870-INS-60
chrX-108397872-INS-654
chrX-108435831-INS-862
chrX-108459982-INS-8230
chrX-108469311-INS-6689
chrX-108500191-INS-321
chrX-108515438-INS-86
chrX-108528514-INS-61
chrX-108568520-INS-527
chrX-108574738-INS-110
chrX-108579230-INS-204
chrX-108611585-INS-894
chrX-108627823-INS-52
chrX-108630182-INS-270
chrX-108642959-INS-204
chrX-108645688-INS-8205
chrX-108667336-INS-639
chrX-108692947-INS-69
chrX-108698133-INS-420
chrX-108703264-INS-59
chrX-108741970-INS-65
chrX-108744108-INS-5357
chrX-108746445-INS-117
chrX-108771037-INS-88
chrX-108774673-INS-183
chrX-108777589-INS-56

chrX-108791740-INS-58
chrX-108793694-INS-7159
chrX-108796607-INS-424
chrX-108820507-INS-218
chrX-108851978-INS-804
chrX-108868516-INS-702
chrX-108884829-INS-777
chrX-108893759-INS-66
chrX-108922191-INS-983
chrX-108926070-INS-307
chrX-108930690-INS-1693
chrX-108953625-INS-624
chrX-108957483-INS-115
chrX-108994329-INS-116
chrX-109017879-INS-534
chrX-109093060-INS-51
chrX-109174928-INS-60
chrX-109177844-INS-184
chrX-109187125-INS-158
chrX-109193168-INS-86
chrX-109232369-INS-166
chrX-109233996-INS-618
chrX-109243921-INS-77
chrX-109251308-INS-4844
chrX-109268525-INS-330
chrX-109276443-INS-92
chrX-109290846-INS-118
chrX-109295513-INS-895
chrX-109304362-INS-391
chrX-109328442-INS-3022
chrX-109335411-INS-73
chrX-109344895-INS-79
chrX-109346095-INS-195
chrX-109368913-INS-690
chrX-109377696-INS-8986
chrX-109395717-INS-84
chrX-109400613-INS-163
chrX-109429956-INS-132
chrX-109507232-INS-231
chrX-109511681-INS-3160
chrX-109517921-INS-57
chrX-109546472-INS-498
chrX-109584461-INS-68
chrX-109586048-INS-71
chrX-109586073-INS-59
chrX-109613052-INS-323
chrX-109622367-INS-4149
chrX-109625369-INS-2845
chrX-109642847-INS-9330
chrX-109689129-INS-230
chrX-109706613-INS-135
chrX-109718453-INS-343
chrX-109781945-INS-133
chrX-109783592-INS-6159
chrX-109787008-INS-775
chrX-109789852-INS-487
chrX-109806290-INS-317
chrX-109808841-INS-6740
chrX-109824452-INS-83
chrX-109870790-INS-74
chrX-109898694-INS-93
chrX-109967874-INS-181
chrX-109972924-INS-110
chrX-109976275-INS-313
chrX-110012679-INS-6600
chrX-110018072-INS-6571
chrX-110021907-INS-205
chrX-110041944-INS-819
chrX-110042845-INS-115
chrX-110066936-INS-52
chrX-110070231-INS-385
chrX-110125124-INS-869
chrX-110128258-INS-187
chrX-110142658-INS-207
chrX-110200654-INS-4791
chrX-110206768-INS-344
chrX-110208654-INS-311

chrX-110234382-INS-591
chrX-110238521-INS-525
chrX-110256253-INS-467
chrX-110268455-INS-55
chrX-110285944-INS-395
chrX-110304932-INS-210
chrX-110308031-INS-52
chrX-110380546-INS-1062
chrX-110425042-INS-233
chrX-110466622-INS-3017
chrX-110516350-INS-438
chrX-110518059-INS-2904
chrX-110533855-INS-1214
chrX-110534646-INS-180
chrX-110539864-INS-7484
chrX-110585947-INS-2236
chrX-110640249-INS-3028
chrX-110648002-INS-359
chrX-110663712-INS-539
chrX-110717547-INS-202
chrX-110758045-INS-58
chrX-110785699-INS-2351
chrX-110788416-INS-93
chrX-110830126-INS-125
chrX-110840497-INS-108
chrX-110857899-INS-124
chrX-110859861-INS-4650
chrX-110861018-INS-78
chrX-110878574-INS-7966
chrX-110887854-INS-501
chrX-110911099-INS-72
chrX-110982102-INS-76
chrX-110996953-INS-1017
chrX-110997754-INS-4667
chrX-111016551-INS-691
chrX-111043956-INS-311
chrX-111044590-INS-480
chrX-111045845-INS-790
chrX-111097671-INS-142
chrX-111122903-INS-6619
chrX-111149932-INS-372
chrX-111240748-INS-238
chrX-111272436-INS-118
chrX-111300231-INS-150
chrX-111300526-INS-5653
chrX-111307926-INS-98
chrX-111407746-INS-77
chrX-111409767-INS-6204
chrX-111418725-INS-66
chrX-111463222-INS-140
chrX-111487193-INS-119
chrX-111546730-INS-7202
chrX-111560771-INS-53
chrX-111594912-INS-279
chrX-111618320-INS-78
chrX-111644228-INS-104
chrX-111693055-INS-516
chrX-111700831-INS-727
chrX-111716272-INS-179
chrX-111735125-INS-55
chrX-111762771-INS-373
chrX-111767300-INS-536
chrX-111772354-INS-93
chrX-111789149-INS-105
chrX-111793648-INS-56
chrX-111811165-INS-3803
chrX-111812649-INS-1889
chrX-111818992-INS-339
chrX-111840593-INS-7421
chrX-111865359-INS-329
chrX-111874215-INS-702
chrX-111875425-INS-6259
chrX-111885408-INS-60
chrX-111908268-INS-237
chrX-111947804-INS-676
chrX-111951008-INS-764
chrX-111993194-INS-728

chrX-112013733-INS-474
chrX-112068433-INS-210
chrX-112086842-INS-205
chrX-112116416-INS-60
chrX-112121053-INS-328
chrX-112124323-INS-60
chrX-112173154-INS-146
chrX-112178321-INS-182
chrX-112218423-INS-726
chrX-112232482-INS-112
chrX-112235082-INS-576
chrX-112275212-INS-68
chrX-112294417-INS-156
chrX-112301978-INS-902
chrX-112316918-INS-1008
chrX-112321519-INS-68
chrX-112340204-INS-6714
chrX-112342883-INS-966
chrX-112342895-INS-661
chrX-112342914-INS-1005
chrX-112400826-INS-128
chrX-112404147-INS-53
chrX-112434359-INS-6392
chrX-112461530-INS-127
chrX-112467652-INS-209
chrX-112469286-INS-52
chrX-112478965-INS-60
chrX-112484275-INS-62
chrX-112485094-INS-1562
chrX-112488498-INS-456
chrX-112495647-INS-6198
chrX-112543961-INS-79
chrX-112617015-INS-52
chrX-112626393-INS-357
chrX-112676171-INS-71
chrX-112734947-INS-51
chrX-112752763-INS-211
chrX-112760113-INS-7503
chrX-112784527-INS-54
chrX-112802059-INS-148
chrX-112804409-INS-58
chrX-112807653-INS-100
chrX-112817608-INS-111
chrX-112845301-INS-102
chrX-112886004-INS-527
chrX-112888407-INS-312
chrX-112896676-INS-79
chrX-112900868-INS-77
chrX-112906141-INS-499
chrX-112963617-INS-325
chrX-112985312-INS-65
chrX-113006417-INS-647
chrX-113035544-INS-2172
chrX-113144391-INS-328
chrX-113153461-INS-117
chrX-113154307-INS-401
chrX-113156149-INS-117
chrX-113160455-INS-277
chrX-113167789-INS-88
chrX-113168281-INS-694
chrX-113195032-INS-2550
chrX-113202256-INS-119
chrX-113221119-INS-122
chrX-113258273-INS-566
chrX-113332204-INS-136
chrX-113382088-INS-713
chrX-113394825-INS-302
chrX-113406387-INS-60
chrX-113414078-INS-344
chrX-113454653-INS-198
chrX-113475676-INS-107
chrX-113544490-INS-230
chrX-113587196-INS-421
chrX-113604773-INS-113
chrX-113622095-INS-141
chrX-113682673-INS-289
chrX-113728182-INS-57

chrX-113748059-INS-433
chrX-113761046-INS-175
chrX-113765249-INS-104
chrX-113791783-INS-76
chrX-113812987-INS-64
chrX-113814921-INS-393
chrX-113817524-INS-125
chrX-113826877-INS-207
chrX-113888853-INS-423
chrX-113895066-INS-467
chrX-113909458-INS-55
chrX-113921859-INS-122
chrX-113933964-INS-794
chrX-113934486-INS-57
chrX-113940845-INS-94
chrX-113959770-INS-84
chrX-113960929-INS-209
chrX-113962847-INS-623
chrX-113964227-INS-607
chrX-113967537-INS-472
chrX-113980926-INS-52
chrX-114034529-INS-57
chrX-114034855-INS-264
chrX-114100009-INS-495
chrX-114101893-INS-56
chrX-114114862-INS-83
chrX-114127789-INS-75
chrX-114154003-INS-589
chrX-114165538-INS-608
chrX-114172986-INS-73
chrX-114201008-INS-62
chrX-114204398-INS-105
chrX-114214116-INS-498
chrX-114239299-INS-392
chrX-114253103-INS-53
chrX-114266320-INS-102
chrX-114277884-INS-4177
chrX-114302547-INS-302
chrX-114307367-INS-581
chrX-114323439-INS-7090
chrX-114343626-INS-58
chrX-114344874-INS-272
chrX-114350094-INS-734
chrX-114364398-INS-6069
chrX-114366221-INS-141
chrX-114409368-INS-51
chrX-114413571-INS-6467
chrX-114433887-INS-6883
chrX-114451324-INS-942
chrX-114490748-INS-200
chrX-114493864-INS-80
chrX-114536444-INS-62
chrX-114537924-INS-364
chrX-114541581-INS-2503
chrX-114550406-INS-444
chrX-114563616-INS-2614
chrX-114597944-INS-66
chrX-114613543-INS-142
chrX-114617517-INS-66
chrX-114621034-INS-70
chrX-114699448-INS-447
chrX-114709267-INS-1206
chrX-114732351-INS-175
chrX-114740672-INS-78
chrX-114782797-INS-2697
chrX-114814366-INS-2094
chrX-114829597-INS-82
chrX-114840757-INS-206
chrX-114847228-INS-132
chrX-114858488-INS-1506
chrX-114866508-INS-52
chrX-114873265-INS-115
chrX-114895786-INS-999
chrX-114904714-INS-177
chrX-114917038-INS-55
chrX-114921447-INS-7118
chrX-114926516-INS-78

chrX-114931887-INS-66
chrX-114947080-INS-210
chrX-114960960-INS-72
chrX-114964974-INS-64
chrX-114988877-INS-57
chrX-114989191-INS-7081
chrX-114994360-INS-8816
chrX-115006406-INS-124
chrX-115015232-INS-74
chrX-115028219-INS-117
chrX-115035117-INS-2067
chrX-115039033-INS-611
chrX-115041381-INS-59
chrX-115047244-INS-8945
chrX-115049085-INS-457
chrX-115062126-INS-4826
chrX-115066138-INS-564
chrX-115078557-INS-323
chrX-115084133-INS-147
chrX-115100300-INS-193
chrX-115106477-INS-60
chrX-115144937-INS-416
chrX-115146294-INS-54
chrX-115176936-INS-2433
chrX-115188503-INS-54
chrX-115194397-INS-976
chrX-115229930-INS-153
chrX-115245160-INS-51
chrX-115254338-INS-66
chrX-115278329-INS-51
chrX-115314031-INS-71
chrX-115380484-INS-438
chrX-115391508-INS-1827
chrX-115403912-INS-8305
chrX-115406077-INS-221
chrX-115446025-INS-289
chrX-115477588-INS-160
chrX-115523131-INS-176
chrX-115524185-INS-58
chrX-115530613-INS-472
chrX-115600416-INS-390
chrX-115618820-INS-77
chrX-115629320-INS-200
chrX-115675169-INS-56
chrX-115676124-INS-390
chrX-115680457-INS-6390
chrX-115698707-INS-6171
chrX-115709926-INS-404
chrX-115730599-INS-150
chrX-115735148-INS-2873
chrX-115737729-INS-254
chrX-115746868-INS-6597
chrX-115760000-INS-438
chrX-115773651-INS-6595
chrX-115778720-INS-4389
chrX-115782879-INS-72
chrX-115789877-INS-6231
chrX-115798684-INS-81
chrX-115804471-INS-354
chrX-115811929-INS-358
chrX-115821812-INS-402
chrX-115823938-INS-322
chrX-115824885-INS-51
chrX-115831423-INS-204
chrX-115840119-INS-117
chrX-115861559-INS-409
chrX-115905441-INS-561
chrX-115932204-INS-430
chrX-115940139-INS-190
chrX-115946350-INS-52
chrX-115957624-INS-55
chrX-115964364-INS-5996
chrX-115989546-INS-126
chrX-116132880-INS-232
chrX-116133476-INS-68
chrX-116137069-INS-134
chrX-116159433-INS-825

chrX-116166310-INS-272
chrX-116176009-INS-713
chrX-116181163-INS-74
chrX-116190553-INS-59
chrX-116206386-INS-65
chrX-116222489-INS-359
chrX-116244944-INS-1211
chrX-116251505-INS-153
chrX-116258272-INS-528
chrX-116273229-INS-116
chrX-116275192-INS-5218
chrX-116298343-INS-439
chrX-116306559-INS-1905
chrX-116314641-INS-444
chrX-116331576-INS-56
chrX-116398854-INS-168
chrX-116440552-INS-247
chrX-116518202-INS-368
chrX-116527438-INS-67
chrX-116571789-INS-8340
chrX-116577514-INS-645
chrX-116624605-INS-143
chrX-116662953-INS-87
chrX-116706445-INS-389
chrX-116742652-INS-83
chrX-116794647-INS-103
chrX-116814003-INS-67
chrX-116820788-INS-169
chrX-116822504-INS-84
chrX-116843583-INS-82
chrX-116845368-INS-354
chrX-116855393-INS-6166
chrX-116858612-INS-722
chrX-116871047-INS-7077
chrX-116876943-INS-122
chrX-116888234-INS-329
chrX-116901124-INS-475
chrX-116923186-INS-401
chrX-116925402-INS-243
chrX-116926482-INS-344
chrX-116928870-INS-344
chrX-116947521-INS-644
chrX-116951359-INS-78
chrX-116967934-INS-55
chrX-116982654-INS-99
chrX-117011177-INS-128
chrX-117020488-INS-118
chrX-117023037-INS-655
chrX-117024749-INS-291
chrX-117031417-INS-3112
chrX-117083628-INS-65
chrX-117085986-INS-688
chrX-117087907-INS-75
chrX-117090344-INS-199
chrX-117092647-INS-985
chrX-117093193-INS-281
chrX-117106882-INS-59
chrX-117129028-INS-2807
chrX-117130207-INS-589
chrX-117136616-INS-7155
chrX-117146148-INS-498
chrX-117146348-INS-6566
chrX-117180152-INS-139
chrX-117190001-INS-61
chrX-117212833-INS-57
chrX-117235966-INS-655
chrX-117249101-INS-64
chrX-117250451-INS-506
chrX-117277315-INS-890
chrX-117284405-INS-359
chrX-117298616-INS-231
chrX-117299911-INS-547
chrX-117314264-INS-985
chrX-117343432-INS-604
chrX-117355130-INS-61
chrX-117375036-INS-60
chrX-117402225-INS-2350

chrX-117409133-INS-112
chrX-117423394-INS-57
chrX-117439342-INS-81
chrX-117442246-INS-64
chrX-117515622-INS-248
chrX-117529034-INS-6187
chrX-117542972-INS-2721
chrX-117561846-INS-78
chrX-117579030-INS-53
chrX-117589545-INS-472
chrX-117596749-INS-72
chrX-117665159-INS-98
chrX-117680166-INS-918
chrX-117685013-INS-68
chrX-117685615-INS-1820
chrX-117738472-INS-6825
chrX-117769626-INS-171
chrX-117850262-INS-64
chrX-117866430-INS-104
chrX-117867794-INS-384
chrX-117868521-INS-2300
chrX-117875896-INS-5383
chrX-117884650-INS-2393
chrX-117902899-INS-56
chrX-117971441-INS-945
chrX-117985284-INS-191
chrX-118133755-INS-269
chrX-118148646-INS-69
chrX-118156906-INS-410
chrX-118159152-INS-142
chrX-118211909-INS-524
chrX-118257200-INS-67
chrX-118273851-INS-199
chrX-118366141-INS-358
chrX-118375696-INS-73
chrX-118437513-INS-127
chrX-118446268-INS-62
chrX-118455102-INS-365
chrX-118467630-INS-82
chrX-118540526-INS-831
chrX-118547614-INS-60
chrX-118576267-INS-438
chrX-118578059-INS-3083
chrX-118581392-INS-8951
chrX-118581548-INS-198
chrX-118606634-INS-468
chrX-118619859-INS-56
chrX-118621748-INS-383
chrX-118633328-INS-122
chrX-118664171-INS-112
chrX-118682132-INS-51
chrX-118736832-INS-52
chrX-118756002-INS-478
chrX-118776990-INS-206
chrX-118782972-INS-922
chrX-118783903-INS-66
chrX-118831307-INS-288
chrX-118845643-INS-6388
chrX-118850125-INS-55
chrX-118879996-INS-152
chrX-118908792-INS-315
chrX-118927618-INS-462
chrX-118981244-INS-6489
chrX-119017751-INS-385
chrX-119022092-INS-518
chrX-119038348-INS-151
chrX-119042291-INS-80
chrX-119056752-INS-120
chrX-119097518-INS-54
chrX-119116184-INS-162
chrX-119148674-INS-6869
chrX-119155668-INS-315
chrX-119172173-INS-84
chrX-119177893-INS-1951
chrX-119217275-INS-54
chrX-119231080-INS-447
chrX-119248136-INS-81

chrX-119293560-INS-525
chrX-119326658-INS-272
chrX-119330953-INS-151
chrX-119360884-INS-51
chrX-119376893-INS-108
chrX-119390469-INS-60
chrX-119403850-INS-136
chrX-119405849-INS-1071
chrX-119432796-INS-546
chrX-119434157-INS-180
chrX-119457004-INS-153
chrX-119480770-INS-7717
chrX-119484421-INS-62
chrX-119489021-INS-215
chrX-119493932-INS-130
chrX-119501112-INS-158
chrX-119518134-INS-156
chrX-119527478-INS-358
chrX-119538068-INS-76
chrX-119591687-INS-74
chrX-119596473-INS-6794
chrX-119600452-INS-52
chrX-119619746-INS-620
chrX-119685243-INS-448
chrX-119702565-INS-208
chrX-119760385-INS-732
chrX-119767924-INS-52
chrX-119769289-INS-60
chrX-119793893-INS-52
chrX-119858527-INS-323
chrX-119860041-INS-5040
chrX-119875169-INS-212
chrX-119922956-INS-1869
chrX-119935189-INS-961
chrX-119940071-INS-79
chrX-119941198-INS-60
chrX-119945780-INS-238
chrX-119952536-INS-76
chrX-119986262-INS-2523
chrX-119997075-INS-697
chrX-120001597-INS-669
chrX-120003723-INS-528
chrX-120018530-INS-982
chrX-120046366-INS-331
chrX-120061230-INS-789
chrX-120069027-INS-4390
chrX-120115761-INS-401
chrX-120174426-INS-72
chrX-120193123-INS-53
chrX-120211938-INS-810
chrX-120332209-INS-807
chrX-120335440-INS-236
chrX-120389735-INS-422
chrX-120396595-INS-322
chrX-120408282-INS-103
chrX-120411080-INS-6401
chrX-120416456-INS-72
chrX-120476759-INS-650
chrX-120539296-INS-55
chrX-120548001-INS-106
chrX-120571401-INS-217
chrX-120589787-INS-5549
chrX-120634511-INS-6931
chrX-120636002-INS-61
chrX-120689872-INS-986
chrX-120708192-INS-475
chrX-120730176-INS-111
chrX-120850168-INS-5767
chrX-120850866-INS-770
chrX-120888498-INS-55
chrX-120925918-INS-117
chrX-120939777-INS-490
chrX-120944686-INS-561
chrX-120958557-INS-243
chrX-120960561-INS-621
chrX-120981338-INS-111
chrX-121001084-INS-55

chrX-121063437-INS-434
chrX-121078631-INS-774
chrX-121088231-INS-293
chrX-121121479-INS-57
chrX-121221317-INS-59
chrX-121229660-INS-336
chrX-121264898-INS-358
chrX-121273952-INS-822
chrX-121282954-INS-113
chrX-121286442-INS-485
chrX-121349191-INS-6655
chrX-121357261-INS-401
chrX-121386760-INS-98
chrX-121438475-INS-5472
chrX-121469938-INS-140
chrX-121540446-INS-788
chrX-121573112-INS-135
chrX-121578522-INS-3643
chrX-121614478-INS-66
chrX-121617819-INS-91
chrX-121631966-INS-204
chrX-121668997-INS-132
chrX-121675077-INS-66
chrX-121676736-INS-174
chrX-121685835-INS-504
chrX-121710763-INS-220
chrX-121713686-INS-67
chrX-121741131-INS-105
chrX-121791579-INS-2428
chrX-121799945-INS-54
chrX-121811333-INS-416
chrX-121916463-INS-57
chrX-121941286-INS-55
chrX-121958023-INS-91
chrX-121974657-INS-2406
chrX-121979470-INS-66
chrX-121983125-INS-72
chrX-121984257-INS-59
chrX-121998403-INS-6494
chrX-122066423-INS-2483
chrX-122150795-INS-382
chrX-122159773-INS-88
chrX-122218418-INS-77
chrX-122337591-INS-7071
chrX-122398541-INS-139
chrX-122577649-INS-1289
chrX-122614998-INS-87
chrX-122619772-INS-157
chrX-122619839-INS-202
chrX-122835304-INS-5154
chrX-122841570-INS-120
chrX-122907290-INS-69
chrX-122908245-INS-100
chrX-122914205-INS-6540
chrX-123031641-INS-81
chrX-123034727-INS-357
chrX-123096735-INS-304
chrX-123122637-INS-124
chrX-123123819-INS-111
chrX-123145803-INS-127
chrX-123161106-INS-925
chrX-123163031-INS-355
chrX-123173266-INS-239
chrX-123288723-INS-66
chrX-123395511-INS-284
chrX-123397534-INS-57
chrX-123404359-INS-99
chrX-123405320-INS-146
chrX-123536620-INS-805
chrX-124055208-INS-173
chrX-124059588-INS-95
chrX-124190402-INS-540
chrX-124196253-INS-83
chrX-124197387-INS-60
chrX-124266870-INS-101
chrX-124271703-INS-57
chrX-124279163-INS-71

chrX-124380794-INS-63
chrX-124574951-INS-53
chrX-124606974-INS-158
chrX-124608130-INS-127
chrX-124611369-INS-58
chrX-124627472-INS-1149
chrX-124769671-INS-284
chrX-124783523-INS-63
chrX-124789822-INS-153
chrX-124790987-INS-131
chrX-124794506-INS-69
chrX-124801900-INS-382
chrX-124948756-INS-88
chrX-124989962-INS-319
chrX-125049093-INS-214
chrX-125050558-INS-121
chrX-125092426-INS-6952
chrX-125291383-INS-83
chrX-125310792-INS-5564
chrX-125322856-INS-52
chrX-125379221-INS-5286
chrX-125400233-INS-521
chrX-125432600-INS-4363
chrX-125480158-INS-160
chrX-125503198-INS-6616
chrX-125582218-INS-54
chrX-125682510-INS-399
chrX-125728967-INS-832
chrX-125755682-INS-2548
chrX-125767572-INS-5483
chrX-125806154-INS-219
chrX-125811568-INS-70
chrX-125840329-INS-54
chrX-125921948-INS-325
chrX-125935410-INS-3253
chrX-125955563-INS-138
chrX-126005926-INS-6511
chrX-126018676-INS-1893
chrX-126041167-INS-7401
chrX-126059887-INS-1348
chrX-126085219-INS-6232
chrX-126120142-INS-60
chrX-126130525-INS-106
chrX-126164333-INS-72
chrX-126189811-INS-122
chrX-126191265-INS-293
chrX-126272720-INS-87
chrX-126295558-INS-59
chrX-126298815-INS-7005
chrX-126304122-INS-122
chrX-126308609-INS-6876
chrX-126311527-INS-6216
chrX-126314050-INS-6783
chrX-126323926-INS-306
chrX-126342758-INS-1351
chrX-126350780-INS-276
chrX-126360639-INS-6953
chrX-126363497-INS-6238
chrX-126380129-INS-4062
chrX-126391213-INS-432
chrX-126409605-INS-53
chrX-126458554-INS-558
chrX-126460318-INS-999
chrX-126460933-INS-394
chrX-126467241-INS-110
chrX-126473994-INS-6444
chrX-126510011-INS-64
chrX-126520336-INS-70
chrX-126605284-INS-498
chrX-126609940-INS-400
chrX-126616205-INS-485
chrX-126619444-INS-68
chrX-126645061-INS-191
chrX-126648962-INS-86
chrX-126693715-INS-119
chrX-126753806-INS-373
chrX-126800961-INS-881

chrX-126827932-INS-52
chrX-126849565-INS-54
chrX-126903506-INS-101
chrX-126940072-INS-189
chrX-126981805-INS-381
chrX-127042335-INS-198
chrX-127063941-INS-97
chrX-127090478-INS-253
chrX-127146787-INS-333
chrX-127149332-INS-386
chrX-127157608-INS-2813
chrX-127217475-INS-112
chrX-127269037-INS-102
chrX-127285506-INS-808
chrX-127290523-INS-83
chrX-127305163-INS-149
chrX-127318463-INS-54
chrX-127330281-INS-52
chrX-127354600-INS-273
chrX-127413226-INS-5215
chrX-127423033-INS-9044
chrX-127445908-INS-248
chrX-127465436-INS-2905
chrX-127490852-INS-51
chrX-127598556-INS-1981
chrX-127620850-INS-6407
chrX-127623722-INS-218
chrX-127624964-INS-358
chrX-127639394-INS-5677
chrX-127647534-INS-600
chrX-127671240-INS-116
chrX-127729110-INS-4780
chrX-127777965-INS-102
chrX-127789109-INS-64
chrX-127829666-INS-774
chrX-127851936-INS-177
chrX-127866857-INS-71
chrX-127867895-INS-153
chrX-127920850-INS-135
chrX-127973857-INS-83
chrX-127981280-INS-59
chrX-127983224-INS-60
chrX-128014746-INS-292
chrX-128031093-INS-335
chrX-128057733-INS-127
chrX-128069734-INS-150
chrX-128086518-INS-54
chrX-128124701-INS-407
chrX-128131829-INS-422
chrX-128147504-INS-487
chrX-128151759-INS-6905
chrX-128172246-INS-6790
chrX-128192638-INS-229
chrX-128214875-INS-4398
chrX-128233141-INS-338
chrX-128250078-INS-102
chrX-128294505-INS-330
chrX-128325413-INS-78
chrX-128360848-INS-535
chrX-128391456-INS-149
chrX-128401362-INS-56
chrX-128433196-INS-554
chrX-128437953-INS-766
chrX-128439702-INS-535
chrX-128442239-INS-273
chrX-128452150-INS-107
chrX-128465025-INS-2448
chrX-128470453-INS-146
chrX-128490731-INS-418
chrX-128493584-INS-214
chrX-128516556-INS-5543
chrX-128518814-INS-372
chrX-128537727-INS-6942
chrX-128542195-INS-208
chrX-128555235-INS-320
chrX-128566102-INS-142
chrX-128580918-INS-6644

chrX-128590938-INS-2527
chrX-128635519-INS-72
chrX-128653823-INS-164
chrX-128655630-INS-505
chrX-128664742-INS-210
chrX-128683451-INS-52
chrX-128727557-INS-64
chrX-128734339-INS-108
chrX-128737939-INS-303
chrX-128752451-INS-94
chrX-128765947-INS-266
chrX-128766262-INS-2589
chrX-128802872-INS-345
chrX-128833887-INS-4393
chrX-128856942-INS-466
chrX-128906833-INS-781
chrX-128929401-INS-2274
chrX-128929564-INS-842
chrX-128941413-INS-467
chrX-128944421-INS-251
chrX-128992795-INS-260
chrX-129007762-INS-292
chrX-129008405-INS-1194
chrX-129021676-INS-121
chrX-129042844-INS-502
chrX-129045322-INS-5781
chrX-129075504-INS-284
chrX-129083678-INS-281
chrX-129087320-INS-256
chrX-129089557-INS-114
chrX-129105998-INS-994
chrX-129107485-INS-66
chrX-129110662-INS-7156
chrX-129129082-INS-696
chrX-129144014-INS-1419
chrX-129158356-INS-68
chrX-129167941-INS-2552
chrX-129181456-INS-215
chrX-129184034-INS-195
chrX-129219772-INS-472
chrX-129221680-INS-186
chrX-129235558-INS-114
chrX-129249756-INS-72
chrX-129256453-INS-52
chrX-129273803-INS-61
chrX-129324244-INS-62
chrX-129370192-INS-58
chrX-129407128-INS-463
chrX-129412082-INS-6913
chrX-129425154-INS-68
chrX-129432853-INS-6749
chrX-129455974-INS-2649
chrX-129458729-INS-1083
chrX-129470159-INS-390
chrX-129491752-INS-470
chrX-129527185-INS-106
chrX-129531367-INS-98
chrX-129577688-INS-81
chrX-129604523-INS-192
chrX-129612100-INS-401
chrX-129656307-INS-111
chrX-129664431-INS-609
chrX-129725312-INS-90
chrX-129728293-INS-98
chrX-129757692-INS-82
chrX-129782163-INS-229
chrX-129794650-INS-58
chrX-129821113-INS-52
chrX-129824910-INS-282
chrX-129866060-INS-3123
chrX-129866365-INS-104
chrX-129928208-INS-51
chrX-129948780-INS-138
chrX-129958069-INS-88
chrX-129964799-INS-57
chrX-129974414-INS-91
chrX-129984361-INS-70

chrX-129994166-INS-2456
chrX-129998755-INS-66
chrX-130002542-INS-188
chrX-130007555-INS-197
chrX-130025879-INS-625
chrX-130026522-INS-52
chrX-130051294-INS-2258
chrX-130056634-INS-6548
chrX-130087174-INS-282
chrX-130093209-INS-498
chrX-130099595-INS-617
chrX-130136275-INS-719
chrX-130144212-INS-678
chrX-130163521-INS-275
chrX-130173764-INS-68
chrX-130248225-INS-2967
chrX-130249288-INS-6435
chrX-130251786-INS-1471
chrX-130315055-INS-88
chrX-130358723-INS-272
chrX-130362218-INS-4719
chrX-130368738-INS-70
chrX-130380020-INS-200
chrX-130382433-INS-56
chrX-130394188-INS-67
chrX-130409246-INS-6177
chrX-130409392-INS-860
chrX-130417345-INS-949
chrX-130418302-INS-101
chrX-130441622-INS-135
chrX-130445519-INS-6473
chrX-130488664-INS-433
chrX-130490864-INS-78
chrX-130504228-INS-135
chrX-130524067-INS-890
chrX-130545761-INS-7266
chrX-130550548-INS-2125
chrX-130551973-INS-201
chrX-130566575-INS-144
chrX-130605839-INS-504
chrX-130667020-INS-8012
chrX-130677945-INS-1409
chrX-130688919-INS-539
chrX-130689970-INS-67
chrX-130707102-INS-2393
chrX-130723283-INS-6169
chrX-130752956-INS-90
chrX-130777070-INS-1773
chrX-130779217-INS-6071
chrX-130788370-INS-3206
chrX-130832424-INS-100
chrX-130863015-INS-201
chrX-130908930-INS-6442
chrX-130921498-INS-87
chrX-130950737-INS-157
chrX-130965669-INS-95
chrX-131003982-INS-1656
chrX-131011631-INS-7540
chrX-131016577-INS-491
chrX-131032293-INS-5688
chrX-131032857-INS-6162
chrX-131035957-INS-5289
chrX-131049352-INS-434
chrX-131067921-INS-136
chrX-131075186-INS-85
chrX-131096141-INS-86
chrX-131123692-INS-278
chrX-131130446-INS-56
chrX-131197100-INS-199
chrX-131235703-INS-6513
chrX-131289408-INS-76
chrX-131293981-INS-60
chrX-131317191-INS-7521
chrX-131323238-INS-97
chrX-131325269-INS-61
chrX-131351168-INS-57
chrX-131363234-INS-2300

chrX-131367851-INS-115
chrX-131378463-INS-88
chrX-131385623-INS-238
chrX-131388872-INS-6692
chrX-131443133-INS-657
chrX-131460878-INS-350
chrX-131501772-INS-1134
chrX-131509675-INS-1060
chrX-131541807-INS-267
chrX-131558368-INS-116
chrX-131561266-INS-74
chrX-131563496-INS-326
chrX-131569555-INS-2808
chrX-131581736-INS-224
chrX-131582105-INS-6694
chrX-131592632-INS-77
chrX-131636893-INS-283
chrX-131646041-INS-215
chrX-131654852-INS-419
chrX-131669674-INS-138
chrX-131670598-INS-4542
chrX-131683306-INS-72
chrX-131705253-INS-313
chrX-131715425-INS-75
chrX-131716889-INS-112
chrX-131732092-INS-235
chrX-131742900-INS-64
chrX-131743850-INS-6282
chrX-131772224-INS-196
chrX-131774499-INS-231
chrX-131782329-INS-579
chrX-131791217-INS-975
chrX-131838156-INS-412
chrX-131860553-INS-105
chrX-131866799-INS-482
chrX-131889251-INS-83
chrX-131961308-INS-255
chrX-131979747-INS-264
chrX-132022437-INS-447
chrX-132030681-INS-311
chrX-132032858-INS-72
chrX-132033924-INS-204
chrX-132058788-INS-200
chrX-132061429-INS-52
chrX-132063762-INS-525
chrX-132068578-INS-91
chrX-132078474-INS-52
chrX-132099614-INS-814
chrX-132110644-INS-6164
chrX-132116974-INS-338
chrX-132123494-INS-75
chrX-132125702-INS-175
chrX-132138442-INS-656
chrX-132184909-INS-135
chrX-132189320-INS-90
chrX-132198023-INS-193
chrX-132217472-INS-466
chrX-132219965-INS-88
chrX-132248662-INS-399
chrX-132252099-INS-97
chrX-132273804-INS-819
chrX-132298133-INS-165
chrX-132355769-INS-189
chrX-132356936-INS-84
chrX-132381201-INS-498
chrX-132393821-INS-309
chrX-132397801-INS-129
chrX-132438900-INS-61
chrX-132458391-INS-6224
chrX-132465756-INS-4161
chrX-132480635-INS-6871
chrX-132496825-INS-2737
chrX-132508843-INS-214
chrX-132537065-INS-891
chrX-132555511-INS-96
chrX-132595651-INS-420
chrX-132637297-INS-399

chrX-132641781-INS-60
chrX-132649818-INS-63
chrX-132692356-INS-129
chrX-132723658-INS-419
chrX-132729676-INS-52
chrX-132750560-INS-5332
chrX-132750812-INS-119
chrX-132762979-INS-714
chrX-132804210-INS-231
chrX-132826499-INS-165
chrX-132886196-INS-381
chrX-132921026-INS-62
chrX-132944630-INS-115
chrX-132947931-INS-63
chrX-132948968-INS-400
chrX-132979290-INS-258
chrX-133002757-INS-56
chrX-133006336-INS-66
chrX-133044353-INS-170
chrX-133059989-INS-329
chrX-133085984-INS-174
chrX-133097788-INS-275
chrX-133114477-INS-191
chrX-133184345-INS-180
chrX-133189622-INS-58
chrX-133203544-INS-78
chrX-133219478-INS-52
chrX-133223334-INS-60
chrX-133230442-INS-198
chrX-133237809-INS-236
chrX-133251745-INS-56
chrX-133264416-INS-69
chrX-133269574-INS-899
chrX-133272487-INS-120
chrX-133277847-INS-160
chrX-133317708-INS-475
chrX-133334094-INS-432
chrX-133388095-INS-765
chrX-133413834-INS-51
chrX-133432950-INS-271
chrX-133441302-INS-53
chrX-133443232-INS-7304
chrX-133448502-INS-178
chrX-133473288-INS-60
chrX-133476684-INS-236
chrX-133482972-INS-474
chrX-133508590-INS-60
chrX-133511062-INS-201
chrX-133512651-INS-140
chrX-133516132-INS-94
chrX-133519620-INS-52
chrX-133526492-INS-177
chrX-133535366-INS-219
chrX-133554146-INS-451
chrX-133560619-INS-124
chrX-133577251-INS-118
chrX-133579948-INS-376
chrX-133607261-INS-749
chrX-133636733-INS-180
chrX-133648238-INS-66
chrX-133654742-INS-80
chrX-133688236-INS-151
chrX-133703562-INS-55
chrX-133710308-INS-110
chrX-133721418-INS-65
chrX-133725256-INS-79
chrX-133727390-INS-180
chrX-133803242-INS-83
chrX-133821174-INS-174
chrX-133846435-INS-57
chrX-133852903-INS-62
chrX-133864113-INS-131
chrX-133871223-INS-957
chrX-133885360-INS-128
chrX-133901675-INS-76
chrX-133920303-INS-56
chrX-133942902-INS-1520

chrX-133948446-INS-1348
chrX-133948992-INS-1518
chrX-133959757-INS-63
chrX-133971281-INS-511
chrX-133987974-INS-161
chrX-133993892-INS-64
chrX-134014184-INS-173
chrX-134047893-INS-6225
chrX-134060732-INS-6440
chrX-134061006-INS-2092
chrX-134083983-INS-74
chrX-134095795-INS-588
chrX-134097396-INS-68
chrX-134150521-INS-419
chrX-134221419-INS-55
chrX-134291283-INS-2887
chrX-134316677-INS-200
chrX-134320855-INS-258
chrX-134381638-INS-85
chrX-134459304-INS-129
chrX-134467153-INS-285
chrX-134523698-INS-206
chrX-134539601-INS-177
chrX-134574640-INS-74
chrX-134576629-INS-57
chrX-134577296-INS-84
chrX-134578834-INS-72
chrX-134586371-INS-82
chrX-134615184-INS-556
chrX-134626271-INS-70
chrX-134641993-INS-51
chrX-134698459-INS-217
chrX-134714759-INS-359
chrX-134724207-INS-150
chrX-134729663-INS-185
chrX-134741316-INS-461
chrX-134741426-INS-235
chrX-134763217-INS-505
chrX-134793563-INS-59
chrX-134801402-INS-345
chrX-134807420-INS-217
chrX-134835619-INS-59
chrX-134852780-INS-127
chrX-134861452-INS-144
chrX-134868255-INS-199
chrX-134879752-INS-74
chrX-134899024-INS-51
chrX-134917829-INS-362
chrX-134923402-INS-938
chrX-134925711-INS-54
chrX-134940192-INS-423
chrX-134951338-INS-210
chrX-134955159-INS-162
chrX-134957675-INS-94
chrX-134971893-INS-67
chrX-135004604-INS-929
chrX-135015338-INS-323
chrX-135034508-INS-56
chrX-135042580-INS-166
chrX-135064106-INS-472
chrX-135074639-INS-54
chrX-135079326-INS-60
chrX-135089157-INS-194
chrX-135096413-INS-57
chrX-135110101-INS-350
chrX-135161665-INS-899
chrX-135191058-INS-191
chrX-135199623-INS-76
chrX-135205750-INS-87
chrX-135210083-INS-198
chrX-135216465-INS-68
chrX-135223727-INS-108
chrX-135243012-INS-333
chrX-135310679-INS-4456
chrX-135325569-INS-6706
chrX-135340653-INS-308
chrX-135405768-INS-6378

chrX-135412343-INS-6670
chrX-135420910-INS-112
chrX-135478396-INS-68
chrX-135480363-INS-79
chrX-135497879-INS-386
chrX-135505749-INS-151
chrX-135548339-INS-58
chrX-135549881-INS-280
chrX-135571373-INS-598
chrX-135576917-INS-51
chrX-135606908-INS-192
chrX-135619204-INS-140
chrX-135622083-INS-76
chrX-135674842-INS-177
chrX-135705767-INS-60
chrX-135709086-INS-868
chrX-135735435-INS-110
chrX-135747930-INS-193
chrX-135763522-INS-1894
chrX-135777199-INS-210
chrX-135780952-INS-52
chrX-135805823-INS-400
chrX-135840392-INS-66
chrX-135879981-INS-196
chrX-135936952-INS-186
chrX-135941715-INS-60
chrX-135953617-INS-66
chrX-135958069-INS-52
chrX-136006867-INS-59
chrX-136019804-INS-119
chrX-136047261-INS-53
chrX-136051129-INS-210
chrX-136057447-INS-125
chrX-136072185-INS-546
chrX-136086653-INS-79
chrX-136096800-INS-217
chrX-136109337-INS-60
chrX-136127360-INS-154
chrX-136149274-INS-940
chrX-136160055-INS-52
chrX-136165388-INS-92
chrX-136173108-INS-64
chrX-136176826-INS-6364
chrX-136185289-INS-934
chrX-136209212-INS-6589
chrX-136229006-INS-998
chrX-136243547-INS-56
chrX-136271421-INS-417
chrX-136279244-INS-250
chrX-136311614-INS-52
chrX-136321821-INS-155
chrX-136337789-INS-54
chrX-136341333-INS-3097
chrX-136418437-INS-703
chrX-136424561-INS-63
chrX-136427290-INS-159
chrX-136433148-INS-184
chrX-136445556-INS-81
chrX-136450897-INS-6903
chrX-136452738-INS-143
chrX-136481393-INS-61
chrX-136505014-INS-1039
chrX-136546871-INS-83
chrX-136552053-INS-4052
chrX-136560063-INS-53
chrX-136565511-INS-1396
chrX-136565513-INS-994
chrX-136576864-INS-4447
chrX-136589374-INS-125
chrX-136603277-INS-60
chrX-136722181-INS-327
chrX-136724162-INS-1947
chrX-136737547-INS-107
chrX-136762476-INS-1094
chrX-136769907-INS-6501
chrX-136790613-INS-932
chrX-136823603-INS-88

chrX-136831364-INS-168
chrX-136833386-INS-6776
chrX-136836616-INS-68
chrX-136844411-INS-303
chrX-136855896-INS-347
chrX-136873350-INS-3484
chrX-136873361-INS-69
chrX-136898609-INS-99
chrX-136924886-INS-443
chrX-136937446-INS-312
chrX-136970432-INS-396
chrX-137002368-INS-6599
chrX-137014394-INS-102
chrX-137023676-INS-108
chrX-137051264-INS-551
chrX-137058613-INS-6387
chrX-137065037-INS-61
chrX-137096439-INS-113
chrX-137098650-INS-7015
chrX-137157462-INS-159
chrX-137160371-INS-428
chrX-137228788-INS-140
chrX-137229851-INS-54
chrX-137269701-INS-130
chrX-137310506-INS-58
chrX-137316955-INS-74
chrX-137318300-INS-692
chrX-137369109-INS-6374
chrX-137400796-INS-2567
chrX-137422627-INS-174
chrX-137450081-INS-294
chrX-137571264-INS-208
chrX-137574953-INS-125
chrX-137575097-INS-6333
chrX-137593492-INS-482
chrX-137645765-INS-106
chrX-137741398-INS-55
chrX-137747853-INS-137
chrX-137760164-INS-62
chrX-137769616-INS-78
chrX-137772476-INS-204
chrX-137782792-INS-511
chrX-137843282-INS-139
chrX-137845773-INS-378
chrX-137852043-INS-64
chrX-137854814-INS-198
chrX-137935202-INS-897
chrX-137943326-INS-58
chrX-137969050-INS-59
chrX-137971913-INS-52
chrX-138043697-INS-58
chrX-138047084-INS-2525
chrX-138053211-INS-52
chrX-138093655-INS-166
chrX-138099114-INS-141
chrX-138149891-INS-911
chrX-138151739-INS-78
chrX-138157627-INS-178
chrX-138180411-INS-205
chrX-138184822-INS-390
chrX-138193876-INS-52
chrX-138260758-INS-342
chrX-138313704-INS-398
chrX-138328717-INS-428
chrX-138334422-INS-100
chrX-138337016-INS-970
chrX-138342218-INS-659
chrX-138347653-INS-7251
chrX-138351021-INS-256
chrX-138383873-INS-201
chrX-138397446-INS-84
chrX-138459069-INS-233
chrX-138474622-INS-81
chrX-138479060-INS-515
chrX-138522808-INS-97
chrX-138540428-INS-918
chrX-138553688-INS-56

chrX-138576401-INS-193
chrX-138590012-INS-176
chrX-138599322-INS-74
chrX-138602406-INS-90
chrX-138625225-INS-54
chrX-138636336-INS-66
chrX-138665413-INS-72
chrX-138667849-INS-341
chrX-138670421-INS-202
chrX-138697709-INS-98
chrX-138713603-INS-119
chrX-138783474-INS-246
chrX-138788623-INS-195
chrX-138801428-INS-72
chrX-138824995-INS-6195
chrX-138847350-INS-6900
chrX-138866273-INS-639
chrX-138898222-INS-213
chrX-138923075-INS-6310
chrX-138951179-INS-1274
chrX-138954568-INS-210
chrX-138964272-INS-171
chrX-138981842-INS-114
chrX-138989009-INS-54
chrX-139005467-INS-52
chrX-139012853-INS-60
chrX-139101216-INS-2982
chrX-139119075-INS-198
chrX-139127281-INS-6224
chrX-139129239-INS-72
chrX-139143539-INS-76
chrX-139156967-INS-6750
chrX-139170800-INS-53
chrX-139174925-INS-75
chrX-139191986-INS-64
chrX-139211421-INS-60
chrX-139275273-INS-892
chrX-139278271-INS-567
chrX-139312683-INS-110
chrX-139348308-INS-189
chrX-139386944-INS-62
chrX-139433921-INS-195
chrX-139447592-INS-155
chrX-139485528-INS-229
chrX-139515753-INS-197
chrX-139545745-INS-101
chrX-139547514-INS-213
chrX-139551223-INS-59
chrX-139570106-INS-88
chrX-139620676-INS-571
chrX-139638660-INS-388
chrX-139684327-INS-2884
chrX-139691138-INS-160
chrX-139695554-INS-4890
chrX-139696524-INS-59
chrX-139713184-INS-164
chrX-139714840-INS-64
chrX-139752539-INS-56
chrX-139786281-INS-605
chrX-139823996-INS-129
chrX-139894830-INS-61
chrX-139904175-INS-974
chrX-139915450-INS-336
chrX-139918355-INS-54
chrX-139964145-INS-116
chrX-139969151-INS-329
chrX-139981477-INS-6481
chrX-139994663-INS-122
chrX-140016224-INS-910
chrX-140036253-INS-561
chrX-140040887-INS-6593
chrX-140081995-INS-54
chrX-140095833-INS-67
chrX-140116864-INS-54
chrX-140145110-INS-52
chrX-140160288-INS-148
chrX-140188683-INS-1063

chrX-140216678-INS-6571
chrX-140220878-INS-391
chrX-140237188-INS-59
chrX-140243327-INS-52
chrX-140250415-INS-108
chrX-140254365-INS-6397
chrX-140270360-INS-662
chrX-140287639-INS-399
chrX-140290396-INS-52
chrX-140320944-INS-107
chrX-140336300-INS-325
chrX-140355300-INS-500
chrX-140371579-INS-66
chrX-140399062-INS-336
chrX-140409210-INS-6381
chrX-140437033-INS-619
chrX-140444086-INS-233
chrX-140454337-INS-457
chrX-140460023-INS-99
chrX-140468275-INS-119
chrX-140515028-INS-358
chrX-140520011-INS-866
chrX-140522310-INS-73
chrX-140550968-INS-2270
chrX-140558834-INS-107
chrX-140562491-INS-923
chrX-140571164-INS-56
chrX-140614875-INS-6924
chrX-140621006-INS-122
chrX-140633198-INS-75
chrX-140633508-INS-954
chrX-140637388-INS-96
chrX-140648586-INS-102
chrX-140666352-INS-250
chrX-140753958-INS-54
chrX-140773579-INS-64
chrX-140781415-INS-364
chrX-140787453-INS-1729
chrX-140792949-INS-68
chrX-140811774-INS-637
chrX-140822726-INS-433
chrX-140824725-INS-2868
chrX-140829803-INS-3128
chrX-140855019-INS-52
chrX-140857616-INS-563
chrX-140861548-INS-174
chrX-140871351-INS-5894
chrX-140876976-INS-100
chrX-140882530-INS-201
chrX-140892802-INS-129
chrX-140973663-INS-165
chrX-140984241-INS-65
chrX-140997938-INS-6702
chrX-141027740-INS-135
chrX-141056063-INS-130
chrX-141058793-INS-55
chrX-141078200-INS-6905
chrX-141098048-INS-200
chrX-141100664-INS-102
chrX-141108321-INS-388
chrX-141112254-INS-62
chrX-141145709-INS-211
chrX-141155490-INS-223
chrX-141168552-INS-63
chrX-141172254-INS-369
chrX-141180977-INS-235
chrX-141182344-INS-430
chrX-141192003-INS-58
chrX-141199168-INS-1916
chrX-141204713-INS-198
chrX-141207453-INS-60
chrX-141249905-INS-181
chrX-141275753-INS-59
chrX-141276832-INS-208
chrX-141280081-INS-396
chrX-141291791-INS-197
chrX-141301721-INS-211

chrX-141309878-INS-51
chrX-141326092-INS-60
chrX-141348122-INS-626
chrX-141351851-INS-235
chrX-141354284-INS-76
chrX-141374398-INS-59
chrX-141399789-INS-87
chrX-141406316-INS-52
chrX-141410307-INS-167
chrX-141417219-INS-68
chrX-141424402-INS-347
chrX-141425303-INS-89
chrX-141431718-INS-324
chrX-141451788-INS-58
chrX-141458638-INS-72
chrX-141462036-INS-198
chrX-141464283-INS-142
chrX-141507864-INS-58
chrX-141508356-INS-81
chrX-141526466-INS-103
chrX-141530028-INS-57
chrX-141547001-INS-60
chrX-141565313-INS-149
chrX-141574902-INS-56
chrX-141583245-INS-331
chrX-141585034-INS-60
chrX-141587443-INS-827
chrX-141593032-INS-81
chrX-141605319-INS-75
chrX-141712250-INS-143
chrX-141723542-INS-81
chrX-141774253-INS-207
chrX-141775985-INS-897
chrX-141798913-INS-78
chrX-141814444-INS-5549
chrX-141896730-INS-70
chrX-141910811-INS-6440
chrX-141954182-INS-400
chrX-141970867-INS-60
chrX-141976448-INS-114
chrX-142007927-INS-6663
chrX-142031095-INS-447
chrX-142054238-INS-212
chrX-142055145-INS-222
chrX-142060550-INS-6307
chrX-142067077-INS-1160
chrX-142123955-INS-6808
chrX-142126805-INS-175
chrX-142184847-INS-234
chrX-142227738-INS-384
chrX-142229042-INS-7351
chrX-142251377-INS-353
chrX-142268295-INS-102
chrX-142361250-INS-776
chrX-142369889-INS-637
chrX-142429918-INS-68
chrX-142434790-INS-6625
chrX-142474871-INS-96
chrX-142475144-INS-228
chrX-142492478-INS-116
chrX-142532627-INS-94
chrX-142555625-INS-693
chrX-142601784-INS-60
chrX-142619439-INS-52
chrX-142655597-INS-52
chrX-142672512-INS-103
chrX-142676820-INS-3226
chrX-142703028-INS-80
chrX-142734811-INS-60
chrX-142738522-INS-6440
chrX-142750288-INS-302
chrX-142751005-INS-71
chrX-142778785-INS-83
chrX-142820699-INS-105
chrX-142871210-INS-7006
chrX-142878290-INS-276
chrX-142879645-INS-6923

chrX-142883699-INS-503
chrX-142886429-INS-197
chrX-142899304-INS-624
chrX-142923656-INS-438
chrX-142934010-INS-121
chrX-142961410-INS-59
chrX-142986069-INS-66
chrX-142993931-INS-182
chrX-142997833-INS-90
chrX-142998606-INS-6893
chrX-142998848-INS-132
chrX-143005038-INS-715
chrX-143016090-INS-89
chrX-143024965-INS-328
chrX-143028535-INS-74
chrX-143037147-INS-96
chrX-143047958-INS-221
chrX-143049432-INS-196
chrX-143085171-INS-315
chrX-143091099-INS-51
chrX-143092661-INS-412
chrX-143100402-INS-52
chrX-143153766-INS-56
chrX-143176061-INS-2076
chrX-143183892-INS-614
chrX-143199813-INS-52
chrX-143206471-INS-7118
chrX-143215309-INS-77
chrX-143216846-INS-137
chrX-143221038-INS-215
chrX-143233314-INS-220
chrX-143246179-INS-69
chrX-143248659-INS-248
chrX-143249775-INS-6580
chrX-143282936-INS-62
chrX-143296109-INS-69
chrX-143335100-INS-112
chrX-143352558-INS-55
chrX-143353834-INS-5154
chrX-143365665-INS-64
chrX-143392996-INS-88
chrX-143396109-INS-148
chrX-143411836-INS-172
chrX-143428124-INS-116
chrX-143454427-INS-180
chrX-143500927-INS-153
chrX-143505540-INS-57
chrX-143507025-INS-58
chrX-143525536-INS-75
chrX-143529519-INS-181
chrX-143546712-INS-606
chrX-143569648-INS-52
chrX-143597954-INS-2410
chrX-143619937-INS-371
chrX-143625718-INS-153
chrX-143626955-INS-427
chrX-143673181-INS-54
chrX-143686482-INS-109
chrX-143740385-INS-227
chrX-143756346-INS-7127
chrX-143803514-INS-466
chrX-143843480-INS-194
chrX-143870429-INS-60
chrX-143884381-INS-58
chrX-143896360-INS-54
chrX-143910440-INS-200
chrX-143927088-INS-257
chrX-144000444-INS-7007
chrX-144046292-INS-55
chrX-144084102-INS-60
chrX-144086392-INS-618
chrX-144095526-INS-7315
chrX-144096367-INS-197
chrX-144097611-INS-6654
chrX-144152289-INS-200
chrX-144162341-INS-6821
chrX-144162583-INS-468

chrX-144168598-INS-52
chrX-144203428-INS-2969
chrX-144205192-INS-124
chrX-144213177-INS-80
chrX-144214196-INS-222
chrX-144220002-INS-101
chrX-144243267-INS-132
chrX-144248571-INS-54
chrX-144283031-INS-81
chrX-144297631-INS-5972
chrX-144310628-INS-100
chrX-144316492-INS-110
chrX-144316670-INS-163
chrX-144346488-INS-872
chrX-144351473-INS-140
chrX-144354956-INS-753
chrX-144360073-INS-68
chrX-144387843-INS-6467
chrX-144388635-INS-7198
chrX-144392916-INS-393
chrX-144412653-INS-55
chrX-144417043-INS-68
chrX-144436904-INS-70
chrX-144442724-INS-156
chrX-144444018-INS-402
chrX-144448337-INS-6701
chrX-144450151-INS-302
chrX-144476349-INS-281
chrX-144489625-INS-6890
chrX-144504369-INS-455
chrX-144506076-INS-68
chrX-144571497-INS-618
chrX-144574661-INS-7174
chrX-144596640-INS-240
chrX-144597319-INS-6660
chrX-144603387-INS-489
chrX-144623235-INS-99
chrX-144638396-INS-119
chrX-144647809-INS-7268
chrX-144656044-INS-60
chrX-144658692-INS-150
chrX-144674624-INS-6398
chrX-144692740-INS-227
chrX-144708963-INS-93
chrX-144721667-INS-7185
chrX-144745610-INS-6659
chrX-144748202-INS-57
chrX-144763071-INS-12097
chrX-144767736-INS-54
chrX-144791337-INS-88
chrX-144807395-INS-7108
chrX-144837175-INS-51
chrX-144842742-INS-78
chrX-144866241-INS-445
chrX-144870281-INS-112
chrX-144885151-INS-233
chrX-144892654-INS-105
chrX-144902427-INS-123
chrX-144909293-INS-76
chrX-144913517-INS-484
chrX-144938130-INS-176
chrX-144952091-INS-7195
chrX-145008882-INS-91
chrX-145017033-INS-6897
chrX-145031956-INS-6424
chrX-145038327-INS-295
chrX-145038892-INS-1376
chrX-145054905-INS-354
chrX-145104095-INS-60
chrX-145115491-INS-76
chrX-145145397-INS-53
chrX-145160329-INS-420
chrX-145174249-INS-70
chrX-145185349-INS-53
chrX-145190555-INS-441
chrX-145197889-INS-520
chrX-145206402-INS-78

chrX-145306163-INS-106
chrX-145319738-INS-1873
chrX-145326839-INS-1475
chrX-145348258-INS-270
chrX-145353442-INS-62
chrX-145376025-INS-60
chrX-145379510-INS-746
chrX-145406814-INS-85
chrX-145408363-INS-397
chrX-145421120-INS-783
chrX-145447484-INS-101
chrX-145472588-INS-6433
chrX-145486902-INS-124
chrX-145487228-INS-7079
chrX-145489618-INS-774
chrX-145506957-INS-237
chrX-145513461-INS-425
chrX-145516911-INS-52
chrX-145537924-INS-737
chrX-145555492-INS-62
chrX-145567867-INS-7193
chrX-145583782-INS-174
chrX-145593748-INS-344
chrX-145627329-INS-392
chrX-145678252-INS-1282
chrX-145689873-INS-200
chrX-145713122-INS-226
chrX-145753580-INS-72
chrX-145767845-INS-65
chrX-145772181-INS-6354
chrX-145773111-INS-56
chrX-145790028-INS-550
chrX-145798829-INS-52
chrX-145802659-INS-52
chrX-145811882-INS-268
chrX-145824328-INS-81
chrX-145842432-INS-641
chrX-145850133-INS-100
chrX-145890536-INS-257
chrX-145905150-INS-111
chrX-145910306-INS-6674
chrX-145951613-INS-499
chrX-145963815-INS-65
chrX-145967220-INS-294
chrX-146006037-INS-601
chrX-146019814-INS-492
chrX-146033578-INS-476
chrX-146067622-INS-214
chrX-146082207-INS-121
chrX-146113120-INS-6935
chrX-146126941-INS-430
chrX-146147605-INS-440
chrX-146155588-INS-113
chrX-146190620-INS-59
chrX-146195174-INS-397
chrX-146225815-INS-54
chrX-146287077-INS-70
chrX-146321264-INS-56
chrX-146368514-INS-161
chrX-146380446-INS-98
chrX-146429568-INS-53
chrX-146543588-INS-3213
chrX-146640564-INS-106
chrX-146655834-INS-115
chrX-146806737-INS-56
chrX-147002374-INS-58
chrX-147019741-INS-105
chrX-147081231-INS-6077
chrX-147141488-INS-6519
chrX-147308181-INS-6519
chrX-147532609-INS-7104
chrX-147674171-INS-5922
chrX-147753160-INS-226
chrX-147753919-INS-4118
chrX-147911202-INS-2935
chrX-148154084-INS-177
chrX-148193396-INS-106

chrX-148200449-INS-6483
chrX-148463322-INS-5062
chrX-148502379-INS-1512
chrX-148508273-INS-366
chrX-148511563-INS-57
chrX-148521423-INS-435
chrX-148680861-INS-115
chrX-148719134-INS-6404
chrX-148766876-INS-123
chrX-148769721-INS-56
chrX-148806652-INS-7058
chrX-148946155-INS-6452
chrX-148992245-INS-82
chrX-149020250-INS-444
chrX-149024287-INS-765
chrX-149050161-INS-740
chrX-149111566-INS-2881
chrX-149144739-INS-3166
chrX-149174123-INS-74
chrX-149180016-INS-122
chrX-149215549-INS-61
chrX-149238621-INS-209
chrX-149248974-INS-99
chrX-149290877-INS-357
chrX-149302184-INS-6401
chrX-149364375-INS-70
chrX-149403439-INS-208
chrX-149423363-INS-80
chrX-149478214-INS-75
chrX-149478751-INS-5899
chrX-149551515-INS-7314
chrX-149557918-INS-6387
chrX-149560463-INS-5472
chrX-149569213-INS-6426
chrX-149571649-INS-255
chrX-149618750-INS-432
chrX-149633558-INS-368
chrX-149640549-INS-274
chrX-149642118-INS-6708
chrX-149655475-INS-118
chrX-149663235-INS-58
chrX-149678157-INS-324
chrX-149680062-INS-767
chrX-149732440-INS-2663
chrX-149734149-INS-198
chrX-149762158-INS-52
chrX-149764380-INS-176
chrX-149777927-INS-57
chrX-149791874-INS-140
chrX-149809524-INS-621
chrX-149818522-INS-121
chrX-149888394-INS-230
chrX-149915118-INS-102
chrX-149928057-INS-112
chrX-149947190-INS-127
chrX-149950636-INS-102
chrX-149952969-INS-60
chrX-149977229-INS-60
chrX-150051432-INS-489
chrX-150072550-INS-72
chrX-150080806-INS-90
chrX-150155535-INS-625
chrX-150164875-INS-51
chrX-150176839-INS-56
chrX-150190613-INS-52
chrX-150193392-INS-595
chrX-150199221-INS-57
chrX-150204548-INS-1474
chrX-150300833-INS-399
chrX-150333455-INS-6278
chrX-150338792-INS-80
chrX-150359332-INS-56
chrX-150378057-INS-76
chrX-150435140-INS-3674
chrX-150436351-INS-3840
chrX-150437190-INS-99
chrX-150449208-INS-422

chrX-150450205-INS-506
chrX-150463384-INS-607
chrX-150494802-INS-4007
chrX-150510919-INS-6550
chrX-150515007-INS-4982
chrX-150515070-INS-6413
chrX-150536157-INS-57
chrX-150544175-INS-401
chrX-150552098-INS-672
chrX-150566546-INS-130
chrX-150594837-INS-81
chrX-150622667-INS-80
chrX-150725313-INS-56
chrX-150759525-INS-95
chrX-150763661-INS-180
chrX-150766392-INS-244
chrX-150772485-INS-51
chrX-150781637-INS-7085
chrX-150791912-INS-345
chrX-150880494-INS-181
chrX-150894344-INS-362
chrX-150936817-INS-83
chrX-150969505-INS-75
chrX-151012351-INS-164
chrX-151027372-INS-199
chrX-151040507-INS-101
chrX-151067471-INS-89
chrX-151165793-INS-1893
chrX-151215341-INS-134
chrX-151216010-INS-972
chrX-151240713-INS-1149
chrX-151253038-INS-51
chrX-151320772-INS-81
chrX-151323488-INS-6299
chrX-151327047-INS-330
chrX-151328658-INS-579
chrX-151334326-INS-52
chrX-151370109-INS-228
chrX-151376989-INS-55
chrX-151390775-INS-399
chrX-151401571-INS-389
chrX-151409396-INS-108
chrX-151429269-INS-363
chrX-151438972-INS-52
chrX-151476495-INS-313
chrX-151510202-INS-51
chrX-151555099-INS-199
chrX-151588523-INS-170
chrX-151673308-INS-3000
chrX-151715021-INS-52
chrX-151754289-INS-395
chrX-151763014-INS-436
chrX-151838866-INS-104
chrX-151841072-INS-88
chrX-151868292-INS-120
chrX-151882106-INS-113
chrX-151886529-INS-94
chrX-151982085-INS-317
chrX-151989941-INS-235
chrX-152008624-INS-68
chrX-152039902-INS-100
chrX-152068606-INS-221
chrX-152082843-INS-102
chrX-152136265-INS-92
chrX-152139155-INS-432
chrX-152146139-INS-607
chrX-152187914-INS-661
chrX-152215118-INS-5236
chrX-152233720-INS-52
chrX-152242266-INS-200
chrX-152295253-INS-7107
chrX-152295276-INS-117
chrX-152302561-INS-58
chrX-152325833-INS-51
chrX-152332000-INS-199
chrX-152362466-INS-85
chrX-152383886-INS-7294

chrX-152390460-INS-6153
chrX-152392076-INS-130
chrX-152399347-INS-6168
chrX-152401251-INS-136
chrX-152432463-INS-192
chrX-152445523-INS-206
chrX-152449311-INS-58
chrX-152450221-INS-5648
chrX-152457645-INS-768
chrX-152479354-INS-165
chrX-152514820-INS-1561
chrX-152531084-INS-53
chrX-152538908-INS-534
chrX-152543863-INS-6933
chrX-152553291-INS-55
chrX-152554315-INS-205
chrX-152590025-INS-72
chrX-152626315-INS-6750
chrX-152636593-INS-77
chrX-152642225-INS-11035
chrX-152673945-INS-330
chrX-152675438-INS-540
chrX-152689942-INS-7551
chrX-152690848-INS-58
chrX-152700887-INS-200
chrX-152734553-INS-383
chrX-152769637-INS-133
chrX-152782858-INS-259
chrX-152843607-INS-5395
chrX-152949908-INS-1173
chrX-152972943-INS-60
chrX-152987158-INS-3350
chrX-152987594-INS-73
chrX-152998569-INS-1473
chrX-153042107-INS-1235
chrX-153086281-INS-5296
chrX-153088944-INS-314
chrX-153097535-INS-4989
chrX-153111951-INS-7130
chrX-153121297-INS-7578
chrX-153127317-INS-1599
chrX-153129238-INS-2840
chrX-153209865-INS-171
chrX-153316231-INS-54
chrX-153318048-INS-376
chrX-153417332-INS-117
chrX-153431615-INS-55
chrX-153446206-INS-6495
chrX-153455099-INS-7290
chrX-153455755-INS-2967
chrX-153461176-INS-63
chrX-153484502-INS-64
chrX-153513775-INS-64
chrX-153541015-INS-210
chrX-153560367-INS-529
chrX-153567896-INS-4038
chrX-153573499-INS-568
chrX-153580963-INS-6184
chrX-153582773-INS-1485
chrX-153584258-INS-81
chrX-153599837-INS-323
chrX-153645876-INS-363
chrX-153667046-INS-944
chrX-153674939-INS-101
chrX-153713620-INS-401
chrX-153862655-INS-114
chrX-153875435-INS-102
chrX-153881710-INS-340
chrX-153888022-INS-115
chrX-153891167-INS-197
chrX-153891961-INS-108
chrX-153898716-INS-2230
chrX-153911585-INS-179
chrX-153982830-INS-67
chrX-154021562-INS-85
chrX-154025688-INS-92
chrX-154026941-INS-603

chrX-154060961-INS-216
chrX-154063108-INS-1047
chrX-154066389-INS-153
chrX-154088970-INS-112
chrX-154089284-INS-6723
chrX-154103610-INS-399
chrX-154155649-INS-777
chrX-154198485-INS-3180
chrX-154198597-INS-62
chrX-154223506-INS-782
chrX-154226347-INS-97
chrX-154238726-INS-6214
chrX-154273582-INS-51
chrX-154298038-INS-692
chrX-154303731-INS-321
chrX-154328208-INS-957
chrX-154358670-INS-299
chrX-154373120-INS-64
chrX-154376677-INS-6473
chrX-154402261-INS-106
chrX-154436002-INS-6395
chrX-154448394-INS-83
chrX-154450370-INS-1204
chrX-154496314-INS-201
chrX-154539281-INS-307
chrX-154593859-INS-7813
chrX-154595355-INS-70
chrX-154609248-INS-103
chrX-154620068-INS-323
chrX-154627261-INS-102
chrX-154673193-INS-275
chrX-154695750-INS-964
chrX-154701398-INS-1470
chrX-154703832-INS-402
chrX-154719814-INS-452
chrX-154749680-INS-169
chrX-154765199-INS-7258
chrX-154805245-INS-83
chrX-154874015-INS-408
chrX-154874831-INS-70
chrX-154881215-INS-250
chrX-154898284-INS-134
chrX-154921267-INS-500
chrX-154930805-INS-881
chrX-154946329-INS-63
chrX-154952358-INS-2779
chrX-154955186-INS-6957
chrX-154970181-INS-220
chrX-154976206-INS-538
chrX-154985948-INS-760
chrX-155021238-INS-108
chrX-155041734-INS-78
chrX-155061916-INS-118
chrX-155072335-INS-464
chrX-155115158-INS-224
chrX-155140618-INS-4968
chrX-155147622-INS-6702
chrX-155163872-INS-5399
chrX-155170341-INS-141
chrX-155205258-INS-75
chrX-155215823-INS-74
chrX-155229609-INS-1864
chrX-155309690-INS-307
chrX-155377077-INS-360
chrX-155389168-INS-221
chrX-155397680-INS-6449
chrX-155433149-INS-7412
chrX-155433483-INS-343
chrX-155436149-INS-55
chrX-155455075-INS-219
chrX-155455669-INS-397
chrX-155457401-INS-2739
chrX-155488064-INS-72
chrX-155490849-INS-172
chrX-155491697-INS-368
chrX-155495325-INS-72
chrX-155507075-INS-353

chrX-155560879-INS-623
chrX-155571531-INS-73
chrX-155574684-INS-142
chrX-155596674-INS-746
chrX-155599778-INS-76
chrX-155617026-INS-6956
chrX-155622316-INS-171
chrX-155629157-INS-89
chrX-155637166-INS-128
chrX-155670454-INS-68
chrX-155698924-INS-518
chrX-155740658-INS-3249
chrX-155753189-INS-56
chrX-155783214-INS-433
chrX-155796469-INS-133
chrX-155803826-INS-256
chrX-155808581-INS-93
chrX-155816149-INS-506
chrX-155829990-INS-1109
chrX-155832886-INS-188
chrX-155845452-INS-6940
chrX-155861783-INS-132
chrX-155868749-INS-1806
chrX-155902051-INS-71
chrX-155920449-INS-6183
chrX-155928200-INS-58
chrX-155937103-INS-78
chrX-155948322-INS-494
chrX-155950455-INS-607
chrX-155981276-INS-231
chrX-155990623-INS-236
chrX-156040711-INS-189
chrX-156043159-INS-834
chrX-156043584-INS-60
chrX-156043663-INS-1260
chrX-156062627-INS-110
chrX-156079346-INS-525
chrX-156081955-INS-61
chrX-156119212-INS-279
chrX-156122464-INS-76
chrX-156124038-INS-6439
chrX-156124172-INS-60
chrX-156153924-INS-202
chrX-156156517-INS-472
chrX-156167946-INS-62
chrX-156177807-INS-196
chrX-156199100-INS-61
chrX-156202305-INS-69
chrX-156211774-INS-52
chrX-156292716-INS-75
chrX-156315792-INS-162
chrX-156335061-INS-575
chrX-156358195-INS-301
chrX-156395878-INS-100
chrX-156397122-INS-390
chrX-156430537-INS-1378
chrX-156432345-INS-148
chrX-156432741-INS-75
chrX-156436963-INS-151
chrX-156437946-INS-186
chrX-156443882-INS-276
chrX-156448029-INS-84
chrX-156473681-INS-52
chrX-156482718-INS-53
chrX-156502649-INS-80
chrX-156569750-INS-7058
chrX-156576672-INS-99
chrX-156578030-INS-54
chrX-156635330-INS-86
chrX-156691467-INS-83
chrX-156697257-INS-93
chrX-156705013-INS-888
chrX-156708244-INS-665
chrX-156717388-INS-84
chrX-156724519-INS-328
chrX-156732298-INS-74
chrX-156734052-INS-240

chrX-156763832-INS-153
chrX-156775380-INS-671
chrX-156778736-INS-423
chrX-156781228-INS-349
chrX-156784249-INS-239
chrX-156809981-INS-56
chrX-156827940-INS-179
chrX-156841128-INS-53
chrX-156869152-INS-957
chrX-156879798-INS-66
chrX-156918167-INS-268
chrX-156920352-INS-72
chrX-156922685-INS-53
chrX-156932407-INS-125
chrX-156958548-INS-76
chrX-156983045-INS-330
chrX-156996233-INS-6144
chrX-156999581-INS-569
chrX-157016538-INS-446
chrX-157066767-INS-465
chrX-157172551-INS-131
chrX-157188681-INS-400
chrX-157194812-INS-401
chrX-157204809-INS-6355
chrX-157219448-INS-831
chrX-157271883-INS-170
chrX-157272590-INS-52
chrX-157273012-INS-7000
chrX-157287725-INS-14858
chrX-157290671-INS-6376
chrX-157302713-INS-6436
chrX-157327893-INS-66
chrX-157337803-INS-85
chrX-157341154-INS-456
chrX-157364791-INS-204
chrX-157366803-INS-116
chrX-157373031-INS-298
chrX-157445324-INS-740
chrX-157445481-INS-2799
chrX-157448085-INS-58
chrX-157510019-INS-311
chrX-157515752-INS-64
chrX-157537343-INS-104
chrX-157538903-INS-335
chrX-157556995-INS-145
chrX-157559190-INS-61
chrX-157563146-INS-73
chrX-157565829-INS-276
chrX-157579240-INS-783
chrX-157618370-INS-63
chrX-157651853-INS-319
chrX-157698521-INS-52
chrX-157730973-INS-462
chrX-157771645-INS-442
chrX-157784527-INS-124
chrX-157787620-INS-84
chrX-157838327-INS-72
chrX-157845215-INS-7101
chrX-157851205-INS-150
chrX-157865141-INS-56
chrX-157870545-INS-52
chrX-157885830-INS-331
chrX-157890885-INS-400
chrX-157901846-INS-233
chrX-157925691-INS-117
chrX-157975832-INS-62
chrX-157978815-INS-54
chrX-157986518-INS-162
chrX-157990724-INS-452
chrX-157991798-INS-850
chrX-158001922-INS-54
chrX-158025458-INS-149
chrX-158051459-INS-274
chrX-158053216-INS-112
chrX-158088648-INS-161
chrX-158129076-INS-79
chrX-158173101-INS-663

chrX-158173928-INS-546
chrX-158179375-INS-277
chrX-158205002-INS-82
chrX-158234136-INS-265
chrX-158239806-INS-190
chrX-158287914-INS-579
chrX-158295370-INS-198
chrX-158300626-INS-73
chrX-158317113-INS-204
chrX-158319699-INS-216
chrX-158339260-INS-200
chrX-158396301-INS-201
chrX-158420149-INS-95
chrX-158420415-INS-941
chrX-158439706-INS-6457
chrX-158455229-INS-117
chrX-158494992-INS-78
chrX-158507767-INS-198
chrX-158513587-INS-621
chrX-158537028-INS-104
chrX-158545906-INS-93
chrX-158588798-INS-70
chrX-158605844-INS-454
chrX-158609446-INS-70
chrX-158619372-INS-6510
chrX-158639125-INS-63
chrX-158640274-INS-410
chrX-158674133-INS-466
chrX-158675256-INS-394
chrX-158692022-INS-848
chrX-158749809-INS-311
chrX-158787594-INS-215
chrX-158821284-INS-108
chrX-158835256-INS-140
chrX-158840851-INS-140
chrX-158886829-INS-60
chrX-158908508-INS-62
chrX-158933430-INS-128
chrX-158945417-INS-56
chrX-158957518-INS-704
chrX-158999090-INS-75
chrX-159010950-INS-520
chrX-159023058-INS-183
chrX-159066675-INS-54
chrX-159096390-INS-145
chrX-159108296-INS-354
chrX-159109973-INS-192
chrX-159136265-INS-1004
chrX-159143546-INS-349
chrX-159160404-INS-185
chrX-159215032-INS-5739
chrX-159223079-INS-200
chrX-159245799-INS-464
chrX-159250953-INS-76
chrX-159259344-INS-56
chrX-159268024-INS-978
chrX-159277716-INS-313
chrX-159298661-INS-415
chrX-159362656-INS-201
chrX-159405841-INS-507
chrX-159452603-INS-6587
chrX-159463167-INS-121
chrX-159495756-INS-2387
chrX-159497321-INS-275
chrX-159518646-INS-68
chrX-159619427-INS-737
chrX-159630545-INS-732
chrX-159664630-INS-348
chrX-159685942-INS-483
chrX-159705844-INS-143
chrX-159708258-INS-52
chrX-159718203-INS-98
chrX-159735164-INS-229
chrX-159751967-INS-76
chrX-159753553-INS-122
chrX-159791270-INS-105
chrX-159799609-INS-1583

chrX-159818706-INS-802
chrX-159819482-INS-56
chrX-159824339-INS-144
chrX-159874844-INS-102
chrX-159906203-INS-75
chrX-159934120-INS-226
chrX-159959589-INS-54
chrX-159995995-INS-6180
chrX-160007410-INS-566
chrX-160019012-INS-72
chrX-160028747-INS-117
chrX-160048817-INS-6544
chrX-160063541-INS-198
chrX-160069857-INS-1060
chrX-160070007-INS-500
chrX-160073004-INS-186
chrX-160083047-INS-176
chrX-160085746-INS-97
chrX-160147085-INS-223
chrX-160158144-INS-392
chrX-160226784-INS-458
chrX-160272749-INS-716
chrX-160309170-INS-642
chrX-160314035-INS-213
chrX-160319505-INS-78
chrX-160341892-INS-132
chrX-160352272-INS-741
chrX-160354344-INS-69
chrX-160363744-INS-538
chrX-160370848-INS-73
chrX-160386156-INS-52
chrX-160390337-INS-71
chrX-160397943-INS-904
chrX-160418716-INS-142
chrX-160461576-INS-70
chrX-160470338-INS-117
chrX-160475513-INS-216
chrX-160477990-INS-6331
chrX-160487134-INS-60
chrX-160495338-INS-208
chrX-160526681-INS-181
chrX-160529603-INS-54
chrX-160543207-INS-333
chrX-160577270-INS-200
chrX-160614353-INS-99
chrX-160632057-INS-146
chrX-160649567-INS-871
chrX-160649580-INS-310
chrX-160659962-INS-117
chrX-160671694-INS-230
chrX-160738325-INS-55
chrX-160744170-INS-1041
chrX-160766198-INS-68
chrX-160771466-INS-98
chrX-160801115-INS-147
chrX-160801992-INS-6840
chrX-160826413-INS-149
chrX-160835314-INS-64
chrX-160838464-INS-54
chrX-160840598-INS-72
chrX-160858778-INS-130
chrX-160867841-INS-1751
chrX-160885061-INS-78
chrX-160951939-INS-61
chrX-160956682-INS-75
chrX-160976027-INS-54
chrX-161052751-INS-128
chrX-161058860-INS-157
chrX-161075050-INS-69
chrX-161080949-INS-91
chrX-161088353-INS-6189
chrX-161090418-INS-6486
chrX-161091628-INS-73
chrX-161125532-INS-6845
chrX-161149546-INS-3966
chrX-161157083-INS-2883
chrX-161169318-INS-73

chrX-161177759-INS-1555
chrX-161226592-INS-179
chrX-161228524-INS-1890
chrX-161240351-INS-370
chrX-161251893-INS-73
chrX-161260864-INS-56
chrX-161265355-INS-571
chrX-161281058-INS-135
chrX-161295126-INS-166
chrX-161319921-INS-55
chrX-161325284-INS-6716
chrX-161331318-INS-325
chrX-161336339-INS-69
chrX-161342950-INS-92
chrX-161358360-INS-212
chrX-161380650-INS-130
chrX-161412899-INS-8701
chrX-161416309-INS-1494
chrX-161421499-INS-65
chrX-161425189-INS-60
chrX-161433016-INS-186
chrX-161447592-INS-59
chrX-161464968-INS-141
chrX-161488478-INS-73
chrX-161527982-INS-220
chrX-161533933-INS-88
chrX-161547902-INS-198
chrX-161561233-INS-76
chrX-161575459-INS-200
chrX-161580671-INS-96
chrX-161585841-INS-214
chrX-161598598-INS-51
chrX-161603248-INS-233
chrX-161605491-INS-187
chrX-161614712-INS-430
chrX-161616478-INS-64
chrX-161623950-INS-104
chrX-161643341-INS-201
chrX-161670099-INS-220
chrX-161675226-INS-205
chrX-161689048-INS-204
chrX-161691767-INS-502
chrX-161699940-INS-53
chrX-161699940-INS-92
chrX-161710160-INS-494
chrX-161716302-INS-78
chrX-161720378-INS-131
chrX-161720463-INS-557
chrX-161754179-INS-116
chrX-161787274-INS-167
chrX-161812259-INS-52
chrX-161841247-INS-193
chrX-161850825-INS-58
chrX-161851419-INS-513
chrX-161858533-INS-343
chrX-161872702-INS-6462
chrX-161872755-INS-1019
chrX-161876617-INS-325
chrX-161885865-INS-6612
chrX-161890206-INS-615
chrX-161913828-INS-129
chrX-161916685-INS-825
chrX-161918749-INS-52
chrX-161932936-INS-169
chrX-161942469-INS-635
chrX-161943789-INS-332
chrX-161952377-INS-5177
chrX-161962308-INS-1759
chrX-161968002-INS-60
chrX-161989611-INS-269
chrX-162021146-INS-67
chrX-162022620-INS-546
chrX-162030243-INS-4000
chrX-162059496-INS-466
chrX-162072471-INS-167
chrX-162100022-INS-54
chrX-162102161-INS-70

chrX-162184367-INS-634
chrX-162195190-INS-324
chrX-162213670-INS-51
chrX-162262396-INS-107
chrX-162273216-INS-1882
chrX-162312338-INS-7242
chrX-162312958-INS-79
chrX-162326536-INS-200
chrX-162326539-INS-6685
chrX-162361432-INS-130
chrX-162363026-INS-335
chrX-162385172-INS-262
chrX-162397352-INS-77
chrX-162398795-INS-70
chrX-162400834-INS-154
chrX-162418160-INS-428
chrX-162421103-INS-98
chrX-162422405-INS-204
chrX-162438837-INS-52
chrX-162470656-INS-60
chrX-162477678-INS-60
chrX-162520171-INS-135
chrX-162556068-INS-86
chrX-162561465-INS-593
chrX-162610831-INS-207
chrX-162653599-INS-167
chrX-162674834-INS-93
chrX-162674910-INS-451
chrX-162681334-INS-97
chrX-162729338-INS-54
chrX-162748535-INS-389
chrX-162774395-INS-576
chrX-162786880-INS-533
chrX-162787216-INS-69
chrX-162789873-INS-53
chrX-162802921-INS-58
chrX-162808460-INS-166
chrX-162812011-INS-53
chrX-162844219-INS-1616
chrX-162869599-INS-66
chrX-162888753-INS-216
chrX-162891017-INS-121
chrX-162918280-INS-58
chrX-162930667-INS-356
chrX-162939735-INS-81
chrX-162944940-INS-104
chrX-162985638-INS-72
chrX-163014489-INS-82
chrX-163016740-INS-60
chrX-163024963-INS-651
chrX-163049017-INS-68
chrX-163055755-INS-107
chrX-163068156-INS-757
chrX-163075573-INS-4027
chrX-163083767-INS-209
chrX-163096771-INS-75
chrX-163114438-INS-6455
chrX-163134106-INS-201
chrX-163154290-INS-148
chrX-163172680-INS-56
chrX-163183114-INS-67
chrX-163194947-INS-197
chrX-163224207-INS-2242
chrX-163238015-INS-334
chrX-163307712-INS-77
chrX-163321162-INS-100
chrX-163336640-INS-72
chrX-163337661-INS-3385
chrX-163338162-INS-6786
chrX-163355569-INS-59
chrX-163357195-INS-54
chrX-163360301-INS-122
chrX-163389451-INS-6957
chrX-163417156-INS-426
chrX-163432227-INS-52
chrX-163444591-INS-72
chrX-163463428-INS-348

chrX-163472081-INS-50
chrX-163479129-INS-76
chrX-163479659-INS-1483
chrX-163500151-INS-7060
chrX-163500239-INS-6454
chrX-163502710-INS-100
chrX-163531145-INS-379
chrX-163533009-INS-2242
chrX-163548482-INS-6425
chrX-163554977-INS-6419
chrX-163558955-INS-279
chrX-163571264-INS-314
chrX-163572303-INS-1223
chrX-163582881-INS-65
chrX-163586883-INS-74
chrX-163597586-INS-63
chrX-163616001-INS-121
chrX-163628783-INS-354
chrX-163660700-INS-6683
chrX-163663590-INS-666
chrX-163678690-INS-204
chrX-163692210-INS-55
chrX-163693284-INS-55
chrX-163705272-INS-74
chrX-163747551-INS-1431
chrX-163795068-INS-329
chrX-163799857-INS-6620
chrX-163803788-INS-55
chrX-163849176-INS-66
chrX-163868044-INS-150
chrX-163888458-INS-144
chrX-163907988-INS-417
chrX-163926868-INS-359
chrX-163942783-INS-76
chrX-163944685-INS-514
chrX-163952478-INS-857
chrX-163968871-INS-57
chrX-163980445-INS-57
chrX-164005636-INS-126
chrX-164006281-INS-574
chrX-164007937-INS-52
chrX-164015909-INS-865
chrX-164044227-INS-135
chrX-164050596-INS-6651
chrX-164052815-INS-139
chrX-164053965-INS-6719
chrX-164092791-INS-3207
chrX-164126077-INS-479
chrX-164142775-INS-52
chrX-164144485-INS-360
chrX-164151586-INS-54
chrX-164171101-INS-66
chrX-164203298-INS-440
chrX-164208079-INS-56
chrX-164220682-INS-169
chrX-164228345-INS-472
chrX-164235393-INS-321
chrX-164239987-INS-66
chrX-164266890-INS-250
chrX-164290710-INS-1092
chrX-164300168-INS-64
chrX-164304041-INS-65
chrX-164308871-INS-666
chrX-164324073-INS-6340
chrX-164331767-INS-72
chrX-164345724-INS-120
chrX-164368155-INS-3888
chrX-164370810-INS-400
chrX-164387070-INS-166
chrX-164442042-INS-209
chrX-164455035-INS-70
chrX-164501768-INS-4327
chrX-164533192-INS-7259
chrX-164548354-INS-426
chrX-164555993-INS-89
chrX-164566301-INS-428
chrX-164579745-INS-203

chrX-164585417-INS-75
chrX-164640817-INS-386
chrX-164648198-INS-470
chrX-164664530-INS-70
chrX-164665684-INS-82
chrX-164678568-INS-58
chrX-164709080-INS-854
chrX-164710377-INS-298
chrX-164759115-INS-52
chrX-164764417-INS-97
chrX-164783346-INS-6792
chrX-164794927-INS-64
chrX-164801193-INS-2839
chrX-164816463-INS-75
chrX-164821095-INS-6339
chrX-164826691-INS-735
chrX-164829117-INS-6206
chrX-164864246-INS-908
chrX-164870536-INS-135
chrX-164883540-INS-80
chrX-164891485-INS-518
chrX-164894845-INS-129
chrX-164898992-INS-6773
chrX-164991152-INS-121
chrX-165039438-INS-92
chrX-165052436-INS-363
chrX-165072812-INS-139
chrX-165078352-INS-56
chrX-165115416-INS-106
chrX-165118471-INS-68
chrX-165191983-INS-6382
chrX-165219077-INS-529
chrX-165221870-INS-750
chrX-165227230-INS-5548
chrX-165239288-INS-411
chrX-165264591-INS-923
chrX-165269457-INS-219
chrX-165282123-INS-95
chrX-165296711-INS-294
chrX-165308234-INS-394
chrX-165340630-INS-51
chrX-165362255-INS-56
chrX-165381306-INS-54
chrX-165428006-INS-727
chrX-165443014-INS-57
chrX-165457681-INS-500
chrX-165463995-INS-4066
chrX-165476793-INS-7002
chrX-165493045-INS-98
chrX-165506013-INS-6164
chrX-165525600-INS-71
chrX-165554620-INS-719
chrX-165561170-INS-56
chrX-165562210-INS-99
chrX-165586104-INS-70
chrX-165604991-INS-9663
chrX-165665964-INS-4087
chrX-165685013-INS-54
chrX-165688501-INS-224
chrX-165691276-INS-453
chrX-165696956-INS-697
chrX-165718362-INS-466
chrX-165725633-INS-358
chrX-165727978-INS-705
chrX-165733025-INS-87
chrX-165736944-INS-129
chrX-165753202-INS-401
chrX-165756683-INS-52
chrX-165777939-INS-64
chrX-165785406-INS-129
chrX-165790222-INS-72
chrX-165819980-INS-102
chrX-165837656-INS-76
chrX-165843817-INS-115
chrX-165849981-INS-83
chrX-165856551-INS-108
chrX-165887712-INS-79

chrX-165921912-INS-332
chrX-165938353-INS-129
chrX-165958345-INS-86
chrX-165960671-INS-73
chrX-165965075-INS-56
chrX-166029154-INS-214
chrX-166066035-INS-115
chrX-166074347-INS-60
chrX-166078554-INS-75
chrX-166095288-INS-188
chrX-166122918-INS-255
chrX-166125959-INS-54
chrX-166167053-INS-89
chrX-166179019-INS-198
chrX-166189565-INS-162
chrX-166228108-INS-77
chrX-166244404-INS-166
chrX-166247463-INS-427
chrX-166262609-INS-61
chrX-166272946-INS-55
chrX-166304447-INS-960
chrX-166309445-INS-447
chrX-166339586-INS-6389
chrX-166348649-INS-140
chrX-166381168-INS-161
chrX-166405284-INS-309
chrX-166410931-INS-7091
chrX-166411066-INS-80
chrX-166418568-INS-1538
chrX-166431670-INS-191
chrX-166445223-INS-99
chrX-166491552-INS-367
chrX-166492238-INS-723
chrX-166542338-INS-6399
chrX-166558932-INS-92
chrX-166566895-INS-52
chrX-166578843-INS-51
chrX-166616214-INS-449
chrX-166620224-INS-52
chrX-166621864-INS-65
chrX-166632681-INS-68
chrX-166643728-INS-59
chrX-166679469-INS-64
chrX-166680002-INS-70
chrX-166693423-INS-95
chrX-166738316-INS-2979
chrX-166756480-INS-175
chrX-166758682-INS-344
chrX-166792462-INS-140
chrX-166807189-INS-110
chrX-166891226-INS-102
chrX-166916296-INS-140
chrX-166928630-INS-102
chrX-166939836-INS-831
chrX-166978352-INS-56
chrX-167076959-INS-62
chrX-167087520-INS-268
chrX-167117696-INS-68
chrX-167162942-INS-9233
chrX-167168320-INS-6479
chrX-167173497-INS-57
chrX-167210512-INS-120
chrX-167213462-INS-80
chrX-167226485-INS-84
chrX-167238552-INS-67
chrX-167249851-INS-96
chrX-167259846-INS-70
chrX-167268828-INS-6570
chrX-167289196-INS-160
chrX-167344576-INS-60
chrX-167367894-INS-78
chrX-167372637-INS-155
chrX-167379971-INS-63
chrX-167386984-INS-51
chrX-167412679-INS-6291
chrX-167413816-INS-369
chrX-167423387-INS-202

chrX-167426757-INS-188
chrX-167443121-INS-72
chrX-167461260-INS-203
chrX-167499680-INS-58
chrX-167551686-INS-198
chrX-167557647-INS-59
chrX-167560347-INS-61
chrX-167572170-INS-3356
chrX-167621110-INS-89
chrX-167652709-INS-51
chrX-167675549-INS-1031
chrX-167694960-INS-407
chrX-167698687-INS-64
chrX-167703232-INS-4000
chrX-167704491-INS-628
chrX-167751686-INS-58
chrX-167758341-INS-1916
chrX-167769239-INS-332
chrX-167770099-INS-157
chrX-167783440-INS-6188
chrX-167793641-INS-267
chrX-167861446-INS-66
chrX-167865953-INS-62
chrX-167873394-INS-55
chrX-167917957-INS-211
chrX-167933715-INS-324
chrX-167938990-INS-82
chrX-167954668-INS-941
chrX-167979122-INS-120
chrX-168008873-INS-2065
chrX-168025337-INS-52
chrX-168043775-INS-253
chrX-168047431-INS-90
chrX-168061680-INS-95
chrX-168078903-INS-6264
chrX-168116741-INS-117
chrX-168131974-INS-2157
chrX-168152029-INS-119
chrX-168159701-INS-1127
chrX-168191432-INS-1486
chrX-168207707-INS-424
chrX-168238650-INS-135
chrX-168239053-INS-336
chrX-168290609-INS-58
chrX-168303883-INS-5360
chrX-168304768-INS-208
chrX-168325077-INS-6692
chrX-168328859-INS-63
chrX-168339292-INS-52
chrX-168372640-INS-281
chrX-168375452-INS-370
chrX-168423596-INS-4361
chrX-168436079-INS-200
chrX-168445520-INS-72
chrX-168446992-INS-258
chrX-168458971-INS-56
chrX-168478166-INS-98
chrX-168503702-INS-136
chrX-168523800-INS-212
chrX-168557058-INS-1598
chrX-168588017-INS-168
chrX-168600632-INS-63
chrX-168609372-INS-855
chrX-168640619-INS-96
chrX-168652305-INS-69
chrX-168660253-INS-848
chrX-168685560-INS-879
chrX-168688717-INS-75
chrX-168704839-INS-58
chrX-168714721-INS-59
chrX-168715862-INS-161
chrX-168728384-INS-160
chrX-168733085-INS-198
chrX-168739942-INS-90
chrX-168742142-INS-54
chrX-168754924-INS-868
chrX-168764478-INS-212

chrX-168769136-INS-93
chrX-168769607-INS-226
chrX-168772563-INS-93
chrX-168773971-INS-955
chrX-168775466-INS-151
chrX-168779459-INS-126
chrX-168781032-INS-150
chrX-168785049-INS-647
chrX-168803497-INS-581
chrX-168804113-INS-217
chrX-169015174-INS-102
chrX-169072099-INS-188
chrX-169077257-INS-158
chrX-169077827-INS-186
chrX-169083535-INS-186
chrX-169086597-INS-83
chrX-169088723-INS-68
chrX-169091942-INS-92
chrX-169096565-INS-899
chrX-169097199-INS-55
chrX-169106568-INS-94
chrX-169107181-INS-186
chrX-169110581-INS-71
chrX-169111105-INS-72
chrX-169118411-INS-61
chrX-169118858-INS-123
chrX-169120120-INS-60
chrX-169121707-INS-62
chrX-169186609-INS-62
chrX-169286418-INS-4996
chrX-169295839-INS-3616
chrX-169305149-INS-1586
chrX-169321063-INS-54

**Identified_By_JAX**
chr1-3090436-INS-2801
chr1-3092238-INS-7104
chr1-3099521-INS-52
chr1-3114752-INS-93
chr1-3120650-INS-162
chr1-3134350-INS-7269
chr1-3149754-INS-51
chr1-3174319-INS-94
chr1-3175032-INS-112
chr1-3241046-INS-4934
chr1-3282080-INS-122
chr1-3320604-INS-893
chr1-3321708-INS-592
chr1-3336930-INS-54
chr1-3448400-INS-84
chr1-3465186-INS-81
chr1-3472093-INS-185
chr1-3482932-INS-142
chr1-3553803-INS-76
chr1-3600450-INS-120
chr1-3627988-INS-7100
chr1-3667582-INS-6544
chr1-3778685-INS-53
chr1-3788760-INS-725
chr1-3860942-INS-462
chr1-3871664-INS-4957
chr1-3890712-INS-55
chr1-3920081-INS-8858
chr1-3924252-INS-2865
chr1-3948259-INS-7199
chr1-3965280-INS-4231
chr1-3984669-INS-194
chr1-3990607-INS-6589
chr1-3998473-INS-126
chr1-4016241-INS-11504
chr1-4020821-INS-1221
chr1-4025598-INS-1837
chr1-4064741-INS-52
chr1-4097516-INS-6192
chr1-4156951-INS-160
chr1-4172668-INS-130
chr1-4172997-INS-164
chr1-4173188-INS-1618
chr1-4180699-INS-8841
chr1-4194111-INS-52
chr1-4221947-INS-6403
chr1-4230915-INS-50
chr1-4244849-INS-4372
chr1-4258014-INS-199
chr1-4259522-INS-6534
chr1-4265416-INS-328
chr1-4278595-INS-909
chr1-4286906-INS-7761
chr1-4298316-INS-126
chr1-4326861-INS-72
chr1-4328160-INS-183
chr1-4355251-INS-80
chr1-4355540-INS-85
chr1-4368748-INS-56
chr1-4369045-INS-544
chr1-4399887-INS-92
chr1-4400128-INS-176
chr1-4400201-INS-70
chr1-4448502-INS-6332
chr1-4455195-INS-198
chr1-4468460-INS-572
chr1-4519972-INS-395
chr1-4520350-INS-199
chr1-4531711-INS-70
chr1-4546566-INS-207
chr1-4550963-INS-684
chr1-4584145-INS-67
chr1-4597361-INS-100
chr1-4649500-INS-107
chr1-4707762-INS-66
chr1-4733566-INS-190

chr1-4746007-INS-165
chr1-4807835-INS-1252
chr1-4810319-INS-164
chr1-4810347-INS-50
chr1-4826051-INS-128
chr1-4834980-INS-1864
chr1-4851908-INS-238
chr1-4860721-INS-66
chr1-4865220-INS-116
chr1-4904178-INS-50
chr1-4904275-INS-400
chr1-4905105-INS-723
chr1-4905789-INS-129
chr1-4906250-INS-105
chr1-4944902-INS-270
chr1-4992622-INS-75
chr1-5014143-INS-118
chr1-5044772-INS-231
chr1-5074396-INS-80
chr1-5074410-INS-226
chr1-5082230-INS-190
chr1-5092647-INS-51
chr1-5094335-INS-6267
chr1-5142828-INS-947
chr1-5143191-INS-1891
chr1-5182608-INS-11544
chr1-5242978-INS-6359
chr1-5366413-INS-7880
chr1-5376697-INS-1157
chr1-5401793-INS-4365
chr1-5512966-INS-525
chr1-5540251-INS-138
chr1-5541847-INS-2148
chr1-5555045-INS-1811
chr1-5564832-INS-6364
chr1-5590872-INS-83
chr1-5649536-INS-580
chr1-5778890-INS-5079
chr1-5813778-INS-500
chr1-5839996-INS-50
chr1-5856978-INS-852
chr1-5862028-INS-5517
chr1-5900965-INS-161
chr1-5901373-INS-307
chr1-5903668-INS-414
chr1-5923658-INS-56
chr1-5930108-INS-249
chr1-5935426-INS-663
chr1-5946888-INS-394
chr1-5963399-INS-104
chr1-6011386-INS-62
chr1-6027111-INS-6445
chr1-6087250-INS-86
chr1-6097307-INS-199
chr1-6117497-INS-449
chr1-6121597-INS-6483
chr1-6129764-INS-1136
chr1-6147571-INS-780
chr1-6152709-INS-53
chr1-6169258-INS-92
chr1-6177000-INS-377
chr1-6186778-INS-121
chr1-6223323-INS-404
chr1-6246566-INS-122
chr1-6257217-INS-9632
chr1-6259719-INS-57
chr1-6288731-INS-2000
chr1-6310626-INS-66
chr1-6323671-INS-718
chr1-6337345-INS-206
chr1-6391872-INS-2896
chr1-6424729-INS-118
chr1-6425187-INS-50
chr1-6453225-INS-52
chr1-6459432-INS-2051
chr1-6471027-INS-100
chr1-6471223-INS-61

chr1-6593636-INS-997
chr1-6661968-INS-50
chr1-6661971-INS-308
chr1-6662134-INS-133
chr1-6676825-INS-328
chr1-6719946-INS-1335
chr1-6732314-INS-380
chr1-6751236-INS-1636
chr1-6797011-INS-155
chr1-6821418-INS-144
chr1-6836702-INS-50
chr1-6848095-INS-100
chr1-6868637-INS-75
chr1-6942086-INS-7418
chr1-6949052-INS-5552
chr1-6990796-INS-5508
chr1-6999862-INS-50
chr1-7004258-INS-193
chr1-7069642-INS-274
chr1-7078405-INS-325
chr1-7092795-INS-545
chr1-7093536-INS-51
chr1-7103276-INS-7117
chr1-7108303-INS-6561
chr1-7128094-INS-199
chr1-7128681-INS-214
chr1-7205322-INS-50
chr1-7209202-INS-174
chr1-7234461-INS-78
chr1-7253931-INS-607
chr1-7254033-INS-547
chr1-7328945-INS-7087
chr1-7342149-INS-214
chr1-7342270-INS-185
chr1-7380932-INS-56
chr1-7392941-INS-403
chr1-7428636-INS-80
chr1-7440101-INS-61
chr1-7447825-INS-52
chr1-7453839-INS-6209
chr1-7456372-INS-54
chr1-7457471-INS-4854
chr1-7457937-INS-972
chr1-7463999-INS-67
chr1-7467401-INS-497
chr1-7471484-INS-54
chr1-7485757-INS-79
chr1-7506448-INS-8792
chr1-7506997-INS-4504
chr1-7519832-INS-98
chr1-7531264-INS-245
chr1-7562627-INS-203
chr1-7568235-INS-4838
chr1-7588359-INS-6473
chr1-7591865-INS-6879
chr1-7593206-INS-52
chr1-7594736-INS-51
chr1-7598344-INS-50
chr1-7627014-INS-7136
chr1-7642883-INS-6273
chr1-7662013-INS-505
chr1-7703082-INS-60
chr1-7725064-INS-11312
chr1-7731220-INS-80
chr1-7749833-INS-2172
chr1-7755459-INS-572
chr1-7778784-INS-76
chr1-7784367-INS-6180
chr1-7814251-INS-1132
chr1-7820877-INS-66
chr1-7842917-INS-7658
chr1-7845122-INS-69
chr1-7880766-INS-412
chr1-7881882-INS-7646
chr1-7896061-INS-353
chr1-7909462-INS-4461
chr1-8071085-INS-269

chr1-8112053-INS-6168
chr1-8164534-INS-1614
chr1-8166113-INS-104
chr1-8167580-INS-13526
chr1-8182484-INS-1798
chr1-8184376-INS-203
chr1-8220930-INS-75
chr1-8245771-INS-70
chr1-8256257-INS-52
chr1-8348974-INS-2746
chr1-8379970-INS-6409
chr1-8387777-INS-439
chr1-8411562-INS-8176
chr1-8436417-INS-50
chr1-8492681-INS-1157
chr1-8496164-INS-58
chr1-8504145-INS-50
chr1-8519101-INS-460
chr1-8526516-INS-98
chr1-8534732-INS-51
chr1-8544679-INS-200
chr1-8550447-INS-52
chr1-8550487-INS-171
chr1-8571440-INS-3055
chr1-8609878-INS-6474
chr1-8614045-INS-90
chr1-8621291-INS-709
chr1-8625105-INS-78
chr1-8676308-INS-55
chr1-8676719-INS-6163
chr1-8698633-INS-3023
chr1-8699114-INS-9255
chr1-8713268-INS-185
chr1-8800535-INS-8584
chr1-8815714-INS-95
chr1-8826601-INS-1289
chr1-8833297-INS-180
chr1-8835287-INS-6380
chr1-8836761-INS-1546
chr1-8937338-INS-5047
chr1-8941478-INS-96
chr1-8971301-INS-189
chr1-8971448-INS-1034
chr1-8998085-INS-7106
chr1-9011605-INS-12575
chr1-9019843-INS-196
chr1-9058178-INS-6882
chr1-9098954-INS-511
chr1-9106171-INS-62
chr1-9106298-INS-311
chr1-9109148-INS-637
chr1-9114630-INS-50
chr1-9150260-INS-8523
chr1-9163652-INS-5065
chr1-9177272-INS-7436
chr1-9181811-INS-109
chr1-9212778-INS-58
chr1-9263075-INS-81
chr1-9328570-INS-65
chr1-9362282-INS-1029
chr1-9366333-INS-51
chr1-9372345-INS-68
chr1-9373369-INS-3588
chr1-9395568-INS-2724
chr1-9445891-INS-6323
chr1-9522152-INS-7118
chr1-9539131-INS-63
chr1-9544765-INS-520
chr1-9551326-INS-436
chr1-9569820-INS-755
chr1-9578697-INS-390
chr1-9579232-INS-227
chr1-9584354-INS-248
chr1-9600726-INS-838
chr1-9641257-INS-85
chr1-9660589-INS-2845
chr1-9661984-INS-50

chr1-9675796-INS-94
chr1-9687146-INS-2786
chr1-9701213-INS-52
chr1-9703896-INS-62
chr1-9706041-INS-91
chr1-9708366-INS-54
chr1-9725370-INS-4755
chr1-9727663-INS-108
chr1-9734309-INS-73
chr1-9773964-INS-195
chr1-9822889-INS-122
chr1-9853317-INS-57
chr1-9854856-INS-666
chr1-9856572-INS-52
chr1-9880074-INS-116
chr1-9896813-INS-544
chr1-9933777-INS-209
chr1-9934820-INS-52
chr1-9944710-INS-451
chr1-9979555-INS-50
chr1-10045252-INS-64
chr1-10050566-INS-1908
chr1-10056105-INS-140
chr1-10074966-INS-167
chr1-10110760-INS-53
chr1-10121289-INS-60
chr1-10122451-INS-422
chr1-10132140-INS-1841
chr1-10132168-INS-59
chr1-10162550-INS-55
chr1-10186778-INS-137
chr1-10310083-INS-66
chr1-10310308-INS-6435
chr1-10311633-INS-50
chr1-10314011-INS-124
chr1-10325361-INS-267
chr1-10341302-INS-7161
chr1-10349176-INS-78
chr1-10378332-INS-59
chr1-10378351-INS-267
chr1-10378451-INS-90
chr1-10416968-INS-66
chr1-10446405-INS-120
chr1-10456977-INS-460
chr1-10458625-INS-704
chr1-10576880-INS-1924
chr1-10577010-INS-59
chr1-10617414-INS-64
chr1-10620821-INS-92
chr1-10621095-INS-70
chr1-10626853-INS-1818
chr1-10660673-INS-85
chr1-10766075-INS-61
chr1-10829344-INS-70
chr1-10931323-INS-196
chr1-10936014-INS-542
chr1-10936943-INS-64
chr1-10937777-INS-8107
chr1-10938214-INS-66
chr1-10939353-INS-123
chr1-11000756-INS-84
chr1-11004454-INS-350
chr1-11036275-INS-66
chr1-11058962-INS-2089
chr1-11086428-INS-220
chr1-11153587-INS-498
chr1-11172606-INS-77
chr1-11183554-INS-6707
chr1-11190597-INS-2565
chr1-11199459-INS-1070
chr1-11212442-INS-64
chr1-11231689-INS-207
chr1-11242289-INS-1321
chr1-11260581-INS-5982
chr1-11273186-INS-50
chr1-11282742-INS-5350
chr1-11315239-INS-63

chr1-11347032-INS-341
chr1-11365813-INS-414
chr1-11387457-INS-2808
chr1-11413681-INS-5147
chr1-11420523-INS-1489
chr1-11451126-INS-7297
chr1-11468383-INS-202
chr1-11468513-INS-756
chr1-11472888-INS-223
chr1-11521145-INS-3541
chr1-11531545-INS-6298
chr1-11549200-INS-6989
chr1-11550226-INS-58
chr1-11550520-INS-64
chr1-11553018-INS-490
chr1-11581513-INS-1666
chr1-11626775-INS-1777
chr1-11652497-INS-124
chr1-11652575-INS-50
chr1-11674841-INS-323
chr1-11720084-INS-7412
chr1-11734150-INS-118
chr1-11805560-INS-1443
chr1-11813347-INS-50
chr1-11826723-INS-142
chr1-11859377-INS-50
chr1-11874179-INS-440
chr1-11878204-INS-6252
chr1-11900219-INS-54
chr1-11923946-INS-400
chr1-11988060-INS-51
chr1-12002725-INS-4139
chr1-12034025-INS-6639
chr1-12036633-INS-1574
chr1-12069105-INS-80
chr1-12078980-INS-58
chr1-12080974-INS-4839
chr1-12086232-INS-6454
chr1-12112912-INS-1697
chr1-12136355-INS-369
chr1-12136426-INS-1472
chr1-12151342-INS-51
chr1-12206951-INS-52
chr1-12232944-INS-86
chr1-12246429-INS-90
chr1-12268741-INS-487
chr1-12319242-INS-2709
chr1-12350187-INS-50
chr1-12355117-INS-63
chr1-12355118-INS-178
chr1-12383307-INS-134
chr1-12383307-INS-54
chr1-12395403-INS-1489
chr1-12396501-INS-1291
chr1-12500855-INS-56
chr1-12543587-INS-54
chr1-12557209-INS-243
chr1-12592512-INS-140
chr1-12592918-INS-406
chr1-12697248-INS-749
chr1-12700255-INS-6303
chr1-12717061-INS-1881
chr1-12722261-INS-65
chr1-12722607-INS-350
chr1-12727784-INS-702
chr1-12727919-INS-1755
chr1-12754710-INS-52
chr1-12763045-INS-54
chr1-12911378-INS-215
chr1-12948800-INS-54
chr1-12972737-INS-798
chr1-13021047-INS-50
chr1-13022210-INS-62
chr1-13022222-INS-186
chr1-13022284-INS-620
chr1-13028349-INS-225
chr1-13032707-INS-56

chr1-13033668-INS-260
chr1-13038258-INS-235
chr1-13041461-INS-62
chr1-13054394-INS-273
chr1-13063880-INS-3746
chr1-13066828-INS-54
chr1-13080147-INS-277
chr1-13101689-INS-54
chr1-13122974-INS-3337
chr1-13140449-INS-206
chr1-13147427-INS-386
chr1-13150454-INS-192
chr1-13157125-INS-56
chr1-13162516-INS-50
chr1-13179505-INS-2490
chr1-13209052-INS-221
chr1-13224523-INS-52
chr1-13230925-INS-215
chr1-13273633-INS-140
chr1-13343212-INS-108
chr1-13389023-INS-648
chr1-13455408-INS-92
chr1-13472689-INS-62
chr1-13502784-INS-180
chr1-13542455-INS-1484
chr1-13545896-INS-462
chr1-13551692-INS-206
chr1-13552986-INS-2190
chr1-13556254-INS-212
chr1-13580400-INS-7260
chr1-13592469-INS-76
chr1-13602735-INS-133
chr1-13641215-INS-330
chr1-13652014-INS-351
chr1-13655755-INS-468
chr1-13662685-INS-56
chr1-13674996-INS-468
chr1-13691664-INS-504
chr1-13700494-INS-51
chr1-13702535-INS-118
chr1-13725113-INS-195
chr1-13732514-INS-5671
chr1-13746569-INS-5643
chr1-13757024-INS-209
chr1-13779129-INS-57
chr1-13779266-INS-233
chr1-13779351-INS-270
chr1-13779355-INS-549
chr1-13784601-INS-213
chr1-13788449-INS-132
chr1-13793639-INS-211
chr1-13794175-INS-326
chr1-13841917-INS-53
chr1-13871593-INS-328
chr1-13873948-INS-59
chr1-13903021-INS-150
chr1-13919444-INS-74
chr1-13923168-INS-60
chr1-13993829-INS-12016
chr1-13993829-INS-5525
chr1-14017491-INS-1894
chr1-14067130-INS-388
chr1-14067163-INS-140
chr1-14075597-INS-71
chr1-14078914-INS-77
chr1-14096602-INS-5069
chr1-14096842-INS-1160
chr1-14120910-INS-95
chr1-14122113-INS-720
chr1-14122308-INS-1761
chr1-14135462-INS-1559
chr1-14141221-INS-226
chr1-14186130-INS-1033
chr1-14226189-INS-54
chr1-14244829-INS-2502
chr1-14267473-INS-223
chr1-14267792-INS-227

chr1-14298387-INS-167
chr1-14308734-INS-1074
chr1-14317164-INS-66
chr1-14336201-INS-116
chr1-14396014-INS-162
chr1-14420105-INS-51
chr1-14470880-INS-749
chr1-14517336-INS-500
chr1-14517800-INS-204
chr1-14554638-INS-184
chr1-14599391-INS-6504
chr1-14643123-INS-190
chr1-14643161-INS-53
chr1-14643235-INS-491
chr1-14668519-INS-118
chr1-14683884-INS-336
chr1-14715523-INS-141
chr1-14717190-INS-448
chr1-14728165-INS-111
chr1-14754657-INS-91
chr1-14769353-INS-5164
chr1-14769549-INS-218
chr1-14781541-INS-1838
chr1-14846684-INS-1046
chr1-14890082-INS-996
chr1-14898492-INS-407
chr1-14923779-INS-1010
chr1-14934165-INS-114
chr1-14952665-INS-64
chr1-14986813-INS-2406
chr1-14990545-INS-65
chr1-15005612-INS-82
chr1-15011979-INS-701
chr1-15105085-INS-52
chr1-15119853-INS-6216
chr1-15167014-INS-639
chr1-15173786-INS-170
chr1-15211963-INS-5701
chr1-15215228-INS-2791
chr1-15243928-INS-62
chr1-15250273-INS-1718
chr1-15250857-INS-62
chr1-15256869-INS-4927
chr1-15268081-INS-2574
chr1-15298670-INS-401
chr1-15347050-INS-167
chr1-15395425-INS-264
chr1-15395425-INS-99
chr1-15411003-INS-874
chr1-15414417-INS-7102
chr1-15450024-INS-50
chr1-15526781-INS-94
chr1-15542287-INS-146
chr1-15542609-INS-52
chr1-15578683-INS-102
chr1-15695126-INS-329
chr1-15712127-INS-51
chr1-15712190-INS-241
chr1-15725420-INS-78
chr1-15752084-INS-740
chr1-15753524-INS-2225
chr1-15753524-INS-476
chr1-15758160-INS-538
chr1-15772651-INS-94
chr1-15800723-INS-52
chr1-15803201-INS-54
chr1-15816905-INS-161
chr1-15829225-INS-1456
chr1-15842518-INS-1037
chr1-15843420-INS-58
chr1-15851554-INS-50
chr1-15855921-INS-396
chr1-15856875-INS-188
chr1-15856875-INS-70
chr1-15858663-INS-1024
chr1-15863724-INS-134
chr1-15871255-INS-3096

chr1-15940276-INS-101
chr1-15941041-INS-198
chr1-16034681-INS-383
chr1-16047461-INS-6305
chr1-16050760-INS-1739
chr1-16078438-INS-134
chr1-16080524-INS-50
chr1-16095470-INS-122
chr1-16103497-INS-3372
chr1-16104742-INS-261
chr1-16113373-INS-51
chr1-16125160-INS-52
chr1-16130981-INS-74
chr1-16133794-INS-140
chr1-16142664-INS-1282
chr1-16146675-INS-579
chr1-16152670-INS-55
chr1-16160289-INS-923
chr1-16164492-INS-221
chr1-16165359-INS-51
chr1-16203214-INS-760
chr1-16218185-INS-60
chr1-16244039-INS-62
chr1-16247533-INS-56
chr1-16300784-INS-104
chr1-16318282-INS-1125
chr1-16327494-INS-50
chr1-16413123-INS-106
chr1-16413123-INS-1354
chr1-16413123-INS-636
chr1-16413344-INS-146
chr1-16429441-INS-57
chr1-16429467-INS-292
chr1-16485072-INS-243
chr1-16547508-INS-1023
chr1-16554544-INS-96
chr1-16582858-INS-176
chr1-16592131-INS-406
chr1-16624720-INS-125
chr1-16673143-INS-882
chr1-16673483-INS-2262
chr1-16709027-INS-116
chr1-16744129-INS-62
chr1-16744228-INS-145
chr1-16745143-INS-7103
chr1-16756145-INS-108
chr1-16759023-INS-71
chr1-16792976-INS-2731
chr1-16805992-INS-52
chr1-16822644-INS-212
chr1-16850013-INS-1893
chr1-16853945-INS-907
chr1-16860766-INS-516
chr1-16881063-INS-846
chr1-16914387-INS-1144
chr1-16928337-INS-52
chr1-16931242-INS-12563
chr1-16931242-INS-5243
chr1-16944108-INS-52
chr1-16946420-INS-306
chr1-16950023-INS-5167
chr1-16981959-INS-52
chr1-17004307-INS-84
chr1-17035843-INS-426
chr1-17060904-INS-88
chr1-17090978-INS-180
chr1-17121667-INS-504
chr1-17146628-INS-396
chr1-17151139-INS-2775
chr1-17213385-INS-1939
chr1-17222930-INS-2560
chr1-17254934-INS-213
chr1-17259391-INS-117
chr1-17275889-INS-81
chr1-17282361-INS-3500
chr1-17302162-INS-187
chr1-17320346-INS-3229

chr1-17324015-INS-6654
chr1-17331438-INS-429
chr1-17335253-INS-1125
chr1-17368206-INS-52
chr1-17375899-INS-102
chr1-17387482-INS-510
chr1-17406588-INS-150
chr1-17417303-INS-864
chr1-17420418-INS-3679
chr1-17445627-INS-1278
chr1-17482740-INS-604
chr1-17492199-INS-60
chr1-17512305-INS-6767
chr1-17592041-INS-60
chr1-17710332-INS-3625
chr1-17711711-INS-8813
chr1-17718712-INS-260
chr1-17725805-INS-52
chr1-17775674-INS-208
chr1-17783478-INS-70
chr1-17783695-INS-55
chr1-17807460-INS-50
chr1-17808559-INS-214
chr1-17819544-INS-60
chr1-17820110-INS-139
chr1-17858922-INS-50
chr1-17864137-INS-7213
chr1-17899643-INS-1854
chr1-17909916-INS-71
chr1-17925704-INS-10059
chr1-17952677-INS-231
chr1-17983988-INS-468
chr1-18039539-INS-84
chr1-18047222-INS-1878
chr1-18103333-INS-5050
chr1-18103430-INS-1558
chr1-18140404-INS-1877
chr1-18189406-INS-6484
chr1-18260016-INS-51
chr1-18269969-INS-4730
chr1-18328198-INS-6195
chr1-18348675-INS-1380
chr1-18351604-INS-9062
chr1-18367321-INS-52
chr1-18386778-INS-7731
chr1-18402402-INS-4708
chr1-18425273-INS-71
chr1-18437864-INS-208
chr1-18441146-INS-65
chr1-18441585-INS-117
chr1-18467384-INS-57
chr1-18626055-INS-91
chr1-18650598-INS-1513
chr1-18660234-INS-50
chr1-18671903-INS-142
chr1-18678392-INS-62
chr1-18695893-INS-78
chr1-18750174-INS-71
chr1-18750434-INS-157
chr1-18750781-INS-58
chr1-18750841-INS-200
chr1-18759203-INS-6300
chr1-18759799-INS-6796
chr1-18775129-INS-55
chr1-18797318-INS-706
chr1-19018665-INS-504
chr1-19019894-INS-8584
chr1-19023935-INS-2061
chr1-19067677-INS-52
chr1-19070746-INS-2189
chr1-19108645-INS-51
chr1-19120277-INS-6463
chr1-19147434-INS-182
chr1-19149564-INS-356
chr1-19168164-INS-78
chr1-19210276-INS-62
chr1-19226511-INS-61

chr1-19235535-INS-63
chr1-19252148-INS-80
chr1-19330936-INS-980
chr1-19331158-INS-72
chr1-19367503-INS-998
chr1-19385162-INS-6421
chr1-19389891-INS-326
chr1-19403740-INS-54
chr1-19404157-INS-59
chr1-19426168-INS-9808
chr1-19442514-INS-834
chr1-19551332-INS-140
chr1-19552540-INS-50
chr1-19566790-INS-7340
chr1-19571542-INS-6615
chr1-19591114-INS-2945
chr1-19662461-INS-133
chr1-19663132-INS-52
chr1-19675646-INS-1436
chr1-19700200-INS-618
chr1-19700739-INS-60
chr1-19765038-INS-143
chr1-19800453-INS-2826
chr1-19820481-INS-112
chr1-19820621-INS-137
chr1-20071776-INS-64
chr1-20073804-INS-305
chr1-20095966-INS-1452
chr1-20099763-INS-50
chr1-20124923-INS-62
chr1-20125559-INS-205
chr1-20211126-INS-50
chr1-20228980-INS-6154
chr1-20236506-INS-1995
chr1-20314249-INS-82
chr1-20342770-INS-52
chr1-20381693-INS-146
chr1-20391584-INS-208
chr1-20434891-INS-499
chr1-20438675-INS-4176
chr1-20440392-INS-241
chr1-20440443-INS-689
chr1-20442379-INS-4195
chr1-20442457-INS-52
chr1-20471783-INS-60
chr1-20542098-INS-93
chr1-20546061-INS-972
chr1-20546197-INS-76
chr1-20551235-INS-1018
chr1-20584378-INS-7353
chr1-20585920-INS-1976
chr1-20634978-INS-64
chr1-20637823-INS-266
chr1-20640191-INS-146
chr1-20654686-INS-1386
chr1-20658150-INS-194
chr1-20658364-INS-52
chr1-20658408-INS-116
chr1-20674610-INS-479
chr1-20718570-INS-201
chr1-20735436-INS-56
chr1-20748705-INS-195
chr1-20763062-INS-52
chr1-20771563-INS-50
chr1-20788647-INS-51
chr1-20789222-INS-215
chr1-20793554-INS-1543
chr1-20793997-INS-91
chr1-20794620-INS-52
chr1-20805047-INS-69
chr1-20832848-INS-83
chr1-20849823-INS-116
chr1-20860617-INS-145
chr1-20870450-INS-200
chr1-20890143-INS-94
chr1-20897036-INS-113
chr1-20897449-INS-4937

chr1-20897569-INS-60
chr1-20918292-INS-128
chr1-20984179-INS-74
chr1-20984203-INS-843
chr1-21014129-INS-186
chr1-21026839-INS-88
chr1-21041638-INS-60
chr1-21083617-INS-57
chr1-21095121-INS-408
chr1-21100771-INS-58
chr1-21213104-INS-5451
chr1-21235247-INS-111
chr1-21236085-INS-60
chr1-21238312-INS-5414
chr1-21253517-INS-60
chr1-21281161-INS-995
chr1-21290972-INS-112
chr1-21290972-INS-528
chr1-21291417-INS-648
chr1-21294036-INS-156
chr1-21301290-INS-71
chr1-21310992-INS-52
chr1-21316191-INS-62
chr1-21321133-INS-153
chr1-21339510-INS-158
chr1-21342019-INS-4453
chr1-21342282-INS-50
chr1-21350904-INS-4539
chr1-21354001-INS-79
chr1-21362187-INS-4312
chr1-21381987-INS-83
chr1-21398510-INS-147
chr1-21422463-INS-194
chr1-21426964-INS-484
chr1-21436903-INS-62
chr1-21437304-INS-186
chr1-21442211-INS-188
chr1-21445651-INS-484
chr1-21457485-INS-183
chr1-21474209-INS-51
chr1-21474373-INS-206
chr1-21489161-INS-155
chr1-21497972-INS-1431
chr1-21500215-INS-61
chr1-21500241-INS-386
chr1-21510463-INS-52
chr1-21513992-INS-196
chr1-21519224-INS-1014
chr1-21565713-INS-433
chr1-21565735-INS-117
chr1-21572031-INS-6482
chr1-21578815-INS-51
chr1-21658920-INS-201
chr1-21672087-INS-64
chr1-21673117-INS-99
chr1-21702478-INS-50
chr1-21702729-INS-199
chr1-21706664-INS-1894
chr1-21710583-INS-202
chr1-21740934-INS-74
chr1-21758825-INS-50
chr1-21772672-INS-52
chr1-21773515-INS-54
chr1-21774449-INS-6458
chr1-21787887-INS-56
chr1-21800669-INS-6881
chr1-21810588-INS-70
chr1-21816057-INS-56
chr1-21837649-INS-60
chr1-21843736-INS-52
chr1-21904542-INS-62
chr1-21933108-INS-70
chr1-21933384-INS-467
chr1-21937208-INS-6739
chr1-21968999-INS-52
chr1-21978052-INS-53
chr1-21986434-INS-6685

chr1-22004133-INS-181
chr1-22060734-INS-3155
chr1-22061540-INS-140
chr1-22155049-INS-4335
chr1-22172266-INS-6372
chr1-22175479-INS-50
chr1-22221957-INS-2792
chr1-22267052-INS-6616
chr1-22269395-INS-2612
chr1-22270161-INS-61
chr1-22271042-INS-78
chr1-22284954-INS-7365
chr1-22295464-INS-208
chr1-22299969-INS-168
chr1-22300753-INS-582
chr1-22306968-INS-199
chr1-22311403-INS-79
chr1-22331212-INS-144
chr1-22351732-INS-65
chr1-22359736-INS-105
chr1-22369275-INS-103
chr1-22391143-INS-499
chr1-22431639-INS-58
chr1-22447308-INS-135
chr1-22447883-INS-66
chr1-22532401-INS-2076
chr1-22532401-INS-861
chr1-22567362-INS-52
chr1-22589140-INS-56
chr1-22608932-INS-64
chr1-22613974-INS-60
chr1-22618232-INS-50
chr1-22627235-INS-54
chr1-22628374-INS-1156
chr1-22648812-INS-60
chr1-22696790-INS-500
chr1-22715636-INS-6405
chr1-22788275-INS-52
chr1-22807248-INS-150
chr1-22827263-INS-53
chr1-22849309-INS-4689
chr1-22877992-INS-60
chr1-22878132-INS-133
chr1-22926807-INS-143
chr1-22936849-INS-68
chr1-22953214-INS-50
chr1-22971278-INS-1488
chr1-23053723-INS-60
chr1-23067443-INS-207
chr1-23072014-INS-242
chr1-23072545-INS-202
chr1-23090330-INS-75
chr1-23098968-INS-141
chr1-23120907-INS-829
chr1-23122121-INS-560
chr1-23127801-INS-162
chr1-23128858-INS-524
chr1-23129174-INS-824
chr1-23179886-INS-98
chr1-23179949-INS-210
chr1-23206440-INS-232
chr1-23227727-INS-197
chr1-23228228-INS-64
chr1-23230141-INS-6685
chr1-23233722-INS-120
chr1-23235295-INS-84
chr1-23266023-INS-99
chr1-23399455-INS-300
chr1-23410138-INS-54
chr1-23448684-INS-1590
chr1-23465891-INS-54
chr1-23470926-INS-53
chr1-23490764-INS-6199
chr1-23514599-INS-52
chr1-23522012-INS-65
chr1-23535034-INS-611
chr1-23535359-INS-1052

chr1-23575821-INS-1515
chr1-23576224-INS-80
chr1-23576597-INS-66
chr1-23615911-INS-108
chr1-23631304-INS-85
chr1-23637914-INS-56
chr1-23638913-INS-2662
chr1-23679860-INS-236
chr1-23720248-INS-249
chr1-23720248-INS-75
chr1-23737660-INS-1607
chr1-23742908-INS-10123
chr1-23751503-INS-186
chr1-23752365-INS-66
chr1-23752940-INS-2501
chr1-23782553-INS-1534
chr1-23829245-INS-2645
chr1-23835790-INS-1176
chr1-23838712-INS-78
chr1-23846600-INS-62
chr1-23848031-INS-120
chr1-23857223-INS-177
chr1-23877433-INS-124
chr1-23915709-INS-51
chr1-23938053-INS-1935
chr1-23947219-INS-78
chr1-24002216-INS-424
chr1-24147860-INS-127
chr1-24156690-INS-1837
chr1-24168029-INS-118
chr1-24168445-INS-52
chr1-24181448-INS-1684
chr1-24189062-INS-951
chr1-24190094-INS-3171
chr1-24224729-INS-72
chr1-24224807-INS-198
chr1-24247602-INS-75
chr1-24248294-INS-51
chr1-24274735-INS-566
chr1-24322811-INS-52
chr1-24344984-INS-62
chr1-24347326-INS-4350
chr1-24347851-INS-54
chr1-24358037-INS-80
chr1-24364004-INS-103
chr1-24370323-INS-387
chr1-24370602-INS-788
chr1-24372264-INS-106
chr1-24397183-INS-1288
chr1-24398894-INS-70
chr1-24398968-INS-141
chr1-24450850-INS-7756
chr1-24461417-INS-67
chr1-24475071-INS-54
chr1-24475852-INS-990
chr1-24475854-INS-160
chr1-24495960-INS-542
chr1-24515238-INS-68
chr1-24542311-INS-259
chr1-24542311-INS-65
chr1-24550509-INS-89
chr1-24591305-INS-201
chr1-24601565-INS-139
chr1-24628093-INS-80
chr1-24632995-INS-166
chr1-24635935-INS-244
chr1-24649688-INS-474
chr1-24656696-INS-394
chr1-24672547-INS-114
chr1-24706358-INS-223
chr1-24714558-INS-54
chr1-24715631-INS-167
chr1-24723127-INS-276
chr1-24723127-INS-3250
chr1-24723223-INS-115
chr1-24723309-INS-666
chr1-24728325-INS-147

chr1-24775641-INS-57
chr1-24821176-INS-50
chr1-24823240-INS-1105
chr1-24831966-INS-2979
chr1-24860314-INS-4739
chr1-24865380-INS-62
chr1-24892168-INS-140
chr1-24899356-INS-5218
chr1-24916892-INS-84
chr1-24947230-INS-58
chr1-24970272-INS-238
chr1-24976481-INS-50
chr1-25044082-INS-4887
chr1-25045571-INS-51
chr1-25091448-INS-66
chr1-25119814-INS-62
chr1-25137584-INS-82
chr1-25142051-INS-50
chr1-25227759-INS-78
chr1-25245567-INS-141
chr1-25272734-INS-6883
chr1-25291970-INS-130
chr1-25410738-INS-112
chr1-25415383-INS-228
chr1-25614364-INS-130
chr1-25615851-INS-927
chr1-25618607-INS-1633
chr1-25685809-INS-436
chr1-25724695-INS-1489
chr1-25764691-INS-1276
chr1-25775482-INS-165
chr1-25775687-INS-1571
chr1-25788578-INS-7108
chr1-25841112-INS-2816
chr1-25881635-INS-66
chr1-25895445-INS-58
chr1-25902038-INS-6079
chr1-25907511-INS-58
chr1-25911313-INS-1343
chr1-25923710-INS-5324
chr1-25940885-INS-192
chr1-25941119-INS-72
chr1-25944277-INS-906
chr1-25969157-INS-1206
chr1-25983013-INS-196
chr1-25995533-INS-149
chr1-25995818-INS-149
chr1-26022734-INS-249
chr1-26022970-INS-271
chr1-26034992-INS-3928
chr1-26055743-INS-205
chr1-26056342-INS-51
chr1-26086031-INS-1936
chr1-26096706-INS-68
chr1-26134155-INS-1310
chr1-26154883-INS-613
chr1-26156984-INS-3146
chr1-26158831-INS-379
chr1-26212211-INS-2588
chr1-26229266-INS-79
chr1-26230880-INS-196
chr1-26280333-INS-81
chr1-26313721-INS-864
chr1-26313900-INS-3168
chr1-26323405-INS-3058
chr1-26341117-INS-56
chr1-26343535-INS-234
chr1-26383580-INS-66
chr1-26463616-INS-852
chr1-26483519-INS-22967
chr1-26493850-INS-140
chr1-26498225-INS-436
chr1-26503878-INS-1446
chr1-26567037-INS-1708
chr1-26579645-INS-52
chr1-26583187-INS-350
chr1-26600786-INS-6202

chr1-26602985-INS-64
chr1-26613596-INS-220
chr1-26619573-INS-67
chr1-26635112-INS-92
chr1-26645388-INS-358
chr1-26647994-INS-54
chr1-26648849-INS-50
chr1-26660068-INS-84
chr1-26669450-INS-115
chr1-26690543-INS-6202
chr1-26703720-INS-233
chr1-26712465-INS-1708
chr1-26715185-INS-6382
chr1-26725896-INS-17946
chr1-26725948-INS-7681
chr1-26725969-INS-25663
chr1-26788670-INS-3301
chr1-26807056-INS-791
chr1-26808465-INS-413
chr1-26809210-INS-75
chr1-26818352-INS-138
chr1-26829610-INS-50
chr1-26885067-INS-2040
chr1-26887786-INS-58
chr1-26922066-INS-54
chr1-26933895-INS-162
chr1-26945905-INS-330
chr1-27043054-INS-130
chr1-27084775-INS-346
chr1-27084894-INS-1643
chr1-27131316-INS-1077
chr1-27151437-INS-3464
chr1-27153430-INS-2521
chr1-27154304-INS-7151
chr1-27217977-INS-60
chr1-27233474-INS-208
chr1-27246984-INS-50
chr1-27261310-INS-71
chr1-27262012-INS-118
chr1-27296034-INS-2879
chr1-27299983-INS-67
chr1-27301044-INS-350
chr1-27332447-INS-194
chr1-27332982-INS-75
chr1-27334524-INS-50
chr1-27342025-INS-5262
chr1-27375716-INS-1992
chr1-27406641-INS-6281
chr1-27407362-INS-2874
chr1-27466961-INS-52
chr1-27497048-INS-533
chr1-27501174-INS-50
chr1-27526203-INS-59
chr1-27559958-INS-172
chr1-27621614-INS-466
chr1-27625463-INS-61
chr1-27628993-INS-880
chr1-27631188-INS-6472
chr1-27635948-INS-366
chr1-27646539-INS-52
chr1-27689888-INS-54
chr1-27751753-INS-1528
chr1-27807795-INS-116
chr1-27807860-INS-263
chr1-27881597-INS-99
chr1-27926784-INS-164
chr1-28002377-INS-587
chr1-28023067-INS-449
chr1-28046882-INS-96
chr1-28079264-INS-83
chr1-28088698-INS-200
chr1-28095142-INS-56
chr1-28106534-INS-1438
chr1-28129211-INS-248
chr1-28131667-INS-6482
chr1-28220728-INS-54
chr1-28237727-INS-437

chr1-28243231-INS-760
chr1-28249066-INS-627
chr1-28256049-INS-52
chr1-28265782-INS-382
chr1-28267755-INS-6667
chr1-28341813-INS-1260
chr1-28361306-INS-89
chr1-28367604-INS-220
chr1-28411160-INS-6397
chr1-28423721-INS-200
chr1-28430083-INS-58
chr1-28431168-INS-1641
chr1-28458797-INS-200
chr1-28493812-INS-6515
chr1-28514186-INS-50
chr1-28571839-INS-2118
chr1-28585231-INS-2316
chr1-28586376-INS-1324
chr1-28596632-INS-4701
chr1-28597507-INS-1008
chr1-28621933-INS-6194
chr1-28628755-INS-498
chr1-28631436-INS-715
chr1-28631568-INS-82
chr1-28666418-INS-56
chr1-28693496-INS-810
chr1-28708561-INS-6487
chr1-28864878-INS-7229
chr1-28890202-INS-56
chr1-28894599-INS-1513
chr1-28895013-INS-600
chr1-28949479-INS-81
chr1-28953660-INS-237
chr1-28960572-INS-320
chr1-28962974-INS-50
chr1-28968438-INS-6175
chr1-28993180-INS-2948
chr1-29035264-INS-50
chr1-29046688-INS-105
chr1-29064283-INS-2119
chr1-29145466-INS-6632
chr1-29159952-INS-1014
chr1-29173918-INS-52
chr1-29179625-INS-171
chr1-29188917-INS-210
chr1-29261298-INS-105
chr1-29308648-INS-51
chr1-29315219-INS-425
chr1-29319559-INS-1113
chr1-29324559-INS-59135
chr1-29340425-INS-1193
chr1-29345252-INS-6218
chr1-29371941-INS-397
chr1-29440311-INS-56
chr1-29495343-INS-6579
chr1-29506912-INS-75
chr1-29513622-INS-523
chr1-29551230-INS-6611
chr1-29571850-INS-451
chr1-29642087-INS-52
chr1-29653979-INS-1424
chr1-29654712-INS-70
chr1-29673618-INS-2751
chr1-29732596-INS-231
chr1-29916178-INS-4499
chr1-29939905-INS-184
chr1-29968534-INS-87
chr1-29995917-INS-1763
chr1-30092656-INS-1126
chr1-30093112-INS-207
chr1-30134522-INS-52
chr1-30143083-INS-438
chr1-30153667-INS-253
chr1-30172684-INS-59
chr1-30224463-INS-399
chr1-30254748-INS-529
chr1-30266148-INS-60

chr1-30276535-INS-57
chr1-30315234-INS-58
chr1-30340940-INS-341
chr1-30341285-INS-102
chr1-30367794-INS-524
chr1-30394402-INS-6591
chr1-30403681-INS-112
chr1-30450486-INS-65
chr1-30471825-INS-75
chr1-30490643-INS-52
chr1-30523223-INS-54
chr1-30558334-INS-282
chr1-30563767-INS-3342
chr1-30669565-INS-1019
chr1-30744827-INS-2286
chr1-30811496-INS-89
chr1-30822888-INS-1616
chr1-30854402-INS-229
chr1-30892985-INS-424
chr1-30926759-INS-101
chr1-30926981-INS-81
chr1-30927103-INS-203
chr1-30951457-INS-6403
chr1-31029052-INS-94
chr1-31057037-INS-56
chr1-31066744-INS-63
chr1-31093223-INS-446
chr1-31094231-INS-138
chr1-31177931-INS-7790
chr1-31184773-INS-6797
chr1-31209698-INS-6594
chr1-31214757-INS-2830
chr1-31311198-INS-14194
chr1-31413656-INS-50
chr1-31443791-INS-2755
chr1-31552117-INS-2176
chr1-31553473-INS-320
chr1-31679290-INS-7195
chr1-31689561-INS-4521
chr1-31756114-INS-208
chr1-31775073-INS-257
chr1-31781462-INS-326
chr1-31805662-INS-6437
chr1-31809889-INS-52
chr1-31832279-INS-288
chr1-31832279-INS-9133
chr1-31847944-INS-55
chr1-31914733-INS-195
chr1-31917697-INS-100
chr1-31938774-INS-248
chr1-31946138-INS-63
chr1-31946586-INS-152
chr1-31951380-INS-6337
chr1-31959341-INS-3214
chr1-31988546-INS-198
chr1-31996087-INS-80
chr1-32004049-INS-4477
chr1-32021215-INS-66
chr1-32073548-INS-164
chr1-32171947-INS-50
chr1-32217014-INS-50
chr1-32240281-INS-4110
chr1-32321927-INS-130
chr1-32435459-INS-450
chr1-32512922-INS-498
chr1-32514651-INS-58
chr1-32595902-INS-54
chr1-32620182-INS-173
chr1-32656988-INS-50
chr1-32661009-INS-92
chr1-32666773-INS-6341
chr1-32682942-INS-254
chr1-32682942-INS-84
chr1-32684365-INS-156
chr1-32717487-INS-121
chr1-32743665-INS-50
chr1-32829106-INS-3580

chr1-32829442-INS-2591
chr1-32860628-INS-584
chr1-32862612-INS-6408
chr1-32879780-INS-54
chr1-32880043-INS-369
chr1-32907037-INS-60
chr1-32923563-INS-90
chr1-33004169-INS-2939
chr1-33045682-INS-50
chr1-33059063-INS-848
chr1-33066706-INS-56
chr1-33113047-INS-6136
chr1-33113119-INS-144
chr1-33113669-INS-119
chr1-33138405-INS-259
chr1-33174767-INS-3536
chr1-33209402-INS-56
chr1-33222990-INS-134
chr1-33272647-INS-553
chr1-33281646-INS-1087
chr1-33295859-INS-617
chr1-33317518-INS-362
chr1-33325071-INS-137
chr1-33408295-INS-441
chr1-33416449-INS-1012
chr1-33488345-INS-8765
chr1-33491777-INS-2925
chr1-33492130-INS-166
chr1-33505657-INS-433
chr1-33508609-INS-727
chr1-33515972-INS-6336
chr1-33525477-INS-114
chr1-33525802-INS-264
chr1-33525877-INS-124
chr1-33532447-INS-55
chr1-33543104-INS-6878
chr1-33570998-INS-6441
chr1-33580495-INS-179
chr1-33609813-INS-2223
chr1-33640184-INS-5194
chr1-33640850-INS-81
chr1-33664018-INS-405
chr1-33670052-INS-1113
chr1-33685529-INS-850
chr1-33706459-INS-179
chr1-33720715-INS-410
chr1-33726815-INS-6674
chr1-33729746-INS-60
chr1-33730250-INS-204
chr1-33730635-INS-215
chr1-33744799-INS-629
chr1-33755510-INS-112
chr1-33781891-INS-128
chr1-33791321-INS-56
chr1-33882172-INS-1074
chr1-33918093-INS-203
chr1-33927174-INS-2477
chr1-33938879-INS-50
chr1-33962872-INS-59
chr1-33963155-INS-108
chr1-33963155-INS-396
chr1-33973099-INS-53
chr1-33994041-INS-2397
chr1-33998826-INS-125
chr1-34004731-INS-404
chr1-34015869-INS-213
chr1-34041205-INS-78
chr1-34094676-INS-144
chr1-34134648-INS-66
chr1-34135943-INS-230
chr1-34146147-INS-1518
chr1-34172874-INS-70
chr1-34395417-INS-65
chr1-34395448-INS-247
chr1-34406791-INS-2452
chr1-34420013-INS-52
chr1-34423558-INS-88

chr1-34451289-INS-126
chr1-34536591-INS-93
chr1-34536869-INS-100
chr1-34548494-INS-1303
chr1-34554289-INS-147
chr1-34559730-INS-1821
chr1-34561171-INS-649
chr1-34570807-INS-112
chr1-34570930-INS-70
chr1-34572094-INS-1549
chr1-34584740-INS-90
chr1-34589677-INS-87
chr1-34590349-INS-74
chr1-34615849-INS-512
chr1-34616178-INS-96
chr1-34641782-INS-5254
chr1-34644503-INS-10164
chr1-34644503-INS-5075
chr1-34662451-INS-1844
chr1-34665978-INS-53
chr1-34682355-INS-90
chr1-34708712-INS-1596
chr1-34760363-INS-79
chr1-34769213-INS-84
chr1-34773098-INS-152
chr1-34782125-INS-1528
chr1-34836994-INS-282
chr1-34898366-INS-220
chr1-34899534-INS-6912
chr1-34919603-INS-117
chr1-34920744-INS-51
chr1-34951680-INS-60
chr1-34951809-INS-63
chr1-34990912-INS-50
chr1-35023346-INS-50
chr1-35038745-INS-77
chr1-35040810-INS-1168
chr1-35062016-INS-1252
chr1-35075917-INS-113
chr1-35100697-INS-67
chr1-35145778-INS-202
chr1-35145890-INS-784
chr1-35170335-INS-159
chr1-35193407-INS-78
chr1-35226696-INS-128
chr1-35238404-INS-59
chr1-35312165-INS-6475
chr1-35316466-INS-6587
chr1-35335816-INS-2670
chr1-35366989-INS-226
chr1-35386095-INS-234
chr1-35395666-INS-134
chr1-35407698-INS-210
chr1-35409888-INS-54
chr1-35434030-INS-180
chr1-35434175-INS-438
chr1-35455862-INS-1243
chr1-35517114-INS-6684
chr1-35536885-INS-66
chr1-35611689-INS-303
chr1-35624121-INS-60
chr1-35656074-INS-73
chr1-35673242-INS-6475
chr1-35678886-INS-1283
chr1-35679013-INS-84
chr1-35679014-INS-165
chr1-35706845-INS-1556
chr1-35733986-INS-498
chr1-35744582-INS-171
chr1-35785791-INS-53
chr1-35838271-INS-75
chr1-35846978-INS-90
chr1-35847731-INS-114
chr1-35847947-INS-200
chr1-35855072-INS-720
chr1-35875639-INS-806
chr1-35875744-INS-75

chr1-35926078-INS-6455
chr1-35958560-INS-88
chr1-35961945-INS-194
chr1-35980003-INS-64
chr1-35986394-INS-74
chr1-35988567-INS-207
chr1-35997950-INS-195
chr1-36013947-INS-92
chr1-36014639-INS-60
chr1-36026764-INS-206
chr1-36049025-INS-590
chr1-36061176-INS-96
chr1-36062658-INS-66
chr1-36062717-INS-138
chr1-36068051-INS-96
chr1-36068546-INS-120
chr1-36068767-INS-530
chr1-36080878-INS-190
chr1-36163625-INS-288
chr1-36168879-INS-1762
chr1-36196739-INS-161
chr1-36262012-INS-128
chr1-36298085-INS-207
chr1-36301259-INS-90
chr1-36322453-INS-4804
chr1-36325332-INS-5955
chr1-36326495-INS-88
chr1-36326825-INS-154
chr1-36398800-INS-196
chr1-36413189-INS-50
chr1-36428349-INS-195
chr1-36455225-INS-64
chr1-36477686-INS-61
chr1-36502054-INS-100
chr1-36553532-INS-53
chr1-36590418-INS-53
chr1-36633378-INS-158
chr1-36658598-INS-275
chr1-36668027-INS-141
chr1-36676884-INS-51
chr1-36702412-INS-60
chr1-36743488-INS-226
chr1-36753475-INS-7142
chr1-36760522-INS-222
chr1-36760942-INS-5274
chr1-36765968-INS-96
chr1-36766031-INS-546
chr1-36768494-INS-71
chr1-36780937-INS-249
chr1-36783045-INS-205
chr1-36793598-INS-531
chr1-36808317-INS-50
chr1-36808798-INS-52
chr1-36822916-INS-58
chr1-36837698-INS-1889
chr1-36841653-INS-1857
chr1-36841769-INS-204
chr1-36887447-INS-406
chr1-36903075-INS-74
chr1-36984724-INS-168
chr1-37003200-INS-486
chr1-37023492-INS-76
chr1-37075220-INS-4550
chr1-37107715-INS-5525
chr1-37122531-INS-7279
chr1-37146265-INS-126
chr1-37150780-INS-518
chr1-37162932-INS-81
chr1-37170421-INS-280
chr1-37170565-INS-112
chr1-37217765-INS-53
chr1-37223953-INS-2595
chr1-37234879-INS-53
chr1-37243298-INS-70
chr1-37262293-INS-61
chr1-37262306-INS-287
chr1-37271935-INS-1114

chr1-37271998-INS-6551
chr1-37272223-INS-78
chr1-37277847-INS-260
chr1-37289255-INS-167
chr1-37289510-INS-62
chr1-37334479-INS-50
chr1-37335219-INS-110
chr1-37352513-INS-69
chr1-37352513-INS-913
chr1-37352530-INS-182
chr1-37352803-INS-320
chr1-37352822-INS-69
chr1-37358507-INS-492
chr1-37374755-INS-56
chr1-37435486-INS-67
chr1-37473540-INS-175
chr1-37554317-INS-61
chr1-37563442-INS-450
chr1-37563517-INS-154
chr1-37606571-INS-430
chr1-37611441-INS-59
chr1-37685553-INS-60
chr1-37803495-INS-69
chr1-37840586-INS-1291
chr1-37868048-INS-1701
chr1-37868048-INS-3601
chr1-37868662-INS-146
chr1-37868905-INS-575
chr1-37883286-INS-392
chr1-37892299-INS-98
chr1-37922597-INS-94
chr1-37945794-INS-119
chr1-37953209-INS-93
chr1-37953352-INS-66
chr1-37972562-INS-132
chr1-37972970-INS-58
chr1-37973062-INS-1878
chr1-37973062-INS-918
chr1-37978492-INS-4271
chr1-38019525-INS-5080
chr1-38069507-INS-464
chr1-38179528-INS-1892
chr1-38213965-INS-68
chr1-38273333-INS-268
chr1-38286973-INS-50
chr1-38293857-INS-52
chr1-38308162-INS-135
chr1-38315905-INS-522
chr1-38317250-INS-61
chr1-38394857-INS-74
chr1-38469476-INS-237
chr1-38499605-INS-63
chr1-38518531-INS-50
chr1-38528269-INS-60
chr1-38572506-INS-62
chr1-38575103-INS-210
chr1-38605493-INS-60
chr1-38605679-INS-336
chr1-38605845-INS-158
chr1-38625533-INS-180
chr1-38649829-INS-148
chr1-38692555-INS-393
chr1-38732835-INS-65
chr1-38732908-INS-170
chr1-38740135-INS-495
chr1-38761959-INS-481
chr1-38848025-INS-363
chr1-38906434-INS-78
chr1-38906826-INS-78
chr1-38907767-INS-65
chr1-38925651-INS-69
chr1-38931719-INS-173
chr1-38955016-INS-80
chr1-38971053-INS-121
chr1-38987181-INS-72
chr1-38999954-INS-5375
chr1-39045962-INS-98

chr1-39092114-INS-50
chr1-39128536-INS-50
chr1-39132086-INS-93
chr1-39150002-INS-6630
chr1-39168177-INS-1074
chr1-39185573-INS-108
chr1-39185664-INS-84
chr1-39185667-INS-272
chr1-39195009-INS-64
chr1-39213393-INS-62
chr1-39312841-INS-64
chr1-39321514-INS-356
chr1-39334014-INS-103
chr1-39392889-INS-53
chr1-39482580-INS-130
chr1-39505727-INS-2794
chr1-39538942-INS-197
chr1-39538958-INS-70
chr1-39572440-INS-99
chr1-39592355-INS-84
chr1-39592467-INS-201
chr1-39619476-INS-60
chr1-39626242-INS-202
chr1-39642568-INS-115
chr1-39642752-INS-56
chr1-39648853-INS-414
chr1-39650888-INS-348
chr1-39650914-INS-84
chr1-39654745-INS-3753
chr1-39671980-INS-252
chr1-39672546-INS-283
chr1-39676551-INS-545
chr1-39703944-INS-4940
chr1-39727985-INS-70
chr1-39728428-INS-113
chr1-39767324-INS-211
chr1-39785241-INS-4375
chr1-39815065-INS-288
chr1-39878347-INS-50
chr1-39947694-INS-235
chr1-39975371-INS-99
chr1-40000579-INS-200
chr1-40001334-INS-54
chr1-40004461-INS-51
chr1-40007100-INS-1756
chr1-40025611-INS-504
chr1-40047142-INS-52
chr1-40052605-INS-305
chr1-40055311-INS-777
chr1-40090139-INS-5126
chr1-40108085-INS-142
chr1-40192448-INS-2081
chr1-40249462-INS-60
chr1-40257696-INS-52
chr1-40284055-INS-52
chr1-40290302-INS-126
chr1-40297790-INS-219
chr1-40369774-INS-219
chr1-40371870-INS-3054
chr1-40383634-INS-73
chr1-40383739-INS-184
chr1-40383758-INS-396
chr1-40386977-INS-52
chr1-40413289-INS-98
chr1-40435655-INS-490
chr1-40460071-INS-1106
chr1-40463770-INS-179
chr1-40521411-INS-75
chr1-40521412-INS-248
chr1-40557163-INS-64
chr1-40575832-INS-53
chr1-40587208-INS-2248
chr1-40598701-INS-3090
chr1-40695865-INS-88
chr1-40758803-INS-4896
chr1-40764200-INS-832
chr1-40821484-INS-68

chr1-40831514-INS-286
chr1-40833700-INS-461
chr1-40833883-INS-60
chr1-40835638-INS-7162
chr1-40912825-INS-186
chr1-40921624-INS-201
chr1-40924498-INS-348
chr1-40938821-INS-76
chr1-40953724-INS-89
chr1-40958908-INS-84
chr1-40992112-INS-103
chr1-40999118-INS-52
chr1-41000986-INS-157
chr1-41022818-INS-50
chr1-41028831-INS-2873
chr1-41033373-INS-288
chr1-41045128-INS-7102
chr1-41061881-INS-983
chr1-41065653-INS-74
chr1-41075330-INS-54
chr1-41075391-INS-128
chr1-41077387-INS-5165
chr1-41086997-INS-373
chr1-41138011-INS-68
chr1-41180839-INS-164
chr1-41185292-INS-96
chr1-41224445-INS-198
chr1-41230679-INS-58
chr1-41290914-INS-1143
chr1-41309422-INS-282
chr1-41317294-INS-72
chr1-41322847-INS-129
chr1-41330882-INS-1351
chr1-41335477-INS-116
chr1-41339965-INS-1045
chr1-41352676-INS-68
chr1-41353425-INS-171
chr1-41360505-INS-968
chr1-41360878-INS-242
chr1-41374312-INS-111
chr1-41432824-INS-2333
chr1-41477603-INS-109
chr1-41493065-INS-58
chr1-41494203-INS-50
chr1-41494819-INS-640
chr1-41523138-INS-58
chr1-41537331-INS-153
chr1-41544250-INS-874
chr1-41569298-INS-852
chr1-41572740-INS-6211
chr1-41585288-INS-54
chr1-41590119-INS-6271
chr1-41616009-INS-1223
chr1-41653223-INS-50
chr1-41677504-INS-2847
chr1-41695140-INS-64
chr1-41715439-INS-168
chr1-41773381-INS-266
chr1-41795325-INS-178
chr1-41806727-INS-2864
chr1-41809354-INS-1104
chr1-41820097-INS-78
chr1-41874251-INS-386
chr1-41874261-INS-64
chr1-41877758-INS-136
chr1-41878310-INS-108
chr1-41878582-INS-62
chr1-41886470-INS-50
chr1-41893884-INS-539
chr1-41902533-INS-1282
chr1-41902786-INS-76
chr1-41905145-INS-14422
chr1-41950342-INS-115
chr1-41950342-INS-244
chr1-41983895-INS-51
chr1-41985355-INS-612
chr1-42015128-INS-117

chr1-42016828-INS-5554
chr1-42059910-INS-54
chr1-42060538-INS-320
chr1-42077139-INS-407
chr1-42081820-INS-947
chr1-42091555-INS-7093
chr1-42121126-INS-87
chr1-42129410-INS-112
chr1-42155126-INS-121
chr1-42197055-INS-52
chr1-42222853-INS-56
chr1-42239578-INS-60
chr1-42250608-INS-128
chr1-42336542-INS-50
chr1-42338300-INS-74
chr1-42345962-INS-114
chr1-42409415-INS-246
chr1-42449270-INS-64
chr1-42459096-INS-1958
chr1-42464775-INS-56
chr1-42465185-INS-2716
chr1-42495526-INS-296
chr1-42546139-INS-158
chr1-42551960-INS-6464
chr1-42553333-INS-2195
chr1-42555027-INS-98
chr1-42616839-INS-114
chr1-42635265-INS-74
chr1-42674658-INS-50
chr1-42678889-INS-65
chr1-42679079-INS-135
chr1-42710130-INS-56
chr1-42766153-INS-3133
chr1-42773466-INS-68
chr1-42777782-INS-208
chr1-42802653-INS-51
chr1-42803127-INS-414
chr1-42804395-INS-841
chr1-42813829-INS-78
chr1-42835435-INS-1806
chr1-42869256-INS-128
chr1-42869572-INS-122
chr1-42879543-INS-4482
chr1-42879543-INS-490
chr1-42976880-INS-6719
chr1-42980903-INS-2892
chr1-42982294-INS-165
chr1-42982428-INS-104
chr1-42982497-INS-239
chr1-43024586-INS-108
chr1-43053769-INS-113
chr1-43158293-INS-66
chr1-43159378-INS-500
chr1-43165848-INS-95
chr1-43226657-INS-74
chr1-43273777-INS-336
chr1-43280433-INS-215
chr1-43293256-INS-52
chr1-43300944-INS-470
chr1-43305411-INS-101
chr1-43332286-INS-209
chr1-43345419-INS-73
chr1-43356964-INS-105
chr1-43378171-INS-58
chr1-43378172-INS-128
chr1-43392896-INS-7110
chr1-43428386-INS-93
chr1-43447323-INS-55
chr1-43471755-INS-60
chr1-43479014-INS-64
chr1-43492941-INS-78
chr1-43502203-INS-228
chr1-43530017-INS-94
chr1-43558809-INS-130
chr1-43573042-INS-1513
chr1-43638246-INS-89
chr1-43638729-INS-194

chr1-43639788-INS-68
chr1-43685423-INS-6526
chr1-43694371-INS-50
chr1-43697723-INS-85
chr1-43729571-INS-1023
chr1-43793776-INS-140
chr1-43806532-INS-3751
chr1-43817607-INS-3973
chr1-43838347-INS-4923
chr1-43854645-INS-52
chr1-43878265-INS-395
chr1-43878337-INS-1192
chr1-43904476-INS-498
chr1-43921578-INS-53
chr1-43929000-INS-1731
chr1-43931074-INS-9965
chr1-43952636-INS-7270
chr1-43955618-INS-168
chr1-43979627-INS-210
chr1-44026835-INS-200
chr1-44077186-INS-6921
chr1-44094418-INS-2899
chr1-44172608-INS-187
chr1-44172611-INS-402
chr1-44172676-INS-58
chr1-44172818-INS-60
chr1-44179037-INS-65
chr1-44185672-INS-506
chr1-44185990-INS-65
chr1-44193732-INS-528
chr1-44198314-INS-216
chr1-44202624-INS-6464
chr1-44209912-INS-620
chr1-44215187-INS-88
chr1-44226531-INS-90
chr1-44227201-INS-6306
chr1-44227667-INS-1858
chr1-44240782-INS-1824
chr1-44249524-INS-136
chr1-44254464-INS-353
chr1-44275494-INS-340
chr1-44324743-INS-416
chr1-44367434-INS-1160
chr1-44478422-INS-560
chr1-44479635-INS-84
chr1-44489404-INS-1179
chr1-44510405-INS-52
chr1-44510885-INS-2335
chr1-44537363-INS-4579
chr1-44551040-INS-572
chr1-44563508-INS-7186
chr1-44590391-INS-69
chr1-44595829-INS-1074
chr1-44621438-INS-283
chr1-44637127-INS-216
chr1-44637127-INS-55
chr1-44662843-INS-141
chr1-44662951-INS-366
chr1-44666033-INS-396
chr1-44676335-INS-50
chr1-44694012-INS-419
chr1-44786090-INS-72
chr1-44788758-INS-1270
chr1-44828537-INS-51
chr1-44878880-INS-88
chr1-44879246-INS-100
chr1-44882997-INS-6419
chr1-44914003-INS-215
chr1-44998735-INS-636
chr1-45022030-INS-56
chr1-45038341-INS-7356
chr1-45047448-INS-2847
chr1-45066404-INS-224
chr1-45106369-INS-2117
chr1-45106577-INS-878
chr1-45122285-INS-3460
chr1-45130100-INS-2828

chr1-45135001-INS-3008
chr1-45140992-INS-460
chr1-45183734-INS-217
chr1-45236041-INS-59
chr1-45247761-INS-54
chr1-45280997-INS-121
chr1-45281048-INS-470
chr1-45342521-INS-260
chr1-45405640-INS-6245
chr1-45480228-INS-64
chr1-45511386-INS-53
chr1-45520810-INS-58
chr1-45523393-INS-428
chr1-45524323-INS-68
chr1-45524370-INS-277
chr1-45548931-INS-96
chr1-45620600-INS-197
chr1-45664206-INS-359
chr1-45728010-INS-2712
chr1-45734843-INS-183
chr1-45739746-INS-326
chr1-45766644-INS-50
chr1-45834004-INS-103
chr1-45858478-INS-2835
chr1-45877318-INS-50
chr1-45902162-INS-61
chr1-45918800-INS-64
chr1-45981073-INS-4715
chr1-45992511-INS-1185
chr1-46001290-INS-105
chr1-46036187-INS-52
chr1-46038692-INS-5140
chr1-46046874-INS-60
chr1-46071902-INS-2747
chr1-46128200-INS-57
chr1-46128288-INS-233
chr1-46128313-INS-1146
chr1-46128313-INS-524
chr1-46142660-INS-3363
chr1-46151397-INS-1330
chr1-46176914-INS-126
chr1-46176914-INS-278
chr1-46247156-INS-471
chr1-46286954-INS-686
chr1-46372025-INS-266
chr1-46445709-INS-5452
chr1-46447136-INS-9282
chr1-46451270-INS-85
chr1-46452838-INS-1031
chr1-46463271-INS-6222
chr1-46542414-INS-57
chr1-46575497-INS-152
chr1-46580430-INS-7632
chr1-46597304-INS-50
chr1-46629423-INS-69
chr1-46691055-INS-6205
chr1-46722188-INS-512
chr1-46729101-INS-200
chr1-46736092-INS-930
chr1-46766892-INS-54
chr1-46839112-INS-763
chr1-46872901-INS-64
chr1-46957152-INS-200
chr1-47009387-INS-56
chr1-47026667-INS-94
chr1-47061290-INS-691
chr1-47062020-INS-1422
chr1-47067243-INS-6305
chr1-47089602-INS-84
chr1-47141307-INS-150
chr1-47159321-INS-79
chr1-47206005-INS-192
chr1-47221090-INS-388
chr1-47227620-INS-461
chr1-47292993-INS-180
chr1-47292993-INS-56
chr1-47335666-INS-451

chr1-47343024-INS-84
chr1-47373716-INS-51
chr1-47432573-INS-60
chr1-47456791-INS-88
chr1-47460156-INS-51
chr1-47496362-INS-88
chr1-47497535-INS-596
chr1-47513967-INS-335
chr1-47648801-INS-709
chr1-47649265-INS-490
chr1-47659102-INS-51
chr1-47666747-INS-2422
chr1-47683272-INS-4544
chr1-47698191-INS-599
chr1-47768748-INS-385
chr1-47781228-INS-347
chr1-47797988-INS-798
chr1-47866402-INS-652
chr1-47881165-INS-99
chr1-47894278-INS-2429
chr1-47964201-INS-340
chr1-47980865-INS-907
chr1-48023038-INS-482
chr1-48041554-INS-2468
chr1-48067134-INS-370
chr1-48151638-INS-1161
chr1-48199590-INS-136
chr1-48211927-INS-50
chr1-48293481-INS-54
chr1-48294369-INS-291
chr1-48294634-INS-1229
chr1-48314502-INS-434
chr1-48463414-INS-208
chr1-48546435-INS-5976
chr1-48548638-INS-102
chr1-48652169-INS-2273
chr1-48661587-INS-3231
chr1-48674867-INS-52
chr1-48688149-INS-6782
chr1-48703092-INS-75
chr1-48703114-INS-184
chr1-48703799-INS-196
chr1-48797852-INS-83
chr1-48808979-INS-50
chr1-48868746-INS-206
chr1-48949699-INS-140
chr1-49040718-INS-875
chr1-49042174-INS-4891
chr1-49059573-INS-127
chr1-49072108-INS-3220
chr1-49083386-INS-90
chr1-49091918-INS-1421
chr1-49151846-INS-54
chr1-49157090-INS-6676
chr1-49169140-INS-79
chr1-49195630-INS-2802
chr1-49215578-INS-1891
chr1-49229264-INS-6228
chr1-49255861-INS-68
chr1-49259597-INS-63
chr1-49264069-INS-2034
chr1-49274509-INS-329
chr1-49274632-INS-149
chr1-49274770-INS-538
chr1-49275165-INS-845
chr1-49278682-INS-50
chr1-49295964-INS-201
chr1-49320072-INS-839
chr1-49350092-INS-100
chr1-49407706-INS-205
chr1-49448304-INS-54
chr1-49472104-INS-74
chr1-49522644-INS-96
chr1-49522926-INS-72
chr1-49548406-INS-343
chr1-49552262-INS-66
chr1-49592186-INS-472

chr1-49614887-INS-628
chr1-49621579-INS-51
chr1-49665237-INS-52
chr1-49671333-INS-50
chr1-49681953-INS-3019
chr1-49705599-INS-101
chr1-49721741-INS-124
chr1-49723084-INS-7044
chr1-49736274-INS-3080
chr1-49738278-INS-7092
chr1-49766478-INS-1348
chr1-49775034-INS-68
chr1-49785004-INS-1490
chr1-49787056-INS-2550
chr1-49796202-INS-1023
chr1-49816831-INS-187
chr1-49822029-INS-6511
chr1-49851819-INS-1383
chr1-49851890-INS-403
chr1-49857552-INS-64
chr1-49892583-INS-515
chr1-49895631-INS-212
chr1-49895640-INS-92
chr1-49905414-INS-115
chr1-49913424-INS-1655
chr1-49925764-INS-151
chr1-49934999-INS-2159
chr1-49935645-INS-54
chr1-49945346-INS-6257
chr1-49970326-INS-188
chr1-49972102-INS-6309
chr1-49972304-INS-62
chr1-49983523-INS-1015
chr1-49992639-INS-5456
chr1-50001788-INS-1492
chr1-50011013-INS-1492
chr1-50037881-INS-134
chr1-50110615-INS-591
chr1-50110841-INS-2390
chr1-50133243-INS-143
chr1-50133247-INS-625
chr1-50184882-INS-50
chr1-50196708-INS-56
chr1-50197113-INS-153
chr1-50214747-INS-170
chr1-50233588-INS-1326
chr1-50261809-INS-2048
chr1-50273275-INS-66
chr1-50276138-INS-4547
chr1-50289124-INS-75
chr1-50299975-INS-65
chr1-50319608-INS-291
chr1-50389773-INS-54
chr1-50433503-INS-53
chr1-50434310-INS-50
chr1-50460445-INS-202
chr1-50475585-INS-1714
chr1-50562554-INS-59
chr1-50569203-INS-90
chr1-50569969-INS-283
chr1-50618812-INS-78
chr1-50623234-INS-1439
chr1-50637692-INS-74
chr1-50638202-INS-157
chr1-50679129-INS-1068
chr1-50685726-INS-60
chr1-50688888-INS-389
chr1-50718024-INS-2130
chr1-50758746-INS-6473
chr1-50767500-INS-111
chr1-50770333-INS-60
chr1-50774181-INS-272
chr1-50856430-INS-6229
chr1-50862294-INS-52
chr1-50923328-INS-332
chr1-50932063-INS-1396
chr1-50935151-INS-80

chr1-51012770-INS-114
chr1-51087547-INS-3061
chr1-51094882-INS-77
chr1-51136676-INS-64
chr1-51151423-INS-140
chr1-51156451-INS-58
chr1-51161630-INS-154
chr1-51174231-INS-79
chr1-51269323-INS-72
chr1-51353272-INS-6213
chr1-51358626-INS-238
chr1-51361715-INS-57
chr1-51381289-INS-81
chr1-51445170-INS-7099
chr1-51462414-INS-64
chr1-51521148-INS-148
chr1-51610718-INS-2969
chr1-51644058-INS-216
chr1-51644066-INS-56
chr1-51664707-INS-86
chr1-51685788-INS-139
chr1-51694781-INS-1380
chr1-51701748-INS-1026
chr1-51713303-INS-5545
chr1-51723077-INS-177
chr1-51736172-INS-429
chr1-51810945-INS-629
chr1-51862263-INS-57
chr1-51938078-INS-69
chr1-51946092-INS-52
chr1-51960835-INS-189
chr1-51962421-INS-217
chr1-51967631-INS-209
chr1-52003153-INS-72
chr1-52023619-INS-261
chr1-52040794-INS-60
chr1-52064975-INS-387
chr1-52135410-INS-66
chr1-52304575-INS-50
chr1-52305918-INS-3930
chr1-52419709-INS-7532
chr1-52421092-INS-58
chr1-52486496-INS-3112
chr1-52495425-INS-13670
chr1-52522379-INS-155
chr1-52549895-INS-3782
chr1-52559936-INS-149
chr1-52571211-INS-154
chr1-52589259-INS-66
chr1-52665496-INS-50
chr1-52707374-INS-599
chr1-52717620-INS-6438
chr1-52741586-INS-1511
chr1-52767714-INS-67
chr1-52790050-INS-185
chr1-52790772-INS-51
chr1-52791200-INS-197
chr1-52791539-INS-60
chr1-52791574-INS-175
chr1-52791831-INS-61
chr1-52794884-INS-156
chr1-52804379-INS-194
chr1-52808616-INS-189
chr1-52828438-INS-69
chr1-52879697-INS-52
chr1-52890741-INS-79
chr1-52891809-INS-1032
chr1-52913319-INS-2764
chr1-52953737-INS-1748
chr1-52969253-INS-132
chr1-52969879-INS-58
chr1-52972669-INS-213
chr1-53048978-INS-2774
chr1-53060298-INS-795
chr1-53060393-INS-76
chr1-53074063-INS-1322
chr1-53096333-INS-91

chr1-53142977-INS-1859
chr1-53218929-INS-56
chr1-53219143-INS-235
chr1-53219911-INS-1503
chr1-53229994-INS-4360
chr1-53235055-INS-434
chr1-53244073-INS-52
chr1-53249678-INS-466
chr1-53266239-INS-2973
chr1-53279449-INS-6417
chr1-53282697-INS-3819
chr1-53289346-INS-226
chr1-53368321-INS-89
chr1-53368529-INS-273
chr1-53372103-INS-4672
chr1-53416077-INS-7562
chr1-53422096-INS-238
chr1-53426614-INS-397
chr1-53426731-INS-1222
chr1-53431223-INS-64
chr1-53432489-INS-2988
chr1-53457793-INS-77
chr1-53458272-INS-54
chr1-53459815-INS-51
chr1-53460062-INS-54
chr1-53510036-INS-64
chr1-53577531-INS-223
chr1-53596399-INS-50
chr1-53611694-INS-68
chr1-53618829-INS-70
chr1-53634319-INS-6420
chr1-53684606-INS-50
chr1-53712901-INS-2224
chr1-53724614-INS-70
chr1-53743252-INS-253
chr1-53763663-INS-3090
chr1-53774962-INS-6440
chr1-53839691-INS-57
chr1-53856543-INS-1335
chr1-53901317-INS-60
chr1-53907332-INS-80
chr1-53916645-INS-66
chr1-53936475-INS-579
chr1-53972868-INS-208
chr1-53978242-INS-303
chr1-53979430-INS-51
chr1-53983449-INS-51
chr1-54002245-INS-510
chr1-54006685-INS-763
chr1-54013630-INS-52
chr1-54018744-INS-50
chr1-54019855-INS-398
chr1-54057384-INS-525
chr1-54059980-INS-60
chr1-54060026-INS-180
chr1-54060780-INS-115
chr1-54083172-INS-50
chr1-54125789-INS-70
chr1-54126105-INS-72
chr1-54194210-INS-59
chr1-54225877-INS-1289
chr1-54252609-INS-7088
chr1-54270812-INS-6276
chr1-54327501-INS-3058
chr1-54381527-INS-6292
chr1-54443231-INS-1031
chr1-54474423-INS-225
chr1-54502449-INS-402
chr1-54537794-INS-489
chr1-54542557-INS-2879
chr1-54555449-INS-988
chr1-54573854-INS-50
chr1-54599625-INS-60
chr1-54623088-INS-64
chr1-54623491-INS-56
chr1-54625180-INS-90
chr1-54634095-INS-6469

chr1-54652804-INS-165
chr1-54661726-INS-400
chr1-54720587-INS-6384
chr1-54725746-INS-500
chr1-54742063-INS-236
chr1-54771168-INS-132
chr1-54790952-INS-156
chr1-54819586-INS-2348
chr1-54840874-INS-267
chr1-54840963-INS-71
chr1-54858583-INS-83
chr1-54873926-INS-59
chr1-54898877-INS-198
chr1-54921585-INS-1026
chr1-54972589-INS-6325
chr1-54996270-INS-52
chr1-54996653-INS-208
chr1-55005362-INS-1895
chr1-55016980-INS-206
chr1-55033239-INS-402
chr1-55047851-INS-78
chr1-55049988-INS-1145
chr1-55050157-INS-366
chr1-55050178-INS-51
chr1-55050441-INS-50
chr1-55064265-INS-1986
chr1-55064268-INS-152
chr1-55073014-INS-241
chr1-55094924-INS-5555
chr1-55101571-INS-235
chr1-55107732-INS-129
chr1-55139578-INS-6461
chr1-55143518-INS-7100
chr1-55171601-INS-84
chr1-55172182-INS-201
chr1-55174150-INS-220
chr1-55186371-INS-810
chr1-55186400-INS-378
chr1-55204506-INS-65
chr1-55204629-INS-62
chr1-55228233-INS-132
chr1-55228382-INS-398
chr1-55239457-INS-92
chr1-55274850-INS-116
chr1-55356906-INS-6475
chr1-55363120-INS-6272
chr1-55373084-INS-79
chr1-55405654-INS-404
chr1-55437733-INS-74
chr1-55453345-INS-50
chr1-55478719-INS-87
chr1-55530115-INS-52
chr1-55533478-INS-84
chr1-55536440-INS-55
chr1-55552077-INS-328
chr1-55555340-INS-209
chr1-55563527-INS-125
chr1-55582816-INS-6555
chr1-55614025-INS-80
chr1-55638911-INS-424
chr1-55641414-INS-103
chr1-55641699-INS-4466
chr1-55643692-INS-405
chr1-55649866-INS-62
chr1-55666695-INS-1134
chr1-55667241-INS-50
chr1-55669270-INS-50
chr1-55679573-INS-4857
chr1-55751039-INS-486
chr1-55751461-INS-953
chr1-55832515-INS-257
chr1-55832667-INS-80
chr1-55853507-INS-6890
chr1-55866951-INS-82
chr1-55882523-INS-114
chr1-55886217-INS-5151
chr1-55904562-INS-128

chr1-55941117-INS-58
chr1-55960176-INS-50
chr1-55961968-INS-53
chr1-55975677-INS-128
chr1-55991563-INS-82
chr1-56003894-INS-1169
chr1-56051141-INS-108
chr1-56078744-INS-124
chr1-56094312-INS-60
chr1-56098208-INS-192
chr1-56098414-INS-3059
chr1-56141710-INS-76
chr1-56256463-INS-50
chr1-56263241-INS-6459
chr1-56280607-INS-58
chr1-56281132-INS-87
chr1-56355997-INS-134
chr1-56396321-INS-51
chr1-56510741-INS-55
chr1-56544609-INS-96
chr1-56544610-INS-217
chr1-56577897-INS-68
chr1-56581813-INS-62
chr1-56581869-INS-7273
chr1-56606018-INS-198
chr1-56620386-INS-251
chr1-56649233-INS-169
chr1-56662127-INS-104
chr1-56698439-INS-63
chr1-56706525-INS-6835
chr1-56709463-INS-267
chr1-56712688-INS-7078
chr1-56713139-INS-471
chr1-56717886-INS-70
chr1-56767783-INS-56
chr1-56795467-INS-6449
chr1-56799631-INS-218
chr1-56907078-INS-62
chr1-56909833-INS-370
chr1-56912724-INS-52
chr1-56920079-INS-98
chr1-56962709-INS-76
chr1-57022253-INS-60
chr1-57074495-INS-97
chr1-57085631-INS-225
chr1-57085650-INS-550
chr1-57156083-INS-52
chr1-57164671-INS-117
chr1-57179819-INS-6238
chr1-57184634-INS-64
chr1-57185378-INS-104
chr1-57208997-INS-90
chr1-57224489-INS-3142
chr1-57261775-INS-421
chr1-57285265-INS-1094
chr1-57293271-INS-3024
chr1-57294314-INS-160
chr1-57310297-INS-125
chr1-57310361-INS-302
chr1-57342968-INS-54
chr1-57347504-INS-2616
chr1-57373564-INS-710
chr1-57382224-INS-68
chr1-57419860-INS-1312
chr1-57419895-INS-256
chr1-57458080-INS-58
chr1-57462268-INS-213
chr1-57494875-INS-4990
chr1-57512788-INS-84
chr1-57538649-INS-56
chr1-57550923-INS-6184
chr1-57587881-INS-88
chr1-57605996-INS-201
chr1-57687312-INS-6779
chr1-57697296-INS-52
chr1-57718457-INS-54
chr1-57798642-INS-87

chr1-57834923-INS-72
chr1-57909567-INS-390
chr1-57909616-INS-105
chr1-57944716-INS-2741
chr1-57984886-INS-126
chr1-57995776-INS-93
chr1-58019436-INS-701
chr1-58038501-INS-201
chr1-58066905-INS-6695
chr1-58125568-INS-94
chr1-58129246-INS-102
chr1-58176526-INS-415
chr1-58188759-INS-309
chr1-58224825-INS-50
chr1-58228964-INS-67
chr1-58265092-INS-104
chr1-58269743-INS-65
chr1-58310190-INS-205
chr1-58362247-INS-199
chr1-58370583-INS-429
chr1-58376137-INS-154
chr1-58383266-INS-92
chr1-58383565-INS-100
chr1-58388820-INS-84
chr1-58388838-INS-348
chr1-58397210-INS-50
chr1-58414150-INS-1230
chr1-58414209-INS-270
chr1-58423570-INS-169
chr1-58424279-INS-203
chr1-58454798-INS-297
chr1-58456099-INS-200
chr1-58465658-INS-530
chr1-58465705-INS-201
chr1-58482322-INS-220
chr1-58670090-INS-70
chr1-58715352-INS-50
chr1-58716996-INS-56
chr1-58742340-INS-5432
chr1-58751277-INS-68
chr1-58753844-INS-52
chr1-58778344-INS-81
chr1-58811202-INS-1076
chr1-58845746-INS-417
chr1-58895792-INS-6336
chr1-58903535-INS-58
chr1-58935976-INS-147
chr1-58969420-INS-869
chr1-59046114-INS-1224
chr1-59049123-INS-454
chr1-59120194-INS-102
chr1-59188386-INS-140
chr1-59192132-INS-68
chr1-59192472-INS-209
chr1-59206252-INS-52
chr1-59238918-INS-391
chr1-59252483-INS-187
chr1-59263164-INS-210
chr1-59324824-INS-50
chr1-59331323-INS-84
chr1-59333034-INS-163
chr1-59338232-INS-79
chr1-59343982-INS-105
chr1-59375085-INS-63
chr1-59377098-INS-516
chr1-59377401-INS-604
chr1-59447411-INS-452
chr1-59456337-INS-54
chr1-59507527-INS-174
chr1-59518739-INS-2090
chr1-59518739-INS-908
chr1-59537975-INS-60
chr1-59586406-INS-126
chr1-59635484-INS-235
chr1-59635489-INS-1543
chr1-59639879-INS-176
chr1-59640753-INS-88

chr1-59702443-INS-110
chr1-59766106-INS-215
chr1-59766666-INS-54
chr1-59766783-INS-137
chr1-59788958-INS-187
chr1-59789961-INS-64
chr1-59792444-INS-230
chr1-59825493-INS-64
chr1-59826198-INS-229
chr1-59827722-INS-233
chr1-59827782-INS-54
chr1-59832877-INS-4639
chr1-59891943-INS-209
chr1-59892729-INS-183
chr1-59921410-INS-199
chr1-59923972-INS-210
chr1-59927130-INS-60
chr1-59928520-INS-80
chr1-59930524-INS-58
chr1-59930960-INS-215
chr1-59937019-INS-537
chr1-59937239-INS-56
chr1-59958140-INS-223
chr1-59969542-INS-114
chr1-59974997-INS-177
chr1-59974997-INS-402
chr1-59991180-INS-167
chr1-60037459-INS-62
chr1-60051703-INS-234
chr1-60077531-INS-1279
chr1-60141983-INS-51
chr1-60142025-INS-112
chr1-60142192-INS-2673
chr1-60156411-INS-2907
chr1-60171907-INS-280
chr1-60187416-INS-595
chr1-60213769-INS-51
chr1-60226272-INS-98
chr1-60242260-INS-202
chr1-60289236-INS-115
chr1-60299027-INS-1492
chr1-60349108-INS-225
chr1-60400789-INS-223
chr1-60406701-INS-50
chr1-60419221-INS-156
chr1-60578906-INS-67
chr1-60628977-INS-51
chr1-60662995-INS-51
chr1-60691353-INS-1088
chr1-60700587-INS-58
chr1-60708217-INS-62
chr1-60710467-INS-6569
chr1-60737424-INS-54
chr1-60748769-INS-5549
chr1-60759313-INS-220
chr1-60759411-INS-71
chr1-60781752-INS-55
chr1-60781981-INS-78
chr1-60819575-INS-6877
chr1-60858027-INS-97
chr1-60895563-INS-52
chr1-60910389-INS-64
chr1-60910925-INS-70
chr1-60913712-INS-59
chr1-60929508-INS-6527
chr1-60944280-INS-2813
chr1-60960295-INS-74
chr1-60960404-INS-270
chr1-60968014-INS-97
chr1-60986664-INS-6376
chr1-60988739-INS-3273
chr1-60995316-INS-469
chr1-61003062-INS-294
chr1-61007542-INS-54
chr1-61009703-INS-75
chr1-61030870-INS-1180
chr1-61033300-INS-51

chr1-61104700-INS-52
chr1-61163897-INS-358
chr1-61232570-INS-308
chr1-61272022-INS-126
chr1-61290990-INS-54
chr1-61291208-INS-309
chr1-61334993-INS-76
chr1-61335633-INS-58
chr1-61395755-INS-1183
chr1-61407512-INS-1809
chr1-61407547-INS-603
chr1-61426321-INS-7219
chr1-61438048-INS-305
chr1-61438386-INS-62
chr1-61439035-INS-56
chr1-61451232-INS-54
chr1-61475085-INS-2242
chr1-61506590-INS-74
chr1-61522084-INS-81
chr1-61529505-INS-96
chr1-61541666-INS-218
chr1-61583411-INS-68
chr1-61584572-INS-6289
chr1-61641770-INS-3054
chr1-61641771-INS-6588
chr1-61664051-INS-6600
chr1-61669654-INS-1123
chr1-61767452-INS-186
chr1-61820422-INS-434
chr1-61895456-INS-6641
chr1-61971176-INS-295
chr1-61990628-INS-2045
chr1-61991904-INS-92
chr1-62012123-INS-138
chr1-62054613-INS-67
chr1-62072993-INS-167
chr1-62123571-INS-6451
chr1-62156124-INS-128
chr1-62185751-INS-60
chr1-62213328-INS-343
chr1-62280199-INS-235
chr1-62280837-INS-68
chr1-62329764-INS-54
chr1-62359033-INS-60
chr1-62389412-INS-54
chr1-62392148-INS-50
chr1-62429842-INS-50
chr1-62438296-INS-7196
chr1-62527330-INS-3868
chr1-62537689-INS-55
chr1-62546176-INS-62
chr1-62595808-INS-76
chr1-62641952-INS-338
chr1-62729361-INS-152
chr1-62729586-INS-108
chr1-62794056-INS-56
chr1-62811194-INS-160
chr1-62903250-INS-55
chr1-62963454-INS-395
chr1-62963555-INS-79
chr1-62963887-INS-54
chr1-62991441-INS-238
chr1-63013940-INS-813
chr1-63014140-INS-6310
chr1-63025544-INS-56
chr1-63136230-INS-55
chr1-63145386-INS-87
chr1-63145977-INS-193
chr1-63146184-INS-99
chr1-63164940-INS-197
chr1-63187319-INS-88
chr1-63234510-INS-202
chr1-63241914-INS-173
chr1-63278857-INS-60
chr1-63278868-INS-270
chr1-63289179-INS-90
chr1-63289523-INS-60

chr1-63320388-INS-50
chr1-63360452-INS-590
chr1-63365736-INS-62
chr1-63368682-INS-1389
chr1-63369118-INS-54
chr1-63405126-INS-2114
chr1-63417523-INS-73
chr1-63417570-INS-347
chr1-63448190-INS-54
chr1-63490371-INS-340
chr1-63501253-INS-58
chr1-63502160-INS-204
chr1-63515865-INS-259
chr1-63523394-INS-50
chr1-63556403-INS-207
chr1-63560303-INS-140
chr1-63560444-INS-680
chr1-63560446-INS-2733
chr1-63566280-INS-64
chr1-63596972-INS-264
chr1-63599828-INS-188
chr1-63669554-INS-138
chr1-63669968-INS-920
chr1-63674832-INS-339
chr1-63705921-INS-99
chr1-63726959-INS-119
chr1-63768632-INS-465
chr1-63794323-INS-7202
chr1-63806496-INS-52
chr1-63827286-INS-128
chr1-63843156-INS-205
chr1-63904986-INS-60
chr1-63906217-INS-62
chr1-63924390-INS-50
chr1-64005557-INS-179
chr1-64010089-INS-132
chr1-64020516-INS-84
chr1-64027572-INS-158
chr1-64049193-INS-63
chr1-64049311-INS-792
chr1-64076699-INS-98
chr1-64215138-INS-85
chr1-64309856-INS-229
chr1-64333758-INS-56
chr1-64335540-INS-84
chr1-64335913-INS-242
chr1-64385880-INS-75
chr1-64440049-INS-1881
chr1-64474497-INS-648
chr1-64506252-INS-104
chr1-64506533-INS-6563
chr1-64531151-INS-70
chr1-64541822-INS-52
chr1-64595789-INS-199
chr1-64602898-INS-206
chr1-64605142-INS-100
chr1-64626463-INS-2260
chr1-64641189-INS-114
chr1-64671202-INS-55
chr1-64681602-INS-152
chr1-64686528-INS-102
chr1-64698711-INS-172
chr1-64705398-INS-51
chr1-64756109-INS-60
chr1-64816101-INS-197
chr1-64852355-INS-3129
chr1-64887691-INS-164
chr1-64956794-INS-2778
chr1-64982181-INS-108
chr1-65028236-INS-2591
chr1-65028468-INS-201
chr1-65043294-INS-1304
chr1-65043330-INS-480
chr1-65076931-INS-222
chr1-65098738-INS-152
chr1-65114285-INS-128
chr1-65139333-INS-58

chr1-65180404-INS-52
chr1-65271669-INS-50
chr1-65273299-INS-1724
chr1-65319441-INS-128
chr1-65340444-INS-68
chr1-65344442-INS-839
chr1-65379758-INS-67
chr1-65431953-INS-6690
chr1-65441589-INS-1059
chr1-65451997-INS-1750
chr1-65478319-INS-11990
chr1-65507197-INS-50
chr1-65507646-INS-177
chr1-65517385-INS-67
chr1-65517853-INS-136
chr1-65528854-INS-1696
chr1-65536167-INS-6594
chr1-65542287-INS-92
chr1-65546278-INS-2527
chr1-65557750-INS-1167
chr1-65602107-INS-254
chr1-65609450-INS-6437
chr1-65688181-INS-2472
chr1-65717572-INS-60
chr1-65728398-INS-15945
chr1-65738594-INS-1274
chr1-65738594-INS-3813
chr1-65750087-INS-486
chr1-65775490-INS-54
chr1-65861970-INS-50
chr1-65889076-INS-52
chr1-65902040-INS-117
chr1-65902231-INS-327
chr1-65903366-INS-2388
chr1-65925024-INS-6181
chr1-66027279-INS-55
chr1-66056488-INS-414
chr1-66087723-INS-201
chr1-66170213-INS-324
chr1-66227553-INS-378
chr1-66243011-INS-51
chr1-66243443-INS-86
chr1-66299934-INS-132
chr1-66348034-INS-6549
chr1-66355123-INS-50
chr1-66356231-INS-50
chr1-66379752-INS-103
chr1-66436447-INS-229
chr1-66461574-INS-1562
chr1-66476652-INS-68
chr1-66498471-INS-64
chr1-66525412-INS-3999
chr1-66531880-INS-50
chr1-66536120-INS-391
chr1-66538569-INS-101
chr1-66538667-INS-211
chr1-66637170-INS-4000
chr1-66786324-INS-100
chr1-66795974-INS-100
chr1-66807772-INS-1941
chr1-66848573-INS-76
chr1-66896202-INS-1534
chr1-66910703-INS-3638
chr1-66912191-INS-6517
chr1-66923924-INS-444
chr1-66924231-INS-470
chr1-66942378-INS-1198
chr1-66947452-INS-7115
chr1-66967760-INS-216
chr1-66986805-INS-58
chr1-67032786-INS-582
chr1-67050919-INS-87
chr1-67089651-INS-2667
chr1-67102195-INS-116
chr1-67102904-INS-406
chr1-67108766-INS-239
chr1-67112367-INS-72

chr1-67120187-INS-458
chr1-67216509-INS-1687
chr1-67259544-INS-4522
chr1-67277605-INS-51
chr1-67372573-INS-56
chr1-67409206-INS-6479
chr1-67425717-INS-99
chr1-67456102-INS-997
chr1-67467679-INS-57
chr1-67478404-INS-245
chr1-67506739-INS-220
chr1-67574312-INS-5787
chr1-67619976-INS-755
chr1-67626592-INS-208
chr1-67633793-INS-4069
chr1-67647142-INS-150
chr1-67647142-INS-60
chr1-67654323-INS-241
chr1-67668620-INS-1710
chr1-67676755-INS-1094
chr1-67697240-INS-502
chr1-67755051-INS-4699
chr1-67767624-INS-6128
chr1-67819465-INS-6548
chr1-67826175-INS-6209
chr1-67854003-INS-1576
chr1-67875080-INS-7640
chr1-67882305-INS-167
chr1-67905342-INS-297
chr1-67905896-INS-88
chr1-67906138-INS-138
chr1-67913412-INS-2663
chr1-67957337-INS-3336
chr1-67966707-INS-466
chr1-68019802-INS-5545
chr1-68050548-INS-5549
chr1-68200453-INS-872
chr1-68221663-INS-8576
chr1-68252834-INS-4169
chr1-68262255-INS-1318
chr1-68269947-INS-2850
chr1-68271741-INS-1254
chr1-68344076-INS-204
chr1-68346413-INS-2854
chr1-68376211-INS-66
chr1-68392991-INS-1269
chr1-68393041-INS-483
chr1-68428115-INS-50
chr1-68466179-INS-6221
chr1-68529579-INS-1492
chr1-68587835-INS-892
chr1-68592653-INS-81
chr1-68694546-INS-55
chr1-68700183-INS-289
chr1-68711009-INS-219
chr1-68746747-INS-56
chr1-68781955-INS-51
chr1-68797512-INS-66
chr1-68847002-INS-50
chr1-68929748-INS-12792
chr1-68944678-INS-2631
chr1-68956463-INS-112
chr1-68975041-INS-912
chr1-69005807-INS-408
chr1-69042290-INS-237
chr1-69048543-INS-6192
chr1-69145987-INS-76
chr1-69180323-INS-52
chr1-69185541-INS-220
chr1-69363963-INS-6538
chr1-69380915-INS-106
chr1-69414981-INS-202
chr1-69425536-INS-58
chr1-69566793-INS-80
chr1-69578868-INS-3236
chr1-69579703-INS-53
chr1-69654152-INS-195

chr1-69654152-INS-87
chr1-69735498-INS-169
chr1-69735498-INS-727
chr1-69743687-INS-200
chr1-69771073-INS-3124
chr1-69777418-INS-50
chr1-69846107-INS-2155
chr1-69854572-INS-434
chr1-69858915-INS-141
chr1-69876412-INS-1433
chr1-69880487-INS-6662
chr1-69881318-INS-1548
chr1-69953311-INS-655
chr1-69956369-INS-366
chr1-69991493-INS-374
chr1-70003747-INS-397
chr1-70023605-INS-203
chr1-70024920-INS-1020
chr1-70092743-INS-240
chr1-70113499-INS-5162
chr1-70215448-INS-63
chr1-70235964-INS-144
chr1-70242355-INS-356
chr1-70246714-INS-251
chr1-70258513-INS-208
chr1-70262786-INS-481
chr1-70286875-INS-2220
chr1-70292947-INS-79
chr1-70295131-INS-60
chr1-70338633-INS-151
chr1-70340649-INS-6875
chr1-70350300-INS-7167
chr1-70355205-INS-183
chr1-70368458-INS-6888
chr1-70368894-INS-3319
chr1-70391578-INS-3425
chr1-70402913-INS-108
chr1-70450523-INS-2152
chr1-70454559-INS-3468
chr1-70544423-INS-5845
chr1-70545897-INS-2316
chr1-70583543-INS-70
chr1-70626821-INS-678
chr1-70627615-INS-4183
chr1-70651758-INS-1516
chr1-70651758-INS-685
chr1-70659485-INS-116
chr1-70660059-INS-53
chr1-70688423-INS-1583
chr1-70690292-INS-6391
chr1-70703121-INS-1338
chr1-70714373-INS-55
chr1-70714476-INS-918
chr1-70714616-INS-96
chr1-70725818-INS-445
chr1-70798105-INS-6398
chr1-70821250-INS-104
chr1-70865615-INS-56
chr1-70865840-INS-564
chr1-70909990-INS-139
chr1-70911189-INS-51
chr1-70943334-INS-604
chr1-70975285-INS-8138
chr1-70988505-INS-188
chr1-71015970-INS-61
chr1-71068704-INS-60
chr1-71096144-INS-163
chr1-71163305-INS-7052
chr1-71202933-INS-84
chr1-71208534-INS-65
chr1-71288912-INS-50
chr1-71310471-INS-100
chr1-71322976-INS-50
chr1-71345357-INS-50
chr1-71373394-INS-166
chr1-71396032-INS-995
chr1-71403498-INS-238

chr1-71464354-INS-6702
chr1-71539668-INS-57
chr1-71584806-INS-61
chr1-71594715-INS-9100
chr1-71612915-INS-179
chr1-71621725-INS-50
chr1-71672922-INS-172
chr1-71673368-INS-307
chr1-71673376-INS-76
chr1-71707706-INS-2319
chr1-71725969-INS-6876
chr1-71732921-INS-61
chr1-71733739-INS-207
chr1-71738330-INS-938
chr1-71751472-INS-6385
chr1-71764986-INS-9049
chr1-71783275-INS-90
chr1-71806747-INS-104
chr1-71807196-INS-70
chr1-71813408-INS-50
chr1-71817738-INS-6439
chr1-71820207-INS-52
chr1-71825892-INS-197
chr1-71831248-INS-144
chr1-71848959-INS-59
chr1-71898073-INS-542
chr1-71904985-INS-492
chr1-71947776-INS-2489
chr1-71990393-INS-64
chr1-71992086-INS-6708
chr1-72011873-INS-66
chr1-72032301-INS-69
chr1-72088559-INS-147
chr1-72119856-INS-1692
chr1-72120371-INS-72
chr1-72130499-INS-135
chr1-72200821-INS-130
chr1-72204553-INS-209
chr1-72210188-INS-528
chr1-72228279-INS-196
chr1-72229173-INS-204
chr1-72231456-INS-195
chr1-72231812-INS-560
chr1-72245633-INS-195
chr1-72257147-INS-219
chr1-72269802-INS-209
chr1-72281516-INS-117
chr1-72282119-INS-505
chr1-72285892-INS-2098
chr1-72287519-INS-201
chr1-72293024-INS-232
chr1-72314969-INS-52
chr1-72319743-INS-3179
chr1-72343258-INS-222
chr1-72361328-INS-78
chr1-72381665-INS-54
chr1-72420311-INS-58
chr1-72420363-INS-136
chr1-72420891-INS-129
chr1-72428570-INS-161
chr1-72428932-INS-69
chr1-72459085-INS-196
chr1-72644727-INS-82
chr1-72661829-INS-54
chr1-72661883-INS-158
chr1-72677364-INS-2630
chr1-72689110-INS-107
chr1-72722869-INS-6653
chr1-72753106-INS-219
chr1-72760668-INS-73
chr1-72764103-INS-272
chr1-72767134-INS-50
chr1-72773499-INS-437
chr1-72810363-INS-80
chr1-72810366-INS-2028
chr1-72842094-INS-518
chr1-72842149-INS-9807

chr1-72846825-INS-233
chr1-72852933-INS-54
chr1-72854367-INS-208
chr1-72860437-INS-3515
chr1-72865111-INS-53
chr1-72885632-INS-131
chr1-72885858-INS-84
chr1-72925734-INS-55
chr1-72934747-INS-62
chr1-72936500-INS-256
chr1-72936911-INS-203
chr1-72945355-INS-434
chr1-72945358-INS-953
chr1-72957385-INS-7306
chr1-72998047-INS-280
chr1-73003598-INS-94
chr1-73022088-INS-204
chr1-73047002-INS-63
chr1-73082185-INS-89
chr1-73109002-INS-194
chr1-73130463-INS-73
chr1-73136427-INS-6261
chr1-73144041-INS-166
chr1-73144139-INS-443
chr1-73160310-INS-88
chr1-73175811-INS-50
chr1-73179310-INS-348
chr1-73179423-INS-169
chr1-73186796-INS-2482
chr1-73243437-INS-619
chr1-73260358-INS-84
chr1-73272147-INS-65
chr1-73287629-INS-192
chr1-73339762-INS-108
chr1-73343703-INS-6711
chr1-73392800-INS-71
chr1-73501629-INS-1313
chr1-73504602-INS-7137
chr1-73529824-INS-1096
chr1-73533999-INS-6690
chr1-73552693-INS-2984
chr1-73605954-INS-184
chr1-73614290-INS-113
chr1-73614741-INS-72
chr1-73626571-INS-165
chr1-73632022-INS-180
chr1-73671368-INS-879
chr1-73695026-INS-3049
chr1-73705392-INS-68
chr1-73713586-INS-60
chr1-73728394-INS-170
chr1-73743036-INS-474
chr1-73801265-INS-1837
chr1-73821928-INS-512
chr1-73821928-INS-5710
chr1-73821992-INS-1113
chr1-73822021-INS-12996
chr1-73877391-INS-84
chr1-73889276-INS-2215
chr1-73916020-INS-95
chr1-73942947-INS-992
chr1-74033914-INS-62
chr1-74116840-INS-50
chr1-74119834-INS-202
chr1-74147113-INS-78
chr1-74154638-INS-204
chr1-74155490-INS-212
chr1-74179877-INS-80
chr1-74204328-INS-642
chr1-74218503-INS-50
chr1-74223599-INS-214
chr1-74226234-INS-72
chr1-74230598-INS-191
chr1-74247373-INS-50
chr1-74258275-INS-50
chr1-74261097-INS-118
chr1-74264827-INS-202

chr1-74299228-INS-200
chr1-74311229-INS-216
chr1-74313237-INS-230
chr1-74328241-INS-52
chr1-74331352-INS-205
chr1-74348174-INS-50
chr1-74360031-INS-66
chr1-74360241-INS-1113
chr1-74384270-INS-64
chr1-74400271-INS-432
chr1-74442706-INS-52
chr1-74478064-INS-50
chr1-74490514-INS-215
chr1-74504325-INS-89
chr1-74540744-INS-96
chr1-74578783-INS-73
chr1-74585454-INS-80
chr1-74592972-INS-69
chr1-74645024-INS-68
chr1-74661027-INS-54
chr1-74689584-INS-104
chr1-74691714-INS-213
chr1-74778516-INS-72
chr1-74803995-INS-1501
chr1-74814427-INS-50
chr1-74874629-INS-290
chr1-74883928-INS-246
chr1-74884221-INS-50
chr1-74900954-INS-148
chr1-74934379-INS-54
chr1-74995425-INS-54
chr1-75015548-INS-189
chr1-75024825-INS-1121
chr1-75058570-INS-96
chr1-75085038-INS-143
chr1-75095377-INS-764
chr1-75128847-INS-169
chr1-75128867-INS-61
chr1-75143493-INS-54
chr1-75179724-INS-327
chr1-75256697-INS-141
chr1-75265255-INS-505
chr1-75281843-INS-68
chr1-75349621-INS-435
chr1-75390439-INS-50
chr1-75391432-INS-108
chr1-75437648-INS-303
chr1-75549276-INS-70
chr1-75560435-INS-228
chr1-75561002-INS-61
chr1-75577627-INS-53
chr1-75604822-INS-62
chr1-75609832-INS-1324
chr1-75619290-INS-105
chr1-75639258-INS-6694
chr1-75678435-INS-62
chr1-75684040-INS-59
chr1-75702685-INS-60
chr1-75709681-INS-72
chr1-75727386-INS-3079
chr1-75743962-INS-6532
chr1-75799815-INS-52
chr1-75814191-INS-51
chr1-75814479-INS-72
chr1-75840022-INS-7420
chr1-75863328-INS-124
chr1-75887561-INS-54
chr1-75912970-INS-57
chr1-75948370-INS-894
chr1-76029395-INS-161
chr1-76044400-INS-5553
chr1-76065699-INS-54
chr1-76096950-INS-140
chr1-76108077-INS-211
chr1-76108596-INS-70
chr1-76115374-INS-72
chr1-76147954-INS-170

chr1-76147974-INS-641
chr1-76170834-INS-89
chr1-76206306-INS-225
chr1-76216504-INS-280
chr1-76235723-INS-196
chr1-76376356-INS-70
chr1-76388116-INS-132
chr1-76418992-INS-146
chr1-76462066-INS-51
chr1-76464504-INS-52
chr1-76491011-INS-124
chr1-76522455-INS-117
chr1-76546891-INS-391
chr1-76547386-INS-726
chr1-76634811-INS-63
chr1-76644241-INS-58
chr1-76709963-INS-51
chr1-76810262-INS-288
chr1-76810334-INS-124
chr1-76825114-INS-215
chr1-76895485-INS-1568
chr1-76906724-INS-800
chr1-76907113-INS-1062
chr1-76911037-INS-112
chr1-76930003-INS-6412
chr1-76940201-INS-731
chr1-77015491-INS-7009
chr1-77037331-INS-6735
chr1-77083446-INS-74
chr1-77085941-INS-253
chr1-77116953-INS-82
chr1-77156929-INS-2805
chr1-77210719-INS-55
chr1-77215224-INS-7723
chr1-77307265-INS-148
chr1-77328362-INS-86
chr1-77331101-INS-88
chr1-77349509-INS-70
chr1-77415912-INS-52
chr1-77448859-INS-138
chr1-77451058-INS-188
chr1-77550609-INS-7416
chr1-77555838-INS-54
chr1-77562012-INS-510
chr1-77579205-INS-1446
chr1-77580157-INS-108
chr1-77582813-INS-120
chr1-77614958-INS-6460
chr1-77670938-INS-321
chr1-77677401-INS-6464
chr1-77680905-INS-4829
chr1-77700083-INS-6756
chr1-77747019-INS-78
chr1-77748376-INS-129
chr1-77757755-INS-56
chr1-77782865-INS-90
chr1-77783002-INS-202
chr1-77798649-INS-68
chr1-77835304-INS-63
chr1-77845421-INS-52
chr1-77854509-INS-3868
chr1-77854933-INS-185
chr1-77868260-INS-58
chr1-77898725-INS-2102
chr1-77948138-INS-200
chr1-77952623-INS-1170
chr1-77965014-INS-2908
chr1-77965672-INS-174
chr1-78022811-INS-3503
chr1-78065921-INS-224
chr1-78321270-INS-219
chr1-78480778-INS-290
chr1-78481339-INS-470
chr1-78487106-INS-50
chr1-78488913-INS-437
chr1-78532200-INS-200
chr1-78532423-INS-388

chr1-78540581-INS-60
chr1-78551178-INS-6772
chr1-78555795-INS-3253
chr1-78559839-INS-150
chr1-78574044-INS-56
chr1-78579687-INS-1404
chr1-78594404-INS-76
chr1-78604438-INS-84
chr1-78621102-INS-62
chr1-78624708-INS-84
chr1-78633146-INS-4934
chr1-78633150-INS-50
chr1-78668214-INS-50
chr1-78691535-INS-78
chr1-78703616-INS-112
chr1-78716127-INS-96
chr1-78786049-INS-125
chr1-78786064-INS-376
chr1-78786498-INS-287
chr1-78801105-INS-212
chr1-78820463-INS-1548
chr1-78854297-INS-55
chr1-78854304-INS-130
chr1-78855063-INS-60
chr1-78857930-INS-6513
chr1-78870837-INS-7235
chr1-78950779-INS-1961
chr1-78974458-INS-2750
chr1-79047494-INS-5650
chr1-79066572-INS-6561
chr1-79129484-INS-1535
chr1-79143700-INS-6706
chr1-79181452-INS-1441
chr1-79185381-INS-50
chr1-79259915-INS-651
chr1-79270155-INS-2334
chr1-79299931-INS-673
chr1-79300503-INS-1074
chr1-79351387-INS-1364
chr1-79351398-INS-437
chr1-79396824-INS-60
chr1-79442207-INS-131
chr1-79475571-INS-1217
chr1-79492294-INS-91
chr1-79504813-INS-54
chr1-79505727-INS-99
chr1-79540707-INS-62
chr1-79562426-INS-72
chr1-79563282-INS-146
chr1-79563282-INS-51
chr1-79569728-INS-261
chr1-79639922-INS-50
chr1-79663263-INS-255
chr1-79663780-INS-230
chr1-79692534-INS-53
chr1-79705255-INS-291
chr1-79727790-INS-875
chr1-79727815-INS-400
chr1-79741811-INS-7116
chr1-79822509-INS-109
chr1-79843936-INS-128
chr1-79854015-INS-209
chr1-79895053-INS-72
chr1-79967207-INS-98
chr1-79972207-INS-1160
chr1-80027329-INS-421
chr1-80133091-INS-131
chr1-80147900-INS-62
chr1-80148284-INS-56
chr1-80149875-INS-56
chr1-80156225-INS-84
chr1-80378845-INS-1489
chr1-80402833-INS-6639
chr1-80419843-INS-161
chr1-80420617-INS-287
chr1-80463306-INS-328
chr1-80465621-INS-308

chr1-80465838-INS-393
chr1-80466076-INS-59
chr1-80484575-INS-51
chr1-80487098-INS-6993
chr1-80495348-INS-844
chr1-80514943-INS-826
chr1-80525978-INS-146
chr1-80526055-INS-51
chr1-80540667-INS-163
chr1-80565800-INS-197
chr1-80593936-INS-2712
chr1-80595429-INS-222
chr1-80605416-INS-55
chr1-80650301-INS-143
chr1-80660375-INS-66
chr1-80660445-INS-89
chr1-80683152-INS-201
chr1-80684435-INS-92
chr1-80685092-INS-51
chr1-80692825-INS-78
chr1-80692852-INS-235
chr1-80695825-INS-271
chr1-80728470-INS-98
chr1-80802459-INS-100
chr1-80819687-INS-1286
chr1-80828341-INS-1588
chr1-80916325-INS-4047
chr1-80958049-INS-50
chr1-81026320-INS-146
chr1-81106173-INS-379
chr1-81106417-INS-64
chr1-81129829-INS-152
chr1-81150205-INS-120
chr1-81177558-INS-1535
chr1-81181720-INS-171
chr1-81229646-INS-1595
chr1-81286537-INS-5304
chr1-81288106-INS-60
chr1-81291140-INS-144
chr1-81291143-INS-53
chr1-81351106-INS-6632
chr1-81387803-INS-139
chr1-81389276-INS-104
chr1-81503810-INS-1309
chr1-81508293-INS-1492
chr1-81522712-INS-165
chr1-81582902-INS-96
chr1-81584328-INS-254
chr1-81611955-INS-50
chr1-81660180-INS-6994
chr1-81720611-INS-2812
chr1-81743996-INS-74
chr1-81760090-INS-92
chr1-81763442-INS-6146
chr1-81811951-INS-4374
chr1-81830946-INS-72
chr1-81841690-INS-435
chr1-81879229-INS-212
chr1-82009905-INS-58
chr1-82014011-INS-53
chr1-82025999-INS-98
chr1-82035431-INS-51
chr1-82037990-INS-99
chr1-82052574-INS-290
chr1-82052806-INS-58
chr1-82100054-INS-7264
chr1-82117138-INS-108
chr1-82130158-INS-69
chr1-82322311-INS-6564
chr1-82489973-INS-876
chr1-82503052-INS-311
chr1-82545487-INS-56
chr1-82601654-INS-1573
chr1-82625451-INS-313
chr1-82662950-INS-6283
chr1-82808324-INS-63
chr1-82827806-INS-3988

chr1-82829600-INS-237
chr1-82835238-INS-377
chr1-82835360-INS-174
chr1-82853176-INS-1635
chr1-82867556-INS-54
chr1-82886260-INS-62
chr1-82910035-INS-58
chr1-82923179-INS-367
chr1-82938044-INS-72
chr1-83021636-INS-141
chr1-83022742-INS-6323
chr1-83059578-INS-7264
chr1-83092664-INS-90
chr1-83104708-INS-540
chr1-83148740-INS-220
chr1-83149007-INS-59
chr1-83169632-INS-1277
chr1-83186715-INS-54
chr1-83219323-INS-50
chr1-83276969-INS-5950
chr1-83302924-INS-50
chr1-83322739-INS-514
chr1-83330683-INS-52
chr1-83360261-INS-132
chr1-83410868-INS-476
chr1-83442896-INS-58
chr1-83444271-INS-1888
chr1-83455771-INS-1056
chr1-83471574-INS-750
chr1-83480095-INS-3534
chr1-83500268-INS-1140
chr1-83516902-INS-6180
chr1-83518532-INS-9520
chr1-83668705-INS-342
chr1-83689978-INS-91
chr1-83709046-INS-502
chr1-83773938-INS-104
chr1-83792212-INS-3657
chr1-83808843-INS-611
chr1-83823086-INS-75
chr1-83823146-INS-91
chr1-83972133-INS-56
chr1-83996408-INS-78
chr1-84068049-INS-129
chr1-84071898-INS-1558
chr1-84081678-INS-1200
chr1-84192978-INS-138
chr1-84278566-INS-51
chr1-84286729-INS-92
chr1-84287273-INS-136
chr1-84292904-INS-202
chr1-84310465-INS-7021
chr1-84357504-INS-212
chr1-84394993-INS-5563
chr1-84405285-INS-60
chr1-84421733-INS-1983
chr1-84438787-INS-59
chr1-84486625-INS-205
chr1-84498155-INS-345
chr1-84498163-INS-109
chr1-84504650-INS-442
chr1-84530378-INS-3620
chr1-84546939-INS-195
chr1-84554572-INS-117
chr1-84597977-INS-50
chr1-84598699-INS-58
chr1-84636219-INS-54
chr1-84638435-INS-100
chr1-84638435-INS-579
chr1-84653789-INS-74
chr1-84653923-INS-60
chr1-84653926-INS-614
chr1-84655483-INS-79
chr1-84665373-INS-2548
chr1-84705031-INS-93
chr1-84768354-INS-2468
chr1-84776207-INS-221

chr1-84778315-INS-123
chr1-84810617-INS-54
chr1-84837199-INS-102
chr1-84843424-INS-258
chr1-84856305-INS-4826
chr1-84856925-INS-209
chr1-84862336-INS-222
chr1-84866460-INS-201
chr1-84893957-INS-222
chr1-84910123-INS-53
chr1-84932455-INS-56
chr1-84934755-INS-51
chr1-84945554-INS-203
chr1-84968254-INS-50
chr1-84998750-INS-617
chr1-85013168-INS-53
chr1-85044797-INS-201
chr1-85049780-INS-162
chr1-85084126-INS-53
chr1-85096584-INS-235
chr1-85096584-INS-75
chr1-85096663-INS-972
chr1-85275307-INS-4973
chr1-85303528-INS-50
chr1-85427310-INS-498
chr1-85475659-INS-98
chr1-85539994-INS-1269
chr1-85540675-INS-50
chr1-85549167-INS-974
chr1-85557752-INS-601
chr1-85595733-INS-990
chr1-85650877-INS-88
chr1-85654585-INS-127
chr1-85696107-INS-165
chr1-85696107-INS-53
chr1-85702312-INS-17520
chr1-85708915-INS-1087
chr1-85735909-INS-152
chr1-85742200-INS-67
chr1-85770097-INS-106
chr1-85853461-INS-99
chr1-85874470-INS-121
chr1-85879316-INS-64
chr1-85884118-INS-52
chr1-85899429-INS-76
chr1-85923208-INS-88
chr1-85942880-INS-168
chr1-85990982-INS-206
chr1-86051628-INS-214
chr1-86102803-INS-62
chr1-86106767-INS-50
chr1-86302026-INS-2672
chr1-86321713-INS-67
chr1-86352412-INS-55
chr1-86378985-INS-2000
chr1-86383727-INS-55
chr1-86383793-INS-206
chr1-86389098-INS-168
chr1-86409231-INS-76
chr1-86416857-INS-52
chr1-86419454-INS-83
chr1-86504820-INS-210
chr1-86541599-INS-52
chr1-86557028-INS-56
chr1-86561126-INS-169
chr1-86613357-INS-101
chr1-86618641-INS-80
chr1-86724548-INS-68
chr1-86758730-INS-391
chr1-86820092-INS-262
chr1-86902705-INS-191
chr1-86916657-INS-68
chr1-86917010-INS-60
chr1-86967569-INS-238
chr1-86988182-INS-50
chr1-87013524-INS-55
chr1-87043222-INS-52

chr1-87060987-INS-50
chr1-87072695-INS-80
chr1-87174420-INS-51
chr1-87182881-INS-88
chr1-87196901-INS-133
chr1-87362246-INS-60
chr1-87367276-INS-89
chr1-87382662-INS-74
chr1-87397443-INS-50
chr1-87447053-INS-51
chr1-87468999-INS-55
chr1-87529009-INS-165
chr1-87536027-INS-6369
chr1-87567286-INS-205
chr1-87573383-INS-3000
chr1-87646062-INS-58
chr1-87658834-INS-102
chr1-87700604-INS-70
chr1-87738363-INS-114
chr1-87749433-INS-60
chr1-87763481-INS-54
chr1-87763594-INS-72
chr1-87801492-INS-106
chr1-87884703-INS-108
chr1-87974037-INS-110
chr1-87974104-INS-4473
chr1-87984575-INS-54
chr1-87991500-INS-17266
chr1-87992287-INS-3533
chr1-88017367-INS-109
chr1-88070941-INS-88
chr1-88091100-INS-3333
chr1-88096049-INS-204
chr1-88152528-INS-3124
chr1-88157308-INS-325
chr1-88175087-INS-553
chr1-88203908-INS-438
chr1-88219237-INS-591
chr1-88250363-INS-50
chr1-88254962-INS-286
chr1-88297939-INS-70
chr1-88297943-INS-195
chr1-88298147-INS-77
chr1-88308586-INS-50
chr1-88313689-INS-420
chr1-88313743-INS-120
chr1-88355269-INS-6475
chr1-88376917-INS-4747
chr1-88385297-INS-54
chr1-88466443-INS-416
chr1-88517580-INS-209
chr1-88521012-INS-2836
chr1-88554599-INS-3341
chr1-88560574-INS-52
chr1-88586025-INS-68
chr1-88653862-INS-58
chr1-88690714-INS-1736
chr1-88768520-INS-71
chr1-88778346-INS-1050
chr1-88788544-INS-88
chr1-88790580-INS-51
chr1-88794216-INS-68
chr1-88816616-INS-6608
chr1-88824590-INS-51
chr1-88842433-INS-6403
chr1-88857837-INS-96
chr1-88857982-INS-115
chr1-88900987-INS-134
chr1-88913511-INS-210
chr1-88926249-INS-1468
chr1-88929070-INS-2506
chr1-88949997-INS-4458
chr1-88960128-INS-2235
chr1-88986869-INS-109
chr1-89109321-INS-90
chr1-89165715-INS-53
chr1-89204546-INS-473

chr1-89222700-INS-2830
chr1-89257599-INS-486
chr1-89270055-INS-207
chr1-89270409-INS-468
chr1-89287981-INS-135
chr1-89289821-INS-2735
chr1-89298269-INS-684
chr1-89310801-INS-84
chr1-89423517-INS-56
chr1-89516573-INS-51
chr1-89517459-INS-805
chr1-89544558-INS-4990
chr1-89549129-INS-153
chr1-89551240-INS-54
chr1-89559294-INS-88
chr1-89607326-INS-156
chr1-90052172-INS-128
chr1-90056070-INS-81
chr1-90061070-INS-186
chr1-90061382-INS-749
chr1-90108344-INS-72
chr1-90128398-INS-88
chr1-90145088-INS-81
chr1-90168378-INS-50
chr1-90180069-INS-50
chr1-90189416-INS-51
chr1-90193629-INS-64
chr1-90238339-INS-342
chr1-90245355-INS-50
chr1-90249586-INS-76
chr1-90250265-INS-999
chr1-90255747-INS-51
chr1-90268957-INS-54
chr1-90284697-INS-3590
chr1-90284749-INS-304
chr1-90303411-INS-90
chr1-90441770-INS-64
chr1-90444276-INS-587
chr1-90469820-INS-399
chr1-90479040-INS-282
chr1-90486254-INS-118
chr1-90549314-INS-491
chr1-90550728-INS-5077
chr1-90571225-INS-87
chr1-90619372-INS-2873
chr1-90625559-INS-951
chr1-90664820-INS-64
chr1-90684226-INS-144
chr1-90728749-INS-87
chr1-90736424-INS-81
chr1-90736628-INS-160
chr1-90736672-INS-74
chr1-90764210-INS-50
chr1-90772594-INS-77
chr1-90775850-INS-5223
chr1-90818232-INS-70
chr1-90821707-INS-84
chr1-90841841-INS-136
chr1-90877755-INS-276
chr1-90892542-INS-85
chr1-90959187-INS-3633
chr1-91052889-INS-50
chr1-91066044-INS-160
chr1-91140540-INS-73
chr1-91142440-INS-161
chr1-91199825-INS-58
chr1-91365344-INS-106
chr1-91369928-INS-205
chr1-91369928-INS-82
chr1-91466156-INS-107
chr1-91481147-INS-196
chr1-91536589-INS-90
chr1-91633805-INS-382
chr1-91675207-INS-168
chr1-91747656-INS-188
chr1-91763216-INS-211
chr1-91804310-INS-64

chr1-91815356-INS-50
chr1-91830564-INS-204
chr1-91836906-INS-100
chr1-91939638-INS-56
chr1-92068844-INS-52
chr1-92080414-INS-335
chr1-92117944-INS-92
chr1-92166873-INS-52
chr1-92170181-INS-54
chr1-92193945-INS-5366
chr1-92259066-INS-2362
chr1-92298391-INS-60
chr1-92307703-INS-53
chr1-92434908-INS-18491
chr1-92452217-INS-153
chr1-92477468-INS-2783
chr1-92478355-INS-1876
chr1-92521403-INS-572
chr1-92521671-INS-986
chr1-92540811-INS-181
chr1-92544109-INS-546
chr1-92567271-INS-52
chr1-92584556-INS-1394
chr1-92588583-INS-64
chr1-92594048-INS-6474
chr1-92610612-INS-268
chr1-92673595-INS-233
chr1-92741996-INS-112
chr1-92742001-INS-606
chr1-92743182-INS-55
chr1-92792060-INS-12880
chr1-92792060-INS-6383
chr1-92818746-INS-96
chr1-92819215-INS-191
chr1-92822226-INS-60
chr1-92853237-INS-200
chr1-92947066-INS-167
chr1-92973186-INS-72
chr1-92990704-INS-120
chr1-93040945-INS-118
chr1-93047725-INS-150
chr1-93049000-INS-855
chr1-93168602-INS-175
chr1-93172519-INS-54
chr1-93217504-INS-1082
chr1-93217504-INS-500
chr1-93235421-INS-963
chr1-93307479-INS-79
chr1-93348723-INS-51
chr1-93492341-INS-52
chr1-93574435-INS-6748
chr1-93580188-INS-396
chr1-93610644-INS-227
chr1-93619333-INS-3114
chr1-93640551-INS-8942
chr1-93659961-INS-135
chr1-93670411-INS-58
chr1-93745033-INS-120
chr1-93764210-INS-95
chr1-93764952-INS-52
chr1-93805885-INS-304
chr1-93813520-INS-50
chr1-93848940-INS-470
chr1-93852606-INS-362
chr1-93863463-INS-205
chr1-93907129-INS-275
chr1-93909075-INS-91
chr1-93958960-INS-112
chr1-93972170-INS-2580
chr1-93992642-INS-1239
chr1-94003421-INS-4358
chr1-94071535-INS-54
chr1-94087335-INS-52
chr1-94173237-INS-9719
chr1-94174490-INS-1770
chr1-94188996-INS-101
chr1-94214321-INS-91

chr1-94239013-INS-50
chr1-94354646-INS-74
chr1-94365397-INS-174
chr1-94431007-INS-50
chr1-94452298-INS-70
chr1-94472472-INS-6308
chr1-94485301-INS-105
chr1-94572954-INS-52
chr1-94595638-INS-52
chr1-94595734-INS-53
chr1-94624653-INS-2147
chr1-94658901-INS-65
chr1-94697046-INS-6530
chr1-94759933-INS-2344
chr1-94800235-INS-150
chr1-94817520-INS-142
chr1-94856266-INS-52
chr1-94894714-INS-4496
chr1-94990703-INS-76
chr1-95020609-INS-1293
chr1-95074934-INS-6604
chr1-95078346-INS-58
chr1-95126159-INS-2073
chr1-95168512-INS-125
chr1-95179595-INS-161
chr1-95183419-INS-64
chr1-95272664-INS-51
chr1-95306573-INS-6683
chr1-95320709-INS-51
chr1-95340475-INS-16655
chr1-95393602-INS-194
chr1-95401986-INS-173
chr1-95401990-INS-73
chr1-95402216-INS-839
chr1-95403236-INS-55
chr1-95415688-INS-263
chr1-95422890-INS-72
chr1-95445577-INS-206
chr1-95537404-INS-318
chr1-95548831-INS-1668
chr1-95616154-INS-1500
chr1-95636129-INS-173
chr1-95670265-INS-208
chr1-95672250-INS-6252
chr1-95703169-INS-510
chr1-95706235-INS-90
chr1-95722508-INS-107
chr1-95726669-INS-5205
chr1-95770504-INS-1855
chr1-95786663-INS-795
chr1-95819095-INS-69
chr1-95892471-INS-569
chr1-95927020-INS-6327
chr1-95991972-INS-244
chr1-96034926-INS-69
chr1-96038184-INS-281
chr1-96055646-INS-179
chr1-96079100-INS-102
chr1-96106136-INS-449
chr1-96119089-INS-208
chr1-96180780-INS-132
chr1-96185369-INS-2377
chr1-96206779-INS-805
chr1-96236607-INS-254
chr1-96275904-INS-247
chr1-96298573-INS-124
chr1-96306289-INS-5537
chr1-96313441-INS-85
chr1-96315794-INS-6186
chr1-96376173-INS-134
chr1-96422213-INS-64
chr1-96436122-INS-2495
chr1-96444369-INS-1396
chr1-96524579-INS-7452
chr1-96529146-INS-52
chr1-96583238-INS-52
chr1-96586823-INS-223

chr1-96664633-INS-175
chr1-96685388-INS-80
chr1-96686258-INS-1470
chr1-96687375-INS-443
chr1-96708515-INS-58
chr1-96777681-INS-51
chr1-96777688-INS-408
chr1-96854693-INS-6252
chr1-96883961-INS-6493
chr1-96886880-INS-136
chr1-96918195-INS-57
chr1-97002737-INS-200
chr1-97305786-INS-50
chr1-97366007-INS-386
chr1-97405234-INS-672
chr1-97421421-INS-700
chr1-97440574-INS-201
chr1-97457864-INS-2168
chr1-97508438-INS-50
chr1-97548840-INS-54
chr1-97595841-INS-6308
chr1-97604965-INS-1044
chr1-97714985-INS-412
chr1-97741172-INS-136
chr1-97741175-INS-66
chr1-97810534-INS-793
chr1-97814028-INS-412
chr1-97820522-INS-8456
chr1-97833852-INS-202
chr1-97842927-INS-85
chr1-97847233-INS-62
chr1-97847326-INS-279
chr1-97862738-INS-807
chr1-97862745-INS-3885
chr1-97909523-INS-140
chr1-97934899-INS-2790
chr1-97974587-INS-6710
chr1-97980564-INS-112
chr1-97988571-INS-6493
chr1-97998397-INS-60
chr1-98044510-INS-62
chr1-98048680-INS-54
chr1-98055767-INS-486
chr1-98070618-INS-56
chr1-98075749-INS-58
chr1-98078019-INS-6331
chr1-98091858-INS-164
chr1-98094625-INS-55
chr1-98108717-INS-155
chr1-98114001-INS-136
chr1-98114001-INS-51
chr1-98116635-INS-3688
chr1-98122811-INS-210
chr1-98136849-INS-73
chr1-98146095-INS-239
chr1-98174046-INS-100
chr1-98178636-INS-2398
chr1-98199285-INS-76
chr1-98233162-INS-50
chr1-98235141-INS-2114
chr1-98258387-INS-1414
chr1-98259715-INS-1452
chr1-98280516-INS-604
chr1-98294169-INS-424
chr1-98302914-INS-98
chr1-98307236-INS-393
chr1-98360284-INS-10382
chr1-98369925-INS-625
chr1-98414542-INS-86
chr1-98433414-INS-3268
chr1-98445165-INS-69
chr1-98567270-INS-1042
chr1-98591077-INS-265
chr1-98608246-INS-68
chr1-98620425-INS-9357
chr1-98640460-INS-166
chr1-98641243-INS-132

chr1-98646194-INS-5453
chr1-98668111-INS-75
chr1-98735669-INS-6291
chr1-98743395-INS-296
chr1-98758856-INS-92
chr1-98808006-INS-58
chr1-98886950-INS-2742
chr1-98909699-INS-922
chr1-98915894-INS-1745
chr1-98921010-INS-88
chr1-98926050-INS-2826
chr1-98929631-INS-8790
chr1-98958895-INS-1545
chr1-99008181-INS-52
chr1-99010962-INS-62
chr1-99015282-INS-78
chr1-99019825-INS-1808
chr1-99051454-INS-56
chr1-99095130-INS-1051
chr1-99138895-INS-3472
chr1-99154924-INS-6914
chr1-99212763-INS-70
chr1-99228176-INS-86
chr1-99228530-INS-665
chr1-99246138-INS-2795
chr1-99268688-INS-57
chr1-99269591-INS-179
chr1-99269967-INS-435
chr1-99306813-INS-1435
chr1-99306932-INS-56
chr1-99310273-INS-85
chr1-99346988-INS-4279
chr1-99351967-INS-106
chr1-99354431-INS-425
chr1-99355416-INS-186
chr1-99370084-INS-63
chr1-99379064-INS-442
chr1-99389247-INS-6604
chr1-99430737-INS-328
chr1-99449000-INS-950
chr1-99462191-INS-106
chr1-99468242-INS-2186
chr1-99478527-INS-100
chr1-99482633-INS-368
chr1-99503420-INS-53
chr1-99524180-INS-52
chr1-99563660-INS-535
chr1-99564358-INS-1894
chr1-99574390-INS-1999
chr1-99592615-INS-437
chr1-99599681-INS-7136
chr1-99623696-INS-150
chr1-99653802-INS-6416
chr1-99725364-INS-60
chr1-99773963-INS-54
chr1-99786146-INS-58
chr1-99789583-INS-324
chr1-99792283-INS-351
chr1-99828544-INS-3003
chr1-99848761-INS-5045
chr1-99852470-INS-597
chr1-99883088-INS-500
chr1-99885186-INS-52
chr1-99888723-INS-84
chr1-99896526-INS-215
chr1-99946586-INS-53
chr1-99989520-INS-257
chr1-99998532-INS-50
chr1-100005170-INS-207
chr1-100005673-INS-78
chr1-100026796-INS-3484
chr1-100048858-INS-7074
chr1-100067536-INS-77
chr1-100168472-INS-296
chr1-100174906-INS-299
chr1-100248973-INS-115
chr1-100288127-INS-89

chr1-100328229-INS-62
chr1-100353997-INS-78
chr1-100361581-INS-4147
chr1-100420682-INS-72
chr1-100426669-INS-3073
chr1-100459067-INS-1618
chr1-100487457-INS-201
chr1-100507786-INS-308
chr1-100507788-INS-747
chr1-100665384-INS-76
chr1-100699398-INS-52
chr1-100727106-INS-60
chr1-100743983-INS-72
chr1-100804802-INS-390
chr1-100830644-INS-1847
chr1-100846942-INS-3005
chr1-100849565-INS-187
chr1-100850192-INS-87
chr1-100859633-INS-606
chr1-100871931-INS-2781
chr1-100879205-INS-2237
chr1-100882059-INS-6470
chr1-100905025-INS-62
chr1-100999660-INS-70
chr1-101014690-INS-1291
chr1-101032813-INS-2602
chr1-101032813-INS-6503
chr1-101040772-INS-1839
chr1-101042858-INS-1050
chr1-101082256-INS-50
chr1-101167552-INS-1687
chr1-101278537-INS-135
chr1-101285110-INS-77
chr1-101285121-INS-458
chr1-101300053-INS-3889
chr1-101315293-INS-50
chr1-101332384-INS-102
chr1-101335756-INS-50
chr1-101337817-INS-100
chr1-101362391-INS-62
chr1-101366679-INS-1100
chr1-101392974-INS-326
chr1-101494406-INS-6491
chr1-101526679-INS-1382
chr1-101538501-INS-55
chr1-101539992-INS-50
chr1-101589752-INS-6234
chr1-101624120-INS-6371
chr1-101674775-INS-66
chr1-101735269-INS-50
chr1-101791504-INS-122
chr1-101812930-INS-1494
chr1-101879653-INS-137
chr1-101928483-INS-880
chr1-101958819-INS-70
chr1-101962385-INS-264
chr1-102009477-INS-195
chr1-102027792-INS-1019
chr1-102028013-INS-3207
chr1-102034118-INS-78
chr1-102060581-INS-56
chr1-102091214-INS-310
chr1-102108620-INS-2414
chr1-102201243-INS-414
chr1-102222284-INS-78
chr1-102245055-INS-440
chr1-102261954-INS-669
chr1-102282495-INS-96
chr1-102301956-INS-89
chr1-102367781-INS-143
chr1-102368739-INS-155
chr1-102374445-INS-200
chr1-102377031-INS-4798
chr1-102378583-INS-102
chr1-102425053-INS-1881
chr1-102430629-INS-102
chr1-102431496-INS-304

chr1-102432096-INS-1313
chr1-102433570-INS-6392
chr1-102467694-INS-138
chr1-102485569-INS-54
chr1-102485576-INS-207
chr1-102514944-INS-136
chr1-102538013-INS-74
chr1-102538450-INS-578
chr1-102582388-INS-100
chr1-102603698-INS-120
chr1-102628950-INS-67
chr1-102635050-INS-85
chr1-102635573-INS-50
chr1-102665211-INS-55
chr1-102694452-INS-1314
chr1-102695712-INS-54
chr1-102720338-INS-412
chr1-102725939-INS-168
chr1-102726617-INS-68
chr1-102729563-INS-1634
chr1-102777419-INS-173
chr1-102781906-INS-1222
chr1-102785224-INS-6894
chr1-102789941-INS-64
chr1-102849960-INS-50
chr1-102872924-INS-61
chr1-102938006-INS-420
chr1-102941728-INS-70
chr1-102950448-INS-1061
chr1-102952492-INS-2904
chr1-102983007-INS-1341
chr1-102987601-INS-50
chr1-102989870-INS-3214
chr1-102992591-INS-6479
chr1-103003040-INS-1026
chr1-103009178-INS-4687
chr1-103019129-INS-59
chr1-103019569-INS-50
chr1-103019791-INS-387
chr1-103019953-INS-1556
chr1-103061960-INS-108
chr1-103074304-INS-80
chr1-103087608-INS-400
chr1-103122350-INS-177
chr1-103152859-INS-1324
chr1-103207789-INS-64
chr1-103208999-INS-196
chr1-103230019-INS-78
chr1-103255816-INS-1093
chr1-103273583-INS-1492
chr1-103293395-INS-332
chr1-103308781-INS-51
chr1-103362061-INS-77
chr1-103362460-INS-1618
chr1-103363181-INS-70
chr1-103368120-INS-221
chr1-103376328-INS-1417
chr1-103384159-INS-1484
chr1-103390347-INS-58
chr1-103416375-INS-6269
chr1-103423300-INS-316
chr1-103428186-INS-387
chr1-103430256-INS-79
chr1-103507403-INS-50
chr1-103527313-INS-664
chr1-103539074-INS-826
chr1-103546587-INS-1307
chr1-103547290-INS-2479
chr1-103552554-INS-99
chr1-103553387-INS-6176
chr1-103556030-INS-1440
chr1-103577071-INS-14332
chr1-103586077-INS-70
chr1-103594144-INS-99
chr1-103602095-INS-7575
chr1-103671781-INS-454
chr1-103673680-INS-5703

chr1-103691582-INS-58
chr1-103709885-INS-6266
chr1-103719170-INS-158
chr1-103727183-INS-156
chr1-103740418-INS-112
chr1-103752327-INS-191
chr1-103780315-INS-8795
chr1-103816106-INS-96
chr1-103824527-INS-13973
chr1-103825886-INS-6591
chr1-103830220-INS-50
chr1-103835085-INS-144
chr1-103844006-INS-50
chr1-103874758-INS-54
chr1-103879177-INS-11308
chr1-103894590-INS-64
chr1-103918543-INS-105
chr1-103935897-INS-36304
chr1-103977908-INS-116
chr1-103988437-INS-892
chr1-103988504-INS-245
chr1-103989182-INS-2391
chr1-103996646-INS-23965
chr1-103996647-INS-10842
chr1-104018063-INS-88
chr1-104047503-INS-4979
chr1-104128581-INS-7133
chr1-104262677-INS-50
chr1-104270910-INS-3096
chr1-104289444-INS-57
chr1-104326203-INS-69
chr1-104357762-INS-50
chr1-104406437-INS-60
chr1-104410110-INS-66
chr1-104457274-INS-129
chr1-104480051-INS-7151
chr1-104496735-INS-116
chr1-104525977-INS-50
chr1-104527228-INS-462
chr1-104528082-INS-588
chr1-104554410-INS-60
chr1-104574131-INS-68
chr1-104589787-INS-4792
chr1-104635857-INS-214
chr1-104639839-INS-62
chr1-104651212-INS-6740
chr1-104656374-INS-54
chr1-104664664-INS-2779
chr1-104665295-INS-50
chr1-104701654-INS-180
chr1-104709551-INS-123
chr1-104709558-INS-60
chr1-104768666-INS-52
chr1-104851549-INS-2840
chr1-104887215-INS-60
chr1-104913534-INS-7112
chr1-104931553-INS-3096
chr1-104977587-INS-104
chr1-104982775-INS-2881
chr1-105001119-INS-76
chr1-105001697-INS-243
chr1-105058007-INS-160
chr1-105152555-INS-1503
chr1-105176518-INS-60
chr1-105185899-INS-6825
chr1-105191357-INS-5996
chr1-105252059-INS-2722
chr1-105275416-INS-7071
chr1-105328608-INS-197
chr1-105342037-INS-6894
chr1-105413325-INS-7116
chr1-105413600-INS-52
chr1-105413843-INS-57
chr1-105418256-INS-66
chr1-105431716-INS-52
chr1-105470762-INS-433
chr1-105609096-INS-51

chr1-105617352-INS-196
chr1-105748160-INS-69
chr1-105773115-INS-382
chr1-105780637-INS-52
chr1-105795540-INS-92
chr1-105795544-INS-254
chr1-105817891-INS-96
chr1-105818752-INS-82
chr1-105822386-INS-767
chr1-105833283-INS-52
chr1-105835014-INS-18326
chr1-105874145-INS-51
chr1-105875615-INS-61
chr1-105975828-INS-5336
chr1-105993115-INS-50
chr1-106011891-INS-3237
chr1-106012624-INS-595
chr1-106016790-INS-78
chr1-106077233-INS-52
chr1-106219155-INS-175
chr1-106260303-INS-6280
chr1-106271854-INS-212
chr1-106289841-INS-109
chr1-106337922-INS-91
chr1-106338196-INS-367
chr1-106338203-INS-143
chr1-106365266-INS-6767
chr1-106397234-INS-60
chr1-106399545-INS-55
chr1-106409195-INS-231
chr1-106413415-INS-5729
chr1-106425672-INS-6379
chr1-106427993-INS-86
chr1-106435174-INS-50
chr1-106508085-INS-70
chr1-106517959-INS-70
chr1-106521001-INS-76
chr1-106619098-INS-91
chr1-106645642-INS-449
chr1-106721808-INS-66
chr1-106760611-INS-6243
chr1-106762415-INS-155
chr1-106762415-INS-55
chr1-106857795-INS-2211
chr1-106868386-INS-166
chr1-107004759-INS-710
chr1-107017984-INS-570
chr1-107058088-INS-130
chr1-107067956-INS-66
chr1-107069037-INS-506
chr1-107073040-INS-1632
chr1-107079217-INS-486
chr1-107113198-INS-856
chr1-107125303-INS-66
chr1-107173796-INS-84
chr1-107212229-INS-446
chr1-107225627-INS-4470
chr1-107418486-INS-72
chr1-107421423-INS-51
chr1-107422915-INS-71
chr1-107457042-INS-75
chr1-107499511-INS-56
chr1-107501894-INS-7144
chr1-107507803-INS-4608
chr1-107517284-INS-68
chr1-107536103-INS-371
chr1-107572897-INS-62
chr1-107577343-INS-55
chr1-107601258-INS-498
chr1-107615217-INS-145
chr1-107615391-INS-54
chr1-107623724-INS-423
chr1-107624381-INS-234
chr1-107625935-INS-196
chr1-107646478-INS-2558
chr1-107672437-INS-324
chr1-107690215-INS-1513

chr1-107704423-INS-118
chr1-107732042-INS-52
chr1-107742355-INS-240
chr1-107771073-INS-1620
chr1-107872347-INS-2140
chr1-107873525-INS-228
chr1-107895338-INS-6643
chr1-107943502-INS-5386
chr1-107985328-INS-5542
chr1-108037041-INS-50
chr1-108040481-INS-519
chr1-108041036-INS-6845
chr1-108053864-INS-55
chr1-108054468-INS-307
chr1-108059410-INS-6416
chr1-108168924-INS-1745
chr1-108169774-INS-1663
chr1-108173980-INS-210
chr1-108187517-INS-50
chr1-108196203-INS-75
chr1-108368102-INS-6260
chr1-108368524-INS-50
chr1-108369696-INS-64
chr1-108416796-INS-490
chr1-108422976-INS-50
chr1-108439227-INS-70
chr1-108585806-INS-808
chr1-108604414-INS-225
chr1-108650321-INS-69
chr1-108671937-INS-1106
chr1-108682391-INS-50
chr1-108684355-INS-52
chr1-108697579-INS-224
chr1-108715665-INS-4471
chr1-108723473-INS-52
chr1-108723800-INS-876
chr1-108723829-INS-259
chr1-108733087-INS-526
chr1-108747536-INS-52
chr1-108753123-INS-6396
chr1-108759552-INS-64
chr1-108798799-INS-138
chr1-108805146-INS-4600
chr1-108815406-INS-512
chr1-108815436-INS-86
chr1-108815682-INS-159
chr1-108816884-INS-548
chr1-108820116-INS-1768
chr1-108853058-INS-6382
chr1-108871314-INS-62
chr1-108884758-INS-7003
chr1-108952741-INS-90
chr1-109020981-INS-6452
chr1-109037858-INS-2101
chr1-109072856-INS-973
chr1-109084880-INS-52
chr1-109158874-INS-72
chr1-109186905-INS-3184
chr1-109215402-INS-5162
chr1-109231447-INS-732
chr1-109231508-INS-60
chr1-109236781-INS-196
chr1-109287356-INS-50
chr1-109287561-INS-100
chr1-109304879-INS-254
chr1-109322871-INS-2205
chr1-109403121-INS-6905
chr1-109444065-INS-612
chr1-109447866-INS-90
chr1-109456381-INS-5705
chr1-109492375-INS-1025
chr1-109511374-INS-6357
chr1-109564061-INS-68
chr1-109642327-INS-2753
chr1-109644933-INS-4944
chr1-109695000-INS-72
chr1-109733553-INS-5646

chr1-109749817-INS-898
chr1-109769531-INS-437
chr1-109875857-INS-54
chr1-109938363-INS-6584
chr1-109952271-INS-2036
chr1-109961806-INS-1047
chr1-109968781-INS-1206
chr1-110077219-INS-1892
chr1-110087024-INS-1824
chr1-110112554-INS-90
chr1-110202969-INS-50
chr1-110301949-INS-50
chr1-110361963-INS-3647
chr1-110398239-INS-4379
chr1-110420583-INS-4485
chr1-110425325-INS-6391
chr1-110449432-INS-6354
chr1-110483859-INS-52
chr1-110509585-INS-6671
chr1-110539964-INS-1505
chr1-110540725-INS-4678
chr1-110542435-INS-497
chr1-110552988-INS-51
chr1-110565274-INS-58
chr1-110577858-INS-54
chr1-110589191-INS-54
chr1-110590577-INS-6433
chr1-110698400-INS-74
chr1-110731877-INS-6649
chr1-110742189-INS-301
chr1-110744159-INS-6552
chr1-110747564-INS-6658
chr1-110757381-INS-50
chr1-110759540-INS-52
chr1-110761766-INS-50
chr1-110810468-INS-3621
chr1-110917507-INS-1135
chr1-111048345-INS-2999
chr1-111054092-INS-261
chr1-111096408-INS-80
chr1-111111348-INS-136
chr1-111152005-INS-500
chr1-111246643-INS-50
chr1-111270431-INS-50
chr1-111273030-INS-608
chr1-111273365-INS-108
chr1-111301416-INS-66
chr1-111349590-INS-73
chr1-111426945-INS-419
chr1-111448615-INS-176
chr1-111470471-INS-86
chr1-111492609-INS-3532
chr1-111535903-INS-1300
chr1-111542969-INS-61
chr1-111554479-INS-64
chr1-111591653-INS-67
chr1-111656523-INS-1296
chr1-111660194-INS-437
chr1-111671889-INS-6175
chr1-111682017-INS-74
chr1-111682868-INS-143
chr1-111691624-INS-593
chr1-111720764-INS-7192
chr1-111721816-INS-88
chr1-111737354-INS-76
chr1-111743847-INS-3223
chr1-111746624-INS-4183
chr1-111778516-INS-2364
chr1-111783106-INS-66
chr1-111821396-INS-6652
chr1-111850150-INS-4116
chr1-111868878-INS-7337
chr1-111880630-INS-3017
chr1-111900939-INS-56
chr1-111903574-INS-1487
chr1-111906519-INS-704
chr1-111921511-INS-2730

chr1-111982887-INS-2983
chr1-112006975-INS-1169
chr1-112021645-INS-13325
chr1-112022412-INS-2086
chr1-112046721-INS-7296
chr1-112074371-INS-84
chr1-112081172-INS-6453
chr1-112090306-INS-1695
chr1-112112052-INS-52
chr1-112119738-INS-332
chr1-112186422-INS-613
chr1-112213739-INS-874
chr1-112239389-INS-50
chr1-112265932-INS-83
chr1-112273064-INS-69
chr1-112289954-INS-3613
chr1-112320486-INS-158
chr1-112321391-INS-52
chr1-112321745-INS-116
chr1-112341791-INS-3035
chr1-112350170-INS-133
chr1-112352779-INS-953
chr1-112355192-INS-2791
chr1-112362839-INS-110
chr1-112363054-INS-50
chr1-112376383-INS-1894
chr1-112392657-INS-6495
chr1-112416238-INS-52
chr1-112420501-INS-7115
chr1-112424633-INS-6270
chr1-112434558-INS-50
chr1-112443794-INS-6256
chr1-112467158-INS-50
chr1-112545674-INS-85
chr1-112555802-INS-176
chr1-112563934-INS-6442
chr1-112622640-INS-436
chr1-112653816-INS-3364
chr1-112674733-INS-2146
chr1-112685926-INS-151
chr1-112699091-INS-50
chr1-112768325-INS-1122
chr1-112770130-INS-527
chr1-112840100-INS-57
chr1-112853897-INS-699
chr1-112880845-INS-1948
chr1-112940927-INS-2933
chr1-112953328-INS-70
chr1-113034217-INS-68
chr1-113070631-INS-446
chr1-113071378-INS-50
chr1-113072066-INS-208
chr1-113077495-INS-421
chr1-113082366-INS-781
chr1-113088713-INS-60
chr1-113101780-INS-50
chr1-113133047-INS-6662
chr1-113148900-INS-50
chr1-113151924-INS-3245
chr1-113164144-INS-117
chr1-113166168-INS-259
chr1-113181547-INS-1320
chr1-113232655-INS-216
chr1-113234722-INS-6798
chr1-113235404-INS-100
chr1-113246699-INS-50
chr1-113268401-INS-157
chr1-113285235-INS-288
chr1-113322114-INS-50
chr1-113331844-INS-1561
chr1-113332686-INS-83
chr1-113356338-INS-2340
chr1-113374272-INS-288
chr1-113375850-INS-2480
chr1-113383505-INS-6191
chr1-113386922-INS-55
chr1-113406528-INS-152

chr1-113408668-INS-54
chr1-113439649-INS-6380
chr1-113448400-INS-4357
chr1-113453968-INS-8015
chr1-113458781-INS-50
chr1-113463398-INS-5132
chr1-113463534-INS-55
chr1-113484292-INS-69
chr1-113486998-INS-6898
chr1-113513574-INS-58
chr1-113515817-INS-169
chr1-113525820-INS-8771
chr1-113576862-INS-350
chr1-113581916-INS-58
chr1-113600327-INS-120
chr1-113604636-INS-2767
chr1-113616754-INS-54
chr1-113618695-INS-82
chr1-113692624-INS-192
chr1-113743599-INS-412
chr1-113749528-INS-120
chr1-113861490-INS-1234
chr1-113864459-INS-52
chr1-113868547-INS-391
chr1-113895209-INS-481
chr1-113899019-INS-50
chr1-113909157-INS-62
chr1-113925750-INS-80
chr1-113972773-INS-55
chr1-113977645-INS-2384
chr1-113988004-INS-52
chr1-113997888-INS-6369
chr1-114014612-INS-50
chr1-114027760-INS-6000
chr1-114138271-INS-192
chr1-114185192-INS-68
chr1-114197408-INS-294
chr1-114204843-INS-703
chr1-114223731-INS-385
chr1-114225915-INS-7240
chr1-114241443-INS-2803
chr1-114285104-INS-2375
chr1-114324443-INS-662
chr1-114327326-INS-66
chr1-114327358-INS-392
chr1-114327473-INS-168
chr1-114327522-INS-60
chr1-114360099-INS-7081
chr1-114379162-INS-66
chr1-114453430-INS-70
chr1-114453615-INS-86
chr1-114460717-INS-2821
chr1-114492133-INS-7141
chr1-114544555-INS-1495
chr1-114565293-INS-54
chr1-114574595-INS-71
chr1-114601305-INS-56
chr1-114614803-INS-6358
chr1-114619689-INS-50
chr1-114700869-INS-53
chr1-114725033-INS-337
chr1-114725082-INS-163
chr1-114756137-INS-62
chr1-114756262-INS-195
chr1-114756557-INS-181
chr1-114760951-INS-885
chr1-114851795-INS-3957
chr1-114875309-INS-2991
chr1-114892196-INS-2866
chr1-114931762-INS-527
chr1-114937070-INS-869
chr1-114938759-INS-2175
chr1-114939290-INS-6534
chr1-114941204-INS-4080
chr1-114998069-INS-173
chr1-115052382-INS-52
chr1-115157334-INS-50

chr1-115169347-INS-56
chr1-115179671-INS-42201
chr1-115192434-INS-299
chr1-115197286-INS-7446
chr1-115207242-INS-5827
chr1-115210482-INS-50
chr1-115236488-INS-18887
chr1-115244349-INS-205
chr1-115249276-INS-316
chr1-115250429-INS-53
chr1-115255581-INS-370
chr1-115261430-INS-1324
chr1-115270888-INS-6203
chr1-115275504-INS-546
chr1-115296874-INS-3526
chr1-115309287-INS-2284
chr1-115315203-INS-642
chr1-115331004-INS-1282
chr1-115338089-INS-50
chr1-115457531-INS-7491
chr1-115485647-INS-275
chr1-115536624-INS-214
chr1-115544092-INS-2080
chr1-115544199-INS-60
chr1-115568672-INS-6836
chr1-115624209-INS-54
chr1-115656009-INS-303
chr1-115699487-INS-52
chr1-115709390-INS-1537
chr1-115719636-INS-61
chr1-115773003-INS-1536
chr1-115779681-INS-397
chr1-115828519-INS-51
chr1-115851039-INS-444
chr1-115858006-INS-467
chr1-115866671-INS-6227
chr1-115907937-INS-6368
chr1-115927522-INS-100
chr1-115966344-INS-6372
chr1-115983885-INS-325
chr1-116001265-INS-124
chr1-116005644-INS-6217
chr1-116010227-INS-2651
chr1-116018276-INS-81
chr1-116050147-INS-101
chr1-116052396-INS-1024
chr1-116052627-INS-72
chr1-116062063-INS-2322
chr1-116085423-INS-185
chr1-116154831-INS-193
chr1-116155260-INS-84
chr1-116236037-INS-404
chr1-116245585-INS-1664
chr1-116262158-INS-68
chr1-116322845-INS-1393
chr1-116358331-INS-6658
chr1-116394660-INS-84
chr1-116437744-INS-6825
chr1-116464427-INS-84
chr1-116496969-INS-6508
chr1-116517261-INS-2430
chr1-116570237-INS-1429
chr1-116609220-INS-50
chr1-116622889-INS-1693
chr1-116632151-INS-52
chr1-116633059-INS-51
chr1-116634585-INS-1146
chr1-116652979-INS-84
chr1-116657194-INS-6458
chr1-116661981-INS-6027
chr1-116661992-INS-2627
chr1-116663368-INS-1197
chr1-116670858-INS-2450
chr1-116714936-INS-54
chr1-116767126-INS-547
chr1-116826081-INS-1168
chr1-116842088-INS-2443

chr1-116844047-INS-6277
chr1-116861697-INS-872
chr1-116877340-INS-119
chr1-116908317-INS-5909
chr1-116941333-INS-76
chr1-116941343-INS-325
chr1-116975350-INS-3045
chr1-117006339-INS-6631
chr1-117014118-INS-142
chr1-117027622-INS-162
chr1-117035256-INS-461
chr1-117044606-INS-383
chr1-117059308-INS-11468
chr1-117060927-INS-2436
chr1-117100203-INS-61
chr1-117102123-INS-3715
chr1-117127080-INS-6367
chr1-117199679-INS-55
chr1-117212077-INS-73
chr1-117220400-INS-199
chr1-117251677-INS-3831
chr1-117260310-INS-6653
chr1-117314561-INS-1820
chr1-117316643-INS-50
chr1-117317830-INS-3529
chr1-117385847-INS-5565
chr1-117391062-INS-105
chr1-117393185-INS-1616
chr1-117431362-INS-1006
chr1-117444345-INS-1950
chr1-117455040-INS-52
chr1-117457045-INS-64
chr1-117462527-INS-52
chr1-117470735-INS-6520
chr1-117478504-INS-6924
chr1-117556267-INS-352
chr1-117781660-INS-1809
chr1-117854639-INS-2220
chr1-117864877-INS-72
chr1-117871331-INS-64
chr1-117871797-INS-2082
chr1-117885161-INS-165
chr1-117904108-INS-1131
chr1-117909944-INS-504
chr1-117970135-INS-3761
chr1-117972806-INS-59
chr1-117974884-INS-82
chr1-118056001-INS-424
chr1-118127034-INS-924
chr1-118131581-INS-689
chr1-118144883-INS-4419
chr1-118187865-INS-6567
chr1-118209561-INS-817
chr1-118213664-INS-495
chr1-118268932-INS-84
chr1-118269394-INS-1400
chr1-118270538-INS-217
chr1-118278257-INS-52
chr1-118394272-INS-120
chr1-118394404-INS-412
chr1-118394486-INS-166
chr1-118423057-INS-58
chr1-118478186-INS-143
chr1-118486568-INS-694
chr1-118486586-INS-202
chr1-118501157-INS-205
chr1-118540926-INS-1766
chr1-118562754-INS-60
chr1-118563006-INS-281
chr1-118580939-INS-52
chr1-118582219-INS-50
chr1-118593288-INS-66
chr1-118596657-INS-270
chr1-118604905-INS-50
chr1-118616096-INS-103
chr1-118620065-INS-70
chr1-118630604-INS-86

chr1-118644980-INS-206
chr1-118655974-INS-157
chr1-118676951-INS-105
chr1-118718780-INS-50
chr1-118721471-INS-891
chr1-118736172-INS-57
chr1-118755224-INS-80
chr1-118760741-INS-5005
chr1-118846776-INS-112
chr1-118856036-INS-204
chr1-118866126-INS-50
chr1-118868478-INS-62
chr1-118931496-INS-157
chr1-118967284-INS-59
chr1-118975125-INS-112
chr1-118977098-INS-115
chr1-118991315-INS-102
chr1-118997104-INS-50
chr1-118999757-INS-84
chr1-119001225-INS-108
chr1-119046163-INS-195
chr1-119154603-INS-62
chr1-119208441-INS-66
chr1-119209216-INS-182
chr1-119209232-INS-76
chr1-119380067-INS-53
chr1-119458438-INS-91
chr1-119459518-INS-2345
chr1-119484459-INS-240
chr1-119567533-INS-52
chr1-119575003-INS-99
chr1-119674062-INS-1633
chr1-119676022-INS-2826
chr1-119741283-INS-50
chr1-119757320-INS-8457
chr1-119776144-INS-1908
chr1-119798956-INS-7174
chr1-119832689-INS-126
chr1-119833276-INS-104
chr1-119860104-INS-150
chr1-119860854-INS-196
chr1-119878191-INS-203
chr1-119911325-INS-60
chr1-119927779-INS-72
chr1-119946063-INS-434
chr1-119958528-INS-50
chr1-119964688-INS-147
chr1-119968140-INS-144
chr1-119968209-INS-62
chr1-119969816-INS-107
chr1-119975076-INS-557
chr1-119997718-INS-52
chr1-120024900-INS-820
chr1-120024964-INS-2099
chr1-120082409-INS-6218
chr1-120132577-INS-3052
chr1-120144384-INS-7003
chr1-120152063-INS-58
chr1-120159087-INS-3850
chr1-120161024-INS-175
chr1-120210141-INS-51
chr1-120245555-INS-207
chr1-120245779-INS-440
chr1-120247002-INS-344
chr1-120255211-INS-60
chr1-120262129-INS-64
chr1-120263211-INS-155
chr1-120263404-INS-96
chr1-120340431-INS-285
chr1-120340789-INS-125
chr1-120345102-INS-80
chr1-120415440-INS-50
chr1-120417188-INS-51
chr1-120433548-INS-1325
chr1-120441601-INS-135
chr1-120487775-INS-1002
chr1-120506989-INS-66

chr1-120513279-INS-50
chr1-120557469-INS-394
chr1-120557869-INS-448
chr1-120562338-INS-51
chr1-120567368-INS-2297
chr1-120585836-INS-74
chr1-120585860-INS-210
chr1-120594032-INS-59
chr1-120628106-INS-1151
chr1-120634437-INS-52
chr1-120703600-INS-142
chr1-120747062-INS-848
chr1-120762376-INS-76
chr1-120808472-INS-80
chr1-120808631-INS-98
chr1-120849195-INS-552
chr1-120989177-INS-534
chr1-121000286-INS-132
chr1-121034672-INS-52
chr1-121036839-INS-66
chr1-121067536-INS-983
chr1-121093598-INS-68
chr1-121115780-INS-4093
chr1-121121421-INS-50
chr1-121126514-INS-213
chr1-121144122-INS-82
chr1-121156206-INS-7112
chr1-121210161-INS-76
chr1-121210452-INS-404
chr1-121239539-INS-62
chr1-121272763-INS-54
chr1-121286124-INS-81
chr1-121286830-INS-166
chr1-121288698-INS-1195
chr1-121297418-INS-101
chr1-121329278-INS-220
chr1-121331938-INS-228
chr1-121347277-INS-59
chr1-121347295-INS-182
chr1-121384602-INS-50
chr1-121389905-INS-144
chr1-121440272-INS-1723
chr1-121448866-INS-560
chr1-121496373-INS-126
chr1-121496660-INS-80
chr1-121506026-INS-386
chr1-121512407-INS-679
chr1-121569722-INS-6755
chr1-121571084-INS-74
chr1-121584634-INS-81
chr1-121598475-INS-212
chr1-121599560-INS-365
chr1-121627662-INS-85
chr1-121632117-INS-3524
chr1-121649174-INS-50
chr1-121676546-INS-120
chr1-121800139-INS-52
chr1-121813463-INS-435
chr1-121822525-INS-7246
chr1-121843604-INS-1443
chr1-121878668-INS-52
chr1-121900909-INS-56
chr1-121901870-INS-52
chr1-121906225-INS-54
chr1-122000691-INS-1494
chr1-122121257-INS-60
chr1-122182807-INS-7049
chr1-122210024-INS-52
chr1-122249406-INS-100
chr1-122250330-INS-6384
chr1-122273719-INS-69
chr1-122283430-INS-95
chr1-122300592-INS-238
chr1-122303831-INS-4776
chr1-122311027-INS-6209
chr1-122376063-INS-1462
chr1-122384632-INS-604

chr1-122415796-INS-2915
chr1-122458233-INS-1256
chr1-122487796-INS-52
chr1-122505542-INS-2791
chr1-122540772-INS-6445
chr1-122543317-INS-126
chr1-122554114-INS-78
chr1-122554429-INS-95
chr1-122586456-INS-60
chr1-122589328-INS-50
chr1-122634070-INS-145
chr1-122673491-INS-4755
chr1-122678968-INS-6687
chr1-122729112-INS-2592
chr1-122749633-INS-1370
chr1-122775720-INS-6387
chr1-122787099-INS-56
chr1-122850799-INS-45357
chr1-122880100-INS-105
chr1-122880103-INS-249
chr1-122931637-INS-6895
chr1-122941649-INS-2610
chr1-122947537-INS-4975
chr1-122995405-INS-50
chr1-123032061-INS-6354
chr1-123038441-INS-71
chr1-123057226-INS-50
chr1-123058408-INS-157
chr1-123076262-INS-54
chr1-123100403-INS-14005
chr1-123115855-INS-197
chr1-123120749-INS-359
chr1-123121802-INS-2512
chr1-123132994-INS-491
chr1-123156455-INS-7164
chr1-123217832-INS-2282
chr1-123231443-INS-1242
chr1-123231443-INS-594
chr1-123231648-INS-2693
chr1-123241349-INS-50
chr1-123306148-INS-6393
chr1-123388646-INS-50
chr1-123399864-INS-3149
chr1-123401017-INS-110
chr1-123485225-INS-73
chr1-123558442-INS-1339
chr1-123562534-INS-114
chr1-123596965-INS-94
chr1-123597527-INS-653
chr1-123600599-INS-94
chr1-123601916-INS-7412
chr1-123606288-INS-6685
chr1-123669375-INS-54
chr1-123709429-INS-1212
chr1-123709546-INS-87
chr1-123829566-INS-106
chr1-123866986-INS-54
chr1-123884767-INS-52
chr1-123885939-INS-52
chr1-123932907-INS-79
chr1-123944318-INS-134
chr1-123951399-INS-87
chr1-124001367-INS-58
chr1-124017190-INS-8175
chr1-124056835-INS-54
chr1-124057400-INS-53
chr1-124143711-INS-131
chr1-124157730-INS-51
chr1-124201721-INS-62
chr1-124203204-INS-7124
chr1-124213435-INS-1253
chr1-124243241-INS-54
chr1-124243791-INS-174
chr1-124256013-INS-6226
chr1-124264856-INS-95
chr1-124266056-INS-200
chr1-124266348-INS-84

chr1-124299285-INS-1242
chr1-124322016-INS-50
chr1-124351102-INS-220
chr1-124377740-INS-54
chr1-124405622-INS-490
chr1-124435248-INS-87
chr1-124466485-INS-166
chr1-124514487-INS-136
chr1-124527143-INS-68
chr1-124561334-INS-1190
chr1-124576136-INS-50
chr1-124645481-INS-1558
chr1-124645481-INS-3427
chr1-124678742-INS-56
chr1-124694848-INS-7008
chr1-124695189-INS-68
chr1-124701101-INS-107
chr1-124711304-INS-139
chr1-124711344-INS-281
chr1-124740084-INS-81
chr1-124759804-INS-100
chr1-124764726-INS-6272
chr1-124835222-INS-59
chr1-124838796-INS-8405
chr1-124901447-INS-54
chr1-124918923-INS-1659
chr1-124932758-INS-1794
chr1-124997512-INS-94
chr1-125046405-INS-50
chr1-125079520-INS-225
chr1-125084463-INS-54
chr1-125100242-INS-66
chr1-125157708-INS-7084
chr1-125289462-INS-208
chr1-125301808-INS-1296
chr1-125342181-INS-75
chr1-125362721-INS-52
chr1-125418231-INS-4365
chr1-125435084-INS-286
chr1-125435281-INS-80
chr1-125443673-INS-1659
chr1-125443673-INS-5142
chr1-125443913-INS-370
chr1-125446299-INS-63
chr1-125450570-INS-5087
chr1-125452296-INS-6684
chr1-125466154-INS-192
chr1-125468835-INS-56
chr1-125486036-INS-6689
chr1-125507204-INS-106
chr1-125511457-INS-58
chr1-125518898-INS-6729
chr1-125526516-INS-50
chr1-125528003-INS-7248
chr1-125569434-INS-2039
chr1-125571017-INS-4970
chr1-125619142-INS-79
chr1-125619573-INS-134
chr1-125641812-INS-63
chr1-125675733-INS-84
chr1-125698074-INS-50
chr1-125698304-INS-60
chr1-125700256-INS-169
chr1-125713983-INS-50
chr1-125757526-INS-180
chr1-125794619-INS-150
chr1-125885923-INS-50
chr1-126006448-INS-68
chr1-126009741-INS-55
chr1-126015981-INS-195
chr1-126020833-INS-62
chr1-126023170-INS-6112
chr1-126058342-INS-58
chr1-126087724-INS-68
chr1-126094869-INS-50
chr1-126115891-INS-6299
chr1-126120448-INS-7092

chr1-126126009-INS-66
chr1-126158410-INS-84
chr1-126221823-INS-7520
chr1-126249674-INS-51
chr1-126316498-INS-233
chr1-126387849-INS-671
chr1-126421095-INS-52
chr1-126448048-INS-59
chr1-126488452-INS-7080
chr1-126494730-INS-60
chr1-126495392-INS-55
chr1-126517104-INS-88
chr1-126517135-INS-284
chr1-126548787-INS-60
chr1-126550496-INS-60
chr1-126550654-INS-58
chr1-126551199-INS-52
chr1-126613612-INS-51
chr1-126627495-INS-52
chr1-126637181-INS-50
chr1-126653054-INS-54
chr1-126684356-INS-80
chr1-126686404-INS-145
chr1-126703156-INS-1229
chr1-126709512-INS-155
chr1-126726267-INS-54
chr1-126836350-INS-1235
chr1-126898864-INS-496
chr1-126968981-INS-102
chr1-127056547-INS-117
chr1-127077534-INS-256
chr1-127077695-INS-57
chr1-127085461-INS-96
chr1-127138432-INS-182
chr1-127138432-INS-58
chr1-127143903-INS-78
chr1-127171205-INS-4854
chr1-127191162-INS-51
chr1-127191370-INS-197
chr1-127259763-INS-262
chr1-127274383-INS-513
chr1-127320251-INS-53
chr1-127385488-INS-52
chr1-127422079-INS-1339
chr1-127447959-INS-72
chr1-127452285-INS-93
chr1-127498917-INS-68
chr1-127502288-INS-196
chr1-127502332-INS-76
chr1-127535956-INS-53
chr1-127586419-INS-52
chr1-127625945-INS-52
chr1-127665167-INS-114
chr1-127709646-INS-576
chr1-127709666-INS-72
chr1-127727588-INS-56
chr1-127739947-INS-163
chr1-127746620-INS-62
chr1-127755713-INS-63
chr1-127761302-INS-192
chr1-127834692-INS-178
chr1-127891660-INS-2700
chr1-127932872-INS-80
chr1-127936727-INS-1008
chr1-127950497-INS-190
chr1-127963769-INS-4989
chr1-127993918-INS-596
chr1-128189810-INS-66
chr1-128194863-INS-5198
chr1-128217041-INS-50
chr1-128217046-INS-112
chr1-128217866-INS-245
chr1-128224070-INS-206
chr1-128227198-INS-292
chr1-128238848-INS-76
chr1-128251414-INS-111
chr1-128291540-INS-170

chr1-128354524-INS-200
chr1-128376792-INS-513
chr1-128377109-INS-720
chr1-128377109-INS-75
chr1-128386042-INS-4812
chr1-128393852-INS-6930
chr1-128394584-INS-207
chr1-128419201-INS-6259
chr1-128430413-INS-2206
chr1-128444803-INS-172
chr1-128462150-INS-143
chr1-128462333-INS-599
chr1-128488482-INS-161
chr1-128585455-INS-412
chr1-128624595-INS-4363
chr1-128711445-INS-53
chr1-128752809-INS-61
chr1-128774133-INS-88
chr1-128801502-INS-685
chr1-128804220-INS-6588
chr1-128823072-INS-172
chr1-128830591-INS-2582
chr1-128834109-INS-2723
chr1-128849213-INS-6132
chr1-128893238-INS-66
chr1-128964162-INS-7970
chr1-128966774-INS-57
chr1-129028144-INS-7535
chr1-129053183-INS-210
chr1-129057071-INS-500
chr1-129082908-INS-87
chr1-129097108-INS-84
chr1-129144232-INS-264
chr1-129174999-INS-199
chr1-129186621-INS-93
chr1-129189687-INS-96
chr1-129196793-INS-78
chr1-129242325-INS-2866
chr1-129265302-INS-90
chr1-129288832-INS-172
chr1-129288903-INS-607
chr1-129383059-INS-197
chr1-129383804-INS-262
chr1-129428363-INS-493
chr1-129475355-INS-339
chr1-129525527-INS-70
chr1-129540737-INS-52
chr1-129568760-INS-50
chr1-129689186-INS-53
chr1-129689194-INS-265
chr1-129783408-INS-384
chr1-129784594-INS-92
chr1-129821416-INS-10489
chr1-129907180-INS-204
chr1-129939692-INS-63
chr1-130007437-INS-376
chr1-130033024-INS-168
chr1-130036068-INS-144
chr1-130086621-INS-53
chr1-130116523-INS-144
chr1-130134502-INS-192
chr1-130137259-INS-109
chr1-130137878-INS-90
chr1-130170266-INS-482
chr1-130194208-INS-146
chr1-130217956-INS-6464
chr1-130305583-INS-54
chr1-130338965-INS-51
chr1-130435131-INS-88
chr1-130518345-INS-201
chr1-130540155-INS-301
chr1-130540731-INS-72
chr1-130560953-INS-6610
chr1-130596163-INS-180
chr1-130596403-INS-193
chr1-130596528-INS-60
chr1-130600781-INS-50

chr1-130625258-INS-60
chr1-130654355-INS-6174
chr1-130671803-INS-60
chr1-130693307-INS-1891
chr1-130701864-INS-332
chr1-130705532-INS-6582
chr1-130712368-INS-83
chr1-130724103-INS-6645
chr1-130783897-INS-344
chr1-130783930-INS-57
chr1-130909288-INS-608
chr1-130911077-INS-78
chr1-131008508-INS-64
chr1-131018578-INS-538
chr1-131030813-INS-54
chr1-131059136-INS-67
chr1-131064647-INS-457
chr1-131176719-INS-2895
chr1-131213120-INS-50
chr1-131213773-INS-65
chr1-131281670-INS-52
chr1-131288343-INS-50
chr1-131288905-INS-102
chr1-131439314-INS-67
chr1-131442086-INS-202
chr1-131459681-INS-56
chr1-131554523-INS-71
chr1-131663266-INS-315
chr1-131803276-INS-203
chr1-131803887-INS-105
chr1-131806075-INS-368
chr1-131813259-INS-200
chr1-131819404-INS-1907
chr1-131827639-INS-181
chr1-131871659-INS-90
chr1-131876102-INS-7088
chr1-131955167-INS-105
chr1-131955500-INS-183
chr1-131958570-INS-345
chr1-132006439-INS-53
chr1-132015152-INS-237
chr1-132033516-INS-61
chr1-132039512-INS-2838
chr1-132059112-INS-1017
chr1-132059214-INS-2261
chr1-132059316-INS-375
chr1-132082785-INS-53
chr1-132101668-INS-201
chr1-132144622-INS-244
chr1-132147598-INS-50
chr1-132147682-INS-63
chr1-132166934-INS-84
chr1-132188439-INS-52
chr1-132201519-INS-176
chr1-132201979-INS-416
chr1-132213630-INS-60
chr1-132239626-INS-252
chr1-132257277-INS-175
chr1-132275105-INS-208
chr1-132275649-INS-393
chr1-132293003-INS-313
chr1-132326875-INS-59
chr1-132327386-INS-6436
chr1-132342583-INS-51
chr1-132352891-INS-62
chr1-132396445-INS-244
chr1-132399337-INS-50
chr1-132437534-INS-56
chr1-132455043-INS-72
chr1-132455129-INS-214
chr1-132481173-INS-208
chr1-132509049-INS-206
chr1-132535266-INS-55
chr1-132539502-INS-156
chr1-132598567-INS-52
chr1-132606979-INS-142
chr1-132640678-INS-344

chr1-132678206-INS-123
chr1-132679128-INS-51
chr1-132691237-INS-184
chr1-132704758-INS-52
chr1-132739757-INS-7131
chr1-132767173-INS-78
chr1-132788449-INS-229
chr1-132792200-INS-197
chr1-132802466-INS-54
chr1-132856039-INS-83
chr1-132877489-INS-50
chr1-132889897-INS-69
chr1-132902916-INS-399
chr1-132904301-INS-60
chr1-132904507-INS-735
chr1-132933905-INS-225
chr1-132937919-INS-56
chr1-132937932-INS-389
chr1-132950796-INS-238
chr1-132965788-INS-2108
chr1-132998209-INS-172
chr1-133020999-INS-185
chr1-133025526-INS-201
chr1-133044111-INS-218
chr1-133068866-INS-1272
chr1-133093901-INS-216
chr1-133173049-INS-233
chr1-133240318-INS-64
chr1-133258868-INS-500
chr1-133274793-INS-31118
chr1-133275908-INS-52
chr1-133289748-INS-242
chr1-133320210-INS-84
chr1-133333689-INS-52
chr1-133369288-INS-119
chr1-133370705-INS-201
chr1-133374723-INS-84
chr1-133383620-INS-243
chr1-133403164-INS-200
chr1-133406656-INS-65
chr1-133438949-INS-135
chr1-133455047-INS-223
chr1-133490574-INS-63
chr1-133514095-INS-1129
chr1-133523310-INS-11548
chr1-133553743-INS-97
chr1-133593175-INS-6253
chr1-133629538-INS-187
chr1-133683883-INS-51
chr1-133692546-INS-202
chr1-133734056-INS-67
chr1-133773419-INS-956
chr1-133786501-INS-57
chr1-133821775-INS-220
chr1-133868217-INS-166
chr1-133875087-INS-55
chr1-133880769-INS-50
chr1-133883936-INS-4757
chr1-133888970-INS-80
chr1-133894681-INS-171
chr1-133916167-INS-200
chr1-133926366-INS-93
chr1-133950278-INS-304
chr1-133950574-INS-106
chr1-133977469-INS-440
chr1-134047096-INS-125
chr1-134047286-INS-151
chr1-134050123-INS-1944
chr1-134058903-INS-54
chr1-134074956-INS-398
chr1-134090901-INS-286
chr1-134098481-INS-144
chr1-134148631-INS-527
chr1-134148665-INS-155
chr1-134231285-INS-57
chr1-134231307-INS-300
chr1-134284508-INS-534

chr1-134300679-INS-54
chr1-134318869-INS-484
chr1-134319024-INS-138
chr1-134355445-INS-5000
chr1-134366196-INS-486
chr1-134374585-INS-213
chr1-134374585-INS-2472
chr1-134440743-INS-143
chr1-134462241-INS-56
chr1-134493970-INS-9047
chr1-134511060-INS-55
chr1-134595804-INS-176
chr1-134605814-INS-6473
chr1-134622408-INS-6442
chr1-134663775-INS-324
chr1-134663775-INS-891
chr1-134663998-INS-364
chr1-134684455-INS-60
chr1-134697480-INS-637
chr1-134726928-INS-141
chr1-134727097-INS-140
chr1-134727472-INS-76
chr1-134795672-INS-52
chr1-134862159-INS-55
chr1-134962737-INS-50
chr1-135101931-INS-181
chr1-135133081-INS-76
chr1-135158074-INS-796
chr1-135173141-INS-178
chr1-135216240-INS-7169
chr1-135233433-INS-131
chr1-135235329-INS-182
chr1-135235562-INS-799
chr1-135329132-INS-223
chr1-135336799-INS-52
chr1-135363185-INS-54
chr1-135371911-INS-2855
chr1-135389160-INS-51
chr1-135422365-INS-246
chr1-135467262-INS-201
chr1-135582557-INS-51
chr1-135588875-INS-54
chr1-135634469-INS-212
chr1-135634854-INS-70
chr1-135638545-INS-4715
chr1-135642816-INS-12386
chr1-135666776-INS-51
chr1-135670156-INS-133
chr1-135752944-INS-220
chr1-135756817-INS-60
chr1-135784533-INS-59
chr1-135810855-INS-50
chr1-135813781-INS-62
chr1-135849215-INS-192
chr1-135869773-INS-62
chr1-135927142-INS-140
chr1-135927365-INS-102
chr1-135927413-INS-316
chr1-135948919-INS-231
chr1-135961375-INS-127
chr1-135984286-INS-50
chr1-136000745-INS-54
chr1-136001661-INS-116
chr1-136011604-INS-80
chr1-136018243-INS-276
chr1-136041522-INS-142
chr1-136041528-INS-63
chr1-136120629-INS-85
chr1-136215515-INS-54
chr1-136263018-INS-52
chr1-136306850-INS-1255
chr1-136314366-INS-217
chr1-136314671-INS-114
chr1-136321618-INS-95
chr1-136321859-INS-547
chr1-136331583-INS-73
chr1-136334481-INS-192

chr1-136335153-INS-58
chr1-136365909-INS-5249
chr1-136366880-INS-189
chr1-136387606-INS-84
chr1-136393254-INS-6430
chr1-136413018-INS-260
chr1-136413303-INS-61
chr1-136421486-INS-1089
chr1-136437766-INS-2809
chr1-136472697-INS-1472
chr1-136488841-INS-1044
chr1-136501924-INS-52
chr1-136503739-INS-87
chr1-136594908-INS-145
chr1-136607118-INS-96
chr1-136607613-INS-379
chr1-136619349-INS-52
chr1-136627395-INS-227
chr1-136680060-INS-178
chr1-136729971-INS-205
chr1-136730574-INS-204
chr1-136799444-INS-232
chr1-136854482-INS-306
chr1-136913128-INS-68
chr1-136925207-INS-116
chr1-136926042-INS-82
chr1-136932990-INS-4460
chr1-136952612-INS-128
chr1-136973201-INS-394
chr1-137015396-INS-75
chr1-137025948-INS-53
chr1-137054771-INS-1110
chr1-137067859-INS-54
chr1-137097554-INS-50
chr1-137097812-INS-52
chr1-137098240-INS-4740
chr1-137108493-INS-52
chr1-137111347-INS-6617
chr1-137225742-INS-79
chr1-137226748-INS-65
chr1-137255170-INS-72
chr1-137294636-INS-391
chr1-137294669-INS-170
chr1-137302263-INS-70
chr1-137309447-INS-50
chr1-137320478-INS-863
chr1-137320777-INS-532
chr1-137323151-INS-518
chr1-137343710-INS-90
chr1-137349755-INS-1240
chr1-137356889-INS-61
chr1-137377536-INS-291
chr1-137412386-INS-4907
chr1-137434117-INS-2370
chr1-137445117-INS-168
chr1-137537271-INS-50
chr1-137568345-INS-566
chr1-137668467-INS-57
chr1-137682249-INS-951
chr1-137682576-INS-4253
chr1-137692047-INS-695
chr1-137710429-INS-62
chr1-137758781-INS-6697
chr1-137769106-INS-6400
chr1-137794035-INS-51
chr1-137811283-INS-61
chr1-137836994-INS-70
chr1-137898861-INS-68
chr1-137899703-INS-152
chr1-137935288-INS-52
chr1-137956584-INS-55
chr1-137968654-INS-50
chr1-137968727-INS-105
chr1-137992974-INS-6389
chr1-138023734-INS-56
chr1-138024858-INS-81
chr1-138025178-INS-177

chr1-138075556-INS-214
chr1-138090973-INS-74
chr1-138126754-INS-6383
chr1-138147904-INS-204
chr1-138164910-INS-95
chr1-138182021-INS-168
chr1-138239867-INS-424
chr1-138348168-INS-124
chr1-138391477-INS-547
chr1-138525714-INS-598
chr1-138526698-INS-168
chr1-138533182-INS-162
chr1-138534202-INS-182
chr1-138534378-INS-286
chr1-138559191-INS-56
chr1-138619914-INS-338
chr1-138630396-INS-175
chr1-138634116-INS-557
chr1-138686348-INS-87
chr1-138686386-INS-413
chr1-138777141-INS-250
chr1-138777242-INS-58
chr1-138786893-INS-200
chr1-138790916-INS-14538
chr1-138794145-INS-33926
chr1-138805062-INS-58
chr1-138808643-INS-80
chr1-138812143-INS-144
chr1-138856192-INS-168
chr1-138924827-INS-55
chr1-139068106-INS-6951
chr1-139139958-INS-193
chr1-139146355-INS-479
chr1-139182236-INS-186
chr1-139182318-INS-124
chr1-139213335-INS-73
chr1-139261632-INS-833
chr1-139307803-INS-202
chr1-139332208-INS-2999
chr1-139335438-INS-1954
chr1-139339664-INS-57
chr1-139339928-INS-181
chr1-139342969-INS-296
chr1-139420436-INS-70
chr1-139432470-INS-6301
chr1-139493673-INS-3408
chr1-139496573-INS-5992
chr1-139507688-INS-3583
chr1-139562419-INS-333
chr1-139562830-INS-909
chr1-139563702-INS-1063
chr1-139616439-INS-69
chr1-139616750-INS-1491
chr1-139685502-INS-6206
chr1-139747189-INS-20357
chr1-139762177-INS-1545
chr1-139787782-INS-4541
chr1-139890205-INS-414
chr1-139925470-INS-50
chr1-139930332-INS-4952
chr1-139945434-INS-180
chr1-139949139-INS-122
chr1-139949139-INS-323
chr1-139949681-INS-312
chr1-139990404-INS-51
chr1-140015913-INS-54
chr1-140084427-INS-87
chr1-140108499-INS-52
chr1-140152433-INS-55
chr1-140198182-INS-68
chr1-140208694-INS-61
chr1-140214694-INS-6183
chr1-140223809-INS-711
chr1-140227089-INS-50
chr1-140227134-INS-117
chr1-140227141-INS-50
chr1-140233480-INS-1475

chr1-140255444-INS-50
chr1-140267799-INS-72
chr1-140314523-INS-60
chr1-140357245-INS-1349
chr1-140369683-INS-50
chr1-140468571-INS-54
chr1-140474865-INS-1136
chr1-140484681-INS-510
chr1-140506741-INS-1687
chr1-140507188-INS-288
chr1-140562687-INS-1238
chr1-140604857-INS-6528
chr1-140648778-INS-53
chr1-140737108-INS-50
chr1-140737215-INS-112
chr1-140767596-INS-6610
chr1-140771935-INS-76
chr1-140772687-INS-268
chr1-140774997-INS-98
chr1-140794819-INS-402
chr1-140796065-INS-6679
chr1-140800438-INS-6416
chr1-140820861-INS-60
chr1-140821062-INS-325
chr1-140893676-INS-16256
chr1-140941864-INS-1566
chr1-140961004-INS-4088
chr1-141020929-INS-6215
chr1-141028690-INS-4155
chr1-141034721-INS-82
chr1-141106710-INS-80
chr1-141130163-INS-80
chr1-141145139-INS-414
chr1-141163946-INS-2700
chr1-141225782-INS-396
chr1-141226159-INS-75
chr1-141240740-INS-141
chr1-141240740-INS-54
chr1-141326971-INS-2730
chr1-141382351-INS-6204
chr1-141402198-INS-947
chr1-141403006-INS-74
chr1-141421241-INS-54
chr1-141441022-INS-50
chr1-141522597-INS-4881
chr1-141532315-INS-7150
chr1-141575733-INS-51
chr1-141604999-INS-68
chr1-141621113-INS-1147
chr1-141631133-INS-424
chr1-141633697-INS-268
chr1-141634362-INS-531
chr1-141657268-INS-54
chr1-141661169-INS-2330
chr1-141701823-INS-172
chr1-141721275-INS-6137
chr1-141722731-INS-6918
chr1-141734618-INS-1619
chr1-141749439-INS-63
chr1-141762250-INS-6162
chr1-141770915-INS-58
chr1-141830466-INS-213
chr1-141840574-INS-3676
chr1-141862931-INS-89
chr1-141895307-INS-2808
chr1-141920360-INS-1031
chr1-141944463-INS-7750
chr1-141947121-INS-1498
chr1-142008688-INS-107
chr1-142089074-INS-384
chr1-142123347-INS-4548
chr1-142138935-INS-60
chr1-142139052-INS-168
chr1-142148068-INS-6668
chr1-142176257-INS-88
chr1-142178358-INS-63
chr1-142211466-INS-70

chr1-142237103-INS-7312
chr1-142263491-INS-195
chr1-142303058-INS-174
chr1-142303060-INS-56
chr1-142316596-INS-209
chr1-142321269-INS-974
chr1-142367106-INS-283
chr1-142401074-INS-214
chr1-142445410-INS-72
chr1-142464656-INS-4469
chr1-142500087-INS-68
chr1-142510501-INS-72
chr1-142554778-INS-10101
chr1-142561465-INS-52
chr1-142585751-INS-13330
chr1-142678802-INS-562
chr1-142680257-INS-6404
chr1-142713433-INS-2823
chr1-142735846-INS-6220
chr1-142765187-INS-50
chr1-142873154-INS-2499
chr1-142875120-INS-780
chr1-142897775-INS-131
chr1-142926322-INS-199
chr1-142955524-INS-240
chr1-143014391-INS-274
chr1-143019727-INS-342
chr1-143019760-INS-96
chr1-143033990-INS-575
chr1-143139486-INS-203
chr1-143144635-INS-198
chr1-143148511-INS-6639
chr1-143181659-INS-52
chr1-143185910-INS-161
chr1-143220957-INS-155
chr1-143250721-INS-166
chr1-143300697-INS-57
chr1-143326762-INS-6583
chr1-143359973-INS-550
chr1-143366165-INS-2888
chr1-143376972-INS-8571
chr1-143391947-INS-54
chr1-143405451-INS-52
chr1-143405808-INS-82
chr1-143406465-INS-65
chr1-143418544-INS-139
chr1-143423023-INS-1290
chr1-143431054-INS-196
chr1-143435316-INS-537
chr1-143445378-INS-405
chr1-143449410-INS-84
chr1-143502840-INS-245
chr1-143580173-INS-204
chr1-143599168-INS-155
chr1-143603817-INS-245
chr1-143604123-INS-1050
chr1-143643333-INS-204
chr1-143663007-INS-1730
chr1-143673252-INS-207
chr1-143803331-INS-50
chr1-143837089-INS-197
chr1-143859164-INS-82
chr1-143864215-INS-993
chr1-143864252-INS-190
chr1-143865387-INS-57
chr1-143865533-INS-754
chr1-143867571-INS-60
chr1-143895608-INS-50
chr1-143939580-INS-725
chr1-143990778-INS-59
chr1-144008206-INS-1235
chr1-144009618-INS-1425
chr1-144012563-INS-50
chr1-144018401-INS-51
chr1-144038517-INS-400
chr1-144039183-INS-180
chr1-144137727-INS-7185

chr1-144148409-INS-6441
chr1-144216709-INS-56
chr1-144256790-INS-1146
chr1-144293862-INS-53
chr1-144356144-INS-6863
chr1-144359887-INS-52
chr1-144388099-INS-394
chr1-144445835-INS-154
chr1-144474130-INS-54
chr1-144488600-INS-162
chr1-144488770-INS-6618
chr1-144494412-INS-50
chr1-144501266-INS-52
chr1-144539848-INS-54
chr1-144591436-INS-428
chr1-144592017-INS-3198
chr1-144605694-INS-478
chr1-144605810-INS-60
chr1-144625559-INS-59
chr1-144626358-INS-52
chr1-144667315-INS-186
chr1-144676372-INS-58
chr1-144715046-INS-1439
chr1-144716082-INS-96
chr1-144722563-INS-54
chr1-144722910-INS-52
chr1-144740768-INS-565
chr1-144745599-INS-50
chr1-144760569-INS-123
chr1-144783235-INS-52
chr1-144819388-INS-376
chr1-144849322-INS-7175
chr1-144911989-INS-3164
chr1-144915484-INS-1308
chr1-144931606-INS-51
chr1-144946710-INS-543
chr1-144951837-INS-191
chr1-144960049-INS-1023
chr1-144960931-INS-1023
chr1-145011689-INS-195
chr1-145012077-INS-407
chr1-145207107-INS-68
chr1-145211242-INS-413
chr1-145291056-INS-85
chr1-145297116-INS-7977
chr1-145315606-INS-70
chr1-145334679-INS-58
chr1-145335176-INS-50
chr1-145353390-INS-3140
chr1-145353622-INS-191
chr1-145353646-INS-54
chr1-145359648-INS-1066
chr1-145387667-INS-6942
chr1-145411704-INS-72
chr1-145417477-INS-66
chr1-145429890-INS-4509
chr1-145443577-INS-88
chr1-145455037-INS-134
chr1-145460700-INS-52
chr1-145487898-INS-2669
chr1-145500323-INS-470
chr1-145511137-INS-50
chr1-145545234-INS-60
chr1-145552637-INS-2850
chr1-145564154-INS-1084
chr1-145573548-INS-1045
chr1-145586317-INS-4363
chr1-145592499-INS-65
chr1-145592919-INS-493
chr1-145594307-INS-6286
chr1-145594845-INS-834
chr1-145645588-INS-2831
chr1-145710626-INS-1486
chr1-145714271-INS-52
chr1-145724375-INS-52
chr1-145724872-INS-448
chr1-145733035-INS-96

chr1-145733117-INS-56
chr1-145740128-INS-58
chr1-145771605-INS-6318
chr1-145796718-INS-6693
chr1-145822131-INS-6151
chr1-145832879-INS-3408
chr1-145838695-INS-361
chr1-145839352-INS-227
chr1-145910800-INS-144
chr1-145916947-INS-3807
chr1-145922752-INS-112
chr1-145926029-INS-2962
chr1-145926686-INS-203
chr1-145944366-INS-3148
chr1-145978151-INS-104
chr1-145978663-INS-100
chr1-145978777-INS-332
chr1-145979030-INS-56
chr1-145996041-INS-370
chr1-146061056-INS-1245
chr1-146171683-INS-214
chr1-146193538-INS-87
chr1-146308174-INS-56
chr1-146453663-INS-6337
chr1-146501575-INS-57
chr1-146503569-INS-50
chr1-146512366-INS-795
chr1-146521605-INS-278
chr1-146570817-INS-584
chr1-146572463-INS-6243
chr1-146736686-INS-269
chr1-146736830-INS-183
chr1-146736833-INS-84
chr1-146803910-INS-50
chr1-146821807-INS-5145
chr1-146855158-INS-6109
chr1-146861819-INS-1060
chr1-146864156-INS-88
chr1-146877633-INS-666
chr1-146937349-INS-52
chr1-146938177-INS-7989
chr1-147001295-INS-62
chr1-147096436-INS-555
chr1-147102768-INS-77
chr1-147103100-INS-118
chr1-147132673-INS-128
chr1-147148335-INS-188
chr1-147151756-INS-1082
chr1-147169803-INS-52
chr1-147191642-INS-589
chr1-147234077-INS-76
chr1-147279120-INS-53
chr1-147309610-INS-7084
chr1-147341210-INS-60
chr1-147428753-INS-197
chr1-147443239-INS-681
chr1-147443243-INS-235
chr1-147656188-INS-303
chr1-147660242-INS-6457
chr1-147680202-INS-820
chr1-147718585-INS-9766
chr1-147746370-INS-3912
chr1-147855828-INS-1481
chr1-147901037-INS-108
chr1-147915315-INS-50
chr1-147915891-INS-7047
chr1-147928728-INS-547
chr1-147959211-INS-55
chr1-147994932-INS-70
chr1-148009432-INS-50
chr1-148054399-INS-1203
chr1-148080366-INS-3213
chr1-148082369-INS-50
chr1-148101407-INS-52
chr1-148106294-INS-52
chr1-148120137-INS-1618
chr1-148124202-INS-50

chr1-148295915-INS-111
chr1-148300337-INS-50
chr1-148319251-INS-148
chr1-148319424-INS-78
chr1-148341710-INS-1220
chr1-148344192-INS-66
chr1-148385399-INS-84
chr1-148445473-INS-52
chr1-148447225-INS-4412
chr1-148516470-INS-1337
chr1-148557693-INS-1115
chr1-148576050-INS-6667
chr1-148598550-INS-667
chr1-148613695-INS-352
chr1-148624170-INS-13831
chr1-148675407-INS-547
chr1-148680437-INS-67
chr1-148680641-INS-72
chr1-148698821-INS-1688
chr1-148724392-INS-58
chr1-148776247-INS-108
chr1-148787551-INS-798
chr1-148799680-INS-6261
chr1-148805791-INS-149
chr1-148811477-INS-50
chr1-148830623-INS-2839
chr1-148916256-INS-12472
chr1-148930460-INS-124
chr1-148930754-INS-6370
chr1-148977526-INS-6976
chr1-149045362-INS-6264
chr1-149077222-INS-1157
chr1-149115472-INS-2664
chr1-149138776-INS-8266
chr1-149142359-INS-198
chr1-149216363-INS-7350
chr1-149265417-INS-75
chr1-149268063-INS-85
chr1-149277670-INS-198
chr1-149293692-INS-392
chr1-149297886-INS-55
chr1-149312663-INS-1951
chr1-149358766-INS-89
chr1-149398619-INS-112
chr1-149430450-INS-79
chr1-149441331-INS-3436
chr1-149562213-INS-478
chr1-149569565-INS-2878
chr1-149577188-INS-51
chr1-149578242-INS-1549
chr1-149602054-INS-57
chr1-149649825-INS-6467
chr1-149706053-INS-79
chr1-149713120-INS-50
chr1-149723409-INS-6382
chr1-149736048-INS-56
chr1-149759545-INS-50
chr1-149798772-INS-104
chr1-149800366-INS-84
chr1-149803833-INS-63
chr1-149803942-INS-466
chr1-149850406-INS-68
chr1-149867888-INS-2395
chr1-149884146-INS-60
chr1-149884169-INS-210
chr1-149890680-INS-1141
chr1-149902315-INS-321
chr1-149939553-INS-690
chr1-149948277-INS-52
chr1-149955422-INS-142
chr1-149958526-INS-5627
chr1-149965431-INS-90
chr1-149984954-INS-89
chr1-149984971-INS-410
chr1-149999828-INS-72
chr1-150000281-INS-50
chr1-150003051-INS-58

chr1-150003719-INS-2537
chr1-150012313-INS-6302
chr1-150025430-INS-53
chr1-150031914-INS-1241
chr1-150056909-INS-7175
chr1-150137702-INS-62
chr1-150138547-INS-56
chr1-150227750-INS-1950
chr1-150252572-INS-336
chr1-150320415-INS-54
chr1-150353496-INS-5018
chr1-150362062-INS-52
chr1-150372245-INS-109
chr1-150372446-INS-77
chr1-150393357-INS-6371
chr1-150395039-INS-60
chr1-150395860-INS-53
chr1-150505347-INS-116
chr1-150541049-INS-1045
chr1-150548899-INS-6424
chr1-150558838-INS-72
chr1-150562130-INS-95
chr1-150771276-INS-71
chr1-150782813-INS-52
chr1-150819799-INS-51
chr1-150844162-INS-632
chr1-150917936-INS-2161
chr1-150920352-INS-51
chr1-150920626-INS-146
chr1-150943906-INS-54
chr1-150998769-INS-163
chr1-151008874-INS-97
chr1-151066158-INS-13415
chr1-151078360-INS-56
chr1-151084082-INS-156
chr1-151087741-INS-1002
chr1-151142124-INS-232
chr1-151143345-INS-88
chr1-151143411-INS-6413
chr1-151144044-INS-59
chr1-151161792-INS-51
chr1-151175731-INS-4707
chr1-151189558-INS-58
chr1-151192839-INS-129
chr1-151202605-INS-6412
chr1-151241690-INS-50
chr1-151251637-INS-82
chr1-151301681-INS-72
chr1-151321559-INS-285
chr1-151323802-INS-112
chr1-151340380-INS-50
chr1-151342828-INS-4116
chr1-151400521-INS-308
chr1-151400779-INS-63
chr1-151403685-INS-171
chr1-151407909-INS-75
chr1-151435942-INS-2741
chr1-151440587-INS-202
chr1-151450479-INS-464
chr1-151450582-INS-58
chr1-151465071-INS-50
chr1-151473139-INS-57
chr1-151479146-INS-52
chr1-151526925-INS-68
chr1-151527847-INS-105
chr1-151533820-INS-3163
chr1-151564291-INS-66
chr1-151587746-INS-51
chr1-151597998-INS-62
chr1-151598556-INS-135
chr1-151609898-INS-50
chr1-151672692-INS-364
chr1-151672692-INS-52
chr1-151716262-INS-117
chr1-151716262-INS-320
chr1-151741239-INS-192
chr1-151749722-INS-358

chr1-151803872-INS-52
chr1-151861683-INS-74
chr1-151862043-INS-607
chr1-151862194-INS-65
chr1-151862557-INS-203
chr1-151892517-INS-65
chr1-151897857-INS-311
chr1-151898080-INS-578
chr1-151898594-INS-56
chr1-151920086-INS-59
chr1-151947749-INS-128
chr1-151975191-INS-230
chr1-151981234-INS-643
chr1-152028832-INS-72
chr1-152067309-INS-148
chr1-152084421-INS-54
chr1-152154717-INS-68
chr1-152156008-INS-66
chr1-152159680-INS-111
chr1-152161207-INS-107
chr1-152167363-INS-6911
chr1-152189704-INS-3719
chr1-152212371-INS-9649
chr1-152244194-INS-2903
chr1-152317985-INS-6253
chr1-152324065-INS-5560
chr1-152337646-INS-3425
chr1-152367457-INS-54
chr1-152387171-INS-144
chr1-152396250-INS-213
chr1-152423392-INS-84
chr1-152511875-INS-72
chr1-152512120-INS-76
chr1-152530178-INS-138
chr1-152549530-INS-406
chr1-152581063-INS-66
chr1-152592280-INS-60
chr1-152594443-INS-209
chr1-152600842-INS-293
chr1-152600858-INS-138
chr1-152611084-INS-264
chr1-152630027-INS-92
chr1-152639006-INS-153
chr1-152639148-INS-611
chr1-152688333-INS-71
chr1-152688870-INS-197
chr1-152728876-INS-234
chr1-152810144-INS-50
chr1-152867885-INS-191
chr1-152884306-INS-7410
chr1-152932331-INS-52
chr1-152979314-INS-252
chr1-152979372-INS-122
chr1-152979400-INS-731
chr1-153038459-INS-1100
chr1-153088944-INS-318
chr1-153147556-INS-2500
chr1-153169575-INS-280
chr1-153177581-INS-132
chr1-153272293-INS-7674
chr1-153275396-INS-232
chr1-153275956-INS-245
chr1-153276455-INS-439
chr1-153289613-INS-306
chr1-153289641-INS-102
chr1-153292595-INS-390
chr1-153367966-INS-487
chr1-153371109-INS-55
chr1-153373934-INS-507
chr1-153382313-INS-156
chr1-153388600-INS-1864
chr1-153390072-INS-190
chr1-153396623-INS-1945
chr1-153408317-INS-209
chr1-153419962-INS-132
chr1-153421495-INS-197
chr1-153425846-INS-222

chr1-153429847-INS-178
chr1-153436035-INS-196
chr1-153441486-INS-5240
chr1-153482577-INS-1890
chr1-153584233-INS-51
chr1-153606125-INS-152
chr1-153615856-INS-60
chr1-153647308-INS-52
chr1-153650829-INS-5081
chr1-153659333-INS-167
chr1-153758137-INS-101
chr1-153758184-INS-703
chr1-153808638-INS-54
chr1-153833999-INS-108
chr1-153846337-INS-55
chr1-153875322-INS-398
chr1-153879703-INS-6414
chr1-153897359-INS-156
chr1-153898293-INS-7232
chr1-153915579-INS-52
chr1-153929163-INS-132
chr1-153929223-INS-548
chr1-153975166-INS-58
chr1-154047835-INS-1888
chr1-154055729-INS-3024
chr1-154055879-INS-301
chr1-154106389-INS-410
chr1-154227895-INS-86
chr1-154237904-INS-7452
chr1-154322722-INS-601
chr1-154322882-INS-87
chr1-154322888-INS-258
chr1-154340224-INS-316
chr1-154380829-INS-1554
chr1-154472295-INS-68
chr1-154508228-INS-51
chr1-154657109-INS-323
chr1-154728640-INS-262
chr1-154787097-INS-94
chr1-154868965-INS-131
chr1-155038156-INS-97
chr1-155038195-INS-515
chr1-155048773-INS-93
chr1-155108014-INS-103
chr1-155124549-INS-159
chr1-155167750-INS-442
chr1-155167893-INS-50
chr1-155191417-INS-1484
chr1-155277043-INS-129
chr1-155349509-INS-50
chr1-155360739-INS-54
chr1-155404293-INS-4044
chr1-155410896-INS-2298
chr1-155419800-INS-8191
chr1-155423332-INS-1851
chr1-155423789-INS-60
chr1-155432432-INS-152
chr1-155446093-INS-7097
chr1-155477529-INS-58
chr1-155485400-INS-54
chr1-155589554-INS-54
chr1-155693829-INS-62
chr1-155694497-INS-235
chr1-155844212-INS-157
chr1-155867704-INS-56
chr1-155962141-INS-327
chr1-155967651-INS-1340
chr1-155996742-INS-249
chr1-156055996-INS-3385
chr1-156074969-INS-7494
chr1-156080810-INS-1493
chr1-156086145-INS-54
chr1-156086363-INS-182
chr1-156089997-INS-5273
chr1-156164342-INS-716
chr1-156169051-INS-54
chr1-156211693-INS-56

chr1-156277507-INS-264
chr1-156288301-INS-445
chr1-156303158-INS-67
chr1-156364235-INS-50
chr1-156374307-INS-536
chr1-156376629-INS-120
chr1-156382027-INS-52
chr1-156387933-INS-66
chr1-156390624-INS-205
chr1-156448173-INS-102
chr1-156448173-INS-1472
chr1-156448173-INS-236
chr1-156448173-INS-552
chr1-156477724-INS-1541
chr1-156477969-INS-184
chr1-156504210-INS-212
chr1-156504426-INS-141
chr1-156739650-INS-411
chr1-156740252-INS-482
chr1-156785442-INS-268
chr1-156803266-INS-166
chr1-156811551-INS-58
chr1-156829507-INS-211
chr1-156858577-INS-60
chr1-156868592-INS-58
chr1-156899790-INS-50
chr1-156903625-INS-65
chr1-156908167-INS-194
chr1-156910325-INS-208
chr1-156916162-INS-60
chr1-156962410-INS-354
chr1-157011500-INS-51
chr1-157021430-INS-51
chr1-157060923-INS-52
chr1-157077780-INS-70
chr1-157096900-INS-51
chr1-157122018-INS-60
chr1-157122380-INS-124
chr1-157124490-INS-50
chr1-157128366-INS-50
chr1-157164384-INS-3039
chr1-157174714-INS-50
chr1-157195207-INS-210
chr1-157291433-INS-841
chr1-157335161-INS-108
chr1-157356355-INS-405
chr1-157356362-INS-103
chr1-157357174-INS-144
chr1-157357248-INS-216
chr1-157369235-INS-66
chr1-157375863-INS-60
chr1-157386556-INS-459
chr1-157387364-INS-52
chr1-157391113-INS-150
chr1-157416266-INS-70
chr1-157420347-INS-50
chr1-157424566-INS-124
chr1-157455102-INS-927
chr1-157465703-INS-300
chr1-157503781-INS-203
chr1-157539210-INS-156
chr1-157539523-INS-652
chr1-157542121-INS-139
chr1-157575367-INS-56
chr1-157619805-INS-6877
chr1-157663552-INS-6835
chr1-157749580-INS-52
chr1-157873204-INS-97
chr1-157873351-INS-161
chr1-157873351-INS-62
chr1-157884837-INS-71
chr1-157976500-INS-99
chr1-158002466-INS-84
chr1-158075856-INS-1517
chr1-158101632-INS-491
chr1-158104210-INS-112
chr1-158130509-INS-55

chr1-158139907-INS-105
chr1-158165684-INS-4459
chr1-158219971-INS-6564
chr1-158239172-INS-244
chr1-158256677-INS-7099
chr1-158257188-INS-216
chr1-158319178-INS-63
chr1-158345976-INS-500
chr1-158404180-INS-76
chr1-158501778-INS-67
chr1-158507026-INS-51
chr1-158530272-INS-4037
chr1-158574275-INS-134
chr1-158594280-INS-6377
chr1-158679871-INS-169
chr1-158679871-INS-80
chr1-158729882-INS-207
chr1-158735523-INS-55
chr1-158753926-INS-6242
chr1-158839405-INS-98
chr1-158850113-INS-428
chr1-158905799-INS-64
chr1-158925244-INS-213
chr1-158927362-INS-6215
chr1-158930962-INS-56
chr1-158941727-INS-6438
chr1-159003764-INS-17451
chr1-159003819-INS-5507
chr1-159009155-INS-86
chr1-159041502-INS-68
chr1-159070440-INS-209
chr1-159078656-INS-64
chr1-159080311-INS-65
chr1-159171311-INS-120
chr1-159218076-INS-243
chr1-159225649-INS-61
chr1-159225750-INS-790
chr1-159242119-INS-7005
chr1-159243733-INS-51
chr1-159271469-INS-678
chr1-159302136-INS-50
chr1-159302263-INS-233
chr1-159401540-INS-175
chr1-159404578-INS-54
chr1-159464782-INS-64
chr1-159487621-INS-315
chr1-159543272-INS-176
chr1-159586162-INS-6395
chr1-159599195-INS-9052
chr1-159606142-INS-6228
chr1-159671177-INS-52
chr1-159672326-INS-73
chr1-159713412-INS-195
chr1-159812134-INS-50
chr1-159842240-INS-64
chr1-159842279-INS-313
chr1-159881149-INS-89
chr1-159970356-INS-6647
chr1-159993208-INS-189
chr1-160133980-INS-54
chr1-160190963-INS-221
chr1-160227908-INS-81
chr1-160239367-INS-176
chr1-160301251-INS-50
chr1-160401390-INS-54
chr1-160472019-INS-243
chr1-160481234-INS-3422
chr1-160493929-INS-648
chr1-160497707-INS-7227
chr1-160543332-INS-187
chr1-160545757-INS-54
chr1-160556367-INS-1143
chr1-160556367-INS-2707
chr1-160598748-INS-356
chr1-160608692-INS-51
chr1-160721232-INS-56
chr1-160728621-INS-141

chr1-160788860-INS-126
chr1-160880748-INS-51
chr1-160903534-INS-62
chr1-160935054-INS-213
chr1-160951598-INS-172
chr1-161014929-INS-174
chr1-161049833-INS-50
chr1-161086544-INS-323
chr1-161086682-INS-56
chr1-161099826-INS-5887
chr1-161101089-INS-68
chr1-161102364-INS-56
chr1-161261153-INS-87
chr1-161261188-INS-207
chr1-161298864-INS-483
chr1-161380601-INS-6201
chr1-161396164-INS-187
chr1-161463967-INS-916
chr1-161466741-INS-6728
chr1-161529476-INS-1346
chr1-161576930-INS-59
chr1-161613068-INS-351
chr1-161641938-INS-56
chr1-161641980-INS-179
chr1-161676813-INS-914
chr1-161706133-INS-2407
chr1-161712828-INS-50
chr1-161742983-INS-58
chr1-161746147-INS-52
chr1-161756582-INS-1707
chr1-161789880-INS-197
chr1-161790575-INS-52
chr1-161825300-INS-160
chr1-161828293-INS-66
chr1-161849665-INS-91
chr1-161861477-INS-62
chr1-161867140-INS-71
chr1-161903919-INS-50
chr1-161974647-INS-55
chr1-161995159-INS-150
chr1-161995166-INS-471
chr1-162104787-INS-58
chr1-162106334-INS-1999
chr1-162259613-INS-2737
chr1-162259922-INS-1056
chr1-162312447-INS-55
chr1-162312501-INS-135
chr1-162312616-INS-792
chr1-162340626-INS-62
chr1-162341321-INS-80
chr1-162434591-INS-1046
chr1-162436421-INS-6591
chr1-162446179-INS-200
chr1-162449781-INS-50
chr1-162458960-INS-1511
chr1-162491379-INS-87
chr1-162616091-INS-1580
chr1-162617196-INS-6171
chr1-162620208-INS-1073
chr1-162684741-INS-330
chr1-162735310-INS-325
chr1-162735380-INS-50
chr1-162740261-INS-70
chr1-162743036-INS-6716
chr1-162801676-INS-68
chr1-162805109-INS-56
chr1-162805616-INS-302
chr1-162912566-INS-1845
chr1-162932978-INS-6359
chr1-162951274-INS-52
chr1-162964419-INS-53
chr1-162964472-INS-139
chr1-163003190-INS-100
chr1-163003840-INS-159
chr1-163003840-INS-50
chr1-163057001-INS-76
chr1-163144118-INS-90

chr1-163170839-INS-1201
chr1-163177707-INS-6382
chr1-163270986-INS-6569
chr1-163336395-INS-6396
chr1-163364184-INS-135
chr1-163471321-INS-7409
chr1-163502518-INS-241
chr1-163512533-INS-50
chr1-163514538-INS-50
chr1-163665092-INS-689
chr1-163679817-INS-1261
chr1-163699742-INS-130
chr1-163718378-INS-3350
chr1-163732189-INS-208
chr1-163748042-INS-1899
chr1-163764300-INS-700
chr1-163770563-INS-95
chr1-163842653-INS-194
chr1-163842666-INS-53
chr1-163858102-INS-1591
chr1-163871788-INS-97
chr1-163872077-INS-55
chr1-163883550-INS-180
chr1-163886211-INS-205
chr1-163920493-INS-6362
chr1-163952103-INS-113
chr1-164010131-INS-131
chr1-164014655-INS-75
chr1-164021192-INS-213
chr1-164022697-INS-326
chr1-164037796-INS-1003
chr1-164068651-INS-51
chr1-164068698-INS-127
chr1-164292179-INS-61
chr1-164302727-INS-102
chr1-164314911-INS-138
chr1-164317031-INS-55
chr1-164320044-INS-222
chr1-164320421-INS-8754
chr1-164366823-INS-844
chr1-164367384-INS-176
chr1-164367780-INS-376
chr1-164368248-INS-6482
chr1-164389019-INS-112
chr1-164420317-INS-7227
chr1-164495922-INS-52
chr1-164496253-INS-110
chr1-164513812-INS-1038
chr1-164578441-INS-76
chr1-164583064-INS-228
chr1-164585472-INS-50
chr1-164603500-INS-2734
chr1-164616295-INS-398
chr1-164640710-INS-76
chr1-164664707-INS-6347
chr1-164668255-INS-6451
chr1-164694139-INS-354
chr1-164705954-INS-138
chr1-164708999-INS-155
chr1-164710490-INS-50
chr1-164713744-INS-6499
chr1-164714457-INS-3034
chr1-164731891-INS-52
chr1-164738650-INS-1892
chr1-164739691-INS-70
chr1-164770303-INS-6188
chr1-164780668-INS-70
chr1-164810657-INS-496
chr1-164822271-INS-221
chr1-164824445-INS-207
chr1-164937704-INS-102
chr1-164957823-INS-51
chr1-164974239-INS-139
chr1-164992466-INS-76
chr1-165026467-INS-96
chr1-165033692-INS-162
chr1-165075482-INS-5562

chr1-165093533-INS-83
chr1-165112085-INS-76
chr1-165112294-INS-54
chr1-165156433-INS-164
chr1-165156493-INS-715
chr1-165178692-INS-5422
chr1-165258305-INS-104
chr1-165267183-INS-131
chr1-165278710-INS-214
chr1-165349560-INS-54
chr1-165367049-INS-68
chr1-165399766-INS-2470
chr1-165402738-INS-100
chr1-165413896-INS-570
chr1-165414002-INS-837
chr1-165474101-INS-56
chr1-165486812-INS-6923
chr1-165499999-INS-195
chr1-165548410-INS-173
chr1-165560211-INS-54
chr1-165567169-INS-434
chr1-165589981-INS-57
chr1-165610882-INS-106
chr1-165639369-INS-180
chr1-165841600-INS-198
chr1-165857884-INS-158
chr1-165891275-INS-240
chr1-165899155-INS-68
chr1-165931035-INS-205
chr1-165954249-INS-527
chr1-165986246-INS-343
chr1-165986308-INS-717
chr1-165998240-INS-65
chr1-165999079-INS-119
chr1-166108026-INS-52
chr1-166238250-INS-2879
chr1-166268218-INS-50
chr1-166321478-INS-82
chr1-166396364-INS-846
chr1-166546482-INS-83
chr1-166572567-INS-146
chr1-166598857-INS-79
chr1-166604535-INS-2085
chr1-166638271-INS-171
chr1-166648984-INS-228
chr1-166649362-INS-644
chr1-166651127-INS-59
chr1-166657423-INS-6511
chr1-166664693-INS-208
chr1-166717736-INS-395
chr1-166723314-INS-1411
chr1-166749791-INS-64
chr1-166758871-INS-519
chr1-166759134-INS-1220
chr1-166762163-INS-6721
chr1-166772541-INS-66
chr1-166836578-INS-53
chr1-166839445-INS-90
chr1-166869475-INS-6586
chr1-166895002-INS-215
chr1-166937513-INS-50
chr1-166941867-INS-116
chr1-166959997-INS-80
chr1-166976597-INS-746
chr1-167039526-INS-77
chr1-167080560-INS-216
chr1-167122839-INS-212
chr1-167255851-INS-100
chr1-167277205-INS-2317
chr1-167292180-INS-51
chr1-167329414-INS-111
chr1-167329473-INS-848
chr1-167356151-INS-228
chr1-167358796-INS-6242
chr1-167406038-INS-79
chr1-167428195-INS-197
chr1-167474198-INS-61

chr1-167483920-INS-52
chr1-167513060-INS-437
chr1-167529492-INS-55
chr1-167554026-INS-150
chr1-167591386-INS-255
chr1-167636845-INS-132
chr1-167686994-INS-7538
chr1-167692661-INS-1213
chr1-167743700-INS-56
chr1-167744711-INS-5392
chr1-167768176-INS-5601
chr1-167796949-INS-50
chr1-167822179-INS-1778
chr1-167865983-INS-6218
chr1-167952138-INS-277
chr1-168042181-INS-50
chr1-168085611-INS-386
chr1-168087588-INS-222
chr1-168088781-INS-76
chr1-168088797-INS-362
chr1-168207236-INS-474
chr1-168226020-INS-90
chr1-168233476-INS-51
chr1-168252702-INS-61
chr1-168285580-INS-170
chr1-168285580-INS-55
chr1-168290554-INS-57
chr1-168291372-INS-70
chr1-168296002-INS-78
chr1-168386601-INS-6434
chr1-168494667-INS-68
chr1-168515147-INS-67
chr1-168573052-INS-5607
chr1-168638459-INS-51
chr1-168658366-INS-6493
chr1-168665401-INS-68
chr1-168667852-INS-164
chr1-168698907-INS-6362
chr1-168727984-INS-7394
chr1-168772728-INS-100
chr1-168807815-INS-50
chr1-168878542-INS-85
chr1-168883114-INS-6587
chr1-168907351-INS-153
chr1-168933674-INS-418
chr1-168996514-INS-50
chr1-169098235-INS-140
chr1-169135488-INS-64
chr1-169210889-INS-3324
chr1-169211213-INS-52
chr1-169211936-INS-52
chr1-169224008-INS-2782
chr1-169237742-INS-7093
chr1-169242841-INS-363
chr1-169249683-INS-140
chr1-169287438-INS-52
chr1-169345474-INS-522
chr1-169345768-INS-559
chr1-169376678-INS-1490
chr1-169378254-INS-58
chr1-169388125-INS-207
chr1-169422697-INS-1167
chr1-169425086-INS-6007
chr1-169442574-INS-246
chr1-169452092-INS-91
chr1-169516606-INS-7295
chr1-169536886-INS-80
chr1-169615812-INS-1290
chr1-169619806-INS-6779
chr1-169692513-INS-136
chr1-169758345-INS-51
chr1-169764083-INS-4399
chr1-169877450-INS-188
chr1-169910019-INS-2764
chr1-169930116-INS-100
chr1-169988458-INS-50
chr1-169996019-INS-7063

chr1-170004295-INS-170
chr1-170043767-INS-2816
chr1-170050040-INS-6198
chr1-170072629-INS-52
chr1-170163069-INS-126
chr1-170206294-INS-81
chr1-170225241-INS-497
chr1-170310128-INS-67
chr1-170314469-INS-51
chr1-170334052-INS-5979
chr1-170352435-INS-210
chr1-170382961-INS-54
chr1-170635861-INS-2829
chr1-170662376-INS-179
chr1-170717300-INS-50
chr1-170765298-INS-227
chr1-170809647-INS-222
chr1-170811564-INS-208
chr1-170855112-INS-4923
chr1-170884782-INS-207
chr1-170888304-INS-4295
chr1-170893890-INS-52
chr1-170913990-INS-1104
chr1-170913990-INS-2594
chr1-170927431-INS-50
chr1-170927550-INS-5377
chr1-170935049-INS-240
chr1-170936340-INS-2665
chr1-170945870-INS-114
chr1-170951774-INS-171
chr1-170962280-INS-79
chr1-171071790-INS-64
chr1-171071941-INS-208
chr1-171088591-INS-1864
chr1-171092168-INS-52
chr1-171132107-INS-395
chr1-171145851-INS-267
chr1-171150849-INS-225
chr1-171208929-INS-213
chr1-171208975-INS-60
chr1-171234158-INS-51
chr1-171248893-INS-54
chr1-171299919-INS-200
chr1-171318029-INS-120
chr1-171326171-INS-182
chr1-171330253-INS-198
chr1-171330387-INS-50
chr1-171343441-INS-223
chr1-171358391-INS-166
chr1-171360181-INS-92
chr1-171362294-INS-245
chr1-171366589-INS-57
chr1-171375671-INS-156
chr1-171375671-INS-58
chr1-171390113-INS-1822
chr1-171395057-INS-56
chr1-171398964-INS-204
chr1-171469948-INS-50
chr1-171639808-INS-7247
chr1-171649605-INS-50
chr1-171650401-INS-1890
chr1-171650401-INS-400
chr1-171666881-INS-162
chr1-171823385-INS-4728
chr1-171869361-INS-182
chr1-171891929-INS-54
chr1-171902084-INS-64
chr1-171936705-INS-50
chr1-172020560-INS-52
chr1-172067092-INS-1621
chr1-172092180-INS-7067
chr1-172101563-INS-54
chr1-172166572-INS-57
chr1-172278251-INS-56
chr1-172419235-INS-2239
chr1-172437977-INS-108
chr1-172544278-INS-416

chr1-172587686-INS-50
chr1-172596234-INS-343
chr1-172614004-INS-56
chr1-172615744-INS-3234
chr1-172621966-INS-440
chr1-172668146-INS-5171
chr1-172672660-INS-7336
chr1-172694055-INS-2614
chr1-172742735-INS-1068
chr1-172821559-INS-60
chr1-172822853-INS-230
chr1-172835971-INS-945
chr1-173030879-INS-1218
chr1-173062352-INS-1225
chr1-173099728-INS-58
chr1-173099816-INS-543
chr1-173108789-INS-3579
chr1-173157251-INS-50
chr1-173322213-INS-74
chr1-173334806-INS-200
chr1-173375572-INS-1003
chr1-173381739-INS-3130
chr1-173439106-INS-1881
chr1-173481343-INS-197
chr1-173498903-INS-60
chr1-173503637-INS-52
chr1-173535930-INS-1019
chr1-173539531-INS-1147
chr1-173544054-INS-6206
chr1-173568191-INS-74
chr1-173581228-INS-4990
chr1-173603119-INS-61
chr1-173741418-INS-66
chr1-173745675-INS-105
chr1-173779386-INS-439
chr1-173848944-INS-7088
chr1-173859168-INS-263
chr1-173923197-INS-1561
chr1-173933631-INS-318
chr1-173933631-INS-55
chr1-173938908-INS-2926
chr1-173984145-INS-70
chr1-173991041-INS-863
chr1-173995151-INS-1884
chr1-174033497-INS-6442
chr1-174041803-INS-122
chr1-174064235-INS-1037
chr1-174064235-INS-360
chr1-174068171-INS-53
chr1-174163151-INS-56
chr1-174163852-INS-1542
chr1-174186984-INS-2067
chr1-174273060-INS-64
chr1-174274638-INS-345
chr1-174274816-INS-54
chr1-174297316-INS-80
chr1-174298096-INS-90
chr1-174303256-INS-54
chr1-174325784-INS-1282
chr1-174345888-INS-7303
chr1-174352053-INS-1703
chr1-174367036-INS-67
chr1-174367969-INS-170
chr1-174368038-INS-56
chr1-174396397-INS-1889
chr1-174403228-INS-1259
chr1-174409024-INS-2148
chr1-174430297-INS-50
chr1-174523424-INS-50
chr1-174635827-INS-62
chr1-174637209-INS-14319
chr1-174637209-INS-6855
chr1-174643543-INS-73
chr1-174753980-INS-58
chr1-174770081-INS-7767
chr1-174848424-INS-220
chr1-174855636-INS-202

chr1-174860696-INS-122
chr1-174956840-INS-7326
chr1-174994188-INS-50
chr1-174994251-INS-187
chr1-175059665-INS-220
chr1-175061729-INS-12789
chr1-175077184-INS-158
chr1-175094983-INS-7143
chr1-175132609-INS-91
chr1-175142514-INS-1385
chr1-175186601-INS-4446
chr1-175193113-INS-52
chr1-175215502-INS-101
chr1-175280256-INS-50
chr1-175353073-INS-50
chr1-175409540-INS-422
chr1-175410093-INS-77
chr1-175429421-INS-50
chr1-175430602-INS-64
chr1-175454905-INS-864
chr1-175525697-INS-50
chr1-175538062-INS-191
chr1-175546524-INS-98
chr1-175574368-INS-50
chr1-175638912-INS-1020
chr1-175683028-INS-70
chr1-175714718-INS-96
chr1-175747482-INS-50
chr1-175771844-INS-2838
chr1-175782959-INS-85
chr1-175794988-INS-163
chr1-175797911-INS-1218
chr1-175825795-INS-5189
chr1-175832109-INS-1862
chr1-175837089-INS-6441
chr1-175853798-INS-71
chr1-175891340-INS-76
chr1-175924917-INS-56
chr1-175953143-INS-52
chr1-175958915-INS-12951
chr1-175960692-INS-66
chr1-175988957-INS-1440
chr1-175989865-INS-52
chr1-176057716-INS-98
chr1-176058221-INS-52
chr1-176065958-INS-7525
chr1-176113472-INS-3000
chr1-176178194-INS-215
chr1-176217493-INS-310
chr1-176220216-INS-872
chr1-176233826-INS-1127
chr1-176240051-INS-2059
chr1-176246255-INS-50
chr1-176249978-INS-52
chr1-176266544-INS-146
chr1-176273299-INS-57
chr1-176286988-INS-2655
chr1-176324094-INS-5663
chr1-176334762-INS-3529
chr1-176415525-INS-52
chr1-176442957-INS-498
chr1-176480848-INS-114
chr1-176545930-INS-3437
chr1-176549729-INS-210
chr1-176566145-INS-70
chr1-176769754-INS-4004
chr1-176837337-INS-3074
chr1-176891620-INS-221
chr1-176892126-INS-518
chr1-176966795-INS-224
chr1-177157066-INS-85
chr1-177157589-INS-278
chr1-177339742-INS-51
chr1-177349966-INS-352
chr1-177437816-INS-50
chr1-177460764-INS-2965
chr1-177550116-INS-7228

chr1-177566173-INS-175
chr1-177643357-INS-205
chr1-177668278-INS-260
chr1-177685735-INS-149
chr1-177685735-INS-301
chr1-177686161-INS-1769
chr1-177720089-INS-332
chr1-177764491-INS-131
chr1-177782578-INS-207
chr1-177782742-INS-197
chr1-177785065-INS-297
chr1-177805755-INS-467
chr1-177806017-INS-4692
chr1-177809913-INS-386
chr1-177880855-INS-3004
chr1-177966648-INS-6397
chr1-178106957-INS-52
chr1-178108656-INS-221
chr1-178124127-INS-6453
chr1-178190573-INS-5413
chr1-178214947-INS-349
chr1-178269420-INS-7067
chr1-178271197-INS-82
chr1-178281676-INS-234
chr1-178293960-INS-88
chr1-178307701-INS-3031
chr1-178366186-INS-80
chr1-178376737-INS-60
chr1-178376741-INS-252
chr1-178376880-INS-1244
chr1-178402123-INS-180
chr1-178560527-INS-4837
chr1-178569674-INS-6400
chr1-178600919-INS-267
chr1-178600928-INS-78
chr1-178603920-INS-50
chr1-178604696-INS-72
chr1-178614904-INS-50
chr1-178646166-INS-342
chr1-178707157-INS-169
chr1-178733712-INS-97
chr1-178820708-INS-408
chr1-178844316-INS-6394
chr1-179023510-INS-4114
chr1-179049233-INS-447
chr1-179049470-INS-210
chr1-179167604-INS-413
chr1-179298489-INS-317
chr1-179404682-INS-133
chr1-179449977-INS-54
chr1-179472412-INS-494
chr1-179540636-INS-50
chr1-179659660-INS-1439
chr1-179665228-INS-1175
chr1-179700282-INS-67
chr1-179715326-INS-161
chr1-179744884-INS-215
chr1-179752872-INS-6729
chr1-179868501-INS-120
chr1-179868501-INS-55
chr1-179869988-INS-78
chr1-179948592-INS-148
chr1-179949122-INS-75
chr1-179949155-INS-200
chr1-180056239-INS-403
chr1-180058435-INS-2758
chr1-180080101-INS-1865
chr1-180151814-INS-50
chr1-180178892-INS-126
chr1-180269627-INS-4848
chr1-180278593-INS-85
chr1-180287992-INS-164
chr1-180296962-INS-63
chr1-180336613-INS-188
chr1-180373771-INS-204
chr1-180380316-INS-1272
chr1-180385472-INS-58

chr1-180386398-INS-210
chr1-180388560-INS-93
chr1-180390651-INS-168
chr1-180402411-INS-156
chr1-180418745-INS-333
chr1-180548398-INS-208
chr1-180577936-INS-134
chr1-180594134-INS-458
chr1-180598129-INS-51
chr1-180618398-INS-503
chr1-180622961-INS-181
chr1-180644567-INS-58
chr1-180662672-INS-351
chr1-180668838-INS-51
chr1-180669448-INS-216
chr1-180689382-INS-56
chr1-180700352-INS-524
chr1-180700358-INS-1752
chr1-180700385-INS-219
chr1-180747722-INS-129
chr1-180768049-INS-80
chr1-180797691-INS-111
chr1-180858609-INS-219
chr1-180864148-INS-649
chr1-180898076-INS-78
chr1-180902955-INS-52
chr1-180942762-INS-326
chr1-181007754-INS-209
chr1-181010592-INS-55
chr1-181088674-INS-210
chr1-181099282-INS-52
chr1-181122854-INS-51
chr1-181132406-INS-50
chr1-181159003-INS-88
chr1-181182532-INS-103
chr1-181182955-INS-50
chr1-181198542-INS-62
chr1-181205805-INS-95
chr1-181229861-INS-181
chr1-181229981-INS-532
chr1-181273807-INS-58
chr1-181348221-INS-364
chr1-181389615-INS-64
chr1-181391761-INS-218
chr1-181410937-INS-2534
chr1-181525206-INS-58
chr1-181529422-INS-50
chr1-181531473-INS-52
chr1-181541456-INS-1490
chr1-181546018-INS-6100
chr1-181571204-INS-4160
chr1-181580268-INS-500
chr1-181619664-INS-711
chr1-181626747-INS-110
chr1-181678873-INS-4981
chr1-181684545-INS-50
chr1-181702873-INS-55
chr1-181703434-INS-174
chr1-181759378-INS-113
chr1-181812106-INS-162
chr1-181812325-INS-691
chr1-181862416-INS-52
chr1-181891881-INS-133
chr1-181896507-INS-215
chr1-181904388-INS-147
chr1-181930965-INS-215
chr1-181935525-INS-189
chr1-181962246-INS-135
chr1-181985922-INS-105
chr1-182015623-INS-4845
chr1-182025369-INS-86
chr1-182080991-INS-60
chr1-182113440-INS-64
chr1-182124181-INS-2105
chr1-182154422-INS-104
chr1-182169738-INS-52
chr1-182193144-INS-60

chr1-182193308-INS-739
chr1-182201263-INS-298
chr1-182311178-INS-981
chr1-182381153-INS-52
chr1-182386331-INS-196
chr1-182386734-INS-88
chr1-182446194-INS-52
chr1-182454232-INS-56
chr1-182458684-INS-427
chr1-182466250-INS-55
chr1-182519900-INS-996
chr1-182649868-INS-180
chr1-182705615-INS-518
chr1-182727606-INS-268
chr1-182812517-INS-87
chr1-182838139-INS-6222
chr1-182838616-INS-67
chr1-182839413-INS-202
chr1-182962262-INS-142
chr1-182968590-INS-53
chr1-182995103-INS-162
chr1-183073820-INS-218
chr1-183080930-INS-76
chr1-183123171-INS-54
chr1-183195169-INS-8671
chr1-183201976-INS-68
chr1-183210789-INS-50
chr1-183266293-INS-312
chr1-183267887-INS-150
chr1-183305570-INS-110
chr1-183322938-INS-108
chr1-183328493-INS-50
chr1-183332467-INS-6443
chr1-183339166-INS-50
chr1-183349485-INS-300
chr1-183360897-INS-88
chr1-183384668-INS-168
chr1-183410181-INS-75
chr1-183414098-INS-10067
chr1-183429559-INS-80
chr1-183480526-INS-169
chr1-183635714-INS-2752
chr1-183667782-INS-52
chr1-183703463-INS-1492
chr1-183742576-INS-68
chr1-183751291-INS-87
chr1-183935404-INS-2241
chr1-183976249-INS-56
chr1-184001104-INS-1064
chr1-184059022-INS-77
chr1-184068484-INS-98
chr1-184069807-INS-123
chr1-184081723-INS-66
chr1-184120142-INS-160
chr1-184122040-INS-56
chr1-184148680-INS-88
chr1-184152351-INS-104
chr1-184191149-INS-68
chr1-184197863-INS-56
chr1-184206497-INS-1508
chr1-184237988-INS-54
chr1-184280694-INS-140
chr1-184401351-INS-58
chr1-184495627-INS-90
chr1-184526610-INS-120
chr1-184526610-INS-257
chr1-184594783-INS-129
chr1-184644905-INS-2158
chr1-184684717-INS-130
chr1-184751849-INS-150
chr1-184766453-INS-236
chr1-184767551-INS-7224
chr1-184810756-INS-186
chr1-184815913-INS-267
chr1-184815913-INS-58
chr1-184832503-INS-52
chr1-184861631-INS-52

chr1-184861707-INS-936
chr1-184933246-INS-2744
chr1-184947152-INS-495
chr1-184951032-INS-604
chr1-184986903-INS-112
chr1-184987130-INS-104
chr1-185010193-INS-256
chr1-185092819-INS-78
chr1-185124871-INS-72
chr1-185125478-INS-71
chr1-185194118-INS-253
chr1-185250091-INS-121
chr1-185250091-INS-50
chr1-185271065-INS-1838
chr1-185272632-INS-87
chr1-185283856-INS-65
chr1-185283899-INS-231
chr1-185314625-INS-1751
chr1-185322843-INS-152
chr1-185378818-INS-60
chr1-185405830-INS-73
chr1-185406171-INS-102
chr1-185410482-INS-94
chr1-185453005-INS-288
chr1-185472982-INS-1832
chr1-185484372-INS-10263
chr1-185706096-INS-52
chr1-185707788-INS-200
chr1-185752099-INS-1326
chr1-185770537-INS-154
chr1-185792423-INS-1519
chr1-185816963-INS-1060
chr1-185883099-INS-6796
chr1-185938719-INS-92
chr1-185955369-INS-161
chr1-185977115-INS-502
chr1-186000544-INS-121
chr1-186034827-INS-3390
chr1-186109147-INS-84
chr1-186121649-INS-612
chr1-186128530-INS-257
chr1-186153071-INS-68
chr1-186153778-INS-67
chr1-186164871-INS-159
chr1-186164916-INS-732
chr1-186165034-INS-241
chr1-186180797-INS-13083
chr1-186186454-INS-270
chr1-186216439-INS-78
chr1-186230558-INS-58
chr1-186285352-INS-168
chr1-186354004-INS-51
chr1-186365994-INS-187
chr1-186382695-INS-101
chr1-186421456-INS-204
chr1-186464168-INS-1705
chr1-186467410-INS-68
chr1-186467939-INS-124
chr1-186521609-INS-50
chr1-186553591-INS-256
chr1-186596189-INS-1264
chr1-186600494-INS-2514
chr1-186618359-INS-2070
chr1-186635269-INS-791
chr1-186637826-INS-50
chr1-186706993-INS-82
chr1-186708082-INS-74
chr1-186714630-INS-1305
chr1-186734370-INS-60
chr1-186735117-INS-94
chr1-186767506-INS-297
chr1-186767749-INS-844
chr1-186819625-INS-173
chr1-186831171-INS-498
chr1-186889020-INS-403
chr1-186898403-INS-7090
chr1-186903572-INS-80

chr1-186922519-INS-59
chr1-186932469-INS-201
chr1-186988612-INS-202
chr1-187101072-INS-2591
chr1-187105698-INS-51
chr1-187119443-INS-296
chr1-187121399-INS-50
chr1-187149068-INS-81
chr1-187166337-INS-452
chr1-187200278-INS-54
chr1-187200689-INS-174
chr1-187268206-INS-52
chr1-187313709-INS-54
chr1-187323498-INS-444
chr1-187324917-INS-52
chr1-187351958-INS-75
chr1-187439181-INS-52
chr1-187440717-INS-1889
chr1-187619444-INS-132
chr1-187623331-INS-76
chr1-187758485-INS-136
chr1-187803291-INS-239
chr1-187805612-INS-81
chr1-187862853-INS-202
chr1-187863027-INS-92
chr1-187895584-INS-68
chr1-187932371-INS-5543
chr1-187934145-INS-654
chr1-188038866-INS-150
chr1-188044989-INS-168
chr1-188078157-INS-51
chr1-188156125-INS-5244
chr1-188178217-INS-181
chr1-188331087-INS-90
chr1-188331789-INS-52
chr1-188386057-INS-72
chr1-188522371-INS-170
chr1-188603845-INS-6723
chr1-188606010-INS-176
chr1-188606707-INS-62
chr1-188619926-INS-352
chr1-188633509-INS-94
chr1-188663188-INS-1248
chr1-188690122-INS-708
chr1-188721200-INS-607
chr1-188742299-INS-389
chr1-188765237-INS-671
chr1-188804484-INS-240
chr1-188830436-INS-52
chr1-188846120-INS-8393
chr1-188846190-INS-152
chr1-188846828-INS-328
chr1-188862920-INS-76
chr1-188878230-INS-286
chr1-188893836-INS-232
chr1-188898403-INS-3281
chr1-188901517-INS-239
chr1-188908374-INS-63
chr1-188911949-INS-142
chr1-188914274-INS-923
chr1-188933359-INS-100
chr1-188981122-INS-1889
chr1-188983335-INS-7165
chr1-188999619-INS-2861
chr1-189013537-INS-87
chr1-189032969-INS-52
chr1-189033425-INS-140
chr1-189052055-INS-102
chr1-189103010-INS-84
chr1-189234023-INS-115
chr1-189244667-INS-88
chr1-189254317-INS-100
chr1-189254567-INS-50
chr1-189255179-INS-68
chr1-189271964-INS-664
chr1-189328107-INS-1892
chr1-189331080-INS-148

chr1-189427647-INS-200
chr1-189427687-INS-560
chr1-189433437-INS-151
chr1-189458184-INS-68
chr1-189478774-INS-294
chr1-189483408-INS-108
chr1-189497092-INS-51
chr1-189546110-INS-289
chr1-189601570-INS-191
chr1-189637932-INS-78
chr1-189638312-INS-580
chr1-189641276-INS-1311
chr1-189673621-INS-50
chr1-189712995-INS-118
chr1-189716162-INS-136
chr1-189723466-INS-89
chr1-189761689-INS-55
chr1-189790460-INS-123
chr1-189790460-INS-353
chr1-189790790-INS-80
chr1-189791136-INS-345
chr1-189916397-INS-552
chr1-189920352-INS-145
chr1-189944340-INS-175
chr1-189944613-INS-145
chr1-190034203-INS-7699
chr1-190034936-INS-2350
chr1-190046246-INS-148
chr1-190054543-INS-56
chr1-190073696-INS-577
chr1-190111437-INS-52
chr1-190121054-INS-360
chr1-190147453-INS-106
chr1-190162512-INS-60
chr1-190245051-INS-215
chr1-190254401-INS-191
chr1-190254468-INS-1504
chr1-190254490-INS-193
chr1-190259500-INS-240
chr1-190275977-INS-1536
chr1-190289864-INS-741
chr1-190339473-INS-91
chr1-190340090-INS-225
chr1-190416809-INS-72
chr1-190475989-INS-50
chr1-190513079-INS-1583
chr1-190517102-INS-535
chr1-190571759-INS-4770
chr1-190572334-INS-501
chr1-190583744-INS-239
chr1-190594548-INS-1266
chr1-190595879-INS-510
chr1-190604731-INS-460
chr1-190610684-INS-103
chr1-190610804-INS-148
chr1-190624857-INS-1102
chr1-190624858-INS-232
chr1-190634894-INS-370
chr1-190635330-INS-840
chr1-190650180-INS-55
chr1-190689351-INS-67
chr1-190697828-INS-116
chr1-190712208-INS-252
chr1-190719791-INS-90
chr1-190727091-INS-71
chr1-190770186-INS-52
chr1-190776648-INS-4839
chr1-190782890-INS-60
chr1-190783925-INS-289
chr1-190784344-INS-223
chr1-190784784-INS-679
chr1-190811275-INS-347
chr1-190888638-INS-62
chr1-190908674-INS-52
chr1-190945317-INS-7095
chr1-191006847-INS-120
chr1-191032872-INS-272

chr1-191038386-INS-4824
chr1-191045936-INS-65
chr1-191050915-INS-61
chr1-191090018-INS-1011
chr1-191092245-INS-1503
chr1-191151039-INS-52
chr1-191152171-INS-102
chr1-191183431-INS-100
chr1-191183718-INS-140
chr1-191221031-INS-7160
chr1-191239525-INS-52
chr1-191245201-INS-72
chr1-191288100-INS-1076
chr1-191403109-INS-1782
chr1-191476312-INS-88
chr1-191486137-INS-62
chr1-191522966-INS-535
chr1-191588245-INS-238
chr1-191588777-INS-149
chr1-191614651-INS-144
chr1-191614651-INS-5427
chr1-191628369-INS-62
chr1-191906382-INS-50
chr1-191934125-INS-136
chr1-191948013-INS-50
chr1-191956564-INS-50
chr1-191962121-INS-7440
chr1-191972390-INS-4223
chr1-191981315-INS-185
chr1-192071012-INS-339
chr1-192153376-INS-2681
chr1-192291205-INS-6216
chr1-192347436-INS-106
chr1-192347609-INS-56
chr1-192348566-INS-1098
chr1-192350779-INS-51
chr1-192352511-INS-58
chr1-192400071-INS-71
chr1-192452835-INS-227
chr1-192510908-INS-7185
chr1-192681915-INS-3382
chr1-192714922-INS-218
chr1-192715155-INS-138
chr1-192728372-INS-90
chr1-192730750-INS-65
chr1-192780552-INS-137
chr1-193058358-INS-112
chr1-193093732-INS-75
chr1-193105421-INS-83
chr1-193170787-INS-90
chr1-193171559-INS-80
chr1-193230464-INS-222
chr1-193239834-INS-57
chr1-193245566-INS-99
chr1-193342075-INS-5249
chr1-193389594-INS-145
chr1-193492203-INS-201
chr1-193557706-INS-114
chr1-193657936-INS-50
chr1-193764850-INS-1386
chr1-193832104-INS-513
chr1-193838931-INS-6905
chr1-193885684-INS-122
chr1-193917967-INS-89
chr1-194007508-INS-132
chr1-194074974-INS-9478
chr1-194123690-INS-442
chr1-194155010-INS-6782
chr1-194234387-INS-70
chr1-194301070-INS-67
chr1-194649480-INS-50
chr1-194844142-INS-1023
chr1-194895121-INS-1222
chr1-194907087-INS-4474
chr1-194907346-INS-188
chr10-3055305-INS-431
chr10-3100157-INS-57

chr10-3155498-INS-52
chr10-3205432-INS-422
chr10-3297937-INS-55
chr10-3330871-INS-75
chr10-3469139-INS-68
chr10-3469552-INS-91
chr10-3469644-INS-345
chr10-3506238-INS-126
chr10-3579879-INS-96
chr10-3706808-INS-1838
chr10-3713955-INS-65
chr10-3727976-INS-50
chr10-3736835-INS-1752
chr10-3755354-INS-1754
chr10-3759897-INS-76
chr10-3805322-INS-315
chr10-3817323-INS-504
chr10-3820688-INS-66
chr10-3868150-INS-225
chr10-3878313-INS-70
chr10-3898038-INS-52
chr10-3938695-INS-80
chr10-3962492-INS-219
chr10-3996556-INS-73
chr10-4005363-INS-5778
chr10-4011934-INS-97
chr10-4015034-INS-157
chr10-4015135-INS-60
chr10-4018665-INS-362
chr10-4026337-INS-54
chr10-4063073-INS-203
chr10-4101534-INS-8877
chr10-4112013-INS-198
chr10-4115233-INS-52
chr10-4151513-INS-70
chr10-4156024-INS-4869
chr10-4175365-INS-225
chr10-4175422-INS-55
chr10-4175865-INS-203
chr10-4182119-INS-277
chr10-4191607-INS-306
chr10-4191920-INS-618
chr10-4191926-INS-254
chr10-4195730-INS-98
chr10-4198948-INS-55
chr10-4199846-INS-137
chr10-4200485-INS-76
chr10-4200855-INS-240
chr10-4202265-INS-215
chr10-4206001-INS-139
chr10-4225676-INS-117
chr10-4238162-INS-53
chr10-4238240-INS-107
chr10-4249141-INS-195
chr10-4265102-INS-115
chr10-4290000-INS-374
chr10-4390122-INS-185
chr10-4423068-INS-61
chr10-4426412-INS-1087
chr10-4428595-INS-75
chr10-4429265-INS-64
chr10-4436649-INS-174
chr10-4451326-INS-79
chr10-4474161-INS-6359
chr10-4477207-INS-55
chr10-4493432-INS-52
chr10-4500949-INS-86
chr10-4501088-INS-645
chr10-4501168-INS-200
chr10-4513534-INS-1267
chr10-4527556-INS-64
chr10-4529634-INS-498
chr10-4537499-INS-50
chr10-4550071-INS-160
chr10-4581748-INS-310
chr10-4581973-INS-228
chr10-4594728-INS-1865

chr10-4594830-INS-51
chr10-4594993-INS-661
chr10-4600737-INS-1829
chr10-4618224-INS-60
chr10-4619022-INS-161
chr10-4668749-INS-55
chr10-4678287-INS-134
chr10-4678287-INS-54
chr10-4694685-INS-52
chr10-4719733-INS-50
chr10-4730083-INS-6413
chr10-4771070-INS-370
chr10-4816231-INS-55
chr10-4866185-INS-1492
chr10-4943186-INS-62
chr10-4985191-INS-130
chr10-5012232-INS-147
chr10-5036531-INS-132
chr10-5080409-INS-52
chr10-5097568-INS-119
chr10-5162589-INS-50
chr10-5180396-INS-58
chr10-5202148-INS-88
chr10-5206542-INS-6083
chr10-5218433-INS-298
chr10-5222497-INS-2110
chr10-5260169-INS-72
chr10-5286475-INS-100
chr10-5291168-INS-377
chr10-5326975-INS-1210
chr10-5333879-INS-56
chr10-5334022-INS-59
chr10-5432313-INS-50
chr10-5441572-INS-312
chr10-5445062-INS-242
chr10-5449114-INS-56
chr10-5489417-INS-1188
chr10-5489482-INS-216
chr10-5497046-INS-50
chr10-5541346-INS-93
chr10-5630826-INS-204
chr10-5645164-INS-372
chr10-5735695-INS-3161
chr10-5756399-INS-184
chr10-5757499-INS-260
chr10-5757499-INS-53
chr10-5779089-INS-82
chr10-5791675-INS-236
chr10-5791858-INS-98
chr10-5792910-INS-191
chr10-5797078-INS-217
chr10-5801608-INS-210
chr10-5808741-INS-56
chr10-5820774-INS-84
chr10-5836122-INS-1158
chr10-5836330-INS-275
chr10-5839016-INS-207
chr10-5844175-INS-102
chr10-5849134-INS-96
chr10-5849765-INS-50
chr10-5884831-INS-6330
chr10-5901572-INS-205
chr10-5918758-INS-78
chr10-5933027-INS-419
chr10-5959679-INS-92
chr10-5973747-INS-50
chr10-5974020-INS-6448
chr10-5975402-INS-5519
chr10-5997859-INS-407
chr10-6000710-INS-180
chr10-6013334-INS-65
chr10-6016116-INS-5502
chr10-6059581-INS-516
chr10-6083776-INS-62
chr10-6092089-INS-6309
chr10-6138382-INS-1581
chr10-6200109-INS-53

chr10-6207605-INS-6487
chr10-6212062-INS-2940
chr10-6214101-INS-6618
chr10-6220817-INS-2915
chr10-6259137-INS-6413
chr10-6261573-INS-68
chr10-6360835-INS-92
chr10-6395518-INS-54
chr10-6395975-INS-90
chr10-6396847-INS-981
chr10-6398649-INS-1166
chr10-6413895-INS-51
chr10-6435034-INS-6390
chr10-6464534-INS-60
chr10-6465000-INS-98
chr10-6482586-INS-6575
chr10-6515339-INS-117
chr10-6530143-INS-50
chr10-6548051-INS-6508
chr10-6549611-INS-6235
chr10-6555544-INS-6488
chr10-6580462-INS-170
chr10-6580618-INS-66
chr10-6580668-INS-616
chr10-6583246-INS-54
chr10-6604814-INS-208
chr10-6634434-INS-3164
chr10-6661277-INS-50
chr10-6716689-INS-106
chr10-6720708-INS-3182
chr10-6729235-INS-2311
chr10-6752087-INS-80
chr10-6796088-INS-527
chr10-6828631-INS-51
chr10-6838592-INS-180
chr10-6841993-INS-6429
chr10-6855509-INS-50
chr10-6875692-INS-76
chr10-6880037-INS-162
chr10-6936926-INS-6637
chr10-6939863-INS-50
chr10-6941003-INS-74
chr10-6954005-INS-280
chr10-6973722-INS-60
chr10-6986596-INS-231
chr10-6999014-INS-55
chr10-7028107-INS-70
chr10-7052976-INS-60
chr10-7089956-INS-2004
chr10-7099606-INS-283
chr10-7118206-INS-221
chr10-7167268-INS-55
chr10-7175774-INS-474
chr10-7219428-INS-4419
chr10-7224276-INS-143
chr10-7273028-INS-5892
chr10-7292265-INS-6255
chr10-7312999-INS-57
chr10-7321445-INS-512
chr10-7323827-INS-5098
chr10-7344481-INS-141
chr10-7405534-INS-4001
chr10-7433567-INS-60
chr10-7441232-INS-1894
chr10-7452780-INS-552
chr10-7468676-INS-156
chr10-7496669-INS-101
chr10-7526264-INS-54
chr10-7532503-INS-74
chr10-7534118-INS-477
chr10-7573159-INS-1418
chr10-7573159-INS-426
chr10-7602818-INS-106
chr10-7603232-INS-52
chr10-7617467-INS-13133
chr10-7642305-INS-485
chr10-7650036-INS-112

chr10-7677021-INS-2512
chr10-7695209-INS-471
chr10-7700999-INS-191
chr10-7778721-INS-402
chr10-7778727-INS-2109
chr10-7850306-INS-50
chr10-7896632-INS-72
chr10-7985718-INS-12964
chr10-7988150-INS-500
chr10-7988699-INS-858
chr10-8011096-INS-54
chr10-8020276-INS-98
chr10-8027476-INS-2720
chr10-8044675-INS-231
chr10-8060084-INS-58
chr10-8073819-INS-185
chr10-8073862-INS-78
chr10-8074144-INS-121
chr10-8085265-INS-2122
chr10-8090860-INS-498
chr10-8091401-INS-92
chr10-8093863-INS-5358
chr10-8109946-INS-3591
chr10-8135376-INS-111
chr10-8177473-INS-241
chr10-8224475-INS-111
chr10-8283074-INS-472
chr10-8296426-INS-253
chr10-8299325-INS-138
chr10-8304758-INS-50
chr10-8325831-INS-65
chr10-8328412-INS-50
chr10-8328415-INS-205
chr10-8329844-INS-567
chr10-8329948-INS-1656
chr10-8353205-INS-244
chr10-8358186-INS-72
chr10-8358312-INS-468
chr10-8366409-INS-7219
chr10-8385379-INS-65
chr10-8416148-INS-72
chr10-8448410-INS-447
chr10-8451121-INS-50
chr10-8462250-INS-207
chr10-8471536-INS-98
chr10-8472976-INS-1413
chr10-8483167-INS-407
chr10-8487756-INS-152
chr10-8494929-INS-509
chr10-8505800-INS-3654
chr10-8508841-INS-129
chr10-8514921-INS-461
chr10-8531690-INS-71
chr10-8540378-INS-506
chr10-8554083-INS-4686
chr10-8571486-INS-397
chr10-8576766-INS-396
chr10-8576998-INS-198
chr10-8639737-INS-354
chr10-8648886-INS-57
chr10-8651269-INS-468
chr10-8666624-INS-151
chr10-8673692-INS-6683
chr10-8685103-INS-1890
chr10-8691989-INS-307
chr10-8699523-INS-4189
chr10-8705156-INS-195
chr10-8717990-INS-50
chr10-8725809-INS-384
chr10-8726379-INS-73
chr10-8731422-INS-62
chr10-8748408-INS-242
chr10-8765377-INS-218
chr10-8772337-INS-55
chr10-8773019-INS-502
chr10-8773059-INS-60
chr10-8773718-INS-149

chr10-8781579-INS-56
chr10-8790683-INS-396
chr10-8790707-INS-153
chr10-8801696-INS-259
chr10-8838102-INS-134
chr10-8870831-INS-6454
chr10-8882564-INS-76
chr10-8891401-INS-227
chr10-8897993-INS-86
chr10-8919504-INS-144
chr10-8925401-INS-58
chr10-8944038-INS-136
chr10-8952178-INS-77
chr10-9059658-INS-56
chr10-9071001-INS-51
chr10-9071181-INS-69
chr10-9071941-INS-84
chr10-9072669-INS-404
chr10-9091298-INS-58
chr10-9091351-INS-321
chr10-9119928-INS-1495
chr10-9120019-INS-210
chr10-9121394-INS-3210
chr10-9131078-INS-6799
chr10-9134781-INS-140
chr10-9143197-INS-114
chr10-9143307-INS-266
chr10-9187589-INS-73
chr10-9220052-INS-64
chr10-9257508-INS-96
chr10-9258161-INS-80
chr10-9258443-INS-366
chr10-9271277-INS-1070
chr10-9271472-INS-214
chr10-9271494-INS-459
chr10-9300590-INS-7230
chr10-9357426-INS-86
chr10-9358336-INS-59
chr10-9358430-INS-64
chr10-9358517-INS-144
chr10-9373384-INS-122
chr10-9389869-INS-888
chr10-9391372-INS-56
chr10-9395717-INS-50
chr10-9402783-INS-1598
chr10-9403040-INS-88
chr10-9407186-INS-79
chr10-9407251-INS-554
chr10-9408072-INS-62
chr10-9457091-INS-432
chr10-9472897-INS-276
chr10-9473181-INS-242
chr10-9492982-INS-122
chr10-9508275-INS-6392
chr10-9543749-INS-3126
chr10-9544204-INS-50
chr10-9552498-INS-58
chr10-9575156-INS-13658
chr10-9593774-INS-136
chr10-9600325-INS-6432
chr10-9624084-INS-54
chr10-9652820-INS-50
chr10-9687563-INS-3435
chr10-9729218-INS-612
chr10-9730485-INS-72
chr10-9733192-INS-70
chr10-9760609-INS-55
chr10-9831414-INS-6314
chr10-9836912-INS-60
chr10-9854165-INS-2788
chr10-9896349-INS-2141
chr10-9927254-INS-135
chr10-9945800-INS-526
chr10-9951516-INS-471
chr10-9973752-INS-55
chr10-9988792-INS-59
chr10-10036813-INS-58

chr10-10054905-INS-77
chr10-10203638-INS-1732
chr10-10211085-INS-156
chr10-10224655-INS-54
chr10-10232167-INS-204
chr10-10309070-INS-100
chr10-10332205-INS-50
chr10-10347118-INS-90
chr10-10423591-INS-108
chr10-10423634-INS-576
chr10-10537268-INS-507
chr10-10568735-INS-84
chr10-10609891-INS-58
chr10-10613825-INS-7240
chr10-10703392-INS-54
chr10-10722897-INS-52
chr10-10771850-INS-7660
chr10-10781396-INS-69
chr10-10781604-INS-67
chr10-10809398-INS-385
chr10-10839772-INS-501
chr10-10887381-INS-648
chr10-10891921-INS-67
chr10-10900555-INS-872
chr10-10909201-INS-744
chr10-10965313-INS-433
chr10-10982114-INS-1116
chr10-10996742-INS-66
chr10-11014489-INS-53
chr10-11033314-INS-307
chr10-11106692-INS-101
chr10-11163503-INS-215
chr10-11164343-INS-200
chr10-11202995-INS-3375
chr10-11222094-INS-3166
chr10-11242836-INS-60
chr10-11243613-INS-176
chr10-11263815-INS-835
chr10-11309358-INS-12741
chr10-11337582-INS-3112
chr10-11344341-INS-120
chr10-11437079-INS-147
chr10-11452803-INS-301
chr10-11501707-INS-3325
chr10-11600681-INS-254
chr10-11618024-INS-65
chr10-11618480-INS-100
chr10-11683807-INS-187
chr10-11683807-INS-55
chr10-11712286-INS-207
chr10-11739446-INS-292
chr10-11748939-INS-518
chr10-11752789-INS-54
chr10-11774184-INS-80
chr10-11804234-INS-3697
chr10-11804263-INS-54
chr10-11804709-INS-955
chr10-11866074-INS-579
chr10-11866399-INS-3248
chr10-11870316-INS-63
chr10-11898945-INS-175
chr10-11949506-INS-1362
chr10-11952541-INS-531
chr10-11952542-INS-199
chr10-11958303-INS-50
chr10-12016159-INS-55
chr10-12127523-INS-50
chr10-12137912-INS-216
chr10-12179319-INS-2517
chr10-12202989-INS-2231
chr10-12203101-INS-56
chr10-12222018-INS-2879
chr10-12245049-INS-4841
chr10-12246899-INS-1487
chr10-12298313-INS-155
chr10-12344563-INS-1141
chr10-12390583-INS-4870

chr10-12414257-INS-192
chr10-12414475-INS-50
chr10-12424268-INS-240
chr10-12430762-INS-186
chr10-12433695-INS-130
chr10-12433918-INS-80
chr10-12513223-INS-56
chr10-12514721-INS-106
chr10-12520995-INS-120
chr10-12520995-INS-54
chr10-12559620-INS-188
chr10-12570390-INS-54
chr10-12637388-INS-209
chr10-12654720-INS-56
chr10-12721621-INS-68
chr10-12742186-INS-2617
chr10-12750452-INS-71
chr10-12753020-INS-72
chr10-12756884-INS-6396
chr10-12789131-INS-135
chr10-12794338-INS-51
chr10-12806579-INS-64
chr10-12812013-INS-297
chr10-12812500-INS-92
chr10-12825712-INS-6475
chr10-12859333-INS-90
chr10-12868380-INS-2645
chr10-12868380-INS-385
chr10-12880348-INS-52
chr10-12901825-INS-7468
chr10-12904389-INS-164
chr10-12929590-INS-1889
chr10-12940511-INS-159
chr10-12945383-INS-98
chr10-12959393-INS-56
chr10-12962768-INS-50
chr10-12977776-INS-51
chr10-13018328-INS-76
chr10-13024013-INS-327
chr10-13075918-INS-1742
chr10-13083046-INS-52
chr10-13138154-INS-3231
chr10-13141861-INS-65
chr10-13156175-INS-82
chr10-13196099-INS-56
chr10-13196228-INS-187
chr10-13271132-INS-293
chr10-13271132-INS-51
chr10-13303089-INS-55
chr10-13306126-INS-90
chr10-13306762-INS-68
chr10-13326660-INS-102
chr10-13361509-INS-76
chr10-13364601-INS-201
chr10-13390850-INS-188
chr10-13489487-INS-130
chr10-13494805-INS-231
chr10-13544291-INS-57
chr10-13609199-INS-4808
chr10-13619458-INS-72
chr10-13674277-INS-7812
chr10-13706337-INS-82
chr10-13723190-INS-50
chr10-13725368-INS-80
chr10-13739213-INS-6229
chr10-13748391-INS-478
chr10-13790829-INS-116
chr10-13816965-INS-106
chr10-13844399-INS-198
chr10-13921297-INS-60
chr10-13927924-INS-711
chr10-13937299-INS-899
chr10-13942647-INS-50
chr10-13956628-INS-55
chr10-13978640-INS-1287
chr10-13995540-INS-76
chr10-14035327-INS-213

chr10-14054494-INS-178
chr10-14081322-INS-1252
chr10-14116995-INS-3636
chr10-14161498-INS-2489
chr10-14184665-INS-525
chr10-14192812-INS-1040
chr10-14198587-INS-6201
chr10-14232029-INS-52
chr10-14235011-INS-50
chr10-14270751-INS-136
chr10-14271279-INS-272
chr10-14293334-INS-54
chr10-14413296-INS-68
chr10-14413661-INS-56
chr10-14421664-INS-97
chr10-14440324-INS-421
chr10-14464699-INS-85
chr10-14491829-INS-773
chr10-14494466-INS-256
chr10-14536571-INS-50
chr10-14627051-INS-212
chr10-14679482-INS-50
chr10-14686366-INS-123
chr10-14686719-INS-86
chr10-14694223-INS-381
chr10-14720783-INS-52
chr10-14721967-INS-2806
chr10-14732985-INS-2024
chr10-14749698-INS-3050
chr10-14780015-INS-7479
chr10-14787419-INS-1036
chr10-14935193-INS-1201
chr10-14936910-INS-57
chr10-14939255-INS-71
chr10-14976918-INS-204
chr10-14977033-INS-369
chr10-14977033-INS-837
chr10-15012702-INS-265
chr10-15012852-INS-6205
chr10-15014153-INS-50
chr10-15049977-INS-50
chr10-15084753-INS-50
chr10-15101858-INS-6214
chr10-15125147-INS-52
chr10-15196403-INS-1210
chr10-15198505-INS-6644
chr10-15215620-INS-1627
chr10-15244443-INS-371
chr10-15283329-INS-5237
chr10-15289864-INS-115
chr10-15297235-INS-97
chr10-15316319-INS-51
chr10-15321665-INS-1374
chr10-15335931-INS-58
chr10-15336122-INS-194
chr10-15362101-INS-940
chr10-15465547-INS-893
chr10-15527226-INS-115
chr10-15535496-INS-215
chr10-15562254-INS-4662
chr10-15588640-INS-6526
chr10-15610965-INS-6695
chr10-15630812-INS-1015
chr10-15631101-INS-3073
chr10-15637816-INS-122
chr10-15646297-INS-1632
chr10-15704001-INS-337
chr10-15730470-INS-325
chr10-15832139-INS-6221
chr10-15860189-INS-207
chr10-15904408-INS-174
chr10-15907225-INS-208
chr10-15921691-INS-56
chr10-15992681-INS-398
chr10-16003564-INS-92
chr10-16019909-INS-63
chr10-16043775-INS-2579

chr10-16053924-INS-1152
chr10-16073985-INS-8325
chr10-16083755-INS-68
chr10-16091732-INS-90
chr10-16093497-INS-523
chr10-16111167-INS-5895
chr10-16118836-INS-1488
chr10-16119604-INS-50
chr10-16139530-INS-54
chr10-16198789-INS-52
chr10-16280255-INS-68
chr10-16301685-INS-5172
chr10-16301688-INS-15100
chr10-16403468-INS-79
chr10-16403532-INS-6398
chr10-16425802-INS-7125
chr10-16429377-INS-6366
chr10-16509496-INS-6589
chr10-16565564-INS-80
chr10-16599566-INS-4825
chr10-16612572-INS-57
chr10-16655497-INS-2740
chr10-16662876-INS-2181
chr10-16677175-INS-88
chr10-16738377-INS-60
chr10-16742130-INS-261
chr10-16753084-INS-1223
chr10-16779466-INS-816
chr10-16793883-INS-3110
chr10-16852033-INS-357
chr10-16863997-INS-2475
chr10-16904699-INS-5985
chr10-17003671-INS-65
chr10-17044806-INS-117
chr10-17066980-INS-400
chr10-17091901-INS-55
chr10-17098691-INS-339
chr10-17186195-INS-80
chr10-17249309-INS-86
chr10-17260701-INS-90
chr10-17266038-INS-6292
chr10-17268657-INS-97
chr10-17323602-INS-58
chr10-17370807-INS-232
chr10-17434826-INS-60
chr10-17489895-INS-51
chr10-17591033-INS-400
chr10-17665354-INS-62
chr10-17665933-INS-78
chr10-17699249-INS-178
chr10-17866304-INS-449
chr10-17947300-INS-68
chr10-17947303-INS-182
chr10-17948033-INS-74
chr10-17953012-INS-176
chr10-17976527-INS-240
chr10-17976897-INS-82
chr10-18060889-INS-3700
chr10-18076715-INS-154
chr10-18077276-INS-86
chr10-18082404-INS-52
chr10-18134367-INS-7292
chr10-18148816-INS-189
chr10-18184480-INS-90
chr10-18225284-INS-50
chr10-18249413-INS-1390
chr10-18257414-INS-54
chr10-18263787-INS-1122
chr10-18300184-INS-1673
chr10-18304072-INS-154
chr10-18328339-INS-545
chr10-18362152-INS-62
chr10-18370702-INS-96
chr10-18411792-INS-6350
chr10-18469901-INS-351
chr10-18529344-INS-406
chr10-18532435-INS-52

chr10-18559326-INS-2591
chr10-18566176-INS-78
chr10-18581347-INS-120
chr10-18585308-INS-112
chr10-18633876-INS-4677
chr10-18688673-INS-259
chr10-18711455-INS-116
chr10-18723923-INS-149
chr10-18738002-INS-433
chr10-18742556-INS-3869
chr10-18784447-INS-7153
chr10-18785193-INS-191
chr10-18791831-INS-95
chr10-18799921-INS-524
chr10-18806776-INS-67
chr10-18818917-INS-6341
chr10-18848251-INS-80
chr10-18920167-INS-173
chr10-18963144-INS-72
chr10-18974151-INS-114
chr10-18999350-INS-67
chr10-19010741-INS-280
chr10-19045560-INS-50
chr10-19065730-INS-79
chr10-19073152-INS-58
chr10-19074504-INS-195
chr10-19077396-INS-109
chr10-19081528-INS-6770
chr10-19110216-INS-1163
chr10-19121898-INS-7087
chr10-19138731-INS-1475
chr10-19156136-INS-66
chr10-19170004-INS-60
chr10-19199812-INS-6751
chr10-19219902-INS-104
chr10-19220389-INS-175
chr10-19220759-INS-169
chr10-19222411-INS-1890
chr10-19233693-INS-76
chr10-19266908-INS-580
chr10-19269226-INS-1343
chr10-19371538-INS-126
chr10-19374736-INS-58
chr10-19379444-INS-60
chr10-19381491-INS-283
chr10-19390261-INS-329
chr10-19391299-INS-117
chr10-19394126-INS-584
chr10-19397800-INS-52
chr10-19454775-INS-125
chr10-19460138-INS-2860
chr10-19460138-INS-464
chr10-19517805-INS-257
chr10-19524443-INS-6777
chr10-19536589-INS-1885
chr10-19557303-INS-177
chr10-19569297-INS-1881
chr10-19574288-INS-79
chr10-19581983-INS-437
chr10-19615606-INS-53
chr10-19618595-INS-207
chr10-19627810-INS-151
chr10-19628015-INS-820
chr10-19643746-INS-770
chr10-19706765-INS-64
chr10-19794181-INS-183
chr10-19794196-INS-66
chr10-19797271-INS-150
chr10-19848906-INS-51
chr10-19883289-INS-440
chr10-19905503-INS-312
chr10-19921504-INS-72
chr10-19961810-INS-4919
chr10-19987323-INS-60
chr10-19988862-INS-91
chr10-20039258-INS-4792
chr10-20041727-INS-143

chr10-20062959-INS-165
chr10-20065826-INS-82
chr10-20109945-INS-64
chr10-20113488-INS-217
chr10-20178339-INS-323
chr10-20199734-INS-209
chr10-20212055-INS-131
chr10-20233683-INS-136
chr10-20244972-INS-905
chr10-20247722-INS-175
chr10-20261170-INS-1675
chr10-20266426-INS-195
chr10-20314636-INS-63
chr10-20315171-INS-6605
chr10-20337892-INS-154
chr10-20342586-INS-68
chr10-20370190-INS-215
chr10-20385123-INS-78
chr10-20486576-INS-50
chr10-20499526-INS-172
chr10-20501148-INS-6474
chr10-20573104-INS-366
chr10-20573210-INS-78
chr10-20602687-INS-96
chr10-20611298-INS-70
chr10-20616509-INS-156
chr10-20660602-INS-91
chr10-20661333-INS-196
chr10-20698719-INS-63
chr10-20706352-INS-6903
chr10-20720840-INS-239
chr10-20720990-INS-60
chr10-20751320-INS-150
chr10-20753167-INS-3049
chr10-20768075-INS-231
chr10-20790343-INS-520
chr10-20805440-INS-219
chr10-20805730-INS-307
chr10-20806159-INS-81
chr10-20824289-INS-60
chr10-20836429-INS-234
chr10-20881952-INS-820
chr10-20890501-INS-50
chr10-20912449-INS-53
chr10-20913503-INS-272
chr10-20922416-INS-675
chr10-20922450-INS-228
chr10-20922512-INS-1439
chr10-20927366-INS-173
chr10-20927571-INS-559
chr10-20984447-INS-50
chr10-20993545-INS-79
chr10-21006647-INS-120
chr10-21010507-INS-58
chr10-21026000-INS-144
chr10-21063608-INS-56
chr10-21112003-INS-58
chr10-21123914-INS-94
chr10-21137270-INS-454
chr10-21137339-INS-1338
chr10-21138738-INS-116
chr10-21138738-INS-53
chr10-21200160-INS-89
chr10-21200224-INS-537
chr10-21200238-INS-182
chr10-21240054-INS-52
chr10-21274682-INS-57
chr10-21274922-INS-95
chr10-21303528-INS-102
chr10-21330798-INS-177
chr10-21330897-INS-59
chr10-21343263-INS-107
chr10-21376337-INS-59
chr10-21409279-INS-84
chr10-21427683-INS-82
chr10-21475881-INS-3674
chr10-21503083-INS-50

chr10-21514550-INS-215
chr10-21556302-INS-144
chr10-21571887-INS-296
chr10-21572029-INS-66
chr10-21584396-INS-1319
chr10-21637023-INS-82
chr10-21641616-INS-287
chr10-21641719-INS-824
chr10-21644817-INS-1863
chr10-21654780-INS-168
chr10-21669936-INS-6505
chr10-21679667-INS-328
chr10-21684614-INS-66
chr10-21694316-INS-50
chr10-21696723-INS-55
chr10-21696845-INS-52
chr10-21702661-INS-189
chr10-21707184-INS-2873
chr10-21748051-INS-607
chr10-21762754-INS-327
chr10-21780658-INS-66
chr10-21783371-INS-58
chr10-21811229-INS-166
chr10-21812175-INS-1008
chr10-21815171-INS-50
chr10-21818093-INS-160
chr10-21818093-INS-64
chr10-21845925-INS-264
chr10-21854216-INS-222
chr10-21906714-INS-402
chr10-21906739-INS-168
chr10-21914655-INS-2540
chr10-21914655-INS-381
chr10-21917290-INS-197
chr10-21938328-INS-66
chr10-22048714-INS-72
chr10-22059318-INS-1892
chr10-22073847-INS-170
chr10-22073905-INS-66
chr10-22093126-INS-766
chr10-22094037-INS-392
chr10-22098684-INS-205
chr10-22129805-INS-106
chr10-22201886-INS-3101
chr10-22205933-INS-575
chr10-22210312-INS-3355
chr10-22231876-INS-1880
chr10-22231876-INS-400
chr10-22237244-INS-98
chr10-22237246-INS-1128
chr10-22264024-INS-254
chr10-22265512-INS-177
chr10-22324230-INS-55
chr10-22336176-INS-59
chr10-22337794-INS-82
chr10-22347372-INS-132
chr10-22350305-INS-1884
chr10-22360428-INS-84
chr10-22425204-INS-728
chr10-22435121-INS-111
chr10-22440616-INS-56
chr10-22441484-INS-131
chr10-22447103-INS-178
chr10-22458601-INS-62
chr10-22463206-INS-749
chr10-22483901-INS-104
chr10-22491829-INS-54
chr10-22491983-INS-389
chr10-22508949-INS-186
chr10-22526288-INS-5337
chr10-22533785-INS-78
chr10-22538297-INS-54
chr10-22538316-INS-671
chr10-22542766-INS-8327
chr10-22661981-INS-54
chr10-22668075-INS-1266
chr10-22682052-INS-3286

chr10-22711127-INS-52
chr10-22712492-INS-93
chr10-22732617-INS-56
chr10-22778675-INS-68
chr10-22800939-INS-859
chr10-22839396-INS-3889
chr10-22934808-INS-180
chr10-22934826-INS-56
chr10-22944319-INS-84
chr10-23020403-INS-51
chr10-23072661-INS-393
chr10-23105718-INS-72
chr10-23150463-INS-92
chr10-23178351-INS-7286
chr10-23187716-INS-4489
chr10-23194175-INS-6899
chr10-23219039-INS-51
chr10-23250899-INS-59
chr10-23301073-INS-1307
chr10-23316326-INS-50
chr10-23351527-INS-424
chr10-23399951-INS-102
chr10-23401999-INS-51
chr10-23415932-INS-86
chr10-23444775-INS-63
chr10-23444858-INS-134
chr10-23452807-INS-4771
chr10-23455246-INS-50
chr10-23475585-INS-108
chr10-23491788-INS-2824
chr10-23521989-INS-59
chr10-23540138-INS-311
chr10-23544693-INS-1138
chr10-23544904-INS-5517
chr10-23551805-INS-50
chr10-23576574-INS-311
chr10-23648375-INS-327
chr10-23648632-INS-314
chr10-23648908-INS-253
chr10-23659886-INS-193
chr10-23671808-INS-198
chr10-23676680-INS-285
chr10-23685055-INS-95
chr10-23727760-INS-202
chr10-23758802-INS-55
chr10-23768072-INS-428
chr10-23798041-INS-66
chr10-23809591-INS-235
chr10-23844792-INS-6941
chr10-23852601-INS-50
chr10-23855319-INS-80
chr10-23880766-INS-2922
chr10-23886590-INS-451
chr10-23912298-INS-564
chr10-23945882-INS-57
chr10-23949023-INS-7295
chr10-23950913-INS-526
chr10-23964089-INS-168
chr10-23975279-INS-316
chr10-23975280-INS-153
chr10-23975757-INS-197
chr10-23976415-INS-6392
chr10-23977876-INS-482
chr10-24014897-INS-2406
chr10-24016298-INS-55
chr10-24037109-INS-175
chr10-24037702-INS-57
chr10-24087915-INS-83
chr10-24097600-INS-1919
chr10-24116438-INS-380
chr10-24128592-INS-86
chr10-24177069-INS-194
chr10-24235411-INS-60
chr10-24243827-INS-98
chr10-24246185-INS-4692
chr10-24287900-INS-52
chr10-24299347-INS-65

chr10-24314615-INS-68
chr10-24388995-INS-212
chr10-24430119-INS-51
chr10-24434605-INS-6628
chr10-24459754-INS-604
chr10-24460469-INS-6536
chr10-24464563-INS-92
chr10-24476690-INS-432
chr10-24477283-INS-576
chr10-24496236-INS-549
chr10-24502108-INS-187
chr10-24521571-INS-56
chr10-24531794-INS-217
chr10-24581891-INS-103
chr10-24596576-INS-5503
chr10-24599998-INS-72
chr10-24614274-INS-56
chr10-24629436-INS-150
chr10-24638574-INS-114
chr10-24642775-INS-188
chr10-24662174-INS-60
chr10-24674211-INS-214
chr10-24707400-INS-60
chr10-24716523-INS-51
chr10-24724921-INS-152
chr10-24725939-INS-50
chr10-24753598-INS-82
chr10-24755366-INS-277
chr10-24780027-INS-50
chr10-24795007-INS-462
chr10-24795007-INS-66
chr10-24800211-INS-50
chr10-24816909-INS-56
chr10-24845462-INS-268
chr10-24850966-INS-2693
chr10-24855275-INS-2878
chr10-24879080-INS-1610
chr10-24900459-INS-1024
chr10-24916879-INS-369
chr10-24926350-INS-1890
chr10-24935169-INS-52
chr10-24938676-INS-2498
chr10-24945282-INS-68
chr10-24947340-INS-6176
chr10-24968138-INS-551
chr10-25084765-INS-101
chr10-25090052-INS-56
chr10-25112912-INS-250
chr10-25120019-INS-80
chr10-25123101-INS-877
chr10-25125899-INS-55
chr10-25128093-INS-55
chr10-25134253-INS-64
chr10-25152265-INS-221
chr10-25167138-INS-370
chr10-25198458-INS-92
chr10-25198468-INS-206
chr10-25205038-INS-5040
chr10-25210505-INS-3981
chr10-25232607-INS-60
chr10-25236978-INS-72
chr10-25264017-INS-132
chr10-25312234-INS-63
chr10-25323954-INS-62
chr10-25334427-INS-377
chr10-25334991-INS-237
chr10-25357974-INS-260
chr10-25357978-INS-126
chr10-25358178-INS-384
chr10-25360959-INS-1171
chr10-25394970-INS-308
chr10-25394974-INS-105
chr10-25433079-INS-50
chr10-25434660-INS-1182
chr10-25467664-INS-7082
chr10-25480983-INS-88
chr10-25493174-INS-106

chr10-25529927-INS-58
chr10-25546477-INS-575
chr10-25557410-INS-3630
chr10-25561308-INS-54
chr10-25587915-INS-3621
chr10-25599263-INS-68
chr10-25621729-INS-79
chr10-25662531-INS-4949
chr10-25725390-INS-6490
chr10-25752317-INS-7401
chr10-25785166-INS-200
chr10-25799450-INS-87
chr10-25836839-INS-50
chr10-25889619-INS-52
chr10-25906438-INS-192
chr10-25917670-INS-57
chr10-25961628-INS-60
chr10-26011832-INS-50
chr10-26013518-INS-84
chr10-26031014-INS-112
chr10-26049371-INS-446
chr10-26051403-INS-6350
chr10-26056370-INS-6352
chr10-26068802-INS-447
chr10-26080057-INS-1899
chr10-26088917-INS-8832
chr10-26090578-INS-273
chr10-26092636-INS-70
chr10-26098448-INS-178
chr10-26145135-INS-127
chr10-26180580-INS-382
chr10-26190916-INS-204
chr10-26232777-INS-58
chr10-26250862-INS-50
chr10-26256589-INS-246
chr10-26267488-INS-68
chr10-26307405-INS-68
chr10-26308741-INS-377
chr10-26311746-INS-1873
chr10-26326135-INS-6201
chr10-26379680-INS-7120
chr10-26389806-INS-2223
chr10-26389834-INS-60
chr10-26389840-INS-280
chr10-26416873-INS-126
chr10-26426182-INS-6132
chr10-26460234-INS-1911
chr10-26503275-INS-161
chr10-26506511-INS-5225
chr10-26578844-INS-6454
chr10-26622461-INS-88
chr10-26638536-INS-50
chr10-26668603-INS-6442
chr10-26757191-INS-78
chr10-26766247-INS-106
chr10-26767739-INS-384
chr10-26768106-INS-152
chr10-26769074-INS-77
chr10-26770589-INS-1460
chr10-26770716-INS-191
chr10-26770896-INS-81
chr10-26846601-INS-3287
chr10-26909410-INS-56
chr10-26954676-INS-1129
chr10-26965708-INS-146
chr10-26985000-INS-52
chr10-26986223-INS-70
chr10-26995394-INS-281
chr10-27008273-INS-92
chr10-27008277-INS-196
chr10-27023663-INS-51
chr10-27057167-INS-6347
chr10-27128220-INS-407
chr10-27177435-INS-50
chr10-27190787-INS-54
chr10-27196607-INS-293
chr10-27201549-INS-874

chr10-27253689-INS-60
chr10-27273313-INS-2083
chr10-27273696-INS-2165
chr10-27297277-INS-1032
chr10-27306571-INS-384
chr10-27314407-INS-68
chr10-27323223-INS-6591
chr10-27347899-INS-348
chr10-27356307-INS-642
chr10-27385455-INS-676
chr10-27424188-INS-50
chr10-27426728-INS-6433
chr10-27480496-INS-199
chr10-27502914-INS-330
chr10-27515278-INS-77
chr10-27538401-INS-88
chr10-27650105-INS-6257
chr10-27650804-INS-55
chr10-27686851-INS-710
chr10-27710544-INS-1056
chr10-27710876-INS-70
chr10-27721311-INS-1828
chr10-27725174-INS-911
chr10-27760888-INS-5976
chr10-27826139-INS-6851
chr10-27882884-INS-207
chr10-27891641-INS-316
chr10-27895265-INS-54
chr10-27905133-INS-52
chr10-27908516-INS-1835
chr10-27908687-INS-64
chr10-27910931-INS-2851
chr10-27956668-INS-165
chr10-27961393-INS-207
chr10-27970596-INS-2611
chr10-27970845-INS-94
chr10-27976633-INS-84
chr10-27992894-INS-50
chr10-27999714-INS-155
chr10-28005064-INS-7074
chr10-28019151-INS-193
chr10-28065073-INS-82
chr10-28068610-INS-71
chr10-28069421-INS-1862
chr10-28096383-INS-60
chr10-28115100-INS-564
chr10-28135941-INS-1329
chr10-28212242-INS-438
chr10-28212861-INS-81
chr10-28216233-INS-66
chr10-28229215-INS-72
chr10-28293047-INS-230
chr10-28293047-INS-90
chr10-28310422-INS-420
chr10-28319002-INS-85
chr10-28324516-INS-56
chr10-28344983-INS-120
chr10-28395287-INS-82
chr10-28411644-INS-1698
chr10-28450027-INS-55
chr10-28453686-INS-100
chr10-28492589-INS-1007
chr10-28496221-INS-999
chr10-28516426-INS-7012
chr10-28531211-INS-2117
chr10-28538352-INS-1281
chr10-28540175-INS-6335
chr10-28588146-INS-76
chr10-28605189-INS-1020
chr10-28605633-INS-4498
chr10-28606394-INS-67
chr10-28606952-INS-212
chr10-28617252-INS-59
chr10-28632577-INS-5899
chr10-28650928-INS-54
chr10-28676984-INS-241
chr10-28687693-INS-454

chr10-28690683-INS-588
chr10-28751850-INS-1349
chr10-28766553-INS-6253
chr10-28782277-INS-2658
chr10-28820085-INS-1417
chr10-28839583-INS-6396
chr10-28914694-INS-5802
chr10-28923239-INS-100
chr10-28982905-INS-3772
chr10-28983801-INS-7546
chr10-29047654-INS-2936
chr10-29058842-INS-6509
chr10-29075710-INS-5544
chr10-29081412-INS-83
chr10-29128803-INS-66
chr10-29166421-INS-1373
chr10-29262933-INS-5398
chr10-29367522-INS-52
chr10-29378293-INS-60
chr10-29423357-INS-60
chr10-29481526-INS-1892
chr10-29483304-INS-1892
chr10-29494589-INS-52
chr10-29526411-INS-58
chr10-29576145-INS-839
chr10-29585329-INS-84
chr10-29589943-INS-3680
chr10-29600443-INS-5015
chr10-29604988-INS-4356
chr10-29623804-INS-3949
chr10-29630099-INS-2830
chr10-29640910-INS-261
chr10-29681546-INS-5550
chr10-29725259-INS-1081
chr10-29726590-INS-1124
chr10-29822942-INS-291
chr10-29829754-INS-88
chr10-29880378-INS-50
chr10-29883565-INS-4991
chr10-29893408-INS-2421
chr10-29897035-INS-2833
chr10-29934344-INS-1023
chr10-29968056-INS-50
chr10-30005559-INS-72
chr10-30009045-INS-2838
chr10-30025966-INS-737
chr10-30058063-INS-66
chr10-30061012-INS-2186
chr10-30085037-INS-586
chr10-30137476-INS-3011
chr10-30233033-INS-1624
chr10-30313566-INS-60
chr10-30337949-INS-96
chr10-30363711-INS-181
chr10-30383886-INS-149
chr10-30445996-INS-105
chr10-30486189-INS-130
chr10-30510104-INS-480
chr10-30537843-INS-183
chr10-30608131-INS-7118
chr10-30661944-INS-132
chr10-30682792-INS-5549
chr10-30723197-INS-132
chr10-30741436-INS-2060
chr10-30746721-INS-3984
chr10-30749109-INS-1196
chr10-30752087-INS-3025
chr10-30813368-INS-83
chr10-30879962-INS-6194
chr10-30889949-INS-231
chr10-30902095-INS-3096
chr10-30910777-INS-134
chr10-30938240-INS-62
chr10-30944449-INS-718
chr10-30944774-INS-119
chr10-30984996-INS-56
chr10-31016745-INS-319

chr10-31017470-INS-2142
chr10-31020085-INS-52
chr10-31032797-INS-4360
chr10-31055293-INS-66
chr10-31106869-INS-695
chr10-31230834-INS-4422
chr10-31234136-INS-377
chr10-31250656-INS-2102
chr10-31340732-INS-86
chr10-31354100-INS-920
chr10-31357275-INS-479
chr10-31357538-INS-125
chr10-31378935-INS-215
chr10-31448222-INS-2768
chr10-31495073-INS-120
chr10-31500518-INS-3864
chr10-31534130-INS-205
chr10-31573447-INS-56
chr10-31588359-INS-6386
chr10-31669867-INS-484
chr10-31674548-INS-6199
chr10-31713762-INS-5823
chr10-31731072-INS-6297
chr10-31750831-INS-76
chr10-31763350-INS-56
chr10-31786488-INS-56
chr10-31929662-INS-99
chr10-31948341-INS-2412
chr10-31966053-INS-5225
chr10-31967352-INS-87
chr10-31993987-INS-178
chr10-31993998-INS-57
chr10-32034367-INS-225
chr10-32067686-INS-116
chr10-32068657-INS-118
chr10-32068803-INS-50
chr10-32092295-INS-64
chr10-32113521-INS-60
chr10-32131874-INS-51
chr10-32137780-INS-491
chr10-32137846-INS-120
chr10-32172707-INS-302
chr10-32185769-INS-6274
chr10-32208106-INS-915
chr10-32235357-INS-55
chr10-32251121-INS-833
chr10-32367468-INS-1194
chr10-32453275-INS-908
chr10-32455640-INS-498
chr10-32456169-INS-74
chr10-32504142-INS-6393
chr10-32601055-INS-481
chr10-32616895-INS-50
chr10-32632612-INS-207
chr10-32651183-INS-469
chr10-32686219-INS-84
chr10-32777052-INS-2601
chr10-32830823-INS-54
chr10-32837719-INS-68
chr10-32851770-INS-1418
chr10-32858330-INS-65
chr10-32872841-INS-1367
chr10-32899504-INS-1531
chr10-32905390-INS-3514
chr10-32906027-INS-10129
chr10-32918651-INS-1226
chr10-32924327-INS-207
chr10-32942826-INS-6303
chr10-32973384-INS-72
chr10-32988513-INS-82
chr10-33005847-INS-1873
chr10-33008058-INS-5343
chr10-33068863-INS-637
chr10-33093095-INS-2555
chr10-33093899-INS-504
chr10-33104941-INS-1969
chr10-33110615-INS-286

chr10-33118425-INS-1601
chr10-33205349-INS-121
chr10-33215234-INS-83
chr10-33222906-INS-50
chr10-33237075-INS-1949
chr10-33253691-INS-66
chr10-33277729-INS-1604
chr10-33317877-INS-1069
chr10-33330936-INS-5537
chr10-33343804-INS-923
chr10-33416626-INS-1225
chr10-33421713-INS-5979
chr10-33424037-INS-52
chr10-33469698-INS-51
chr10-33494628-INS-56
chr10-33499289-INS-54
chr10-33499429-INS-155
chr10-33505028-INS-52
chr10-33578111-INS-1337
chr10-33578858-INS-70
chr10-33590708-INS-1251
chr10-33601340-INS-587
chr10-33627167-INS-6253
chr10-33628910-INS-2655
chr10-33634912-INS-56
chr10-33652969-INS-6380
chr10-33808977-INS-97
chr10-33829209-INS-63
chr10-33851549-INS-3212
chr10-33887764-INS-159
chr10-33928492-INS-50
chr10-33929434-INS-289
chr10-33935289-INS-239
chr10-33935322-INS-59
chr10-33938338-INS-53
chr10-33941683-INS-3708
chr10-33950934-INS-1082
chr10-33951504-INS-72
chr10-33983302-INS-69
chr10-33991615-INS-6476
chr10-33998339-INS-55
chr10-34042342-INS-2686
chr10-34048299-INS-57
chr10-34050167-INS-7348
chr10-34051617-INS-417
chr10-34054232-INS-140
chr10-34055465-INS-297
chr10-34081524-INS-210
chr10-34115852-INS-206
chr10-34115981-INS-52
chr10-34125119-INS-60
chr10-34192175-INS-176
chr10-34222538-INS-143
chr10-34226654-INS-6480
chr10-34269433-INS-51
chr10-34285311-INS-62
chr10-34341872-INS-119
chr10-34344945-INS-53
chr10-34347181-INS-7006
chr10-34373656-INS-60
chr10-34385156-INS-6168
chr10-34408158-INS-96
chr10-34419314-INS-582
chr10-34420110-INS-5998
chr10-34427892-INS-53
chr10-34433537-INS-149
chr10-34441387-INS-230
chr10-34441913-INS-327
chr10-34449714-INS-424
chr10-34451865-INS-3035
chr10-34457360-INS-152
chr10-34465336-INS-442
chr10-34474044-INS-243
chr10-34475087-INS-79
chr10-34535006-INS-60
chr10-34538756-INS-915
chr10-34574246-INS-6184

chr10-34583724-INS-85
chr10-34613087-INS-120
chr10-34634131-INS-1264
chr10-34634521-INS-170
chr10-34672797-INS-9064
chr10-34715918-INS-7087
chr10-34726404-INS-66
chr10-34736585-INS-881
chr10-34738786-INS-67
chr10-34744037-INS-202
chr10-34746341-INS-104
chr10-34791825-INS-2059
chr10-34795097-INS-108
chr10-34815446-INS-4715
chr10-34819301-INS-2814
chr10-34828704-INS-1190
chr10-34903601-INS-15712
chr10-34903601-INS-7517
chr10-34955909-INS-205
chr10-34966378-INS-171
chr10-34969809-INS-7437
chr10-34985020-INS-3747
chr10-35076829-INS-61
chr10-35113418-INS-6477
chr10-35116367-INS-56
chr10-35120234-INS-61
chr10-35120242-INS-153
chr10-35122832-INS-6541
chr10-35141260-INS-5321
chr10-35160739-INS-54
chr10-35246244-INS-54
chr10-35283220-INS-218
chr10-35283363-INS-295
chr10-35294166-INS-61
chr10-35305540-INS-60
chr10-35408420-INS-1230
chr10-35448421-INS-6289
chr10-35479481-INS-56
chr10-35489026-INS-56
chr10-35532166-INS-60
chr10-35584201-INS-50
chr10-35592583-INS-142
chr10-35633364-INS-11642
chr10-35649759-INS-2811
chr10-35678922-INS-7636
chr10-35680423-INS-1762
chr10-35688807-INS-1085
chr10-35700826-INS-8487
chr10-35741259-INS-68
chr10-35763760-INS-6203
chr10-35840432-INS-1475
chr10-35903702-INS-1387
chr10-35906877-INS-1574
chr10-35916227-INS-55
chr10-35943130-INS-193
chr10-35944308-INS-1249
chr10-36008774-INS-5220
chr10-36022561-INS-1550
chr10-36030896-INS-64
chr10-36065838-INS-52
chr10-36129484-INS-67
chr10-36142986-INS-1405
chr10-36174291-INS-6631
chr10-36177017-INS-149
chr10-36229405-INS-203
chr10-36263693-INS-2264
chr10-36309060-INS-268
chr10-36316180-INS-1194
chr10-36320196-INS-411
chr10-36331568-INS-293
chr10-36343000-INS-6738
chr10-36355640-INS-104
chr10-36429945-INS-6995
chr10-36432616-INS-226
chr10-36465198-INS-6280
chr10-36488697-INS-6385
chr10-36511736-INS-2990

chr10-36719014-INS-114
chr10-36720916-INS-6712
chr10-36743160-INS-7080
chr10-36753150-INS-2878
chr10-36782072-INS-50
chr10-36794065-INS-2936
chr10-36794156-INS-131
chr10-36806033-INS-50
chr10-36807608-INS-1713
chr10-36856150-INS-84
chr10-36876433-INS-115
chr10-36898475-INS-1007
chr10-36898475-INS-6666
chr10-36904115-INS-194
chr10-36904279-INS-64
chr10-36978169-INS-50
chr10-36985077-INS-69
chr10-37003770-INS-101
chr10-37113849-INS-294
chr10-37121218-INS-399
chr10-37157395-INS-66
chr10-37242575-INS-6898
chr10-37284954-INS-296
chr10-37353167-INS-57
chr10-37353739-INS-114
chr10-37364327-INS-3086
chr10-37463873-INS-59
chr10-37487685-INS-6829
chr10-37655594-INS-2209
chr10-37688077-INS-7340
chr10-37694162-INS-58
chr10-37696334-INS-1220
chr10-37698344-INS-73
chr10-37732343-INS-116
chr10-37769953-INS-72
chr10-37840539-INS-1410
chr10-37850582-INS-55
chr10-37868448-INS-1514
chr10-37899920-INS-445
chr10-37916985-INS-50
chr10-37917825-INS-7215
chr10-37939762-INS-2217
chr10-37947309-INS-2483
chr10-37953255-INS-192
chr10-38045206-INS-92
chr10-38059361-INS-6516
chr10-38082736-INS-84
chr10-38118359-INS-1008
chr10-38176254-INS-5542
chr10-38181407-INS-5376
chr10-38196756-INS-4993
chr10-38203953-INS-8357
chr10-38217438-INS-56
chr10-38228949-INS-6375
chr10-38264694-INS-174
chr10-38276048-INS-50
chr10-38288271-INS-53
chr10-38289500-INS-1666
chr10-38294071-INS-53
chr10-38306883-INS-382
chr10-38312727-INS-7831
chr10-38340563-INS-58
chr10-38362099-INS-68
chr10-38382124-INS-172
chr10-38423855-INS-54
chr10-38508921-INS-450
chr10-38524855-INS-50
chr10-38571429-INS-123
chr10-38571434-INS-51
chr10-38578901-INS-476
chr10-38581633-INS-736
chr10-38659881-INS-500
chr10-38691142-INS-220
chr10-38757816-INS-59
chr10-38770500-INS-77
chr10-38849849-INS-1150
chr10-38849849-INS-230

chr10-38868656-INS-694
chr10-38899139-INS-67
chr10-38952538-INS-1129
chr10-38952643-INS-72
chr10-38996916-INS-72
chr10-38997502-INS-56
chr10-39019630-INS-52
chr10-39021051-INS-400
chr10-39023251-INS-136
chr10-39035945-INS-826
chr10-39036711-INS-112
chr10-39043147-INS-2284
chr10-39043147-INS-451
chr10-39082347-INS-103
chr10-39088899-INS-161
chr10-39119159-INS-208
chr10-39132600-INS-316
chr10-39148018-INS-509
chr10-39169320-INS-2753
chr10-39175202-INS-6391
chr10-39193733-INS-261
chr10-39195355-INS-2737
chr10-39209286-INS-185
chr10-39237877-INS-161
chr10-39271917-INS-252
chr10-39271923-INS-60
chr10-39281283-INS-156
chr10-39282075-INS-2116
chr10-39308577-INS-241
chr10-39325047-INS-369
chr10-39358998-INS-549
chr10-39400590-INS-60
chr10-39422253-INS-321
chr10-39437514-INS-210
chr10-39474723-INS-241
chr10-39493540-INS-55
chr10-39512959-INS-340
chr10-39540828-INS-50
chr10-39545946-INS-1038
chr10-39571221-INS-105
chr10-39632097-INS-1370
chr10-39667435-INS-238
chr10-39667437-INS-98
chr10-39677814-INS-604
chr10-39715768-INS-306
chr10-39715843-INS-127
chr10-39731503-INS-120
chr10-39769251-INS-231
chr10-39772071-INS-180
chr10-39772148-INS-7091
chr10-39819688-INS-114
chr10-39871977-INS-59
chr10-39873967-INS-4786
chr10-39874876-INS-52
chr10-39875136-INS-1581
chr10-39884440-INS-288
chr10-39943532-INS-73
chr10-40011733-INS-57
chr10-40027590-INS-554
chr10-40035234-INS-63
chr10-40046824-INS-52
chr10-40053326-INS-56
chr10-40057161-INS-5372
chr10-40068302-INS-85
chr10-40068353-INS-511
chr10-40112595-INS-72
chr10-40117186-INS-540
chr10-40126524-INS-211
chr10-40131390-INS-298
chr10-40131393-INS-835
chr10-40149332-INS-54
chr10-40156664-INS-13879
chr10-40156664-INS-5097
chr10-40181466-INS-268
chr10-40217492-INS-4089
chr10-40272019-INS-186
chr10-40295924-INS-56

chr10-40328703-INS-266
chr10-40342170-INS-1210
chr10-40372597-INS-60
chr10-40384201-INS-295
chr10-40389467-INS-191
chr10-40437813-INS-3151
chr10-40468521-INS-4134
chr10-40471045-INS-203
chr10-40471456-INS-223
chr10-40484876-INS-51
chr10-40512706-INS-207
chr10-40522018-INS-1896
chr10-40522980-INS-2152
chr10-40525941-INS-207
chr10-40529760-INS-108
chr10-40529827-INS-60
chr10-40531486-INS-54
chr10-40566737-INS-462
chr10-40576128-INS-196
chr10-40594277-INS-262
chr10-40609247-INS-54
chr10-40633553-INS-7075
chr10-40647391-INS-1888
chr10-40649465-INS-1622
chr10-40653398-INS-70
chr10-40679071-INS-1907
chr10-40708602-INS-101
chr10-40715560-INS-240
chr10-40744517-INS-861
chr10-40744528-INS-237
chr10-40831113-INS-107
chr10-40832048-INS-84
chr10-40880775-INS-80
chr10-40881156-INS-225
chr10-40888371-INS-7099
chr10-40936354-INS-6279
chr10-40939077-INS-207
chr10-41032317-INS-64
chr10-41040068-INS-178
chr10-41065351-INS-469
chr10-41102165-INS-1078
chr10-41109703-INS-315
chr10-41216596-INS-305
chr10-41242198-INS-58
chr10-41289153-INS-278
chr10-41298548-INS-204
chr10-41303029-INS-210
chr10-41311941-INS-109
chr10-41333053-INS-212
chr10-41358344-INS-105
chr10-41359008-INS-74
chr10-41461051-INS-198
chr10-41493062-INS-4871
chr10-41540257-INS-2671
chr10-41552814-INS-60
chr10-41553390-INS-249
chr10-41559790-INS-1413
chr10-41583574-INS-51
chr10-41605722-INS-135
chr10-41605810-INS-440
chr10-41613025-INS-95
chr10-41617516-INS-50
chr10-41625239-INS-90
chr10-41652349-INS-1951
chr10-41671667-INS-8684
chr10-41684743-INS-148
chr10-41695054-INS-50
chr10-41701657-INS-558
chr10-41743210-INS-174
chr10-41767061-INS-110
chr10-41771533-INS-132
chr10-41805580-INS-226
chr10-41805612-INS-54
chr10-41806431-INS-81
chr10-41848232-INS-162
chr10-41889187-INS-80
chr10-41935593-INS-154

chr10-41940601-INS-68
chr10-41960220-INS-101
chr10-41960750-INS-55
chr10-41971152-INS-69
chr10-42013357-INS-418
chr10-42026825-INS-3272
chr10-42052119-INS-62
chr10-42053609-INS-195
chr10-42067554-INS-97
chr10-42087662-INS-351
chr10-42102854-INS-267
chr10-42114905-INS-60
chr10-42175862-INS-103
chr10-42186245-INS-204
chr10-42189404-INS-130
chr10-42204386-INS-1170
chr10-42269191-INS-50
chr10-42288635-INS-94
chr10-42305525-INS-177
chr10-42306565-INS-6403
chr10-42315730-INS-1795
chr10-42322649-INS-68
chr10-42326078-INS-269
chr10-42366622-INS-8457
chr10-42425874-INS-89
chr10-42488676-INS-106
chr10-42502121-INS-196
chr10-42525573-INS-119
chr10-42532023-INS-180
chr10-42551093-INS-116
chr10-42557556-INS-228
chr10-42565573-INS-54
chr10-42610194-INS-60
chr10-42660367-INS-137
chr10-42707900-INS-69
chr10-42716546-INS-1874
chr10-42729487-INS-50
chr10-42769886-INS-351
chr10-42791004-INS-56
chr10-42801174-INS-54
chr10-42815623-INS-70
chr10-42816207-INS-205
chr10-42830224-INS-80
chr10-42832561-INS-60
chr10-42843269-INS-211
chr10-42874355-INS-1563
chr10-42874805-INS-471
chr10-42884851-INS-136
chr10-42920167-INS-152
chr10-42937396-INS-6442
chr10-43034632-INS-50
chr10-43059314-INS-164
chr10-43069911-INS-84
chr10-43106090-INS-4763
chr10-43112837-INS-96
chr10-43113678-INS-168
chr10-43122554-INS-76
chr10-43140567-INS-60
chr10-43140625-INS-149
chr10-43143139-INS-484
chr10-43157200-INS-99
chr10-43183421-INS-64
chr10-43254342-INS-70
chr10-43262940-INS-111
chr10-43291152-INS-1604
chr10-43291160-INS-796
chr10-43291225-INS-358
chr10-43307192-INS-112
chr10-43323282-INS-68
chr10-43327485-INS-59
chr10-43335177-INS-191
chr10-43347601-INS-50
chr10-43396462-INS-50
chr10-43398758-INS-50
chr10-43421028-INS-176
chr10-43422536-INS-248
chr10-43425202-INS-52

chr10-43433404-INS-64
chr10-43438578-INS-5515
chr10-43445844-INS-64
chr10-43464350-INS-50
chr10-43477150-INS-160
chr10-43477153-INS-399
chr10-43477998-INS-52
chr10-43479743-INS-3799
chr10-43487468-INS-62
chr10-43490275-INS-6637
chr10-43491159-INS-3240
chr10-43494041-INS-50
chr10-43496678-INS-6913
chr10-43523547-INS-205
chr10-43574619-INS-4617
chr10-43596033-INS-136
chr10-43604676-INS-61
chr10-43623345-INS-113
chr10-43623904-INS-60
chr10-43635821-INS-52
chr10-43643633-INS-1021
chr10-43653061-INS-282
chr10-43663866-INS-66
chr10-43696227-INS-631
chr10-43737629-INS-1012
chr10-43739414-INS-407
chr10-43763565-INS-322
chr10-43765574-INS-99
chr10-43785613-INS-66
chr10-43807557-INS-1633
chr10-43814379-INS-140
chr10-43844257-INS-84
chr10-43920587-INS-80
chr10-43935603-INS-58
chr10-43952462-INS-312
chr10-43955951-INS-56
chr10-43960891-INS-108
chr10-43990638-INS-78
chr10-43991373-INS-985
chr10-43998081-INS-70
chr10-44018363-INS-1923
chr10-44068793-INS-235
chr10-44095755-INS-718
chr10-44106666-INS-60
chr10-44106717-INS-552
chr10-44111548-INS-1752
chr10-44172713-INS-60
chr10-44189578-INS-64
chr10-44207404-INS-52
chr10-44271332-INS-922
chr10-44271435-INS-115
chr10-44309923-INS-63
chr10-44352303-INS-262
chr10-44358445-INS-116
chr10-44366034-INS-54
chr10-44374142-INS-1938
chr10-44374142-INS-608
chr10-44385706-INS-126
chr10-44428490-INS-207
chr10-44428975-INS-731
chr10-44439528-INS-5920
chr10-44484429-INS-52
chr10-44513584-INS-88
chr10-44629219-INS-63
chr10-44665715-INS-96
chr10-44689418-INS-268
chr10-44729390-INS-64
chr10-44737051-INS-52
chr10-44765045-INS-282
chr10-44838093-INS-6260
chr10-44838422-INS-54
chr10-44856068-INS-194
chr10-44859347-INS-55
chr10-44874741-INS-59
chr10-44875726-INS-51
chr10-44895957-INS-4710
chr10-44907947-INS-57

chr10-44910690-INS-100
chr10-44915746-INS-78
chr10-44986961-INS-364
chr10-45050453-INS-1792
chr10-45060694-INS-136
chr10-45066174-INS-58
chr10-45110543-INS-6108
chr10-45156017-INS-52
chr10-45164211-INS-508
chr10-45167441-INS-60
chr10-45168597-INS-215
chr10-45170002-INS-57
chr10-45184222-INS-72
chr10-45225391-INS-65
chr10-45229527-INS-344
chr10-45236716-INS-450
chr10-45236767-INS-50
chr10-45251790-INS-96
chr10-45302027-INS-98
chr10-45390364-INS-62
chr10-45482973-INS-59
chr10-45579573-INS-78
chr10-45590981-INS-440
chr10-45660548-INS-390
chr10-45722165-INS-55
chr10-45764483-INS-76
chr10-45768270-INS-72
chr10-45795487-INS-132
chr10-45809783-INS-277
chr10-45853161-INS-3647
chr10-45861654-INS-90
chr10-45863790-INS-52
chr10-45914393-INS-72
chr10-45926767-INS-74
chr10-46031930-INS-6488
chr10-46045001-INS-170
chr10-46135833-INS-163
chr10-46142493-INS-55
chr10-46204879-INS-4723
chr10-46238870-INS-82
chr10-46280099-INS-50
chr10-46311230-INS-235
chr10-46335350-INS-60
chr10-46350980-INS-793
chr10-46362632-INS-75
chr10-46368873-INS-169
chr10-46388582-INS-109
chr10-46427367-INS-2311
chr10-46465302-INS-166
chr10-46472543-INS-3299
chr10-46514975-INS-50
chr10-46653292-INS-5557
chr10-46666851-INS-70
chr10-46734982-INS-207
chr10-46765354-INS-141
chr10-46770557-INS-6258
chr10-46817469-INS-200
chr10-46828800-INS-931
chr10-46830985-INS-2717
chr10-46895525-INS-52
chr10-46971350-INS-3206
chr10-46974697-INS-1390
chr10-46987531-INS-123
chr10-47004441-INS-1444
chr10-47044183-INS-51
chr10-47046024-INS-131
chr10-47046608-INS-490
chr10-47066502-INS-327
chr10-47066934-INS-761
chr10-47082421-INS-128
chr10-47172784-INS-57
chr10-47190203-INS-168
chr10-47222578-INS-95
chr10-47260373-INS-6304
chr10-47309339-INS-111
chr10-47324018-INS-168
chr10-47350589-INS-406

chr10-47371798-INS-58
chr10-47372393-INS-465
chr10-47375934-INS-112
chr10-47506661-INS-434
chr10-47526083-INS-60
chr10-47528035-INS-92
chr10-47528958-INS-106
chr10-47538721-INS-84
chr10-47579591-INS-774
chr10-47585300-INS-438
chr10-47600820-INS-497
chr10-47696826-INS-72
chr10-47811762-INS-55
chr10-47855603-INS-220
chr10-48022679-INS-67
chr10-48032555-INS-51
chr10-48036603-INS-3238
chr10-48040397-INS-1888
chr10-48043054-INS-52
chr10-48065440-INS-6692
chr10-48166645-INS-58
chr10-48166771-INS-251
chr10-48198672-INS-1171
chr10-48208000-INS-234
chr10-48209715-INS-5891
chr10-48224816-INS-6163
chr10-48318849-INS-6563
chr10-48321787-INS-50
chr10-48389395-INS-76
chr10-48402408-INS-10773
chr10-48412701-INS-64
chr10-48414188-INS-373
chr10-48424670-INS-60
chr10-48531484-INS-176
chr10-48571757-INS-88
chr10-48607808-INS-558
chr10-48608323-INS-68
chr10-48608401-INS-468
chr10-48609677-INS-274
chr10-48611882-INS-4898
chr10-48625567-INS-1339
chr10-48641152-INS-4828
chr10-48684386-INS-6617
chr10-48759349-INS-50
chr10-48771781-INS-56
chr10-48771805-INS-126
chr10-48804726-INS-6207
chr10-48839868-INS-62
chr10-48883789-INS-3541
chr10-48891368-INS-4782
chr10-48955150-INS-70
chr10-48972592-INS-54
chr10-49008531-INS-1179
chr10-49014481-INS-1798
chr10-49045147-INS-70
chr10-49050531-INS-105
chr10-49083929-INS-1246
chr10-49086385-INS-1889
chr10-49088470-INS-64
chr10-49095356-INS-107
chr10-49215785-INS-2856
chr10-49216095-INS-116
chr10-49227657-INS-8290
chr10-49239368-INS-3490
chr10-49247188-INS-135
chr10-49252950-INS-56
chr10-49253575-INS-370
chr10-49269073-INS-860
chr10-49345340-INS-50
chr10-49373789-INS-4305
chr10-49471553-INS-6651
chr10-49521329-INS-797
chr10-49536824-INS-217
chr10-49638337-INS-1865
chr10-49669623-INS-984
chr10-49673550-INS-664
chr10-49695456-INS-1432

chr10-49720129-INS-73
chr10-49723883-INS-204
chr10-49758397-INS-124
chr10-49770508-INS-850
chr10-49770924-INS-2287
chr10-49770924-INS-5348
chr10-49779634-INS-545
chr10-49785925-INS-1074
chr10-49799421-INS-4225
chr10-49801288-INS-6258
chr10-49804646-INS-457
chr10-49822597-INS-2936
chr10-49847742-INS-50
chr10-49848507-INS-52
chr10-49913007-INS-121
chr10-49991895-INS-52
chr10-50076518-INS-2635
chr10-50081170-INS-84
chr10-50084379-INS-1406
chr10-50108387-INS-9713
chr10-50129770-INS-201
chr10-50156296-INS-137
chr10-50198958-INS-56
chr10-50218561-INS-125
chr10-50234348-INS-178
chr10-50247613-INS-185
chr10-50249125-INS-309
chr10-50251457-INS-101
chr10-50254610-INS-1089
chr10-50259916-INS-2613
chr10-50308407-INS-348
chr10-50309317-INS-57
chr10-50384034-INS-70
chr10-50395140-INS-136
chr10-50422022-INS-120
chr10-50457288-INS-90
chr10-50457678-INS-328
chr10-50475630-INS-54
chr10-50573262-INS-1523
chr10-50574115-INS-1460
chr10-50574504-INS-52
chr10-50668568-INS-85
chr10-50673348-INS-202
chr10-50792690-INS-252
chr10-50921530-INS-175
chr10-50942389-INS-236
chr10-50957049-INS-1445
chr10-50975353-INS-52
chr10-51077622-INS-76
chr10-51136354-INS-6762
chr10-51289240-INS-51
chr10-51366750-INS-54
chr10-51373092-INS-3865
chr10-51407535-INS-118
chr10-51410141-INS-136
chr10-51468747-INS-80
chr10-51476166-INS-113
chr10-51493059-INS-55
chr10-51535655-INS-52
chr10-51538219-INS-50
chr10-51570579-INS-184
chr10-51580871-INS-2051
chr10-51592489-INS-54
chr10-51619048-INS-6698
chr10-51625867-INS-1209
chr10-51655315-INS-58
chr10-51658836-INS-6666
chr10-51673236-INS-56
chr10-51674682-INS-1858
chr10-51681037-INS-449
chr10-51739251-INS-559
chr10-51750065-INS-182
chr10-51750342-INS-3113
chr10-51785948-INS-88
chr10-51803943-INS-318
chr10-51811045-INS-1517
chr10-51823771-INS-50

chr10-51826642-INS-67
chr10-51829886-INS-479
chr10-51848963-INS-3488
chr10-51867394-INS-214
chr10-51879028-INS-1498
chr10-51941895-INS-251
chr10-51964578-INS-554
chr10-51971711-INS-124
chr10-52018967-INS-220
chr10-52035944-INS-60
chr10-52048503-INS-52
chr10-52073387-INS-4821
chr10-52075039-INS-1072
chr10-52075039-INS-425
chr10-52076662-INS-81
chr10-52076671-INS-275
chr10-52097499-INS-66
chr10-52115595-INS-70
chr10-52119529-INS-50
chr10-52119775-INS-211
chr10-52124709-INS-6442
chr10-52125682-INS-1245
chr10-52143233-INS-198
chr10-52149579-INS-59
chr10-52180977-INS-232
chr10-52217049-INS-217
chr10-52222830-INS-215
chr10-52251182-INS-211
chr10-52278544-INS-399
chr10-52285065-INS-69
chr10-52371729-INS-1847
chr10-52390117-INS-60
chr10-52444765-INS-2884
chr10-52458689-INS-6954
chr10-52480947-INS-449
chr10-52490870-INS-4943
chr10-52496535-INS-172
chr10-52496753-INS-68
chr10-52581587-INS-114
chr10-52592811-INS-173
chr10-52620626-INS-1193
chr10-52631888-INS-2683
chr10-52680411-INS-55
chr10-52700703-INS-52
chr10-52768827-INS-148
chr10-52790235-INS-316
chr10-52792382-INS-3166
chr10-52878103-INS-6925
chr10-52918164-INS-59
chr10-52930542-INS-2580
chr10-52956337-INS-55
chr10-52998207-INS-65
chr10-53022804-INS-6240
chr10-53034323-INS-60
chr10-53070647-INS-6492
chr10-53100579-INS-810
chr10-53102340-INS-7188
chr10-53109915-INS-126
chr10-53111832-INS-1510
chr10-53147704-INS-123
chr10-53191165-INS-207
chr10-53196531-INS-367
chr10-53251842-INS-1357
chr10-53262548-INS-461
chr10-53263192-INS-7317
chr10-53266109-INS-50
chr10-53293612-INS-60
chr10-53350423-INS-410
chr10-53377358-INS-1612
chr10-53409858-INS-51
chr10-53422943-INS-51
chr10-53440144-INS-220
chr10-53537378-INS-54
chr10-53549553-INS-324
chr10-53570407-INS-56
chr10-53588354-INS-6493
chr10-53597842-INS-50

chr10-53632966-INS-4556
chr10-53646852-INS-185
chr10-53648226-INS-52
chr10-53731496-INS-176
chr10-53732648-INS-141
chr10-53788837-INS-215
chr10-53846493-INS-1015
chr10-53853586-INS-1448
chr10-53896096-INS-59
chr10-53921158-INS-78
chr10-53943444-INS-61
chr10-53965394-INS-52
chr10-54024776-INS-208
chr10-54029431-INS-52
chr10-54074801-INS-80
chr10-54084121-INS-50
chr10-54119814-INS-57
chr10-54133030-INS-624
chr10-54133934-INS-764
chr10-54134576-INS-6312
chr10-54247461-INS-6488
chr10-54274617-INS-2444
chr10-54275688-INS-55
chr10-54305759-INS-178
chr10-54306746-INS-100
chr10-54389283-INS-1213
chr10-54396503-INS-64
chr10-54457407-INS-202
chr10-54599795-INS-53
chr10-54636091-INS-834
chr10-54666025-INS-52
chr10-54687285-INS-548
chr10-54690038-INS-66
chr10-54710614-INS-123
chr10-54722410-INS-198
chr10-54722410-INS-80
chr10-54739657-INS-51
chr10-54752138-INS-343
chr10-54797516-INS-1984
chr10-54808866-INS-52
chr10-54841596-INS-1625
chr10-54881398-INS-136
chr10-54893386-INS-498
chr10-54923621-INS-50
chr10-54944135-INS-815
chr10-54952269-INS-50
chr10-54967393-INS-2192
chr10-54982651-INS-435
chr10-54994919-INS-110
chr10-55034742-INS-1686
chr10-55100692-INS-836
chr10-55144545-INS-445
chr10-55180786-INS-111
chr10-55189810-INS-6472
chr10-55233650-INS-50
chr10-55284013-INS-1199
chr10-55308555-INS-270
chr10-55309141-INS-1530
chr10-55392942-INS-162
chr10-55415054-INS-1230
chr10-55437050-INS-507
chr10-55440854-INS-736
chr10-55444004-INS-206
chr10-55486398-INS-667
chr10-55490523-INS-186
chr10-55492798-INS-2834
chr10-55547994-INS-2329
chr10-55563885-INS-72
chr10-55605430-INS-108
chr10-55626480-INS-942
chr10-55659149-INS-6177
chr10-55687214-INS-8357
chr10-55700037-INS-82
chr10-55704281-INS-56
chr10-55717965-INS-856
chr10-55770767-INS-214
chr10-55790573-INS-133

chr10-55806641-INS-9553
chr10-55846930-INS-180
chr10-55846930-INS-60
chr10-55862078-INS-590
chr10-55862373-INS-689
chr10-55875798-INS-154
chr10-55896419-INS-900
chr10-55901346-INS-56
chr10-55934361-INS-201
chr10-55997913-INS-1478
chr10-56016063-INS-1499
chr10-56020977-INS-408
chr10-56037913-INS-199
chr10-56063804-INS-53
chr10-56073634-INS-5702
chr10-56092593-INS-60
chr10-56108132-INS-62
chr10-56123298-INS-551
chr10-56123417-INS-94
chr10-56137962-INS-76
chr10-56138808-INS-117
chr10-56175183-INS-52
chr10-56199689-INS-6371
chr10-56212649-INS-1177
chr10-56223123-INS-6583
chr10-56238098-INS-107
chr10-56263005-INS-84
chr10-56276296-INS-156
chr10-56276784-INS-59
chr10-56351536-INS-174
chr10-56352488-INS-60
chr10-56370970-INS-84
chr10-56385554-INS-859
chr10-56421574-INS-55
chr10-56435005-INS-63
chr10-56436872-INS-33571
chr10-56451006-INS-56
chr10-56453765-INS-1424
chr10-56518460-INS-63
chr10-56522754-INS-1847
chr10-56535421-INS-478
chr10-56538677-INS-1199
chr10-56568099-INS-211
chr10-56586011-INS-63
chr10-56618564-INS-77
chr10-56668492-INS-640
chr10-56684690-INS-72
chr10-56700915-INS-58
chr10-56718036-INS-52
chr10-56741816-INS-4870
chr10-56753812-INS-1312
chr10-56755042-INS-2006
chr10-56755042-INS-538
chr10-56758443-INS-51
chr10-56760368-INS-9088
chr10-56760726-INS-5564
chr10-56823347-INS-50
chr10-56863232-INS-1334
chr10-56898170-INS-628
chr10-56908331-INS-50
chr10-56944014-INS-52
chr10-57043622-INS-80
chr10-57060079-INS-84
chr10-57092236-INS-99
chr10-57115654-INS-601
chr10-57122140-INS-484
chr10-57138025-INS-320
chr10-57155805-INS-66
chr10-57160524-INS-6068
chr10-57170917-INS-71
chr10-57170977-INS-870
chr10-57175215-INS-485
chr10-57189180-INS-128
chr10-57222812-INS-461
chr10-57234941-INS-5113
chr10-57239228-INS-60
chr10-57241668-INS-55

chr10-57247689-INS-2203
chr10-57270430-INS-6809
chr10-57313744-INS-160
chr10-57318126-INS-1952
chr10-57324964-INS-2263
chr10-57327131-INS-142
chr10-57351651-INS-186
chr10-57386788-INS-176
chr10-57430179-INS-51
chr10-57431501-INS-193
chr10-57434159-INS-50
chr10-57444374-INS-50
chr10-57444448-INS-105
chr10-57490598-INS-145
chr10-57539366-INS-98
chr10-57572759-INS-69
chr10-57614147-INS-50
chr10-57619410-INS-301
chr10-57633521-INS-1979
chr10-57665467-INS-3042
chr10-58000107-INS-108
chr10-58037302-INS-74
chr10-58079645-INS-57
chr10-58103211-INS-51
chr10-58208116-INS-50
chr10-58209131-INS-2900
chr10-58273331-INS-1466
chr10-58309019-INS-208
chr10-58324188-INS-271
chr10-58330209-INS-54
chr10-58330465-INS-77
chr10-58363556-INS-180
chr10-58399924-INS-365
chr10-58423875-INS-1053
chr10-58451917-INS-54
chr10-58478027-INS-225
chr10-58500298-INS-78
chr10-58525591-INS-190
chr10-58554034-INS-60
chr10-58560016-INS-6448
chr10-58563859-INS-2131
chr10-58566072-INS-1425
chr10-58613117-INS-315
chr10-58616981-INS-60
chr10-58620490-INS-3700
chr10-58645813-INS-189
chr10-58662878-INS-1884
chr10-58667263-INS-83
chr10-58677465-INS-404
chr10-58696785-INS-66
chr10-58715205-INS-52
chr10-58730642-INS-3353
chr10-58734433-INS-136
chr10-58734593-INS-63
chr10-58794126-INS-374
chr10-58838240-INS-14154
chr10-58842252-INS-124
chr10-58898350-INS-201
chr10-58917339-INS-58
chr10-58928435-INS-73
chr10-58953207-INS-64
chr10-58972910-INS-795
chr10-58993678-INS-52
chr10-59034127-INS-3890
chr10-59038990-INS-1048
chr10-59046681-INS-6439
chr10-59067500-INS-2824
chr10-59079222-INS-123
chr10-59097766-INS-109
chr10-59148572-INS-50
chr10-59182186-INS-692
chr10-59202151-INS-271
chr10-59285058-INS-91
chr10-59295544-INS-386
chr10-59295547-INS-59
chr10-59324586-INS-7036
chr10-59358481-INS-117

chr10-59358738-INS-58
chr10-59389212-INS-181
chr10-59414924-INS-507
chr10-59454449-INS-54
chr10-59455859-INS-157
chr10-59456950-INS-56
chr10-59461258-INS-60
chr10-59461374-INS-251
chr10-59486584-INS-61
chr10-59492164-INS-105
chr10-59508139-INS-6313
chr10-59530326-INS-5032
chr10-59573290-INS-1492
chr10-59589552-INS-104
chr10-59672835-INS-150
chr10-59672835-INS-450
chr10-59683899-INS-197
chr10-59712733-INS-205
chr10-59745794-INS-118
chr10-59746140-INS-208
chr10-59769925-INS-531
chr10-59775399-INS-52
chr10-59777330-INS-144
chr10-59798359-INS-64
chr10-59811599-INS-59
chr10-59912532-INS-61
chr10-59915762-INS-50
chr10-59918536-INS-50
chr10-59930974-INS-717
chr10-59950705-INS-117
chr10-59980129-INS-244
chr10-59986625-INS-72
chr10-60003894-INS-738
chr10-60012416-INS-152
chr10-60014940-INS-3014
chr10-60020860-INS-2455
chr10-60031744-INS-1864
chr10-60032491-INS-202
chr10-60059754-INS-53
chr10-60073043-INS-100
chr10-60094597-INS-67
chr10-60100954-INS-52
chr10-60126740-INS-55
chr10-60127597-INS-51
chr10-60142837-INS-286
chr10-60164608-INS-504
chr10-60282902-INS-237
chr10-60313098-INS-2208
chr10-60316321-INS-8673
chr10-60340169-INS-60
chr10-60350924-INS-77
chr10-60414418-INS-76
chr10-60437399-INS-6403
chr10-60487635-INS-596
chr10-60513070-INS-50
chr10-60543475-INS-199
chr10-60642924-INS-160
chr10-60675216-INS-62
chr10-60684704-INS-476
chr10-60689792-INS-112
chr10-60691266-INS-80
chr10-60697886-INS-68
chr10-60793525-INS-54
chr10-60841386-INS-447
chr10-60868777-INS-111
chr10-60878863-INS-70
chr10-60913728-INS-252
chr10-60913743-INS-553
chr10-60919777-INS-420
chr10-60920958-INS-109
chr10-60949703-INS-1507
chr10-60962936-INS-212
chr10-60968604-INS-111
chr10-60970394-INS-319
chr10-60990293-INS-542
chr10-61032605-INS-198
chr10-61049996-INS-56

chr10-61057919-INS-120
chr10-61066551-INS-138
chr10-61066553-INS-580
chr10-61086098-INS-205
chr10-61144302-INS-4801
chr10-61165721-INS-208
chr10-61170418-INS-227
chr10-61185243-INS-223
chr10-61198499-INS-80
chr10-61214383-INS-50
chr10-61231030-INS-173
chr10-61233183-INS-82
chr10-61233224-INS-116
chr10-61296592-INS-3568
chr10-61298248-INS-855
chr10-61298363-INS-104
chr10-61299449-INS-66
chr10-61300127-INS-349
chr10-61409101-INS-50
chr10-61438782-INS-8683
chr10-61442866-INS-51
chr10-61448022-INS-252
chr10-61448155-INS-86
chr10-61474748-INS-75
chr10-61540302-INS-211
chr10-61637272-INS-130
chr10-61640567-INS-203
chr10-61833734-INS-50
chr10-61871424-INS-165
chr10-61957736-INS-183
chr10-61974187-INS-84
chr10-61974286-INS-258
chr10-61979414-INS-264
chr10-61984566-INS-296
chr10-61987987-INS-57
chr10-62010286-INS-96
chr10-62038149-INS-51
chr10-62042499-INS-62
chr10-62051491-INS-92
chr10-62059541-INS-184
chr10-62070463-INS-1657
chr10-62070463-INS-8896
chr10-62087427-INS-66
chr10-62186938-INS-208
chr10-62220562-INS-95
chr10-62232239-INS-51
chr10-62234299-INS-51
chr10-62241745-INS-193
chr10-62270589-INS-80
chr10-62296215-INS-63
chr10-62326750-INS-4573
chr10-62327236-INS-51
chr10-62331906-INS-4429
chr10-62383299-INS-88
chr10-62388953-INS-1951
chr10-62689670-INS-52
chr10-62717474-INS-202
chr10-62717561-INS-406
chr10-62728532-INS-50
chr10-62737588-INS-84
chr10-62741712-INS-56
chr10-62746194-INS-176
chr10-62750720-INS-3186
chr10-62750742-INS-6602
chr10-62753293-INS-1466
chr10-62778083-INS-383
chr10-62778610-INS-207
chr10-62783547-INS-73
chr10-62786303-INS-50
chr10-62795944-INS-119
chr10-62796443-INS-180
chr10-62800207-INS-380
chr10-62803107-INS-233
chr10-62825432-INS-199
chr10-62872065-INS-68
chr10-62887382-INS-223
chr10-62894673-INS-166

chr10-62919222-INS-70
chr10-62920179-INS-2820
chr10-62953663-INS-133
chr10-62956828-INS-1125
chr10-62964890-INS-84
chr10-62969982-INS-111
chr10-62986255-INS-67
chr10-62986505-INS-52
chr10-62994745-INS-56
chr10-63002692-INS-65
chr10-63002855-INS-120
chr10-63019683-INS-173
chr10-63020223-INS-372
chr10-63050791-INS-680
chr10-63061288-INS-64
chr10-63075951-INS-76
chr10-63147859-INS-185
chr10-63167862-INS-204
chr10-63178068-INS-74
chr10-63206381-INS-175
chr10-63215646-INS-62
chr10-63216127-INS-96
chr10-63250503-INS-180
chr10-63269625-INS-222
chr10-63295725-INS-1653
chr10-63322625-INS-57
chr10-63322860-INS-235
chr10-63323169-INS-206
chr10-63333249-INS-198
chr10-63337332-INS-2050
chr10-63347322-INS-122
chr10-63357530-INS-60
chr10-63358394-INS-638
chr10-63374432-INS-52
chr10-63381365-INS-158
chr10-63382283-INS-279
chr10-63382843-INS-181
chr10-63382855-INS-56
chr10-63407745-INS-511
chr10-63419623-INS-85
chr10-63457668-INS-393
chr10-63518348-INS-76
chr10-63521463-INS-332
chr10-63593605-INS-2297
chr10-63735277-INS-498
chr10-63813951-INS-6392
chr10-63824608-INS-80
chr10-63834407-INS-223
chr10-63880812-INS-257
chr10-63880814-INS-831
chr10-63934457-INS-65
chr10-63942437-INS-76
chr10-63989647-INS-236
chr10-63991664-INS-63
chr10-64023157-INS-120
chr10-64030458-INS-66
chr10-64109540-INS-83
chr10-64114061-INS-56
chr10-64120359-INS-174
chr10-64168444-INS-58
chr10-64184069-INS-1007
chr10-64291150-INS-118
chr10-64291162-INS-758
chr10-64300931-INS-136
chr10-64313253-INS-63
chr10-64329196-INS-57
chr10-64457435-INS-50
chr10-64468040-INS-2237
chr10-64475958-INS-6382
chr10-64509791-INS-154
chr10-64510577-INS-390
chr10-64551775-INS-7083
chr10-64613815-INS-12177
chr10-64635057-INS-52
chr10-64643686-INS-2353
chr10-64644992-INS-4324
chr10-64656378-INS-3632

chr10-64670928-INS-59
chr10-64687375-INS-6418
chr10-64713298-INS-52
chr10-64735184-INS-6766
chr10-64755040-INS-56
chr10-64773341-INS-54
chr10-64788415-INS-5756
chr10-64937628-INS-50
chr10-64960135-INS-136
chr10-64964594-INS-52
chr10-64974731-INS-17879
chr10-64980833-INS-54
chr10-65109668-INS-7093
chr10-65136693-INS-132
chr10-65168763-INS-68
chr10-65181837-INS-3183
chr10-65207525-INS-50
chr10-65253519-INS-50
chr10-65253648-INS-1112
chr10-65276654-INS-51
chr10-65280051-INS-66
chr10-65280509-INS-215
chr10-65281674-INS-60
chr10-65333571-INS-441
chr10-65341621-INS-4074
chr10-65412399-INS-50
chr10-65447978-INS-481
chr10-65455567-INS-1093
chr10-65459601-INS-449
chr10-65534877-INS-6477
chr10-65591171-INS-70
chr10-65596159-INS-143
chr10-65607527-INS-435
chr10-65659089-INS-716
chr10-65672509-INS-73
chr10-65686345-INS-50
chr10-65703575-INS-1287
chr10-65709042-INS-6389
chr10-65758182-INS-115
chr10-65827044-INS-159
chr10-65852518-INS-129
chr10-65853019-INS-80
chr10-65870833-INS-60
chr10-65877740-INS-424
chr10-65894329-INS-261
chr10-65980761-INS-6898
chr10-65987538-INS-446
chr10-65987933-INS-737
chr10-65989407-INS-4714
chr10-66053612-INS-215
chr10-66053636-INS-594
chr10-66115452-INS-547
chr10-66188565-INS-96
chr10-66261180-INS-14165
chr10-66326508-INS-326
chr10-66346286-INS-260
chr10-66371191-INS-100
chr10-66377942-INS-162
chr10-66390439-INS-81
chr10-66413376-INS-50
chr10-66489384-INS-72
chr10-66508924-INS-216
chr10-66582634-INS-142
chr10-66585359-INS-63
chr10-66607125-INS-94
chr10-66627966-INS-4265
chr10-66634181-INS-67
chr10-66665526-INS-278
chr10-66666253-INS-239
chr10-66668612-INS-1542
chr10-66703513-INS-199
chr10-66727706-INS-71
chr10-66732004-INS-207
chr10-66752062-INS-76
chr10-66752272-INS-208
chr10-66780596-INS-141
chr10-66784962-INS-175

chr10-66788196-INS-76
chr10-66824925-INS-7101
chr10-66842320-INS-608
chr10-66842606-INS-246
chr10-66844374-INS-50
chr10-66848266-INS-54
chr10-66916087-INS-284
chr10-66916087-INS-84
chr10-66917314-INS-202
chr10-66917851-INS-59
chr10-66921647-INS-254
chr10-66927092-INS-62
chr10-66965769-INS-64
chr10-66966429-INS-50
chr10-67043330-INS-117
chr10-67110192-INS-70
chr10-67138183-INS-331
chr10-67143479-INS-94
chr10-67161053-INS-2395
chr10-67175096-INS-160
chr10-67175665-INS-66
chr10-67191154-INS-217
chr10-67204655-INS-236
chr10-67225420-INS-113
chr10-67246734-INS-169
chr10-67264044-INS-78
chr10-67278236-INS-179
chr10-67331087-INS-80
chr10-67337598-INS-110
chr10-67338621-INS-229
chr10-67402714-INS-7300
chr10-67430147-INS-356
chr10-67590250-INS-69
chr10-67633166-INS-343
chr10-67646056-INS-7256
chr10-67660993-INS-56
chr10-67690304-INS-56
chr10-67703376-INS-96
chr10-67707007-INS-52
chr10-67710723-INS-224
chr10-67754999-INS-118
chr10-67772730-INS-4660
chr10-67778738-INS-159
chr10-67779415-INS-151
chr10-67802745-INS-84
chr10-67814221-INS-120
chr10-67866197-INS-1862
chr10-67869428-INS-290
chr10-67869652-INS-309
chr10-67873515-INS-118
chr10-67947655-INS-94
chr10-67947858-INS-134
chr10-68084465-INS-50
chr10-68094703-INS-60
chr10-68135349-INS-54
chr10-68211320-INS-63
chr10-68330120-INS-214
chr10-68332075-INS-109
chr10-68454394-INS-110
chr10-68454539-INS-144
chr10-68454731-INS-77
chr10-68478776-INS-1548
chr10-68482684-INS-122
chr10-68522941-INS-56
chr10-68529622-INS-50
chr10-68529624-INS-329
chr10-68529966-INS-185
chr10-68531301-INS-57
chr10-68531873-INS-73
chr10-68533877-INS-2750
chr10-68553554-INS-6370
chr10-68616691-INS-116
chr10-68651025-INS-389
chr10-68660888-INS-2924
chr10-68697273-INS-1269
chr10-68697830-INS-756
chr10-68705101-INS-50

chr10-68719065-INS-106
chr10-68735683-INS-202
chr10-68854535-INS-60
chr10-68855560-INS-144
chr10-68859376-INS-87
chr10-68918316-INS-202
chr10-68976020-INS-233
chr10-69045804-INS-50
chr10-69055563-INS-56
chr10-69058861-INS-78
chr10-69063244-INS-686
chr10-69064595-INS-568
chr10-69079655-INS-103
chr10-69133721-INS-50
chr10-69138405-INS-64
chr10-69193376-INS-195
chr10-69197919-INS-226
chr10-69198542-INS-80
chr10-69202660-INS-63
chr10-69217583-INS-56
chr10-69219711-INS-204
chr10-69224404-INS-174
chr10-69225292-INS-2091
chr10-69233837-INS-68
chr10-69258071-INS-206
chr10-69259781-INS-66
chr10-69261803-INS-1103
chr10-69299364-INS-258
chr10-69312378-INS-167
chr10-69326090-INS-185
chr10-69383424-INS-163
chr10-69387232-INS-3818
chr10-69401463-INS-83
chr10-69411832-INS-89
chr10-69440189-INS-74
chr10-69478705-INS-193
chr10-69478737-INS-81
chr10-69506223-INS-528
chr10-69508891-INS-64
chr10-69509251-INS-663
chr10-69584318-INS-50
chr10-69584443-INS-480
chr10-69598809-INS-198
chr10-69599518-INS-118
chr10-69690485-INS-81
chr10-69720252-INS-198
chr10-69732593-INS-62
chr10-69745881-INS-73
chr10-69776317-INS-431
chr10-69783910-INS-50
chr10-69799586-INS-108
chr10-69856786-INS-60
chr10-69868336-INS-57
chr10-69868339-INS-220
chr10-69884084-INS-130
chr10-69910531-INS-128
chr10-69915407-INS-52
chr10-69958358-INS-105
chr10-69996279-INS-173
chr10-69998767-INS-90
chr10-70016157-INS-62
chr10-70056621-INS-924
chr10-70063693-INS-57
chr10-70077762-INS-293
chr10-70082535-INS-212
chr10-70135232-INS-411
chr10-70142290-INS-50
chr10-70142393-INS-266
chr10-70161981-INS-1895
chr10-70194336-INS-5080
chr10-70201480-INS-6603
chr10-70202237-INS-98
chr10-70206381-INS-1190
chr10-70243925-INS-207
chr10-70256508-INS-668
chr10-70260548-INS-2475
chr10-70261930-INS-152

chr10-70263749-INS-216
chr10-70284861-INS-101
chr10-70303115-INS-68
chr10-70305569-INS-1450
chr10-70343732-INS-64
chr10-70344225-INS-86
chr10-70344245-INS-201
chr10-70448565-INS-123
chr10-70480918-INS-238
chr10-70513327-INS-4192
chr10-70608695-INS-171
chr10-70624596-INS-323
chr10-70659459-INS-7408
chr10-70667704-INS-52
chr10-70712747-INS-4807
chr10-70751698-INS-50
chr10-70782257-INS-105
chr10-70805697-INS-132
chr10-70858675-INS-70
chr10-70861703-INS-434
chr10-70869772-INS-50
chr10-70893841-INS-6274
chr10-70907509-INS-50
chr10-70920820-INS-50
chr10-70932014-INS-196
chr10-70936264-INS-872
chr10-71000422-INS-325
chr10-71035482-INS-470
chr10-71035719-INS-2750
chr10-71035907-INS-191
chr10-71039417-INS-68
chr10-71039891-INS-274
chr10-71061206-INS-160
chr10-71062118-INS-55
chr10-71062164-INS-127
chr10-71094848-INS-326
chr10-71121586-INS-74
chr10-71129675-INS-69
chr10-71130163-INS-7249
chr10-71130228-INS-191
chr10-71203678-INS-112
chr10-71213793-INS-76
chr10-71494187-INS-66
chr10-71503902-INS-6260
chr10-71548300-INS-209
chr10-71571262-INS-790
chr10-71583861-INS-7565
chr10-71584116-INS-50
chr10-71594206-INS-65
chr10-71615647-INS-1081
chr10-71622738-INS-1760
chr10-71720936-INS-76
chr10-71748970-INS-2528
chr10-71775188-INS-138
chr10-71835613-INS-148
chr10-71867283-INS-56
chr10-71873963-INS-167
chr10-71888228-INS-89
chr10-71922184-INS-1615
chr10-71930074-INS-132
chr10-71946138-INS-1165
chr10-71947302-INS-51
chr10-71961787-INS-53
chr10-71968961-INS-543
chr10-71990787-INS-4575
chr10-72013664-INS-6524
chr10-72018372-INS-50
chr10-72023653-INS-64
chr10-72116128-INS-59
chr10-72169361-INS-70
chr10-72172129-INS-225
chr10-72180928-INS-230
chr10-72180928-INS-79
chr10-72190304-INS-466
chr10-72217952-INS-6462
chr10-72394256-INS-2719
chr10-72436804-INS-60

chr10-72648954-INS-6159
chr10-72685456-INS-293
chr10-72685460-INS-76
chr10-72686100-INS-164
chr10-72686263-INS-888
chr10-72689786-INS-80
chr10-72699452-INS-1866
chr10-72702025-INS-116
chr10-72734861-INS-2497
chr10-72766494-INS-1171
chr10-72769692-INS-1429
chr10-72779814-INS-1544
chr10-72879881-INS-54
chr10-72895518-INS-7376
chr10-72937371-INS-184
chr10-72961144-INS-76
chr10-72976648-INS-122
chr10-72976664-INS-549
chr10-73007071-INS-6596
chr10-73007254-INS-118
chr10-73007254-INS-640
chr10-73133892-INS-1397
chr10-73166682-INS-50
chr10-73178480-INS-65
chr10-73186217-INS-2300
chr10-73230086-INS-1775
chr10-73231929-INS-1041
chr10-73238764-INS-328
chr10-73308566-INS-7102
chr10-73382890-INS-294
chr10-73394329-INS-90
chr10-73394563-INS-698
chr10-73394718-INS-239
chr10-73394718-INS-97
chr10-73396268-INS-120
chr10-73402148-INS-4701
chr10-73407299-INS-1249
chr10-73455227-INS-78
chr10-73654856-INS-90
chr10-73674980-INS-75
chr10-73740076-INS-61
chr10-73779776-INS-2197
chr10-73793405-INS-7635
chr10-73798466-INS-1701
chr10-73847646-INS-51
chr10-73898206-INS-415
chr10-73969317-INS-167
chr10-73969392-INS-266
chr10-74005644-INS-56
chr10-74014889-INS-6401
chr10-74035512-INS-72
chr10-74067171-INS-152
chr10-74101464-INS-122
chr10-74170013-INS-120
chr10-74172472-INS-827
chr10-74194594-INS-1499
chr10-74205608-INS-88
chr10-74224036-INS-50
chr10-74229468-INS-118
chr10-74274945-INS-1477
chr10-74279047-INS-2429
chr10-74421700-INS-58
chr10-74421755-INS-364
chr10-74425887-INS-95
chr10-74444478-INS-1057
chr10-74476474-INS-81
chr10-74531468-INS-152
chr10-74535144-INS-6853
chr10-74641407-INS-6517
chr10-74660086-INS-84
chr10-74665618-INS-6490
chr10-74674195-INS-63
chr10-74704087-INS-65
chr10-74746341-INS-61
chr10-74772133-INS-1870
chr10-74792261-INS-215
chr10-74841256-INS-309

chr10-74841256-INS-833
chr10-74843972-INS-190
chr10-74844212-INS-225
chr10-74844212-INS-767
chr10-74869519-INS-135
chr10-74870387-INS-66
chr10-74872404-INS-56
chr10-74876884-INS-50
chr10-74891260-INS-56
chr10-74903682-INS-165
chr10-74941196-INS-262
chr10-74948021-INS-284
chr10-74968017-INS-214
chr10-75000186-INS-363
chr10-75069281-INS-261
chr10-75069347-INS-891
chr10-75079686-INS-220
chr10-75121331-INS-51
chr10-75129034-INS-55
chr10-75218075-INS-79
chr10-75269684-INS-342
chr10-75269850-INS-1001
chr10-75270740-INS-200
chr10-75310028-INS-58
chr10-75358288-INS-214
chr10-75358852-INS-72
chr10-75406576-INS-50
chr10-75421689-INS-64
chr10-75547770-INS-138
chr10-75596607-INS-788
chr10-75605624-INS-186
chr10-75642594-INS-225
chr10-75661310-INS-124
chr10-75666158-INS-1366
chr10-75710102-INS-197
chr10-75745380-INS-110
chr10-75748080-INS-99
chr10-75755918-INS-109
chr10-75799890-INS-1219
chr10-75800708-INS-119
chr10-75823425-INS-201
chr10-75832709-INS-72
chr10-75834519-INS-7399
chr10-75837839-INS-68
chr10-75838716-INS-211
chr10-75840442-INS-108
chr10-75855032-INS-720
chr10-75855147-INS-348
chr10-75858865-INS-775
chr10-75887380-INS-60
chr10-75911059-INS-3748
chr10-75913429-INS-3153
chr10-75934734-INS-102
chr10-75972289-INS-1838
chr10-76003116-INS-62
chr10-76082455-INS-87
chr10-76166917-INS-210
chr10-76170216-INS-214
chr10-76175193-INS-74
chr10-76209694-INS-207
chr10-76238551-INS-108
chr10-76246407-INS-68
chr10-76261451-INS-59
chr10-76296174-INS-1455
chr10-76309214-INS-385
chr10-76314677-INS-131
chr10-76317069-INS-103
chr10-76317069-INS-328
chr10-76320495-INS-56
chr10-76338238-INS-220
chr10-76350063-INS-269
chr10-76417686-INS-746
chr10-76417785-INS-85
chr10-76418044-INS-81
chr10-76467906-INS-96
chr10-76488274-INS-81
chr10-76493554-INS-100

chr10-76502000-INS-1858
chr10-76568739-INS-52
chr10-76581381-INS-189
chr10-76581519-INS-353
chr10-76581802-INS-242
chr10-76594125-INS-70
chr10-76662987-INS-235
chr10-76662987-INS-79
chr10-76666613-INS-74
chr10-76679316-INS-88
chr10-76686510-INS-131
chr10-76732158-INS-106
chr10-76739150-INS-52
chr10-76753880-INS-88
chr10-76834945-INS-72
chr10-76940080-INS-202
chr10-76965406-INS-56
chr10-76985665-INS-105
chr10-76985834-INS-51
chr10-77010553-INS-71
chr10-77018608-INS-61
chr10-77018839-INS-72
chr10-77019386-INS-178
chr10-77019745-INS-1542
chr10-77024136-INS-4344
chr10-77037434-INS-80
chr10-77068836-INS-68
chr10-77084576-INS-64
chr10-77086296-INS-56
chr10-77124243-INS-700
chr10-77165725-INS-3059
chr10-77180352-INS-1314
chr10-77211669-INS-65
chr10-77259436-INS-2575
chr10-77341803-INS-4824
chr10-77389951-INS-54
chr10-77421169-INS-56
chr10-77421274-INS-742
chr10-77474317-INS-185
chr10-77539343-INS-311
chr10-77568359-INS-200
chr10-77605126-INS-2859
chr10-77639083-INS-167
chr10-77657568-INS-2217
chr10-77693277-INS-206
chr10-77696189-INS-52
chr10-77846166-INS-67
chr10-77868800-INS-66
chr10-77881282-INS-264
chr10-77889331-INS-55
chr10-77901293-INS-161
chr10-77902417-INS-1903
chr10-77904146-INS-74
chr10-77945837-INS-86
chr10-77945839-INS-218
chr10-77949624-INS-120
chr10-77964187-INS-170
chr10-77967936-INS-226
chr10-77968241-INS-100
chr10-77992342-INS-53
chr10-77993601-INS-148
chr10-78027871-INS-140
chr10-78077310-INS-218
chr10-78085625-INS-53
chr10-78115347-INS-336
chr10-78195382-INS-51
chr10-78210698-INS-56
chr10-78291102-INS-143
chr10-78312982-INS-63
chr10-78317125-INS-134
chr10-78330852-INS-50
chr10-78331185-INS-235
chr10-78331540-INS-200
chr10-78332668-INS-50
chr10-78334805-INS-521
chr10-78341749-INS-200
chr10-78342860-INS-192

chr10-78374034-INS-9058
chr10-78405499-INS-68
chr10-78407962-INS-70
chr10-78423835-INS-60
chr10-78494125-INS-471
chr10-78494125-INS-95
chr10-78501786-INS-108
chr10-78506464-INS-71
chr10-78512482-INS-3409
chr10-78517739-INS-6453
chr10-78563366-INS-52
chr10-78563380-INS-215
chr10-78573242-INS-319
chr10-78583351-INS-9800
chr10-78594531-INS-52
chr10-78628528-INS-66
chr10-78638034-INS-50
chr10-78638270-INS-55
chr10-78638367-INS-76
chr10-78640400-INS-2001
chr10-78788074-INS-122
chr10-78849251-INS-54
chr10-78852920-INS-7441
chr10-78935018-INS-57
chr10-78949972-INS-6082
chr10-79014883-INS-2884
chr10-79049687-INS-61
chr10-79051000-INS-1716
chr10-79103757-INS-502
chr10-79141934-INS-197
chr10-79167875-INS-4045
chr10-79237240-INS-2758
chr10-79361172-INS-67
chr10-79425507-INS-222
chr10-79425613-INS-64
chr10-79434439-INS-58
chr10-79477552-INS-404
chr10-79566326-INS-233
chr10-79605272-INS-203
chr10-79619813-INS-55
chr10-79624293-INS-394
chr10-79664268-INS-50
chr10-79761765-INS-89
chr10-79793383-INS-55
chr10-79795055-INS-54
chr10-79883551-INS-519
chr10-79885628-INS-210
chr10-79904934-INS-132
chr10-79946845-INS-108
chr10-80027242-INS-72
chr10-80099952-INS-122
chr10-80122156-INS-187
chr10-80132180-INS-125
chr10-80132180-INS-50
chr10-80193931-INS-74
chr10-80198244-INS-843
chr10-80209931-INS-174
chr10-80273122-INS-75
chr10-80300141-INS-60
chr10-80394666-INS-66
chr10-80598150-INS-76
chr10-80638755-INS-62
chr10-80652052-INS-857
chr10-80652175-INS-7218
chr10-80690082-INS-111
chr10-80747899-INS-61
chr10-80759298-INS-55
chr10-80810708-INS-62
chr10-80820583-INS-215
chr10-80827303-INS-51
chr10-80867909-INS-2128
chr10-80899196-INS-216
chr10-81045019-INS-184
chr10-81267974-INS-209
chr10-81315673-INS-53
chr10-81319178-INS-201
chr10-81383854-INS-76

chr10-81387363-INS-50
chr10-81446926-INS-207
chr10-81466565-INS-94
chr10-81473409-INS-225
chr10-81492905-INS-2897
chr10-81493535-INS-510
chr10-81503682-INS-205
chr10-81503935-INS-233
chr10-81585650-INS-497
chr10-81643382-INS-62
chr10-81644193-INS-216
chr10-81875079-INS-169
chr10-81875404-INS-310
chr10-81912397-INS-2649
chr10-81973318-INS-123
chr10-82195411-INS-6438
chr10-82213665-INS-252
chr10-82398115-INS-1427
chr10-82416702-INS-1430
chr10-82442580-INS-129
chr10-82443375-INS-204
chr10-82482086-INS-108
chr10-82512823-INS-160
chr10-82515875-INS-167
chr10-82546081-INS-318
chr10-82608274-INS-2596
chr10-82632114-INS-89
chr10-82651282-INS-55
chr10-82655678-INS-214
chr10-82675809-INS-551
chr10-82733216-INS-170
chr10-82756597-INS-158
chr10-82761332-INS-1864
chr10-82763896-INS-169
chr10-82766618-INS-74
chr10-82800975-INS-81
chr10-82801362-INS-304
chr10-82822046-INS-55
chr10-82848946-INS-54
chr10-82886151-INS-1379
chr10-82986943-INS-113
chr10-82989518-INS-6619
chr10-82996900-INS-51
chr10-83004773-INS-70
chr10-83064288-INS-448
chr10-83101931-INS-54
chr10-83120947-INS-172
chr10-83153746-INS-51
chr10-83360528-INS-79
chr10-83364207-INS-50
chr10-83442712-INS-51
chr10-83462974-INS-1850
chr10-83505106-INS-59
chr10-83525774-INS-971
chr10-83533792-INS-50
chr10-83535443-INS-1119
chr10-83551284-INS-50
chr10-83589554-INS-70
chr10-83691197-INS-552
chr10-83691317-INS-112
chr10-83716033-INS-1547
chr10-83716243-INS-52
chr10-83735599-INS-124
chr10-83737712-INS-179
chr10-83741099-INS-82
chr10-83743625-INS-105
chr10-83763953-INS-52
chr10-83770116-INS-3084
chr10-83807242-INS-89
chr10-83809276-INS-63
chr10-83815840-INS-110
chr10-83821966-INS-62
chr10-83838950-INS-186
chr10-83918232-INS-147
chr10-84020442-INS-63
chr10-84023821-INS-89
chr10-84039071-INS-5514

chr10-84068651-INS-50
chr10-84108569-INS-263
chr10-84127084-INS-722
chr10-84189135-INS-412
chr10-84211408-INS-272
chr10-84295896-INS-4133
chr10-84321779-INS-116
chr10-84342864-INS-17342
chr10-84351792-INS-61
chr10-84382832-INS-177
chr10-84385019-INS-50
chr10-84390825-INS-105
chr10-84411126-INS-6989
chr10-84457577-INS-6330
chr10-84532325-INS-1197
chr10-84551894-INS-7249
chr10-84553444-INS-340
chr10-84574372-INS-240
chr10-84574372-INS-960
chr10-84575845-INS-50
chr10-84590174-INS-75
chr10-84641311-INS-76
chr10-84642792-INS-387
chr10-84644773-INS-217
chr10-84644911-INS-284
chr10-84716437-INS-292
chr10-84716441-INS-140
chr10-84866554-INS-50
chr10-84885388-INS-114
chr10-84885894-INS-220
chr10-84902167-INS-241
chr10-84927383-INS-128
chr10-84981491-INS-199
chr10-84986531-INS-128
chr10-85006382-INS-74
chr10-85086051-INS-75
chr10-85087426-INS-5342
chr10-85094440-INS-115
chr10-85102284-INS-60
chr10-85116559-INS-1676
chr10-85117187-INS-199
chr10-85117311-INS-92
chr10-85181965-INS-6118
chr10-85339389-INS-64
chr10-85351691-INS-197
chr10-85352283-INS-62
chr10-85356389-INS-134
chr10-85357193-INS-116
chr10-85372210-INS-121
chr10-85383223-INS-648
chr10-85383447-INS-186
chr10-85383548-INS-52
chr10-85423590-INS-210
chr10-85445457-INS-66
chr10-85482796-INS-50
chr10-85489183-INS-79
chr10-85579581-INS-6530
chr10-85648387-INS-57
chr10-85673624-INS-55
chr10-85686258-INS-3567
chr10-85756349-INS-2519
chr10-85767495-INS-50
chr10-85794014-INS-88
chr10-85794191-INS-52
chr10-85803362-INS-50
chr10-85809708-INS-202
chr10-85836102-INS-8575
chr10-85845662-INS-64
chr10-85857283-INS-2019
chr10-85928705-INS-90
chr10-85954496-INS-2173
chr10-85969081-INS-84
chr10-85976068-INS-6396
chr10-85980247-INS-103
chr10-86204076-INS-50
chr10-86248719-INS-209
chr10-86285507-INS-118

chr10-86286924-INS-346
chr10-86287987-INS-50
chr10-86458294-INS-1282
chr10-86483524-INS-4210
chr10-86519655-INS-8702
chr10-86558131-INS-135
chr10-86558139-INS-62
chr10-86558876-INS-123
chr10-86562191-INS-62
chr10-86580725-INS-2891
chr10-86590740-INS-348
chr10-86595368-INS-164
chr10-86602983-INS-207
chr10-86603783-INS-231
chr10-86624006-INS-210
chr10-86649008-INS-497
chr10-86681266-INS-210
chr10-86741028-INS-1132
chr10-86742733-INS-60
chr10-86743769-INS-54
chr10-86764856-INS-100
chr10-86765011-INS-450
chr10-86774388-INS-550
chr10-86795124-INS-6547
chr10-86798349-INS-129
chr10-86832922-INS-59
chr10-86843960-INS-5336
chr10-86853264-INS-6231
chr10-86857709-INS-144
chr10-86860722-INS-238
chr10-86918280-INS-4049
chr10-86920046-INS-1396
chr10-86935617-INS-78
chr10-87031664-INS-50
chr10-87109056-INS-1520
chr10-87121975-INS-6748
chr10-87222565-INS-6693
chr10-87251318-INS-1213
chr10-87280258-INS-549
chr10-87295882-INS-6476
chr10-87303939-INS-171
chr10-87335466-INS-383
chr10-87348686-INS-51
chr10-87372020-INS-185
chr10-87391724-INS-50
chr10-87428903-INS-93
chr10-87471646-INS-810
chr10-87628193-INS-267
chr10-87795062-INS-311
chr10-87804158-INS-141
chr10-87805494-INS-76
chr10-87878616-INS-50
chr10-87891391-INS-575
chr10-87939945-INS-257
chr10-87948501-INS-216
chr10-88028070-INS-92
chr10-88032685-INS-1787
chr10-88046591-INS-342
chr10-88046640-INS-77
chr10-88051299-INS-55
chr10-88081509-INS-148
chr10-88082202-INS-64
chr10-88082398-INS-162
chr10-88119865-INS-1236
chr10-88155353-INS-221
chr10-88155381-INS-649
chr10-88174133-INS-78
chr10-88193934-INS-1735
chr10-88198019-INS-50
chr10-88202707-INS-300
chr10-88204709-INS-60
chr10-88204985-INS-53
chr10-88214470-INS-230
chr10-88223506-INS-195
chr10-88230158-INS-174
chr10-88232418-INS-50
chr10-88260267-INS-248

chr10-88278066-INS-202
chr10-88278184-INS-4506
chr10-88285943-INS-60
chr10-88287466-INS-6399
chr10-88318295-INS-79
chr10-88323955-INS-481
chr10-88334875-INS-4797
chr10-88367799-INS-102
chr10-88393111-INS-205
chr10-88435105-INS-204
chr10-88447586-INS-192
chr10-88448351-INS-6375
chr10-88466418-INS-492
chr10-88469356-INS-3472
chr10-88529253-INS-126
chr10-88532470-INS-144
chr10-88610019-INS-144
chr10-88610019-INS-432
chr10-88617523-INS-733
chr10-88706051-INS-119
chr10-88706054-INS-596
chr10-88724215-INS-2154
chr10-88748570-INS-88
chr10-88755862-INS-54
chr10-88756142-INS-84
chr10-88758611-INS-111
chr10-88761065-INS-99
chr10-88822175-INS-256
chr10-88824497-INS-75
chr10-88842902-INS-998
chr10-88851410-INS-85
chr10-88909057-INS-55
chr10-88960223-INS-54
chr10-88980034-INS-52
chr10-89000358-INS-290
chr10-89003155-INS-1831
chr10-89014560-INS-6634
chr10-89038677-INS-5054
chr10-89052333-INS-50
chr10-89061754-INS-318
chr10-89106345-INS-377
chr10-89141625-INS-1986
chr10-89145372-INS-15078
chr10-89213894-INS-2307
chr10-89222977-INS-175
chr10-89366574-INS-57
chr10-89373248-INS-5606
chr10-89397648-INS-215
chr10-89424463-INS-50
chr10-89424838-INS-103
chr10-89448011-INS-138
chr10-89448237-INS-577
chr10-89448348-INS-2293
chr10-89453772-INS-150
chr10-89506512-INS-431
chr10-89518481-INS-166
chr10-89592071-INS-213
chr10-89600205-INS-52
chr10-89678578-INS-58
chr10-89684331-INS-118
chr10-89703584-INS-236
chr10-89703962-INS-130
chr10-89706859-INS-80
chr10-89715912-INS-171
chr10-89735984-INS-68
chr10-89736956-INS-67
chr10-89746382-INS-4860
chr10-89767294-INS-1134
chr10-89819982-INS-62
chr10-89823878-INS-6591
chr10-89824600-INS-81
chr10-89828962-INS-62
chr10-89858406-INS-5184
chr10-89872892-INS-150
chr10-89874428-INS-551
chr10-89941207-INS-50
chr10-89955032-INS-4170

chr10-90007511-INS-6564
chr10-90020197-INS-2853
chr10-90071350-INS-1040
chr10-90092123-INS-58
chr10-90122381-INS-90
chr10-90123636-INS-1489
chr10-90174547-INS-510
chr10-90182464-INS-6231
chr10-90185892-INS-2354
chr10-90198901-INS-251
chr10-90205249-INS-6644
chr10-90216033-INS-443
chr10-90229978-INS-173
chr10-90272577-INS-6380
chr10-90328408-INS-1840
chr10-90368865-INS-52
chr10-90392693-INS-313
chr10-90402179-INS-55
chr10-90421630-INS-83
chr10-90454404-INS-7043
chr10-90464899-INS-253
chr10-90544473-INS-50
chr10-90812077-INS-10572
chr10-90819971-INS-52
chr10-90822795-INS-113
chr10-90826338-INS-174
chr10-90842844-INS-780
chr10-90843345-INS-217
chr10-90852445-INS-50
chr10-90856798-INS-387
chr10-90856910-INS-60
chr10-90906227-INS-219
chr10-90921871-INS-405
chr10-90926629-INS-66
chr10-90942325-INS-224
chr10-90967664-INS-5435
chr10-91031335-INS-54
chr10-91036735-INS-7121
chr10-91046396-INS-1693
chr10-91056198-INS-239
chr10-91062025-INS-211
chr10-91128046-INS-65
chr10-91134682-INS-306
chr10-91134856-INS-61
chr10-91135088-INS-115
chr10-91135088-INS-305
chr10-91211050-INS-3096
chr10-91212904-INS-70
chr10-91295239-INS-68
chr10-91318478-INS-67
chr10-91347511-INS-773
chr10-91361441-INS-60
chr10-91368389-INS-94
chr10-91401786-INS-92
chr10-91429505-INS-6555
chr10-91435316-INS-852
chr10-91440098-INS-55
chr10-91461144-INS-489
chr10-91498611-INS-61
chr10-91500687-INS-6146
chr10-91573009-INS-56
chr10-91656797-INS-51
chr10-91663829-INS-4766
chr10-91680395-INS-88
chr10-91689061-INS-66
chr10-91700813-INS-51
chr10-91710316-INS-1947
chr10-91813050-INS-4867
chr10-91821387-INS-62
chr10-91852080-INS-108
chr10-91863495-INS-66
chr10-91895048-INS-528
chr10-91899513-INS-82
chr10-91962209-INS-54
chr10-91972292-INS-150
chr10-92015300-INS-166
chr10-92018956-INS-4847

chr10-92040538-INS-132
chr10-92088732-INS-2242
chr10-92136359-INS-1133
chr10-92208588-INS-6584
chr10-92213914-INS-62
chr10-92223879-INS-452
chr10-92224148-INS-123
chr10-92280934-INS-6520
chr10-92290391-INS-6924
chr10-92295378-INS-60
chr10-92303366-INS-6834
chr10-92344662-INS-6151
chr10-92354202-INS-94
chr10-92423248-INS-1588
chr10-92457101-INS-52
chr10-92469770-INS-5439
chr10-92480929-INS-66
chr10-92597281-INS-78
chr10-92604098-INS-6259
chr10-92642671-INS-2001
chr10-92731121-INS-156
chr10-92755727-INS-74
chr10-92824321-INS-955
chr10-92841998-INS-75
chr10-92887726-INS-184
chr10-92887974-INS-229
chr10-92888023-INS-657
chr10-92934930-INS-126
chr10-92935036-INS-1606
chr10-92945062-INS-1864
chr10-92985850-INS-241
chr10-92989939-INS-70
chr10-93023293-INS-53
chr10-93031141-INS-65
chr10-93031520-INS-97
chr10-93034252-INS-100
chr10-93058796-INS-277
chr10-93058829-INS-53
chr10-93070209-INS-176
chr10-93070209-INS-71
chr10-93082951-INS-544
chr10-93083036-INS-264
chr10-93170286-INS-140
chr10-93188923-INS-50
chr10-93251742-INS-400
chr10-93251786-INS-4661
chr10-93263831-INS-59
chr10-93302128-INS-98
chr10-93304008-INS-234
chr10-93304039-INS-1470
chr10-93306082-INS-76
chr10-93306299-INS-276
chr10-93324071-INS-50
chr10-93326293-INS-325
chr10-93353366-INS-125
chr10-93384679-INS-84
chr10-93433560-INS-3678
chr10-93473266-INS-50
chr10-93491743-INS-60
chr10-93494927-INS-365
chr10-93563409-INS-330
chr10-93629147-INS-50
chr10-93634016-INS-50
chr10-93656954-INS-56
chr10-93657966-INS-75
chr10-93673163-INS-199
chr10-93745894-INS-2832
chr10-93764019-INS-1220
chr10-93764349-INS-7142
chr10-93765995-INS-50
chr10-93772390-INS-4489
chr10-93783736-INS-95
chr10-93909454-INS-109
chr10-93964857-INS-72
chr10-93988572-INS-51
chr10-94075227-INS-208
chr10-94075461-INS-101

chr10-94095922-INS-6449
chr10-94175205-INS-64
chr10-94460368-INS-76
chr10-94477114-INS-55
chr10-94477612-INS-228
chr10-94502383-INS-122
chr10-94563945-INS-111
chr10-94669913-INS-154
chr10-94675337-INS-56
chr10-94679253-INS-62
chr10-94722399-INS-67
chr10-94729859-INS-213
chr10-94744727-INS-126
chr10-94750956-INS-106
chr10-94817192-INS-65
chr10-94889707-INS-82
chr10-94912608-INS-5553
chr10-94925001-INS-2714
chr10-94930937-INS-500
chr10-94992762-INS-119
chr10-94992840-INS-51
chr10-95004874-INS-68
chr10-95062477-INS-7229
chr10-95083669-INS-456
chr10-95106884-INS-62
chr10-95110298-INS-57
chr10-95124644-INS-8457
chr10-95148704-INS-78
chr10-95234810-INS-50
chr10-95267990-INS-59
chr10-95307800-INS-118
chr10-95347683-INS-534
chr10-95371972-INS-241
chr10-95382581-INS-50
chr10-95404934-INS-50
chr10-95405372-INS-341
chr10-95413069-INS-128
chr10-95418457-INS-1894
chr10-95420503-INS-255
chr10-95451437-INS-166
chr10-95511782-INS-78
chr10-95533006-INS-5039
chr10-95545974-INS-169
chr10-95585045-INS-293
chr10-95599837-INS-50
chr10-95620793-INS-55
chr10-95636067-INS-66
chr10-95648589-INS-65
chr10-95651267-INS-55
chr10-95693156-INS-1331
chr10-95696601-INS-147
chr10-95697031-INS-124
chr10-95699959-INS-118
chr10-95701487-INS-50
chr10-95711353-INS-86
chr10-95718671-INS-50
chr10-95729142-INS-480
chr10-95729668-INS-1559
chr10-95745449-INS-104
chr10-95827784-INS-171
chr10-95872024-INS-2140
chr10-95899333-INS-52
chr10-95931788-INS-6346
chr10-95954593-INS-555
chr10-95962853-INS-153
chr10-96025867-INS-55
chr10-96038303-INS-54
chr10-96040931-INS-344
chr10-96041844-INS-70
chr10-96044741-INS-599
chr10-96054140-INS-386
chr10-96086287-INS-2848
chr10-96113774-INS-7068
chr10-96124219-INS-75
chr10-96124949-INS-54
chr10-96136501-INS-102
chr10-96152198-INS-2303

chr10-96172768-INS-129
chr10-96173931-INS-2730
chr10-96176459-INS-443
chr10-96183192-INS-76
chr10-96183310-INS-144
chr10-96186180-INS-69
chr10-96190278-INS-69
chr10-96200339-INS-52
chr10-96236219-INS-1266
chr10-96250659-INS-114
chr10-96256244-INS-58
chr10-96295056-INS-6589
chr10-96323646-INS-1033
chr10-96349075-INS-68
chr10-96387180-INS-257
chr10-96392186-INS-88
chr10-96404603-INS-63
chr10-96411667-INS-105
chr10-96412171-INS-360
chr10-96480263-INS-65
chr10-96480708-INS-1836
chr10-96487481-INS-61
chr10-96511732-INS-140
chr10-96525308-INS-54
chr10-96549221-INS-126
chr10-96554826-INS-80
chr10-96555160-INS-56
chr10-96611985-INS-50
chr10-96623968-INS-68
chr10-96637253-INS-495
chr10-96670792-INS-65
chr10-96688636-INS-55
chr10-96750073-INS-2809
chr10-96763833-INS-57
chr10-96769483-INS-116
chr10-96795718-INS-50
chr10-96799623-INS-50
chr10-96807356-INS-6204
chr10-96807575-INS-72
chr10-96820846-INS-168
chr10-96834618-INS-93
chr10-96848319-INS-88
chr10-96872915-INS-116
chr10-96927542-INS-703
chr10-96976342-INS-56
chr10-97022707-INS-541
chr10-97040288-INS-1204
chr10-97051507-INS-6528
chr10-97052190-INS-656
chr10-97068369-INS-3629
chr10-97109213-INS-133
chr10-97115072-INS-9553
chr10-97122371-INS-72
chr10-97153890-INS-50
chr10-97202294-INS-8457
chr10-97211267-INS-237
chr10-97221427-INS-598
chr10-97237831-INS-1406
chr10-97248210-INS-4373
chr10-97266962-INS-58
chr10-97274020-INS-92
chr10-97313228-INS-52
chr10-97348591-INS-983
chr10-97372617-INS-105
chr10-97411412-INS-6401
chr10-97412887-INS-498
chr10-97515329-INS-60
chr10-97552625-INS-65
chr10-97556145-INS-6723
chr10-97572517-INS-83
chr10-97581383-INS-3079
chr10-97623629-INS-58
chr10-97640501-INS-98
chr10-97697368-INS-64
chr10-97724861-INS-1130
chr10-97766839-INS-164
chr10-97855716-INS-120

chr10-97900304-INS-52
chr10-97900916-INS-859
chr10-98020053-INS-52
chr10-98055006-INS-1767
chr10-98056634-INS-276
chr10-98063902-INS-1516
chr10-98078571-INS-127
chr10-98088183-INS-237
chr10-98096892-INS-168
chr10-98105220-INS-691
chr10-98105255-INS-98
chr10-98107809-INS-487
chr10-98123322-INS-1930
chr10-98176685-INS-1010
chr10-98225623-INS-56
chr10-98229915-INS-110
chr10-98252435-INS-106
chr10-98282477-INS-85
chr10-98338224-INS-53
chr10-98357814-INS-51
chr10-98399186-INS-54
chr10-98412009-INS-333
chr10-98412009-INS-90
chr10-98456382-INS-669
chr10-98466225-INS-50
chr10-98493105-INS-6278
chr10-98517265-INS-190
chr10-98525095-INS-56
chr10-98534453-INS-81
chr10-98572852-INS-73
chr10-98600836-INS-359
chr10-98632567-INS-135
chr10-98641653-INS-89
chr10-98648641-INS-76
chr10-98648904-INS-201
chr10-98662617-INS-176
chr10-98670600-INS-199
chr10-98670693-INS-59
chr10-98672394-INS-56
chr10-98687353-INS-84
chr10-98698147-INS-50
chr10-98713597-INS-50
chr10-98802554-INS-60
chr10-98811061-INS-393
chr10-98877714-INS-75
chr10-98918052-INS-56
chr10-98928267-INS-1301
chr10-98952357-INS-215
chr10-98966198-INS-131
chr10-98967049-INS-1736
chr10-98971052-INS-2127
chr10-98983275-INS-224
chr10-98983338-INS-56
chr10-99019786-INS-2954
chr10-99020795-INS-3211
chr10-99021420-INS-81
chr10-99033396-INS-178
chr10-99041383-INS-1191
chr10-99042061-INS-50
chr10-99042763-INS-55
chr10-99049029-INS-54
chr10-99049967-INS-68
chr10-99049975-INS-240
chr10-99068230-INS-590
chr10-99128613-INS-190
chr10-99137726-INS-61
chr10-99181877-INS-1997
chr10-99209979-INS-6263
chr10-99219596-INS-52
chr10-99242062-INS-56
chr10-99242562-INS-138
chr10-99330547-INS-66
chr10-99330655-INS-182
chr10-99332210-INS-491
chr10-99332219-INS-50
chr10-99353074-INS-5188
chr10-99378856-INS-60

chr10-99387837-INS-158
chr10-99394278-INS-389
chr10-99413681-INS-7077
chr10-99457793-INS-1325
chr10-99496184-INS-52
chr10-99504056-INS-214
chr10-99505222-INS-73
chr10-99511683-INS-60
chr10-99603083-INS-6486
chr10-99615701-INS-1442
chr10-99619603-INS-1236
chr10-99639160-INS-791
chr10-99769963-INS-6648
chr10-99773499-INS-426
chr10-99812666-INS-101
chr10-99852708-INS-51
chr10-99862985-INS-106
chr10-99883945-INS-721
chr10-99908981-INS-180
chr10-99967326-INS-3388
chr10-100009700-INS-86
chr10-100011318-INS-75
chr10-100073352-INS-184
chr10-100083105-INS-456
chr10-100139106-INS-57
chr10-100139752-INS-140
chr10-100163809-INS-50
chr10-100203058-INS-417
chr10-100221434-INS-1079
chr10-100278899-INS-52
chr10-100313505-INS-2683
chr10-100363869-INS-50
chr10-100431622-INS-79
chr10-100476458-INS-58
chr10-100521319-INS-552
chr10-100524174-INS-994
chr10-100527317-INS-124
chr10-100528015-INS-52
chr10-100561242-INS-244
chr10-100561309-INS-433
chr10-100589691-INS-1032
chr10-100596854-INS-3959
chr10-100604286-INS-6196
chr10-100627454-INS-542
chr10-100755433-INS-52
chr10-100769210-INS-528
chr10-100789865-INS-50
chr10-100888588-INS-468
chr10-100888750-INS-78
chr10-100938175-INS-331
chr10-100979672-INS-103
chr10-100991396-INS-50
chr10-101006369-INS-6739
chr10-101052588-INS-6189
chr10-101071700-INS-3000
chr10-101100346-INS-2272
chr10-101136037-INS-2827
chr10-101164900-INS-59
chr10-101173758-INS-209
chr10-101222563-INS-254
chr10-101223202-INS-55
chr10-101233358-INS-206
chr10-101254776-INS-62
chr10-101256329-INS-354
chr10-101256329-INS-86
chr10-101334158-INS-1091
chr10-101339387-INS-1107
chr10-101342673-INS-6599
chr10-101385082-INS-57
chr10-101389322-INS-3419
chr10-101418897-INS-100
chr10-101441166-INS-177
chr10-101442929-INS-55
chr10-101448922-INS-50
chr10-101449369-INS-3226
chr10-101449675-INS-410
chr10-101493176-INS-1950

chr10-101495246-INS-4724
chr10-101506744-INS-814
chr10-101526619-INS-61
chr10-101539996-INS-300
chr10-101580790-INS-63
chr10-101595962-INS-55
chr10-101602233-INS-55
chr10-101624022-INS-69
chr10-101640428-INS-50
chr10-101647518-INS-1399
chr10-101653728-INS-106
chr10-101664301-INS-70
chr10-101675636-INS-110
chr10-101696169-INS-6308
chr10-101731694-INS-105
chr10-101763379-INS-6410
chr10-101799169-INS-432
chr10-101799608-INS-110
chr10-101813673-INS-1130
chr10-101818733-INS-2879
chr10-101836451-INS-59
chr10-101850746-INS-654
chr10-101861455-INS-50
chr10-101870424-INS-1886
chr10-101897700-INS-57
chr10-101905768-INS-90
chr10-101915530-INS-6124
chr10-101916168-INS-68
chr10-101922465-INS-56
chr10-101941957-INS-1201
chr10-101955378-INS-291
chr10-102007252-INS-528
chr10-102047457-INS-4531
chr10-102068121-INS-6686
chr10-102102859-INS-7100
chr10-102124581-INS-6599
chr10-102148339-INS-53
chr10-102169544-INS-1891
chr10-102195103-INS-52
chr10-102195664-INS-1761
chr10-102205921-INS-54
chr10-102215989-INS-127
chr10-102226340-INS-92
chr10-102238692-INS-2277
chr10-102302880-INS-96
chr10-102312343-INS-52
chr10-102312976-INS-549
chr10-102313617-INS-317
chr10-102405448-INS-133
chr10-102432378-INS-2148
chr10-102433214-INS-50
chr10-102452279-INS-3134
chr10-102454502-INS-72
chr10-102474403-INS-376
chr10-102481716-INS-3762
chr10-102521349-INS-1459
chr10-102531820-INS-1514
chr10-102534035-INS-1090
chr10-102552487-INS-6657
chr10-102566843-INS-77
chr10-102603392-INS-62
chr10-102609752-INS-100
chr10-102643723-INS-6679
chr10-102649607-INS-54
chr10-102672946-INS-329
chr10-102680532-INS-108
chr10-102685237-INS-5267
chr10-102700193-INS-59
chr10-102721578-INS-211
chr10-102732703-INS-87
chr10-102735874-INS-6137
chr10-102748746-INS-306
chr10-102770205-INS-140
chr10-102771519-INS-2670
chr10-102791252-INS-55
chr10-102798026-INS-209
chr10-102799920-INS-1798

chr10-102820456-INS-61
chr10-102838232-INS-79
chr10-102849276-INS-257
chr10-102994931-INS-1625
chr10-102998488-INS-53
chr10-103057616-INS-329
chr10-103096836-INS-74
chr10-103097352-INS-72
chr10-103097998-INS-134
chr10-103102214-INS-4013
chr10-103107773-INS-116
chr10-103111713-INS-72
chr10-103128775-INS-50
chr10-103134185-INS-136
chr10-103134890-INS-848
chr10-103140976-INS-52
chr10-103153998-INS-6504
chr10-103166847-INS-205
chr10-103198475-INS-50
chr10-103199272-INS-6586
chr10-103226507-INS-100
chr10-103235418-INS-54
chr10-103241939-INS-80
chr10-103246174-INS-60
chr10-103284977-INS-7086
chr10-103300551-INS-6392
chr10-103308088-INS-455
chr10-103315183-INS-3229
chr10-103318020-INS-62
chr10-103321174-INS-6277
chr10-103379949-INS-55
chr10-103382800-INS-8386
chr10-103384067-INS-1697
chr10-103387237-INS-132
chr10-103411167-INS-199
chr10-103522357-INS-214
chr10-103522358-INS-76
chr10-103545179-INS-1514
chr10-103590122-INS-212
chr10-103614702-INS-3151
chr10-103615631-INS-50
chr10-103678376-INS-51
chr10-103734370-INS-322
chr10-103749163-INS-319
chr10-103780606-INS-198
chr10-103783730-INS-92
chr10-103789652-INS-68
chr10-103867942-INS-7187
chr10-103888475-INS-2598
chr10-103904289-INS-98
chr10-103906149-INS-6656
chr10-103944876-INS-7299
chr10-103959300-INS-80
chr10-103975485-INS-3497
chr10-104051707-INS-3372
chr10-104105395-INS-50
chr10-104180678-INS-81
chr10-104182818-INS-573
chr10-104188191-INS-7089
chr10-104221173-INS-3047
chr10-104243714-INS-208
chr10-104252934-INS-476
chr10-104260911-INS-1038
chr10-104267438-INS-3248
chr10-104279637-INS-55
chr10-104281765-INS-58
chr10-104345417-INS-50
chr10-104348679-INS-56
chr10-104352841-INS-50
chr10-104417149-INS-6233
chr10-104490848-INS-6546
chr10-104502252-INS-52
chr10-104502350-INS-68
chr10-104518152-INS-65
chr10-104560472-INS-50
chr10-104619863-INS-60
chr10-104619942-INS-152

chr10-104620000-INS-74
chr10-104625952-INS-56
chr10-104655315-INS-242
chr10-104819092-INS-50
chr10-104831029-INS-62
chr10-104837945-INS-593
chr10-104844385-INS-523
chr10-104846038-INS-95
chr10-104852385-INS-70
chr10-104853615-INS-156
chr10-104869373-INS-888
chr10-104869685-INS-94
chr10-104870054-INS-201
chr10-104895277-INS-222
chr10-104964246-INS-1696
chr10-104965566-INS-1709
chr10-104967329-INS-529
chr10-104969064-INS-52
chr10-104970049-INS-237
chr10-104974028-INS-13793
chr10-104974028-INS-6595
chr10-104991206-INS-606
chr10-105048662-INS-50
chr10-105053860-INS-361
chr10-105061000-INS-6800
chr10-105170012-INS-89
chr10-105183999-INS-72
chr10-105196720-INS-78
chr10-105251586-INS-69
chr10-105263040-INS-57
chr10-105292375-INS-57
chr10-105295775-INS-72
chr10-105324440-INS-7179
chr10-105325549-INS-134
chr10-105357244-INS-231
chr10-105398850-INS-234
chr10-105422014-INS-56
chr10-105449268-INS-432
chr10-105453448-INS-51
chr10-105460004-INS-50
chr10-105496275-INS-52
chr10-105539586-INS-91
chr10-105562629-INS-3540
chr10-105564697-INS-1022
chr10-105592358-INS-1176
chr10-105606305-INS-235
chr10-105637743-INS-346
chr10-105637752-INS-82
chr10-105696004-INS-69
chr10-105702820-INS-71
chr10-105708091-INS-174
chr10-105708161-INS-517
chr10-105735397-INS-461
chr10-105754868-INS-879
chr10-105785819-INS-169
chr10-105787539-INS-2924
chr10-105821034-INS-7184
chr10-105846393-INS-4819
chr10-105868796-INS-53
chr10-105875956-INS-2989
chr10-105896778-INS-7017
chr10-105931542-INS-177
chr10-105939101-INS-1275
chr10-105978235-INS-1891
chr10-105981938-INS-50
chr10-106007810-INS-3167
chr10-106043219-INS-212
chr10-106061938-INS-106
chr10-106062296-INS-444
chr10-106134186-INS-1041
chr10-106135280-INS-2071
chr10-106141004-INS-67
chr10-106208643-INS-85
chr10-106243466-INS-132
chr10-106247964-INS-1692
chr10-106300752-INS-1927
chr10-106405527-INS-4830

chr10-106474182-INS-2756
chr10-106504798-INS-4454
chr10-106510444-INS-253
chr10-106511205-INS-108
chr10-106515383-INS-50
chr10-106546161-INS-1069
chr10-106548667-INS-88
chr10-106554674-INS-76
chr10-106582544-INS-52
chr10-106583359-INS-84
chr10-106585590-INS-195
chr10-106619730-INS-6801
chr10-106700144-INS-1181
chr10-106704642-INS-177
chr10-106720411-INS-82
chr10-106722979-INS-264
chr10-106779378-INS-52
chr10-106789776-INS-7141
chr10-106817785-INS-195
chr10-106838633-INS-164
chr10-106841813-INS-7090
chr10-106878210-INS-2478
chr10-106884339-INS-589
chr10-106898033-INS-252
chr10-106932749-INS-498
chr10-106997327-INS-1415
chr10-107042633-INS-56
chr10-107053142-INS-145
chr10-107135710-INS-200
chr10-107178615-INS-1220
chr10-107178907-INS-360
chr10-107212640-INS-431
chr10-107227686-INS-2908
chr10-107235489-INS-64
chr10-107236652-INS-50
chr10-107239508-INS-51
chr10-107243189-INS-1360
chr10-107243189-INS-289
chr10-107272370-INS-6666
chr10-107290230-INS-75
chr10-107294672-INS-2172
chr10-107308184-INS-6724
chr10-107324894-INS-81
chr10-107337770-INS-449
chr10-107340327-INS-3888
chr10-107408050-INS-637
chr10-107426336-INS-113
chr10-107453253-INS-2727
chr10-107467918-INS-90
chr10-107487817-INS-52
chr10-107510132-INS-112
chr10-107513615-INS-1402
chr10-107530575-INS-297
chr10-107545225-INS-6399
chr10-107567101-INS-1799
chr10-107587367-INS-66
chr10-107587437-INS-174
chr10-107589258-INS-129
chr10-107594467-INS-14000
chr10-107626943-INS-63
chr10-107647118-INS-66
chr10-107648201-INS-6526
chr10-107653379-INS-146
chr10-107675368-INS-108
chr10-107749604-INS-1887
chr10-107763214-INS-341
chr10-107780523-INS-107
chr10-107789119-INS-6813
chr10-107790734-INS-50
chr10-107813862-INS-13832
chr10-107818921-INS-76
chr10-107826364-INS-639
chr10-107860718-INS-3379
chr10-107863289-INS-6356
chr10-107868302-INS-603
chr10-107877542-INS-89
chr10-107878678-INS-55

chr10-107896221-INS-16181
chr10-107900050-INS-1291
chr10-107900589-INS-405
chr10-107900592-INS-185
chr10-107901086-INS-562
chr10-107912149-INS-70
chr10-107913488-INS-158
chr10-107930141-INS-2748
chr10-107945713-INS-55
chr10-107948828-INS-51
chr10-107952551-INS-136
chr10-108004183-INS-137
chr10-108010003-INS-1417
chr10-108024944-INS-50
chr10-108072309-INS-50
chr10-108106538-INS-205
chr10-108125461-INS-88
chr10-108138296-INS-235
chr10-108148210-INS-86
chr10-108148971-INS-86
chr10-108153139-INS-126
chr10-108154706-INS-219
chr10-108154753-INS-76
chr10-108159502-INS-6436
chr10-108207436-INS-63
chr10-108227068-INS-100
chr10-108227698-INS-222
chr10-108301704-INS-396
chr10-108322368-INS-90
chr10-108326473-INS-130
chr10-108339157-INS-138
chr10-108364103-INS-215
chr10-108370594-INS-261
chr10-108406237-INS-220
chr10-108453199-INS-56
chr10-108518545-INS-469
chr10-108598511-INS-82
chr10-108664416-INS-65
chr10-108696514-INS-120
chr10-108718450-INS-76
chr10-108721152-INS-63
chr10-108725907-INS-60
chr10-108763142-INS-70
chr10-108763956-INS-164
chr10-108771305-INS-66
chr10-108795764-INS-570
chr10-108795940-INS-155
chr10-108798176-INS-6167
chr10-108802146-INS-2499
chr10-108813583-INS-90
chr10-108882095-INS-1022
chr10-108892971-INS-135
chr10-108897275-INS-534
chr10-108902618-INS-66
chr10-108911169-INS-64
chr10-108934051-INS-381
chr10-108942029-INS-50
chr10-108984840-INS-271
chr10-108987271-INS-6464
chr10-109014858-INS-189
chr10-109016880-INS-182
chr10-109110264-INS-135
chr10-109136663-INS-177
chr10-109176136-INS-52
chr10-109176377-INS-604
chr10-109200811-INS-76
chr10-109206489-INS-6205
chr10-109210171-INS-2953
chr10-109268433-INS-145
chr10-109354775-INS-387
chr10-109358138-INS-50
chr10-109389452-INS-66
chr10-109477323-INS-7099
chr10-109501890-INS-2883
chr10-109535053-INS-100
chr10-109535443-INS-108
chr10-109552636-INS-101

chr10-109586267-INS-6372
chr10-109634824-INS-67
chr10-109756319-INS-6522
chr10-109784223-INS-64
chr10-109807712-INS-72
chr10-109808170-INS-3117
chr10-109815328-INS-816
chr10-109817510-INS-54
chr10-109819859-INS-679
chr10-109822732-INS-1035
chr10-109862720-INS-60
chr10-109916328-INS-761
chr10-109919929-INS-98
chr10-109948592-INS-50
chr10-109949439-INS-100
chr10-109955896-INS-54
chr10-109983221-INS-6323
chr10-109985215-INS-204
chr10-110022370-INS-98
chr10-110024675-INS-60
chr10-110024829-INS-388
chr10-110024889-INS-60
chr10-110035744-INS-76
chr10-110174392-INS-112
chr10-110197969-INS-506
chr10-110197975-INS-72
chr10-110224313-INS-52
chr10-110224648-INS-112
chr10-110258250-INS-50
chr10-110270648-INS-124
chr10-110281882-INS-60
chr10-110282461-INS-50
chr10-110309633-INS-54
chr10-110312721-INS-116
chr10-110368752-INS-871
chr10-110385763-INS-301
chr10-110392007-INS-4585
chr10-110396123-INS-8924
chr10-110401236-INS-50
chr10-110406071-INS-52
chr10-110449148-INS-4729
chr10-110505074-INS-90
chr10-110531718-INS-183
chr10-110540942-INS-61
chr10-110544996-INS-98
chr10-110549456-INS-52
chr10-110565763-INS-68
chr10-110568262-INS-5396
chr10-110571865-INS-134
chr10-110571876-INS-456
chr10-110572388-INS-415
chr10-110661723-INS-208
chr10-110678716-INS-719
chr10-110678844-INS-132
chr10-110679150-INS-949
chr10-110738900-INS-7460
chr10-110745841-INS-4982
chr10-110749259-INS-52
chr10-110773936-INS-978
chr10-110787937-INS-74
chr10-110795147-INS-595
chr10-110854235-INS-273
chr10-110871902-INS-59
chr10-110892058-INS-50
chr10-110893602-INS-217
chr10-110904488-INS-174
chr10-110957882-INS-251
chr10-111044316-INS-5218
chr10-111147436-INS-50
chr10-111160859-INS-896
chr10-111187951-INS-939
chr10-111263882-INS-4039
chr10-111304573-INS-79
chr10-111354789-INS-52
chr10-111355404-INS-234
chr10-111372032-INS-50
chr10-111382002-INS-440

chr10-111382238-INS-67
chr10-111427080-INS-1056
chr10-111428920-INS-185
chr10-111437980-INS-148
chr10-111443259-INS-68
chr10-111445710-INS-74
chr10-111455506-INS-5218
chr10-111458323-INS-50
chr10-111495259-INS-482
chr10-111500921-INS-849
chr10-111500925-INS-374
chr10-111523524-INS-204
chr10-111610782-INS-1224
chr10-111632530-INS-239
chr10-111696256-INS-53
chr10-111732429-INS-50
chr10-111775333-INS-87
chr10-111791378-INS-51
chr10-111799187-INS-199
chr10-111806352-INS-242
chr10-111814883-INS-64
chr10-111821118-INS-98
chr10-111841567-INS-837
chr10-111860183-INS-50
chr10-111863752-INS-121
chr10-111883979-INS-60
chr10-111886115-INS-561
chr10-111887918-INS-134
chr10-111926107-INS-57
chr10-112031159-INS-6646
chr10-112045911-INS-64
chr10-112046840-INS-99
chr10-112052742-INS-51
chr10-112055431-INS-52
chr10-112060641-INS-62
chr10-112197715-INS-141
chr10-112201319-INS-173
chr10-112251102-INS-50
chr10-112261366-INS-212
chr10-112279596-INS-53
chr10-112318911-INS-60
chr10-112367087-INS-162
chr10-112419341-INS-1237
chr10-112419709-INS-175
chr10-112456094-INS-2184
chr10-112460304-INS-300
chr10-112467196-INS-70
chr10-112492461-INS-50
chr10-112508230-INS-126
chr10-112529329-INS-1139
chr10-112530221-INS-342
chr10-112531094-INS-190
chr10-112545487-INS-210
chr10-112560528-INS-8960
chr10-112575917-INS-152
chr10-112602464-INS-56
chr10-112656992-INS-55
chr10-112657622-INS-1996
chr10-112658786-INS-75
chr10-112659320-INS-228
chr10-112660462-INS-75
chr10-112698466-INS-72
chr10-112750893-INS-1363
chr10-112770244-INS-89
chr10-112781615-INS-125
chr10-112781615-INS-50
chr10-112784398-INS-2324
chr10-112789106-INS-168
chr10-112801037-INS-445
chr10-112835964-INS-61
chr10-112843214-INS-2228
chr10-112843214-INS-8721
chr10-112856378-INS-89
chr10-112871247-INS-100
chr10-112872147-INS-56
chr10-112891991-INS-64
chr10-112916383-INS-3006

chr10-112925835-INS-216
chr10-112926033-INS-92
chr10-112972592-INS-64
chr10-112999966-INS-54
chr10-113014154-INS-54
chr10-113044835-INS-2794
chr10-113078740-INS-116
chr10-113085311-INS-2518
chr10-113101347-INS-273
chr10-113109437-INS-156
chr10-113109501-INS-61
chr10-113125845-INS-6495
chr10-113178857-INS-50
chr10-113180893-INS-50
chr10-113215658-INS-2650
chr10-113219340-INS-1499
chr10-113273172-INS-57
chr10-113274894-INS-108
chr10-113302611-INS-442
chr10-113304042-INS-82
chr10-113304680-INS-96
chr10-113372730-INS-2711
chr10-113374748-INS-54
chr10-113462649-INS-4376
chr10-113477971-INS-51
chr10-113512006-INS-1763
chr10-113512596-INS-2561
chr10-113535975-INS-75
chr10-113543984-INS-2325
chr10-113590192-INS-6311
chr10-113662174-INS-86
chr10-113682787-INS-52
chr10-113701734-INS-2955
chr10-113756191-INS-56
chr10-113765533-INS-75
chr10-113827237-INS-1186
chr10-113931806-INS-1152
chr10-113936349-INS-120
chr10-113937421-INS-1053
chr10-113939044-INS-767
chr10-113949947-INS-393
chr10-113963620-INS-642
chr10-113967808-INS-57
chr10-113972887-INS-56
chr10-114068354-INS-84
chr10-114118010-INS-3070
chr10-114153817-INS-4036
chr10-114220922-INS-68
chr10-114279796-INS-64
chr10-114321022-INS-50
chr10-114407648-INS-50
chr10-114410395-INS-6796
chr10-114425721-INS-124
chr10-114451241-INS-100
chr10-114582584-INS-97
chr10-114587493-INS-2879
chr10-114639415-INS-56
chr10-114651645-INS-179
chr10-114651788-INS-86
chr10-114662381-INS-283
chr10-114662393-INS-115
chr10-114662469-INS-96
chr10-114664380-INS-65
chr10-114664449-INS-7062
chr10-114684177-INS-54
chr10-114696256-INS-106
chr10-114720324-INS-204
chr10-114735180-INS-2805
chr10-114784134-INS-90
chr10-114799003-INS-58
chr10-114800138-INS-6477
chr10-114934095-INS-2637
chr10-115030298-INS-7237
chr10-115063521-INS-132
chr10-115079431-INS-906
chr10-115102418-INS-62
chr10-115109507-INS-103

chr10-115109993-INS-190
chr10-115111699-INS-5767
chr10-115165672-INS-4278
chr10-115210446-INS-2101
chr10-115233635-INS-135
chr10-115247289-INS-50
chr10-115271430-INS-228
chr10-115296133-INS-112
chr10-115296208-INS-320
chr10-115336999-INS-258
chr10-115343702-INS-1891
chr10-115367816-INS-74
chr10-115368341-INS-161
chr10-115368403-INS-1300
chr10-115408897-INS-195
chr10-115432395-INS-159
chr10-115470736-INS-5417
chr10-115484194-INS-119
chr10-115484916-INS-62
chr10-115497712-INS-484
chr10-115616516-INS-328
chr10-115616664-INS-336
chr10-115622463-INS-6428
chr10-115648347-INS-50
chr10-115677347-INS-52
chr10-115692787-INS-135
chr10-115700758-INS-277
chr10-115721382-INS-61
chr10-115743020-INS-68
chr10-115752954-INS-1563
chr10-115773430-INS-389
chr10-115829986-INS-52
chr10-115839659-INS-58
chr10-115844095-INS-61
chr10-115882054-INS-60
chr10-115995470-INS-50
chr10-116031723-INS-270
chr10-116047319-INS-159
chr10-116071538-INS-60
chr10-116118489-INS-54
chr10-116127000-INS-52
chr10-116127060-INS-257
chr10-116129139-INS-54
chr10-116147908-INS-140
chr10-116178409-INS-782
chr10-116226503-INS-52
chr10-116237029-INS-195
chr10-116251672-INS-343
chr10-116281948-INS-389
chr10-116295660-INS-55
chr10-116314542-INS-89
chr10-116328115-INS-53
chr10-116360862-INS-131
chr10-116400020-INS-50
chr10-116430657-INS-208
chr10-116437269-INS-1477
chr10-116454237-INS-74
chr10-116463121-INS-51
chr10-116472393-INS-279
chr10-116484424-INS-64
chr10-116503324-INS-563
chr10-116523991-INS-2435
chr10-116538891-INS-52
chr10-116552289-INS-627
chr10-116619241-INS-64
chr10-116623631-INS-74
chr10-116625492-INS-81
chr10-116650657-INS-154
chr10-116662759-INS-1891
chr10-116669730-INS-81
chr10-116714320-INS-76
chr10-116759360-INS-312
chr10-116817534-INS-4745
chr10-116842048-INS-215
chr10-116842417-INS-210
chr10-116846218-INS-59
chr10-116861562-INS-687

chr10-116876831-INS-288
chr10-116882736-INS-50
chr10-116892592-INS-88
chr10-116897534-INS-227
chr10-116898324-INS-195
chr10-116933852-INS-1098
chr10-117000172-INS-235
chr10-117035970-INS-121
chr10-117037474-INS-72
chr10-117037762-INS-50
chr10-117045858-INS-172
chr10-117049621-INS-248
chr10-117064994-INS-5421
chr10-117068384-INS-180
chr10-117068411-INS-72
chr10-117081213-INS-61
chr10-117081574-INS-150
chr10-117126429-INS-106
chr10-117156657-INS-221
chr10-117162899-INS-5965
chr10-117235497-INS-114
chr10-117260700-INS-55
chr10-117294068-INS-50
chr10-117298622-INS-424
chr10-117300000-INS-1056
chr10-117331624-INS-51
chr10-117332953-INS-320
chr10-117332966-INS-135
chr10-117359692-INS-65
chr10-117391879-INS-52
chr10-117485030-INS-66
chr10-117506758-INS-182
chr10-117529386-INS-1973
chr10-117659845-INS-54
chr10-117722313-INS-68
chr10-117727875-INS-80
chr10-117732312-INS-71
chr10-117768652-INS-51
chr10-117771634-INS-129
chr10-117785663-INS-111
chr10-117794401-INS-65
chr10-117794993-INS-429
chr10-117848160-INS-64
chr10-117852431-INS-185
chr10-117868993-INS-2456
chr10-117871233-INS-52
chr10-117881598-INS-179
chr10-117882150-INS-578
chr10-117894987-INS-1167
chr10-117895127-INS-174
chr10-117900431-INS-62
chr10-117922255-INS-90
chr10-117923354-INS-166
chr10-117938550-INS-50
chr10-117989126-INS-68
chr10-118033907-INS-456
chr10-118126839-INS-274
chr10-118143167-INS-100
chr10-118238746-INS-4721
chr10-118251101-INS-72
chr10-118257497-INS-111
chr10-118302315-INS-100
chr10-118345347-INS-135
chr10-118356542-INS-188
chr10-118376432-INS-166
chr10-118402481-INS-186
chr10-118449723-INS-173
chr10-118449731-INS-77
chr10-118493624-INS-134
chr10-118495208-INS-51
chr10-118573895-INS-170
chr10-118603404-INS-127
chr10-118628126-INS-50
chr10-118646646-INS-72
chr10-118648256-INS-53
chr10-118650934-INS-63
chr10-118654973-INS-549

chr10-118661839-INS-10489
chr10-118710670-INS-183
chr10-118717346-INS-50
chr10-118739896-INS-198
chr10-118814798-INS-115
chr10-118821739-INS-128
chr10-118828032-INS-166
chr10-118838069-INS-1105
chr10-118843187-INS-50
chr10-118859878-INS-1615
chr10-118886689-INS-73
chr10-118919192-INS-156
chr10-118935451-INS-76
chr10-118935559-INS-51
chr10-118940354-INS-54
chr10-118943372-INS-337
chr10-118975264-INS-78
chr10-118976731-INS-13386
chr10-118990879-INS-183
chr10-119087616-INS-124
chr10-119102247-INS-53
chr10-119193173-INS-69
chr10-119209665-INS-564
chr10-119213130-INS-1989
chr10-119253371-INS-139
chr10-119258831-INS-114
chr10-119259025-INS-259
chr10-119318337-INS-204
chr10-119361974-INS-6366
chr10-119375602-INS-1282
chr10-119377093-INS-54
chr10-119381933-INS-64
chr10-119446643-INS-50
chr10-119501293-INS-69
chr10-119505831-INS-287
chr10-119509066-INS-336
chr10-119509341-INS-129
chr10-119514747-INS-6430
chr10-119531517-INS-1265
chr10-119588307-INS-2482
chr10-119588801-INS-140
chr10-119591352-INS-174
chr10-119626584-INS-6988
chr10-119650254-INS-6701
chr10-119695810-INS-231
chr10-119713689-INS-75
chr10-119715069-INS-74
chr10-119722412-INS-142
chr10-119723026-INS-135
chr10-119723026-INS-54
chr10-119763541-INS-64
chr10-119798453-INS-73
chr10-119801258-INS-388
chr10-119842963-INS-52
chr10-119857062-INS-92
chr10-119864655-INS-292
chr10-119885090-INS-832
chr10-119934524-INS-80
chr10-119934564-INS-88
chr10-120036719-INS-64
chr10-120051290-INS-52
chr10-120052745-INS-2000
chr10-120052986-INS-300
chr10-120055386-INS-160
chr10-120059096-INS-305
chr10-120063704-INS-442
chr10-120070560-INS-50
chr10-120097955-INS-89
chr10-120146237-INS-56
chr10-120148952-INS-116
chr10-120221310-INS-836
chr10-120301978-INS-56
chr10-120317762-INS-83
chr10-120354204-INS-324
chr10-120417943-INS-159
chr10-120438056-INS-445
chr10-120450643-INS-96

chr10-120502006-INS-210
chr10-120519383-INS-55
chr10-120521595-INS-54
chr10-120522872-INS-8457
chr10-120574765-INS-50
chr10-120589402-INS-74
chr10-120590186-INS-176
chr10-120597680-INS-50
chr10-120609623-INS-56
chr10-120639303-INS-52
chr10-120641127-INS-296
chr10-120641135-INS-62
chr10-120674032-INS-106
chr10-120683373-INS-669
chr10-120701691-INS-489
chr10-120720710-INS-56
chr10-120730404-INS-60
chr10-120733249-INS-52
chr10-120733934-INS-116
chr10-120737465-INS-3027
chr10-120739963-INS-250
chr10-120744247-INS-56
chr10-120776103-INS-167
chr10-120790226-INS-58
chr10-120804355-INS-560
chr10-120804790-INS-208
chr10-120811425-INS-74
chr10-120812061-INS-208
chr10-120832934-INS-1068
chr10-120836265-INS-131
chr10-120836265-INS-411
chr10-120842577-INS-837
chr10-120869503-INS-142
chr10-120891189-INS-66
chr10-120901111-INS-164
chr10-120901402-INS-222
chr10-120903032-INS-50
chr10-120905848-INS-401
chr10-120939591-INS-50
chr10-120946136-INS-225
chr10-120949989-INS-50
chr10-120964417-INS-115
chr10-120964793-INS-172
chr10-121020135-INS-468
chr10-121020761-INS-86
chr10-121032110-INS-4864
chr10-121041456-INS-200
chr10-121049065-INS-205
chr10-121069832-INS-9913
chr10-121083161-INS-94
chr10-121088257-INS-60
chr10-121105179-INS-54
chr10-121115179-INS-239
chr10-121135182-INS-4013
chr10-121176200-INS-6606
chr10-121189701-INS-78
chr10-121191998-INS-5284
chr10-121217989-INS-186
chr10-121220372-INS-58
chr10-121286012-INS-278
chr10-121321075-INS-90
chr10-121321241-INS-186
chr10-121354572-INS-163
chr10-121376560-INS-220
chr10-121416108-INS-224
chr10-121426283-INS-198
chr10-121439717-INS-188
chr10-121455349-INS-1039
chr10-121484853-INS-224
chr10-121491129-INS-213
chr10-121498326-INS-105
chr10-121504137-INS-75
chr10-121508761-INS-8756
chr10-121531910-INS-172
chr10-121553250-INS-171
chr10-121572871-INS-7105
chr10-121579071-INS-217

chr10-121587402-INS-54
chr10-121593629-INS-64
chr10-121610540-INS-177
chr10-121710366-INS-85
chr10-121733244-INS-71
chr10-121738010-INS-2828
chr10-121754451-INS-3315
chr10-121760139-INS-407
chr10-121812377-INS-64
chr10-121812431-INS-300
chr10-121825794-INS-2188
chr10-121838334-INS-70
chr10-121838338-INS-1236
chr10-121896710-INS-29110
chr10-121909996-INS-955
chr10-121961734-INS-1012
chr10-121963470-INS-1045
chr10-121966367-INS-144
chr10-121981555-INS-143
chr10-121983493-INS-51
chr10-121983630-INS-267
chr10-122028452-INS-62
chr10-122105999-INS-1493
chr10-122125612-INS-53
chr10-122147510-INS-172
chr10-122155381-INS-64
chr10-122170944-INS-61
chr10-122171062-INS-57
chr10-122171190-INS-195
chr10-122235756-INS-66
chr10-122267415-INS-2199
chr10-122269309-INS-115
chr10-122293714-INS-7553
chr10-122310975-INS-2856
chr10-122314967-INS-6253
chr10-122324632-INS-144
chr10-122373375-INS-174
chr10-122389472-INS-60
chr10-122410815-INS-4602
chr10-122424179-INS-112
chr10-122427072-INS-7011
chr10-122455597-INS-3235
chr10-122467484-INS-201
chr10-122467811-INS-201
chr10-122474437-INS-218
chr10-122481063-INS-4402
chr10-122504068-INS-316
chr10-122504068-INS-63
chr10-122529825-INS-54
chr10-122587269-INS-8422
chr10-122597885-INS-1372
chr10-122622073-INS-590
chr10-122622110-INS-51
chr10-122627976-INS-55
chr10-122631753-INS-4754
chr10-122689394-INS-62
chr10-122690726-INS-55
chr10-122696387-INS-112
chr10-122696415-INS-52
chr10-122696559-INS-229
chr10-122719064-INS-96
chr10-122722158-INS-158
chr10-122745386-INS-54
chr10-122774346-INS-80
chr10-122788862-INS-112
chr10-122791198-INS-61
chr10-122825456-INS-170
chr10-122857599-INS-429
chr10-122862653-INS-102
chr10-122875291-INS-412
chr10-122897854-INS-359
chr10-122910341-INS-497
chr10-122914209-INS-232
chr10-122916267-INS-400
chr10-122928363-INS-65
chr10-122938583-INS-57
chr10-122940269-INS-106

chr10-122971480-INS-360
chr10-122971578-INS-135
chr10-123045264-INS-63
chr10-123055709-INS-6269
chr10-123068462-INS-72
chr10-123130868-INS-50
chr10-123133810-INS-294
chr10-123165755-INS-130
chr10-123166625-INS-347
chr10-123169318-INS-4702
chr10-123170681-INS-51
chr10-123180696-INS-227
chr10-123180728-INS-63
chr10-123181581-INS-214
chr10-123207681-INS-224
chr10-123207681-INS-96
chr10-123247755-INS-188
chr10-123262490-INS-178
chr10-123262490-INS-376
chr10-123280650-INS-51
chr10-123288927-INS-51
chr10-123320276-INS-1112
chr10-123321071-INS-65
chr10-123341255-INS-51
chr10-123358481-INS-54
chr10-123372253-INS-1206
chr10-123372253-INS-570
chr10-123372253-INS-78
chr10-123383724-INS-645
chr10-123384952-INS-587
chr10-123387294-INS-2777
chr10-123392549-INS-54
chr10-123402928-INS-518
chr10-123428736-INS-60
chr10-123447493-INS-7130
chr10-123458406-INS-1006
chr10-123504344-INS-97
chr10-123534544-INS-57
chr10-123546134-INS-6196
chr10-123552719-INS-340
chr10-123584845-INS-7098
chr10-123611561-INS-173
chr10-123631170-INS-2126
chr10-123633662-INS-1213
chr10-123644632-INS-6266
chr10-123645677-INS-74
chr10-123671566-INS-6222
chr10-123678731-INS-284
chr10-123683204-INS-109
chr10-123683681-INS-112
chr10-123724828-INS-182
chr10-123760990-INS-3662
chr10-123784377-INS-1863
chr10-123830929-INS-136
chr10-123843537-INS-318
chr10-123846985-INS-4990
chr10-123905020-INS-52
chr10-123911178-INS-62
chr10-123911362-INS-374
chr10-123950898-INS-395
chr10-123955169-INS-6440
chr10-123971211-INS-100
chr10-123993994-INS-2000
chr10-123997948-INS-63
chr10-124007013-INS-139
chr10-124007099-INS-56
chr10-124010910-INS-73
chr10-124028438-INS-312
chr10-124121777-INS-69
chr10-124128706-INS-172
chr10-124128768-INS-59
chr10-124187941-INS-68
chr10-124209763-INS-64
chr10-124231067-INS-2773
chr10-124282617-INS-7173
chr10-124284863-INS-178
chr10-124361862-INS-2362

chr10-124362728-INS-7136
chr10-124375372-INS-54
chr10-124378472-INS-2902
chr10-124384557-INS-2295
chr10-124453009-INS-109
chr10-124468100-INS-61
chr10-124482147-INS-50
chr10-124563027-INS-176
chr10-124587952-INS-6399
chr10-124593002-INS-65
chr10-124643709-INS-63
chr10-124649136-INS-68
chr10-124684815-INS-111
chr10-124749932-INS-437
chr10-124767893-INS-119
chr10-124772820-INS-142
chr10-124772950-INS-96
chr10-124782599-INS-2000
chr10-124784053-INS-95
chr10-124801231-INS-82
chr10-124805900-INS-182
chr10-124805947-INS-2923
chr10-124807838-INS-92
chr10-124809563-INS-606
chr10-124817512-INS-6473
chr10-124846998-INS-3437
chr10-124858131-INS-55
chr10-124870934-INS-54
chr10-124887230-INS-72
chr10-124901642-INS-210
chr10-124918581-INS-91
chr10-124923319-INS-114
chr10-124959178-INS-2238
chr10-124962729-INS-199
chr10-124962932-INS-123
chr10-124963555-INS-117
chr10-124963586-INS-1142
chr10-124966157-INS-141
chr10-124967025-INS-138
chr10-124997541-INS-542
chr10-124997572-INS-1125
chr10-124997595-INS-156
chr10-125012845-INS-74
chr10-125026902-INS-52
chr10-125061807-INS-58
chr10-125062260-INS-52
chr10-125110540-INS-331
chr10-125116983-INS-317
chr10-125121179-INS-54
chr10-125182479-INS-253
chr10-125222894-INS-62
chr10-125230508-INS-6242
chr10-125230815-INS-481
chr10-125237043-INS-279
chr10-125282618-INS-110
chr10-125315823-INS-52
chr10-125318434-INS-80
chr10-125323585-INS-96
chr10-125330217-INS-115
chr10-125345215-INS-51
chr10-125359787-INS-283
chr10-125374452-INS-2833
chr10-125388994-INS-76
chr10-125433736-INS-1026
chr10-125434523-INS-430
chr10-125449623-INS-498
chr10-125456258-INS-6407
chr10-125461617-INS-96
chr10-125498957-INS-106
chr10-125510504-INS-467
chr10-125513447-INS-80
chr10-125528131-INS-141
chr10-125537138-INS-197
chr10-125537826-INS-224
chr10-125537991-INS-94
chr10-125567386-INS-660
chr10-125648493-INS-244

chr10-125688218-INS-215
chr10-125694277-INS-62
chr10-125724162-INS-52
chr10-125729721-INS-1033
chr10-125739306-INS-85
chr10-125772972-INS-81
chr10-125773139-INS-1000
chr10-125800136-INS-50
chr10-125812185-INS-168
chr10-125813723-INS-70
chr10-125832264-INS-50
chr10-125837464-INS-124
chr10-125862158-INS-50
chr10-125893590-INS-284
chr10-125901137-INS-60
chr10-125912883-INS-56
chr10-125940374-INS-178
chr10-125942846-INS-54
chr10-125962408-INS-236
chr10-125968199-INS-130
chr10-125971494-INS-51
chr10-125976263-INS-78
chr10-125976372-INS-159
chr10-126045807-INS-78
chr10-126069680-INS-354
chr10-126116107-INS-106
chr10-126121460-INS-599
chr10-126179497-INS-13619
chr10-126189773-INS-64
chr10-126206583-INS-81
chr10-126228451-INS-411
chr10-126276775-INS-1903
chr10-126277131-INS-75
chr10-126292089-INS-207
chr10-126300088-INS-56
chr10-126308073-INS-112
chr10-126308334-INS-68
chr10-126310859-INS-77
chr10-126347501-INS-14873
chr10-126350030-INS-88
chr10-126350872-INS-2838
chr10-126351373-INS-64
chr10-126363998-INS-189
chr10-126383317-INS-138
chr10-126385242-INS-66
chr10-126385790-INS-72
chr10-126409501-INS-7186
chr10-126423168-INS-6655
chr10-126428820-INS-50
chr10-126450325-INS-90
chr10-126451110-INS-66
chr10-126474752-INS-1655
chr10-126476958-INS-111
chr10-126484116-INS-64
chr10-126485169-INS-1544
chr10-126496905-INS-50
chr10-126498224-INS-76
chr10-126503370-INS-498
chr10-126518477-INS-208
chr10-126616452-INS-75
chr10-126617100-INS-78
chr10-126632925-INS-214
chr10-126653271-INS-139
chr10-126658623-INS-7144
chr10-126702663-INS-52
chr10-126707193-INS-184
chr10-126707419-INS-2551
chr10-126723000-INS-61
chr10-126743876-INS-478
chr10-126744706-INS-502
chr10-126747565-INS-1706
chr10-126752863-INS-214
chr10-126757955-INS-98
chr10-126758090-INS-108
chr10-126766976-INS-66
chr10-126819450-INS-102
chr10-126819646-INS-459

chr10-126853462-INS-344
chr10-126909895-INS-50
chr10-126938782-INS-1391
chr10-126938782-INS-3481
chr10-126942470-INS-50
chr10-126983903-INS-6532
chr10-127042163-INS-52
chr10-127043973-INS-159
chr10-127054697-INS-198
chr10-127071089-INS-140
chr10-127114494-INS-220
chr10-127135644-INS-156
chr10-127174697-INS-170
chr10-127211187-INS-232
chr10-127213028-INS-60
chr10-127255933-INS-237
chr10-127262095-INS-192
chr10-127291862-INS-1016
chr10-127329092-INS-1004
chr10-127329376-INS-264
chr10-127352253-INS-54
chr10-127383473-INS-85
chr10-127403873-INS-50
chr10-127485053-INS-167
chr10-127552366-INS-183
chr10-127552599-INS-224
chr10-127566116-INS-50
chr10-127581270-INS-110
chr10-127685331-INS-218
chr10-127713132-INS-8665
chr10-127728220-INS-4743
chr10-127744299-INS-312
chr10-127756909-INS-7083
chr10-127782465-INS-257
chr10-127793867-INS-55
chr10-127794527-INS-58
chr10-127795654-INS-60
chr10-127799709-INS-609
chr10-127799782-INS-100
chr10-127802467-INS-310
chr10-127838301-INS-189
chr10-127846198-INS-323
chr10-127860375-INS-147
chr10-127864439-INS-105
chr10-127870327-INS-52
chr10-127877995-INS-192
chr10-127878240-INS-138
chr10-127972075-INS-66
chr10-127985413-INS-150
chr10-128006830-INS-2867
chr10-128008869-INS-7732
chr10-128020649-INS-66
chr10-128033800-INS-354
chr10-128033869-INS-174
chr10-128055944-INS-201
chr10-128058971-INS-64
chr10-128063258-INS-172
chr10-128069230-INS-212
chr10-128072956-INS-144
chr10-128110139-INS-52
chr10-128166357-INS-52
chr10-128257562-INS-52
chr10-128277819-INS-4365
chr10-128283937-INS-191
chr10-128293481-INS-50
chr10-128309836-INS-203
chr10-128313109-INS-412
chr10-128313930-INS-78
chr10-128324451-INS-105
chr10-128393515-INS-136
chr10-128449282-INS-221
chr10-128449862-INS-84
chr10-128453893-INS-106
chr10-128457254-INS-54
chr10-128457739-INS-110
chr10-128501088-INS-195
chr10-128546359-INS-124

chr10-128574632-INS-202
chr10-128598159-INS-52
chr10-128621898-INS-170
chr10-128622577-INS-136
chr10-128691524-INS-1483
chr10-128729440-INS-62
chr10-128735291-INS-60
chr10-128743431-INS-60
chr10-128760132-INS-54
chr10-128760734-INS-56
chr10-128788691-INS-63
chr10-128795223-INS-62
chr10-128796010-INS-202
chr10-128798487-INS-76
chr10-128798795-INS-225
chr10-128798796-INS-459
chr10-128807452-INS-218
chr10-128815624-INS-54
chr10-128828370-INS-66
chr10-128838796-INS-207
chr10-128841667-INS-50
chr10-128851467-INS-54
chr10-128866496-INS-6545
chr10-128878358-INS-158
chr10-128878461-INS-1011
chr10-128878593-INS-77
chr10-128889904-INS-10753
chr10-128897563-INS-54
chr10-128924621-INS-6440
chr10-128946923-INS-6595
chr10-128949452-INS-59
chr10-128950412-INS-145
chr10-128952008-INS-1313
chr10-128961848-INS-6251
chr10-129008086-INS-13500
chr10-129047227-INS-146
chr10-129057441-INS-117
chr10-129069222-INS-109
chr10-129080674-INS-72
chr10-129140588-INS-52
chr10-129189524-INS-147
chr10-129292433-INS-413
chr10-129452994-INS-6472
chr10-129479751-INS-6592
chr10-129484667-INS-849
chr10-129532630-INS-6591
chr10-129570478-INS-1639
chr10-129576485-INS-64
chr10-129643671-INS-393
chr10-129669881-INS-65
chr10-129670178-INS-155
chr10-129671748-INS-52
chr10-129683700-INS-1766
chr10-129684693-INS-7985
chr10-129687310-INS-68
chr10-129701282-INS-4715
chr10-129743203-INS-5544
chr10-129793542-INS-4352
chr10-129829045-INS-70
chr10-129909398-INS-85
chr10-130043117-INS-7477
chr10-130071415-INS-50
chr10-130220451-INS-52
chr10-130272706-INS-1495
chr10-130273653-INS-71
chr10-130321710-INS-2491
chr10-130345658-INS-86
chr10-130372261-INS-11980
chr10-130376571-INS-60
chr11-3064426-INS-270
chr11-3068840-INS-1451
chr11-3070994-INS-267
chr11-3072803-INS-152
chr11-3087134-INS-79
chr11-3090376-INS-50
chr11-3118935-INS-205
chr11-3130185-INS-108

chr11-3141229-INS-211
chr11-3144461-INS-140
chr11-3147278-INS-162
chr11-3158221-INS-212
chr11-3299683-INS-59
chr11-3299874-INS-395
chr11-3299985-INS-60
chr11-3311221-INS-78
chr11-3330054-INS-219
chr11-3340551-INS-604
chr11-3395175-INS-432
chr11-3421830-INS-66
chr11-3473651-INS-56
chr11-3473975-INS-309
chr11-3499428-INS-108
chr11-3539803-INS-83
chr11-3547102-INS-188
chr11-3547897-INS-247
chr11-3563796-INS-816
chr11-3613418-INS-206
chr11-3650764-INS-280
chr11-3737287-INS-105
chr11-3737484-INS-315
chr11-3737882-INS-140
chr11-3738022-INS-500
chr11-3782215-INS-56
chr11-3904014-INS-1431
chr11-3922115-INS-56
chr11-3930886-INS-290
chr11-4035796-INS-6442
chr11-4076128-INS-78
chr11-4203580-INS-363
chr11-4239807-INS-94
chr11-4272646-INS-146
chr11-4306222-INS-959
chr11-4358366-INS-216
chr11-4420787-INS-3901
chr11-4510153-INS-97
chr11-4510300-INS-143
chr11-4510668-INS-89
chr11-4517590-INS-97
chr11-4523401-INS-662
chr11-4529893-INS-180
chr11-4547957-INS-779
chr11-4583977-INS-50
chr11-4602481-INS-1240
chr11-4641719-INS-56
chr11-4642973-INS-203
chr11-4818707-INS-50
chr11-4829902-INS-50
chr11-4901224-INS-99
chr11-4906132-INS-75
chr11-4929997-INS-51
chr11-5093304-INS-197
chr11-5134769-INS-366
chr11-5136229-INS-1350
chr11-5172415-INS-50
chr11-5172874-INS-50
chr11-5239547-INS-149
chr11-5249279-INS-404
chr11-5249393-INS-168
chr11-5318933-INS-64
chr11-5385193-INS-56
chr11-5385438-INS-68
chr11-5405171-INS-50
chr11-5416969-INS-87
chr11-5421038-INS-131
chr11-5426240-INS-125
chr11-5442985-INS-696
chr11-5443978-INS-204
chr11-5538801-INS-66
chr11-5547133-INS-287
chr11-5547923-INS-274
chr11-5638665-INS-173
chr11-5668678-INS-50
chr11-5692946-INS-3965
chr11-5725283-INS-1719

chr11-5855913-INS-245
chr11-5890187-INS-166
chr11-5894583-INS-422
chr11-5909805-INS-128
chr11-5955381-INS-69
chr11-5962137-INS-62
chr11-5962517-INS-177
chr11-6022090-INS-135
chr11-6041315-INS-124
chr11-6042084-INS-113
chr11-6159683-INS-115
chr11-6184084-INS-577
chr11-6227536-INS-279
chr11-6332625-INS-1226
chr11-6337185-INS-76
chr11-6433992-INS-208
chr11-6444882-INS-234
chr11-6447345-INS-10691
chr11-6514669-INS-156
chr11-6531961-INS-308
chr11-6588072-INS-66
chr11-6592493-INS-386
chr11-6594973-INS-253
chr11-6615259-INS-127
chr11-6675899-INS-80
chr11-6679988-INS-50
chr11-6722689-INS-116
chr11-6769553-INS-1619
chr11-6778738-INS-230
chr11-6780720-INS-60
chr11-6788099-INS-210
chr11-6838084-INS-181
chr11-6838339-INS-243
chr11-6888404-INS-51
chr11-6900923-INS-68
chr11-6909863-INS-55
chr11-6935941-INS-54
chr11-6958699-INS-108
chr11-6959008-INS-481
chr11-6963920-INS-1346
chr11-6976833-INS-62
chr11-6977520-INS-440
chr11-7031239-INS-520
chr11-7240980-INS-50
chr11-7242684-INS-1203
chr11-7271644-INS-51
chr11-7289863-INS-569
chr11-7296183-INS-6228
chr11-7311632-INS-268
chr11-7347308-INS-5554
chr11-7348798-INS-268
chr11-7420781-INS-160
chr11-7450241-INS-299
chr11-7484377-INS-562
chr11-7491248-INS-52
chr11-7499630-INS-162
chr11-7506170-INS-54
chr11-7588208-INS-441
chr11-7588422-INS-50
chr11-7588755-INS-66
chr11-7590974-INS-104
chr11-7663947-INS-137
chr11-7671690-INS-918
chr11-7680716-INS-156
chr11-7837296-INS-50
chr11-7867878-INS-63
chr11-7881176-INS-948
chr11-8002313-INS-90
chr11-8002741-INS-841
chr11-8065004-INS-65
chr11-8080713-INS-5095
chr11-8109687-INS-274
chr11-8249815-INS-514
chr11-8249825-INS-71
chr11-8250081-INS-56
chr11-8250105-INS-136
chr11-8273633-INS-9533

chr11-8280557-INS-929
chr11-8352451-INS-7080
chr11-8353761-INS-176
chr11-8394484-INS-56
chr11-8404778-INS-52
chr11-8404959-INS-171
chr11-8460937-INS-121
chr11-8490485-INS-92
chr11-8502868-INS-50
chr11-8581342-INS-366
chr11-8639581-INS-51
chr11-8640093-INS-77
chr11-8650568-INS-51
chr11-8656137-INS-188
chr11-8663859-INS-241
chr11-8670416-INS-103
chr11-8681166-INS-214
chr11-8681410-INS-574
chr11-8699743-INS-64
chr11-8705459-INS-51
chr11-8719566-INS-1888
chr11-8722303-INS-58
chr11-8731377-INS-67
chr11-8735781-INS-2307
chr11-8760078-INS-123
chr11-8765439-INS-184
chr11-8802643-INS-3793
chr11-8812413-INS-83
chr11-8823028-INS-57
chr11-8838738-INS-178
chr11-8885638-INS-185
chr11-8905404-INS-50
chr11-8915941-INS-5471
chr11-8917705-INS-76
chr11-8917947-INS-63
chr11-8977150-INS-6679
chr11-8982344-INS-147
chr11-9011603-INS-51
chr11-9017376-INS-714
chr11-9027311-INS-65
chr11-9027395-INS-122
chr11-9138189-INS-1565
chr11-9139785-INS-735
chr11-9151296-INS-2336
chr11-9152684-INS-6486
chr11-9169691-INS-648
chr11-9190480-INS-72
chr11-9192327-INS-105
chr11-9193749-INS-406
chr11-9194179-INS-445
chr11-9200267-INS-729
chr11-9203811-INS-60
chr11-9208277-INS-2832
chr11-9225702-INS-6971
chr11-9320608-INS-1851
chr11-9344782-INS-56
chr11-9365914-INS-7994
chr11-9376991-INS-2825
chr11-9441665-INS-421
chr11-9462611-INS-65
chr11-9481496-INS-5080
chr11-9540648-INS-100
chr11-9572281-INS-6576
chr11-9574791-INS-77
chr11-9587277-INS-74
chr11-9596530-INS-6390
chr11-9623260-INS-1667
chr11-9629328-INS-1164
chr11-9742817-INS-98
chr11-9767936-INS-73
chr11-9784171-INS-6225
chr11-9786158-INS-57
chr11-9812867-INS-52
chr11-9812869-INS-251
chr11-9823860-INS-50
chr11-9825510-INS-6392
chr11-9834560-INS-130

chr11-9843746-INS-52
chr11-9856631-INS-6394
chr11-9868534-INS-526
chr11-9872336-INS-6576
chr11-9877879-INS-1383
chr11-9878602-INS-2898
chr11-9886859-INS-2230
chr11-9899411-INS-50
chr11-9906323-INS-85
chr11-9907528-INS-2786
chr11-9916131-INS-51
chr11-9930300-INS-2610
chr11-9954840-INS-51
chr11-9981984-INS-54
chr11-10052421-INS-147
chr11-10085626-INS-250
chr11-10098092-INS-504
chr11-10101781-INS-243
chr11-10152484-INS-1476
chr11-10167095-INS-300
chr11-10178123-INS-2374
chr11-10178185-INS-82
chr11-10198663-INS-120
chr11-10198663-INS-59
chr11-10201464-INS-579
chr11-10248321-INS-95
chr11-10251281-INS-72
chr11-10251373-INS-50
chr11-10251713-INS-57
chr11-10256987-INS-6418
chr11-10298973-INS-1179
chr11-10311583-INS-50
chr11-10342694-INS-539
chr11-10342930-INS-54
chr11-10344525-INS-300
chr11-10357394-INS-207
chr11-10427867-INS-55
chr11-10435354-INS-233
chr11-10456026-INS-337
chr11-10457221-INS-597
chr11-10487287-INS-283
chr11-10492175-INS-1455
chr11-10519320-INS-1295
chr11-10548171-INS-7125
chr11-10553316-INS-4206
chr11-10602038-INS-841
chr11-10633491-INS-2968
chr11-10647618-INS-148
chr11-10697966-INS-52
chr11-10743126-INS-50
chr11-10762287-INS-55
chr11-10770218-INS-1102
chr11-10775038-INS-1420
chr11-10776572-INS-129
chr11-10787191-INS-52
chr11-10795444-INS-587
chr11-10816927-INS-55
chr11-10820630-INS-159
chr11-10854191-INS-6260
chr11-10857339-INS-615
chr11-10859871-INS-337
chr11-10863192-INS-76
chr11-10889767-INS-2939
chr11-10902576-INS-144
chr11-10907615-INS-6590
chr11-11088438-INS-56
chr11-11186971-INS-188
chr11-11196802-INS-6003
chr11-11291136-INS-94
chr11-11291463-INS-457
chr11-11304167-INS-6010
chr11-11320300-INS-327
chr11-11394169-INS-496
chr11-11406111-INS-346
chr11-11408490-INS-162
chr11-11418407-INS-52
chr11-11418453-INS-149

chr11-11427698-INS-52
chr11-11431332-INS-6223
chr11-11468825-INS-54
chr11-11495999-INS-2829
chr11-11515183-INS-118
chr11-11521673-INS-246
chr11-11522923-INS-991
chr11-11522938-INS-76
chr11-11523129-INS-191
chr11-11541536-INS-6392
chr11-11628945-INS-68
chr11-11638473-INS-68
chr11-11743870-INS-204
chr11-11753711-INS-903
chr11-11777947-INS-244
chr11-11777988-INS-970
chr11-11836284-INS-123
chr11-11849186-INS-481
chr11-11860784-INS-279
chr11-11861752-INS-325
chr11-11861754-INS-160
chr11-11868130-INS-198
chr11-11871937-INS-204
chr11-11872891-INS-205
chr11-11880242-INS-210
chr11-11890549-INS-212
chr11-11899460-INS-134
chr11-11954849-INS-96
chr11-11974546-INS-109
chr11-11974582-INS-240
chr11-12030729-INS-258
chr11-12032460-INS-712
chr11-12054387-INS-1499
chr11-12062507-INS-6270
chr11-12115637-INS-310
chr11-12122975-INS-432
chr11-12122979-INS-116
chr11-12131101-INS-55
chr11-12131347-INS-252
chr11-12160164-INS-92
chr11-12178474-INS-96
chr11-12178585-INS-58
chr11-12261614-INS-59
chr11-12262405-INS-171
chr11-12263843-INS-812
chr11-12266674-INS-110
chr11-12294288-INS-60
chr11-12322668-INS-50
chr11-12325023-INS-4007
chr11-12325173-INS-1496
chr11-12337040-INS-248
chr11-12351095-INS-59
chr11-12431198-INS-51
chr11-12432650-INS-1459
chr11-12453756-INS-204
chr11-12466126-INS-57
chr11-12467450-INS-201
chr11-12507483-INS-173
chr11-12527803-INS-57
chr11-12558954-INS-93
chr11-12558972-INS-393
chr11-12598412-INS-2835
chr11-12610540-INS-860
chr11-12627831-INS-1394
chr11-12658240-INS-911
chr11-12658367-INS-565
chr11-12661887-INS-407
chr11-12730807-INS-195
chr11-12785523-INS-543
chr11-12837782-INS-59
chr11-12842935-INS-57
chr11-12856250-INS-399
chr11-12872154-INS-91
chr11-12983335-INS-108
chr11-12991592-INS-6580
chr11-13022927-INS-75
chr11-13046318-INS-56

chr11-13058816-INS-964
chr11-13058906-INS-71
chr11-13078527-INS-241
chr11-13132962-INS-245
chr11-13145300-INS-416
chr11-13162943-INS-6511
chr11-13165931-INS-2264
chr11-13194990-INS-857
chr11-13232813-INS-84
chr11-13245991-INS-101
chr11-13246103-INS-64
chr11-13311106-INS-66
chr11-13327625-INS-86
chr11-13376664-INS-102
chr11-13378607-INS-1418
chr11-13379670-INS-342
chr11-13415789-INS-53
chr11-13416396-INS-80
chr11-13431660-INS-62
chr11-13431686-INS-2744
chr11-13458465-INS-6534
chr11-13458536-INS-76
chr11-13493913-INS-52
chr11-13503741-INS-2505
chr11-13511789-INS-56
chr11-13513504-INS-1719
chr11-13541809-INS-1214
chr11-13544801-INS-2669
chr11-13550371-INS-1055
chr11-13563694-INS-51
chr11-13597471-INS-2602
chr11-13609230-INS-10097
chr11-13609230-INS-4912
chr11-13738074-INS-8795
chr11-13896893-INS-204
chr11-13922885-INS-6092
chr11-13949617-INS-12309
chr11-13956920-INS-56
chr11-13956970-INS-324
chr11-13958836-INS-6906
chr11-13960795-INS-6652
chr11-13994794-INS-1405
chr11-13998563-INS-1369
chr11-14022540-INS-1312
chr11-14076355-INS-4950
chr11-14110840-INS-77
chr11-14110875-INS-367
chr11-14138754-INS-1173
chr11-14220172-INS-2979
chr11-14240079-INS-187
chr11-14240901-INS-107
chr11-14248851-INS-5113
chr11-14271308-INS-6438
chr11-14280993-INS-99
chr11-14282750-INS-116
chr11-14297941-INS-140
chr11-14299274-INS-282
chr11-14310836-INS-68
chr11-14312472-INS-62
chr11-14366821-INS-6407
chr11-14409826-INS-197
chr11-14420543-INS-72
chr11-14430168-INS-64
chr11-14431058-INS-53
chr11-14433448-INS-109
chr11-14448378-INS-4120
chr11-14552557-INS-7249
chr11-14598604-INS-1635
chr11-14660200-INS-2836
chr11-14661333-INS-6333
chr11-14681910-INS-3836
chr11-14684796-INS-299
chr11-14685485-INS-598
chr11-14765388-INS-568
chr11-14766029-INS-3569
chr11-14806285-INS-55
chr11-14813932-INS-50

chr11-14820071-INS-115
chr11-14820225-INS-217
chr11-14836136-INS-4817
chr11-14842719-INS-6901
chr11-14854311-INS-1032
chr11-14929183-INS-50
chr11-14935066-INS-198
chr11-14945689-INS-524
chr11-14945764-INS-102
chr11-15044757-INS-68
chr11-15071642-INS-7135
chr11-15071668-INS-78
chr11-15202990-INS-199
chr11-15220512-INS-87
chr11-15303357-INS-1068
chr11-15319905-INS-185
chr11-15320099-INS-523
chr11-15353670-INS-6124
chr11-15356849-INS-3433
chr11-15379410-INS-284
chr11-15382668-INS-100
chr11-15393967-INS-1889
chr11-15404393-INS-99
chr11-15418608-INS-7066
chr11-15432187-INS-204
chr11-15453542-INS-7035
chr11-15460198-INS-3098
chr11-15473859-INS-1443
chr11-15475820-INS-118
chr11-15497194-INS-6219
chr11-15525510-INS-1802
chr11-15534994-INS-1476
chr11-15540894-INS-7231
chr11-15548523-INS-1062
chr11-15550498-INS-6710
chr11-15577743-INS-6302
chr11-15665569-INS-76
chr11-15667654-INS-60
chr11-15668205-INS-2424
chr11-15671997-INS-1487
chr11-15724509-INS-169
chr11-15735028-INS-662
chr11-15742715-INS-389
chr11-15774773-INS-50
chr11-15774855-INS-578
chr11-15864080-INS-4549
chr11-15869702-INS-639
chr11-15880105-INS-54
chr11-15882727-INS-965
chr11-15923278-INS-1427
chr11-15925146-INS-88
chr11-15925812-INS-60
chr11-15951812-INS-190
chr11-15971908-INS-193
chr11-16012414-INS-196
chr11-16069311-INS-4960
chr11-16109694-INS-54
chr11-16109701-INS-122
chr11-16146872-INS-6201
chr11-16156385-INS-67
chr11-16163229-INS-2036
chr11-16252377-INS-62
chr11-16270681-INS-2814
chr11-16286850-INS-1196
chr11-16288436-INS-109
chr11-16377322-INS-75
chr11-16378787-INS-2864
chr11-16434025-INS-1405
chr11-16549459-INS-4784
chr11-16566555-INS-4784
chr11-16605067-INS-145
chr11-16611296-INS-426
chr11-16621281-INS-156
chr11-16633971-INS-1116
chr11-16634787-INS-2930
chr11-16654672-INS-105
chr11-16676374-INS-1895

chr11-16676390-INS-13461
chr11-16708663-INS-50
chr11-16761056-INS-316
chr11-16784042-INS-929
chr11-16805862-INS-56
chr11-16829210-INS-106
chr11-16834241-INS-81
chr11-16842429-INS-56
chr11-16888122-INS-56
chr11-16981046-INS-55
chr11-16993234-INS-6424
chr11-17055333-INS-56
chr11-17091707-INS-75
chr11-17102320-INS-70
chr11-17142909-INS-388
chr11-17164804-INS-72
chr11-17201763-INS-6177
chr11-17207465-INS-116
chr11-17265465-INS-98
chr11-17319444-INS-73
chr11-17350847-INS-64
chr11-17365206-INS-194
chr11-17371808-INS-61
chr11-17402220-INS-1052
chr11-17436217-INS-2507
chr11-17436282-INS-120
chr11-17436282-INS-528
chr11-17457390-INS-940
chr11-17524608-INS-63
chr11-17570458-INS-6578
chr11-17607220-INS-1927
chr11-17662522-INS-76
chr11-17668492-INS-399
chr11-17668667-INS-104
chr11-17668744-INS-319
chr11-17678546-INS-54
chr11-17708562-INS-6766
chr11-17708858-INS-233
chr11-17743937-INS-5066
chr11-17753196-INS-3715
chr11-17768801-INS-3771
chr11-17774182-INS-6316
chr11-17840369-INS-7179
chr11-17848799-INS-52
chr11-17909021-INS-2000
chr11-17935413-INS-195
chr11-17950124-INS-120
chr11-17950327-INS-390
chr11-18022612-INS-1521
chr11-18110773-INS-434
chr11-18119999-INS-245
chr11-18146254-INS-198
chr11-18162409-INS-397
chr11-18202884-INS-4999
chr11-18243042-INS-6809
chr11-18258331-INS-4357
chr11-18281671-INS-240
chr11-18345001-INS-194
chr11-18345004-INS-407
chr11-18407320-INS-6392
chr11-18432084-INS-54
chr11-18458394-INS-53
chr11-18461693-INS-485
chr11-18461789-INS-200
chr11-18473621-INS-1894
chr11-18477392-INS-196
chr11-18494327-INS-5936
chr11-18515674-INS-236
chr11-18569337-INS-164
chr11-18648111-INS-6549
chr11-18673325-INS-2832
chr11-18750556-INS-3087
chr11-18776777-INS-1508
chr11-18790660-INS-7300
chr11-18808020-INS-197
chr11-18872199-INS-2048
chr11-18874208-INS-3694

chr11-18876027-INS-60
chr11-18888522-INS-63
chr11-18903535-INS-203
chr11-19004962-INS-50
chr11-19019602-INS-52
chr11-19022919-INS-64
chr11-19055934-INS-52
chr11-19113914-INS-59
chr11-19185832-INS-356
chr11-19185921-INS-52
chr11-19264060-INS-50
chr11-19341605-INS-184
chr11-19351675-INS-630
chr11-19382690-INS-169
chr11-19413172-INS-1173
chr11-19423279-INS-67
chr11-19433184-INS-60
chr11-19450692-INS-55
chr11-19475685-INS-171
chr11-19478462-INS-58
chr11-19503715-INS-178
chr11-19544373-INS-115
chr11-19570791-INS-142
chr11-19597705-INS-178
chr11-19652968-INS-253
chr11-19685901-INS-209
chr11-19692921-INS-87
chr11-19702163-INS-370
chr11-19734647-INS-327
chr11-19761344-INS-90
chr11-19762621-INS-199
chr11-19770558-INS-80
chr11-19776693-INS-201
chr11-19776975-INS-90
chr11-19782258-INS-202
chr11-19801399-INS-72
chr11-19822641-INS-84
chr11-19823222-INS-207
chr11-19839045-INS-322
chr11-19839045-INS-84
chr11-19863734-INS-177
chr11-19873483-INS-91
chr11-19873544-INS-1651
chr11-19873547-INS-269
chr11-19912938-INS-80
chr11-19913594-INS-169
chr11-19913819-INS-50
chr11-19940697-INS-171
chr11-19980297-INS-62
chr11-19980947-INS-434
chr11-19990607-INS-1045
chr11-20021911-INS-128
chr11-20026579-INS-7174
chr11-20032732-INS-216
chr11-20033922-INS-134
chr11-20033923-INS-1568
chr11-20034274-INS-215
chr11-20063818-INS-205
chr11-20064568-INS-210
chr11-20065661-INS-7473
chr11-20069673-INS-668
chr11-20082574-INS-92
chr11-20085181-INS-221
chr11-20090235-INS-50
chr11-20098424-INS-116
chr11-20098468-INS-71
chr11-20098699-INS-50
chr11-20098704-INS-115
chr11-20099833-INS-193
chr11-20109237-INS-199
chr11-20110020-INS-184
chr11-20112119-INS-61
chr11-20112527-INS-61
chr11-20128588-INS-218
chr11-20131904-INS-124
chr11-20166988-INS-135
chr11-20167420-INS-184

chr11-20168349-INS-185
chr11-20222474-INS-78
chr11-20222821-INS-195
chr11-20224736-INS-56
chr11-20309765-INS-195
chr11-20315190-INS-2781
chr11-20315503-INS-8545
chr11-20315557-INS-80
chr11-20315710-INS-360
chr11-20326336-INS-7252
chr11-20329703-INS-903
chr11-20360912-INS-120
chr11-20379307-INS-54
chr11-20380287-INS-180
chr11-20380448-INS-136
chr11-20390420-INS-232
chr11-20402626-INS-445
chr11-20408661-INS-59
chr11-20413488-INS-92
chr11-20414918-INS-57
chr11-20414939-INS-402
chr11-20414994-INS-146
chr11-20447782-INS-66
chr11-20560348-INS-60
chr11-20560349-INS-224
chr11-20596824-INS-86
chr11-20619576-INS-101
chr11-20627251-INS-499
chr11-20701522-INS-338
chr11-20746645-INS-73
chr11-20749942-INS-153
chr11-20761084-INS-239
chr11-20784458-INS-142
chr11-20807036-INS-399
chr11-20810114-INS-3118
chr11-20879078-INS-83
chr11-20886541-INS-230
chr11-20901365-INS-52
chr11-20908702-INS-88
chr11-20920449-INS-230
chr11-20933194-INS-210
chr11-20957935-INS-2470
chr11-20961509-INS-2573
chr11-21117324-INS-64
chr11-21130576-INS-50
chr11-21136267-INS-173
chr11-21139876-INS-829
chr11-21141922-INS-59
chr11-21179005-INS-348
chr11-21191897-INS-207
chr11-21220906-INS-58
chr11-21262558-INS-6671
chr11-21285210-INS-3598
chr11-21296400-INS-272
chr11-21300681-INS-1219
chr11-21305021-INS-72
chr11-21327588-INS-90
chr11-21341206-INS-2906
chr11-21385952-INS-185
chr11-21416570-INS-54
chr11-21426269-INS-274
chr11-21430703-INS-213
chr11-21451124-INS-100
chr11-21473030-INS-6023
chr11-21477344-INS-616
chr11-21491877-INS-144
chr11-21531821-INS-1127
chr11-21565546-INS-56
chr11-21584916-INS-74
chr11-21585071-INS-4370
chr11-21619686-INS-1117
chr11-21620078-INS-93
chr11-21620286-INS-567
chr11-21626794-INS-6226
chr11-21660591-INS-370
chr11-21660675-INS-178
chr11-21678393-INS-147

chr11-21697036-INS-82
chr11-21704898-INS-726
chr11-21734480-INS-4014
chr11-21768275-INS-126
chr11-21822110-INS-63
chr11-21871426-INS-52
chr11-21907813-INS-1562
chr11-21917957-INS-65
chr11-21918690-INS-206
chr11-21929670-INS-180
chr11-21930723-INS-84
chr11-21942616-INS-52
chr11-21952371-INS-56
chr11-21964261-INS-96
chr11-21978768-INS-1863
chr11-22078319-INS-821
chr11-22177131-INS-530
chr11-22177214-INS-81
chr11-22179996-INS-74
chr11-22200923-INS-117
chr11-22201246-INS-511
chr11-22246835-INS-3622
chr11-22255635-INS-52
chr11-22256146-INS-90
chr11-22263848-INS-219
chr11-22303336-INS-6473
chr11-22305573-INS-473
chr11-22340289-INS-161
chr11-22374032-INS-75
chr11-22403275-INS-2500
chr11-22408767-INS-647
chr11-22410182-INS-8743
chr11-22423260-INS-191
chr11-22424095-INS-6673
chr11-22427056-INS-4212
chr11-22438502-INS-202
chr11-22442580-INS-58
chr11-22451224-INS-52
chr11-22470653-INS-328
chr11-22471366-INS-66
chr11-22488903-INS-371
chr11-22492598-INS-851
chr11-22497722-INS-6441
chr11-22503464-INS-3144
chr11-22522250-INS-204
chr11-22546566-INS-61
chr11-22550339-INS-436
chr11-22566185-INS-265
chr11-22574807-INS-70
chr11-22578508-INS-144
chr11-22589748-INS-211
chr11-22596747-INS-129
chr11-22627124-INS-209
chr11-22641859-INS-6661
chr11-22646327-INS-143
chr11-22674778-INS-150
chr11-22688323-INS-2160
chr11-22695102-INS-8734
chr11-22711315-INS-109
chr11-22741603-INS-135
chr11-22742076-INS-55
chr11-22747904-INS-1374
chr11-22753096-INS-58
chr11-22754056-INS-212
chr11-22785692-INS-50
chr11-22824534-INS-56
chr11-22872691-INS-191
chr11-22873538-INS-62
chr11-22888858-INS-487
chr11-22961058-INS-88
chr11-22961178-INS-224
chr11-22971752-INS-235
chr11-22986311-INS-118
chr11-22987228-INS-171
chr11-22997268-INS-177
chr11-23005899-INS-210
chr11-23015641-INS-167

chr11-23022947-INS-840
chr11-23031553-INS-177
chr11-23039064-INS-328
chr11-23039310-INS-469
chr11-23044962-INS-50
chr11-23060804-INS-1018
chr11-23065152-INS-52
chr11-23065376-INS-198
chr11-23077194-INS-52
chr11-23078829-INS-211
chr11-23081282-INS-6475
chr11-23109867-INS-1476
chr11-23110832-INS-8459
chr11-23127355-INS-76
chr11-23129093-INS-4947
chr11-23130205-INS-76
chr11-23134838-INS-96
chr11-23141613-INS-228
chr11-23142698-INS-201
chr11-23158429-INS-50
chr11-23177004-INS-428
chr11-23183261-INS-194
chr11-23186828-INS-75
chr11-23221542-INS-71
chr11-23249346-INS-63
chr11-23264073-INS-212
chr11-23286349-INS-145
chr11-23309509-INS-55
chr11-23328237-INS-109
chr11-23343085-INS-70
chr11-23373845-INS-201
chr11-23385806-INS-159
chr11-23434902-INS-260
chr11-23472103-INS-54
chr11-23473574-INS-645
chr11-23482233-INS-4846
chr11-23493018-INS-487
chr11-23495142-INS-50
chr11-23521058-INS-178
chr11-23521135-INS-73
chr11-23522265-INS-764
chr11-23546884-INS-200
chr11-23578765-INS-300
chr11-23656219-INS-60
chr11-23672581-INS-874
chr11-23679383-INS-509
chr11-23690345-INS-175
chr11-23694204-INS-162
chr11-23695367-INS-52
chr11-23711974-INS-88
chr11-23723691-INS-215
chr11-23727941-INS-55
chr11-23768613-INS-190
chr11-23809185-INS-1122
chr11-23879299-INS-1863
chr11-23902915-INS-194
chr11-23905555-INS-94
chr11-23947206-INS-199
chr11-23960449-INS-612
chr11-23968157-INS-316
chr11-23970081-INS-58
chr11-23977398-INS-50
chr11-23978310-INS-330
chr11-24011475-INS-142
chr11-24011475-INS-60
chr11-24012230-INS-105
chr11-24017402-INS-61
chr11-24022125-INS-63
chr11-24064020-INS-50
chr11-24081827-INS-193
chr11-24108525-INS-51
chr11-24150137-INS-175
chr11-24173987-INS-57
chr11-24202469-INS-74
chr11-24239907-INS-242
chr11-24286146-INS-1037
chr11-24286634-INS-105

chr11-24307450-INS-57
chr11-24307947-INS-107
chr11-24307987-INS-277
chr11-24329817-INS-77
chr11-24334523-INS-205
chr11-24362628-INS-131
chr11-24450994-INS-115
chr11-24487400-INS-125
chr11-24497945-INS-876
chr11-24518750-INS-76
chr11-24532421-INS-1442
chr11-24558499-INS-50
chr11-24572794-INS-6442
chr11-24577992-INS-3316
chr11-24636757-INS-53
chr11-24692357-INS-56
chr11-24766760-INS-1477
chr11-24821156-INS-58
chr11-24860610-INS-50
chr11-24872992-INS-87
chr11-24951262-INS-100
chr11-25033799-INS-54
chr11-25078835-INS-6624
chr11-25126880-INS-716
chr11-25162361-INS-1878
chr11-25170201-INS-50
chr11-25246176-INS-6339
chr11-25434009-INS-60
chr11-25486968-INS-54
chr11-25489186-INS-157
chr11-25503662-INS-1444
chr11-25522003-INS-641
chr11-25542815-INS-269
chr11-25575581-INS-187
chr11-25575686-INS-52
chr11-25583395-INS-231
chr11-25615420-INS-52
chr11-25629048-INS-61
chr11-25676307-INS-58
chr11-25687058-INS-50
chr11-25734021-INS-60
chr11-25772924-INS-6468
chr11-25796987-INS-3000
chr11-25844377-INS-60
chr11-25849552-INS-50
chr11-25850018-INS-52
chr11-25969024-INS-3136
chr11-26095077-INS-89
chr11-26194423-INS-2277
chr11-26205094-INS-435
chr11-26218422-INS-1068
chr11-26237204-INS-3319
chr11-26249651-INS-72
chr11-26255193-INS-201
chr11-26298458-INS-2187
chr11-26350661-INS-631
chr11-26350870-INS-2847
chr11-26363208-INS-154
chr11-26363208-INS-50
chr11-26391433-INS-52
chr11-26472209-INS-551
chr11-26472238-INS-81
chr11-26478661-INS-432
chr11-26494933-INS-105
chr11-26529342-INS-3063
chr11-26596124-INS-846
chr11-26664385-INS-6834
chr11-26706017-INS-3420
chr11-26755711-INS-123
chr11-26807400-INS-52
chr11-26895871-INS-160
chr11-27034105-INS-212
chr11-27034286-INS-83
chr11-27039251-INS-92
chr11-27144823-INS-124
chr11-27190885-INS-505
chr11-27216024-INS-64

chr11-27220943-INS-3912
chr11-27223123-INS-6556
chr11-27226351-INS-8377
chr11-27249621-INS-1779
chr11-27252278-INS-2645
chr11-27258171-INS-7465
chr11-27318644-INS-129
chr11-27318747-INS-139
chr11-27324226-INS-70
chr11-27330061-INS-58
chr11-27371052-INS-135
chr11-27375718-INS-1708
chr11-27397376-INS-2875
chr11-27399431-INS-5938
chr11-27406250-INS-4118
chr11-27427801-INS-140
chr11-27503479-INS-3146
chr11-27513646-INS-3259
chr11-27533226-INS-80
chr11-27613151-INS-2891
chr11-27621479-INS-82
chr11-27671414-INS-551
chr11-27729740-INS-179
chr11-27730024-INS-636
chr11-27731315-INS-169
chr11-27731358-INS-103
chr11-27744968-INS-153
chr11-27745598-INS-371
chr11-27817945-INS-68
chr11-27889726-INS-74
chr11-27890034-INS-997
chr11-27914920-INS-156
chr11-27915736-INS-50
chr11-27929930-INS-3186
chr11-27950296-INS-1252
chr11-27994290-INS-1911
chr11-28078714-INS-104
chr11-28080712-INS-391
chr11-28093083-INS-212
chr11-28103508-INS-1785
chr11-28110704-INS-510
chr11-28143965-INS-61
chr11-28144067-INS-161
chr11-28150254-INS-68
chr11-28165570-INS-2971
chr11-28233438-INS-125
chr11-28234072-INS-362
chr11-28238681-INS-92
chr11-28239279-INS-110
chr11-28270085-INS-58
chr11-28271999-INS-2289
chr11-28319478-INS-155
chr11-28356893-INS-55
chr11-28368462-INS-1995
chr11-28383245-INS-75
chr11-28383308-INS-328
chr11-28387771-INS-54
chr11-28431867-INS-52
chr11-28487248-INS-56
chr11-28508332-INS-119
chr11-28520794-INS-2954
chr11-28572336-INS-2093
chr11-28579262-INS-184
chr11-28629642-INS-1773
chr11-28677102-INS-6600
chr11-28745465-INS-4354
chr11-28787458-INS-7988
chr11-28994310-INS-9046
chr11-29029908-INS-56
chr11-29055019-INS-54
chr11-29080111-INS-52
chr11-29190599-INS-50
chr11-29245077-INS-62
chr11-29332541-INS-215
chr11-29336388-INS-202
chr11-29359401-INS-70
chr11-29362926-INS-491

chr11-29396071-INS-56
chr11-29544158-INS-50
chr11-29552747-INS-180
chr11-29552747-INS-463
chr11-29553608-INS-189
chr11-29560700-INS-66
chr11-29566202-INS-55
chr11-29593196-INS-123
chr11-29598789-INS-171
chr11-29599556-INS-459
chr11-29603356-INS-54
chr11-29609059-INS-190
chr11-29619506-INS-219
chr11-29623704-INS-54
chr11-29633415-INS-213
chr11-29672533-INS-170
chr11-29698356-INS-308
chr11-29708778-INS-417
chr11-29730789-INS-50
chr11-29732271-INS-682
chr11-29791753-INS-1193
chr11-29795714-INS-1712
chr11-29797911-INS-88
chr11-29797953-INS-303
chr11-29804190-INS-93
chr11-29807179-INS-6217
chr11-29807609-INS-1352
chr11-29809840-INS-54
chr11-29810601-INS-56
chr11-29813159-INS-146
chr11-29862271-INS-56
chr11-29905173-INS-120
chr11-29922508-INS-3445
chr11-29923086-INS-108
chr11-29973965-INS-64
chr11-29992882-INS-2689
chr11-29993030-INS-522
chr11-30187856-INS-3346
chr11-30209266-INS-70
chr11-30211189-INS-130
chr11-30245624-INS-1839
chr11-30273422-INS-1030
chr11-30281132-INS-3200
chr11-30326620-INS-6296
chr11-30350227-INS-447
chr11-30358000-INS-604
chr11-30383090-INS-74
chr11-30426834-INS-1812
chr11-30428358-INS-1312
chr11-30455447-INS-21420
chr11-30465436-INS-1449
chr11-30499410-INS-6878
chr11-30508625-INS-72
chr11-30520971-INS-2892
chr11-30561347-INS-6001
chr11-30575617-INS-193
chr11-30578269-INS-2887
chr11-30583972-INS-231
chr11-30585078-INS-1445
chr11-30590110-INS-2517
chr11-30598348-INS-57
chr11-30604087-INS-6649
chr11-30615763-INS-199
chr11-30703546-INS-3727
chr11-30831584-INS-5301
chr11-30856591-INS-70
chr11-30861723-INS-2374
chr11-30919205-INS-52
chr11-31025120-INS-3567
chr11-31119666-INS-58
chr11-31120432-INS-50
chr11-31122224-INS-1019
chr11-31177055-INS-1860
chr11-31195951-INS-6525
chr11-31204606-INS-6796
chr11-31230346-INS-139
chr11-31264230-INS-3267

chr11-31285463-INS-53
chr11-31288197-INS-7408
chr11-31341056-INS-77
chr11-31366935-INS-651
chr11-31406714-INS-50
chr11-31413695-INS-1231
chr11-31436997-INS-7162
chr11-31459440-INS-305
chr11-31488346-INS-1860
chr11-31500213-INS-1229
chr11-31525717-INS-276
chr11-31528135-INS-6211
chr11-31659681-INS-64
chr11-31678508-INS-213
chr11-31681499-INS-1199
chr11-31700517-INS-7163
chr11-31704703-INS-5049
chr11-31723853-INS-50
chr11-31787139-INS-7465
chr11-31799127-INS-50
chr11-31909679-INS-487
chr11-31989229-INS-120
chr11-31989499-INS-402
chr11-32043100-INS-7224
chr11-32053106-INS-143
chr11-32089211-INS-4806
chr11-32105554-INS-57
chr11-32117844-INS-216
chr11-32124391-INS-50
chr11-32127369-INS-78
chr11-32154757-INS-50
chr11-32238118-INS-150
chr11-32269532-INS-2086
chr11-32322390-INS-68
chr11-32325832-INS-694
chr11-32332768-INS-66
chr11-32334970-INS-102
chr11-32339278-INS-53
chr11-32416635-INS-215
chr11-32519872-INS-73
chr11-32534124-INS-214
chr11-32544796-INS-50
chr11-32559726-INS-3137
chr11-32565295-INS-80
chr11-32577910-INS-239
chr11-32578828-INS-206
chr11-32703087-INS-279
chr11-32711542-INS-50
chr11-32716715-INS-1080
chr11-32747821-INS-129
chr11-32747821-INS-59
chr11-32764097-INS-257
chr11-32770875-INS-130
chr11-32770882-INS-50
chr11-32789852-INS-5009
chr11-32807298-INS-52
chr11-32810976-INS-148
chr11-32820369-INS-543
chr11-32820605-INS-226
chr11-32820705-INS-84
chr11-32839732-INS-425
chr11-32854132-INS-394
chr11-32886602-INS-351
chr11-32892284-INS-51
chr11-32896083-INS-50
chr11-32904261-INS-231
chr11-32906614-INS-432
chr11-32906619-INS-50
chr11-32920705-INS-55
chr11-32935545-INS-67
chr11-32967059-INS-50
chr11-32986124-INS-76
chr11-32988264-INS-4866
chr11-33021949-INS-54
chr11-33046248-INS-359
chr11-33046677-INS-84
chr11-33056253-INS-64

chr11-33077907-INS-106
chr11-33095858-INS-60
chr11-33193605-INS-7598
chr11-33304109-INS-78
chr11-33329798-INS-108
chr11-33354919-INS-211
chr11-33368778-INS-6464
chr11-33369442-INS-57
chr11-33385920-INS-239
chr11-33410808-INS-64
chr11-33451300-INS-89
chr11-33466128-INS-752
chr11-33488731-INS-93
chr11-33504806-INS-240
chr11-33509778-INS-52
chr11-33554098-INS-320
chr11-33554101-INS-77
chr11-33555952-INS-279
chr11-33555952-INS-711
chr11-33556233-INS-1282
chr11-33565218-INS-987
chr11-33595903-INS-296
chr11-33605657-INS-248
chr11-33605764-INS-72
chr11-33617149-INS-165
chr11-33642751-INS-78
chr11-33643414-INS-50
chr11-33643858-INS-196
chr11-33644704-INS-97
chr11-33658909-INS-75
chr11-33660363-INS-63
chr11-33660643-INS-138
chr11-33660771-INS-202
chr11-33673076-INS-374
chr11-33674780-INS-170
chr11-33688301-INS-84
chr11-33690970-INS-95
chr11-33719610-INS-311
chr11-33733666-INS-494
chr11-33761595-INS-50
chr11-33776657-INS-220
chr11-33818185-INS-433
chr11-33857972-INS-50
chr11-33884410-INS-50
chr11-33908788-INS-335
chr11-33951617-INS-54
chr11-33955568-INS-55
chr11-33980406-INS-266
chr11-33986045-INS-4282
chr11-33994951-INS-58
chr11-33995490-INS-56
chr11-33998818-INS-413
chr11-34004735-INS-50
chr11-34057662-INS-50
chr11-34073597-INS-63
chr11-34123777-INS-281
chr11-34138600-INS-108
chr11-34332713-INS-252
chr11-34332719-INS-96
chr11-34365135-INS-81
chr11-34394374-INS-59
chr11-34457841-INS-80
chr11-34462569-INS-179
chr11-34466093-INS-51
chr11-34518460-INS-96
chr11-34544770-INS-860
chr11-34548798-INS-860
chr11-34666209-INS-54
chr11-34667352-INS-50
chr11-34680724-INS-68
chr11-34703491-INS-548
chr11-34703663-INS-163
chr11-34763508-INS-180
chr11-34790136-INS-55
chr11-34836113-INS-51
chr11-34836972-INS-50
chr11-34849465-INS-99

chr11-34901255-INS-977
chr11-34901387-INS-203
chr11-34912330-INS-50
chr11-35036436-INS-275
chr11-35043741-INS-198
chr11-35050695-INS-54
chr11-35071368-INS-52
chr11-35118012-INS-1054
chr11-35173103-INS-214
chr11-35178816-INS-165
chr11-35221287-INS-175
chr11-35225031-INS-283
chr11-35248721-INS-60
chr11-35316105-INS-303
chr11-35319382-INS-55
chr11-35320052-INS-56
chr11-35363802-INS-1150
chr11-35522340-INS-6466
chr11-35622674-INS-2466
chr11-35623350-INS-2500
chr11-35734078-INS-737
chr11-35742515-INS-2847
chr11-35832920-INS-349
chr11-35856286-INS-129
chr11-35856431-INS-115
chr11-35885016-INS-6898
chr11-35896535-INS-141
chr11-35913027-INS-348
chr11-35929931-INS-50
chr11-35961430-INS-182
chr11-36036808-INS-209
chr11-36036822-INS-1522
chr11-36040486-INS-62
chr11-36063348-INS-6709
chr11-36084365-INS-95
chr11-36231551-INS-82
chr11-36271821-INS-127
chr11-36415974-INS-401
chr11-36426625-INS-353
chr11-36485607-INS-52
chr11-36519387-INS-1100
chr11-36551936-INS-138
chr11-36585174-INS-95
chr11-36617206-INS-54
chr11-36635240-INS-378
chr11-36644719-INS-52
chr11-36771934-INS-149
chr11-36788672-INS-1241
chr11-36789568-INS-6628
chr11-36808223-INS-440
chr11-36866258-INS-117
chr11-36970744-INS-480
chr11-36992336-INS-205
chr11-37058443-INS-50
chr11-37080376-INS-50
chr11-37159214-INS-891
chr11-37179197-INS-2154
chr11-37182242-INS-1131
chr11-37189072-INS-82
chr11-37335443-INS-50
chr11-37372210-INS-53
chr11-37386021-INS-8245
chr11-37389633-INS-64
chr11-37451269-INS-340
chr11-37455399-INS-543
chr11-37456633-INS-58
chr11-37482212-INS-90
chr11-37519211-INS-1495
chr11-37555374-INS-50
chr11-37723969-INS-50
chr11-37730537-INS-255
chr11-37843179-INS-84
chr11-37878873-INS-58
chr11-37914669-INS-3994
chr11-37961680-INS-506
chr11-37994345-INS-138
chr11-38013931-INS-51

chr11-38044723-INS-65
chr11-38099646-INS-109
chr11-38112447-INS-54
chr11-38135138-INS-282
chr11-38135382-INS-145
chr11-38149368-INS-72
chr11-38202721-INS-52
chr11-38206437-INS-4276
chr11-38325878-INS-6888
chr11-38397748-INS-50
chr11-38407591-INS-53
chr11-38407912-INS-56
chr11-38429985-INS-100
chr11-38430741-INS-197
chr11-38433611-INS-120
chr11-38448568-INS-53
chr11-38488994-INS-137
chr11-38532477-INS-50
chr11-38543008-INS-1697
chr11-38567926-INS-69
chr11-38569285-INS-54
chr11-38627832-INS-1972
chr11-38652246-INS-6249
chr11-38660459-INS-97
chr11-38676526-INS-268
chr11-38687505-INS-6443
chr11-38764319-INS-2750
chr11-38800324-INS-416
chr11-38854180-INS-6579
chr11-38963029-INS-147
chr11-39029329-INS-65
chr11-39112155-INS-132
chr11-39112555-INS-88
chr11-39160948-INS-295
chr11-39229098-INS-51
chr11-39238659-INS-50
chr11-39245406-INS-221
chr11-39251149-INS-76
chr11-39261488-INS-6411
chr11-39315972-INS-6692
chr11-39400986-INS-60
chr11-39401232-INS-73
chr11-39425498-INS-231
chr11-39450356-INS-100
chr11-39493991-INS-57
chr11-39542636-INS-50
chr11-39551739-INS-122
chr11-39573841-INS-80
chr11-39695416-INS-6821
chr11-39702336-INS-146
chr11-39768600-INS-68
chr11-39846500-INS-69
chr11-39861239-INS-105
chr11-39887580-INS-2836
chr11-39901719-INS-102
chr11-39927597-INS-1039
chr11-39939440-INS-859
chr11-39945493-INS-117
chr11-39964791-INS-76
chr11-40031669-INS-58
chr11-40031691-INS-474
chr11-40077494-INS-1493
chr11-40100574-INS-3185
chr11-40114351-INS-50
chr11-40166933-INS-7076
chr11-40172401-INS-66
chr11-40172406-INS-149
chr11-40263624-INS-64
chr11-40405075-INS-407
chr11-40414429-INS-204
chr11-40543452-INS-440
chr11-40586227-INS-1514
chr11-40586227-INS-739
chr11-40614040-INS-60
chr11-40614838-INS-50
chr11-40620308-INS-863
chr11-40670297-INS-54

chr11-40675400-INS-8381
chr11-40680793-INS-2836
chr11-40706917-INS-59
chr11-40736214-INS-2025
chr11-40779893-INS-3601
chr11-40807639-INS-5847
chr11-40830695-INS-843
chr11-40831486-INS-55
chr11-40942869-INS-6601
chr11-40957287-INS-91
chr11-40965250-INS-140
chr11-41039972-INS-382
chr11-41193138-INS-80
chr11-41194695-INS-120
chr11-41212071-INS-164
chr11-41215896-INS-69
chr11-41252406-INS-6439
chr11-41257898-INS-6236
chr11-41265214-INS-144
chr11-41308033-INS-11651
chr11-41320038-INS-86
chr11-41320653-INS-395
chr11-41330778-INS-174
chr11-41334578-INS-6246
chr11-41338497-INS-67
chr11-41348446-INS-2750
chr11-41356800-INS-469
chr11-41386340-INS-60
chr11-41521004-INS-138
chr11-41523503-INS-658
chr11-41555398-INS-3075
chr11-41557337-INS-50
chr11-41683644-INS-402
chr11-41685695-INS-4403
chr11-41728830-INS-1930
chr11-41800393-INS-52
chr11-41819732-INS-4548
chr11-41819732-INS-9210
chr11-41840338-INS-117
chr11-41904166-INS-385
chr11-41912041-INS-1620
chr11-41929728-INS-52
chr11-41930970-INS-108
chr11-41946973-INS-3801
chr11-41986299-INS-1012
chr11-42095931-INS-643
chr11-42096396-INS-682
chr11-42106674-INS-156
chr11-42111765-INS-3302
chr11-42128814-INS-6185
chr11-42129190-INS-1511
chr11-42175537-INS-3171
chr11-42196133-INS-98
chr11-42198668-INS-77
chr11-42256127-INS-102
chr11-42261683-INS-376
chr11-42265483-INS-604
chr11-42389141-INS-926
chr11-42389798-INS-8055
chr11-42450872-INS-545
chr11-42457480-INS-2346
chr11-42526216-INS-50
chr11-42611241-INS-6677
chr11-42649026-INS-176
chr11-42686773-INS-1441
chr11-42729870-INS-1381
chr11-42739539-INS-60
chr11-42793447-INS-54
chr11-42806806-INS-61
chr11-42858074-INS-2539
chr11-42881580-INS-52
chr11-42914368-INS-2046
chr11-42920626-INS-559
chr11-42958360-INS-50
chr11-42975886-INS-703
chr11-42980552-INS-105
chr11-42988802-INS-51

chr11-42988918-INS-52
chr11-43016422-INS-94
chr11-43016636-INS-92
chr11-43036576-INS-6222
chr11-43050299-INS-80
chr11-43064369-INS-78
chr11-43082522-INS-1003
chr11-43132631-INS-1402
chr11-43173358-INS-66
chr11-43177220-INS-1293
chr11-43185945-INS-235
chr11-43248581-INS-123
chr11-43282300-INS-55
chr11-43298185-INS-202
chr11-43330465-INS-185
chr11-43340846-INS-94
chr11-43354770-INS-133
chr11-43357564-INS-201
chr11-43360348-INS-62
chr11-43362246-INS-100
chr11-43368685-INS-118
chr11-43390395-INS-216
chr11-43391524-INS-156
chr11-43405587-INS-1143
chr11-43411578-INS-139
chr11-43432615-INS-118
chr11-43474637-INS-154
chr11-43488805-INS-223
chr11-43493325-INS-103
chr11-43503069-INS-60
chr11-43510444-INS-2061
chr11-43520455-INS-5388
chr11-43532655-INS-203
chr11-43547706-INS-218
chr11-43566913-INS-196
chr11-43594316-INS-408
chr11-43594316-INS-72
chr11-43607501-INS-237
chr11-43647237-INS-307
chr11-43649840-INS-102
chr11-43656183-INS-120
chr11-43661724-INS-709
chr11-43703436-INS-222
chr11-43709616-INS-1393
chr11-43729194-INS-120
chr11-43743119-INS-70
chr11-43757524-INS-1171
chr11-43781938-INS-77
chr11-43825118-INS-274
chr11-43825120-INS-607
chr11-43829212-INS-826
chr11-43829974-INS-64
chr11-43830093-INS-70
chr11-43830373-INS-88
chr11-43853395-INS-52
chr11-43870212-INS-69
chr11-43889805-INS-67
chr11-43915013-INS-138
chr11-43933024-INS-1912
chr11-43936284-INS-75
chr11-43968104-INS-52
chr11-44071260-INS-66
chr11-44114191-INS-174
chr11-44114956-INS-7498
chr11-44136124-INS-70
chr11-44144994-INS-130
chr11-44157886-INS-58
chr11-44158984-INS-119
chr11-44158984-INS-325
chr11-44169723-INS-62
chr11-44170358-INS-50
chr11-44170526-INS-350
chr11-44211311-INS-79
chr11-44234186-INS-143
chr11-44239660-INS-346
chr11-44331593-INS-59
chr11-44335440-INS-205

chr11-44353499-INS-301
chr11-44353701-INS-101
chr11-44353730-INS-725
chr11-44391161-INS-123
chr11-44393615-INS-2136
chr11-44393615-INS-4595
chr11-44463300-INS-50
chr11-44463863-INS-212
chr11-44467895-INS-1360
chr11-44476108-INS-64
chr11-44526641-INS-174
chr11-44528841-INS-330
chr11-44660826-INS-55
chr11-44720267-INS-278
chr11-44732968-INS-351
chr11-44803580-INS-85
chr11-44844516-INS-50
chr11-44849049-INS-142
chr11-44849073-INS-312
chr11-44929543-INS-258
chr11-44953818-INS-1250
chr11-44974493-INS-67
chr11-45096494-INS-64
chr11-45107702-INS-80
chr11-45193794-INS-96
chr11-45252728-INS-366
chr11-45273412-INS-3482
chr11-45313669-INS-6478
chr11-45328054-INS-76
chr11-45383596-INS-129
chr11-45408709-INS-342
chr11-45412178-INS-7186
chr11-45469223-INS-78
chr11-45473947-INS-601
chr11-45487405-INS-400
chr11-45549654-INS-56
chr11-45567938-INS-60
chr11-45604099-INS-50
chr11-45637134-INS-51
chr11-45650335-INS-2658
chr11-45679815-INS-108
chr11-45695439-INS-115
chr11-45700166-INS-54
chr11-45723437-INS-200
chr11-45727328-INS-98
chr11-45747439-INS-225
chr11-45814230-INS-330
chr11-45851809-INS-506
chr11-45925729-INS-177
chr11-45926241-INS-406
chr11-45952234-INS-110
chr11-45953692-INS-63
chr11-45978984-INS-50
chr11-46027734-INS-64
chr11-46051928-INS-134
chr11-46057893-INS-66
chr11-46058223-INS-343
chr11-46077537-INS-64
chr11-46088092-INS-66
chr11-46101695-INS-59
chr11-46153364-INS-63
chr11-46153468-INS-166
chr11-46175947-INS-132
chr11-46228236-INS-74
chr11-46228481-INS-52
chr11-46235441-INS-62
chr11-46347819-INS-52
chr11-46406535-INS-54
chr11-46436234-INS-98
chr11-46464380-INS-120
chr11-46470346-INS-58
chr11-46476021-INS-200
chr11-46488791-INS-77
chr11-46556067-INS-176
chr11-46556072-INS-509
chr11-46559212-INS-5061
chr11-46587047-INS-170

chr11-46591796-INS-93
chr11-46598264-INS-52
chr11-46609902-INS-99
chr11-46634073-INS-405
chr11-46639624-INS-208
chr11-46679345-INS-204
chr11-46709782-INS-72
chr11-46718494-INS-50
chr11-46735500-INS-1896
chr11-46745827-INS-2065
chr11-46756952-INS-715
chr11-46759450-INS-1461
chr11-46760526-INS-55
chr11-46760838-INS-615
chr11-46760838-INS-89
chr11-46767976-INS-7081
chr11-46781414-INS-199
chr11-46787039-INS-826
chr11-46799630-INS-180
chr11-46802820-INS-609
chr11-46805876-INS-243
chr11-46846647-INS-116
chr11-46847178-INS-1891
chr11-46857806-INS-65
chr11-46895435-INS-152
chr11-46913467-INS-52
chr11-46918343-INS-50
chr11-46933165-INS-68
chr11-46963739-INS-60
chr11-47022593-INS-83
chr11-47022632-INS-195
chr11-47037567-INS-60
chr11-47046364-INS-64
chr11-47272123-INS-155
chr11-47344663-INS-53
chr11-47420838-INS-52
chr11-47421156-INS-120
chr11-47468548-INS-92
chr11-47513232-INS-997
chr11-47531733-INS-343
chr11-47540230-INS-6395
chr11-47555560-INS-2481
chr11-47591250-INS-73
chr11-47591385-INS-680
chr11-47622293-INS-1287
chr11-47700438-INS-1909
chr11-47752546-INS-186
chr11-47795562-INS-120
chr11-47806789-INS-51
chr11-47860741-INS-675
chr11-47906086-INS-6532
chr11-47942992-INS-186
chr11-47970018-INS-3436
chr11-48009967-INS-54
chr11-48082825-INS-59
chr11-48088591-INS-332
chr11-48088608-INS-804
chr11-48099218-INS-60
chr11-48132661-INS-50
chr11-48188785-INS-6752
chr11-48190147-INS-4245
chr11-48192602-INS-131
chr11-48193303-INS-3632
chr11-48201082-INS-55
chr11-48238272-INS-453
chr11-48245694-INS-74
chr11-48251459-INS-76
chr11-48252188-INS-129
chr11-48298218-INS-60
chr11-48332392-INS-358
chr11-48355524-INS-183
chr11-48370516-INS-367
chr11-48410844-INS-60
chr11-48477659-INS-84
chr11-48496772-INS-50
chr11-48548569-INS-268
chr11-48567605-INS-1415

chr11-48635033-INS-218
chr11-48645575-INS-187
chr11-48645575-INS-75
chr11-48658346-INS-2454
chr11-48658346-INS-721
chr11-48680981-INS-116
chr11-48681767-INS-691
chr11-48710581-INS-400
chr11-48719344-INS-213
chr11-48719534-INS-51
chr11-48751437-INS-110
chr11-48771578-INS-2278
chr11-48781636-INS-72
chr11-48782622-INS-68
chr11-48789072-INS-7765
chr11-48793854-INS-423
chr11-48809187-INS-52
chr11-48809282-INS-1509
chr11-48883970-INS-69
chr11-48955575-INS-122
chr11-48972096-INS-362
chr11-48988419-INS-422
chr11-48991778-INS-536
chr11-48996502-INS-114
chr11-49030824-INS-1221
chr11-49155285-INS-217
chr11-49183032-INS-6661
chr11-49274334-INS-1412
chr11-49322935-INS-84
chr11-49354969-INS-481
chr11-49358294-INS-24070
chr11-49394645-INS-1482
chr11-49427044-INS-612
chr11-49461982-INS-497
chr11-49463033-INS-60
chr11-49463612-INS-66
chr11-49521911-INS-99
chr11-49530209-INS-119
chr11-49552829-INS-193
chr11-49552982-INS-359
chr11-49555505-INS-56
chr11-49586741-INS-57
chr11-49609326-INS-1024
chr11-49655961-INS-1846
chr11-49655961-INS-78
chr11-49666285-INS-269
chr11-49670341-INS-535
chr11-49678266-INS-54
chr11-49864324-INS-256
chr11-49874243-INS-10159
chr11-49879523-INS-759
chr11-49880349-INS-113
chr11-49880454-INS-56
chr11-49893927-INS-1177
chr11-49905362-INS-108
chr11-49939530-INS-90
chr11-49954683-INS-63
chr11-49955411-INS-186
chr11-49961094-INS-325
chr11-49969439-INS-57
chr11-49969462-INS-201
chr11-50002067-INS-58
chr11-50047228-INS-81
chr11-50077319-INS-214
chr11-50086466-INS-71
chr11-50158884-INS-1032
chr11-50175550-INS-207
chr11-50197098-INS-59
chr11-50197482-INS-80
chr11-50227694-INS-228
chr11-50252560-INS-97
chr11-50259389-INS-132
chr11-50286728-INS-61
chr11-50306558-INS-191
chr11-50310128-INS-108
chr11-50351285-INS-52
chr11-50376236-INS-76

chr11-50377402-INS-249
chr11-50377685-INS-183
chr11-50381731-INS-6412
chr11-50413466-INS-110
chr11-50413572-INS-50
chr11-50498995-INS-2242
chr11-50505443-INS-89
chr11-50515131-INS-60
chr11-50572071-INS-4255
chr11-50574078-INS-54
chr11-50586121-INS-197
chr11-50588684-INS-53
chr11-50607964-INS-72
chr11-50616694-INS-51
chr11-50680951-INS-94
chr11-50755185-INS-64
chr11-50757178-INS-95
chr11-50772119-INS-269
chr11-50818614-INS-72
chr11-50824568-INS-582
chr11-50827735-INS-5380
chr11-50836305-INS-51
chr11-50855735-INS-6185
chr11-50872033-INS-10319
chr11-50881902-INS-851
chr11-50912442-INS-1181
chr11-50926114-INS-519
chr11-50927475-INS-210
chr11-50927482-INS-65
chr11-50970486-INS-228
chr11-50971006-INS-94
chr11-50990615-INS-86
chr11-50991673-INS-1583
chr11-51040214-INS-7844
chr11-51066305-INS-7520
chr11-51066506-INS-460
chr11-51070614-INS-499
chr11-51077193-INS-1346
chr11-51092808-INS-52
chr11-51117867-INS-165
chr11-51151411-INS-60
chr11-51193425-INS-52
chr11-51215306-INS-186
chr11-51224315-INS-112
chr11-51271386-INS-52
chr11-51271486-INS-202
chr11-51281804-INS-50
chr11-51286764-INS-51
chr11-51291016-INS-688
chr11-51302971-INS-50
chr11-51313024-INS-133
chr11-51333244-INS-210
chr11-51346912-INS-6231
chr11-51390275-INS-173
chr11-51395009-INS-1944
chr11-51423653-INS-419
chr11-51454838-INS-8620
chr11-51461771-INS-116
chr11-51462107-INS-99
chr11-51462490-INS-58
chr11-51479901-INS-62
chr11-51503399-INS-62
chr11-51503539-INS-130
chr11-51528029-INS-55
chr11-51550958-INS-50
chr11-51638757-INS-62
chr11-51639494-INS-123
chr11-51653578-INS-455
chr11-51694305-INS-339
chr11-51694318-INS-76
chr11-51702782-INS-212
chr11-51782395-INS-65
chr11-51782889-INS-250
chr11-51796880-INS-81
chr11-51802229-INS-122
chr11-51811601-INS-3042
chr11-51849297-INS-55

chr11-51883202-INS-211
chr11-51906507-INS-67
chr11-51939714-INS-1372
chr11-51939738-INS-408
chr11-51959474-INS-57
chr11-52001045-INS-125
chr11-52022184-INS-206
chr11-52034073-INS-198
chr11-52066558-INS-238
chr11-52098073-INS-499
chr11-52102453-INS-64
chr11-52103167-INS-397
chr11-52105175-INS-131
chr11-52192435-INS-180
chr11-52193920-INS-189
chr11-52197194-INS-55
chr11-52216478-INS-498
chr11-52256229-INS-50
chr11-52259324-INS-94
chr11-52294766-INS-6279
chr11-52340799-INS-55
chr11-52340819-INS-247
chr11-52349981-INS-1240
chr11-52350125-INS-466
chr11-52350248-INS-112
chr11-52350475-INS-475
chr11-52389663-INS-128
chr11-52398137-INS-569
chr11-52409872-INS-140
chr11-52409888-INS-60
chr11-52534019-INS-138
chr11-52548713-INS-50
chr11-52554049-INS-80
chr11-52554210-INS-80
chr11-52554210-INS-860
chr11-52602937-INS-106
chr11-52627266-INS-272
chr11-52630152-INS-180
chr11-52654809-INS-95
chr11-52678521-INS-156
chr11-52933246-INS-91
chr11-52997153-INS-209
chr11-53003254-INS-52
chr11-53017865-INS-52
chr11-53058429-INS-3014
chr11-53104740-INS-76
chr11-53104794-INS-74
chr11-53116414-INS-55
chr11-53133105-INS-343
chr11-53209502-INS-7110
chr11-53227443-INS-51
chr11-53317682-INS-60
chr11-53404451-INS-64
chr11-53413879-INS-7232
chr11-53451741-INS-216
chr11-53451953-INS-284
chr11-53469119-INS-80
chr11-53510806-INS-164
chr11-53528307-INS-167
chr11-53530642-INS-51
chr11-53573320-INS-249
chr11-53612419-INS-52
chr11-53616979-INS-72
chr11-53625122-INS-246
chr11-53636021-INS-52
chr11-53636452-INS-197
chr11-53642310-INS-57
chr11-53725054-INS-209
chr11-53759347-INS-76
chr11-53768450-INS-63
chr11-53769298-INS-3031
chr11-53800971-INS-88
chr11-53815720-INS-68
chr11-53856981-INS-5717
chr11-53873392-INS-106
chr11-53874982-INS-175
chr11-53907622-INS-78

chr11-53908481-INS-57
chr11-53921975-INS-397
chr11-53923423-INS-54
chr11-53923726-INS-84
chr11-53927318-INS-2435
chr11-53941237-INS-205
chr11-53967431-INS-61
chr11-53967601-INS-1834
chr11-53968343-INS-204
chr11-53980978-INS-1890
chr11-53986290-INS-80
chr11-53990783-INS-105
chr11-54019011-INS-54
chr11-54126165-INS-57
chr11-54143740-INS-56
chr11-54217174-INS-517
chr11-54298289-INS-60
chr11-54317867-INS-112
chr11-54326814-INS-60
chr11-54327223-INS-67
chr11-54333444-INS-211
chr11-54375571-INS-296
chr11-54385980-INS-55
chr11-54419359-INS-197
chr11-54480864-INS-64
chr11-54566123-INS-244
chr11-54573393-INS-750
chr11-54599805-INS-7818
chr11-54699100-INS-2080
chr11-54727125-INS-1260
chr11-54782136-INS-299
chr11-54791171-INS-269
chr11-54791218-INS-105
chr11-54892228-INS-184
chr11-54927380-INS-164
chr11-54963045-INS-71
chr11-54979514-INS-4524
chr11-54979516-INS-142
chr11-54993997-INS-52
chr11-55014161-INS-6470
chr11-55034653-INS-54
chr11-55039692-INS-124
chr11-55066410-INS-50
chr11-55091744-INS-80
chr11-55095898-INS-90
chr11-55121867-INS-104
chr11-55145613-INS-55
chr11-55209749-INS-50
chr11-55265030-INS-138
chr11-55265107-INS-590
chr11-55268803-INS-557
chr11-55294771-INS-460
chr11-55294846-INS-172
chr11-55318487-INS-61
chr11-55332347-INS-63
chr11-55412981-INS-58
chr11-55417116-INS-53
chr11-55420851-INS-250
chr11-55420925-INS-61
chr11-55427836-INS-188
chr11-55450806-INS-60
chr11-55489855-INS-187
chr11-55509639-INS-121
chr11-55538169-INS-67
chr11-55538371-INS-669
chr11-55549166-INS-4352
chr11-55567588-INS-261
chr11-55568745-INS-152
chr11-55577426-INS-7365
chr11-55707276-INS-1205
chr11-55747836-INS-4292
chr11-55786622-INS-57
chr11-55816630-INS-110
chr11-55842427-INS-50
chr11-55874845-INS-242
chr11-55892333-INS-1044
chr11-55892345-INS-470

chr11-55899737-INS-149
chr11-55909186-INS-4253
chr11-55950172-INS-365
chr11-55974615-INS-141
chr11-55980059-INS-74
chr11-55995860-INS-434
chr11-56046582-INS-6607
chr11-56052223-INS-1610
chr11-56062119-INS-50
chr11-56075473-INS-2704
chr11-56078157-INS-4457
chr11-56086587-INS-93
chr11-56111702-INS-50
chr11-56157216-INS-50
chr11-56157365-INS-138
chr11-56160482-INS-123
chr11-56219073-INS-5640
chr11-56220846-INS-520
chr11-56236864-INS-61
chr11-56245887-INS-51
chr11-56399932-INS-1321
chr11-56411648-INS-50
chr11-56442454-INS-67
chr11-56565141-INS-50
chr11-56631803-INS-57
chr11-56689680-INS-63
chr11-56711296-INS-6211
chr11-56734910-INS-62
chr11-56807180-INS-2722
chr11-56809953-INS-1257
chr11-56811505-INS-6254
chr11-56865711-INS-54
chr11-56884135-INS-56
chr11-56959718-INS-1548
chr11-56975079-INS-141
chr11-57010384-INS-192
chr11-57055763-INS-51
chr11-57065819-INS-52
chr11-57126798-INS-60
chr11-57227550-INS-305
chr11-57272236-INS-245
chr11-57279541-INS-220
chr11-57301673-INS-4142
chr11-57312984-INS-366
chr11-57322024-INS-4727
chr11-57344799-INS-2188
chr11-57355056-INS-1892
chr11-57356857-INS-2929
chr11-57361330-INS-124
chr11-57362256-INS-90
chr11-57363385-INS-2332
chr11-57370422-INS-50
chr11-57381892-INS-52
chr11-57419409-INS-80
chr11-57420172-INS-212
chr11-57513644-INS-161
chr11-57537175-INS-228
chr11-57546405-INS-51
chr11-57586909-INS-55
chr11-57707644-INS-66
chr11-57707647-INS-357
chr11-57756855-INS-50
chr11-57776212-INS-1173
chr11-57778204-INS-56
chr11-57778344-INS-418
chr11-57794291-INS-212
chr11-57866061-INS-217
chr11-57876633-INS-256
chr11-57879385-INS-209
chr11-57997504-INS-113
chr11-58040799-INS-4205
chr11-58125815-INS-221
chr11-58128277-INS-997
chr11-58162928-INS-104
chr11-58165157-INS-86
chr11-58173209-INS-57
chr11-58235187-INS-55

chr11-58235275-INS-1066
chr11-58246804-INS-6524
chr11-58247114-INS-3983
chr11-58263289-INS-933
chr11-58272568-INS-198
chr11-58284619-INS-327
chr11-58290329-INS-188
chr11-58290623-INS-316
chr11-58303203-INS-121
chr11-58333715-INS-144
chr11-58348071-INS-1475
chr11-58429978-INS-2831
chr11-58473254-INS-3233
chr11-58519496-INS-132
chr11-58534377-INS-593
chr11-58551839-INS-1888
chr11-58576539-INS-68
chr11-58698090-INS-1488
chr11-58704597-INS-58
chr11-58731125-INS-65
chr11-58731807-INS-1237
chr11-58748745-INS-147
chr11-58768597-INS-245
chr11-58784940-INS-260
chr11-58806452-INS-132
chr11-58806662-INS-492
chr11-58825200-INS-258
chr11-58853331-INS-161
chr11-58896535-INS-510
chr11-58957406-INS-55
chr11-58957624-INS-136
chr11-58983071-INS-1352
chr11-59013728-INS-6415
chr11-59093241-INS-53
chr11-59120412-INS-200
chr11-59121001-INS-372
chr11-59162697-INS-50
chr11-59191615-INS-2136
chr11-59228791-INS-124
chr11-59232177-INS-5040
chr11-59332306-INS-64
chr11-59386369-INS-71
chr11-59441639-INS-270
chr11-59448422-INS-134
chr11-59458619-INS-218
chr11-59476766-INS-50
chr11-59513613-INS-2429
chr11-59517560-INS-2343
chr11-59521751-INS-103
chr11-59522482-INS-219
chr11-59695011-INS-50
chr11-59809734-INS-559
chr11-59814983-INS-323
chr11-59908494-INS-73
chr11-59915857-INS-52
chr11-59918163-INS-204
chr11-59932168-INS-54
chr11-59948769-INS-341
chr11-59953886-INS-2978
chr11-59954620-INS-112
chr11-60001693-INS-59
chr11-60011942-INS-55
chr11-60215167-INS-400
chr11-60215528-INS-106
chr11-60256790-INS-56
chr11-60267491-INS-57
chr11-60301362-INS-213
chr11-60342277-INS-1681
chr11-60406727-INS-90
chr11-60450393-INS-2875
chr11-60496196-INS-76
chr11-60505486-INS-70
chr11-60512269-INS-173
chr11-60515999-INS-51
chr11-60537151-INS-63
chr11-60540446-INS-486
chr11-60642538-INS-175

chr11-60686638-INS-460
chr11-60732066-INS-191
chr11-60749771-INS-216
chr11-60750492-INS-58
chr11-60788235-INS-70
chr11-60825562-INS-78
chr11-60830064-INS-224
chr11-60836133-INS-2080
chr11-60866336-INS-2664
chr11-60885832-INS-100
chr11-60931539-INS-98
chr11-60949955-INS-50
chr11-60969056-INS-148
chr11-60970515-INS-57
chr11-60979174-INS-231
chr11-60984506-INS-112
chr11-60986029-INS-60
chr11-60996520-INS-50
chr11-61009685-INS-172
chr11-61074375-INS-167
chr11-61074397-INS-83
chr11-61097979-INS-145
chr11-61151904-INS-408
chr11-61166852-INS-90
chr11-61170148-INS-483
chr11-61198798-INS-62
chr11-61207708-INS-50
chr11-61210125-INS-5546
chr11-61264903-INS-116
chr11-61278378-INS-50
chr11-61299416-INS-50
chr11-61363523-INS-100
chr11-61451128-INS-250
chr11-61459023-INS-207
chr11-61481017-INS-51
chr11-61524718-INS-70
chr11-61538051-INS-2364
chr11-61607314-INS-50
chr11-61656059-INS-182
chr11-61696055-INS-513
chr11-61698148-INS-210
chr11-61714439-INS-204
chr11-61730132-INS-481
chr11-61749685-INS-53
chr11-61753241-INS-5127
chr11-61769595-INS-10931
chr11-61778641-INS-70
chr11-61798289-INS-3167
chr11-61804970-INS-116
chr11-61832344-INS-1149
chr11-61843995-INS-310
chr11-61847188-INS-56
chr11-61864396-INS-125
chr11-61875702-INS-92
chr11-61885092-INS-2713
chr11-61887572-INS-78
chr11-61911731-INS-900
chr11-61924940-INS-5058
chr11-61947615-INS-66
chr11-61955900-INS-206
chr11-61989549-INS-1054
chr11-62015851-INS-200
chr11-62029895-INS-50
chr11-62040236-INS-199
chr11-62042674-INS-649
chr11-62055633-INS-688
chr11-62067444-INS-132
chr11-62067653-INS-56
chr11-62090750-INS-68
chr11-62130817-INS-52
chr11-62151483-INS-201
chr11-62252681-INS-72
chr11-62263207-INS-190
chr11-62301148-INS-166
chr11-62301150-INS-533
chr11-62311146-INS-96
chr11-62320788-INS-54

chr11-62359196-INS-165
chr11-62372916-INS-183
chr11-62375990-INS-75
chr11-62379878-INS-563
chr11-62388539-INS-201
chr11-62401825-INS-186
chr11-62435406-INS-67
chr11-62436147-INS-198
chr11-62438259-INS-223
chr11-62439451-INS-185
chr11-62453015-INS-108
chr11-62457573-INS-415
chr11-62457753-INS-192
chr11-62464980-INS-171
chr11-62480860-INS-54
chr11-62481316-INS-247
chr11-62511516-INS-8497
chr11-62511995-INS-52
chr11-62533741-INS-52
chr11-62543626-INS-205
chr11-62570776-INS-1283
chr11-62570838-INS-103
chr11-62576240-INS-180
chr11-62637707-INS-265
chr11-62653474-INS-68
chr11-62663890-INS-79
chr11-62703637-INS-139
chr11-62704594-INS-102
chr11-62733964-INS-150
chr11-62734065-INS-90
chr11-62734403-INS-51
chr11-62734497-INS-152
chr11-62740833-INS-54
chr11-62746099-INS-1892
chr11-62748988-INS-901
chr11-62763043-INS-192
chr11-62806118-INS-162
chr11-62865259-INS-624
chr11-62979889-INS-52
chr11-62984132-INS-5389
chr11-62985090-INS-6440
chr11-62985210-INS-50
chr11-63013476-INS-168
chr11-63013476-INS-64
chr11-63029549-INS-134
chr11-63070142-INS-82
chr11-63142059-INS-60
chr11-63155279-INS-6622
chr11-63179981-INS-122
chr11-63248189-INS-50
chr11-63253107-INS-2465
chr11-63262651-INS-88
chr11-63285546-INS-791
chr11-63298013-INS-6841
chr11-63315904-INS-50
chr11-63347010-INS-2838
chr11-63381581-INS-7081
chr11-63385363-INS-79
chr11-63450060-INS-62
chr11-63465142-INS-83
chr11-63479411-INS-55
chr11-63483334-INS-7110
chr11-63491441-INS-62
chr11-63522390-INS-141
chr11-63526646-INS-60
chr11-63530584-INS-424
chr11-63530584-INS-53
chr11-63567062-INS-51
chr11-63612483-INS-80
chr11-63614842-INS-7230
chr11-63624755-INS-54
chr11-63625338-INS-173
chr11-63643398-INS-285
chr11-63665529-INS-291
chr11-63689301-INS-62
chr11-63703355-INS-2522
chr11-63711102-INS-1434

chr11-63724788-INS-50
chr11-63732373-INS-6930
chr11-63753527-INS-78
chr11-63757649-INS-136
chr11-63757748-INS-59
chr11-63824969-INS-72
chr11-63835249-INS-94
chr11-63836002-INS-80
chr11-63860377-INS-435
chr11-63877437-INS-77
chr11-63902958-INS-727
chr11-63948872-INS-52
chr11-63960749-INS-1354
chr11-64014679-INS-55
chr11-64028647-INS-54
chr11-64129505-INS-4409
chr11-64141285-INS-51
chr11-64194621-INS-563
chr11-64236818-INS-52
chr11-64242018-INS-394
chr11-64277587-INS-2191
chr11-64299369-INS-108
chr11-64333434-INS-52
chr11-64362479-INS-83
chr11-64362592-INS-78
chr11-64365829-INS-57
chr11-64367392-INS-64
chr11-64399128-INS-58
chr11-64399159-INS-5517
chr11-64422546-INS-305
chr11-64439443-INS-50
chr11-64442824-INS-63
chr11-64510375-INS-402
chr11-64510375-INS-54
chr11-64510898-INS-770
chr11-64556108-INS-6439
chr11-64557451-INS-51
chr11-64570194-INS-1144
chr11-64573239-INS-56
chr11-64604441-INS-99
chr11-64608971-INS-127
chr11-64653939-INS-1769
chr11-64676445-INS-109
chr11-64676451-INS-53
chr11-64677299-INS-64
chr11-64677939-INS-121
chr11-64702347-INS-113
chr11-64708803-INS-6178
chr11-64732765-INS-121
chr11-64732985-INS-51
chr11-64740668-INS-399
chr11-64789312-INS-58
chr11-64802129-INS-517
chr11-64805718-INS-168
chr11-64827448-INS-148
chr11-64838044-INS-142
chr11-64859025-INS-598
chr11-64881009-INS-368
chr11-64972827-INS-56
chr11-65063776-INS-7224
chr11-65082633-INS-193
chr11-65094269-INS-323
chr11-65123582-INS-50
chr11-65127720-INS-72
chr11-65133774-INS-201
chr11-65134242-INS-216
chr11-65143183-INS-173
chr11-65192997-INS-158
chr11-65204192-INS-50
chr11-65232950-INS-50
chr11-65237853-INS-91
chr11-65246636-INS-6657
chr11-65258473-INS-285
chr11-65291950-INS-307
chr11-65333407-INS-215
chr11-65338085-INS-4676
chr11-65369595-INS-69

chr11-65379075-INS-380
chr11-65380380-INS-99
chr11-65404476-INS-5860
chr11-65440676-INS-810
chr11-65471131-INS-50
chr11-65536232-INS-50
chr11-65540173-INS-92
chr11-65560719-INS-55
chr11-65564552-INS-107
chr11-65601507-INS-168
chr11-65662197-INS-3383
chr11-65677457-INS-76
chr11-65683154-INS-3129
chr11-65688140-INS-176
chr11-65714603-INS-108
chr11-65736898-INS-256
chr11-65775804-INS-625
chr11-65775812-INS-2753
chr11-65782095-INS-1276
chr11-65782214-INS-61
chr11-65792684-INS-3211
chr11-65798322-INS-53
chr11-65893495-INS-6122
chr11-65898178-INS-197
chr11-65934100-INS-104
chr11-65948886-INS-660
chr11-66014058-INS-1976
chr11-66121976-INS-399
chr11-66198514-INS-67
chr11-66230832-INS-138
chr11-66246377-INS-73
chr11-66267314-INS-51
chr11-66317566-INS-160
chr11-66317595-INS-61
chr11-66328272-INS-53
chr11-66390198-INS-57
chr11-66455651-INS-205
chr11-66456137-INS-713
chr11-66466488-INS-52
chr11-66467230-INS-175
chr11-66534200-INS-1783
chr11-66536783-INS-71
chr11-66539783-INS-6574
chr11-66540937-INS-55
chr11-66550781-INS-75
chr11-66595591-INS-52
chr11-66657957-INS-176
chr11-66668244-INS-61
chr11-66724720-INS-3694
chr11-66732317-INS-290
chr11-66752008-INS-3360
chr11-66762854-INS-54
chr11-66769430-INS-108
chr11-66769859-INS-112
chr11-66769944-INS-475
chr11-66774053-INS-240
chr11-66788418-INS-194
chr11-66810977-INS-228
chr11-66811951-INS-5573
chr11-66824695-INS-62
chr11-66857089-INS-74
chr11-66898323-INS-350
chr11-66926724-INS-7768
chr11-66981545-INS-99
chr11-66983389-INS-697
chr11-66983871-INS-722
chr11-67020103-INS-181
chr11-67043834-INS-156
chr11-67164378-INS-74
chr11-67178527-INS-65
chr11-67191171-INS-201
chr11-67198816-INS-78
chr11-67243774-INS-221
chr11-67299085-INS-90
chr11-67383804-INS-165
chr11-67437149-INS-60
chr11-67508719-INS-242

chr11-67516761-INS-366
chr11-67517832-INS-1019
chr11-67564749-INS-80
chr11-67619411-INS-253
chr11-67658830-INS-54
chr11-67700002-INS-92
chr11-67709483-INS-184
chr11-67719931-INS-80
chr11-67734528-INS-57
chr11-67755417-INS-58
chr11-67828088-INS-52
chr11-67844203-INS-50
chr11-67853494-INS-50
chr11-67855468-INS-6402
chr11-67864908-INS-53
chr11-67872690-INS-51
chr11-67873933-INS-72
chr11-67874007-INS-52
chr11-67874077-INS-67
chr11-67893339-INS-4065
chr11-67947632-INS-94
chr11-67951030-INS-912
chr11-67962120-INS-156
chr11-67964954-INS-257
chr11-67993099-INS-209
chr11-67993998-INS-202
chr11-68006457-INS-153
chr11-68017513-INS-81
chr11-68018128-INS-367
chr11-68021940-INS-95
chr11-68052584-INS-461
chr11-68114010-INS-65
chr11-68149110-INS-529
chr11-68187241-INS-96
chr11-68190910-INS-1932
chr11-68227096-INS-153
chr11-68268996-INS-80
chr11-68293350-INS-63
chr11-68338576-INS-88
chr11-68435240-INS-53
chr11-68440547-INS-69
chr11-68464046-INS-56
chr11-68464726-INS-771
chr11-68482234-INS-52
chr11-68496932-INS-163
chr11-68500161-INS-50
chr11-68523588-INS-1464
chr11-68547202-INS-108
chr11-68594684-INS-210
chr11-68641039-INS-52
chr11-68641115-INS-204
chr11-68657313-INS-519
chr11-68677783-INS-137
chr11-68677880-INS-66
chr11-68719955-INS-183
chr11-68732290-INS-174
chr11-68791265-INS-60
chr11-68796306-INS-67
chr11-68814329-INS-63
chr11-68838294-INS-76
chr11-68839026-INS-248
chr11-68850015-INS-212
chr11-68853859-INS-198
chr11-68856781-INS-195
chr11-68867712-INS-230
chr11-68867930-INS-175
chr11-68896717-INS-221
chr11-68957280-INS-58
chr11-68960558-INS-120
chr11-69005811-INS-59
chr11-69032007-INS-61
chr11-69052446-INS-178
chr11-69053237-INS-132
chr11-69062833-INS-74
chr11-69072128-INS-153
chr11-69104176-INS-1116
chr11-69104372-INS-252

chr11-69174699-INS-268
chr11-69207379-INS-521
chr11-69210420-INS-160
chr11-69242294-INS-50
chr11-69245496-INS-209
chr11-69272538-INS-207
chr11-69340866-INS-226
chr11-69353980-INS-152
chr11-69373339-INS-4680
chr11-69419036-INS-1707
chr11-69426514-INS-92
chr11-69436657-INS-184
chr11-69441477-INS-651
chr11-69456787-INS-210
chr11-69517042-INS-203
chr11-69580324-INS-55
chr11-69595685-INS-80
chr11-69605073-INS-117
chr11-69605260-INS-208
chr11-69615404-INS-186
chr11-69747108-INS-64
chr11-69756281-INS-72
chr11-69824351-INS-176
chr11-69845862-INS-1656
chr11-69951622-INS-194
chr11-69962521-INS-136
chr11-69969850-INS-204
chr11-69982148-INS-56
chr11-69989305-INS-383
chr11-70014460-INS-1191
chr11-70015992-INS-2645
chr11-70093327-INS-70
chr11-70093372-INS-260
chr11-70122121-INS-232
chr11-70147667-INS-542
chr11-70148638-INS-99
chr11-70159338-INS-145
chr11-70164276-INS-85
chr11-70174454-INS-161
chr11-70174523-INS-479
chr11-70196876-INS-4809
chr11-70214705-INS-108
chr11-70252929-INS-42845
chr11-70261316-INS-444
chr11-70262781-INS-260
chr11-70264553-INS-57
chr11-70284300-INS-74
chr11-70295864-INS-205
chr11-70320991-INS-197
chr11-70334127-INS-199
chr11-70338816-INS-60
chr11-70392245-INS-219
chr11-70392674-INS-406
chr11-70419502-INS-200
chr11-70461708-INS-1893
chr11-70491560-INS-62
chr11-70492016-INS-80
chr11-70500650-INS-80
chr11-70637118-INS-217
chr11-70639383-INS-62
chr11-70641765-INS-1042
chr11-70665120-INS-975
chr11-70667679-INS-63
chr11-70669959-INS-60
chr11-70675314-INS-204
chr11-70734417-INS-52
chr11-70806933-INS-80
chr11-70807388-INS-105
chr11-70914155-INS-4897
chr11-70941582-INS-58
chr11-71022547-INS-60
chr11-71036880-INS-17962
chr11-71036880-INS-8319
chr11-71076214-INS-112
chr11-71080959-INS-555
chr11-71117254-INS-342
chr11-71187589-INS-228

chr11-71206785-INS-1595
chr11-71275089-INS-208
chr11-71291197-INS-9637
chr11-71404053-INS-312
chr11-71411572-INS-52
chr11-71428129-INS-139
chr11-71455289-INS-50
chr11-71497884-INS-6932
chr11-71522131-INS-715
chr11-71525517-INS-7087
chr11-71545336-INS-54
chr11-71558802-INS-6876
chr11-71560119-INS-211
chr11-71580871-INS-90
chr11-71585957-INS-181
chr11-71600421-INS-463
chr11-71605006-INS-321
chr11-71605657-INS-69
chr11-71736122-INS-882
chr11-71774304-INS-242
chr11-71784724-INS-1933
chr11-71859144-INS-58
chr11-71861998-INS-175
chr11-71862123-INS-75
chr11-71873427-INS-195
chr11-71873596-INS-52
chr11-71880083-INS-636
chr11-71880487-INS-192
chr11-71880611-INS-54
chr11-71881024-INS-350
chr11-71898469-INS-117
chr11-71917883-INS-50
chr11-71998417-INS-240
chr11-72000512-INS-283
chr11-72000540-INS-126
chr11-72140764-INS-136
chr11-72164982-INS-120
chr11-72166473-INS-68
chr11-72166871-INS-184
chr11-72264033-INS-213
chr11-72321915-INS-64
chr11-72322473-INS-156
chr11-72322493-INS-66
chr11-72325238-INS-221
chr11-72390045-INS-60
chr11-72412053-INS-7145
chr11-72423282-INS-164
chr11-72511715-INS-52
chr11-72624237-INS-56
chr11-72689680-INS-50
chr11-72808231-INS-2133
chr11-72839761-INS-52
chr11-72956936-INS-692
chr11-73004857-INS-66
chr11-73044112-INS-58
chr11-73053459-INS-193
chr11-73147158-INS-6482
chr11-73167954-INS-78
chr11-73183192-INS-81
chr11-73183988-INS-196
chr11-73201321-INS-102
chr11-73205955-INS-60
chr11-73206550-INS-151
chr11-73240410-INS-870
chr11-73296122-INS-1915
chr11-73350662-INS-1065
chr11-73350899-INS-95
chr11-73399252-INS-8442
chr11-73400442-INS-65
chr11-73443903-INS-28193
chr11-73480681-INS-3762
chr11-73495968-INS-105
chr11-73500762-INS-132
chr11-73523406-INS-123
chr11-73523485-INS-760
chr11-73682456-INS-497
chr11-73683079-INS-318

chr11-73710695-INS-441
chr11-73711634-INS-501
chr11-73789257-INS-1320
chr11-73825523-INS-106
chr11-73825561-INS-1370
chr11-73882217-INS-216
chr11-73882856-INS-6344
chr11-73907136-INS-84
chr11-73933171-INS-935
chr11-73956131-INS-163
chr11-74060675-INS-3544
chr11-74256640-INS-13279
chr11-74280509-INS-1262
chr11-74290085-INS-177
chr11-74295508-INS-1162
chr11-74314272-INS-50
chr11-74318954-INS-52
chr11-74338298-INS-332
chr11-74351171-INS-54
chr11-74396664-INS-52
chr11-74437432-INS-65
chr11-74469060-INS-7204
chr11-74499370-INS-1875
chr11-74516354-INS-217
chr11-74622882-INS-3305
chr11-74639912-INS-5636
chr11-74640974-INS-312
chr11-74649737-INS-161
chr11-74668619-INS-150
chr11-74668972-INS-103
chr11-74671934-INS-954
chr11-74673681-INS-1999
chr11-74769305-INS-228
chr11-74769469-INS-193
chr11-74770588-INS-204
chr11-74808222-INS-239
chr11-74813090-INS-199
chr11-74838435-INS-74
chr11-74839568-INS-168
chr11-74849683-INS-70
chr11-74850096-INS-206
chr11-74851841-INS-54
chr11-74853520-INS-206
chr11-74854839-INS-224
chr11-74860240-INS-372
chr11-74891251-INS-207
chr11-74941921-INS-207
chr11-74972904-INS-206
chr11-75088884-INS-88
chr11-75111405-INS-204
chr11-75114519-INS-86
chr11-75200141-INS-183
chr11-75205609-INS-56
chr11-75207399-INS-220
chr11-75269834-INS-69
chr11-75277765-INS-211
chr11-75284798-INS-79
chr11-75296458-INS-204
chr11-75301481-INS-52
chr11-75309981-INS-137
chr11-75317303-INS-89
chr11-75317441-INS-57
chr11-75348309-INS-171
chr11-75374681-INS-232
chr11-75375622-INS-69
chr11-75395772-INS-229
chr11-75406717-INS-100
chr11-75533732-INS-175
chr11-75550634-INS-219
chr11-75576232-INS-88
chr11-75605784-INS-178
chr11-75610831-INS-9184
chr11-75638739-INS-179
chr11-75650598-INS-193
chr11-75669825-INS-140
chr11-75694952-INS-50
chr11-75708401-INS-52

chr11-75754155-INS-610
chr11-75777662-INS-132
chr11-75823998-INS-388
chr11-75842642-INS-56
chr11-75861253-INS-7242
chr11-75875610-INS-52
chr11-75882265-INS-230
chr11-75888351-INS-177
chr11-75896709-INS-200
chr11-75897022-INS-52
chr11-75943808-INS-72
chr11-75986099-INS-222
chr11-75989610-INS-201
chr11-75995188-INS-372
chr11-76034474-INS-103
chr11-76050323-INS-50
chr11-76056305-INS-218
chr11-76075005-INS-250
chr11-76085540-INS-91
chr11-76122583-INS-228
chr11-76126483-INS-154
chr11-76128415-INS-221
chr11-76129215-INS-183
chr11-76137204-INS-67
chr11-76165891-INS-362
chr11-76165903-INS-126
chr11-76172165-INS-201
chr11-76172352-INS-76
chr11-76187751-INS-400
chr11-76207853-INS-207
chr11-76276737-INS-206
chr11-76282284-INS-190
chr11-76311676-INS-102
chr11-76337528-INS-125
chr11-76337528-INS-55
chr11-76352530-INS-194
chr11-76425745-INS-242
chr11-76431562-INS-123
chr11-76471334-INS-202
chr11-76487099-INS-207
chr11-76541340-INS-156
chr11-76541615-INS-165
chr11-76573844-INS-94
chr11-76579784-INS-220
chr11-76610201-INS-79
chr11-76636141-INS-71
chr11-76673639-INS-527
chr11-76744579-INS-215
chr11-76764012-INS-199
chr11-76788872-INS-523
chr11-76809803-INS-772
chr11-76919434-INS-300
chr11-76936391-INS-54
chr11-76950804-INS-339
chr11-76974049-INS-50
chr11-77002944-INS-288
chr11-77010782-INS-236
chr11-77028407-INS-180
chr11-77037405-INS-141
chr11-77058681-INS-463
chr11-77211534-INS-1493
chr11-77220941-INS-83
chr11-77226802-INS-5387
chr11-77238362-INS-545
chr11-77448108-INS-222
chr11-77463193-INS-93
chr11-77480888-INS-210
chr11-77503308-INS-324
chr11-77506877-INS-203
chr11-77528911-INS-155
chr11-77528991-INS-293
chr11-77599834-INS-388
chr11-77636740-INS-269
chr11-77697376-INS-327
chr11-77780884-INS-190
chr11-77782310-INS-66
chr11-77832418-INS-50

chr11-77834004-INS-193
chr11-77878741-INS-72
chr11-77883606-INS-58
chr11-77978317-INS-200
chr11-78003054-INS-203
chr11-78003890-INS-205
chr11-78035293-INS-254
chr11-78073486-INS-203
chr11-78081772-INS-132
chr11-78133141-INS-230
chr11-78139870-INS-344
chr11-78140697-INS-206
chr11-78151403-INS-288
chr11-78172194-INS-207
chr11-78209943-INS-128
chr11-78231605-INS-238
chr11-78231606-INS-57
chr11-78243898-INS-57
chr11-78245056-INS-5101
chr11-78247430-INS-52
chr11-78323692-INS-164
chr11-78323702-INS-53
chr11-78372820-INS-51
chr11-78465766-INS-606
chr11-78486858-INS-58
chr11-78498595-INS-204
chr11-78510098-INS-240
chr11-78540216-INS-220
chr11-78549759-INS-61
chr11-78631601-INS-387
chr11-78645505-INS-56
chr11-78651167-INS-52
chr11-78677993-INS-1277
chr11-78737070-INS-340
chr11-78745436-INS-196
chr11-78748277-INS-50
chr11-78766178-INS-462
chr11-78777011-INS-200
chr11-78777894-INS-55
chr11-78782652-INS-210
chr11-78782948-INS-54
chr11-78784396-INS-6768
chr11-78785560-INS-54
chr11-78785586-INS-137
chr11-78817996-INS-68
chr11-78829299-INS-53
chr11-78879273-INS-50
chr11-78885388-INS-210
chr11-78931075-INS-70
chr11-78941035-INS-174
chr11-78981379-INS-72
chr11-78988707-INS-288
chr11-78990696-INS-52
chr11-78991128-INS-53
chr11-79001877-INS-1075
chr11-79001877-INS-255
chr11-79015675-INS-3216
chr11-79022754-INS-89
chr11-79103091-INS-50
chr11-79119559-INS-240
chr11-79135920-INS-187
chr11-79155115-INS-315
chr11-79159916-INS-167
chr11-79166774-INS-172
chr11-79171832-INS-70
chr11-79176335-INS-203
chr11-79225480-INS-214
chr11-79225860-INS-52
chr11-79243755-INS-63
chr11-79243895-INS-130
chr11-79281515-INS-84
chr11-79295279-INS-110
chr11-79342514-INS-105
chr11-79342535-INS-252
chr11-79351000-INS-770
chr11-79474041-INS-239
chr11-79474573-INS-54

chr11-79493734-INS-386
chr11-79521068-INS-62
chr11-79563611-INS-112
chr11-79593221-INS-86
chr11-79605625-INS-232
chr11-79625234-INS-558
chr11-79641601-INS-5467
chr11-79644377-INS-3168
chr11-79665656-INS-56
chr11-79682169-INS-288
chr11-79685801-INS-50
chr11-79689225-INS-463
chr11-79703337-INS-99
chr11-79726046-INS-215
chr11-79773189-INS-396
chr11-79773414-INS-207
chr11-79794703-INS-405
chr11-79804502-INS-126
chr11-79812688-INS-114
chr11-79868053-INS-221
chr11-79886819-INS-1105
chr11-79887333-INS-94
chr11-79911214-INS-208
chr11-79957942-INS-62
chr11-79974096-INS-56
chr11-80026623-INS-198
chr11-80066034-INS-232
chr11-80072905-INS-130
chr11-80088196-INS-524
chr11-80097056-INS-210
chr11-80127436-INS-234
chr11-80158178-INS-3186
chr11-80160150-INS-174
chr11-80167245-INS-1258
chr11-80172121-INS-171
chr11-80172985-INS-197
chr11-80225931-INS-52
chr11-80248867-INS-152
chr11-80250233-INS-1892
chr11-80258660-INS-220
chr11-80276663-INS-148
chr11-80312707-INS-216
chr11-80328212-INS-200
chr11-80341496-INS-216
chr11-80356394-INS-52
chr11-80418743-INS-631
chr11-80440123-INS-92
chr11-80444312-INS-123
chr11-80448547-INS-198
chr11-80490810-INS-197
chr11-80511441-INS-185
chr11-80512098-INS-358
chr11-80522327-INS-136
chr11-80524197-INS-7705
chr11-80623047-INS-70
chr11-80696558-INS-62
chr11-80696912-INS-69
chr11-80732764-INS-62
chr11-80737612-INS-196
chr11-80740630-INS-1955
chr11-80751978-INS-221
chr11-80752835-INS-83
chr11-80802441-INS-462
chr11-80820723-INS-1311
chr11-80842473-INS-50
chr11-80847392-INS-376
chr11-80892391-INS-66
chr11-80939122-INS-402
chr11-80972327-INS-50
chr11-81007658-INS-52
chr11-81082555-INS-2844
chr11-81126088-INS-61
chr11-81131576-INS-727
chr11-81143482-INS-52
chr11-81150902-INS-1187
chr11-81195332-INS-82
chr11-81195448-INS-109

chr11-81283460-INS-162
chr11-81302597-INS-7225
chr11-81354761-INS-334
chr11-81363232-INS-52
chr11-81370763-INS-50
chr11-81393652-INS-51
chr11-81417401-INS-55
chr11-81463169-INS-60
chr11-81496903-INS-1889
chr11-81537328-INS-88
chr11-81590530-INS-112
chr11-81619422-INS-118
chr11-81619497-INS-377
chr11-81661730-INS-256
chr11-81666670-INS-52
chr11-81672435-INS-153
chr11-81775965-INS-4458
chr11-81824025-INS-114
chr11-81834215-INS-6889
chr11-81875989-INS-571
chr11-81881474-INS-52
chr11-81905536-INS-1089
chr11-81905760-INS-107
chr11-81943862-INS-586
chr11-81984569-INS-51
chr11-82010189-INS-474
chr11-82022751-INS-436
chr11-82088385-INS-60
chr11-82107887-INS-113
chr11-82161578-INS-63
chr11-82180075-INS-112
chr11-82212666-INS-72
chr11-82513571-INS-50
chr11-82563654-INS-55
chr11-82613426-INS-1040
chr11-82620745-INS-1184
chr11-82621488-INS-134
chr11-82664812-INS-123
chr11-82687360-INS-176
chr11-82699859-INS-50
chr11-82706668-INS-262
chr11-82711640-INS-81
chr11-82755335-INS-202
chr11-82759309-INS-99
chr11-82777029-INS-50
chr11-82778303-INS-56
chr11-82778386-INS-660
chr11-82821098-INS-84
chr11-82821457-INS-249
chr11-82821545-INS-646
chr11-82821965-INS-659
chr11-82823083-INS-569
chr11-82823309-INS-51
chr11-82826227-INS-65
chr11-82854181-INS-55
chr11-82863130-INS-136
chr11-82889732-INS-161
chr11-82889734-INS-67
chr11-82891056-INS-155
chr11-82896892-INS-409
chr11-82898417-INS-327
chr11-82902122-INS-130
chr11-82913832-INS-201
chr11-82914078-INS-312
chr11-82926409-INS-5255
chr11-82932782-INS-74
chr11-83063423-INS-896
chr11-83135481-INS-3429
chr11-83153042-INS-1651
chr11-83191007-INS-72
chr11-83207763-INS-125
chr11-83208527-INS-72
chr11-83432844-INS-479
chr11-83446368-INS-56
chr11-83446560-INS-482
chr11-83470083-INS-50
chr11-83470998-INS-201

chr11-83482669-INS-51
chr11-83543476-INS-52
chr11-83572483-INS-2853
chr11-83590628-INS-106
chr11-83611266-INS-6440
chr11-83633861-INS-74
chr11-83720339-INS-56
chr11-83721582-INS-56
chr11-83730668-INS-126
chr11-83774519-INS-80
chr11-83775465-INS-84
chr11-83779129-INS-100
chr11-83827734-INS-67
chr11-83874562-INS-118
chr11-83936205-INS-626
chr11-83936331-INS-238
chr11-83936331-INS-79
chr11-83936560-INS-503
chr11-83976170-INS-218
chr11-83977432-INS-5336
chr11-84001525-INS-205
chr11-84002150-INS-232
chr11-84009654-INS-209
chr11-84028632-INS-55
chr11-84042964-INS-326
chr11-84043264-INS-116
chr11-84043264-INS-54
chr11-84059124-INS-54
chr11-84123408-INS-130
chr11-84145171-INS-206
chr11-84212137-INS-193
chr11-84223248-INS-337
chr11-84255924-INS-744
chr11-84294660-INS-97
chr11-84302118-INS-73
chr11-84302199-INS-217
chr11-84313352-INS-62
chr11-84390838-INS-347
chr11-84407108-INS-190
chr11-84483601-INS-208
chr11-84519847-INS-59
chr11-84565127-INS-52
chr11-84586806-INS-58
chr11-84595233-INS-830
chr11-84606574-INS-50
chr11-84660722-INS-50
chr11-84695751-INS-146
chr11-84717390-INS-59
chr11-84718891-INS-66
chr11-84739750-INS-225
chr11-84766766-INS-128
chr11-84772624-INS-129
chr11-84807597-INS-98
chr11-84822659-INS-209
chr11-85020980-INS-286
chr11-85024823-INS-287
chr11-85041753-INS-54
chr11-85072618-INS-208
chr11-85140753-INS-99
chr11-85141861-INS-77
chr11-85153787-INS-50
chr11-85156442-INS-50
chr11-85197017-INS-169
chr11-85216020-INS-13759
chr11-85217568-INS-198
chr11-85239922-INS-303
chr11-85310301-INS-1953
chr11-85315991-INS-52
chr11-85525817-INS-91
chr11-85565054-INS-692
chr11-85810738-INS-99
chr11-85816760-INS-74
chr11-85859877-INS-57
chr11-85872940-INS-325
chr11-85879709-INS-196
chr11-85891746-INS-147
chr11-85924137-INS-64

chr11-85927043-INS-4959
chr11-85959833-INS-1458
chr11-85967264-INS-391
chr11-86027086-INS-235
chr11-86033426-INS-80
chr11-86044698-INS-89
chr11-86045440-INS-211
chr11-86065754-INS-103
chr11-86065904-INS-833
chr11-86093230-INS-200
chr11-86126959-INS-192
chr11-86205136-INS-68
chr11-86230441-INS-445
chr11-86235970-INS-122
chr11-86274270-INS-53
chr11-86301209-INS-7087
chr11-86306059-INS-211
chr11-86330564-INS-95
chr11-86342612-INS-58
chr11-86365023-INS-950
chr11-86370627-INS-173
chr11-86458702-INS-226
chr11-86458849-INS-82
chr11-86486715-INS-172
chr11-86499305-INS-149
chr11-86513848-INS-267
chr11-86539290-INS-172
chr11-86626482-INS-116
chr11-86671967-INS-226
chr11-86708126-INS-50
chr11-86721330-INS-1325
chr11-86817583-INS-253
chr11-86826725-INS-377
chr11-86861436-INS-519
chr11-86891972-INS-51
chr11-86892586-INS-52
chr11-86895771-INS-228
chr11-86902766-INS-2946
chr11-86910635-INS-197
chr11-86919922-INS-779
chr11-86956033-INS-52
chr11-87075178-INS-65
chr11-87253859-INS-143
chr11-87309468-INS-299
chr11-87374687-INS-210
chr11-87382454-INS-6337
chr11-87398043-INS-66
chr11-87582660-INS-116
chr11-87586302-INS-60
chr11-87724217-INS-563
chr11-87790102-INS-248
chr11-87795865-INS-52
chr11-87807607-INS-262
chr11-87807774-INS-100
chr11-87815804-INS-66
chr11-87822404-INS-63
chr11-87826817-INS-1675
chr11-87863581-INS-96
chr11-87901576-INS-156
chr11-87902042-INS-219
chr11-87905262-INS-200
chr11-87919893-INS-1269
chr11-88012474-INS-254
chr11-88053264-INS-70
chr11-88072546-INS-52
chr11-88073116-INS-147
chr11-88083753-INS-68
chr11-88096265-INS-165
chr11-88096818-INS-104
chr11-88120421-INS-234
chr11-88190183-INS-71
chr11-88223810-INS-3955
chr11-88461388-INS-209
chr11-88507726-INS-55
chr11-88583739-INS-282
chr11-88630073-INS-226
chr11-88758600-INS-74

chr11-88769993-INS-179
chr11-88809890-INS-1705
chr11-88850411-INS-370
chr11-88859124-INS-6524
chr11-88860477-INS-15134
chr11-88959579-INS-97
chr11-88968122-INS-193
chr11-88968203-INS-391
chr11-88970745-INS-66
chr11-88978595-INS-176
chr11-88978595-INS-61
chr11-89001878-INS-66
chr11-89023907-INS-3053
chr11-89024601-INS-50
chr11-89030299-INS-249
chr11-89031792-INS-50
chr11-89032094-INS-212
chr11-89041978-INS-168
chr11-89081400-INS-174
chr11-89099125-INS-221
chr11-89153110-INS-1160
chr11-89180160-INS-210
chr11-89183831-INS-151
chr11-89194505-INS-50
chr11-89210343-INS-190
chr11-89234757-INS-202
chr11-89242696-INS-2606
chr11-89252103-INS-67
chr11-89392358-INS-63
chr11-89408477-INS-216
chr11-89445078-INS-208
chr11-89467990-INS-1893
chr11-89476264-INS-96
chr11-89508720-INS-207
chr11-89520772-INS-72
chr11-89548275-INS-121
chr11-89592755-INS-145
chr11-89593411-INS-203
chr11-89620369-INS-7156
chr11-89624927-INS-78
chr11-89699608-INS-126
chr11-89748379-INS-105
chr11-89748670-INS-67
chr11-89752343-INS-258
chr11-89771085-INS-52
chr11-89780324-INS-65
chr11-89783909-INS-54
chr11-89824987-INS-52
chr11-89897606-INS-50
chr11-89909985-INS-1319
chr11-89914584-INS-50
chr11-89914842-INS-328
chr11-89929622-INS-170
chr11-89982137-INS-96
chr11-89995270-INS-50
chr11-90065017-INS-67
chr11-90077802-INS-136
chr11-90096257-INS-84
chr11-90107917-INS-124
chr11-90165885-INS-62
chr11-90180434-INS-50
chr11-90286873-INS-60
chr11-90342125-INS-52
chr11-90346489-INS-403
chr11-90356200-INS-218
chr11-90426099-INS-137
chr11-90427061-INS-150
chr11-90507002-INS-60
chr11-90518067-INS-145
chr11-90535497-INS-116
chr11-90539205-INS-2207
chr11-90558848-INS-200
chr11-90562959-INS-498
chr11-90563096-INS-78
chr11-90583231-INS-1220
chr11-90583910-INS-55
chr11-90584870-INS-56

chr11-90594467-INS-172
chr11-90599063-INS-167
chr11-90602076-INS-159
chr11-90621411-INS-50
chr11-90627612-INS-190
chr11-90704579-INS-138
chr11-90709940-INS-138
chr11-90726923-INS-135
chr11-90764427-INS-490
chr11-90805401-INS-235
chr11-90809304-INS-247
chr11-90824857-INS-105
chr11-90847158-INS-50
chr11-90867325-INS-325
chr11-90881173-INS-106
chr11-90923946-INS-362
chr11-90979698-INS-730
chr11-91002374-INS-521
chr11-91029876-INS-194
chr11-91033159-INS-11269
chr11-91043546-INS-3992
chr11-91048654-INS-166
chr11-91069701-INS-50
chr11-91079459-INS-7129
chr11-91084153-INS-376
chr11-91098534-INS-52
chr11-91122631-INS-247
chr11-91123301-INS-179
chr11-91133364-INS-155
chr11-91135252-INS-1089
chr11-91151681-INS-6360
chr11-91164849-INS-50
chr11-91165471-INS-1669
chr11-91211857-INS-87
chr11-91215117-INS-1392
chr11-91229893-INS-6464
chr11-91245278-INS-64
chr11-91245741-INS-2680
chr11-91249876-INS-108
chr11-91263572-INS-7568
chr11-91276340-INS-4553
chr11-91280694-INS-52
chr11-91280725-INS-160
chr11-91327367-INS-97
chr11-91332376-INS-214
chr11-91391660-INS-63
chr11-91394883-INS-1532
chr11-91412134-INS-1307
chr11-91436789-INS-4719
chr11-91440873-INS-6620
chr11-91449247-INS-58
chr11-91449686-INS-6722
chr11-91458257-INS-1539
chr11-91458680-INS-143
chr11-91458732-INS-71
chr11-91487857-INS-6316
chr11-91491601-INS-199
chr11-91598160-INS-54
chr11-91606187-INS-13997
chr11-91606187-INS-6187
chr11-91625384-INS-2345
chr11-91645900-INS-6022
chr11-91658925-INS-178
chr11-91674712-INS-143
chr11-91674712-INS-59
chr11-91676501-INS-60
chr11-91695470-INS-56
chr11-91698221-INS-6461
chr11-91703897-INS-73
chr11-91705607-INS-4233
chr11-91717058-INS-924
chr11-91725808-INS-2969
chr11-91726864-INS-54
chr11-91730633-INS-73
chr11-91807565-INS-216
chr11-91846313-INS-170
chr11-91860011-INS-5042

chr11-91876220-INS-6137
chr11-91941627-INS-538
chr11-91989323-INS-184
chr11-92016722-INS-7091
chr11-92025359-INS-841
chr11-92035704-INS-64
chr11-92129377-INS-174
chr11-92213314-INS-934
chr11-92218363-INS-328
chr11-92218544-INS-736
chr11-92248024-INS-184
chr11-92248024-INS-76
chr11-92262667-INS-460
chr11-92284435-INS-50
chr11-92411247-INS-200
chr11-92415646-INS-56
chr11-92432026-INS-328
chr11-92499450-INS-52
chr11-92531478-INS-75
chr11-92544555-INS-169
chr11-92544561-INS-55
chr11-92573100-INS-52
chr11-92577687-INS-232
chr11-92644062-INS-252
chr11-92686378-INS-9086
chr11-92703415-INS-1680
chr11-92725715-INS-207
chr11-92731988-INS-56
chr11-92732178-INS-4918
chr11-92772826-INS-670
chr11-92784152-INS-581
chr11-92803281-INS-441
chr11-92810722-INS-4201
chr11-92875420-INS-182
chr11-92948030-INS-6799
chr11-93009795-INS-3376
chr11-93043698-INS-6435
chr11-93054005-INS-58
chr11-93064240-INS-194
chr11-93098587-INS-50
chr11-93100276-INS-50
chr11-93131739-INS-81
chr11-93156271-INS-1235
chr11-93162256-INS-1161
chr11-93180294-INS-1442
chr11-93208128-INS-366
chr11-93219375-INS-7583
chr11-93300461-INS-366
chr11-93356246-INS-352
chr11-93361517-INS-62
chr11-93399619-INS-150
chr11-93399641-INS-58
chr11-93408189-INS-52
chr11-93408413-INS-72
chr11-93425918-INS-54
chr11-93452344-INS-139
chr11-93495682-INS-167
chr11-93505116-INS-11337
chr11-93505678-INS-50
chr11-93521538-INS-51
chr11-93538112-INS-6649
chr11-93559329-INS-50
chr11-93577900-INS-74
chr11-93578086-INS-6632
chr11-93588459-INS-50
chr11-93593190-INS-112
chr11-93630592-INS-6100
chr11-93657813-INS-465
chr11-93661124-INS-1460
chr11-93718550-INS-6701
chr11-93747387-INS-6407
chr11-93780047-INS-270
chr11-93845449-INS-57
chr11-93852343-INS-84
chr11-93852720-INS-211
chr11-93871372-INS-199
chr11-93877042-INS-209

chr11-93893269-INS-123
chr11-93982468-INS-50
chr11-94036926-INS-960
chr11-94039031-INS-54
chr11-94061123-INS-55
chr11-94082995-INS-52
chr11-94126086-INS-213
chr11-94195996-INS-401
chr11-94196795-INS-222
chr11-94200952-INS-406
chr11-94244896-INS-74
chr11-94282499-INS-124
chr11-94296637-INS-50
chr11-94298121-INS-52
chr11-94326530-INS-96
chr11-94396234-INS-209
chr11-94426958-INS-136
chr11-94448825-INS-271
chr11-94474393-INS-210
chr11-94475239-INS-66
chr11-94491035-INS-64
chr11-94497974-INS-213
chr11-94505000-INS-407
chr11-94599596-INS-1569
chr11-94605517-INS-442
chr11-94605666-INS-566
chr11-94627218-INS-192
chr11-94627249-INS-403
chr11-94627265-INS-66
chr11-94631014-INS-50
chr11-94631049-INS-109
chr11-94655283-INS-213
chr11-94680832-INS-208
chr11-94681263-INS-377
chr11-94681273-INS-107
chr11-94781279-INS-89
chr11-94782971-INS-78
chr11-94873517-INS-83
chr11-94873775-INS-65
chr11-94874102-INS-135
chr11-94874204-INS-52
chr11-94901065-INS-52
chr11-94904380-INS-600
chr11-94904487-INS-77
chr11-94915072-INS-67
chr11-94940648-INS-106
chr11-95018244-INS-68
chr11-95055289-INS-58
chr11-95067869-INS-184
chr11-95079712-INS-50
chr11-95108803-INS-186
chr11-95147307-INS-224
chr11-95198188-INS-180
chr11-95314522-INS-106
chr11-95427926-INS-61
chr11-95444516-INS-160
chr11-95452145-INS-175
chr11-95456971-INS-70
chr11-95482728-INS-54
chr11-95528680-INS-75
chr11-95580969-INS-231
chr11-95594693-INS-71
chr11-95597731-INS-74
chr11-95598201-INS-186
chr11-95623274-INS-1197
chr11-95681081-INS-56
chr11-95801715-INS-1034
chr11-95805005-INS-50
chr11-95808782-INS-161
chr11-95811808-INS-203
chr11-95812602-INS-73
chr11-95830185-INS-64
chr11-95830193-INS-181
chr11-95835364-INS-55
chr11-95838465-INS-203
chr11-95838502-INS-167
chr11-95845858-INS-181

chr11-95863068-INS-250
chr11-95863096-INS-50
chr11-95913765-INS-60
chr11-95953552-INS-204
chr11-95990821-INS-147
chr11-96023757-INS-606
chr11-96057445-INS-202
chr11-96125991-INS-81
chr11-96131482-INS-74
chr11-96189670-INS-58
chr11-96278160-INS-177
chr11-96283938-INS-200
chr11-96293935-INS-81
chr11-96309563-INS-200
chr11-96322198-INS-3224
chr11-96363691-INS-76
chr11-96364200-INS-66
chr11-96393092-INS-84
chr11-96412912-INS-2146
chr11-96452868-INS-116
chr11-96453034-INS-52
chr11-96503228-INS-114
chr11-96509222-INS-68
chr11-96554262-INS-50
chr11-96570520-INS-197
chr11-96593556-INS-60
chr11-96594072-INS-116
chr11-96594072-INS-54
chr11-96619719-INS-198
chr11-96625143-INS-159
chr11-96625213-INS-64
chr11-96642812-INS-225
chr11-96642826-INS-103
chr11-96654369-INS-5219
chr11-96670615-INS-447
chr11-96672715-INS-64
chr11-96672950-INS-177
chr11-96674032-INS-64
chr11-96737527-INS-50
chr11-96746581-INS-6475
chr11-96757247-INS-1186
chr11-96757309-INS-51
chr11-96760923-INS-51
chr11-96770804-INS-316
chr11-96774070-INS-50
chr11-96795421-INS-74
chr11-96822156-INS-95
chr11-96822222-INS-195
chr11-96842134-INS-8689
chr11-96897749-INS-60
chr11-96906856-INS-1999
chr11-96964551-INS-86
chr11-96990416-INS-106
chr11-96992215-INS-65
chr11-97087106-INS-1430
chr11-97091524-INS-3385
chr11-97092472-INS-207
chr11-97110171-INS-107
chr11-97195103-INS-181
chr11-97221790-INS-1767
chr11-97270439-INS-213
chr11-97279253-INS-214
chr11-97316061-INS-248
chr11-97323870-INS-206
chr11-97345667-INS-66
chr11-97398034-INS-151
chr11-97418236-INS-94
chr11-97484027-INS-197
chr11-97492972-INS-51
chr11-97510537-INS-82
chr11-97511815-INS-650
chr11-97512001-INS-60
chr11-97523127-INS-200
chr11-97524779-INS-67
chr11-97642947-INS-207
chr11-97652574-INS-205
chr11-97653234-INS-204

chr11-97679507-INS-63
chr11-97687428-INS-52
chr11-97745350-INS-230
chr11-97777080-INS-198
chr11-97777695-INS-537
chr11-97781486-INS-170
chr11-97785285-INS-194
chr11-97799636-INS-50
chr11-97803605-INS-81
chr11-97882426-INS-228
chr11-97970621-INS-250
chr11-97976973-INS-210
chr11-97978889-INS-109
chr11-97997610-INS-94
chr11-98015685-INS-232
chr11-98016372-INS-174
chr11-98033892-INS-51
chr11-98037960-INS-60
chr11-98085325-INS-207
chr11-98087561-INS-218
chr11-98088236-INS-205
chr11-98110240-INS-237
chr11-98192954-INS-208
chr11-98204337-INS-52
chr11-98272333-INS-126
chr11-98292464-INS-146
chr11-98389247-INS-248
chr11-98452118-INS-66
chr11-98496607-INS-66
chr11-98498338-INS-70
chr11-98501688-INS-436
chr11-98514550-INS-3459
chr11-98536803-INS-222
chr11-98551159-INS-50
chr11-98710190-INS-198
chr11-98739567-INS-66
chr11-98747902-INS-208
chr11-98753621-INS-5673
chr11-98882562-INS-170
chr11-98923254-INS-233
chr11-98940762-INS-4362
chr11-98985157-INS-60
chr11-99006134-INS-204
chr11-99006134-INS-456
chr11-99086628-INS-3224
chr11-99090007-INS-151
chr11-99122588-INS-80
chr11-99132704-INS-983
chr11-99151299-INS-50
chr11-99156910-INS-3503
chr11-99157709-INS-57
chr11-99160970-INS-152
chr11-99171324-INS-288
chr11-99184927-INS-184
chr11-99185093-INS-980
chr11-99186874-INS-70
chr11-99195141-INS-4441
chr11-99245496-INS-249
chr11-99259642-INS-226
chr11-99297650-INS-168
chr11-99343246-INS-188
chr11-99349371-INS-84
chr11-99390158-INS-67
chr11-99411477-INS-77
chr11-99418124-INS-251
chr11-99424407-INS-72
chr11-99440208-INS-120
chr11-99451477-INS-953
chr11-99454251-INS-2699
chr11-99491703-INS-60
chr11-99527819-INS-50
chr11-99536497-INS-4797
chr11-99551076-INS-51
chr11-99589776-INS-72
chr11-99611714-INS-52
chr11-99625083-INS-5225
chr11-99667709-INS-143

chr11-99668021-INS-127
chr11-99693127-INS-233
chr11-99698660-INS-216
chr11-99842842-INS-71
chr11-99845629-INS-5484
chr11-99856339-INS-60
chr11-99856582-INS-121
chr11-99875485-INS-6017
chr11-99932958-INS-200
chr11-99981504-INS-152
chr11-99981528-INS-64
chr11-100024189-INS-60
chr11-100050272-INS-99
chr11-100051825-INS-129
chr11-100052075-INS-65
chr11-100063479-INS-170
chr11-100106051-INS-204
chr11-100128509-INS-127
chr11-100131296-INS-64
chr11-100135189-INS-58
chr11-100154339-INS-70
chr11-100163649-INS-80
chr11-100183904-INS-1140
chr11-100183988-INS-3774
chr11-100188269-INS-1000
chr11-100195313-INS-2140
chr11-100200054-INS-54
chr11-100202380-INS-4807
chr11-100205046-INS-172
chr11-100226495-INS-115
chr11-100227163-INS-70
chr11-100237833-INS-78
chr11-100255822-INS-75
chr11-100292172-INS-88
chr11-100310909-INS-53
chr11-100323387-INS-165
chr11-100398196-INS-213
chr11-100428163-INS-181
chr11-100432114-INS-150
chr11-100432169-INS-60
chr11-100432914-INS-88
chr11-100503510-INS-70
chr11-100505244-INS-248
chr11-100511780-INS-68
chr11-100527248-INS-729
chr11-100546181-INS-51
chr11-100572482-INS-203
chr11-100651035-INS-66
chr11-100657067-INS-142
chr11-100658450-INS-54
chr11-100667854-INS-763
chr11-100674148-INS-216
chr11-100682280-INS-66
chr11-100698367-INS-55
chr11-100698400-INS-220
chr11-100704381-INS-50
chr11-100748286-INS-62
chr11-100748299-INS-143
chr11-100786152-INS-191
chr11-100795391-INS-90
chr11-100809710-INS-62
chr11-100810608-INS-53
chr11-100826140-INS-115
chr11-100827329-INS-210
chr11-100877810-INS-108
chr11-100896093-INS-120
chr11-100896095-INS-313
chr11-100927268-INS-50
chr11-101016520-INS-72
chr11-101029562-INS-181
chr11-101049815-INS-298
chr11-101065601-INS-56
chr11-101103420-INS-76
chr11-101103448-INS-92
chr11-101127552-INS-144
chr11-101163782-INS-66
chr11-101195566-INS-128

chr11-101265138-INS-52
chr11-101277202-INS-168
chr11-101300083-INS-59
chr11-101303497-INS-50
chr11-101304093-INS-67
chr11-101309320-INS-103
chr11-101318523-INS-179
chr11-101326743-INS-182
chr11-101343983-INS-100
chr11-101389437-INS-64
chr11-101438513-INS-289
chr11-101439398-INS-212
chr11-101448785-INS-54
chr11-101450858-INS-171
chr11-101479268-INS-324
chr11-101494244-INS-4757
chr11-101495905-INS-348
chr11-101496155-INS-193
chr11-101544771-INS-5837
chr11-101584921-INS-184
chr11-101592779-INS-53
chr11-101631093-INS-262
chr11-101631373-INS-1516
chr11-101643929-INS-59
chr11-101650428-INS-207
chr11-101651817-INS-58
chr11-101651877-INS-194
chr11-101715501-INS-78
chr11-101734351-INS-210
chr11-101744020-INS-102
chr11-101744366-INS-50
chr11-101785454-INS-52
chr11-101790138-INS-67
chr11-101790910-INS-637
chr11-101808091-INS-51
chr11-101832414-INS-55
chr11-101847720-INS-12302
chr11-101870076-INS-68
chr11-101885040-INS-58
chr11-101903116-INS-128
chr11-101955711-INS-182
chr11-101955733-INS-64
chr11-101963315-INS-86
chr11-101974053-INS-186
chr11-101985375-INS-204
chr11-102028853-INS-504
chr11-102067536-INS-61
chr11-102067542-INS-128
chr11-102067836-INS-201
chr11-102078921-INS-208
chr11-102104006-INS-80
chr11-102127750-INS-147
chr11-102128397-INS-227
chr11-102161097-INS-62
chr11-102173079-INS-230
chr11-102198419-INS-227
chr11-102220145-INS-201
chr11-102223769-INS-175
chr11-102237557-INS-342
chr11-102262092-INS-50
chr11-102279478-INS-54
chr11-102384787-INS-254
chr11-102423076-INS-137
chr11-102423079-INS-62
chr11-102453647-INS-352
chr11-102480749-INS-82
chr11-102490039-INS-212
chr11-102491538-INS-197
chr11-102492308-INS-68
chr11-102504292-INS-115
chr11-102510088-INS-197
chr11-102525961-INS-165
chr11-102532340-INS-536
chr11-102532945-INS-235
chr11-102537771-INS-231
chr11-102541650-INS-193
chr11-102541722-INS-52

chr11-102599050-INS-575
chr11-102639750-INS-121
chr11-102680309-INS-60
chr11-102699065-INS-67
chr11-102723956-INS-221
chr11-102738477-INS-490
chr11-102740479-INS-213
chr11-102754129-INS-159
chr11-102755197-INS-211
chr11-102758314-INS-157
chr11-102763700-INS-295
chr11-102764262-INS-3351
chr11-102782042-INS-392
chr11-102807649-INS-118
chr11-102840636-INS-55
chr11-102871230-INS-50
chr11-102876416-INS-64
chr11-102876898-INS-465
chr11-102904882-INS-700
chr11-102932459-INS-163
chr11-102950395-INS-1005
chr11-103045717-INS-98
chr11-103046524-INS-172
chr11-103055943-INS-115
chr11-103055943-INS-290
chr11-103055946-INS-3620
chr11-103141935-INS-223
chr11-103145953-INS-52
chr11-103146658-INS-207
chr11-103166004-INS-115
chr11-103177062-INS-299
chr11-103184070-INS-6145
chr11-103205549-INS-132
chr11-103228727-INS-53
chr11-103233644-INS-104
chr11-103248198-INS-5267
chr11-103279364-INS-390
chr11-103311662-INS-515
chr11-103344189-INS-206
chr11-103398836-INS-6401
chr11-103399054-INS-50
chr11-103409917-INS-58
chr11-103413619-INS-5554
chr11-103416074-INS-410
chr11-103417842-INS-1743
chr11-103485721-INS-7085
chr11-103530442-INS-452
chr11-103558605-INS-179
chr11-103563985-INS-52
chr11-103565973-INS-52
chr11-103566348-INS-145
chr11-103572649-INS-139
chr11-103657042-INS-196
chr11-103657281-INS-184
chr11-103728091-INS-88
chr11-103734074-INS-89
chr11-103734897-INS-252
chr11-103768458-INS-52
chr11-103778137-INS-524
chr11-103805297-INS-901
chr11-103811744-INS-432
chr11-103862197-INS-145
chr11-103886595-INS-1892
chr11-103887634-INS-194
chr11-103915104-INS-56
chr11-103934266-INS-56
chr11-103935381-INS-229
chr11-103942375-INS-51
chr11-103949857-INS-207
chr11-103960575-INS-86
chr11-103983120-INS-54
chr11-103985851-INS-52
chr11-104012164-INS-105
chr11-104051435-INS-58
chr11-104076906-INS-1455
chr11-104092245-INS-179
chr11-104095454-INS-63

chr11-104116454-INS-50
chr11-104154920-INS-64
chr11-104172211-INS-2999
chr11-104205561-INS-77
chr11-104206806-INS-212
chr11-104214708-INS-251
chr11-104214904-INS-182
chr11-104231798-INS-74
chr11-104231842-INS-194
chr11-104242835-INS-181
chr11-104266645-INS-206
chr11-104280947-INS-129
chr11-104309235-INS-208
chr11-104321795-INS-126
chr11-104375467-INS-242
chr11-104377294-INS-158
chr11-104380375-INS-148
chr11-104380383-INS-65
chr11-104400329-INS-220
chr11-104479131-INS-220
chr11-104483979-INS-88
chr11-104496296-INS-51
chr11-104496553-INS-135
chr11-104504086-INS-108
chr11-104520397-INS-106
chr11-104520447-INS-332
chr11-104527736-INS-180
chr11-104566358-INS-340
chr11-104572445-INS-6469
chr11-104599060-INS-90
chr11-104611881-INS-99
chr11-104611958-INS-594
chr11-104616241-INS-211
chr11-104618670-INS-99
chr11-104618947-INS-120
chr11-104618951-INS-765
chr11-104622446-INS-212
chr11-104632988-INS-50
chr11-104638679-INS-210
chr11-104665560-INS-6415
chr11-104670966-INS-60
chr11-104680581-INS-6325
chr11-104685780-INS-50
chr11-104698042-INS-128
chr11-104705569-INS-270
chr11-104716860-INS-1727
chr11-104757364-INS-82
chr11-104770489-INS-431
chr11-104825759-INS-93
chr11-104859224-INS-235
chr11-104879914-INS-1893
chr11-104901898-INS-5162
chr11-104929628-INS-76
chr11-104954641-INS-51
chr11-104954721-INS-235
chr11-104965143-INS-66
chr11-105001432-INS-1188
chr11-105004075-INS-3034
chr11-105007269-INS-108
chr11-105035633-INS-164
chr11-105037536-INS-170
chr11-105051287-INS-72
chr11-105060068-INS-146
chr11-105060990-INS-370
chr11-105084631-INS-201
chr11-105085934-INS-66
chr11-105161648-INS-676
chr11-105199369-INS-741
chr11-105226549-INS-89
chr11-105230710-INS-149
chr11-105233292-INS-56
chr11-105268266-INS-66
chr11-105327684-INS-377
chr11-105377457-INS-169
chr11-105383613-INS-120
chr11-105391700-INS-83
chr11-105393874-INS-64

chr11-105400336-INS-50
chr11-105505849-INS-205
chr11-105568681-INS-52
chr11-105602621-INS-428
chr11-105636973-INS-6608
chr11-105646530-INS-126
chr11-105688100-INS-6186
chr11-105699940-INS-6877
chr11-105704958-INS-1860
chr11-105735052-INS-229
chr11-105803209-INS-81
chr11-105819430-INS-60
chr11-105842185-INS-58
chr11-105845916-INS-189
chr11-105848806-INS-324
chr11-105869070-INS-626
chr11-105916918-INS-66
chr11-105930055-INS-210
chr11-105935269-INS-1224
chr11-105940368-INS-192
chr11-105943119-INS-174
chr11-105943991-INS-101
chr11-105967362-INS-104
chr11-106016209-INS-236
chr11-106016328-INS-203
chr11-106076259-INS-169
chr11-106118572-INS-216
chr11-106193537-INS-192
chr11-106196707-INS-235
chr11-106213237-INS-107
chr11-106214202-INS-146
chr11-106229686-INS-50
chr11-106231710-INS-88
chr11-106246407-INS-2977
chr11-106250536-INS-385
chr11-106260481-INS-198
chr11-106263967-INS-204
chr11-106267406-INS-50
chr11-106319207-INS-210
chr11-106338906-INS-196
chr11-106365054-INS-252
chr11-106365399-INS-72
chr11-106391276-INS-215
chr11-106399476-INS-1088
chr11-106399635-INS-433
chr11-106439061-INS-208
chr11-106462406-INS-228
chr11-106480810-INS-261
chr11-106482463-INS-224
chr11-106500052-INS-217
chr11-106501768-INS-202
chr11-106506097-INS-126
chr11-106507494-INS-7088
chr11-106511583-INS-120
chr11-106531195-INS-68
chr11-106551998-INS-65
chr11-106558684-INS-57
chr11-106585552-INS-202
chr11-106586056-INS-199
chr11-106635000-INS-200
chr11-106646176-INS-78
chr11-106663231-INS-71
chr11-106682431-INS-58
chr11-106685138-INS-203
chr11-106697356-INS-1359
chr11-106697428-INS-246
chr11-106727504-INS-209
chr11-106732981-INS-204
chr11-106756562-INS-104
chr11-106767296-INS-52
chr11-106767399-INS-5095
chr11-106790493-INS-149
chr11-106820957-INS-176
chr11-106846055-INS-92
chr11-106855298-INS-152
chr11-106859308-INS-272
chr11-106904665-INS-213

chr11-106907615-INS-136
chr11-106917240-INS-58
chr11-106930650-INS-51
chr11-106939513-INS-105
chr11-106940064-INS-1581
chr11-106940064-INS-212
chr11-106950673-INS-728
chr11-107000412-INS-148
chr11-107006247-INS-50
chr11-107027468-INS-1191
chr11-107029876-INS-2936
chr11-107034570-INS-136
chr11-107034858-INS-52
chr11-107037068-INS-526
chr11-107045760-INS-85
chr11-107045779-INS-204
chr11-107045963-INS-789
chr11-107048179-INS-53
chr11-107051284-INS-1888
chr11-107096067-INS-203
chr11-107122477-INS-198
chr11-107161341-INS-55
chr11-107164218-INS-352
chr11-107172192-INS-177
chr11-107191635-INS-197
chr11-107216062-INS-237
chr11-107231507-INS-208
chr11-107272551-INS-107
chr11-107300213-INS-63
chr11-107304975-INS-162
chr11-107305767-INS-173
chr11-107312090-INS-530
chr11-107319758-INS-490
chr11-107358697-INS-91
chr11-107362116-INS-54
chr11-107368210-INS-241
chr11-107371821-INS-213
chr11-107372605-INS-210
chr11-107399296-INS-523
chr11-107405485-INS-50
chr11-107406956-INS-141
chr11-107410158-INS-211
chr11-107418343-INS-287
chr11-107419852-INS-325
chr11-107419853-INS-104
chr11-107423242-INS-214
chr11-107425834-INS-102
chr11-107428007-INS-210
chr11-107428646-INS-50
chr11-107432484-INS-52
chr11-107442036-INS-214
chr11-107450466-INS-193
chr11-107451259-INS-59
chr11-107452362-INS-112
chr11-107453115-INS-217
chr11-107470824-INS-177
chr11-107497544-INS-260
chr11-107501099-INS-87
chr11-107508454-INS-95
chr11-107508864-INS-52
chr11-107513482-INS-214
chr11-107523515-INS-193
chr11-107567654-INS-223
chr11-107570533-INS-158
chr11-107591952-INS-270
chr11-107597509-INS-58
chr11-107697554-INS-190
chr11-107698005-INS-483
chr11-107741474-INS-78
chr11-107743303-INS-197
chr11-107770588-INS-396
chr11-107779872-INS-128
chr11-107845362-INS-80
chr11-107845551-INS-920
chr11-107881563-INS-368
chr11-107904442-INS-115
chr11-107904442-INS-57

chr11-107974258-INS-701
chr11-107980439-INS-52
chr11-107983242-INS-348
chr11-107985572-INS-63
chr11-108058614-INS-64
chr11-108063306-INS-60
chr11-108149507-INS-84
chr11-108210167-INS-72
chr11-108213719-INS-51
chr11-108254127-INS-124
chr11-108311758-INS-80
chr11-108333395-INS-102
chr11-108351372-INS-168
chr11-108353707-INS-2927
chr11-108359136-INS-6559
chr11-108364888-INS-125
chr11-108368737-INS-6547
chr11-108379651-INS-7153
chr11-108394991-INS-2635
chr11-108413499-INS-6748
chr11-108442750-INS-379
chr11-108470952-INS-65
chr11-108471947-INS-198
chr11-108480624-INS-204
chr11-108485351-INS-62
chr11-108547003-INS-203
chr11-108555281-INS-2070
chr11-108569863-INS-826
chr11-108650592-INS-52
chr11-108652729-INS-5904
chr11-108660965-INS-360
chr11-108704195-INS-6998
chr11-108708484-INS-102
chr11-108708517-INS-186
chr11-108770262-INS-159
chr11-108770535-INS-114
chr11-108770677-INS-491
chr11-108859696-INS-130
chr11-108898339-INS-50
chr11-108898715-INS-72
chr11-108898873-INS-346
chr11-108920647-INS-53
chr11-108921318-INS-2936
chr11-108923012-INS-1592
chr11-108924355-INS-316
chr11-108939096-INS-50
chr11-108967074-INS-50
chr11-109000128-INS-102
chr11-109030043-INS-60
chr11-109033845-INS-358
chr11-109056120-INS-1028
chr11-109057109-INS-1890
chr11-109067980-INS-64
chr11-109087975-INS-56
chr11-109088569-INS-60
chr11-109108636-INS-77
chr11-109127517-INS-157
chr11-109147065-INS-280
chr11-109173542-INS-66
chr11-109179500-INS-162
chr11-109206257-INS-8121
chr11-109212733-INS-89
chr11-109232217-INS-88
chr11-109245574-INS-460
chr11-109248383-INS-108
chr11-109248961-INS-67
chr11-109257931-INS-92
chr11-109290470-INS-201
chr11-109315116-INS-228
chr11-109322168-INS-90
chr11-109333431-INS-4834
chr11-109338721-INS-569
chr11-109344883-INS-230
chr11-109356829-INS-66
chr11-109407038-INS-81
chr11-109407149-INS-526
chr11-109430651-INS-50

chr11-109438449-INS-52
chr11-109439723-INS-50
chr11-109442389-INS-168
chr11-109445871-INS-126
chr11-109446532-INS-120
chr11-109450997-INS-3695
chr11-109471629-INS-212
chr11-109484334-INS-110
chr11-109508054-INS-140
chr11-109536124-INS-64
chr11-109552570-INS-89
chr11-109579599-INS-3441
chr11-109582112-INS-2886
chr11-109598296-INS-56
chr11-109635771-INS-484
chr11-109635829-INS-204
chr11-109640363-INS-108
chr11-109671068-INS-50
chr11-109705291-INS-56
chr11-109717915-INS-2743
chr11-109719361-INS-192
chr11-109746377-INS-499
chr11-109752322-INS-116
chr11-109756095-INS-451
chr11-109760525-INS-212
chr11-109761803-INS-53
chr11-109768838-INS-98
chr11-109769233-INS-144
chr11-109770786-INS-352
chr11-109771890-INS-60
chr11-109805756-INS-96
chr11-109826252-INS-185
chr11-109830072-INS-83
chr11-109878871-INS-5609
chr11-109894663-INS-4095
chr11-109959380-INS-50
chr11-109961393-INS-126
chr11-109961764-INS-57
chr11-109981544-INS-139
chr11-109981564-INS-55
chr11-109987631-INS-60
chr11-110001910-INS-6431
chr11-110004280-INS-50
chr11-110008451-INS-56
chr11-110008458-INS-168
chr11-110009127-INS-84
chr11-110015334-INS-1456
chr11-110025489-INS-845
chr11-110031016-INS-111
chr11-110037643-INS-55
chr11-110050655-INS-4716
chr11-110052471-INS-923
chr11-110052499-INS-181
chr11-110070788-INS-363
chr11-110092575-INS-1430
chr11-110092577-INS-503
chr11-110098732-INS-62
chr11-110111459-INS-51
chr11-110119682-INS-560
chr11-110139988-INS-254
chr11-110163470-INS-52
chr11-110195458-INS-498
chr11-110215459-INS-4644
chr11-110229366-INS-328
chr11-110307870-INS-54
chr11-110334561-INS-55
chr11-110344684-INS-86
chr11-110354451-INS-418
chr11-110366318-INS-188
chr11-110408068-INS-355
chr11-110418729-INS-141
chr11-110418729-INS-307
chr11-110429631-INS-249
chr11-110450211-INS-6374
chr11-110458636-INS-556
chr11-110466739-INS-7660
chr11-110475167-INS-306

chr11-110481604-INS-321
chr11-110495372-INS-67
chr11-110497608-INS-112
chr11-110502446-INS-58
chr11-110502694-INS-50
chr11-110516078-INS-50
chr11-110525144-INS-455
chr11-110525213-INS-94
chr11-110557172-INS-866
chr11-110561399-INS-412
chr11-110567795-INS-300
chr11-110574060-INS-339
chr11-110584516-INS-213
chr11-110596453-INS-60
chr11-110605255-INS-104
chr11-110613203-INS-636
chr11-110613757-INS-197
chr11-110613866-INS-243
chr11-110648321-INS-469
chr11-110668871-INS-52
chr11-110712848-INS-417
chr11-110734766-INS-486
chr11-110741816-INS-143
chr11-110755933-INS-170
chr11-110756795-INS-6092
chr11-110839349-INS-1887
chr11-110840139-INS-50
chr11-110860129-INS-50
chr11-110861789-INS-1203
chr11-110937425-INS-110
chr11-110941188-INS-7701
chr11-110983994-INS-5318
chr11-110988648-INS-70
chr11-111098257-INS-56
chr11-111117366-INS-354
chr11-111121062-INS-88
chr11-111146023-INS-479
chr11-111159906-INS-3207
chr11-111185070-INS-270
chr11-111196920-INS-75
chr11-111213930-INS-50
chr11-111273021-INS-87
chr11-111273440-INS-66
chr11-111273690-INS-76
chr11-111307949-INS-208
chr11-111327166-INS-78
chr11-111331814-INS-60
chr11-111332423-INS-310
chr11-111333156-INS-280
chr11-111366110-INS-57
chr11-111391072-INS-626
chr11-111403027-INS-51
chr11-111419282-INS-437
chr11-111445687-INS-144
chr11-111466609-INS-1831
chr11-111467641-INS-1069
chr11-111487357-INS-52
chr11-111493454-INS-351
chr11-111513091-INS-70
chr11-111532842-INS-120
chr11-111551642-INS-84
chr11-111552676-INS-56
chr11-111566507-INS-61
chr11-111566930-INS-60
chr11-111567848-INS-204
chr11-111569056-INS-53
chr11-111580275-INS-54
chr11-111603315-INS-60
chr11-111628979-INS-589
chr11-111636780-INS-1030
chr11-111648811-INS-6471
chr11-111653291-INS-58
chr11-111671724-INS-1053
chr11-111682668-INS-1335
chr11-111699854-INS-72
chr11-111710216-INS-50
chr11-111728462-INS-64

chr11-111740943-INS-50
chr11-111768542-INS-50
chr11-111784064-INS-110
chr11-111796932-INS-52
chr11-111844852-INS-6887
chr11-111889171-INS-3560
chr11-111898334-INS-60
chr11-111899684-INS-50
chr11-111913170-INS-58
chr11-111913586-INS-54
chr11-111922894-INS-6149
chr11-111923335-INS-416
chr11-111967139-INS-79
chr11-111992781-INS-219
chr11-111992955-INS-60
chr11-111995263-INS-7553
chr11-112056229-INS-1429
chr11-112087852-INS-66
chr11-112106532-INS-54
chr11-112107433-INS-185
chr11-112111007-INS-1997
chr11-112111186-INS-127
chr11-112111401-INS-60
chr11-112114640-INS-78
chr11-112172596-INS-121
chr11-112173159-INS-54
chr11-112174125-INS-68
chr11-112175181-INS-84
chr11-112178898-INS-86
chr11-112194625-INS-56
chr11-112203236-INS-75
chr11-112231287-INS-64
chr11-112231361-INS-122
chr11-112234186-INS-1114
chr11-112270936-INS-220
chr11-112270939-INS-70
chr11-112288421-INS-92
chr11-112307555-INS-62
chr11-112309035-INS-491
chr11-112335983-INS-86
chr11-112336508-INS-148
chr11-112336581-INS-62
chr11-112342285-INS-60
chr11-112355836-INS-52
chr11-112456848-INS-306
chr11-112482528-INS-72
chr11-112511510-INS-55
chr11-112512292-INS-108
chr11-112528594-INS-75
chr11-112528601-INS-166
chr11-112538440-INS-60
chr11-112540959-INS-93
chr11-112565413-INS-56
chr11-112585247-INS-633
chr11-112585252-INS-259
chr11-112596103-INS-1203
chr11-112614875-INS-92
chr11-112618757-INS-50
chr11-112631953-INS-499
chr11-112660321-INS-54
chr11-112677151-INS-52
chr11-112679693-INS-57
chr11-112680064-INS-50
chr11-112728206-INS-51
chr11-112744086-INS-2627
chr11-112749968-INS-51
chr11-112752773-INS-65
chr11-112753530-INS-59
chr11-112778708-INS-7252
chr11-112787543-INS-185
chr11-112809194-INS-149
chr11-112809238-INS-61
chr11-112868208-INS-57
chr11-112885621-INS-59
chr11-112893709-INS-386
chr11-112898646-INS-54
chr11-112929230-INS-72

chr11-112952792-INS-50
chr11-113025108-INS-75
chr11-113064441-INS-102
chr11-113078240-INS-83
chr11-113101555-INS-76
chr11-113110997-INS-4959
chr11-113163340-INS-56
chr11-113197162-INS-126
chr11-113197276-INS-287
chr11-113210735-INS-71
chr11-113230280-INS-3705
chr11-113231343-INS-377
chr11-113236977-INS-50
chr11-113312880-INS-126
chr11-113341752-INS-80
chr11-113382302-INS-142
chr11-113434662-INS-252
chr11-113434774-INS-112
chr11-113436072-INS-206
chr11-113466794-INS-60
chr11-113469414-INS-193
chr11-113469709-INS-4516
chr11-113507120-INS-116
chr11-113525243-INS-64
chr11-113525245-INS-315
chr11-113527999-INS-6685
chr11-113542204-INS-315
chr11-113560796-INS-128
chr11-113582347-INS-394
chr11-113603385-INS-50
chr11-113623830-INS-209
chr11-113681502-INS-504
chr11-113681608-INS-1408
chr11-113682065-INS-52
chr11-113692064-INS-2838
chr11-113705789-INS-147
chr11-113759112-INS-6848
chr11-113762628-INS-4050
chr11-113811065-INS-54
chr11-113994962-INS-52
chr11-114024869-INS-57
chr11-114032885-INS-121
chr11-114084494-INS-5971
chr11-114156585-INS-55
chr11-114195566-INS-109
chr11-114195726-INS-474
chr11-114210292-INS-1046
chr11-114216881-INS-54
chr11-114225647-INS-50
chr11-114226128-INS-532
chr11-114239824-INS-50
chr11-114260369-INS-101
chr11-114262778-INS-63
chr11-114280114-INS-1490
chr11-114287879-INS-50
chr11-114303042-INS-858
chr11-114325724-INS-50
chr11-114331501-INS-110
chr11-114365001-INS-227
chr11-114366983-INS-52
chr11-114373161-INS-2422
chr11-114374009-INS-60
chr11-114387942-INS-84
chr11-114395384-INS-204
chr11-114404725-INS-99
chr11-114431341-INS-358
chr11-114456919-INS-736
chr11-114470465-INS-58
chr11-114479637-INS-156
chr11-114480938-INS-480
chr11-114502012-INS-5528
chr11-114561561-INS-1175
chr11-114563646-INS-3962
chr11-114617273-INS-56
chr11-114622657-INS-68
chr11-114674395-INS-178
chr11-114686101-INS-52

chr11-114693530-INS-52
chr11-114713032-INS-124
chr11-114718465-INS-54
chr11-114723219-INS-200
chr11-114725278-INS-756
chr11-114730438-INS-58
chr11-114738501-INS-52
chr11-114749615-INS-6741
chr11-114750850-INS-199
chr11-114751540-INS-92
chr11-114779252-INS-90
chr11-114818953-INS-111
chr11-114829779-INS-528
chr11-114829792-INS-193
chr11-114847141-INS-7165
chr11-114848577-INS-51
chr11-114849090-INS-511
chr11-114857905-INS-346
chr11-114866247-INS-3794
chr11-114879237-INS-684
chr11-114889954-INS-768
chr11-114890045-INS-60
chr11-114899434-INS-489
chr11-114911528-INS-444
chr11-114922211-INS-258
chr11-114939348-INS-429
chr11-114951991-INS-52
chr11-114963376-INS-1449
chr11-114963392-INS-96
chr11-114977598-INS-76
chr11-114978860-INS-52
chr11-115005055-INS-198
chr11-115014681-INS-50
chr11-115086614-INS-72
chr11-115091358-INS-125
chr11-115185827-INS-55
chr11-115191898-INS-200
chr11-115198888-INS-97
chr11-115201911-INS-56
chr11-115202277-INS-136
chr11-115202354-INS-56
chr11-115206744-INS-60
chr11-115326061-INS-210
chr11-115335388-INS-76
chr11-115335444-INS-101
chr11-115343638-INS-50
chr11-115352281-INS-218
chr11-115369422-INS-200
chr11-115384086-INS-196
chr11-115416032-INS-131
chr11-115416810-INS-262
chr11-115434460-INS-182
chr11-115434495-INS-186
chr11-115442521-INS-8776
chr11-115471019-INS-106
chr11-115524501-INS-54
chr11-115546549-INS-218
chr11-115547311-INS-93
chr11-115549584-INS-66
chr11-115550472-INS-262
chr11-115559031-INS-100
chr11-115569085-INS-473
chr11-115597093-INS-175
chr11-115615882-INS-110
chr11-115621588-INS-2812
chr11-115626852-INS-212
chr11-115631612-INS-205
chr11-115632270-INS-1451
chr11-115659681-INS-187
chr11-115677888-INS-50
chr11-115677953-INS-350
chr11-115681494-INS-155
chr11-115734843-INS-9097
chr11-115736698-INS-2373
chr11-115737159-INS-142
chr11-115751727-INS-261
chr11-115766742-INS-151

chr11-115814568-INS-63
chr11-115814910-INS-108
chr11-115833981-INS-51
chr11-115847255-INS-76
chr11-115860368-INS-58
chr11-115878077-INS-175
chr11-115883643-INS-74
chr11-115887114-INS-114
chr11-115896695-INS-55
chr11-115907840-INS-53
chr11-115937758-INS-174
chr11-115985450-INS-232
chr11-116015190-INS-483
chr11-116016435-INS-198
chr11-116053488-INS-320
chr11-116053663-INS-72
chr11-116055085-INS-244
chr11-116060015-INS-58
chr11-116060332-INS-977
chr11-116126794-INS-201
chr11-116167424-INS-81
chr11-116170481-INS-52
chr11-116177521-INS-369
chr11-116185815-INS-480
chr11-116190497-INS-110
chr11-116251815-INS-62
chr11-116252060-INS-217
chr11-116265245-INS-91
chr11-116265390-INS-209
chr11-116287440-INS-50
chr11-116292562-INS-204
chr11-116348158-INS-60
chr11-116348328-INS-120
chr11-116358622-INS-59
chr11-116359183-INS-51
chr11-116366132-INS-205
chr11-116376280-INS-58
chr11-116376394-INS-203
chr11-116384382-INS-50
chr11-116393539-INS-387
chr11-116405672-INS-255
chr11-116412421-INS-218
chr11-116460642-INS-525
chr11-116588160-INS-152
chr11-116597105-INS-7070
chr11-116606820-INS-5250
chr11-116629814-INS-1466
chr11-116636222-INS-421
chr11-116636818-INS-139
chr11-116692834-INS-63
chr11-116698247-INS-100
chr11-116698307-INS-204
chr11-116781845-INS-377
chr11-116782128-INS-194
chr11-116783771-INS-201
chr11-116786645-INS-54
chr11-116868091-INS-60
chr11-116879000-INS-214
chr11-116883866-INS-5174
chr11-116893823-INS-509
chr11-116907151-INS-172
chr11-116914017-INS-2876
chr11-116923554-INS-132
chr11-116957980-INS-1748
chr11-116960590-INS-492
chr11-116977928-INS-97
chr11-116983982-INS-56
chr11-116986748-INS-480
chr11-117001219-INS-215
chr11-117002531-INS-65
chr11-117009011-INS-245
chr11-117015877-INS-206
chr11-117022679-INS-84
chr11-117031807-INS-96
chr11-117084661-INS-95
chr11-117112662-INS-190
chr11-117123722-INS-67

chr11-117128449-INS-138
chr11-117128793-INS-130
chr11-117128793-INS-267
chr11-117149399-INS-52
chr11-117171823-INS-54
chr11-117184373-INS-88
chr11-117192149-INS-141
chr11-117232152-INS-213
chr11-117238568-INS-2733
chr11-117260361-INS-209
chr11-117263943-INS-74
chr11-117267815-INS-52
chr11-117269254-INS-263
chr11-117270145-INS-3264
chr11-117292534-INS-128
chr11-117310079-INS-55
chr11-117313668-INS-206
chr11-117315838-INS-120
chr11-117343141-INS-120
chr11-117343256-INS-3833
chr11-117354655-INS-109
chr11-117358978-INS-197
chr11-117365800-INS-128
chr11-117366186-INS-198
chr11-117380982-INS-159
chr11-117383376-INS-50
chr11-117425491-INS-61
chr11-117425510-INS-165
chr11-117437225-INS-93
chr11-117438833-INS-127
chr11-117442839-INS-327
chr11-117446388-INS-176
chr11-117449331-INS-56
chr11-117467505-INS-154
chr11-117467543-INS-331
chr11-117483363-INS-203
chr11-117484850-INS-128
chr11-117528885-INS-1055
chr11-117568531-INS-60
chr11-117648130-INS-83
chr11-117707037-INS-109
chr11-117707212-INS-198
chr11-117711587-INS-230
chr11-117727945-INS-1160
chr11-117733827-INS-51
chr11-117744788-INS-170
chr11-117817580-INS-1444
chr11-117820026-INS-218
chr11-117835117-INS-1080
chr11-117835564-INS-107
chr11-117944405-INS-152
chr11-117959396-INS-135
chr11-117959396-INS-411
chr11-117968442-INS-1275
chr11-117968496-INS-225
chr11-118108796-INS-76
chr11-118140932-INS-1147
chr11-118182320-INS-181
chr11-118187459-INS-50
chr11-118203725-INS-52
chr11-118226239-INS-68
chr11-118226490-INS-72
chr11-118236215-INS-2688
chr11-118251213-INS-210
chr11-118257122-INS-120
chr11-118267369-INS-74
chr11-118271696-INS-52
chr11-118349213-INS-210
chr11-118403303-INS-57
chr11-118451594-INS-87
chr11-118466075-INS-84
chr11-118470291-INS-54
chr11-118471104-INS-68
chr11-118578798-INS-57
chr11-118591460-INS-150
chr11-118632545-INS-50
chr11-118697899-INS-158

chr11-118738717-INS-66
chr11-118773358-INS-253
chr11-118778107-INS-152
chr11-118778228-INS-104
chr11-118884261-INS-50
chr11-118951822-INS-54
chr11-118995317-INS-50
chr11-119007480-INS-101
chr11-119132051-INS-1121
chr11-119139248-INS-2498
chr11-119200189-INS-263
chr11-119225133-INS-50
chr11-119239912-INS-237
chr11-119281696-INS-212
chr11-119292599-INS-4473
chr11-119294407-INS-221
chr11-119317503-INS-54
chr11-119319670-INS-138
chr11-119402687-INS-174
chr11-119418735-INS-279
chr11-119420604-INS-98
chr11-119456462-INS-82
chr11-119473826-INS-437
chr11-119474114-INS-216
chr11-119504240-INS-802
chr11-119519416-INS-141
chr11-119519698-INS-114
chr11-119519896-INS-166
chr11-119519896-INS-335
chr11-119537664-INS-57
chr11-119545220-INS-981
chr11-119556160-INS-126
chr11-119564258-INS-50
chr11-119597571-INS-343
chr11-119606937-INS-238
chr11-119617400-INS-50
chr11-119626404-INS-68
chr11-119626572-INS-174
chr11-119652024-INS-52
chr11-119692747-INS-58
chr11-119720635-INS-209
chr11-119933958-INS-80
chr11-120027136-INS-85
chr11-120049365-INS-58
chr11-120050089-INS-172
chr11-120053588-INS-72
chr11-120059750-INS-7472
chr11-120089527-INS-50
chr11-120105482-INS-253
chr11-120106148-INS-53
chr11-120137192-INS-135
chr11-120191744-INS-75
chr11-120268639-INS-54
chr11-120305451-INS-217
chr11-120322800-INS-75
chr11-120322819-INS-1390
chr11-120323097-INS-72
chr11-120330880-INS-58
chr11-120331082-INS-216
chr11-120331930-INS-7172
chr11-120337858-INS-206
chr11-120342419-INS-350
chr11-120448849-INS-899
chr11-120529767-INS-227
chr11-120529804-INS-59
chr11-120559219-INS-194
chr11-120629680-INS-222
chr11-120668900-INS-853
chr11-120689665-INS-397
chr11-120689733-INS-111
chr11-120689947-INS-190
chr11-120754754-INS-191
chr11-120781551-INS-205
chr11-120820492-INS-75
chr11-120894254-INS-202
chr11-120906967-INS-2890
chr11-120908093-INS-102

chr11-120920503-INS-679
chr11-120920510-INS-286
chr11-120951857-INS-50
chr11-120969075-INS-202
chr11-120987033-INS-168
chr11-121072404-INS-156
chr11-121072604-INS-611
chr11-121075579-INS-182
chr11-121125590-INS-201
chr11-121167321-INS-194
chr11-121170359-INS-323
chr11-121170658-INS-63
chr11-121176951-INS-52
chr11-121187408-INS-177
chr11-121188203-INS-197
chr11-121196266-INS-147
chr11-121202324-INS-5695
chr11-121255087-INS-210
chr11-121257310-INS-68
chr11-121260148-INS-80
chr11-121295738-INS-50
chr11-121321763-INS-74
chr11-121336744-INS-81
chr11-121393621-INS-260
chr11-121633293-INS-1308
chr11-121656014-INS-157
chr11-121704379-INS-50
chr11-121723534-INS-204
chr11-121739844-INS-2701
chr11-121769531-INS-6232
chr11-121833760-INS-504
chr11-121841970-INS-4310
chr12-3142862-INS-60
chr12-3172836-INS-52
chr12-3443708-INS-456
chr12-3490592-INS-87
chr12-3543868-INS-94
chr12-3561618-INS-208
chr12-3566408-INS-71
chr12-3582389-INS-171
chr12-3593327-INS-194
chr12-3606785-INS-8816
chr12-3611941-INS-111
chr12-3612468-INS-236
chr12-3625202-INS-50
chr12-3639596-INS-75
chr12-3816709-INS-432
chr12-3840811-INS-77
chr12-3840876-INS-236
chr12-3876682-INS-54
chr12-3892842-INS-185
chr12-3922470-INS-794
chr12-3926541-INS-85
chr12-3979894-INS-59
chr12-3980564-INS-70
chr12-4039755-INS-7085
chr12-4043729-INS-235
chr12-4070208-INS-54
chr12-4101854-INS-92
chr12-4112862-INS-2042
chr12-4114153-INS-70
chr12-4142532-INS-56
chr12-4159507-INS-109
chr12-4163106-INS-72
chr12-4167335-INS-3826
chr12-4173259-INS-66
chr12-4174192-INS-92
chr12-4190764-INS-218
chr12-4212484-INS-506
chr12-4239221-INS-50
chr12-4241392-INS-64
chr12-4271648-INS-5778
chr12-4271711-INS-2761
chr12-4306256-INS-96
chr12-4315536-INS-204
chr12-4317431-INS-212
chr12-4327017-INS-56

chr12-4335050-INS-102
chr12-4368108-INS-66
chr12-4398076-INS-498
chr12-4398641-INS-51
chr12-4398648-INS-255
chr12-4405754-INS-1529
chr12-4414305-INS-189
chr12-4418125-INS-95
chr12-4444180-INS-1039
chr12-4458911-INS-1431
chr12-4469063-INS-136
chr12-4474551-INS-1716
chr12-4477704-INS-106
chr12-4494510-INS-54
chr12-4494578-INS-498
chr12-4500350-INS-140
chr12-4516755-INS-170
chr12-4517212-INS-680
chr12-4559129-INS-51
chr12-4559129-INS-786
chr12-4564628-INS-115
chr12-4571239-INS-207
chr12-4580309-INS-150
chr12-4580413-INS-638
chr12-4580605-INS-462
chr12-4581068-INS-1005
chr12-4601583-INS-3847
chr12-4616992-INS-151
chr12-4627213-INS-2881
chr12-4628542-INS-243
chr12-4753412-INS-445
chr12-4784086-INS-124
chr12-4784371-INS-169
chr12-4819132-INS-68
chr12-4848088-INS-370
chr12-4866302-INS-199
chr12-4873195-INS-161
chr12-4877758-INS-948
chr12-4880442-INS-457
chr12-4893461-INS-84
chr12-4913997-INS-108
chr12-4926840-INS-1000
chr12-4943403-INS-252
chr12-4950122-INS-201
chr12-4961476-INS-215
chr12-4972272-INS-181
chr12-5079048-INS-136
chr12-5083098-INS-238
chr12-5097655-INS-175
chr12-5101837-INS-5286
chr12-5102046-INS-2492
chr12-5113282-INS-7095
chr12-5115903-INS-184
chr12-5174243-INS-246
chr12-5218138-INS-64
chr12-5357307-INS-370
chr12-5411975-INS-435
chr12-5495783-INS-771
chr12-5505226-INS-438
chr12-5565311-INS-62
chr12-5579961-INS-205
chr12-5624769-INS-159
chr12-5630586-INS-71
chr12-5638586-INS-66
chr12-5642419-INS-769
chr12-5648904-INS-54
chr12-5676980-INS-482
chr12-5708058-INS-4735
chr12-5713141-INS-130
chr12-5729047-INS-6798
chr12-5790923-INS-70
chr12-5790948-INS-6395
chr12-5808501-INS-50
chr12-5830049-INS-54
chr12-5842722-INS-100
chr12-5847241-INS-399
chr12-5866018-INS-58

chr12-5915062-INS-3213
chr12-5947098-INS-84
chr12-5986588-INS-985
chr12-6013476-INS-3587
chr12-6083429-INS-54
chr12-6110387-INS-206
chr12-6125755-INS-52
chr12-6155939-INS-1890
chr12-6157734-INS-412
chr12-6184996-INS-234
chr12-6223790-INS-165
chr12-6269935-INS-322
chr12-6269979-INS-8583
chr12-6276737-INS-486
chr12-6327027-INS-2660
chr12-6402413-INS-60
chr12-6403712-INS-60
chr12-6403855-INS-218
chr12-6444886-INS-122
chr12-6448867-INS-5405
chr12-6464576-INS-1469
chr12-6478732-INS-14281
chr12-6487635-INS-428
chr12-6499213-INS-98
chr12-6499299-INS-153
chr12-6515619-INS-1391
chr12-6550477-INS-504
chr12-6589222-INS-56
chr12-6616777-INS-274
chr12-6674685-INS-52
chr12-6676444-INS-135
chr12-6787993-INS-3019
chr12-6793353-INS-1035
chr12-6886008-INS-6702
chr12-6907312-INS-199
chr12-6924886-INS-80
chr12-6953627-INS-52
chr12-6963616-INS-56
chr12-6964005-INS-7100
chr12-6978290-INS-56
chr12-7023670-INS-6272
chr12-7050987-INS-1737
chr12-7083551-INS-64
chr12-7122058-INS-6350
chr12-7138320-INS-58
chr12-7182291-INS-52
chr12-7188162-INS-132
chr12-7220427-INS-181
chr12-7232676-INS-1331
chr12-7235106-INS-2726
chr12-7236446-INS-2996
chr12-7256756-INS-96
chr12-7273936-INS-2569
chr12-7304728-INS-6670
chr12-7309152-INS-65
chr12-7311189-INS-206
chr12-7314962-INS-88
chr12-7374357-INS-54
chr12-7379614-INS-50
chr12-7401230-INS-74
chr12-7405360-INS-1922
chr12-7415033-INS-120
chr12-7425451-INS-66
chr12-7460219-INS-63
chr12-7489522-INS-82
chr12-7511931-INS-144
chr12-7584267-INS-395
chr12-7618594-INS-120
chr12-7651060-INS-137
chr12-7761587-INS-50
chr12-7763423-INS-251
chr12-7773171-INS-270
chr12-7796114-INS-163
chr12-7801721-INS-6347
chr12-7850006-INS-60
chr12-7996485-INS-373
chr12-8073134-INS-7561

chr12-8084275-INS-175
chr12-8120692-INS-18205
chr12-8132415-INS-50
chr12-8144449-INS-50
chr12-8189721-INS-1265
chr12-8195756-INS-74
chr12-8222266-INS-499
chr12-8224726-INS-789
chr12-8229856-INS-50
chr12-8238752-INS-1502
chr12-8275246-INS-116
chr12-8281135-INS-109
chr12-8357949-INS-182
chr12-8360347-INS-4817
chr12-8361975-INS-88
chr12-8424744-INS-201
chr12-8426878-INS-50
chr12-8437974-INS-115
chr12-8463700-INS-50
chr12-8471875-INS-180
chr12-8483698-INS-181
chr12-8540347-INS-52
chr12-8556473-INS-199
chr12-8563038-INS-92
chr12-8580814-INS-154
chr12-8580817-INS-532
chr12-8609322-INS-260
chr12-8618820-INS-176
chr12-8646087-INS-177
chr12-8671029-INS-6678
chr12-8720140-INS-8648
chr12-8921675-INS-50
chr12-8922250-INS-50
chr12-8951608-INS-50
chr12-9005849-INS-165
chr12-9005849-INS-66
chr12-9018583-INS-190
chr12-9033645-INS-597
chr12-9042283-INS-805
chr12-9057240-INS-864
chr12-9057490-INS-199
chr12-9075619-INS-199
chr12-9087302-INS-90
chr12-9136646-INS-56
chr12-9172976-INS-9046
chr12-9269509-INS-52
chr12-9280317-INS-513
chr12-9280443-INS-251
chr12-9298476-INS-158
chr12-9304908-INS-74
chr12-9322561-INS-65
chr12-9371306-INS-395
chr12-9460156-INS-2676
chr12-9476007-INS-946
chr12-9493767-INS-247
chr12-9536023-INS-174
chr12-9552320-INS-58
chr12-9569063-INS-90
chr12-9570853-INS-234
chr12-9622485-INS-70
chr12-9690558-INS-1521
chr12-9742823-INS-60
chr12-9749669-INS-1233
chr12-9751406-INS-705
chr12-9778752-INS-52
chr12-9799042-INS-4464
chr12-9801829-INS-91
chr12-9803075-INS-190
chr12-9803231-INS-54
chr12-9826252-INS-2141
chr12-9836749-INS-60
chr12-9851381-INS-158
chr12-9865664-INS-6199
chr12-9880524-INS-1039
chr12-9884512-INS-3454
chr12-9928992-INS-316
chr12-9939732-INS-64

chr12-9962403-INS-80
chr12-9964615-INS-284
chr12-9964999-INS-80
chr12-10005635-INS-225
chr12-10017940-INS-6158
chr12-10030041-INS-80
chr12-10056791-INS-104
chr12-10152630-INS-52
chr12-10173990-INS-1580
chr12-10247463-INS-242
chr12-10298676-INS-50
chr12-10388138-INS-5825
chr12-10397104-INS-7509
chr12-10421353-INS-61
chr12-10426537-INS-50
chr12-10427793-INS-50
chr12-10457862-INS-194
chr12-10458750-INS-50
chr12-10464851-INS-885
chr12-10466608-INS-65
chr12-10482043-INS-489
chr12-10501089-INS-66
chr12-10503602-INS-66
chr12-10518837-INS-302
chr12-10573267-INS-3022
chr12-10628906-INS-50
chr12-10662496-INS-291
chr12-10662626-INS-114
chr12-10662829-INS-584
chr12-10665761-INS-471
chr12-10667357-INS-448
chr12-10695875-INS-8908
chr12-10701644-INS-180
chr12-10740796-INS-184
chr12-10742874-INS-1237
chr12-10785674-INS-50
chr12-10786592-INS-5748
chr12-10815246-INS-237
chr12-10815288-INS-663
chr12-10816834-INS-6914
chr12-10821479-INS-52
chr12-10821646-INS-864
chr12-10833368-INS-76
chr12-10889248-INS-52
chr12-10895887-INS-1297
chr12-10908496-INS-172
chr12-10999892-INS-316
chr12-10999892-INS-652
chr12-11011875-INS-1398
chr12-11012261-INS-4032
chr12-11012384-INS-3150
chr12-11012742-INS-1350
chr12-11012895-INS-126
chr12-11046032-INS-54
chr12-11068563-INS-248
chr12-11071652-INS-223
chr12-11079958-INS-2500
chr12-11081434-INS-612
chr12-11106067-INS-6694
chr12-11113961-INS-86
chr12-11125994-INS-100
chr12-11164554-INS-612
chr12-11207553-INS-1102
chr12-11223323-INS-362
chr12-11229935-INS-156
chr12-11249872-INS-51
chr12-11269939-INS-68
chr12-11302104-INS-50
chr12-11374215-INS-57
chr12-11379032-INS-629
chr12-11384167-INS-130
chr12-11391842-INS-85
chr12-11392404-INS-60
chr12-11394879-INS-56
chr12-11395489-INS-185
chr12-11420070-INS-59
chr12-11427889-INS-52

chr12-11454165-INS-84
chr12-11492194-INS-389
chr12-11495548-INS-50
chr12-11523744-INS-6163
chr12-11585243-INS-124
chr12-11631887-INS-65
chr12-11642135-INS-7196
chr12-11651630-INS-1181
chr12-11662276-INS-64
chr12-11685716-INS-1142
chr12-11690575-INS-50
chr12-11712091-INS-146
chr12-11728366-INS-423
chr12-11730722-INS-399
chr12-11734669-INS-339
chr12-11751244-INS-8171
chr12-11764672-INS-787
chr12-11776442-INS-54
chr12-11807902-INS-6161
chr12-11821284-INS-60
chr12-11877549-INS-125
chr12-11889621-INS-55
chr12-11897939-INS-290
chr12-11900768-INS-90
chr12-11918096-INS-168
chr12-11923035-INS-75
chr12-11941389-INS-66
chr12-11960051-INS-67
chr12-11960900-INS-9368
chr12-11998472-INS-2109
chr12-12013716-INS-50
chr12-12015812-INS-577
chr12-12029314-INS-56
chr12-12058249-INS-50
chr12-12065947-INS-202
chr12-12081769-INS-268
chr12-12200914-INS-158
chr12-12212788-INS-595
chr12-12213158-INS-50
chr12-12226308-INS-52
chr12-12237175-INS-75
chr12-12240797-INS-6238
chr12-12244303-INS-120
chr12-12302342-INS-64
chr12-12327444-INS-55
chr12-12336277-INS-386
chr12-12336277-INS-924
chr12-12413341-INS-406
chr12-12432987-INS-507
chr12-12500731-INS-6007
chr12-12596721-INS-50
chr12-12623416-INS-1044
chr12-12623418-INS-166
chr12-12623429-INS-2372
chr12-12668722-INS-51
chr12-12685320-INS-75
chr12-12741525-INS-55
chr12-12758197-INS-100
chr12-12764408-INS-94
chr12-12764661-INS-967
chr12-12800323-INS-292
chr12-12813617-INS-56
chr12-12814048-INS-81
chr12-12844989-INS-239
chr12-12845023-INS-50
chr12-12845529-INS-96
chr12-12845567-INS-200
chr12-12865881-INS-50
chr12-12870979-INS-70
chr12-12887639-INS-83
chr12-12887799-INS-61
chr12-12979471-INS-218
chr12-12979702-INS-64
chr12-13021989-INS-72
chr12-13032714-INS-274
chr12-13033813-INS-75
chr12-13062984-INS-367

chr12-13124479-INS-171
chr12-13187087-INS-82
chr12-13227495-INS-421
chr12-13227553-INS-1221
chr12-13229105-INS-1958
chr12-13239886-INS-100
chr12-13246578-INS-66
chr12-13332975-INS-144
chr12-13410346-INS-2026
chr12-13411554-INS-172
chr12-13414089-INS-1588
chr12-13418243-INS-208
chr12-13418996-INS-194
chr12-13489124-INS-112
chr12-13549791-INS-6733
chr12-13555538-INS-320
chr12-13617628-INS-258
chr12-13622380-INS-5152
chr12-13639032-INS-117
chr12-13647851-INS-56
chr12-13650696-INS-50
chr12-13671085-INS-69
chr12-13701418-INS-629
chr12-13726795-INS-122
chr12-13751977-INS-6283
chr12-13769730-INS-3636
chr12-13781828-INS-64
chr12-13840353-INS-72
chr12-13840891-INS-55
chr12-13906659-INS-4899
chr12-13909760-INS-6149
chr12-13981693-INS-59
chr12-14006963-INS-5051
chr12-14015602-INS-60
chr12-14029101-INS-368
chr12-14029101-INS-54
chr12-14088511-INS-6132
chr12-14091939-INS-2774
chr12-14095463-INS-76
chr12-14110928-INS-50
chr12-14141493-INS-1951
chr12-14171393-INS-108
chr12-14171815-INS-6506
chr12-14232540-INS-58
chr12-14267173-INS-4741
chr12-14268943-INS-6362
chr12-14302810-INS-6390
chr12-14326535-INS-6398
chr12-14335310-INS-6023
chr12-14341436-INS-1006
chr12-14410494-INS-3143
chr12-14428636-INS-2806
chr12-14454788-INS-59
chr12-14464469-INS-57
chr12-14531180-INS-3390
chr12-14596080-INS-128
chr12-14702521-INS-9847
chr12-14761611-INS-6472
chr12-14795837-INS-50
chr12-14813079-INS-477
chr12-14840388-INS-214
chr12-14871338-INS-7088
chr12-14937154-INS-6310
chr12-14940511-INS-57
chr12-14942237-INS-6497
chr12-14943044-INS-145
chr12-14946999-INS-7094
chr12-15021489-INS-69
chr12-15049685-INS-6381
chr12-15064913-INS-74
chr12-15068907-INS-50
chr12-15072956-INS-777
chr12-15121565-INS-1034
chr12-15127520-INS-74
chr12-15151603-INS-50
chr12-15226139-INS-50
chr12-15230127-INS-50

chr12-15234840-INS-244
chr12-15241510-INS-208
chr12-15267806-INS-7449
chr12-15277135-INS-116
chr12-15323188-INS-76
chr12-15339150-INS-84
chr12-15343571-INS-780
chr12-15343630-INS-272
chr12-15355331-INS-647
chr12-15404416-INS-2312
chr12-15408622-INS-51
chr12-15436979-INS-2618
chr12-15465327-INS-123
chr12-15473424-INS-421
chr12-15476832-INS-6100
chr12-15496472-INS-56
chr12-15496610-INS-231
chr12-15506207-INS-5414
chr12-15514726-INS-6611
chr12-15555853-INS-1994
chr12-15578297-INS-95
chr12-15650508-INS-65
chr12-15650519-INS-423
chr12-15742947-INS-92
chr12-15756459-INS-2122
chr12-15773407-INS-6393
chr12-15831725-INS-80
chr12-15831770-INS-209
chr12-15921854-INS-406
chr12-15922314-INS-208
chr12-15945331-INS-2267
chr12-15963908-INS-2659
chr12-15974360-INS-2045
chr12-16003519-INS-75
chr12-16011589-INS-52
chr12-16051246-INS-435
chr12-16100679-INS-6988
chr12-16110468-INS-50
chr12-16117422-INS-1125
chr12-16117931-INS-50
chr12-16145555-INS-6535
chr12-16179611-INS-7177
chr12-16185434-INS-53
chr12-16206644-INS-168
chr12-16266940-INS-51
chr12-16312325-INS-50
chr12-16315430-INS-50
chr12-16354295-INS-50
chr12-16408558-INS-72
chr12-16423169-INS-5977
chr12-16484079-INS-52
chr12-16497523-INS-163
chr12-16515670-INS-1916
chr12-16550134-INS-144
chr12-16624248-INS-273
chr12-16626587-INS-495
chr12-16636916-INS-80
chr12-16641911-INS-2355
chr12-16682721-INS-256
chr12-16687005-INS-264
chr12-16687021-INS-74
chr12-16812441-INS-50
chr12-16812776-INS-92
chr12-16837886-INS-192
chr12-16892389-INS-125
chr12-16953902-INS-210
chr12-17016290-INS-52
chr12-17067840-INS-4984
chr12-17105893-INS-63
chr12-17115589-INS-84
chr12-17141988-INS-1890
chr12-17181926-INS-59
chr12-17204198-INS-4291
chr12-17219710-INS-989
chr12-17230289-INS-216
chr12-17232978-INS-56
chr12-17234231-INS-7110

chr12-17245425-INS-1734
chr12-17259192-INS-52
chr12-17271552-INS-499
chr12-17280368-INS-52
chr12-17282235-INS-50
chr12-17307715-INS-58
chr12-17453448-INS-54
chr12-17478545-INS-115
chr12-17478849-INS-554
chr12-17491276-INS-50
chr12-17505823-INS-53
chr12-17507975-INS-231
chr12-17508376-INS-368
chr12-17527678-INS-72
chr12-17552988-INS-182
chr12-17580960-INS-80
chr12-17581597-INS-110
chr12-17666411-INS-208
chr12-17724829-INS-299
chr12-17727998-INS-6900
chr12-17745439-INS-51
chr12-17756693-INS-7165
chr12-17810223-INS-449
chr12-17832298-INS-53
chr12-17834070-INS-60
chr12-17842548-INS-1889
chr12-17844689-INS-216
chr12-17968787-INS-1013
chr12-17974431-INS-60
chr12-18148095-INS-50
chr12-18172030-INS-80
chr12-18322675-INS-466
chr12-18335281-INS-64
chr12-18432824-INS-484
chr12-18435801-INS-246
chr12-18443023-INS-57
chr12-18455647-INS-84
chr12-18572301-INS-60
chr12-18646205-INS-1893
chr12-18652455-INS-2894
chr12-18771807-INS-945
chr12-18814234-INS-215
chr12-18984271-INS-77
chr12-18987228-INS-434
chr12-19073200-INS-1708
chr12-19136630-INS-107
chr12-19190997-INS-435
chr12-19223276-INS-172
chr12-19368243-INS-104
chr12-19370403-INS-154
chr12-19482063-INS-74
chr12-19535131-INS-55
chr12-19774663-INS-381
chr12-19823745-INS-204
chr12-19872445-INS-55
chr12-19931463-INS-2557
chr12-19932426-INS-101
chr12-19934648-INS-192
chr12-19941300-INS-70
chr12-19953512-INS-174
chr12-19955772-INS-76
chr12-19971592-INS-98
chr12-19972792-INS-114
chr12-20135164-INS-168
chr12-20301478-INS-6803
chr12-20332266-INS-174
chr12-20334479-INS-67
chr12-20343982-INS-96
chr12-20364137-INS-3500
chr12-20372244-INS-252
chr12-20435550-INS-4831
chr12-20440770-INS-75
chr12-20471213-INS-57
chr12-20486219-INS-84
chr12-20510288-INS-220
chr12-20522346-INS-1886
chr12-20545699-INS-4919

chr12-20654114-INS-86
chr12-20669864-INS-114
chr12-20685997-INS-83
chr12-20686225-INS-338
chr12-20699845-INS-56
chr12-20719671-INS-60
chr12-20793694-INS-62
chr12-20853613-INS-71
chr12-20877263-INS-3288
chr12-20883246-INS-6433
chr12-20892146-INS-68
chr12-20909638-INS-202
chr12-20910473-INS-52
chr12-20988677-INS-93
chr12-21115230-INS-2090
chr12-21127829-INS-5120
chr12-21142086-INS-171
chr12-21143932-INS-1997
chr12-21143932-INS-503
chr12-21149704-INS-3148
chr12-21166378-INS-347
chr12-21191693-INS-122
chr12-21265177-INS-285
chr12-21276247-INS-396
chr12-21300123-INS-65
chr12-21365222-INS-203
chr12-21365997-INS-422
chr12-21382990-INS-70
chr12-21391127-INS-259
chr12-21435822-INS-216
chr12-21455812-INS-1611
chr12-21506004-INS-2322
chr12-21515298-INS-81
chr12-21515909-INS-319
chr12-21528222-INS-1812
chr12-21530788-INS-126
chr12-21536680-INS-50
chr12-21538324-INS-195
chr12-21559299-INS-109
chr12-21567570-INS-342
chr12-21568388-INS-472
chr12-21625626-INS-6263
chr12-21626594-INS-162
chr12-21808714-INS-84
chr12-21930635-INS-169
chr12-22062110-INS-1046
chr12-22093984-INS-229
chr12-22781791-INS-163
chr12-22785970-INS-55
chr12-23048202-INS-373
chr12-23055502-INS-231
chr12-23117909-INS-2180
chr12-23151737-INS-65
chr12-23152990-INS-3470
chr12-23161519-INS-92
chr12-23169289-INS-69
chr12-23194880-INS-995
chr12-23235077-INS-172
chr12-23460618-INS-58
chr12-23665619-INS-84
chr12-23725734-INS-60
chr12-23831578-INS-420
chr12-23831905-INS-335
chr12-23846064-INS-115
chr12-23967228-INS-193
chr12-24121608-INS-168
chr12-24122481-INS-336
chr12-24142413-INS-87
chr12-24179897-INS-59
chr12-24182029-INS-2393
chr12-24342103-INS-57
chr12-24405679-INS-114
chr12-24538423-INS-118
chr12-24693421-INS-51
chr12-24696749-INS-50
chr12-24697346-INS-205
chr12-24710696-INS-50

chr12-24725104-INS-2872
chr12-24726663-INS-50
chr12-24727335-INS-54
chr12-24769751-INS-254
chr12-24770176-INS-456
chr12-24778425-INS-166
chr12-24803614-INS-54
chr12-24813223-INS-877
chr12-24818611-INS-54
chr12-24826848-INS-202
chr12-24859362-INS-122
chr12-24946454-INS-642
chr12-24948041-INS-286
chr12-24972826-INS-275
chr12-25006851-INS-347
chr12-25115453-INS-480
chr12-25123277-INS-1592
chr12-25123277-INS-656
chr12-25153348-INS-104
chr12-25187946-INS-577
chr12-25266257-INS-59
chr12-25293204-INS-408
chr12-25293377-INS-170
chr12-25293377-INS-57
chr12-25297770-INS-2907
chr12-25338934-INS-1030
chr12-25338949-INS-1739
chr12-25407379-INS-54
chr12-25418731-INS-220
chr12-25418788-INS-609
chr12-25464667-INS-104
chr12-25467182-INS-169
chr12-25476432-INS-347
chr12-25476951-INS-84
chr12-25486007-INS-50
chr12-25512993-INS-6319
chr12-25635686-INS-6764
chr12-25685818-INS-99
chr12-25711134-INS-190
chr12-25757127-INS-6658
chr12-25782167-INS-1142
chr12-25785645-INS-96
chr12-25897519-INS-6489
chr12-26048218-INS-206
chr12-26061340-INS-2578
chr12-26065008-INS-6654
chr12-26106422-INS-252
chr12-26119421-INS-64
chr12-26131454-INS-108
chr12-26235869-INS-50
chr12-26267589-INS-60
chr12-26267693-INS-248
chr12-26384356-INS-78
chr12-26446661-INS-250
chr12-26479334-INS-481
chr12-26503579-INS-63
chr12-26580429-INS-1804
chr12-26591959-INS-100
chr12-26593017-INS-56
chr12-26602223-INS-483
chr12-26630461-INS-4668
chr12-26635347-INS-1033
chr12-26648599-INS-644
chr12-26701025-INS-1875
chr12-26799681-INS-6379
chr12-26818111-INS-83
chr12-26848887-INS-66
chr12-26858250-INS-108
chr12-26872674-INS-13449
chr12-26919032-INS-91
chr12-26990801-INS-112
chr12-26994858-INS-54
chr12-27003190-INS-50
chr12-27008073-INS-143
chr12-27045535-INS-8779
chr12-27055333-INS-3027
chr12-27106946-INS-51

chr12-27120683-INS-610
chr12-27214308-INS-53
chr12-27265063-INS-235
chr12-27265587-INS-163
chr12-27448234-INS-1305
chr12-27538029-INS-294
chr12-27538037-INS-770
chr12-27548605-INS-126
chr12-27552225-INS-52
chr12-27646654-INS-50
chr12-27665010-INS-6605
chr12-27725963-INS-109
chr12-27731809-INS-13022
chr12-27748697-INS-515
chr12-27759803-INS-8265
chr12-27761949-INS-149
chr12-27768883-INS-400
chr12-27846356-INS-15891
chr12-27848554-INS-2606
chr12-27971958-INS-237
chr12-27972664-INS-336
chr12-28040414-INS-6222
chr12-28063080-INS-3054
chr12-28075290-INS-6401
chr12-28085077-INS-93
chr12-28131075-INS-54
chr12-28167335-INS-5361
chr12-28177620-INS-85
chr12-28225752-INS-4060
chr12-28305399-INS-4999
chr12-28377380-INS-83
chr12-28414013-INS-6169
chr12-28440679-INS-85
chr12-28440706-INS-185
chr12-28446371-INS-68
chr12-28455094-INS-4385
chr12-28582769-INS-6778
chr12-28616380-INS-207
chr12-28629845-INS-6020
chr12-28642743-INS-79
chr12-28692864-INS-1882
chr12-28694442-INS-55
chr12-28706605-INS-206
chr12-28730302-INS-1885
chr12-28820164-INS-981
chr12-28820199-INS-2115
chr12-28908457-INS-3149
chr12-28931236-INS-1884
chr12-28961812-INS-1890
chr12-28976790-INS-100
chr12-29046728-INS-50
chr12-29066310-INS-2001
chr12-29082669-INS-121
chr12-29086629-INS-6726
chr12-29098171-INS-71
chr12-29121123-INS-1312
chr12-29156567-INS-159
chr12-29208314-INS-4523
chr12-29312725-INS-83
chr12-29316274-INS-312
chr12-29404400-INS-62
chr12-29457588-INS-282
chr12-29600154-INS-66
chr12-29683865-INS-62
chr12-29737920-INS-63
chr12-29738661-INS-86
chr12-29914437-INS-56
chr12-29921357-INS-1067
chr12-30215136-INS-1137
chr12-30220404-INS-1171
chr12-30230165-INS-69
chr12-30275155-INS-210
chr12-30330666-INS-155
chr12-30405612-INS-1310
chr12-30412827-INS-52
chr12-30469845-INS-406
chr12-30470122-INS-160

chr12-30491961-INS-6602
chr12-30495581-INS-492
chr12-30500209-INS-74
chr12-30524180-INS-441
chr12-30584456-INS-646
chr12-30628615-INS-1690
chr12-30628935-INS-6187
chr12-30736803-INS-214
chr12-30785433-INS-287
chr12-30785525-INS-64
chr12-30792330-INS-100
chr12-30796870-INS-55
chr12-30809287-INS-6446
chr12-30942688-INS-60
chr12-30957396-INS-118
chr12-30972964-INS-1060
chr12-31007363-INS-1315
chr12-31034934-INS-77
chr12-31048863-INS-211
chr12-31051998-INS-156
chr12-31078111-INS-208
chr12-31196972-INS-50
chr12-31253372-INS-165
chr12-31284763-INS-348
chr12-31299989-INS-116
chr12-31309322-INS-202
chr12-31383116-INS-211
chr12-31466415-INS-180
chr12-31481336-INS-166
chr12-31491177-INS-54
chr12-31494160-INS-580
chr12-31510256-INS-76
chr12-31597741-INS-52
chr12-31599834-INS-116
chr12-31611064-INS-112
chr12-31613694-INS-132
chr12-31619005-INS-194
chr12-31622013-INS-93
chr12-31726970-INS-201
chr12-31756494-INS-207
chr12-31820585-INS-176
chr12-31820794-INS-424
chr12-31842798-INS-6443
chr12-31853682-INS-158
chr12-31863525-INS-124
chr12-31887547-INS-52
chr12-31911566-INS-205
chr12-31926348-INS-554
chr12-31950587-INS-185
chr12-31952423-INS-53
chr12-31994236-INS-206
chr12-31996740-INS-1254
chr12-32024362-INS-1559
chr12-32028150-INS-6163
chr12-32047725-INS-203
chr12-32057746-INS-195
chr12-32059252-INS-183
chr12-32060139-INS-241
chr12-32060741-INS-397
chr12-32066084-INS-937
chr12-32161304-INS-54
chr12-32185007-INS-662
chr12-32185516-INS-5125
chr12-32187788-INS-3701
chr12-32191464-INS-89
chr12-32206437-INS-100
chr12-32206437-INS-312
chr12-32222408-INS-6445
chr12-32227354-INS-88
chr12-32270673-INS-10168
chr12-32294783-INS-75
chr12-32330914-INS-370
chr12-32339634-INS-6511
chr12-32368802-INS-1084
chr12-32378672-INS-1912
chr12-32407922-INS-68
chr12-32456920-INS-50

chr12-32556701-INS-217
chr12-32562333-INS-114
chr12-32575569-INS-62
chr12-32576450-INS-60
chr12-32666457-INS-6644
chr12-32774118-INS-198
chr12-32836356-INS-3689
chr12-32898195-INS-352
chr12-32903587-INS-51
chr12-32909793-INS-549
chr12-32914813-INS-120
chr12-32963023-INS-1095
chr12-32971637-INS-219
chr12-32971729-INS-361
chr12-32972470-INS-834
chr12-33036709-INS-445
chr12-33043662-INS-91
chr12-33043824-INS-357
chr12-33047929-INS-88
chr12-33048045-INS-171
chr12-33049727-INS-250
chr12-33049850-INS-506
chr12-33056226-INS-64
chr12-33056866-INS-58
chr12-33095415-INS-5706
chr12-33110918-INS-65
chr12-33133210-INS-56
chr12-33150488-INS-54
chr12-33150497-INS-120
chr12-33154994-INS-2291
chr12-33251172-INS-126
chr12-33289747-INS-4736
chr12-33309744-INS-52
chr12-33310088-INS-151
chr12-33310399-INS-58
chr12-33344150-INS-66
chr12-33356662-INS-51
chr12-33363786-INS-56
chr12-33395182-INS-54
chr12-33431067-INS-60
chr12-33438744-INS-127
chr12-33500352-INS-1984
chr12-33501624-INS-60
chr12-33506767-INS-215
chr12-33506925-INS-101
chr12-33517729-INS-1421
chr12-33527047-INS-506
chr12-33545235-INS-7116
chr12-33550182-INS-52
chr12-33561785-INS-228
chr12-33571702-INS-238
chr12-33600285-INS-1147
chr12-33601371-INS-218
chr12-33611795-INS-14366
chr12-33626861-INS-7476
chr12-33663702-INS-50
chr12-33677393-INS-1578
chr12-33679495-INS-675
chr12-33700149-INS-52
chr12-33791510-INS-51
chr12-33810414-INS-96
chr12-33824538-INS-344
chr12-33875381-INS-108
chr12-33937620-INS-141
chr12-33937620-INS-701
chr12-33937630-INS-341
chr12-33938105-INS-883
chr12-33940875-INS-50
chr12-33956628-INS-50
chr12-33966715-INS-56
chr12-34050072-INS-208
chr12-34133599-INS-226
chr12-34232211-INS-52
chr12-34284050-INS-279
chr12-34288621-INS-57
chr12-34310134-INS-148
chr12-34374593-INS-279

chr12-34390470-INS-495
chr12-34463541-INS-112
chr12-34497526-INS-70
chr12-34508229-INS-53
chr12-34519862-INS-55
chr12-34520095-INS-216
chr12-34620721-INS-75
chr12-34636188-INS-74
chr12-34649372-INS-76
chr12-34649685-INS-1552
chr12-34659343-INS-62
chr12-34661656-INS-251
chr12-34680251-INS-1634
chr12-34703209-INS-123
chr12-34716702-INS-196
chr12-34729865-INS-545
chr12-34729909-INS-153
chr12-34729909-INS-51
chr12-34753688-INS-699
chr12-34753992-INS-336
chr12-34754357-INS-461
chr12-34759671-INS-147
chr12-34767289-INS-6293
chr12-34769928-INS-120
chr12-34795436-INS-593
chr12-34795547-INS-3231
chr12-34824515-INS-7005
chr12-34829993-INS-1283
chr12-34897425-INS-3709
chr12-34898791-INS-60
chr12-34905154-INS-1423
chr12-34940594-INS-71
chr12-34940688-INS-147
chr12-34948781-INS-168
chr12-34969290-INS-68
chr12-34969309-INS-544
chr12-34973534-INS-6626
chr12-35103880-INS-57
chr12-35125012-INS-297
chr12-35177375-INS-2745
chr12-35178253-INS-872
chr12-35186281-INS-200
chr12-35186614-INS-161
chr12-35238709-INS-6414
chr12-35240560-INS-499
chr12-35241828-INS-5014
chr12-35248573-INS-2972
chr12-35258016-INS-311
chr12-35271553-INS-1576
chr12-35310685-INS-92
chr12-35382589-INS-50
chr12-35431391-INS-1751
chr12-35438877-INS-1625
chr12-35517084-INS-6272
chr12-35542267-INS-67
chr12-35598928-INS-66
chr12-35685276-INS-185
chr12-35686880-INS-52
chr12-35707294-INS-3034
chr12-35831566-INS-50
chr12-35846583-INS-15063
chr12-35888344-INS-1892
chr12-35891787-INS-136
chr12-35895958-INS-556
chr12-35937856-INS-1047
chr12-35963137-INS-1230
chr12-35996801-INS-444
chr12-36021331-INS-55
chr12-36023439-INS-4771
chr12-36099437-INS-268
chr12-36112059-INS-50
chr12-36130317-INS-50
chr12-36142372-INS-480
chr12-36192742-INS-231
chr12-36252937-INS-1000
chr12-36263618-INS-50
chr12-36263648-INS-264

chr12-36264026-INS-188
chr12-36464756-INS-89
chr12-36520158-INS-6179
chr12-36576883-INS-797
chr12-36591831-INS-547
chr12-36594017-INS-50
chr12-36636752-INS-6436
chr12-36647290-INS-50
chr12-36649667-INS-1260
chr12-36664176-INS-185
chr12-36732672-INS-646
chr12-36743027-INS-335
chr12-36744114-INS-2714
chr12-36784683-INS-56
chr12-36785480-INS-239
chr12-36785958-INS-1405
chr12-36801732-INS-121
chr12-36808531-INS-125
chr12-36811920-INS-341
chr12-36818759-INS-7306
chr12-36826629-INS-58
chr12-36830026-INS-54
chr12-36855583-INS-6288
chr12-36861233-INS-969
chr12-36892636-INS-13069
chr12-36928704-INS-52
chr12-36937084-INS-80
chr12-36937233-INS-60
chr12-36952618-INS-222
chr12-36955532-INS-71
chr12-36955609-INS-274
chr12-36975860-INS-8249
chr12-36991943-INS-1960
chr12-37017540-INS-53
chr12-37026367-INS-1177
chr12-37038528-INS-185
chr12-37093905-INS-58
chr12-37097775-INS-356
chr12-37282283-INS-1237
chr12-37296877-INS-90
chr12-37340406-INS-50
chr12-37433088-INS-4069
chr12-37451585-INS-72
chr12-37480726-INS-576
chr12-37481621-INS-210
chr12-37489109-INS-312
chr12-37489438-INS-329
chr12-37489539-INS-50
chr12-37501323-INS-248
chr12-37501327-INS-709
chr12-37501327-INS-76
chr12-37501896-INS-1512
chr12-37540365-INS-134
chr12-37562493-INS-76
chr12-37590636-INS-52
chr12-37627674-INS-2147
chr12-37640278-INS-291
chr12-37733688-INS-2330
chr12-37758398-INS-106
chr12-37816957-INS-387
chr12-37823669-INS-167
chr12-37844949-INS-6357
chr12-37858463-INS-54
chr12-37859569-INS-512
chr12-37883731-INS-459
chr12-37886873-INS-158
chr12-37907255-INS-1177
chr12-37907753-INS-100
chr12-37915293-INS-6221
chr12-37928809-INS-1077
chr12-37941268-INS-138
chr12-37946698-INS-78
chr12-37957592-INS-104
chr12-38013101-INS-123
chr12-38028080-INS-6383
chr12-38035987-INS-97
chr12-38051218-INS-76

chr12-38068434-INS-50
chr12-38089327-INS-2014
chr12-38105937-INS-1052
chr12-38118225-INS-1879
chr12-38174075-INS-54
chr12-38259336-INS-72
chr12-38282167-INS-420
chr12-38303670-INS-7585
chr12-38306412-INS-798
chr12-38308204-INS-6213
chr12-38311314-INS-4076
chr12-38321417-INS-50
chr12-38322797-INS-70
chr12-38329079-INS-52
chr12-38338813-INS-1245
chr12-38440842-INS-178
chr12-38475562-INS-7298
chr12-38478701-INS-52
chr12-38487810-INS-58
chr12-38541502-INS-172
chr12-38617707-INS-72
chr12-38625836-INS-390
chr12-38643121-INS-668
chr12-38651468-INS-87
chr12-38651493-INS-229
chr12-38726627-INS-63
chr12-38736064-INS-3054
chr12-38736064-INS-6667
chr12-38757626-INS-78
chr12-38758753-INS-60
chr12-38775993-INS-6156
chr12-38788950-INS-120
chr12-38933583-INS-191
chr12-38936409-INS-2822
chr12-38938056-INS-1113
chr12-39005903-INS-400
chr12-39042931-INS-50
chr12-39057168-INS-669
chr12-39066149-INS-2910
chr12-39081482-INS-6574
chr12-39100206-INS-52
chr12-39112106-INS-50
chr12-39143900-INS-1616
chr12-39144722-INS-548
chr12-39231181-INS-205
chr12-39264557-INS-3070
chr12-39282430-INS-6785
chr12-39326648-INS-6801
chr12-39328608-INS-7054
chr12-39416088-INS-457
chr12-39481439-INS-56
chr12-39503875-INS-68
chr12-39503920-INS-208
chr12-39511408-INS-6516
chr12-39525454-INS-537
chr12-39633153-INS-52
chr12-39662942-INS-6407
chr12-39689773-INS-248
chr12-39689777-INS-110
chr12-39698114-INS-2499
chr12-39708252-INS-6468
chr12-39720029-INS-200
chr12-39734899-INS-6795
chr12-39761912-INS-1368
chr12-39776262-INS-158
chr12-39782976-INS-2793
chr12-39847403-INS-488
chr12-39847737-INS-254
chr12-39898944-INS-300
chr12-39910318-INS-86
chr12-39922089-INS-1756
chr12-39930126-INS-625
chr12-39930143-INS-172
chr12-39937439-INS-58
chr12-39941570-INS-440
chr12-39969133-INS-570
chr12-39969979-INS-508

chr12-39980255-INS-2492
chr12-40005234-INS-185
chr12-40059006-INS-50
chr12-40091025-INS-330
chr12-40091479-INS-354
chr12-40124135-INS-3098
chr12-40130593-INS-444
chr12-40159534-INS-3472
chr12-40163686-INS-1502
chr12-40178920-INS-453
chr12-40220444-INS-294
chr12-40222636-INS-70
chr12-40223552-INS-50
chr12-40235270-INS-64
chr12-40306901-INS-96
chr12-40314588-INS-116
chr12-40334770-INS-124
chr12-40361836-INS-76
chr12-40385738-INS-78
chr12-40388042-INS-328
chr12-40443822-INS-60
chr12-40451748-INS-50
chr12-40505940-INS-205
chr12-40506021-INS-82
chr12-40582138-INS-343
chr12-40593190-INS-63
chr12-40609801-INS-1367
chr12-40645046-INS-1433
chr12-40697776-INS-546
chr12-40697791-INS-2637
chr12-40734667-INS-463
chr12-40781186-INS-136
chr12-40799748-INS-50
chr12-40818725-INS-50
chr12-40852686-INS-1497
chr12-40921032-INS-342
chr12-40938512-INS-270
chr12-40943021-INS-58
chr12-40949387-INS-4502
chr12-40960394-INS-77
chr12-40981580-INS-1097
chr12-40985394-INS-50
chr12-41039867-INS-126
chr12-41041109-INS-63
chr12-41043443-INS-56
chr12-41048229-INS-92
chr12-41051035-INS-6296
chr12-41079511-INS-178
chr12-41080424-INS-1293
chr12-41117236-INS-577
chr12-41159614-INS-2605
chr12-41175308-INS-163
chr12-41197677-INS-52
chr12-41208656-INS-3205
chr12-41221851-INS-76
chr12-41253819-INS-187
chr12-41271545-INS-77
chr12-41305236-INS-60
chr12-41305475-INS-125
chr12-41328747-INS-3033
chr12-41345332-INS-1421
chr12-41378370-INS-80
chr12-41379162-INS-64
chr12-41379243-INS-162
chr12-41396938-INS-60
chr12-41437265-INS-92
chr12-41445324-INS-246
chr12-41445746-INS-438
chr12-41536547-INS-58
chr12-41687294-INS-8950
chr12-41687936-INS-50
chr12-41720269-INS-54
chr12-41729476-INS-492
chr12-41791254-INS-104
chr12-41915391-INS-4230
chr12-41918267-INS-6927
chr12-41969631-INS-55

chr12-41983239-INS-6437
chr12-42119118-INS-498
chr12-42130770-INS-6401
chr12-42140363-INS-1855
chr12-42180140-INS-2998
chr12-42208019-INS-174
chr12-42226636-INS-190
chr12-42266902-INS-440
chr12-42276562-INS-64
chr12-42276876-INS-51
chr12-42297934-INS-479
chr12-42328896-INS-6243
chr12-42330539-INS-420
chr12-42341519-INS-62
chr12-42349274-INS-124
chr12-42426745-INS-6405
chr12-42434977-INS-6314
chr12-42453500-INS-310
chr12-42470204-INS-56
chr12-42479351-INS-1114
chr12-42523507-INS-58
chr12-42532620-INS-3048
chr12-42555658-INS-52
chr12-42582884-INS-64
chr12-42626376-INS-95
chr12-42626514-INS-166
chr12-42626524-INS-75
chr12-42679000-INS-122
chr12-42679000-INS-50
chr12-42702220-INS-1892
chr12-42720573-INS-50
chr12-42753962-INS-194
chr12-42767877-INS-376
chr12-42772543-INS-100
chr12-42784481-INS-96
chr12-42796527-INS-1327
chr12-42846735-INS-56
chr12-42847392-INS-2765
chr12-42862279-INS-987
chr12-42868627-INS-207
chr12-42935180-INS-50
chr12-42955646-INS-52
chr12-42957112-INS-3071
chr12-42964486-INS-6859
chr12-42992770-INS-50
chr12-42993014-INS-114
chr12-42996652-INS-53
chr12-43074481-INS-2975
chr12-43085673-INS-3221
chr12-43104223-INS-2508
chr12-43107119-INS-532
chr12-43173218-INS-66
chr12-43210543-INS-122
chr12-43236370-INS-219
chr12-43243199-INS-75
chr12-43248602-INS-2836
chr12-43260032-INS-770
chr12-43282532-INS-118
chr12-43284843-INS-417
chr12-43300592-INS-55
chr12-43302044-INS-1627
chr12-43303384-INS-129
chr12-43331271-INS-6367
chr12-43381376-INS-50
chr12-43401805-INS-50
chr12-43416310-INS-52
chr12-43470312-INS-6922
chr12-43484523-INS-68
chr12-43498280-INS-60
chr12-43507637-INS-182
chr12-43559627-INS-6789
chr12-43596112-INS-52
chr12-43606030-INS-56
chr12-43612821-INS-562
chr12-43620638-INS-73
chr12-43644817-INS-2817
chr12-43646549-INS-11446

chr12-43650421-INS-2356
chr12-43664233-INS-1616
chr12-43678312-INS-2634
chr12-43715104-INS-52
chr12-43728579-INS-806
chr12-43731010-INS-862
chr12-43735931-INS-55
chr12-43736137-INS-200
chr12-43740463-INS-90
chr12-43747568-INS-6090
chr12-43807569-INS-52
chr12-43827012-INS-64
chr12-43839161-INS-12257
chr12-43853135-INS-312
chr12-43854537-INS-54
chr12-43883546-INS-335
chr12-43928655-INS-2500
chr12-43945892-INS-50
chr12-43947983-INS-180
chr12-43956390-INS-492
chr12-43970338-INS-1620
chr12-44086471-INS-134
chr12-44114963-INS-2917
chr12-44139579-INS-321
chr12-44167809-INS-6347
chr12-44171774-INS-828
chr12-44213005-INS-252
chr12-44243818-INS-1151
chr12-44243882-INS-113
chr12-44263841-INS-69
chr12-44268673-INS-84
chr12-44388189-INS-467
chr12-44404064-INS-100
chr12-44407879-INS-53
chr12-44426727-INS-340
chr12-44460436-INS-50
chr12-44460710-INS-130
chr12-44470389-INS-70
chr12-44608090-INS-601
chr12-44611754-INS-547
chr12-44622060-INS-4686
chr12-44635069-INS-205
chr12-44638360-INS-50
chr12-44641917-INS-1019
chr12-44652779-INS-65
chr12-44652939-INS-52
chr12-44670304-INS-1351
chr12-44670791-INS-1422
chr12-44673932-INS-661
chr12-44675149-INS-198
chr12-44845823-INS-2499
chr12-44860116-INS-6801
chr12-44892491-INS-62
chr12-44903263-INS-2027
chr12-44904781-INS-51
chr12-44932742-INS-155
chr12-45028605-INS-1460
chr12-45030182-INS-168
chr12-45056136-INS-88
chr12-45056697-INS-64
chr12-45125937-INS-66
chr12-45160178-INS-7113
chr12-45165008-INS-51
chr12-45165481-INS-66
chr12-45165495-INS-4906
chr12-45181851-INS-160
chr12-45195239-INS-1861
chr12-45209123-INS-6364
chr12-45212021-INS-60
chr12-45216851-INS-1889
chr12-45224047-INS-4509
chr12-45248593-INS-1122
chr12-45253855-INS-6386
chr12-45259458-INS-6328
chr12-45297337-INS-294
chr12-45368906-INS-52
chr12-45430979-INS-91

chr12-45446804-INS-156
chr12-45447474-INS-115
chr12-45449210-INS-518
chr12-45477993-INS-90
chr12-45515739-INS-56
chr12-45520577-INS-4607
chr12-45553016-INS-68
chr12-45597055-INS-323
chr12-45664698-INS-6182
chr12-45677669-INS-426
chr12-45749530-INS-64
chr12-45765965-INS-463
chr12-45804352-INS-195
chr12-45842937-INS-190
chr12-45845274-INS-585
chr12-45859924-INS-200
chr12-45902952-INS-1936
chr12-45920731-INS-720
chr12-45938904-INS-641
chr12-45939351-INS-86
chr12-45940938-INS-4629
chr12-46086309-INS-171
chr12-46139424-INS-1748
chr12-46169848-INS-210
chr12-46177411-INS-66
chr12-46182905-INS-60
chr12-46206777-INS-3188
chr12-46211053-INS-52
chr12-46236209-INS-59
chr12-46237097-INS-194
chr12-46237945-INS-58
chr12-46272223-INS-56
chr12-46307052-INS-50
chr12-46409346-INS-62
chr12-46417498-INS-205
chr12-46452580-INS-345
chr12-46511268-INS-13181
chr12-46560855-INS-1543
chr12-46599005-INS-105
chr12-46601181-INS-58
chr12-46661990-INS-374
chr12-46726613-INS-148
chr12-46829652-INS-75
chr12-46831128-INS-106
chr12-46832067-INS-193
chr12-46834289-INS-1662
chr12-46873942-INS-781
chr12-46906300-INS-1162
chr12-46909637-INS-221
chr12-46925293-INS-355
chr12-46925465-INS-114
chr12-46925670-INS-62
chr12-46958666-INS-119
chr12-46987864-INS-3016
chr12-46988625-INS-6131
chr12-47037513-INS-55
chr12-47043895-INS-180
chr12-47043895-INS-402
chr12-47044119-INS-743
chr12-47051874-INS-229
chr12-47089407-INS-51
chr12-47135562-INS-108
chr12-47179511-INS-335
chr12-47230660-INS-57
chr12-47230784-INS-6362
chr12-47351657-INS-55
chr12-47353715-INS-5526
chr12-47373364-INS-53
chr12-47373567-INS-64
chr12-47419293-INS-933
chr12-47437559-INS-54
chr12-47444280-INS-66
chr12-47456649-INS-57
chr12-47463115-INS-1992
chr12-47465253-INS-2151
chr12-47498420-INS-747
chr12-47498864-INS-868

chr12-47545209-INS-192
chr12-47549556-INS-694
chr12-47595406-INS-59
chr12-47636278-INS-64
chr12-47701688-INS-1296
chr12-47733045-INS-5936
chr12-47746511-INS-2881
chr12-47815552-INS-7211
chr12-47820691-INS-4136
chr12-47820698-INS-1737
chr12-47872030-INS-106
chr12-47893552-INS-501
chr12-47935330-INS-6650
chr12-47940438-INS-3024
chr12-47997758-INS-144
chr12-48062070-INS-3133
chr12-48070795-INS-66
chr12-48075400-INS-60
chr12-48078559-INS-418
chr12-48144219-INS-6762
chr12-48161713-INS-52
chr12-48218164-INS-138
chr12-48241721-INS-50
chr12-48243539-INS-108
chr12-48349395-INS-3671
chr12-48429598-INS-974
chr12-48459452-INS-694
chr12-48472147-INS-203
chr12-48473132-INS-191
chr12-48481739-INS-62
chr12-48507191-INS-50
chr12-48538822-INS-54
chr12-48629283-INS-74
chr12-48666012-INS-140
chr12-48680195-INS-65
chr12-48703622-INS-6628
chr12-48800531-INS-3004
chr12-48831524-INS-543
chr12-48853137-INS-1089
chr12-48921564-INS-104
chr12-48941947-INS-7063
chr12-49009560-INS-1978
chr12-49010805-INS-76
chr12-49062739-INS-1986
chr12-49155136-INS-204
chr12-49169817-INS-198
chr12-49169817-INS-96
chr12-49217994-INS-441
chr12-49270742-INS-839
chr12-49285681-INS-1631
chr12-49333120-INS-52
chr12-49365249-INS-397
chr12-49394673-INS-433
chr12-49490849-INS-60
chr12-49495601-INS-1253
chr12-49545521-INS-108
chr12-49616449-INS-432
chr12-49687846-INS-135
chr12-49694615-INS-6681
chr12-49702049-INS-1017
chr12-49715389-INS-6018
chr12-49718770-INS-401
chr12-49750200-INS-3042
chr12-49772541-INS-160
chr12-49774209-INS-65
chr12-49799652-INS-6385
chr12-49862666-INS-445
chr12-49896541-INS-8291
chr12-49917414-INS-220
chr12-50017066-INS-72
chr12-50056217-INS-54
chr12-50061044-INS-319
chr12-50148304-INS-1581
chr12-50239854-INS-85
chr12-50240662-INS-82
chr12-50244028-INS-100
chr12-50252361-INS-5463

chr12-50267569-INS-4671
chr12-50368139-INS-2301
chr12-50396318-INS-540
chr12-50398282-INS-2876
chr12-50421199-INS-60
chr12-50437552-INS-449
chr12-50511459-INS-50
chr12-50540073-INS-838
chr12-50598753-INS-204
chr12-50657335-INS-2075
chr12-50666826-INS-1349
chr12-50769305-INS-349
chr12-50775058-INS-148
chr12-50838489-INS-55
chr12-50866734-INS-5561
chr12-50954236-INS-5473
chr12-50978331-INS-2418
chr12-50981014-INS-423
chr12-51138483-INS-87
chr12-51150905-INS-168
chr12-51179333-INS-223
chr12-51179400-INS-550
chr12-51188110-INS-1213
chr12-51198174-INS-906
chr12-51231287-INS-3559
chr12-51260834-INS-475
chr12-51308178-INS-71
chr12-51334831-INS-157
chr12-51344924-INS-76
chr12-51382927-INS-196
chr12-51438396-INS-215
chr12-51449869-INS-92
chr12-51450703-INS-1003
chr12-51490573-INS-1045
chr12-51516609-INS-307
chr12-51516613-INS-64
chr12-51530680-INS-120
chr12-51538774-INS-874
chr12-51545915-INS-1738
chr12-51550345-INS-214
chr12-51603221-INS-151
chr12-51603239-INS-66
chr12-51603516-INS-116
chr12-51603770-INS-348
chr12-51607572-INS-300
chr12-51607788-INS-74
chr12-51611960-INS-1540
chr12-51621661-INS-68
chr12-51660678-INS-209
chr12-51746854-INS-429
chr12-51754620-INS-53
chr12-51754721-INS-3362
chr12-51757978-INS-190
chr12-51758524-INS-183
chr12-51770552-INS-765
chr12-51771585-INS-128
chr12-51810429-INS-206
chr12-51845288-INS-58
chr12-51917509-INS-405
chr12-51934719-INS-132
chr12-51938839-INS-56
chr12-51947941-INS-164
chr12-51947952-INS-1226
chr12-51948657-INS-92
chr12-52035275-INS-423
chr12-52066498-INS-414
chr12-52099590-INS-1102
chr12-52108498-INS-58
chr12-52120785-INS-622
chr12-52219579-INS-3550
chr12-52273207-INS-64
chr12-52293124-INS-1205
chr12-52303905-INS-942
chr12-52323575-INS-2206
chr12-52379150-INS-50
chr12-52396018-INS-315
chr12-52419191-INS-60

chr12-52438425-INS-78
chr12-52458288-INS-183
chr12-52664407-INS-178
chr12-52667692-INS-1626
chr12-52678897-INS-109
chr12-52725472-INS-202
chr12-52771778-INS-66
chr12-52814759-INS-51
chr12-52885144-INS-150
chr12-52907765-INS-106
chr12-52913835-INS-1108
chr12-52943305-INS-740
chr12-52953758-INS-149
chr12-52954089-INS-301
chr12-52954183-INS-726
chr12-53046804-INS-6724
chr12-53135217-INS-130
chr12-53178163-INS-74
chr12-53178775-INS-61
chr12-53182455-INS-61
chr12-53182634-INS-168
chr12-53204700-INS-6327
chr12-53279312-INS-59
chr12-53316835-INS-207
chr12-53317580-INS-148
chr12-53326123-INS-99
chr12-53414516-INS-55
chr12-53538313-INS-162
chr12-53538364-INS-69
chr12-53586422-INS-52
chr12-53659109-INS-210
chr12-53691250-INS-50
chr12-53710063-INS-77
chr12-53820196-INS-6705
chr12-53991706-INS-286
chr12-54142972-INS-51
chr12-54192353-INS-109
chr12-54193033-INS-6432
chr12-54202044-INS-1557
chr12-54202104-INS-242
chr12-54208093-INS-52
chr12-54208139-INS-145
chr12-54208519-INS-50
chr12-54213844-INS-1766
chr12-54219272-INS-56
chr12-54237260-INS-426
chr12-54262040-INS-54
chr12-54264418-INS-77
chr12-54270996-INS-76
chr12-54317762-INS-467
chr12-54320299-INS-165
chr12-54320359-INS-200
chr12-54339552-INS-3215
chr12-54364498-INS-74
chr12-54373598-INS-84
chr12-54426078-INS-50
chr12-54458241-INS-50
chr12-54462041-INS-68
chr12-54462076-INS-800
chr12-54466715-INS-53
chr12-54470822-INS-1035
chr12-54501081-INS-415
chr12-54523230-INS-119
chr12-54557124-INS-56
chr12-54568065-INS-130
chr12-54611975-INS-87
chr12-54634409-INS-256
chr12-54637229-INS-70
chr12-54655556-INS-161
chr12-54730051-INS-2352
chr12-54735083-INS-205
chr12-54758097-INS-105
chr12-54764144-INS-54
chr12-54766786-INS-73
chr12-54772696-INS-5525
chr12-54783893-INS-71
chr12-54795929-INS-80

chr12-54811494-INS-343
chr12-54838122-INS-56
chr12-54848367-INS-155
chr12-54878394-INS-50
chr12-54878779-INS-309
chr12-54887402-INS-232
chr12-54922473-INS-50
chr12-54944012-INS-233
chr12-54946588-INS-1300
chr12-55014806-INS-135
chr12-55025189-INS-432
chr12-55108064-INS-64
chr12-55114077-INS-91
chr12-55121037-INS-497
chr12-55155263-INS-217
chr12-55320108-INS-72
chr12-55324513-INS-99
chr12-55325032-INS-56
chr12-55385139-INS-345
chr12-55386076-INS-78
chr12-55402440-INS-285
chr12-55418300-INS-600
chr12-55477405-INS-252
chr12-55482283-INS-101
chr12-55496495-INS-233
chr12-55498830-INS-52
chr12-55585443-INS-384
chr12-55603319-INS-55
chr12-55603501-INS-55
chr12-55626490-INS-76
chr12-55627319-INS-60
chr12-55705450-INS-81
chr12-55710947-INS-50
chr12-55713270-INS-67
chr12-55803810-INS-91
chr12-55820461-INS-60
chr12-55862379-INS-248
chr12-55879383-INS-164
chr12-55886602-INS-51
chr12-55893123-INS-56
chr12-55893162-INS-139
chr12-55943881-INS-449
chr12-55955263-INS-521
chr12-56090651-INS-68
chr12-56098933-INS-100
chr12-56108964-INS-1624
chr12-56109180-INS-84
chr12-56115352-INS-1829
chr12-56118252-INS-498
chr12-56194652-INS-2998
chr12-56200585-INS-54
chr12-56247611-INS-102
chr12-56301709-INS-310
chr12-56309633-INS-85
chr12-56310372-INS-162
chr12-56327721-INS-788
chr12-56337492-INS-76
chr12-56489278-INS-50
chr12-56491606-INS-1922
chr12-56544046-INS-70
chr12-56563290-INS-147
chr12-56589939-INS-226
chr12-56597724-INS-6878
chr12-56642854-INS-124
chr12-56650219-INS-204
chr12-56663940-INS-188
chr12-56804886-INS-50
chr12-56805061-INS-60
chr12-56808983-INS-70
chr12-56839975-INS-112
chr12-56845279-INS-60
chr12-56893580-INS-119
chr12-56906084-INS-224
chr12-56927473-INS-157
chr12-56973748-INS-548
chr12-56979698-INS-176
chr12-56983805-INS-6594

chr12-57061922-INS-236
chr12-57179366-INS-181
chr12-57179702-INS-61
chr12-57198912-INS-4829
chr12-57233459-INS-223
chr12-57233628-INS-92
chr12-57271322-INS-8079
chr12-57292777-INS-247
chr12-57313457-INS-66
chr12-57346858-INS-3028
chr12-57365580-INS-7190
chr12-57380861-INS-59
chr12-57390266-INS-3070
chr12-57412115-INS-316
chr12-57463040-INS-50
chr12-57474612-INS-579
chr12-57478961-INS-759
chr12-57481709-INS-160
chr12-57485789-INS-235
chr12-57568380-INS-83
chr12-57571900-INS-563
chr12-57609220-INS-1873
chr12-57610279-INS-1873
chr12-57678701-INS-98
chr12-57689992-INS-90
chr12-57709667-INS-432
chr12-57735982-INS-63
chr12-57755668-INS-495
chr12-57757093-INS-50
chr12-57760633-INS-3512
chr12-57800643-INS-420
chr12-57826613-INS-128
chr12-57826661-INS-329
chr12-57838383-INS-1469
chr12-57840993-INS-13314
chr12-57874617-INS-56
chr12-57916930-INS-1500
chr12-57925786-INS-6433
chr12-57927350-INS-3021
chr12-57930471-INS-1631
chr12-57940829-INS-1043
chr12-58001559-INS-7188
chr12-58008427-INS-74
chr12-58022438-INS-63
chr12-58052153-INS-109
chr12-58062799-INS-3321
chr12-58081240-INS-1883
chr12-58091670-INS-136
chr12-58119744-INS-1647
chr12-58191534-INS-56
chr12-58203667-INS-208
chr12-58266061-INS-164
chr12-58266076-INS-745
chr12-58282164-INS-59
chr12-58283296-INS-50
chr12-58375670-INS-2113
chr12-58438242-INS-171
chr12-58438608-INS-13426
chr12-58451430-INS-56
chr12-58462859-INS-2836
chr12-58481005-INS-1329
chr12-58514142-INS-6200
chr12-58546427-INS-199
chr12-58550177-INS-139
chr12-58581764-INS-6480
chr12-58588186-INS-146
chr12-58635930-INS-198
chr12-58706664-INS-201
chr12-58719478-INS-115
chr12-58719501-INS-449
chr12-58746919-INS-796
chr12-58756452-INS-262
chr12-58792154-INS-246
chr12-58792258-INS-112
chr12-58792318-INS-108
chr12-58811619-INS-442
chr12-58828245-INS-324

chr12-58847075-INS-291
chr12-58875143-INS-6250
chr12-58890219-INS-118
chr12-58909093-INS-2255
chr12-58977174-INS-54
chr12-58988053-INS-108
chr12-59000283-INS-134
chr12-59000329-INS-54
chr12-59045952-INS-398
chr12-59049660-INS-68
chr12-59073710-INS-198
chr12-59078670-INS-222
chr12-59111569-INS-201
chr12-59112533-INS-78
chr12-59144406-INS-203
chr12-59167786-INS-54
chr12-59187699-INS-53
chr12-59197793-INS-975
chr12-59259813-INS-6171
chr12-59288410-INS-1565
chr12-59296711-INS-1624
chr12-59375170-INS-60
chr12-59396104-INS-5999
chr12-59432431-INS-119
chr12-59432474-INS-59
chr12-59484225-INS-51
chr12-59509669-INS-313
chr12-59523738-INS-53
chr12-59523785-INS-282
chr12-59538147-INS-116
chr12-59556497-INS-64
chr12-59594820-INS-1010
chr12-59594820-INS-94
chr12-59709145-INS-326
chr12-59717493-INS-60
chr12-59717529-INS-143
chr12-59717533-INS-543
chr12-59754895-INS-56
chr12-59780296-INS-177
chr12-59795660-INS-95
chr12-59830054-INS-591
chr12-59847134-INS-1419
chr12-59884609-INS-3001
chr12-59952608-INS-6301
chr12-59971034-INS-615
chr12-59972468-INS-6808
chr12-59994196-INS-199
chr12-60031572-INS-6228
chr12-60032874-INS-94
chr12-60063311-INS-213
chr12-60074774-INS-112
chr12-60088787-INS-52
chr12-60090448-INS-2902
chr12-60106303-INS-423
chr12-60146174-INS-6180
chr12-60184691-INS-297
chr12-60224714-INS-6197
chr12-60226658-INS-142
chr12-60380687-INS-216
chr12-60397683-INS-421
chr12-60397719-INS-54
chr12-60397964-INS-147
chr12-60404150-INS-1453
chr12-60413344-INS-86
chr12-60475221-INS-214
chr12-60524295-INS-3158
chr12-60542947-INS-1445
chr12-60632010-INS-4041
chr12-60643043-INS-54
chr12-60657561-INS-9319
chr12-60662979-INS-100
chr12-60738332-INS-6455
chr12-60769771-INS-3135
chr12-60813357-INS-50
chr12-60814030-INS-811
chr12-60827185-INS-3962
chr12-60828662-INS-6482

chr12-60904170-INS-2262
chr12-60908675-INS-2473
chr12-60913146-INS-5241
chr12-60943369-INS-1412
chr12-60972438-INS-4249
chr12-60980606-INS-52
chr12-60999381-INS-2938
chr12-61003428-INS-68
chr12-61033828-INS-6239
chr12-61037132-INS-72
chr12-61039171-INS-52
chr12-61039586-INS-4440
chr12-61045374-INS-967
chr12-61087603-INS-3017
chr12-61095271-INS-84
chr12-61117690-INS-52
chr12-61233129-INS-103
chr12-61235563-INS-1411
chr12-61277977-INS-2011
chr12-61284271-INS-51
chr12-61365823-INS-88
chr12-61379588-INS-1866
chr12-61383955-INS-8801
chr12-61392522-INS-50
chr12-61407987-INS-7243
chr12-61427128-INS-105
chr12-61427250-INS-368
chr12-61427406-INS-70
chr12-61448425-INS-180
chr12-61521626-INS-6365
chr12-61524401-INS-2304
chr12-61531753-INS-2379
chr12-61602840-INS-990
chr12-61606917-INS-4465
chr12-61619632-INS-75
chr12-61620154-INS-69
chr12-61623786-INS-2418
chr12-61662701-INS-475
chr12-61679799-INS-95
chr12-61680019-INS-212
chr12-61852847-INS-59
chr12-61912030-INS-72
chr12-61943428-INS-162
chr12-61949027-INS-1948
chr12-61957049-INS-6217
chr12-61983835-INS-160
chr12-61999353-INS-1944
chr12-62068634-INS-7106
chr12-62071558-INS-50
chr12-62163077-INS-1636
chr12-62180179-INS-184
chr12-62190703-INS-51
chr12-62202648-INS-1196
chr12-62217130-INS-3676
chr12-62245643-INS-1662
chr12-62251616-INS-67
chr12-62262657-INS-54
chr12-62285208-INS-56
chr12-62285497-INS-54
chr12-62288902-INS-3526
chr12-62385207-INS-6239
chr12-62386213-INS-2180
chr12-62412012-INS-5026
chr12-62419582-INS-2684
chr12-62442824-INS-7080
chr12-62467096-INS-6895
chr12-62522423-INS-110
chr12-62568061-INS-84
chr12-62574881-INS-259
chr12-62595993-INS-212
chr12-62596810-INS-58
chr12-62627899-INS-233
chr12-62732539-INS-116
chr12-62751358-INS-6676
chr12-62751927-INS-102
chr12-62759014-INS-1829
chr12-62792878-INS-432

chr12-62822399-INS-2483
chr12-62857913-INS-70
chr12-62865070-INS-1417
chr12-62877141-INS-7260
chr12-62923091-INS-6495
chr12-62928519-INS-199
chr12-62939610-INS-6916
chr12-62947052-INS-6492
chr12-62950883-INS-56
chr12-62952259-INS-6386
chr12-62982608-INS-114
chr12-63049663-INS-1578
chr12-63126056-INS-790
chr12-63149037-INS-664
chr12-63152235-INS-401
chr12-63188220-INS-1054
chr12-63188229-INS-2222
chr12-63192750-INS-893
chr12-63238456-INS-2799
chr12-63247531-INS-7286
chr12-63248563-INS-313
chr12-63249277-INS-177
chr12-63249277-INS-70
chr12-63265382-INS-9659
chr12-63276694-INS-103
chr12-63282355-INS-281
chr12-63287717-INS-1154
chr12-63297881-INS-3790
chr12-63326165-INS-6302
chr12-63328674-INS-521
chr12-63355511-INS-1954
chr12-63356294-INS-3678
chr12-63357016-INS-322
chr12-63392251-INS-1885
chr12-63420230-INS-56
chr12-63474411-INS-52
chr12-63541330-INS-3191
chr12-63542393-INS-6617
chr12-63572717-INS-351
chr12-63594397-INS-76
chr12-63597082-INS-83
chr12-63607976-INS-1990
chr12-63713116-INS-195
chr12-63714691-INS-4804
chr12-63806014-INS-53
chr12-63809646-INS-6391
chr12-63825635-INS-164
chr12-63872680-INS-84
chr12-63872792-INS-112
chr12-63931199-INS-74
chr12-63938155-INS-1844
chr12-63974585-INS-1423
chr12-63977281-INS-2455
chr12-63992660-INS-3561
chr12-63996196-INS-2399
chr12-63996971-INS-2993
chr12-64006113-INS-56
chr12-64013533-INS-1220
chr12-64063849-INS-60
chr12-64069691-INS-545
chr12-64090303-INS-2440
chr12-64120649-INS-116
chr12-64125852-INS-1764
chr12-64168823-INS-6209
chr12-64211642-INS-2944
chr12-64234441-INS-157
chr12-64249747-INS-5062
chr12-64270675-INS-468
chr12-64274569-INS-240
chr12-64314662-INS-53
chr12-64318605-INS-53
chr12-64319169-INS-57
chr12-64330596-INS-50
chr12-64359057-INS-1935
chr12-64361117-INS-4742
chr12-64383572-INS-6238
chr12-64421689-INS-62

chr12-64422086-INS-63
chr12-64422217-INS-309
chr12-64422294-INS-148
chr12-64422940-INS-934
chr12-64471611-INS-350
chr12-64526556-INS-6853
chr12-64551617-INS-92
chr12-64552526-INS-1354
chr12-64552732-INS-6363
chr12-64603581-INS-1158
chr12-64646003-INS-58
chr12-64656249-INS-3957
chr12-64735197-INS-6886
chr12-64737135-INS-58
chr12-64740399-INS-130
chr12-64751018-INS-6530
chr12-64801208-INS-4747
chr12-64841034-INS-50
chr12-64865467-INS-52
chr12-64874927-INS-501
chr12-64883602-INS-381
chr12-64909325-INS-100
chr12-64913178-INS-8639
chr12-64925069-INS-1932
chr12-64967229-INS-129
chr12-64982608-INS-88
chr12-64982736-INS-182
chr12-64982757-INS-88
chr12-64990398-INS-50
chr12-65036547-INS-229
chr12-65037281-INS-68
chr12-65061734-INS-200
chr12-65063941-INS-62
chr12-65067664-INS-218
chr12-65073176-INS-72
chr12-65092717-INS-122
chr12-65132011-INS-218
chr12-65138743-INS-65
chr12-65240593-INS-93
chr12-65253694-INS-95
chr12-65262337-INS-706
chr12-65264706-INS-78
chr12-65349200-INS-249
chr12-65359751-INS-6423
chr12-65405320-INS-1893
chr12-65406941-INS-289
chr12-65415402-INS-4047
chr12-65449663-INS-7418
chr12-65472412-INS-8048
chr12-65522683-INS-1293
chr12-65561046-INS-111
chr12-65561914-INS-92
chr12-65587722-INS-50
chr12-65590064-INS-6339
chr12-65596942-INS-199
chr12-65627621-INS-6465
chr12-65635138-INS-135
chr12-65648246-INS-57
chr12-65694564-INS-2920
chr12-65695073-INS-59
chr12-65724814-INS-1227
chr12-65751458-INS-57
chr12-65763177-INS-4387
chr12-65843998-INS-1352
chr12-65931337-INS-54
chr12-65931701-INS-166
chr12-65971549-INS-163
chr12-65997338-INS-415
chr12-66024502-INS-5074
chr12-66070449-INS-50
chr12-66091933-INS-7098
chr12-66219016-INS-156
chr12-66287770-INS-209
chr12-66307621-INS-208
chr12-66320184-INS-92
chr12-66337607-INS-120
chr12-66337681-INS-248

chr12-66350467-INS-56
chr12-66397376-INS-265
chr12-66406444-INS-1685
chr12-66557585-INS-73
chr12-66603638-INS-62
chr12-66628846-INS-858
chr12-66629184-INS-297
chr12-66640965-INS-80
chr12-66651671-INS-1111
chr12-66664332-INS-130
chr12-66672464-INS-65
chr12-66684975-INS-52
chr12-66713584-INS-50
chr12-66716637-INS-581
chr12-66758561-INS-72
chr12-66798372-INS-1209
chr12-66807665-INS-5543
chr12-66879121-INS-106
chr12-66895364-INS-66
chr12-66919126-INS-327
chr12-66957384-INS-652
chr12-66964284-INS-1493
chr12-67038967-INS-222
chr12-67038967-INS-84
chr12-67042119-INS-735
chr12-67223604-INS-86
chr12-67223605-INS-277
chr12-67234580-INS-234
chr12-67240632-INS-324
chr12-67262550-INS-80
chr12-67287583-INS-1594
chr12-67304892-INS-55
chr12-67304912-INS-130
chr12-67328586-INS-624
chr12-67394953-INS-83
chr12-67402181-INS-74
chr12-67402610-INS-276
chr12-67444708-INS-1918
chr12-67469771-INS-13215
chr12-67502601-INS-6703
chr12-67524490-INS-127
chr12-67525497-INS-61
chr12-67542439-INS-224
chr12-67542503-INS-84
chr12-67549602-INS-330
chr12-67607757-INS-208
chr12-67740197-INS-95
chr12-67797887-INS-1528
chr12-67822426-INS-61
chr12-67840074-INS-146
chr12-67848467-INS-111
chr12-67936749-INS-120
chr12-67945622-INS-312
chr12-67962075-INS-190
chr12-67991958-INS-7883
chr12-68008421-INS-70
chr12-68008448-INS-244
chr12-68029350-INS-6202
chr12-68055577-INS-53
chr12-68075383-INS-3857
chr12-68112851-INS-1465
chr12-68116587-INS-6251
chr12-68145182-INS-663
chr12-68145872-INS-59
chr12-68180050-INS-9013
chr12-68279774-INS-101
chr12-68280078-INS-431
chr12-68289017-INS-6486
chr12-68357352-INS-1002
chr12-68367029-INS-6172
chr12-68370340-INS-54
chr12-68370383-INS-149
chr12-68390144-INS-4046
chr12-68390697-INS-128
chr12-68394647-INS-52
chr12-68398051-INS-56
chr12-68411332-INS-336

chr12-68417062-INS-52
chr12-68440455-INS-54
chr12-68485846-INS-2734
chr12-68527109-INS-54
chr12-68558885-INS-528
chr12-68562694-INS-70
chr12-68662870-INS-131
chr12-68720590-INS-6991
chr12-68810876-INS-51
chr12-68811722-INS-121
chr12-68817787-INS-287
chr12-68875704-INS-7421
chr12-68880709-INS-60
chr12-68882817-INS-56
chr12-68882860-INS-182
chr12-69015528-INS-300
chr12-69022052-INS-206
chr12-69092703-INS-108
chr12-69138715-INS-6610
chr12-69139254-INS-126
chr12-69181796-INS-61
chr12-69192368-INS-116
chr12-69225153-INS-208
chr12-69242087-INS-111
chr12-69250258-INS-93
chr12-69252218-INS-156
chr12-69282325-INS-7083
chr12-69301131-INS-50
chr12-69324107-INS-260
chr12-69340604-INS-120
chr12-69409041-INS-217
chr12-69477918-INS-103
chr12-69478811-INS-132
chr12-69489790-INS-285
chr12-69490735-INS-178
chr12-69495607-INS-1160
chr12-69547346-INS-85
chr12-69551826-INS-100
chr12-69551900-INS-340
chr12-69556660-INS-1107
chr12-69556732-INS-333
chr12-69571252-INS-171
chr12-69574730-INS-168
chr12-69619626-INS-200
chr12-69634796-INS-61
chr12-69636392-INS-52
chr12-69661760-INS-55
chr12-69671017-INS-624
chr12-69697321-INS-215
chr12-69700867-INS-4881
chr12-69700880-INS-1749
chr12-69713233-INS-118
chr12-69714816-INS-3499
chr12-69779082-INS-5507
chr12-69811654-INS-74
chr12-69812189-INS-192
chr12-69817405-INS-3704
chr12-69849795-INS-206
chr12-69902421-INS-50
chr12-69916315-INS-108
chr12-69968683-INS-54
chr12-69988444-INS-50
chr12-70015592-INS-55
chr12-70016481-INS-217
chr12-70063666-INS-199
chr12-70111263-INS-50
chr12-70150867-INS-50
chr12-70168949-INS-81
chr12-70207152-INS-225
chr12-70240301-INS-54
chr12-70245651-INS-567
chr12-70285090-INS-56
chr12-70285816-INS-158
chr12-70318495-INS-56
chr12-70333929-INS-56
chr12-70414919-INS-86
chr12-70424192-INS-474

chr12-70424301-INS-57
chr12-70434933-INS-64
chr12-70455797-INS-2016
chr12-70475958-INS-220
chr12-70478252-INS-162
chr12-70487955-INS-1332
chr12-70494787-INS-62
chr12-70569571-INS-198
chr12-70578694-INS-3065
chr12-70618403-INS-6648
chr12-70624183-INS-2818
chr12-70625406-INS-6236
chr12-70628605-INS-1860
chr12-70663700-INS-100
chr12-70666143-INS-128
chr12-70677291-INS-124
chr12-70685673-INS-7176
chr12-70692302-INS-506
chr12-70693751-INS-57
chr12-70693891-INS-584
chr12-70730439-INS-124
chr12-70736185-INS-608
chr12-70794095-INS-68
chr12-70794873-INS-56
chr12-70811804-INS-107
chr12-70819174-INS-324
chr12-70820148-INS-8985
chr12-70889227-INS-76
chr12-70916424-INS-69
chr12-70924517-INS-200
chr12-70924695-INS-556
chr12-70952383-INS-60
chr12-70995759-INS-195
chr12-71028759-INS-4081
chr12-71041498-INS-496
chr12-71041824-INS-497
chr12-71064701-INS-64
chr12-71065214-INS-52
chr12-71075030-INS-98
chr12-71075299-INS-343
chr12-71162896-INS-56
chr12-71208050-INS-53
chr12-71208329-INS-836
chr12-71215052-INS-166
chr12-71226528-INS-2012
chr12-71236495-INS-212
chr12-71332951-INS-51
chr12-71369915-INS-209
chr12-71375681-INS-111
chr12-71404423-INS-50
chr12-71409708-INS-294
chr12-71409747-INS-54
chr12-71422640-INS-229
chr12-71439373-INS-286
chr12-71462352-INS-84
chr12-71472340-INS-597
chr12-71495774-INS-66
chr12-71508305-INS-50
chr12-71565526-INS-57
chr12-71566851-INS-167
chr12-71566928-INS-73
chr12-71578590-INS-3161
chr12-71583936-INS-5633
chr12-71592556-INS-54
chr12-71593160-INS-7085
chr12-71604044-INS-2114
chr12-71677067-INS-60
chr12-71691585-INS-4972
chr12-71715451-INS-1074
chr12-71729362-INS-79
chr12-71732659-INS-1053
chr12-71734991-INS-9001
chr12-71780299-INS-101
chr12-71785973-INS-105
chr12-71796500-INS-76
chr12-71809375-INS-107
chr12-71817780-INS-87

chr12-71818811-INS-55
chr12-71820383-INS-181
chr12-71820388-INS-51
chr12-71829018-INS-320
chr12-71829048-INS-74
chr12-71864543-INS-181
chr12-71934208-INS-105
chr12-71952339-INS-52
chr12-72044944-INS-62
chr12-72053065-INS-138
chr12-72084355-INS-209
chr12-72106492-INS-139
chr12-72106492-INS-61
chr12-72132649-INS-206
chr12-72134484-INS-242
chr12-72162075-INS-435
chr12-72162326-INS-530
chr12-72186158-INS-129
chr12-72186982-INS-52
chr12-72242976-INS-76
chr12-72246530-INS-160
chr12-72246574-INS-79
chr12-72246610-INS-492
chr12-72311238-INS-84
chr12-72312801-INS-84
chr12-72341802-INS-7026
chr12-72343149-INS-62
chr12-72347621-INS-50
chr12-72354139-INS-165
chr12-72363680-INS-52
chr12-72380390-INS-69
chr12-72380501-INS-414
chr12-72441559-INS-65
chr12-72466546-INS-68
chr12-72479761-INS-54
chr12-72539229-INS-2289
chr12-72542840-INS-63
chr12-72643759-INS-8871
chr12-72674182-INS-50
chr12-72682691-INS-116
chr12-72683060-INS-57
chr12-72694558-INS-80
chr12-72703675-INS-466
chr12-72719598-INS-50
chr12-72725152-INS-58
chr12-72733444-INS-5715
chr12-72738260-INS-189
chr12-72743433-INS-942
chr12-72747564-INS-216
chr12-72747566-INS-63
chr12-72751517-INS-3239
chr12-72755504-INS-196
chr12-72755939-INS-173
chr12-72766094-INS-50
chr12-72777957-INS-147
chr12-72779413-INS-200
chr12-72783004-INS-52
chr12-72784686-INS-177
chr12-72803012-INS-126
chr12-72814197-INS-218
chr12-72826844-INS-66
chr12-72866134-INS-74
chr12-72867438-INS-66
chr12-72870889-INS-56
chr12-72873802-INS-54
chr12-72883720-INS-50
chr12-72898640-INS-60
chr12-72925605-INS-2714
chr12-72938068-INS-63
chr12-72973064-INS-72
chr12-72973314-INS-340
chr12-73039886-INS-2907
chr12-73056736-INS-127
chr12-73068202-INS-50
chr12-73118296-INS-797
chr12-73118473-INS-73
chr12-73140935-INS-227

chr12-73161600-INS-53
chr12-73172917-INS-226
chr12-73179275-INS-290
chr12-73201473-INS-652
chr12-73201961-INS-200
chr12-73253726-INS-80
chr12-73266607-INS-1150
chr12-73354423-INS-86
chr12-73369677-INS-50
chr12-73411466-INS-3089
chr12-73424412-INS-72
chr12-73425236-INS-190
chr12-73431377-INS-145
chr12-73445220-INS-5094
chr12-73446263-INS-2336
chr12-73475799-INS-1327
chr12-73532632-INS-515
chr12-73541512-INS-55
chr12-73603438-INS-1522
chr12-73609785-INS-56
chr12-73612137-INS-54
chr12-73615329-INS-84
chr12-73622624-INS-96
chr12-73623462-INS-199
chr12-73628053-INS-64
chr12-73633688-INS-67
chr12-73689551-INS-80
chr12-73729487-INS-62
chr12-73782454-INS-78
chr12-73789063-INS-77
chr12-73791841-INS-64
chr12-73813512-INS-261
chr12-73819067-INS-138
chr12-73820030-INS-513
chr12-73947464-INS-152
chr12-73953008-INS-81
chr12-74017963-INS-1107
chr12-74018170-INS-168
chr12-74040778-INS-105
chr12-74086737-INS-1246
chr12-74087098-INS-533
chr12-74097531-INS-9574
chr12-74103019-INS-187
chr12-74103019-INS-423
chr12-74104821-INS-6606
chr12-74274790-INS-276
chr12-74321804-INS-62
chr12-74358026-INS-375
chr12-74360972-INS-111
chr12-74466401-INS-58
chr12-74478036-INS-1255
chr12-74552455-INS-4628
chr12-74677169-INS-1832
chr12-74680720-INS-6686
chr12-74738610-INS-1485
chr12-74786129-INS-1552
chr12-74795200-INS-52
chr12-74799205-INS-7318
chr12-74810433-INS-210
chr12-74827681-INS-109
chr12-74856063-INS-2355
chr12-74865618-INS-2008
chr12-74914911-INS-58
chr12-75025218-INS-474
chr12-75031289-INS-3153
chr12-75035054-INS-86
chr12-75074778-INS-58
chr12-75120537-INS-50
chr12-75150716-INS-2227
chr12-75165482-INS-51
chr12-75166547-INS-115
chr12-75182124-INS-157
chr12-75186910-INS-200
chr12-75208831-INS-71
chr12-75225284-INS-56
chr12-75233050-INS-484
chr12-75247354-INS-98

chr12-75263205-INS-138
chr12-75291953-INS-6983
chr12-75316623-INS-230
chr12-75322540-INS-113
chr12-75334267-INS-1634
chr12-75372303-INS-103
chr12-75398511-INS-755
chr12-75421597-INS-76
chr12-75449937-INS-3134
chr12-75487259-INS-2392
chr12-75487273-INS-9487
chr12-75601750-INS-152
chr12-75670034-INS-82
chr12-75702060-INS-52
chr12-75707107-INS-335
chr12-75718426-INS-182
chr12-75741531-INS-230
chr12-75764037-INS-281
chr12-75807529-INS-207
chr12-75807976-INS-122
chr12-75808028-INS-696
chr12-75840020-INS-200
chr12-75849883-INS-228
chr12-75903613-INS-144
chr12-75995163-INS-88
chr12-75996710-INS-68
chr12-76008918-INS-480
chr12-76018947-INS-67
chr12-76019415-INS-92
chr12-76031102-INS-213
chr12-76034810-INS-87
chr12-76035362-INS-50
chr12-76140157-INS-201
chr12-76148149-INS-164
chr12-76211284-INS-1285
chr12-76230225-INS-80
chr12-76237984-INS-218
chr12-76251631-INS-4318
chr12-76279320-INS-71
chr12-76280545-INS-100
chr12-76280956-INS-282
chr12-76280985-INS-1471
chr12-76402319-INS-52
chr12-76431359-INS-111
chr12-76502797-INS-215
chr12-76514699-INS-243
chr12-76558616-INS-376
chr12-76716129-INS-73
chr12-76762356-INS-50
chr12-76828140-INS-87
chr12-76852519-INS-2878
chr12-76872824-INS-655
chr12-76905930-INS-202
chr12-76906036-INS-216
chr12-76913716-INS-50
chr12-76937048-INS-295
chr12-76948699-INS-91
chr12-76949519-INS-122
chr12-76953206-INS-101
chr12-76953359-INS-223
chr12-76965137-INS-621
chr12-76975971-INS-234
chr12-77034377-INS-63
chr12-77048163-INS-56
chr12-77065714-INS-202
chr12-77077277-INS-675
chr12-77079549-INS-243
chr12-77079549-INS-99
chr12-77083657-INS-54
chr12-77103360-INS-89
chr12-77135661-INS-55
chr12-77144357-INS-2395
chr12-77216679-INS-80
chr12-77218664-INS-50
chr12-77228442-INS-70
chr12-77258425-INS-203
chr12-77293681-INS-1455

chr12-77379917-INS-247
chr12-77451624-INS-766
chr12-77468863-INS-55
chr12-77498864-INS-74
chr12-77547785-INS-60
chr12-77559534-INS-58
chr12-77590170-INS-1348
chr12-77624345-INS-63
chr12-77649727-INS-114
chr12-77708116-INS-55
chr12-77720722-INS-84
chr12-77895626-INS-52
chr12-77955272-INS-309
chr12-77959436-INS-296
chr12-77993330-INS-551
chr12-77995575-INS-188
chr12-78016424-INS-6407
chr12-78052856-INS-215
chr12-78109539-INS-108
chr12-78160008-INS-6609
chr12-78200250-INS-50
chr12-78207983-INS-50
chr12-78261000-INS-216
chr12-78305039-INS-692
chr12-78324930-INS-189
chr12-78325797-INS-197
chr12-78325937-INS-264
chr12-78359927-INS-50
chr12-78585433-INS-88
chr12-78700924-INS-108
chr12-78708892-INS-261
chr12-78859935-INS-56
chr12-78894987-INS-60
chr12-78895957-INS-256
chr12-78903865-INS-67
chr12-78917677-INS-197
chr12-78952960-INS-54
chr12-78979819-INS-83
chr12-78993743-INS-172
chr12-79018454-INS-1971
chr12-79020630-INS-185
chr12-79067442-INS-68
chr12-79074355-INS-218
chr12-79075340-INS-52
chr12-79113830-INS-51
chr12-79129444-INS-50
chr12-79181169-INS-64
chr12-79228995-INS-224
chr12-79284214-INS-1200
chr12-79310283-INS-538
chr12-79328670-INS-59
chr12-79366158-INS-51
chr12-79400903-INS-205
chr12-79451769-INS-6647
chr12-79492009-INS-56
chr12-79492420-INS-137
chr12-79537881-INS-197
chr12-79618100-INS-53
chr12-79641833-INS-94
chr12-79646851-INS-96
chr12-79698389-INS-328
chr12-79715161-INS-60
chr12-79724767-INS-136
chr12-79728152-INS-50
chr12-79735473-INS-121
chr12-79785465-INS-53
chr12-79866408-INS-159
chr12-79875319-INS-84
chr12-79976290-INS-84
chr12-79985755-INS-104
chr12-79991385-INS-2822
chr12-80029874-INS-214
chr12-80077473-INS-64
chr12-80165086-INS-65
chr12-80176111-INS-206
chr12-80193301-INS-52
chr12-80203353-INS-254

chr12-80209613-INS-84
chr12-80261429-INS-106
chr12-80298453-INS-60
chr12-80312162-INS-84
chr12-80314143-INS-541
chr12-80322015-INS-379
chr12-80341385-INS-6442
chr12-80408420-INS-249
chr12-80411415-INS-227
chr12-80547240-INS-217
chr12-80576084-INS-180
chr12-80640034-INS-107
chr12-80672515-INS-130
chr12-80699084-INS-334
chr12-80735541-INS-80
chr12-80783028-INS-203
chr12-80792461-INS-70
chr12-80794290-INS-57
chr12-80828498-INS-209
chr12-80889096-INS-326
chr12-80930515-INS-203
chr12-80931068-INS-179
chr12-80935837-INS-59
chr12-80968531-INS-207
chr12-80986272-INS-424
chr12-81023776-INS-3127
chr12-81066671-INS-50
chr12-81114557-INS-145
chr12-81119036-INS-58
chr12-81119191-INS-1515
chr12-81160501-INS-203
chr12-81293109-INS-52
chr12-81313930-INS-50
chr12-81345902-INS-919
chr12-81544513-INS-66
chr12-81550698-INS-56
chr12-81584609-INS-4908
chr12-81655740-INS-59
chr12-81673533-INS-3063
chr12-81701155-INS-88
chr12-81708255-INS-60
chr12-81776705-INS-74
chr12-81788393-INS-187
chr12-81800951-INS-922
chr12-81803978-INS-130
chr12-81811046-INS-50
chr12-81818142-INS-172
chr12-81818327-INS-54
chr12-81821749-INS-52
chr12-81826111-INS-501
chr12-81838191-INS-2495
chr12-81845793-INS-107
chr12-81850446-INS-219
chr12-81855842-INS-392
chr12-81855899-INS-2268
chr12-81877253-INS-108
chr12-81898597-INS-61
chr12-81903187-INS-224
chr12-81908355-INS-50
chr12-81917388-INS-199
chr12-81927610-INS-89
chr12-81931859-INS-76
chr12-81932513-INS-89
chr12-81936904-INS-99
chr12-81968281-INS-60
chr12-81982512-INS-58
chr12-81986096-INS-124
chr12-82001509-INS-362
chr12-82029612-INS-224
chr12-82032296-INS-1680
chr12-82053149-INS-123
chr12-82144753-INS-54
chr12-82156117-INS-313
chr12-82156199-INS-68
chr12-82177952-INS-98
chr12-82196420-INS-219
chr12-82214092-INS-52

chr12-82236804-INS-124
chr12-82241055-INS-156
chr12-82291192-INS-206
chr12-82328460-INS-912
chr12-82337382-INS-300
chr12-82340913-INS-155
chr12-82344831-INS-150
chr12-82350514-INS-4876
chr12-82354195-INS-149
chr12-82366088-INS-3729
chr12-82377318-INS-50
chr12-82415617-INS-237
chr12-82492277-INS-1696
chr12-82498134-INS-51
chr12-82514634-INS-68
chr12-82515786-INS-382
chr12-82539480-INS-51
chr12-82585677-INS-4019
chr12-82596018-INS-2891
chr12-82596018-INS-458
chr12-82608393-INS-6471
chr12-82654438-INS-259
chr12-82692995-INS-1174
chr12-82705932-INS-61
chr12-82726121-INS-451
chr12-82753906-INS-208
chr12-82772658-INS-1887
chr12-82818156-INS-51
chr12-82828936-INS-76
chr12-82881836-INS-578
chr12-82895369-INS-1357
chr12-82909128-INS-2264
chr12-82930285-INS-62
chr12-82988256-INS-166
chr12-82992234-INS-126
chr12-83015574-INS-55
chr12-83061681-INS-167
chr12-83067255-INS-163
chr12-83123799-INS-50
chr12-83130291-INS-98
chr12-83159218-INS-84
chr12-83165626-INS-248
chr12-83243125-INS-656
chr12-83243155-INS-88
chr12-83244913-INS-1152
chr12-83244913-INS-62
chr12-83245325-INS-53
chr12-83334885-INS-1565
chr12-83380843-INS-70
chr12-83396983-INS-1866
chr12-83413850-INS-285
chr12-83413886-INS-51
chr12-83453805-INS-65
chr12-83487641-INS-1119
chr12-83509492-INS-134
chr12-83510630-INS-263
chr12-83517572-INS-88
chr12-83525853-INS-442
chr12-83526656-INS-327
chr12-83548517-INS-673
chr12-83561553-INS-3959
chr12-83564963-INS-188
chr12-83583585-INS-434
chr12-83596876-INS-193
chr12-83606993-INS-198
chr12-83612328-INS-158
chr12-83616610-INS-229
chr12-83620610-INS-87
chr12-83663772-INS-58
chr12-83669641-INS-69
chr12-83705440-INS-202
chr12-83726146-INS-203
chr12-83726529-INS-491
chr12-83751819-INS-183
chr12-83755894-INS-127
chr12-83885718-INS-7097
chr12-83913741-INS-6587

chr12-83916914-INS-212
chr12-83917752-INS-101
chr12-83930918-INS-52
chr12-83957051-INS-50
chr12-83979233-INS-3208
chr12-83980152-INS-96
chr12-83989319-INS-2000
chr12-84049801-INS-2579
chr12-84068152-INS-1701
chr12-84114171-INS-86
chr12-84114197-INS-212
chr12-84119100-INS-50
chr12-84119415-INS-50
chr12-84153058-INS-141
chr12-84154749-INS-4804
chr12-84158443-INS-2019
chr12-84163919-INS-81
chr12-84186711-INS-60
chr12-84187386-INS-78
chr12-84274434-INS-8457
chr12-84284310-INS-232
chr12-84303886-INS-198
chr12-84304390-INS-7218
chr12-84306727-INS-137
chr12-84321682-INS-200
chr12-84329547-INS-2430
chr12-84329684-INS-6404
chr12-84337574-INS-432
chr12-84427567-INS-58
chr12-84466141-INS-53
chr12-84478060-INS-234
chr12-84502425-INS-3059
chr12-84543213-INS-243
chr12-84562438-INS-100
chr12-84562438-INS-590
chr12-84572541-INS-86
chr12-84590043-INS-160
chr12-84594246-INS-498
chr12-84611969-INS-60
chr12-84631279-INS-221
chr12-84642295-INS-166
chr12-84642493-INS-118
chr12-84669331-INS-108
chr12-84680335-INS-216
chr12-84686961-INS-61
chr12-84713268-INS-249
chr12-84716829-INS-84
chr12-84734000-INS-65
chr12-84865894-INS-52
chr12-84880366-INS-50
chr12-84902330-INS-263
chr12-84930038-INS-436
chr12-85066385-INS-50
chr12-85099533-INS-81
chr12-85121686-INS-379
chr12-85175876-INS-165
chr12-85177583-INS-346
chr12-85179079-INS-137
chr12-85229600-INS-60
chr12-85230361-INS-383
chr12-85233798-INS-214
chr12-85233798-INS-66
chr12-85245645-INS-50
chr12-85263604-INS-426
chr12-85286126-INS-66
chr12-85321325-INS-131
chr12-85321398-INS-310
chr12-85386526-INS-263
chr12-85404354-INS-1248
chr12-85495390-INS-7118
chr12-85510272-INS-64
chr12-85551873-INS-50
chr12-85570437-INS-2730
chr12-85659984-INS-239
chr12-85684153-INS-62
chr12-85695961-INS-57
chr12-85715203-INS-64

chr12-85718649-INS-56
chr12-85719481-INS-105
chr12-85749540-INS-50
chr12-85759835-INS-250
chr12-85764125-INS-96
chr12-85779673-INS-136
chr12-85780078-INS-1698
chr12-85787017-INS-60
chr12-85892608-INS-2084
chr12-85893428-INS-5817
chr12-85894186-INS-925
chr12-85896733-INS-124
chr12-85922530-INS-287
chr12-85952827-INS-108
chr12-86040627-INS-51
chr12-86041368-INS-205
chr12-86057831-INS-276
chr12-86063704-INS-125
chr12-86137514-INS-109
chr12-86165208-INS-160
chr12-86172457-INS-162
chr12-86230589-INS-76
chr12-86231234-INS-204
chr12-86266298-INS-74
chr12-86322450-INS-82
chr12-86347159-INS-2102
chr12-86405303-INS-136
chr12-86430277-INS-272
chr12-86430277-INS-88
chr12-86459965-INS-55
chr12-86528576-INS-150
chr12-86549787-INS-116
chr12-86599171-INS-111
chr12-86613860-INS-588
chr12-86653787-INS-176
chr12-86654605-INS-58
chr12-86654717-INS-256
chr12-86662932-INS-2896
chr12-86679292-INS-59
chr12-86689633-INS-361
chr12-86690391-INS-72
chr12-86707671-INS-106
chr12-86712789-INS-198
chr12-86717806-INS-54
chr12-86721818-INS-51
chr12-86732806-INS-116
chr12-86746480-INS-80
chr12-86755898-INS-1806
chr12-86786649-INS-163
chr12-86786672-INS-636
chr12-86799799-INS-140
chr12-86812132-INS-477
chr12-86812570-INS-52
chr12-86832049-INS-204
chr12-86850241-INS-62
chr12-86850749-INS-107
chr12-86866437-INS-195
chr12-86944394-INS-52
chr12-86956367-INS-233
chr12-86956880-INS-50
chr12-86983292-INS-329
chr12-87018304-INS-7232
chr12-87026434-INS-2270
chr12-87050841-INS-52
chr12-87056628-INS-126
chr12-87056830-INS-58
chr12-87081601-INS-58
chr12-87087319-INS-126
chr12-87093858-INS-52
chr12-87120231-INS-215
chr12-87120985-INS-78
chr12-87147374-INS-194
chr12-87235881-INS-197
chr12-87256646-INS-230
chr12-87344697-INS-69
chr12-87383615-INS-2373
chr12-87423431-INS-359

chr12-87444232-INS-148
chr12-87444899-INS-104
chr12-87445425-INS-209
chr12-87449983-INS-62
chr12-87467618-INS-203
chr12-87471809-INS-151
chr12-87484616-INS-50
chr12-87489971-INS-217
chr12-87522133-INS-1892
chr12-87568493-INS-3167
chr12-87588433-INS-202
chr12-87601746-INS-6181
chr12-87817353-INS-1245
chr12-87828870-INS-498
chr12-87857339-INS-2835
chr12-88051125-INS-60
chr12-88154468-INS-498
chr12-88174377-INS-208
chr12-88187931-INS-50
chr12-88190235-INS-205
chr12-88219203-INS-64
chr12-88231563-INS-52
chr12-88249776-INS-76
chr12-88252748-INS-84
chr12-88253088-INS-497
chr12-88256910-INS-5426
chr12-88270958-INS-2659
chr12-88297274-INS-6886
chr12-88321376-INS-189
chr12-88334215-INS-5001
chr12-88347027-INS-136
chr12-88368566-INS-310
chr12-88464444-INS-156
chr12-88477711-INS-70
chr12-88484773-INS-104
chr12-88484828-INS-54
chr12-88507578-INS-94
chr12-88510207-INS-53
chr12-88565144-INS-52
chr12-88583411-INS-3856
chr12-88623895-INS-173
chr12-88641599-INS-1710
chr12-88643719-INS-5804
chr12-88646616-INS-79
chr12-88667601-INS-66
chr12-88667609-INS-134
chr12-88679184-INS-55
chr12-88739692-INS-564
chr12-88811222-INS-84
chr12-88826202-INS-50
chr12-88842780-INS-584
chr12-88851954-INS-204
chr12-88932292-INS-69
chr12-88940502-INS-1397
chr12-89083648-INS-158
chr12-89133217-INS-54
chr12-89168757-INS-1578
chr12-89241501-INS-202
chr12-89265723-INS-52
chr12-89274610-INS-202
chr12-89317026-INS-80
chr12-89347046-INS-1425
chr12-89379047-INS-52
chr12-89416792-INS-739
chr12-89417217-INS-457
chr12-89417496-INS-190
chr12-89430865-INS-76
chr12-89582758-INS-974
chr12-89596456-INS-50
chr12-89677046-INS-237
chr12-89685497-INS-2131
chr12-89692314-INS-50
chr12-89815389-INS-342
chr12-89843412-INS-628
chr12-89877961-INS-52
chr12-89888488-INS-53
chr12-90148153-INS-74

chr12-90161283-INS-50
chr12-90204007-INS-80
chr12-90211197-INS-6462
chr12-90227053-INS-555
chr12-90240314-INS-54
chr12-90261588-INS-51
chr12-90261628-INS-367
chr12-90295271-INS-115
chr12-90295403-INS-161
chr12-90313528-INS-632
chr12-90342568-INS-752
chr12-90397034-INS-3144
chr12-90397034-INS-429
chr12-90397448-INS-198
chr12-90494731-INS-58
chr12-90495195-INS-64
chr12-90496841-INS-633
chr12-90496907-INS-316
chr12-90545058-INS-589
chr12-90549938-INS-153
chr12-90563401-INS-50
chr12-90566923-INS-186
chr12-90568076-INS-6373
chr12-90581957-INS-50
chr12-90647404-INS-1231
chr12-90669765-INS-92
chr12-90688823-INS-100
chr12-90709261-INS-57
chr12-90757888-INS-165
chr12-90758028-INS-549
chr12-90782670-INS-54
chr12-90782748-INS-117
chr12-90831298-INS-113
chr12-90837697-INS-2709
chr12-90839262-INS-1295
chr12-90845730-INS-548
chr12-90903828-INS-284
chr12-90903887-INS-76
chr12-90931487-INS-3347
chr12-90956861-INS-166
chr12-90969976-INS-96
chr12-91026684-INS-2739
chr12-91065940-INS-64
chr12-91088936-INS-1213
chr12-91099388-INS-86
chr12-91114573-INS-278
chr12-91114740-INS-82
chr12-91150118-INS-138
chr12-91177605-INS-104
chr12-91181943-INS-86
chr12-91213630-INS-247
chr12-91221896-INS-61
chr12-91259853-INS-125
chr12-91260127-INS-250
chr12-91297756-INS-357
chr12-91303327-INS-61
chr12-91305681-INS-336
chr12-91359462-INS-209
chr12-91397490-INS-1036
chr12-91411326-INS-50
chr12-91467710-INS-85
chr12-91469046-INS-134
chr12-91513009-INS-736
chr12-91513145-INS-349
chr12-91519778-INS-638
chr12-91529143-INS-6531
chr12-91560298-INS-840
chr12-91583101-INS-14576
chr12-91588734-INS-57
chr12-91629281-INS-160
chr12-91630329-INS-1041
chr12-91630329-INS-3862
chr12-91696916-INS-99
chr12-91736111-INS-60
chr12-91829058-INS-6197
chr12-91841759-INS-198
chr12-91942074-INS-2382

chr12-91943057-INS-50
chr12-91970766-INS-62
chr12-91993713-INS-1652
chr12-92032488-INS-156
chr12-92033029-INS-52
chr12-92056408-INS-63
chr12-92101195-INS-63
chr12-92115680-INS-387
chr12-92118248-INS-6698
chr12-92122502-INS-177
chr12-92122502-INS-65
chr12-92131931-INS-74
chr12-92145150-INS-52
chr12-92190844-INS-58
chr12-92215444-INS-94
chr12-92252879-INS-51
chr12-92304983-INS-50
chr12-92305943-INS-969
chr12-92370952-INS-116
chr12-92388270-INS-56
chr12-92395200-INS-1392
chr12-92416245-INS-922
chr12-92416270-INS-76
chr12-92428033-INS-6438
chr12-92463415-INS-216
chr12-92494197-INS-55
chr12-92586423-INS-7885
chr12-92635032-INS-2982
chr12-92652117-INS-2889
chr12-92668042-INS-52
chr12-92674968-INS-509
chr12-92677404-INS-52
chr12-92703623-INS-292
chr12-92715246-INS-128
chr12-92715852-INS-126
chr12-92719219-INS-76
chr12-92764411-INS-6924
chr12-92792025-INS-1862
chr12-92895893-INS-108
chr12-92949775-INS-6260
chr12-93051464-INS-1009
chr12-93097079-INS-549
chr12-93128038-INS-52
chr12-93170799-INS-112
chr12-93187075-INS-6630
chr12-93194419-INS-50
chr12-93222977-INS-1203
chr12-93309244-INS-1898
chr12-93336003-INS-2929
chr12-93349535-INS-1879
chr12-93352067-INS-1583
chr12-93360834-INS-503
chr12-93361833-INS-4311
chr12-93363006-INS-2343
chr12-93364433-INS-75
chr12-93392481-INS-6248
chr12-93414142-INS-7242
chr12-93433539-INS-1987
chr12-93434198-INS-76
chr12-93435962-INS-50
chr12-93437433-INS-52
chr12-93490819-INS-1022
chr12-93514279-INS-1886
chr12-93518266-INS-6384
chr12-93550638-INS-479
chr12-93596185-INS-2583
chr12-93619400-INS-60
chr12-93722307-INS-56
chr12-93725679-INS-918
chr12-93738849-INS-813
chr12-93753034-INS-50
chr12-93753343-INS-363
chr12-93795765-INS-4544
chr12-93802776-INS-6461
chr12-93817510-INS-50
chr12-93819472-INS-72
chr12-93826825-INS-1126

chr12-93830083-INS-610
chr12-93844638-INS-50
chr12-93899707-INS-50
chr12-93904874-INS-289
chr12-93922806-INS-98
chr12-93934076-INS-108
chr12-93978516-INS-1129
chr12-93979586-INS-61
chr12-94026834-INS-93
chr12-94034045-INS-61
chr12-94117882-INS-70
chr12-94118034-INS-609
chr12-94257176-INS-68
chr12-94344498-INS-290
chr12-94346131-INS-78
chr12-94358819-INS-2245
chr12-94360788-INS-116
chr12-94376947-INS-4793
chr12-94462058-INS-64
chr12-94489560-INS-598
chr12-94559227-INS-6444
chr12-94566583-INS-456
chr12-94594177-INS-390
chr12-94614035-INS-900
chr12-94659437-INS-53
chr12-94675006-INS-157
chr12-94681422-INS-96
chr12-94685675-INS-1075
chr12-94694238-INS-547
chr12-94799140-INS-50
chr12-94840022-INS-5107
chr12-94841330-INS-72
chr12-94844428-INS-50
chr12-94848998-INS-57
chr12-94862302-INS-4678
chr12-94875084-INS-66
chr12-94896209-INS-6293
chr12-94983145-INS-477
chr12-94994899-INS-1049
chr12-95006350-INS-433
chr12-95006469-INS-3116
chr12-95015279-INS-50
chr12-95043873-INS-52
chr12-95060831-INS-444
chr12-95065719-INS-189
chr12-95079886-INS-80
chr12-95082935-INS-575
chr12-95118031-INS-2537
chr12-95164346-INS-7085
chr12-95185592-INS-11159
chr12-95198990-INS-55
chr12-95223270-INS-136
chr12-95263279-INS-111
chr12-95274936-INS-6670
chr12-95290043-INS-51
chr12-95293985-INS-375
chr12-95309838-INS-1379
chr12-95373259-INS-736
chr12-95397740-INS-323
chr12-95415472-INS-55
chr12-95425315-INS-268
chr12-95461157-INS-6432
chr12-95485228-INS-50
chr12-95521199-INS-51
chr12-95521395-INS-443
chr12-95522901-INS-50
chr12-95530030-INS-64
chr12-95562975-INS-232
chr12-95563656-INS-55
chr12-95577696-INS-75
chr12-95621783-INS-58
chr12-95628148-INS-3932
chr12-95656579-INS-54
chr12-95727545-INS-52
chr12-95809182-INS-750
chr12-95815132-INS-52
chr12-95837538-INS-52

chr12-95856815-INS-52
chr12-95866634-INS-128
chr12-95868692-INS-245
chr12-95880427-INS-52
chr12-95893075-INS-337
chr12-96000844-INS-61
chr12-96055819-INS-885
chr12-96087592-INS-60
chr12-96089526-INS-92
chr12-96099720-INS-66
chr12-96102832-INS-62
chr12-96112076-INS-75
chr12-96125138-INS-1038
chr12-96131190-INS-135
chr12-96139931-INS-4823
chr12-96161840-INS-513
chr12-96268865-INS-2233
chr12-96269020-INS-68
chr12-96269780-INS-359
chr12-96323454-INS-6394
chr12-96335467-INS-6213
chr12-96341001-INS-228
chr12-96362901-INS-108
chr12-96373294-INS-6514
chr12-96385624-INS-62
chr12-96393150-INS-50
chr12-96405348-INS-975
chr12-96465461-INS-100
chr12-96469164-INS-137
chr12-96519199-INS-58
chr12-96545894-INS-10168
chr12-96556277-INS-192
chr12-96562326-INS-1373
chr12-96590025-INS-4899
chr12-96626697-INS-462
chr12-96648716-INS-1038
chr12-96648848-INS-102
chr12-96660495-INS-135
chr12-96746569-INS-56
chr12-96770849-INS-116
chr12-96773706-INS-80
chr12-96827305-INS-80
chr12-96865073-INS-50
chr12-96912300-INS-66
chr12-96924817-INS-177
chr12-96951143-INS-6525
chr12-97052520-INS-251
chr12-97090161-INS-6449
chr12-97108657-INS-56
chr12-97137664-INS-4759
chr12-97158599-INS-6173
chr12-97163924-INS-53
chr12-97169846-INS-218
chr12-97234421-INS-52
chr12-97277003-INS-672
chr12-97288915-INS-52
chr12-97313744-INS-2518
chr12-97364993-INS-114
chr12-97370808-INS-207
chr12-97376605-INS-2654
chr12-97404955-INS-2745
chr12-97429160-INS-6222
chr12-97451102-INS-115
chr12-97572441-INS-54
chr12-97578644-INS-80
chr12-97603586-INS-3296
chr12-97615617-INS-3500
chr12-97631219-INS-56
chr12-97673663-INS-447
chr12-97685997-INS-1377
chr12-97708355-INS-253
chr12-97736219-INS-5558
chr12-97738198-INS-367
chr12-97782759-INS-1195
chr12-97805135-INS-120
chr12-97830641-INS-84
chr12-97842736-INS-52

chr12-97871390-INS-470
chr12-97880469-INS-201
chr12-97901531-INS-1837
chr12-97992226-INS-72
chr12-97993693-INS-308
chr12-98021887-INS-51
chr12-98027742-INS-179
chr12-98037731-INS-110
chr12-98042647-INS-6447
chr12-98045303-INS-1452
chr12-98069652-INS-1112
chr12-98073047-INS-105
chr12-98102807-INS-520
chr12-98110560-INS-939
chr12-98123257-INS-1288
chr12-98196220-INS-2655
chr12-98198113-INS-220
chr12-98231770-INS-6175
chr12-98239554-INS-101
chr12-98246864-INS-51
chr12-98275361-INS-1911
chr12-98296302-INS-67
chr12-98296984-INS-75
chr12-98296985-INS-156
chr12-98304559-INS-6285
chr12-98317416-INS-825
chr12-98433847-INS-53
chr12-98461670-INS-1035
chr12-98595685-INS-217
chr12-98649051-INS-217
chr12-98674554-INS-282
chr12-98741057-INS-288
chr12-98794158-INS-138
chr12-98796319-INS-210
chr12-98811475-INS-7308
chr12-98852562-INS-232
chr12-98856310-INS-196
chr12-98863831-INS-200
chr12-98864009-INS-144
chr12-98864510-INS-104
chr12-98888558-INS-52
chr12-99041232-INS-939
chr12-99048867-INS-211
chr12-99081741-INS-158
chr12-99081747-INS-500
chr12-99150252-INS-95
chr12-99153908-INS-207
chr12-99187958-INS-214
chr12-99239376-INS-123
chr12-99311349-INS-54
chr12-99339777-INS-104
chr12-99339777-INS-50
chr12-99360931-INS-210
chr12-99362277-INS-171
chr12-99402849-INS-59
chr12-99544480-INS-484
chr12-99545533-INS-2882
chr12-99550945-INS-55
chr12-99558022-INS-174
chr12-99561882-INS-50
chr12-99563640-INS-50
chr12-99564829-INS-1802
chr12-99580151-INS-66
chr12-99602957-INS-84
chr12-99614709-INS-50
chr12-99641023-INS-51
chr12-99671590-INS-282
chr12-99709719-INS-168
chr12-99892873-INS-260
chr12-99920530-INS-52
chr12-99927611-INS-52
chr12-100001476-INS-57
chr12-100011386-INS-199
chr12-100011998-INS-7140
chr12-100014200-INS-54
chr12-100065398-INS-194
chr12-100116881-INS-221

chr12-100119684-INS-337
chr12-100124693-INS-103
chr12-100130059-INS-405
chr12-100130136-INS-162
chr12-100130998-INS-205
chr12-100136954-INS-2105
chr12-100161607-INS-6885
chr12-100186936-INS-89
chr12-100206746-INS-323
chr12-100217746-INS-190
chr12-100221656-INS-56
chr12-100263717-INS-114
chr12-100309045-INS-309
chr12-100317334-INS-210
chr12-100317853-INS-592
chr12-100332550-INS-106
chr12-100343877-INS-54
chr12-100384236-INS-80
chr12-100407832-INS-77
chr12-100443052-INS-359
chr12-100452797-INS-179
chr12-100498776-INS-129
chr12-100500127-INS-9098
chr12-100504170-INS-81
chr12-100513295-INS-186
chr12-100533490-INS-53
chr12-100539387-INS-217
chr12-100539597-INS-262
chr12-100550449-INS-133
chr12-100550657-INS-60
chr12-100554273-INS-84
chr12-100557644-INS-864
chr12-100567363-INS-222
chr12-100576254-INS-3029
chr12-100578387-INS-53
chr12-100610997-INS-3271
chr12-100667237-INS-72
chr12-100694655-INS-143
chr12-100697187-INS-4770
chr12-100700567-INS-413
chr12-100700671-INS-70
chr12-100705962-INS-151
chr12-100717950-INS-123
chr12-100733111-INS-7164
chr12-100764883-INS-236
chr12-100785364-INS-223
chr12-100842726-INS-405
chr12-100850210-INS-560
chr12-100915773-INS-86
chr12-100929690-INS-66
chr12-100933770-INS-135
chr12-100933813-INS-50
chr12-100934377-INS-207
chr12-100941217-INS-56
chr12-100941560-INS-70
chr12-100953326-INS-277
chr12-101001297-INS-155
chr12-101079370-INS-282
chr12-101103092-INS-6055
chr12-101279501-INS-5548
chr12-101280776-INS-50
chr12-101285866-INS-1058
chr12-101578655-INS-193
chr12-101589639-INS-113
chr12-101624771-INS-4939
chr12-101692026-INS-1143
chr12-101698662-INS-343
chr12-101710363-INS-132
chr12-101718817-INS-170
chr12-101737951-INS-2094
chr12-101743671-INS-112
chr12-101807346-INS-321
chr12-101820396-INS-78
chr12-101824137-INS-688
chr12-101833857-INS-194
chr12-101858824-INS-181
chr12-101866571-INS-188

chr12-101866645-INS-69
chr12-101882358-INS-6344
chr12-101887101-INS-571
chr12-101904372-INS-199
chr12-101910075-INS-80
chr12-101910077-INS-268
chr12-101963901-INS-2002
chr12-102007223-INS-72
chr12-102035613-INS-64
chr12-102064281-INS-1220
chr12-102158539-INS-407
chr12-102267101-INS-464
chr12-102321423-INS-213
chr12-102336690-INS-55
chr12-102366846-INS-477
chr12-102401388-INS-51
chr12-102404137-INS-50
chr12-102418328-INS-392
chr12-102442281-INS-245
chr12-102454392-INS-999
chr12-102477173-INS-7113
chr12-102514376-INS-64
chr12-102515372-INS-133
chr12-102522876-INS-52
chr12-102533558-INS-228
chr12-102552522-INS-58
chr12-102585800-INS-66
chr12-102601093-INS-260
chr12-102625380-INS-306
chr12-102675100-INS-75
chr12-102679502-INS-61
chr12-102680068-INS-187
chr12-102680981-INS-164
chr12-102682273-INS-164
chr12-102689963-INS-135
chr12-102718581-INS-219
chr12-102727879-INS-219
chr12-102764324-INS-131
chr12-102864489-INS-128
chr12-102887044-INS-211
chr12-102952098-INS-157
chr12-102955324-INS-56
chr12-103042593-INS-1366
chr12-103045453-INS-72
chr12-103064432-INS-7174
chr12-103066823-INS-330
chr12-103117614-INS-182
chr12-103144425-INS-70
chr12-103213719-INS-187
chr12-103230835-INS-50
chr12-103239012-INS-100
chr12-103259577-INS-163
chr12-103289926-INS-76
chr12-103312846-INS-299
chr12-103330633-INS-84
chr12-103410837-INS-139
chr12-103459813-INS-56
chr12-103583885-INS-201
chr12-103603826-INS-112
chr12-103604589-INS-8327
chr12-103630866-INS-72
chr12-103689234-INS-3236
chr12-103724657-INS-3310
chr12-103836571-INS-1417
chr12-103953012-INS-56
chr12-103990397-INS-202
chr12-104020895-INS-51
chr12-104065038-INS-1192
chr12-104074521-INS-394
chr12-104075577-INS-355
chr12-104077471-INS-97
chr12-104165361-INS-257
chr12-104166089-INS-115
chr12-104192058-INS-6906
chr12-104197017-INS-2342
chr12-104208788-INS-926
chr12-104213462-INS-2485

chr12-104223781-INS-144
chr12-104224820-INS-50
chr12-104335743-INS-256
chr12-104346640-INS-6425
chr12-104357148-INS-7011
chr12-104371590-INS-179
chr12-104387952-INS-139
chr12-104390719-INS-7253
chr12-104408984-INS-175
chr12-104408984-INS-390
chr12-104419612-INS-84
chr12-104448244-INS-258
chr12-104449892-INS-7262
chr12-104466568-INS-90
chr12-104478331-INS-93
chr12-104505056-INS-188
chr12-104532227-INS-60
chr12-104575669-INS-4095
chr12-104577978-INS-62
chr12-104584135-INS-275
chr12-104611128-INS-87
chr12-104611160-INS-348
chr12-104633585-INS-119
chr12-104672996-INS-72
chr12-104699647-INS-90
chr12-104706931-INS-948
chr12-104801314-INS-58
chr12-104801740-INS-128
chr12-104847604-INS-4438
chr12-104853051-INS-169
chr12-104865924-INS-3063
chr12-104871416-INS-132
chr12-104871416-INS-64
chr12-104877602-INS-126
chr12-104881544-INS-65
chr12-104903974-INS-112
chr12-104903984-INS-371
chr12-104909518-INS-1309
chr12-104915639-INS-646
chr12-104933051-INS-84
chr12-104946872-INS-576
chr12-104955726-INS-6259
chr12-104960261-INS-139
chr12-105004974-INS-349
chr12-105016456-INS-216
chr12-105043402-INS-54
chr12-105061015-INS-3624
chr12-105074622-INS-67
chr12-105074851-INS-53
chr12-105075516-INS-234
chr12-105098110-INS-123
chr12-105105236-INS-210
chr12-105173904-INS-111
chr12-105179941-INS-68
chr12-105195903-INS-176
chr12-105220254-INS-123
chr12-105245078-INS-50
chr12-105251696-INS-438
chr12-105290978-INS-78
chr12-105338887-INS-289
chr12-105361688-INS-1091
chr12-105378878-INS-714
chr12-105382990-INS-160
chr12-105402364-INS-764
chr12-105446627-INS-207
chr12-105462163-INS-506
chr12-105491062-INS-60
chr12-105494753-INS-194
chr12-105516270-INS-353
chr12-105546104-INS-54
chr12-105597530-INS-430
chr12-105621902-INS-630
chr12-105638741-INS-1082
chr12-105638743-INS-170
chr12-105701513-INS-57
chr12-105710111-INS-187
chr12-105713091-INS-72

chr12-105749555-INS-113
chr12-105761776-INS-204
chr12-105801749-INS-210
chr12-105849490-INS-259
chr12-105853686-INS-474
chr12-105888829-INS-108
chr12-105906191-INS-216
chr12-105906640-INS-52
chr12-105909800-INS-58
chr12-105923224-INS-246
chr12-105925699-INS-227
chr12-105925789-INS-723
chr12-105950193-INS-52
chr12-105966964-INS-50
chr12-105984205-INS-62
chr12-106072307-INS-198
chr12-106072903-INS-104
chr12-106086249-INS-258
chr12-106086877-INS-192
chr12-106088687-INS-99
chr12-106124947-INS-70
chr12-106139857-INS-82
chr12-106143271-INS-70
chr12-106168111-INS-92
chr12-106194346-INS-172
chr12-106233638-INS-52
chr12-106304587-INS-166
chr12-106321195-INS-360
chr12-106328809-INS-52
chr12-106331648-INS-60
chr12-106395068-INS-379
chr12-106400259-INS-203
chr12-106410692-INS-72
chr12-106469081-INS-112
chr12-106472885-INS-64
chr12-106472957-INS-108
chr12-106479089-INS-95
chr12-106479544-INS-336
chr12-106479730-INS-1230
chr12-106488972-INS-78
chr12-106489455-INS-60
chr12-106503033-INS-188
chr12-106537691-INS-66
chr12-106569092-INS-60
chr12-106603994-INS-276
chr12-106604006-INS-99
chr12-106604342-INS-53
chr12-106612751-INS-90
chr12-106630693-INS-68
chr12-106648290-INS-59
chr12-106651044-INS-56
chr12-106655311-INS-157
chr12-106655350-INS-73
chr12-106692263-INS-340
chr12-106692263-INS-65
chr12-106706313-INS-3223
chr12-106734391-INS-51
chr12-106741668-INS-218
chr12-106741668-INS-76
chr12-106741997-INS-140
chr12-106745080-INS-382
chr12-106767897-INS-61
chr12-106840091-INS-760
chr12-106847481-INS-70
chr12-106850033-INS-6723
chr12-106854504-INS-750
chr12-106885489-INS-188
chr12-106885977-INS-55
chr12-106945792-INS-65
chr12-107018839-INS-176
chr12-107030706-INS-52
chr12-107040298-INS-50
chr12-107055903-INS-51
chr12-107067223-INS-50
chr12-107069430-INS-50
chr12-107124326-INS-99
chr12-107124594-INS-52

chr12-107168229-INS-197
chr12-107174025-INS-179
chr12-107262542-INS-62
chr12-107297421-INS-201
chr12-107303180-INS-80
chr12-107332611-INS-97
chr12-107344005-INS-120
chr12-107359125-INS-50
chr12-107377480-INS-63
chr12-107418387-INS-1566
chr12-107419405-INS-72
chr12-107422442-INS-214
chr12-107456316-INS-84
chr12-107489107-INS-273
chr12-107489133-INS-63
chr12-107499686-INS-242
chr12-107506853-INS-112
chr12-107522345-INS-79
chr12-107566475-INS-60
chr12-107575062-INS-597
chr12-107600401-INS-156
chr12-107646989-INS-93
chr12-107681390-INS-75
chr12-107701799-INS-168
chr12-107704958-INS-112
chr12-107753836-INS-1500
chr12-107758545-INS-502
chr12-107764161-INS-190
chr12-107784308-INS-188
chr12-107812480-INS-237
chr12-107873853-INS-364
chr12-107951463-INS-54
chr12-107952775-INS-52
chr12-107988406-INS-364
chr12-108003381-INS-59
chr12-108021852-INS-1006
chr12-108035498-INS-54
chr12-108111190-INS-72
chr12-108195107-INS-147
chr12-108212312-INS-285
chr12-108258229-INS-3892
chr12-108290703-INS-92
chr12-108303897-INS-217
chr12-108304093-INS-62
chr12-108315686-INS-699
chr12-108316073-INS-798
chr12-108325965-INS-117
chr12-108349800-INS-6975
chr12-108363602-INS-50
chr12-108434948-INS-120
chr12-108435133-INS-363
chr12-108437662-INS-304
chr12-108452006-INS-110
chr12-108452006-INS-260
chr12-108465237-INS-323
chr12-108498969-INS-50
chr12-108507680-INS-142
chr12-108508492-INS-247
chr12-108509228-INS-204
chr12-108524990-INS-387
chr12-108562619-INS-85
chr12-108632082-INS-215
chr12-108648615-INS-165
chr12-108664397-INS-311
chr12-108692841-INS-68
chr12-108728550-INS-63
chr12-108728554-INS-198
chr12-108806064-INS-53
chr12-108875665-INS-672
chr12-108899047-INS-216
chr12-108901391-INS-59
chr12-108908028-INS-50
chr12-108932499-INS-197
chr12-108932569-INS-794
chr12-108968569-INS-66
chr12-108969053-INS-90
chr12-108984265-INS-189

chr12-109028737-INS-128
chr12-109032699-INS-169
chr12-109048758-INS-55
chr12-109048766-INS-150
chr12-109056875-INS-51
chr12-109067581-INS-234
chr12-109071498-INS-83
chr12-109085558-INS-116
chr12-109085744-INS-702
chr12-109096443-INS-180
chr12-109106124-INS-448
chr12-109110060-INS-202
chr12-109126567-INS-6411
chr12-109127802-INS-512
chr12-109134013-INS-1361
chr12-109157753-INS-91
chr12-109160427-INS-6389
chr12-109219470-INS-51
chr12-109221704-INS-244
chr12-109244566-INS-161
chr12-109328682-INS-203
chr12-109346988-INS-50
chr12-109352459-INS-146
chr12-109441470-INS-114
chr12-109452901-INS-52
chr12-109540819-INS-142
chr12-109720240-INS-76
chr12-109757096-INS-62
chr12-109804215-INS-97
chr12-109816549-INS-153
chr12-109818273-INS-4857
chr12-109823435-INS-710
chr12-109823469-INS-288
chr12-109823688-INS-150
chr12-109854926-INS-59
chr12-109893549-INS-53
chr12-109898313-INS-225
chr12-109905802-INS-70
chr12-109930966-INS-1153
chr12-109931905-INS-477
chr12-109932175-INS-585
chr12-109953596-INS-5311
chr12-109954038-INS-174
chr12-109970720-INS-112
chr12-109989066-INS-79
chr12-110021044-INS-121
chr12-110076107-INS-1476
chr12-110076248-INS-2023
chr12-110081780-INS-54
chr12-110094764-INS-1232
chr12-110100087-INS-353
chr12-110100707-INS-329
chr12-110159633-INS-190
chr12-110161812-INS-3510
chr12-110227002-INS-51
chr12-110227151-INS-313
chr12-110264546-INS-202
chr12-110269573-INS-67
chr12-110281799-INS-96
chr12-110335304-INS-489
chr12-110335652-INS-101
chr12-110348566-INS-99
chr12-110358626-INS-149
chr12-110361966-INS-60
chr12-110362038-INS-398
chr12-110366170-INS-221
chr12-110380670-INS-228
chr12-110381325-INS-118
chr12-110384736-INS-52
chr12-110399483-INS-478
chr12-110457362-INS-88
chr12-110467719-INS-109
chr12-110468327-INS-50
chr12-110476693-INS-198
chr12-110487099-INS-723
chr12-110589139-INS-220
chr12-110616532-INS-60

chr12-110617454-INS-658
chr12-110639696-INS-423
chr12-110640014-INS-364
chr12-110646993-INS-91
chr12-110647236-INS-90
chr12-110668034-INS-57
chr12-110670907-INS-193
chr12-110686401-INS-72
chr12-110689101-INS-131
chr12-110689108-INS-265
chr12-110689369-INS-202
chr12-110690628-INS-192
chr12-110707221-INS-210
chr12-110816114-INS-50
chr12-110842610-INS-205
chr12-110843105-INS-448
chr12-110862962-INS-50
chr12-110970346-INS-2503
chr12-111039244-INS-161
chr12-111039654-INS-82
chr12-111044358-INS-52
chr12-111149950-INS-7319
chr12-111165617-INS-224
chr12-111172868-INS-198
chr12-111173071-INS-72
chr12-111174267-INS-200
chr12-111185853-INS-212
chr12-111194714-INS-51
chr12-111216817-INS-64
chr12-111239675-INS-62
chr12-111247576-INS-52
chr12-111251177-INS-2890
chr12-111279185-INS-608
chr12-111307677-INS-288
chr12-111308122-INS-78
chr12-111314695-INS-72
chr12-111327176-INS-89
chr12-111329268-INS-192
chr12-111329836-INS-50
chr12-111330030-INS-155
chr12-111345689-INS-140
chr12-111374875-INS-51
chr12-111377232-INS-282
chr12-111400095-INS-194
chr12-111419056-INS-112
chr12-111442339-INS-2272
chr12-111467958-INS-480
chr12-111473495-INS-200
chr12-111495745-INS-95
chr12-111498416-INS-52
chr12-111515265-INS-311
chr12-111527983-INS-205
chr12-111552889-INS-198
chr12-111580189-INS-216
chr12-111611860-INS-57
chr12-111633806-INS-3073
chr12-111645942-INS-215
chr12-111646204-INS-526
chr12-111678739-INS-210
chr12-111703326-INS-1034
chr12-111703326-INS-2416
chr12-111708427-INS-125
chr12-111708518-INS-51
chr12-111720267-INS-163
chr12-111751401-INS-168
chr12-111751426-INS-56
chr12-111751786-INS-530
chr12-111751793-INS-225
chr12-111753847-INS-4896
chr12-111757161-INS-133
chr12-111757161-INS-66
chr12-111804498-INS-4374
chr12-111816679-INS-210
chr12-111869169-INS-6440
chr12-111889421-INS-1868
chr12-112002004-INS-151
chr12-112002154-INS-377

chr12-112002601-INS-1866
chr12-112014325-INS-262
chr12-112014354-INS-643
chr12-112039930-INS-4690
chr12-112103734-INS-200
chr12-112146422-INS-75
chr12-112146559-INS-280
chr12-112182944-INS-123
chr12-112213457-INS-102
chr12-112225836-INS-533
chr12-112235086-INS-6440
chr12-112261318-INS-50
chr12-112263956-INS-274
chr12-112264215-INS-259
chr12-112292289-INS-90
chr12-112292336-INS-270
chr12-112316073-INS-4005
chr12-112320390-INS-68
chr12-112324838-INS-2863
chr12-112390556-INS-50
chr12-112423472-INS-51
chr12-112490975-INS-7130
chr12-112548539-INS-120
chr12-112548628-INS-570
chr12-112597674-INS-72
chr12-112609241-INS-50
chr12-112666170-INS-184
chr12-112675156-INS-76
chr12-112677253-INS-280
chr12-112713734-INS-89
chr12-112729209-INS-70
chr12-112826315-INS-118
chr12-112835758-INS-5254
chr12-112862711-INS-6401
chr12-112949807-INS-171
chr12-112967191-INS-461
chr12-112990166-INS-58
chr12-113007637-INS-311
chr12-113041351-INS-196
chr12-113046791-INS-132
chr12-113079354-INS-455
chr12-113129057-INS-231
chr12-113129846-INS-112
chr12-113165789-INS-337
chr12-113169603-INS-3016
chr12-113183154-INS-1272
chr12-113183545-INS-125
chr12-113184427-INS-62
chr12-113199119-INS-104
chr12-113202183-INS-104
chr12-113207744-INS-186
chr12-113216373-INS-1168
chr12-113224276-INS-7875
chr12-113225838-INS-528
chr12-113226029-INS-503
chr12-113226377-INS-538
chr12-113226484-INS-160
chr12-113246690-INS-3449
chr12-113247753-INS-219
chr12-113249487-INS-17042
chr12-113257946-INS-52
chr12-113258311-INS-416
chr12-113258548-INS-78
chr12-113274654-INS-2396
chr12-113276327-INS-98
chr12-113287640-INS-3387
chr12-113298854-INS-3670
chr12-113298978-INS-244
chr12-113327907-INS-1386
chr12-113327940-INS-184
chr12-113388033-INS-1332
chr12-113397119-INS-6187
chr12-113412051-INS-54
chr12-113412535-INS-51
chr12-113422379-INS-52
chr12-113425996-INS-327
chr12-113436705-INS-8675

chr12-113443412-INS-165
chr12-113484310-INS-6341
chr12-113539804-INS-15203
chr12-113554681-INS-105
chr12-113554813-INS-51
chr12-113576466-INS-52
chr12-113586952-INS-221
chr12-113614954-INS-55
chr12-113615448-INS-108
chr12-113650546-INS-500
chr12-113652059-INS-54
chr12-113683274-INS-1098
chr12-113686637-INS-2019
chr12-113688787-INS-64
chr12-113689716-INS-93
chr12-113691758-INS-109
chr12-113691843-INS-55
chr12-113701848-INS-292
chr12-113702120-INS-780
chr12-113702805-INS-589
chr12-113712383-INS-63
chr12-113724947-INS-183
chr12-113735061-INS-64
chr12-113753114-INS-4372
chr12-113760654-INS-50
chr12-113766489-INS-741
chr12-113785464-INS-5077
chr12-113828597-INS-6391
chr12-113838280-INS-1414
chr12-113860438-INS-84
chr12-113895440-INS-62
chr12-113901354-INS-175
chr12-113903914-INS-1802
chr12-113904696-INS-75
chr12-113906259-INS-681
chr12-113907270-INS-424
chr12-113913043-INS-522
chr12-113913495-INS-129
chr12-113918898-INS-241
chr12-113921182-INS-4451
chr12-113949666-INS-578
chr12-113968874-INS-148
chr12-113978360-INS-325
chr12-113981245-INS-104
chr12-114003220-INS-215
chr12-114015396-INS-217
chr12-114036103-INS-165
chr12-114042370-INS-504
chr12-114042698-INS-2916
chr12-114094501-INS-191
chr12-114099386-INS-223
chr12-114103336-INS-834
chr12-114126780-INS-1874
chr12-114145823-INS-314
chr12-114153858-INS-10701
chr12-114336406-INS-69
chr12-114351749-INS-79
chr12-114352681-INS-104
chr12-114354353-INS-1354
chr12-114361550-INS-441
chr12-114361766-INS-477
chr12-114370163-INS-6642
chr12-114377900-INS-3935
chr12-114378562-INS-4294
chr12-114382346-INS-317
chr12-114383622-INS-65
chr12-114387721-INS-75
chr12-114390715-INS-103
chr12-114392735-INS-233
chr12-114394815-INS-9942
chr12-114396064-INS-6422
chr12-114406963-INS-53
chr12-114410316-INS-173
chr12-114427863-INS-4675
chr12-114428230-INS-471
chr12-114434629-INS-6272
chr12-114436721-INS-750

chr12-114437801-INS-64
chr12-114442164-INS-64
chr12-114449105-INS-328
chr12-114450078-INS-498
chr12-114466005-INS-768
chr12-114478198-INS-946
chr12-114481445-INS-254
chr12-114483890-INS-3438
chr12-114486605-INS-100
chr12-114503833-INS-87
chr12-114505696-INS-106
chr12-114513736-INS-80
chr12-114521485-INS-82
chr12-114523820-INS-132
chr12-114533118-INS-126
chr12-114535006-INS-1566
chr12-114546093-INS-4647
chr12-114549320-INS-177
chr12-114553863-INS-68
chr12-114560656-INS-214
chr12-114584235-INS-50
chr12-114607937-INS-1459
chr12-114612291-INS-50
chr12-114613085-INS-77
chr12-114622719-INS-131
chr12-114634210-INS-53
chr12-114642672-INS-11828
chr12-114643649-INS-69
chr12-114653489-INS-60
chr12-114660021-INS-58
chr12-114680158-INS-54
chr12-114734783-INS-3381
chr12-114738532-INS-70
chr12-114765710-INS-56
chr12-114776495-INS-175
chr12-114781933-INS-124
chr12-114782795-INS-199
chr12-114802409-INS-436
chr12-114803300-INS-65
chr12-114863055-INS-82
chr12-114864019-INS-1535
chr12-114869448-INS-198
chr12-114875996-INS-7118
chr12-114882336-INS-8734
chr12-114909307-INS-318
chr12-114910369-INS-3489
chr12-114914089-INS-119
chr12-114914620-INS-194
chr12-114919415-INS-213
chr12-114927436-INS-79
chr12-114928675-INS-104
chr12-114929616-INS-4887
chr12-114930851-INS-354
chr12-114932607-INS-63
chr12-114946372-INS-71
chr12-114954035-INS-1282
chr12-114956296-INS-2080
chr12-114964741-INS-151
chr12-114965353-INS-51
chr12-114973442-INS-7279
chr12-114985712-INS-185
chr12-114998853-INS-7815
chr12-115005201-INS-1230
chr12-115019968-INS-85
chr12-115026948-INS-208
chr12-115131269-INS-305
chr12-115140028-INS-84
chr12-115145806-INS-320
chr12-115149630-INS-73
chr12-115164365-INS-50
chr12-115168454-INS-121
chr12-115177619-INS-76
chr12-115179351-INS-78
chr12-115228409-INS-184
chr12-115248112-INS-251
chr12-115268157-INS-254
chr12-115274463-INS-56

chr12-115289131-INS-387
chr12-115304055-INS-486
chr12-115318954-INS-62
chr12-115341538-INS-74
chr12-115415382-INS-1170
chr12-115415928-INS-62
chr12-115449136-INS-567
chr12-115462076-INS-262
chr12-115467849-INS-53
chr12-115470855-INS-4912
chr12-115479600-INS-837
chr12-115494506-INS-93
chr12-115506538-INS-52
chr12-115514752-INS-100
chr12-115515252-INS-300
chr12-115515278-INS-117
chr12-115515890-INS-506
chr12-115560211-INS-72
chr12-115564219-INS-189
chr12-115566080-INS-466
chr12-115568474-INS-63
chr12-115572193-INS-154
chr12-115577964-INS-436
chr12-115605496-INS-51
chr12-115606001-INS-88
chr12-115606002-INS-222
chr12-115608090-INS-830
chr12-115659333-INS-1061
chr12-115660447-INS-199
chr12-115674526-INS-4819
chr12-115679735-INS-1418
chr12-115738607-INS-54
chr12-115744410-INS-320
chr12-115747785-INS-1770
chr12-115755235-INS-499
chr12-115759455-INS-1929
chr12-115763082-INS-84
chr12-115798451-INS-68
chr12-115798978-INS-192
chr12-115826778-INS-72
chr12-115855761-INS-218
chr12-115880155-INS-4366
chr12-115924508-INS-75
chr12-115925494-INS-72
chr12-115931415-INS-4293
chr12-115956510-INS-481
chr12-115960101-INS-101
chr12-115960709-INS-67
chr12-115960931-INS-440
chr12-115966324-INS-200
chr12-115974267-INS-126
chr12-116029237-INS-2843
chr12-116037785-INS-208
chr12-116055871-INS-224
chr12-116071370-INS-126
chr12-116127545-INS-3677
chr12-116128975-INS-86
chr12-116157496-INS-2173
chr12-116166236-INS-6461
chr12-116177594-INS-62
chr12-116190649-INS-1938
chr12-116262413-INS-95
chr12-116323451-INS-732
chr12-116329725-INS-1073
chr12-116338078-INS-4847
chr12-116345638-INS-5052
chr12-116354526-INS-213
chr12-116359613-INS-205
chr12-116417998-INS-139
chr12-116477101-INS-64
chr12-116554066-INS-104
chr12-116588948-INS-7633
chr12-116597920-INS-7150
chr12-116604319-INS-4199
chr12-116622846-INS-408
chr12-116641884-INS-134
chr12-116642478-INS-455

chr12-116644635-INS-2080
chr12-116648548-INS-5376
chr12-116765972-INS-62
chr12-116774285-INS-6616
chr12-116791250-INS-6640
chr12-116833072-INS-6458
chr12-116869546-INS-110
chr12-116870503-INS-433
chr12-116879765-INS-50
chr12-116881266-INS-140
chr12-116886187-INS-501
chr12-116970828-INS-128
chr12-116975704-INS-4103
chr12-116996998-INS-86
chr12-117016403-INS-68
chr12-117040746-INS-69
chr12-117158718-INS-50
chr12-117189553-INS-50
chr12-117251080-INS-133
chr12-117258144-INS-132
chr12-117258556-INS-173
chr12-117286925-INS-100
chr12-117303076-INS-59
chr12-117329853-INS-19654
chr12-117349329-INS-65
chr12-117367534-INS-64
chr12-117379463-INS-282
chr12-117381223-INS-817
chr12-117403670-INS-198
chr12-117403670-INS-726
chr12-117404272-INS-54
chr12-117434191-INS-188
chr12-117450779-INS-7085
chr12-117480470-INS-7227
chr12-117481830-INS-51
chr12-117489422-INS-100
chr12-117553183-INS-255
chr12-117578205-INS-170
chr12-117637699-INS-179
chr12-117648890-INS-444
chr12-117680567-INS-87
chr12-117687135-INS-182
chr12-117700096-INS-6399
chr12-117711176-INS-171
chr12-117724126-INS-236
chr12-117728948-INS-55
chr12-117741745-INS-52
chr12-117779035-INS-335
chr12-117780396-INS-50
chr12-117788967-INS-2948
chr12-117800895-INS-58
chr12-117824991-INS-986
chr12-117826001-INS-355
chr12-117840296-INS-50
chr12-117869708-INS-443
chr12-117891426-INS-6265
chr12-117895636-INS-468
chr12-117913154-INS-118
chr12-117920645-INS-54
chr12-117929558-INS-65
chr12-117933527-INS-77
chr12-117950508-INS-81
chr12-117951622-INS-174
chr12-117965444-INS-166
chr12-117970889-INS-264
chr12-117970891-INS-75
chr12-118004603-INS-207
chr12-118017166-INS-591
chr12-118066742-INS-64
chr12-118071029-INS-54
chr12-118133790-INS-1521
chr12-118226620-INS-60
chr12-118258281-INS-52
chr12-118313113-INS-50
chr12-118384569-INS-1174
chr12-118384569-INS-3239
chr12-118397292-INS-126

chr12-118416237-INS-52
chr12-118461754-INS-436
chr12-118481585-INS-3179
chr12-118536322-INS-66
chr12-118605111-INS-55
chr12-118668389-INS-100
chr12-118721662-INS-64
chr12-118733211-INS-3073
chr12-118740323-INS-199
chr12-118744425-INS-1895
chr12-118758791-INS-50
chr12-118761120-INS-354
chr12-118825712-INS-60
chr12-118829071-INS-50
chr12-118903285-INS-225
chr12-118908007-INS-513
chr12-118915285-INS-6743
chr12-118936115-INS-6394
chr12-119054669-INS-1440
chr12-119235251-INS-55
chr12-119239463-INS-2870
chr12-119281076-INS-53
chr12-119384709-INS-698
chr12-119420199-INS-50
chr12-119424989-INS-78
chr12-119425224-INS-484
chr12-119460537-INS-66
chr12-119549718-INS-100
chr12-119580501-INS-1857
chr12-119581841-INS-267
chr12-119586891-INS-6252
chr12-119603728-INS-1350
chr12-119607976-INS-104
chr12-119608047-INS-51
chr12-119805844-INS-601
chr12-119822722-INS-186
chr12-119830368-INS-11620
chr12-119853599-INS-138
chr12-119923310-INS-50
chr12-119972534-INS-7445
chr13-3086031-INS-1062
chr13-3097859-INS-853
chr13-3113497-INS-3451
chr13-3268553-INS-50
chr13-3289147-INS-199
chr13-3305784-INS-2847
chr13-3335057-INS-5088
chr13-3339068-INS-54
chr13-3461068-INS-700
chr13-3523653-INS-83
chr13-3523741-INS-566
chr13-3547494-INS-4610
chr13-3571625-INS-70
chr13-3698199-INS-7083
chr13-3794758-INS-189
chr13-3802025-INS-390
chr13-3802230-INS-64
chr13-3828287-INS-50
chr13-3910441-INS-68
chr13-3947774-INS-203
chr13-3948254-INS-124
chr13-3952110-INS-2041
chr13-3956111-INS-4871
chr13-4043487-INS-50
chr13-4048474-INS-75
chr13-4052098-INS-50
chr13-4061211-INS-2069
chr13-4086292-INS-99
chr13-4087068-INS-177
chr13-4153421-INS-50
chr13-4159826-INS-6223
chr13-4165719-INS-4438
chr13-4204781-INS-67
chr13-4256919-INS-105
chr13-4263107-INS-5615
chr13-4302691-INS-1147
chr13-4339737-INS-1028

chr13-4373386-INS-3089
chr13-4500346-INS-61
chr13-4500532-INS-217
chr13-4500984-INS-57
chr13-4542042-INS-2985
chr13-4544992-INS-50
chr13-4550284-INS-4474
chr13-4556100-INS-6128
chr13-4613149-INS-7046
chr13-4631085-INS-6427
chr13-4650926-INS-787
chr13-4656369-INS-298
chr13-4656759-INS-197
chr13-4676748-INS-8757
chr13-4713274-INS-411
chr13-4766432-INS-182
chr13-4779605-INS-69
chr13-4779693-INS-243
chr13-4780315-INS-104
chr13-4802734-INS-78
chr13-4828566-INS-25271
chr13-4843270-INS-999
chr13-4890125-INS-4524
chr13-4978900-INS-6633
chr13-5040804-INS-76
chr13-5084123-INS-3026
chr13-5089453-INS-91
chr13-5091721-INS-7851
chr13-5100931-INS-208
chr13-5110978-INS-470
chr13-5212719-INS-4354
chr13-5276798-INS-6260
chr13-5590334-INS-60
chr13-5619621-INS-179
chr13-5619666-INS-57
chr13-5641650-INS-50
chr13-5652925-INS-54
chr13-5660137-INS-6714
chr13-5674266-INS-1139
chr13-5675388-INS-3673
chr13-5715050-INS-858
chr13-5829761-INS-7524
chr13-5878848-INS-3035
chr13-5902739-INS-212
chr13-5902739-INS-487
chr13-5906311-INS-50
chr13-5920966-INS-73
chr13-5976834-INS-54
chr13-5997466-INS-88
chr13-5998255-INS-68
chr13-6005131-INS-60
chr13-6005193-INS-780
chr13-6010545-INS-1812
chr13-6086011-INS-3076
chr13-6099626-INS-415
chr13-6173635-INS-6549
chr13-6178383-INS-317
chr13-6185450-INS-72
chr13-6185522-INS-252
chr13-6189248-INS-205
chr13-6202890-INS-3066
chr13-6213338-INS-75
chr13-6221944-INS-1856
chr13-6244860-INS-57
chr13-6267551-INS-265
chr13-6334619-INS-2600
chr13-6337302-INS-70
chr13-6365469-INS-203
chr13-6365469-INS-551
chr13-6377563-INS-158
chr13-6432147-INS-6305
chr13-6446860-INS-52
chr13-6503110-INS-71
chr13-6503329-INS-505
chr13-6507659-INS-50
chr13-6515318-INS-515
chr13-6550344-INS-166

chr13-6584772-INS-6299
chr13-6590871-INS-2274
chr13-6592455-INS-1088
chr13-6630144-INS-212
chr13-6630708-INS-421
chr13-6681172-INS-735
chr13-6706271-INS-162
chr13-6726200-INS-410
chr13-6800937-INS-52
chr13-6838087-INS-1815
chr13-6839459-INS-1030
chr13-6859799-INS-476
chr13-6865603-INS-60
chr13-6887995-INS-112
chr13-6899042-INS-1467
chr13-6908212-INS-6924
chr13-6918729-INS-388
chr13-6919391-INS-159
chr13-6937571-INS-524
chr13-6943650-INS-248
chr13-6946566-INS-59
chr13-6953060-INS-132
chr13-6961274-INS-6592
chr13-6972831-INS-216
chr13-6997070-INS-573
chr13-7027382-INS-6373
chr13-7031506-INS-185
chr13-7056472-INS-58
chr13-7187287-INS-52
chr13-7218040-INS-55
chr13-7230019-INS-50
chr13-7294597-INS-60
chr13-7302618-INS-1264
chr13-7310434-INS-6482
chr13-7343880-INS-395
chr13-7382013-INS-50
chr13-7464521-INS-50
chr13-7493118-INS-341
chr13-7515715-INS-50
chr13-7578536-INS-160
chr13-7621376-INS-202
chr13-7624645-INS-50
chr13-7646659-INS-50
chr13-7651058-INS-50
chr13-7658474-INS-150
chr13-7658611-INS-388
chr13-7666222-INS-1738
chr13-7679860-INS-6337
chr13-7723202-INS-7628
chr13-7750104-INS-52
chr13-7785132-INS-55
chr13-7856196-INS-1130
chr13-7918091-INS-70
chr13-7921321-INS-119
chr13-7945693-INS-2662
chr13-7945779-INS-7593
chr13-7948161-INS-400
chr13-8011330-INS-114
chr13-8011372-INS-296
chr13-8011874-INS-66
chr13-8011882-INS-158
chr13-8016601-INS-72
chr13-8079511-INS-68
chr13-8081390-INS-1260
chr13-8095756-INS-82
chr13-8155600-INS-58
chr13-8172731-INS-1000
chr13-8215455-INS-130
chr13-8216004-INS-82
chr13-8272692-INS-50
chr13-8282922-INS-58
chr13-8375573-INS-55
chr13-8380955-INS-59
chr13-8404268-INS-136
chr13-8407488-INS-80
chr13-8413329-INS-7105
chr13-8426171-INS-1501

chr13-8430490-INS-1417
chr13-8441779-INS-1857
chr13-8446853-INS-171
chr13-8465641-INS-6178
chr13-8501341-INS-60
chr13-8504402-INS-1390
chr13-8545114-INS-144
chr13-8545114-INS-53
chr13-8545492-INS-108
chr13-8610423-INS-80
chr13-8612646-INS-730
chr13-8612938-INS-2596
chr13-8627786-INS-6360
chr13-8668043-INS-6483
chr13-8705263-INS-2287
chr13-8711006-INS-279
chr13-8741211-INS-50
chr13-8741458-INS-646
chr13-8781376-INS-60
chr13-8798416-INS-6389
chr13-8802171-INS-141
chr13-8804571-INS-88
chr13-8851759-INS-126
chr13-8851759-INS-414
chr13-8877520-INS-63
chr13-8883303-INS-242
chr13-8887861-INS-109
chr13-8938986-INS-60
chr13-8981883-INS-173
chr13-8993031-INS-252
chr13-9004822-INS-516
chr13-9008222-INS-216
chr13-9019616-INS-56
chr13-9040257-INS-56
chr13-9055011-INS-7202
chr13-9056973-INS-1974
chr13-9062091-INS-122
chr13-9062100-INS-444
chr13-9062344-INS-50
chr13-9070174-INS-423
chr13-9084970-INS-55
chr13-9135166-INS-50
chr13-9139910-INS-50
chr13-9164232-INS-202
chr13-9240787-INS-70
chr13-9267552-INS-412
chr13-9268694-INS-182
chr13-9283015-INS-6424
chr13-9288893-INS-124
chr13-9308461-INS-60
chr13-9310843-INS-50
chr13-9318535-INS-52
chr13-9335915-INS-86
chr13-9369254-INS-115
chr13-9371211-INS-433
chr13-9371337-INS-145
chr13-9371453-INS-752
chr13-9414599-INS-85
chr13-9458527-INS-163
chr13-9523094-INS-64
chr13-9582485-INS-58
chr13-9693416-INS-253
chr13-9703044-INS-182
chr13-9783193-INS-94
chr13-9865339-INS-1228
chr13-9875853-INS-164
chr13-9924102-INS-72
chr13-9933591-INS-310
chr13-9950351-INS-84
chr13-9950922-INS-232
chr13-9950922-INS-96
chr13-9956937-INS-731
chr13-9957047-INS-138
chr13-9966801-INS-2856
chr13-10021323-INS-304
chr13-10038519-INS-52
chr13-10042064-INS-50

chr13-10064323-INS-255
chr13-10064916-INS-76
chr13-10073476-INS-50
chr13-10082288-INS-6158
chr13-10107007-INS-50
chr13-10120778-INS-1176
chr13-10149271-INS-536
chr13-10169509-INS-205
chr13-10209594-INS-1351
chr13-10224814-INS-11673
chr13-10232201-INS-190
chr13-10232201-INS-74
chr13-10232284-INS-64
chr13-10236619-INS-70
chr13-10271041-INS-105
chr13-10271501-INS-96
chr13-10327799-INS-200
chr13-10346665-INS-89
chr13-10431799-INS-974
chr13-10493121-INS-62
chr13-10530077-INS-201
chr13-10544125-INS-6543
chr13-10552148-INS-2904
chr13-10558841-INS-566
chr13-10559260-INS-79
chr13-10670247-INS-503
chr13-10680480-INS-80
chr13-10690503-INS-12445
chr13-10781583-INS-97
chr13-10786687-INS-58
chr13-10814181-INS-52
chr13-10879587-INS-6222
chr13-10922419-INS-6317
chr13-10924047-INS-2439
chr13-10985419-INS-892
chr13-11039691-INS-525
chr13-11055683-INS-52
chr13-11061160-INS-213
chr13-11078452-INS-574
chr13-11105542-INS-192
chr13-11108910-INS-3870
chr13-11119639-INS-254
chr13-11133259-INS-50
chr13-11139712-INS-52
chr13-11141702-INS-80
chr13-11162982-INS-88
chr13-11199722-INS-66
chr13-11209230-INS-108
chr13-11215136-INS-1517
chr13-11224227-INS-51
chr13-11282728-INS-72
chr13-11295436-INS-4768
chr13-11298613-INS-63
chr13-11299106-INS-342
chr13-11306628-INS-402
chr13-11307134-INS-2490
chr13-11307276-INS-57
chr13-11307469-INS-6533
chr13-11333759-INS-52
chr13-11339395-INS-111
chr13-11361928-INS-80
chr13-11362879-INS-128
chr13-11442168-INS-94
chr13-11464062-INS-6411
chr13-11500242-INS-662
chr13-11531316-INS-54
chr13-11531876-INS-319
chr13-11609487-INS-53
chr13-11647027-INS-472
chr13-11654602-INS-447
chr13-11666631-INS-58
chr13-11717546-INS-94
chr13-11748419-INS-807
chr13-11748760-INS-303
chr13-11775028-INS-58
chr13-11824679-INS-159
chr13-11881540-INS-70

chr13-11901855-INS-346
chr13-11902471-INS-131
chr13-11948240-INS-121
chr13-11975049-INS-216
chr13-12006545-INS-54
chr13-12018956-INS-56
chr13-12066031-INS-50
chr13-12097236-INS-320
chr13-12097276-INS-56
chr13-12124041-INS-264
chr13-12124055-INS-86
chr13-12182208-INS-108
chr13-12183233-INS-180
chr13-12183255-INS-444
chr13-12183258-INS-54
chr13-12185003-INS-61
chr13-12252375-INS-115
chr13-12263726-INS-77
chr13-12263732-INS-473
chr13-12266221-INS-52
chr13-12274775-INS-6751
chr13-12317950-INS-68
chr13-12317962-INS-254
chr13-12322385-INS-208
chr13-12325288-INS-148
chr13-12344011-INS-128
chr13-12367140-INS-3201
chr13-12380796-INS-206
chr13-12407666-INS-55
chr13-12458450-INS-207
chr13-12467764-INS-164
chr13-12513160-INS-51
chr13-12521101-INS-121
chr13-12538185-INS-50
chr13-12550310-INS-211
chr13-12585433-INS-213
chr13-12586279-INS-94
chr13-12611603-INS-64
chr13-12619709-INS-56
chr13-12633769-INS-348
chr13-12644505-INS-971
chr13-12748779-INS-4274
chr13-12752529-INS-3037
chr13-12898860-INS-1998
chr13-13267897-INS-364
chr13-13509196-INS-206
chr13-13523960-INS-314
chr13-13531575-INS-1344
chr13-13537139-INS-348
chr13-13543358-INS-218
chr13-13607816-INS-208
chr13-13639010-INS-213
chr13-13639264-INS-572
chr13-13669241-INS-783
chr13-13683696-INS-3134
chr13-13696312-INS-4044
chr13-13715687-INS-7080
chr13-13750325-INS-6476
chr13-13769797-INS-280
chr13-13774548-INS-77
chr13-13777173-INS-6429
chr13-13782061-INS-50
chr13-13834581-INS-564
chr13-13870267-INS-7147
chr13-13873364-INS-191
chr13-13915483-INS-6430
chr13-13928390-INS-824
chr13-13929073-INS-7886
chr13-13976837-INS-1496
chr13-14004147-INS-56
chr13-14004786-INS-549
chr13-14020338-INS-6167
chr13-14029300-INS-78
chr13-14031957-INS-620
chr13-14035358-INS-60
chr13-14035644-INS-54
chr13-14035879-INS-174

chr13-14037339-INS-52
chr13-14056120-INS-781
chr13-14156532-INS-74
chr13-14171319-INS-1181
chr13-14187089-INS-64
chr13-14239551-INS-88
chr13-14249648-INS-148
chr13-14263153-INS-197
chr13-14267340-INS-253
chr13-14268587-INS-292
chr13-14268675-INS-66
chr13-14280861-INS-52
chr13-14411664-INS-52
chr13-14413841-INS-353
chr13-14418545-INS-170
chr13-14517286-INS-60
chr13-14547470-INS-63
chr13-14554949-INS-50
chr13-14599252-INS-645
chr13-14632714-INS-231
chr13-14632718-INS-504
chr13-14670311-INS-1698
chr13-14707145-INS-78
chr13-14707223-INS-2184
chr13-14759670-INS-260
chr13-14759818-INS-69
chr13-14816661-INS-183
chr13-14884771-INS-758
chr13-14900724-INS-2031
chr13-14901718-INS-479
chr13-15011470-INS-149
chr13-15077130-INS-63
chr13-15120674-INS-210
chr13-15131604-INS-5312
chr13-15139877-INS-536
chr13-15168188-INS-6257
chr13-15197926-INS-1410
chr13-15206827-INS-1580
chr13-15295397-INS-533
chr13-15306901-INS-58
chr13-15356182-INS-2149
chr13-15400612-INS-58
chr13-15437072-INS-138
chr13-15479395-INS-191
chr13-15484694-INS-58
chr13-15485712-INS-248
chr13-15508900-INS-7101
chr13-15515533-INS-2721
chr13-15556977-INS-979
chr13-15559822-INS-6754
chr13-15627998-INS-6387
chr13-15640087-INS-87
chr13-15735599-INS-84
chr13-15750681-INS-84
chr13-15760002-INS-73
chr13-15884033-INS-105
chr13-15929490-INS-7278
chr13-15958960-INS-6153
chr13-15972852-INS-8780
chr13-15996482-INS-286
chr13-16006246-INS-4925
chr13-16107217-INS-50
chr13-16129670-INS-4402
chr13-16183299-INS-50
chr13-16202408-INS-221
chr13-16211305-INS-6497
chr13-16216573-INS-256
chr13-16242633-INS-3534
chr13-16264057-INS-7230
chr13-16278387-INS-7254
chr13-16355071-INS-7194
chr13-16389938-INS-126
chr13-16406421-INS-108
chr13-16424356-INS-6894
chr13-16509428-INS-544
chr13-16556904-INS-56
chr13-16566774-INS-755

chr13-16578567-INS-73
chr13-16602300-INS-2445
chr13-16634798-INS-152
chr13-16661028-INS-59
chr13-16703025-INS-125
chr13-16890793-INS-109
chr13-16896831-INS-58
chr13-16958321-INS-72
chr13-16973621-INS-152
chr13-16973993-INS-577
chr13-16981283-INS-72
chr13-16982744-INS-540
chr13-16993493-INS-86
chr13-17024887-INS-54
chr13-17064492-INS-64
chr13-17091364-INS-1445
chr13-17114579-INS-212
chr13-17157885-INS-1026
chr13-17161071-INS-7154
chr13-17171070-INS-54
chr13-17181657-INS-6442
chr13-17245775-INS-85
chr13-17245977-INS-368
chr13-17281975-INS-6629
chr13-17344086-INS-58
chr13-17404830-INS-3815
chr13-17453255-INS-3875
chr13-17463889-INS-1099
chr13-17522309-INS-348
chr13-17524566-INS-870
chr13-17540839-INS-203
chr13-17672750-INS-6769
chr13-17688733-INS-50
chr13-17707668-INS-6386
chr13-17765300-INS-180
chr13-17772922-INS-68
chr13-17776174-INS-6567
chr13-17827766-INS-6322
chr13-17996360-INS-2524
chr13-18008727-INS-444
chr13-18021509-INS-4826
chr13-18032528-INS-843
chr13-18084813-INS-2000
chr13-18166534-INS-1570
chr13-18174122-INS-202
chr13-18185985-INS-78
chr13-18191843-INS-134
chr13-18201292-INS-60
chr13-18214136-INS-57
chr13-18237280-INS-4828
chr13-18252299-INS-242
chr13-18272091-INS-61
chr13-18279166-INS-2824
chr13-18314070-INS-50
chr13-18377272-INS-127
chr13-18466509-INS-6288
chr13-18469371-INS-69
chr13-18484240-INS-1712
chr13-18496888-INS-53
chr13-18496956-INS-128
chr13-18518491-INS-81
chr13-18537010-INS-90
chr13-18558185-INS-479
chr13-18634008-INS-54
chr13-18644545-INS-105
chr13-18644545-INS-245
chr13-18653009-INS-155
chr13-18705887-INS-50
chr13-18739909-INS-6306
chr13-18754272-INS-532
chr13-18762146-INS-62
chr13-18762726-INS-55
chr13-18776093-INS-1040
chr13-18803057-INS-1268
chr13-18810853-INS-50
chr13-18911918-INS-5311
chr13-18951000-INS-65

chr13-18953662-INS-343
chr13-18959486-INS-294
chr13-18969066-INS-178
chr13-18990141-INS-90
chr13-19001955-INS-194
chr13-19043876-INS-110
chr13-19063141-INS-548
chr13-19063165-INS-90
chr13-19087659-INS-672
chr13-19087732-INS-319
chr13-19199859-INS-56
chr13-19211818-INS-135
chr13-19264083-INS-54
chr13-19324928-INS-1246
chr13-19327798-INS-52
chr13-19464829-INS-2793
chr13-19582929-INS-2518
chr13-19690259-INS-207
chr13-19771115-INS-212
chr13-19851231-INS-4894
chr13-19908590-INS-6996
chr13-19919919-INS-762
chr13-19959034-INS-1082
chr13-19962374-INS-78
chr13-19962404-INS-244
chr13-20057844-INS-5735
chr13-20102266-INS-74
chr13-20103716-INS-597
chr13-20147504-INS-614
chr13-20177680-INS-6485
chr13-20192639-INS-64
chr13-20212489-INS-6459
chr13-20243972-INS-2883
chr13-20292969-INS-151
chr13-20326565-INS-57
chr13-20329632-INS-72
chr13-20329778-INS-106
chr13-20334665-INS-148
chr13-20353945-INS-70
chr13-20356354-INS-177
chr13-20403285-INS-5343
chr13-20420506-INS-137
chr13-20420761-INS-172
chr13-20463133-INS-2312
chr13-20463313-INS-149
chr13-20464781-INS-365
chr13-20484172-INS-52
chr13-20530953-INS-95
chr13-20615554-INS-124
chr13-20616303-INS-87
chr13-20672030-INS-74
chr13-20686922-INS-532
chr13-20715317-INS-1294
chr13-20759115-INS-122
chr13-20768109-INS-1406
chr13-20879548-INS-4428
chr13-20922254-INS-6359
chr13-20977356-INS-50
chr13-20990619-INS-7393
chr13-21069720-INS-267
chr13-21069977-INS-195
chr13-21070269-INS-145
chr13-21074582-INS-2713
chr13-21083845-INS-57
chr13-21118902-INS-191
chr13-21119130-INS-224
chr13-21119682-INS-53
chr13-21126677-INS-7341
chr13-21155785-INS-6886
chr13-21216632-INS-1354
chr13-21224352-INS-6393
chr13-21261926-INS-6493
chr13-21271578-INS-6225
chr13-21287331-INS-5427
chr13-21306877-INS-208
chr13-21309196-INS-54
chr13-21314895-INS-1581

chr13-21332775-INS-50
chr13-21362798-INS-1000
chr13-21386892-INS-4592
chr13-21403701-INS-1743
chr13-21413400-INS-173
chr13-21413633-INS-497
chr13-21452371-INS-50
chr13-21453367-INS-50
chr13-21463045-INS-359
chr13-21499579-INS-333
chr13-21539568-INS-83
chr13-21561143-INS-1409
chr13-21611430-INS-313
chr13-21611430-INS-59
chr13-21762259-INS-481
chr13-21770589-INS-1452
chr13-21779446-INS-6379
chr13-21786358-INS-65
chr13-21837308-INS-1871
chr13-21885708-INS-129
chr13-21946674-INS-1317
chr13-21946674-INS-652
chr13-21984677-INS-70
chr13-22028648-INS-4526
chr13-22066069-INS-7021
chr13-22106462-INS-237
chr13-22106513-INS-109
chr13-22116521-INS-104
chr13-22171441-INS-116
chr13-22260312-INS-1306
chr13-22280454-INS-283
chr13-22364383-INS-545
chr13-22367344-INS-6450
chr13-22401354-INS-3484
chr13-22405057-INS-419
chr13-22510650-INS-52
chr13-22517561-INS-7339
chr13-22589131-INS-1499
chr13-22666198-INS-83
chr13-22744031-INS-157
chr13-22818745-INS-401
chr13-22819251-INS-196
chr13-22849202-INS-101
chr13-22909570-INS-5504
chr13-22943693-INS-52
chr13-22948899-INS-4729
chr13-22972938-INS-997
chr13-22979202-INS-254
chr13-23054173-INS-195
chr13-23055650-INS-7188
chr13-23130870-INS-199
chr13-23147835-INS-57
chr13-23159611-INS-126
chr13-23165257-INS-3108
chr13-23168019-INS-11775
chr13-23178424-INS-556
chr13-23178424-INS-8739
chr13-23217869-INS-129
chr13-23225777-INS-348
chr13-23408839-INS-1150
chr13-23595355-INS-12100
chr13-23601757-INS-583
chr13-23621618-INS-75
chr13-23657741-INS-201
chr13-23811185-INS-56
chr13-23870017-INS-51
chr13-23886832-INS-1475
chr13-23984391-INS-50
chr13-23984714-INS-235
chr13-24030529-INS-50
chr13-24151146-INS-561
chr13-24157219-INS-1412
chr13-24180764-INS-8413
chr13-24196020-INS-52
chr13-24288722-INS-486
chr13-24338887-INS-62
chr13-24340206-INS-132

chr13-24364864-INS-1673
chr13-24369136-INS-449
chr13-24407491-INS-64
chr13-24418198-INS-50
chr13-24500987-INS-334
chr13-24530626-INS-6499
chr13-24544591-INS-267
chr13-24545420-INS-51
chr13-24566043-INS-2272
chr13-24590762-INS-56
chr13-24605891-INS-72
chr13-24627973-INS-55
chr13-24632165-INS-208
chr13-24641873-INS-70
chr13-24661655-INS-69
chr13-24721072-INS-50
chr13-24767895-INS-65
chr13-24788489-INS-1706
chr13-24851013-INS-4806
chr13-24870185-INS-102
chr13-24922876-INS-52
chr13-24933147-INS-4882
chr13-24955652-INS-106
chr13-24979434-INS-1029
chr13-25050252-INS-292
chr13-25058031-INS-461
chr13-25081185-INS-162
chr13-25120986-INS-598
chr13-25145789-INS-280
chr13-25148180-INS-50
chr13-25176455-INS-207
chr13-25205395-INS-243
chr13-25300638-INS-62
chr13-25358691-INS-68
chr13-25367629-INS-7090
chr13-25381229-INS-70
chr13-25505694-INS-2578
chr13-25524298-INS-52
chr13-25533605-INS-104
chr13-25573419-INS-123
chr13-25586223-INS-2747
chr13-25619114-INS-4089
chr13-25634389-INS-359
chr13-25649701-INS-70
chr13-25653633-INS-316
chr13-25653865-INS-236
chr13-25653897-INS-72
chr13-25661327-INS-6592
chr13-25670849-INS-115
chr13-25701820-INS-2844
chr13-25727015-INS-208
chr13-25753421-INS-50
chr13-25817607-INS-376
chr13-25822381-INS-882
chr13-25860151-INS-62
chr13-25861101-INS-126
chr13-25901467-INS-134
chr13-25916303-INS-52
chr13-25938570-INS-160
chr13-25955866-INS-54
chr13-25957663-INS-6164
chr13-25997012-INS-6184
chr13-26038963-INS-242
chr13-26039026-INS-96
chr13-26045803-INS-3442
chr13-26056009-INS-1993
chr13-26062968-INS-314
chr13-26079976-INS-7083
chr13-26091431-INS-3206
chr13-26147419-INS-84
chr13-26162392-INS-2488
chr13-26169684-INS-77
chr13-26169910-INS-249
chr13-26188488-INS-1427
chr13-26190254-INS-50
chr13-26200093-INS-1832
chr13-26243707-INS-60

chr13-26251335-INS-87
chr13-26274435-INS-452
chr13-26276452-INS-411
chr13-26313376-INS-50
chr13-26328888-INS-74
chr13-26361890-INS-3297
chr13-26464750-INS-6545
chr13-26506548-INS-65
chr13-26506639-INS-186
chr13-26515395-INS-142
chr13-26567842-INS-476
chr13-26595386-INS-133
chr13-26611136-INS-407
chr13-26640061-INS-96
chr13-26640097-INS-562
chr13-26675532-INS-1226
chr13-26739369-INS-92
chr13-26743462-INS-201
chr13-26767257-INS-63
chr13-26826248-INS-121
chr13-26826433-INS-6206
chr13-26826846-INS-129
chr13-26852836-INS-65
chr13-26918988-INS-13403
chr13-26918989-INS-6166
chr13-26942304-INS-415
chr13-27072130-INS-1889
chr13-27085722-INS-2419
chr13-27086573-INS-60
chr13-27088312-INS-132
chr13-27089700-INS-6161
chr13-27093428-INS-1748
chr13-27109951-INS-4480
chr13-27219066-INS-236
chr13-27236719-INS-50
chr13-27274982-INS-251
chr13-27325709-INS-51
chr13-27336213-INS-5028
chr13-27361423-INS-873
chr13-27390142-INS-103
chr13-27410332-INS-127
chr13-27483455-INS-72
chr13-27528426-INS-90
chr13-27639465-INS-70
chr13-27642001-INS-100
chr13-27654606-INS-8413
chr13-27677795-INS-64
chr13-27678963-INS-1368
chr13-27686797-INS-7086
chr13-27720263-INS-169
chr13-27829873-INS-56
chr13-27841874-INS-87
chr13-28017923-INS-1268
chr13-28056506-INS-236
chr13-28091713-INS-8978
chr13-28095418-INS-1212
chr13-28095697-INS-506
chr13-28140009-INS-231
chr13-28140450-INS-125
chr13-28232339-INS-373
chr13-28284716-INS-6700
chr13-28323106-INS-50
chr13-28334134-INS-50
chr13-28349511-INS-58
chr13-28353552-INS-152
chr13-28363085-INS-1668
chr13-28413702-INS-10982
chr13-28413702-INS-2516
chr13-28414798-INS-3523
chr13-28477708-INS-90
chr13-28514069-INS-2999
chr13-28537993-INS-6335
chr13-28541786-INS-1552
chr13-28548061-INS-6383
chr13-28550406-INS-824
chr13-28550844-INS-1583
chr13-28602884-INS-62

chr13-28679839-INS-6491
chr13-28703278-INS-50
chr13-28750429-INS-186
chr13-28815749-INS-72
chr13-28861761-INS-233
chr13-28937578-INS-297
chr13-28945349-INS-268
chr13-28961170-INS-57
chr13-29023438-INS-388
chr13-29027172-INS-105
chr13-29034284-INS-112
chr13-29037486-INS-157
chr13-29075970-INS-60
chr13-29123430-INS-15922
chr13-29178552-INS-62
chr13-29211878-INS-51
chr13-29213960-INS-90
chr13-29222637-INS-189
chr13-29222810-INS-1057
chr13-29227378-INS-134
chr13-29276643-INS-175
chr13-29352017-INS-51
chr13-29447150-INS-52
chr13-29447430-INS-341
chr13-29447592-INS-72
chr13-29483421-INS-6463
chr13-29483925-INS-56
chr13-29495650-INS-54
chr13-29513564-INS-607
chr13-29579263-INS-60
chr13-29678953-INS-77
chr13-29708448-INS-196
chr13-29711776-INS-89
chr13-29749434-INS-51
chr13-29760123-INS-51
chr13-29760135-INS-391
chr13-29760392-INS-535
chr13-29829368-INS-76
chr13-29866996-INS-431
chr13-29876109-INS-1856
chr13-29886672-INS-85
chr13-29934495-INS-335
chr13-29934495-INS-801
chr13-29936593-INS-60
chr13-29963178-INS-54
chr13-29972696-INS-60
chr13-30043324-INS-520
chr13-30043409-INS-198
chr13-30046359-INS-207
chr13-30047829-INS-77
chr13-30177413-INS-104
chr13-30215419-INS-54
chr13-30235492-INS-52
chr13-30258236-INS-4520
chr13-30283053-INS-62
chr13-30288089-INS-54
chr13-30294012-INS-301
chr13-30314548-INS-77
chr13-30322748-INS-50
chr13-30329680-INS-66
chr13-30346005-INS-3659
chr13-30348622-INS-152
chr13-30353031-INS-69
chr13-30383606-INS-153
chr13-30384421-INS-275
chr13-30395947-INS-160
chr13-30398733-INS-102
chr13-30411425-INS-60
chr13-30418853-INS-236
chr13-30451028-INS-6532
chr13-30471336-INS-204
chr13-30518942-INS-59
chr13-30519657-INS-334
chr13-30529950-INS-336
chr13-30531165-INS-1356
chr13-30550690-INS-123
chr13-30670075-INS-585

chr13-30680526-INS-447
chr13-30696803-INS-128
chr13-30752556-INS-72
chr13-30781959-INS-111
chr13-30782360-INS-600
chr13-30804133-INS-306
chr13-30877089-INS-117
chr13-30902799-INS-136
chr13-30964586-INS-105
chr13-30997643-INS-56
chr13-31077031-INS-57
chr13-31230186-INS-85
chr13-31241956-INS-88
chr13-31269246-INS-362
chr13-31269374-INS-112
chr13-31303205-INS-226
chr13-31311809-INS-401
chr13-31386376-INS-102
chr13-31386616-INS-84
chr13-31399446-INS-65
chr13-31405817-INS-60
chr13-31406375-INS-52
chr13-31429625-INS-4666
chr13-31438584-INS-576
chr13-31457524-INS-1740
chr13-31484529-INS-6347
chr13-31538543-INS-618
chr13-31588021-INS-7254
chr13-31616489-INS-71
chr13-31737075-INS-273
chr13-31747620-INS-51
chr13-31758927-INS-1547
chr13-31760044-INS-1047
chr13-31760440-INS-241
chr13-31766783-INS-6847
chr13-31768693-INS-154
chr13-31772950-INS-1076
chr13-31864625-INS-1669
chr13-31900204-INS-63
chr13-31911738-INS-5804
chr13-31947474-INS-56
chr13-31958046-INS-200
chr13-31958328-INS-90
chr13-31967058-INS-56
chr13-31967108-INS-90
chr13-31984372-INS-50
chr13-32030678-INS-185
chr13-32031418-INS-215
chr13-32039779-INS-60
chr13-32119196-INS-1892
chr13-32450147-INS-51
chr13-32453974-INS-678
chr13-32521353-INS-211
chr13-32616331-INS-354
chr13-32681409-INS-1018
chr13-32835393-INS-213
chr13-32835393-INS-543
chr13-32880078-INS-994
chr13-32904261-INS-2917
chr13-32993644-INS-86
chr13-33062719-INS-120
chr13-33088072-INS-197
chr13-33088092-INS-56
chr13-33155653-INS-1189
chr13-33267704-INS-63
chr13-33270745-INS-6583
chr13-33272992-INS-354
chr13-33291799-INS-6595
chr13-33359180-INS-66
chr13-33368466-INS-80
chr13-33756727-INS-256
chr13-33756820-INS-881
chr13-33767741-INS-12529
chr13-33773357-INS-78
chr13-33776068-INS-51
chr13-33818347-INS-432
chr13-33860597-INS-70

chr13-33861296-INS-135
chr13-33952602-INS-98
chr13-34011739-INS-1451
chr13-34014825-INS-162
chr13-34028225-INS-81
chr13-34093097-INS-6921
chr13-34110990-INS-6666
chr13-34122352-INS-317
chr13-34178135-INS-157
chr13-34182763-INS-79
chr13-34184211-INS-420
chr13-34275524-INS-2775
chr13-34343609-INS-178
chr13-34473165-INS-50
chr13-34592186-INS-498
chr13-34622983-INS-454
chr13-34714876-INS-146
chr13-34719381-INS-98
chr13-34720971-INS-110
chr13-34726527-INS-58
chr13-34757923-INS-498
chr13-34759189-INS-3781
chr13-34774236-INS-585
chr13-34841388-INS-285
chr13-34871366-INS-494
chr13-34874780-INS-75
chr13-34896002-INS-72
chr13-34896492-INS-59
chr13-34901151-INS-414
chr13-34933030-INS-52
chr13-34943577-INS-444
chr13-34956557-INS-4196
chr13-34969444-INS-62
chr13-35026673-INS-404
chr13-35027463-INS-486
chr13-35029862-INS-598
chr13-35045604-INS-6442
chr13-35068564-INS-78
chr13-35133276-INS-196
chr13-35153610-INS-6254
chr13-35159528-INS-62
chr13-35169046-INS-80
chr13-35181203-INS-565
chr13-35196201-INS-326
chr13-35224791-INS-68
chr13-35225127-INS-147
chr13-35280772-INS-54
chr13-35295051-INS-65
chr13-35295106-INS-134
chr13-35372017-INS-66
chr13-35389375-INS-6191
chr13-35394311-INS-50
chr13-35442229-INS-80
chr13-35527530-INS-171
chr13-35545905-INS-242
chr13-35562780-INS-1800
chr13-35572153-INS-3662
chr13-35592932-INS-2154
chr13-35625518-INS-6123
chr13-35645880-INS-69
chr13-35654003-INS-145
chr13-35670149-INS-56
chr13-35734671-INS-56
chr13-35840709-INS-51
chr13-35889207-INS-120
chr13-35889987-INS-64
chr13-35890247-INS-194
chr13-35890247-INS-56
chr13-35907371-INS-1653
chr13-35921443-INS-424
chr13-35974377-INS-203
chr13-36006045-INS-258
chr13-36067916-INS-184
chr13-36067972-INS-471
chr13-36068226-INS-69
chr13-36099422-INS-107
chr13-36099908-INS-240

chr13-36196284-INS-74
chr13-36196349-INS-638
chr13-36203369-INS-1096
chr13-36218391-INS-6251
chr13-36223933-INS-50
chr13-36262620-INS-51
chr13-36263787-INS-70
chr13-36289267-INS-172
chr13-36315698-INS-5634
chr13-36343658-INS-96
chr13-36390340-INS-294
chr13-36401580-INS-409
chr13-36422299-INS-1951
chr13-36524632-INS-54
chr13-36547015-INS-104
chr13-36568378-INS-2563
chr13-36581728-INS-250
chr13-36583393-INS-612
chr13-36588303-INS-267
chr13-36622785-INS-55
chr13-36653297-INS-168
chr13-36659315-INS-200
chr13-36687248-INS-369
chr13-36718272-INS-88
chr13-36718584-INS-661
chr13-36735002-INS-50
chr13-36742885-INS-191
chr13-36748315-INS-240
chr13-36756650-INS-164
chr13-36765108-INS-2732
chr13-36806473-INS-63
chr13-36811101-INS-311
chr13-36813414-INS-224
chr13-36819162-INS-6394
chr13-36886173-INS-111
chr13-36935007-INS-60
chr13-36939561-INS-51
chr13-37000156-INS-6350
chr13-37006830-INS-325
chr13-37017950-INS-52
chr13-37021697-INS-400
chr13-37075215-INS-952
chr13-37100513-INS-51
chr13-37101251-INS-52
chr13-37114476-INS-116
chr13-37132888-INS-112
chr13-37160935-INS-1225
chr13-37162735-INS-516
chr13-37198261-INS-62
chr13-37273794-INS-8671
chr13-37280924-INS-56
chr13-37282560-INS-6440
chr13-37325600-INS-87
chr13-37383564-INS-2104
chr13-37438219-INS-54
chr13-37518341-INS-525
chr13-37554403-INS-50
chr13-37559295-INS-59
chr13-37568429-INS-1724
chr13-37584197-INS-50
chr13-37617568-INS-147
chr13-37621252-INS-70
chr13-37666464-INS-130
chr13-37680013-INS-88
chr13-37680678-INS-140
chr13-37698446-INS-55
chr13-37704684-INS-126
chr13-37705473-INS-199
chr13-37760980-INS-147
chr13-37790995-INS-211
chr13-37839262-INS-109
chr13-37851304-INS-280
chr13-37855153-INS-138
chr13-37857824-INS-66
chr13-37914690-INS-61
chr13-37928578-INS-84
chr13-37999784-INS-66

chr13-38016705-INS-288
chr13-38020278-INS-130
chr13-38038549-INS-68
chr13-38047932-INS-58
chr13-38047992-INS-134
chr13-38109806-INS-186
chr13-38138154-INS-64
chr13-38138736-INS-498
chr13-38138736-INS-6475
chr13-38144552-INS-56
chr13-38148453-INS-223
chr13-38168100-INS-1900
chr13-38173719-INS-868
chr13-38256223-INS-198
chr13-38261577-INS-1340
chr13-38270764-INS-1889
chr13-38276109-INS-6474
chr13-38283064-INS-460
chr13-38319276-INS-81
chr13-38324495-INS-7119
chr13-38343189-INS-55
chr13-38344217-INS-312
chr13-38344364-INS-667
chr13-38356380-INS-355
chr13-38387891-INS-56
chr13-38439148-INS-275
chr13-38477038-INS-90
chr13-38508299-INS-52
chr13-38518202-INS-195
chr13-38522171-INS-132
chr13-38522474-INS-520
chr13-38554926-INS-102
chr13-38599643-INS-99
chr13-38660145-INS-54
chr13-38679130-INS-68
chr13-38714926-INS-82
chr13-38773144-INS-146
chr13-38775807-INS-76
chr13-38857588-INS-88
chr13-38867954-INS-210
chr13-38869259-INS-208
chr13-38938543-INS-55
chr13-38955946-INS-386
chr13-38956224-INS-52
chr13-38960048-INS-534
chr13-38969895-INS-288
chr13-38970138-INS-663
chr13-39001238-INS-1078
chr13-39015646-INS-7246
chr13-39023577-INS-1818
chr13-39027915-INS-1404
chr13-39037424-INS-178
chr13-39044082-INS-484
chr13-39052751-INS-50
chr13-39079803-INS-802
chr13-39080123-INS-162
chr13-39081009-INS-78
chr13-39107653-INS-7384
chr13-39118060-INS-56
chr13-39118175-INS-478
chr13-39169127-INS-196
chr13-39202247-INS-52
chr13-39243421-INS-59
chr13-39244327-INS-2809
chr13-39260015-INS-1489
chr13-39319086-INS-131
chr13-39326013-INS-70
chr13-39338563-INS-123
chr13-39344348-INS-1164
chr13-39347573-INS-434
chr13-39373128-INS-201
chr13-39389383-INS-57
chr13-39423462-INS-6363
chr13-39440534-INS-382
chr13-39441792-INS-1752
chr13-39462794-INS-67
chr13-39478458-INS-1055

chr13-39484513-INS-1788
chr13-39513024-INS-2689
chr13-39519252-INS-56
chr13-39520822-INS-6456
chr13-39526024-INS-240
chr13-39540813-INS-56
chr13-39545262-INS-66
chr13-39547373-INS-52
chr13-39623071-INS-211
chr13-39632357-INS-1120
chr13-39633095-INS-98
chr13-39671843-INS-52
chr13-39679819-INS-50
chr13-39702075-INS-62
chr13-39720793-INS-58
chr13-39727617-INS-120
chr13-39806950-INS-355
chr13-39808044-INS-67
chr13-39808264-INS-59
chr13-39837220-INS-236
chr13-39874872-INS-158
chr13-39899044-INS-1989
chr13-40066805-INS-71
chr13-40229729-INS-104
chr13-40261718-INS-198
chr13-40284030-INS-66
chr13-40327048-INS-1242
chr13-40327048-INS-253
chr13-40359402-INS-52
chr13-40359712-INS-59
chr13-40377139-INS-132
chr13-40417993-INS-60
chr13-40419154-INS-104
chr13-40452452-INS-52
chr13-40534193-INS-80
chr13-40593867-INS-153
chr13-40635216-INS-84
chr13-40635896-INS-159
chr13-40645165-INS-6215
chr13-40668829-INS-72
chr13-40701603-INS-381
chr13-40716313-INS-72
chr13-40718314-INS-56
chr13-40741535-INS-212
chr13-40754139-INS-695
chr13-40775392-INS-175
chr13-40776886-INS-443
chr13-40810880-INS-1417
chr13-40852768-INS-95
chr13-40872156-INS-66
chr13-40894115-INS-54
chr13-40898937-INS-59
chr13-40909916-INS-1482
chr13-40917511-INS-1138
chr13-40917511-INS-146
chr13-40938955-INS-94
chr13-40978749-INS-190
chr13-40981198-INS-7407
chr13-40985105-INS-201
chr13-40998750-INS-3601
chr13-41001172-INS-62
chr13-41006787-INS-167
chr13-41025107-INS-72
chr13-41025673-INS-785
chr13-41033827-INS-72
chr13-41085567-INS-126
chr13-41194859-INS-342
chr13-41238452-INS-62
chr13-41276033-INS-96
chr13-41346992-INS-61
chr13-41443166-INS-50
chr13-41611962-INS-258
chr13-41612623-INS-1010
chr13-41642988-INS-54
chr13-41673093-INS-81
chr13-41685298-INS-3410
chr13-41720967-INS-1321

chr13-41735354-INS-60
chr13-41757315-INS-4092
chr13-41775877-INS-196
chr13-41787197-INS-51
chr13-41802358-INS-51
chr13-41805791-INS-81
chr13-41811309-INS-88
chr13-41825493-INS-79
chr13-41868412-INS-65
chr13-41884865-INS-6725
chr13-41937620-INS-52
chr13-41955642-INS-5381
chr13-42074628-INS-58
chr13-42075609-INS-65
chr13-42129108-INS-427
chr13-42153883-INS-210
chr13-42157515-INS-162
chr13-42177745-INS-327
chr13-42236618-INS-1321
chr13-42246973-INS-234
chr13-42266536-INS-54
chr13-42325003-INS-338
chr13-42349703-INS-51
chr13-42357071-INS-3667
chr13-42364307-INS-60
chr13-42366143-INS-72
chr13-42384451-INS-488
chr13-42386590-INS-65
chr13-42392514-INS-124
chr13-42402158-INS-127
chr13-42405034-INS-194
chr13-42406568-INS-52
chr13-42417980-INS-52
chr13-42420395-INS-82
chr13-42421788-INS-72
chr13-42445997-INS-81
chr13-42446001-INS-172
chr13-42454323-INS-50
chr13-42459914-INS-380
chr13-42462776-INS-162
chr13-42470592-INS-86
chr13-42494097-INS-390
chr13-42505612-INS-353
chr13-42618056-INS-147
chr13-42625577-INS-396
chr13-42638012-INS-100
chr13-42730142-INS-86
chr13-42737376-INS-728
chr13-42762937-INS-60
chr13-42771780-INS-1825
chr13-42785997-INS-1888
chr13-42791222-INS-60
chr13-42869034-INS-257
chr13-42869756-INS-98
chr13-42884419-INS-94
chr13-42932214-INS-2609
chr13-42933015-INS-61
chr13-42937965-INS-57
chr13-43184330-INS-221
chr13-43189563-INS-136
chr13-43201486-INS-64
chr13-43224903-INS-382
chr13-43256081-INS-166
chr13-43313790-INS-52
chr13-43328487-INS-65
chr13-43509047-INS-51
chr13-43509068-INS-177
chr13-43515227-INS-220
chr13-43522793-INS-771
chr13-43522922-INS-275
chr13-43726386-INS-150
chr13-43766508-INS-1042
chr13-43858501-INS-2359
chr13-43911022-INS-167
chr13-43915841-INS-54
chr13-43941107-INS-108
chr13-43984491-INS-3911

chr13-44038186-INS-57
chr13-44038582-INS-112
chr13-44102754-INS-70
chr13-44134072-INS-1350
chr13-44194152-INS-66
chr13-44207277-INS-90
chr13-44245816-INS-222
chr13-44450498-INS-54
chr13-44508708-INS-124
chr13-44517862-INS-56
chr13-44617103-INS-248
chr13-44624271-INS-74
chr13-44629755-INS-60
chr13-44635187-INS-3500
chr13-44681549-INS-207
chr13-44743127-INS-52
chr13-44744271-INS-129
chr13-44747866-INS-81
chr13-44768725-INS-52
chr13-44821495-INS-74
chr13-44821686-INS-102
chr13-44898738-INS-68
chr13-44920655-INS-203
chr13-45023626-INS-68
chr13-45082213-INS-386
chr13-45085997-INS-204
chr13-45128690-INS-894
chr13-45195911-INS-3516
chr13-45284686-INS-201
chr13-45336835-INS-295
chr13-45347453-INS-359
chr13-45457118-INS-214
chr13-45481618-INS-412
chr13-45540860-INS-87
chr13-45580498-INS-55
chr13-45624645-INS-280
chr13-45631553-INS-228
chr13-45673575-INS-62
chr13-45679459-INS-67
chr13-45936845-INS-50
chr13-45993022-INS-7131
chr13-46009203-INS-208
chr13-46053485-INS-56
chr13-46056833-INS-69
chr13-46073152-INS-177
chr13-46128868-INS-245
chr13-46301280-INS-307
chr13-46312276-INS-64
chr13-46320134-INS-72
chr13-46320258-INS-586
chr13-46320338-INS-271
chr13-46396616-INS-3537
chr13-46414354-INS-170
chr13-46420348-INS-4138
chr13-46423769-INS-112
chr13-46510450-INS-436
chr13-46538272-INS-87
chr13-46592056-INS-61
chr13-46598784-INS-57
chr13-46606859-INS-7183
chr13-46672346-INS-155
chr13-46680525-INS-208
chr13-46692932-INS-121
chr13-46729723-INS-201
chr13-46757341-INS-50
chr13-46798858-INS-52
chr13-46940779-INS-1103
chr13-46951706-INS-50
chr13-46978141-INS-96
chr13-46992468-INS-103
chr13-47189179-INS-162
chr13-47216726-INS-141
chr13-47282178-INS-92
chr13-47376538-INS-50
chr13-47409310-INS-112
chr13-47421282-INS-6390
chr13-47423598-INS-96

chr13-47445195-INS-51
chr13-47524677-INS-917
chr13-47526321-INS-397
chr13-47535277-INS-6168
chr13-47579961-INS-181
chr13-47580766-INS-64
chr13-47639796-INS-260
chr13-47693579-INS-555
chr13-47735236-INS-56
chr13-47745617-INS-68
chr13-47750974-INS-71
chr13-47786493-INS-202
chr13-47857398-INS-2310
chr13-47869186-INS-52
chr13-47881766-INS-7500
chr13-47908816-INS-53
chr13-47969530-INS-60
chr13-47983155-INS-56
chr13-47999799-INS-59
chr13-48058762-INS-66
chr13-48120723-INS-213
chr13-48156213-INS-155
chr13-48161238-INS-55
chr13-48337876-INS-7756
chr13-48400037-INS-62
chr13-48400789-INS-68
chr13-48426942-INS-54
chr13-48427710-INS-62
chr13-48451189-INS-212
chr13-48484143-INS-2833
chr13-48491910-INS-764
chr13-48519703-INS-87
chr13-48520682-INS-2816
chr13-48529599-INS-76
chr13-48607100-INS-125
chr13-48620950-INS-69
chr13-48623107-INS-51
chr13-48680948-INS-125
chr13-48786984-INS-255
chr13-48824145-INS-68
chr13-48830482-INS-86
chr13-48837476-INS-58
chr13-48942600-INS-96
chr13-48944321-INS-400
chr13-48962615-INS-11647
chr13-48999734-INS-185
chr13-49159824-INS-1884
chr13-49179759-INS-209
chr13-49186145-INS-1176
chr13-49316576-INS-51
chr13-49350924-INS-54
chr13-49431118-INS-236
chr13-49466596-INS-56
chr13-49472678-INS-85
chr13-49524202-INS-133
chr13-49526698-INS-127
chr13-49546468-INS-204
chr13-49564140-INS-184
chr13-49746506-INS-69
chr13-49746520-INS-503
chr13-49768524-INS-204
chr13-49861381-INS-62
chr13-49893670-INS-179
chr13-49893979-INS-54
chr13-49907133-INS-162
chr13-49907133-INS-349
chr13-49915100-INS-2051
chr13-49935397-INS-53
chr13-49947066-INS-2126
chr13-49951925-INS-6211
chr13-49969453-INS-345
chr13-49969552-INS-50
chr13-50017882-INS-53
chr13-50113299-INS-1765
chr13-50121446-INS-384
chr13-50127450-INS-120
chr13-50196830-INS-2718

chr13-50224467-INS-4594
chr13-50238174-INS-574
chr13-50250754-INS-110
chr13-50342586-INS-1044
chr13-50485065-INS-5562
chr13-50519892-INS-1054
chr13-50520026-INS-3550
chr13-50547031-INS-6965
chr13-50579539-INS-104
chr13-50600783-INS-4795
chr13-50618010-INS-1506
chr13-50630912-INS-80
chr13-50704089-INS-1880
chr13-50758244-INS-727
chr13-50776641-INS-1044
chr13-50805058-INS-80
chr13-50806983-INS-342
chr13-50835845-INS-1937
chr13-50855947-INS-497
chr13-50917288-INS-510
chr13-50940104-INS-62
chr13-50966763-INS-289
chr13-50968785-INS-121
chr13-51016605-INS-66
chr13-51028799-INS-1779
chr13-51124906-INS-53
chr13-51154133-INS-7109
chr13-51156810-INS-175
chr13-51169160-INS-51
chr13-51190492-INS-212
chr13-51224358-INS-203
chr13-51236431-INS-72
chr13-51339627-INS-1508
chr13-51344656-INS-240
chr13-51344672-INS-57
chr13-51405835-INS-90
chr13-51483629-INS-164
chr13-51509158-INS-86
chr13-51527559-INS-277
chr13-51546663-INS-63
chr13-51554229-INS-86
chr13-51633439-INS-625
chr13-51709281-INS-89
chr13-51709441-INS-204
chr13-51763728-INS-165
chr13-51798463-INS-59
chr13-51818248-INS-277
chr13-51847163-INS-211
chr13-51860387-INS-59
chr13-51983724-INS-308
chr13-51983775-INS-64
chr13-51984384-INS-498
chr13-51998592-INS-60
chr13-52029941-INS-61
chr13-52089740-INS-71
chr13-52187283-INS-200
chr13-52187839-INS-52
chr13-52195098-INS-131
chr13-52211556-INS-2782
chr13-52256655-INS-155
chr13-52260681-INS-116
chr13-52294292-INS-52
chr13-52335926-INS-136
chr13-52339157-INS-144
chr13-52339792-INS-364
chr13-52340102-INS-832
chr13-52345578-INS-50
chr13-52346151-INS-430
chr13-52397853-INS-125
chr13-52397875-INS-50
chr13-52431940-INS-608
chr13-52432082-INS-245
chr13-52469359-INS-101
chr13-52490735-INS-5517
chr13-52492222-INS-498
chr13-52495908-INS-52
chr13-52545473-INS-192

chr13-52545697-INS-322
chr13-52551316-INS-63
chr13-52556971-INS-4669
chr13-52561046-INS-68
chr13-52606912-INS-125
chr13-52637100-INS-7438
chr13-52641581-INS-87
chr13-52644815-INS-63
chr13-52702789-INS-3558
chr13-52742460-INS-284
chr13-52742460-INS-91
chr13-52752244-INS-126
chr13-52759765-INS-7049
chr13-52782681-INS-54
chr13-52789644-INS-130
chr13-52805378-INS-106
chr13-52828811-INS-220
chr13-52899791-INS-445
chr13-52918813-INS-1581
chr13-52942431-INS-72
chr13-52968287-INS-58
chr13-52978578-INS-78
chr13-52987197-INS-50
chr13-53111348-INS-52
chr13-53114230-INS-52
chr13-53128479-INS-50
chr13-53140654-INS-52
chr13-53150004-INS-108
chr13-53203608-INS-68
chr13-53204173-INS-56
chr13-53283332-INS-181
chr13-53354149-INS-496
chr13-53357394-INS-83
chr13-53394009-INS-85
chr13-53406145-INS-267
chr13-53409988-INS-231
chr13-53429382-INS-92
chr13-53438581-INS-211
chr13-53457310-INS-2452
chr13-53515790-INS-118
chr13-53515790-INS-826
chr13-53605974-INS-66
chr13-53620645-INS-70
chr13-53656300-INS-52
chr13-53720408-INS-3520
chr13-53754002-INS-361
chr13-53777511-INS-115
chr13-53777609-INS-330
chr13-53793046-INS-56
chr13-53816498-INS-52
chr13-53846593-INS-6390
chr13-53858242-INS-3100
chr13-53859307-INS-294
chr13-53864641-INS-1889
chr13-53887501-INS-55
chr13-53896639-INS-456
chr13-53896693-INS-654
chr13-53960830-INS-2599
chr13-53988088-INS-5605
chr13-53989854-INS-206
chr13-53998287-INS-5233
chr13-54025773-INS-127
chr13-54030962-INS-60
chr13-54084521-INS-323
chr13-54088530-INS-191
chr13-54091838-INS-70
chr13-54146030-INS-72
chr13-54148655-INS-259
chr13-54173725-INS-72
chr13-54174186-INS-421
chr13-54244375-INS-55
chr13-54283385-INS-64
chr13-54308140-INS-301
chr13-54316746-INS-89
chr13-54327369-INS-180
chr13-54327369-INS-575
chr13-54348804-INS-66

chr13-54434054-INS-72
chr13-54450663-INS-50
chr13-54488870-INS-53
chr13-54497064-INS-51
chr13-54580257-INS-61
chr13-54599566-INS-50
chr13-54599795-INS-5369
chr13-54636425-INS-60
chr13-54636481-INS-260
chr13-54637112-INS-75
chr13-54653806-INS-231
chr13-54656303-INS-213
chr13-54656455-INS-202
chr13-54684052-INS-1034
chr13-54686515-INS-222
chr13-54697738-INS-497
chr13-54721072-INS-686
chr13-54802378-INS-4352
chr13-54832528-INS-176
chr13-54845109-INS-64
chr13-54854488-INS-201
chr13-54928179-INS-7691
chr13-54958683-INS-279
chr13-54967164-INS-52
chr13-54978776-INS-238
chr13-54979539-INS-50
chr13-55172631-INS-922
chr13-55202209-INS-170
chr13-55202864-INS-81
chr13-55203023-INS-4139
chr13-55215766-INS-344
chr13-55218330-INS-230
chr13-55225109-INS-208
chr13-55234963-INS-68
chr13-55237638-INS-786
chr13-55245977-INS-87
chr13-55286942-INS-5675
chr13-55287213-INS-50
chr13-55295170-INS-121
chr13-55298441-INS-54
chr13-55329430-INS-167
chr13-55329975-INS-8712
chr13-55365450-INS-180
chr13-55367741-INS-206
chr13-55368341-INS-64
chr13-55398068-INS-50
chr13-55429442-INS-196
chr13-55432678-INS-217
chr13-55450023-INS-223
chr13-55500736-INS-65
chr13-55514069-INS-176
chr13-55525026-INS-56
chr13-55543534-INS-111
chr13-55572687-INS-342
chr13-55586763-INS-54
chr13-55589528-INS-107
chr13-55626459-INS-179
chr13-55636900-INS-60
chr13-55638563-INS-68
chr13-55651892-INS-203
chr13-55652678-INS-67
chr13-55678722-INS-108
chr13-55678730-INS-756
chr13-55739160-INS-110
chr13-55769364-INS-193
chr13-55792320-INS-53
chr13-55808749-INS-56
chr13-55839437-INS-254
chr13-55844334-INS-50
chr13-55845076-INS-201
chr13-55870264-INS-159
chr13-55878477-INS-59
chr13-55942151-INS-296
chr13-55967280-INS-108
chr13-55967612-INS-136
chr13-56009568-INS-52
chr13-56086749-INS-54

chr13-56190384-INS-530
chr13-56247216-INS-648
chr13-56314678-INS-741
chr13-56382907-INS-122
chr13-56387273-INS-448
chr13-56415284-INS-53
chr13-56457335-INS-302
chr13-56526846-INS-385
chr13-56526879-INS-148
chr13-56557681-INS-224
chr13-56562783-INS-6546
chr13-56624396-INS-162
chr13-56662374-INS-50
chr13-56665040-INS-91
chr13-56666024-INS-318
chr13-56677709-INS-128
chr13-56678490-INS-77
chr13-56714793-INS-290
chr13-56725113-INS-54
chr13-56732781-INS-57
chr13-56738117-INS-54
chr13-56821107-INS-58
chr13-56823307-INS-1534
chr13-56838811-INS-86
chr13-56857467-INS-244
chr13-56892802-INS-94
chr13-56892966-INS-250
chr13-56893425-INS-50
chr13-56915923-INS-57
chr13-56919276-INS-66
chr13-56948506-INS-97
chr13-56989142-INS-199
chr13-57024339-INS-54
chr13-57081277-INS-416
chr13-57096120-INS-264
chr13-57096140-INS-103
chr13-57206630-INS-94
chr13-57264354-INS-124
chr13-57265144-INS-58
chr13-57277384-INS-6969
chr13-57316046-INS-260
chr13-57331747-INS-154
chr13-57341056-INS-2549
chr13-57375612-INS-167
chr13-57376196-INS-396
chr13-57376208-INS-2385
chr13-57381351-INS-54
chr13-57381799-INS-58
chr13-57381899-INS-7156
chr13-57478854-INS-54
chr13-57497821-INS-53
chr13-57540278-INS-50
chr13-57615796-INS-68
chr13-57635142-INS-88
chr13-57663209-INS-171
chr13-57696471-INS-66
chr13-57707900-INS-95
chr13-57723606-INS-5732
chr13-57801123-INS-731
chr13-57806459-INS-1624
chr13-57806637-INS-266
chr13-57825968-INS-56
chr13-57830962-INS-119
chr13-57859856-INS-2498
chr13-57881623-INS-51
chr13-57883556-INS-164
chr13-57890116-INS-64
chr13-57895084-INS-164
chr13-57895184-INS-56
chr13-57911767-INS-70
chr13-57928557-INS-60
chr13-57995617-INS-3105
chr13-58001416-INS-133
chr13-58007617-INS-2339
chr13-58007680-INS-111
chr13-58044442-INS-76
chr13-58058069-INS-138

chr13-58058069-INS-672
chr13-58084687-INS-51
chr13-58101979-INS-70
chr13-58102165-INS-361
chr13-58120255-INS-1510
chr13-58125974-INS-108
chr13-58134608-INS-51
chr13-58142912-INS-64
chr13-58145604-INS-54
chr13-58146399-INS-114
chr13-58146498-INS-117
chr13-58149975-INS-25563
chr13-58155171-INS-82
chr13-58191617-INS-52
chr13-58195655-INS-211
chr13-58229461-INS-208
chr13-58234302-INS-86
chr13-58244769-INS-97
chr13-58308768-INS-176
chr13-58370461-INS-507
chr13-58409901-INS-410
chr13-58416814-INS-199
chr13-58425268-INS-138
chr13-58432780-INS-80
chr13-58504024-INS-53
chr13-58508666-INS-321
chr13-58566523-INS-88
chr13-58567612-INS-64
chr13-58584095-INS-65
chr13-58640503-INS-114
chr13-58724880-INS-123
chr13-58725774-INS-166
chr13-58783562-INS-60
chr13-58788249-INS-78
chr13-58808537-INS-179
chr13-58808560-INS-89
chr13-58825836-INS-190
chr13-58857807-INS-89
chr13-58857819-INS-182
chr13-58874429-INS-198
chr13-58888168-INS-72
chr13-58888283-INS-378
chr13-58888288-INS-72
chr13-58892760-INS-102
chr13-58912793-INS-248
chr13-58952084-INS-67
chr13-58961041-INS-50
chr13-58974224-INS-54
chr13-58994140-INS-60
chr13-59033779-INS-174
chr13-59035557-INS-142
chr13-59080504-INS-111
chr13-59100850-INS-112
chr13-59121174-INS-55
chr13-59131500-INS-102
chr13-59161420-INS-57
chr13-59174464-INS-3021
chr13-59192235-INS-50
chr13-59198171-INS-102
chr13-59202869-INS-72
chr13-59213506-INS-50
chr13-59285977-INS-53
chr13-59374299-INS-8974
chr13-59387945-INS-133
chr13-59400672-INS-100
chr13-59410573-INS-6490
chr13-59482242-INS-259
chr13-59482242-INS-55
chr13-59490523-INS-61
chr13-59552798-INS-498
chr13-59569570-INS-50
chr13-59575510-INS-341
chr13-59611404-INS-208
chr13-59611407-INS-564
chr13-59625733-INS-786
chr13-59644560-INS-7582
chr13-59670564-INS-163

chr13-59673989-INS-294
chr13-59682894-INS-102
chr13-59686540-INS-58
chr13-59698083-INS-208
chr13-59708107-INS-3414
chr13-59715682-INS-62
chr13-59716492-INS-80
chr13-59729029-INS-788
chr13-59744349-INS-54
chr13-59752788-INS-387
chr13-59767710-INS-258
chr13-59804846-INS-486
chr13-59817278-INS-50
chr13-59821997-INS-60
chr13-59853094-INS-441
chr13-59854987-INS-1318
chr13-59932284-INS-108
chr13-59934768-INS-344
chr13-60025481-INS-54
chr13-60026126-INS-132
chr13-60046740-INS-238
chr13-60046847-INS-62
chr13-60056966-INS-14739
chr13-60061866-INS-51
chr13-60080957-INS-60
chr13-60131188-INS-75
chr13-60151478-INS-1046
chr13-60157644-INS-60
chr13-60190169-INS-56
chr13-60210817-INS-477
chr13-60221064-INS-4374
chr13-60227651-INS-3096
chr13-60238065-INS-68
chr13-60238258-INS-187
chr13-60240196-INS-3509
chr13-60244685-INS-694
chr13-60244797-INS-69
chr13-60245134-INS-243
chr13-60308079-INS-284
chr13-60318664-INS-52
chr13-60327262-INS-120
chr13-60337242-INS-50
chr13-60338298-INS-83
chr13-60338383-INS-204
chr13-60342131-INS-1888
chr13-60387902-INS-70
chr13-60405648-INS-51
chr13-60419402-INS-50
chr13-60435263-INS-216
chr13-60446167-INS-508
chr13-60494556-INS-456
chr13-60508240-INS-92
chr13-60523977-INS-55
chr13-60560040-INS-55
chr13-60590166-INS-1012
chr13-60590607-INS-7277
chr13-60609806-INS-2986
chr13-60617382-INS-4676
chr13-60664199-INS-81
chr13-60725920-INS-328
chr13-60732461-INS-240
chr13-60740053-INS-237
chr13-60747916-INS-5082
chr13-60747928-INS-67
chr13-60748373-INS-210
chr13-60788968-INS-200
chr13-60801106-INS-6579
chr13-60803501-INS-172
chr13-60824274-INS-406
chr13-60871828-INS-58
chr13-60903584-INS-74
chr13-60928115-INS-96
chr13-60929315-INS-498
chr13-60954150-INS-123
chr13-60970679-INS-175
chr13-60976300-INS-3595
chr13-60985794-INS-124

chr13-60985911-INS-54
chr13-61015204-INS-61
chr13-61086827-INS-199
chr13-61092344-INS-54
chr13-61109329-INS-80
chr13-61178534-INS-3742
chr13-61225285-INS-5575
chr13-61228654-INS-11155
chr13-61271381-INS-2932
chr13-61292847-INS-1150
chr13-61304813-INS-66
chr13-61311706-INS-180
chr13-61314241-INS-1482
chr13-61321027-INS-50
chr13-61326201-INS-526
chr13-61386811-INS-3229
chr13-61398026-INS-3304
chr13-61405387-INS-400
chr13-61414340-INS-79
chr13-61417892-INS-1344
chr13-61419588-INS-1564
chr13-61420154-INS-428
chr13-61454968-INS-6224
chr13-61460134-INS-3673
chr13-61476856-INS-1878
chr13-61527547-INS-199
chr13-61527945-INS-88
chr13-61590744-INS-6609
chr13-61641302-INS-6558
chr13-61681075-INS-142
chr13-61696032-INS-1473
chr13-61730487-INS-8259
chr13-61773795-INS-1888
chr13-61790653-INS-114
chr13-61805130-INS-84
chr13-61805476-INS-174
chr13-61809375-INS-401
chr13-61810273-INS-331
chr13-61811825-INS-169
chr13-61889464-INS-217
chr13-61895489-INS-66
chr13-61902388-INS-209
chr13-61907977-INS-80
chr13-61908147-INS-191
chr13-61908157-INS-60
chr13-61909024-INS-123
chr13-61910218-INS-169
chr13-61911170-INS-162
chr13-61926458-INS-174
chr13-61962594-INS-252
chr13-61989726-INS-166
chr13-61992959-INS-195
chr13-62244021-INS-210
chr13-62246056-INS-202
chr13-62253556-INS-210
chr13-62254698-INS-180
chr13-62272088-INS-1496
chr13-62324851-INS-524
chr13-62325645-INS-4013
chr13-62332004-INS-198
chr13-62372163-INS-1681
chr13-62412924-INS-213
chr13-62534685-INS-196
chr13-62582891-INS-50
chr13-62651110-INS-50
chr13-62672719-INS-3750
chr13-62675060-INS-500
chr13-62714283-INS-178
chr13-62726386-INS-50
chr13-62728076-INS-4813
chr13-62737102-INS-1403
chr13-62785514-INS-54
chr13-62831532-INS-229
chr13-62840961-INS-408
chr13-62841438-INS-439
chr13-62858852-INS-70
chr13-62894645-INS-221

chr13-62947923-INS-190
chr13-62960569-INS-4365
chr13-62971959-INS-2515
chr13-62979340-INS-87
chr13-63101775-INS-1738
chr13-63109364-INS-200
chr13-63109985-INS-216
chr13-63199421-INS-6403
chr13-63201280-INS-200
chr13-63203310-INS-118
chr13-63311837-INS-164
chr13-63312408-INS-104
chr13-63327654-INS-3200
chr13-63327654-INS-56
chr13-63339565-INS-58
chr13-63348845-INS-509
chr13-63348949-INS-366
chr13-63382345-INS-50
chr13-63399807-INS-210
chr13-63412057-INS-68
chr13-63457946-INS-50
chr13-63487112-INS-160
chr13-63495122-INS-52
chr13-63512077-INS-288
chr13-63512544-INS-162
chr13-63518977-INS-2810
chr13-63527415-INS-266
chr13-63553289-INS-50
chr13-63553403-INS-7134
chr13-63570250-INS-88
chr13-63581182-INS-60
chr13-63654292-INS-59
chr13-63726824-INS-210
chr13-63748380-INS-152
chr13-63748606-INS-853
chr13-63770115-INS-372
chr13-63834840-INS-239
chr13-63834840-INS-65
chr13-63846905-INS-1001
chr13-63869457-INS-125
chr13-63887253-INS-405
chr13-63897515-INS-615
chr13-63902509-INS-149
chr13-63936731-INS-6431
chr13-63955922-INS-6580
chr13-63960802-INS-7203
chr13-64011297-INS-1948
chr13-64012285-INS-210
chr13-64120271-INS-52
chr13-64143903-INS-1767
chr13-64144357-INS-99
chr13-64145248-INS-536
chr13-64184301-INS-311
chr13-64184310-INS-142
chr13-64199350-INS-213
chr13-64203978-INS-6360
chr13-64240321-INS-1990
chr13-64242128-INS-162
chr13-64242128-INS-73
chr13-64248973-INS-54
chr13-64256850-INS-6440
chr13-64315101-INS-88
chr13-64322214-INS-105
chr13-64323957-INS-52
chr13-64329353-INS-203
chr13-64332561-INS-56
chr13-64338705-INS-208
chr13-64339155-INS-830
chr13-64339271-INS-370
chr13-64364616-INS-215
chr13-64476256-INS-139
chr13-64503690-INS-5318
chr13-64534653-INS-138
chr13-64535297-INS-171
chr13-64554617-INS-157
chr13-64556099-INS-171
chr13-64557976-INS-2928

chr13-64574676-INS-1886
chr13-64575975-INS-2993
chr13-64581476-INS-57
chr13-64594504-INS-5154
chr13-64605508-INS-1983
chr13-64610184-INS-59
chr13-64624901-INS-205
chr13-64628501-INS-1253
chr13-64641547-INS-221
chr13-64643788-INS-126
chr13-64643834-INS-822
chr13-64648358-INS-285
chr13-64655949-INS-205
chr13-64678280-INS-2346
chr13-64683830-INS-1107
chr13-64701600-INS-85
chr13-64706972-INS-67
chr13-64707264-INS-649
chr13-64710899-INS-60
chr13-64711774-INS-2986
chr13-64748343-INS-114
chr13-64833973-INS-7129
chr13-64841693-INS-567
chr13-64852265-INS-80
chr13-64853936-INS-54
chr13-64857357-INS-10712
chr13-64859882-INS-4235
chr13-64882541-INS-70
chr13-64888086-INS-6441
chr13-64906314-INS-1237
chr13-64948061-INS-338
chr13-64956175-INS-202
chr13-64961053-INS-7436
chr13-64991698-INS-52
chr13-65089792-INS-132
chr13-65089792-INS-347
chr13-65099959-INS-7414
chr13-65114531-INS-2696
chr13-65170271-INS-1592
chr13-65172079-INS-1954
chr13-65173236-INS-200
chr13-65209899-INS-54
chr13-65224426-INS-84
chr13-65224433-INS-456
chr13-65294484-INS-6273
chr13-65314450-INS-58
chr13-65357498-INS-52
chr13-65360973-INS-168
chr13-65374513-INS-210
chr13-65381767-INS-2684
chr13-65396629-INS-215
chr13-65510739-INS-2002
chr13-65522757-INS-77
chr13-65647875-INS-1860
chr13-65660298-INS-252
chr13-65670812-INS-978
chr13-65706135-INS-716
chr13-65838341-INS-84
chr13-65839134-INS-84
chr13-65839343-INS-381
chr13-65893331-INS-93
chr13-65897684-INS-104
chr13-65899607-INS-826
chr13-66020000-INS-220
chr13-66024941-INS-418
chr13-66056138-INS-83
chr13-66060406-INS-654
chr13-66082184-INS-67
chr13-66269718-INS-520
chr13-66384649-INS-52
chr13-66392330-INS-468
chr13-66392410-INS-351
chr13-66414601-INS-93
chr13-66471565-INS-197
chr13-66477810-INS-200
chr13-66579716-INS-252
chr13-66589995-INS-58

chr13-66690618-INS-307
chr13-66692003-INS-69
chr13-66746179-INS-104
chr13-66769182-INS-170
chr13-66770942-INS-213
chr13-66818173-INS-336
chr13-66824911-INS-260
chr13-66952895-INS-166
chr13-67010709-INS-231
chr13-67060916-INS-465
chr13-67122286-INS-60
chr13-67174082-INS-64
chr13-67177437-INS-6679
chr13-67212256-INS-6415
chr13-67213475-INS-3065
chr13-67222055-INS-4865
chr13-67324027-INS-78
chr13-67324103-INS-351
chr13-67346315-INS-234
chr13-67361672-INS-161
chr13-67444341-INS-244
chr13-67456533-INS-261
chr13-67476319-INS-158
chr13-67502655-INS-732
chr13-67553108-INS-208
chr13-67563432-INS-319
chr13-67572075-INS-208
chr13-67573811-INS-168
chr13-67613127-INS-2582
chr13-67620145-INS-6441
chr13-67625635-INS-3612
chr13-67737283-INS-6512
chr13-67851433-INS-201
chr13-67863853-INS-204
chr13-67880957-INS-211
chr13-67947963-INS-488
chr13-67970602-INS-54
chr13-68004974-INS-111
chr13-68054176-INS-160
chr13-68136982-INS-381
chr13-68147915-INS-135
chr13-68181724-INS-152
chr13-68204204-INS-510
chr13-68309850-INS-510
chr13-68337778-INS-78
chr13-68342326-INS-152
chr13-68463899-INS-118
chr13-68478086-INS-3537
chr13-68516678-INS-255
chr13-68518445-INS-7174
chr13-68524360-INS-55
chr13-68527442-INS-50
chr13-68529772-INS-64
chr13-68538821-INS-195
chr13-68565324-INS-498
chr13-68585398-INS-51
chr13-68598869-INS-192
chr13-68620366-INS-206
chr13-68663218-INS-214
chr13-68694276-INS-316
chr13-68699133-INS-302
chr13-68858569-INS-6227
chr13-68866816-INS-67
chr13-68884202-INS-6251
chr13-68895965-INS-6523
chr13-68953957-INS-167
chr13-68953976-INS-69
chr13-68960838-INS-63
chr13-68969758-INS-1020
chr13-68993660-INS-6832
chr13-69001944-INS-195
chr13-69006297-INS-134
chr13-69013427-INS-102
chr13-69064459-INS-54
chr13-69180968-INS-182
chr13-69183849-INS-50
chr13-69241994-INS-99

chr13-69266058-INS-69
chr13-69283596-INS-6401
chr13-69330082-INS-1052
chr13-69339262-INS-123
chr13-69340072-INS-50
chr13-69344541-INS-104
chr13-69350417-INS-6371
chr13-69351581-INS-53
chr13-69392313-INS-478
chr13-69456206-INS-58
chr13-69467719-INS-258
chr13-69485046-INS-410
chr13-69485300-INS-113
chr13-69494715-INS-5555
chr13-69499980-INS-7026
chr13-69517752-INS-208
chr13-69518606-INS-55
chr13-69574899-INS-666
chr13-69585517-INS-74
chr13-69594568-INS-161
chr13-69613768-INS-57
chr13-69620987-INS-6592
chr13-69628781-INS-1009
chr13-69656619-INS-64
chr13-69659300-INS-104
chr13-69664289-INS-337
chr13-69705414-INS-93
chr13-69708918-INS-126
chr13-69717969-INS-1174
chr13-69723448-INS-80
chr13-69742909-INS-76
chr13-69825591-INS-184
chr13-69875277-INS-198
chr13-69909050-INS-159
chr13-69920887-INS-512
chr13-69921079-INS-126
chr13-69921148-INS-59
chr13-69921226-INS-326
chr13-70030375-INS-402
chr13-70151559-INS-50
chr13-70178731-INS-1864
chr13-70208535-INS-1844
chr13-70308468-INS-56
chr13-70335051-INS-143
chr13-70357207-INS-6855
chr13-70357208-INS-84
chr13-70357369-INS-140
chr13-70368447-INS-66
chr13-70394835-INS-209
chr13-70425876-INS-368
chr13-70436395-INS-51
chr13-70449360-INS-71
chr13-70450019-INS-60
chr13-70477090-INS-7112
chr13-70491483-INS-569
chr13-70557323-INS-66
chr13-70569063-INS-1684
chr13-70605377-INS-52
chr13-70622884-INS-8847
chr13-70627440-INS-374
chr13-70689654-INS-114
chr13-70771709-INS-781
chr13-70820958-INS-161
chr13-70860944-INS-80
chr13-70897742-INS-6920
chr13-70934317-INS-72
chr13-70957220-INS-1365
chr13-71001719-INS-112
chr13-71004172-INS-51
chr13-71063909-INS-1138
chr13-71068516-INS-58
chr13-71090547-INS-148
chr13-71123763-INS-6469
chr13-71168363-INS-183
chr13-71180548-INS-52
chr13-71302223-INS-278
chr13-71310616-INS-50

chr13-71331188-INS-151
chr13-71347354-INS-467
chr13-71363288-INS-62
chr13-71602849-INS-92
chr13-71619924-INS-166
chr13-71620079-INS-513
chr13-71741881-INS-212
chr13-71747901-INS-58
chr13-71827935-INS-6389
chr13-71866172-INS-67
chr13-71890415-INS-1176
chr13-71903502-INS-56
chr13-71982134-INS-52
chr13-72051951-INS-3620
chr13-72057091-INS-8486
chr13-72072570-INS-6426
chr13-72077574-INS-377
chr13-72092639-INS-214
chr13-72130055-INS-2604
chr13-72133920-INS-50
chr13-72146114-INS-62
chr13-72146370-INS-14636
chr13-72151117-INS-160
chr13-72160712-INS-139
chr13-72320431-INS-50
chr13-72407258-INS-93
chr13-72459867-INS-396
chr13-72460099-INS-78
chr13-72487582-INS-161
chr13-72499543-INS-299
chr13-72596235-INS-86
chr13-72619700-INS-280
chr13-72642465-INS-2861
chr13-72653740-INS-6238
chr13-72704348-INS-7944
chr13-72714275-INS-50
chr13-72726886-INS-90
chr13-72727689-INS-183
chr13-72756649-INS-60
chr13-72760675-INS-194
chr13-72760906-INS-86
chr13-72799275-INS-50
chr13-72846228-INS-56
chr13-72867599-INS-66
chr13-72867651-INS-150
chr13-72870174-INS-55
chr13-72891622-INS-6266
chr13-72908292-INS-74
chr13-72942761-INS-3024
chr13-72952144-INS-64
chr13-72952207-INS-148
chr13-72957354-INS-474
chr13-72969321-INS-145
chr13-73003503-INS-86
chr13-73074659-INS-88
chr13-73081078-INS-50
chr13-73089103-INS-56
chr13-73115057-INS-1889
chr13-73119845-INS-63
chr13-73121574-INS-554
chr13-73123474-INS-116
chr13-73148647-INS-2045
chr13-73166673-INS-7203
chr13-73172418-INS-2432
chr13-73191490-INS-6309
chr13-73196103-INS-73
chr13-73209687-INS-6300
chr13-73268952-INS-94
chr13-73288041-INS-72
chr13-73291096-INS-3066
chr13-73313885-INS-55
chr13-73313914-INS-165
chr13-73320609-INS-110
chr13-73344777-INS-479
chr13-73346795-INS-323
chr13-73365449-INS-497
chr13-73414234-INS-215

chr13-73462971-INS-356
chr13-73463195-INS-69
chr13-73491625-INS-68
chr13-73492933-INS-1169
chr13-73507730-INS-321
chr13-73513193-INS-72
chr13-73560833-INS-584
chr13-73569362-INS-50
chr13-73578827-INS-56
chr13-73627984-INS-103
chr13-73652108-INS-207
chr13-73660390-INS-219
chr13-73661306-INS-537
chr13-73678065-INS-106
chr13-73678076-INS-52
chr13-73678826-INS-60
chr13-73798401-INS-104
chr13-73807851-INS-112
chr13-73807881-INS-50
chr13-73846064-INS-56
chr13-73846568-INS-63
chr13-73853052-INS-145
chr13-73862008-INS-57
chr13-73909292-INS-50
chr13-73982388-INS-58
chr13-73989587-INS-342
chr13-74165833-INS-245
chr13-74166614-INS-190
chr13-74187913-INS-756
chr13-74217069-INS-1246
chr13-74277510-INS-60
chr13-74371840-INS-65
chr13-74372354-INS-72
chr13-74403058-INS-52
chr13-74415137-INS-62
chr13-74434923-INS-56
chr13-74446964-INS-178
chr13-74524280-INS-1892
chr13-74528080-INS-79
chr13-74535042-INS-50
chr13-74542782-INS-1841
chr13-74544682-INS-1092
chr13-74628391-INS-1469
chr13-74630774-INS-496
chr13-74665354-INS-877
chr13-74677404-INS-497
chr13-74698394-INS-198
chr13-74741949-INS-50
chr13-74777903-INS-7088
chr13-74809826-INS-60
chr13-74910659-INS-211
chr13-74914806-INS-57
chr13-74915285-INS-116
chr13-74971801-INS-57
chr13-74993268-INS-160
chr13-75064080-INS-473
chr13-75099987-INS-6380
chr13-75106067-INS-80
chr13-75127087-INS-206
chr13-75158193-INS-94
chr13-75205298-INS-1108
chr13-75350653-INS-2067
chr13-75364099-INS-148
chr13-75371395-INS-1306
chr13-75390352-INS-52
chr13-75400381-INS-215
chr13-75407387-INS-100
chr13-75411121-INS-54
chr13-75414336-INS-6418
chr13-75418444-INS-50
chr13-75436527-INS-2326
chr13-75490432-INS-3619
chr13-75490542-INS-8674
chr13-75499823-INS-1101
chr13-75503409-INS-1384
chr13-75528927-INS-398
chr13-75531140-INS-356

chr13-75537310-INS-123
chr13-75537749-INS-622
chr13-75563465-INS-56
chr13-75620283-INS-68
chr13-75620489-INS-7532
chr13-75654125-INS-98
chr13-75667958-INS-252
chr13-75668024-INS-68
chr13-75700924-INS-68
chr13-75701513-INS-6917
chr13-75707420-INS-106
chr13-75708036-INS-6352
chr13-75735948-INS-2910
chr13-75850744-INS-114
chr13-75852130-INS-82
chr13-75953466-INS-94
chr13-75953521-INS-522
chr13-75971124-INS-5157
chr13-75992148-INS-426
chr13-76053875-INS-232
chr13-76071214-INS-2973
chr13-76099190-INS-5262
chr13-76142067-INS-507
chr13-76165170-INS-22015
chr13-76172460-INS-287
chr13-76210109-INS-1889
chr13-76215788-INS-55
chr13-76229367-INS-7475
chr13-76261250-INS-6501
chr13-76265319-INS-54
chr13-76295508-INS-6374
chr13-76302854-INS-162
chr13-76309578-INS-7102
chr13-76358812-INS-64
chr13-76360444-INS-90
chr13-76374556-INS-1111
chr13-76414511-INS-6383
chr13-76422021-INS-2953
chr13-76453198-INS-1039
chr13-76458207-INS-50
chr13-76523167-INS-54
chr13-76544821-INS-1229
chr13-76572829-INS-2712
chr13-76591605-INS-161
chr13-76593813-INS-50
chr13-76657206-INS-1458
chr13-76661868-INS-1825
chr13-76674605-INS-1390
chr13-76685060-INS-5087
chr13-76724505-INS-1237
chr13-76725725-INS-334
chr13-76738333-INS-160
chr13-76739196-INS-526
chr13-76920826-INS-480
chr13-76943204-INS-2757
chr13-76945712-INS-1040
chr13-76980102-INS-302
chr13-77084033-INS-225
chr13-77088588-INS-594
chr13-77088769-INS-162
chr13-77168649-INS-78
chr13-77257864-INS-322
chr13-77484228-INS-4119
chr13-77573135-INS-1030
chr13-77631095-INS-1421
chr13-77662560-INS-6610
chr13-77675401-INS-252
chr13-77675997-INS-595
chr13-77710488-INS-56
chr13-77868690-INS-155
chr13-77922762-INS-89
chr13-77996499-INS-1845
chr13-78014729-INS-87
chr13-78014744-INS-231
chr13-78093453-INS-1129
chr13-78111811-INS-61
chr13-78113817-INS-139

chr13-78286820-INS-204
chr13-78348373-INS-50
chr13-78373453-INS-2921
chr13-78407510-INS-62
chr13-78417686-INS-6917
chr13-78432618-INS-4315
chr13-78480198-INS-56
chr13-78545126-INS-51
chr13-78558122-INS-50
chr13-78567878-INS-7171
chr13-78578125-INS-198
chr13-78653632-INS-7914
chr13-78657125-INS-68
chr13-78687045-INS-6795
chr13-78716559-INS-82
chr13-78761311-INS-50
chr13-78871826-INS-205
chr13-78974705-INS-627
chr13-78979804-INS-7077
chr13-78988518-INS-6480
chr13-79126493-INS-163
chr13-79126743-INS-187
chr13-79130886-INS-1015
chr13-79161021-INS-2771
chr13-79171950-INS-169
chr13-79197844-INS-50
chr13-79198058-INS-135
chr13-79209375-INS-399
chr13-79228870-INS-52
chr13-79292370-INS-414
chr13-79396871-INS-72
chr13-79515242-INS-70
chr13-79554985-INS-669
chr13-79587913-INS-326
chr13-79606345-INS-409
chr13-79609979-INS-51
chr13-79683923-INS-207
chr13-79692001-INS-877
chr13-79710764-INS-1190
chr13-79920119-INS-122
chr13-79920215-INS-58
chr13-79924519-INS-66
chr13-79935503-INS-187
chr13-79996810-INS-8136
chr13-80020008-INS-59
chr13-80120602-INS-58
chr13-80155265-INS-55
chr13-80159683-INS-132
chr13-80197702-INS-64
chr13-80223924-INS-343
chr13-80241017-INS-54
chr13-80248472-INS-6875
chr13-80281287-INS-775
chr13-80283817-INS-1046
chr13-80286475-INS-120
chr13-80289840-INS-204
chr13-80291353-INS-64
chr13-80292917-INS-284
chr13-80337417-INS-191
chr13-80349373-INS-2507
chr13-80373573-INS-7726
chr13-80404821-INS-6539
chr13-80542172-INS-3941
chr13-80563100-INS-148
chr13-80598259-INS-66
chr13-80605781-INS-6074
chr13-80660231-INS-6365
chr13-80696027-INS-2477
chr13-80712331-INS-407
chr13-80744328-INS-4644
chr13-80750997-INS-396
chr13-80756261-INS-52
chr13-80808006-INS-4804
chr13-80816389-INS-56
chr13-80927311-INS-55
chr13-80927528-INS-86
chr13-80934853-INS-6600

chr13-81081328-INS-80
chr13-81113363-INS-1888
chr13-81120441-INS-5544
chr13-81136528-INS-66
chr13-81303590-INS-660
chr13-81377541-INS-6493
chr13-81382171-INS-60
chr13-81420499-INS-3177
chr13-81420812-INS-133
chr13-81434894-INS-1294
chr13-81442809-INS-7041
chr13-81453229-INS-64
chr13-81519700-INS-2135
chr13-81561884-INS-52
chr13-81561923-INS-259
chr13-81561938-INS-5695
chr13-81608723-INS-52
chr13-81617508-INS-1169
chr13-81618148-INS-1144
chr13-81621776-INS-539
chr13-81663682-INS-82
chr13-81664475-INS-71
chr13-81675078-INS-6717
chr13-81679087-INS-54
chr13-81679252-INS-126
chr13-81679411-INS-1122
chr13-81679532-INS-1151
chr13-81696187-INS-283
chr13-81700295-INS-4210
chr13-81714687-INS-67
chr13-81720937-INS-144
chr13-81765994-INS-206
chr13-81766155-INS-1618
chr13-81768562-INS-226
chr13-81783777-INS-1373
chr13-81783797-INS-3682
chr13-81823257-INS-81
chr13-81865363-INS-100
chr13-81906093-INS-233
chr13-81913150-INS-252
chr13-81957043-INS-147
chr13-81958929-INS-1200
chr13-81962165-INS-136
chr13-81964324-INS-153
chr13-81985908-INS-1062
chr13-81993704-INS-401
chr13-82005901-INS-1027
chr13-82014111-INS-50
chr13-82028676-INS-116
chr13-82028771-INS-180
chr13-82074636-INS-6196
chr13-82080040-INS-147
chr13-82091481-INS-56
chr13-82096434-INS-75
chr13-82106613-INS-177
chr13-82111494-INS-6317
chr13-82111945-INS-210
chr13-82125839-INS-120
chr13-82131083-INS-293
chr13-82194821-INS-1023
chr13-82208745-INS-230
chr13-82263728-INS-63
chr13-82279600-INS-6230
chr13-82304130-INS-119
chr13-82304245-INS-90
chr13-82305302-INS-60
chr13-82306514-INS-143
chr13-82308427-INS-54
chr13-82333892-INS-120
chr13-82363041-INS-146
chr13-82375495-INS-540
chr13-82432444-INS-53
chr13-82444381-INS-51
chr13-82508183-INS-1866
chr13-82540084-INS-2048
chr13-82569700-INS-1111
chr13-82598031-INS-5443

chr13-82689175-INS-128
chr13-82713953-INS-109
chr13-82721274-INS-142
chr13-82727495-INS-54
chr13-82735245-INS-1608
chr13-82781571-INS-458
chr13-82830733-INS-78
chr13-82868460-INS-54
chr13-82874780-INS-2561
chr13-82987032-INS-76
chr13-83007000-INS-120
chr13-83017015-INS-6497
chr13-83018775-INS-56
chr13-83044905-INS-52
chr13-83069734-INS-256
chr13-83136457-INS-3142
chr13-83141300-INS-68
chr13-83171410-INS-55
chr13-83174777-INS-6570
chr13-83256109-INS-78
chr13-83323297-INS-53
chr13-83367783-INS-6838
chr13-83397929-INS-102
chr13-83419098-INS-68
chr13-83487059-INS-78
chr13-83529468-INS-396
chr13-83532285-INS-72
chr13-83534123-INS-228
chr13-83541402-INS-68
chr13-83541810-INS-64
chr13-83583647-INS-54
chr13-83583907-INS-157
chr13-83583907-INS-57
chr13-83612256-INS-408
chr13-83663722-INS-76
chr13-83699505-INS-50
chr13-83701560-INS-770
chr13-83707508-INS-62
chr13-83709396-INS-325
chr13-83713561-INS-50
chr13-83739096-INS-858
chr13-83924506-INS-2774
chr13-83941084-INS-1521
chr13-83941764-INS-389
chr13-83942034-INS-54
chr13-83948924-INS-96
chr13-84009860-INS-191
chr13-84068552-INS-6170
chr13-84068893-INS-50
chr13-84115647-INS-1150
chr13-84181482-INS-6046
chr13-84190222-INS-150
chr13-84276599-INS-211
chr13-84282866-INS-72
chr13-84304747-INS-4961
chr13-84339409-INS-6268
chr13-84342415-INS-50
chr13-84345577-INS-52
chr13-84482323-INS-1063
chr13-84510938-INS-5029
chr13-84548306-INS-61
chr13-84560559-INS-55
chr13-84576340-INS-285
chr13-84615862-INS-299
chr13-84648554-INS-387
chr13-84696225-INS-2694
chr13-84796923-INS-54
chr13-84796981-INS-64
chr13-84797303-INS-3739
chr13-84802019-INS-6090
chr13-84802459-INS-80
chr13-84815965-INS-144
chr13-84830603-INS-76
chr13-84865320-INS-4362
chr13-84882368-INS-6580
chr13-84894011-INS-76
chr13-84903701-INS-62

chr13-84907901-INS-108
chr13-84935053-INS-1299
chr13-85051006-INS-124
chr13-85084046-INS-6214
chr13-85116486-INS-187
chr13-85190789-INS-9917
chr13-85261497-INS-96
chr13-85290201-INS-202
chr13-85411832-INS-50
chr13-85455045-INS-1217
chr13-85455270-INS-50
chr13-85464289-INS-519
chr13-85479335-INS-303
chr13-85493932-INS-1568
chr13-85538999-INS-72
chr13-85540381-INS-1492
chr13-85557075-INS-1886
chr13-85561313-INS-534
chr13-85567776-INS-65
chr13-85647699-INS-118
chr13-85661785-INS-60
chr13-85680099-INS-2388
chr13-85746264-INS-276
chr13-85764012-INS-287
chr13-85789608-INS-6393
chr13-85790469-INS-567
chr13-85795283-INS-246
chr13-85796182-INS-65
chr13-85807985-INS-6602
chr13-85820687-INS-208
chr13-85856572-INS-2678
chr13-85862570-INS-8974
chr13-85889026-INS-201
chr13-85892717-INS-50
chr13-85915218-INS-243
chr13-85979410-INS-54
chr13-86013468-INS-87
chr13-86036018-INS-80
chr13-86047583-INS-1155
chr13-86054770-INS-837
chr13-86099631-INS-529
chr13-86162312-INS-232
chr13-86165070-INS-61
chr13-86166264-INS-136
chr13-86206548-INS-52
chr13-86206551-INS-106
chr13-86238916-INS-4780
chr13-86249994-INS-268
chr13-86251755-INS-170
chr13-86283395-INS-328
chr13-86314931-INS-1322
chr13-86371060-INS-6554
chr13-86377106-INS-211
chr13-86392652-INS-664
chr13-86415753-INS-1594
chr13-86530301-INS-72
chr13-86536450-INS-1098
chr13-86536625-INS-203
chr13-86564063-INS-58
chr13-86586273-INS-2932
chr13-86594481-INS-1052
chr13-86607240-INS-6675
chr13-86610901-INS-68
chr13-86612928-INS-78
chr13-86680615-INS-7235
chr13-86715325-INS-3564
chr13-86726479-INS-65
chr13-86746468-INS-60
chr13-86757139-INS-25711
chr13-86757140-INS-6869
chr13-86763149-INS-1636
chr13-86803310-INS-352
chr13-86807732-INS-3010
chr13-86935860-INS-3044
chr13-86944482-INS-6607
chr13-86952456-INS-454
chr13-86956382-INS-143

chr13-86956954-INS-2675
chr13-86961664-INS-90
chr13-86961740-INS-320
chr13-86973856-INS-50
chr13-86974116-INS-54
chr13-86975382-INS-86
chr13-87145424-INS-6591
chr13-87148100-INS-84
chr13-87204178-INS-1293
chr13-87223425-INS-3924
chr13-87239716-INS-56
chr13-87240680-INS-2786
chr13-87265451-INS-70
chr13-87266288-INS-1201
chr13-87272290-INS-6531
chr13-87303856-INS-3230
chr13-87329136-INS-538
chr13-87359110-INS-3764
chr13-87361047-INS-6166
chr13-87365573-INS-2876
chr13-87400569-INS-140
chr13-87401342-INS-70
chr13-87421050-INS-168
chr13-87454104-INS-78
chr13-87467434-INS-50
chr13-87477345-INS-3055
chr13-87484533-INS-1420
chr13-87490078-INS-66
chr13-87554520-INS-188
chr13-87554520-INS-571
chr13-87576481-INS-427
chr13-87576862-INS-366
chr13-87637458-INS-50
chr13-87783241-INS-7667
chr13-87800413-INS-50
chr13-87936614-INS-6446
chr13-87976068-INS-348
chr13-88005408-INS-55
chr13-88010851-INS-3357
chr13-88010851-INS-606
chr13-88081805-INS-53
chr13-88087986-INS-214
chr13-88105807-INS-51
chr13-88148893-INS-55
chr13-88155636-INS-6106
chr13-88180078-INS-703
chr13-88237764-INS-63
chr13-88240579-INS-342
chr13-88296367-INS-50
chr13-88497538-INS-58
chr13-88531124-INS-551
chr13-88597884-INS-51
chr13-88601592-INS-84
chr13-88611219-INS-6797
chr13-88637500-INS-348
chr13-88657261-INS-70
chr13-88696792-INS-150
chr13-88747257-INS-6898
chr13-88892689-INS-1939
chr13-88994674-INS-8437
chr13-89057590-INS-85
chr13-89099970-INS-1588
chr13-89166076-INS-2162
chr13-89239163-INS-6848
chr13-89248095-INS-608
chr13-89370539-INS-640
chr13-89372186-INS-78
chr13-89479975-INS-1323
chr13-89480329-INS-7151
chr13-89495557-INS-7217
chr13-89502097-INS-3618
chr13-89507222-INS-240
chr13-89522836-INS-6382
chr13-89532130-INS-1391
chr13-89532276-INS-58
chr13-89534877-INS-7507
chr13-89559340-INS-4731

chr13-89595774-INS-52
chr13-89628932-INS-107
chr13-89645911-INS-78
chr13-89671286-INS-54
chr13-89698077-INS-50
chr13-89699096-INS-928
chr13-89699224-INS-80
chr13-89716018-INS-95
chr13-89721497-INS-175
chr13-89721497-INS-602
chr13-89771968-INS-478
chr13-89772203-INS-655
chr13-89772269-INS-165
chr13-89790412-INS-940
chr13-89898718-INS-891
chr13-89898762-INS-425
chr13-89910354-INS-77
chr13-89954135-INS-2618
chr13-89956574-INS-83
chr13-89963492-INS-206
chr13-89974590-INS-168
chr13-89983813-INS-2103
chr13-90090939-INS-57
chr13-90092825-INS-119
chr13-90180975-INS-82
chr13-90199390-INS-615
chr13-90223473-INS-240
chr13-90278560-INS-88
chr13-90348878-INS-633
chr13-90359419-INS-6705
chr13-90360884-INS-4458
chr13-90385118-INS-52
chr13-90389172-INS-372
chr13-90425777-INS-425
chr13-90457446-INS-499
chr13-90471363-INS-58
chr13-90504272-INS-1267
chr13-90571704-INS-78
chr13-90576664-INS-1189
chr13-90663022-INS-3385
chr13-90717733-INS-6623
chr13-90767857-INS-119
chr13-90771595-INS-53
chr13-90783106-INS-68
chr13-90838540-INS-236
chr13-90847205-INS-1143
chr13-90917404-INS-6723
chr13-90929779-INS-794
chr13-90978634-INS-4076
chr13-91016962-INS-5337
chr13-91040981-INS-437
chr13-91041418-INS-61
chr13-91057113-INS-5264
chr13-91074170-INS-221
chr13-91090585-INS-3209
chr13-91091278-INS-498
chr13-91098383-INS-291
chr13-91108114-INS-1102
chr13-91117507-INS-54
chr13-91140719-INS-52
chr13-91155594-INS-6535
chr13-91164964-INS-555
chr13-91165195-INS-111
chr13-91219566-INS-604
chr13-91225929-INS-1779
chr13-91316850-INS-150
chr13-91322790-INS-390
chr13-91325678-INS-122
chr13-91364688-INS-114
chr13-91382743-INS-55
chr13-91393110-INS-4393
chr13-91438623-INS-7088
chr13-91452600-INS-71
chr13-91466378-INS-235
chr13-91466492-INS-57
chr13-91488445-INS-764
chr13-91530578-INS-280

chr13-91551151-INS-126
chr13-91615691-INS-208
chr13-91637233-INS-55
chr13-91637797-INS-66
chr13-91639830-INS-58
chr13-91654454-INS-2098
chr13-91675054-INS-50
chr13-91678599-INS-56
chr13-91684902-INS-3040
chr13-91701531-INS-1369
chr13-91783089-INS-199
chr13-91786526-INS-918
chr13-91786950-INS-204
chr13-91802135-INS-208
chr13-91821548-INS-72
chr13-91896290-INS-160
chr13-91896372-INS-64
chr13-91941750-INS-52
chr13-92033406-INS-51
chr13-92040388-INS-66
chr13-92047373-INS-234
chr13-92102644-INS-147
chr13-92198316-INS-51
chr13-92207655-INS-7065
chr13-92212447-INS-50
chr13-92320497-INS-50
chr13-92329241-INS-4400
chr13-92338182-INS-1075
chr13-92339573-INS-52
chr13-92346490-INS-1074
chr13-92366176-INS-440
chr13-92366963-INS-156
chr13-92385322-INS-279
chr13-92385420-INS-110
chr13-92396542-INS-614
chr13-92396641-INS-52
chr13-92448415-INS-349
chr13-92448447-INS-153
chr13-92457275-INS-796
chr13-92468811-INS-2299
chr13-92473848-INS-451
chr13-92533516-INS-68
chr13-92542911-INS-74
chr13-92549543-INS-209
chr13-92550012-INS-344
chr13-92551721-INS-61
chr13-92587317-INS-525
chr13-92626079-INS-1005
chr13-92662206-INS-180
chr13-92697651-INS-58
chr13-92750084-INS-54
chr13-92773825-INS-96
chr13-92779389-INS-745
chr13-92869301-INS-202
chr13-92872300-INS-54
chr13-92872857-INS-64
chr13-92906862-INS-148
chr13-92907051-INS-147
chr13-93067533-INS-90
chr13-93134780-INS-85
chr13-93225072-INS-105
chr13-93233147-INS-183
chr13-93233546-INS-120
chr13-93302064-INS-772
chr13-93315929-INS-657
chr13-93324692-INS-941
chr13-93325855-INS-356
chr13-93346147-INS-193
chr13-93368962-INS-56
chr13-93377303-INS-111
chr13-93426657-INS-1876
chr13-93434066-INS-103
chr13-93477098-INS-138
chr13-93563662-INS-94
chr13-93576873-INS-617
chr13-93576933-INS-234
chr13-93656047-INS-52

chr13-93664507-INS-55
chr13-93708289-INS-209
chr13-93711129-INS-72
chr13-93715951-INS-648
chr13-93717905-INS-80
chr13-93723278-INS-70
chr13-93725840-INS-55
chr13-93741946-INS-581
chr13-93764009-INS-388
chr13-93768537-INS-65
chr13-93778240-INS-171
chr13-93778240-INS-441
chr13-93778825-INS-76
chr13-93820341-INS-96
chr13-93826252-INS-4782
chr13-93841495-INS-63
chr13-93852862-INS-372
chr13-93863088-INS-220
chr13-93865414-INS-84
chr13-93879777-INS-219
chr13-93894084-INS-430
chr13-93894084-INS-86
chr13-93897623-INS-188
chr13-93920780-INS-52
chr13-93936301-INS-1223
chr13-93944959-INS-55
chr13-93945483-INS-76
chr13-93953915-INS-536
chr13-93954029-INS-237
chr13-93954194-INS-511
chr13-94008956-INS-54
chr13-94024152-INS-3923
chr13-94031344-INS-161
chr13-94053015-INS-51
chr13-94064719-INS-74
chr13-94064951-INS-121
chr13-94065012-INS-63
chr13-94071295-INS-150
chr13-94085492-INS-200
chr13-94085634-INS-98
chr13-94099973-INS-150
chr13-94120687-INS-72
chr13-94175114-INS-66
chr13-94175934-INS-52
chr13-94187575-INS-98
chr13-94232413-INS-131
chr13-94232706-INS-75
chr13-94245205-INS-5563
chr13-94298310-INS-50
chr13-94307052-INS-3293
chr13-94336534-INS-66
chr13-94359016-INS-529
chr13-94359135-INS-175
chr13-94365341-INS-280
chr13-94365434-INS-1062
chr13-94377704-INS-442
chr13-94379937-INS-52
chr13-94387744-INS-783
chr13-94390924-INS-55
chr13-94442237-INS-7227
chr13-94445834-INS-202
chr13-94483864-INS-50
chr13-94539010-INS-80
chr13-94546469-INS-72
chr13-94561441-INS-178
chr13-94573997-INS-1288
chr13-94573997-INS-92
chr13-94719904-INS-78
chr13-94752609-INS-238
chr13-94758257-INS-75
chr13-94758453-INS-221
chr13-94771689-INS-501
chr13-94786687-INS-87
chr13-94787445-INS-54
chr13-94787517-INS-161
chr13-94792788-INS-68
chr13-94798782-INS-50

chr13-94825945-INS-71
chr13-94855320-INS-129
chr13-94867646-INS-393
chr13-94879408-INS-1328
chr13-94895877-INS-298
chr13-94947572-INS-54
chr13-94979599-INS-815
chr13-94979798-INS-233
chr13-95053511-INS-470
chr13-95053974-INS-54
chr13-95079279-INS-89
chr13-95082670-INS-1156
chr13-95090774-INS-1516
chr13-95100289-INS-7027
chr13-95106264-INS-1865
chr13-95114839-INS-3509
chr13-95114839-INS-463
chr13-95134923-INS-157
chr13-95148990-INS-72
chr13-95149318-INS-126
chr13-95151977-INS-54
chr13-95186401-INS-574
chr13-95265410-INS-286
chr13-95296033-INS-142
chr13-95306560-INS-93
chr13-95314799-INS-54
chr13-95349526-INS-52
chr13-95382048-INS-385
chr13-95382090-INS-191
chr13-95410554-INS-501
chr13-95437492-INS-69
chr13-95445732-INS-575
chr13-95448094-INS-56
chr13-95450721-INS-89
chr13-95488871-INS-51
chr13-95495819-INS-1449
chr13-95532693-INS-123
chr13-95543522-INS-84
chr13-95548850-INS-87
chr13-95565302-INS-68
chr13-95590290-INS-6401
chr13-95606217-INS-218
chr13-95611038-INS-66
chr13-95623003-INS-124
chr13-95632845-INS-183
chr13-95639759-INS-2502
chr13-95681335-INS-50
chr13-95681955-INS-78
chr13-95692002-INS-3251
chr13-95696773-INS-542
chr13-95696847-INS-61
chr13-95696851-INS-250
chr13-95697086-INS-1349
chr13-95697102-INS-401
chr13-95731530-INS-318
chr13-95732486-INS-708
chr13-95811211-INS-96
chr13-95830549-INS-381
chr13-95830647-INS-100
chr13-95831251-INS-56
chr13-95831317-INS-60
chr13-95854828-INS-50
chr13-95944120-INS-6544
chr13-95988635-INS-498
chr13-95988874-INS-681
chr13-96004739-INS-90
chr13-96005363-INS-378
chr13-96015988-INS-464
chr13-96074493-INS-52
chr13-96077924-INS-1586
chr13-96099241-INS-84
chr13-96169141-INS-513
chr13-96205754-INS-63
chr13-96208472-INS-2585
chr13-96226561-INS-52
chr13-96236979-INS-99
chr13-96259136-INS-54

chr13-96260395-INS-80
chr13-96261095-INS-55
chr13-96286550-INS-76
chr13-96287142-INS-2846
chr13-96287253-INS-647
chr13-96304286-INS-103
chr13-96304710-INS-226
chr13-96348280-INS-54
chr13-96351120-INS-507
chr13-96352037-INS-74
chr13-96381649-INS-3732
chr13-96421168-INS-100
chr13-96425333-INS-133
chr13-96462082-INS-6606
chr13-96473735-INS-62
chr13-96500331-INS-416
chr13-96502639-INS-2511
chr13-96518610-INS-2238
chr13-96518610-INS-277
chr13-96518610-INS-76
chr13-96582639-INS-6495
chr13-96585306-INS-176
chr13-96602565-INS-54
chr13-96609377-INS-77
chr13-96635345-INS-59
chr13-96687831-INS-227
chr13-96708642-INS-641
chr13-96709205-INS-196
chr13-96716406-INS-328
chr13-96731579-INS-164
chr13-96737726-INS-52
chr13-96758438-INS-60
chr13-96760532-INS-69
chr13-96777623-INS-238
chr13-96802866-INS-464
chr13-96806418-INS-80
chr13-96838886-INS-52
chr13-96862806-INS-66
chr13-96881334-INS-205
chr13-96882116-INS-80
chr13-96890279-INS-60
chr13-96898372-INS-99
chr13-96916938-INS-373
chr13-96918951-INS-202
chr13-96935711-INS-379
chr13-96936572-INS-68
chr13-96971433-INS-52
chr13-96971482-INS-197
chr13-97037463-INS-585
chr13-97050191-INS-56
chr13-97050811-INS-92
chr13-97079703-INS-50
chr13-97098834-INS-206
chr13-97113343-INS-68
chr13-97116833-INS-51
chr13-97128872-INS-175
chr13-97132906-INS-3001
chr13-97133256-INS-50
chr13-97140191-INS-80
chr13-97145618-INS-538
chr13-97183331-INS-189
chr13-97192504-INS-62
chr13-97204042-INS-64
chr13-97217607-INS-7295
chr13-97222378-INS-203
chr13-97230959-INS-210
chr13-97239199-INS-203
chr13-97244137-INS-370
chr13-97246058-INS-385
chr13-97256629-INS-219
chr13-97270921-INS-213
chr13-97278873-INS-50
chr13-97280568-INS-209
chr13-97281692-INS-201
chr13-97283239-INS-60
chr13-97284170-INS-211
chr13-97292774-INS-198

chr13-97295067-INS-176
chr13-97302383-INS-63
chr13-97303273-INS-196
chr13-97303454-INS-168
chr13-97304131-INS-3000
chr13-97307243-INS-185
chr13-97320794-INS-214
chr13-97328013-INS-58
chr13-97331372-INS-92
chr13-97341358-INS-5649
chr13-97345588-INS-54
chr13-97349589-INS-192
chr13-97350042-INS-206
chr13-97355548-INS-50
chr13-97391013-INS-101
chr13-97394919-INS-76
chr13-97432703-INS-50
chr13-97453403-INS-433
chr13-97464149-INS-385
chr13-97478746-INS-53
chr13-97505732-INS-89
chr13-97508814-INS-72
chr13-97508871-INS-256
chr13-97526888-INS-50
chr13-97534539-INS-7118
chr13-97543141-INS-54
chr13-97607759-INS-52
chr13-97652210-INS-168
chr13-97668192-INS-76
chr13-97687968-INS-519
chr13-97689753-INS-5168
chr13-97693238-INS-118
chr13-97706811-INS-4803
chr13-97716173-INS-158
chr13-97745276-INS-58
chr13-97776317-INS-77
chr13-97778872-INS-54
chr13-97794727-INS-62
chr13-97794733-INS-164
chr13-97804911-INS-5542
chr13-97846792-INS-108
chr13-97875918-INS-91
chr13-97889953-INS-201
chr13-97890071-INS-85
chr13-97918181-INS-160
chr13-97937772-INS-227
chr13-97958210-INS-72
chr13-97958646-INS-59
chr13-97960543-INS-82
chr13-97982164-INS-45582
chr13-98005966-INS-52
chr13-98030697-INS-180
chr13-98045010-INS-64
chr13-98045997-INS-403
chr13-98055984-INS-104
chr13-98065344-INS-51
chr13-98131551-INS-64
chr13-98168545-INS-52
chr13-98174726-INS-54
chr13-98213007-INS-225
chr13-98229491-INS-61
chr13-98239863-INS-52
chr13-98300657-INS-63
chr13-98301030-INS-518
chr13-98303179-INS-70
chr13-98303883-INS-56
chr13-98321280-INS-192
chr13-98324901-INS-56
chr13-98353527-INS-194
chr13-98358975-INS-211
chr13-98397058-INS-228
chr13-98397769-INS-72
chr13-98415036-INS-4986
chr13-98419657-INS-167
chr13-98426817-INS-57
chr13-98435885-INS-102
chr13-98435917-INS-481

chr13-98439268-INS-64
chr13-98441465-INS-52
chr13-98468033-INS-201
chr13-98469617-INS-71
chr13-98473020-INS-54
chr13-98475597-INS-509
chr13-98482796-INS-127
chr13-98483220-INS-75
chr13-98516745-INS-437
chr13-98522019-INS-60
chr13-98526669-INS-65
chr13-98538093-INS-116
chr13-98546430-INS-268
chr13-98546515-INS-132
chr13-98546860-INS-2965
chr13-98631564-INS-58
chr13-98632200-INS-586
chr13-98645095-INS-6479
chr13-98680075-INS-80
chr13-98710874-INS-103
chr13-98733124-INS-194
chr13-98743812-INS-54
chr13-98757559-INS-382
chr13-98757971-INS-330
chr13-98762312-INS-84
chr13-98762443-INS-148
chr13-98764265-INS-183
chr13-98777946-INS-2841
chr13-98793863-INS-64
chr13-98813035-INS-238
chr13-98828352-INS-65
chr13-98852700-INS-79
chr13-98853762-INS-282
chr13-98859445-INS-1096
chr13-98866458-INS-133
chr13-98873836-INS-96
chr13-98874733-INS-581
chr13-98890636-INS-52
chr13-98890716-INS-189
chr13-98948780-INS-62
chr13-98996829-INS-173
chr13-99031456-INS-199
chr13-99039972-INS-174
chr13-99043867-INS-61
chr13-99067600-INS-421
chr13-99068703-INS-110
chr13-99070440-INS-1995
chr13-99176019-INS-125
chr13-99180194-INS-162
chr13-99204602-INS-379
chr13-99206814-INS-173
chr13-99210407-INS-2515
chr13-99220863-INS-1403
chr13-99224080-INS-3476
chr13-99225110-INS-207
chr13-99225539-INS-6469
chr13-99238112-INS-317
chr13-99278511-INS-50
chr13-99297324-INS-336
chr13-99301813-INS-50
chr13-99353427-INS-120
chr13-99354813-INS-69
chr13-99390465-INS-416
chr13-99432979-INS-59
chr13-99455516-INS-50
chr13-99471636-INS-1608
chr13-99485954-INS-50
chr13-99492211-INS-344
chr13-99497453-INS-479
chr13-99508882-INS-104
chr13-99605837-INS-152
chr13-99659429-INS-72
chr13-99667921-INS-1889
chr13-99680398-INS-7097
chr13-99701312-INS-52
chr13-99707589-INS-136
chr13-99712318-INS-83

chr13-99722900-INS-109
chr13-99743139-INS-80
chr13-99781844-INS-268
chr13-99796566-INS-2972
chr13-99800971-INS-54
chr13-99808127-INS-221
chr13-99818046-INS-76
chr13-99822408-INS-455
chr13-99864047-INS-276
chr13-99873323-INS-3098
chr13-99889315-INS-868
chr13-99912493-INS-1760
chr13-99912589-INS-748
chr13-99924522-INS-98
chr13-99982433-INS-456
chr13-100008578-INS-60
chr13-100018326-INS-184
chr13-100030902-INS-500
chr13-100045898-INS-69
chr13-100046022-INS-86
chr13-100075195-INS-82
chr13-100076768-INS-73
chr13-100084910-INS-56
chr13-100086059-INS-203
chr13-100106903-INS-3501
chr13-100106938-INS-167
chr13-100147969-INS-222
chr13-100149034-INS-50
chr13-100194476-INS-371
chr13-100194476-INS-795
chr13-100221289-INS-247
chr13-100306461-INS-51
chr13-100313471-INS-59
chr13-100331490-INS-742
chr13-100332871-INS-66
chr13-100346605-INS-1096
chr13-100370418-INS-624
chr13-100400285-INS-9524
chr13-100453821-INS-248
chr13-100467819-INS-1351
chr13-100479950-INS-203
chr13-100497133-INS-3608
chr13-100535832-INS-74
chr13-100537463-INS-162
chr13-100537464-INS-67
chr13-100541478-INS-506
chr13-100546064-INS-308
chr13-100572770-INS-177
chr13-100582733-INS-151
chr13-100590139-INS-207
chr13-100606331-INS-234
chr13-100636348-INS-133
chr13-100636559-INS-191
chr13-100640574-INS-104
chr13-100647183-INS-957
chr13-100657657-INS-199
chr13-100661236-INS-199
chr13-100662117-INS-276
chr13-100662801-INS-65
chr13-100695546-INS-157
chr13-100744152-INS-206
chr13-100799187-INS-369
chr13-100810392-INS-1466
chr13-100822938-INS-558
chr13-100823095-INS-180
chr13-100869917-INS-203
chr13-100873869-INS-486
chr13-100874099-INS-890
chr13-100874237-INS-320
chr13-100884672-INS-168
chr13-100909384-INS-1733
chr13-100926432-INS-234
chr13-100930723-INS-182
chr13-100932353-INS-253
chr13-100932514-INS-84
chr13-100933222-INS-358
chr13-100974817-INS-50

chr13-101002003-INS-1843
chr13-101003025-INS-206
chr13-101007520-INS-52
chr13-101012263-INS-1863
chr13-101014635-INS-76
chr13-101031874-INS-521
chr13-101033617-INS-398
chr13-101090319-INS-120
chr13-101090335-INS-312
chr13-101109140-INS-98
chr13-101109437-INS-221
chr13-101115988-INS-223
chr13-101116076-INS-458
chr13-101127294-INS-637
chr13-101127377-INS-65
chr13-101210656-INS-1867
chr13-101211511-INS-6261
chr13-101289516-INS-69
chr13-101294321-INS-493
chr13-101304695-INS-50
chr13-101306535-INS-79
chr13-101306656-INS-60
chr13-101306792-INS-442
chr13-101314674-INS-188
chr13-101318653-INS-6156
chr13-101325038-INS-50
chr13-101336797-INS-7179
chr13-101399871-INS-80
chr13-101407025-INS-154
chr13-101513351-INS-55
chr13-101531835-INS-208
chr13-101537321-INS-57
chr13-101680308-INS-6511
chr13-101697977-INS-119
chr13-101703727-INS-7102
chr13-101725969-INS-194
chr13-101776407-INS-60
chr13-101813672-INS-72
chr13-101821171-INS-52
chr13-101912923-INS-199
chr13-101913110-INS-1045
chr13-101931587-INS-1480
chr13-102019614-INS-138
chr13-102046875-INS-54
chr13-102120316-INS-53
chr13-102182654-INS-7152
chr13-102440347-INS-56
chr13-102550362-INS-198
chr13-102565456-INS-80
chr13-102584925-INS-7078
chr13-102585145-INS-126
chr13-102605643-INS-52
chr13-102628514-INS-399
chr13-102657879-INS-1468
chr13-102692003-INS-613
chr13-102692686-INS-1252
chr13-102707768-INS-154
chr13-102722778-INS-69
chr13-102760686-INS-121
chr13-102768088-INS-61
chr13-102768212-INS-224
chr13-102768281-INS-75
chr13-102777048-INS-52
chr13-102779674-INS-6153
chr13-102799029-INS-1260
chr13-102849492-INS-55
chr13-102996057-INS-56
chr13-103059530-INS-832
chr13-103061356-INS-56
chr13-103066111-INS-56
chr13-103074493-INS-66
chr13-103077943-INS-209
chr13-103099224-INS-2713
chr13-103109142-INS-50
chr13-103109772-INS-6179
chr13-103115652-INS-236
chr13-103169327-INS-56

chr13-103195415-INS-56
chr13-103221009-INS-193
chr13-103221011-INS-77
chr13-103293054-INS-106
chr13-103319961-INS-80
chr13-103337407-INS-114
chr13-103343393-INS-1085
chr13-103371182-INS-56
chr13-103434996-INS-96
chr13-103436589-INS-112
chr13-103483295-INS-1446
chr13-103508361-INS-76
chr13-103527488-INS-84
chr13-103528528-INS-3561
chr13-103638532-INS-785
chr13-103646763-INS-753
chr13-103656825-INS-1179
chr13-103656859-INS-112
chr13-103661769-INS-50
chr13-103680755-INS-79
chr13-103730082-INS-50
chr13-103737356-INS-52
chr13-103745554-INS-108
chr13-103745679-INS-53
chr13-103776074-INS-108
chr13-103798146-INS-186
chr13-103798791-INS-64
chr13-103810130-INS-50
chr13-103819024-INS-3073
chr13-103892686-INS-1076
chr13-103893014-INS-136
chr13-103893050-INS-508
chr13-103913439-INS-52
chr13-103918739-INS-6655
chr13-103932436-INS-6569
chr13-103996410-INS-364
chr13-104005098-INS-1205
chr13-104031738-INS-6620
chr13-104046659-INS-54
chr13-104078089-INS-80
chr13-104078576-INS-78
chr13-104079654-INS-57
chr13-104107489-INS-263
chr13-104119522-INS-6669
chr13-104131444-INS-285
chr13-104185698-INS-52
chr13-104187709-INS-140
chr13-104193214-INS-1536
chr13-104194003-INS-106
chr13-104194003-INS-52
chr13-104202801-INS-50
chr13-104220835-INS-376
chr13-104220859-INS-135
chr13-104274117-INS-66
chr13-104274199-INS-70
chr13-104285732-INS-496
chr13-104312680-INS-66
chr13-104313017-INS-204
chr13-104334577-INS-5424
chr13-104370169-INS-50
chr13-104390744-INS-5525
chr13-104395148-INS-50
chr13-104464784-INS-185
chr13-104471343-INS-4283
chr13-104473693-INS-1546
chr13-104498954-INS-87
chr13-104536693-INS-1999
chr13-104545035-INS-52
chr13-104550717-INS-126
chr13-104573273-INS-158
chr13-104579130-INS-91
chr13-104639741-INS-52
chr13-104692336-INS-92
chr13-104709667-INS-50
chr13-104712615-INS-52
chr13-104879507-INS-310
chr13-104880101-INS-66

chr13-104972768-INS-132
chr13-104992142-INS-50
chr13-105092235-INS-150
chr13-105117339-INS-54
chr13-105196929-INS-794
chr13-105201370-INS-56
chr13-105211764-INS-3714
chr13-105234560-INS-51
chr13-105291337-INS-50
chr13-105309691-INS-1714
chr13-105336143-INS-4796
chr13-105372691-INS-196
chr13-105401043-INS-6636
chr13-105418317-INS-98
chr13-105436879-INS-124
chr13-105444066-INS-50
chr13-105469510-INS-59
chr13-105478038-INS-212
chr13-105491567-INS-6246
chr13-105527402-INS-1897
chr13-105539250-INS-6248
chr13-105560089-INS-217
chr13-105562073-INS-67
chr13-105585190-INS-116
chr13-105586640-INS-498
chr13-105592370-INS-349
chr13-105656839-INS-70
chr13-105657769-INS-114
chr13-105667556-INS-5843
chr13-105689905-INS-4527
chr13-105730147-INS-876
chr13-105733630-INS-1663
chr13-105764570-INS-5217
chr13-105782637-INS-700
chr13-105814575-INS-190
chr13-105815081-INS-64
chr13-105824645-INS-154
chr13-105834963-INS-429
chr13-105869198-INS-842
chr13-105906785-INS-83
chr13-105929051-INS-408
chr13-105962679-INS-1401
chr13-105963500-INS-93
chr13-105963678-INS-901
chr13-105975005-INS-238
chr13-105975005-INS-68
chr13-105978972-INS-1257
chr13-105993320-INS-6385
chr13-105998437-INS-1510
chr13-106010411-INS-490
chr13-106018090-INS-55
chr13-106020778-INS-144
chr13-106029940-INS-102
chr13-106064823-INS-5635
chr13-106079982-INS-53
chr13-106084201-INS-2626
chr13-106130946-INS-8061
chr13-106150041-INS-199
chr13-106162125-INS-400
chr13-106189095-INS-61
chr13-106226711-INS-3070
chr13-106262357-INS-60
chr13-106399834-INS-135
chr13-106415402-INS-7167
chr13-106490548-INS-56
chr13-106538986-INS-52
chr13-106549206-INS-198
chr13-106549415-INS-299
chr13-106604720-INS-2407
chr13-106607083-INS-190
chr13-106610042-INS-7102
chr13-106636255-INS-6898
chr13-106637059-INS-252
chr13-106649550-INS-162
chr13-106703872-INS-186
chr13-106798784-INS-2012
chr13-106817116-INS-950

chr13-106826461-INS-491
chr13-106843255-INS-472
chr13-107002619-INS-114
chr13-107064487-INS-114
chr13-107070159-INS-221
chr13-107070169-INS-667
chr13-107074016-INS-487
chr13-107079984-INS-70
chr13-107090267-INS-777
chr13-107090268-INS-185
chr13-107107122-INS-58
chr13-107153343-INS-466
chr13-107190938-INS-155
chr13-107197470-INS-95
chr13-107200534-INS-56
chr13-107206427-INS-4292
chr13-107212793-INS-69
chr13-107212964-INS-168
chr13-107213098-INS-482
chr13-107213239-INS-57
chr13-107220774-INS-148
chr13-107225839-INS-6476
chr13-107239774-INS-4351
chr13-107254486-INS-7196
chr13-107290582-INS-250
chr13-107333147-INS-457
chr13-107333947-INS-66
chr13-107340820-INS-69
chr13-107362286-INS-6268
chr13-107378225-INS-55
chr13-107383956-INS-58
chr13-107402066-INS-106
chr13-107411738-INS-350
chr13-107411738-INS-7239
chr13-107421142-INS-129
chr13-107424903-INS-2538
chr13-107425400-INS-90
chr13-107501022-INS-100
chr13-107511601-INS-202
chr13-107603836-INS-166
chr13-107619454-INS-114
chr13-107668514-INS-339
chr13-107686453-INS-84
chr13-107703047-INS-52
chr13-107709510-INS-6329
chr13-107709940-INS-60
chr13-107716143-INS-60
chr13-107791683-INS-104
chr13-107798632-INS-130
chr13-107816520-INS-54
chr13-107817101-INS-207
chr13-107921194-INS-80
chr13-107956138-INS-239
chr13-107995218-INS-626
chr13-108037313-INS-158
chr13-108049945-INS-254
chr13-108087666-INS-140
chr13-108118192-INS-168
chr13-108119902-INS-51
chr13-108123572-INS-477
chr13-108152932-INS-88
chr13-108156442-INS-80
chr13-108164021-INS-342
chr13-108202679-INS-51
chr13-108222352-INS-5311
chr13-108225598-INS-56
chr13-108249438-INS-646
chr13-108274894-INS-53
chr13-108322693-INS-83
chr13-108339896-INS-121
chr13-108343888-INS-348
chr13-108349651-INS-199
chr13-108396452-INS-5111
chr13-108409169-INS-1985
chr13-108414952-INS-51
chr13-108444381-INS-7294
chr13-108444637-INS-256

chr13-108466828-INS-61
chr13-108471618-INS-52
chr13-108500967-INS-90
chr13-108574367-INS-117
chr13-108575291-INS-7132
chr13-108585625-INS-61
chr13-108590426-INS-54
chr13-108591099-INS-6277
chr13-108659030-INS-189
chr13-108666169-INS-50
chr13-108700126-INS-52
chr13-108701906-INS-68
chr13-108721522-INS-7123
chr13-108771711-INS-6161
chr13-108802333-INS-94
chr13-108804764-INS-50
chr13-108850544-INS-156
chr13-108938430-INS-349
chr13-108938650-INS-374
chr13-108972147-INS-77
chr13-108985814-INS-52
chr13-108989435-INS-54
chr13-109008854-INS-64
chr13-109027511-INS-2733
chr13-109032580-INS-4118
chr13-109034046-INS-418
chr13-109055789-INS-472
chr13-109056028-INS-284
chr13-109110164-INS-154
chr13-109110963-INS-200
chr13-109111994-INS-6450
chr13-109122522-INS-968
chr13-109122585-INS-2577
chr13-109122602-INS-285
chr13-109122630-INS-78
chr13-109139789-INS-685
chr13-109150079-INS-5763
chr13-109152257-INS-2052
chr13-109185125-INS-51
chr13-109239574-INS-52
chr13-109335435-INS-204
chr13-109340547-INS-214
chr13-109355458-INS-92
chr13-109371592-INS-52
chr13-109391327-INS-67
chr13-109391778-INS-50
chr13-109414206-INS-53
chr13-109435911-INS-302
chr13-109646358-INS-59
chr13-109650783-INS-204
chr13-109665198-INS-104
chr13-109722194-INS-265
chr13-109730567-INS-125
chr13-109767006-INS-54
chr13-109785933-INS-52
chr13-109786005-INS-81
chr13-109796747-INS-55
chr13-109814023-INS-1033
chr13-109817783-INS-53
chr13-109838169-INS-50
chr13-109843018-INS-54
chr13-109846536-INS-1059
chr13-109856161-INS-1808
chr13-110024307-INS-71
chr13-110033995-INS-163
chr13-110099024-INS-454
chr13-110159209-INS-82
chr13-110266875-INS-50
chr13-110267406-INS-224
chr13-110286566-INS-577
chr13-110317889-INS-60
chr13-110319886-INS-50
chr13-110332519-INS-52
chr13-110351050-INS-100
chr13-110353493-INS-261
chr13-110422572-INS-2530
chr13-110444209-INS-2117

chr13-110485521-INS-60
chr13-110485731-INS-120
chr13-110495852-INS-128
chr13-110519896-INS-1197
chr13-110547799-INS-93
chr13-110589170-INS-53
chr13-110645237-INS-75
chr13-110669107-INS-4717
chr13-110773889-INS-1614
chr13-110829776-INS-6363
chr13-110837611-INS-135
chr13-110845257-INS-86
chr13-110861853-INS-8744
chr13-110862762-INS-86
chr13-110879513-INS-6388
chr13-110885431-INS-195
chr13-110901490-INS-2813
chr13-110931758-INS-2168
chr13-110955717-INS-7098
chr13-111070465-INS-56
chr13-111070807-INS-183
chr13-111077434-INS-82
chr13-111096463-INS-288
chr13-111101151-INS-51
chr13-111119514-INS-475
chr13-111137175-INS-189
chr13-111141361-INS-2589
chr13-111166517-INS-64
chr13-111174087-INS-7241
chr13-111183202-INS-93
chr13-111185805-INS-2320
chr13-111308975-INS-1814
chr13-111349909-INS-1433
chr13-111362730-INS-1611
chr13-111362746-INS-378
chr13-111374087-INS-60
chr13-111410231-INS-228
chr13-111433502-INS-1270
chr13-111433846-INS-344
chr13-111433891-INS-5225
chr13-111469237-INS-1873
chr13-111493898-INS-3457
chr13-111494838-INS-6230
chr13-111499510-INS-92
chr13-111511295-INS-211
chr13-111516042-INS-3487
chr13-111516042-INS-822
chr13-111517266-INS-553
chr13-111535764-INS-266
chr13-111545030-INS-88
chr13-111545865-INS-236
chr13-111554440-INS-223
chr13-111556024-INS-98
chr13-111559751-INS-186
chr13-111559891-INS-78
chr13-111650298-INS-55
chr13-111653641-INS-62
chr13-111656508-INS-164
chr13-111658369-INS-60
chr13-111666592-INS-147
chr13-111666601-INS-759
chr13-111667813-INS-330
chr13-111670182-INS-185
chr13-111681285-INS-2816
chr13-111685972-INS-68
chr13-111685978-INS-370
chr13-111686746-INS-617
chr13-111687991-INS-2149
chr13-111692753-INS-180
chr13-111693000-INS-153
chr13-111693194-INS-70
chr13-111693397-INS-242
chr13-111704450-INS-86
chr13-111709457-INS-2862
chr13-111714622-INS-50
chr13-111732665-INS-867
chr13-111761027-INS-50

chr13-111769926-INS-71
chr13-111770241-INS-56
chr13-111770437-INS-60
chr13-111774407-INS-76
chr13-111777910-INS-60
chr13-111911187-INS-88
chr13-111930330-INS-70
chr13-111935256-INS-218
chr13-111955560-INS-60
chr13-111962415-INS-272
chr13-112004204-INS-54
chr13-112029568-INS-84
chr13-112037117-INS-74
chr13-112069581-INS-50
chr13-112083672-INS-2633
chr13-112103527-INS-50
chr13-112103559-INS-321
chr13-112106593-INS-89
chr13-112137386-INS-99
chr13-112161125-INS-51
chr13-112176988-INS-52
chr13-112178044-INS-3641
chr13-112211097-INS-63
chr13-112213471-INS-88
chr13-112213603-INS-426
chr13-112241668-INS-50
chr13-112246954-INS-1064
chr13-112258650-INS-50
chr13-112286091-INS-476
chr13-112345115-INS-64
chr13-112348108-INS-279
chr13-112356404-INS-191
chr13-112391539-INS-82
chr13-112392059-INS-198
chr13-112392123-INS-68
chr13-112400253-INS-3473
chr13-112421327-INS-64
chr13-112422084-INS-62
chr13-112446939-INS-400
chr13-112510743-INS-56
chr13-112535067-INS-138
chr13-112535072-INS-66
chr13-112551542-INS-50
chr13-112564025-INS-50
chr13-112565442-INS-70
chr13-112571899-INS-50
chr13-112580614-INS-141
chr13-112580676-INS-63
chr13-112583064-INS-68
chr13-112583462-INS-118
chr13-112628991-INS-416
chr13-112628991-INS-960
chr13-112645694-INS-120
chr13-112679759-INS-50
chr13-112681366-INS-1501
chr13-112690483-INS-69
chr13-112703140-INS-980
chr13-112703167-INS-76
chr13-112728254-INS-204
chr13-112729073-INS-84
chr13-112768772-INS-326
chr13-112835054-INS-81
chr13-112835198-INS-221
chr13-112836435-INS-141
chr13-112848776-INS-104
chr13-112848997-INS-234
chr13-112854509-INS-159
chr13-112861467-INS-327
chr13-112866021-INS-208
chr13-112866492-INS-179
chr13-112903046-INS-216
chr13-112921175-INS-2919
chr13-112925369-INS-175
chr13-112939110-INS-155
chr13-112949143-INS-539
chr13-112949244-INS-232
chr13-112949249-INS-58

chr13-112956752-INS-124
chr13-112962606-INS-2596
chr13-113031005-INS-200
chr13-113044277-INS-53
chr13-113068696-INS-6478
chr13-113119756-INS-218
chr13-113129557-INS-160
chr13-113164463-INS-1380
chr13-113176340-INS-7196
chr13-113184737-INS-99
chr13-113189231-INS-1140
chr13-113193337-INS-50
chr13-113216574-INS-54
chr13-113230099-INS-29035
chr13-113230261-INS-9974
chr13-113241939-INS-94
chr13-113242414-INS-590
chr13-113255623-INS-102
chr13-113277287-INS-111
chr13-113310195-INS-500
chr13-113312183-INS-50
chr13-113340025-INS-60
chr13-113354985-INS-4132
chr13-113354985-INS-64
chr13-113369655-INS-211
chr13-113370013-INS-276
chr13-113399882-INS-1614
chr13-113440543-INS-198
chr13-113523283-INS-80
chr13-113592650-INS-108
chr13-113592746-INS-292
chr13-113592782-INS-813
chr13-113639586-INS-168
chr13-113650420-INS-137
chr13-113772044-INS-8420
chr13-113793826-INS-744
chr13-113928313-INS-1176
chr13-113935115-INS-216
chr13-113941562-INS-352
chr13-113951849-INS-100
chr13-113978541-INS-247
chr13-114013317-INS-79
chr13-114020544-INS-190
chr13-114056004-INS-52
chr13-114069055-INS-58
chr13-114069131-INS-146
chr13-114069762-INS-58
chr13-114115647-INS-1564
chr13-114117818-INS-52
chr13-114165187-INS-220
chr13-114252931-INS-51
chr13-114299198-INS-128
chr13-114364239-INS-8758
chr13-114372326-INS-52
chr13-114399709-INS-159
chr13-114447774-INS-1500
chr13-114525505-INS-50
chr13-114525845-INS-398
chr13-114571771-INS-51
chr13-114636207-INS-3383
chr13-114672264-INS-354
chr13-114672270-INS-823
chr13-114672768-INS-113
chr13-114895184-INS-59
chr13-114895448-INS-56
chr13-114905824-INS-6452
chr13-114935549-INS-81
chr13-114987572-INS-52
chr13-114988757-INS-64
chr13-115017688-INS-52
chr13-115034270-INS-1155
chr13-115047272-INS-52
chr13-115087692-INS-4173
chr13-115107964-INS-556
chr13-115149817-INS-50
chr13-115151198-INS-149
chr13-115179703-INS-1288

chr13-115206101-INS-1223
chr13-115239286-INS-96
chr13-115294464-INS-202
chr13-115304882-INS-117
chr13-115320911-INS-1058
chr13-115332991-INS-6442
chr13-115360978-INS-6195
chr13-115377975-INS-190
chr13-115384105-INS-298
chr13-115384221-INS-68
chr13-115441773-INS-66
chr13-115446172-INS-67
chr13-115491025-INS-6132
chr13-115515063-INS-1000
chr13-115532612-INS-1932
chr13-115577199-INS-7256
chr13-115613722-INS-51
chr13-115643798-INS-60
chr13-115665535-INS-2845
chr13-115674346-INS-5000
chr13-115701877-INS-213
chr13-115778360-INS-63
chr13-115779651-INS-189
chr13-115849563-INS-3097
chr13-115855092-INS-81
chr13-115870074-INS-1004
chr13-115870074-INS-408
chr13-115901010-INS-88
chr13-115970450-INS-77
chr13-115971272-INS-60
chr13-116061343-INS-3045
chr13-116062440-INS-4946
chr13-116139833-INS-779
chr13-116166260-INS-6255
chr13-116210270-INS-3182
chr13-116213834-INS-104
chr13-116215377-INS-3088
chr13-116261193-INS-50
chr13-116278124-INS-247
chr13-116329013-INS-440
chr13-116349663-INS-50
chr13-116355322-INS-55
chr13-116389591-INS-221
chr13-116408754-INS-560
chr13-116409115-INS-421
chr13-116425830-INS-1163
chr13-116449985-INS-56
chr13-116460935-INS-2360
chr13-116464701-INS-2952
chr13-116479089-INS-2232
chr13-116518734-INS-51
chr13-116546273-INS-110
chr13-116546613-INS-403
chr13-116555272-INS-105
chr13-116556009-INS-6390
chr13-116583200-INS-51
chr13-116599934-INS-4817
chr13-116616582-INS-50
chr13-116657422-INS-50
chr13-116780827-INS-1867
chr13-116781007-INS-64
chr13-116807579-INS-1206
chr13-116814742-INS-111
chr13-116891847-INS-599
chr13-116938182-INS-50
chr13-117001928-INS-2687
chr13-117009033-INS-92
chr13-117012844-INS-5822
chr13-117038841-INS-4955
chr13-117065029-INS-4104
chr13-117065399-INS-2744
chr13-117069345-INS-1645
chr13-117101461-INS-151
chr13-117140142-INS-1947
chr13-117144494-INS-220
chr13-117146358-INS-166
chr13-117147303-INS-58

chr13-117211712-INS-229
chr13-117229410-INS-143
chr13-117255443-INS-565
chr13-117315169-INS-167
chr13-117315620-INS-134
chr13-117337664-INS-100
chr13-117363158-INS-181
chr13-117466789-INS-359
chr13-117488550-INS-69
chr13-117612072-INS-66
chr13-117627934-INS-994
chr13-117638102-INS-505
chr13-117642732-INS-7300
chr13-117657061-INS-126
chr13-117657269-INS-901
chr13-117666485-INS-6810
chr13-117696955-INS-6408
chr13-117701318-INS-145
chr13-117706178-INS-6908
chr13-117753255-INS-217
chr13-117766948-INS-95
chr13-117775041-INS-205
chr13-117800659-INS-54
chr13-117801449-INS-56
chr13-117865767-INS-100
chr13-117916901-INS-54
chr13-117925911-INS-1058
chr13-118014913-INS-50
chr13-118021966-INS-3089
chr13-118031345-INS-6328
chr13-118092431-INS-220
chr13-118127020-INS-7518
chr13-118220538-INS-60
chr13-118245799-INS-3307
chr13-118261577-INS-66
chr13-118263166-INS-717
chr13-118266513-INS-6354
chr13-118280811-INS-300
chr13-118281599-INS-197
chr13-118291204-INS-6332
chr13-118335734-INS-2211
chr13-118351813-INS-197
chr13-118389511-INS-96
chr13-118407282-INS-7419
chr13-118438363-INS-78
chr13-118494037-INS-3167
chr13-118525114-INS-5155
chr13-118530564-INS-688
chr13-118534494-INS-205
chr13-118544087-INS-1073
chr13-118575715-INS-635
chr13-118689199-INS-222
chr13-118699050-INS-121
chr13-118738155-INS-63
chr13-118746093-INS-7126
chr13-118752995-INS-59
chr13-118762081-INS-50
chr13-118764076-INS-423
chr13-118770201-INS-110
chr13-118770984-INS-6438
chr13-118825374-INS-609
chr13-118833782-INS-206
chr13-118859500-INS-1197
chr13-118896944-INS-76
chr13-118973512-INS-54
chr13-118998082-INS-181
chr13-119018487-INS-8114
chr13-119094534-INS-66
chr13-119151602-INS-66
chr13-119160068-INS-63
chr13-119174430-INS-52
chr13-119177815-INS-6655
chr13-119211597-INS-54
chr13-119244352-INS-138
chr13-119244639-INS-117
chr13-119244768-INS-315
chr13-119248720-INS-198

chr13-119252618-INS-1229
chr13-119290476-INS-1263
chr13-119418340-INS-4788
chr13-119519582-INS-199
chr13-119557772-INS-1290
chr13-119560297-INS-52
chr13-119564467-INS-458
chr13-119564805-INS-392
chr13-119566329-INS-910
chr13-119576436-INS-938
chr13-119614059-INS-181
chr13-119625183-INS-172
chr13-119625843-INS-84
chr13-120071171-INS-109
chr13-120093086-INS-227
chr13-120093254-INS-50
chr13-120093429-INS-58
chr13-120136856-INS-50
chr13-120194077-INS-60
chr13-120194455-INS-440
chr13-120264585-INS-1460
chr13-120280648-INS-462
chr13-120281018-INS-60
chr13-120292549-INS-107
chr13-120342792-INS-346
chr13-120343423-INS-217
chr13-120345195-INS-490
chr13-120352540-INS-1890
chr13-120363364-INS-279
chr13-120364043-INS-6273
chr13-120407798-INS-193
chr13-120492434-INS-3238
chr13-120504397-INS-81
chr13-120510996-INS-206
chr13-120526180-INS-177
chr13-120626906-INS-182
chr13-120646487-INS-2993
chr13-120736283-INS-58
chr14-3112389-INS-486
chr14-3155999-INS-50
chr14-3293692-INS-52
chr14-3329944-INS-7221
chr14-3375180-INS-449
chr14-3410040-INS-72
chr14-3478090-INS-59
chr14-3554399-INS-460
chr14-3663199-INS-4365
chr14-3810513-INS-343
chr14-3879019-INS-73
chr14-3923069-INS-4163
chr14-3962046-INS-66
chr14-4026382-INS-3000
chr14-4056479-INS-6486
chr14-4067556-INS-210
chr14-4068859-INS-4239
chr14-4103268-INS-88
chr14-4107975-INS-2070
chr14-4121094-INS-54
chr14-4137450-INS-330
chr14-4137457-INS-150
chr14-4159605-INS-1876
chr14-4225245-INS-2477
chr14-4248700-INS-591
chr14-4248730-INS-1887
chr14-4253647-INS-78
chr14-4284125-INS-124
chr14-4294182-INS-506
chr14-4335601-INS-50
chr14-4414768-INS-1520
chr14-4434791-INS-62
chr14-4454699-INS-52
chr14-4493126-INS-54
chr14-4524463-INS-1146
chr14-4526099-INS-1276
chr14-4526522-INS-2447
chr14-4533498-INS-1769
chr14-4545253-INS-77

chr14-4559205-INS-168
chr14-4559365-INS-56
chr14-4574026-INS-211
chr14-4603176-INS-363
chr14-4603292-INS-172
chr14-4604474-INS-5003
chr14-4632077-INS-73
chr14-4649949-INS-147
chr14-4687342-INS-1912
chr14-4694178-INS-6962
chr14-4729685-INS-51
chr14-4805192-INS-173
chr14-4838237-INS-60
chr14-4843874-INS-53
chr14-4844199-INS-85
chr14-4881164-INS-119
chr14-4956611-INS-4998
chr14-4964499-INS-284
chr14-4969614-INS-110
chr14-5008659-INS-4788
chr14-5122156-INS-180
chr14-5231845-INS-54
chr14-5256583-INS-6752
chr14-5270639-INS-50
chr14-5273631-INS-69
chr14-5384467-INS-794
chr14-5407631-INS-6563
chr14-5456383-INS-69
chr14-5524655-INS-56
chr14-5528547-INS-2687
chr14-5533987-INS-6345
chr14-5547922-INS-2650
chr14-5574764-INS-50
chr14-5593225-INS-7595
chr14-5595352-INS-196
chr14-5595923-INS-56
chr14-5612266-INS-77
chr14-5624115-INS-1450
chr14-5700458-INS-142
chr14-5785390-INS-108
chr14-5926650-INS-63
chr14-5926716-INS-211
chr14-5942616-INS-80
chr14-5962643-INS-51
chr14-5966013-INS-229
chr14-5979110-INS-336
chr14-6004250-INS-56
chr14-6021068-INS-397
chr14-6058375-INS-50
chr14-6131255-INS-192
chr14-6140185-INS-336
chr14-6174236-INS-141
chr14-6174236-INS-70
chr14-6244312-INS-524
chr14-6244312-INS-8632
chr14-6264882-INS-3258
chr14-6264882-INS-517
chr14-6301517-INS-54
chr14-6320931-INS-1979
chr14-6375976-INS-105
chr14-6405951-INS-1253
chr14-6429185-INS-3110
chr14-6459348-INS-415
chr14-6490672-INS-5653
chr14-6536980-INS-1661
chr14-6560364-INS-57
chr14-6583041-INS-256
chr14-6589353-INS-6506
chr14-6598613-INS-1896
chr14-6613482-INS-3319
chr14-6613482-INS-6639
chr14-6668498-INS-385
chr14-6685043-INS-210
chr14-6703572-INS-55
chr14-6710400-INS-60
chr14-6711246-INS-140
chr14-6736940-INS-130

chr14-6763452-INS-1006
chr14-6788419-INS-118
chr14-6822972-INS-728
chr14-6883353-INS-54
chr14-6935766-INS-50
chr14-6968198-INS-139
chr14-6968235-INS-477
chr14-7089131-INS-554
chr14-7093692-INS-7107
chr14-7104032-INS-1639
chr14-7186177-INS-331
chr14-7186181-INS-134
chr14-7186591-INS-248
chr14-7186838-INS-329
chr14-7232934-INS-284
chr14-7297958-INS-55
chr14-7300514-INS-6544
chr14-7324265-INS-2544
chr14-7337659-INS-1181
chr14-7395515-INS-50
chr14-7414578-INS-7092
chr14-7432467-INS-706
chr14-7454617-INS-6989
chr14-7471999-INS-2835
chr14-7485849-INS-264
chr14-7495605-INS-50
chr14-7503329-INS-1010
chr14-7505381-INS-221
chr14-7507100-INS-1284
chr14-7530007-INS-315
chr14-7541117-INS-1096
chr14-7547059-INS-4706
chr14-7560702-INS-58
chr14-7560760-INS-133
chr14-7561660-INS-1075
chr14-7572512-INS-1387
chr14-7593995-INS-6462
chr14-7595837-INS-117
chr14-7662537-INS-6444
chr14-7662908-INS-101
chr14-7666612-INS-52
chr14-7714160-INS-7462
chr14-7732221-INS-2812
chr14-7762488-INS-218
chr14-7763930-INS-58
chr14-7764610-INS-79
chr14-7764645-INS-211
chr14-7793908-INS-519
chr14-7793908-INS-60
chr14-7826602-INS-465
chr14-7849553-INS-55
chr14-7895741-INS-66
chr14-7909374-INS-1493
chr14-7952039-INS-60
chr14-7964638-INS-69
chr14-7964962-INS-3700
chr14-7982366-INS-1616
chr14-8067658-INS-56
chr14-8167101-INS-939
chr14-8191695-INS-90
chr14-8224960-INS-165
chr14-8227411-INS-191
chr14-8431828-INS-298
chr14-8432079-INS-94
chr14-8445882-INS-370
chr14-8472690-INS-50
chr14-8472832-INS-195
chr14-8477521-INS-794
chr14-8479083-INS-180
chr14-8496151-INS-68
chr14-8496689-INS-51
chr14-8513611-INS-66
chr14-8526825-INS-188
chr14-8538759-INS-54
chr14-8541460-INS-76
chr14-8591574-INS-205
chr14-8623104-INS-108

chr14-8645895-INS-51
chr14-8648773-INS-65
chr14-8651598-INS-6267
chr14-8680613-INS-50
chr14-8741401-INS-54
chr14-8743782-INS-624
chr14-8768045-INS-234
chr14-8789960-INS-50
chr14-8856618-INS-5905
chr14-8857079-INS-56
chr14-8868849-INS-2295
chr14-9021923-INS-4200
chr14-9065688-INS-58
chr14-9093379-INS-4809
chr14-9110537-INS-1447
chr14-9112544-INS-77
chr14-9113898-INS-185
chr14-9113898-INS-52
chr14-9114377-INS-66
chr14-9160083-INS-3581
chr14-9170717-INS-6710
chr14-9180582-INS-54
chr14-9183400-INS-50
chr14-9188900-INS-89
chr14-9188905-INS-202
chr14-9196110-INS-90
chr14-9221041-INS-66
chr14-9233995-INS-56
chr14-9240662-INS-6047
chr14-9268946-INS-306
chr14-9273953-INS-963
chr14-9278708-INS-69
chr14-9333665-INS-130
chr14-9338992-INS-50
chr14-9358571-INS-60
chr14-9400713-INS-50
chr14-9421404-INS-50
chr14-9435151-INS-87
chr14-9497740-INS-111
chr14-9509622-INS-52
chr14-9509763-INS-3152
chr14-9511071-INS-491
chr14-9517306-INS-50
chr14-9538440-INS-2955
chr14-9548720-INS-209
chr14-9548822-INS-549
chr14-9585610-INS-51
chr14-9600039-INS-6788
chr14-9665653-INS-222
chr14-9691494-INS-76
chr14-9720829-INS-60
chr14-9721937-INS-174
chr14-9792786-INS-195
chr14-9793154-INS-6358
chr14-9811028-INS-1014
chr14-9814574-INS-85
chr14-9835274-INS-52
chr14-9864914-INS-6880
chr14-9871714-INS-50
chr14-9920554-INS-64
chr14-9920586-INS-130
chr14-9930852-INS-50
chr14-9937022-INS-50
chr14-9973711-INS-132
chr14-9974332-INS-166
chr14-9974563-INS-569
chr14-9974608-INS-100
chr14-10134113-INS-64
chr14-10154894-INS-132
chr14-10155255-INS-146
chr14-10155303-INS-52
chr14-10179522-INS-525
chr14-10331272-INS-78
chr14-10484315-INS-102
chr14-10488405-INS-6442
chr14-10500770-INS-106
chr14-10526917-INS-98

chr14-10533415-INS-201
chr14-10559065-INS-157
chr14-10579885-INS-175
chr14-10587514-INS-53
chr14-10615227-INS-53
chr14-10639021-INS-63
chr14-10647329-INS-52
chr14-10660046-INS-50
chr14-10695890-INS-166
chr14-10705093-INS-58
chr14-10707937-INS-4897
chr14-10758281-INS-144
chr14-10769221-INS-1892
chr14-10770489-INS-150
chr14-10808680-INS-88
chr14-10815429-INS-64
chr14-10821932-INS-84
chr14-10822916-INS-55
chr14-10836132-INS-203
chr14-10862708-INS-59
chr14-10866181-INS-82
chr14-10880848-INS-50
chr14-10884048-INS-299
chr14-10884160-INS-3498
chr14-10891954-INS-6442
chr14-10905985-INS-105
chr14-10923702-INS-53
chr14-10956255-INS-50
chr14-10960912-INS-50
chr14-10989177-INS-50
chr14-10999293-INS-57
chr14-11047726-INS-1262
chr14-11069074-INS-3686
chr14-11069675-INS-52
chr14-11097017-INS-6765
chr14-11097783-INS-250
chr14-11099452-INS-1518
chr14-11104051-INS-76
chr14-11181453-INS-6454
chr14-11185720-INS-8821
chr14-11209939-INS-1585
chr14-11209939-INS-3174
chr14-11217607-INS-935
chr14-11217899-INS-161
chr14-11218039-INS-55
chr14-11275884-INS-174
chr14-11283767-INS-325
chr14-11284797-INS-34110
chr14-11290091-INS-850
chr14-11386331-INS-54
chr14-11423389-INS-612
chr14-11449727-INS-363
chr14-11449771-INS-64
chr14-11492455-INS-573
chr14-11492608-INS-183
chr14-11503967-INS-268
chr14-11549611-INS-2162
chr14-11577288-INS-6111
chr14-11580311-INS-477
chr14-11581601-INS-6197
chr14-11595981-INS-50
chr14-11598850-INS-2652
chr14-11609944-INS-62
chr14-11625417-INS-84
chr14-11712618-INS-102
chr14-11733518-INS-119
chr14-11789340-INS-60
chr14-11802651-INS-50
chr14-11837996-INS-184
chr14-11866345-INS-2876
chr14-11911190-INS-879
chr14-11915962-INS-206
chr14-11916660-INS-50
chr14-11952177-INS-204
chr14-11956310-INS-52
chr14-12005697-INS-52
chr14-12034478-INS-121

chr14-12070257-INS-6509
chr14-12088723-INS-1211
chr14-12117643-INS-55
chr14-12138088-INS-115
chr14-12138088-INS-50
chr14-12138152-INS-249
chr14-12241256-INS-58
chr14-12246301-INS-1518
chr14-12275768-INS-52
chr14-12495247-INS-6225
chr14-12507995-INS-52
chr14-12508590-INS-99
chr14-12509454-INS-78
chr14-12577626-INS-59
chr14-12703364-INS-6407
chr14-12733377-INS-50
chr14-12783987-INS-366
chr14-12785023-INS-52
chr14-12817860-INS-362
chr14-12844539-INS-570
chr14-12873777-INS-1524
chr14-12909198-INS-55
chr14-12920013-INS-413
chr14-12928147-INS-1506
chr14-12929125-INS-6628
chr14-12932600-INS-164
chr14-12953478-INS-2939
chr14-12973517-INS-745
chr14-13022917-INS-532
chr14-13028076-INS-270
chr14-13081247-INS-111
chr14-13082191-INS-84
chr14-13084087-INS-50
chr14-13126176-INS-64
chr14-13147997-INS-208
chr14-13147997-INS-78
chr14-13171345-INS-191
chr14-13266258-INS-381
chr14-13310313-INS-74
chr14-13480193-INS-1813
chr14-13485812-INS-69
chr14-13491520-INS-6801
chr14-13518304-INS-50
chr14-13529814-INS-1430
chr14-13572877-INS-145
chr14-13573241-INS-677
chr14-13589279-INS-214
chr14-13646369-INS-2027
chr14-13649686-INS-56
chr14-13671953-INS-70
chr14-13739661-INS-3796
chr14-13764341-INS-298
chr14-13772723-INS-7253
chr14-13832719-INS-628
chr14-13850446-INS-198
chr14-13853224-INS-897
chr14-13855085-INS-216
chr14-13884952-INS-191
chr14-13885794-INS-58
chr14-13911960-INS-50
chr14-13971314-INS-1657
chr14-14013166-INS-159
chr14-14022282-INS-2200
chr14-14074632-INS-77
chr14-14087121-INS-65
chr14-14109564-INS-910
chr14-14171217-INS-52
chr14-14181535-INS-531
chr14-14232587-INS-205
chr14-14268229-INS-52
chr14-14276074-INS-114
chr14-14280164-INS-5107
chr14-14356655-INS-52
chr14-14362925-INS-211
chr14-14363536-INS-530
chr14-14426426-INS-126
chr14-14430803-INS-152

chr14-14462609-INS-387
chr14-14490175-INS-168
chr14-14598141-INS-1654
chr14-14599223-INS-2339
chr14-14627175-INS-58
chr14-14662821-INS-240
chr14-14680610-INS-7087
chr14-14691100-INS-59
chr14-14703694-INS-73
chr14-14719268-INS-501
chr14-14896748-INS-125
chr14-15095754-INS-224
chr14-15483516-INS-87
chr14-15550452-INS-200
chr14-18122398-INS-68
chr14-18378015-INS-489
chr14-18400113-INS-61
chr14-18571659-INS-444
chr14-18937745-INS-331
chr14-18937783-INS-64
chr14-18958154-INS-104
chr14-19636907-INS-101
chr14-19660506-INS-620
chr14-19660507-INS-235
chr14-19660892-INS-360
chr14-19681358-INS-330
chr14-19725851-INS-794
chr14-19744892-INS-274
chr14-19762179-INS-238
chr14-19766094-INS-71
chr14-19787926-INS-184
chr14-19789480-INS-224
chr14-19831708-INS-884
chr14-19853884-INS-69
chr14-19884512-INS-2851
chr14-19926964-INS-147
chr14-19935707-INS-446
chr14-19940431-INS-982
chr14-19958415-INS-50
chr14-19962445-INS-260
chr14-20004886-INS-203
chr14-20006745-INS-386
chr14-20007385-INS-54
chr14-20018712-INS-210
chr14-20022867-INS-1114
chr14-20031015-INS-118
chr14-20092835-INS-591
chr14-20104001-INS-4864
chr14-20115054-INS-66
chr14-20134523-INS-52
chr14-20158424-INS-156
chr14-20173788-INS-65
chr14-20176204-INS-154
chr14-20212137-INS-90
chr14-20259143-INS-117
chr14-20289739-INS-51
chr14-20303782-INS-896
chr14-20306730-INS-534
chr14-20315638-INS-98
chr14-20316105-INS-85
chr14-20339132-INS-64
chr14-20342928-INS-121
chr14-20343157-INS-215
chr14-20349659-INS-171
chr14-20349686-INS-71
chr14-20352716-INS-79
chr14-20356863-INS-58
chr14-20361271-INS-510
chr14-20383237-INS-52
chr14-20383343-INS-52
chr14-20383807-INS-6774
chr14-20453589-INS-66
chr14-20478632-INS-7331
chr14-20499913-INS-58
chr14-20525196-INS-1680
chr14-20618903-INS-425
chr14-20622010-INS-152

chr14-20662466-INS-52
chr14-20666389-INS-56
chr14-20752672-INS-94
chr14-20762709-INS-384
chr14-20857603-INS-94
chr14-20860749-INS-96
chr14-20872403-INS-461
chr14-20906087-INS-4240
chr14-20924821-INS-230
chr14-20955528-INS-160
chr14-20957381-INS-126
chr14-20968049-INS-206
chr14-20968281-INS-190
chr14-21063834-INS-82
chr14-21066197-INS-19531
chr14-21107833-INS-71
chr14-21163344-INS-145
chr14-21198304-INS-205
chr14-21223802-INS-5313
chr14-21254200-INS-3289
chr14-21277029-INS-58
chr14-21292469-INS-115
chr14-21298239-INS-50
chr14-21304007-INS-64
chr14-21361841-INS-3294
chr14-21363515-INS-3300
chr14-21371055-INS-72
chr14-21402343-INS-53
chr14-21429990-INS-1093
chr14-21443138-INS-2688
chr14-21465218-INS-1476
chr14-21538261-INS-567
chr14-21558760-INS-56
chr14-21571957-INS-92
chr14-21587565-INS-301
chr14-21598627-INS-78
chr14-21598744-INS-169
chr14-21612308-INS-81
chr14-21639039-INS-2003
chr14-21642241-INS-197
chr14-21668550-INS-86
chr14-21668566-INS-174
chr14-21680152-INS-99
chr14-21685978-INS-52
chr14-21692675-INS-372
chr14-21693851-INS-205
chr14-21744500-INS-826
chr14-21745181-INS-932
chr14-21768121-INS-85
chr14-21772094-INS-119
chr14-21788403-INS-272
chr14-21799372-INS-202
chr14-21806862-INS-357
chr14-21827155-INS-50
chr14-21827960-INS-149
chr14-21831950-INS-79
chr14-21871788-INS-51
chr14-21893537-INS-96
chr14-21924714-INS-1564
chr14-21940936-INS-76
chr14-21948738-INS-76
chr14-21949424-INS-1102
chr14-21949652-INS-157
chr14-21970482-INS-155
chr14-22014519-INS-120
chr14-22031381-INS-53
chr14-22060049-INS-313
chr14-22060109-INS-649
chr14-22060245-INS-145
chr14-22103559-INS-314
chr14-22130057-INS-276
chr14-22152769-INS-6753
chr14-22159134-INS-92
chr14-22209263-INS-57
chr14-22252481-INS-63
chr14-22252521-INS-52
chr14-22264436-INS-78

chr14-22273707-INS-228
chr14-22286993-INS-194
chr14-22349469-INS-110
chr14-22349670-INS-81
chr14-22350020-INS-192
chr14-22371308-INS-50
chr14-22402382-INS-1916
chr14-22406995-INS-56
chr14-22447150-INS-1055
chr14-22453229-INS-2188
chr14-22524064-INS-55
chr14-22624677-INS-674
chr14-22624679-INS-191
chr14-22657785-INS-1585
chr14-22660938-INS-58
chr14-22663545-INS-6693
chr14-22666709-INS-132
chr14-22668682-INS-60
chr14-22670138-INS-400
chr14-22678463-INS-1476
chr14-22687238-INS-66
chr14-22687312-INS-63
chr14-22747071-INS-64
chr14-22762987-INS-50
chr14-22799147-INS-161
chr14-22863357-INS-56
chr14-22888109-INS-4073
chr14-23027123-INS-1195
chr14-23037265-INS-62
chr14-23042169-INS-50
chr14-23090590-INS-53
chr14-23165752-INS-98
chr14-23227146-INS-6468
chr14-23245568-INS-70
chr14-23368804-INS-490
chr14-23377771-INS-5591
chr14-23403706-INS-100
chr14-23403706-INS-280
chr14-23418992-INS-72
chr14-23423434-INS-500
chr14-23458633-INS-64
chr14-23468674-INS-2633
chr14-23508431-INS-93
chr14-23513572-INS-98
chr14-23566725-INS-82
chr14-23566735-INS-169
chr14-23606521-INS-1402
chr14-23608124-INS-3702
chr14-23793823-INS-51
chr14-23901769-INS-52
chr14-23922979-INS-496
chr14-23937858-INS-153
chr14-23943420-INS-314
chr14-23960077-INS-3038
chr14-24089028-INS-60
chr14-24089186-INS-133
chr14-24125802-INS-7213
chr14-24126230-INS-14313
chr14-24182924-INS-206
chr14-24197914-INS-52
chr14-24246118-INS-1700
chr14-24299583-INS-98
chr14-24322113-INS-107
chr14-24323062-INS-121
chr14-24330049-INS-2251
chr14-24338259-INS-106
chr14-24368261-INS-2133
chr14-24373571-INS-636
chr14-24375457-INS-123
chr14-24375564-INS-537
chr14-24419014-INS-60
chr14-24453517-INS-218
chr14-24454229-INS-65
chr14-24485636-INS-6228
chr14-24547333-INS-223
chr14-24570941-INS-78
chr14-24590428-INS-207

chr14-24590939-INS-96
chr14-24593829-INS-54
chr14-24631541-INS-1006
chr14-24632409-INS-319
chr14-24647235-INS-1595
chr14-24674967-INS-72
chr14-24744568-INS-129
chr14-24830803-INS-1491
chr14-24883850-INS-50
chr14-24889875-INS-50
chr14-24898985-INS-507
chr14-24913000-INS-198
chr14-24914099-INS-70
chr14-24916842-INS-58
chr14-24958240-INS-92
chr14-24959022-INS-122
chr14-24988218-INS-54
chr14-24989318-INS-203
chr14-24999023-INS-52
chr14-25016292-INS-62
chr14-25046202-INS-52
chr14-25051077-INS-52
chr14-25148559-INS-51
chr14-25153835-INS-130
chr14-25184781-INS-143
chr14-25248719-INS-265
chr14-25286223-INS-1097
chr14-25303249-INS-91
chr14-25323663-INS-58
chr14-25380186-INS-144
chr14-25396036-INS-5551
chr14-25589086-INS-182
chr14-25623589-INS-50
chr14-25703003-INS-535
chr14-25792000-INS-468
chr14-25810295-INS-1510
chr14-25822383-INS-1189
chr14-25834527-INS-291
chr14-25836300-INS-265
chr14-25890602-INS-420
chr14-25906892-INS-549
chr14-25912255-INS-1781
chr14-25922195-INS-111
chr14-25933926-INS-1892
chr14-25972265-INS-57
chr14-26025322-INS-114
chr14-26026614-INS-91
chr14-26026854-INS-253
chr14-26030979-INS-5778
chr14-26096873-INS-77
chr14-26110531-INS-3101
chr14-26119148-INS-414
chr14-26122357-INS-1085
chr14-26208365-INS-168
chr14-26257846-INS-200
chr14-26279901-INS-540
chr14-26304589-INS-218
chr14-26306985-INS-72
chr14-26311926-INS-375
chr14-26340035-INS-73
chr14-26398357-INS-101
chr14-26408564-INS-1158
chr14-26408669-INS-2853
chr14-26452914-INS-1340
chr14-26467787-INS-4808
chr14-26474621-INS-104
chr14-26499273-INS-1210
chr14-26509231-INS-6909
chr14-26523841-INS-6281
chr14-26546709-INS-893
chr14-26568944-INS-204
chr14-26573832-INS-83
chr14-26587754-INS-101
chr14-26609904-INS-84
chr14-26620855-INS-302
chr14-26621025-INS-59
chr14-26628772-INS-5294

chr14-26652993-INS-2562
chr14-26652993-INS-5897
chr14-26652993-INS-630
chr14-26655529-INS-50
chr14-26661286-INS-54
chr14-26686092-INS-209
chr14-26749207-INS-73
chr14-26750390-INS-202
chr14-26753110-INS-182
chr14-26756948-INS-221
chr14-26777203-INS-1124
chr14-26777276-INS-5862
chr14-26778177-INS-148
chr14-26780600-INS-2060
chr14-26808647-INS-393
chr14-26830449-INS-58
chr14-26854764-INS-151
chr14-26854914-INS-52
chr14-26858801-INS-4282
chr14-26878595-INS-320
chr14-26970653-INS-122
chr14-26977576-INS-84
chr14-26977774-INS-419
chr14-27030463-INS-231
chr14-27045602-INS-218
chr14-27050139-INS-75
chr14-27055003-INS-235
chr14-27066164-INS-137
chr14-27080630-INS-130
chr14-27089069-INS-448
chr14-27113669-INS-225
chr14-27129772-INS-564
chr14-27168976-INS-473
chr14-27239166-INS-4820
chr14-27240436-INS-8753
chr14-27243896-INS-2162
chr14-27381088-INS-60
chr14-27387197-INS-97
chr14-27460839-INS-70
chr14-27461190-INS-172
chr14-27469404-INS-2809
chr14-27483081-INS-405
chr14-27483543-INS-99
chr14-27496525-INS-170
chr14-27505448-INS-51
chr14-27505450-INS-109
chr14-27506120-INS-116
chr14-27551069-INS-235
chr14-27551647-INS-6205
chr14-27583499-INS-124
chr14-27583529-INS-62
chr14-27585032-INS-51
chr14-27672127-INS-335
chr14-27672807-INS-98
chr14-27676098-INS-52
chr14-27732302-INS-222
chr14-27753512-INS-160
chr14-27753512-INS-56
chr14-27768954-INS-395
chr14-27768966-INS-143
chr14-27807867-INS-415
chr14-27807869-INS-200
chr14-27866319-INS-88
chr14-27867196-INS-60
chr14-27885782-INS-50
chr14-27910556-INS-206
chr14-27949710-INS-6853
chr14-27961903-INS-74
chr14-27966636-INS-180
chr14-27999852-INS-51
chr14-28088086-INS-188
chr14-28250649-INS-5405
chr14-28275414-INS-52
chr14-28292239-INS-181
chr14-28304235-INS-3491
chr14-28358165-INS-1844
chr14-28358165-INS-3731

chr14-28360622-INS-50
chr14-28423418-INS-6355
chr14-28462503-INS-92
chr14-28514632-INS-6402
chr14-28522980-INS-316
chr14-28578022-INS-93
chr14-28626149-INS-726
chr14-28672252-INS-318
chr14-28693607-INS-176
chr14-28735574-INS-200
chr14-28825476-INS-169
chr14-28840470-INS-52
chr14-28861302-INS-51
chr14-28922223-INS-54
chr14-28933597-INS-102
chr14-28940088-INS-6265
chr14-28949959-INS-1462
chr14-28959628-INS-527
chr14-28979923-INS-76
chr14-28985427-INS-184
chr14-28985427-INS-76
chr14-29000732-INS-69
chr14-29054190-INS-58
chr14-29106436-INS-657
chr14-29132401-INS-519
chr14-29133585-INS-110
chr14-29135687-INS-107
chr14-29161193-INS-57
chr14-29168262-INS-205
chr14-29230751-INS-54
chr14-29233394-INS-50
chr14-29249925-INS-53
chr14-29292625-INS-7275
chr14-29360779-INS-82
chr14-29360858-INS-282
chr14-29360929-INS-600
chr14-29385349-INS-6513
chr14-29389706-INS-350
chr14-29414833-INS-440
chr14-29414985-INS-4028
chr14-29423657-INS-78
chr14-29475687-INS-222
chr14-29502053-INS-60
chr14-29584038-INS-74
chr14-29612516-INS-93
chr14-29676296-INS-2068
chr14-29684878-INS-1928
chr14-29746733-INS-78
chr14-29749160-INS-246
chr14-29749389-INS-112
chr14-29781848-INS-68
chr14-29867854-INS-234
chr14-29867921-INS-86
chr14-29882842-INS-145
chr14-29898757-INS-196
chr14-30094077-INS-52
chr14-30101081-INS-1925
chr14-30106632-INS-221
chr14-30117803-INS-52
chr14-30119735-INS-52
chr14-30134974-INS-94
chr14-30160557-INS-161
chr14-30173860-INS-60
chr14-30261249-INS-191
chr14-30261470-INS-198
chr14-30263241-INS-69
chr14-30263287-INS-299
chr14-30268095-INS-116
chr14-30304289-INS-144
chr14-30304860-INS-52
chr14-30334532-INS-198
chr14-30381859-INS-186
chr14-30433443-INS-861
chr14-30446456-INS-219
chr14-30447620-INS-217
chr14-30534521-INS-4350
chr14-30559634-INS-57

chr14-30607288-INS-76
chr14-30621078-INS-78
chr14-30628646-INS-319
chr14-30628694-INS-1740
chr14-30684593-INS-1462
chr14-30685202-INS-55
chr14-30685293-INS-82
chr14-30692885-INS-203
chr14-30693959-INS-199
chr14-30702549-INS-7418
chr14-30766020-INS-52
chr14-30792998-INS-94
chr14-30808290-INS-210
chr14-30849985-INS-124
chr14-30908187-INS-207
chr14-30932958-INS-76
chr14-31031327-INS-1409
chr14-31092500-INS-239
chr14-31093205-INS-240
chr14-31109186-INS-190
chr14-31135552-INS-50
chr14-31145307-INS-197
chr14-31169369-INS-119
chr14-31240952-INS-63
chr14-31248956-INS-116
chr14-31268166-INS-199
chr14-31297780-INS-112
chr14-31302575-INS-5028
chr14-31304081-INS-227
chr14-31314554-INS-199
chr14-31354115-INS-206
chr14-31367333-INS-54
chr14-31380932-INS-138
chr14-31459984-INS-52
chr14-31507793-INS-4179
chr14-31529515-INS-68
chr14-31531455-INS-50
chr14-31543930-INS-65
chr14-31630042-INS-2119
chr14-31635376-INS-62
chr14-31674733-INS-51
chr14-31674861-INS-119
chr14-31677826-INS-67
chr14-31692913-INS-52
chr14-31694889-INS-6604
chr14-31703605-INS-135
chr14-31704852-INS-68
chr14-31708374-INS-174
chr14-31708406-INS-54
chr14-31717080-INS-889
chr14-31717160-INS-62
chr14-31721951-INS-115
chr14-31722708-INS-223
chr14-31763745-INS-317
chr14-31768755-INS-2158
chr14-31785404-INS-50
chr14-31793394-INS-332
chr14-31802271-INS-76
chr14-31804588-INS-223
chr14-31806370-INS-110
chr14-31839801-INS-125
chr14-31853473-INS-55
chr14-31855284-INS-215
chr14-31865581-INS-195
chr14-31866982-INS-84
chr14-31876829-INS-72
chr14-31899195-INS-420
chr14-32096279-INS-976
chr14-32129083-INS-226
chr14-32134546-INS-76
chr14-32204189-INS-473
chr14-32214035-INS-5206
chr14-32214187-INS-337
chr14-32232417-INS-144
chr14-32284981-INS-2750
chr14-32296563-INS-363
chr14-32305182-INS-2974

chr14-32491837-INS-242
chr14-32497251-INS-50
chr14-32497714-INS-558
chr14-32542816-INS-59
chr14-32564329-INS-52
chr14-32589107-INS-172
chr14-32622312-INS-62
chr14-32655075-INS-2697
chr14-32694530-INS-50
chr14-32762339-INS-58
chr14-32762609-INS-195
chr14-32762614-INS-57
chr14-32769972-INS-134
chr14-32770853-INS-55
chr14-32772641-INS-60
chr14-32807336-INS-476
chr14-32837235-INS-425
chr14-32853684-INS-50
chr14-32856026-INS-51
chr14-32898568-INS-52
chr14-32898966-INS-185
chr14-32899923-INS-1502
chr14-32916672-INS-125
chr14-32921880-INS-204
chr14-32938117-INS-50
chr14-32965744-INS-184
chr14-32966267-INS-60
chr14-33044509-INS-50
chr14-33076654-INS-50
chr14-33077765-INS-52
chr14-33079160-INS-903
chr14-33192098-INS-52
chr14-33201400-INS-66
chr14-33220372-INS-144
chr14-33223688-INS-55
chr14-33224125-INS-50
chr14-33252236-INS-55
chr14-33291043-INS-6217
chr14-33321907-INS-64
chr14-33356516-INS-1341
chr14-33403626-INS-674
chr14-33404667-INS-118
chr14-33404714-INS-295
chr14-33406358-INS-50
chr14-33435096-INS-8418
chr14-33492283-INS-617
chr14-33493575-INS-52
chr14-33500458-INS-6787
chr14-33536047-INS-10409
chr14-33536047-INS-461
chr14-33536481-INS-506
chr14-33545541-INS-64
chr14-33584899-INS-54
chr14-33592372-INS-50
chr14-33609844-INS-51
chr14-33661986-INS-197
chr14-33683126-INS-103
chr14-33732017-INS-64
chr14-33734305-INS-4054
chr14-33786557-INS-171
chr14-33786705-INS-302
chr14-33830857-INS-240
chr14-33830913-INS-96
chr14-33834378-INS-2053
chr14-33849834-INS-7218
chr14-33890626-INS-171
chr14-33901046-INS-732
chr14-33934555-INS-212
chr14-33978097-INS-56
chr14-33978349-INS-230
chr14-33980966-INS-7114
chr14-33999677-INS-104
chr14-34002192-INS-62
chr14-34023970-INS-368
chr14-34061239-INS-211
chr14-34085730-INS-10764
chr14-34122427-INS-102

chr14-34122825-INS-141
chr14-34200062-INS-4358
chr14-34229719-INS-154
chr14-34254889-INS-362
chr14-34255294-INS-54
chr14-34273148-INS-101
chr14-34368112-INS-74
chr14-34386739-INS-51
chr14-34410624-INS-95
chr14-34437223-INS-62
chr14-34437472-INS-143
chr14-34437492-INS-871
chr14-34445050-INS-478
chr14-34449268-INS-60
chr14-34487985-INS-52
chr14-34502798-INS-277
chr14-34503396-INS-1480
chr14-34509037-INS-90
chr14-34516968-INS-149
chr14-34530464-INS-98
chr14-34562614-INS-80
chr14-34562702-INS-76
chr14-34574707-INS-88
chr14-34603662-INS-205
chr14-34607856-INS-52
chr14-34634970-INS-80
chr14-34694051-INS-164
chr14-34698502-INS-64
chr14-34779663-INS-115
chr14-34796040-INS-88
chr14-34800363-INS-50
chr14-34864027-INS-1431
chr14-34898975-INS-91
chr14-34950610-INS-50
chr14-34952894-INS-3799
chr14-34954828-INS-70
chr14-35104163-INS-6174
chr14-35171049-INS-421
chr14-35174946-INS-50
chr14-35191133-INS-85
chr14-35217194-INS-406
chr14-35325990-INS-2909
chr14-35347091-INS-8219
chr14-35394334-INS-6889
chr14-35449942-INS-2568
chr14-35491548-INS-51
chr14-35508514-INS-1089
chr14-35568429-INS-51
chr14-35569468-INS-569
chr14-35585595-INS-6531
chr14-35590563-INS-1886
chr14-35634800-INS-1530
chr14-35634813-INS-111
chr14-35687489-INS-510
chr14-35687574-INS-134
chr14-35710939-INS-66
chr14-35725801-INS-2682
chr14-35728428-INS-8030
chr14-35764193-INS-1978
chr14-35778947-INS-6950
chr14-35870406-INS-2892
chr14-35903912-INS-59
chr14-35926491-INS-62
chr14-35937591-INS-2420
chr14-35997972-INS-50
chr14-36020591-INS-1920
chr14-36043350-INS-4641
chr14-36055272-INS-1255
chr14-36060078-INS-62
chr14-36077803-INS-50
chr14-36183143-INS-257
chr14-36183890-INS-99
chr14-36183892-INS-470
chr14-36184213-INS-489
chr14-36216373-INS-1200
chr14-36230774-INS-56
chr14-36283371-INS-315

chr14-36283395-INS-737
chr14-36283628-INS-1058
chr14-36284057-INS-6579
chr14-36284207-INS-58
chr14-36291179-INS-970
chr14-36301264-INS-607
chr14-36307481-INS-314
chr14-36327724-INS-516
chr14-36332786-INS-51
chr14-36369464-INS-52
chr14-36425561-INS-430
chr14-36432781-INS-6734
chr14-36439637-INS-56
chr14-36452494-INS-72
chr14-36550924-INS-75
chr14-36627434-INS-50
chr14-36632505-INS-15343
chr14-36632687-INS-146
chr14-36632924-INS-7380
chr14-36730372-INS-602
chr14-36748368-INS-209
chr14-36762154-INS-62
chr14-36777882-INS-140
chr14-36786826-INS-7124
chr14-36834138-INS-205
chr14-36854570-INS-7095
chr14-36865920-INS-168
chr14-36874929-INS-105
chr14-36878396-INS-177
chr14-36892507-INS-50
chr14-36913655-INS-2387
chr14-36919919-INS-212
chr14-36942940-INS-406
chr14-36972948-INS-52
chr14-37105703-INS-57
chr14-37106546-INS-51
chr14-37178539-INS-82
chr14-37184862-INS-7019
chr14-37245171-INS-64
chr14-37283240-INS-433
chr14-37362414-INS-10188
chr14-37399703-INS-470
chr14-37410629-INS-415
chr14-37423465-INS-130
chr14-37433110-INS-238
chr14-37457603-INS-7382
chr14-37481749-INS-4791
chr14-37483806-INS-66
chr14-37484096-INS-1894
chr14-37506469-INS-208
chr14-37526320-INS-3005
chr14-37544184-INS-50
chr14-37652433-INS-1974
chr14-37708483-INS-51
chr14-37717521-INS-3973
chr14-37730379-INS-268
chr14-37730910-INS-121
chr14-37799621-INS-8710
chr14-37806500-INS-6290
chr14-37839774-INS-4079
chr14-37860814-INS-783
chr14-37883872-INS-3102
chr14-37971269-INS-11804
chr14-37993912-INS-6131
chr14-38019020-INS-5969
chr14-38024509-INS-67
chr14-38024827-INS-105
chr14-38024950-INS-66
chr14-38028767-INS-3046
chr14-38050995-INS-52
chr14-38057335-INS-9279
chr14-38058143-INS-650
chr14-38123627-INS-75
chr14-38198932-INS-64
chr14-38242648-INS-50
chr14-38246432-INS-1666
chr14-38271124-INS-170

chr14-38275783-INS-539
chr14-38316352-INS-530
chr14-38344894-INS-1105
chr14-38356162-INS-60
chr14-38379487-INS-52
chr14-38392652-INS-7767
chr14-38404281-INS-56
chr14-38469795-INS-383
chr14-38470332-INS-97
chr14-38470679-INS-86
chr14-38485424-INS-1293
chr14-38510458-INS-927
chr14-38621270-INS-202
chr14-38628082-INS-1156
chr14-38660464-INS-57
chr14-38661187-INS-222
chr14-38710639-INS-1798
chr14-38737766-INS-147
chr14-38737842-INS-51
chr14-38780644-INS-56
chr14-38824781-INS-56
chr14-38851186-INS-6482
chr14-38863146-INS-88
chr14-38878338-INS-216
chr14-38887518-INS-5011
chr14-38895670-INS-52
chr14-38961739-INS-628
chr14-38962097-INS-1210
chr14-38974522-INS-129
chr14-39015767-INS-1145
chr14-39056293-INS-146
chr14-39056847-INS-132
chr14-39077916-INS-53
chr14-39133029-INS-60
chr14-39137717-INS-347
chr14-39149079-INS-1264
chr14-39233225-INS-3058
chr14-39241971-INS-54
chr14-39260844-INS-393
chr14-39318387-INS-56
chr14-39338130-INS-1695
chr14-39359494-INS-171
chr14-39359494-INS-56
chr14-39365773-INS-642
chr14-39395515-INS-1002
chr14-39413579-INS-6522
chr14-39421466-INS-56
chr14-39427611-INS-1390
chr14-39489750-INS-56
chr14-39552673-INS-56
chr14-39559925-INS-162
chr14-39578751-INS-7153
chr14-39589099-INS-106
chr14-39598051-INS-50
chr14-39600691-INS-1679
chr14-39619382-INS-52
chr14-39662905-INS-2963
chr14-39680375-INS-256
chr14-39680414-INS-80
chr14-39684837-INS-504
chr14-39695417-INS-50
chr14-39758643-INS-50
chr14-39775259-INS-56
chr14-39826733-INS-204
chr14-39894000-INS-51
chr14-39927711-INS-374
chr14-39987132-INS-494
chr14-39988455-INS-2146
chr14-40002411-INS-6486
chr14-40031633-INS-1320
chr14-40032067-INS-7315
chr14-40047243-INS-1744
chr14-40055025-INS-6378
chr14-40084871-INS-144
chr14-40110318-INS-63
chr14-40147513-INS-7354
chr14-40152577-INS-1209

chr14-40158605-INS-50
chr14-40266188-INS-94
chr14-40319875-INS-665
chr14-40338599-INS-207
chr14-40382476-INS-5351
chr14-40401181-INS-78
chr14-40406789-INS-65
chr14-40406872-INS-158
chr14-40407747-INS-691
chr14-40417937-INS-6403
chr14-40418676-INS-2376
chr14-40421316-INS-1293
chr14-40426205-INS-922
chr14-40440597-INS-4039
chr14-40444044-INS-2529
chr14-40448706-INS-553
chr14-40450032-INS-56
chr14-40459789-INS-6658
chr14-40493111-INS-66
chr14-40504409-INS-6313
chr14-40507662-INS-2158
chr14-40516607-INS-64
chr14-40540498-INS-57
chr14-40550051-INS-4968
chr14-40593418-INS-567
chr14-40600649-INS-190
chr14-40606935-INS-6680
chr14-40622245-INS-92
chr14-40634461-INS-187
chr14-40634647-INS-63
chr14-40634647-INS-942
chr14-40693835-INS-200
chr14-40770462-INS-50
chr14-40920763-INS-4535
chr14-40923037-INS-4934
chr14-40940647-INS-6391
chr14-41200778-INS-3215
chr14-41207945-INS-51
chr14-41325923-INS-1515
chr14-41329451-INS-369
chr14-41454064-INS-106
chr14-41490917-INS-268
chr14-41743916-INS-71
chr14-41748293-INS-50
chr14-42054632-INS-77
chr14-42223415-INS-315
chr14-42946242-INS-66
chr14-42946417-INS-156
chr14-42977285-INS-90
chr14-42977377-INS-210
chr14-42994836-INS-56
chr14-43189813-INS-686
chr14-43766534-INS-165
chr14-43785614-INS-230
chr14-43791082-INS-504
chr14-43994628-INS-287
chr14-44031874-INS-341
chr14-44115841-INS-220
chr14-44136698-INS-498
chr14-44218317-INS-425
chr14-44220550-INS-362
chr14-44221397-INS-406
chr14-44323467-INS-529
chr14-44330804-INS-53
chr14-44335752-INS-174
chr14-44336036-INS-56
chr14-44715615-INS-394
chr14-44945492-INS-1146
chr14-44964691-INS-156
chr14-45031752-INS-80
chr14-45070113-INS-4803
chr14-45070925-INS-2494
chr14-45077973-INS-118
chr14-45125702-INS-3876
chr14-45151508-INS-73
chr14-45163638-INS-479
chr14-45169199-INS-274

chr14-45211062-INS-2859
chr14-45214128-INS-314
chr14-45222933-INS-80
chr14-45240869-INS-67
chr14-45333238-INS-98
chr14-45337916-INS-8512
chr14-45337918-INS-1446
chr14-45359077-INS-77
chr14-45359367-INS-190
chr14-45361822-INS-1862
chr14-45366668-INS-1878
chr14-45398541-INS-185
chr14-45440930-INS-6805
chr14-45509929-INS-390
chr14-45552329-INS-78
chr14-45653623-INS-1100
chr14-45732568-INS-179
chr14-45805463-INS-94
chr14-45806907-INS-2411
chr14-45852130-INS-209
chr14-45910399-INS-205
chr14-45931494-INS-462
chr14-45958293-INS-89
chr14-46009092-INS-1278
chr14-46009118-INS-406
chr14-46022752-INS-6440
chr14-46055096-INS-400
chr14-46091223-INS-2318
chr14-46162207-INS-192
chr14-46165272-INS-55
chr14-46177509-INS-78
chr14-46186355-INS-460
chr14-46195710-INS-2311
chr14-46199375-INS-7087
chr14-46218531-INS-1626
chr14-46245306-INS-96
chr14-46285067-INS-51
chr14-46300503-INS-58
chr14-46320027-INS-6535
chr14-46330599-INS-236
chr14-46330599-INS-621
chr14-46346652-INS-152
chr14-46351234-INS-200
chr14-46353122-INS-420
chr14-46353122-INS-72
chr14-46509540-INS-3712
chr14-46531195-INS-7617
chr14-46535951-INS-50
chr14-46550576-INS-57
chr14-46602125-INS-165
chr14-46693834-INS-2870
chr14-46791062-INS-98
chr14-46791455-INS-404
chr14-46921144-INS-6831
chr14-46928130-INS-64
chr14-46971170-INS-99
chr14-46976929-INS-52
chr14-46981583-INS-2862
chr14-47040031-INS-154
chr14-47063469-INS-4325
chr14-47067724-INS-382
chr14-47135128-INS-50
chr14-47137218-INS-138
chr14-47137487-INS-81
chr14-47168992-INS-168
chr14-47175028-INS-367
chr14-47175095-INS-171
chr14-47201411-INS-165
chr14-47201574-INS-415
chr14-47202224-INS-86
chr14-47224730-INS-52
chr14-47260778-INS-224
chr14-47299548-INS-226
chr14-47309521-INS-5218
chr14-47323858-INS-52
chr14-47351699-INS-50
chr14-47412923-INS-90

chr14-47440364-INS-7084
chr14-47476644-INS-4151
chr14-47498297-INS-168
chr14-47529935-INS-60
chr14-47591436-INS-202
chr14-47606272-INS-7317
chr14-47632509-INS-408
chr14-47636154-INS-179
chr14-47666612-INS-372
chr14-47670263-INS-163
chr14-47675042-INS-204
chr14-47684964-INS-480
chr14-47691928-INS-123
chr14-47701050-INS-6595
chr14-47725484-INS-352
chr14-47769790-INS-208
chr14-47791680-INS-94
chr14-47819350-INS-5383
chr14-47836838-INS-4736
chr14-47840768-INS-233
chr14-47840967-INS-127
chr14-47867839-INS-8701
chr14-47868751-INS-190
chr14-47884542-INS-52
chr14-47890228-INS-54
chr14-47892000-INS-263
chr14-47923264-INS-207
chr14-47955841-INS-54
chr14-47959495-INS-885
chr14-48095701-INS-50
chr14-48167375-INS-73
chr14-48167480-INS-151
chr14-48241876-INS-196
chr14-48268138-INS-1865
chr14-48301210-INS-102
chr14-48322999-INS-170
chr14-48332625-INS-53
chr14-48341378-INS-240
chr14-48504489-INS-196
chr14-48504489-INS-84
chr14-48514382-INS-5140
chr14-48604338-INS-491
chr14-48612824-INS-51
chr14-48618471-INS-3000
chr14-48653208-INS-181
chr14-48739112-INS-2668
chr14-48798374-INS-71
chr14-48833367-INS-84
chr14-48985421-INS-172
chr14-49045762-INS-134
chr14-49079513-INS-153
chr14-49079723-INS-693
chr14-49094570-INS-690
chr14-49094788-INS-1626
chr14-49186827-INS-66
chr14-49349003-INS-266
chr14-49451728-INS-8470
chr14-49453114-INS-1813
chr14-49453210-INS-802
chr14-49458890-INS-50
chr14-49475946-INS-50
chr14-49560224-INS-6213
chr14-49577454-INS-497
chr14-49633995-INS-66
chr14-49652969-INS-6171
chr14-49705031-INS-195
chr14-49721997-INS-189
chr14-49729168-INS-60
chr14-49729208-INS-150
chr14-49797876-INS-4552
chr14-49934713-INS-3952
chr14-49959061-INS-7179
chr14-49977757-INS-50
chr14-49995959-INS-6222
chr14-50030674-INS-50
chr14-50052807-INS-499
chr14-50171851-INS-293

chr14-50190565-INS-1790
chr14-50366325-INS-68
chr14-50384079-INS-50
chr14-50408095-INS-4389
chr14-50593747-INS-89
chr14-50605071-INS-6396
chr14-50666472-INS-13204
chr14-50708010-INS-250
chr14-50708732-INS-599
chr14-50735232-INS-3543
chr14-50753091-INS-6594
chr14-50795383-INS-64
chr14-50818527-INS-206
chr14-50857488-INS-508
chr14-50857552-INS-249
chr14-50936843-INS-6296
chr14-50940370-INS-3388
chr14-50969718-INS-1306
chr14-50969718-INS-3899
chr14-50978565-INS-54
chr14-51006649-INS-171
chr14-51040514-INS-217
chr14-51042710-INS-1965
chr14-51060315-INS-2092
chr14-51072688-INS-84
chr14-51072898-INS-66
chr14-51186638-INS-2997
chr14-51211478-INS-1288
chr14-51217270-INS-7447
chr14-51250035-INS-4944
chr14-51262365-INS-52
chr14-51345768-INS-50
chr14-51358985-INS-52
chr14-51419979-INS-3979
chr14-51470650-INS-12527
chr14-51479324-INS-50
chr14-51525764-INS-615
chr14-51525891-INS-1589
chr14-51535240-INS-362
chr14-51547018-INS-3040
chr14-51547018-INS-465
chr14-51561965-INS-7482
chr14-51674347-INS-172
chr14-51675680-INS-113
chr14-51809854-INS-50
chr14-51813472-INS-438
chr14-51826707-INS-418
chr14-51844948-INS-51
chr14-51856887-INS-132
chr14-51870131-INS-498
chr14-51871372-INS-70
chr14-51871657-INS-172
chr14-51979298-INS-500
chr14-51982134-INS-151
chr14-52011696-INS-56
chr14-52021484-INS-562
chr14-52028483-INS-273
chr14-52031288-INS-4336
chr14-52040809-INS-6419
chr14-52043745-INS-2009
chr14-52142545-INS-54
chr14-52168323-INS-50
chr14-52173291-INS-78
chr14-52177208-INS-209
chr14-52179812-INS-52
chr14-52181280-INS-906
chr14-52181340-INS-348
chr14-52184714-INS-2467
chr14-52266777-INS-219
chr14-52305025-INS-164
chr14-52305264-INS-110
chr14-52331357-INS-209
chr14-52444142-INS-66
chr14-52478128-INS-88
chr14-52522334-INS-193
chr14-52526784-INS-64
chr14-52636004-INS-1093

chr14-52639934-INS-2998
chr14-52642127-INS-1983
chr14-52653965-INS-1846
chr14-52663069-INS-124
chr14-52663243-INS-409
chr14-52663330-INS-80
chr14-52676829-INS-9201
chr14-52708329-INS-6647
chr14-52717489-INS-144
chr14-52723696-INS-1862
chr14-52734417-INS-188
chr14-52801158-INS-226
chr14-52801780-INS-50
chr14-52856372-INS-8327
chr14-52929762-INS-2932
chr14-52946370-INS-204
chr14-53039542-INS-264
chr14-53095630-INS-59
chr14-53529632-INS-1057
chr14-53539189-INS-78
chr14-53566221-INS-109
chr14-53570634-INS-323
chr14-53573378-INS-445
chr14-53597973-INS-3876
chr14-53691589-INS-1915
chr14-53732214-INS-187
chr14-53732760-INS-56
chr14-53732764-INS-134
chr14-53745868-INS-6190
chr14-53748074-INS-474
chr14-53757839-INS-353
chr14-53758483-INS-173
chr14-53773365-INS-846
chr14-53782127-INS-52
chr14-53793845-INS-218
chr14-53793845-INS-57
chr14-53820705-INS-3009
chr14-53857090-INS-831
chr14-53986516-INS-83
chr14-53994160-INS-498
chr14-53996435-INS-298
chr14-54002319-INS-483
chr14-54012902-INS-6200
chr14-54024547-INS-1496
chr14-54024553-INS-3664
chr14-54036339-INS-3367
chr14-54042540-INS-3250
chr14-54057848-INS-6235
chr14-54070330-INS-7734
chr14-54074414-INS-792
chr14-54088051-INS-668
chr14-54105009-INS-1360
chr14-54130169-INS-3763
chr14-54132098-INS-59
chr14-54141915-INS-6363
chr14-54161914-INS-2225
chr14-54210473-INS-91
chr14-54263618-INS-7005
chr14-54338980-INS-51
chr14-54423981-INS-261
chr14-54469120-INS-380
chr14-54560432-INS-214
chr14-54571528-INS-52
chr14-54603019-INS-219
chr14-54603750-INS-76
chr14-54726790-INS-460
chr14-54733695-INS-2129
chr14-54758456-INS-731
chr14-54758477-INS-139
chr14-54761709-INS-156
chr14-54829919-INS-50
chr14-54871703-INS-64
chr14-54964936-INS-199
chr14-55026877-INS-58
chr14-55031958-INS-50
chr14-55046088-INS-57
chr14-55047864-INS-505

chr14-55068835-INS-4877
chr14-55085330-INS-202
chr14-55160507-INS-334
chr14-55161050-INS-64
chr14-55178509-INS-50
chr14-55257489-INS-387
chr14-55399578-INS-218
chr14-55403442-INS-230
chr14-55403442-INS-69
chr14-55444368-INS-61
chr14-55461087-INS-100
chr14-55488163-INS-54
chr14-55489539-INS-6234
chr14-55503397-INS-140
chr14-55547907-INS-171
chr14-55572688-INS-52
chr14-55572949-INS-58
chr14-55620162-INS-50
chr14-55625472-INS-20153
chr14-55682947-INS-104
chr14-55714077-INS-104
chr14-55763688-INS-55
chr14-55820337-INS-79
chr14-55961656-INS-2071
chr14-55997029-INS-420
chr14-56197729-INS-4801
chr14-56212817-INS-1961
chr14-56238004-INS-50
chr14-56306710-INS-1890
chr14-56398180-INS-73
chr14-56518233-INS-1490
chr14-56541026-INS-451
chr14-56542735-INS-59
chr14-56542834-INS-596
chr14-56542887-INS-93
chr14-56562373-INS-58
chr14-56579661-INS-654
chr14-56585235-INS-158
chr14-56612758-INS-253
chr14-56642912-INS-118
chr14-56820392-INS-297
chr14-56834765-INS-52
chr14-56840505-INS-68
chr14-56857627-INS-74
chr14-56896263-INS-54
chr14-57001122-INS-60
chr14-57061679-INS-225
chr14-57073652-INS-171
chr14-57166795-INS-52
chr14-57218179-INS-56
chr14-57254558-INS-7240
chr14-57254850-INS-14611
chr14-57383876-INS-60
chr14-57399299-INS-52
chr14-57409473-INS-60
chr14-57586472-INS-526
chr14-57595499-INS-99
chr14-57738324-INS-320
chr14-57759429-INS-709
chr14-57921580-INS-166
chr14-57960384-INS-56
chr14-57962302-INS-5450
chr14-58018229-INS-424
chr14-58026378-INS-50
chr14-58026425-INS-557
chr14-58059219-INS-51
chr14-58072440-INS-876
chr14-58072497-INS-103
chr14-58098659-INS-171
chr14-58122869-INS-143
chr14-58139867-INS-52
chr14-58142472-INS-50
chr14-58149532-INS-86
chr14-58165298-INS-580
chr14-58168209-INS-340
chr14-58183753-INS-216
chr14-58184723-INS-114

chr14-58224850-INS-60
chr14-58266763-INS-114
chr14-58303599-INS-50
chr14-58319760-INS-96
chr14-58361119-INS-86
chr14-58361320-INS-123
chr14-58408995-INS-50
chr14-58481334-INS-137
chr14-58495420-INS-50
chr14-58499079-INS-508
chr14-58513218-INS-60
chr14-58519097-INS-500
chr14-58614041-INS-86
chr14-58614117-INS-215
chr14-58714875-INS-105
chr14-58725880-INS-6443
chr14-58863203-INS-55
chr14-58946128-INS-221
chr14-58974761-INS-232
chr14-58975461-INS-61
chr14-58989996-INS-200
chr14-58990076-INS-76
chr14-59123098-INS-1887
chr14-59246485-INS-71
chr14-59289813-INS-6649
chr14-59293754-INS-197
chr14-59297266-INS-51
chr14-59643418-INS-4041
chr14-59683485-INS-2118
chr14-59683485-INS-683
chr14-59709115-INS-80
chr14-59849195-INS-1912
chr14-60112232-INS-129
chr14-60192771-INS-66
chr14-60215916-INS-150
chr14-60261441-INS-519
chr14-60278352-INS-770
chr14-60318207-INS-145
chr14-60343071-INS-86
chr14-60347805-INS-570
chr14-60359684-INS-133
chr14-60383299-INS-104
chr14-60383652-INS-67
chr14-60423901-INS-69
chr14-60535806-INS-50
chr14-60719456-INS-1731
chr14-60732282-INS-64
chr14-60823637-INS-234
chr14-60841686-INS-110
chr14-60905453-INS-356
chr14-60954185-INS-53
chr14-61075413-INS-141
chr14-61232212-INS-255
chr14-61297356-INS-58
chr14-61335413-INS-94
chr14-61348716-INS-76
chr14-61459494-INS-294
chr14-61483852-INS-105
chr14-61494647-INS-58
chr14-61503198-INS-52
chr14-61515131-INS-133
chr14-61536443-INS-113
chr14-61585751-INS-62
chr14-61627458-INS-101
chr14-61666315-INS-1288
chr14-61991089-INS-6677
chr14-62040227-INS-84
chr14-62096674-INS-101
chr14-62130236-INS-50
chr14-62139568-INS-223
chr14-62144103-INS-4200
chr14-62144991-INS-1385
chr14-62173456-INS-6215
chr14-62175311-INS-1257
chr14-62237356-INS-477
chr14-62307910-INS-50
chr14-62323037-INS-52

chr14-62328199-INS-90
chr14-62334108-INS-60
chr14-62418000-INS-134
chr14-62463226-INS-52
chr14-62468414-INS-317
chr14-62468700-INS-356
chr14-62470022-INS-149
chr14-62520101-INS-94
chr14-62522417-INS-1689
chr14-62559622-INS-193
chr14-62566622-INS-510
chr14-62617952-INS-416
chr14-62636884-INS-92
chr14-62640755-INS-2382
chr14-62642773-INS-110
chr14-62655391-INS-1623
chr14-62668639-INS-152
chr14-62670679-INS-450
chr14-62674498-INS-60
chr14-62682440-INS-56
chr14-62694831-INS-84
chr14-62703494-INS-150
chr14-62765207-INS-109
chr14-62798167-INS-312
chr14-62818676-INS-2263
chr14-62839083-INS-6660
chr14-62849521-INS-97
chr14-62867941-INS-124
chr14-62907731-INS-1680
chr14-63104548-INS-545
chr14-63108154-INS-207
chr14-63131738-INS-400
chr14-63170942-INS-2244
chr14-63170982-INS-814
chr14-63204412-INS-687
chr14-63204756-INS-569
chr14-63242626-INS-902
chr14-63250969-INS-90
chr14-63257351-INS-116
chr14-63260947-INS-50
chr14-63308875-INS-120
chr14-63312090-INS-99
chr14-63314881-INS-136
chr14-63319161-INS-351
chr14-63334759-INS-52
chr14-63342078-INS-87
chr14-63376784-INS-66
chr14-63385922-INS-3000
chr14-63392636-INS-52
chr14-63427538-INS-96
chr14-63443350-INS-492
chr14-63515869-INS-216
chr14-63518045-INS-6726
chr14-63531196-INS-53
chr14-63565118-INS-67
chr14-63590143-INS-502
chr14-63639172-INS-141
chr14-63674362-INS-89
chr14-63699163-INS-209
chr14-63708912-INS-95
chr14-63784465-INS-52
chr14-63800220-INS-151
chr14-63805351-INS-264
chr14-63841526-INS-207
chr14-63919067-INS-66
chr14-64002734-INS-248
chr14-64104763-INS-112
chr14-64272365-INS-1170
chr14-64335897-INS-177
chr14-64383381-INS-50
chr14-64400182-INS-56
chr14-64454349-INS-4800
chr14-64478303-INS-265
chr14-64498485-INS-52
chr14-64524359-INS-192
chr14-64611329-INS-52
chr14-64646534-INS-110

chr14-64648335-INS-109
chr14-64659159-INS-407
chr14-64740859-INS-1755
chr14-64753012-INS-212
chr14-64764870-INS-183
chr14-64803100-INS-89
chr14-64865452-INS-50
chr14-64869324-INS-800
chr14-64874230-INS-1721
chr14-64877923-INS-199
chr14-64877923-INS-55
chr14-64880108-INS-54
chr14-64880433-INS-137
chr14-64884444-INS-730
chr14-64900422-INS-229
chr14-65167289-INS-202
chr14-65174490-INS-65
chr14-65218381-INS-215
chr14-65284430-INS-108
chr14-65330534-INS-118
chr14-65405976-INS-109
chr14-65421533-INS-217
chr14-65421969-INS-400
chr14-65422041-INS-50
chr14-65429268-INS-87
chr14-65543939-INS-190
chr14-65544663-INS-124
chr14-65577580-INS-210
chr14-65605759-INS-56
chr14-65607307-INS-999
chr14-65678384-INS-838
chr14-65689204-INS-53
chr14-65710712-INS-68
chr14-65739799-INS-237
chr14-65791965-INS-638
chr14-65865258-INS-341
chr14-65866918-INS-712
chr14-65867397-INS-96
chr14-65867576-INS-1090
chr14-65880913-INS-81
chr14-65906653-INS-60
chr14-65914844-INS-68
chr14-65920947-INS-204
chr14-65924929-INS-207
chr14-65930365-INS-52
chr14-65939170-INS-6713
chr14-65942286-INS-53
chr14-66004545-INS-3614
chr14-66023578-INS-120
chr14-66026147-INS-50
chr14-66035396-INS-498
chr14-66039043-INS-50
chr14-66041649-INS-117
chr14-66044516-INS-94
chr14-66105965-INS-167
chr14-66124112-INS-54
chr14-66130760-INS-6548
chr14-66135317-INS-211
chr14-66139699-INS-106
chr14-66140857-INS-50
chr14-66174478-INS-230
chr14-66258206-INS-66
chr14-66291005-INS-50
chr14-66338033-INS-2010
chr14-66369726-INS-50
chr14-66425290-INS-375
chr14-66432832-INS-70
chr14-66462052-INS-54
chr14-66462229-INS-202
chr14-66488917-INS-50
chr14-66508015-INS-78
chr14-66514712-INS-52
chr14-66607877-INS-779
chr14-66629046-INS-1487
chr14-66629420-INS-3806
chr14-66644865-INS-52
chr14-66673530-INS-125

chr14-66711153-INS-351
chr14-66755749-INS-66
chr14-66803036-INS-151
chr14-66816195-INS-1877
chr14-66857354-INS-1736
chr14-66872149-INS-55
chr14-66943682-INS-52
chr14-66944483-INS-69
chr14-66976017-INS-52
chr14-66994540-INS-13665
chr14-67007231-INS-58
chr14-67013536-INS-50
chr14-67022307-INS-58
chr14-67036647-INS-306
chr14-67068681-INS-360
chr14-67068804-INS-272
chr14-67069007-INS-642
chr14-67084296-INS-50
chr14-67119769-INS-50
chr14-67209213-INS-54
chr14-67209364-INS-348
chr14-67266165-INS-189
chr14-67280425-INS-195
chr14-67325259-INS-84
chr14-67331854-INS-124
chr14-67347154-INS-792
chr14-67349831-INS-52
chr14-67350076-INS-688
chr14-67368352-INS-201
chr14-67371494-INS-92
chr14-67380636-INS-54
chr14-67383437-INS-118
chr14-67402770-INS-133
chr14-67420528-INS-1890
chr14-67428949-INS-4508
chr14-67433837-INS-642
chr14-67446892-INS-110
chr14-67456659-INS-234
chr14-67476178-INS-52
chr14-67523300-INS-62
chr14-67575294-INS-135
chr14-67589365-INS-270
chr14-67609295-INS-52
chr14-67617748-INS-114
chr14-67629716-INS-74
chr14-67636618-INS-104
chr14-67676416-INS-114
chr14-67763630-INS-144
chr14-67777178-INS-190
chr14-67794171-INS-124
chr14-67828274-INS-102
chr14-67829003-INS-2943
chr14-67839841-INS-167
chr14-67991625-INS-367
chr14-68088634-INS-435
chr14-68174836-INS-74
chr14-68174984-INS-141
chr14-68176719-INS-82
chr14-68191943-INS-622
chr14-68192167-INS-121
chr14-68228538-INS-413
chr14-68254586-INS-3877
chr14-68356783-INS-116
chr14-68409502-INS-4346
chr14-68425717-INS-1032
chr14-68518429-INS-222
chr14-68530249-INS-116
chr14-68537308-INS-62
chr14-68568820-INS-6947
chr14-68572520-INS-59
chr14-68591132-INS-50
chr14-68603999-INS-204
chr14-68640367-INS-387
chr14-68678170-INS-1042
chr14-68684775-INS-7271
chr14-68727613-INS-3027
chr14-68736766-INS-145

chr14-68736766-INS-69
chr14-68741412-INS-151
chr14-68826457-INS-579
chr14-68826653-INS-210
chr14-68832428-INS-61
chr14-68835550-INS-2352
chr14-68839898-INS-7478
chr14-68845669-INS-2383
chr14-68846718-INS-129
chr14-68925015-INS-502
chr14-69019222-INS-50
chr14-69034793-INS-107
chr14-69034793-INS-278
chr14-69042886-INS-3407
chr14-69079487-INS-52
chr14-69098356-INS-521
chr14-69120043-INS-6731
chr14-69195215-INS-62
chr14-69231045-INS-118
chr14-69289695-INS-56
chr14-69514091-INS-158
chr14-69514437-INS-99
chr14-69561075-INS-2835
chr14-69732687-INS-96
chr14-69796925-INS-93
chr14-69807487-INS-198
chr14-69875577-INS-733
chr14-69915909-INS-58
chr14-69930741-INS-187
chr14-69942283-INS-82
chr14-70039407-INS-206
chr14-70039407-INS-55
chr14-70044801-INS-210
chr14-70079500-INS-61
chr14-70102265-INS-333
chr14-70102721-INS-198
chr14-70162543-INS-6478
chr14-70207970-INS-116
chr14-70249631-INS-60
chr14-70288842-INS-88
chr14-70456135-INS-117
chr14-70473704-INS-1170
chr14-70660717-INS-128
chr14-70850954-INS-72
chr14-70851665-INS-163
chr14-70891583-INS-210
chr14-70943479-INS-208
chr14-70964216-INS-399
chr14-71019529-INS-71
chr14-71054210-INS-70
chr14-71063819-INS-6414
chr14-71069023-INS-6240
chr14-71124325-INS-50
chr14-71140609-INS-72
chr14-71140909-INS-58
chr14-71158694-INS-2849
chr14-71263254-INS-178
chr14-71279168-INS-6917
chr14-71287372-INS-144
chr14-71402670-INS-13060
chr14-71428547-INS-1551
chr14-71478439-INS-4397
chr14-71509702-INS-84
chr14-71544626-INS-124
chr14-71553544-INS-464
chr14-71563520-INS-178
chr14-71563850-INS-160
chr14-71921089-INS-625
chr14-71921461-INS-2253
chr14-71956429-INS-2092
chr14-71957306-INS-413
chr14-71967856-INS-274
chr14-71988918-INS-8137
chr14-72052636-INS-6529
chr14-72061433-INS-113
chr14-72068618-INS-50
chr14-72096449-INS-429

chr14-72100899-INS-77
chr14-72101194-INS-78
chr14-72143322-INS-10198
chr14-72187278-INS-1607
chr14-72220135-INS-89
chr14-72220842-INS-2382
chr14-72272284-INS-3117
chr14-72295474-INS-58
chr14-72346785-INS-1450
chr14-72360360-INS-2844
chr14-72393383-INS-268
chr14-72403013-INS-51
chr14-72423953-INS-50
chr14-72435771-INS-3285
chr14-72502778-INS-50
chr14-72507860-INS-427
chr14-72518532-INS-6634
chr14-72523889-INS-174
chr14-72523895-INS-456
chr14-72542166-INS-3900
chr14-72586451-INS-514
chr14-72614573-INS-76
chr14-72625047-INS-371
chr14-72625047-INS-50
chr14-72666542-INS-2725
chr14-72686536-INS-112
chr14-72747415-INS-66
chr14-72806027-INS-451
chr14-72863342-INS-359
chr14-72901871-INS-1432
chr14-72912284-INS-816
chr14-72912507-INS-2046
chr14-72987558-INS-78
chr14-73012704-INS-131
chr14-73012704-INS-425
chr14-73052513-INS-126
chr14-73059222-INS-66
chr14-73069476-INS-52
chr14-73069662-INS-154
chr14-73096675-INS-62
chr14-73099915-INS-53
chr14-73138736-INS-54
chr14-73164010-INS-91
chr14-73205517-INS-213
chr14-73211840-INS-58
chr14-73214220-INS-56
chr14-73262881-INS-6237
chr14-73300638-INS-3195
chr14-73321632-INS-790
chr14-73321972-INS-174
chr14-73328118-INS-1894
chr14-73342460-INS-1206
chr14-73384328-INS-203
chr14-73392971-INS-189
chr14-73396502-INS-79
chr14-73396827-INS-7153
chr14-73412737-INS-72
chr14-73454969-INS-58
chr14-73480862-INS-172
chr14-73521055-INS-251
chr14-73678968-INS-129
chr14-73679905-INS-53
chr14-73717768-INS-115
chr14-73731402-INS-190
chr14-73731624-INS-160
chr14-73732437-INS-641
chr14-73733176-INS-94
chr14-73742544-INS-52
chr14-73770259-INS-7262
chr14-73809089-INS-72
chr14-73830266-INS-498
chr14-73877616-INS-86
chr14-73935910-INS-3952
chr14-73936614-INS-90
chr14-73936627-INS-309
chr14-73937229-INS-80
chr14-73946943-INS-7501

chr14-74036226-INS-50
chr14-74038161-INS-405
chr14-74126990-INS-177
chr14-74221498-INS-55
chr14-74226312-INS-6307
chr14-74266234-INS-160
chr14-74284712-INS-2102
chr14-74301588-INS-1712
chr14-74313744-INS-198
chr14-74360263-INS-206
chr14-74465081-INS-197
chr14-74465247-INS-72
chr14-74470443-INS-7135
chr14-74477696-INS-212
chr14-74478883-INS-50
chr14-74515589-INS-3669
chr14-74546119-INS-444
chr14-74546558-INS-4797
chr14-74571704-INS-50
chr14-74667449-INS-286
chr14-74763554-INS-6363
chr14-74798622-INS-335
chr14-74799796-INS-6700
chr14-74802244-INS-498
chr14-74807040-INS-5994
chr14-74814394-INS-4569
chr14-74815024-INS-1981
chr14-74839808-INS-50
chr14-74886599-INS-54
chr14-74893933-INS-168
chr14-74895339-INS-277
chr14-74903952-INS-6483
chr14-74915857-INS-1142
chr14-74956268-INS-1294
chr14-74957583-INS-6185
chr14-75045031-INS-468
chr14-75074578-INS-60
chr14-75095781-INS-87
chr14-75168556-INS-910
chr14-75214266-INS-50
chr14-75240230-INS-2961
chr14-75245395-INS-77
chr14-75245555-INS-6773
chr14-75247292-INS-2704
chr14-75262233-INS-5716
chr14-75283652-INS-100
chr14-75284166-INS-50
chr14-75331202-INS-7053
chr14-75396324-INS-221
chr14-75400615-INS-66
chr14-75438001-INS-70
chr14-75442969-INS-164
chr14-75444818-INS-124
chr14-75462568-INS-88
chr14-75492166-INS-6439
chr14-75494542-INS-118
chr14-75520298-INS-241
chr14-75602234-INS-64
chr14-75614982-INS-52
chr14-75627374-INS-52
chr14-75653420-INS-4510
chr14-75656749-INS-56
chr14-75679681-INS-53
chr14-75682603-INS-159
chr14-75704214-INS-1499
chr14-75704481-INS-96
chr14-75706516-INS-140
chr14-75735439-INS-6473
chr14-75762475-INS-55
chr14-75824461-INS-87
chr14-75845687-INS-349
chr14-75869301-INS-3418
chr14-75893352-INS-52
chr14-75896457-INS-374
chr14-75911543-INS-204
chr14-75911663-INS-68
chr14-75938165-INS-8024

chr14-76020801-INS-126
chr14-76116629-INS-249
chr14-76122211-INS-114
chr14-76125504-INS-5047
chr14-76146989-INS-60
chr14-76147065-INS-440
chr14-76147215-INS-210
chr14-76206977-INS-402
chr14-76283998-INS-114
chr14-76320816-INS-90
chr14-76377558-INS-56
chr14-76397003-INS-220
chr14-76397006-INS-96
chr14-76430253-INS-74
chr14-76430840-INS-338
chr14-76431675-INS-103
chr14-76438925-INS-50
chr14-76452620-INS-64
chr14-76452761-INS-68
chr14-76510837-INS-165
chr14-76523033-INS-117
chr14-76526809-INS-117
chr14-76530000-INS-554
chr14-76530205-INS-82
chr14-76558617-INS-106
chr14-76620015-INS-586
chr14-76631335-INS-64
chr14-76632156-INS-50
chr14-76826246-INS-2519
chr14-76880711-INS-1096
chr14-76901594-INS-136
chr14-76972879-INS-52
chr14-77196031-INS-5003
chr14-77199445-INS-185
chr14-77211106-INS-468
chr14-77254589-INS-184
chr14-77268501-INS-1484
chr14-77273467-INS-52
chr14-77314670-INS-208
chr14-77333150-INS-6286
chr14-77361386-INS-222
chr14-77377133-INS-89
chr14-77457397-INS-82
chr14-77533471-INS-565
chr14-77556448-INS-94
chr14-77556982-INS-54
chr14-77563852-INS-1073
chr14-77598804-INS-51
chr14-77681597-INS-186
chr14-78005032-INS-52
chr14-78171447-INS-2586
chr14-78173590-INS-217
chr14-78251665-INS-277
chr14-78283113-INS-341
chr14-78317634-INS-2067
chr14-78345119-INS-108
chr14-78388445-INS-6710
chr14-78465018-INS-99
chr14-78472077-INS-1215
chr14-78491132-INS-2280
chr14-78518051-INS-161
chr14-78519417-INS-4783
chr14-78532689-INS-56
chr14-78541499-INS-54
chr14-78603436-INS-6408
chr14-78661638-INS-6843
chr14-78710996-INS-50
chr14-78787172-INS-62
chr14-78832234-INS-193
chr14-78894337-INS-1181
chr14-78915070-INS-99
chr14-78946543-INS-674
chr14-78947925-INS-219
chr14-78954479-INS-50
chr14-78999202-INS-317
chr14-79066014-INS-150
chr14-79097564-INS-7113

chr14-79132713-INS-420
chr14-79224282-INS-50
chr14-79232955-INS-336
chr14-79295979-INS-211
chr14-79459650-INS-6204
chr14-79524428-INS-240
chr14-79547717-INS-51
chr14-79558017-INS-50
chr14-79615712-INS-55
chr14-79653023-INS-1494
chr14-79747294-INS-103
chr14-79830732-INS-265
chr14-79833009-INS-152
chr14-79968462-INS-135
chr14-80030581-INS-262
chr14-80061985-INS-804
chr14-80097593-INS-73
chr14-80101212-INS-50
chr14-80106708-INS-124
chr14-80124320-INS-66
chr14-80137375-INS-500
chr14-80169912-INS-50
chr14-80228890-INS-100
chr14-80257981-INS-169
chr14-80264186-INS-104
chr14-80316405-INS-1482
chr14-80367875-INS-64
chr14-80478365-INS-2107
chr14-80540384-INS-6681
chr14-80652504-INS-210
chr14-80685607-INS-145
chr14-80790293-INS-122
chr14-80919384-INS-161
chr14-81037622-INS-116
chr14-81063905-INS-6804
chr14-81102394-INS-1592
chr14-81118954-INS-179
chr14-81246582-INS-78
chr14-81250606-INS-1295
chr14-81394304-INS-72
chr14-81420395-INS-205
chr14-81449093-INS-53
chr14-81449738-INS-321
chr14-81453685-INS-82
chr14-81517734-INS-359
chr14-81585523-INS-6795
chr14-81647051-INS-60
chr14-81691116-INS-218
chr14-81705212-INS-119
chr14-81776215-INS-6003
chr14-81929516-INS-4153
chr14-81963254-INS-3771
chr14-81964437-INS-72
chr14-81973112-INS-240
chr14-81991703-INS-167
chr14-81994323-INS-6176
chr14-82011945-INS-86
chr14-82044592-INS-82
chr14-82067415-INS-87
chr14-82067417-INS-175
chr14-82071387-INS-576
chr14-82184217-INS-5105
chr14-82213127-INS-6599
chr14-82225657-INS-75
chr14-82228171-INS-99
chr14-82231350-INS-6467
chr14-82240074-INS-98
chr14-82261202-INS-3980
chr14-82281880-INS-167
chr14-82283603-INS-50
chr14-82286138-INS-93
chr14-82291582-INS-2503
chr14-82300123-INS-5572
chr14-82338874-INS-2505
chr14-82495126-INS-214
chr14-82547628-INS-122
chr14-82579933-INS-6151

chr14-82614028-INS-4919
chr14-82624458-INS-83
chr14-82624716-INS-248
chr14-82727756-INS-56
chr14-82828332-INS-3409
chr14-82866118-INS-9902
chr14-82955589-INS-11442
chr14-82960806-INS-82
chr14-82997957-INS-7116
chr14-83004950-INS-200
chr14-83004950-INS-56
chr14-83057537-INS-2524
chr14-83121274-INS-60
chr14-83123419-INS-105
chr14-83123905-INS-225
chr14-83203090-INS-1310
chr14-83294759-INS-88
chr14-83295359-INS-6432
chr14-83338894-INS-166
chr14-83392624-INS-59
chr14-83490302-INS-1999
chr14-83505164-INS-607
chr14-83557095-INS-50
chr14-83565607-INS-76
chr14-83566574-INS-50
chr14-83615180-INS-58
chr14-83618873-INS-429
chr14-83674387-INS-1414
chr14-83684287-INS-1079
chr14-83685871-INS-11997
chr14-83685871-INS-4932
chr14-83697128-INS-8623
chr14-83705783-INS-4700
chr14-83821817-INS-32731
chr14-83854917-INS-3832
chr14-83856334-INS-729
chr14-83879897-INS-53
chr14-83886099-INS-1495
chr14-83890329-INS-141
chr14-83942144-INS-1319
chr14-83960771-INS-60
chr14-84036370-INS-133
chr14-84041478-INS-1931
chr14-84124724-INS-78
chr14-84155935-INS-8921
chr14-84177778-INS-6471
chr14-84225883-INS-374
chr14-84226620-INS-492
chr14-84249677-INS-1833
chr14-84270696-INS-4154
chr14-84351719-INS-1001
chr14-84354059-INS-7521
chr14-84358350-INS-2585
chr14-84375062-INS-6221
chr14-84410738-INS-50
chr14-84411752-INS-52
chr14-84420954-INS-7026
chr14-84459385-INS-63
chr14-84468243-INS-76
chr14-84503049-INS-85
chr14-84513163-INS-6872
chr14-84560018-INS-50
chr14-84565683-INS-6586
chr14-84568983-INS-215
chr14-84584532-INS-1751
chr14-84663668-INS-72
chr14-84672681-INS-235
chr14-84751464-INS-222
chr14-84791994-INS-224
chr14-84799103-INS-2525
chr14-84820298-INS-6438
chr14-84878600-INS-50
chr14-84880090-INS-6215
chr14-84964460-INS-605
chr14-84975430-INS-73
chr14-84975532-INS-414
chr14-84976507-INS-78

chr14-84983114-INS-176
chr14-84983165-INS-433
chr14-84983212-INS-2463
chr14-84992123-INS-987
chr14-85049409-INS-296
chr14-85049495-INS-132
chr14-85143255-INS-4405
chr14-85143744-INS-1971
chr14-85185027-INS-353
chr14-85330356-INS-52
chr14-85331756-INS-80
chr14-85425564-INS-107
chr14-85432487-INS-59
chr14-85451635-INS-557
chr14-85519146-INS-1716
chr14-85534169-INS-362
chr14-85534453-INS-370
chr14-85654003-INS-779
chr14-85774402-INS-54
chr14-85812303-INS-934
chr14-85842853-INS-6667
chr14-85959810-INS-88
chr14-85997145-INS-140
chr14-86209964-INS-50
chr14-86280467-INS-683
chr14-86287159-INS-61
chr14-86289125-INS-75
chr14-86347748-INS-1297
chr14-86348825-INS-3856
chr14-86518865-INS-7052
chr14-86560518-INS-54
chr14-86579602-INS-3101
chr14-86599309-INS-7770
chr14-86627992-INS-126
chr14-86679056-INS-1778
chr14-86716496-INS-6180
chr14-86727312-INS-1500
chr14-86755498-INS-2190
chr14-86781020-INS-129
chr14-86807470-INS-224
chr14-86807471-INS-56
chr14-86807728-INS-183
chr14-87033820-INS-7517
chr14-87061717-INS-50
chr14-87062175-INS-92
chr14-87078712-INS-6479
chr14-87166503-INS-1820
chr14-87166503-INS-5012
chr14-87200156-INS-2972
chr14-87215995-INS-236
chr14-87305498-INS-160
chr14-87319241-INS-83
chr14-87351648-INS-6512
chr14-87361180-INS-2597
chr14-87423732-INS-186
chr14-87441205-INS-6317
chr14-87478135-INS-6718
chr14-87487637-INS-330
chr14-87531838-INS-368
chr14-87540881-INS-7117
chr14-87562216-INS-197
chr14-87576694-INS-86
chr14-87604385-INS-106
chr14-87605929-INS-56
chr14-87606219-INS-461
chr14-87677276-INS-53
chr14-87677630-INS-367
chr14-87690403-INS-66
chr14-87700576-INS-70
chr14-87707688-INS-66
chr14-87708483-INS-210
chr14-87742204-INS-108
chr14-87817668-INS-6902
chr14-87828082-INS-1940
chr14-87860140-INS-771
chr14-87870483-INS-6482
chr14-87970898-INS-1016

chr14-87975120-INS-5149
chr14-87983055-INS-7089
chr14-88073038-INS-138
chr14-88086261-INS-50
chr14-88123350-INS-206
chr14-88131469-INS-6359
chr14-88139267-INS-4041
chr14-88163455-INS-668
chr14-88311710-INS-1746
chr14-88340370-INS-58
chr14-88343933-INS-629
chr14-88346697-INS-6514
chr14-88365520-INS-52
chr14-88371470-INS-50
chr14-88378077-INS-6269
chr14-88384827-INS-50
chr14-88399364-INS-1000
chr14-88413376-INS-157
chr14-88442339-INS-6181
chr14-88479458-INS-51
chr14-88482301-INS-1168
chr14-88509026-INS-89
chr14-88528006-INS-1254
chr14-88590524-INS-54
chr14-88661620-INS-6294
chr14-88678733-INS-2997
chr14-88683576-INS-54
chr14-88826362-INS-178
chr14-88847266-INS-84
chr14-88935173-INS-6426
chr14-88940275-INS-605
chr14-88942457-INS-3746
chr14-88997786-INS-1405
chr14-89019943-INS-290
chr14-89022940-INS-51
chr14-89032788-INS-1518
chr14-89067512-INS-3211
chr14-89076667-INS-6266
chr14-89129627-INS-1320
chr14-89158060-INS-70
chr14-89206470-INS-808
chr14-89211312-INS-982
chr14-89211740-INS-109
chr14-89234215-INS-7539
chr14-89252322-INS-1071
chr14-89348738-INS-72
chr14-89363601-INS-51
chr14-89404231-INS-1006
chr14-89516583-INS-3270
chr14-89527650-INS-8341
chr14-89534056-INS-2304
chr14-89553658-INS-235
chr14-89663449-INS-4919
chr14-89664561-INS-50
chr14-89665054-INS-6445
chr14-89766652-INS-78
chr14-89808501-INS-82
chr14-89926473-INS-3013
chr14-89947365-INS-7492
chr14-89953482-INS-359
chr14-89956987-INS-1241
chr14-89995892-INS-510
chr14-90121153-INS-157
chr14-90121183-INS-56
chr14-90143343-INS-301
chr14-90186919-INS-352
chr14-90194494-INS-75
chr14-90195193-INS-66
chr14-90225213-INS-4161
chr14-90273233-INS-223
chr14-90339404-INS-132
chr14-90348868-INS-500
chr14-90374944-INS-6482
chr14-90385838-INS-51
chr14-90394995-INS-50
chr14-90404409-INS-6192
chr14-90446894-INS-1055

chr14-90463585-INS-862
chr14-90569799-INS-54
chr14-90592911-INS-66
chr14-90614480-INS-50
chr14-90641918-INS-211
chr14-90661617-INS-5370
chr14-90665609-INS-53
chr14-90693022-INS-1140
chr14-90702389-INS-59
chr14-90705777-INS-1242
chr14-90724390-INS-4658
chr14-90737058-INS-200
chr14-90737121-INS-1357
chr14-90737567-INS-153
chr14-90769662-INS-50
chr14-90857632-INS-1936
chr14-90899214-INS-51
chr14-90978439-INS-2664
chr14-90979772-INS-1575
chr14-90998146-INS-2647
chr14-91036825-INS-3999
chr14-91085354-INS-52
chr14-91163545-INS-1648
chr14-91181266-INS-1894
chr14-91237246-INS-50
chr14-91279990-INS-57
chr14-91339199-INS-500
chr14-91362461-INS-1161
chr14-91362470-INS-225
chr14-91466066-INS-7101
chr14-91472337-INS-3213
chr14-91483333-INS-1329
chr14-91499102-INS-50
chr14-91558518-INS-3239
chr14-91565614-INS-6386
chr14-91599030-INS-374
chr14-91703580-INS-6786
chr14-91713668-INS-414
chr14-91714411-INS-118
chr14-91743932-INS-56
chr14-91872766-INS-50
chr14-91882305-INS-1899
chr14-91885915-INS-139
chr14-91934071-INS-1887
chr14-91945589-INS-4423
chr14-91980986-INS-76
chr14-92001451-INS-640
chr14-92052907-INS-78
chr14-92142049-INS-60
chr14-92159320-INS-50
chr14-92187470-INS-1430
chr14-92220104-INS-55
chr14-92248802-INS-1160
chr14-92257412-INS-50
chr14-92319468-INS-3156
chr14-92384049-INS-1202
chr14-92392824-INS-72
chr14-92408053-INS-6321
chr14-92419735-INS-57
chr14-92458805-INS-3194
chr14-92639214-INS-61
chr14-92680507-INS-2789
chr14-92692237-INS-367
chr14-92693657-INS-78
chr14-92696830-INS-6035
chr14-92746137-INS-67
chr14-92755085-INS-146
chr14-92802934-INS-6587
chr14-92865546-INS-3221
chr14-92875254-INS-6227
chr14-92930422-INS-6608
chr14-92946967-INS-6442
chr14-92969421-INS-9038
chr14-93016871-INS-2950
chr14-93029987-INS-1581
chr14-93030080-INS-64
chr14-93088844-INS-3491

chr14-93106441-INS-5310
chr14-93120919-INS-2900
chr14-93129751-INS-51
chr14-93164899-INS-994
chr14-93185374-INS-1439
chr14-93204081-INS-6403
chr14-93320943-INS-226
chr14-93359614-INS-290
chr14-93387669-INS-334
chr14-93514604-INS-344
chr14-93515791-INS-56
chr14-93516648-INS-136
chr14-93527010-INS-50
chr14-93589942-INS-76
chr14-93626504-INS-1579
chr14-93627276-INS-78
chr14-93647566-INS-295
chr14-93686795-INS-463
chr14-93730294-INS-50
chr14-93814990-INS-54
chr14-93845656-INS-6560
chr14-93998320-INS-56
chr14-94014994-INS-1122
chr14-94049579-INS-166
chr14-94158421-INS-1702
chr14-94190460-INS-50
chr14-94352436-INS-455
chr14-94352712-INS-1453
chr14-94352816-INS-162
chr14-94364431-INS-77
chr14-94375460-INS-6270
chr14-94388194-INS-99
chr14-94409672-INS-317
chr14-94439677-INS-1202
chr14-94458197-INS-50
chr14-94464777-INS-6332
chr14-94490182-INS-6045
chr14-94494477-INS-1686
chr14-94531361-INS-60
chr14-94533211-INS-7155
chr14-94543725-INS-8676
chr14-94603391-INS-2493
chr14-94604841-INS-130
chr14-94604841-INS-280
chr14-94609591-INS-120
chr14-94610345-INS-60
chr14-94702464-INS-2213
chr14-94781723-INS-6485
chr14-94829919-INS-50
chr14-94839008-INS-398
chr14-94875653-INS-6864
chr14-94934734-INS-222
chr14-94995229-INS-3393
chr14-95005811-INS-4860
chr14-95008269-INS-1648
chr14-95025941-INS-4465
chr14-95053145-INS-1927
chr14-95055772-INS-2015
chr14-95060994-INS-1834
chr14-95061311-INS-66
chr14-95120992-INS-160
chr14-95172874-INS-282
chr14-95173092-INS-1446
chr14-95225557-INS-96
chr14-95360603-INS-200
chr14-95360844-INS-163
chr14-95506924-INS-244
chr14-95532115-INS-6579
chr14-95549766-INS-794
chr14-95566682-INS-2982
chr14-95588406-INS-63
chr14-95638193-INS-54
chr14-95662890-INS-2050
chr14-95667848-INS-313
chr14-95690426-INS-79
chr14-95719972-INS-6584
chr14-95742648-INS-2888

chr14-95824533-INS-53
chr14-95835659-INS-6315
chr14-95851880-INS-4997
chr14-95939145-INS-6745
chr14-95945980-INS-1514
chr14-95955004-INS-58
chr14-95966252-INS-3709
chr14-95971813-INS-98
chr14-96170241-INS-4516
chr14-96244585-INS-51
chr14-96251332-INS-50
chr14-96304297-INS-3685
chr14-96317662-INS-65
chr14-96386864-INS-3241
chr14-96397419-INS-6294
chr14-96411266-INS-3470
chr14-96418715-INS-1629
chr14-96418715-INS-3346
chr14-96421123-INS-1690
chr14-96443606-INS-66
chr14-96462070-INS-116
chr14-96490009-INS-390
chr14-96508675-INS-1619
chr14-96551867-INS-4526
chr14-96572810-INS-50
chr14-96676460-INS-211
chr14-96691838-INS-54
chr14-96749070-INS-6153
chr14-96783437-INS-1195
chr14-96846346-INS-230
chr14-96866539-INS-72
chr14-96877626-INS-6453
chr14-96880233-INS-1509
chr14-96894339-INS-83
chr14-97004792-INS-7403
chr14-97007763-INS-3540
chr14-97132580-INS-1760
chr14-97143318-INS-60
chr14-97144040-INS-516
chr14-97189181-INS-2725
chr14-97326222-INS-59
chr14-97526079-INS-208
chr14-97528787-INS-54
chr14-97529700-INS-54
chr14-97599698-INS-6162
chr14-97631335-INS-114
chr14-97736131-INS-589
chr14-97736238-INS-159
chr14-97762080-INS-508
chr14-97762891-INS-668
chr14-97881754-INS-879
chr14-98055537-INS-1014
chr14-98119198-INS-91
chr14-98411770-INS-1452
chr14-98413992-INS-2995
chr14-98429473-INS-1433
chr14-98438434-INS-61
chr14-98491574-INS-3284
chr14-98502784-INS-2681
chr14-98560302-INS-2341
chr14-98565065-INS-761
chr14-98567980-INS-131
chr14-98577525-INS-2593
chr14-98589272-INS-2875
chr14-98601123-INS-1713
chr14-98610622-INS-66
chr14-98612638-INS-7139
chr14-98612844-INS-50
chr14-98666439-INS-1720
chr14-98714192-INS-93
chr14-98756689-INS-52
chr14-98781601-INS-160
chr14-98785342-INS-324
chr14-98805137-INS-7101
chr14-98815869-INS-74
chr14-98815917-INS-1640
chr14-98860178-INS-50

chr14-98866708-INS-1498
chr14-98914538-INS-544
chr14-98916023-INS-717
chr14-98958309-INS-442
chr14-99052236-INS-110
chr14-99204775-INS-219
chr14-99226287-INS-110
chr14-99264035-INS-2938
chr14-99293556-INS-140
chr14-99293786-INS-488
chr14-99297733-INS-120
chr14-99299999-INS-1465
chr14-99307943-INS-330
chr14-99308097-INS-146
chr14-99453330-INS-50
chr14-99454010-INS-1378
chr14-99469961-INS-93
chr14-99493291-INS-113
chr14-99502745-INS-98
chr14-99510084-INS-181
chr14-99525688-INS-77
chr14-99559283-INS-202
chr14-99573601-INS-370
chr14-99609208-INS-52
chr14-99690915-INS-453
chr14-99695962-INS-50
chr14-99699243-INS-466
chr14-99705581-INS-316
chr14-99705701-INS-58
chr14-99757585-INS-91
chr14-99784696-INS-234
chr14-99861061-INS-50
chr14-99866801-INS-62
chr14-99916257-INS-4010
chr14-99941552-INS-6401
chr14-99950911-INS-6663
chr14-99956958-INS-52
chr14-99997679-INS-69
chr14-100001695-INS-6378
chr14-100028651-INS-52
chr14-100033537-INS-96
chr14-100036372-INS-265
chr14-100036372-INS-531
chr14-100059514-INS-8173
chr14-100067382-INS-5080
chr14-100154161-INS-367
chr14-100197397-INS-322
chr14-100207215-INS-355
chr14-100208308-INS-50
chr14-100348572-INS-113
chr14-100349810-INS-3061
chr14-100360683-INS-195
chr14-100426047-INS-84
chr14-100469373-INS-65
chr14-100479617-INS-55
chr14-100514008-INS-105
chr14-100534096-INS-63
chr14-100567126-INS-50
chr14-100648161-INS-2069
chr14-100651722-INS-1759
chr14-100674554-INS-2145
chr14-100677776-INS-56
chr14-100769611-INS-154
chr14-100809606-INS-195
chr14-100822976-INS-6485
chr14-100827978-INS-77
chr14-100865617-INS-403
chr14-100906510-INS-301
chr14-100907284-INS-106
chr14-100974854-INS-208
chr14-100986214-INS-236
chr14-100986466-INS-314
chr14-101028816-INS-132
chr14-101042187-INS-181
chr14-101168432-INS-8457
chr14-101169003-INS-717
chr14-101176387-INS-222

chr14-101214548-INS-6466
chr14-101222223-INS-98
chr14-101225987-INS-7102
chr14-101262437-INS-6838
chr14-101272157-INS-2703
chr14-101372208-INS-52
chr14-101441477-INS-1562
chr14-101448548-INS-500
chr14-101449108-INS-403
chr14-101455573-INS-458
chr14-101455599-INS-171
chr14-101455695-INS-203
chr14-101516122-INS-608
chr14-101571942-INS-386
chr14-101572175-INS-374
chr14-101572289-INS-58
chr14-101593389-INS-231
chr14-101670508-INS-836
chr14-101697056-INS-84
chr14-101704617-INS-56
chr14-101723240-INS-63
chr14-101732734-INS-1954
chr14-101750235-INS-177
chr14-101770936-INS-184
chr14-101789846-INS-268
chr14-101794827-INS-7253
chr14-101806870-INS-177
chr14-101812529-INS-147
chr14-101823789-INS-1500
chr14-101852227-INS-1430
chr14-101928644-INS-52
chr14-101945694-INS-50
chr14-101948147-INS-262
chr14-102056485-INS-2881
chr14-102093065-INS-1899
chr14-102177293-INS-6937
chr14-102187039-INS-52
chr14-102190940-INS-220
chr14-102190940-INS-648
chr14-102190943-INS-82
chr14-102194178-INS-451
chr14-102230347-INS-2369
chr14-102235732-INS-448
chr14-102312467-INS-6210
chr14-102314732-INS-225
chr14-102322334-INS-337
chr14-102322561-INS-52
chr14-102342793-INS-6326
chr14-102350919-INS-4812
chr14-102390777-INS-929
chr14-102391594-INS-1328
chr14-102438864-INS-206
chr14-102527016-INS-6167
chr14-102580768-INS-154
chr14-102596230-INS-50
chr14-102645755-INS-6348
chr14-102656686-INS-1981
chr14-102713672-INS-7176
chr14-102736107-INS-51
chr14-102771551-INS-206
chr14-102772113-INS-174
chr14-102884489-INS-625
chr14-102931968-INS-60
chr14-102965446-INS-315
chr14-103022452-INS-2161
chr14-103048871-INS-50
chr14-103061511-INS-130
chr14-103103102-INS-500
chr14-103128111-INS-78
chr14-103192099-INS-89
chr14-103226040-INS-161
chr14-103248705-INS-54
chr14-103248916-INS-62
chr14-103264424-INS-54
chr14-103283152-INS-276
chr14-103283152-INS-732
chr14-103310003-INS-6431

chr14-103344117-INS-93
chr14-103344512-INS-51
chr14-103390264-INS-1391
chr14-103499113-INS-60
chr14-103499363-INS-60
chr14-103570558-INS-5603
chr14-103575387-INS-816
chr14-103575482-INS-7560
chr14-103586669-INS-100
chr14-103589382-INS-2788
chr14-103687304-INS-3414
chr14-103701956-INS-159
chr14-103743143-INS-470
chr14-103787247-INS-54
chr14-103787677-INS-292
chr14-103787893-INS-6385
chr14-103823794-INS-100
chr14-103853252-INS-7158
chr14-103873208-INS-81
chr14-103911953-INS-1659
chr14-103914656-INS-1148
chr14-103915001-INS-3313
chr14-103950473-INS-350
chr14-104009050-INS-95
chr14-104029466-INS-108
chr14-104160344-INS-209
chr14-104200637-INS-200
chr14-104228541-INS-270
chr14-104239663-INS-455
chr14-104241336-INS-62
chr14-104308963-INS-212
chr14-104404662-INS-6315
chr14-104405362-INS-198
chr14-104414629-INS-365
chr14-104484701-INS-53
chr14-104508808-INS-66
chr14-104549033-INS-88
chr14-104558736-INS-6136
chr14-104564066-INS-54
chr14-104601957-INS-402
chr14-104606010-INS-161
chr14-104611420-INS-50
chr14-104674589-INS-111
chr14-104691389-INS-54
chr14-104730820-INS-115
chr14-104730820-INS-263
chr14-104753320-INS-50
chr14-104784536-INS-51
chr14-104812950-INS-52
chr14-104832262-INS-56
chr14-104885222-INS-246
chr14-104934524-INS-607
chr14-105089498-INS-85
chr14-105089823-INS-367
chr14-105090980-INS-108
chr14-105110392-INS-74
chr14-105133276-INS-84
chr14-105179786-INS-1374
chr14-105223262-INS-152
chr14-105249543-INS-4879
chr14-105290917-INS-177
chr14-105317950-INS-61
chr14-105341418-INS-64
chr14-105342106-INS-3332
chr14-105460779-INS-516
chr14-105484890-INS-182
chr14-105502399-INS-499
chr14-105533955-INS-75
chr14-105553014-INS-231
chr14-105612976-INS-142
chr14-105630114-INS-1500
chr14-105659288-INS-1718
chr14-105670866-INS-63
chr14-105714317-INS-75
chr14-105715554-INS-106
chr14-105737672-INS-634
chr14-105752626-INS-50

chr14-105782551-INS-54
chr14-105876901-INS-288
chr14-105891199-INS-65
chr14-105934182-INS-6603
chr14-105975185-INS-240
chr14-105981270-INS-54
chr14-105987416-INS-200
chr14-106054357-INS-56
chr14-106063900-INS-7104
chr14-106117236-INS-5852
chr14-106117781-INS-192
chr14-106286707-INS-201
chr14-106320659-INS-1049
chr14-106322613-INS-11077
chr14-106419325-INS-209
chr14-106422776-INS-52
chr14-106450416-INS-126
chr14-106676733-INS-6052
chr14-106804018-INS-2108
chr14-106843309-INS-138
chr14-106863829-INS-56
chr14-106863841-INS-304
chr14-106865187-INS-480
chr14-106913202-INS-52
chr14-106920651-INS-50
chr14-106933778-INS-1007
chr14-106955781-INS-52
chr14-107066195-INS-101
chr14-107130558-INS-250
chr14-107185265-INS-1631
chr14-107217923-INS-1847
chr14-107218501-INS-3465
chr14-107312799-INS-97
chr14-107336814-INS-2774
chr14-107372461-INS-6424
chr14-107418680-INS-88
chr14-107440039-INS-52
chr14-107492179-INS-3936
chr14-107492867-INS-114
chr14-107520312-INS-7858
chr14-107543095-INS-274
chr14-107601070-INS-591
chr14-107657040-INS-208
chr14-107660961-INS-2657
chr14-107684475-INS-6258
chr14-107748319-INS-50
chr14-107779968-INS-60
chr14-107792676-INS-487
chr14-107794429-INS-352
chr14-107794474-INS-533
chr14-107799885-INS-219
chr14-107800600-INS-55
chr14-107809427-INS-6605
chr14-107877032-INS-57
chr14-107893223-INS-126
chr14-107904627-INS-6240
chr14-107989374-INS-50
chr14-108027323-INS-201
chr14-108027926-INS-263
chr14-108112143-INS-1290
chr14-108114319-INS-1029
chr14-108116118-INS-1259
chr14-108123707-INS-4555
chr14-108125493-INS-1389
chr14-108159270-INS-51
chr14-108241188-INS-176
chr14-108248553-INS-5520
chr14-108347735-INS-120
chr14-108371321-INS-1680
chr14-108375259-INS-1065
chr14-108381061-INS-60
chr14-108435104-INS-56
chr14-108560859-INS-211
chr14-108609710-INS-124
chr14-108610091-INS-389
chr14-108663743-INS-405
chr14-108719464-INS-10272

chr14-108751726-INS-2123
chr14-108753357-INS-108
chr14-108754848-INS-52
chr14-108770440-INS-2905
chr14-108830386-INS-2245
chr14-108879583-INS-1045
chr14-108979249-INS-402
chr14-109009372-INS-1135
chr14-109021120-INS-124
chr14-109044280-INS-7218
chr14-109061585-INS-234
chr14-109081501-INS-52
chr14-109159913-INS-51
chr14-109183801-INS-81
chr14-109194510-INS-51
chr14-109214912-INS-7058
chr14-109226512-INS-60
chr14-109278416-INS-51
chr14-109305469-INS-658
chr14-109305659-INS-256
chr14-109316078-INS-6403
chr14-109329143-INS-62
chr14-109330848-INS-114
chr14-109339604-INS-1589
chr14-109353751-INS-4426
chr14-109362671-INS-1279
chr14-109370686-INS-450
chr14-109381139-INS-57
chr14-109390752-INS-6190
chr14-109438713-INS-6159
chr14-109444558-INS-262
chr14-109462226-INS-2922
chr14-109502676-INS-5216
chr14-109510296-INS-54
chr14-109542795-INS-186
chr14-109579325-INS-928
chr14-109609164-INS-3243
chr14-109651634-INS-93
chr14-109669212-INS-59
chr14-109690890-INS-70
chr14-109715250-INS-413
chr14-109716145-INS-460
chr14-109721073-INS-52
chr14-109727868-INS-3247
chr14-109728185-INS-4353
chr14-109731422-INS-76
chr14-109740739-INS-801
chr14-109838260-INS-52
chr14-109838654-INS-62
chr14-109857104-INS-1895
chr14-109913930-INS-100
chr14-109915466-INS-124
chr14-109922051-INS-4710
chr14-109923569-INS-747
chr14-109923720-INS-230
chr14-109930877-INS-70
chr14-109934740-INS-271
chr14-109948814-INS-2489
chr14-109958682-INS-7174
chr14-109999237-INS-519
chr14-110002454-INS-1548
chr14-110006114-INS-7053
chr14-110014300-INS-4160
chr14-110021457-INS-52
chr14-110022547-INS-51
chr14-110056248-INS-89
chr14-110083255-INS-308
chr14-110106326-INS-331
chr14-110220412-INS-766
chr14-110243004-INS-57
chr14-110248107-INS-50
chr14-110276421-INS-1165
chr14-110356295-INS-332
chr14-110363104-INS-138
chr14-110387345-INS-112
chr14-110414765-INS-2867
chr14-110418739-INS-50

chr14-110427623-INS-6401
chr14-110435564-INS-1165
chr14-110435564-INS-2945
chr14-110445300-INS-50
chr14-110457655-INS-6391
chr14-110475923-INS-601
chr14-110502043-INS-2478
chr14-110515047-INS-3504
chr14-110519693-INS-6468
chr14-110537194-INS-886
chr14-110545194-INS-96
chr14-110572145-INS-2119
chr14-110585101-INS-3200
chr14-110670027-INS-2263
chr14-110670055-INS-123
chr14-110718704-INS-1225
chr14-110766295-INS-110
chr14-110800461-INS-197
chr14-110814934-INS-1072
chr14-110820714-INS-108
chr14-110820882-INS-395
chr14-110821013-INS-194
chr14-110903703-INS-53
chr14-110923035-INS-52
chr14-110976510-INS-103
chr14-110995965-INS-1272
chr14-111002679-INS-313
chr14-111022188-INS-62
chr14-111134428-INS-401
chr14-111142095-INS-214
chr14-111213508-INS-2037
chr14-111220053-INS-278
chr14-111261800-INS-6999
chr14-111268654-INS-7979
chr14-111268729-INS-107
chr14-111269173-INS-250
chr14-111287264-INS-2687
chr14-111317481-INS-413
chr14-111320876-INS-167
chr14-111341367-INS-8783
chr14-111346012-INS-364
chr14-111352387-INS-214
chr14-111507124-INS-118
chr14-111526233-INS-210
chr14-111541445-INS-6402
chr14-111546443-INS-68
chr14-111546599-INS-102
chr14-111546827-INS-53
chr14-111640004-INS-299
chr14-111672403-INS-50
chr14-111681234-INS-408
chr14-111681992-INS-242
chr14-111690988-INS-6559
chr14-111702761-INS-72
chr14-111704536-INS-50
chr14-111719198-INS-50
chr14-111806196-INS-4430
chr14-111830849-INS-50
chr14-111879876-INS-13875
chr14-111879876-INS-6205
chr14-111979872-INS-59
chr14-111996679-INS-113
chr14-112008952-INS-63
chr14-112010995-INS-1988
chr14-112024615-INS-76
chr14-112056034-INS-193
chr14-112108439-INS-197
chr14-112118291-INS-6258
chr14-112129499-INS-72
chr14-112139511-INS-632
chr14-112145232-INS-4048
chr14-112174866-INS-50
chr14-112202981-INS-58
chr14-112204567-INS-6986
chr14-112207331-INS-67
chr14-112250539-INS-581
chr14-112268550-INS-6204

chr14-112278031-INS-2925
chr14-112278499-INS-68
chr14-112281619-INS-200
chr14-112312147-INS-1503
chr14-112316760-INS-110
chr14-112320657-INS-5024
chr14-112320969-INS-1401
chr14-112331721-INS-6605
chr14-112336920-INS-1480
chr14-112353437-INS-3041
chr14-112381542-INS-56
chr14-112386582-INS-50
chr14-112409876-INS-72
chr14-112421756-INS-147
chr14-112440857-INS-116
chr14-112494812-INS-2842
chr14-112519850-INS-135
chr14-112591766-INS-147
chr14-112615584-INS-52
chr14-112616893-INS-1156
chr14-112712934-INS-79
chr14-112732331-INS-60
chr14-112752684-INS-51
chr14-112757769-INS-2322
chr14-112777806-INS-50
chr14-112789016-INS-7192
chr14-112813939-INS-878
chr14-112833065-INS-488
chr14-112833413-INS-562
chr14-112837347-INS-346
chr14-113059140-INS-1471
chr14-113070264-INS-612
chr14-113090320-INS-3351
chr14-113101282-INS-649
chr14-113101528-INS-53
chr14-113116845-INS-114
chr14-113116904-INS-56
chr14-113124668-INS-1195
chr14-113153287-INS-7570
chr14-113194621-INS-2565
chr14-113196744-INS-8630
chr14-113197171-INS-1888
chr14-113271576-INS-56
chr14-113286544-INS-391
chr14-113422143-INS-51
chr14-113471312-INS-6437
chr14-113474767-INS-587
chr14-113507817-INS-207
chr14-113516011-INS-132
chr14-113554563-INS-309
chr14-113555615-INS-448
chr14-113572764-INS-1488
chr14-113594434-INS-445
chr14-113637704-INS-1512
chr14-113662929-INS-52
chr14-113675880-INS-2855
chr14-113685876-INS-821
chr14-113693781-INS-111
chr14-113694432-INS-955
chr14-113710576-INS-167
chr14-113727384-INS-410
chr14-113749925-INS-179
chr14-113760381-INS-5961
chr14-113814752-INS-1570
chr14-113913559-INS-1914
chr14-113938398-INS-2015
chr14-113943961-INS-50
chr14-113983330-INS-1111
chr14-113985365-INS-1752
chr14-113988398-INS-4800
chr14-114060105-INS-126
chr14-114071568-INS-4308
chr14-114092763-INS-2029
chr14-114097627-INS-3951
chr14-114104622-INS-6384
chr14-114157596-INS-66
chr14-114193469-INS-82

chr14-114255755-INS-128
chr14-114257004-INS-78
chr14-114305068-INS-6906
chr14-114311018-INS-325
chr14-114327202-INS-359
chr14-114399125-INS-59
chr14-114408767-INS-1443
chr14-114409373-INS-166
chr14-114507837-INS-417
chr14-114511789-INS-7251
chr14-114548421-INS-84
chr14-114556947-INS-1060
chr14-114608593-INS-979
chr14-114665187-INS-67
chr14-114666111-INS-6481
chr14-114753335-INS-57
chr14-114760407-INS-153
chr14-114760478-INS-311
chr14-114822067-INS-368
chr14-114848782-INS-114
chr14-114858493-INS-1024
chr14-114864933-INS-130
chr14-114922226-INS-65
chr14-114935772-INS-60
chr14-114967950-INS-1890
chr14-114979957-INS-4704
chr14-115022155-INS-65
chr14-115029930-INS-79
chr14-115049944-INS-51
chr14-115050142-INS-82
chr14-115074897-INS-58
chr14-115114425-INS-64
chr14-115118483-INS-52
chr14-115133047-INS-120
chr14-115167678-INS-51
chr14-115178892-INS-96
chr14-115179563-INS-1172
chr14-115182664-INS-51
chr14-115182674-INS-140
chr14-115197370-INS-99
chr14-115215615-INS-869
chr14-115250669-INS-210
chr14-115253957-INS-321
chr14-115355182-INS-115
chr14-115365304-INS-2885
chr14-115378221-INS-1599
chr14-115383797-INS-52
chr14-115393870-INS-50
chr14-115410322-INS-90
chr14-115435598-INS-169
chr14-115459364-INS-52
chr14-115576641-INS-2571
chr14-115581327-INS-173
chr14-115581498-INS-795
chr14-115603180-INS-67
chr14-115620705-INS-8407
chr14-115629430-INS-67
chr14-115630373-INS-57
chr14-115637404-INS-110
chr14-115643269-INS-220
chr14-115665190-INS-86
chr14-115761892-INS-115
chr14-115857589-INS-50
chr14-115869334-INS-55
chr14-115909367-INS-59
chr14-116051387-INS-165
chr14-116051637-INS-56
chr14-116072215-INS-250
chr14-116138128-INS-545
chr14-116175123-INS-50
chr14-116242822-INS-50
chr14-116302813-INS-82
chr14-116311197-INS-295
chr14-116318576-INS-669
chr14-116325451-INS-4893
chr14-116332058-INS-582
chr14-116389521-INS-50

chr14-116421679-INS-108
chr14-116459393-INS-50
chr14-116484455-INS-352
chr14-116500967-INS-275
chr14-116535652-INS-409
chr14-116535799-INS-140
chr14-116589925-INS-6550
chr14-116600042-INS-217
chr14-116622504-INS-53
chr14-116623386-INS-1954
chr14-116657619-INS-58
chr14-116657748-INS-370
chr14-116663042-INS-387
chr14-116680564-INS-54
chr14-116703039-INS-205
chr14-116712969-INS-120
chr14-116721973-INS-1625
chr14-116732472-INS-54
chr14-116738130-INS-100
chr14-116738416-INS-420
chr14-116801623-INS-64
chr14-116821016-INS-74
chr14-116902831-INS-81
chr14-116907682-INS-304
chr14-116909368-INS-160
chr14-116923318-INS-1181
chr14-116933655-INS-316
chr14-116933973-INS-353
chr14-116934185-INS-1379
chr14-116982735-INS-123
chr14-116985972-INS-6549
chr14-117007623-INS-52
chr14-117049745-INS-50
chr14-117052588-INS-429
chr14-117052933-INS-208
chr14-117072977-INS-2170
chr14-117091382-INS-136
chr14-117098266-INS-52
chr14-117141290-INS-1281
chr14-117162088-INS-138
chr14-117172174-INS-100
chr14-117177497-INS-68
chr14-117177566-INS-66
chr14-117185889-INS-3554
chr14-117199509-INS-311
chr14-117206975-INS-62
chr14-117221807-INS-61
chr14-117228971-INS-82
chr14-117254006-INS-112
chr14-117278241-INS-975
chr14-117322661-INS-242
chr14-117342308-INS-6622
chr14-117359765-INS-209
chr14-117365461-INS-96
chr14-117440172-INS-60
chr14-117460279-INS-88
chr14-117489094-INS-140
chr14-117491124-INS-363
chr14-117506617-INS-240
chr14-117506683-INS-92
chr14-117576080-INS-2149
chr14-117576469-INS-62
chr14-117579012-INS-789
chr14-117588496-INS-1183
chr14-117591538-INS-82
chr14-117592397-INS-104
chr14-117597133-INS-6474
chr14-117606331-INS-50
chr14-117608596-INS-258
chr14-117623824-INS-76
chr14-117667303-INS-278
chr14-117678922-INS-126
chr14-117706470-INS-119
chr14-117732747-INS-685
chr14-117741458-INS-6633
chr14-117765313-INS-120
chr14-117790058-INS-130

chr14-117817892-INS-1214
chr14-117835739-INS-6383
chr14-117855787-INS-56
chr14-117880759-INS-475
chr14-117901891-INS-392
chr14-117958290-INS-138
chr14-117958873-INS-61
chr14-117959719-INS-134
chr14-117976324-INS-223
chr14-117977146-INS-162
chr14-117985816-INS-91
chr14-117997756-INS-194
chr14-118049354-INS-445
chr14-118082376-INS-118
chr14-118142631-INS-55
chr14-118145514-INS-50
chr14-118147590-INS-522
chr14-118158477-INS-56
chr14-118166726-INS-95
chr14-118167275-INS-92
chr14-118178758-INS-1521
chr14-118183159-INS-99
chr14-118234132-INS-223
chr14-118249339-INS-222
chr14-118261174-INS-393
chr14-118262567-INS-1315
chr14-118279105-INS-591
chr14-118296274-INS-104
chr14-118352429-INS-528
chr14-118355962-INS-106
chr14-118356747-INS-130
chr14-118357250-INS-181
chr14-118359806-INS-223
chr14-118389409-INS-315
chr14-118389409-INS-81
chr14-118393674-INS-378
chr14-118436545-INS-148
chr14-118443614-INS-231
chr14-118443682-INS-66
chr14-118445442-INS-842
chr14-118466630-INS-57
chr14-118495257-INS-7079
chr14-118519591-INS-416
chr14-118530809-INS-61
chr14-118530959-INS-406
chr14-118571703-INS-150
chr14-118573420-INS-146
chr14-118578004-INS-72
chr14-118596144-INS-4979
chr14-118598307-INS-52
chr14-118626241-INS-99
chr14-118627320-INS-239
chr14-118667082-INS-253
chr14-118689362-INS-187
chr14-118705983-INS-316
chr14-118709082-INS-150
chr14-118729897-INS-184
chr14-118835736-INS-204
chr14-118844428-INS-165
chr14-118878155-INS-168
chr14-118957971-INS-54
chr14-118958579-INS-210
chr14-118973247-INS-54
chr14-118980867-INS-1904
chr14-118989572-INS-72
chr14-118989680-INS-276
chr14-119006376-INS-153
chr14-119008482-INS-199
chr14-119010101-INS-90
chr14-119014440-INS-3350
chr14-119014809-INS-8796
chr14-119015870-INS-209
chr14-119022252-INS-3074
chr14-119039365-INS-3230
chr14-119039600-INS-245
chr14-119040144-INS-5091
chr14-119053998-INS-103

chr14-119065521-INS-498
chr14-119084791-INS-50
chr14-119118736-INS-667
chr14-119118762-INS-207
chr14-119119091-INS-299
chr14-119141008-INS-203
chr14-119141420-INS-50
chr14-119158861-INS-144
chr14-119180225-INS-291
chr14-119234573-INS-1330
chr14-119254919-INS-84
chr14-119265629-INS-272
chr14-119271461-INS-172
chr14-119271714-INS-52
chr14-119272780-INS-156
chr14-119283595-INS-5261
chr14-119309667-INS-7122
chr14-119323816-INS-50
chr14-119328373-INS-374
chr14-119347567-INS-776
chr14-119365050-INS-83
chr14-119365701-INS-159
chr14-119373273-INS-60
chr14-119407026-INS-82
chr14-119418273-INS-125
chr14-119443168-INS-7180
chr14-119476593-INS-1907
chr14-119495094-INS-58
chr14-119513567-INS-396
chr14-119533361-INS-6591
chr14-119552500-INS-1291
chr14-119571704-INS-52
chr14-119585531-INS-2804
chr14-119669369-INS-308
chr14-119669956-INS-2149
chr14-119696311-INS-6365
chr14-119696558-INS-8644
chr14-119711978-INS-66
chr14-119734564-INS-4119
chr14-119759318-INS-547
chr14-119860426-INS-52
chr14-119862238-INS-6801
chr14-119905504-INS-68
chr14-119905740-INS-68
chr14-119933773-INS-3069
chr14-119955222-INS-50
chr14-119959927-INS-99
chr14-120005526-INS-260
chr14-120010249-INS-6336
chr14-120017376-INS-114
chr14-120041733-INS-64
chr14-120051279-INS-52
chr14-120066649-INS-10695
chr14-120130218-INS-241
chr14-120155867-INS-94
chr14-120158684-INS-4532
chr14-120178070-INS-450
chr14-120183762-INS-190
chr14-120187191-INS-123
chr14-120187625-INS-55
chr14-120201279-INS-221
chr14-120232157-INS-6192
chr14-120265046-INS-50
chr14-120269726-INS-1923
chr14-120272009-INS-50
chr14-120286986-INS-69
chr14-120288108-INS-62
chr14-120323523-INS-100
chr14-120346254-INS-152
chr14-120358081-INS-6262
chr14-120366246-INS-82
chr14-120370934-INS-178
chr14-120380458-INS-6433
chr14-120406078-INS-68
chr14-120450595-INS-3319
chr14-120457393-INS-141
chr14-120546098-INS-134

chr14-120554469-INS-183
chr14-120601693-INS-94
chr14-120722385-INS-2142
chr14-120740136-INS-406
chr14-120740136-INS-71
chr14-120756528-INS-2854
chr14-120762753-INS-3313
chr14-120821745-INS-56
chr14-120876677-INS-1200
chr14-120887624-INS-116
chr14-120889564-INS-74
chr14-120894681-INS-72
chr14-120953706-INS-64
chr14-120975468-INS-1657
chr14-121030297-INS-1477
chr14-121052674-INS-466
chr14-121068116-INS-332
chr14-121068119-INS-795
chr14-121074093-INS-446
chr14-121095442-INS-183
chr14-121121991-INS-64
chr14-121122503-INS-4168
chr14-121130856-INS-215
chr14-121135889-INS-167
chr14-121140944-INS-116
chr14-121194292-INS-2259
chr14-121207488-INS-128
chr14-121210527-INS-79
chr14-121211993-INS-58
chr14-121235760-INS-669
chr14-121235856-INS-72
chr14-121243777-INS-53
chr14-121253271-INS-284
chr14-121253342-INS-101
chr14-121265331-INS-60
chr14-121276353-INS-215
chr14-121278949-INS-458
chr14-121293020-INS-52
chr14-121310975-INS-69
chr14-121341657-INS-432
chr14-121342987-INS-276
chr14-121350597-INS-153
chr14-121350856-INS-67
chr14-121369036-INS-95
chr14-121369055-INS-314
chr14-121422026-INS-60
chr14-121449579-INS-426
chr14-121449897-INS-106
chr14-121461009-INS-178
chr14-121463714-INS-60
chr14-121465392-INS-153
chr14-121471371-INS-1536
chr14-121471442-INS-262
chr14-121555324-INS-277
chr14-121558629-INS-83
chr14-121561785-INS-50
chr14-121623622-INS-3124
chr14-121626289-INS-2419
chr14-121630044-INS-64
chr14-121630532-INS-1383
chr14-121653964-INS-143
chr14-121661572-INS-50
chr14-121662692-INS-619
chr14-121662856-INS-77
chr14-121677866-INS-208
chr14-121685987-INS-102
chr14-121697618-INS-66
chr14-121701381-INS-69
chr14-121724231-INS-224
chr14-121756173-INS-72
chr14-121759784-INS-191
chr14-121761305-INS-114
chr14-121763899-INS-52
chr14-121818166-INS-77
chr14-121876360-INS-70
chr14-121887367-INS-68
chr14-121898940-INS-154

chr14-121926797-INS-94
chr14-121952301-INS-69
chr14-121962774-INS-209
chr14-121962783-INS-50
chr14-121962943-INS-3136
chr14-121966784-INS-350
chr14-121972298-INS-89
chr14-121975124-INS-174
chr14-122002830-INS-217
chr14-122005557-INS-340
chr14-122010722-INS-310
chr14-122015785-INS-51
chr14-122027928-INS-201
chr14-122046474-INS-530
chr14-122046540-INS-261
chr14-122064319-INS-736
chr14-122068749-INS-284
chr14-122093489-INS-6572
chr14-122099242-INS-2864
chr14-122132652-INS-52
chr14-122148462-INS-64
chr14-122148495-INS-262
chr14-122157969-INS-136
chr14-122158038-INS-358
chr14-122158070-INS-748
chr14-122217250-INS-396
chr14-122227042-INS-222
chr14-122240017-INS-945
chr14-122245567-INS-74
chr14-122295728-INS-60
chr14-122332699-INS-1034
chr14-122350137-INS-63
chr14-122350154-INS-231
chr14-122350214-INS-850
chr14-122430163-INS-58
chr14-122430882-INS-61
chr14-122430893-INS-314
chr14-122479881-INS-58
chr14-122481885-INS-71
chr14-122515948-INS-50
chr14-122523294-INS-201
chr14-122534718-INS-52
chr14-122617121-INS-60
chr14-122638235-INS-1459
chr14-122680379-INS-152
chr14-122790606-INS-90
chr14-122818039-INS-498
chr14-122821430-INS-54
chr14-122821556-INS-2381
chr14-122823400-INS-317
chr14-122843337-INS-236
chr14-122849538-INS-471
chr14-122858334-INS-799
chr14-122876313-INS-50
chr14-122882469-INS-71
chr14-122938970-INS-8796
chr14-123160729-INS-319
chr14-123174813-INS-167
chr14-123228296-INS-6998
chr14-123232215-INS-181
chr14-123242362-INS-229
chr14-123267287-INS-6201
chr14-123291909-INS-1551
chr14-123386675-INS-90
chr14-123395907-INS-1391
chr14-123406313-INS-58
chr14-123413519-INS-56
chr14-123414067-INS-197
chr14-123434444-INS-64
chr14-123471087-INS-6282
chr14-123502639-INS-622
chr14-123511561-INS-60
chr14-123593198-INS-105
chr14-123700541-INS-246
chr14-123741347-INS-69
chr14-123746122-INS-333
chr14-123746138-INS-6105

chr14-123765256-INS-51
chr14-123796662-INS-232
chr14-123872265-INS-6374
chr14-123874768-INS-64
chr14-123903128-INS-216
chr14-123994178-INS-193
chr14-124018841-INS-98
chr14-124068107-INS-199
chr14-124105555-INS-3233
chr14-124109434-INS-70
chr14-124118087-INS-208
chr14-124140172-INS-9320
chr14-124241338-INS-95
chr14-124269707-INS-755
chr14-124286283-INS-5308
chr14-124294923-INS-72
chr14-124356888-INS-150
chr14-124357195-INS-78
chr14-124382671-INS-6849
chr14-124433812-INS-50
chr14-124531640-INS-73
chr14-124581031-INS-51
chr14-124591107-INS-1294
chr14-124627520-INS-56
chr14-124652422-INS-410
chr14-124653043-INS-409
chr14-124688492-INS-6230
chr14-124693885-INS-4668
chr14-124736279-INS-98
chr14-124876910-INS-52
chr14-124918615-INS-54
chr14-124962184-INS-6713
chr14-124969093-INS-195
chr14-124977820-INS-6176
chr14-124980827-INS-83
chr15-3124068-INS-78
chr15-3263507-INS-65
chr15-3442868-INS-467
chr15-3553269-INS-566
chr15-3553535-INS-736
chr15-3570259-INS-588
chr15-3732006-INS-50
chr15-3745635-INS-196
chr15-3791179-INS-98
chr15-3830869-INS-2148
chr15-3831568-INS-54
chr15-3930345-INS-338
chr15-3960588-INS-336
chr15-3960597-INS-71
chr15-3962893-INS-212
chr15-3972788-INS-199
chr15-3996778-INS-114
chr15-4033146-INS-84
chr15-4052068-INS-255
chr15-4142713-INS-242
chr15-4187022-INS-2194
chr15-4243878-INS-356
chr15-4250045-INS-52
chr15-4254399-INS-52
chr15-4274788-INS-99
chr15-4284056-INS-6399
chr15-4315914-INS-1000
chr15-4321821-INS-205
chr15-4447103-INS-3078
chr15-4462589-INS-412
chr15-4467332-INS-90
chr15-4545483-INS-86
chr15-4549922-INS-1096
chr15-4633976-INS-50
chr15-4637542-INS-270
chr15-4642269-INS-6440
chr15-4686820-INS-6268
chr15-4734496-INS-140
chr15-4763432-INS-114
chr15-4842318-INS-260
chr15-4895362-INS-8296
chr15-4961068-INS-427

chr15-4961679-INS-105
chr15-4961679-INS-52
chr15-5045191-INS-180
chr15-5045934-INS-155
chr15-5063126-INS-220
chr15-5115020-INS-95
chr15-5120284-INS-428
chr15-5120493-INS-87
chr15-5131780-INS-1440
chr15-5151267-INS-199
chr15-5151804-INS-109
chr15-5155621-INS-100
chr15-5170679-INS-207
chr15-5170913-INS-164
chr15-5183460-INS-144
chr15-5197615-INS-123
chr15-5206853-INS-56
chr15-5257233-INS-52
chr15-5353174-INS-52
chr15-5360489-INS-320
chr15-5360639-INS-400
chr15-5360953-INS-268
chr15-5528132-INS-2525
chr15-5532464-INS-56
chr15-5616767-INS-62
chr15-5631173-INS-1494
chr15-5646428-INS-56
chr15-5664665-INS-56
chr15-5665141-INS-52
chr15-5702679-INS-60
chr15-5723655-INS-130
chr15-5775887-INS-1072
chr15-5790873-INS-194
chr15-5890708-INS-1209
chr15-5897811-INS-6351
chr15-5939588-INS-4048
chr15-5947302-INS-1982
chr15-5957658-INS-54
chr15-5962507-INS-3943
chr15-5996857-INS-7464
chr15-6053802-INS-1889
chr15-6053880-INS-209
chr15-6055875-INS-1500
chr15-6063399-INS-1038
chr15-6088356-INS-1660
chr15-6091087-INS-6552
chr15-6130496-INS-152
chr15-6131192-INS-2830
chr15-6133576-INS-6489
chr15-6222241-INS-72
chr15-6285124-INS-64
chr15-6326102-INS-52
chr15-6336593-INS-120
chr15-6336595-INS-50
chr15-6369804-INS-52
chr15-6372585-INS-53
chr15-6381503-INS-881
chr15-6387192-INS-50
chr15-6397411-INS-51
chr15-6404358-INS-98
chr15-6410093-INS-476
chr15-6416505-INS-50
chr15-6426766-INS-190
chr15-6439921-INS-318
chr15-6449750-INS-101
chr15-6486490-INS-54
chr15-6488730-INS-54
chr15-6524136-INS-261
chr15-6577366-INS-57
chr15-6579511-INS-6139
chr15-6627054-INS-164
chr15-6660307-INS-600
chr15-6660324-INS-290
chr15-6662576-INS-168
chr15-6682317-INS-138
chr15-6697964-INS-2054
chr15-6700649-INS-69

chr15-6907132-INS-5508
chr15-7015213-INS-101
chr15-7015251-INS-50
chr15-7015895-INS-80
chr15-7036877-INS-120
chr15-7082161-INS-92
chr15-7087831-INS-199
chr15-7151359-INS-146
chr15-7209145-INS-194
chr15-7243984-INS-104
chr15-7250992-INS-338
chr15-7261682-INS-3789
chr15-7274061-INS-146
chr15-7325346-INS-78
chr15-7356235-INS-50
chr15-7359758-INS-106
chr15-7418374-INS-74
chr15-7418711-INS-484
chr15-7447942-INS-78
chr15-7449779-INS-2636
chr15-7470516-INS-76
chr15-7472935-INS-60
chr15-7473914-INS-202
chr15-7475219-INS-442
chr15-7475261-INS-159
chr15-7482568-INS-50
chr15-7530869-INS-164
chr15-7587088-INS-169
chr15-7613857-INS-230
chr15-7664047-INS-6243
chr15-7861361-INS-190
chr15-7875694-INS-184
chr15-7933018-INS-605
chr15-7948537-INS-239
chr15-7978019-INS-79
chr15-8007587-INS-79
chr15-8021441-INS-512
chr15-8025287-INS-7962
chr15-8027434-INS-194
chr15-8059007-INS-50
chr15-8084143-INS-63
chr15-8116510-INS-279
chr15-8116511-INS-67
chr15-8120608-INS-106
chr15-8149867-INS-318
chr15-8174299-INS-157
chr15-8174735-INS-107
chr15-8185100-INS-412
chr15-8188859-INS-55
chr15-8206045-INS-540
chr15-8209868-INS-86
chr15-8257477-INS-763
chr15-8262125-INS-3667
chr15-8293198-INS-368
chr15-8303748-INS-1894
chr15-8333780-INS-66
chr15-8486057-INS-199
chr15-8493087-INS-172
chr15-8521154-INS-86
chr15-8536829-INS-52
chr15-8572686-INS-862
chr15-8575754-INS-222
chr15-8619650-INS-217
chr15-8634951-INS-54
chr15-8672606-INS-287
chr15-8699293-INS-77
chr15-8701579-INS-50
chr15-8705103-INS-230
chr15-8705621-INS-160
chr15-8717721-INS-148
chr15-8730058-INS-63
chr15-8751123-INS-82
chr15-8757513-INS-377
chr15-8768820-INS-148
chr15-8770289-INS-206
chr15-8770426-INS-81
chr15-8782935-INS-60

chr15-8807056-INS-64
chr15-8814518-INS-204
chr15-8824478-INS-189
chr15-8824510-INS-84
chr15-8834675-INS-1697
chr15-8834943-INS-13419
chr15-8870923-INS-179
chr15-8876797-INS-1888
chr15-8899079-INS-1222
chr15-8916611-INS-4051
chr15-8920587-INS-112
chr15-8944449-INS-6489
chr15-8973510-INS-431
chr15-8975615-INS-2128
chr15-8978407-INS-117
chr15-8978861-INS-6276
chr15-8994587-INS-70
chr15-9004277-INS-187
chr15-9011246-INS-125
chr15-9026770-INS-60
chr15-9035849-INS-108
chr15-9085375-INS-18099
chr15-9085407-INS-95
chr15-9085540-INS-551
chr15-9105072-INS-92
chr15-9154348-INS-90
chr15-9203025-INS-106
chr15-9229421-INS-341
chr15-9237074-INS-245
chr15-9246240-INS-82
chr15-9253730-INS-6274
chr15-9290634-INS-6583
chr15-9292984-INS-87
chr15-9293012-INS-203
chr15-9350426-INS-71
chr15-9352140-INS-87
chr15-9358612-INS-485
chr15-9364353-INS-615
chr15-9458160-INS-211
chr15-9458654-INS-7115
chr15-9494788-INS-10161
chr15-9515868-INS-68
chr15-9551286-INS-314
chr15-9569220-INS-50
chr15-9623527-INS-741
chr15-9657534-INS-88
chr15-9738310-INS-70
chr15-9740329-INS-64
chr15-9746381-INS-51
chr15-9808506-INS-145
chr15-9814418-INS-10644
chr15-9814752-INS-852
chr15-9863981-INS-80
chr15-9866943-INS-100
chr15-9866963-INS-259
chr15-9867307-INS-716
chr15-9924163-INS-98
chr15-9937536-INS-6351
chr15-9942132-INS-50
chr15-9970470-INS-14801
chr15-10004603-INS-677
chr15-10009447-INS-397
chr15-10044798-INS-1609
chr15-10052920-INS-102
chr15-10053010-INS-90
chr15-10053036-INS-205
chr15-10091151-INS-61
chr15-10165165-INS-115
chr15-10165165-INS-57
chr15-10166959-INS-6883
chr15-10174740-INS-54
chr15-10202344-INS-281
chr15-10221209-INS-58
chr15-10244613-INS-55
chr15-10273725-INS-2568
chr15-10300360-INS-136
chr15-10370806-INS-94

chr15-10445874-INS-225
chr15-10493843-INS-7100
chr15-10497009-INS-119
chr15-10504771-INS-58
chr15-10515730-INS-75
chr15-10524381-INS-157
chr15-10534665-INS-158
chr15-10538128-INS-92
chr15-10568939-INS-195
chr15-10590329-INS-61
chr15-10591033-INS-528
chr15-10663821-INS-52
chr15-10686891-INS-528
chr15-10704049-INS-57
chr15-10707053-INS-4526
chr15-10755087-INS-6006
chr15-10767561-INS-2836
chr15-10796030-INS-80
chr15-10800928-INS-54
chr15-10801390-INS-212
chr15-10832049-INS-131
chr15-10838065-INS-3650
chr15-10922232-INS-224
chr15-10924493-INS-169
chr15-10939681-INS-6822
chr15-11049590-INS-1889
chr15-11073418-INS-141
chr15-11073636-INS-427
chr15-11073637-INS-146
chr15-11081185-INS-7154
chr15-11112486-INS-124
chr15-11209631-INS-54
chr15-11273698-INS-160
chr15-11284358-INS-1488
chr15-11294385-INS-52
chr15-11294581-INS-6807
chr15-11334769-INS-353
chr15-11341558-INS-72
chr15-11350319-INS-138
chr15-11368899-INS-379
chr15-11389333-INS-895
chr15-11413266-INS-56
chr15-11429037-INS-907
chr15-11438268-INS-3225
chr15-11453213-INS-70
chr15-11462804-INS-1187
chr15-11480720-INS-117
chr15-11528101-INS-129
chr15-11535899-INS-132
chr15-11536045-INS-581
chr15-11544318-INS-60
chr15-11549716-INS-6713
chr15-11559302-INS-264
chr15-11563566-INS-377
chr15-11593113-INS-54
chr15-11594422-INS-64
chr15-11665711-INS-230
chr15-11666427-INS-54
chr15-11671950-INS-69
chr15-11673502-INS-59
chr15-11676913-INS-202
chr15-11681107-INS-189
chr15-11762968-INS-3337
chr15-11794458-INS-8454
chr15-11814981-INS-157
chr15-11853530-INS-190
chr15-11872901-INS-414
chr15-11879594-INS-50
chr15-11885558-INS-114
chr15-11898583-INS-60
chr15-12013515-INS-104
chr15-12041045-INS-513
chr15-12075347-INS-315
chr15-12099162-INS-555
chr15-12099462-INS-217
chr15-12122681-INS-171
chr15-12250356-INS-689

chr15-12253958-INS-737
chr15-12260082-INS-1494
chr15-12275071-INS-208
chr15-12288438-INS-140
chr15-12351936-INS-56
chr15-12355324-INS-58
chr15-12420777-INS-574
chr15-12421173-INS-134
chr15-12470106-INS-190
chr15-12516704-INS-52
chr15-12594845-INS-79
chr15-12604027-INS-2869
chr15-12625197-INS-72
chr15-12632132-INS-184
chr15-12642399-INS-188
chr15-12673334-INS-77
chr15-12678449-INS-2208
chr15-12686598-INS-1202
chr15-12690865-INS-353
chr15-12698864-INS-2822
chr15-12708856-INS-52
chr15-12715051-INS-1892
chr15-12743235-INS-3241
chr15-12748940-INS-143
chr15-12788583-INS-230
chr15-12789058-INS-162
chr15-12801351-INS-53
chr15-12801362-INS-124
chr15-12869141-INS-6934
chr15-12873126-INS-6068
chr15-12873163-INS-999
chr15-12885360-INS-92
chr15-12901731-INS-336
chr15-12921570-INS-50
chr15-12926171-INS-51
chr15-13063213-INS-51
chr15-13080242-INS-105
chr15-13127597-INS-102
chr15-13131706-INS-567
chr15-13143608-INS-76
chr15-13143752-INS-171
chr15-13147429-INS-326
chr15-13166049-INS-57
chr15-13180626-INS-69
chr15-13215114-INS-68
chr15-13220345-INS-567
chr15-13220484-INS-61
chr15-13227879-INS-50
chr15-13240066-INS-2743
chr15-13268308-INS-6173
chr15-13292736-INS-248
chr15-13310118-INS-68
chr15-13341914-INS-13112
chr15-13382871-INS-239
chr15-13388383-INS-6631
chr15-13448666-INS-278
chr15-13485131-INS-50
chr15-13503360-INS-343
chr15-13510197-INS-121
chr15-13541638-INS-6365
chr15-13543886-INS-118
chr15-13545208-INS-418
chr15-13565451-INS-265
chr15-13602667-INS-398
chr15-13603314-INS-2776
chr15-13619422-INS-132
chr15-13635209-INS-96
chr15-13652193-INS-62
chr15-13738064-INS-61
chr15-13738064-INS-879
chr15-13764521-INS-76
chr15-13773703-INS-79
chr15-13778020-INS-50
chr15-13799200-INS-237
chr15-13834121-INS-50
chr15-13848094-INS-2772
chr15-13957366-INS-54

chr15-14025804-INS-5758
chr15-14026243-INS-6338
chr15-14040191-INS-86
chr15-14044567-INS-6445
chr15-14063746-INS-460
chr15-14150793-INS-79
chr15-14252345-INS-58
chr15-14252797-INS-196
chr15-14335470-INS-112
chr15-14336715-INS-76
chr15-14340814-INS-6208
chr15-14383725-INS-58
chr15-14384836-INS-120
chr15-14409550-INS-1866
chr15-14418288-INS-178
chr15-14425121-INS-125
chr15-14449902-INS-434
chr15-14466907-INS-402
chr15-14467714-INS-303
chr15-14510580-INS-52
chr15-14524411-INS-545
chr15-14524415-INS-52
chr15-14564393-INS-52
chr15-14660399-INS-2785
chr15-14693924-INS-2001
chr15-14720633-INS-55
chr15-14735436-INS-119
chr15-14735436-INS-54
chr15-14775052-INS-50
chr15-14802545-INS-946
chr15-14804267-INS-345
chr15-14819585-INS-58
chr15-14912421-INS-3284
chr15-14940111-INS-1980
chr15-15115242-INS-50
chr15-15143808-INS-7732
chr15-15170015-INS-532
chr15-15174885-INS-4668
chr15-15193313-INS-6319
chr15-15218265-INS-4092
chr15-15247321-INS-6585
chr15-15265564-INS-214
chr15-15293369-INS-1337
chr15-15328092-INS-2798
chr15-15364531-INS-204
chr15-15369209-INS-67
chr15-15422203-INS-2820
chr15-15426172-INS-3244
chr15-15433174-INS-238
chr15-15570815-INS-219
chr15-15675509-INS-79
chr15-15676398-INS-60
chr15-15698794-INS-62
chr15-15699792-INS-120
chr15-15710038-INS-72
chr15-15742830-INS-1487
chr15-15743297-INS-2547
chr15-15761230-INS-6186
chr15-15762540-INS-99
chr15-15775393-INS-516
chr15-15847488-INS-235
chr15-15893288-INS-988
chr15-15933481-INS-200
chr15-16041515-INS-72
chr15-16076907-INS-2837
chr15-16100484-INS-50
chr15-16108578-INS-70
chr15-16110599-INS-588
chr15-16135852-INS-4700
chr15-16160831-INS-1110
chr15-16164840-INS-6593
chr15-16218257-INS-95
chr15-16225558-INS-7066
chr15-16226958-INS-118
chr15-16241038-INS-98
chr15-16271036-INS-162
chr15-16271108-INS-69

chr15-16271881-INS-214
chr15-16273525-INS-673
chr15-16298609-INS-169
chr15-16302613-INS-154
chr15-16311950-INS-60
chr15-16323150-INS-777
chr15-16345431-INS-6188
chr15-16371621-INS-192
chr15-16373993-INS-50
chr15-16397782-INS-252
chr15-16415153-INS-2028
chr15-16456523-INS-81
chr15-16464531-INS-5975
chr15-16518885-INS-215
chr15-16561182-INS-4994
chr15-16562914-INS-1034
chr15-16577661-INS-83
chr15-16596127-INS-821
chr15-16603387-INS-6746
chr15-16648953-INS-92
chr15-16671214-INS-6725
chr15-16689956-INS-6276
chr15-16739531-INS-209
chr15-16739654-INS-98
chr15-16804167-INS-60
chr15-16860314-INS-107
chr15-16860487-INS-77
chr15-16922500-INS-174
chr15-16927650-INS-6559
chr15-16960574-INS-52
chr15-16966282-INS-103
chr15-16967128-INS-50
chr15-17100109-INS-52
chr15-17139517-INS-1349
chr15-17149532-INS-239
chr15-17189108-INS-3479
chr15-17208646-INS-2346
chr15-17226402-INS-6442
chr15-17228914-INS-3846
chr15-17232617-INS-198
chr15-17365549-INS-54
chr15-17445272-INS-8934
chr15-17515200-INS-3341
chr15-17532259-INS-50
chr15-17538728-INS-1892
chr15-17543083-INS-6255
chr15-17555680-INS-1302
chr15-17587957-INS-55
chr15-17644344-INS-372
chr15-17644884-INS-105
chr15-17662583-INS-65
chr15-17675815-INS-13563
chr15-17680389-INS-3487
chr15-17697659-INS-64
chr15-17701110-INS-120
chr15-17702234-INS-1380
chr15-17716691-INS-297
chr15-17716697-INS-644
chr15-17721006-INS-60
chr15-17773906-INS-1031
chr15-17774014-INS-110
chr15-17781933-INS-6385
chr15-17836684-INS-375
chr15-17865562-INS-4213
chr15-17875127-INS-1588
chr15-17883869-INS-434
chr15-17915467-INS-54
chr15-17928199-INS-210
chr15-17937866-INS-53
chr15-17965500-INS-62
chr15-17978700-INS-3250
chr15-18048955-INS-1138
chr15-18098374-INS-6419
chr15-18111206-INS-6460
chr15-18139137-INS-54
chr15-18158103-INS-675
chr15-18163966-INS-304

chr15-18183800-INS-205
chr15-18188150-INS-52
chr15-18223390-INS-50
chr15-18231723-INS-18572
chr15-18327621-INS-149
chr15-18345699-INS-198
chr15-18364620-INS-1005
chr15-18370892-INS-641
chr15-18375393-INS-207
chr15-18391181-INS-70
chr15-18424551-INS-1162
chr15-18453880-INS-1890
chr15-18461852-INS-232
chr15-18493699-INS-2506
chr15-18548812-INS-62
chr15-18613062-INS-63
chr15-18675237-INS-68
chr15-18683913-INS-187
chr15-18684032-INS-72
chr15-18714364-INS-70
chr15-18831805-INS-6396
chr15-18863998-INS-54
chr15-18871255-INS-2727
chr15-18980862-INS-61
chr15-18981171-INS-54
chr15-18982558-INS-1807
chr15-18982811-INS-1421
chr15-18999427-INS-179
chr15-19009878-INS-50
chr15-19034378-INS-897
chr15-19034919-INS-390
chr15-19040761-INS-488
chr15-19084442-INS-82
chr15-19091801-INS-328
chr15-19092149-INS-494
chr15-19142943-INS-8736
chr15-19144836-INS-54
chr15-19146040-INS-52
chr15-19172609-INS-1505
chr15-19196581-INS-352
chr15-19203491-INS-4158
chr15-19226794-INS-124
chr15-19226981-INS-667
chr15-19229666-INS-2063
chr15-19271048-INS-195
chr15-19281620-INS-13516
chr15-19292006-INS-3169
chr15-19324742-INS-646
chr15-19325161-INS-1677
chr15-19343118-INS-1160
chr15-19349815-INS-188
chr15-19355924-INS-6373
chr15-19382116-INS-52
chr15-19395265-INS-136
chr15-19407876-INS-54
chr15-19426735-INS-208
chr15-19426990-INS-281
chr15-19427173-INS-465
chr15-19427540-INS-516
chr15-19492390-INS-1988
chr15-19655920-INS-2380
chr15-19662299-INS-147
chr15-19667184-INS-76
chr15-19676394-INS-70
chr15-19718274-INS-66
chr15-19723467-INS-58
chr15-19774117-INS-6287
chr15-19783377-INS-6219
chr15-19836831-INS-7716
chr15-19845775-INS-7096
chr15-19865018-INS-5567
chr15-19914629-INS-1661
chr15-19920775-INS-7081
chr15-19921842-INS-2000
chr15-19935584-INS-180
chr15-19960815-INS-103
chr15-19961653-INS-1979

chr15-20214753-INS-81
chr15-20385139-INS-1224
chr15-20393315-INS-56
chr15-20397797-INS-2258
chr15-20418657-INS-54
chr15-20538767-INS-184
chr15-20556720-INS-1037
chr15-20590867-INS-3331
chr15-20595498-INS-4335
chr15-20605209-INS-52
chr15-20624650-INS-981
chr15-20640754-INS-86
chr15-20641344-INS-53
chr15-20687917-INS-6482
chr15-20746501-INS-6179
chr15-20770314-INS-7764
chr15-20859002-INS-60
chr15-20867651-INS-691
chr15-20879107-INS-7516
chr15-20887952-INS-951
chr15-20892116-INS-1175
chr15-20930346-INS-2108
chr15-20930346-INS-766
chr15-21021187-INS-100
chr15-21031765-INS-6382
chr15-21036686-INS-52
chr15-21062072-INS-52
chr15-21101837-INS-594
chr15-21188484-INS-65
chr15-21188626-INS-84
chr15-21248092-INS-537
chr15-21293041-INS-72
chr15-21293694-INS-85
chr15-21315961-INS-8716
chr15-21498242-INS-50
chr15-21767853-INS-50
chr15-21804046-INS-2054
chr15-21828753-INS-6780
chr15-21829842-INS-5930
chr15-21829868-INS-53
chr15-21861093-INS-200
chr15-21971256-INS-51
chr15-21986340-INS-335
chr15-21986340-INS-713
chr15-21989956-INS-7161
chr15-22003704-INS-3042
chr15-22082949-INS-54
chr15-22113412-INS-62
chr15-22146806-INS-7519
chr15-22249268-INS-50
chr15-22318648-INS-2878
chr15-22396100-INS-2505
chr15-22478887-INS-6667
chr15-22539814-INS-1730
chr15-22541913-INS-1552
chr15-22578641-INS-121
chr15-22611797-INS-6802
chr15-22618190-INS-6643
chr15-22631905-INS-7235
chr15-22674274-INS-63
chr15-22707453-INS-52
chr15-22793730-INS-4439
chr15-22799718-INS-715
chr15-22816035-INS-80
chr15-22841027-INS-2900
chr15-22845268-INS-62
chr15-22848857-INS-215
chr15-22861907-INS-1936
chr15-22915305-INS-103
chr15-22943567-INS-50
chr15-22957603-INS-126
chr15-22957603-INS-59
chr15-22968488-INS-3358
chr15-23001109-INS-252
chr15-23008337-INS-54
chr15-23015833-INS-54
chr15-23056118-INS-50

chr15-23115486-INS-55
chr15-23119102-INS-98
chr15-23184387-INS-500
chr15-23206895-INS-133
chr15-23247932-INS-57
chr15-23247982-INS-277
chr15-23290672-INS-933
chr15-23371154-INS-50
chr15-23395440-INS-197
chr15-23412094-INS-235
chr15-23441700-INS-3499
chr15-23454403-INS-52
chr15-23461383-INS-213
chr15-23505035-INS-5891
chr15-23547887-INS-86
chr15-23626158-INS-69
chr15-23627450-INS-486
chr15-23635619-INS-126
chr15-23686190-INS-647
chr15-23686198-INS-139
chr15-23686309-INS-282
chr15-23686313-INS-133
chr15-23690250-INS-50
chr15-23725474-INS-480
chr15-23793803-INS-763
chr15-23794035-INS-76
chr15-23877451-INS-8902
chr15-23908490-INS-52
chr15-23942930-INS-53
chr15-23988494-INS-92
chr15-23989035-INS-109
chr15-24126287-INS-84
chr15-24129262-INS-182
chr15-24145864-INS-96
chr15-24146020-INS-685
chr15-24232765-INS-6435
chr15-24249032-INS-2419
chr15-24262038-INS-56
chr15-24268979-INS-50
chr15-24270218-INS-70
chr15-24302886-INS-116
chr15-24337079-INS-8593
chr15-24388090-INS-66
chr15-24530840-INS-64
chr15-24577573-INS-55
chr15-24585580-INS-341
chr15-24595316-INS-138
chr15-24602936-INS-1389
chr15-24604278-INS-186
chr15-24605142-INS-3284
chr15-24687716-INS-615
chr15-24688082-INS-385
chr15-24714975-INS-7388
chr15-24785195-INS-68
chr15-24785396-INS-51
chr15-24816054-INS-57
chr15-24886351-INS-358
chr15-24905671-INS-186
chr15-24910078-INS-138
chr15-24978527-INS-56
chr15-25015923-INS-2065
chr15-25025389-INS-54
chr15-25025445-INS-56
chr15-25050917-INS-206
chr15-25050920-INS-484
chr15-25057119-INS-340
chr15-25057124-INS-50
chr15-25060933-INS-1195
chr15-25072695-INS-52
chr15-25146339-INS-84
chr15-25155995-INS-94
chr15-25156706-INS-152
chr15-25202217-INS-74
chr15-25217110-INS-62
chr15-25217936-INS-50
chr15-25217990-INS-175
chr15-25328164-INS-78

chr15-25329122-INS-84
chr15-25380631-INS-60
chr15-25405476-INS-177
chr15-25417750-INS-102
chr15-25436403-INS-154
chr15-25436697-INS-334
chr15-25437038-INS-692
chr15-25447077-INS-290
chr15-25453292-INS-116
chr15-25500037-INS-589
chr15-25504373-INS-88
chr15-25521392-INS-76
chr15-25525173-INS-62
chr15-25538644-INS-68
chr15-25548067-INS-170
chr15-25549039-INS-98
chr15-25570189-INS-300
chr15-25597724-INS-59
chr15-25644161-INS-182
chr15-25662030-INS-89
chr15-25662043-INS-593
chr15-25677389-INS-86
chr15-25698889-INS-66
chr15-25710456-INS-4448
chr15-25736457-INS-203
chr15-25739152-INS-380
chr15-25752282-INS-115
chr15-25771574-INS-123
chr15-25791365-INS-50
chr15-25827291-INS-106
chr15-25890006-INS-71
chr15-25894470-INS-280
chr15-25960712-INS-58
chr15-25998573-INS-181
chr15-25999678-INS-2111
chr15-26002171-INS-145
chr15-26042094-INS-6134
chr15-26078139-INS-387
chr15-26081657-INS-2128
chr15-26142697-INS-8619
chr15-26220694-INS-6221
chr15-26251413-INS-514
chr15-26269697-INS-112
chr15-26302664-INS-1224
chr15-26307526-INS-358
chr15-26330038-INS-6426
chr15-26338407-INS-1890
chr15-26397267-INS-322
chr15-26398700-INS-92
chr15-26401892-INS-53
chr15-26422570-INS-1468
chr15-26453975-INS-1290
chr15-26468042-INS-117
chr15-26479019-INS-1419
chr15-26502807-INS-52
chr15-26573867-INS-76
chr15-26607083-INS-64
chr15-26630337-INS-392
chr15-26650613-INS-6553
chr15-26662498-INS-3615
chr15-26665204-INS-561
chr15-26691447-INS-357
chr15-26709972-INS-874
chr15-26732935-INS-2744
chr15-26745694-INS-50
chr15-26785221-INS-52
chr15-26828458-INS-205
chr15-26831806-INS-722
chr15-26836056-INS-56
chr15-26837787-INS-721
chr15-26848857-INS-67
chr15-26854185-INS-1368
chr15-26855478-INS-733
chr15-26869094-INS-54
chr15-26880045-INS-102
chr15-26889756-INS-57
chr15-26899748-INS-171

chr15-26901186-INS-810
chr15-26918472-INS-1178
chr15-26918562-INS-2845
chr15-26920930-INS-159
chr15-26920969-INS-51
chr15-26929485-INS-486
chr15-26937885-INS-194
chr15-26955985-INS-54
chr15-26961921-INS-164
chr15-26971346-INS-378
chr15-26986227-INS-56
chr15-26993671-INS-6396
chr15-27015365-INS-7044
chr15-27019407-INS-1485
chr15-27105271-INS-62
chr15-27116173-INS-1276
chr15-27139402-INS-7587
chr15-27156563-INS-2057
chr15-27156578-INS-517
chr15-27156819-INS-56
chr15-27233307-INS-141
chr15-27239692-INS-6733
chr15-27350655-INS-52
chr15-27351208-INS-365
chr15-27356737-INS-88
chr15-27407157-INS-1196
chr15-27420735-INS-89
chr15-27427586-INS-2733
chr15-27431406-INS-73
chr15-27456708-INS-164
chr15-27532795-INS-170
chr15-27561113-INS-76
chr15-27612529-INS-944
chr15-27622444-INS-72
chr15-27674077-INS-293
chr15-27682452-INS-51
chr15-27686566-INS-642
chr15-27695933-INS-110
chr15-27702227-INS-93
chr15-27704070-INS-401
chr15-27704255-INS-501
chr15-27708409-INS-1207
chr15-27742264-INS-544
chr15-27758124-INS-76
chr15-27761928-INS-1500
chr15-27867954-INS-53
chr15-27917721-INS-59
chr15-27946760-INS-57
chr15-28011528-INS-86
chr15-28011622-INS-173
chr15-28039130-INS-2793
chr15-28055563-INS-155
chr15-28074191-INS-2823
chr15-28080931-INS-206
chr15-28090541-INS-247
chr15-28119529-INS-1407
chr15-28122937-INS-67
chr15-28134821-INS-59
chr15-28137393-INS-104
chr15-28143948-INS-1063
chr15-28147248-INS-134
chr15-28149795-INS-178
chr15-28163220-INS-476
chr15-28169774-INS-50
chr15-28243886-INS-57
chr15-28272777-INS-1719
chr15-28286690-INS-214
chr15-28291802-INS-86
chr15-28332986-INS-6879
chr15-28373667-INS-56
chr15-28384883-INS-134
chr15-28396978-INS-65
chr15-28422375-INS-1146
chr15-28435231-INS-358
chr15-28447670-INS-469
chr15-28476874-INS-177
chr15-28500657-INS-121

chr15-28502113-INS-51
chr15-28504665-INS-61
chr15-28504728-INS-276
chr15-28506865-INS-928
chr15-28534365-INS-75
chr15-28556387-INS-63
chr15-28563398-INS-53
chr15-28570321-INS-141
chr15-28586537-INS-56
chr15-28594656-INS-122
chr15-28605100-INS-50
chr15-28605141-INS-1880
chr15-28609317-INS-50
chr15-28613299-INS-493
chr15-28614147-INS-52
chr15-28632169-INS-2361
chr15-28640202-INS-5463
chr15-28656962-INS-501
chr15-28657634-INS-106
chr15-28661298-INS-148
chr15-28662391-INS-90
chr15-28700788-INS-231
chr15-28708193-INS-70
chr15-28712403-INS-129
chr15-28712443-INS-1276
chr15-28734975-INS-50
chr15-28747720-INS-1918
chr15-28767731-INS-204
chr15-28768138-INS-84
chr15-28768209-INS-396
chr15-28772398-INS-86
chr15-28790830-INS-80
chr15-28813738-INS-80
chr15-28813927-INS-836
chr15-28832308-INS-561
chr15-28839645-INS-6279
chr15-28887485-INS-162
chr15-28889474-INS-235
chr15-28890880-INS-6485
chr15-28895568-INS-1026
chr15-28927426-INS-6297
chr15-28937893-INS-3846
chr15-28954877-INS-1590
chr15-28957127-INS-3873
chr15-28986759-INS-3021
chr15-28999371-INS-148
chr15-28999371-INS-436
chr15-29003182-INS-500
chr15-29033727-INS-94
chr15-29044369-INS-340
chr15-29070312-INS-376
chr15-29071196-INS-5987
chr15-29074266-INS-52
chr15-29111452-INS-16585
chr15-29149264-INS-81
chr15-29209665-INS-199
chr15-29224786-INS-74
chr15-29251728-INS-105
chr15-29258125-INS-2786
chr15-29263469-INS-88
chr15-29276498-INS-88
chr15-29330978-INS-472
chr15-29331797-INS-2318
chr15-29363497-INS-2795
chr15-29386937-INS-52
chr15-29407662-INS-453
chr15-29440569-INS-80
chr15-29525420-INS-52
chr15-29527022-INS-479
chr15-29606093-INS-2519
chr15-29610221-INS-149
chr15-29621784-INS-173
chr15-29629384-INS-110
chr15-29671640-INS-160
chr15-29675025-INS-2339
chr15-29682322-INS-3155
chr15-29692277-INS-1458

chr15-29723420-INS-6452
chr15-29785177-INS-1891
chr15-29785177-INS-404
chr15-29788024-INS-6484
chr15-29816861-INS-6425
chr15-29842800-INS-2625
chr15-29845457-INS-52
chr15-29859540-INS-7549
chr15-29872918-INS-510
chr15-29873137-INS-3488
chr15-29874571-INS-5278
chr15-29878716-INS-568
chr15-29887071-INS-56
chr15-29894809-INS-1024
chr15-29921026-INS-132
chr15-29958347-INS-90
chr15-29958539-INS-120
chr15-29969505-INS-588
chr15-30018276-INS-52
chr15-30085810-INS-1682
chr15-30114918-INS-298
chr15-30117673-INS-56
chr15-30144166-INS-6357
chr15-30149991-INS-235
chr15-30160509-INS-847
chr15-30165341-INS-56
chr15-30166665-INS-172
chr15-30175244-INS-1390
chr15-30197996-INS-130
chr15-30201277-INS-6797
chr15-30205037-INS-194
chr15-30250819-INS-162
chr15-30251677-INS-214
chr15-30258281-INS-234
chr15-30443267-INS-3770
chr15-30505017-INS-96
chr15-30511308-INS-3280
chr15-30514397-INS-52
chr15-30532649-INS-1793
chr15-30540023-INS-696
chr15-30547470-INS-5136
chr15-30649251-INS-120
chr15-30649294-INS-318
chr15-30649384-INS-732
chr15-30657306-INS-57
chr15-30664531-INS-659
chr15-30677743-INS-487
chr15-30677900-INS-781
chr15-30787206-INS-227
chr15-30836620-INS-50
chr15-30841055-INS-235
chr15-30841244-INS-73
chr15-30850242-INS-210
chr15-30901383-INS-1116
chr15-30904772-INS-97
chr15-30919054-INS-1891
chr15-30946383-INS-58
chr15-30949035-INS-175
chr15-30955826-INS-82
chr15-30994987-INS-147
chr15-30995188-INS-234
chr15-31040624-INS-50
chr15-31046403-INS-6667
chr15-31055972-INS-206
chr15-31056949-INS-434
chr15-31076911-INS-1000
chr15-31123655-INS-66
chr15-31181650-INS-3245
chr15-31208607-INS-65
chr15-31208710-INS-198
chr15-31230136-INS-149
chr15-31308574-INS-163
chr15-31319156-INS-157
chr15-31330209-INS-56
chr15-31405894-INS-300
chr15-31414049-INS-56
chr15-31415205-INS-52

chr15-31424033-INS-52
chr15-31470511-INS-212
chr15-31520469-INS-3112
chr15-31570573-INS-105
chr15-31573150-INS-179
chr15-31573888-INS-85
chr15-31605833-INS-728
chr15-31605889-INS-84
chr15-31634435-INS-96
chr15-31653777-INS-6526
chr15-31672936-INS-1887
chr15-31691748-INS-161
chr15-31701568-INS-2453
chr15-31703217-INS-416
chr15-31714717-INS-396
chr15-31715380-INS-7329
chr15-31721899-INS-68
chr15-31722314-INS-6581
chr15-31802097-INS-6416
chr15-31809924-INS-2320
chr15-31820143-INS-7079
chr15-31865094-INS-2264
chr15-31885510-INS-1044
chr15-31889433-INS-6363
chr15-31903098-INS-7095
chr15-31915221-INS-81
chr15-32007707-INS-50
chr15-32110535-INS-80
chr15-32110973-INS-203
chr15-32110973-INS-58
chr15-32111255-INS-108
chr15-32196650-INS-50
chr15-32228634-INS-84
chr15-32306599-INS-1101
chr15-32367298-INS-7475
chr15-32384948-INS-83
chr15-32405561-INS-113
chr15-32414769-INS-141
chr15-32417320-INS-6209
chr15-32518832-INS-1492
chr15-32536073-INS-126
chr15-32561033-INS-6410
chr15-32615051-INS-1600
chr15-32719737-INS-50
chr15-32766849-INS-163
chr15-32812276-INS-609
chr15-32880843-INS-76
chr15-32880948-INS-128
chr15-32933100-INS-50
chr15-32994154-INS-6332
chr15-33129280-INS-6394
chr15-33161330-INS-998
chr15-33168761-INS-184
chr15-33178894-INS-145
chr15-33207491-INS-1971
chr15-33285939-INS-192
chr15-33297514-INS-6188
chr15-33338589-INS-1546
chr15-33386339-INS-277
chr15-33427955-INS-56
chr15-33428342-INS-308
chr15-33452094-INS-6788
chr15-33461429-INS-117
chr15-33504879-INS-2579
chr15-33511835-INS-50
chr15-33512451-INS-99
chr15-33527017-INS-6324
chr15-33532707-INS-996
chr15-33538276-INS-6577
chr15-33582220-INS-72
chr15-33650542-INS-2392
chr15-33655688-INS-307
chr15-33767848-INS-6434
chr15-33846754-INS-514
chr15-33887407-INS-14026
chr15-33935043-INS-106
chr15-33995876-INS-67

chr15-34009507-INS-445
chr15-34035694-INS-6215
chr15-34057812-INS-213
chr15-34057987-INS-316
chr15-34063072-INS-572
chr15-34176111-INS-108
chr15-34176375-INS-50
chr15-34226754-INS-2850
chr15-34302321-INS-81
chr15-34302554-INS-208
chr15-34303149-INS-203
chr15-34320078-INS-5256
chr15-34340837-INS-208
chr15-34354115-INS-50
chr15-34366435-INS-6493
chr15-34369240-INS-70
chr15-34509250-INS-386
chr15-34570783-INS-214
chr15-34576616-INS-218
chr15-34591008-INS-60
chr15-34591314-INS-412
chr15-34608540-INS-50
chr15-34610482-INS-450
chr15-34630121-INS-320
chr15-34688768-INS-3797
chr15-34698972-INS-900
chr15-34775025-INS-1520
chr15-34862556-INS-52
chr15-34934307-INS-192
chr15-35003360-INS-3228
chr15-35130133-INS-440
chr15-35130288-INS-892
chr15-35213378-INS-4472
chr15-35253051-INS-887
chr15-35330049-INS-215
chr15-35332141-INS-1288
chr15-35344709-INS-119
chr15-35355644-INS-611
chr15-35404795-INS-675
chr15-35456208-INS-128
chr15-35457012-INS-241
chr15-35571840-INS-9600
chr15-35590215-INS-424
chr15-35626286-INS-70
chr15-35664996-INS-70
chr15-35719723-INS-201
chr15-35785041-INS-489
chr15-35820035-INS-50
chr15-35858389-INS-199
chr15-35881677-INS-50
chr15-35892177-INS-59
chr15-35892323-INS-53
chr15-35892437-INS-172
chr15-35909477-INS-78
chr15-35953453-INS-6247
chr15-35962184-INS-200
chr15-35968254-INS-114
chr15-35969059-INS-2881
chr15-36012280-INS-213
chr15-36043973-INS-244
chr15-36056361-INS-51
chr15-36104761-INS-1002
chr15-36104761-INS-2058
chr15-36106428-INS-92
chr15-36125515-INS-1534
chr15-36180067-INS-1390
chr15-36223424-INS-94
chr15-36223924-INS-62
chr15-36228295-INS-5538
chr15-36324194-INS-437
chr15-36332548-INS-64
chr15-36414521-INS-60
chr15-36453625-INS-190
chr15-36457961-INS-66
chr15-36458016-INS-268
chr15-36458199-INS-52
chr15-36537102-INS-82

chr15-36543043-INS-1550
chr15-36568668-INS-81
chr15-36607254-INS-263
chr15-36630961-INS-218
chr15-36747769-INS-2767
chr15-36767556-INS-78
chr15-36778678-INS-50
chr15-36821232-INS-51
chr15-36876841-INS-54
chr15-36909366-INS-220
chr15-36909406-INS-2354
chr15-36916049-INS-325
chr15-36916052-INS-162
chr15-36921070-INS-220
chr15-36921583-INS-64
chr15-36964119-INS-52
chr15-36972587-INS-385
chr15-36977032-INS-258
chr15-36995825-INS-1409
chr15-37001362-INS-206
chr15-37006343-INS-213
chr15-37017631-INS-178
chr15-37049813-INS-200
chr15-37067208-INS-150
chr15-37086713-INS-122
chr15-37106381-INS-226
chr15-37142793-INS-345
chr15-37165708-INS-95
chr15-37170867-INS-142
chr15-37170869-INS-54
chr15-37203471-INS-2680
chr15-37222087-INS-52
chr15-37225330-INS-85
chr15-37225940-INS-76
chr15-37228439-INS-53
chr15-37229834-INS-7087
chr15-37263143-INS-96
chr15-37263171-INS-1513
chr15-37265689-INS-460
chr15-37265747-INS-55
chr15-37309599-INS-123
chr15-37309772-INS-665
chr15-37317061-INS-54
chr15-37329255-INS-6538
chr15-37349807-INS-261
chr15-37388880-INS-179
chr15-37396551-INS-50
chr15-37430794-INS-72
chr15-37446357-INS-208
chr15-37474234-INS-124
chr15-37481991-INS-70
chr15-37488582-INS-50
chr15-37535588-INS-1940
chr15-37552976-INS-64
chr15-37566518-INS-161
chr15-37566518-INS-351
chr15-37567923-INS-525
chr15-37574191-INS-73
chr15-37629510-INS-66
chr15-37751745-INS-52
chr15-37795634-INS-59
chr15-37935297-INS-201
chr15-37936140-INS-209
chr15-37966610-INS-172
chr15-37982010-INS-97
chr15-37986905-INS-157
chr15-37988013-INS-89
chr15-38024078-INS-61
chr15-38090343-INS-93
chr15-38090354-INS-386
chr15-38097309-INS-52
chr15-38105189-INS-1636
chr15-38112732-INS-50
chr15-38119911-INS-364
chr15-38121761-INS-1140
chr15-38124224-INS-82
chr15-38138682-INS-66

chr15-38181668-INS-55
chr15-38188705-INS-322
chr15-38207097-INS-50
chr15-38210770-INS-386
chr15-38241524-INS-473
chr15-38250455-INS-87
chr15-38254070-INS-80
chr15-38265529-INS-2766
chr15-38281661-INS-128
chr15-38286877-INS-52
chr15-38330922-INS-53
chr15-38331935-INS-125
chr15-38370166-INS-216
chr15-38383360-INS-167
chr15-38383659-INS-150
chr15-38394832-INS-126
chr15-38396995-INS-232
chr15-38404740-INS-50
chr15-38411164-INS-100
chr15-38444099-INS-181
chr15-38449267-INS-62
chr15-38459827-INS-50
chr15-38487315-INS-50
chr15-38510198-INS-54
chr15-38581519-INS-210
chr15-38595578-INS-110
chr15-38649805-INS-64
chr15-38699988-INS-198
chr15-38701951-INS-210
chr15-38712227-INS-207
chr15-38712910-INS-138
chr15-38729929-INS-76
chr15-38732394-INS-54
chr15-38734716-INS-80
chr15-38735493-INS-76
chr15-38761329-INS-77
chr15-38767481-INS-5228
chr15-38803048-INS-1520
chr15-38817301-INS-70
chr15-38823214-INS-450
chr15-38847442-INS-95
chr15-38992877-INS-210
chr15-39034886-INS-2639
chr15-39052280-INS-5571
chr15-39088223-INS-97
chr15-39097471-INS-145
chr15-39098583-INS-3552
chr15-39102875-INS-208
chr15-39125766-INS-214
chr15-39166420-INS-194
chr15-39188009-INS-2126
chr15-39221630-INS-1039
chr15-39232476-INS-1890
chr15-39263934-INS-58
chr15-39274747-INS-56
chr15-39286627-INS-556
chr15-39337741-INS-52
chr15-39337810-INS-5315
chr15-39340739-INS-333
chr15-39343269-INS-2282
chr15-39378251-INS-649
chr15-39422183-INS-148
chr15-39463331-INS-79
chr15-39464021-INS-356
chr15-39481847-INS-5634
chr15-39613368-INS-69
chr15-39622203-INS-7446
chr15-39623231-INS-219
chr15-39666299-INS-66
chr15-39666373-INS-265
chr15-39678552-INS-118
chr15-39678722-INS-183
chr15-39684062-INS-240
chr15-39715473-INS-57
chr15-39723932-INS-155
chr15-39723932-INS-544
chr15-39727227-INS-127

chr15-39727373-INS-384
chr15-39777913-INS-588
chr15-39809583-INS-51
chr15-39860026-INS-97
chr15-39870200-INS-306
chr15-39870242-INS-2392
chr15-39892969-INS-781
chr15-39893538-INS-116
chr15-39904471-INS-65
chr15-39925678-INS-84
chr15-39925697-INS-195
chr15-39949507-INS-59
chr15-40015593-INS-89
chr15-40075665-INS-360
chr15-40087877-INS-52
chr15-40088848-INS-54
chr15-40189140-INS-184
chr15-40327203-INS-140
chr15-40331342-INS-74
chr15-40331607-INS-106
chr15-40335218-INS-6225
chr15-40359229-INS-440
chr15-40419128-INS-573
chr15-40459117-INS-1057
chr15-40460019-INS-1170
chr15-40637713-INS-60
chr15-40705409-INS-378
chr15-40773196-INS-1714
chr15-40792741-INS-56
chr15-40811661-INS-253
chr15-40910869-INS-59
chr15-40914031-INS-3313
chr15-40983227-INS-6802
chr15-40986321-INS-123
chr15-40987382-INS-203
chr15-40989076-INS-69
chr15-40989243-INS-350
chr15-41019809-INS-50
chr15-41021648-INS-192
chr15-41031251-INS-441
chr15-41056655-INS-215
chr15-41094285-INS-1118
chr15-41097454-INS-2810
chr15-41126102-INS-62
chr15-41154147-INS-50
chr15-41267836-INS-56
chr15-41271446-INS-6145
chr15-41289145-INS-6881
chr15-41295991-INS-58
chr15-41304658-INS-64
chr15-41305326-INS-6930
chr15-41328708-INS-522
chr15-41328916-INS-122
chr15-41328916-INS-792
chr15-41429433-INS-389
chr15-41459932-INS-1811
chr15-41511023-INS-106
chr15-41511527-INS-50
chr15-41521254-INS-101
chr15-41577867-INS-75
chr15-41578766-INS-604
chr15-41600828-INS-1999
chr15-41624315-INS-811
chr15-41636260-INS-442
chr15-41640348-INS-147
chr15-41675982-INS-204
chr15-41681477-INS-50
chr15-41697331-INS-185
chr15-41697331-INS-78
chr15-41697671-INS-736
chr15-41748650-INS-3035
chr15-41748958-INS-116
chr15-41760455-INS-221
chr15-41760627-INS-53
chr15-41771533-INS-77
chr15-41776419-INS-978
chr15-41802707-INS-56

chr15-41810221-INS-2842
chr15-41814404-INS-6607
chr15-41838698-INS-6226
chr15-41853528-INS-65
chr15-41853664-INS-223
chr15-41862852-INS-145
chr15-41872256-INS-56
chr15-41877410-INS-60
chr15-41880859-INS-1072
chr15-41898818-INS-70
chr15-41905989-INS-5055
chr15-41912901-INS-6038
chr15-41934399-INS-106
chr15-41982366-INS-168
chr15-42012140-INS-7118
chr15-42052244-INS-6689
chr15-42121001-INS-6179
chr15-42141637-INS-2761
chr15-42153664-INS-6287
chr15-42175684-INS-648
chr15-42185604-INS-6225
chr15-42185865-INS-13005
chr15-42204752-INS-1358
chr15-42232235-INS-528
chr15-42246023-INS-1249
chr15-42255012-INS-7254
chr15-42324549-INS-112
chr15-42336488-INS-50
chr15-42345659-INS-1240
chr15-42395990-INS-440
chr15-42431494-INS-54
chr15-42435329-INS-100
chr15-42620127-INS-70
chr15-42624137-INS-53
chr15-42635074-INS-1558
chr15-42637567-INS-401
chr15-42662844-INS-3173
chr15-42694546-INS-2788
chr15-42755115-INS-2066
chr15-42755744-INS-321
chr15-42932803-INS-61
chr15-43049942-INS-52
chr15-43050007-INS-58
chr15-43090231-INS-2008
chr15-43147037-INS-52
chr15-43242122-INS-236
chr15-43245039-INS-1997
chr15-43253921-INS-142
chr15-43305208-INS-1045
chr15-43507949-INS-386
chr15-43525014-INS-50
chr15-43536220-INS-52
chr15-43618984-INS-95
chr15-43638076-INS-50
chr15-43639598-INS-4298
chr15-43705848-INS-6736
chr15-43754766-INS-55
chr15-43792178-INS-5743
chr15-43808425-INS-159
chr15-43880680-INS-383
chr15-43880896-INS-327
chr15-43910090-INS-87
chr15-43971089-INS-990
chr15-44114042-INS-112
chr15-44122438-INS-7547
chr15-44140759-INS-7861
chr15-44169081-INS-141
chr15-44198095-INS-60
chr15-44231185-INS-8463
chr15-44245637-INS-251
chr15-44250386-INS-6454
chr15-44319525-INS-376
chr15-44356176-INS-187
chr15-44436781-INS-604
chr15-44448734-INS-50
chr15-44557511-INS-50
chr15-44669844-INS-87

chr15-44840831-INS-2066
chr15-44860425-INS-158
chr15-44899447-INS-874
chr15-44901530-INS-1922
chr15-44921721-INS-360
chr15-44942391-INS-50
chr15-44951437-INS-6496
chr15-44967916-INS-4808
chr15-44989965-INS-947
chr15-44995656-INS-60
chr15-45050065-INS-577
chr15-45065323-INS-4509
chr15-45083550-INS-1298
chr15-45090014-INS-3239
chr15-45091419-INS-428
chr15-45104198-INS-86
chr15-45116157-INS-179
chr15-45143008-INS-383
chr15-45153678-INS-7604
chr15-45209019-INS-6223
chr15-45338559-INS-1163
chr15-45363378-INS-184
chr15-45390725-INS-68
chr15-45435543-INS-6475
chr15-45440269-INS-2637
chr15-45453185-INS-848
chr15-45477358-INS-129
chr15-45479467-INS-50
chr15-45485050-INS-2788
chr15-45540154-INS-198
chr15-45543186-INS-179
chr15-45543194-INS-374
chr15-45566856-INS-264
chr15-45585081-INS-51
chr15-45612189-INS-66
chr15-45617929-INS-56
chr15-45636235-INS-2773
chr15-45650219-INS-2024
chr15-45652330-INS-1674
chr15-45712158-INS-300
chr15-45767235-INS-15327
chr15-45847945-INS-1496
chr15-45943032-INS-50
chr15-45954549-INS-1010
chr15-45981501-INS-6180
chr15-45983892-INS-64
chr15-45985636-INS-4652
chr15-46017945-INS-91
chr15-46032120-INS-1773
chr15-46046731-INS-140
chr15-46102460-INS-664
chr15-46134174-INS-2887
chr15-46158540-INS-6650
chr15-46168362-INS-2719
chr15-46170033-INS-73
chr15-46170970-INS-1217
chr15-46190481-INS-72
chr15-46191655-INS-7717
chr15-46209719-INS-329
chr15-46215299-INS-52
chr15-46227097-INS-214
chr15-46237346-INS-52
chr15-46265276-INS-74
chr15-46289212-INS-735
chr15-46385944-INS-6151
chr15-46403038-INS-4930
chr15-46457875-INS-50
chr15-46481211-INS-52
chr15-46509467-INS-3908
chr15-46510182-INS-466
chr15-46587839-INS-82
chr15-46609315-INS-215
chr15-46620641-INS-265
chr15-46623205-INS-2227
chr15-46633181-INS-244
chr15-46671499-INS-68
chr15-46691622-INS-1486

chr15-46711725-INS-6437
chr15-46803325-INS-9823
chr15-46817338-INS-6445
chr15-46856016-INS-50
chr15-46858963-INS-50
chr15-46863255-INS-440
chr15-46873823-INS-2392
chr15-46897595-INS-60
chr15-46912724-INS-380
chr15-46968226-INS-59
chr15-46968364-INS-6473
chr15-46977078-INS-72
chr15-46984843-INS-199
chr15-47001259-INS-134
chr15-47022450-INS-273
chr15-47068813-INS-50
chr15-47142787-INS-138
chr15-47205443-INS-50
chr15-47250994-INS-52
chr15-47257559-INS-6590
chr15-47312703-INS-13714
chr15-47312703-INS-6452
chr15-47325364-INS-264
chr15-47325521-INS-612
chr15-47340250-INS-119
chr15-47340250-INS-54
chr15-47352379-INS-80
chr15-47364742-INS-6147
chr15-47379526-INS-640
chr15-47383612-INS-59
chr15-47397261-INS-506
chr15-47397549-INS-878
chr15-47397878-INS-1324
chr15-47402925-INS-74
chr15-47427827-INS-116
chr15-47441331-INS-75
chr15-47441352-INS-945
chr15-47443977-INS-1559
chr15-47478232-INS-6816
chr15-47489524-INS-195
chr15-47489909-INS-60
chr15-47489999-INS-812
chr15-47523210-INS-6601
chr15-47525463-INS-89
chr15-47549981-INS-110
chr15-47552142-INS-558
chr15-47573324-INS-88
chr15-47609176-INS-52
chr15-47644976-INS-67
chr15-47645092-INS-293
chr15-47650880-INS-188
chr15-47655713-INS-7196
chr15-47661509-INS-3331
chr15-47720174-INS-147
chr15-47720315-INS-321
chr15-47731178-INS-1965
chr15-47778680-INS-215
chr15-47778680-INS-68
chr15-47836875-INS-224
chr15-47840645-INS-207
chr15-47845870-INS-2352
chr15-47846098-INS-52
chr15-47846503-INS-64
chr15-47847219-INS-72
chr15-47871361-INS-8941
chr15-47930761-INS-50
chr15-47952056-INS-55
chr15-47984246-INS-1755
chr15-48095343-INS-297
chr15-48118681-INS-51
chr15-48132311-INS-1959
chr15-48164175-INS-56
chr15-48232605-INS-6186
chr15-48254892-INS-201
chr15-48278534-INS-132
chr15-48282390-INS-1775
chr15-48285885-INS-98

chr15-48305994-INS-7213
chr15-48325648-INS-51
chr15-48369703-INS-1130
chr15-48409694-INS-104
chr15-48455170-INS-76
chr15-48520584-INS-63
chr15-48537190-INS-3099
chr15-48551472-INS-400
chr15-48612289-INS-406
chr15-48643554-INS-105
chr15-48738678-INS-6722
chr15-48751973-INS-76
chr15-48760182-INS-6611
chr15-48806289-INS-6734
chr15-48822002-INS-71
chr15-48826636-INS-5952
chr15-48839475-INS-6458
chr15-48919051-INS-1007
chr15-48923105-INS-7335
chr15-48934645-INS-114
chr15-48938925-INS-106
chr15-48941321-INS-88
chr15-48945225-INS-212
chr15-48987544-INS-1474
chr15-48998931-INS-452
chr15-49012737-INS-549
chr15-49019220-INS-59
chr15-49021865-INS-1612
chr15-49030987-INS-51
chr15-49060337-INS-6234
chr15-49060675-INS-52
chr15-49068815-INS-1735
chr15-49081907-INS-109
chr15-49083293-INS-4119
chr15-49177685-INS-1272
chr15-49220413-INS-153
chr15-49220459-INS-54
chr15-49245009-INS-67
chr15-49253586-INS-900
chr15-49255827-INS-1243
chr15-49259721-INS-178
chr15-49278206-INS-99
chr15-49309434-INS-158
chr15-49309814-INS-140
chr15-49329806-INS-6059
chr15-49365618-INS-366
chr15-49369728-INS-110
chr15-49387634-INS-50
chr15-49399441-INS-50
chr15-49412054-INS-490
chr15-49414732-INS-60
chr15-49428621-INS-366
chr15-49430398-INS-68
chr15-49430638-INS-701
chr15-49441665-INS-4149
chr15-49449642-INS-5956
chr15-49481988-INS-638
chr15-49485190-INS-54
chr15-49513559-INS-1175
chr15-49556802-INS-1964
chr15-49567055-INS-54
chr15-49568013-INS-64
chr15-49577498-INS-66
chr15-49586496-INS-4004
chr15-49614224-INS-70
chr15-49614980-INS-54
chr15-49617140-INS-368
chr15-49617453-INS-80
chr15-49617545-INS-1774
chr15-49617769-INS-160
chr15-49621266-INS-75
chr15-49692223-INS-2920
chr15-49730653-INS-10385
chr15-49744835-INS-56
chr15-49747845-INS-309
chr15-49771773-INS-7351
chr15-49780128-INS-642

chr15-49797088-INS-2588
chr15-49807670-INS-866
chr15-49808092-INS-653
chr15-49837598-INS-6233
chr15-49878484-INS-6239
chr15-49882323-INS-1821
chr15-49916720-INS-3101
chr15-49919714-INS-152
chr15-49930693-INS-133
chr15-49943212-INS-7048
chr15-49960007-INS-76
chr15-49963996-INS-6381
chr15-49974824-INS-57
chr15-50015521-INS-64
chr15-50041545-INS-5924
chr15-50061740-INS-1805
chr15-50072118-INS-168
chr15-50075940-INS-2976
chr15-50092464-INS-75
chr15-50118481-INS-1028
chr15-50121748-INS-999
chr15-50130140-INS-58
chr15-50130451-INS-281
chr15-50145664-INS-6334
chr15-50200122-INS-54
chr15-50210057-INS-429
chr15-50212209-INS-2739
chr15-50236332-INS-50
chr15-50239049-INS-182
chr15-50239049-INS-76
chr15-50259737-INS-55
chr15-50273437-INS-7261
chr15-50283765-INS-104
chr15-50289295-INS-84
chr15-50308424-INS-52
chr15-50346519-INS-638
chr15-50410499-INS-50
chr15-50415703-INS-13548
chr15-50415703-INS-6106
chr15-50485659-INS-1688
chr15-50509968-INS-52
chr15-50550559-INS-60
chr15-50583548-INS-62
chr15-50584394-INS-140
chr15-50658959-INS-7177
chr15-50711383-INS-4360
chr15-50761724-INS-260
chr15-50761724-INS-86
chr15-50771181-INS-1726
chr15-50783684-INS-66
chr15-50806484-INS-200
chr15-50806519-INS-507
chr15-50863959-INS-6637
chr15-50878968-INS-109
chr15-50928810-INS-58
chr15-51012874-INS-1074
chr15-51015369-INS-1168
chr15-51018710-INS-111
chr15-51020807-INS-114
chr15-51030298-INS-50
chr15-51031085-INS-6195
chr15-51058649-INS-87
chr15-51058749-INS-261
chr15-51061648-INS-461
chr15-51061648-INS-76
chr15-51164453-INS-2894
chr15-51176254-INS-2781
chr15-51177065-INS-111
chr15-51258498-INS-377
chr15-51258516-INS-75
chr15-51281919-INS-255
chr15-51283702-INS-62
chr15-51316461-INS-10104
chr15-51317300-INS-150
chr15-51331516-INS-294
chr15-51353259-INS-69
chr15-51391764-INS-52

chr15-51401140-INS-58
chr15-51402809-INS-50
chr15-51403702-INS-6334
chr15-51519388-INS-121
chr15-51543789-INS-54
chr15-51543870-INS-138
chr15-51571878-INS-76
chr15-51626906-INS-64
chr15-51710838-INS-181
chr15-51795761-INS-94
chr15-51796654-INS-70
chr15-51858992-INS-212
chr15-51873207-INS-8952
chr15-51905940-INS-205
chr15-51925467-INS-145
chr15-51925748-INS-174
chr15-51929140-INS-90
chr15-51952597-INS-2699
chr15-51982285-INS-84
chr15-52037528-INS-1552
chr15-52037719-INS-121
chr15-52037915-INS-633
chr15-52070020-INS-53
chr15-52070737-INS-6212
chr15-52079683-INS-65
chr15-52080087-INS-160
chr15-52090753-INS-3716
chr15-52106353-INS-51
chr15-52201219-INS-332
chr15-52208857-INS-1245
chr15-52221695-INS-4673
chr15-52235807-INS-122
chr15-52240995-INS-4876
chr15-52284400-INS-14994
chr15-52359335-INS-229
chr15-52367546-INS-3060
chr15-52384795-INS-2484
chr15-52385725-INS-11413
chr15-52390860-INS-62
chr15-52397933-INS-206
chr15-52402162-INS-7067
chr15-52432434-INS-6353
chr15-52460071-INS-485
chr15-52467742-INS-6456
chr15-52569687-INS-80
chr15-52632834-INS-67
chr15-52645135-INS-1122
chr15-52683068-INS-3291
chr15-52684164-INS-484
chr15-52721980-INS-391
chr15-52732623-INS-160
chr15-52758397-INS-6188
chr15-52769401-INS-432
chr15-52790148-INS-50
chr15-52790756-INS-252
chr15-52798737-INS-2826
chr15-52819498-INS-3373
chr15-52824071-INS-110
chr15-52841537-INS-451
chr15-52849090-INS-8783
chr15-52859921-INS-6280
chr15-52910024-INS-256
chr15-52915412-INS-78
chr15-52916126-INS-662
chr15-52943751-INS-141
chr15-52984802-INS-60
chr15-53011654-INS-69
chr15-53081198-INS-157
chr15-53094693-INS-208
chr15-53116455-INS-1616
chr15-53134108-INS-88
chr15-53150651-INS-82
chr15-53151263-INS-193
chr15-53164478-INS-60
chr15-53181405-INS-491
chr15-53201267-INS-196
chr15-53201505-INS-367

chr15-53230027-INS-152
chr15-53231469-INS-101
chr15-53246006-INS-73
chr15-53309904-INS-6518
chr15-53362903-INS-7801
chr15-53373713-INS-6419
chr15-53392500-INS-72
chr15-53482337-INS-54
chr15-53505821-INS-50
chr15-53523762-INS-50
chr15-53536608-INS-72
chr15-53552417-INS-4175
chr15-53573019-INS-290
chr15-53583360-INS-54
chr15-53584857-INS-284
chr15-53589746-INS-90
chr15-53590508-INS-56
chr15-53594998-INS-7299
chr15-53622915-INS-68
chr15-53650125-INS-50
chr15-53717274-INS-7242
chr15-53742746-INS-1611
chr15-53777887-INS-895
chr15-53804845-INS-935
chr15-53807760-INS-1892
chr15-53854760-INS-264
chr15-53886557-INS-200
chr15-53909998-INS-60
chr15-53922220-INS-2990
chr15-54024438-INS-56
chr15-54038625-INS-66
chr15-54040372-INS-100
chr15-54042380-INS-100
chr15-54058240-INS-52
chr15-54148286-INS-65
chr15-54186283-INS-86
chr15-54211488-INS-434
chr15-54370134-INS-64
chr15-54374162-INS-105
chr15-54375021-INS-52
chr15-54375037-INS-120
chr15-54428101-INS-240
chr15-54446418-INS-69
chr15-54487273-INS-182
chr15-54499630-INS-1938
chr15-54517711-INS-2617
chr15-54526519-INS-69
chr15-54537169-INS-6292
chr15-54558642-INS-60
chr15-54564825-INS-1964
chr15-54577961-INS-4662
chr15-54605734-INS-1065
chr15-54610497-INS-112
chr15-54611654-INS-93
chr15-54642554-INS-52
chr15-54664138-INS-55
chr15-54667049-INS-1237
chr15-54706470-INS-363
chr15-54717425-INS-52
chr15-54788833-INS-421
chr15-54842196-INS-99
chr15-54845437-INS-972
chr15-54845470-INS-140
chr15-54848922-INS-477
chr15-54864372-INS-159
chr15-54944567-INS-113
chr15-54946222-INS-315
chr15-54973448-INS-184
chr15-54974205-INS-500
chr15-55062064-INS-50
chr15-55072909-INS-51
chr15-55073601-INS-58
chr15-55087581-INS-78
chr15-55098815-INS-52
chr15-55125121-INS-8982
chr15-55157218-INS-256
chr15-55161933-INS-172

chr15-55162169-INS-271
chr15-55179535-INS-1809
chr15-55184137-INS-451
chr15-55190958-INS-1046
chr15-55217704-INS-67
chr15-55243719-INS-2023
chr15-55251592-INS-56
chr15-55321621-INS-87
chr15-55429969-INS-1083
chr15-55476155-INS-1520
chr15-55476977-INS-307
chr15-55478833-INS-1785
chr15-55507897-INS-2830
chr15-55516324-INS-557
chr15-55547238-INS-212
chr15-55556893-INS-89
chr15-55561339-INS-328
chr15-55566129-INS-56
chr15-55587588-INS-777
chr15-55618989-INS-50
chr15-55620672-INS-54
chr15-55626933-INS-1988
chr15-55705485-INS-6211
chr15-55751497-INS-108
chr15-55809972-INS-72
chr15-55836980-INS-377
chr15-55876705-INS-6576
chr15-55888400-INS-54
chr15-55910551-INS-3113
chr15-55916714-INS-3874
chr15-56083841-INS-6795
chr15-56098401-INS-278
chr15-56106736-INS-104
chr15-56130068-INS-2643
chr15-56171560-INS-4768
chr15-56172704-INS-50
chr15-56176469-INS-52
chr15-56203584-INS-74
chr15-56238970-INS-60
chr15-56287408-INS-13949
chr15-56357123-INS-120
chr15-56368855-INS-6488
chr15-56405015-INS-51
chr15-56405102-INS-498
chr15-56431834-INS-6565
chr15-56447989-INS-2729
chr15-56453153-INS-860
chr15-56454268-INS-448
chr15-56481043-INS-198
chr15-56485205-INS-2235
chr15-56494090-INS-72
chr15-56520434-INS-110
chr15-56550027-INS-337
chr15-56550106-INS-199
chr15-56568655-INS-71
chr15-56582233-INS-54
chr15-56596802-INS-77
chr15-56629172-INS-98
chr15-56688910-INS-6617
chr15-56692769-INS-72
chr15-56716405-INS-92
chr15-56730120-INS-251
chr15-56731857-INS-1178
chr15-56828550-INS-386
chr15-56884266-INS-1489
chr15-56901548-INS-54
chr15-56904610-INS-54
chr15-56905488-INS-147
chr15-56905488-INS-70
chr15-56912051-INS-756
chr15-56971626-INS-57
chr15-56973488-INS-68
chr15-56987838-INS-200
chr15-57001516-INS-117
chr15-57031703-INS-6254
chr15-57099029-INS-6380
chr15-57119387-INS-406

chr15-57150957-INS-51
chr15-57174760-INS-5621
chr15-57179529-INS-303
chr15-57191712-INS-183
chr15-57192550-INS-50
chr15-57206361-INS-1855
chr15-57206476-INS-169
chr15-57209204-INS-6342
chr15-57240549-INS-54
chr15-57255338-INS-60
chr15-57270997-INS-1700
chr15-57354723-INS-6190
chr15-57357868-INS-3142
chr15-57378837-INS-76
chr15-57405368-INS-75
chr15-57405554-INS-52
chr15-57453230-INS-78
chr15-57461793-INS-6369
chr15-57488095-INS-1320
chr15-57488606-INS-118
chr15-57488618-INS-51
chr15-57499881-INS-58
chr15-57544357-INS-1109
chr15-57545928-INS-135
chr15-57610768-INS-52
chr15-57705256-INS-1299
chr15-57707028-INS-70
chr15-57707095-INS-315
chr15-57716503-INS-122
chr15-57716547-INS-51
chr15-57732850-INS-114
chr15-57777058-INS-282
chr15-57777118-INS-578
chr15-57794675-INS-1380
chr15-57796373-INS-207
chr15-57807069-INS-586
chr15-57817141-INS-62
chr15-57820897-INS-315
chr15-57832091-INS-92
chr15-57850882-INS-50
chr15-57851818-INS-110
chr15-57883266-INS-50
chr15-57905423-INS-56
chr15-57926762-INS-343
chr15-57926809-INS-7824
chr15-58009257-INS-139
chr15-58034773-INS-555
chr15-58038648-INS-1197
chr15-58038648-INS-460
chr15-58065585-INS-113
chr15-58084787-INS-54
chr15-58108402-INS-205
chr15-58112599-INS-60
chr15-58112747-INS-277
chr15-58193309-INS-132
chr15-58194541-INS-54
chr15-58212618-INS-50
chr15-58236074-INS-7040
chr15-58310143-INS-1824
chr15-58311391-INS-3359
chr15-58334583-INS-130
chr15-58361519-INS-105
chr15-58362547-INS-54
chr15-58368742-INS-2205
chr15-58386284-INS-112
chr15-58422167-INS-66
chr15-58527776-INS-73
chr15-58534061-INS-51
chr15-58596123-INS-216
chr15-58611201-INS-233
chr15-58620124-INS-78
chr15-58620484-INS-1925
chr15-58686065-INS-50
chr15-58705955-INS-560
chr15-58787554-INS-246
chr15-58840181-INS-147
chr15-58848144-INS-448

chr15-58849698-INS-67
chr15-58856463-INS-52
chr15-58884724-INS-6698
chr15-58911127-INS-799
chr15-58922828-INS-585
chr15-58923126-INS-2200
chr15-58934932-INS-856
chr15-58988873-INS-82
chr15-59016241-INS-164
chr15-59017449-INS-138
chr15-59032597-INS-186
chr15-59044286-INS-80
chr15-59052489-INS-1386
chr15-59080269-INS-103
chr15-59080500-INS-164
chr15-59080504-INS-53
chr15-59114477-INS-52
chr15-59114561-INS-382
chr15-59137094-INS-82
chr15-59176973-INS-259
chr15-59177031-INS-80
chr15-59185040-INS-207
chr15-59251918-INS-76
chr15-59251986-INS-182
chr15-59261426-INS-670
chr15-59339638-INS-60
chr15-59422896-INS-225
chr15-59454078-INS-206
chr15-59456575-INS-50
chr15-59495006-INS-63
chr15-59503181-INS-176
chr15-59512858-INS-108
chr15-59536279-INS-59
chr15-59536698-INS-60
chr15-59543326-INS-102
chr15-59550499-INS-321
chr15-59553248-INS-50
chr15-59555796-INS-52
chr15-59557796-INS-691
chr15-59562725-INS-56
chr15-59575705-INS-1651
chr15-59576023-INS-72
chr15-59601804-INS-279
chr15-59601804-INS-50
chr15-59624464-INS-60
chr15-59628842-INS-60
chr15-59629595-INS-132
chr15-59634745-INS-1822
chr15-59661685-INS-52
chr15-59670578-INS-1886
chr15-59684555-INS-71
chr15-59752016-INS-59
chr15-59760811-INS-68
chr15-59762173-INS-478
chr15-59790765-INS-133
chr15-59807173-INS-2465
chr15-59831427-INS-60
chr15-59834016-INS-214
chr15-59867485-INS-162
chr15-59928993-INS-108
chr15-59932703-INS-855
chr15-59953179-INS-196
chr15-59970313-INS-601
chr15-59990730-INS-7092
chr15-60005218-INS-52
chr15-60017227-INS-2489
chr15-60020858-INS-72
chr15-60125601-INS-68
chr15-60167340-INS-2735
chr15-60171876-INS-52
chr15-60184744-INS-72
chr15-60185470-INS-126
chr15-60197356-INS-191
chr15-60237040-INS-600
chr15-60237474-INS-50
chr15-60249380-INS-630
chr15-60257911-INS-78

chr15-60360030-INS-61
chr15-60392855-INS-215
chr15-60393221-INS-212
chr15-60393221-INS-83
chr15-60431199-INS-166
chr15-60439377-INS-61
chr15-60440395-INS-72
chr15-60443644-INS-7197
chr15-60459638-INS-60
chr15-60506100-INS-217
chr15-60531169-INS-2943
chr15-60535195-INS-6430
chr15-60568164-INS-218
chr15-60572806-INS-55
chr15-60596760-INS-70
chr15-60629349-INS-50
chr15-60629528-INS-113
chr15-60640646-INS-338
chr15-60643624-INS-555
chr15-60643644-INS-183
chr15-60644148-INS-6436
chr15-60698580-INS-52
chr15-60715845-INS-92
chr15-60722517-INS-89
chr15-60852727-INS-489
chr15-60852893-INS-136
chr15-60860177-INS-57
chr15-60877506-INS-102
chr15-60904262-INS-1238
chr15-60912448-INS-90
chr15-60929509-INS-76
chr15-60930751-INS-81
chr15-60933568-INS-6225
chr15-60973456-INS-4760
chr15-60988182-INS-50
chr15-61009866-INS-425
chr15-61009984-INS-113
chr15-61060009-INS-236
chr15-61090334-INS-2647
chr15-61106711-INS-1107
chr15-61122061-INS-501
chr15-61126007-INS-6587
chr15-61192451-INS-6233
chr15-61194021-INS-55
chr15-61231294-INS-54
chr15-61260763-INS-6271
chr15-61290807-INS-6487
chr15-61311153-INS-50
chr15-61347862-INS-2389
chr15-61348622-INS-7090
chr15-61385859-INS-7110
chr15-61396952-INS-97
chr15-61412067-INS-55
chr15-61477172-INS-2661
chr15-61509505-INS-84
chr15-61524587-INS-60
chr15-61546939-INS-1188
chr15-61573377-INS-493
chr15-61592624-INS-594
chr15-61613163-INS-735
chr15-61636996-INS-6362
chr15-61681622-INS-628
chr15-61685419-INS-1803
chr15-61685419-INS-831
chr15-61694025-INS-140
chr15-61711049-INS-278
chr15-61729843-INS-203
chr15-61746250-INS-50
chr15-61770526-INS-2889
chr15-61778238-INS-192
chr15-61786210-INS-482
chr15-61789307-INS-6693
chr15-61815751-INS-309
chr15-61971848-INS-51
chr15-61992397-INS-60
chr15-62023026-INS-87
chr15-62030058-INS-74

chr15-62059340-INS-52
chr15-62100716-INS-390
chr15-62172871-INS-58
chr15-62200746-INS-88
chr15-62232086-INS-50
chr15-62239860-INS-6491
chr15-62248475-INS-1914
chr15-62265003-INS-50
chr15-62292378-INS-6492
chr15-62314820-INS-297
chr15-62340928-INS-50
chr15-62348696-INS-50
chr15-62408728-INS-1186
chr15-62415734-INS-288
chr15-62433626-INS-62
chr15-62445729-INS-6398
chr15-62451305-INS-50
chr15-62457765-INS-76
chr15-62469034-INS-6374
chr15-62481540-INS-1776
chr15-62510839-INS-6709
chr15-62513863-INS-171
chr15-62514068-INS-68
chr15-62541893-INS-3602
chr15-62666849-INS-55
chr15-62677142-INS-58
chr15-62756494-INS-3453
chr15-62762317-INS-80
chr15-62781277-INS-95
chr15-62790829-INS-55
chr15-62851049-INS-8971
chr15-62855383-INS-2756
chr15-62878242-INS-207
chr15-62882794-INS-51
chr15-62898739-INS-2088
chr15-62996486-INS-2818
chr15-63022839-INS-6016
chr15-63035224-INS-52
chr15-63084447-INS-56
chr15-63157288-INS-329
chr15-63234797-INS-6949
chr15-63246539-INS-454
chr15-63262152-INS-63
chr15-63278031-INS-63
chr15-63278652-INS-493
chr15-63281581-INS-938
chr15-63281609-INS-369
chr15-63315748-INS-65
chr15-63336760-INS-52
chr15-63343996-INS-68
chr15-63383656-INS-1732
chr15-63394785-INS-99
chr15-63444982-INS-102
chr15-63458746-INS-133
chr15-63460403-INS-6250
chr15-63506784-INS-115
chr15-63523661-INS-75
chr15-63537902-INS-2053
chr15-63538963-INS-1053
chr15-63540716-INS-1268
chr15-63567470-INS-228
chr15-63583216-INS-1722
chr15-63624381-INS-110
chr15-63624609-INS-96
chr15-63631057-INS-54
chr15-63691601-INS-1142
chr15-63712155-INS-94
chr15-63721194-INS-189
chr15-63750874-INS-67
chr15-63754802-INS-33693
chr15-63804034-INS-384
chr15-63820244-INS-50
chr15-63860215-INS-142
chr15-63884012-INS-133
chr15-63885883-INS-61
chr15-63900778-INS-53
chr15-63908722-INS-139

chr15-63927970-INS-60
chr15-64080071-INS-116
chr15-64098439-INS-666
chr15-64098447-INS-87
chr15-64135085-INS-449
chr15-64147644-INS-114
chr15-64168715-INS-411
chr15-64168715-INS-84
chr15-64227246-INS-66
chr15-64238954-INS-207
chr15-64277747-INS-119
chr15-64294880-INS-126
chr15-64298999-INS-2996
chr15-64328788-INS-439
chr15-64334345-INS-52
chr15-64366896-INS-7041
chr15-64417553-INS-517
chr15-64433362-INS-170
chr15-64437469-INS-277
chr15-64437964-INS-1889
chr15-64445388-INS-51
chr15-64460542-INS-111
chr15-64506392-INS-1490
chr15-64519421-INS-72
chr15-64521147-INS-6305
chr15-64526972-INS-144
chr15-64556128-INS-122
chr15-64557811-INS-1895
chr15-64575053-INS-67
chr15-64575467-INS-56
chr15-64593943-INS-853
chr15-64610632-INS-6326
chr15-64611134-INS-2368
chr15-64621797-INS-84
chr15-64680614-INS-56
chr15-64712205-INS-84
chr15-64724807-INS-50
chr15-64732088-INS-94
chr15-64738537-INS-5251
chr15-64760958-INS-50
chr15-64761323-INS-112
chr15-64784538-INS-959
chr15-64811555-INS-75
chr15-64811963-INS-380
chr15-64811963-INS-64
chr15-64817256-INS-244
chr15-64817256-INS-70
chr15-64820863-INS-10830
chr15-64841591-INS-304
chr15-64842034-INS-1281
chr15-64850068-INS-93
chr15-64879605-INS-1758
chr15-64911191-INS-75
chr15-64933406-INS-90
chr15-64963287-INS-2154
chr15-64980727-INS-68
chr15-65007795-INS-50
chr15-65065581-INS-144
chr15-65104448-INS-6564
chr15-65132790-INS-84
chr15-65167161-INS-63
chr15-65171623-INS-266
chr15-65199140-INS-1770
chr15-65200058-INS-379
chr15-65203381-INS-2229
chr15-65262667-INS-8854
chr15-65266933-INS-206
chr15-65267919-INS-50
chr15-65308480-INS-6237
chr15-65344593-INS-1387
chr15-65352168-INS-52
chr15-65355034-INS-728
chr15-65364087-INS-79
chr15-65394455-INS-64
chr15-65396383-INS-70
chr15-65410506-INS-76
chr15-65421320-INS-1102

chr15-65446162-INS-3742
chr15-65452041-INS-6870
chr15-65535017-INS-96
chr15-65594544-INS-957
chr15-65618245-INS-203
chr15-65618756-INS-54
chr15-65623522-INS-68
chr15-65762002-INS-53
chr15-65839113-INS-58
chr15-65879396-INS-111
chr15-65879439-INS-53
chr15-66030313-INS-2070
chr15-66077421-INS-224
chr15-66138399-INS-50
chr15-66198886-INS-74
chr15-66199041-INS-61
chr15-66226833-INS-54
chr15-66258458-INS-6669
chr15-66262153-INS-2547
chr15-66307231-INS-397
chr15-66314893-INS-5733
chr15-66332736-INS-506
chr15-66339313-INS-2790
chr15-66371040-INS-72
chr15-66373323-INS-61
chr15-66443446-INS-2475
chr15-66451305-INS-125
chr15-66520492-INS-6269
chr15-66546406-INS-406
chr15-66682353-INS-225
chr15-66693776-INS-374
chr15-66715572-INS-62
chr15-66726738-INS-68
chr15-66744624-INS-2585
chr15-66744655-INS-85
chr15-66787352-INS-72
chr15-66842690-INS-50
chr15-66945696-INS-58
chr15-66955835-INS-461
chr15-67032310-INS-80
chr15-67034152-INS-100
chr15-67064749-INS-2094
chr15-67066554-INS-178
chr15-67067109-INS-52
chr15-67083693-INS-593
chr15-67084383-INS-50
chr15-67112049-INS-52
chr15-67125389-INS-4202
chr15-67157371-INS-449
chr15-67175011-INS-1131
chr15-67189639-INS-84
chr15-67195747-INS-4915
chr15-67216042-INS-81
chr15-67218231-INS-63
chr15-67222921-INS-64
chr15-67240536-INS-65
chr15-67305062-INS-50
chr15-67358872-INS-60
chr15-67369744-INS-1112
chr15-67398945-INS-1084
chr15-67481890-INS-94
chr15-67495477-INS-413
chr15-67496280-INS-178
chr15-67527807-INS-2273
chr15-67543462-INS-58
chr15-67581709-INS-755
chr15-67594309-INS-160
chr15-67603727-INS-70
chr15-67604986-INS-1504
chr15-67613254-INS-207
chr15-67618969-INS-6273
chr15-67631477-INS-3138
chr15-67641411-INS-150
chr15-67660475-INS-494
chr15-67695383-INS-2323
chr15-67735614-INS-60
chr15-67852781-INS-340

chr15-67852781-INS-68
chr15-67924272-INS-50
chr15-67933448-INS-203
chr15-67934993-INS-74
chr15-67987388-INS-67
chr15-67989424-INS-78
chr15-68019027-INS-63
chr15-68027214-INS-312
chr15-68042768-INS-54
chr15-68090786-INS-50
chr15-68131517-INS-5541
chr15-68145315-INS-54
chr15-68156937-INS-50
chr15-68158355-INS-54
chr15-68170577-INS-6169
chr15-68253027-INS-50
chr15-68262902-INS-2323
chr15-68274118-INS-60
chr15-68293300-INS-74
chr15-68412983-INS-4081
chr15-68417243-INS-6369
chr15-68434557-INS-54
chr15-68443360-INS-2364
chr15-68487599-INS-113
chr15-68525059-INS-2724
chr15-68527366-INS-7190
chr15-68534452-INS-65
chr15-68569230-INS-6498
chr15-68569832-INS-520
chr15-68570071-INS-87
chr15-68618852-INS-1199
chr15-68656036-INS-147
chr15-68657382-INS-56
chr15-68731242-INS-51
chr15-68769916-INS-83
chr15-68772950-INS-106
chr15-68772957-INS-265
chr15-68822510-INS-1209
chr15-68830528-INS-75
chr15-68887648-INS-64
chr15-68887833-INS-120
chr15-68892651-INS-493
chr15-68897198-INS-278
chr15-69028862-INS-435
chr15-69031132-INS-226
chr15-69126541-INS-113
chr15-69152141-INS-52
chr15-69178400-INS-205
chr15-69212549-INS-206
chr15-69213755-INS-152
chr15-69217462-INS-390
chr15-69253437-INS-547
chr15-69276293-INS-110
chr15-69276681-INS-54
chr15-69278286-INS-1404
chr15-69312056-INS-204
chr15-69316898-INS-5998
chr15-69331194-INS-203
chr15-69334278-INS-163
chr15-69341914-INS-127
chr15-69345764-INS-156
chr15-69374919-INS-58
chr15-69417321-INS-69
chr15-69426670-INS-140
chr15-69427187-INS-4333
chr15-69496930-INS-54
chr15-69565329-INS-6266
chr15-69566259-INS-1730
chr15-69574547-INS-74
chr15-69578463-INS-6661
chr15-69593434-INS-442
chr15-69620849-INS-6265
chr15-69622601-INS-51
chr15-69650067-INS-582
chr15-69659133-INS-273
chr15-69663076-INS-668
chr15-69667717-INS-139

chr15-69676593-INS-5619
chr15-69690612-INS-1162
chr15-69709407-INS-878
chr15-69719380-INS-409
chr15-69719724-INS-70
chr15-69738347-INS-60
chr15-69739507-INS-60
chr15-69766003-INS-51
chr15-69767032-INS-6390
chr15-69776719-INS-3841
chr15-69799213-INS-64
chr15-69818517-INS-336
chr15-69855070-INS-72
chr15-69932036-INS-68
chr15-69972586-INS-50
chr15-70006283-INS-1068
chr15-70009743-INS-1263
chr15-70099894-INS-68
chr15-70115254-INS-3470
chr15-70126103-INS-2721
chr15-70132621-INS-110
chr15-70140390-INS-62
chr15-70141971-INS-1000
chr15-70143373-INS-72
chr15-70152378-INS-6524
chr15-70164791-INS-2231
chr15-70190050-INS-92
chr15-70201420-INS-213
chr15-70201897-INS-109
chr15-70218684-INS-88
chr15-70234901-INS-71
chr15-70251361-INS-1894
chr15-70262149-INS-114
chr15-70290622-INS-55
chr15-70322723-INS-3043
chr15-70328417-INS-1264
chr15-70331258-INS-1122
chr15-70331808-INS-6621
chr15-70332960-INS-196
chr15-70332995-INS-53
chr15-70366525-INS-52
chr15-70382384-INS-68
chr15-70465064-INS-81
chr15-70467373-INS-88
chr15-70490882-INS-3094
chr15-70497292-INS-3401
chr15-70535902-INS-1676
chr15-70542311-INS-124
chr15-70542311-INS-258
chr15-70549921-INS-504
chr15-70602946-INS-216
chr15-70605384-INS-251
chr15-70606170-INS-431
chr15-70695577-INS-471
chr15-70707325-INS-50
chr15-70741873-INS-514
chr15-70743150-INS-1294
chr15-70751139-INS-50
chr15-70792012-INS-70
chr15-70819141-INS-82
chr15-70837429-INS-375
chr15-70837635-INS-74
chr15-70855672-INS-57
chr15-70859893-INS-6260
chr15-70891829-INS-106
chr15-70962553-INS-60
chr15-70963089-INS-56
chr15-70966405-INS-343
chr15-70987663-INS-68
chr15-71003960-INS-60
chr15-71006857-INS-578
chr15-71031863-INS-50
chr15-71035147-INS-4610
chr15-71060380-INS-855
chr15-71082774-INS-710
chr15-71130468-INS-6214
chr15-71134676-INS-108

chr15-71159979-INS-6433
chr15-71178900-INS-56
chr15-71190956-INS-52
chr15-71194116-INS-52
chr15-71196318-INS-1224
chr15-71198967-INS-57
chr15-71208937-INS-930
chr15-71218799-INS-332
chr15-71237436-INS-1575
chr15-71238506-INS-88
chr15-71238709-INS-65
chr15-71251155-INS-57
chr15-71254427-INS-758
chr15-71258743-INS-80
chr15-71258787-INS-209
chr15-71348672-INS-64
chr15-71360054-INS-81
chr15-71360319-INS-54
chr15-71381384-INS-6654
chr15-71392367-INS-63
chr15-71409909-INS-7043
chr15-71419980-INS-62
chr15-71445878-INS-263
chr15-71494209-INS-92
chr15-71499857-INS-440
chr15-71508552-INS-72
chr15-71528706-INS-1105
chr15-71586592-INS-84
chr15-71633582-INS-196
chr15-71636195-INS-1254
chr15-71655470-INS-60
chr15-71660274-INS-66
chr15-71736191-INS-93
chr15-71745140-INS-56
chr15-71767236-INS-6457
chr15-71829185-INS-112
chr15-71845081-INS-76
chr15-71861159-INS-107
chr15-71862385-INS-419
chr15-71892849-INS-7304
chr15-71893874-INS-127
chr15-71910982-INS-52
chr15-71918149-INS-151
chr15-71918169-INS-56
chr15-71918268-INS-384
chr15-71928953-INS-62
chr15-71938539-INS-115
chr15-72018345-INS-50
chr15-72043747-INS-6262
chr15-72043829-INS-88
chr15-72088311-INS-1196
chr15-72105604-INS-1246
chr15-72280429-INS-6915
chr15-72307372-INS-2190
chr15-72307380-INS-4381
chr15-72322968-INS-128
chr15-72322995-INS-60
chr15-72334268-INS-2428
chr15-72361855-INS-62
chr15-72373220-INS-243
chr15-72448995-INS-50
chr15-72455149-INS-51
chr15-72482567-INS-284
chr15-72597850-INS-55
chr15-72645729-INS-52
chr15-72685890-INS-512
chr15-72722370-INS-564
chr15-72722463-INS-68
chr15-72778382-INS-56
chr15-72800390-INS-3374
chr15-72835215-INS-85
chr15-72835722-INS-283
chr15-72865500-INS-52
chr15-72892113-INS-213
chr15-72940405-INS-50
chr15-73009486-INS-54
chr15-73009512-INS-230

chr15-73048131-INS-125
chr15-73078474-INS-50
chr15-73092265-INS-200
chr15-73137192-INS-1849
chr15-73222464-INS-54
chr15-73258189-INS-210
chr15-73319387-INS-52
chr15-73344682-INS-1197
chr15-73344962-INS-92
chr15-73345329-INS-204
chr15-73348036-INS-237
chr15-73357198-INS-197
chr15-73357865-INS-227
chr15-73367381-INS-84
chr15-73374911-INS-92
chr15-73375507-INS-204
chr15-73427280-INS-50
chr15-73430486-INS-181
chr15-73549716-INS-70
chr15-73549756-INS-1675
chr15-73559131-INS-50
chr15-73572788-INS-70
chr15-73590240-INS-50
chr15-73646672-INS-118
chr15-73646922-INS-118
chr15-73656038-INS-66
chr15-73656498-INS-64
chr15-73656548-INS-240
chr15-73712757-INS-689
chr15-73732166-INS-104
chr15-73763651-INS-63
chr15-73850190-INS-460
chr15-73888780-INS-202
chr15-73888991-INS-109
chr15-73892902-INS-52
chr15-73904285-INS-75
chr15-73977371-INS-561
chr15-73977737-INS-72
chr15-73979793-INS-54
chr15-73998553-INS-54
chr15-74017326-INS-54
chr15-74084129-INS-60
chr15-74119800-INS-2340
chr15-74159869-INS-585
chr15-74162605-INS-65
chr15-74239842-INS-410
chr15-74298857-INS-27306
chr15-74309756-INS-50
chr15-74377339-INS-64
chr15-74424651-INS-1281
chr15-74496200-INS-58
chr15-74562238-INS-440
chr15-74630740-INS-130
chr15-74630748-INS-338
chr15-74631103-INS-380
chr15-74673365-INS-61
chr15-74703063-INS-462
chr15-74706304-INS-54
chr15-74759868-INS-266
chr15-74780055-INS-105
chr15-74792236-INS-67
chr15-74802012-INS-233
chr15-74879723-INS-72
chr15-74880753-INS-3252
chr15-74928298-INS-251
chr15-74943818-INS-498
chr15-75015092-INS-85
chr15-75062561-INS-104
chr15-75120316-INS-2215
chr15-75162988-INS-50
chr15-75179236-INS-179
chr15-75195581-INS-50
chr15-75244006-INS-5560
chr15-75311742-INS-2253
chr15-75313978-INS-135
chr15-75338091-INS-66
chr15-75376853-INS-24444

chr15-75395809-INS-6453
chr15-75516618-INS-308
chr15-75516696-INS-112
chr15-75519878-INS-118
chr15-75652446-INS-147
chr15-75652508-INS-336
chr15-75657922-INS-610
chr15-75750103-INS-60
chr15-75830021-INS-3905
chr15-75893770-INS-74
chr15-75901297-INS-60
chr15-75984019-INS-4636
chr15-76038921-INS-151
chr15-76193659-INS-217
chr15-76208416-INS-217
chr15-76208416-INS-5847
chr15-76208650-INS-75
chr15-76275965-INS-51
chr15-76307605-INS-209
chr15-76309457-INS-2794
chr15-76309457-INS-694
chr15-76335014-INS-106
chr15-76343782-INS-101
chr15-76357920-INS-88
chr15-76388294-INS-965
chr15-76466914-INS-211
chr15-76570798-INS-602
chr15-76603602-INS-68
chr15-76612503-INS-239
chr15-76633645-INS-241
chr15-76675622-INS-226
chr15-76683080-INS-50
chr15-76697358-INS-332
chr15-76722132-INS-253
chr15-76746298-INS-171
chr15-76747730-INS-74
chr15-76748093-INS-320
chr15-76750776-INS-382
chr15-76750962-INS-434
chr15-76760563-INS-6385
chr15-76779949-INS-107
chr15-76794062-INS-1096
chr15-76798911-INS-174
chr15-76807021-INS-52
chr15-76836272-INS-419
chr15-76836304-INS-987
chr15-76860415-INS-54
chr15-76877891-INS-4438
chr15-76879567-INS-511
chr15-76886841-INS-351
chr15-76887165-INS-432
chr15-76917525-INS-162
chr15-76923882-INS-594
chr15-76950993-INS-66
chr15-76984642-INS-72
chr15-77029310-INS-57
chr15-77060088-INS-211
chr15-77073743-INS-215
chr15-77088495-INS-243
chr15-77110114-INS-84
chr15-77149891-INS-76
chr15-77198838-INS-492
chr15-77246001-INS-1776
chr15-77250425-INS-54
chr15-77299976-INS-781
chr15-77304998-INS-2827
chr15-77332493-INS-1206
chr15-77335294-INS-826
chr15-77354284-INS-233
chr15-77354648-INS-119
chr15-77390023-INS-51
chr15-77394261-INS-633
chr15-77421340-INS-3182
chr15-77428827-INS-52
chr15-77448175-INS-58
chr15-77462112-INS-78
chr15-77462608-INS-56

chr15-77515955-INS-50
chr15-77529220-INS-214
chr15-77535731-INS-419
chr15-77633393-INS-6428
chr15-77636072-INS-686
chr15-77636879-INS-91
chr15-77638097-INS-111
chr15-77639707-INS-76
chr15-77677548-INS-112
chr15-77703818-INS-1460
chr15-77753430-INS-182
chr15-77810591-INS-89
chr15-77839421-INS-56
chr15-77868058-INS-63
chr15-77882926-INS-85
chr15-77923336-INS-52
chr15-77944553-INS-52
chr15-77951112-INS-63
chr15-77961754-INS-117
chr15-77962311-INS-52
chr15-77972623-INS-56
chr15-78022271-INS-57
chr15-78046554-INS-140
chr15-78098799-INS-98
chr15-78140675-INS-1958
chr15-78140913-INS-653
chr15-78140995-INS-784
chr15-78212914-INS-50
chr15-78249999-INS-322
chr15-78262410-INS-1000
chr15-78263951-INS-50
chr15-78267258-INS-384
chr15-78267354-INS-104
chr15-78277000-INS-74
chr15-78277821-INS-181
chr15-78362638-INS-438
chr15-78447514-INS-1371
chr15-78477643-INS-52
chr15-78593750-INS-79
chr15-78612322-INS-60
chr15-78617780-INS-77
chr15-78632011-INS-60
chr15-78637312-INS-2670
chr15-78637402-INS-5864
chr15-78648828-INS-408
chr15-78714649-INS-385
chr15-78748946-INS-168
chr15-78804875-INS-212
chr15-78805795-INS-185
chr15-78822672-INS-213
chr15-78833995-INS-106
chr15-78865052-INS-102
chr15-78892466-INS-251
chr15-78917850-INS-66
chr15-78956753-INS-139
chr15-78977196-INS-59
chr15-78977410-INS-151
chr15-79066760-INS-64
chr15-79068070-INS-658
chr15-79068935-INS-266
chr15-79117678-INS-95
chr15-79121290-INS-112
chr15-79121398-INS-255
chr15-79121412-INS-94
chr15-79172149-INS-594
chr15-79221043-INS-50
chr15-79222942-INS-433
chr15-79267507-INS-238
chr15-79311753-INS-396
chr15-79341620-INS-75
chr15-79346127-INS-662
chr15-79347617-INS-63
chr15-79377577-INS-63
chr15-79436005-INS-216
chr15-79436050-INS-329
chr15-79436641-INS-215
chr15-79445024-INS-224

chr15-79453996-INS-195
chr15-79479973-INS-55
chr15-79491086-INS-208
chr15-79503763-INS-173
chr15-79573024-INS-183
chr15-79615727-INS-214
chr15-79627520-INS-66
chr15-79643486-INS-54
chr15-79661125-INS-4806
chr15-79666828-INS-124
chr15-79667419-INS-72
chr15-79667638-INS-75
chr15-79707815-INS-192
chr15-79737377-INS-205
chr15-79749292-INS-69
chr15-79749554-INS-1331
chr15-79751173-INS-148
chr15-79770033-INS-209
chr15-79829819-INS-187
chr15-79839286-INS-231
chr15-79871418-INS-113
chr15-79871477-INS-53
chr15-79928062-INS-500
chr15-79952891-INS-100
chr15-80014738-INS-203
chr15-80147764-INS-246
chr15-80155437-INS-66
chr15-80188179-INS-126
chr15-80193345-INS-230
chr15-80234526-INS-7163
chr15-80296896-INS-94
chr15-80328634-INS-108
chr15-80412757-INS-78
chr15-80474847-INS-6466
chr15-80481046-INS-117
chr15-80482434-INS-89
chr15-80496858-INS-138
chr15-80554688-INS-54
chr15-80569977-INS-52
chr15-80589656-INS-231
chr15-80625149-INS-457
chr15-80677301-INS-205
chr15-80677313-INS-642
chr15-80750285-INS-68
chr15-80832185-INS-847
chr15-80832385-INS-1120
chr15-80832501-INS-220
chr15-80847349-INS-206
chr15-80866026-INS-73
chr15-80866211-INS-179
chr15-80868323-INS-916
chr15-80886469-INS-84
chr15-80912244-INS-701
chr15-80917648-INS-82
chr15-80924985-INS-84
chr15-80925716-INS-194
chr15-80955760-INS-430
chr15-81014960-INS-183
chr15-81068889-INS-240
chr15-81069108-INS-384
chr15-81092809-INS-250
chr15-81093288-INS-52
chr15-81100592-INS-4115
chr15-81138312-INS-6442
chr15-81150376-INS-1348
chr15-81172904-INS-6606
chr15-81183289-INS-136
chr15-81191249-INS-202
chr15-81191941-INS-50
chr15-81193232-INS-245
chr15-81193996-INS-68
chr15-81212446-INS-227
chr15-81216415-INS-588
chr15-81224504-INS-176
chr15-81267853-INS-55
chr15-81331459-INS-59
chr15-81331741-INS-166

chr15-81332047-INS-124
chr15-81342696-INS-219
chr15-81343639-INS-189
chr15-81361809-INS-442
chr15-81362354-INS-68
chr15-81364713-INS-60
chr15-81373031-INS-122
chr15-81399835-INS-53
chr15-81403609-INS-89
chr15-81441396-INS-382
chr15-81463125-INS-73
chr15-81463596-INS-162
chr15-81475187-INS-140
chr15-81476098-INS-208
chr15-81523414-INS-62
chr15-81531259-INS-76
chr15-81534944-INS-499
chr15-81641761-INS-52
chr15-81671636-INS-172
chr15-81745697-INS-185
chr15-81760341-INS-171
chr15-81761000-INS-163
chr15-81775283-INS-236
chr15-81793428-INS-60
chr15-81803636-INS-997
chr15-81812394-INS-65
chr15-81815475-INS-1110
chr15-81815475-INS-96
chr15-81815574-INS-421
chr15-81816352-INS-58
chr15-81827105-INS-69
chr15-81828451-INS-205
chr15-81846894-INS-180
chr15-81881110-INS-146
chr15-81904589-INS-74
chr15-81909179-INS-59
chr15-81914571-INS-207
chr15-81925030-INS-134
chr15-81944819-INS-84
chr15-81960371-INS-210
chr15-81960902-INS-1024
chr15-81963762-INS-75
chr15-81981653-INS-50
chr15-81985930-INS-68
chr15-81986034-INS-203
chr15-81992268-INS-115
chr15-82098691-INS-165
chr15-82098708-INS-79
chr15-82115540-INS-77
chr15-82115544-INS-381
chr15-82132750-INS-197
chr15-82134096-INS-79
chr15-82134425-INS-50
chr15-82144946-INS-2026
chr15-82147327-INS-712
chr15-82184994-INS-70
chr15-82191903-INS-58
chr15-82196940-INS-185
chr15-82201433-INS-94
chr15-82204247-INS-109
chr15-82246303-INS-403
chr15-82263487-INS-64
chr15-82300345-INS-53
chr15-82304921-INS-174
chr15-82337104-INS-168
chr15-82347068-INS-1977
chr15-82473054-INS-943
chr15-82568190-INS-2075
chr15-82597347-INS-125
chr15-82597571-INS-125
chr15-82597874-INS-313
chr15-82608017-INS-56
chr15-82629662-INS-51
chr15-82660946-INS-3784
chr15-82746986-INS-50
chr15-82770822-INS-219
chr15-82875836-INS-205

chr15-82889369-INS-92
chr15-82959537-INS-226
chr15-82965985-INS-4899
chr15-83013343-INS-800
chr15-83058235-INS-145
chr15-83058881-INS-197
chr15-83059957-INS-205
chr15-83068721-INS-233
chr15-83139462-INS-50
chr15-83144844-INS-186
chr15-83181806-INS-214
chr15-83193295-INS-116
chr15-83193961-INS-60
chr15-83195999-INS-2976
chr15-83199131-INS-206
chr15-83213014-INS-207
chr15-83223852-INS-54
chr15-83236253-INS-574
chr15-83252870-INS-88
chr15-83266989-INS-468
chr15-83312452-INS-150
chr15-83357380-INS-72
chr15-83364350-INS-5560
chr15-83386066-INS-100
chr15-83392713-INS-208
chr15-83441314-INS-197
chr15-83441434-INS-430
chr15-83443857-INS-423
chr15-83449041-INS-55
chr15-83450720-INS-147
chr15-83450829-INS-50
chr15-83516413-INS-619
chr15-83521385-INS-214
chr15-83558939-INS-80
chr15-83560165-INS-65
chr15-83601457-INS-142
chr15-83624237-INS-114
chr15-83626758-INS-75
chr15-83645738-INS-66
chr15-83686204-INS-52
chr15-83700135-INS-500
chr15-83710752-INS-154
chr15-83714854-INS-98
chr15-83718698-INS-50
chr15-83760771-INS-2291
chr15-83769169-INS-96
chr15-83783385-INS-58
chr15-83804434-INS-1218
chr15-83804434-INS-294
chr15-83828760-INS-60
chr15-83833242-INS-83
chr15-83833346-INS-430
chr15-83848436-INS-646
chr15-83878650-INS-162
chr15-83883902-INS-1864
chr15-83905470-INS-726
chr15-83915438-INS-296
chr15-83925081-INS-594
chr15-83937060-INS-83
chr15-84005211-INS-72
chr15-84027794-INS-102
chr15-84040936-INS-255
chr15-84041996-INS-171
chr15-84048930-INS-108
chr15-84049628-INS-70
chr15-84071453-INS-178
chr15-84085753-INS-100
chr15-84085805-INS-520
chr15-84106645-INS-100
chr15-84107618-INS-74
chr15-84114238-INS-4840
chr15-84172734-INS-217
chr15-84211446-INS-77
chr15-84224229-INS-3158
chr15-84242247-INS-2321
chr15-84268762-INS-264
chr15-84281234-INS-553

chr15-84305841-INS-52
chr15-84319821-INS-62
chr15-84334082-INS-130
chr15-84364735-INS-52
chr15-84370086-INS-484
chr15-84378826-INS-56
chr15-84387100-INS-54
chr15-84394808-INS-57
chr15-84398746-INS-255
chr15-84399656-INS-60
chr15-84428707-INS-312
chr15-84453328-INS-4887
chr15-84459573-INS-53
chr15-84459861-INS-195
chr15-84490391-INS-131
chr15-84490431-INS-63
chr15-84520739-INS-1411
chr15-84521376-INS-2032
chr15-84524500-INS-128
chr15-84589350-INS-165
chr15-84675002-INS-144
chr15-84675213-INS-167
chr15-84677883-INS-90
chr15-84723022-INS-52
chr15-84769731-INS-50
chr15-84789716-INS-178
chr15-84839124-INS-4707
chr15-84855003-INS-120
chr15-84855518-INS-80
chr15-84982496-INS-359
chr15-84982819-INS-359
chr15-84993704-INS-5438
chr15-85013753-INS-273
chr15-85013753-INS-725
chr15-85027964-INS-60
chr15-85045963-INS-204
chr15-85053729-INS-100
chr15-85070359-INS-816
chr15-85119747-INS-705
chr15-85133083-INS-87
chr15-85133839-INS-434
chr15-85167831-INS-85
chr15-85170516-INS-7116
chr15-85177601-INS-330
chr15-85184350-INS-296
chr15-85191016-INS-175
chr15-85191038-INS-52
chr15-85202175-INS-620
chr15-85214222-INS-72
chr15-85216574-INS-52
chr15-85223318-INS-52
chr15-85241657-INS-84
chr15-85256077-INS-967
chr15-85284036-INS-73
chr15-85327680-INS-58
chr15-85355543-INS-89
chr15-85380214-INS-108
chr15-85380785-INS-285
chr15-85380802-INS-53
chr15-85382105-INS-244
chr15-85386412-INS-68
chr15-85423220-INS-131
chr15-85428451-INS-58
chr15-85533913-INS-50
chr15-85572898-INS-102
chr15-85630080-INS-71
chr15-85640008-INS-471
chr15-85649327-INS-1910
chr15-85662039-INS-50
chr15-85662866-INS-64
chr15-85669876-INS-271
chr15-85669980-INS-812
chr15-85698413-INS-78
chr15-85698473-INS-157
chr15-85724020-INS-52
chr15-85732270-INS-95
chr15-85757164-INS-209

chr15-85788569-INS-175
chr15-85797856-INS-53
chr15-85836247-INS-322
chr15-85865935-INS-169
chr15-85882078-INS-246
chr15-85882078-INS-56
chr15-85928691-INS-130
chr15-85969731-INS-88
chr15-85972965-INS-477
chr15-85980024-INS-129
chr15-85996197-INS-65
chr15-86012575-INS-229
chr15-86045122-INS-100
chr15-86106076-INS-72
chr15-86107981-INS-94
chr15-86117772-INS-83
chr15-86131684-INS-54
chr15-86133527-INS-229
chr15-86135662-INS-180
chr15-86150647-INS-100
chr15-86151693-INS-392
chr15-86163056-INS-65
chr15-86172595-INS-68
chr15-86183423-INS-3536
chr15-86192001-INS-76
chr15-86246146-INS-144
chr15-86298862-INS-270
chr15-86320343-INS-60
chr15-86405928-INS-116
chr15-86406877-INS-68
chr15-86439480-INS-54
chr15-86459597-INS-56
chr15-86500823-INS-2822
chr15-86506954-INS-176
chr15-86545364-INS-60
chr15-86545715-INS-96
chr15-86545865-INS-121
chr15-86623443-INS-5990
chr15-86628176-INS-51
chr15-86632960-INS-56
chr15-86633198-INS-90
chr15-86639047-INS-127
chr15-86642012-INS-55
chr15-86676536-INS-128
chr15-86677409-INS-60
chr15-86681675-INS-395
chr15-86688899-INS-227
chr15-86689654-INS-84
chr15-86703089-INS-51
chr15-86710596-INS-55
chr15-86740194-INS-66
chr15-86769614-INS-50
chr15-86770654-INS-63
chr15-86772997-INS-62
chr15-86796280-INS-401
chr15-86802770-INS-381
chr15-86835396-INS-79
chr15-86857066-INS-80
chr15-86873554-INS-120
chr15-86898274-INS-58
chr15-86967827-INS-188
chr15-86971011-INS-2071
chr15-86977491-INS-295
chr15-86977549-INS-761
chr15-87010560-INS-92
chr15-87023952-INS-52
chr15-87065957-INS-54
chr15-87070698-INS-223
chr15-87087247-INS-78
chr15-87122628-INS-151
chr15-87149800-INS-62
chr15-87151222-INS-65
chr15-87212671-INS-84
chr15-87237549-INS-72
chr15-87242549-INS-176
chr15-87246352-INS-112
chr15-87280801-INS-72

chr15-87280903-INS-177
chr15-87303174-INS-72
chr15-87328231-INS-58
chr15-87336902-INS-73
chr15-87341713-INS-66
chr15-87342228-INS-140
chr15-87378974-INS-56
chr15-87457280-INS-977
chr15-87469570-INS-1000
chr15-87599136-INS-206
chr15-87606068-INS-172
chr15-87664650-INS-50
chr15-87684023-INS-179
chr15-87690539-INS-72
chr15-87716185-INS-129
chr15-87716284-INS-50
chr15-87735204-INS-2102
chr15-87735230-INS-209
chr15-87735258-INS-516
chr15-87757895-INS-63
chr15-87779445-INS-70
chr15-87791136-INS-69
chr15-87800452-INS-105
chr15-87806204-INS-141
chr15-87807775-INS-78
chr15-87836119-INS-761
chr15-87837256-INS-189
chr15-87846641-INS-1305
chr15-87852122-INS-85
chr15-87857654-INS-113
chr15-87860256-INS-2921
chr15-87889266-INS-1829
chr15-87894210-INS-52
chr15-87915984-INS-86
chr15-87917323-INS-52
chr15-87951053-INS-102
chr15-87976838-INS-65
chr15-88009586-INS-60
chr15-88011212-INS-50
chr15-88031158-INS-448
chr15-88163090-INS-2070
chr15-88163688-INS-203
chr15-88169343-INS-110
chr15-88193815-INS-212
chr15-88210796-INS-58
chr15-88213494-INS-54
chr15-88217369-INS-6413
chr15-88226832-INS-1972
chr15-88226856-INS-640
chr15-88248285-INS-2377
chr15-88249186-INS-1244
chr15-88257553-INS-82
chr15-88263584-INS-2320
chr15-88287096-INS-594
chr15-88287555-INS-203
chr15-88333560-INS-106
chr15-88348470-INS-56
chr15-88369997-INS-197
chr15-88387995-INS-56
chr15-88397215-INS-102
chr15-88460815-INS-585
chr15-88550710-INS-55
chr15-88569259-INS-52
chr15-88603826-INS-233
chr15-88621600-INS-223
chr15-88622622-INS-208
chr15-88637947-INS-57
chr15-88638553-INS-213
chr15-88643543-INS-55
chr15-88669032-INS-205
chr15-88676687-INS-177
chr15-88722816-INS-496
chr15-88725417-INS-54
chr15-88738212-INS-69
chr15-88750245-INS-116
chr15-88782825-INS-104
chr15-88795342-INS-72

chr15-88815513-INS-588
chr15-88840187-INS-60
chr15-88987460-INS-81
chr15-89032511-INS-197
chr15-89082795-INS-200
chr15-89090862-INS-95
chr15-89103660-INS-420
chr15-89104878-INS-65
chr15-89108341-INS-96
chr15-89125106-INS-193
chr15-89125968-INS-60
chr15-89133090-INS-280
chr15-89144917-INS-80
chr15-89182915-INS-100
chr15-89183081-INS-431
chr15-89214259-INS-132
chr15-89228518-INS-98
chr15-89233674-INS-138
chr15-89290408-INS-52
chr15-89291288-INS-210
chr15-89296084-INS-1076
chr15-89296210-INS-261
chr15-89296498-INS-99
chr15-89297452-INS-57
chr15-89321508-INS-60
chr15-89321978-INS-234
chr15-89329054-INS-80
chr15-89370226-INS-66
chr15-89394869-INS-96
chr15-89401074-INS-85
chr15-89473314-INS-243
chr15-89473332-INS-2617
chr15-89485994-INS-530
chr15-89510142-INS-1899
chr15-89519753-INS-3572
chr15-89562727-INS-6262
chr15-89568387-INS-6812
chr15-89582998-INS-726
chr15-89583081-INS-200
chr15-89625740-INS-6168
chr15-89644394-INS-1793
chr15-89645768-INS-2938
chr15-89653881-INS-66
chr15-89659074-INS-2410
chr15-89659074-INS-314
chr15-89672044-INS-300
chr15-89672273-INS-248
chr15-89680789-INS-6245
chr15-89703686-INS-177
chr15-89705242-INS-92
chr15-89706172-INS-116
chr15-89709741-INS-78
chr15-89742926-INS-1155
chr15-89753677-INS-6267
chr15-89754085-INS-50
chr15-89773127-INS-1492
chr15-89785629-INS-289
chr15-89800607-INS-6811
chr15-89889071-INS-3079
chr15-89898685-INS-120
chr15-89908168-INS-12870
chr15-89908168-INS-6228
chr15-89929642-INS-74
chr15-89945500-INS-3068
chr15-89967690-INS-51
chr15-89973518-INS-224
chr15-89994467-INS-50
chr15-89995579-INS-4092
chr15-89998161-INS-1677
chr15-90003458-INS-657
chr15-90009213-INS-50
chr15-90027212-INS-6455
chr15-90027976-INS-2730
chr15-90030188-INS-407
chr15-90061339-INS-6084
chr15-90072577-INS-1012
chr15-90162881-INS-86

chr15-90208198-INS-98
chr15-90210603-INS-393
chr15-90213011-INS-6484
chr15-90215165-INS-259
chr15-90216014-INS-321
chr15-90254802-INS-280
chr15-90322041-INS-106
chr15-90373476-INS-164
chr15-90416899-INS-181
chr15-90506200-INS-60
chr15-90583728-INS-50
chr15-90586928-INS-693
chr15-90597852-INS-133
chr15-90608453-INS-56
chr15-90621200-INS-650
chr15-90630986-INS-6192
chr15-90696236-INS-206
chr15-90780124-INS-52
chr15-90798614-INS-395
chr15-90807930-INS-78
chr15-90807932-INS-176
chr15-90809163-INS-70
chr15-90816511-INS-166
chr15-90827486-INS-390
chr15-90832439-INS-187
chr15-90882509-INS-125
chr15-90911931-INS-234
chr15-90976159-INS-238
chr15-90990933-INS-94
chr15-90996121-INS-198
chr15-91084131-INS-50
chr15-91100318-INS-6292
chr15-91186859-INS-75
chr15-91211111-INS-777
chr15-91219464-INS-1340
chr15-91252532-INS-533
chr15-91252532-INS-96
chr15-91252533-INS-1429
chr15-91269897-INS-4875
chr15-91280976-INS-23634
chr15-91296928-INS-61
chr15-91328763-INS-151
chr15-91343576-INS-60
chr15-91344507-INS-1726
chr15-91392355-INS-515
chr15-91396944-INS-104
chr15-91410171-INS-57
chr15-91412140-INS-82
chr15-91462669-INS-66
chr15-91480449-INS-7365
chr15-91505941-INS-93
chr15-91532416-INS-50
chr15-91552531-INS-54
chr15-91591618-INS-56
chr15-91606264-INS-1472
chr15-91642729-INS-190
chr15-91669460-INS-199
chr15-91677236-INS-336
chr15-91702583-INS-320
chr15-91709580-INS-216
chr15-91733903-INS-66
chr15-91733915-INS-195
chr15-91823345-INS-62
chr15-91833138-INS-51
chr15-91865281-INS-629
chr15-91880465-INS-86
chr15-91881073-INS-424
chr15-91881115-INS-73
chr15-91931029-INS-52
chr15-91939508-INS-52
chr15-92021345-INS-57
chr15-92023394-INS-98
chr15-92046073-INS-779
chr15-92065704-INS-74
chr15-92067442-INS-208
chr15-92068039-INS-90
chr15-92083556-INS-54

chr15-92101335-INS-3219
chr15-92103158-INS-64
chr15-92115083-INS-190
chr15-92119898-INS-116
chr15-92119925-INS-56
chr15-92124672-INS-7853
chr15-92176640-INS-1090
chr15-92222951-INS-212
chr15-92261804-INS-2365
chr15-92276062-INS-54
chr15-92276104-INS-157
chr15-92283946-INS-60
chr15-92310037-INS-54
chr15-92317426-INS-55
chr15-92338354-INS-78
chr15-92341921-INS-92
chr15-92355390-INS-3165
chr15-92491833-INS-195
chr15-92557900-INS-80
chr15-92558398-INS-92
chr15-92558934-INS-491
chr15-92591875-INS-6447
chr15-92606689-INS-70
chr15-92664260-INS-175
chr15-92671639-INS-461
chr15-92677665-INS-1093
chr15-92680113-INS-50
chr15-92680871-INS-1104
chr15-92735136-INS-98
chr15-92751248-INS-288
chr15-92751282-INS-169
chr15-92751289-INS-66
chr15-92836479-INS-1052
chr15-92864707-INS-58
chr15-92867738-INS-129
chr15-92879014-INS-50
chr15-92905047-INS-60
chr15-92934620-INS-791
chr15-92940991-INS-213
chr15-92955748-INS-542
chr15-92972858-INS-50
chr15-92977358-INS-57
chr15-92980619-INS-55
chr15-92991315-INS-443
chr15-93021876-INS-422
chr15-93039637-INS-455
chr15-93045423-INS-78
chr15-93053465-INS-386
chr15-93067625-INS-212
chr15-93102168-INS-95
chr15-93127948-INS-84
chr15-93128834-INS-69
chr15-93131320-INS-143
chr15-93161222-INS-232
chr15-93161244-INS-58
chr15-93204997-INS-481
chr15-93222858-INS-7193
chr15-93229018-INS-64
chr15-93246948-INS-104
chr15-93275284-INS-175
chr15-93276684-INS-1060
chr15-93285021-INS-98
chr15-93309465-INS-196
chr15-93334540-INS-213
chr15-93365755-INS-2909
chr15-93366897-INS-1452
chr15-93379331-INS-68
chr15-93399706-INS-470
chr15-93424898-INS-335
chr15-93424983-INS-107
chr15-93430057-INS-50
chr15-93454830-INS-108
chr15-93465974-INS-236
chr15-93469367-INS-645
chr15-93505207-INS-5543
chr15-93505533-INS-146
chr15-93522509-INS-257

chr15-93555557-INS-204
chr15-93573741-INS-78
chr15-93583208-INS-279
chr15-93591217-INS-995
chr15-93676096-INS-174
chr15-93719916-INS-56
chr15-93760033-INS-96
chr15-93776672-INS-54
chr15-93780023-INS-140
chr15-93811580-INS-84
chr15-93811860-INS-191
chr15-93816476-INS-60
chr15-93821250-INS-50
chr15-93829118-INS-429
chr15-93900063-INS-115
chr15-93935370-INS-4712
chr15-93982412-INS-52
chr15-94029501-INS-247
chr15-94059783-INS-129
chr15-94105783-INS-50
chr15-94140068-INS-1927
chr15-94160784-INS-65
chr15-94184550-INS-88
chr15-94223010-INS-394
chr15-94260749-INS-204
chr15-94287319-INS-182
chr15-94325701-INS-1968
chr15-94339077-INS-52
chr15-94379579-INS-3135
chr15-94408506-INS-1898
chr15-94419278-INS-5531
chr15-94425122-INS-455
chr15-94442357-INS-1429
chr15-94467126-INS-7096
chr15-94467326-INS-68
chr15-94471424-INS-56
chr15-94517698-INS-204
chr15-94582970-INS-2349
chr15-94583808-INS-1088
chr15-94605859-INS-6423
chr15-94613075-INS-80
chr15-94621468-INS-6233
chr15-94635292-INS-50
chr15-94635464-INS-1152
chr15-94637399-INS-6999
chr15-94644496-INS-90
chr15-94674298-INS-66
chr15-94733480-INS-6576
chr15-94741492-INS-80
chr15-94803790-INS-236
chr15-94817901-INS-112
chr15-94818238-INS-440
chr15-94829058-INS-64
chr15-94832514-INS-150
chr15-94832514-INS-50
chr15-94878509-INS-76
chr15-94892009-INS-68
chr15-94908532-INS-1991
chr15-94923832-INS-50
chr15-94928673-INS-549
chr15-94929090-INS-64
chr15-94935331-INS-1018
chr15-94940967-INS-52
chr15-94969418-INS-391
chr15-95005430-INS-6386
chr15-95011690-INS-70
chr15-95060236-INS-157
chr15-95086768-INS-64
chr15-95088252-INS-157
chr15-95088300-INS-65
chr15-95118688-INS-53
chr15-95121333-INS-90
chr15-95154786-INS-789
chr15-95166394-INS-101
chr15-95273022-INS-64
chr15-95276136-INS-748
chr15-95368348-INS-6285

chr15-95382786-INS-1506
chr15-95420501-INS-76
chr15-95449582-INS-3467
chr15-95455963-INS-144
chr15-95459566-INS-65
chr15-95478994-INS-324
chr15-95529785-INS-51
chr15-95533780-INS-51
chr15-95545641-INS-70
chr15-95553879-INS-590
chr15-95559237-INS-94
chr15-95577041-INS-54
chr15-95613218-INS-2972
chr15-95616630-INS-53
chr15-95664484-INS-82
chr15-95665411-INS-56
chr15-95706756-INS-52
chr15-95716938-INS-6333
chr15-95719055-INS-50
chr15-95770546-INS-332
chr15-95789270-INS-64
chr15-95827327-INS-6288
chr15-95839669-INS-114
chr15-95915510-INS-61
chr15-95961842-INS-408
chr15-95975382-INS-52
chr15-95986132-INS-522
chr15-95999253-INS-378
chr15-96018460-INS-27211
chr15-96038792-INS-160
chr15-96120768-INS-197
chr15-96125950-INS-50
chr15-96128524-INS-200
chr15-96146496-INS-199
chr15-96147018-INS-507
chr15-96147890-INS-7134
chr15-96149649-INS-429
chr15-96216908-INS-68
chr15-96278606-INS-52
chr15-96282012-INS-52
chr15-96347045-INS-70
chr15-96378484-INS-452
chr15-96382084-INS-2541
chr15-96402057-INS-60
chr15-96430401-INS-50
chr15-96480853-INS-2484
chr15-96507245-INS-63
chr15-96510986-INS-196
chr15-96524107-INS-258
chr15-96524766-INS-84
chr15-96547441-INS-68
chr15-96555940-INS-110
chr15-96562566-INS-1246
chr15-96574913-INS-142
chr15-96614509-INS-507
chr15-96692745-INS-410
chr15-96695524-INS-207
chr15-96716066-INS-72
chr15-96721659-INS-162
chr15-96721782-INS-63
chr15-96747573-INS-122
chr15-96761763-INS-108
chr15-96772543-INS-60
chr15-96806869-INS-52
chr15-96851043-INS-8986
chr15-96903542-INS-141
chr15-96944014-INS-198
chr15-96944321-INS-400
chr15-96976907-INS-172
chr15-97052821-INS-432
chr15-97081687-INS-68
chr15-97129218-INS-52
chr15-97166498-INS-83
chr15-97169433-INS-54
chr15-97171995-INS-104
chr15-97179973-INS-182
chr15-97202390-INS-396

chr15-97233019-INS-7692
chr15-97233366-INS-360
chr15-97237880-INS-88
chr15-97237963-INS-271
chr15-97249615-INS-56
chr15-97251293-INS-54
chr15-97257060-INS-65
chr15-97257584-INS-51
chr15-97293112-INS-168
chr15-97297619-INS-252
chr15-97303305-INS-50
chr15-97309860-INS-3093
chr15-97311767-INS-673
chr15-97347583-INS-3374
chr15-97355175-INS-595
chr15-97364597-INS-179
chr15-97402704-INS-58
chr15-97419452-INS-193
chr15-97426121-INS-54
chr15-97430902-INS-50
chr15-97438013-INS-779
chr15-97455638-INS-67
chr15-97461238-INS-56
chr15-97474320-INS-50
chr15-97500454-INS-69
chr15-97502629-INS-124
chr15-97515011-INS-142
chr15-97515581-INS-7083
chr15-97515639-INS-444
chr15-97519767-INS-201
chr15-97535999-INS-89
chr15-97536064-INS-568
chr15-97545016-INS-130
chr15-97549270-INS-764
chr15-97550273-INS-52
chr15-97552530-INS-217
chr15-97552918-INS-372
chr15-97566919-INS-250
chr15-97577411-INS-262
chr15-97580346-INS-60
chr15-97583759-INS-89
chr15-97583959-INS-62
chr15-97584264-INS-231
chr15-97590528-INS-60
chr15-97592678-INS-194
chr15-97613251-INS-315
chr15-97613383-INS-74
chr15-97662144-INS-192
chr15-97715693-INS-445
chr15-97747966-INS-294
chr15-97818646-INS-320
chr15-97845176-INS-459
chr15-97865619-INS-66
chr15-97887910-INS-116
chr15-97907916-INS-216
chr15-97952791-INS-192
chr15-97953186-INS-126
chr15-97956435-INS-148
chr15-98027562-INS-165
chr15-98031758-INS-269
chr15-98032251-INS-280
chr15-98039426-INS-68
chr15-98081891-INS-55
chr15-98097015-INS-20457
chr15-98104073-INS-381
chr15-98106095-INS-7657
chr15-98112634-INS-709
chr15-98121122-INS-1280
chr15-98138262-INS-62
chr15-98145782-INS-55
chr15-98146030-INS-495
chr15-98158705-INS-59
chr15-98160057-INS-72
chr15-98174024-INS-414
chr15-98177021-INS-6673
chr15-98181704-INS-94
chr15-98194792-INS-50

chr15-98208468-INS-26789
chr15-98245219-INS-105
chr15-98269176-INS-65
chr15-98293299-INS-1009
chr15-98314706-INS-92
chr15-98314948-INS-233
chr15-98317373-INS-2752
chr15-98333178-INS-194
chr15-98333399-INS-149
chr15-98338952-INS-6263
chr15-98355643-INS-1035
chr15-98394607-INS-81
chr15-98394834-INS-445
chr15-98407956-INS-65
chr15-98414488-INS-383
chr15-98454076-INS-223
chr15-98482444-INS-82
chr15-98484696-INS-58
chr15-98513915-INS-190
chr15-98517340-INS-423
chr15-98517340-INS-68
chr15-98517571-INS-204
chr15-98538450-INS-126
chr15-98563491-INS-505
chr15-98569791-INS-209
chr15-98583068-INS-75
chr15-98585767-INS-180
chr15-98599804-INS-166
chr15-98599882-INS-59
chr15-98641963-INS-58
chr15-98695342-INS-106
chr15-98699026-INS-79
chr15-98719634-INS-50
chr15-98749244-INS-204
chr15-98754086-INS-50
chr15-98821936-INS-92
chr15-98825407-INS-232
chr15-98888659-INS-5626
chr15-98900917-INS-13883
chr15-98906365-INS-223
chr15-98916843-INS-195
chr15-98916926-INS-1528
chr15-98917794-INS-54
chr15-98964709-INS-396
chr15-98967510-INS-82
chr15-99000196-INS-195
chr15-99002832-INS-112
chr15-99019108-INS-54
chr15-99055048-INS-595
chr15-99056631-INS-300
chr15-99063740-INS-50
chr15-99083122-INS-500
chr15-99093528-INS-66
chr15-99098839-INS-60
chr15-99106405-INS-76
chr15-99161807-INS-57
chr15-99174898-INS-13538
chr15-99223920-INS-72
chr15-99271926-INS-1700
chr15-99272011-INS-179
chr15-99279660-INS-112
chr15-99335400-INS-203
chr15-99343983-INS-1056
chr15-99345825-INS-56
chr15-99345951-INS-68
chr15-99374504-INS-329
chr15-99382106-INS-56
chr15-99387800-INS-404
chr15-99432179-INS-568
chr15-99434495-INS-120
chr15-99436861-INS-112
chr15-99500077-INS-50
chr15-99516421-INS-132
chr15-99516936-INS-224
chr15-99539050-INS-60
chr15-99588449-INS-64
chr15-99606174-INS-1224

chr15-99633215-INS-118
chr15-99637837-INS-50
chr15-99644412-INS-204
chr15-99660706-INS-112
chr15-99761160-INS-51
chr15-99763236-INS-58
chr15-99763620-INS-139
chr15-99814699-INS-51
chr15-99834971-INS-50
chr15-99841036-INS-309
chr15-99841329-INS-501
chr15-99886161-INS-54
chr15-99934619-INS-110
chr15-99972386-INS-59
chr15-99973088-INS-155
chr15-99992137-INS-68
chr15-100002893-INS-165
chr15-100063736-INS-1003
chr15-100063962-INS-3148
chr15-100071918-INS-2102
chr15-100074248-INS-3766
chr15-100087636-INS-75
chr15-100090696-INS-64
chr15-100091367-INS-212
chr15-100096471-INS-279
chr15-100176062-INS-54
chr15-100219967-INS-1507
chr15-100222040-INS-126
chr15-100230910-INS-66
chr15-100325713-INS-954
chr15-100331233-INS-1893
chr15-100340601-INS-168
chr15-100340752-INS-2411
chr15-100341467-INS-269
chr15-100344049-INS-183
chr15-100355434-INS-76
chr15-100356090-INS-51
chr15-100385510-INS-181
chr15-100386075-INS-82
chr15-100394704-INS-187
chr15-100395497-INS-160
chr15-100443584-INS-154
chr15-100444288-INS-56
chr15-100449796-INS-13017
chr15-100495952-INS-123
chr15-100495961-INS-272
chr15-100500913-INS-185
chr15-100543923-INS-140
chr15-100546358-INS-735
chr15-100600400-INS-202
chr15-100605082-INS-208
chr15-100608073-INS-190
chr15-100610163-INS-96
chr15-100642643-INS-610
chr15-100657403-INS-5155
chr15-100678860-INS-8326
chr15-100690567-INS-66
chr15-100690581-INS-148
chr15-100747067-INS-145
chr15-100748165-INS-65
chr15-100815339-INS-157
chr15-100815635-INS-812
chr15-100818482-INS-92
chr15-100818814-INS-138
chr15-100886752-INS-2218
chr15-101216975-INS-3059
chr15-101244524-INS-52
chr15-101257319-INS-64
chr15-101337418-INS-238
chr15-101346402-INS-150
chr15-101425496-INS-80
chr15-101489910-INS-132
chr15-101506754-INS-67
chr15-101515940-INS-12642
chr15-101517781-INS-2971
chr15-101528919-INS-628
chr15-101564093-INS-424

chr15-101618088-INS-55
chr15-101659114-INS-92
chr15-101685235-INS-173
chr15-101685533-INS-782
chr15-101738265-INS-50
chr15-101745984-INS-96
chr15-101750914-INS-56
chr15-101753360-INS-246
chr15-101787992-INS-72
chr15-101814988-INS-3210
chr15-101815573-INS-63
chr15-101837494-INS-58
chr15-101853024-INS-2755
chr15-101856287-INS-1644
chr15-101868532-INS-60
chr15-101871974-INS-178
chr15-101878401-INS-22035
chr15-101918084-INS-56
chr15-101918189-INS-119
chr15-101921554-INS-276
chr15-101922021-INS-56
chr15-101935842-INS-66
chr15-101954759-INS-54
chr15-102016185-INS-52
chr15-102049132-INS-62
chr15-102050575-INS-201
chr15-102076802-INS-50
chr15-102077443-INS-52
chr15-102080963-INS-152
chr15-102082923-INS-92
chr15-102119795-INS-50
chr15-102204426-INS-219
chr15-102211563-INS-53
chr15-102269225-INS-72
chr15-102270365-INS-81
chr15-102278237-INS-65
chr15-102294045-INS-134
chr15-102294130-INS-214
chr15-102299071-INS-2071
chr15-102323261-INS-63
chr15-102323765-INS-204
chr15-102331923-INS-54
chr15-102332429-INS-197
chr15-102346704-INS-207
chr15-102389601-INS-268
chr15-102462524-INS-214
chr15-102478464-INS-3521
chr15-102483822-INS-82
chr15-102484103-INS-201
chr15-102484791-INS-203
chr15-102516803-INS-202
chr15-102568245-INS-50
chr15-102587188-INS-196
chr15-102590440-INS-95
chr15-102673273-INS-468
chr15-102673676-INS-63
chr15-102728312-INS-205
chr15-102775483-INS-825
chr15-102779902-INS-50
chr15-102804135-INS-429
chr15-102839451-INS-54
chr15-102942071-INS-56
chr15-102953531-INS-999
chr15-102959681-INS-7836
chr15-102961285-INS-517
chr15-102970392-INS-99
chr15-102993990-INS-200
chr15-103026777-INS-175
chr15-103027414-INS-50
chr15-103027547-INS-267
chr15-103082350-INS-209
chr15-103140136-INS-52
chr15-103142999-INS-68
chr15-103189339-INS-204
chr15-103282729-INS-2961
chr15-103319376-INS-424
chr15-103340038-INS-227

chr15-103342076-INS-52
chr15-103356080-INS-203
chr15-103361547-INS-224
chr15-103422998-INS-102
chr15-103469249-INS-652
chr15-103482843-INS-770
chr15-103505320-INS-6432
chr15-103546054-INS-983
chr15-103606353-INS-20884
chr15-103636231-INS-211
chr15-103731370-INS-382
chr15-103825059-INS-2784
chr15-103834762-INS-1223
chr15-103835515-INS-1920
chr15-103909531-INS-212
chr15-103913057-INS-4439
chr15-103933181-INS-7078
chr15-103933451-INS-259
chr16-3054680-INS-325
chr16-3076505-INS-1165
chr16-3080193-INS-2657
chr16-3105409-INS-58
chr16-3106501-INS-50
chr16-3133844-INS-64
chr16-3133898-INS-214
chr16-3156391-INS-197
chr16-3168158-INS-2122
chr16-3169634-INS-68
chr16-3187774-INS-51
chr16-3219079-INS-401
chr16-3225390-INS-66
chr16-3225802-INS-89
chr16-3263464-INS-662
chr16-3317903-INS-236
chr16-3421516-INS-60
chr16-3469459-INS-269
chr16-3507660-INS-1177
chr16-3518646-INS-50
chr16-3537032-INS-296
chr16-3556489-INS-228
chr16-3598393-INS-80
chr16-3615318-INS-53
chr16-3631882-INS-130
chr16-3631894-INS-52
chr16-3648444-INS-124
chr16-3648444-INS-52
chr16-3649667-INS-160
chr16-3675929-INS-215
chr16-3688073-INS-52
chr16-3779198-INS-430
chr16-3780820-INS-186
chr16-3784670-INS-136
chr16-3789948-INS-248
chr16-3829792-INS-209
chr16-3837703-INS-200
chr16-3858578-INS-216
chr16-3879702-INS-208
chr16-3882757-INS-163
chr16-3913533-INS-215
chr16-3969600-INS-218
chr16-3989920-INS-294
chr16-4035527-INS-434
chr16-4066479-INS-1581
chr16-4067799-INS-212
chr16-4068090-INS-105
chr16-4081015-INS-195
chr16-4082711-INS-206
chr16-4118782-INS-220
chr16-4143130-INS-74
chr16-4146808-INS-730
chr16-4157218-INS-1800
chr16-4185669-INS-418
chr16-4197514-INS-429
chr16-4202779-INS-50
chr16-4203617-INS-72
chr16-4219779-INS-397
chr16-4220510-INS-584

chr16-4220576-INS-165
chr16-4222225-INS-195
chr16-4227307-INS-208
chr16-4242526-INS-256
chr16-4260922-INS-209
chr16-4280615-INS-54
chr16-4286028-INS-1500
chr16-4291734-INS-178
chr16-4350690-INS-248
chr16-4353482-INS-2841
chr16-4353942-INS-216
chr16-4354751-INS-173
chr16-4357861-INS-60
chr16-4357877-INS-319
chr16-4359232-INS-181
chr16-4364567-INS-450
chr16-4364639-INS-186
chr16-4366743-INS-256
chr16-4377592-INS-50
chr16-4385304-INS-211
chr16-4390465-INS-200
chr16-4390820-INS-225
chr16-4400075-INS-177
chr16-4406203-INS-59
chr16-4509900-INS-51
chr16-4536159-INS-1511
chr16-4547302-INS-196
chr16-4579248-INS-877
chr16-4613186-INS-90
chr16-4633540-INS-195
chr16-4633880-INS-576
chr16-4645230-INS-50
chr16-4679886-INS-265
chr16-4679990-INS-102
chr16-4680003-INS-296
chr16-4685073-INS-316
chr16-4706606-INS-222
chr16-4735503-INS-120
chr16-4753587-INS-502
chr16-4772474-INS-342
chr16-4800006-INS-95
chr16-4812781-INS-108
chr16-4813474-INS-455
chr16-4813620-INS-110
chr16-4860263-INS-246
chr16-4917338-INS-449
chr16-4974089-INS-165
chr16-4984394-INS-220
chr16-4997533-INS-61
chr16-5005594-INS-259
chr16-5006367-INS-85
chr16-5010149-INS-50
chr16-5026765-INS-72
chr16-5027578-INS-219
chr16-5078488-INS-3926
chr16-5087139-INS-81
chr16-5099078-INS-259
chr16-5102084-INS-153
chr16-5124040-INS-58
chr16-5138292-INS-90
chr16-5152506-INS-74
chr16-5178422-INS-2133
chr16-5185341-INS-168
chr16-5188681-INS-64
chr16-5211559-INS-6642
chr16-5217086-INS-406
chr16-5221402-INS-89
chr16-5223845-INS-351
chr16-5223916-INS-63
chr16-5283537-INS-190
chr16-5332181-INS-110
chr16-5347196-INS-92
chr16-5361862-INS-52
chr16-5362887-INS-110
chr16-5400347-INS-142
chr16-5407942-INS-195
chr16-5449613-INS-2908

chr16-5461146-INS-157
chr16-5483726-INS-55
chr16-5496246-INS-3794
chr16-5588880-INS-123
chr16-5621626-INS-219
chr16-5623782-INS-1890
chr16-5625867-INS-112
chr16-5836223-INS-169
chr16-5836226-INS-80
chr16-5886619-INS-230
chr16-5887055-INS-664
chr16-5903389-INS-126
chr16-5905280-INS-262
chr16-5905298-INS-62
chr16-5919483-INS-399
chr16-5928448-INS-60
chr16-5961608-INS-205
chr16-5997613-INS-80
chr16-6133727-INS-50
chr16-6134586-INS-114
chr16-6378053-INS-423
chr16-6426148-INS-187
chr16-6426348-INS-65
chr16-6435871-INS-236
chr16-6564041-INS-6648
chr16-6575040-INS-152
chr16-6589911-INS-50
chr16-6611130-INS-6386
chr16-6614743-INS-558
chr16-6626366-INS-6211
chr16-6714700-INS-114
chr16-6718625-INS-1931
chr16-6731694-INS-76
chr16-6732342-INS-51
chr16-6737376-INS-66
chr16-6738852-INS-50
chr16-6778059-INS-64
chr16-6778864-INS-194
chr16-6925601-INS-6712
chr16-7102339-INS-80
chr16-7102359-INS-234
chr16-7126623-INS-50
chr16-7143513-INS-52
chr16-7143539-INS-118
chr16-7186564-INS-65
chr16-7190389-INS-1055
chr16-7251868-INS-76
chr16-7314758-INS-471
chr16-7325988-INS-50
chr16-7353598-INS-53
chr16-7356475-INS-193
chr16-7374786-INS-6417
chr16-7383565-INS-52
chr16-7483812-INS-60
chr16-7505294-INS-80
chr16-7509935-INS-56
chr16-7513173-INS-294
chr16-7513249-INS-91
chr16-7543954-INS-54
chr16-7600216-INS-54
chr16-7606794-INS-80
chr16-7755792-INS-290
chr16-7755842-INS-754
chr16-7835444-INS-68
chr16-7911280-INS-6677
chr16-7929177-INS-102
chr16-7960763-INS-1433
chr16-8060916-INS-306
chr16-8088105-INS-72
chr16-8105394-INS-214
chr16-8112240-INS-6337
chr16-8113807-INS-50
chr16-8139336-INS-51
chr16-8149295-INS-93
chr16-8171621-INS-220
chr16-8209813-INS-125
chr16-8235786-INS-64

chr16-8269572-INS-311
chr16-8334181-INS-72
chr16-8334776-INS-106
chr16-8345262-INS-82
chr16-8348950-INS-120
chr16-8382550-INS-80
chr16-8412015-INS-809
chr16-8419674-INS-114
chr16-8422513-INS-81
chr16-8423057-INS-69
chr16-8443010-INS-162
chr16-8497972-INS-181
chr16-8576503-INS-8734
chr16-8591722-INS-144
chr16-8610955-INS-197
chr16-8612319-INS-7477
chr16-8615980-INS-1012
chr16-8637675-INS-150
chr16-8662703-INS-249
chr16-8669616-INS-192
chr16-8696663-INS-308
chr16-8706937-INS-52
chr16-8707362-INS-222
chr16-8707624-INS-84
chr16-8710765-INS-112
chr16-8717071-INS-570
chr16-8722996-INS-97
chr16-8742358-INS-4357
chr16-8742799-INS-130
chr16-8779995-INS-52
chr16-8840670-INS-50
chr16-8855210-INS-99
chr16-8870826-INS-1523
chr16-8968340-INS-64
chr16-8968523-INS-231
chr16-8968994-INS-7396
chr16-8982325-INS-79
chr16-9084144-INS-173
chr16-9088907-INS-7043
chr16-9112706-INS-406
chr16-9128210-INS-50
chr16-9129557-INS-52
chr16-9147219-INS-3341
chr16-9167122-INS-70
chr16-9268914-INS-6137
chr16-9279967-INS-1294
chr16-9283347-INS-554
chr16-9293963-INS-6774
chr16-9342551-INS-375
chr16-9342734-INS-378
chr16-9370914-INS-66
chr16-9404352-INS-72
chr16-9435762-INS-79
chr16-9456388-INS-54
chr16-9464822-INS-60
chr16-9481616-INS-54
chr16-9555550-INS-55
chr16-9603021-INS-50
chr16-9638590-INS-66
chr16-9737481-INS-140
chr16-9740130-INS-165
chr16-9765802-INS-50
chr16-9778271-INS-75
chr16-9817683-INS-558
chr16-9817725-INS-80
chr16-9894038-INS-260
chr16-9894284-INS-208
chr16-9904377-INS-84
chr16-9904808-INS-78
chr16-9905812-INS-312
chr16-9911666-INS-131
chr16-9999758-INS-458
chr16-10148345-INS-53
chr16-10154379-INS-8626
chr16-10157394-INS-336
chr16-10195072-INS-187
chr16-10218058-INS-214

chr16-10246897-INS-440
chr16-10285977-INS-60
chr16-10287564-INS-1462
chr16-10322677-INS-50
chr16-10331980-INS-56
chr16-10332743-INS-613
chr16-10341444-INS-471
chr16-10434212-INS-7575
chr16-10464291-INS-102
chr16-10500855-INS-245
chr16-10564075-INS-150
chr16-10588576-INS-52
chr16-10629504-INS-2780
chr16-10674424-INS-357
chr16-10679294-INS-228
chr16-10683211-INS-3192
chr16-10691043-INS-260
chr16-10700523-INS-1777
chr16-10704244-INS-132
chr16-10704244-INS-484
chr16-10705379-INS-51
chr16-10709088-INS-5505
chr16-10712690-INS-162
chr16-10717256-INS-99
chr16-10717614-INS-128
chr16-10771018-INS-3005
chr16-10772793-INS-114
chr16-10833550-INS-53
chr16-10848580-INS-1682
chr16-10850992-INS-52
chr16-10861580-INS-144
chr16-10865996-INS-568
chr16-10878698-INS-263
chr16-10941889-INS-75
chr16-10961271-INS-97
chr16-10973502-INS-108
chr16-10979981-INS-63
chr16-10981502-INS-618
chr16-10995009-INS-52
chr16-10997274-INS-6262
chr16-11010775-INS-421
chr16-11026633-INS-92
chr16-11066719-INS-288
chr16-11080112-INS-241
chr16-11091380-INS-221
chr16-11115912-INS-54
chr16-11122205-INS-234
chr16-11129462-INS-82
chr16-11134955-INS-216
chr16-11138894-INS-86
chr16-11161991-INS-70
chr16-11191873-INS-56
chr16-11199849-INS-338
chr16-11201692-INS-201
chr16-11219153-INS-368
chr16-11229723-INS-82
chr16-11243077-INS-2479
chr16-11255082-INS-400
chr16-11284181-INS-897
chr16-11294802-INS-159
chr16-11370498-INS-400
chr16-11371070-INS-370
chr16-11372977-INS-145
chr16-11474587-INS-214
chr16-11505024-INS-108
chr16-11505523-INS-99
chr16-11519718-INS-1511
chr16-11521692-INS-194
chr16-11549757-INS-55
chr16-11616562-INS-64
chr16-11657395-INS-1948
chr16-11669910-INS-327
chr16-11669946-INS-841
chr16-11677783-INS-424
chr16-11742129-INS-104
chr16-11796484-INS-54
chr16-11799337-INS-5195

chr16-11881395-INS-645
chr16-11888439-INS-100
chr16-11917661-INS-113
chr16-11962316-INS-64
chr16-11963113-INS-50
chr16-11973566-INS-52
chr16-11995674-INS-91
chr16-11996441-INS-214
chr16-12031843-INS-154
chr16-12038273-INS-90
chr16-12049374-INS-6438
chr16-12110828-INS-53
chr16-12171907-INS-1865
chr16-12187581-INS-6199
chr16-12196753-INS-50
chr16-12213046-INS-53
chr16-12222725-INS-4162
chr16-12254153-INS-54
chr16-12255777-INS-1680
chr16-12353375-INS-2561
chr16-12354615-INS-1874
chr16-12358006-INS-1153
chr16-12366430-INS-54
chr16-12483272-INS-50
chr16-12497989-INS-62
chr16-12504039-INS-5794
chr16-12647950-INS-470
chr16-12663937-INS-2588
chr16-12675559-INS-58
chr16-12707837-INS-134
chr16-12729246-INS-54
chr16-12749344-INS-60
chr16-12782360-INS-986
chr16-12807921-INS-3703
chr16-12840731-INS-120
chr16-12845804-INS-61
chr16-12846007-INS-82
chr16-12846565-INS-403
chr16-12846670-INS-127
chr16-12862693-INS-91
chr16-12895364-INS-2983
chr16-12975185-INS-2866
chr16-13024094-INS-506
chr16-13081663-INS-52
chr16-13081686-INS-116
chr16-13120097-INS-873
chr16-13130372-INS-761
chr16-13130668-INS-126
chr16-13130713-INS-1120
chr16-13131745-INS-107
chr16-13181473-INS-559
chr16-13223297-INS-181
chr16-13253445-INS-338
chr16-13259564-INS-88
chr16-13272924-INS-364
chr16-13310477-INS-64
chr16-13359130-INS-63
chr16-13381178-INS-92
chr16-13385355-INS-188
chr16-13385598-INS-414
chr16-13404456-INS-550
chr16-13409291-INS-453
chr16-13409585-INS-861
chr16-13417004-INS-1762
chr16-13424601-INS-171
chr16-13474289-INS-233
chr16-13475786-INS-50
chr16-13498260-INS-206
chr16-13500248-INS-188
chr16-13504110-INS-175
chr16-13505836-INS-50
chr16-13510712-INS-112
chr16-13518498-INS-184
chr16-13559448-INS-194
chr16-13564655-INS-52
chr16-13565627-INS-257
chr16-13602505-INS-204

chr16-13609996-INS-66
chr16-13610178-INS-181
chr16-13610372-INS-592
chr16-13684895-INS-72
chr16-13726737-INS-8567
chr16-13753223-INS-198
chr16-13761863-INS-172
chr16-13790411-INS-74
chr16-13863719-INS-64
chr16-13890926-INS-93
chr16-14006718-INS-104
chr16-14006953-INS-544
chr16-14076047-INS-50
chr16-14103548-INS-389
chr16-14114106-INS-137
chr16-14147323-INS-58
chr16-14205253-INS-88
chr16-14320721-INS-111
chr16-14340043-INS-150
chr16-14340078-INS-55
chr16-14340383-INS-8590
chr16-14425788-INS-122
chr16-14436945-INS-6235
chr16-14437519-INS-195
chr16-14526329-INS-54
chr16-14661159-INS-50
chr16-14681609-INS-3219
chr16-14772454-INS-153
chr16-14781070-INS-122
chr16-14818919-INS-52
chr16-14838727-INS-6428
chr16-14921718-INS-77
chr16-14921817-INS-218
chr16-14950799-INS-6489
chr16-15028693-INS-2216
chr16-15054780-INS-84
chr16-15162589-INS-4351
chr16-15266253-INS-17235
chr16-15286256-INS-6502
chr16-15299413-INS-60
chr16-15334046-INS-50
chr16-15363422-INS-1460
chr16-15421407-INS-6369
chr16-15422353-INS-116
chr16-15437336-INS-64
chr16-15489061-INS-1866
chr16-15489061-INS-306
chr16-15489061-INS-4140
chr16-15489061-INS-8460
chr16-15503468-INS-211
chr16-15504196-INS-77
chr16-15505368-INS-315
chr16-15513512-INS-456
chr16-15513640-INS-56
chr16-15518074-INS-54
chr16-15578828-INS-677
chr16-15588341-INS-63
chr16-15615739-INS-196
chr16-15635825-INS-618
chr16-15647832-INS-64
chr16-15655272-INS-198
chr16-15659160-INS-222
chr16-15670953-INS-201
chr16-15732208-INS-111
chr16-15763693-INS-347
chr16-15881809-INS-238
chr16-15914376-INS-193
chr16-15935438-INS-53
chr16-15935949-INS-325
chr16-15943515-INS-224
chr16-16112119-INS-5369
chr16-16118014-INS-50
chr16-16120375-INS-50
chr16-16224488-INS-1391
chr16-16247271-INS-1129
chr16-16272075-INS-109
chr16-16272794-INS-300

chr16-16311237-INS-159
chr16-16343087-INS-58
chr16-16358320-INS-61
chr16-16432051-INS-8145
chr16-16475843-INS-6445
chr16-16501003-INS-791
chr16-16502864-INS-7274
chr16-16541900-INS-62
chr16-16556940-INS-132
chr16-16569635-INS-6569
chr16-16583556-INS-6344
chr16-16589214-INS-6071
chr16-16614761-INS-207
chr16-16616975-INS-198
chr16-16624760-INS-52
chr16-16646245-INS-118
chr16-16646849-INS-63
chr16-16687732-INS-96
chr16-16687754-INS-295
chr16-16758247-INS-101
chr16-16774249-INS-328
chr16-16775422-INS-661
chr16-16849032-INS-50
chr16-16991583-INS-158
chr16-17072422-INS-804
chr16-17072523-INS-1651
chr16-17139846-INS-1500
chr16-17145950-INS-196
chr16-17266731-INS-50
chr16-17315106-INS-88
chr16-17374595-INS-77
chr16-17382390-INS-66
chr16-17403630-INS-211
chr16-17403961-INS-70
chr16-17439825-INS-2719
chr16-17439850-INS-225
chr16-17458542-INS-138
chr16-17486806-INS-216
chr16-17524379-INS-220
chr16-17530052-INS-492
chr16-17551180-INS-3355
chr16-17591003-INS-68
chr16-17762225-INS-153
chr16-17786306-INS-1053
chr16-17793904-INS-5305
chr16-17814023-INS-1796
chr16-17816549-INS-55
chr16-17842352-INS-201
chr16-17848341-INS-56
chr16-17854824-INS-213
chr16-17856732-INS-62
chr16-17868007-INS-211
chr16-17893682-INS-66
chr16-17914420-INS-7261
chr16-17940091-INS-198
chr16-17980407-INS-50
chr16-18108586-INS-104
chr16-18117337-INS-6952
chr16-18133453-INS-86
chr16-18139573-INS-226
chr16-18182683-INS-206
chr16-18241633-INS-135
chr16-18281332-INS-82
chr16-18311062-INS-197
chr16-18334584-INS-214
chr16-18372284-INS-72
chr16-18378925-INS-65
chr16-18379207-INS-1212
chr16-18379207-INS-51
chr16-18385020-INS-207
chr16-18389474-INS-72
chr16-18390025-INS-58
chr16-18411055-INS-235
chr16-18421846-INS-98
chr16-18461306-INS-4696
chr16-18490622-INS-195
chr16-18501368-INS-519

chr16-18504153-INS-1008
chr16-18504158-INS-54
chr16-18507891-INS-927
chr16-18563172-INS-78
chr16-18573054-INS-123
chr16-18577740-INS-150
chr16-18602070-INS-69
chr16-18644143-INS-203
chr16-18656999-INS-149
chr16-18703485-INS-286
chr16-18731761-INS-134
chr16-18733268-INS-77
chr16-18749401-INS-69
chr16-18762300-INS-129
chr16-18771425-INS-60
chr16-18783366-INS-70
chr16-18786063-INS-168
chr16-18797596-INS-6059
chr16-18858020-INS-87
chr16-18869974-INS-5374
chr16-19000744-INS-2526
chr16-19065064-INS-237
chr16-19130503-INS-4226
chr16-19133438-INS-2445
chr16-19141899-INS-5255
chr16-19177039-INS-170
chr16-19195010-INS-199
chr16-19201253-INS-3006
chr16-19264464-INS-5546
chr16-19338495-INS-443
chr16-19342208-INS-1116
chr16-19386265-INS-56
chr16-19389150-INS-110
chr16-19435784-INS-16058
chr16-19514332-INS-3577
chr16-19563168-INS-2143
chr16-19584732-INS-171
chr16-19596930-INS-4768
chr16-19612836-INS-3064
chr16-19617408-INS-50
chr16-19701640-INS-137
chr16-19781153-INS-108
chr16-19781593-INS-7304
chr16-19788662-INS-255
chr16-19816002-INS-202
chr16-19820466-INS-260
chr16-19829298-INS-4621
chr16-19833328-INS-100
chr16-19838816-INS-210
chr16-19890648-INS-129
chr16-19891962-INS-59
chr16-19939491-INS-197
chr16-19940081-INS-65
chr16-20007216-INS-325
chr16-20007630-INS-520
chr16-20011767-INS-52
chr16-20013910-INS-476
chr16-20066498-INS-208
chr16-20082277-INS-52
chr16-20115726-INS-50
chr16-20135998-INS-343
chr16-20147821-INS-60
chr16-20166068-INS-7910
chr16-20188508-INS-62
chr16-20217144-INS-559
chr16-20217458-INS-112
chr16-20225635-INS-1995
chr16-20228540-INS-206
chr16-20265860-INS-159
chr16-20282733-INS-50
chr16-20283758-INS-6403
chr16-20285106-INS-401
chr16-20292263-INS-51
chr16-20316493-INS-5075
chr16-20333187-INS-82
chr16-20335142-INS-133
chr16-20338640-INS-184

chr16-20385876-INS-56
chr16-20406416-INS-122
chr16-20517124-INS-88
chr16-20556564-INS-205
chr16-20559079-INS-90
chr16-20567311-INS-208
chr16-20615818-INS-151
chr16-20628635-INS-93
chr16-20643559-INS-201
chr16-20703789-INS-556
chr16-20721594-INS-2732
chr16-20727131-INS-160
chr16-20731991-INS-60
chr16-20756497-INS-9104
chr16-20783316-INS-7182
chr16-20788112-INS-4724
chr16-20814555-INS-98
chr16-20833362-INS-1896
chr16-20845527-INS-1545
chr16-20859891-INS-341
chr16-20861407-INS-51
chr16-20871740-INS-2824
chr16-20885019-INS-8339
chr16-20936468-INS-84
chr16-20937473-INS-9079
chr16-20959578-INS-195
chr16-21016336-INS-52
chr16-21098006-INS-70
chr16-21146106-INS-3496
chr16-21164787-INS-113
chr16-21166348-INS-534
chr16-21166559-INS-240
chr16-21172834-INS-63
chr16-21173734-INS-161
chr16-21178252-INS-55
chr16-21187732-INS-160
chr16-21199886-INS-54
chr16-21202784-INS-1095
chr16-21228446-INS-323
chr16-21288005-INS-593
chr16-21310436-INS-2779
chr16-21323692-INS-61
chr16-21324461-INS-162
chr16-21326048-INS-90
chr16-21330392-INS-52
chr16-21330507-INS-521
chr16-21371327-INS-122
chr16-21413539-INS-149
chr16-21431971-INS-76
chr16-21443713-INS-56
chr16-21477422-INS-94
chr16-21481726-INS-191
chr16-21487413-INS-69
chr16-21487710-INS-150
chr16-21487710-INS-54
chr16-21488590-INS-51
chr16-21502940-INS-4558
chr16-21521424-INS-8423
chr16-21540868-INS-89
chr16-21545090-INS-247
chr16-21547360-INS-1977
chr16-21552665-INS-166
chr16-21585967-INS-263
chr16-21612668-INS-207
chr16-21616274-INS-52
chr16-21622322-INS-176
chr16-21630903-INS-163
chr16-21666216-INS-138
chr16-21695304-INS-246
chr16-21703803-INS-74
chr16-21706904-INS-51
chr16-21745564-INS-62
chr16-21779053-INS-290
chr16-21881032-INS-122
chr16-21881099-INS-486
chr16-21912952-INS-411
chr16-21939635-INS-68

chr16-21940868-INS-102
chr16-21947079-INS-1110
chr16-21973844-INS-226
chr16-22016091-INS-220
chr16-22027960-INS-6454
chr16-22032181-INS-218
chr16-22042840-INS-228
chr16-22050587-INS-7087
chr16-22111030-INS-4524
chr16-22137044-INS-364
chr16-22195747-INS-443
chr16-22196987-INS-70
chr16-22269504-INS-1194
chr16-22269562-INS-398
chr16-22270345-INS-50
chr16-22289764-INS-158
chr16-22325393-INS-52
chr16-22379736-INS-86
chr16-22408979-INS-158
chr16-22408979-INS-65
chr16-22497849-INS-65
chr16-22553067-INS-508
chr16-22617990-INS-296
chr16-22635343-INS-232
chr16-22640390-INS-135
chr16-22647803-INS-120
chr16-22650656-INS-6471
chr16-22653206-INS-2010
chr16-22656999-INS-54
chr16-22735091-INS-58
chr16-22784911-INS-172
chr16-22793149-INS-58
chr16-22813539-INS-1883
chr16-22835922-INS-774
chr16-22905577-INS-226
chr16-22919601-INS-100
chr16-22937534-INS-217
chr16-22938015-INS-122
chr16-22938237-INS-208
chr16-22940791-INS-162
chr16-22970138-INS-85
chr16-22982570-INS-206
chr16-23007236-INS-391
chr16-23030616-INS-52
chr16-23052016-INS-64
chr16-23072829-INS-499
chr16-23143991-INS-50
chr16-23184976-INS-158
chr16-23200117-INS-958
chr16-23210023-INS-104
chr16-23226511-INS-63
chr16-23233843-INS-67
chr16-23255268-INS-836
chr16-23296879-INS-140
chr16-23299721-INS-56
chr16-23324194-INS-55
chr16-23403053-INS-360
chr16-23439878-INS-500
chr16-23542199-INS-6601
chr16-23680484-INS-56
chr16-23762144-INS-124
chr16-23762661-INS-53
chr16-23768482-INS-57
chr16-23768876-INS-109
chr16-23811834-INS-434
chr16-23859078-INS-60
chr16-23868106-INS-50
chr16-23899248-INS-56
chr16-23906436-INS-2544
chr16-23972404-INS-77
chr16-23975229-INS-68
chr16-24036908-INS-62
chr16-24085337-INS-6775
chr16-24111403-INS-6794
chr16-24130716-INS-7093
chr16-24220125-INS-75
chr16-24224758-INS-113

chr16-24224769-INS-261
chr16-24276729-INS-364
chr16-24276866-INS-108
chr16-24311385-INS-60
chr16-24402530-INS-266
chr16-24405950-INS-56
chr16-24430359-INS-544
chr16-24459027-INS-55
chr16-24551285-INS-218
chr16-24602841-INS-1422
chr16-24617395-INS-127
chr16-24629614-INS-139
chr16-24672264-INS-1185
chr16-24733890-INS-6366
chr16-24759556-INS-51
chr16-24838102-INS-79
chr16-24850856-INS-150
chr16-24851177-INS-333
chr16-24935619-INS-307
chr16-24936002-INS-144
chr16-24961813-INS-58
chr16-24965868-INS-246
chr16-24974749-INS-4881
chr16-24974798-INS-205
chr16-24978432-INS-374
chr16-24998725-INS-86
chr16-24999083-INS-549
chr16-25002428-INS-60
chr16-25016725-INS-834
chr16-25018116-INS-286
chr16-25018162-INS-78
chr16-25044351-INS-2787
chr16-25051150-INS-208
chr16-25051150-INS-58
chr16-25089114-INS-2037
chr16-25094192-INS-154
chr16-25097619-INS-125
chr16-25169618-INS-160
chr16-25169655-INS-66
chr16-25175505-INS-116
chr16-25198368-INS-213
chr16-25238549-INS-448
chr16-25246208-INS-81
chr16-25253054-INS-11756
chr16-25318681-INS-65
chr16-25364701-INS-6197
chr16-25367806-INS-52
chr16-25382560-INS-143
chr16-25408584-INS-6173
chr16-25416736-INS-323
chr16-25417411-INS-193
chr16-25472054-INS-245
chr16-25490861-INS-288
chr16-25579158-INS-52
chr16-25677826-INS-115
chr16-25733446-INS-1651
chr16-25745736-INS-110
chr16-25782075-INS-102
chr16-25808831-INS-7200
chr16-25834502-INS-57
chr16-25885793-INS-1384
chr16-25906833-INS-76
chr16-25961149-INS-315
chr16-25961511-INS-200
chr16-26025323-INS-2832
chr16-26037877-INS-5500
chr16-26052625-INS-50
chr16-26079522-INS-503
chr16-26090457-INS-376
chr16-26098509-INS-70
chr16-26098705-INS-144
chr16-26169167-INS-84
chr16-26219164-INS-60
chr16-26240305-INS-59
chr16-26243011-INS-2798
chr16-26256734-INS-1532
chr16-26260906-INS-51

chr16-26260986-INS-92
chr16-26283613-INS-58
chr16-26285684-INS-76
chr16-26299888-INS-7194
chr16-26312814-INS-96
chr16-26340036-INS-4165
chr16-26350194-INS-344
chr16-26360738-INS-500
chr16-26375083-INS-485
chr16-26397832-INS-54
chr16-26415601-INS-219
chr16-26417862-INS-821
chr16-26418111-INS-276
chr16-26460815-INS-133
chr16-26468198-INS-50
chr16-26517200-INS-57
chr16-26549804-INS-50
chr16-26557592-INS-88
chr16-26560298-INS-2894
chr16-26575899-INS-1527
chr16-26596140-INS-136
chr16-26597765-INS-378
chr16-26613815-INS-132
chr16-26618969-INS-667
chr16-26672961-INS-1319
chr16-26676878-INS-1785
chr16-26681051-INS-1340
chr16-26739216-INS-255
chr16-26765641-INS-60
chr16-26769536-INS-177
chr16-26789011-INS-180
chr16-26804666-INS-78
chr16-26816183-INS-72
chr16-26844369-INS-58
chr16-26844727-INS-78
chr16-26857929-INS-388
chr16-26879274-INS-1003
chr16-26922248-INS-58
chr16-26929272-INS-74
chr16-26979829-INS-129
chr16-26997730-INS-2186
chr16-27024800-INS-785
chr16-27067432-INS-84
chr16-27073795-INS-1420
chr16-27076820-INS-6448
chr16-27080650-INS-84
chr16-27089832-INS-6712
chr16-27112455-INS-370
chr16-27169090-INS-2496
chr16-27194620-INS-51
chr16-27194730-INS-50
chr16-27206634-INS-3207
chr16-27207694-INS-54
chr16-27308571-INS-6447
chr16-27314536-INS-590
chr16-27319042-INS-70
chr16-27325230-INS-2718
chr16-27326363-INS-76
chr16-27344749-INS-160
chr16-27350483-INS-1313
chr16-27360782-INS-204
chr16-27376895-INS-65
chr16-27452295-INS-6486
chr16-27468802-INS-98
chr16-27503521-INS-52
chr16-27509995-INS-177
chr16-27510079-INS-440
chr16-27594065-INS-6394
chr16-27637158-INS-6220
chr16-27664853-INS-6466
chr16-27700830-INS-7348
chr16-27715986-INS-6057
chr16-27743026-INS-6145
chr16-27759179-INS-1004
chr16-27853561-INS-160
chr16-27876294-INS-277
chr16-28079434-INS-3727

chr16-28132345-INS-3438
chr16-28160862-INS-103
chr16-28161645-INS-79
chr16-28187151-INS-521
chr16-28198855-INS-76
chr16-28246170-INS-70
chr16-28283660-INS-50
chr16-28294013-INS-54
chr16-28311527-INS-52
chr16-28333710-INS-410
chr16-28387915-INS-356
chr16-28412259-INS-60
chr16-28435714-INS-50
chr16-28487415-INS-50
chr16-28541622-INS-70
chr16-28573460-INS-740
chr16-28604175-INS-52
chr16-28627528-INS-1113
chr16-28685475-INS-51
chr16-28775638-INS-70
chr16-28776530-INS-58
chr16-28917265-INS-61
chr16-29010922-INS-50
chr16-29016054-INS-50
chr16-29072174-INS-54
chr16-29098960-INS-181
chr16-29122586-INS-181
chr16-29157405-INS-170
chr16-29180844-INS-52
chr16-29207672-INS-52
chr16-29219941-INS-78
chr16-29220777-INS-78
chr16-29227564-INS-248
chr16-29236859-INS-130
chr16-29237023-INS-326
chr16-29276601-INS-62
chr16-29296473-INS-56
chr16-29358385-INS-84
chr16-29368327-INS-56
chr16-29368810-INS-78
chr16-29414151-INS-55
chr16-29503978-INS-184
chr16-29504939-INS-81
chr16-29540171-INS-82
chr16-29543027-INS-52
chr16-29547147-INS-155
chr16-29551494-INS-64
chr16-29556629-INS-182
chr16-29571502-INS-81
chr16-29623172-INS-4134
chr16-29626013-INS-60
chr16-29635377-INS-178
chr16-29645859-INS-7062
chr16-29700882-INS-61
chr16-29717669-INS-102
chr16-29720596-INS-76
chr16-29736718-INS-309
chr16-29774223-INS-65
chr16-29820920-INS-55
chr16-29850946-INS-52
chr16-29940308-INS-715
chr16-29943906-INS-569
chr16-29951520-INS-179
chr16-29954666-INS-198
chr16-29956436-INS-130
chr16-29963384-INS-379
chr16-30066195-INS-56
chr16-30081353-INS-168
chr16-30157088-INS-212
chr16-30164697-INS-209
chr16-30192478-INS-132
chr16-30193240-INS-208
chr16-30217009-INS-75
chr16-30220700-INS-74
chr16-30226546-INS-328
chr16-30238698-INS-211
chr16-30240748-INS-191

chr16-30240748-INS-75
chr16-30241310-INS-70
chr16-30245285-INS-51
chr16-30255789-INS-50
chr16-30271826-INS-235
chr16-30290739-INS-218
chr16-30300239-INS-215
chr16-30373809-INS-1866
chr16-30379256-INS-190
chr16-30386847-INS-133
chr16-30463192-INS-328
chr16-30466163-INS-462
chr16-30536793-INS-1334
chr16-30537796-INS-3275
chr16-30574589-INS-50
chr16-30575294-INS-159
chr16-30604235-INS-54
chr16-30636973-INS-3716
chr16-30644143-INS-56
chr16-30645671-INS-191
chr16-30721485-INS-132
chr16-30759388-INS-63
chr16-30768890-INS-333
chr16-30794781-INS-5612
chr16-30795420-INS-71
chr16-30807645-INS-197
chr16-30842103-INS-96
chr16-30891945-INS-248
chr16-30969226-INS-63
chr16-30981576-INS-468
chr16-30981815-INS-144
chr16-30997951-INS-317
chr16-31000631-INS-287
chr16-31000990-INS-164
chr16-31017805-INS-284
chr16-31048662-INS-624
chr16-31101597-INS-64
chr16-31139220-INS-64
chr16-31237282-INS-342
chr16-31241521-INS-3902
chr16-31243732-INS-3917
chr16-31307057-INS-193
chr16-31311083-INS-55
chr16-31321198-INS-60
chr16-31331147-INS-70
chr16-31331308-INS-210
chr16-31335088-INS-190
chr16-31338209-INS-78
chr16-31393638-INS-2929
chr16-31464116-INS-376
chr16-31579544-INS-50
chr16-31630215-INS-495
chr16-31734085-INS-205
chr16-31742923-INS-72
chr16-31801843-INS-213
chr16-31812737-INS-148
chr16-31866405-INS-123
chr16-31868995-INS-160
chr16-31980847-INS-218
chr16-31999790-INS-422
chr16-32031049-INS-74
chr16-32077605-INS-61
chr16-32079442-INS-108
chr16-32127069-INS-176
chr16-32137559-INS-206
chr16-32137918-INS-171
chr16-32157259-INS-272
chr16-32158854-INS-211
chr16-32231748-INS-65
chr16-32242119-INS-157
chr16-32285670-INS-257
chr16-32301153-INS-75
chr16-32301183-INS-438
chr16-32305780-INS-316
chr16-32311359-INS-3204
chr16-32349649-INS-71
chr16-32358244-INS-50

chr16-32366408-INS-51
chr16-32367051-INS-206
chr16-32381081-INS-1883
chr16-32393639-INS-142
chr16-32418384-INS-276
chr16-32428737-INS-63
chr16-32513096-INS-50
chr16-32516818-INS-150
chr16-32521482-INS-246
chr16-32521482-INS-82
chr16-32539913-INS-428
chr16-32566699-INS-942
chr16-32576267-INS-372
chr16-32631987-INS-1326
chr16-32638642-INS-3500
chr16-32660839-INS-372
chr16-32664794-INS-64
chr16-32671003-INS-190
chr16-32686464-INS-233
chr16-32705386-INS-58
chr16-32716530-INS-296
chr16-32795837-INS-3223
chr16-32838830-INS-268
chr16-32848337-INS-89
chr16-32851830-INS-97
chr16-32854045-INS-128
chr16-32868701-INS-3195
chr16-32868701-INS-645
chr16-32908442-INS-174
chr16-32913654-INS-125
chr16-32924283-INS-257
chr16-32927323-INS-50
chr16-32959272-INS-51
chr16-32967396-INS-237
chr16-32967529-INS-59
chr16-32967749-INS-231
chr16-32974880-INS-1000
chr16-32975276-INS-78
chr16-32978926-INS-208
chr16-32990848-INS-480
chr16-32998768-INS-8954
chr16-33020405-INS-111
chr16-33091869-INS-237
chr16-33118383-INS-52
chr16-33133498-INS-4887
chr16-33133814-INS-1887
chr16-33147418-INS-52
chr16-33152500-INS-4745
chr16-33211374-INS-206
chr16-33329505-INS-65
chr16-33347835-INS-186
chr16-33363966-INS-513
chr16-33371759-INS-79
chr16-33419948-INS-150
chr16-33420293-INS-50
chr16-33437993-INS-357
chr16-33532135-INS-52
chr16-33604531-INS-72
chr16-33608241-INS-57
chr16-33638446-INS-52
chr16-33646416-INS-214
chr16-33674286-INS-774
chr16-33675397-INS-1444
chr16-33685424-INS-84
chr16-33687444-INS-209
chr16-33721495-INS-63
chr16-33729711-INS-163
chr16-33734645-INS-104
chr16-33734838-INS-868
chr16-33745747-INS-116
chr16-33745747-INS-56
chr16-33747562-INS-66
chr16-33789333-INS-449
chr16-33798769-INS-80
chr16-33800909-INS-124
chr16-33804308-INS-135
chr16-33805220-INS-84

chr16-33811856-INS-84
chr16-33919333-INS-75
chr16-34011360-INS-461
chr16-34054969-INS-80
chr16-34059319-INS-52
chr16-34071501-INS-1965
chr16-34180790-INS-13169
chr16-34195521-INS-262
chr16-34205137-INS-6865
chr16-34211309-INS-201
chr16-34211329-INS-465
chr16-34211329-INS-65
chr16-34260208-INS-56
chr16-34341257-INS-59
chr16-34355138-INS-140
chr16-34392050-INS-51
chr16-34395573-INS-53
chr16-34396481-INS-86
chr16-34430018-INS-166
chr16-34439999-INS-54
chr16-34443994-INS-169
chr16-34444003-INS-65
chr16-34475421-INS-100
chr16-34483260-INS-116
chr16-34483292-INS-247
chr16-34506387-INS-205
chr16-34506590-INS-68
chr16-34722927-INS-68
chr16-34757678-INS-78
chr16-34874278-INS-230
chr16-34935893-INS-186
chr16-34936481-INS-60
chr16-34956549-INS-76
chr16-35005601-INS-53
chr16-35018329-INS-81
chr16-35020007-INS-52
chr16-35029946-INS-50
chr16-35064493-INS-878
chr16-35157247-INS-67
chr16-35157472-INS-121
chr16-35269358-INS-198
chr16-35320226-INS-180
chr16-35332617-INS-95
chr16-35332780-INS-329
chr16-35363842-INS-50
chr16-35378700-INS-181
chr16-35415918-INS-60
chr16-35416184-INS-59
chr16-35440134-INS-187
chr16-35450176-INS-54
chr16-35456459-INS-60
chr16-35457132-INS-127
chr16-35464552-INS-313
chr16-35464806-INS-595
chr16-35489330-INS-199
chr16-35509495-INS-4924
chr16-35516776-INS-150
chr16-35524080-INS-203
chr16-35547968-INS-526
chr16-35562490-INS-212
chr16-35569896-INS-203
chr16-35585733-INS-176
chr16-35659899-INS-202
chr16-35663267-INS-179
chr16-35664953-INS-225
chr16-35675531-INS-395
chr16-35686142-INS-64
chr16-35697065-INS-6350
chr16-35697180-INS-216
chr16-35700395-INS-51
chr16-35704298-INS-203
chr16-35720268-INS-202
chr16-35720907-INS-213
chr16-35728142-INS-242
chr16-35786420-INS-98
chr16-35786635-INS-520
chr16-35790962-INS-182

chr16-35828967-INS-212
chr16-35890973-INS-205
chr16-35898311-INS-234
chr16-35900730-INS-53
chr16-35960343-INS-7435
chr16-35963357-INS-2927
chr16-35981157-INS-1307
chr16-35993651-INS-524
chr16-35993760-INS-549
chr16-36021907-INS-61
chr16-36129878-INS-6355
chr16-36144721-INS-6765
chr16-36192814-INS-50
chr16-36224730-INS-64
chr16-36229319-INS-500
chr16-36250042-INS-1424
chr16-36271151-INS-111
chr16-36290427-INS-547
chr16-36298764-INS-140
chr16-36299208-INS-173
chr16-36304647-INS-72
chr16-36339846-INS-433
chr16-36391832-INS-51
chr16-36476969-INS-114
chr16-36491691-INS-96
chr16-36511193-INS-52
chr16-36551388-INS-6230
chr16-36568758-INS-6276
chr16-36574529-INS-169
chr16-36575117-INS-56
chr16-36608826-INS-6475
chr16-36610098-INS-65
chr16-36742251-INS-304
chr16-36777860-INS-120
chr16-36929459-INS-122
chr16-36985879-INS-214
chr16-37000501-INS-6278
chr16-37020139-INS-52
chr16-37100097-INS-7132
chr16-37107733-INS-205
chr16-37108338-INS-208
chr16-37123920-INS-64
chr16-37237548-INS-224
chr16-37275792-INS-1941
chr16-37290029-INS-205
chr16-37318311-INS-6246
chr16-37394924-INS-2719
chr16-37431086-INS-50
chr16-37453840-INS-92
chr16-37481861-INS-4905
chr16-37488257-INS-50
chr16-37494318-INS-53
chr16-37645075-INS-364
chr16-37655853-INS-99
chr16-37669945-INS-487
chr16-37793915-INS-80
chr16-37849846-INS-58
chr16-37957000-INS-92
chr16-37993547-INS-64
chr16-38027594-INS-66
chr16-38052646-INS-731
chr16-38089121-INS-51
chr16-38105982-INS-245
chr16-38105982-INS-70
chr16-38106050-INS-595
chr16-38106650-INS-6586
chr16-38137589-INS-72
chr16-38162755-INS-52
chr16-38175559-INS-5059
chr16-38213592-INS-106
chr16-38298166-INS-525
chr16-38306589-INS-54
chr16-38324565-INS-65
chr16-38324771-INS-540
chr16-38339923-INS-152
chr16-38364259-INS-184
chr16-38398001-INS-237

chr16-38426921-INS-53
chr16-38464883-INS-52
chr16-38503865-INS-60
chr16-38517412-INS-1055
chr16-38647682-INS-54
chr16-38904430-INS-522
chr16-39001705-INS-471
chr16-39001709-INS-180
chr16-39001848-INS-85
chr16-39034583-INS-116
chr16-39161437-INS-685
chr16-39200684-INS-1856
chr16-39207805-INS-1894
chr16-39213792-INS-50
chr16-39246273-INS-52
chr16-39293693-INS-1742
chr16-39307679-INS-82
chr16-39467135-INS-182
chr16-39467135-INS-50
chr16-39467135-INS-650
chr16-39471545-INS-227
chr16-39471555-INS-85
chr16-39471848-INS-2219
chr16-39648929-INS-1967
chr16-39654190-INS-96
chr16-39666425-INS-145
chr16-39856364-INS-1743
chr16-39903608-INS-54
chr16-39903673-INS-210
chr16-39921687-INS-78
chr16-40101208-INS-228
chr16-40118609-INS-66
chr16-40120660-INS-674
chr16-40146829-INS-500
chr16-40147400-INS-500
chr16-40156811-INS-80
chr16-40191909-INS-109
chr16-40220712-INS-377
chr16-40282370-INS-93
chr16-40495664-INS-50
chr16-40496945-INS-2500
chr16-40634817-INS-214
chr16-40793689-INS-64
chr16-40840778-INS-1287
chr16-40925295-INS-2375
chr16-40927241-INS-69
chr16-41000943-INS-126
chr16-41068242-INS-185
chr16-41089808-INS-7505
chr16-41097424-INS-82
chr16-41164086-INS-98
chr16-41194086-INS-137
chr16-41284616-INS-102
chr16-41299438-INS-1478
chr16-41336627-INS-52
chr16-41477635-INS-54
chr16-41547022-INS-56
chr16-41615850-INS-887
chr16-41654038-INS-6377
chr16-41808918-INS-52
chr16-41809365-INS-186
chr16-41823636-INS-3486
chr16-41922704-INS-6510
chr16-41936394-INS-186
chr16-41946270-INS-76
chr16-42011704-INS-50
chr16-42015003-INS-62
chr16-42027074-INS-98
chr16-42046749-INS-500
chr16-42049814-INS-55
chr16-42067646-INS-82
chr16-42105946-INS-147
chr16-42161017-INS-70
chr16-42173140-INS-2554
chr16-42233360-INS-70
chr16-42295616-INS-62
chr16-42311003-INS-77

chr16-42320019-INS-2447
chr16-42388166-INS-1086
chr16-42391475-INS-810
chr16-42393479-INS-1170
chr16-42472215-INS-58
chr16-42477752-INS-393
chr16-42477776-INS-85
chr16-42505789-INS-64
chr16-42575777-INS-62
chr16-42577463-INS-6193
chr16-42585174-INS-66
chr16-42616831-INS-51
chr16-42620716-INS-80
chr16-42777169-INS-109
chr16-42842838-INS-6806
chr16-42861929-INS-6651
chr16-42867569-INS-1807
chr16-42999114-INS-50
chr16-43000594-INS-50
chr16-43129948-INS-60
chr16-43266083-INS-209
chr16-43555256-INS-206
chr16-43582420-INS-719
chr16-43614033-INS-69
chr16-43614941-INS-1670
chr16-43623498-INS-68
chr16-43817102-INS-7402
chr16-43890282-INS-140
chr16-43937417-INS-62
chr16-43967778-INS-62
chr16-43988395-INS-724
chr16-44014122-INS-179
chr16-44015208-INS-76
chr16-44015646-INS-78
chr16-44016428-INS-22966
chr16-44123149-INS-1102
chr16-44133042-INS-198
chr16-44137886-INS-5034
chr16-44198286-INS-110
chr16-44263305-INS-400
chr16-44270670-INS-104
chr16-44295879-INS-51
chr16-44317133-INS-60
chr16-44343070-INS-71
chr16-44343530-INS-66
chr16-44345832-INS-63
chr16-44346031-INS-57
chr16-44375579-INS-172
chr16-44407740-INS-664
chr16-44468156-INS-80
chr16-44472129-INS-86
chr16-44491030-INS-65
chr16-44491127-INS-192
chr16-44506265-INS-282
chr16-44517252-INS-993
chr16-44522438-INS-866
chr16-44551188-INS-271
chr16-44626963-INS-383
chr16-44629946-INS-4644
chr16-44636864-INS-457
chr16-44641754-INS-152
chr16-44688788-INS-56
chr16-44696672-INS-53
chr16-44742951-INS-507
chr16-44765336-INS-50
chr16-44765872-INS-6475
chr16-44767698-INS-1494
chr16-44774128-INS-83
chr16-44797183-INS-84
chr16-44834405-INS-3189
chr16-44860067-INS-324
chr16-44897141-INS-2975
chr16-44938745-INS-55
chr16-45019845-INS-1893
chr16-45020601-INS-53
chr16-45112114-INS-1162
chr16-45137124-INS-201

chr16-45145628-INS-3183
chr16-45153621-INS-84
chr16-45153902-INS-69
chr16-45160052-INS-6166
chr16-45169289-INS-3021
chr16-45173418-INS-364
chr16-45173498-INS-3096
chr16-45173722-INS-358
chr16-45173865-INS-62
chr16-45185443-INS-7349
chr16-45225711-INS-206
chr16-45246015-INS-168
chr16-45255075-INS-50
chr16-45289426-INS-297
chr16-45360929-INS-284
chr16-45363811-INS-84
chr16-45364022-INS-168
chr16-45378925-INS-1866
chr16-45445167-INS-108
chr16-45508401-INS-137
chr16-45519112-INS-76
chr16-45521787-INS-77
chr16-45529507-INS-167
chr16-45610205-INS-64
chr16-45665258-INS-51
chr16-45706689-INS-382
chr16-45721578-INS-6464
chr16-45733161-INS-92
chr16-45741740-INS-210
chr16-45810135-INS-1002
chr16-45826724-INS-148
chr16-45835589-INS-104
chr16-45835778-INS-246
chr16-45845933-INS-58
chr16-45992753-INS-74
chr16-46003726-INS-244
chr16-46004370-INS-199
chr16-46030116-INS-7957
chr16-46089436-INS-90
chr16-46166505-INS-93
chr16-46283992-INS-100
chr16-46323883-INS-6483
chr16-46350130-INS-243
chr16-46356439-INS-2363
chr16-46359764-INS-207
chr16-46359978-INS-168
chr16-46363799-INS-3614
chr16-46376400-INS-221
chr16-46426704-INS-76
chr16-46442046-INS-165
chr16-46487623-INS-58
chr16-46501716-INS-116
chr16-46539710-INS-727
chr16-46594633-INS-111
chr16-46628666-INS-135
chr16-46645958-INS-58
chr16-46653336-INS-6475
chr16-46676616-INS-68
chr16-46687059-INS-84
chr16-46687610-INS-55
chr16-46700911-INS-69
chr16-46707066-INS-6454
chr16-46731277-INS-3509
chr16-46751325-INS-6632
chr16-46833973-INS-6691
chr16-46841036-INS-125
chr16-46878225-INS-84
chr16-46878983-INS-52
chr16-46891316-INS-138
chr16-46891377-INS-303
chr16-46893076-INS-64
chr16-46980513-INS-56
chr16-46994591-INS-1108
chr16-47034447-INS-165
chr16-47100281-INS-104
chr16-47151208-INS-6193
chr16-47172472-INS-2220

chr16-47180423-INS-50
chr16-47201265-INS-3977
chr16-47224459-INS-4374
chr16-47233610-INS-117
chr16-47233692-INS-50
chr16-47272820-INS-6519
chr16-47275846-INS-7050
chr16-47358971-INS-1366
chr16-47400106-INS-58
chr16-47417310-INS-5954
chr16-47462636-INS-50
chr16-47467397-INS-58
chr16-47487462-INS-7134
chr16-47498873-INS-6970
chr16-47516845-INS-53
chr16-47522707-INS-5506
chr16-47528277-INS-3049
chr16-47547394-INS-6689
chr16-47566156-INS-2024
chr16-47579400-INS-64
chr16-47579731-INS-81
chr16-47596214-INS-245
chr16-47606070-INS-175
chr16-47636431-INS-69
chr16-47656602-INS-118
chr16-47656602-INS-388
chr16-47657210-INS-192
chr16-47665343-INS-1083
chr16-47702112-INS-2000
chr16-47714829-INS-155
chr16-47737630-INS-142
chr16-47741523-INS-2441
chr16-47749397-INS-2159
chr16-47861402-INS-4607
chr16-47867398-INS-2099
chr16-47898515-INS-1894
chr16-47939856-INS-1820
chr16-47973953-INS-50
chr16-48038639-INS-66
chr16-48066102-INS-165
chr16-48069897-INS-82
chr16-48070730-INS-51
chr16-48080076-INS-1079
chr16-48099887-INS-84
chr16-48125573-INS-50
chr16-48141128-INS-53
chr16-48142492-INS-4769
chr16-48150642-INS-52
chr16-48151884-INS-639
chr16-48165354-INS-6251
chr16-48175278-INS-2054
chr16-48175278-INS-498
chr16-48175713-INS-216
chr16-48175981-INS-88
chr16-48176433-INS-79
chr16-48176775-INS-57
chr16-48217299-INS-81
chr16-48236262-INS-7370
chr16-48243576-INS-381
chr16-48288806-INS-6425
chr16-48333557-INS-143
chr16-48347670-INS-1711
chr16-48409920-INS-1046
chr16-48439321-INS-6646
chr16-48472253-INS-52
chr16-48493763-INS-66
chr16-48522933-INS-85
chr16-48523523-INS-51
chr16-48524219-INS-2745
chr16-48543752-INS-70
chr16-48560277-INS-54
chr16-48609714-INS-6232
chr16-48617246-INS-2869
chr16-48625532-INS-809
chr16-48627958-INS-808
chr16-48629764-INS-6798
chr16-48630785-INS-1858

chr16-48674278-INS-2539
chr16-48735073-INS-161
chr16-48808683-INS-83
chr16-48875717-INS-50
chr16-48881712-INS-51
chr16-48906157-INS-174
chr16-48906157-INS-60
chr16-49083494-INS-2681
chr16-49086349-INS-61
chr16-49086412-INS-234
chr16-49092629-INS-1310
chr16-49104048-INS-486
chr16-49108389-INS-131
chr16-49116797-INS-1230
chr16-49155246-INS-1890
chr16-49161189-INS-15427
chr16-49163022-INS-3392
chr16-49163406-INS-66
chr16-49199203-INS-90
chr16-49199865-INS-74
chr16-49239168-INS-58
chr16-49268114-INS-415
chr16-49362526-INS-147
chr16-49370305-INS-174
chr16-49381125-INS-203
chr16-49381173-INS-75
chr16-49402096-INS-222
chr16-49403999-INS-2420
chr16-49413073-INS-102
chr16-49431657-INS-50
chr16-49452603-INS-7190
chr16-49484926-INS-56
chr16-49535472-INS-56
chr16-49545255-INS-101
chr16-49564210-INS-51
chr16-49615071-INS-1208
chr16-49726627-INS-52
chr16-49758809-INS-64
chr16-49803181-INS-52
chr16-49834097-INS-51
chr16-49851105-INS-257
chr16-49851119-INS-779
chr16-49899223-INS-476
chr16-49903848-INS-1290
chr16-49922018-INS-54
chr16-49922148-INS-66
chr16-49939205-INS-410
chr16-49944521-INS-173
chr16-49963918-INS-1456
chr16-49973574-INS-71
chr16-49973801-INS-1041
chr16-49987472-INS-116
chr16-49992217-INS-2001
chr16-50054850-INS-376
chr16-50088356-INS-50
chr16-50134944-INS-62
chr16-50139762-INS-50
chr16-50169724-INS-161
chr16-50178081-INS-235
chr16-50212171-INS-535
chr16-50251642-INS-54
chr16-50286161-INS-6507
chr16-50293798-INS-202
chr16-50315413-INS-57
chr16-50319948-INS-67
chr16-50327925-INS-54
chr16-50364865-INS-104
chr16-50433280-INS-68
chr16-50433967-INS-189
chr16-50435244-INS-55
chr16-50458811-INS-159
chr16-50459105-INS-221
chr16-50466473-INS-538
chr16-50466515-INS-105
chr16-50468923-INS-6609
chr16-50470021-INS-1885
chr16-50473149-INS-402

chr16-50490272-INS-4595
chr16-50679142-INS-1700
chr16-50699029-INS-236
chr16-50699460-INS-6592
chr16-50711002-INS-10158
chr16-50717687-INS-135
chr16-50717733-INS-59
chr16-50721055-INS-2927
chr16-50728637-INS-2390
chr16-50745633-INS-96
chr16-50770326-INS-90
chr16-50788141-INS-198
chr16-50807117-INS-1582
chr16-50819826-INS-167
chr16-50933950-INS-52
chr16-50946959-INS-1157
chr16-50952332-INS-4277
chr16-50984169-INS-72
chr16-50985355-INS-6233
chr16-50988971-INS-3188
chr16-51024526-INS-2857
chr16-51047367-INS-69
chr16-51052773-INS-3980
chr16-51115248-INS-1092
chr16-51124911-INS-6212
chr16-51197134-INS-202
chr16-51321662-INS-16572
chr16-51525903-INS-252
chr16-51526071-INS-50
chr16-51538226-INS-50
chr16-51604501-INS-56
chr16-51751404-INS-210
chr16-51860769-INS-56
chr16-51864603-INS-354
chr16-51877445-INS-241
chr16-51996169-INS-104
chr16-52043200-INS-138
chr16-52060025-INS-58
chr16-52060531-INS-149
chr16-52120355-INS-166
chr16-52124527-INS-50
chr16-52185575-INS-2792
chr16-52205965-INS-67
chr16-52206307-INS-401
chr16-52369632-INS-3096
chr16-52413876-INS-66
chr16-52414319-INS-270
chr16-52418229-INS-2881
chr16-52429327-INS-480
chr16-52432591-INS-778
chr16-52456188-INS-96
chr16-52477882-INS-683
chr16-52491087-INS-1535
chr16-52512107-INS-2914
chr16-52521152-INS-554
chr16-52629357-INS-52
chr16-52631330-INS-1695
chr16-52668904-INS-123
chr16-52701035-INS-88
chr16-52701786-INS-1914
chr16-52772916-INS-717
chr16-52786017-INS-2782
chr16-52804783-INS-3586
chr16-52847427-INS-1539
chr16-52857854-INS-903
chr16-52893555-INS-267
chr16-52948159-INS-6189
chr16-52967692-INS-6694
chr16-52976498-INS-4002
chr16-52998243-INS-50
chr16-53036805-INS-74
chr16-53060162-INS-296
chr16-53122179-INS-468
chr16-53125185-INS-60
chr16-53166082-INS-546
chr16-53201003-INS-6434
chr16-53220817-INS-6616

chr16-53252660-INS-228
chr16-53294133-INS-357
chr16-53334784-INS-54
chr16-53341044-INS-84
chr16-53388912-INS-74
chr16-53389078-INS-52
chr16-53418879-INS-50
chr16-53423415-INS-2867
chr16-53441935-INS-7320
chr16-53447523-INS-476
chr16-53459998-INS-68
chr16-53465527-INS-3665
chr16-53480294-INS-592
chr16-53491258-INS-867
chr16-53512394-INS-1188
chr16-53555405-INS-3203
chr16-53564712-INS-345
chr16-53580773-INS-66
chr16-53588858-INS-105
chr16-53624685-INS-997
chr16-53627320-INS-2500
chr16-53694929-INS-54
chr16-53704842-INS-214
chr16-53705641-INS-160
chr16-53724537-INS-1225
chr16-53726884-INS-1216
chr16-53736803-INS-1145
chr16-53772324-INS-476
chr16-53799382-INS-536
chr16-53869897-INS-52
chr16-53875117-INS-170
chr16-53898326-INS-1315
chr16-53992842-INS-1290
chr16-53999821-INS-52
chr16-54008128-INS-6687
chr16-54032265-INS-6191
chr16-54032560-INS-1182
chr16-54034608-INS-113
chr16-54035610-INS-999
chr16-54069104-INS-2274
chr16-54086813-INS-63
chr16-54156635-INS-388
chr16-54170935-INS-50
chr16-54177503-INS-6443
chr16-54196974-INS-94
chr16-54264059-INS-1028
chr16-54283600-INS-90
chr16-54297478-INS-1327
chr16-54394788-INS-14719
chr16-54397287-INS-3586
chr16-54405337-INS-369
chr16-54454639-INS-50
chr16-54536327-INS-1120
chr16-54544145-INS-112
chr16-54586328-INS-198
chr16-54590314-INS-5765
chr16-54590965-INS-512
chr16-54660562-INS-54
chr16-54677484-INS-1163
chr16-54738059-INS-58
chr16-54742310-INS-7080
chr16-54742918-INS-158
chr16-54745786-INS-78
chr16-54819380-INS-74
chr16-54829690-INS-80
chr16-54861432-INS-339
chr16-54861512-INS-138
chr16-54870245-INS-3362
chr16-54880451-INS-8856
chr16-54900947-INS-81
chr16-54934979-INS-3298
chr16-54957705-INS-53
chr16-54986067-INS-68
chr16-54992074-INS-145
chr16-55068138-INS-57
chr16-55068409-INS-336
chr16-55098313-INS-84

chr16-55126115-INS-126
chr16-55127431-INS-2679
chr16-55129729-INS-1807
chr16-55131194-INS-524
chr16-55134579-INS-144
chr16-55137563-INS-79
chr16-55180579-INS-70
chr16-55197680-INS-280
chr16-55215779-INS-72
chr16-55279573-INS-6606
chr16-55290274-INS-138
chr16-55290398-INS-993
chr16-55293256-INS-1152
chr16-55305550-INS-63
chr16-55312407-INS-574
chr16-55315259-INS-80
chr16-55318495-INS-6341
chr16-55327972-INS-1245
chr16-55342133-INS-496
chr16-55359140-INS-60
chr16-55385945-INS-319
chr16-55406973-INS-148
chr16-55424349-INS-506
chr16-55452997-INS-62
chr16-55453357-INS-585
chr16-55514957-INS-480
chr16-55526382-INS-106
chr16-55530286-INS-222
chr16-55533704-INS-239
chr16-55533794-INS-56
chr16-55534278-INS-479
chr16-55563553-INS-58
chr16-55563797-INS-255
chr16-55575600-INS-88
chr16-55589547-INS-66
chr16-55594663-INS-58
chr16-55598176-INS-210
chr16-55680258-INS-297
chr16-55685124-INS-77
chr16-55688700-INS-216
chr16-55689101-INS-117
chr16-55707932-INS-71
chr16-55708242-INS-392
chr16-55719896-INS-855
chr16-55724192-INS-219
chr16-55738649-INS-57
chr16-55743951-INS-105
chr16-55744627-INS-94
chr16-55750028-INS-207
chr16-55758183-INS-198
chr16-55767379-INS-78
chr16-55769958-INS-200
chr16-55887079-INS-210
chr16-55892375-INS-203
chr16-55985280-INS-472
chr16-55994342-INS-108
chr16-56043350-INS-6663
chr16-56047688-INS-73
chr16-56052893-INS-120
chr16-56061555-INS-51
chr16-56063711-INS-59
chr16-56063808-INS-273
chr16-56065525-INS-443
chr16-56065532-INS-110
chr16-56065610-INS-222
chr16-56066125-INS-217
chr16-56102081-INS-94
chr16-56133380-INS-63
chr16-56133634-INS-283
chr16-56168204-INS-4683
chr16-56178621-INS-53
chr16-56206729-INS-52
chr16-56213630-INS-6465
chr16-56233862-INS-7309
chr16-56277753-INS-102
chr16-56309060-INS-118
chr16-56309060-INS-345

chr16-56396290-INS-1180
chr16-56425283-INS-51
chr16-56425790-INS-57
chr16-56460826-INS-82
chr16-56595557-INS-6417
chr16-56596505-INS-2000
chr16-56623823-INS-52
chr16-56658691-INS-2653
chr16-56659151-INS-5316
chr16-56698981-INS-62
chr16-56729095-INS-406
chr16-56751899-INS-211
chr16-56780107-INS-1887
chr16-56787793-INS-139
chr16-56792429-INS-6331
chr16-56840780-INS-2847
chr16-56861694-INS-6514
chr16-56865967-INS-332
chr16-56923675-INS-6582
chr16-56947088-INS-205
chr16-56994073-INS-71
chr16-57013453-INS-116
chr16-57013572-INS-147
chr16-57076681-INS-515
chr16-57076775-INS-187
chr16-57082170-INS-257
chr16-57091910-INS-2760
chr16-57112583-INS-50
chr16-57113019-INS-292
chr16-57195255-INS-54
chr16-57228853-INS-122
chr16-57230237-INS-6658
chr16-57458498-INS-477
chr16-57471133-INS-50
chr16-57541927-INS-90
chr16-57604876-INS-56
chr16-57653351-INS-278
chr16-57680538-INS-128
chr16-57726911-INS-50
chr16-57766287-INS-5495
chr16-57769417-INS-50
chr16-57808142-INS-50
chr16-57815465-INS-55
chr16-57847221-INS-1582
chr16-57854459-INS-54
chr16-57909303-INS-56
chr16-57909371-INS-226
chr16-57976329-INS-120
chr16-58077827-INS-256
chr16-58083779-INS-328
chr16-58091855-INS-548
chr16-58091962-INS-1989
chr16-58091962-INS-883
chr16-58095314-INS-4220
chr16-58116330-INS-52
chr16-58117464-INS-2214
chr16-58176482-INS-195
chr16-58188754-INS-206
chr16-58242837-INS-279
chr16-58264970-INS-791
chr16-58299618-INS-58
chr16-58328666-INS-79
chr16-58331086-INS-188
chr16-58345288-INS-50
chr16-58444493-INS-58
chr16-58482725-INS-2258
chr16-58511573-INS-1153
chr16-58522966-INS-243
chr16-58543464-INS-1208
chr16-58543464-INS-429
chr16-58601590-INS-103
chr16-58613489-INS-50
chr16-58614018-INS-6016
chr16-58619142-INS-439
chr16-58629735-INS-1587
chr16-58631729-INS-2701
chr16-58636255-INS-1140

chr16-58645980-INS-50
chr16-58647778-INS-676
chr16-58651496-INS-2441
chr16-58667229-INS-7107
chr16-58804627-INS-366
chr16-58809124-INS-107
chr16-58829561-INS-6635
chr16-58837720-INS-4066
chr16-58854405-INS-227
chr16-58902189-INS-54
chr16-58958526-INS-50
chr16-58966000-INS-306
chr16-59158154-INS-6359
chr16-59161102-INS-7122
chr16-59172044-INS-132
chr16-59177817-INS-14227
chr16-59187204-INS-268
chr16-59209162-INS-786
chr16-59209659-INS-4735
chr16-59222327-INS-3050
chr16-59224179-INS-7282
chr16-59234940-INS-1208
chr16-59239890-INS-6499
chr16-59257621-INS-199
chr16-59314250-INS-547
chr16-59337211-INS-116
chr16-59352385-INS-215
chr16-59385112-INS-80
chr16-59413055-INS-93
chr16-59445309-INS-1056
chr16-59476365-INS-1677
chr16-59529380-INS-2042
chr16-59529380-INS-379
chr16-59542473-INS-6377
chr16-59587196-INS-68
chr16-59633390-INS-1568
chr16-59641906-INS-3419
chr16-59669653-INS-3353
chr16-59700169-INS-6193
chr16-59704129-INS-379
chr16-59723521-INS-1877
chr16-59730221-INS-949
chr16-59786846-INS-8309
chr16-59804121-INS-941
chr16-59847959-INS-1240
chr16-59855280-INS-6432
chr16-59896875-INS-58
chr16-59906302-INS-63
chr16-59937343-INS-6621
chr16-59958579-INS-1799
chr16-59977179-INS-6169
chr16-59993050-INS-55
chr16-59993151-INS-206
chr16-60000358-INS-52
chr16-60012262-INS-405
chr16-60043637-INS-6592
chr16-60053250-INS-60
chr16-60065121-INS-66
chr16-60069955-INS-75
chr16-60135116-INS-1336
chr16-60159429-INS-1654
chr16-60236997-INS-92
chr16-60357737-INS-50
chr16-60368086-INS-237
chr16-60397881-INS-60
chr16-60403423-INS-2409
chr16-60455786-INS-1533
chr16-60469035-INS-28769
chr16-60478836-INS-87
chr16-60523577-INS-65
chr16-60524232-INS-390
chr16-60640861-INS-1021
chr16-60640960-INS-154
chr16-60644667-INS-70
chr16-60667131-INS-4328
chr16-60687021-INS-10524
chr16-60704813-INS-602

chr16-60768865-INS-373
chr16-60775236-INS-100
chr16-60785834-INS-64
chr16-60837336-INS-99
chr16-60841881-INS-376
chr16-60876685-INS-626
chr16-60902495-INS-54
chr16-60942654-INS-2097
chr16-60961247-INS-2048
chr16-61000444-INS-52
chr16-61035748-INS-2850
chr16-61066596-INS-1410
chr16-61098836-INS-653
chr16-61105781-INS-1318
chr16-61143835-INS-172
chr16-61171825-INS-2392
chr16-61177684-INS-371
chr16-61195591-INS-163
chr16-61195708-INS-145
chr16-61207590-INS-56
chr16-61250607-INS-6168
chr16-61261133-INS-2698
chr16-61273597-INS-56
chr16-61289507-INS-88
chr16-61321936-INS-719
chr16-61391227-INS-334
chr16-61392263-INS-686
chr16-61395479-INS-78
chr16-61438558-INS-1431
chr16-61442831-INS-56
chr16-61445779-INS-2548
chr16-61450363-INS-52
chr16-61464852-INS-1726
chr16-61504015-INS-59
chr16-61556028-INS-3570
chr16-61578012-INS-6400
chr16-61580742-INS-51
chr16-61581833-INS-64
chr16-61583757-INS-210
chr16-61621765-INS-423
chr16-61623737-INS-52
chr16-61637988-INS-6174
chr16-61644537-INS-3921
chr16-61665235-INS-62
chr16-61665236-INS-167
chr16-61668703-INS-2743
chr16-61681489-INS-50
chr16-61689729-INS-74
chr16-61701059-INS-75
chr16-61701168-INS-195
chr16-61728387-INS-4542
chr16-61737966-INS-886
chr16-61752827-INS-2540
chr16-61756069-INS-3031
chr16-61769962-INS-2396
chr16-61791467-INS-52
chr16-61877357-INS-1093
chr16-61879384-INS-6205
chr16-61892880-INS-50
chr16-61944742-INS-50
chr16-61945448-INS-266
chr16-61948370-INS-648
chr16-61995462-INS-56
chr16-62022442-INS-2462
chr16-62046895-INS-261
chr16-62068919-INS-293
chr16-62085188-INS-232
chr16-62085325-INS-477
chr16-62096556-INS-6486
chr16-62147499-INS-81
chr16-62275721-INS-58
chr16-62308355-INS-97
chr16-62320186-INS-134
chr16-62333189-INS-50
chr16-62360095-INS-1184
chr16-62371356-INS-433
chr16-62417126-INS-2136

chr16-62434518-INS-75
chr16-62474793-INS-60
chr16-62483210-INS-261
chr16-62505674-INS-6390
chr16-62514144-INS-141
chr16-62514166-INS-70
chr16-62536136-INS-72
chr16-62561439-INS-116
chr16-62567869-INS-332
chr16-62641912-INS-56
chr16-62655423-INS-3731
chr16-62656501-INS-68
chr16-62718407-INS-56
chr16-62735791-INS-62
chr16-62783563-INS-52
chr16-62790588-INS-76
chr16-62795780-INS-178
chr16-62796605-INS-3701
chr16-62800194-INS-6334
chr16-62830082-INS-2810
chr16-62840445-INS-54
chr16-62870432-INS-1246
chr16-62881919-INS-210
chr16-62909880-INS-1078
chr16-62918799-INS-1574
chr16-63037679-INS-65
chr16-63080017-INS-536
chr16-63100471-INS-4997
chr16-63102662-INS-182
chr16-63145093-INS-52
chr16-63149571-INS-53
chr16-63157801-INS-8521
chr16-63252956-INS-85
chr16-63270822-INS-1643
chr16-63283170-INS-344
chr16-63290856-INS-54
chr16-63349006-INS-52
chr16-63369140-INS-118
chr16-63369434-INS-74
chr16-63369728-INS-75
chr16-63389040-INS-6591
chr16-63396631-INS-6442
chr16-63413136-INS-146
chr16-63503954-INS-88
chr16-63589428-INS-132
chr16-63643441-INS-271
chr16-63648602-INS-382
chr16-63654170-INS-498
chr16-63686622-INS-55
chr16-63702692-INS-274
chr16-63713773-INS-762
chr16-63769980-INS-52
chr16-63935428-INS-50
chr16-63937131-INS-52
chr16-63972684-INS-52
chr16-63981275-INS-891
chr16-63983355-INS-1422
chr16-64073641-INS-56
chr16-64122800-INS-125
chr16-64154670-INS-78
chr16-64196389-INS-166
chr16-64200308-INS-64
chr16-64200904-INS-150
chr16-64325399-INS-4513
chr16-64333660-INS-50
chr16-64346588-INS-52
chr16-64364085-INS-5583
chr16-64376385-INS-1189
chr16-64378798-INS-173
chr16-64378969-INS-277
chr16-64380438-INS-346
chr16-64401775-INS-480
chr16-64409163-INS-84
chr16-64419068-INS-52
chr16-64421967-INS-3152
chr16-64430253-INS-6269
chr16-64466083-INS-1622

chr16-64493959-INS-51
chr16-64494608-INS-231
chr16-64507301-INS-1326
chr16-64523496-INS-88
chr16-64543398-INS-189
chr16-64562090-INS-56
chr16-64562106-INS-132
chr16-64574766-INS-52
chr16-64619973-INS-57
chr16-64627283-INS-832
chr16-64648685-INS-145
chr16-64665754-INS-4919
chr16-64701937-INS-217
chr16-64702527-INS-7112
chr16-64715881-INS-3356
chr16-64717818-INS-430
chr16-64721358-INS-382
chr16-64755184-INS-252
chr16-64832923-INS-157
chr16-64859732-INS-243
chr16-64902235-INS-1222
chr16-64936613-INS-52
chr16-64981365-INS-1396
chr16-64988686-INS-50
chr16-65165793-INS-68
chr16-65178708-INS-6387
chr16-65190936-INS-7101
chr16-65192698-INS-6798
chr16-65207941-INS-514
chr16-65208327-INS-993
chr16-65224857-INS-6010
chr16-65307706-INS-2003
chr16-65313350-INS-7233
chr16-65378172-INS-2294
chr16-65422953-INS-58
chr16-65445344-INS-7222
chr16-65480127-INS-1439
chr16-65493148-INS-140
chr16-65560619-INS-1699
chr16-65577687-INS-3561
chr16-65589867-INS-1274
chr16-65620494-INS-175
chr16-65663125-INS-659
chr16-65707521-INS-177
chr16-65708926-INS-283
chr16-65741660-INS-1045
chr16-65746525-INS-55
chr16-65815707-INS-1835
chr16-65874071-INS-60
chr16-65881321-INS-2278
chr16-65896578-INS-290
chr16-65896669-INS-82
chr16-65941058-INS-7895
chr16-65968669-INS-7193
chr16-66009186-INS-53
chr16-66013290-INS-65
chr16-66059202-INS-64
chr16-66076874-INS-64
chr16-66076901-INS-456
chr16-66085273-INS-6383
chr16-66148735-INS-84
chr16-66148925-INS-195
chr16-66181239-INS-770
chr16-66185128-INS-1509
chr16-66192899-INS-776
chr16-66204295-INS-6635
chr16-66214905-INS-204
chr16-66215637-INS-83
chr16-66225654-INS-3674
chr16-66229155-INS-64
chr16-66251000-INS-6631
chr16-66386842-INS-2203
chr16-66388667-INS-71
chr16-66419524-INS-1599
chr16-66422030-INS-6193
chr16-66429124-INS-51
chr16-66494866-INS-111

chr16-66495248-INS-587
chr16-66570067-INS-2595
chr16-66627127-INS-1709
chr16-66640468-INS-104
chr16-66708653-INS-55
chr16-66756511-INS-59
chr16-66777683-INS-7442
chr16-66789934-INS-364
chr16-66789999-INS-68
chr16-66791715-INS-3034
chr16-66861883-INS-480
chr16-66874916-INS-6413
chr16-66913025-INS-212
chr16-66933298-INS-52
chr16-66941066-INS-60
chr16-66964233-INS-3095
chr16-67001786-INS-62
chr16-67002222-INS-50
chr16-67004496-INS-124
chr16-67005586-INS-6832
chr16-67006817-INS-61
chr16-67031720-INS-567
chr16-67046806-INS-6488
chr16-67084845-INS-6649
chr16-67111039-INS-6654
chr16-67114243-INS-543
chr16-67240938-INS-1174
chr16-67242035-INS-96
chr16-67242307-INS-121
chr16-67302569-INS-487
chr16-67322951-INS-53
chr16-67402518-INS-52
chr16-67421657-INS-824
chr16-67457179-INS-492
chr16-67465675-INS-761
chr16-67479958-INS-7179
chr16-67499513-INS-126
chr16-67506800-INS-1039
chr16-67518635-INS-318
chr16-67518979-INS-796
chr16-67519834-INS-220
chr16-67521107-INS-52
chr16-67543090-INS-168
chr16-67550613-INS-2995
chr16-67574097-INS-52
chr16-67576201-INS-214
chr16-67576201-INS-75
chr16-67600540-INS-6239
chr16-67657660-INS-212
chr16-67678155-INS-99
chr16-67680047-INS-249
chr16-67685407-INS-6361
chr16-67694553-INS-52
chr16-67758036-INS-1560
chr16-67761439-INS-88
chr16-67780079-INS-304
chr16-67786257-INS-3500
chr16-67795190-INS-57
chr16-67795871-INS-60
chr16-67838675-INS-1281
chr16-67849666-INS-415
chr16-67872920-INS-458
chr16-67886750-INS-106
chr16-67888470-INS-57
chr16-67890565-INS-598
chr16-67954963-INS-50
chr16-67967043-INS-128
chr16-67967637-INS-73
chr16-68013422-INS-960
chr16-68029431-INS-465
chr16-68029440-INS-140
chr16-68043369-INS-52
chr16-68046896-INS-6173
chr16-68056613-INS-4638
chr16-68080291-INS-124
chr16-68086000-INS-1965
chr16-68095557-INS-6281

chr16-68104358-INS-99
chr16-68114473-INS-501
chr16-68133904-INS-4891
chr16-68152901-INS-421
chr16-68159414-INS-102
chr16-68187529-INS-1686
chr16-68234396-INS-3163
chr16-68243635-INS-2203
chr16-68244776-INS-865
chr16-68252949-INS-238
chr16-68266349-INS-75
chr16-68303422-INS-5006
chr16-68399406-INS-6250
chr16-68412098-INS-221
chr16-68444425-INS-108
chr16-68512270-INS-55
chr16-68514772-INS-202
chr16-68526408-INS-56
chr16-68526840-INS-80
chr16-68558117-INS-108
chr16-68599061-INS-642
chr16-68625456-INS-1015
chr16-68641259-INS-6533
chr16-68673843-INS-532
chr16-68690782-INS-7224
chr16-68702964-INS-3077
chr16-68730886-INS-115
chr16-68749711-INS-4025
chr16-68790879-INS-4368
chr16-68821958-INS-8110
chr16-68845987-INS-1235
chr16-68915208-INS-6238
chr16-68919047-INS-1452
chr16-68927812-INS-445
chr16-68928320-INS-2806
chr16-68931746-INS-8981
chr16-68939658-INS-565
chr16-68959368-INS-768
chr16-68989716-INS-697
chr16-68990112-INS-3702
chr16-68997577-INS-104
chr16-68999326-INS-181
chr16-68999598-INS-68
chr16-69000008-INS-550
chr16-69032283-INS-67
chr16-69042690-INS-3049
chr16-69056141-INS-473
chr16-69065530-INS-1021
chr16-69075644-INS-1537
chr16-69115404-INS-170
chr16-69184498-INS-50
chr16-69224465-INS-58
chr16-69241555-INS-619
chr16-69277316-INS-1248
chr16-69341011-INS-286
chr16-69341369-INS-387
chr16-69351112-INS-6130
chr16-69429158-INS-6453
chr16-69493642-INS-1092
chr16-69507120-INS-349
chr16-69507665-INS-77
chr16-69542715-INS-3168
chr16-69556201-INS-176
chr16-69620100-INS-2884
chr16-69708209-INS-3008
chr16-69737251-INS-50
chr16-69785133-INS-63
chr16-69785344-INS-52
chr16-69820588-INS-300
chr16-69850214-INS-55
chr16-69860364-INS-1653
chr16-69871197-INS-6675
chr16-69907924-INS-264
chr16-69939792-INS-6760
chr16-69979625-INS-426
chr16-69986657-INS-510
chr16-69994601-INS-880

chr16-70022462-INS-6524
chr16-70098815-INS-69
chr16-70099599-INS-50
chr16-70128837-INS-6249
chr16-70170196-INS-824
chr16-70214268-INS-1579
chr16-70296816-INS-53
chr16-70297533-INS-120
chr16-70348804-INS-795
chr16-70365109-INS-198
chr16-70457345-INS-104
chr16-70457759-INS-376
chr16-70492609-INS-197
chr16-70496443-INS-478
chr16-70498676-INS-522
chr16-70505386-INS-1005
chr16-70505829-INS-60
chr16-70600557-INS-215
chr16-70630980-INS-129
chr16-70633008-INS-87
chr16-70645494-INS-718
chr16-70712326-INS-1391
chr16-70714471-INS-56
chr16-70786122-INS-104
chr16-70815337-INS-206
chr16-70815340-INS-600
chr16-70864656-INS-2293
chr16-70879060-INS-195
chr16-70903493-INS-328
chr16-70904145-INS-1047
chr16-70958636-INS-66
chr16-70965177-INS-1237
chr16-70983118-INS-1451
chr16-70994700-INS-53
chr16-70996776-INS-130
chr16-70996872-INS-292
chr16-71003778-INS-2387
chr16-71005422-INS-1112
chr16-71021675-INS-4050
chr16-71040668-INS-1913
chr16-71041764-INS-6182
chr16-71041765-INS-2944
chr16-71042240-INS-257
chr16-71042241-INS-568
chr16-71109930-INS-94
chr16-71134547-INS-164
chr16-71174679-INS-54
chr16-71174701-INS-128
chr16-71181358-INS-986
chr16-71239635-INS-61
chr16-71327130-INS-7009
chr16-71335122-INS-1201
chr16-71354629-INS-2866
chr16-71375508-INS-1823
chr16-71389435-INS-6218
chr16-71393473-INS-1492
chr16-71407731-INS-4160
chr16-71412095-INS-115
chr16-71412133-INS-664
chr16-71437160-INS-1240
chr16-71448164-INS-6335
chr16-71463789-INS-1871
chr16-71463789-INS-7790
chr16-71464477-INS-895
chr16-71466850-INS-69
chr16-71467927-INS-2482
chr16-71468771-INS-1097
chr16-71518949-INS-14938
chr16-71518949-INS-2650
chr16-71554424-INS-54
chr16-71590068-INS-498
chr16-71614281-INS-6310
chr16-71621381-INS-418
chr16-71635937-INS-1555
chr16-71637147-INS-76
chr16-71639361-INS-1167
chr16-71670165-INS-6317

chr16-71708556-INS-220
chr16-71708556-INS-65
chr16-71743859-INS-991
chr16-71749279-INS-51
chr16-71800554-INS-743
chr16-71802021-INS-50
chr16-71813902-INS-7350
chr16-71823302-INS-56
chr16-71837823-INS-171
chr16-71839438-INS-397
chr16-71881217-INS-2438
chr16-71921795-INS-50
chr16-71954023-INS-55
chr16-71963005-INS-154
chr16-71994242-INS-50
chr16-72054425-INS-718
chr16-72067650-INS-126
chr16-72088488-INS-1375
chr16-72093485-INS-55
chr16-72100348-INS-6446
chr16-72146628-INS-51
chr16-72157944-INS-957
chr16-72158051-INS-175
chr16-72167919-INS-279
chr16-72168409-INS-290
chr16-72189101-INS-57
chr16-72189450-INS-172
chr16-72206653-INS-70
chr16-72226584-INS-79
chr16-72238726-INS-4776
chr16-72279333-INS-1190
chr16-72342706-INS-69
chr16-72343579-INS-6225
chr16-72369003-INS-3955
chr16-72387966-INS-759
chr16-72392211-INS-51
chr16-72410347-INS-55
chr16-72495640-INS-104
chr16-72498302-INS-6357
chr16-72602374-INS-195
chr16-72616343-INS-3037
chr16-72657556-INS-54
chr16-72678819-INS-118
chr16-72693505-INS-2451
chr16-72695867-INS-54
chr16-72777791-INS-54
chr16-72794741-INS-124
chr16-72836880-INS-52
chr16-72873794-INS-92
chr16-72880567-INS-7267
chr16-72880704-INS-59
chr16-72903019-INS-3248
chr16-72911011-INS-1251
chr16-72913438-INS-6401
chr16-72925142-INS-94
chr16-72926038-INS-385
chr16-72931815-INS-288
chr16-72941665-INS-56
chr16-72967935-INS-3076
chr16-73005110-INS-74
chr16-73013715-INS-106
chr16-73034882-INS-7151
chr16-73088495-INS-144
chr16-73094594-INS-429
chr16-73094941-INS-543
chr16-73095126-INS-126
chr16-73104611-INS-115
chr16-73111074-INS-86
chr16-73140013-INS-58
chr16-73148954-INS-1946
chr16-73176589-INS-51
chr16-73206782-INS-572
chr16-73319809-INS-54
chr16-73320290-INS-158
chr16-73383070-INS-3673
chr16-73398193-INS-9015
chr16-73404541-INS-1200

chr16-73422765-INS-70
chr16-73423035-INS-204
chr16-73510590-INS-220
chr16-73547977-INS-6383
chr16-73558216-INS-3154
chr16-73593047-INS-60
chr16-73632671-INS-64
chr16-73904160-INS-139
chr16-73904361-INS-96
chr16-73909160-INS-95
chr16-73924954-INS-224
chr16-74025811-INS-210
chr16-74030346-INS-54
chr16-74031229-INS-62
chr16-74059146-INS-70
chr16-74067857-INS-87
chr16-74078714-INS-71
chr16-74145070-INS-50
chr16-74153662-INS-60
chr16-74160419-INS-201
chr16-74199462-INS-58
chr16-74210561-INS-80
chr16-74211911-INS-79
chr16-74214076-INS-170
chr16-74331975-INS-158
chr16-74336775-INS-160
chr16-74403295-INS-490
chr16-74403576-INS-174
chr16-74406221-INS-362
chr16-74417903-INS-77
chr16-74426837-INS-218
chr16-74439730-INS-200
chr16-74446891-INS-1307
chr16-74462686-INS-682
chr16-74468968-INS-132
chr16-74522585-INS-80
chr16-74522599-INS-332
chr16-74525627-INS-1035
chr16-74621707-INS-6873
chr16-74625382-INS-56
chr16-74657374-INS-549
chr16-74658559-INS-315
chr16-74790134-INS-50
chr16-74790940-INS-64
chr16-74923102-INS-6450
chr16-74983685-INS-190
chr16-75020950-INS-7195
chr16-75023473-INS-54
chr16-75047752-INS-498
chr16-75060450-INS-211
chr16-75074306-INS-55
chr16-75075720-INS-390
chr16-75075771-INS-194
chr16-75078839-INS-344
chr16-75085807-INS-258
chr16-75097380-INS-214
chr16-75105134-INS-51
chr16-75108552-INS-226
chr16-75109816-INS-68
chr16-75111197-INS-54
chr16-75256474-INS-209
chr16-75273983-INS-12100
chr16-75274626-INS-5522
chr16-75282253-INS-64
chr16-75287138-INS-4549
chr16-75332176-INS-1577
chr16-75333309-INS-58
chr16-75346178-INS-55
chr16-75384087-INS-749
chr16-75439079-INS-272
chr16-75443672-INS-127
chr16-75470913-INS-356
chr16-75483941-INS-1728
chr16-75489388-INS-431
chr16-75523707-INS-52
chr16-75569961-INS-50
chr16-75586628-INS-89

chr16-75623249-INS-53
chr16-75647953-INS-6442
chr16-75675247-INS-67
chr16-75679131-INS-766
chr16-75679997-INS-6596
chr16-75691203-INS-6146
chr16-75719909-INS-127
chr16-75740341-INS-74
chr16-75764707-INS-56
chr16-75800110-INS-124
chr16-75832295-INS-60
chr16-75841249-INS-51
chr16-75894796-INS-116
chr16-75956270-INS-56
chr16-76003547-INS-1894
chr16-76003710-INS-50
chr16-76032632-INS-1261
chr16-76033744-INS-51
chr16-76145793-INS-52
chr16-76246514-INS-318
chr16-76253458-INS-89
chr16-76253563-INS-58
chr16-76309297-INS-57
chr16-76348206-INS-1898
chr16-76357722-INS-827
chr16-76357835-INS-1974
chr16-76371986-INS-54
chr16-76395668-INS-54
chr16-76442171-INS-261
chr16-76469022-INS-7714
chr16-76488039-INS-888
chr16-76563203-INS-84
chr16-76586068-INS-424
chr16-76590403-INS-5918
chr16-76596823-INS-236
chr16-76623852-INS-56
chr16-76637294-INS-90
chr16-76656638-INS-80
chr16-76659472-INS-1755
chr16-76697129-INS-1647
chr16-76719359-INS-4640
chr16-76719711-INS-600
chr16-76765712-INS-60
chr16-76776259-INS-444
chr16-76776485-INS-151
chr16-76793519-INS-325
chr16-76802025-INS-67
chr16-76804318-INS-76
chr16-76819790-INS-59
chr16-76857263-INS-416
chr16-76877797-INS-58
chr16-76882759-INS-130
chr16-76909028-INS-51
chr16-76909830-INS-96
chr16-76981013-INS-2199
chr16-77007612-INS-5040
chr16-77008969-INS-56
chr16-77009710-INS-993
chr16-77106750-INS-55
chr16-77135034-INS-382
chr16-77135095-INS-95
chr16-77135606-INS-64
chr16-77162193-INS-5783
chr16-77183924-INS-88
chr16-77185924-INS-546
chr16-77191714-INS-54
chr16-77196600-INS-53
chr16-77226335-INS-57
chr16-77322588-INS-82
chr16-77322914-INS-271
chr16-77390300-INS-2522
chr16-77395388-INS-133
chr16-77398569-INS-94
chr16-77448963-INS-68
chr16-77485116-INS-2871
chr16-77491852-INS-63
chr16-77491917-INS-344

chr16-77527831-INS-209
chr16-77659348-INS-223
chr16-77732097-INS-267
chr16-77744892-INS-812
chr16-77788442-INS-121
chr16-77788671-INS-149
chr16-77822371-INS-1980
chr16-77847633-INS-52
chr16-77857002-INS-116
chr16-77890428-INS-3213
chr16-77900413-INS-204
chr16-77902394-INS-3913
chr16-77904796-INS-143
chr16-77936100-INS-581
chr16-77978784-INS-53
chr16-77991646-INS-93
chr16-78058984-INS-414
chr16-78059336-INS-1345
chr16-78067843-INS-66
chr16-78075520-INS-124
chr16-78081576-INS-60
chr16-78203259-INS-68
chr16-78209213-INS-575
chr16-78214158-INS-62
chr16-78250384-INS-50
chr16-78253949-INS-604
chr16-78262878-INS-178
chr16-78265447-INS-53
chr16-78326771-INS-124
chr16-78330085-INS-110
chr16-78332437-INS-3474
chr16-78332636-INS-399
chr16-78370525-INS-302
chr16-78381356-INS-96
chr16-78387081-INS-129
chr16-78392421-INS-3899
chr16-78399997-INS-110
chr16-78415773-INS-81
chr16-78426305-INS-1579
chr16-78440048-INS-199
chr16-78440106-INS-537
chr16-78457892-INS-1290
chr16-78492932-INS-53
chr16-78495557-INS-85
chr16-78498269-INS-107
chr16-78515512-INS-1468
chr16-78541563-INS-78
chr16-78556564-INS-3507
chr16-78621825-INS-160
chr16-78622303-INS-646
chr16-78623122-INS-52
chr16-78634202-INS-6361
chr16-78643266-INS-257
chr16-78669212-INS-228
chr16-78670500-INS-2819
chr16-78695968-INS-2789
chr16-78706150-INS-393
chr16-78715097-INS-6153
chr16-78862120-INS-207
chr16-78862120-INS-555
chr16-78891009-INS-56
chr16-78930408-INS-286
chr16-78950344-INS-56
chr16-78986345-INS-68
chr16-78991486-INS-52
chr16-79052285-INS-5510
chr16-79100329-INS-6307
chr16-79100710-INS-50
chr16-79129888-INS-58
chr16-79182776-INS-3891
chr16-79190510-INS-3030
chr16-79207407-INS-194
chr16-79214101-INS-54
chr16-79232113-INS-117
chr16-79232302-INS-95
chr16-79245249-INS-57
chr16-79248834-INS-6404

chr16-79253973-INS-5283
chr16-79266515-INS-57
chr16-79273204-INS-2450
chr16-79273997-INS-175
chr16-79280629-INS-6389
chr16-79307457-INS-1335
chr16-79317927-INS-4037
chr16-79349894-INS-56
chr16-79369811-INS-134
chr16-79395252-INS-3071
chr16-79398746-INS-140
chr16-79419878-INS-66
chr16-79500939-INS-80
chr16-79507179-INS-67
chr16-79512984-INS-126
chr16-79551544-INS-168
chr16-79566625-INS-96
chr16-79652723-INS-56
chr16-79659127-INS-5272
chr16-79756933-INS-115
chr16-79771876-INS-2865
chr16-79821982-INS-2358
chr16-79853157-INS-142
chr16-79910455-INS-429
chr16-79971190-INS-8858
chr16-79997594-INS-231
chr16-79997654-INS-586
chr16-79997701-INS-76
chr16-80021873-INS-1117
chr16-80049350-INS-1236
chr16-80052731-INS-68
chr16-80060671-INS-58
chr16-80060839-INS-54
chr16-80075678-INS-79
chr16-80125476-INS-50
chr16-80130124-INS-9068
chr16-80130145-INS-393
chr16-80294308-INS-904
chr16-80319212-INS-105
chr16-80326644-INS-55
chr16-80335809-INS-50
chr16-80381245-INS-84
chr16-80385992-INS-1619
chr16-80399319-INS-6406
chr16-80414372-INS-320
chr16-80451974-INS-76
chr16-80465255-INS-76
chr16-80468417-INS-1680
chr16-80471073-INS-50
chr16-80504229-INS-52
chr16-80551100-INS-63
chr16-80564510-INS-72
chr16-80565286-INS-52
chr16-80565413-INS-105
chr16-80579687-INS-1547
chr16-80665255-INS-52
chr16-80744407-INS-77
chr16-80745245-INS-367
chr16-80745247-INS-73
chr16-80752704-INS-676
chr16-80769707-INS-1396
chr16-80800459-INS-101
chr16-80811268-INS-498
chr16-80842470-INS-60
chr16-80858259-INS-562
chr16-80858446-INS-1283
chr16-80864270-INS-1993
chr16-80896455-INS-784
chr16-80899580-INS-1722
chr16-80907243-INS-132
chr16-80918698-INS-2901
chr16-80927898-INS-1064
chr16-80940259-INS-132
chr16-80941167-INS-85
chr16-80962679-INS-7627
chr16-80963599-INS-94
chr16-80978574-INS-70

chr16-81114673-INS-196
chr16-81129246-INS-372
chr16-81137397-INS-2017
chr16-81146890-INS-2441
chr16-81164441-INS-6155
chr16-81225471-INS-283
chr16-81260822-INS-59
chr16-81269055-INS-204
chr16-81287585-INS-206
chr16-81287587-INS-53
chr16-81310732-INS-126
chr16-81368659-INS-1890
chr16-81382856-INS-89
chr16-81383326-INS-50
chr16-81387672-INS-1489
chr16-81415511-INS-180
chr16-81503820-INS-108
chr16-81561873-INS-12054
chr16-81598353-INS-5856
chr16-81613267-INS-1204
chr16-81616584-INS-3445
chr16-81632073-INS-189
chr16-81656341-INS-212
chr16-81695509-INS-308
chr16-81696296-INS-181
chr16-81707016-INS-50
chr16-81715205-INS-52
chr16-81751773-INS-2339
chr16-81753560-INS-4360
chr16-81776849-INS-7085
chr16-81792998-INS-147
chr16-81797278-INS-461
chr16-81917842-INS-2304
chr16-81922193-INS-2443
chr16-81967243-INS-107
chr16-82044345-INS-3077
chr16-82082871-INS-4490
chr16-82112140-INS-547
chr16-82116627-INS-2931
chr16-82137268-INS-2733
chr16-82237057-INS-6277
chr16-82243009-INS-52
chr16-82291193-INS-6176
chr16-82299378-INS-1373
chr16-82303414-INS-62
chr16-82316925-INS-50
chr16-82323833-INS-68
chr16-82333619-INS-64
chr16-82354960-INS-97
chr16-82357678-INS-324
chr16-82482401-INS-69
chr16-82482544-INS-947
chr16-82561912-INS-123
chr16-82562059-INS-6220
chr16-82592678-INS-164
chr16-82651649-INS-6377
chr16-82670409-INS-102
chr16-82670444-INS-110
chr16-82732313-INS-164
chr16-82736907-INS-54
chr16-82737894-INS-72
chr16-82764722-INS-983
chr16-82847940-INS-112
chr16-82858978-INS-50
chr16-82862567-INS-58
chr16-82863075-INS-470
chr16-82933460-INS-2694
chr16-82965471-INS-51
chr16-82965822-INS-411
chr16-82974808-INS-6532
chr16-82991152-INS-56
chr16-82992789-INS-249
chr16-83092188-INS-50
chr16-83104724-INS-110
chr16-83129177-INS-1477
chr16-83132070-INS-61
chr16-83132145-INS-112

chr16-83142342-INS-51
chr16-83154448-INS-2501
chr16-83212580-INS-55
chr16-83227178-INS-203
chr16-83227275-INS-71
chr16-83227859-INS-1293
chr16-83242541-INS-1253
chr16-83302471-INS-61
chr16-83315136-INS-51
chr16-83322985-INS-108
chr16-83387446-INS-1092
chr16-83409750-INS-127
chr16-83412885-INS-5630
chr16-83431125-INS-56
chr16-83452849-INS-2133
chr16-83462586-INS-7261
chr16-83467498-INS-1244
chr16-83470735-INS-1440
chr16-83504619-INS-4763
chr16-83512267-INS-1632
chr16-83522302-INS-2842
chr16-83532702-INS-6297
chr16-83556285-INS-1229
chr16-83579558-INS-1531
chr16-83609303-INS-302
chr16-83668885-INS-1050
chr16-83701776-INS-4511
chr16-83704912-INS-1059
chr16-83705122-INS-52
chr16-83708701-INS-1203
chr16-83715360-INS-50
chr16-83716571-INS-458
chr16-83768780-INS-470
chr16-83771068-INS-190
chr16-83787617-INS-332
chr16-83790901-INS-909
chr16-83802411-INS-6419
chr16-83806628-INS-78
chr16-83807205-INS-51
chr16-83809168-INS-873
chr16-83817248-INS-127
chr16-83845244-INS-109
chr16-83846380-INS-6202
chr16-83903819-INS-254
chr16-83934487-INS-54
chr16-83948378-INS-697
chr16-84064622-INS-94
chr16-84085178-INS-2320
chr16-84087809-INS-117
chr16-84087827-INS-1077
chr16-84113746-INS-6657
chr16-84138382-INS-1380
chr16-84161915-INS-52
chr16-84165196-INS-324
chr16-84199405-INS-181
chr16-84202639-INS-3418
chr16-84203936-INS-266
chr16-84204106-INS-449
chr16-84208205-INS-5800
chr16-84211190-INS-152
chr16-84216583-INS-90
chr16-84235743-INS-52
chr16-84259897-INS-1183
chr16-84284424-INS-6968
chr16-84384389-INS-644
chr16-84384704-INS-366
chr16-84412673-INS-186
chr16-84417843-INS-318
chr16-84423131-INS-5046
chr16-84470751-INS-1893
chr16-84507754-INS-111
chr16-84524713-INS-174
chr16-84540142-INS-50
chr16-84541229-INS-141
chr16-84560017-INS-60
chr16-84588033-INS-51
chr16-84590743-INS-188

chr16-84618398-INS-1210
chr16-84619319-INS-255
chr16-84619319-INS-51
chr16-84673969-INS-5759
chr16-84677594-INS-1253
chr16-84697592-INS-6458
chr16-84711685-INS-305
chr16-84722375-INS-200
chr16-84722579-INS-4176
chr16-84727757-INS-201
chr16-84738921-INS-212
chr16-84743655-INS-242
chr16-84825336-INS-193
chr16-84842766-INS-1992
chr16-84870734-INS-157
chr16-84880080-INS-7331
chr16-84894422-INS-98
chr16-84895382-INS-6716
chr16-84902147-INS-55
chr16-84905559-INS-80
chr16-84932785-INS-1797
chr16-84947356-INS-210
chr16-84959595-INS-221
chr16-84961642-INS-105
chr16-84981808-INS-298
chr16-85001223-INS-1006
chr16-85009053-INS-62
chr16-85012452-INS-70
chr16-85014184-INS-55
chr16-85030603-INS-96
chr16-85052036-INS-90
chr16-85068990-INS-6353
chr16-85072998-INS-50
chr16-85083232-INS-6969
chr16-85106069-INS-117
chr16-85128306-INS-927
chr16-85223685-INS-486
chr16-85232302-INS-51
chr16-85232955-INS-55
chr16-85244526-INS-52
chr16-85265288-INS-6419
chr16-85275242-INS-234
chr16-85294624-INS-1879
chr16-85399628-INS-1895
chr16-85421954-INS-56
chr16-85444556-INS-70
chr16-85460766-INS-52
chr16-85460991-INS-6673
chr16-85476058-INS-51
chr16-85476959-INS-88
chr16-85481438-INS-1701
chr16-85483474-INS-6312
chr16-85491601-INS-369
chr16-85497447-INS-84
chr16-85505454-INS-660
chr16-85527850-INS-140
chr16-85531267-INS-50
chr16-85562576-INS-53
chr16-85568602-INS-1097
chr16-85585202-INS-58
chr16-85601650-INS-324
chr16-85642026-INS-255
chr16-85650010-INS-3140
chr16-85678079-INS-51
chr16-85678758-INS-106
chr16-85703561-INS-72
chr16-85716000-INS-180
chr16-85739406-INS-380
chr16-85740097-INS-52
chr16-85740328-INS-560
chr16-85751612-INS-1220
chr16-85800601-INS-50
chr16-85822417-INS-2285
chr16-85824446-INS-4613
chr16-85849830-INS-147
chr16-85868592-INS-357
chr16-85895211-INS-112

chr16-85910110-INS-52
chr16-85916945-INS-1455
chr16-85961811-INS-1142
chr16-85999323-INS-7228
chr16-86027910-INS-6227
chr16-86120914-INS-54
chr16-86156619-INS-50
chr16-86164937-INS-473
chr16-86173419-INS-2026
chr16-86212291-INS-6242
chr16-86219424-INS-122
chr16-86229281-INS-12855
chr16-86235723-INS-1653
chr16-86294900-INS-108
chr16-86508111-INS-54
chr16-86643325-INS-54
chr16-86672468-INS-6187
chr16-86682443-INS-4654
chr16-86707217-INS-301
chr16-86707227-INS-7546
chr16-86731673-INS-143
chr16-86735234-INS-231
chr16-86783987-INS-87
chr16-86784119-INS-56
chr16-86799689-INS-308
chr16-86844543-INS-63
chr16-86857561-INS-51
chr16-86925070-INS-1786
chr16-86977068-INS-1350
chr16-86984642-INS-170
chr16-86984896-INS-113
chr16-86986882-INS-582
chr16-87010395-INS-129
chr16-87053242-INS-80
chr16-87079758-INS-58
chr16-87092259-INS-3025
chr16-87100113-INS-146
chr16-87111280-INS-229
chr16-87113232-INS-284
chr16-87114213-INS-226
chr16-87168171-INS-94
chr16-87169274-INS-217
chr16-87173752-INS-205
chr16-87183269-INS-870
chr16-87183719-INS-504
chr16-87189520-INS-6495
chr16-87237047-INS-119
chr16-87237799-INS-54
chr16-87238938-INS-50
chr16-87239691-INS-126
chr16-87240698-INS-7081
chr16-87241417-INS-52
chr16-87246629-INS-131
chr16-87251311-INS-537
chr16-87252793-INS-66
chr16-87266544-INS-220
chr16-87268472-INS-214
chr16-87276915-INS-64
chr16-87281018-INS-178
chr16-87296512-INS-114
chr16-87302418-INS-7202
chr16-87319671-INS-246
chr16-87323114-INS-725
chr16-87323220-INS-97
chr16-87340779-INS-186
chr16-87340857-INS-450
chr16-87342537-INS-9728
chr16-87353900-INS-62
chr16-87354199-INS-209
chr16-87355027-INS-773
chr16-87373032-INS-120
chr16-87389402-INS-78
chr16-87389900-INS-65
chr16-87403114-INS-159
chr16-87405857-INS-118
chr16-87413366-INS-457
chr16-87416666-INS-2305

chr16-87431523-INS-56
chr16-87431751-INS-50
chr16-87445808-INS-71
chr16-87446415-INS-187
chr16-87481934-INS-112
chr16-87483048-INS-276
chr16-87553017-INS-6505
chr16-87599726-INS-95
chr16-87633703-INS-5550
chr16-87650784-INS-2859
chr16-87651680-INS-83
chr16-87681608-INS-140
chr16-87697929-INS-197
chr16-87698788-INS-358
chr16-87711767-INS-246
chr16-87711863-INS-138
chr16-87730204-INS-58
chr16-87765724-INS-6784
chr16-87765930-INS-459
chr16-87765995-INS-115
chr16-87791960-INS-189
chr16-87802416-INS-560
chr16-87849834-INS-50
chr16-87862596-INS-1029
chr16-87880812-INS-64
chr16-87886601-INS-614
chr16-87887929-INS-54
chr16-87895842-INS-211
chr16-87896756-INS-1111
chr16-87967317-INS-3195
chr16-87992744-INS-1122
chr16-87996659-INS-110
chr16-87996950-INS-460
chr16-88000018-INS-148
chr16-88078860-INS-65
chr16-88135491-INS-99
chr16-88146776-INS-6546
chr16-88159629-INS-445
chr16-88210582-INS-62
chr16-88235864-INS-1259
chr16-88266990-INS-258
chr16-88268763-INS-128
chr16-88281531-INS-1444
chr16-88286486-INS-58
chr16-88286860-INS-1139
chr16-88293655-INS-402
chr16-88295616-INS-2247
chr16-88316465-INS-52
chr16-88335824-INS-3139
chr16-88340096-INS-56
chr16-88341718-INS-965
chr16-88372188-INS-174
chr16-88402933-INS-7967
chr16-88445090-INS-52
chr16-88462399-INS-933
chr16-88505578-INS-54
chr16-88535070-INS-75
chr16-88557655-INS-51
chr16-88559689-INS-308
chr16-88566290-INS-1514
chr16-88570251-INS-90
chr16-88570751-INS-75
chr16-88626743-INS-506
chr16-88670924-INS-4478
chr16-88671970-INS-92
chr16-88697538-INS-102
chr16-88707563-INS-2908
chr16-88729814-INS-50
chr16-88746335-INS-1461
chr16-88750572-INS-569
chr16-88772561-INS-60
chr16-88843587-INS-531
chr16-88857534-INS-8269
chr16-88874138-INS-50
chr16-88986241-INS-67
chr16-89002207-INS-113
chr16-89020436-INS-6411

chr16-89065720-INS-2033
chr16-89085828-INS-3380
chr16-89122858-INS-100
chr16-89138870-INS-61
chr16-89159031-INS-1078
chr16-89207126-INS-62
chr16-89210885-INS-122
chr16-89211023-INS-321
chr16-89313561-INS-102
chr16-89316486-INS-96
chr16-89323174-INS-316
chr16-89340831-INS-126
chr16-89350274-INS-176
chr16-89369862-INS-50
chr16-89386303-INS-74
chr16-89388554-INS-55
chr16-89442284-INS-213
chr16-89456322-INS-455
chr16-89456353-INS-1565
chr16-89458172-INS-6195
chr16-89458889-INS-7102
chr16-89490867-INS-58
chr16-89496023-INS-112
chr16-89504954-INS-137
chr16-89505872-INS-110
chr16-89528596-INS-77
chr16-89535161-INS-60
chr16-89538951-INS-6617
chr16-89548688-INS-750
chr16-89555027-INS-168
chr16-89555212-INS-69
chr16-89555233-INS-1283
chr16-89571868-INS-2984
chr16-89571980-INS-6217
chr16-89701962-INS-2945
chr16-89702625-INS-843
chr16-89746182-INS-236
chr16-89849992-INS-120
chr16-89868138-INS-1053
chr16-89868246-INS-3176
chr16-89903761-INS-608
chr16-89907936-INS-306
chr16-89930283-INS-94
chr16-89951329-INS-56
chr16-89953483-INS-54
chr16-89958489-INS-78
chr16-89958587-INS-289
chr16-89965468-INS-202
chr16-89995715-INS-13516
chr16-90003918-INS-169
chr16-90004922-INS-1701
chr16-90009985-INS-162
chr16-90017052-INS-330
chr16-90094532-INS-51
chr16-90098881-INS-1398
chr16-90103641-INS-60
chr16-90125443-INS-50
chr16-90137520-INS-1113
chr16-90152594-INS-336
chr16-90152727-INS-278
chr16-90165364-INS-1535
chr16-90165832-INS-6401
chr16-90172511-INS-81
chr16-90172713-INS-185
chr16-90212422-INS-76
chr16-90224987-INS-130
chr16-90227095-INS-52
chr16-90232088-INS-5537
chr16-90257839-INS-127
chr16-90262692-INS-380
chr16-90286460-INS-50
chr16-90310645-INS-401
chr16-90311950-INS-180
chr16-90313476-INS-122
chr16-90333365-INS-106
chr16-90334211-INS-4320
chr16-90355052-INS-50

chr16-90387820-INS-92
chr16-90440454-INS-6904
chr16-90453259-INS-50
chr16-90454624-INS-52
chr16-90455170-INS-64
chr16-90458658-INS-50
chr16-90481654-INS-7150
chr16-90486702-INS-748
chr16-90486744-INS-201
chr16-90486866-INS-60
chr16-90488242-INS-132
chr16-90488860-INS-308
chr16-90528437-INS-108
chr16-90529124-INS-2092
chr16-90531188-INS-155
chr16-90560194-INS-592
chr16-90613838-INS-68
chr16-90616026-INS-96
chr16-90638491-INS-50
chr16-90655924-INS-1452
chr16-90662301-INS-64
chr16-90662331-INS-165
chr16-90676528-INS-54
chr16-90686011-INS-5536
chr16-90689606-INS-265
chr16-90707302-INS-56
chr16-90737563-INS-196
chr16-90738190-INS-73
chr16-90771960-INS-78
chr16-90772537-INS-210
chr16-90782662-INS-778
chr16-90782693-INS-232
chr16-90826407-INS-200
chr16-90864086-INS-80
chr16-90872846-INS-58
chr16-90885329-INS-324
chr16-90888925-INS-82
chr16-90889271-INS-489
chr16-90934170-INS-7541
chr16-91000775-INS-52
chr16-91005067-INS-1725
chr16-91010670-INS-143
chr16-91078120-INS-54
chr16-91120461-INS-144
chr16-91120461-INS-3015
chr16-91123007-INS-79
chr16-91139606-INS-512
chr16-91139784-INS-192
chr16-91140232-INS-608
chr16-91140328-INS-96
chr16-91142784-INS-142
chr16-91155168-INS-1417
chr16-91155168-INS-588
chr16-91235648-INS-79
chr16-91236024-INS-1129
chr16-91236372-INS-201
chr16-91240966-INS-196
chr16-91251785-INS-250
chr16-91252100-INS-2078
chr16-91252264-INS-52
chr16-91260247-INS-84
chr16-91270777-INS-3391
chr16-91293299-INS-480
chr16-91317132-INS-3005
chr16-91322955-INS-180
chr16-91329407-INS-1209
chr16-91377508-INS-279
chr16-91400333-INS-394
chr16-91403738-INS-50
chr16-91446718-INS-179
chr16-91465632-INS-762
chr16-91518347-INS-276
chr16-91521138-INS-244
chr16-91536378-INS-150
chr16-91555040-INS-368
chr16-91571493-INS-823
chr16-91595222-INS-337

chr16-91670637-INS-50
chr16-91685851-INS-206
chr16-91687857-INS-50
chr16-91691096-INS-76
chr16-91709614-INS-50
chr16-91717795-INS-67
chr16-91732664-INS-178
chr16-91734517-INS-112
chr16-91737329-INS-1370
chr16-91737331-INS-5220
chr16-91776249-INS-202
chr16-91787364-INS-64
chr16-91788435-INS-446
chr16-91808127-INS-191
chr16-91825853-INS-55
chr16-91872664-INS-193
chr16-91891101-INS-391
chr16-91909484-INS-77
chr16-91919173-INS-139
chr16-91933505-INS-149
chr16-91938348-INS-62
chr16-91951129-INS-76
chr16-91984860-INS-267
chr16-92017560-INS-1038
chr16-92047483-INS-440
chr16-92068398-INS-52
chr16-92134904-INS-131
chr16-92135569-INS-390
chr16-92139093-INS-84
chr16-92146841-INS-52
chr16-92166076-INS-401
chr16-92166288-INS-250
chr16-92172986-INS-172
chr16-92175894-INS-483
chr16-92181107-INS-7219
chr16-92188239-INS-51
chr16-92198596-INS-7137
chr16-92204769-INS-1625
chr16-92211714-INS-1711
chr16-92310345-INS-294
chr16-92326023-INS-144
chr16-92327944-INS-73
chr16-92371424-INS-288
chr16-92382472-INS-117
chr16-92382472-INS-576
chr16-92448251-INS-108
chr16-92457837-INS-290
chr16-92503026-INS-50
chr16-92584401-INS-690
chr16-92584470-INS-92
chr16-92595784-INS-132
chr16-92619947-INS-54
chr16-92645169-INS-89
chr16-92662623-INS-61
chr16-92669934-INS-96
chr16-92700121-INS-86
chr16-92730780-INS-104
chr16-92730903-INS-338
chr16-92731415-INS-55
chr16-92733285-INS-298
chr16-92744426-INS-875
chr16-92749891-INS-2824
chr16-92751939-INS-74
chr16-92752038-INS-217
chr16-92752148-INS-62
chr16-92761464-INS-160
chr16-92761899-INS-69
chr16-92841380-INS-165
chr16-92878988-INS-194
chr16-92881121-INS-1918
chr16-92891303-INS-111
chr16-92943558-INS-74
chr16-92950796-INS-1050
chr16-92961691-INS-198
chr16-92967541-INS-52
chr16-92969041-INS-1213
chr16-93008898-INS-50

chr16-93079855-INS-211
chr16-93086535-INS-140
chr16-93105436-INS-1060
chr16-93145492-INS-360
chr16-93161767-INS-6460
chr16-93177392-INS-88
chr16-93204204-INS-216
chr16-93205630-INS-224
chr16-93205653-INS-1152
chr16-93249016-INS-498
chr16-93253106-INS-292
chr16-93266172-INS-52
chr16-93280462-INS-53
chr16-93296715-INS-170
chr16-93302959-INS-4384
chr16-93308499-INS-50
chr16-93335093-INS-203
chr16-93364325-INS-410
chr16-93369093-INS-74
chr16-93391715-INS-2237
chr16-93393078-INS-112
chr16-93404333-INS-51
chr16-93410838-INS-73
chr16-93451503-INS-312
chr16-93451987-INS-708
chr16-93453444-INS-51
chr16-93454543-INS-4953
chr16-93455366-INS-117
chr16-93457264-INS-56
chr16-93473922-INS-3408
chr16-93473925-INS-102
chr16-93474027-INS-512
chr16-93570607-INS-498
chr16-93575229-INS-442
chr16-93575469-INS-242
chr16-93595561-INS-497
chr16-93597789-INS-1107
chr16-93599900-INS-1340
chr16-93600888-INS-112
chr16-93643960-INS-50
chr16-93692339-INS-208
chr16-93710212-INS-480
chr16-93712728-INS-239
chr16-93724379-INS-168
chr16-93732119-INS-52
chr16-93734454-INS-201
chr16-93782743-INS-223
chr16-93797705-INS-3096
chr16-93805015-INS-57
chr16-93817035-INS-2775
chr16-93838627-INS-160
chr16-93858282-INS-120
chr16-93866215-INS-1141
chr16-93883683-INS-76
chr16-93917712-INS-284
chr16-93937344-INS-10333
chr16-93946270-INS-215
chr16-93970810-INS-1078
chr16-94002086-INS-51
chr16-94022238-INS-233
chr16-94032966-INS-163
chr16-94038537-INS-113
chr16-94057452-INS-120
chr16-94074826-INS-65
chr16-94100590-INS-125
chr16-94112276-INS-543
chr16-94117364-INS-159
chr16-94123916-INS-201
chr16-94146532-INS-50
chr16-94150216-INS-52
chr16-94151987-INS-7100
chr16-94159509-INS-64
chr16-94190414-INS-305
chr16-94245689-INS-209
chr16-94264972-INS-298
chr16-94281242-INS-56
chr16-94290532-INS-776

chr16-94296155-INS-122
chr16-94302682-INS-132
chr16-94307146-INS-51
chr16-94327466-INS-151
chr16-94415798-INS-110
chr16-94423494-INS-208
chr16-94429734-INS-451
chr16-94476752-INS-96
chr16-94477907-INS-1104
chr16-94529113-INS-4751
chr16-94539826-INS-2520
chr16-94539911-INS-504
chr16-94560647-INS-731
chr16-94578745-INS-7116
chr16-94586162-INS-70
chr16-94599431-INS-286
chr16-94599484-INS-573
chr16-94641202-INS-275
chr16-94680383-INS-59
chr16-94743093-INS-74
chr16-94743149-INS-50
chr16-94746267-INS-1080
chr16-94746273-INS-282
chr16-94833558-INS-78
chr16-94849644-INS-52
chr16-94867702-INS-4552
chr16-94871950-INS-50
chr16-94876483-INS-64
chr16-94898007-INS-72
chr16-94901316-INS-4803
chr16-94923515-INS-2810
chr16-94926968-INS-206
chr16-94940570-INS-1081
chr16-94960393-INS-63
chr16-94966513-INS-192
chr16-94966977-INS-138
chr16-94990888-INS-6388
chr16-94991558-INS-50
chr16-95065947-INS-78
chr16-95121009-INS-145
chr16-95127270-INS-90
chr16-95133158-INS-6962
chr16-95145247-INS-61
chr16-95157023-INS-80
chr16-95192405-INS-52
chr16-95203639-INS-50
chr16-95219855-INS-64
chr16-95287834-INS-433
chr16-95302226-INS-598
chr16-95337464-INS-102
chr16-95456486-INS-2885
chr16-95475999-INS-55
chr16-95494309-INS-148
chr16-95593902-INS-50
chr16-95605339-INS-240
chr16-95608960-INS-333
chr16-95614442-INS-90
chr16-95638317-INS-216
chr16-95638317-INS-504
chr16-95675222-INS-148
chr16-95687770-INS-213
chr16-95732015-INS-84
chr16-95760182-INS-3811
chr16-95774938-INS-3219
chr16-95780749-INS-817
chr16-95780950-INS-2098
chr16-95781043-INS-199
chr16-95901584-INS-178
chr16-95907858-INS-177
chr16-95915371-INS-55
chr16-95933867-INS-118
chr16-95934106-INS-86
chr16-95938000-INS-50
chr16-95942778-INS-68
chr16-95951972-INS-194
chr16-95955835-INS-668
chr16-95969088-INS-609

chr16-95978199-INS-54
chr16-96037481-INS-306
chr16-96094804-INS-128
chr16-96095223-INS-240
chr16-96100944-INS-2027
chr16-96106819-INS-82
chr16-96138187-INS-1077
chr16-96139598-INS-71
chr16-96148560-INS-51
chr16-96165507-INS-108
chr16-96165507-INS-243
chr16-96206394-INS-445
chr16-96232440-INS-54
chr16-96258308-INS-107
chr16-96263816-INS-472
chr16-96279243-INS-1177
chr16-96281745-INS-55
chr16-96311760-INS-60
chr16-96318655-INS-64
chr16-96343970-INS-209
chr16-96377670-INS-3023
chr16-96411013-INS-1519
chr16-96427661-INS-52
chr16-96436725-INS-58
chr16-96565267-INS-207
chr16-96565804-INS-61
chr16-96587870-INS-225
chr16-96599903-INS-6419
chr16-96615344-INS-52
chr16-96684928-INS-2993
chr16-96762411-INS-102
chr16-96763152-INS-66
chr16-96764473-INS-122
chr16-96853424-INS-58
chr16-96854244-INS-50
chr16-96871083-INS-404
chr16-96874687-INS-68
chr16-96875251-INS-133
chr16-96879586-INS-58
chr16-96882196-INS-117
chr16-96935354-INS-67
chr16-97006863-INS-116
chr16-97010383-INS-50
chr16-97015310-INS-105
chr16-97024795-INS-7075
chr16-97027911-INS-64
chr16-97043937-INS-1112
chr16-97067073-INS-152
chr16-97073997-INS-60
chr16-97091379-INS-164
chr16-97131149-INS-354
chr16-97197268-INS-50
chr16-97253671-INS-3446
chr16-97269579-INS-50
chr16-97279046-INS-54
chr16-97286970-INS-140
chr16-97287092-INS-320
chr16-97300096-INS-6573
chr16-97311028-INS-70
chr16-97331270-INS-221
chr16-97372732-INS-50
chr16-97379293-INS-135
chr16-97379294-INS-50
chr16-97381042-INS-186
chr16-97439466-INS-63
chr16-97439891-INS-6755
chr16-97442593-INS-208
chr16-97461318-INS-246
chr16-97483737-INS-52
chr16-97711787-INS-50
chr16-97733740-INS-54
chr16-97735775-INS-936
chr16-97786081-INS-4880
chr16-97812743-INS-501
chr16-97813088-INS-588
chr16-97871563-INS-62
chr16-97873358-INS-1977

chr16-97873358-INS-5768
chr16-97873358-INS-928
chr16-97873432-INS-311
chr17-3053093-INS-97
chr17-3053575-INS-116
chr17-3077085-INS-376
chr17-3078311-INS-4915
chr17-3105000-INS-8743
chr17-3123942-INS-55
chr17-3127501-INS-9099
chr17-3183162-INS-55
chr17-3183180-INS-395
chr17-3224290-INS-50
chr17-3324645-INS-108
chr17-3328119-INS-50
chr17-3339473-INS-205
chr17-3351236-INS-58
chr17-3355080-INS-472
chr17-3375349-INS-203
chr17-3403261-INS-353
chr17-3403476-INS-1215
chr17-3411552-INS-186
chr17-3433361-INS-102
chr17-3461417-INS-122
chr17-3514216-INS-78
chr17-3583825-INS-1136
chr17-3588584-INS-286
chr17-3602242-INS-440
chr17-3658557-INS-66
chr17-3694595-INS-54
chr17-3704352-INS-62
chr17-3713240-INS-64
chr17-3735482-INS-507
chr17-3790179-INS-515
chr17-3825202-INS-122
chr17-3847141-INS-176
chr17-3888911-INS-336
chr17-3916573-INS-250
chr17-3930107-INS-193
chr17-4090183-INS-93
chr17-4091843-INS-56
chr17-4101319-INS-54
chr17-4114830-INS-2106
chr17-4157884-INS-7083
chr17-4161133-INS-202
chr17-4186074-INS-148
chr17-4186467-INS-167
chr17-4208434-INS-378
chr17-4275908-INS-856
chr17-4298479-INS-686
chr17-4321541-INS-60
chr17-4322606-INS-141
chr17-4356650-INS-69
chr17-4366706-INS-55
chr17-4383001-INS-1956
chr17-4386134-INS-1214
chr17-4402715-INS-131
chr17-4427151-INS-334
chr17-4446597-INS-6499
chr17-4456844-INS-3858
chr17-4482834-INS-52
chr17-4492143-INS-50
chr17-4504322-INS-6268
chr17-4519540-INS-104
chr17-4531521-INS-372
chr17-4549489-INS-98
chr17-4557145-INS-100
chr17-4572553-INS-174
chr17-4583841-INS-96
chr17-4609278-INS-315
chr17-4652571-INS-171
chr17-4656132-INS-68
chr17-4790478-INS-152
chr17-4791694-INS-230
chr17-4823005-INS-1008
chr17-4843142-INS-1891
chr17-4843584-INS-215

chr17-4846102-INS-78
chr17-4849394-INS-220
chr17-4858834-INS-655
chr17-4865551-INS-586
chr17-4867340-INS-66
chr17-4872063-INS-97
chr17-4897531-INS-915
chr17-4912159-INS-1582
chr17-4914149-INS-94
chr17-4997658-INS-57
chr17-5012195-INS-202
chr17-5018692-INS-302
chr17-5024534-INS-66
chr17-5027007-INS-205
chr17-5055973-INS-308
chr17-5061898-INS-1162
chr17-5062030-INS-106
chr17-5080779-INS-67
chr17-5084689-INS-144
chr17-5106600-INS-196
chr17-5140014-INS-58
chr17-5182887-INS-213
chr17-5190733-INS-170
chr17-5191997-INS-53
chr17-5194841-INS-60
chr17-5194901-INS-150
chr17-5224801-INS-384
chr17-5228117-INS-440
chr17-5242974-INS-216
chr17-5253416-INS-210
chr17-5291311-INS-594
chr17-5291481-INS-99
chr17-5291957-INS-99
chr17-5364015-INS-849
chr17-5365181-INS-139
chr17-5366886-INS-61
chr17-5374968-INS-50
chr17-5435221-INS-54
chr17-5440655-INS-51
chr17-5443555-INS-388
chr17-5444113-INS-99
chr17-5475976-INS-1978
chr17-5497880-INS-213
chr17-5501546-INS-52
chr17-5512697-INS-88
chr17-5549927-INS-60
chr17-5551329-INS-6443
chr17-5622007-INS-52
chr17-5632612-INS-60
chr17-5666425-INS-5771
chr17-5678679-INS-113
chr17-5691360-INS-60
chr17-5706207-INS-1422
chr17-5710232-INS-1495
chr17-5763193-INS-56
chr17-5791771-INS-270
chr17-5833405-INS-66
chr17-5841410-INS-400
chr17-5842558-INS-288
chr17-5853294-INS-481
chr17-5857372-INS-56
chr17-5864485-INS-140
chr17-5864500-INS-450
chr17-5931264-INS-121
chr17-5938464-INS-95
chr17-5943943-INS-156
chr17-5951633-INS-65
chr17-5967443-INS-52
chr17-5978640-INS-166
chr17-5978940-INS-656
chr17-6025369-INS-62
chr17-6064764-INS-102
chr17-6107411-INS-86
chr17-6111075-INS-66
chr17-6145249-INS-775
chr17-6192364-INS-504
chr17-6206624-INS-221

chr17-6232559-INS-560
chr17-6234893-INS-112
chr17-6241399-INS-50
chr17-6259519-INS-670
chr17-6259746-INS-80
chr17-6269955-INS-172
chr17-6300738-INS-56
chr17-6351837-INS-158
chr17-6375266-INS-3478
chr17-6441298-INS-416
chr17-6784946-INS-88
chr17-6797827-INS-209
chr17-6894996-INS-63
chr17-6958131-INS-498
chr17-6958541-INS-408
chr17-6961099-INS-932
chr17-6961406-INS-254
chr17-6976047-INS-6414
chr17-6991237-INS-5496
chr17-6997824-INS-352
chr17-6997915-INS-95
chr17-7016602-INS-135
chr17-7031185-INS-71
chr17-7054834-INS-213
chr17-7058530-INS-50
chr17-7081409-INS-80
chr17-7083004-INS-260
chr17-7171496-INS-596
chr17-7171591-INS-12481
chr17-7212335-INS-412
chr17-7217646-INS-7097
chr17-7219456-INS-828
chr17-7318826-INS-62
chr17-7321140-INS-219
chr17-7329245-INS-399
chr17-7337269-INS-59
chr17-7349948-INS-71
chr17-7359447-INS-120
chr17-7361055-INS-574
chr17-7382708-INS-200
chr17-7393776-INS-4232
chr17-7407740-INS-57
chr17-7408060-INS-73
chr17-7409678-INS-170
chr17-7460418-INS-1306
chr17-7460538-INS-4046
chr17-7467954-INS-57
chr17-7539609-INS-267
chr17-7539636-INS-104
chr17-7541930-INS-581
chr17-7545285-INS-51
chr17-7585922-INS-1062
chr17-7646008-INS-100
chr17-7705834-INS-204
chr17-7727192-INS-339
chr17-7727237-INS-64
chr17-7735394-INS-2851
chr17-7741225-INS-7083
chr17-7757207-INS-482
chr17-7782549-INS-174
chr17-7782561-INS-490
chr17-7787524-INS-1946
chr17-7807299-INS-89
chr17-7824100-INS-718
chr17-7824355-INS-50
chr17-7824925-INS-52
chr17-7829027-INS-57
chr17-7842608-INS-103
chr17-7856806-INS-66
chr17-7857664-INS-91
chr17-7911305-INS-2821
chr17-7911305-INS-404
chr17-7922375-INS-6293
chr17-7934935-INS-347
chr17-7937179-INS-6435
chr17-7962276-INS-226
chr17-7962450-INS-94

chr17-7970973-INS-117
chr17-7971104-INS-54
chr17-8004439-INS-297
chr17-8021688-INS-268
chr17-8047177-INS-55
chr17-8054273-INS-71
chr17-8084194-INS-3121
chr17-8088440-INS-4826
chr17-8114056-INS-198
chr17-8123507-INS-78
chr17-8123651-INS-90
chr17-8125581-INS-54
chr17-8126388-INS-1716
chr17-8139854-INS-2918
chr17-8168239-INS-412
chr17-8175310-INS-925
chr17-8195181-INS-210
chr17-8199751-INS-197
chr17-8200912-INS-376
chr17-8202873-INS-104
chr17-8252723-INS-109
chr17-8293970-INS-92
chr17-8339181-INS-1890
chr17-8341059-INS-956
chr17-8342457-INS-434
chr17-8358646-INS-131
chr17-8361857-INS-399
chr17-8396477-INS-174
chr17-8396590-INS-406
chr17-8402722-INS-205
chr17-8427069-INS-537
chr17-8427148-INS-219
chr17-8427423-INS-84
chr17-8427562-INS-1313
chr17-8427581-INS-412
chr17-8428118-INS-57
chr17-8446303-INS-191
chr17-8446616-INS-407
chr17-8448539-INS-399
chr17-8455765-INS-66
chr17-8455947-INS-60
chr17-8470470-INS-256
chr17-8503767-INS-199
chr17-8525718-INS-58
chr17-8525746-INS-815
chr17-8526844-INS-89
chr17-8548125-INS-879
chr17-8566106-INS-75
chr17-8566258-INS-195
chr17-8570515-INS-1896
chr17-8596166-INS-178
chr17-8604708-INS-191
chr17-8620822-INS-56
chr17-8646344-INS-59
chr17-8649732-INS-79
chr17-8665256-INS-96
chr17-8685149-INS-56
chr17-8719923-INS-148
chr17-8789192-INS-152
chr17-8796383-INS-59
chr17-8807780-INS-120
chr17-8824977-INS-1500
chr17-8847786-INS-72
chr17-8885417-INS-58
chr17-8889965-INS-594
chr17-8923336-INS-176
chr17-8923659-INS-64
chr17-8932821-INS-1147
chr17-8939230-INS-93
chr17-8959702-INS-124
chr17-8979751-INS-119
chr17-9012555-INS-1696
chr17-9044075-INS-70
chr17-9046173-INS-120
chr17-9067812-INS-205
chr17-9071617-INS-84
chr17-9071921-INS-112

chr17-9072252-INS-66
chr17-9096827-INS-876
chr17-9109710-INS-60
chr17-9154947-INS-297
chr17-9155598-INS-502
chr17-9234173-INS-303
chr17-9234176-INS-86
chr17-9285737-INS-110
chr17-9285819-INS-227
chr17-9310698-INS-53
chr17-9316216-INS-52
chr17-9319682-INS-1169
chr17-9339155-INS-52
chr17-9396011-INS-368
chr17-9419877-INS-122
chr17-9419877-INS-350
chr17-9432787-INS-4406
chr17-9434265-INS-958
chr17-9436470-INS-140
chr17-9486027-INS-197
chr17-9488643-INS-6597
chr17-9496585-INS-56
chr17-9497312-INS-57
chr17-9506709-INS-355
chr17-9508146-INS-493
chr17-9508152-INS-230
chr17-9542315-INS-68
chr17-9593297-INS-76
chr17-9681233-INS-251
chr17-9681249-INS-108
chr17-9720155-INS-194
chr17-9742095-INS-498
chr17-9793688-INS-72
chr17-9850520-INS-270
chr17-9857895-INS-52
chr17-9900980-INS-50
chr17-9908406-INS-50
chr17-9924804-INS-697
chr17-9947326-INS-6906
chr17-9963892-INS-506
chr17-9964028-INS-116
chr17-9970025-INS-50
chr17-9983497-INS-56
chr17-9987166-INS-495
chr17-10057100-INS-272
chr17-10102578-INS-50
chr17-10122085-INS-58
chr17-10128382-INS-127
chr17-10133661-INS-442
chr17-10133665-INS-103
chr17-10147880-INS-152
chr17-10184786-INS-56
chr17-10233844-INS-52
chr17-10242424-INS-3048
chr17-10244072-INS-104
chr17-10337478-INS-90
chr17-10345524-INS-126
chr17-10345524-INS-393
chr17-10453324-INS-241
chr17-10473630-INS-166
chr17-10475797-INS-50
chr17-10503458-INS-205
chr17-10608522-INS-52
chr17-10610170-INS-54
chr17-10628398-INS-327
chr17-10657421-INS-156
chr17-10664189-INS-164
chr17-10685067-INS-108
chr17-10708651-INS-82
chr17-10815028-INS-309
chr17-10816770-INS-7317
chr17-10826185-INS-116
chr17-10831319-INS-2378
chr17-10834738-INS-60
chr17-10841276-INS-6924
chr17-10865974-INS-6457
chr17-10891532-INS-145

chr17-10910883-INS-56
chr17-10944885-INS-178
chr17-10980738-INS-90
chr17-11004212-INS-354
chr17-11054240-INS-108
chr17-11101094-INS-7118
chr17-11149805-INS-266
chr17-11166025-INS-141
chr17-11166096-INS-377
chr17-11185222-INS-60
chr17-11227650-INS-253
chr17-11268017-INS-315
chr17-11282646-INS-56
chr17-11326145-INS-210
chr17-11326160-INS-63
chr17-11350031-INS-217
chr17-11477668-INS-52
chr17-11478418-INS-6344
chr17-11500504-INS-30592
chr17-11511898-INS-648
chr17-11536494-INS-129
chr17-11537198-INS-69
chr17-11603820-INS-69
chr17-11608608-INS-1632
chr17-11620174-INS-116
chr17-11621111-INS-68
chr17-11653902-INS-68
chr17-11680563-INS-1933
chr17-11682906-INS-66
chr17-11741157-INS-72
chr17-11743720-INS-110
chr17-11773307-INS-6436
chr17-11874020-INS-62
chr17-11905863-INS-434
chr17-11906033-INS-54
chr17-11972521-INS-350
chr17-11992480-INS-54
chr17-12000615-INS-463
chr17-12026528-INS-67
chr17-12068115-INS-62
chr17-12157048-INS-50
chr17-12194895-INS-50
chr17-12262313-INS-6337
chr17-12324643-INS-5641
chr17-12360561-INS-214
chr17-12509223-INS-97
chr17-12513117-INS-57
chr17-12572182-INS-76
chr17-12575894-INS-162
chr17-12584080-INS-132
chr17-12612647-INS-71
chr17-12612725-INS-996
chr17-12616877-INS-52
chr17-12655380-INS-60
chr17-12655688-INS-102
chr17-12676303-INS-206
chr17-12681447-INS-1874
chr17-12825418-INS-52
chr17-12848112-INS-74
chr17-12913409-INS-66
chr17-12947429-INS-216
chr17-13024553-INS-64
chr17-13054149-INS-89
chr17-13112634-INS-206
chr17-13172804-INS-442
chr17-13176368-INS-220
chr17-13220711-INS-50
chr17-13248932-INS-208
chr17-13273892-INS-499
chr17-13285597-INS-1044
chr17-13343872-INS-79
chr17-13390120-INS-1286
chr17-13411734-INS-198
chr17-13418267-INS-112
chr17-13426922-INS-90
chr17-13536110-INS-141
chr17-13581491-INS-58

chr17-13736407-INS-64
chr17-13748016-INS-84
chr17-13770354-INS-945
chr17-13771179-INS-75
chr17-13775498-INS-6380
chr17-13802018-INS-392
chr17-13807330-INS-437
chr17-13963145-INS-71
chr17-14031347-INS-58
chr17-14107363-INS-225
chr17-14107363-INS-90
chr17-14137304-INS-122
chr17-14137687-INS-212
chr17-14142138-INS-59
chr17-14181093-INS-201
chr17-14182145-INS-542
chr17-14192348-INS-1939
chr17-14267669-INS-203
chr17-14277364-INS-54
chr17-14282643-INS-79
chr17-14320004-INS-67
chr17-14348096-INS-3068
chr17-14355689-INS-69
chr17-14368697-INS-75
chr17-14412158-INS-50
chr17-14478082-INS-3489
chr17-14511480-INS-858
chr17-14511582-INS-379
chr17-14511848-INS-363
chr17-14561301-INS-339
chr17-14575932-INS-88
chr17-14581790-INS-54
chr17-14647906-INS-89
chr17-14654479-INS-2481
chr17-14682133-INS-80
chr17-14712672-INS-434
chr17-14747364-INS-1971
chr17-14771254-INS-134
chr17-14797736-INS-103
chr17-14797995-INS-172
chr17-14808000-INS-880
chr17-14848311-INS-56
chr17-14863783-INS-215
chr17-14922865-INS-50
chr17-14949823-INS-6344
chr17-14957746-INS-116
chr17-14959961-INS-7301
chr17-14977578-INS-500
chr17-15013461-INS-57
chr17-15015730-INS-74
chr17-15027030-INS-214
chr17-15043027-INS-1243
chr17-15051953-INS-1982
chr17-15061890-INS-56
chr17-15062669-INS-205
chr17-15081995-INS-741
chr17-15082296-INS-350
chr17-15154010-INS-68
chr17-15168543-INS-3473
chr17-15221487-INS-205
chr17-15224721-INS-220
chr17-15252854-INS-205
chr17-15307198-INS-8419
chr17-15332279-INS-63
chr17-15349212-INS-90
chr17-15351335-INS-128
chr17-15402834-INS-62
chr17-15442374-INS-53
chr17-15473602-INS-97
chr17-15481330-INS-70
chr17-15481433-INS-249
chr17-15487514-INS-63
chr17-15534741-INS-56
chr17-15580573-INS-80
chr17-15625743-INS-446
chr17-15630568-INS-54
chr17-15632115-INS-124

chr17-15632298-INS-128
chr17-15634742-INS-238
chr17-15643227-INS-1000
chr17-15647380-INS-136
chr17-15662112-INS-209
chr17-15666381-INS-291
chr17-15678153-INS-2251
chr17-15695556-INS-64
chr17-15700566-INS-64
chr17-15700584-INS-134
chr17-15750757-INS-127
chr17-15751057-INS-199
chr17-15753957-INS-213
chr17-15754281-INS-273
chr17-15758126-INS-68
chr17-15770892-INS-84
chr17-15779857-INS-998
chr17-15790122-INS-4155
chr17-15810298-INS-639
chr17-15811336-INS-82
chr17-15812910-INS-4199
chr17-15814763-INS-56
chr17-15822400-INS-50
chr17-15866908-INS-586
chr17-15869203-INS-70
chr17-15875712-INS-72
chr17-15881309-INS-219
chr17-15905953-INS-188
chr17-15943319-INS-115
chr17-15966579-INS-211
chr17-15973688-INS-108
chr17-15983590-INS-68
chr17-15994016-INS-88
chr17-16010203-INS-104
chr17-16013375-INS-189
chr17-16023211-INS-130
chr17-16032059-INS-50
chr17-16052483-INS-60
chr17-16053208-INS-64
chr17-16061123-INS-7002
chr17-16073600-INS-51
chr17-16107059-INS-155
chr17-16120405-INS-169
chr17-16189638-INS-55
chr17-16212270-INS-54
chr17-16237220-INS-116
chr17-16238199-INS-442
chr17-16273435-INS-221
chr17-16312041-INS-6191
chr17-16312306-INS-57
chr17-16313962-INS-148
chr17-16387280-INS-5007
chr17-16424721-INS-528
chr17-16489144-INS-6025
chr17-16532776-INS-497
chr17-16532833-INS-914
chr17-16544112-INS-66
chr17-16550422-INS-6218
chr17-16582091-INS-4574
chr17-16667864-INS-4793
chr17-16674573-INS-3109
chr17-16692231-INS-7155
chr17-16696806-INS-62
chr17-16702558-INS-2763
chr17-16709373-INS-935
chr17-16711972-INS-319
chr17-16740743-INS-298
chr17-16740993-INS-166
chr17-16760712-INS-5278
chr17-16820404-INS-61
chr17-16828510-INS-175
chr17-16896863-INS-212
chr17-16943728-INS-54
chr17-16965150-INS-10646
chr17-16992615-INS-470
chr17-17111222-INS-183
chr17-17213526-INS-68

chr17-17230562-INS-208
chr17-17230643-INS-833
chr17-17248163-INS-84
chr17-17273767-INS-199
chr17-17331483-INS-171
chr17-17360389-INS-63
chr17-17374948-INS-723
chr17-17389404-INS-188
chr17-17422427-INS-1114
chr17-17430860-INS-68
chr17-17436589-INS-5507
chr17-17437661-INS-205
chr17-17461198-INS-57
chr17-17490796-INS-50
chr17-17512536-INS-780
chr17-17563769-INS-6491
chr17-17616907-INS-216
chr17-17635052-INS-123
chr17-17635052-INS-58
chr17-17643015-INS-58
chr17-17683105-INS-60
chr17-17702719-INS-258
chr17-17718568-INS-68
chr17-17721811-INS-156
chr17-17731267-INS-54
chr17-17796969-INS-60
chr17-17837613-INS-240
chr17-17873949-INS-2862
chr17-17891159-INS-345
chr17-17891165-INS-54
chr17-17939770-INS-426
chr17-18003120-INS-11494
chr17-18007345-INS-4920
chr17-18026406-INS-4499
chr17-18064929-INS-288
chr17-18076486-INS-54
chr17-18078237-INS-51
chr17-18167682-INS-96
chr17-18187462-INS-455
chr17-18192297-INS-721
chr17-18196540-INS-11890
chr17-18208213-INS-6979
chr17-18220436-INS-176
chr17-18227245-INS-108
chr17-18243680-INS-7852
chr17-18265503-INS-9377
chr17-18306429-INS-78
chr17-18343024-INS-106
chr17-18351740-INS-83
chr17-18393857-INS-185
chr17-18397799-INS-1348
chr17-18457025-INS-2147
chr17-18506698-INS-78
chr17-18519453-INS-6613
chr17-18581381-INS-92
chr17-18581819-INS-103
chr17-18610078-INS-165
chr17-18617671-INS-55
chr17-18657625-INS-179
chr17-18658455-INS-355
chr17-18718724-INS-1873
chr17-18731109-INS-7713
chr17-18764848-INS-1745
chr17-18800689-INS-3080
chr17-19057922-INS-1270
chr17-19079265-INS-413
chr17-19135673-INS-60
chr17-19290086-INS-6643
chr17-19292625-INS-50
chr17-19294454-INS-257
chr17-19339036-INS-157
chr17-19354129-INS-14074
chr17-19665776-INS-556
chr17-19668984-INS-86
chr17-19672729-INS-199
chr17-19682967-INS-351
chr17-19693298-INS-142

chr17-19712670-INS-58
chr17-19837605-INS-327
chr17-19907314-INS-56
chr17-19921442-INS-498
chr17-19926993-INS-54
chr17-19962281-INS-399
chr17-19970361-INS-1855
chr17-19989512-INS-144
chr17-19992880-INS-69
chr17-20007557-INS-6746
chr17-20050418-INS-2984
chr17-20054944-INS-466
chr17-20089867-INS-4185
chr17-20110257-INS-169
chr17-20130051-INS-1934
chr17-20154482-INS-55
chr17-20202460-INS-6432
chr17-20312033-INS-4770
chr17-20348120-INS-64
chr17-20386480-INS-53
chr17-20391297-INS-50
chr17-20417498-INS-54
chr17-20421800-INS-63
chr17-20422253-INS-60
chr17-20432972-INS-177
chr17-20456085-INS-123
chr17-20456085-INS-58
chr17-20479924-INS-1417
chr17-20481194-INS-2848
chr17-20509793-INS-6603
chr17-20529442-INS-2630
chr17-20533303-INS-1720
chr17-20538373-INS-572
chr17-20542742-INS-3951
chr17-20545932-INS-4920
chr17-20546354-INS-4708
chr17-20549513-INS-789
chr17-20582612-INS-180
chr17-20586409-INS-63
chr17-20586621-INS-2045
chr17-20594888-INS-285
chr17-20594965-INS-87
chr17-20649692-INS-50
chr17-20669276-INS-202
chr17-20677717-INS-6280
chr17-20717930-INS-394
chr17-20737727-INS-5457
chr17-20743133-INS-716
chr17-20830655-INS-6165
chr17-20867107-INS-74
chr17-20878263-INS-2097
chr17-20885201-INS-52
chr17-20886699-INS-126
chr17-20898393-INS-6371
chr17-21045456-INS-51
chr17-21046765-INS-2927
chr17-21118226-INS-134
chr17-21148186-INS-50
chr17-21194270-INS-99
chr17-21243077-INS-52
chr17-21256657-INS-1863
chr17-21268197-INS-68
chr17-21274588-INS-328
chr17-21284346-INS-751
chr17-21309677-INS-6788
chr17-21315055-INS-6049
chr17-21424424-INS-428
chr17-21427648-INS-3142
chr17-21438965-INS-233
chr17-21467582-INS-199
chr17-21473096-INS-15652
chr17-21473096-INS-5744
chr17-21503704-INS-25978
chr17-21551495-INS-112
chr17-21557927-INS-889
chr17-21589986-INS-2064
chr17-21594896-INS-9156

chr17-21665948-INS-6540
chr17-21679693-INS-198
chr17-21697476-INS-498
chr17-21737188-INS-90
chr17-21737786-INS-1411
chr17-21737786-INS-256
chr17-21740767-INS-1428
chr17-21742949-INS-77
chr17-21750529-INS-432
chr17-21762912-INS-3291
chr17-21817199-INS-178
chr17-21852549-INS-83
chr17-21857936-INS-1405
chr17-21889039-INS-7231
chr17-21891427-INS-335
chr17-21894468-INS-6448
chr17-21949441-INS-4119
chr17-21958924-INS-67
chr17-21981964-INS-117
chr17-21992986-INS-2086
chr17-22004636-INS-567
chr17-22018981-INS-841
chr17-22067266-INS-230
chr17-22077452-INS-236
chr17-22079412-INS-6416
chr17-22127641-INS-388
chr17-22132213-INS-12371
chr17-22161761-INS-2308
chr17-22176237-INS-422
chr17-22188190-INS-72
chr17-22268700-INS-24123
chr17-22298690-INS-472
chr17-22346292-INS-653
chr17-22362134-INS-2853
chr17-22371517-INS-634
chr17-22376926-INS-71
chr17-22383550-INS-80
chr17-22394115-INS-3030
chr17-22401467-INS-98
chr17-22401564-INS-263
chr17-22409535-INS-53
chr17-22413020-INS-102
chr17-22416885-INS-4378
chr17-22459025-INS-604
chr17-22466694-INS-441
chr17-22500316-INS-58
chr17-22512752-INS-6566
chr17-22610881-INS-354
chr17-22766797-INS-996
chr17-22800071-INS-77
chr17-22823221-INS-54
chr17-22823239-INS-195
chr17-22847699-INS-178
chr17-22860459-INS-1000
chr17-22895984-INS-1500
chr17-22999851-INS-50
chr17-23006866-INS-6826
chr17-23026097-INS-200
chr17-23027779-INS-2836
chr17-23086910-INS-50
chr17-23103544-INS-1515
chr17-23140017-INS-88
chr17-23151566-INS-57
chr17-23179437-INS-3097
chr17-23270742-INS-100
chr17-23277962-INS-78
chr17-23292352-INS-156
chr17-23298062-INS-3097
chr17-23376974-INS-1079
chr17-23381275-INS-6018
chr17-23418016-INS-7295
chr17-23598454-INS-3036
chr17-23639888-INS-5549
chr17-23666916-INS-208
chr17-23686242-INS-3137
chr17-23692116-INS-207
chr17-23708826-INS-61

chr17-23750080-INS-52
chr17-23750432-INS-106
chr17-23756169-INS-154
chr17-23767706-INS-52
chr17-23770850-INS-76
chr17-23776149-INS-196
chr17-23789348-INS-644
chr17-23802449-INS-133
chr17-23822463-INS-60
chr17-23923193-INS-81
chr17-23972985-INS-69
chr17-23973501-INS-199
chr17-23999252-INS-219
chr17-24000492-INS-103
chr17-24003072-INS-62
chr17-24030204-INS-1219
chr17-24030204-INS-384
chr17-24062118-INS-320
chr17-24097104-INS-5271
chr17-24116661-INS-206
chr17-24120712-INS-205
chr17-24124317-INS-14853
chr17-24147116-INS-189
chr17-24175267-INS-59
chr17-24219855-INS-211
chr17-24305626-INS-124
chr17-24432959-INS-216
chr17-24446455-INS-58
chr17-24459224-INS-224
chr17-24519901-INS-214
chr17-24526295-INS-164
chr17-24529585-INS-382
chr17-24538436-INS-205
chr17-24561772-INS-210
chr17-24561917-INS-6333
chr17-24575631-INS-148
chr17-24592560-INS-211
chr17-24614883-INS-149
chr17-24614883-INS-56
chr17-24629831-INS-209
chr17-24650976-INS-99
chr17-24702161-INS-183
chr17-24703410-INS-232
chr17-24716717-INS-92
chr17-24760563-INS-51
chr17-24766271-INS-400
chr17-24828241-INS-68
chr17-24836112-INS-135
chr17-24836234-INS-3475
chr17-24843864-INS-109
chr17-24844058-INS-769
chr17-24844197-INS-247
chr17-24844222-INS-61
chr17-24845215-INS-6399
chr17-24881955-INS-53
chr17-24882385-INS-212
chr17-24892745-INS-50
chr17-24899580-INS-3419
chr17-24901897-INS-189
chr17-24905976-INS-91
chr17-24956460-INS-202
chr17-24974378-INS-6915
chr17-24988471-INS-126
chr17-24995568-INS-118
chr17-25014329-INS-5255
chr17-25014329-INS-628
chr17-25071265-INS-200
chr17-25261516-INS-696
chr17-25310437-INS-340
chr17-25328474-INS-50
chr17-25344424-INS-96
chr17-25362686-INS-187
chr17-25414692-INS-124
chr17-25415293-INS-61
chr17-25420775-INS-383
chr17-25478721-INS-208
chr17-25494786-INS-206

chr17-25502152-INS-56
chr17-25510327-INS-3097
chr17-25522581-INS-154
chr17-25526058-INS-202
chr17-25540511-INS-57
chr17-25544980-INS-66
chr17-25554504-INS-337
chr17-25554890-INS-54
chr17-25657292-INS-207
chr17-25705680-INS-484
chr17-25726278-INS-158
chr17-25772963-INS-50
chr17-25839806-INS-54
chr17-25839814-INS-112
chr17-25848752-INS-63
chr17-25848879-INS-2855
chr17-25849226-INS-6626
chr17-25877082-INS-378
chr17-25878199-INS-543
chr17-25878432-INS-371
chr17-25889952-INS-192
chr17-25890652-INS-202
chr17-25919016-INS-661
chr17-25923778-INS-52
chr17-25926107-INS-187
chr17-25948487-INS-60
chr17-25961113-INS-62
chr17-25979741-INS-66
chr17-26089708-INS-116
chr17-26090441-INS-212
chr17-26124260-INS-84
chr17-26125449-INS-52
chr17-26129903-INS-210
chr17-26180299-INS-69
chr17-26205577-INS-88
chr17-26257066-INS-70
chr17-26290841-INS-100
chr17-26292077-INS-1700
chr17-26295782-INS-74
chr17-26301794-INS-227
chr17-26318312-INS-10309
chr17-26318873-INS-4959
chr17-26320077-INS-478
chr17-26326676-INS-155
chr17-26327474-INS-63
chr17-26373979-INS-198
chr17-26378069-INS-198
chr17-26386561-INS-75
chr17-26386601-INS-388
chr17-26386698-INS-329
chr17-26398654-INS-199
chr17-26398975-INS-99
chr17-26399025-INS-477
chr17-26433897-INS-80
chr17-26465012-INS-81
chr17-26474273-INS-99
chr17-26474831-INS-195
chr17-26483960-INS-73
chr17-26484746-INS-177
chr17-26496277-INS-3246
chr17-26522161-INS-201
chr17-26561037-INS-64
chr17-26581399-INS-92
chr17-26616132-INS-66
chr17-26679854-INS-53
chr17-26680432-INS-72
chr17-26690310-INS-60
chr17-26730931-INS-190
chr17-26757472-INS-1207
chr17-26777277-INS-51
chr17-26797923-INS-62
chr17-26798588-INS-202
chr17-26805245-INS-104
chr17-26811173-INS-594
chr17-26852033-INS-203
chr17-26899941-INS-68
chr17-26900644-INS-169

chr17-26902829-INS-199
chr17-26902913-INS-490
chr17-26910624-INS-69
chr17-26983590-INS-437
chr17-26983838-INS-4470
chr17-27006920-INS-65
chr17-27021238-INS-58
chr17-27023417-INS-215
chr17-27050567-INS-5319
chr17-27062306-INS-50
chr17-27065948-INS-424
chr17-27070481-INS-126
chr17-27095919-INS-226
chr17-27110506-INS-198
chr17-27152904-INS-60
chr17-27200083-INS-143
chr17-27213801-INS-94
chr17-27214723-INS-165
chr17-27278233-INS-74
chr17-27292527-INS-97
chr17-27298874-INS-57
chr17-27311111-INS-120
chr17-27381106-INS-250
chr17-27381410-INS-100
chr17-27416318-INS-69
chr17-27422130-INS-504
chr17-27425762-INS-478
chr17-27448458-INS-52
chr17-27527851-INS-52
chr17-27541104-INS-76
chr17-27610770-INS-88
chr17-27680838-INS-210
chr17-27689447-INS-81
chr17-27790617-INS-158
chr17-27800397-INS-2572
chr17-27809255-INS-595
chr17-27826531-INS-62
chr17-27834789-INS-203
chr17-27834791-INS-75
chr17-27835020-INS-66
chr17-27844562-INS-7633
chr17-27857135-INS-60
chr17-27870729-INS-221
chr17-27886891-INS-195
chr17-27910859-INS-208
chr17-28047599-INS-1003
chr17-28057666-INS-54
chr17-28082827-INS-211
chr17-28107754-INS-199
chr17-28107895-INS-65
chr17-28138524-INS-124
chr17-28146700-INS-600
chr17-28206572-INS-6481
chr17-28231737-INS-208
chr17-28240774-INS-353
chr17-28295377-INS-84
chr17-28403580-INS-88
chr17-28565615-INS-50
chr17-28580650-INS-65
chr17-28585463-INS-52
chr17-28604134-INS-75
chr17-28615201-INS-208
chr17-28622140-INS-953
chr17-28627716-INS-212
chr17-28646950-INS-84
chr17-28673247-INS-224
chr17-28673471-INS-112
chr17-28692098-INS-174
chr17-28711638-INS-163
chr17-28752020-INS-76
chr17-28766379-INS-5632
chr17-28776100-INS-178
chr17-28777033-INS-210
chr17-28781861-INS-55
chr17-28782279-INS-119
chr17-28784994-INS-179
chr17-28785638-INS-196

chr17-28786978-INS-206
chr17-28788119-INS-114
chr17-28788305-INS-104
chr17-28789190-INS-63
chr17-28789248-INS-171
chr17-28828297-INS-111
chr17-28848600-INS-88
chr17-28849963-INS-64
chr17-28853497-INS-1218
chr17-28853503-INS-58
chr17-28864378-INS-459
chr17-28951781-INS-120
chr17-28987734-INS-56
chr17-29003740-INS-149
chr17-29003770-INS-56
chr17-29008484-INS-1176
chr17-29034492-INS-576
chr17-29103720-INS-63
chr17-29105125-INS-167
chr17-29136897-INS-4974
chr17-29162437-INS-161
chr17-29199776-INS-579
chr17-29223744-INS-417
chr17-29229484-INS-219
chr17-29235648-INS-1955
chr17-29247443-INS-973
chr17-29248722-INS-98
chr17-29282061-INS-62
chr17-29306136-INS-191
chr17-29306398-INS-58
chr17-29326101-INS-182
chr17-29337626-INS-62
chr17-29351900-INS-182
chr17-29410742-INS-143
chr17-29423142-INS-175
chr17-29423880-INS-67
chr17-29424172-INS-215
chr17-29430766-INS-50
chr17-29442585-INS-53
chr17-29478347-INS-382
chr17-29503805-INS-230
chr17-29505688-INS-967
chr17-29505829-INS-50
chr17-29506035-INS-277
chr17-29506546-INS-198
chr17-29508736-INS-77
chr17-29554663-INS-59
chr17-29555345-INS-160
chr17-29555349-INS-60
chr17-29563996-INS-161
chr17-29578696-INS-177
chr17-29578696-INS-370
chr17-29578844-INS-53
chr17-29589828-INS-407
chr17-29651916-INS-205
chr17-29654061-INS-86
chr17-29667559-INS-56
chr17-29671787-INS-92
chr17-29697993-INS-523
chr17-29716835-INS-76
chr17-29753560-INS-82
chr17-29754527-INS-231
chr17-29761392-INS-138
chr17-29764326-INS-327
chr17-29797196-INS-58
chr17-29869849-INS-50
chr17-29871598-INS-238
chr17-29875200-INS-238
chr17-29879396-INS-208
chr17-29883194-INS-60
chr17-29954921-INS-66
chr17-29963881-INS-7088
chr17-29980347-INS-50
chr17-30003726-INS-167
chr17-30035807-INS-200
chr17-30044805-INS-85
chr17-30051599-INS-261

chr17-30081679-INS-306
chr17-30082235-INS-2514
chr17-30133572-INS-84
chr17-30198010-INS-76
chr17-30205138-INS-654
chr17-30302341-INS-377
chr17-30556477-INS-1782
chr17-30665908-INS-63
chr17-30667910-INS-50
chr17-30702430-INS-3400
chr17-30702463-INS-604
chr17-30702704-INS-181
chr17-30752406-INS-6403
chr17-30792804-INS-58
chr17-30793519-INS-201
chr17-30814885-INS-366
chr17-30869568-INS-84
chr17-30887996-INS-426
chr17-30937021-INS-110
chr17-30953225-INS-190
chr17-30958040-INS-54
chr17-30958126-INS-162
chr17-30962548-INS-92
chr17-30969533-INS-135
chr17-30974272-INS-160
chr17-30982059-INS-56
chr17-31017974-INS-864
chr17-31030186-INS-64
chr17-31041920-INS-1152
chr17-31059799-INS-104
chr17-31108311-INS-54
chr17-31110193-INS-1001
chr17-31111529-INS-189
chr17-31114969-INS-757
chr17-31115217-INS-100
chr17-31115222-INS-1249
chr17-31117283-INS-63
chr17-31132106-INS-143
chr17-31136645-INS-100
chr17-31175117-INS-254
chr17-31175604-INS-51
chr17-31177322-INS-50
chr17-31182413-INS-103
chr17-31193832-INS-277
chr17-31203834-INS-4580
chr17-31209216-INS-78
chr17-31362465-INS-100
chr17-31362742-INS-96
chr17-31426124-INS-160
chr17-31434592-INS-56
chr17-31441585-INS-50
chr17-31449582-INS-85
chr17-31453342-INS-64
chr17-31453567-INS-84
chr17-31465280-INS-205
chr17-31467254-INS-50
chr17-31482005-INS-223
chr17-31495113-INS-85
chr17-31510467-INS-165
chr17-31561005-INS-57
chr17-31575800-INS-4796
chr17-31578242-INS-219
chr17-31579726-INS-35426
chr17-31580890-INS-319
chr17-31592192-INS-462
chr17-31615284-INS-130
chr17-31651365-INS-248
chr17-31654417-INS-256
chr17-31659653-INS-135
chr17-31659793-INS-165
chr17-31675540-INS-100
chr17-31698686-INS-15546
chr17-31719779-INS-54
chr17-31749723-INS-176
chr17-31765196-INS-50
chr17-31766751-INS-52
chr17-31780020-INS-202

chr17-31843081-INS-56
chr17-31901626-INS-7088
chr17-31908238-INS-2751
chr17-31908349-INS-118
chr17-31908385-INS-50
chr17-31942810-INS-2867
chr17-31961171-INS-151
chr17-32009122-INS-52
chr17-32075427-INS-115
chr17-32085749-INS-108
chr17-32130614-INS-57
chr17-32139968-INS-326
chr17-32140757-INS-55
chr17-32160743-INS-50
chr17-32207012-INS-252
chr17-32207012-INS-575
chr17-32232812-INS-292
chr17-32261609-INS-163
chr17-32295053-INS-100
chr17-32301794-INS-72
chr17-32304483-INS-204
chr17-32307750-INS-66
chr17-32321329-INS-210
chr17-32364894-INS-77
chr17-32402089-INS-205
chr17-32403260-INS-96
chr17-32430568-INS-129
chr17-32434322-INS-60
chr17-32434760-INS-89
chr17-32496613-INS-208
chr17-32508158-INS-198
chr17-32510706-INS-166
chr17-32511583-INS-198
chr17-32521036-INS-201
chr17-32529232-INS-407
chr17-32617728-INS-82
chr17-32619462-INS-138
chr17-32640595-INS-8457
chr17-32641134-INS-79
chr17-32674794-INS-55
chr17-32694815-INS-56
chr17-32703969-INS-397
chr17-32723677-INS-68
chr17-32729946-INS-7115
chr17-32742720-INS-157
chr17-32744465-INS-130
chr17-32745516-INS-500
chr17-32831713-INS-2788
chr17-32852842-INS-5946
chr17-32856645-INS-4788
chr17-32863293-INS-3658
chr17-32863630-INS-132
chr17-32874355-INS-6635
chr17-32880869-INS-80
chr17-32896919-INS-56
chr17-32927026-INS-52
chr17-32927422-INS-50
chr17-32931346-INS-1347
chr17-32943142-INS-109
chr17-32943181-INS-54
chr17-32950306-INS-3786
chr17-32963381-INS-2878
chr17-32975456-INS-88
chr17-32976139-INS-4801
chr17-33024481-INS-62
chr17-33034968-INS-183
chr17-33047413-INS-240
chr17-33047782-INS-288
chr17-33059228-INS-95
chr17-33081462-INS-50
chr17-33084756-INS-384
chr17-33090358-INS-60
chr17-33118596-INS-61
chr17-33119329-INS-60
chr17-33123443-INS-200
chr17-33133373-INS-81
chr17-33151279-INS-939

chr17-33180044-INS-69
chr17-33185307-INS-54
chr17-33188236-INS-113
chr17-33199695-INS-185
chr17-33226757-INS-498
chr17-33229736-INS-171
chr17-33229976-INS-232
chr17-33239581-INS-255
chr17-33342521-INS-3702
chr17-33381592-INS-189
chr17-33433554-INS-454
chr17-33514632-INS-85
chr17-33633429-INS-214
chr17-33683861-INS-281
chr17-33691030-INS-82
chr17-33760758-INS-93
chr17-33765999-INS-186
chr17-33766237-INS-202
chr17-33776421-INS-56
chr17-33807596-INS-132
chr17-33810920-INS-51
chr17-33822742-INS-192
chr17-33824116-INS-64
chr17-34005663-INS-3245
chr17-34051293-INS-300
chr17-34055988-INS-7398
chr17-34057470-INS-172
chr17-34069066-INS-86
chr17-34079227-INS-81
chr17-34114763-INS-211
chr17-34115354-INS-226
chr17-34117974-INS-622
chr17-34163437-INS-69
chr17-34163437-INS-690
chr17-34182758-INS-119
chr17-34183824-INS-70
chr17-34189725-INS-1992
chr17-34192097-INS-196
chr17-34201725-INS-1573
chr17-34203145-INS-2896
chr17-34224248-INS-506
chr17-34266353-INS-64
chr17-34326486-INS-264
chr17-34326604-INS-85
chr17-34362859-INS-191
chr17-34368812-INS-4832
chr17-34371171-INS-1266
chr17-34376911-INS-63
chr17-34398756-INS-6400
chr17-34432935-INS-84
chr17-34468730-INS-96
chr17-34473081-INS-237
chr17-34484488-INS-186
chr17-34485037-INS-527
chr17-34491276-INS-548
chr17-34492064-INS-2484
chr17-34492742-INS-253
chr17-34494968-INS-6176
chr17-34498728-INS-6252
chr17-34499901-INS-72
chr17-34503797-INS-92
chr17-34506168-INS-3490
chr17-34517946-INS-100
chr17-34518612-INS-73
chr17-34549558-INS-158
chr17-34549678-INS-333
chr17-34572514-INS-60
chr17-34572854-INS-140
chr17-34591519-INS-127
chr17-34599080-INS-50
chr17-34615067-INS-54
chr17-34615435-INS-50
chr17-34626455-INS-578
chr17-34645163-INS-995
chr17-34669256-INS-185
chr17-34669775-INS-71
chr17-34697808-INS-286

chr17-34720789-INS-343
chr17-34721312-INS-212
chr17-34722518-INS-211
chr17-34726025-INS-74
chr17-34740390-INS-60
chr17-34740876-INS-1766
chr17-34754065-INS-60
chr17-34777569-INS-183
chr17-34952745-INS-467
chr17-35024474-INS-1215
chr17-35027013-INS-533
chr17-35046162-INS-7732
chr17-35055693-INS-95
chr17-35301333-INS-70
chr17-35311114-INS-113
chr17-35328400-INS-204
chr17-35416510-INS-55
chr17-35474643-INS-5813
chr17-35483696-INS-52
chr17-35488812-INS-54
chr17-35490865-INS-279
chr17-35494819-INS-72
chr17-35496750-INS-3525
chr17-35504000-INS-155
chr17-35533670-INS-388
chr17-35540201-INS-123
chr17-35621332-INS-89
chr17-35648316-INS-319
chr17-35755725-INS-200
chr17-35767315-INS-1922
chr17-35773445-INS-1880
chr17-35778774-INS-64
chr17-35782778-INS-241
chr17-35798108-INS-162
chr17-35799132-INS-264
chr17-35804057-INS-56
chr17-35863765-INS-65
chr17-35898601-INS-161
chr17-35899384-INS-50
chr17-35905270-INS-169
chr17-35909759-INS-51
chr17-35915057-INS-7096
chr17-35923406-INS-58
chr17-35930787-INS-135
chr17-35931530-INS-315
chr17-35933868-INS-90
chr17-35935562-INS-195
chr17-35946647-INS-216
chr17-35989522-INS-217
chr17-36020482-INS-159
chr17-36151419-INS-207
chr17-36167428-INS-227
chr17-36244092-INS-206
chr17-36309592-INS-108
chr17-36332690-INS-2665
chr17-36337838-INS-186
chr17-36358551-INS-218
chr17-36426587-INS-203
chr17-36427135-INS-86
chr17-36427448-INS-4647
chr17-36433623-INS-145
chr17-36433624-INS-55
chr17-36434111-INS-193
chr17-36445626-INS-72
chr17-36478680-INS-190
chr17-36494065-INS-60
chr17-36511031-INS-133
chr17-36511629-INS-160
chr17-36511790-INS-310
chr17-36527624-INS-582
chr17-36539459-INS-52
chr17-36541015-INS-7118
chr17-36544824-INS-56
chr17-36553948-INS-205
chr17-36610078-INS-2939
chr17-36671177-INS-29533
chr17-36707876-INS-8975

chr17-36733389-INS-348
chr17-36812567-INS-10350
chr17-36856025-INS-6245
chr17-36863609-INS-108
chr17-36874529-INS-1365
chr17-36902153-INS-2480
chr17-36907742-INS-1152
chr17-36917500-INS-510
chr17-36956082-INS-6362
chr17-36989404-INS-6867
chr17-37035994-INS-4760
chr17-37037651-INS-677
chr17-37048276-INS-60
chr17-37053094-INS-669
chr17-37054651-INS-1314
chr17-37105202-INS-106
chr17-37118526-INS-92
chr17-37140547-INS-6292
chr17-37143212-INS-65
chr17-37200668-INS-54
chr17-37205279-INS-5280
chr17-37205517-INS-55
chr17-37211272-INS-50
chr17-37282942-INS-50
chr17-37294152-INS-2878
chr17-37301496-INS-87
chr17-37342604-INS-256
chr17-37367512-INS-59
chr17-37367574-INS-210
chr17-37393890-INS-100
chr17-37406944-INS-7187
chr17-37433485-INS-229
chr17-37452777-INS-971
chr17-37452927-INS-180
chr17-37459291-INS-2810
chr17-37461800-INS-51
chr17-37625608-INS-11357
chr17-37660965-INS-623
chr17-37684213-INS-167
chr17-37738236-INS-991
chr17-37783592-INS-204
chr17-37832264-INS-50
chr17-38045219-INS-2843
chr17-38164581-INS-831
chr17-38219787-INS-200
chr17-38228918-INS-1568
chr17-38281995-INS-50
chr17-38451152-INS-336
chr17-38491288-INS-14457
chr17-38542005-INS-737
chr17-38799406-INS-2159
chr17-38808283-INS-86
chr17-38816150-INS-74
chr17-38819188-INS-87
chr17-38824987-INS-78
chr17-38859918-INS-1749
chr17-38861193-INS-100
chr17-38874434-INS-52
chr17-38876650-INS-62
chr17-38884433-INS-194
chr17-38890923-INS-2997
chr17-38896058-INS-961
chr17-38904941-INS-252
chr17-38905418-INS-6657
chr17-38907343-INS-58
chr17-39047614-INS-62
chr17-39181691-INS-217
chr17-39242455-INS-6830
chr17-39465693-INS-80
chr17-39512876-INS-414
chr17-39577585-INS-365
chr17-39646590-INS-214
chr17-39663715-INS-508
chr17-39760654-INS-623
chr17-39879665-INS-1999
chr17-39947494-INS-4492
chr17-39959389-INS-102

chr17-40025728-INS-3148
chr17-40031872-INS-513
chr17-40079943-INS-214
chr17-40126763-INS-549
chr17-40171681-INS-52
chr17-40225131-INS-52
chr17-40264944-INS-1134
chr17-40390939-INS-463
chr17-40417536-INS-5497
chr17-40485823-INS-7202
chr17-40492097-INS-7129
chr17-40528676-INS-208
chr17-40588745-INS-68041
chr17-40604147-INS-51
chr17-40604205-INS-64
chr17-40613944-INS-60
chr17-40614263-INS-878
chr17-40614286-INS-52
chr17-40691988-INS-143
chr17-40732849-INS-3572
chr17-40747421-INS-60
chr17-40814694-INS-70
chr17-40991455-INS-154
chr17-40991828-INS-69
chr17-41020527-INS-135
chr17-41020708-INS-649
chr17-41097023-INS-703
chr17-41108522-INS-1181
chr17-41204853-INS-53
chr17-41205674-INS-50
chr17-41206010-INS-1889
chr17-41206317-INS-59
chr17-41213824-INS-6171
chr17-41218581-INS-75
chr17-41225324-INS-4597
chr17-41226307-INS-95
chr17-41316643-INS-51
chr17-41384622-INS-3715
chr17-41415325-INS-1951
chr17-41418786-INS-7635
chr17-41445614-INS-6285
chr17-41472898-INS-1702
chr17-41473180-INS-54
chr17-41499293-INS-6645
chr17-41508703-INS-86
chr17-41524707-INS-7482
chr17-41526280-INS-1081
chr17-41566802-INS-351
chr17-41570024-INS-7082
chr17-41677790-INS-2097
chr17-41697139-INS-480
chr17-41710744-INS-50
chr17-41859857-INS-1521
chr17-41924012-INS-121
chr17-41929252-INS-1158
chr17-41954884-INS-223
chr17-41991364-INS-50
chr17-42015299-INS-120
chr17-42032194-INS-471
chr17-42151103-INS-5729
chr17-42164151-INS-186
chr17-42171769-INS-977
chr17-42172251-INS-2285
chr17-42213872-INS-3210
chr17-42270180-INS-2027
chr17-42276468-INS-6492
chr17-42317884-INS-208
chr17-42325787-INS-201
chr17-42334850-INS-195
chr17-42348911-INS-88
chr17-42349496-INS-129
chr17-42352021-INS-705
chr17-42389745-INS-1785
chr17-42392882-INS-58
chr17-42438474-INS-94
chr17-42504372-INS-72
chr17-42505594-INS-2406

chr17-42620234-INS-51
chr17-42628680-INS-64
chr17-42639208-INS-50
chr17-42648209-INS-114
chr17-42662347-INS-246
chr17-42714578-INS-225
chr17-42732637-INS-107
chr17-42765886-INS-52
chr17-42781990-INS-2465
chr17-42791871-INS-72
chr17-42793901-INS-7511
chr17-42816215-INS-88
chr17-42822776-INS-527
chr17-42833688-INS-51
chr17-42844805-INS-72
chr17-42861986-INS-57
chr17-42872489-INS-50
chr17-42915182-INS-285
chr17-42915200-INS-58
chr17-42915530-INS-210
chr17-42917483-INS-73
chr17-42942848-INS-7086
chr17-42947890-INS-1013
chr17-42984520-INS-101
chr17-42999805-INS-1125
chr17-42999810-INS-78
chr17-43028722-INS-756
chr17-43054386-INS-1229
chr17-43055230-INS-96
chr17-43055327-INS-2566
chr17-43152097-INS-412
chr17-43153822-INS-207
chr17-43155015-INS-212
chr17-43163807-INS-3190
chr17-43167135-INS-91
chr17-43191026-INS-1476
chr17-43216632-INS-59
chr17-43216656-INS-950
chr17-43216775-INS-436
chr17-43242210-INS-50
chr17-43254892-INS-60
chr17-43265557-INS-4373
chr17-43266610-INS-53
chr17-43289105-INS-754
chr17-43289917-INS-197
chr17-43370853-INS-104
chr17-43379614-INS-105
chr17-43522514-INS-266
chr17-43522650-INS-117
chr17-43534609-INS-102
chr17-43536577-INS-50
chr17-43549467-INS-3580
chr17-43626544-INS-63
chr17-43630182-INS-150
chr17-43640969-INS-79
chr17-43642830-INS-756
chr17-43647013-INS-2879
chr17-43648911-INS-53
chr17-43651720-INS-86
chr17-43653265-INS-7421
chr17-43667571-INS-6037
chr17-43668816-INS-96
chr17-43685207-INS-4729
chr17-43703610-INS-53
chr17-43724841-INS-68
chr17-43763553-INS-75
chr17-43764620-INS-162
chr17-43785149-INS-50
chr17-43794063-INS-155
chr17-43796608-INS-61
chr17-43804789-INS-378
chr17-43811803-INS-82
chr17-43815097-INS-51
chr17-43819753-INS-1301
chr17-43830639-INS-549
chr17-43831421-INS-52
chr17-43833650-INS-54

chr17-43840706-INS-200
chr17-43856479-INS-179
chr17-43865232-INS-4055
chr17-43889386-INS-231
chr17-43889587-INS-172
chr17-43890659-INS-3124
chr17-43912258-INS-58
chr17-43921346-INS-276
chr17-44008220-INS-76
chr17-44008232-INS-480
chr17-44015330-INS-240
chr17-44015860-INS-62
chr17-44066527-INS-966
chr17-44066689-INS-126
chr17-44066893-INS-364
chr17-44066898-INS-126
chr17-44096929-INS-72
chr17-44101230-INS-121
chr17-44101461-INS-132
chr17-44103502-INS-1068
chr17-44107961-INS-1917
chr17-44136340-INS-109
chr17-44138670-INS-55
chr17-44141528-INS-117
chr17-44142145-INS-232
chr17-44168974-INS-54
chr17-44177383-INS-84
chr17-44185489-INS-388
chr17-44185498-INS-105
chr17-44192727-INS-6293
chr17-44193079-INS-80
chr17-44210237-INS-5881
chr17-44232878-INS-124
chr17-44234683-INS-1339
chr17-44265321-INS-54
chr17-44267311-INS-54
chr17-44315369-INS-6378
chr17-44366269-INS-1124
chr17-44375118-INS-955
chr17-44418012-INS-2818
chr17-44418363-INS-72
chr17-44418693-INS-159
chr17-44504993-INS-58
chr17-44518805-INS-94
chr17-44531573-INS-91
chr17-44560184-INS-1917
chr17-44563473-INS-435
chr17-44563492-INS-1232
chr17-44563804-INS-113
chr17-44564756-INS-116
chr17-44568362-INS-1000
chr17-44631591-INS-208
chr17-44631672-INS-62
chr17-44631843-INS-523
chr17-44661722-INS-136
chr17-44668582-INS-7224
chr17-44696839-INS-204
chr17-44765645-INS-207
chr17-44778843-INS-143
chr17-44778869-INS-59
chr17-44792995-INS-6936
chr17-44801583-INS-53
chr17-44820337-INS-52
chr17-44860489-INS-108
chr17-44901828-INS-3424
chr17-44906876-INS-82
chr17-44913308-INS-189
chr17-44913739-INS-210
chr17-44951674-INS-51
chr17-45003184-INS-72
chr17-45004795-INS-57
chr17-45058665-INS-1500
chr17-45073509-INS-2084
chr17-45091611-INS-1221
chr17-45148913-INS-53
chr17-45228890-INS-326
chr17-45229294-INS-438

chr17-45257724-INS-1457
chr17-45345800-INS-86
chr17-45420859-INS-207
chr17-45434830-INS-627
chr17-45438062-INS-206
chr17-45453843-INS-1099
chr17-45483302-INS-2050
chr17-45524636-INS-340
chr17-45534814-INS-110
chr17-45539090-INS-368
chr17-45542342-INS-56
chr17-45548798-INS-174
chr17-45559735-INS-443
chr17-45559735-INS-68
chr17-45559921-INS-117
chr17-45594979-INS-63
chr17-45676776-INS-198
chr17-45778260-INS-310
chr17-45782691-INS-66
chr17-45790262-INS-214
chr17-45791185-INS-200
chr17-45794130-INS-200
chr17-45795820-INS-202
chr17-45798878-INS-807
chr17-45816034-INS-73
chr17-45831665-INS-71
chr17-45837903-INS-212
chr17-45870429-INS-50
chr17-45891726-INS-78
chr17-45898255-INS-227
chr17-45913633-INS-50
chr17-45914825-INS-217
chr17-45932188-INS-204
chr17-45940686-INS-354
chr17-45954025-INS-58
chr17-45954860-INS-133
chr17-45983109-INS-52
chr17-46019847-INS-73
chr17-46050786-INS-54
chr17-46082032-INS-157
chr17-46088888-INS-117
chr17-46088940-INS-53
chr17-46110393-INS-156
chr17-46110419-INS-68
chr17-46120006-INS-464
chr17-46131771-INS-56
chr17-46194795-INS-204
chr17-46194795-INS-534
chr17-46204994-INS-54
chr17-46252968-INS-103
chr17-46261364-INS-121
chr17-46273898-INS-341
chr17-46300572-INS-270
chr17-46323013-INS-464
chr17-46353719-INS-208
chr17-46355495-INS-85
chr17-46367010-INS-52
chr17-46406322-INS-2899
chr17-46443974-INS-487
chr17-46444146-INS-84
chr17-46448063-INS-206
chr17-46487915-INS-69
chr17-46501406-INS-1092
chr17-46516185-INS-50
chr17-46546294-INS-164
chr17-46586151-INS-212
chr17-46601330-INS-84
chr17-46611603-INS-114
chr17-46648911-INS-706
chr17-46670532-INS-53
chr17-46706054-INS-68
chr17-46789160-INS-171
chr17-46803006-INS-60
chr17-46840548-INS-213
chr17-46840679-INS-199
chr17-46879139-INS-52
chr17-46894603-INS-71

chr17-46927761-INS-65
chr17-46949472-INS-167
chr17-47001598-INS-161
chr17-47043247-INS-219
chr17-47072314-INS-198
chr17-47073094-INS-205
chr17-47096753-INS-209
chr17-47103231-INS-79
chr17-47130843-INS-275
chr17-47145977-INS-1150
chr17-47156217-INS-196
chr17-47179835-INS-76
chr17-47189965-INS-52
chr17-47190315-INS-190
chr17-47190315-INS-64
chr17-47191928-INS-112
chr17-47195387-INS-125
chr17-47210882-INS-161
chr17-47211094-INS-55
chr17-47216501-INS-106
chr17-47226573-INS-82
chr17-47256715-INS-215
chr17-47363175-INS-1909
chr17-47471227-INS-233
chr17-47471227-INS-489
chr17-47509598-INS-208
chr17-47524555-INS-236
chr17-47603958-INS-8740
chr17-47616131-INS-51
chr17-47624037-INS-1104
chr17-47640403-INS-93
chr17-47640478-INS-214
chr17-47640789-INS-57
chr17-47650215-INS-183
chr17-47695462-INS-77
chr17-47732158-INS-58
chr17-47755712-INS-204
chr17-47792955-INS-88
chr17-47820768-INS-60
chr17-47849073-INS-64
chr17-47855190-INS-216
chr17-47866255-INS-220
chr17-47872170-INS-115
chr17-47876150-INS-186
chr17-47885593-INS-120
chr17-47914095-INS-207
chr17-47939948-INS-136
chr17-47953443-INS-63
chr17-47957563-INS-180
chr17-48109623-INS-57
chr17-48112840-INS-59
chr17-48117080-INS-240
chr17-48129821-INS-50
chr17-48137478-INS-259
chr17-48142507-INS-50
chr17-48160540-INS-1196
chr17-48179845-INS-102
chr17-48182648-INS-220
chr17-48204200-INS-86
chr17-48213030-INS-54
chr17-48258853-INS-64
chr17-48268231-INS-85
chr17-48293782-INS-50
chr17-48311732-INS-52
chr17-48316548-INS-50
chr17-48364302-INS-286
chr17-48364302-INS-52
chr17-48372621-INS-84
chr17-48414404-INS-59
chr17-48435120-INS-93
chr17-48443106-INS-137
chr17-48463961-INS-4599
chr17-48503348-INS-91
chr17-48527929-INS-77
chr17-48555880-INS-263
chr17-48556175-INS-59
chr17-48559770-INS-293

chr17-48560420-INS-380
chr17-48562975-INS-200
chr17-48600392-INS-66
chr17-48604353-INS-776
chr17-48604486-INS-127
chr17-48626626-INS-101
chr17-48663692-INS-135
chr17-48664589-INS-122
chr17-48677246-INS-8420
chr17-48679126-INS-1756
chr17-48683620-INS-266
chr17-48712038-INS-211
chr17-48733607-INS-52
chr17-48737864-INS-70
chr17-48752099-INS-240
chr17-48752742-INS-403
chr17-48759865-INS-200
chr17-48760049-INS-77
chr17-48792671-INS-968
chr17-48800697-INS-1399
chr17-48809379-INS-279
chr17-48822481-INS-162
chr17-48822776-INS-72
chr17-48827958-INS-7061
chr17-48830822-INS-2052
chr17-48843657-INS-196
chr17-48847124-INS-205
chr17-48895257-INS-124
chr17-49026617-INS-50
chr17-49043209-INS-50
chr17-49046516-INS-4496
chr17-49059235-INS-52
chr17-49073631-INS-2791
chr17-49079929-INS-84
chr17-49107276-INS-91
chr17-49170589-INS-105
chr17-49203382-INS-146
chr17-49308088-INS-108
chr17-49320111-INS-73
chr17-49375517-INS-62
chr17-49382592-INS-116
chr17-49422437-INS-121
chr17-49491864-INS-3288
chr17-49521773-INS-51
chr17-49551508-INS-298
chr17-49560308-INS-210
chr17-49570920-INS-50
chr17-49575736-INS-102
chr17-49580471-INS-62
chr17-49585077-INS-54
chr17-49585117-INS-154
chr17-49588888-INS-111
chr17-49593073-INS-63
chr17-49683315-INS-100
chr17-49689950-INS-368
chr17-49694036-INS-92
chr17-49718692-INS-159
chr17-49758563-INS-134
chr17-49834234-INS-416
chr17-49852165-INS-649
chr17-49867928-INS-998
chr17-49924434-INS-67
chr17-49971830-INS-50
chr17-50071792-INS-328
chr17-50125273-INS-128
chr17-50161270-INS-363
chr17-50198000-INS-96
chr17-50228712-INS-132
chr17-50249031-INS-109
chr17-50252818-INS-7162
chr17-50350957-INS-198
chr17-50394268-INS-154
chr17-50395607-INS-175
chr17-50401052-INS-72
chr17-50417503-INS-125
chr17-50439807-INS-58
chr17-50448764-INS-162

chr17-50451264-INS-524
chr17-50467324-INS-58
chr17-50482866-INS-1175
chr17-50514837-INS-88
chr17-50534346-INS-310
chr17-50536205-INS-94
chr17-50539320-INS-117
chr17-50540707-INS-2647
chr17-50541535-INS-402
chr17-50621769-INS-62
chr17-50635051-INS-575
chr17-50701929-INS-164
chr17-50706288-INS-54
chr17-50707628-INS-1008
chr17-50743883-INS-67
chr17-50769195-INS-383
chr17-50791359-INS-60
chr17-50836333-INS-406
chr17-50876877-INS-246
chr17-50883221-INS-3470
chr17-50891198-INS-539
chr17-50903743-INS-104
chr17-50911175-INS-149
chr17-50938666-INS-50
chr17-50942346-INS-1074
chr17-50945009-INS-6498
chr17-50960285-INS-82
chr17-50960318-INS-345
chr17-50960576-INS-271
chr17-51257942-INS-4501
chr17-51322059-INS-674
chr17-51373025-INS-7576
chr17-51375650-INS-375
chr17-51375925-INS-150
chr17-51396765-INS-72
chr17-51406388-INS-7254
chr17-51466161-INS-6825
chr17-51513922-INS-60
chr17-51536865-INS-6513
chr17-51556194-INS-2947
chr17-51651995-INS-99
chr17-51654052-INS-55
chr17-51737968-INS-72
chr17-51803887-INS-6517
chr17-51818861-INS-50
chr17-51860841-INS-53
chr17-51888128-INS-52
chr17-51888319-INS-683
chr17-51930282-INS-52
chr17-51934079-INS-2910
chr17-51938741-INS-113
chr17-51940567-INS-1764
chr17-51944743-INS-60
chr17-51951013-INS-50
chr17-51970044-INS-200
chr17-52021118-INS-145
chr17-52021649-INS-132
chr17-52021721-INS-414
chr17-52023341-INS-7602
chr17-52196466-INS-4677
chr17-52262543-INS-119
chr17-52264683-INS-189
chr17-52289316-INS-138
chr17-52336297-INS-69
chr17-52340925-INS-297
chr17-52363155-INS-156
chr17-52396184-INS-66
chr17-52417689-INS-66
chr17-52430583-INS-58
chr17-52430689-INS-149
chr17-52431875-INS-1247
chr17-52449902-INS-1971
chr17-52469028-INS-54
chr17-52513616-INS-6440
chr17-52537131-INS-226
chr17-52570248-INS-2342
chr17-52708034-INS-758

chr17-52722001-INS-307
chr17-52722424-INS-76
chr17-52740177-INS-50
chr17-52765479-INS-75
chr17-52774343-INS-122
chr17-52808617-INS-1891
chr17-52819455-INS-145
chr17-52823924-INS-1779
chr17-52823924-INS-6263
chr17-52894010-INS-696
chr17-52895874-INS-100
chr17-52902983-INS-63
chr17-52911755-INS-70
chr17-52937377-INS-94
chr17-52952428-INS-59
chr17-52990442-INS-125
chr17-52990470-INS-264
chr17-53001483-INS-60
chr17-53038545-INS-610
chr17-53152254-INS-6531
chr17-53177873-INS-6255
chr17-53192114-INS-134
chr17-53251891-INS-51
chr17-53264276-INS-60
chr17-53273014-INS-55
chr17-53307314-INS-951
chr17-53387288-INS-60
chr17-53393771-INS-89
chr17-53400307-INS-54
chr17-53443035-INS-7510
chr17-53452123-INS-7003
chr17-53457448-INS-1978
chr17-53472246-INS-3687
chr17-53473307-INS-439
chr17-53495121-INS-1429
chr17-53497046-INS-4748
chr17-53502736-INS-65
chr17-53528294-INS-4721
chr17-53533797-INS-10291
chr17-53540328-INS-5509
chr17-53549432-INS-2606
chr17-53549711-INS-195
chr17-53549711-INS-400
chr17-53553887-INS-757
chr17-53554669-INS-92
chr17-53562794-INS-4916
chr17-53602227-INS-163
chr17-53616869-INS-126
chr17-53631434-INS-50
chr17-53639325-INS-5510
chr17-53654428-INS-168
chr17-53654550-INS-50
chr17-53714096-INS-185
chr17-53757043-INS-160
chr17-53761869-INS-235
chr17-53775427-INS-8793
chr17-53782679-INS-68
chr17-53783478-INS-53
chr17-53812900-INS-193
chr17-53813278-INS-202
chr17-53842956-INS-196
chr17-53847698-INS-95
chr17-53860627-INS-1205
chr17-53865452-INS-56
chr17-53865701-INS-294
chr17-53868138-INS-205
chr17-53870016-INS-301
chr17-53871539-INS-293
chr17-53877976-INS-417
chr17-53883412-INS-230
chr17-53887403-INS-278
chr17-53888174-INS-200
chr17-53896576-INS-7211
chr17-53904937-INS-81
chr17-53905044-INS-232
chr17-53905157-INS-510
chr17-53906855-INS-63

chr17-53930935-INS-120
chr17-53947349-INS-3080
chr17-53947352-INS-194
chr17-53956861-INS-135
chr17-53964815-INS-762
chr17-53999021-INS-615
chr17-54005626-INS-434
chr17-54008474-INS-790
chr17-54008475-INS-253
chr17-54009997-INS-6191
chr17-54014410-INS-1358
chr17-54034833-INS-2500
chr17-54094403-INS-2907
chr17-54096744-INS-4739
chr17-54150022-INS-87
chr17-54154581-INS-56
chr17-54163056-INS-102
chr17-54171464-INS-147
chr17-54191136-INS-75
chr17-54208270-INS-156
chr17-54218096-INS-3329
chr17-54234265-INS-448
chr17-54234288-INS-116
chr17-54243423-INS-212
chr17-54288236-INS-71
chr17-54341325-INS-3871
chr17-54341376-INS-354
chr17-54345764-INS-1160
chr17-54363165-INS-4086
chr17-54387106-INS-5059
chr17-54482888-INS-1966
chr17-54505048-INS-1580
chr17-54509896-INS-128
chr17-54523858-INS-5601
chr17-54528433-INS-120
chr17-54534732-INS-132
chr17-54547479-INS-95
chr17-54548605-INS-118
chr17-54604734-INS-52
chr17-54625374-INS-76
chr17-54634452-INS-1082
chr17-54639869-INS-1051
chr17-54649346-INS-66
chr17-54672436-INS-2327
chr17-54690333-INS-1430
chr17-54744460-INS-52
chr17-54751990-INS-178
chr17-54782436-INS-75
chr17-54789418-INS-2324
chr17-54801720-INS-2663
chr17-54801792-INS-1808
chr17-54803158-INS-8675
chr17-54820259-INS-477
chr17-54820358-INS-214
chr17-54820450-INS-106
chr17-54827242-INS-6013
chr17-54828814-INS-70
chr17-54831907-INS-2116
chr17-54837125-INS-6453
chr17-54861123-INS-142
chr17-54872736-INS-60
chr17-54886897-INS-54
chr17-54911332-INS-120
chr17-54911771-INS-499
chr17-54914520-INS-2313
chr17-54916823-INS-333
chr17-54919915-INS-6400
chr17-54930764-INS-1898
chr17-54965681-INS-126
chr17-54966812-INS-74
chr17-54968161-INS-2662
chr17-54987838-INS-6199
chr17-55007144-INS-519
chr17-55018594-INS-110
chr17-55046591-INS-6440
chr17-55064753-INS-1135
chr17-55068114-INS-9876

chr17-55074226-INS-4597
chr17-55108896-INS-171
chr17-55115929-INS-5381
chr17-55118563-INS-1429
chr17-55123297-INS-3495
chr17-55124260-INS-175
chr17-55142010-INS-469
chr17-55142160-INS-943
chr17-55146259-INS-75
chr17-55150167-INS-52
chr17-55188178-INS-3313
chr17-55243024-INS-1963
chr17-55267620-INS-581
chr17-55272556-INS-83
chr17-55278758-INS-536
chr17-55292124-INS-189
chr17-55325422-INS-680
chr17-55346818-INS-1873
chr17-55481158-INS-416
chr17-55488218-INS-291
chr17-55523131-INS-5348
chr17-55545315-INS-1750
chr17-55556597-INS-68
chr17-55564969-INS-171
chr17-55581425-INS-1008
chr17-55617765-INS-4165
chr17-55650354-INS-1031
chr17-55659787-INS-3502
chr17-55665338-INS-5370
chr17-55727808-INS-2674
chr17-55730736-INS-6575
chr17-55777447-INS-50
chr17-55783699-INS-144
chr17-55822293-INS-103
chr17-55906984-INS-955
chr17-55941534-INS-6421
chr17-55949064-INS-1169
chr17-55979658-INS-58
chr17-55997720-INS-6284
chr17-56041185-INS-568
chr17-56082132-INS-4231
chr17-56103662-INS-116
chr17-56146187-INS-479
chr17-56195770-INS-147
chr17-56228375-INS-184
chr17-56228925-INS-492
chr17-56239279-INS-118
chr17-56273044-INS-120
chr17-56282573-INS-102
chr17-56308018-INS-50
chr17-56348420-INS-138
chr17-56413589-INS-99
chr17-56474159-INS-331
chr17-56543569-INS-91
chr17-56562834-INS-207
chr17-56630813-INS-206
chr17-56741999-INS-6033
chr17-56774383-INS-357
chr17-56787876-INS-213
chr17-56791462-INS-588
chr17-56831344-INS-121
chr17-56874171-INS-999
chr17-56887031-INS-866
chr17-56953402-INS-202
chr17-56976625-INS-73
chr17-57032798-INS-343
chr17-57101816-INS-61
chr17-57124918-INS-216
chr17-57163565-INS-50
chr17-57167523-INS-214
chr17-57223547-INS-56
chr17-57235390-INS-321
chr17-57238644-INS-79
chr17-57249026-INS-218
chr17-57266896-INS-110
chr17-57267503-INS-124
chr17-57269027-INS-171

chr17-57324352-INS-343
chr17-57357456-INS-168
chr17-57382177-INS-267
chr17-57437492-INS-61
chr17-57452660-INS-69
chr17-57463977-INS-1292
chr17-57487325-INS-113
chr17-57507499-INS-259
chr17-57613946-INS-66
chr17-57623486-INS-348
chr17-57626356-INS-141
chr17-57634042-INS-59
chr17-57635608-INS-152
chr17-57643831-INS-201
chr17-57643880-INS-58
chr17-57671362-INS-817
chr17-57726517-INS-321
chr17-57784539-INS-199
chr17-57805281-INS-209
chr17-57817856-INS-7021
chr17-57823078-INS-101
chr17-57841660-INS-10113
chr17-57861374-INS-7099
chr17-57929216-INS-704
chr17-57994079-INS-200
chr17-58021560-INS-573
chr17-58037389-INS-1141
chr17-58051700-INS-74
chr17-58056391-INS-243
chr17-58067163-INS-102
chr17-58101327-INS-177
chr17-58109756-INS-6365
chr17-58110063-INS-96
chr17-58136593-INS-1554
chr17-58139059-INS-56
chr17-58156801-INS-2191
chr17-58184248-INS-90
chr17-58217976-INS-580
chr17-58227761-INS-72
chr17-58237703-INS-175
chr17-58254357-INS-200
chr17-58262944-INS-533
chr17-58283396-INS-80
chr17-58295531-INS-153
chr17-58307146-INS-202
chr17-58311446-INS-838
chr17-58345278-INS-72
chr17-58350357-INS-191
chr17-58350498-INS-50
chr17-58365483-INS-6683
chr17-58381033-INS-710
chr17-58422041-INS-1409
chr17-58426367-INS-50
chr17-58432240-INS-50
chr17-58432972-INS-1912
chr17-58448256-INS-1218
chr17-58454339-INS-6200
chr17-58471556-INS-76
chr17-58471580-INS-216
chr17-58475178-INS-60
chr17-58578824-INS-201
chr17-58580115-INS-67
chr17-58653704-INS-2987
chr17-58658711-INS-1908
chr17-58664691-INS-72
chr17-58665620-INS-69
chr17-58677387-INS-786
chr17-58682597-INS-336
chr17-58685462-INS-55
chr17-58732074-INS-81
chr17-58744920-INS-1152
chr17-58768697-INS-1296
chr17-58769951-INS-6517
chr17-58806245-INS-6417
chr17-58810886-INS-4371
chr17-58816634-INS-54
chr17-58859714-INS-1787

chr17-58906429-INS-70
chr17-58906912-INS-105
chr17-58916269-INS-2070
chr17-58923297-INS-56
chr17-58924083-INS-79
chr17-58934560-INS-417
chr17-58937808-INS-486
chr17-58942370-INS-90
chr17-58942504-INS-1779
chr17-58944928-INS-1579
chr17-58949699-INS-931
chr17-58956872-INS-7658
chr17-58962230-INS-731
chr17-58978207-INS-1115
chr17-58978319-INS-365
chr17-58981148-INS-60
chr17-58993638-INS-1586
chr17-58994066-INS-130
chr17-59006422-INS-4435
chr17-59022730-INS-202
chr17-59031706-INS-58
chr17-59050835-INS-1570
chr17-59053208-INS-1590
chr17-59079657-INS-158
chr17-59082292-INS-995
chr17-59083043-INS-923
chr17-59084691-INS-7504
chr17-59087104-INS-347
chr17-59089016-INS-1726
chr17-59112383-INS-1160
chr17-59116397-INS-687
chr17-59119822-INS-224
chr17-59119822-INS-52
chr17-59131337-INS-74
chr17-59133573-INS-336
chr17-59139162-INS-276
chr17-59159790-INS-154
chr17-59217750-INS-124
chr17-59223595-INS-52
chr17-59225200-INS-336
chr17-59233863-INS-62
chr17-59233937-INS-298
chr17-59245812-INS-7158
chr17-59276780-INS-904
chr17-59330906-INS-120
chr17-59343568-INS-121
chr17-59353160-INS-416
chr17-59358203-INS-1132
chr17-59364092-INS-120
chr17-59372035-INS-73
chr17-59380621-INS-185
chr17-59401046-INS-60
chr17-59416019-INS-7230
chr17-59429567-INS-1768
chr17-59508690-INS-370
chr17-59567738-INS-412
chr17-59611174-INS-221
chr17-59662620-INS-59
chr17-59674997-INS-2489
chr17-59698986-INS-50
chr17-59793204-INS-452
chr17-59816838-INS-60
chr17-59825413-INS-57
chr17-59866614-INS-299
chr17-59950850-INS-4425
chr17-59956866-INS-72
chr17-59957462-INS-60
chr17-59964057-INS-6199
chr17-59964659-INS-59
chr17-59965440-INS-108
chr17-59975731-INS-130
chr17-60007363-INS-2455
chr17-60020368-INS-694
chr17-60021636-INS-720
chr17-60026292-INS-380
chr17-60028929-INS-59
chr17-60035697-INS-2036

chr17-60040166-INS-50
chr17-60041922-INS-6257
chr17-60066203-INS-6866
chr17-60074578-INS-142
chr17-60076471-INS-50
chr17-60088535-INS-85
chr17-60094052-INS-298
chr17-60102677-INS-1078
chr17-60113577-INS-1893
chr17-60181877-INS-69
chr17-60209032-INS-82
chr17-60221446-INS-52
chr17-60223006-INS-6209
chr17-60241993-INS-486
chr17-60306490-INS-6193
chr17-60375491-INS-204
chr17-60387762-INS-124
chr17-60392981-INS-144
chr17-60424794-INS-974
chr17-60434411-INS-78
chr17-60445899-INS-56
chr17-60454026-INS-1889
chr17-60475557-INS-140
chr17-60476124-INS-941
chr17-60477701-INS-64
chr17-60513477-INS-1028
chr17-60540869-INS-70
chr17-60559532-INS-1465
chr17-60566293-INS-60
chr17-60633318-INS-2854
chr17-60662125-INS-1718
chr17-60688160-INS-53
chr17-60690264-INS-506
chr17-60710836-INS-53
chr17-60728105-INS-431
chr17-60743546-INS-486
chr17-60794119-INS-7058
chr17-60805567-INS-6401
chr17-60821567-INS-426
chr17-60822114-INS-114
chr17-60838056-INS-1135
chr17-60865865-INS-3129
chr17-60974414-INS-280
chr17-60975160-INS-361
chr17-60981575-INS-2713
chr17-61010627-INS-55
chr17-61061988-INS-96
chr17-61069771-INS-646
chr17-61086048-INS-100
chr17-61095022-INS-2178
chr17-61121386-INS-6364
chr17-61135023-INS-210
chr17-61152488-INS-1848
chr17-61152575-INS-206
chr17-61152648-INS-89
chr17-61155672-INS-1171
chr17-61177529-INS-6606
chr17-61193988-INS-2221
chr17-61212295-INS-691
chr17-61213635-INS-1528
chr17-61222106-INS-54
chr17-61294611-INS-14025
chr17-61302572-INS-3975
chr17-61309237-INS-6554
chr17-61317617-INS-167
chr17-61358481-INS-2872
chr17-61379679-INS-56
chr17-61387793-INS-52
chr17-61409125-INS-130
chr17-61421640-INS-204
chr17-61437579-INS-2138
chr17-61441694-INS-403
chr17-61458839-INS-61
chr17-61477166-INS-69
chr17-61585870-INS-13329
chr17-61585870-INS-6190
chr17-61613789-INS-3339

chr17-61616068-INS-435
chr17-61620786-INS-2908
chr17-61641545-INS-187
chr17-61676540-INS-1274
chr17-61723157-INS-204
chr17-61771821-INS-331
chr17-61791608-INS-52
chr17-61813468-INS-4807
chr17-61880254-INS-122
chr17-61936198-INS-90
chr17-61946737-INS-6234
chr17-61966378-INS-132
chr17-61966994-INS-1130
chr17-62032123-INS-6430
chr17-62045192-INS-5338
chr17-62052865-INS-1535
chr17-62064526-INS-1511
chr17-62109304-INS-2965
chr17-62147844-INS-6235
chr17-62161644-INS-52
chr17-62177757-INS-120
chr17-62183472-INS-64
chr17-62234800-INS-84
chr17-62240804-INS-7103
chr17-62305461-INS-125
chr17-62305631-INS-105
chr17-62320021-INS-64
chr17-62361935-INS-4711
chr17-62401478-INS-158
chr17-62491248-INS-52
chr17-62515928-INS-50
chr17-62519855-INS-96
chr17-62553454-INS-80
chr17-62568555-INS-96
chr17-62591375-INS-56
chr17-62608492-INS-70
chr17-62619964-INS-758
chr17-62620590-INS-1695
chr17-62621373-INS-63
chr17-62709715-INS-201
chr17-62713004-INS-92
chr17-62713224-INS-66
chr17-62716998-INS-4378
chr17-62725068-INS-50
chr17-62744290-INS-159
chr17-62757375-INS-219
chr17-62786764-INS-1637
chr17-62792817-INS-52
chr17-62798923-INS-196
chr17-62830678-INS-178
chr17-62830750-INS-56
chr17-62853169-INS-198
chr17-62862954-INS-2207
chr17-62872448-INS-288
chr17-62872587-INS-117
chr17-62878902-INS-56
chr17-62889846-INS-55
chr17-62903951-INS-216
chr17-62903951-INS-65
chr17-62933529-INS-52
chr17-62997210-INS-297
chr17-63014200-INS-709
chr17-63019776-INS-53
chr17-63113216-INS-209
chr17-63273716-INS-188
chr17-63284858-INS-52
chr17-63330113-INS-137
chr17-63343972-INS-112
chr17-63455845-INS-1342
chr17-63587335-INS-107
chr17-63587481-INS-2017
chr17-63601447-INS-68
chr17-63690545-INS-295
chr17-63720608-INS-1862
chr17-63720662-INS-812
chr17-63730328-INS-288
chr17-63735982-INS-184

chr17-63761238-INS-116
chr17-63788902-INS-206
chr17-63814566-INS-54
chr17-63819790-INS-177
chr17-63845088-INS-143
chr17-63870569-INS-68
chr17-63881185-INS-215
chr17-63889511-INS-96
chr17-63907600-INS-370
chr17-63907616-INS-74
chr17-63930588-INS-81
chr17-63936962-INS-1744
chr17-63947667-INS-451
chr17-63957786-INS-56
chr17-63962619-INS-65
chr17-63962679-INS-144
chr17-63994832-INS-1288
chr17-64027884-INS-1177
chr17-64034638-INS-345
chr17-64061025-INS-113
chr17-64067347-INS-2019
chr17-64084856-INS-6198
chr17-64085742-INS-1032
chr17-64107129-INS-7238
chr17-64119989-INS-272
chr17-64121306-INS-53
chr17-64142185-INS-90
chr17-64173525-INS-80
chr17-64232542-INS-76
chr17-64256699-INS-96
chr17-64354376-INS-505
chr17-64389569-INS-6235
chr17-64422385-INS-249
chr17-64430376-INS-66
chr17-64434788-INS-9394
chr17-64458406-INS-1515
chr17-64497161-INS-348
chr17-64498387-INS-6591
chr17-64552743-INS-50
chr17-64562236-INS-3212
chr17-64565855-INS-508
chr17-64689841-INS-273
chr17-64770574-INS-1305
chr17-64781981-INS-6716
chr17-64802049-INS-75
chr17-64813452-INS-1861
chr17-64814414-INS-52
chr17-64815313-INS-56
chr17-64835220-INS-2769
chr17-64850983-INS-62
chr17-64851647-INS-70
chr17-64895101-INS-560
chr17-64895360-INS-716
chr17-64923694-INS-320
chr17-64923715-INS-1445
chr17-64979465-INS-212
chr17-64979671-INS-306
chr17-65043559-INS-7245
chr17-65044156-INS-68
chr17-65086308-INS-295
chr17-65089065-INS-118
chr17-65089139-INS-50
chr17-65106376-INS-161
chr17-65110460-INS-74
chr17-65120605-INS-4161
chr17-65138989-INS-4876
chr17-65140731-INS-51
chr17-65160332-INS-2492
chr17-65171907-INS-50
chr17-65252960-INS-54
chr17-65255226-INS-52
chr17-65266480-INS-88
chr17-65291562-INS-1163
chr17-65334160-INS-51
chr17-65345459-INS-178
chr17-65350720-INS-249
chr17-65374740-INS-120

chr17-65430882-INS-6386
chr17-65504789-INS-50
chr17-65539377-INS-10071
chr17-65539377-INS-2654
chr17-65551493-INS-465
chr17-65585688-INS-122
chr17-65614645-INS-62
chr17-65629715-INS-3926
chr17-65636390-INS-50
chr17-65664688-INS-50
chr17-65684658-INS-6123
chr17-65696569-INS-6428
chr17-65697877-INS-56
chr17-65719159-INS-3223
chr17-65725256-INS-715
chr17-65731407-INS-88
chr17-65733882-INS-1916
chr17-65736721-INS-55
chr17-65766279-INS-2365
chr17-65793033-INS-72
chr17-65793063-INS-162
chr17-65801657-INS-184
chr17-65821483-INS-180
chr17-65833160-INS-510
chr17-65849010-INS-1751
chr17-65849093-INS-181
chr17-65849310-INS-92
chr17-65850128-INS-52
chr17-65869362-INS-382
chr17-65896294-INS-286
chr17-65899528-INS-50
chr17-65923527-INS-60
chr17-65957899-INS-73
chr17-65978208-INS-72
chr17-65980820-INS-77
chr17-65982697-INS-3054
chr17-66029331-INS-2737
chr17-66038017-INS-50
chr17-66054619-INS-217
chr17-66059382-INS-334
chr17-66063976-INS-140
chr17-66077774-INS-91
chr17-66118031-INS-372
chr17-66154460-INS-436
chr17-66163821-INS-167
chr17-66195122-INS-207
chr17-66196067-INS-391
chr17-66201687-INS-2292
chr17-66215785-INS-79
chr17-66216284-INS-234
chr17-66222210-INS-7184
chr17-66241075-INS-2811
chr17-66245251-INS-246
chr17-66253885-INS-2631
chr17-66269083-INS-52
chr17-66269978-INS-195
chr17-66284103-INS-180
chr17-66300075-INS-200
chr17-66335087-INS-193
chr17-66346477-INS-160
chr17-66378543-INS-1001
chr17-66387919-INS-82
chr17-66414252-INS-100
chr17-66414558-INS-1066
chr17-66426822-INS-51
chr17-66429388-INS-64
chr17-66449752-INS-195
chr17-66449845-INS-271
chr17-66449919-INS-420
chr17-66494102-INS-61
chr17-66516613-INS-102
chr17-66517575-INS-83
chr17-66538190-INS-514
chr17-66571157-INS-179
chr17-66581483-INS-203
chr17-66594624-INS-105
chr17-66620168-INS-56

chr17-66621428-INS-206
chr17-66636774-INS-303
chr17-66639418-INS-6464
chr17-66727382-INS-117
chr17-66735798-INS-124
chr17-66742832-INS-50
chr17-66748069-INS-119
chr17-66748900-INS-88
chr17-66761211-INS-87
chr17-66761253-INS-209
chr17-66772573-INS-415
chr17-66780742-INS-209
chr17-66792191-INS-8419
chr17-66795802-INS-93
chr17-66795966-INS-437
chr17-66797728-INS-60
chr17-66799923-INS-126
chr17-66823607-INS-173
chr17-66832059-INS-69
chr17-66840584-INS-208
chr17-66845088-INS-142
chr17-66857969-INS-116
chr17-66880151-INS-80
chr17-66902568-INS-382
chr17-66933449-INS-78
chr17-66933482-INS-187
chr17-66952006-INS-72
chr17-66973616-INS-3284
chr17-66985907-INS-89
chr17-67030364-INS-57
chr17-67038721-INS-52
chr17-67174551-INS-50
chr17-67253955-INS-102
chr17-67280557-INS-279
chr17-67289496-INS-51
chr17-67304502-INS-1273
chr17-67346972-INS-50
chr17-67358069-INS-103
chr17-67396059-INS-264
chr17-67419546-INS-289
chr17-67446607-INS-411
chr17-67469506-INS-1586
chr17-67475627-INS-85
chr17-67496430-INS-6489
chr17-67512540-INS-90
chr17-67522492-INS-80
chr17-67523023-INS-4167
chr17-67532501-INS-308
chr17-67532509-INS-68
chr17-67532767-INS-76
chr17-67555480-INS-78
chr17-67555728-INS-52
chr17-67564054-INS-1194
chr17-67618391-INS-191
chr17-67638596-INS-62
chr17-67638806-INS-328
chr17-67658510-INS-64
chr17-67664051-INS-56
chr17-67721940-INS-2913
chr17-67747467-INS-210
chr17-67750094-INS-64
chr17-67793543-INS-311
chr17-67793544-INS-77
chr17-67800804-INS-108
chr17-67810985-INS-151
chr17-67850757-INS-56
chr17-67874591-INS-440
chr17-67884745-INS-198
chr17-67952944-INS-6420
chr17-67964672-INS-254
chr17-67966587-INS-330
chr17-67974691-INS-56
chr17-67991283-INS-52
chr17-68011115-INS-312
chr17-68027262-INS-204
chr17-68027299-INS-80
chr17-68027683-INS-241

chr17-68027728-INS-119
chr17-68036655-INS-142
chr17-68042528-INS-114
chr17-68048405-INS-228
chr17-68048408-INS-96
chr17-68057316-INS-606
chr17-68066669-INS-649
chr17-68067604-INS-240
chr17-68080155-INS-6380
chr17-68147186-INS-1784
chr17-68153113-INS-274
chr17-68153348-INS-148
chr17-68153364-INS-52
chr17-68153590-INS-614
chr17-68163923-INS-54
chr17-68178442-INS-96
chr17-68183592-INS-619
chr17-68184066-INS-581
chr17-68210499-INS-6616
chr17-68231913-INS-50
chr17-68237213-INS-57
chr17-68239366-INS-7164
chr17-68275586-INS-56
chr17-68331951-INS-6490
chr17-68363894-INS-341
chr17-68403956-INS-91
chr17-68403975-INS-275
chr17-68428331-INS-59
chr17-68429864-INS-3082
chr17-68439382-INS-6664
chr17-68444188-INS-108
chr17-68444192-INS-303
chr17-68447815-INS-395
chr17-68455618-INS-1347
chr17-68457080-INS-6514
chr17-68461829-INS-72
chr17-68466360-INS-56
chr17-68468224-INS-78
chr17-68469115-INS-59
chr17-68495408-INS-7230
chr17-68521463-INS-260
chr17-68543270-INS-66
chr17-68551083-INS-70
chr17-68597977-INS-50
chr17-68626677-INS-499
chr17-68626738-INS-1214
chr17-68634475-INS-1555
chr17-68650336-INS-79
chr17-68670106-INS-255
chr17-68757119-INS-2132
chr17-68763964-INS-146
chr17-68819538-INS-90
chr17-68828633-INS-56
chr17-68844639-INS-6372
chr17-68847601-INS-148
chr17-68858234-INS-413
chr17-68879309-INS-77
chr17-68918759-INS-52
chr17-68926044-INS-6687
chr17-68957351-INS-479
chr17-69023090-INS-313
chr17-69025667-INS-392
chr17-69025667-INS-84
chr17-69070116-INS-137
chr17-69122560-INS-92
chr17-69157087-INS-104
chr17-69162039-INS-60
chr17-69173113-INS-84
chr17-69175463-INS-221
chr17-69300226-INS-54
chr17-69327468-INS-51
chr17-69392033-INS-128
chr17-69398493-INS-67
chr17-69401756-INS-69
chr17-69402622-INS-253
chr17-69411465-INS-56
chr17-69430889-INS-6402

chr17-69452453-INS-50
chr17-69460360-INS-312
chr17-69492846-INS-949
chr17-69498322-INS-180
chr17-69542031-INS-104
chr17-69542358-INS-178
chr17-69545966-INS-265
chr17-69546239-INS-190
chr17-69606059-INS-142
chr17-69630418-INS-253
chr17-69665839-INS-148
chr17-69713033-INS-4340
chr17-69718132-INS-72
chr17-69732257-INS-353
chr17-69744344-INS-73
chr17-69752380-INS-67
chr17-69752484-INS-295
chr17-69768074-INS-58
chr17-69782545-INS-6400
chr17-69801635-INS-66
chr17-69802543-INS-198
chr17-69850795-INS-114
chr17-69921425-INS-7152
chr17-69950372-INS-1516
chr17-69976672-INS-2853
chr17-70024175-INS-75
chr17-70043053-INS-56
chr17-70045713-INS-1547
chr17-70049181-INS-204
chr17-70053539-INS-4463
chr17-70090027-INS-94
chr17-70090242-INS-63
chr17-70092422-INS-6596
chr17-70147511-INS-63
chr17-70152041-INS-279
chr17-70152082-INS-56
chr17-70156129-INS-50
chr17-70209926-INS-392
chr17-70214704-INS-413
chr17-70215357-INS-68
chr17-70234923-INS-3183
chr17-70237599-INS-107
chr17-70245146-INS-1888
chr17-70338597-INS-207
chr17-70377259-INS-276
chr17-70449751-INS-90
chr17-70449831-INS-281
chr17-70475042-INS-78
chr17-70546218-INS-51
chr17-70551366-INS-1084
chr17-70597535-INS-1424
chr17-70605420-INS-689
chr17-70649765-INS-79
chr17-70671792-INS-52
chr17-70689254-INS-100
chr17-70689312-INS-228
chr17-70689611-INS-261
chr17-70698711-INS-185
chr17-70715287-INS-1246
chr17-70727321-INS-318
chr17-70735833-INS-50
chr17-70752226-INS-233
chr17-70760908-INS-163
chr17-70773301-INS-93
chr17-70816083-INS-87
chr17-70836429-INS-87
chr17-70857571-INS-141
chr17-70863051-INS-65
chr17-70866999-INS-52
chr17-70875633-INS-2650
chr17-70926564-INS-77
chr17-70927353-INS-51
chr17-70939840-INS-4664
chr17-70944654-INS-50
chr17-70959520-INS-81
chr17-70959841-INS-384
chr17-70960179-INS-57

chr17-70968250-INS-242
chr17-70969019-INS-80
chr17-70987030-INS-356
chr17-70997703-INS-6720
chr17-71005002-INS-231
chr17-71005090-INS-345
chr17-71010016-INS-54
chr17-71012352-INS-316
chr17-71012371-INS-86
chr17-71012580-INS-84
chr17-71014394-INS-78
chr17-71040855-INS-336
chr17-71053791-INS-186
chr17-71056295-INS-201
chr17-71074248-INS-238
chr17-71092678-INS-194
chr17-71099032-INS-3411
chr17-71110542-INS-141
chr17-71121157-INS-202
chr17-71163449-INS-50
chr17-71185012-INS-61
chr17-71201182-INS-1106
chr17-71212552-INS-242
chr17-71213384-INS-7194
chr17-71220891-INS-5160
chr17-71221579-INS-56
chr17-71225025-INS-5542
chr17-71237292-INS-1892
chr17-71243153-INS-98
chr17-71245919-INS-60
chr17-71245953-INS-234
chr17-71246556-INS-209
chr17-71247841-INS-60
chr17-71247957-INS-6159
chr17-71249334-INS-358
chr17-71252823-INS-1269
chr17-71255242-INS-68
chr17-71262807-INS-582
chr17-71266431-INS-56
chr17-71267939-INS-197
chr17-71268563-INS-193
chr17-71268710-INS-68
chr17-71315799-INS-68
chr17-71315873-INS-201
chr17-71321977-INS-4344
chr17-71323023-INS-71
chr17-71335862-INS-408
chr17-71377687-INS-100
chr17-71427550-INS-128
chr17-71444502-INS-54
chr17-71445588-INS-50
chr17-71449661-INS-51
chr17-71520702-INS-1968
chr17-71535386-INS-265
chr17-71552336-INS-155
chr17-71582561-INS-5853
chr17-71583140-INS-65
chr17-71609520-INS-138
chr17-71609891-INS-422
chr17-71611106-INS-83
chr17-71611489-INS-122
chr17-71628837-INS-54
chr17-71643066-INS-53
chr17-71646927-INS-313
chr17-71646927-INS-645
chr17-71658963-INS-176
chr17-71675687-INS-201
chr17-71687588-INS-56
chr17-71706945-INS-155
chr17-71739301-INS-64
chr17-71744959-INS-202
chr17-71795496-INS-56
chr17-71798932-INS-100
chr17-71805275-INS-714
chr17-71817732-INS-208
chr17-71840579-INS-202
chr17-71841235-INS-470

chr17-71843873-INS-714
chr17-71930981-INS-50
chr17-71945057-INS-652
chr17-71951854-INS-199
chr17-71960923-INS-477
chr17-72008617-INS-214
chr17-72032648-INS-52
chr17-72038141-INS-56
chr17-72075673-INS-4177
chr17-72077731-INS-120
chr17-72083980-INS-155
chr17-72111225-INS-51
chr17-72127537-INS-79
chr17-72127581-INS-420
chr17-72127600-INS-1094
chr17-72172467-INS-1497
chr17-72177249-INS-138
chr17-72182349-INS-52
chr17-72182842-INS-54
chr17-72206784-INS-178
chr17-72290209-INS-132
chr17-72290209-INS-442
chr17-72299859-INS-114
chr17-72339239-INS-76
chr17-72348891-INS-284
chr17-72384276-INS-54
chr17-72406486-INS-1938
chr17-72421361-INS-54
chr17-72421393-INS-501
chr17-72442386-INS-87
chr17-72448201-INS-435
chr17-72482178-INS-51
chr17-72482279-INS-84
chr17-72499680-INS-6469
chr17-72515984-INS-759
chr17-72515990-INS-213
chr17-72536952-INS-56
chr17-72560128-INS-144
chr17-72626881-INS-423
chr17-72639274-INS-52
chr17-72662805-INS-2266
chr17-72663465-INS-50
chr17-72732166-INS-68
chr17-72732177-INS-150
chr17-72746678-INS-612
chr17-72746763-INS-204
chr17-72779339-INS-6323
chr17-72790043-INS-60
chr17-72803288-INS-81
chr17-72808377-INS-75
chr17-72846445-INS-95
chr17-72857254-INS-52
chr17-72890842-INS-178
chr17-72905895-INS-6453
chr17-72921496-INS-56
chr17-72947905-INS-1583
chr17-72980753-INS-5313
chr17-72986550-INS-83
chr17-72991880-INS-52
chr17-72992495-INS-247
chr17-73011713-INS-64
chr17-73013203-INS-90
chr17-73022119-INS-6978
chr17-73025437-INS-70
chr17-73047635-INS-4758
chr17-73055551-INS-6922
chr17-73111785-INS-244
chr17-73111929-INS-53
chr17-73125975-INS-76
chr17-73128935-INS-134
chr17-73130080-INS-3176
chr17-73149105-INS-7722
chr17-73173879-INS-794
chr17-73173882-INS-132
chr17-73173938-INS-296
chr17-73206392-INS-58
chr17-73206918-INS-565

chr17-73207007-INS-125
chr17-73246161-INS-54
chr17-73308414-INS-202
chr17-73310526-INS-92
chr17-73317525-INS-64
chr17-73333175-INS-1601
chr17-73359845-INS-52
chr17-73385885-INS-56
chr17-73405452-INS-579
chr17-73437239-INS-788
chr17-73516421-INS-65
chr17-73578105-INS-1265
chr17-73590927-INS-92
chr17-73590944-INS-372
chr17-73613311-INS-53
chr17-73625257-INS-211
chr17-73652748-INS-66
chr17-73652749-INS-150
chr17-73666480-INS-55
chr17-73705681-INS-90
chr17-73763048-INS-96
chr17-73763125-INS-3005
chr17-73772931-INS-154
chr17-73788436-INS-82
chr17-73791342-INS-5710
chr17-73831414-INS-208
chr17-73834413-INS-6998
chr17-73838463-INS-162
chr17-73838975-INS-9317
chr17-73893830-INS-50
chr17-73911437-INS-1134
chr17-73914163-INS-76
chr17-73915185-INS-54
chr17-73917982-INS-6391
chr17-73918570-INS-51
chr17-73920064-INS-81
chr17-73958411-INS-119
chr17-73958492-INS-406
chr17-73958503-INS-170
chr17-74039102-INS-174
chr17-74060956-INS-2699
chr17-74062335-INS-7600
chr17-74084707-INS-6382
chr17-74106334-INS-7103
chr17-74122175-INS-62
chr17-74142925-INS-768
chr17-74146382-INS-69
chr17-74147235-INS-75
chr17-74159193-INS-254
chr17-74174517-INS-72
chr17-74215850-INS-146
chr17-74220225-INS-525
chr17-74236352-INS-59
chr17-74252765-INS-50
chr17-74263261-INS-90
chr17-74273607-INS-2611
chr17-74281501-INS-7168
chr17-74377084-INS-114
chr17-74377196-INS-267
chr17-74379098-INS-279
chr17-74380518-INS-7078
chr17-74393099-INS-1586
chr17-74393390-INS-121
chr17-74405558-INS-60
chr17-74405752-INS-180
chr17-74416775-INS-56
chr17-74421554-INS-998
chr17-74456262-INS-417
chr17-74459427-INS-6531
chr17-74463698-INS-158
chr17-74465597-INS-58
chr17-74508961-INS-345
chr17-74516751-INS-77
chr17-74519435-INS-120
chr17-74590076-INS-207
chr17-74590706-INS-84
chr17-74609753-INS-202

chr17-74610178-INS-443
chr17-74624706-INS-141
chr17-74626177-INS-60
chr17-74635427-INS-224
chr17-74641632-INS-6820
chr17-74642448-INS-491
chr17-74655833-INS-590
chr17-74655833-INS-94
chr17-74689776-INS-52
chr17-74691267-INS-249
chr17-74691976-INS-180
chr17-74692453-INS-262
chr17-74715460-INS-350
chr17-74720295-INS-224
chr17-74720639-INS-639
chr17-74722470-INS-158
chr17-74735295-INS-55
chr17-74735714-INS-177
chr17-74760452-INS-54
chr17-74776814-INS-1622
chr17-74778034-INS-5442
chr17-74803115-INS-199
chr17-74811056-INS-1375
chr17-74816383-INS-195
chr17-74817234-INS-358
chr17-74847913-INS-211
chr17-74848587-INS-498
chr17-74854877-INS-56
chr17-74854964-INS-224
chr17-74856344-INS-50
chr17-74865284-INS-135
chr17-74877940-INS-1002
chr17-74878942-INS-120
chr17-74965744-INS-159
chr17-74966043-INS-213
chr17-74998659-INS-206
chr17-75009150-INS-161
chr17-75036767-INS-169
chr17-75037369-INS-3805
chr17-75037369-INS-498
chr17-75037880-INS-5603
chr17-75077175-INS-3225
chr17-75087386-INS-251
chr17-75097755-INS-3388
chr17-75173875-INS-50
chr17-75192838-INS-668
chr17-75205232-INS-624
chr17-75222615-INS-165
chr17-75245687-INS-203
chr17-75257579-INS-95
chr17-75258034-INS-54
chr17-75277072-INS-86
chr17-75309517-INS-50
chr17-75311104-INS-54
chr17-75376968-INS-54
chr17-75395510-INS-6502
chr17-75421097-INS-2843
chr17-75438113-INS-66
chr17-75438675-INS-1193
chr17-75441148-INS-52
chr17-75511641-INS-2777
chr17-75525244-INS-51
chr17-75576624-INS-151
chr17-75591227-INS-53
chr17-75670111-INS-50
chr17-75721223-INS-2183
chr17-75749951-INS-232
chr17-75770507-INS-80
chr17-75820563-INS-1272
chr17-75824727-INS-59
chr17-75883718-INS-78
chr17-75891708-INS-465
chr17-75896995-INS-68
chr17-75901144-INS-374
chr17-75908937-INS-431
chr17-75913157-INS-8284
chr17-75949973-INS-839

chr17-75951754-INS-401
chr17-75963297-INS-2829
chr17-75999719-INS-1188
chr17-76002458-INS-6447
chr17-76008028-INS-1277
chr17-76026199-INS-422
chr17-76034380-INS-1296
chr17-76043760-INS-2737
chr17-76084876-INS-50
chr17-76093345-INS-6459
chr17-76129188-INS-166
chr17-76130090-INS-55
chr17-76153386-INS-6189
chr17-76156296-INS-1086
chr17-76167780-INS-162
chr17-76182707-INS-88
chr17-76228073-INS-7308
chr17-76310842-INS-64
chr17-76318723-INS-50
chr17-76334751-INS-101
chr17-76338889-INS-159
chr17-76338980-INS-64
chr17-76368861-INS-50
chr17-76372918-INS-1038
chr17-76393257-INS-84
chr17-76430164-INS-6258
chr17-76446578-INS-51
chr17-76466112-INS-409
chr17-76517267-INS-3947
chr17-76530605-INS-56
chr17-76572075-INS-1467
chr17-76592080-INS-173
chr17-76630267-INS-246
chr17-76645394-INS-56
chr17-76652131-INS-6176
chr17-76668437-INS-2744
chr17-76671947-INS-6170
chr17-76695829-INS-52
chr17-76757268-INS-2108
chr17-76806847-INS-1077
chr17-76809307-INS-60
chr17-76833546-INS-343
chr17-76841455-INS-90
chr17-76860121-INS-193
chr17-76866632-INS-723
chr17-76868788-INS-6226
chr17-76887315-INS-58
chr17-76887336-INS-171
chr17-76915491-INS-214
chr17-76934370-INS-7185
chr17-76940444-INS-6357
chr17-76947225-INS-62
chr17-76959211-INS-77
chr17-77012430-INS-52
chr17-77020522-INS-6235
chr17-77051301-INS-1804
chr17-77057406-INS-98
chr17-77084217-INS-694
chr17-77095935-INS-56
chr17-77100432-INS-56
chr17-77111458-INS-2485
chr17-77129289-INS-469
chr17-77143957-INS-136
chr17-77164484-INS-95
chr17-77168711-INS-733
chr17-77187552-INS-64
chr17-77197612-INS-56
chr17-77202597-INS-2153
chr17-77288834-INS-50
chr17-77392820-INS-3019
chr17-77394129-INS-5034
chr17-77407396-INS-57
chr17-77407436-INS-182
chr17-77428744-INS-284
chr17-77439604-INS-52
chr17-77439999-INS-183
chr17-77447027-INS-92

chr17-77450248-INS-78
chr17-77491982-INS-57
chr17-77512189-INS-6680
chr17-77552574-INS-52
chr17-77566689-INS-157
chr17-77584142-INS-72
chr17-77767161-INS-51
chr17-77775114-INS-5757
chr17-77819680-INS-489
chr17-77819960-INS-52
chr17-77876855-INS-52
chr17-77885342-INS-68
chr17-77902042-INS-293
chr17-77911045-INS-56
chr17-77930339-INS-764
chr17-77942677-INS-690
chr17-77943026-INS-324
chr17-77943033-INS-806
chr17-77979227-INS-60
chr17-78017938-INS-356
chr17-78037380-INS-82
chr17-78037681-INS-2691
chr17-78094358-INS-1226
chr17-78113009-INS-87
chr17-78125076-INS-64
chr17-78157968-INS-1511
chr17-78163806-INS-88
chr17-78177008-INS-123
chr17-78205423-INS-126
chr17-78210853-INS-58
chr17-78225812-INS-85
chr17-78245059-INS-1160
chr17-78263866-INS-407
chr17-78315053-INS-6209
chr17-78315146-INS-95
chr17-78328886-INS-90
chr17-78359845-INS-322
chr17-78364448-INS-219
chr17-78397007-INS-58
chr17-78427213-INS-7168
chr17-78441851-INS-6168
chr17-78443170-INS-123
chr17-78447989-INS-1171
chr17-78483731-INS-183
chr17-78483811-INS-451
chr17-78494230-INS-680
chr17-78559569-INS-126
chr17-78735498-INS-3327
chr17-78737781-INS-207
chr17-78745849-INS-52
chr17-78761336-INS-88
chr17-78765453-INS-64
chr17-78766532-INS-77
chr17-78766953-INS-152
chr17-78778027-INS-52
chr17-78833147-INS-133
chr17-78843895-INS-1562
chr17-78854080-INS-1234
chr17-78872123-INS-63
chr17-78873096-INS-56
chr17-78906448-INS-54
chr17-78933115-INS-72
chr17-78958449-INS-50
chr17-78968984-INS-244
chr17-79021427-INS-126
chr17-79058293-INS-234
chr17-79058293-INS-54
chr17-79058293-INS-558
chr17-79079194-INS-71
chr17-79079625-INS-209
chr17-79098113-INS-60
chr17-79117044-INS-57
chr17-79123941-INS-125
chr17-79162396-INS-303
chr17-79198959-INS-693
chr17-79202584-INS-105
chr17-79236395-INS-2545

chr17-79248551-INS-55
chr17-79253005-INS-118
chr17-79265067-INS-213
chr17-79316422-INS-55
chr17-79326736-INS-77
chr17-79329628-INS-74
chr17-79329708-INS-382
chr17-79370042-INS-953
chr17-79379366-INS-3128
chr17-79390458-INS-72
chr17-79403442-INS-1342
chr17-79424301-INS-60
chr17-79431387-INS-62
chr17-79431652-INS-139
chr17-79516330-INS-348
chr17-79530358-INS-124
chr17-79536070-INS-174
chr17-79543352-INS-6875
chr17-79621584-INS-86
chr17-79635430-INS-176
chr17-79646005-INS-3594
chr17-79724014-INS-65
chr17-79761736-INS-7499
chr17-79807396-INS-70
chr17-79836609-INS-1685
chr17-79839444-INS-100
chr17-79839564-INS-322
chr17-79946178-INS-129
chr17-79976836-INS-120
chr17-79998859-INS-204
chr17-80065502-INS-52
chr17-80087095-INS-102
chr17-80098940-INS-5499
chr17-80102181-INS-504
chr17-80106897-INS-67
chr17-80114581-INS-58
chr17-80124010-INS-257
chr17-80134875-INS-2270
chr17-80135079-INS-162
chr17-80173398-INS-68
chr17-80187838-INS-94
chr17-80191884-INS-193
chr17-80212044-INS-1895
chr17-80266693-INS-207
chr17-80267335-INS-226
chr17-80267432-INS-54
chr17-80270189-INS-251
chr17-80271064-INS-64
chr17-80271143-INS-210
chr17-80277153-INS-231
chr17-80286411-INS-52
chr17-80305283-INS-51
chr17-80306985-INS-50
chr17-80319258-INS-7400
chr17-80408946-INS-222
chr17-80409225-INS-82
chr17-80410703-INS-1387
chr17-80429131-INS-144
chr17-80450959-INS-207
chr17-80458285-INS-103
chr17-80458751-INS-616
chr17-80458898-INS-1245
chr17-80461002-INS-573
chr17-80494325-INS-218
chr17-80501700-INS-61
chr17-80538070-INS-60
chr17-80544887-INS-7086
chr17-80567545-INS-117
chr17-80571494-INS-56
chr17-80571500-INS-122
chr17-80596273-INS-50
chr17-80624034-INS-240
chr17-80654155-INS-50
chr17-80743668-INS-332
chr17-80744368-INS-80
chr17-80764511-INS-57
chr17-80798746-INS-184

chr17-80801016-INS-208
chr17-80817448-INS-210
chr17-80826614-INS-1754
chr17-80848355-INS-500
chr17-80876486-INS-64
chr17-80906486-INS-588
chr17-80907500-INS-3182
chr17-80911659-INS-55
chr17-80912517-INS-110
chr17-80928389-INS-2920
chr17-80950348-INS-183
chr17-80956316-INS-68
chr17-81003134-INS-7325
chr17-81068714-INS-66
chr17-81121466-INS-50
chr17-81126966-INS-1042
chr17-81132118-INS-2875
chr17-81177332-INS-509
chr17-81201291-INS-157
chr17-81201396-INS-194
chr17-81223983-INS-1602
chr17-81224982-INS-55
chr17-81260439-INS-72
chr17-81262214-INS-382
chr17-81262362-INS-690
chr17-81272392-INS-6415
chr17-81273925-INS-1239
chr17-81296108-INS-248
chr17-81340521-INS-200
chr17-81340521-INS-75
chr17-81341087-INS-72
chr17-81341099-INS-184
chr17-81355921-INS-323
chr17-81388474-INS-7081
chr17-81406867-INS-66
chr17-81407038-INS-368
chr17-81411334-INS-140
chr17-81475444-INS-95
chr17-81478356-INS-479
chr17-81493114-INS-1501
chr17-81495145-INS-79
chr17-81495608-INS-178
chr17-81517668-INS-1258
chr17-81517775-INS-286
chr17-81522849-INS-223
chr17-81582063-INS-905
chr17-81585293-INS-3176
chr17-81642729-INS-63
chr17-81646896-INS-322
chr17-81646896-INS-598
chr17-81646969-INS-138
chr17-81650054-INS-69
chr17-81738556-INS-87
chr17-81744150-INS-63
chr17-81803350-INS-50
chr17-81806606-INS-1886
chr17-81819960-INS-178
chr17-81839012-INS-52
chr17-81879275-INS-270
chr17-81915549-INS-6501
chr17-82023974-INS-424
chr17-82107710-INS-55
chr17-82156935-INS-825
chr17-82217153-INS-7260
chr17-82254081-INS-7120
chr17-82293767-INS-1346
chr17-82325474-INS-50
chr17-82332705-INS-1000
chr17-82344486-INS-60
chr17-82345398-INS-62
chr17-82349083-INS-357
chr17-82367926-INS-3450
chr17-82383321-INS-118
chr17-82401565-INS-6191
chr17-82423030-INS-51
chr17-82463108-INS-434
chr17-82477976-INS-68

chr17-82501228-INS-2257
chr17-82505396-INS-440
chr17-82542854-INS-203
chr17-82561310-INS-6460
chr17-82561909-INS-315
chr17-82574693-INS-1858
chr17-82593076-INS-2124
chr17-82613301-INS-111
chr17-82619976-INS-1020
chr17-82623616-INS-99
chr17-82632979-INS-6179
chr17-82666612-INS-57
chr17-82681568-INS-6396
chr17-82691641-INS-50
chr17-82706064-INS-55
chr17-82706363-INS-123
chr17-82859887-INS-124
chr17-82912288-INS-68
chr17-82922716-INS-199
chr17-82956772-INS-179
chr17-82957524-INS-66
chr17-82970651-INS-595
chr17-82975534-INS-63
chr17-82988349-INS-1000
chr17-82989053-INS-544
chr17-83012578-INS-52
chr17-83013624-INS-56
chr17-83018009-INS-406
chr17-83024457-INS-2007
chr17-83061901-INS-191
chr17-83069923-INS-52
chr17-83076642-INS-64
chr17-83094613-INS-2769
chr17-83112079-INS-1375
chr17-83113891-INS-177
chr17-83135564-INS-178
chr17-83139014-INS-2514
chr17-83141404-INS-146
chr17-83141411-INS-411
chr17-83148613-INS-172
chr17-83170603-INS-4428
chr17-83178474-INS-70
chr17-83189619-INS-63
chr17-83189652-INS-6683
chr17-83191010-INS-1110
chr17-83213536-INS-1890
chr17-83224420-INS-333
chr17-83224525-INS-7458
chr17-83241179-INS-79
chr17-83241218-INS-383
chr17-83264891-INS-137
chr17-83264891-INS-61
chr17-83265261-INS-480
chr17-83327652-INS-995
chr17-83364397-INS-251
chr17-83368573-INS-5125
chr17-83415955-INS-92
chr17-83465478-INS-224
chr17-83465920-INS-175
chr17-83475459-INS-68
chr17-83488111-INS-171
chr17-83500148-INS-231
chr17-83510716-INS-307
chr17-83539884-INS-212
chr17-83540260-INS-240
chr17-83540528-INS-71
chr17-83549127-INS-615
chr17-83551539-INS-141
chr17-83578716-INS-50
chr17-83599588-INS-201
chr17-83600191-INS-84
chr17-83607457-INS-118
chr17-83607617-INS-297
chr17-83611327-INS-53
chr17-83614030-INS-62
chr17-83626317-INS-90
chr17-83664248-INS-117

chr17-83699601-INS-7111
chr17-83728958-INS-628
chr17-83738598-INS-57
chr17-83752198-INS-981
chr17-83796643-INS-229
chr17-83797869-INS-169
chr17-83798127-INS-538
chr17-83838725-INS-206
chr17-83866236-INS-98
chr17-83873450-INS-268
chr17-83875965-INS-173
chr17-83876178-INS-100
chr17-83876290-INS-168
chr17-83876732-INS-99
chr17-83915038-INS-1263
chr17-83916349-INS-214
chr17-83916910-INS-197
chr17-83916949-INS-69
chr17-83924359-INS-148
chr17-83926697-INS-1856
chr17-83928495-INS-60
chr17-83932877-INS-195
chr17-83954531-INS-74
chr17-83974177-INS-73
chr17-84003178-INS-363
chr17-84003248-INS-874
chr17-84005529-INS-53
chr17-84022521-INS-54
chr17-84032536-INS-298
chr17-84032536-INS-84
chr17-84039066-INS-126
chr17-84039066-INS-505
chr17-84065220-INS-76
chr17-84080008-INS-184
chr17-84118706-INS-58
chr17-84135040-INS-152
chr17-84143651-INS-243
chr17-84143651-INS-81
chr17-84150010-INS-108
chr17-84159750-INS-78
chr17-84161264-INS-63
chr17-84161564-INS-344
chr17-84174348-INS-8673
chr17-84177805-INS-3209
chr17-84205791-INS-421
chr17-84210510-INS-50
chr17-84260404-INS-1908
chr17-84280207-INS-462
chr17-84296148-INS-416
chr17-84313399-INS-4672
chr17-84380861-INS-104
chr17-84392063-INS-5545
chr17-84431010-INS-244
chr17-84466371-INS-590
chr17-84467275-INS-56
chr17-84503661-INS-7188
chr17-84529472-INS-67
chr17-84576186-INS-666
chr17-84623517-INS-138
chr17-84635104-INS-117
chr17-84636247-INS-1743
chr17-84658349-INS-50
chr17-84771417-INS-230
chr17-84809190-INS-1472
chr17-84813796-INS-1220
chr17-84813796-INS-192
chr17-84822041-INS-72
chr17-84846657-INS-1123
chr17-84846694-INS-243
chr17-84854066-INS-158
chr17-84929566-INS-9103
chr17-84937167-INS-144
chr17-84937924-INS-96
chr17-84941292-INS-52
chr17-84946112-INS-194
chr17-84960950-INS-52
chr17-84961800-INS-52

chr17-84963596-INS-56
chr17-84963909-INS-195
chr17-84964841-INS-452
chr17-84973379-INS-788
chr17-84973784-INS-104
chr17-84974639-INS-186
chr17-84974721-INS-386
chr17-84981630-INS-53
chr17-85002892-INS-120
chr17-85005550-INS-481
chr17-85047777-INS-206
chr17-85054509-INS-60
chr17-85055493-INS-153
chr17-85077147-INS-153
chr17-85110267-INS-170
chr17-85118140-INS-87
chr17-85118231-INS-272
chr17-85119215-INS-481
chr17-85138591-INS-105
chr17-85147189-INS-125
chr17-85147189-INS-395
chr17-85169297-INS-52
chr17-85177196-INS-56
chr17-85179370-INS-204
chr17-85217058-INS-70
chr17-85219259-INS-1101
chr17-85222536-INS-84
chr17-85223963-INS-187
chr17-85225737-INS-2966
chr17-85237972-INS-334
chr17-85238516-INS-390
chr17-85250884-INS-194
chr17-85253232-INS-207
chr17-85254554-INS-833
chr17-85308857-INS-181
chr17-85336986-INS-110
chr17-85347083-INS-94
chr17-85347269-INS-56
chr17-85362109-INS-52
chr17-85428039-INS-104
chr17-85440029-INS-1171
chr17-85440436-INS-1672
chr17-85448231-INS-129
chr17-85458279-INS-59
chr17-85471347-INS-64
chr17-85486992-INS-60
chr17-85504677-INS-133
chr17-85504740-INS-60
chr17-85545804-INS-50
chr17-85547877-INS-2979
chr17-85552069-INS-213
chr17-85560677-INS-675
chr17-85585642-INS-56
chr17-85604501-INS-65
chr17-85642224-INS-56
chr17-85646373-INS-50
chr17-85664904-INS-329
chr17-85672655-INS-54
chr17-85691309-INS-158
chr17-85839170-INS-1222
chr17-85859910-INS-258
chr17-85896885-INS-123
chr17-85911545-INS-159
chr17-85945655-INS-1629
chr17-85945705-INS-648
chr17-85946086-INS-1019
chr17-85954850-INS-372
chr17-85962950-INS-322
chr17-85990677-INS-4921
chr17-86013434-INS-62
chr17-86014884-INS-72
chr17-86034966-INS-69
chr17-86035104-INS-82
chr17-86046512-INS-64
chr17-86052449-INS-7233
chr17-86064716-INS-86
chr17-86084565-INS-152

chr17-86093954-INS-3892
chr17-86184752-INS-200
chr17-86288183-INS-76
chr17-86402102-INS-50
chr17-86422793-INS-1500
chr17-86431738-INS-56
chr17-86434070-INS-58
chr17-86458503-INS-52
chr17-86491100-INS-96
chr17-86496221-INS-93
chr17-86498508-INS-329
chr17-86524779-INS-120
chr17-86524826-INS-50
chr17-86539825-INS-6906
chr17-86565351-INS-58
chr17-86603550-INS-68
chr17-86644468-INS-55
chr17-86668089-INS-144
chr17-86668248-INS-63
chr17-86709916-INS-312
chr17-86721517-INS-54
chr17-86734449-INS-4828
chr17-86775375-INS-62
chr17-86776020-INS-207
chr17-86810949-INS-2608
chr17-86814236-INS-4455
chr17-86827195-INS-84
chr17-86846158-INS-60
chr17-86885489-INS-285
chr17-86893812-INS-50
chr17-86910738-INS-111
chr17-86910910-INS-77
chr17-86913621-INS-156
chr17-86940410-INS-184
chr17-86970298-INS-57
chr17-86996774-INS-138
chr17-86998152-INS-132
chr17-87000989-INS-61
chr17-87027718-INS-77
chr17-87069561-INS-154
chr17-87111089-INS-223
chr17-87171383-INS-268
chr17-87218505-INS-58
chr17-87219204-INS-105
chr17-87278388-INS-108
chr17-87278598-INS-94
chr17-87379068-INS-6226
chr17-87405820-INS-9584
chr17-87460292-INS-228
chr17-87460292-INS-93
chr17-87465544-INS-7169
chr17-87518144-INS-54
chr17-87524809-INS-85
chr17-87539793-INS-647
chr17-87555935-INS-1314
chr17-87555935-INS-357
chr17-87565929-INS-59
chr17-87604451-INS-593
chr17-87681912-INS-214
chr17-87702500-INS-58
chr17-87709404-INS-1561
chr17-87714052-INS-73
chr17-87714431-INS-1727
chr17-87714431-INS-353
chr17-87729123-INS-290
chr17-87823105-INS-440
chr17-87882854-INS-144
chr17-87892502-INS-1888
chr17-87908399-INS-5455
chr17-87913542-INS-136
chr17-87914142-INS-208
chr17-87940854-INS-403
chr17-87941375-INS-249
chr17-87964175-INS-1822
chr17-87984501-INS-83
chr17-87995711-INS-50
chr17-88062077-INS-287

chr17-88126604-INS-118
chr17-88127450-INS-79
chr17-88136463-INS-338
chr17-88137199-INS-104
chr17-88153024-INS-266
chr17-88169703-INS-138
chr17-88173362-INS-284
chr17-88213626-INS-86
chr17-88227520-INS-88
chr17-88238209-INS-599
chr17-88259191-INS-7205
chr17-88331321-INS-2035
chr17-88394454-INS-663
chr17-88407470-INS-144
chr17-88411268-INS-53
chr17-88413334-INS-508
chr17-88416535-INS-52
chr17-88423020-INS-62
chr17-88423143-INS-150
chr17-88425024-INS-56
chr17-88452618-INS-128
chr17-88453421-INS-56
chr17-88458434-INS-856
chr17-88474047-INS-279
chr17-88484071-INS-64
chr17-88484415-INS-95
chr17-88486293-INS-7140
chr17-88488024-INS-291
chr17-88497407-INS-388
chr17-88497726-INS-78
chr17-88504971-INS-2885
chr17-88508360-INS-55
chr17-88535668-INS-76
chr17-88552187-INS-344
chr17-88569169-INS-478
chr17-88601245-INS-467
chr17-88617623-INS-73
chr17-88617963-INS-222
chr17-88626830-INS-68
chr17-88627008-INS-108
chr17-88628202-INS-82
chr17-88630940-INS-154
chr17-88631790-INS-102
chr17-88644072-INS-8969
chr17-88664204-INS-107
chr17-88691397-INS-822
chr17-88697014-INS-384
chr17-88716053-INS-55
chr17-88731432-INS-508
chr17-88741306-INS-69
chr17-88741422-INS-156
chr17-88742360-INS-110
chr17-88802731-INS-315
chr17-88823920-INS-2808
chr17-88841066-INS-100
chr17-88862130-INS-207
chr17-88865495-INS-209
chr17-88876741-INS-252
chr17-88881120-INS-134
chr17-88883248-INS-831
chr17-88891873-INS-426
chr17-88892328-INS-869
chr17-88898446-INS-358
chr17-88902420-INS-7013
chr17-88907605-INS-62
chr17-88907687-INS-299
chr17-88914690-INS-165
chr17-88915174-INS-115
chr17-88921958-INS-51
chr17-88944770-INS-90
chr17-88951020-INS-478
chr17-88964212-INS-335
chr17-88964231-INS-55
chr17-88965766-INS-628
chr17-88965886-INS-70
chr17-88993495-INS-655
chr17-88995023-INS-845

chr17-88999036-INS-164
chr17-89011034-INS-140
chr17-89015506-INS-2728
chr17-89017806-INS-7917
chr17-89025393-INS-78
chr17-89082626-INS-78
chr17-89085972-INS-74
chr17-89124760-INS-771
chr17-89137956-INS-51
chr17-89144134-INS-131
chr17-89173148-INS-116
chr17-89213970-INS-938
chr17-89230395-INS-102
chr17-89265050-INS-59
chr17-89281362-INS-73
chr17-89308594-INS-62
chr17-89317779-INS-80
chr17-89321450-INS-1769
chr17-89360840-INS-114
chr17-89429511-INS-71
chr17-89429587-INS-162
chr17-89429596-INS-509
chr17-89473854-INS-50
chr17-89475355-INS-120
chr17-89484076-INS-79
chr17-89484426-INS-80
chr17-89533917-INS-78
chr17-89556916-INS-504
chr17-89562325-INS-193
chr17-89587902-INS-6483
chr17-89590550-INS-50
chr17-89626054-INS-110
chr17-89660040-INS-581
chr17-89742547-INS-221
chr17-89751486-INS-1204
chr17-89758376-INS-78
chr17-89762220-INS-2787
chr17-89763306-INS-53
chr17-89778524-INS-61
chr17-89797054-INS-50
chr17-89827594-INS-6371
chr17-89849848-INS-66
chr17-89882913-INS-57
chr17-89888184-INS-720
chr17-89894103-INS-3726
chr17-89898600-INS-6842
chr17-89934740-INS-54
chr17-89973961-INS-1023
chr17-89987971-INS-62
chr17-90024540-INS-159
chr17-90042429-INS-7125
chr17-90049799-INS-2635
chr17-90084580-INS-58
chr17-90092824-INS-216
chr17-90165482-INS-54
chr17-90174478-INS-8407
chr17-90214819-INS-50
chr17-90230039-INS-63
chr17-90260999-INS-2604
chr17-90265400-INS-50
chr17-90271365-INS-549
chr17-90307817-INS-627
chr17-90316200-INS-3011
chr17-90362494-INS-54
chr17-90543963-INS-52
chr17-90565576-INS-64
chr17-90571543-INS-434
chr17-90574617-INS-53
chr17-90589736-INS-291
chr17-90595561-INS-102
chr17-90609609-INS-1650
chr17-90657506-INS-610
chr17-90671536-INS-51
chr17-90825940-INS-398
chr17-90900192-INS-1854
chr17-90900771-INS-624
chr17-90900910-INS-60

chr17-90943033-INS-50
chr17-90968957-INS-58
chr17-90986210-INS-224
chr17-91051770-INS-7162
chr17-91120968-INS-109
chr17-91192147-INS-8774
chr17-91204364-INS-375
chr17-91379319-INS-202
chr17-91447666-INS-657
chr17-91449534-INS-320
chr17-91453695-INS-222
chr17-91500158-INS-55
chr17-91508890-INS-52
chr17-91572948-INS-1603
chr17-91608076-INS-6305
chr17-91632756-INS-52
chr17-91636989-INS-116
chr17-91637799-INS-126
chr17-91645991-INS-6046
chr17-91695777-INS-54
chr17-91793012-INS-282
chr17-91842602-INS-211
chr17-91843571-INS-2495
chr17-91937353-INS-1756
chr17-91981525-INS-750
chr17-91981915-INS-60
chr17-91982213-INS-915
chr17-91985527-INS-50
chr17-92066674-INS-7011
chr17-92077825-INS-3126
chr17-92110090-INS-2748
chr17-92119406-INS-163
chr17-92120782-INS-50
chr17-92131065-INS-1368
chr17-92230067-INS-58
chr17-92230262-INS-390
chr17-92231770-INS-4521
chr17-92235811-INS-2369
chr17-92239402-INS-2699
chr17-92335861-INS-1107
chr17-92337069-INS-3162
chr17-92359362-INS-215
chr17-92363872-INS-54
chr17-92388692-INS-69
chr17-92400489-INS-58
chr17-92401087-INS-479
chr17-92408211-INS-55
chr17-92500286-INS-51
chr17-92508848-INS-58
chr17-92517136-INS-1058
chr17-92550711-INS-481
chr17-92584304-INS-449
chr17-92670299-INS-405
chr17-92700762-INS-189
chr17-92758698-INS-62
chr17-92764369-INS-1208
chr17-92787355-INS-51
chr17-92807295-INS-65
chr17-92808492-INS-94
chr17-92950758-INS-6513
chr17-93049368-INS-554
chr17-93051674-INS-50
chr17-93069164-INS-107
chr17-93112200-INS-7903
chr17-93137104-INS-56
chr17-93146225-INS-453
chr17-93149736-INS-61
chr17-93149928-INS-1537
chr17-93152998-INS-13955
chr17-93157098-INS-57
chr17-93161362-INS-57
chr17-93229683-INS-88
chr17-93266811-INS-146
chr17-93282041-INS-1008
chr17-93352277-INS-65
chr17-93356463-INS-2882
chr17-93372823-INS-6782

chr17-93383386-INS-978
chr17-93388672-INS-6705
chr17-93424498-INS-50
chr17-93482893-INS-1620
chr17-93491986-INS-389
chr17-93556438-INS-4807
chr17-93644015-INS-5434
chr17-93682604-INS-1235
chr17-93799919-INS-823
chr17-93838002-INS-1164
chr17-93845954-INS-358
chr17-93846507-INS-171
chr17-93867210-INS-202
chr17-93893968-INS-57
chr17-93937170-INS-2137
chr17-93976429-INS-128
chr17-93976429-INS-56
chr17-94042067-INS-16093
chr17-94059987-INS-203
chr17-94086469-INS-50
chr17-94161271-INS-95
chr17-94182531-INS-50
chr17-94228355-INS-56
chr17-94264434-INS-1783
chr17-94324802-INS-71
chr17-94395447-INS-3285
chr17-94418127-INS-2605
chr17-94448762-INS-52
chr17-94507535-INS-7166
chr17-94547832-INS-50
chr17-94576105-INS-6394
chr17-94578333-INS-57
chr17-94623460-INS-151
chr17-94639348-INS-66
chr17-94762845-INS-59
chr17-94833782-INS-66
chr17-94833848-INS-140
chr17-94834534-INS-91
chr17-94866360-INS-2845
chr17-94876626-INS-52
chr17-94888348-INS-1132
chr17-94898501-INS-2587
chr17-94901688-INS-50
chr17-94937014-INS-50
chr17-94946978-INS-151
chr17-94950300-INS-3515
chr17-95000037-INS-60
chr17-95044892-INS-435
chr17-95107300-INS-370
chr17-95177450-INS-302
chr18-3007217-INS-51
chr18-3007542-INS-469
chr18-3007874-INS-187
chr18-3011540-INS-53
chr18-3098716-INS-1099
chr18-3098716-INS-494
chr18-3303525-INS-50
chr18-3328868-INS-93
chr18-3332243-INS-50
chr18-3354317-INS-5184
chr18-3355690-INS-60
chr18-3400731-INS-6918
chr18-3473976-INS-203
chr18-3485360-INS-193
chr18-3499758-INS-110
chr18-3545895-INS-322
chr18-3548903-INS-52
chr18-3553144-INS-138
chr18-3812490-INS-3515
chr18-3876787-INS-86
chr18-3894984-INS-2969
chr18-3908005-INS-258
chr18-3909571-INS-936
chr18-3911576-INS-73
chr18-3934596-INS-352
chr18-3938071-INS-454
chr18-3975574-INS-96

chr18-3999479-INS-218
chr18-4054367-INS-420
chr18-4057814-INS-98
chr18-4109164-INS-68
chr18-4147318-INS-416
chr18-4165528-INS-116
chr18-4213423-INS-152
chr18-4213432-INS-494
chr18-4240479-INS-2052
chr18-4251231-INS-106
chr18-4290746-INS-84
chr18-4292070-INS-107
chr18-4301567-INS-172
chr18-4309928-INS-153
chr18-4316247-INS-113
chr18-4367626-INS-234
chr18-4384168-INS-240
chr18-4432732-INS-582
chr18-4546656-INS-186
chr18-4550770-INS-178
chr18-4554577-INS-62
chr18-4562574-INS-3395
chr18-4576462-INS-176
chr18-4618505-INS-6412
chr18-4676779-INS-116
chr18-4676907-INS-68
chr18-4725593-INS-9233
chr18-4742330-INS-55
chr18-4755726-INS-126
chr18-4817130-INS-5240
chr18-4826051-INS-65
chr18-4862120-INS-60
chr18-4880235-INS-51
chr18-4881563-INS-55
chr18-4916549-INS-2250
chr18-4937706-INS-221
chr18-4976748-INS-54
chr18-4982170-INS-384
chr18-5016574-INS-56
chr18-5037529-INS-140
chr18-5057967-INS-52
chr18-5061219-INS-93
chr18-5062430-INS-223
chr18-5090589-INS-566
chr18-5102117-INS-139
chr18-5113903-INS-922
chr18-5141883-INS-360
chr18-5170855-INS-2874
chr18-5170861-INS-1077
chr18-5191199-INS-60
chr18-5267441-INS-301
chr18-5347855-INS-60
chr18-5363209-INS-562
chr18-5377530-INS-201
chr18-5440488-INS-972
chr18-5456760-INS-50
chr18-5460267-INS-154
chr18-5480071-INS-9793
chr18-5484461-INS-352
chr18-5527992-INS-104
chr18-5636034-INS-6506
chr18-5681271-INS-177
chr18-5681664-INS-350
chr18-5742848-INS-62
chr18-5742960-INS-274
chr18-5748127-INS-82
chr18-5813674-INS-445
chr18-5840823-INS-106
chr18-5841824-INS-58
chr18-5845729-INS-4292
chr18-5906188-INS-6348
chr18-5921588-INS-6531
chr18-5962302-INS-514
chr18-5962411-INS-158
chr18-5972940-INS-505
chr18-5982063-INS-808
chr18-6019227-INS-177

chr18-6034097-INS-315
chr18-6074239-INS-1200
chr18-6082252-INS-1509
chr18-6082348-INS-180
chr18-6107925-INS-71
chr18-6124790-INS-135
chr18-6127098-INS-125
chr18-6127114-INS-50
chr18-6153962-INS-618
chr18-6185376-INS-7101
chr18-6210884-INS-189
chr18-6242668-INS-212
chr18-6265953-INS-89
chr18-6299930-INS-206
chr18-6387401-INS-574
chr18-6389794-INS-53
chr18-6411896-INS-1561
chr18-6462037-INS-740
chr18-6523697-INS-322
chr18-6527089-INS-3833
chr18-6559108-INS-88
chr18-6563729-INS-204
chr18-6577279-INS-8686
chr18-6595861-INS-6295
chr18-6650803-INS-3000
chr18-6654088-INS-2063
chr18-6675770-INS-469
chr18-6675808-INS-228
chr18-6683020-INS-1182
chr18-6683770-INS-80
chr18-6686664-INS-64
chr18-6719590-INS-2608
chr18-6740046-INS-98
chr18-6755331-INS-78
chr18-6759646-INS-58
chr18-6780774-INS-199
chr18-6787721-INS-196
chr18-6845619-INS-267
chr18-6845653-INS-76
chr18-6894209-INS-7348
chr18-7027214-INS-247
chr18-7113145-INS-473
chr18-7113899-INS-57
chr18-7134172-INS-363
chr18-7215470-INS-58
chr18-7255566-INS-228
chr18-7261699-INS-6504
chr18-7275692-INS-825
chr18-7336459-INS-6878
chr18-7379681-INS-221
chr18-7384375-INS-52
chr18-7410658-INS-61
chr18-7490592-INS-1351
chr18-7490630-INS-144
chr18-7491108-INS-60
chr18-7504773-INS-353
chr18-7518863-INS-1713
chr18-7523110-INS-89
chr18-7524256-INS-53
chr18-7546160-INS-966
chr18-7546269-INS-210
chr18-7556837-INS-5442
chr18-7574127-INS-257
chr18-7591318-INS-6810
chr18-7608117-INS-162
chr18-7609701-INS-270
chr18-7609737-INS-599
chr18-7609867-INS-53
chr18-7623470-INS-135
chr18-7628613-INS-91
chr18-7628694-INS-61
chr18-7628698-INS-301
chr18-7628954-INS-487
chr18-7652492-INS-3108
chr18-7662542-INS-400
chr18-7669875-INS-205
chr18-7698866-INS-206

chr18-7721453-INS-6403
chr18-7751132-INS-400
chr18-7795619-INS-2728
chr18-7806187-INS-350
chr18-7831342-INS-1862
chr18-7836135-INS-418
chr18-7909670-INS-180
chr18-7933805-INS-50
chr18-7953798-INS-143
chr18-7975706-INS-136
chr18-7975748-INS-61
chr18-7975802-INS-399
chr18-7977708-INS-868
chr18-7987474-INS-4749
chr18-8011209-INS-72
chr18-8031601-INS-84
chr18-8039086-INS-6564
chr18-8041499-INS-241
chr18-8046921-INS-220
chr18-8059632-INS-62
chr18-8061947-INS-82
chr18-8104678-INS-2932
chr18-8105797-INS-6608
chr18-8115251-INS-1597
chr18-8135583-INS-483
chr18-8147655-INS-119
chr18-8149234-INS-889
chr18-8154343-INS-852
chr18-8193457-INS-60
chr18-8260954-INS-1675
chr18-8283963-INS-56
chr18-8290080-INS-1672
chr18-8299658-INS-7097
chr18-8327150-INS-123
chr18-8344767-INS-68
chr18-8345533-INS-4358
chr18-8373549-INS-60
chr18-8412755-INS-508
chr18-8431945-INS-1001
chr18-8435067-INS-216
chr18-8442984-INS-3010
chr18-8566559-INS-6158
chr18-8575354-INS-72
chr18-8606515-INS-54
chr18-8618154-INS-7106
chr18-8642841-INS-7069
chr18-8656734-INS-56
chr18-8700272-INS-81
chr18-8700581-INS-7212
chr18-8707421-INS-186
chr18-8757406-INS-7188
chr18-8813259-INS-2801
chr18-8837910-INS-424
chr18-8838759-INS-1045
chr18-8846559-INS-1009
chr18-8877664-INS-2260
chr18-8884491-INS-93
chr18-8909363-INS-1175
chr18-8970552-INS-3917
chr18-8975638-INS-263
chr18-8976937-INS-95
chr18-8979772-INS-85
chr18-9010191-INS-413
chr18-9014775-INS-1496
chr18-9035263-INS-6662
chr18-9177787-INS-10380
chr18-9199032-INS-2268
chr18-9283026-INS-233
chr18-9291680-INS-50
chr18-9296154-INS-50
chr18-9296837-INS-115
chr18-9414065-INS-110
chr18-9417978-INS-940
chr18-9442187-INS-208
chr18-9442794-INS-52
chr18-9443387-INS-169
chr18-9451605-INS-116

chr18-9465492-INS-140
chr18-9475839-INS-97
chr18-9515337-INS-3311
chr18-9530795-INS-342
chr18-9530797-INS-113
chr18-9531164-INS-408
chr18-9554661-INS-112
chr18-9568641-INS-50
chr18-9598706-INS-1010
chr18-9633218-INS-6277
chr18-9635224-INS-4425
chr18-9676083-INS-156
chr18-9723622-INS-50
chr18-9812600-INS-6135
chr18-9839015-INS-76
chr18-9842038-INS-51
chr18-9868799-INS-50
chr18-9942990-INS-1635
chr18-9949881-INS-50
chr18-9980586-INS-801
chr18-9985257-INS-208
chr18-10059897-INS-267
chr18-10076091-INS-135
chr18-10265490-INS-124
chr18-10276640-INS-7846
chr18-10288149-INS-106
chr18-10302201-INS-922
chr18-10316848-INS-221
chr18-10351448-INS-104
chr18-10355033-INS-68
chr18-10361113-INS-300
chr18-10370089-INS-1415
chr18-10412344-INS-386
chr18-10440898-INS-50
chr18-10508400-INS-124
chr18-10531207-INS-80
chr18-10565802-INS-5349
chr18-10581142-INS-171
chr18-10642591-INS-208
chr18-10642591-INS-70
chr18-10653228-INS-289
chr18-10653362-INS-116
chr18-10684249-INS-360
chr18-10823536-INS-1946
chr18-10853494-INS-187
chr18-10898828-INS-1531
chr18-10921152-INS-56
chr18-10938826-INS-180
chr18-10939116-INS-68
chr18-10973709-INS-88
chr18-11011522-INS-229
chr18-11012182-INS-50
chr18-11032916-INS-174
chr18-11034008-INS-161
chr18-11037162-INS-104
chr18-11038061-INS-57
chr18-11038444-INS-119
chr18-11043004-INS-55
chr18-11043469-INS-130
chr18-11067816-INS-50
chr18-11080504-INS-54
chr18-11084868-INS-59
chr18-11087264-INS-97
chr18-11112669-INS-85
chr18-11132083-INS-64
chr18-11160305-INS-94
chr18-11171530-INS-88
chr18-11193521-INS-51
chr18-11193689-INS-136
chr18-11208352-INS-174
chr18-11216708-INS-72
chr18-11220103-INS-50
chr18-11231487-INS-169
chr18-11236203-INS-224
chr18-11278084-INS-52
chr18-11288664-INS-19809
chr18-11288664-INS-2574

chr18-11288664-INS-7826
chr18-11288765-INS-153
chr18-11289876-INS-67
chr18-11303068-INS-188
chr18-11303283-INS-329
chr18-11303571-INS-80
chr18-11303576-INS-205
chr18-11312511-INS-227
chr18-11318848-INS-60
chr18-11332673-INS-164
chr18-11332875-INS-156
chr18-11332877-INS-318
chr18-11334562-INS-120
chr18-11349781-INS-120
chr18-11352892-INS-6585
chr18-11361717-INS-436
chr18-11361803-INS-161
chr18-11394940-INS-409
chr18-11408961-INS-3351
chr18-11413906-INS-1010
chr18-11415518-INS-96
chr18-11416725-INS-3719
chr18-11436645-INS-50
chr18-11464312-INS-242
chr18-11548029-INS-106
chr18-11555337-INS-93
chr18-11582779-INS-72
chr18-11582859-INS-204
chr18-11622980-INS-100
chr18-11639208-INS-50
chr18-11639226-INS-1230
chr18-11644575-INS-52
chr18-11681269-INS-7254
chr18-11709110-INS-61
chr18-11737021-INS-52
chr18-11737627-INS-52
chr18-11743915-INS-1365
chr18-11751593-INS-64
chr18-11782742-INS-64
chr18-11811831-INS-234
chr18-11815878-INS-95
chr18-11822349-INS-419
chr18-11863991-INS-2725
chr18-11865849-INS-2461
chr18-11895930-INS-1633
chr18-11901785-INS-568
chr18-11917810-INS-81
chr18-11918299-INS-241
chr18-11948945-INS-86
chr18-11960563-INS-2138
chr18-12012833-INS-6427
chr18-12033316-INS-120
chr18-12057060-INS-180
chr18-12083304-INS-3769
chr18-12096409-INS-1439
chr18-12120674-INS-328
chr18-12120799-INS-56
chr18-12123170-INS-209
chr18-12128812-INS-338
chr18-12139338-INS-238
chr18-12143633-INS-139
chr18-12143900-INS-208
chr18-12145676-INS-62
chr18-12145969-INS-434
chr18-12151033-INS-189
chr18-12153178-INS-207
chr18-12161953-INS-54
chr18-12167433-INS-70
chr18-12175616-INS-60
chr18-12175681-INS-53
chr18-12187112-INS-137
chr18-12187933-INS-208
chr18-12190505-INS-76
chr18-12200435-INS-4710
chr18-12224651-INS-915
chr18-12231301-INS-212
chr18-12231631-INS-190

chr18-12290243-INS-70
chr18-12292055-INS-5086
chr18-12302921-INS-50
chr18-12305708-INS-416
chr18-12310644-INS-76
chr18-12310797-INS-304
chr18-12311416-INS-140
chr18-12329759-INS-1865
chr18-12346682-INS-50
chr18-12348316-INS-211
chr18-12350727-INS-288
chr18-12352207-INS-203
chr18-12361207-INS-80
chr18-12366096-INS-209
chr18-12450673-INS-68
chr18-12465870-INS-68
chr18-12498938-INS-162
chr18-12507208-INS-140
chr18-12507217-INS-61
chr18-12514588-INS-66
chr18-12552422-INS-199
chr18-12555452-INS-1885
chr18-12556981-INS-56
chr18-12564893-INS-2030
chr18-12585888-INS-101
chr18-12603632-INS-127
chr18-12603874-INS-135
chr18-12604722-INS-128
chr18-12604722-INS-480
chr18-12609669-INS-412
chr18-12623089-INS-369
chr18-12651332-INS-303
chr18-12651930-INS-62
chr18-12655479-INS-167
chr18-12668439-INS-886
chr18-12668838-INS-434
chr18-12684041-INS-123
chr18-12694188-INS-370
chr18-12694214-INS-81
chr18-12708282-INS-6684
chr18-12741858-INS-1000
chr18-12750446-INS-53
chr18-12818520-INS-200
chr18-12823749-INS-111
chr18-12843385-INS-4514
chr18-12850698-INS-126
chr18-12863841-INS-58
chr18-12866981-INS-207
chr18-12873036-INS-63
chr18-12900796-INS-713
chr18-12919360-INS-169
chr18-12928867-INS-68
chr18-12939716-INS-264
chr18-12989249-INS-841
chr18-12989254-INS-171
chr18-12993126-INS-786
chr18-12997343-INS-102
chr18-13010653-INS-3542
chr18-13015047-INS-482
chr18-13020896-INS-853
chr18-13021587-INS-197
chr18-13044402-INS-698
chr18-13058904-INS-675
chr18-13058909-INS-135
chr18-13061635-INS-203
chr18-13068393-INS-3319
chr18-13094715-INS-78
chr18-13129733-INS-204
chr18-13135591-INS-64
chr18-13167022-INS-64
chr18-13169624-INS-52
chr18-13172292-INS-138
chr18-13225468-INS-210
chr18-13234660-INS-6759
chr18-13243291-INS-193
chr18-13255661-INS-342
chr18-13255670-INS-124

chr18-13257293-INS-77
chr18-13258433-INS-66
chr18-13324781-INS-88
chr18-13324799-INS-461
chr18-13393288-INS-6253
chr18-13420781-INS-1000
chr18-13429678-INS-160
chr18-13432636-INS-150
chr18-13458803-INS-4124
chr18-13467184-INS-704
chr18-13495706-INS-52
chr18-13524409-INS-86
chr18-13524760-INS-280
chr18-13611619-INS-50
chr18-13625018-INS-84
chr18-13670741-INS-832
chr18-13687032-INS-940
chr18-13687154-INS-210
chr18-13687325-INS-2870
chr18-13701704-INS-54
chr18-13749401-INS-174
chr18-13763395-INS-88
chr18-13766887-INS-228
chr18-13767003-INS-6386
chr18-13866365-INS-105
chr18-13883841-INS-169
chr18-13887086-INS-6180
chr18-14090358-INS-115
chr18-14120758-INS-50
chr18-14195803-INS-60
chr18-14238979-INS-717
chr18-14239510-INS-1132
chr18-14239510-INS-2328
chr18-14261889-INS-8745
chr18-14265580-INS-56
chr18-14266177-INS-96
chr18-14268789-INS-1493
chr18-14313546-INS-57
chr18-14329983-INS-54
chr18-14375246-INS-11658
chr18-14375246-INS-2711
chr18-14402577-INS-393
chr18-14422164-INS-68
chr18-14458913-INS-82
chr18-14475176-INS-6468
chr18-14480546-INS-292
chr18-14497171-INS-457
chr18-14511720-INS-6289
chr18-14544513-INS-56
chr18-14551107-INS-223
chr18-14551569-INS-436
chr18-14564759-INS-1306
chr18-14596088-INS-998
chr18-14604464-INS-372
chr18-14609067-INS-54
chr18-14639419-INS-50
chr18-14644855-INS-59
chr18-14664674-INS-201
chr18-14687846-INS-52
chr18-14714556-INS-2802
chr18-14725290-INS-7016
chr18-14742158-INS-94
chr18-14743923-INS-6825
chr18-14791338-INS-108
chr18-14798459-INS-74
chr18-14819990-INS-5679
chr18-14827405-INS-51
chr18-14835160-INS-52
chr18-14901843-INS-1573
chr18-15015107-INS-58
chr18-15048599-INS-149
chr18-15057806-INS-1481
chr18-15063006-INS-6321
chr18-15192021-INS-213
chr18-15220351-INS-78
chr18-15254503-INS-102
chr18-15254660-INS-571

chr18-15259897-INS-56
chr18-15305885-INS-72
chr18-15357220-INS-60
chr18-15370872-INS-58
chr18-15377612-INS-1900
chr18-15378197-INS-135
chr18-15389393-INS-1041
chr18-15404544-INS-62
chr18-15422631-INS-5203
chr18-15461480-INS-6139
chr18-15482066-INS-52
chr18-15491066-INS-56
chr18-15491066-INS-584
chr18-15555073-INS-60
chr18-15595944-INS-130
chr18-15636468-INS-188
chr18-15647692-INS-64
chr18-15702441-INS-175
chr18-15806443-INS-3074
chr18-15811076-INS-6720
chr18-15828242-INS-1501
chr18-15850262-INS-51
chr18-15936019-INS-6484
chr18-15984048-INS-50
chr18-15989455-INS-85
chr18-16136058-INS-309
chr18-16152487-INS-2219
chr18-16157478-INS-121
chr18-16175935-INS-431
chr18-16225091-INS-507
chr18-16250373-INS-56
chr18-16255418-INS-3951
chr18-16292328-INS-62
chr18-16307602-INS-189
chr18-16307603-INS-54
chr18-16322539-INS-440
chr18-16355594-INS-698
chr18-16377135-INS-52
chr18-16439556-INS-9807
chr18-16496759-INS-7252
chr18-16513782-INS-6598
chr18-16546427-INS-202
chr18-16621183-INS-56
chr18-16635812-INS-93
chr18-16636517-INS-381
chr18-16676892-INS-1796
chr18-16705174-INS-51
chr18-16776220-INS-108
chr18-16847680-INS-66
chr18-16922306-INS-52
chr18-16948097-INS-73
chr18-16967153-INS-640
chr18-16967153-INS-9162
chr18-17011926-INS-2822
chr18-17024514-INS-1779
chr18-17024514-INS-3642
chr18-17050018-INS-50
chr18-17058099-INS-345
chr18-17059457-INS-135
chr18-17063527-INS-196
chr18-17094422-INS-341
chr18-17095329-INS-212
chr18-17103296-INS-2013
chr18-17117790-INS-54
chr18-17127869-INS-212
chr18-17171010-INS-128
chr18-17206006-INS-6088
chr18-17215419-INS-447
chr18-17230550-INS-57
chr18-17241067-INS-1019
chr18-17242691-INS-54
chr18-17255839-INS-8497
chr18-17257568-INS-99
chr18-17257670-INS-56
chr18-17291453-INS-6985
chr18-17358165-INS-335
chr18-17437738-INS-56

chr18-17466716-INS-442
chr18-17490223-INS-88
chr18-17506007-INS-141
chr18-17527853-INS-54
chr18-17576955-INS-108
chr18-17606556-INS-58
chr18-17638371-INS-105
chr18-17641476-INS-90
chr18-17642031-INS-188
chr18-17724806-INS-3056
chr18-17768841-INS-1955
chr18-17778255-INS-58
chr18-17807807-INS-1178
chr18-17856500-INS-7087
chr18-17896687-INS-62
chr18-17925850-INS-5864
chr18-17926453-INS-222
chr18-17961780-INS-50
chr18-17995624-INS-397
chr18-18108095-INS-5720
chr18-18113890-INS-9589
chr18-18131382-INS-1796
chr18-18160739-INS-73
chr18-18175973-INS-496
chr18-18211904-INS-451
chr18-18251123-INS-2661
chr18-18254766-INS-98
chr18-18294463-INS-2274
chr18-18295986-INS-1666
chr18-18297555-INS-54
chr18-18331922-INS-65
chr18-18337246-INS-74
chr18-18381526-INS-2776
chr18-18408150-INS-6569
chr18-18419813-INS-108
chr18-18422421-INS-8309
chr18-18569861-INS-2999
chr18-18570230-INS-50
chr18-18637815-INS-56
chr18-18658782-INS-413
chr18-18680011-INS-5566
chr18-18774410-INS-50
chr18-18791087-INS-481
chr18-18810635-INS-2785
chr18-18848420-INS-2896
chr18-18873572-INS-1888
chr18-18936613-INS-6478
chr18-19045234-INS-58
chr18-19070246-INS-54
chr18-19084580-INS-261
chr18-19103908-INS-65
chr18-19171302-INS-458
chr18-19230017-INS-50
chr18-19246930-INS-8838
chr18-19248921-INS-2919
chr18-19249832-INS-59
chr18-19250146-INS-401
chr18-19302490-INS-212
chr18-19323826-INS-151
chr18-19433440-INS-295
chr18-19433496-INS-111
chr18-19446117-INS-476
chr18-19525814-INS-50
chr18-19545845-INS-1862
chr18-19579254-INS-754
chr18-19595533-INS-199
chr18-19607327-INS-217
chr18-19665868-INS-675
chr18-19669079-INS-2857
chr18-19677871-INS-100
chr18-19703058-INS-56
chr18-19763155-INS-136
chr18-19837815-INS-990
chr18-19880241-INS-65
chr18-19968688-INS-123
chr18-20087502-INS-147
chr18-20091183-INS-2094

chr18-20091183-INS-8389
chr18-20111690-INS-76
chr18-20230439-INS-54
chr18-20278896-INS-50
chr18-20311185-INS-54
chr18-20316819-INS-70
chr18-20317914-INS-1871
chr18-20333450-INS-341
chr18-20333498-INS-1315
chr18-20336002-INS-1474
chr18-20339288-INS-10152
chr18-20349136-INS-64
chr18-20366835-INS-81
chr18-20373891-INS-77
chr18-20382778-INS-117
chr18-20439623-INS-81
chr18-20452237-INS-50
chr18-20498938-INS-6590
chr18-20521486-INS-168
chr18-20528398-INS-6423
chr18-20542925-INS-52
chr18-20580739-INS-206
chr18-20646756-INS-179
chr18-20652483-INS-50
chr18-20659177-INS-205
chr18-20659550-INS-106
chr18-20659842-INS-2725
chr18-20684596-INS-6695
chr18-20711737-INS-100
chr18-20720788-INS-2416
chr18-20728391-INS-1989
chr18-20728408-INS-289
chr18-20735919-INS-122
chr18-20766963-INS-8709
chr18-20787377-INS-52
chr18-20838676-INS-50
chr18-20852388-INS-171
chr18-20889178-INS-100
chr18-20913932-INS-175
chr18-20916857-INS-3576
chr18-20936947-INS-60
chr18-20962291-INS-88
chr18-20994140-INS-394
chr18-21056546-INS-1037
chr18-21058971-INS-3442
chr18-21071573-INS-1242
chr18-21080525-INS-92
chr18-21112861-INS-60
chr18-21121190-INS-2719
chr18-21122538-INS-205
chr18-21127871-INS-194
chr18-21130633-INS-68
chr18-21140697-INS-140
chr18-21148948-INS-196
chr18-21151618-INS-191
chr18-21181724-INS-114
chr18-21191500-INS-69
chr18-21196392-INS-183
chr18-21251560-INS-54
chr18-21279833-INS-54
chr18-21280311-INS-144
chr18-21315402-INS-87
chr18-21346262-INS-56
chr18-21445246-INS-70
chr18-21461335-INS-68
chr18-21472287-INS-6379
chr18-21498292-INS-98
chr18-21512736-INS-2412
chr18-21541514-INS-71
chr18-21550742-INS-2978
chr18-21558826-INS-235
chr18-21568627-INS-8824
chr18-21578763-INS-54
chr18-21588679-INS-1128
chr18-21601657-INS-1863
chr18-21645538-INS-1163
chr18-21665901-INS-1633

chr18-21666809-INS-164
chr18-21680968-INS-1305
chr18-21804988-INS-74
chr18-21814021-INS-58
chr18-21896514-INS-60
chr18-21899067-INS-56
chr18-21907363-INS-453
chr18-21941550-INS-172
chr18-21952552-INS-96
chr18-21956796-INS-305
chr18-21971474-INS-138
chr18-21976270-INS-65
chr18-22000017-INS-51
chr18-22016128-INS-60
chr18-22025168-INS-62
chr18-22025223-INS-58
chr18-22057344-INS-99
chr18-22121529-INS-55
chr18-22271086-INS-89
chr18-22293701-INS-435
chr18-22298799-INS-50
chr18-22331263-INS-1584
chr18-22356055-INS-491
chr18-22411327-INS-68
chr18-22459726-INS-1541
chr18-22462218-INS-73
chr18-22462254-INS-1111
chr18-22565817-INS-56
chr18-22568379-INS-500
chr18-22569774-INS-135
chr18-22585832-INS-54
chr18-22585905-INS-534
chr18-22606190-INS-61
chr18-22610116-INS-211
chr18-22736807-INS-6265
chr18-22747189-INS-1012
chr18-22756577-INS-362
chr18-22756801-INS-54
chr18-22798946-INS-3125
chr18-22805675-INS-7462
chr18-22807237-INS-1176
chr18-22818085-INS-571
chr18-22830802-INS-54
chr18-22831496-INS-197
chr18-22842479-INS-269
chr18-22916249-INS-8629
chr18-23015465-INS-967
chr18-23016021-INS-1029
chr18-23020146-INS-9946
chr18-23039296-INS-315
chr18-23087837-INS-76
chr18-23146883-INS-341
chr18-23148449-INS-3981
chr18-23157720-INS-7074
chr18-23200868-INS-50
chr18-23202107-INS-53
chr18-23205870-INS-1475
chr18-23230695-INS-56
chr18-23231496-INS-88
chr18-23233945-INS-2447
chr18-23237226-INS-1140
chr18-23239286-INS-649
chr18-23263587-INS-66
chr18-23270831-INS-195
chr18-23283140-INS-56
chr18-23285948-INS-144
chr18-23286024-INS-1658
chr18-23289355-INS-54
chr18-23333050-INS-52
chr18-23333073-INS-3551
chr18-23336493-INS-97
chr18-23429821-INS-51
chr18-23447718-INS-266
chr18-23469584-INS-52
chr18-23498136-INS-442
chr18-23518018-INS-50
chr18-23561122-INS-1101

chr18-23607178-INS-100
chr18-23658387-INS-156
chr18-23658543-INS-78
chr18-23660508-INS-52
chr18-23836738-INS-434
chr18-23846046-INS-6621
chr18-23872768-INS-6325
chr18-23912406-INS-108
chr18-23913048-INS-61
chr18-24022706-INS-6466
chr18-24024286-INS-51
chr18-24043141-INS-56
chr18-24048454-INS-118
chr18-24049851-INS-334
chr18-24105016-INS-6672
chr18-24171344-INS-1630
chr18-24173322-INS-58
chr18-24182176-INS-7254
chr18-24224878-INS-335
chr18-24224880-INS-50
chr18-24224975-INS-108
chr18-24238714-INS-353
chr18-24248872-INS-251
chr18-24279414-INS-60
chr18-24279765-INS-210
chr18-24280293-INS-420
chr18-24280583-INS-390
chr18-24283879-INS-198
chr18-24293390-INS-60
chr18-24324886-INS-7163
chr18-24329627-INS-206
chr18-24357183-INS-214
chr18-24368859-INS-691
chr18-24381282-INS-70
chr18-24392263-INS-110
chr18-24400044-INS-68
chr18-24426686-INS-375
chr18-24439157-INS-107
chr18-24441635-INS-90
chr18-24442037-INS-105
chr18-24443619-INS-3227
chr18-24445137-INS-195
chr18-24458426-INS-216
chr18-24459148-INS-121
chr18-24459148-INS-58
chr18-24533696-INS-7449
chr18-24560647-INS-342
chr18-24588327-INS-210
chr18-24598005-INS-79
chr18-24601845-INS-64
chr18-24602264-INS-307
chr18-24656412-INS-116
chr18-24659066-INS-50
chr18-24716185-INS-223
chr18-24763772-INS-213
chr18-24765964-INS-212
chr18-24801770-INS-59
chr18-24805109-INS-51
chr18-24806063-INS-50
chr18-24810000-INS-128
chr18-24814614-INS-936
chr18-24834242-INS-1106
chr18-24859211-INS-268
chr18-24876764-INS-862
chr18-24877052-INS-1213
chr18-24880068-INS-196
chr18-24894667-INS-96
chr18-24912088-INS-195
chr18-24913381-INS-50
chr18-24918516-INS-50
chr18-24925980-INS-164
chr18-24926348-INS-210
chr18-24926565-INS-76
chr18-24934435-INS-114
chr18-24935160-INS-335
chr18-24968580-INS-61
chr18-24987320-INS-94

chr18-24987529-INS-57
chr18-25079461-INS-54
chr18-25097208-INS-152
chr18-25119536-INS-292
chr18-25167008-INS-52
chr18-25182889-INS-52
chr18-25186325-INS-58
chr18-25191510-INS-51
chr18-25216750-INS-308
chr18-25225640-INS-54
chr18-25225794-INS-90
chr18-25264649-INS-50
chr18-25264653-INS-230
chr18-25320512-INS-72
chr18-25353796-INS-403
chr18-25379573-INS-1571
chr18-25384362-INS-254
chr18-25385556-INS-1494
chr18-25404982-INS-140
chr18-25412628-INS-1974
chr18-25413316-INS-189
chr18-25416854-INS-1116
chr18-25433360-INS-1208
chr18-25434636-INS-84
chr18-25434843-INS-1042
chr18-25505932-INS-5633
chr18-25559260-INS-472
chr18-25570152-INS-1613
chr18-25585094-INS-53
chr18-25590418-INS-52
chr18-25679303-INS-51
chr18-25682335-INS-64
chr18-25687764-INS-5708
chr18-25740885-INS-262
chr18-25757784-INS-344
chr18-25759846-INS-291
chr18-25760110-INS-80
chr18-25771314-INS-3200
chr18-25771790-INS-54
chr18-25804454-INS-55
chr18-25843082-INS-218
chr18-25846636-INS-80
chr18-25846647-INS-299
chr18-25883470-INS-52
chr18-25935295-INS-50
chr18-25950104-INS-998
chr18-25996045-INS-109
chr18-26001489-INS-243
chr18-26002554-INS-83
chr18-26002758-INS-166
chr18-26002998-INS-50
chr18-26012910-INS-94
chr18-26035552-INS-60
chr18-26055124-INS-211
chr18-26068367-INS-2981
chr18-26068779-INS-117
chr18-26074276-INS-60
chr18-26086117-INS-5223
chr18-26087939-INS-234
chr18-26155497-INS-2939
chr18-26168656-INS-3464
chr18-26176068-INS-52
chr18-26181484-INS-50
chr18-26200582-INS-401
chr18-26206840-INS-51
chr18-26208073-INS-131
chr18-26282230-INS-1913
chr18-26329138-INS-56
chr18-26359609-INS-5975
chr18-26364320-INS-158
chr18-26364531-INS-60
chr18-26365065-INS-850
chr18-26365115-INS-66
chr18-26370259-INS-125
chr18-26435485-INS-53
chr18-26470244-INS-1006
chr18-26481826-INS-6052

chr18-26507722-INS-3110
chr18-26579786-INS-7484
chr18-26587332-INS-6458
chr18-26635559-INS-50
chr18-26692773-INS-457
chr18-26767983-INS-62
chr18-26769857-INS-60
chr18-26790756-INS-63
chr18-26854654-INS-56
chr18-26959168-INS-779
chr18-26975977-INS-54
chr18-26985411-INS-1398
chr18-26985599-INS-544
chr18-26989944-INS-50
chr18-26998342-INS-54
chr18-26998346-INS-112
chr18-27012656-INS-56
chr18-27067947-INS-50
chr18-27068684-INS-207
chr18-27069135-INS-103
chr18-27069854-INS-108
chr18-27087393-INS-2159
chr18-27087398-INS-6348
chr18-27209582-INS-72
chr18-27249742-INS-53
chr18-27263407-INS-60
chr18-27263995-INS-104
chr18-27334353-INS-148
chr18-27362727-INS-68
chr18-27461109-INS-62
chr18-27464624-INS-124
chr18-27465345-INS-76
chr18-27480980-INS-93
chr18-27496022-INS-2805
chr18-27513435-INS-67
chr18-27518355-INS-2926
chr18-27531700-INS-2363
chr18-27561378-INS-4767
chr18-27562169-INS-50
chr18-27639390-INS-1281
chr18-27693920-INS-2314
chr18-27697151-INS-1024
chr18-27775733-INS-260
chr18-27783481-INS-6274
chr18-27886959-INS-56
chr18-27891052-INS-58
chr18-27895815-INS-52
chr18-27917371-INS-82
chr18-27953538-INS-688
chr18-27966581-INS-2005
chr18-27966848-INS-62
chr18-27967153-INS-52
chr18-27969219-INS-120
chr18-28108317-INS-970
chr18-28111670-INS-103
chr18-28129047-INS-361
chr18-28162011-INS-6293
chr18-28216056-INS-6576
chr18-28282430-INS-3081
chr18-28291628-INS-206
chr18-28293745-INS-70
chr18-28327789-INS-6713
chr18-28335225-INS-489
chr18-28335747-INS-181
chr18-28349951-INS-104
chr18-28389369-INS-50
chr18-28449833-INS-3064
chr18-28511428-INS-82
chr18-28526741-INS-105
chr18-28546703-INS-59
chr18-28582823-INS-524
chr18-28582851-INS-129
chr18-28624516-INS-64
chr18-28633871-INS-393
chr18-28639332-INS-56
chr18-28661514-INS-56
chr18-28669213-INS-4011

chr18-28679905-INS-128
chr18-28679922-INS-63
chr18-28681300-INS-83
chr18-28686626-INS-66
chr18-28690576-INS-56
chr18-28693317-INS-60
chr18-28697794-INS-2058
chr18-28705494-INS-54
chr18-28707680-INS-1962
chr18-28707680-INS-4118
chr18-28713802-INS-349
chr18-28772059-INS-72
chr18-28816528-INS-98
chr18-28834054-INS-56
chr18-28856297-INS-113
chr18-28920643-INS-51
chr18-28934614-INS-6301
chr18-28943282-INS-76
chr18-28982505-INS-124
chr18-29056467-INS-1192
chr18-29108926-INS-1527
chr18-29127565-INS-6173
chr18-29135000-INS-96
chr18-29141430-INS-1466
chr18-29154932-INS-22361
chr18-29176417-INS-138
chr18-29193923-INS-54
chr18-29209565-INS-1370
chr18-29211596-INS-77
chr18-29226796-INS-85
chr18-29227430-INS-113
chr18-29232514-INS-1534
chr18-29241435-INS-6005
chr18-29242291-INS-51
chr18-29242352-INS-123
chr18-29244640-INS-64
chr18-29245027-INS-80
chr18-29247194-INS-6588
chr18-29262777-INS-236
chr18-29286057-INS-2828
chr18-29313556-INS-63
chr18-29346502-INS-113
chr18-29346610-INS-65
chr18-29355937-INS-59
chr18-29377883-INS-6187
chr18-29379008-INS-56
chr18-29379524-INS-1868
chr18-29414052-INS-2660
chr18-29440426-INS-6361
chr18-29452781-INS-120
chr18-29516289-INS-152
chr18-29532959-INS-190
chr18-29552537-INS-79
chr18-29608694-INS-60
chr18-29628037-INS-58
chr18-29631270-INS-100
chr18-29631649-INS-1875
chr18-29729965-INS-55
chr18-29747795-INS-146
chr18-29763228-INS-146
chr18-29763764-INS-348
chr18-29790895-INS-1116
chr18-29829990-INS-138
chr18-29829990-INS-58
chr18-29860586-INS-64
chr18-29976437-INS-51
chr18-29976461-INS-303
chr18-29976467-INS-612
chr18-29991034-INS-56
chr18-29998405-INS-51
chr18-29998483-INS-54
chr18-30006484-INS-77
chr18-30047710-INS-400
chr18-30048282-INS-122
chr18-30050653-INS-66
chr18-30139900-INS-112
chr18-30152841-INS-64

chr18-30164628-INS-6797
chr18-30165237-INS-64
chr18-30195415-INS-94
chr18-30202582-INS-77
chr18-30255383-INS-509
chr18-30294695-INS-6192
chr18-30295041-INS-439
chr18-30295139-INS-177
chr18-30302829-INS-1122
chr18-30387153-INS-69
chr18-30458194-INS-87
chr18-30459094-INS-1733
chr18-30460038-INS-477
chr18-30491883-INS-149
chr18-30500672-INS-72
chr18-30501202-INS-190
chr18-30527267-INS-2888
chr18-30534756-INS-70
chr18-30541912-INS-3205
chr18-30561315-INS-2038
chr18-30608531-INS-128
chr18-30613935-INS-62
chr18-30677131-INS-57
chr18-30684018-INS-13605
chr18-30684018-INS-6452
chr18-30704335-INS-173
chr18-30707825-INS-161
chr18-30731934-INS-6442
chr18-30732148-INS-191
chr18-30758131-INS-543
chr18-30821847-INS-663
chr18-30826710-INS-73
chr18-30829190-INS-7444
chr18-30871118-INS-94
chr18-30871120-INS-218
chr18-30872932-INS-408
chr18-30875647-INS-8843
chr18-30950457-INS-1000
chr18-30968389-INS-1214
chr18-30982119-INS-205
chr18-31015437-INS-56
chr18-31021159-INS-52
chr18-31029063-INS-1436
chr18-31036099-INS-68
chr18-31042807-INS-6698
chr18-31050910-INS-215
chr18-31065223-INS-2358
chr18-31065712-INS-122
chr18-31067130-INS-267
chr18-31067378-INS-56
chr18-31077153-INS-3119
chr18-31090908-INS-84
chr18-31110741-INS-6363
chr18-31190752-INS-490
chr18-31225520-INS-12388
chr18-31225520-INS-6013
chr18-31253092-INS-52
chr18-31315068-INS-1539
chr18-31328740-INS-212
chr18-31331321-INS-3676
chr18-31427069-INS-215
chr18-31445547-INS-56
chr18-31531286-INS-6325
chr18-31532725-INS-1964
chr18-31534796-INS-312
chr18-31541376-INS-1434
chr18-31546154-INS-6648
chr18-31629572-INS-3016
chr18-31648292-INS-7163
chr18-31668800-INS-55
chr18-31675742-INS-412
chr18-31771362-INS-8445
chr18-31865053-INS-343
chr18-31871767-INS-69
chr18-31919512-INS-53
chr18-31931989-INS-69
chr18-31943633-INS-204

chr18-31949653-INS-3276
chr18-31949682-INS-661
chr18-31961948-INS-205
chr18-32015004-INS-60
chr18-32047367-INS-80
chr18-32052571-INS-514
chr18-32061407-INS-188
chr18-32061407-INS-421
chr18-32088100-INS-4947
chr18-32096399-INS-101
chr18-32120767-INS-54
chr18-32132636-INS-50
chr18-32166943-INS-239
chr18-32190426-INS-50
chr18-32228838-INS-1533
chr18-32240650-INS-205
chr18-32261220-INS-274
chr18-32320249-INS-6358
chr18-32323370-INS-198
chr18-32323873-INS-331
chr18-32327184-INS-417
chr18-32327271-INS-210
chr18-32347779-INS-398
chr18-32357568-INS-566
chr18-32377767-INS-59
chr18-32390838-INS-5642
chr18-32415221-INS-784
chr18-32419898-INS-90
chr18-32437527-INS-216
chr18-32439930-INS-202
chr18-32514161-INS-175
chr18-32520005-INS-188
chr18-32628040-INS-321
chr18-32653322-INS-1033
chr18-32695976-INS-81
chr18-32699722-INS-110
chr18-32706262-INS-173
chr18-32730964-INS-456
chr18-32731741-INS-72
chr18-32760553-INS-2696
chr18-32764992-INS-52
chr18-32777444-INS-117
chr18-32789802-INS-1535
chr18-32800567-INS-7105
chr18-32832383-INS-1266
chr18-32832561-INS-267
chr18-32833050-INS-1640
chr18-32843366-INS-176
chr18-32928601-INS-159
chr18-32959485-INS-6771
chr18-33008789-INS-4461
chr18-33054209-INS-247
chr18-33081118-INS-1338
chr18-33107644-INS-3150
chr18-33131492-INS-161
chr18-33132846-INS-3138
chr18-33184371-INS-78
chr18-33203499-INS-1757
chr18-33220709-INS-6014
chr18-33245970-INS-1749
chr18-33245982-INS-873
chr18-33276535-INS-72
chr18-33373370-INS-50
chr18-33375512-INS-484
chr18-33383157-INS-423
chr18-33411823-INS-1161
chr18-33446681-INS-64
chr18-33466421-INS-1592
chr18-33555866-INS-220
chr18-33556086-INS-339
chr18-33605187-INS-181
chr18-33605187-INS-72
chr18-33605387-INS-77
chr18-33641911-INS-141
chr18-33657862-INS-75
chr18-33663293-INS-423
chr18-33675790-INS-151

chr18-33675797-INS-54
chr18-33696956-INS-54
chr18-33712279-INS-6733
chr18-33741660-INS-50
chr18-33744048-INS-393
chr18-33744139-INS-1279
chr18-33777856-INS-605
chr18-33791149-INS-1887
chr18-33793968-INS-66
chr18-33799094-INS-66
chr18-33812087-INS-6477
chr18-33857990-INS-96
chr18-33869895-INS-50
chr18-33883721-INS-158
chr18-33898159-INS-70
chr18-33926171-INS-58
chr18-33932305-INS-78
chr18-33943527-INS-462
chr18-33945023-INS-101
chr18-33954903-INS-210
chr18-33958153-INS-82
chr18-33985436-INS-105
chr18-34003650-INS-223
chr18-34003664-INS-461
chr18-34017431-INS-212
chr18-34034593-INS-8695
chr18-34041674-INS-57
chr18-34044035-INS-199
chr18-34072708-INS-62
chr18-34085208-INS-150
chr18-34085225-INS-360
chr18-34094835-INS-55
chr18-34104915-INS-914
chr18-34180496-INS-110
chr18-34205626-INS-51
chr18-34225022-INS-5658
chr18-34246691-INS-4788
chr18-34255590-INS-180
chr18-34269217-INS-60
chr18-34294914-INS-116
chr18-34332073-INS-80
chr18-34336167-INS-194
chr18-34338467-INS-126
chr18-34373588-INS-429
chr18-34417368-INS-1553
chr18-34457085-INS-71
chr18-34457333-INS-223
chr18-34459359-INS-66
chr18-34461163-INS-52
chr18-34495468-INS-65
chr18-34496147-INS-174
chr18-34497005-INS-168
chr18-34497079-INS-338
chr18-34523007-INS-54
chr18-34563592-INS-141
chr18-34584870-INS-50
chr18-34645083-INS-260
chr18-34654814-INS-132
chr18-34662463-INS-1646
chr18-34663884-INS-121
chr18-34668065-INS-5392
chr18-34691241-INS-50
chr18-34699420-INS-140
chr18-34710871-INS-179
chr18-34779084-INS-219
chr18-34783497-INS-166
chr18-34783885-INS-585
chr18-34785311-INS-345
chr18-34785564-INS-53
chr18-34785721-INS-107
chr18-34790412-INS-52
chr18-34794357-INS-488
chr18-34799333-INS-6800
chr18-34832766-INS-127
chr18-34838383-INS-72
chr18-34864967-INS-69
chr18-34869878-INS-66

chr18-34875067-INS-56
chr18-34875154-INS-215
chr18-34886843-INS-186
chr18-34918048-INS-56
chr18-34919865-INS-289
chr18-34923440-INS-143
chr18-34924853-INS-60
chr18-34968632-INS-138
chr18-34981046-INS-63
chr18-34981507-INS-242
chr18-35006969-INS-204
chr18-35007679-INS-72
chr18-35017611-INS-56
chr18-35024965-INS-154
chr18-35028744-INS-123
chr18-35042601-INS-200
chr18-35095410-INS-199
chr18-35114394-INS-326
chr18-35149103-INS-7169
chr18-35170340-INS-191
chr18-35190497-INS-2399
chr18-35190617-INS-148
chr18-35199884-INS-168
chr18-35235895-INS-281
chr18-35306095-INS-52
chr18-35388649-INS-5374
chr18-35459202-INS-4013
chr18-35488412-INS-304
chr18-35560485-INS-252
chr18-35578996-INS-56
chr18-35586587-INS-1563
chr18-35588362-INS-218
chr18-35592559-INS-179
chr18-35593669-INS-62
chr18-35599649-INS-160
chr18-35671205-INS-74
chr18-35673741-INS-207
chr18-35719842-INS-192
chr18-35728812-INS-207
chr18-35737591-INS-202
chr18-35817106-INS-216
chr18-35819369-INS-227
chr18-35850100-INS-216
chr18-35858028-INS-214
chr18-35883368-INS-854
chr18-35912278-INS-219
chr18-35928893-INS-243
chr18-36030072-INS-196
chr18-36040692-INS-51
chr18-36044216-INS-1226
chr18-36056598-INS-564
chr18-36064320-INS-113
chr18-36097068-INS-1061
chr18-36103022-INS-107
chr18-36145395-INS-165
chr18-36183898-INS-212
chr18-36199166-INS-184
chr18-36255125-INS-194
chr18-36285837-INS-128
chr18-36297927-INS-53
chr18-36324841-INS-88
chr18-36338421-INS-292
chr18-36344253-INS-100
chr18-36348993-INS-70
chr18-36355715-INS-192
chr18-36356447-INS-75
chr18-36373798-INS-312
chr18-36380447-INS-300
chr18-36394162-INS-50
chr18-36470345-INS-104
chr18-36492354-INS-5349
chr18-36511799-INS-1115
chr18-36619790-INS-52
chr18-36660624-INS-50
chr18-36675941-INS-63
chr18-36681696-INS-232
chr18-36681996-INS-72

chr18-36692233-INS-168
chr18-36806812-INS-68
chr18-36806948-INS-404
chr18-36820723-INS-60
chr18-36823010-INS-4028
chr18-36831918-INS-1353
chr18-36851676-INS-4968
chr18-36864341-INS-204
chr18-36905925-INS-53
chr18-36907795-INS-214
chr18-36909385-INS-209
chr18-36923023-INS-218
chr18-36955042-INS-208
chr18-37021393-INS-54
chr18-37128572-INS-114
chr18-37141453-INS-76
chr18-37152878-INS-62
chr18-37304899-INS-3581
chr18-37337722-INS-7234
chr18-37421616-INS-6387
chr18-37446788-INS-68
chr18-37457153-INS-154
chr18-37593491-INS-57
chr18-37730074-INS-7699
chr18-37773240-INS-397
chr18-37843737-INS-54
chr18-37910244-INS-228
chr18-37926133-INS-196
chr18-38032889-INS-310
chr18-38050030-INS-57
chr18-38167859-INS-51
chr18-38198045-INS-154
chr18-38206999-INS-251
chr18-38212488-INS-62
chr18-38232711-INS-50
chr18-38256182-INS-52
chr18-38290222-INS-215
chr18-38306269-INS-64
chr18-38339622-INS-70
chr18-38357333-INS-446
chr18-38359238-INS-50
chr18-38369381-INS-186
chr18-38404998-INS-90
chr18-38437451-INS-339
chr18-38447029-INS-656
chr18-38463316-INS-239
chr18-38475152-INS-209
chr18-38490867-INS-211
chr18-38492235-INS-3268
chr18-38513431-INS-6883
chr18-38515509-INS-64
chr18-38553964-INS-70
chr18-38556360-INS-215
chr18-38556919-INS-117
chr18-38563708-INS-86
chr18-38572613-INS-52
chr18-38578047-INS-50
chr18-38604728-INS-5268
chr18-38609841-INS-768
chr18-38610410-INS-286
chr18-38645169-INS-70
chr18-38663906-INS-186
chr18-38666848-INS-111
chr18-38759618-INS-90
chr18-38791864-INS-155
chr18-38792290-INS-193
chr18-38794967-INS-215
chr18-38804143-INS-78
chr18-38804914-INS-52
chr18-38841813-INS-140
chr18-38841840-INS-317
chr18-38855938-INS-487
chr18-38864174-INS-215
chr18-38919296-INS-205
chr18-38920068-INS-140
chr18-39027339-INS-50
chr18-39049257-INS-161

chr18-39069309-INS-477
chr18-39069313-INS-106
chr18-39093184-INS-97
chr18-39170843-INS-78
chr18-39204173-INS-360
chr18-39256878-INS-76
chr18-39284480-INS-72
chr18-39308161-INS-296
chr18-39308298-INS-64
chr18-39308924-INS-50
chr18-39312514-INS-82
chr18-39317127-INS-6475
chr18-39364787-INS-173
chr18-39368063-INS-234
chr18-39454803-INS-2932
chr18-39484517-INS-127
chr18-39484563-INS-559
chr18-39511996-INS-2876
chr18-39644526-INS-449
chr18-39651817-INS-50
chr18-39699119-INS-52
chr18-39711663-INS-233
chr18-39744053-INS-54
chr18-39763613-INS-90
chr18-39855491-INS-259
chr18-39985803-INS-5062
chr18-40006077-INS-6445
chr18-40045361-INS-65
chr18-40063681-INS-61
chr18-40064773-INS-51
chr18-40069146-INS-54
chr18-40087522-INS-221
chr18-40107900-INS-452
chr18-40108175-INS-599
chr18-40123255-INS-1914
chr18-40130459-INS-6146
chr18-40191412-INS-66
chr18-40194263-INS-6453
chr18-40212828-INS-2104
chr18-40239065-INS-6498
chr18-40241015-INS-2518
chr18-40241069-INS-90
chr18-40247703-INS-320
chr18-40249849-INS-50
chr18-40252023-INS-188
chr18-40263271-INS-910
chr18-40333800-INS-485
chr18-40439333-INS-53
chr18-40553161-INS-85
chr18-40567690-INS-7286
chr18-40598678-INS-199
chr18-40630623-INS-6023
chr18-40630674-INS-572
chr18-40650105-INS-340
chr18-40650675-INS-124
chr18-40700799-INS-135
chr18-40767904-INS-236
chr18-40818193-INS-202
chr18-40819225-INS-63
chr18-40825227-INS-662
chr18-40862739-INS-730
chr18-40907423-INS-75
chr18-40932189-INS-132
chr18-40949970-INS-124
chr18-40985333-INS-6337
chr18-40988352-INS-62
chr18-41003413-INS-6455
chr18-41008002-INS-1430
chr18-41113901-INS-485
chr18-41140711-INS-1378
chr18-41155407-INS-66
chr18-41158832-INS-1575
chr18-41245727-INS-166
chr18-41246460-INS-354
chr18-41253256-INS-5506
chr18-41267107-INS-8222
chr18-41274723-INS-64

chr18-41284577-INS-385
chr18-41296171-INS-5743
chr18-41314807-INS-568
chr18-41314911-INS-758
chr18-41315458-INS-6205
chr18-41328467-INS-344
chr18-41346463-INS-187
chr18-41347927-INS-303
chr18-41355624-INS-6392
chr18-41360403-INS-96
chr18-41444887-INS-69
chr18-41462506-INS-104
chr18-41548766-INS-197
chr18-41617425-INS-1066
chr18-41617780-INS-794
chr18-41625849-INS-53
chr18-41632935-INS-1343
chr18-41677416-INS-272
chr18-41677482-INS-69
chr18-41725594-INS-252
chr18-41753498-INS-831
chr18-41753777-INS-54
chr18-41805349-INS-66
chr18-41805478-INS-438
chr18-41859803-INS-50
chr18-41906686-INS-7640
chr18-41932977-INS-360
chr18-41944489-INS-82
chr18-41973653-INS-105
chr18-41973694-INS-67
chr18-41974970-INS-2985
chr18-41976629-INS-127
chr18-42025343-INS-194
chr18-42042895-INS-51
chr18-42079104-INS-76
chr18-42117555-INS-199
chr18-42126772-INS-172
chr18-42198873-INS-62
chr18-42199413-INS-63
chr18-42206530-INS-68
chr18-42210371-INS-6886
chr18-42225849-INS-53
chr18-42245894-INS-52
chr18-42246515-INS-50
chr18-42249643-INS-159
chr18-42270657-INS-144
chr18-42270657-INS-54
chr18-42270747-INS-486
chr18-42283095-INS-2751
chr18-42302338-INS-3516
chr18-42363935-INS-151
chr18-42376618-INS-204
chr18-42377055-INS-505
chr18-42500193-INS-220
chr18-42506255-INS-83
chr18-42518036-INS-70
chr18-42540011-INS-50
chr18-42622724-INS-1888
chr18-42630111-INS-54
chr18-42642105-INS-58
chr18-42691000-INS-348
chr18-42691242-INS-278
chr18-42692034-INS-667
chr18-42734212-INS-52
chr18-42744195-INS-1869
chr18-42746910-INS-56
chr18-42747142-INS-68
chr18-42796810-INS-64
chr18-42820438-INS-184
chr18-42820438-INS-91
chr18-42840486-INS-50
chr18-42907186-INS-94
chr18-43049171-INS-56
chr18-43049368-INS-71
chr18-43055326-INS-59
chr18-43108909-INS-104
chr18-43122655-INS-3018

chr18-43143810-INS-161
chr18-43177466-INS-192
chr18-43216406-INS-1351
chr18-43216927-INS-2686
chr18-43232705-INS-597
chr18-43233627-INS-528
chr18-43336913-INS-54
chr18-43384998-INS-81
chr18-43399201-INS-68
chr18-43401744-INS-6753
chr18-43417443-INS-2738
chr18-43419240-INS-4158
chr18-43460772-INS-55
chr18-43461736-INS-63
chr18-43478618-INS-1000
chr18-43485197-INS-318
chr18-43518129-INS-3381
chr18-43534738-INS-74
chr18-43539070-INS-2411
chr18-43589178-INS-68
chr18-43591256-INS-249
chr18-43603278-INS-2606
chr18-43727999-INS-325
chr18-43728512-INS-78
chr18-43766044-INS-78
chr18-43769087-INS-1978
chr18-43780580-INS-457
chr18-43826493-INS-64
chr18-43853337-INS-2166
chr18-43869143-INS-112
chr18-43894795-INS-50
chr18-43936625-INS-76
chr18-43937082-INS-408
chr18-43938928-INS-1319
chr18-43946478-INS-1149
chr18-43947956-INS-3074
chr18-43960267-INS-157
chr18-43986987-INS-122
chr18-44090856-INS-210
chr18-44210359-INS-66
chr18-44242979-INS-116
chr18-44465080-INS-60
chr18-44545032-INS-600
chr18-44552542-INS-50
chr18-44586492-INS-51
chr18-44609517-INS-6732
chr18-44660651-INS-525
chr18-44685535-INS-71
chr18-44716580-INS-7353
chr18-44732134-INS-182
chr18-44766519-INS-53
chr18-44773866-INS-53
chr18-44835288-INS-1792
chr18-44865483-INS-61
chr18-44867129-INS-116
chr18-44867929-INS-78
chr18-44885808-INS-161
chr18-44889929-INS-207
chr18-44912952-INS-144
chr18-44914689-INS-78
chr18-44935952-INS-70
chr18-44955829-INS-146
chr18-45025747-INS-2201
chr18-45043760-INS-5448
chr18-45051126-INS-114
chr18-45051126-INS-240
chr18-45058445-INS-7136
chr18-45086188-INS-54
chr18-45101099-INS-56
chr18-45114113-INS-5313
chr18-45192447-INS-68
chr18-45226734-INS-4428
chr18-45242973-INS-114
chr18-45257850-INS-93
chr18-45333269-INS-92
chr18-45401222-INS-111
chr18-45500817-INS-56

chr18-45527137-INS-3703
chr18-45528148-INS-7405
chr18-45611502-INS-6263
chr18-45648639-INS-696
chr18-45665826-INS-195
chr18-45779660-INS-56
chr18-45819987-INS-115
chr18-45850802-INS-1877
chr18-45864917-INS-112
chr18-45910951-INS-6707
chr18-45963594-INS-2079
chr18-45978388-INS-117
chr18-45993236-INS-163
chr18-46047473-INS-6469
chr18-46132817-INS-197
chr18-46145392-INS-1000
chr18-46174743-INS-2121
chr18-46183855-INS-487
chr18-46244452-INS-150
chr18-46266488-INS-2244
chr18-46289279-INS-63
chr18-46326860-INS-1065
chr18-46409710-INS-259
chr18-46417436-INS-269
chr18-46427818-INS-1596
chr18-46466361-INS-90
chr18-46471523-INS-55
chr18-46518726-INS-354
chr18-46524423-INS-99
chr18-46532896-INS-74
chr18-46539539-INS-51
chr18-46586681-INS-66
chr18-46596689-INS-80
chr18-46614718-INS-2024
chr18-46644962-INS-204
chr18-46652370-INS-199
chr18-46688301-INS-2822
chr18-46688302-INS-7460
chr18-46689865-INS-74
chr18-46699664-INS-190
chr18-46758353-INS-220
chr18-46801136-INS-253
chr18-46801631-INS-60
chr18-46828825-INS-203
chr18-46835308-INS-412
chr18-46935244-INS-151
chr18-46942156-INS-7611
chr18-46964837-INS-101
chr18-46971022-INS-7191
chr18-47012143-INS-142
chr18-47037605-INS-203
chr18-47037869-INS-341
chr18-47050722-INS-6504
chr18-47071161-INS-324
chr18-47123049-INS-372
chr18-47131194-INS-5223
chr18-47150181-INS-54
chr18-47155598-INS-53
chr18-47216394-INS-427
chr18-47232976-INS-1917
chr18-47266232-INS-129
chr18-47266232-INS-283
chr18-47306744-INS-397
chr18-47326776-INS-88
chr18-47354033-INS-60
chr18-47373151-INS-52
chr18-47444579-INS-210
chr18-47456353-INS-77
chr18-47467087-INS-64
chr18-47512447-INS-5975
chr18-47557488-INS-66
chr18-47572461-INS-1954
chr18-47578928-INS-52
chr18-47588626-INS-206
chr18-47634904-INS-198
chr18-47686858-INS-110
chr18-47712178-INS-401

chr18-47727920-INS-508
chr18-47743374-INS-52
chr18-47763407-INS-61
chr18-47763922-INS-787
chr18-47765294-INS-245
chr18-47765453-INS-59
chr18-47793427-INS-52
chr18-47797582-INS-1044
chr18-47816891-INS-312
chr18-47816988-INS-56
chr18-47819705-INS-1860
chr18-47823485-INS-56
chr18-47914120-INS-1249
chr18-47922726-INS-2737
chr18-47950653-INS-1815
chr18-47955005-INS-68
chr18-47983959-INS-625
chr18-48086174-INS-188
chr18-48086174-INS-53
chr18-48107380-INS-101
chr18-48185959-INS-183
chr18-48205471-INS-1075
chr18-48216805-INS-671
chr18-48231646-INS-137
chr18-48273022-INS-68
chr18-48287351-INS-56
chr18-48305926-INS-6458
chr18-48310811-INS-52
chr18-48357067-INS-52
chr18-48357601-INS-224
chr18-48371770-INS-6193
chr18-48375155-INS-50
chr18-48413942-INS-3249
chr18-48415271-INS-172
chr18-48421646-INS-4491
chr18-48444606-INS-413
chr18-48449478-INS-220
chr18-48464619-INS-58
chr18-48508957-INS-524
chr18-48519941-INS-6369
chr18-48520954-INS-126
chr18-48528143-INS-1104
chr18-48610741-INS-1665
chr18-48621387-INS-1177
chr18-48655815-INS-72
chr18-48696908-INS-453
chr18-48713428-INS-6414
chr18-48715666-INS-187
chr18-48724894-INS-1337
chr18-48727638-INS-4849
chr18-48732541-INS-52
chr18-48746778-INS-54
chr18-48758431-INS-53
chr18-48774010-INS-50
chr18-48793955-INS-407
chr18-48830700-INS-95
chr18-48884783-INS-50
chr18-48901767-INS-7155
chr18-48910004-INS-999
chr18-48932918-INS-4746
chr18-48933516-INS-1358
chr18-48936604-INS-65
chr18-48950384-INS-100
chr18-49008304-INS-52
chr18-49022310-INS-351
chr18-49022310-INS-4103
chr18-49034090-INS-541
chr18-49047278-INS-1254
chr18-49085686-INS-164
chr18-49157544-INS-3069
chr18-49167357-INS-63
chr18-49226930-INS-131
chr18-49245111-INS-766
chr18-49249721-INS-484
chr18-49305707-INS-215
chr18-49321953-INS-84
chr18-49342666-INS-78

chr18-49347862-INS-50
chr18-49357514-INS-84
chr18-49414127-INS-6644
chr18-49434701-INS-105
chr18-49469147-INS-1607
chr18-49471109-INS-7316
chr18-49488408-INS-54
chr18-49522072-INS-5441
chr18-49536220-INS-51
chr18-49541232-INS-55
chr18-49568866-INS-129
chr18-49575752-INS-998
chr18-49641206-INS-444
chr18-49658157-INS-105
chr18-49662479-INS-52
chr18-49704653-INS-66
chr18-49711335-INS-54
chr18-49712107-INS-56
chr18-49713443-INS-511
chr18-49741751-INS-7928
chr18-49762515-INS-6389
chr18-49772939-INS-303
chr18-49843230-INS-306
chr18-49843230-INS-828
chr18-49880433-INS-570
chr18-49898391-INS-6348
chr18-49908812-INS-112
chr18-49913471-INS-204
chr18-49929485-INS-169
chr18-49942042-INS-4426
chr18-49953536-INS-6185
chr18-49962226-INS-191
chr18-49985678-INS-531
chr18-50089022-INS-215
chr18-50107117-INS-69
chr18-50146402-INS-200
chr18-50229017-INS-217
chr18-50248008-INS-11494
chr18-50248470-INS-208
chr18-50253266-INS-570
chr18-50269696-INS-61
chr18-50276106-INS-246
chr18-50276193-INS-542
chr18-50287085-INS-54
chr18-50301651-INS-5261
chr18-50323117-INS-145
chr18-50323119-INS-50
chr18-50323907-INS-187
chr18-50379178-INS-551
chr18-50416307-INS-2903
chr18-50423605-INS-495
chr18-50435966-INS-6213
chr18-50499416-INS-1491
chr18-50516826-INS-65
chr18-50540428-INS-53
chr18-50562067-INS-1154
chr18-50572052-INS-769
chr18-50613538-INS-359
chr18-50618643-INS-64
chr18-50623715-INS-6691
chr18-50630379-INS-65
chr18-50637081-INS-3493
chr18-50653407-INS-1266
chr18-50680549-INS-87
chr18-50817436-INS-134
chr18-50866881-INS-8665
chr18-50943921-INS-7422
chr18-50974329-INS-175
chr18-50974330-INS-445
chr18-50974509-INS-57
chr18-50998711-INS-303
chr18-51003460-INS-60
chr18-51018635-INS-59
chr18-51020414-INS-893
chr18-51085038-INS-50
chr18-51085400-INS-526
chr18-51152790-INS-50

chr18-51198248-INS-917
chr18-51239804-INS-6507
chr18-51262935-INS-50
chr18-51283204-INS-81
chr18-51291428-INS-52
chr18-51298990-INS-65
chr18-51348857-INS-100
chr18-51348905-INS-272
chr18-51349020-INS-126
chr18-51349141-INS-60
chr18-51349195-INS-278
chr18-51349598-INS-86
chr18-51350153-INS-836
chr18-51369591-INS-54
chr18-51369655-INS-134
chr18-51376415-INS-54
chr18-51402323-INS-60
chr18-51404779-INS-7351
chr18-51425096-INS-177
chr18-51431743-INS-1970
chr18-51443977-INS-288
chr18-51444286-INS-81
chr18-51457758-INS-52
chr18-51460191-INS-62
chr18-51464275-INS-122
chr18-51483570-INS-68
chr18-51497359-INS-61
chr18-51533966-INS-52
chr18-51534558-INS-50
chr18-51563308-INS-4106
chr18-51574153-INS-64
chr18-51584694-INS-6376
chr18-51613097-INS-962
chr18-51626366-INS-63
chr18-51645168-INS-1417
chr18-51653956-INS-646
chr18-51654609-INS-328
chr18-51663240-INS-1448
chr18-51714672-INS-101
chr18-51732949-INS-77
chr18-51740087-INS-6033
chr18-51741613-INS-89
chr18-51747403-INS-1065
chr18-51762282-INS-6271
chr18-51771837-INS-3241
chr18-51808362-INS-189
chr18-51836810-INS-50
chr18-51841759-INS-94
chr18-51850620-INS-50
chr18-51858438-INS-498
chr18-51882323-INS-1618
chr18-51883098-INS-100
chr18-51920361-INS-3320
chr18-51932229-INS-50
chr18-51967523-INS-85
chr18-51975673-INS-51
chr18-51982221-INS-7146
chr18-52125939-INS-66
chr18-52127288-INS-6152
chr18-52131752-INS-92
chr18-52135111-INS-50
chr18-52204860-INS-50
chr18-52209489-INS-3190
chr18-52228963-INS-12487
chr18-52238890-INS-84
chr18-52250325-INS-57
chr18-52250391-INS-148
chr18-52271984-INS-50
chr18-52284090-INS-4123
chr18-52287309-INS-535
chr18-52325438-INS-1372
chr18-52331396-INS-50
chr18-52425117-INS-6323
chr18-52453048-INS-558
chr18-52489093-INS-60
chr18-52499894-INS-528
chr18-52514678-INS-351

chr18-52538470-INS-571
chr18-52547375-INS-1466
chr18-52556965-INS-545
chr18-52585134-INS-8958
chr18-52630206-INS-61
chr18-52697806-INS-109
chr18-52716941-INS-6364
chr18-52833341-INS-63
chr18-52852544-INS-64
chr18-52853006-INS-56
chr18-52853010-INS-211
chr18-52869374-INS-78
chr18-52889417-INS-7099
chr18-52922710-INS-53
chr18-52977928-INS-98
chr18-52987786-INS-259
chr18-53006426-INS-980
chr18-53052840-INS-2468
chr18-53059492-INS-532
chr18-53064413-INS-55
chr18-53134662-INS-50
chr18-53153392-INS-75
chr18-53162011-INS-54
chr18-53224759-INS-76
chr18-53227126-INS-329
chr18-53246111-INS-64
chr18-53252776-INS-117
chr18-53315756-INS-76
chr18-53315880-INS-72
chr18-53346348-INS-2084
chr18-53355436-INS-76
chr18-53356042-INS-203
chr18-53362059-INS-4993
chr18-53370572-INS-114
chr18-53477133-INS-987
chr18-53488696-INS-80
chr18-53531832-INS-3147
chr18-53563589-INS-58
chr18-53573701-INS-60
chr18-53575082-INS-5975
chr18-53580014-INS-54
chr18-53649801-INS-6821
chr18-53651683-INS-54
chr18-53715197-INS-198
chr18-53722527-INS-186
chr18-53753920-INS-411
chr18-53791177-INS-6216
chr18-53799994-INS-86
chr18-53810492-INS-100
chr18-53839779-INS-214
chr18-53855831-INS-231
chr18-53879151-INS-74
chr18-53910231-INS-52
chr18-53920135-INS-178
chr18-53920383-INS-221
chr18-53951807-INS-60
chr18-54028204-INS-382
chr18-54047143-INS-50
chr18-54123971-INS-3265
chr18-54135947-INS-1965
chr18-54137173-INS-5545
chr18-54163572-INS-384
chr18-54171844-INS-51
chr18-54219429-INS-111
chr18-54236800-INS-71
chr18-54238971-INS-6444
chr18-54270645-INS-1154
chr18-54430491-INS-56
chr18-54431058-INS-391
chr18-54431539-INS-132
chr18-54435114-INS-6917
chr18-54457032-INS-99
chr18-54555138-INS-58
chr18-54649486-INS-62
chr18-54649548-INS-113
chr18-54700304-INS-51
chr18-54723087-INS-50

chr18-54724800-INS-65
chr18-54728252-INS-68
chr18-54738410-INS-440
chr18-54783460-INS-142
chr18-54815018-INS-102
chr18-54816910-INS-321
chr18-54843152-INS-69
chr18-54886139-INS-144
chr18-54960360-INS-4107
chr18-54984071-INS-74
chr18-54993509-INS-3099
chr18-55106736-INS-56
chr18-55113872-INS-142
chr18-55144108-INS-261
chr18-55164551-INS-858
chr18-55170000-INS-52
chr18-55201069-INS-2743
chr18-55201069-INS-7335
chr18-55233563-INS-144
chr18-55292968-INS-118
chr18-55305305-INS-75
chr18-55325379-INS-76
chr18-55385603-INS-1017
chr18-55393293-INS-64
chr18-55429868-INS-6477
chr18-55430231-INS-648
chr18-55487446-INS-53
chr18-55492120-INS-489
chr18-55504672-INS-412
chr18-55528783-INS-54
chr18-55533035-INS-244
chr18-55571412-INS-414
chr18-55727559-INS-1865
chr18-55768143-INS-51
chr18-55778595-INS-74
chr18-55795648-INS-52
chr18-55806721-INS-64
chr18-55808546-INS-6212
chr18-55840909-INS-51
chr18-55870778-INS-80
chr18-55878152-INS-3225
chr18-55879468-INS-50
chr18-55908085-INS-5183
chr18-55930934-INS-6177
chr18-55969019-INS-7024
chr18-55990530-INS-53
chr18-56052261-INS-1620
chr18-56062512-INS-4625
chr18-56089679-INS-6828
chr18-56096165-INS-324
chr18-56096213-INS-3028
chr18-56102454-INS-278
chr18-56110798-INS-332
chr18-56116211-INS-7013
chr18-56265537-INS-6401
chr18-56281186-INS-1050
chr18-56289328-INS-82
chr18-56294374-INS-88
chr18-56295252-INS-2552
chr18-56301288-INS-5998
chr18-56306206-INS-213
chr18-56336937-INS-171
chr18-56342930-INS-99
chr18-56363983-INS-425
chr18-56380087-INS-50
chr18-56380305-INS-59
chr18-56396580-INS-104
chr18-56439860-INS-50
chr18-56453940-INS-54
chr18-56459913-INS-776
chr18-56493337-INS-6593
chr18-56513392-INS-52
chr18-56546551-INS-280
chr18-56551316-INS-159
chr18-56580793-INS-62
chr18-56605381-INS-68
chr18-56605399-INS-376

chr18-56607891-INS-4942
chr18-56648805-INS-98
chr18-56650645-INS-1062
chr18-56657683-INS-104
chr18-56680990-INS-246
chr18-56733641-INS-85
chr18-56747107-INS-184
chr18-56747758-INS-73
chr18-56751052-INS-55
chr18-56753160-INS-130
chr18-56756566-INS-206
chr18-56759788-INS-52
chr18-56761571-INS-84
chr18-56766613-INS-83
chr18-56772823-INS-211
chr18-56776724-INS-54
chr18-56778221-INS-52
chr18-56789151-INS-213
chr18-56794658-INS-57
chr18-56797628-INS-52
chr18-56806590-INS-54
chr18-56809947-INS-8755
chr18-56823018-INS-184
chr18-56834478-INS-194
chr18-56835078-INS-73
chr18-56837332-INS-260
chr18-56854923-INS-52
chr18-56855453-INS-220
chr18-56866814-INS-371
chr18-56899479-INS-176
chr18-56937494-INS-619
chr18-56943497-INS-214
chr18-56991273-INS-105
chr18-57027446-INS-63
chr18-57071300-INS-56
chr18-57071589-INS-263
chr18-57201990-INS-83
chr18-57208995-INS-70
chr18-57233493-INS-111
chr18-57233923-INS-7220
chr18-57236955-INS-80
chr18-57242857-INS-1183
chr18-57299509-INS-50
chr18-57357283-INS-224
chr18-57377299-INS-99
chr18-57377368-INS-199
chr18-57452622-INS-174
chr18-57482393-INS-52
chr18-57491594-INS-80
chr18-57494995-INS-87
chr18-57497703-INS-112
chr18-57512469-INS-396
chr18-57516575-INS-199
chr18-57517642-INS-293
chr18-57550569-INS-2589
chr18-57568060-INS-54
chr18-57586578-INS-207
chr18-57607165-INS-50
chr18-57611535-INS-228
chr18-57613157-INS-6476
chr18-57616785-INS-58
chr18-57623759-INS-60
chr18-57638842-INS-65
chr18-57703386-INS-61
chr18-57708583-INS-56
chr18-57709429-INS-3422
chr18-57737224-INS-52
chr18-57757407-INS-122
chr18-57771426-INS-80
chr18-57778664-INS-60
chr18-57814718-INS-248
chr18-57817945-INS-87
chr18-57818206-INS-341
chr18-57818452-INS-716
chr18-57818820-INS-252
chr18-57818820-INS-80
chr18-57818897-INS-760

chr18-57830198-INS-82
chr18-57834895-INS-60
chr18-57867847-INS-77
chr18-57872320-INS-369
chr18-57875791-INS-2034
chr18-57883526-INS-87
chr18-57889412-INS-7091
chr18-57893266-INS-221
chr18-57916341-INS-78
chr18-57963083-INS-405
chr18-57972700-INS-588
chr18-58001796-INS-169
chr18-58004517-INS-214
chr18-58005186-INS-91
chr18-58022062-INS-231
chr18-58029011-INS-737
chr18-58089028-INS-86
chr18-58111989-INS-60
chr18-58113027-INS-434
chr18-58175429-INS-51
chr18-58221954-INS-85
chr18-58230799-INS-105
chr18-58340371-INS-64
chr18-58340409-INS-155
chr18-58358002-INS-62
chr18-58359643-INS-66
chr18-58367879-INS-80
chr18-58376850-INS-1512
chr18-58383070-INS-50
chr18-58401454-INS-6902
chr18-58412344-INS-1228
chr18-58415570-INS-260
chr18-58470138-INS-53
chr18-58473327-INS-180
chr18-58653440-INS-78
chr18-58662385-INS-83
chr18-58677550-INS-131
chr18-58682777-INS-56
chr18-58734201-INS-6452
chr18-58744219-INS-968
chr18-58757610-INS-364
chr18-58757618-INS-97
chr18-58758871-INS-2290
chr18-58770685-INS-107
chr18-58780565-INS-2979
chr18-58828321-INS-193
chr18-58860061-INS-52
chr18-58860191-INS-198
chr18-58860287-INS-73
chr18-58870071-INS-174
chr18-58940520-INS-614
chr18-58947685-INS-4703
chr18-58986956-INS-199
chr18-59037569-INS-95
chr18-59038972-INS-265
chr18-59043850-INS-302
chr18-59045828-INS-64
chr18-59048333-INS-354
chr18-59168948-INS-487
chr18-59183220-INS-762
chr18-59262139-INS-56
chr18-59287440-INS-3230
chr18-59298329-INS-53
chr18-59323776-INS-56
chr18-59372776-INS-149
chr18-59373620-INS-498
chr18-59393140-INS-4982
chr18-59436580-INS-456
chr18-59438598-INS-84
chr18-59524811-INS-56
chr18-59528281-INS-78
chr18-59551231-INS-83
chr18-59565206-INS-51
chr18-59573455-INS-2063
chr18-59603032-INS-395
chr18-59608693-INS-1298
chr18-59650357-INS-54

chr18-59660432-INS-401
chr18-59660450-INS-90
chr18-59662929-INS-778
chr18-59697150-INS-307
chr18-59702918-INS-1457
chr18-59719987-INS-76
chr18-59762573-INS-6192
chr18-59768118-INS-312
chr18-59769738-INS-2526
chr18-59777905-INS-555
chr18-59785959-INS-66
chr18-59794281-INS-137
chr18-59922403-INS-744
chr18-59942035-INS-369
chr18-59951745-INS-2937
chr18-59952192-INS-172
chr18-59954688-INS-5474
chr18-59962757-INS-861
chr18-59968466-INS-6287
chr18-59982643-INS-99
chr18-60007273-INS-964
chr18-60020271-INS-6575
chr18-60129758-INS-338
chr18-60141994-INS-52
chr18-60152349-INS-56
chr18-60156131-INS-6305
chr18-60194479-INS-57
chr18-60196440-INS-100
chr18-60215683-INS-82
chr18-60229012-INS-155
chr18-60249404-INS-70
chr18-60253277-INS-1381
chr18-60272804-INS-6436
chr18-60303135-INS-70
chr18-60307133-INS-75
chr18-60324521-INS-125
chr18-60370417-INS-6464
chr18-60381230-INS-2020
chr18-60481779-INS-918
chr18-60573187-INS-145
chr18-60593659-INS-1651
chr18-60594175-INS-113
chr18-60597780-INS-62
chr18-60668316-INS-127
chr18-60795138-INS-50
chr18-60811777-INS-52
chr18-60831407-INS-52
chr18-60858739-INS-208
chr18-60892477-INS-7241
chr18-60911860-INS-1864
chr18-60920911-INS-172
chr18-60931912-INS-198
chr18-60939976-INS-977
chr18-60966240-INS-268
chr18-60966400-INS-54
chr18-60981695-INS-2598
chr18-61001680-INS-2309
chr18-61007589-INS-56
chr18-61030794-INS-190
chr18-61060549-INS-68
chr18-61088101-INS-52
chr18-61145882-INS-58
chr18-61167119-INS-70
chr18-61207097-INS-104
chr18-61207331-INS-305
chr18-61207436-INS-110
chr18-61207844-INS-65
chr18-61244419-INS-51
chr18-61315494-INS-415
chr18-61315565-INS-135
chr18-61337702-INS-215
chr18-61338206-INS-519
chr18-61376398-INS-50
chr18-61418580-INS-210
chr18-61480696-INS-141
chr18-61484788-INS-52
chr18-61513855-INS-62

chr18-61516059-INS-144
chr18-61551859-INS-72
chr18-61552118-INS-166
chr18-61592171-INS-90
chr18-61641243-INS-95
chr18-61650657-INS-86
chr18-61650680-INS-609
chr18-61715456-INS-203
chr18-61728817-INS-113
chr18-61746028-INS-751
chr18-61758605-INS-85
chr18-61760810-INS-228
chr18-61795593-INS-59
chr18-61812033-INS-60
chr18-61833766-INS-171
chr18-61833766-INS-70
chr18-61833771-INS-394
chr18-61833879-INS-79
chr18-61834424-INS-537
chr18-61855432-INS-183
chr18-61869032-INS-50
chr18-61908059-INS-60
chr18-61946170-INS-56
chr18-61964546-INS-248
chr18-61981803-INS-232
chr18-61997433-INS-466
chr18-62031558-INS-1529
chr18-62033364-INS-660
chr18-62059701-INS-55
chr18-62078960-INS-89
chr18-62081320-INS-966
chr18-62092315-INS-617
chr18-62101790-INS-7214
chr18-62140385-INS-13664
chr18-62141938-INS-200
chr18-62193592-INS-128
chr18-62193690-INS-352
chr18-62212626-INS-135
chr18-62213156-INS-3563
chr18-62213156-INS-454
chr18-62220339-INS-116
chr18-62226115-INS-119
chr18-62268150-INS-449
chr18-62273176-INS-52
chr18-62341322-INS-50
chr18-62385862-INS-50
chr18-62397585-INS-58
chr18-62405449-INS-111
chr18-62406212-INS-70
chr18-62406228-INS-587
chr18-62406364-INS-210
chr18-62429443-INS-53
chr18-62440642-INS-74
chr18-62481795-INS-58
chr18-62522197-INS-56
chr18-62524764-INS-3227
chr18-62576205-INS-60
chr18-62626684-INS-728
chr18-62631229-INS-71
chr18-62690141-INS-447
chr18-62694507-INS-84
chr18-62694767-INS-440
chr18-62696679-INS-72
chr18-62714846-INS-6180
chr18-62724934-INS-55
chr18-62730637-INS-132
chr18-62769307-INS-111
chr18-62799917-INS-131
chr18-62802624-INS-6541
chr18-62819204-INS-140
chr18-62839457-INS-140
chr18-62872461-INS-185
chr18-62875120-INS-100
chr18-62886564-INS-374
chr18-62887260-INS-724
chr18-62891741-INS-290
chr18-62894440-INS-50

chr18-62901569-INS-56
chr18-62916647-INS-75
chr18-62939543-INS-56
chr18-62967910-INS-5060
chr18-62985787-INS-824
chr18-63090132-INS-52
chr18-63090542-INS-401
chr18-63099508-INS-66
chr18-63105907-INS-1332
chr18-63134580-INS-6207
chr18-63204798-INS-222
chr18-63205022-INS-154
chr18-63240667-INS-346
chr18-63240798-INS-700
chr18-63245021-INS-136
chr18-63306336-INS-54
chr18-63319471-INS-159
chr18-63321966-INS-248
chr18-63322011-INS-104
chr18-63346978-INS-108
chr18-63382517-INS-715
chr18-63388147-INS-55
chr18-63388213-INS-150
chr18-63402678-INS-51
chr18-63434725-INS-6153
chr18-63435927-INS-542
chr18-63436395-INS-173
chr18-63437634-INS-54
chr18-63460866-INS-736
chr18-63461040-INS-2648
chr18-63463041-INS-6442
chr18-63490846-INS-436
chr18-63553249-INS-191
chr18-63618244-INS-1831
chr18-63622190-INS-326
chr18-63632429-INS-120
chr18-63635973-INS-1468
chr18-63654676-INS-5186
chr18-63673837-INS-416
chr18-63681737-INS-54
chr18-63716832-INS-6259
chr18-63744890-INS-191
chr18-63748974-INS-2554
chr18-63774437-INS-958
chr18-63845579-INS-60
chr18-63955234-INS-150
chr18-63962433-INS-62
chr18-64029508-INS-54
chr18-64030035-INS-50
chr18-64030143-INS-206
chr18-64086793-INS-52
chr18-64097627-INS-93
chr18-64203150-INS-84
chr18-64203948-INS-500
chr18-64216515-INS-142
chr18-64243455-INS-88
chr18-64289315-INS-966
chr18-64308904-INS-6243
chr18-64321328-INS-590
chr18-64329722-INS-50
chr18-64332512-INS-1986
chr18-64363084-INS-52
chr18-64374840-INS-1575
chr18-64454732-INS-134
chr18-64496432-INS-56
chr18-64497458-INS-85
chr18-64502129-INS-54
chr18-64528851-INS-196
chr18-64544635-INS-112
chr18-64560527-INS-50
chr18-64607326-INS-130
chr18-64636320-INS-63
chr18-64636323-INS-129
chr18-64639354-INS-61
chr18-64644042-INS-958
chr18-64690236-INS-95
chr18-64703525-INS-743

chr18-64715809-INS-183
chr18-64791205-INS-52
chr18-64795166-INS-59
chr18-64798638-INS-217
chr18-64806057-INS-78
chr18-64810031-INS-453
chr18-64810080-INS-190
chr18-64816253-INS-52
chr18-64830991-INS-5047
chr18-64843498-INS-87
chr18-64878549-INS-3614
chr18-64893096-INS-252
chr18-64925251-INS-221
chr18-64935872-INS-1908
chr18-64936294-INS-50
chr18-64961294-INS-54
chr18-64963630-INS-7194
chr18-64973508-INS-51
chr18-65011034-INS-64
chr18-65044736-INS-66
chr18-65049200-INS-78
chr18-65094422-INS-2129
chr18-65129892-INS-139
chr18-65129892-INS-461
chr18-65147941-INS-80
chr18-65179732-INS-81
chr18-65183070-INS-70
chr18-65200290-INS-318
chr18-65208317-INS-120
chr18-65224213-INS-174
chr18-65295445-INS-72
chr18-65380451-INS-402
chr18-65381501-INS-185
chr18-65390279-INS-132
chr18-65405262-INS-50
chr18-65406041-INS-148
chr18-65436199-INS-52
chr18-65490390-INS-137
chr18-65506855-INS-159
chr18-65519661-INS-84
chr18-65544784-INS-347
chr18-65572267-INS-4349
chr18-65594888-INS-195
chr18-65602027-INS-63
chr18-65629513-INS-639
chr18-65647478-INS-279
chr18-65648356-INS-103
chr18-65648356-INS-217
chr18-65660905-INS-991
chr18-65662483-INS-262
chr18-65662818-INS-491
chr18-65668728-INS-391
chr18-65669168-INS-283
chr18-65677365-INS-83
chr18-65707010-INS-221
chr18-65709087-INS-5642
chr18-65750325-INS-80
chr18-65755229-INS-270
chr18-65755357-INS-106
chr18-65755357-INS-753
chr18-65755789-INS-247
chr18-65765576-INS-270
chr18-65775813-INS-136
chr18-65776861-INS-112
chr18-65799058-INS-268
chr18-65799323-INS-117
chr18-65856408-INS-70
chr18-65981934-INS-66
chr18-66000589-INS-92
chr18-66000681-INS-76
chr18-66055051-INS-149
chr18-66061686-INS-103
chr18-66105899-INS-5114
chr18-66119366-INS-60
chr18-66138980-INS-449
chr18-66142871-INS-6387
chr18-66163326-INS-132

chr18-66168994-INS-129
chr18-66169075-INS-57
chr18-66216117-INS-174
chr18-66220076-INS-58
chr18-66224668-INS-192
chr18-66234975-INS-2532
chr18-66324344-INS-6176
chr18-66378887-INS-63
chr18-66432200-INS-54
chr18-66439912-INS-5434
chr18-66475805-INS-533
chr18-66484023-INS-50
chr18-66484044-INS-128
chr18-66486256-INS-6381
chr18-66487772-INS-53
chr18-66487977-INS-75
chr18-66498268-INS-83
chr18-66498969-INS-52
chr18-66499130-INS-142
chr18-66501685-INS-278
chr18-66507687-INS-62
chr18-66525135-INS-7103
chr18-66546688-INS-51
chr18-66549358-INS-67
chr18-66556594-INS-546
chr18-66600545-INS-52
chr18-66604863-INS-4388
chr18-66628189-INS-50
chr18-66645135-INS-328
chr18-66645358-INS-717
chr18-66697427-INS-64
chr18-66701536-INS-54
chr18-66726615-INS-204
chr18-66727216-INS-1373
chr18-66729051-INS-174
chr18-66747603-INS-289
chr18-66774007-INS-120
chr18-66781619-INS-64
chr18-66782104-INS-3730
chr18-66809728-INS-70
chr18-66810127-INS-167
chr18-66818979-INS-114
chr18-66835235-INS-56
chr18-66842872-INS-252
chr18-66850888-INS-55
chr18-66850918-INS-512
chr18-66853489-INS-60
chr18-66853590-INS-66
chr18-66892287-INS-390
chr18-66905629-INS-224
chr18-66912642-INS-115
chr18-66919212-INS-1121
chr18-66927844-INS-7080
chr18-66928242-INS-188
chr18-67033640-INS-265
chr18-67055481-INS-449
chr18-67066334-INS-2538
chr18-67093569-INS-52
chr18-67097687-INS-61
chr18-67139052-INS-6197
chr18-67166862-INS-80
chr18-67207827-INS-153
chr18-67207877-INS-629
chr18-67242782-INS-201
chr18-67256845-INS-132
chr18-67261988-INS-50
chr18-67262489-INS-124
chr18-67280862-INS-108
chr18-67286477-INS-76
chr18-67345206-INS-111
chr18-67376368-INS-56
chr18-67388746-INS-515
chr18-67391460-INS-6794
chr18-67416362-INS-60
chr18-67437987-INS-1278
chr18-67449108-INS-209
chr18-67472573-INS-54

chr18-67474895-INS-212
chr18-67475184-INS-8306
chr18-67487355-INS-212
chr18-67491174-INS-66
chr18-67492324-INS-70
chr18-67496293-INS-57
chr18-67496660-INS-2109
chr18-67512387-INS-5566
chr18-67516090-INS-51
chr18-67526543-INS-208
chr18-67530330-INS-142
chr18-67532199-INS-201
chr18-67532573-INS-212
chr18-67574811-INS-226
chr18-67587675-INS-82
chr18-67629205-INS-247
chr18-67647010-INS-177
chr18-67652606-INS-507
chr18-67652752-INS-201
chr18-67670367-INS-1248
chr18-67683132-INS-56
chr18-67683957-INS-195
chr18-67698697-INS-168
chr18-67699201-INS-57
chr18-67701680-INS-226
chr18-67701962-INS-66
chr18-67702340-INS-88
chr18-67707283-INS-1062
chr18-67713045-INS-169
chr18-67726878-INS-205
chr18-67745390-INS-1286
chr18-67756931-INS-57
chr18-67758519-INS-176
chr18-67760506-INS-98
chr18-67765502-INS-207
chr18-67766972-INS-771
chr18-67767541-INS-86
chr18-67776870-INS-94
chr18-67799979-INS-140
chr18-67825257-INS-278
chr18-67842616-INS-50
chr18-67844896-INS-136
chr18-67852606-INS-168
chr18-67868719-INS-197
chr18-67876178-INS-178
chr18-67877912-INS-466
chr18-67880903-INS-58
chr18-67889821-INS-1883
chr18-67892391-INS-106
chr18-67893321-INS-122
chr18-67893597-INS-149
chr18-67896494-INS-267
chr18-67906426-INS-172
chr18-67946364-INS-60
chr18-67964449-INS-6442
chr18-67966951-INS-204
chr18-67969949-INS-444
chr18-67969973-INS-114
chr18-67991638-INS-180
chr18-67991662-INS-60
chr18-68010057-INS-210
chr18-68012586-INS-221
chr18-68015331-INS-4436
chr18-68019770-INS-2861
chr18-68023465-INS-50
chr18-68032138-INS-86
chr18-68034447-INS-968
chr18-68043563-INS-54
chr18-68077973-INS-60
chr18-68078214-INS-76
chr18-68091010-INS-208
chr18-68092234-INS-203
chr18-68099466-INS-63
chr18-68105690-INS-94
chr18-68142084-INS-63
chr18-68175200-INS-475
chr18-68194660-INS-62

chr18-68195663-INS-1139
chr18-68229014-INS-56
chr18-68245984-INS-51
chr18-68246299-INS-85
chr18-68247403-INS-84
chr18-68299846-INS-203
chr18-68300130-INS-176
chr18-68304340-INS-199
chr18-68352571-INS-73
chr18-68360961-INS-56
chr18-68365390-INS-50
chr18-68403369-INS-2512
chr18-68410395-INS-704
chr18-68412158-INS-364
chr18-68412816-INS-4867
chr18-68418126-INS-2830
chr18-68418300-INS-319
chr18-68418492-INS-422
chr18-68441605-INS-632
chr18-68448064-INS-550
chr18-68461631-INS-203
chr18-68467419-INS-7815
chr18-68516266-INS-34900
chr18-68538637-INS-276
chr18-68539145-INS-60
chr18-68543443-INS-233
chr18-68570236-INS-78
chr18-68579332-INS-81
chr18-68579476-INS-104
chr18-68581930-INS-56
chr18-68587771-INS-583
chr18-68605174-INS-784
chr18-68605641-INS-486
chr18-68652048-INS-158
chr18-68653402-INS-193
chr18-68654065-INS-65
chr18-68680331-INS-82
chr18-68682981-INS-198
chr18-68684544-INS-65
chr18-68736995-INS-71
chr18-68797264-INS-65
chr18-68805245-INS-446
chr18-68842414-INS-425
chr18-68892136-INS-76
chr18-68897006-INS-328
chr18-68907251-INS-52
chr18-68916510-INS-66
chr18-68928309-INS-204
chr18-68944004-INS-194
chr18-68949224-INS-1859
chr18-68950902-INS-52
chr18-68981196-INS-84
chr18-69019959-INS-61
chr18-69019962-INS-140
chr18-69026999-INS-153
chr18-69043410-INS-53
chr18-69048497-INS-5342
chr18-69067444-INS-120
chr18-69075188-INS-79
chr18-69085656-INS-219
chr18-69103897-INS-52
chr18-69113694-INS-70
chr18-69133086-INS-84
chr18-69159648-INS-50
chr18-69163248-INS-155
chr18-69173056-INS-2402
chr18-69197045-INS-50
chr18-69257149-INS-50
chr18-69265146-INS-50
chr18-69324223-INS-81
chr18-69324265-INS-295
chr18-69324463-INS-52
chr18-69324518-INS-278
chr18-69361910-INS-89
chr18-69363659-INS-264
chr18-69421240-INS-54
chr18-69422335-INS-6285

chr18-69465236-INS-50
chr18-69492730-INS-74
chr18-69515663-INS-197
chr18-69552075-INS-191
chr18-69602580-INS-195
chr18-69635984-INS-63
chr18-69644636-INS-80
chr18-69877718-INS-285
chr18-69901950-INS-6621
chr18-69933738-INS-1915
chr18-69936537-INS-60
chr18-69936647-INS-212
chr18-70042760-INS-6067
chr18-70112944-INS-2486
chr18-70134806-INS-236
chr18-70149793-INS-299
chr18-70218085-INS-50
chr18-70289988-INS-885
chr18-70299916-INS-6685
chr18-70329030-INS-77
chr18-70332378-INS-63
chr18-70336775-INS-100
chr18-70400571-INS-246
chr18-70474134-INS-54
chr18-70484046-INS-51
chr18-70484176-INS-7201
chr18-70555486-INS-380
chr18-70555608-INS-62
chr18-70555795-INS-87
chr18-70562915-INS-3231
chr18-70615038-INS-186
chr18-70616221-INS-7126
chr18-70667100-INS-186
chr18-70667475-INS-7254
chr18-70668735-INS-7084
chr18-70671162-INS-112
chr18-70779228-INS-73
chr18-70795162-INS-6484
chr18-70795586-INS-119
chr18-70796182-INS-57
chr18-70849257-INS-6166
chr18-70891304-INS-377
chr18-70896089-INS-55
chr18-70907567-INS-89
chr18-70945061-INS-55
chr18-70952351-INS-203
chr18-70952432-INS-362
chr18-70998743-INS-50
chr18-71018643-INS-5020
chr18-71025508-INS-54
chr18-71033218-INS-2350
chr18-71035866-INS-50
chr18-71035915-INS-527
chr18-71043911-INS-195
chr18-71043919-INS-71
chr18-71052761-INS-168
chr18-71126327-INS-93
chr18-71126524-INS-176
chr18-71133666-INS-1892
chr18-71140660-INS-125
chr18-71141101-INS-309
chr18-71186822-INS-544
chr18-71253549-INS-50
chr18-71268613-INS-498
chr18-71283135-INS-50
chr18-71295484-INS-108
chr18-71313197-INS-445
chr18-71313508-INS-443
chr18-71318102-INS-445
chr18-71324477-INS-2606
chr18-71368251-INS-2303
chr18-71374092-INS-1226
chr18-71424959-INS-500
chr18-71484205-INS-60
chr18-71518868-INS-6405
chr18-71547439-INS-1178
chr18-71559458-INS-6925

chr18-71623332-INS-52
chr18-71627026-INS-6054
chr18-71647746-INS-214
chr18-71691953-INS-96
chr18-71736943-INS-148
chr18-71848873-INS-60
chr18-71851789-INS-120
chr18-71881776-INS-56
chr18-71892203-INS-144
chr18-71906125-INS-6219
chr18-71997416-INS-6636
chr18-72014500-INS-6297
chr18-72071947-INS-230
chr18-72260316-INS-189
chr18-72270653-INS-1750
chr18-72296628-INS-6676
chr18-72297014-INS-364
chr18-72297043-INS-68
chr18-72299385-INS-50
chr18-72352598-INS-70
chr18-72384537-INS-1413
chr18-72404456-INS-50
chr18-72412812-INS-497
chr18-72430553-INS-68
chr18-72469909-INS-77
chr18-72481992-INS-50
chr18-72493380-INS-2456
chr18-72510403-INS-95
chr18-72513206-INS-51
chr18-72522961-INS-248
chr18-72530183-INS-171
chr18-72531694-INS-146
chr18-72545002-INS-352
chr18-72545273-INS-51
chr18-72548112-INS-71
chr18-72550128-INS-52
chr18-72574199-INS-1188
chr18-72582947-INS-2327
chr18-72585117-INS-90
chr18-72585206-INS-342
chr18-72601633-INS-82
chr18-72609459-INS-1552
chr18-72617143-INS-252
chr18-72650572-INS-66
chr18-72689942-INS-227
chr18-72728997-INS-1199
chr18-72745619-INS-50
chr18-72749031-INS-58
chr18-72765727-INS-980
chr18-72768706-INS-101
chr18-72768884-INS-90
chr18-72793902-INS-162
chr18-72793957-INS-51
chr18-72809906-INS-54
chr18-72862986-INS-5743
chr18-72875082-INS-276
chr18-72976161-INS-6204
chr18-72979230-INS-103
chr18-72979412-INS-132
chr18-72994171-INS-6237
chr18-73023025-INS-117
chr18-73046542-INS-108
chr18-73065891-INS-150
chr18-73108910-INS-4806
chr18-73120948-INS-56
chr18-73120980-INS-136
chr18-73136118-INS-220
chr18-73139742-INS-6363
chr18-73144088-INS-178
chr18-73171996-INS-439
chr18-73191766-INS-251
chr18-73222987-INS-146
chr18-73256739-INS-462
chr18-73264286-INS-50
chr18-73307371-INS-7711
chr18-73390108-INS-720
chr18-73401196-INS-1086

chr18-73422277-INS-1392
chr18-73423778-INS-199
chr18-73424925-INS-210
chr18-73429044-INS-101
chr18-73447365-INS-6468
chr18-73487680-INS-526
chr18-73493105-INS-2431
chr18-73511679-INS-4237
chr18-73553201-INS-121
chr18-73577411-INS-6459
chr18-73580782-INS-529
chr18-73580920-INS-2229
chr18-73582349-INS-237
chr18-73621044-INS-132
chr18-73668057-INS-72
chr18-73674027-INS-6843
chr18-73683210-INS-128
chr18-73692660-INS-66
chr18-73695069-INS-56
chr18-73754054-INS-84
chr18-73767688-INS-52
chr18-73785574-INS-1302
chr18-73829249-INS-191
chr18-73853300-INS-52
chr18-73878306-INS-52
chr18-73895467-INS-83
chr18-73908926-INS-156
chr18-73978636-INS-1187
chr18-74022959-INS-50
chr18-74032844-INS-1160
chr18-74071203-INS-587
chr18-74074622-INS-56
chr18-74121710-INS-60
chr18-74130518-INS-86
chr18-74134920-INS-70
chr18-74168472-INS-121
chr18-74183032-INS-56
chr18-74207829-INS-68
chr18-74233073-INS-132
chr18-74236246-INS-50
chr18-74236825-INS-162
chr18-74286454-INS-3458
chr18-74331578-INS-226
chr18-74337113-INS-238
chr18-74384988-INS-2262
chr18-74433954-INS-228
chr18-74441865-INS-1743
chr18-74443672-INS-4300
chr18-74450359-INS-72
chr18-74507358-INS-85
chr18-74521642-INS-50
chr18-74593042-INS-226
chr18-74593337-INS-193
chr18-74617447-INS-1082
chr18-74617585-INS-528
chr18-74623117-INS-130
chr18-74639580-INS-1229
chr18-74662279-INS-155
chr18-74689397-INS-402
chr18-74714072-INS-70
chr18-74714907-INS-124
chr18-74715340-INS-363
chr18-74721177-INS-99
chr18-74743935-INS-388
chr18-74757106-INS-58
chr18-74778854-INS-6368
chr18-74806491-INS-535
chr18-74819442-INS-100
chr18-74819514-INS-250
chr18-74839234-INS-1085
chr18-74895886-INS-156
chr18-74962341-INS-589
chr18-74969194-INS-302
chr18-74969627-INS-50
chr18-74972416-INS-70
chr18-74977849-INS-463
chr18-74980075-INS-89

chr18-74980147-INS-1104
chr18-74996509-INS-96
chr18-75002207-INS-208
chr18-75023137-INS-124
chr18-75023359-INS-485
chr18-75029511-INS-1531
chr18-75050320-INS-314
chr18-75065941-INS-826
chr18-75128407-INS-395
chr18-75128407-INS-678
chr18-75129406-INS-210
chr18-75144705-INS-191
chr18-75150380-INS-1114
chr18-75161011-INS-62
chr18-75168970-INS-211
chr18-75218890-INS-176
chr18-75219500-INS-84
chr18-75231699-INS-2038
chr18-75250602-INS-53
chr18-75270525-INS-208
chr18-75276188-INS-488
chr18-75333263-INS-315
chr18-75339410-INS-207
chr18-75371737-INS-2319
chr18-75385111-INS-825
chr18-75397498-INS-5635
chr18-75415459-INS-1870
chr18-75415459-INS-820
chr18-75415515-INS-304
chr18-75433506-INS-290
chr18-75446630-INS-793
chr18-75449335-INS-72
chr18-75494792-INS-50
chr18-75539737-INS-272
chr18-75549272-INS-51
chr18-75594616-INS-50
chr18-75610493-INS-192
chr18-75616907-INS-92
chr18-75630622-INS-52
chr18-75633949-INS-221
chr18-75646277-INS-51
chr18-75656393-INS-76
chr18-75701641-INS-58
chr18-75709503-INS-173
chr18-75733847-INS-58
chr18-75748606-INS-122
chr18-75761361-INS-100
chr18-75762284-INS-65
chr18-75768000-INS-222
chr18-75785527-INS-50
chr18-75814145-INS-51
chr18-75834134-INS-1851
chr18-75834134-INS-876
chr18-75839702-INS-1864
chr18-75890046-INS-51
chr18-75908127-INS-122
chr18-75909981-INS-2583
chr18-75914485-INS-2573
chr18-75931979-INS-130
chr18-75946887-INS-214
chr18-75951723-INS-216
chr18-76029714-INS-56
chr18-76086504-INS-52
chr18-76135475-INS-50
chr18-76220312-INS-6467
chr18-76287084-INS-52
chr18-76327470-INS-51
chr18-76340161-INS-840
chr18-76345118-INS-4995
chr18-76359504-INS-51
chr18-76401822-INS-566
chr18-76402028-INS-1056
chr18-76492768-INS-481
chr18-76504070-INS-94
chr18-76510485-INS-84
chr18-76520685-INS-51
chr18-76545003-INS-53

chr18-76549104-INS-324
chr18-76590055-INS-52
chr18-76590122-INS-836
chr18-76611232-INS-571
chr18-76614767-INS-162
chr18-76614888-INS-327
chr18-76615022-INS-129
chr18-76638487-INS-52
chr18-76656799-INS-231
chr18-76681794-INS-96
chr18-76682750-INS-50
chr18-76689974-INS-56
chr18-76720189-INS-7325
chr18-76759972-INS-92
chr18-76768409-INS-1890
chr18-76781358-INS-325
chr18-76781655-INS-981
chr18-76786938-INS-316
chr18-76788481-INS-203
chr18-76814826-INS-63
chr18-76815254-INS-399
chr18-76819558-INS-6249
chr18-76849905-INS-279
chr18-76865954-INS-1249
chr18-76883033-INS-57
chr18-76995096-INS-102
chr18-77013353-INS-1797
chr18-77029192-INS-206
chr18-77072781-INS-54
chr18-77088515-INS-2791
chr18-77090733-INS-62
chr18-77106385-INS-109
chr18-77142333-INS-109
chr18-77147545-INS-99
chr18-77166517-INS-306
chr18-77204470-INS-107
chr18-77262007-INS-208
chr18-77270129-INS-295
chr18-77337100-INS-207
chr18-77348671-INS-525
chr18-77420931-INS-113
chr18-77436919-INS-54
chr18-77457585-INS-7377
chr18-77471260-INS-50
chr18-77674940-INS-96
chr18-77675618-INS-259
chr18-77710146-INS-50
chr18-77711304-INS-195
chr18-77725418-INS-4345
chr18-77784411-INS-114
chr18-77795527-INS-236
chr18-77821649-INS-65
chr18-77821968-INS-228
chr18-77836787-INS-7100
chr18-77844416-INS-50
chr18-77859033-INS-1031
chr18-77859084-INS-107
chr18-77871046-INS-187
chr18-77893646-INS-52
chr18-77895678-INS-93
chr18-77939475-INS-80
chr18-77939907-INS-93
chr18-77945316-INS-70
chr18-77956808-INS-231
chr18-77967500-INS-152
chr18-77973329-INS-66
chr18-77977712-INS-202
chr18-77994072-INS-84
chr18-78000272-INS-60
chr18-78019880-INS-116
chr18-78020766-INS-56
chr18-78020869-INS-370
chr18-78030608-INS-150
chr18-78060462-INS-690
chr18-78074324-INS-502
chr18-78087992-INS-185
chr18-78124090-INS-197

chr18-78131832-INS-1031
chr18-78170874-INS-372
chr18-78180577-INS-305
chr18-78180597-INS-727
chr18-78180733-INS-146
chr18-78187032-INS-123
chr18-78208838-INS-184
chr18-78209669-INS-62
chr18-78216381-INS-60
chr18-78270215-INS-101
chr18-78351504-INS-50
chr18-78362411-INS-50
chr18-78378559-INS-160
chr18-78404626-INS-51
chr18-78417699-INS-75
chr18-78541876-INS-83
chr18-78541940-INS-328
chr18-78542405-INS-118
chr18-78542465-INS-252
chr18-78545387-INS-3884
chr18-78605206-INS-66
chr18-78629673-INS-82
chr18-78629798-INS-808
chr18-78633066-INS-50
chr18-78638895-INS-194
chr18-78678139-INS-282
chr18-78697524-INS-232
chr18-78736892-INS-320
chr18-78772104-INS-50
chr18-78786905-INS-282
chr18-78812659-INS-68
chr18-78882098-INS-68
chr18-78916299-INS-50
chr18-78937719-INS-110
chr18-78937976-INS-208
chr18-78945511-INS-54
chr18-78976525-INS-94
chr18-78985019-INS-270
chr18-79053972-INS-56
chr18-79057956-INS-56
chr18-79058187-INS-125
chr18-79058578-INS-409
chr18-79070694-INS-79
chr18-79089016-INS-52
chr18-79150015-INS-82
chr18-79174894-INS-137
chr18-79258327-INS-1074
chr18-79268177-INS-132
chr18-79268255-INS-387
chr18-79280516-INS-52
chr18-79300527-INS-78
chr18-79358782-INS-52
chr18-79362379-INS-11250
chr18-79442735-INS-6189
chr18-79457434-INS-174
chr18-79535263-INS-3539
chr18-79542460-INS-1006
chr18-79564692-INS-50
chr18-79572267-INS-67
chr18-79574454-INS-71
chr18-79595665-INS-50
chr18-79606001-INS-474
chr18-79659340-INS-15116
chr18-79664672-INS-3660
chr18-79670010-INS-209
chr18-79714576-INS-1092
chr18-79719544-INS-59
chr18-79736826-INS-101
chr18-79775527-INS-70
chr18-79800444-INS-255
chr18-79857170-INS-2192
chr18-79871252-INS-7068
chr18-79877973-INS-6123
chr18-79902791-INS-255
chr18-79928960-INS-75
chr18-79941561-INS-52
chr18-79942066-INS-167

chr18-79943672-INS-95
chr18-79989892-INS-55
chr18-79998436-INS-98
chr18-80008261-INS-202
chr18-80008958-INS-437
chr18-80010383-INS-3107
chr18-80030129-INS-433
chr18-80035891-INS-180
chr18-80035891-INS-80
chr18-80053042-INS-535
chr18-80055159-INS-389
chr18-80062343-INS-379
chr18-80068882-INS-379
chr18-80125518-INS-1110
chr18-80133056-INS-188
chr18-80165190-INS-179
chr18-80178189-INS-115
chr18-80178248-INS-251
chr18-80178261-INS-54
chr18-80187612-INS-119
chr18-80201600-INS-1703
chr18-80214399-INS-87
chr18-80226030-INS-1854
chr18-80239464-INS-176
chr18-80248194-INS-4358
chr18-80249238-INS-205
chr18-80249713-INS-122
chr18-80259590-INS-129
chr18-80285740-INS-642
chr18-80303027-INS-52
chr18-80303089-INS-105
chr18-80348205-INS-1754
chr18-80375848-INS-104
chr18-80376645-INS-116
chr18-80385591-INS-195
chr18-80385733-INS-393
chr18-80396482-INS-102
chr18-80396704-INS-64
chr18-80428403-INS-498
chr18-80428494-INS-222
chr18-80436085-INS-190
chr18-80479663-INS-58
chr18-80521310-INS-7144
chr18-80537131-INS-161
chr18-80581151-INS-180
chr18-80701882-INS-95
chr18-80771435-INS-250
chr18-80793785-INS-74
chr18-80832596-INS-51
chr18-80837181-INS-152
chr18-80843821-INS-235
chr18-80850946-INS-927
chr18-80851077-INS-338
chr18-80886502-INS-1147
chr18-80941478-INS-161
chr18-80969007-INS-1923
chr18-80971502-INS-64
chr18-80999403-INS-61
chr18-81006475-INS-60
chr18-81099165-INS-108
chr18-81107658-INS-54
chr18-81117059-INS-480
chr18-81136293-INS-76
chr18-81139474-INS-157
chr18-81198097-INS-64
chr18-81242950-INS-461
chr18-81252563-INS-50
chr18-81253764-INS-60
chr18-81257067-INS-180
chr18-81257081-INS-392
chr18-81262816-INS-120
chr18-81289571-INS-299
chr18-81310453-INS-330
chr18-81315639-INS-285
chr18-81316023-INS-297
chr18-81316437-INS-125
chr18-81317902-INS-111

chr18-81351246-INS-178
chr18-81399602-INS-2238
chr18-81418498-INS-72
chr18-81421115-INS-50
chr18-81434481-INS-6394
chr18-81444677-INS-156
chr18-81480315-INS-78
chr18-81489599-INS-60
chr18-81513842-INS-675
chr18-81563910-INS-104
chr18-81647352-INS-146
chr18-81657746-INS-136
chr18-81659217-INS-397
chr18-81679202-INS-229
chr18-81684407-INS-60
chr18-81706700-INS-662
chr18-81707406-INS-54
chr18-81731701-INS-54
chr18-81733205-INS-54
chr18-81742506-INS-396
chr18-81755993-INS-973
chr18-81756516-INS-64
chr18-81765095-INS-174
chr18-81765201-INS-390
chr18-81778818-INS-378
chr18-81779516-INS-196
chr18-81793376-INS-52
chr18-81838509-INS-680
chr18-81846289-INS-105
chr18-81846610-INS-52
chr18-81856373-INS-110
chr18-81864861-INS-58
chr18-81886866-INS-64
chr18-81901220-INS-56
chr18-81947326-INS-575
chr18-81984390-INS-298
chr18-81984405-INS-75
chr18-81984569-INS-50
chr18-81992798-INS-74
chr18-82011502-INS-74
chr18-82022953-INS-265
chr18-82024047-INS-89
chr18-82028994-INS-1961
chr18-82036403-INS-80
chr18-82043359-INS-6175
chr18-82063320-INS-60
chr18-82084172-INS-6639
chr18-82095340-INS-402
chr18-82140565-INS-77
chr18-82142903-INS-6421
chr18-82154031-INS-179
chr18-82176021-INS-54
chr18-82177943-INS-58
chr18-82178144-INS-51
chr18-82178328-INS-316
chr18-82182442-INS-6442
chr18-82186556-INS-58
chr18-82204195-INS-81
chr18-82230405-INS-164
chr18-82239706-INS-392
chr18-82271346-INS-54
chr18-82271347-INS-336
chr18-82280651-INS-82
chr18-82298096-INS-496
chr18-82304483-INS-129
chr18-82329959-INS-243
chr18-82339745-INS-4418
chr18-82397928-INS-384
chr18-82405750-INS-114
chr18-82453487-INS-120
chr18-82454201-INS-531
chr18-82480372-INS-52
chr18-82490190-INS-420
chr18-82501525-INS-51
chr18-82502375-INS-61
chr18-82514104-INS-82
chr18-82514380-INS-130

chr18-82522528-INS-7178
chr18-82562483-INS-250
chr18-82562720-INS-202
chr18-82604888-INS-426
chr18-82622694-INS-66
chr18-82627776-INS-945
chr18-82633669-INS-399
chr18-82633669-INS-856
chr18-82637884-INS-66
chr18-82649981-INS-108
chr18-82650277-INS-504
chr18-82665273-INS-66
chr18-82690346-INS-488
chr18-82694976-INS-702
chr18-82738041-INS-114
chr18-82751998-INS-192
chr18-82766975-INS-1093
chr18-82766975-INS-233
chr18-82779742-INS-58
chr18-82806398-INS-165
chr18-82833001-INS-237
chr18-82859079-INS-174
chr18-82883442-INS-158
chr18-82942115-INS-168
chr18-82952974-INS-102
chr18-82969796-INS-118
chr18-83003796-INS-188
chr18-83008933-INS-75
chr18-83011648-INS-53
chr18-83013179-INS-6413
chr18-83035332-INS-268
chr18-83035817-INS-107
chr18-83075952-INS-126
chr18-83083993-INS-77
chr18-83101328-INS-78
chr18-83120782-INS-452
chr18-83127214-INS-493
chr18-83137328-INS-197
chr18-83146723-INS-50
chr18-83175839-INS-180
chr18-83176928-INS-209
chr18-83214905-INS-71
chr18-83238619-INS-214
chr18-83269164-INS-52
chr18-83269625-INS-134
chr18-83273371-INS-166
chr18-83361874-INS-52
chr18-83376332-INS-52
chr18-83376390-INS-115
chr18-83392897-INS-101
chr18-83393300-INS-381
chr18-83436789-INS-4737
chr18-83449755-INS-208
chr18-83460004-INS-113
chr18-83463450-INS-134
chr18-83503369-INS-116
chr18-83503464-INS-56
chr18-83503565-INS-4433
chr18-83560148-INS-72
chr18-83588513-INS-58
chr18-83621029-INS-52
chr18-83715655-INS-462
chr18-83716105-INS-70
chr18-83749870-INS-50
chr18-83810992-INS-6393
chr18-83866459-INS-63
chr18-83869554-INS-869
chr18-83908987-INS-349
chr18-83937324-INS-51
chr18-83943873-INS-146
chr18-83953923-INS-63
chr18-84000666-INS-236
chr18-84048550-INS-150
chr18-84057707-INS-106
chr18-84083925-INS-54
chr18-84084114-INS-54
chr18-84122776-INS-51

chr18-84149560-INS-162
chr18-84214229-INS-138
chr18-84221251-INS-516
chr18-84294476-INS-343
chr18-84294956-INS-76
chr18-84326406-INS-112
chr18-84376268-INS-237
chr18-84402906-INS-6288
chr18-84418302-INS-86
chr18-84443064-INS-79
chr18-84503949-INS-51
chr18-84504048-INS-50
chr18-84538991-INS-143
chr18-84540972-INS-6360
chr18-84546121-INS-131
chr18-84582386-INS-322
chr18-84620068-INS-60
chr18-84621105-INS-76
chr18-84653691-INS-72
chr18-84654555-INS-112
chr18-84659502-INS-58
chr18-84672983-INS-1409
chr18-84680322-INS-6258
chr18-84701496-INS-140
chr18-84707869-INS-1049
chr18-84707869-INS-371
chr18-84710307-INS-180
chr18-84715113-INS-127
chr18-84721800-INS-2251
chr18-84741770-INS-58
chr18-84743161-INS-7867
chr18-84750307-INS-69
chr18-84751251-INS-108
chr18-84751862-INS-423
chr18-84752015-INS-137
chr18-84753373-INS-467
chr18-84761175-INS-630
chr18-84785975-INS-55
chr18-84788086-INS-706
chr18-84794386-INS-64
chr18-84797579-INS-1178
chr18-84826715-INS-791
chr18-84872451-INS-246
chr18-84878287-INS-488
chr18-84881466-INS-246
chr18-84893090-INS-221
chr18-84921326-INS-187
chr18-84934696-INS-50
chr18-84936268-INS-8955
chr18-84954490-INS-172
chr18-84997113-INS-50
chr18-85001607-INS-60
chr18-85009641-INS-4088
chr18-85018886-INS-94
chr18-85026904-INS-314
chr18-85034500-INS-341
chr18-85034642-INS-844
chr18-85041883-INS-1486
chr18-85041883-INS-475
chr18-85060709-INS-4889
chr18-85070493-INS-73
chr18-85101879-INS-139
chr18-85102780-INS-314
chr18-85103122-INS-324
chr18-85117992-INS-60
chr18-85145137-INS-333
chr18-85228787-INS-2768
chr18-85229388-INS-6540
chr18-85255999-INS-84
chr18-85256581-INS-62
chr18-85309750-INS-816
chr18-85332036-INS-493
chr18-85342242-INS-500
chr18-85386482-INS-139
chr18-85388392-INS-182
chr18-85452247-INS-164
chr18-85518737-INS-121

chr18-85547608-INS-255
chr18-85564289-INS-6389
chr18-85588999-INS-307
chr18-85618098-INS-50
chr18-85660738-INS-1724
chr18-85669478-INS-52
chr18-85677752-INS-238
chr18-85692029-INS-71
chr18-85717544-INS-1043
chr18-85731685-INS-2660
chr18-85731685-INS-324
chr18-85762786-INS-6213
chr18-85763491-INS-982
chr18-85781741-INS-3197
chr18-85820093-INS-3464
chr18-85987432-INS-187
chr18-85993459-INS-51
chr18-86013945-INS-6437
chr18-86041708-INS-6162
chr18-86049758-INS-184
chr18-86052874-INS-144
chr18-86060754-INS-913
chr18-86081332-INS-1086
chr18-86081756-INS-1490
chr18-86086463-INS-136
chr18-86108862-INS-1297
chr18-86148988-INS-6483
chr18-86155631-INS-50
chr18-86193846-INS-2980
chr18-86196203-INS-194
chr18-86233605-INS-147
chr18-86242150-INS-58
chr18-86247466-INS-1576
chr18-86257045-INS-53
chr18-86264355-INS-6680
chr18-86273444-INS-344
chr18-86287394-INS-341
chr18-86291176-INS-182
chr18-86303498-INS-53
chr18-86320160-INS-118
chr18-86337043-INS-2037
chr18-86382593-INS-52
chr18-86410063-INS-60
chr18-86459174-INS-124
chr18-86564911-INS-50
chr18-86565712-INS-1185
chr18-86572334-INS-6194
chr18-86576703-INS-108
chr18-86650631-INS-12256
chr18-86718786-INS-306
chr18-86724725-INS-499
chr18-86755842-INS-214
chr18-86818246-INS-1317
chr18-86913465-INS-1483
chr18-86914449-INS-3329
chr18-86929417-INS-6114
chr18-86953692-INS-529
chr18-86955325-INS-1752
chr18-86980619-INS-1749
chr18-86980804-INS-56
chr18-87018616-INS-1110
chr18-87059230-INS-172
chr18-87062283-INS-834
chr18-87089208-INS-486
chr18-87120691-INS-165
chr18-87131841-INS-529
chr18-87155410-INS-291
chr18-87155691-INS-59
chr18-87187704-INS-101
chr18-87188091-INS-118
chr18-87266122-INS-574
chr18-87323277-INS-50
chr18-87335839-INS-2830
chr18-87361677-INS-164
chr18-87409870-INS-226
chr18-87415662-INS-50
chr18-87435351-INS-2279

chr18-87436109-INS-3616
chr18-87481463-INS-6445
chr18-87500022-INS-108
chr18-87502352-INS-6242
chr18-87503504-INS-1214
chr18-87536590-INS-56
chr18-87592210-INS-402
chr18-87639977-INS-157
chr18-87690461-INS-2758
chr18-87739909-INS-50
chr18-87752369-INS-289
chr18-87770459-INS-422
chr18-87776468-INS-215
chr18-87947309-INS-2297
chr18-87948615-INS-76
chr18-87998391-INS-103
chr18-88106203-INS-2852
chr18-88229927-INS-186
chr18-88240470-INS-76
chr18-88263903-INS-1791
chr18-88290682-INS-6500
chr18-88337463-INS-64
chr18-88364361-INS-4151
chr18-88392420-INS-74
chr18-88460102-INS-7079
chr18-88519263-INS-137
chr18-88538107-INS-883
chr18-88563587-INS-80
chr18-88573296-INS-454
chr18-88573368-INS-1229
chr18-88574282-INS-50
chr18-88579820-INS-164
chr18-88602744-INS-471
chr18-88635767-INS-3205
chr18-88741861-INS-6404
chr18-88746166-INS-1477
chr18-88780294-INS-6662
chr18-88788733-INS-57
chr18-88830761-INS-8982
chr18-88847376-INS-6335
chr18-88964509-INS-60
chr18-88974207-INS-1097
chr18-89007176-INS-58
chr18-89023917-INS-148
chr18-89023917-INS-58
chr18-89047932-INS-5811
chr18-89081800-INS-6002
chr18-89249227-INS-55
chr18-89280712-INS-2205
chr18-89313106-INS-56
chr18-89352614-INS-2791
chr18-89356118-INS-277
chr18-89356122-INS-133
chr18-89357396-INS-6482
chr18-89358084-INS-294
chr18-89396205-INS-2450
chr18-89410086-INS-10321
chr18-89413551-INS-1132
chr18-89438892-INS-18396
chr18-89446607-INS-158
chr18-89474071-INS-52
chr18-89475377-INS-57
chr18-89527131-INS-723
chr18-89570060-INS-160
chr18-89573392-INS-4207
chr18-89592102-INS-1465
chr18-89599137-INS-875
chr18-89633644-INS-312
chr18-89643165-INS-122
chr18-89739982-INS-993
chr18-89759277-INS-13824
chr18-89790560-INS-6177
chr18-89831720-INS-62
chr18-89899440-INS-6560
chr18-90034227-INS-474
chr18-90047171-INS-3327
chr18-90071121-INS-111

chr18-90104179-INS-213
chr18-90169532-INS-3627
chr18-90177944-INS-494
chr18-90260635-INS-70
chr18-90260708-INS-6502
chr18-90268705-INS-50
chr18-90274171-INS-3161
chr18-90310306-INS-480
chr18-90321030-INS-3555
chr18-90386836-INS-88
chr18-90409652-INS-2000
chr18-90445275-INS-400
chr18-90462823-INS-201
chr18-90468854-INS-110
chr18-90475026-INS-316
chr18-90495437-INS-179
chr18-90513054-INS-2318
chr18-90561725-INS-1889
chr18-90583395-INS-249
chr18-90620373-INS-137
chr19-3181954-INS-118
chr19-3182682-INS-101
chr19-3255655-INS-56
chr19-3257268-INS-50
chr19-3292285-INS-11200
chr19-3403354-INS-1053
chr19-3419579-INS-70
chr19-3422349-INS-371
chr19-3422579-INS-282
chr19-3424870-INS-118
chr19-3501861-INS-152
chr19-3502232-INS-68
chr19-3517734-INS-66
chr19-3559946-INS-1223
chr19-3590876-INS-199
chr19-3591708-INS-200
chr19-3633412-INS-343
chr19-3643511-INS-462
chr19-3645078-INS-216
chr19-3657391-INS-449
chr19-3684285-INS-156
chr19-3687035-INS-55
chr19-3702546-INS-51
chr19-3710899-INS-426
chr19-3710966-INS-652
chr19-3741170-INS-205
chr19-3759718-INS-2806
chr19-3763993-INS-182
chr19-3787302-INS-168
chr19-3794500-INS-76
chr19-3795279-INS-191
chr19-3911526-INS-74
chr19-3983139-INS-201
chr19-3988137-INS-132
chr19-4007076-INS-161
chr19-4010996-INS-88
chr19-4012711-INS-59
chr19-4041559-INS-111
chr19-4056517-INS-216
chr19-4068937-INS-62
chr19-4072682-INS-368
chr19-4078130-INS-5673
chr19-4099363-INS-993
chr19-4123918-INS-50
chr19-4159418-INS-144
chr19-4175570-INS-100
chr19-4230364-INS-213
chr19-4255262-INS-60
chr19-4258532-INS-80
chr19-4265977-INS-53
chr19-4266009-INS-184
chr19-4292139-INS-187
chr19-4311032-INS-84
chr19-4338239-INS-306
chr19-4364196-INS-284
chr19-4403604-INS-67
chr19-4427939-INS-218

chr19-4481602-INS-130
chr19-4504247-INS-138
chr19-4528973-INS-249
chr19-4528973-INS-53
chr19-4535644-INS-51
chr19-4544279-INS-190
chr19-4608154-INS-105
chr19-4625052-INS-1120
chr19-4630116-INS-304
chr19-4645829-INS-52
chr19-4703916-INS-6467
chr19-4707515-INS-256
chr19-4725680-INS-2487
chr19-4739163-INS-5613
chr19-4801230-INS-66
chr19-4866247-INS-56
chr19-4866982-INS-159
chr19-4871783-INS-7465
chr19-4919188-INS-64
chr19-4925366-INS-65
chr19-4931422-INS-245
chr19-4932279-INS-228
chr19-4975706-INS-791
chr19-5007278-INS-6468
chr19-5010815-INS-504
chr19-5012808-INS-70
chr19-5015331-INS-2802
chr19-5016683-INS-180
chr19-5021224-INS-67
chr19-5023572-INS-5398
chr19-5032820-INS-139
chr19-5110408-INS-432
chr19-5126735-INS-54
chr19-5197272-INS-390
chr19-5199051-INS-105
chr19-5199106-INS-335
chr19-5199750-INS-622
chr19-5205470-INS-205
chr19-5218379-INS-50
chr19-5230543-INS-50
chr19-5247847-INS-1253
chr19-5255161-INS-74
chr19-5309943-INS-55
chr19-5310573-INS-401
chr19-5330371-INS-100
chr19-5335185-INS-90
chr19-5338263-INS-92
chr19-5347865-INS-997
chr19-5358482-INS-51
chr19-5359231-INS-1532
chr19-5359255-INS-3118
chr19-5413593-INS-65
chr19-5447115-INS-52
chr19-5456391-INS-4676
chr19-5457434-INS-76
chr19-5508185-INS-50
chr19-5573022-INS-106
chr19-5573669-INS-458
chr19-5623652-INS-176
chr19-5636096-INS-192
chr19-5650653-INS-284
chr19-5656294-INS-65
chr19-5656587-INS-141
chr19-5666976-INS-85
chr19-5728311-INS-1004
chr19-5736993-INS-336
chr19-5736993-INS-840
chr19-5788424-INS-82
chr19-5825721-INS-105
chr19-5830454-INS-211
chr19-5830968-INS-180
chr19-5838656-INS-80
chr19-5859356-INS-142
chr19-5863043-INS-200
chr19-5869617-INS-132
chr19-5873928-INS-95
chr19-5950070-INS-229

chr19-5987050-INS-122
chr19-6015873-INS-242
chr19-6016332-INS-60
chr19-6032316-INS-52
chr19-6061450-INS-224
chr19-6072121-INS-172
chr19-6072160-INS-4535
chr19-6072636-INS-1155
chr19-6081041-INS-215
chr19-6083556-INS-328
chr19-6109911-INS-76
chr19-6110278-INS-172
chr19-6115343-INS-58
chr19-6204147-INS-178
chr19-6228609-INS-52
chr19-6265025-INS-204
chr19-6299064-INS-670
chr19-6379529-INS-152
chr19-6379599-INS-588
chr19-6381612-INS-211
chr19-6486883-INS-107
chr19-6497655-INS-124
chr19-6515921-INS-67
chr19-6558935-INS-177
chr19-6565645-INS-86
chr19-6605024-INS-62
chr19-6605058-INS-1480
chr19-6614051-INS-5447
chr19-6615975-INS-51
chr19-6618791-INS-201
chr19-6627681-INS-54
chr19-6676028-INS-84
chr19-6683886-INS-60
chr19-6684365-INS-80
chr19-6684655-INS-1218
chr19-6686328-INS-56
chr19-6690452-INS-50
chr19-6705859-INS-50
chr19-6706394-INS-84
chr19-6721812-INS-816
chr19-6721824-INS-276
chr19-6726803-INS-60
chr19-6821161-INS-66
chr19-6843681-INS-68
chr19-6859537-INS-100
chr19-6880794-INS-5981
chr19-6909174-INS-64
chr19-6938961-INS-50
chr19-6985645-INS-202
chr19-7010680-INS-64
chr19-7011221-INS-210
chr19-7022667-INS-200
chr19-7089838-INS-107
chr19-7090195-INS-926
chr19-7132520-INS-53
chr19-7160559-INS-86
chr19-7179349-INS-841
chr19-7179351-INS-141
chr19-7228566-INS-195
chr19-7232608-INS-184
chr19-7270873-INS-198
chr19-7281939-INS-86
chr19-7300957-INS-52
chr19-7320000-INS-69
chr19-7320763-INS-200
chr19-7321416-INS-150
chr19-7324416-INS-380
chr19-7355875-INS-205
chr19-7362827-INS-50
chr19-7373190-INS-52
chr19-7374195-INS-211
chr19-7374807-INS-689
chr19-7385520-INS-4939
chr19-7390709-INS-55
chr19-7432130-INS-220
chr19-7437768-INS-62
chr19-7481150-INS-138

chr19-7494401-INS-246
chr19-7507124-INS-203
chr19-7542813-INS-51
chr19-7548026-INS-85
chr19-7548266-INS-65
chr19-7577970-INS-89
chr19-7594989-INS-264
chr19-7595052-INS-59
chr19-7606686-INS-6796
chr19-7630132-INS-4327
chr19-7658119-INS-609
chr19-7678655-INS-2346
chr19-7680081-INS-6473
chr19-7730610-INS-1268
chr19-7827796-INS-69
chr19-7969978-INS-1930
chr19-8021886-INS-62
chr19-8033011-INS-422
chr19-8041283-INS-73
chr19-8050153-INS-387
chr19-8210466-INS-919
chr19-8287328-INS-1943
chr19-8331002-INS-497
chr19-8334073-INS-327
chr19-8346040-INS-6615
chr19-8390711-INS-2941
chr19-8443998-INS-56
chr19-8479334-INS-4355
chr19-8526199-INS-58
chr19-8593507-INS-160
chr19-8600476-INS-50
chr19-8630816-INS-96
chr19-8647530-INS-151
chr19-8651374-INS-68
chr19-8674424-INS-2816
chr19-8681970-INS-577
chr19-8682182-INS-129
chr19-8682242-INS-54
chr19-8708529-INS-60
chr19-8709351-INS-237
chr19-8738203-INS-115
chr19-8738203-INS-56
chr19-8839518-INS-198
chr19-8877074-INS-180
chr19-8877896-INS-143
chr19-8882786-INS-64
chr19-8912564-INS-213
chr19-8924236-INS-126
chr19-9031886-INS-210
chr19-9041776-INS-205
chr19-9064473-INS-224
chr19-9083889-INS-66
chr19-9394471-INS-197
chr19-9607708-INS-3218
chr19-9631455-INS-1287
chr19-9638623-INS-282
chr19-9719491-INS-6902
chr19-9884640-INS-186
chr19-9911159-INS-52
chr19-9911583-INS-50
chr19-9937958-INS-227
chr19-9941192-INS-263
chr19-9970895-INS-64
chr19-9974259-INS-133
chr19-9976025-INS-68
chr19-9977231-INS-180
chr19-9984976-INS-217
chr19-9999124-INS-50
chr19-10054358-INS-1086
chr19-10086949-INS-4955
chr19-10103797-INS-52
chr19-10104105-INS-234
chr19-10105444-INS-2504
chr19-10138302-INS-70
chr19-10142657-INS-56
chr19-10159314-INS-222
chr19-10243121-INS-54

chr19-10252367-INS-804
chr19-10295153-INS-3369
chr19-10295171-INS-165
chr19-10295177-INS-66
chr19-10344235-INS-129
chr19-10410027-INS-52
chr19-10442897-INS-54
chr19-10444505-INS-77
chr19-10468159-INS-1370
chr19-10489058-INS-56
chr19-10504156-INS-204
chr19-10538535-INS-180
chr19-10538980-INS-174
chr19-10543182-INS-56
chr19-10550417-INS-66
chr19-10551607-INS-4087
chr19-10566954-INS-56
chr19-10587636-INS-65
chr19-10636233-INS-199
chr19-10639746-INS-236
chr19-10640452-INS-218
chr19-10692496-INS-222
chr19-10692496-INS-82
chr19-10695225-INS-108
chr19-10717708-INS-90
chr19-10722224-INS-56
chr19-10727275-INS-81
chr19-10728138-INS-1001
chr19-10728222-INS-54
chr19-10745712-INS-187
chr19-10745856-INS-73
chr19-10749923-INS-92
chr19-10749974-INS-409
chr19-10798251-INS-50
chr19-10798483-INS-76
chr19-10810916-INS-89
chr19-10825672-INS-59
chr19-10825824-INS-146
chr19-10843488-INS-75
chr19-10854998-INS-66
chr19-10856403-INS-55
chr19-10903383-INS-121
chr19-10909900-INS-50
chr19-10976569-INS-157
chr19-10998060-INS-70
chr19-11042343-INS-3143
chr19-11044237-INS-2135
chr19-11050778-INS-54
chr19-11050975-INS-62
chr19-11084588-INS-7039
chr19-11085888-INS-60
chr19-11112098-INS-52
chr19-11113291-INS-598
chr19-11120166-INS-6503
chr19-11156901-INS-4164
chr19-11251963-INS-1605
chr19-11251994-INS-278
chr19-11309642-INS-7012
chr19-11353414-INS-941
chr19-11370319-INS-290
chr19-11421823-INS-106
chr19-11435701-INS-7290
chr19-11443079-INS-66
chr19-11451564-INS-4931
chr19-11462496-INS-7144
chr19-11474409-INS-6825
chr19-11478639-INS-1629
chr19-11514609-INS-303
chr19-11521096-INS-2021
chr19-11594732-INS-219
chr19-11650160-INS-4128
chr19-11657730-INS-275
chr19-11657730-INS-558
chr19-11669355-INS-202
chr19-11670171-INS-78
chr19-11679024-INS-340
chr19-11689671-INS-3296

chr19-11695678-INS-4564
chr19-11720096-INS-58
chr19-11720417-INS-6730
chr19-11723612-INS-2529
chr19-11735041-INS-54
chr19-11744726-INS-291
chr19-11760998-INS-299
chr19-11769504-INS-188
chr19-11792893-INS-162
chr19-11793216-INS-510
chr19-11801854-INS-5620
chr19-11810745-INS-2121
chr19-11811912-INS-200
chr19-11817864-INS-223
chr19-11858387-INS-172
chr19-11871215-INS-54
chr19-11876057-INS-7085
chr19-11908372-INS-322
chr19-11931167-INS-1810
chr19-11990605-INS-203
chr19-11993631-INS-212
chr19-12010131-INS-126
chr19-12015388-INS-66
chr19-12017911-INS-56
chr19-12060421-INS-60
chr19-12062182-INS-51
chr19-12110799-INS-7486
chr19-12182453-INS-97
chr19-12190019-INS-51
chr19-12241581-INS-54
chr19-12257595-INS-6194
chr19-12271731-INS-6157
chr19-12290383-INS-1513
chr19-12304131-INS-2848
chr19-12304805-INS-1458
chr19-12307303-INS-131
chr19-12318792-INS-70
chr19-12328781-INS-50
chr19-12336406-INS-51
chr19-12349210-INS-416
chr19-12394256-INS-60
chr19-12456762-INS-5825
chr19-12459112-INS-53
chr19-12459430-INS-6442
chr19-12471763-INS-62
chr19-12479312-INS-70
chr19-12504654-INS-120
chr19-12623350-INS-217
chr19-12643618-INS-6359
chr19-12703146-INS-366
chr19-12713397-INS-56
chr19-12718115-INS-144
chr19-12719511-INS-54
chr19-12811279-INS-72
chr19-12811301-INS-168
chr19-12811811-INS-135
chr19-12827517-INS-72
chr19-12829149-INS-60
chr19-12834177-INS-2069
chr19-12848253-INS-610
chr19-12855063-INS-526
chr19-12920177-INS-81
chr19-12928114-INS-182
chr19-12935854-INS-497
chr19-12936084-INS-185
chr19-12936084-INS-50
chr19-12979651-INS-342
chr19-12983073-INS-160
chr19-12994669-INS-85
chr19-13075756-INS-221
chr19-13076058-INS-307
chr19-13127519-INS-140
chr19-13134066-INS-448
chr19-13146637-INS-56
chr19-13159162-INS-1484
chr19-13166368-INS-1329
chr19-13190850-INS-21878

chr19-13191002-INS-1807
chr19-13214008-INS-204
chr19-13255784-INS-52
chr19-13302993-INS-6678
chr19-13410483-INS-2249
chr19-13419856-INS-50
chr19-13445854-INS-56
chr19-13456811-INS-6920
chr19-13488274-INS-64
chr19-13522701-INS-5695
chr19-13528965-INS-313
chr19-13536758-INS-71
chr19-13540939-INS-102
chr19-13565121-INS-68
chr19-13567000-INS-200
chr19-13573728-INS-850
chr19-13578849-INS-52
chr19-13586563-INS-177
chr19-13631231-INS-104
chr19-13650121-INS-177
chr19-13650121-INS-51
chr19-13662048-INS-204
chr19-13664260-INS-3251
chr19-13732158-INS-179
chr19-13815157-INS-3099
chr19-13821589-INS-231
chr19-13855296-INS-390
chr19-13855819-INS-4714
chr19-13921041-INS-50
chr19-13921716-INS-53
chr19-13934515-INS-6467
chr19-13936624-INS-87
chr19-13956869-INS-6204
chr19-13974925-INS-173
chr19-14011658-INS-185
chr19-14021645-INS-537
chr19-14026956-INS-185
chr19-14031304-INS-4456
chr19-14043036-INS-186
chr19-14050543-INS-6204
chr19-14050907-INS-1798
chr19-14063331-INS-65
chr19-14076471-INS-141
chr19-14078831-INS-400
chr19-14110459-INS-1045
chr19-14124191-INS-7229
chr19-14153331-INS-401
chr19-14191828-INS-192
chr19-14276445-INS-2121
chr19-14290490-INS-6242
chr19-14343484-INS-51
chr19-14349735-INS-64
chr19-14349897-INS-2040
chr19-14350047-INS-6307
chr19-14374512-INS-2322
chr19-14378490-INS-71
chr19-14379271-INS-331
chr19-14505641-INS-72
chr19-14513761-INS-227
chr19-14519880-INS-343
chr19-14535504-INS-431
chr19-14536364-INS-124
chr19-14557547-INS-203
chr19-14559841-INS-204
chr19-14603503-INS-64
chr19-14618018-INS-50
chr19-14668191-INS-209
chr19-14669145-INS-200
chr19-14673217-INS-4381
chr19-14697034-INS-1301
chr19-14726197-INS-641
chr19-14793054-INS-1315
chr19-14871004-INS-108
chr19-14962721-INS-208
chr19-14962813-INS-358
chr19-14993351-INS-1489
chr19-15050199-INS-261

chr19-15050265-INS-120
chr19-15077174-INS-54
chr19-15118856-INS-5905
chr19-15164356-INS-6461
chr19-15176630-INS-832
chr19-15225298-INS-380
chr19-15226620-INS-256
chr19-15243337-INS-101
chr19-15243493-INS-445
chr19-15275385-INS-204
chr19-15351793-INS-538
chr19-15384099-INS-6203
chr19-15412927-INS-2500
chr19-15418652-INS-94
chr19-15456958-INS-184
chr19-15487493-INS-1307
chr19-15530899-INS-56
chr19-15586412-INS-50
chr19-15604776-INS-112
chr19-15640903-INS-96
chr19-15641413-INS-180
chr19-15642175-INS-60
chr19-15643176-INS-129
chr19-15652937-INS-351
chr19-15665623-INS-123
chr19-15721240-INS-70
chr19-15742415-INS-6566
chr19-15744147-INS-1981
chr19-15763536-INS-52
chr19-15763551-INS-354
chr19-15780302-INS-59
chr19-15787842-INS-50
chr19-15796589-INS-105
chr19-15796613-INS-220
chr19-15834114-INS-68
chr19-15842022-INS-62
chr19-15856362-INS-58
chr19-15856377-INS-236
chr19-15871751-INS-52
chr19-15879224-INS-904
chr19-15879678-INS-65
chr19-15881101-INS-8327
chr19-15893553-INS-52
chr19-15904219-INS-135
chr19-15917898-INS-1093
chr19-15918001-INS-54
chr19-15966181-INS-84
chr19-15991026-INS-212
chr19-16006085-INS-107
chr19-16023512-INS-51
chr19-16065874-INS-4432
chr19-16095174-INS-55
chr19-16095421-INS-125
chr19-16102699-INS-72
chr19-16103836-INS-60
chr19-16132539-INS-139
chr19-16171912-INS-70
chr19-16190364-INS-237
chr19-16195811-INS-52
chr19-16250955-INS-112
chr19-16251725-INS-171
chr19-16311300-INS-114
chr19-16333396-INS-52
chr19-16383169-INS-4990
chr19-16424513-INS-55
chr19-16451575-INS-208
chr19-16461503-INS-64
chr19-16541763-INS-97
chr19-16565232-INS-55
chr19-16574054-INS-150
chr19-16637258-INS-145
chr19-16637301-INS-64
chr19-16679636-INS-148
chr19-16680132-INS-3711
chr19-16741322-INS-1421
chr19-16748578-INS-76
chr19-16773415-INS-69

chr19-16779063-INS-148
chr19-16795782-INS-128
chr19-16837699-INS-78
chr19-16867856-INS-58
chr19-16874516-INS-105
chr19-16884093-INS-50
chr19-16889901-INS-67
chr19-16902498-INS-198
chr19-16916476-INS-6428
chr19-16923893-INS-4088
chr19-16929871-INS-50
chr19-16962703-INS-58
chr19-16962909-INS-218
chr19-16988452-INS-54
chr19-17015953-INS-6253
chr19-17069640-INS-52
chr19-17090251-INS-86
chr19-17138946-INS-54
chr19-17139516-INS-2310
chr19-17204847-INS-64
chr19-17243082-INS-125
chr19-17243163-INS-129
chr19-17254302-INS-2805
chr19-17270236-INS-367
chr19-17273206-INS-60
chr19-17293069-INS-131
chr19-17295829-INS-64
chr19-17304348-INS-57
chr19-17304406-INS-175
chr19-17312798-INS-1622
chr19-17318822-INS-3133
chr19-17318926-INS-55
chr19-17319024-INS-119
chr19-17322483-INS-50
chr19-17346911-INS-214
chr19-17354876-INS-510
chr19-17359729-INS-4864
chr19-17399273-INS-8735
chr19-17410069-INS-64
chr19-17501604-INS-84
chr19-17501914-INS-285
chr19-17511503-INS-50
chr19-17526380-INS-64
chr19-17526639-INS-69
chr19-17561401-INS-129
chr19-17572442-INS-824
chr19-17577002-INS-378
chr19-17577377-INS-52
chr19-17661374-INS-54
chr19-17786799-INS-66
chr19-17791554-INS-56
chr19-17932188-INS-50
chr19-17949124-INS-7440
chr19-17973132-INS-60
chr19-17978443-INS-64
chr19-17995723-INS-196
chr19-18021350-INS-56
chr19-18040911-INS-52
chr19-18047117-INS-51
chr19-18052600-INS-62
chr19-18053305-INS-1312
chr19-18057594-INS-57
chr19-18078390-INS-373
chr19-18079950-INS-68
chr19-18112757-INS-255
chr19-18128565-INS-212
chr19-18129696-INS-50
chr19-18227793-INS-66
chr19-18277197-INS-191
chr19-18288286-INS-2684
chr19-18291365-INS-6242
chr19-18307918-INS-6308
chr19-18311357-INS-60
chr19-18332685-INS-1494
chr19-18359326-INS-335
chr19-18363215-INS-200
chr19-18387992-INS-64

chr19-18443467-INS-6219
chr19-18443981-INS-77
chr19-18469844-INS-189
chr19-18526411-INS-60
chr19-18536516-INS-658
chr19-18536518-INS-122
chr19-18543366-INS-52
chr19-18543785-INS-7087
chr19-18547861-INS-208
chr19-18548353-INS-53
chr19-18592957-INS-2015
chr19-18620366-INS-431
chr19-18639534-INS-346
chr19-18639903-INS-180
chr19-18686690-INS-1859
chr19-18741532-INS-1264
chr19-18742917-INS-7879
chr19-18765375-INS-205
chr19-18765520-INS-574
chr19-18770854-INS-1120
chr19-18770984-INS-63
chr19-18835728-INS-1959
chr19-18867312-INS-58
chr19-18875434-INS-144
chr19-18890287-INS-56
chr19-18908292-INS-51
chr19-19009103-INS-155
chr19-19049794-INS-52
chr19-19113990-INS-2634
chr19-19117308-INS-2090
chr19-19140778-INS-68
chr19-19160757-INS-64
chr19-19175081-INS-7218
chr19-19191963-INS-216
chr19-19192090-INS-251
chr19-19221270-INS-6680
chr19-19250755-INS-7116
chr19-19268945-INS-7297
chr19-19274204-INS-125
chr19-19292126-INS-135
chr19-19316237-INS-52
chr19-19338894-INS-54
chr19-19349657-INS-6840
chr19-19417167-INS-151
chr19-19419801-INS-55
chr19-19452465-INS-1676
chr19-19549310-INS-2470
chr19-19549563-INS-193
chr19-19554897-INS-602
chr19-19569268-INS-3533
chr19-19603167-INS-186
chr19-19656191-INS-6352
chr19-19678992-INS-134
chr19-19679303-INS-51
chr19-19702282-INS-1565
chr19-19713445-INS-3417
chr19-19731618-INS-7275
chr19-19846930-INS-138
chr19-19940926-INS-1028
chr19-19945907-INS-506
chr19-19948387-INS-337
chr19-19961397-INS-572
chr19-19966035-INS-52
chr19-19972679-INS-2895
chr19-19993435-INS-1412
chr19-20003869-INS-321
chr19-20004050-INS-6123
chr19-20011901-INS-52
chr19-20020130-INS-54
chr19-20032299-INS-6369
chr19-20038729-INS-4807
chr19-20056087-INS-68
chr19-20056631-INS-124
chr19-20071316-INS-7291
chr19-20145615-INS-495
chr19-20181416-INS-54
chr19-20181488-INS-126

chr19-20181817-INS-87
chr19-20198464-INS-6596
chr19-20206895-INS-108
chr19-20249007-INS-2755
chr19-20274216-INS-1595
chr19-20288626-INS-84
chr19-20289219-INS-58
chr19-20302541-INS-92
chr19-20302747-INS-159
chr19-20304077-INS-6420
chr19-20369223-INS-87
chr19-20377453-INS-54
chr19-20400881-INS-817
chr19-20414964-INS-285
chr19-20415076-INS-4903
chr19-20419748-INS-3908
chr19-20465465-INS-6513
chr19-20473472-INS-417
chr19-20477991-INS-162
chr19-20479657-INS-72
chr19-20550784-INS-212
chr19-20551032-INS-212
chr19-20580259-INS-54
chr19-20583247-INS-50
chr19-20646462-INS-50
chr19-20660534-INS-6097
chr19-20664568-INS-2576
chr19-20726293-INS-1577
chr19-20755515-INS-207
chr19-20756259-INS-7125
chr19-20757793-INS-6374
chr19-20851523-INS-143
chr19-20866147-INS-6615
chr19-20881584-INS-64
chr19-20978636-INS-6244
chr19-21004722-INS-6409
chr19-21019835-INS-202
chr19-21047608-INS-3747
chr19-21055040-INS-4808
chr19-21067635-INS-102
chr19-21079241-INS-106
chr19-21084202-INS-111
chr19-21091532-INS-202
chr19-21103096-INS-1192
chr19-21106286-INS-58
chr19-21110013-INS-62
chr19-21119112-INS-50
chr19-21151175-INS-135
chr19-21195720-INS-427
chr19-21214363-INS-84
chr19-21231728-INS-58
chr19-21283720-INS-2975
chr19-21283925-INS-54
chr19-21292330-INS-3156
chr19-21310298-INS-52
chr19-21321702-INS-3055
chr19-21344285-INS-240
chr19-21367393-INS-206
chr19-21378872-INS-326
chr19-21384653-INS-50
chr19-21384661-INS-315
chr19-21409671-INS-6538
chr19-21411312-INS-278
chr19-21436423-INS-95
chr19-21440432-INS-184
chr19-21440432-INS-86
chr19-21455742-INS-7275
chr19-21476604-INS-50
chr19-21481121-INS-702
chr19-21495972-INS-256
chr19-21526833-INS-76
chr19-21531922-INS-173
chr19-21580577-INS-56
chr19-21580731-INS-323
chr19-21580961-INS-1164
chr19-21581181-INS-309
chr19-21651770-INS-77

chr19-21684034-INS-2828
chr19-21686422-INS-372
chr19-21713334-INS-250
chr19-21745411-INS-1864
chr19-21751893-INS-66
chr19-21781459-INS-66
chr19-21796199-INS-374
chr19-21798810-INS-50
chr19-21837342-INS-58
chr19-21845305-INS-500
chr19-21877705-INS-242
chr19-21879766-INS-1945
chr19-21889032-INS-568
chr19-21889092-INS-164
chr19-21916831-INS-50
chr19-21977770-INS-132
chr19-21977835-INS-112
chr19-21989016-INS-654
chr19-21995159-INS-2819
chr19-22013003-INS-144
chr19-22014034-INS-6178
chr19-22027115-INS-77
chr19-22038027-INS-263
chr19-22056139-INS-69
chr19-22056139-INS-831
chr19-22056827-INS-2713
chr19-22131048-INS-4779
chr19-22143033-INS-56
chr19-22145832-INS-1311
chr19-22261402-INS-50
chr19-22282997-INS-74
chr19-22284895-INS-2740
chr19-22286617-INS-1889
chr19-22303755-INS-463
chr19-22340349-INS-343
chr19-22367667-INS-7252
chr19-22371542-INS-7131
chr19-22379914-INS-197
chr19-22379991-INS-399
chr19-22487316-INS-58
chr19-22493491-INS-4776
chr19-22499630-INS-88
chr19-22523721-INS-92
chr19-22556401-INS-1196
chr19-22565488-INS-54
chr19-22565777-INS-172
chr19-22573297-INS-60
chr19-22583594-INS-58
chr19-22617939-INS-90
chr19-22632444-INS-1312
chr19-22658259-INS-1656
chr19-22663448-INS-6788
chr19-22664587-INS-70
chr19-22664782-INS-199
chr19-22669232-INS-58
chr19-22684589-INS-516
chr19-22730267-INS-149
chr19-22756947-INS-260
chr19-22792604-INS-51
chr19-22817509-INS-201
chr19-22831349-INS-598
chr19-22831378-INS-125
chr19-22831484-INS-169
chr19-22831682-INS-433
chr19-22849726-INS-209
chr19-22880367-INS-483
chr19-22886169-INS-56
chr19-22897591-INS-129
chr19-22897591-INS-367
chr19-22911849-INS-1871
chr19-22930173-INS-156
chr19-22931668-INS-454
chr19-22963561-INS-54
chr19-22982146-INS-111
chr19-22995092-INS-3180
chr19-23003232-INS-3896
chr19-23008919-INS-99

chr19-23020601-INS-72
chr19-23031224-INS-123
chr19-23031563-INS-540
chr19-23033046-INS-156
chr19-23042411-INS-222
chr19-23059886-INS-178
chr19-23070625-INS-2394
chr19-23082280-INS-230
chr19-23088987-INS-72
chr19-23100596-INS-207
chr19-23134196-INS-202
chr19-23154702-INS-211
chr19-23164174-INS-58
chr19-23178654-INS-57
chr19-23216017-INS-1352
chr19-23217588-INS-233
chr19-23217630-INS-109
chr19-23254497-INS-1039
chr19-23256268-INS-60
chr19-23315516-INS-81
chr19-23347359-INS-57
chr19-23347360-INS-136
chr19-23353132-INS-210
chr19-23423207-INS-142
chr19-23425912-INS-86
chr19-23432110-INS-54
chr19-23433912-INS-54
chr19-23454921-INS-50
chr19-23481768-INS-8454
chr19-23506232-INS-252
chr19-23531646-INS-126
chr19-23532006-INS-295
chr19-23549120-INS-63
chr19-23592547-INS-56
chr19-23626836-INS-685
chr19-23643282-INS-50
chr19-23682387-INS-70
chr19-23745410-INS-67
chr19-23795666-INS-122
chr19-23820732-INS-158
chr19-23857623-INS-76
chr19-23867504-INS-203
chr19-23923361-INS-56
chr19-23966098-INS-115
chr19-23971436-INS-94
chr19-23992466-INS-261
chr19-24020396-INS-921
chr19-24020571-INS-192
chr19-24043319-INS-50
chr19-24046842-INS-480
chr19-24063708-INS-629
chr19-24079815-INS-140
chr19-24132021-INS-326
chr19-24177189-INS-64
chr19-24177199-INS-151
chr19-24178406-INS-216
chr19-24185360-INS-72
chr19-24197224-INS-90
chr19-24197246-INS-250
chr19-24217814-INS-1490
chr19-24229916-INS-562
chr19-24231470-INS-228
chr19-24232205-INS-64
chr19-24256964-INS-4851
chr19-24271545-INS-322
chr19-24272059-INS-57
chr19-24274147-INS-105
chr19-24323421-INS-208
chr19-24334610-INS-55
chr19-24360983-INS-218
chr19-24510934-INS-56
chr19-24542613-INS-144
chr19-24568639-INS-203
chr19-24600958-INS-60
chr19-24602235-INS-60
chr19-24604378-INS-528
chr19-24604383-INS-3144

chr19-24640422-INS-84
chr19-24718826-INS-88
chr19-24722934-INS-50
chr19-24756830-INS-52
chr19-24831920-INS-123
chr19-24831921-INS-248
chr19-24835586-INS-239
chr19-24835718-INS-115
chr19-24843297-INS-99
chr19-24849409-INS-84
chr19-24869312-INS-50
chr19-24875342-INS-52
chr19-24875393-INS-151
chr19-24949746-INS-7127
chr19-24959959-INS-56
chr19-24970048-INS-54
chr19-24998116-INS-7176
chr19-25000622-INS-50
chr19-25003331-INS-56
chr19-25004007-INS-70
chr19-25004486-INS-51
chr19-25022033-INS-259
chr19-25022181-INS-53
chr19-25071941-INS-6383
chr19-25092966-INS-23223
chr19-25102303-INS-54
chr19-25102310-INS-422
chr19-25176448-INS-50
chr19-25195873-INS-94
chr19-25200598-INS-60
chr19-25250915-INS-56
chr19-25274137-INS-164
chr19-25349500-INS-460
chr19-25353917-INS-235
chr19-25354229-INS-248
chr19-25422969-INS-197
chr19-25425252-INS-56
chr19-25431562-INS-1663
chr19-25433771-INS-80
chr19-25433827-INS-719
chr19-25442278-INS-96
chr19-25449173-INS-153
chr19-25452156-INS-203
chr19-25485869-INS-257
chr19-25493085-INS-58
chr19-25506421-INS-1953
chr19-25508973-INS-202
chr19-25531521-INS-78
chr19-25536017-INS-402
chr19-25540707-INS-207
chr19-25542721-INS-78
chr19-25598639-INS-55
chr19-25656604-INS-99
chr19-25684569-INS-80
chr19-25685139-INS-124
chr19-25696851-INS-6494
chr19-25715603-INS-578
chr19-25720323-INS-1289
chr19-25741804-INS-10608
chr19-25756562-INS-182
chr19-25763136-INS-712
chr19-25763232-INS-182
chr19-25767539-INS-58
chr19-25821019-INS-8191
chr19-25891332-INS-2599
chr19-25893401-INS-1088
chr19-25897450-INS-3460
chr19-25927689-INS-413
chr19-25930012-INS-154
chr19-25966482-INS-12829
chr19-25980073-INS-494
chr19-26039490-INS-345
chr19-26076869-INS-199
chr19-26081498-INS-2835
chr19-26086882-INS-93
chr19-26147824-INS-247
chr19-26157423-INS-6358

chr19-26167584-INS-63
chr19-26181710-INS-52
chr19-26267694-INS-3693
chr19-26272679-INS-6616
chr19-26277049-INS-54
chr19-26350813-INS-523
chr19-26370482-INS-94
chr19-26497285-INS-96
chr19-26513809-INS-1530
chr19-26514343-INS-75
chr19-26568092-INS-81
chr19-26615065-INS-58
chr19-26658695-INS-51
chr19-26670661-INS-97
chr19-26692052-INS-73
chr19-26706858-INS-69
chr19-26707117-INS-153
chr19-26748084-INS-80
chr19-26786116-INS-144
chr19-26789665-INS-58
chr19-26794838-INS-75
chr19-26795171-INS-1690
chr19-26796589-INS-61
chr19-26823202-INS-104
chr19-26828734-INS-211
chr19-26829261-INS-211
chr19-26835755-INS-60
chr19-26915017-INS-6205
chr19-26939673-INS-83
chr19-26939830-INS-332
chr19-26939838-INS-150
chr19-26979888-INS-51
chr19-26994535-INS-6194
chr19-27008068-INS-50
chr19-27021974-INS-115
chr19-27024862-INS-98
chr19-27058461-INS-94
chr19-27060127-INS-71
chr19-27072088-INS-732
chr19-27074355-INS-7215
chr19-27098714-INS-54
chr19-27184788-INS-111
chr19-27185117-INS-746
chr19-27219328-INS-66
chr19-27219676-INS-804
chr19-27426152-INS-66
chr19-27429990-INS-384
chr19-27429994-INS-172
chr19-27429994-INS-72
chr19-27475414-INS-70
chr19-27599409-INS-80
chr19-27603131-INS-180
chr19-27628781-INS-486
chr19-27636864-INS-139
chr19-27657215-INS-50
chr19-27667858-INS-2493
chr19-27681748-INS-1240
chr19-27685116-INS-4843
chr19-27699482-INS-70
chr19-27719829-INS-1557
chr19-27747092-INS-357
chr19-27749490-INS-57
chr19-27768735-INS-6330
chr19-27817192-INS-4978
chr19-27913662-INS-52
chr19-27956231-INS-50
chr19-27957921-INS-60
chr19-28135630-INS-2230
chr19-28140213-INS-57
chr19-28176742-INS-58
chr19-28194809-INS-1333
chr19-28246549-INS-7847
chr19-28255285-INS-68
chr19-28255342-INS-90
chr19-28285252-INS-89
chr19-28288540-INS-1766
chr19-28288540-INS-8169

chr19-28348227-INS-280
chr19-28349577-INS-64
chr19-28389365-INS-70
chr19-28465925-INS-60
chr19-28483902-INS-2530
chr19-28507788-INS-51
chr19-28575239-INS-1098
chr19-28614766-INS-160
chr19-28715598-INS-333
chr19-28738477-INS-72
chr19-28738560-INS-174
chr19-28762385-INS-72
chr19-28773994-INS-3376
chr19-28778875-INS-54
chr19-28788474-INS-366
chr19-28811118-INS-54
chr19-28825913-INS-50
chr19-28910760-INS-125
chr19-28911146-INS-76
chr19-28911664-INS-72
chr19-28954416-INS-7218
chr19-28980442-INS-108
chr19-28980558-INS-1080
chr19-29035252-INS-154
chr19-29074482-INS-7406
chr19-29075567-INS-408
chr19-29130982-INS-3221
chr19-29133725-INS-1575
chr19-29148106-INS-675
chr19-29160448-INS-8729
chr19-29169574-INS-209
chr19-29181218-INS-60
chr19-29181295-INS-88
chr19-29203043-INS-5950
chr19-29206966-INS-444
chr19-29207254-INS-332
chr19-29291538-INS-50
chr19-29322369-INS-164
chr19-29324490-INS-65
chr19-29335239-INS-631
chr19-29452010-INS-62
chr19-29458819-INS-51
chr19-29459276-INS-318
chr19-29474546-INS-52
chr19-29479561-INS-65
chr19-29488995-INS-76
chr19-29509033-INS-53
chr19-29516328-INS-7143
chr19-29606676-INS-461
chr19-29627653-INS-398
chr19-29643049-INS-205
chr19-29643272-INS-367
chr19-29663906-INS-76
chr19-29740745-INS-116
chr19-29752931-INS-73
chr19-29811508-INS-203
chr19-29815205-INS-453
chr19-29863910-INS-189
chr19-29870912-INS-50
chr19-29875818-INS-106
chr19-29885689-INS-102
chr19-29889808-INS-92
chr19-29947453-INS-119
chr19-29961270-INS-538
chr19-30088034-INS-1248
chr19-30118156-INS-56
chr19-30159470-INS-189
chr19-30161447-INS-500
chr19-30170800-INS-338
chr19-30175864-INS-137
chr19-30193347-INS-131
chr19-30194259-INS-6226
chr19-30222095-INS-2721
chr19-30236732-INS-7263
chr19-30254434-INS-1107
chr19-30270563-INS-413
chr19-30273989-INS-86

chr19-30274275-INS-76
chr19-30365025-INS-54
chr19-30382162-INS-6154
chr19-30396960-INS-76
chr19-30427970-INS-91
chr19-30442942-INS-543
chr19-30459564-INS-53
chr19-30475283-INS-6504
chr19-30476523-INS-243
chr19-30477151-INS-301
chr19-30483199-INS-61
chr19-30517445-INS-51
chr19-30544233-INS-182
chr19-30555766-INS-51
chr19-30576642-INS-326
chr19-30598890-INS-52
chr19-30622954-INS-132
chr19-30674752-INS-1498
chr19-30684205-INS-6812
chr19-30722713-INS-2808
chr19-30740791-INS-212
chr19-30741610-INS-96
chr19-30747114-INS-109
chr19-30784401-INS-76
chr19-30787645-INS-182
chr19-30801918-INS-93
chr19-30817956-INS-184
chr19-30849101-INS-3203
chr19-30862939-INS-197
chr19-30905686-INS-68
chr19-30907887-INS-386
chr19-31002354-INS-165
chr19-31029558-INS-52
chr19-31086988-INS-50
chr19-31090039-INS-900
chr19-31114560-INS-176
chr19-31133385-INS-775
chr19-31139318-INS-6390
chr19-31142692-INS-71
chr19-31143440-INS-227
chr19-31189888-INS-1094
chr19-31201969-INS-10007
chr19-31209415-INS-742
chr19-31224703-INS-1604
chr19-31255184-INS-50
chr19-31260963-INS-312
chr19-31266714-INS-1152
chr19-31303124-INS-54
chr19-31317261-INS-163
chr19-31319283-INS-58
chr19-31321645-INS-66
chr19-31396901-INS-66
chr19-31514494-INS-3756
chr19-31586223-INS-159
chr19-31591032-INS-148
chr19-31672824-INS-54
chr19-31728158-INS-73
chr19-31728294-INS-625
chr19-31777206-INS-163
chr19-31798084-INS-106
chr19-31802563-INS-122
chr19-31808757-INS-387
chr19-31812619-INS-1500
chr19-31939994-INS-4312
chr19-31941597-INS-7248
chr19-31967675-INS-1861
chr19-32031713-INS-372
chr19-32039296-INS-205
chr19-32039827-INS-54
chr19-32041413-INS-731
chr19-32064376-INS-209
chr19-32106302-INS-59
chr19-32141803-INS-249
chr19-32189822-INS-58
chr19-32204167-INS-67
chr19-32220266-INS-209
chr19-32306687-INS-1282

chr19-32339650-INS-82
chr19-32339717-INS-380
chr19-32396131-INS-60
chr19-32430681-INS-143
chr19-32448590-INS-977
chr19-32479923-INS-448
chr19-32489999-INS-1734
chr19-32515267-INS-215
chr19-32616495-INS-132
chr19-32627579-INS-112
chr19-32663876-INS-338
chr19-32664019-INS-65
chr19-32678598-INS-239
chr19-32682297-INS-7148
chr19-32699374-INS-153
chr19-32706795-INS-54
chr19-32709398-INS-205
chr19-32710049-INS-514
chr19-32725739-INS-206
chr19-32728374-INS-383
chr19-32728967-INS-453
chr19-32767557-INS-80
chr19-32816324-INS-202
chr19-32825922-INS-193
chr19-32829313-INS-211
chr19-32835760-INS-178
chr19-32837278-INS-63
chr19-32842919-INS-122
chr19-32852550-INS-142
chr19-32857203-INS-474
chr19-32902832-INS-65
chr19-32948426-INS-142
chr19-32960745-INS-50
chr19-32972663-INS-129
chr19-32977742-INS-100
chr19-32984531-INS-54
chr19-32995252-INS-207
chr19-33073791-INS-58
chr19-33073801-INS-222
chr19-33075973-INS-1253
chr19-33098189-INS-577
chr19-33134949-INS-50
chr19-33208904-INS-59
chr19-33348206-INS-4757
chr19-33356086-INS-367
chr19-33359971-INS-101
chr19-33366691-INS-174
chr19-33400262-INS-61
chr19-33409588-INS-95
chr19-33468449-INS-6194
chr19-33496759-INS-111
chr19-33500038-INS-148
chr19-33507523-INS-121
chr19-33543208-INS-6442
chr19-33588889-INS-80
chr19-33705470-INS-3128
chr19-33797519-INS-1091
chr19-33797947-INS-336
chr19-33855568-INS-150
chr19-33901929-INS-1566
chr19-33914602-INS-82
chr19-33968358-INS-326
chr19-33984538-INS-57
chr19-34002379-INS-51
chr19-34020134-INS-879
chr19-34060758-INS-156
chr19-34071577-INS-581
chr19-34071751-INS-1218
chr19-34120444-INS-2354
chr19-34189033-INS-50
chr19-34261890-INS-52
chr19-34264981-INS-50
chr19-34274585-INS-54
chr19-34277037-INS-50
chr19-34292419-INS-84
chr19-34329503-INS-105
chr19-34329870-INS-75

chr19-34374904-INS-60
chr19-34377591-INS-79
chr19-34377640-INS-468
chr19-34377696-INS-175
chr19-34386565-INS-3713
chr19-34397153-INS-74
chr19-34401900-INS-50
chr19-34426807-INS-64
chr19-34428932-INS-1285
chr19-34430625-INS-50
chr19-34430721-INS-2568
chr19-34430721-INS-6159
chr19-34455734-INS-62
chr19-34466317-INS-388
chr19-34487314-INS-70
chr19-34538318-INS-56
chr19-34595218-INS-6009
chr19-34662290-INS-103
chr19-34721202-INS-66
chr19-34727352-INS-216
chr19-34727406-INS-96
chr19-34755383-INS-56
chr19-34755410-INS-114
chr19-34772696-INS-279
chr19-34776321-INS-575
chr19-34777400-INS-70
chr19-34837130-INS-335
chr19-34864740-INS-7098
chr19-34896487-INS-333
chr19-34920226-INS-52
chr19-34952421-INS-50
chr19-34956618-INS-1512
chr19-34985081-INS-52
chr19-34997310-INS-380
chr19-34997603-INS-91
chr19-35016731-INS-56
chr19-35020086-INS-6146
chr19-35049498-INS-4817
chr19-35053202-INS-62
chr19-35068480-INS-2689
chr19-35079854-INS-68
chr19-35100245-INS-401
chr19-35104864-INS-6511
chr19-35107983-INS-656
chr19-35118843-INS-6176
chr19-35131482-INS-50
chr19-35178185-INS-100
chr19-35186561-INS-6440
chr19-35189199-INS-58
chr19-35223162-INS-4651
chr19-35235947-INS-384
chr19-35236001-INS-64
chr19-35248618-INS-6411
chr19-35317846-INS-156
chr19-35326661-INS-1847
chr19-35334662-INS-100
chr19-35381691-INS-1838
chr19-35391124-INS-88
chr19-35391797-INS-55
chr19-35391803-INS-126
chr19-35427456-INS-6593
chr19-35449268-INS-59
chr19-35479350-INS-130
chr19-35486907-INS-392
chr19-35579883-INS-2644
chr19-35582437-INS-77
chr19-35611089-INS-6256
chr19-35625696-INS-78
chr19-35626187-INS-72
chr19-35627753-INS-7248
chr19-35644481-INS-204
chr19-35651369-INS-223
chr19-35651821-INS-218
chr19-35717100-INS-62
chr19-35727368-INS-170
chr19-35784569-INS-57
chr19-35847433-INS-7203

chr19-35856045-INS-2848
chr19-35918133-INS-674
chr19-35918159-INS-105
chr19-35928255-INS-2100
chr19-35964667-INS-430
chr19-35970677-INS-93
chr19-35971144-INS-1173
chr19-36040601-INS-52
chr19-36041417-INS-56
chr19-36079488-INS-144
chr19-36095097-INS-116
chr19-36127087-INS-58
chr19-36130606-INS-1205
chr19-36170664-INS-6495
chr19-36198698-INS-3458
chr19-36209109-INS-199
chr19-36249867-INS-51
chr19-36277539-INS-145
chr19-36278255-INS-1892
chr19-36280555-INS-79
chr19-36285126-INS-7455
chr19-36332306-INS-211
chr19-36337602-INS-6376
chr19-36355269-INS-55
chr19-36359309-INS-7607
chr19-36375682-INS-66
chr19-36469329-INS-92
chr19-36487566-INS-150
chr19-36489749-INS-1491
chr19-36495199-INS-690
chr19-36505940-INS-6220
chr19-36541085-INS-4823
chr19-36625822-INS-133
chr19-36631235-INS-543
chr19-36633755-INS-1237
chr19-36636392-INS-86
chr19-36659452-INS-60
chr19-36684254-INS-73
chr19-36807468-INS-6582
chr19-36893874-INS-172
chr19-36913561-INS-1038
chr19-36979689-INS-115
chr19-37005687-INS-60
chr19-37005802-INS-1224
chr19-37015617-INS-144
chr19-37016278-INS-60
chr19-37054590-INS-105
chr19-37071297-INS-103
chr19-37091094-INS-1895
chr19-37109992-INS-388
chr19-37120701-INS-87
chr19-37170996-INS-1176
chr19-37174402-INS-94
chr19-37174537-INS-1243
chr19-37176637-INS-448
chr19-37176676-INS-126
chr19-37195043-INS-3084
chr19-37205314-INS-202
chr19-37210638-INS-480
chr19-37217944-INS-701
chr19-37243270-INS-211
chr19-37250863-INS-61
chr19-37278338-INS-231
chr19-37307209-INS-6442
chr19-37315118-INS-121
chr19-37332696-INS-90
chr19-37349463-INS-172
chr19-37365783-INS-264
chr19-37370205-INS-193
chr19-37371463-INS-217
chr19-37374384-INS-59
chr19-37403575-INS-90
chr19-37408339-INS-209
chr19-37494430-INS-92
chr19-37545150-INS-160
chr19-37548061-INS-210
chr19-37550524-INS-53

chr19-37553766-INS-87
chr19-37557783-INS-64
chr19-37560125-INS-900
chr19-37563731-INS-1995
chr19-37603122-INS-184
chr19-37616167-INS-54
chr19-37635774-INS-214
chr19-37831795-INS-72
chr19-37831819-INS-176
chr19-37846742-INS-194
chr19-37913412-INS-66
chr19-37991673-INS-120
chr19-37994949-INS-427
chr19-38017798-INS-51
chr19-38032221-INS-168
chr19-38040843-INS-207
chr19-38041360-INS-54
chr19-38049483-INS-206
chr19-38073416-INS-62
chr19-38092786-INS-50
chr19-38131017-INS-113
chr19-38131017-INS-576
chr19-38209331-INS-76
chr19-38209697-INS-1000
chr19-38221774-INS-52
chr19-38259360-INS-219
chr19-38267143-INS-232
chr19-38306429-INS-70
chr19-38324326-INS-140
chr19-38331099-INS-208
chr19-38332250-INS-7527
chr19-38341826-INS-206
chr19-38343969-INS-200
chr19-38349862-INS-208
chr19-38356637-INS-186
chr19-38408233-INS-84
chr19-38408475-INS-72
chr19-38410554-INS-52
chr19-38413002-INS-64
chr19-38421252-INS-182
chr19-38440393-INS-63
chr19-38448405-INS-3151
chr19-38463307-INS-2214
chr19-38486328-INS-192
chr19-38507156-INS-1060
chr19-38539031-INS-194
chr19-38552292-INS-166
chr19-38554159-INS-50
chr19-38567930-INS-60
chr19-38570108-INS-227
chr19-38600347-INS-205
chr19-38601256-INS-524
chr19-38631007-INS-50
chr19-38652521-INS-54
chr19-38673758-INS-416
chr19-38689432-INS-52
chr19-38699074-INS-138
chr19-38739544-INS-100
chr19-38768836-INS-75
chr19-38791957-INS-411
chr19-38795961-INS-54
chr19-38810472-INS-990
chr19-38810855-INS-64
chr19-38839073-INS-206
chr19-38840794-INS-765
chr19-38840986-INS-72
chr19-38871763-INS-50
chr19-38878874-INS-99
chr19-38903436-INS-80
chr19-38926144-INS-87
chr19-38930327-INS-421
chr19-38988335-INS-267
chr19-38990272-INS-6460
chr19-38992167-INS-536
chr19-38994279-INS-673
chr19-39040268-INS-485
chr19-39051768-INS-60

chr19-39084191-INS-68
chr19-39100869-INS-150
chr19-39101039-INS-114
chr19-39189746-INS-6626
chr19-39214723-INS-5121
chr19-39244183-INS-600
chr19-39341416-INS-3984
chr19-39343882-INS-3963
chr19-39365456-INS-106
chr19-39386906-INS-214
chr19-39452690-INS-2785
chr19-39676136-INS-61
chr19-39727010-INS-56
chr19-39779226-INS-6162
chr19-39795205-INS-195
chr19-39795261-INS-80
chr19-39795320-INS-82
chr19-39833945-INS-79
chr19-39869009-INS-62
chr19-40020249-INS-3629
chr19-40026271-INS-1110
chr19-40031766-INS-1141
chr19-40042915-INS-7187
chr19-40070782-INS-8499
chr19-40077827-INS-201
chr19-40104187-INS-203
chr19-40119438-INS-108
chr19-40119899-INS-104
chr19-40122314-INS-802
chr19-40123301-INS-78
chr19-40126572-INS-2585
chr19-40138094-INS-3678
chr19-40144502-INS-52
chr19-40157519-INS-1755
chr19-40197116-INS-1309
chr19-40200055-INS-216
chr19-40200240-INS-1123
chr19-40215627-INS-56
chr19-40268684-INS-52
chr19-40271859-INS-289
chr19-40271859-INS-66
chr19-40301682-INS-62
chr19-40317357-INS-246
chr19-40322225-INS-98
chr19-40339459-INS-50
chr19-40383939-INS-52
chr19-40402035-INS-157
chr19-40402414-INS-409
chr19-40420243-INS-72
chr19-40467625-INS-120
chr19-40497155-INS-310
chr19-40553184-INS-66
chr19-40554004-INS-145
chr19-40554902-INS-8340
chr19-40562956-INS-82
chr19-40605361-INS-558
chr19-40605422-INS-187
chr19-40608932-INS-179
chr19-40616028-INS-220
chr19-40617608-INS-54
chr19-40658074-INS-111
chr19-40697500-INS-54
chr19-40697649-INS-207
chr19-40702099-INS-51
chr19-40735233-INS-1283
chr19-40746944-INS-203
chr19-40749035-INS-132
chr19-40754850-INS-306
chr19-40787417-INS-60
chr19-40787515-INS-135
chr19-40805447-INS-115
chr19-40805657-INS-164
chr19-40840600-INS-63
chr19-40842493-INS-143
chr19-40856257-INS-172
chr19-40858689-INS-107
chr19-40858689-INS-52

chr19-40867124-INS-6684
chr19-40867486-INS-218
chr19-40915205-INS-91
chr19-40933074-INS-68
chr19-40972366-INS-437
chr19-40974783-INS-199
chr19-40977281-INS-99
chr19-41016177-INS-56
chr19-41039337-INS-649
chr19-41124898-INS-150
chr19-41127910-INS-212
chr19-41154486-INS-1384
chr19-41156257-INS-1443
chr19-41158269-INS-430
chr19-41160761-INS-66
chr19-41257591-INS-72
chr19-41260077-INS-143
chr19-41350898-INS-56
chr19-41377786-INS-69
chr19-41403506-INS-173
chr19-41412837-INS-245
chr19-41425471-INS-1033
chr19-41434026-INS-218
chr19-41439894-INS-1077
chr19-41445624-INS-3370
chr19-41475262-INS-210
chr19-41477321-INS-169
chr19-41489756-INS-201
chr19-41490933-INS-3300
chr19-41524313-INS-62
chr19-41528939-INS-54
chr19-41621653-INS-148
chr19-41629915-INS-50
chr19-41674616-INS-50
chr19-41690926-INS-62
chr19-41750025-INS-150
chr19-41778496-INS-160
chr19-41788257-INS-192
chr19-41817374-INS-72
chr19-41840275-INS-320
chr19-41872323-INS-462
chr19-41872383-INS-77
chr19-41888060-INS-168
chr19-41895148-INS-50
chr19-41949984-INS-60
chr19-41957487-INS-63
chr19-41957974-INS-339
chr19-41960752-INS-249
chr19-41983223-INS-234
chr19-42031800-INS-181
chr19-42061756-INS-2977
chr19-42091998-INS-87
chr19-42093173-INS-111
chr19-42096968-INS-662
chr19-42097008-INS-200
chr19-42101133-INS-189
chr19-42101364-INS-86
chr19-42120883-INS-52
chr19-42121898-INS-251
chr19-42122027-INS-604
chr19-42124508-INS-592
chr19-42175756-INS-1353
chr19-42206370-INS-134
chr19-42210406-INS-195
chr19-42252472-INS-349
chr19-42252983-INS-78
chr19-42274654-INS-50
chr19-42309264-INS-190
chr19-42391900-INS-339
chr19-42430041-INS-104
chr19-42430790-INS-88
chr19-42442126-INS-59
chr19-42460545-INS-441
chr19-42537131-INS-604
chr19-42597674-INS-376
chr19-42609741-INS-107
chr19-42610490-INS-54

chr19-42610988-INS-64
chr19-42690398-INS-58
chr19-42701151-INS-100
chr19-42703949-INS-141
chr19-42717335-INS-115
chr19-42717335-INS-56
chr19-42717686-INS-408
chr19-42726141-INS-144
chr19-42733386-INS-445
chr19-42762137-INS-129
chr19-42772984-INS-54
chr19-42856051-INS-50
chr19-42876137-INS-136
chr19-42892154-INS-6301
chr19-42930355-INS-6668
chr19-42938986-INS-62
chr19-43018216-INS-369
chr19-43083858-INS-75
chr19-43084098-INS-126
chr19-43101752-INS-4763
chr19-43103194-INS-1383
chr19-43119715-INS-683
chr19-43131287-INS-254
chr19-43146945-INS-1899
chr19-43148307-INS-1041
chr19-43239125-INS-50
chr19-43289134-INS-1852
chr19-43292614-INS-2212
chr19-43292614-INS-310
chr19-43382202-INS-390
chr19-43436445-INS-87
chr19-43461135-INS-61
chr19-43471751-INS-67
chr19-43498555-INS-147
chr19-43499029-INS-70
chr19-43547889-INS-2987
chr19-43561456-INS-66
chr19-43579350-INS-94
chr19-43597138-INS-52
chr19-43599371-INS-91
chr19-43610947-INS-56
chr19-43619349-INS-56
chr19-43619859-INS-111
chr19-43622175-INS-72
chr19-43622846-INS-132
chr19-43637343-INS-76
chr19-43639143-INS-201
chr19-43676058-INS-66
chr19-43696504-INS-98
chr19-43696665-INS-212
chr19-43704949-INS-58
chr19-43730033-INS-53
chr19-43764933-INS-121
chr19-43783553-INS-219
chr19-43808834-INS-93
chr19-43881065-INS-67
chr19-43882987-INS-180
chr19-43883243-INS-56
chr19-43892221-INS-174
chr19-43924219-INS-173
chr19-43933997-INS-338
chr19-43934442-INS-183
chr19-43987573-INS-86
chr19-44077783-INS-165
chr19-44148885-INS-208
chr19-44174925-INS-218
chr19-44175328-INS-86
chr19-44175360-INS-224
chr19-44187306-INS-50
chr19-44193332-INS-4835
chr19-44195226-INS-1801
chr19-44221308-INS-54
chr19-44228071-INS-201
chr19-44231101-INS-196
chr19-44240291-INS-70
chr19-44247335-INS-2896
chr19-44253247-INS-208

chr19-44254206-INS-112
chr19-44257481-INS-712
chr19-44307099-INS-108
chr19-44307621-INS-128
chr19-44307621-INS-317
chr19-44319739-INS-116
chr19-44341914-INS-3381
chr19-44353843-INS-143
chr19-44372217-INS-117
chr19-44392304-INS-195
chr19-44422777-INS-189
chr19-44437463-INS-267
chr19-44443643-INS-171
chr19-44467396-INS-3335
chr19-44475835-INS-4433
chr19-44476327-INS-1866
chr19-44486459-INS-210
chr19-44486582-INS-57
chr19-44486680-INS-306
chr19-44515403-INS-111
chr19-44516907-INS-78
chr19-44524320-INS-287
chr19-44526777-INS-6436
chr19-44532449-INS-78
chr19-44533082-INS-83
chr19-44548055-INS-71
chr19-44560842-INS-72
chr19-44595527-INS-55
chr19-44599208-INS-130
chr19-44602853-INS-52
chr19-44635661-INS-52
chr19-44636231-INS-146
chr19-44641115-INS-203
chr19-44641859-INS-86
chr19-44841929-INS-127
chr19-44868384-INS-120
chr19-44883480-INS-155
chr19-44901653-INS-52
chr19-44908722-INS-1248
chr19-44940775-INS-207
chr19-44978791-INS-56
chr19-44984466-INS-167
chr19-45006182-INS-99
chr19-45023736-INS-190
chr19-45108033-INS-60
chr19-45119015-INS-1240
chr19-45119015-INS-2920
chr19-45126420-INS-240
chr19-45174008-INS-3502
chr19-45180945-INS-223
chr19-45181012-INS-106
chr19-45182823-INS-498
chr19-45185189-INS-207
chr19-45187080-INS-244
chr19-45187610-INS-61
chr19-45187723-INS-149
chr19-45198870-INS-5970
chr19-45249529-INS-222
chr19-45249575-INS-77
chr19-45258130-INS-183
chr19-45274518-INS-1650
chr19-45274518-INS-3372
chr19-45274533-INS-66
chr19-45312537-INS-66
chr19-45314484-INS-134
chr19-45347961-INS-114
chr19-45361877-INS-214
chr19-45363293-INS-2322
chr19-45363335-INS-149
chr19-45363540-INS-1073
chr19-45379811-INS-1079
chr19-45381952-INS-211
chr19-45388494-INS-88
chr19-45389978-INS-153
chr19-45389978-INS-51
chr19-45401500-INS-813
chr19-45425405-INS-64

chr19-45509186-INS-328
chr19-45510009-INS-208
chr19-45520573-INS-50
chr19-45536829-INS-110
chr19-45582030-INS-256
chr19-45607231-INS-561
chr19-45627358-INS-206
chr19-45654931-INS-53
chr19-45718002-INS-60
chr19-45777388-INS-72
chr19-45779797-INS-918
chr19-45816531-INS-52
chr19-45835149-INS-73
chr19-45838394-INS-845
chr19-45838439-INS-187
chr19-45838661-INS-201
chr19-45842471-INS-132
chr19-45844267-INS-3238
chr19-45885684-INS-139
chr19-45889423-INS-922
chr19-45948869-INS-3191
chr19-45961007-INS-179
chr19-45973435-INS-136
chr19-45973873-INS-67
chr19-45984572-INS-175
chr19-45997830-INS-57
chr19-46009726-INS-722
chr19-46009912-INS-314
chr19-46010283-INS-52
chr19-46013901-INS-206
chr19-46015590-INS-207
chr19-46038341-INS-659
chr19-46041608-INS-175
chr19-46042259-INS-203
chr19-46069689-INS-169
chr19-46107810-INS-4738
chr19-46109003-INS-60
chr19-46110282-INS-178
chr19-46118544-INS-87
chr19-46119057-INS-167
chr19-46121035-INS-62
chr19-46196942-INS-52
chr19-46217486-INS-2146
chr19-46218152-INS-183
chr19-46242545-INS-411
chr19-46262861-INS-127
chr19-46284511-INS-5333
chr19-46284972-INS-50
chr19-46331763-INS-72
chr19-46334639-INS-60
chr19-46343141-INS-64
chr19-46343186-INS-669
chr19-46374072-INS-206
chr19-46374233-INS-200
chr19-46393323-INS-102
chr19-46399830-INS-7076
chr19-46404168-INS-210
chr19-46418789-INS-2974
chr19-46472144-INS-56
chr19-46488001-INS-364
chr19-46510245-INS-5599
chr19-46527407-INS-336
chr19-46527727-INS-64
chr19-46533075-INS-74
chr19-46535218-INS-5531
chr19-46539018-INS-1497
chr19-46540712-INS-206
chr19-46568482-INS-308
chr19-46568614-INS-792
chr19-46572548-INS-85
chr19-46594036-INS-313
chr19-46599859-INS-55
chr19-46628995-INS-96
chr19-46693427-INS-51
chr19-46701184-INS-124
chr19-46738085-INS-103
chr19-46742366-INS-471

chr19-46749166-INS-60
chr19-46765300-INS-145
chr19-46766142-INS-162
chr19-46775630-INS-262
chr19-46799781-INS-50
chr19-46821459-INS-143
chr19-46860671-INS-60
chr19-46915325-INS-65
chr19-46926850-INS-116
chr19-46932263-INS-137
chr19-46932321-INS-50
chr19-46993799-INS-70
chr19-46997813-INS-98
chr19-47020781-INS-188
chr19-47048957-INS-465
chr19-47055676-INS-58
chr19-47080093-INS-55
chr19-47090130-INS-84
chr19-47095916-INS-115
chr19-47104503-INS-210
chr19-47111569-INS-206
chr19-47133800-INS-1495
chr19-47138345-INS-63
chr19-47138770-INS-72
chr19-47147568-INS-1963
chr19-47153094-INS-152
chr19-47221281-INS-82
chr19-47223308-INS-109
chr19-47315753-INS-120
chr19-47438689-INS-52
chr19-47444395-INS-66
chr19-47451072-INS-284
chr19-47451072-INS-3026
chr19-47451072-INS-914
chr19-47458282-INS-64
chr19-47466788-INS-218
chr19-47467759-INS-164
chr19-47471008-INS-208
chr19-47471552-INS-218
chr19-47474490-INS-62
chr19-47514225-INS-566
chr19-47527771-INS-52
chr19-47533864-INS-74
chr19-47534122-INS-165
chr19-47535007-INS-202
chr19-47549957-INS-217
chr19-47559299-INS-83
chr19-47614339-INS-219
chr19-47624044-INS-861
chr19-47643515-INS-174
chr19-47689051-INS-953
chr19-47692783-INS-206
chr19-47741542-INS-732
chr19-47743257-INS-133
chr19-47755934-INS-50
chr19-47756190-INS-182
chr19-47771885-INS-145
chr19-47782759-INS-113
chr19-47791791-INS-6054
chr19-47802722-INS-55
chr19-47830393-INS-7786
chr19-47836275-INS-4987
chr19-47841304-INS-57
chr19-47849771-INS-1510
chr19-47849771-INS-241
chr19-47958418-INS-136
chr19-47962780-INS-7507
chr19-47968365-INS-140
chr19-47976830-INS-2511
chr19-47978986-INS-197
chr19-47994317-INS-142
chr19-48020224-INS-56
chr19-48021960-INS-551
chr19-48082683-INS-4721
chr19-48084703-INS-213
chr19-48109665-INS-50
chr19-48154919-INS-61

chr19-48162479-INS-88
chr19-48165207-INS-1929
chr19-48212448-INS-100
chr19-48212480-INS-273
chr19-48212848-INS-151
chr19-48317185-INS-146
chr19-48338325-INS-55
chr19-48384407-INS-4378
chr19-48409077-INS-115
chr19-48419653-INS-1161
chr19-48420790-INS-204
chr19-48421006-INS-140
chr19-48421843-INS-124
chr19-48428436-INS-50
chr19-48494867-INS-59
chr19-48495400-INS-1759
chr19-48503267-INS-1126
chr19-48507648-INS-50
chr19-48568429-INS-474
chr19-48568440-INS-62
chr19-48602310-INS-132
chr19-48618917-INS-80
chr19-48662139-INS-58
chr19-48696964-INS-55
chr19-48743087-INS-99
chr19-48761521-INS-55
chr19-48761741-INS-387
chr19-48767257-INS-180
chr19-48767314-INS-411
chr19-48802808-INS-180
chr19-48820160-INS-52
chr19-48823418-INS-1468
chr19-48825370-INS-56
chr19-48857467-INS-110
chr19-48862988-INS-1756
chr19-48906186-INS-51
chr19-48943830-INS-388
chr19-48965911-INS-199
chr19-48983148-INS-57
chr19-49021521-INS-75
chr19-49021617-INS-324
chr19-49051494-INS-112
chr19-49071815-INS-59
chr19-49086921-INS-58
chr19-49101370-INS-124
chr19-49117677-INS-2951
chr19-49128675-INS-79
chr19-49149607-INS-64
chr19-49237118-INS-71
chr19-49254972-INS-260
chr19-49306811-INS-57
chr19-49313004-INS-50
chr19-49332709-INS-540
chr19-49361011-INS-2500
chr19-49371324-INS-210
chr19-49371328-INS-63
chr19-49375820-INS-180
chr19-49384258-INS-199
chr19-49400681-INS-1551
chr19-49419028-INS-162
chr19-49419028-INS-342
chr19-49434277-INS-100
chr19-49435012-INS-688
chr19-49490853-INS-420
chr19-49512060-INS-212
chr19-49530891-INS-1266
chr19-49562397-INS-134
chr19-49598798-INS-52
chr19-49629257-INS-72
chr19-49646497-INS-308
chr19-49678944-INS-64
chr19-49697931-INS-107
chr19-49702503-INS-54
chr19-49703403-INS-2925
chr19-49709156-INS-60
chr19-49710545-INS-99
chr19-49712635-INS-71

chr19-49712731-INS-6448
chr19-49765765-INS-3204
chr19-49778166-INS-97
chr19-49791823-INS-253
chr19-49793699-INS-391
chr19-49800324-INS-3527
chr19-49830686-INS-2932
chr19-49867643-INS-72
chr19-49900203-INS-55
chr19-49918959-INS-75
chr19-49945049-INS-56
chr19-49946465-INS-51
chr19-49954173-INS-3924
chr19-49962596-INS-50
chr19-49999933-INS-718
chr19-50000402-INS-71
chr19-50004926-INS-76
chr19-50017106-INS-134
chr19-50021206-INS-56
chr19-50026064-INS-4935
chr19-50046852-INS-116
chr19-50046852-INS-56
chr19-50047393-INS-162
chr19-50072275-INS-78
chr19-50079218-INS-54
chr19-50118091-INS-58
chr19-50143374-INS-51
chr19-50150789-INS-225
chr19-50150913-INS-65
chr19-50160180-INS-52
chr19-50172883-INS-139
chr19-50172883-INS-58
chr19-50175548-INS-72
chr19-50192395-INS-105
chr19-50197112-INS-55
chr19-50233678-INS-50
chr19-50243360-INS-143
chr19-50243438-INS-159
chr19-50243773-INS-89
chr19-50299116-INS-54
chr19-50307364-INS-641
chr19-50307411-INS-222
chr19-50307688-INS-140
chr19-50372460-INS-66
chr19-50405303-INS-62
chr19-50422711-INS-66
chr19-50451451-INS-1204
chr19-50462392-INS-66
chr19-50462868-INS-243
chr19-50469512-INS-138
chr19-50472417-INS-209
chr19-50488838-INS-69
chr19-50511115-INS-339
chr19-50511245-INS-140
chr19-50569382-INS-1196
chr19-50598088-INS-160
chr19-50689789-INS-50
chr19-50791530-INS-421
chr19-50794972-INS-3895
chr19-50795815-INS-1429
chr19-50851826-INS-1588
chr19-50872500-INS-222
chr19-50898452-INS-1963
chr19-50900091-INS-54
chr19-50910567-INS-6208
chr19-50910855-INS-172
chr19-50914132-INS-52
chr19-50954376-INS-77
chr19-50954996-INS-55
chr19-50969500-INS-60
chr19-50976231-INS-1673
chr19-50977881-INS-93
chr19-50986428-INS-50
chr19-50995651-INS-2093
chr19-50999489-INS-50
chr19-51009027-INS-54
chr19-51010749-INS-80

chr19-51011538-INS-523
chr19-51043383-INS-197
chr19-51095029-INS-78
chr19-51110050-INS-1312
chr19-51140491-INS-94
chr19-51142602-INS-70
chr19-51169044-INS-6236
chr19-51175057-INS-343
chr19-51209270-INS-193
chr19-51211241-INS-50
chr19-51273988-INS-180
chr19-51274543-INS-246
chr19-51274842-INS-293
chr19-51280241-INS-108
chr19-51280258-INS-261
chr19-51288964-INS-332
chr19-51288994-INS-159
chr19-51290841-INS-99
chr19-51291688-INS-68
chr19-51299201-INS-951
chr19-51312721-INS-7281
chr19-51317744-INS-114
chr19-51325758-INS-56
chr19-51326286-INS-74
chr19-51341922-INS-6468
chr19-51353608-INS-380
chr19-51368524-INS-66
chr19-51387826-INS-457
chr19-51408958-INS-1166
chr19-51409529-INS-59
chr19-51476601-INS-7191
chr19-51492866-INS-6376
chr19-51523064-INS-601
chr19-51555425-INS-83
chr19-51611846-INS-63
chr19-51627492-INS-54
chr19-51690102-INS-199
chr19-51690510-INS-54
chr19-51730212-INS-79
chr19-51738225-INS-7264
chr19-51740605-INS-56
chr19-51750832-INS-73
chr19-51760937-INS-64
chr19-51761401-INS-108
chr19-51795267-INS-2718
chr19-51798619-INS-295
chr19-51804034-INS-5566
chr19-51860023-INS-108
chr19-51868049-INS-3355
chr19-51870752-INS-50
chr19-51879797-INS-5768
chr19-51884338-INS-85
chr19-51897336-INS-119
chr19-51897509-INS-102
chr19-51919864-INS-1904
chr19-51943949-INS-636
chr19-52009630-INS-4168
chr19-52025474-INS-2688
chr19-52076019-INS-7218
chr19-52115873-INS-82
chr19-52142244-INS-500
chr19-52148377-INS-54
chr19-52156947-INS-690
chr19-52181243-INS-157
chr19-52243072-INS-60
chr19-52254030-INS-1292
chr19-52263793-INS-53
chr19-52315748-INS-3446
chr19-52330729-INS-128
chr19-52335429-INS-3770
chr19-52382469-INS-3292
chr19-52412722-INS-6552
chr19-52444640-INS-74
chr19-52456976-INS-6168
chr19-52476373-INS-217
chr19-52476397-INS-84
chr19-52476398-INS-658

chr19-52482348-INS-2847
chr19-52484291-INS-1917
chr19-52497223-INS-60
chr19-52503574-INS-1469
chr19-52505209-INS-66
chr19-52510927-INS-59
chr19-52513841-INS-2393
chr19-52517422-INS-842
chr19-52521028-INS-2820
chr19-52543404-INS-66
chr19-52552733-INS-53
chr19-52578045-INS-2974
chr19-52590933-INS-56
chr19-52644888-INS-50
chr19-52651263-INS-4029
chr19-52651408-INS-90
chr19-52652701-INS-312
chr19-52671042-INS-6184
chr19-52679426-INS-54
chr19-52679858-INS-317
chr19-52685791-INS-105
chr19-52694811-INS-3476
chr19-52707547-INS-320
chr19-52709167-INS-318
chr19-52715141-INS-58
chr19-52721360-INS-52
chr19-52732790-INS-7338
chr19-52738571-INS-1385
chr19-52754524-INS-64
chr19-52769885-INS-6404
chr19-52775732-INS-6504
chr19-52802651-INS-217
chr19-52802651-INS-53
chr19-52843369-INS-1320
chr19-52844709-INS-432
chr19-52886242-INS-70
chr19-52900607-INS-66
chr19-52956367-INS-1490
chr19-52983693-INS-195
chr19-53009384-INS-200
chr19-53034371-INS-158
chr19-53037390-INS-70
chr19-53055785-INS-207
chr19-53059704-INS-76
chr19-53077221-INS-112
chr19-53077287-INS-267
chr19-53098402-INS-704
chr19-53203711-INS-53
chr19-53203840-INS-347
chr19-53214312-INS-95
chr19-53256519-INS-55
chr19-53294573-INS-60
chr19-53408655-INS-68
chr19-53414882-INS-63
chr19-53415576-INS-57
chr19-53448188-INS-192
chr19-53460274-INS-80
chr19-53462785-INS-59
chr19-53478168-INS-4333
chr19-53485976-INS-84
chr19-53494658-INS-206
chr19-53561569-INS-103
chr19-53590547-INS-50
chr19-53631713-INS-188
chr19-53651339-INS-50
chr19-53651560-INS-239
chr19-53656568-INS-75
chr19-53673189-INS-189
chr19-53729818-INS-122
chr19-53730284-INS-152
chr19-53745009-INS-2459
chr19-53767154-INS-132
chr19-53776635-INS-211
chr19-53795463-INS-168
chr19-53895091-INS-316
chr19-53928523-INS-144
chr19-53928523-INS-384

chr19-53939396-INS-50
chr19-53988874-INS-54
chr19-53997080-INS-76
chr19-54043669-INS-50
chr19-54046947-INS-50
chr19-54064427-INS-50
chr19-54070588-INS-50
chr19-54096566-INS-129
chr19-54096718-INS-60
chr19-54139259-INS-550
chr19-54139781-INS-242
chr19-54149461-INS-6909
chr19-54170248-INS-56
chr19-54170406-INS-136
chr19-54174124-INS-205
chr19-54174163-INS-631
chr19-54175618-INS-201
chr19-54182652-INS-142
chr19-54197449-INS-60
chr19-54202876-INS-50
chr19-54209367-INS-78
chr19-54238978-INS-60
chr19-54259317-INS-66
chr19-54260076-INS-54
chr19-54262090-INS-71
chr19-54371186-INS-81
chr19-54371278-INS-334
chr19-54378323-INS-1877
chr19-54391090-INS-88
chr19-54397641-INS-147
chr19-54408711-INS-78
chr19-54431402-INS-874
chr19-54433084-INS-525
chr19-54477875-INS-58
chr19-54489118-INS-76
chr19-54489146-INS-162
chr19-54489173-INS-53
chr19-54570964-INS-50
chr19-54602273-INS-480
chr19-54620091-INS-6718
chr19-54630560-INS-68
chr19-54637602-INS-112
chr19-54648894-INS-6504
chr19-54651446-INS-402
chr19-54652158-INS-2988
chr19-54661016-INS-1578
chr19-54685676-INS-2388
chr19-54703438-INS-200
chr19-54704300-INS-6760
chr19-54707408-INS-64
chr19-54711631-INS-428
chr19-54727255-INS-127
chr19-54761624-INS-59
chr19-54788675-INS-62
chr19-54791238-INS-65
chr19-54806446-INS-2984
chr19-54840898-INS-6201
chr19-54843447-INS-218
chr19-54849341-INS-253
chr19-54887572-INS-77
chr19-54889985-INS-68
chr19-54890053-INS-74
chr19-54890150-INS-88
chr19-54898368-INS-80
chr19-54901975-INS-497
chr19-54914133-INS-108
chr19-54946549-INS-288
chr19-54949992-INS-110
chr19-54950030-INS-233
chr19-54965630-INS-242
chr19-55027313-INS-56
chr19-55027434-INS-203
chr19-55029681-INS-627
chr19-55043481-INS-50
chr19-55047106-INS-78
chr19-55047261-INS-130
chr19-55050327-INS-601

chr19-55100886-INS-128
chr19-55131088-INS-60
chr19-55135285-INS-120
chr19-55138580-INS-60
chr19-55151373-INS-102
chr19-55188286-INS-63
chr19-55188898-INS-390
chr19-55228799-INS-116
chr19-55235335-INS-109
chr19-55235637-INS-205
chr19-55287688-INS-68
chr19-55317326-INS-64
chr19-55322456-INS-181
chr19-55324261-INS-571
chr19-55325054-INS-92
chr19-55337074-INS-100
chr19-55352177-INS-230
chr19-55389512-INS-480
chr19-55405289-INS-102
chr19-55516937-INS-500
chr19-55517262-INS-1321
chr19-55559247-INS-60
chr19-55625020-INS-235
chr19-55641001-INS-74
chr19-55650394-INS-75
chr19-55653821-INS-165
chr19-55673537-INS-50
chr19-55701871-INS-64
chr19-55719882-INS-288
chr19-55724147-INS-206
chr19-55778889-INS-59
chr19-55802370-INS-92
chr19-55824356-INS-52
chr19-55841814-INS-50
chr19-55849109-INS-266
chr19-55858486-INS-50
chr19-55874769-INS-54
chr19-55906936-INS-137
chr19-55933347-INS-73
chr19-55958253-INS-56
chr19-55976568-INS-510
chr19-56020078-INS-175
chr19-56020078-INS-85
chr19-56025552-INS-80
chr19-56079019-INS-1346
chr19-56099756-INS-213
chr19-56112879-INS-217
chr19-56112894-INS-62
chr19-56114606-INS-117
chr19-56114645-INS-457
chr19-56141746-INS-144
chr19-56157394-INS-171
chr19-56162321-INS-52
chr19-56168385-INS-356
chr19-56184219-INS-66
chr19-56199530-INS-54
chr19-56263611-INS-2726
chr19-56285295-INS-429
chr19-56290559-INS-171
chr19-56290841-INS-427
chr19-56291414-INS-108
chr19-56291462-INS-220
chr19-56298888-INS-130
chr19-56304313-INS-108
chr19-56311359-INS-56
chr19-56311884-INS-140
chr19-56317607-INS-123
chr19-56374882-INS-230
chr19-56388846-INS-139
chr19-56410253-INS-1062
chr19-56478815-INS-3118
chr19-56490770-INS-68
chr19-56571465-INS-50
chr19-56622575-INS-132
chr19-56631275-INS-2770
chr19-56631944-INS-192
chr19-56639730-INS-214

chr19-56646604-INS-7078
chr19-56708273-INS-64
chr19-56767217-INS-72
chr19-56836553-INS-899
chr19-56852236-INS-382
chr19-56876949-INS-75
chr19-56883800-INS-74
chr19-56917125-INS-69
chr19-56922963-INS-147
chr19-56934780-INS-2985
chr19-56940113-INS-126
chr19-56940279-INS-81
chr19-56941878-INS-76
chr19-56961839-INS-205
chr19-56965118-INS-98
chr19-57101841-INS-913
chr19-57101861-INS-3104
chr19-57109888-INS-68
chr19-57123756-INS-2379
chr19-57128010-INS-51
chr19-57128437-INS-138
chr19-57129266-INS-80
chr19-57141911-INS-255
chr19-57165475-INS-78
chr19-57191400-INS-68
chr19-57193084-INS-583
chr19-57215550-INS-98
chr19-57220490-INS-60
chr19-57233201-INS-87
chr19-57276702-INS-52
chr19-57286743-INS-103
chr19-57307512-INS-57
chr19-57331713-INS-1474
chr19-57334356-INS-2471
chr19-57345207-INS-176
chr19-57345245-INS-394
chr19-57370834-INS-94
chr19-57383469-INS-213
chr19-57403972-INS-90
chr19-57438022-INS-3044
chr19-57446793-INS-147
chr19-57466878-INS-316
chr19-57467085-INS-1409
chr19-57481988-INS-50
chr19-57495989-INS-60
chr19-57496316-INS-60
chr19-57537634-INS-215
chr19-57537770-INS-55
chr19-57539305-INS-51
chr19-57548133-INS-92
chr19-57605073-INS-56
chr19-57650455-INS-1304
chr19-57670179-INS-114
chr19-57753692-INS-1806
chr19-57783781-INS-144
chr19-57807820-INS-57
chr19-57833479-INS-52
chr19-57838425-INS-83
chr19-57888969-INS-1931
chr19-57904439-INS-64
chr19-57917799-INS-78
chr19-57917923-INS-469
chr19-57955803-INS-1097
chr19-57968889-INS-71
chr19-57970861-INS-50
chr19-57978064-INS-60
chr19-57985288-INS-52
chr19-58025944-INS-92
chr19-58105028-INS-140
chr19-58130454-INS-2157
chr19-58142742-INS-86
chr19-58192994-INS-508
chr19-58251126-INS-53
chr19-58251434-INS-245
chr19-58258136-INS-50
chr19-58280347-INS-431
chr19-58291724-INS-1352

chr19-58307508-INS-64
chr19-58307548-INS-170
chr19-58312596-INS-162
chr19-58321519-INS-369
chr19-58406507-INS-51
chr19-58406628-INS-186
chr19-58478086-INS-113
chr19-58505630-INS-6565
chr19-58506908-INS-851
chr19-58509394-INS-2756
chr19-58514585-INS-86
chr19-58540997-INS-72
chr19-58601945-INS-64
chr19-58656015-INS-418
chr19-58698592-INS-5016
chr19-58699342-INS-1026
chr19-58758818-INS-51
chr19-58820988-INS-346
chr19-58821025-INS-172
chr19-58821352-INS-127
chr19-58976012-INS-97
chr19-59034407-INS-52
chr19-59070897-INS-50
chr19-59092030-INS-75
chr19-59130410-INS-51
chr19-59170008-INS-67
chr19-59175161-INS-212
chr19-59187609-INS-2925
chr19-59342856-INS-84
chr19-59343070-INS-272
chr19-59452791-INS-50
chr19-59488515-INS-122
chr19-59537284-INS-1259
chr19-59541375-INS-69
chr19-59581724-INS-118
chr19-59622509-INS-109
chr19-59654495-INS-56
chr19-59672039-INS-56
chr19-59724416-INS-158
chr19-59724876-INS-312
chr19-59839968-INS-76
chr19-59840833-INS-61
chr19-59968292-INS-6499
chr19-59995692-INS-81
chr19-59996168-INS-517
chr19-60044786-INS-70
chr19-60058708-INS-172
chr19-60093969-INS-5203
chr19-60220096-INS-1862
chr19-60236607-INS-400
chr19-60260294-INS-6989
chr19-60282417-INS-55
chr19-60397751-INS-1160
chr19-60411220-INS-1715
chr19-60432938-INS-2541
chr19-60539352-INS-201
chr19-60583843-INS-215
chr19-60657151-INS-51
chr19-60664604-INS-61
chr19-60668717-INS-1462
chr19-60673737-INS-1539
chr19-60736114-INS-107
chr19-60777010-INS-209
chr19-60784848-INS-50
chr19-60809938-INS-154
chr19-60815538-INS-195
chr19-60815751-INS-76
chr19-61022065-INS-52
chr19-61047766-INS-127
chr19-61052575-INS-56
chr19-61139494-INS-365
chr2-3050698-INS-80
chr2-3051442-INS-72
chr2-3051877-INS-126
chr2-3058851-INS-75
chr2-3092377-INS-55
chr2-3130075-INS-3057

chr2-3150836-INS-922
chr2-3153918-INS-105
chr2-3165551-INS-178
chr2-3192388-INS-79
chr2-3213318-INS-1704
chr2-3235577-INS-4819
chr2-3259956-INS-97
chr2-3281582-INS-4365
chr2-3307052-INS-206
chr2-3315930-INS-61
chr2-3358475-INS-65
chr2-3366997-INS-61
chr2-3368660-INS-66
chr2-3394964-INS-82
chr2-3411451-INS-64
chr2-3429788-INS-66
chr2-3434434-INS-192
chr2-3434721-INS-342
chr2-3440555-INS-1655
chr2-3440780-INS-720
chr2-3530403-INS-3349
chr2-3545475-INS-185
chr2-3568520-INS-59
chr2-3592171-INS-2399
chr2-3630808-INS-74
chr2-3656568-INS-51
chr2-3662780-INS-89
chr2-3668968-INS-644
chr2-3685446-INS-88
chr2-3688365-INS-7295
chr2-3704221-INS-147
chr2-3711545-INS-54
chr2-3754048-INS-100
chr2-3786496-INS-367
chr2-3813679-INS-159
chr2-3830528-INS-437
chr2-3870420-INS-433
chr2-3880970-INS-91
chr2-3904534-INS-56
chr2-3909855-INS-54
chr2-3921549-INS-764
chr2-3959153-INS-52
chr2-3967204-INS-294
chr2-3967986-INS-11030
chr2-3993301-INS-96
chr2-4021178-INS-87
chr2-4044023-INS-462
chr2-4050405-INS-112
chr2-4050855-INS-272
chr2-4062137-INS-409
chr2-4070466-INS-6436
chr2-4071115-INS-86
chr2-4123760-INS-67
chr2-4137794-INS-4553
chr2-4141676-INS-1196
chr2-4141754-INS-3538
chr2-4150874-INS-2187
chr2-4179113-INS-73
chr2-4227646-INS-296
chr2-4321438-INS-60
chr2-4354747-INS-88
chr2-4360511-INS-65
chr2-4392029-INS-60
chr2-4394065-INS-81
chr2-4394110-INS-206
chr2-4399244-INS-56
chr2-4425440-INS-60
chr2-4438363-INS-100
chr2-4462446-INS-52
chr2-4520761-INS-50
chr2-4523386-INS-3167
chr2-4524325-INS-71
chr2-4610241-INS-197
chr2-4635504-INS-121
chr2-4637413-INS-487
chr2-4644758-INS-283
chr2-4649113-INS-1139

chr2-4666096-INS-192
chr2-4671114-INS-252
chr2-4684104-INS-644
chr2-4701285-INS-20317
chr2-4719231-INS-387
chr2-4721066-INS-100
chr2-4733540-INS-112
chr2-4734235-INS-52
chr2-4742204-INS-304
chr2-4743891-INS-210
chr2-4748240-INS-6239
chr2-4760026-INS-138
chr2-4761452-INS-1471
chr2-4771438-INS-1525
chr2-4818711-INS-203
chr2-4832391-INS-77
chr2-4850034-INS-147
chr2-4850355-INS-259
chr2-4853348-INS-66
chr2-4874062-INS-531
chr2-4875250-INS-259
chr2-4875567-INS-300
chr2-4896734-INS-53
chr2-4927835-INS-180
chr2-4930753-INS-201
chr2-4935272-INS-116
chr2-4968074-INS-52
chr2-4984303-INS-52
chr2-5020259-INS-193
chr2-5030943-INS-150
chr2-5042864-INS-172
chr2-5042882-INS-442
chr2-5053219-INS-2648
chr2-5072875-INS-54
chr2-5073233-INS-651
chr2-5079925-INS-84
chr2-5087444-INS-72
chr2-5088337-INS-220
chr2-5088541-INS-79
chr2-5096279-INS-330
chr2-5100707-INS-165
chr2-5104395-INS-114
chr2-5111040-INS-180
chr2-5114659-INS-88
chr2-5119378-INS-6726
chr2-5123641-INS-2380
chr2-5123767-INS-58
chr2-5124375-INS-63
chr2-5135914-INS-60
chr2-5149969-INS-409
chr2-5151870-INS-82
chr2-5164084-INS-491
chr2-5166692-INS-54
chr2-5174465-INS-209
chr2-5178657-INS-137
chr2-5179290-INS-130
chr2-5230569-INS-2739
chr2-5231419-INS-503
chr2-5231439-INS-60
chr2-5239714-INS-228
chr2-5244877-INS-86
chr2-5250462-INS-6438
chr2-5272195-INS-55
chr2-5319121-INS-2791
chr2-5323490-INS-172
chr2-5323490-INS-989
chr2-5344365-INS-124
chr2-5363989-INS-266
chr2-5364526-INS-224
chr2-5377400-INS-50
chr2-5382713-INS-50
chr2-5393428-INS-790
chr2-5404755-INS-548
chr2-5417283-INS-332
chr2-5429686-INS-82
chr2-5430411-INS-58
chr2-5457972-INS-56

chr2-5462107-INS-55
chr2-5496072-INS-172
chr2-5510146-INS-1573
chr2-5518268-INS-64
chr2-5529942-INS-50
chr2-5530576-INS-72
chr2-5530577-INS-259
chr2-5572205-INS-146
chr2-5587351-INS-774
chr2-5606924-INS-67
chr2-5622056-INS-52
chr2-5684488-INS-222
chr2-5684488-INS-74
chr2-5702553-INS-135
chr2-5707299-INS-51
chr2-5726228-INS-50
chr2-5726475-INS-240
chr2-5727035-INS-53
chr2-5728203-INS-104
chr2-5742257-INS-66
chr2-5749266-INS-56
chr2-5767197-INS-176
chr2-5819435-INS-76
chr2-5844594-INS-366
chr2-5847307-INS-51
chr2-5851092-INS-54
chr2-5905934-INS-69
chr2-5912720-INS-318
chr2-5912762-INS-1113
chr2-5919887-INS-2939
chr2-5924164-INS-974
chr2-5933616-INS-204
chr2-5938982-INS-212
chr2-5945765-INS-96
chr2-5962694-INS-3551
chr2-5962705-INS-95
chr2-5998822-INS-204
chr2-6022200-INS-66
chr2-6026082-INS-202
chr2-6028884-INS-463
chr2-6041267-INS-204
chr2-6074838-INS-69
chr2-6088359-INS-98
chr2-6094143-INS-199
chr2-6108060-INS-210
chr2-6109756-INS-60
chr2-6117362-INS-223
chr2-6170453-INS-202
chr2-6177763-INS-521
chr2-6188978-INS-586
chr2-6206149-INS-148
chr2-6206149-INS-312
chr2-6240888-INS-52
chr2-6240932-INS-424
chr2-6249176-INS-134
chr2-6307988-INS-84
chr2-6326471-INS-52
chr2-6332109-INS-124
chr2-6333164-INS-311
chr2-6333164-INS-93
chr2-6350334-INS-73
chr2-6364830-INS-208
chr2-6401069-INS-378
chr2-6408402-INS-71
chr2-6418529-INS-102
chr2-6418917-INS-126
chr2-6443738-INS-6610
chr2-6463227-INS-198
chr2-6474708-INS-329
chr2-6484997-INS-175
chr2-6494094-INS-82
chr2-6494410-INS-104
chr2-6503773-INS-178
chr2-6505685-INS-54
chr2-6517991-INS-7340
chr2-6536029-INS-278
chr2-6598282-INS-103

chr2-6622013-INS-242
chr2-6633381-INS-52
chr2-6641329-INS-644
chr2-6642728-INS-145
chr2-6650339-INS-2272
chr2-6679166-INS-130
chr2-6791057-INS-65
chr2-6804843-INS-55
chr2-6805804-INS-50
chr2-6818145-INS-52
chr2-6890108-INS-56
chr2-6893312-INS-117
chr2-6906074-INS-6687
chr2-6986653-INS-76
chr2-6992586-INS-57
chr2-7055744-INS-186
chr2-7082742-INS-50
chr2-7117896-INS-213
chr2-7131234-INS-60
chr2-7133132-INS-52
chr2-7136856-INS-58
chr2-7156447-INS-204
chr2-7169845-INS-128
chr2-7179965-INS-217
chr2-7205298-INS-111
chr2-7205300-INS-534
chr2-7205690-INS-84
chr2-7208822-INS-1719
chr2-7209314-INS-397
chr2-7210919-INS-134
chr2-7213383-INS-69
chr2-7264842-INS-54
chr2-7267734-INS-1100
chr2-7283529-INS-105
chr2-7309371-INS-132
chr2-7395740-INS-2835
chr2-7429723-INS-56
chr2-7432587-INS-610
chr2-7441121-INS-369
chr2-7484700-INS-228
chr2-7507111-INS-115
chr2-7517218-INS-60
chr2-7531199-INS-74
chr2-7540856-INS-50
chr2-7552924-INS-6867
chr2-7699326-INS-89
chr2-7791326-INS-7462
chr2-7791913-INS-54
chr2-7813422-INS-196
chr2-7827884-INS-80
chr2-7852670-INS-314
chr2-7853782-INS-76
chr2-7856256-INS-234
chr2-7859622-INS-126
chr2-7859676-INS-1074
chr2-7881183-INS-87
chr2-7885600-INS-130
chr2-7895449-INS-50
chr2-7900410-INS-6935
chr2-7900737-INS-94
chr2-7978324-INS-1546
chr2-8011661-INS-8368
chr2-8022880-INS-218
chr2-8048668-INS-452
chr2-8057240-INS-111
chr2-8064362-INS-54
chr2-8092500-INS-1973
chr2-8110956-INS-50
chr2-8116825-INS-1320
chr2-8119991-INS-282
chr2-8143931-INS-541
chr2-8177446-INS-57
chr2-8211762-INS-3517
chr2-8235744-INS-489
chr2-8235750-INS-201
chr2-8238916-INS-71
chr2-8338241-INS-52

chr2-8368798-INS-3244
chr2-8459591-INS-102
chr2-8464938-INS-176
chr2-8495620-INS-485
chr2-8513708-INS-921
chr2-8565835-INS-299
chr2-8566083-INS-249
chr2-8571281-INS-54
chr2-8573764-INS-60
chr2-8653938-INS-6386
chr2-8664639-INS-130
chr2-8678509-INS-235
chr2-8678531-INS-96
chr2-8695010-INS-59
chr2-8707899-INS-6158
chr2-8772193-INS-50
chr2-8801539-INS-138
chr2-8832801-INS-1728
chr2-8850910-INS-328
chr2-8941029-INS-72
chr2-8986482-INS-154
chr2-9054174-INS-50
chr2-9076107-INS-219
chr2-9196691-INS-6443
chr2-9201344-INS-60
chr2-9244826-INS-54
chr2-9319968-INS-92
chr2-9320045-INS-825
chr2-9323851-INS-2317
chr2-9324924-INS-50
chr2-9354795-INS-63
chr2-9384446-INS-3646
chr2-9417808-INS-74
chr2-9445410-INS-6402
chr2-9507152-INS-56
chr2-9507694-INS-394
chr2-9528085-INS-1142
chr2-9538309-INS-72
chr2-9558004-INS-52
chr2-9571158-INS-134
chr2-9613669-INS-123
chr2-9618153-INS-84
chr2-9707779-INS-56
chr2-9766808-INS-168
chr2-9778246-INS-52
chr2-9785363-INS-223
chr2-9802485-INS-52
chr2-9802763-INS-452
chr2-9802763-INS-98
chr2-9804361-INS-127
chr2-9804423-INS-142
chr2-9858638-INS-76
chr2-9905410-INS-50
chr2-9909694-INS-84
chr2-9930785-INS-171
chr2-9930837-INS-65
chr2-9935573-INS-344
chr2-10022167-INS-175
chr2-10086917-INS-193
chr2-10093505-INS-80
chr2-10095721-INS-231
chr2-10098139-INS-380
chr2-10141204-INS-6910
chr2-10151663-INS-797
chr2-10153086-INS-84
chr2-10163323-INS-96
chr2-10176637-INS-98
chr2-10190905-INS-133
chr2-10191222-INS-1218
chr2-10203991-INS-80
chr2-10229074-INS-209
chr2-10234929-INS-52
chr2-10236439-INS-218
chr2-10247726-INS-51
chr2-10252329-INS-517
chr2-10281905-INS-148
chr2-10282279-INS-132

chr2-10324427-INS-2751
chr2-10326625-INS-158
chr2-10330445-INS-582
chr2-10330665-INS-52
chr2-10333559-INS-97
chr2-10334412-INS-92
chr2-10337564-INS-56
chr2-10362366-INS-73
chr2-10462949-INS-517
chr2-10540196-INS-474
chr2-10622899-INS-5316
chr2-10720261-INS-418
chr2-10757378-INS-6619
chr2-10758709-INS-2125
chr2-10763601-INS-122
chr2-10772944-INS-50
chr2-10786904-INS-6014
chr2-10814265-INS-52
chr2-10829336-INS-7121
chr2-10860747-INS-66
chr2-10860846-INS-198
chr2-10888894-INS-56
chr2-10897771-INS-60
chr2-10904994-INS-116
chr2-10957173-INS-2291
chr2-10985126-INS-4862
chr2-10987628-INS-2488
chr2-11011068-INS-1479
chr2-11031473-INS-312
chr2-11054188-INS-51
chr2-11065307-INS-340
chr2-11067117-INS-60
chr2-11067147-INS-1230
chr2-11072817-INS-6316
chr2-11094155-INS-352
chr2-11114203-INS-4770
chr2-11141955-INS-2988
chr2-11159569-INS-132
chr2-11159960-INS-59
chr2-11168356-INS-76
chr2-11185323-INS-54
chr2-11185912-INS-174
chr2-11196578-INS-50
chr2-11222457-INS-135
chr2-11222902-INS-196
chr2-11232299-INS-141
chr2-11242354-INS-64
chr2-11258764-INS-53
chr2-11284208-INS-226
chr2-11289830-INS-1486
chr2-11308255-INS-113
chr2-11308543-INS-56
chr2-11328333-INS-386
chr2-11330501-INS-605
chr2-11339914-INS-1041
chr2-11347769-INS-1053
chr2-11378751-INS-4358
chr2-11399658-INS-2011
chr2-11424086-INS-7097
chr2-11450383-INS-354
chr2-11451182-INS-1730
chr2-11454888-INS-57
chr2-11459499-INS-92
chr2-11460623-INS-206
chr2-11466728-INS-94
chr2-11544131-INS-73
chr2-11544397-INS-188
chr2-11545065-INS-375
chr2-11577795-INS-336
chr2-11583532-INS-1693
chr2-11625962-INS-90
chr2-11631171-INS-67
chr2-11631686-INS-225
chr2-11657449-INS-90
chr2-11691164-INS-235
chr2-11712272-INS-1100
chr2-11778606-INS-124

chr2-11902319-INS-61
chr2-11910581-INS-523
chr2-11928913-INS-72
chr2-11941210-INS-6477
chr2-11948173-INS-201
chr2-12005743-INS-85
chr2-12012855-INS-15402
chr2-12030937-INS-588
chr2-12073964-INS-702
chr2-12074345-INS-140
chr2-12085163-INS-6394
chr2-12106138-INS-12268
chr2-12106682-INS-1596
chr2-12143187-INS-7294
chr2-12147358-INS-456
chr2-12173147-INS-226
chr2-12175350-INS-97
chr2-12175489-INS-118
chr2-12176949-INS-57
chr2-12179115-INS-50
chr2-12181029-INS-50
chr2-12230463-INS-54
chr2-12235206-INS-120
chr2-12307725-INS-56
chr2-12324010-INS-81
chr2-12437962-INS-137
chr2-12458267-INS-4789
chr2-12469114-INS-84
chr2-12501438-INS-56
chr2-12506202-INS-2504
chr2-12518296-INS-181
chr2-12518346-INS-90
chr2-12522587-INS-52
chr2-12553323-INS-68
chr2-12559609-INS-335
chr2-12660481-INS-964
chr2-12663167-INS-1998
chr2-12663172-INS-57
chr2-12669898-INS-363
chr2-12682315-INS-173
chr2-12694597-INS-108
chr2-12696408-INS-109
chr2-12706782-INS-180
chr2-12712867-INS-2103
chr2-12741004-INS-1612
chr2-12761270-INS-720
chr2-12761519-INS-3013
chr2-12765459-INS-168
chr2-12775709-INS-64
chr2-12796566-INS-6349
chr2-12809346-INS-7200
chr2-12837163-INS-7113
chr2-12847110-INS-65
chr2-12913369-INS-817
chr2-12919850-INS-1208
chr2-12948409-INS-1000
chr2-12999486-INS-105
chr2-13034409-INS-442
chr2-13036318-INS-132
chr2-13055079-INS-81
chr2-13095637-INS-144
chr2-13115517-INS-257
chr2-13170164-INS-101
chr2-13229914-INS-52
chr2-13235228-INS-400
chr2-13273292-INS-59
chr2-13342509-INS-93
chr2-13417434-INS-93
chr2-13417488-INS-217
chr2-13440080-INS-2500
chr2-13476701-INS-942
chr2-13486172-INS-66
chr2-13500408-INS-182
chr2-13538531-INS-60
chr2-13540667-INS-344
chr2-13602279-INS-126
chr2-13609337-INS-1062

chr2-13626050-INS-3048
chr2-13709572-INS-6576
chr2-13717731-INS-63
chr2-13756656-INS-53
chr2-13767671-INS-705
chr2-13803159-INS-84
chr2-13844945-INS-105
chr2-13865604-INS-74
chr2-13875496-INS-6580
chr2-13898169-INS-7095
chr2-13898489-INS-3091
chr2-13928006-INS-919
chr2-13928604-INS-2187
chr2-13938988-INS-3946
chr2-13979788-INS-1211
chr2-13979996-INS-1451
chr2-13998080-INS-6219
chr2-14046280-INS-60
chr2-14155862-INS-1009
chr2-14180424-INS-188
chr2-14182304-INS-96
chr2-14201152-INS-2308
chr2-14201466-INS-5991
chr2-14208750-INS-207
chr2-14212131-INS-54
chr2-14215219-INS-207
chr2-14229530-INS-51
chr2-14229547-INS-144
chr2-14233739-INS-167
chr2-14254403-INS-498
chr2-14254699-INS-166
chr2-14282500-INS-50
chr2-14339821-INS-70
chr2-14344373-INS-139
chr2-14344487-INS-240
chr2-14396669-INS-54
chr2-14396980-INS-2650
chr2-14427289-INS-98
chr2-14441820-INS-5841
chr2-14467604-INS-615
chr2-14471576-INS-64
chr2-14475167-INS-162
chr2-14478808-INS-50
chr2-14479329-INS-5519
chr2-14506841-INS-1164
chr2-14549158-INS-64
chr2-14569444-INS-382
chr2-14569444-INS-74
chr2-14575001-INS-100
chr2-14663081-INS-52
chr2-14762904-INS-83
chr2-14792947-INS-331
chr2-14814967-INS-50
chr2-14818922-INS-50
chr2-14839469-INS-80
chr2-14874021-INS-177
chr2-14880922-INS-112
chr2-14904293-INS-150
chr2-14974748-INS-50
chr2-14982856-INS-1309
chr2-15118113-INS-347
chr2-15127811-INS-88
chr2-15155494-INS-7102
chr2-15165815-INS-1494
chr2-15183839-INS-6724
chr2-15341921-INS-3491
chr2-15365096-INS-2490
chr2-15375709-INS-295
chr2-15454805-INS-56
chr2-15454884-INS-181
chr2-15470873-INS-822
chr2-15546390-INS-2097
chr2-15574586-INS-1494
chr2-15634162-INS-6255
chr2-15719196-INS-479
chr2-15787685-INS-2112
chr2-15791734-INS-389

chr2-15797948-INS-80
chr2-15815392-INS-1445
chr2-15860676-INS-6964
chr2-15915702-INS-4325
chr2-15927077-INS-934
chr2-15955728-INS-116
chr2-15956908-INS-6084
chr2-15981152-INS-62
chr2-16002294-INS-67
chr2-16008640-INS-640
chr2-16119411-INS-279
chr2-16119411-INS-56
chr2-16137094-INS-4792
chr2-16140721-INS-6206
chr2-16140724-INS-1790
chr2-16165633-INS-52
chr2-16167776-INS-481
chr2-16168616-INS-50
chr2-16173222-INS-1943
chr2-16188776-INS-2266
chr2-16277798-INS-3957
chr2-16311796-INS-135
chr2-16339230-INS-74
chr2-16339310-INS-855
chr2-16379405-INS-202
chr2-16407077-INS-336
chr2-16463623-INS-2035
chr2-16486470-INS-64
chr2-16524346-INS-6443
chr2-16546650-INS-6329
chr2-16548051-INS-1492
chr2-16616730-INS-52
chr2-16619036-INS-93
chr2-16727688-INS-1175
chr2-16752798-INS-602
chr2-16752981-INS-297
chr2-16753200-INS-59
chr2-16766198-INS-50
chr2-16840256-INS-1254
chr2-16870612-INS-60
chr2-16920437-INS-1561
chr2-16977308-INS-460
chr2-16983030-INS-67
chr2-16992297-INS-1097
chr2-17044402-INS-468
chr2-17147052-INS-313
chr2-17228936-INS-1242
chr2-17234052-INS-1890
chr2-17280717-INS-1378
chr2-17342232-INS-54
chr2-17480865-INS-51
chr2-17498222-INS-196
chr2-17547087-INS-239
chr2-17547676-INS-2061
chr2-17571803-INS-676
chr2-17602950-INS-1864
chr2-17671994-INS-86
chr2-17750336-INS-67
chr2-17769741-INS-178
chr2-17861647-INS-132
chr2-17861763-INS-52
chr2-17875009-INS-165
chr2-17879083-INS-93
chr2-17890321-INS-112
chr2-17905537-INS-201
chr2-17906396-INS-505
chr2-17946816-INS-88
chr2-17956104-INS-7389
chr2-17966498-INS-288
chr2-17992039-INS-305
chr2-18020869-INS-943
chr2-18027611-INS-209
chr2-18042933-INS-54
chr2-18075538-INS-65
chr2-18137738-INS-54
chr2-18149207-INS-1106
chr2-18161847-INS-3337

chr2-18161921-INS-311
chr2-18161931-INS-50
chr2-18221937-INS-88
chr2-18261525-INS-840
chr2-18294639-INS-7516
chr2-18298578-INS-210
chr2-18373729-INS-1446
chr2-18418884-INS-4810
chr2-18420914-INS-51
chr2-18421795-INS-12947
chr2-18456187-INS-1894
chr2-18501038-INS-471
chr2-18514045-INS-1706
chr2-18514409-INS-914
chr2-18516648-INS-4800
chr2-18570820-INS-120
chr2-18574682-INS-1026
chr2-18576974-INS-77
chr2-18586783-INS-52
chr2-18619517-INS-86
chr2-18647032-INS-96
chr2-18651820-INS-629
chr2-18653277-INS-217
chr2-18666852-INS-199
chr2-18672012-INS-52
chr2-18740585-INS-157
chr2-18772597-INS-145
chr2-18780216-INS-169
chr2-18783593-INS-78
chr2-18793673-INS-98
chr2-18809905-INS-264
chr2-18892283-INS-62
chr2-18898257-INS-561
chr2-18898379-INS-62
chr2-18909127-INS-215
chr2-19024445-INS-54
chr2-19026894-INS-508
chr2-19070718-INS-76
chr2-19087105-INS-1974
chr2-19092346-INS-1883
chr2-19096640-INS-70
chr2-19148755-INS-68
chr2-19163892-INS-118
chr2-19199875-INS-141
chr2-19249073-INS-52
chr2-19256996-INS-52
chr2-19272895-INS-1054
chr2-19274253-INS-216
chr2-19274703-INS-52
chr2-19275526-INS-112
chr2-19288915-INS-124
chr2-19289005-INS-60
chr2-19345873-INS-65
chr2-19348299-INS-3680
chr2-19372006-INS-78
chr2-19408388-INS-185
chr2-19412434-INS-6182
chr2-19421869-INS-73
chr2-19433415-INS-75
chr2-19512662-INS-60
chr2-19519950-INS-175
chr2-19520010-INS-75
chr2-19522000-INS-70
chr2-19544789-INS-2631
chr2-19574689-INS-50
chr2-19579439-INS-180
chr2-19598557-INS-52
chr2-19600751-INS-52
chr2-19657932-INS-56
chr2-19692008-INS-299
chr2-19692068-INS-125
chr2-19700267-INS-157
chr2-19702045-INS-256
chr2-19702096-INS-102
chr2-19766362-INS-174
chr2-19804819-INS-71
chr2-19820519-INS-5629

chr2-19827500-INS-112
chr2-19843282-INS-332
chr2-19843480-INS-5962
chr2-19861032-INS-7281
chr2-19862482-INS-1891
chr2-19876990-INS-1781
chr2-19880296-INS-4150
chr2-19885582-INS-2411
chr2-19889009-INS-359
chr2-19903028-INS-162
chr2-19905383-INS-50
chr2-19908626-INS-216
chr2-19910192-INS-6569
chr2-19920433-INS-988
chr2-19943719-INS-6286
chr2-19967334-INS-2148
chr2-20038015-INS-161
chr2-20052252-INS-260
chr2-20087057-INS-1020
chr2-20087316-INS-325
chr2-20100102-INS-4409
chr2-20104643-INS-293
chr2-20141300-INS-270
chr2-20141348-INS-758
chr2-20201073-INS-6700
chr2-20212253-INS-13509
chr2-20267668-INS-57
chr2-20280132-INS-700
chr2-20280394-INS-50
chr2-20280512-INS-205
chr2-20347118-INS-122
chr2-20365242-INS-420
chr2-20372702-INS-186
chr2-20469017-INS-73
chr2-20469058-INS-207
chr2-20470631-INS-238
chr2-20479271-INS-196
chr2-20507833-INS-6879
chr2-20510812-INS-186
chr2-20511685-INS-78
chr2-20515429-INS-156
chr2-20516905-INS-243
chr2-20521316-INS-191
chr2-20579956-INS-224
chr2-20589426-INS-204
chr2-20589928-INS-55
chr2-20593488-INS-147
chr2-20605709-INS-50
chr2-20611919-INS-50
chr2-20641515-INS-89
chr2-20641861-INS-190
chr2-20671778-INS-72
chr2-20675699-INS-212
chr2-20716401-INS-209
chr2-20725562-INS-100
chr2-20799000-INS-212
chr2-20832882-INS-6162
chr2-20835489-INS-174
chr2-20889292-INS-224
chr2-20889886-INS-97
chr2-20900026-INS-54
chr2-20912707-INS-202
chr2-20915297-INS-480
chr2-20939532-INS-52
chr2-20942229-INS-672
chr2-20960756-INS-188
chr2-20979034-INS-225
chr2-20986285-INS-50
chr2-20992278-INS-129
chr2-20992443-INS-74
chr2-21004954-INS-105
chr2-21011496-INS-186
chr2-21018368-INS-56
chr2-21022105-INS-162
chr2-21030238-INS-60
chr2-21030253-INS-187
chr2-21032470-INS-66

chr2-21034209-INS-210
chr2-21036887-INS-1188
chr2-21050570-INS-171
chr2-21073326-INS-445
chr2-21105598-INS-556
chr2-21118852-INS-54
chr2-21120425-INS-1047
chr2-21122253-INS-199
chr2-21122277-INS-56
chr2-21135387-INS-58
chr2-21136772-INS-343
chr2-21145392-INS-2394
chr2-21157252-INS-6124
chr2-21166360-INS-62
chr2-21199639-INS-459
chr2-21223241-INS-1181
chr2-21260729-INS-63
chr2-21290530-INS-2748
chr2-21299916-INS-115
chr2-21319159-INS-100
chr2-21331597-INS-110
chr2-21339405-INS-1187
chr2-21342391-INS-314
chr2-21343427-INS-379
chr2-21465039-INS-143
chr2-21469747-INS-165
chr2-21469747-INS-53
chr2-21477440-INS-1286
chr2-21486872-INS-50
chr2-21499072-INS-235
chr2-21517636-INS-4332
chr2-21525128-INS-3011
chr2-21557158-INS-274
chr2-21557170-INS-81
chr2-21655443-INS-503
chr2-21705864-INS-409
chr2-21741917-INS-13132
chr2-21741917-INS-6464
chr2-21757844-INS-3215
chr2-21825741-INS-6726
chr2-21854642-INS-1306
chr2-21858116-INS-7240
chr2-21904763-INS-2353
chr2-21940153-INS-73
chr2-21954955-INS-2685
chr2-21983100-INS-751
chr2-22002811-INS-375
chr2-22024586-INS-1978
chr2-22046086-INS-526
chr2-22051546-INS-64
chr2-22074342-INS-93
chr2-22132088-INS-145
chr2-22178166-INS-187
chr2-22210632-INS-224
chr2-22238282-INS-5436
chr2-22283212-INS-133
chr2-22283513-INS-221
chr2-22283806-INS-1964
chr2-22293078-INS-2007
chr2-22293487-INS-3892
chr2-22364712-INS-323
chr2-22367148-INS-170
chr2-22435031-INS-79
chr2-22442199-INS-2776
chr2-22460911-INS-2198
chr2-22483446-INS-152
chr2-22488609-INS-3599
chr2-22493513-INS-2837
chr2-22537004-INS-68
chr2-22552508-INS-71
chr2-22575742-INS-149
chr2-22575990-INS-146
chr2-22588009-INS-398
chr2-22630336-INS-319
chr2-22632423-INS-2145
chr2-22646768-INS-123
chr2-22650400-INS-72

chr2-22653448-INS-407
chr2-22698696-INS-216
chr2-22732412-INS-52
chr2-22762263-INS-65
chr2-22768109-INS-122
chr2-22777325-INS-5329
chr2-22783212-INS-1199
chr2-22920321-INS-156
chr2-22931213-INS-366
chr2-22931476-INS-4923
chr2-22940972-INS-1780
chr2-22943513-INS-114
chr2-22961934-INS-55
chr2-22969311-INS-201
chr2-22981133-INS-118
chr2-23059310-INS-415
chr2-23059310-INS-85
chr2-23060165-INS-79
chr2-23078923-INS-203
chr2-23080116-INS-68
chr2-23125350-INS-6385
chr2-23133510-INS-5311
chr2-23146712-INS-4382
chr2-23165528-INS-2356
chr2-23171280-INS-15739
chr2-23181346-INS-5026
chr2-23190926-INS-1767
chr2-23213511-INS-109
chr2-23255971-INS-6818
chr2-23288378-INS-199
chr2-23301306-INS-63
chr2-23323779-INS-503
chr2-23328441-INS-230
chr2-23339208-INS-56
chr2-23339287-INS-66
chr2-23360892-INS-52
chr2-23388057-INS-216
chr2-23470378-INS-693
chr2-23476327-INS-205
chr2-23479990-INS-254
chr2-23511466-INS-1754
chr2-23520632-INS-4995
chr2-23555169-INS-548
chr2-23571639-INS-83
chr2-23573326-INS-3326
chr2-23576128-INS-856
chr2-23576287-INS-220
chr2-23624047-INS-50
chr2-23636640-INS-2953
chr2-23692473-INS-1073
chr2-23712666-INS-161
chr2-23721678-INS-257
chr2-23722796-INS-6469
chr2-23742309-INS-156
chr2-23769947-INS-6566
chr2-23777203-INS-60
chr2-23793199-INS-334
chr2-23857237-INS-1626
chr2-23860035-INS-52
chr2-23867466-INS-231
chr2-23888896-INS-50
chr2-23897454-INS-6237
chr2-23900068-INS-1159
chr2-23903595-INS-205
chr2-23958208-INS-1268
chr2-24057660-INS-75
chr2-24068766-INS-4246
chr2-24071582-INS-71
chr2-24081940-INS-446
chr2-24102503-INS-121
chr2-24170211-INS-50
chr2-24181008-INS-1266
chr2-24268301-INS-439
chr2-24322028-INS-104
chr2-24356791-INS-131
chr2-24404849-INS-111
chr2-24408839-INS-51

chr2-24417319-INS-1237
chr2-24436258-INS-161
chr2-24436286-INS-62
chr2-24436782-INS-55
chr2-24467939-INS-9923
chr2-24473810-INS-5157
chr2-24484522-INS-544
chr2-24522940-INS-428
chr2-24524817-INS-218
chr2-24524847-INS-957
chr2-24556951-INS-88
chr2-24557045-INS-1196
chr2-24557665-INS-54
chr2-24600142-INS-498
chr2-24613602-INS-72
chr2-24624973-INS-116
chr2-24624973-INS-532
chr2-24631371-INS-203
chr2-24640783-INS-167
chr2-24660251-INS-985
chr2-24724965-INS-66
chr2-24725022-INS-78
chr2-24754249-INS-54
chr2-24769333-INS-92
chr2-24828431-INS-102
chr2-24844275-INS-174
chr2-24844479-INS-208
chr2-24846248-INS-54
chr2-24880866-INS-650
chr2-24881683-INS-346
chr2-24917076-INS-210
chr2-24917955-INS-275
chr2-24934739-INS-1650
chr2-24999336-INS-66
chr2-24999811-INS-566
chr2-25014142-INS-3189
chr2-25045867-INS-205
chr2-25078688-INS-216
chr2-25081420-INS-204
chr2-25082218-INS-62
chr2-25142604-INS-206
chr2-25171673-INS-207
chr2-25196482-INS-191
chr2-25198679-INS-98
chr2-25199317-INS-204
chr2-25213570-INS-8457
chr2-25219011-INS-112
chr2-25276353-INS-162
chr2-25310271-INS-79
chr2-25310837-INS-294
chr2-25342141-INS-54
chr2-25352415-INS-112
chr2-25402173-INS-4514
chr2-25443498-INS-138
chr2-25448617-INS-499
chr2-25462676-INS-226
chr2-25512988-INS-73
chr2-25556248-INS-60
chr2-25561685-INS-505
chr2-25654448-INS-138
chr2-25708417-INS-599
chr2-25709045-INS-848
chr2-25709767-INS-508
chr2-25753667-INS-54
chr2-25843309-INS-204
chr2-25877392-INS-120
chr2-25959888-INS-1280
chr2-26045643-INS-55
chr2-26071077-INS-579
chr2-26128867-INS-126
chr2-26176903-INS-110
chr2-26191330-INS-186
chr2-26414045-INS-2473
chr2-26418982-INS-50
chr2-26455598-INS-104
chr2-26464097-INS-96
chr2-26475151-INS-203

chr2-26488475-INS-177
chr2-26506907-INS-206
chr2-26540684-INS-73
chr2-26548846-INS-208
chr2-26564707-INS-100
chr2-26620977-INS-60
chr2-26659403-INS-823
chr2-26663976-INS-214
chr2-26816180-INS-51
chr2-26855404-INS-311
chr2-26920584-INS-225
chr2-26954573-INS-404
chr2-26954734-INS-388
chr2-26955010-INS-902
chr2-27020250-INS-264
chr2-27020695-INS-60
chr2-27080361-INS-204
chr2-27080714-INS-66
chr2-27099022-INS-60
chr2-27135885-INS-202
chr2-27165868-INS-324
chr2-27166428-INS-483
chr2-27166471-INS-7709
chr2-27170166-INS-57
chr2-27251522-INS-56
chr2-27264676-INS-225
chr2-27269641-INS-52
chr2-27288128-INS-60
chr2-27324454-INS-7105
chr2-27324814-INS-88
chr2-27328130-INS-189
chr2-27372653-INS-204
chr2-27373286-INS-166
chr2-27381874-INS-1191
chr2-27392379-INS-205
chr2-27406765-INS-177
chr2-27440813-INS-210
chr2-27476172-INS-129
chr2-27484598-INS-65
chr2-27494972-INS-50
chr2-27545843-INS-50
chr2-27549472-INS-74
chr2-27565828-INS-202
chr2-27784137-INS-82
chr2-27831612-INS-245
chr2-27839090-INS-52
chr2-27930947-INS-165
chr2-27963598-INS-4383
chr2-27992922-INS-96
chr2-28157440-INS-78
chr2-28186794-INS-320
chr2-28187946-INS-56
chr2-28188585-INS-57
chr2-28196472-INS-60
chr2-28226144-INS-62
chr2-28281218-INS-460
chr2-28281218-INS-972
chr2-28316903-INS-142
chr2-28331020-INS-66
chr2-28331478-INS-199
chr2-28349359-INS-2825
chr2-28391367-INS-60
chr2-28406926-INS-50
chr2-28437923-INS-343
chr2-28443690-INS-222
chr2-28453830-INS-87
chr2-28472374-INS-202
chr2-28476222-INS-75
chr2-28476386-INS-183
chr2-28482036-INS-50
chr2-28485441-INS-86
chr2-28485938-INS-118
chr2-28485994-INS-496
chr2-28499835-INS-3297
chr2-28504174-INS-51
chr2-28511529-INS-52
chr2-28520927-INS-208

chr2-28522139-INS-63
chr2-28549644-INS-216
chr2-28572888-INS-250
chr2-28608729-INS-111
chr2-28622848-INS-61
chr2-28638482-INS-68
chr2-28659745-INS-51
chr2-28689316-INS-59
chr2-28709733-INS-50
chr2-28728035-INS-416
chr2-28772457-INS-390
chr2-28831188-INS-60
chr2-28842558-INS-51
chr2-28866033-INS-51
chr2-28867661-INS-79
chr2-28905518-INS-168
chr2-28906137-INS-83
chr2-28927106-INS-348
chr2-28933229-INS-608
chr2-28945342-INS-694
chr2-28945658-INS-210
chr2-28982744-INS-397
chr2-29111254-INS-58
chr2-29111894-INS-78
chr2-29121746-INS-71
chr2-29123425-INS-226
chr2-29216068-INS-129
chr2-29235848-INS-210
chr2-29263092-INS-50
chr2-29270573-INS-67
chr2-29280939-INS-179
chr2-29304917-INS-152
chr2-29333774-INS-120
chr2-29347659-INS-215
chr2-29357972-INS-60
chr2-29358043-INS-192
chr2-29430828-INS-74
chr2-29441188-INS-152
chr2-29446544-INS-149
chr2-29461468-INS-72
chr2-29479212-INS-76
chr2-29479751-INS-1371
chr2-29486173-INS-83
chr2-29506141-INS-2027
chr2-29550707-INS-66
chr2-29614512-INS-168
chr2-29630541-INS-1501
chr2-29642157-INS-207
chr2-29642785-INS-56
chr2-29713746-INS-302
chr2-29767524-INS-122
chr2-29767614-INS-444
chr2-29778875-INS-51
chr2-29822970-INS-133
chr2-29859597-INS-188
chr2-29896793-INS-189
chr2-29943691-INS-197
chr2-29965015-INS-147
chr2-30003037-INS-50
chr2-30019623-INS-149
chr2-30022117-INS-1475
chr2-30049131-INS-198
chr2-30088428-INS-113
chr2-30133280-INS-1153
chr2-30178927-INS-261
chr2-30253434-INS-64
chr2-30339796-INS-340
chr2-30342779-INS-718
chr2-30345120-INS-50
chr2-30353121-INS-448
chr2-30381460-INS-59
chr2-30383227-INS-294
chr2-30383487-INS-359
chr2-30447302-INS-492
chr2-30448350-INS-1261
chr2-30453091-INS-105
chr2-30454509-INS-99

chr2-30475406-INS-144
chr2-30475406-INS-333
chr2-30511577-INS-64
chr2-30517680-INS-52
chr2-30533951-INS-699
chr2-30533951-INS-84
chr2-30558262-INS-561
chr2-30576131-INS-102
chr2-30593226-INS-231
chr2-30595612-INS-114
chr2-30601295-INS-1861
chr2-30621278-INS-106
chr2-30653496-INS-3829
chr2-30657080-INS-138
chr2-30657832-INS-5550
chr2-30667199-INS-268
chr2-30669950-INS-57
chr2-30711389-INS-64
chr2-30736602-INS-149
chr2-30770125-INS-540
chr2-30785496-INS-119
chr2-30794606-INS-50
chr2-30802752-INS-232
chr2-30802769-INS-76
chr2-30812813-INS-87
chr2-30814734-INS-1016
chr2-30822810-INS-235
chr2-30823323-INS-211
chr2-30831212-INS-4414
chr2-30862298-INS-107
chr2-30908037-INS-102
chr2-30923598-INS-206
chr2-30968906-INS-242
chr2-30992971-INS-115
chr2-30993837-INS-500
chr2-30998954-INS-232
chr2-31006956-INS-98
chr2-31035777-INS-205
chr2-31038412-INS-218
chr2-31073496-INS-51
chr2-31101082-INS-54
chr2-31120293-INS-232
chr2-31120293-INS-58
chr2-31166406-INS-51
chr2-31248774-INS-242
chr2-31265254-INS-56
chr2-31265804-INS-68
chr2-31299850-INS-54
chr2-31326316-INS-171
chr2-31326634-INS-140
chr2-31331065-INS-3500
chr2-31355596-INS-52
chr2-31366751-INS-7145
chr2-31368418-INS-1490
chr2-31369431-INS-51
chr2-31458199-INS-445
chr2-31458414-INS-212
chr2-31499657-INS-58
chr2-31580056-INS-219
chr2-31583672-INS-96
chr2-31584460-INS-155
chr2-31591523-INS-403
chr2-31597350-INS-497
chr2-31759113-INS-76
chr2-31759966-INS-80
chr2-31826493-INS-232
chr2-31827053-INS-68
chr2-31831700-INS-204
chr2-31833863-INS-54
chr2-31837744-INS-70
chr2-31971003-INS-480
chr2-31974797-INS-94
chr2-32017635-INS-379
chr2-32025094-INS-95
chr2-32025256-INS-91
chr2-32026725-INS-211
chr2-32031974-INS-61

chr2-32034695-INS-533
chr2-32066807-INS-556
chr2-32157223-INS-169
chr2-32160886-INS-231
chr2-32211474-INS-54
chr2-32284692-INS-213
chr2-32298310-INS-67
chr2-32360246-INS-78
chr2-32387063-INS-56
chr2-32388603-INS-6799
chr2-32493114-INS-867
chr2-32493501-INS-203
chr2-32562447-INS-61
chr2-32588938-INS-228
chr2-32588938-INS-52
chr2-32654128-INS-56
chr2-32675598-INS-85
chr2-32715070-INS-112
chr2-32723522-INS-175
chr2-32743546-INS-52
chr2-32751808-INS-76
chr2-32752206-INS-60
chr2-32754132-INS-191
chr2-32762718-INS-69
chr2-32768381-INS-50
chr2-32773651-INS-203
chr2-32777488-INS-62
chr2-32836854-INS-144
chr2-32841236-INS-94
chr2-32841443-INS-185
chr2-32860538-INS-126
chr2-32886889-INS-60
chr2-32890528-INS-230
chr2-32890707-INS-205
chr2-32900224-INS-493
chr2-32905523-INS-209
chr2-32910039-INS-69
chr2-32938152-INS-64
chr2-32959791-INS-407
chr2-32976055-INS-56
chr2-32997915-INS-143
chr2-33045811-INS-204
chr2-33048031-INS-96
chr2-33054597-INS-171
chr2-33126609-INS-66
chr2-33159211-INS-185
chr2-33206114-INS-50
chr2-33232557-INS-1454
chr2-33243916-INS-1174
chr2-33271348-INS-9555
chr2-33307224-INS-51
chr2-33323227-INS-808
chr2-33327833-INS-95
chr2-33336142-INS-101
chr2-33357846-INS-50
chr2-33363148-INS-239
chr2-33384589-INS-210
chr2-33386351-INS-6120
chr2-33416102-INS-168
chr2-33446470-INS-70
chr2-33454281-INS-97
chr2-33479857-INS-122
chr2-33602292-INS-194
chr2-33631149-INS-297
chr2-33632159-INS-69
chr2-33668176-INS-55
chr2-33714840-INS-205
chr2-33729159-INS-90
chr2-33746164-INS-152
chr2-33752207-INS-60
chr2-33791173-INS-318
chr2-33824111-INS-3179
chr2-33824206-INS-6690
chr2-33879614-INS-56
chr2-33952064-INS-181
chr2-33973105-INS-58
chr2-33979314-INS-80

chr2-33985025-INS-201
chr2-34049375-INS-182
chr2-34051691-INS-4239
chr2-34163624-INS-86
chr2-34185925-INS-63
chr2-34190745-INS-328
chr2-34197524-INS-51
chr2-34208559-INS-170
chr2-34324457-INS-124
chr2-34328913-INS-60
chr2-34341614-INS-202
chr2-34358933-INS-98
chr2-34363697-INS-205
chr2-34401358-INS-204
chr2-34426919-INS-51
chr2-34451971-INS-986
chr2-34476549-INS-223
chr2-34544239-INS-50
chr2-34558872-INS-76
chr2-34584553-INS-51
chr2-34601991-INS-230
chr2-34631963-INS-5994
chr2-34640085-INS-220
chr2-34642574-INS-57
chr2-34647027-INS-190
chr2-34654487-INS-800
chr2-34674109-INS-214
chr2-34726430-INS-208
chr2-34730292-INS-213
chr2-34732974-INS-567
chr2-34751426-INS-278
chr2-34905964-INS-64
chr2-34931265-INS-50
chr2-34939252-INS-12405
chr2-35004474-INS-57
chr2-35016993-INS-1344
chr2-35024993-INS-68
chr2-35053644-INS-90
chr2-35056650-INS-1663
chr2-35105397-INS-280
chr2-35134110-INS-232
chr2-35134290-INS-106
chr2-35268321-INS-63
chr2-35283819-INS-1109
chr2-35318316-INS-98
chr2-35394578-INS-4442
chr2-35407704-INS-232
chr2-35408385-INS-80
chr2-35408819-INS-178
chr2-35427148-INS-210
chr2-35531768-INS-72
chr2-35541968-INS-113
chr2-35544428-INS-56
chr2-35615405-INS-81
chr2-35628142-INS-54
chr2-35639716-INS-233
chr2-35681688-INS-52
chr2-35726453-INS-116
chr2-35733267-INS-191
chr2-35748939-INS-214
chr2-35798375-INS-50
chr2-35946214-INS-60
chr2-35986283-INS-54
chr2-36052105-INS-290
chr2-36068953-INS-232
chr2-36090102-INS-272
chr2-36111648-INS-60
chr2-36177186-INS-156
chr2-36212079-INS-364
chr2-36224463-INS-1116
chr2-36225057-INS-64
chr2-36268770-INS-191
chr2-36283550-INS-53
chr2-36333528-INS-208
chr2-36351508-INS-1838
chr2-36357177-INS-13921
chr2-36443188-INS-65

chr2-36493977-INS-317
chr2-36508757-INS-233
chr2-36514076-INS-7083
chr2-36538267-INS-115
chr2-36584203-INS-861
chr2-36601367-INS-4985
chr2-36712713-INS-9181
chr2-36713020-INS-6371
chr2-36736599-INS-360
chr2-36777895-INS-6262
chr2-36783972-INS-1318
chr2-36786085-INS-136
chr2-36861235-INS-6417
chr2-36862237-INS-1538
chr2-36881091-INS-98
chr2-36891741-INS-152
chr2-36904976-INS-2597
chr2-37020191-INS-4017
chr2-37054186-INS-57
chr2-37060970-INS-415
chr2-37088712-INS-7232
chr2-37111139-INS-52
chr2-37148454-INS-1196
chr2-37150048-INS-194
chr2-37156479-INS-58
chr2-37162374-INS-116
chr2-37165626-INS-70
chr2-37238022-INS-81
chr2-37252081-INS-5573
chr2-37263066-INS-76
chr2-37289114-INS-165
chr2-37364236-INS-208
chr2-37554916-INS-51
chr2-37596422-INS-237
chr2-37703485-INS-68
chr2-37867131-INS-148
chr2-37877767-INS-106
chr2-37894323-INS-133
chr2-37993223-INS-56
chr2-38095761-INS-1766
chr2-38122357-INS-165
chr2-38122357-INS-55
chr2-38130070-INS-103
chr2-38144078-INS-199
chr2-38144771-INS-59
chr2-38217095-INS-50
chr2-38221353-INS-68
chr2-38237275-INS-77
chr2-38324169-INS-52
chr2-38324723-INS-3025
chr2-38358076-INS-50
chr2-38361059-INS-78
chr2-38373743-INS-56
chr2-38432769-INS-515
chr2-38489472-INS-658
chr2-38489628-INS-1449
chr2-38522452-INS-203
chr2-38525793-INS-179
chr2-38549676-INS-88
chr2-38554368-INS-337
chr2-38568963-INS-6461
chr2-38623811-INS-79
chr2-38637795-INS-52
chr2-38645776-INS-1370
chr2-38646021-INS-50
chr2-38647853-INS-102
chr2-38648740-INS-96
chr2-38666280-INS-171
chr2-38675655-INS-54
chr2-38694430-INS-428
chr2-38711224-INS-112
chr2-38774040-INS-468
chr2-38784227-INS-215
chr2-38804208-INS-708
chr2-38824460-INS-126
chr2-38864239-INS-60
chr2-38882286-INS-62

chr2-38988577-INS-113
chr2-39022742-INS-52
chr2-39074967-INS-60
chr2-39118818-INS-78
chr2-39157970-INS-2499
chr2-39165263-INS-1422
chr2-39175437-INS-7289
chr2-39186722-INS-203
chr2-39191397-INS-102
chr2-39209177-INS-6601
chr2-39221485-INS-85
chr2-39250542-INS-82
chr2-39282206-INS-497
chr2-39297575-INS-1495
chr2-39303242-INS-6814
chr2-39320700-INS-341
chr2-39356453-INS-336
chr2-39363204-INS-1604
chr2-39371326-INS-13303
chr2-39385478-INS-4703
chr2-39448484-INS-8315
chr2-39480241-INS-6460
chr2-39524945-INS-51
chr2-39642017-INS-6403
chr2-39668184-INS-60
chr2-39675879-INS-199
chr2-39763568-INS-68
chr2-39781221-INS-747
chr2-39781797-INS-87
chr2-39800914-INS-452
chr2-39868251-INS-2831
chr2-39871049-INS-416
chr2-39897331-INS-135
chr2-39928093-INS-437
chr2-39928773-INS-216
chr2-39946359-INS-83
chr2-39990017-INS-62
chr2-39992244-INS-233
chr2-40017916-INS-2550
chr2-40046018-INS-1863
chr2-40137546-INS-3051
chr2-40203000-INS-6564
chr2-40271164-INS-102
chr2-40325758-INS-4709
chr2-40380241-INS-4131
chr2-40398962-INS-358
chr2-40400280-INS-105
chr2-40400347-INS-356
chr2-40498985-INS-253
chr2-40511536-INS-772
chr2-40527271-INS-2547
chr2-40532119-INS-144
chr2-40556737-INS-141
chr2-40575373-INS-60
chr2-40576119-INS-50
chr2-40595618-INS-69
chr2-40709068-INS-50
chr2-40762510-INS-1862
chr2-40879818-INS-54
chr2-40894292-INS-57
chr2-40898897-INS-250
chr2-40904854-INS-3769
chr2-40941611-INS-491
chr2-40996193-INS-3433
chr2-40996259-INS-152
chr2-41039510-INS-1273
chr2-41083907-INS-4897
chr2-41129476-INS-7134
chr2-41177503-INS-2771
chr2-41178088-INS-58
chr2-41203479-INS-159
chr2-41227208-INS-76
chr2-41244329-INS-1378
chr2-41267378-INS-775
chr2-41393396-INS-122
chr2-41418333-INS-296
chr2-41418719-INS-660

chr2-41449125-INS-6155
chr2-41454912-INS-50
chr2-41458018-INS-54
chr2-41471181-INS-1161
chr2-41475444-INS-6972
chr2-41486832-INS-1319
chr2-41506447-INS-96
chr2-41532555-INS-466
chr2-41533734-INS-54
chr2-41561193-INS-50
chr2-41587307-INS-448
chr2-41591832-INS-63
chr2-41604524-INS-1411
chr2-41692521-INS-219
chr2-41701410-INS-3642
chr2-41710698-INS-113
chr2-41919381-INS-1515
chr2-41956321-INS-53
chr2-41981165-INS-65
chr2-42009826-INS-52
chr2-42010087-INS-80
chr2-42012362-INS-525
chr2-42035168-INS-74
chr2-42038200-INS-558
chr2-42043335-INS-115
chr2-42052160-INS-56
chr2-42150917-INS-79
chr2-42201520-INS-52
chr2-42249818-INS-1247
chr2-42254834-INS-51
chr2-42257993-INS-104
chr2-42310885-INS-6109
chr2-42336807-INS-461
chr2-42447777-INS-7354
chr2-42459758-INS-214
chr2-42502971-INS-3195
chr2-42589230-INS-13972
chr2-42603987-INS-1961
chr2-42613385-INS-54
chr2-42662577-INS-5401
chr2-42680414-INS-79
chr2-42700569-INS-52
chr2-42704922-INS-212
chr2-42717541-INS-5554
chr2-42789173-INS-7088
chr2-42792233-INS-316
chr2-42792969-INS-93
chr2-42831808-INS-119
chr2-42839733-INS-60
chr2-42864184-INS-488
chr2-42873155-INS-4822
chr2-42883424-INS-260
chr2-42888230-INS-73
chr2-42892230-INS-66
chr2-42909587-INS-52
chr2-42910741-INS-6158
chr2-42935130-INS-70
chr2-42992095-INS-1267
chr2-43031775-INS-405
chr2-43044117-INS-308
chr2-43094243-INS-4607
chr2-43115593-INS-1160
chr2-43146981-INS-123
chr2-43149357-INS-1243
chr2-43150977-INS-6621
chr2-43177739-INS-1089
chr2-43200069-INS-596
chr2-43203094-INS-1725
chr2-43354666-INS-69
chr2-43370845-INS-3164
chr2-43377824-INS-54
chr2-43410513-INS-3957
chr2-43456561-INS-56
chr2-43534920-INS-51
chr2-43573733-INS-86
chr2-43636353-INS-82
chr2-43636507-INS-142

chr2-43648119-INS-6464
chr2-43655235-INS-118
chr2-43655822-INS-50
chr2-43657646-INS-3788
chr2-43708248-INS-268
chr2-43740381-INS-60
chr2-43762347-INS-1311
chr2-43790584-INS-128
chr2-43969237-INS-50
chr2-44002981-INS-2330
chr2-44064576-INS-214
chr2-44126540-INS-207
chr2-44189257-INS-51
chr2-44199659-INS-62
chr2-44224459-INS-53
chr2-44226197-INS-470
chr2-44246911-INS-54
chr2-44248187-INS-268
chr2-44260408-INS-100
chr2-44291209-INS-202
chr2-44393701-INS-337
chr2-44398824-INS-125
chr2-44427015-INS-469
chr2-44494695-INS-68
chr2-44501663-INS-162
chr2-44562770-INS-7025
chr2-44641813-INS-56
chr2-44738276-INS-698
chr2-44758857-INS-151
chr2-44766324-INS-54
chr2-44798166-INS-155
chr2-44848579-INS-240
chr2-44872927-INS-300
chr2-44894342-INS-174
chr2-44937428-INS-58
chr2-45030152-INS-164
chr2-45055769-INS-326
chr2-45080784-INS-114
chr2-45081320-INS-387
chr2-45081345-INS-102
chr2-45097334-INS-52
chr2-45105501-INS-54
chr2-45159931-INS-52
chr2-45310750-INS-168
chr2-45315896-INS-68
chr2-45358220-INS-164
chr2-45431837-INS-6411
chr2-45506324-INS-193
chr2-45541936-INS-7071
chr2-45766242-INS-1365
chr2-45773079-INS-1009
chr2-45796595-INS-161
chr2-45803218-INS-61
chr2-45817071-INS-78
chr2-45817170-INS-50
chr2-45869413-INS-3230
chr2-45906236-INS-72
chr2-45956801-INS-102
chr2-46063615-INS-2496
chr2-46096715-INS-51
chr2-46096911-INS-132
chr2-46171832-INS-220
chr2-46256892-INS-1428
chr2-46298874-INS-96
chr2-46328378-INS-52
chr2-46346567-INS-6375
chr2-46350892-INS-4374
chr2-46351845-INS-471
chr2-46584903-INS-1013
chr2-46609098-INS-2681
chr2-46632493-INS-2968
chr2-46648903-INS-1397
chr2-46661545-INS-3936
chr2-46716297-INS-580
chr2-46721486-INS-166
chr2-46731327-INS-50
chr2-46770397-INS-1313

chr2-46786836-INS-93
chr2-46862257-INS-6421
chr2-46899126-INS-624
chr2-46928593-INS-54
chr2-46932414-INS-810
chr2-46950824-INS-2133
chr2-46970118-INS-108
chr2-47053006-INS-58
chr2-47081737-INS-50
chr2-47093914-INS-6431
chr2-47167716-INS-121
chr2-47168022-INS-130
chr2-47230828-INS-108
chr2-47231537-INS-326
chr2-47239631-INS-54
chr2-47377482-INS-63
chr2-47377694-INS-86
chr2-47401365-INS-6659
chr2-47500182-INS-277
chr2-47509520-INS-207
chr2-47549212-INS-64
chr2-47598623-INS-59
chr2-47754170-INS-51
chr2-47757648-INS-6380
chr2-47786650-INS-6350
chr2-47793363-INS-3149
chr2-47793448-INS-551
chr2-47811439-INS-66
chr2-47814903-INS-2835
chr2-47815474-INS-6579
chr2-47826920-INS-4739
chr2-47856552-INS-83
chr2-47925858-INS-6496
chr2-47996322-INS-353
chr2-48025952-INS-129
chr2-48168051-INS-1348
chr2-48173425-INS-60
chr2-48235270-INS-98
chr2-48255101-INS-6619
chr2-48261604-INS-360
chr2-48291902-INS-427
chr2-48294111-INS-1003
chr2-48326781-INS-57
chr2-48327214-INS-120
chr2-48351569-INS-54
chr2-48356640-INS-62
chr2-48402584-INS-65
chr2-48403425-INS-6331
chr2-48414269-INS-68
chr2-48417269-INS-59
chr2-48443935-INS-420
chr2-48501634-INS-468
chr2-48593674-INS-56
chr2-48654493-INS-68
chr2-48700655-INS-228
chr2-48740938-INS-144
chr2-48746169-INS-6919
chr2-48796399-INS-205
chr2-48796601-INS-768
chr2-48829561-INS-52
chr2-48880884-INS-173
chr2-48883040-INS-52
chr2-48899127-INS-876
chr2-48919385-INS-553
chr2-48928283-INS-76
chr2-48962059-INS-4843
chr2-48973637-INS-889
chr2-48986685-INS-1627
chr2-49015149-INS-210
chr2-49124836-INS-90
chr2-49167963-INS-52
chr2-49174592-INS-147
chr2-49251027-INS-65
chr2-49253523-INS-56
chr2-49288340-INS-251
chr2-49356268-INS-500
chr2-49357482-INS-1001

chr2-49358070-INS-140
chr2-49358547-INS-999
chr2-49361543-INS-103
chr2-49374941-INS-1302
chr2-49391890-INS-185
chr2-49470218-INS-7550
chr2-49495208-INS-6401
chr2-49524995-INS-76
chr2-49578357-INS-3116
chr2-49587535-INS-196
chr2-49608005-INS-50
chr2-49669236-INS-168
chr2-49669482-INS-286
chr2-49683769-INS-104
chr2-49684272-INS-53
chr2-49684524-INS-117
chr2-49686426-INS-88
chr2-49686522-INS-583
chr2-49722820-INS-51
chr2-49723012-INS-277
chr2-49858238-INS-72
chr2-49894891-INS-217
chr2-49964920-INS-54
chr2-50022652-INS-60
chr2-50054313-INS-1171
chr2-50062701-INS-67
chr2-50099252-INS-52
chr2-50109372-INS-139
chr2-50190218-INS-92
chr2-50208662-INS-88
chr2-50218169-INS-56
chr2-50253266-INS-96
chr2-50268544-INS-168
chr2-50277521-INS-50
chr2-50291792-INS-60
chr2-50293433-INS-1006
chr2-50298542-INS-2469
chr2-50347681-INS-50
chr2-50351151-INS-353
chr2-50352862-INS-468
chr2-50367695-INS-1459
chr2-50382166-INS-1840
chr2-50388928-INS-216
chr2-50389238-INS-2592
chr2-50426819-INS-78
chr2-50438967-INS-7215
chr2-50454167-INS-84
chr2-50490145-INS-364
chr2-50508318-INS-65
chr2-50544272-INS-2780
chr2-50599955-INS-2932
chr2-50630919-INS-559
chr2-50726058-INS-256
chr2-50778357-INS-4616
chr2-50778357-INS-9913
chr2-50783481-INS-114
chr2-50789936-INS-1261
chr2-50798604-INS-1903
chr2-50800000-INS-6421
chr2-50803477-INS-2239
chr2-50811687-INS-131
chr2-50893086-INS-1346
chr2-50896341-INS-60
chr2-50994037-INS-177
chr2-51042017-INS-203
chr2-51066150-INS-1698
chr2-51086593-INS-254
chr2-51163767-INS-1459
chr2-51163767-INS-6682
chr2-51188056-INS-118
chr2-51233205-INS-86
chr2-51253339-INS-52
chr2-51254373-INS-6229
chr2-51293629-INS-68
chr2-51314421-INS-394
chr2-51386080-INS-424
chr2-51438999-INS-6946

chr2-51442997-INS-279
chr2-51444488-INS-2664
chr2-51450418-INS-52
chr2-51460437-INS-6374
chr2-51468984-INS-427
chr2-51524695-INS-306
chr2-51529534-INS-232
chr2-51552339-INS-2839
chr2-51552909-INS-202
chr2-51618102-INS-3425
chr2-51661230-INS-375
chr2-51661460-INS-700
chr2-51661665-INS-1144
chr2-51679365-INS-782
chr2-51747718-INS-6326
chr2-51807096-INS-70
chr2-51836416-INS-108
chr2-51836538-INS-1338
chr2-51851851-INS-180
chr2-51872896-INS-5430
chr2-51919151-INS-102
chr2-51922087-INS-876
chr2-51957540-INS-1082
chr2-51957540-INS-3026
chr2-51975946-INS-842
chr2-52129724-INS-326
chr2-52191834-INS-156
chr2-52208886-INS-183
chr2-52250109-INS-72
chr2-52279949-INS-126
chr2-52328939-INS-675
chr2-52339725-INS-288
chr2-52345434-INS-249
chr2-52369159-INS-50
chr2-52392786-INS-84
chr2-52392831-INS-472
chr2-52394434-INS-480
chr2-52415247-INS-54
chr2-52415526-INS-169
chr2-52421664-INS-1000
chr2-52425516-INS-82
chr2-52425712-INS-73
chr2-52462870-INS-78
chr2-52463256-INS-152
chr2-52503529-INS-285
chr2-52515512-INS-2490
chr2-52519477-INS-90
chr2-52526699-INS-54
chr2-52586968-INS-3835
chr2-52608482-INS-50
chr2-52611041-INS-737
chr2-52612418-INS-52
chr2-52634614-INS-212
chr2-52650081-INS-60
chr2-52650561-INS-179
chr2-52661743-INS-886
chr2-52661785-INS-149
chr2-52666857-INS-844
chr2-52667013-INS-332
chr2-52692279-INS-56
chr2-52745267-INS-56
chr2-52745308-INS-116
chr2-52750025-INS-52
chr2-52762609-INS-205
chr2-52791094-INS-130
chr2-52847072-INS-54
chr2-52847176-INS-3615
chr2-52893835-INS-60
chr2-52916944-INS-167
chr2-52948149-INS-64
chr2-53005974-INS-130
chr2-53006060-INS-912
chr2-53014329-INS-150
chr2-53040329-INS-222
chr2-53056460-INS-82
chr2-53142256-INS-90
chr2-53154998-INS-3187

chr2-53157658-INS-152
chr2-53220736-INS-62
chr2-53230757-INS-977
chr2-53262223-INS-50
chr2-53290451-INS-1000
chr2-53299424-INS-145
chr2-53354590-INS-185
chr2-53363875-INS-1318
chr2-53384609-INS-59
chr2-53404900-INS-50
chr2-53420213-INS-6415
chr2-53449931-INS-64
chr2-53451026-INS-60
chr2-53471225-INS-220
chr2-53506945-INS-6681
chr2-53510281-INS-56
chr2-53547044-INS-128
chr2-53547044-INS-50
chr2-53559380-INS-54
chr2-53587031-INS-170
chr2-53588883-INS-4947
chr2-53704122-INS-287
chr2-53747042-INS-6149
chr2-53754012-INS-68
chr2-53763101-INS-57
chr2-53804128-INS-274
chr2-53806322-INS-2851
chr2-53855490-INS-6117
chr2-53910531-INS-168
chr2-53941158-INS-178
chr2-53972727-INS-61
chr2-53991749-INS-328
chr2-53997861-INS-1458
chr2-54001452-INS-768
chr2-54013796-INS-76
chr2-54046390-INS-6385
chr2-54085394-INS-6430
chr2-54091202-INS-176
chr2-54130818-INS-448
chr2-54133862-INS-1490
chr2-54209178-INS-146
chr2-54209238-INS-54
chr2-54237306-INS-361
chr2-54257371-INS-6938
chr2-54260974-INS-2664
chr2-54274133-INS-1069
chr2-54333614-INS-118
chr2-54351607-INS-7101
chr2-54355835-INS-80
chr2-54368896-INS-865
chr2-54393399-INS-50
chr2-54429888-INS-66
chr2-54561520-INS-3431
chr2-54565475-INS-1990
chr2-54618593-INS-165
chr2-54640184-INS-438
chr2-54644549-INS-516
chr2-54675647-INS-6263
chr2-54687263-INS-573
chr2-54698271-INS-67
chr2-54805610-INS-1189
chr2-54971584-INS-6756
chr2-54980673-INS-400
chr2-54981060-INS-158
chr2-55129760-INS-82
chr2-55204487-INS-201
chr2-55233855-INS-52
chr2-55242822-INS-1450
chr2-55261497-INS-50
chr2-55328980-INS-189
chr2-55352184-INS-2103
chr2-55357500-INS-6804
chr2-55457567-INS-66
chr2-55538503-INS-132
chr2-55551919-INS-675
chr2-55594693-INS-308
chr2-55596587-INS-10595

chr2-55602391-INS-470
chr2-55666852-INS-70
chr2-55858343-INS-117
chr2-55858683-INS-74
chr2-55859269-INS-163
chr2-55862366-INS-74
chr2-55912467-INS-6108
chr2-56014855-INS-4490
chr2-56031879-INS-79
chr2-56054699-INS-6868
chr2-56087096-INS-80
chr2-56106068-INS-1677
chr2-56112938-INS-500
chr2-56116277-INS-80
chr2-56137010-INS-104
chr2-56151315-INS-245
chr2-56181101-INS-50
chr2-56183211-INS-117
chr2-56266637-INS-6507
chr2-56283894-INS-4058
chr2-56297855-INS-56
chr2-56324572-INS-6591
chr2-56374255-INS-57
chr2-56419119-INS-228
chr2-56420145-INS-2166
chr2-56606440-INS-269
chr2-56615997-INS-64
chr2-56678553-INS-51
chr2-56679513-INS-217
chr2-56680013-INS-95
chr2-56715677-INS-57
chr2-56795428-INS-1703
chr2-56820048-INS-6914
chr2-56822999-INS-1987
chr2-56840230-INS-63
chr2-56868446-INS-509
chr2-56873983-INS-2671
chr2-56946692-INS-777
chr2-56960999-INS-6448
chr2-56962726-INS-2724
chr2-56980786-INS-56
chr2-56994697-INS-70
chr2-57043742-INS-50
chr2-57065247-INS-195
chr2-57104154-INS-265
chr2-57116265-INS-638
chr2-57136500-INS-54
chr2-57182153-INS-60
chr2-57220265-INS-724
chr2-57220298-INS-167
chr2-57261066-INS-56
chr2-57270403-INS-54
chr2-57279037-INS-60
chr2-57298586-INS-497
chr2-57302046-INS-665
chr2-57346751-INS-330
chr2-57346979-INS-136
chr2-57346979-INS-343
chr2-57347031-INS-59
chr2-57351445-INS-2687
chr2-57367115-INS-50
chr2-57368288-INS-50
chr2-57395212-INS-376
chr2-57397738-INS-1130
chr2-57461823-INS-75
chr2-57479005-INS-5673
chr2-57487476-INS-89
chr2-57545211-INS-54
chr2-57549221-INS-3128
chr2-57549944-INS-4818
chr2-57566392-INS-134
chr2-57573435-INS-5575
chr2-57602164-INS-50
chr2-57667535-INS-607
chr2-57778751-INS-107
chr2-57786723-INS-163
chr2-57786868-INS-50

chr2-57813068-INS-80
chr2-57814054-INS-54
chr2-57818586-INS-751
chr2-57831217-INS-50
chr2-57836079-INS-513
chr2-57837282-INS-54
chr2-57847193-INS-468
chr2-57969254-INS-82
chr2-58026777-INS-490
chr2-58041429-INS-56
chr2-58041624-INS-280
chr2-58042257-INS-1122
chr2-58062904-INS-6127
chr2-58145014-INS-58
chr2-58153798-INS-122
chr2-58203171-INS-60
chr2-58326674-INS-54
chr2-58328143-INS-56
chr2-58330224-INS-387
chr2-58330225-INS-165
chr2-58342649-INS-96
chr2-58342866-INS-90
chr2-58343060-INS-243
chr2-58357303-INS-61
chr2-58382876-INS-175
chr2-58384661-INS-104
chr2-58387558-INS-58
chr2-58418914-INS-80
chr2-58461286-INS-1173
chr2-58467534-INS-222
chr2-58516142-INS-227
chr2-58525381-INS-240
chr2-58525949-INS-60
chr2-58527935-INS-2690
chr2-58534671-INS-170
chr2-58572001-INS-60
chr2-58616661-INS-207
chr2-58618469-INS-51
chr2-58619063-INS-96
chr2-58651888-INS-1721
chr2-58671585-INS-160
chr2-58685940-INS-53
chr2-58708962-INS-548
chr2-58728399-INS-6432
chr2-58883321-INS-102
chr2-58883486-INS-57
chr2-58897382-INS-122
chr2-58939013-INS-190
chr2-58952849-INS-74
chr2-58984284-INS-132
chr2-59006836-INS-10415
chr2-59027344-INS-50
chr2-59083349-INS-711
chr2-59083350-INS-1432
chr2-59083497-INS-222
chr2-59189637-INS-261
chr2-59220417-INS-6180
chr2-59221292-INS-966
chr2-59224172-INS-6151
chr2-59224986-INS-937
chr2-59228808-INS-155
chr2-59295235-INS-93
chr2-59296618-INS-53
chr2-59299041-INS-140
chr2-59318133-INS-74
chr2-59326672-INS-85
chr2-59354852-INS-195
chr2-59360894-INS-245
chr2-59379380-INS-207
chr2-59380100-INS-210
chr2-59381324-INS-66
chr2-59390581-INS-50
chr2-59427043-INS-50
chr2-59438621-INS-631
chr2-59514914-INS-91
chr2-59534962-INS-82
chr2-59535905-INS-418

chr2-59545794-INS-60
chr2-59579494-INS-4505
chr2-59580665-INS-175
chr2-59580902-INS-63
chr2-59607159-INS-183
chr2-59616638-INS-192
chr2-59649583-INS-86
chr2-59761063-INS-6259
chr2-59801836-INS-954
chr2-59937088-INS-50
chr2-59939700-INS-236
chr2-59991483-INS-191
chr2-60016880-INS-64
chr2-60019501-INS-58
chr2-60020546-INS-304
chr2-60090304-INS-370
chr2-60100747-INS-300
chr2-60103136-INS-154
chr2-60128002-INS-235
chr2-60130363-INS-1453
chr2-60138736-INS-2318
chr2-60164571-INS-116
chr2-60193746-INS-119
chr2-60214895-INS-68
chr2-60236180-INS-885
chr2-60236181-INS-245
chr2-60239272-INS-216
chr2-60255151-INS-50
chr2-60259221-INS-71
chr2-60259787-INS-449
chr2-60353668-INS-7225
chr2-60411055-INS-182
chr2-60511378-INS-6494
chr2-60623365-INS-162
chr2-60625904-INS-51
chr2-60630540-INS-1999
chr2-60707823-INS-200
chr2-60727826-INS-61
chr2-60738986-INS-7011
chr2-60750285-INS-56
chr2-60761871-INS-1829
chr2-60803531-INS-52
chr2-60814543-INS-165
chr2-60821825-INS-110
chr2-60843345-INS-476
chr2-60881148-INS-712
chr2-60950212-INS-2344
chr2-60952379-INS-150
chr2-60952379-INS-440
chr2-60952673-INS-113
chr2-60968269-INS-7115
chr2-60968554-INS-182
chr2-60983045-INS-300
chr2-60985559-INS-94
chr2-61014557-INS-51
chr2-61042146-INS-56
chr2-61048587-INS-6485
chr2-61048818-INS-1896
chr2-61064188-INS-84
chr2-61064290-INS-184
chr2-61092953-INS-186
chr2-61110315-INS-7267
chr2-61122405-INS-52
chr2-61128564-INS-66
chr2-61358603-INS-58
chr2-61358642-INS-1707
chr2-61371701-INS-6745
chr2-61375921-INS-171
chr2-61424700-INS-136
chr2-61448663-INS-1343
chr2-61449539-INS-2795
chr2-61463511-INS-1215
chr2-61562712-INS-193
chr2-61571115-INS-828
chr2-61577938-INS-434
chr2-61583654-INS-483
chr2-61655559-INS-492

chr2-61668347-INS-2187
chr2-61721176-INS-6413
chr2-61779221-INS-151
chr2-61796238-INS-8751
chr2-61809485-INS-517
chr2-61857248-INS-160
chr2-61862450-INS-72
chr2-61898564-INS-78
chr2-61935387-INS-16458
chr2-61948633-INS-489
chr2-62113714-INS-50
chr2-62150417-INS-58
chr2-62182412-INS-64
chr2-62302854-INS-2416
chr2-62373924-INS-59
chr2-62374207-INS-211
chr2-62401163-INS-56
chr2-62433042-INS-322
chr2-62433082-INS-75
chr2-62517455-INS-3046
chr2-62521067-INS-72
chr2-62556947-INS-68
chr2-62557319-INS-260
chr2-62578018-INS-1315
chr2-62649071-INS-6482
chr2-62720415-INS-1489
chr2-62723304-INS-95
chr2-62730660-INS-655
chr2-62738258-INS-834
chr2-62767692-INS-100
chr2-62768382-INS-1153
chr2-62782711-INS-74
chr2-62800132-INS-6603
chr2-62843340-INS-472
chr2-62915811-INS-68
chr2-62921172-INS-50
chr2-62942142-INS-171
chr2-63115560-INS-288
chr2-63144113-INS-77
chr2-63146221-INS-52
chr2-63163161-INS-343
chr2-63206871-INS-2274
chr2-63255763-INS-1001
chr2-63260315-INS-1079
chr2-63380423-INS-113
chr2-63438582-INS-1632
chr2-63445377-INS-1679
chr2-63484441-INS-50
chr2-63579409-INS-152
chr2-63627134-INS-285
chr2-63642899-INS-63
chr2-63642906-INS-134
chr2-63647581-INS-80
chr2-63656686-INS-102
chr2-63667970-INS-92
chr2-63719922-INS-50
chr2-63767469-INS-431
chr2-63767469-INS-52
chr2-63796728-INS-121
chr2-63889335-INS-6248
chr2-63955529-INS-157
chr2-63975530-INS-832
chr2-64014965-INS-358
chr2-64059800-INS-2436
chr2-64063547-INS-2891
chr2-64095465-INS-570
chr2-64105909-INS-238
chr2-64117313-INS-1469
chr2-64145418-INS-3483
chr2-64243956-INS-96
chr2-64251967-INS-82
chr2-64300924-INS-475
chr2-64303694-INS-5229
chr2-64305911-INS-68
chr2-64338747-INS-100
chr2-64339633-INS-64
chr2-64358445-INS-50

chr2-64384074-INS-2479
chr2-64394162-INS-790
chr2-64394196-INS-348
chr2-64446941-INS-58
chr2-64524470-INS-60
chr2-64546989-INS-6390
chr2-64560736-INS-5974
chr2-64739045-INS-3066
chr2-64760021-INS-1742
chr2-64814884-INS-50
chr2-64860641-INS-201
chr2-64861154-INS-404
chr2-64863841-INS-6069
chr2-64867929-INS-178
chr2-64935833-INS-145
chr2-65010930-INS-52
chr2-65031549-INS-64
chr2-65082978-INS-62
chr2-65089201-INS-1844
chr2-65090378-INS-739
chr2-65110412-INS-70
chr2-65130031-INS-205
chr2-65132028-INS-162
chr2-65132028-INS-73
chr2-65141591-INS-6403
chr2-65146877-INS-4883
chr2-65179185-INS-409
chr2-65280109-INS-50
chr2-65282067-INS-7065
chr2-65285060-INS-246
chr2-65292883-INS-50
chr2-65304932-INS-66
chr2-65305568-INS-103
chr2-65311941-INS-399
chr2-65332410-INS-205
chr2-65523896-INS-58
chr2-65587247-INS-4906
chr2-65600133-INS-1412
chr2-65602016-INS-72
chr2-65615108-INS-91
chr2-65626206-INS-81
chr2-65634982-INS-56
chr2-65643519-INS-176
chr2-65646903-INS-3073
chr2-65658044-INS-6926
chr2-65671939-INS-93
chr2-65744819-INS-119
chr2-65843489-INS-993
chr2-65843566-INS-209
chr2-65845014-INS-50
chr2-65872287-INS-58
chr2-65881368-INS-70
chr2-65886494-INS-200
chr2-66011411-INS-209
chr2-66019618-INS-66
chr2-66128806-INS-100
chr2-66135775-INS-117
chr2-66176027-INS-196
chr2-66182715-INS-397
chr2-66192165-INS-58
chr2-66208447-INS-6409
chr2-66242682-INS-72
chr2-66247737-INS-107
chr2-66305945-INS-6437
chr2-66315602-INS-169
chr2-66334832-INS-590
chr2-66335099-INS-804
chr2-66337617-INS-59
chr2-66377071-INS-256
chr2-66380312-INS-50
chr2-66380489-INS-412
chr2-66420933-INS-401
chr2-66422558-INS-208
chr2-66454525-INS-50
chr2-66479665-INS-62
chr2-66584400-INS-1451
chr2-66592849-INS-77

chr2-66629588-INS-515
chr2-66650311-INS-50
chr2-66654291-INS-6465
chr2-66657543-INS-6772
chr2-66662312-INS-356
chr2-66667035-INS-66
chr2-66715960-INS-111
chr2-66716465-INS-326
chr2-66775524-INS-6442
chr2-66809469-INS-56
chr2-66818066-INS-111
chr2-66821166-INS-193
chr2-66864092-INS-52
chr2-66901030-INS-1312
chr2-66909928-INS-156
chr2-66933118-INS-616
chr2-66933830-INS-84
chr2-66943672-INS-186
chr2-66945664-INS-1893
chr2-66975283-INS-6669
chr2-66983568-INS-1616
chr2-66984440-INS-88
chr2-66998382-INS-7098
chr2-67011832-INS-1014
chr2-67033097-INS-235
chr2-67056338-INS-599
chr2-67078995-INS-8781
chr2-67136369-INS-78
chr2-67143357-INS-58
chr2-67146255-INS-1753
chr2-67166507-INS-387
chr2-67186585-INS-1392
chr2-67197443-INS-2837
chr2-67218421-INS-7098
chr2-67267264-INS-3446
chr2-67278226-INS-112
chr2-67281080-INS-9128
chr2-67310772-INS-196
chr2-67399457-INS-364
chr2-67418996-INS-284
chr2-67419772-INS-137
chr2-67430073-INS-58
chr2-67439899-INS-60
chr2-67447537-INS-6543
chr2-67462493-INS-52
chr2-67510579-INS-220
chr2-67518861-INS-72
chr2-67526850-INS-84
chr2-67550425-INS-6914
chr2-67555212-INS-205
chr2-67572693-INS-322
chr2-67594911-INS-310
chr2-67605826-INS-9396
chr2-67693114-INS-6400
chr2-67705110-INS-6586
chr2-67717776-INS-194
chr2-67718427-INS-56
chr2-67738944-INS-825
chr2-67825347-INS-70
chr2-67844987-INS-1562
chr2-67847603-INS-245
chr2-67863456-INS-5772
chr2-67930656-INS-6419
chr2-67963741-INS-7190
chr2-67967714-INS-176
chr2-67969711-INS-74
chr2-67989722-INS-6258
chr2-68008969-INS-7467
chr2-68011151-INS-11205
chr2-68017553-INS-52
chr2-68032375-INS-399
chr2-68036989-INS-1938
chr2-68055713-INS-70
chr2-68099023-INS-52
chr2-68099852-INS-50
chr2-68204051-INS-1030
chr2-68213955-INS-196

chr2-68279404-INS-260
chr2-68280368-INS-90
chr2-68290270-INS-61
chr2-68298375-INS-214
chr2-68437585-INS-6392
chr2-68463196-INS-6589
chr2-68463312-INS-65
chr2-68467066-INS-56
chr2-68525509-INS-560
chr2-68553705-INS-4081
chr2-68555857-INS-924
chr2-68577119-INS-3947
chr2-68711854-INS-89
chr2-68718335-INS-99
chr2-68718368-INS-264
chr2-68740804-INS-72
chr2-68783115-INS-446
chr2-68787168-INS-71
chr2-68838441-INS-56
chr2-68862602-INS-3215
chr2-68932065-INS-50
chr2-68957165-INS-7158
chr2-68988594-INS-768
chr2-68991340-INS-62
chr2-69004383-INS-68
chr2-69031809-INS-252
chr2-69081522-INS-4222
chr2-69096912-INS-54
chr2-69146372-INS-6196
chr2-69155051-INS-113
chr2-69202550-INS-6482
chr2-69204533-INS-64
chr2-69207158-INS-50
chr2-69221342-INS-139
chr2-69250323-INS-78
chr2-69272935-INS-55
chr2-69301324-INS-6390
chr2-69301864-INS-80
chr2-69320194-INS-222
chr2-69327389-INS-50
chr2-69381129-INS-282
chr2-69392297-INS-62
chr2-69392776-INS-80
chr2-69404583-INS-50
chr2-69442387-INS-54
chr2-69449127-INS-73
chr2-69456955-INS-108
chr2-69457684-INS-69
chr2-69472040-INS-51
chr2-69486900-INS-108
chr2-69554036-INS-2072
chr2-69555976-INS-4880
chr2-69661998-INS-240
chr2-69668720-INS-61
chr2-69669780-INS-105
chr2-69716889-INS-7109
chr2-69735535-INS-211
chr2-69740118-INS-1875
chr2-69743585-INS-51
chr2-69746093-INS-233
chr2-69747308-INS-197
chr2-69748228-INS-210
chr2-69847108-INS-236
chr2-69865649-INS-1178
chr2-69866592-INS-84
chr2-69878744-INS-532
chr2-69924401-INS-73
chr2-69941909-INS-152
chr2-69942646-INS-344
chr2-69954968-INS-98
chr2-69963089-INS-210
chr2-69985163-INS-173
chr2-69989633-INS-55
chr2-69995335-INS-60
chr2-69996014-INS-76
chr2-69996995-INS-827
chr2-70014439-INS-79

chr2-70024745-INS-84
chr2-70066651-INS-62
chr2-70106919-INS-132
chr2-70108592-INS-529
chr2-70126989-INS-6197
chr2-70130554-INS-57
chr2-70181300-INS-131
chr2-70208871-INS-104
chr2-70209655-INS-55
chr2-70211483-INS-50
chr2-70240809-INS-52
chr2-70262320-INS-430
chr2-70263016-INS-59
chr2-70268002-INS-8688
chr2-70290655-INS-80
chr2-70295621-INS-364
chr2-70298606-INS-246
chr2-70323957-INS-51
chr2-70363644-INS-51
chr2-70367580-INS-67
chr2-70368638-INS-110
chr2-70368780-INS-62
chr2-70372380-INS-167
chr2-70419096-INS-74
chr2-70419299-INS-309
chr2-70503192-INS-221
chr2-70522540-INS-184
chr2-70560351-INS-6475
chr2-70584295-INS-265
chr2-70661635-INS-50
chr2-70668686-INS-66
chr2-70689618-INS-119
chr2-70705291-INS-52
chr2-70716051-INS-66
chr2-70718397-INS-51
chr2-70732108-INS-461
chr2-70761658-INS-54
chr2-70778913-INS-55
chr2-70803798-INS-50
chr2-70811281-INS-106
chr2-70840106-INS-72
chr2-70840595-INS-129
chr2-70856358-INS-9936
chr2-70928594-INS-7302
chr2-70943902-INS-60
chr2-71039044-INS-291
chr2-71039116-INS-136
chr2-71100802-INS-3075
chr2-71118686-INS-672
chr2-71136781-INS-148
chr2-71147504-INS-69
chr2-71147762-INS-2957
chr2-71180983-INS-79
chr2-71181031-INS-53
chr2-71181041-INS-149
chr2-71190343-INS-50
chr2-71193095-INS-50
chr2-71206168-INS-56
chr2-71206182-INS-203
chr2-71281514-INS-2470
chr2-71281514-INS-6536
chr2-71299699-INS-996
chr2-71299993-INS-406
chr2-71309175-INS-94
chr2-71309832-INS-181
chr2-71311785-INS-71
chr2-71322422-INS-83
chr2-71406512-INS-137
chr2-71420659-INS-120
chr2-71427082-INS-6386
chr2-71434954-INS-109
chr2-71491616-INS-187
chr2-71491818-INS-372
chr2-71496392-INS-12670
chr2-71511158-INS-50
chr2-71518485-INS-221
chr2-71539067-INS-198

chr2-71547893-INS-152
chr2-71789118-INS-82
chr2-71837341-INS-56
chr2-71857264-INS-66
chr2-71867584-INS-1000
chr2-71916846-INS-217
chr2-71926656-INS-90
chr2-71940181-INS-224
chr2-72057339-INS-69
chr2-72057971-INS-61
chr2-72121609-INS-434
chr2-72226166-INS-211
chr2-72244521-INS-397
chr2-72244892-INS-524
chr2-72525227-INS-50
chr2-72532793-INS-285
chr2-72541169-INS-143
chr2-72565192-INS-80
chr2-72570497-INS-510
chr2-72604120-INS-900
chr2-72665165-INS-242
chr2-72673162-INS-232
chr2-72676042-INS-222
chr2-72781341-INS-75
chr2-72797714-INS-398
chr2-72862862-INS-7076
chr2-72869364-INS-56
chr2-72880930-INS-6156
chr2-72886104-INS-2284
chr2-72899473-INS-1911
chr2-73194418-INS-66
chr2-73309920-INS-293
chr2-73372454-INS-1021
chr2-73383231-INS-137
chr2-73513292-INS-469
chr2-73528155-INS-6380
chr2-73539444-INS-70
chr2-73587828-INS-75
chr2-73596190-INS-79
chr2-73596220-INS-230
chr2-73767631-INS-1870
chr2-73776106-INS-83
chr2-73792258-INS-2019
chr2-73794102-INS-76
chr2-73828537-INS-217
chr2-73842971-INS-420
chr2-73843853-INS-67
chr2-73888186-INS-120
chr2-73900621-INS-52
chr2-74027572-INS-144
chr2-74062436-INS-50
chr2-74110504-INS-52
chr2-74126948-INS-112
chr2-74157976-INS-93
chr2-74182180-INS-116
chr2-74292936-INS-310
chr2-74293455-INS-498
chr2-74295146-INS-170
chr2-74375500-INS-959
chr2-74430437-INS-6378
chr2-74441496-INS-3489
chr2-74450521-INS-54
chr2-74451791-INS-129
chr2-74452313-INS-276
chr2-74478209-INS-52
chr2-74497327-INS-70
chr2-74625443-INS-198
chr2-74682239-INS-5086
chr2-74742788-INS-165
chr2-74786412-INS-116
chr2-74789566-INS-164
chr2-74802609-INS-64
chr2-74814345-INS-65
chr2-74827465-INS-115
chr2-74829606-INS-231
chr2-74876058-INS-6586
chr2-74894739-INS-66

chr2-74995383-INS-268
chr2-75137103-INS-50
chr2-75231237-INS-504
chr2-75299778-INS-732
chr2-75309057-INS-5975
chr2-75332189-INS-929
chr2-75332297-INS-945
chr2-75374514-INS-232
chr2-75443924-INS-744
chr2-75453152-INS-360
chr2-75453157-INS-1748
chr2-75457009-INS-209
chr2-75457697-INS-75
chr2-75465088-INS-740
chr2-75573184-INS-219
chr2-75573995-INS-66
chr2-75578406-INS-3440
chr2-75578406-INS-52
chr2-75604674-INS-66
chr2-75604861-INS-116
chr2-75610271-INS-87
chr2-75610514-INS-276
chr2-75617819-INS-87
chr2-75619418-INS-180
chr2-75771552-INS-5721
chr2-75798838-INS-60
chr2-75802034-INS-268
chr2-75805479-INS-327
chr2-75805684-INS-94
chr2-75857918-INS-168
chr2-75921417-INS-297
chr2-75958038-INS-306
chr2-75975401-INS-200
chr2-75985975-INS-106
chr2-75990112-INS-54
chr2-76011640-INS-126
chr2-76012797-INS-320
chr2-76015051-INS-5460
chr2-76021348-INS-1923
chr2-76038957-INS-1736
chr2-76038957-INS-3476
chr2-76041056-INS-50
chr2-76071687-INS-6085
chr2-76077899-INS-517
chr2-76078704-INS-54
chr2-76135252-INS-52
chr2-76135863-INS-1947
chr2-76141586-INS-688
chr2-76149341-INS-66
chr2-76184934-INS-70
chr2-76186927-INS-215
chr2-76230533-INS-51
chr2-76233837-INS-148
chr2-76256516-INS-209
chr2-76299293-INS-147
chr2-76342787-INS-72
chr2-76348775-INS-98
chr2-76349889-INS-104
chr2-76349917-INS-780
chr2-76350422-INS-56
chr2-76352815-INS-88
chr2-76487467-INS-162
chr2-76521152-INS-69
chr2-76901157-INS-134
chr2-76922810-INS-55
chr2-76942227-INS-114
chr2-76946439-INS-117
chr2-76970818-INS-228
chr2-77001375-INS-504
chr2-77123967-INS-52
chr2-77125688-INS-150
chr2-77141130-INS-211
chr2-77141371-INS-140
chr2-77149492-INS-1254
chr2-77151143-INS-56
chr2-77166898-INS-204
chr2-77169067-INS-75

chr2-77196060-INS-54
chr2-77203722-INS-124
chr2-77203838-INS-223
chr2-77222465-INS-54
chr2-77272113-INS-100
chr2-77336297-INS-93
chr2-77355255-INS-108
chr2-77355611-INS-50
chr2-77429269-INS-123
chr2-77479420-INS-198
chr2-77480186-INS-74
chr2-77501422-INS-62
chr2-77507199-INS-58
chr2-77530966-INS-50
chr2-77558868-INS-2229
chr2-77567112-INS-4346
chr2-77581937-INS-66
chr2-77582269-INS-345
chr2-77653284-INS-3154
chr2-77678331-INS-6164
chr2-77701331-INS-4859
chr2-77710352-INS-3127
chr2-77710755-INS-50
chr2-77734800-INS-396
chr2-77766658-INS-183
chr2-77771050-INS-363
chr2-77777352-INS-65
chr2-77777839-INS-52
chr2-77802151-INS-310
chr2-77802705-INS-56
chr2-77869084-INS-77
chr2-77885380-INS-52
chr2-77933840-INS-27313
chr2-77963152-INS-80
chr2-78016716-INS-69
chr2-78092943-INS-54
chr2-78099466-INS-50
chr2-78100607-INS-3781
chr2-78110047-INS-50
chr2-78192305-INS-108
chr2-78233822-INS-4965
chr2-78242140-INS-6422
chr2-78256262-INS-710
chr2-78273964-INS-6679
chr2-78324513-INS-446
chr2-78349296-INS-3006
chr2-78362284-INS-6969
chr2-78419942-INS-136
chr2-78425906-INS-220
chr2-78463437-INS-56
chr2-78516580-INS-1235
chr2-78517765-INS-92
chr2-78520360-INS-90
chr2-78520992-INS-50
chr2-78538066-INS-388
chr2-78538604-INS-51
chr2-78542230-INS-10709
chr2-78628753-INS-6130
chr2-78643410-INS-85
chr2-78692304-INS-179
chr2-78712224-INS-141
chr2-78712376-INS-138
chr2-78770481-INS-50
chr2-78787644-INS-561
chr2-78789715-INS-6505
chr2-78805729-INS-68
chr2-78839705-INS-114
chr2-78839936-INS-419
chr2-78852519-INS-105
chr2-78939620-INS-50
chr2-78950672-INS-125
chr2-78973707-INS-480
chr2-78984163-INS-76
chr2-78985740-INS-253
chr2-78987357-INS-190
chr2-78987357-INS-800
chr2-78987410-INS-70

chr2-79000826-INS-2093
chr2-79000826-INS-756
chr2-79062465-INS-80
chr2-79062937-INS-56
chr2-79082200-INS-96
chr2-79134670-INS-5061
chr2-79201105-INS-61
chr2-79266707-INS-163
chr2-79273469-INS-211
chr2-79334622-INS-70
chr2-79436113-INS-58
chr2-79451326-INS-124
chr2-79496777-INS-50
chr2-79525580-INS-13220
chr2-79526606-INS-52
chr2-79557570-INS-158
chr2-79557799-INS-78
chr2-79558191-INS-93
chr2-79569643-INS-516
chr2-79569831-INS-88
chr2-79583179-INS-54
chr2-79589331-INS-88
chr2-79613428-INS-89
chr2-79619861-INS-208
chr2-79623640-INS-2943
chr2-79627827-INS-63
chr2-79730976-INS-277
chr2-79730993-INS-95
chr2-79747383-INS-82
chr2-79814623-INS-51
chr2-79848595-INS-172
chr2-79848625-INS-901
chr2-79915822-INS-6413
chr2-79929759-INS-8432
chr2-79986210-INS-329
chr2-79987406-INS-97
chr2-80029462-INS-2953
chr2-80094553-INS-526
chr2-80094613-INS-50
chr2-80105105-INS-6449
chr2-80131916-INS-7228
chr2-80168919-INS-260
chr2-80169687-INS-148
chr2-80213823-INS-1276
chr2-80227503-INS-213
chr2-80241414-INS-51
chr2-80359052-INS-219
chr2-80369222-INS-60
chr2-80369228-INS-123
chr2-80388631-INS-255
chr2-80405863-INS-222
chr2-80408962-INS-194
chr2-80419820-INS-726
chr2-80435438-INS-2001
chr2-80458377-INS-157
chr2-80482075-INS-2330
chr2-80519451-INS-3251
chr2-80553188-INS-7043
chr2-80558228-INS-276
chr2-80587748-INS-6282
chr2-80591986-INS-97
chr2-80644962-INS-624
chr2-80688560-INS-7349
chr2-80706510-INS-301
chr2-80707190-INS-57
chr2-80767518-INS-928
chr2-80767530-INS-351
chr2-80824044-INS-165
chr2-80850107-INS-6356
chr2-80898838-INS-103
chr2-80918958-INS-7105
chr2-80932678-INS-124
chr2-80965099-INS-2190
chr2-80985352-INS-1217
chr2-81090569-INS-64
chr2-81120687-INS-262
chr2-81144875-INS-134

chr2-81196821-INS-1820
chr2-81231159-INS-179
chr2-81325486-INS-93
chr2-81328244-INS-1115
chr2-81458681-INS-51
chr2-81466823-INS-517
chr2-81467947-INS-52
chr2-81524452-INS-52
chr2-81526312-INS-84
chr2-81531592-INS-390
chr2-81542210-INS-50
chr2-81559069-INS-6477
chr2-81578705-INS-5098
chr2-81601303-INS-50
chr2-81601522-INS-248
chr2-81627066-INS-1933
chr2-81693290-INS-616
chr2-81693570-INS-131
chr2-81758543-INS-6206
chr2-81775486-INS-4349
chr2-81805790-INS-50
chr2-81891933-INS-941
chr2-81898249-INS-124
chr2-81903716-INS-422
chr2-81927531-INS-84
chr2-81949878-INS-64
chr2-81950647-INS-51
chr2-82010566-INS-2243
chr2-82065287-INS-667
chr2-82098628-INS-1780
chr2-82134297-INS-55
chr2-82177223-INS-770
chr2-82212998-INS-342
chr2-82304170-INS-200
chr2-82310157-INS-3534
chr2-82358208-INS-96
chr2-82397652-INS-216
chr2-82425307-INS-6513
chr2-82427693-INS-50
chr2-82430744-INS-67
chr2-82437740-INS-288
chr2-82449673-INS-847
chr2-82462008-INS-83
chr2-82561777-INS-73
chr2-82562147-INS-51
chr2-82666923-INS-57
chr2-82667057-INS-187
chr2-82676939-INS-204
chr2-82733051-INS-1455
chr2-82842091-INS-492
chr2-82851571-INS-97
chr2-82925043-INS-2003
chr2-82925832-INS-121
chr2-82943619-INS-523
chr2-83017957-INS-525
chr2-83062183-INS-787
chr2-83063118-INS-171
chr2-83063221-INS-169
chr2-83069144-INS-50
chr2-83090840-INS-112
chr2-83107956-INS-155
chr2-83110238-INS-198
chr2-83156139-INS-368
chr2-83225793-INS-205
chr2-83271920-INS-194
chr2-83271944-INS-84
chr2-83272231-INS-132
chr2-83272237-INS-57
chr2-83311859-INS-50
chr2-83369188-INS-631
chr2-83426844-INS-66
chr2-83454839-INS-116
chr2-83493604-INS-372
chr2-83563211-INS-214
chr2-83570148-INS-120
chr2-83592786-INS-50
chr2-83619447-INS-509

chr2-83675679-INS-50
chr2-83743153-INS-1724
chr2-83752066-INS-62
chr2-83753496-INS-53
chr2-83753818-INS-129
chr2-83862485-INS-100
chr2-83935901-INS-461
chr2-83958745-INS-1491
chr2-83991904-INS-66
chr2-84057793-INS-192
chr2-84100131-INS-667
chr2-84117972-INS-411
chr2-84127196-INS-50
chr2-84128111-INS-70
chr2-84165432-INS-8841
chr2-84203432-INS-120
chr2-84233841-INS-129
chr2-84285586-INS-77
chr2-84285603-INS-184
chr2-84317570-INS-6401
chr2-84362422-INS-53
chr2-84362430-INS-184
chr2-84367911-INS-76
chr2-84387315-INS-241
chr2-84394098-INS-202
chr2-84460245-INS-189
chr2-84476897-INS-271
chr2-84488976-INS-188
chr2-84510183-INS-205
chr2-84523641-INS-61
chr2-84526794-INS-220
chr2-84529169-INS-206
chr2-84548136-INS-54
chr2-84637852-INS-74
chr2-84658402-INS-157
chr2-84678843-INS-1275
chr2-84694905-INS-56
chr2-84695928-INS-152
chr2-84745335-INS-64
chr2-84769562-INS-706
chr2-84769993-INS-55
chr2-84783470-INS-209
chr2-84792946-INS-123
chr2-84801044-INS-190
chr2-84811754-INS-54
chr2-84838481-INS-264
chr2-84916025-INS-398
chr2-84939956-INS-2000
chr2-85000896-INS-81
chr2-85001763-INS-1403
chr2-85001765-INS-106
chr2-85016537-INS-220
chr2-85030245-INS-50
chr2-85030255-INS-302
chr2-85032661-INS-72
chr2-85048589-INS-50
chr2-85052365-INS-102
chr2-85069133-INS-213
chr2-85074314-INS-1766
chr2-85090070-INS-112
chr2-85100954-INS-304
chr2-85130675-INS-206
chr2-85133880-INS-897
chr2-85242368-INS-161
chr2-85256011-INS-525
chr2-85261866-INS-84
chr2-85387196-INS-1275
chr2-85394554-INS-428
chr2-85506267-INS-58
chr2-85646240-INS-51
chr2-85669367-INS-6339
chr2-85686997-INS-6900
chr2-85747632-INS-162
chr2-85843295-INS-237
chr2-85843465-INS-58
chr2-85921635-INS-730
chr2-85925124-INS-232

chr2-85927715-INS-61
chr2-85960173-INS-184
chr2-85985197-INS-108
chr2-86030760-INS-4759
chr2-86039960-INS-986
chr2-86055566-INS-289
chr2-86063053-INS-311
chr2-86092874-INS-499
chr2-86111152-INS-57
chr2-86112602-INS-55
chr2-86115565-INS-56
chr2-86137133-INS-669
chr2-86220085-INS-5982
chr2-86249493-INS-203
chr2-86324065-INS-84
chr2-86330687-INS-1037
chr2-86357809-INS-1510
chr2-86374112-INS-193
chr2-86378728-INS-853
chr2-86406186-INS-70
chr2-86481059-INS-1500
chr2-86552816-INS-2976
chr2-86690912-INS-68
chr2-86709051-INS-498
chr2-86752205-INS-7644
chr2-86842063-INS-10584
chr2-86859654-INS-142
chr2-86860283-INS-139
chr2-86863285-INS-125
chr2-86892850-INS-75
chr2-86940357-INS-1483
chr2-86945552-INS-1892
chr2-86970042-INS-6916
chr2-86985572-INS-131
chr2-86994293-INS-255
chr2-87017511-INS-3702
chr2-87077225-INS-465
chr2-87090507-INS-967
chr2-87093890-INS-1318
chr2-87156953-INS-2971
chr2-87182278-INS-138
chr2-87183337-INS-60
chr2-87188882-INS-92
chr2-87195011-INS-408
chr2-87199607-INS-2433
chr2-87229114-INS-109
chr2-87241654-INS-71
chr2-87255610-INS-58
chr2-87256536-INS-198
chr2-87259223-INS-192
chr2-87266702-INS-210
chr2-87272903-INS-5225
chr2-87277672-INS-243
chr2-87347313-INS-798
chr2-87360185-INS-209
chr2-87363989-INS-185
chr2-87364133-INS-495
chr2-87370509-INS-198
chr2-87373607-INS-567
chr2-87383961-INS-573
chr2-87390258-INS-2225
chr2-87401451-INS-206
chr2-87450184-INS-6335
chr2-87537167-INS-201
chr2-87614117-INS-70
chr2-87626286-INS-612
chr2-87651683-INS-6424
chr2-87672763-INS-1616
chr2-87722558-INS-203
chr2-87729782-INS-294
chr2-87802244-INS-6725
chr2-87821272-INS-1478
chr2-87825176-INS-6998
chr2-87831683-INS-1842
chr2-87831914-INS-140
chr2-87868945-INS-6698
chr2-87884809-INS-84

chr2-87898837-INS-489
chr2-87927909-INS-51
chr2-87935592-INS-1500
chr2-87973550-INS-498
chr2-88012820-INS-169
chr2-88051667-INS-2932
chr2-88094507-INS-15848
chr2-88100206-INS-1501
chr2-88230453-INS-86
chr2-88231389-INS-111
chr2-88252620-INS-50
chr2-88283804-INS-198
chr2-88326748-INS-604
chr2-88349082-INS-207
chr2-88351922-INS-88
chr2-88432969-INS-65
chr2-88444109-INS-53
chr2-88465092-INS-68
chr2-88484079-INS-935
chr2-88487026-INS-505
chr2-88493363-INS-64
chr2-88505155-INS-56
chr2-88508702-INS-2977
chr2-88594193-INS-1705
chr2-88621551-INS-1499
chr2-88691152-INS-91
chr2-88774399-INS-64
chr2-88830284-INS-52
chr2-88856248-INS-210
chr2-88869411-INS-620
chr2-89008140-INS-72
chr2-89014868-INS-204
chr2-89015515-INS-86
chr2-89033508-INS-191
chr2-89033793-INS-243
chr2-89035890-INS-98
chr2-89075473-INS-52
chr2-89093252-INS-61
chr2-89094712-INS-85
chr2-89143051-INS-866
chr2-89185603-INS-195
chr2-89189797-INS-60
chr2-89215225-INS-165
chr2-89217967-INS-662
chr2-89220002-INS-61
chr2-89338729-INS-695
chr2-89338760-INS-2195
chr2-89338834-INS-297
chr2-89339037-INS-1069
chr2-89412813-INS-777
chr2-89446400-INS-6612
chr2-89460331-INS-4265
chr2-89468696-INS-59
chr2-89590849-INS-2361
chr2-89604349-INS-1130
chr2-89698247-INS-218
chr2-89785487-INS-50
chr2-89818341-INS-6234
chr2-89822014-INS-60
chr2-89838173-INS-6438
chr2-89914692-INS-51
chr2-90232233-INS-499
chr2-90236100-INS-79
chr2-90240109-INS-500
chr2-90280873-INS-50
chr2-90287337-INS-820
chr2-90334236-INS-1543
chr2-90347583-INS-213
chr2-90393422-INS-61
chr2-90420800-INS-1183
chr2-90438321-INS-7368
chr2-90444074-INS-179
chr2-90455126-INS-51
chr2-90456133-INS-51
chr2-90505178-INS-80
chr2-90660428-INS-79
chr2-90742913-INS-212

chr2-90750548-INS-178
chr2-90813899-INS-588
chr2-90832020-INS-1798
chr2-90864896-INS-60
chr2-90864900-INS-255
chr2-90872945-INS-106
chr2-90880026-INS-52
chr2-90923814-INS-109
chr2-90925838-INS-104
chr2-90936363-INS-56
chr2-90947179-INS-128
chr2-90956481-INS-55
chr2-91026215-INS-54
chr2-91050519-INS-54
chr2-91069524-INS-413
chr2-91123658-INS-369
chr2-91125630-INS-52
chr2-91140856-INS-517
chr2-91188094-INS-52
chr2-91339460-INS-187
chr2-91371892-INS-210
chr2-91372716-INS-213
chr2-91413724-INS-209
chr2-91414621-INS-177
chr2-91520223-INS-55
chr2-91554740-INS-205
chr2-91577500-INS-63
chr2-91590551-INS-360
chr2-91781929-INS-904
chr2-91781938-INS-70
chr2-91786868-INS-225
chr2-91811052-INS-97
chr2-91823242-INS-92
chr2-91855253-INS-50
chr2-91875094-INS-198
chr2-91893400-INS-90
chr2-91910195-INS-376
chr2-91914648-INS-6974
chr2-91934007-INS-369
chr2-91934802-INS-110
chr2-91939403-INS-50
chr2-91966362-INS-62
chr2-91980328-INS-56
chr2-92034517-INS-228
chr2-92049331-INS-61
chr2-92165084-INS-658
chr2-92165088-INS-112
chr2-92177090-INS-50
chr2-92183038-INS-120
chr2-92203720-INS-51
chr2-92241484-INS-50
chr2-92304915-INS-220
chr2-92304915-INS-50
chr2-92309657-INS-428
chr2-92310059-INS-591
chr2-92354188-INS-334
chr2-92379621-INS-51
chr2-92430177-INS-50
chr2-92458839-INS-100
chr2-92479501-INS-88
chr2-92487461-INS-189
chr2-92491282-INS-138
chr2-92491316-INS-396
chr2-92498982-INS-1625
chr2-92506753-INS-54
chr2-92536273-INS-51
chr2-92536891-INS-70
chr2-92580393-INS-761
chr2-92587407-INS-296
chr2-92621953-INS-688
chr2-92671364-INS-73
chr2-92684536-INS-140
chr2-92700158-INS-112
chr2-92700158-INS-54
chr2-92708238-INS-253
chr2-92714437-INS-155
chr2-92753567-INS-102

chr2-92790607-INS-63
chr2-92840296-INS-51
chr2-92864339-INS-62
chr2-92887382-INS-76
chr2-92912253-INS-66
chr2-92916549-INS-85
chr2-92940761-INS-95
chr2-92947212-INS-50
chr2-92951366-INS-132
chr2-92959906-INS-78
chr2-92970334-INS-223
chr2-93049765-INS-206
chr2-93136867-INS-268
chr2-93179491-INS-78
chr2-93183042-INS-209
chr2-93212070-INS-1315
chr2-93408987-INS-50
chr2-93411251-INS-208
chr2-93469493-INS-52
chr2-93489993-INS-51
chr2-93496869-INS-94
chr2-93515364-INS-81
chr2-93543484-INS-54
chr2-93572960-INS-260
chr2-93603780-INS-300
chr2-93604473-INS-192
chr2-93606630-INS-110
chr2-93634164-INS-57
chr2-93645176-INS-279
chr2-93675802-INS-50
chr2-93682503-INS-58
chr2-93682505-INS-201
chr2-93690623-INS-269
chr2-93700641-INS-134
chr2-93707697-INS-172
chr2-93708244-INS-4847
chr2-93710324-INS-291
chr2-93710692-INS-728
chr2-93801829-INS-180
chr2-93824754-INS-5136
chr2-93842556-INS-77
chr2-93862025-INS-6522
chr2-93947509-INS-51
chr2-93960631-INS-204
chr2-93972046-INS-355
chr2-93975855-INS-166
chr2-93989955-INS-539
chr2-94003980-INS-54
chr2-94012513-INS-50
chr2-94122617-INS-191
chr2-94141448-INS-320
chr2-94161934-INS-91
chr2-94175756-INS-292
chr2-94175760-INS-71
chr2-94179384-INS-207
chr2-94179456-INS-441
chr2-94220359-INS-594
chr2-94220384-INS-92
chr2-94221292-INS-264
chr2-94295665-INS-57
chr2-94299448-INS-6175
chr2-94306988-INS-1066
chr2-94321598-INS-56
chr2-94323446-INS-220
chr2-94336516-INS-149
chr2-94340917-INS-7426
chr2-94386988-INS-92
chr2-94392281-INS-3724
chr2-94413305-INS-5033
chr2-94493051-INS-66
chr2-94518473-INS-118
chr2-94544808-INS-7938
chr2-94569611-INS-1691
chr2-94573715-INS-1872
chr2-94785317-INS-7000
chr2-94813285-INS-8151
chr2-94814995-INS-50

chr2-94860591-INS-2374
chr2-94922625-INS-4958
chr2-94956982-INS-343
chr2-95018181-INS-865
chr2-95037639-INS-4613
chr2-95210591-INS-56
chr2-95215944-INS-7058
chr2-95232895-INS-66
chr2-95244612-INS-212
chr2-95295113-INS-140
chr2-95332171-INS-210
chr2-95335981-INS-4745
chr2-95414322-INS-114
chr2-95417058-INS-8761
chr2-95490405-INS-88
chr2-95688848-INS-71
chr2-95706668-INS-126
chr2-95765531-INS-72
chr2-95789684-INS-494
chr2-95796232-INS-1899
chr2-95820322-INS-56
chr2-95848025-INS-66
chr2-95879694-INS-120
chr2-95926204-INS-201
chr2-95950728-INS-7538
chr2-95979212-INS-54
chr2-96031337-INS-625
chr2-96068382-INS-65
chr2-96069089-INS-78
chr2-96126552-INS-6589
chr2-96153452-INS-2781
chr2-96170536-INS-64
chr2-96194609-INS-68
chr2-96288730-INS-1760
chr2-96407853-INS-52
chr2-96426129-INS-50
chr2-96474912-INS-126
chr2-96500064-INS-449
chr2-96509939-INS-373
chr2-96515528-INS-53
chr2-96528453-INS-2119
chr2-96561526-INS-487
chr2-96577174-INS-7007
chr2-96577576-INS-309
chr2-96579639-INS-988
chr2-96599182-INS-6774
chr2-96616608-INS-57
chr2-96641824-INS-50
chr2-96648945-INS-83
chr2-96661403-INS-56
chr2-96682447-INS-52
chr2-96703329-INS-158
chr2-96709388-INS-50
chr2-96713689-INS-50
chr2-96730844-INS-1251
chr2-96750954-INS-3080
chr2-96761783-INS-7544
chr2-96776902-INS-747
chr2-96779505-INS-284
chr2-96783604-INS-97
chr2-96792860-INS-57
chr2-96795940-INS-342
chr2-96810654-INS-50
chr2-96823283-INS-133
chr2-96823283-INS-302
chr2-96880632-INS-278
chr2-96885598-INS-58
chr2-96932076-INS-2339
chr2-96985935-INS-342
chr2-97075010-INS-510
chr2-97078190-INS-52
chr2-97083298-INS-52
chr2-97109931-INS-82
chr2-97115379-INS-50
chr2-97144286-INS-201
chr2-97180501-INS-551
chr2-97185078-INS-81

chr2-97200322-INS-60
chr2-97204823-INS-144
chr2-97266523-INS-6189
chr2-97271037-INS-1241
chr2-97313921-INS-749
chr2-97316684-INS-54
chr2-97357002-INS-412
chr2-97370330-INS-189
chr2-97417978-INS-98
chr2-97427729-INS-135
chr2-97452411-INS-77
chr2-97476304-INS-6396
chr2-97483036-INS-595
chr2-97522006-INS-52
chr2-97529530-INS-50
chr2-97547539-INS-516
chr2-97554578-INS-82
chr2-97612858-INS-762
chr2-97616720-INS-2515
chr2-97685846-INS-52
chr2-97685919-INS-110
chr2-97720471-INS-54
chr2-97747916-INS-390
chr2-97757401-INS-375
chr2-97803183-INS-6733
chr2-97891218-INS-102
chr2-97922543-INS-1640
chr2-97949673-INS-1633
chr2-98001377-INS-1507
chr2-98018497-INS-52
chr2-98042623-INS-54
chr2-98068114-INS-68
chr2-98139710-INS-469
chr2-98148464-INS-435
chr2-98148686-INS-153
chr2-98176340-INS-6501
chr2-98177906-INS-64
chr2-98275912-INS-1697
chr2-98295145-INS-1386
chr2-98301607-INS-123
chr2-98316401-INS-68
chr2-98317167-INS-136
chr2-98337034-INS-331
chr2-98372247-INS-4315
chr2-98382681-INS-866
chr2-98384055-INS-3359
chr2-98418697-INS-730
chr2-98473459-INS-91
chr2-98480565-INS-50
chr2-98520234-INS-170
chr2-98536153-INS-367
chr2-98554516-INS-515
chr2-98578367-INS-342
chr2-98639825-INS-53
chr2-98649699-INS-2217
chr2-98658852-INS-70
chr2-98711182-INS-5917
chr2-98755126-INS-231
chr2-98770963-INS-606
chr2-98797831-INS-1643
chr2-98803063-INS-5891
chr2-98812549-INS-2968
chr2-98818732-INS-2273
chr2-98820522-INS-4391
chr2-98829596-INS-56
chr2-98844889-INS-1448
chr2-98903392-INS-56
chr2-98904248-INS-112
chr2-98906786-INS-318
chr2-98961766-INS-152
chr2-99017033-INS-7041
chr2-99026328-INS-70
chr2-99097091-INS-136
chr2-99102009-INS-2791
chr2-99102972-INS-55
chr2-99103087-INS-112
chr2-99161421-INS-6466

chr2-99178439-INS-117
chr2-99203775-INS-7281
chr2-99208405-INS-6753
chr2-99224493-INS-84
chr2-99238004-INS-78
chr2-99241359-INS-3332
chr2-99256135-INS-7285
chr2-99257259-INS-2851
chr2-99277214-INS-354
chr2-99302126-INS-51
chr2-99303688-INS-255
chr2-99317864-INS-6564
chr2-99321874-INS-688
chr2-99338144-INS-987
chr2-99344806-INS-3424
chr2-99348131-INS-4907
chr2-99356547-INS-187
chr2-99362397-INS-7083
chr2-99458917-INS-86
chr2-99462428-INS-52
chr2-99480918-INS-417
chr2-99499211-INS-287
chr2-99503578-INS-808
chr2-99504214-INS-930
chr2-99542904-INS-6382
chr2-99554107-INS-85
chr2-99573495-INS-6275
chr2-99653674-INS-7448
chr2-99714278-INS-6449
chr2-99720548-INS-161
chr2-99725345-INS-299
chr2-99727624-INS-58
chr2-99777118-INS-3092
chr2-99785173-INS-195
chr2-99797818-INS-1443
chr2-99809148-INS-50
chr2-99825991-INS-60
chr2-99826809-INS-58
chr2-99864873-INS-6323
chr2-99904306-INS-2185
chr2-99948504-INS-116
chr2-99995129-INS-1518
chr2-99996825-INS-290
chr2-100017332-INS-150
chr2-100021686-INS-6380
chr2-100037897-INS-51
chr2-100063130-INS-1997
chr2-100064698-INS-7103
chr2-100076982-INS-102
chr2-100077218-INS-714
chr2-100077489-INS-2224
chr2-100082182-INS-73
chr2-100082184-INS-154
chr2-100083553-INS-4927
chr2-100188823-INS-56
chr2-100188937-INS-120
chr2-100189092-INS-60
chr2-100236998-INS-115
chr2-100258958-INS-132
chr2-100278644-INS-6592
chr2-100345991-INS-2852
chr2-100392350-INS-6250
chr2-100426909-INS-177
chr2-100552990-INS-815
chr2-100620771-INS-593
chr2-100624375-INS-203
chr2-100645997-INS-1119
chr2-100694124-INS-56
chr2-100702062-INS-677
chr2-100719904-INS-1480
chr2-100797698-INS-457
chr2-100801143-INS-112
chr2-100849157-INS-5995
chr2-100858837-INS-1916
chr2-101078268-INS-405
chr2-101083271-INS-6379
chr2-101097813-INS-5214

chr2-101099144-INS-58
chr2-101217090-INS-7552
chr2-101221310-INS-6178
chr2-101313709-INS-1222
chr2-101337250-INS-440
chr2-101352700-INS-171
chr2-101357227-INS-187
chr2-101370413-INS-6207
chr2-101379106-INS-131
chr2-101379684-INS-194
chr2-101398927-INS-406
chr2-101403189-INS-145
chr2-101450722-INS-280
chr2-101504901-INS-80
chr2-101539146-INS-82
chr2-101539657-INS-199
chr2-101619461-INS-132
chr2-101676320-INS-50
chr2-101699593-INS-113
chr2-101837567-INS-55
chr2-101856729-INS-312
chr2-101857117-INS-369
chr2-101857129-INS-102
chr2-101962547-INS-553
chr2-101984647-INS-50
chr2-102042994-INS-54
chr2-102055716-INS-835
chr2-102082204-INS-208
chr2-102160309-INS-943
chr2-102198994-INS-64
chr2-102267141-INS-88
chr2-102273917-INS-207
chr2-102303234-INS-499
chr2-102329213-INS-2500
chr2-102348093-INS-52
chr2-102366588-INS-59
chr2-102406574-INS-6226
chr2-102425330-INS-109
chr2-102449880-INS-399
chr2-102450341-INS-51
chr2-102484040-INS-177
chr2-102495405-INS-138
chr2-102596695-INS-7227
chr2-102605897-INS-285
chr2-102630944-INS-55
chr2-102665834-INS-62
chr2-102732338-INS-320
chr2-102741968-INS-150
chr2-102795955-INS-116
chr2-102804195-INS-104
chr2-102827357-INS-108
chr2-102884745-INS-1492
chr2-102890271-INS-6462
chr2-102925249-INS-55
chr2-102925290-INS-116
chr2-102959886-INS-494
chr2-103050780-INS-84
chr2-103054080-INS-288
chr2-103055569-INS-79
chr2-103071711-INS-182
chr2-103126822-INS-75
chr2-103126822-INS-933
chr2-103144975-INS-776
chr2-103144992-INS-143
chr2-103145509-INS-98
chr2-103202592-INS-75
chr2-103212451-INS-74
chr2-103236235-INS-50
chr2-103261116-INS-126
chr2-103264516-INS-245
chr2-103316878-INS-1220
chr2-103362912-INS-878
chr2-103401710-INS-68
chr2-103453858-INS-81
chr2-103464387-INS-124
chr2-103510556-INS-984
chr2-103636841-INS-128

chr2-103650179-INS-194
chr2-103672584-INS-231
chr2-103738102-INS-215
chr2-103814975-INS-3290
chr2-103873180-INS-124
chr2-103873236-INS-279
chr2-103904392-INS-1375
chr2-103913990-INS-3465
chr2-103916066-INS-4800
chr2-103921991-INS-57
chr2-103957306-INS-1549
chr2-103957314-INS-912
chr2-103989583-INS-50
chr2-103997904-INS-200
chr2-104050060-INS-156
chr2-104050130-INS-72
chr2-104093359-INS-498
chr2-104110809-INS-52
chr2-104126525-INS-134
chr2-104126525-INS-50
chr2-104132410-INS-54
chr2-104135111-INS-75
chr2-104156551-INS-66
chr2-104175241-INS-63
chr2-104211017-INS-63
chr2-104211057-INS-185
chr2-104217571-INS-6181
chr2-104237651-INS-204
chr2-104241934-INS-2843
chr2-104261825-INS-202
chr2-104278619-INS-82
chr2-104317357-INS-1401
chr2-104317480-INS-174
chr2-104317630-INS-55
chr2-104317687-INS-322
chr2-104317855-INS-373
chr2-104351497-INS-88
chr2-104353124-INS-7395
chr2-104360009-INS-70
chr2-104372435-INS-474
chr2-104377864-INS-243
chr2-104393671-INS-50
chr2-104405032-INS-599
chr2-104405567-INS-5545
chr2-104416710-INS-68
chr2-104438194-INS-56
chr2-104471235-INS-200
chr2-104471416-INS-67
chr2-104502329-INS-212
chr2-104530920-INS-64
chr2-104540865-INS-202
chr2-104554545-INS-54
chr2-104576154-INS-212
chr2-104652550-INS-56
chr2-104655123-INS-131
chr2-104687618-INS-197
chr2-104705231-INS-92
chr2-104715116-INS-77
chr2-104715639-INS-235
chr2-104738467-INS-1727
chr2-104738632-INS-83
chr2-104738678-INS-395
chr2-104759000-INS-54
chr2-104780241-INS-393
chr2-104790249-INS-75
chr2-104834723-INS-276
chr2-104835972-INS-50
chr2-104955227-INS-52
chr2-104966308-INS-83
chr2-104984264-INS-108
chr2-104985061-INS-213
chr2-105015145-INS-10930
chr2-105079068-INS-217
chr2-105083305-INS-726
chr2-105083319-INS-148
chr2-105108833-INS-116
chr2-105168774-INS-78

chr2-105193355-INS-159
chr2-105235230-INS-73
chr2-105236540-INS-265
chr2-105240029-INS-179
chr2-105240029-INS-494
chr2-105250999-INS-68
chr2-105258209-INS-1291
chr2-105266210-INS-5548
chr2-105292528-INS-160
chr2-105292528-INS-420
chr2-105306040-INS-114
chr2-105311679-INS-84
chr2-105317029-INS-1819
chr2-105322810-INS-5791
chr2-105327254-INS-301
chr2-105327438-INS-507
chr2-105340297-INS-1152
chr2-105366814-INS-268
chr2-105407471-INS-72
chr2-105434582-INS-100
chr2-105465358-INS-1000
chr2-105514019-INS-52
chr2-105596312-INS-50
chr2-105599656-INS-57
chr2-105601640-INS-58
chr2-105736831-INS-3056
chr2-105762187-INS-54
chr2-105823224-INS-99
chr2-105844317-INS-13371
chr2-105957524-INS-585
chr2-105971729-INS-104
chr2-105972984-INS-6592
chr2-106026777-INS-50
chr2-106045141-INS-84
chr2-106046479-INS-3167
chr2-106050153-INS-2000
chr2-106053183-INS-86
chr2-106067714-INS-84
chr2-106098505-INS-228
chr2-106146116-INS-172
chr2-106167813-INS-72
chr2-106168644-INS-1318
chr2-106173711-INS-52
chr2-106240871-INS-52
chr2-106253996-INS-1371
chr2-106295965-INS-6440
chr2-106324481-INS-124
chr2-106333428-INS-400
chr2-106333992-INS-6391
chr2-106338853-INS-3088
chr2-106386116-INS-60
chr2-106437456-INS-343
chr2-106441629-INS-7104
chr2-106443708-INS-344
chr2-106538133-INS-214
chr2-106559450-INS-62
chr2-106608010-INS-81
chr2-106633390-INS-72
chr2-106638384-INS-58
chr2-106646235-INS-134
chr2-106654596-INS-454
chr2-106672406-INS-55
chr2-106744486-INS-53
chr2-106745364-INS-105
chr2-106745364-INS-260
chr2-106750989-INS-117
chr2-106758620-INS-153
chr2-106784808-INS-2715
chr2-106786580-INS-74
chr2-106788739-INS-211
chr2-106817118-INS-5846
chr2-106856507-INS-2967
chr2-106864264-INS-3189
chr2-106912700-INS-7686
chr2-106918060-INS-1126
chr2-106956548-INS-540
chr2-106958026-INS-1505

chr2-107004554-INS-62
chr2-107034597-INS-58
chr2-107057314-INS-6576
chr2-107240639-INS-60
chr2-107267788-INS-2010
chr2-107278186-INS-4126
chr2-107282678-INS-52
chr2-107302549-INS-6384
chr2-107331178-INS-68
chr2-107339912-INS-58
chr2-107352597-INS-5060
chr2-107357817-INS-54
chr2-107378508-INS-2529
chr2-107423759-INS-120
chr2-107455544-INS-1102
chr2-107507366-INS-2445
chr2-107512797-INS-235
chr2-107512797-INS-57
chr2-107516603-INS-2355
chr2-107546256-INS-130
chr2-107546275-INS-52
chr2-107546916-INS-336
chr2-107594774-INS-54
chr2-107636172-INS-50
chr2-107772211-INS-7542
chr2-107851033-INS-52
chr2-107859515-INS-346
chr2-107859517-INS-122
chr2-107870903-INS-6437
chr2-107882411-INS-1190
chr2-107891925-INS-7104
chr2-107938840-INS-50
chr2-107973303-INS-227
chr2-107976694-INS-6205
chr2-108036093-INS-58
chr2-108100192-INS-6153
chr2-108140718-INS-530
chr2-108157050-INS-114
chr2-108168246-INS-54
chr2-108189914-INS-50
chr2-108263242-INS-5672
chr2-108282519-INS-55
chr2-108299258-INS-3000
chr2-108304016-INS-6016
chr2-108337619-INS-253
chr2-108394408-INS-81
chr2-108429091-INS-14032
chr2-108433916-INS-181
chr2-108500563-INS-86
chr2-108583640-INS-106
chr2-108638237-INS-50
chr2-108648397-INS-2023
chr2-108680752-INS-6612
chr2-108700233-INS-410
chr2-108704193-INS-1561
chr2-108727569-INS-6844
chr2-108767342-INS-13665
chr2-108770048-INS-140
chr2-108793046-INS-169
chr2-108806647-INS-265
chr2-108862107-INS-75
chr2-108862364-INS-108
chr2-108935083-INS-997
chr2-108945845-INS-437
chr2-108947294-INS-592
chr2-108955641-INS-77
chr2-109055045-INS-154
chr2-109064079-INS-407
chr2-109064159-INS-117
chr2-109192789-INS-106
chr2-109194719-INS-326
chr2-109199084-INS-6386
chr2-109208189-INS-6739
chr2-109232332-INS-112
chr2-109252866-INS-215
chr2-109333379-INS-50
chr2-109405646-INS-58

chr2-109406247-INS-72
chr2-109418048-INS-60
chr2-109418921-INS-133
chr2-109440229-INS-51
chr2-109488416-INS-52
chr2-109580134-INS-416
chr2-109618101-INS-1020
chr2-109712065-INS-9923
chr2-109715969-INS-76
chr2-109782112-INS-120
chr2-109844913-INS-67
chr2-109904389-INS-1855
chr2-109964485-INS-149
chr2-109964990-INS-104
chr2-109965182-INS-1876
chr2-109983456-INS-123
chr2-110052584-INS-2152
chr2-110059951-INS-850
chr2-110127520-INS-1109
chr2-110128313-INS-5648
chr2-110164195-INS-2876
chr2-110282494-INS-195
chr2-110312827-INS-6262
chr2-110348799-INS-6157
chr2-110365646-INS-56
chr2-110365884-INS-1136
chr2-110384003-INS-1019
chr2-110480559-INS-216
chr2-110495514-INS-50
chr2-110519531-INS-76
chr2-110652538-INS-73
chr2-110655804-INS-997
chr2-110685362-INS-50
chr2-110685605-INS-54
chr2-110702858-INS-204
chr2-110709361-INS-66
chr2-110734652-INS-6267
chr2-110761260-INS-1332
chr2-110762446-INS-80
chr2-110788367-INS-1156
chr2-110789140-INS-6894
chr2-110832404-INS-61
chr2-110882390-INS-1326
chr2-110892137-INS-3203
chr2-110898122-INS-466
chr2-110898444-INS-5951
chr2-110919746-INS-3579
chr2-110921821-INS-2414
chr2-111103875-INS-50
chr2-111106192-INS-80
chr2-111124424-INS-636
chr2-111192396-INS-1011
chr2-111194517-INS-52
chr2-111231342-INS-60
chr2-111238720-INS-68
chr2-111269230-INS-555
chr2-111275640-INS-11942
chr2-111336010-INS-3069
chr2-111406116-INS-60
chr2-111439022-INS-54
chr2-111567488-INS-112
chr2-111567488-INS-50
chr2-111589476-INS-6521
chr2-111604012-INS-296
chr2-111622202-INS-115
chr2-111622281-INS-491
chr2-111624929-INS-157
chr2-111634010-INS-649
chr2-111641152-INS-11766
chr2-111654199-INS-302
chr2-111701399-INS-52
chr2-111776006-INS-145
chr2-111776006-INS-291
chr2-111780282-INS-5668
chr2-111853393-INS-517
chr2-111928373-INS-2972
chr2-111939417-INS-76

chr2-111942914-INS-3075
chr2-111975063-INS-134
chr2-112157715-INS-390
chr2-112169930-INS-182
chr2-112206965-INS-157
chr2-112213634-INS-143
chr2-112284114-INS-5843
chr2-112314711-INS-124
chr2-112324640-INS-52
chr2-112331637-INS-144
chr2-112353822-INS-407
chr2-112421137-INS-367
chr2-112434478-INS-902
chr2-112437920-INS-7377
chr2-112442653-INS-7103
chr2-112464132-INS-89
chr2-112464237-INS-56
chr2-112506848-INS-551
chr2-112512975-INS-252
chr2-112554203-INS-50
chr2-112580792-INS-6592
chr2-112605423-INS-6505
chr2-112655881-INS-140
chr2-112693490-INS-70
chr2-112717052-INS-6239
chr2-112786095-INS-6568
chr2-112833424-INS-52
chr2-112835577-INS-1477
chr2-112847156-INS-488
chr2-112973405-INS-2193
chr2-112980640-INS-6212
chr2-112997415-INS-6219
chr2-113012397-INS-416
chr2-113073914-INS-5059
chr2-113112646-INS-5391
chr2-113119728-INS-63
chr2-113128985-INS-2105
chr2-113138834-INS-287
chr2-113153151-INS-306
chr2-113153284-INS-1452
chr2-113159404-INS-64
chr2-113201044-INS-56
chr2-113201673-INS-120
chr2-113240207-INS-60
chr2-113286012-INS-50
chr2-113286293-INS-417
chr2-113298602-INS-54
chr2-113298960-INS-238
chr2-113315901-INS-1134
chr2-113398460-INS-50
chr2-113405401-INS-54
chr2-113501346-INS-424
chr2-113501346-INS-53
chr2-113501533-INS-2229
chr2-113503639-INS-56
chr2-113589770-INS-52
chr2-113607652-INS-2710
chr2-113649189-INS-464
chr2-113650588-INS-633
chr2-113674617-INS-204
chr2-113674867-INS-118
chr2-113679904-INS-62
chr2-113715200-INS-82
chr2-113741716-INS-52
chr2-113763277-INS-6253
chr2-113772077-INS-55
chr2-113773015-INS-1193
chr2-113886522-INS-69
chr2-113900693-INS-5014
chr2-113908126-INS-1228
chr2-113908644-INS-222
chr2-113973390-INS-52
chr2-114038635-INS-70
chr2-114044355-INS-62
chr2-114049934-INS-176
chr2-114059000-INS-147
chr2-114066266-INS-87

chr2-114123433-INS-52
chr2-114162690-INS-218
chr2-114168741-INS-653
chr2-114213881-INS-51
chr2-114344821-INS-52
chr2-114407686-INS-6722
chr2-114580274-INS-2917
chr2-114632370-INS-61
chr2-114686709-INS-10547
chr2-114695507-INS-281
chr2-114742149-INS-50
chr2-114786588-INS-7522
chr2-114824414-INS-7412
chr2-114831853-INS-4144
chr2-114854148-INS-1071
chr2-114888897-INS-2673
chr2-114999053-INS-116
chr2-115072255-INS-3241
chr2-115110794-INS-50
chr2-115132770-INS-2756
chr2-115133582-INS-6206
chr2-115136991-INS-51
chr2-115149644-INS-68
chr2-115174480-INS-6190
chr2-115248929-INS-51
chr2-115350275-INS-401
chr2-115371936-INS-57
chr2-115371936-INS-955
chr2-115485465-INS-1108
chr2-115509450-INS-1226
chr2-115527782-INS-211
chr2-115542839-INS-69
chr2-115630640-INS-2058
chr2-115630640-INS-854
chr2-115744717-INS-60
chr2-115804385-INS-123
chr2-115811492-INS-80
chr2-115881371-INS-53
chr2-115907231-INS-52
chr2-115911430-INS-81
chr2-115962641-INS-1240
chr2-115963109-INS-54
chr2-115967638-INS-1094
chr2-115972651-INS-112
chr2-116006750-INS-6367
chr2-116020202-INS-54
chr2-116022118-INS-1579
chr2-116030322-INS-225
chr2-116060649-INS-297
chr2-116075541-INS-5411
chr2-116089970-INS-50
chr2-116106492-INS-581
chr2-116186439-INS-50
chr2-116222486-INS-52
chr2-116306883-INS-92
chr2-116385429-INS-58
chr2-116388851-INS-3000
chr2-116403811-INS-274
chr2-116442615-INS-2287
chr2-116466171-INS-6178
chr2-116466945-INS-106
chr2-116469722-INS-128
chr2-116496303-INS-380
chr2-116534008-INS-52
chr2-116541498-INS-54
chr2-116542135-INS-75
chr2-116612098-INS-63
chr2-116615602-INS-7121
chr2-116685729-INS-330
chr2-116698700-INS-7240
chr2-116703576-INS-481
chr2-116728141-INS-790
chr2-116733249-INS-100
chr2-116764769-INS-50
chr2-116764939-INS-773
chr2-116786216-INS-50
chr2-116865534-INS-1363

chr2-116885947-INS-132
chr2-116896885-INS-1794
chr2-116906264-INS-180
chr2-117001174-INS-219
chr2-117037357-INS-2097
chr2-117067704-INS-90
chr2-117079365-INS-93
chr2-117128471-INS-78
chr2-117128550-INS-189
chr2-117200812-INS-157
chr2-117205988-INS-54
chr2-117262382-INS-211
chr2-117306844-INS-58
chr2-117330739-INS-50
chr2-117399721-INS-69
chr2-117415108-INS-6803
chr2-117445842-INS-96
chr2-117484519-INS-84
chr2-117518792-INS-85
chr2-117525402-INS-360
chr2-117529534-INS-52
chr2-117610178-INS-80
chr2-117610212-INS-194
chr2-117610224-INS-481
chr2-117610419-INS-152
chr2-117657566-INS-3974
chr2-117668742-INS-92
chr2-117747510-INS-50
chr2-117791807-INS-95
chr2-117800241-INS-6588
chr2-117804062-INS-83
chr2-117806007-INS-1499
chr2-117809238-INS-124
chr2-117824195-INS-60
chr2-117829301-INS-496
chr2-117833412-INS-376
chr2-117853172-INS-94
chr2-117897670-INS-96
chr2-117976228-INS-168
chr2-118005617-INS-2603
chr2-118031597-INS-726
chr2-118054026-INS-2207
chr2-118058990-INS-177
chr2-118064345-INS-123
chr2-118083534-INS-89
chr2-118156372-INS-77
chr2-118158300-INS-840
chr2-118168021-INS-230
chr2-118196994-INS-126
chr2-118210751-INS-200
chr2-118257303-INS-153
chr2-118275031-INS-1708
chr2-118275241-INS-666
chr2-118284101-INS-52
chr2-118332587-INS-74
chr2-118333233-INS-429
chr2-118347643-INS-62
chr2-118482359-INS-225
chr2-118485808-INS-54
chr2-118486556-INS-457
chr2-118576197-INS-1504
chr2-118604453-INS-50
chr2-118633854-INS-4988
chr2-118638629-INS-210
chr2-118767088-INS-78
chr2-118822195-INS-50
chr2-118858287-INS-167
chr2-118858472-INS-52
chr2-118874941-INS-72
chr2-118893436-INS-174
chr2-118988304-INS-189
chr2-119044891-INS-98
chr2-119057781-INS-237
chr2-119061212-INS-98
chr2-119149539-INS-177
chr2-119149607-INS-66
chr2-119149957-INS-62

chr2-119209141-INS-52
chr2-119219925-INS-274
chr2-119227427-INS-1032
chr2-119270931-INS-64
chr2-119285014-INS-344
chr2-119287672-INS-911
chr2-119287831-INS-322
chr2-119298031-INS-209
chr2-119302570-INS-202
chr2-119305593-INS-184
chr2-119316379-INS-266
chr2-119354869-INS-182
chr2-119355282-INS-58
chr2-119374489-INS-57
chr2-119398092-INS-194
chr2-119413514-INS-83
chr2-119419043-INS-7138
chr2-119420110-INS-280
chr2-119432381-INS-124
chr2-119465710-INS-239
chr2-119478846-INS-213
chr2-119480982-INS-212
chr2-119487131-INS-612
chr2-119487483-INS-601
chr2-119489616-INS-876
chr2-119489728-INS-187
chr2-119517844-INS-4713
chr2-119533007-INS-510
chr2-119535414-INS-577
chr2-119541768-INS-273
chr2-119543725-INS-222
chr2-119544455-INS-204
chr2-119561430-INS-220
chr2-119599218-INS-176
chr2-119652877-INS-137
chr2-119670713-INS-195
chr2-119699608-INS-7225
chr2-119710837-INS-58
chr2-119735956-INS-657
chr2-119753691-INS-6295
chr2-119815498-INS-60
chr2-120041498-INS-95
chr2-120062024-INS-70
chr2-120063655-INS-372
chr2-120086104-INS-3080
chr2-120097922-INS-96
chr2-120098206-INS-53
chr2-120098464-INS-114
chr2-120127765-INS-348
chr2-120128900-INS-210
chr2-120134722-INS-202
chr2-120260799-INS-188
chr2-120266581-INS-140
chr2-120266611-INS-50
chr2-120310656-INS-1706
chr2-120317599-INS-458
chr2-120394622-INS-209
chr2-120403510-INS-201
chr2-120403945-INS-1932
chr2-120408383-INS-215
chr2-120445225-INS-80
chr2-120473126-INS-196
chr2-120484637-INS-74
chr2-120490088-INS-114
chr2-120490295-INS-206
chr2-120498035-INS-208
chr2-120498363-INS-259
chr2-120510047-INS-202
chr2-120521937-INS-95
chr2-120522649-INS-200
chr2-120661806-INS-196
chr2-120783104-INS-51
chr2-120792595-INS-1132
chr2-120899999-INS-51
chr2-120914391-INS-765
chr2-120918576-INS-55
chr2-120923070-INS-73

chr2-120977036-INS-113
chr2-121096540-INS-54
chr2-121210813-INS-50
chr2-121229408-INS-268
chr2-121236612-INS-202
chr2-121256616-INS-544
chr2-121262940-INS-217
chr2-121285419-INS-83
chr2-121289975-INS-203
chr2-121295975-INS-407
chr2-121308543-INS-213
chr2-121309282-INS-208
chr2-121320901-INS-4380
chr2-121321652-INS-50
chr2-121338782-INS-53
chr2-121355175-INS-373
chr2-121382614-INS-276
chr2-121386782-INS-282
chr2-121405399-INS-118
chr2-121410994-INS-55
chr2-121411516-INS-94
chr2-121429710-INS-156
chr2-121590924-INS-115
chr2-121626075-INS-334
chr2-121665407-INS-56
chr2-121679423-INS-132
chr2-121720924-INS-62
chr2-121723360-INS-481
chr2-121728068-INS-210
chr2-121739015-INS-102
chr2-121739606-INS-236
chr2-121747794-INS-182
chr2-121764472-INS-147
chr2-121765004-INS-226
chr2-121804144-INS-179
chr2-121804996-INS-199
chr2-121812599-INS-229
chr2-121813098-INS-256
chr2-121832914-INS-777
chr2-121836685-INS-52
chr2-121887371-INS-214
chr2-121906877-INS-538
chr2-121928554-INS-775
chr2-121994005-INS-1236
chr2-121999014-INS-216
chr2-122002966-INS-389
chr2-122017687-INS-217
chr2-122042229-INS-476
chr2-122056974-INS-3783
chr2-122106200-INS-5316
chr2-122207496-INS-66
chr2-122216245-INS-236
chr2-122216567-INS-363
chr2-122222217-INS-3105
chr2-122233536-INS-72
chr2-122268698-INS-1903
chr2-122294450-INS-6566
chr2-122311316-INS-5871
chr2-122326418-INS-3465
chr2-122353631-INS-4830
chr2-122406694-INS-102
chr2-122411591-INS-50
chr2-122453171-INS-3210
chr2-122460633-INS-5401
chr2-122590745-INS-240
chr2-122656191-INS-112
chr2-122691438-INS-66
chr2-122727433-INS-2072
chr2-122727435-INS-1003
chr2-122752277-INS-126
chr2-122798248-INS-208
chr2-122841308-INS-214
chr2-122845339-INS-1490
chr2-122849442-INS-82
chr2-122853846-INS-517
chr2-122908526-INS-166
chr2-122913612-INS-55

chr2-122916730-INS-182
chr2-122936958-INS-6485
chr2-122941017-INS-450
chr2-122941094-INS-6389
chr2-123030959-INS-2761
chr2-123030959-INS-6312
chr2-123046385-INS-51
chr2-123048768-INS-64338
chr2-123079329-INS-62
chr2-123170707-INS-82
chr2-123180786-INS-498
chr2-123205849-INS-6388
chr2-123219215-INS-4888
chr2-123258349-INS-340
chr2-123379082-INS-55
chr2-123380017-INS-1093
chr2-123507148-INS-4624
chr2-123586866-INS-10149
chr2-123668820-INS-87
chr2-123701801-INS-7090
chr2-123776367-INS-442
chr2-123779769-INS-498
chr2-123879733-INS-67
chr2-123894954-INS-4093
chr2-123914099-INS-3129
chr2-123988607-INS-254
chr2-123997997-INS-77
chr2-124037137-INS-134
chr2-124123537-INS-1637
chr2-124176887-INS-107
chr2-124185927-INS-76
chr2-124208726-INS-7532
chr2-124216949-INS-59
chr2-124237642-INS-52
chr2-124258179-INS-58
chr2-124298538-INS-128
chr2-124308673-INS-558
chr2-124390220-INS-55
chr2-124520771-INS-52
chr2-124521756-INS-4699
chr2-124531732-INS-1057
chr2-124536437-INS-78
chr2-124537684-INS-203
chr2-124555803-INS-50
chr2-124556741-INS-164
chr2-124667182-INS-194
chr2-124693336-INS-70
chr2-124693377-INS-870
chr2-124787518-INS-168
chr2-124808345-INS-78
chr2-124832632-INS-113
chr2-124844602-INS-208
chr2-124876502-INS-51
chr2-124891039-INS-797
chr2-124940312-INS-174
chr2-125040183-INS-163
chr2-125080728-INS-1081
chr2-125126489-INS-50
chr2-125275319-INS-2003
chr2-125307111-INS-54
chr2-125336772-INS-68
chr2-125343556-INS-106
chr2-125349847-INS-594
chr2-125354907-INS-68
chr2-125404318-INS-67
chr2-125429472-INS-81
chr2-125444029-INS-286
chr2-125446031-INS-875
chr2-125459585-INS-224
chr2-125464024-INS-50
chr2-125478983-INS-119
chr2-125482331-INS-51
chr2-125483007-INS-73
chr2-125488142-INS-77
chr2-125510576-INS-56
chr2-125521627-INS-167
chr2-125525541-INS-84

chr2-125526245-INS-160
chr2-125568792-INS-50
chr2-125608103-INS-153
chr2-125608950-INS-102
chr2-125644801-INS-89
chr2-125659510-INS-56
chr2-125659793-INS-180
chr2-125740842-INS-105
chr2-125781915-INS-64
chr2-125782497-INS-136
chr2-125822070-INS-54
chr2-125846771-INS-78
chr2-125859133-INS-68
chr2-125868700-INS-426
chr2-125893543-INS-74
chr2-125972223-INS-18037
chr2-125972223-INS-7915
chr2-125993193-INS-65
chr2-125993261-INS-577
chr2-126009823-INS-6165
chr2-126042050-INS-6272
chr2-126050745-INS-6595
chr2-126123651-INS-6791
chr2-126126573-INS-249
chr2-126142274-INS-6268
chr2-126169342-INS-5016
chr2-126197083-INS-2246
chr2-126212484-INS-62
chr2-126227741-INS-6901
chr2-126257294-INS-80
chr2-126265625-INS-85
chr2-126325998-INS-228
chr2-126326326-INS-84
chr2-126357337-INS-6552
chr2-126377382-INS-1417
chr2-126441093-INS-54
chr2-126509323-INS-50
chr2-126509501-INS-1231
chr2-126527157-INS-95
chr2-126537739-INS-7658
chr2-126538873-INS-755
chr2-126545823-INS-75
chr2-126558677-INS-122
chr2-126558796-INS-201
chr2-126586508-INS-280
chr2-126595363-INS-53
chr2-126615249-INS-70
chr2-126650368-INS-88
chr2-126721257-INS-901
chr2-126723442-INS-120
chr2-126723990-INS-50
chr2-126741199-INS-76
chr2-126753891-INS-1377
chr2-126763954-INS-50
chr2-126826714-INS-1249
chr2-126834316-INS-186
chr2-126854013-INS-60
chr2-126873266-INS-50
chr2-126873747-INS-86
chr2-126882689-INS-52
chr2-126893011-INS-375
chr2-126893011-INS-68
chr2-126919570-INS-81
chr2-126932495-INS-323
chr2-126942695-INS-832
chr2-126965197-INS-55
chr2-126998860-INS-82
chr2-126998879-INS-4986
chr2-126998995-INS-183
chr2-127002417-INS-8441
chr2-127011068-INS-1858
chr2-127012037-INS-175
chr2-127012743-INS-95
chr2-127015640-INS-1493
chr2-127021319-INS-51
chr2-127026841-INS-5289
chr2-127063183-INS-270

chr2-127163985-INS-2286
chr2-127297084-INS-121
chr2-127297141-INS-542
chr2-127408389-INS-87
chr2-127448292-INS-54
chr2-127462840-INS-63
chr2-127477047-INS-441
chr2-127478506-INS-68
chr2-127509140-INS-203
chr2-127522798-INS-52
chr2-127528665-INS-119
chr2-127552761-INS-392
chr2-127557477-INS-139
chr2-127577256-INS-126
chr2-127579578-INS-191
chr2-127579782-INS-290
chr2-127579788-INS-90
chr2-127593581-INS-444
chr2-127610881-INS-411
chr2-127685353-INS-50
chr2-127691916-INS-1217
chr2-127738489-INS-69
chr2-127819141-INS-64
chr2-127851315-INS-317
chr2-127913300-INS-60
chr2-127916927-INS-56
chr2-127989301-INS-187
chr2-128042075-INS-205
chr2-128054706-INS-102
chr2-128060768-INS-116
chr2-128074015-INS-7086
chr2-128115062-INS-404
chr2-128133255-INS-6475
chr2-128192008-INS-130
chr2-128249009-INS-50
chr2-128251577-INS-252
chr2-128291963-INS-202
chr2-128296086-INS-132
chr2-128296095-INS-63
chr2-128350667-INS-50
chr2-128370684-INS-60
chr2-128377581-INS-93
chr2-128404210-INS-52
chr2-128404540-INS-497
chr2-128406568-INS-125
chr2-128451369-INS-1520
chr2-128519153-INS-176
chr2-128575142-INS-328
chr2-128583382-INS-54
chr2-128583501-INS-162
chr2-128640104-INS-64
chr2-128656767-INS-113
chr2-128666833-INS-362
chr2-128720418-INS-134
chr2-128728268-INS-135
chr2-128753836-INS-145
chr2-128753838-INS-1826
chr2-128754353-INS-116
chr2-128837678-INS-1889
chr2-128861186-INS-6482
chr2-128906469-INS-56
chr2-128935304-INS-705
chr2-128956906-INS-193
chr2-129015201-INS-149
chr2-129064707-INS-53
chr2-129156163-INS-50
chr2-129173966-INS-276
chr2-129212774-INS-160
chr2-129212932-INS-58
chr2-129218758-INS-152
chr2-129260188-INS-224
chr2-129262319-INS-452
chr2-129267545-INS-7279
chr2-129284148-INS-689
chr2-129284297-INS-1554
chr2-129296154-INS-428
chr2-129372984-INS-58

chr2-129380536-INS-50
chr2-129445561-INS-773
chr2-129495179-INS-124
chr2-129495252-INS-52
chr2-129536812-INS-50
chr2-129568647-INS-318
chr2-129572471-INS-7094
chr2-129583917-INS-69
chr2-129585803-INS-4864
chr2-129613578-INS-50
chr2-129620102-INS-166
chr2-129655860-INS-2977
chr2-129657247-INS-240
chr2-129691284-INS-189
chr2-129782785-INS-160
chr2-129789829-INS-2902
chr2-129789829-INS-451
chr2-129940503-INS-8120
chr2-129971339-INS-87
chr2-129972363-INS-370
chr2-129972381-INS-56
chr2-129972440-INS-120
chr2-129980069-INS-7276
chr2-129989020-INS-124
chr2-130000607-INS-105
chr2-130005205-INS-464
chr2-130072353-INS-6632
chr2-130193085-INS-50
chr2-130214733-INS-300
chr2-130291719-INS-62
chr2-130316070-INS-50
chr2-130320696-INS-180
chr2-130367785-INS-230
chr2-130435911-INS-686
chr2-130438630-INS-4393
chr2-130449642-INS-60
chr2-130463345-INS-99
chr2-130469478-INS-8751
chr2-130487907-INS-84
chr2-130523698-INS-58
chr2-130547877-INS-224
chr2-130605337-INS-193
chr2-130630735-INS-492
chr2-130635590-INS-54
chr2-130637710-INS-201
chr2-130638488-INS-58
chr2-130655518-INS-219
chr2-130675937-INS-70
chr2-130711697-INS-6635
chr2-130715790-INS-57
chr2-130728044-INS-413
chr2-130730505-INS-70
chr2-130846836-INS-231
chr2-130847394-INS-76
chr2-130938981-INS-1215
chr2-130972366-INS-248
chr2-131061158-INS-11141
chr2-131065128-INS-112
chr2-131066102-INS-97
chr2-131120559-INS-58
chr2-131146927-INS-374
chr2-131165341-INS-60
chr2-131165536-INS-135
chr2-131166842-INS-62
chr2-131182672-INS-208
chr2-131221771-INS-3828
chr2-131236820-INS-56
chr2-131256311-INS-115
chr2-131264821-INS-1028
chr2-131269899-INS-92
chr2-131270420-INS-1600
chr2-131292472-INS-2250
chr2-131293034-INS-52
chr2-131311721-INS-186
chr2-131312486-INS-62
chr2-131323548-INS-53
chr2-131338667-INS-236

chr2-131424672-INS-263
chr2-131446188-INS-3558
chr2-131473063-INS-72
chr2-131491180-INS-80
chr2-131514526-INS-193
chr2-131515619-INS-9879
chr2-131528170-INS-105
chr2-131561071-INS-6124
chr2-131576377-INS-2366
chr2-131584320-INS-360
chr2-131602324-INS-50
chr2-131608771-INS-54
chr2-131644456-INS-6338
chr2-131670264-INS-60
chr2-131699361-INS-191
chr2-131716858-INS-142
chr2-131758529-INS-50
chr2-131786095-INS-1832
chr2-131828604-INS-54
chr2-131968753-INS-590
chr2-131969019-INS-4335
chr2-131979675-INS-331
chr2-131979675-INS-81
chr2-131991325-INS-50
chr2-132000617-INS-60
chr2-132000819-INS-50
chr2-132111750-INS-125
chr2-132123186-INS-68
chr2-132123235-INS-383
chr2-132123244-INS-832
chr2-132123736-INS-249
chr2-132141121-INS-224
chr2-132159075-INS-174
chr2-132174129-INS-53
chr2-132182200-INS-61
chr2-132216408-INS-212
chr2-132239029-INS-62
chr2-132239030-INS-186
chr2-132256556-INS-1074
chr2-132263702-INS-61
chr2-132272871-INS-2795
chr2-132281359-INS-84
chr2-132285624-INS-50
chr2-132317436-INS-478
chr2-132319175-INS-708
chr2-132334067-INS-6437
chr2-132350849-INS-231
chr2-132353539-INS-52
chr2-132353596-INS-127
chr2-132436709-INS-7172
chr2-132440619-INS-55
chr2-132440671-INS-154
chr2-132475862-INS-2799
chr2-132510942-INS-457
chr2-132521031-INS-192
chr2-132542177-INS-6244
chr2-132567851-INS-82
chr2-132570089-INS-74
chr2-132575722-INS-60
chr2-132591156-INS-51
chr2-132603071-INS-792
chr2-132617337-INS-184
chr2-132617836-INS-176
chr2-132631536-INS-57
chr2-132633477-INS-64
chr2-132674617-INS-1499
chr2-132674617-INS-499
chr2-132681805-INS-75
chr2-132722347-INS-213
chr2-132765321-INS-59
chr2-132769802-INS-54
chr2-132799653-INS-66
chr2-132859876-INS-82
chr2-132860030-INS-52
chr2-132916114-INS-78
chr2-132921476-INS-130
chr2-132953877-INS-50

chr2-132954533-INS-179
chr2-132971968-INS-2865
chr2-132990744-INS-1187
chr2-133074472-INS-339
chr2-133074992-INS-1893
chr2-133081177-INS-428
chr2-133100633-INS-762
chr2-133162188-INS-50
chr2-133205726-INS-1344
chr2-133205768-INS-3685
chr2-133205792-INS-216
chr2-133429201-INS-58
chr2-133430786-INS-281
chr2-133433319-INS-50
chr2-133444661-INS-72
chr2-133466811-INS-56
chr2-133489171-INS-84
chr2-133495691-INS-118
chr2-133513028-INS-81
chr2-133560113-INS-1261
chr2-133613615-INS-63
chr2-133628540-INS-372
chr2-133633266-INS-76
chr2-133652416-INS-1114
chr2-133726993-INS-9992
chr2-133727620-INS-82
chr2-133783897-INS-1307
chr2-133803108-INS-61
chr2-133861840-INS-1243
chr2-133873471-INS-116
chr2-133928552-INS-1265
chr2-133966522-INS-6886
chr2-134020059-INS-160
chr2-134025931-INS-6454
chr2-134077881-INS-52
chr2-134118935-INS-73
chr2-134129667-INS-478
chr2-134142590-INS-1121
chr2-134168542-INS-50
chr2-134172973-INS-128
chr2-134228066-INS-6836
chr2-134292522-INS-6182
chr2-134315974-INS-2869
chr2-134356645-INS-196
chr2-134373784-INS-112
chr2-134375996-INS-1506
chr2-134384761-INS-62
chr2-134412665-INS-15126
chr2-134425569-INS-83
chr2-134429596-INS-52
chr2-134477507-INS-219
chr2-134478701-INS-50
chr2-134491005-INS-69
chr2-134526751-INS-1492
chr2-134532788-INS-2167
chr2-134541103-INS-1665
chr2-134543897-INS-74
chr2-134556942-INS-60
chr2-134579878-INS-134
chr2-134580049-INS-410
chr2-134607457-INS-6614
chr2-134739329-INS-139
chr2-134762002-INS-121
chr2-134784851-INS-633
chr2-134798956-INS-316
chr2-134801657-INS-100
chr2-134838101-INS-2587
chr2-134846880-INS-122
chr2-134900573-INS-56
chr2-135020628-INS-355
chr2-135060566-INS-106
chr2-135060585-INS-402
chr2-135099926-INS-104
chr2-135142792-INS-6598
chr2-135192993-INS-1549
chr2-135198217-INS-6066
chr2-135220418-INS-682

chr2-135250911-INS-6382
chr2-135258972-INS-66
chr2-135326802-INS-1927
chr2-135335444-INS-50
chr2-135335451-INS-140
chr2-135416638-INS-52
chr2-135417890-INS-107
chr2-135447252-INS-50
chr2-135480998-INS-52
chr2-135488909-INS-72
chr2-135504669-INS-14067
chr2-135516297-INS-117
chr2-135561773-INS-224
chr2-135599800-INS-6774
chr2-135600621-INS-213
chr2-135600662-INS-3075
chr2-135647547-INS-52
chr2-135719739-INS-837
chr2-135728406-INS-905
chr2-135830550-INS-1798
chr2-135867628-INS-1957
chr2-135867672-INS-54
chr2-135881782-INS-6435
chr2-135881783-INS-16652
chr2-135936772-INS-72
chr2-135974783-INS-56
chr2-136001807-INS-6599
chr2-136011136-INS-62
chr2-136012059-INS-165
chr2-136029617-INS-62
chr2-136060150-INS-1555
chr2-136062189-INS-4471
chr2-136124502-INS-108
chr2-136143996-INS-56
chr2-136154949-INS-144
chr2-136163051-INS-6242
chr2-136175901-INS-185
chr2-136226237-INS-56
chr2-136242789-INS-5043
chr2-136268068-INS-249
chr2-136320728-INS-58
chr2-136332152-INS-87
chr2-136348683-INS-202
chr2-136372409-INS-66
chr2-136373035-INS-59
chr2-136413045-INS-3918
chr2-136468045-INS-404
chr2-136468164-INS-952
chr2-136487113-INS-212
chr2-136547064-INS-6408
chr2-136561560-INS-144
chr2-136665369-INS-90
chr2-136731407-INS-392
chr2-136778226-INS-180
chr2-136782363-INS-52
chr2-136867880-INS-50
chr2-136902417-INS-500
chr2-136914095-INS-65
chr2-136967249-INS-6610
chr2-137035415-INS-667
chr2-137077434-INS-50
chr2-137105477-INS-131
chr2-137186823-INS-50
chr2-137371390-INS-57
chr2-137486512-INS-1755
chr2-137512322-INS-52
chr2-137517599-INS-61
chr2-137523984-INS-6531
chr2-137540624-INS-2471
chr2-137592754-INS-350
chr2-137593531-INS-350
chr2-137654098-INS-93
chr2-137762552-INS-2752
chr2-137770336-INS-57
chr2-137777040-INS-108
chr2-137802991-INS-6050
chr2-137827873-INS-6537

chr2-137903204-INS-6333
chr2-137918928-INS-99
chr2-137928326-INS-72
chr2-137953000-INS-383
chr2-137961681-INS-54
chr2-138000404-INS-261
chr2-138030200-INS-164
chr2-138087946-INS-50
chr2-138110512-INS-160
chr2-138185088-INS-6498
chr2-138209682-INS-76
chr2-138225035-INS-122
chr2-138277333-INS-569
chr2-138277860-INS-203
chr2-138373713-INS-55
chr2-138373716-INS-401
chr2-138467397-INS-1010
chr2-138499812-INS-50
chr2-138557529-INS-54
chr2-138591276-INS-2750
chr2-138601010-INS-6458
chr2-138612156-INS-102
chr2-138612162-INS-50
chr2-138623993-INS-95
chr2-138626070-INS-50
chr2-138639065-INS-76
chr2-138639129-INS-54
chr2-138673247-INS-50
chr2-138702626-INS-498
chr2-138706064-INS-55
chr2-138706086-INS-160
chr2-138706341-INS-75
chr2-138709670-INS-75
chr2-138713522-INS-58
chr2-138726771-INS-7094
chr2-138741043-INS-2183
chr2-138784381-INS-50
chr2-138790265-INS-7340
chr2-138798154-INS-52
chr2-138826472-INS-6206
chr2-138866962-INS-1170
chr2-138887066-INS-54
chr2-138931935-INS-50
chr2-138946081-INS-50
chr2-138946865-INS-74
chr2-139005792-INS-365
chr2-139013821-INS-244
chr2-139030766-INS-3042
chr2-139038389-INS-52
chr2-139043926-INS-2000
chr2-139048131-INS-82
chr2-139077247-INS-292
chr2-139147131-INS-57
chr2-139171780-INS-53
chr2-139189272-INS-6349
chr2-139192383-INS-207
chr2-139194890-INS-1463
chr2-139230428-INS-3393
chr2-139322029-INS-1712
chr2-139343406-INS-51
chr2-139375719-INS-3564
chr2-139417842-INS-6008
chr2-139419457-INS-2439
chr2-139438414-INS-184
chr2-139464548-INS-53
chr2-139464553-INS-158
chr2-139505056-INS-58
chr2-139531763-INS-1098
chr2-139536641-INS-56
chr2-139585342-INS-7104
chr2-139596838-INS-145
chr2-139601777-INS-7176
chr2-139673530-INS-207
chr2-139701660-INS-68
chr2-139734293-INS-7560
chr2-139811983-INS-2632
chr2-139824260-INS-54

chr2-139858176-INS-7118
chr2-139860888-INS-281
chr2-139860889-INS-812
chr2-139953414-INS-124
chr2-140046538-INS-160
chr2-140048816-INS-6579
chr2-140049065-INS-93
chr2-140050276-INS-1501
chr2-140069529-INS-4109
chr2-140100965-INS-6904
chr2-140150969-INS-54
chr2-140223998-INS-785
chr2-140278110-INS-2175
chr2-140529498-INS-50
chr2-140560200-INS-140
chr2-140572599-INS-52
chr2-140633651-INS-6380
chr2-140652146-INS-282
chr2-140652162-INS-117
chr2-140660612-INS-141
chr2-140689684-INS-1890
chr2-140739293-INS-5276
chr2-140762876-INS-102
chr2-140819817-INS-191
chr2-140819818-INS-86
chr2-140851428-INS-139
chr2-140851930-INS-1430
chr2-140861967-INS-6240
chr2-140876498-INS-50
chr2-140897497-INS-148
chr2-140898367-INS-64
chr2-140900738-INS-4759
chr2-140978658-INS-2837
chr2-140995677-INS-54
chr2-140998288-INS-51
chr2-141043185-INS-6505
chr2-141090015-INS-87
chr2-141148374-INS-64
chr2-141184007-INS-62
chr2-141195829-INS-768
chr2-141208503-INS-6891
chr2-141227391-INS-62
chr2-141308850-INS-4835
chr2-141313674-INS-6375
chr2-141320375-INS-1281
chr2-141355529-INS-412
chr2-141461059-INS-56
chr2-141462629-INS-735
chr2-141479303-INS-513
chr2-141488358-INS-923
chr2-141549301-INS-56
chr2-141550532-INS-51
chr2-141556509-INS-6616
chr2-141564033-INS-55
chr2-141568726-INS-50
chr2-141574281-INS-69
chr2-141591116-INS-74
chr2-141591301-INS-332
chr2-141608264-INS-50
chr2-141620317-INS-83
chr2-141625739-INS-2076
chr2-141640787-INS-84
chr2-141641007-INS-185
chr2-141652465-INS-6508
chr2-141655703-INS-277
chr2-141657090-INS-78
chr2-141661285-INS-52
chr2-141682971-INS-203
chr2-141765916-INS-3021
chr2-141799050-INS-400
chr2-141805704-INS-58
chr2-141875388-INS-395
chr2-141907961-INS-285
chr2-141976926-INS-90
chr2-142028402-INS-441
chr2-142069505-INS-98
chr2-142070172-INS-50

chr2-142079553-INS-720
chr2-142181606-INS-496
chr2-142216463-INS-89
chr2-142254034-INS-984
chr2-142258053-INS-74
chr2-142292151-INS-126
chr2-142304884-INS-139
chr2-142352529-INS-1713
chr2-142394836-INS-531
chr2-142412929-INS-171
chr2-142424408-INS-6625
chr2-142440603-INS-52
chr2-142450257-INS-6497
chr2-142481359-INS-90
chr2-142513758-INS-91
chr2-142530370-INS-76
chr2-142530507-INS-68
chr2-142574198-INS-60
chr2-142613504-INS-139
chr2-142620130-INS-118
chr2-142620175-INS-272
chr2-142664227-INS-204
chr2-142668305-INS-54
chr2-142691931-INS-468
chr2-142696794-INS-167
chr2-142703246-INS-136
chr2-142719849-INS-97
chr2-142729608-INS-80
chr2-142735324-INS-88
chr2-142740501-INS-187
chr2-142757390-INS-7175
chr2-142761184-INS-8985
chr2-142763700-INS-74
chr2-142767954-INS-66
chr2-142809730-INS-65
chr2-142824267-INS-56
chr2-142905849-INS-165
chr2-142906386-INS-204
chr2-142914291-INS-412
chr2-142946773-INS-144
chr2-142963727-INS-61
chr2-143013401-INS-72
chr2-143014258-INS-1016
chr2-143015710-INS-6233
chr2-143058107-INS-52
chr2-143064181-INS-8064
chr2-143070923-INS-121
chr2-143077115-INS-3475
chr2-143129754-INS-54
chr2-143171747-INS-1747
chr2-143172183-INS-65
chr2-143179434-INS-108
chr2-143187545-INS-88
chr2-143187639-INS-2989
chr2-143188415-INS-93
chr2-143193432-INS-5680
chr2-143237600-INS-90
chr2-143237634-INS-290
chr2-143306902-INS-6848
chr2-143313221-INS-304
chr2-143313312-INS-68
chr2-143368630-INS-60
chr2-143400835-INS-6403
chr2-143411621-INS-1236
chr2-143464516-INS-160
chr2-143474678-INS-67
chr2-143476379-INS-86
chr2-143537164-INS-154
chr2-143555548-INS-55
chr2-143606140-INS-3193
chr2-143606807-INS-68
chr2-143610761-INS-116
chr2-143667592-INS-52
chr2-143671542-INS-62
chr2-143671845-INS-85
chr2-143727257-INS-1367
chr2-143739263-INS-1309

chr2-143775347-INS-3128
chr2-143778825-INS-197
chr2-143805132-INS-241
chr2-143861517-INS-597
chr2-143880602-INS-91
chr2-143903291-INS-50
chr2-143908614-INS-54
chr2-143913612-INS-67
chr2-143923808-INS-234
chr2-143933630-INS-56
chr2-143967069-INS-115
chr2-144020468-INS-293
chr2-144056441-INS-50
chr2-144074792-INS-204
chr2-144080998-INS-51
chr2-144083326-INS-88
chr2-144083644-INS-216
chr2-144086448-INS-52
chr2-144124536-INS-197
chr2-144239728-INS-194
chr2-144276006-INS-52
chr2-144281447-INS-233
chr2-144299228-INS-517
chr2-144352569-INS-56
chr2-144424299-INS-391
chr2-144424299-INS-53
chr2-144442928-INS-320
chr2-144555198-INS-335
chr2-144560013-INS-80
chr2-144621305-INS-58
chr2-144682746-INS-6470
chr2-144690077-INS-85
chr2-144700645-INS-54
chr2-144714422-INS-138
chr2-144737475-INS-1998
chr2-144772516-INS-84
chr2-144804888-INS-106
chr2-144835055-INS-50
chr2-144929557-INS-88
chr2-144940254-INS-336
chr2-145063154-INS-6422
chr2-145096757-INS-2564
chr2-145166242-INS-554
chr2-145184174-INS-132
chr2-145198656-INS-2950
chr2-145327586-INS-126
chr2-145327651-INS-374
chr2-145379818-INS-52
chr2-145435480-INS-54
chr2-145440866-INS-104
chr2-145490072-INS-88
chr2-145490131-INS-103
chr2-145496788-INS-58
chr2-145497181-INS-1036
chr2-145536297-INS-2053
chr2-145537470-INS-8860
chr2-145540891-INS-182
chr2-145555337-INS-1767
chr2-145562670-INS-54
chr2-145593323-INS-1485
chr2-145704104-INS-52
chr2-145725881-INS-2561
chr2-145733305-INS-422
chr2-145854595-INS-543
chr2-145868269-INS-98
chr2-145882128-INS-6674
chr2-145918738-INS-52
chr2-145945819-INS-6440
chr2-145964529-INS-328
chr2-145964531-INS-698
chr2-146378133-INS-6261
chr2-146382613-INS-56
chr2-146394568-INS-52
chr2-146446380-INS-727
chr2-146450371-INS-331
chr2-146557898-INS-50
chr2-146571681-INS-6870

chr2-146601296-INS-2978
chr2-146622501-INS-569
chr2-146633578-INS-6829
chr2-146650095-INS-71
chr2-146685920-INS-112
chr2-146818969-INS-7071
chr2-146939866-INS-51
chr2-147008743-INS-5372
chr2-147018199-INS-120
chr2-147019042-INS-1711
chr2-147074851-INS-208
chr2-147111183-INS-248
chr2-147131310-INS-117
chr2-147135162-INS-1867
chr2-147135373-INS-2658
chr2-147164697-INS-140
chr2-147164795-INS-55
chr2-147171253-INS-120
chr2-147224970-INS-163
chr2-147315054-INS-92
chr2-147351305-INS-87
chr2-147351466-INS-485
chr2-147361863-INS-64
chr2-147364505-INS-343
chr2-147409605-INS-104
chr2-147529647-INS-62
chr2-147534431-INS-80
chr2-147541601-INS-5356
chr2-147567076-INS-54
chr2-147639660-INS-229
chr2-147639698-INS-56
chr2-147709653-INS-3849
chr2-147718018-INS-54
chr2-147744864-INS-67
chr2-147824936-INS-5856
chr2-147853307-INS-261
chr2-147854401-INS-212
chr2-147866969-INS-51
chr2-148014501-INS-289
chr2-148016567-INS-52
chr2-148155821-INS-54
chr2-148174240-INS-168
chr2-148354068-INS-1892
chr2-148382948-INS-62
chr2-148488890-INS-213
chr2-148583948-INS-52
chr2-148586412-INS-52
chr2-148607435-INS-1718
chr2-148611281-INS-72
chr2-148611999-INS-70
chr2-148616192-INS-60
chr2-148617239-INS-3295
chr2-148617914-INS-140
chr2-148629580-INS-325
chr2-148641631-INS-57
chr2-148698457-INS-3100
chr2-148803564-INS-70
chr2-148861952-INS-6467
chr2-148865022-INS-91
chr2-148870979-INS-1132
chr2-148903486-INS-6572
chr2-148933034-INS-70
chr2-148955467-INS-1607
chr2-148961230-INS-444
chr2-148971456-INS-2806
chr2-148972518-INS-962
chr2-148978603-INS-199
chr2-148995003-INS-368
chr2-149004240-INS-85
chr2-149006694-INS-56
chr2-149017059-INS-77
chr2-149020939-INS-3845
chr2-149021651-INS-752
chr2-149161597-INS-78
chr2-149163956-INS-58
chr2-149169454-INS-5681
chr2-149179719-INS-131

chr2-149192708-INS-52
chr2-149196040-INS-2406
chr2-149255791-INS-218
chr2-149260204-INS-977
chr2-149336213-INS-204
chr2-149336683-INS-60
chr2-149394721-INS-236
chr2-149466800-INS-3647
chr2-149483565-INS-60
chr2-149494878-INS-6587
chr2-149533441-INS-64
chr2-149575443-INS-5010
chr2-149697809-INS-6502
chr2-149744164-INS-4432
chr2-149758293-INS-200
chr2-149821842-INS-6597
chr2-149948562-INS-88
chr2-149990426-INS-5604
chr2-149995284-INS-3021
chr2-149996166-INS-8687
chr2-150011076-INS-5177
chr2-150047265-INS-1596
chr2-150058231-INS-58
chr2-150076985-INS-4107
chr2-150110338-INS-253
chr2-150110356-INS-84
chr2-150176507-INS-3534
chr2-150179970-INS-1876
chr2-150179970-INS-399
chr2-150188237-INS-4113
chr2-150192537-INS-1931
chr2-150207090-INS-270
chr2-150217502-INS-1285
chr2-150237506-INS-511
chr2-150248467-INS-52
chr2-150255762-INS-4037
chr2-150263807-INS-1541
chr2-150301614-INS-4848
chr2-150405175-INS-89
chr2-150405257-INS-185
chr2-150483132-INS-378
chr2-150619620-INS-84
chr2-150643771-INS-244
chr2-150663779-INS-325
chr2-151014682-INS-484
chr2-151130577-INS-2841
chr2-151168994-INS-1014
chr2-151185027-INS-201
chr2-151329741-INS-50
chr2-151367420-INS-225
chr2-151481977-INS-50
chr2-151485429-INS-1885
chr2-151506062-INS-413
chr2-151588574-INS-82
chr2-151588637-INS-394
chr2-151623692-INS-64
chr2-151623699-INS-320
chr2-151648533-INS-56
chr2-151667335-INS-88
chr2-151697856-INS-60
chr2-151704389-INS-324
chr2-151787536-INS-130
chr2-151789059-INS-85
chr2-151865605-INS-96
chr2-151883552-INS-118
chr2-151884863-INS-183
chr2-151885356-INS-112
chr2-151901468-INS-67
chr2-151976298-INS-163
chr2-152148789-INS-312
chr2-152158754-INS-212
chr2-152182877-INS-103
chr2-152185599-INS-96
chr2-152193540-INS-199
chr2-152221280-INS-206
chr2-152231395-INS-166
chr2-152231778-INS-190

chr2-152234148-INS-218
chr2-152235026-INS-70
chr2-152262496-INS-176
chr2-152267973-INS-165
chr2-152289215-INS-232
chr2-152296637-INS-57
chr2-152302998-INS-770
chr2-152392013-INS-656
chr2-152398080-INS-694
chr2-152419840-INS-2822
chr2-152494195-INS-50
chr2-152610244-INS-131
chr2-152628527-INS-98
chr2-152665203-INS-68
chr2-152707881-INS-108
chr2-152743109-INS-58
chr2-152744642-INS-273
chr2-152754930-INS-160
chr2-152757715-INS-56
chr2-152761596-INS-52
chr2-152771524-INS-136
chr2-152827533-INS-72
chr2-152840095-INS-75
chr2-152863814-INS-50
chr2-152911301-INS-54
chr2-152921681-INS-51
chr2-152924801-INS-6475
chr2-152938273-INS-550
chr2-152964180-INS-58
chr2-152967395-INS-61
chr2-152997441-INS-235
chr2-153099055-INS-5549
chr2-153114950-INS-50
chr2-153117885-INS-484
chr2-153123896-INS-299
chr2-153140003-INS-207
chr2-153140950-INS-213
chr2-153152940-INS-257
chr2-153153314-INS-459
chr2-153177347-INS-2319
chr2-153178011-INS-214
chr2-153222399-INS-119
chr2-153238120-INS-541
chr2-153268665-INS-52
chr2-153376645-INS-213
chr2-153430231-INS-181
chr2-153430231-INS-70
chr2-153432479-INS-80
chr2-153440227-INS-54
chr2-153441059-INS-7442
chr2-153466798-INS-59
chr2-153477670-INS-201
chr2-153501872-INS-187
chr2-153549799-INS-68
chr2-153593165-INS-133
chr2-153676168-INS-200
chr2-153677559-INS-91
chr2-153699404-INS-140
chr2-153699893-INS-1866
chr2-153781114-INS-76
chr2-153781174-INS-431
chr2-153793637-INS-7091
chr2-153809717-INS-6857
chr2-153820872-INS-210
chr2-153907920-INS-4322
chr2-153918709-INS-54
chr2-153970305-INS-1564
chr2-154015967-INS-346
chr2-154028782-INS-166
chr2-154049630-INS-147
chr2-154078927-INS-168
chr2-154079327-INS-64
chr2-154079450-INS-287
chr2-154089849-INS-2254
chr2-154135070-INS-56
chr2-154147714-INS-7198
chr2-154172860-INS-7434

chr2-154174766-INS-116
chr2-154174766-INS-392
chr2-154204761-INS-210
chr2-154205428-INS-481
chr2-154229950-INS-200
chr2-154233257-INS-64
chr2-154236083-INS-78
chr2-154254891-INS-75
chr2-154260790-INS-215
chr2-154265584-INS-68
chr2-154266421-INS-140
chr2-154283502-INS-59
chr2-154288249-INS-213
chr2-154291256-INS-363
chr2-154291270-INS-68
chr2-154312557-INS-525
chr2-154317328-INS-106
chr2-154320261-INS-426
chr2-154320261-INS-59
chr2-154320281-INS-177
chr2-154453678-INS-195
chr2-154458459-INS-79
chr2-154458879-INS-260
chr2-154484180-INS-87
chr2-154502075-INS-63
chr2-154506837-INS-60
chr2-154517064-INS-58
chr2-154522402-INS-1750
chr2-154535734-INS-104
chr2-154555151-INS-193
chr2-154562557-INS-90
chr2-154584251-INS-156
chr2-154593321-INS-54
chr2-154636687-INS-225
chr2-154668512-INS-221
chr2-154676930-INS-174
chr2-154682542-INS-200
chr2-154693301-INS-214
chr2-154753085-INS-64
chr2-154761672-INS-62
chr2-154769933-INS-129
chr2-154791856-INS-213
chr2-154800667-INS-50
chr2-154818733-INS-114
chr2-154818960-INS-587
chr2-154881023-INS-301
chr2-154891878-INS-191
chr2-154927153-INS-192
chr2-154927927-INS-212
chr2-154989232-INS-204
chr2-155000781-INS-54
chr2-155009616-INS-231
chr2-155022853-INS-203
chr2-155042833-INS-190
chr2-155085122-INS-694
chr2-155101155-INS-191
chr2-155152390-INS-226
chr2-155216434-INS-84
chr2-155269801-INS-817
chr2-155387977-INS-134
chr2-155494688-INS-84
chr2-155600093-INS-4600
chr2-155681837-INS-97
chr2-155709120-INS-448
chr2-155762494-INS-212
chr2-155795587-INS-729
chr2-155797481-INS-81
chr2-155797892-INS-221
chr2-155839677-INS-67
chr2-155863904-INS-214
chr2-155892804-INS-705
chr2-155899504-INS-810
chr2-155900154-INS-238
chr2-155998621-INS-62
chr2-156007835-INS-76
chr2-156031883-INS-75
chr2-156034925-INS-179

chr2-156046440-INS-50
chr2-156165307-INS-580
chr2-156170893-INS-176
chr2-156190079-INS-50
chr2-156196861-INS-68
chr2-156196957-INS-138
chr2-156239316-INS-179
chr2-156252061-INS-125
chr2-156252111-INS-300
chr2-156326158-INS-294
chr2-156358816-INS-314
chr2-156358818-INS-154
chr2-156401267-INS-55
chr2-156444623-INS-78
chr2-156498811-INS-3420
chr2-156552141-INS-496
chr2-156560538-INS-50
chr2-156561426-INS-61
chr2-156640005-INS-1774
chr2-156646183-INS-201
chr2-156654036-INS-75
chr2-156668688-INS-212
chr2-156679336-INS-180
chr2-156709876-INS-66
chr2-156731335-INS-56
chr2-156734410-INS-52
chr2-156742605-INS-108
chr2-156755331-INS-7186
chr2-156756752-INS-2596
chr2-156762264-INS-173
chr2-156794962-INS-1877
chr2-156807432-INS-104
chr2-156874600-INS-56
chr2-156874694-INS-186
chr2-156913000-INS-64
chr2-156932528-INS-222
chr2-156952163-INS-52
chr2-156960283-INS-876
chr2-156999078-INS-137
chr2-157066405-INS-643
chr2-157073816-INS-243
chr2-157073816-INS-51
chr2-157123367-INS-220
chr2-157134673-INS-108
chr2-157140741-INS-122
chr2-157153845-INS-333
chr2-157168444-INS-72
chr2-157169709-INS-116
chr2-157180217-INS-346
chr2-157181140-INS-129
chr2-157225073-INS-216
chr2-157226510-INS-210
chr2-157236708-INS-190
chr2-157323415-INS-50
chr2-157365764-INS-84
chr2-157385627-INS-63
chr2-157386213-INS-1728
chr2-157409881-INS-92
chr2-157411078-INS-455
chr2-157440627-INS-90
chr2-157469055-INS-54
chr2-157483971-INS-13247
chr2-157490524-INS-106
chr2-157531140-INS-128
chr2-157571982-INS-516
chr2-157622292-INS-114
chr2-157679074-INS-1301
chr2-157711165-INS-56
chr2-157801896-INS-67
chr2-157807813-INS-74
chr2-157809045-INS-87
chr2-157809073-INS-435
chr2-157818946-INS-52
chr2-157867988-INS-73
chr2-157893014-INS-150
chr2-157943446-INS-161
chr2-157997392-INS-64

chr2-158005807-INS-182
chr2-158031975-INS-69
chr2-158049015-INS-294
chr2-158049030-INS-129
chr2-158066897-INS-70
chr2-158084787-INS-50
chr2-158092940-INS-57
chr2-158112464-INS-7481
chr2-158115435-INS-74
chr2-158171891-INS-68
chr2-158175461-INS-174
chr2-158180111-INS-1750
chr2-158182378-INS-214
chr2-158194999-INS-284
chr2-158225122-INS-56
chr2-158225940-INS-221
chr2-158236788-INS-205
chr2-158242916-INS-105
chr2-158243927-INS-409
chr2-158288425-INS-226
chr2-158371917-INS-50
chr2-158382540-INS-114
chr2-158387722-INS-208
chr2-158406785-INS-106
chr2-158409282-INS-703
chr2-158419950-INS-131
chr2-158482600-INS-7236
chr2-158487649-INS-51
chr2-158501982-INS-112
chr2-158516431-INS-50
chr2-158546026-INS-51
chr2-158546178-INS-87
chr2-158588772-INS-98
chr2-158589362-INS-168
chr2-158620230-INS-271
chr2-158644131-INS-3608
chr2-158644596-INS-360
chr2-158650123-INS-181
chr2-158650609-INS-60
chr2-158650716-INS-391
chr2-158683029-INS-306
chr2-158707573-INS-229
chr2-158738923-INS-50
chr2-158738965-INS-428
chr2-158766527-INS-58
chr2-158789093-INS-126
chr2-158835963-INS-94
chr2-158893543-INS-64
chr2-158901450-INS-610
chr2-158941887-INS-71
chr2-159018232-INS-6606
chr2-159018598-INS-69
chr2-159064962-INS-58
chr2-159067183-INS-5085
chr2-159139145-INS-1796
chr2-159268193-INS-6446
chr2-159282052-INS-52
chr2-159320095-INS-174
chr2-159322094-INS-56
chr2-159325392-INS-58
chr2-159387490-INS-211
chr2-159513977-INS-1010
chr2-159682142-INS-60
chr2-159682336-INS-200
chr2-159685805-INS-81
chr2-159695957-INS-7084
chr2-159705502-INS-54
chr2-159721389-INS-56
chr2-159721439-INS-134
chr2-159723394-INS-96
chr2-159723434-INS-221
chr2-159755184-INS-61
chr2-159789493-INS-206
chr2-159789689-INS-266
chr2-159821028-INS-1894
chr2-159864821-INS-382
chr2-159885179-INS-188

chr2-159925683-INS-6798
chr2-159930873-INS-2134
chr2-159933955-INS-76
chr2-159944616-INS-80
chr2-159981481-INS-55
chr2-159981852-INS-80
chr2-160001644-INS-87
chr2-160023075-INS-80
chr2-160101519-INS-210
chr2-160113069-INS-100
chr2-160115401-INS-6341
chr2-160119024-INS-134
chr2-160204709-INS-56
chr2-160244202-INS-65
chr2-160291663-INS-152
chr2-160296172-INS-3020
chr2-160309348-INS-116
chr2-160314331-INS-948
chr2-160360580-INS-105
chr2-160419805-INS-78
chr2-160435545-INS-1117
chr2-160449344-INS-116
chr2-160453371-INS-50
chr2-160471938-INS-52
chr2-160484824-INS-66
chr2-160501202-INS-195
chr2-160653554-INS-80
chr2-160654350-INS-59
chr2-160763552-INS-62
chr2-160769833-INS-117
chr2-160777687-INS-81
chr2-160781089-INS-56
chr2-160799550-INS-303
chr2-160932506-INS-77
chr2-160961464-INS-11461
chr2-160984842-INS-54
chr2-160998137-INS-176
chr2-161020882-INS-1153
chr2-161041437-INS-4325
chr2-161069917-INS-1133
chr2-161116347-INS-151
chr2-161121540-INS-498
chr2-161123737-INS-220
chr2-161251382-INS-87
chr2-161267762-INS-6780
chr2-161320096-INS-92
chr2-161356680-INS-658
chr2-161438974-INS-192
chr2-161477642-INS-6676
chr2-161505066-INS-74
chr2-161585456-INS-6242
chr2-161591227-INS-55
chr2-161613579-INS-285
chr2-161613579-INS-77
chr2-161636038-INS-56
chr2-161666676-INS-100
chr2-161716316-INS-134
chr2-161751196-INS-75
chr2-161761904-INS-135
chr2-161762174-INS-745
chr2-161767552-INS-400
chr2-161792797-INS-1632
chr2-161835706-INS-1115
chr2-161839051-INS-213
chr2-161946123-INS-127
chr2-162071044-INS-299
chr2-162071060-INS-678
chr2-162089732-INS-60
chr2-162123818-INS-7099
chr2-162150475-INS-50
chr2-162188459-INS-63
chr2-162200544-INS-1206
chr2-162241868-INS-1368
chr2-162285792-INS-166
chr2-162291630-INS-78
chr2-162293174-INS-1331
chr2-162400427-INS-5092

chr2-162461401-INS-52
chr2-162475782-INS-106
chr2-162488643-INS-176
chr2-162514008-INS-52
chr2-162514890-INS-59
chr2-162662006-INS-158
chr2-162687854-INS-6196
chr2-162694443-INS-6285
chr2-162695324-INS-61
chr2-162705761-INS-53
chr2-162725783-INS-51
chr2-162736430-INS-52
chr2-162769581-INS-291
chr2-162797700-INS-252
chr2-162875469-INS-8018
chr2-162879712-INS-334
chr2-162909719-INS-8585
chr2-162940334-INS-199
chr2-162960106-INS-160
chr2-162960371-INS-68
chr2-162992841-INS-98
chr2-163068952-INS-88
chr2-163098283-INS-51
chr2-163144785-INS-531
chr2-163154782-INS-147
chr2-163172460-INS-63
chr2-163186089-INS-51
chr2-163186867-INS-52
chr2-163219767-INS-59
chr2-163241058-INS-52
chr2-163241117-INS-113
chr2-163251728-INS-1055
chr2-163271915-INS-228
chr2-163308427-INS-50
chr2-163308978-INS-222
chr2-163320947-INS-499
chr2-163339575-INS-168
chr2-163370118-INS-87
chr2-163439617-INS-258
chr2-163440236-INS-50
chr2-163440481-INS-167
chr2-163441442-INS-548
chr2-163480823-INS-98
chr2-163484572-INS-54
chr2-163524126-INS-180
chr2-163559974-INS-56
chr2-163601397-INS-222
chr2-163601418-INS-68
chr2-163601719-INS-96
chr2-163604015-INS-53
chr2-163604101-INS-123
chr2-163608680-INS-129
chr2-163615768-INS-96
chr2-163652113-INS-52
chr2-163699292-INS-396
chr2-163765613-INS-50
chr2-163813596-INS-268
chr2-163814008-INS-145
chr2-163827983-INS-75
chr2-163832797-INS-5622
chr2-163848002-INS-50
chr2-163852197-INS-1737
chr2-163877308-INS-60
chr2-163882557-INS-89
chr2-163888836-INS-129
chr2-163889872-INS-174
chr2-163890607-INS-136
chr2-163899624-INS-115
chr2-163899674-INS-264
chr2-163940161-INS-1748
chr2-164036562-INS-1653
chr2-164046313-INS-6239
chr2-164053422-INS-4275
chr2-164056213-INS-476
chr2-164070644-INS-6329
chr2-164170397-INS-2184
chr2-164216675-INS-78

chr2-164218592-INS-2927
chr2-164229780-INS-50
chr2-164240425-INS-199
chr2-164240431-INS-1454
chr2-164240611-INS-73
chr2-164241936-INS-7112
chr2-164341590-INS-282
chr2-164341590-INS-94
chr2-164362007-INS-50
chr2-164363304-INS-376
chr2-164400052-INS-101
chr2-164402426-INS-57
chr2-164495478-INS-54
chr2-164496461-INS-52
chr2-164531201-INS-874
chr2-164539482-INS-52
chr2-164545169-INS-58
chr2-164546948-INS-302
chr2-164547213-INS-1872
chr2-164547253-INS-928
chr2-164556083-INS-59
chr2-164568598-INS-292
chr2-164568863-INS-358
chr2-164578445-INS-195
chr2-164583379-INS-336
chr2-164591894-INS-50
chr2-164620824-INS-244
chr2-164681340-INS-1000
chr2-164702074-INS-454
chr2-164708491-INS-52
chr2-164717884-INS-201
chr2-164742731-INS-256
chr2-164766994-INS-208
chr2-164909359-INS-76
chr2-164915294-INS-56
chr2-164917620-INS-50
chr2-164924005-INS-106
chr2-164937293-INS-59
chr2-164937564-INS-267
chr2-164977445-INS-54
chr2-164995850-INS-249
chr2-164996470-INS-52
chr2-165021790-INS-170
chr2-165021981-INS-74
chr2-165073079-INS-50
chr2-165107608-INS-274
chr2-165107763-INS-107
chr2-165114426-INS-1897
chr2-165126714-INS-215
chr2-165132746-INS-53
chr2-165182566-INS-74
chr2-165184692-INS-84
chr2-165188183-INS-372
chr2-165197768-INS-1112
chr2-165226910-INS-1356
chr2-165230858-INS-206
chr2-165265532-INS-350
chr2-165277254-INS-357
chr2-165307477-INS-210
chr2-165321683-INS-62
chr2-165336383-INS-77
chr2-165355961-INS-508
chr2-165361696-INS-50
chr2-165382795-INS-357
chr2-165384395-INS-200
chr2-165408469-INS-66
chr2-165422056-INS-329
chr2-165422590-INS-113
chr2-165450868-INS-54
chr2-165461521-INS-151
chr2-165469546-INS-81
chr2-165469776-INS-118
chr2-165502472-INS-827
chr2-165502933-INS-59
chr2-165551954-INS-492
chr2-165577822-INS-175
chr2-165586130-INS-106

chr2-165604454-INS-75
chr2-165618973-INS-510
chr2-165619084-INS-76
chr2-165638365-INS-332
chr2-165638591-INS-1577
chr2-165644001-INS-408
chr2-165675114-INS-56
chr2-165688777-INS-84
chr2-165689777-INS-188
chr2-165734032-INS-781
chr2-165749939-INS-58
chr2-165754242-INS-186
chr2-165754321-INS-199
chr2-165766841-INS-231
chr2-165766939-INS-61
chr2-165777039-INS-134
chr2-165780838-INS-52
chr2-165784222-INS-197
chr2-165784569-INS-504
chr2-165784595-INS-114
chr2-165789691-INS-119
chr2-165791341-INS-65
chr2-165797633-INS-218
chr2-165803296-INS-207
chr2-165816878-INS-78
chr2-165832160-INS-206
chr2-165930661-INS-174
chr2-166003793-INS-94
chr2-166013016-INS-52
chr2-166077891-INS-54
chr2-166106149-INS-50
chr2-166114463-INS-150
chr2-166130856-INS-7096
chr2-166193357-INS-358
chr2-166200503-INS-493
chr2-166209862-INS-96
chr2-166218106-INS-50
chr2-166224711-INS-390
chr2-166244662-INS-149
chr2-166257828-INS-255
chr2-166266232-INS-470
chr2-166270660-INS-117
chr2-166289547-INS-440
chr2-166290088-INS-292
chr2-166309090-INS-72
chr2-166312596-INS-1123
chr2-166328385-INS-291
chr2-166335395-INS-92
chr2-166347099-INS-50
chr2-166353783-INS-1384
chr2-166390916-INS-86
chr2-166402403-INS-74
chr2-166404629-INS-69
chr2-166421061-INS-139
chr2-166449719-INS-66
chr2-166475715-INS-54
chr2-166490842-INS-52
chr2-166501404-INS-71
chr2-166510636-INS-210
chr2-166544645-INS-270
chr2-166557717-INS-254
chr2-166580765-INS-106
chr2-166586127-INS-67
chr2-166626995-INS-2867
chr2-166629388-INS-102
chr2-166632350-INS-209
chr2-166665147-INS-4035
chr2-166689520-INS-145
chr2-166698937-INS-59
chr2-166721796-INS-129
chr2-166725853-INS-174
chr2-166752149-INS-179
chr2-166770308-INS-225
chr2-166785885-INS-333
chr2-166842048-INS-92
chr2-166856549-INS-2155
chr2-166874526-INS-69

chr2-166919909-INS-57
chr2-166922685-INS-50
chr2-166926830-INS-226
chr2-166932915-INS-132
chr2-166948710-INS-66
chr2-166999169-INS-72
chr2-167042604-INS-248
chr2-167081162-INS-148
chr2-167082781-INS-51
chr2-167106672-INS-52
chr2-167111329-INS-795
chr2-167123235-INS-64
chr2-167164461-INS-154
chr2-167206040-INS-75
chr2-167212644-INS-65
chr2-167213100-INS-62
chr2-167220952-INS-433
chr2-167240656-INS-127
chr2-167240937-INS-367
chr2-167248325-INS-1268
chr2-167270391-INS-238
chr2-167297226-INS-302
chr2-167299981-INS-64
chr2-167348436-INS-200
chr2-167369624-INS-96
chr2-167370050-INS-228
chr2-167432053-INS-155
chr2-167440810-INS-402
chr2-167505135-INS-183
chr2-167511784-INS-72
chr2-167524836-INS-244
chr2-167543780-INS-116
chr2-167549612-INS-497
chr2-167555229-INS-56
chr2-167573839-INS-72
chr2-167581962-INS-76
chr2-167583298-INS-383
chr2-167632554-INS-70
chr2-167633252-INS-171
chr2-167634080-INS-216
chr2-167658258-INS-200
chr2-167658708-INS-56
chr2-167672026-INS-454
chr2-167718894-INS-608
chr2-167738016-INS-2886
chr2-167746615-INS-94
chr2-167755646-INS-130
chr2-167757750-INS-1060
chr2-167805144-INS-61
chr2-167810215-INS-169
chr2-167846767-INS-382
chr2-167856776-INS-561
chr2-167860732-INS-52
chr2-167872168-INS-319
chr2-167929810-INS-50
chr2-167954747-INS-52
chr2-167954994-INS-168
chr2-167979969-INS-96
chr2-167992961-INS-209
chr2-167995325-INS-72
chr2-167996789-INS-62
chr2-167998679-INS-310
chr2-168004949-INS-331
chr2-168028965-INS-208
chr2-168086646-INS-60
chr2-168126667-INS-62
chr2-168126908-INS-95
chr2-168142935-INS-169
chr2-168146412-INS-164
chr2-168165785-INS-511
chr2-168168002-INS-92
chr2-168177382-INS-65
chr2-168193779-INS-116
chr2-168215454-INS-6692
chr2-168252152-INS-105
chr2-168252564-INS-288
chr2-168344868-INS-141

chr2-168349526-INS-299
chr2-168385478-INS-100
chr2-168410603-INS-162
chr2-168439206-INS-68
chr2-168461321-INS-297
chr2-168480638-INS-58
chr2-168487142-INS-239
chr2-168503507-INS-50
chr2-168507603-INS-212
chr2-168508436-INS-5544
chr2-168522186-INS-161
chr2-168533442-INS-74
chr2-168556029-INS-4328
chr2-168565970-INS-458
chr2-168572124-INS-58
chr2-168578740-INS-217
chr2-168585595-INS-8985
chr2-168589084-INS-152
chr2-168601250-INS-148
chr2-168603858-INS-194
chr2-168644089-INS-150
chr2-168648439-INS-78
chr2-168665871-INS-50
chr2-168674716-INS-108
chr2-168674997-INS-70
chr2-168675548-INS-70
chr2-168676551-INS-1795
chr2-168677786-INS-135
chr2-168690386-INS-459
chr2-168691807-INS-51
chr2-168697075-INS-118
chr2-168705097-INS-215
chr2-168715651-INS-58
chr2-168728742-INS-54
chr2-168731976-INS-236
chr2-168734105-INS-7080
chr2-168738191-INS-51
chr2-168755953-INS-151
chr2-168792820-INS-306
chr2-168792820-INS-51
chr2-168813314-INS-50
chr2-168844106-INS-52
chr2-168855784-INS-64
chr2-168862785-INS-226
chr2-168881655-INS-56
chr2-168884628-INS-1495
chr2-168923925-INS-104
chr2-168954807-INS-622
chr2-168957521-INS-72
chr2-168986291-INS-162
chr2-168986977-INS-82
chr2-168993017-INS-77
chr2-169004822-INS-406
chr2-169005069-INS-457
chr2-169016169-INS-99
chr2-169025712-INS-50
chr2-169041985-INS-1000
chr2-169091066-INS-177
chr2-169091505-INS-290
chr2-169105317-INS-103
chr2-169112449-INS-118
chr2-169127126-INS-132
chr2-169141088-INS-255
chr2-169163152-INS-6292
chr2-169177447-INS-75
chr2-169218400-INS-50
chr2-169222867-INS-57
chr2-169225197-INS-50
chr2-169249330-INS-128
chr2-169257335-INS-326
chr2-169321101-INS-1153
chr2-169321862-INS-80
chr2-169364304-INS-157
chr2-169369314-INS-50
chr2-169382085-INS-66
chr2-169382757-INS-96
chr2-169411385-INS-82

chr2-169437073-INS-79
chr2-169440853-INS-146
chr2-169449625-INS-66
chr2-169472440-INS-121
chr2-169472440-INS-270
chr2-169485902-INS-155
chr2-169486967-INS-687
chr2-169487206-INS-220
chr2-169487217-INS-87
chr2-169521303-INS-6695
chr2-169573853-INS-78
chr2-169598749-INS-214
chr2-169619036-INS-52
chr2-169648930-INS-52
chr2-169654952-INS-1898
chr2-169658756-INS-186
chr2-169699328-INS-220
chr2-169700897-INS-218
chr2-169700979-INS-103
chr2-169717011-INS-269
chr2-169731060-INS-50
chr2-169746444-INS-197
chr2-169746613-INS-409
chr2-169781965-INS-67
chr2-169789597-INS-69
chr2-169789731-INS-242
chr2-169795581-INS-260
chr2-169806541-INS-155
chr2-169817698-INS-51
chr2-169832771-INS-815
chr2-169846483-INS-50
chr2-169850129-INS-94
chr2-169869015-INS-216
chr2-169873411-INS-52
chr2-169873542-INS-2008
chr2-169880195-INS-100
chr2-169880408-INS-89
chr2-169896980-INS-267
chr2-169900374-INS-66
chr2-169926034-INS-96
chr2-169965687-INS-50
chr2-169982174-INS-213
chr2-170010812-INS-158
chr2-170010873-INS-50
chr2-170025300-INS-51
chr2-170037961-INS-80
chr2-170038935-INS-54
chr2-170046306-INS-404
chr2-170067207-INS-7214
chr2-170068701-INS-376
chr2-170073916-INS-177
chr2-170085778-INS-947
chr2-170106631-INS-279
chr2-170113226-INS-202
chr2-170118558-INS-1034
chr2-170121502-INS-650
chr2-170132347-INS-288
chr2-170140759-INS-117
chr2-170168689-INS-56
chr2-170173069-INS-78
chr2-170179869-INS-6473
chr2-170191786-INS-60
chr2-170193971-INS-178
chr2-170221405-INS-442
chr2-170261367-INS-78
chr2-170273868-INS-232
chr2-170275498-INS-96
chr2-170276267-INS-120
chr2-170294540-INS-1993
chr2-170300620-INS-773
chr2-170320121-INS-167
chr2-170335349-INS-50
chr2-170346366-INS-711
chr2-170346443-INS-113
chr2-170347074-INS-2544
chr2-170374472-INS-8417
chr2-170389221-INS-76

chr2-170399022-INS-5151
chr2-170403491-INS-56
chr2-170407746-INS-126
chr2-170413496-INS-546
chr2-170414596-INS-131
chr2-170426698-INS-336
chr2-170457206-INS-407
chr2-170475295-INS-93
chr2-170476862-INS-82
chr2-170501926-INS-52
chr2-170522416-INS-204
chr2-170541698-INS-240
chr2-170548768-INS-3759
chr2-170563259-INS-72
chr2-170601048-INS-114
chr2-170601176-INS-69
chr2-170604802-INS-101
chr2-170605275-INS-228
chr2-170605288-INS-106
chr2-170606418-INS-4741
chr2-170651646-INS-743
chr2-170656577-INS-74
chr2-170666648-INS-66
chr2-170667114-INS-132
chr2-170684373-INS-110
chr2-170714261-INS-71
chr2-170714417-INS-64
chr2-170714730-INS-617
chr2-170728022-INS-62
chr2-170732750-INS-1188
chr2-170744078-INS-137
chr2-170781450-INS-6618
chr2-170795825-INS-56
chr2-170814230-INS-197
chr2-170815636-INS-90
chr2-170846193-INS-68
chr2-170847095-INS-63
chr2-170867399-INS-91
chr2-170876695-INS-116
chr2-170877170-INS-184
chr2-170878259-INS-63
chr2-170884220-INS-333
chr2-170884931-INS-160
chr2-170891845-INS-4794
chr2-170901220-INS-92
chr2-170920293-INS-392
chr2-170920524-INS-198
chr2-170932916-INS-67
chr2-170936445-INS-144
chr2-170966867-INS-50
chr2-171005992-INS-240
chr2-171010591-INS-323
chr2-171035363-INS-1310
chr2-171086324-INS-1231
chr2-171093864-INS-124
chr2-171096166-INS-87
chr2-171103756-INS-84
chr2-171114086-INS-54
chr2-171137670-INS-54
chr2-171138384-INS-58
chr2-171139992-INS-449
chr2-171141616-INS-98
chr2-171157639-INS-1144
chr2-171182158-INS-194
chr2-171199034-INS-97
chr2-171220292-INS-207
chr2-171236299-INS-2089
chr2-171236587-INS-1015
chr2-171236926-INS-4298
chr2-171240995-INS-64
chr2-171253674-INS-69
chr2-171262222-INS-323
chr2-171265562-INS-673
chr2-171272179-INS-7019
chr2-171274610-INS-507
chr2-171276224-INS-56
chr2-171280814-INS-52

chr2-171285381-INS-109
chr2-171289437-INS-52
chr2-171290270-INS-56
chr2-171291474-INS-50
chr2-171303113-INS-585
chr2-171311243-INS-891
chr2-171340423-INS-110
chr2-171347944-INS-165
chr2-171359865-INS-196
chr2-171368217-INS-341
chr2-171368597-INS-184
chr2-171368605-INS-91
chr2-171410064-INS-380
chr2-171410207-INS-100
chr2-171463254-INS-60
chr2-171463541-INS-689
chr2-171507127-INS-63
chr2-171507249-INS-405
chr2-171513625-INS-620
chr2-171526063-INS-134
chr2-171526155-INS-666
chr2-171531330-INS-51
chr2-171565868-INS-64
chr2-171567982-INS-6698
chr2-171575797-INS-80
chr2-171582026-INS-60
chr2-171597823-INS-62
chr2-171645350-INS-186
chr2-171655683-INS-129
chr2-171657732-INS-1220
chr2-171658600-INS-54
chr2-171659039-INS-55
chr2-171667491-INS-4002
chr2-171674829-INS-210
chr2-171692138-INS-6379
chr2-171728290-INS-4910
chr2-171749426-INS-50
chr2-171751213-INS-593
chr2-171833272-INS-399
chr2-171847492-INS-164
chr2-171865745-INS-196
chr2-171893350-INS-77
chr2-171908336-INS-1840
chr2-171913455-INS-82
chr2-171988825-INS-149
chr2-171988886-INS-566
chr2-171994717-INS-196
chr2-172017176-INS-240
chr2-172045883-INS-50
chr2-172046205-INS-9354
chr2-172048113-INS-147
chr2-172048113-INS-60
chr2-172051832-INS-281
chr2-172064524-INS-97
chr2-172074735-INS-1372
chr2-172099385-INS-1441
chr2-172104952-INS-190
chr2-172120796-INS-52
chr2-172136558-INS-89
chr2-172149713-INS-1254
chr2-172151255-INS-50
chr2-172168052-INS-276
chr2-172168052-INS-62
chr2-172168554-INS-62
chr2-172169416-INS-68
chr2-172172019-INS-155
chr2-172176309-INS-184
chr2-172208716-INS-67
chr2-172284973-INS-52
chr2-172286270-INS-1096
chr2-172286278-INS-3285
chr2-172286503-INS-385
chr2-172302063-INS-52329
chr2-172309056-INS-95
chr2-172312438-INS-200
chr2-172318451-INS-80
chr2-172325004-INS-51

chr2-172325100-INS-148
chr2-172325611-INS-269
chr2-172328299-INS-240
chr2-172335593-INS-448
chr2-172342392-INS-102
chr2-172343332-INS-731
chr2-172366026-INS-52
chr2-172374821-INS-194
chr2-172374882-INS-429
chr2-172381096-INS-50
chr2-172384499-INS-142
chr2-172424881-INS-76
chr2-172430066-INS-68
chr2-172434382-INS-188
chr2-172466834-INS-61
chr2-172474930-INS-213
chr2-172475492-INS-311
chr2-172480053-INS-234
chr2-172483541-INS-63
chr2-172490345-INS-56
chr2-172506697-INS-219
chr2-172515006-INS-143
chr2-172516012-INS-63
chr2-172516128-INS-348
chr2-172518382-INS-62
chr2-172518766-INS-64
chr2-172525550-INS-254
chr2-172553807-INS-2600
chr2-172559389-INS-412
chr2-172565269-INS-124
chr2-172575214-INS-114
chr2-172594116-INS-4161
chr2-172595266-INS-66
chr2-172604656-INS-383
chr2-172604893-INS-51
chr2-172644688-INS-394
chr2-172677021-INS-140
chr2-172677137-INS-60
chr2-172683742-INS-162
chr2-172692695-INS-6499
chr2-172698406-INS-248
chr2-172762844-INS-72
chr2-172763540-INS-144
chr2-172763566-INS-52
chr2-172777946-INS-117
chr2-172790125-INS-144
chr2-172790149-INS-1686
chr2-172790339-INS-201
chr2-172808803-INS-89
chr2-172818225-INS-59
chr2-172818234-INS-133
chr2-172837232-INS-190
chr2-172920325-INS-68
chr2-172924156-INS-106
chr2-172924520-INS-104
chr2-172924621-INS-91
chr2-172925813-INS-280
chr2-172937774-INS-50
chr2-172962135-INS-210
chr2-172990234-INS-62
chr2-172999598-INS-59
chr2-173004970-INS-189
chr2-173006155-INS-95
chr2-173044200-INS-9892
chr2-173055391-INS-759
chr2-173061781-INS-441
chr2-173062321-INS-210
chr2-173065166-INS-3065
chr2-173085275-INS-56
chr2-173154409-INS-96
chr2-173156476-INS-52
chr2-173171766-INS-90
chr2-173187538-INS-68
chr2-173188638-INS-67
chr2-173190318-INS-60
chr2-173198051-INS-199
chr2-173212403-INS-79

chr2-173212428-INS-1989
chr2-173220553-INS-51
chr2-173223009-INS-84
chr2-173237996-INS-52
chr2-173246327-INS-50
chr2-173250380-INS-2279
chr2-173257655-INS-51
chr2-173292398-INS-70
chr2-173320238-INS-102
chr2-173327960-INS-187
chr2-173365306-INS-316
chr2-173368096-INS-151
chr2-173368099-INS-55
chr2-173379342-INS-5020
chr2-173384232-INS-360
chr2-173386968-INS-50
chr2-173462933-INS-66
chr2-173477827-INS-296
chr2-173541940-INS-105
chr2-173542189-INS-140
chr2-173549497-INS-2550
chr2-173565483-INS-825
chr2-173566309-INS-256
chr2-173575320-INS-120
chr2-173593797-INS-56
chr2-173598020-INS-72
chr2-173603640-INS-68
chr2-173612119-INS-64
chr2-173680980-INS-66
chr2-173703300-INS-288
chr2-173809828-INS-61
chr2-173993456-INS-312
chr2-174037128-INS-57
chr2-174089189-INS-75
chr2-174157286-INS-172
chr2-174190449-INS-125
chr2-174202168-INS-68
chr2-174207751-INS-50
chr2-174229232-INS-62
chr2-174253616-INS-52
chr2-174272920-INS-413
chr2-174273630-INS-53
chr2-174293490-INS-60
chr2-174329791-INS-264
chr2-174340496-INS-55
chr2-174343952-INS-70
chr2-174425913-INS-54
chr2-174428687-INS-94
chr2-174475043-INS-259
chr2-174480154-INS-302
chr2-174503999-INS-582
chr2-174525870-INS-362
chr2-174581303-INS-80
chr2-174617260-INS-165
chr2-174654549-INS-138
chr2-174671939-INS-127
chr2-174679587-INS-59
chr2-174679656-INS-200
chr2-174729144-INS-56
chr2-174750580-INS-409
chr2-174759723-INS-73
chr2-175011021-INS-84
chr2-176758469-INS-147
chr2-176817815-INS-84
chr2-176915884-INS-345
chr2-177062090-INS-202
chr2-177070401-INS-98
chr2-177213195-INS-4749
chr2-177587571-INS-84
chr2-177610548-INS-274
chr2-177727847-INS-1001
chr2-177735647-INS-291
chr2-177735827-INS-64
chr2-177750868-INS-4584
chr2-177771917-INS-58
chr2-177778818-INS-366
chr2-177808048-INS-63

chr2-177817820-INS-218
chr2-177856564-INS-62
chr2-177865055-INS-55
chr2-177900499-INS-53
chr2-177980006-INS-86
chr2-177989772-INS-63
chr2-178003244-INS-328
chr2-178095043-INS-80
chr2-178104130-INS-147
chr2-178104213-INS-463
chr2-178134473-INS-120
chr2-178170115-INS-580
chr2-178172787-INS-103
chr2-178175483-INS-3071
chr2-178200444-INS-121
chr2-178281133-INS-392
chr2-178282532-INS-54
chr2-178377861-INS-174
chr2-178388480-INS-6198
chr2-178395120-INS-96
chr2-178428823-INS-503
chr2-178442387-INS-104
chr2-178504277-INS-116
chr2-178505172-INS-60
chr2-178546666-INS-2255
chr2-178575833-INS-60
chr2-178644706-INS-110
chr2-178645433-INS-615
chr2-178657408-INS-1887
chr2-178660082-INS-102
chr2-178672631-INS-322
chr2-178697221-INS-126
chr2-178785428-INS-117
chr2-178841632-INS-1892
chr2-178858141-INS-84
chr2-178863135-INS-50
chr2-178885765-INS-179
chr2-178897566-INS-96
chr2-178898682-INS-50
chr2-178952294-INS-1056
chr2-178964206-INS-131
chr2-178971113-INS-277
chr2-178992794-INS-231
chr2-178993104-INS-52
chr2-179087863-INS-73
chr2-179108910-INS-71
chr2-179134190-INS-428
chr2-179159134-INS-210
chr2-179159134-INS-70
chr2-179229693-INS-94
chr2-179230535-INS-234
chr2-179294279-INS-448
chr2-179317650-INS-50
chr2-179320996-INS-6071
chr2-179329669-INS-52
chr2-179341452-INS-210
chr2-179345703-INS-286
chr2-179345898-INS-66
chr2-179361205-INS-2865
chr2-179361310-INS-255
chr2-179379663-INS-140
chr2-179408984-INS-50
chr2-179430025-INS-230
chr2-179430096-INS-659
chr2-179437286-INS-52
chr2-179443876-INS-56
chr2-179445143-INS-55
chr2-179537272-INS-173
chr2-179605920-INS-209
chr2-179614996-INS-213
chr2-179643223-INS-50
chr2-179645522-INS-225
chr2-179708424-INS-416
chr2-179716797-INS-52
chr2-179717683-INS-50
chr2-179778233-INS-84
chr2-179879996-INS-189

chr2-179880749-INS-73
chr2-179889671-INS-211
chr2-179898127-INS-228
chr2-179949573-INS-51
chr2-179986595-INS-92
chr2-179989601-INS-55
chr2-180096424-INS-50
chr2-180116973-INS-168
chr2-180187565-INS-58
chr2-180201989-INS-138
chr2-180327594-INS-372
chr2-180362150-INS-305
chr2-180394060-INS-217
chr2-180422552-INS-50
chr2-180424681-INS-8864
chr2-180469196-INS-50
chr2-180511511-INS-64
chr2-180512847-INS-50
chr2-180545352-INS-5453
chr2-180631718-INS-79
chr2-180748601-INS-62
chr2-180768694-INS-193
chr2-180784426-INS-145
chr2-180851977-INS-64
chr2-180898628-INS-64
chr2-181013628-INS-50
chr2-181045711-INS-257
chr2-181055713-INS-133
chr2-181056085-INS-201
chr2-181081486-INS-103
chr2-181159934-INS-410
chr2-181182940-INS-137
chr2-181249426-INS-75
chr2-181265667-INS-55
chr2-181265690-INS-277
chr2-181268691-INS-55
chr2-181306151-INS-206
chr2-181341386-INS-198
chr2-181352700-INS-201
chr2-181424103-INS-71
chr2-181485292-INS-72
chr2-181491162-INS-98
chr2-181491589-INS-162
chr2-181529963-INS-52
chr2-181530128-INS-400
chr2-181531446-INS-3451
chr2-181559116-INS-52
chr2-181571184-INS-184
chr3-3035380-INS-2587
chr3-3119431-INS-93
chr3-3123910-INS-72
chr3-3148785-INS-6645
chr3-3185381-INS-2868
chr3-3196635-INS-181
chr3-3196635-INS-72
chr3-3210661-INS-80
chr3-3373399-INS-4655
chr3-3414763-INS-5336
chr3-3465976-INS-358
chr3-3469175-INS-7928
chr3-3489760-INS-6266
chr3-3506093-INS-3555
chr3-3521679-INS-99
chr3-3521781-INS-120
chr3-3568254-INS-63
chr3-3568429-INS-57
chr3-3614824-INS-2567
chr3-3615606-INS-52
chr3-3628653-INS-75
chr3-3628783-INS-228
chr3-3632777-INS-7906
chr3-3658883-INS-50
chr3-3719321-INS-173
chr3-3729184-INS-319
chr3-3778841-INS-457
chr3-3822775-INS-52
chr3-3833959-INS-134

chr3-3852085-INS-186
chr3-3872264-INS-73
chr3-3898610-INS-6365
chr3-3919872-INS-56
chr3-3944158-INS-50
chr3-4023071-INS-152
chr3-4023201-INS-64
chr3-4095194-INS-265
chr3-4103714-INS-6400
chr3-4119695-INS-820
chr3-4193445-INS-67
chr3-4232002-INS-2622
chr3-4268743-INS-1509
chr3-4309619-INS-6602
chr3-4346977-INS-62
chr3-4347095-INS-51
chr3-4357655-INS-91
chr3-4361771-INS-152
chr3-4367410-INS-59
chr3-4527002-INS-52
chr3-4546800-INS-1365
chr3-4547215-INS-194
chr3-4578381-INS-54
chr3-4590534-INS-266
chr3-4598967-INS-65
chr3-4599302-INS-330
chr3-4599377-INS-129
chr3-4608142-INS-6799
chr3-4632947-INS-12883
chr3-4721796-INS-3266
chr3-4738492-INS-54
chr3-4745128-INS-79
chr3-4762403-INS-53
chr3-4805777-INS-436
chr3-4841907-INS-7217
chr3-4850598-INS-56
chr3-4867560-INS-77
chr3-4869422-INS-7230
chr3-4942537-INS-6525
chr3-4951455-INS-704
chr3-5003103-INS-50
chr3-5004713-INS-914
chr3-5011745-INS-6215
chr3-5016667-INS-81
chr3-5016785-INS-1103
chr3-5016785-INS-199
chr3-5016785-INS-98
chr3-5095666-INS-4536
chr3-5162071-INS-655
chr3-5171597-INS-3637
chr3-5188459-INS-1250
chr3-5197163-INS-827
chr3-5202161-INS-61
chr3-5224118-INS-52
chr3-5254146-INS-68
chr3-5271586-INS-58
chr3-5284948-INS-951
chr3-5285509-INS-95
chr3-5286422-INS-52
chr3-5328207-INS-489
chr3-5340921-INS-147
chr3-5340921-INS-446
chr3-5413978-INS-52
chr3-5444632-INS-64
chr3-5455131-INS-74
chr3-5495162-INS-1154
chr3-5537332-INS-6456
chr3-5745226-INS-87
chr3-5750594-INS-247
chr3-5794817-INS-76
chr3-5808721-INS-3584
chr3-5877451-INS-69
chr3-5969134-INS-4584
chr3-5978288-INS-5875
chr3-6003513-INS-724
chr3-6003812-INS-120
chr3-6027458-INS-2975

chr3-6028675-INS-120
chr3-6033691-INS-100
chr3-6034114-INS-52
chr3-6034505-INS-66
chr3-6043387-INS-4141
chr3-6045053-INS-75
chr3-6052873-INS-72
chr3-6056412-INS-52
chr3-6090540-INS-60
chr3-6096886-INS-117
chr3-6118371-INS-1086
chr3-6136829-INS-52
chr3-6282925-INS-144
chr3-6327053-INS-6518
chr3-6360139-INS-3294
chr3-6363412-INS-624
chr3-6364065-INS-416
chr3-6368997-INS-1872
chr3-6369337-INS-6449
chr3-6416022-INS-3052
chr3-6438589-INS-200
chr3-6501943-INS-60
chr3-6545681-INS-104
chr3-6551016-INS-62
chr3-6559009-INS-590
chr3-6588011-INS-178
chr3-6588171-INS-56
chr3-6603816-INS-6958
chr3-6611644-INS-539
chr3-6720324-INS-3050
chr3-6780514-INS-1000
chr3-6787121-INS-50
chr3-6815599-INS-77
chr3-6829998-INS-4249
chr3-6834667-INS-3907
chr3-6859115-INS-6378
chr3-6875066-INS-93
chr3-6959728-INS-65
chr3-7012000-INS-740
chr3-7066673-INS-351
chr3-7093856-INS-90
chr3-7099491-INS-193
chr3-7111350-INS-51
chr3-7125291-INS-135
chr3-7165576-INS-119
chr3-7185078-INS-4526
chr3-7205633-INS-112
chr3-7209881-INS-7083
chr3-7219049-INS-171
chr3-7224349-INS-148
chr3-7257278-INS-7718
chr3-7270250-INS-1101
chr3-7281587-INS-84
chr3-7306912-INS-66
chr3-7342643-INS-6255
chr3-7359327-INS-58
chr3-7380259-INS-582
chr3-7386673-INS-69
chr3-7421496-INS-165
chr3-7438069-INS-62
chr3-7441892-INS-198
chr3-7474131-INS-1306
chr3-7475923-INS-1255
chr3-7477686-INS-1057
chr3-7479138-INS-2033
chr3-7492962-INS-1651
chr3-7500897-INS-66
chr3-7501122-INS-231
chr3-7501897-INS-165
chr3-7501910-INS-429
chr3-7512792-INS-56
chr3-7522950-INS-6800
chr3-7573412-INS-67
chr3-7573602-INS-534
chr3-7598316-INS-54
chr3-7625684-INS-318
chr3-7643439-INS-1058

chr3-7666997-INS-340
chr3-7670389-INS-52
chr3-7675904-INS-50
chr3-7692082-INS-175
chr3-7713953-INS-220
chr3-7741255-INS-1104
chr3-7768802-INS-52
chr3-7769283-INS-7437
chr3-7775326-INS-1730
chr3-7794245-INS-220
chr3-7821397-INS-412
chr3-7821407-INS-58
chr3-7882424-INS-3165
chr3-7946137-INS-109
chr3-7983473-INS-115
chr3-7990448-INS-189
chr3-7990480-INS-591
chr3-7991772-INS-52
chr3-8017458-INS-7202
chr3-8033003-INS-162
chr3-8054105-INS-3522
chr3-8072100-INS-51
chr3-8074045-INS-5412
chr3-8092120-INS-111
chr3-8097526-INS-52
chr3-8106033-INS-9483
chr3-8126548-INS-6436
chr3-8157550-INS-85
chr3-8158008-INS-50
chr3-8165840-INS-52
chr3-8177871-INS-531
chr3-8178617-INS-2178
chr3-8194162-INS-58
chr3-8212272-INS-1411
chr3-8228703-INS-55
chr3-8232534-INS-381
chr3-8255846-INS-96
chr3-8258839-INS-72
chr3-8272174-INS-54
chr3-8286180-INS-6675
chr3-8313328-INS-2375
chr3-8321970-INS-6167
chr3-8333122-INS-162
chr3-8333122-INS-54
chr3-8347072-INS-7100
chr3-8356643-INS-6887
chr3-8368706-INS-3812
chr3-8377371-INS-84
chr3-8380280-INS-155
chr3-8380311-INS-322
chr3-8381335-INS-3364
chr3-8391961-INS-7478
chr3-8399879-INS-2733
chr3-8409859-INS-55
chr3-8434840-INS-1485
chr3-8435641-INS-1893
chr3-8482858-INS-68
chr3-8525597-INS-53
chr3-8526123-INS-299
chr3-8539842-INS-77
chr3-8540136-INS-71
chr3-8558139-INS-5589
chr3-8580037-INS-513
chr3-8592246-INS-1036
chr3-8600091-INS-78
chr3-8672943-INS-93
chr3-8673068-INS-573
chr3-8680566-INS-2769
chr3-8692490-INS-51
chr3-8698837-INS-74
chr3-8748134-INS-55
chr3-8754895-INS-76
chr3-8768250-INS-58
chr3-8771049-INS-530
chr3-8771373-INS-54
chr3-8788569-INS-7475
chr3-8794301-INS-134

chr3-8834419-INS-1034
chr3-8841665-INS-108
chr3-8848217-INS-2693
chr3-8858122-INS-52
chr3-8859320-INS-366
chr3-8872603-INS-144
chr3-8928899-INS-64
chr3-8946962-INS-52
chr3-8956299-INS-1004
chr3-8956299-INS-451
chr3-8960859-INS-340
chr3-8967975-INS-59
chr3-8975881-INS-3224
chr3-8980823-INS-331
chr3-8984023-INS-486
chr3-8984023-INS-7492
chr3-9005378-INS-60
chr3-9013283-INS-66
chr3-9044672-INS-2340
chr3-9045382-INS-5558
chr3-9051414-INS-350
chr3-9071933-INS-115
chr3-9079849-INS-575
chr3-9088612-INS-54
chr3-9089127-INS-87
chr3-9098936-INS-56
chr3-9112266-INS-369
chr3-9112442-INS-210
chr3-9115845-INS-204
chr3-9117559-INS-183
chr3-9118746-INS-82
chr3-9124186-INS-222
chr3-9126191-INS-190
chr3-9135273-INS-210
chr3-9149037-INS-134
chr3-9153271-INS-58
chr3-9180036-INS-52
chr3-9184597-INS-105
chr3-9193748-INS-86
chr3-9199894-INS-78
chr3-9200217-INS-184
chr3-9204881-INS-145
chr3-9205706-INS-198
chr3-9237597-INS-161
chr3-9243926-INS-96
chr3-9296058-INS-54
chr3-9339679-INS-212
chr3-9369798-INS-52
chr3-9370009-INS-54
chr3-9392827-INS-227
chr3-9392968-INS-139
chr3-9425568-INS-423
chr3-9427353-INS-128
chr3-9432837-INS-7086
chr3-9463331-INS-3707
chr3-9471901-INS-311
chr3-9519261-INS-198
chr3-9531398-INS-7305
chr3-9544699-INS-1113
chr3-9544959-INS-60
chr3-9572826-INS-302
chr3-9573015-INS-111
chr3-9595838-INS-50
chr3-9610859-INS-64
chr3-9618291-INS-161
chr3-9630891-INS-203
chr3-9666408-INS-50
chr3-9683184-INS-50
chr3-9683933-INS-158
chr3-9686364-INS-157
chr3-9689766-INS-52
chr3-9696918-INS-240
chr3-9696924-INS-106
chr3-9702093-INS-186
chr3-9721708-INS-121
chr3-9732978-INS-4350
chr3-9748082-INS-828

chr3-9748537-INS-116
chr3-9789230-INS-54
chr3-9789658-INS-229
chr3-9852108-INS-362
chr3-9854616-INS-6443
chr3-9864018-INS-216
chr3-9864308-INS-201
chr3-9867599-INS-172
chr3-9867854-INS-1259
chr3-9868663-INS-4456
chr3-9920661-INS-2436
chr3-9926739-INS-223
chr3-9926753-INS-1076
chr3-9927077-INS-80
chr3-9983064-INS-143
chr3-9985006-INS-3586
chr3-9991320-INS-57
chr3-9994769-INS-2901
chr3-10001632-INS-58
chr3-10005849-INS-128
chr3-10006577-INS-2153
chr3-10022430-INS-56
chr3-10023138-INS-96
chr3-10024468-INS-637
chr3-10033240-INS-212
chr3-10038032-INS-216
chr3-10050874-INS-136
chr3-10061229-INS-775
chr3-10074403-INS-54
chr3-10085147-INS-68
chr3-10097995-INS-6711
chr3-10100704-INS-507
chr3-10103916-INS-100
chr3-10116520-INS-90
chr3-10117183-INS-357
chr3-10124080-INS-411
chr3-10139712-INS-3972
chr3-10142889-INS-127
chr3-10150222-INS-51
chr3-10185083-INS-55
chr3-10233737-INS-50
chr3-10236030-INS-1891
chr3-10249716-INS-64
chr3-10270031-INS-115
chr3-10275304-INS-61
chr3-10303699-INS-80
chr3-10308284-INS-120
chr3-10317572-INS-452
chr3-10322488-INS-76
chr3-10324835-INS-6200
chr3-10329279-INS-64
chr3-10332116-INS-400
chr3-10370354-INS-178
chr3-10393157-INS-97
chr3-10423775-INS-82
chr3-10429759-INS-108
chr3-10429763-INS-52
chr3-10432713-INS-7584
chr3-10438990-INS-204
chr3-10455684-INS-212
chr3-10458902-INS-166
chr3-10461993-INS-1860
chr3-10471655-INS-63
chr3-10498714-INS-158
chr3-10517885-INS-4942
chr3-10579255-INS-955
chr3-10601153-INS-1336
chr3-10617003-INS-7007
chr3-10626102-INS-318
chr3-10634797-INS-503
chr3-10635876-INS-194
chr3-10639694-INS-50
chr3-10639863-INS-211
chr3-10656009-INS-4665
chr3-10664121-INS-1872
chr3-10668337-INS-208
chr3-10673104-INS-62

chr3-10714086-INS-2445
chr3-10729647-INS-50
chr3-10729932-INS-133
chr3-10729987-INS-62
chr3-10744274-INS-131
chr3-10747132-INS-273
chr3-10787772-INS-1171
chr3-10796062-INS-58
chr3-10812473-INS-70
chr3-10855526-INS-3089
chr3-10865778-INS-400
chr3-10866429-INS-128
chr3-10878522-INS-161
chr3-10888811-INS-89
chr3-10891025-INS-56
chr3-10898858-INS-3230
chr3-10939131-INS-3399
chr3-10996886-INS-94
chr3-10998215-INS-2847
chr3-11003280-INS-101
chr3-11003494-INS-461
chr3-11029988-INS-992
chr3-11034685-INS-160
chr3-11047606-INS-3221
chr3-11050536-INS-6312
chr3-11159694-INS-54
chr3-11183713-INS-138
chr3-11266962-INS-107
chr3-11285547-INS-3171
chr3-11288693-INS-195
chr3-11329032-INS-54
chr3-11378696-INS-63
chr3-11403195-INS-2731
chr3-11413505-INS-3195
chr3-11415335-INS-4946
chr3-11453272-INS-552
chr3-11510158-INS-95
chr3-11513168-INS-116
chr3-11525884-INS-80
chr3-11558003-INS-3465
chr3-11564791-INS-65
chr3-11583250-INS-737
chr3-11619758-INS-824
chr3-11670603-INS-68
chr3-11693148-INS-1920
chr3-11694310-INS-3186
chr3-11708380-INS-7782
chr3-11734627-INS-51
chr3-11745198-INS-4477
chr3-11820864-INS-1306
chr3-11865703-INS-62
chr3-11924290-INS-56
chr3-11924702-INS-2510
chr3-11926286-INS-58
chr3-11941238-INS-1197
chr3-11962477-INS-206
chr3-11962517-INS-72
chr3-12058507-INS-75
chr3-12065088-INS-1499
chr3-12136950-INS-54
chr3-12144062-INS-172
chr3-12147640-INS-416
chr3-12208386-INS-1368
chr3-12208411-INS-9322
chr3-12224928-INS-1217
chr3-12232148-INS-60
chr3-12284412-INS-68
chr3-12374208-INS-53
chr3-12375569-INS-74
chr3-12465085-INS-661
chr3-12565662-INS-481
chr3-12565672-INS-170
chr3-12576808-INS-52
chr3-12634380-INS-1896
chr3-12636462-INS-1230
chr3-12650871-INS-3795
chr3-12690590-INS-5324

chr3-12698487-INS-201
chr3-12751706-INS-1950
chr3-12758188-INS-98
chr3-12767221-INS-1862
chr3-12771338-INS-88
chr3-12773047-INS-561
chr3-12798621-INS-6212
chr3-12802508-INS-173
chr3-12816789-INS-5060
chr3-12824874-INS-2079
chr3-12837345-INS-59
chr3-12864269-INS-322
chr3-12940339-INS-52
chr3-12955622-INS-154
chr3-12973191-INS-363
chr3-13007651-INS-4441
chr3-13108817-INS-62
chr3-13116432-INS-7079
chr3-13161004-INS-6591
chr3-13190337-INS-6461
chr3-13195953-INS-78
chr3-13283135-INS-57
chr3-13300985-INS-60
chr3-13325896-INS-98
chr3-13325944-INS-6163
chr3-13352710-INS-194
chr3-13352904-INS-393
chr3-13369188-INS-52
chr3-13383059-INS-115
chr3-13383226-INS-137
chr3-13383383-INS-128
chr3-13447921-INS-264
chr3-13455597-INS-99
chr3-13481064-INS-115
chr3-13481163-INS-272
chr3-13483137-INS-65
chr3-13489885-INS-2919
chr3-13512683-INS-2496
chr3-13515546-INS-1419
chr3-13528234-INS-72
chr3-13558712-INS-133
chr3-13621977-INS-8022
chr3-13622212-INS-50
chr3-13627187-INS-1887
chr3-13636723-INS-52
chr3-13641311-INS-5313
chr3-13664470-INS-6349
chr3-13710350-INS-316
chr3-13726988-INS-1023
chr3-13729921-INS-64
chr3-13760146-INS-50
chr3-13801696-INS-255
chr3-13844365-INS-264
chr3-13845112-INS-153
chr3-13845234-INS-1083
chr3-13884528-INS-163
chr3-13944024-INS-172
chr3-13944024-INS-66
chr3-13998653-INS-6212
chr3-14075858-INS-184
chr3-14076566-INS-88
chr3-14086470-INS-1197
chr3-14148571-INS-5504
chr3-14169732-INS-4324
chr3-14174136-INS-2722
chr3-14184771-INS-56
chr3-14185123-INS-65
chr3-14197518-INS-74
chr3-14197539-INS-204
chr3-14200347-INS-58
chr3-14280241-INS-175
chr3-14311640-INS-1058
chr3-14366206-INS-1046
chr3-14417582-INS-458
chr3-14453312-INS-5566
chr3-14466788-INS-4612
chr3-14496915-INS-1525

chr3-14526891-INS-199
chr3-14635958-INS-50
chr3-14649726-INS-180
chr3-14659532-INS-73
chr3-14683316-INS-65
chr3-14686715-INS-7126
chr3-14752256-INS-6209
chr3-14849921-INS-76
chr3-14858785-INS-564
chr3-14882159-INS-300
chr3-14889225-INS-405
chr3-14893888-INS-140
chr3-14907088-INS-357
chr3-14947851-INS-355
chr3-14974118-INS-2210
chr3-15008905-INS-837
chr3-15035740-INS-165
chr3-15066743-INS-4690
chr3-15100804-INS-53
chr3-15114554-INS-74
chr3-15114830-INS-120
chr3-15182480-INS-74
chr3-15189294-INS-205
chr3-15203291-INS-417
chr3-15205782-INS-86
chr3-15214854-INS-73
chr3-15222609-INS-6375
chr3-15236247-INS-2500
chr3-15238500-INS-50
chr3-15241065-INS-3354
chr3-15262233-INS-3161
chr3-15273232-INS-362
chr3-15273502-INS-1673
chr3-15282108-INS-656
chr3-15282610-INS-1891
chr3-15299442-INS-2536
chr3-15311946-INS-52
chr3-15312102-INS-164
chr3-15324494-INS-2841
chr3-15346607-INS-187
chr3-15356566-INS-6175
chr3-15365361-INS-1711
chr3-15379502-INS-328
chr3-15395032-INS-1519
chr3-15400071-INS-3026
chr3-15410596-INS-56
chr3-15412948-INS-64
chr3-15420472-INS-135
chr3-15433969-INS-6182
chr3-15447683-INS-199
chr3-15458611-INS-6590
chr3-15470641-INS-169
chr3-15477252-INS-331
chr3-15482318-INS-58
chr3-15544563-INS-452
chr3-15602329-INS-169
chr3-15602671-INS-2636
chr3-15606673-INS-106
chr3-15614080-INS-58
chr3-15638294-INS-258
chr3-15638639-INS-56
chr3-15644267-INS-144
chr3-15663066-INS-6412
chr3-15665380-INS-70
chr3-15679962-INS-90
chr3-15694128-INS-59
chr3-15697589-INS-545
chr3-15825356-INS-64
chr3-15832892-INS-84
chr3-15874030-INS-462
chr3-15890524-INS-392
chr3-15908230-INS-50
chr3-15941095-INS-126
chr3-15950241-INS-355
chr3-15993750-INS-2332
chr3-16004662-INS-96
chr3-16007189-INS-51

chr3-16023455-INS-62
chr3-16025405-INS-68
chr3-16025482-INS-116
chr3-16025505-INS-299
chr3-16033891-INS-52
chr3-16046292-INS-72
chr3-16053982-INS-149
chr3-16058917-INS-389
chr3-16058920-INS-52
chr3-16059061-INS-150
chr3-16059296-INS-172
chr3-16059296-INS-84
chr3-16102708-INS-60
chr3-16105796-INS-333
chr3-16105865-INS-196
chr3-16116965-INS-396
chr3-16139341-INS-113
chr3-16154343-INS-105
chr3-16163241-INS-128
chr3-16180369-INS-5056
chr3-16207878-INS-186
chr3-16225331-INS-216
chr3-16229028-INS-3035
chr3-16229028-INS-403
chr3-16263772-INS-802
chr3-16300919-INS-365
chr3-16310946-INS-6383
chr3-16321807-INS-176
chr3-16321826-INS-66
chr3-16332737-INS-416
chr3-16332754-INS-173
chr3-16359637-INS-94
chr3-16391600-INS-165
chr3-16391738-INS-68
chr3-16403114-INS-1062
chr3-16450999-INS-328
chr3-16465656-INS-5988
chr3-16489860-INS-5329
chr3-16501322-INS-65
chr3-16506356-INS-52
chr3-16552544-INS-201
chr3-16554689-INS-52
chr3-16583081-INS-7561
chr3-16613829-INS-6271
chr3-16666525-INS-318
chr3-16681553-INS-2410
chr3-16683857-INS-12056
chr3-16683857-INS-4823
chr3-16730662-INS-50
chr3-16735481-INS-2094
chr3-16757315-INS-1753
chr3-16780502-INS-52
chr3-16815634-INS-55
chr3-16824936-INS-4730
chr3-16895360-INS-479
chr3-16928305-INS-80
chr3-17011219-INS-3850
chr3-17064239-INS-6530
chr3-17122979-INS-2886
chr3-17123094-INS-70
chr3-17136107-INS-56
chr3-17143799-INS-67
chr3-17184927-INS-81
chr3-17186657-INS-1264
chr3-17194739-INS-2207
chr3-17288286-INS-63
chr3-17294010-INS-2731
chr3-17413322-INS-56
chr3-17492983-INS-119
chr3-17530319-INS-104
chr3-17539257-INS-6272
chr3-17540686-INS-303
chr3-17540686-INS-780
chr3-17553958-INS-6899
chr3-17578213-INS-1601
chr3-17581740-INS-836
chr3-17600851-INS-52

chr3-17623775-INS-6382
chr3-17626277-INS-1718
chr3-17630987-INS-3156
chr3-17653814-INS-6384
chr3-17698700-INS-80
chr3-17717248-INS-52
chr3-17740768-INS-1194
chr3-17788665-INS-411
chr3-17797091-INS-56
chr3-17826408-INS-938
chr3-17894038-INS-3025
chr3-17894038-INS-9875
chr3-17907616-INS-60
chr3-17951984-INS-1730
chr3-17980808-INS-145
chr3-17992574-INS-62
chr3-18062201-INS-416
chr3-18070406-INS-789
chr3-18088626-INS-2858
chr3-18130785-INS-6443
chr3-18170791-INS-88
chr3-18170961-INS-84
chr3-18229717-INS-230
chr3-18237489-INS-893
chr3-18247737-INS-57
chr3-18277078-INS-154
chr3-18277769-INS-120
chr3-18279582-INS-54
chr3-18313841-INS-7126
chr3-18417846-INS-51
chr3-18426628-INS-75
chr3-18438961-INS-64
chr3-18445302-INS-165
chr3-18447043-INS-59
chr3-18538791-INS-54
chr3-18545031-INS-654
chr3-18551052-INS-1209
chr3-18565574-INS-348
chr3-18571738-INS-54
chr3-18572587-INS-2104
chr3-18573985-INS-1382
chr3-18580719-INS-480
chr3-18671560-INS-54
chr3-18736084-INS-197
chr3-18750682-INS-5124
chr3-18817759-INS-589
chr3-18819924-INS-172
chr3-18830023-INS-1271
chr3-18847016-INS-68
chr3-18866493-INS-60
chr3-18943467-INS-50
chr3-18952111-INS-3049
chr3-18971523-INS-265
chr3-19006343-INS-62
chr3-19011437-INS-2211
chr3-19013154-INS-278
chr3-19017555-INS-117
chr3-19018274-INS-52
chr3-19020345-INS-426
chr3-19028137-INS-51
chr3-19042825-INS-180
chr3-19042825-INS-533
chr3-19052382-INS-6390
chr3-19100431-INS-84
chr3-19104055-INS-6191
chr3-19126740-INS-8764
chr3-19133778-INS-64
chr3-19134043-INS-130
chr3-19225663-INS-63
chr3-19261289-INS-124
chr3-19307568-INS-160
chr3-19307733-INS-557
chr3-19316134-INS-2291
chr3-19320397-INS-168
chr3-19344130-INS-141
chr3-19379439-INS-214
chr3-19384459-INS-50

chr3-19389129-INS-1523
chr3-19393278-INS-374
chr3-19395126-INS-50
chr3-19428262-INS-50
chr3-19447073-INS-2897
chr3-19468491-INS-53
chr3-19492100-INS-423
chr3-19498389-INS-8542
chr3-19538932-INS-51
chr3-19542872-INS-6189
chr3-19592910-INS-7094
chr3-19597852-INS-6408
chr3-19618000-INS-176
chr3-19679717-INS-50
chr3-19690160-INS-60
chr3-19713540-INS-75
chr3-19721389-INS-293
chr3-19724202-INS-84
chr3-19736685-INS-120
chr3-19737267-INS-150
chr3-19737710-INS-60
chr3-19782700-INS-2881
chr3-19826523-INS-6135
chr3-19827464-INS-636
chr3-19842480-INS-480
chr3-19880166-INS-196
chr3-19909184-INS-98
chr3-19914683-INS-2221
chr3-19920809-INS-178
chr3-19921235-INS-176
chr3-19922714-INS-201
chr3-19925128-INS-5480
chr3-19930481-INS-6205
chr3-19931923-INS-1729
chr3-19934800-INS-59
chr3-19938452-INS-96
chr3-19959847-INS-320
chr3-19960841-INS-1435
chr3-19989012-INS-84
chr3-19989242-INS-167
chr3-20010502-INS-53
chr3-20017745-INS-1094
chr3-20031801-INS-69
chr3-20052466-INS-96
chr3-20055217-INS-92
chr3-20066105-INS-50
chr3-20069726-INS-6241
chr3-20077798-INS-257
chr3-20082659-INS-159
chr3-20087652-INS-2884
chr3-20090756-INS-1705
chr3-20113652-INS-62
chr3-20145320-INS-383
chr3-20176030-INS-324
chr3-20191787-INS-51
chr3-20197409-INS-210
chr3-20231002-INS-2276
chr3-20237141-INS-63
chr3-20253351-INS-4773
chr3-20263653-INS-588
chr3-20279076-INS-102
chr3-20280551-INS-160
chr3-20285256-INS-132
chr3-20297312-INS-78
chr3-20324228-INS-212
chr3-20332245-INS-202
chr3-20336354-INS-3115
chr3-20339247-INS-326
chr3-20353438-INS-60
chr3-20365485-INS-192
chr3-20366214-INS-413
chr3-20377181-INS-2468
chr3-20382721-INS-54
chr3-20385452-INS-50
chr3-20389386-INS-72
chr3-20389437-INS-71
chr3-20404645-INS-3055

chr3-20406540-INS-58
chr3-20433196-INS-2516
chr3-20447646-INS-341
chr3-20455401-INS-484
chr3-20490315-INS-1566
chr3-20506437-INS-1311
chr3-20508996-INS-8669
chr3-20552426-INS-805
chr3-20553027-INS-2380
chr3-20570667-INS-105
chr3-20572264-INS-68
chr3-20583773-INS-8669
chr3-20591886-INS-8804
chr3-20623369-INS-3627
chr3-20632700-INS-380
chr3-20686977-INS-311
chr3-20708737-INS-6253
chr3-20767559-INS-126
chr3-20775056-INS-50
chr3-20783545-INS-1541
chr3-20786571-INS-588
chr3-20821285-INS-1790
chr3-20860954-INS-83
chr3-20958792-INS-357
chr3-20974133-INS-58
chr3-20982545-INS-51
chr3-21007354-INS-1199
chr3-21007373-INS-7689
chr3-21014425-INS-1080
chr3-21029056-INS-431
chr3-21065046-INS-2990
chr3-21065535-INS-1118
chr3-21100056-INS-2481
chr3-21139293-INS-2040
chr3-21181061-INS-65
chr3-21191089-INS-493
chr3-21201580-INS-58
chr3-21295105-INS-293
chr3-21315850-INS-812
chr3-21327189-INS-90
chr3-21358814-INS-56
chr3-21413603-INS-50
chr3-21421912-INS-6178
chr3-21422846-INS-11945
chr3-21446159-INS-227
chr3-21453334-INS-114
chr3-21463910-INS-404
chr3-21472781-INS-93
chr3-21513513-INS-159
chr3-21533533-INS-58
chr3-21666258-INS-7628
chr3-21672971-INS-6352
chr3-21686436-INS-60
chr3-21686455-INS-130
chr3-21786094-INS-93
chr3-21817567-INS-78
chr3-21822565-INS-196
chr3-21927102-INS-3510
chr3-21932762-INS-601
chr3-21953641-INS-1320
chr3-21966418-INS-195
chr3-22103251-INS-76
chr3-22109149-INS-64
chr3-22215699-INS-56
chr3-22275142-INS-105
chr3-22296216-INS-1369
chr3-22349317-INS-213
chr3-22360448-INS-50
chr3-22383096-INS-84
chr3-22397139-INS-449
chr3-22503661-INS-499
chr3-22514306-INS-1028
chr3-22596910-INS-326
chr3-22618273-INS-3347
chr3-22633967-INS-240
chr3-22633971-INS-116
chr3-22634077-INS-672

chr3-22658862-INS-50
chr3-22760861-INS-78
chr3-22824615-INS-4843
chr3-22844566-INS-5318
chr3-22858939-INS-7349
chr3-22898751-INS-56
chr3-22950929-INS-2272
chr3-22979448-INS-56
chr3-23050492-INS-90
chr3-23052257-INS-612
chr3-23052812-INS-56
chr3-23069071-INS-208
chr3-23138707-INS-56
chr3-23147311-INS-62
chr3-23157514-INS-7523
chr3-23253971-INS-167
chr3-23268376-INS-50
chr3-23297394-INS-58
chr3-23297904-INS-315
chr3-23302185-INS-88
chr3-23302225-INS-290
chr3-23306486-INS-51
chr3-23334071-INS-80
chr3-23340459-INS-188
chr3-23341161-INS-84
chr3-23385297-INS-517
chr3-23397776-INS-4169
chr3-23417970-INS-238
chr3-23464678-INS-6169
chr3-23477187-INS-7145
chr3-23523450-INS-198
chr3-23535892-INS-625
chr3-23536325-INS-58
chr3-23545939-INS-58
chr3-23546065-INS-269
chr3-23685115-INS-56
chr3-23685322-INS-98
chr3-23707826-INS-54
chr3-23708717-INS-58
chr3-23718799-INS-155
chr3-23719071-INS-52
chr3-23749593-INS-58
chr3-23749654-INS-493
chr3-23750173-INS-2387
chr3-23751260-INS-56
chr3-23758815-INS-70
chr3-23760380-INS-950
chr3-23774445-INS-55
chr3-23776147-INS-6344
chr3-23788590-INS-5138
chr3-23802325-INS-8846
chr3-23809387-INS-358
chr3-23841447-INS-7232
chr3-23841897-INS-190
chr3-23869188-INS-58
chr3-23881143-INS-5279
chr3-23888792-INS-7725
chr3-23954153-INS-475
chr3-23961056-INS-59
chr3-23986046-INS-452
chr3-24131235-INS-6055
chr3-24149119-INS-60
chr3-24153989-INS-50
chr3-24172709-INS-59
chr3-24173668-INS-54
chr3-24198438-INS-68
chr3-24212285-INS-56
chr3-24306733-INS-962
chr3-24324913-INS-128
chr3-24394429-INS-74
chr3-24395953-INS-100
chr3-24397233-INS-83
chr3-24413393-INS-911
chr3-24419184-INS-52
chr3-24419809-INS-66
chr3-24503949-INS-54
chr3-24588867-INS-58

chr3-24609703-INS-50
chr3-24610006-INS-241
chr3-24639912-INS-72
chr3-24648293-INS-137
chr3-24720639-INS-1800
chr3-24733612-INS-60
chr3-24734942-INS-1188
chr3-24740978-INS-60
chr3-24781667-INS-341
chr3-24825177-INS-72
chr3-24843709-INS-131
chr3-24855318-INS-50
chr3-24869571-INS-60
chr3-24884306-INS-6581
chr3-24896151-INS-3371
chr3-24924643-INS-310
chr3-24940353-INS-69
chr3-24941371-INS-6767
chr3-24948968-INS-52
chr3-24958868-INS-446
chr3-24968580-INS-240
chr3-24993335-INS-1065
chr3-24993562-INS-135
chr3-25062572-INS-228
chr3-25121853-INS-54
chr3-25152949-INS-55
chr3-25187336-INS-3101
chr3-25190218-INS-50
chr3-25223342-INS-63
chr3-25239684-INS-192
chr3-25248726-INS-3722
chr3-25255097-INS-252
chr3-25259208-INS-456
chr3-25275868-INS-64
chr3-25294106-INS-297
chr3-25304455-INS-428
chr3-25305692-INS-2500
chr3-25307446-INS-1500
chr3-25318502-INS-1180
chr3-25372239-INS-294
chr3-25385394-INS-58
chr3-25392057-INS-1999
chr3-25392628-INS-6399
chr3-25404461-INS-473
chr3-25423065-INS-4828
chr3-25457619-INS-55
chr3-25462776-INS-197
chr3-25482935-INS-461
chr3-25511165-INS-50
chr3-25538027-INS-54
chr3-25539050-INS-64
chr3-25681694-INS-2836
chr3-25716643-INS-64
chr3-25844353-INS-155
chr3-25844353-INS-50
chr3-25874699-INS-116
chr3-25895992-INS-7694
chr3-25936329-INS-54
chr3-25937941-INS-65
chr3-25941383-INS-2109
chr3-25952161-INS-68
chr3-25989774-INS-137
chr3-25990028-INS-520
chr3-26003620-INS-2029
chr3-26024166-INS-1239
chr3-26063425-INS-74
chr3-26064038-INS-59
chr3-26100139-INS-126
chr3-26104233-INS-4878
chr3-26105565-INS-7267
chr3-26113748-INS-836
chr3-26146922-INS-70
chr3-26166952-INS-424
chr3-26201866-INS-53
chr3-26255985-INS-3539
chr3-26283489-INS-117
chr3-26302846-INS-6471

chr3-26307953-INS-4253
chr3-26308134-INS-52
chr3-26322916-INS-1596
chr3-26337828-INS-2938
chr3-26362127-INS-203
chr3-26398320-INS-3167
chr3-26399993-INS-111
chr3-26400085-INS-52
chr3-26438585-INS-3764
chr3-26505531-INS-579
chr3-26515265-INS-55
chr3-26542734-INS-62
chr3-26572846-INS-72
chr3-26579963-INS-6375
chr3-26580510-INS-80
chr3-26590881-INS-195
chr3-26656028-INS-52
chr3-26656157-INS-624
chr3-26656191-INS-286
chr3-26671079-INS-584
chr3-26725649-INS-113
chr3-26747690-INS-54
chr3-26778375-INS-61
chr3-26779068-INS-113
chr3-26779293-INS-74
chr3-26779708-INS-557
chr3-26803539-INS-2010
chr3-26814560-INS-2154
chr3-26928555-INS-52
chr3-26936857-INS-84
chr3-26985360-INS-2546
chr3-27051607-INS-917
chr3-27051618-INS-166
chr3-27051850-INS-292
chr3-27104254-INS-80
chr3-27106884-INS-6708
chr3-27121293-INS-50
chr3-27129492-INS-2842
chr3-27171067-INS-128
chr3-27332713-INS-112
chr3-27333784-INS-14384
chr3-27340110-INS-253
chr3-27342446-INS-78
chr3-27355608-INS-188
chr3-27400527-INS-179
chr3-27408146-INS-221
chr3-27408765-INS-132
chr3-27465736-INS-87
chr3-27469637-INS-98
chr3-27480761-INS-68
chr3-27490412-INS-2755
chr3-27494897-INS-138
chr3-27571542-INS-142
chr3-27699957-INS-326
chr3-27700980-INS-57
chr3-27727594-INS-367
chr3-27735783-INS-69
chr3-27788877-INS-50
chr3-27792633-INS-73
chr3-27795121-INS-228
chr3-27804890-INS-1236
chr3-27811716-INS-60
chr3-27818680-INS-53
chr3-27835938-INS-581
chr3-27839922-INS-74
chr3-27871578-INS-74
chr3-27875047-INS-127
chr3-27924599-INS-124
chr3-27932748-INS-74
chr3-27949850-INS-105
chr3-27981711-INS-69
chr3-28013968-INS-338
chr3-28065527-INS-153
chr3-28097995-INS-1477
chr3-28152034-INS-69
chr3-28182640-INS-7053
chr3-28256709-INS-253

chr3-28278295-INS-141
chr3-28296384-INS-155
chr3-28311305-INS-389
chr3-28403825-INS-255
chr3-28404117-INS-9469
chr3-28447517-INS-50
chr3-28463469-INS-84
chr3-28532739-INS-3280
chr3-28538689-INS-7018
chr3-28539158-INS-130
chr3-28550971-INS-4050
chr3-28618142-INS-70
chr3-28660691-INS-176
chr3-28695391-INS-123
chr3-28759666-INS-55
chr3-28787877-INS-188
chr3-28788954-INS-213
chr3-28801751-INS-173
chr3-28820946-INS-4011
chr3-28839087-INS-50
chr3-28841487-INS-86
chr3-28885961-INS-6976
chr3-28902905-INS-79
chr3-28903073-INS-296
chr3-28903082-INS-1439
chr3-28910682-INS-5492
chr3-28936766-INS-1209
chr3-28960377-INS-7095
chr3-29000859-INS-436
chr3-29022911-INS-60
chr3-29050736-INS-2003
chr3-29077096-INS-3082
chr3-29102636-INS-6902
chr3-29111252-INS-58
chr3-29113625-INS-218
chr3-29115470-INS-6369
chr3-29122050-INS-6666
chr3-29150656-INS-313
chr3-29166715-INS-52
chr3-29184459-INS-6303
chr3-29221662-INS-50
chr3-29228602-INS-125
chr3-29258968-INS-8943
chr3-29329255-INS-84
chr3-29329533-INS-318
chr3-29398121-INS-5889
chr3-29437239-INS-174
chr3-29445529-INS-3462
chr3-29467889-INS-115
chr3-29468801-INS-51
chr3-29535598-INS-148
chr3-29558967-INS-365
chr3-29624130-INS-7086
chr3-29727271-INS-1331
chr3-29805302-INS-122
chr3-29834147-INS-6541
chr3-29856236-INS-55
chr3-29896670-INS-363
chr3-29896711-INS-57
chr3-29897709-INS-108
chr3-29898063-INS-297
chr3-29898574-INS-1975
chr3-29904089-INS-2159
chr3-29951567-INS-122
chr3-29959753-INS-269
chr3-29996138-INS-7136
chr3-30040432-INS-54
chr3-30040511-INS-58
chr3-30164755-INS-66
chr3-30223281-INS-1150
chr3-30247705-INS-662
chr3-30331226-INS-52
chr3-30387927-INS-6442
chr3-30417864-INS-81
chr3-30418624-INS-58
chr3-30418637-INS-145
chr3-30443790-INS-60

chr3-30458809-INS-50
chr3-30473610-INS-62
chr3-30475593-INS-250
chr3-30475701-INS-742
chr3-30518761-INS-2931
chr3-30525900-INS-105
chr3-30549921-INS-68
chr3-30586032-INS-5113
chr3-30616875-INS-231
chr3-30623685-INS-1131
chr3-30626302-INS-54
chr3-30650471-INS-598
chr3-30656263-INS-742
chr3-30674905-INS-478
chr3-30695454-INS-50
chr3-30712970-INS-211
chr3-30730763-INS-2003
chr3-30734408-INS-2727
chr3-30757306-INS-187
chr3-30768790-INS-6654
chr3-30781988-INS-52
chr3-30787520-INS-2685
chr3-30802020-INS-529
chr3-30808354-INS-60
chr3-30833801-INS-281
chr3-30835045-INS-3034
chr3-30919718-INS-1229
chr3-30938656-INS-68
chr3-30984654-INS-1240
chr3-30992131-INS-52
chr3-31043076-INS-58
chr3-31075792-INS-165
chr3-31078962-INS-51
chr3-31101066-INS-74
chr3-31102580-INS-5501
chr3-31110806-INS-50
chr3-31124236-INS-2603
chr3-31124236-INS-6081
chr3-31133564-INS-376
chr3-31181672-INS-72
chr3-31192902-INS-211
chr3-31209571-INS-161
chr3-31230476-INS-1476
chr3-31232732-INS-1477
chr3-31234627-INS-1060
chr3-31254043-INS-78
chr3-31279247-INS-50
chr3-31344442-INS-100
chr3-31346541-INS-238
chr3-31348933-INS-86
chr3-31378874-INS-50
chr3-31383317-INS-250
chr3-31389260-INS-62
chr3-31413011-INS-197
chr3-31418078-INS-725
chr3-31435034-INS-668
chr3-31435967-INS-591
chr3-31461854-INS-603
chr3-31463973-INS-578
chr3-31464755-INS-5457
chr3-31476546-INS-1389
chr3-31482035-INS-5816
chr3-31521224-INS-52
chr3-31548056-INS-2242
chr3-31559989-INS-66
chr3-31583985-INS-1156
chr3-31588000-INS-295
chr3-31618998-INS-64
chr3-31621049-INS-2984
chr3-31631034-INS-1966
chr3-31642813-INS-54
chr3-31655516-INS-156
chr3-31664733-INS-444
chr3-31703817-INS-65
chr3-31741178-INS-51
chr3-31773487-INS-126
chr3-31795886-INS-4740

chr3-31801636-INS-16994
chr3-31821214-INS-260
chr3-31849020-INS-181
chr3-31910355-INS-386
chr3-31920149-INS-6288
chr3-31924091-INS-4093
chr3-31932023-INS-60
chr3-31992926-INS-6618
chr3-32050074-INS-1891
chr3-32243209-INS-206
chr3-32273917-INS-1838
chr3-32292562-INS-186
chr3-32311859-INS-3015
chr3-32318346-INS-6554
chr3-32318650-INS-7120
chr3-32319200-INS-210
chr3-32346609-INS-8309
chr3-32381485-INS-6176
chr3-32394611-INS-50
chr3-32407713-INS-1034
chr3-32409171-INS-301
chr3-32413823-INS-91
chr3-32443657-INS-7440
chr3-32445103-INS-456
chr3-32505024-INS-104
chr3-32543456-INS-93
chr3-32544113-INS-84
chr3-32547694-INS-70
chr3-32552088-INS-261
chr3-32585009-INS-177
chr3-32597248-INS-66
chr3-32613886-INS-197
chr3-32622126-INS-53
chr3-32626709-INS-58
chr3-32643822-INS-780
chr3-32650947-INS-58
chr3-32680005-INS-208
chr3-32711426-INS-384
chr3-32717754-INS-65
chr3-32725211-INS-50
chr3-32771186-INS-316
chr3-32773673-INS-74
chr3-32822234-INS-51
chr3-32822377-INS-72
chr3-32856860-INS-2977
chr3-32872386-INS-58
chr3-32914047-INS-53
chr3-32928420-INS-204
chr3-32931939-INS-492
chr3-32945813-INS-256
chr3-32945813-INS-84
chr3-32976696-INS-64
chr3-32984747-INS-2780
chr3-32999560-INS-109
chr3-33039902-INS-394
chr3-33091164-INS-1500
chr3-33103925-INS-54
chr3-33114156-INS-74
chr3-33129300-INS-1287
chr3-33154687-INS-50
chr3-33159829-INS-694
chr3-33168140-INS-815
chr3-33209845-INS-101
chr3-33213828-INS-182
chr3-33213828-INS-88
chr3-33244177-INS-60
chr3-33317339-INS-50
chr3-33317401-INS-62
chr3-33320372-INS-486
chr3-33345973-INS-752
chr3-33353832-INS-7081
chr3-33364195-INS-114
chr3-33406209-INS-3088
chr3-33406503-INS-52
chr3-33422388-INS-75
chr3-33436011-INS-57
chr3-33456788-INS-227

chr3-33509423-INS-2115
chr3-33554179-INS-2594
chr3-33554459-INS-60
chr3-33562279-INS-75
chr3-33607227-INS-1178
chr3-33608355-INS-90
chr3-33614161-INS-6583
chr3-33637215-INS-54
chr3-33647941-INS-7521
chr3-33677016-INS-3217
chr3-33694648-INS-1039
chr3-33718206-INS-1923
chr3-33745862-INS-2990
chr3-33751501-INS-18995
chr3-33765022-INS-2116
chr3-33798154-INS-1182
chr3-33849084-INS-6555
chr3-33932582-INS-144
chr3-33994023-INS-837
chr3-34024383-INS-1136
chr3-34037983-INS-52
chr3-34049943-INS-328
chr3-34050665-INS-599
chr3-34063568-INS-219
chr3-34111627-INS-50
chr3-34164370-INS-19791
chr3-34164883-INS-967
chr3-34179079-INS-84
chr3-34187778-INS-1369
chr3-34205599-INS-165
chr3-34248647-INS-55
chr3-34265926-INS-6484
chr3-34290394-INS-140
chr3-34350769-INS-82
chr3-34433941-INS-50
chr3-34445688-INS-6351
chr3-34476130-INS-261
chr3-34493053-INS-111
chr3-34493225-INS-50
chr3-34494496-INS-369
chr3-34525937-INS-53
chr3-34534331-INS-60
chr3-34616906-INS-84
chr3-34624023-INS-415
chr3-34670912-INS-214
chr3-34685499-INS-112
chr3-34687328-INS-420
chr3-34747569-INS-275
chr3-34769356-INS-267
chr3-34799182-INS-8755
chr3-34826547-INS-78
chr3-34827726-INS-52
chr3-34831413-INS-507
chr3-34838080-INS-140
chr3-34857968-INS-676
chr3-34874970-INS-50
chr3-34909643-INS-237
chr3-34920218-INS-220
chr3-34964990-INS-50
chr3-34999101-INS-345
chr3-34999216-INS-132
chr3-35003807-INS-137
chr3-35061694-INS-102
chr3-35061780-INS-164
chr3-35080005-INS-116
chr3-35224409-INS-59
chr3-35232677-INS-64
chr3-35245319-INS-3201
chr3-35256299-INS-1289
chr3-35330304-INS-54
chr3-35349600-INS-50
chr3-35354975-INS-123
chr3-35545574-INS-57
chr3-35547429-INS-52
chr3-35577785-INS-58
chr3-35588878-INS-56
chr3-35596297-INS-429

chr3-35601739-INS-6207
chr3-35608319-INS-178
chr3-35608581-INS-92
chr3-35698701-INS-1459
chr3-35715624-INS-6454
chr3-35719875-INS-6746
chr3-35720469-INS-6670
chr3-35720629-INS-2067
chr3-35770598-INS-207
chr3-35771294-INS-371
chr3-35775538-INS-67
chr3-35794773-INS-206
chr3-35816657-INS-369
chr3-35820139-INS-1193
chr3-35850534-INS-184
chr3-35950543-INS-57
chr3-35957075-INS-54
chr3-35983147-INS-96
chr3-35998266-INS-2849
chr3-36009974-INS-50
chr3-36050296-INS-174
chr3-36082631-INS-3795
chr3-36082631-INS-439
chr3-36085597-INS-291
chr3-36099107-INS-142
chr3-36128385-INS-221
chr3-36128563-INS-1220
chr3-36156548-INS-6716
chr3-36228876-INS-60
chr3-36266880-INS-123
chr3-36276983-INS-58
chr3-36308613-INS-764
chr3-36348501-INS-464
chr3-36361639-INS-196
chr3-36522585-INS-59
chr3-36522932-INS-114
chr3-36539971-INS-494
chr3-36542556-INS-134
chr3-36565385-INS-76
chr3-36603612-INS-1886
chr3-36634397-INS-50
chr3-36634398-INS-135
chr3-36726438-INS-60
chr3-36783933-INS-1864
chr3-36792792-INS-1300
chr3-36792864-INS-68
chr3-36814212-INS-358
chr3-36821979-INS-1895
chr3-36826771-INS-3750
chr3-36828897-INS-1351
chr3-36921970-INS-7016
chr3-36922920-INS-88
chr3-36930151-INS-1889
chr3-36932013-INS-3848
chr3-36936116-INS-6916
chr3-36959602-INS-188
chr3-37003438-INS-771
chr3-37021515-INS-8420
chr3-37158639-INS-608
chr3-37184318-INS-89
chr3-37200527-INS-141
chr3-37200527-INS-51
chr3-37238125-INS-2873
chr3-37294198-INS-682
chr3-37302570-INS-4856
chr3-37304975-INS-323
chr3-37328624-INS-507
chr3-37328624-INS-78
chr3-37333843-INS-859
chr3-37342813-INS-52
chr3-37404192-INS-120
chr3-37420001-INS-242
chr3-37501094-INS-69
chr3-37516133-INS-2453
chr3-37689208-INS-58
chr3-37699940-INS-108
chr3-37709111-INS-72

chr3-37735325-INS-494
chr3-37765339-INS-234
chr3-37774613-INS-50
chr3-37888016-INS-69
chr3-37913870-INS-86
chr3-37924602-INS-136
chr3-37944664-INS-132
chr3-37944700-INS-51
chr3-38105538-INS-4458
chr3-38124823-INS-52
chr3-38138373-INS-4544
chr3-38209916-INS-176
chr3-38210865-INS-191
chr3-38244046-INS-198
chr3-38246334-INS-72
chr3-38246430-INS-265
chr3-38246433-INS-87
chr3-38251501-INS-6488
chr3-38324251-INS-59
chr3-38328969-INS-2605
chr3-38348087-INS-62
chr3-38365459-INS-419
chr3-38408838-INS-6243
chr3-38430740-INS-6604
chr3-38442551-INS-133
chr3-38467224-INS-144
chr3-38496523-INS-62
chr3-38530418-INS-392
chr3-38552137-INS-204
chr3-38552137-INS-68
chr3-38554656-INS-270
chr3-38554889-INS-340
chr3-38561469-INS-1129
chr3-38579756-INS-66
chr3-38599751-INS-399
chr3-38604631-INS-6211
chr3-38629700-INS-51
chr3-38629833-INS-613
chr3-38633959-INS-376
chr3-38733638-INS-246
chr3-38753725-INS-80
chr3-38792398-INS-728
chr3-38807592-INS-50
chr3-38830015-INS-1485
chr3-38836616-INS-448
chr3-38869517-INS-7665
chr3-38881442-INS-5041
chr3-38905876-INS-1017
chr3-38909970-INS-50
chr3-39018906-INS-225
chr3-39027499-INS-94
chr3-39031841-INS-62
chr3-39052858-INS-54
chr3-39073421-INS-51
chr3-39089841-INS-5750
chr3-39101523-INS-1194
chr3-39108335-INS-6172
chr3-39115795-INS-50
chr3-39134047-INS-3916
chr3-39152853-INS-50
chr3-39161274-INS-50
chr3-39192749-INS-659
chr3-39194388-INS-13933
chr3-39194432-INS-6199
chr3-39198615-INS-64
chr3-39202656-INS-50
chr3-39206007-INS-2089
chr3-39286127-INS-123
chr3-39313310-INS-54
chr3-39340329-INS-60
chr3-39348688-INS-50
chr3-39385107-INS-1091
chr3-39404802-INS-99
chr3-39406531-INS-144
chr3-39443264-INS-50
chr3-39461551-INS-112
chr3-39498703-INS-528

chr3-39511322-INS-67
chr3-39513750-INS-50
chr3-39516381-INS-6360
chr3-39539071-INS-2602
chr3-39545506-INS-5239
chr3-39548562-INS-3054
chr3-39621705-INS-52
chr3-39622382-INS-247
chr3-39626695-INS-69
chr3-39627365-INS-201
chr3-39629867-INS-268
chr3-39638800-INS-500
chr3-39658817-INS-3488
chr3-39693956-INS-60
chr3-39713405-INS-62
chr3-39714503-INS-522
chr3-39723082-INS-58
chr3-39723133-INS-6253
chr3-39769307-INS-52
chr3-39783655-INS-64
chr3-39809024-INS-6804
chr3-39852389-INS-2814
chr3-39877106-INS-1140
chr3-39986945-INS-6363
chr3-39994368-INS-54
chr3-40035657-INS-2795
chr3-40063026-INS-64
chr3-40073769-INS-2017
chr3-40122752-INS-54
chr3-40135976-INS-50
chr3-40289831-INS-1330
chr3-40338455-INS-68
chr3-40379386-INS-52
chr3-40381637-INS-1089
chr3-40423208-INS-782
chr3-40576350-INS-444
chr3-40604738-INS-1033
chr3-40605051-INS-486
chr3-40605539-INS-199
chr3-40605711-INS-57
chr3-40623219-INS-52
chr3-40624727-INS-197
chr3-40630966-INS-4839
chr3-40689150-INS-4866
chr3-40705038-INS-224
chr3-40770549-INS-229
chr3-40780774-INS-70
chr3-40873578-INS-822
chr3-40973081-INS-467
chr3-40980601-INS-3390
chr3-41076685-INS-7079
chr3-41080390-INS-116
chr3-41085509-INS-958
chr3-41099222-INS-215
chr3-41130022-INS-67
chr3-41131654-INS-1395
chr3-41148238-INS-241
chr3-41229223-INS-6457
chr3-41248629-INS-54
chr3-41269167-INS-2243
chr3-41286113-INS-1274
chr3-41288336-INS-51
chr3-41289828-INS-112
chr3-41337015-INS-830
chr3-41350272-INS-75
chr3-41356069-INS-52
chr3-41356779-INS-284
chr3-41356847-INS-118
chr3-41357613-INS-234
chr3-41357613-INS-78
chr3-41357677-INS-2028
chr3-41394590-INS-7070
chr3-41399629-INS-74
chr3-41450404-INS-205
chr3-41456180-INS-400
chr3-41481669-INS-492
chr3-41482198-INS-52

chr3-41515213-INS-2170
chr3-41549574-INS-265
chr3-41675523-INS-1020
chr3-41769462-INS-90
chr3-41785988-INS-388
chr3-41822845-INS-185
chr3-41831821-INS-9953
chr3-41840301-INS-9958
chr3-41872077-INS-60
chr3-41877796-INS-171
chr3-41904661-INS-4501
chr3-41906068-INS-1391
chr3-41975580-INS-96
chr3-41990288-INS-6796
chr3-42002266-INS-6882
chr3-42024468-INS-6938
chr3-42057445-INS-337
chr3-42070577-INS-832
chr3-42095188-INS-60
chr3-42103649-INS-434
chr3-42119295-INS-2001
chr3-42162603-INS-74
chr3-42180906-INS-71
chr3-42206327-INS-6638
chr3-42339757-INS-6376
chr3-42345076-INS-11825
chr3-42354664-INS-6333
chr3-42380645-INS-6369
chr3-42426073-INS-410
chr3-42432031-INS-1752
chr3-42449394-INS-50
chr3-42451198-INS-358
chr3-42490737-INS-6719
chr3-42535948-INS-524
chr3-42559032-INS-72
chr3-42565824-INS-65
chr3-42570906-INS-1492
chr3-42590005-INS-1104
chr3-42613839-INS-7898
chr3-42633074-INS-319
chr3-42654157-INS-50
chr3-42716146-INS-70
chr3-42739078-INS-247
chr3-42745689-INS-50
chr3-42786017-INS-52
chr3-42796602-INS-58
chr3-42823327-INS-64
chr3-42840278-INS-1890
chr3-42853379-INS-79
chr3-42868273-INS-76
chr3-42881312-INS-5203
chr3-42920733-INS-814
chr3-42948896-INS-2254
chr3-42951678-INS-328
chr3-42952709-INS-520
chr3-43013783-INS-2586
chr3-43067859-INS-500
chr3-43069337-INS-78
chr3-43124969-INS-50
chr3-43154734-INS-907
chr3-43195151-INS-2968
chr3-43244796-INS-50
chr3-43467694-INS-966
chr3-43469790-INS-350
chr3-43478435-INS-50
chr3-43481936-INS-62
chr3-43483761-INS-498
chr3-43486165-INS-1865
chr3-43517249-INS-1815
chr3-43545433-INS-8138
chr3-43608888-INS-6382
chr3-43610043-INS-108
chr3-43694198-INS-60
chr3-43777238-INS-2170
chr3-43785981-INS-66
chr3-43921369-INS-89
chr3-43944947-INS-1951

chr3-43958648-INS-60
chr3-43992422-INS-120
chr3-44013171-INS-97
chr3-44084216-INS-6506
chr3-44146849-INS-1284
chr3-44177922-INS-52
chr3-44240400-INS-211
chr3-44240969-INS-66
chr3-44315312-INS-5035
chr3-44346336-INS-3108
chr3-44383174-INS-52
chr3-44421288-INS-780
chr3-44432553-INS-2180
chr3-44434516-INS-85
chr3-44448357-INS-323
chr3-44456301-INS-6404
chr3-44517832-INS-36663
chr3-44549388-INS-6182
chr3-44552581-INS-59
chr3-44601687-INS-145
chr3-44632682-INS-172
chr3-44649396-INS-839
chr3-44694606-INS-54
chr3-44694669-INS-152
chr3-44702858-INS-105
chr3-44713985-INS-3326
chr3-44727537-INS-84
chr3-44835913-INS-6496
chr3-44837512-INS-56
chr3-44871068-INS-132
chr3-44887246-INS-4732
chr3-44896936-INS-2018
chr3-44923211-INS-1626
chr3-44926230-INS-13699
chr3-44940278-INS-81
chr3-44940424-INS-181
chr3-44956502-INS-4731
chr3-44959905-INS-410
chr3-44960721-INS-396
chr3-44973385-INS-280
chr3-44973521-INS-842
chr3-44987879-INS-52
chr3-45025545-INS-6382
chr3-45043418-INS-6206
chr3-45044169-INS-57
chr3-45088421-INS-417
chr3-45103592-INS-461
chr3-45167401-INS-111
chr3-45167436-INS-290
chr3-45167545-INS-89
chr3-45167707-INS-447
chr3-45207721-INS-50
chr3-45271949-INS-114
chr3-45272211-INS-1239
chr3-45279960-INS-9054
chr3-45279980-INS-656
chr3-45287210-INS-157
chr3-45392592-INS-101
chr3-45398718-INS-2008
chr3-45412630-INS-397
chr3-45413040-INS-1379
chr3-45444077-INS-6226
chr3-45472067-INS-904
chr3-45482749-INS-2368
chr3-45551157-INS-1093
chr3-45624404-INS-351
chr3-45634198-INS-104
chr3-45660940-INS-58
chr3-45668601-INS-185
chr3-45709731-INS-221
chr3-45727121-INS-6099
chr3-45728785-INS-2474
chr3-45734722-INS-1981
chr3-45751659-INS-1499
chr3-45756659-INS-72
chr3-45766534-INS-4937
chr3-45770817-INS-1190

chr3-45775669-INS-3588
chr3-45841076-INS-3220
chr3-45886593-INS-7060
chr3-45917828-INS-3796
chr3-45946519-INS-428
chr3-45978946-INS-6815
chr3-46005183-INS-66
chr3-46005241-INS-793
chr3-46011103-INS-50
chr3-46020754-INS-80
chr3-46025816-INS-62
chr3-46095474-INS-208
chr3-46136649-INS-7095
chr3-46165905-INS-165
chr3-46199746-INS-235
chr3-46224396-INS-54
chr3-46226002-INS-71
chr3-46269133-INS-4651
chr3-46280929-INS-50
chr3-46281243-INS-199
chr3-46325401-INS-437
chr3-46349611-INS-221
chr3-46372276-INS-57
chr3-46407199-INS-70
chr3-46426132-INS-67
chr3-46440363-INS-299
chr3-46484949-INS-1161
chr3-46501140-INS-96
chr3-46536879-INS-3412
chr3-46550123-INS-592
chr3-46576315-INS-81
chr3-46630183-INS-60
chr3-46640214-INS-58
chr3-46648210-INS-1576
chr3-46652349-INS-449
chr3-46708236-INS-2119
chr3-46711371-INS-445
chr3-46712070-INS-72
chr3-46717706-INS-1661
chr3-46729097-INS-50
chr3-46739152-INS-4087
chr3-46742901-INS-3980
chr3-46780749-INS-2996
chr3-46781669-INS-6171
chr3-46787717-INS-72
chr3-46788321-INS-6575
chr3-46821283-INS-101
chr3-46823839-INS-52
chr3-46843875-INS-634
chr3-46854716-INS-168
chr3-46873118-INS-57
chr3-46898446-INS-56
chr3-46901784-INS-50
chr3-46945955-INS-387
chr3-46945955-INS-869
chr3-46982156-INS-133
chr3-47017086-INS-124
chr3-47025330-INS-40266
chr3-47063606-INS-93
chr3-47131626-INS-146
chr3-47152277-INS-236
chr3-47186546-INS-54
chr3-47205144-INS-1382
chr3-47249076-INS-1096
chr3-47281506-INS-87
chr3-47299515-INS-51
chr3-47301136-INS-169
chr3-47321798-INS-66
chr3-47323935-INS-172
chr3-47351358-INS-1053
chr3-47351534-INS-67
chr3-47354445-INS-100
chr3-47405709-INS-96
chr3-47406044-INS-424
chr3-47415005-INS-476
chr3-47455227-INS-1588
chr3-47464673-INS-6848

chr3-47472295-INS-255
chr3-47507759-INS-9004
chr3-47509562-INS-4087
chr3-47521693-INS-1179
chr3-47527525-INS-9462
chr3-47554715-INS-56
chr3-47585078-INS-270
chr3-47585321-INS-54
chr3-47588927-INS-753
chr3-47620229-INS-88
chr3-47640444-INS-2417
chr3-47655346-INS-70
chr3-47670891-INS-6222
chr3-47691394-INS-6180
chr3-47692009-INS-69
chr3-47706832-INS-50
chr3-47722909-INS-1117
chr3-47734559-INS-3535
chr3-47756465-INS-52
chr3-47817877-INS-65
chr3-47886790-INS-6391
chr3-47894527-INS-7080
chr3-47899045-INS-6248
chr3-47909498-INS-1584
chr3-47927665-INS-65
chr3-47994682-INS-870
chr3-48000155-INS-6339
chr3-48016726-INS-6256
chr3-48056102-INS-698
chr3-48078759-INS-524
chr3-48082440-INS-146
chr3-48117119-INS-78
chr3-48164185-INS-199
chr3-48188727-INS-84
chr3-48215058-INS-53
chr3-48216391-INS-72
chr3-48284802-INS-52
chr3-48315990-INS-1751
chr3-48329611-INS-996
chr3-48333263-INS-5516
chr3-48420828-INS-120
chr3-48435589-INS-64
chr3-48447470-INS-2202
chr3-48453186-INS-3067
chr3-48463143-INS-6246
chr3-48472972-INS-87
chr3-48477415-INS-6442
chr3-48531504-INS-145
chr3-48550051-INS-50
chr3-48550599-INS-7132
chr3-48640189-INS-386
chr3-48640223-INS-50
chr3-48701318-INS-4428
chr3-48720494-INS-51
chr3-48772819-INS-88
chr3-48784902-INS-4344
chr3-48814599-INS-60
chr3-48816554-INS-58
chr3-48839827-INS-55
chr3-48873981-INS-2916
chr3-48903742-INS-545
chr3-48925443-INS-5841
chr3-48940519-INS-102
chr3-48954224-INS-78
chr3-48955158-INS-132
chr3-48976935-INS-2490
chr3-48995666-INS-54
chr3-49004667-INS-6834
chr3-49012513-INS-200
chr3-49036438-INS-118
chr3-49077954-INS-960
chr3-49119822-INS-3103
chr3-49122478-INS-6769
chr3-49166789-INS-4581
chr3-49192023-INS-6307
chr3-49221384-INS-203
chr3-49238383-INS-122

chr3-49253590-INS-1742
chr3-49261946-INS-4580
chr3-49281424-INS-2896
chr3-49318270-INS-2789
chr3-49350316-INS-7862
chr3-49373214-INS-466
chr3-49443927-INS-66
chr3-49453786-INS-241
chr3-49474984-INS-63
chr3-49484336-INS-562
chr3-49486827-INS-4164
chr3-49497777-INS-6343
chr3-49507984-INS-193
chr3-49535849-INS-94
chr3-49582224-INS-54
chr3-49590743-INS-7071
chr3-49659078-INS-123
chr3-49791210-INS-50
chr3-49876784-INS-2794
chr3-49904811-INS-6396
chr3-49905413-INS-140
chr3-49952958-INS-118
chr3-50017581-INS-891
chr3-50039964-INS-71
chr3-50082556-INS-60
chr3-50083241-INS-250
chr3-50116830-INS-138
chr3-50136524-INS-6415
chr3-50183411-INS-1227
chr3-50208337-INS-1093
chr3-50210854-INS-231
chr3-50221915-INS-50
chr3-50227995-INS-675
chr3-50227995-INS-84
chr3-50235662-INS-78
chr3-50246389-INS-62
chr3-50402316-INS-222
chr3-50407113-INS-63
chr3-50491851-INS-63
chr3-50501462-INS-208
chr3-50551088-INS-1039
chr3-50594243-INS-1245
chr3-50597661-INS-7103
chr3-50620787-INS-50
chr3-50666857-INS-126
chr3-50692797-INS-310
chr3-50721006-INS-66
chr3-50753260-INS-2836
chr3-50754863-INS-7309
chr3-50768809-INS-1241
chr3-50818448-INS-917
chr3-50826645-INS-50
chr3-50829229-INS-65
chr3-50844293-INS-50
chr3-50859240-INS-56
chr3-50886761-INS-222
chr3-50896405-INS-69
chr3-50896415-INS-414
chr3-50900879-INS-72
chr3-50907870-INS-127
chr3-50911312-INS-192
chr3-50916296-INS-6559
chr3-50917925-INS-56
chr3-50919243-INS-158
chr3-50962765-INS-320
chr3-50971792-INS-2640
chr3-50977708-INS-50
chr3-50998646-INS-723
chr3-50998723-INS-75
chr3-51006910-INS-121
chr3-51008475-INS-3782
chr3-51028663-INS-114
chr3-51030678-INS-255
chr3-51055759-INS-128
chr3-51060446-INS-53
chr3-51100766-INS-467
chr3-51120965-INS-5560

chr3-51158806-INS-219
chr3-51177442-INS-87
chr3-51180593-INS-78
chr3-51194799-INS-64
chr3-51198230-INS-50
chr3-51200075-INS-82
chr3-51200804-INS-134
chr3-51201602-INS-144
chr3-51209474-INS-174
chr3-51209508-INS-82
chr3-51209678-INS-486
chr3-51212200-INS-438
chr3-51240250-INS-216
chr3-51252527-INS-4979
chr3-51253065-INS-294
chr3-51263863-INS-63
chr3-51271491-INS-373
chr3-51283945-INS-995
chr3-51289394-INS-50
chr3-51300607-INS-194
chr3-51335873-INS-69
chr3-51340082-INS-212
chr3-51375661-INS-394
chr3-51387357-INS-960
chr3-51393693-INS-195
chr3-51460592-INS-69
chr3-51475252-INS-68
chr3-51475531-INS-131
chr3-51488173-INS-218
chr3-51522436-INS-74
chr3-51532518-INS-3423
chr3-51532972-INS-78
chr3-51535666-INS-114
chr3-51619647-INS-151
chr3-51621689-INS-83
chr3-51673755-INS-1522
chr3-51673990-INS-2934
chr3-51678796-INS-860
chr3-51686297-INS-67
chr3-51703902-INS-60
chr3-51719436-INS-56
chr3-51727715-INS-50
chr3-51741792-INS-290
chr3-51741799-INS-725
chr3-51765165-INS-490
chr3-51765195-INS-74
chr3-51864251-INS-66
chr3-51874600-INS-86
chr3-51900170-INS-3394
chr3-51920986-INS-62
chr3-51946056-INS-201
chr3-52016742-INS-50
chr3-52017479-INS-116
chr3-52068182-INS-52
chr3-52087570-INS-63
chr3-52096325-INS-164
chr3-52097108-INS-173
chr3-52158205-INS-227
chr3-52164124-INS-171
chr3-52208257-INS-925
chr3-52208267-INS-242
chr3-52247025-INS-279
chr3-52261487-INS-80
chr3-52262546-INS-80
chr3-52269300-INS-111
chr3-52309797-INS-193
chr3-52319761-INS-196
chr3-52323631-INS-52
chr3-52361871-INS-341
chr3-52387484-INS-83
chr3-52412827-INS-232
chr3-52425250-INS-61
chr3-52427562-INS-106
chr3-52492742-INS-126
chr3-52504516-INS-134
chr3-52509909-INS-51
chr3-52527302-INS-55

chr3-52573980-INS-196
chr3-52575080-INS-56
chr3-52581214-INS-6361
chr3-52642656-INS-93
chr3-52644085-INS-64
chr3-52693862-INS-50
chr3-52706755-INS-441
chr3-52718358-INS-806
chr3-52734464-INS-62
chr3-52739081-INS-80
chr3-52775832-INS-79
chr3-52825184-INS-416
chr3-52833330-INS-120
chr3-52845638-INS-190
chr3-52885352-INS-53
chr3-52901013-INS-50
chr3-52902766-INS-707
chr3-52902779-INS-343
chr3-52917404-INS-56
chr3-52969997-INS-259
chr3-53081925-INS-122
chr3-53082865-INS-261
chr3-53103719-INS-4746
chr3-53121843-INS-371
chr3-53137481-INS-50
chr3-53161763-INS-4113
chr3-53173094-INS-3136
chr3-53188207-INS-4281
chr3-53203846-INS-6335
chr3-53230978-INS-64
chr3-53231087-INS-144
chr3-53282852-INS-6439
chr3-53288110-INS-60
chr3-53290168-INS-2499
chr3-53306010-INS-94
chr3-53332986-INS-485
chr3-53354094-INS-60
chr3-53363514-INS-65
chr3-53406401-INS-189
chr3-53407354-INS-60
chr3-53409227-INS-56
chr3-53409521-INS-1240
chr3-53413006-INS-751
chr3-53428392-INS-277
chr3-53439132-INS-1381
chr3-53442363-INS-65
chr3-53517593-INS-108
chr3-53558983-INS-370
chr3-53559070-INS-177
chr3-53588454-INS-3560
chr3-53590906-INS-374
chr3-53612611-INS-59
chr3-53617342-INS-137
chr3-53618857-INS-181
chr3-53630058-INS-257
chr3-53633781-INS-68
chr3-53676578-INS-193
chr3-53682990-INS-144
chr3-53683014-INS-50
chr3-53683284-INS-789
chr3-53692511-INS-6907
chr3-53707940-INS-229
chr3-53717222-INS-68
chr3-53720225-INS-184
chr3-53720455-INS-110
chr3-53741158-INS-50
chr3-53743463-INS-90
chr3-53794501-INS-64
chr3-53800878-INS-63
chr3-53817030-INS-206
chr3-53823452-INS-7116
chr3-53829555-INS-6564
chr3-53862641-INS-852
chr3-53880461-INS-54
chr3-53991604-INS-100
chr3-54005623-INS-6835
chr3-54011998-INS-198

chr3-54012325-INS-124
chr3-54014241-INS-52
chr3-54015340-INS-500
chr3-54044980-INS-394
chr3-54102590-INS-258
chr3-54113342-INS-56
chr3-54177109-INS-56
chr3-54183934-INS-3305
chr3-54208203-INS-52
chr3-54209084-INS-60
chr3-54322989-INS-1202
chr3-54326886-INS-113
chr3-54346886-INS-231
chr3-54368650-INS-218
chr3-54380436-INS-52
chr3-54391166-INS-1162
chr3-54397239-INS-6590
chr3-54401649-INS-51
chr3-54401704-INS-166
chr3-54433793-INS-1074
chr3-54441013-INS-100
chr3-54456483-INS-1583
chr3-54457944-INS-51
chr3-54461123-INS-125
chr3-54479885-INS-108
chr3-54496484-INS-58
chr3-54500379-INS-7080
chr3-54503762-INS-567
chr3-54503985-INS-2792
chr3-54504031-INS-180
chr3-54527641-INS-198
chr3-54543233-INS-1658
chr3-54556936-INS-2052
chr3-54660818-INS-63
chr3-54690527-INS-55
chr3-54697771-INS-62
chr3-54740882-INS-206
chr3-54769595-INS-56
chr3-54783813-INS-78
chr3-54791744-INS-84
chr3-54811722-INS-152
chr3-54855069-INS-60
chr3-54915718-INS-228
chr3-54915766-INS-479
chr3-54916013-INS-55
chr3-54929533-INS-6217
chr3-54931084-INS-78
chr3-54937586-INS-282
chr3-54965401-INS-549
chr3-54975493-INS-241
chr3-54976545-INS-732
chr3-54988215-INS-4438
chr3-54996878-INS-50
chr3-55007249-INS-74
chr3-55014353-INS-268
chr3-55017699-INS-88
chr3-55039497-INS-730
chr3-55137599-INS-1036
chr3-55146158-INS-195
chr3-55146160-INS-66
chr3-55202688-INS-384
chr3-55236128-INS-56
chr3-55281066-INS-74
chr3-55323648-INS-250
chr3-55323648-INS-50
chr3-55343718-INS-820
chr3-55381465-INS-62
chr3-55383088-INS-50
chr3-55408739-INS-691
chr3-55460913-INS-353
chr3-55461656-INS-140
chr3-55470231-INS-3002
chr3-55487679-INS-475
chr3-55506463-INS-150
chr3-55518990-INS-54
chr3-55519892-INS-50
chr3-55521350-INS-60

chr3-55541370-INS-358
chr3-55541977-INS-270
chr3-55568813-INS-64
chr3-55577713-INS-56
chr3-55659779-INS-3557
chr3-55710560-INS-331
chr3-55710582-INS-678
chr3-55716424-INS-300
chr3-55746061-INS-52
chr3-55757286-INS-234
chr3-55757320-INS-66
chr3-55788425-INS-9511
chr3-55790595-INS-1085
chr3-55801089-INS-134
chr3-55823115-INS-55
chr3-55902060-INS-366
chr3-55902150-INS-122
chr3-55902803-INS-128
chr3-56037809-INS-143
chr3-56037927-INS-857
chr3-56066841-INS-456
chr3-56071230-INS-281
chr3-56084455-INS-1134
chr3-56084586-INS-116
chr3-56119506-INS-1118
chr3-56162545-INS-114
chr3-56169286-INS-3944
chr3-56189326-INS-68
chr3-56189580-INS-101
chr3-56215793-INS-50
chr3-56224750-INS-53
chr3-56227575-INS-62
chr3-56228461-INS-65
chr3-56244345-INS-7151
chr3-56324170-INS-705
chr3-56324369-INS-71
chr3-56324507-INS-244
chr3-56329840-INS-1307
chr3-56376185-INS-60
chr3-56376207-INS-157
chr3-56478436-INS-266
chr3-56483202-INS-50
chr3-56569595-INS-5092
chr3-56584946-INS-3253
chr3-56596610-INS-56
chr3-56607314-INS-186
chr3-56609103-INS-6439
chr3-56621714-INS-210
chr3-56653125-INS-498
chr3-56744455-INS-62
chr3-56745034-INS-148
chr3-56785493-INS-1476
chr3-56874347-INS-56
chr3-56886433-INS-6536
chr3-56950692-INS-3282
chr3-56962881-INS-68
chr3-56964038-INS-480
chr3-56976965-INS-1411
chr3-56996542-INS-50
chr3-57022406-INS-6490
chr3-57039602-INS-53
chr3-57046780-INS-53
chr3-57069692-INS-1596
chr3-57079117-INS-118
chr3-57118819-INS-6713
chr3-57127272-INS-80
chr3-57188955-INS-3086
chr3-57193898-INS-1145
chr3-57216030-INS-4860
chr3-57226273-INS-285
chr3-57299132-INS-52
chr3-57321933-INS-6228
chr3-57324807-INS-1280
chr3-57346609-INS-210
chr3-57346947-INS-54
chr3-57352037-INS-6612
chr3-57371647-INS-2932

chr3-57405568-INS-64
chr3-57409014-INS-500
chr3-57444559-INS-6273
chr3-57471571-INS-55
chr3-57493153-INS-187
chr3-57523696-INS-50
chr3-57532689-INS-219
chr3-57534324-INS-125
chr3-57601370-INS-154
chr3-57621193-INS-250
chr3-57628192-INS-502
chr3-57628606-INS-1041
chr3-57653202-INS-3037
chr3-57672707-INS-137
chr3-57709571-INS-180
chr3-57725181-INS-1060
chr3-57735519-INS-60
chr3-57824847-INS-72
chr3-57825616-INS-50
chr3-57832198-INS-1910
chr3-57846639-INS-79
chr3-57897549-INS-76
chr3-57905130-INS-1864
chr3-57920160-INS-201
chr3-57920312-INS-202
chr3-57924835-INS-1453
chr3-57956416-INS-63
chr3-57977574-INS-96
chr3-58027344-INS-50
chr3-58036925-INS-1265
chr3-58065503-INS-7543
chr3-58078746-INS-215
chr3-58115322-INS-735
chr3-58121681-INS-5038
chr3-58134271-INS-13872
chr3-58154452-INS-1926
chr3-58155909-INS-51
chr3-58164488-INS-921
chr3-58180377-INS-500
chr3-58190703-INS-53
chr3-58194810-INS-192
chr3-58211030-INS-287
chr3-58239422-INS-68
chr3-58380047-INS-98
chr3-58408356-INS-56
chr3-58458923-INS-54
chr3-58522227-INS-132
chr3-58529161-INS-610
chr3-58530723-INS-6004
chr3-58567931-INS-62
chr3-58714056-INS-50
chr3-58723516-INS-54
chr3-58742249-INS-197
chr3-58793648-INS-120
chr3-58832122-INS-8658
chr3-58838951-INS-4751
chr3-58838951-INS-759
chr3-58887444-INS-143
chr3-58892062-INS-71
chr3-58907537-INS-7126
chr3-58940000-INS-105
chr3-59016355-INS-68
chr3-59045531-INS-1111
chr3-59097634-INS-1024
chr3-59097640-INS-342
chr3-59128337-INS-240
chr3-59161722-INS-208
chr3-59179181-INS-81
chr3-59200778-INS-51
chr3-59216025-INS-1493
chr3-59332388-INS-457
chr3-59359653-INS-619
chr3-59360655-INS-1231
chr3-59364899-INS-1164
chr3-59366391-INS-66
chr3-59385114-INS-1333
chr3-59388176-INS-6466

chr3-59393398-INS-1387
chr3-59407764-INS-1125
chr3-59467588-INS-76
chr3-59602488-INS-6892
chr3-59611874-INS-365
chr3-59612009-INS-52
chr3-59612346-INS-79
chr3-59612489-INS-184
chr3-59721147-INS-58
chr3-59724450-INS-56
chr3-59724616-INS-64
chr3-59738022-INS-4138
chr3-59754951-INS-80
chr3-59782404-INS-150
chr3-59782404-INS-60
chr3-59796859-INS-2895
chr3-59808957-INS-6414
chr3-59810458-INS-226
chr3-59853376-INS-372
chr3-59865470-INS-70
chr3-59952936-INS-263
chr3-59956893-INS-80
chr3-59966964-INS-6325
chr3-59991635-INS-2193
chr3-60006371-INS-5871
chr3-60008960-INS-50
chr3-60111890-INS-64
chr3-60158456-INS-1204
chr3-60233418-INS-60
chr3-60241497-INS-1118
chr3-60250324-INS-1500
chr3-60252631-INS-56
chr3-60266130-INS-6266
chr3-60278443-INS-6553
chr3-60294640-INS-64
chr3-60304009-INS-3798
chr3-60314504-INS-5729
chr3-60367804-INS-2696
chr3-60439530-INS-11849
chr3-60542926-INS-74
chr3-60579235-INS-96
chr3-60600020-INS-813
chr3-60605550-INS-80
chr3-60615829-INS-212
chr3-60615835-INS-50
chr3-60625622-INS-351
chr3-60693390-INS-74
chr3-60722119-INS-16242
chr3-60745629-INS-6364
chr3-60825752-INS-56
chr3-60829211-INS-1794
chr3-60921104-INS-65
chr3-60937661-INS-2671
chr3-60993430-INS-123
chr3-61024392-INS-1266
chr3-61130874-INS-52
chr3-61134342-INS-6840
chr3-61189720-INS-165
chr3-61219698-INS-364
chr3-61266681-INS-503
chr3-61276196-INS-56
chr3-61286663-INS-100
chr3-61365447-INS-82
chr3-61368557-INS-559
chr3-61397717-INS-54
chr3-61462529-INS-525
chr3-61511349-INS-53
chr3-61605382-INS-1909
chr3-61659927-INS-52
chr3-61661333-INS-6795
chr3-61730361-INS-51
chr3-61730364-INS-130
chr3-61738091-INS-60
chr3-61777886-INS-50
chr3-61786660-INS-656
chr3-61791119-INS-69
chr3-61814456-INS-6311

chr3-61816878-INS-54
chr3-61834485-INS-620
chr3-61850649-INS-3098
chr3-61856278-INS-6212
chr3-61924223-INS-1091
chr3-61931920-INS-96
chr3-61974181-INS-7334
chr3-61978031-INS-485
chr3-61995856-INS-70
chr3-62029046-INS-3132
chr3-62086173-INS-72
chr3-62091123-INS-50
chr3-62091784-INS-119
chr3-62123136-INS-102
chr3-62132173-INS-1170
chr3-62182175-INS-7068
chr3-62199787-INS-884
chr3-62202219-INS-156
chr3-62218780-INS-122
chr3-62315536-INS-55
chr3-62356991-INS-205
chr3-62357812-INS-58
chr3-62446758-INS-68
chr3-62455119-INS-1545
chr3-62461216-INS-577
chr3-62504701-INS-131
chr3-62521169-INS-6575
chr3-62547506-INS-60
chr3-62664400-INS-66
chr3-62665075-INS-203
chr3-62701101-INS-3662
chr3-62716299-INS-6299
chr3-62745692-INS-480
chr3-62771682-INS-68
chr3-62819848-INS-527
chr3-62826903-INS-188
chr3-62908031-INS-2358
chr3-62924338-INS-65
chr3-62925816-INS-50
chr3-62931700-INS-144
chr3-62949350-INS-188
chr3-62949354-INS-83
chr3-62974014-INS-66
chr3-63125555-INS-65
chr3-63128406-INS-360
chr3-63144085-INS-68
chr3-63146342-INS-7725
chr3-63236249-INS-66
chr3-63308062-INS-306
chr3-63308088-INS-129
chr3-63337997-INS-2580
chr3-63374720-INS-6290
chr3-63395775-INS-6141
chr3-63437354-INS-200
chr3-63440071-INS-64
chr3-63455302-INS-51
chr3-63466413-INS-54
chr3-63573710-INS-127
chr3-63573929-INS-245
chr3-63677144-INS-65
chr3-63781819-INS-63
chr3-63820998-INS-163
chr3-63843245-INS-289
chr3-63917307-INS-226
chr3-64017161-INS-9426
chr3-64079364-INS-320
chr3-64152777-INS-9866
chr3-64318537-INS-118
chr3-64319042-INS-55
chr3-64319819-INS-86
chr3-64323376-INS-52
chr3-64409678-INS-66
chr3-64410001-INS-52
chr3-64629504-INS-62
chr3-64642187-INS-52
chr3-64647747-INS-72
chr3-64768682-INS-8296

chr3-64776295-INS-288
chr3-64787118-INS-52
chr3-64807822-INS-382
chr3-64814525-INS-6802
chr3-64853822-INS-661
chr3-64874099-INS-56
chr3-64924233-INS-1693
chr3-64924937-INS-138
chr3-64925071-INS-149
chr3-64958916-INS-6402
chr3-64964678-INS-62
chr3-64965070-INS-355
chr3-64967894-INS-112
chr3-65009452-INS-52
chr3-65048209-INS-52
chr3-65116359-INS-127
chr3-65138379-INS-57
chr3-65142204-INS-50
chr3-65142324-INS-89
chr3-65143561-INS-1076
chr3-65144478-INS-376
chr3-65146853-INS-66
chr3-65165607-INS-116
chr3-65168352-INS-84
chr3-65175556-INS-174
chr3-65177079-INS-54
chr3-65182016-INS-375
chr3-65182016-INS-50
chr3-65182669-INS-60
chr3-65187335-INS-90
chr3-65189183-INS-5379
chr3-65197027-INS-1891
chr3-65221718-INS-97
chr3-65258808-INS-219
chr3-65281499-INS-66
chr3-65304319-INS-1291
chr3-65307886-INS-7385
chr3-65328972-INS-119
chr3-65329573-INS-99
chr3-65374370-INS-123
chr3-65388526-INS-4794
chr3-65424948-INS-209
chr3-65492124-INS-63
chr3-65514509-INS-254
chr3-65514509-INS-59
chr3-65538152-INS-200
chr3-65592185-INS-50
chr3-65602883-INS-51
chr3-65654829-INS-159
chr3-65714860-INS-88
chr3-65715236-INS-60
chr3-65814345-INS-50
chr3-65832312-INS-162
chr3-65849955-INS-138
chr3-65885984-INS-2920
chr3-65895818-INS-64
chr3-65904124-INS-1888
chr3-65907296-INS-69
chr3-65930446-INS-479
chr3-65932560-INS-4933
chr3-65939921-INS-66
chr3-65949664-INS-486
chr3-65951518-INS-3290
chr3-65971387-INS-3236
chr3-65979299-INS-208
chr3-66015156-INS-6394
chr3-66017106-INS-76
chr3-66022974-INS-50
chr3-66023007-INS-346
chr3-66059071-INS-6370
chr3-66072375-INS-940
chr3-66075672-INS-1740
chr3-66112872-INS-72
chr3-66116164-INS-52
chr3-66156575-INS-56
chr3-66166504-INS-68
chr3-66176396-INS-166

chr3-66188120-INS-814
chr3-66237114-INS-208
chr3-66244210-INS-50
chr3-66256369-INS-562
chr3-66257264-INS-2955
chr3-66261559-INS-543
chr3-66265289-INS-6294
chr3-66265790-INS-130
chr3-66269245-INS-306
chr3-66291451-INS-61
chr3-66293442-INS-358
chr3-66301139-INS-2936
chr3-66426449-INS-260
chr3-66426608-INS-159
chr3-66466439-INS-142
chr3-66514094-INS-73
chr3-66514511-INS-1801
chr3-66515848-INS-53
chr3-66549693-INS-498
chr3-66600525-INS-641
chr3-66635648-INS-6556
chr3-66636404-INS-62
chr3-66670292-INS-338
chr3-66716918-INS-126
chr3-66755094-INS-60
chr3-66789372-INS-66
chr3-66789621-INS-112
chr3-66789626-INS-1003
chr3-66790470-INS-1243
chr3-66803532-INS-927
chr3-66892431-INS-52
chr3-66894708-INS-53
chr3-66900570-INS-68
chr3-66904262-INS-214
chr3-66953551-INS-1739
chr3-66953569-INS-252
chr3-66953591-INS-116
chr3-66956877-INS-52
chr3-66980982-INS-146
chr3-66980982-INS-66
chr3-66981109-INS-406
chr3-67011958-INS-1751
chr3-67027307-INS-6765
chr3-67033843-INS-6482
chr3-67048532-INS-6289
chr3-67116839-INS-490
chr3-67164166-INS-221
chr3-67194618-INS-63
chr3-67195198-INS-483
chr3-67297678-INS-163
chr3-67307374-INS-125
chr3-67336415-INS-1048
chr3-67346656-INS-55
chr3-67359071-INS-50
chr3-67378210-INS-50
chr3-67381517-INS-70
chr3-67417236-INS-207
chr3-67449779-INS-492
chr3-67472889-INS-50
chr3-67476766-INS-145
chr3-67480743-INS-283
chr3-67481445-INS-260
chr3-67495984-INS-200
chr3-67497792-INS-60
chr3-67511619-INS-4142
chr3-67525531-INS-58
chr3-67526075-INS-483
chr3-67535844-INS-2050
chr3-67539721-INS-50
chr3-67541036-INS-6465
chr3-67544684-INS-78
chr3-67549469-INS-1849
chr3-67604185-INS-177
chr3-67613515-INS-2844
chr3-67623357-INS-304
chr3-67624001-INS-507
chr3-67624790-INS-104

chr3-67627252-INS-65
chr3-67632839-INS-171
chr3-67637704-INS-6037
chr3-67643130-INS-50
chr3-67668743-INS-88
chr3-67668914-INS-78
chr3-67674612-INS-99
chr3-67726855-INS-79
chr3-67745790-INS-203
chr3-67763680-INS-195
chr3-67763680-INS-425
chr3-67786879-INS-57
chr3-67790861-INS-70
chr3-67794428-INS-85
chr3-67833215-INS-50
chr3-67856382-INS-6399
chr3-67875265-INS-460
chr3-67885502-INS-66
chr3-67951282-INS-54
chr3-67991146-INS-178
chr3-67992344-INS-7157
chr3-68014963-INS-63
chr3-68033239-INS-141
chr3-68098385-INS-7419
chr3-68121481-INS-52
chr3-68123176-INS-1271
chr3-68124893-INS-1880
chr3-68131916-INS-2141
chr3-68136346-INS-2387
chr3-68155997-INS-50
chr3-68181081-INS-6187
chr3-68196120-INS-2114
chr3-68208717-INS-5476
chr3-68211247-INS-4041
chr3-68225334-INS-102
chr3-68226193-INS-587
chr3-68234938-INS-168
chr3-68235538-INS-269
chr3-68250675-INS-1067
chr3-68268571-INS-104
chr3-68268765-INS-152
chr3-68286527-INS-50
chr3-68297869-INS-89
chr3-68300553-INS-379
chr3-68347361-INS-135
chr3-68409909-INS-186
chr3-68409998-INS-392
chr3-68414588-INS-204
chr3-68485249-INS-122
chr3-68500906-INS-132
chr3-68501313-INS-123
chr3-68502024-INS-52
chr3-68505283-INS-51
chr3-68521117-INS-142
chr3-68527220-INS-5259
chr3-68566784-INS-180
chr3-68570917-INS-138
chr3-68606737-INS-100
chr3-68664434-INS-469
chr3-68691575-INS-1649
chr3-68691612-INS-426
chr3-68755294-INS-96
chr3-68765518-INS-444
chr3-68766654-INS-54
chr3-68855784-INS-108
chr3-68870174-INS-221
chr3-68896127-INS-375
chr3-68901744-INS-310
chr3-68901744-INS-62
chr3-68950527-INS-96
chr3-69012006-INS-492
chr3-69054981-INS-196
chr3-69059553-INS-159
chr3-69061555-INS-262
chr3-69142730-INS-6374
chr3-69146073-INS-1143
chr3-69159610-INS-64

chr3-69162812-INS-103
chr3-69176718-INS-1028
chr3-69178435-INS-2595
chr3-69191516-INS-129
chr3-69204284-INS-422
chr3-69220317-INS-90
chr3-69242061-INS-623
chr3-69273652-INS-344
chr3-69283031-INS-1149
chr3-69283861-INS-1766
chr3-69284179-INS-120
chr3-69284429-INS-207
chr3-69305542-INS-56
chr3-69310832-INS-77
chr3-69331256-INS-580
chr3-69331660-INS-52
chr3-69336897-INS-56
chr3-69344388-INS-62
chr3-69348603-INS-3167
chr3-69369662-INS-57
chr3-69412259-INS-9601
chr3-69421921-INS-205
chr3-69434271-INS-56
chr3-69440596-INS-2865
chr3-69448028-INS-1341
chr3-69457749-INS-183
chr3-69508076-INS-50
chr3-69513691-INS-375
chr3-69539315-INS-845
chr3-69542314-INS-107
chr3-69552878-INS-214
chr3-69580744-INS-10500
chr3-69580749-INS-3473
chr3-69588207-INS-3239
chr3-69595068-INS-781
chr3-69606969-INS-52
chr3-69617531-INS-1178
chr3-69627811-INS-233
chr3-69683278-INS-553
chr3-69705738-INS-6411
chr3-69752058-INS-64
chr3-69773019-INS-69
chr3-69792145-INS-154
chr3-69796168-INS-140
chr3-69848907-INS-5983
chr3-69853503-INS-1847
chr3-69880063-INS-6598
chr3-69887715-INS-50
chr3-69891737-INS-639
chr3-69899153-INS-1281
chr3-69903677-INS-50
chr3-69950971-INS-51
chr3-69974661-INS-1999
chr3-70008268-INS-52
chr3-70021455-INS-102
chr3-70021787-INS-354
chr3-70029180-INS-67
chr3-70033408-INS-259
chr3-70034327-INS-6888
chr3-70065924-INS-76
chr3-70082966-INS-2535
chr3-70087619-INS-50
chr3-70112123-INS-131
chr3-70114411-INS-5161
chr3-70124780-INS-4616
chr3-70127324-INS-69
chr3-70130008-INS-58
chr3-70130010-INS-338
chr3-70133610-INS-3599
chr3-70135029-INS-4520
chr3-70141917-INS-64
chr3-70176528-INS-50
chr3-70182358-INS-1864
chr3-70206744-INS-3126
chr3-70226823-INS-1129
chr3-70230824-INS-96
chr3-70246469-INS-7067

chr3-70302138-INS-66
chr3-70333534-INS-54
chr3-70333867-INS-720
chr3-70346880-INS-54
chr3-70354613-INS-1491
chr3-70355555-INS-119
chr3-70372968-INS-2273
chr3-70394442-INS-55
chr3-70396839-INS-126
chr3-70396923-INS-395
chr3-70398609-INS-82
chr3-70430263-INS-101
chr3-70430550-INS-128
chr3-70567415-INS-382
chr3-70608274-INS-68
chr3-70608949-INS-239
chr3-70609012-INS-58
chr3-70609401-INS-1102
chr3-70623155-INS-50
chr3-70638426-INS-1890
chr3-70650132-INS-1201
chr3-70655377-INS-550
chr3-70670639-INS-186
chr3-70710432-INS-58
chr3-70728007-INS-5625
chr3-70751746-INS-113
chr3-70758746-INS-158
chr3-70766146-INS-154
chr3-70777064-INS-60
chr3-70778580-INS-4495
chr3-70788013-INS-100
chr3-70801336-INS-59
chr3-70811016-INS-2879
chr3-70816703-INS-2782
chr3-70827053-INS-160
chr3-70827053-INS-64
chr3-70864352-INS-6369
chr3-70876450-INS-1259
chr3-70909106-INS-54
chr3-70909887-INS-208
chr3-70917536-INS-6387
chr3-70927714-INS-2678
chr3-70943836-INS-1024
chr3-70974026-INS-498
chr3-70993144-INS-227
chr3-70997608-INS-254
chr3-71007005-INS-371
chr3-71007413-INS-445
chr3-71014981-INS-482
chr3-71039543-INS-727
chr3-71084316-INS-6530
chr3-71191070-INS-9684
chr3-71197579-INS-245
chr3-71239821-INS-219
chr3-71247393-INS-2774
chr3-71250041-INS-67
chr3-71250412-INS-651
chr3-71254163-INS-4792
chr3-71287062-INS-469
chr3-71344888-INS-201
chr3-71373238-INS-396
chr3-71381926-INS-537
chr3-71383963-INS-213
chr3-71388080-INS-1448
chr3-71400244-INS-6458
chr3-71402939-INS-1493
chr3-71461199-INS-50
chr3-71473822-INS-6607
chr3-71530601-INS-7017
chr3-71574044-INS-1057
chr3-71576819-INS-66
chr3-71630903-INS-60
chr3-71654159-INS-6340
chr3-71665646-INS-4613
chr3-71689562-INS-7208
chr3-71710273-INS-60
chr3-71730219-INS-71

chr3-71740728-INS-78
chr3-71756176-INS-207
chr3-71765587-INS-319
chr3-71776110-INS-295
chr3-71823761-INS-50
chr3-71838727-INS-60
chr3-71839075-INS-93
chr3-71870282-INS-2688
chr3-71892490-INS-50
chr3-71931602-INS-50
chr3-71973123-INS-114
chr3-72058618-INS-1986
chr3-72077904-INS-7123
chr3-72132895-INS-113
chr3-72161153-INS-1862
chr3-72162241-INS-68
chr3-72162419-INS-273
chr3-72260768-INS-600
chr3-72275640-INS-390
chr3-72282266-INS-72
chr3-72287587-INS-64
chr3-72293983-INS-6423
chr3-72316271-INS-51
chr3-72336676-INS-3464
chr3-72336816-INS-57
chr3-72340139-INS-239
chr3-72340500-INS-294
chr3-72351734-INS-1499
chr3-72353288-INS-269
chr3-72361242-INS-64
chr3-72366718-INS-6369
chr3-72415224-INS-816
chr3-72631623-INS-54
chr3-72746542-INS-50
chr3-72756748-INS-5976
chr3-72870122-INS-998
chr3-72879842-INS-496
chr3-72893216-INS-6581
chr3-72912336-INS-434
chr3-72942759-INS-477
chr3-72944710-INS-52
chr3-72951312-INS-52
chr3-72987310-INS-1361
chr3-73007038-INS-66
chr3-73023421-INS-554
chr3-73033802-INS-604
chr3-73056386-INS-249
chr3-73060211-INS-76
chr3-73087889-INS-64
chr3-73112709-INS-6433
chr3-73134612-INS-112
chr3-73204032-INS-1023
chr3-73204032-INS-415
chr3-73214126-INS-2149
chr3-73226692-INS-6328
chr3-73244589-INS-7812
chr3-73284177-INS-50
chr3-73286791-INS-2778
chr3-73289244-INS-6269
chr3-73296935-INS-118
chr3-73321071-INS-3738
chr3-73323902-INS-54
chr3-73331471-INS-70
chr3-73349133-INS-617
chr3-73382081-INS-52
chr3-73383195-INS-76
chr3-73395668-INS-1489
chr3-73399849-INS-6035
chr3-73402496-INS-65
chr3-73420663-INS-198
chr3-73431365-INS-64
chr3-73478930-INS-94
chr3-73499184-INS-70
chr3-73553842-INS-140
chr3-73577740-INS-65
chr3-73626376-INS-17879
chr3-73662432-INS-677

chr3-73663288-INS-6246
chr3-73706422-INS-6472
chr3-73745263-INS-111
chr3-73787885-INS-54
chr3-73792754-INS-50
chr3-73794081-INS-1511
chr3-73796961-INS-50
chr3-73800555-INS-55
chr3-73801627-INS-1379
chr3-73803628-INS-272
chr3-73816226-INS-1491
chr3-73866839-INS-7076
chr3-73876784-INS-6297
chr3-73892982-INS-52
chr3-73900736-INS-439
chr3-73903915-INS-2888
chr3-73906562-INS-271
chr3-73925170-INS-1004
chr3-73984039-INS-309
chr3-74002861-INS-60
chr3-74019093-INS-52
chr3-74044488-INS-564
chr3-74044543-INS-259
chr3-74068509-INS-6540
chr3-74138110-INS-51
chr3-74168257-INS-58
chr3-74189822-INS-1443
chr3-74215634-INS-9325
chr3-74231920-INS-362
chr3-74236623-INS-2911
chr3-74245051-INS-1671
chr3-74252704-INS-3022
chr3-74256365-INS-66
chr3-74257033-INS-3797
chr3-74270062-INS-105
chr3-74278721-INS-6164
chr3-74305473-INS-1086
chr3-74336496-INS-119
chr3-74388593-INS-50
chr3-74416770-INS-133
chr3-74434560-INS-1101
chr3-74450250-INS-139
chr3-74488868-INS-201
chr3-74539562-INS-183
chr3-74572190-INS-584
chr3-74572199-INS-1192
chr3-74594103-INS-50
chr3-74597065-INS-374
chr3-74609900-INS-276
chr3-74610730-INS-51
chr3-74680740-INS-6777
chr3-74732036-INS-72
chr3-74811322-INS-51
chr3-74833695-INS-76
chr3-74856922-INS-1053
chr3-74859089-INS-229
chr3-74888773-INS-227
chr3-74917671-INS-63
chr3-74974584-INS-7515
chr3-75023941-INS-183
chr3-75040677-INS-1129
chr3-75040679-INS-346
chr3-75069971-INS-6838
chr3-75077411-INS-635
chr3-75083208-INS-357
chr3-75086805-INS-830
chr3-75092794-INS-652
chr3-75104566-INS-63
chr3-75108585-INS-414
chr3-75121165-INS-50
chr3-75181938-INS-6351
chr3-75208778-INS-6443
chr3-75209893-INS-788
chr3-75310148-INS-64
chr3-75335045-INS-50
chr3-75360647-INS-5343
chr3-75396324-INS-2201

chr3-75408284-INS-51
chr3-75438932-INS-167
chr3-75462449-INS-203
chr3-75541042-INS-1579
chr3-75574798-INS-196
chr3-75620327-INS-1065
chr3-75629874-INS-6656
chr3-75631287-INS-226
chr3-75666719-INS-679
chr3-75668766-INS-319
chr3-75677516-INS-51
chr3-75695044-INS-2904
chr3-75727960-INS-208
chr3-75735520-INS-205
chr3-75735644-INS-1075
chr3-75772627-INS-6225
chr3-75789208-INS-2475
chr3-75820901-INS-121
chr3-75887847-INS-896
chr3-75891990-INS-56
chr3-75907825-INS-6462
chr3-75933675-INS-6521
chr3-75951206-INS-6860
chr3-75963960-INS-6126
chr3-75967846-INS-431
chr3-75979792-INS-77
chr3-76004079-INS-787
chr3-76010933-INS-105
chr3-76015475-INS-6261
chr3-76107250-INS-6672
chr3-76108624-INS-125
chr3-76111551-INS-60
chr3-76115605-INS-7083
chr3-76138660-INS-2203
chr3-76149906-INS-6595
chr3-76154139-INS-6773
chr3-76234181-INS-200
chr3-76234925-INS-1025
chr3-76253550-INS-6164
chr3-76297843-INS-261
chr3-76321691-INS-1275
chr3-76331134-INS-1075
chr3-76349416-INS-467
chr3-76487645-INS-5379
chr3-76490990-INS-56
chr3-76610009-INS-141
chr3-76626199-INS-2962
chr3-76629962-INS-6575
chr3-76630785-INS-52
chr3-76662425-INS-50
chr3-76723869-INS-55
chr3-76730666-INS-63
chr3-76732561-INS-1929
chr3-76753822-INS-147
chr3-76771792-INS-464
chr3-76835540-INS-58
chr3-76874134-INS-5279
chr3-76938330-INS-52
chr3-76944167-INS-83
chr3-76975154-INS-51
chr3-77005184-INS-2538
chr3-77014566-INS-160
chr3-77022521-INS-103
chr3-77027454-INS-1373
chr3-77028905-INS-72
chr3-77035068-INS-54
chr3-77043502-INS-1135
chr3-77046766-INS-54
chr3-77047362-INS-6241
chr3-77047912-INS-259
chr3-77087935-INS-1488
chr3-77126033-INS-1400
chr3-77153673-INS-1000
chr3-77176347-INS-61
chr3-77182501-INS-56
chr3-77188437-INS-92
chr3-77198056-INS-6256

chr3-77292342-INS-3647
chr3-77363501-INS-68
chr3-77402553-INS-6729
chr3-77411704-INS-993
chr3-77440825-INS-6428
chr3-77467462-INS-1468
chr3-77467786-INS-6196
chr3-77486826-INS-5618
chr3-77502947-INS-50
chr3-77514121-INS-50
chr3-77530922-INS-1281
chr3-77608579-INS-51
chr3-77712317-INS-50
chr3-77751546-INS-820
chr3-77778792-INS-69
chr3-77779052-INS-5990
chr3-77829599-INS-260
chr3-77900637-INS-6919
chr3-77930923-INS-56
chr3-77961809-INS-50
chr3-77986245-INS-61
chr3-77988054-INS-6578
chr3-78056804-INS-391
chr3-78063270-INS-70
chr3-78091236-INS-142
chr3-78143683-INS-1954
chr3-78163404-INS-52
chr3-78179266-INS-3739
chr3-78180126-INS-575
chr3-78183186-INS-1539
chr3-78186458-INS-3159
chr3-78194749-INS-13853
chr3-78196285-INS-75
chr3-78196327-INS-59
chr3-78199616-INS-1956
chr3-78222138-INS-6348
chr3-78230482-INS-50
chr3-78249386-INS-97
chr3-78249393-INS-576
chr3-78250785-INS-1203
chr3-78258134-INS-71
chr3-78260824-INS-72
chr3-78287728-INS-6221
chr3-78294247-INS-3193
chr3-78319387-INS-156
chr3-78326596-INS-61
chr3-78328391-INS-861
chr3-78330683-INS-6396
chr3-78360979-INS-687
chr3-78408819-INS-895
chr3-78532269-INS-53
chr3-78534081-INS-52
chr3-78573559-INS-81
chr3-78589077-INS-8084
chr3-78625998-INS-52
chr3-78633976-INS-4837
chr3-78694113-INS-498
chr3-78700970-INS-126
chr3-78703149-INS-57
chr3-78716504-INS-64
chr3-78770438-INS-1910
chr3-78775949-INS-2864
chr3-78779881-INS-1860
chr3-78784000-INS-1002
chr3-78790802-INS-6776
chr3-78845040-INS-256
chr3-78845707-INS-58
chr3-78847637-INS-60
chr3-78848414-INS-6307
chr3-78857000-INS-106
chr3-78857001-INS-276
chr3-78857166-INS-115
chr3-78869013-INS-3381
chr3-78884434-INS-5570
chr3-78905140-INS-4801
chr3-78913085-INS-1388
chr3-78921668-INS-1696

chr3-78974894-INS-207
chr3-78983248-INS-993
chr3-78983326-INS-3702
chr3-79025526-INS-274
chr3-79094787-INS-148
chr3-79169927-INS-160
chr3-79273245-INS-152
chr3-79273327-INS-464
chr3-79304043-INS-56
chr3-79347963-INS-108
chr3-79348071-INS-288
chr3-79360529-INS-130
chr3-79374042-INS-110
chr3-79381326-INS-468
chr3-79381396-INS-195
chr3-79418335-INS-56
chr3-79454107-INS-98
chr3-79463162-INS-1231
chr3-79463399-INS-75
chr3-79481699-INS-70
chr3-79486191-INS-204
chr3-79490002-INS-60
chr3-79493010-INS-264
chr3-79493010-INS-624
chr3-79570447-INS-176
chr3-79588548-INS-54
chr3-79593621-INS-50
chr3-79597072-INS-405
chr3-79598363-INS-107
chr3-79608007-INS-8040
chr3-79612250-INS-356
chr3-79651693-INS-427
chr3-79661138-INS-74
chr3-79663528-INS-56
chr3-79717025-INS-284
chr3-79717649-INS-740
chr3-79751368-INS-78
chr3-79753514-INS-5090
chr3-79762917-INS-50
chr3-79763941-INS-198
chr3-79764509-INS-107
chr3-79769137-INS-6185
chr3-79775586-INS-54
chr3-79828564-INS-66
chr3-79838088-INS-97
chr3-79843391-INS-2576
chr3-79893552-INS-231
chr3-79920489-INS-2274
chr3-79920964-INS-11361
chr3-79932177-INS-69
chr3-79932357-INS-153
chr3-79942868-INS-237
chr3-79977758-INS-1148
chr3-79986719-INS-71
chr3-80015326-INS-52
chr3-80016088-INS-84
chr3-80066538-INS-54
chr3-80067370-INS-92
chr3-80071499-INS-6397
chr3-80087032-INS-66
chr3-80101465-INS-131
chr3-80104700-INS-56
chr3-80108627-INS-156
chr3-80147475-INS-1268
chr3-80177179-INS-447
chr3-80209670-INS-128
chr3-80223019-INS-200
chr3-80223073-INS-90
chr3-80234686-INS-390
chr3-80243806-INS-232
chr3-80243825-INS-98
chr3-80277418-INS-6730
chr3-80279292-INS-54
chr3-80318496-INS-6115
chr3-80318765-INS-58
chr3-80329321-INS-72
chr3-80394078-INS-295

chr3-80394296-INS-353
chr3-80417240-INS-403
chr3-80419528-INS-211
chr3-80434251-INS-6420
chr3-80485747-INS-7084
chr3-80521073-INS-236
chr3-80525921-INS-162
chr3-80547049-INS-583
chr3-80549470-INS-50
chr3-80563758-INS-65
chr3-80567604-INS-151
chr3-80598359-INS-98
chr3-80608814-INS-1483
chr3-80611908-INS-51
chr3-80734183-INS-222
chr3-80734473-INS-102
chr3-80742329-INS-60
chr3-80743679-INS-4280
chr3-80764541-INS-5050
chr3-80774499-INS-360
chr3-80778565-INS-286
chr3-80799545-INS-76
chr3-80836686-INS-68
chr3-80880025-INS-1890
chr3-80881365-INS-6484
chr3-80904797-INS-6962
chr3-80941271-INS-56
chr3-80951628-INS-408
chr3-81018782-INS-6363
chr3-81075868-INS-113
chr3-81126148-INS-5647
chr3-81134112-INS-494
chr3-81142327-INS-257
chr3-81153638-INS-50
chr3-81154054-INS-1136
chr3-81167528-INS-62
chr3-81170314-INS-186
chr3-81219136-INS-4244
chr3-81242530-INS-57
chr3-81293751-INS-50
chr3-81298462-INS-8174
chr3-81306359-INS-82
chr3-81307575-INS-1434
chr3-81323839-INS-1070
chr3-81324162-INS-711
chr3-81331380-INS-6468
chr3-81357087-INS-7090
chr3-81413604-INS-2390
chr3-81435040-INS-50
chr3-81463104-INS-55
chr3-81467502-INS-52
chr3-81481177-INS-6305
chr3-81495187-INS-148
chr3-81495438-INS-414
chr3-81505141-INS-1212
chr3-81529252-INS-1494
chr3-81538640-INS-110
chr3-81640926-INS-138
chr3-81650387-INS-60
chr3-81673799-INS-6467
chr3-81681463-INS-67
chr3-81689014-INS-500
chr3-81689309-INS-56
chr3-81708752-INS-1213
chr3-81715126-INS-3278
chr3-81760752-INS-62
chr3-81778053-INS-128
chr3-81778054-INS-54
chr3-81889989-INS-52
chr3-81917501-INS-54
chr3-81918952-INS-326
chr3-81983268-INS-52
chr3-82062553-INS-110
chr3-82093281-INS-7669
chr3-82210998-INS-1748
chr3-82213651-INS-72
chr3-82224824-INS-6824

chr3-82244103-INS-50
chr3-82254252-INS-120
chr3-82287750-INS-5086
chr3-82288316-INS-716
chr3-82326211-INS-6156
chr3-82336356-INS-76
chr3-82364542-INS-68
chr3-82375728-INS-328
chr3-82375849-INS-549
chr3-82399941-INS-1065
chr3-82413400-INS-1000
chr3-82417693-INS-294
chr3-82423742-INS-2168
chr3-82458269-INS-1022
chr3-82458289-INS-66
chr3-82487634-INS-1345
chr3-82496542-INS-3916
chr3-82503734-INS-51
chr3-82530163-INS-306
chr3-82530235-INS-142
chr3-82544233-INS-74
chr3-82571734-INS-125
chr3-82571793-INS-258
chr3-82585941-INS-6707
chr3-82613025-INS-3094
chr3-82648498-INS-50
chr3-82650865-INS-3024
chr3-82665885-INS-120
chr3-82680541-INS-963
chr3-82698231-INS-1941
chr3-82711440-INS-215
chr3-82714843-INS-478
chr3-82895083-INS-97
chr3-82971638-INS-294
chr3-82971997-INS-682
chr3-83008861-INS-117
chr3-83031414-INS-751
chr3-83056316-INS-138
chr3-83070435-INS-51
chr3-83109304-INS-86
chr3-83121315-INS-270
chr3-83152297-INS-6653
chr3-83185219-INS-285
chr3-83194847-INS-68
chr3-83228456-INS-50
chr3-83307312-INS-50
chr3-83335942-INS-122
chr3-83343484-INS-451
chr3-83375427-INS-7125
chr3-83423069-INS-117
chr3-83446592-INS-53
chr3-83460517-INS-1259
chr3-83472400-INS-76
chr3-83477259-INS-369
chr3-83477263-INS-69
chr3-83477339-INS-80
chr3-83478905-INS-724
chr3-83530610-INS-2989
chr3-83575357-INS-2583
chr3-83597371-INS-52
chr3-83615253-INS-2685
chr3-83689942-INS-76
chr3-83713526-INS-54
chr3-83714144-INS-234
chr3-83735359-INS-52
chr3-83765046-INS-69
chr3-83827432-INS-129
chr3-83848838-INS-62
chr3-83865324-INS-51
chr3-83905732-INS-116
chr3-83905761-INS-290
chr3-83937044-INS-182
chr3-83937152-INS-76
chr3-83951057-INS-191
chr3-83951710-INS-78
chr3-83959594-INS-6473
chr3-83979512-INS-51

chr3-83979543-INS-103
chr3-83998732-INS-6600
chr3-84021827-INS-418
chr3-84042607-INS-2847
chr3-84089059-INS-57
chr3-84107186-INS-50
chr3-84202104-INS-83
chr3-84217887-INS-2420
chr3-84288675-INS-222
chr3-84308645-INS-550
chr3-84310210-INS-7186
chr3-84318503-INS-152
chr3-84325286-INS-67
chr3-84348542-INS-50
chr3-84496201-INS-77
chr3-84516460-INS-6575
chr3-84531419-INS-104
chr3-84531648-INS-313
chr3-84532512-INS-224
chr3-84533938-INS-213
chr3-84540308-INS-142
chr3-84582419-INS-70
chr3-84590729-INS-62
chr3-84734317-INS-6327
chr3-84750531-INS-287
chr3-84978606-INS-138
chr3-84998917-INS-157
chr3-85036898-INS-521
chr3-85036898-INS-80
chr3-85037143-INS-202
chr3-85161835-INS-899
chr3-85170512-INS-2792
chr3-85189131-INS-148
chr3-85203708-INS-3808
chr3-85219046-INS-102
chr3-85231312-INS-50
chr3-85308648-INS-403
chr3-85308716-INS-62
chr3-85332495-INS-60
chr3-85335958-INS-6905
chr3-85384105-INS-63
chr3-85409501-INS-75
chr3-85420968-INS-51
chr3-85465151-INS-1637
chr3-85536875-INS-163
chr3-85713432-INS-56
chr3-85735912-INS-6344
chr3-85759639-INS-309
chr3-85763930-INS-98
chr3-85773077-INS-6410
chr3-85790159-INS-68
chr3-85811397-INS-2932
chr3-85848534-INS-268
chr3-85848615-INS-87
chr3-85848988-INS-104
chr3-85857062-INS-212
chr3-85865427-INS-124
chr3-85875278-INS-8747
chr3-85905902-INS-69
chr3-85994856-INS-266
chr3-85998473-INS-203
chr3-86009181-INS-54
chr3-86027332-INS-52
chr3-86039286-INS-104
chr3-86143071-INS-216
chr3-86143537-INS-222
chr3-86154030-INS-418
chr3-86184684-INS-477
chr3-86185246-INS-206
chr3-86276935-INS-174
chr3-86508386-INS-4845
chr3-86528545-INS-6389
chr3-86544696-INS-527
chr3-86645595-INS-232
chr3-86671308-INS-106
chr3-86672183-INS-155
chr3-86723968-INS-200

chr3-86724308-INS-407
chr3-86750688-INS-51
chr3-86773440-INS-4967
chr3-86797221-INS-74
chr3-86817652-INS-8671
chr3-86904200-INS-64
chr3-86932728-INS-1843
chr3-86936769-INS-823
chr3-86941679-INS-64
chr3-86947763-INS-106
chr3-86959213-INS-82
chr3-87041931-INS-1000
chr3-87055145-INS-56
chr3-87090049-INS-3237
chr3-87116473-INS-5801
chr3-87153273-INS-508
chr3-87153866-INS-184
chr3-87179298-INS-60
chr3-87212942-INS-403
chr3-87232959-INS-66
chr3-87278540-INS-1938
chr3-87318999-INS-7327
chr3-87339506-INS-108
chr3-87478140-INS-90
chr3-87482689-INS-376
chr3-87531136-INS-94
chr3-87727791-INS-217
chr3-87741225-INS-62
chr3-87748139-INS-203
chr3-87748552-INS-61
chr3-87770965-INS-202
chr3-87819262-INS-220
chr3-87845756-INS-206
chr3-87914022-INS-480
chr3-87914354-INS-211
chr3-87988621-INS-204
chr3-88102415-INS-116
chr3-88126362-INS-50
chr3-88147308-INS-141
chr3-88173503-INS-55
chr3-88173678-INS-203
chr3-88177708-INS-60
chr3-88178402-INS-2363
chr3-88180465-INS-940
chr3-88181006-INS-95
chr3-88186748-INS-209
chr3-88202871-INS-177
chr3-88235727-INS-390
chr3-88236146-INS-201
chr3-88237577-INS-212
chr3-88239127-INS-74
chr3-88246261-INS-212
chr3-88309482-INS-116
chr3-88332726-INS-70
chr3-88342503-INS-202
chr3-88349100-INS-202
chr3-88351626-INS-91
chr3-88373989-INS-62
chr3-88377326-INS-208
chr3-88391714-INS-93
chr3-88401477-INS-194
chr3-88401945-INS-86
chr3-88418177-INS-217
chr3-88432134-INS-206
chr3-88435606-INS-50
chr3-88492713-INS-243
chr3-88513077-INS-84
chr3-88519260-INS-198
chr3-88519506-INS-52
chr3-88519950-INS-201
chr3-88566459-INS-203
chr3-88581140-INS-209
chr3-88626866-INS-449
chr3-88632075-INS-384
chr3-88686095-INS-153
chr3-88686394-INS-212
chr3-88688580-INS-198

chr3-88698034-INS-212
chr3-88759538-INS-61
chr3-88790451-INS-176
chr3-88791261-INS-998
chr3-88794246-INS-496
chr3-88839218-INS-54
chr3-88924044-INS-498
chr3-88950192-INS-164
chr3-89019330-INS-88
chr3-89043031-INS-113
chr3-89046182-INS-52
chr3-89105128-INS-52
chr3-89105665-INS-210
chr3-89109724-INS-59
chr3-89116343-INS-554
chr3-89155735-INS-196
chr3-89190570-INS-203
chr3-89305038-INS-82
chr3-89349731-INS-52
chr3-89377754-INS-343
chr3-89379978-INS-187
chr3-89387604-INS-211
chr3-89393148-INS-113
chr3-89399262-INS-54
chr3-89406757-INS-207
chr3-89413084-INS-214
chr3-89414446-INS-461
chr3-89453042-INS-130
chr3-89470510-INS-54
chr3-89537522-INS-431
chr3-89546188-INS-52
chr3-89552332-INS-56
chr3-89553530-INS-2497
chr3-89557121-INS-579
chr3-89589102-INS-3017
chr3-89609878-INS-1137
chr3-89673433-INS-225
chr3-89703393-INS-50
chr3-89710795-INS-54
chr3-89733970-INS-178
chr3-89751123-INS-182
chr3-89755846-INS-53
chr3-89770054-INS-66
chr3-89807986-INS-144
chr3-89809999-INS-66
chr3-89832233-INS-105
chr3-89832248-INS-50
chr3-89885112-INS-63
chr3-89889626-INS-176
chr3-89894103-INS-1079
chr3-89894235-INS-80
chr3-89929826-INS-243
chr3-89936549-INS-68
chr3-89950872-INS-200
chr3-89971936-INS-50
chr3-90012978-INS-569
chr3-90012978-INS-60
chr3-90013388-INS-593
chr3-90017672-INS-1230
chr3-90024607-INS-92
chr3-90066578-INS-136
chr3-90078841-INS-3067
chr3-90114224-INS-181
chr3-90137992-INS-187
chr3-90142224-INS-207
chr3-90195060-INS-53
chr3-90195240-INS-189
chr3-90197781-INS-206
chr3-90242925-INS-175
chr3-90275515-INS-96
chr3-90275938-INS-515
chr3-90287724-INS-371
chr3-90290641-INS-345
chr3-90389231-INS-207
chr3-90398905-INS-126
chr3-90415392-INS-219
chr3-90423026-INS-240

chr3-90431365-INS-100
chr3-90443393-INS-54
chr3-90457347-INS-3460
chr3-90474012-INS-306
chr3-90505567-INS-105
chr3-90511251-INS-50
chr3-90604188-INS-262
chr3-90702165-INS-2292
chr3-90702361-INS-343
chr3-90876770-INS-212
chr3-91003041-INS-208
chr3-91110985-INS-54
chr3-91158386-INS-675
chr3-91193398-INS-75
chr3-91244456-INS-1210
chr3-91246757-INS-90
chr3-91347816-INS-7157
chr3-91395056-INS-841
chr3-91397356-INS-7195
chr3-91408062-INS-1896
chr3-91415215-INS-458
chr3-91534187-INS-482
chr3-91625013-INS-2537
chr3-91675015-INS-617
chr3-91709501-INS-476
chr3-91770600-INS-146
chr3-91782704-INS-53
chr3-91949388-INS-51
chr3-91950323-INS-62
chr3-92053079-INS-72
chr3-92246871-INS-100
chr3-92247301-INS-60
chr3-92351442-INS-76
chr3-92358342-INS-50
chr3-92374451-INS-6516
chr3-92380760-INS-367
chr3-92479264-INS-312
chr3-92484953-INS-51
chr3-92489394-INS-89
chr3-92494964-INS-64
chr3-92503270-INS-6364
chr3-92516866-INS-1400
chr3-92524955-INS-4341
chr3-92535095-INS-55
chr3-92565889-INS-135
chr3-92566119-INS-260
chr3-92591976-INS-202
chr3-92622062-INS-62
chr3-92622722-INS-100
chr3-92624410-INS-144
chr3-92638979-INS-1444
chr3-92667294-INS-6473
chr3-92692639-INS-58
chr3-92694321-INS-246
chr3-92715701-INS-579
chr3-92722764-INS-5169
chr3-92742928-INS-858
chr3-92800318-INS-110
chr3-92811776-INS-80
chr3-92813452-INS-74
chr3-92813988-INS-6713
chr3-92859653-INS-144
chr3-92861766-INS-6452
chr3-92879158-INS-349
chr3-92885830-INS-124
chr3-92939056-INS-1064
chr3-92966422-INS-196
chr3-92974533-INS-54
chr3-92976696-INS-124
chr3-92977758-INS-52
chr3-93003669-INS-6571
chr3-93007039-INS-56
chr3-93007044-INS-130
chr3-93033328-INS-3941
chr3-93209776-INS-50
chr3-93229682-INS-50
chr3-93236187-INS-1575

chr3-93237578-INS-513
chr3-93241751-INS-171
chr3-93243623-INS-60
chr3-93258353-INS-372
chr3-93269013-INS-6415
chr3-93386536-INS-171
chr3-93386761-INS-407
chr3-93402506-INS-58
chr3-93424567-INS-196
chr3-93429684-INS-378
chr3-93432664-INS-75
chr3-93436719-INS-5537
chr3-93462247-INS-64
chr3-93480611-INS-6533
chr3-93495135-INS-80
chr3-93513576-INS-175
chr3-93548962-INS-564
chr3-93564012-INS-305
chr3-93623157-INS-7625
chr3-93650460-INS-55
chr3-93665880-INS-205
chr3-93670219-INS-56
chr3-93670979-INS-96
chr3-93671472-INS-629
chr3-93684503-INS-1671
chr3-93689391-INS-58
chr3-93691169-INS-497
chr3-93711426-INS-1497
chr3-93713538-INS-2387
chr3-93714226-INS-198
chr3-93746937-INS-108
chr3-93848013-INS-76
chr3-93946792-INS-185
chr3-93968146-INS-210
chr3-94037563-INS-302
chr3-94119069-INS-216
chr3-94122247-INS-1509
chr3-94127483-INS-3039
chr3-94177607-INS-273
chr3-94234635-INS-58
chr3-94234966-INS-58
chr3-94314871-INS-205
chr3-94339863-INS-272
chr3-94358474-INS-151
chr3-94359758-INS-50
chr3-94360330-INS-129
chr3-94361459-INS-76
chr3-94374272-INS-76
chr3-94375176-INS-60
chr3-94376026-INS-112
chr3-94443411-INS-76
chr3-94464462-INS-112
chr3-94496701-INS-312
chr3-94509391-INS-58
chr3-94512628-INS-202
chr3-94543938-INS-207
chr3-94548742-INS-179
chr3-94550964-INS-217
chr3-94551011-INS-153
chr3-94574231-INS-72
chr3-94574240-INS-251
chr3-94596325-INS-102
chr3-94613492-INS-7626
chr3-94623702-INS-1045
chr3-94656813-INS-737
chr3-94663610-INS-236
chr3-94663862-INS-147
chr3-94688205-INS-228
chr3-94700772-INS-212
chr3-94707506-INS-1258
chr3-94710292-INS-55
chr3-94722924-INS-56
chr3-94722979-INS-162
chr3-94765427-INS-113
chr3-94783670-INS-627
chr3-94800024-INS-131
chr3-94813361-INS-4924

chr3-94817559-INS-208
chr3-94818784-INS-55
chr3-94841372-INS-235
chr3-94853500-INS-199
chr3-94895735-INS-200
chr3-94946868-INS-100
chr3-94947282-INS-204
chr3-94951118-INS-164
chr3-94957325-INS-197
chr3-94957879-INS-99
chr3-94959561-INS-266
chr3-94970388-INS-60
chr3-95016369-INS-217
chr3-95057412-INS-219
chr3-95085678-INS-406
chr3-95149434-INS-211
chr3-95183377-INS-218
chr3-95186343-INS-56
chr3-95204272-INS-52
chr3-95211119-INS-363
chr3-95258493-INS-190
chr3-95287122-INS-113
chr3-95287249-INS-191
chr3-95352622-INS-1080
chr3-95379457-INS-816
chr3-95417922-INS-228
chr3-95425252-INS-100
chr3-95431859-INS-63
chr3-95432167-INS-102
chr3-95449698-INS-192
chr3-95449722-INS-564
chr3-95519161-INS-226
chr3-95519522-INS-205
chr3-95546225-INS-210
chr3-95546925-INS-406
chr3-95612733-INS-165
chr3-95618335-INS-177
chr3-95636015-INS-112
chr3-95648211-INS-65
chr3-95699658-INS-91
chr3-95734546-INS-198
chr3-95734605-INS-794
chr3-95745189-INS-266
chr3-95749602-INS-60
chr3-95820491-INS-78
chr3-95835402-INS-1270
chr3-95933459-INS-270
chr3-95951825-INS-136
chr3-96000435-INS-58
chr3-96048449-INS-366
chr3-96123736-INS-56
chr3-96136647-INS-2603
chr3-96189693-INS-335
chr3-96198441-INS-129
chr3-96198726-INS-160
chr3-96265540-INS-197
chr3-96283127-INS-214
chr3-96306862-INS-203
chr3-96313928-INS-54
chr3-96328711-INS-58
chr3-96345589-INS-377
chr3-96351723-INS-52
chr3-96383627-INS-66
chr3-96455969-INS-50
chr3-96501937-INS-65
chr3-96573589-INS-55
chr3-96645795-INS-211
chr3-96658790-INS-122
chr3-96658832-INS-58
chr3-96756124-INS-374
chr3-96781262-INS-451
chr3-96795335-INS-52
chr3-96797405-INS-81
chr3-96814085-INS-6475
chr3-96873484-INS-215
chr3-96893209-INS-334
chr3-96893553-INS-219

chr3-96900710-INS-134
chr3-96954722-INS-55
chr3-96974676-INS-14203
chr3-96986994-INS-269
chr3-96987873-INS-75
chr3-97009409-INS-52
chr3-97032292-INS-255
chr3-97032518-INS-535
chr3-97044284-INS-335
chr3-97064941-INS-216
chr3-97090616-INS-2851
chr3-97160826-INS-205
chr3-97222824-INS-140
chr3-97222913-INS-121
chr3-97227330-INS-64
chr3-97235079-INS-449
chr3-97235083-INS-134
chr3-97283902-INS-5538
chr3-97286454-INS-207
chr3-97308318-INS-1793
chr3-97311021-INS-50
chr3-97318171-INS-81919
chr3-97326624-INS-4806
chr3-97369547-INS-445
chr3-97369796-INS-381
chr3-97441861-INS-161
chr3-97442289-INS-75
chr3-97449503-INS-4169
chr3-97458208-INS-222
chr3-97458714-INS-137
chr3-97460189-INS-144
chr3-97460279-INS-453
chr3-97490799-INS-65
chr3-97497217-INS-297
chr3-97498995-INS-120
chr3-97499131-INS-74
chr3-97584096-INS-130
chr3-97651747-INS-320
chr3-97680632-INS-201
chr3-97714507-INS-55
chr3-97725656-INS-78
chr3-97733971-INS-7394
chr3-97743932-INS-2296
chr3-97793834-INS-270
chr3-97887350-INS-199
chr3-97890846-INS-205
chr3-97905229-INS-51
chr3-97924096-INS-646
chr3-97932189-INS-105
chr3-97939535-INS-226
chr3-97976030-INS-88
chr3-98039189-INS-619
chr3-98039258-INS-74
chr3-98102535-INS-116
chr3-98113347-INS-260
chr3-98115290-INS-72
chr3-98138796-INS-3321
chr3-98159397-INS-64
chr3-98159419-INS-248
chr3-98165604-INS-410
chr3-98181021-INS-570
chr3-98296587-INS-58
chr3-98301261-INS-116
chr3-98330672-INS-50
chr3-98340121-INS-147
chr3-98484056-INS-189
chr3-98484084-INS-64
chr3-98484329-INS-969
chr3-98488580-INS-541
chr3-98513032-INS-52
chr3-98519107-INS-82
chr3-98519407-INS-287
chr3-98521391-INS-74
chr3-98532307-INS-97
chr3-98550848-INS-416
chr3-98554513-INS-307
chr3-98569574-INS-210

chr3-98588664-INS-1806
chr3-98593150-INS-6167
chr3-98630275-INS-2793
chr3-98645313-INS-6667
chr3-98650151-INS-7198
chr3-98691727-INS-342
chr3-98692204-INS-392
chr3-98698309-INS-376
chr3-98732962-INS-51
chr3-98801745-INS-75
chr3-98822765-INS-128
chr3-98824848-INS-170
chr3-98824869-INS-445
chr3-98824980-INS-56
chr3-98871927-INS-6295
chr3-98875977-INS-50
chr3-98901573-INS-53
chr3-98910118-INS-93
chr3-98918177-INS-4639
chr3-98964925-INS-6258
chr3-98975831-INS-2820
chr3-99010854-INS-6314
chr3-99070627-INS-394
chr3-99070643-INS-93
chr3-99102150-INS-114
chr3-99133335-INS-175
chr3-99134645-INS-285
chr3-99166808-INS-173
chr3-99296840-INS-270
chr3-99300757-INS-6009
chr3-99310665-INS-1997
chr3-99349836-INS-3797
chr3-99380173-INS-4915
chr3-99387131-INS-174
chr3-99445571-INS-1620
chr3-99445600-INS-535
chr3-99445850-INS-656
chr3-99465466-INS-410
chr3-99492018-INS-75
chr3-99502011-INS-3256
chr3-99502173-INS-294
chr3-99587875-INS-82
chr3-99692662-INS-88
chr3-99694675-INS-66
chr3-99749326-INS-50
chr3-99753244-INS-139
chr3-99887540-INS-51
chr3-99892392-INS-60
chr3-99905686-INS-220
chr3-99982145-INS-88
chr3-99991963-INS-64
chr3-99993719-INS-6421
chr3-99994444-INS-371
chr3-99998352-INS-89
chr3-99998488-INS-167
chr3-100026596-INS-678
chr3-100067461-INS-273
chr3-100134145-INS-334
chr3-100156847-INS-180
chr3-100189835-INS-83
chr3-100229220-INS-56
chr3-100351144-INS-501
chr3-100361174-INS-1063
chr3-100365040-INS-420
chr3-100423608-INS-4741
chr3-100431956-INS-56
chr3-100448279-INS-56
chr3-100448892-INS-6701
chr3-100453230-INS-72
chr3-100456905-INS-2064
chr3-100620115-INS-300
chr3-100651515-INS-148
chr3-100668141-INS-6479
chr3-100686311-INS-109
chr3-100739060-INS-2823
chr3-100800727-INS-7095
chr3-100849531-INS-111

chr3-100856150-INS-97
chr3-100877812-INS-446
chr3-100887088-INS-164
chr3-100961872-INS-320
chr3-101039721-INS-357
chr3-101060481-INS-50
chr3-101080148-INS-65
chr3-101082733-INS-269
chr3-101171979-INS-565
chr3-101217854-INS-152
chr3-101227016-INS-106
chr3-101321635-INS-240
chr3-101376582-INS-63
chr3-101379673-INS-67
chr3-101436953-INS-50
chr3-101450848-INS-97
chr3-101457471-INS-202
chr3-101521015-INS-144
chr3-101538398-INS-53
chr3-101557685-INS-62
chr3-101575893-INS-50
chr3-101584632-INS-59
chr3-101602768-INS-80
chr3-101637095-INS-174
chr3-101719325-INS-6364
chr3-101728701-INS-172
chr3-101814835-INS-652
chr3-101840362-INS-499
chr3-101886444-INS-123
chr3-101913303-INS-376
chr3-101913886-INS-66
chr3-101968731-INS-116
chr3-101996403-INS-51
chr3-102040505-INS-355
chr3-102139807-INS-7057
chr3-102163133-INS-60
chr3-102171485-INS-61
chr3-102195205-INS-6131
chr3-102209011-INS-66
chr3-102253049-INS-169
chr3-102269806-INS-105
chr3-102274525-INS-380
chr3-102280752-INS-2094
chr3-102318546-INS-186
chr3-102343934-INS-404
chr3-102350554-INS-198
chr3-102350609-INS-594
chr3-102359442-INS-61
chr3-102399385-INS-148
chr3-102399950-INS-184
chr3-102399981-INS-76
chr3-102431926-INS-60
chr3-102445712-INS-160
chr3-102460020-INS-50
chr3-102466087-INS-462
chr3-102474606-INS-4232
chr3-102485881-INS-129
chr3-102614617-INS-66
chr3-102621623-INS-7151
chr3-102636099-INS-145
chr3-102684540-INS-664
chr3-102723912-INS-1630
chr3-102821350-INS-50
chr3-102884282-INS-54
chr3-102917172-INS-88
chr3-102923710-INS-104
chr3-102959745-INS-60
chr3-102962224-INS-60
chr3-103003262-INS-172
chr3-103024958-INS-4078
chr3-103027426-INS-50
chr3-103029349-INS-440
chr3-103041218-INS-1802
chr3-103043180-INS-4791
chr3-103045168-INS-91
chr3-103048093-INS-642
chr3-103048313-INS-285

chr3-103077230-INS-216
chr3-103079813-INS-198
chr3-103124764-INS-193
chr3-103130801-INS-236
chr3-103130870-INS-50
chr3-103141654-INS-155
chr3-103144361-INS-209
chr3-103175762-INS-4855
chr3-103204959-INS-229
chr3-103235761-INS-144
chr3-103235767-INS-62
chr3-103311050-INS-202
chr3-103324012-INS-184
chr3-103351887-INS-696
chr3-103354893-INS-84
chr3-103406724-INS-135
chr3-103406724-INS-460
chr3-103460382-INS-52
chr3-103465288-INS-52
chr3-103515658-INS-66
chr3-103525498-INS-403
chr3-103581005-INS-74
chr3-103588907-INS-7101
chr3-103598274-INS-108
chr3-103598955-INS-210
chr3-103601519-INS-250
chr3-103664938-INS-54
chr3-103668741-INS-87
chr3-103678716-INS-194
chr3-103702923-INS-4715
chr3-103725186-INS-110
chr3-103736930-INS-64
chr3-103744644-INS-224
chr3-103745189-INS-423
chr3-103780318-INS-135
chr3-103782758-INS-50
chr3-103789631-INS-183
chr3-103790326-INS-50
chr3-103791954-INS-184
chr3-103798070-INS-150
chr3-103813606-INS-6228
chr3-104110360-INS-1214
chr3-104120556-INS-50
chr3-104137115-INS-50
chr3-104176885-INS-117
chr3-104400246-INS-254
chr3-104400274-INS-56
chr3-104446462-INS-590
chr3-104529773-INS-220
chr3-104589166-INS-256
chr3-104597372-INS-56
chr3-104613148-INS-1314
chr3-104633029-INS-61
chr3-104638790-INS-79
chr3-104652849-INS-203
chr3-104664671-INS-184
chr3-104705879-INS-136
chr3-104706105-INS-107
chr3-104722699-INS-122
chr3-104737329-INS-64
chr3-104866871-INS-73
chr3-104893963-INS-91
chr3-104894893-INS-52
chr3-104929452-INS-766
chr3-104935959-INS-80
chr3-104936731-INS-340
chr3-104991824-INS-205
chr3-105064640-INS-216
chr3-105064648-INS-51
chr3-105078223-INS-68
chr3-105097632-INS-1805
chr3-105102556-INS-72
chr3-105127599-INS-50
chr3-105133882-INS-771
chr3-105134062-INS-102
chr3-105141388-INS-50
chr3-105166141-INS-108

chr3-105169050-INS-143
chr3-105178505-INS-555
chr3-105191144-INS-126
chr3-105191415-INS-205
chr3-105218492-INS-52
chr3-105239109-INS-245
chr3-105255092-INS-97
chr3-105265580-INS-5397
chr3-105299999-INS-54
chr3-105300230-INS-105
chr3-105317033-INS-6978
chr3-105324839-INS-60
chr3-105330321-INS-116
chr3-105417352-INS-105
chr3-105427595-INS-1844
chr3-105427604-INS-159
chr3-105504646-INS-100
chr3-105525744-INS-320
chr3-105525825-INS-155
chr3-105544462-INS-50
chr3-105608622-INS-64
chr3-105626846-INS-281
chr3-105731254-INS-6814
chr3-105731508-INS-146
chr3-105810625-INS-200
chr3-105811454-INS-148
chr3-105819275-INS-54
chr3-105828185-INS-152
chr3-105893273-INS-57
chr3-105893744-INS-210
chr3-105952174-INS-85
chr3-105952193-INS-134
chr3-105969023-INS-276
chr3-105987978-INS-417
chr3-106003160-INS-186
chr3-106039520-INS-52
chr3-106045171-INS-4757
chr3-106120089-INS-3214
chr3-106174219-INS-208
chr3-106238651-INS-178
chr3-106249610-INS-1230
chr3-106301040-INS-50
chr3-106313146-INS-52
chr3-106313165-INS-106
chr3-106315553-INS-478
chr3-106350686-INS-2861
chr3-106377440-INS-61
chr3-106384448-INS-51
chr3-106402655-INS-54
chr3-106423333-INS-214
chr3-106429110-INS-1534
chr3-106429627-INS-238
chr3-106550722-INS-185
chr3-106550722-INS-90
chr3-106654954-INS-5810
chr3-106655408-INS-2606
chr3-106754179-INS-54
chr3-106754799-INS-3054
chr3-106754799-INS-459
chr3-106755391-INS-79
chr3-106782096-INS-6410
chr3-106815537-INS-544
chr3-106864085-INS-52
chr3-106891637-INS-50
chr3-107105130-INS-842
chr3-107133843-INS-80
chr3-107136486-INS-68
chr3-107175141-INS-273
chr3-107180867-INS-64
chr3-107195119-INS-56
chr3-107225761-INS-153
chr3-107225826-INS-233
chr3-107267748-INS-50
chr3-107274115-INS-666
chr3-107290278-INS-276
chr3-107298525-INS-135
chr3-107309750-INS-50

chr3-107356158-INS-138
chr3-107386684-INS-168
chr3-107414546-INS-118
chr3-107430577-INS-2849
chr3-107475312-INS-224
chr3-107531205-INS-158
chr3-107552633-INS-51
chr3-107642671-INS-50
chr3-107755188-INS-138
chr3-107764924-INS-111
chr3-107843073-INS-52
chr3-107857337-INS-1876
chr3-107886484-INS-6750
chr3-107904636-INS-434
chr3-107922659-INS-308
chr3-107944195-INS-201
chr3-107988815-INS-52
chr3-108038246-INS-56
chr3-108073715-INS-57
chr3-108091277-INS-7191
chr3-108091349-INS-54
chr3-108125953-INS-120
chr3-108138172-INS-56
chr3-108140168-INS-209
chr3-108168703-INS-200
chr3-108182454-INS-195
chr3-108183029-INS-61
chr3-108201396-INS-810
chr3-108206968-INS-210
chr3-108236664-INS-63
chr3-108279270-INS-57
chr3-108293439-INS-60
chr3-108297699-INS-72
chr3-108297879-INS-756
chr3-108298503-INS-121
chr3-108329680-INS-175
chr3-108399665-INS-50
chr3-108412876-INS-418
chr3-108419032-INS-7323
chr3-108419040-INS-64
chr3-108421365-INS-119
chr3-108434680-INS-99
chr3-108448436-INS-175
chr3-108448836-INS-177
chr3-108458450-INS-341
chr3-108487227-INS-209
chr3-108497566-INS-206
chr3-108511669-INS-213
chr3-108535432-INS-208
chr3-108544094-INS-66
chr3-108560942-INS-52
chr3-108597239-INS-259
chr3-108673355-INS-61
chr3-108674541-INS-50
chr3-108714887-INS-76
chr3-108715533-INS-124
chr3-108725138-INS-385
chr3-108774749-INS-665
chr3-108808839-INS-5114
chr3-108863964-INS-52
chr3-108906974-INS-2900
chr3-108923468-INS-128
chr3-108948496-INS-2786
chr3-108962916-INS-7251
chr3-108964399-INS-58
chr3-109001146-INS-191
chr3-109005754-INS-6485
chr3-109013410-INS-96
chr3-109094187-INS-104
chr3-109095880-INS-3006
chr3-109151304-INS-6153
chr3-109165232-INS-390
chr3-109166868-INS-471
chr3-109169724-INS-4883
chr3-109201695-INS-210
chr3-109218082-INS-174
chr3-109224822-INS-54

chr3-109231101-INS-68
chr3-109265956-INS-84
chr3-109278938-INS-503
chr3-109288472-INS-315
chr3-109294251-INS-984
chr3-109310913-INS-6414
chr3-109316662-INS-6188
chr3-109329340-INS-3040
chr3-109333248-INS-276
chr3-109358201-INS-60
chr3-109460031-INS-50
chr3-109480248-INS-144
chr3-109532600-INS-52
chr3-109606537-INS-217
chr3-109645509-INS-7191
chr3-109664752-INS-4908
chr3-109685741-INS-52
chr3-109698599-INS-50
chr3-109703528-INS-1981
chr3-109703556-INS-438
chr3-109779444-INS-92
chr3-109780855-INS-6636
chr3-109829259-INS-108
chr3-109829848-INS-324
chr3-109833206-INS-52
chr3-109841284-INS-50
chr3-109862384-INS-50
chr3-109885985-INS-1030
chr3-110057870-INS-67
chr3-110062655-INS-216
chr3-110063566-INS-55
chr3-110086975-INS-343
chr3-110166740-INS-51
chr3-110205098-INS-3302
chr3-110231350-INS-1000
chr3-110331369-INS-114
chr3-110343022-INS-246
chr3-110380103-INS-2179
chr3-110495108-INS-2241
chr3-110508651-INS-4837
chr3-110667406-INS-76
chr3-110693972-INS-272
chr3-110788292-INS-50
chr3-110813638-INS-60
chr3-110837893-INS-116
chr3-110852870-INS-128
chr3-110857056-INS-50
chr3-110870333-INS-346
chr3-110877382-INS-6615
chr3-110885171-INS-90
chr3-110896273-INS-1194
chr3-110907099-INS-68
chr3-110927158-INS-3631
chr3-110963427-INS-58
chr3-110967036-INS-2219
chr3-111009666-INS-1872
chr3-111067995-INS-56
chr3-111068004-INS-195
chr3-111085647-INS-617
chr3-111087233-INS-6795
chr3-111105675-INS-6226
chr3-111117065-INS-194
chr3-111139692-INS-172
chr3-111145128-INS-175
chr3-111262336-INS-55
chr3-111270344-INS-93
chr3-111288725-INS-210
chr3-111298711-INS-174
chr3-111317000-INS-56
chr3-111321922-INS-90
chr3-111328875-INS-72
chr3-111377506-INS-7242
chr3-111404792-INS-63
chr3-111405532-INS-2835
chr3-111453459-INS-50
chr3-111464796-INS-1653
chr3-111544271-INS-306

chr3-111544271-INS-56
chr3-111545650-INS-54
chr3-111560806-INS-108
chr3-111568287-INS-401
chr3-111581896-INS-62
chr3-111595303-INS-5545
chr3-111628898-INS-56
chr3-111629542-INS-345
chr3-111662345-INS-107
chr3-111679467-INS-55
chr3-111690712-INS-50
chr3-111713493-INS-59
chr3-111713497-INS-382
chr3-111719263-INS-4540
chr3-111762737-INS-53
chr3-111788444-INS-58
chr3-111850131-INS-6806
chr3-111893811-INS-62
chr3-111901721-INS-1104
chr3-111923846-INS-124
chr3-111924178-INS-996
chr3-111972602-INS-4415
chr3-111987214-INS-50
chr3-112008795-INS-1084
chr3-112014781-INS-6378
chr3-112028766-INS-52
chr3-112099842-INS-214
chr3-112182279-INS-14580
chr3-112207028-INS-745
chr3-112264653-INS-1925
chr3-112281260-INS-62
chr3-112317524-INS-509
chr3-112370285-INS-217
chr3-112441482-INS-6686
chr3-112596792-INS-6164
chr3-112604641-INS-495
chr3-112622908-INS-558
chr3-112626621-INS-100
chr3-112647300-INS-5791
chr3-112647932-INS-498
chr3-112778288-INS-60
chr3-112778710-INS-209
chr3-112778906-INS-59
chr3-112779030-INS-325
chr3-112820336-INS-112
chr3-112856062-INS-55
chr3-112856064-INS-155
chr3-112928954-INS-6195
chr3-112971926-INS-77
chr3-112971979-INS-322
chr3-113038446-INS-87
chr3-113038509-INS-390
chr3-113090668-INS-1621
chr3-113127834-INS-55
chr3-113149617-INS-181
chr3-113317367-INS-52
chr3-113342069-INS-3542
chr3-113372580-INS-972
chr3-113453664-INS-221
chr3-113482267-INS-80
chr3-113547507-INS-61
chr3-113559203-INS-6205
chr3-113564277-INS-2089
chr3-113589135-INS-1831
chr3-113595975-INS-6380
chr3-113610109-INS-55
chr3-113668157-INS-76
chr3-113692323-INS-55
chr3-113723119-INS-8307
chr3-113837176-INS-422
chr3-113839824-INS-7091
chr3-113851055-INS-194
chr3-114004416-INS-112
chr3-114126065-INS-1163
chr3-114132096-INS-6273
chr3-114170755-INS-6695
chr3-114188274-INS-4363

chr3-114196576-INS-1738
chr3-114197823-INS-453
chr3-114209605-INS-5019
chr3-114256432-INS-81
chr3-114317068-INS-6195
chr3-114332228-INS-81
chr3-114332742-INS-50
chr3-114346960-INS-1584
chr3-114349084-INS-6166
chr3-114475682-INS-65
chr3-114564189-INS-84
chr3-114579073-INS-341
chr3-114594949-INS-4332
chr3-114613043-INS-56
chr3-114613646-INS-88
chr3-114623129-INS-88
chr3-114623698-INS-3723
chr3-114627687-INS-1180
chr3-114751913-INS-60
chr3-114752778-INS-88
chr3-114774541-INS-2059
chr3-114805762-INS-589
chr3-114808758-INS-236
chr3-114901291-INS-92
chr3-114910499-INS-75
chr3-114920557-INS-52
chr3-114966240-INS-604
chr3-114976699-INS-5715
chr3-114993317-INS-1806
chr3-114993659-INS-52
chr3-115002252-INS-352
chr3-115037615-INS-1341
chr3-115037776-INS-70
chr3-115097586-INS-2973
chr3-115098021-INS-6207
chr3-115105351-INS-3568
chr3-115114980-INS-50
chr3-115186176-INS-51
chr3-115196091-INS-158
chr3-115264108-INS-170
chr3-115264666-INS-6598
chr3-115266269-INS-503
chr3-115272142-INS-322
chr3-115299094-INS-50
chr3-115307627-INS-4898
chr3-115358561-INS-1535
chr3-115358876-INS-578
chr3-115364192-INS-54
chr3-115372644-INS-151
chr3-115373225-INS-300
chr3-115378894-INS-60
chr3-115400120-INS-10859
chr3-115405005-INS-552
chr3-115419284-INS-54
chr3-115434145-INS-304
chr3-115446862-INS-421
chr3-115454491-INS-1336
chr3-115475885-INS-1288
chr3-115497427-INS-85
chr3-115517322-INS-144
chr3-115517334-INS-314
chr3-115530966-INS-415
chr3-115537570-INS-127
chr3-115582665-INS-2317
chr3-115614975-INS-369
chr3-115615009-INS-90
chr3-115624976-INS-209
chr3-115624976-INS-59
chr3-115646221-INS-66
chr3-115662679-INS-6230
chr3-115676048-INS-136
chr3-115676060-INS-592
chr3-115676177-INS-66
chr3-115747561-INS-184
chr3-115748953-INS-54
chr3-115756920-INS-52
chr3-115823903-INS-141

chr3-115827897-INS-6475
chr3-115853212-INS-452
chr3-115857332-INS-198
chr3-115870060-INS-1181
chr3-115880659-INS-52
chr3-115953834-INS-175
chr3-115954139-INS-701
chr3-116018912-INS-6398
chr3-116029828-INS-60
chr3-116056774-INS-6424
chr3-116065273-INS-177
chr3-116081355-INS-953
chr3-116107448-INS-6510
chr3-116116758-INS-181
chr3-116124703-INS-53
chr3-116206613-INS-92
chr3-116207513-INS-88
chr3-116211463-INS-60
chr3-116212082-INS-221
chr3-116237994-INS-4023
chr3-116263454-INS-199
chr3-116263707-INS-70
chr3-116266094-INS-6653
chr3-116266858-INS-70
chr3-116279260-INS-943
chr3-116285104-INS-80
chr3-116286518-INS-258
chr3-116286518-INS-86
chr3-116312167-INS-200
chr3-116313183-INS-189
chr3-116313366-INS-386
chr3-116325409-INS-3242
chr3-116332296-INS-79
chr3-116357432-INS-97
chr3-116363092-INS-52
chr3-116370690-INS-6382
chr3-116402565-INS-392
chr3-116486292-INS-63
chr3-116486551-INS-63
chr3-116502984-INS-152
chr3-116512753-INS-178
chr3-116516456-INS-602
chr3-116518399-INS-50
chr3-116522608-INS-95
chr3-116522782-INS-181
chr3-116528352-INS-2629
chr3-116530951-INS-2489
chr3-116530951-INS-970
chr3-116552730-INS-211
chr3-116553476-INS-213
chr3-116591886-INS-230
chr3-116604599-INS-160
chr3-116617023-INS-422
chr3-116632861-INS-7222
chr3-116635385-INS-96
chr3-116636660-INS-898
chr3-116637505-INS-666
chr3-116642235-INS-66
chr3-116655963-INS-142
chr3-116655963-INS-324
chr3-116656964-INS-153
chr3-116659461-INS-311
chr3-116659461-INS-68
chr3-116664138-INS-6443
chr3-116670486-INS-4796
chr3-116674037-INS-7116
chr3-116685318-INS-62
chr3-116692052-INS-51
chr3-116692358-INS-205
chr3-116710290-INS-211
chr3-116743566-INS-63
chr3-116757270-INS-50
chr3-116768367-INS-2764
chr3-116771023-INS-78
chr3-116773447-INS-54
chr3-116774591-INS-144
chr3-116776480-INS-193

chr3-116780178-INS-5360
chr3-116783063-INS-1892
chr3-116805160-INS-3640
chr3-116842998-INS-6257
chr3-116844718-INS-279
chr3-116845828-INS-189
chr3-116849229-INS-171
chr3-116859603-INS-6280
chr3-116880634-INS-4972
chr3-116881005-INS-1218
chr3-116927233-INS-117
chr3-116929527-INS-158
chr3-116943915-INS-13802
chr3-116978108-INS-50
chr3-116978389-INS-77
chr3-117041132-INS-50
chr3-117064057-INS-497
chr3-117119147-INS-53
chr3-117127200-INS-1401
chr3-117157576-INS-1759
chr3-117186780-INS-381
chr3-117187277-INS-1882
chr3-117204871-INS-53
chr3-117237262-INS-283
chr3-117251599-INS-52
chr3-117287160-INS-6215
chr3-117412003-INS-204
chr3-117416192-INS-58
chr3-117438228-INS-95
chr3-117450269-INS-87
chr3-117452092-INS-163
chr3-117467340-INS-6197
chr3-117472380-INS-58
chr3-117488021-INS-52
chr3-117514616-INS-7102
chr3-117523620-INS-50
chr3-117531304-INS-72
chr3-117592628-INS-57
chr3-117596432-INS-138
chr3-117603877-INS-637
chr3-117610092-INS-54
chr3-117617983-INS-6562
chr3-117676863-INS-50
chr3-117696941-INS-84
chr3-117732158-INS-2498
chr3-117746734-INS-50
chr3-117754037-INS-6687
chr3-117839866-INS-56
chr3-117895310-INS-371
chr3-117905153-INS-55
chr3-117941271-INS-52
chr3-117953381-INS-156
chr3-117979111-INS-201
chr3-117979781-INS-614
chr3-118021163-INS-176
chr3-118022087-INS-126
chr3-118022139-INS-50
chr3-118047184-INS-57
chr3-118066491-INS-50
chr3-118140215-INS-50
chr3-118214721-INS-60
chr3-118236936-INS-317
chr3-118279976-INS-7124
chr3-118330956-INS-56
chr3-118334890-INS-56
chr3-118335141-INS-64
chr3-118366004-INS-1186
chr3-118420892-INS-616
chr3-118421366-INS-88
chr3-118421717-INS-50
chr3-118441211-INS-58
chr3-118443488-INS-3143
chr3-118448947-INS-1336
chr3-118475660-INS-56
chr3-118492144-INS-52
chr3-118501255-INS-402
chr3-118520097-INS-51

chr3-118556440-INS-50
chr3-118577258-INS-102
chr3-118589025-INS-52
chr3-118626525-INS-58
chr3-118752943-INS-2860
chr3-118790920-INS-139
chr3-118873805-INS-52
chr3-118879501-INS-53
chr3-118879565-INS-312
chr3-118886203-INS-57
chr3-118888407-INS-6380
chr3-118931303-INS-603
chr3-118931369-INS-52
chr3-118969408-INS-8358
chr3-118984747-INS-225
chr3-118991512-INS-66
chr3-119093784-INS-114
chr3-119110804-INS-52
chr3-119132486-INS-76
chr3-119206791-INS-66
chr3-119246998-INS-1366
chr3-119296168-INS-462
chr3-119301240-INS-185
chr3-119301240-INS-64
chr3-119312730-INS-231
chr3-119313072-INS-90
chr3-119317977-INS-56
chr3-119319618-INS-7089
chr3-119319814-INS-52
chr3-119330282-INS-52
chr3-119419776-INS-543
chr3-119425558-INS-255
chr3-119501695-INS-135
chr3-119592318-INS-171
chr3-119641984-INS-96
chr3-119660660-INS-52
chr3-119665322-INS-72
chr3-119673261-INS-1319
chr3-119675573-INS-448
chr3-119746053-INS-311
chr3-119761541-INS-177
chr3-119816714-INS-84
chr3-119821232-INS-2279
chr3-119836869-INS-6394
chr3-119862768-INS-87
chr3-119935100-INS-52
chr3-119943231-INS-58
chr3-119945568-INS-68
chr3-119967371-INS-458
chr3-119967974-INS-316
chr3-119990995-INS-306
chr3-120020234-INS-90
chr3-120020528-INS-88
chr3-120023285-INS-125
chr3-120023352-INS-51
chr3-120026750-INS-212
chr3-120035373-INS-706
chr3-120045155-INS-106
chr3-120051661-INS-7175
chr3-120133271-INS-50
chr3-120178227-INS-51
chr3-120190477-INS-2878
chr3-120211758-INS-603
chr3-120322948-INS-84
chr3-120333797-INS-1095
chr3-120342735-INS-2603
chr3-120343762-INS-1854
chr3-120379070-INS-94
chr3-120433870-INS-1206
chr3-120433871-INS-559
chr3-120489668-INS-50
chr3-120491709-INS-52
chr3-120538896-INS-2179
chr3-120565736-INS-7969
chr3-120567510-INS-60
chr3-120572285-INS-106
chr3-120670938-INS-150

chr3-120684561-INS-85
chr3-120702467-INS-52
chr3-120710371-INS-54
chr3-120751586-INS-51
chr3-120752137-INS-342
chr3-120765887-INS-6301
chr3-120769774-INS-50
chr3-120775248-INS-160
chr3-120786733-INS-114
chr3-120791611-INS-57
chr3-120792381-INS-1366
chr3-120805729-INS-116
chr3-120833738-INS-140
chr3-120929999-INS-605
chr3-120935182-INS-736
chr3-120939850-INS-155
chr3-120946468-INS-184
chr3-120946649-INS-365
chr3-120946745-INS-113
chr3-120961321-INS-54
chr3-120973779-INS-2877
chr3-121010418-INS-187
chr3-121016713-INS-342
chr3-121043191-INS-368
chr3-121043243-INS-164
chr3-121051839-INS-198
chr3-121066867-INS-198
chr3-121066922-INS-84
chr3-121097160-INS-256
chr3-121112462-INS-96
chr3-121118302-INS-119
chr3-121119573-INS-3926
chr3-121119573-INS-418
chr3-121144552-INS-1227
chr3-121168335-INS-52
chr3-121307963-INS-56
chr3-121336075-INS-276
chr3-121361429-INS-60
chr3-121407423-INS-134
chr3-121410386-INS-205
chr3-121454453-INS-69
chr3-121548850-INS-5642
chr3-121584081-INS-459
chr3-121617440-INS-216
chr3-121618288-INS-218
chr3-121619062-INS-69
chr3-121623448-INS-125
chr3-121630150-INS-499
chr3-121634716-INS-51
chr3-121645353-INS-64
chr3-121649145-INS-391
chr3-121649145-INS-5365
chr3-121659803-INS-63
chr3-121672240-INS-119
chr3-121732033-INS-50
chr3-121755663-INS-52
chr3-121784442-INS-174
chr3-121816804-INS-265
chr3-121840837-INS-68
chr3-121851385-INS-79
chr3-121873735-INS-1471
chr3-121889456-INS-109
chr3-121907642-INS-7080
chr3-121929140-INS-495
chr3-121947610-INS-278
chr3-121955698-INS-1758
chr3-121955698-INS-320
chr3-121959880-INS-4396
chr3-121961486-INS-9021
chr3-121991927-INS-70
chr3-122092557-INS-7067
chr3-122096148-INS-188
chr3-122101792-INS-874
chr3-122102052-INS-2286
chr3-122102102-INS-554
chr3-122102532-INS-11655
chr3-122104480-INS-2226

chr3-122122886-INS-720
chr3-122123686-INS-224
chr3-122129760-INS-936
chr3-122133878-INS-229
chr3-122137603-INS-51
chr3-122159358-INS-167
chr3-122168315-INS-66
chr3-122182188-INS-1670
chr3-122196499-INS-50
chr3-122207324-INS-57
chr3-122228703-INS-229
chr3-122241879-INS-100
chr3-122253841-INS-241
chr3-122312234-INS-120
chr3-122318445-INS-120
chr3-122327234-INS-6334
chr3-122367796-INS-55
chr3-122398396-INS-104
chr3-122403598-INS-619
chr3-122408990-INS-141
chr3-122426037-INS-273
chr3-122444851-INS-139
chr3-122447544-INS-245
chr3-122464635-INS-68
chr3-122509119-INS-4231
chr3-122540174-INS-116
chr3-122542795-INS-370
chr3-122564401-INS-85
chr3-122576510-INS-6356
chr3-122628638-INS-162
chr3-122628765-INS-56
chr3-122629752-INS-57
chr3-122669486-INS-5859
chr3-122684242-INS-100
chr3-122699026-INS-198
chr3-122703501-INS-5071
chr3-122735056-INS-1292
chr3-122764436-INS-251
chr3-122764436-INS-69
chr3-122837529-INS-776
chr3-122857063-INS-572
chr3-122857376-INS-219
chr3-122878313-INS-99
chr3-122879141-INS-58
chr3-122996804-INS-2000
chr3-123005981-INS-53
chr3-123007381-INS-240
chr3-123009976-INS-63
chr3-123010057-INS-498
chr3-123022277-INS-52
chr3-123033119-INS-7261
chr3-123048551-INS-4735
chr3-123049630-INS-434
chr3-123050021-INS-8413
chr3-123094176-INS-95
chr3-123111330-INS-157
chr3-123121866-INS-64
chr3-123139392-INS-6558
chr3-123193615-INS-96
chr3-123210526-INS-51
chr3-123273033-INS-90
chr3-123305090-INS-298
chr3-123305090-INS-950
chr3-123317232-INS-58
chr3-123326855-INS-81
chr3-123338653-INS-73
chr3-123368616-INS-143
chr3-123369086-INS-61
chr3-123369155-INS-823
chr3-123369786-INS-320
chr3-123397168-INS-2902
chr3-123421983-INS-84
chr3-123422081-INS-176
chr3-123425383-INS-75
chr3-123434786-INS-152
chr3-123470113-INS-4841
chr3-123471793-INS-91

chr3-123489863-INS-2896
chr3-123508043-INS-6806
chr3-123537426-INS-192
chr3-123545017-INS-206
chr3-123552752-INS-84
chr3-123558006-INS-7366
chr3-123580295-INS-808
chr3-123588585-INS-70
chr3-123638696-INS-6206
chr3-123708152-INS-94
chr3-123753466-INS-3469
chr3-123797543-INS-54
chr3-123803903-INS-1319
chr3-123823193-INS-53
chr3-123826742-INS-208
chr3-123879235-INS-6222
chr3-123887506-INS-135
chr3-123897780-INS-72
chr3-123918873-INS-51
chr3-123948772-INS-2235
chr3-124027468-INS-7083
chr3-124043986-INS-56
chr3-124052139-INS-90
chr3-124063858-INS-77
chr3-124105496-INS-1749
chr3-124131756-INS-2004
chr3-124138164-INS-368
chr3-124138189-INS-922
chr3-124139455-INS-319
chr3-124152623-INS-742
chr3-124196410-INS-50
chr3-124208928-INS-140
chr3-124252826-INS-132
chr3-124284295-INS-50
chr3-124319622-INS-2999
chr3-124338422-INS-2027
chr3-124346568-INS-583
chr3-124347420-INS-626
chr3-124382022-INS-226
chr3-124385953-INS-128
chr3-124393499-INS-50
chr3-124393885-INS-122
chr3-124425491-INS-195
chr3-124450120-INS-6488
chr3-124470262-INS-84
chr3-124482010-INS-4028
chr3-124539651-INS-8948
chr3-124585014-INS-2821
chr3-124620723-INS-102
chr3-124620888-INS-166
chr3-124632856-INS-2151
chr3-124646044-INS-2445
chr3-124648353-INS-52
chr3-124655607-INS-97
chr3-124695659-INS-62
chr3-124731228-INS-85
chr3-124763939-INS-212
chr3-124769970-INS-3985
chr3-124773066-INS-66
chr3-124795481-INS-6330
chr3-124828592-INS-6917
chr3-124842594-INS-222
chr3-124846022-INS-2843
chr3-124853997-INS-57
chr3-124882217-INS-1899
chr3-124961620-INS-2710
chr3-125030218-INS-270
chr3-125039101-INS-64
chr3-125068328-INS-170
chr3-125085742-INS-50
chr3-125159583-INS-164
chr3-125241218-INS-5651
chr3-125322030-INS-243
chr3-125367554-INS-62
chr3-125592916-INS-52
chr3-125640885-INS-1619
chr3-125642767-INS-1834

chr3-125654285-INS-68
chr3-125654875-INS-59
chr3-125672332-INS-4782
chr3-125682354-INS-54
chr3-125815165-INS-4202
chr3-125844250-INS-2304
chr3-125862036-INS-60
chr3-125878299-INS-51
chr3-125961094-INS-64
chr3-125973345-INS-78
chr3-126014096-INS-54
chr3-126024094-INS-62
chr3-126037845-INS-4960
chr3-126040895-INS-54
chr3-126050244-INS-7064
chr3-126090627-INS-3107
chr3-126105424-INS-8914
chr3-126143387-INS-2672
chr3-126269120-INS-1628
chr3-126307178-INS-58
chr3-126313226-INS-341
chr3-126331137-INS-72
chr3-126334750-INS-52
chr3-126368108-INS-124
chr3-126373437-INS-80
chr3-126401001-INS-55
chr3-126480028-INS-72
chr3-126514378-INS-152
chr3-126516484-INS-6182
chr3-126525271-INS-546
chr3-126536052-INS-224
chr3-126540723-INS-202
chr3-126540923-INS-1466
chr3-126560466-INS-55
chr3-126626307-INS-56
chr3-126650561-INS-6381
chr3-126698889-INS-1845
chr3-126711707-INS-116
chr3-126820833-INS-315
chr3-126821145-INS-185
chr3-126834945-INS-50
chr3-126863012-INS-6474
chr3-126875824-INS-54
chr3-126876969-INS-113
chr3-126890386-INS-62
chr3-126894603-INS-2876
chr3-126899667-INS-60
chr3-126925635-INS-52
chr3-126966172-INS-56
chr3-126972070-INS-62
chr3-127012115-INS-8986
chr3-127055616-INS-50
chr3-127057969-INS-156
chr3-127060433-INS-306
chr3-127081747-INS-128
chr3-127081996-INS-520
chr3-127121398-INS-1032
chr3-127133934-INS-108
chr3-127153334-INS-2067
chr3-127154102-INS-480
chr3-127164395-INS-107
chr3-127197847-INS-68
chr3-127197873-INS-167
chr3-127212224-INS-51
chr3-127214507-INS-1431
chr3-127222435-INS-196
chr3-127224225-INS-168
chr3-127225200-INS-1841
chr3-127226304-INS-215
chr3-127227403-INS-74
chr3-127230793-INS-97
chr3-127234809-INS-50
chr3-127254985-INS-50
chr3-127256523-INS-205
chr3-127266509-INS-60
chr3-127266905-INS-198
chr3-127271749-INS-200

chr3-127295448-INS-208
chr3-127309052-INS-105
chr3-127309345-INS-1298
chr3-127315616-INS-218
chr3-127324710-INS-345
chr3-127359106-INS-209
chr3-127369915-INS-60
chr3-127375930-INS-86
chr3-127412432-INS-61
chr3-127430006-INS-67
chr3-127441365-INS-320
chr3-127442513-INS-4817
chr3-127453662-INS-6491
chr3-127453968-INS-205
chr3-127462477-INS-427
chr3-127481956-INS-51
chr3-127483306-INS-819
chr3-127492160-INS-446
chr3-127583894-INS-54
chr3-127602730-INS-120
chr3-127608357-INS-389
chr3-127608371-INS-187
chr3-127640428-INS-68
chr3-127661187-INS-328
chr3-127661232-INS-64
chr3-127694599-INS-356
chr3-127697153-INS-117
chr3-127698166-INS-227
chr3-127698338-INS-75
chr3-127721410-INS-78
chr3-127748878-INS-52
chr3-127749447-INS-404
chr3-127764187-INS-165
chr3-127814379-INS-201
chr3-127814490-INS-92
chr3-127826191-INS-989
chr3-127826242-INS-2042
chr3-127883490-INS-235
chr3-127921882-INS-84
chr3-127929512-INS-65
chr3-128001937-INS-379
chr3-128001938-INS-55
chr3-128003703-INS-132
chr3-128047409-INS-52
chr3-128055188-INS-6742
chr3-128067303-INS-58
chr3-128109444-INS-88
chr3-128114410-INS-100
chr3-128136161-INS-6389
chr3-128152782-INS-56
chr3-128156735-INS-6804
chr3-128159295-INS-51
chr3-128159347-INS-312
chr3-128211128-INS-6586
chr3-128233660-INS-56
chr3-128253297-INS-50
chr3-128262695-INS-50
chr3-128265941-INS-966
chr3-128312223-INS-1164
chr3-128316037-INS-108
chr3-128322922-INS-456
chr3-128356256-INS-5518
chr3-128368795-INS-52
chr3-128380697-INS-777
chr3-128382281-INS-4717
chr3-128388583-INS-2624
chr3-128545852-INS-162
chr3-128556760-INS-1767
chr3-128609218-INS-65
chr3-128609509-INS-147
chr3-128635937-INS-7733
chr3-128660431-INS-252
chr3-128739164-INS-63
chr3-128741706-INS-50
chr3-128777484-INS-1127
chr3-128792131-INS-640
chr3-128803744-INS-70

chr3-128810413-INS-120
chr3-128810567-INS-209
chr3-128836360-INS-67
chr3-128865132-INS-65
chr3-128892904-INS-198
chr3-128896651-INS-115
chr3-128905790-INS-54
chr3-128936100-INS-955
chr3-128971598-INS-78
chr3-129004042-INS-306
chr3-129004839-INS-50
chr3-129036469-INS-58
chr3-129052361-INS-118
chr3-129073857-INS-52
chr3-129079002-INS-423
chr3-129084923-INS-82
chr3-129086164-INS-73
chr3-129089995-INS-1622
chr3-129118889-INS-261
chr3-129213457-INS-1962
chr3-129285391-INS-64
chr3-129295166-INS-52
chr3-129322499-INS-56
chr3-129343706-INS-90
chr3-129347223-INS-286
chr3-129347223-INS-58
chr3-129354364-INS-209
chr3-129354521-INS-393
chr3-129357949-INS-89
chr3-129373761-INS-218
chr3-129403092-INS-63
chr3-129410005-INS-232
chr3-129424506-INS-52
chr3-129439522-INS-4991
chr3-129455032-INS-56
chr3-129513499-INS-183
chr3-129543554-INS-222
chr3-129543554-INS-74
chr3-129572402-INS-60
chr3-129575278-INS-69
chr3-129575309-INS-230
chr3-129599305-INS-842
chr3-129637505-INS-50
chr3-129644863-INS-58
chr3-129645514-INS-60
chr3-129648924-INS-208
chr3-129664716-INS-68
chr3-129664941-INS-194
chr3-129670153-INS-3710
chr3-129701190-INS-60
chr3-129701419-INS-60
chr3-129746839-INS-4402
chr3-129750902-INS-387
chr3-129751580-INS-216
chr3-129760922-INS-64
chr3-129836466-INS-56
chr3-129844543-INS-161
chr3-129867289-INS-123
chr3-129869508-INS-76
chr3-129870278-INS-165
chr3-129900503-INS-121
chr3-129906830-INS-137
chr3-129907722-INS-280
chr3-129920126-INS-996
chr3-129923804-INS-4796
chr3-129949600-INS-397
chr3-129993238-INS-136
chr3-129998850-INS-50
chr3-130032810-INS-6273
chr3-130061735-INS-66
chr3-130061773-INS-231
chr3-130077777-INS-92
chr3-130120606-INS-64
chr3-130122102-INS-176
chr3-130129673-INS-59
chr3-130153315-INS-66
chr3-130153347-INS-298

chr3-130166064-INS-362
chr3-130170590-INS-78
chr3-130173283-INS-217
chr3-130173564-INS-114
chr3-130180652-INS-1900
chr3-130204348-INS-80
chr3-130205114-INS-68
chr3-130215133-INS-58
chr3-130223155-INS-79
chr3-130236962-INS-65
chr3-130237276-INS-56
chr3-130237317-INS-242
chr3-130247563-INS-7438
chr3-130274504-INS-55
chr3-130296509-INS-171
chr3-130325294-INS-99
chr3-130338688-INS-76
chr3-130402824-INS-60
chr3-130425324-INS-149
chr3-130433531-INS-1767
chr3-130436803-INS-1036
chr3-130481919-INS-275
chr3-130515898-INS-192
chr3-130528146-INS-50
chr3-130594868-INS-205
chr3-130595003-INS-398
chr3-130606100-INS-248
chr3-130614075-INS-1110
chr3-130645934-INS-380
chr3-130646125-INS-108
chr3-130687653-INS-6689
chr3-130702379-INS-72
chr3-130719689-INS-104
chr3-130731954-INS-184
chr3-130740169-INS-51
chr3-130762499-INS-7058
chr3-130792702-INS-75
chr3-130820741-INS-80
chr3-130854576-INS-199
chr3-130858495-INS-123
chr3-130865056-INS-225
chr3-130875943-INS-50
chr3-130883201-INS-62
chr3-130924328-INS-80
chr3-130924541-INS-65
chr3-130924542-INS-140
chr3-130933861-INS-209
chr3-130934267-INS-68
chr3-130935587-INS-229
chr3-130964433-INS-165
chr3-130966120-INS-50
chr3-130988533-INS-50
chr3-131003093-INS-54
chr3-131086527-INS-181
chr3-131109638-INS-96
chr3-131111392-INS-1249
chr3-131111723-INS-537
chr3-131111739-INS-221
chr3-131145092-INS-264
chr3-131145660-INS-56
chr3-131152229-INS-1500
chr3-131155294-INS-63
chr3-131194637-INS-159
chr3-131239707-INS-2102
chr3-131247655-INS-72
chr3-131248022-INS-119
chr3-131269311-INS-56
chr3-131277233-INS-86
chr3-131277299-INS-313
chr3-131277417-INS-115
chr3-131288664-INS-62
chr3-131295348-INS-2745
chr3-131297288-INS-52
chr3-131303991-INS-2101
chr3-131314366-INS-256
chr3-131315778-INS-75
chr3-131328773-INS-380

chr3-131328773-INS-940
chr3-131333890-INS-207
chr3-131387377-INS-95
chr3-131405655-INS-52
chr3-131406258-INS-263
chr3-131438605-INS-118
chr3-131441571-INS-60
chr3-131463415-INS-91
chr3-131475300-INS-75
chr3-131488959-INS-66
chr3-131493812-INS-6998
chr3-131506773-INS-50
chr3-131531450-INS-265
chr3-131532291-INS-135
chr3-131534715-INS-596
chr3-131535095-INS-430
chr3-131552801-INS-1028
chr3-131552801-INS-173
chr3-131572925-INS-64
chr3-131596402-INS-123
chr3-131611084-INS-5536
chr3-131630790-INS-449
chr3-131630816-INS-66
chr3-131681617-INS-56
chr3-131687693-INS-1898
chr3-131698214-INS-91
chr3-131708123-INS-56
chr3-131736138-INS-175
chr3-131777884-INS-3030
chr3-131816629-INS-1540
chr3-131826069-INS-52
chr3-131839760-INS-119
chr3-131875174-INS-123
chr3-131885423-INS-60
chr3-131903014-INS-6168
chr3-131923398-INS-12472
chr3-131959828-INS-7202
chr3-131967991-INS-52
chr3-131972054-INS-184
chr3-131993992-INS-4149
chr3-131994131-INS-80
chr3-131994196-INS-302
chr3-132001844-INS-6594
chr3-132041811-INS-56
chr3-132086077-INS-2278
chr3-132097629-INS-59
chr3-132109326-INS-617
chr3-132110599-INS-203
chr3-132117868-INS-411
chr3-132132990-INS-51
chr3-132160345-INS-137
chr3-132160385-INS-53
chr3-132160553-INS-6595
chr3-132181636-INS-2960
chr3-132187214-INS-7089
chr3-132214579-INS-206
chr3-132215532-INS-6437
chr3-132237764-INS-52
chr3-132258901-INS-240
chr3-132294400-INS-64
chr3-132297648-INS-689
chr3-132305750-INS-206
chr3-132441467-INS-160
chr3-132455058-INS-1102
chr3-132505219-INS-498
chr3-132521997-INS-62
chr3-132567347-INS-84
chr3-132568690-INS-64
chr3-132661736-INS-1624
chr3-132683793-INS-65
chr3-132693621-INS-58
chr3-132706943-INS-4850
chr3-132712675-INS-1486
chr3-132749874-INS-169
chr3-132759526-INS-96
chr3-132841565-INS-126
chr3-132842000-INS-50

chr3-132846370-INS-68
chr3-132890476-INS-3484
chr3-132894663-INS-60
chr3-132899855-INS-1610
chr3-132903259-INS-54
chr3-132907820-INS-7567
chr3-132916796-INS-1099
chr3-132930849-INS-67
chr3-132934650-INS-50
chr3-132947367-INS-231
chr3-132969965-INS-200
chr3-132976504-INS-4371
chr3-132988362-INS-6181
chr3-132996115-INS-1058
chr3-133012553-INS-488
chr3-133012630-INS-59
chr3-133012805-INS-154
chr3-133012961-INS-188
chr3-133021762-INS-126
chr3-133022071-INS-184
chr3-133058610-INS-528
chr3-133099597-INS-211
chr3-133118019-INS-6543
chr3-133141461-INS-8783
chr3-133146586-INS-1418
chr3-133147099-INS-196
chr3-133150964-INS-6637
chr3-133151866-INS-1997
chr3-133241580-INS-94
chr3-133250359-INS-84
chr3-133260317-INS-212
chr3-133262349-INS-164
chr3-133263215-INS-73
chr3-133271003-INS-216
chr3-133276214-INS-2921
chr3-133278925-INS-76
chr3-133280645-INS-6583
chr3-133290566-INS-50
chr3-133308046-INS-484
chr3-133312801-INS-1173
chr3-133336634-INS-50
chr3-133359787-INS-478
chr3-133361327-INS-1382
chr3-133376373-INS-50
chr3-133377560-INS-727
chr3-133377981-INS-7110
chr3-133436972-INS-79
chr3-133473451-INS-76
chr3-133480609-INS-50
chr3-133481136-INS-52
chr3-133494580-INS-477
chr3-133495363-INS-77
chr3-133498745-INS-215
chr3-133507374-INS-2065
chr3-133509356-INS-1195
chr3-133522049-INS-69
chr3-133586481-INS-509
chr3-133604018-INS-53
chr3-133619221-INS-104
chr3-133643694-INS-147
chr3-133659672-INS-60
chr3-133665861-INS-57
chr3-133747435-INS-140
chr3-133748145-INS-163
chr3-133749122-INS-172
chr3-133763670-INS-305
chr3-133763673-INS-70
chr3-133854988-INS-445
chr3-133875827-INS-537
chr3-133877163-INS-177
chr3-133887435-INS-50
chr3-133938242-INS-129
chr3-133977983-INS-673
chr3-133983053-INS-90
chr3-133986661-INS-56
chr3-134034945-INS-432
chr3-134045101-INS-756

chr3-134203215-INS-52
chr3-134205221-INS-106
chr3-134235173-INS-289
chr3-134251674-INS-623
chr3-134278630-INS-76
chr3-134284513-INS-198
chr3-134313512-INS-52
chr3-134357480-INS-2372
chr3-134366772-INS-2641
chr3-134375735-INS-589
chr3-134395363-INS-1251
chr3-134584878-INS-74
chr3-134601101-INS-748
chr3-134657099-INS-67
chr3-134691762-INS-1340
chr3-134738716-INS-60
chr3-134757370-INS-52
chr3-134775009-INS-925
chr3-134821961-INS-7596
chr3-134823073-INS-1260
chr3-134826840-INS-50
chr3-134827435-INS-80
chr3-134832384-INS-117
chr3-134832602-INS-199
chr3-134832853-INS-116
chr3-134834896-INS-230
chr3-134858843-INS-172
chr3-134886421-INS-288
chr3-134900067-INS-278
chr3-134906529-INS-2045
chr3-134990194-INS-4535
chr3-135013633-INS-51
chr3-135014810-INS-68
chr3-135015091-INS-747
chr3-135168041-INS-158
chr3-135180153-INS-569
chr3-135194258-INS-200
chr3-135212847-INS-250
chr3-135213851-INS-133
chr3-135214440-INS-204
chr3-135236185-INS-224
chr3-135236185-INS-56
chr3-135256110-INS-746
chr3-135256484-INS-534
chr3-135323730-INS-60
chr3-135357725-INS-66
chr3-135363310-INS-73
chr3-135369623-INS-137
chr3-135386260-INS-722
chr3-135419336-INS-1072
chr3-135447253-INS-120
chr3-135452307-INS-357
chr3-135489271-INS-525
chr3-135581111-INS-280
chr3-135616904-INS-64
chr3-135617827-INS-6362
chr3-135618812-INS-3070
chr3-135661189-INS-624
chr3-135690121-INS-6392
chr3-135740502-INS-136
chr3-135753729-INS-1348
chr3-135762828-INS-1807
chr3-135766343-INS-6289
chr3-135854076-INS-2833
chr3-135859437-INS-199
chr3-135862943-INS-96
chr3-135913636-INS-1292
chr3-135934784-INS-56
chr3-135950889-INS-1860
chr3-135969192-INS-6624
chr3-135978300-INS-236
chr3-135992930-INS-77
chr3-136023257-INS-5810
chr3-136026983-INS-159
chr3-136043157-INS-4661
chr3-136052922-INS-6406
chr3-136053822-INS-52

chr3-136166865-INS-6268
chr3-136224055-INS-3124
chr3-136264313-INS-4873
chr3-136270980-INS-76
chr3-136273185-INS-96
chr3-136273783-INS-52
chr3-136318787-INS-95
chr3-136334051-INS-56
chr3-136383099-INS-72
chr3-136389092-INS-153
chr3-136544509-INS-1952
chr3-136548049-INS-52
chr3-136550945-INS-72
chr3-136637114-INS-66
chr3-136649486-INS-76
chr3-136654953-INS-470
chr3-136670056-INS-53
chr3-136678578-INS-1293
chr3-136678578-INS-4193
chr3-136710726-INS-1606
chr3-136768259-INS-75
chr3-136782836-INS-402
chr3-136785531-INS-51
chr3-136805571-INS-513
chr3-136810839-INS-6418
chr3-136847522-INS-3880
chr3-136857934-INS-514
chr3-136881183-INS-276
chr3-136882780-INS-122
chr3-136934876-INS-15041
chr3-136940042-INS-225
chr3-136940358-INS-57
chr3-136941690-INS-7212
chr3-137005905-INS-113
chr3-137013549-INS-2289
chr3-137016990-INS-81
chr3-137019357-INS-1392
chr3-137039331-INS-134
chr3-137039331-INS-272
chr3-137074978-INS-54
chr3-137075625-INS-57
chr3-137101288-INS-124
chr3-137127348-INS-82
chr3-137128129-INS-62
chr3-137187172-INS-66
chr3-137195054-INS-2093
chr3-137253487-INS-59
chr3-137275093-INS-141
chr3-137297691-INS-64
chr3-137298935-INS-51
chr3-137332627-INS-64
chr3-137335429-INS-353
chr3-137341537-INS-2743
chr3-137382983-INS-779
chr3-137428884-INS-76
chr3-137431455-INS-51
chr3-137440246-INS-1062
chr3-137456951-INS-1138
chr3-137461483-INS-6648
chr3-137483280-INS-201
chr3-137483280-INS-65
chr3-137495873-INS-128
chr3-137503765-INS-76
chr3-137540108-INS-241
chr3-137558714-INS-574
chr3-137562957-INS-74
chr3-137615581-INS-52
chr3-137646238-INS-429
chr3-137646343-INS-69
chr3-137652088-INS-7098
chr3-137658431-INS-213
chr3-137689154-INS-60
chr3-137695940-INS-2708
chr3-137751236-INS-188
chr3-137761990-INS-4666
chr3-137768477-INS-88
chr3-137812095-INS-132

chr3-137820023-INS-80
chr3-137820808-INS-96
chr3-137868690-INS-374
chr3-137985300-INS-160
chr3-138091766-INS-60
chr3-138122468-INS-564
chr3-138126821-INS-56
chr3-138128985-INS-527
chr3-138146987-INS-56
chr3-138154851-INS-189
chr3-138199105-INS-512
chr3-138208646-INS-2637
chr3-138240682-INS-64
chr3-138240988-INS-159
chr3-138275943-INS-194
chr3-138276589-INS-508
chr3-138294561-INS-128
chr3-138294561-INS-60
chr3-138307584-INS-8821
chr3-138313465-INS-50
chr3-138340029-INS-693
chr3-138384519-INS-6701
chr3-138387652-INS-60
chr3-138427420-INS-193
chr3-138459829-INS-832
chr3-138464567-INS-145
chr3-138475014-INS-83
chr3-138486177-INS-1974
chr3-138486268-INS-200
chr3-138486296-INS-754
chr3-138603215-INS-6753
chr3-138698821-INS-228
chr3-138723345-INS-118
chr3-138729163-INS-2818
chr3-138753551-INS-6196
chr3-138795011-INS-64
chr3-138863866-INS-51
chr3-138922046-INS-175
chr3-138927285-INS-52
chr3-138962394-INS-7121
chr3-139015789-INS-6477
chr3-139026132-INS-68
chr3-139062874-INS-76
chr3-139063588-INS-4656
chr3-139099458-INS-2774
chr3-139102506-INS-320
chr3-139102506-INS-96
chr3-139108941-INS-1314
chr3-139126675-INS-100
chr3-139129919-INS-111
chr3-139146300-INS-1481
chr3-139150292-INS-566
chr3-139330123-INS-84
chr3-139397591-INS-4555
chr3-139420045-INS-110
chr3-139421799-INS-50
chr3-139518298-INS-6160
chr3-139518817-INS-12020
chr3-139533088-INS-1036
chr3-139578989-INS-967
chr3-139639730-INS-7080
chr3-139717062-INS-52
chr3-139723624-INS-1205
chr3-139736798-INS-118
chr3-139768408-INS-444
chr3-139772271-INS-50
chr3-139804695-INS-54
chr3-139811600-INS-958
chr3-139818136-INS-112
chr3-139820290-INS-93
chr3-139845005-INS-117
chr3-139871563-INS-436
chr3-139896949-INS-171
chr3-139902342-INS-762
chr3-139971208-INS-54
chr3-139983011-INS-1692
chr3-140021134-INS-369

chr3-140052271-INS-6211
chr3-140052618-INS-50
chr3-140053149-INS-60
chr3-140053260-INS-248
chr3-140073262-INS-973
chr3-140073748-INS-71
chr3-140099243-INS-2336
chr3-140099604-INS-207
chr3-140111632-INS-1409
chr3-140137071-INS-73
chr3-140234136-INS-3997
chr3-140244542-INS-5142
chr3-140257438-INS-2511
chr3-140257726-INS-1593
chr3-140264813-INS-1430
chr3-140275430-INS-62
chr3-140316900-INS-112
chr3-140360564-INS-1292
chr3-140365061-INS-64
chr3-140389606-INS-6218
chr3-140395829-INS-6150
chr3-140405529-INS-698
chr3-140431993-INS-195
chr3-140442361-INS-2926
chr3-140448304-INS-3017
chr3-140448304-INS-402
chr3-140452602-INS-65
chr3-140478274-INS-624
chr3-140479219-INS-2708
chr3-140482014-INS-1147
chr3-140493272-INS-1012
chr3-140541966-INS-56
chr3-140579861-INS-1224
chr3-140608200-INS-6900
chr3-140621308-INS-4583
chr3-140621308-INS-498
chr3-140640408-INS-390
chr3-140654755-INS-7084
chr3-140682669-INS-50
chr3-140689136-INS-6470
chr3-140736008-INS-7207
chr3-140769475-INS-1657
chr3-140776225-INS-483
chr3-140805387-INS-319
chr3-140806646-INS-122
chr3-140806646-INS-302
chr3-140822529-INS-78
chr3-140839525-INS-737
chr3-140843150-INS-50
chr3-140887824-INS-1826
chr3-140952322-INS-70
chr3-140959798-INS-2841
chr3-140976539-INS-3583
chr3-141031499-INS-595
chr3-141067978-INS-52
chr3-141090845-INS-1329
chr3-141112804-INS-2228
chr3-141206165-INS-1092
chr3-141249017-INS-67
chr3-141259785-INS-94
chr3-141273312-INS-64
chr3-141287641-INS-8461
chr3-141288969-INS-127
chr3-141297150-INS-293
chr3-141297186-INS-128
chr3-141336407-INS-145
chr3-141353684-INS-2314
chr3-141362760-INS-7206
chr3-141391766-INS-208
chr3-141395682-INS-376
chr3-141402981-INS-6644
chr3-141403557-INS-6491
chr3-141422580-INS-96
chr3-141459638-INS-2974
chr3-141498027-INS-288
chr3-141557376-INS-971
chr3-141599010-INS-129

chr3-141613537-INS-2318
chr3-141619668-INS-60
chr3-141619713-INS-210
chr3-141768797-INS-1284
chr3-141805625-INS-3571
chr3-141880032-INS-68
chr3-141896799-INS-349
chr3-141897391-INS-4022
chr3-141897478-INS-20492
chr3-141919161-INS-60
chr3-141946998-INS-82
chr3-141967276-INS-6433
chr3-141991084-INS-97
chr3-142000621-INS-769
chr3-142000721-INS-60
chr3-142077938-INS-58
chr3-142077939-INS-621
chr3-142078044-INS-2424
chr3-142085371-INS-1178
chr3-142109612-INS-1846
chr3-142115542-INS-130
chr3-142118444-INS-474
chr3-142174768-INS-2649
chr3-142227901-INS-246
chr3-142228028-INS-72
chr3-142230464-INS-70
chr3-142268592-INS-78
chr3-142275589-INS-460
chr3-142295884-INS-64431
chr3-142299083-INS-67
chr3-142307427-INS-31458
chr3-142316044-INS-204
chr3-142316558-INS-6016
chr3-142432927-INS-112
chr3-142448732-INS-1199
chr3-142475311-INS-3000
chr3-142482108-INS-209
chr3-142495601-INS-4939
chr3-142526041-INS-51
chr3-142558677-INS-60
chr3-142586723-INS-52
chr3-142619676-INS-100
chr3-142619720-INS-615
chr3-142636324-INS-4406
chr3-142643832-INS-56
chr3-142680583-INS-6196
chr3-142834128-INS-116
chr3-142863297-INS-75
chr3-142863349-INS-162
chr3-142863767-INS-183
chr3-142885679-INS-52
chr3-142893361-INS-1834
chr3-142894605-INS-1396
chr3-142953842-INS-228
chr3-142953888-INS-112
chr3-143006288-INS-58
chr3-143015321-INS-180
chr3-143016845-INS-99
chr3-143016881-INS-561
chr3-143029763-INS-85
chr3-143074770-INS-72
chr3-143138077-INS-80
chr3-143146906-INS-146
chr3-143147217-INS-51
chr3-143147369-INS-51
chr3-143274003-INS-64
chr3-143350408-INS-188
chr3-143360152-INS-52
chr3-143367370-INS-74
chr3-143405811-INS-1668
chr3-143421080-INS-50
chr3-143467579-INS-66
chr3-143489721-INS-70
chr3-143507293-INS-7112
chr3-143577104-INS-165
chr3-143577141-INS-369
chr3-143577159-INS-60

chr3-143580317-INS-83
chr3-143633649-INS-659
chr3-143667980-INS-55
chr3-143681895-INS-492
chr3-143711383-INS-6201
chr3-143807414-INS-391
chr3-143808787-INS-245
chr3-143883073-INS-66
chr3-144108837-INS-288
chr3-144118173-INS-83
chr3-144136486-INS-4566
chr3-144196137-INS-154
chr3-144196386-INS-1248
chr3-144196420-INS-208
chr3-144196961-INS-110
chr3-144218126-INS-1074
chr3-144218553-INS-58
chr3-144232678-INS-58
chr3-144237842-INS-215
chr3-144252908-INS-836
chr3-144268033-INS-54
chr3-144289091-INS-69
chr3-144304905-INS-731
chr3-144317150-INS-212
chr3-144367051-INS-402
chr3-144390146-INS-60
chr3-144449282-INS-440
chr3-144451085-INS-990
chr3-144451772-INS-1353
chr3-144481503-INS-1338
chr3-144597537-INS-5701
chr3-144628511-INS-50
chr3-144633339-INS-83
chr3-144649663-INS-1950
chr3-144689064-INS-92
chr3-144689342-INS-80
chr3-144716983-INS-182
chr3-144717823-INS-112
chr3-144752045-INS-214
chr3-144843434-INS-153
chr3-144844049-INS-421
chr3-144857318-INS-6429
chr3-144868937-INS-250
chr3-144869311-INS-1508
chr3-144991306-INS-210
chr3-145126949-INS-96
chr3-145161514-INS-303
chr3-145186214-INS-56
chr3-145220440-INS-296
chr3-145243838-INS-276
chr3-145249550-INS-126
chr3-145255070-INS-156
chr3-145272019-INS-53
chr3-145277449-INS-222
chr3-145277449-INS-74
chr3-145279160-INS-5244
chr3-145292816-INS-2153
chr3-145361331-INS-134
chr3-145362394-INS-225
chr3-145376281-INS-140
chr3-145384772-INS-1992
chr3-145396678-INS-60
chr3-145398854-INS-259
chr3-145398854-INS-63
chr3-145408855-INS-4425
chr3-145428182-INS-50
chr3-145455833-INS-62
chr3-145484626-INS-56
chr3-145505068-INS-1066
chr3-145505240-INS-98
chr3-145537268-INS-1898
chr3-145539216-INS-170
chr3-145547449-INS-112
chr3-145549961-INS-244
chr3-145579533-INS-280
chr3-145604492-INS-1338
chr3-145618783-INS-50

chr3-145650882-INS-7276
chr3-145652905-INS-191
chr3-145665087-INS-280
chr3-145688709-INS-1887
chr3-145691132-INS-72
chr3-145697201-INS-847
chr3-145712179-INS-56
chr3-145735696-INS-60
chr3-145743241-INS-3397
chr3-145746424-INS-136
chr3-145747768-INS-381
chr3-145749518-INS-60
chr3-145778182-INS-143
chr3-145785771-INS-985
chr3-145785856-INS-96
chr3-145800330-INS-80
chr3-145807243-INS-1807
chr3-145807381-INS-843
chr3-145807886-INS-182
chr3-145820892-INS-91
chr3-145829806-INS-994
chr3-145844764-INS-50
chr3-145868056-INS-225
chr3-145886290-INS-100
chr3-145899663-INS-623
chr3-145904410-INS-1177
chr3-145912751-INS-316
chr3-145912878-INS-76
chr3-145950962-INS-62
chr3-145998981-INS-3715
chr3-145999638-INS-214
chr3-146031646-INS-563
chr3-146034828-INS-2305
chr3-146039571-INS-124
chr3-146040834-INS-98
chr3-146074880-INS-520
chr3-146080798-INS-3101
chr3-146100345-INS-183
chr3-146100995-INS-51
chr3-146147541-INS-272
chr3-146147656-INS-84
chr3-146160948-INS-7073
chr3-146236467-INS-638
chr3-146266177-INS-362
chr3-146277409-INS-55
chr3-146277496-INS-1844
chr3-146286672-INS-152
chr3-146294586-INS-2929
chr3-146300517-INS-446
chr3-146307411-INS-132
chr3-146329617-INS-252
chr3-146363133-INS-1444
chr3-146373050-INS-106
chr3-146373250-INS-485
chr3-146402658-INS-6399
chr3-146410379-INS-497
chr3-146446139-INS-187
chr3-146565766-INS-92
chr3-146573433-INS-957
chr3-146601380-INS-6466
chr3-146614075-INS-1455
chr3-146616563-INS-114
chr3-146632175-INS-60
chr3-146647805-INS-204
chr3-146647916-INS-63
chr3-146654329-INS-115
chr3-146728720-INS-164
chr3-146771000-INS-178
chr3-146775375-INS-50
chr3-146775500-INS-51
chr3-146778433-INS-6384
chr3-146828916-INS-50
chr3-146834573-INS-78
chr3-146893143-INS-168
chr3-146911953-INS-1989
chr3-146930901-INS-212
chr3-146931153-INS-424

chr3-146981017-INS-4813
chr3-147027248-INS-500
chr3-147040733-INS-86
chr3-147080538-INS-468
chr3-147080761-INS-520
chr3-147081118-INS-341
chr3-147085247-INS-3429
chr3-147111893-INS-124
chr3-147120205-INS-93
chr3-147139895-INS-50
chr3-147149585-INS-52
chr3-147185917-INS-2792
chr3-147215597-INS-6211
chr3-147224805-INS-72
chr3-147241492-INS-5278
chr3-147253931-INS-51
chr3-147288029-INS-6419
chr3-147292674-INS-80
chr3-147319699-INS-150
chr3-147353493-INS-205
chr3-147369223-INS-1888
chr3-147387999-INS-52
chr3-147424752-INS-6186
chr3-147445648-INS-459
chr3-147483568-INS-2083
chr3-147487958-INS-50
chr3-147513883-INS-694
chr3-147515232-INS-149
chr3-147515416-INS-1810
chr3-147516048-INS-588
chr3-147556568-INS-6292
chr3-147561812-INS-258
chr3-147561812-INS-90
chr3-147564804-INS-54
chr3-147610872-INS-255
chr3-147652273-INS-6363
chr3-147733064-INS-1562
chr3-147769377-INS-126
chr3-147839429-INS-130
chr3-147848524-INS-168
chr3-147861721-INS-76
chr3-147868567-INS-64
chr3-147912797-INS-50
chr3-147934827-INS-74
chr3-147935681-INS-135
chr3-147935711-INS-64
chr3-148017771-INS-96
chr3-148052928-INS-98
chr3-148056365-INS-192
chr3-148061347-INS-4432
chr3-148144278-INS-6172
chr3-148168352-INS-104
chr3-148169443-INS-56
chr3-148190316-INS-71
chr3-148218099-INS-60
chr3-148283774-INS-62
chr3-148299046-INS-1070
chr3-148328715-INS-65
chr3-148328805-INS-190
chr3-148332901-INS-6467
chr3-148345411-INS-212
chr3-148345782-INS-530
chr3-148357214-INS-1293
chr3-148391430-INS-159
chr3-148391631-INS-177
chr3-148393945-INS-50
chr3-148403363-INS-51
chr3-148404627-INS-221
chr3-148474146-INS-50
chr3-148494977-INS-131
chr3-148507039-INS-92
chr3-148541263-INS-50
chr3-148620670-INS-54
chr3-148633159-INS-245
chr3-148735707-INS-1289
chr3-148738987-INS-146
chr3-148745775-INS-262

chr3-148745775-INS-89
chr3-148763633-INS-147
chr3-148769133-INS-54
chr3-148775094-INS-53
chr3-148778939-INS-50
chr3-148792897-INS-740
chr3-148841553-INS-116
chr3-148843610-INS-199
chr3-148853013-INS-231
chr3-148855779-INS-480
chr3-148878937-INS-64
chr3-148880712-INS-200
chr3-148881719-INS-76
chr3-148909921-INS-54
chr3-148912119-INS-88
chr3-148912525-INS-60
chr3-148933954-INS-65
chr3-148934462-INS-54
chr3-148956396-INS-50
chr3-148985907-INS-156
chr3-149033617-INS-63
chr3-149078258-INS-116
chr3-149078370-INS-280
chr3-149104048-INS-515
chr3-149133772-INS-126
chr3-149151029-INS-50
chr3-149166048-INS-64
chr3-149244972-INS-70
chr3-149284201-INS-130
chr3-149299804-INS-65
chr3-149306906-INS-78
chr3-149311225-INS-120
chr3-149323519-INS-54
chr3-149374755-INS-2598
chr3-149405515-INS-70
chr3-149408689-INS-85
chr3-149420484-INS-96
chr3-149423882-INS-6261
chr3-149446095-INS-60
chr3-149450488-INS-88
chr3-149451040-INS-81
chr3-149451132-INS-84
chr3-149451768-INS-155
chr3-149452595-INS-134
chr3-149462332-INS-1729
chr3-149488043-INS-52
chr3-149506984-INS-3326
chr3-149514669-INS-204
chr3-149536132-INS-456
chr3-149556806-INS-1878
chr3-149557456-INS-220
chr3-149618507-INS-56
chr3-149663980-INS-395
chr3-149726378-INS-184
chr3-149773398-INS-80
chr3-149800717-INS-61
chr3-149800872-INS-162
chr3-149818807-INS-425
chr3-149819554-INS-6523
chr3-149823466-INS-130
chr3-149837427-INS-2500
chr3-149844089-INS-888
chr3-149846992-INS-64
chr3-149847037-INS-129
chr3-149874390-INS-497
chr3-149901423-INS-84
chr3-149931309-INS-51
chr3-149999152-INS-142
chr3-150028672-INS-6401
chr3-150063027-INS-1889
chr3-150065893-INS-64
chr3-150069216-INS-123
chr3-150080331-INS-6712
chr3-150120753-INS-311
chr3-150124541-INS-337
chr3-150128677-INS-88
chr3-150137276-INS-3143

chr3-150144489-INS-1293
chr3-150155309-INS-3308
chr3-150159130-INS-360
chr3-150327847-INS-82
chr3-150328223-INS-61
chr3-150328231-INS-2934
chr3-150336405-INS-106
chr3-150336405-INS-275
chr3-150352735-INS-218
chr3-150386287-INS-68
chr3-150398752-INS-2791
chr3-150411610-INS-2255
chr3-150422755-INS-67
chr3-150434069-INS-1957
chr3-150447347-INS-56
chr3-150454995-INS-52
chr3-150469643-INS-88
chr3-150469849-INS-78
chr3-150486629-INS-58
chr3-150503341-INS-95
chr3-150515183-INS-62
chr3-150573924-INS-330
chr3-150583500-INS-73
chr3-150607957-INS-51
chr3-150613101-INS-425
chr3-150619925-INS-2946
chr3-150664810-INS-50
chr3-150683507-INS-1781
chr3-150683885-INS-193
chr3-150710452-INS-198
chr3-150718715-INS-104
chr3-150750401-INS-4108
chr3-150775417-INS-1661
chr3-150785530-INS-2849
chr3-150795255-INS-62
chr3-150864238-INS-61
chr3-150869245-INS-219
chr3-150870461-INS-52
chr3-150895064-INS-54
chr3-150901694-INS-228
chr3-150915305-INS-54
chr3-150927224-INS-76
chr3-150947853-INS-50
chr3-150975041-INS-1871
chr3-150985180-INS-146
chr3-151010853-INS-672
chr3-151052254-INS-92
chr3-151061983-INS-55
chr3-151065096-INS-1443
chr3-151080232-INS-116
chr3-151082986-INS-363
chr3-151095419-INS-7130
chr3-151108674-INS-578
chr3-151121115-INS-4100
chr3-151132591-INS-64
chr3-151135724-INS-146
chr3-151190674-INS-55
chr3-151215194-INS-55
chr3-151253658-INS-1493
chr3-151257573-INS-3067
chr3-151258965-INS-6564
chr3-151286521-INS-53
chr3-151288582-INS-57
chr3-151318240-INS-96
chr3-151355067-INS-206
chr3-151439358-INS-1385
chr3-151444600-INS-4800
chr3-151447770-INS-6121
chr3-151469089-INS-456
chr3-151469194-INS-1810
chr3-151469376-INS-541
chr3-151470036-INS-11999
chr3-151491968-INS-91
chr3-151495588-INS-5540
chr3-151555218-INS-1232
chr3-151588380-INS-72
chr3-151604024-INS-1389

chr3-151607690-INS-69
chr3-151623052-INS-755
chr3-151625587-INS-923
chr3-151627767-INS-1067
chr3-151643131-INS-6729
chr3-151656177-INS-100
chr3-151672909-INS-70
chr3-151673766-INS-126
chr3-151678363-INS-60
chr3-151685962-INS-6392
chr3-151693866-INS-7261
chr3-151702526-INS-453
chr3-151732185-INS-70
chr3-151733342-INS-232
chr3-151740637-INS-6476
chr3-151761710-INS-50
chr3-151769457-INS-971
chr3-151793655-INS-270
chr3-151793682-INS-699
chr3-151821008-INS-202
chr3-151830999-INS-263
chr3-151831071-INS-105
chr3-151832356-INS-183
chr3-151837185-INS-322
chr3-151842993-INS-68
chr3-151845208-INS-126
chr3-151851817-INS-93
chr3-151851971-INS-537
chr3-151852004-INS-109
chr3-151866776-INS-57
chr3-151878990-INS-72
chr3-151879689-INS-74
chr3-151902660-INS-73
chr3-151935249-INS-210
chr3-151941407-INS-68
chr3-151950237-INS-185
chr3-152001182-INS-150
chr3-152020272-INS-586
chr3-152022184-INS-151
chr3-152022655-INS-126
chr3-152022755-INS-1480
chr3-152046974-INS-165
chr3-152046988-INS-54
chr3-152047560-INS-112
chr3-152060851-INS-225
chr3-152075609-INS-320
chr3-152107476-INS-64
chr3-152124339-INS-50
chr3-152184641-INS-212
chr3-152191921-INS-96
chr3-152217285-INS-7202
chr3-152239764-INS-423
chr3-152267580-INS-489
chr3-152269484-INS-99
chr3-152293437-INS-214
chr3-152295273-INS-1235
chr3-152297358-INS-1256
chr3-152304737-INS-103
chr3-152311075-INS-56
chr3-152311369-INS-1037
chr3-152313507-INS-86
chr3-152320363-INS-1408
chr3-152328896-INS-2043
chr3-152334719-INS-72
chr3-152358531-INS-82
chr3-152359879-INS-89
chr3-152387592-INS-102
chr3-152407296-INS-106
chr3-152418081-INS-4308
chr3-152436475-INS-1145
chr3-152446036-INS-79
chr3-152447088-INS-181
chr3-152449731-INS-104
chr3-152454096-INS-175
chr3-152461290-INS-157
chr3-152531739-INS-94
chr3-152532539-INS-88

chr3-152532545-INS-182
chr3-152556933-INS-99
chr3-152654754-INS-1556
chr3-152709219-INS-91
chr3-152712406-INS-63
chr3-152713570-INS-1550
chr3-152726444-INS-172
chr3-152738598-INS-1950
chr3-152748429-INS-63
chr3-152776277-INS-74
chr3-152776314-INS-178
chr3-152776610-INS-54
chr3-152784483-INS-170
chr3-152813836-INS-208
chr3-152817080-INS-56
chr3-152832807-INS-55
chr3-152833150-INS-208
chr3-152860769-INS-58
chr3-152866472-INS-76
chr3-152944772-INS-205
chr3-152963925-INS-66
chr3-152965886-INS-6218
chr3-152977559-INS-499
chr3-152987234-INS-349
chr3-153020331-INS-68
chr3-153021142-INS-325
chr3-153035300-INS-268
chr3-153048548-INS-66
chr3-153048567-INS-237
chr3-153074665-INS-68
chr3-153087955-INS-160
chr3-153119738-INS-79
chr3-153134194-INS-504
chr3-153158256-INS-111
chr3-153192510-INS-50
chr3-153241751-INS-2536
chr3-153253740-INS-180
chr3-153278306-INS-54
chr3-153296290-INS-52
chr3-153302531-INS-50
chr3-153313933-INS-100
chr3-153314236-INS-54
chr3-153320570-INS-3105
chr3-153320798-INS-196
chr3-153338712-INS-66
chr3-153338976-INS-6514
chr3-153342358-INS-53
chr3-153348476-INS-6914
chr3-153364165-INS-2105
chr3-153380952-INS-96
chr3-153409190-INS-209
chr3-153409190-INS-56
chr3-153415929-INS-166
chr3-153433143-INS-93
chr3-153433224-INS-95
chr3-153461198-INS-554
chr3-153472441-INS-5541
chr3-153475879-INS-95
chr3-153500072-INS-101
chr3-153512883-INS-215
chr3-153518266-INS-6693
chr3-153520736-INS-487
chr3-153524676-INS-56
chr3-153543894-INS-1218
chr3-153557341-INS-697
chr3-153557476-INS-164
chr3-153605441-INS-352
chr3-153605515-INS-120
chr3-153608048-INS-203
chr3-153627054-INS-204
chr3-153627507-INS-198
chr3-153627522-INS-68
chr3-153627718-INS-152
chr3-153632343-INS-90
chr3-153641112-INS-1863
chr3-153645673-INS-226
chr3-153669237-INS-94

chr3-153672245-INS-56
chr3-153683727-INS-2233
chr3-153694916-INS-1161
chr3-153708036-INS-194
chr3-153717469-INS-230
chr3-153718251-INS-64
chr3-153723932-INS-1997
chr3-153765061-INS-161
chr3-153767781-INS-59
chr3-153780025-INS-128
chr3-153784797-INS-1065
chr3-153792127-INS-888
chr3-153827328-INS-7155
chr3-153840183-INS-54
chr3-153881170-INS-113
chr3-153881272-INS-235
chr3-153884958-INS-57
chr3-153884962-INS-255
chr3-153914135-INS-1568
chr3-153923932-INS-493
chr3-153924432-INS-4204
chr3-153925256-INS-58
chr3-153933215-INS-60
chr3-153959962-INS-320
chr3-153961050-INS-282
chr3-154055107-INS-120
chr3-154075499-INS-402
chr3-154078920-INS-105
chr3-154123948-INS-246
chr3-154124485-INS-733
chr3-154135021-INS-587
chr3-154160755-INS-96
chr3-154176324-INS-1597
chr3-154176504-INS-90
chr3-154178582-INS-129
chr3-154230454-INS-55
chr3-154241433-INS-93
chr3-154242517-INS-82
chr3-154246817-INS-78
chr3-154249766-INS-437
chr3-154266748-INS-5568
chr3-154277550-INS-54
chr3-154281587-INS-214
chr3-154283843-INS-2220
chr3-154283843-INS-296
chr3-154286376-INS-77
chr3-154286846-INS-50
chr3-154292885-INS-312
chr3-154298937-INS-7087
chr3-154314708-INS-432
chr3-154315838-INS-82
chr3-154322543-INS-318
chr3-154323590-INS-258
chr3-154333524-INS-1504
chr3-154333964-INS-268
chr3-154334894-INS-400
chr3-154356334-INS-54
chr3-154364231-INS-3108
chr3-154371660-INS-701
chr3-154379927-INS-901
chr3-154385635-INS-4836
chr3-154392695-INS-60
chr3-154433632-INS-54
chr3-154439615-INS-1166
chr3-154442815-INS-336
chr3-154459180-INS-95
chr3-154483355-INS-70
chr3-154604465-INS-912
chr3-154608452-INS-6402
chr3-154616182-INS-50
chr3-154630736-INS-145
chr3-154630736-INS-333
chr3-154642145-INS-182
chr3-154648510-INS-367
chr3-154663039-INS-205
chr3-154680847-INS-7194
chr3-154716442-INS-50

chr3-154739104-INS-194
chr3-154739113-INS-79
chr3-154774418-INS-172
chr3-154851590-INS-244
chr3-154922490-INS-5490
chr3-154960680-INS-679
chr3-154961008-INS-1079
chr3-154992408-INS-4998
chr3-155050242-INS-54
chr3-155083157-INS-3416
chr3-155088953-INS-68
chr3-155091912-INS-499
chr3-155094742-INS-50
chr3-155124367-INS-140
chr3-155156639-INS-208
chr3-155156684-INS-494
chr3-155156718-INS-78
chr3-155192199-INS-50
chr3-155236477-INS-13694
chr3-155256126-INS-120
chr3-155256226-INS-216
chr3-155421568-INS-58
chr3-155422126-INS-88
chr3-155449343-INS-968
chr3-155452225-INS-173
chr3-155522313-INS-80
chr3-155537454-INS-3242
chr3-155549995-INS-80
chr3-155550242-INS-114
chr3-155595881-INS-6399
chr3-155638161-INS-93
chr3-155689712-INS-6314
chr3-155693758-INS-4064
chr3-155703345-INS-302
chr3-155704372-INS-117
chr3-155707498-INS-1898
chr3-155719679-INS-6729
chr3-155789353-INS-2790
chr3-155841044-INS-175
chr3-155901266-INS-204
chr3-155925183-INS-52
chr3-155959266-INS-257
chr3-155982653-INS-50
chr3-155990387-INS-201
chr3-156111254-INS-2705
chr3-156119597-INS-1784
chr3-156166861-INS-2799
chr3-156182554-INS-166
chr3-156184048-INS-2806
chr3-156306421-INS-319
chr3-156318996-INS-62
chr3-156348291-INS-6900
chr3-156365635-INS-76
chr3-156432372-INS-1583
chr3-156440382-INS-1068
chr3-156472900-INS-166
chr3-156503722-INS-57
chr3-156503750-INS-2600
chr3-156556770-INS-6670
chr3-156580788-INS-135
chr3-156599671-INS-368
chr3-156599711-INS-52
chr3-156655477-INS-6482
chr3-156661060-INS-50
chr3-156691820-INS-1178
chr3-156727132-INS-66
chr3-156737867-INS-70
chr3-156758566-INS-1485
chr3-156795975-INS-255
chr3-156932938-INS-60
chr3-156956655-INS-139
chr3-157066969-INS-5624
chr3-157067752-INS-54
chr3-157085061-INS-106
chr3-157085525-INS-64
chr3-157087395-INS-414
chr3-157146661-INS-637

chr3-157148054-INS-144
chr3-157184233-INS-585
chr3-157192131-INS-153
chr3-157226880-INS-82
chr3-157228146-INS-78
chr3-157229713-INS-6245
chr3-157260983-INS-866
chr3-157274438-INS-50
chr3-157302005-INS-2448
chr3-157309895-INS-133
chr3-157315932-INS-52
chr3-157334748-INS-79
chr3-157339791-INS-202
chr3-157361566-INS-55
chr3-157367097-INS-2776
chr3-157373964-INS-88
chr3-157397739-INS-57
chr3-157427697-INS-1895
chr3-157447977-INS-1360
chr3-157455129-INS-103
chr3-157457741-INS-206
chr3-157471524-INS-6313
chr3-157476971-INS-7456
chr3-157478752-INS-405
chr3-157498104-INS-76
chr3-157518357-INS-81
chr3-157533729-INS-2236
chr3-157538142-INS-1312
chr3-157540743-INS-2640
chr3-157591797-INS-198
chr3-157595613-INS-52
chr3-157596390-INS-61
chr3-157687574-INS-223
chr3-157891679-INS-54
chr3-157894010-INS-50
chr3-157916075-INS-51
chr3-157952727-INS-60
chr3-157969642-INS-120
chr3-158034752-INS-54
chr3-158083394-INS-219
chr3-158158113-INS-122
chr3-158210457-INS-61
chr3-158248632-INS-203
chr3-158272059-INS-6493
chr3-158285383-INS-92
chr3-158321230-INS-4690
chr3-158324187-INS-485
chr3-158347602-INS-64
chr3-158391189-INS-74
chr3-158395980-INS-95
chr3-158443507-INS-832
chr3-158514672-INS-573
chr3-158518033-INS-72
chr3-158545251-INS-390
chr3-158550834-INS-552
chr3-158683932-INS-328
chr3-158788242-INS-66
chr3-158815922-INS-706
chr3-158905187-INS-80
chr3-158946551-INS-2339
chr3-158978583-INS-86
chr3-159041919-INS-1185
chr3-159048484-INS-6303
chr3-159076573-INS-2001
chr3-159082310-INS-7082
chr3-159104359-INS-69
chr3-159105892-INS-6523
chr3-159130536-INS-3047
chr3-159140641-INS-60
chr3-159162803-INS-504
chr3-159163092-INS-161
chr3-159167090-INS-556
chr3-159219135-INS-81
chr3-159230716-INS-7085
chr3-159268985-INS-1731
chr3-159351220-INS-397
chr3-159398967-INS-324

chr3-159429104-INS-16496
chr3-159441882-INS-421
chr3-159472789-INS-412
chr3-159483771-INS-1899
chr3-159516271-INS-498
chr3-159528958-INS-50
chr3-159591935-INS-52
chr3-159637234-INS-50
chr4-3069274-INS-82
chr4-3156709-INS-2970
chr4-3170206-INS-60
chr4-3175377-INS-1228
chr4-3180977-INS-66
chr4-3219277-INS-58
chr4-3219440-INS-1360
chr4-3222290-INS-568
chr4-3223366-INS-2073
chr4-3227361-INS-136
chr4-3231256-INS-188
chr4-3231740-INS-57
chr4-3232851-INS-68
chr4-3234855-INS-342
chr4-3236901-INS-52
chr4-3244385-INS-445
chr4-3342542-INS-72
chr4-3403901-INS-7514
chr4-3412204-INS-133
chr4-3416610-INS-191
chr4-3416610-INS-75
chr4-3422995-INS-135
chr4-3426109-INS-5366
chr4-3430020-INS-414
chr4-3430563-INS-415
chr4-3453883-INS-186
chr4-3471105-INS-114
chr4-3512485-INS-66
chr4-3516280-INS-50
chr4-3606985-INS-71
chr4-3661529-INS-712
chr4-3669189-INS-238
chr4-3672560-INS-183
chr4-3700682-INS-50
chr4-3706162-INS-3248
chr4-3717723-INS-70
chr4-3732870-INS-6327
chr4-3764956-INS-250
chr4-3812369-INS-56
chr4-3846781-INS-66
chr4-3864612-INS-3102
chr4-3894541-INS-212
chr4-3907823-INS-58
chr4-3914578-INS-62
chr4-3926877-INS-75
chr4-3927083-INS-75
chr4-3967135-INS-65
chr4-3984293-INS-4406
chr4-4096839-INS-202
chr4-4140086-INS-56
chr4-4176959-INS-180
chr4-4177006-INS-62
chr4-4179303-INS-2535
chr4-4182440-INS-80
chr4-4205188-INS-464
chr4-4247396-INS-56
chr4-4334879-INS-242
chr4-4387993-INS-1336
chr4-4403037-INS-3170
chr4-4420275-INS-177
chr4-4425870-INS-50
chr4-4432708-INS-6760
chr4-4433506-INS-72
chr4-4447079-INS-87
chr4-4454533-INS-591
chr4-4459631-INS-6475
chr4-4481480-INS-2280
chr4-4556975-INS-167
chr4-4558817-INS-6175

chr4-4559383-INS-173
chr4-4562299-INS-59
chr4-4584936-INS-355
chr4-4601460-INS-6182
chr4-4632728-INS-59
chr4-4679013-INS-7148
chr4-4691821-INS-330
chr4-4723265-INS-138
chr4-4725216-INS-76
chr4-4730295-INS-401
chr4-4734028-INS-65
chr4-4768691-INS-6722
chr4-4824198-INS-58
chr4-4879092-INS-55
chr4-4881465-INS-166
chr4-4881592-INS-227
chr4-4930041-INS-141
chr4-4942965-INS-92
chr4-4966305-INS-5113
chr4-4999225-INS-76
chr4-5074684-INS-236
chr4-5075973-INS-420
chr4-5138292-INS-125
chr4-5211649-INS-85
chr4-5317389-INS-549
chr4-5345185-INS-69
chr4-5386073-INS-199
chr4-5409567-INS-282
chr4-5411415-INS-150
chr4-5421590-INS-465
chr4-5446391-INS-5798
chr4-5479891-INS-2498
chr4-5485015-INS-67
chr4-5541367-INS-6434
chr4-5554000-INS-4659
chr4-5567399-INS-138
chr4-5583770-INS-701
chr4-5621553-INS-6068
chr4-5626448-INS-70
chr4-5626611-INS-607
chr4-5638851-INS-51
chr4-5697455-INS-92
chr4-5704931-INS-105
chr4-5748660-INS-910
chr4-5784003-INS-197
chr4-5790491-INS-251
chr4-5790642-INS-1060
chr4-5817778-INS-290
chr4-5829183-INS-1891
chr4-5879443-INS-328
chr4-5900478-INS-349
chr4-5902870-INS-70
chr4-5923662-INS-2031
chr4-5927769-INS-422
chr4-5929758-INS-181
chr4-5931379-INS-112
chr4-5931579-INS-191
chr4-5932769-INS-118
chr4-5963796-INS-85
chr4-5966874-INS-288
chr4-5978669-INS-164
chr4-6051620-INS-658
chr4-6077850-INS-192
chr4-6101939-INS-148
chr4-6163175-INS-78
chr4-6170114-INS-56
chr4-6197677-INS-60
chr4-6259041-INS-87
chr4-6314847-INS-1350
chr4-6335917-INS-7237
chr4-6352894-INS-143
chr4-6377575-INS-114
chr4-6399406-INS-50
chr4-6431904-INS-935
chr4-6433584-INS-2497
chr4-6439614-INS-68
chr4-6470283-INS-52

chr4-6472293-INS-199
chr4-6498220-INS-424
chr4-6505678-INS-104
chr4-6507568-INS-79
chr4-6507641-INS-162
chr4-6510681-INS-104
chr4-6515483-INS-308
chr4-6520316-INS-8409
chr4-6533483-INS-116
chr4-6533731-INS-211
chr4-6539591-INS-1142
chr4-6556372-INS-2761
chr4-6556687-INS-70
chr4-6560891-INS-77
chr4-6566184-INS-68
chr4-6608869-INS-2786
chr4-6618773-INS-2213
chr4-6629566-INS-52
chr4-6629608-INS-172
chr4-6663551-INS-140
chr4-6683486-INS-92
chr4-6683835-INS-58
chr4-6690922-INS-589
chr4-6745556-INS-335
chr4-6757782-INS-87
chr4-6793939-INS-50
chr4-6825292-INS-209
chr4-6837689-INS-60
chr4-6880068-INS-52
chr4-6937670-INS-1907
chr4-7010763-INS-168
chr4-7024306-INS-330
chr4-7024317-INS-164
chr4-7024510-INS-55
chr4-7024875-INS-205
chr4-7024875-INS-439
chr4-7024875-INS-61
chr4-7033603-INS-120
chr4-7058444-INS-6436
chr4-7113282-INS-56
chr4-7114030-INS-75
chr4-7138892-INS-282
chr4-7246049-INS-2772
chr4-7315924-INS-327
chr4-7327124-INS-356
chr4-7330248-INS-466
chr4-7380463-INS-6717
chr4-7381590-INS-58
chr4-7403793-INS-54
chr4-7411492-INS-6481
chr4-7447472-INS-109
chr4-7447569-INS-52
chr4-7507970-INS-1894
chr4-7624590-INS-416
chr4-7628545-INS-408
chr4-7684888-INS-6475
chr4-7748445-INS-67
chr4-7752256-INS-50
chr4-7778910-INS-1337
chr4-7782596-INS-6239
chr4-7783090-INS-335
chr4-7784244-INS-2828
chr4-7787855-INS-6264
chr4-7798127-INS-56
chr4-7812357-INS-52
chr4-7830311-INS-51
chr4-7862739-INS-84
chr4-7879282-INS-204
chr4-7912496-INS-6623
chr4-7914599-INS-1723
chr4-7947464-INS-3726
chr4-7957138-INS-168
chr4-7978975-INS-6487
chr4-7991056-INS-395
chr4-8018974-INS-140
chr4-8027890-INS-5628
chr4-8095828-INS-6254

chr4-8102033-INS-82
chr4-8115721-INS-451
chr4-8152162-INS-110
chr4-8152162-INS-256
chr4-8159825-INS-284
chr4-8174521-INS-7194
chr4-8212841-INS-58
chr4-8216193-INS-7093
chr4-8218435-INS-359
chr4-8230135-INS-266
chr4-8236321-INS-95
chr4-8244189-INS-192
chr4-8249903-INS-60
chr4-8262053-INS-1002
chr4-8263125-INS-322
chr4-8273840-INS-52
chr4-8274038-INS-105
chr4-8279290-INS-132
chr4-8283612-INS-252
chr4-8283896-INS-587
chr4-8308554-INS-3497
chr4-8330854-INS-1185
chr4-8333684-INS-81
chr4-8334922-INS-60
chr4-8338653-INS-84
chr4-8360286-INS-81
chr4-8383342-INS-88
chr4-8392261-INS-408
chr4-8431734-INS-276
chr4-8447731-INS-105
chr4-8447988-INS-275
chr4-8498039-INS-50
chr4-8509722-INS-50
chr4-8512226-INS-192
chr4-8629735-INS-168
chr4-8630206-INS-50
chr4-8630550-INS-718
chr4-8640826-INS-180
chr4-8643475-INS-450
chr4-8652145-INS-132
chr4-8652423-INS-50
chr4-8663907-INS-64
chr4-8680464-INS-86
chr4-8731165-INS-1000
chr4-8740506-INS-316
chr4-8786692-INS-50
chr4-8792090-INS-50
chr4-8817383-INS-546
chr4-8829202-INS-134
chr4-8842615-INS-65
chr4-8858960-INS-439
chr4-8895600-INS-302
chr4-8901348-INS-81
chr4-8903376-INS-267
chr4-8914951-INS-418
chr4-8917771-INS-180
chr4-8919228-INS-201
chr4-8931464-INS-50
chr4-8947104-INS-130
chr4-8960520-INS-60
chr4-8968373-INS-85
chr4-8978135-INS-50
chr4-8991593-INS-50
chr4-8997422-INS-1140
chr4-9004295-INS-7237
chr4-9014174-INS-129
chr4-9029389-INS-379
chr4-9031213-INS-136
chr4-9036384-INS-444
chr4-9063785-INS-65
chr4-9063980-INS-390
chr4-9084625-INS-148
chr4-9086587-INS-140
chr4-9123888-INS-112
chr4-9135144-INS-6270
chr4-9143541-INS-82
chr4-9146856-INS-60

chr4-9146882-INS-168
chr4-9207144-INS-50
chr4-9209852-INS-86
chr4-9249847-INS-68
chr4-9249870-INS-698
chr4-9299374-INS-236
chr4-9301426-INS-6415
chr4-9325915-INS-1307
chr4-9334663-INS-3048
chr4-9364314-INS-1373
chr4-9385886-INS-58
chr4-9386058-INS-60
chr4-9388775-INS-7310
chr4-9407584-INS-108
chr4-9442445-INS-51
chr4-9470016-INS-1506
chr4-9493857-INS-428
chr4-9516950-INS-52
chr4-9540644-INS-65
chr4-9573957-INS-98
chr4-9609207-INS-98
chr4-9620675-INS-51
chr4-9654427-INS-309
chr4-9663097-INS-166
chr4-9692759-INS-52
chr4-9703734-INS-63
chr4-9703808-INS-206
chr4-9749035-INS-157
chr4-9761381-INS-2892
chr4-9769606-INS-1198
chr4-9772079-INS-56
chr4-9784152-INS-261
chr4-9790373-INS-386
chr4-9814061-INS-1926
chr4-9827713-INS-84
chr4-9830747-INS-1153
chr4-9834700-INS-6197
chr4-9878642-INS-68
chr4-9912636-INS-2811
chr4-9952638-INS-105
chr4-9954354-INS-119
chr4-9954918-INS-56
chr4-9958013-INS-231
chr4-9969863-INS-6418
chr4-9989991-INS-198
chr4-10011726-INS-56
chr4-10020461-INS-1614
chr4-10030833-INS-94
chr4-10030861-INS-230
chr4-10032205-INS-104
chr4-10048789-INS-175
chr4-10126397-INS-2327
chr4-10126397-INS-939
chr4-10135316-INS-51
chr4-10137523-INS-2473
chr4-10150348-INS-191
chr4-10157688-INS-54
chr4-10157772-INS-111
chr4-10161629-INS-211
chr4-10164122-INS-406
chr4-10181684-INS-59
chr4-10210936-INS-410
chr4-10225114-INS-132
chr4-10226813-INS-417
chr4-10247123-INS-552
chr4-10249789-INS-1103
chr4-10273216-INS-68
chr4-10290923-INS-56
chr4-10292904-INS-1973
chr4-10298522-INS-50
chr4-10311378-INS-7103
chr4-10347054-INS-6509
chr4-10360684-INS-191
chr4-10361589-INS-96
chr4-10383897-INS-51
chr4-10466779-INS-238
chr4-10470332-INS-370

chr4-10517165-INS-88
chr4-10518200-INS-64
chr4-10532238-INS-148
chr4-10551718-INS-239
chr4-10551794-INS-1795
chr4-10591963-INS-6236
chr4-10593362-INS-478
chr4-10599380-INS-154
chr4-10621428-INS-261
chr4-10650696-INS-122
chr4-10656247-INS-170
chr4-10694100-INS-505
chr4-10694135-INS-91
chr4-10694610-INS-202
chr4-10694610-INS-95
chr4-10708889-INS-1749
chr4-10716918-INS-1699
chr4-10779813-INS-68
chr4-10842294-INS-199
chr4-10845572-INS-975
chr4-10889474-INS-95
chr4-10889871-INS-59
chr4-10915341-INS-52
chr4-10917298-INS-1140
chr4-10917894-INS-56
chr4-10939650-INS-126
chr4-10939696-INS-567
chr4-10950853-INS-4421
chr4-10970774-INS-162
chr4-10996306-INS-167
chr4-10996839-INS-113
chr4-10997730-INS-182
chr4-11024803-INS-206
chr4-11031734-INS-63
chr4-11035115-INS-218
chr4-11035644-INS-164
chr4-11044201-INS-175
chr4-11049107-INS-180
chr4-11053801-INS-202
chr4-11054876-INS-90
chr4-11059169-INS-182
chr4-11066291-INS-193
chr4-11127508-INS-127
chr4-11177976-INS-99
chr4-11217257-INS-2710
chr4-11222977-INS-231
chr4-11269393-INS-112
chr4-11287074-INS-175
chr4-11287303-INS-211
chr4-11288861-INS-266
chr4-11300447-INS-64
chr4-11436368-INS-222
chr4-11453629-INS-52
chr4-11453664-INS-284
chr4-11463600-INS-178
chr4-11466248-INS-111
chr4-11520909-INS-51
chr4-11535834-INS-250
chr4-11571207-INS-655
chr4-11571215-INS-157
chr4-11586721-INS-6470
chr4-11605481-INS-54
chr4-11605539-INS-180
chr4-11608981-INS-87
chr4-11617975-INS-222
chr4-11646982-INS-1535
chr4-11656712-INS-6509
chr4-11657912-INS-454
chr4-11676276-INS-223
chr4-11680328-INS-189
chr4-11695005-INS-1377
chr4-11695067-INS-237
chr4-11710472-INS-66
chr4-11717027-INS-204
chr4-11786450-INS-64
chr4-11802253-INS-75
chr4-11806029-INS-313

chr4-11849766-INS-50
chr4-11867868-INS-1528
chr4-11874342-INS-6258
chr4-11903884-INS-52
chr4-11906978-INS-52
chr4-11913585-INS-52
chr4-11915535-INS-901
chr4-11947080-INS-294
chr4-11947104-INS-74
chr4-11949770-INS-178
chr4-12029386-INS-879
chr4-12037350-INS-6495
chr4-12037470-INS-55
chr4-12037473-INS-205
chr4-12037473-INS-2447
chr4-12037473-INS-564
chr4-12037695-INS-85
chr4-12048373-INS-182
chr4-12050492-INS-351
chr4-12072100-INS-450
chr4-12113549-INS-10771
chr4-12121298-INS-70
chr4-12133153-INS-4434
chr4-12139554-INS-264
chr4-12158469-INS-202
chr4-12221010-INS-56
chr4-12363285-INS-55
chr4-12370418-INS-115
chr4-12387204-INS-192
chr4-12387404-INS-608
chr4-12389236-INS-2196
chr4-12409697-INS-2909
chr4-12414358-INS-6690
chr4-12428179-INS-1634
chr4-12501017-INS-6506
chr4-12517581-INS-59
chr4-12541570-INS-334
chr4-12592676-INS-226
chr4-12633547-INS-129
chr4-12706191-INS-258
chr4-12713229-INS-208
chr4-12723056-INS-944
chr4-12740583-INS-54
chr4-12740932-INS-120
chr4-12746851-INS-1635
chr4-12762042-INS-72
chr4-12809241-INS-77
chr4-12830209-INS-4797
chr4-12836810-INS-51
chr4-12863376-INS-6344
chr4-12875085-INS-74
chr4-12891531-INS-150
chr4-12894508-INS-17130
chr4-12911844-INS-60
chr4-12911884-INS-547
chr4-12945989-INS-96
chr4-12989505-INS-4919
chr4-13003361-INS-64
chr4-13003425-INS-334
chr4-13074585-INS-76
chr4-13095242-INS-59
chr4-13096144-INS-2412
chr4-13140676-INS-96
chr4-13175386-INS-1080
chr4-13307655-INS-609
chr4-13350225-INS-1343
chr4-13353074-INS-6169
chr4-13380973-INS-5543
chr4-13400920-INS-52
chr4-13417067-INS-513
chr4-13442944-INS-94
chr4-13454664-INS-852
chr4-13556720-INS-1951
chr4-13562118-INS-3353
chr4-13570602-INS-7116
chr4-13632605-INS-74
chr4-13636214-INS-331

chr4-13656921-INS-86
chr4-13656957-INS-288
chr4-13664222-INS-164
chr4-13664452-INS-309
chr4-13710289-INS-409
chr4-13760906-INS-169
chr4-13964388-INS-66
chr4-13965013-INS-156
chr4-13986182-INS-2501
chr4-14006512-INS-287
chr4-14033260-INS-1835
chr4-14068416-INS-1838
chr4-14077904-INS-2872
chr4-14137573-INS-75
chr4-14198853-INS-397
chr4-14269816-INS-68
chr4-14287794-INS-6876
chr4-14387635-INS-52
chr4-14394791-INS-3878
chr4-14438728-INS-994
chr4-14460266-INS-252
chr4-14505715-INS-6421
chr4-14533877-INS-392
chr4-14572488-INS-264
chr4-14597998-INS-62
chr4-14604166-INS-76
chr4-14610723-INS-133
chr4-14624392-INS-1316
chr4-14655145-INS-1290
chr4-14662850-INS-65
chr4-14695176-INS-494
chr4-14695176-INS-98
chr4-14711329-INS-601
chr4-14711873-INS-476
chr4-14736072-INS-341
chr4-14755194-INS-2278
chr4-14758877-INS-211
chr4-14820669-INS-50
chr4-14833520-INS-72
chr4-14843576-INS-2594
chr4-14843895-INS-7158
chr4-14844158-INS-72
chr4-14871661-INS-189
chr4-14879748-INS-1113
chr4-14887392-INS-207
chr4-14888059-INS-152
chr4-14920361-INS-99
chr4-14924209-INS-155
chr4-14977610-INS-50
chr4-14986376-INS-1184
chr4-15010126-INS-60
chr4-15056148-INS-7014
chr4-15088420-INS-2588
chr4-15092955-INS-10157
chr4-15098441-INS-2307
chr4-15122595-INS-400
chr4-15176393-INS-67
chr4-15222386-INS-1833
chr4-15262287-INS-370
chr4-15303014-INS-1492
chr4-15306855-INS-466
chr4-15316272-INS-156
chr4-15317176-INS-86
chr4-15348545-INS-6370
chr4-15353803-INS-199
chr4-15365470-INS-314
chr4-15365470-INS-85
chr4-15365918-INS-8888
chr4-15367359-INS-58
chr4-15369052-INS-1228
chr4-15369468-INS-1165
chr4-15419514-INS-6195
chr4-15422288-INS-1336
chr4-15481473-INS-9712
chr4-15500946-INS-56
chr4-15518930-INS-323
chr4-15519808-INS-793

chr4-15521933-INS-77
chr4-15523678-INS-6163
chr4-15529019-INS-191
chr4-15529019-INS-72
chr4-15573031-INS-64
chr4-15629746-INS-1356
chr4-15642000-INS-60
chr4-15670350-INS-66
chr4-15711130-INS-6573
chr4-15733937-INS-326
chr4-15754294-INS-1851
chr4-15767059-INS-51
chr4-15796076-INS-1468
chr4-15799449-INS-5184
chr4-15800605-INS-520
chr4-15819886-INS-53
chr4-15825799-INS-8752
chr4-15902653-INS-50
chr4-15903536-INS-301
chr4-15921179-INS-7090
chr4-15956543-INS-50
chr4-15992785-INS-72
chr4-15993611-INS-1552
chr4-16025654-INS-588
chr4-16026703-INS-2777
chr4-16040410-INS-382
chr4-16054112-INS-2857
chr4-16065973-INS-65
chr4-16120635-INS-6223
chr4-16177265-INS-199
chr4-16194262-INS-60
chr4-16203650-INS-151
chr4-16203927-INS-483
chr4-16252169-INS-2626
chr4-16263376-INS-50
chr4-16279673-INS-2223
chr4-16309982-INS-409
chr4-16337535-INS-110
chr4-16341690-INS-413
chr4-16394466-INS-3320
chr4-16394466-INS-9573
chr4-16451875-INS-6279
chr4-16467631-INS-696
chr4-16500978-INS-2047
chr4-16593871-INS-64
chr4-16605797-INS-214
chr4-16608520-INS-6261
chr4-16628693-INS-75
chr4-16629229-INS-118
chr4-16633145-INS-54
chr4-16637453-INS-74
chr4-16688514-INS-226
chr4-16727987-INS-6682
chr4-16734319-INS-117
chr4-16751068-INS-6207
chr4-16764668-INS-78
chr4-16805688-INS-1934
chr4-16826264-INS-2863
chr4-16833019-INS-5331
chr4-16870987-INS-222
chr4-16871294-INS-302
chr4-16929084-INS-6164
chr4-16960379-INS-2914
chr4-17045414-INS-1219
chr4-17048811-INS-63
chr4-17076178-INS-1448
chr4-17085481-INS-62
chr4-17105393-INS-84
chr4-17108617-INS-64
chr4-17117276-INS-507
chr4-17117524-INS-83
chr4-17188241-INS-136
chr4-17230087-INS-4337
chr4-17264078-INS-50
chr4-17309589-INS-214
chr4-17365651-INS-199
chr4-17379639-INS-50

chr4-17450806-INS-1078
chr4-17474914-INS-60
chr4-17477057-INS-1099
chr4-17512346-INS-131
chr4-17533431-INS-407
chr4-17701525-INS-67
chr4-17701592-INS-3434
chr4-17701602-INS-72
chr4-17720194-INS-146
chr4-17739575-INS-5147
chr4-17748604-INS-2190
chr4-17766229-INS-76
chr4-17823211-INS-6187
chr4-17824296-INS-50
chr4-17965251-INS-209
chr4-17975962-INS-50
chr4-17995423-INS-6319
chr4-18050305-INS-5544
chr4-18058669-INS-678
chr4-18079234-INS-1740
chr4-18090974-INS-1947
chr4-18115592-INS-160
chr4-18139799-INS-825
chr4-18152436-INS-1378
chr4-18163126-INS-657
chr4-18200596-INS-874
chr4-18322108-INS-6245
chr4-18329413-INS-653
chr4-18375783-INS-52
chr4-18415907-INS-794
chr4-18457053-INS-124
chr4-18460671-INS-790
chr4-18492466-INS-2295
chr4-18506403-INS-1585
chr4-18512975-INS-489
chr4-18555506-INS-199
chr4-18574275-INS-367
chr4-18582941-INS-6259
chr4-18627435-INS-647
chr4-18631502-INS-556
chr4-18654538-INS-6401
chr4-18704938-INS-201
chr4-18739327-INS-51
chr4-18742902-INS-6138
chr4-18764209-INS-74
chr4-18784666-INS-417
chr4-18788272-INS-1414
chr4-18826911-INS-60
chr4-18829179-INS-51
chr4-18832252-INS-436
chr4-18832302-INS-60
chr4-18850747-INS-245
chr4-18852576-INS-53
chr4-18853146-INS-1092
chr4-18868314-INS-6518
chr4-18888284-INS-6355
chr4-18899314-INS-2358
chr4-18902214-INS-440
chr4-18913882-INS-126
chr4-18913882-INS-58
chr4-18914603-INS-62
chr4-18919601-INS-106
chr4-18925848-INS-7451
chr4-18968754-INS-7471
chr4-18978709-INS-6466
chr4-18984838-INS-6348
chr4-19000091-INS-1143
chr4-19005840-INS-51
chr4-19008248-INS-778
chr4-19044063-INS-9696
chr4-19081629-INS-268
chr4-19084459-INS-110
chr4-19104748-INS-212
chr4-19177467-INS-2111
chr4-19177814-INS-92
chr4-19208098-INS-97
chr4-19254758-INS-1892

chr4-19260698-INS-6245
chr4-19278585-INS-203
chr4-19347540-INS-1079
chr4-19360600-INS-172
chr4-19386894-INS-228
chr4-19406190-INS-1802
chr4-19411363-INS-1956
chr4-19422514-INS-1134
chr4-19430282-INS-1090
chr4-19445781-INS-139
chr4-19446084-INS-2794
chr4-19449807-INS-1355
chr4-19462143-INS-7064
chr4-19527456-INS-159
chr4-19550502-INS-2705
chr4-19564604-INS-50
chr4-19590127-INS-52
chr4-19592220-INS-13707
chr4-19662679-INS-59
chr4-19666184-INS-1198
chr4-19702369-INS-50
chr4-19733504-INS-160
chr4-19794987-INS-6398
chr4-19821536-INS-256
chr4-19828943-INS-186
chr4-19845646-INS-878
chr4-19885745-INS-985
chr4-19899301-INS-245
chr4-19899770-INS-52
chr4-19937387-INS-7084
chr4-19938990-INS-96
chr4-19958717-INS-191
chr4-19964047-INS-2823
chr4-19999583-INS-130
chr4-20005950-INS-138
chr4-20013879-INS-111
chr4-20016744-INS-6525
chr4-20024733-INS-185
chr4-20032830-INS-1545
chr4-20035521-INS-1059
chr4-20048036-INS-75
chr4-20051633-INS-50
chr4-20075371-INS-1019
chr4-20087713-INS-1527
chr4-20127068-INS-193
chr4-20174992-INS-208
chr4-20191915-INS-5096
chr4-20195448-INS-150
chr4-20222246-INS-52
chr4-20248500-INS-13701
chr4-20248500-INS-6769
chr4-20272207-INS-105
chr4-20333185-INS-114
chr4-20400854-INS-234
chr4-20420761-INS-212
chr4-20489717-INS-322
chr4-20489954-INS-3447
chr4-20545525-INS-52
chr4-20546060-INS-333
chr4-20547900-INS-179
chr4-20548464-INS-251
chr4-20551665-INS-74
chr4-20571707-INS-6895
chr4-20581849-INS-2392
chr4-20647084-INS-695
chr4-20648200-INS-1488
chr4-20663570-INS-55
chr4-20702263-INS-422
chr4-20741840-INS-57
chr4-20748667-INS-188
chr4-20753056-INS-6646
chr4-20851248-INS-1747
chr4-20888917-INS-5197
chr4-20958292-INS-1392
chr4-20977135-INS-66
chr4-21003042-INS-71
chr4-21050480-INS-4948

chr4-21051985-INS-82
chr4-21068105-INS-85
chr4-21100832-INS-62
chr4-21107838-INS-5162
chr4-21116719-INS-174
chr4-21145951-INS-358
chr4-21151478-INS-567
chr4-21213823-INS-1545
chr4-21265286-INS-240
chr4-21269681-INS-55
chr4-21326642-INS-3338
chr4-21330433-INS-515
chr4-21343792-INS-8369
chr4-21347049-INS-68
chr4-21360454-INS-56
chr4-21361422-INS-1886
chr4-21433599-INS-66
chr4-21446401-INS-2875
chr4-21446401-INS-461
chr4-21474908-INS-1595
chr4-21552331-INS-5097
chr4-21572048-INS-196
chr4-21573489-INS-209
chr4-21588894-INS-125
chr4-21608679-INS-440
chr4-21609161-INS-75
chr4-21625213-INS-1563
chr4-21633379-INS-1888
chr4-21638188-INS-105
chr4-21639060-INS-214
chr4-21681871-INS-50
chr4-21683550-INS-59
chr4-21692745-INS-2536
chr4-21698006-INS-69
chr4-21725404-INS-166
chr4-21754243-INS-51
chr4-21754320-INS-129
chr4-21758766-INS-224
chr4-21759203-INS-216
chr4-21767467-INS-158
chr4-21768309-INS-192
chr4-21777169-INS-6605
chr4-21791728-INS-1207
chr4-21793583-INS-208
chr4-21798743-INS-438
chr4-21812820-INS-3255
chr4-21834072-INS-198
chr4-21838059-INS-855
chr4-21885598-INS-215
chr4-21946026-INS-379
chr4-21954049-INS-52
chr4-22001876-INS-199
chr4-22003861-INS-64
chr4-22011464-INS-83
chr4-22034126-INS-1139
chr4-22038644-INS-59
chr4-22041657-INS-56
chr4-22042748-INS-6457
chr4-22043717-INS-204
chr4-22100566-INS-99
chr4-22121478-INS-138
chr4-22123236-INS-90
chr4-22123897-INS-400
chr4-22141203-INS-354
chr4-22160395-INS-6181
chr4-22162038-INS-52
chr4-22194211-INS-6576
chr4-22197909-INS-2107
chr4-22200252-INS-1209
chr4-22204183-INS-209
chr4-22218114-INS-435
chr4-22219717-INS-119
chr4-22221383-INS-60
chr4-22224972-INS-120
chr4-22259679-INS-105
chr4-22275371-INS-50
chr4-22306769-INS-3136

chr4-22330829-INS-473
chr4-22364440-INS-82
chr4-22375376-INS-390
chr4-22390463-INS-242
chr4-22459231-INS-209
chr4-22509470-INS-207
chr4-22562076-INS-6501
chr4-22583651-INS-1326
chr4-22627721-INS-11478
chr4-22654434-INS-337
chr4-22660222-INS-135
chr4-22663641-INS-55
chr4-22664393-INS-150
chr4-22667599-INS-80
chr4-22714512-INS-60
chr4-22715307-INS-731
chr4-22738467-INS-1863
chr4-22757133-INS-293
chr4-22761581-INS-107
chr4-22780056-INS-74
chr4-22790634-INS-122
chr4-22805006-INS-1171
chr4-22814771-INS-130
chr4-22868766-INS-6047
chr4-22872304-INS-3693
chr4-22899551-INS-368
chr4-22918286-INS-59
chr4-22920987-INS-159
chr4-22922934-INS-154
chr4-22954469-INS-144
chr4-22982595-INS-666
chr4-23007864-INS-5880
chr4-23032003-INS-50
chr4-23034535-INS-68
chr4-23049726-INS-90
chr4-23070657-INS-776
chr4-23090525-INS-1868
chr4-23096674-INS-14181
chr4-23125787-INS-1005
chr4-23132403-INS-50
chr4-23132478-INS-104
chr4-23165795-INS-6250
chr4-23167249-INS-6637
chr4-23177723-INS-202
chr4-23182105-INS-6472
chr4-23212408-INS-166
chr4-23238279-INS-89
chr4-23302042-INS-59
chr4-23313365-INS-50
chr4-23330642-INS-1439
chr4-23332164-INS-54
chr4-23408644-INS-75
chr4-23435757-INS-130
chr4-23475887-INS-3845
chr4-23478048-INS-2987
chr4-23516692-INS-81
chr4-23561167-INS-64
chr4-23566085-INS-147
chr4-23568546-INS-76
chr4-23622678-INS-50
chr4-23656231-INS-62
chr4-23664736-INS-67
chr4-23675869-INS-107
chr4-23685910-INS-2466
chr4-23692399-INS-50
chr4-23709054-INS-6794
chr4-23746353-INS-149
chr4-23766997-INS-382
chr4-23793358-INS-111
chr4-23836535-INS-79
chr4-23885913-INS-64
chr4-23889459-INS-148
chr4-23902447-INS-153
chr4-23915130-INS-219
chr4-23934414-INS-571
chr4-23954435-INS-409
chr4-23976277-INS-7236

chr4-23993811-INS-56
chr4-24027543-INS-5150
chr4-24051113-INS-1131
chr4-24087237-INS-6466
chr4-24169139-INS-6606
chr4-24177201-INS-199
chr4-24177201-INS-65
chr4-24191810-INS-245
chr4-24245684-INS-174
chr4-24249864-INS-50
chr4-24258743-INS-1250
chr4-24260141-INS-221
chr4-24315509-INS-83
chr4-24345418-INS-8515
chr4-24346135-INS-233
chr4-24369540-INS-74
chr4-24385549-INS-6289
chr4-24389071-INS-50
chr4-24392042-INS-4299
chr4-24407472-INS-128
chr4-24427164-INS-364
chr4-24453072-INS-1656
chr4-24493556-INS-208
chr4-24570154-INS-167
chr4-24625029-INS-403
chr4-24638634-INS-114
chr4-24704222-INS-3740
chr4-24717135-INS-2500
chr4-24717556-INS-50
chr4-24731514-INS-302
chr4-24740832-INS-6594
chr4-24760808-INS-6179
chr4-24761411-INS-468
chr4-24781671-INS-6100
chr4-24800566-INS-78
chr4-24818118-INS-2139
chr4-24832450-INS-198
chr4-24834466-INS-52
chr4-24854637-INS-56
chr4-24854761-INS-500
chr4-24873111-INS-5610
chr4-24906693-INS-7178
chr4-24937795-INS-132
chr4-24937795-INS-632
chr4-24945947-INS-72
chr4-24954206-INS-95
chr4-24991126-INS-240
chr4-24993862-INS-6545
chr4-25004904-INS-224
chr4-25006971-INS-199
chr4-25032523-INS-8271
chr4-25042168-INS-50
chr4-25073810-INS-322
chr4-25138248-INS-402
chr4-25206037-INS-1085
chr4-25222395-INS-1186
chr4-25238150-INS-193
chr4-25275563-INS-254
chr4-25343034-INS-200
chr4-25372493-INS-78
chr4-25391978-INS-404
chr4-25425541-INS-52
chr4-25435277-INS-6357
chr4-25443558-INS-706
chr4-25446753-INS-110
chr4-25518620-INS-1148
chr4-25597133-INS-15330
chr4-25597133-INS-7220
chr4-25603458-INS-3686
chr4-25621959-INS-172
chr4-25642553-INS-156
chr4-25658495-INS-206
chr4-25677931-INS-82
chr4-25728057-INS-208
chr4-25748751-INS-325
chr4-25765664-INS-215
chr4-25776736-INS-231

chr4-25777008-INS-82
chr4-25871678-INS-1264
chr4-25889449-INS-551
chr4-25891536-INS-94
chr4-25902899-INS-243
chr4-25912498-INS-53
chr4-25954402-INS-53
chr4-25955576-INS-609
chr4-26067868-INS-85
chr4-26096261-INS-243
chr4-26106925-INS-70
chr4-26231609-INS-51
chr4-26238277-INS-797
chr4-26240280-INS-55
chr4-26249379-INS-67
chr4-26259101-INS-53
chr4-26273862-INS-919
chr4-26274857-INS-3416
chr4-26324720-INS-722
chr4-26341914-INS-50
chr4-26356794-INS-409
chr4-26358209-INS-2470
chr4-26407143-INS-82
chr4-26444069-INS-57
chr4-26456734-INS-978
chr4-26481338-INS-65
chr4-26516178-INS-1875
chr4-26522515-INS-1190
chr4-26523028-INS-226
chr4-26538833-INS-56
chr4-26541161-INS-2738
chr4-26579059-INS-2141
chr4-26625792-INS-5804
chr4-26644441-INS-212
chr4-26694342-INS-390
chr4-26712381-INS-212
chr4-26729760-INS-1055
chr4-26780694-INS-64
chr4-26802167-INS-6280
chr4-26813689-INS-4801
chr4-26822134-INS-101
chr4-26835071-INS-504
chr4-26842258-INS-564
chr4-26863270-INS-1340
chr4-26873098-INS-60
chr4-26876970-INS-50
chr4-26888327-INS-6334
chr4-26912126-INS-5245
chr4-26995264-INS-113
chr4-26995316-INS-50
chr4-27024173-INS-2838
chr4-27030982-INS-1914
chr4-27049232-INS-1365
chr4-27056246-INS-107
chr4-27095324-INS-60
chr4-27119054-INS-611
chr4-27153306-INS-50
chr4-27184873-INS-6225
chr4-27203793-INS-81
chr4-27206117-INS-13647
chr4-27257925-INS-224
chr4-27261296-INS-888
chr4-27263042-INS-122
chr4-27274601-INS-95
chr4-27325496-INS-3021
chr4-27338297-INS-60
chr4-27342935-INS-330
chr4-27350351-INS-6239
chr4-27351024-INS-2676
chr4-27375434-INS-13823
chr4-27376881-INS-1493
chr4-27384135-INS-1451
chr4-27394309-INS-539
chr4-27409486-INS-51
chr4-27423363-INS-148
chr4-27431353-INS-12596
chr4-27433592-INS-2631

chr4-27439025-INS-52
chr4-27464467-INS-3734
chr4-27474461-INS-1175
chr4-27485180-INS-6229
chr4-27493942-INS-617
chr4-27502753-INS-186
chr4-27511814-INS-84
chr4-27538594-INS-290
chr4-27567373-INS-50
chr4-27574339-INS-371
chr4-27574672-INS-82
chr4-27576623-INS-6287
chr4-27578649-INS-1307
chr4-27588384-INS-5863
chr4-27595021-INS-64
chr4-27606304-INS-2465
chr4-27654610-INS-156
chr4-27656373-INS-2529
chr4-27661896-INS-148
chr4-27705821-INS-6360
chr4-27725054-INS-50
chr4-27725716-INS-118
chr4-27771535-INS-82
chr4-27841780-INS-150
chr4-27848862-INS-160
chr4-27879840-INS-2150
chr4-27881523-INS-512
chr4-27895186-INS-60
chr4-27912960-INS-205
chr4-27926922-INS-2621
chr4-27940398-INS-194
chr4-27957483-INS-78
chr4-27992667-INS-2319
chr4-28003520-INS-3166
chr4-28006800-INS-1292
chr4-28012385-INS-60
chr4-28041300-INS-294
chr4-28066812-INS-1173
chr4-28081582-INS-6382
chr4-28105662-INS-492
chr4-28108682-INS-58
chr4-28128492-INS-52
chr4-28128928-INS-89
chr4-28163329-INS-2722
chr4-28170251-INS-6185
chr4-28182706-INS-6004
chr4-28183508-INS-1641
chr4-28183939-INS-4530
chr4-28199036-INS-3282
chr4-28218778-INS-2420
chr4-28225504-INS-56
chr4-28233551-INS-2374
chr4-28274276-INS-56
chr4-28280535-INS-52
chr4-28323355-INS-304
chr4-28331325-INS-1911
chr4-28369668-INS-5175
chr4-28382246-INS-120
chr4-28382246-INS-305
chr4-28400993-INS-68
chr4-28439914-INS-152
chr4-28455420-INS-92
chr4-28475025-INS-88
chr4-28485668-INS-64
chr4-28487457-INS-6185
chr4-28539183-INS-264
chr4-28539236-INS-1147
chr4-28557739-INS-323
chr4-28582689-INS-50
chr4-28640128-INS-149
chr4-28658760-INS-1048
chr4-28689004-INS-1481
chr4-28695992-INS-272
chr4-28697668-INS-2920
chr4-28715422-INS-68
chr4-28719481-INS-2299
chr4-28758531-INS-56

chr4-28774244-INS-1062
chr4-28829900-INS-72
chr4-28842216-INS-64
chr4-28929400-INS-414
chr4-28929899-INS-1296
chr4-28959615-INS-66
chr4-28982851-INS-74
chr4-28984083-INS-7258
chr4-28992366-INS-50
chr4-28994868-INS-201
chr4-29002622-INS-95
chr4-29027054-INS-584
chr4-29029224-INS-83
chr4-29032506-INS-550
chr4-29073496-INS-363
chr4-29085392-INS-100
chr4-29112844-INS-482
chr4-29140221-INS-3164
chr4-29288878-INS-293
chr4-29319737-INS-204
chr4-29337750-INS-143
chr4-29374271-INS-237
chr4-29374372-INS-94
chr4-29381841-INS-6125
chr4-29385047-INS-2385
chr4-29401486-INS-201
chr4-29460311-INS-54
chr4-29489757-INS-50
chr4-29549496-INS-3762
chr4-29550777-INS-456
chr4-29550777-INS-76
chr4-29617114-INS-72
chr4-29619009-INS-590
chr4-29631497-INS-6238
chr4-29644748-INS-60
chr4-29646297-INS-3379
chr4-29656931-INS-84
chr4-29662779-INS-5324
chr4-29679054-INS-6877
chr4-29688496-INS-124
chr4-29693529-INS-72
chr4-29722754-INS-72
chr4-29758827-INS-177
chr4-29763995-INS-8430
chr4-29764421-INS-117
chr4-29774463-INS-146
chr4-29817824-INS-103
chr4-29824998-INS-125
chr4-29834795-INS-68
chr4-29846047-INS-149
chr4-29865906-INS-4088
chr4-29911566-INS-6220
chr4-29919887-INS-56
chr4-29933269-INS-579
chr4-29934058-INS-78
chr4-29952227-INS-50
chr4-30003093-INS-6274
chr4-30017389-INS-6423
chr4-30044993-INS-4890
chr4-30114680-INS-3234
chr4-30114781-INS-60
chr4-30141739-INS-8804
chr4-30142746-INS-5709
chr4-30149764-INS-795
chr4-30166545-INS-1519
chr4-30207542-INS-106
chr4-30207580-INS-214
chr4-30209216-INS-1339
chr4-30217717-INS-94
chr4-30238170-INS-95
chr4-30241832-INS-3575
chr4-30264225-INS-7509
chr4-30286355-INS-84
chr4-30292342-INS-2466
chr4-30305425-INS-72
chr4-30318193-INS-180
chr4-30336188-INS-358

chr4-30339750-INS-3587
chr4-30356305-INS-109
chr4-30384552-INS-3050
chr4-30390730-INS-1595
chr4-30405131-INS-2943
chr4-30415503-INS-2278
chr4-30424565-INS-52
chr4-30433606-INS-3210
chr4-30469080-INS-130
chr4-30471029-INS-1646
chr4-30484594-INS-52
chr4-30493866-INS-70
chr4-30600257-INS-262
chr4-30704738-INS-70
chr4-30733893-INS-644
chr4-30734642-INS-157
chr4-30734642-INS-64
chr4-30739401-INS-13835
chr4-30749962-INS-76
chr4-30771459-INS-1424
chr4-30786246-INS-6126
chr4-30794715-INS-1972
chr4-30821145-INS-114
chr4-30852219-INS-204
chr4-30930554-INS-374
chr4-30930566-INS-58
chr4-30982950-INS-52
chr4-30984053-INS-3532
chr4-31008108-INS-81
chr4-31008138-INS-393
chr4-31147874-INS-6214
chr4-31151532-INS-82
chr4-31202941-INS-99
chr4-31207001-INS-6685
chr4-31229110-INS-2969
chr4-31252651-INS-452
chr4-31253301-INS-99
chr4-31271796-INS-1637
chr4-31277173-INS-84
chr4-31277724-INS-269
chr4-31279827-INS-65
chr4-31303633-INS-99
chr4-31340624-INS-234
chr4-31355266-INS-1568
chr4-31372277-INS-102
chr4-31433035-INS-94
chr4-31471666-INS-6801
chr4-31472409-INS-1600
chr4-31486458-INS-1113
chr4-31489123-INS-6874
chr4-31552949-INS-242
chr4-31552957-INS-74
chr4-31567723-INS-396
chr4-31595264-INS-1023
chr4-31595264-INS-8244
chr4-31619018-INS-52
chr4-31654751-INS-128
chr4-31666588-INS-573
chr4-31675125-INS-403
chr4-31683930-INS-72
chr4-31690626-INS-2659
chr4-31693176-INS-1177
chr4-31706261-INS-4402
chr4-31719413-INS-227
chr4-31777699-INS-208
chr4-31804626-INS-184
chr4-31814953-INS-130
chr4-31840743-INS-84
chr4-31841138-INS-497
chr4-31855104-INS-124
chr4-31943222-INS-134
chr4-31946866-INS-52
chr4-31986957-INS-91
chr4-31986975-INS-273
chr4-32053579-INS-50
chr4-32083129-INS-3519
chr4-32088614-INS-739

chr4-32089866-INS-63
chr4-32103741-INS-52
chr4-32165626-INS-111
chr4-32167041-INS-7121
chr4-32194117-INS-50
chr4-32196241-INS-122
chr4-32263882-INS-245
chr4-32279936-INS-1075
chr4-32295046-INS-53
chr4-32302774-INS-162
chr4-32303595-INS-50
chr4-32356718-INS-3511
chr4-32428360-INS-52
chr4-32430580-INS-204
chr4-32443467-INS-237
chr4-32451299-INS-469
chr4-32456095-INS-203
chr4-32497071-INS-52
chr4-32588493-INS-401
chr4-32588739-INS-160
chr4-32592651-INS-173
chr4-32594049-INS-734
chr4-32611548-INS-52
chr4-32628875-INS-64
chr4-32659597-INS-1956
chr4-32673640-INS-1845
chr4-32673738-INS-60
chr4-32750678-INS-1305
chr4-32786425-INS-190
chr4-32821636-INS-204
chr4-32830123-INS-54
chr4-32850624-INS-206
chr4-32901894-INS-62
chr4-32903843-INS-196
chr4-32943662-INS-66
chr4-32957038-INS-219
chr4-32958138-INS-489
chr4-32966361-INS-157
chr4-32967022-INS-219
chr4-32967165-INS-178
chr4-33000415-INS-204
chr4-33010454-INS-67
chr4-33010793-INS-84
chr4-33028505-INS-1184
chr4-33054834-INS-483
chr4-33057565-INS-73
chr4-33058714-INS-84
chr4-33073670-INS-641
chr4-33097318-INS-7482
chr4-33102152-INS-148
chr4-33107230-INS-1047
chr4-33112189-INS-467
chr4-33141448-INS-216
chr4-33148171-INS-53
chr4-33153562-INS-51
chr4-33166279-INS-166
chr4-33171424-INS-1052
chr4-33200008-INS-6403
chr4-33240985-INS-7549
chr4-33244235-INS-85
chr4-33259697-INS-67
chr4-33263629-INS-104
chr4-33292207-INS-6090
chr4-33296291-INS-571
chr4-33299698-INS-51
chr4-33344833-INS-52
chr4-33358481-INS-564
chr4-33540433-INS-52
chr4-33540865-INS-73
chr4-33541199-INS-112
chr4-33551677-INS-206
chr4-33606462-INS-21210
chr4-33622004-INS-1436
chr4-33633476-INS-1017
chr4-33653506-INS-150
chr4-33671933-INS-121
chr4-33678388-INS-343

chr4-33680042-INS-181
chr4-33681148-INS-477
chr4-33682717-INS-4868
chr4-33695775-INS-326
chr4-33701543-INS-54
chr4-33729885-INS-60
chr4-33813464-INS-3215
chr4-33844016-INS-271
chr4-33845394-INS-106
chr4-33926439-INS-54
chr4-34057761-INS-6663
chr4-34143915-INS-54
chr4-34197412-INS-60
chr4-34238332-INS-50
chr4-34260280-INS-2942
chr4-34380300-INS-1153
chr4-34388607-INS-764
chr4-34443016-INS-3337
chr4-34450667-INS-1701
chr4-34458235-INS-192
chr4-34461853-INS-2709
chr4-34489738-INS-209
chr4-34490005-INS-392
chr4-34494177-INS-380
chr4-34519819-INS-77
chr4-34531168-INS-1263
chr4-34579043-INS-60
chr4-34708661-INS-3728
chr4-34718068-INS-74
chr4-34736821-INS-5305
chr4-34738178-INS-165
chr4-34741419-INS-50
chr4-34762537-INS-203
chr4-34782490-INS-68
chr4-34783698-INS-72
chr4-34783925-INS-54
chr4-34786344-INS-222
chr4-34786344-INS-71
chr4-34798925-INS-203
chr4-34802445-INS-213
chr4-34812187-INS-101
chr4-34862141-INS-58
chr4-34862729-INS-203
chr4-34878988-INS-403
chr4-34928531-INS-6847
chr4-34945659-INS-64
chr4-34966682-INS-304
chr4-34990751-INS-459
chr4-34994689-INS-239
chr4-35010214-INS-465
chr4-35028766-INS-212
chr4-35051293-INS-1290
chr4-35113818-INS-55
chr4-35166651-INS-6829
chr4-35190792-INS-1473
chr4-35200296-INS-665
chr4-35207020-INS-2668
chr4-35215319-INS-50
chr4-35257898-INS-1649
chr4-35285524-INS-101
chr4-35345327-INS-65
chr4-35360864-INS-125
chr4-35384516-INS-1263
chr4-35400301-INS-52
chr4-35450045-INS-847
chr4-35470446-INS-7021
chr4-35506953-INS-662
chr4-35569163-INS-2410
chr4-35647128-INS-132
chr4-35717449-INS-62
chr4-35857240-INS-66
chr4-35875375-INS-127
chr4-35875748-INS-4443
chr4-35948346-INS-52
chr4-35960465-INS-161
chr4-36015169-INS-3476
chr4-36105508-INS-160

chr4-36154604-INS-50
chr4-36199510-INS-500
chr4-36205176-INS-500
chr4-36285210-INS-164
chr4-36325902-INS-76
chr4-36403177-INS-60
chr4-36421411-INS-2715
chr4-36515783-INS-1651
chr4-36518374-INS-88
chr4-36538266-INS-54
chr4-36554077-INS-156
chr4-36564907-INS-349
chr4-36598572-INS-50
chr4-36624641-INS-1637
chr4-36626371-INS-873
chr4-36698545-INS-6177
chr4-36756977-INS-58
chr4-36809251-INS-66
chr4-36859081-INS-54
chr4-36911733-INS-50
chr4-36920091-INS-106
chr4-37011656-INS-2723
chr4-37066758-INS-542
chr4-37067102-INS-66
chr4-37132110-INS-7191
chr4-37186711-INS-9092
chr4-37193078-INS-344
chr4-37396657-INS-762
chr4-37428949-INS-6256
chr4-37478652-INS-313
chr4-37508771-INS-2741
chr4-37534498-INS-6869
chr4-37559874-INS-56
chr4-37623740-INS-176
chr4-37675087-INS-52
chr4-37711952-INS-51
chr4-37712200-INS-6373
chr4-37732782-INS-81
chr4-37795801-INS-164
chr4-37800845-INS-52
chr4-37827253-INS-54
chr4-37897180-INS-1679
chr4-37940101-INS-64
chr4-37979075-INS-2371
chr4-38008150-INS-298
chr4-38013298-INS-6782
chr4-38042885-INS-641
chr4-38101447-INS-631
chr4-38154650-INS-1748
chr4-38218447-INS-2424
chr4-38241633-INS-1059
chr4-38268659-INS-6431
chr4-38283740-INS-7341
chr4-38296089-INS-1288
chr4-38296659-INS-51
chr4-38297154-INS-67
chr4-38339961-INS-680
chr4-38340570-INS-2243
chr4-38341031-INS-9135
chr4-38447669-INS-150
chr4-38490929-INS-7156
chr4-38528393-INS-15835
chr4-38620598-INS-55
chr4-38658278-INS-50
chr4-38659050-INS-64
chr4-38660795-INS-6990
chr4-38666891-INS-1065
chr4-38668640-INS-1445
chr4-38678964-INS-4594
chr4-38687743-INS-12318
chr4-38699445-INS-2275
chr4-38707610-INS-391
chr4-38719646-INS-1741
chr4-38722751-INS-65
chr4-38757531-INS-195
chr4-38759073-INS-693
chr4-38761891-INS-2857

chr4-38852263-INS-50
chr4-38878775-INS-178
chr4-38919502-INS-66
chr4-38933489-INS-8353
chr4-38934330-INS-802
chr4-38937345-INS-1930
chr4-38938284-INS-1133
chr4-38950411-INS-1296
chr4-38952826-INS-400
chr4-38962018-INS-4452
chr4-39030712-INS-54
chr4-39070839-INS-236
chr4-39146462-INS-193
chr4-39150323-INS-7778
chr4-39151322-INS-52
chr4-39174037-INS-171
chr4-39176797-INS-87
chr4-39214339-INS-80
chr4-39248267-INS-153
chr4-39248505-INS-344
chr4-39274152-INS-67
chr4-39284115-INS-62
chr4-39318077-INS-54
chr4-39321496-INS-605
chr4-39430349-INS-51
chr4-39487812-INS-1037
chr4-39499649-INS-84
chr4-39546281-INS-50
chr4-39597232-INS-6369
chr4-39614517-INS-2489
chr4-39615438-INS-3935
chr4-39634975-INS-787
chr4-39636731-INS-268
chr4-39641249-INS-54
chr4-39683837-INS-1351
chr4-39687256-INS-62
chr4-39712852-INS-341
chr4-39797480-INS-185
chr4-39805466-INS-456
chr4-39847831-INS-779
chr4-39856093-INS-720
chr4-39929105-INS-131
chr4-39944333-INS-52
chr4-40011702-INS-176
chr4-40069311-INS-1890
chr4-40131962-INS-199
chr4-40151336-INS-1087
chr4-40161376-INS-7156
chr4-40218243-INS-3749
chr4-40291423-INS-778
chr4-40315230-INS-184
chr4-40319971-INS-79
chr4-40323165-INS-128
chr4-40323364-INS-192
chr4-40352471-INS-60
chr4-40364863-INS-68
chr4-40382693-INS-6282
chr4-40390338-INS-1214
chr4-40407142-INS-2946
chr4-40422282-INS-374
chr4-40422810-INS-172
chr4-40461484-INS-3827
chr4-40505359-INS-66
chr4-40505383-INS-264
chr4-40522516-INS-258
chr4-40528950-INS-1335
chr4-40537892-INS-6381
chr4-40548291-INS-198
chr4-40562325-INS-1997
chr4-40633407-INS-127
chr4-40637843-INS-122
chr4-40638078-INS-354
chr4-40641886-INS-103
chr4-40647588-INS-1058
chr4-40674179-INS-6662
chr4-40677743-INS-437
chr4-40685484-INS-802

chr4-40690033-INS-82
chr4-40703726-INS-110
chr4-40710429-INS-52
chr4-40711318-INS-80
chr4-40765065-INS-190
chr4-40766714-INS-1634
chr4-40773141-INS-106
chr4-40778498-INS-55
chr4-40782674-INS-2957
chr4-40785449-INS-120
chr4-40831550-INS-87
chr4-40839138-INS-200
chr4-40850159-INS-66
chr4-40850408-INS-192
chr4-40861729-INS-52
chr4-40887014-INS-2601
chr4-40907357-INS-522
chr4-40913962-INS-673
chr4-40923890-INS-195
chr4-41037809-INS-225
chr4-41042540-INS-54
chr4-41050484-INS-50
chr4-41050632-INS-105
chr4-41057757-INS-158
chr4-41058790-INS-475
chr4-41059101-INS-2003
chr4-41079433-INS-71
chr4-41099914-INS-52
chr4-41100674-INS-55
chr4-41104968-INS-82
chr4-41144015-INS-215
chr4-41153095-INS-200
chr4-41153513-INS-134
chr4-41174366-INS-58
chr4-41215281-INS-185
chr4-41246295-INS-191
chr4-41249229-INS-262
chr4-41283189-INS-75
chr4-41303633-INS-188
chr4-41303681-INS-204
chr4-41322587-INS-56
chr4-41330255-INS-4636
chr4-41332332-INS-208
chr4-41337280-INS-59
chr4-41348056-INS-59
chr4-41364418-INS-139
chr4-41370921-INS-122
chr4-41372649-INS-52
chr4-41460603-INS-55
chr4-41475058-INS-80
chr4-41560082-INS-120
chr4-41565226-INS-64
chr4-41577659-INS-68
chr4-41595438-INS-204
chr4-41599410-INS-50
chr4-41666233-INS-69
chr4-41721548-INS-62
chr4-41732317-INS-60
chr4-41776018-INS-171
chr4-41855911-INS-88
chr4-41856828-INS-192
chr4-41857162-INS-322
chr4-42140409-INS-72
chr4-42206263-INS-271
chr4-42856144-INS-498
chr4-42897011-INS-221
chr4-42904416-INS-1400
chr4-42928490-INS-200
chr4-42964699-INS-229
chr4-43026972-INS-50
chr4-43054400-INS-50
chr4-43056956-INS-210
chr4-43057678-INS-203
chr4-43063278-INS-53
chr4-43066166-INS-50
chr4-43082694-INS-207
chr4-43090077-INS-90

chr4-43108896-INS-63
chr4-43113352-INS-1114
chr4-43130757-INS-1558
chr4-43211809-INS-125
chr4-43284050-INS-64
chr4-43291894-INS-489
chr4-43295065-INS-111
chr4-43295244-INS-5255
chr4-43310618-INS-52
chr4-43326136-INS-2963
chr4-43326575-INS-195
chr4-43329453-INS-6620
chr4-43354238-INS-317
chr4-43365227-INS-169
chr4-43410820-INS-72
chr4-43411250-INS-121
chr4-43429467-INS-149
chr4-43429497-INS-66
chr4-43431123-INS-1329
chr4-43432195-INS-54
chr4-43438821-INS-7073
chr4-43440237-INS-239
chr4-43463527-INS-937
chr4-43477892-INS-60
chr4-43541272-INS-82
chr4-43596360-INS-6475
chr4-43658558-INS-68
chr4-43681341-INS-173
chr4-43714750-INS-71
chr4-43717408-INS-78
chr4-43746084-INS-177
chr4-43756892-INS-360
chr4-43758064-INS-4454
chr4-43758190-INS-268
chr4-43763932-INS-54
chr4-43780476-INS-2251
chr4-43803134-INS-53
chr4-43803340-INS-72
chr4-43806852-INS-188
chr4-43810721-INS-207
chr4-43825014-INS-63
chr4-43829751-INS-3602
chr4-43830470-INS-136
chr4-43830470-INS-64
chr4-43887999-INS-1890
chr4-43951817-INS-126
chr4-43969084-INS-205
chr4-43971088-INS-128
chr4-43974354-INS-60
chr4-43991607-INS-72
chr4-43997521-INS-208
chr4-44000806-INS-219
chr4-44004431-INS-6389
chr4-44011818-INS-93
chr4-44035984-INS-58
chr4-44037959-INS-394
chr4-44038280-INS-334
chr4-44097819-INS-137
chr4-44199140-INS-8678
chr4-44213875-INS-121
chr4-44220411-INS-65
chr4-44220437-INS-453
chr4-44247057-INS-64
chr4-44248890-INS-393
chr4-44255221-INS-59
chr4-44296330-INS-62
chr4-44333871-INS-131
chr4-44358103-INS-403
chr4-44375100-INS-161
chr4-44375100-INS-64
chr4-44437382-INS-56
chr4-44484312-INS-50
chr4-44499352-INS-64
chr4-44499412-INS-188
chr4-44507692-INS-50
chr4-44597544-INS-198
chr4-44600609-INS-113

chr4-44627752-INS-116
chr4-44679896-INS-52
chr4-44683042-INS-102
chr4-44721762-INS-327
chr4-44722064-INS-52
chr4-44747880-INS-56
chr4-44807640-INS-50
chr4-44838705-INS-396
chr4-44872394-INS-72
chr4-44889932-INS-241
chr4-44890553-INS-75
chr4-44898355-INS-962
chr4-44947386-INS-55
chr4-44951418-INS-222
chr4-44966796-INS-429
chr4-44974925-INS-407
chr4-44986436-INS-100
chr4-45042412-INS-184
chr4-45050197-INS-157
chr4-45073403-INS-1328
chr4-45102481-INS-181
chr4-45102525-INS-50
chr4-45130934-INS-88
chr4-45145356-INS-56
chr4-45146356-INS-65
chr4-45150634-INS-58
chr4-45151253-INS-80
chr4-45155133-INS-126
chr4-45165576-INS-226
chr4-45174677-INS-118
chr4-45193282-INS-2877
chr4-45193889-INS-905
chr4-45220823-INS-72
chr4-45232108-INS-250
chr4-45257680-INS-62
chr4-45276772-INS-116
chr4-45276772-INS-318
chr4-45321460-INS-216
chr4-45404393-INS-81
chr4-45404911-INS-150
chr4-45446985-INS-243
chr4-45447771-INS-69
chr4-45486696-INS-197
chr4-45561467-INS-212
chr4-45583726-INS-66
chr4-45595023-INS-55
chr4-45600259-INS-2744
chr4-45613732-INS-126
chr4-45613732-INS-504
chr4-45615030-INS-169
chr4-45615899-INS-192
chr4-45637991-INS-440
chr4-45641106-INS-112
chr4-45641859-INS-327
chr4-45660079-INS-1487
chr4-45686821-INS-51
chr4-45707054-INS-2294
chr4-45728458-INS-258
chr4-45740172-INS-75
chr4-45774264-INS-171
chr4-45785240-INS-7098
chr4-45799672-INS-54
chr4-45805046-INS-482
chr4-45847019-INS-207
chr4-45847336-INS-72
chr4-45847441-INS-176
chr4-45856694-INS-72
chr4-45858553-INS-61
chr4-45881557-INS-2503
chr4-45883782-INS-215
chr4-45891703-INS-150
chr4-45892095-INS-69
chr4-45918951-INS-838
chr4-45942157-INS-56
chr4-45958123-INS-8944
chr4-46075565-INS-102
chr4-46082403-INS-84

chr4-46104381-INS-70
chr4-46108864-INS-68
chr4-46178939-INS-76
chr4-46217244-INS-239
chr4-46223539-INS-87
chr4-46224415-INS-52
chr4-46246303-INS-9606
chr4-46252883-INS-204
chr4-46269564-INS-258
chr4-46281346-INS-1240
chr4-46289355-INS-64
chr4-46300547-INS-160
chr4-46314070-INS-2171
chr4-46321013-INS-50
chr4-46357418-INS-135
chr4-46372358-INS-118
chr4-46373274-INS-1382
chr4-46399688-INS-71
chr4-46409990-INS-58
chr4-46455941-INS-221
chr4-46457860-INS-6442
chr4-46479600-INS-108
chr4-46480050-INS-139
chr4-46481983-INS-112
chr4-46482946-INS-201
chr4-46486390-INS-3947
chr4-46507750-INS-100
chr4-46508529-INS-231
chr4-46517903-INS-189
chr4-46521182-INS-78
chr4-46525942-INS-56
chr4-46527314-INS-566
chr4-46538104-INS-2867
chr4-46553215-INS-52
chr4-46587783-INS-50
chr4-46612192-INS-64
chr4-46658531-INS-59
chr4-46661271-INS-103
chr4-46670983-INS-52
chr4-46745911-INS-55
chr4-46750167-INS-64
chr4-46757128-INS-381
chr4-46760271-INS-60
chr4-46767712-INS-286
chr4-46768344-INS-50
chr4-46768394-INS-250
chr4-46769740-INS-91
chr4-46773107-INS-51
chr4-46777678-INS-359
chr4-46777837-INS-5289
chr4-46914632-INS-52
chr4-46940994-INS-124
chr4-46962519-INS-604
chr4-46986667-INS-51
chr4-46994875-INS-204
chr4-47005462-INS-6806
chr4-47074458-INS-195
chr4-47077579-INS-6339
chr4-47094050-INS-3339
chr4-47096237-INS-4845
chr4-47099063-INS-192
chr4-47134797-INS-56
chr4-47164949-INS-191
chr4-47188030-INS-2780
chr4-47280227-INS-953
chr4-47339069-INS-52
chr4-47387804-INS-6468
chr4-47397696-INS-70
chr4-47441951-INS-80
chr4-47502590-INS-2607
chr4-47507913-INS-6390
chr4-47518499-INS-184
chr4-47518827-INS-64
chr4-47569024-INS-2644
chr4-47571504-INS-118
chr4-47582085-INS-105
chr4-47606569-INS-64

chr4-47687814-INS-58
chr4-47723984-INS-57
chr4-47766396-INS-84
chr4-47805096-INS-58
chr4-47828706-INS-173
chr4-47868723-INS-52
chr4-47915139-INS-50
chr4-47942198-INS-83
chr4-47967934-INS-6583
chr4-47968243-INS-56
chr4-47992386-INS-68
chr4-48029934-INS-60
chr4-48137400-INS-52
chr4-48243862-INS-230
chr4-48300493-INS-223
chr4-48340674-INS-3076
chr4-48370411-INS-50
chr4-48391799-INS-60
chr4-48398971-INS-258
chr4-48409603-INS-764
chr4-48450887-INS-682
chr4-48502420-INS-5565
chr4-48510010-INS-506
chr4-48524766-INS-56
chr4-48558024-INS-60
chr4-48572368-INS-57
chr4-48575078-INS-58
chr4-48591449-INS-54
chr4-48685045-INS-64
chr4-48685847-INS-74
chr4-48717725-INS-6478
chr4-48730123-INS-5227
chr4-48784022-INS-212
chr4-48830395-INS-99
chr4-48831076-INS-232
chr4-48856079-INS-104
chr4-48935672-INS-6774
chr4-48941762-INS-50
chr4-48962500-INS-6563
chr4-48964453-INS-140
chr4-48973950-INS-116
chr4-48986922-INS-3441
chr4-48989698-INS-2984
chr4-49040551-INS-504
chr4-49089589-INS-126
chr4-49116981-INS-6540
chr4-49124853-INS-221
chr4-49163431-INS-2852
chr4-49164572-INS-6283
chr4-49178335-INS-1281
chr4-49191981-INS-1711
chr4-49199643-INS-2543
chr4-49200216-INS-6416
chr4-49227179-INS-585
chr4-49237144-INS-1080
chr4-49240421-INS-113
chr4-49279365-INS-56
chr4-49323739-INS-1359
chr4-49336017-INS-101
chr4-49396848-INS-170
chr4-49446698-INS-199
chr4-49505841-INS-84
chr4-49629093-INS-5683
chr4-49684779-INS-4147
chr4-49687069-INS-212
chr4-49707372-INS-1309
chr4-49759580-INS-72
chr4-49763669-INS-6832
chr4-49781109-INS-158
chr4-49813365-INS-76
chr4-49816410-INS-1386
chr4-49883248-INS-201
chr4-49883321-INS-107
chr4-49885765-INS-6212
chr4-49920681-INS-72
chr4-49930358-INS-63
chr4-49973509-INS-50

chr4-50004037-INS-944
chr4-50068968-INS-68
chr4-50070551-INS-64
chr4-50088200-INS-2813
chr4-50158910-INS-1320
chr4-50184215-INS-182
chr4-50191024-INS-159
chr4-50217140-INS-6688
chr4-50221977-INS-85
chr4-50285824-INS-396
chr4-50357984-INS-76
chr4-50429761-INS-6296
chr4-50447155-INS-2650
chr4-50573059-INS-132
chr4-50574231-INS-50
chr4-50621314-INS-3676
chr4-50635134-INS-50
chr4-50662480-INS-136
chr4-50733986-INS-2723
chr4-50737855-INS-3138
chr4-50793831-INS-6707
chr4-50815938-INS-84
chr4-50824266-INS-879
chr4-50834535-INS-2882
chr4-50843074-INS-85
chr4-50850466-INS-2710
chr4-50864435-INS-1165
chr4-50872903-INS-2677
chr4-50922376-INS-83
chr4-50925234-INS-6211
chr4-50958132-INS-52
chr4-50980982-INS-1353
chr4-50997963-INS-6379
chr4-51052837-INS-54
chr4-51081880-INS-53
chr4-51087496-INS-210
chr4-51109267-INS-562
chr4-51126513-INS-6464
chr4-51151687-INS-2425
chr4-51173827-INS-6398
chr4-51207546-INS-3556
chr4-51215488-INS-80
chr4-51229432-INS-81
chr4-51268453-INS-70
chr4-51276677-INS-88
chr4-51282059-INS-2394
chr4-51325228-INS-2638
chr4-51345642-INS-2408
chr4-51357608-INS-6672
chr4-51442726-INS-3057
chr4-51465929-INS-1631
chr4-51485397-INS-1533
chr4-51488956-INS-321
chr4-51522269-INS-4822
chr4-51581382-INS-51
chr4-51610793-INS-403
chr4-51614261-INS-54
chr4-51615977-INS-2475
chr4-51621507-INS-2800
chr4-51669323-INS-1229
chr4-51817868-INS-2953
chr4-51844336-INS-2917
chr4-51963992-INS-50
chr4-51964881-INS-92
chr4-51968534-INS-3230
chr4-51977658-INS-1688
chr4-52026949-INS-50
chr4-52245777-INS-360
chr4-52246481-INS-1291
chr4-52253345-INS-61
chr4-52274205-INS-6610
chr4-52275839-INS-111
chr4-52318070-INS-100
chr4-52318801-INS-108
chr4-52342256-INS-2788
chr4-52403930-INS-88
chr4-52404047-INS-235

chr4-52406198-INS-888
chr4-52406651-INS-157
chr4-52406777-INS-60
chr4-52419383-INS-6426
chr4-52453570-INS-102
chr4-52495427-INS-7464
chr4-52507345-INS-84
chr4-52518609-INS-5113
chr4-52551501-INS-302
chr4-52600569-INS-1426
chr4-52600691-INS-409
chr4-52609694-INS-58
chr4-52616769-INS-74
chr4-52629623-INS-5085
chr4-52629906-INS-1331
chr4-52640029-INS-60
chr4-52654758-INS-481
chr4-52683165-INS-2809
chr4-52708279-INS-107
chr4-52714994-INS-1230
chr4-52718060-INS-90
chr4-52748834-INS-6209
chr4-52749794-INS-56
chr4-52774864-INS-2602
chr4-52802625-INS-2111
chr4-52810657-INS-55
chr4-52815244-INS-61
chr4-52833021-INS-54
chr4-52880285-INS-60
chr4-52886974-INS-196
chr4-52942511-INS-634
chr4-52960376-INS-109
chr4-52965776-INS-50
chr4-52969789-INS-81
chr4-52990666-INS-1441
chr4-53028810-INS-50
chr4-53040999-INS-52
chr4-53094683-INS-64
chr4-53103920-INS-1397
chr4-53110000-INS-7168
chr4-53210695-INS-79
chr4-53226705-INS-80
chr4-53226769-INS-171
chr4-53237706-INS-173
chr4-53277366-INS-56
chr4-53346212-INS-248
chr4-53346475-INS-96
chr4-53363391-INS-208
chr4-53364138-INS-140
chr4-53412969-INS-192
chr4-53416378-INS-166
chr4-53557614-INS-64
chr4-53557880-INS-211
chr4-53597790-INS-2495
chr4-53773211-INS-87
chr4-53791832-INS-6995
chr4-53794166-INS-172
chr4-53817519-INS-1217
chr4-53838536-INS-72
chr4-53838601-INS-1069
chr4-53847154-INS-199
chr4-53879138-INS-50
chr4-53880949-INS-50
chr4-53928112-INS-66
chr4-53962138-INS-6484
chr4-53964247-INS-1107
chr4-53992779-INS-56
chr4-54015523-INS-54
chr4-54017794-INS-63
chr4-54042940-INS-189
chr4-54117270-INS-3153
chr4-54148795-INS-124
chr4-54166404-INS-60
chr4-54166741-INS-54
chr4-54176220-INS-9902
chr4-54184105-INS-168
chr4-54193244-INS-1786

chr4-54303596-INS-1071
chr4-54321890-INS-109
chr4-54338304-INS-74
chr4-54423633-INS-9273
chr4-54424306-INS-1805
chr4-54544729-INS-2550
chr4-54575348-INS-60
chr4-54620385-INS-116
chr4-54706729-INS-78
chr4-54706806-INS-786
chr4-54748774-INS-210
chr4-54793501-INS-3163
chr4-54816045-INS-305
chr4-54828422-INS-54
chr4-54854314-INS-136
chr4-54877899-INS-66
chr4-54882501-INS-2803
chr4-54962855-INS-4741
chr4-54988321-INS-50
chr4-55155958-INS-82
chr4-55198151-INS-50
chr4-55226205-INS-288
chr4-55283807-INS-222
chr4-55285524-INS-140
chr4-55297981-INS-184
chr4-55388520-INS-165
chr4-55446237-INS-89
chr4-55463717-INS-175
chr4-55509769-INS-50
chr4-55561805-INS-6433
chr4-55582997-INS-109
chr4-55588510-INS-72
chr4-55591997-INS-54
chr4-55625786-INS-2135
chr4-55717881-INS-86
chr4-55761075-INS-187
chr4-55849584-INS-129
chr4-55908521-INS-339
chr4-55934960-INS-845
chr4-55940067-INS-226
chr4-55986233-INS-6620
chr4-56048220-INS-99
chr4-56096459-INS-305
chr4-56103871-INS-50
chr4-56199617-INS-56
chr4-56225710-INS-50
chr4-56239779-INS-113
chr4-56312623-INS-60
chr4-56330070-INS-6274
chr4-56336082-INS-51
chr4-56424674-INS-1227
chr4-56428977-INS-66
chr4-56444317-INS-274
chr4-56447852-INS-2056
chr4-56497660-INS-52
chr4-56501391-INS-989
chr4-56529063-INS-6418
chr4-56582394-INS-76
chr4-56625283-INS-127
chr4-56625351-INS-129
chr4-56652112-INS-52
chr4-56652117-INS-359
chr4-56692974-INS-1619
chr4-56693514-INS-50
chr4-56698041-INS-71
chr4-56698113-INS-256
chr4-56712522-INS-2334
chr4-56727999-INS-666
chr4-56728215-INS-54
chr4-56826875-INS-207
chr4-56828600-INS-111
chr4-56840345-INS-79
chr4-56842912-INS-242
chr4-56911169-INS-50
chr4-56939285-INS-229
chr4-56939883-INS-242
chr4-56960012-INS-57

chr4-56970654-INS-65
chr4-56975151-INS-805
chr4-57015879-INS-55
chr4-57034057-INS-83
chr4-57048291-INS-86
chr4-57112377-INS-4846
chr4-57182018-INS-59
chr4-57183461-INS-205
chr4-57199355-INS-151
chr4-57222914-INS-203
chr4-57283351-INS-137
chr4-57304816-INS-90
chr4-57312017-INS-60
chr4-57315029-INS-3074
chr4-57315194-INS-81
chr4-57337305-INS-50
chr4-57352056-INS-54
chr4-57365588-INS-231
chr4-57372107-INS-52
chr4-57376004-INS-204
chr4-57384197-INS-2985
chr4-57385012-INS-89
chr4-57387649-INS-1889
chr4-57410741-INS-52
chr4-57425147-INS-1484
chr4-57463914-INS-297
chr4-57523506-INS-732
chr4-57555727-INS-79
chr4-57555784-INS-179
chr4-57559894-INS-413
chr4-57611061-INS-79
chr4-57615302-INS-116
chr4-57619989-INS-119
chr4-57728364-INS-54
chr4-57773568-INS-66
chr4-57774488-INS-113
chr4-57774494-INS-56
chr4-57774495-INS-557
chr4-57779649-INS-215
chr4-57780359-INS-89
chr4-57799955-INS-99
chr4-57811521-INS-362
chr4-57811813-INS-425
chr4-57839042-INS-59
chr4-57846498-INS-53
chr4-57851615-INS-120
chr4-57864823-INS-211
chr4-57906441-INS-68
chr4-57920825-INS-52
chr4-57927518-INS-78
chr4-57927539-INS-157
chr4-57928465-INS-96
chr4-57936471-INS-66
chr4-57937019-INS-59
chr4-57997957-INS-497
chr4-58005421-INS-53
chr4-58026168-INS-115
chr4-58026507-INS-61
chr4-58030261-INS-583
chr4-58035372-INS-211
chr4-58036465-INS-6245
chr4-58061570-INS-223
chr4-58082351-INS-118
chr4-58097215-INS-108
chr4-58115233-INS-316
chr4-58148594-INS-3126
chr4-58151780-INS-172
chr4-58198330-INS-323
chr4-58199061-INS-395
chr4-58222936-INS-102
chr4-58223923-INS-50
chr4-58228304-INS-2595
chr4-58246205-INS-70
chr4-58251844-INS-188
chr4-58256940-INS-7066
chr4-58270549-INS-548
chr4-58274041-INS-131

chr4-58274041-INS-305
chr4-58275103-INS-6237
chr4-58276694-INS-462
chr4-58279361-INS-9718
chr4-58348597-INS-57
chr4-58380149-INS-51
chr4-58381812-INS-78
chr4-58385296-INS-66
chr4-58427003-INS-50
chr4-58484777-INS-77
chr4-58492297-INS-60
chr4-58505001-INS-507
chr4-58505017-INS-118
chr4-58579753-INS-66
chr4-58585398-INS-5006
chr4-58596912-INS-2337
chr4-58606400-INS-222
chr4-58616050-INS-193
chr4-58639823-INS-220
chr4-58659729-INS-377
chr4-58662752-INS-64
chr4-58677617-INS-51
chr4-58681836-INS-50
chr4-58763185-INS-4270
chr4-58765159-INS-50
chr4-58767388-INS-279
chr4-58780285-INS-432
chr4-58789278-INS-136
chr4-58906653-INS-84
chr4-58929358-INS-196
chr4-58997763-INS-105
chr4-59020768-INS-276
chr4-59067262-INS-135
chr4-59068117-INS-52
chr4-59107618-INS-507
chr4-59111815-INS-1151
chr4-59114638-INS-75
chr4-59127469-INS-1444
chr4-59127953-INS-203
chr4-59156906-INS-227
chr4-59163602-INS-194
chr4-59173901-INS-54
chr4-59180337-INS-2763
chr4-59198925-INS-83
chr4-59203226-INS-219
chr4-59207177-INS-79
chr4-59229510-INS-50
chr4-59242937-INS-12386
chr4-59270338-INS-1412
chr4-59280240-INS-201
chr4-59287871-INS-1671
chr4-59288112-INS-60
chr4-59291799-INS-56
chr4-59293075-INS-979
chr4-59301821-INS-197
chr4-59367225-INS-200
chr4-59397984-INS-118
chr4-59412738-INS-72
chr4-59414519-INS-60
chr4-59427047-INS-395
chr4-59431861-INS-6693
chr4-59445577-INS-50
chr4-59514973-INS-380
chr4-59538177-INS-505
chr4-59573352-INS-57
chr4-59573978-INS-96
chr4-59586014-INS-165
chr4-59586565-INS-202
chr4-59608395-INS-1532
chr4-59615901-INS-123
chr4-59638174-INS-52
chr4-59668611-INS-731
chr4-59687935-INS-92
chr4-59693372-INS-182
chr4-59712870-INS-208
chr4-59772814-INS-192
chr4-59773024-INS-286

chr4-59784639-INS-62
chr4-59788426-INS-1447
chr4-59790761-INS-223
chr4-59794970-INS-175
chr4-59827947-INS-153
chr4-59850874-INS-50
chr4-59872013-INS-50
chr4-59915778-INS-58
chr4-59974818-INS-4788
chr4-60001674-INS-2401
chr4-61226262-INS-50
chr4-61442120-INS-562
chr4-61627751-INS-1350
chr4-61646685-INS-54
chr4-61669165-INS-1002
chr4-61701346-INS-175
chr4-61743197-INS-220
chr4-61747132-INS-6396
chr4-62008955-INS-71
chr4-62014239-INS-1168
chr4-62030380-INS-358
chr4-62031424-INS-80
chr4-62171539-INS-54
chr4-62195716-INS-128
chr4-62196510-INS-363
chr4-62197314-INS-249
chr4-62266701-INS-6454
chr4-62285158-INS-352
chr4-62300668-INS-60
chr4-62329338-INS-217
chr4-62389536-INS-229
chr4-62413148-INS-16989
chr4-62413148-INS-7844
chr4-62432834-INS-24495
chr4-62434664-INS-4036
chr4-62434738-INS-50
chr4-62480731-INS-2930
chr4-62490502-INS-210
chr4-62505065-INS-223
chr4-62507795-INS-60
chr4-62592434-INS-77
chr4-62619925-INS-170
chr4-62703563-INS-236
chr4-62703811-INS-7085
chr4-62730030-INS-130
chr4-62765418-INS-480
chr4-62801212-INS-99
chr4-62806285-INS-512
chr4-62821460-INS-52
chr4-62823417-INS-84
chr4-62824313-INS-6676
chr4-62837591-INS-171
chr4-62851463-INS-67
chr4-62876256-INS-64
chr4-62933916-INS-106
chr4-62959714-INS-64
chr4-63025623-INS-4494
chr4-63049617-INS-459
chr4-63071613-INS-79
chr4-63076187-INS-178
chr4-63091382-INS-2004
chr4-63104115-INS-221
chr4-63107557-INS-72
chr4-63107611-INS-79
chr4-63127712-INS-68
chr4-63128522-INS-52
chr4-63171140-INS-70
chr4-63196262-INS-52
chr4-63273675-INS-11349
chr4-63277738-INS-5551
chr4-63304361-INS-393
chr4-63431702-INS-64
chr4-63442984-INS-8117
chr4-63467755-INS-1134
chr4-63481334-INS-188
chr4-63509304-INS-6336
chr4-63520133-INS-172

chr4-63538011-INS-6161
chr4-63553588-INS-50
chr4-63573177-INS-1466
chr4-63577866-INS-4053
chr4-63577866-INS-680
chr4-63581898-INS-152
chr4-63582619-INS-934
chr4-63596829-INS-6182
chr4-63610065-INS-2046
chr4-63612729-INS-206
chr4-63612839-INS-1759
chr4-63619974-INS-80
chr4-63625293-INS-2535
chr4-63699809-INS-758
chr4-63802848-INS-6683
chr4-63841021-INS-64
chr4-63913720-INS-54
chr4-63928126-INS-150
chr4-63928160-INS-74
chr4-64052309-INS-50
chr4-64070460-INS-6566
chr4-64080494-INS-54
chr4-64093748-INS-5545
chr4-64120883-INS-1180
chr4-64133692-INS-183
chr4-64155233-INS-50
chr4-64241387-INS-645
chr4-64253356-INS-50
chr4-64267818-INS-4491
chr4-64267967-INS-50
chr4-64319284-INS-135
chr4-64422636-INS-6824
chr4-64468577-INS-74
chr4-64480816-INS-84
chr4-64484823-INS-132
chr4-64497699-INS-4150
chr4-64562145-INS-6415
chr4-64610250-INS-299
chr4-64627800-INS-7083
chr4-64704244-INS-50
chr4-64717791-INS-190
chr4-64717818-INS-388
chr4-64721070-INS-300
chr4-64721126-INS-99
chr4-64724803-INS-180
chr4-64749812-INS-54
chr4-64778583-INS-121
chr4-64895975-INS-728
chr4-64897203-INS-1474
chr4-64915715-INS-78
chr4-64920256-INS-175
chr4-64930123-INS-1204
chr4-64930123-INS-2612
chr4-64941067-INS-532
chr4-64955875-INS-846
chr4-64981952-INS-6356
chr4-65013289-INS-50
chr4-65044646-INS-50
chr4-65145551-INS-6810
chr4-65155648-INS-103
chr4-65155720-INS-188
chr4-65190373-INS-211
chr4-65299177-INS-52
chr4-65405053-INS-96
chr4-65435254-INS-60
chr4-65491926-INS-137
chr4-65511278-INS-287
chr4-65573403-INS-62
chr4-65580828-INS-613
chr4-65605967-INS-79
chr4-65616907-INS-1713
chr4-65623169-INS-53
chr4-65629368-INS-51
chr4-65630327-INS-58
chr4-65633202-INS-74
chr4-65663394-INS-498
chr4-65675313-INS-6171

chr4-65689984-INS-157
chr4-65692979-INS-120
chr4-65709700-INS-70
chr4-65738247-INS-278
chr4-65817155-INS-6662
chr4-65843243-INS-78
chr4-65867090-INS-7955
chr4-65874067-INS-13466
chr4-65927784-INS-6426
chr4-66055509-INS-88
chr4-66077471-INS-382
chr4-66084650-INS-230
chr4-66105635-INS-54
chr4-66110305-INS-6173
chr4-66305919-INS-90
chr4-66330975-INS-144
chr4-66334466-INS-50
chr4-66343737-INS-54
chr4-66415968-INS-206
chr4-66415981-INS-6626
chr4-66501679-INS-611
chr4-66528342-INS-58
chr4-66633905-INS-28630
chr4-66662660-INS-58
chr4-66685152-INS-67
chr4-66713572-INS-3016
chr4-66725730-INS-6535
chr4-66814864-INS-163
chr4-66819712-INS-50
chr4-66907864-INS-6453
chr4-66935627-INS-160
chr4-67001615-INS-96
chr4-67021528-INS-110
chr4-67045940-INS-52
chr4-67082644-INS-392
chr4-67111181-INS-7097
chr4-67150484-INS-110
chr4-67196562-INS-13990
chr4-67315680-INS-258
chr4-67320679-INS-6751
chr4-67332593-INS-96
chr4-67340494-INS-1164
chr4-67389897-INS-72
chr4-67417423-INS-223
chr4-67473013-INS-2820
chr4-67478281-INS-157
chr4-67514545-INS-52
chr4-67517424-INS-1584
chr4-67542929-INS-60
chr4-67576520-INS-409
chr4-67607671-INS-58
chr4-67620736-INS-62
chr4-67620754-INS-137
chr4-67625903-INS-918
chr4-67638821-INS-299
chr4-67688275-INS-205
chr4-67780199-INS-219
chr4-67823714-INS-8337
chr4-67892264-INS-64
chr4-67894684-INS-54
chr4-68042687-INS-83
chr4-68070670-INS-6225
chr4-68078945-INS-6695
chr4-68144690-INS-3500
chr4-68152289-INS-6576
chr4-68156395-INS-2592
chr4-68170291-INS-3218
chr4-68202357-INS-52
chr4-68202964-INS-200
chr4-68247159-INS-6338
chr4-68321598-INS-302
chr4-68397973-INS-6656
chr4-68406147-INS-72
chr4-68428643-INS-2348
chr4-68434842-INS-50
chr4-68457937-INS-6309
chr4-68498346-INS-777

chr4-68498511-INS-1709
chr4-68688314-INS-82
chr4-68722022-INS-76
chr4-68763677-INS-63
chr4-68800098-INS-56
chr4-68813126-INS-515
chr4-68822620-INS-348
chr4-68893641-INS-728
chr4-68923512-INS-6283
chr4-68950373-INS-51
chr4-69010463-INS-609
chr4-69042486-INS-228
chr4-69074393-INS-50
chr4-69124218-INS-6253
chr4-69210186-INS-60
chr4-69249876-INS-50
chr4-69277590-INS-3911
chr4-69323351-INS-60
chr4-69555128-INS-7003
chr4-69564637-INS-3123
chr4-69683172-INS-72
chr4-69686781-INS-131
chr4-69759460-INS-50
chr4-69791676-INS-64
chr4-69874455-INS-6404
chr4-69879108-INS-527
chr4-69881542-INS-90
chr4-69932798-INS-834
chr4-69936980-INS-58
chr4-69973845-INS-1408
chr4-70139311-INS-52
chr4-70151160-INS-1566
chr4-70305441-INS-240
chr4-70311980-INS-7467
chr4-70322888-INS-1735
chr4-70383537-INS-55
chr4-70436661-INS-52
chr4-70482260-INS-84
chr4-70492078-INS-1181
chr4-70497714-INS-8206
chr4-70502714-INS-1299
chr4-70511693-INS-50
chr4-70524783-INS-2802
chr4-70542152-INS-14177
chr4-70549375-INS-3777
chr4-70567960-INS-195
chr4-70572599-INS-7270
chr4-70576449-INS-325
chr4-70589106-INS-2266
chr4-70601241-INS-4449
chr4-70627200-INS-195
chr4-70665249-INS-2971
chr4-70780244-INS-423
chr4-70784557-INS-499
chr4-70805167-INS-4808
chr4-70877270-INS-560
chr4-70888573-INS-63
chr4-70941147-INS-50
chr4-70957898-INS-54
chr4-71003659-INS-78
chr4-71003984-INS-206
chr4-71016275-INS-53
chr4-71019535-INS-1152
chr4-71038669-INS-119
chr4-71048512-INS-50
chr4-71054858-INS-3142
chr4-71130251-INS-4074
chr4-71165835-INS-4704
chr4-71309593-INS-142
chr4-71319266-INS-6522
chr4-71370036-INS-91
chr4-71379634-INS-3446
chr4-71434372-INS-1643
chr4-71607752-INS-1034
chr4-71616258-INS-2825
chr4-71638739-INS-64
chr4-71646088-INS-6679

chr4-71667063-INS-5878
chr4-71723418-INS-88
chr4-71723743-INS-625
chr4-71723748-INS-304
chr4-71738323-INS-799
chr4-71905976-INS-50
chr4-71906106-INS-115
chr4-71930603-INS-5289
chr4-71969047-INS-78
chr4-71974899-INS-4196
chr4-71975379-INS-6644
chr4-72023549-INS-3232
chr4-72145618-INS-7844
chr4-72163701-INS-4777
chr4-72164843-INS-1265
chr4-72216892-INS-136
chr4-72236605-INS-2516
chr4-72236605-INS-9329
chr4-72298839-INS-100
chr4-72405344-INS-2791
chr4-72461350-INS-201
chr4-72478951-INS-50
chr4-72573824-INS-400
chr4-72585312-INS-1265
chr4-72586644-INS-74
chr4-72671832-INS-50
chr4-72737968-INS-8603
chr4-72747969-INS-258
chr4-72748151-INS-87
chr4-72811533-INS-1437
chr4-72815364-INS-941
chr4-72828196-INS-108
chr4-72839681-INS-56
chr4-72923194-INS-3982
chr4-72925175-INS-886
chr4-72927958-INS-177
chr4-72928117-INS-358
chr4-72929350-INS-6307
chr4-72971786-INS-165
chr4-72983470-INS-6648
chr4-72997113-INS-63
chr4-73003573-INS-874
chr4-73032894-INS-1891
chr4-73039294-INS-6519
chr4-73077877-INS-374
chr4-73147319-INS-58
chr4-73147617-INS-4390
chr4-73192451-INS-199
chr4-73199846-INS-416
chr4-73265022-INS-2765
chr4-73268396-INS-2128
chr4-73325242-INS-721
chr4-73343686-INS-921
chr4-73365099-INS-145
chr4-73446216-INS-4542
chr4-73551881-INS-52
chr4-73567007-INS-56
chr4-73582107-INS-56
chr4-73705969-INS-123
chr4-73858782-INS-92
chr4-73882959-INS-835
chr4-73883405-INS-2312
chr4-73900937-INS-6660
chr4-73930519-INS-57
chr4-74044165-INS-221
chr4-74044971-INS-56
chr4-74058318-INS-1495
chr4-74063968-INS-381
chr4-74065324-INS-2146
chr4-74067340-INS-5974
chr4-74127841-INS-74
chr4-74137230-INS-778
chr4-74139141-INS-5980
chr4-74142821-INS-178
chr4-74148613-INS-119
chr4-74150969-INS-180
chr4-74152193-INS-1550

chr4-74179027-INS-214
chr4-74179041-INS-2967
chr4-74214225-INS-87
chr4-74266950-INS-198
chr4-74273021-INS-9513
chr4-74310190-INS-225
chr4-74389345-INS-55
chr4-74405586-INS-491
chr4-74431527-INS-80
chr4-74486489-INS-6781
chr4-74493939-INS-6799
chr4-74499399-INS-120
chr4-74504201-INS-52
chr4-74508813-INS-62
chr4-74542437-INS-63
chr4-74581859-INS-2384
chr4-74590670-INS-124
chr4-74679452-INS-50
chr4-74773579-INS-93
chr4-74773636-INS-446
chr4-74826615-INS-52
chr4-74827563-INS-56
chr4-74838444-INS-80
chr4-74886660-INS-50
chr4-74940027-INS-76
chr4-74944568-INS-1116
chr4-74954104-INS-4942
chr4-74974797-INS-211
chr4-75034862-INS-1382
chr4-75067428-INS-73
chr4-75101829-INS-1443
chr4-75142590-INS-62
chr4-75160026-INS-53
chr4-75162734-INS-482
chr4-75164069-INS-233
chr4-75186662-INS-296
chr4-75186699-INS-5311
chr4-75232726-INS-710
chr4-75272934-INS-201
chr4-75273818-INS-201
chr4-75342494-INS-50
chr4-75372543-INS-59
chr4-75376322-INS-6569
chr4-75387015-INS-102
chr4-75401309-INS-178
chr4-75434545-INS-92
chr4-75437016-INS-3157
chr4-75455765-INS-228
chr4-75467731-INS-60
chr4-75467822-INS-130
chr4-75487777-INS-88
chr4-75495946-INS-6995
chr4-75512846-INS-3031
chr4-75556801-INS-72
chr4-75567329-INS-104
chr4-75573583-INS-1494
chr4-75608710-INS-2079
chr4-75698366-INS-87
chr4-75713877-INS-261
chr4-75725186-INS-99
chr4-75725596-INS-936
chr4-75762988-INS-174
chr4-75811311-INS-79
chr4-76091391-INS-72
chr4-76154309-INS-247
chr4-76199376-INS-6238
chr4-76290324-INS-199
chr4-76290960-INS-61
chr4-76307392-INS-50
chr4-76317114-INS-84
chr4-76357975-INS-575
chr4-76392235-INS-50
chr4-76401277-INS-74
chr4-76410474-INS-199
chr4-76418168-INS-6243
chr4-76534634-INS-246
chr4-76584339-INS-50

chr4-76584382-INS-58
chr4-76585485-INS-1012
chr4-76618245-INS-148
chr4-76619883-INS-108
chr4-76638454-INS-50
chr4-76644144-INS-2754
chr4-76666700-INS-340
chr4-76679954-INS-6443
chr4-76729020-INS-62
chr4-76762480-INS-296
chr4-76779746-INS-56
chr4-76780461-INS-56
chr4-76786045-INS-1599
chr4-76893800-INS-60
chr4-76900651-INS-100
chr4-76913572-INS-252
chr4-76923843-INS-8574
chr4-76949116-INS-3646
chr4-77001770-INS-61
chr4-77018291-INS-64
chr4-77052351-INS-59
chr4-77059520-INS-6675
chr4-77074823-INS-2054
chr4-77127227-INS-58
chr4-77131703-INS-50
chr4-77146184-INS-55
chr4-77152044-INS-85
chr4-77153217-INS-148
chr4-77187728-INS-6801
chr4-77225522-INS-213
chr4-77263342-INS-501
chr4-77355553-INS-70
chr4-77424727-INS-176
chr4-77476455-INS-146
chr4-77539840-INS-3971
chr4-77588880-INS-1261
chr4-77614412-INS-364
chr4-77617036-INS-54
chr4-77647191-INS-349
chr4-77647191-INS-92
chr4-77709617-INS-6202
chr4-77721225-INS-132
chr4-77721509-INS-124
chr4-77727235-INS-248
chr4-77797728-INS-50
chr4-77809782-INS-1820
chr4-77812083-INS-308
chr4-77824428-INS-510
chr4-77829081-INS-50
chr4-77852580-INS-67
chr4-77856947-INS-88
chr4-77877936-INS-82
chr4-77885520-INS-54
chr4-77885720-INS-64
chr4-77924602-INS-234
chr4-77968686-INS-498
chr4-77970229-INS-1276
chr4-77975507-INS-245
chr4-78052596-INS-5182
chr4-78070187-INS-50
chr4-78146681-INS-51
chr4-78160983-INS-66
chr4-78174968-INS-339
chr4-78184427-INS-60
chr4-78184637-INS-256
chr4-78273670-INS-6230
chr4-78347983-INS-51
chr4-78375682-INS-180
chr4-78444698-INS-72
chr4-78454616-INS-6615
chr4-78467474-INS-276
chr4-78468341-INS-61
chr4-78477174-INS-100
chr4-78481444-INS-5370
chr4-78492405-INS-6307
chr4-78524787-INS-107
chr4-78538528-INS-50

chr4-78553675-INS-98
chr4-78563439-INS-58
chr4-78678181-INS-332
chr4-78678374-INS-6178
chr4-78707572-INS-174
chr4-78727686-INS-6449
chr4-78749159-INS-158
chr4-78840658-INS-6999
chr4-78861592-INS-50
chr4-78924936-INS-981
chr4-78939560-INS-145
chr4-79012920-INS-54
chr4-79047311-INS-59
chr4-79111317-INS-60
chr4-79209660-INS-1187
chr4-79213524-INS-50
chr4-79229452-INS-2248
chr4-79260479-INS-2931
chr4-79276481-INS-1287
chr4-79293578-INS-1050
chr4-79293578-INS-7733
chr4-79315690-INS-6828
chr4-79321276-INS-114
chr4-79357418-INS-56
chr4-79373663-INS-484
chr4-79395632-INS-803
chr4-79409107-INS-400
chr4-79521344-INS-134
chr4-79620603-INS-3018
chr4-79721306-INS-89
chr4-79727871-INS-4729
chr4-79741778-INS-57
chr4-79741802-INS-1491
chr4-79778337-INS-1256
chr4-79778337-INS-5551
chr4-79794207-INS-202
chr4-79823343-INS-385
chr4-79882935-INS-56
chr4-79895568-INS-6707
chr4-79907724-INS-209
chr4-79940994-INS-50
chr4-79957683-INS-1480
chr4-79981010-INS-204
chr4-79991781-INS-1535
chr4-80007635-INS-8865
chr4-80029604-INS-242
chr4-80040954-INS-80
chr4-80075211-INS-782
chr4-80096621-INS-72
chr4-80111432-INS-194
chr4-80139606-INS-329
chr4-80172369-INS-288
chr4-80189890-INS-53
chr4-80202573-INS-60
chr4-80222156-INS-221
chr4-80293562-INS-6451
chr4-80357769-INS-6626
chr4-80362475-INS-52
chr4-80436730-INS-6985
chr4-80459988-INS-3086
chr4-80462581-INS-84
chr4-80463082-INS-1338
chr4-80469951-INS-68
chr4-80469963-INS-1004
chr4-80523299-INS-52
chr4-80533820-INS-5939
chr4-80564295-INS-6401
chr4-80594861-INS-196
chr4-80606779-INS-104
chr4-80613245-INS-642
chr4-80613816-INS-50
chr4-80705470-INS-6394
chr4-80742361-INS-929
chr4-80797728-INS-74
chr4-80828080-INS-52
chr4-80828332-INS-90
chr4-80836918-INS-3118

chr4-80903803-INS-82
chr4-80925331-INS-182
chr4-80925748-INS-248
chr4-80936801-INS-6376
chr4-80954538-INS-144
chr4-80954538-INS-66
chr4-80963500-INS-56
chr4-81015060-INS-1940
chr4-81042930-INS-58
chr4-81046413-INS-6367
chr4-81057015-INS-113
chr4-81074693-INS-4185
chr4-81144199-INS-224
chr4-81187849-INS-388
chr4-81220843-INS-689
chr4-81284276-INS-742
chr4-81318259-INS-195
chr4-81367186-INS-1393
chr4-81385833-INS-104
chr4-81385833-INS-50
chr4-81389229-INS-296
chr4-81389684-INS-210
chr4-81419308-INS-163
chr4-81424346-INS-162
chr4-81442172-INS-464
chr4-81444697-INS-180
chr4-81445567-INS-50
chr4-81468658-INS-229
chr4-81501549-INS-265
chr4-81516117-INS-68
chr4-81548840-INS-5919
chr4-81601766-INS-498
chr4-81606536-INS-79
chr4-81627035-INS-55
chr4-81643021-INS-66
chr4-81647717-INS-52
chr4-81650625-INS-146
chr4-81650989-INS-239
chr4-81662726-INS-88
chr4-81753827-INS-201
chr4-81770454-INS-2732
chr4-81788533-INS-1553
chr4-81805921-INS-1275
chr4-81865192-INS-6441
chr4-81868536-INS-6880
chr4-81889678-INS-62
chr4-81890996-INS-6407
chr4-82001693-INS-7819
chr4-82022733-INS-204
chr4-82028565-INS-56
chr4-82045051-INS-161
chr4-82063774-INS-51
chr4-82129605-INS-66
chr4-82163686-INS-103
chr4-82171263-INS-84
chr4-82174422-INS-6440
chr4-82279793-INS-84
chr4-82368177-INS-50
chr4-82482473-INS-72
chr4-82493719-INS-50
chr4-82526690-INS-5138
chr4-82534624-INS-1830
chr4-82564306-INS-143
chr4-82574458-INS-6240
chr4-82600795-INS-90
chr4-82630342-INS-190
chr4-82635387-INS-143
chr4-82636565-INS-50
chr4-82687320-INS-296
chr4-82755038-INS-651
chr4-82842209-INS-273
chr4-82893106-INS-401
chr4-82893310-INS-549
chr4-82903514-INS-3022
chr4-82904954-INS-50
chr4-82911947-INS-51
chr4-82951776-INS-144

chr4-82981941-INS-82
chr4-83005602-INS-73
chr4-83025694-INS-5600
chr4-83065913-INS-725
chr4-83084043-INS-4030
chr4-83095333-INS-440
chr4-83129774-INS-233
chr4-83132037-INS-136
chr4-83170729-INS-393
chr4-83207899-INS-69
chr4-83208074-INS-596
chr4-83208139-INS-99
chr4-83208285-INS-1017
chr4-83239508-INS-120
chr4-83250891-INS-51
chr4-83277796-INS-236
chr4-83283555-INS-50
chr4-83317302-INS-7280
chr4-83328844-INS-252
chr4-83333744-INS-227
chr4-83334029-INS-52
chr4-83345648-INS-50
chr4-83415678-INS-216
chr4-83436289-INS-1751
chr4-83438055-INS-109
chr4-83462025-INS-319
chr4-83462053-INS-1425
chr4-83474027-INS-700
chr4-83485219-INS-508
chr4-83497981-INS-1019
chr4-83500225-INS-62
chr4-83500865-INS-76
chr4-83510229-INS-4938
chr4-83514856-INS-60
chr4-83523648-INS-140
chr4-83534409-INS-50
chr4-83582704-INS-53
chr4-83589895-INS-6201
chr4-83593749-INS-60
chr4-83628703-INS-4715
chr4-83631707-INS-2722
chr4-83643654-INS-1476
chr4-83692527-INS-308
chr4-83692568-INS-74
chr4-83715179-INS-54
chr4-83762568-INS-6468
chr4-83815462-INS-50
chr4-83858384-INS-50
chr4-83896780-INS-50
chr4-83899182-INS-76
chr4-83977095-INS-91
chr4-84029607-INS-237
chr4-84062946-INS-105
chr4-84063172-INS-192
chr4-84063172-INS-87
chr4-84138836-INS-52
chr4-84158497-INS-197
chr4-84251903-INS-90
chr4-84295979-INS-72
chr4-84337466-INS-2366
chr4-84391412-INS-250
chr4-84407850-INS-68
chr4-84428462-INS-92
chr4-84531934-INS-647
chr4-84537367-INS-56
chr4-84562447-INS-50
chr4-84628851-INS-88
chr4-84635310-INS-98
chr4-84659078-INS-503
chr4-84690019-INS-291
chr4-84698607-INS-228
chr4-84880351-INS-50
chr4-84880396-INS-130
chr4-84960196-INS-214
chr4-84976079-INS-714
chr4-85002263-INS-2976
chr4-85007960-INS-115

chr4-85099736-INS-6284
chr4-85129010-INS-87
chr4-85129215-INS-160
chr4-85140234-INS-213
chr4-85148755-INS-196
chr4-85151739-INS-68
chr4-85170244-INS-50
chr4-85171172-INS-54
chr4-85177397-INS-93
chr4-85177622-INS-204
chr4-85274511-INS-60
chr4-85303393-INS-110
chr4-85312095-INS-158
chr4-85319557-INS-1556
chr4-85324970-INS-60
chr4-85420421-INS-6074
chr4-85458863-INS-69
chr4-85463582-INS-1135
chr4-85470456-INS-7406
chr4-85490583-INS-2934
chr4-85503150-INS-3924
chr4-85514942-INS-1181
chr4-85524107-INS-180
chr4-85560365-INS-137
chr4-85562068-INS-689
chr4-85562204-INS-11218
chr4-85590215-INS-55
chr4-85603655-INS-1892
chr4-85619337-INS-88
chr4-85626256-INS-6677
chr4-85636451-INS-526
chr4-85698857-INS-1625
chr4-85707772-INS-6372
chr4-85751134-INS-462
chr4-85766817-INS-181
chr4-85814166-INS-2172
chr4-85839644-INS-2308
chr4-85862243-INS-88
chr4-85908766-INS-1524
chr4-85927577-INS-451
chr4-85927775-INS-62
chr4-85951766-INS-54
chr4-85966287-INS-115
chr4-86000941-INS-76
chr4-86012905-INS-52
chr4-86098640-INS-69
chr4-86117007-INS-54
chr4-86230409-INS-50
chr4-86265357-INS-427
chr4-86265637-INS-52
chr4-86356039-INS-88
chr4-86425342-INS-148
chr4-86455866-INS-404
chr4-86456758-INS-550
chr4-86481280-INS-74
chr4-86590055-INS-240
chr4-86596657-INS-104
chr4-86598570-INS-50
chr4-86608767-INS-60
chr4-86610644-INS-108
chr4-86610781-INS-50
chr4-86613604-INS-1010
chr4-86621054-INS-8772
chr4-86645145-INS-62
chr4-86652167-INS-73
chr4-86653995-INS-216
chr4-86654399-INS-166
chr4-86665744-INS-169
chr4-86729429-INS-70
chr4-86751043-INS-573
chr4-86783617-INS-9085
chr4-86799994-INS-54
chr4-86808081-INS-153
chr4-86816583-INS-160
chr4-86832127-INS-62
chr4-86838023-INS-2809
chr4-86861699-INS-218

chr4-86863630-INS-3418
chr4-86879328-INS-85
chr4-86886201-INS-3447
chr4-86891294-INS-6729
chr4-86892489-INS-4745
chr4-86896780-INS-1837
chr4-86899979-INS-62
chr4-86917898-INS-52
chr4-86921595-INS-70
chr4-86944599-INS-4788
chr4-86944858-INS-72
chr4-86970782-INS-51
chr4-86996941-INS-779
chr4-87048958-INS-6203
chr4-87057795-INS-1536
chr4-87062586-INS-935
chr4-87071737-INS-72
chr4-87087823-INS-50
chr4-87103798-INS-343
chr4-87135626-INS-54
chr4-87139725-INS-6412
chr4-87166125-INS-6214
chr4-87179104-INS-3260
chr4-87283384-INS-50
chr4-87284346-INS-62
chr4-87307237-INS-54
chr4-87321556-INS-164
chr4-87331915-INS-2581
chr4-87361929-INS-54
chr4-87524874-INS-54
chr4-87544987-INS-203
chr4-87548263-INS-84
chr4-87561800-INS-224
chr4-87565152-INS-109
chr4-87619191-INS-134
chr4-87647710-INS-76
chr4-87648610-INS-72
chr4-87733651-INS-55
chr4-87767462-INS-101
chr4-87878763-INS-50
chr4-87880692-INS-1163
chr4-87889868-INS-240
chr4-87889932-INS-80
chr4-87916126-INS-104
chr4-87952784-INS-117
chr4-87971847-INS-325
chr4-88048443-INS-144
chr4-88066525-INS-2417
chr4-88068627-INS-69
chr4-88091393-INS-80
chr4-88091819-INS-77
chr4-88168477-INS-11734
chr4-88179003-INS-154
chr4-88257879-INS-6477
chr4-88263146-INS-174
chr4-88308544-INS-103
chr4-88321204-INS-50
chr4-88417182-INS-1494
chr4-88417182-INS-3003
chr4-88505064-INS-7080
chr4-88516173-INS-144
chr4-88520489-INS-91
chr4-88550023-INS-96
chr4-88605138-INS-187
chr4-88605233-INS-431
chr4-88616721-INS-76
chr4-88665154-INS-50
chr4-88665400-INS-51
chr4-88725312-INS-502
chr4-88733024-INS-64
chr4-88771703-INS-55
chr4-88773711-INS-4478
chr4-88779011-INS-189
chr4-88792379-INS-3276
chr4-88800882-INS-310
chr4-88819121-INS-214
chr4-88911705-INS-115

chr4-88931525-INS-126
chr4-88968829-INS-4348
chr4-88981829-INS-80
chr4-88985038-INS-54
chr4-88985094-INS-6669
chr4-89046170-INS-57
chr4-89167100-INS-52
chr4-89175659-INS-64
chr4-89182937-INS-4947
chr4-89183144-INS-1523
chr4-89217532-INS-500
chr4-89222032-INS-56
chr4-89235037-INS-10645
chr4-89235037-INS-3541
chr4-89236620-INS-59
chr4-89237117-INS-204
chr4-89241198-INS-54
chr4-89253568-INS-389
chr4-89275979-INS-50
chr4-89276089-INS-122
chr4-89276117-INS-290
chr4-89297142-INS-66
chr4-89297996-INS-84
chr4-89305846-INS-68
chr4-89325262-INS-1892
chr4-89328482-INS-364
chr4-89348894-INS-318
chr4-89366246-INS-6731
chr4-89408513-INS-90
chr4-89434084-INS-112
chr4-89449880-INS-112
chr4-89452010-INS-51
chr4-89452926-INS-356
chr4-89461038-INS-216
chr4-89470336-INS-50
chr4-89472221-INS-102
chr4-89477743-INS-1901
chr4-89525839-INS-20746
chr4-89603468-INS-162
chr4-89628385-INS-184
chr4-89729478-INS-6224
chr4-89740239-INS-67
chr4-89776852-INS-62
chr4-89789565-INS-7057
chr4-89862457-INS-77
chr4-89918227-INS-7581
chr4-89963528-INS-6454
chr4-90047227-INS-139
chr4-90081207-INS-105
chr4-90092628-INS-6517
chr4-90121401-INS-343
chr4-90137860-INS-66
chr4-90167883-INS-1263
chr4-90198217-INS-52
chr4-90204711-INS-66
chr4-90238343-INS-6443
chr4-90257886-INS-1887
chr4-90268557-INS-54
chr4-90280499-INS-3482
chr4-90294638-INS-7415
chr4-90302199-INS-8149
chr4-90319270-INS-70
chr4-90319332-INS-182
chr4-90354253-INS-1349
chr4-90376867-INS-247
chr4-90378621-INS-93
chr4-90382315-INS-50
chr4-90394211-INS-1493
chr4-90418639-INS-1934
chr4-90419588-INS-643
chr4-90421925-INS-6486
chr4-90521114-INS-262
chr4-90570682-INS-6419
chr4-90641446-INS-6367
chr4-90669189-INS-54
chr4-90690122-INS-148
chr4-90696013-INS-5186

chr4-90776716-INS-116
chr4-90802353-INS-6585
chr4-90809787-INS-57
chr4-90827499-INS-4618
chr4-90835151-INS-50
chr4-90835906-INS-50
chr4-90867872-INS-636
chr4-90960188-INS-1388
chr4-91007945-INS-56
chr4-91096707-INS-500
chr4-91111287-INS-71
chr4-91167561-INS-161
chr4-91340968-INS-1864
chr4-91408217-INS-52
chr4-91414886-INS-6231
chr4-91429937-INS-50
chr4-91439959-INS-1874
chr4-91450721-INS-189
chr4-91480101-INS-235
chr4-91526435-INS-500
chr4-91599811-INS-184
chr4-91609465-INS-191
chr4-91625318-INS-3868
chr4-91651792-INS-50
chr4-91660001-INS-58
chr4-91805395-INS-72
chr4-91902853-INS-101
chr4-91904629-INS-525
chr4-91933347-INS-6365
chr4-91965091-INS-50
chr4-91996196-INS-1083
chr4-92047458-INS-2295
chr4-92047458-INS-861
chr4-92054912-INS-10069
chr4-92204167-INS-50
chr4-92207666-INS-50
chr4-92241795-INS-188
chr4-92267975-INS-847
chr4-92317911-INS-3675
chr4-92389298-INS-64
chr4-92405232-INS-412
chr4-92422155-INS-1516
chr4-92442615-INS-6874
chr4-92494163-INS-60
chr4-92494343-INS-88
chr4-92506900-INS-194
chr4-92574542-INS-55
chr4-92580386-INS-82
chr4-92600759-INS-236
chr4-92625605-INS-2026
chr4-92781868-INS-330
chr4-92854014-INS-229
chr4-92938101-INS-414
chr4-92970137-INS-1189
chr4-92970142-INS-128
chr4-93061773-INS-79
chr4-93112549-INS-7263
chr4-93113363-INS-114
chr4-93362931-INS-4504
chr4-93370687-INS-214
chr4-93371074-INS-120
chr4-93378324-INS-703
chr4-93397409-INS-64
chr4-93400036-INS-167
chr4-93426513-INS-111
chr4-93485714-INS-499
chr4-93490743-INS-54
chr4-93495587-INS-6571
chr4-93684240-INS-423
chr4-93684450-INS-80
chr4-93684996-INS-54
chr4-93715225-INS-51
chr4-93722707-INS-4176
chr4-93723028-INS-168
chr4-93757936-INS-2509
chr4-93765217-INS-194
chr4-93823855-INS-1587

chr4-93839399-INS-76
chr4-93890342-INS-72
chr4-93932653-INS-90
chr4-94018423-INS-414
chr4-94026661-INS-6612
chr4-94134269-INS-141
chr4-94134283-INS-501
chr4-94160083-INS-8609
chr4-94161705-INS-460
chr4-94203071-INS-1304
chr4-94228029-INS-6509
chr4-94243608-INS-1873
chr4-94276387-INS-922
chr4-94322114-INS-51
chr4-94352563-INS-210
chr4-94372322-INS-7249
chr4-94522549-INS-1189
chr4-94525414-INS-111
chr4-94539463-INS-60
chr4-94554415-INS-1706
chr4-94573225-INS-385
chr4-94573947-INS-480
chr4-94611234-INS-50
chr4-94634352-INS-556
chr4-94637289-INS-90
chr4-94670614-INS-1964
chr4-94821468-INS-131
chr4-94833037-INS-52
chr4-94858683-INS-183
chr4-94865351-INS-143
chr4-94883213-INS-7107
chr4-94940963-INS-60
chr4-94965568-INS-129
chr4-95082921-INS-73
chr4-95085923-INS-120
chr4-95111946-INS-251
chr4-95115875-INS-1887
chr4-95129412-INS-123
chr4-95171073-INS-52
chr4-95171441-INS-532
chr4-95174463-INS-6032
chr4-95199443-INS-96
chr4-95289864-INS-184
chr4-95362371-INS-50
chr4-95369112-INS-121
chr4-95387120-INS-1800
chr4-95448806-INS-6703
chr4-95457093-INS-6852
chr4-95505041-INS-165
chr4-95512958-INS-239
chr4-95536178-INS-58
chr4-95551720-INS-172
chr4-95562023-INS-210
chr4-95646352-INS-151
chr4-95646447-INS-54
chr4-95668137-INS-341
chr4-95767866-INS-7835
chr4-95813615-INS-316
chr4-95844918-INS-6184
chr4-95901946-INS-2163
chr4-95960369-INS-72
chr4-96059647-INS-450
chr4-96123259-INS-222
chr4-96133266-INS-76
chr4-96221253-INS-14943
chr4-96230431-INS-2347
chr4-96234316-INS-50
chr4-96245369-INS-60
chr4-96280531-INS-343
chr4-96282594-INS-445
chr4-96282746-INS-100
chr4-96388922-INS-52
chr4-96399521-INS-870
chr4-96399940-INS-1024
chr4-96449458-INS-72
chr4-96477571-INS-106
chr4-96477572-INS-235

chr4-96509232-INS-1529
chr4-96509389-INS-143
chr4-96551006-INS-105
chr4-96583022-INS-10756
chr4-96595288-INS-112
chr4-96776008-INS-56
chr4-96851651-INS-60
chr4-96908812-INS-492
chr4-96921670-INS-6371
chr4-96924203-INS-102
chr4-96953011-INS-1258
chr4-97023717-INS-5014
chr4-97041386-INS-84
chr4-97042497-INS-50
chr4-97079831-INS-7152
chr4-97082489-INS-76
chr4-97083146-INS-69
chr4-97098846-INS-72
chr4-97191943-INS-115
chr4-97265033-INS-5258
chr4-97277585-INS-18944
chr4-97280558-INS-6821
chr4-97282580-INS-278
chr4-97291334-INS-76
chr4-97331464-INS-334
chr4-97361626-INS-2916
chr4-97386469-INS-84
chr4-97386702-INS-266
chr4-97386702-INS-3338
chr4-97386735-INS-646
chr4-97400469-INS-85
chr4-97437256-INS-1264
chr4-97459826-INS-67
chr4-97507834-INS-93
chr4-97511864-INS-817
chr4-97556070-INS-87
chr4-97622228-INS-2860
chr4-97625848-INS-92
chr4-97748525-INS-50
chr4-97781788-INS-400
chr4-97808593-INS-50
chr4-97866396-INS-75
chr4-97908658-INS-462
chr4-97975837-INS-572
chr4-97977182-INS-260
chr4-97984721-INS-86
chr4-97998057-INS-579
chr4-97998143-INS-290
chr4-98011284-INS-56
chr4-98041708-INS-146
chr4-98115687-INS-3380
chr4-98130253-INS-128
chr4-98135341-INS-200
chr4-98138831-INS-50
chr4-98141001-INS-1142
chr4-98157904-INS-50
chr4-98192557-INS-1503
chr4-98222941-INS-200
chr4-98223896-INS-68
chr4-98223897-INS-415
chr4-98268251-INS-148
chr4-98308591-INS-90
chr4-98310693-INS-130
chr4-98366743-INS-74
chr4-98375582-INS-680
chr4-98421592-INS-340
chr4-98439983-INS-54
chr4-98469356-INS-386
chr4-98471040-INS-75
chr4-98492788-INS-639
chr4-98502433-INS-1198
chr4-98511091-INS-953
chr4-98517665-INS-261
chr4-98538798-INS-55
chr4-98569601-INS-56
chr4-98597007-INS-198
chr4-98602173-INS-6473

chr4-98609813-INS-141
chr4-98614089-INS-108
chr4-98631999-INS-56
chr4-98633196-INS-3601
chr4-98642032-INS-199
chr4-98649012-INS-207
chr4-98681350-INS-201
chr4-98687114-INS-1534
chr4-98687129-INS-211
chr4-98709062-INS-70
chr4-98710140-INS-186
chr4-98711678-INS-57
chr4-98714628-INS-144
chr4-98716033-INS-160
chr4-98749930-INS-60
chr4-98750476-INS-173
chr4-98756293-INS-52
chr4-98784283-INS-80
chr4-98797518-INS-209
chr4-98808105-INS-197
chr4-98832279-INS-199
chr4-98843808-INS-255
chr4-98853335-INS-68
chr4-98866381-INS-1536
chr4-98900203-INS-72
chr4-98949969-INS-66
chr4-98980507-INS-113
chr4-98986699-INS-1296
chr4-99004399-INS-61
chr4-99025509-INS-211
chr4-99031704-INS-103
chr4-99053666-INS-128
chr4-99054211-INS-88
chr4-99059261-INS-56
chr4-99059410-INS-132
chr4-99077813-INS-333
chr4-99086375-INS-205
chr4-99089571-INS-83
chr4-99091747-INS-65
chr4-99094240-INS-479
chr4-99169241-INS-53
chr4-99232183-INS-156
chr4-99235152-INS-140
chr4-99235238-INS-72
chr4-99246406-INS-50
chr4-99296552-INS-234
chr4-99298183-INS-516
chr4-99307554-INS-164
chr4-99308393-INS-59
chr4-99312517-INS-83
chr4-99343770-INS-115
chr4-99413744-INS-76
chr4-99420662-INS-228
chr4-99470891-INS-908
chr4-99478382-INS-2358
chr4-99493139-INS-220
chr4-99511498-INS-66
chr4-99532565-INS-57
chr4-99558276-INS-72
chr4-99565444-INS-8669
chr4-99595733-INS-61
chr4-99596539-INS-92
chr4-99597064-INS-1018
chr4-99629387-INS-88
chr4-99633916-INS-61
chr4-99644289-INS-225
chr4-99670511-INS-219
chr4-99691707-INS-143
chr4-99693318-INS-1149
chr4-99713256-INS-61
chr4-99729382-INS-214
chr4-99752774-INS-172
chr4-99761045-INS-202
chr4-99796140-INS-50
chr4-99841649-INS-54
chr4-99856316-INS-1698
chr4-99904023-INS-300

chr4-99948432-INS-661
chr4-100009177-INS-59
chr4-100010564-INS-104
chr4-100036993-INS-53
chr4-100167399-INS-175
chr4-100172252-INS-75
chr4-100176807-INS-192
chr4-100199087-INS-201
chr4-100213265-INS-7230
chr4-100216370-INS-52
chr4-100244379-INS-273
chr4-100281475-INS-6312
chr4-100297944-INS-401
chr4-100317195-INS-3041
chr4-100358945-INS-58
chr4-100443084-INS-3199
chr4-100447163-INS-94
chr4-100447881-INS-54
chr4-100460149-INS-198
chr4-100519928-INS-94
chr4-100551993-INS-127
chr4-100558867-INS-8677
chr4-100565117-INS-52
chr4-100570434-INS-60
chr4-100572715-INS-518
chr4-100575582-INS-61
chr4-100584956-INS-209
chr4-100648904-INS-202
chr4-100663196-INS-208
chr4-100717006-INS-7839
chr4-100726726-INS-50
chr4-100732031-INS-66
chr4-100774661-INS-56
chr4-100778654-INS-3135
chr4-100805416-INS-50
chr4-100839894-INS-100
chr4-100846861-INS-52
chr4-100882162-INS-51
chr4-100893098-INS-3137
chr4-100898114-INS-154
chr4-100911878-INS-201
chr4-100947189-INS-204
chr4-100998399-INS-72
chr4-101092376-INS-354
chr4-101122840-INS-209
chr4-101145102-INS-192
chr4-101169813-INS-1890
chr4-101171430-INS-92
chr4-101172566-INS-55
chr4-101220428-INS-60
chr4-101220791-INS-199
chr4-101235201-INS-616
chr4-101241408-INS-228
chr4-101243672-INS-76
chr4-101243960-INS-300
chr4-101244068-INS-68
chr4-101261648-INS-205
chr4-101289942-INS-206
chr4-101290967-INS-56
chr4-101311918-INS-280
chr4-101336582-INS-4653
chr4-101337613-INS-201
chr4-101342168-INS-174
chr4-101350573-INS-113
chr4-101392664-INS-51
chr4-101397373-INS-85
chr4-101487909-INS-63
chr4-101517917-INS-468
chr4-101555817-INS-119
chr4-101570306-INS-126
chr4-101570509-INS-98
chr4-101572487-INS-50
chr4-101573473-INS-60
chr4-101578063-INS-2799
chr4-101609629-INS-1200
chr4-101609641-INS-325
chr4-101627477-INS-2088

chr4-101636068-INS-50
chr4-101688512-INS-50
chr4-101697507-INS-51
chr4-101754875-INS-14282
chr4-101788036-INS-269
chr4-101801757-INS-231
chr4-101803347-INS-16221
chr4-101849688-INS-2982
chr4-101879635-INS-6779
chr4-101912962-INS-6677
chr4-101980743-INS-2063
chr4-101986740-INS-50
chr4-102055711-INS-1552
chr4-102061431-INS-56
chr4-102063041-INS-78
chr4-102063185-INS-179
chr4-102069390-INS-6175
chr4-102072078-INS-377
chr4-102092356-INS-6587
chr4-102237888-INS-964
chr4-102238293-INS-477
chr4-102281250-INS-238
chr4-102312919-INS-106
chr4-102318175-INS-108
chr4-102322921-INS-50
chr4-102351308-INS-122
chr4-102351537-INS-603
chr4-102490117-INS-243
chr4-102499479-INS-67
chr4-102522169-INS-117
chr4-102576080-INS-6179
chr4-102591251-INS-111
chr4-102595821-INS-75
chr4-102634359-INS-225
chr4-102654377-INS-66
chr4-102667858-INS-487
chr4-102676619-INS-110
chr4-102684844-INS-6454
chr4-102692436-INS-105
chr4-102698735-INS-171
chr4-102722624-INS-193
chr4-102818145-INS-504
chr4-102874695-INS-252
chr4-102934173-INS-818
chr4-102949159-INS-2738
chr4-103019116-INS-54
chr4-103050963-INS-450
chr4-103059432-INS-96
chr4-103078174-INS-60
chr4-103123343-INS-86
chr4-103175910-INS-55
chr4-103181575-INS-237
chr4-103197433-INS-92
chr4-103199790-INS-118
chr4-103241498-INS-124
chr4-103289658-INS-92
chr4-103401959-INS-459
chr4-103422590-INS-56
chr4-103422610-INS-133
chr4-103551953-INS-50
chr4-103554214-INS-52
chr4-103628177-INS-133
chr4-103628177-INS-61
chr4-103655208-INS-204
chr4-103657770-INS-133
chr4-103657830-INS-55
chr4-103673954-INS-909
chr4-103702408-INS-248
chr4-103741056-INS-68
chr4-103758187-INS-941
chr4-103802879-INS-110
chr4-103860320-INS-503
chr4-103881780-INS-54
chr4-103885878-INS-214
chr4-104032480-INS-88
chr4-104046095-INS-6386
chr4-104060931-INS-60

chr4-104080942-INS-338
chr4-104103377-INS-6456
chr4-104114275-INS-50
chr4-104182699-INS-2451
chr4-104191067-INS-117
chr4-104322139-INS-123
chr4-104325673-INS-63
chr4-104344134-INS-245
chr4-104363958-INS-54
chr4-104505944-INS-173
chr4-104505944-INS-83
chr4-104506090-INS-480
chr4-104532356-INS-1102
chr4-104629432-INS-6428
chr4-104738082-INS-108
chr4-104746403-INS-54
chr4-104763467-INS-85
chr4-104823370-INS-408
chr4-104831152-INS-1327
chr4-104863558-INS-276
chr4-104863571-INS-52
chr4-104880239-INS-82
chr4-104884183-INS-62
chr4-104900173-INS-185
chr4-104900175-INS-76
chr4-104939926-INS-83
chr4-104940507-INS-188
chr4-104999161-INS-7254
chr4-105004174-INS-55
chr4-105109765-INS-56
chr4-105110636-INS-186
chr4-105160514-INS-82
chr4-105194195-INS-50
chr4-105200851-INS-118
chr4-105208395-INS-74
chr4-105213637-INS-56
chr4-105221192-INS-550
chr4-105230388-INS-92
chr4-105259639-INS-366
chr4-105268240-INS-124
chr4-105268991-INS-102
chr4-105282890-INS-1025
chr4-105305972-INS-51
chr4-105423620-INS-3824
chr4-105444812-INS-3489
chr4-105526439-INS-6500
chr4-105528555-INS-52
chr4-105550032-INS-400
chr4-105645221-INS-1980
chr4-105649787-INS-54
chr4-105702315-INS-50
chr4-105714094-INS-59
chr4-105775418-INS-276
chr4-105814152-INS-125
chr4-105863549-INS-54
chr4-105910274-INS-6388
chr4-105949519-INS-5895
chr4-105969761-INS-53
chr4-106003187-INS-76
chr4-106038007-INS-2221
chr4-106059097-INS-403
chr4-106059145-INS-121
chr4-106074767-INS-112
chr4-106074992-INS-326
chr4-106076035-INS-974
chr4-106103815-INS-61
chr4-106107247-INS-69
chr4-106119015-INS-66
chr4-106143972-INS-540
chr4-106145659-INS-87
chr4-106160911-INS-167
chr4-106235636-INS-240
chr4-106277314-INS-76
chr4-106317268-INS-113
chr4-106317350-INS-271
chr4-106317561-INS-228
chr4-106322939-INS-89

chr4-106351103-INS-1892
chr4-106408857-INS-64
chr4-106441521-INS-90
chr4-106456604-INS-112
chr4-106483585-INS-124
chr4-106483647-INS-59
chr4-106541084-INS-50
chr4-106547162-INS-251
chr4-106575517-INS-383
chr4-106583855-INS-199
chr4-106599013-INS-104
chr4-106607467-INS-290
chr4-106607467-INS-676
chr4-106631656-INS-84
chr4-106631727-INS-571
chr4-106634162-INS-128
chr4-106679932-INS-122
chr4-106683295-INS-247
chr4-106688949-INS-70
chr4-106725859-INS-2941
chr4-106732946-INS-63
chr4-106750824-INS-241
chr4-106753834-INS-87
chr4-106759092-INS-464
chr4-106797568-INS-52
chr4-106828662-INS-64
chr4-106917185-INS-925
chr4-106947835-INS-116
chr4-107028893-INS-372
chr4-107028976-INS-1115
chr4-107056312-INS-57
chr4-107062913-INS-78
chr4-107099228-INS-420
chr4-107102190-INS-2833
chr4-107107430-INS-223
chr4-107107761-INS-203
chr4-107127880-INS-6475
chr4-107139650-INS-119
chr4-107140485-INS-363
chr4-107153056-INS-72
chr4-107177674-INS-2888
chr4-107214404-INS-202
chr4-107215330-INS-204
chr4-107220658-INS-202
chr4-107234501-INS-621
chr4-107242440-INS-176
chr4-107253115-INS-90
chr4-107289571-INS-71
chr4-107337407-INS-868
chr4-107419026-INS-88
chr4-107432408-INS-51
chr4-107432436-INS-357
chr4-107432510-INS-153
chr4-107441711-INS-106
chr4-107450587-INS-481
chr4-107463819-INS-68
chr4-107498353-INS-440
chr4-107514421-INS-72
chr4-107553643-INS-68
chr4-107606638-INS-3281
chr4-107619505-INS-7152
chr4-107740412-INS-204
chr4-107760906-INS-52
chr4-107772190-INS-52
chr4-107773777-INS-224
chr4-107793328-INS-67
chr4-107793545-INS-82
chr4-107796089-INS-2459
chr4-107831014-INS-95
chr4-107834880-INS-297
chr4-107841356-INS-50
chr4-107854854-INS-56
chr4-107911755-INS-206
chr4-107921963-INS-68
chr4-107929008-INS-132
chr4-107932939-INS-55
chr4-107933153-INS-291

chr4-107937911-INS-520
chr4-107937913-INS-78
chr4-107937968-INS-206
chr4-107959810-INS-209
chr4-107960303-INS-528
chr4-107967157-INS-50
chr4-107974584-INS-190
chr4-107977285-INS-200
chr4-108020247-INS-213
chr4-108026429-INS-116
chr4-108059516-INS-234
chr4-108060421-INS-225
chr4-108076820-INS-195
chr4-108085141-INS-209
chr4-108100046-INS-76
chr4-108109638-INS-205
chr4-108114068-INS-137
chr4-108119110-INS-508
chr4-108155882-INS-77
chr4-108173222-INS-84
chr4-108176798-INS-63
chr4-108177262-INS-199
chr4-108224240-INS-5721
chr4-108224252-INS-52
chr4-108228139-INS-202
chr4-108229497-INS-214
chr4-108236010-INS-54
chr4-108252453-INS-68
chr4-108303031-INS-379
chr4-108322908-INS-185
chr4-108388353-INS-259
chr4-108442017-INS-704
chr4-108444972-INS-52
chr4-108451405-INS-50
chr4-108518216-INS-214
chr4-108541358-INS-209
chr4-108543069-INS-201
chr4-108543763-INS-105
chr4-108629687-INS-212
chr4-108634560-INS-60
chr4-108634961-INS-209
chr4-108646080-INS-162
chr4-108653534-INS-414
chr4-108656177-INS-169
chr4-108728658-INS-7155
chr4-108744601-INS-76
chr4-108751570-INS-203
chr4-108824517-INS-84
chr4-108832813-INS-1145
chr4-108834973-INS-354
chr4-108835658-INS-456
chr4-108839416-INS-53
chr4-108847563-INS-201
chr4-108855461-INS-13426
chr4-108856678-INS-201
chr4-108878897-INS-201
chr4-108882505-INS-57
chr4-108882595-INS-50
chr4-108886921-INS-152
chr4-108908569-INS-93
chr4-108953669-INS-179
chr4-109015383-INS-61
chr4-109060185-INS-214
chr4-109080548-INS-8740
chr4-109081736-INS-144
chr4-109092935-INS-51
chr4-109144631-INS-320
chr4-109148534-INS-213
chr4-109175893-INS-703
chr4-109238237-INS-212
chr4-109254915-INS-96
chr4-109286509-INS-257
chr4-109289993-INS-107
chr4-109352388-INS-83
chr4-109362051-INS-202
chr4-109368307-INS-194
chr4-109373277-INS-4298

chr4-109374499-INS-110
chr4-109382934-INS-122
chr4-109395088-INS-252
chr4-109405029-INS-134
chr4-109428648-INS-55
chr4-109446654-INS-156
chr4-109448041-INS-71
chr4-109486075-INS-57
chr4-109508131-INS-490
chr4-109508529-INS-549
chr4-109553331-INS-9208
chr4-109558536-INS-167
chr4-109576541-INS-207
chr4-109580962-INS-254
chr4-109595064-INS-945
chr4-109656190-INS-52
chr4-109753387-INS-64
chr4-109803121-INS-483
chr4-109848998-INS-61
chr4-109862638-INS-3001
chr4-109863258-INS-6591
chr4-109886185-INS-59
chr4-109901939-INS-84
chr4-109979553-INS-50
chr4-109989616-INS-133
chr4-110023261-INS-6000
chr4-110080869-INS-261
chr4-110109962-INS-50
chr4-110152949-INS-655
chr4-110233416-INS-1994
chr4-110238876-INS-6371
chr4-110290529-INS-50
chr4-110377705-INS-800
chr4-110378351-INS-1077
chr4-110390213-INS-56
chr4-110409356-INS-113
chr4-110447156-INS-3638
chr4-110469613-INS-201
chr4-110493394-INS-463
chr4-110532004-INS-2565
chr4-110532669-INS-430
chr4-110533313-INS-446
chr4-110539676-INS-188
chr4-110574964-INS-214
chr4-110617000-INS-62
chr4-110624739-INS-268
chr4-110659438-INS-551
chr4-110659760-INS-722
chr4-110764697-INS-466
chr4-110764777-INS-186
chr4-110782521-INS-1477
chr4-110853212-INS-8707
chr4-110892385-INS-4826
chr4-110931627-INS-70
chr4-110988753-INS-73
chr4-111037247-INS-2182
chr4-111045692-INS-484
chr4-111083723-INS-552
chr4-111093876-INS-273
chr4-111175963-INS-50
chr4-111286086-INS-51
chr4-111296449-INS-536
chr4-111319115-INS-57
chr4-111328222-INS-431
chr4-111351815-INS-4424
chr4-111421681-INS-751
chr4-111427735-INS-56
chr4-111461817-INS-82
chr4-111467613-INS-7310
chr4-111606317-INS-157
chr4-111698672-INS-1127
chr4-111745582-INS-170
chr4-111777570-INS-572
chr4-111777882-INS-90
chr4-111777940-INS-677
chr4-111877596-INS-50
chr4-111912126-INS-588

chr4-112473073-INS-119
chr4-112474126-INS-217
chr4-112557194-INS-991
chr4-112590964-INS-60
chr4-112628361-INS-346
chr4-112638676-INS-9771
chr4-112641365-INS-2111
chr4-112641673-INS-64
chr4-112655884-INS-54
chr4-112666811-INS-346
chr4-113283268-INS-294
chr4-113288329-INS-100
chr4-113299436-INS-199
chr4-113317097-INS-52
chr4-113485525-INS-580
chr4-113486783-INS-500
chr4-113490387-INS-613
chr4-113494995-INS-52
chr4-113499421-INS-298
chr4-113502517-INS-156
chr4-113511324-INS-349
chr4-113513484-INS-87
chr4-113518529-INS-70
chr4-113523859-INS-65
chr4-113524097-INS-188
chr4-113530026-INS-64
chr4-113532906-INS-186
chr4-113535018-INS-900
chr4-113549020-INS-86
chr4-113570866-INS-156
chr4-113579209-INS-56
chr4-113579219-INS-187
chr4-113624761-INS-52
chr4-113626726-INS-62
chr4-113635708-INS-56
chr4-113637380-INS-347
chr4-113682071-INS-163
chr4-113758353-INS-120
chr4-113761554-INS-107
chr4-113762678-INS-351
chr4-113798684-INS-145
chr4-113805183-INS-118
chr4-113811919-INS-1004
chr4-113814949-INS-64
chr4-113831829-INS-3166
chr4-113851482-INS-3712
chr4-113862593-INS-1666
chr4-113875144-INS-2625
chr4-113885955-INS-1804
chr4-113898500-INS-2772
chr4-113922278-INS-287
chr4-113984518-INS-50
chr4-114015761-INS-58
chr4-114024854-INS-60
chr4-114047225-INS-334
chr4-114047440-INS-55
chr4-114056219-INS-235
chr4-114159763-INS-699
chr4-114165342-INS-59
chr4-114169624-INS-197
chr4-114182614-INS-1121
chr4-114215080-INS-53
chr4-114250027-INS-112
chr4-114262034-INS-60
chr4-114272171-INS-118
chr4-114342637-INS-50
chr4-114352390-INS-53
chr4-114494906-INS-53
chr4-114527431-INS-152
chr4-114584390-INS-50
chr4-114625315-INS-50
chr4-114664452-INS-106
chr4-114664655-INS-53
chr4-114665195-INS-148
chr4-114674072-INS-7242
chr4-114811157-INS-95
chr4-114832477-INS-64

chr4-114879721-INS-363
chr4-114910590-INS-70
chr4-114962756-INS-811
chr4-114972310-INS-150
chr4-114973713-INS-192
chr4-114973806-INS-78
chr4-115043549-INS-6298
chr4-115169068-INS-16327
chr4-115180207-INS-640
chr4-115181784-INS-62
chr4-115241862-INS-395
chr4-115319045-INS-3105
chr4-115366789-INS-874
chr4-115399894-INS-396
chr4-115434314-INS-485
chr4-115440984-INS-124
chr4-115451049-INS-183
chr4-115485733-INS-4481
chr4-115510819-INS-53
chr4-115510865-INS-139
chr4-115525521-INS-6395
chr4-115541088-INS-6109
chr4-115558355-INS-281
chr4-115588260-INS-50
chr4-115724583-INS-494
chr4-115770076-INS-52
chr4-115776687-INS-154
chr4-115972514-INS-181
chr4-115973214-INS-57
chr4-116057472-INS-213
chr4-116065414-INS-88
chr4-116095062-INS-72
chr4-116095127-INS-113
chr4-116129313-INS-73
chr4-116154690-INS-689
chr4-116202350-INS-227
chr4-116296594-INS-301
chr4-116296627-INS-120
chr4-116317714-INS-886
chr4-116375380-INS-86
chr4-116387119-INS-2349
chr4-116446619-INS-197
chr4-116471672-INS-203
chr4-116472275-INS-100
chr4-116515813-INS-2932
chr4-116519332-INS-211
chr4-116535354-INS-206
chr4-116545557-INS-318
chr4-116598242-INS-131
chr4-116601031-INS-198
chr4-116601831-INS-386
chr4-116604413-INS-56
chr4-116605768-INS-177
chr4-116636230-INS-123
chr4-116655474-INS-241
chr4-116668177-INS-1990
chr4-116672255-INS-420
chr4-116672378-INS-60
chr4-116705373-INS-1036
chr4-116715097-INS-54
chr4-116715423-INS-145
chr4-116743445-INS-244
chr4-116751700-INS-249
chr4-116751700-INS-52
chr4-116765901-INS-135
chr4-116795172-INS-308
chr4-116808609-INS-82
chr4-116813229-INS-1815
chr4-116826866-INS-192
chr4-116834845-INS-237
chr4-116835679-INS-121
chr4-116897065-INS-210
chr4-116901484-INS-56
chr4-116903513-INS-51
chr4-116927163-INS-217
chr4-116941328-INS-4462
chr4-116964495-INS-301

chr4-116970198-INS-183
chr4-116970933-INS-1247
chr4-117001047-INS-173
chr4-117014628-INS-442
chr4-117026850-INS-238
chr4-117028778-INS-53
chr4-117041762-INS-230
chr4-117045479-INS-184
chr4-117082301-INS-90
chr4-117088450-INS-52
chr4-117102194-INS-167
chr4-117189345-INS-94
chr4-117287757-INS-61
chr4-117313892-INS-52
chr4-117317139-INS-84
chr4-117317933-INS-117
chr4-117326431-INS-68
chr4-117364602-INS-70
chr4-117364717-INS-63
chr4-117366249-INS-120
chr4-117370070-INS-82
chr4-117372420-INS-64
chr4-117373356-INS-54
chr4-117374595-INS-189
chr4-117434638-INS-102
chr4-117627702-INS-195
chr4-117652174-INS-70
chr4-117654712-INS-5863
chr4-117661652-INS-55
chr4-117829447-INS-52
chr4-117830207-INS-143
chr4-117869898-INS-325
chr4-117870418-INS-606
chr4-117876421-INS-1368
chr4-117888024-INS-230
chr4-117918037-INS-103
chr4-117919072-INS-299
chr4-117959322-INS-412
chr4-117959933-INS-143
chr4-117962019-INS-50
chr4-118051652-INS-160
chr4-118052348-INS-196
chr4-118261028-INS-93
chr4-118261887-INS-291
chr4-118306840-INS-148
chr4-118370026-INS-50
chr4-118393255-INS-6457
chr4-118406683-INS-1214
chr4-118416540-INS-2567
chr4-118493999-INS-2913
chr4-118499195-INS-178
chr4-118499229-INS-866
chr4-118499238-INS-68
chr4-118509305-INS-202
chr4-118518317-INS-83
chr4-118550066-INS-8496
chr4-118659356-INS-3737
chr4-118676169-INS-160
chr4-118690544-INS-51
chr4-118710611-INS-1893
chr4-118738008-INS-60
chr4-118767111-INS-5483
chr4-118787727-INS-5010
chr4-118797660-INS-403
chr4-118810757-INS-158
chr4-118827941-INS-222
chr4-118832707-INS-125
chr4-118835734-INS-52
chr4-118846452-INS-1069
chr4-118849109-INS-684
chr4-118873220-INS-216
chr4-118873822-INS-74
chr4-118900054-INS-145
chr4-118913469-INS-94
chr4-118913519-INS-208
chr4-118920006-INS-299
chr4-118920192-INS-223

chr4-119060046-INS-238
chr4-119121727-INS-50
chr4-119191867-INS-64
chr4-119213618-INS-120
chr4-119238640-INS-199
chr4-119238819-INS-113
chr4-119273299-INS-96
chr4-119341217-INS-50
chr4-119371310-INS-359
chr4-119486963-INS-75
chr4-119563604-INS-211
chr4-119570887-INS-132
chr4-119571548-INS-110
chr4-119613892-INS-1461
chr4-119614256-INS-249
chr4-119758379-INS-51
chr4-119760591-INS-66
chr4-119837060-INS-52
chr4-119842637-INS-50
chr4-119861072-INS-245
chr4-119931940-INS-293
chr4-119932056-INS-108
chr4-119936751-INS-116
chr4-119950357-INS-124
chr4-119958171-INS-68
chr4-119964197-INS-54
chr4-120008615-INS-464
chr4-120033462-INS-64
chr4-120072708-INS-610
chr4-120159055-INS-57
chr4-120165364-INS-60
chr4-120299971-INS-4000
chr4-120359822-INS-219
chr4-120380448-INS-234
chr4-120395818-INS-60
chr4-120404813-INS-79
chr4-120454697-INS-118
chr4-120484448-INS-50
chr4-120510770-INS-158
chr4-120513111-INS-70
chr4-120534847-INS-428
chr4-120534947-INS-207
chr4-120537986-INS-832
chr4-120623843-INS-117
chr4-120719364-INS-274
chr4-120758517-INS-756
chr4-120759878-INS-166
chr4-120773034-INS-82
chr4-120773761-INS-72
chr4-120801406-INS-116
chr4-120815943-INS-203
chr4-120833786-INS-117
chr4-120847969-INS-5543
chr4-120850562-INS-389
chr4-120862768-INS-58
chr4-121146810-INS-795
chr4-121162678-INS-813
chr4-121266534-INS-981
chr4-121272902-INS-633
chr4-121274964-INS-1630
chr4-121279210-INS-172
chr4-121301465-INS-234
chr4-121310897-INS-64
chr4-121313395-INS-56
chr4-121362524-INS-2200
chr4-121364534-INS-50
chr4-121376396-INS-645
chr4-121378722-INS-1889
chr4-121391093-INS-59
chr4-121431551-INS-6689
chr4-121435597-INS-79
chr4-121444882-INS-50
chr4-121451898-INS-3847
chr4-121453354-INS-50
chr4-121472675-INS-1604
chr4-121483684-INS-179
chr4-121492836-INS-779

chr4-121509805-INS-63
chr4-121511160-INS-2717
chr4-121522311-INS-50
chr4-121523921-INS-1966
chr4-121536190-INS-3324
chr4-121539063-INS-6451
chr4-121546939-INS-56
chr4-121548933-INS-1543
chr4-121549556-INS-64
chr4-121550701-INS-50
chr4-121574284-INS-62
chr4-121576742-INS-60
chr4-121578029-INS-106
chr4-121584667-INS-1075
chr4-121584878-INS-278
chr4-121585378-INS-91
chr4-121630915-INS-96
chr4-122412437-INS-660
chr4-122438358-INS-273
chr4-122458816-INS-84
chr4-122459422-INS-72
chr4-122460985-INS-72
chr4-122461536-INS-448
chr4-122466632-INS-654
chr4-122478952-INS-63
chr4-122491515-INS-2339
chr4-122623859-INS-114
chr4-122637233-INS-66
chr4-122637493-INS-160
chr4-122662720-INS-71
chr4-122697616-INS-6180
chr4-122701398-INS-1077
chr4-122709556-INS-95
chr4-122716372-INS-2502
chr4-122728368-INS-214
chr4-122788455-INS-69
chr4-122793404-INS-277
chr4-122805893-INS-78
chr4-122812961-INS-198
chr4-122819243-INS-107
chr4-122835519-INS-309
chr4-122836123-INS-99
chr4-122855854-INS-97
chr4-122913627-INS-213
chr4-122954313-INS-187
chr4-123040041-INS-304
chr4-123046438-INS-64
chr4-123061317-INS-165
chr4-123061318-INS-339
chr4-123097384-INS-212
chr4-123102963-INS-2498
chr4-123214742-INS-54
chr4-123329819-INS-219
chr4-123336794-INS-74
chr4-123347007-INS-57
chr4-123347495-INS-112
chr4-123442510-INS-181
chr4-123494079-INS-57
chr4-123522652-INS-684
chr4-123572241-INS-78
chr4-123589970-INS-230
chr4-123595254-INS-1414
chr4-123595715-INS-381
chr4-123596073-INS-435
chr4-123605018-INS-85
chr4-123607539-INS-206
chr4-123615486-INS-196
chr4-123632429-INS-218
chr4-123647283-INS-336
chr4-123647482-INS-72
chr4-123647937-INS-72
chr4-123682133-INS-60
chr4-123690929-INS-1418
chr4-123692695-INS-56
chr4-123697915-INS-5349
chr4-123706001-INS-50
chr4-123711437-INS-68

chr4-123713834-INS-9459
chr4-123720797-INS-5510
chr4-123770731-INS-289
chr4-123793435-INS-198
chr4-123834717-INS-58
chr4-123854603-INS-63
chr4-123877134-INS-52
chr4-123877164-INS-142
chr4-123877164-INS-380
chr4-123906791-INS-1315
chr4-123918095-INS-194
chr4-123955528-INS-171
chr4-123969036-INS-6440
chr4-123972531-INS-450
chr4-123989970-INS-234
chr4-124005365-INS-50
chr4-124039279-INS-234
chr4-124039361-INS-53
chr4-124039542-INS-168
chr4-124049082-INS-225
chr4-124053116-INS-52
chr4-124053128-INS-154
chr4-124053764-INS-50
chr4-124088085-INS-270
chr4-124091698-INS-50
chr4-124166638-INS-65
chr4-124189674-INS-102
chr4-124267142-INS-69
chr4-124272821-INS-84
chr4-124286055-INS-92
chr4-124310491-INS-52
chr4-124318757-INS-76
chr4-124351940-INS-61
chr4-124378931-INS-134
chr4-124386882-INS-120
chr4-124428885-INS-60
chr4-124444514-INS-86
chr4-124461588-INS-232
chr4-124466457-INS-314
chr4-124505100-INS-207
chr4-124590493-INS-65
chr4-124611237-INS-71
chr4-124617874-INS-444
chr4-124638309-INS-50
chr4-124663766-INS-1154
chr4-124683415-INS-2742
chr4-124689305-INS-216
chr4-124690095-INS-114
chr4-124712952-INS-394
chr4-124716674-INS-88
chr4-124758193-INS-204
chr4-124759351-INS-1491
chr4-124775048-INS-58
chr4-124802495-INS-57
chr4-124804285-INS-57
chr4-124834411-INS-975
chr4-124834461-INS-2325
chr4-124834470-INS-475
chr4-124834672-INS-150
chr4-124836823-INS-81
chr4-124853541-INS-218
chr4-124904674-INS-112
chr4-124906032-INS-3084
chr4-124913786-INS-58
chr4-124929951-INS-128
chr4-124933308-INS-50
chr4-125025253-INS-4772
chr4-125031893-INS-66
chr4-125033954-INS-63
chr4-125035553-INS-314
chr4-125071408-INS-262
chr4-125075919-INS-91
chr4-125077164-INS-76
chr4-125077350-INS-512
chr4-125096294-INS-168
chr4-125108773-INS-168
chr4-125140809-INS-3299

chr4-125166431-INS-198
chr4-125184261-INS-160
chr4-125197634-INS-124
chr4-125218881-INS-316
chr4-125228515-INS-311
chr4-125244968-INS-51
chr4-125273468-INS-53
chr4-125282439-INS-96
chr4-125289516-INS-3536
chr4-125422502-INS-50
chr4-125462124-INS-2498
chr4-125465968-INS-145
chr4-125466913-INS-263
chr4-125527110-INS-114
chr4-125603837-INS-50
chr4-125619152-INS-82
chr4-125669991-INS-62
chr4-125693902-INS-150
chr4-125712101-INS-117
chr4-125714891-INS-54
chr4-125722791-INS-182
chr4-125758821-INS-54
chr4-125761975-INS-199
chr4-125761979-INS-56
chr4-125777248-INS-52
chr4-125785627-INS-60
chr4-125790530-INS-52
chr4-125809764-INS-249
chr4-125828477-INS-569
chr4-125840080-INS-80
chr4-125849004-INS-76
chr4-125870293-INS-90
chr4-125872336-INS-78
chr4-125872430-INS-5753
chr4-125875745-INS-84
chr4-125904337-INS-206
chr4-125907843-INS-1424
chr4-125934162-INS-62
chr4-125955512-INS-243
chr4-125967047-INS-220
chr4-125967109-INS-55
chr4-125973638-INS-54
chr4-125974575-INS-50
chr4-125983392-INS-180
chr4-125990197-INS-104
chr4-126014770-INS-203
chr4-126037022-INS-2707
chr4-126102302-INS-627
chr4-126102410-INS-52
chr4-126104494-INS-203
chr4-126108958-INS-346
chr4-126112780-INS-193
chr4-126119760-INS-206
chr4-126123020-INS-213
chr4-126130613-INS-1222
chr4-126147101-INS-208
chr4-126151863-INS-54
chr4-126165413-INS-126
chr4-126166251-INS-231
chr4-126168202-INS-889
chr4-126168455-INS-428
chr4-126168771-INS-153
chr4-126173604-INS-75
chr4-126183082-INS-120
chr4-126225917-INS-56
chr4-126226244-INS-202
chr4-126231736-INS-208
chr4-126257885-INS-160
chr4-126264590-INS-207
chr4-126308363-INS-76
chr4-126334722-INS-178
chr4-126335485-INS-217
chr4-126345452-INS-119
chr4-126363756-INS-245
chr4-126401629-INS-653
chr4-126401629-INS-68
chr4-126407074-INS-117

chr4-126414468-INS-1130
chr4-126415478-INS-235
chr4-126443075-INS-202
chr4-126470983-INS-179
chr4-126478892-INS-308
chr4-126483565-INS-729
chr4-126489313-INS-217
chr4-126517016-INS-66
chr4-126543304-INS-78
chr4-126604335-INS-1280
chr4-126650001-INS-74
chr4-126651220-INS-54
chr4-126651362-INS-228
chr4-126660563-INS-2319
chr4-126675851-INS-53
chr4-126727028-INS-50
chr4-126731975-INS-1473
chr4-126732359-INS-60
chr4-126783744-INS-94
chr4-126793005-INS-93
chr4-126827687-INS-68
chr4-126872304-INS-1347
chr4-126877158-INS-60
chr4-126883764-INS-219
chr4-126884452-INS-209
chr4-126891335-INS-516
chr4-126891524-INS-205
chr4-126967427-INS-73
chr4-126967598-INS-212
chr4-126969815-INS-70
chr4-126975574-INS-210
chr4-127038908-INS-58
chr4-127039686-INS-202
chr4-127050950-INS-120
chr4-127054092-INS-538
chr4-127054248-INS-184
chr4-127065756-INS-126
chr4-127146190-INS-52
chr4-127159675-INS-341
chr4-127188537-INS-58
chr4-127188823-INS-213
chr4-127227448-INS-82
chr4-127229317-INS-143
chr4-127229785-INS-474
chr4-127229902-INS-618
chr4-127244764-INS-52
chr4-127256316-INS-150
chr4-127270565-INS-218
chr4-127322753-INS-574
chr4-127326495-INS-50
chr4-127351714-INS-54
chr4-127363885-INS-5586
chr4-127391657-INS-78
chr4-127393343-INS-75
chr4-127393351-INS-192
chr4-127434144-INS-142
chr4-127442388-INS-71
chr4-127451228-INS-109
chr4-127451257-INS-52
chr4-127468091-INS-226
chr4-127482895-INS-100
chr4-127489653-INS-92
chr4-127515225-INS-96
chr4-127525892-INS-130
chr4-127525990-INS-643
chr4-127526181-INS-226
chr4-127544047-INS-105
chr4-127548156-INS-445
chr4-127591102-INS-52
chr4-127629947-INS-96
chr4-127633127-INS-1378
chr4-127657406-INS-69
chr4-127688518-INS-5567
chr4-127699426-INS-64
chr4-127699894-INS-112
chr4-127725833-INS-64
chr4-127778322-INS-223

chr4-127792837-INS-1404
chr4-127797729-INS-541
chr4-127801002-INS-68
chr4-127918680-INS-54
chr4-127927947-INS-1535
chr4-128012608-INS-50
chr4-128019228-INS-100
chr4-128022189-INS-694
chr4-128059590-INS-115
chr4-128060225-INS-322
chr4-128083700-INS-56
chr4-128083863-INS-116
chr4-128106237-INS-217
chr4-128115362-INS-2841
chr4-128134274-INS-55
chr4-128153831-INS-126
chr4-128178665-INS-132
chr4-128254685-INS-56
chr4-128271750-INS-55
chr4-128317573-INS-155
chr4-128353544-INS-304
chr4-128398918-INS-62
chr4-128441622-INS-2377
chr4-128467136-INS-262
chr4-128467737-INS-87
chr4-128475691-INS-400
chr4-128488294-INS-52
chr4-128496555-INS-69
chr4-128520469-INS-5466
chr4-128524531-INS-74
chr4-128607899-INS-70
chr4-128646891-INS-94
chr4-128715895-INS-58
chr4-128729551-INS-428
chr4-128752087-INS-80
chr4-128766683-INS-64
chr4-128802778-INS-3257
chr4-128816177-INS-485
chr4-128816320-INS-839
chr4-128822242-INS-1489
chr4-128837849-INS-68
chr4-128850353-INS-174
chr4-128866033-INS-75
chr4-128866167-INS-63
chr4-128904057-INS-205
chr4-128923005-INS-1302
chr4-128937670-INS-227
chr4-128993252-INS-110
chr4-129010391-INS-57
chr4-129039582-INS-87
chr4-129058017-INS-268
chr4-129073066-INS-117
chr4-129073285-INS-216
chr4-129079198-INS-290
chr4-129086259-INS-207
chr4-129092805-INS-68
chr4-129104025-INS-204
chr4-129127699-INS-287
chr4-129127758-INS-68
chr4-129148480-INS-462
chr4-129163231-INS-244
chr4-129171101-INS-191
chr4-129171560-INS-64
chr4-129179239-INS-358
chr4-129185066-INS-191
chr4-129264504-INS-194
chr4-129266040-INS-205
chr4-129306253-INS-211
chr4-129306983-INS-202
chr4-129307991-INS-219
chr4-129357588-INS-199
chr4-129367029-INS-204
chr4-129419593-INS-6665
chr4-129439688-INS-206
chr4-129445902-INS-51
chr4-129456360-INS-58
chr4-129457555-INS-206

chr4-129526774-INS-285
chr4-129529336-INS-189
chr4-129621533-INS-1888
chr4-129638764-INS-96
chr4-129639093-INS-72
chr4-129659122-INS-328
chr4-129668080-INS-62
chr4-129673322-INS-199
chr4-129710629-INS-218
chr4-129777557-INS-332
chr4-129777653-INS-110
chr4-129788233-INS-68
chr4-129813640-INS-65
chr4-129831768-INS-119
chr4-129890718-INS-546
chr4-129890718-INS-75
chr4-129890924-INS-75
chr4-129973867-INS-227
chr4-129996545-INS-76
chr4-130046694-INS-1085
chr4-130046869-INS-431
chr4-130090316-INS-111
chr4-130097821-INS-1725
chr4-130097821-INS-4056
chr4-130098194-INS-112
chr4-130139480-INS-84
chr4-130153261-INS-70
chr4-130179553-INS-6429
chr4-130180019-INS-52
chr4-130187162-INS-147
chr4-130187618-INS-91
chr4-130196967-INS-72
chr4-130215318-INS-63
chr4-130235003-INS-54
chr4-130242523-INS-437
chr4-130250095-INS-214
chr4-130260137-INS-50
chr4-130276959-INS-331
chr4-130283022-INS-126
chr4-130288554-INS-52
chr4-130290203-INS-1910
chr4-130291450-INS-273
chr4-130301362-INS-61
chr4-130344429-INS-66
chr4-130386906-INS-224
chr4-130439337-INS-50
chr4-130569883-INS-123
chr4-130582937-INS-51
chr4-130596563-INS-50
chr4-130599196-INS-74
chr4-130603057-INS-1123
chr4-130603169-INS-519
chr4-130613955-INS-60
chr4-130631101-INS-276
chr4-130657895-INS-62
chr4-130659980-INS-925
chr4-130667563-INS-87
chr4-130667810-INS-192
chr4-130678475-INS-52
chr4-130697526-INS-516
chr4-130697988-INS-540
chr4-130700484-INS-54
chr4-130736548-INS-438
chr4-130745320-INS-134
chr4-130757091-INS-342
chr4-130758537-INS-202
chr4-130775250-INS-2428
chr4-130782244-INS-63
chr4-130809753-INS-585
chr4-130815730-INS-390
chr4-130826408-INS-60
chr4-130827223-INS-149
chr4-130827309-INS-59
chr4-130836653-INS-384
chr4-130859031-INS-95
chr4-130872975-INS-51
chr4-130873493-INS-118

chr4-130897022-INS-106
chr4-130900227-INS-76
chr4-130977741-INS-1895
chr4-131002909-INS-2882
chr4-131024707-INS-75
chr4-131068899-INS-464
chr4-131075397-INS-54
chr4-131079283-INS-452
chr4-131100424-INS-60
chr4-131117951-INS-489
chr4-131123453-INS-7187
chr4-131141227-INS-54
chr4-131232177-INS-5429
chr4-131275213-INS-7247
chr4-131318937-INS-64
chr4-131382612-INS-1429
chr4-131420592-INS-59
chr4-131473241-INS-170
chr4-131531969-INS-108
chr4-131532065-INS-344
chr4-131536371-INS-117
chr4-131536672-INS-251
chr4-131541817-INS-139
chr4-131556135-INS-77
chr4-131786042-INS-288
chr4-131829694-INS-112
chr4-131831437-INS-384
chr4-132016545-INS-592
chr4-132017656-INS-69
chr4-132017970-INS-410
chr4-132018157-INS-85
chr4-132020873-INS-58
chr4-132027116-INS-4842
chr4-132135774-INS-72
chr4-132164973-INS-76
chr4-132174597-INS-94
chr4-132185291-INS-210
chr4-132195320-INS-55
chr4-132275495-INS-62
chr4-132278590-INS-582
chr4-132301943-INS-87
chr4-132349108-INS-157
chr4-132414156-INS-59
chr4-132479146-INS-433
chr4-132479562-INS-227
chr4-132482186-INS-183
chr4-132491340-INS-218
chr4-132513911-INS-195
chr4-132617404-INS-53
chr4-132659787-INS-72
chr4-132679419-INS-55
chr4-132681311-INS-58
chr4-132723914-INS-215
chr4-132890626-INS-190
chr4-133048678-INS-160
chr4-133052567-INS-210
chr4-133220510-INS-123
chr4-133278267-INS-5284
chr4-133288043-INS-189
chr4-133304303-INS-206
chr4-133313029-INS-638
chr4-133321207-INS-81
chr4-133323818-INS-1017
chr4-133382075-INS-67
chr4-133498962-INS-226
chr4-133508278-INS-274
chr4-133522123-INS-62
chr4-133527775-INS-804
chr4-133527785-INS-6924
chr4-133527790-INS-3113
chr4-133562577-INS-177
chr4-133566062-INS-396
chr4-133566431-INS-87
chr4-133631083-INS-2945
chr4-133643799-INS-67
chr4-133643833-INS-269
chr4-133659756-INS-8419

chr4-133670061-INS-1811
chr4-133725780-INS-218
chr4-133726602-INS-212
chr4-133752928-INS-473
chr4-133772954-INS-3159
chr4-133813682-INS-208
chr4-133968146-INS-1920
chr4-133994827-INS-434
chr4-134000605-INS-103
chr4-134105004-INS-54
chr4-134240016-INS-4690
chr4-134251344-INS-50
chr4-134301502-INS-172
chr4-134322236-INS-11623
chr4-134376593-INS-219
chr4-134407756-INS-59
chr4-134409425-INS-85
chr4-134443841-INS-5058
chr4-134446392-INS-55
chr4-134511584-INS-262
chr4-134572046-INS-225
chr4-134572151-INS-72
chr4-134591297-INS-60
chr4-134593547-INS-136
chr4-134594394-INS-51
chr4-134598607-INS-96
chr4-134632794-INS-220
chr4-134663790-INS-1182
chr4-134667614-INS-210
chr4-134675333-INS-142
chr4-134678869-INS-76
chr4-134680720-INS-57
chr4-134683080-INS-52
chr4-134693602-INS-299
chr4-134711063-INS-268
chr4-134712796-INS-3178
chr4-134719783-INS-4806
chr4-134733196-INS-2538
chr4-134739115-INS-59
chr4-134770293-INS-98
chr4-134779316-INS-210
chr4-134827837-INS-88
chr4-134828539-INS-52
chr4-134906400-INS-51
chr4-134910596-INS-60
chr4-134929929-INS-162
chr4-134929930-INS-624
chr4-134948282-INS-240
chr4-134979713-INS-4377
chr4-134992493-INS-5534
chr4-134993092-INS-68
chr4-135009297-INS-4105
chr4-135020401-INS-55
chr4-135038421-INS-192
chr4-135038681-INS-558
chr4-135044728-INS-50
chr4-135086327-INS-216
chr4-135090160-INS-211
chr4-135091513-INS-96
chr4-135092244-INS-82
chr4-135094569-INS-2237
chr4-135096681-INS-186
chr4-135097107-INS-240
chr4-135102588-INS-171
chr4-135134427-INS-380
chr4-135134652-INS-204
chr4-135165965-INS-793
chr4-135166147-INS-228
chr4-135167922-INS-66
chr4-135214714-INS-81
chr4-135214993-INS-50
chr4-135228880-INS-240
chr4-135308357-INS-108
chr4-135378338-INS-7070
chr4-135387492-INS-50
chr4-135393239-INS-1995
chr4-135407768-INS-3855

chr4-135416242-INS-209
chr4-135429218-INS-61
chr4-135442912-INS-150
chr4-135451273-INS-64
chr4-135471825-INS-184
chr4-135485990-INS-610
chr4-135486090-INS-230
chr4-135503454-INS-239
chr4-135504693-INS-205
chr4-135511683-INS-94
chr4-135513634-INS-122
chr4-135544084-INS-202
chr4-135568613-INS-72
chr4-135606271-INS-3200
chr4-135615369-INS-208
chr4-135621399-INS-67
chr4-135652654-INS-198
chr4-135707885-INS-214
chr4-135729007-INS-197
chr4-135729481-INS-227
chr4-135766053-INS-68
chr4-135797117-INS-78
chr4-135815340-INS-120
chr4-135815425-INS-172
chr4-135824950-INS-56
chr4-135831545-INS-56
chr4-135836230-INS-160
chr4-135836426-INS-70
chr4-135845370-INS-207
chr4-135851671-INS-151
chr4-135880739-INS-50
chr4-135886113-INS-207
chr4-135916995-INS-72
chr4-135917247-INS-73
chr4-135918209-INS-210
chr4-135927615-INS-98
chr4-135933152-INS-218
chr4-135933369-INS-150
chr4-135984892-INS-60
chr4-135989928-INS-106
chr4-135993970-INS-452
chr4-135996022-INS-162
chr4-135997149-INS-73
chr4-136028777-INS-1503
chr4-136034295-INS-207
chr4-136078916-INS-57
chr4-136089455-INS-66
chr4-136094370-INS-128
chr4-136097423-INS-521
chr4-136100277-INS-50
chr4-136125543-INS-137
chr4-136136815-INS-230
chr4-136159784-INS-305
chr4-136162741-INS-209
chr4-136167134-INS-664
chr4-136167139-INS-92
chr4-136207154-INS-194
chr4-136316222-INS-2520
chr4-136333075-INS-138
chr4-136337347-INS-480
chr4-136338302-INS-60
chr4-136338981-INS-399
chr4-136346320-INS-50
chr4-136352050-INS-330
chr4-136357683-INS-439
chr4-136361293-INS-206
chr4-136366018-INS-4470
chr4-136366921-INS-51
chr4-136422866-INS-102
chr4-136431085-INS-54
chr4-136432927-INS-71
chr4-136443246-INS-95
chr4-136462404-INS-96
chr4-136529104-INS-212
chr4-136546116-INS-50
chr4-136546753-INS-91
chr4-136581683-INS-84

chr4-136599771-INS-6350
chr4-136610437-INS-56
chr4-136648299-INS-54
chr4-136801255-INS-63
chr4-136801601-INS-150
chr4-136807474-INS-52
chr4-136807514-INS-109
chr4-136808176-INS-264
chr4-136868656-INS-2717
chr4-136876715-INS-2681
chr4-137132544-INS-60
chr4-137140282-INS-65
chr4-137161103-INS-124
chr4-137174071-INS-368
chr4-137174094-INS-74
chr4-137291225-INS-62
chr4-137306612-INS-293
chr4-137308723-INS-181
chr4-137314263-INS-374
chr4-137314266-INS-93
chr4-137340538-INS-1582
chr4-137461399-INS-70
chr4-137533987-INS-75
chr4-137555540-INS-80
chr4-137558893-INS-202
chr4-137651965-INS-377
chr4-137665420-INS-53
chr4-137738106-INS-50
chr4-137745760-INS-184
chr4-137759800-INS-71
chr4-137769363-INS-243
chr4-137777112-INS-248
chr4-137781534-INS-1244
chr4-137790027-INS-348
chr4-137796563-INS-213
chr4-137840270-INS-167
chr4-137897465-INS-270
chr4-137921673-INS-60
chr4-137924818-INS-156
chr4-137958433-INS-544
chr4-137958485-INS-96
chr4-138024999-INS-54
chr4-138058173-INS-5086
chr4-138089040-INS-50
chr4-138090506-INS-1889
chr4-138113047-INS-3899
chr4-138193651-INS-186
chr4-138204526-INS-70
chr4-138205149-INS-53
chr4-138229995-INS-473
chr4-138230889-INS-51
chr4-138258366-INS-175
chr4-138258486-INS-65
chr4-138275134-INS-141
chr4-138322683-INS-68
chr4-138322791-INS-448
chr4-138328059-INS-50
chr4-138365980-INS-2091
chr4-138374322-INS-1824
chr4-138388451-INS-62
chr4-138389991-INS-67
chr4-138408579-INS-5259
chr4-138409564-INS-675
chr4-138409580-INS-94
chr4-138416065-INS-51
chr4-138425146-INS-64
chr4-138463918-INS-74
chr4-138591075-INS-5401
chr4-138611243-INS-144
chr4-138624533-INS-140
chr4-138635608-INS-256
chr4-138702363-INS-55
chr4-138716375-INS-216
chr4-138739117-INS-56
chr4-138748709-INS-396
chr4-138757556-INS-1475
chr4-138805020-INS-172

chr4-138825390-INS-52
chr4-138833342-INS-205
chr4-138864868-INS-62
chr4-138865846-INS-174
chr4-138869586-INS-6452
chr4-138871761-INS-209
chr4-138872428-INS-50
chr4-138876750-INS-66
chr4-138934146-INS-1020
chr4-138934146-INS-2125
chr4-138934194-INS-425
chr4-139047731-INS-199
chr4-139048375-INS-391
chr4-139052346-INS-246
chr4-139052560-INS-427
chr4-139052828-INS-201
chr4-139053274-INS-250
chr4-139053457-INS-97
chr4-139061519-INS-1856
chr4-139068533-INS-115
chr4-139069437-INS-4976
chr4-139082523-INS-94
chr4-139083334-INS-180
chr4-139090074-INS-126
chr4-139090649-INS-87
chr4-139116011-INS-53
chr4-139230869-INS-53
chr4-139230881-INS-589
chr4-139260692-INS-128
chr4-139263037-INS-121
chr4-139286894-INS-55
chr4-139316467-INS-734
chr4-139316495-INS-231
chr4-139416067-INS-186
chr4-139446126-INS-543
chr4-139448858-INS-52
chr4-139457455-INS-595
chr4-139492612-INS-211
chr4-139584441-INS-352
chr4-139593054-INS-1596
chr4-139622510-INS-360
chr4-139723829-INS-184
chr4-139724559-INS-377
chr4-139747941-INS-86
chr4-139757780-INS-151
chr4-139770850-INS-254
chr4-139824914-INS-313
chr4-139825034-INS-60
chr4-139833262-INS-212
chr4-139881664-INS-1444
chr4-139898914-INS-102
chr4-139914307-INS-50
chr4-139981416-INS-50
chr4-140034575-INS-182
chr4-140038081-INS-50
chr4-140042155-INS-212
chr4-140114133-INS-50
chr4-140120552-INS-123
chr4-140121225-INS-136
chr4-140158564-INS-53
chr4-140175249-INS-87
chr4-140244368-INS-877
chr4-140284729-INS-407
chr4-140287773-INS-128
chr4-140298472-INS-705
chr4-140298604-INS-50
chr4-140298640-INS-328
chr4-140301543-INS-76
chr4-140378985-INS-114
chr4-140398176-INS-575
chr4-140410474-INS-198
chr4-140423307-INS-5512
chr4-140450560-INS-860
chr4-140494168-INS-99
chr4-140531432-INS-80
chr4-140531439-INS-5180
chr4-140531575-INS-56

chr4-140531672-INS-287
chr4-140535487-INS-51
chr4-140567482-INS-6444
chr4-140573333-INS-4737
chr4-140585856-INS-4871
chr4-140639843-INS-5402
chr4-140765021-INS-175
chr4-140789605-INS-206
chr4-140829703-INS-67
chr4-140856120-INS-133
chr4-140950853-INS-6042
chr4-140989564-INS-169
chr4-140992656-INS-4368
chr4-141029828-INS-170
chr4-141032896-INS-52
chr4-141146361-INS-52
chr4-141164150-INS-1887
chr4-141168683-INS-94
chr4-141254987-INS-55
chr4-141277995-INS-216
chr4-141278853-INS-197
chr4-141289348-INS-197
chr4-141311695-INS-50
chr4-141312358-INS-84
chr4-141334194-INS-60
chr4-141393622-INS-778
chr4-141395073-INS-52
chr4-141446692-INS-60
chr4-141468709-INS-77
chr4-141477460-INS-1188
chr4-141478124-INS-1460
chr4-141478124-INS-728
chr4-141500628-INS-100
chr4-141545061-INS-89
chr4-141556764-INS-54
chr4-141561053-INS-58
chr4-141564304-INS-70
chr4-141565691-INS-60
chr4-141576530-INS-816
chr4-141615791-INS-160
chr4-141615906-INS-573
chr4-141616334-INS-725
chr4-141616377-INS-71
chr4-141715790-INS-557
chr4-141721326-INS-114
chr4-141725419-INS-1065
chr4-141739584-INS-157
chr4-141762365-INS-137
chr4-141801653-INS-73
chr4-141824388-INS-94
chr4-141824608-INS-124
chr4-141826979-INS-1355
chr4-141827004-INS-534
chr4-141927302-INS-159
chr4-141967923-INS-79
chr4-141972944-INS-271
chr4-141973227-INS-50
chr4-141999887-INS-270
chr4-142065535-INS-64
chr4-142120157-INS-50
chr4-142150818-INS-198
chr4-142177631-INS-108
chr4-142181256-INS-62
chr4-142251443-INS-343
chr4-142309472-INS-1865
chr4-142317106-INS-768
chr4-142318756-INS-529
chr4-142319278-INS-60
chr4-142325320-INS-96
chr4-142341087-INS-70
chr4-142347784-INS-111
chr4-142394189-INS-60
chr4-142417648-INS-61
chr4-142447433-INS-65
chr4-142447448-INS-410
chr4-142507604-INS-4831
chr4-142537662-INS-54

chr4-142559951-INS-99
chr4-142567861-INS-428
chr4-142574193-INS-6207
chr4-142598495-INS-223
chr4-142624686-INS-6359
chr4-142702858-INS-152
chr4-142736868-INS-423
chr4-142753005-INS-52
chr4-142755982-INS-510
chr4-142807947-INS-51
chr4-142817680-INS-320
chr4-142817694-INS-50
chr4-142850438-INS-330
chr4-142917615-INS-421
chr4-142929961-INS-117
chr4-142930336-INS-507
chr4-142937412-INS-218
chr4-142946998-INS-244
chr4-142954581-INS-440
chr4-142958024-INS-134
chr4-142992791-INS-7187
chr4-143034756-INS-52
chr4-143036414-INS-169
chr4-143044040-INS-57
chr4-143044325-INS-190
chr4-143070245-INS-58
chr4-143090915-INS-136
chr4-143109530-INS-50
chr4-143242505-INS-3060
chr4-143262216-INS-175
chr4-143389109-INS-1402
chr4-143403347-INS-7310
chr4-143421832-INS-130
chr4-143471774-INS-342
chr4-143636822-INS-212
chr4-143765422-INS-305
chr4-143774201-INS-2054
chr4-143779883-INS-1060
chr4-143875506-INS-201
chr4-143931589-INS-1864
chr4-143990368-INS-161
chr4-144028287-INS-183
chr4-144028527-INS-139
chr4-144032803-INS-221
chr4-144036367-INS-4001
chr4-144048971-INS-120
chr4-144061511-INS-196
chr4-144091511-INS-223
chr4-144156319-INS-159
chr4-144156319-INS-423
chr4-144202058-INS-6561
chr4-144332463-INS-1498
chr4-144345592-INS-52
chr4-144410407-INS-54
chr4-144441356-INS-56
chr4-144456585-INS-115
chr4-144482090-INS-14071
chr4-144557556-INS-1358
chr4-144605265-INS-52
chr4-144622579-INS-496
chr4-144686905-INS-417
chr4-144696976-INS-89
chr4-144765150-INS-52
chr4-144772643-INS-51
chr4-144772810-INS-57
chr4-144849628-INS-7180
chr4-144869574-INS-315
chr4-145021330-INS-1364
chr4-145046452-INS-1350
chr4-145046452-INS-495
chr4-145086028-INS-14597
chr4-145086028-INS-7021
chr4-145088223-INS-58
chr4-145088361-INS-376
chr4-145088463-INS-57
chr4-145088540-INS-105
chr4-145136958-INS-141

chr4-145140880-INS-281
chr4-145161684-INS-70
chr4-145164835-INS-444
chr4-145166302-INS-567
chr4-145168743-INS-272
chr4-145172369-INS-422
chr4-145177206-INS-167
chr4-145222525-INS-169
chr4-145254241-INS-685
chr4-145278143-INS-1885
chr4-145278143-INS-400
chr4-145278143-INS-9310
chr4-145282477-INS-1892
chr4-145286808-INS-81
chr4-145301833-INS-199
chr4-145312031-INS-178
chr4-145313885-INS-240
chr4-145316197-INS-200
chr4-145319979-INS-463
chr4-145405472-INS-54
chr4-145412899-INS-1184
chr4-145455079-INS-67
chr4-145570092-INS-214
chr4-145991259-INS-222
chr4-146091538-INS-56
chr4-146132300-INS-251
chr4-146133320-INS-84
chr4-146220034-INS-444
chr4-146224156-INS-381
chr4-146268996-INS-214
chr4-146306843-INS-2913
chr4-146432473-INS-523
chr4-146477369-INS-168
chr4-146490398-INS-6223
chr4-146491727-INS-338
chr4-146527858-INS-4168
chr4-146532979-INS-204
chr4-146534045-INS-194
chr4-146556710-INS-192
chr4-146577648-INS-197
chr4-146577980-INS-199
chr4-146604103-INS-1297
chr4-146620028-INS-346
chr4-146622265-INS-84
chr4-146624014-INS-109
chr4-146635894-INS-202
chr4-146995966-INS-389
chr4-147008721-INS-351
chr4-147017827-INS-65
chr4-147041818-INS-178
chr4-147214137-INS-69
chr4-147406597-INS-276
chr4-147416914-INS-86
chr4-147417453-INS-84
chr4-147431593-INS-193
chr4-147451060-INS-476
chr4-147475400-INS-420
chr4-147475797-INS-84
chr4-147511098-INS-163
chr4-147514413-INS-358
chr4-147520265-INS-157
chr4-147521313-INS-1001
chr4-147522603-INS-246
chr4-147525824-INS-492
chr4-147544776-INS-227
chr4-147577097-INS-812
chr4-147594664-INS-207
chr4-147599105-INS-84
chr4-147600067-INS-2235
chr4-147601005-INS-183
chr4-147637348-INS-225
chr4-147637600-INS-251
chr4-147639105-INS-67
chr4-147639701-INS-227
chr4-147640891-INS-125
chr4-147642815-INS-196
chr4-147679284-INS-2796

chr4-147762868-INS-204
chr4-147769172-INS-86
chr4-147888059-INS-209
chr4-147890967-INS-233
chr4-147891757-INS-233
chr4-147894469-INS-195
chr4-147898593-INS-212
chr4-147899605-INS-64
chr4-147901237-INS-54
chr4-147905958-INS-1079
chr4-147915553-INS-2251
chr4-147928790-INS-211
chr4-147936761-INS-1824
chr4-147956486-INS-7092
chr4-148012163-INS-500
chr4-148016488-INS-4052
chr4-148075275-INS-441
chr4-148129786-INS-57
chr4-148179817-INS-963
chr4-148198236-INS-2677
chr4-148203829-INS-214
chr4-148204337-INS-126
chr4-148206970-INS-56
chr4-148211045-INS-3672
chr4-148253710-INS-53
chr4-148286431-INS-50
chr4-148301450-INS-312
chr4-148301450-INS-62
chr4-148301450-INS-800
chr4-148309925-INS-190
chr4-148310419-INS-51
chr4-148326224-INS-62
chr4-148408142-INS-62
chr4-148408343-INS-59
chr4-148423709-INS-150
chr4-148466277-INS-5542
chr4-148472052-INS-239
chr4-148487172-INS-997
chr4-148505736-INS-282
chr4-148505751-INS-1160
chr4-148524158-INS-62
chr4-148625659-INS-208
chr4-148629541-INS-52
chr4-148682372-INS-9597
chr4-148767723-INS-307
chr4-148774661-INS-208
chr4-148818140-INS-50
chr4-149067215-INS-54
chr4-149094915-INS-382
chr4-149127491-INS-95
chr4-149127493-INS-396
chr4-149165335-INS-396
chr4-149165636-INS-1353
chr4-149165759-INS-429
chr4-149172998-INS-147
chr4-149173002-INS-364
chr4-149177734-INS-186
chr4-149178243-INS-125
chr4-149203394-INS-809
chr4-149205433-INS-209
chr4-149221754-INS-51
chr4-149237223-INS-751
chr4-149260659-INS-51
chr4-149274006-INS-199
chr4-149298960-INS-51
chr4-149360151-INS-328
chr4-149465101-INS-192
chr4-149481021-INS-704
chr4-149496677-INS-169
chr4-149497148-INS-465
chr4-149525575-INS-210
chr4-149526283-INS-60
chr4-149551952-INS-5546
chr4-149557461-INS-149
chr4-149581710-INS-56
chr4-149616353-INS-359
chr4-149677597-INS-52

chr4-149684009-INS-1515
chr4-149702479-INS-219
chr4-149768880-INS-51
chr4-149771571-INS-371
chr4-149789446-INS-280
chr4-149790165-INS-761
chr4-149790360-INS-1920
chr4-149844301-INS-52
chr4-149850835-INS-456
chr4-149895213-INS-1857
chr4-149895235-INS-16533
chr4-149896201-INS-1893
chr4-149909258-INS-83
chr4-149913623-INS-627
chr4-150031558-INS-50
chr4-150037126-INS-98
chr4-150052561-INS-59
chr4-150058320-INS-204
chr4-150067398-INS-57
chr4-150117406-INS-78
chr4-150143837-INS-375
chr4-150152231-INS-138
chr4-150166161-INS-225
chr4-150183628-INS-204
chr4-150240702-INS-328
chr4-150251601-INS-1268
chr4-150254355-INS-59
chr4-150258189-INS-50
chr4-150265239-INS-62
chr4-150287694-INS-225
chr4-150300252-INS-90
chr4-150313821-INS-166
chr4-150336016-INS-50
chr4-150361562-INS-208
chr4-150381585-INS-229
chr4-150398679-INS-170
chr4-150409214-INS-327
chr4-150411934-INS-210
chr4-150420834-INS-66
chr4-150421656-INS-158
chr4-150421658-INS-74
chr4-150429848-INS-197
chr4-150432991-INS-197
chr4-150435845-INS-200
chr4-150441533-INS-52
chr4-150444023-INS-52
chr4-150455276-INS-204
chr4-150529714-INS-252
chr4-150550757-INS-358
chr4-150555528-INS-87
chr4-150556339-INS-510
chr4-150565215-INS-51
chr4-150570266-INS-152
chr4-150570351-INS-606
chr4-150582878-INS-236
chr4-150588495-INS-111
chr4-150608926-INS-1549
chr4-150682422-INS-52
chr4-150695389-INS-615
chr4-150723607-INS-51
chr4-150723648-INS-207
chr4-150738162-INS-57
chr4-150756825-INS-50
chr4-150762809-INS-344
chr4-150778729-INS-80
chr4-150788861-INS-60
chr4-150795010-INS-52
chr4-150799932-INS-90
chr4-150809027-INS-7117
chr4-150811843-INS-416
chr4-150811867-INS-204
chr4-150829286-INS-108
chr4-150830805-INS-50
chr4-150836254-INS-51
chr4-150859347-INS-75
chr4-150886860-INS-441
chr4-150902626-INS-337

chr4-150966613-INS-126
chr4-150966621-INS-469
chr4-150966973-INS-154
chr4-151033373-INS-207
chr4-151055905-INS-1377
chr4-151071602-INS-52
chr4-151072319-INS-52
chr4-151077637-INS-54
chr4-151165795-INS-72
chr4-151208292-INS-193
chr4-151238315-INS-440
chr4-151238581-INS-374
chr4-151313555-INS-68
chr4-151317470-INS-64
chr4-151336073-INS-128
chr4-151360983-INS-170
chr4-151367539-INS-96
chr4-151378352-INS-62
chr4-151393480-INS-128
chr4-151393741-INS-120
chr4-151459130-INS-4839
chr4-151460072-INS-56
chr4-151474991-INS-64
chr4-151481650-INS-63
chr4-151488878-INS-54
chr4-151493575-INS-50
chr4-151501935-INS-71
chr4-151516609-INS-96
chr4-151533356-INS-201
chr4-151553280-INS-128
chr4-151569548-INS-175
chr4-151569576-INS-400
chr4-151584376-INS-70
chr4-151600731-INS-62
chr4-151605435-INS-52
chr4-151616985-INS-1012
chr4-151683153-INS-60
chr4-151708793-INS-68
chr4-151790186-INS-106
chr4-151818442-INS-339
chr4-151819068-INS-11840
chr4-151822619-INS-779
chr4-151822629-INS-384
chr4-151822822-INS-95
chr4-151829482-INS-87
chr4-151880363-INS-60
chr4-151885575-INS-353
chr4-151887176-INS-132
chr4-151895856-INS-66
chr4-151929902-INS-405
chr4-151930060-INS-171
chr4-151950361-INS-277
chr4-151955522-INS-60
chr4-151970737-INS-700
chr4-151973417-INS-6559
chr4-151975987-INS-102
chr4-151979369-INS-205
chr4-151979374-INS-71
chr4-151982918-INS-907
chr4-152000070-INS-2000
chr4-152008678-INS-72
chr4-152017917-INS-55
chr4-152085676-INS-1090
chr4-152101398-INS-124
chr4-152165240-INS-114
chr4-152166729-INS-52
chr4-152192354-INS-173
chr4-152206327-INS-204
chr4-152222441-INS-82
chr4-152234657-INS-223
chr4-152256338-INS-118
chr4-152256466-INS-197
chr4-152280785-INS-1220
chr4-152283179-INS-231
chr4-152294005-INS-72
chr4-152322817-INS-199
chr4-152323549-INS-81

chr4-152323651-INS-186
chr4-152330218-INS-200
chr4-152369237-INS-4678
chr4-152374073-INS-81
chr4-152448375-INS-132
chr4-152451617-INS-500
chr4-152491838-INS-7217
chr4-152528446-INS-92
chr4-152529977-INS-54
chr4-152535098-INS-169
chr4-152550499-INS-50
chr4-152553269-INS-154
chr4-152574450-INS-1642
chr4-152595894-INS-60
chr4-152597465-INS-108
chr4-152612059-INS-100
chr4-152612541-INS-53
chr4-152613085-INS-676
chr4-152613095-INS-146
chr4-152640295-INS-953
chr4-152640379-INS-96
chr4-152642083-INS-76
chr4-152651755-INS-116
chr4-152654622-INS-51
chr4-152654812-INS-209
chr4-152654997-INS-485
chr4-152659464-INS-76
chr4-152668841-INS-76
chr4-152669606-INS-60
chr4-152689855-INS-620
chr4-152693874-INS-190
chr4-152746730-INS-63
chr4-152752602-INS-52
chr4-152752770-INS-90
chr4-152753329-INS-50
chr4-152760289-INS-62
chr4-152781460-INS-52
chr4-152787134-INS-401
chr4-152797184-INS-7080
chr4-152814916-INS-260
chr4-152827011-INS-74
chr4-152843527-INS-75
chr4-152847041-INS-68
chr4-152861361-INS-1933
chr4-152898204-INS-312
chr4-152900202-INS-50
chr4-152900225-INS-114
chr4-152908778-INS-103
chr4-152965114-INS-92
chr4-152978564-INS-70
chr4-152982259-INS-326
chr4-152989631-INS-50
chr4-152991127-INS-55
chr4-152999090-INS-162
chr4-153016369-INS-105
chr4-153026710-INS-1927
chr4-153035945-INS-209
chr4-153043621-INS-52
chr4-153045206-INS-394
chr4-153085770-INS-50
chr4-153109900-INS-66
chr4-153143151-INS-100
chr4-153179571-INS-100
chr4-153181118-INS-64
chr4-153184343-INS-2822
chr4-153215309-INS-116
chr4-153215932-INS-2888
chr4-153224937-INS-57
chr4-153225077-INS-428
chr4-153238130-INS-54
chr4-153242323-INS-50
chr4-153258387-INS-78
chr4-153258881-INS-56
chr4-153300682-INS-52
chr4-153338030-INS-138
chr4-153352916-INS-148
chr4-153368110-INS-90

chr4-153395506-INS-118
chr4-153398068-INS-68
chr4-153434755-INS-56
chr4-153437084-INS-62
chr4-153467143-INS-50
chr4-153467266-INS-357
chr4-153467502-INS-306
chr4-153481706-INS-357
chr4-153481868-INS-117
chr4-153486228-INS-60
chr4-153488607-INS-280
chr4-153502769-INS-54
chr4-153510023-INS-50
chr4-153525978-INS-140
chr4-153570836-INS-1000
chr4-153593605-INS-225
chr4-153600928-INS-54
chr4-153601230-INS-258
chr4-153601380-INS-710
chr4-153636497-INS-152
chr4-153636497-INS-72
chr4-153651802-INS-150
chr4-153652660-INS-82
chr4-153663842-INS-62
chr4-153665209-INS-253
chr4-153669629-INS-86
chr4-153702399-INS-50
chr4-153756453-INS-114
chr4-153756453-INS-54
chr4-153758387-INS-144
chr4-153765823-INS-195
chr4-153782237-INS-50
chr4-153795856-INS-337
chr4-153826868-INS-65
chr4-153884314-INS-96
chr4-153890209-INS-208
chr4-153894790-INS-350
chr4-153922930-INS-333
chr4-153944285-INS-59
chr4-153954903-INS-2497
chr4-153955008-INS-50
chr4-153977172-INS-158
chr4-153981286-INS-61
chr4-153981892-INS-68
chr4-153995792-INS-50
chr4-154004051-INS-72
chr4-154004211-INS-127
chr4-154015714-INS-2106
chr4-154070645-INS-288
chr4-154144392-INS-111
chr4-154150299-INS-78
chr4-154157586-INS-54
chr4-154158454-INS-1190
chr4-154158517-INS-574
chr4-154177243-INS-144
chr4-154178223-INS-302
chr4-154186597-INS-57
chr4-154192853-INS-89
chr4-154214093-INS-611
chr4-154232620-INS-99
chr4-154233606-INS-322
chr4-154233925-INS-63
chr4-154237970-INS-164
chr4-154373347-INS-163
chr4-154374211-INS-116
chr4-154396476-INS-343
chr4-154446176-INS-80
chr4-154464319-INS-60
chr4-154516053-INS-195
chr4-154516165-INS-397
chr4-154518694-INS-52
chr4-154560548-INS-56
chr4-154575760-INS-63
chr4-154610637-INS-129
chr4-154610754-INS-64
chr4-154622628-INS-335
chr4-154675404-INS-124

chr4-154738341-INS-82
chr4-154749207-INS-164
chr4-154757069-INS-204
chr4-154764042-INS-61
chr4-154765206-INS-492
chr4-154765820-INS-119
chr4-154869731-INS-50
chr4-154901909-INS-50
chr4-154903909-INS-196
chr4-154918823-INS-97
chr4-154946889-INS-51
chr4-154957379-INS-149
chr4-154966073-INS-7250
chr4-154969733-INS-662
chr4-154973213-INS-67
chr4-154996918-INS-183
chr4-155004370-INS-50
chr4-155026642-INS-258
chr4-155029981-INS-60
chr4-155078348-INS-3749
chr4-155110935-INS-50
chr4-155111039-INS-130
chr4-155112545-INS-98
chr4-155123705-INS-54
chr4-155135258-INS-209
chr4-155139778-INS-349
chr4-155221246-INS-582
chr4-155253046-INS-54
chr4-155313419-INS-209
chr4-155313761-INS-174
chr4-155358196-INS-120
chr4-155358421-INS-380
chr4-155358635-INS-1239
chr4-155405857-INS-50
chr4-155434019-INS-199
chr4-155439834-INS-223
chr4-155458176-INS-404
chr4-155458189-INS-1464
chr4-155461461-INS-5071
chr4-155478705-INS-9096
chr4-155497631-INS-58
chr4-155527740-INS-181
chr4-155527784-INS-67
chr4-155535507-INS-106
chr4-155549005-INS-238
chr4-155555963-INS-930
chr4-155564145-INS-124
chr4-155568808-INS-54
chr4-155594963-INS-211
chr4-155599463-INS-327
chr4-155603498-INS-198
chr4-155607492-INS-194
chr4-155631599-INS-166
chr4-155698931-INS-408
chr4-155698931-INS-841
chr4-155729802-INS-100
chr4-155729803-INS-302
chr4-155747691-INS-1063
chr4-155756044-INS-209
chr4-155769709-INS-179
chr4-155785150-INS-116
chr4-155787250-INS-237
chr4-155849265-INS-68
chr4-155852704-INS-185
chr4-155865124-INS-144
chr4-155918106-INS-223
chr4-155958681-INS-58
chr4-155959277-INS-209
chr4-155982270-INS-336
chr4-155996064-INS-53
chr4-156010496-INS-343
chr4-156043815-INS-248
chr4-156052784-INS-51
chr4-156067956-INS-191
chr4-156077275-INS-76
chr4-156077321-INS-220
chr4-156086196-INS-420

chr4-156086455-INS-156
chr4-156104609-INS-2843
chr4-156116408-INS-52
chr4-156120870-INS-205
chr4-156121355-INS-60
chr4-156171381-INS-552
chr4-156171590-INS-82
chr4-156173836-INS-52
chr4-156180608-INS-216
chr4-156189274-INS-6376
chr4-156194838-INS-54
chr4-156194939-INS-205
chr4-156242285-INS-94
chr4-156245237-INS-191
chr4-156267362-INS-50
chr4-156268865-INS-73
chr4-156296554-INS-371
chr4-156302625-INS-792
chr4-156321337-INS-550
chr4-156322593-INS-209
chr4-156322842-INS-378
chr4-156334140-INS-50
chr4-156337941-INS-260
chr4-156346719-INS-213
chr4-156346847-INS-64
chr4-156371300-INS-62
chr4-156373836-INS-232
chr4-156379641-INS-545
chr4-156379700-INS-1110
chr4-156380077-INS-187
chr4-156380793-INS-260
chr4-156396672-INS-2394
chr4-156404221-INS-200
chr4-156408141-INS-1079
chr4-156476260-INS-264
chr4-156477751-INS-595
chr4-156569545-INS-956
chr5-3052893-INS-106
chr5-3052931-INS-51
chr5-3053673-INS-61
chr5-3138725-INS-3197
chr5-3267260-INS-66
chr5-3267503-INS-68
chr5-3271582-INS-422
chr5-3291634-INS-54
chr5-3291977-INS-66
chr5-3333860-INS-87
chr5-3369123-INS-7116
chr5-3382944-INS-64
chr5-3470814-INS-415
chr5-3475441-INS-972
chr5-3495159-INS-201
chr5-3501060-INS-2004
chr5-3512566-INS-118
chr5-3512895-INS-212
chr5-3526492-INS-51
chr5-3539145-INS-70
chr5-3545061-INS-66
chr5-3545081-INS-165
chr5-3575979-INS-221
chr5-3584081-INS-81
chr5-3584913-INS-356
chr5-3585031-INS-780
chr5-3663194-INS-104
chr5-3663744-INS-64
chr5-3727494-INS-92
chr5-3771665-INS-180
chr5-3917456-INS-125
chr5-3954923-INS-66
chr5-3983832-INS-346
chr5-4004220-INS-51
chr5-4012735-INS-356
chr5-4013033-INS-414
chr5-4052135-INS-102
chr5-4102260-INS-196
chr5-4104388-INS-396
chr5-4107494-INS-154

chr5-4108129-INS-144
chr5-4116940-INS-88
chr5-4119641-INS-248
chr5-4138358-INS-310
chr5-4199358-INS-205
chr5-4205942-INS-72
chr5-4226173-INS-914
chr5-4282258-INS-176
chr5-4350536-INS-6199
chr5-4351579-INS-140
chr5-4413786-INS-324
chr5-4447111-INS-1274
chr5-4447199-INS-593
chr5-4487746-INS-1035
chr5-4489350-INS-401
chr5-4521784-INS-118
chr5-4546072-INS-54
chr5-4550245-INS-116
chr5-4587884-INS-443
chr5-4597880-INS-237
chr5-4597880-INS-522
chr5-4613504-INS-768
chr5-4688100-INS-2254
chr5-4733196-INS-62
chr5-4733323-INS-72
chr5-4773683-INS-344
chr5-4784747-INS-7095
chr5-4800958-INS-1801
chr5-4834786-INS-6375
chr5-4845446-INS-408
chr5-4873388-INS-76
chr5-4897834-INS-6280
chr5-4916616-INS-84
chr5-4935501-INS-6394
chr5-4962845-INS-151
chr5-4997069-INS-65
chr5-5005917-INS-54
chr5-5037827-INS-169
chr5-5058792-INS-445
chr5-5121239-INS-1254
chr5-5136042-INS-101
chr5-5223729-INS-6236
chr5-5239656-INS-50
chr5-5258996-INS-72
chr5-5269997-INS-11914
chr5-5272543-INS-210
chr5-5278652-INS-835
chr5-5325826-INS-3319
chr5-5361624-INS-212
chr5-5361708-INS-89
chr5-5367039-INS-61
chr5-5378505-INS-224
chr5-5382350-INS-140
chr5-5387280-INS-1575
chr5-5413535-INS-98
chr5-5432236-INS-202
chr5-5442656-INS-544
chr5-5448336-INS-54
chr5-5458063-INS-1630
chr5-5463743-INS-236
chr5-5472709-INS-1177
chr5-5491917-INS-52
chr5-5496511-INS-1884
chr5-5532230-INS-52
chr5-5533342-INS-3410
chr5-5534611-INS-50
chr5-5546163-INS-52
chr5-5548094-INS-198
chr5-5562779-INS-1000
chr5-5592229-INS-64
chr5-5623593-INS-51
chr5-5680762-INS-201
chr5-5770244-INS-3053
chr5-5778684-INS-169
chr5-5797793-INS-7061
chr5-5801047-INS-52
chr5-5802033-INS-55

chr5-5835992-INS-159
chr5-5846283-INS-6416
chr5-5861561-INS-51
chr5-5866001-INS-50
chr5-5876924-INS-2304
chr5-5893547-INS-6647
chr5-5903469-INS-1521
chr5-5930329-INS-132
chr5-5968410-INS-526
chr5-5969471-INS-50
chr5-6010645-INS-104
chr5-6012232-INS-70
chr5-6048258-INS-7185
chr5-6071578-INS-92
chr5-6071882-INS-432
chr5-6120947-INS-7213
chr5-6155194-INS-150
chr5-6157816-INS-3309
chr5-6159130-INS-64
chr5-6165792-INS-126
chr5-6192724-INS-1234
chr5-6220279-INS-51
chr5-6329760-INS-245
chr5-6337780-INS-52
chr5-6338184-INS-385
chr5-6521950-INS-64
chr5-6529936-INS-460
chr5-6530551-INS-250
chr5-6540938-INS-499
chr5-6590484-INS-1415
chr5-6621268-INS-2367
chr5-6678429-INS-322
chr5-6683968-INS-6708
chr5-6686146-INS-2856
chr5-6692106-INS-1892
chr5-6695321-INS-806
chr5-6718515-INS-65
chr5-6753048-INS-2856
chr5-6758145-INS-3409
chr5-6758145-INS-8050
chr5-6765427-INS-200
chr5-6798768-INS-52
chr5-6888523-INS-7104
chr5-6918183-INS-7817
chr5-6928518-INS-282
chr5-6928606-INS-120
chr5-6971263-INS-6483
chr5-7036142-INS-104
chr5-7039066-INS-50
chr5-7070338-INS-123
chr5-7076636-INS-50
chr5-7093230-INS-6233
chr5-7103839-INS-72
chr5-7116484-INS-1476
chr5-7163659-INS-2793
chr5-7170741-INS-112
chr5-7183267-INS-54
chr5-7190969-INS-57
chr5-7262692-INS-54
chr5-7283836-INS-88
chr5-7285751-INS-55
chr5-7287213-INS-66
chr5-7308065-INS-55
chr5-7312742-INS-6516
chr5-7571492-INS-70
chr5-7601718-INS-6356
chr5-7623918-INS-1214
chr5-7626070-INS-60
chr5-7628197-INS-57
chr5-7630206-INS-56
chr5-7635043-INS-50
chr5-7650277-INS-11616
chr5-7696765-INS-1255
chr5-7754163-INS-5737
chr5-7899608-INS-51
chr5-7950956-INS-50
chr5-7952692-INS-5009

chr5-7958548-INS-50
chr5-7979090-INS-66
chr5-8001515-INS-3494
chr5-8032365-INS-736
chr5-8057884-INS-2013
chr5-8058165-INS-531
chr5-8061883-INS-6729
chr5-8133717-INS-658
chr5-8147281-INS-355
chr5-8164066-INS-6413
chr5-8164530-INS-1505
chr5-8171681-INS-50
chr5-8210529-INS-304
chr5-8210547-INS-54
chr5-8230926-INS-3128
chr5-8230960-INS-99
chr5-8235351-INS-159
chr5-8239391-INS-84
chr5-8242940-INS-8270
chr5-8282935-INS-80
chr5-8283206-INS-328
chr5-8304816-INS-3553
chr5-8316602-INS-211
chr5-8530599-INS-117
chr5-8534920-INS-6703
chr5-8590364-INS-201
chr5-8614258-INS-2486
chr5-8741773-INS-212
chr5-8761989-INS-848
chr5-8764183-INS-291
chr5-8781661-INS-112
chr5-8793682-INS-243
chr5-8822836-INS-178
chr5-8826530-INS-5197
chr5-8839470-INS-2790
chr5-8839909-INS-2929
chr5-8882668-INS-252
chr5-8882668-INS-63
chr5-8924283-INS-2854
chr5-8951597-INS-1214
chr5-8976307-INS-60
chr5-9002298-INS-1188
chr5-9024524-INS-223
chr5-9091027-INS-50
chr5-9140830-INS-2231
chr5-9296663-INS-52
chr5-9303618-INS-632
chr5-9313043-INS-185
chr5-9325967-INS-1336
chr5-9433142-INS-6274
chr5-9440482-INS-6981
chr5-9453346-INS-4361
chr5-9477226-INS-4330
chr5-9482421-INS-6438
chr5-9502651-INS-1461
chr5-9502651-INS-364
chr5-9534290-INS-144
chr5-9580033-INS-90
chr5-9590271-INS-69
chr5-9593772-INS-1750
chr5-9594277-INS-4306
chr5-9610310-INS-160
chr5-9643249-INS-5196
chr5-9722496-INS-52
chr5-9738466-INS-731
chr5-9773524-INS-527
chr5-9790367-INS-119
chr5-9809125-INS-69
chr5-9809128-INS-158
chr5-9861298-INS-645
chr5-9959842-INS-54
chr5-9965836-INS-77
chr5-10084490-INS-2828
chr5-10091606-INS-390
chr5-10104052-INS-108
chr5-10129332-INS-50
chr5-10157400-INS-54

chr5-10161442-INS-10171
chr5-10194446-INS-161
chr5-10208554-INS-1096
chr5-10208798-INS-1400
chr5-10215034-INS-8839
chr5-10239273-INS-4474
chr5-10332624-INS-80
chr5-10405164-INS-2627
chr5-10433953-INS-588
chr5-10437499-INS-50
chr5-10438763-INS-89
chr5-10465093-INS-1584
chr5-10470235-INS-58
chr5-10472359-INS-6244
chr5-10477145-INS-1795
chr5-10569213-INS-4496
chr5-10594864-INS-6514
chr5-10611355-INS-193
chr5-10708387-INS-192
chr5-10711973-INS-62
chr5-10736897-INS-6486
chr5-10824763-INS-207
chr5-10824900-INS-3568
chr5-10866273-INS-54
chr5-11995279-INS-3114
chr5-12040432-INS-2999
chr5-12083877-INS-189
chr5-12091557-INS-195
chr5-12099182-INS-1887
chr5-12121460-INS-168
chr5-12148783-INS-617
chr5-12178887-INS-68
chr5-12191837-INS-1289
chr5-12227591-INS-62
chr5-12227656-INS-699
chr5-12255823-INS-202
chr5-12274583-INS-321
chr5-12282089-INS-64
chr5-12287159-INS-71
chr5-12289961-INS-458
chr5-12293893-INS-50
chr5-12339206-INS-389
chr5-12339325-INS-123
chr5-12362023-INS-86
chr5-12372730-INS-6891
chr5-12403792-INS-100
chr5-12431080-INS-62
chr5-12439481-INS-3235
chr5-12509467-INS-6575
chr5-12534175-INS-887
chr5-12565828-INS-52
chr5-12567069-INS-3720
chr5-12581982-INS-245
chr5-12591687-INS-67
chr5-12604063-INS-66
chr5-12617273-INS-50
chr5-12667974-INS-81
chr5-12689811-INS-1407
chr5-12694173-INS-50
chr5-12694807-INS-103
chr5-12695210-INS-140
chr5-12696139-INS-106
chr5-12725372-INS-2937
chr5-12738707-INS-7266
chr5-12777219-INS-344
chr5-12777813-INS-185
chr5-12800549-INS-98
chr5-12858115-INS-1819
chr5-12885920-INS-6390
chr5-12892823-INS-1021
chr5-12915346-INS-7123
chr5-12930216-INS-54
chr5-12934204-INS-88
chr5-12944880-INS-3061
chr5-12947925-INS-58
chr5-12948896-INS-60
chr5-12956646-INS-6288

chr5-12959305-INS-3804
chr5-12970419-INS-6480
chr5-12997476-INS-265
chr5-12997498-INS-81
chr5-12997743-INS-52
chr5-13062760-INS-2055
chr5-13064078-INS-166
chr5-13077372-INS-2933
chr5-13077949-INS-168
chr5-13078234-INS-4788
chr5-13082069-INS-52
chr5-13087033-INS-50
chr5-13099298-INS-61
chr5-13108707-INS-50
chr5-13128658-INS-1196
chr5-13140194-INS-204
chr5-13157176-INS-809
chr5-13189575-INS-5009
chr5-13215147-INS-56
chr5-13215625-INS-1518
chr5-13215735-INS-305
chr5-13215791-INS-51
chr5-13223160-INS-180
chr5-13225426-INS-6211
chr5-13233829-INS-6447
chr5-13241863-INS-71
chr5-13327673-INS-1688
chr5-13331826-INS-6641
chr5-13364750-INS-52
chr5-13509190-INS-1100
chr5-13509695-INS-67
chr5-13573911-INS-58
chr5-13575202-INS-98
chr5-13602845-INS-6426
chr5-13603351-INS-1657
chr5-13623152-INS-197
chr5-13631979-INS-63
chr5-13694731-INS-52
chr5-13698485-INS-6587
chr5-13729966-INS-6275
chr5-13744270-INS-6387
chr5-13787727-INS-2173
chr5-13810291-INS-6371
chr5-13824587-INS-278
chr5-13826520-INS-1614
chr5-13827170-INS-22854
chr5-13857362-INS-2802
chr5-13857731-INS-2618
chr5-13863206-INS-1227
chr5-13945902-INS-6453
chr5-13964729-INS-160
chr5-13974476-INS-7216
chr5-13987085-INS-6997
chr5-13995510-INS-55
chr5-14000391-INS-6494
chr5-14013891-INS-317
chr5-14055187-INS-55
chr5-14056315-INS-58
chr5-14070058-INS-145
chr5-14187295-INS-74
chr5-14230470-INS-636
chr5-14238139-INS-59
chr5-14249832-INS-148
chr5-14252820-INS-90
chr5-14253382-INS-57
chr5-14286560-INS-210
chr5-14308929-INS-6274
chr5-14320634-INS-554
chr5-14330661-INS-3122
chr5-14335987-INS-1486
chr5-14339236-INS-437
chr5-14345694-INS-7136
chr5-14378276-INS-246
chr5-14425670-INS-86
chr5-14427269-INS-499
chr5-14457852-INS-497
chr5-14466019-INS-97

chr5-14468637-INS-937
chr5-14519283-INS-164
chr5-14522832-INS-63
chr5-14537201-INS-3551
chr5-14596509-INS-200
chr5-14619416-INS-242
chr5-14703245-INS-72
chr5-14706583-INS-50
chr5-14707750-INS-58
chr5-14708833-INS-477
chr5-14721534-INS-56
chr5-14737273-INS-199
chr5-14747585-INS-22668
chr5-14783093-INS-6399
chr5-14790086-INS-94
chr5-14790183-INS-254
chr5-14830050-INS-1034
chr5-14843538-INS-361
chr5-14843635-INS-154
chr5-14858089-INS-55
chr5-14877072-INS-1388
chr5-14920751-INS-54
chr5-14932049-INS-1194
chr5-15000723-INS-800
chr5-15038425-INS-1122
chr5-15054590-INS-731
chr5-15076881-INS-167
chr5-15666770-INS-1446
chr5-15702240-INS-815
chr5-15735270-INS-81
chr5-15741492-INS-610
chr5-15823067-INS-68
chr5-15869368-INS-1286
chr5-15904920-INS-269
chr5-15907849-INS-1000
chr5-16028438-INS-230
chr5-16065023-INS-51
chr5-16088992-INS-2187
chr5-16096070-INS-2770
chr5-16121745-INS-72
chr5-16129173-INS-6420
chr5-16168918-INS-72
chr5-16184847-INS-87
chr5-16194186-INS-86
chr5-16227676-INS-1437
chr5-16232423-INS-253
chr5-16299016-INS-153
chr5-16299856-INS-50
chr5-16339839-INS-52
chr5-16446416-INS-168
chr5-16501603-INS-631
chr5-16501793-INS-1326
chr5-16501796-INS-56
chr5-16502201-INS-135
chr5-16543020-INS-520
chr5-16556883-INS-55
chr5-16609364-INS-54
chr5-16634651-INS-127
chr5-16635242-INS-50
chr5-16635758-INS-88
chr5-16642043-INS-6993
chr5-16699014-INS-62
chr5-16699302-INS-97
chr5-16699987-INS-81
chr5-16745496-INS-132
chr5-16810991-INS-122
chr5-16860231-INS-74
chr5-16884181-INS-1124
chr5-16908839-INS-148
chr5-16917980-INS-571
chr5-16937967-INS-63
chr5-16938934-INS-58
chr5-16974183-INS-245
chr5-17055671-INS-11701
chr5-17057882-INS-50
chr5-17077823-INS-6205
chr5-17088038-INS-58

chr5-17273715-INS-6589
chr5-17284304-INS-52
chr5-17326943-INS-4557
chr5-17354577-INS-681
chr5-17354965-INS-80
chr5-17359408-INS-6408
chr5-17377559-INS-690
chr5-17397812-INS-5972
chr5-17405703-INS-191
chr5-17433678-INS-224
chr5-17447108-INS-3623
chr5-17454209-INS-2677
chr5-17459635-INS-1421
chr5-17480069-INS-62
chr5-17506358-INS-54
chr5-17588004-INS-145
chr5-17601076-INS-56
chr5-17612594-INS-3572
chr5-17628007-INS-1263
chr5-17654047-INS-185
chr5-17657459-INS-6463
chr5-17673065-INS-64
chr5-17676857-INS-6493
chr5-17716093-INS-146
chr5-17790781-INS-13861
chr5-17805601-INS-65
chr5-17813446-INS-636
chr5-17906414-INS-50
chr5-17938167-INS-53
chr5-17975433-INS-72
chr5-17991331-INS-153
chr5-17992072-INS-63
chr5-18021409-INS-2590
chr5-18054939-INS-80
chr5-18058080-INS-175
chr5-18095621-INS-5219
chr5-18157851-INS-6813
chr5-18174545-INS-60
chr5-18230612-INS-6958
chr5-18244589-INS-50
chr5-18247921-INS-229
chr5-18248492-INS-50
chr5-18278423-INS-7070
chr5-18281631-INS-2678
chr5-18288662-INS-50
chr5-18341854-INS-434
chr5-18415587-INS-491
chr5-18421224-INS-52
chr5-18431467-INS-228
chr5-18431469-INS-109
chr5-18431916-INS-496
chr5-18447939-INS-2127
chr5-18457541-INS-92
chr5-18461712-INS-51
chr5-18518212-INS-460
chr5-18519351-INS-1884
chr5-18534317-INS-56
chr5-18534321-INS-253
chr5-18542437-INS-51
chr5-18546971-INS-63
chr5-18547741-INS-165
chr5-18547751-INS-70
chr5-18612839-INS-147
chr5-18623991-INS-850
chr5-18637422-INS-3511
chr5-18657787-INS-10569
chr5-18699890-INS-84
chr5-18703792-INS-6440
chr5-18776468-INS-307
chr5-18777103-INS-80
chr5-18787128-INS-6210
chr5-18791190-INS-6278
chr5-18802476-INS-6264
chr5-18810114-INS-384
chr5-18824666-INS-146
chr5-18836743-INS-519
chr5-18836947-INS-469

chr5-18857117-INS-64
chr5-18888425-INS-324
chr5-18917808-INS-262
chr5-18927085-INS-1053
chr5-18971171-INS-1524
chr5-18987466-INS-2198
chr5-18994304-INS-50
chr5-18999674-INS-3500
chr5-19001815-INS-126
chr5-19001992-INS-923
chr5-19010336-INS-1181
chr5-19069412-INS-195
chr5-19069491-INS-498
chr5-19128161-INS-60
chr5-19159177-INS-2617
chr5-19203323-INS-1128
chr5-19248105-INS-3840
chr5-19248566-INS-63
chr5-19277035-INS-437
chr5-19310753-INS-6160
chr5-19332558-INS-199
chr5-19379713-INS-837
chr5-19380576-INS-853
chr5-19383158-INS-550
chr5-19413688-INS-97
chr5-19413858-INS-286
chr5-19414054-INS-618
chr5-19414306-INS-809
chr5-19440883-INS-81
chr5-19484467-INS-69
chr5-19510332-INS-3273
chr5-19512399-INS-6219
chr5-19513119-INS-107
chr5-19523906-INS-400
chr5-19541408-INS-55
chr5-19559352-INS-59
chr5-19561046-INS-1949
chr5-19567848-INS-3006
chr5-19570592-INS-156
chr5-19582353-INS-69
chr5-19582773-INS-3175
chr5-19584191-INS-50
chr5-19607589-INS-205
chr5-19692389-INS-3286
chr5-19709527-INS-60
chr5-19710017-INS-66
chr5-19714033-INS-1178
chr5-19775926-INS-52
chr5-19851074-INS-69
chr5-19866335-INS-76
chr5-19898834-INS-64
chr5-19981450-INS-75
chr5-20045810-INS-60
chr5-20049892-INS-6393
chr5-20064258-INS-6382
chr5-20075999-INS-70
chr5-20080851-INS-74
chr5-20081562-INS-908
chr5-20159755-INS-594
chr5-20192602-INS-120
chr5-20193523-INS-54
chr5-20199156-INS-1481
chr5-20232066-INS-229
chr5-20265135-INS-649
chr5-20301729-INS-6502
chr5-20325995-INS-381
chr5-20410727-INS-70
chr5-20425813-INS-50
chr5-20451460-INS-80
chr5-20463315-INS-56
chr5-20469672-INS-2154
chr5-20593309-INS-50
chr5-20609480-INS-51
chr5-20655371-INS-764
chr5-20678587-INS-68
chr5-20719279-INS-272
chr5-20720097-INS-166

chr5-20735084-INS-3132
chr5-20750872-INS-197
chr5-20779766-INS-52
chr5-20814566-INS-80
chr5-20843649-INS-453
chr5-20877805-INS-193
chr5-20878166-INS-478
chr5-20900248-INS-178
chr5-20929154-INS-323
chr5-20943974-INS-105
chr5-20943974-INS-289
chr5-20944601-INS-208
chr5-20987051-INS-205
chr5-20988703-INS-55
chr5-20991714-INS-135
chr5-20991925-INS-315
chr5-20999211-INS-2736
chr5-21008110-INS-434
chr5-21040189-INS-122
chr5-21040271-INS-122
chr5-21063013-INS-494
chr5-21174413-INS-229
chr5-21175085-INS-52
chr5-21175094-INS-165
chr5-21179481-INS-530
chr5-21181331-INS-811
chr5-21193501-INS-462
chr5-21212443-INS-217
chr5-21290265-INS-295
chr5-21331128-INS-6559
chr5-21346292-INS-375
chr5-21351995-INS-156
chr5-21352723-INS-1254
chr5-21365665-INS-70
chr5-21384117-INS-50
chr5-21385772-INS-105
chr5-21392316-INS-78
chr5-21392364-INS-2275
chr5-21402939-INS-1148
chr5-21429011-INS-141
chr5-21471323-INS-270
chr5-21471793-INS-165
chr5-21529658-INS-153
chr5-21533384-INS-114
chr5-21620140-INS-6363
chr5-21634097-INS-54
chr5-21670695-INS-324
chr5-21691720-INS-72
chr5-21706682-INS-3062
chr5-21720350-INS-58
chr5-21720690-INS-56
chr5-21737127-INS-106
chr5-21744424-INS-1408
chr5-21791750-INS-378
chr5-21792391-INS-66
chr5-21793829-INS-203
chr5-21805946-INS-240
chr5-21843885-INS-1011
chr5-21844070-INS-70
chr5-21844177-INS-239
chr5-21846775-INS-190
chr5-21861741-INS-4989
chr5-21865663-INS-106
chr5-21888985-INS-63
chr5-21889572-INS-195
chr5-21890471-INS-203
chr5-21897967-INS-73
chr5-21898190-INS-514
chr5-21916444-INS-262
chr5-21921018-INS-181
chr5-21921431-INS-572
chr5-21922880-INS-5918
chr5-21924632-INS-1018
chr5-21932359-INS-68
chr5-21932806-INS-2710
chr5-21935453-INS-370
chr5-21935709-INS-724

chr5-21957162-INS-137
chr5-21960628-INS-93
chr5-21961280-INS-393
chr5-21985899-INS-211
chr5-22009897-INS-186
chr5-22027404-INS-217
chr5-22033604-INS-296
chr5-22033691-INS-103
chr5-22060478-INS-78
chr5-22062543-INS-283
chr5-22100391-INS-50
chr5-22111669-INS-198
chr5-22114090-INS-195
chr5-22136988-INS-644
chr5-22174726-INS-90
chr5-22329535-INS-68
chr5-22352824-INS-84
chr5-22467796-INS-222
chr5-22468402-INS-51
chr5-22469214-INS-50
chr5-22520649-INS-58
chr5-22551412-INS-52
chr5-22589312-INS-186
chr5-22612587-INS-7705
chr5-22621749-INS-74
chr5-22648978-INS-655
chr5-22648990-INS-297
chr5-22654242-INS-240
chr5-22673918-INS-59
chr5-22698631-INS-709
chr5-22707605-INS-7214
chr5-22723542-INS-52
chr5-22724068-INS-121
chr5-22833274-INS-3438
chr5-22839772-INS-165
chr5-22840358-INS-668
chr5-22864825-INS-6260
chr5-22874702-INS-157
chr5-22908719-INS-60
chr5-23005442-INS-1569
chr5-23009884-INS-60
chr5-23017008-INS-122
chr5-23019525-INS-6013
chr5-23020282-INS-596
chr5-23038809-INS-7001
chr5-23067418-INS-182
chr5-23067470-INS-52
chr5-23075962-INS-7595
chr5-23095624-INS-1861
chr5-23097229-INS-6213
chr5-23112914-INS-2803
chr5-23125804-INS-6845
chr5-23158906-INS-2767
chr5-23201703-INS-2012
chr5-23210861-INS-6258
chr5-23222740-INS-59
chr5-23227851-INS-92
chr5-23242831-INS-324
chr5-23242990-INS-682
chr5-23262021-INS-224
chr5-23262026-INS-677
chr5-23270228-INS-462
chr5-23270308-INS-184
chr5-23290709-INS-54
chr5-23293926-INS-6256
chr5-23311064-INS-166
chr5-23321502-INS-6442
chr5-23328230-INS-573
chr5-23331778-INS-696
chr5-23361289-INS-50
chr5-23364708-INS-85
chr5-23385905-INS-3045
chr5-23388856-INS-543
chr5-23404988-INS-57
chr5-23409266-INS-2580
chr5-23446988-INS-68
chr5-23447090-INS-317

chr5-23454281-INS-6947
chr5-23465947-INS-166
chr5-23496358-INS-3063
chr5-23534015-INS-6295
chr5-23555992-INS-191
chr5-23556109-INS-1226
chr5-23567759-INS-174
chr5-23587677-INS-186
chr5-23590104-INS-3810
chr5-23751569-INS-197
chr5-23755061-INS-98
chr5-23758416-INS-156
chr5-23786989-INS-542
chr5-23842094-INS-60
chr5-23844357-INS-189
chr5-23844489-INS-546
chr5-23866575-INS-129
chr5-23895399-INS-705
chr5-23939201-INS-203
chr5-23940416-INS-450
chr5-23947455-INS-50
chr5-23972915-INS-504
chr5-23986504-INS-208
chr5-24017245-INS-72
chr5-24017965-INS-222
chr5-24020012-INS-146
chr5-24023971-INS-58
chr5-24050625-INS-244
chr5-24052195-INS-390
chr5-24084241-INS-2913
chr5-24090328-INS-200
chr5-24094494-INS-140
chr5-24094494-INS-56
chr5-24094498-INS-476
chr5-24110216-INS-126
chr5-24149552-INS-204
chr5-24155817-INS-3302
chr5-24157536-INS-99
chr5-24157564-INS-205
chr5-24200747-INS-147
chr5-24219618-INS-260
chr5-24232078-INS-982
chr5-24250938-INS-62
chr5-24286414-INS-691
chr5-24397838-INS-4561
chr5-24409931-INS-2397
chr5-24410798-INS-300
chr5-24418672-INS-228
chr5-24428503-INS-334
chr5-24428813-INS-638
chr5-24428837-INS-56
chr5-24465063-INS-51
chr5-24487086-INS-611
chr5-24496615-INS-2821
chr5-24525238-INS-83
chr5-24575387-INS-50
chr5-24712101-INS-186
chr5-24819331-INS-411
chr5-24837466-INS-50
chr5-24837550-INS-103
chr5-24848689-INS-60
chr5-24851875-INS-66
chr5-24860429-INS-8692
chr5-24866822-INS-624
chr5-24873840-INS-344
chr5-24885083-INS-9062
chr5-24889289-INS-58
chr5-24912456-INS-627
chr5-24922528-INS-54
chr5-24927219-INS-138
chr5-24950435-INS-398
chr5-24950948-INS-101
chr5-24951831-INS-508
chr5-24982259-INS-84
chr5-24983127-INS-205
chr5-24993004-INS-152
chr5-24993183-INS-1819

chr5-24994965-INS-124
chr5-24996848-INS-59
chr5-25070592-INS-199
chr5-25130348-INS-61
chr5-25162561-INS-238
chr5-25169585-INS-127
chr5-25178081-INS-273
chr5-25180079-INS-397
chr5-25192457-INS-70
chr5-25192471-INS-320
chr5-25192471-INS-734
chr5-25194219-INS-52
chr5-25220405-INS-336
chr5-25236606-INS-54
chr5-25236879-INS-208
chr5-25282414-INS-150
chr5-25295136-INS-68
chr5-25343463-INS-58
chr5-25346429-INS-54
chr5-25366375-INS-2102
chr5-25369828-INS-51
chr5-25369861-INS-199
chr5-25417486-INS-5466
chr5-25420538-INS-374
chr5-25420718-INS-445
chr5-25421086-INS-338
chr5-25421151-INS-230
chr5-25422718-INS-738
chr5-25447851-INS-160
chr5-25564050-INS-376
chr5-25598046-INS-233
chr5-25611231-INS-1331
chr5-25623269-INS-198
chr5-25623351-INS-174
chr5-25642152-INS-90
chr5-25642154-INS-295
chr5-25647659-INS-50
chr5-25654815-INS-135
chr5-25656135-INS-74
chr5-25660393-INS-102
chr5-25689246-INS-255
chr5-25700069-INS-116
chr5-25717655-INS-58
chr5-25724753-INS-56
chr5-25753347-INS-214
chr5-25754629-INS-2086
chr5-25757865-INS-52
chr5-25758703-INS-52
chr5-25789007-INS-1038
chr5-25803810-INS-100
chr5-25824197-INS-168
chr5-25824349-INS-495
chr5-25831282-INS-208
chr5-25835010-INS-195
chr5-25846218-INS-4397
chr5-25856502-INS-493
chr5-25872373-INS-448
chr5-25935079-INS-317
chr5-25952951-INS-62
chr5-25954376-INS-1188
chr5-26018503-INS-288
chr5-26020218-INS-120
chr5-26020260-INS-560
chr5-26024025-INS-220
chr5-26030544-INS-207
chr5-26030562-INS-73
chr5-26049214-INS-78
chr5-26060076-INS-50
chr5-26087417-INS-5190
chr5-26094053-INS-467
chr5-26102215-INS-2721
chr5-26114951-INS-719
chr5-26135581-INS-2935
chr5-26261609-INS-60
chr5-26334402-INS-404
chr5-26384735-INS-1276
chr5-26425333-INS-50

chr5-26466494-INS-18417
chr5-26470813-INS-1597
chr5-26537745-INS-50
chr5-26563707-INS-2900
chr5-26580658-INS-70
chr5-26595975-INS-139
chr5-26636943-INS-2462
chr5-26669035-INS-56
chr5-26675097-INS-185
chr5-26681166-INS-6177
chr5-26711432-INS-477
chr5-26718838-INS-81
chr5-26718860-INS-570
chr5-26743091-INS-7165
chr5-26780422-INS-4404
chr5-26801944-INS-98
chr5-26802290-INS-52
chr5-26806471-INS-6465
chr5-26846893-INS-3082
chr5-26912260-INS-6425
chr5-26920601-INS-428
chr5-26974527-INS-522
chr5-26981204-INS-1289
chr5-26986493-INS-4669
chr5-27012449-INS-76
chr5-27036808-INS-108
chr5-27084001-INS-272
chr5-27084001-INS-7567
chr5-27115230-INS-926
chr5-27222242-INS-61
chr5-27303529-INS-1544
chr5-27303814-INS-318
chr5-27304013-INS-106
chr5-27324334-INS-54
chr5-27376640-INS-75
chr5-27379504-INS-54
chr5-27380163-INS-52
chr5-27387263-INS-6168
chr5-27451273-INS-54
chr5-27469329-INS-72
chr5-27479529-INS-54
chr5-27510415-INS-51
chr5-27578169-INS-85
chr5-27598468-INS-105
chr5-27602184-INS-75
chr5-27647838-INS-1861
chr5-27671096-INS-173
chr5-27674631-INS-60
chr5-27681810-INS-50
chr5-27700662-INS-3861
chr5-27860372-INS-54
chr5-27860632-INS-514
chr5-27864819-INS-58
chr5-27872070-INS-115
chr5-27878619-INS-50
chr5-27881998-INS-180
chr5-27927835-INS-112
chr5-27930006-INS-126
chr5-27949901-INS-242
chr5-27949934-INS-3374
chr5-27962803-INS-284
chr5-27997524-INS-201
chr5-27998364-INS-54
chr5-28036193-INS-542
chr5-28044636-INS-3033
chr5-28045108-INS-5327
chr5-28049871-INS-62
chr5-28052255-INS-6936
chr5-28107553-INS-2816
chr5-28220824-INS-87
chr5-28252813-INS-426
chr5-28253611-INS-1649
chr5-28259064-INS-3178
chr5-28261614-INS-72
chr5-28274201-INS-257
chr5-28289791-INS-1744
chr5-28323938-INS-130

chr5-28333901-INS-59
chr5-28353132-INS-62
chr5-28369836-INS-62
chr5-28445094-INS-200
chr5-28462095-INS-153
chr5-28462286-INS-424
chr5-28477064-INS-2977
chr5-28496941-INS-164
chr5-28516197-INS-76
chr5-28568710-INS-220
chr5-28633366-INS-195
chr5-28656910-INS-50
chr5-28684698-INS-70
chr5-28684808-INS-90
chr5-28685848-INS-54
chr5-28697883-INS-496
chr5-28742103-INS-476
chr5-28839678-INS-6602
chr5-28858276-INS-6594
chr5-28877016-INS-11100
chr5-28934611-INS-57
chr5-29023035-INS-131
chr5-29057607-INS-54
chr5-29075601-INS-157
chr5-29091453-INS-6131
chr5-29099972-INS-96
chr5-29114887-INS-1566
chr5-29133477-INS-210
chr5-29279795-INS-50
chr5-29311454-INS-3254
chr5-29357668-INS-166
chr5-29360802-INS-3153
chr5-29361648-INS-548
chr5-29372965-INS-56
chr5-29415900-INS-115
chr5-29466959-INS-66
chr5-29466959-INS-7463
chr5-29578319-INS-60
chr5-29587469-INS-60
chr5-29592976-INS-293
chr5-29593040-INS-65
chr5-29634297-INS-529
chr5-29717764-INS-261
chr5-29766055-INS-295
chr5-29766741-INS-66
chr5-29767043-INS-538
chr5-29768991-INS-76
chr5-29769058-INS-1233
chr5-29795404-INS-219
chr5-29903310-INS-2843
chr5-29910401-INS-76
chr5-29922391-INS-1064
chr5-29929062-INS-242
chr5-29929540-INS-267
chr5-29930881-INS-78
chr5-29931000-INS-240
chr5-29931613-INS-190
chr5-30009410-INS-50
chr5-30019269-INS-191
chr5-30027432-INS-58
chr5-30035863-INS-153
chr5-30040832-INS-149
chr5-30107704-INS-92
chr5-30126553-INS-85
chr5-30163988-INS-252
chr5-30167631-INS-3205
chr5-30169157-INS-346
chr5-30187336-INS-6460
chr5-30192415-INS-73
chr5-30194978-INS-70
chr5-30231675-INS-1264
chr5-30283157-INS-410
chr5-30283298-INS-900
chr5-30336815-INS-64
chr5-30403134-INS-76
chr5-30434645-INS-81
chr5-30460382-INS-68

chr5-30463887-INS-498
chr5-30519161-INS-484
chr5-30519237-INS-105
chr5-30525826-INS-212
chr5-30544768-INS-72
chr5-30562999-INS-59
chr5-30590827-INS-180
chr5-30594474-INS-85
chr5-30604083-INS-466
chr5-30649693-INS-247
chr5-30651624-INS-242
chr5-30674916-INS-50
chr5-30696245-INS-172
chr5-30699560-INS-1490
chr5-30723901-INS-96
chr5-30727503-INS-206
chr5-30758388-INS-117
chr5-30767501-INS-3939
chr5-30769243-INS-50
chr5-30796638-INS-223
chr5-30827293-INS-75
chr5-30875852-INS-60
chr5-30958988-INS-80
chr5-31155090-INS-69
chr5-31155308-INS-105
chr5-31170470-INS-236
chr5-31215658-INS-272
chr5-31262175-INS-1205
chr5-31264694-INS-165
chr5-31273429-INS-72
chr5-31283876-INS-82
chr5-31373042-INS-199
chr5-31387611-INS-211
chr5-31479642-INS-56
chr5-31487385-INS-6547
chr5-31545546-INS-219
chr5-31599948-INS-63
chr5-31621213-INS-62
chr5-31648740-INS-140
chr5-31665705-INS-495
chr5-31667644-INS-2000
chr5-31691100-INS-51
chr5-31696553-INS-2462
chr5-31711547-INS-78
chr5-31726825-INS-243
chr5-31729546-INS-236
chr5-31730420-INS-114
chr5-31764341-INS-2403
chr5-31771051-INS-51
chr5-31783151-INS-5766
chr5-31809284-INS-161
chr5-31832887-INS-218
chr5-31854197-INS-223
chr5-31881437-INS-1783
chr5-31899758-INS-88
chr5-31899850-INS-287
chr5-31922160-INS-1438
chr5-31982484-INS-6406
chr5-32089377-INS-53
chr5-32126065-INS-50
chr5-32169571-INS-89
chr5-32186434-INS-617
chr5-32186520-INS-441
chr5-32202991-INS-134
chr5-32327002-INS-54
chr5-32338291-INS-65
chr5-32361232-INS-56
chr5-32361895-INS-94
chr5-32376056-INS-1832
chr5-32395482-INS-104
chr5-32407598-INS-70
chr5-32445675-INS-2355
chr5-32491955-INS-1153
chr5-32547741-INS-1502
chr5-32573740-INS-63
chr5-32676421-INS-56
chr5-32676796-INS-534

chr5-32683428-INS-4240
chr5-32694400-INS-67
chr5-32738434-INS-2097
chr5-32743400-INS-69
chr5-32871984-INS-61
chr5-32874111-INS-1298
chr5-32874268-INS-4568
chr5-32874425-INS-2058
chr5-32938590-INS-64
chr5-32945647-INS-1560
chr5-32982237-INS-57
chr5-33012530-INS-50
chr5-33038153-INS-50
chr5-33043448-INS-313
chr5-33043448-INS-80
chr5-33076431-INS-477
chr5-33179793-INS-209
chr5-33194791-INS-128
chr5-33198533-INS-80
chr5-33215089-INS-120
chr5-33230664-INS-123
chr5-33231226-INS-213
chr5-33251243-INS-133
chr5-33251405-INS-380
chr5-33257143-INS-116
chr5-33287968-INS-81
chr5-33339728-INS-6895
chr5-33346493-INS-659
chr5-33364042-INS-110
chr5-33385281-INS-51
chr5-33506982-INS-176
chr5-33550602-INS-50
chr5-33580148-INS-60
chr5-33603248-INS-64
chr5-33623740-INS-73
chr5-33687997-INS-258
chr5-33710187-INS-168
chr5-33734435-INS-153
chr5-33744559-INS-66
chr5-33751885-INS-96
chr5-33788527-INS-3126
chr5-33821686-INS-144
chr5-33822094-INS-72
chr5-33838088-INS-215
chr5-33868552-INS-202
chr5-33945301-INS-183
chr5-33981602-INS-78
chr5-33989477-INS-70
chr5-33992719-INS-254
chr5-34170621-INS-3106
chr5-34251368-INS-280
chr5-34264389-INS-52
chr5-34285494-INS-166
chr5-34287736-INS-2078
chr5-34358231-INS-196
chr5-34368579-INS-50
chr5-34404377-INS-78
chr5-34405875-INS-64
chr5-34472656-INS-50
chr5-34474152-INS-51
chr5-34498389-INS-235
chr5-34535797-INS-594
chr5-34548336-INS-72
chr5-34554193-INS-60
chr5-34576793-INS-50
chr5-34647474-INS-198
chr5-34657296-INS-105
chr5-34661835-INS-58
chr5-34683756-INS-6454
chr5-34724325-INS-176
chr5-34745452-INS-219
chr5-34748226-INS-2823
chr5-34753061-INS-174
chr5-34805199-INS-1088
chr5-34812900-INS-100
chr5-34836242-INS-82
chr5-34872426-INS-332

chr5-34874797-INS-75
chr5-34906343-INS-191
chr5-34925069-INS-103
chr5-35030601-INS-203
chr5-35061614-INS-215
chr5-35091734-INS-52
chr5-35161480-INS-54
chr5-35285454-INS-239
chr5-35337126-INS-60
chr5-35343852-INS-58
chr5-35366586-INS-50
chr5-35382306-INS-270
chr5-35456267-INS-50
chr5-35473334-INS-168
chr5-35473586-INS-84
chr5-35479349-INS-58
chr5-35487647-INS-252
chr5-35533742-INS-82
chr5-35569224-INS-160
chr5-35590593-INS-162
chr5-35633905-INS-78
chr5-35641799-INS-75
chr5-35641980-INS-142
chr5-35718361-INS-346
chr5-35733734-INS-6587
chr5-35760584-INS-260
chr5-35810641-INS-50
chr5-35871127-INS-1366
chr5-35909611-INS-2757
chr5-35981022-INS-180
chr5-35984916-INS-727
chr5-35999858-INS-144
chr5-36017081-INS-85
chr5-36018324-INS-104
chr5-36018590-INS-128
chr5-36028895-INS-64
chr5-36091769-INS-80
chr5-36097958-INS-1354
chr5-36132307-INS-88
chr5-36265357-INS-78
chr5-36265554-INS-286
chr5-36277324-INS-53
chr5-36345544-INS-374
chr5-36354738-INS-85
chr5-36373540-INS-133
chr5-36380245-INS-2498
chr5-36396751-INS-212
chr5-36397145-INS-212
chr5-36449994-INS-55
chr5-36461022-INS-52
chr5-36461527-INS-54
chr5-36476070-INS-88
chr5-36482393-INS-1499
chr5-36614781-INS-420
chr5-36697052-INS-983
chr5-36698648-INS-52
chr5-36711057-INS-260
chr5-36859555-INS-166
chr5-36898818-INS-210
chr5-36936744-INS-59
chr5-36960581-INS-51
chr5-36961945-INS-96
chr5-37058707-INS-52
chr5-37105719-INS-1870
chr5-37163901-INS-400
chr5-37220571-INS-54
chr5-37251766-INS-100
chr5-37326483-INS-71
chr5-37359678-INS-145
chr5-37453367-INS-50
chr5-37507916-INS-150
chr5-37518740-INS-1034
chr5-37554750-INS-330
chr5-37564360-INS-107
chr5-37564409-INS-81
chr5-37564636-INS-150
chr5-37640932-INS-54

chr5-37669680-INS-124
chr5-37692597-INS-59
chr5-37704396-INS-423
chr5-37753610-INS-560
chr5-37753805-INS-144
chr5-37753805-INS-1728
chr5-37787834-INS-342
chr5-37790336-INS-50
chr5-37881659-INS-62
chr5-37882113-INS-1878
chr5-37890451-INS-6229
chr5-37935905-INS-97
chr5-37968321-INS-84
chr5-37969192-INS-72
chr5-37973957-INS-67
chr5-38002309-INS-3137
chr5-38019214-INS-1122
chr5-38024458-INS-66
chr5-38067899-INS-7198
chr5-38090953-INS-52
chr5-38160996-INS-74
chr5-38272591-INS-234
chr5-38450101-INS-72
chr5-38517813-INS-394
chr5-38531008-INS-182
chr5-38553848-INS-1180
chr5-38587018-INS-2901
chr5-38636112-INS-150
chr5-38643663-INS-1598
chr5-38713864-INS-52
chr5-38754912-INS-50
chr5-38758450-INS-197
chr5-38780508-INS-12755
chr5-38788580-INS-360
chr5-38797440-INS-221
chr5-38816935-INS-630
chr5-38821627-INS-1526
chr5-38856222-INS-53
chr5-38867532-INS-58
chr5-38879470-INS-2836
chr5-38889520-INS-175
chr5-38916497-INS-2267
chr5-38938577-INS-50
chr5-38955340-INS-81
chr5-38987104-INS-400
chr5-38987753-INS-95
chr5-39016425-INS-50
chr5-39048478-INS-279
chr5-39084781-INS-446
chr5-39166620-INS-130
chr5-39201407-INS-836
chr5-39224476-INS-309
chr5-39225778-INS-8154
chr5-39233316-INS-95
chr5-39238837-INS-62
chr5-39262737-INS-52
chr5-39281538-INS-6732
chr5-39360571-INS-2845
chr5-39536882-INS-136
chr5-39543475-INS-343
chr5-39543660-INS-678
chr5-39563965-INS-1709
chr5-39597902-INS-2500
chr5-39650088-INS-122
chr5-39652776-INS-54
chr5-39652842-INS-187
chr5-39687634-INS-10098
chr5-39825567-INS-50
chr5-39875353-INS-50
chr5-39911007-INS-6646
chr5-39912870-INS-464
chr5-39969298-INS-199
chr5-39973907-INS-428
chr5-39976898-INS-4195
chr5-40039912-INS-1035
chr5-40045325-INS-54
chr5-40069877-INS-440

chr5-40098621-INS-62
chr5-40115484-INS-218
chr5-40119196-INS-1673
chr5-40125321-INS-6034
chr5-40242935-INS-922
chr5-40260886-INS-92
chr5-40264139-INS-218
chr5-40295567-INS-611
chr5-40325759-INS-398
chr5-40383777-INS-62
chr5-40425856-INS-64
chr5-40441173-INS-620
chr5-40446143-INS-82
chr5-40484582-INS-6979
chr5-40501191-INS-106
chr5-40528281-INS-916
chr5-40599604-INS-72
chr5-40609619-INS-55
chr5-40609619-INS-848
chr5-40697915-INS-1889
chr5-40719441-INS-90
chr5-40745572-INS-1341
chr5-40748400-INS-183
chr5-40821348-INS-1954
chr5-40847482-INS-64
chr5-40854100-INS-2501
chr5-40855685-INS-50
chr5-40942294-INS-921
chr5-40990279-INS-50
chr5-41047555-INS-52
chr5-41057466-INS-64
chr5-41059596-INS-66
chr5-41109570-INS-1499
chr5-41110161-INS-56
chr5-41116324-INS-6379
chr5-41153168-INS-3060
chr5-41153607-INS-9015
chr5-41189287-INS-60
chr5-41199478-INS-1955
chr5-41200143-INS-6394
chr5-41216281-INS-761
chr5-41237385-INS-56
chr5-41266738-INS-3852
chr5-41330017-INS-1244
chr5-41387306-INS-51
chr5-41430752-INS-726
chr5-41431263-INS-623
chr5-41434350-INS-1519
chr5-41467813-INS-6211
chr5-41493199-INS-53
chr5-41493336-INS-1491
chr5-41493678-INS-608
chr5-41495639-INS-2019
chr5-41496706-INS-52
chr5-41509521-INS-50
chr5-41514245-INS-6993
chr5-41551922-INS-1934
chr5-41565086-INS-60
chr5-41573948-INS-3202
chr5-41650164-INS-2034
chr5-41672500-INS-1089
chr5-41712673-INS-88
chr5-41738647-INS-5500
chr5-41813488-INS-54
chr5-41813556-INS-995
chr5-41839608-INS-859
chr5-41871557-INS-255
chr5-41872876-INS-609
chr5-41872891-INS-151
chr5-41873064-INS-431
chr5-41873116-INS-81
chr5-41873459-INS-1515
chr5-41873956-INS-7869
chr5-41891365-INS-736
chr5-41917233-INS-53
chr5-41953101-INS-488
chr5-41981779-INS-344

chr5-42006369-INS-219
chr5-42036234-INS-7082
chr5-42045221-INS-780
chr5-42056652-INS-7117
chr5-42070420-INS-54
chr5-42100370-INS-754
chr5-42117114-INS-2358
chr5-42118026-INS-69
chr5-42120006-INS-173
chr5-42125281-INS-298
chr5-42202186-INS-60
chr5-42238234-INS-436
chr5-42238292-INS-69
chr5-42238542-INS-102
chr5-42238683-INS-6405
chr5-42258057-INS-5075
chr5-42293173-INS-57
chr5-42321866-INS-52
chr5-42350501-INS-2182
chr5-42368981-INS-150
chr5-42390519-INS-6450
chr5-42408430-INS-396
chr5-42408601-INS-54
chr5-42436770-INS-2851
chr5-42473158-INS-508
chr5-42501197-INS-50
chr5-42525820-INS-4773
chr5-42541325-INS-148
chr5-42561165-INS-3442
chr5-42568483-INS-190
chr5-42568491-INS-91
chr5-42573145-INS-1949
chr5-42592923-INS-6407
chr5-42594102-INS-587
chr5-42692605-INS-168
chr5-42707930-INS-6331
chr5-42729672-INS-6172
chr5-42734893-INS-6748
chr5-42743247-INS-488
chr5-42747072-INS-6379
chr5-42759067-INS-307
chr5-42767842-INS-7206
chr5-42825384-INS-249
chr5-42825954-INS-111
chr5-42924122-INS-847
chr5-42946167-INS-2868
chr5-43017975-INS-186
chr5-43054721-INS-93
chr5-43067136-INS-373
chr5-43086854-INS-2184
chr5-43097681-INS-128
chr5-43112806-INS-1802
chr5-43124755-INS-66
chr5-43179441-INS-50
chr5-43191790-INS-195
chr5-43235811-INS-60
chr5-43237192-INS-176
chr5-43303817-INS-393
chr5-43312257-INS-237
chr5-43313183-INS-552
chr5-43315743-INS-1182
chr5-43320901-INS-913
chr5-43343943-INS-127
chr5-43376709-INS-77
chr5-43409412-INS-86
chr5-43453103-INS-203
chr5-43454566-INS-200
chr5-43470256-INS-74
chr5-43472127-INS-450
chr5-43511665-INS-88
chr5-43513133-INS-433
chr5-43514908-INS-7649
chr5-43521665-INS-6384
chr5-43563788-INS-52
chr5-43657643-INS-102
chr5-43657931-INS-157
chr5-43690874-INS-408

chr5-43698749-INS-151
chr5-43786539-INS-6767
chr5-43798331-INS-6481
chr5-43800832-INS-1595
chr5-43817900-INS-481
chr5-43819253-INS-2456
chr5-43851050-INS-254
chr5-43851142-INS-64
chr5-43883885-INS-879
chr5-43976031-INS-104
chr5-44004474-INS-103
chr5-44013512-INS-1604
chr5-44013620-INS-216
chr5-44038858-INS-3109
chr5-44074604-INS-6780
chr5-44099822-INS-157
chr5-44099822-INS-70
chr5-44116359-INS-2597
chr5-44145331-INS-168
chr5-44145675-INS-175
chr5-44178317-INS-119
chr5-44190707-INS-256
chr5-44227968-INS-425
chr5-44229372-INS-125
chr5-44267594-INS-216
chr5-44288351-INS-272
chr5-44323973-INS-5413
chr5-44416342-INS-76
chr5-44421575-INS-148
chr5-44426661-INS-144
chr5-44434430-INS-1954
chr5-44438113-INS-159
chr5-44438164-INS-77
chr5-44440119-INS-6578
chr5-44468766-INS-58
chr5-44501960-INS-1855
chr5-44503304-INS-5347
chr5-44516334-INS-8434
chr5-44571231-INS-6846
chr5-44575199-INS-7009
chr5-44578266-INS-2810
chr5-44592313-INS-201
chr5-44611559-INS-1897
chr5-44646308-INS-6481
chr5-44655883-INS-3677
chr5-44668804-INS-69
chr5-44677614-INS-6327
chr5-44727218-INS-186
chr5-44779719-INS-226
chr5-44780349-INS-90
chr5-44846014-INS-54
chr5-44867000-INS-58
chr5-44882836-INS-318
chr5-44882894-INS-112
chr5-44941244-INS-60
chr5-45057646-INS-7186
chr5-45068438-INS-6479
chr5-45092886-INS-1436
chr5-45139259-INS-2588
chr5-45187979-INS-6580
chr5-45194917-INS-52
chr5-45195562-INS-54
chr5-45235555-INS-7485
chr5-45245327-INS-863
chr5-45286693-INS-394
chr5-45350927-INS-1355
chr5-45352291-INS-54
chr5-45359678-INS-50
chr5-45369846-INS-7189
chr5-45372490-INS-254
chr5-45374299-INS-6481
chr5-45392768-INS-69
chr5-45421003-INS-1469
chr5-45485366-INS-6310
chr5-45641571-INS-1265
chr5-45744732-INS-51
chr5-45745704-INS-52

chr5-45752539-INS-50
chr5-45778215-INS-58
chr5-45790964-INS-232
chr5-45831386-INS-165
chr5-45838002-INS-1730
chr5-45838016-INS-50
chr5-45886310-INS-6454
chr5-45986322-INS-3062
chr5-46044799-INS-83
chr5-46072161-INS-83
chr5-46078578-INS-342
chr5-46083975-INS-754
chr5-46097262-INS-1198
chr5-46103049-INS-505
chr5-46112294-INS-5073
chr5-46213494-INS-7788
chr5-46232967-INS-71
chr5-46241847-INS-7276
chr5-46245794-INS-172
chr5-46252393-INS-338
chr5-46355913-INS-358
chr5-46356204-INS-441
chr5-46384576-INS-57
chr5-46399954-INS-6354
chr5-46460818-INS-433
chr5-46471650-INS-4216
chr5-46483539-INS-53
chr5-46487531-INS-3636
chr5-46509527-INS-66
chr5-46514778-INS-194
chr5-46529230-INS-6246
chr5-46558541-INS-212
chr5-46582405-INS-6467
chr5-46627154-INS-53
chr5-46671190-INS-92
chr5-46688076-INS-6491
chr5-46726270-INS-4608
chr5-46766357-INS-52
chr5-46774276-INS-1635
chr5-46780883-INS-6709
chr5-46787353-INS-1045
chr5-46791397-INS-1162
chr5-46868350-INS-56
chr5-46880358-INS-60
chr5-46901514-INS-6407
chr5-46904094-INS-2815
chr5-46905111-INS-56
chr5-46962990-INS-3008
chr5-46971708-INS-4629
chr5-46995665-INS-68
chr5-46996468-INS-118
chr5-47026659-INS-159
chr5-47127606-INS-210
chr5-47140780-INS-50
chr5-47174702-INS-67
chr5-47196990-INS-56
chr5-47232972-INS-1781
chr5-47246922-INS-615
chr5-47297955-INS-377
chr5-47338340-INS-62
chr5-47363305-INS-74
chr5-47374955-INS-60
chr5-47396200-INS-10806
chr5-47403732-INS-608
chr5-47420394-INS-51
chr5-47446740-INS-64
chr5-47532278-INS-2816
chr5-47594950-INS-184
chr5-47621568-INS-814
chr5-47631988-INS-1897
chr5-47641087-INS-52
chr5-47654681-INS-62
chr5-47685904-INS-50
chr5-47751919-INS-237
chr5-47761339-INS-265
chr5-47780843-INS-51
chr5-47800586-INS-5997

chr5-47802631-INS-112
chr5-47815067-INS-1628
chr5-47843741-INS-131
chr5-47930867-INS-2846
chr5-47964893-INS-6238
chr5-47981354-INS-1892
chr5-48007214-INS-6021
chr5-48029128-INS-54
chr5-48071352-INS-52
chr5-48071419-INS-82
chr5-48074541-INS-68
chr5-48084005-INS-6419
chr5-48183672-INS-178
chr5-48183900-INS-174
chr5-48189543-INS-50
chr5-48255896-INS-63
chr5-48346458-INS-55
chr5-48408044-INS-8363
chr5-48487844-INS-154
chr5-48487844-INS-50
chr5-48502323-INS-368
chr5-48504921-INS-102
chr5-48505103-INS-272
chr5-48505804-INS-198
chr5-48523225-INS-56
chr5-48575272-INS-140
chr5-48629324-INS-266
chr5-48629892-INS-60
chr5-48672113-INS-68
chr5-48703709-INS-174
chr5-48706758-INS-504
chr5-48727790-INS-255
chr5-48769958-INS-110
chr5-48770027-INS-484
chr5-48823084-INS-60
chr5-48861451-INS-182
chr5-48862569-INS-1987
chr5-48868020-INS-1323
chr5-48869109-INS-4210
chr5-48874065-INS-1979
chr5-48912194-INS-1561
chr5-48938244-INS-437
chr5-48949036-INS-933
chr5-48967752-INS-6163
chr5-49071616-INS-74
chr5-49086938-INS-7632
chr5-49089920-INS-53
chr5-49314453-INS-1190
chr5-49326253-INS-1812
chr5-49357825-INS-526
chr5-49438900-INS-584
chr5-49495003-INS-2695
chr5-49503355-INS-1293
chr5-49521372-INS-417
chr5-49521392-INS-853
chr5-49557599-INS-90
chr5-49589978-INS-6237
chr5-49626650-INS-6490
chr5-49742279-INS-65
chr5-49742911-INS-1047
chr5-49743816-INS-428
chr5-49744674-INS-264
chr5-49747806-INS-6377
chr5-49754553-INS-2826
chr5-49756020-INS-117
chr5-49756448-INS-425
chr5-49768021-INS-2844
chr5-49783338-INS-135
chr5-49790923-INS-52
chr5-49796479-INS-6048
chr5-49800199-INS-170
chr5-49800541-INS-273
chr5-49844912-INS-272
chr5-49859699-INS-405
chr5-49948403-INS-266
chr5-49948499-INS-642
chr5-49977707-INS-97

chr5-49977854-INS-775
chr5-49988025-INS-2325
chr5-49989532-INS-829
chr5-50022742-INS-7219
chr5-50029624-INS-109
chr5-50046935-INS-1266
chr5-50046935-INS-591
chr5-50049032-INS-6220
chr5-50073781-INS-58
chr5-50084504-INS-9523
chr5-50165956-INS-6428
chr5-50183578-INS-50
chr5-50197366-INS-50
chr5-50264737-INS-303
chr5-50343134-INS-54
chr5-50344778-INS-91
chr5-50345953-INS-53
chr5-50361703-INS-1875
chr5-50388150-INS-92
chr5-50436937-INS-61
chr5-50507771-INS-1076
chr5-50520952-INS-3249
chr5-50541688-INS-58
chr5-50546069-INS-6367
chr5-50582737-INS-60
chr5-50582950-INS-308
chr5-50583204-INS-100
chr5-50651821-INS-1687
chr5-50661752-INS-327
chr5-50717924-INS-3977
chr5-50742978-INS-58
chr5-50808100-INS-120
chr5-50825918-INS-64
chr5-50897721-INS-50
chr5-50897799-INS-1401
chr5-50927363-INS-1007
chr5-50931586-INS-52
chr5-50941995-INS-1893
chr5-50955825-INS-6218
chr5-50978403-INS-1511
chr5-50985253-INS-2906
chr5-51045746-INS-1914
chr5-51059632-INS-3003
chr5-51100112-INS-1287
chr5-51109186-INS-56
chr5-51214471-INS-13432
chr5-51225630-INS-7900
chr5-51250493-INS-4395
chr5-51251523-INS-87
chr5-51255153-INS-76
chr5-51259722-INS-2233
chr5-51277462-INS-52
chr5-51300079-INS-402
chr5-51307989-INS-76
chr5-51316794-INS-117
chr5-51324894-INS-6298
chr5-51344119-INS-143
chr5-51384389-INS-195
chr5-51559367-INS-349
chr5-51573088-INS-996
chr5-51577442-INS-547
chr5-51602473-INS-219
chr5-51603013-INS-548
chr5-51664826-INS-400
chr5-51793449-INS-152
chr5-51815005-INS-114
chr5-51832342-INS-4624
chr5-51846575-INS-1116
chr5-51846575-INS-121
chr5-51850853-INS-6071
chr5-51901732-INS-76
chr5-51911543-INS-142
chr5-51933212-INS-360
chr5-51943561-INS-64
chr5-51945489-INS-64
chr5-51970981-INS-87
chr5-52143465-INS-88

chr5-52195600-INS-50
chr5-52248514-INS-296
chr5-52428203-INS-69
chr5-52450480-INS-72
chr5-52458764-INS-6388
chr5-52468773-INS-399
chr5-52495448-INS-162
chr5-52507367-INS-54
chr5-52536194-INS-56
chr5-52562178-INS-190
chr5-52580011-INS-176
chr5-52591029-INS-160
chr5-52621586-INS-65
chr5-52660121-INS-247
chr5-52663209-INS-100
chr5-52679019-INS-115
chr5-52688452-INS-173
chr5-52750015-INS-222
chr5-52750583-INS-54
chr5-52750726-INS-496
chr5-52787327-INS-72
chr5-52841745-INS-2999
chr5-52851476-INS-64
chr5-52878845-INS-58
chr5-52897756-INS-220
chr5-52922458-INS-50
chr5-52930845-INS-7162
chr5-52933870-INS-1469
chr5-52967905-INS-517
chr5-52998987-INS-198
chr5-53065291-INS-327
chr5-53096721-INS-305
chr5-53106075-INS-231
chr5-53108535-INS-52
chr5-53110041-INS-1578
chr5-53123051-INS-392
chr5-53127758-INS-52
chr5-53127950-INS-53
chr5-53160827-INS-2133
chr5-53164321-INS-75
chr5-53167320-INS-57
chr5-53185615-INS-135
chr5-53185615-INS-60
chr5-53333159-INS-50
chr5-53342592-INS-70
chr5-53349232-INS-60
chr5-53387478-INS-55
chr5-53402446-INS-89
chr5-53412737-INS-228
chr5-53426518-INS-86
chr5-53474759-INS-1016
chr5-53474759-INS-299
chr5-53496583-INS-199
chr5-53501139-INS-10775
chr5-53507260-INS-1063
chr5-53507263-INS-2771
chr5-53513023-INS-60
chr5-53548096-INS-181
chr5-53563143-INS-50
chr5-53571762-INS-90
chr5-53598067-INS-58
chr5-53619237-INS-50
chr5-53627979-INS-109
chr5-53632944-INS-85
chr5-53636514-INS-205
chr5-53654580-INS-52
chr5-53657010-INS-50
chr5-53673611-INS-65
chr5-53686134-INS-113
chr5-53686167-INS-84
chr5-53732371-INS-55
chr5-53758449-INS-203
chr5-53809277-INS-1290
chr5-53833628-INS-832
chr5-53841129-INS-368
chr5-53898963-INS-3316
chr5-53922369-INS-102

chr5-53931227-INS-1513
chr5-54016522-INS-124
chr5-54055820-INS-70
chr5-54117096-INS-52
chr5-54120679-INS-111
chr5-54122869-INS-965
chr5-54177625-INS-224
chr5-54221655-INS-210
chr5-54273912-INS-205
chr5-54307714-INS-120
chr5-54315996-INS-78
chr5-54327914-INS-7007
chr5-54329075-INS-65
chr5-54351988-INS-50
chr5-54367867-INS-6422
chr5-54372757-INS-66
chr5-54466951-INS-9077
chr5-54498195-INS-50
chr5-54499142-INS-63
chr5-54512616-INS-60
chr5-54538390-INS-166
chr5-54599591-INS-148
chr5-54632181-INS-8504
chr5-54686464-INS-757
chr5-54693741-INS-1522
chr5-54735848-INS-2916
chr5-54805785-INS-216
chr5-54864916-INS-64
chr5-54921226-INS-51
chr5-54959762-INS-72
chr5-54964407-INS-7043
chr5-54977850-INS-86
chr5-54987052-INS-591
chr5-54992298-INS-76
chr5-55001721-INS-135
chr5-55048317-INS-2628
chr5-55091474-INS-127
chr5-55112602-INS-1381
chr5-55130637-INS-4218
chr5-55145250-INS-360
chr5-55163645-INS-414
chr5-55163970-INS-3129
chr5-55209638-INS-51
chr5-55290946-INS-50
chr5-55357995-INS-4075
chr5-55422409-INS-6601
chr5-55426589-INS-1499
chr5-55475216-INS-76
chr5-55499567-INS-307
chr5-55516318-INS-1722
chr5-55523435-INS-481
chr5-55542052-INS-1271
chr5-55551485-INS-3052
chr5-55571337-INS-228
chr5-55591283-INS-234
chr5-55593553-INS-144
chr5-55598444-INS-1479
chr5-55613669-INS-58
chr5-55614384-INS-2205
chr5-55624575-INS-2779
chr5-55640141-INS-504
chr5-55652242-INS-4157
chr5-55682978-INS-5134
chr5-55693467-INS-56
chr5-55756028-INS-1738
chr5-55772599-INS-6735
chr5-55796006-INS-116
chr5-55799198-INS-4455
chr5-55825924-INS-56
chr5-55826025-INS-70
chr5-55843265-INS-72
chr5-55843347-INS-615
chr5-55865339-INS-100
chr5-55900012-INS-1332
chr5-55943890-INS-6242
chr5-55946931-INS-1326
chr5-55970383-INS-2987

chr5-55975795-INS-70
chr5-56007713-INS-275
chr5-56010264-INS-565
chr5-56010912-INS-166
chr5-56014074-INS-251
chr5-56022937-INS-381
chr5-56023049-INS-64
chr5-56033683-INS-60
chr5-56064831-INS-88
chr5-56085779-INS-55
chr5-56132195-INS-78
chr5-56138433-INS-156
chr5-56222305-INS-108
chr5-56317438-INS-70
chr5-56333678-INS-554
chr5-56375123-INS-6366
chr5-56400645-INS-63
chr5-56442680-INS-5098
chr5-56476442-INS-116
chr5-56494697-INS-2071
chr5-56541456-INS-1693
chr5-56545100-INS-50
chr5-56586087-INS-126
chr5-56602234-INS-86
chr5-56608480-INS-52
chr5-56608837-INS-1892
chr5-56649950-INS-161
chr5-56730184-INS-372
chr5-56748426-INS-4171
chr5-56749662-INS-82
chr5-56760356-INS-76
chr5-56761776-INS-1050
chr5-56792531-INS-2964
chr5-56799517-INS-3032
chr5-56816810-INS-52
chr5-56839419-INS-1728
chr5-56853097-INS-75
chr5-56874246-INS-1000
chr5-56879570-INS-7214
chr5-56906101-INS-56
chr5-56915701-INS-218
chr5-56968450-INS-6378
chr5-57017570-INS-55
chr5-57038937-INS-70
chr5-57068123-INS-57
chr5-57069718-INS-105
chr5-57192064-INS-935
chr5-57252050-INS-216
chr5-57261143-INS-7089
chr5-57288884-INS-50
chr5-57302399-INS-176
chr5-57346527-INS-174
chr5-57349996-INS-68
chr5-57352031-INS-556
chr5-57381045-INS-851
chr5-57382732-INS-107
chr5-57384343-INS-6240
chr5-57404340-INS-71
chr5-57461827-INS-6261
chr5-57469233-INS-95
chr5-57473198-INS-57
chr5-57492316-INS-54
chr5-57512756-INS-1179
chr5-57520448-INS-2826
chr5-57536227-INS-77
chr5-57769562-INS-1395
chr5-57775417-INS-2978
chr5-57857548-INS-500
chr5-57866099-INS-2927
chr5-57882861-INS-52
chr5-57980446-INS-581
chr5-57982981-INS-68
chr5-57987924-INS-6171
chr5-58048149-INS-51
chr5-58139063-INS-56
chr5-58356967-INS-52
chr5-58367573-INS-133

chr5-58367678-INS-4603
chr5-58368137-INS-56
chr5-58460078-INS-93
chr5-58477514-INS-329
chr5-58482699-INS-76
chr5-58506773-INS-131
chr5-58587532-INS-1529
chr5-58602658-INS-2985
chr5-58623375-INS-92
chr5-58637970-INS-2685
chr5-58643177-INS-2152
chr5-58650041-INS-7529
chr5-58772209-INS-88
chr5-58777260-INS-5446
chr5-58802458-INS-60
chr5-58848733-INS-6304
chr5-58860083-INS-54
chr5-58904258-INS-68
chr5-58926863-INS-475
chr5-58956601-INS-731
chr5-58968526-INS-70
chr5-59018770-INS-651
chr5-59054770-INS-1887
chr5-59077121-INS-1993
chr5-59110684-INS-4904
chr5-59116438-INS-256
chr5-59209720-INS-50
chr5-59253156-INS-86
chr5-59258281-INS-55
chr5-59324587-INS-481
chr5-59334242-INS-468
chr5-59396501-INS-6818
chr5-59426048-INS-143
chr5-59442249-INS-6213
chr5-59453699-INS-218
chr5-59511306-INS-3062
chr5-59515466-INS-1557
chr5-59542576-INS-1892
chr5-59547864-INS-6421
chr5-59556648-INS-6213
chr5-59564423-INS-1807
chr5-59631370-INS-61
chr5-59645251-INS-180
chr5-59645692-INS-388
chr5-59659168-INS-6360
chr5-59667709-INS-52
chr5-59691265-INS-84
chr5-59697071-INS-52
chr5-59700101-INS-2835
chr5-59721799-INS-209
chr5-59725383-INS-2701
chr5-59745146-INS-87
chr5-59772218-INS-175
chr5-59781586-INS-149
chr5-59789529-INS-996
chr5-59815835-INS-68
chr5-59845650-INS-286
chr5-59946282-INS-1914
chr5-59952788-INS-138
chr5-59958448-INS-530
chr5-59959277-INS-138
chr5-59961093-INS-80
chr5-59967160-INS-3057
chr5-59971748-INS-63
chr5-59972709-INS-6452
chr5-59973381-INS-130
chr5-59974355-INS-51
chr5-59980093-INS-68
chr5-59985243-INS-1022
chr5-59985913-INS-6600
chr5-60015566-INS-272
chr5-60038456-INS-60
chr5-60050266-INS-3527
chr5-60053734-INS-52
chr5-60064122-INS-270
chr5-60072428-INS-4813
chr5-60160150-INS-110

chr5-60200702-INS-117
chr5-60239594-INS-56
chr5-60280004-INS-7119
chr5-60290335-INS-4819
chr5-60353699-INS-54
chr5-60356309-INS-72
chr5-60376880-INS-52
chr5-60389605-INS-2934
chr5-60398560-INS-99
chr5-60398690-INS-229
chr5-60406947-INS-54
chr5-60414436-INS-2837
chr5-60419769-INS-213
chr5-60434465-INS-1320
chr5-60460632-INS-380
chr5-60460634-INS-1700
chr5-60473164-INS-100
chr5-60603357-INS-416
chr5-60694081-INS-214
chr5-60725512-INS-3096
chr5-60737528-INS-188
chr5-60782151-INS-3751
chr5-60809137-INS-212
chr5-60821082-INS-2562
chr5-60835269-INS-570
chr5-60847301-INS-779
chr5-60865488-INS-3581
chr5-60870957-INS-50
chr5-60924542-INS-54
chr5-60951401-INS-377
chr5-61015162-INS-60
chr5-61022242-INS-50
chr5-61030641-INS-1395
chr5-61035790-INS-2952
chr5-61049163-INS-873
chr5-61051581-INS-202
chr5-61081682-INS-135
chr5-61094830-INS-6613
chr5-61123708-INS-62
chr5-61131263-INS-54
chr5-61182067-INS-282
chr5-61196721-INS-75
chr5-61208018-INS-5923
chr5-61247328-INS-130
chr5-61250105-INS-186
chr5-61260525-INS-129
chr5-61338914-INS-1517
chr5-61339582-INS-2854
chr5-61341649-INS-82
chr5-61386118-INS-1917
chr5-61434229-INS-6325
chr5-61468848-INS-1453
chr5-61508328-INS-7165
chr5-61522632-INS-611
chr5-61563171-INS-6420
chr5-61598272-INS-187
chr5-61675012-INS-4177
chr5-61695771-INS-708
chr5-61717888-INS-170
chr5-61718039-INS-220
chr5-61718372-INS-2325
chr5-61727861-INS-6387
chr5-61727966-INS-52
chr5-61777752-INS-2462
chr5-61809515-INS-120
chr5-61809813-INS-168
chr5-61820354-INS-4839
chr5-61857102-INS-948
chr5-61882807-INS-50
chr5-61913106-INS-223
chr5-61931789-INS-352
chr5-62022431-INS-6205
chr5-62071697-INS-624
chr5-62259031-INS-68
chr5-62274118-INS-204
chr5-62338715-INS-170
chr5-62412632-INS-454

chr5-62489566-INS-409
chr5-62512225-INS-60
chr5-62540397-INS-2331
chr5-62560476-INS-138
chr5-62595373-INS-68
chr5-62649616-INS-97
chr5-62649743-INS-1212
chr5-62649929-INS-291
chr5-62654562-INS-86
chr5-62654775-INS-78
chr5-62655019-INS-62
chr5-62659950-INS-1552
chr5-62682054-INS-201
chr5-62709667-INS-54
chr5-62756071-INS-182
chr5-62789342-INS-68
chr5-62794730-INS-335
chr5-62893842-INS-65
chr5-62898904-INS-100
chr5-62940544-INS-50
chr5-62964984-INS-57
chr5-63003932-INS-498
chr5-63009745-INS-202
chr5-63025885-INS-2000
chr5-63026519-INS-64
chr5-63026705-INS-332
chr5-63026717-INS-64
chr5-63035325-INS-4191
chr5-63072648-INS-129
chr5-63185320-INS-6172
chr5-63213258-INS-476
chr5-63227746-INS-219
chr5-63230488-INS-50
chr5-63266397-INS-6681
chr5-63296544-INS-56
chr5-63307526-INS-203
chr5-63349801-INS-52
chr5-63350194-INS-54
chr5-63369516-INS-52
chr5-63398023-INS-165
chr5-63399692-INS-199
chr5-63401121-INS-55
chr5-63444445-INS-368
chr5-63447808-INS-1447
chr5-63466447-INS-2057
chr5-63502288-INS-1336
chr5-63513236-INS-110
chr5-63546123-INS-7103
chr5-63553882-INS-1743
chr5-63616605-INS-3471
chr5-63638028-INS-55
chr5-63638942-INS-76
chr5-63639966-INS-82
chr5-63691964-INS-60
chr5-63700145-INS-106
chr5-63720164-INS-183
chr5-63744609-INS-3040
chr5-63749592-INS-2268
chr5-63754381-INS-616
chr5-63770959-INS-8690
chr5-63784807-INS-68
chr5-63808421-INS-61
chr5-63817557-INS-2231
chr5-63822343-INS-81
chr5-63850903-INS-221
chr5-63860207-INS-53
chr5-63863950-INS-6656
chr5-63873024-INS-873
chr5-63929576-INS-120
chr5-63931548-INS-126
chr5-63932684-INS-266
chr5-63934774-INS-483
chr5-63945027-INS-62
chr5-63980981-INS-86
chr5-63987783-INS-51
chr5-63994891-INS-570
chr5-64026425-INS-59

chr5-64135866-INS-57
chr5-64169642-INS-54
chr5-64226458-INS-327
chr5-64235276-INS-2007
chr5-64357915-INS-146
chr5-64405048-INS-96
chr5-64448041-INS-1626
chr5-64448041-INS-4795
chr5-64456762-INS-4832
chr5-64468326-INS-50
chr5-64479684-INS-96
chr5-64497101-INS-99
chr5-64501609-INS-153
chr5-64510671-INS-51
chr5-64613598-INS-671
chr5-64726043-INS-108
chr5-64815669-INS-7128
chr5-64850998-INS-87
chr5-64901013-INS-50
chr5-64916482-INS-51
chr5-64930635-INS-72
chr5-64953950-INS-72
chr5-64975565-INS-114
chr5-65037057-INS-65
chr5-65037284-INS-102
chr5-65041357-INS-1206
chr5-65067895-INS-140
chr5-65201933-INS-923
chr5-65202220-INS-151
chr5-65213670-INS-132
chr5-65251505-INS-74
chr5-65305640-INS-61
chr5-65334265-INS-480
chr5-65389333-INS-70
chr5-65389690-INS-588
chr5-65397924-INS-177
chr5-65402263-INS-181
chr5-65406483-INS-233
chr5-65406540-INS-676
chr5-65432974-INS-215
chr5-65469293-INS-1476
chr5-65504508-INS-80
chr5-65522458-INS-208
chr5-65523236-INS-495
chr5-65523236-INS-53
chr5-65523611-INS-211
chr5-65561043-INS-55
chr5-65562910-INS-206
chr5-65564806-INS-3151
chr5-65586803-INS-203
chr5-65601970-INS-54
chr5-65610851-INS-50
chr5-65612972-INS-78
chr5-65635637-INS-198
chr5-65659092-INS-50
chr5-65675827-INS-483
chr5-65728573-INS-552
chr5-65729864-INS-213
chr5-65747325-INS-52
chr5-65747441-INS-151
chr5-65769556-INS-77
chr5-65809640-INS-157
chr5-65818327-INS-108
chr5-65823933-INS-224
chr5-65862592-INS-75
chr5-65865146-INS-92
chr5-65890648-INS-68
chr5-65902787-INS-56
chr5-65910599-INS-209
chr5-65926274-INS-226
chr5-65953781-INS-465
chr5-65993356-INS-195
chr5-65993606-INS-580
chr5-65994098-INS-163
chr5-66000111-INS-56
chr5-66009448-INS-806
chr5-66009612-INS-1260

chr5-66015763-INS-50
chr5-66063382-INS-106
chr5-66064679-INS-630
chr5-66066707-INS-60
chr5-66072437-INS-245
chr5-66082198-INS-88
chr5-66118388-INS-5245
chr5-66125851-INS-218
chr5-66136870-INS-173
chr5-66140940-INS-338
chr5-66141050-INS-51
chr5-66142040-INS-239
chr5-66145069-INS-205
chr5-66150936-INS-90
chr5-66192874-INS-51
chr5-66198084-INS-81
chr5-66209280-INS-79
chr5-66256771-INS-201
chr5-66276590-INS-460
chr5-66276784-INS-60
chr5-66277407-INS-211
chr5-66318273-INS-56
chr5-66318654-INS-147
chr5-66340971-INS-130
chr5-66342363-INS-4836
chr5-66347471-INS-54
chr5-66361582-INS-51
chr5-66378440-INS-68
chr5-66391099-INS-53
chr5-66391731-INS-59
chr5-66398126-INS-151
chr5-66405880-INS-1369
chr5-66407559-INS-86
chr5-66408275-INS-158
chr5-66408276-INS-320
chr5-66413159-INS-66
chr5-66418050-INS-310
chr5-66418126-INS-736
chr5-66426112-INS-67
chr5-66447735-INS-345
chr5-66502113-INS-248
chr5-66524634-INS-72
chr5-66540141-INS-189
chr5-66586231-INS-318
chr5-66586254-INS-690
chr5-66597386-INS-137
chr5-66635715-INS-64
chr5-66684008-INS-160
chr5-66686620-INS-1945
chr5-66702208-INS-404
chr5-66714272-INS-173
chr5-66720393-INS-6264
chr5-66735523-INS-170
chr5-66740581-INS-66
chr5-66741296-INS-1610
chr5-66800646-INS-66
chr5-66815347-INS-767
chr5-66818845-INS-284
chr5-66822825-INS-170
chr5-66857972-INS-54
chr5-66869918-INS-139
chr5-66869921-INS-65
chr5-66870741-INS-340
chr5-66870741-INS-80
chr5-66877642-INS-11315
chr5-66884212-INS-52
chr5-66892071-INS-209
chr5-66896600-INS-205
chr5-66899913-INS-197
chr5-66915510-INS-81
chr5-66915629-INS-308
chr5-66961356-INS-6580
chr5-66986229-INS-73
chr5-66986366-INS-201
chr5-66986366-INS-921
chr5-67010760-INS-51
chr5-67073107-INS-7086

chr5-67095721-INS-125
chr5-67135721-INS-116
chr5-67166499-INS-70
chr5-67174303-INS-620
chr5-67177411-INS-407
chr5-67181287-INS-100
chr5-67205512-INS-1065
chr5-67208456-INS-72
chr5-67229711-INS-590
chr5-67233355-INS-106
chr5-67276800-INS-51
chr5-67324178-INS-69
chr5-67337281-INS-525
chr5-67352800-INS-141
chr5-67355799-INS-472
chr5-67374758-INS-54
chr5-67393666-INS-709
chr5-67403750-INS-4757
chr5-67409723-INS-3607
chr5-67426658-INS-1542
chr5-67477479-INS-86
chr5-67481520-INS-198
chr5-67498574-INS-98
chr5-67537258-INS-126
chr5-67579478-INS-320
chr5-67595204-INS-73
chr5-67612655-INS-400
chr5-67615263-INS-69
chr5-67629616-INS-6442
chr5-67638381-INS-1368
chr5-67691265-INS-50
chr5-67739716-INS-404
chr5-67773494-INS-1658
chr5-67792529-INS-51
chr5-67815473-INS-63
chr5-67852429-INS-1672
chr5-67918976-INS-87
chr5-67920499-INS-1902
chr5-67942754-INS-112
chr5-67949417-INS-616
chr5-67968077-INS-56
chr5-67985303-INS-63
chr5-68012049-INS-56
chr5-68012887-INS-348
chr5-68013113-INS-601
chr5-68034686-INS-153
chr5-68047367-INS-1875
chr5-68113339-INS-50
chr5-68136201-INS-474
chr5-68231725-INS-81
chr5-68238369-INS-6401
chr5-68239960-INS-657
chr5-68264597-INS-122
chr5-68346200-INS-75
chr5-68354485-INS-262
chr5-68394625-INS-65
chr5-68407690-INS-1615
chr5-68413170-INS-1387
chr5-68440777-INS-52
chr5-68472817-INS-2437
chr5-68488323-INS-461
chr5-68513486-INS-270
chr5-68513495-INS-546
chr5-68515052-INS-64
chr5-68538493-INS-3142
chr5-68554663-INS-187
chr5-68565388-INS-53
chr5-68565692-INS-50
chr5-68612045-INS-603
chr5-68620889-INS-9115
chr5-68641075-INS-2284
chr5-68659578-INS-1886
chr5-68661942-INS-80
chr5-68662080-INS-86
chr5-68678178-INS-59
chr5-68689509-INS-1485
chr5-68719356-INS-151

chr5-68772530-INS-328
chr5-68791130-INS-924
chr5-68794587-INS-3234
chr5-68798622-INS-6412
chr5-68809037-INS-4644
chr5-68823832-INS-161
chr5-68824043-INS-82
chr5-68835687-INS-802
chr5-68859081-INS-752
chr5-68860761-INS-104
chr5-68894348-INS-103
chr5-68918625-INS-55
chr5-68918999-INS-192
chr5-68919515-INS-218
chr5-68934793-INS-169
chr5-68939273-INS-487
chr5-68939415-INS-209
chr5-68952642-INS-58
chr5-68976682-INS-92
chr5-69016287-INS-104
chr5-69038183-INS-63
chr5-69141335-INS-6397
chr5-69158495-INS-950
chr5-69169295-INS-59
chr5-69170434-INS-509
chr5-69173393-INS-100
chr5-69183854-INS-116
chr5-69249138-INS-54
chr5-69299988-INS-306
chr5-69300659-INS-85
chr5-69301566-INS-53
chr5-69322648-INS-144
chr5-69343360-INS-2959
chr5-69382656-INS-289
chr5-69394003-INS-2028
chr5-69395825-INS-100
chr5-69411059-INS-294
chr5-69421290-INS-174
chr5-69439442-INS-56
chr5-69472675-INS-6921
chr5-69491354-INS-57
chr5-69520996-INS-1650
chr5-69557956-INS-66
chr5-69558756-INS-80
chr5-69559265-INS-50
chr5-69561598-INS-525
chr5-69625358-INS-158
chr5-69625771-INS-631
chr5-69849535-INS-290
chr5-69894835-INS-828
chr5-69922306-INS-150
chr5-69966471-INS-92
chr5-69968823-INS-68
chr5-69974196-INS-2895
chr5-70004050-INS-73
chr5-70004190-INS-402
chr5-70006045-INS-197
chr5-70033773-INS-50
chr5-70083686-INS-52
chr5-70086853-INS-123
chr5-70099602-INS-189
chr5-70100054-INS-62
chr5-70139442-INS-110
chr5-70167269-INS-69
chr5-70252726-INS-128
chr5-70261302-INS-517
chr5-70349426-INS-115
chr5-70373298-INS-1304
chr5-70384796-INS-1042
chr5-70385915-INS-159
chr5-70386825-INS-995
chr5-70404228-INS-132
chr5-70422133-INS-2141
chr5-70422938-INS-58
chr5-70441049-INS-223
chr5-70457055-INS-1438
chr5-70490517-INS-171

chr5-70496034-INS-405
chr5-70514795-INS-53
chr5-70516005-INS-1339
chr5-70565281-INS-1914
chr5-70566961-INS-566
chr5-70596754-INS-1129
chr5-70607552-INS-1494
chr5-70637370-INS-1706
chr5-70651113-INS-6041
chr5-70763126-INS-266
chr5-70834435-INS-7159
chr5-70843996-INS-52
chr5-70851888-INS-96
chr5-70918842-INS-412
chr5-70933617-INS-4611
chr5-70951156-INS-203
chr5-70962310-INS-108
chr5-70962525-INS-250
chr5-70962538-INS-70
chr5-70994708-INS-50
chr5-71007068-INS-88
chr5-71036196-INS-52
chr5-71060618-INS-90
chr5-71061771-INS-57
chr5-71118259-INS-235
chr5-71135586-INS-72
chr5-71190690-INS-164
chr5-71231581-INS-157
chr5-71252985-INS-56
chr5-71312718-INS-1197
chr5-71339115-INS-1287
chr5-71419240-INS-500
chr5-71458811-INS-68
chr5-71458849-INS-194
chr5-71469933-INS-2930
chr5-71476940-INS-214
chr5-71614056-INS-1082
chr5-71729846-INS-72
chr5-71741716-INS-57
chr5-71743515-INS-50
chr5-71787989-INS-311
chr5-71788113-INS-150
chr5-71831345-INS-526
chr5-71832303-INS-500
chr5-71842576-INS-50
chr5-71866247-INS-159
chr5-71930871-INS-64
chr5-72013804-INS-4613
chr5-72039927-INS-70
chr5-72040962-INS-197
chr5-72133352-INS-280
chr5-72144149-INS-50
chr5-72151480-INS-52
chr5-72152065-INS-133
chr5-72162249-INS-4195
chr5-72179508-INS-56
chr5-72182664-INS-445
chr5-72189163-INS-280
chr5-72202261-INS-630
chr5-72204291-INS-233
chr5-72220007-INS-3194
chr5-72281875-INS-161
chr5-72282982-INS-642
chr5-72308135-INS-374
chr5-72312114-INS-50
chr5-72315016-INS-150
chr5-72336215-INS-198
chr5-72350617-INS-9101
chr5-72356700-INS-136
chr5-72374794-INS-51
chr5-72381753-INS-1116
chr5-72384898-INS-63
chr5-72424962-INS-3135
chr5-72431600-INS-58
chr5-72526132-INS-232
chr5-72528142-INS-1658
chr5-72540217-INS-340

chr5-72540221-INS-141
chr5-72540319-INS-119
chr5-72559424-INS-1311
chr5-72570454-INS-114
chr5-72597824-INS-102
chr5-72692250-INS-311
chr5-72721206-INS-212
chr5-72727990-INS-2169
chr5-72741490-INS-72
chr5-72755718-INS-68
chr5-72760068-INS-66
chr5-72843636-INS-76
chr5-72843726-INS-2592
chr5-72871227-INS-78
chr5-72880681-INS-69
chr5-72890264-INS-104
chr5-72890934-INS-194
chr5-72890934-INS-66
chr5-72903404-INS-7183
chr5-72923712-INS-179
chr5-72939151-INS-252
chr5-72964697-INS-223
chr5-72974181-INS-79
chr5-72981762-INS-56
chr5-72982097-INS-662
chr5-72983782-INS-55
chr5-73004788-INS-80
chr5-73014980-INS-51
chr5-73026583-INS-211
chr5-73030234-INS-80
chr5-73033276-INS-55
chr5-73033329-INS-2480
chr5-73040992-INS-246
chr5-73041584-INS-200
chr5-73047249-INS-202
chr5-73112247-INS-200
chr5-73132261-INS-132
chr5-73143943-INS-90
chr5-73146045-INS-158
chr5-73146586-INS-3003
chr5-73146894-INS-94
chr5-73151017-INS-134
chr5-73215769-INS-155
chr5-73215771-INS-332
chr5-73235541-INS-318
chr5-73265163-INS-76
chr5-73299945-INS-218
chr5-73305735-INS-106
chr5-73396502-INS-64
chr5-73452744-INS-215
chr5-73453315-INS-204
chr5-73477059-INS-178
chr5-73479864-INS-116
chr5-73514137-INS-65
chr5-73551993-INS-502
chr5-73552552-INS-983
chr5-73556506-INS-222
chr5-73556578-INS-907
chr5-73556949-INS-402
chr5-73566691-INS-208
chr5-73579422-INS-211
chr5-73590402-INS-76
chr5-73600285-INS-244
chr5-73605707-INS-75
chr5-73624519-INS-1894
chr5-73652930-INS-342
chr5-73652935-INS-86
chr5-73656790-INS-66
chr5-73709613-INS-5458
chr5-73732561-INS-52
chr5-73733123-INS-55
chr5-73842095-INS-200
chr5-73845920-INS-1006
chr5-73856790-INS-596
chr5-73857903-INS-209
chr5-73883044-INS-1766
chr5-73883170-INS-114

chr5-73897788-INS-113
chr5-73918463-INS-185
chr5-73970645-INS-111
chr5-73980623-INS-5914
chr5-74004959-INS-52
chr5-74037689-INS-5474
chr5-74053489-INS-66
chr5-74091373-INS-9381
chr5-74110183-INS-53
chr5-74116121-INS-60
chr5-74116358-INS-147
chr5-74137739-INS-289
chr5-74155064-INS-178
chr5-74156501-INS-94
chr5-74168277-INS-520
chr5-74168292-INS-1190
chr5-74202252-INS-80
chr5-74218469-INS-165
chr5-74244368-INS-200
chr5-74295915-INS-68
chr5-74313339-INS-56
chr5-74325999-INS-53
chr5-74331975-INS-58
chr5-74432560-INS-54
chr5-74449141-INS-1900
chr5-74467996-INS-123
chr5-74469036-INS-516
chr5-74469275-INS-52
chr5-74469690-INS-680
chr5-74470696-INS-118
chr5-74504319-INS-213
chr5-74506691-INS-91
chr5-74510785-INS-167
chr5-74511597-INS-6385
chr5-74569754-INS-55
chr5-74597411-INS-205
chr5-74635002-INS-206
chr5-74647968-INS-5253
chr5-74660921-INS-165
chr5-74666752-INS-276
chr5-74686070-INS-58
chr5-74687832-INS-4909
chr5-74687835-INS-107
chr5-74705186-INS-208
chr5-74711009-INS-3852
chr5-74711365-INS-84
chr5-74715136-INS-50
chr5-74742226-INS-1864
chr5-74799056-INS-108
chr5-74817815-INS-191
chr5-74884923-INS-78
chr5-74884969-INS-1252
chr5-74895887-INS-103
chr5-74905437-INS-328
chr5-74922003-INS-249
chr5-74962607-INS-78
chr5-74965250-INS-155
chr5-74975070-INS-116
chr5-74994286-INS-55
chr5-74994481-INS-3102
chr5-75025113-INS-50
chr5-75028676-INS-531
chr5-75038689-INS-84
chr5-75044526-INS-384
chr5-75044526-INS-54
chr5-75107687-INS-72
chr5-75110064-INS-2730
chr5-75119336-INS-4958
chr5-75154103-INS-68
chr5-75201676-INS-774
chr5-75239758-INS-122
chr5-75249096-INS-213
chr5-75249573-INS-114
chr5-75249720-INS-110
chr5-75260592-INS-1197
chr5-75261567-INS-247
chr5-75272848-INS-75

chr5-75285670-INS-1332
chr5-75299705-INS-117
chr5-75328240-INS-125
chr5-75336281-INS-152
chr5-75341474-INS-50
chr5-75370422-INS-205
chr5-75373285-INS-149
chr5-75387712-INS-51
chr5-75426148-INS-2784
chr5-75436742-INS-105
chr5-75436991-INS-516
chr5-75458001-INS-66
chr5-75463211-INS-80
chr5-75471564-INS-53
chr5-75514843-INS-88
chr5-75533538-INS-50
chr5-75534057-INS-74
chr5-75566066-INS-54
chr5-75574408-INS-2592
chr5-75610362-INS-72
chr5-75613430-INS-252
chr5-75613430-INS-648
chr5-75622532-INS-242
chr5-75642837-INS-121
chr5-75669129-INS-156
chr5-75680032-INS-1599
chr5-75701969-INS-82
chr5-75702573-INS-53
chr5-75714065-INS-4764
chr5-75821258-INS-64
chr5-75833201-INS-171
chr5-75846165-INS-140
chr5-75851265-INS-2349
chr5-75880330-INS-176
chr5-76049648-INS-52
chr5-76074464-INS-617
chr5-76137950-INS-100
chr5-76137950-INS-210
chr5-76181448-INS-52
chr5-76196237-INS-231
chr5-76234727-INS-360
chr5-76242928-INS-190
chr5-76247833-INS-70
chr5-76260577-INS-1390
chr5-76262407-INS-54
chr5-76282736-INS-7165
chr5-76347827-INS-187
chr5-76351637-INS-57
chr5-76358797-INS-174
chr5-76411510-INS-111
chr5-76447138-INS-199
chr5-76448457-INS-975
chr5-76468913-INS-80
chr5-76490992-INS-64
chr5-76512753-INS-64
chr5-76546791-INS-71
chr5-76551413-INS-68
chr5-76558197-INS-4743
chr5-76651264-INS-458
chr5-76662486-INS-50
chr5-76722082-INS-1071
chr5-76731720-INS-507
chr5-76801581-INS-56
chr5-76823558-INS-72
chr5-76826548-INS-648
chr5-76826553-INS-232
chr5-76826772-INS-196
chr5-76830751-INS-66
chr5-76888711-INS-192
chr5-76958991-INS-106
chr5-76966153-INS-1723
chr5-77008105-INS-106
chr5-77025113-INS-92
chr5-77034942-INS-187
chr5-77056692-INS-183
chr5-77104203-INS-224
chr5-77112228-INS-62

chr5-77116772-INS-86
chr5-77126153-INS-82
chr5-77160687-INS-120
chr5-77180489-INS-50
chr5-77222034-INS-50
chr5-77224437-INS-68
chr5-77240138-INS-252
chr5-77246169-INS-87
chr5-77260181-INS-90
chr5-77275345-INS-206
chr5-77292046-INS-471
chr5-77295070-INS-2914
chr5-77296696-INS-1020
chr5-77297341-INS-410
chr5-77298280-INS-877
chr5-77298603-INS-51
chr5-77307984-INS-208
chr5-77328783-INS-51
chr5-77332460-INS-50
chr5-77345473-INS-852
chr5-77345548-INS-74
chr5-77355725-INS-1184
chr5-77367264-INS-52
chr5-77374952-INS-144
chr5-77391156-INS-277
chr5-77397025-INS-209
chr5-77408742-INS-52
chr5-77488792-INS-56
chr5-77490136-INS-272
chr5-77521106-INS-103
chr5-77526622-INS-327
chr5-77549947-INS-164
chr5-77551381-INS-50
chr5-77587017-INS-6177
chr5-77587284-INS-933
chr5-77591545-INS-282
chr5-77617571-INS-94
chr5-77617581-INS-330
chr5-77661067-INS-634
chr5-77661717-INS-17549
chr5-77667743-INS-65
chr5-77667810-INS-135
chr5-77668024-INS-405
chr5-77691989-INS-80
chr5-77692045-INS-52
chr5-77732641-INS-5391
chr5-77759489-INS-196
chr5-77765878-INS-59
chr5-77800221-INS-507
chr5-77800642-INS-138
chr5-77808732-INS-206
chr5-77808962-INS-265
chr5-77814319-INS-50
chr5-77864547-INS-59
chr5-77880196-INS-502
chr5-77899320-INS-600
chr5-77926123-INS-147
chr5-77934671-INS-1766
chr5-77945793-INS-64
chr5-77982341-INS-88
chr5-78034670-INS-3539
chr5-78044965-INS-82
chr5-78069362-INS-6364
chr5-78072755-INS-191
chr5-78105607-INS-112
chr5-78131830-INS-105
chr5-78196562-INS-205
chr5-78231063-INS-50
chr5-78253408-INS-385
chr5-78273558-INS-50
chr5-78309291-INS-795
chr5-78313099-INS-147
chr5-78340587-INS-110
chr5-78395354-INS-2449
chr5-78454238-INS-53
chr5-78457129-INS-810
chr5-78458346-INS-2818

chr5-78458402-INS-1031
chr5-78462738-INS-540
chr5-78509713-INS-6692
chr5-78510433-INS-150
chr5-78519941-INS-405
chr5-78520451-INS-68
chr5-78521001-INS-74
chr5-78569844-INS-54
chr5-78599518-INS-5058
chr5-78608906-INS-238
chr5-78646598-INS-1562
chr5-78666207-INS-50
chr5-78675754-INS-50
chr5-78694229-INS-129
chr5-78699217-INS-74
chr5-78701468-INS-78
chr5-78705615-INS-54
chr5-78716851-INS-554
chr5-78722919-INS-58
chr5-78759063-INS-431
chr5-78783633-INS-3731
chr5-78784386-INS-170
chr5-78826024-INS-412
chr5-78843972-INS-374
chr5-78891819-INS-17957
chr5-78975008-INS-1728
chr5-78982661-INS-95
chr5-78985940-INS-60
chr5-78995751-INS-1469
chr5-78998218-INS-6561
chr5-79017587-INS-56
chr5-79050077-INS-57
chr5-79058726-INS-562
chr5-79082110-INS-2537
chr5-79088556-INS-2512
chr5-79091054-INS-4971
chr5-79091399-INS-1613
chr5-79110246-INS-50
chr5-79110934-INS-366
chr5-79131130-INS-243
chr5-79131702-INS-297
chr5-79132260-INS-121
chr5-79134662-INS-54
chr5-79161390-INS-2877
chr5-79163511-INS-174
chr5-79207612-INS-6431
chr5-79229728-INS-70
chr5-79250474-INS-5002
chr5-79281028-INS-196
chr5-79287101-INS-1770
chr5-79293564-INS-6477
chr5-79347402-INS-130
chr5-79362447-INS-152
chr5-79362506-INS-70
chr5-79416150-INS-1379
chr5-79432768-INS-52
chr5-79446634-INS-163
chr5-79454756-INS-522
chr5-79527749-INS-266
chr5-79536101-INS-51
chr5-79569558-INS-352
chr5-79576609-INS-1563
chr5-79607760-INS-153
chr5-79612619-INS-339
chr5-79640745-INS-513
chr5-79641250-INS-80
chr5-79692290-INS-52
chr5-79696522-INS-830
chr5-79703799-INS-1427
chr5-79713141-INS-1122
chr5-79743153-INS-164
chr5-79743544-INS-68
chr5-79763136-INS-52
chr5-79765693-INS-284
chr5-79775411-INS-60
chr5-79779359-INS-75
chr5-79788747-INS-237

chr5-79789391-INS-555
chr5-79801723-INS-3004
chr5-79813094-INS-72
chr5-79835962-INS-344
chr5-79841523-INS-3113
chr5-79883173-INS-66
chr5-79908186-INS-62
chr5-79919304-INS-135
chr5-79921519-INS-5945
chr5-79929789-INS-463
chr5-79936029-INS-100
chr5-79979194-INS-3251
chr5-80029985-INS-69
chr5-80037807-INS-2329
chr5-80098849-INS-60
chr5-80143499-INS-3704
chr5-80150539-INS-6255
chr5-80151517-INS-68
chr5-80190547-INS-51
chr5-80190551-INS-270
chr5-80259646-INS-1489
chr5-80272083-INS-660
chr5-80273805-INS-51
chr5-80314875-INS-50
chr5-80375063-INS-3279
chr5-80397924-INS-1000
chr5-80463748-INS-6645
chr5-80581631-INS-60
chr5-80652848-INS-146
chr5-80697330-INS-3149
chr5-80712861-INS-1597
chr5-80730038-INS-53
chr5-80767590-INS-64
chr5-80768342-INS-78
chr5-80820480-INS-70
chr5-80884352-INS-348
chr5-80884722-INS-3078
chr5-80896862-INS-12795
chr5-80898336-INS-2472
chr5-80906972-INS-657
chr5-80910246-INS-50
chr5-80944857-INS-106
chr5-80946221-INS-85
chr5-80962145-INS-10283
chr5-80962312-INS-750
chr5-81033274-INS-108
chr5-81046925-INS-88
chr5-81108426-INS-6465
chr5-81120181-INS-575
chr5-81123910-INS-160
chr5-81129337-INS-3058
chr5-81210766-INS-91
chr5-81218214-INS-59
chr5-81230102-INS-85
chr5-81308834-INS-94
chr5-81309188-INS-630
chr5-81409637-INS-7277
chr5-81592569-INS-202
chr5-81691136-INS-6301
chr5-81743801-INS-73
chr5-81755371-INS-61
chr5-81813249-INS-50
chr5-81867797-INS-68
chr5-81880636-INS-66
chr5-81895134-INS-169
chr5-81895134-INS-52
chr5-81897382-INS-82
chr5-81967180-INS-5292
chr5-81986606-INS-83
chr5-81992117-INS-210
chr5-82029461-INS-64
chr5-82033996-INS-56
chr5-82047473-INS-765
chr5-82062766-INS-64
chr5-82063691-INS-3115
chr5-82093376-INS-503
chr5-82095860-INS-496

chr5-82096340-INS-96
chr5-82106448-INS-110
chr5-82118153-INS-329
chr5-82124290-INS-7135
chr5-82154822-INS-81
chr5-82159975-INS-11788
chr5-82199859-INS-62
chr5-82227644-INS-58
chr5-82254365-INS-70
chr5-82328068-INS-201
chr5-82346763-INS-9172
chr5-82400093-INS-51
chr5-82404292-INS-508
chr5-82473324-INS-6728
chr5-82510853-INS-8781
chr5-82532815-INS-59
chr5-82540476-INS-6388
chr5-82557939-INS-214
chr5-82573932-INS-342
chr5-82598905-INS-1227
chr5-82603348-INS-58
chr5-82605901-INS-50
chr5-82626036-INS-4999
chr5-82652383-INS-151
chr5-82659238-INS-397
chr5-82668166-INS-6401
chr5-82741054-INS-56
chr5-82742181-INS-4560
chr5-82747639-INS-158
chr5-82813185-INS-132
chr5-82814064-INS-54
chr5-82874754-INS-600
chr5-82884885-INS-185
chr5-82909306-INS-2028
chr5-82948330-INS-3079
chr5-82966084-INS-7531
chr5-82971381-INS-79
chr5-82973754-INS-6451
chr5-83117157-INS-66
chr5-83129416-INS-64
chr5-83135221-INS-72
chr5-83139042-INS-101
chr5-83221822-INS-1820
chr5-83246294-INS-349
chr5-83279864-INS-136
chr5-83299977-INS-50
chr5-83312168-INS-90
chr5-83319070-INS-6465
chr5-83329046-INS-110
chr5-83419427-INS-829
chr5-83431724-INS-74
chr5-83439660-INS-142
chr5-83439661-INS-65
chr5-83443634-INS-312
chr5-83443860-INS-161
chr5-83464587-INS-52
chr5-83464658-INS-237
chr5-83492482-INS-67
chr5-83567408-INS-3004
chr5-83572731-INS-1563
chr5-83634500-INS-913
chr5-83636007-INS-338
chr5-83638879-INS-175
chr5-83639175-INS-78
chr5-83645392-INS-142
chr5-83673387-INS-1369
chr5-83708816-INS-78
chr5-83709133-INS-150
chr5-83789001-INS-56
chr5-83829640-INS-50
chr5-83871572-INS-52
chr5-83902140-INS-70
chr5-83925763-INS-204
chr5-83952769-INS-57
chr5-83959876-INS-93
chr5-83975265-INS-446
chr5-83988597-INS-173

chr5-83991149-INS-610
chr5-83994790-INS-280
chr5-84007368-INS-533
chr5-84035725-INS-2849
chr5-84058036-INS-214
chr5-84089513-INS-2967
chr5-84118396-INS-1000
chr5-84305390-INS-122
chr5-84309518-INS-127
chr5-84375721-INS-711
chr5-84394624-INS-6719
chr5-84401889-INS-6369
chr5-84408190-INS-72
chr5-84444333-INS-526
chr5-84480374-INS-54
chr5-84481290-INS-150
chr5-84509931-INS-58
chr5-84533376-INS-397
chr5-84564838-INS-50
chr5-84574425-INS-54
chr5-84586162-INS-52
chr5-84595198-INS-1104
chr5-84647313-INS-62
chr5-84647351-INS-303
chr5-84664616-INS-56
chr5-84691918-INS-111
chr5-84731232-INS-514
chr5-84749064-INS-1315
chr5-84751385-INS-160
chr5-84826486-INS-72
chr5-84849298-INS-676
chr5-84874402-INS-197
chr5-84880542-INS-90
chr5-84880676-INS-84
chr5-84936682-INS-1999
chr5-84938160-INS-120
chr5-84941252-INS-1961
chr5-84952174-INS-6280
chr5-84974054-INS-860
chr5-84974782-INS-3150
chr5-84983324-INS-208
chr5-84986195-INS-50
chr5-85000086-INS-459
chr5-85022851-INS-56
chr5-85061564-INS-175
chr5-85073720-INS-58
chr5-85080466-INS-56
chr5-85095488-INS-189
chr5-85097819-INS-480
chr5-85101776-INS-66
chr5-85115467-INS-1504
chr5-85156763-INS-132
chr5-85189932-INS-101
chr5-85230350-INS-205
chr5-85251226-INS-1110
chr5-85263234-INS-4173
chr5-85392923-INS-6548
chr5-85433212-INS-6245
chr5-85437907-INS-72
chr5-85442609-INS-1203
chr5-85471586-INS-244
chr5-85488979-INS-118
chr5-85493622-INS-1322
chr5-85512885-INS-71
chr5-85521221-INS-363
chr5-85554211-INS-56
chr5-85558444-INS-78
chr5-85558540-INS-121
chr5-85574939-INS-50
chr5-85583669-INS-69
chr5-85584920-INS-172
chr5-85596576-INS-201
chr5-85601714-INS-317
chr5-85654617-INS-2044
chr5-85698671-INS-88
chr5-85728058-INS-55
chr5-85736426-INS-5834

chr5-85747044-INS-6535
chr5-85762483-INS-2345
chr5-85777926-INS-6444
chr5-85788407-INS-465
chr5-85890001-INS-221
chr5-85917458-INS-4328
chr5-85921370-INS-794
chr5-85932256-INS-7011
chr5-85976903-INS-259
chr5-86014046-INS-477
chr5-86022914-INS-645
chr5-86026807-INS-132
chr5-86046767-INS-67
chr5-86046793-INS-256
chr5-86057853-INS-6179
chr5-86101653-INS-52
chr5-86149514-INS-3666
chr5-86181780-INS-716
chr5-86192953-INS-537
chr5-86233918-INS-56
chr5-86339114-INS-8068
chr5-86347069-INS-404
chr5-86375210-INS-449
chr5-86392341-INS-6219
chr5-86425255-INS-79
chr5-86444107-INS-5371
chr5-86449362-INS-196
chr5-86464749-INS-6403
chr5-86477444-INS-78
chr5-86499869-INS-87
chr5-86546735-INS-6508
chr5-86547196-INS-50
chr5-86564099-INS-50
chr5-86590205-INS-82
chr5-86605254-INS-405
chr5-86609881-INS-279
chr5-86612048-INS-428
chr5-86653953-INS-139
chr5-86653953-INS-52
chr5-86686760-INS-132
chr5-86691738-INS-65
chr5-86691742-INS-214
chr5-86693414-INS-55
chr5-86740751-INS-182
chr5-86760125-INS-166
chr5-86761907-INS-548
chr5-86772602-INS-73
chr5-86811475-INS-114
chr5-86823404-INS-106
chr5-86851113-INS-1987
chr5-86857419-INS-226
chr5-86870384-INS-221
chr5-86880687-INS-81
chr5-86900314-INS-226
chr5-86924669-INS-88
chr5-86959829-INS-770
chr5-86967411-INS-194
chr5-86970998-INS-215
chr5-87043206-INS-52
chr5-87049270-INS-6750
chr5-87051311-INS-248
chr5-87106873-INS-12828
chr5-87123524-INS-2084
chr5-87130244-INS-1728
chr5-87134440-INS-460
chr5-87156543-INS-6390
chr5-87162826-INS-4722
chr5-87261444-INS-1567
chr5-87398253-INS-192
chr5-87448650-INS-6590
chr5-87449763-INS-495
chr5-87481018-INS-1255
chr5-87503108-INS-496
chr5-87565310-INS-95
chr5-87617200-INS-487
chr5-87630928-INS-1448
chr5-87648076-INS-97

chr5-87723775-INS-84
chr5-87747002-INS-90
chr5-87790908-INS-108
chr5-87793238-INS-103
chr5-87816644-INS-70
chr5-87823940-INS-259
chr5-87878798-INS-6465
chr5-87892743-INS-124
chr5-87893552-INS-578
chr5-87951737-INS-415
chr5-87969731-INS-52
chr5-87975158-INS-177
chr5-87980086-INS-130
chr5-88002028-INS-354
chr5-88003155-INS-1884
chr5-88009580-INS-6206
chr5-88016752-INS-101
chr5-88038298-INS-56
chr5-88038442-INS-536
chr5-88047214-INS-229
chr5-88069653-INS-70
chr5-88078784-INS-108
chr5-88079007-INS-300
chr5-88083836-INS-861
chr5-88090737-INS-6549
chr5-88137815-INS-1707
chr5-88145180-INS-5696
chr5-88145982-INS-134
chr5-88150005-INS-1333
chr5-88161003-INS-227
chr5-88209522-INS-251
chr5-88246053-INS-2853
chr5-88252867-INS-6521
chr5-88325762-INS-105
chr5-88332328-INS-70
chr5-88378976-INS-1165
chr5-88385184-INS-100
chr5-88422445-INS-6501
chr5-88430009-INS-995
chr5-88453615-INS-1255
chr5-88463100-INS-168
chr5-88470394-INS-120
chr5-88470978-INS-157
chr5-88484714-INS-8289
chr5-88486076-INS-10770
chr5-88504381-INS-6974
chr5-88553130-INS-52
chr5-88568908-INS-3032
chr5-88584129-INS-1342
chr5-88614440-INS-148
chr5-88639907-INS-4300
chr5-88676397-INS-134
chr5-88678429-INS-352
chr5-88685904-INS-74
chr5-88690617-INS-462
chr5-88701104-INS-3188
chr5-88842462-INS-311
chr5-88843235-INS-382
chr5-88869220-INS-64
chr5-88873411-INS-327
chr5-88967910-INS-2323
chr5-89009766-INS-354
chr5-89033724-INS-50
chr5-89097440-INS-56
chr5-89099244-INS-53
chr5-89124097-INS-1595
chr5-89140390-INS-181
chr5-89146326-INS-50
chr5-89236996-INS-69
chr5-89237183-INS-6701
chr5-89250207-INS-50
chr5-89269118-INS-643
chr5-89288277-INS-108
chr5-89335198-INS-6651
chr5-89357564-INS-275
chr5-89388219-INS-5546
chr5-89393133-INS-64

chr5-89422433-INS-51
chr5-89449821-INS-114
chr5-89467044-INS-650
chr5-89508895-INS-60
chr5-89512714-INS-50
chr5-89515460-INS-226
chr5-89518877-INS-52
chr5-89522425-INS-1440
chr5-89531084-INS-126
chr5-89531225-INS-1995
chr5-89531930-INS-585
chr5-89565617-INS-98
chr5-89573396-INS-282
chr5-89615169-INS-527
chr5-89629129-INS-872
chr5-89644260-INS-6590
chr5-89644750-INS-61
chr5-89645537-INS-6912
chr5-89654885-INS-1635
chr5-89670149-INS-114
chr5-89737827-INS-1427
chr5-89739979-INS-84
chr5-89740244-INS-157
chr5-89779074-INS-489
chr5-89798008-INS-6652
chr5-89800866-INS-2719
chr5-89810225-INS-6076
chr5-89817963-INS-166
chr5-89835057-INS-212
chr5-89840642-INS-76
chr5-89880289-INS-53
chr5-89890395-INS-1086
chr5-89898529-INS-6206
chr5-89959589-INS-7196
chr5-89974639-INS-332
chr5-89974980-INS-6304
chr5-89983168-INS-488
chr5-90001188-INS-164
chr5-90005898-INS-855
chr5-90015453-INS-132
chr5-90058433-INS-1360
chr5-90064569-INS-126
chr5-90066758-INS-104
chr5-90067139-INS-250
chr5-90070765-INS-1282
chr5-90084585-INS-10378
chr5-90088326-INS-52
chr5-90101106-INS-204
chr5-90137880-INS-124
chr5-90186907-INS-2415
chr5-90195413-INS-52
chr5-90209440-INS-52
chr5-90212902-INS-78
chr5-90232082-INS-59
chr5-90237949-INS-67
chr5-90242022-INS-2143
chr5-90273452-INS-133
chr5-90274017-INS-130
chr5-90287813-INS-2816
chr5-90292848-INS-601
chr5-90299968-INS-73
chr5-90306536-INS-207
chr5-90333341-INS-225
chr5-90334319-INS-305
chr5-90342137-INS-164
chr5-90357823-INS-297
chr5-90364458-INS-4797
chr5-90366333-INS-202
chr5-90367965-INS-80
chr5-90404592-INS-200
chr5-90424660-INS-53
chr5-90452810-INS-61
chr5-90525548-INS-7220
chr5-90533048-INS-3787
chr5-90537928-INS-566
chr5-90602919-INS-78
chr5-90606420-INS-90

chr5-90657566-INS-52
chr5-90668533-INS-163
chr5-90668909-INS-126
chr5-90697203-INS-480
chr5-90733477-INS-50
chr5-90740305-INS-1096
chr5-90741367-INS-112
chr5-90746021-INS-13568
chr5-90758605-INS-198
chr5-90758614-INS-483
chr5-90790269-INS-50
chr5-90815837-INS-68
chr5-90825119-INS-50
chr5-90857725-INS-65
chr5-90858392-INS-55
chr5-90858843-INS-56
chr5-90877951-INS-301
chr5-90879755-INS-6166
chr5-90892328-INS-1514
chr5-90902821-INS-133
chr5-90943802-INS-52
chr5-91014338-INS-65
chr5-91030084-INS-55
chr5-91030858-INS-117
chr5-91034041-INS-229
chr5-91035672-INS-83
chr5-91054523-INS-320
chr5-91081560-INS-58
chr5-91100780-INS-80
chr5-91121263-INS-372
chr5-91146779-INS-161
chr5-91147197-INS-6380
chr5-91148849-INS-60
chr5-91180508-INS-121
chr5-91198931-INS-60
chr5-91261555-INS-198
chr5-91289811-INS-60
chr5-91325315-INS-59
chr5-91423008-INS-51
chr5-91423496-INS-934
chr5-91438292-INS-52
chr5-91471186-INS-77
chr5-91480595-INS-1078
chr5-91482625-INS-6512
chr5-91483607-INS-80
chr5-91483797-INS-219
chr5-91495804-INS-221
chr5-91543265-INS-56
chr5-91547228-INS-58
chr5-91577301-INS-104
chr5-91578080-INS-54
chr5-91584295-INS-148
chr5-91602005-INS-63
chr5-91607940-INS-51
chr5-91609559-INS-6226
chr5-91631359-INS-145
chr5-91644468-INS-165
chr5-91659423-INS-71
chr5-91720475-INS-50
chr5-91767980-INS-65
chr5-91813187-INS-208
chr5-91822869-INS-59
chr5-91852982-INS-790
chr5-91854760-INS-51
chr5-91867662-INS-78
chr5-91898023-INS-54
chr5-91906230-INS-81
chr5-91920737-INS-247
chr5-91927556-INS-59
chr5-91930509-INS-140
chr5-91938244-INS-50
chr5-91941941-INS-96
chr5-91942283-INS-913
chr5-91942608-INS-396
chr5-91953808-INS-142
chr5-92005491-INS-53
chr5-92018696-INS-74

chr5-92027606-INS-14109
chr5-92037458-INS-259
chr5-92043776-INS-572
chr5-92055862-INS-56
chr5-92132523-INS-130
chr5-92151202-INS-521
chr5-92151251-INS-97
chr5-92160488-INS-52
chr5-92164179-INS-128
chr5-92179883-INS-87
chr5-92184511-INS-178
chr5-92184763-INS-64
chr5-92240824-INS-224
chr5-92246873-INS-166
chr5-92269897-INS-211
chr5-92273214-INS-213
chr5-92274465-INS-204
chr5-92293759-INS-74
chr5-92320962-INS-174
chr5-92343950-INS-112
chr5-92370981-INS-81
chr5-92371362-INS-120
chr5-92381560-INS-54
chr5-92420247-INS-117
chr5-92438694-INS-753
chr5-92483307-INS-50
chr5-92498913-INS-54
chr5-92503122-INS-202
chr5-92510708-INS-60
chr5-92587946-INS-202
chr5-92610717-INS-386
chr5-92639269-INS-215
chr5-92654748-INS-50
chr5-92654902-INS-121
chr5-92683086-INS-523
chr5-92684569-INS-204
chr5-92689688-INS-1335
chr5-92701153-INS-202
chr5-92702143-INS-179
chr5-92705031-INS-533
chr5-92722183-INS-181
chr5-92741079-INS-64
chr5-92741197-INS-90
chr5-92771377-INS-117
chr5-92775676-INS-208
chr5-92776217-INS-51
chr5-92776725-INS-195
chr5-92779853-INS-52
chr5-92792780-INS-289
chr5-92794395-INS-1228
chr5-92794406-INS-605
chr5-92795031-INS-60
chr5-92797367-INS-159
chr5-92797391-INS-6789
chr5-92826694-INS-6391
chr5-92827564-INS-108
chr5-92879768-INS-88
chr5-92911917-INS-50
chr5-92933342-INS-1537
chr5-92933563-INS-97
chr5-92933605-INS-74
chr5-92942368-INS-174
chr5-92975588-INS-140
chr5-92988253-INS-57
chr5-93001691-INS-53
chr5-93007485-INS-54
chr5-93019559-INS-88
chr5-93034654-INS-232
chr5-93050150-INS-71
chr5-93052371-INS-83
chr5-93126881-INS-182
chr5-93127537-INS-64
chr5-93127551-INS-152
chr5-93186122-INS-54
chr5-93186429-INS-126
chr5-93186541-INS-316
chr5-93234546-INS-103

chr5-93259434-INS-96
chr5-93308023-INS-185
chr5-93308253-INS-179
chr5-93314728-INS-106
chr5-93315029-INS-122
chr5-93371958-INS-1637
chr5-93374594-INS-69
chr5-93391565-INS-70
chr5-93426986-INS-80
chr5-93457846-INS-201
chr5-93461354-INS-2002
chr5-93488980-INS-2183
chr5-93512300-INS-1814
chr5-93566922-INS-4606
chr5-93655057-INS-3423
chr5-93788199-INS-73
chr5-93880333-INS-6244
chr5-95622203-INS-161
chr5-95641841-INS-572
chr5-95659655-INS-671
chr5-95666278-INS-165
chr5-95936263-INS-66
chr5-96023790-INS-3013
chr5-96027504-INS-51
chr5-96070960-INS-58
chr5-96089115-INS-50
chr5-96129376-INS-7496
chr5-96155355-INS-3056
chr5-96181180-INS-331
chr5-96204413-INS-66
chr5-96291620-INS-1009
chr5-96352009-INS-156
chr5-96352134-INS-57
chr5-96395965-INS-55
chr5-96431890-INS-4550
chr5-96461843-INS-80
chr5-96490788-INS-60
chr5-96508177-INS-79
chr5-96578141-INS-88
chr5-96579021-INS-95
chr5-96584020-INS-1872
chr5-96584144-INS-120
chr5-96651161-INS-62
chr5-96659232-INS-113
chr5-96727251-INS-86
chr5-96733445-INS-157
chr5-96735391-INS-182
chr5-96814635-INS-54
chr5-96877992-INS-221
chr5-96908000-INS-1208
chr5-96948915-INS-104
chr5-96956462-INS-6466
chr5-96987384-INS-195
chr5-96989511-INS-53
chr5-96992776-INS-568
chr5-96994790-INS-837
chr5-97009941-INS-166
chr5-97051198-INS-56
chr5-97051601-INS-66
chr5-97060136-INS-279
chr5-97094290-INS-63
chr5-97107534-INS-3410
chr5-97127593-INS-53
chr5-97139858-INS-6956
chr5-97167146-INS-233
chr5-97168999-INS-279
chr5-97175144-INS-467
chr5-97183007-INS-62
chr5-97248306-INS-91
chr5-97252718-INS-5026
chr5-97264585-INS-1757
chr5-97277108-INS-1182
chr5-97285650-INS-50
chr5-97317875-INS-555
chr5-97352083-INS-52
chr5-97363065-INS-132
chr5-97364714-INS-1460

chr5-97391951-INS-70
chr5-97412183-INS-196
chr5-97428144-INS-226
chr5-97428204-INS-80
chr5-97431338-INS-7231
chr5-97443224-INS-6276
chr5-97445522-INS-147
chr5-97455998-INS-68
chr5-97472945-INS-204
chr5-97474791-INS-572
chr5-97478549-INS-58
chr5-97505966-INS-50
chr5-97550636-INS-539
chr5-97551445-INS-549
chr5-97592258-INS-925
chr5-97625144-INS-7182
chr5-97634503-INS-7000
chr5-97676529-INS-58
chr5-97677230-INS-342
chr5-97695654-INS-124
chr5-97708939-INS-287
chr5-97711135-INS-94
chr5-97782560-INS-519
chr5-97799179-INS-130
chr5-97829170-INS-54
chr5-97864992-INS-1918
chr5-97865398-INS-54
chr5-97866496-INS-166
chr5-97884565-INS-3134
chr5-97898075-INS-6368
chr5-97912090-INS-50
chr5-97937505-INS-215
chr5-97958064-INS-6794
chr5-97962437-INS-74
chr5-98028325-INS-1120
chr5-98038267-INS-1683
chr5-98043414-INS-1142
chr5-98093843-INS-82
chr5-98106570-INS-2006
chr5-98125119-INS-143
chr5-98132553-INS-2911
chr5-98133262-INS-128
chr5-98165638-INS-354
chr5-98186853-INS-50
chr5-98200749-INS-126
chr5-98239508-INS-1423
chr5-98244286-INS-56
chr5-98277287-INS-67
chr5-98277421-INS-7166
chr5-98338393-INS-165
chr5-98364214-INS-214
chr5-98367411-INS-108
chr5-98387983-INS-197
chr5-98396361-INS-98
chr5-98428951-INS-349
chr5-98467740-INS-2679
chr5-98470606-INS-1263
chr5-98487781-INS-167
chr5-98492328-INS-50
chr5-98543753-INS-1985
chr5-98563870-INS-6285
chr5-98573840-INS-1277
chr5-98606072-INS-3321
chr5-98632520-INS-6601
chr5-98676869-INS-537
chr5-98687541-INS-56
chr5-98688189-INS-219
chr5-98721276-INS-52
chr5-98726057-INS-1229
chr5-98728244-INS-110
chr5-98734594-INS-141
chr5-98745079-INS-152
chr5-98758749-INS-6616
chr5-98760175-INS-1889
chr5-98773132-INS-68
chr5-98777882-INS-1410
chr5-98783887-INS-2797

chr5-98794517-INS-102
chr5-98819937-INS-6226
chr5-98857291-INS-294
chr5-98868186-INS-875
chr5-98887886-INS-674
chr5-98888596-INS-52
chr5-98905160-INS-56
chr5-98905372-INS-84
chr5-98907044-INS-1663
chr5-98907901-INS-11564
chr5-98942284-INS-66
chr5-98978334-INS-153
chr5-99009522-INS-190
chr5-99027781-INS-126
chr5-99038008-INS-243
chr5-99047351-INS-59
chr5-99050909-INS-546
chr5-99063312-INS-424
chr5-99082847-INS-573
chr5-99108159-INS-88
chr5-99117674-INS-50
chr5-99127017-INS-89
chr5-99129842-INS-113
chr5-99171636-INS-83
chr5-99173542-INS-295
chr5-99200770-INS-58
chr5-99211266-INS-50
chr5-99233885-INS-361
chr5-99233888-INS-159
chr5-99246449-INS-647
chr5-99258701-INS-1910
chr5-99285974-INS-107
chr5-99296154-INS-228
chr5-99318364-INS-1158
chr5-99353471-INS-58
chr5-99353748-INS-54
chr5-99362801-INS-87
chr5-99385748-INS-222
chr5-99393052-INS-74
chr5-99396097-INS-114
chr5-99435304-INS-84
chr5-99441283-INS-100
chr5-99516587-INS-57
chr5-99517626-INS-76
chr5-99522910-INS-4945
chr5-99532278-INS-165
chr5-99542534-INS-5874
chr5-99555627-INS-1390
chr5-99573254-INS-50
chr5-99608639-INS-125
chr5-99609418-INS-56
chr5-99611798-INS-6175
chr5-99630622-INS-2286
chr5-99636044-INS-139
chr5-99636053-INS-55
chr5-99638676-INS-50
chr5-99718160-INS-193
chr5-99722085-INS-62
chr5-99724603-INS-86
chr5-99740191-INS-2455
chr5-99741032-INS-52
chr5-99757662-INS-6403
chr5-99764474-INS-1442
chr5-99805214-INS-52
chr5-99815558-INS-128
chr5-99829172-INS-52
chr5-99838562-INS-301
chr5-99844445-INS-260
chr5-99864172-INS-66
chr5-99895086-INS-1005
chr5-99922898-INS-1132
chr5-99941749-INS-458
chr5-99941843-INS-221
chr5-99966132-INS-55
chr5-99970925-INS-110
chr5-99985530-INS-68
chr5-99987129-INS-51

chr5-99990203-INS-5269
chr5-100002772-INS-828
chr5-100037613-INS-74
chr5-100052102-INS-3114
chr5-100054934-INS-6666
chr5-100070260-INS-252
chr5-100075902-INS-51
chr5-100143315-INS-969
chr5-100168816-INS-172
chr5-100169407-INS-52
chr5-100176042-INS-205
chr5-100218072-INS-305
chr5-100236904-INS-62
chr5-100247168-INS-473
chr5-100271733-INS-57
chr5-100271910-INS-192
chr5-100325994-INS-93
chr5-100330166-INS-94
chr5-100342440-INS-50
chr5-100354923-INS-196
chr5-100392433-INS-156
chr5-100398341-INS-156
chr5-100398990-INS-153
chr5-100400503-INS-57
chr5-100404729-INS-51
chr5-100405805-INS-62
chr5-100407022-INS-10204
chr5-100425032-INS-301
chr5-100459397-INS-79
chr5-100465666-INS-130
chr5-100467488-INS-50
chr5-100469808-INS-76
chr5-100472927-INS-54
chr5-100474820-INS-6442
chr5-100495814-INS-75
chr5-100522804-INS-62
chr5-100526244-INS-902
chr5-100559592-INS-171
chr5-100594363-INS-56
chr5-100609225-INS-109
chr5-100613082-INS-50
chr5-100659721-INS-82
chr5-100666004-INS-4983
chr5-100668999-INS-138
chr5-100669193-INS-243
chr5-100678752-INS-209
chr5-100685399-INS-206
chr5-100686400-INS-223
chr5-100700889-INS-552
chr5-100738839-INS-513
chr5-100750281-INS-147
chr5-100782868-INS-57
chr5-100790920-INS-385
chr5-100825886-INS-72
chr5-100844908-INS-166
chr5-100861912-INS-63
chr5-100883584-INS-110
chr5-100896104-INS-90
chr5-100896631-INS-56
chr5-100897592-INS-164
chr5-100900700-INS-67
chr5-100902223-INS-504
chr5-100934212-INS-145
chr5-100950995-INS-212
chr5-100966279-INS-58
chr5-100974858-INS-448
chr5-101025661-INS-166
chr5-101026488-INS-203
chr5-101036036-INS-83
chr5-101043524-INS-214
chr5-101051689-INS-96
chr5-101083853-INS-509
chr5-101089864-INS-50
chr5-101134361-INS-201
chr5-101138071-INS-50
chr5-101183271-INS-51
chr5-101219548-INS-289

chr5-101223199-INS-106
chr5-101233199-INS-229
chr5-101234541-INS-119
chr5-101248207-INS-154
chr5-101262839-INS-54
chr5-101282138-INS-72
chr5-101299696-INS-75
chr5-101301121-INS-227
chr5-101301496-INS-130
chr5-101317699-INS-1104
chr5-101324657-INS-51
chr5-101358537-INS-101
chr5-101398682-INS-60
chr5-101398807-INS-150
chr5-101400234-INS-73
chr5-101409120-INS-113
chr5-101415184-INS-50
chr5-101427193-INS-678
chr5-101452307-INS-1577
chr5-101495809-INS-2543
chr5-101544863-INS-64
chr5-101545102-INS-736
chr5-101545947-INS-60
chr5-101563914-INS-275
chr5-101581189-INS-261
chr5-101581280-INS-51
chr5-101590175-INS-522
chr5-101600099-INS-234
chr5-101611142-INS-6697
chr5-101613654-INS-1442
chr5-101667910-INS-196
chr5-101684964-INS-125
chr5-101730231-INS-60
chr5-101742330-INS-228
chr5-101758446-INS-314
chr5-101785568-INS-167
chr5-101789919-INS-108
chr5-101815295-INS-52
chr5-101817833-INS-84
chr5-101844850-INS-4967
chr5-101847944-INS-120
chr5-101869966-INS-112
chr5-101910645-INS-56
chr5-101924131-INS-372
chr5-101925016-INS-2851
chr5-102017215-INS-58
chr5-102021426-INS-53
chr5-102024867-INS-121
chr5-102027441-INS-124
chr5-102028269-INS-106
chr5-102032396-INS-81
chr5-102046607-INS-126
chr5-102050066-INS-159
chr5-102056062-INS-64
chr5-102056728-INS-110
chr5-102086771-INS-56
chr5-102087494-INS-62
chr5-102090686-INS-95
chr5-102096864-INS-570
chr5-102099550-INS-1534
chr5-102118714-INS-53
chr5-102119268-INS-58
chr5-102136549-INS-69
chr5-102154871-INS-52
chr5-102156895-INS-7142
chr5-102164165-INS-66
chr5-102165478-INS-260
chr5-102205710-INS-155
chr5-102207111-INS-220
chr5-102222214-INS-58
chr5-102244787-INS-210
chr5-102285023-INS-3026
chr5-102348921-INS-608
chr5-102352538-INS-84
chr5-102361772-INS-2457
chr5-102362157-INS-84
chr5-102373271-INS-6211

chr5-102382386-INS-78
chr5-102406583-INS-52
chr5-102410855-INS-700
chr5-102411782-INS-4119
chr5-102421327-INS-65
chr5-102422876-INS-70
chr5-102437017-INS-176
chr5-102446710-INS-150
chr5-102455625-INS-175
chr5-102455675-INS-75
chr5-102467142-INS-452
chr5-102480232-INS-1131
chr5-102497081-INS-59
chr5-102515472-INS-136
chr5-102515542-INS-52
chr5-102525149-INS-316
chr5-102525701-INS-257
chr5-102550542-INS-76
chr5-102582922-INS-50
chr5-102597270-INS-110
chr5-102604115-INS-4803
chr5-102623902-INS-69
chr5-102639078-INS-334
chr5-102656243-INS-137
chr5-102683103-INS-67
chr5-102696304-INS-3627
chr5-102701059-INS-67
chr5-102701419-INS-60
chr5-102707470-INS-82
chr5-102743223-INS-100
chr5-102782716-INS-61
chr5-102783112-INS-85
chr5-102786886-INS-76
chr5-102815649-INS-52
chr5-102847102-INS-2986
chr5-102848130-INS-4479
chr5-102911574-INS-51
chr5-102926806-INS-70
chr5-102956612-INS-58
chr5-102987221-INS-146
chr5-103075919-INS-113
chr5-103089123-INS-6341
chr5-103168489-INS-60
chr5-103171175-INS-2260
chr5-103256600-INS-50
chr5-103259961-INS-80
chr5-103268701-INS-1285
chr5-103276821-INS-208
chr5-103323894-INS-518
chr5-103385158-INS-399
chr5-103386834-INS-7249
chr5-103405075-INS-3268
chr5-103423208-INS-54
chr5-103447224-INS-328
chr5-103459529-INS-512
chr5-103490294-INS-52
chr5-103490470-INS-100
chr5-103493955-INS-76
chr5-103506325-INS-495
chr5-103506559-INS-197
chr5-103511760-INS-109
chr5-103533771-INS-91
chr5-103534848-INS-7089
chr5-103536749-INS-175
chr5-103567473-INS-206
chr5-103578323-INS-58
chr5-103578546-INS-220
chr5-103581892-INS-205
chr5-103618719-INS-190
chr5-103659547-INS-228
chr5-103671927-INS-4529
chr5-103680345-INS-57
chr5-103696275-INS-122
chr5-103705024-INS-99
chr5-103718470-INS-216
chr5-103718583-INS-735
chr5-103727750-INS-161

chr5-103753357-INS-326
chr5-103800521-INS-222
chr5-103817236-INS-132
chr5-103855251-INS-181
chr5-103874511-INS-52
chr5-103891816-INS-598
chr5-103899759-INS-988
chr5-103910606-INS-186
chr5-103942753-INS-96
chr5-104006731-INS-80
chr5-104007397-INS-50
chr5-104030616-INS-67
chr5-104039034-INS-5311
chr5-104040144-INS-507
chr5-104050736-INS-204
chr5-104054842-INS-233
chr5-104087725-INS-70
chr5-104104336-INS-5514
chr5-104106402-INS-56
chr5-104107217-INS-50
chr5-104110337-INS-105
chr5-104115870-INS-158
chr5-104121238-INS-50
chr5-104142109-INS-73
chr5-104144940-INS-2670
chr5-104179502-INS-1257
chr5-104191025-INS-149
chr5-104197259-INS-260
chr5-104250991-INS-672
chr5-104251047-INS-84
chr5-104252321-INS-185
chr5-104300760-INS-100
chr5-104304624-INS-1199
chr5-104326287-INS-57
chr5-104346638-INS-56
chr5-104364181-INS-96
chr5-104401595-INS-170
chr5-104403324-INS-7140
chr5-104460078-INS-238
chr5-104461886-INS-140
chr5-104479241-INS-224
chr5-104479700-INS-629
chr5-104480093-INS-5388
chr5-104496542-INS-274
chr5-104496625-INS-700
chr5-104520238-INS-162
chr5-104520327-INS-66
chr5-104521117-INS-136
chr5-104521129-INS-322
chr5-104521211-INS-88
chr5-104521435-INS-261
chr5-104536918-INS-5522
chr5-104567052-INS-9288
chr5-104570882-INS-205
chr5-104571324-INS-64
chr5-104610428-INS-235
chr5-104707006-INS-68
chr5-104721738-INS-178
chr5-104723078-INS-81
chr5-104723638-INS-261
chr5-104730610-INS-1344
chr5-104730610-INS-504
chr5-104735645-INS-50
chr5-104748467-INS-176
chr5-104749156-INS-191
chr5-104788242-INS-3000
chr5-104789201-INS-80
chr5-104852652-INS-1019
chr5-104853447-INS-827
chr5-104921444-INS-332
chr5-104962364-INS-84
chr5-104966385-INS-217
chr5-104979954-INS-67
chr5-104985422-INS-5014
chr5-104998330-INS-541
chr5-105005443-INS-1545
chr5-105088010-INS-2810

chr5-105097245-INS-55
chr5-105109428-INS-1210
chr5-105112568-INS-182
chr5-105118367-INS-124
chr5-105152213-INS-7241
chr5-105165472-INS-66
chr5-105190990-INS-206
chr5-105241177-INS-2403
chr5-105402096-INS-350
chr5-105429697-INS-330
chr5-105447308-INS-259
chr5-105484922-INS-66
chr5-105545173-INS-63
chr5-105549690-INS-174
chr5-105574576-INS-75
chr5-105574584-INS-200
chr5-105616833-INS-6581
chr5-105655789-INS-815
chr5-105687048-INS-180
chr5-105707143-INS-1316
chr5-105740828-INS-2556
chr5-105765626-INS-724
chr5-105780939-INS-56
chr5-105786110-INS-115
chr5-105787091-INS-56
chr5-105806352-INS-1348
chr5-105807985-INS-1136
chr5-105809371-INS-217
chr5-105814413-INS-336
chr5-105814413-INS-4416
chr5-105838164-INS-50
chr5-105841332-INS-81
chr5-105859049-INS-1180
chr5-105885711-INS-88
chr5-105886950-INS-57
chr5-105917231-INS-72
chr5-105929670-INS-50
chr5-105977720-INS-6870
chr5-105993449-INS-332
chr5-106077232-INS-390
chr5-106086618-INS-444
chr5-106087942-INS-65
chr5-106091266-INS-93
chr5-106093073-INS-84
chr5-106094094-INS-85
chr5-106158795-INS-7152
chr5-106158876-INS-50
chr5-106198456-INS-87
chr5-106207342-INS-74
chr5-106210991-INS-457
chr5-106245452-INS-551
chr5-106252263-INS-338
chr5-106294656-INS-258
chr5-106304174-INS-538
chr5-106304964-INS-94
chr5-106305577-INS-86
chr5-106316774-INS-255
chr5-106347453-INS-98
chr5-106350384-INS-129
chr5-106358669-INS-171
chr5-106393654-INS-234
chr5-106396740-INS-7121
chr5-106401801-INS-61
chr5-106436393-INS-148
chr5-106448217-INS-2499
chr5-106512336-INS-81
chr5-106516699-INS-510
chr5-106516699-INS-57
chr5-106528241-INS-60
chr5-106543172-INS-540
chr5-106556524-INS-60
chr5-106562170-INS-340
chr5-106585123-INS-168
chr5-106610205-INS-72
chr5-106632632-INS-7190
chr5-106667533-INS-58
chr5-106709261-INS-122

chr5-106728633-INS-80
chr5-106849039-INS-228
chr5-106856635-INS-54
chr5-106858294-INS-50
chr5-106878751-INS-152
chr5-106880436-INS-896
chr5-106880462-INS-195
chr5-106897956-INS-78
chr5-106928353-INS-89
chr5-106931474-INS-66
chr5-106938651-INS-73
chr5-106956039-INS-935
chr5-106956377-INS-88
chr5-106956435-INS-690
chr5-106959994-INS-228
chr5-107051545-INS-102
chr5-107051785-INS-416
chr5-107055680-INS-1566
chr5-107079607-INS-2742
chr5-107091098-INS-202
chr5-107091219-INS-359
chr5-107095876-INS-206
chr5-107096893-INS-65
chr5-107096899-INS-141
chr5-107097483-INS-155
chr5-107098275-INS-108
chr5-107117757-INS-209
chr5-107147783-INS-210
chr5-107148911-INS-212
chr5-107155788-INS-199
chr5-107157752-INS-1029
chr5-107158068-INS-1328
chr5-107180364-INS-190
chr5-107239623-INS-51
chr5-107264230-INS-112
chr5-107334141-INS-52
chr5-107344437-INS-76
chr5-107354966-INS-210
chr5-107357403-INS-50
chr5-107401621-INS-441
chr5-107413524-INS-66
chr5-107450574-INS-3758
chr5-107461869-INS-194
chr5-107473473-INS-1599
chr5-107499178-INS-88
chr5-107501484-INS-56
chr5-107573501-INS-59
chr5-107628189-INS-610
chr5-107675791-INS-202
chr5-107697535-INS-51
chr5-107735834-INS-4408
chr5-107739443-INS-4675
chr5-107769605-INS-6405
chr5-107839377-INS-157
chr5-107839510-INS-72
chr5-107850327-INS-146
chr5-107880836-INS-50
chr5-107903818-INS-68
chr5-107903987-INS-163
chr5-107970928-INS-208
chr5-107993840-INS-1832
chr5-107999683-INS-191
chr5-108009850-INS-200
chr5-108019971-INS-56
chr5-108024085-INS-244
chr5-108024977-INS-196
chr5-108035221-INS-90
chr5-108042053-INS-453
chr5-108042630-INS-324
chr5-108042729-INS-132
chr5-108043955-INS-192
chr5-108086101-INS-200
chr5-108096948-INS-168
chr5-108108869-INS-221
chr5-108118087-INS-205
chr5-108132766-INS-191
chr5-108145453-INS-66

chr5-108158716-INS-221
chr5-108264379-INS-227
chr5-108275042-INS-169
chr5-108275381-INS-54
chr5-108308355-INS-189
chr5-108319915-INS-2809
chr5-108343243-INS-66
chr5-108414017-INS-50
chr5-108448935-INS-68
chr5-108495687-INS-270
chr5-108505139-INS-78
chr5-108540920-INS-202
chr5-108541696-INS-111
chr5-108553968-INS-66
chr5-108554715-INS-164
chr5-108588768-INS-194
chr5-108592145-INS-114
chr5-108600969-INS-210
chr5-108624364-INS-208
chr5-108624692-INS-98
chr5-108664281-INS-207
chr5-108665976-INS-170
chr5-108712251-INS-60
chr5-108742450-INS-586
chr5-108742523-INS-85
chr5-108810385-INS-50
chr5-108830113-INS-188
chr5-108830547-INS-373
chr5-108874590-INS-132
chr5-108875758-INS-173
chr5-108946668-INS-6402
chr5-108957517-INS-2658
chr5-108972502-INS-4710
chr5-108987051-INS-6812
chr5-109004453-INS-55
chr5-109279564-INS-527
chr5-109339279-INS-76
chr5-109396710-INS-1308
chr5-109557791-INS-2491
chr5-109586423-INS-5452
chr5-109658272-INS-76
chr5-109660983-INS-50
chr5-109697787-INS-52
chr5-109805063-INS-107
chr5-109853512-INS-91
chr5-109853795-INS-174
chr5-109889579-INS-990
chr5-109904912-INS-323
chr5-109911326-INS-1376
chr5-109913805-INS-617
chr5-109960447-INS-4083
chr5-109965845-INS-139
chr5-109967385-INS-743
chr5-109968795-INS-367
chr5-109995530-INS-201
chr5-110021055-INS-207
chr5-110047131-INS-194
chr5-110062110-INS-202
chr5-110077215-INS-207
chr5-110083991-INS-123
chr5-110084562-INS-75
chr5-110106497-INS-254
chr5-110146549-INS-2157
chr5-110151784-INS-3853
chr5-110156847-INS-756
chr5-110157289-INS-756
chr5-110172902-INS-98
chr5-110175667-INS-236
chr5-110199917-INS-3961
chr5-110211613-INS-5274
chr5-110221543-INS-147
chr5-110262399-INS-196
chr5-110265568-INS-240
chr5-110274748-INS-199
chr5-110280565-INS-4529
chr5-110285736-INS-204
chr5-110293402-INS-196

chr5-110297622-INS-56
chr5-110312453-INS-106
chr5-110312691-INS-204
chr5-110355134-INS-217
chr5-110419865-INS-498
chr5-110438802-INS-206
chr5-110478352-INS-167
chr5-110504798-INS-76
chr5-110546961-INS-50
chr5-110553864-INS-159
chr5-110599648-INS-3047
chr5-110603401-INS-1242
chr5-110612494-INS-207
chr5-110640407-INS-80
chr5-110657585-INS-209
chr5-110665011-INS-50
chr5-110665380-INS-216
chr5-110669540-INS-415
chr5-110672133-INS-53
chr5-110705146-INS-106
chr5-110714260-INS-216
chr5-110714517-INS-177
chr5-110798724-INS-296
chr5-110805962-INS-358
chr5-110872981-INS-56
chr5-110882311-INS-69
chr5-110893407-INS-136
chr5-110898254-INS-72
chr5-110913476-INS-180
chr5-110917625-INS-52
chr5-110961897-INS-388
chr5-110964083-INS-52
chr5-110969420-INS-94
chr5-110970327-INS-179
chr5-110971899-INS-208
chr5-110978095-INS-1047
chr5-110978389-INS-158
chr5-110980298-INS-104
chr5-110991691-INS-81
chr5-110997052-INS-50
chr5-111002533-INS-201
chr5-111023539-INS-2498
chr5-111042418-INS-208
chr5-111043622-INS-326
chr5-111052921-INS-254
chr5-111084371-INS-140
chr5-111084658-INS-143
chr5-111085125-INS-84
chr5-111134875-INS-205
chr5-111136481-INS-60
chr5-111137671-INS-144
chr5-111137699-INS-432
chr5-111224217-INS-52
chr5-111283004-INS-421
chr5-111296035-INS-1029
chr5-111318896-INS-62
chr5-111319023-INS-175
chr5-111337528-INS-58
chr5-111340718-INS-93
chr5-111395110-INS-389
chr5-111445335-INS-52
chr5-111503282-INS-205
chr5-111510666-INS-54
chr5-111513790-INS-58
chr5-111592853-INS-330
chr5-111607463-INS-53
chr5-111676603-INS-136
chr5-111740414-INS-388
chr5-111741036-INS-439
chr5-111776926-INS-1098
chr5-111777345-INS-450
chr5-111787523-INS-72
chr5-111818339-INS-77
chr5-111833671-INS-60
chr5-111874191-INS-78
chr5-111900799-INS-276
chr5-111922804-INS-2715

chr5-112015282-INS-1013
chr5-112015929-INS-78
chr5-112020590-INS-209
chr5-112024701-INS-70
chr5-112038465-INS-51
chr5-112087359-INS-276
chr5-112087565-INS-55
chr5-112147549-INS-50
chr5-112151412-INS-62
chr5-112169970-INS-52
chr5-112174808-INS-254
chr5-112185806-INS-4112
chr5-112191281-INS-324
chr5-112199108-INS-56
chr5-112203978-INS-136
chr5-112203978-INS-52
chr5-112242644-INS-3003
chr5-112292061-INS-151
chr5-112342310-INS-181
chr5-112350810-INS-56
chr5-112391447-INS-59
chr5-112392398-INS-50
chr5-112458079-INS-103
chr5-112458590-INS-50
chr5-112495085-INS-218
chr5-112564034-INS-54
chr5-112610902-INS-1000
chr5-112625282-INS-50
chr5-112647567-INS-1212
chr5-112648806-INS-50
chr5-112663185-INS-64
chr5-112689194-INS-68
chr5-112691197-INS-81
chr5-112770601-INS-240
chr5-112798051-INS-75
chr5-112807355-INS-179
chr5-112814481-INS-99
chr5-112824324-INS-233
chr5-112855309-INS-232
chr5-112855328-INS-99
chr5-112901080-INS-143
chr5-112939371-INS-62
chr5-112949663-INS-50
chr5-113016870-INS-66
chr5-113038651-INS-1425
chr5-113045724-INS-215
chr5-113050558-INS-6401
chr5-113055950-INS-62
chr5-113069759-INS-56
chr5-113114902-INS-87
chr5-113188085-INS-128
chr5-113199342-INS-53
chr5-113256012-INS-132
chr5-113258277-INS-564
chr5-113258606-INS-343
chr5-113294714-INS-52
chr5-113304753-INS-90
chr5-113320454-INS-52
chr5-113334330-INS-192
chr5-113394355-INS-59
chr5-113407765-INS-514
chr5-113413037-INS-202
chr5-113420488-INS-173
chr5-113429827-INS-76
chr5-113461799-INS-52
chr5-113467637-INS-465
chr5-113467637-INS-59
chr5-113487480-INS-174
chr5-113525775-INS-62
chr5-113563656-INS-2743
chr5-113589368-INS-78
chr5-113713543-INS-70
chr5-113713686-INS-360
chr5-113722240-INS-57
chr5-113740112-INS-149
chr5-113742569-INS-260
chr5-113807366-INS-116

chr5-113833604-INS-2698
chr5-113838538-INS-117
chr5-113895155-INS-156
chr5-113946863-INS-306
chr5-114060961-INS-199
chr5-114061540-INS-174
chr5-114085685-INS-232
chr5-114100416-INS-3885
chr5-114144053-INS-62
chr5-114149922-INS-159
chr5-114150182-INS-267
chr5-114178848-INS-109
chr5-114256171-INS-61
chr5-114256524-INS-73
chr5-114280583-INS-963
chr5-114298437-INS-143
chr5-114298441-INS-60
chr5-114315622-INS-8822
chr5-114336734-INS-274
chr5-114356597-INS-6747
chr5-114391162-INS-440
chr5-114391615-INS-676
chr5-114422660-INS-172
chr5-114441038-INS-288
chr5-114441164-INS-308
chr5-114441165-INS-79
chr5-114445786-INS-58
chr5-114459527-INS-72
chr5-114475317-INS-219
chr5-114475328-INS-70
chr5-114491280-INS-162
chr5-114587454-INS-465
chr5-114617569-INS-195
chr5-114617658-INS-177
chr5-114627251-INS-124
chr5-114641772-INS-115
chr5-114643737-INS-2759
chr5-114682391-INS-85
chr5-114715254-INS-81
chr5-114799962-INS-221
chr5-114850511-INS-51
chr5-114891880-INS-245
chr5-114911708-INS-108
chr5-114911737-INS-239
chr5-114993961-INS-211
chr5-114994097-INS-81
chr5-115067590-INS-145
chr5-115094950-INS-7104
chr5-115113752-INS-164
chr5-115153035-INS-192
chr5-115155942-INS-486
chr5-115161156-INS-215
chr5-115187896-INS-162
chr5-115199248-INS-207
chr5-115204763-INS-53
chr5-115204849-INS-208
chr5-115216964-INS-264
chr5-115247382-INS-285
chr5-115247382-INS-84
chr5-115257519-INS-232
chr5-115257872-INS-52
chr5-115271161-INS-53
chr5-115300663-INS-202
chr5-115335420-INS-343
chr5-115337483-INS-69
chr5-115339126-INS-1892
chr5-115351994-INS-54
chr5-115359881-INS-66
chr5-115365342-INS-64
chr5-115401876-INS-374
chr5-115418501-INS-249
chr5-115519442-INS-50
chr5-115526047-INS-95
chr5-115563536-INS-51
chr5-115631568-INS-214
chr5-115636195-INS-94
chr5-115679434-INS-60

chr5-115799828-INS-84
chr5-115814388-INS-201
chr5-115815198-INS-135
chr5-115889304-INS-60
chr5-115901245-INS-54
chr5-115917513-INS-4636
chr5-115919525-INS-54
chr5-115922376-INS-1111
chr5-115938317-INS-56
chr5-115959301-INS-405
chr5-115965198-INS-54
chr5-115969921-INS-549
chr5-116016049-INS-75
chr5-116022348-INS-63
chr5-116052443-INS-88
chr5-116076606-INS-50
chr5-116164761-INS-51
chr5-116175409-INS-77
chr5-116179647-INS-106
chr5-116183526-INS-201
chr5-116187060-INS-54
chr5-116191630-INS-175
chr5-116193537-INS-69
chr5-116209791-INS-700
chr5-116223462-INS-242
chr5-116231532-INS-54
chr5-116233491-INS-126
chr5-116256510-INS-308
chr5-116257814-INS-272
chr5-116277138-INS-681
chr5-116277789-INS-101
chr5-116279548-INS-54
chr5-116279952-INS-600
chr5-116325889-INS-7331
chr5-116337918-INS-92
chr5-116364510-INS-80
chr5-116386863-INS-51
chr5-116474911-INS-229
chr5-116474951-INS-108
chr5-116492086-INS-114
chr5-116501827-INS-90
chr5-116507151-INS-504
chr5-116534898-INS-190
chr5-116559413-INS-141
chr5-116594198-INS-112
chr5-116640591-INS-70
chr5-116652876-INS-157
chr5-116689922-INS-58
chr5-116741265-INS-98
chr5-116749413-INS-208
chr5-116760544-INS-134
chr5-116765493-INS-110
chr5-116776406-INS-60
chr5-116797514-INS-4042
chr5-116822058-INS-88
chr5-116838225-INS-159
chr5-116848418-INS-79
chr5-116857791-INS-6101
chr5-116886668-INS-252
chr5-116903273-INS-57
chr5-116904256-INS-90
chr5-116912652-INS-56
chr5-116922123-INS-251
chr5-116945353-INS-95
chr5-116950231-INS-65
chr5-116994643-INS-61
chr5-117074002-INS-230
chr5-117074002-INS-92
chr5-117074281-INS-204
chr5-117075457-INS-52
chr5-117149134-INS-84
chr5-117150055-INS-50
chr5-117191817-INS-52
chr5-117202039-INS-1291
chr5-117209269-INS-1233
chr5-117209445-INS-3349
chr5-117211897-INS-107

chr5-117211898-INS-497
chr5-117220535-INS-98
chr5-117227434-INS-50
chr5-117305653-INS-58
chr5-117327580-INS-52
chr5-117332644-INS-56
chr5-117344018-INS-214
chr5-117416636-INS-235
chr5-117417078-INS-372
chr5-117476241-INS-5041
chr5-117536053-INS-57
chr5-117571609-INS-203
chr5-117590469-INS-67
chr5-117722425-INS-77
chr5-117751338-INS-287
chr5-117751504-INS-181
chr5-117772445-INS-493
chr5-117835371-INS-175
chr5-117885493-INS-370
chr5-117890671-INS-259
chr5-117916697-INS-50
chr5-117951013-INS-3038
chr5-117973096-INS-469
chr5-118038088-INS-814
chr5-118038269-INS-4888
chr5-118084309-INS-53
chr5-118163323-INS-50
chr5-118355890-INS-58
chr5-118356605-INS-134
chr5-118392117-INS-6508
chr5-118394366-INS-107
chr5-118403244-INS-90
chr5-118408553-INS-133
chr5-118408553-INS-63
chr5-118455170-INS-74
chr5-118460536-INS-2598
chr5-118474131-INS-469
chr5-118474613-INS-232
chr5-118474912-INS-257
chr5-118569931-INS-234
chr5-118587928-INS-60
chr5-118607932-INS-248
chr5-118633830-INS-86
chr5-118687416-INS-1684
chr5-118764406-INS-99
chr5-118801402-INS-437
chr5-118801402-INS-99
chr5-118913320-INS-99
chr5-118982903-INS-228
chr5-118983047-INS-100
chr5-119066862-INS-100
chr5-119092729-INS-188
chr5-119124696-INS-108
chr5-119130630-INS-259
chr5-119130786-INS-64
chr5-119131551-INS-76
chr5-119159220-INS-70
chr5-119219282-INS-64
chr5-119257994-INS-51
chr5-119266869-INS-50
chr5-119284224-INS-177
chr5-119302225-INS-443
chr5-119307295-INS-7053
chr5-119360043-INS-70
chr5-119373229-INS-66
chr5-119449591-INS-50
chr5-119472339-INS-54
chr5-119489513-INS-113
chr5-119538297-INS-57
chr5-119584222-INS-96
chr5-119586160-INS-56
chr5-119632011-INS-103
chr5-119635231-INS-60
chr5-119637314-INS-57
chr5-119638340-INS-56
chr5-119644498-INS-180
chr5-119661544-INS-201

chr5-119686460-INS-523
chr5-119692588-INS-71
chr5-119700329-INS-110
chr5-119700548-INS-95
chr5-119729221-INS-495
chr5-119755793-INS-220
chr5-119843244-INS-50
chr5-119848764-INS-72
chr5-119857091-INS-96
chr5-119866405-INS-51
chr5-119892504-INS-143
chr5-119894868-INS-132
chr5-119921669-INS-500
chr5-119992818-INS-103
chr5-120032866-INS-59
chr5-120032946-INS-300
chr5-120054407-INS-70
chr5-120063173-INS-185
chr5-120063378-INS-176
chr5-120089129-INS-113
chr5-120089325-INS-77
chr5-120099459-INS-72
chr5-120102458-INS-71
chr5-120224571-INS-490
chr5-120245925-INS-120
chr5-120254234-INS-205
chr5-120299580-INS-102
chr5-120346705-INS-54
chr5-120388817-INS-78
chr5-120454058-INS-205
chr5-120465294-INS-52
chr5-120466504-INS-986
chr5-120477086-INS-67
chr5-120477202-INS-60
chr5-120535813-INS-115
chr5-120560392-INS-91
chr5-120564055-INS-90
chr5-120603012-INS-58
chr5-120603061-INS-119
chr5-120603127-INS-416
chr5-120606175-INS-218
chr5-120656796-INS-50
chr5-120687824-INS-348
chr5-120724050-INS-78
chr5-120732357-INS-68
chr5-120799464-INS-184
chr5-120885740-INS-80
chr5-120904936-INS-50
chr5-120945922-INS-62
chr5-121023657-INS-206
chr5-121024428-INS-62
chr5-121026068-INS-105
chr5-121072475-INS-2758
chr5-121096110-INS-205
chr5-121107212-INS-61
chr5-121111265-INS-357
chr5-121111265-INS-54
chr5-121136598-INS-228
chr5-121136695-INS-307
chr5-121157135-INS-141
chr5-121164847-INS-54
chr5-121174670-INS-5066
chr5-121192861-INS-75
chr5-121199748-INS-106
chr5-121218430-INS-504
chr5-121225513-INS-74
chr5-121265258-INS-53
chr5-121286098-INS-69
chr5-121339037-INS-221
chr5-121365719-INS-210
chr5-121378932-INS-83
chr5-121384784-INS-164
chr5-121465545-INS-276
chr5-121465549-INS-631
chr5-121470595-INS-124
chr5-121506676-INS-250
chr5-121512154-INS-358

chr5-121527716-INS-52
chr5-121563643-INS-79
chr5-121621119-INS-2022
chr5-121621249-INS-300
chr5-121655614-INS-3082
chr5-121693403-INS-1608
chr5-121695377-INS-279
chr5-121703046-INS-115
chr5-121713091-INS-7113
chr5-121747141-INS-51
chr5-121755349-INS-8395
chr5-121788259-INS-198
chr5-121826694-INS-131
chr5-121833283-INS-1063
chr5-121833283-INS-484
chr5-121840119-INS-246
chr5-121840153-INS-80
chr5-121877240-INS-395
chr5-121881323-INS-86
chr5-121881398-INS-577
chr5-121925574-INS-628
chr5-122019404-INS-229
chr5-122151842-INS-124
chr5-122193435-INS-194
chr5-122199514-INS-1121
chr5-122201146-INS-339
chr5-122203981-INS-912
chr5-122205017-INS-2965
chr5-122218003-INS-50
chr5-122222062-INS-200
chr5-122228831-INS-57
chr5-122238703-INS-51
chr5-122257182-INS-52
chr5-122278678-INS-694
chr5-122280930-INS-52
chr5-122290340-INS-229
chr5-122291164-INS-170
chr5-122307816-INS-246
chr5-122315177-INS-193
chr5-122328398-INS-83
chr5-122343805-INS-60
chr5-122365319-INS-95
chr5-122417364-INS-71
chr5-122418100-INS-200
chr5-122434289-INS-576
chr5-122486261-INS-98
chr5-122508390-INS-211
chr5-122546240-INS-53
chr5-122564995-INS-76
chr5-122578795-INS-1293
chr5-122631545-INS-71
chr5-122643242-INS-165
chr5-122643255-INS-66
chr5-122656372-INS-910
chr5-122662649-INS-50
chr5-122689842-INS-232
chr5-122700029-INS-116
chr5-122809531-INS-61
chr5-122812540-INS-68
chr5-122828276-INS-145
chr5-122829217-INS-57
chr5-122849541-INS-51
chr5-122883263-INS-1400
chr5-122914322-INS-483
chr5-122932068-INS-201
chr5-122933832-INS-244
chr5-122968331-INS-121
chr5-122968997-INS-199
chr5-122979066-INS-61
chr5-122982264-INS-3655
chr5-122984880-INS-4874
chr5-122992679-INS-50
chr5-122996224-INS-217
chr5-122996514-INS-160
chr5-123119209-INS-203
chr5-123119947-INS-60
chr5-123162658-INS-5671

chr5-123189033-INS-202
chr5-123251317-INS-77
chr5-123258319-INS-60
chr5-123273850-INS-89
chr5-123305772-INS-60
chr5-123321890-INS-63
chr5-123322167-INS-72
chr5-123358463-INS-64
chr5-123359226-INS-55
chr5-123376604-INS-1190
chr5-123444991-INS-156
chr5-123446246-INS-68
chr5-123501906-INS-112
chr5-123513084-INS-54
chr5-123623634-INS-91
chr5-123628952-INS-56
chr5-123643962-INS-59
chr5-123697990-INS-82
chr5-123705683-INS-222
chr5-123707192-INS-1878
chr5-123730631-INS-90
chr5-123730885-INS-89
chr5-123756658-INS-66
chr5-123758404-INS-484
chr5-123775594-INS-104
chr5-123806975-INS-289
chr5-123811371-INS-199
chr5-123817562-INS-120
chr5-123840185-INS-64
chr5-123861512-INS-95
chr5-123861886-INS-96
chr5-123863973-INS-321
chr5-123864119-INS-201
chr5-123908725-INS-203
chr5-123911192-INS-198
chr5-123981285-INS-78
chr5-123993762-INS-272
chr5-124019234-INS-66
chr5-124041239-INS-80
chr5-124043927-INS-116
chr5-124058373-INS-59
chr5-124072794-INS-53
chr5-124075317-INS-57
chr5-124078665-INS-8204
chr5-124091408-INS-66
chr5-124178310-INS-50
chr5-124179000-INS-52
chr5-124180164-INS-50
chr5-124190692-INS-91
chr5-124207297-INS-72
chr5-124223991-INS-229
chr5-124241000-INS-143
chr5-124315727-INS-181
chr5-124376405-INS-54
chr5-124413786-INS-52
chr5-124424079-INS-227
chr5-124482212-INS-203
chr5-124517809-INS-136
chr5-124602462-INS-172
chr5-124609400-INS-57
chr5-124615267-INS-105
chr5-124618718-INS-213
chr5-124651634-INS-185
chr5-124707538-INS-223
chr5-124755095-INS-367
chr5-124757702-INS-175
chr5-124787015-INS-396
chr5-124788273-INS-90
chr5-124804387-INS-727
chr5-124813562-INS-169
chr5-124814477-INS-151
chr5-124822617-INS-68
chr5-124851328-INS-108
chr5-124856413-INS-633
chr5-124856483-INS-276
chr5-124856698-INS-249
chr5-124869271-INS-7121

chr5-124892440-INS-196
chr5-124908588-INS-50
chr5-124914142-INS-50
chr5-124959532-INS-379
chr5-124970035-INS-3124
chr5-125051874-INS-132
chr5-125056036-INS-60
chr5-125056256-INS-50
chr5-125059857-INS-50
chr5-125061458-INS-7169
chr5-125064732-INS-217
chr5-125189606-INS-212
chr5-125212741-INS-444
chr5-125243672-INS-382
chr5-125253226-INS-64
chr5-125270949-INS-444
chr5-125279121-INS-3046
chr5-125288052-INS-72
chr5-125303983-INS-64
chr5-125311365-INS-190
chr5-125318043-INS-236
chr5-125319988-INS-202
chr5-125320538-INS-79
chr5-125322274-INS-53
chr5-125360002-INS-225
chr5-125375091-INS-53
chr5-125422404-INS-214
chr5-125463227-INS-684
chr5-125474817-INS-632
chr5-125485234-INS-56
chr5-125490120-INS-106
chr5-125547701-INS-56
chr5-125563645-INS-187
chr5-125574115-INS-245
chr5-125574527-INS-542
chr5-125583615-INS-1174
chr5-125614306-INS-148
chr5-125618983-INS-51
chr5-125630630-INS-76
chr5-125630643-INS-168
chr5-125672769-INS-100
chr5-125685870-INS-58
chr5-125704191-INS-58
chr5-125708079-INS-447
chr5-125746876-INS-159
chr5-125780096-INS-887
chr5-125830251-INS-168
chr5-125853743-INS-68
chr5-125867256-INS-1388
chr5-125887153-INS-203
chr5-125919846-INS-189
chr5-125979001-INS-52
chr5-126022924-INS-396
chr5-126024637-INS-5300
chr5-126095000-INS-171
chr5-126116279-INS-164
chr5-126157739-INS-927
chr5-126186929-INS-54
chr5-126194388-INS-60
chr5-126222970-INS-52
chr5-126378109-INS-5004
chr5-126390886-INS-52
chr5-126391585-INS-66
chr5-126409249-INS-74
chr5-126411190-INS-441
chr5-126417491-INS-1018
chr5-126417671-INS-444
chr5-126426447-INS-88
chr5-126473661-INS-137
chr5-126483617-INS-77
chr5-126487946-INS-6287
chr5-126489243-INS-56
chr5-126529811-INS-6725
chr5-126543682-INS-573
chr5-126543682-INS-8748
chr5-126549345-INS-70
chr5-126575118-INS-86

chr5-126575144-INS-321
chr5-126575454-INS-103
chr5-126602242-INS-189
chr5-126606749-INS-60
chr5-126610170-INS-51
chr5-126610479-INS-226
chr5-126616805-INS-67
chr5-126629758-INS-64
chr5-126636485-INS-147
chr5-126688350-INS-3479
chr5-126706612-INS-6348
chr5-126707277-INS-65
chr5-126731310-INS-75
chr5-126733097-INS-136
chr5-126740589-INS-1300
chr5-126794732-INS-57
chr5-126794826-INS-350
chr5-126800184-INS-108
chr5-126807415-INS-8015
chr5-126814613-INS-144
chr5-126814627-INS-368
chr5-126827979-INS-68
chr5-126828657-INS-56
chr5-126834297-INS-340
chr5-126844070-INS-56
chr5-126851005-INS-5387
chr5-126853205-INS-6491
chr5-126893447-INS-139
chr5-126906730-INS-1192
chr5-126940629-INS-62
chr5-126942442-INS-87
chr5-126944308-INS-104
chr5-126963629-INS-98
chr5-126995572-INS-3155
chr5-126996314-INS-53
chr5-126996447-INS-229
chr5-126997502-INS-232
chr5-126998873-INS-2627
chr5-127006450-INS-659
chr5-127160882-INS-102
chr5-127204529-INS-486
chr5-127235136-INS-84
chr5-127241222-INS-2429
chr5-127246443-INS-865
chr5-127254220-INS-4259
chr5-127266801-INS-1157
chr5-127274483-INS-387
chr5-127275151-INS-147
chr5-127383335-INS-120
chr5-127387002-INS-159
chr5-127419712-INS-6415
chr5-127421034-INS-54
chr5-127443819-INS-50
chr5-127475057-INS-73
chr5-127475844-INS-51
chr5-127501106-INS-67
chr5-127551083-INS-102
chr5-127576371-INS-4352
chr5-127705699-INS-93
chr5-127713291-INS-61
chr5-127763520-INS-7158
chr5-127846357-INS-285
chr5-127863767-INS-82
chr5-127867806-INS-61
chr5-127879356-INS-284
chr5-127924913-INS-2645
chr5-127935439-INS-68
chr5-127976682-INS-230
chr5-127990226-INS-80
chr5-127991892-INS-54
chr5-127992432-INS-73
chr5-128097070-INS-65
chr5-128107947-INS-120
chr5-128111495-INS-878
chr5-128117765-INS-1018
chr5-128191208-INS-1546
chr5-128278664-INS-2384

chr5-128303709-INS-626
chr5-128344767-INS-51
chr5-128380720-INS-52
chr5-128410130-INS-58
chr5-128434601-INS-704
chr5-128480901-INS-213
chr5-128580925-INS-64
chr5-128584368-INS-50
chr5-128588973-INS-165
chr5-128594774-INS-6442
chr5-128598264-INS-109
chr5-128622078-INS-53
chr5-128645846-INS-255
chr5-128665386-INS-171
chr5-128681304-INS-108
chr5-128751612-INS-52
chr5-128782982-INS-2336
chr5-128805233-INS-129
chr5-128805279-INS-530
chr5-128867682-INS-222
chr5-128871135-INS-482
chr5-128871148-INS-180
chr5-128951700-INS-54
chr5-128954828-INS-50
chr5-128961766-INS-492
chr5-128961883-INS-118
chr5-128963398-INS-60
chr5-128969318-INS-347
chr5-128970241-INS-166
chr5-128972977-INS-52
chr5-128986874-INS-3871
chr5-128989271-INS-79
chr5-129000409-INS-60
chr5-129000451-INS-574
chr5-129000723-INS-177
chr5-129036295-INS-5311
chr5-129050277-INS-75
chr5-129050891-INS-52
chr5-129054541-INS-56
chr5-129056079-INS-5644
chr5-129093596-INS-88
chr5-129121637-INS-51
chr5-129121821-INS-185
chr5-129132769-INS-85
chr5-129136029-INS-129
chr5-129166638-INS-53
chr5-129187401-INS-172
chr5-129202726-INS-240
chr5-129202873-INS-660
chr5-129212857-INS-54
chr5-129251202-INS-407
chr5-129276696-INS-317
chr5-129305316-INS-89
chr5-129331775-INS-286
chr5-129350306-INS-716
chr5-129350731-INS-336
chr5-129365039-INS-124
chr5-129377051-INS-119
chr5-129386178-INS-152
chr5-129403234-INS-80
chr5-129411875-INS-93
chr5-129440923-INS-78
chr5-129460621-INS-52
chr5-129461052-INS-149
chr5-129466422-INS-59
chr5-129495071-INS-64
chr5-129527685-INS-84
chr5-129548895-INS-56
chr5-129658802-INS-56
chr5-129692025-INS-50
chr5-129695013-INS-105
chr5-129698590-INS-4827
chr5-129707231-INS-99
chr5-129707418-INS-775
chr5-129709606-INS-63
chr5-129710271-INS-518
chr5-129775318-INS-51

chr5-129778398-INS-494
chr5-129778710-INS-285
chr5-129778718-INS-108
chr5-129785013-INS-1318
chr5-129789679-INS-10710
chr5-129823934-INS-51
chr5-129847226-INS-81
chr5-129850059-INS-53
chr5-129865595-INS-208
chr5-129871127-INS-213
chr5-129880152-INS-179
chr5-129880288-INS-71
chr5-129888336-INS-205
chr5-129898972-INS-6865
chr5-129901008-INS-105
chr5-129903515-INS-174
chr5-129909767-INS-1109
chr5-129916971-INS-3132
chr5-129925723-INS-50
chr5-129926629-INS-56
chr5-129928964-INS-594
chr5-129933319-INS-14344
chr5-130064762-INS-452
chr5-130076347-INS-110
chr5-130077392-INS-69
chr5-130094539-INS-62
chr5-130095175-INS-416
chr5-130101493-INS-3254
chr5-130157036-INS-220
chr5-130167668-INS-137
chr5-130171199-INS-1301
chr5-130247786-INS-197
chr5-130288501-INS-287
chr5-130297368-INS-53
chr5-130298381-INS-56
chr5-130338108-INS-84
chr5-130338447-INS-56
chr5-130356535-INS-189
chr5-130360447-INS-54
chr5-130371931-INS-110
chr5-130374291-INS-105
chr5-130382527-INS-82
chr5-130410651-INS-58
chr5-130484949-INS-72
chr5-130485277-INS-60
chr5-130486761-INS-62
chr5-130498904-INS-114
chr5-130504929-INS-69
chr5-130509900-INS-211
chr5-130520265-INS-198
chr5-130529823-INS-120
chr5-130545814-INS-64
chr5-130559837-INS-54
chr5-130622058-INS-161
chr5-130628035-INS-92
chr5-130655608-INS-65
chr5-130664109-INS-80
chr5-130667662-INS-308
chr5-130720804-INS-52
chr5-130730355-INS-97
chr5-130737985-INS-318
chr5-130773022-INS-1159
chr5-130778192-INS-105
chr5-130793247-INS-80
chr5-130818415-INS-52
chr5-130825193-INS-556
chr5-130829710-INS-83
chr5-130914933-INS-108
chr5-130924160-INS-1234
chr5-130934542-INS-478
chr5-130942655-INS-50
chr5-130947070-INS-6372
chr5-130950351-INS-88
chr5-130950377-INS-344
chr5-130972522-INS-72
chr5-130989584-INS-60
chr5-130996662-INS-54

chr5-131013326-INS-50
chr5-131076935-INS-106
chr5-131079134-INS-624
chr5-131080147-INS-4875
chr5-131087321-INS-52
chr5-131118556-INS-60
chr5-131138204-INS-52
chr5-131166435-INS-70
chr5-131167205-INS-161
chr5-131204019-INS-2115
chr5-131213096-INS-97
chr5-131248608-INS-265
chr5-131287179-INS-66
chr5-131322438-INS-57
chr5-131350989-INS-72
chr5-131356804-INS-50
chr5-131379722-INS-836
chr5-131397518-INS-2629
chr5-131398308-INS-99
chr5-131458061-INS-76
chr5-131460290-INS-229
chr5-131525274-INS-50
chr5-131525689-INS-330
chr5-131622928-INS-65
chr5-131645846-INS-50
chr5-131650733-INS-76
chr5-131652173-INS-583
chr5-131690405-INS-139
chr5-131690470-INS-7437
chr5-131690771-INS-53
chr5-131701563-INS-106
chr5-131714911-INS-131
chr5-131720768-INS-53
chr5-131721830-INS-141
chr5-131721888-INS-352
chr5-131722135-INS-64
chr5-131724782-INS-50
chr5-131775457-INS-8292
chr5-131784186-INS-135
chr5-131811801-INS-76
chr5-131826609-INS-60
chr5-131840298-INS-254
chr5-131882450-INS-53
chr5-131897802-INS-446
chr5-131935758-INS-1356
chr5-131998186-INS-65
chr5-132034064-INS-1352
chr5-132057048-INS-60
chr5-132077888-INS-74
chr5-132107293-INS-270
chr5-132121296-INS-72
chr5-132174640-INS-50
chr5-132192238-INS-73
chr5-132257919-INS-65
chr5-132281709-INS-155
chr5-132313124-INS-254
chr5-132313541-INS-66
chr5-132354427-INS-73
chr5-132381034-INS-80
chr5-132381339-INS-60
chr5-132419216-INS-670
chr5-132440568-INS-66
chr5-132566847-INS-176
chr5-132581174-INS-80
chr5-132593072-INS-88
chr5-132603191-INS-52
chr5-132624673-INS-106
chr5-132649974-INS-161
chr5-132664280-INS-134
chr5-132680920-INS-167
chr5-132698304-INS-117
chr5-132706579-INS-59
chr5-132728116-INS-106
chr5-132737798-INS-69
chr5-132778945-INS-238
chr5-132797154-INS-2802
chr5-132837700-INS-1113

chr5-132844248-INS-428
chr5-132852169-INS-60
chr5-132899989-INS-58
chr5-132906836-INS-53
chr5-132914952-INS-699
chr5-132922973-INS-56
chr5-132958324-INS-1105
chr5-132980504-INS-74
chr5-132992453-INS-76
chr5-132997726-INS-208
chr5-133013571-INS-106
chr5-133040850-INS-56
chr5-133041505-INS-200
chr5-133109535-INS-52
chr5-133126252-INS-53
chr5-133151900-INS-293
chr5-133163990-INS-4063
chr5-133169344-INS-7156
chr5-133172108-INS-114
chr5-133172314-INS-157
chr5-133195449-INS-86
chr5-133211028-INS-180
chr5-133236751-INS-1345
chr5-133342297-INS-125
chr5-133342297-INS-55
chr5-133411638-INS-149
chr5-133412353-INS-211
chr5-133425225-INS-1647
chr5-133452543-INS-66
chr5-133452634-INS-118
chr5-133521595-INS-63
chr5-133521968-INS-1270
chr5-133539867-INS-50
chr5-133639545-INS-58
chr5-133639564-INS-2226
chr5-133652496-INS-1054
chr5-133653084-INS-6775
chr5-133661166-INS-56
chr5-133664803-INS-204
chr5-133664954-INS-74
chr5-133686381-INS-533
chr5-133687083-INS-347
chr5-133716215-INS-132
chr5-133717040-INS-58
chr5-133722717-INS-63
chr5-133752548-INS-78
chr5-133764242-INS-234
chr5-133769311-INS-1132
chr5-133800991-INS-56
chr5-133820868-INS-288
chr5-133830129-INS-4794
chr5-133849542-INS-105
chr5-133862530-INS-122
chr5-133895717-INS-120
chr5-133929199-INS-213
chr5-133929324-INS-7069
chr5-133938814-INS-5018
chr5-133940280-INS-210
chr5-133957283-INS-632
chr5-134036858-INS-51
chr5-134039131-INS-106
chr5-134044822-INS-96
chr5-134053223-INS-1895
chr5-134070462-INS-127
chr5-134080778-INS-297
chr5-134080842-INS-60
chr5-134086707-INS-60
chr5-134090332-INS-419
chr5-134101054-INS-106
chr5-134112606-INS-336
chr5-134113391-INS-86
chr5-134113453-INS-276
chr5-134114668-INS-58
chr5-134118853-INS-54
chr5-134136176-INS-159
chr5-134158711-INS-159
chr5-134210877-INS-113

chr5-134233786-INS-400
chr5-134241297-INS-493
chr5-134244133-INS-56
chr5-134285693-INS-86
chr5-134292821-INS-214
chr5-134314655-INS-50
chr5-134349641-INS-8673
chr5-134375498-INS-204
chr5-134383581-INS-167
chr5-134383700-INS-354
chr5-134385205-INS-204
chr5-134388228-INS-53
chr5-134389021-INS-205
chr5-134415733-INS-54
chr5-134417732-INS-159
chr5-134417804-INS-53
chr5-134419300-INS-259
chr5-134424036-INS-954
chr5-134424301-INS-313
chr5-134424372-INS-106
chr5-134428934-INS-2847
chr5-134447987-INS-95
chr5-134468511-INS-265
chr5-134494111-INS-55
chr5-134502560-INS-58
chr5-134503253-INS-68
chr5-134515621-INS-174
chr5-134520485-INS-55
chr5-134547267-INS-108
chr5-134564988-INS-197
chr5-134585846-INS-67
chr5-134585965-INS-91
chr5-134591613-INS-200
chr5-134598520-INS-7101
chr5-134605374-INS-199
chr5-134651763-INS-54
chr5-134669320-INS-50
chr5-134672125-INS-68
chr5-134678215-INS-182
chr5-134687055-INS-235
chr5-134709791-INS-7124
chr5-134718590-INS-200
chr5-134732788-INS-56
chr5-134733756-INS-212
chr5-134733756-INS-53
chr5-134735710-INS-53
chr5-134764378-INS-56
chr5-134764431-INS-424
chr5-134781405-INS-470
chr5-134789906-INS-92
chr5-134802371-INS-52
chr5-134808828-INS-261
chr5-134830302-INS-50
chr5-134836351-INS-2006
chr5-134836700-INS-337
chr5-134850884-INS-2169
chr5-134852487-INS-212
chr5-134873051-INS-317
chr5-134879814-INS-7306
chr5-134905122-INS-176
chr5-134913567-INS-81
chr5-134938529-INS-1414
chr5-134938674-INS-56
chr5-134938900-INS-89
chr5-134942056-INS-379
chr5-134960797-INS-53
chr5-134973005-INS-175
chr5-134987237-INS-183
chr5-135005424-INS-53
chr5-135005475-INS-159
chr5-135115363-INS-61
chr5-135126330-INS-68
chr5-135131839-INS-57
chr5-135148780-INS-179
chr5-135171208-INS-53
chr5-135174317-INS-208
chr5-135286547-INS-1042

chr5-135296239-INS-2223
chr5-135296398-INS-59
chr5-135296687-INS-113
chr5-135316407-INS-213
chr5-135316660-INS-61
chr5-135398064-INS-103
chr5-135417357-INS-62
chr5-135417357-INS-742
chr5-135426252-INS-211
chr5-135448116-INS-158
chr5-135467233-INS-184
chr5-135468497-INS-164
chr5-135475302-INS-106
chr5-135477505-INS-228
chr5-135515644-INS-78
chr5-135520062-INS-87
chr5-135529679-INS-53
chr5-135541112-INS-280
chr5-135543392-INS-142
chr5-135556316-INS-56
chr5-135556664-INS-165
chr5-135556703-INS-66
chr5-135587247-INS-858
chr5-135587280-INS-370
chr5-135621740-INS-221
chr5-135642149-INS-1209
chr5-135648755-INS-592
chr5-135648788-INS-1907
chr5-135658367-INS-70
chr5-135690524-INS-205
chr5-135694348-INS-244
chr5-135704011-INS-217
chr5-135728542-INS-228
chr5-135740091-INS-237
chr5-135742297-INS-416
chr5-135766584-INS-212
chr5-135778585-INS-65
chr5-135827074-INS-79
chr5-135827932-INS-226
chr5-135834492-INS-323
chr5-135859491-INS-318
chr5-135859557-INS-54
chr5-135860344-INS-53
chr5-135869874-INS-53
chr5-135882326-INS-80
chr5-135909812-INS-74
chr5-135920780-INS-66
chr5-135931202-INS-231
chr5-135955967-INS-50
chr5-135968853-INS-157
chr5-135970949-INS-174
chr5-135972467-INS-54
chr5-135976495-INS-120
chr5-136004703-INS-480
chr5-136027718-INS-183
chr5-136083265-INS-317
chr5-136084353-INS-280
chr5-136093060-INS-3738
chr5-136096243-INS-2242
chr5-136096362-INS-63
chr5-136096717-INS-51
chr5-136142524-INS-53
chr5-136173048-INS-5195
chr5-136195857-INS-158
chr5-136254714-INS-56
chr5-136374585-INS-84
chr5-136406855-INS-157
chr5-136472578-INS-344
chr5-136486289-INS-105
chr5-136495995-INS-109
chr5-136523518-INS-50
chr5-136530591-INS-117
chr5-136576405-INS-185
chr5-136622077-INS-148
chr5-136626444-INS-7161
chr5-136641423-INS-76
chr5-136661834-INS-53

chr5-136662000-INS-159
chr5-136665725-INS-53
chr5-136668243-INS-62
chr5-136675201-INS-53
chr5-136675456-INS-53
chr5-136678881-INS-53
chr5-136722454-INS-54
chr5-136740785-INS-74
chr5-136741057-INS-110
chr5-136745804-INS-53
chr5-136755019-INS-884
chr5-136779546-INS-212
chr5-136786043-INS-52
chr5-136788185-INS-116
chr5-136808032-INS-366
chr5-136808499-INS-180
chr5-136819356-INS-187
chr5-136823805-INS-216
chr5-136824155-INS-159
chr5-136838914-INS-160
chr5-136839124-INS-86
chr5-136847892-INS-80
chr5-136870826-INS-651
chr5-136895485-INS-192
chr5-136900712-INS-186
chr5-136900923-INS-374
chr5-136900928-INS-170
chr5-136904390-INS-76
chr5-136923125-INS-171
chr5-136927478-INS-51
chr5-136970996-INS-249
chr5-136973101-INS-143
chr5-137003235-INS-60
chr5-137005354-INS-205
chr5-137029041-INS-66
chr5-137045709-INS-242
chr5-137045816-INS-160
chr5-137048576-INS-262
chr5-137048623-INS-120
chr5-137048973-INS-130
chr5-137092612-INS-215
chr5-137097756-INS-61
chr5-137097921-INS-53
chr5-137105518-INS-68
chr5-137109481-INS-75
chr5-137110992-INS-54
chr5-137123813-INS-60
chr5-137153704-INS-106
chr5-137169019-INS-158
chr5-137173596-INS-105
chr5-137176486-INS-50
chr5-137179753-INS-108
chr5-137182526-INS-90
chr5-137189886-INS-54
chr5-137201101-INS-52
chr5-137201982-INS-55
chr5-137205449-INS-94
chr5-137219998-INS-52
chr5-137221385-INS-85
chr5-137233735-INS-105
chr5-137274231-INS-105
chr5-137282663-INS-53
chr5-137317673-INS-106
chr5-137327993-INS-90
chr5-137351714-INS-105
chr5-137361159-INS-317
chr5-137361167-INS-1041
chr5-137365666-INS-691
chr5-137365753-INS-53
chr5-137366561-INS-54
chr5-137366575-INS-319
chr5-137377476-INS-67
chr5-137395417-INS-159
chr5-137409186-INS-133
chr5-137433629-INS-96
chr5-137442492-INS-58
chr5-137445800-INS-1230

chr5-137446454-INS-585
chr5-137450214-INS-106
chr5-137459564-INS-212
chr5-137461977-INS-99
chr5-137467858-INS-122
chr5-137487018-INS-303
chr5-137487253-INS-65
chr5-137493699-INS-60
chr5-137504743-INS-254
chr5-137537356-INS-99
chr5-137539236-INS-83
chr5-137547681-INS-206
chr5-137563587-INS-66
chr5-137574490-INS-88
chr5-137583113-INS-107
chr5-137588732-INS-52
chr5-137592010-INS-115
chr5-137592542-INS-94
chr5-137597160-INS-145
chr5-137597821-INS-245
chr5-137599814-INS-198
chr5-137657892-INS-106
chr5-137663200-INS-527
chr5-137666672-INS-95
chr5-137666911-INS-172
chr5-137696924-INS-197
chr5-137697856-INS-92
chr5-137699781-INS-54
chr5-137703087-INS-211
chr5-137707319-INS-51
chr5-137709243-INS-98
chr5-137713631-INS-264
chr5-137756034-INS-52
chr5-137769266-INS-71
chr5-137769359-INS-2333
chr5-137852738-INS-96
chr5-137920401-INS-64
chr5-137958395-INS-111
chr5-137964294-INS-196
chr5-137975086-INS-52
chr5-137978718-INS-180
chr5-137981183-INS-52
chr5-137981469-INS-337
chr5-137986237-INS-211
chr5-138005178-INS-60
chr5-138008743-INS-224
chr5-138013590-INS-1525
chr5-138022347-INS-126
chr5-138038502-INS-128
chr5-138079177-INS-50
chr5-138089192-INS-59
chr5-138117879-INS-201
chr5-138121811-INS-197
chr5-138202687-INS-367
chr5-138202687-INS-5500
chr5-138207604-INS-53
chr5-138217157-INS-399
chr5-138217401-INS-144
chr5-138228145-INS-58064
chr5-138231764-INS-197
chr5-138242858-INS-80
chr5-138256590-INS-105
chr5-138415259-INS-52
chr5-138433356-INS-117
chr5-138463226-INS-50
chr5-138468030-INS-202
chr5-138509819-INS-392
chr5-138544080-INS-148
chr5-138544293-INS-63
chr5-138559889-INS-5045
chr5-138565066-INS-2153
chr5-138631659-INS-68
chr5-138640217-INS-51
chr5-138640450-INS-106
chr5-138658660-INS-416
chr5-138674067-INS-312
chr5-138674310-INS-292

chr5-138678233-INS-140
chr5-138701985-INS-107
chr5-138715294-INS-647
chr5-138722108-INS-1136
chr5-138765290-INS-125
chr5-138767664-INS-105
chr5-138767734-INS-422
chr5-138818740-INS-1205
chr5-138821944-INS-125
chr5-138889987-INS-111
chr5-138890157-INS-430
chr5-138894311-INS-53
chr5-138895827-INS-181
chr5-138896033-INS-68
chr5-138912259-INS-157
chr5-138921217-INS-53
chr5-138927019-INS-56
chr5-138941131-INS-53
chr5-138976693-INS-60
chr5-138989741-INS-210
chr5-138992383-INS-190
chr5-139026218-INS-104
chr5-139026845-INS-77
chr5-139038343-INS-106
chr5-139058103-INS-53
chr5-139073693-INS-518
chr5-139081115-INS-64
chr5-139086321-INS-158
chr5-139094603-INS-156
chr5-139094673-INS-474
chr5-139119596-INS-315
chr5-139160546-INS-182
chr5-139162361-INS-100
chr5-139175597-INS-412
chr5-139175921-INS-202
chr5-139181476-INS-126
chr5-139181872-INS-142
chr5-139182085-INS-184
chr5-139203144-INS-62
chr5-139242449-INS-63
chr5-139264986-INS-312
chr5-139273425-INS-319
chr5-139285333-INS-785
chr5-139285792-INS-470
chr5-139292092-INS-188
chr5-139294153-INS-52
chr5-139317220-INS-810
chr5-139326043-INS-194
chr5-139326565-INS-52
chr5-139329519-INS-206
chr5-139333683-INS-812
chr5-139362508-INS-1156
chr5-139364141-INS-655
chr5-139381574-INS-454
chr5-139416514-INS-1001
chr5-139424652-INS-297
chr5-139475576-INS-1287
chr5-139479942-INS-70
chr5-139480223-INS-53
chr5-139480241-INS-370
chr5-139480345-INS-121
chr5-139493231-INS-55
chr5-139494207-INS-56
chr5-139508545-INS-157
chr5-139512095-INS-158
chr5-139536595-INS-56
chr5-139553328-INS-158
chr5-139557012-INS-163
chr5-139574279-INS-52
chr5-139625943-INS-104
chr5-139626029-INS-212
chr5-139636895-INS-162
chr5-139637588-INS-60
chr5-139637991-INS-440
chr5-139640007-INS-294
chr5-139640007-INS-65
chr5-139648257-INS-62

chr5-139666182-INS-105
chr5-139675863-INS-68
chr5-139685527-INS-106
chr5-139705969-INS-528
chr5-139719131-INS-70
chr5-139739446-INS-831
chr5-139749474-INS-174
chr5-139749561-INS-401
chr5-139759028-INS-104
chr5-139776274-INS-73
chr5-139776317-INS-173
chr5-139829261-INS-439
chr5-139837082-INS-53
chr5-139838413-INS-157
chr5-139851331-INS-88
chr5-139862726-INS-154
chr5-139863062-INS-53
chr5-139875972-INS-54
chr5-139906160-INS-174
chr5-139934752-INS-370
chr5-139934762-INS-104
chr5-139952809-INS-528
chr5-139967898-INS-1186
chr5-139968825-INS-64
chr5-139969032-INS-90
chr5-140081401-INS-429
chr5-140107231-INS-51
chr5-140151127-INS-50
chr5-140189939-INS-170
chr5-140214523-INS-52
chr5-140259163-INS-344
chr5-140259405-INS-689
chr5-140268010-INS-100
chr5-140282881-INS-200
chr5-140291751-INS-89
chr5-140291909-INS-791
chr5-140313158-INS-128
chr5-140314629-INS-176
chr5-140315639-INS-54
chr5-140352452-INS-209
chr5-140357484-INS-608
chr5-140357998-INS-106
chr5-140379431-INS-73
chr5-140384273-INS-53
chr5-140393957-INS-80
chr5-140436968-INS-5310
chr5-140449833-INS-50
chr5-140460212-INS-92
chr5-140467008-INS-147
chr5-140469478-INS-55
chr5-140469820-INS-52
chr5-140473196-INS-61
chr5-140473339-INS-403
chr5-140473350-INS-145
chr5-140473906-INS-890
chr5-140478531-INS-56
chr5-140486934-INS-106
chr5-140496778-INS-70
chr5-140499190-INS-53
chr5-140506357-INS-174
chr5-140538653-INS-82
chr5-140540380-INS-592
chr5-140562030-INS-75
chr5-140584900-INS-64
chr5-140654093-INS-50
chr5-140658319-INS-263
chr5-140658908-INS-52
chr5-140673318-INS-107
chr5-140676486-INS-157
chr5-140680867-INS-50
chr5-140694054-INS-57
chr5-140725096-INS-159
chr5-140731333-INS-266
chr5-140732046-INS-65
chr5-140826354-INS-52
chr5-140884589-INS-158
chr5-140884685-INS-956

chr5-140905002-INS-159
chr5-140911078-INS-51
chr5-140916981-INS-81
chr5-140951980-INS-159
chr5-140977745-INS-186
chr5-140984405-INS-157
chr5-140984473-INS-370
chr5-141000755-INS-322
chr5-141041579-INS-54
chr5-141049353-INS-103
chr5-141062687-INS-53
chr5-141099375-INS-60
chr5-141128263-INS-210
chr5-141152116-INS-176
chr5-141167565-INS-56
chr5-141259220-INS-62
chr5-141271307-INS-61
chr5-141315011-INS-3711
chr5-141322145-INS-54
chr5-141352839-INS-98
chr5-141379530-INS-3083
chr5-141403871-INS-138
chr5-141491885-INS-52
chr5-141527046-INS-244
chr5-141528075-INS-114
chr5-141537330-INS-112
chr5-141539530-INS-1000
chr5-141561056-INS-191
chr5-141565237-INS-51
chr5-141576052-INS-101
chr5-141622913-INS-166
chr5-141622933-INS-735
chr5-141697289-INS-315
chr5-141697390-INS-116
chr5-141709881-INS-472
chr5-141739518-INS-50
chr5-141740063-INS-58
chr5-141750252-INS-456
chr5-141753761-INS-98
chr5-141813681-INS-56
chr5-141813720-INS-128
chr5-141825517-INS-318
chr5-141825517-INS-52
chr5-141829164-INS-64
chr5-141845081-INS-104
chr5-141853028-INS-72
chr5-141856435-INS-78
chr5-141857423-INS-62
chr5-141873758-INS-67
chr5-141873929-INS-313
chr5-141886526-INS-68
chr5-141929843-INS-74
chr5-141933598-INS-54
chr5-141949139-INS-60
chr5-141981237-INS-309
chr5-141983239-INS-136
chr5-141991331-INS-64
chr5-142001382-INS-66
chr5-142019860-INS-53
chr5-142023345-INS-52
chr5-142081340-INS-106
chr5-142112776-INS-50
chr5-142119302-INS-74
chr5-142137769-INS-213
chr5-142138681-INS-228
chr5-142175903-INS-60
chr5-142209938-INS-66
chr5-142244954-INS-210
chr5-142245947-INS-363
chr5-142246713-INS-172
chr5-142265918-INS-261
chr5-142266048-INS-109
chr5-142307112-INS-52
chr5-142346564-INS-408
chr5-142348636-INS-2320
chr5-142356585-INS-52
chr5-142374888-INS-70

chr5-142379516-INS-55
chr5-142400049-INS-2957
chr5-142429111-INS-54
chr5-142430905-INS-50
chr5-142437573-INS-71
chr5-142437671-INS-196
chr5-142461692-INS-240
chr5-142462179-INS-273
chr5-142503276-INS-315
chr5-142505707-INS-224
chr5-142543171-INS-141
chr5-142549117-INS-56
chr5-142556827-INS-56
chr5-142576501-INS-50
chr5-142601902-INS-408
chr5-142613291-INS-108
chr5-142665418-INS-110
chr5-142670793-INS-55
chr5-142676642-INS-137
chr5-142677522-INS-314
chr5-142697463-INS-60
chr5-142706615-INS-208
chr5-142718974-INS-52
chr5-142744761-INS-112
chr5-142746245-INS-109
chr5-142756549-INS-104
chr5-142768850-INS-126
chr5-142794870-INS-313
chr5-142823155-INS-66
chr5-142823984-INS-198
chr5-142833283-INS-218
chr5-142835087-INS-53
chr5-142840126-INS-196
chr5-142877655-INS-5219
chr5-142924680-INS-165
chr5-142929493-INS-779
chr5-142929940-INS-235
chr5-142952943-INS-146
chr5-142957185-INS-317
chr5-142967116-INS-70
chr5-142968226-INS-194
chr5-142968235-INS-417
chr5-142969260-INS-57
chr5-142969732-INS-53
chr5-142971204-INS-213
chr5-142997988-INS-186
chr5-143050190-INS-117
chr5-143087037-INS-3067
chr5-143113122-INS-8375
chr5-143125759-INS-367
chr5-143132066-INS-100
chr5-143134878-INS-212
chr5-143145844-INS-100
chr5-143168737-INS-368
chr5-143168737-INS-737
chr5-143180259-INS-172
chr5-143181501-INS-50
chr5-143216270-INS-434
chr5-143218990-INS-69
chr5-143242060-INS-72
chr5-143245801-INS-205
chr5-143283075-INS-869
chr5-143332551-INS-76
chr5-143332563-INS-182
chr5-143344202-INS-170
chr5-143363558-INS-53
chr5-143363782-INS-130
chr5-143364497-INS-54
chr5-143394125-INS-329
chr5-143394177-INS-124
chr5-143394710-INS-219
chr5-143404670-INS-84
chr5-143418673-INS-206
chr5-143448426-INS-4358
chr5-143462639-INS-376
chr5-143464920-INS-656
chr5-143480953-INS-421

chr5-143480962-INS-161
chr5-143497062-INS-498
chr5-143497083-INS-56
chr5-143497249-INS-160
chr5-143522227-INS-211
chr5-143544821-INS-144
chr5-143554954-INS-50
chr5-143557701-INS-466
chr5-143561828-INS-82
chr5-143561870-INS-1206
chr5-143561922-INS-89
chr5-143567284-INS-264
chr5-143593129-INS-50
chr5-143596242-INS-89
chr5-143639695-INS-80
chr5-143639750-INS-712
chr5-143692258-INS-66
chr5-143733771-INS-74
chr5-143741243-INS-76
chr5-143747120-INS-296
chr5-143791142-INS-116
chr5-143818346-INS-50
chr5-143819991-INS-299
chr5-143820654-INS-742
chr5-143845024-INS-205
chr5-143879213-INS-68
chr5-143902291-INS-52
chr5-143904700-INS-437
chr5-143906774-INS-249
chr5-143909630-INS-77
chr5-143909817-INS-1078
chr5-143909922-INS-588
chr5-143951939-INS-54
chr5-143954602-INS-50
chr5-143974767-INS-3046
chr5-143977225-INS-62
chr5-143981494-INS-66
chr5-143981745-INS-488
chr5-144004227-INS-1388
chr5-144004227-INS-200
chr5-144007643-INS-166
chr5-144009661-INS-8749
chr5-144031385-INS-188
chr5-144034278-INS-640
chr5-144050313-INS-147
chr5-144070662-INS-87
chr5-144079646-INS-319
chr5-144117417-INS-123
chr5-144153029-INS-818
chr5-144160761-INS-220
chr5-144175308-INS-152
chr5-144206103-INS-218
chr5-144218573-INS-135
chr5-144222742-INS-64
chr5-144226930-INS-797
chr5-144228714-INS-114
chr5-144229058-INS-219
chr5-144238007-INS-101
chr5-144238108-INS-1000
chr5-144245610-INS-53
chr5-144252708-INS-158
chr5-144260039-INS-52
chr5-144304235-INS-2139
chr5-144313108-INS-1345
chr5-144327980-INS-50
chr5-144329719-INS-50
chr5-144333518-INS-106
chr5-144337652-INS-58
chr5-144339898-INS-213
chr5-144340138-INS-470
chr5-144340986-INS-573
chr5-144349694-INS-7452
chr5-144351028-INS-66
chr5-144351424-INS-342
chr5-144360094-INS-52
chr5-144367607-INS-652
chr5-144368825-INS-418

chr5-144377032-INS-227
chr5-144382911-INS-159
chr5-144383091-INS-440
chr5-144396325-INS-50
chr5-144407644-INS-57
chr5-144408959-INS-84
chr5-144426180-INS-240
chr5-144428390-INS-1952
chr5-144431730-INS-53
chr5-144433951-INS-52
chr5-144434213-INS-209
chr5-144434494-INS-114
chr5-144441835-INS-105
chr5-144457289-INS-84
chr5-144494934-INS-88
chr5-144498252-INS-62
chr5-144556089-INS-99
chr5-144561938-INS-261
chr5-144562047-INS-103
chr5-144572547-INS-54
chr5-144572617-INS-262
chr5-144575090-INS-788
chr5-144603382-INS-50
chr5-144634482-INS-204
chr5-144645127-INS-106
chr5-144659884-INS-104
chr5-144660133-INS-574
chr5-144678940-INS-494
chr5-144732784-INS-1194
chr5-144797873-INS-185
chr5-144826884-INS-158
chr5-144832155-INS-460
chr5-144854467-INS-166
chr5-144869471-INS-56
chr5-144870059-INS-106
chr5-144870059-INS-52
chr5-144893925-INS-50
chr5-144906934-INS-308
chr5-144906934-INS-88
chr5-144933505-INS-1097
chr5-144937351-INS-168
chr5-144937688-INS-58
chr5-144958943-INS-231
chr5-144963474-INS-58
chr5-144972978-INS-4389
chr5-144981033-INS-153
chr5-145018788-INS-117
chr5-145019406-INS-62
chr5-145019420-INS-230
chr5-145020150-INS-59
chr5-145039415-INS-57
chr5-145052610-INS-5195
chr5-145069372-INS-52
chr5-145079051-INS-196
chr5-145079056-INS-93
chr5-145093921-INS-149
chr5-145093957-INS-2722
chr5-145102713-INS-206
chr5-145129674-INS-726
chr5-145140398-INS-120
chr5-145143814-INS-896
chr5-145151121-INS-306
chr5-145166665-INS-53
chr5-145229551-INS-64
chr5-145257679-INS-3134
chr5-145274831-INS-2464
chr5-145281112-INS-369
chr5-145300978-INS-120
chr5-145320949-INS-50
chr5-145352967-INS-6668
chr5-145380328-INS-392
chr5-145385722-INS-1962
chr5-145409256-INS-773
chr5-145460786-INS-116
chr5-145508406-INS-64
chr5-145535376-INS-644
chr5-145538653-INS-2336

chr5-145539611-INS-69
chr5-145558918-INS-1325
chr5-145560897-INS-202
chr5-145574110-INS-50
chr5-145665093-INS-56
chr5-145675639-INS-54
chr5-145679248-INS-648
chr5-145687964-INS-54
chr5-145689981-INS-1456
chr5-145700312-INS-52
chr5-145712452-INS-66
chr5-145806399-INS-76
chr5-145846593-INS-66
chr5-145949503-INS-355
chr5-145949724-INS-780
chr5-146008920-INS-1881
chr5-146012582-INS-52
chr5-146019431-INS-125
chr5-146041278-INS-121
chr5-146045385-INS-53
chr5-146113852-INS-72
chr5-146119603-INS-2635
chr5-146204100-INS-265
chr5-146217620-INS-561
chr5-146217752-INS-1556
chr5-146226587-INS-520
chr5-146264806-INS-224
chr5-146265013-INS-64
chr5-146292565-INS-215
chr5-146324901-INS-152
chr5-146336431-INS-50
chr5-146351280-INS-585
chr5-146379927-INS-1557
chr5-146387068-INS-2059
chr5-146458292-INS-190
chr5-146467961-INS-77
chr5-146501393-INS-64
chr5-146511001-INS-1295
chr5-146529412-INS-4814
chr5-146535036-INS-339
chr5-146553187-INS-126
chr5-146579596-INS-3111
chr5-146594809-INS-418
chr5-146602927-INS-59
chr5-146606392-INS-55
chr5-146611847-INS-192
chr5-146612342-INS-230
chr5-146653802-INS-72
chr5-146655164-INS-64
chr5-146675297-INS-237
chr5-146715903-INS-211
chr5-146719422-INS-224
chr5-146751494-INS-8854
chr5-146757923-INS-123
chr5-146757961-INS-714
chr5-146762175-INS-104
chr5-146763054-INS-74
chr5-146792272-INS-130
chr5-146792272-INS-50
chr5-146799574-INS-66
chr5-146813424-INS-351
chr5-146818791-INS-222
chr5-146825119-INS-305
chr5-146846247-INS-70
chr5-146853536-INS-448
chr5-146894273-INS-125
chr5-146912919-INS-213
chr5-146913380-INS-146
chr5-146920088-INS-460
chr5-146920813-INS-377
chr5-146923313-INS-101
chr5-146931588-INS-120
chr5-146982415-INS-50
chr5-146989677-INS-164
chr5-147102998-INS-276
chr5-147130433-INS-50
chr5-147138775-INS-7171

chr5-147167246-INS-268
chr5-147167274-INS-120
chr5-147193755-INS-80
chr5-147198023-INS-50
chr5-147202447-INS-214
chr5-147210251-INS-94
chr5-147257722-INS-64
chr5-147265795-INS-76
chr5-147270074-INS-78
chr5-147272634-INS-80
chr5-147278650-INS-4115
chr5-147283944-INS-74
chr5-147284093-INS-53
chr5-147302829-INS-80
chr5-147307915-INS-537
chr5-147308049-INS-203
chr5-147321006-INS-50
chr5-147333848-INS-334
chr5-147351101-INS-198
chr5-147352523-INS-72
chr5-147383330-INS-1043
chr5-147388022-INS-273
chr5-147405355-INS-214
chr5-147418879-INS-2218
chr5-147418910-INS-180
chr5-147487745-INS-118
chr5-147488477-INS-156
chr5-147509305-INS-54
chr5-147512074-INS-60
chr5-147514237-INS-68
chr5-147516576-INS-8329
chr5-147528026-INS-54
chr5-147548442-INS-304
chr5-147557676-INS-57
chr5-147668401-INS-50
chr5-147678015-INS-85
chr5-147682061-INS-431
chr5-147688994-INS-295
chr5-147688994-INS-68
chr5-147689272-INS-86
chr5-147701372-INS-164
chr5-147722263-INS-58
chr5-147768156-INS-299
chr5-147788931-INS-158
chr5-147793599-INS-66
chr5-147800337-INS-58
chr5-147815752-INS-215
chr5-147815752-INS-53
chr5-147818106-INS-87
chr5-147833012-INS-66
chr5-147833079-INS-76
chr5-147841013-INS-458
chr5-147855968-INS-50
chr5-147864294-INS-155
chr5-147873812-INS-178
chr5-147880394-INS-4346
chr5-147887428-INS-52
chr5-147897483-INS-75
chr5-147929090-INS-439
chr5-147929510-INS-211
chr5-147933998-INS-4985
chr5-147967611-INS-102
chr5-147990774-INS-343
chr5-148003373-INS-61
chr5-148012459-INS-73
chr5-148015484-INS-74
chr5-148030228-INS-7138
chr5-148030241-INS-372
chr5-148046012-INS-60
chr5-148046091-INS-56
chr5-148078216-INS-54
chr5-148119991-INS-4767
chr5-148121262-INS-60
chr5-148126441-INS-1862
chr5-148136339-INS-80
chr5-148139178-INS-50
chr5-148144798-INS-1893

chr5-148154241-INS-84
chr5-148155087-INS-561
chr5-148157590-INS-89
chr5-148204506-INS-62
chr5-148220286-INS-52
chr5-148222825-INS-157
chr5-148271642-INS-127
chr5-148284300-INS-7089
chr5-148292105-INS-103
chr5-148297172-INS-103
chr5-148303167-INS-2148
chr5-148305785-INS-162
chr5-148352451-INS-147
chr5-148358828-INS-2057
chr5-148367675-INS-88
chr5-148367799-INS-84
chr5-148456807-INS-508
chr5-148456870-INS-224
chr5-148485462-INS-88
chr5-148537766-INS-198
chr5-148602088-INS-77
chr5-148603517-INS-66
chr5-148611922-INS-61
chr5-148646682-INS-2514
chr5-148653679-INS-470
chr5-148663631-INS-88
chr5-148717815-INS-62
chr5-148718986-INS-85
chr5-148730958-INS-4870
chr5-148737916-INS-8457
chr5-148755300-INS-215
chr5-148761857-INS-94
chr5-148770588-INS-362
chr5-148778728-INS-1876
chr5-148819585-INS-740
chr5-148819614-INS-311
chr5-148822519-INS-1634
chr5-148840681-INS-80
chr5-148857125-INS-540
chr5-148880388-INS-4807
chr5-148914336-INS-425
chr5-148997156-INS-51
chr5-149001095-INS-514
chr5-149017847-INS-143
chr5-149051330-INS-271
chr5-149077149-INS-72
chr5-149086720-INS-210
chr5-149098894-INS-276
chr5-149147802-INS-360
chr5-149190519-INS-64
chr5-149193012-INS-63
chr5-149193587-INS-169
chr5-149207708-INS-78
chr5-149211026-INS-69
chr5-149215130-INS-72
chr5-149215909-INS-244
chr5-149229854-INS-66
chr5-149234722-INS-73
chr5-149244912-INS-165
chr5-149266179-INS-108
chr5-149280459-INS-64
chr5-149281016-INS-204
chr5-149281170-INS-136
chr5-149290678-INS-4833
chr5-149297293-INS-110
chr5-149351286-INS-217
chr5-149356633-INS-64
chr5-149372785-INS-137
chr5-149374449-INS-117
chr5-149374477-INS-255
chr5-149400948-INS-140
chr5-149418392-INS-3831
chr5-149434581-INS-711
chr5-149435627-INS-294
chr5-149445573-INS-50
chr5-149473690-INS-55
chr5-149483803-INS-102

chr5-149486472-INS-83
chr5-149498650-INS-360
chr5-149516375-INS-379
chr5-149516994-INS-213
chr5-149540146-INS-217
chr5-149582731-INS-69
chr5-149585691-INS-52
chr5-149589643-INS-389
chr5-149601353-INS-7350
chr5-149613743-INS-84
chr5-149649478-INS-66
chr5-149649486-INS-140
chr5-149650557-INS-54
chr5-149659422-INS-188
chr5-149660087-INS-74
chr5-149665346-INS-50
chr5-149676267-INS-201
chr5-149794086-INS-137
chr5-149794119-INS-663
chr5-149807008-INS-128
chr5-149818306-INS-77
chr5-149818359-INS-212
chr5-149835238-INS-80
chr5-149850297-INS-205
chr5-149861517-INS-164
chr5-149866270-INS-69
chr5-149867201-INS-4666
chr5-149883800-INS-66
chr5-149918630-INS-66
chr5-149930237-INS-377
chr5-150004301-INS-62
chr5-150036633-INS-171
chr5-150102708-INS-5360
chr5-150104346-INS-142
chr5-150105568-INS-68
chr5-150131996-INS-89
chr5-150151489-INS-70
chr5-150169885-INS-60
chr5-150191705-INS-126
chr5-150229558-INS-56
chr5-150243208-INS-156
chr5-150248417-INS-90
chr5-150272437-INS-99
chr5-150297205-INS-6414
chr5-150313037-INS-72
chr5-150351835-INS-209
chr5-150406854-INS-60
chr5-150407151-INS-117
chr5-150418057-INS-110
chr5-150431674-INS-54
chr5-150449484-INS-203
chr5-150486992-INS-80
chr5-150512979-INS-101
chr5-150539614-INS-54
chr5-150548063-INS-168
chr5-150579380-INS-211
chr5-150604990-INS-55
chr5-150607413-INS-89
chr5-150615419-INS-88
chr5-150632838-INS-221
chr5-150664035-INS-53
chr5-150775293-INS-189
chr5-150805013-INS-2872
chr5-150808457-INS-325
chr5-150808511-INS-85
chr5-150826171-INS-75
chr5-150837879-INS-51
chr5-150840417-INS-110
chr5-150841919-INS-247
chr5-150843680-INS-6441
chr5-150847002-INS-166
chr5-150848552-INS-4328
chr5-150958453-INS-192
chr5-151010934-INS-58
chr5-151041798-INS-51
chr5-151044297-INS-124
chr5-151046474-INS-4778

chr5-151057112-INS-1125
chr5-151061898-INS-221
chr5-151064560-INS-50
chr5-151088683-INS-4892
chr5-151183568-INS-6233
chr5-151198764-INS-189
chr5-151199079-INS-74
chr5-151215028-INS-2844
chr5-151261530-INS-84
chr5-151377759-INS-1261
chr5-151424901-INS-332
chr5-151502090-INS-462
chr5-151506651-INS-510
chr5-151556911-INS-52
chr5-151594476-INS-70
chr6-3289637-INS-50
chr6-3292307-INS-265
chr6-3319696-INS-60
chr6-3344870-INS-113
chr6-3380346-INS-2923
chr6-3383940-INS-112
chr6-3385383-INS-549
chr6-3407261-INS-4973
chr6-3407915-INS-51
chr6-3429441-INS-1609
chr6-3439043-INS-134
chr6-3439123-INS-290
chr6-3440924-INS-76
chr6-3469486-INS-242
chr6-3493248-INS-52
chr6-3500908-INS-55
chr6-3501551-INS-188
chr6-3514418-INS-52
chr6-3573962-INS-90
chr6-3620128-INS-252
chr6-3621510-INS-52
chr6-3663185-INS-50
chr6-3677411-INS-5652
chr6-3781934-INS-50
chr6-3847816-INS-104
chr6-3852770-INS-938
chr6-3918342-INS-74
chr6-3957595-INS-2975
chr6-3993043-INS-6363
chr6-4039751-INS-70
chr6-4062846-INS-222
chr6-4065578-INS-65
chr6-4076764-INS-2140
chr6-4092091-INS-2163
chr6-4111479-INS-73
chr6-4156407-INS-412
chr6-4156443-INS-94
chr6-4220422-INS-50
chr6-4271925-INS-1085
chr6-4296229-INS-126
chr6-4315638-INS-52
chr6-4322359-INS-4505
chr6-4378102-INS-50
chr6-4391391-INS-413
chr6-4406085-INS-90
chr6-4417781-INS-61
chr6-4464960-INS-500
chr6-4464966-INS-1298
chr6-4496209-INS-210
chr6-4502811-INS-50
chr6-4503901-INS-54
chr6-4515447-INS-244
chr6-4515447-INS-547
chr6-4572983-INS-7105
chr6-4578722-INS-60
chr6-4579119-INS-59
chr6-4586415-INS-78
chr6-4599161-INS-60
chr6-4647951-INS-183
chr6-4735356-INS-115
chr6-4762240-INS-1320
chr6-4762441-INS-2995

chr6-4786715-INS-58
chr6-4814382-INS-204
chr6-4842969-INS-250
chr6-4865307-INS-54
chr6-4867812-INS-2472
chr6-4867812-INS-351
chr6-4868270-INS-4775
chr6-4911331-INS-89
chr6-4911625-INS-480
chr6-4917994-INS-117
chr6-4927935-INS-1720
chr6-4930404-INS-220
chr6-4940290-INS-213
chr6-4946261-INS-50
chr6-4961200-INS-96
chr6-4967657-INS-115
chr6-4993041-INS-154
chr6-5001882-INS-163
chr6-5032603-INS-184
chr6-5051209-INS-50
chr6-5148680-INS-66
chr6-5150906-INS-72
chr6-5177724-INS-272
chr6-5196299-INS-58
chr6-5239432-INS-54
chr6-5243767-INS-62
chr6-5263731-INS-567
chr6-5268828-INS-230
chr6-5269557-INS-56
chr6-5345115-INS-7183
chr6-5355010-INS-150
chr6-5362696-INS-50
chr6-5412451-INS-213
chr6-5550768-INS-110
chr6-5550927-INS-87
chr6-5572692-INS-79
chr6-5578797-INS-538
chr6-5591841-INS-183
chr6-5639483-INS-148
chr6-5640159-INS-197
chr6-5652099-INS-1271
chr6-5662135-INS-69
chr6-5662874-INS-77
chr6-5668851-INS-564
chr6-5674991-INS-2573
chr6-5692896-INS-50
chr6-5705210-INS-2624
chr6-5706790-INS-7059
chr6-5709276-INS-51
chr6-5712722-INS-13383
chr6-5712722-INS-6495
chr6-5714917-INS-56
chr6-5720803-INS-201
chr6-5779498-INS-52
chr6-5794015-INS-1383
chr6-5805560-INS-60
chr6-5814003-INS-56
chr6-5822115-INS-82
chr6-5866658-INS-61
chr6-5897665-INS-57
chr6-5948952-INS-68
chr6-5955124-INS-98
chr6-5966045-INS-50
chr6-5966096-INS-161
chr6-6032453-INS-1050
chr6-6032471-INS-454
chr6-6032892-INS-276
chr6-6105017-INS-909
chr6-6123489-INS-265
chr6-6129939-INS-52
chr6-6130237-INS-179
chr6-6152295-INS-58
chr6-6179381-INS-99
chr6-6192421-INS-143
chr6-6192430-INS-341
chr6-6198106-INS-215
chr6-6231925-INS-282

chr6-6235528-INS-50
chr6-6244484-INS-548
chr6-6252819-INS-2809
chr6-6270069-INS-6328
chr6-6275437-INS-62
chr6-6285875-INS-1216
chr6-6334242-INS-2917
chr6-6341591-INS-69
chr6-6341784-INS-400
chr6-6357708-INS-52
chr6-6380891-INS-1079
chr6-6416467-INS-300
chr6-6416476-INS-108
chr6-6438167-INS-92
chr6-6439338-INS-50
chr6-6446749-INS-188
chr6-6459297-INS-116
chr6-6459374-INS-473
chr6-6469423-INS-205
chr6-6481692-INS-61
chr6-6489354-INS-64
chr6-6497341-INS-56
chr6-6552928-INS-400
chr6-6598135-INS-76
chr6-6611809-INS-194
chr6-6640574-INS-115
chr6-6687393-INS-7341
chr6-6731075-INS-122
chr6-6779971-INS-84
chr6-6805957-INS-82
chr6-6805966-INS-168
chr6-6806617-INS-59
chr6-6840656-INS-6350
chr6-6934874-INS-7217
chr6-6936680-INS-2675
chr6-6961036-INS-3108
chr6-6962876-INS-9722
chr6-6963310-INS-630
chr6-7051000-INS-6352
chr6-7067235-INS-80
chr6-7113745-INS-2562
chr6-7117200-INS-147
chr6-7186496-INS-196
chr6-7205467-INS-6436
chr6-7217083-INS-91
chr6-7222790-INS-1591
chr6-7224784-INS-2941
chr6-7228804-INS-377
chr6-7236760-INS-66
chr6-7247233-INS-73
chr6-7253422-INS-4156
chr6-7256347-INS-50
chr6-7296467-INS-51
chr6-7346412-INS-136
chr6-7448927-INS-5579
chr6-7531313-INS-7087
chr6-7544070-INS-550
chr6-7567473-INS-144
chr6-7574021-INS-50
chr6-7591520-INS-5504
chr6-7596417-INS-6242
chr6-7600054-INS-201
chr6-7615008-INS-66
chr6-7621448-INS-3490
chr6-7629032-INS-387
chr6-7660636-INS-110
chr6-7690569-INS-275
chr6-7758258-INS-1778
chr6-7774856-INS-75
chr6-7804658-INS-8646
chr6-7805349-INS-73
chr6-7806523-INS-532
chr6-7827052-INS-640
chr6-7834152-INS-102
chr6-7834152-INS-50
chr6-7885725-INS-111
chr6-7906861-INS-411

chr6-7958635-INS-6546
chr6-7960562-INS-257
chr6-7984123-INS-117
chr6-8001714-INS-54
chr6-8011087-INS-888
chr6-8024205-INS-7254
chr6-8049071-INS-128
chr6-8064142-INS-7599
chr6-8067883-INS-200
chr6-8165657-INS-145
chr6-8165657-INS-608
chr6-8188782-INS-68
chr6-8191057-INS-84
chr6-8210669-INS-5526
chr6-8221328-INS-2718
chr6-8265417-INS-1813
chr6-8451993-INS-435
chr6-8452009-INS-88
chr6-8490681-INS-57
chr6-8494728-INS-54
chr6-8600064-INS-337
chr6-8619600-INS-50
chr6-8627948-INS-50
chr6-8646682-INS-516
chr6-8681905-INS-69
chr6-8684256-INS-82
chr6-8738407-INS-5156
chr6-8738430-INS-1747
chr6-8819319-INS-1043
chr6-8819983-INS-967
chr6-8882025-INS-6339
chr6-8924228-INS-54
chr6-8999109-INS-50
chr6-9035706-INS-291
chr6-9044109-INS-50
chr6-9055166-INS-471
chr6-9157088-INS-55
chr6-9162094-INS-6397
chr6-9176038-INS-1545
chr6-9178092-INS-58
chr6-9178250-INS-89
chr6-9178395-INS-81
chr6-9210213-INS-403
chr6-9211370-INS-84
chr6-9245810-INS-213
chr6-9273270-INS-4340
chr6-9309516-INS-51
chr6-9359764-INS-6802
chr6-9392515-INS-6567
chr6-9408312-INS-4668
chr6-9494311-INS-336
chr6-9512027-INS-488
chr6-9518572-INS-6381
chr6-9536901-INS-201
chr6-9596725-INS-107
chr6-9631006-INS-1816
chr6-9742756-INS-70
chr6-9762817-INS-6611
chr6-9781591-INS-5218
chr6-9788443-INS-67
chr6-9791499-INS-75
chr6-9801871-INS-50
chr6-9887283-INS-79
chr6-9955087-INS-112
chr6-10064343-INS-3804
chr6-10079023-INS-3006
chr6-10094500-INS-57
chr6-10106428-INS-304
chr6-10132353-INS-919
chr6-10143977-INS-62
chr6-10198679-INS-7006
chr6-10252504-INS-75
chr6-10278478-INS-62
chr6-10298375-INS-2344
chr6-10361991-INS-202
chr6-10368513-INS-63
chr6-10453461-INS-50

chr6-10453983-INS-478
chr6-10602010-INS-3866
chr6-10666769-INS-100
chr6-10667323-INS-167
chr6-10703175-INS-662
chr6-10726043-INS-58
chr6-10752017-INS-2444
chr6-10757059-INS-376
chr6-10757059-INS-754
chr6-10764535-INS-231
chr6-10776549-INS-51
chr6-10850341-INS-106
chr6-10868294-INS-12600
chr6-10896934-INS-2808
chr6-10914995-INS-88
chr6-10966624-INS-65
chr6-10990053-INS-5020
chr6-11005452-INS-1080
chr6-11016412-INS-1467
chr6-11029824-INS-83
chr6-11043858-INS-1734
chr6-11065829-INS-838
chr6-11069523-INS-60
chr6-11127608-INS-1483
chr6-11143970-INS-126
chr6-11200860-INS-1045
chr6-11225295-INS-52
chr6-11233675-INS-56
chr6-11265703-INS-1060
chr6-11319405-INS-370
chr6-11341403-INS-3755
chr6-11354805-INS-6442
chr6-11428734-INS-199
chr6-11491997-INS-133
chr6-11510746-INS-357
chr6-11518950-INS-6261
chr6-11537696-INS-341
chr6-11538306-INS-185
chr6-11550161-INS-66
chr6-11587253-INS-7026
chr6-11734787-INS-54
chr6-11770217-INS-132
chr6-11797552-INS-213
chr6-11799135-INS-64
chr6-11815393-INS-4421
chr6-11899478-INS-58
chr6-11910989-INS-2015
chr6-11995536-INS-94
chr6-12059462-INS-64
chr6-12066116-INS-50
chr6-12068895-INS-716
chr6-12070209-INS-199
chr6-12075197-INS-233
chr6-12097551-INS-50
chr6-12097937-INS-150
chr6-12097959-INS-52
chr6-12125973-INS-182
chr6-12125979-INS-85
chr6-12126225-INS-54
chr6-12126253-INS-2100
chr6-12133701-INS-60
chr6-12149859-INS-2455
chr6-12154146-INS-370
chr6-12159764-INS-53
chr6-12162474-INS-130
chr6-12207263-INS-64
chr6-12210998-INS-1738
chr6-12274319-INS-116
chr6-12311274-INS-1973
chr6-12319161-INS-1501
chr6-12359144-INS-64
chr6-12433456-INS-307
chr6-12435990-INS-860
chr6-12469654-INS-52
chr6-12556986-INS-76
chr6-12599514-INS-69
chr6-12611533-INS-206

chr6-12653673-INS-52
chr6-12673590-INS-2429
chr6-12725978-INS-115
chr6-12752184-INS-74
chr6-12800157-INS-147
chr6-12805046-INS-57
chr6-12821346-INS-7322
chr6-12838557-INS-198
chr6-12846210-INS-207
chr6-12886582-INS-1864
chr6-12900666-INS-198
chr6-12909367-INS-434
chr6-12932170-INS-7072
chr6-12939358-INS-6719
chr6-12963316-INS-241
chr6-12963690-INS-328
chr6-12988934-INS-6397
chr6-13002090-INS-949
chr6-13012045-INS-50
chr6-13025111-INS-140
chr6-13046869-INS-1528
chr6-13115046-INS-154
chr6-13148611-INS-6396
chr6-13154009-INS-420
chr6-13154124-INS-65
chr6-13154333-INS-59
chr6-13154529-INS-93
chr6-13173804-INS-5147
chr6-13235772-INS-1311
chr6-13282394-INS-238
chr6-13320142-INS-270
chr6-13320712-INS-391
chr6-13321507-INS-116
chr6-13324754-INS-52
chr6-13419369-INS-6204
chr6-13492411-INS-497
chr6-13517689-INS-276
chr6-13518157-INS-340
chr6-13521269-INS-412
chr6-13561748-INS-192
chr6-13600409-INS-59
chr6-13600708-INS-2898
chr6-13620541-INS-6265
chr6-13695453-INS-112
chr6-13699724-INS-1328
chr6-13723608-INS-54
chr6-13734297-INS-1075
chr6-13749317-INS-59
chr6-13776757-INS-90
chr6-13795169-INS-65
chr6-13797456-INS-52
chr6-13802808-INS-675
chr6-13815494-INS-268
chr6-13822100-INS-236
chr6-13822170-INS-84
chr6-13852327-INS-50
chr6-13906160-INS-54
chr6-13939359-INS-1861
chr6-13948703-INS-7614
chr6-13950444-INS-1738
chr6-13969140-INS-116
chr6-13970630-INS-66
chr6-13981949-INS-440
chr6-14006506-INS-99
chr6-14029501-INS-917
chr6-14032977-INS-7344
chr6-14066460-INS-78
chr6-14072901-INS-492
chr6-14118060-INS-57
chr6-14124087-INS-8300
chr6-14142982-INS-50
chr6-14149720-INS-412
chr6-14187267-INS-58
chr6-14195423-INS-283
chr6-14221279-INS-65
chr6-14244989-INS-1947
chr6-14261427-INS-81

chr6-14262614-INS-377
chr6-14266426-INS-6436
chr6-14343843-INS-960
chr6-14362474-INS-50
chr6-14404045-INS-121
chr6-14418381-INS-51
chr6-14420152-INS-73
chr6-14447792-INS-6356
chr6-14485245-INS-60
chr6-14487304-INS-58
chr6-14498453-INS-109
chr6-14498994-INS-96
chr6-14511416-INS-1599
chr6-14512674-INS-51
chr6-14520641-INS-119
chr6-14572804-INS-1941
chr6-14576167-INS-1761
chr6-14606125-INS-52
chr6-14679694-INS-2776
chr6-14691011-INS-86
chr6-14704313-INS-283
chr6-14704543-INS-1020
chr6-14745356-INS-367
chr6-14775467-INS-62
chr6-14812004-INS-50
chr6-14824298-INS-727
chr6-14842874-INS-62
chr6-14844203-INS-553
chr6-14846342-INS-88
chr6-14886278-INS-1416
chr6-14902796-INS-54
chr6-14943195-INS-60
chr6-14978226-INS-132
chr6-15023598-INS-213
chr6-15023616-INS-60
chr6-15031083-INS-2099
chr6-15037041-INS-1064
chr6-15052478-INS-6223
chr6-15062902-INS-56
chr6-15103221-INS-58
chr6-15172633-INS-59
chr6-15174946-INS-152
chr6-15175223-INS-106
chr6-15265279-INS-74
chr6-15297432-INS-176
chr6-15336190-INS-118
chr6-15359757-INS-59
chr6-15446784-INS-83
chr6-15472786-INS-1350
chr6-15477522-INS-4830
chr6-15478689-INS-2533
chr6-15497224-INS-191
chr6-15518703-INS-603
chr6-15529985-INS-3088
chr6-15544173-INS-3104
chr6-15616317-INS-174
chr6-15618341-INS-80
chr6-15618345-INS-245
chr6-15618466-INS-63
chr6-15621743-INS-412
chr6-15637735-INS-51
chr6-15641444-INS-6382
chr6-15660723-INS-50
chr6-15682728-INS-1451
chr6-15710561-INS-3188
chr6-15756737-INS-93
chr6-15786637-INS-63
chr6-15794528-INS-673
chr6-15794551-INS-318
chr6-15895512-INS-58
chr6-15930290-INS-752
chr6-15994053-INS-133
chr6-16014043-INS-240
chr6-16014169-INS-512
chr6-16045978-INS-1766
chr6-16050586-INS-2859
chr6-16108171-INS-126

chr6-16117393-INS-58
chr6-16156852-INS-2546
chr6-16239979-INS-311
chr6-16247467-INS-6457
chr6-16269603-INS-6047
chr6-16291141-INS-3374
chr6-16351570-INS-4815
chr6-16391327-INS-213
chr6-16440550-INS-4579
chr6-16454446-INS-3106
chr6-16454667-INS-6383
chr6-16456727-INS-497
chr6-16469441-INS-2970
chr6-16487546-INS-6480
chr6-16496842-INS-84
chr6-16563240-INS-60
chr6-16564959-INS-2301
chr6-16570430-INS-2009
chr6-16588979-INS-60
chr6-16594419-INS-60
chr6-16611719-INS-78
chr6-16690163-INS-7225
chr6-16717005-INS-208
chr6-16721039-INS-2798
chr6-16730625-INS-50
chr6-16773501-INS-67
chr6-16892316-INS-239
chr6-16892962-INS-361
chr6-16950117-INS-125
chr6-17023860-INS-204
chr6-17045839-INS-1540
chr6-17123089-INS-229
chr6-17140057-INS-329
chr6-17145960-INS-93
chr6-17145964-INS-275
chr6-17182953-INS-6192
chr6-17225986-INS-191
chr6-17233530-INS-169
chr6-17326347-INS-114
chr6-17328795-INS-69
chr6-17336276-INS-80
chr6-17345526-INS-1254
chr6-17355454-INS-959
chr6-17355563-INS-3418
chr6-17376235-INS-59
chr6-17379922-INS-64
chr6-17390042-INS-140
chr6-17397993-INS-51
chr6-17398336-INS-778
chr6-17401173-INS-6570
chr6-17429188-INS-60
chr6-17493388-INS-2572
chr6-17569411-INS-92
chr6-17570339-INS-54
chr6-17587390-INS-1864
chr6-17590175-INS-69
chr6-17590250-INS-235
chr6-17591124-INS-86
chr6-17615398-INS-6813
chr6-17681270-INS-6457
chr6-17696384-INS-6190
chr6-17711484-INS-507
chr6-17727854-INS-287
chr6-17753261-INS-212
chr6-17753955-INS-67
chr6-17760542-INS-60
chr6-17766215-INS-124
chr6-17772967-INS-50
chr6-17794454-INS-146
chr6-17805925-INS-209
chr6-17807644-INS-2982
chr6-17823553-INS-60
chr6-17935874-INS-189
chr6-17936899-INS-243
chr6-17988932-INS-587
chr6-18020284-INS-411
chr6-18020284-INS-913

chr6-18038851-INS-155
chr6-18110270-INS-6309
chr6-18132489-INS-616
chr6-18133106-INS-8983
chr6-18138342-INS-4217
chr6-18194847-INS-70
chr6-18195547-INS-250
chr6-18198896-INS-52
chr6-18244035-INS-56
chr6-18244472-INS-107
chr6-18244472-INS-286
chr6-18261872-INS-6453
chr6-18262851-INS-92
chr6-18295630-INS-56
chr6-18315776-INS-653
chr6-18352465-INS-1569
chr6-18374627-INS-140
chr6-18386473-INS-151
chr6-18430537-INS-54
chr6-18449281-INS-56
chr6-18453230-INS-116
chr6-18480074-INS-84
chr6-18505151-INS-50
chr6-18558397-INS-96
chr6-18560049-INS-50
chr6-18575284-INS-1255
chr6-18604232-INS-68
chr6-18617856-INS-4748
chr6-18645608-INS-6630
chr6-18656169-INS-6489
chr6-18665603-INS-4782
chr6-18701217-INS-2884
chr6-18705820-INS-54
chr6-18755126-INS-67
chr6-18802359-INS-142
chr6-18873874-INS-626
chr6-18914943-INS-50
chr6-18927416-INS-7137
chr6-18989722-INS-347
chr6-18992318-INS-92
chr6-19006184-INS-54
chr6-19006375-INS-152
chr6-19020702-INS-3200
chr6-19024874-INS-2502
chr6-19057922-INS-130
chr6-19060844-INS-1248
chr6-19062508-INS-70
chr6-19089446-INS-12932
chr6-19109498-INS-6423
chr6-19132494-INS-54
chr6-19169823-INS-5364
chr6-19171115-INS-52
chr6-19198193-INS-50
chr6-19210164-INS-100
chr6-19251072-INS-6456
chr6-19277782-INS-120
chr6-19280077-INS-50
chr6-19284859-INS-5043
chr6-19302608-INS-57
chr6-19306211-INS-390
chr6-19546287-INS-104
chr6-19548903-INS-1322
chr6-19581347-INS-2100
chr6-19582946-INS-53
chr6-19623466-INS-3173
chr6-19675991-INS-1856
chr6-19734089-INS-484
chr6-19752419-INS-54
chr6-19772235-INS-64
chr6-19772727-INS-6622
chr6-19802366-INS-7466
chr6-19868086-INS-52
chr6-19884786-INS-127
chr6-19917754-INS-6571
chr6-19923021-INS-56
chr6-19924873-INS-1266
chr6-20036702-INS-58

chr6-20051996-INS-80
chr6-20107065-INS-1050
chr6-20116980-INS-6401
chr6-20143026-INS-3345
chr6-20188434-INS-1823
chr6-20281805-INS-102
chr6-20299540-INS-311
chr6-20362754-INS-1001
chr6-20368976-INS-871
chr6-20387470-INS-101
chr6-20473797-INS-51
chr6-20475160-INS-58
chr6-20477033-INS-1694
chr6-20482176-INS-956
chr6-20532283-INS-4704
chr6-20550861-INS-142
chr6-20560188-INS-2726
chr6-20564787-INS-193
chr6-20582003-INS-50
chr6-20603134-INS-55
chr6-20639583-INS-7209
chr6-20654963-INS-498
chr6-20661792-INS-467
chr6-20717178-INS-6516
chr6-20771675-INS-58
chr6-20787380-INS-299
chr6-20806491-INS-397
chr6-20809885-INS-50
chr6-20826273-INS-2261
chr6-20828045-INS-1613
chr6-20855015-INS-780
chr6-20878398-INS-3256
chr6-20901531-INS-165
chr6-20905075-INS-4735
chr6-20913671-INS-381
chr6-20913678-INS-134
chr6-20951900-INS-89
chr6-20954393-INS-52
chr6-20955756-INS-5185
chr6-20969757-INS-500
chr6-21013562-INS-55
chr6-21025178-INS-26479
chr6-21047175-INS-51
chr6-21051344-INS-1536
chr6-21053679-INS-8592
chr6-21069108-INS-2484
chr6-21079853-INS-2831
chr6-21083094-INS-6346
chr6-21092137-INS-50
chr6-21114671-INS-108
chr6-21114768-INS-399
chr6-21121209-INS-1368
chr6-21131337-INS-810
chr6-21131698-INS-2682
chr6-21154789-INS-165
chr6-21176596-INS-199
chr6-21182391-INS-510
chr6-21196025-INS-7558
chr6-21212162-INS-66
chr6-21229601-INS-269
chr6-21257214-INS-6477
chr6-21288534-INS-62
chr6-21318874-INS-133
chr6-21323187-INS-130
chr6-21414426-INS-144
chr6-21448473-INS-56
chr6-21478986-INS-292
chr6-21494353-INS-54
chr6-21505710-INS-6591
chr6-21524628-INS-6368
chr6-21539217-INS-60
chr6-21539904-INS-77
chr6-21575493-INS-929
chr6-21604336-INS-146
chr6-21666695-INS-1041
chr6-21668374-INS-78
chr6-21738064-INS-60

chr6-21750017-INS-3544
chr6-21775781-INS-1580
chr6-21788991-INS-6520
chr6-21821086-INS-71
chr6-21821428-INS-218
chr6-21860544-INS-113
chr6-21868010-INS-386
chr6-21876927-INS-58
chr6-21877540-INS-212
chr6-21880326-INS-76
chr6-21901241-INS-60
chr6-21916702-INS-6658
chr6-21920582-INS-50
chr6-21971435-INS-63
chr6-21976946-INS-84
chr6-22001946-INS-1909
chr6-22025635-INS-68
chr6-22098671-INS-148
chr6-22102981-INS-401
chr6-22136635-INS-50
chr6-22139965-INS-72
chr6-22144957-INS-94
chr6-22151153-INS-327
chr6-22154181-INS-76
chr6-22168220-INS-69
chr6-22209150-INS-74
chr6-22226883-INS-3419
chr6-22235104-INS-114
chr6-22235609-INS-795
chr6-22254045-INS-2774
chr6-22259650-INS-855
chr6-22262444-INS-6713
chr6-22285759-INS-55
chr6-22298732-INS-53
chr6-22332287-INS-222
chr6-22360021-INS-58
chr6-22397480-INS-126
chr6-22397558-INS-57
chr6-22401708-INS-128
chr6-22405312-INS-406
chr6-22438634-INS-431
chr6-22472105-INS-58
chr6-22474413-INS-86
chr6-22498972-INS-992
chr6-22520259-INS-52
chr6-22526641-INS-279
chr6-22527363-INS-160
chr6-22537517-INS-496
chr6-22540689-INS-348
chr6-22540798-INS-64
chr6-22547830-INS-6333
chr6-22558704-INS-1250
chr6-22560345-INS-182
chr6-22583084-INS-6498
chr6-22638011-INS-101
chr6-22638011-INS-50
chr6-22679693-INS-272
chr6-22680036-INS-241
chr6-22701275-INS-4745
chr6-22708456-INS-60
chr6-22773679-INS-2262
chr6-22859504-INS-2666
chr6-22877895-INS-109
chr6-22927638-INS-230
chr6-22927679-INS-56
chr6-22930809-INS-4870
chr6-22947175-INS-102
chr6-22947578-INS-470
chr6-22954853-INS-3114
chr6-22968130-INS-4833
chr6-22969881-INS-852
chr6-22988104-INS-65
chr6-23029586-INS-110
chr6-23042103-INS-52
chr6-23066521-INS-816
chr6-23067672-INS-224
chr6-23068725-INS-67

chr6-23080914-INS-56
chr6-23121904-INS-426
chr6-23136931-INS-2075
chr6-23151156-INS-4355
chr6-23152396-INS-51
chr6-23178466-INS-228
chr6-23238251-INS-1144
chr6-23266514-INS-136
chr6-23296580-INS-66
chr6-23297177-INS-58
chr6-23333667-INS-338
chr6-23334787-INS-51
chr6-23354104-INS-2026
chr6-23399995-INS-412
chr6-23402978-INS-926
chr6-23474178-INS-407
chr6-23485836-INS-1062
chr6-23488910-INS-591
chr6-23495870-INS-1052
chr6-23498081-INS-1062
chr6-23498364-INS-4944
chr6-23541891-INS-907
chr6-23548955-INS-50
chr6-23570901-INS-264
chr6-23570976-INS-282
chr6-23595541-INS-386
chr6-23614759-INS-84
chr6-23615285-INS-123
chr6-23643481-INS-884
chr6-23645890-INS-6630
chr6-23676324-INS-53
chr6-23699923-INS-78
chr6-23736422-INS-6362
chr6-23758030-INS-56
chr6-23762271-INS-1063
chr6-23762857-INS-50
chr6-23919312-INS-199
chr6-23924292-INS-61
chr6-23945970-INS-192
chr6-24024018-INS-4277
chr6-24041428-INS-50
chr6-24048347-INS-2998
chr6-24050664-INS-2000
chr6-24056007-INS-1674
chr6-24085156-INS-6230
chr6-24110612-INS-211
chr6-24185016-INS-50
chr6-24199448-INS-123
chr6-24199448-INS-50
chr6-24225052-INS-3713
chr6-24226738-INS-14218
chr6-24226738-INS-7105
chr6-24246952-INS-7028
chr6-24250433-INS-85
chr6-24261399-INS-220
chr6-24307831-INS-304
chr6-24307832-INS-70
chr6-24323031-INS-390
chr6-24330876-INS-175
chr6-24344587-INS-106
chr6-24369596-INS-220
chr6-24373165-INS-51
chr6-24381660-INS-50
chr6-24399372-INS-4162
chr6-24437258-INS-668
chr6-24455745-INS-118
chr6-24455822-INS-2989
chr6-24472535-INS-202
chr6-24510024-INS-3702
chr6-24565443-INS-1555
chr6-24566672-INS-589
chr6-24568219-INS-184
chr6-24644215-INS-68
chr6-24646605-INS-52
chr6-24708059-INS-683
chr6-24712907-INS-917
chr6-24721295-INS-6715

chr6-24732543-INS-64
chr6-24733613-INS-7729
chr6-24737113-INS-99
chr6-24739091-INS-67
chr6-24759036-INS-6914
chr6-24765390-INS-50
chr6-24769368-INS-64
chr6-24769520-INS-764
chr6-24776174-INS-836
chr6-24776696-INS-208
chr6-24799062-INS-160
chr6-24834436-INS-138
chr6-24839292-INS-52
chr6-24889327-INS-396
chr6-24889757-INS-95
chr6-24906774-INS-2090
chr6-24911303-INS-130
chr6-24911303-INS-54
chr6-24916640-INS-6475
chr6-24963760-INS-50
chr6-24971240-INS-62
chr6-24976914-INS-137
chr6-24982550-INS-251
chr6-24983248-INS-307
chr6-24983755-INS-6387
chr6-25011810-INS-450
chr6-25020983-INS-121
chr6-25027294-INS-169
chr6-25031172-INS-196
chr6-25036456-INS-60
chr6-25083272-INS-85
chr6-25091109-INS-6381
chr6-25170188-INS-94
chr6-25192776-INS-4097
chr6-25307270-INS-200
chr6-25349471-INS-106
chr6-25377463-INS-58
chr6-25388607-INS-120
chr6-25391884-INS-2890
chr6-25410963-INS-326
chr6-25427026-INS-2719
chr6-25433190-INS-58
chr6-25459117-INS-205
chr6-25500770-INS-62
chr6-25518331-INS-6343
chr6-25568444-INS-306
chr6-25604942-INS-61
chr6-25644455-INS-104
chr6-25666422-INS-2440
chr6-25720664-INS-55
chr6-25755261-INS-150
chr6-25756063-INS-68
chr6-25822143-INS-557
chr6-25827101-INS-50
chr6-25856106-INS-54
chr6-25880325-INS-52
chr6-25895450-INS-54
chr6-25902713-INS-54
chr6-25912422-INS-7084
chr6-25918908-INS-206
chr6-25930898-INS-67
chr6-25938159-INS-7199
chr6-25949251-INS-154
chr6-26013327-INS-2654
chr6-26042955-INS-313
chr6-26058802-INS-2892
chr6-26111324-INS-365
chr6-26120078-INS-175
chr6-26131863-INS-1034
chr6-26154041-INS-315
chr6-26172054-INS-113
chr6-26172141-INS-585
chr6-26172268-INS-208
chr6-26182323-INS-67
chr6-26182352-INS-3174
chr6-26185806-INS-1426
chr6-26213923-INS-3143

chr6-26349680-INS-64
chr6-26364723-INS-343
chr6-26391263-INS-234
chr6-26394135-INS-1453
chr6-26399436-INS-1238
chr6-26422432-INS-329
chr6-26450880-INS-160
chr6-26485660-INS-2113
chr6-26546087-INS-2883
chr6-26564173-INS-500
chr6-26564480-INS-566
chr6-26577496-INS-14620
chr6-26631420-INS-6555
chr6-26646505-INS-67
chr6-26668562-INS-1842
chr6-26724533-INS-6308
chr6-26735789-INS-58
chr6-26739938-INS-594
chr6-26777558-INS-6597
chr6-26781136-INS-546
chr6-26781386-INS-177
chr6-26783065-INS-52
chr6-26788743-INS-1364
chr6-26801292-INS-99
chr6-26815991-INS-6901
chr6-26844657-INS-132
chr6-26845916-INS-2506
chr6-26856420-INS-219
chr6-26916828-INS-13431
chr6-26932898-INS-60
chr6-26933059-INS-469
chr6-26940679-INS-4340
chr6-26974422-INS-54
chr6-27051797-INS-61
chr6-27053914-INS-2764
chr6-27054344-INS-50
chr6-27058490-INS-93
chr6-27071527-INS-7890
chr6-27090765-INS-4159
chr6-27155859-INS-80
chr6-27255615-INS-51
chr6-27286229-INS-6471
chr6-27287910-INS-52
chr6-27336627-INS-6159
chr6-27343837-INS-508
chr6-27356679-INS-55
chr6-27369927-INS-6690
chr6-27372350-INS-91
chr6-27406243-INS-2842
chr6-27426875-INS-146
chr6-27438439-INS-3195
chr6-27458022-INS-50
chr6-27515186-INS-93
chr6-27520051-INS-106
chr6-27552113-INS-56
chr6-27628349-INS-66
chr6-27630100-INS-771
chr6-27675517-INS-52
chr6-27675986-INS-2057
chr6-27711512-INS-193
chr6-27711527-INS-82
chr6-27767995-INS-189
chr6-27794688-INS-6783
chr6-27804676-INS-50
chr6-27844304-INS-148
chr6-27895920-INS-315
chr6-27921185-INS-102
chr6-27975860-INS-66
chr6-28027774-INS-118
chr6-28046165-INS-102
chr6-28047348-INS-185
chr6-28047348-INS-716
chr6-28200281-INS-6055
chr6-28227374-INS-50
chr6-28291689-INS-202
chr6-28295657-INS-61
chr6-28326148-INS-170

chr6-28328849-INS-50
chr6-28357683-INS-459
chr6-28404433-INS-6485
chr6-28412445-INS-54
chr6-28428240-INS-60
chr6-28428875-INS-52
chr6-28502310-INS-76
chr6-28531956-INS-360
chr6-28579523-INS-201
chr6-28582068-INS-88
chr6-28597096-INS-176
chr6-28633202-INS-463
chr6-28641090-INS-823
chr6-28655045-INS-476
chr6-28662390-INS-538
chr6-28740941-INS-355
chr6-28788884-INS-65
chr6-28818702-INS-417
chr6-28848797-INS-3621
chr6-28962676-INS-456
chr6-28982929-INS-1872
chr6-28997259-INS-52
chr6-28998559-INS-58
chr6-29085944-INS-1867
chr6-29114040-INS-72
chr6-29150543-INS-214
chr6-29151995-INS-578
chr6-29157047-INS-50
chr6-29175114-INS-102
chr6-29221487-INS-116
chr6-29224824-INS-56
chr6-29233649-INS-202
chr6-29233854-INS-64
chr6-29249215-INS-98
chr6-29249251-INS-254
chr6-29268753-INS-489
chr6-29272004-INS-405
chr6-29280319-INS-428
chr6-29288494-INS-737
chr6-29327102-INS-64
chr6-29407294-INS-208
chr6-29407464-INS-59
chr6-29431127-INS-70
chr6-29432768-INS-72
chr6-29454864-INS-58
chr6-29469715-INS-157
chr6-29490656-INS-215
chr6-29573239-INS-50
chr6-29626698-INS-68
chr6-29640219-INS-57
chr6-29640554-INS-177
chr6-29665543-INS-50
chr6-29720130-INS-51
chr6-29724466-INS-92
chr6-29731182-INS-61
chr6-29748011-INS-902
chr6-29748661-INS-6205
chr6-29762574-INS-7553
chr6-29774704-INS-202
chr6-29810621-INS-207
chr6-29848449-INS-376
chr6-29848596-INS-874
chr6-29873434-INS-62
chr6-29913603-INS-4804
chr6-29925310-INS-159
chr6-29928063-INS-60
chr6-29960235-INS-58
chr6-29992782-INS-74
chr6-30024575-INS-173
chr6-30084080-INS-60
chr6-30146424-INS-56
chr6-30158726-INS-212
chr6-30195095-INS-161
chr6-30195096-INS-54
chr6-30201981-INS-190
chr6-30270354-INS-201
chr6-30277661-INS-978

chr6-30282457-INS-64
chr6-30296469-INS-88
chr6-30302626-INS-58
chr6-30314284-INS-7207
chr6-30321672-INS-89
chr6-30333573-INS-200
chr6-30341822-INS-54
chr6-30348923-INS-117
chr6-30358655-INS-198
chr6-30360588-INS-180
chr6-30363618-INS-4788
chr6-30394156-INS-207
chr6-30394472-INS-56
chr6-30397827-INS-197
chr6-30414026-INS-508
chr6-30415439-INS-463
chr6-30420610-INS-135
chr6-30420610-INS-290
chr6-30445586-INS-110
chr6-30470438-INS-219
chr6-30477059-INS-210
chr6-30524805-INS-61
chr6-30575499-INS-61
chr6-30577791-INS-283
chr6-30588601-INS-101
chr6-30595554-INS-80
chr6-30660253-INS-50
chr6-30717775-INS-63
chr6-30920445-INS-60
chr6-30921817-INS-68
chr6-30921919-INS-149
chr6-30928209-INS-179
chr6-30931642-INS-71
chr6-30953410-INS-75
chr6-31044815-INS-51
chr6-31057004-INS-60
chr6-31100395-INS-129
chr6-31130957-INS-56
chr6-31205099-INS-120
chr6-31211675-INS-63
chr6-31216728-INS-67
chr6-31226599-INS-205
chr6-31245432-INS-76
chr6-31252355-INS-235
chr6-31252499-INS-4803
chr6-31273466-INS-58
chr6-31385246-INS-481
chr6-31405381-INS-84
chr6-31409380-INS-345
chr6-31430441-INS-592
chr6-31433284-INS-100
chr6-31459216-INS-98
chr6-31509207-INS-52
chr6-31562661-INS-58
chr6-31572892-INS-101
chr6-31575528-INS-1090
chr6-31592193-INS-91
chr6-31608772-INS-70
chr6-31608791-INS-201
chr6-31623305-INS-96
chr6-31643464-INS-50
chr6-31664548-INS-506
chr6-31682666-INS-59
chr6-31683042-INS-81
chr6-31702882-INS-7092
chr6-31703267-INS-287
chr6-31703570-INS-54
chr6-31706494-INS-54
chr6-31730329-INS-648
chr6-31756400-INS-5262
chr6-31772362-INS-1629
chr6-31785576-INS-570
chr6-31811398-INS-72
chr6-31849202-INS-50
chr6-31861044-INS-66
chr6-31876616-INS-7098
chr6-31925890-INS-3050

chr6-31932102-INS-52
chr6-31933398-INS-390
chr6-31958938-INS-50
chr6-32079699-INS-208
chr6-32194577-INS-126
chr6-32210783-INS-261
chr6-32220771-INS-69
chr6-32313742-INS-54
chr6-32344108-INS-2745
chr6-32367212-INS-163
chr6-32368268-INS-55
chr6-32395936-INS-84
chr6-32459280-INS-81
chr6-32556505-INS-234
chr6-32588516-INS-153
chr6-32616636-INS-56
chr6-32629716-INS-740
chr6-32694546-INS-155
chr6-32702322-INS-6797
chr6-32724332-INS-104
chr6-32754303-INS-110
chr6-32776115-INS-96
chr6-32776655-INS-239
chr6-32778499-INS-6644
chr6-32784209-INS-131
chr6-32862332-INS-105
chr6-32909753-INS-118
chr6-32922581-INS-6615
chr6-32946737-INS-563
chr6-32965225-INS-56
chr6-32968254-INS-1879
chr6-33064128-INS-72
chr6-33070438-INS-2384
chr6-33084849-INS-136
chr6-33085384-INS-400
chr6-33087109-INS-377
chr6-33112443-INS-364
chr6-33128153-INS-172
chr6-33158972-INS-6453
chr6-33210320-INS-1320
chr6-33215528-INS-62
chr6-33215659-INS-127
chr6-33218802-INS-2742
chr6-33271093-INS-56
chr6-33295803-INS-90
chr6-33352531-INS-194
chr6-33352535-INS-582
chr6-33394878-INS-52
chr6-33410853-INS-276
chr6-33456954-INS-133
chr6-33476495-INS-171
chr6-33485202-INS-52
chr6-33493223-INS-1965
chr6-33595666-INS-74
chr6-33595727-INS-1301
chr6-33595801-INS-3252
chr6-33608046-INS-2600
chr6-33708438-INS-150
chr6-33708447-INS-63
chr6-33768622-INS-983
chr6-33778695-INS-2068
chr6-33779711-INS-341
chr6-33817125-INS-96
chr6-33827775-INS-3651
chr6-33860260-INS-4210
chr6-33884780-INS-1459
chr6-33960302-INS-385
chr6-33986805-INS-3519
chr6-33995002-INS-76
chr6-34003499-INS-76
chr6-34020761-INS-115
chr6-34065492-INS-58
chr6-34094621-INS-3151
chr6-34094656-INS-50
chr6-34096808-INS-50
chr6-34156585-INS-7100
chr6-34170901-INS-1830

chr6-34179566-INS-51
chr6-34191296-INS-131
chr6-34227924-INS-180
chr6-34231458-INS-2781
chr6-34231575-INS-399
chr6-34237104-INS-85
chr6-34239988-INS-93
chr6-34296616-INS-505
chr6-34300162-INS-74
chr6-34359327-INS-2076
chr6-34372709-INS-142
chr6-34411954-INS-173
chr6-34507603-INS-158
chr6-34511498-INS-146
chr6-34536080-INS-60
chr6-34577662-INS-50
chr6-34578434-INS-53
chr6-34587142-INS-76
chr6-34624802-INS-243
chr6-34646106-INS-465
chr6-34657060-INS-121
chr6-34657060-INS-258
chr6-34667111-INS-230
chr6-34667111-INS-580
chr6-34678630-INS-175
chr6-34706423-INS-77
chr6-34832934-INS-70
chr6-34841195-INS-292
chr6-34841289-INS-99
chr6-34860418-INS-891
chr6-34861031-INS-588
chr6-34882358-INS-209
chr6-34888274-INS-1789
chr6-34888341-INS-255
chr6-34912917-INS-234
chr6-34913027-INS-918
chr6-34919887-INS-54
chr6-34920265-INS-192
chr6-34939784-INS-63
chr6-34947013-INS-76
chr6-34947065-INS-154
chr6-34995229-INS-66
chr6-35052427-INS-392
chr6-35071110-INS-89
chr6-35122036-INS-570
chr6-35149462-INS-180
chr6-35154057-INS-75
chr6-35219955-INS-201
chr6-35228241-INS-56
chr6-35239649-INS-218
chr6-35268748-INS-111
chr6-35285803-INS-85
chr6-35343836-INS-171
chr6-35364681-INS-101
chr6-35365476-INS-229
chr6-35394797-INS-206
chr6-35501555-INS-57
chr6-35638217-INS-9632
chr6-35667597-INS-166
chr6-35732701-INS-1797
chr6-35739262-INS-1898
chr6-35741478-INS-149
chr6-35788076-INS-73
chr6-35792307-INS-380
chr6-35792323-INS-125
chr6-35794135-INS-9782
chr6-35818458-INS-62
chr6-35819148-INS-84
chr6-35831585-INS-60
chr6-35867596-INS-6469
chr6-35923172-INS-128
chr6-35947479-INS-61
chr6-35949223-INS-62
chr6-35974708-INS-174
chr6-35990059-INS-386
chr6-35990068-INS-81
chr6-36001931-INS-55

chr6-36004304-INS-371
chr6-36066298-INS-50
chr6-36073374-INS-2886
chr6-36078249-INS-50
chr6-36081638-INS-1385
chr6-36103901-INS-172
chr6-36109045-INS-64
chr6-36113040-INS-5042
chr6-36115610-INS-2222
chr6-36127006-INS-251
chr6-36137129-INS-368
chr6-36144632-INS-113
chr6-36160246-INS-6304
chr6-36164983-INS-401
chr6-36172857-INS-138
chr6-36172873-INS-50
chr6-36190266-INS-137
chr6-36190311-INS-64
chr6-36192747-INS-1892
chr6-36403962-INS-92
chr6-36412004-INS-1179
chr6-36516898-INS-317
chr6-36566147-INS-119
chr6-36568550-INS-606
chr6-36607747-INS-65
chr6-36613905-INS-6372
chr6-36619069-INS-64
chr6-36701395-INS-60
chr6-36747395-INS-194
chr6-36747431-INS-560
chr6-36748276-INS-58
chr6-36769159-INS-192
chr6-36784613-INS-68
chr6-36814237-INS-54
chr6-36866113-INS-134
chr6-36910060-INS-86
chr6-36991070-INS-2567
chr6-37008613-INS-63
chr6-37021743-INS-60
chr6-37048138-INS-300
chr6-37105696-INS-63
chr6-37110342-INS-551
chr6-37155903-INS-70
chr6-37159868-INS-52
chr6-37162979-INS-51
chr6-37184287-INS-1841
chr6-37184815-INS-2679
chr6-37201435-INS-121
chr6-37204076-INS-82
chr6-37204469-INS-3315
chr6-37276397-INS-58
chr6-37291526-INS-2668
chr6-37316369-INS-183
chr6-37316941-INS-498
chr6-37377898-INS-60
chr6-37410855-INS-106
chr6-37481365-INS-466
chr6-37495126-INS-222
chr6-37517107-INS-58
chr6-37563364-INS-279
chr6-37565124-INS-85
chr6-37572314-INS-50
chr6-37575845-INS-89
chr6-37582331-INS-474
chr6-37582406-INS-58
chr6-37583770-INS-64
chr6-37586164-INS-51
chr6-37650140-INS-211
chr6-37652952-INS-51
chr6-37663082-INS-200
chr6-37667599-INS-248
chr6-37669403-INS-9045
chr6-37688434-INS-70
chr6-37710806-INS-384
chr6-37710806-INS-4033
chr6-37766329-INS-1490
chr6-37779231-INS-5771

chr6-37790007-INS-174
chr6-37811832-INS-7083
chr6-37870168-INS-138
chr6-37880308-INS-56
chr6-37883096-INS-408
chr6-37919699-INS-240
chr6-37984191-INS-8859
chr6-37991001-INS-130
chr6-37991001-INS-50
chr6-38022396-INS-7096
chr6-38080502-INS-223
chr6-38121568-INS-58
chr6-38137807-INS-618
chr6-38236881-INS-254
chr6-38281647-INS-167
chr6-38297331-INS-107
chr6-38299194-INS-60
chr6-38307155-INS-58
chr6-38318574-INS-80
chr6-38344703-INS-66
chr6-38344893-INS-910
chr6-38386067-INS-205
chr6-38386190-INS-279
chr6-38387650-INS-173
chr6-38436085-INS-487
chr6-38517980-INS-439
chr6-38616279-INS-92
chr6-38632421-INS-468
chr6-38637159-INS-558
chr6-38663088-INS-256
chr6-38699082-INS-50
chr6-38732566-INS-216
chr6-38751260-INS-145
chr6-38871938-INS-189
chr6-38888728-INS-54
chr6-38937567-INS-3237
chr6-38961228-INS-200
chr6-38973520-INS-317
chr6-39009830-INS-53
chr6-39031241-INS-66
chr6-39084173-INS-50
chr6-39103839-INS-58
chr6-39146925-INS-139
chr6-39176060-INS-187
chr6-39203951-INS-51
chr6-39207966-INS-54
chr6-39212343-INS-345
chr6-39216270-INS-231
chr6-39225648-INS-62
chr6-39228794-INS-401
chr6-39256444-INS-80
chr6-39285390-INS-167
chr6-39304332-INS-139
chr6-39311756-INS-202
chr6-39336320-INS-228
chr6-39338929-INS-161
chr6-39355618-INS-198
chr6-39388720-INS-101
chr6-39390280-INS-203
chr6-39410079-INS-116
chr6-39415492-INS-195
chr6-39428559-INS-197
chr6-39438629-INS-158
chr6-39462336-INS-68
chr6-39464479-INS-64
chr6-39471473-INS-243
chr6-39523655-INS-6434
chr6-39540709-INS-150
chr6-39540765-INS-210
chr6-39574500-INS-527
chr6-39608751-INS-200
chr6-39609500-INS-220
chr6-39653149-INS-188
chr6-39653150-INS-50
chr6-39708082-INS-107
chr6-39708138-INS-630
chr6-39711593-INS-534

chr6-39735237-INS-50
chr6-39735385-INS-177
chr6-39740036-INS-2691
chr6-39745297-INS-227
chr6-39747405-INS-87
chr6-39760873-INS-193
chr6-39765862-INS-228
chr6-39774717-INS-301
chr6-39917159-INS-55
chr6-39921476-INS-58
chr6-39966716-INS-6150
chr6-39975408-INS-367
chr6-39979762-INS-59
chr6-40046346-INS-64
chr6-40121973-INS-327
chr6-40158407-INS-54
chr6-40220575-INS-56
chr6-40235029-INS-1890
chr6-40414476-INS-145
chr6-40447470-INS-53
chr6-40463692-INS-1305
chr6-40489333-INS-8064
chr6-40502884-INS-96
chr6-40503090-INS-222
chr6-40508651-INS-50
chr6-40525835-INS-6714
chr6-40566914-INS-13820
chr6-40595489-INS-50
chr6-40646143-INS-2002
chr6-40681651-INS-972
chr6-40694994-INS-2152
chr6-40729629-INS-182
chr6-40760973-INS-131
chr6-40828796-INS-162
chr6-40844849-INS-480
chr6-40888136-INS-2222
chr6-40894135-INS-6803
chr6-40916160-INS-609
chr6-40932211-INS-2462
chr6-41022831-INS-166
chr6-41025805-INS-967
chr6-41037076-INS-1244
chr6-41058585-INS-1473
chr6-41059218-INS-3343
chr6-41065835-INS-6712
chr6-41090623-INS-51
chr6-41093757-INS-84
chr6-41139632-INS-1120
chr6-41152267-INS-4415
chr6-41168585-INS-380
chr6-41172392-INS-119
chr6-41202358-INS-2492
chr6-41207093-INS-6159
chr6-41213803-INS-74
chr6-41260187-INS-2856
chr6-41318921-INS-217
chr6-41363417-INS-6481
chr6-41369938-INS-1134
chr6-41431444-INS-1242
chr6-41432313-INS-997
chr6-41450148-INS-2210
chr6-41450429-INS-117
chr6-41504202-INS-76
chr6-41507062-INS-72
chr6-41513649-INS-425
chr6-41514827-INS-195
chr6-41548803-INS-54
chr6-41557955-INS-199
chr6-41568588-INS-51
chr6-41651722-INS-78
chr6-41655398-INS-50
chr6-41690338-INS-2905
chr6-41762843-INS-51
chr6-41765492-INS-6285
chr6-41819682-INS-6365
chr6-41863588-INS-1293
chr6-41876290-INS-298

chr6-41884715-INS-7466
chr6-41895986-INS-70
chr6-41937721-INS-8662
chr6-42031938-INS-76
chr6-42037629-INS-1774
chr6-42171492-INS-7083
chr6-42177464-INS-61
chr6-42224191-INS-466
chr6-42224340-INS-116
chr6-42368314-INS-418
chr6-42378604-INS-104
chr6-42378712-INS-98
chr6-42397013-INS-68
chr6-42399173-INS-96
chr6-42404016-INS-6228
chr6-42409347-INS-106
chr6-42424641-INS-263
chr6-42482655-INS-62
chr6-42510660-INS-122
chr6-42527772-INS-6246
chr6-42541674-INS-66
chr6-42567738-INS-64
chr6-42604511-INS-88
chr6-42608132-INS-50
chr6-42612128-INS-1144
chr6-42636734-INS-321
chr6-42764861-INS-1634
chr6-42770928-INS-52
chr6-42840612-INS-76
chr6-42841356-INS-1142
chr6-42875462-INS-210
chr6-42880615-INS-54
chr6-42906828-INS-64
chr6-42972161-INS-58
chr6-42975902-INS-144
chr6-42976508-INS-1126
chr6-42977484-INS-57
chr6-42977836-INS-521
chr6-42985401-INS-376
chr6-42985784-INS-59
chr6-42986996-INS-66
chr6-42987876-INS-50
chr6-42995744-INS-2478
chr6-43006054-INS-168
chr6-43022374-INS-6488
chr6-43039728-INS-7014
chr6-43041145-INS-98
chr6-43112427-INS-199
chr6-43125275-INS-99
chr6-43141931-INS-175
chr6-43142960-INS-58
chr6-43157001-INS-53
chr6-43188133-INS-13432
chr6-43267967-INS-56
chr6-43292060-INS-560
chr6-43351782-INS-6561
chr6-43382944-INS-533
chr6-43423892-INS-10111
chr6-43432175-INS-59
chr6-43502238-INS-352
chr6-43509261-INS-352
chr6-43546039-INS-398
chr6-43583108-INS-2178
chr6-43614286-INS-925
chr6-43634347-INS-2655
chr6-43705852-INS-51
chr6-43729437-INS-56
chr6-43782490-INS-624
chr6-43821254-INS-160
chr6-43871556-INS-11184
chr6-43877059-INS-50
chr6-43883021-INS-594
chr6-43997302-INS-100
chr6-44000083-INS-6466
chr6-44010386-INS-6973
chr6-44017195-INS-65
chr6-44017592-INS-150

chr6-44041411-INS-6288
chr6-44048778-INS-73
chr6-44065448-INS-78
chr6-44128160-INS-499
chr6-44158660-INS-57
chr6-44163388-INS-6215
chr6-44164576-INS-142
chr6-44191238-INS-4771
chr6-44198044-INS-4847
chr6-44259487-INS-1240
chr6-44274968-INS-55
chr6-44301496-INS-1010
chr6-44331155-INS-148
chr6-44339794-INS-84
chr6-44343500-INS-703
chr6-44410122-INS-490
chr6-44413500-INS-601
chr6-44414536-INS-546
chr6-44438433-INS-196
chr6-44442949-INS-208
chr6-44453976-INS-50
chr6-44454071-INS-2628
chr6-44524756-INS-104
chr6-44535531-INS-3043
chr6-44543486-INS-195
chr6-44552293-INS-3945
chr6-44554687-INS-5555
chr6-44596048-INS-2916
chr6-44646575-INS-4664
chr6-44666910-INS-216
chr6-44676716-INS-242
chr6-44739239-INS-149
chr6-44743128-INS-68
chr6-44748125-INS-71
chr6-44749236-INS-486
chr6-44789833-INS-144
chr6-44805640-INS-56
chr6-44829645-INS-6268
chr6-44852742-INS-114
chr6-44874482-INS-251
chr6-44884809-INS-98
chr6-44892996-INS-1867
chr6-44917265-INS-1268
chr6-44918435-INS-3077
chr6-44921298-INS-300
chr6-44926606-INS-712
chr6-44958287-INS-6369
chr6-44958441-INS-2627
chr6-44959202-INS-52
chr6-44978209-INS-1560
chr6-44982656-INS-2934
chr6-45008145-INS-55
chr6-45028659-INS-2690
chr6-45045440-INS-80
chr6-45045968-INS-58
chr6-45116745-INS-68
chr6-45124252-INS-263
chr6-45124732-INS-52
chr6-45156541-INS-99
chr6-45157427-INS-71
chr6-45176472-INS-190
chr6-45209249-INS-142
chr6-45244457-INS-127
chr6-45273254-INS-57
chr6-45312783-INS-706
chr6-45373020-INS-1650
chr6-45390025-INS-477
chr6-45405704-INS-68
chr6-45413776-INS-52
chr6-45425360-INS-1793
chr6-45497617-INS-50
chr6-45541521-INS-71
chr6-45541549-INS-166
chr6-45572083-INS-51
chr6-45589312-INS-1787
chr6-45603631-INS-2450
chr6-45631971-INS-140

chr6-45646283-INS-4861
chr6-45651294-INS-5330
chr6-45665894-INS-1048
chr6-45673207-INS-73
chr6-45676938-INS-149
chr6-45716885-INS-6688
chr6-45719985-INS-120
chr6-45779800-INS-146
chr6-45780502-INS-66
chr6-45792669-INS-80
chr6-45803207-INS-267
chr6-45803309-INS-76
chr6-45852184-INS-209
chr6-45881409-INS-404
chr6-45883276-INS-167
chr6-45903363-INS-50
chr6-45926044-INS-52
chr6-45932850-INS-50
chr6-45942117-INS-2077
chr6-45951330-INS-7114
chr6-45970358-INS-73
chr6-46072803-INS-907
chr6-46093791-INS-551
chr6-46236377-INS-1019
chr6-46283866-INS-505
chr6-46284886-INS-57
chr6-46298704-INS-1216
chr6-46313610-INS-229
chr6-46314525-INS-76
chr6-46345735-INS-1444
chr6-46358090-INS-4982
chr6-46423494-INS-89
chr6-46521485-INS-54
chr6-46596905-INS-66
chr6-46602279-INS-240
chr6-46636832-INS-6439
chr6-46727434-INS-2749
chr6-46759158-INS-458
chr6-46759224-INS-80
chr6-46759819-INS-60
chr6-46817261-INS-72
chr6-46880392-INS-78
chr6-46892385-INS-421
chr6-46914994-INS-8102
chr6-46924318-INS-65
chr6-46934433-INS-62
chr6-46975246-INS-6616
chr6-46983542-INS-6295
chr6-46986003-INS-52
chr6-46988580-INS-644
chr6-47006953-INS-376
chr6-47023544-INS-50
chr6-47045650-INS-60
chr6-47098714-INS-519
chr6-47099101-INS-401
chr6-47136145-INS-58
chr6-47180040-INS-118
chr6-47186638-INS-74
chr6-47194244-INS-185
chr6-47195509-INS-6213
chr6-47260287-INS-77
chr6-47263536-INS-75
chr6-47279281-INS-3477
chr6-47279961-INS-73
chr6-47286925-INS-8680
chr6-47304940-INS-105
chr6-47305728-INS-661
chr6-47391803-INS-100
chr6-47406947-INS-1582
chr6-47412725-INS-206
chr6-47565582-INS-202
chr6-47568576-INS-60
chr6-47593694-INS-78
chr6-47597514-INS-52
chr6-47600232-INS-92
chr6-47720740-INS-100
chr6-47729416-INS-51

chr6-47740713-INS-51
chr6-47744187-INS-100
chr6-47765746-INS-56
chr6-47774896-INS-59
chr6-47820288-INS-166
chr6-47861252-INS-205
chr6-47864789-INS-223
chr6-47892267-INS-185
chr6-47915170-INS-5626
chr6-47922265-INS-5183
chr6-47922265-INS-578
chr6-47929233-INS-52
chr6-47945332-INS-126
chr6-47955844-INS-227
chr6-47962821-INS-4724
chr6-47990716-INS-86
chr6-48003944-INS-58
chr6-48092086-INS-186
chr6-48096264-INS-436
chr6-48158288-INS-406
chr6-48158288-INS-60
chr6-48159632-INS-345
chr6-48172392-INS-361
chr6-48200803-INS-104
chr6-48243090-INS-104
chr6-48245392-INS-1781
chr6-48334411-INS-6021
chr6-48335494-INS-50
chr6-48474916-INS-62
chr6-48485667-INS-171
chr6-48494987-INS-337
chr6-48512814-INS-102
chr6-48512947-INS-204
chr6-48541443-INS-58
chr6-48580473-INS-50
chr6-48725703-INS-140
chr6-48769428-INS-140
chr6-48771457-INS-191
chr6-48794026-INS-351
chr6-48823241-INS-84
chr6-48823986-INS-203
chr6-48897488-INS-124
chr6-48919856-INS-6349
chr6-48935794-INS-88
chr6-48937761-INS-7017
chr6-48949732-INS-84
chr6-48988714-INS-98
chr6-48988822-INS-88
chr6-49004258-INS-149
chr6-49018597-INS-208
chr6-49020712-INS-67
chr6-49030134-INS-50
chr6-49036427-INS-637
chr6-49045922-INS-59
chr6-49115821-INS-108
chr6-49123988-INS-50
chr6-49126507-INS-64
chr6-49134350-INS-56
chr6-49206401-INS-198
chr6-49206981-INS-55
chr6-49209490-INS-543
chr6-49214842-INS-195
chr6-49215611-INS-69
chr6-49232000-INS-151
chr6-49232398-INS-166
chr6-49233823-INS-211
chr6-49246681-INS-97
chr6-49268694-INS-3729
chr6-49278069-INS-335
chr6-49287452-INS-210
chr6-49301923-INS-114
chr6-49302618-INS-180
chr6-49315256-INS-175
chr6-49322171-INS-200
chr6-49327646-INS-327
chr6-49329402-INS-127
chr6-49336445-INS-157

chr6-49370522-INS-439
chr6-49382964-INS-997
chr6-49383008-INS-473
chr6-49383086-INS-52
chr6-49383380-INS-140
chr6-49463724-INS-4129
chr6-49480556-INS-91
chr6-49501670-INS-58
chr6-49528827-INS-50
chr6-49538101-INS-1536
chr6-49556715-INS-63
chr6-49566430-INS-8018
chr6-49573649-INS-50
chr6-49600546-INS-155
chr6-49625180-INS-54
chr6-49643891-INS-59
chr6-49684959-INS-102
chr6-49711036-INS-136
chr6-49725435-INS-553
chr6-49725754-INS-50
chr6-49753554-INS-396
chr6-49764069-INS-53
chr6-49764707-INS-343
chr6-49770863-INS-537
chr6-49770866-INS-252
chr6-49808789-INS-1814
chr6-49855751-INS-52
chr6-49873794-INS-72
chr6-49878476-INS-60
chr6-49979127-INS-89
chr6-49993686-INS-72
chr6-50018669-INS-137
chr6-50023520-INS-69
chr6-50028977-INS-144
chr6-50035565-INS-622
chr6-50035933-INS-195
chr6-50042793-INS-248
chr6-50113461-INS-50
chr6-50115155-INS-458
chr6-50144855-INS-64
chr6-50146182-INS-60
chr6-50146386-INS-64
chr6-50153679-INS-203
chr6-50173863-INS-429
chr6-50174214-INS-429
chr6-50189009-INS-249
chr6-50219546-INS-101
chr6-50227274-INS-58
chr6-50254616-INS-112
chr6-50259522-INS-285
chr6-50259522-INS-50
chr6-50394279-INS-498
chr6-50428406-INS-80
chr6-50431355-INS-82
chr6-50456516-INS-6574
chr6-50469061-INS-136
chr6-50473380-INS-6442
chr6-50485039-INS-191
chr6-50499181-INS-60
chr6-50503394-INS-4373
chr6-50526529-INS-60
chr6-50531545-INS-120
chr6-50535494-INS-119
chr6-50538260-INS-6403
chr6-50612757-INS-292
chr6-50612933-INS-525
chr6-50615983-INS-306
chr6-50689747-INS-124
chr6-50706563-INS-1412
chr6-50708999-INS-52
chr6-50813696-INS-53
chr6-50815220-INS-825
chr6-50832812-INS-68
chr6-50843247-INS-951
chr6-50870152-INS-124
chr6-50881845-INS-65
chr6-50899969-INS-106

chr6-50911886-INS-259
chr6-50930661-INS-72
chr6-50975482-INS-179
chr6-50985944-INS-288
chr6-51030167-INS-52
chr6-51032223-INS-57
chr6-51091187-INS-61
chr6-51145564-INS-97
chr6-51163052-INS-1084
chr6-51163059-INS-226
chr6-51201766-INS-74
chr6-51215692-INS-64
chr6-51232066-INS-1160
chr6-51272290-INS-95
chr6-51272965-INS-92
chr6-51295903-INS-164
chr6-51302772-INS-50
chr6-51328862-INS-70
chr6-51371941-INS-86
chr6-51376970-INS-182
chr6-51385166-INS-2705
chr6-51387149-INS-409
chr6-51408457-INS-60
chr6-51423746-INS-53
chr6-51432699-INS-596
chr6-51447465-INS-60
chr6-51452553-INS-53
chr6-51474047-INS-171
chr6-51479589-INS-194
chr6-51504181-INS-72
chr6-51507810-INS-6319
chr6-51514210-INS-70
chr6-51514735-INS-70
chr6-51525772-INS-112
chr6-51578323-INS-7082
chr6-51579256-INS-99
chr6-51607849-INS-60
chr6-51613034-INS-60
chr6-51629821-INS-131
chr6-51632785-INS-1420
chr6-51640256-INS-53
chr6-51656926-INS-1379
chr6-51656926-INS-8578
chr6-51665410-INS-97
chr6-51683368-INS-132
chr6-51690582-INS-558
chr6-51690804-INS-52
chr6-51690889-INS-564
chr6-51715093-INS-200
chr6-51740187-INS-78
chr6-51741368-INS-213
chr6-51761512-INS-52
chr6-51762296-INS-243
chr6-51797143-INS-7238
chr6-51801141-INS-60
chr6-51900504-INS-50
chr6-51917436-INS-52
chr6-51922656-INS-308
chr6-52003594-INS-81
chr6-52006515-INS-1978
chr6-52047363-INS-112
chr6-52048435-INS-50
chr6-52064504-INS-391
chr6-52071003-INS-1889
chr6-52071498-INS-507
chr6-52071833-INS-3043
chr6-52072336-INS-197
chr6-52108586-INS-80
chr6-52121019-INS-80
chr6-52275037-INS-138
chr6-52275091-INS-60
chr6-52278782-INS-459
chr6-52281703-INS-144
chr6-52282540-INS-199
chr6-52304372-INS-85
chr6-52319150-INS-52
chr6-52319390-INS-203

chr6-52322632-INS-255
chr6-52323178-INS-55
chr6-52396895-INS-112
chr6-52399302-INS-52
chr6-52414677-INS-94
chr6-52431178-INS-60
chr6-52435722-INS-50
chr6-52439406-INS-1176
chr6-52446086-INS-712
chr6-52446124-INS-91
chr6-52456667-INS-208
chr6-52483758-INS-63
chr6-52488264-INS-174
chr6-52489240-INS-50
chr6-52497463-INS-142
chr6-52510066-INS-72
chr6-52518366-INS-66
chr6-52557182-INS-888
chr6-52565540-INS-120
chr6-52602031-INS-473
chr6-52656268-INS-6232
chr6-52656445-INS-272
chr6-52686520-INS-92
chr6-52686618-INS-370
chr6-52688045-INS-205
chr6-52706630-INS-478
chr6-52740248-INS-72
chr6-52839739-INS-6055
chr6-52882732-INS-72
chr6-52893089-INS-2440
chr6-52908086-INS-51
chr6-53006088-INS-885
chr6-53073835-INS-56
chr6-53109023-INS-146
chr6-53110350-INS-278
chr6-53130770-INS-88
chr6-53134321-INS-211
chr6-53138801-INS-69
chr6-53147997-INS-4532
chr6-53164242-INS-1225
chr6-53197000-INS-180
chr6-53225926-INS-55
chr6-53226246-INS-118
chr6-53226322-INS-654
chr6-53226442-INS-310
chr6-53233741-INS-70
chr6-53235107-INS-3166
chr6-53331162-INS-97
chr6-53341741-INS-186
chr6-53388052-INS-78
chr6-53402490-INS-302
chr6-53417854-INS-2734
chr6-53449567-INS-50
chr6-53473255-INS-149
chr6-53492332-INS-270
chr6-53517038-INS-50
chr6-53526430-INS-159
chr6-53536383-INS-696
chr6-53543084-INS-85
chr6-53575924-INS-50
chr6-53656899-INS-128
chr6-53691030-INS-156
chr6-53709352-INS-5507
chr6-53723272-INS-842
chr6-53723554-INS-164
chr6-53723712-INS-1454
chr6-53774181-INS-78
chr6-53774198-INS-173
chr6-53774780-INS-418
chr6-53776908-INS-380
chr6-53828164-INS-531
chr6-53828169-INS-74
chr6-53866598-INS-70
chr6-53890931-INS-158
chr6-53914526-INS-607
chr6-53929625-INS-456
chr6-53965218-INS-1293

chr6-54013120-INS-67
chr6-54031576-INS-55
chr6-54046235-INS-50
chr6-54133113-INS-194
chr6-54141834-INS-9015
chr6-54145712-INS-353
chr6-54146028-INS-352
chr6-54148634-INS-385
chr6-54166059-INS-72
chr6-54183096-INS-100
chr6-54185163-INS-1734
chr6-54190571-INS-99
chr6-54191320-INS-79
chr6-54205972-INS-50
chr6-54290508-INS-84
chr6-54337236-INS-606
chr6-54344988-INS-71
chr6-54350939-INS-107
chr6-54351979-INS-686
chr6-54352659-INS-410
chr6-54366892-INS-51
chr6-54373534-INS-80
chr6-54394495-INS-70
chr6-54424509-INS-82
chr6-54424976-INS-90
chr6-54445441-INS-93
chr6-54445875-INS-211
chr6-54484801-INS-58
chr6-54497316-INS-98
chr6-54498179-INS-3396
chr6-54506629-INS-3413
chr6-54509895-INS-4703
chr6-54556844-INS-64
chr6-54565526-INS-187
chr6-54565662-INS-680
chr6-54566035-INS-238
chr6-54570182-INS-464
chr6-54630636-INS-52
chr6-54706950-INS-67
chr6-54722895-INS-57
chr6-54752267-INS-56
chr6-54759973-INS-1142
chr6-54887873-INS-87
chr6-54925268-INS-94
chr6-54929076-INS-202
chr6-54935482-INS-52
chr6-54977725-INS-2727
chr6-55005490-INS-88
chr6-55009977-INS-60
chr6-55011617-INS-232
chr6-55060724-INS-220
chr6-55125879-INS-60
chr6-55129245-INS-1273
chr6-55169598-INS-74
chr6-55234958-INS-110
chr6-55260251-INS-338
chr6-55317089-INS-68
chr6-55344720-INS-929
chr6-55359841-INS-76
chr6-55370486-INS-183
chr6-55392276-INS-78
chr6-55476146-INS-53
chr6-55497182-INS-68
chr6-55589915-INS-50
chr6-55599558-INS-97
chr6-55607735-INS-261
chr6-55667865-INS-166
chr6-55688238-INS-175
chr6-55744584-INS-52
chr6-55779313-INS-51
chr6-55838273-INS-121
chr6-55861111-INS-50
chr6-55899692-INS-72
chr6-55935170-INS-191
chr6-55941996-INS-214
chr6-55971920-INS-1078
chr6-55972416-INS-59

chr6-55984802-INS-6327
chr6-55991001-INS-78
chr6-56009756-INS-50
chr6-56111868-INS-67
chr6-56170696-INS-68
chr6-56170735-INS-156
chr6-56171729-INS-50
chr6-56172651-INS-50
chr6-56294989-INS-7123
chr6-56351924-INS-81
chr6-56353087-INS-6346
chr6-56397836-INS-89
chr6-56481313-INS-75
chr6-56495635-INS-220
chr6-56514877-INS-130
chr6-56516351-INS-172
chr6-56526985-INS-130
chr6-56547452-INS-86
chr6-56586622-INS-1359
chr6-56588265-INS-454
chr6-56600389-INS-151
chr6-56600471-INS-50
chr6-56602723-INS-1624
chr6-56612657-INS-122
chr6-56662444-INS-205
chr6-56671402-INS-2269
chr6-56713296-INS-435
chr6-56715843-INS-111
chr6-56744816-INS-92
chr6-56760087-INS-205
chr6-56787401-INS-400
chr6-56787727-INS-795
chr6-56797367-INS-885
chr6-56803281-INS-52
chr6-56811458-INS-205
chr6-56839214-INS-2155
chr6-56909828-INS-4018
chr6-56910202-INS-1183
chr6-56910278-INS-7739
chr6-56917341-INS-4834
chr6-56922256-INS-3823
chr6-56922601-INS-221
chr6-56956801-INS-6779
chr6-57022053-INS-196
chr6-57039047-INS-320
chr6-57057028-INS-280
chr6-57057028-INS-82
chr6-57082036-INS-264
chr6-57102069-INS-52
chr6-57106536-INS-5139
chr6-57106907-INS-90
chr6-57107535-INS-172
chr6-57107553-INS-81
chr6-57120100-INS-6487
chr6-57159188-INS-6519
chr6-57183471-INS-74
chr6-57281420-INS-214
chr6-57333517-INS-6679
chr6-57360262-INS-216
chr6-57433044-INS-4126
chr6-57467321-INS-355
chr6-57471338-INS-117
chr6-57471618-INS-77
chr6-57472526-INS-15550
chr6-57643470-INS-54
chr6-57670851-INS-50
chr6-57751286-INS-72
chr6-57799974-INS-95
chr6-57800543-INS-74
chr6-57828155-INS-112
chr6-57828489-INS-199
chr6-57837172-INS-189
chr6-57842985-INS-1376
chr6-57859146-INS-56
chr6-57867187-INS-6374
chr6-57876874-INS-6182
chr6-57888993-INS-947

chr6-57896765-INS-214
chr6-57948051-INS-6232
chr6-57958513-INS-236
chr6-58000681-INS-52
chr6-58069652-INS-2500
chr6-58080017-INS-360
chr6-58098659-INS-3862
chr6-58110857-INS-3335
chr6-58147808-INS-116
chr6-58390613-INS-499
chr6-58445937-INS-1994
chr6-58481763-INS-1070
chr6-58491414-INS-1084
chr6-58507895-INS-1466
chr6-58524972-INS-60
chr6-58550186-INS-3634
chr6-58620979-INS-4372
chr6-58666318-INS-129
chr6-58673388-INS-6201
chr6-58699611-INS-159
chr6-58706775-INS-61
chr6-58709741-INS-242
chr6-58716293-INS-14235
chr6-58727182-INS-224
chr6-58760765-INS-50
chr6-58776039-INS-141
chr6-58792147-INS-508
chr6-58926243-INS-521
chr6-59004135-INS-1119
chr6-59040514-INS-584
chr6-59080732-INS-293
chr6-59097896-INS-500
chr6-59159327-INS-6354
chr6-59161142-INS-59
chr6-59193360-INS-1832
chr6-59195085-INS-672
chr6-59195855-INS-499
chr6-59202787-INS-1498
chr6-59294756-INS-7085
chr6-59306098-INS-6671
chr6-59337482-INS-2727
chr6-59373708-INS-1369
chr6-59373708-INS-9377
chr6-59432166-INS-81
chr6-59635723-INS-134
chr6-59689220-INS-53
chr6-59767901-INS-53
chr6-59771544-INS-4816
chr6-59800468-INS-62
chr6-59811423-INS-66
chr6-59829061-INS-404
chr6-59838185-INS-399
chr6-59848069-INS-97
chr6-59857645-INS-59
chr6-59874795-INS-64
chr6-59874934-INS-439
chr6-59896660-INS-52
chr6-59903500-INS-1746
chr6-59915188-INS-145
chr6-59918769-INS-88
chr6-59920093-INS-68
chr6-59930987-INS-3707
chr6-59935316-INS-57
chr6-59938300-INS-94
chr6-59943377-INS-498
chr6-59944380-INS-575
chr6-59947423-INS-85
chr6-59948790-INS-62
chr6-59955273-INS-54
chr6-59971660-INS-542
chr6-59975149-INS-191
chr6-59980064-INS-7289
chr6-59983104-INS-1833
chr6-59984086-INS-87
chr6-59984094-INS-180
chr6-59990236-INS-302
chr6-59998894-INS-51

chr6-60022796-INS-6551
chr6-60044430-INS-68
chr6-60045656-INS-1569
chr6-60129075-INS-142
chr6-60146924-INS-1050
chr6-60201627-INS-1196
chr6-60209276-INS-161
chr6-60214061-INS-549
chr6-60221460-INS-65
chr6-60223822-INS-400
chr6-60235790-INS-105
chr6-60317149-INS-2334
chr6-60326958-INS-75
chr6-60335240-INS-6165
chr6-60337861-INS-4937
chr6-60427538-INS-84
chr6-60457555-INS-903
chr6-60462177-INS-104
chr6-60481600-INS-80
chr6-60483386-INS-53
chr6-60507946-INS-4344
chr6-60579824-INS-362
chr6-60584530-INS-188
chr6-60625472-INS-6420
chr6-60639368-INS-8852
chr6-60655376-INS-514
chr6-60699799-INS-132
chr6-60802353-INS-1643
chr6-60818196-INS-11338
chr6-60870876-INS-2823
chr6-60871197-INS-217
chr6-60871529-INS-464
chr6-60899647-INS-72
chr6-60908056-INS-60
chr6-60919057-INS-50
chr6-60933407-INS-2030
chr6-60956679-INS-50
chr6-60971160-INS-150
chr6-60977038-INS-117
chr6-60982023-INS-3021
chr6-60983242-INS-6485
chr6-60985871-INS-2227
chr6-60998678-INS-50
chr6-61006722-INS-387
chr6-61016862-INS-467
chr6-61022468-INS-300
chr6-61092050-INS-174
chr6-61104166-INS-4420
chr6-61138390-INS-51
chr6-61148754-INS-120
chr6-61208062-INS-89
chr6-61224464-INS-7245
chr6-61230152-INS-1438
chr6-61241825-INS-390
chr6-61252155-INS-626
chr6-61252593-INS-6020
chr6-61341139-INS-6436
chr6-61349165-INS-54
chr6-61356110-INS-52
chr6-61392790-INS-194
chr6-61393420-INS-6379
chr6-61402320-INS-104
chr6-61407045-INS-434
chr6-61420081-INS-824
chr6-61449352-INS-6468
chr6-61501513-INS-3527
chr6-61614895-INS-94
chr6-61658789-INS-210
chr6-61697591-INS-421
chr6-61718872-INS-1551
chr6-61722636-INS-66
chr6-61726330-INS-68
chr6-61736246-INS-50
chr6-61791338-INS-226
chr6-61800141-INS-2498
chr6-61803364-INS-51
chr6-61824551-INS-50

chr6-61844625-INS-291
chr6-61849281-INS-73
chr6-61853753-INS-50
chr6-61891649-INS-52
chr6-61903627-INS-265
chr6-61936496-INS-50
chr6-61938885-INS-56
chr6-61939242-INS-50
chr6-61942102-INS-1110
chr6-61957016-INS-965
chr6-61969002-INS-329
chr6-61969341-INS-554
chr6-61982664-INS-52
chr6-61992429-INS-72
chr6-62002976-INS-168
chr6-62021399-INS-1312
chr6-62053614-INS-312
chr6-62088098-INS-491
chr6-62233945-INS-60
chr6-62234685-INS-98
chr6-62257515-INS-53
chr6-62286324-INS-595
chr6-62294059-INS-50
chr6-62295455-INS-56
chr6-62298380-INS-302
chr6-62317903-INS-6489
chr6-62332230-INS-75
chr6-62336414-INS-50
chr6-62340635-INS-6384
chr6-62353383-INS-852
chr6-62355417-INS-68
chr6-62374624-INS-390
chr6-62421196-INS-3893
chr6-62429791-INS-1787
chr6-62450701-INS-6341
chr6-62456570-INS-360
chr6-62456686-INS-52
chr6-62461605-INS-51
chr6-62472859-INS-6937
chr6-62504160-INS-1429
chr6-62531498-INS-52
chr6-62549604-INS-54
chr6-62596849-INS-1154
chr6-62601487-INS-50
chr6-62611293-INS-2307
chr6-62629720-INS-1609
chr6-62663622-INS-149
chr6-62724876-INS-138
chr6-62752599-INS-6387
chr6-62767643-INS-6628
chr6-62807667-INS-51
chr6-62854266-INS-175
chr6-62861966-INS-74
chr6-62863903-INS-2825
chr6-62895302-INS-400
chr6-62947769-INS-163
chr6-62958290-INS-449
chr6-62992373-INS-212
chr6-63069156-INS-437
chr6-63083772-INS-2046
chr6-63089924-INS-301
chr6-63107231-INS-188
chr6-63113001-INS-6792
chr6-63136775-INS-50
chr6-63157016-INS-4247
chr6-63223910-INS-96
chr6-63281583-INS-819
chr6-63345298-INS-69
chr6-63376516-INS-95
chr6-63391293-INS-53
chr6-63391627-INS-595
chr6-63404791-INS-512
chr6-63408742-INS-68
chr6-63411489-INS-84
chr6-63427861-INS-6108
chr6-63437084-INS-75
chr6-63441207-INS-5048

chr6-63485435-INS-820
chr6-63542828-INS-1570
chr6-63567017-INS-2858
chr6-63606671-INS-383
chr6-63607910-INS-153
chr6-63622330-INS-1852
chr6-63656402-INS-60
chr6-63665235-INS-3183
chr6-63737876-INS-7533
chr6-63823585-INS-59
chr6-63831151-INS-52
chr6-63871049-INS-118
chr6-63884741-INS-7560
chr6-63886859-INS-155
chr6-63886859-INS-60
chr6-63887068-INS-95
chr6-63897368-INS-50
chr6-63925817-INS-52
chr6-63955815-INS-83
chr6-63985086-INS-611
chr6-64003703-INS-164
chr6-64016382-INS-6366
chr6-64052068-INS-250
chr6-64059923-INS-114
chr6-64077959-INS-90
chr6-64104213-INS-124
chr6-64181506-INS-7400
chr6-64214710-INS-118
chr6-64235435-INS-6396
chr6-64236774-INS-398
chr6-64244563-INS-177
chr6-64289517-INS-1451
chr6-64307520-INS-72
chr6-64427671-INS-1490
chr6-64509618-INS-2361
chr6-64552103-INS-2779
chr6-64557192-INS-1181
chr6-64583284-INS-1000
chr6-64596988-INS-2113
chr6-64628286-INS-66
chr6-64657576-INS-96
chr6-64670251-INS-1172
chr6-64701520-INS-7245
chr6-64734236-INS-105
chr6-64734380-INS-141
chr6-64759346-INS-4107
chr6-64820167-INS-521
chr6-64832902-INS-668
chr6-64833737-INS-966
chr6-64834110-INS-9295
chr6-64849464-INS-204
chr6-64849807-INS-216
chr6-64849858-INS-1932
chr6-64850051-INS-174
chr6-64888445-INS-209
chr6-64923660-INS-120
chr6-64923714-INS-552
chr6-64925714-INS-116
chr6-64944437-INS-267
chr6-64944550-INS-1897
chr6-64945044-INS-92
chr6-64946244-INS-120
chr6-64950866-INS-82
chr6-64950866-INS-821
chr6-64951402-INS-54
chr6-64962725-INS-320
chr6-64977745-INS-187
chr6-64982064-INS-101
chr6-64987482-INS-52
chr6-64988077-INS-140
chr6-64988954-INS-106
chr6-65000729-INS-57
chr6-65006738-INS-189
chr6-65008222-INS-6718
chr6-65070610-INS-207
chr6-65117359-INS-52
chr6-65126057-INS-65

chr6-65177458-INS-1314
chr6-65243599-INS-1499
chr6-65244724-INS-86
chr6-65287238-INS-700
chr6-65298001-INS-9692
chr6-65339533-INS-494
chr6-65343429-INS-1427
chr6-65344740-INS-453
chr6-65355455-INS-1486
chr6-65464750-INS-82
chr6-65475627-INS-104
chr6-65494707-INS-96
chr6-65578821-INS-567
chr6-65580488-INS-62
chr6-65590438-INS-197
chr6-65628133-INS-141
chr6-65723496-INS-7087
chr6-65731356-INS-52
chr6-65789199-INS-191
chr6-65789944-INS-2064
chr6-65790045-INS-87
chr6-65790163-INS-353
chr6-65790367-INS-499
chr6-65802916-INS-52
chr6-65807578-INS-85
chr6-65882257-INS-2849
chr6-65906882-INS-51
chr6-65972490-INS-174
chr6-65998755-INS-2135
chr6-66010052-INS-6442
chr6-66097705-INS-1968
chr6-66100907-INS-1690
chr6-66104515-INS-4219
chr6-66111145-INS-1150
chr6-66113523-INS-805
chr6-66115383-INS-2467
chr6-66126352-INS-1715
chr6-66194283-INS-361
chr6-66265325-INS-1716
chr6-66308515-INS-180
chr6-66316802-INS-512
chr6-66325101-INS-4439
chr6-66378060-INS-90
chr6-66379890-INS-187
chr6-66550758-INS-6585
chr6-66642174-INS-485
chr6-66644122-INS-25554
chr6-66655272-INS-1334
chr6-66727985-INS-6762
chr6-66834390-INS-50
chr6-66864609-INS-68
chr6-66917784-INS-99
chr6-66947033-INS-81
chr6-66960229-INS-50
chr6-66960303-INS-150
chr6-66962278-INS-3249
chr6-67016280-INS-249
chr6-67043121-INS-184
chr6-67043684-INS-116
chr6-67094789-INS-84
chr6-67102509-INS-67
chr6-67184550-INS-159
chr6-67259477-INS-167
chr6-67284234-INS-74
chr6-67284888-INS-273
chr6-67345886-INS-193
chr6-67403698-INS-241
chr6-67464766-INS-291
chr6-67497131-INS-6224
chr6-67521861-INS-491
chr6-67536091-INS-537
chr6-67661394-INS-67
chr6-67664365-INS-509
chr6-67672709-INS-50
chr6-67672784-INS-328
chr6-67672919-INS-64
chr6-67673985-INS-88

chr6-67674827-INS-352
chr6-67699631-INS-63
chr6-67705417-INS-126
chr6-67705429-INS-1322
chr6-67709091-INS-91
chr6-67711046-INS-286
chr6-67711730-INS-105
chr6-67742259-INS-18010
chr6-67769583-INS-2328
chr6-67862313-INS-342
chr6-67874608-INS-50
chr6-67883704-INS-187
chr6-67883704-INS-55
chr6-67957526-INS-56
chr6-67967673-INS-2312
chr6-67980850-INS-5385
chr6-68027804-INS-64
chr6-68028043-INS-52
chr6-68049174-INS-2995
chr6-68050201-INS-2077
chr6-68055621-INS-12523
chr6-68060240-INS-611
chr6-68086728-INS-50
chr6-68145338-INS-2579
chr6-68156813-INS-57
chr6-68179433-INS-143
chr6-68245240-INS-151
chr6-68250026-INS-55
chr6-68259632-INS-58
chr6-68270968-INS-96
chr6-68282124-INS-94
chr6-68289701-INS-223
chr6-68290894-INS-52
chr6-68296214-INS-388
chr6-68299683-INS-386
chr6-68303802-INS-9040
chr6-68308390-INS-891
chr6-68313111-INS-66
chr6-68314502-INS-488
chr6-68315279-INS-68
chr6-68315280-INS-286
chr6-68318927-INS-191
chr6-68330301-INS-230
chr6-68330344-INS-785
chr6-68330364-INS-180
chr6-68370937-INS-831
chr6-68404519-INS-156
chr6-68409985-INS-175
chr6-68415537-INS-436
chr6-68421576-INS-461
chr6-68435872-INS-162
chr6-68444542-INS-138
chr6-68444845-INS-209
chr6-68445037-INS-2859
chr6-68463541-INS-7086
chr6-68475450-INS-2473
chr6-68478975-INS-3035
chr6-68483350-INS-7047
chr6-68496483-INS-5067
chr6-68505783-INS-6752
chr6-68522253-INS-52
chr6-68535841-INS-66
chr6-68553697-INS-6797
chr6-68555289-INS-74
chr6-68559364-INS-1413
chr6-68569292-INS-81
chr6-68579880-INS-1051
chr6-68602586-INS-52
chr6-68680605-INS-59
chr6-68703477-INS-81
chr6-68706259-INS-201
chr6-68706360-INS-441
chr6-68709718-INS-52
chr6-68796079-INS-94
chr6-68812686-INS-741
chr6-68832612-INS-326
chr6-68859994-INS-69

chr6-68886310-INS-443
chr6-68896475-INS-411
chr6-68959053-INS-473
chr6-69152793-INS-774
chr6-69153489-INS-179
chr6-69156549-INS-67
chr6-69166804-INS-566
chr6-69183286-INS-1222
chr6-69187813-INS-51
chr6-69290128-INS-1456
chr6-69300813-INS-50
chr6-69322617-INS-326
chr6-69332336-INS-734
chr6-69336736-INS-58
chr6-69374820-INS-1016
chr6-69478814-INS-2546
chr6-69478944-INS-958
chr6-69494899-INS-1598
chr6-69498587-INS-1025
chr6-69577563-INS-208
chr6-69577635-INS-200
chr6-69603939-INS-1481
chr6-69663492-INS-6662
chr6-69664545-INS-5107
chr6-69672126-INS-2366
chr6-69701769-INS-326
chr6-69706142-INS-69
chr6-69706234-INS-61
chr6-69712477-INS-3226
chr6-69723422-INS-1402
chr6-69734095-INS-52
chr6-69756285-INS-59
chr6-69820983-INS-456
chr6-69837289-INS-1244
chr6-69839578-INS-689
chr6-69844062-INS-739
chr6-69854137-INS-3127
chr6-69855516-INS-158
chr6-69891777-INS-358
chr6-69896647-INS-6107
chr6-69896958-INS-961
chr6-69901076-INS-118
chr6-69909732-INS-6459
chr6-69919665-INS-768
chr6-69938028-INS-2810
chr6-69948276-INS-398
chr6-69950956-INS-2751
chr6-69958349-INS-54
chr6-69959682-INS-86
chr6-69968839-INS-1147
chr6-70006043-INS-464
chr6-70024021-INS-293
chr6-70036778-INS-60
chr6-70039360-INS-749
chr6-70043010-INS-1778
chr6-70050245-INS-113
chr6-70052631-INS-1092
chr6-70080354-INS-1207
chr6-70084210-INS-53
chr6-70093424-INS-6170
chr6-70095819-INS-54
chr6-70096501-INS-58
chr6-70097331-INS-324
chr6-70104534-INS-2049
chr6-70109979-INS-62
chr6-70171346-INS-50
chr6-70186440-INS-419
chr6-70198836-INS-83
chr6-70199128-INS-94
chr6-70205289-INS-78
chr6-70207782-INS-152
chr6-70209351-INS-446
chr6-70209951-INS-52
chr6-70219974-INS-81
chr6-70253753-INS-3554
chr6-70273265-INS-2096
chr6-70338551-INS-3247

chr6-70366248-INS-51
chr6-70389808-INS-453
chr6-70395817-INS-4725
chr6-70397969-INS-116
chr6-70408458-INS-50
chr6-70454512-INS-199
chr6-70489381-INS-79
chr6-70615122-INS-198
chr6-70676595-INS-72
chr6-70692600-INS-50
chr6-70708475-INS-84
chr6-70726179-INS-1077
chr6-70765378-INS-528
chr6-70788171-INS-63
chr6-70815720-INS-76
chr6-70855770-INS-564
chr6-70858718-INS-56
chr6-70870438-INS-108
chr6-70964644-INS-222
chr6-70990981-INS-50
chr6-70993028-INS-6214
chr6-71007589-INS-4573
chr6-71008767-INS-121
chr6-71014491-INS-1890
chr6-71015338-INS-69
chr6-71015527-INS-394
chr6-71024638-INS-7288
chr6-71036433-INS-50
chr6-71037770-INS-84
chr6-71103924-INS-7263
chr6-71145963-INS-5903
chr6-71198466-INS-50
chr6-71210966-INS-202
chr6-71229516-INS-1797
chr6-71235118-INS-99
chr6-71242134-INS-60
chr6-71244107-INS-198
chr6-71267196-INS-64
chr6-71292779-INS-278
chr6-71336364-INS-74
chr6-71429117-INS-6083
chr6-71441067-INS-86
chr6-71449059-INS-7076
chr6-71498542-INS-56
chr6-71510225-INS-51
chr6-71613523-INS-197
chr6-71621775-INS-60
chr6-71628821-INS-6417
chr6-71629971-INS-473
chr6-71640557-INS-164
chr6-71640558-INS-74
chr6-71645992-INS-1063
chr6-71646913-INS-198
chr6-71703219-INS-68
chr6-71706649-INS-68
chr6-71743233-INS-2773
chr6-71774589-INS-92
chr6-71775333-INS-67
chr6-71810099-INS-52
chr6-71820718-INS-1226
chr6-71838784-INS-252
chr6-71839228-INS-117
chr6-71970431-INS-68
chr6-71984116-INS-264
chr6-71995374-INS-6551
chr6-72009332-INS-2816
chr6-72032336-INS-70
chr6-72047783-INS-7205
chr6-72060796-INS-167
chr6-72060826-INS-81
chr6-72067658-INS-153
chr6-72096092-INS-81
chr6-72133715-INS-2846
chr6-72162408-INS-58
chr6-72172368-INS-165
chr6-72226748-INS-249
chr6-72234745-INS-200

chr6-72241332-INS-112
chr6-72241368-INS-55
chr6-72256213-INS-75
chr6-72274375-INS-52
chr6-72276424-INS-202
chr6-72326961-INS-51
chr6-72335945-INS-103
chr6-72407723-INS-72
chr6-72407858-INS-212
chr6-72428966-INS-1195
chr6-72438641-INS-59
chr6-72440727-INS-210
chr6-72441121-INS-953
chr6-72458600-INS-194
chr6-72459734-INS-104
chr6-72465905-INS-857
chr6-72471991-INS-64
chr6-72514362-INS-90
chr6-72601346-INS-52
chr6-72602973-INS-473
chr6-72684336-INS-93
chr6-72703079-INS-1497
chr6-72704233-INS-124
chr6-72727791-INS-188
chr6-72728010-INS-92
chr6-72728030-INS-298
chr6-72735079-INS-237
chr6-72735079-INS-535
chr6-72735200-INS-60
chr6-72752342-INS-1996
chr6-72786193-INS-198
chr6-72804678-INS-64
chr6-72804773-INS-257
chr6-72806740-INS-199
chr6-72808179-INS-206
chr6-72878892-INS-5558
chr6-72889969-INS-5407
chr6-72909061-INS-60
chr6-72919192-INS-1770
chr6-72928444-INS-188
chr6-72955570-INS-50
chr6-72962596-INS-105
chr6-72971191-INS-50
chr6-72974247-INS-4416
chr6-72989347-INS-129
chr6-73027197-INS-170
chr6-73095955-INS-110
chr6-73095982-INS-224
chr6-73109042-INS-62
chr6-73110174-INS-56
chr6-73110409-INS-351
chr6-73110961-INS-13247
chr6-73141601-INS-712
chr6-73152703-INS-76
chr6-73168285-INS-756
chr6-73211682-INS-4853
chr6-73214492-INS-138
chr6-73271658-INS-437
chr6-73316510-INS-541
chr6-73321375-INS-552
chr6-73337723-INS-2450
chr6-73340189-INS-82
chr6-73347019-INS-313
chr6-73355610-INS-53
chr6-73365687-INS-232
chr6-73410193-INS-4724
chr6-73434657-INS-255
chr6-73453383-INS-84
chr6-73453391-INS-225
chr6-73454116-INS-217
chr6-73492378-INS-2699
chr6-73497215-INS-138
chr6-73516599-INS-60
chr6-73541856-INS-204
chr6-73550906-INS-84
chr6-73564381-INS-1594
chr6-73595371-INS-64

chr6-73595407-INS-184
chr6-73611915-INS-56
chr6-73625301-INS-73
chr6-73628299-INS-50
chr6-73640471-INS-63
chr6-73652903-INS-2586
chr6-73663061-INS-6261
chr6-74008155-INS-245
chr6-74008870-INS-1055
chr6-74018534-INS-62
chr6-74029522-INS-1378
chr6-74049452-INS-77
chr6-74056859-INS-2493
chr6-74057474-INS-50
chr6-74067524-INS-6941
chr6-74216802-INS-3244
chr6-74234542-INS-144
chr6-74237136-INS-55
chr6-74238458-INS-106
chr6-74238458-INS-240
chr6-74238493-INS-50
chr6-74290766-INS-6385
chr6-74307796-INS-6804
chr6-74327290-INS-2682
chr6-74351204-INS-55
chr6-74395671-INS-70
chr6-74454269-INS-154
chr6-74460889-INS-4619
chr6-74478361-INS-423
chr6-74492898-INS-117
chr6-74600624-INS-7123
chr6-74604983-INS-1117
chr6-74618135-INS-56
chr6-74625923-INS-5180
chr6-74763343-INS-312
chr6-74837344-INS-6361
chr6-75012806-INS-6564
chr6-75020776-INS-56
chr6-75047227-INS-3731
chr6-75061670-INS-51
chr6-75098122-INS-50
chr6-75157256-INS-645
chr6-75157339-INS-262
chr6-75157684-INS-150
chr6-75157763-INS-98
chr6-75187686-INS-6574
chr6-75196705-INS-3184
chr6-75201956-INS-1108
chr6-75202931-INS-50
chr6-75210267-INS-389
chr6-75210979-INS-66
chr6-75211672-INS-2410
chr6-75284037-INS-715
chr6-75299511-INS-50
chr6-75314122-INS-96
chr6-75316146-INS-164
chr6-75347058-INS-50
chr6-75382496-INS-50
chr6-75386473-INS-2904
chr6-75460769-INS-84
chr6-75475850-INS-2128
chr6-75485673-INS-60
chr6-75521402-INS-54
chr6-75533417-INS-1677
chr6-75580882-INS-56
chr6-75581645-INS-59
chr6-75585568-INS-78
chr6-75610486-INS-1400
chr6-75623920-INS-2833
chr6-75630156-INS-4790
chr6-75679686-INS-92
chr6-75730520-INS-1891
chr6-75741904-INS-6203
chr6-75757739-INS-50
chr6-75781426-INS-52
chr6-75819043-INS-1681
chr6-75821670-INS-80

chr6-75837105-INS-2591
chr6-75895945-INS-6215
chr6-75901852-INS-7092
chr6-75903594-INS-250
chr6-75940633-INS-53
chr6-75948047-INS-2549
chr6-75955014-INS-2813
chr6-75958239-INS-116
chr6-75979403-INS-6290
chr6-75995204-INS-58
chr6-75998394-INS-6360
chr6-76012572-INS-70
chr6-76023037-INS-6163
chr6-76027772-INS-2846
chr6-76028278-INS-759
chr6-76036954-INS-3763
chr6-76064215-INS-3794
chr6-76085985-INS-132
chr6-76086191-INS-98
chr6-76091126-INS-6505
chr6-76139277-INS-3072
chr6-76144844-INS-197
chr6-76158005-INS-1715
chr6-76183903-INS-52
chr6-76184401-INS-62
chr6-76186703-INS-6535
chr6-76191238-INS-1018
chr6-76202488-INS-63
chr6-76258719-INS-5795
chr6-76276640-INS-57
chr6-76282049-INS-2414
chr6-76292945-INS-1000
chr6-76296465-INS-6006
chr6-76299313-INS-144
chr6-76304939-INS-6523
chr6-76332298-INS-56
chr6-76350494-INS-52
chr6-76367354-INS-92
chr6-76373400-INS-7164
chr6-76446062-INS-2933
chr6-76461552-INS-971
chr6-76563816-INS-405
chr6-76592728-INS-6433
chr6-76779914-INS-52
chr6-76849171-INS-9187
chr6-76871912-INS-1434
chr6-76871913-INS-456
chr6-76886500-INS-710
chr6-76888304-INS-58
chr6-76898106-INS-544
chr6-76899309-INS-1890
chr6-76905245-INS-414
chr6-76947931-INS-51
chr6-76963419-INS-592
chr6-76963644-INS-1014
chr6-76963720-INS-60
chr6-77084659-INS-510
chr6-77104911-INS-106
chr6-77113424-INS-57
chr6-77120103-INS-6443
chr6-77164969-INS-54
chr6-77187115-INS-3420
chr6-77219192-INS-193
chr6-77235482-INS-117
chr6-77245151-INS-52
chr6-77300801-INS-62
chr6-77352611-INS-545
chr6-77393449-INS-158
chr6-77413659-INS-112
chr6-77524432-INS-276
chr6-77612791-INS-116
chr6-77696866-INS-586
chr6-77706546-INS-72
chr6-77747499-INS-55
chr6-77798154-INS-5499
chr6-77803306-INS-1351
chr6-77882704-INS-1046

chr6-77900971-INS-292
chr6-77948264-INS-1478
chr6-77982969-INS-640
chr6-77989709-INS-515
chr6-77990229-INS-210
chr6-78020591-INS-89
chr6-78152506-INS-56
chr6-78223850-INS-60
chr6-78258636-INS-51
chr6-78266223-INS-57
chr6-78275184-INS-5630
chr6-78289278-INS-3135
chr6-78356144-INS-1845
chr6-78358516-INS-56
chr6-78366376-INS-207
chr6-78413418-INS-4458
chr6-78450027-INS-1485
chr6-78450797-INS-1484
chr6-78489004-INS-53
chr6-78489364-INS-513
chr6-78500878-INS-6713
chr6-78545660-INS-100
chr6-78590891-INS-78
chr6-78593914-INS-3360
chr6-78637239-INS-4171
chr6-78704069-INS-1362
chr6-78737995-INS-118
chr6-78756525-INS-118
chr6-78841402-INS-207
chr6-79007524-INS-2983
chr6-79019272-INS-149
chr6-79022301-INS-1412
chr6-79047127-INS-1000
chr6-79129793-INS-1306
chr6-79147121-INS-6014
chr6-79154288-INS-60
chr6-79210603-INS-86
chr6-79258214-INS-1921
chr6-79314115-INS-292
chr6-79362832-INS-51
chr6-79373301-INS-54
chr6-79417783-INS-50
chr6-79564295-INS-64
chr6-79573489-INS-496
chr6-79581783-INS-4757
chr6-79582926-INS-54
chr6-79617010-INS-654
chr6-79623411-INS-96
chr6-79623458-INS-448
chr6-79629868-INS-356
chr6-79629878-INS-927
chr6-79630035-INS-461
chr6-79660380-INS-8218
chr6-79756247-INS-3216
chr6-79756757-INS-59
chr6-79779336-INS-7168
chr6-79798298-INS-168
chr6-79875848-INS-57
chr6-79898425-INS-50
chr6-79906891-INS-50
chr6-79973599-INS-50
chr6-79986365-INS-208
chr6-79996912-INS-51
chr6-80118666-INS-63
chr6-80141123-INS-50
chr6-80143905-INS-62
chr6-80144025-INS-237
chr6-80255907-INS-92
chr6-80297533-INS-50
chr6-80394834-INS-488
chr6-80400113-INS-510
chr6-80403607-INS-56
chr6-80415800-INS-56
chr6-80469505-INS-513
chr6-80499833-INS-204
chr6-80554160-INS-85
chr6-80567419-INS-78

chr6-80580288-INS-82
chr6-80647347-INS-1882
chr6-80689912-INS-50
chr6-80699881-INS-665
chr6-80728884-INS-63
chr6-80731025-INS-3304
chr6-80790982-INS-145
chr6-80791028-INS-68
chr6-80827682-INS-66
chr6-80869535-INS-7170
chr6-80875825-INS-583
chr6-80956393-INS-128
chr6-80956569-INS-50
chr6-80973050-INS-78
chr6-80976953-INS-50
chr6-81036244-INS-248
chr6-81071961-INS-631
chr6-81073768-INS-140
chr6-81081789-INS-133
chr6-81081873-INS-64
chr6-81177783-INS-58
chr6-81229164-INS-148
chr6-81271117-INS-79
chr6-81277274-INS-274
chr6-81318258-INS-6309
chr6-81323900-INS-855
chr6-81391401-INS-68
chr6-81615039-INS-58
chr6-81634900-INS-6243
chr6-81679745-INS-283
chr6-81679847-INS-98
chr6-81712310-INS-6692
chr6-81723578-INS-124
chr6-81734749-INS-134
chr6-81747751-INS-8413
chr6-81755359-INS-250
chr6-81759021-INS-96
chr6-81773893-INS-6491
chr6-81795578-INS-232
chr6-81795595-INS-78
chr6-81795819-INS-110
chr6-81795819-INS-273
chr6-81819454-INS-1498
chr6-81861095-INS-66
chr6-81896760-INS-311
chr6-81944813-INS-618
chr6-81951228-INS-1830
chr6-81970443-INS-54
chr6-82052563-INS-50
chr6-82088089-INS-50
chr6-82100672-INS-56
chr6-82102873-INS-1023
chr6-82105485-INS-1956
chr6-82111773-INS-229
chr6-82183665-INS-50
chr6-82206899-INS-303
chr6-82208399-INS-60
chr6-82208964-INS-54
chr6-82246582-INS-65
chr6-82355346-INS-139
chr6-82367335-INS-5288
chr6-82412653-INS-99
chr6-82413212-INS-59
chr6-82482426-INS-6199
chr6-82493585-INS-6871
chr6-82497482-INS-58
chr6-82540234-INS-1142
chr6-82541407-INS-6475
chr6-82586197-INS-64
chr6-82663667-INS-58
chr6-82666621-INS-67
chr6-82666978-INS-3222
chr6-82718377-INS-157
chr6-82726483-INS-53
chr6-82757181-INS-92
chr6-82765051-INS-54
chr6-82793506-INS-557

chr6-82805756-INS-86
chr6-82808025-INS-50
chr6-82813931-INS-50
chr6-82826474-INS-50
chr6-82853033-INS-154
chr6-82866416-INS-170
chr6-82879998-INS-50
chr6-82901357-INS-162
chr6-82902629-INS-295
chr6-82950003-INS-58
chr6-82950469-INS-76
chr6-82989298-INS-7095
chr6-82996032-INS-50
chr6-83117771-INS-180
chr6-83127327-INS-58
chr6-83128152-INS-60
chr6-83132856-INS-205
chr6-83132856-INS-54
chr6-83150820-INS-60
chr6-83216797-INS-77
chr6-83224997-INS-264
chr6-83227785-INS-3117
chr6-83232342-INS-90
chr6-83246743-INS-319
chr6-83246765-INS-142
chr6-83259727-INS-60
chr6-83259752-INS-268
chr6-83284150-INS-208
chr6-83285767-INS-100
chr6-83316197-INS-56
chr6-83378356-INS-353
chr6-83378510-INS-85
chr6-83413496-INS-120
chr6-83413563-INS-265
chr6-83438827-INS-82
chr6-83463385-INS-86
chr6-83472086-INS-50
chr6-83508300-INS-178
chr6-83508592-INS-154
chr6-83508642-INS-66
chr6-83553358-INS-142
chr6-83582369-INS-826
chr6-83582434-INS-159
chr6-83625434-INS-294
chr6-83639746-INS-1278
chr6-83660629-INS-55
chr6-83665268-INS-208
chr6-83677619-INS-53
chr6-83723299-INS-984
chr6-83767720-INS-7120
chr6-83793627-INS-242
chr6-83833603-INS-52
chr6-83852775-INS-357
chr6-83856593-INS-86
chr6-83974545-INS-139
chr6-84019379-INS-50
chr6-84044083-INS-2805
chr6-84105434-INS-227
chr6-84135976-INS-745
chr6-84140383-INS-96
chr6-84166118-INS-78
chr6-84166818-INS-90
chr6-84191729-INS-93
chr6-84209923-INS-124
chr6-84210284-INS-58
chr6-84222186-INS-66
chr6-84222204-INS-198
chr6-84229415-INS-50
chr6-84238885-INS-58
chr6-84240567-INS-320
chr6-84264062-INS-138
chr6-84295328-INS-82
chr6-84303213-INS-60
chr6-84323517-INS-225
chr6-84352106-INS-68
chr6-84365394-INS-123
chr6-84370035-INS-55

chr6-84398239-INS-61
chr6-84433503-INS-51
chr6-84434010-INS-223
chr6-84447598-INS-56
chr6-84448385-INS-98
chr6-84454955-INS-50
chr6-84488696-INS-50
chr6-84491610-INS-283
chr6-84501624-INS-64
chr6-84577426-INS-53
chr6-84589756-INS-51
chr6-84661450-INS-1197
chr6-84680544-INS-2499
chr6-84700545-INS-113
chr6-84794630-INS-1567
chr6-84798412-INS-6481
chr6-84826930-INS-630
chr6-84829393-INS-278
chr6-84840334-INS-1343
chr6-84855842-INS-235
chr6-84904926-INS-53
chr6-84956802-INS-76
chr6-84990387-INS-74
chr6-85031603-INS-182
chr6-85048478-INS-322
chr6-85049093-INS-79
chr6-85061942-INS-498
chr6-85080952-INS-213
chr6-85118073-INS-137
chr6-85118311-INS-56
chr6-85118676-INS-232
chr6-85145032-INS-65
chr6-85373221-INS-63
chr6-85381012-INS-168
chr6-85483323-INS-4375
chr6-85533397-INS-58
chr6-85603627-INS-50
chr6-85772510-INS-57
chr6-85839426-INS-228
chr6-85889131-INS-572
chr6-86012619-INS-106
chr6-86021390-INS-54
chr6-86031306-INS-7181
chr6-86116880-INS-56
chr6-86117121-INS-89
chr6-86136013-INS-85
chr6-86140075-INS-63
chr6-86140110-INS-762
chr6-86140257-INS-334
chr6-86140725-INS-85
chr6-86152685-INS-1530
chr6-86154875-INS-2093
chr6-86160373-INS-166
chr6-86165253-INS-1356
chr6-86193918-INS-76
chr6-86259243-INS-57
chr6-86265646-INS-4748
chr6-86298911-INS-55
chr6-86301959-INS-527
chr6-86308923-INS-1677
chr6-86308923-INS-3771
chr6-86331568-INS-56
chr6-86340204-INS-62
chr6-86423993-INS-346
chr6-86489210-INS-274
chr6-86490142-INS-209
chr6-86574405-INS-52
chr6-86593024-INS-432
chr6-86620427-INS-78
chr6-86685167-INS-1633
chr6-86728614-INS-199
chr6-86733984-INS-484
chr6-86745015-INS-64
chr6-86753411-INS-50
chr6-86783870-INS-86
chr6-86784451-INS-57
chr6-86786335-INS-102

chr6-86794014-INS-132
chr6-86816199-INS-5313
chr6-86860480-INS-268
chr6-86992892-INS-216
chr6-86993274-INS-897
chr6-87007451-INS-4641
chr6-87011241-INS-929
chr6-87025288-INS-92
chr6-87025840-INS-156
chr6-87080361-INS-67
chr6-87080868-INS-72
chr6-87080974-INS-398
chr6-87096475-INS-172
chr6-87115415-INS-62
chr6-87124760-INS-60
chr6-87185080-INS-70
chr6-87228835-INS-183
chr6-87336257-INS-87
chr6-87375226-INS-58
chr6-87377643-INS-74
chr6-87434833-INS-196
chr6-87450933-INS-550
chr6-87451021-INS-100
chr6-87503200-INS-64
chr6-87599431-INS-50
chr6-87611894-INS-52
chr6-87612033-INS-74
chr6-87632084-INS-50
chr6-87650844-INS-361
chr6-87664805-INS-139
chr6-87668831-INS-279
chr6-87685634-INS-116
chr6-87723452-INS-54
chr6-87732889-INS-96
chr6-87744969-INS-69
chr6-87744982-INS-152
chr6-87752681-INS-2566
chr6-87783075-INS-203
chr6-87802215-INS-224
chr6-87810472-INS-147
chr6-87862720-INS-3520
chr6-87904579-INS-183
chr6-87906007-INS-214
chr6-87918511-INS-202
chr6-87974569-INS-212
chr6-88031736-INS-64
chr6-88032095-INS-292
chr6-88034856-INS-1497
chr6-88112169-INS-52
chr6-88112321-INS-142
chr6-88115917-INS-180
chr6-88116715-INS-152
chr6-88164916-INS-91
chr6-88216089-INS-63
chr6-88338072-INS-50
chr6-88407776-INS-243
chr6-88420933-INS-1638
chr6-88448733-INS-228
chr6-88499964-INS-82
chr6-88506875-INS-66
chr6-88510665-INS-1158
chr6-88564880-INS-997
chr6-88591154-INS-50
chr6-88674299-INS-400
chr6-88705855-INS-50
chr6-88723669-INS-78
chr6-88791521-INS-11501
chr6-88792162-INS-1477
chr6-88830494-INS-52
chr6-88845929-INS-204
chr6-88846057-INS-88
chr6-88968626-INS-210
chr6-88968626-INS-50
chr6-88969656-INS-93
chr6-88972450-INS-148
chr6-88972795-INS-60
chr6-88981428-INS-291

chr6-88981481-INS-97
chr6-89045784-INS-62
chr6-89055683-INS-417
chr6-89058386-INS-72
chr6-89069079-INS-54
chr6-89073635-INS-54
chr6-89073719-INS-264
chr6-89142069-INS-56
chr6-89142075-INS-357
chr6-89142162-INS-57
chr6-89142210-INS-164
chr6-89232802-INS-62
chr6-89254172-INS-168
chr6-89567623-INS-59
chr6-89599719-INS-205
chr6-89630265-INS-109
chr6-89657878-INS-416
chr6-89709488-INS-3153
chr6-89824394-INS-2580
chr6-89856170-INS-147
chr6-89864517-INS-77
chr6-89930460-INS-1919
chr6-89945042-INS-63
chr6-89980208-INS-2499
chr6-89996058-INS-52
chr6-90112500-INS-6442
chr6-90186654-INS-123
chr6-90298049-INS-118
chr6-90466099-INS-180
chr6-90494265-INS-2010
chr6-90519933-INS-50
chr6-90522638-INS-127
chr6-90550913-INS-120
chr6-90635733-INS-246
chr6-90824253-INS-123
chr6-90841261-INS-144
chr6-90880874-INS-6455
chr6-91010914-INS-149
chr6-91189343-INS-60
chr6-91282177-INS-110
chr6-91363273-INS-52
chr6-91514338-INS-144
chr6-91574987-INS-53
chr6-91586553-INS-64
chr6-91723634-INS-150
chr6-91771633-INS-50
chr6-91783669-INS-54
chr6-91793582-INS-489
chr6-91797521-INS-58
chr6-91885073-INS-312
chr6-91886946-INS-82
chr6-91890722-INS-52
chr6-92108082-INS-50
chr6-92163010-INS-6121
chr6-92196058-INS-50
chr6-92203618-INS-224
chr6-92223852-INS-68
chr6-92224077-INS-204
chr6-92243557-INS-212
chr6-92260421-INS-7084
chr6-92265682-INS-70
chr6-92274268-INS-76
chr6-92325613-INS-3649
chr6-92346217-INS-115
chr6-92419798-INS-498
chr6-92440395-INS-50
chr6-92452670-INS-64
chr6-92603359-INS-6902
chr6-92618431-INS-6569
chr6-92640708-INS-50
chr6-92725218-INS-54
chr6-92726216-INS-6443
chr6-92742761-INS-58
chr6-92751736-INS-52
chr6-92759278-INS-597
chr6-92847253-INS-63
chr6-92856318-INS-56

chr6-92862382-INS-54
chr6-92863075-INS-62
chr6-92895539-INS-304
chr6-92897866-INS-52
chr6-92926268-INS-676
chr6-92932696-INS-219
chr6-92937523-INS-472
chr6-92951493-INS-52
chr6-92952101-INS-214
chr6-93014361-INS-293
chr6-93033192-INS-7055
chr6-93073980-INS-6847
chr6-93100429-INS-121
chr6-93101981-INS-78
chr6-93138549-INS-115
chr6-93158179-INS-72
chr6-93165218-INS-4533
chr6-93211234-INS-136
chr6-93376111-INS-6847
chr6-93386127-INS-286
chr6-93449520-INS-1025
chr6-93489630-INS-80
chr6-93518339-INS-72
chr6-93521918-INS-11221
chr6-93547270-INS-56
chr6-93547501-INS-243
chr6-93600279-INS-52
chr6-93603689-INS-52
chr6-93669197-INS-67
chr6-93782096-INS-50
chr6-93812566-INS-1175
chr6-93841746-INS-50
chr6-93846451-INS-52
chr6-93860628-INS-516
chr6-93982958-INS-58
chr6-93985159-INS-1602
chr6-93993092-INS-6335
chr6-93993321-INS-96
chr6-93995522-INS-54
chr6-94015621-INS-52
chr6-94031878-INS-126
chr6-94059116-INS-92
chr6-94087795-INS-65
chr6-94152506-INS-51
chr6-94153062-INS-195
chr6-94231110-INS-96
chr6-94262400-INS-51
chr6-94297533-INS-148
chr6-94297810-INS-362
chr6-94306894-INS-93
chr6-94323175-INS-7375
chr6-94332080-INS-64
chr6-94336562-INS-132
chr6-94343328-INS-131
chr6-94343956-INS-52
chr6-94351894-INS-1218
chr6-94373515-INS-6502
chr6-94375178-INS-2824
chr6-94378905-INS-2453
chr6-94415926-INS-50
chr6-94452288-INS-2559
chr6-94460687-INS-52
chr6-94486017-INS-51
chr6-94490073-INS-1879
chr6-94501138-INS-2906
chr6-94502511-INS-50
chr6-94521710-INS-50
chr6-94540104-INS-7116
chr6-94556139-INS-56
chr6-94567635-INS-464
chr6-94567635-INS-87
chr6-94568179-INS-102
chr6-94685716-INS-168
chr6-94694000-INS-76
chr6-94743371-INS-140
chr6-94780362-INS-1714
chr6-94792733-INS-1416

chr6-94800787-INS-54
chr6-94817365-INS-220
chr6-94818831-INS-7056
chr6-94827101-INS-90
chr6-94832882-INS-138
chr6-94846128-INS-116
chr6-94865599-INS-2002
chr6-94906452-INS-130
chr6-94916194-INS-1298
chr6-94920124-INS-50
chr6-94924724-INS-839
chr6-94958032-INS-71
chr6-94958080-INS-235
chr6-94997768-INS-52
chr6-95032989-INS-64
chr6-95073426-INS-1417
chr6-95074058-INS-1749
chr6-95083929-INS-380
chr6-95087663-INS-2217
chr6-95171387-INS-1314
chr6-95183155-INS-352
chr6-95201132-INS-72
chr6-95216662-INS-127
chr6-95216887-INS-121
chr6-95217910-INS-2437
chr6-95251957-INS-62
chr6-95252040-INS-175
chr6-95355894-INS-56
chr6-95362559-INS-7085
chr6-95367388-INS-86
chr6-95429016-INS-62
chr6-95432893-INS-2767
chr6-95462096-INS-50
chr6-95470786-INS-120
chr6-95476249-INS-200
chr6-95565641-INS-51
chr6-95603344-INS-78
chr6-95616315-INS-301
chr6-95665212-INS-5440
chr6-95676005-INS-618
chr6-95677482-INS-55
chr6-95724276-INS-373
chr6-95729043-INS-55
chr6-95753040-INS-6455
chr6-95756975-INS-248
chr6-95766797-INS-51
chr6-95834596-INS-57
chr6-95866440-INS-78
chr6-95921455-INS-51
chr6-96010128-INS-2947
chr6-96015840-INS-6588
chr6-96019831-INS-132
chr6-96044850-INS-140
chr6-96047193-INS-52
chr6-96167356-INS-6452
chr6-96171900-INS-60
chr6-96253589-INS-181
chr6-96273936-INS-848
chr6-96304629-INS-6606
chr6-96305539-INS-780
chr6-96314905-INS-58
chr6-96334017-INS-596
chr6-96364418-INS-216
chr6-96393460-INS-208
chr6-96400034-INS-135
chr6-96433997-INS-52
chr6-96462813-INS-59
chr6-96464591-INS-343
chr6-96530739-INS-78
chr6-96537529-INS-1066
chr6-96541743-INS-525
chr6-96553839-INS-165
chr6-96593300-INS-132
chr6-96665889-INS-214
chr6-96666523-INS-6478
chr6-96697815-INS-54
chr6-96715980-INS-287

chr6-96779854-INS-85
chr6-96824273-INS-556
chr6-96923844-INS-130
chr6-96923964-INS-84
chr6-96946990-INS-212
chr6-96955182-INS-180
chr6-96967716-INS-69
chr6-96967802-INS-156
chr6-96968498-INS-6499
chr6-96987981-INS-576
chr6-97015809-INS-62
chr6-97026030-INS-68
chr6-97026924-INS-222
chr6-97053782-INS-150
chr6-97085162-INS-97
chr6-97085253-INS-613
chr6-97139341-INS-259
chr6-97151761-INS-75
chr6-97249135-INS-105
chr6-97251792-INS-118
chr6-97260017-INS-70
chr6-97278989-INS-62
chr6-97343433-INS-279
chr6-97450724-INS-180
chr6-97452831-INS-461
chr6-97480826-INS-56
chr6-97481667-INS-6635
chr6-97488631-INS-105
chr6-97527827-INS-54
chr6-97545507-INS-571
chr6-97545852-INS-62
chr6-97560540-INS-58
chr6-97562420-INS-330
chr6-97665676-INS-333
chr6-97703669-INS-206
chr6-97742419-INS-573
chr6-97743803-INS-161
chr6-97753036-INS-62
chr6-97851886-INS-399
chr6-97909055-INS-6822
chr6-97910586-INS-52
chr6-97948028-INS-361
chr6-97961374-INS-51
chr6-97975582-INS-75
chr6-98006376-INS-4111
chr6-98040123-INS-68
chr6-98041546-INS-742
chr6-98092691-INS-54
chr6-98113213-INS-134
chr6-98118227-INS-140
chr6-98118890-INS-76
chr6-98193191-INS-72
chr6-98196881-INS-93
chr6-98205244-INS-100
chr6-98222872-INS-190
chr6-98244499-INS-60
chr6-98260724-INS-111
chr6-98412264-INS-80
chr6-98437348-INS-81
chr6-98437595-INS-276
chr6-98575393-INS-6670
chr6-98599640-INS-255
chr6-98625236-INS-140
chr6-98625263-INS-64
chr6-98672899-INS-410
chr6-98708352-INS-54
chr6-98712016-INS-168
chr6-98748191-INS-523
chr6-98763436-INS-52
chr6-98792527-INS-392
chr6-98810112-INS-51
chr6-98833215-INS-743
chr6-98869768-INS-405
chr6-98886539-INS-56
chr6-98910711-INS-247
chr6-98910711-INS-57
chr6-98942740-INS-64

chr6-98970159-INS-74
chr6-98970621-INS-52
chr6-99032405-INS-99
chr6-99231649-INS-204
chr6-99233904-INS-52
chr6-99255170-INS-130
chr6-99319905-INS-217
chr6-99320083-INS-51
chr6-99412717-INS-50
chr6-99423138-INS-51
chr6-99513812-INS-76
chr6-99525639-INS-58
chr6-99542128-INS-52
chr6-99611822-INS-274
chr6-99615814-INS-51
chr6-99657347-INS-72
chr6-99720990-INS-2498
chr6-99746267-INS-128
chr6-99766994-INS-298
chr6-99768008-INS-59
chr6-99783641-INS-68
chr6-99786411-INS-999
chr6-99786923-INS-132
chr6-99787843-INS-60
chr6-99825042-INS-378
chr6-99826946-INS-1820
chr6-99837912-INS-222
chr6-99881371-INS-111
chr6-99942815-INS-476
chr6-99961331-INS-54
chr6-99999942-INS-54
chr6-100035294-INS-59
chr6-100054192-INS-52
chr6-100065477-INS-52
chr6-100145187-INS-130
chr6-100162190-INS-87
chr6-100169062-INS-98
chr6-100177978-INS-80
chr6-100200808-INS-221
chr6-100271316-INS-138
chr6-100321622-INS-58
chr6-100327004-INS-199
chr6-100341860-INS-60
chr6-100388661-INS-51
chr6-100400815-INS-162
chr6-100402918-INS-204
chr6-100411960-INS-412
chr6-100413321-INS-92
chr6-100430748-INS-768
chr6-100430787-INS-153
chr6-100438174-INS-96
chr6-100440157-INS-208
chr6-100440985-INS-63
chr6-100441568-INS-1450
chr6-100444707-INS-261
chr6-100506447-INS-55
chr6-100556232-INS-160
chr6-100587759-INS-390
chr6-100587759-INS-826
chr6-100602258-INS-66
chr6-100612687-INS-6325
chr6-100612848-INS-52
chr6-100618538-INS-4830
chr6-100653439-INS-62
chr6-100672171-INS-1779
chr6-100690883-INS-50
chr6-100706530-INS-82
chr6-100738810-INS-2967
chr6-100815836-INS-100
chr6-100820705-INS-175
chr6-100879739-INS-172
chr6-100891074-INS-459
chr6-100891978-INS-2556
chr6-100905301-INS-52
chr6-100928611-INS-3951
chr6-100932098-INS-77
chr6-100940082-INS-69

chr6-100946608-INS-2535
chr6-101000191-INS-102
chr6-101000433-INS-162
chr6-101017870-INS-1873
chr6-101030669-INS-54
chr6-101046183-INS-253
chr6-101051215-INS-118
chr6-101061122-INS-227
chr6-101061122-INS-62
chr6-101078246-INS-416
chr6-101091329-INS-1865
chr6-101104471-INS-54
chr6-101140601-INS-52
chr6-101188713-INS-403
chr6-101223437-INS-231
chr6-101368671-INS-97
chr6-101383095-INS-81
chr6-101414981-INS-54
chr6-101437329-INS-311
chr6-101437333-INS-95
chr6-101446740-INS-52
chr6-101448128-INS-63
chr6-101448503-INS-102
chr6-101479088-INS-69
chr6-101480503-INS-1725
chr6-101481790-INS-63
chr6-101484569-INS-6239
chr6-101488153-INS-6699
chr6-101491195-INS-50
chr6-101513184-INS-688
chr6-101543801-INS-1303
chr6-101597584-INS-2813
chr6-101599633-INS-1452
chr6-101617503-INS-3503
chr6-101670295-INS-85
chr6-101682218-INS-50
chr6-101695013-INS-7108
chr6-101705278-INS-513
chr6-101759053-INS-16026
chr6-101777862-INS-504
chr6-101781182-INS-2997
chr6-101796252-INS-3370
chr6-101802198-INS-200
chr6-101802927-INS-112
chr6-101808047-INS-50
chr6-101808663-INS-6899
chr6-101831790-INS-171
chr6-101843905-INS-1148
chr6-101854671-INS-105
chr6-101893847-INS-551
chr6-101896157-INS-5875
chr6-101959294-INS-114
chr6-101997728-INS-51
chr6-102009964-INS-60
chr6-102024832-INS-52
chr6-102031676-INS-1246
chr6-102091129-INS-144
chr6-102117317-INS-4023
chr6-102214002-INS-2598
chr6-102255962-INS-84
chr6-102257385-INS-89
chr6-102257615-INS-51
chr6-102260931-INS-69
chr6-102317582-INS-58
chr6-102323841-INS-6159
chr6-102350747-INS-170
chr6-102405024-INS-52
chr6-102426491-INS-50
chr6-102449030-INS-1572
chr6-102510171-INS-74
chr6-102511980-INS-240
chr6-102515579-INS-52
chr6-102521029-INS-5269
chr6-102595831-INS-352
chr6-102671227-INS-50
chr6-102686554-INS-153
chr6-102720804-INS-8406

chr6-102727877-INS-270
chr6-102738665-INS-85
chr6-102738929-INS-229
chr6-102831381-INS-50
chr6-102870564-INS-6329
chr6-102878118-INS-6591
chr6-102879153-INS-438
chr6-102938928-INS-76
chr6-102939312-INS-357
chr6-102952620-INS-2682
chr6-102959470-INS-1222
chr6-102988916-INS-7085
chr6-102994814-INS-50
chr6-103016935-INS-51
chr6-103018118-INS-2930
chr6-103026279-INS-54
chr6-103049140-INS-60
chr6-103062785-INS-115
chr6-103066395-INS-138
chr6-103084217-INS-9917
chr6-103127438-INS-420
chr6-103128106-INS-2560
chr6-103231877-INS-50
chr6-103253269-INS-1404
chr6-103319997-INS-60
chr6-103331487-INS-75
chr6-103346936-INS-1745
chr6-103350883-INS-50
chr6-103388315-INS-18319
chr6-103392160-INS-50
chr6-103393324-INS-108
chr6-103416081-INS-60
chr6-103472833-INS-70
chr6-103479246-INS-188
chr6-103479922-INS-1294
chr6-103482258-INS-96
chr6-103496633-INS-50
chr6-103506060-INS-1311
chr6-103508474-INS-6157
chr6-103590368-INS-1094
chr6-103630167-INS-155
chr6-103630647-INS-55
chr6-103632023-INS-78
chr6-103649373-INS-4740
chr6-103652953-INS-88
chr6-103659926-INS-50
chr6-103777770-INS-205
chr6-103808992-INS-975
chr6-103831516-INS-278
chr6-103866067-INS-167
chr6-103876204-INS-50
chr6-103876759-INS-135
chr6-103902362-INS-6061
chr6-103906222-INS-172
chr6-103926352-INS-68
chr6-103975251-INS-50
chr6-103998417-INS-64
chr6-103998921-INS-56
chr6-104020089-INS-307
chr6-104029677-INS-52
chr6-104043881-INS-2839
chr6-104119999-INS-155
chr6-104158930-INS-56
chr6-104160742-INS-62
chr6-104210753-INS-682
chr6-104210753-INS-92
chr6-104232570-INS-138
chr6-104310096-INS-3729
chr6-104315583-INS-9357
chr6-104323781-INS-1083
chr6-104327958-INS-147
chr6-104344871-INS-105
chr6-104506479-INS-50
chr6-104508419-INS-53
chr6-104591477-INS-297
chr6-104623045-INS-1969
chr6-104627824-INS-52

chr6-104778546-INS-1227
chr6-104828160-INS-87
chr6-104828224-INS-222
chr6-104828249-INS-105
chr6-104888487-INS-1812
chr6-104945432-INS-155
chr6-104946221-INS-54
chr6-104953285-INS-54
chr6-104965045-INS-182
chr6-104965046-INS-52
chr6-104969608-INS-314
chr6-105028099-INS-2595
chr6-105044020-INS-2476
chr6-105084787-INS-55
chr6-105118167-INS-1624
chr6-105118627-INS-6499
chr6-105157502-INS-54
chr6-105213576-INS-107
chr6-105325576-INS-153
chr6-105400262-INS-2217
chr6-105473593-INS-110
chr6-105474193-INS-70
chr6-105474465-INS-59
chr6-105474528-INS-120
chr6-105487130-INS-7168
chr6-105505476-INS-66
chr6-105506098-INS-84
chr6-105520369-INS-50
chr6-105585226-INS-50
chr6-105595447-INS-490
chr6-105629637-INS-199
chr6-105747719-INS-153
chr6-105774639-INS-185
chr6-105795803-INS-6424
chr6-105829462-INS-6463
chr6-105920390-INS-102
chr6-105921501-INS-64
chr6-106049049-INS-77
chr6-106127835-INS-622
chr6-106136641-INS-90
chr6-106206852-INS-54
chr6-106357477-INS-522
chr6-106358300-INS-56
chr6-106382584-INS-265
chr6-106382856-INS-68
chr6-106384997-INS-103
chr6-106433448-INS-757
chr6-106454274-INS-169
chr6-106495908-INS-460
chr6-106496061-INS-50
chr6-106509444-INS-6290
chr6-106509590-INS-114
chr6-106525953-INS-355
chr6-106546815-INS-194
chr6-106553418-INS-50
chr6-106612510-INS-1557
chr6-106624952-INS-52
chr6-106625186-INS-66
chr6-106625337-INS-198
chr6-106742280-INS-7080
chr6-106800519-INS-55
chr6-106827956-INS-1379
chr6-106883583-INS-1375
chr6-106897294-INS-1527
chr6-106958160-INS-4676
chr6-106969254-INS-51
chr6-106969934-INS-212
chr6-106971916-INS-50
chr6-107036785-INS-59
chr6-107053147-INS-7112
chr6-107109986-INS-352
chr6-107110609-INS-109
chr6-107130103-INS-1175
chr6-107130103-INS-7611
chr6-107171756-INS-6484
chr6-107304922-INS-149
chr6-107364752-INS-246

chr6-107366549-INS-60
chr6-107405412-INS-58
chr6-107412205-INS-2395
chr6-107454700-INS-84
chr6-107487495-INS-67
chr6-107516351-INS-264
chr6-107575399-INS-51
chr6-107625254-INS-527
chr6-107666998-INS-727
chr6-107672598-INS-357
chr6-107678476-INS-50
chr6-107715401-INS-52
chr6-107720131-INS-6432
chr6-107741447-INS-200
chr6-107742485-INS-6660
chr6-107744526-INS-55
chr6-107787629-INS-1697
chr6-107799338-INS-1164
chr6-107822244-INS-2983
chr6-107823263-INS-52
chr6-107864956-INS-504
chr6-107872972-INS-146
chr6-107899873-INS-6568
chr6-107912734-INS-7171
chr6-107914310-INS-89
chr6-107915277-INS-2003
chr6-107957246-INS-1179
chr6-107958768-INS-87
chr6-107995252-INS-6971
chr6-107998361-INS-139
chr6-108000131-INS-58
chr6-108007430-INS-55
chr6-108040335-INS-507
chr6-108059335-INS-2147
chr6-108074417-INS-202
chr6-108076109-INS-74
chr6-108104614-INS-538
chr6-108104650-INS-1493
chr6-108110133-INS-59
chr6-108111083-INS-69
chr6-108264889-INS-2544
chr6-108336287-INS-315
chr6-108395981-INS-172
chr6-108536567-INS-56
chr6-108553353-INS-60
chr6-108581358-INS-365
chr6-108581433-INS-749
chr6-108581486-INS-155
chr6-108603538-INS-75
chr6-108603556-INS-187
chr6-108650203-INS-238
chr6-108653487-INS-230
chr6-108660399-INS-6461
chr6-108670655-INS-92
chr6-108676161-INS-167
chr6-108677212-INS-52
chr6-108689828-INS-135
chr6-108693908-INS-53
chr6-108695095-INS-51
chr6-108703973-INS-57
chr6-108738989-INS-196
chr6-108741648-INS-51
chr6-108742481-INS-162
chr6-108746393-INS-4784
chr6-108753343-INS-52
chr6-108808080-INS-62
chr6-108826262-INS-190
chr6-108874695-INS-50
chr6-108878886-INS-1593
chr6-108893095-INS-96
chr6-108893267-INS-1510
chr6-108893267-INS-169
chr6-108895876-INS-169
chr6-108895904-INS-6183
chr6-108905069-INS-99
chr6-108908538-INS-1319
chr6-108912861-INS-8430

chr6-108958562-INS-2636
chr6-108965405-INS-133
chr6-108967245-INS-6747
chr6-108972508-INS-590
chr6-108974969-INS-1101
chr6-108978286-INS-243
chr6-108978571-INS-153
chr6-108978655-INS-464
chr6-109004899-INS-1066
chr6-109036292-INS-1123
chr6-109076113-INS-64
chr6-109180165-INS-483
chr6-109181004-INS-2013
chr6-109208927-INS-3244
chr6-109240129-INS-394
chr6-109243692-INS-100
chr6-109243692-INS-388
chr6-109243831-INS-50
chr6-109248069-INS-2966
chr6-109250633-INS-6335
chr6-109267433-INS-168
chr6-109273517-INS-988
chr6-109314620-INS-52
chr6-109393490-INS-2285
chr6-109429120-INS-50
chr6-109434586-INS-366
chr6-109448866-INS-50
chr6-109456854-INS-1018
chr6-109477510-INS-56
chr6-109511395-INS-62
chr6-109559875-INS-1132
chr6-109669679-INS-413
chr6-109670593-INS-494
chr6-109709616-INS-411
chr6-109712593-INS-2930
chr6-109713008-INS-5173
chr6-109729314-INS-1727
chr6-109739744-INS-52
chr6-109775916-INS-3399
chr6-109787201-INS-52
chr6-109825727-INS-76
chr6-109842477-INS-74
chr6-109842557-INS-310
chr6-109848225-INS-6773
chr6-109848469-INS-3035
chr6-109865419-INS-50
chr6-109867835-INS-52
chr6-109909856-INS-428
chr6-109919267-INS-7068
chr6-109923812-INS-50
chr6-109945837-INS-50
chr6-109986623-INS-504
chr6-110006140-INS-197
chr6-110014661-INS-72
chr6-110038932-INS-902
chr6-110043532-INS-766
chr6-110059551-INS-1331
chr6-110078555-INS-466
chr6-110093800-INS-1967
chr6-110096249-INS-4024
chr6-110160139-INS-2386
chr6-110160646-INS-7982
chr6-110169009-INS-95
chr6-110198105-INS-1117
chr6-110198765-INS-53
chr6-110211519-INS-213
chr6-110317429-INS-7082
chr6-110360153-INS-75
chr6-110377691-INS-171
chr6-110390337-INS-117
chr6-110403333-INS-1654
chr6-110405164-INS-394
chr6-110420197-INS-50
chr6-110441564-INS-112
chr6-110464037-INS-1811
chr6-110535986-INS-129
chr6-110535986-INS-64

chr6-110543385-INS-61
chr6-110582952-INS-801
chr6-110585750-INS-1597
chr6-110593253-INS-1433
chr6-110593643-INS-4681
chr6-110682906-INS-65
chr6-110693284-INS-70
chr6-110713199-INS-720
chr6-110732517-INS-60
chr6-110733742-INS-1810
chr6-110749431-INS-1228
chr6-110764594-INS-6677
chr6-110775683-INS-135
chr6-110776400-INS-85
chr6-110788574-INS-6437
chr6-110797932-INS-223
chr6-110840100-INS-588
chr6-110844874-INS-103
chr6-110844876-INS-50
chr6-110845669-INS-80
chr6-110854972-INS-1516
chr6-110980139-INS-1246
chr6-111015213-INS-1333
chr6-111015988-INS-61
chr6-111023337-INS-6486
chr6-111063167-INS-68
chr6-111106310-INS-919
chr6-111209153-INS-3801
chr6-111231774-INS-1185
chr6-111255326-INS-398
chr6-111285498-INS-169
chr6-111377616-INS-1445
chr6-111382771-INS-595
chr6-111382807-INS-227
chr6-111460489-INS-2015
chr6-111492850-INS-68
chr6-111572179-INS-2921
chr6-111603619-INS-117
chr6-111675739-INS-50
chr6-111683517-INS-53
chr6-111710516-INS-5207
chr6-111780700-INS-216
chr6-111888720-INS-6406
chr6-111928151-INS-50
chr6-111932030-INS-1280
chr6-111935661-INS-57
chr6-111964962-INS-6810
chr6-111983848-INS-4382
chr6-112058549-INS-72
chr6-112092290-INS-6382
chr6-112113348-INS-4278
chr6-112129472-INS-63
chr6-112137787-INS-146
chr6-112203170-INS-276
chr6-112220384-INS-246
chr6-112220384-INS-74
chr6-112220489-INS-72
chr6-112269814-INS-52
chr6-112283268-INS-50
chr6-112326901-INS-388
chr6-112338259-INS-1038
chr6-112340337-INS-280
chr6-112341475-INS-932
chr6-112375560-INS-83
chr6-112402713-INS-79
chr6-112428841-INS-72
chr6-112459060-INS-51
chr6-112470631-INS-64
chr6-112494578-INS-50
chr6-112494965-INS-2828
chr6-112504907-INS-123
chr6-112507842-INS-144
chr6-112508178-INS-756
chr6-112521080-INS-2172
chr6-112555714-INS-50
chr6-112565191-INS-6306
chr6-112605596-INS-315

chr6-112652340-INS-202
chr6-112663454-INS-1305
chr6-112663503-INS-323
chr6-112719886-INS-702
chr6-112748604-INS-692
chr6-112820790-INS-332
chr6-112837087-INS-224
chr6-112863052-INS-85
chr6-112886651-INS-3044
chr6-112911422-INS-87
chr6-112921723-INS-51
chr6-112936826-INS-513
chr6-112937049-INS-627
chr6-112954943-INS-66
chr6-112971339-INS-66
chr6-113016597-INS-147
chr6-113058832-INS-187
chr6-113140678-INS-53
chr6-113204228-INS-102
chr6-113210177-INS-6478
chr6-113218658-INS-918
chr6-113229811-INS-179
chr6-113272902-INS-54
chr6-113323201-INS-154
chr6-113323212-INS-52
chr6-113332173-INS-5221
chr6-113373731-INS-180
chr6-113399931-INS-69
chr6-113400457-INS-137
chr6-113400457-INS-53
chr6-113403695-INS-180
chr6-113454824-INS-80
chr6-113558995-INS-87
chr6-113584669-INS-226
chr6-113598079-INS-62
chr6-113599757-INS-50
chr6-113623056-INS-51
chr6-113634331-INS-234
chr6-113634496-INS-64
chr6-113645225-INS-55
chr6-113647763-INS-64
chr6-113655363-INS-4360
chr6-113662099-INS-103
chr6-113713708-INS-72
chr6-113746752-INS-408
chr6-113777305-INS-172
chr6-113826792-INS-138
chr6-113838422-INS-117
chr6-113869693-INS-90
chr6-113906865-INS-61
chr6-114001221-INS-50
chr6-114010259-INS-88
chr6-114085165-INS-50
chr6-114230689-INS-52
chr6-114245016-INS-136
chr6-114296659-INS-56
chr6-114303766-INS-81
chr6-114314152-INS-102
chr6-114347839-INS-53
chr6-114372689-INS-200
chr6-114388505-INS-50
chr6-114421076-INS-6411
chr6-114430484-INS-84
chr6-114478201-INS-59
chr6-114507012-INS-64
chr6-114656514-INS-180
chr6-114677339-INS-804
chr6-114687526-INS-266
chr6-114749938-INS-114
chr6-114756053-INS-50
chr6-114767515-INS-502
chr6-114767705-INS-352
chr6-114830074-INS-535
chr6-114830291-INS-3691
chr6-114955165-INS-64
chr6-114961741-INS-127
chr6-115037048-INS-72

chr6-115097549-INS-105
chr6-115104558-INS-2494
chr6-115111099-INS-57
chr6-115147835-INS-132
chr6-115159760-INS-4480
chr6-115185084-INS-6351
chr6-115204810-INS-94
chr6-115217519-INS-492
chr6-115232983-INS-56
chr6-115233690-INS-147
chr6-115289942-INS-71
chr6-115290784-INS-52
chr6-115328220-INS-81
chr6-115328230-INS-508
chr6-115408442-INS-50
chr6-115411664-INS-1277
chr6-115411832-INS-50
chr6-115539869-INS-173
chr6-115565258-INS-84
chr6-115619416-INS-209
chr6-115619646-INS-539
chr6-115666110-INS-680
chr6-115666280-INS-210
chr6-115680701-INS-3158
chr6-115695955-INS-1796
chr6-115696078-INS-561
chr6-115730195-INS-54
chr6-115747194-INS-239
chr6-115750968-INS-54
chr6-115751475-INS-450
chr6-115751475-INS-64
chr6-115789311-INS-123
chr6-115809053-INS-208
chr6-115862798-INS-76
chr6-115972179-INS-136
chr6-116024897-INS-55
chr6-116085940-INS-4156
chr6-116101553-INS-218
chr6-116136747-INS-73
chr6-116153284-INS-178
chr6-116153318-INS-581
chr6-116157409-INS-721
chr6-116207987-INS-303
chr6-116219271-INS-1192
chr6-116241278-INS-75
chr6-116268257-INS-60
chr6-116294945-INS-166
chr6-116322218-INS-145
chr6-116323203-INS-100
chr6-116323828-INS-75
chr6-116333072-INS-8419
chr6-116372153-INS-123
chr6-116384482-INS-489
chr6-116429683-INS-314
chr6-116477711-INS-478
chr6-116492171-INS-100
chr6-116492171-INS-237
chr6-116494745-INS-11404
chr6-116538509-INS-1903
chr6-116549500-INS-57
chr6-116597754-INS-422
chr6-116600922-INS-544
chr6-116623614-INS-158
chr6-116792217-INS-1755
chr6-116793371-INS-54
chr6-116814033-INS-5421
chr6-116816811-INS-191
chr6-116903881-INS-424
chr6-116955280-INS-6366
chr6-116960195-INS-55
chr6-116960302-INS-864
chr6-116978309-INS-68
chr6-116999427-INS-100
chr6-117017848-INS-3687
chr6-117043469-INS-428
chr6-117104594-INS-7573
chr6-117118321-INS-52

chr6-117126184-INS-65
chr6-117128274-INS-2588
chr6-117170451-INS-996
chr6-117268186-INS-60
chr6-117298776-INS-119
chr6-117334287-INS-5606
chr6-117363190-INS-2113
chr6-117369703-INS-171
chr6-117378827-INS-62
chr6-117421552-INS-114
chr6-117435581-INS-53
chr6-117441539-INS-20636
chr6-117530777-INS-544
chr6-117542045-INS-6364
chr6-117546075-INS-60
chr6-117599463-INS-337
chr6-117639837-INS-253
chr6-117653646-INS-7136
chr6-117697249-INS-70
chr6-117712968-INS-3332
chr6-117728653-INS-95
chr6-117742190-INS-192
chr6-117749356-INS-970
chr6-117753780-INS-210
chr6-117800964-INS-6473
chr6-117821641-INS-366
chr6-117821747-INS-854
chr6-117837733-INS-63
chr6-117837832-INS-190
chr6-117840867-INS-189
chr6-117844954-INS-54
chr6-117870178-INS-222
chr6-117886730-INS-78
chr6-117918288-INS-78
chr6-117935599-INS-522
chr6-117942185-INS-121
chr6-117958292-INS-63
chr6-117959254-INS-51
chr6-117961011-INS-516
chr6-117962341-INS-50
chr6-117977485-INS-232
chr6-118006899-INS-190
chr6-118007669-INS-133
chr6-118007689-INS-50
chr6-118027666-INS-200
chr6-118078147-INS-1632
chr6-118175766-INS-156
chr6-118328485-INS-62
chr6-118333013-INS-2347
chr6-118367828-INS-209
chr6-118375826-INS-980
chr6-118416789-INS-50
chr6-118423156-INS-66
chr6-118494608-INS-362
chr6-118670916-INS-699
chr6-118676366-INS-7108
chr6-118689630-INS-60
chr6-118703968-INS-275
chr6-118787056-INS-130
chr6-118792827-INS-65
chr6-118841149-INS-64
chr6-118887431-INS-61
chr6-118893651-INS-70
chr6-118894564-INS-81
chr6-118905986-INS-33309
chr6-118911843-INS-8329
chr6-118934562-INS-408
chr6-118944443-INS-60
chr6-118944818-INS-65
chr6-118947068-INS-73
chr6-118949492-INS-50
chr6-119069082-INS-265
chr6-119097770-INS-120
chr6-119175996-INS-562
chr6-119220771-INS-754
chr6-119220953-INS-52
chr6-119221976-INS-50

chr6-119225660-INS-96
chr6-119234769-INS-264
chr6-119238672-INS-64
chr6-119239194-INS-68
chr6-119283216-INS-200
chr6-119362290-INS-210
chr6-119381865-INS-5120
chr6-119394767-INS-72
chr6-119400544-INS-52
chr6-119411409-INS-3148
chr6-119430340-INS-170
chr6-119447136-INS-2828
chr6-119447337-INS-282
chr6-119489136-INS-95
chr6-119572085-INS-52
chr6-119592108-INS-94
chr6-119592415-INS-775
chr6-119613821-INS-1663
chr6-119620133-INS-163
chr6-119682671-INS-75
chr6-119797206-INS-218
chr6-119797452-INS-474
chr6-119835274-INS-55
chr6-119848554-INS-1886
chr6-119872590-INS-52
chr6-119872653-INS-296
chr6-119872664-INS-125
chr6-119994202-INS-1555
chr6-120001213-INS-184
chr6-120018105-INS-60
chr6-120026542-INS-119
chr6-120028783-INS-57
chr6-120038124-INS-210
chr6-120041490-INS-132
chr6-120084198-INS-412
chr6-120091567-INS-55
chr6-120121611-INS-113
chr6-120132338-INS-52
chr6-120133506-INS-96
chr6-120158778-INS-183
chr6-120162290-INS-60
chr6-120172993-INS-94
chr6-120237137-INS-75
chr6-120249869-INS-92
chr6-120280451-INS-54
chr6-120281167-INS-58
chr6-120285766-INS-4913
chr6-120290087-INS-51
chr6-120313499-INS-62
chr6-120324129-INS-212
chr6-120324151-INS-488
chr6-120334852-INS-118
chr6-120341984-INS-72
chr6-120344495-INS-168
chr6-120356009-INS-52
chr6-120356402-INS-195
chr6-120383456-INS-217
chr6-120429090-INS-157
chr6-120437560-INS-52
chr6-120464894-INS-50
chr6-120477184-INS-76
chr6-120479846-INS-50
chr6-120479957-INS-217
chr6-120490534-INS-209
chr6-120491054-INS-337
chr6-120497062-INS-192
chr6-120497814-INS-163
chr6-120502926-INS-54
chr6-120528178-INS-128
chr6-120540958-INS-54
chr6-120587386-INS-56
chr6-120606294-INS-70
chr6-120609337-INS-92
chr6-120609509-INS-322
chr6-120622982-INS-210
chr6-120626553-INS-56
chr6-120627372-INS-77

chr6-120627525-INS-170
chr6-120637243-INS-784
chr6-120665242-INS-52
chr6-120704528-INS-217
chr6-120716166-INS-1212
chr6-120717751-INS-1169
chr6-120724726-INS-5246
chr6-120728994-INS-976
chr6-120778330-INS-548
chr6-120798726-INS-209
chr6-120805994-INS-56
chr6-120806175-INS-193
chr6-120807611-INS-175
chr6-120823179-INS-62
chr6-120825089-INS-212
chr6-120904383-INS-50
chr6-120982853-INS-162
chr6-121036521-INS-63
chr6-121051370-INS-52
chr6-121057078-INS-78
chr6-121074450-INS-56
chr6-121127352-INS-224
chr6-121135197-INS-52
chr6-121160096-INS-14279
chr6-121176469-INS-69
chr6-121177235-INS-125
chr6-121189749-INS-68
chr6-121230119-INS-104
chr6-121234387-INS-66
chr6-121246599-INS-162
chr6-121293493-INS-126
chr6-121343747-INS-617
chr6-121343875-INS-260
chr6-121377083-INS-74
chr6-121377531-INS-74
chr6-121456050-INS-78
chr6-121492372-INS-1491
chr6-121505076-INS-130
chr6-121508621-INS-146
chr6-121586073-INS-7264
chr6-121644440-INS-469
chr6-121700146-INS-110
chr6-121722155-INS-6480
chr6-121762072-INS-3892
chr6-121789445-INS-135
chr6-121809651-INS-56
chr6-121826093-INS-2343
chr6-121843170-INS-2794
chr6-121885203-INS-3109
chr6-121954220-INS-3542
chr6-121958672-INS-62
chr6-121967744-INS-504
chr6-122006583-INS-1951
chr6-122007178-INS-518
chr6-122008281-INS-975
chr6-122115800-INS-52
chr6-122116237-INS-138
chr6-122249533-INS-225
chr6-122249551-INS-96
chr6-122257533-INS-222
chr6-122332508-INS-7130
chr6-122336054-INS-181
chr6-122336571-INS-206
chr6-122340836-INS-59
chr6-122341513-INS-192
chr6-122350696-INS-268
chr6-122356877-INS-65
chr6-122375759-INS-1100
chr6-122382389-INS-75
chr6-122398465-INS-106
chr6-122398768-INS-64
chr6-122403816-INS-6367
chr6-122404178-INS-1054
chr6-122410978-INS-74
chr6-122415007-INS-6457
chr6-122439482-INS-158
chr6-122508992-INS-52

chr6-122521649-INS-93
chr6-122524866-INS-108
chr6-122525077-INS-416
chr6-122529812-INS-114
chr6-122537451-INS-54
chr6-122556398-INS-2000
chr6-122567309-INS-70
chr6-122569778-INS-86
chr6-122573318-INS-51
chr6-122582806-INS-207
chr6-122587771-INS-137
chr6-122587772-INS-52
chr6-122592262-INS-1246
chr6-122592795-INS-16705
chr6-122593068-INS-1730
chr6-122595484-INS-62
chr6-122612102-INS-2604
chr6-122639394-INS-513
chr6-122642365-INS-751
chr6-122653457-INS-53
chr6-122663760-INS-73
chr6-122685951-INS-138
chr6-122686769-INS-66
chr6-122767340-INS-150
chr6-122776766-INS-2472
chr6-122807883-INS-592
chr6-122808779-INS-64
chr6-122863568-INS-500
chr6-122919320-INS-1303
chr6-123044631-INS-58
chr6-123082669-INS-6901
chr6-123097957-INS-1252
chr6-123111631-INS-2751
chr6-123121307-INS-3958
chr6-123128956-INS-97
chr6-123134296-INS-53
chr6-123134369-INS-53
chr6-123134381-INS-111
chr6-123182290-INS-4429
chr6-123192775-INS-4931
chr6-123234250-INS-3194
chr6-123266229-INS-393
chr6-123279828-INS-6657
chr6-123304147-INS-5563
chr6-123308257-INS-52
chr6-123318357-INS-2827
chr6-123375015-INS-4245
chr6-123448774-INS-173
chr6-123485049-INS-1769
chr6-123485049-INS-669
chr6-123495377-INS-154
chr6-123542522-INS-159
chr6-123564126-INS-230
chr6-123602270-INS-669
chr6-123695616-INS-72
chr6-123710436-INS-51
chr6-123754965-INS-1139
chr6-123768281-INS-55
chr6-123813189-INS-92
chr6-123883005-INS-6490
chr6-123884332-INS-1005
chr6-124000678-INS-171
chr6-124006260-INS-257
chr6-124033313-INS-7127
chr6-124054262-INS-51
chr6-124066380-INS-6375
chr6-124068000-INS-225
chr6-124079587-INS-2174
chr6-124106198-INS-63
chr6-124147926-INS-200
chr6-124149041-INS-5194
chr6-124180848-INS-218
chr6-124187904-INS-121
chr6-124205048-INS-2980
chr6-124206677-INS-213
chr6-124218588-INS-1825
chr6-124219241-INS-4198

chr6-124258625-INS-6796
chr6-124269556-INS-3177
chr6-124320183-INS-59
chr6-124337271-INS-125
chr6-124338684-INS-116
chr6-124366958-INS-216
chr6-124419804-INS-300
chr6-124431402-INS-1096
chr6-124469051-INS-390
chr6-124489842-INS-121
chr6-124501079-INS-80
chr6-124506756-INS-225
chr6-124506799-INS-92
chr6-124547608-INS-51
chr6-124549743-INS-169
chr6-124549899-INS-52
chr6-124558182-INS-11473
chr6-124592034-INS-481
chr6-124750106-INS-96
chr6-124750362-INS-92
chr6-124763389-INS-200
chr6-124781203-INS-357
chr6-124781565-INS-215
chr6-124831059-INS-99
chr6-124833783-INS-70
chr6-124861679-INS-60
chr6-124898444-INS-213
chr6-124919064-INS-116
chr6-124928109-INS-155
chr6-124928192-INS-68
chr6-124996945-INS-260
chr6-125082810-INS-469
chr6-125159149-INS-171
chr6-125231674-INS-86
chr6-125269030-INS-149
chr6-125288105-INS-192
chr6-125308315-INS-56
chr6-125313081-INS-105
chr6-125363779-INS-291
chr6-125394991-INS-230
chr6-125396196-INS-70
chr6-125413570-INS-82
chr6-125487188-INS-104
chr6-125500365-INS-129
chr6-125501353-INS-95
chr6-125501433-INS-270
chr6-125501532-INS-570
chr6-125502637-INS-55
chr6-125513299-INS-108
chr6-125516982-INS-198
chr6-125517023-INS-60
chr6-125520066-INS-209
chr6-125520898-INS-203
chr6-125538552-INS-115
chr6-125544590-INS-546
chr6-125558790-INS-138
chr6-125568358-INS-1095
chr6-125571582-INS-156
chr6-125572433-INS-119
chr6-125595139-INS-64
chr6-125606190-INS-50
chr6-125615534-INS-799
chr6-125621818-INS-60
chr6-125658622-INS-148
chr6-125709757-INS-1100
chr6-125726089-INS-126
chr6-125726287-INS-119
chr6-125726443-INS-52
chr6-125759696-INS-52
chr6-125781445-INS-194
chr6-125854480-INS-252
chr6-125923692-INS-71
chr6-125936928-INS-50
chr6-125967108-INS-112
chr6-125969549-INS-68
chr6-125969969-INS-265
chr6-125973044-INS-56

chr6-125973232-INS-113
chr6-125976314-INS-210
chr6-125989939-INS-356
chr6-126006763-INS-7802
chr6-126044920-INS-6653
chr6-126066194-INS-199
chr6-126076697-INS-56
chr6-126082603-INS-56
chr6-126105204-INS-138
chr6-126142311-INS-80
chr6-126164258-INS-108
chr6-126172469-INS-195
chr6-126206485-INS-58
chr6-126276471-INS-74
chr6-126360913-INS-292
chr6-126377279-INS-157
chr6-126445162-INS-1565
chr6-126461228-INS-50
chr6-126469006-INS-58
chr6-126470320-INS-6804
chr6-126494902-INS-55
chr6-126495336-INS-128
chr6-126521907-INS-54
chr6-126523373-INS-117
chr6-126625568-INS-153
chr6-126639935-INS-6139
chr6-126664734-INS-670
chr6-126727773-INS-54
chr6-126728328-INS-162
chr6-126734502-INS-148
chr6-126735262-INS-149
chr6-126750166-INS-64
chr6-126873524-INS-194
chr6-126891403-INS-52
chr6-127026417-INS-55
chr6-127068259-INS-173
chr6-127087107-INS-146
chr6-127110661-INS-591
chr6-127138710-INS-136
chr6-127138940-INS-101
chr6-127139154-INS-422
chr6-127213433-INS-498
chr6-127231485-INS-201
chr6-127283760-INS-291
chr6-127292642-INS-74
chr6-127324679-INS-102
chr6-127324881-INS-82
chr6-127357368-INS-125
chr6-127374670-INS-2104
chr6-127418231-INS-76
chr6-127445572-INS-1731
chr6-127445838-INS-479
chr6-127445937-INS-182
chr6-127483669-INS-52
chr6-127486104-INS-202
chr6-127501638-INS-263
chr6-127507906-INS-102
chr6-127524859-INS-128
chr6-127560430-INS-353
chr6-127566744-INS-74
chr6-127567621-INS-72
chr6-127633407-INS-50
chr6-127658579-INS-57
chr6-127658635-INS-133
chr6-127685004-INS-161
chr6-127719979-INS-54
chr6-127720138-INS-156
chr6-127799063-INS-68
chr6-127857656-INS-79
chr6-127897030-INS-129
chr6-127917293-INS-130
chr6-127917565-INS-128
chr6-127925264-INS-58
chr6-127972973-INS-608
chr6-128012479-INS-64
chr6-128064759-INS-158
chr6-128118353-INS-50

chr6-128251625-INS-78
chr6-128289072-INS-2100
chr6-128289266-INS-307
chr6-128291830-INS-195
chr6-128306237-INS-181
chr6-128308558-INS-105
chr6-128326465-INS-54
chr6-128365408-INS-277
chr6-128365408-INS-58
chr6-128368507-INS-50
chr6-128444829-INS-1014
chr6-128444863-INS-194
chr6-128459499-INS-5544
chr6-128467419-INS-79
chr6-128483580-INS-84
chr6-128513302-INS-454
chr6-128518974-INS-180
chr6-128535095-INS-201
chr6-128537143-INS-718
chr6-128555355-INS-2886
chr6-128562480-INS-70
chr6-128575792-INS-55
chr6-128576056-INS-114
chr6-128582643-INS-144
chr6-128606213-INS-77
chr6-128611317-INS-8320
chr6-128616977-INS-52
chr6-128624325-INS-101
chr6-128632662-INS-78
chr6-128648556-INS-400
chr6-128691963-INS-206
chr6-128699557-INS-1199
chr6-128708997-INS-3029
chr6-128720169-INS-352
chr6-128728617-INS-201
chr6-128731335-INS-79
chr6-128748433-INS-130
chr6-128748448-INS-50
chr6-128748986-INS-190
chr6-128767723-INS-196
chr6-128767723-INS-84
chr6-128795202-INS-1177
chr6-128801139-INS-7099
chr6-128820366-INS-6031
chr6-128826049-INS-198
chr6-128828435-INS-100
chr6-128829354-INS-490
chr6-128830551-INS-213
chr6-128831306-INS-21371
chr6-128868756-INS-498
chr6-128893132-INS-65
chr6-128907170-INS-79
chr6-128907858-INS-159
chr6-128912038-INS-3490
chr6-128945139-INS-4171
chr6-128956217-INS-209
chr6-129023287-INS-295
chr6-129045151-INS-2149
chr6-129090072-INS-7366
chr6-129095024-INS-1798
chr6-129099631-INS-344
chr6-129099631-INS-7084
chr6-129110580-INS-80
chr6-129113455-INS-2207
chr6-129147792-INS-184
chr6-129155223-INS-52
chr6-129157196-INS-4193
chr6-129158845-INS-70
chr6-129177066-INS-53
chr6-129188715-INS-60
chr6-129191371-INS-54
chr6-129191648-INS-1837
chr6-129239465-INS-1304
chr6-129247944-INS-100
chr6-129257148-INS-190
chr6-129271202-INS-160
chr6-129272132-INS-141

chr6-129272227-INS-68
chr6-129340045-INS-218
chr6-129340616-INS-69
chr6-129347875-INS-1953
chr6-129348082-INS-4718
chr6-129348956-INS-81
chr6-129365425-INS-7191
chr6-129380676-INS-3767
chr6-129415006-INS-50
chr6-129416394-INS-50
chr6-129429815-INS-155
chr6-129453897-INS-762
chr6-129469369-INS-161
chr6-129486537-INS-236
chr6-129495018-INS-52
chr6-129526690-INS-6486
chr6-129553126-INS-70
chr6-129553565-INS-56
chr6-129575187-INS-105
chr6-129592709-INS-159
chr6-129608139-INS-132
chr6-129652475-INS-3831
chr6-129681936-INS-1655
chr6-129684255-INS-4284
chr6-129700226-INS-70
chr6-129700236-INS-231
chr6-129702472-INS-52
chr6-129712568-INS-112
chr6-129712672-INS-80
chr6-129717947-INS-6404
chr6-129728848-INS-1066
chr6-129732297-INS-50
chr6-129748864-INS-1322
chr6-129965177-INS-60
chr6-129985612-INS-5242
chr6-129993431-INS-52
chr6-130006700-INS-68
chr6-130012904-INS-50
chr6-130015698-INS-1927
chr6-130020165-INS-162
chr6-130020188-INS-57
chr6-130020543-INS-62
chr6-130031720-INS-2237
chr6-130059336-INS-750
chr6-130059340-INS-281
chr6-130065553-INS-205
chr6-130127356-INS-232
chr6-130157461-INS-56
chr6-130159788-INS-1606
chr6-130161675-INS-205
chr6-130201807-INS-53
chr6-130222003-INS-2041
chr6-130267754-INS-175
chr6-130272425-INS-55
chr6-130281687-INS-153
chr6-130290059-INS-342
chr6-130290178-INS-1880
chr6-130294066-INS-227
chr6-130294481-INS-100
chr6-130297375-INS-80
chr6-130304469-INS-1604
chr6-130310836-INS-51
chr6-130318883-INS-68
chr6-130340345-INS-163
chr6-130341417-INS-6339
chr6-130343773-INS-310
chr6-130346574-INS-1515
chr6-130347389-INS-52
chr6-130348801-INS-190
chr6-130349386-INS-56
chr6-130395174-INS-52
chr6-130411104-INS-52
chr6-130429395-INS-18000
chr6-130536966-INS-64
chr6-130546909-INS-121
chr6-130619613-INS-1297
chr6-130640988-INS-212

chr6-130648166-INS-1592
chr6-130670828-INS-548
chr6-130737604-INS-57
chr6-130748841-INS-6392
chr6-130753285-INS-1926
chr6-130763457-INS-84
chr6-130763672-INS-252
chr6-130766720-INS-58
chr6-130786312-INS-2604
chr6-130789572-INS-5180
chr6-130840485-INS-2346
chr6-130890961-INS-58
chr6-130893585-INS-230
chr6-130915343-INS-168
chr6-130967188-INS-83
chr6-130970466-INS-4356
chr6-131049872-INS-50
chr6-131143415-INS-1176
chr6-131241453-INS-66
chr6-131266279-INS-62
chr6-131285746-INS-244
chr6-131324570-INS-552
chr6-131324824-INS-120
chr6-131393151-INS-3627
chr6-131433456-INS-6167
chr6-131439888-INS-63
chr6-131440485-INS-65
chr6-131452620-INS-6615
chr6-131455953-INS-60
chr6-131460359-INS-198
chr6-131460578-INS-109
chr6-131486884-INS-51
chr6-131496431-INS-316
chr6-131500229-INS-441
chr6-131500823-INS-64
chr6-131503036-INS-956
chr6-131512473-INS-82
chr6-131517496-INS-675
chr6-131631297-INS-50
chr6-131648416-INS-1274
chr6-131716685-INS-6612
chr6-131734469-INS-100
chr6-131746647-INS-1845
chr6-131801643-INS-180
chr6-131908451-INS-832
chr6-131932434-INS-56
chr6-131984633-INS-6759
chr6-131994886-INS-478
chr6-131994956-INS-58
chr6-131995030-INS-549
chr6-132001978-INS-55
chr6-132010738-INS-138
chr6-132036991-INS-198
chr6-132075455-INS-2988
chr6-132091902-INS-54
chr6-132167422-INS-168
chr6-132172203-INS-6240
chr6-132192287-INS-481
chr6-132289137-INS-207
chr6-132367067-INS-50
chr6-132424592-INS-406
chr6-132453175-INS-55
chr6-132485405-INS-53
chr6-132542295-INS-70
chr6-132547737-INS-2010
chr6-132564579-INS-50
chr6-132624050-INS-319
chr6-132640075-INS-6411
chr6-132641900-INS-58
chr6-132692255-INS-3777
chr6-132726831-INS-50
chr6-132752303-INS-1132
chr6-132755932-INS-66
chr6-132847127-INS-215
chr6-132865982-INS-60
chr6-132885059-INS-1153
chr6-132900169-INS-11507

chr6-132905085-INS-51
chr6-132918368-INS-225
chr6-132922251-INS-62
chr6-132962513-INS-68
chr6-132998918-INS-523
chr6-133006853-INS-54
chr6-133027990-INS-4963
chr6-133029614-INS-255
chr6-133029666-INS-75
chr6-133037700-INS-2011
chr6-133085649-INS-7086
chr6-133108375-INS-2255
chr6-133123612-INS-112
chr6-133127899-INS-51
chr6-133139069-INS-52
chr6-133164761-INS-1414
chr6-133165438-INS-195
chr6-133173305-INS-308
chr6-133192026-INS-2859
chr6-133198666-INS-207
chr6-133226730-INS-56
chr6-133282364-INS-52
chr6-133336848-INS-93
chr6-133342030-INS-1380
chr6-133346382-INS-51
chr6-133351089-INS-312
chr6-133352247-INS-104
chr6-133357787-INS-197
chr6-133435777-INS-172
chr6-133448943-INS-52
chr6-133467408-INS-108
chr6-133469154-INS-95
chr6-133469191-INS-1265
chr6-133469194-INS-421
chr6-133469344-INS-90
chr6-133475617-INS-55
chr6-133475626-INS-147
chr6-133481825-INS-6580
chr6-133502904-INS-1999
chr6-133529852-INS-317
chr6-133551202-INS-7472
chr6-133568762-INS-2582
chr6-133600463-INS-50
chr6-133600468-INS-1588
chr6-133686014-INS-1499
chr6-133714801-INS-172
chr6-133753982-INS-84
chr6-133755831-INS-60
chr6-133763762-INS-30877
chr6-133785494-INS-831
chr6-133816827-INS-50
chr6-133822528-INS-196
chr6-133836346-INS-1139
chr6-133842394-INS-690
chr6-133845680-INS-7222
chr6-133879176-INS-202
chr6-133898603-INS-50
chr6-133908189-INS-835
chr6-133929895-INS-76
chr6-133956456-INS-238
chr6-134004015-INS-202
chr6-134007271-INS-203
chr6-134040498-INS-62
chr6-134103592-INS-78
chr6-134136033-INS-50
chr6-134302167-INS-1062
chr6-134305712-INS-197
chr6-134333345-INS-152
chr6-134361452-INS-99
chr6-134371901-INS-76
chr6-134381369-INS-99
chr6-134388975-INS-179
chr6-134391715-INS-8060
chr6-134423234-INS-4434
chr6-134459027-INS-171
chr6-134461447-INS-223
chr6-134465782-INS-406

chr6-134466036-INS-724
chr6-134499206-INS-257
chr6-134523954-INS-50
chr6-134568404-INS-121
chr6-134574500-INS-636
chr6-134579935-INS-75
chr6-134644271-INS-168
chr6-134651128-INS-465
chr6-134651337-INS-6722
chr6-134655424-INS-605
chr6-134752710-INS-1734
chr6-134774921-INS-348
chr6-134788975-INS-1856
chr6-134842650-INS-149
chr6-134872818-INS-104
chr6-134908896-INS-72
chr6-134922841-INS-8686
chr6-134924061-INS-211
chr6-134924453-INS-91
chr6-134947686-INS-86
chr6-134973372-INS-249
chr6-134974420-INS-146
chr6-134976059-INS-66
chr6-134999144-INS-391
chr6-135005357-INS-50
chr6-135010512-INS-1711
chr6-135024638-INS-1538
chr6-135024638-INS-560
chr6-135027211-INS-7106
chr6-135036135-INS-2512
chr6-135039146-INS-1260
chr6-135045802-INS-164
chr6-135060529-INS-8302
chr6-135066059-INS-52
chr6-135068360-INS-149
chr6-135132720-INS-52
chr6-135133230-INS-107
chr6-135136600-INS-233
chr6-135136734-INS-836
chr6-135154473-INS-62
chr6-135159533-INS-594
chr6-135159924-INS-51
chr6-135163823-INS-6295
chr6-135210638-INS-50
chr6-135253090-INS-203
chr6-135277816-INS-378
chr6-135292600-INS-102
chr6-135306645-INS-6192
chr6-135383594-INS-1118
chr6-135383594-INS-494
chr6-135383885-INS-104
chr6-135390440-INS-53
chr6-135393460-INS-142
chr6-135401830-INS-56
chr6-135405659-INS-81
chr6-135405721-INS-165
chr6-135449182-INS-64
chr6-135476607-INS-2186
chr6-135554922-INS-1196
chr6-135571654-INS-366
chr6-135571828-INS-882
chr6-135572821-INS-385
chr6-135573520-INS-6054
chr6-135603261-INS-207
chr6-135603322-INS-60
chr6-135625575-INS-51
chr6-135654329-INS-56
chr6-135669944-INS-8923
chr6-135684073-INS-97
chr6-135698324-INS-54
chr6-135779465-INS-50
chr6-135803726-INS-92
chr6-135855469-INS-335
chr6-135874265-INS-71
chr6-135890496-INS-241
chr6-135896690-INS-87
chr6-135927812-INS-165

chr6-136032268-INS-86
chr6-136056681-INS-94
chr6-136141037-INS-96
chr6-136158569-INS-6401
chr6-136172364-INS-1532
chr6-136193966-INS-50
chr6-136202938-INS-82
chr6-136203021-INS-95
chr6-136225928-INS-162
chr6-136291739-INS-6440
chr6-136299464-INS-3744
chr6-136299592-INS-384
chr6-136302302-INS-61
chr6-136302330-INS-25788
chr6-136302405-INS-403
chr6-136304169-INS-79
chr6-136308210-INS-72
chr6-136310637-INS-152
chr6-136314561-INS-63
chr6-136363831-INS-50
chr6-136369005-INS-546
chr6-136369199-INS-72
chr6-136382486-INS-81
chr6-136382503-INS-1811
chr6-136383944-INS-3519
chr6-136385815-INS-198
chr6-136391703-INS-1455
chr6-136410417-INS-180
chr6-136411128-INS-108
chr6-136431272-INS-2920
chr6-136454168-INS-55
chr6-136463818-INS-5668
chr6-136465733-INS-2364
chr6-136500590-INS-214
chr6-136613957-INS-1146
chr6-136620475-INS-202
chr6-136659307-INS-604
chr6-136660144-INS-96
chr6-136664995-INS-126
chr6-136665127-INS-344
chr6-136679055-INS-864
chr6-136679129-INS-68
chr6-136685575-INS-50
chr6-136687072-INS-67
chr6-136687241-INS-136
chr6-136696118-INS-180
chr6-136732268-INS-5453
chr6-136741026-INS-132
chr6-136776525-INS-5119
chr6-136793810-INS-890
chr6-136802731-INS-1858
chr6-136813010-INS-62
chr6-136828412-INS-438
chr6-136829984-INS-64
chr6-136884223-INS-63
chr6-136894697-INS-50
chr6-136903537-INS-192
chr6-136909566-INS-66
chr6-136916299-INS-581
chr6-136922759-INS-162
chr6-136955312-INS-54
chr6-136959415-INS-379
chr6-136966430-INS-60
chr6-136975545-INS-5305
chr6-136977005-INS-3867
chr6-136979852-INS-52
chr6-136980200-INS-4989
chr6-136993117-INS-88
chr6-137004969-INS-113
chr6-137019513-INS-130
chr6-137020177-INS-6173
chr6-137024728-INS-3749
chr6-137098835-INS-64
chr6-137107110-INS-317
chr6-137112156-INS-685
chr6-137128424-INS-52
chr6-137138219-INS-3097

chr6-137154594-INS-53
chr6-137165096-INS-142
chr6-137194438-INS-70
chr6-137251093-INS-110
chr6-137251143-INS-398
chr6-137261279-INS-63
chr6-137262778-INS-148
chr6-137279499-INS-380
chr6-137279511-INS-153
chr6-137288016-INS-1455
chr6-137297730-INS-67
chr6-137321550-INS-58
chr6-137339836-INS-478
chr6-137353256-INS-976
chr6-137407497-INS-50
chr6-137423178-INS-52
chr6-137452305-INS-50
chr6-137475464-INS-56
chr6-137576463-INS-54
chr6-137590296-INS-99
chr6-137598444-INS-320
chr6-137624547-INS-51
chr6-137636167-INS-185
chr6-137660793-INS-61
chr6-137669560-INS-404
chr6-137673050-INS-124
chr6-137685576-INS-107
chr6-137704517-INS-2847
chr6-137708069-INS-57
chr6-137709341-INS-2598
chr6-137758176-INS-6442
chr6-137788777-INS-129
chr6-137788936-INS-587
chr6-137791007-INS-196
chr6-137802221-INS-113
chr6-137808509-INS-50
chr6-137857721-INS-109
chr6-137860503-INS-5608
chr6-137875933-INS-100
chr6-137919442-INS-228
chr6-137919442-INS-54
chr6-137919620-INS-56
chr6-137934340-INS-112
chr6-137967398-INS-106
chr6-138016055-INS-53
chr6-138039030-INS-363
chr6-138039271-INS-270
chr6-138062632-INS-564
chr6-138063094-INS-688
chr6-138111085-INS-6713
chr6-138128525-INS-157
chr6-138162110-INS-64
chr6-138175829-INS-1891
chr6-138193485-INS-434
chr6-138219495-INS-351
chr6-138267338-INS-105
chr6-138279704-INS-78
chr6-138297987-INS-166
chr6-138329557-INS-73
chr6-138375421-INS-140
chr6-138384422-INS-63
chr6-138409128-INS-336
chr6-138434193-INS-158
chr6-138478452-INS-62
chr6-138486191-INS-54
chr6-138507911-INS-67
chr6-138533275-INS-557
chr6-138597100-INS-76
chr6-138601625-INS-903
chr6-138619046-INS-110
chr6-138622851-INS-575
chr6-138623022-INS-50
chr6-138640839-INS-450
chr6-138640972-INS-84
chr6-138700048-INS-52
chr6-138739858-INS-54
chr6-138739858-INS-729

chr6-138739887-INS-151
chr6-138742942-INS-50
chr6-138793102-INS-60
chr6-138808813-INS-51
chr6-138844926-INS-7116
chr6-138861990-INS-879
chr6-138870392-INS-50
chr6-138885615-INS-10891
chr6-138934135-INS-140
chr6-138934266-INS-442
chr6-138934501-INS-724
chr6-138934994-INS-364
chr6-138945733-INS-63
chr6-138945812-INS-141
chr6-138946035-INS-426
chr6-138946418-INS-878
chr6-138961564-INS-109
chr6-138964068-INS-158
chr6-138981523-INS-50
chr6-138993128-INS-131
chr6-139005857-INS-90
chr6-139011350-INS-50
chr6-139014714-INS-303
chr6-139014918-INS-152
chr6-139014921-INS-2050
chr6-139014930-INS-566
chr6-139075233-INS-162
chr6-139076506-INS-1305
chr6-139202204-INS-4452
chr6-139203867-INS-56
chr6-139216791-INS-4863
chr6-139233199-INS-62
chr6-139250497-INS-108
chr6-139301016-INS-60
chr6-139332364-INS-206
chr6-139332367-INS-80
chr6-139389181-INS-54
chr6-139392767-INS-212
chr6-139407716-INS-6193
chr6-139432691-INS-1235
chr6-139438998-INS-370
chr6-139532220-INS-6508
chr6-139533984-INS-52
chr6-139555667-INS-1192
chr6-139555667-INS-5615
chr6-139575332-INS-53
chr6-139609939-INS-7084
chr6-139615384-INS-6743
chr6-139622567-INS-513
chr6-139625769-INS-211
chr6-139633592-INS-153
chr6-139633761-INS-51
chr6-139638958-INS-5719
chr6-139667879-INS-60
chr6-139667879-INS-698
chr6-139714108-INS-6396
chr6-139800204-INS-126
chr6-139853427-INS-3004
chr6-139860672-INS-607
chr6-139897668-INS-66
chr6-139898020-INS-86
chr6-139907559-INS-64
chr6-139991158-INS-186
chr6-140054306-INS-140
chr6-140055695-INS-74
chr6-140073144-INS-56
chr6-140090528-INS-57
chr6-140106771-INS-70
chr6-140109907-INS-79
chr6-140168837-INS-7101
chr6-140180945-INS-74
chr6-140194700-INS-207
chr6-140211216-INS-267
chr6-140213404-INS-155
chr6-140213782-INS-77
chr6-140213799-INS-232
chr6-140215617-INS-1647

chr6-140236348-INS-165
chr6-140252067-INS-3192
chr6-140273130-INS-56
chr6-140286503-INS-75
chr6-140354084-INS-2732
chr6-140354422-INS-86
chr6-140377263-INS-201
chr6-140384328-INS-50
chr6-140389834-INS-257
chr6-140419058-INS-990
chr6-140427734-INS-533
chr6-140446152-INS-284
chr6-140529173-INS-86
chr6-140529509-INS-181
chr6-140533867-INS-181
chr6-140536853-INS-191
chr6-140537013-INS-92
chr6-140563888-INS-220
chr6-140592789-INS-51
chr6-140605901-INS-555
chr6-140623101-INS-400
chr6-140626074-INS-94
chr6-140661748-INS-296
chr6-140661763-INS-1084
chr6-140662492-INS-206
chr6-140664867-INS-1413
chr6-140704460-INS-204
chr6-140719108-INS-710
chr6-140732669-INS-1067
chr6-140767286-INS-2031
chr6-140773494-INS-88
chr6-140795180-INS-60
chr6-140803920-INS-57
chr6-140815683-INS-205
chr6-140821412-INS-184
chr6-140838879-INS-106
chr6-140843602-INS-1860
chr6-140843773-INS-158
chr6-140857050-INS-3249
chr6-140864241-INS-2234
chr6-140864567-INS-576
chr6-140880253-INS-78
chr6-140884901-INS-189
chr6-140885084-INS-1525
chr6-140979834-INS-814
chr6-141033165-INS-9417
chr6-141033926-INS-443
chr6-141065925-INS-7163
chr6-141093130-INS-4875
chr6-141098463-INS-1076
chr6-141100782-INS-1536
chr6-141108927-INS-2303
chr6-141195104-INS-60
chr6-141212330-INS-2527
chr6-141222048-INS-896
chr6-141367313-INS-64
chr6-141403273-INS-187
chr6-141492117-INS-200
chr6-141590668-INS-1689
chr6-141615711-INS-65
chr6-141615848-INS-62
chr6-141656838-INS-2708
chr6-141688640-INS-158
chr6-141721685-INS-401
chr6-141728979-INS-6397
chr6-141759916-INS-107
chr6-141804690-INS-145
chr6-141814084-INS-78
chr6-141843433-INS-50
chr6-141847242-INS-1485
chr6-141869127-INS-50
chr6-141900974-INS-114
chr6-141922584-INS-7476
chr6-141965478-INS-6291
chr6-141987467-INS-6418
chr6-142140433-INS-136
chr6-142167458-INS-1128

chr6-142195054-INS-65
chr6-142198178-INS-413
chr6-142240637-INS-74
chr6-142255508-INS-152
chr6-142281716-INS-8458
chr6-142326376-INS-456
chr6-142344886-INS-256
chr6-142364112-INS-51
chr6-142374882-INS-74
chr6-142410236-INS-92
chr6-142410767-INS-71
chr6-142411264-INS-153
chr6-142468208-INS-88
chr6-142471401-INS-64
chr6-142516358-INS-51
chr6-142531610-INS-280
chr6-142651285-INS-64
chr6-142687780-INS-320
chr6-142696345-INS-249
chr6-142701068-INS-2931
chr6-142716604-INS-198
chr6-142716604-INS-2343
chr6-142740705-INS-100
chr6-142749616-INS-490
chr6-142833148-INS-347
chr6-142836336-INS-226
chr6-142853827-INS-362
chr6-142858182-INS-182
chr6-142882676-INS-224
chr6-142888375-INS-184
chr6-142902637-INS-387
chr6-142902990-INS-1121
chr6-142922341-INS-5209
chr6-142944620-INS-228
chr6-142955929-INS-1113
chr6-142955929-INS-3170
chr6-143003317-INS-360
chr6-143061864-INS-58
chr6-143171582-INS-1158
chr6-143196672-INS-6222
chr6-143263923-INS-1093
chr6-143273526-INS-265
chr6-143283564-INS-319
chr6-143288983-INS-6564
chr6-143302466-INS-153
chr6-143316548-INS-71
chr6-143333929-INS-64
chr6-143354423-INS-14850
chr6-143391167-INS-103
chr6-143408796-INS-110
chr6-143410487-INS-1412
chr6-143410685-INS-227
chr6-143418797-INS-56
chr6-143426033-INS-562
chr6-143426325-INS-704
chr6-143441163-INS-176
chr6-143506327-INS-53
chr6-143556228-INS-129
chr6-143576234-INS-113
chr6-143583885-INS-52
chr6-143587640-INS-52
chr6-143622300-INS-2751
chr6-143728661-INS-1026
chr6-143760169-INS-131
chr6-143788312-INS-109
chr6-143868604-INS-51
chr6-143872090-INS-64
chr6-143896674-INS-79
chr6-143930714-INS-143
chr6-143937399-INS-903
chr6-143941476-INS-58
chr6-143972802-INS-119
chr6-144123899-INS-61
chr6-144165058-INS-180
chr6-144195992-INS-1831
chr6-144281977-INS-50
chr6-144325573-INS-98

chr6-144335344-INS-674
chr6-144337711-INS-157
chr6-144337711-INS-62
chr6-144337940-INS-2565
chr6-144344972-INS-360
chr6-144362460-INS-156
chr6-144377411-INS-64
chr6-144390448-INS-248
chr6-144390471-INS-2573
chr6-144428091-INS-232
chr6-144435468-INS-70
chr6-144455402-INS-72
chr6-144455570-INS-240
chr6-144457433-INS-50
chr6-144464821-INS-72
chr6-144486419-INS-50
chr6-144488723-INS-194
chr6-144518717-INS-50
chr6-144547125-INS-799
chr6-144558450-INS-214
chr6-144560953-INS-143
chr6-144605264-INS-238
chr6-144605293-INS-510
chr6-144647555-INS-82
chr6-144712055-INS-102
chr6-144712702-INS-63
chr6-144735485-INS-53
chr6-144785800-INS-97
chr6-144818471-INS-500
chr6-144836891-INS-82
chr6-144852858-INS-54
chr6-144856990-INS-108
chr6-144871359-INS-6742
chr6-144876304-INS-70
chr6-144906478-INS-50
chr6-144906583-INS-207
chr6-144929716-INS-460
chr6-144929812-INS-226
chr6-144950171-INS-440
chr6-144950358-INS-88
chr6-144956143-INS-58
chr6-144956405-INS-56
chr6-144973385-INS-3836
chr6-144979640-INS-326
chr6-144989058-INS-299
chr6-145013023-INS-169
chr6-145016308-INS-387
chr6-145030595-INS-3208
chr6-145056410-INS-432
chr6-145060158-INS-96
chr6-145103201-INS-441
chr6-145106517-INS-54
chr6-145114672-INS-68
chr6-145114822-INS-528
chr6-145192178-INS-74
chr6-145201494-INS-64
chr6-145226343-INS-136
chr6-145227215-INS-379
chr6-145227216-INS-83
chr6-145244185-INS-206
chr6-145244829-INS-129
chr6-145266713-INS-62
chr6-145278214-INS-52
chr6-145278310-INS-55
chr6-145283781-INS-60
chr6-145283866-INS-1274
chr6-145317367-INS-50
chr6-145322540-INS-95
chr6-145322793-INS-75
chr6-145330112-INS-121
chr6-145356895-INS-1990
chr6-145357985-INS-970
chr6-145357997-INS-4017
chr6-145367680-INS-4807
chr6-145371520-INS-52
chr6-145386089-INS-826
chr6-145390694-INS-58

chr6-145395116-INS-50
chr6-145399405-INS-7150
chr6-145400143-INS-1897
chr6-145400511-INS-3123
chr6-145404091-INS-104
chr6-145408096-INS-2656
chr6-145411632-INS-2250
chr6-145448476-INS-543
chr6-145453669-INS-234
chr6-145475452-INS-290
chr6-145482593-INS-7137
chr6-145495929-INS-2147
chr6-145502196-INS-300
chr6-145520694-INS-53
chr6-145538242-INS-6421
chr6-145551842-INS-50
chr6-145563728-INS-50
chr6-145570803-INS-5856
chr6-145582874-INS-102
chr6-145583129-INS-100
chr6-145595422-INS-337
chr6-145597315-INS-3090
chr6-145609238-INS-92
chr6-145614213-INS-775
chr6-145614213-INS-92
chr6-145667521-INS-78
chr6-145691464-INS-54
chr6-145699171-INS-50
chr6-145736816-INS-8682
chr6-145749442-INS-193
chr6-145755195-INS-50
chr6-145775746-INS-57
chr6-145784868-INS-6437
chr6-145785697-INS-498
chr6-145791197-INS-66
chr6-145802447-INS-196
chr6-145803444-INS-3481
chr6-145855300-INS-62
chr6-145855476-INS-53
chr6-145920283-INS-56
chr6-145926674-INS-5773
chr6-145929989-INS-11432
chr6-145966337-INS-294
chr6-145980594-INS-260
chr6-145985909-INS-3281
chr6-146014533-INS-50
chr6-146015212-INS-60
chr6-146024450-INS-72
chr6-146104033-INS-50
chr6-146133703-INS-126
chr6-146135303-INS-496
chr6-146155132-INS-7111
chr6-146247219-INS-143
chr6-146252959-INS-240
chr6-146258321-INS-160
chr6-146266781-INS-1313
chr6-146280535-INS-806
chr6-146280541-INS-348
chr6-146281046-INS-50
chr6-146313005-INS-78
chr6-146341595-INS-55
chr6-146353441-INS-58
chr6-146359304-INS-1267
chr6-146401231-INS-1244
chr6-146424938-INS-1424
chr6-146432911-INS-277
chr6-146449319-INS-1003
chr6-146468238-INS-4168
chr6-146480776-INS-50
chr6-146518491-INS-50
chr6-146546361-INS-178
chr6-146588308-INS-204
chr6-146618768-INS-328
chr6-146619717-INS-484
chr6-146630460-INS-191
chr6-146635421-INS-886
chr6-146635766-INS-1090

chr6-146651771-INS-288
chr6-146703683-INS-310
chr6-146710995-INS-169
chr6-146756875-INS-220
chr6-146763298-INS-310
chr6-146771062-INS-51
chr6-146809917-INS-1841
chr6-146839137-INS-72
chr6-146938458-INS-55
chr6-146955119-INS-218
chr6-147007205-INS-72
chr6-147012352-INS-192
chr6-147020496-INS-599
chr6-147020783-INS-799
chr6-147026651-INS-89
chr6-147058846-INS-144
chr6-147064571-INS-1506
chr6-147102590-INS-2074
chr6-147108929-INS-6433
chr6-147128129-INS-211
chr6-147268448-INS-594
chr6-147345852-INS-1993
chr6-147347224-INS-147
chr6-147367115-INS-57
chr6-147398647-INS-68
chr6-147410890-INS-394
chr6-147431092-INS-58
chr6-147473231-INS-50
chr6-147522729-INS-212
chr6-147528113-INS-286
chr6-147578942-INS-496
chr6-147620498-INS-67
chr6-147666347-INS-81
chr6-147666496-INS-182
chr6-147669803-INS-78
chr6-147680436-INS-6854
chr6-147705383-INS-1466
chr6-147769383-INS-6379
chr6-147781544-INS-794
chr6-147783239-INS-50
chr6-147787161-INS-128
chr6-147853542-INS-172
chr6-147865400-INS-6466
chr6-147875840-INS-55
chr6-147895711-INS-77
chr6-147916282-INS-704
chr6-147927720-INS-1539
chr6-147964438-INS-64
chr6-147998125-INS-55
chr6-148045143-INS-58
chr6-148050377-INS-216
chr6-148055322-INS-50
chr6-148171054-INS-5044
chr6-148193410-INS-310
chr6-148195981-INS-331
chr6-148197700-INS-7103
chr6-148198349-INS-1065
chr6-148214286-INS-50
chr6-148214743-INS-311
chr6-148277945-INS-141
chr6-148291142-INS-3492
chr6-148321074-INS-351
chr6-148327852-INS-50
chr6-148359772-INS-481
chr6-148364690-INS-132
chr6-148377763-INS-64
chr6-148392115-INS-86
chr6-148392154-INS-196
chr6-148421294-INS-1626
chr6-148429541-INS-54
chr6-148481091-INS-72
chr6-148501506-INS-127
chr6-148542094-INS-187
chr6-148557471-INS-104
chr6-148615993-INS-4644
chr6-148648210-INS-6464
chr6-148663128-INS-6583

chr6-148670664-INS-117
chr6-148705095-INS-147
chr6-148816665-INS-210
chr6-148879173-INS-56
chr6-148883011-INS-50
chr6-148886144-INS-435
chr6-148887272-INS-58
chr6-148972068-INS-4442
chr6-148982881-INS-161
chr6-149026032-INS-209
chr6-149026032-INS-98
chr6-149027194-INS-212
chr6-149049190-INS-2547
chr6-149070485-INS-76
chr6-149092830-INS-211
chr6-149112841-INS-52
chr6-149153423-INS-204
chr6-149179368-INS-408
chr6-149186434-INS-205
chr6-149189449-INS-5339
chr6-149278982-INS-50
chr6-149303934-INS-626
chr6-149334499-INS-53
chr6-149346247-INS-187
chr6-149378666-INS-58
chr7-3125028-INS-203
chr7-3140034-INS-1311
chr7-3187459-INS-500
chr7-3201161-INS-120
chr7-3209731-INS-196
chr7-3213366-INS-301
chr7-3215564-INS-205
chr7-3223561-INS-209
chr7-3238747-INS-228
chr7-3245029-INS-177
chr7-3245239-INS-196
chr7-3255984-INS-79
chr7-3261366-INS-205
chr7-3264690-INS-64
chr7-3273089-INS-184
chr7-3299921-INS-350
chr7-3307824-INS-166
chr7-3309377-INS-178
chr7-3337196-INS-108
chr7-3381134-INS-63
chr7-3400811-INS-134
chr7-3424035-INS-52
chr7-3433125-INS-586
chr7-3433155-INS-170
chr7-3436703-INS-384
chr7-3451346-INS-96
chr7-3482989-INS-52
chr7-3494163-INS-70
chr7-3511889-INS-3036
chr7-3517643-INS-52
chr7-3551381-INS-82
chr7-3557248-INS-242
chr7-3560029-INS-498
chr7-3565020-INS-107
chr7-3575991-INS-110
chr7-3584416-INS-1762
chr7-3584489-INS-105
chr7-3605925-INS-268
chr7-3606676-INS-112
chr7-3606841-INS-170
chr7-3607472-INS-69
chr7-3630812-INS-172
chr7-3686443-INS-76
chr7-3705834-INS-220
chr7-3726082-INS-111
chr7-3726248-INS-680
chr7-3726263-INS-196
chr7-3777828-INS-175
chr7-3822451-INS-1895
chr7-3831515-INS-875
chr7-3909992-INS-236
chr7-3950072-INS-6679

chr7-3979298-INS-4775
chr7-3993090-INS-965
chr7-3993238-INS-273
chr7-4067067-INS-138
chr7-4077125-INS-293
chr7-4095758-INS-1739
chr7-4113202-INS-456
chr7-4159299-INS-55
chr7-4173906-INS-199
chr7-4213325-INS-638
chr7-4213552-INS-853
chr7-4217350-INS-520
chr7-4232275-INS-7268
chr7-4245696-INS-170
chr7-4256196-INS-221
chr7-4289988-INS-201
chr7-4295303-INS-256
chr7-4317259-INS-112
chr7-4328716-INS-5768
chr7-4330682-INS-434
chr7-4343416-INS-1884
chr7-4357780-INS-87
chr7-4361693-INS-54
chr7-4400383-INS-58
chr7-4410452-INS-196
chr7-4425713-INS-172
chr7-4434093-INS-172
chr7-4442780-INS-133
chr7-4445010-INS-316
chr7-4456361-INS-68
chr7-4456479-INS-56
chr7-4490854-INS-205
chr7-4509300-INS-170
chr7-4509300-INS-892
chr7-4532581-INS-51
chr7-4544762-INS-402
chr7-4545016-INS-53
chr7-4616961-INS-245
chr7-4625652-INS-58
chr7-4630293-INS-67
chr7-4672118-INS-217
chr7-4689844-INS-190
chr7-4702565-INS-215
chr7-4716560-INS-54
chr7-4727700-INS-68
chr7-4730096-INS-56
chr7-4763546-INS-406
chr7-4763594-INS-102
chr7-4766539-INS-183
chr7-4775213-INS-78
chr7-4777236-INS-212
chr7-4794318-INS-206
chr7-4818716-INS-3157
chr7-4822104-INS-217
chr7-4831346-INS-54
chr7-4851774-INS-1885
chr7-4855791-INS-120
chr7-4857432-INS-4192
chr7-4858790-INS-399
chr7-4863952-INS-198
chr7-4884517-INS-206
chr7-4888282-INS-1445
chr7-4891126-INS-252
chr7-4917383-INS-201
chr7-4938531-INS-107
chr7-4958072-INS-82
chr7-4989425-INS-80
chr7-5013389-INS-204
chr7-5013513-INS-270
chr7-5027922-INS-71
chr7-5028017-INS-675
chr7-5058496-INS-2186
chr7-5073744-INS-307
chr7-5079217-INS-103
chr7-5085958-INS-176
chr7-5097915-INS-3164
chr7-5105055-INS-122

chr7-5117714-INS-54
chr7-5138591-INS-2086
chr7-5138918-INS-6111
chr7-5156147-INS-194
chr7-5178836-INS-1294
chr7-5307385-INS-805
chr7-5318845-INS-50
chr7-5320202-INS-121
chr7-5394008-INS-6226
chr7-5418684-INS-3896
chr7-5467404-INS-389
chr7-5520507-INS-3113
chr7-5543524-INS-87
chr7-5569217-INS-83
chr7-5601691-INS-591
chr7-5604188-INS-56
chr7-5609802-INS-183
chr7-5655824-INS-505
chr7-5659896-INS-74
chr7-5660269-INS-896
chr7-5784564-INS-1828
chr7-5785725-INS-899
chr7-5804777-INS-79
chr7-5919441-INS-1644
chr7-5926318-INS-2676
chr7-5930013-INS-79
chr7-5931033-INS-71
chr7-5945812-INS-449
chr7-5955029-INS-4181
chr7-5959871-INS-398
chr7-6032209-INS-244
chr7-6040627-INS-621
chr7-6052086-INS-229
chr7-6084748-INS-66
chr7-6129133-INS-158
chr7-6139143-INS-284
chr7-6144280-INS-217
chr7-6144419-INS-58
chr7-6155046-INS-190
chr7-6183066-INS-96
chr7-6216708-INS-64
chr7-6217637-INS-180
chr7-6219151-INS-150
chr7-6225830-INS-450
chr7-6228662-INS-60
chr7-6228767-INS-209
chr7-6243351-INS-340
chr7-6251049-INS-210
chr7-6259992-INS-278
chr7-6268493-INS-4560
chr7-6269991-INS-199
chr7-6271554-INS-90
chr7-6273354-INS-52
chr7-6292481-INS-393
chr7-6299052-INS-57
chr7-6299423-INS-72
chr7-6305717-INS-2309
chr7-6312168-INS-63
chr7-6312365-INS-1241
chr7-6318427-INS-50
chr7-6321588-INS-156
chr7-6322980-INS-70
chr7-6323001-INS-227
chr7-6323896-INS-327
chr7-6347210-INS-1874
chr7-6354869-INS-180
chr7-6376840-INS-191
chr7-6383055-INS-4746
chr7-6427915-INS-223
chr7-6456060-INS-1733
chr7-6457221-INS-60
chr7-6476104-INS-52
chr7-6486412-INS-50
chr7-6508402-INS-172
chr7-6508408-INS-75
chr7-6520527-INS-219
chr7-6581357-INS-51

chr7-6641324-INS-5338
chr7-6649219-INS-83
chr7-6676635-INS-73
chr7-6705948-INS-143
chr7-6749786-INS-175
chr7-6817166-INS-62
chr7-6818850-INS-288
chr7-6839666-INS-70
chr7-6843544-INS-213
chr7-6851360-INS-315
chr7-6880167-INS-9133
chr7-6899666-INS-60
chr7-6981894-INS-1944
chr7-6984874-INS-13508
chr7-7034815-INS-52
chr7-7047722-INS-4735
chr7-7063292-INS-149
chr7-7084355-INS-183
chr7-7096518-INS-174
chr7-7105034-INS-3043
chr7-7106653-INS-61
chr7-7114286-INS-82
chr7-7126812-INS-55
chr7-7128200-INS-163
chr7-7141452-INS-188
chr7-7192256-INS-256
chr7-7303385-INS-97
chr7-7376492-INS-96
chr7-7522044-INS-7144
chr7-8502112-INS-3116
chr7-8506043-INS-2773
chr7-8570250-INS-55
chr7-8811659-INS-181
chr7-9723635-INS-50
chr7-9980827-INS-38947
chr7-10125673-INS-2126
chr7-10445300-INS-6185
chr7-10446430-INS-150
chr7-10514897-INS-429
chr7-10522737-INS-392
chr7-10527739-INS-506
chr7-10874917-INS-86
chr7-10892883-INS-498
chr7-10936761-INS-6573
chr7-11078396-INS-5490
chr7-11101847-INS-615
chr7-11104103-INS-200
chr7-11124834-INS-1122
chr7-11347544-INS-5693
chr7-11348998-INS-259
chr7-11444235-INS-6198
chr7-11459212-INS-638
chr7-11486005-INS-6895
chr7-11495734-INS-145
chr7-11495776-INS-72
chr7-11523608-INS-11182
chr7-11535444-INS-354
chr7-11548839-INS-54
chr7-11556383-INS-66
chr7-11557311-INS-53
chr7-11610953-INS-1265
chr7-11642585-INS-4561
chr7-11680239-INS-82
chr7-11699404-INS-1870
chr7-11763599-INS-103
chr7-11768071-INS-64
chr7-11857023-INS-2369
chr7-11858534-INS-72
chr7-11866324-INS-5703
chr7-11883249-INS-82
chr7-11908241-INS-128
chr7-11946039-INS-4958
chr7-11951319-INS-1453
chr7-11956200-INS-343
chr7-11965929-INS-5842
chr7-12180136-INS-51
chr7-12233775-INS-186

chr7-12233831-INS-60
chr7-12270350-INS-204
chr7-12334199-INS-52
chr7-12363239-INS-8431
chr7-12389086-INS-110
chr7-12407309-INS-624
chr7-12439142-INS-3284
chr7-12471478-INS-63
chr7-12520532-INS-80
chr7-12559493-INS-323
chr7-12581235-INS-50
chr7-12629552-INS-52
chr7-12655046-INS-73
chr7-12692512-INS-202
chr7-12693331-INS-284
chr7-12694072-INS-191
chr7-12721253-INS-120
chr7-12736460-INS-54
chr7-12745508-INS-56
chr7-12790914-INS-205
chr7-12794104-INS-159
chr7-12794724-INS-1198
chr7-12822225-INS-68
chr7-12854200-INS-50
chr7-13044058-INS-204
chr7-13044058-INS-411
chr7-13085618-INS-591
chr7-13092573-INS-681
chr7-13113602-INS-167
chr7-13127722-INS-1344
chr7-13128416-INS-1301
chr7-13131285-INS-6635
chr7-13158634-INS-1745
chr7-13256356-INS-70
chr7-13266271-INS-4229
chr7-13269136-INS-215
chr7-13281990-INS-1289
chr7-13306905-INS-14989
chr7-13573619-INS-2998
chr7-13596661-INS-13417
chr7-13603650-INS-50
chr7-13654352-INS-89
chr7-13667984-INS-3304
chr7-13673625-INS-466
chr7-13683274-INS-3122
chr7-13687268-INS-68
chr7-13768969-INS-6691
chr7-13777031-INS-100
chr7-13814681-INS-1344
chr7-13857554-INS-493
chr7-13995575-INS-3155
chr7-14000378-INS-480
chr7-14000436-INS-680
chr7-14007948-INS-219
chr7-14008747-INS-205
chr7-14021255-INS-56
chr7-14039123-INS-1379
chr7-14056752-INS-52
chr7-14083561-INS-371
chr7-14210565-INS-54
chr7-14218462-INS-2699
chr7-14237843-INS-50
chr7-14316543-INS-267
chr7-14324139-INS-207
chr7-14364499-INS-126
chr7-14364791-INS-68
chr7-14384658-INS-6891
chr7-14487949-INS-3253
chr7-14505947-INS-4663
chr7-14532597-INS-6374
chr7-14572012-INS-52
chr7-14649729-INS-852
chr7-14672770-INS-112
chr7-14673632-INS-58
chr7-14674049-INS-463
chr7-14678416-INS-1506
chr7-14750696-INS-806

chr7-14845356-INS-214
chr7-14858263-INS-443
chr7-15018617-INS-1305
chr7-15140843-INS-201
chr7-15344822-INS-825
chr7-15356434-INS-147
chr7-15424204-INS-62
chr7-15525588-INS-52
chr7-15638201-INS-11405
chr7-15644389-INS-382
chr7-15650797-INS-62
chr7-15666556-INS-52
chr7-15666605-INS-106
chr7-15666605-INS-261
chr7-15667193-INS-144
chr7-15671212-INS-157
chr7-15742864-INS-106
chr7-15742864-INS-265
chr7-15755418-INS-364
chr7-15770917-INS-50
chr7-15785535-INS-66
chr7-15801898-INS-65
chr7-15885440-INS-73
chr7-15910712-INS-212
chr7-15910712-INS-425
chr7-15926618-INS-56
chr7-15927884-INS-13945
chr7-15928563-INS-158
chr7-15929909-INS-277
chr7-15940646-INS-86
chr7-15974610-INS-55
chr7-15978468-INS-74
chr7-15984076-INS-50
chr7-15994209-INS-637
chr7-15998250-INS-159
chr7-15998250-INS-53
chr7-16000066-INS-566
chr7-16027790-INS-2510
chr7-16035885-INS-53
chr7-16035893-INS-159
chr7-16054274-INS-87
chr7-16055229-INS-55
chr7-16068674-INS-76
chr7-16076898-INS-229
chr7-16123504-INS-222
chr7-16125930-INS-200
chr7-16176163-INS-256
chr7-16190472-INS-108
chr7-16221207-INS-52
chr7-16251697-INS-50
chr7-16268834-INS-60
chr7-16359820-INS-5417
chr7-16360384-INS-58
chr7-16375369-INS-1135
chr7-16383261-INS-2273
chr7-16394288-INS-2364
chr7-16395247-INS-25585
chr7-16395347-INS-212
chr7-16435230-INS-1500
chr7-16439027-INS-210
chr7-16440775-INS-132
chr7-16445995-INS-224
chr7-16447543-INS-4892
chr7-16459518-INS-140
chr7-16461363-INS-806
chr7-16500921-INS-51
chr7-16500996-INS-203
chr7-16501661-INS-107
chr7-16505008-INS-108
chr7-16505107-INS-206
chr7-16509166-INS-52
chr7-16532711-INS-204
chr7-16533817-INS-212
chr7-16536670-INS-442
chr7-16537403-INS-128
chr7-16537732-INS-201
chr7-16598630-INS-215

chr7-16612492-INS-105
chr7-16698829-INS-262
chr7-16698830-INS-53
chr7-16706605-INS-65
chr7-16708753-INS-2567
chr7-16723533-INS-52
chr7-16726878-INS-228
chr7-16727074-INS-81
chr7-16734301-INS-182
chr7-16735174-INS-212
chr7-16774570-INS-53
chr7-16775223-INS-157
chr7-16798186-INS-105
chr7-16798432-INS-264
chr7-16801942-INS-5519
chr7-16833627-INS-4667
chr7-16903502-INS-52
chr7-16904015-INS-201
chr7-16904113-INS-52
chr7-16910598-INS-52
chr7-16930462-INS-1368
chr7-16938331-INS-67
chr7-16947061-INS-3253
chr7-16966861-INS-2019
chr7-16982293-INS-624
chr7-16997555-INS-183
chr7-17041743-INS-1487
chr7-17145361-INS-573
chr7-17165341-INS-1698
chr7-17356810-INS-94
chr7-17519745-INS-6567
chr7-17599501-INS-50
chr7-17630773-INS-999
chr7-17632298-INS-2014
chr7-17665881-INS-262
chr7-17675930-INS-462
chr7-17713059-INS-54
chr7-17717089-INS-2141
chr7-17731776-INS-6524
chr7-17743196-INS-52
chr7-17783376-INS-5651
chr7-17817574-INS-1382
chr7-17877670-INS-6403
chr7-17909535-INS-52
chr7-17921980-INS-1632
chr7-17922121-INS-214
chr7-17922246-INS-603
chr7-17975974-INS-5084
chr7-17988902-INS-51
chr7-18011207-INS-77
chr7-18031770-INS-76
chr7-18047076-INS-1340
chr7-18056654-INS-80
chr7-18089656-INS-468
chr7-18104098-INS-86
chr7-18141819-INS-69
chr7-18142371-INS-204
chr7-18145997-INS-80
chr7-18147028-INS-56
chr7-18183187-INS-3501
chr7-18206485-INS-76
chr7-18265346-INS-71
chr7-18272518-INS-52
chr7-18278165-INS-112
chr7-18349510-INS-155
chr7-18364199-INS-8561
chr7-18367906-INS-70
chr7-18374549-INS-176
chr7-18440617-INS-251
chr7-18466037-INS-2756
chr7-18482805-INS-435
chr7-18540833-INS-80
chr7-18540970-INS-72
chr7-18544205-INS-58
chr7-18557561-INS-8416
chr7-18578756-INS-3041
chr7-18617087-INS-90

chr7-18617292-INS-53
chr7-18637179-INS-83
chr7-18637332-INS-508
chr7-18682808-INS-56
chr7-18683180-INS-53
chr7-18710884-INS-264
chr7-18710884-INS-52
chr7-18715047-INS-166
chr7-18731030-INS-53
chr7-18752921-INS-154
chr7-18790277-INS-54
chr7-18793263-INS-2702
chr7-18901984-INS-106
chr7-18923031-INS-90
chr7-18923247-INS-60
chr7-18937540-INS-106
chr7-18944331-INS-75
chr7-18944349-INS-648
chr7-18945273-INS-188
chr7-18946747-INS-116
chr7-18960351-INS-196
chr7-18971460-INS-201
chr7-19008566-INS-208
chr7-19021713-INS-287
chr7-19022224-INS-831
chr7-19022285-INS-263
chr7-19056266-INS-159
chr7-19070326-INS-220
chr7-19083362-INS-142
chr7-19098930-INS-53
chr7-19169232-INS-204
chr7-19176480-INS-158
chr7-19176480-INS-52
chr7-19176538-INS-424
chr7-19180754-INS-107
chr7-19188445-INS-52
chr7-19196309-INS-8950
chr7-19204618-INS-78
chr7-19219796-INS-212
chr7-19219951-INS-51
chr7-19226318-INS-107
chr7-19279262-INS-51
chr7-19344039-INS-62
chr7-19355168-INS-371
chr7-19374271-INS-51
chr7-19374273-INS-318
chr7-19402309-INS-610
chr7-19418926-INS-105
chr7-19422765-INS-87
chr7-19422926-INS-396
chr7-19453471-INS-162
chr7-19460757-INS-50
chr7-19461432-INS-204
chr7-19464350-INS-137
chr7-19464357-INS-346
chr7-19491473-INS-52
chr7-19494743-INS-105
chr7-19527449-INS-107
chr7-19535512-INS-53
chr7-19536938-INS-147
chr7-19560473-INS-277
chr7-19561820-INS-50
chr7-19571242-INS-159
chr7-19579783-INS-158
chr7-19593152-INS-53
chr7-19614056-INS-236
chr7-19618823-INS-105
chr7-19626228-INS-516
chr7-19627820-INS-1842
chr7-19642937-INS-475
chr7-19654830-INS-133
chr7-19663993-INS-57
chr7-19675999-INS-4514
chr7-19677319-INS-154
chr7-19677341-INS-310
chr7-19704698-INS-311
chr7-19705052-INS-439

chr7-19709122-INS-64
chr7-19712302-INS-546
chr7-19732195-INS-410
chr7-19732690-INS-202
chr7-19756326-INS-184
chr7-19756335-INS-918
chr7-19778554-INS-170
chr7-19781349-INS-1956
chr7-19824439-INS-2813
chr7-19882824-INS-787
chr7-20068469-INS-329
chr7-20641567-INS-60
chr7-20656084-INS-193
chr7-20669734-INS-926
chr7-20695216-INS-158
chr7-20701205-INS-758
chr7-20710717-INS-81
chr7-22882523-INS-1809
chr7-22951663-INS-143
chr7-22967564-INS-72
chr7-22995202-INS-1095
chr7-23037552-INS-242
chr7-23037617-INS-66
chr7-23043943-INS-96
chr7-23130527-INS-135
chr7-23130964-INS-231
chr7-23133544-INS-160
chr7-23154072-INS-190
chr7-23154072-INS-68
chr7-23162701-INS-54
chr7-23234147-INS-7114
chr7-23245224-INS-1477
chr7-23356906-INS-5499
chr7-23378421-INS-135
chr7-23391126-INS-69
chr7-23416055-INS-2667
chr7-23502530-INS-1447
chr7-23507344-INS-68
chr7-23558147-INS-161
chr7-23602678-INS-50
chr7-23610657-INS-6233
chr7-23745198-INS-2465
chr7-23775452-INS-6449
chr7-23789194-INS-567
chr7-23828032-INS-50
chr7-23829303-INS-84
chr7-23832641-INS-7781
chr7-23891704-INS-1340
chr7-23892125-INS-3812
chr7-23929921-INS-106
chr7-23930405-INS-539
chr7-23988875-INS-589
chr7-23989776-INS-94
chr7-24033842-INS-495
chr7-24033857-INS-102
chr7-24033913-INS-50
chr7-24048995-INS-241
chr7-24049185-INS-628
chr7-24061421-INS-80
chr7-24061474-INS-64
chr7-24062313-INS-60
chr7-24066446-INS-53
chr7-24068502-INS-68
chr7-24075559-INS-264
chr7-24091429-INS-140
chr7-24143014-INS-50
chr7-24147291-INS-212
chr7-24147424-INS-53
chr7-24152703-INS-50
chr7-24160767-INS-213
chr7-24181976-INS-88
chr7-24183685-INS-58
chr7-24222005-INS-64
chr7-24251143-INS-206
chr7-24260499-INS-209
chr7-24261819-INS-7305
chr7-24296737-INS-107

chr7-24296970-INS-321
chr7-24311554-INS-70
chr7-24315557-INS-157
chr7-24334810-INS-159
chr7-24368409-INS-114
chr7-24388712-INS-56
chr7-24400954-INS-2041
chr7-24402983-INS-838
chr7-24435860-INS-12797
chr7-24462911-INS-442
chr7-24485837-INS-60
chr7-24488889-INS-1483
chr7-24510258-INS-168
chr7-24558518-INS-207
chr7-24561453-INS-2749
chr7-24563081-INS-859
chr7-24615513-INS-312
chr7-24647776-INS-1501
chr7-24651684-INS-80
chr7-24659573-INS-292
chr7-24659673-INS-5260
chr7-24659722-INS-1559
chr7-24659899-INS-494
chr7-24663202-INS-482
chr7-24663206-INS-76
chr7-24668555-INS-193
chr7-24675095-INS-185
chr7-24675937-INS-202
chr7-24676336-INS-219
chr7-24697441-INS-140
chr7-24715822-INS-65
chr7-24738848-INS-173
chr7-24760897-INS-59
chr7-24761234-INS-90
chr7-24812926-INS-741
chr7-24813032-INS-369
chr7-24813047-INS-158
chr7-24828659-INS-140
chr7-24840691-INS-120
chr7-24843176-INS-102
chr7-24843343-INS-918
chr7-24843354-INS-219
chr7-24884490-INS-50
chr7-24898005-INS-58
chr7-24957259-INS-214
chr7-24995391-INS-210
chr7-25022064-INS-171
chr7-25023504-INS-225
chr7-25028487-INS-121
chr7-25035925-INS-82
chr7-25047159-INS-369
chr7-25061891-INS-585
chr7-25096492-INS-125
chr7-25116171-INS-216
chr7-25180828-INS-52
chr7-25185898-INS-123
chr7-25274431-INS-5574
chr7-25282340-INS-72
chr7-25295584-INS-57
chr7-25342423-INS-172
chr7-25368119-INS-65
chr7-25371161-INS-223
chr7-25371967-INS-56
chr7-25399851-INS-304
chr7-25406018-INS-192
chr7-25464532-INS-106
chr7-25473078-INS-268
chr7-25474934-INS-140
chr7-25493553-INS-316
chr7-25495465-INS-78
chr7-25521172-INS-72
chr7-25521559-INS-148
chr7-25527167-INS-130
chr7-25533820-INS-90
chr7-25572507-INS-448
chr7-25581165-INS-1244
chr7-25581257-INS-91

chr7-25585049-INS-56
chr7-25585862-INS-52
chr7-25594192-INS-66
chr7-25594935-INS-200
chr7-25603009-INS-186
chr7-25609860-INS-1074
chr7-25613242-INS-342
chr7-25620393-INS-760
chr7-25652633-INS-51
chr7-25665144-INS-144
chr7-25682919-INS-1128
chr7-25688401-INS-50
chr7-25697203-INS-53
chr7-25736181-INS-1488
chr7-25788698-INS-58
chr7-25822343-INS-576
chr7-25887478-INS-55
chr7-25945758-INS-7968
chr7-25953093-INS-66
chr7-26118126-INS-160
chr7-26277836-INS-92
chr7-26278074-INS-202
chr7-26278665-INS-890
chr7-26278715-INS-52
chr7-26284313-INS-610
chr7-26284318-INS-250
chr7-26284612-INS-634
chr7-26307055-INS-6780
chr7-26307413-INS-6596
chr7-26316546-INS-1025
chr7-26319575-INS-6151
chr7-26359340-INS-2194
chr7-26367115-INS-60
chr7-26367551-INS-70
chr7-26371839-INS-994
chr7-26373006-INS-54
chr7-26419659-INS-118
chr7-26419665-INS-598
chr7-26443929-INS-213
chr7-26467613-INS-265
chr7-26479836-INS-193
chr7-26495437-INS-83
chr7-26528234-INS-1079
chr7-26528573-INS-335
chr7-26537711-INS-63
chr7-26547953-INS-2985
chr7-26661142-INS-60
chr7-26755535-INS-1888
chr7-26764961-INS-6225
chr7-26777032-INS-3849
chr7-26779885-INS-6447
chr7-26784323-INS-2867
chr7-26793185-INS-53
chr7-26810519-INS-83
chr7-26816954-INS-167
chr7-26823751-INS-77
chr7-26825507-INS-105
chr7-26839425-INS-3165
chr7-26851833-INS-1898
chr7-26852267-INS-1243
chr7-26900251-INS-54
chr7-26901100-INS-851
chr7-26918460-INS-154
chr7-26936150-INS-390
chr7-26965308-INS-265
chr7-26965766-INS-104
chr7-26973152-INS-55
chr7-26996518-INS-318
chr7-26996545-INS-53
chr7-27000715-INS-614
chr7-27011798-INS-156
chr7-27018117-INS-60
chr7-27049543-INS-87
chr7-27062133-INS-71
chr7-27062884-INS-55
chr7-27073074-INS-105
chr7-27077353-INS-60

chr7-27089811-INS-159
chr7-27110502-INS-183
chr7-27113161-INS-206
chr7-27118921-INS-109
chr7-27122612-INS-265
chr7-27135107-INS-86
chr7-27151477-INS-148
chr7-27153348-INS-217
chr7-27187366-INS-88
chr7-27194341-INS-159
chr7-27210309-INS-293
chr7-27229888-INS-106
chr7-27234314-INS-146
chr7-27259081-INS-205
chr7-27262048-INS-202
chr7-27265624-INS-50
chr7-27283982-INS-170
chr7-27285493-INS-52
chr7-27285658-INS-986
chr7-27309939-INS-391
chr7-27311766-INS-619
chr7-27312339-INS-204
chr7-27344024-INS-203
chr7-27351760-INS-205
chr7-27351982-INS-64
chr7-27360348-INS-212
chr7-27363586-INS-201
chr7-27377223-INS-182
chr7-27377499-INS-65
chr7-27383117-INS-53
chr7-27383170-INS-318
chr7-27390595-INS-405
chr7-27396984-INS-206
chr7-27397277-INS-969
chr7-27404788-INS-202
chr7-27419859-INS-4454
chr7-27423924-INS-53
chr7-27454187-INS-62
chr7-27457066-INS-212
chr7-27466236-INS-50
chr7-27469441-INS-125
chr7-27487052-INS-278
chr7-27508706-INS-244
chr7-27528556-INS-909
chr7-27536061-INS-59
chr7-27563431-INS-4855
chr7-27599114-INS-119
chr7-27599239-INS-392
chr7-27599485-INS-60
chr7-27601406-INS-69
chr7-27622865-INS-260
chr7-27622897-INS-99
chr7-27623145-INS-80
chr7-27644426-INS-343
chr7-27648229-INS-52
chr7-27712269-INS-184
chr7-27730418-INS-1746
chr7-27751166-INS-5398
chr7-27759666-INS-236
chr7-27760713-INS-195
chr7-27788073-INS-210
chr7-27788073-INS-84
chr7-27790377-INS-60
chr7-27794493-INS-78
chr7-27796136-INS-306
chr7-27849638-INS-100
chr7-27849870-INS-353
chr7-27903919-INS-56
chr7-27904002-INS-814
chr7-27911847-INS-159
chr7-27943908-INS-132
chr7-27947997-INS-1610
chr7-27970884-INS-55
chr7-27985451-INS-162
chr7-27987500-INS-104
chr7-27994650-INS-65
chr7-28046708-INS-185

chr7-28047992-INS-219
chr7-28065035-INS-84
chr7-28129265-INS-180
chr7-28129957-INS-207
chr7-28172689-INS-1896
chr7-28185170-INS-208
chr7-28185521-INS-86
chr7-28204605-INS-129
chr7-28234913-INS-205
chr7-28236106-INS-55
chr7-28242078-INS-53
chr7-28251621-INS-156
chr7-28252167-INS-228
chr7-28299238-INS-259
chr7-28301976-INS-212
chr7-28302018-INS-53
chr7-28310599-INS-74
chr7-28311260-INS-194
chr7-28311281-INS-90
chr7-28319562-INS-198
chr7-28332966-INS-52
chr7-28333034-INS-138
chr7-28336111-INS-59
chr7-28355228-INS-53
chr7-28355239-INS-110
chr7-28362169-INS-433
chr7-28366733-INS-277
chr7-28379326-INS-6606
chr7-28402744-INS-432
chr7-28404223-INS-188
chr7-28404441-INS-66
chr7-28405427-INS-317
chr7-28406444-INS-5624
chr7-28418541-INS-218
chr7-28471116-INS-60
chr7-28475138-INS-209
chr7-28479975-INS-591
chr7-28489206-INS-201
chr7-28529089-INS-1303
chr7-28536486-INS-215
chr7-28540462-INS-162
chr7-28541526-INS-11160
chr7-28544640-INS-198
chr7-28549727-INS-413
chr7-28562240-INS-56
chr7-28578290-INS-211
chr7-28583429-INS-53
chr7-28583570-INS-212
chr7-28583693-INS-53
chr7-28604197-INS-221
chr7-28654628-INS-217
chr7-28664512-INS-52
chr7-28703622-INS-902
chr7-28720897-INS-61
chr7-28721201-INS-53
chr7-28733894-INS-77
chr7-28734126-INS-63
chr7-28734820-INS-106
chr7-28742128-INS-150
chr7-28751202-INS-57
chr7-28779617-INS-72
chr7-28808593-INS-212
chr7-28827238-INS-52
chr7-28844894-INS-53
chr7-28983187-INS-69
chr7-29001716-INS-56
chr7-29012897-INS-79
chr7-29061834-INS-81
chr7-29075435-INS-634
chr7-29085029-INS-406
chr7-29085387-INS-504
chr7-29089017-INS-202
chr7-29113595-INS-395
chr7-29202056-INS-52
chr7-29211453-INS-134
chr7-29231373-INS-52
chr7-29250737-INS-191

chr7-29265482-INS-60
chr7-29268121-INS-92
chr7-29275703-INS-369
chr7-29349492-INS-161
chr7-29349492-INS-372
chr7-29349514-INS-66
chr7-29459452-INS-1490
chr7-29516633-INS-132
chr7-29516633-INS-308
chr7-29538409-INS-3136
chr7-29542327-INS-52
chr7-29564281-INS-333
chr7-29584025-INS-3858
chr7-29638077-INS-118
chr7-29644639-INS-138
chr7-29644639-INS-3371
chr7-29650184-INS-7496
chr7-29698673-INS-112
chr7-29708401-INS-76
chr7-29735430-INS-542
chr7-29735752-INS-179
chr7-29762473-INS-65
chr7-29763681-INS-219
chr7-29775701-INS-239
chr7-29781235-INS-72
chr7-29813784-INS-1198
chr7-29847160-INS-204
chr7-29865814-INS-220
chr7-29874257-INS-1130
chr7-29876021-INS-206
chr7-29901695-INS-314
chr7-29930961-INS-3370
chr7-29974414-INS-209
chr7-29987898-INS-1890
chr7-30042592-INS-1222
chr7-30064890-INS-68
chr7-30065053-INS-377
chr7-30065322-INS-934
chr7-30065841-INS-343
chr7-30083329-INS-254
chr7-30095527-INS-318
chr7-30096442-INS-200
chr7-30136744-INS-182
chr7-30138943-INS-52
chr7-30177583-INS-173
chr7-30183712-INS-2143
chr7-30187254-INS-209
chr7-30187379-INS-52
chr7-30187390-INS-9007
chr7-30190784-INS-51
chr7-30238354-INS-78
chr7-30238600-INS-120
chr7-30243558-INS-142
chr7-30243824-INS-206
chr7-30244976-INS-199
chr7-30245303-INS-597
chr7-30255611-INS-183
chr7-30310977-INS-52
chr7-30314167-INS-215
chr7-30314603-INS-52
chr7-30314650-INS-105
chr7-30314686-INS-212
chr7-30315967-INS-50
chr7-30320737-INS-61
chr7-30321296-INS-162
chr7-30321427-INS-65
chr7-30403843-INS-700
chr7-30434701-INS-157
chr7-30451450-INS-312
chr7-30452385-INS-54
chr7-30466322-INS-62
chr7-30467338-INS-210
chr7-30483137-INS-83
chr7-30494183-INS-2244
chr7-30526222-INS-85
chr7-30527935-INS-197
chr7-30531304-INS-596

chr7-30558164-INS-53
chr7-30564640-INS-53
chr7-30587841-INS-81
chr7-30761671-INS-192
chr7-30778696-INS-4979
chr7-30787569-INS-263
chr7-30805409-INS-424
chr7-30805543-INS-53
chr7-30986975-INS-5392
chr7-30993116-INS-1375
chr7-31017201-INS-2290
chr7-31042016-INS-7750
chr7-31043374-INS-101
chr7-31043530-INS-826
chr7-31055444-INS-162
chr7-31056117-INS-74
chr7-31082046-INS-4002
chr7-31088202-INS-76140
chr7-31137001-INS-2446
chr7-31170152-INS-112
chr7-31173380-INS-212
chr7-31173723-INS-68
chr7-31177258-INS-87
chr7-31177694-INS-700
chr7-31188502-INS-90
chr7-31219689-INS-148
chr7-31500273-INS-113
chr7-31536008-INS-2090
chr7-31557702-INS-6885
chr7-31730435-INS-2873
chr7-31809809-INS-76
chr7-31815325-INS-309
chr7-32823532-INS-56
chr7-32834447-INS-497
chr7-32909592-INS-71
chr7-33001671-INS-51
chr7-33131627-INS-56
chr7-33160003-INS-113
chr7-33422597-INS-8822
chr7-33490282-INS-64
chr7-33493611-INS-50
chr7-33496615-INS-77
chr7-33540443-INS-2788
chr7-33545356-INS-1110
chr7-33555308-INS-3585
chr7-33579367-INS-52
chr7-33605272-INS-4761
chr7-33606081-INS-1040
chr7-33614099-INS-188
chr7-33614211-INS-70
chr7-33631843-INS-6134
chr7-33703300-INS-50
chr7-33723045-INS-56
chr7-33769858-INS-1892
chr7-33772192-INS-3900
chr7-33789787-INS-358
chr7-33790383-INS-4716
chr7-33808501-INS-86
chr7-33827364-INS-500
chr7-33855837-INS-201
chr7-33877737-INS-3577
chr7-33902768-INS-60
chr7-33903159-INS-56
chr7-34046721-INS-572
chr7-34068901-INS-291
chr7-34069596-INS-59
chr7-34113215-INS-52
chr7-34115626-INS-308
chr7-34115854-INS-195
chr7-34121578-INS-155
chr7-34122660-INS-484
chr7-34135524-INS-263
chr7-34152888-INS-70
chr7-34153045-INS-114
chr7-34171628-INS-94
chr7-34174557-INS-5081
chr7-34268959-INS-53

chr7-34272375-INS-703
chr7-34290522-INS-68
chr7-34335566-INS-62
chr7-34381748-INS-107
chr7-34382644-INS-156
chr7-34413188-INS-287
chr7-34432557-INS-211
chr7-34460822-INS-289
chr7-34460871-INS-139
chr7-34488962-INS-211
chr7-34490917-INS-376
chr7-34515658-INS-159
chr7-34622321-INS-528
chr7-34623720-INS-2808
chr7-34757035-INS-460
chr7-34794392-INS-395
chr7-34794394-INS-189
chr7-34805562-INS-55
chr7-34811176-INS-263
chr7-34832908-INS-100
chr7-34859215-INS-1800
chr7-34860307-INS-1264
chr7-34860307-INS-4485
chr7-34904714-INS-118
chr7-34921329-INS-50
chr7-34937408-INS-57
chr7-34960675-INS-373
chr7-34960675-INS-52
chr7-34978121-INS-2323
chr7-35003699-INS-164
chr7-35011890-INS-1484
chr7-35018933-INS-530
chr7-35030184-INS-53
chr7-35051426-INS-7091
chr7-35051519-INS-236
chr7-35058704-INS-50
chr7-35060361-INS-67
chr7-35071447-INS-53
chr7-35082532-INS-107
chr7-35097792-INS-56
chr7-35114375-INS-54
chr7-35121148-INS-188
chr7-35125256-INS-50
chr7-35126478-INS-196
chr7-35128777-INS-210
chr7-35137202-INS-6449
chr7-35139417-INS-50
chr7-35141281-INS-2003
chr7-35141727-INS-370
chr7-35152236-INS-50
chr7-35244076-INS-137
chr7-35270758-INS-51
chr7-35273450-INS-130
chr7-35293531-INS-51
chr7-35310334-INS-783
chr7-35310760-INS-211
chr7-35313690-INS-160
chr7-35313774-INS-945
chr7-35318485-INS-53
chr7-35320709-INS-218
chr7-35320710-INS-654
chr7-35335259-INS-422
chr7-35356481-INS-6367
chr7-35373161-INS-95
chr7-35378549-INS-174
chr7-35384178-INS-105
chr7-35387315-INS-75
chr7-35397825-INS-114
chr7-35404474-INS-58
chr7-35404599-INS-261
chr7-35410893-INS-50
chr7-35411226-INS-51
chr7-35416658-INS-140
chr7-35425479-INS-164
chr7-35443777-INS-52
chr7-35461384-INS-159
chr7-35465866-INS-54

chr7-35484660-INS-53
chr7-35518109-INS-156
chr7-35548606-INS-112
chr7-35580507-INS-158
chr7-35580559-INS-53
chr7-35607492-INS-57
chr7-35607611-INS-138
chr7-35639387-INS-52
chr7-35653147-INS-110
chr7-35653666-INS-72
chr7-35653708-INS-186
chr7-35655812-INS-264
chr7-35661022-INS-182
chr7-35748141-INS-54
chr7-35783363-INS-82
chr7-35792528-INS-6713
chr7-35821323-INS-210
chr7-35821341-INS-82
chr7-35860494-INS-60
chr7-35893834-INS-79
chr7-35894849-INS-51
chr7-35904636-INS-52
chr7-35933403-INS-1868
chr7-35948020-INS-157
chr7-35979905-INS-77
chr7-36003906-INS-502
chr7-36004329-INS-100
chr7-36065175-INS-218
chr7-36140620-INS-52
chr7-36250708-INS-69
chr7-36250922-INS-76
chr7-36259549-INS-264
chr7-36259646-INS-53
chr7-36300659-INS-148
chr7-36308941-INS-52
chr7-36309463-INS-98
chr7-36314261-INS-6297
chr7-36318260-INS-6620
chr7-36344686-INS-66
chr7-36344970-INS-79
chr7-36365054-INS-54
chr7-36365247-INS-53
chr7-36387040-INS-52
chr7-36429001-INS-1460
chr7-36452530-INS-424
chr7-36463988-INS-58
chr7-36481476-INS-752
chr7-36562593-INS-312
chr7-36643585-INS-50
chr7-36705435-INS-96
chr7-36733262-INS-115
chr7-36762205-INS-1402
chr7-36816938-INS-116
chr7-36894996-INS-190
chr7-36968089-INS-168
chr7-37008849-INS-289
chr7-37052606-INS-316
chr7-37062877-INS-50
chr7-37077957-INS-161
chr7-37093096-INS-186
chr7-37208097-INS-50
chr7-37335329-INS-108
chr7-37481609-INS-50
chr7-37492939-INS-52
chr7-37493699-INS-50
chr7-37577545-INS-328
chr7-37612154-INS-163
chr7-37612155-INS-391
chr7-37614029-INS-98
chr7-37614128-INS-588
chr7-37684890-INS-50
chr7-37724378-INS-208
chr7-37741339-INS-204
chr7-37741728-INS-261
chr7-37745861-INS-120
chr7-37813167-INS-57
chr7-37814089-INS-63

chr7-37819878-INS-328
chr7-37823376-INS-62
chr7-37852172-INS-50
chr7-37860154-INS-298
chr7-37863896-INS-68
chr7-37880874-INS-4835
chr7-37954087-INS-6715
chr7-37971727-INS-73
chr7-38009270-INS-276
chr7-38019106-INS-561
chr7-38028549-INS-375
chr7-38029029-INS-12341
chr7-38035676-INS-1183
chr7-38036666-INS-1180
chr7-38049454-INS-7542
chr7-38067618-INS-6122
chr7-38086928-INS-81
chr7-38098141-INS-92
chr7-38102915-INS-58
chr7-38109531-INS-145
chr7-38165539-INS-9049
chr7-38201051-INS-50
chr7-38206948-INS-81
chr7-38208686-INS-235
chr7-38216290-INS-88
chr7-38258648-INS-62
chr7-38492064-INS-157
chr7-38495696-INS-112
chr7-38516092-INS-642
chr7-38526829-INS-765
chr7-38527619-INS-669
chr7-38533126-INS-62
chr7-38537648-INS-56
chr7-38542813-INS-132
chr7-38543112-INS-1643
chr7-38617933-INS-50
chr7-38631908-INS-80
chr7-38641182-INS-397
chr7-38673958-INS-94
chr7-38817900-INS-76
chr7-38824192-INS-2062
chr7-38830612-INS-73
chr7-38849621-INS-199
chr7-38857947-INS-2801
chr7-38927957-INS-56
chr7-38986071-INS-200
chr7-38995268-INS-52
chr7-38995285-INS-177
chr7-38996574-INS-50
chr7-39000581-INS-1049
chr7-39006504-INS-1614
chr7-39010252-INS-605
chr7-39212860-INS-60
chr7-39212863-INS-294
chr7-39245104-INS-886
chr7-39265127-INS-52
chr7-39278696-INS-2138
chr7-39279625-INS-724
chr7-39299935-INS-70
chr7-39300122-INS-460
chr7-39300696-INS-68
chr7-39321619-INS-50
chr7-39339481-INS-207
chr7-39363244-INS-61
chr7-39382569-INS-259
chr7-39394328-INS-229
chr7-39394349-INS-60
chr7-39413119-INS-50
chr7-39435667-INS-14720
chr7-39463663-INS-67
chr7-39474248-INS-60
chr7-39519501-INS-587
chr7-39627965-INS-59
chr7-39647985-INS-6347
chr7-39689792-INS-7365
chr7-39689793-INS-3188
chr7-39693070-INS-51

chr7-39695435-INS-350
chr7-39721381-INS-132
chr7-39732141-INS-2054
chr7-39741448-INS-6411
chr7-39742264-INS-250
chr7-39749322-INS-2491
chr7-39750553-INS-66
chr7-39781552-INS-81
chr7-39788210-INS-552
chr7-39789185-INS-6522
chr7-39822261-INS-6821
chr7-39830376-INS-296
chr7-39831999-INS-50
chr7-39873044-INS-56
chr7-39873397-INS-355
chr7-39888258-INS-1727
chr7-39913705-INS-50
chr7-39935328-INS-6687
chr7-39943312-INS-69
chr7-40050035-INS-54
chr7-40050811-INS-1350
chr7-40081245-INS-3103
chr7-40148403-INS-213
chr7-40224838-INS-83
chr7-40255600-INS-1809
chr7-40297974-INS-136
chr7-40339876-INS-288
chr7-40419972-INS-78
chr7-40431986-INS-56
chr7-40431997-INS-220
chr7-40436594-INS-1492
chr7-40490882-INS-479
chr7-40494166-INS-79
chr7-40494727-INS-50
chr7-40503229-INS-1031
chr7-40515803-INS-6115
chr7-40521589-INS-52
chr7-40734975-INS-210
chr7-40735133-INS-1255
chr7-40738010-INS-64
chr7-40758323-INS-6429
chr7-40796911-INS-55
chr7-40821954-INS-208
chr7-40851080-INS-11347
chr7-40855143-INS-1253
chr7-40860285-INS-389
chr7-40892435-INS-50
chr7-40896718-INS-128
chr7-40929838-INS-688
chr7-40940377-INS-6672
chr7-40951406-INS-67
chr7-40961080-INS-1199
chr7-40961577-INS-1294
chr7-40977277-INS-7118
chr7-40989071-INS-125
chr7-41085136-INS-80
chr7-41144965-INS-181
chr7-41145656-INS-102
chr7-41145946-INS-164
chr7-41152694-INS-209
chr7-41153671-INS-1509
chr7-41160469-INS-6183
chr7-41177933-INS-50
chr7-41230867-INS-4609
chr7-41236293-INS-645
chr7-41249878-INS-6258
chr7-41270146-INS-68
chr7-41270341-INS-272
chr7-41281421-INS-597
chr7-41289254-INS-217
chr7-41364717-INS-4112
chr7-41383849-INS-56
chr7-41385376-INS-962
chr7-41413161-INS-517
chr7-41424451-INS-254
chr7-41425229-INS-6210
chr7-41514572-INS-996

chr7-41532661-INS-481
chr7-41726953-INS-96
chr7-41734091-INS-108
chr7-41752091-INS-56
chr7-41812772-INS-3390
chr7-41948276-INS-6230
chr7-41978115-INS-69
chr7-41990779-INS-64
chr7-41999725-INS-147
chr7-42000352-INS-70
chr7-42029011-INS-82
chr7-42033633-INS-297
chr7-42110645-INS-8722
chr7-42147850-INS-2479
chr7-42213504-INS-105
chr7-42282351-INS-5123
chr7-42284094-INS-1215
chr7-42301268-INS-91
chr7-42398215-INS-546
chr7-42469529-INS-168
chr7-42502679-INS-5913
chr7-42514079-INS-1999
chr7-42526223-INS-372
chr7-42529603-INS-154
chr7-42559697-INS-57
chr7-42610659-INS-1228
chr7-42613257-INS-1891
chr7-42642579-INS-69
chr7-42644285-INS-204
chr7-42644452-INS-60
chr7-42663806-INS-170
chr7-42668899-INS-69
chr7-42680338-INS-841
chr7-42680994-INS-339
chr7-42689280-INS-56
chr7-42704548-INS-222
chr7-42709861-INS-74
chr7-42711240-INS-52
chr7-42712076-INS-223
chr7-42751200-INS-573
chr7-42771075-INS-1214
chr7-42772411-INS-6662
chr7-42851396-INS-240
chr7-42926250-INS-362
chr7-42927058-INS-7210
chr7-42930337-INS-56
chr7-42932218-INS-96
chr7-42956969-INS-204
chr7-42964568-INS-50
chr7-43003987-INS-57
chr7-43009282-INS-257
chr7-43009282-INS-81
chr7-43018865-INS-223
chr7-43028294-INS-166
chr7-43028591-INS-214
chr7-43079816-INS-61
chr7-43099960-INS-147
chr7-43142077-INS-84
chr7-43146579-INS-53
chr7-43158038-INS-194
chr7-43158038-INS-56
chr7-43198605-INS-50
chr7-43215253-INS-196
chr7-43216616-INS-64
chr7-43218843-INS-52
chr7-43219754-INS-7297
chr7-43252278-INS-500
chr7-43255269-INS-6369
chr7-43268282-INS-181
chr7-43273321-INS-52
chr7-43276475-INS-2664
chr7-43284502-INS-138
chr7-43290602-INS-130
chr7-43290602-INS-60
chr7-43293888-INS-110
chr7-43304757-INS-275
chr7-43322914-INS-8414

chr7-43332222-INS-3459
chr7-43334158-INS-53
chr7-43344545-INS-400
chr7-43404191-INS-209
chr7-43464040-INS-68
chr7-43501989-INS-106
chr7-43502157-INS-4979
chr7-43515545-INS-6063
chr7-43521637-INS-56
chr7-43586317-INS-50
chr7-43586638-INS-1074
chr7-43609901-INS-93
chr7-43623485-INS-3294
chr7-43645806-INS-207
chr7-43646872-INS-97
chr7-43659350-INS-50
chr7-43678417-INS-345
chr7-43684141-INS-204
chr7-43707749-INS-50
chr7-43783927-INS-62
chr7-43783959-INS-137
chr7-43824707-INS-2001
chr7-43858666-INS-74
chr7-43889967-INS-160
chr7-43890112-INS-110
chr7-43903841-INS-50
chr7-43925339-INS-147
chr7-43925479-INS-123
chr7-43948020-INS-102
chr7-43980166-INS-78
chr7-43996584-INS-430
chr7-44029563-INS-65
chr7-44064493-INS-94
chr7-44068806-INS-129
chr7-44124955-INS-2001
chr7-44132517-INS-56
chr7-44133650-INS-175
chr7-44133894-INS-353
chr7-44133907-INS-106
chr7-44142631-INS-1459
chr7-44159423-INS-550
chr7-44159862-INS-880
chr7-44170376-INS-64
chr7-44207949-INS-55
chr7-44208113-INS-7089
chr7-44233025-INS-195
chr7-44277228-INS-104
chr7-44333989-INS-115
chr7-44349459-INS-167
chr7-44357123-INS-172
chr7-44357645-INS-60
chr7-44357802-INS-549
chr7-44359018-INS-109
chr7-44365800-INS-84
chr7-44366024-INS-57
chr7-44373038-INS-168
chr7-44388124-INS-737
chr7-44390439-INS-384
chr7-44471884-INS-94
chr7-44483099-INS-308
chr7-44659313-INS-106
chr7-44747953-INS-60
chr7-44756815-INS-1264
chr7-44810286-INS-50
chr7-44854690-INS-135
chr7-44868752-INS-50
chr7-44887117-INS-50
chr7-44896761-INS-8826
chr7-44906596-INS-54
chr7-44907339-INS-80
chr7-44919626-INS-55
chr7-44919693-INS-199
chr7-44943993-INS-204
chr7-44970662-INS-68
chr7-45004961-INS-216
chr7-45029300-INS-120
chr7-45029667-INS-206

chr7-45046084-INS-205
chr7-45076313-INS-54
chr7-45089285-INS-213
chr7-45089863-INS-284
chr7-45090142-INS-230
chr7-45098929-INS-198
chr7-45127519-INS-52
chr7-45176276-INS-194
chr7-45178150-INS-56
chr7-45191854-INS-223
chr7-45258275-INS-50
chr7-45258795-INS-58
chr7-45265326-INS-68
chr7-45265490-INS-52
chr7-45309517-INS-4808
chr7-45403159-INS-390
chr7-45543824-INS-51
chr7-45606296-INS-4930
chr7-45614963-INS-54
chr7-45628462-INS-590
chr7-45631297-INS-2239
chr7-45652233-INS-50
chr7-45655777-INS-81
chr7-45655887-INS-209
chr7-45660617-INS-470
chr7-45660865-INS-616
chr7-45665265-INS-161
chr7-45669735-INS-324
chr7-45685904-INS-78
chr7-45685922-INS-1417
chr7-45698206-INS-60
chr7-45698692-INS-120
chr7-45711435-INS-50
chr7-45720344-INS-147
chr7-45725930-INS-61
chr7-45741121-INS-123
chr7-45741147-INS-2095
chr7-45779814-INS-60
chr7-45795391-INS-7232
chr7-45801567-INS-220
chr7-45801567-INS-56
chr7-45811451-INS-68
chr7-45822143-INS-50
chr7-45830519-INS-330
chr7-45830585-INS-154
chr7-45907313-INS-1182
chr7-45921889-INS-104
chr7-45985779-INS-190
chr7-46102871-INS-309
chr7-46133370-INS-79
chr7-46138758-INS-1060
chr7-46161635-INS-2320
chr7-46181581-INS-57
chr7-46269357-INS-62
chr7-46273324-INS-669
chr7-46273381-INS-102
chr7-46347402-INS-50
chr7-46360279-INS-249
chr7-46388506-INS-203
chr7-46394133-INS-12960
chr7-46407641-INS-106
chr7-46418888-INS-216
chr7-46419409-INS-215
chr7-46449459-INS-214
chr7-46493949-INS-209
chr7-46510938-INS-210
chr7-46544703-INS-191
chr7-46552574-INS-208
chr7-46562913-INS-226
chr7-46630923-INS-68
chr7-46668681-INS-112
chr7-46676819-INS-100
chr7-46680963-INS-1020
chr7-46704558-INS-2984
chr7-46788341-INS-112
chr7-46807553-INS-1066
chr7-46848150-INS-2781

chr7-46866364-INS-54
chr7-46893822-INS-345
chr7-46953042-INS-6412
chr7-47004841-INS-60
chr7-47006264-INS-261
chr7-47036442-INS-3105
chr7-47045927-INS-400
chr7-47141965-INS-2122
chr7-47155389-INS-1303
chr7-47158628-INS-2743
chr7-47165013-INS-716
chr7-47167964-INS-1015
chr7-47168497-INS-93
chr7-47180616-INS-50
chr7-47180851-INS-64
chr7-47215084-INS-16263
chr7-47247484-INS-1896
chr7-47280168-INS-90
chr7-47284934-INS-70
chr7-47288988-INS-1347
chr7-47295614-INS-745
chr7-47307365-INS-196
chr7-47317928-INS-5995
chr7-47375765-INS-191
chr7-47391846-INS-58
chr7-47395086-INS-140
chr7-47395111-INS-63
chr7-47412581-INS-4273
chr7-47420533-INS-53
chr7-47425474-INS-3447
chr7-47435953-INS-66
chr7-47435989-INS-144
chr7-47449259-INS-76
chr7-47462595-INS-293
chr7-47463152-INS-80
chr7-47465335-INS-6470
chr7-47467979-INS-53
chr7-47474209-INS-448
chr7-47489753-INS-397
chr7-47492192-INS-390
chr7-47494526-INS-543
chr7-47497862-INS-1689
chr7-47499535-INS-1586
chr7-47546030-INS-4983
chr7-47566181-INS-492
chr7-47580506-INS-51
chr7-47580836-INS-156
chr7-47583136-INS-3553
chr7-47587064-INS-74
chr7-47593997-INS-1448
chr7-47599144-INS-50
chr7-47607988-INS-147
chr7-47613225-INS-719
chr7-47613353-INS-266
chr7-47613391-INS-54
chr7-47622879-INS-10128
chr7-47624249-INS-198
chr7-47635086-INS-606
chr7-47639945-INS-62
chr7-47655341-INS-52
chr7-47661111-INS-1968
chr7-47662432-INS-61
chr7-47664713-INS-1880
chr7-47673470-INS-1557
chr7-47685816-INS-50
chr7-47693403-INS-53
chr7-47705133-INS-2266
chr7-47725559-INS-5473
chr7-47760438-INS-205
chr7-47763648-INS-501
chr7-47807919-INS-3368
chr7-47812001-INS-50
chr7-47836112-INS-4226
chr7-47848721-INS-1130
chr7-47849205-INS-1693
chr7-47945970-INS-99
chr7-48015844-INS-315

chr7-48040088-INS-3272
chr7-48053599-INS-3356
chr7-48078772-INS-730
chr7-48089694-INS-672
chr7-48124291-INS-7089
chr7-48164939-INS-54
chr7-48231927-INS-1000
chr7-48247955-INS-1289
chr7-48313286-INS-68
chr7-48339062-INS-340
chr7-48377966-INS-113
chr7-48468625-INS-6704
chr7-48480083-INS-106
chr7-48535363-INS-50
chr7-48572002-INS-310
chr7-48588573-INS-50
chr7-48595808-INS-314
chr7-48643342-INS-83
chr7-48673512-INS-75
chr7-48757718-INS-88
chr7-48765340-INS-220
chr7-48789825-INS-62
chr7-48793127-INS-52
chr7-48862380-INS-52
chr7-48944182-INS-52
chr7-48946958-INS-54
chr7-48985237-INS-147
chr7-49000012-INS-127
chr7-49016407-INS-220
chr7-49044470-INS-69
chr7-49137020-INS-144
chr7-49137044-INS-70
chr7-49192891-INS-52
chr7-49196274-INS-145
chr7-49210419-INS-51
chr7-49220136-INS-209
chr7-49226808-INS-4544
chr7-49240027-INS-207
chr7-49353165-INS-53
chr7-49357630-INS-345
chr7-49378269-INS-2838
chr7-49383392-INS-2847
chr7-49398526-INS-170
chr7-49407941-INS-122
chr7-49452741-INS-204
chr7-49461774-INS-477
chr7-49514363-INS-72
chr7-49578841-INS-66
chr7-49669502-INS-467
chr7-49688722-INS-911
chr7-49688995-INS-159
chr7-49689037-INS-189
chr7-49725362-INS-6535
chr7-49726951-INS-3212
chr7-49734575-INS-286
chr7-49741829-INS-363
chr7-49795231-INS-1105
chr7-49838811-INS-52
chr7-49881678-INS-135
chr7-49893939-INS-60
chr7-49926933-INS-7911
chr7-49966292-INS-72
chr7-49988977-INS-94
chr7-50035116-INS-50
chr7-50050448-INS-1903
chr7-50064718-INS-51
chr7-50073183-INS-80
chr7-50085482-INS-74
chr7-50086374-INS-51
chr7-50087062-INS-67
chr7-50093839-INS-380
chr7-50123312-INS-863
chr7-50143822-INS-1043
chr7-50151104-INS-6596
chr7-50163702-INS-367
chr7-50224863-INS-1252
chr7-50233012-INS-90

chr7-50241438-INS-356
chr7-50241497-INS-155
chr7-50250856-INS-88
chr7-50289019-INS-6303
chr7-50299839-INS-116
chr7-50348942-INS-59
chr7-50372134-INS-184
chr7-50372134-INS-86
chr7-50419297-INS-52
chr7-50428334-INS-180
chr7-50436257-INS-100
chr7-50438613-INS-50
chr7-50486975-INS-6442
chr7-50518024-INS-52
chr7-50518134-INS-363
chr7-50530668-INS-228
chr7-50530774-INS-55
chr7-50596284-INS-142
chr7-50597110-INS-68
chr7-50605849-INS-88
chr7-50613782-INS-107
chr7-50622535-INS-118
chr7-50623349-INS-60
chr7-50631289-INS-99
chr7-50632101-INS-58
chr7-50719005-INS-51
chr7-50721266-INS-126
chr7-50734311-INS-64
chr7-50785010-INS-156
chr7-50789486-INS-1724
chr7-51001328-INS-2920
chr7-51016891-INS-53
chr7-51017870-INS-6202
chr7-51046820-INS-5149
chr7-51077993-INS-743
chr7-51090145-INS-91
chr7-51090622-INS-6230
chr7-51095225-INS-6322
chr7-51099958-INS-1996
chr7-51103261-INS-400
chr7-51128829-INS-64
chr7-51158160-INS-68
chr7-51216453-INS-62
chr7-51251155-INS-61
chr7-51264064-INS-102
chr7-51327685-INS-231
chr7-51336623-INS-72
chr7-51407765-INS-2866
chr7-51410799-INS-964
chr7-51411277-INS-1174
chr7-51439264-INS-73
chr7-51441987-INS-80
chr7-51464092-INS-59
chr7-51547349-INS-245
chr7-51605266-INS-235
chr7-51605583-INS-72
chr7-51605621-INS-25007
chr7-51667592-INS-2381
chr7-51669101-INS-2889
chr7-51680130-INS-51
chr7-51680142-INS-128
chr7-51723271-INS-1563
chr7-51852144-INS-372
chr7-51862820-INS-2795
chr7-51863606-INS-54
chr7-51884055-INS-68
chr7-51884366-INS-401
chr7-51899911-INS-5593
chr7-51901661-INS-2203
chr7-51903003-INS-7746
chr7-51913327-INS-3671
chr7-51988513-INS-54
chr7-52029468-INS-3547
chr7-52029854-INS-636
chr7-52033799-INS-1348
chr7-52052410-INS-7115
chr7-52072586-INS-6989

chr7-52131833-INS-120
chr7-52145938-INS-66
chr7-52147233-INS-6400
chr7-52244906-INS-62
chr7-52283265-INS-100
chr7-52286934-INS-58
chr7-52293123-INS-61
chr7-52313862-INS-6917
chr7-52320692-INS-9472
chr7-52327416-INS-1036
chr7-52348906-INS-400
chr7-52353885-INS-2954
chr7-52362774-INS-1346
chr7-52376633-INS-60
chr7-52407472-INS-1181
chr7-52430731-INS-680
chr7-52530555-INS-72
chr7-52569931-INS-573
chr7-52572627-INS-54
chr7-52583651-INS-417
chr7-52583927-INS-112
chr7-52596444-INS-155
chr7-52620474-INS-58
chr7-52628174-INS-60
chr7-52633029-INS-6442
chr7-52668734-INS-6208
chr7-52696897-INS-51
chr7-52726749-INS-197
chr7-52749262-INS-7066
chr7-52837726-INS-2567
chr7-52852567-INS-239
chr7-52879996-INS-56
chr7-52910699-INS-1464
chr7-52911643-INS-92
chr7-52935907-INS-73
chr7-53001933-INS-50
chr7-53011835-INS-145
chr7-53061372-INS-70
chr7-53068959-INS-62
chr7-53105744-INS-3204
chr7-53136346-INS-1236
chr7-53193274-INS-62
chr7-53230691-INS-50
chr7-53271822-INS-407
chr7-53299864-INS-52
chr7-53301201-INS-405
chr7-53343071-INS-50
chr7-53369649-INS-2102
chr7-53369649-INS-6477
chr7-53370103-INS-302
chr7-53384683-INS-6592
chr7-53395424-INS-1808
chr7-53400919-INS-2939
chr7-53407242-INS-164
chr7-53407589-INS-633
chr7-53460668-INS-317
chr7-53533951-INS-50
chr7-53553048-INS-440
chr7-53578563-INS-1890
chr7-53581744-INS-1607
chr7-53593855-INS-1689
chr7-53601103-INS-7648
chr7-53635614-INS-161
chr7-53651603-INS-549
chr7-53693223-INS-51
chr7-53699553-INS-823
chr7-53735166-INS-1208
chr7-53743090-INS-560
chr7-53750810-INS-123
chr7-53754931-INS-110
chr7-53769886-INS-750
chr7-53776040-INS-134
chr7-53800694-INS-50
chr7-53862556-INS-344
chr7-53863237-INS-151
chr7-53900760-INS-715
chr7-53973895-INS-1114

chr7-54050759-INS-52
chr7-54050912-INS-2851
chr7-54056391-INS-73
chr7-54069820-INS-4669
chr7-54079626-INS-4484
chr7-54096974-INS-7101
chr7-54128311-INS-1093
chr7-54131353-INS-500
chr7-54173211-INS-2353
chr7-54189669-INS-495
chr7-54225624-INS-6671
chr7-54233264-INS-6419
chr7-54270504-INS-526
chr7-54270883-INS-52
chr7-54322584-INS-234
chr7-54331496-INS-1749
chr7-54337733-INS-2466
chr7-54344049-INS-6556
chr7-54357921-INS-141
chr7-54376654-INS-6428
chr7-54386292-INS-886
chr7-54537850-INS-1026
chr7-54653437-INS-69
chr7-54679957-INS-2625
chr7-54688285-INS-225
chr7-54791154-INS-323
chr7-54796702-INS-3108
chr7-54865475-INS-90
chr7-54940103-INS-2188
chr7-54965982-INS-58
chr7-54978953-INS-89
chr7-55038524-INS-60
chr7-55083904-INS-4536
chr7-55095945-INS-148
chr7-55096194-INS-522
chr7-55139492-INS-6445
chr7-55144749-INS-3134
chr7-55173042-INS-1451
chr7-55177382-INS-51
chr7-55185235-INS-8272
chr7-55221716-INS-1689
chr7-55396380-INS-50
chr7-55401515-INS-50
chr7-55402366-INS-413
chr7-55407069-INS-206
chr7-55409869-INS-3498
chr7-55428558-INS-68
chr7-55431597-INS-184
chr7-55455556-INS-52
chr7-55462485-INS-50
chr7-55507221-INS-60
chr7-55541320-INS-55
chr7-55554467-INS-523
chr7-55620461-INS-328
chr7-55671409-INS-187
chr7-55673499-INS-199
chr7-55674201-INS-261
chr7-55765908-INS-610
chr7-55811526-INS-1141
chr7-55812514-INS-1254
chr7-55935352-INS-140
chr7-55938015-INS-50
chr7-55938402-INS-6055
chr7-55945941-INS-231
chr7-55945943-INS-105
chr7-55955087-INS-760
chr7-56015180-INS-11473
chr7-56031755-INS-51
chr7-56034576-INS-7008
chr7-56060396-INS-3435
chr7-56077885-INS-6434
chr7-56094030-INS-6170
chr7-56130528-INS-56
chr7-56132529-INS-6701
chr7-56137677-INS-6130
chr7-56169409-INS-2760
chr7-56184419-INS-6542

chr7-56200689-INS-65
chr7-56228220-INS-348
chr7-56247024-INS-2975
chr7-56248526-INS-6660
chr7-56365250-INS-118
chr7-56440502-INS-2897
chr7-56444183-INS-1074
chr7-56496051-INS-60
chr7-56499913-INS-52
chr7-56522867-INS-54
chr7-56642708-INS-188
chr7-56652793-INS-172
chr7-56697624-INS-1871
chr7-56713415-INS-198
chr7-56724347-INS-6809
chr7-56806203-INS-52
chr7-56808030-INS-61
chr7-56842214-INS-1802
chr7-56935946-INS-7118
chr7-56944070-INS-488
chr7-56953776-INS-214
chr7-56953776-INS-2394
chr7-56956692-INS-2956
chr7-56960711-INS-109
chr7-56963572-INS-6228
chr7-56976670-INS-7025
chr7-56999052-INS-68
chr7-57004235-INS-120
chr7-57084237-INS-7899
chr7-57089302-INS-118
chr7-57096430-INS-393
chr7-57111413-INS-64
chr7-57143608-INS-50
chr7-57176069-INS-211
chr7-57176475-INS-347
chr7-57178333-INS-58
chr7-57186430-INS-52
chr7-57218963-INS-4808
chr7-57294625-INS-51
chr7-57315196-INS-6458
chr7-57349531-INS-474
chr7-57365469-INS-64
chr7-57377642-INS-100
chr7-57403806-INS-600
chr7-57438935-INS-54
chr7-57485616-INS-166
chr7-57485616-INS-51
chr7-57485810-INS-529
chr7-57494517-INS-872
chr7-57524266-INS-1444
chr7-57600851-INS-1199
chr7-57621344-INS-56
chr7-57683922-INS-105
chr7-57738319-INS-511
chr7-57785057-INS-4532
chr7-57853392-INS-74
chr7-57859113-INS-137
chr7-57872350-INS-8850
chr7-57891257-INS-1030
chr7-57900217-INS-50
chr7-57952850-INS-52
chr7-57964878-INS-116
chr7-57982880-INS-103
chr7-58021149-INS-7088
chr7-58079274-INS-1176
chr7-58103018-INS-1903
chr7-58131153-INS-6396
chr7-58149994-INS-1489
chr7-58159501-INS-6472
chr7-58162426-INS-70
chr7-58211068-INS-199
chr7-58211096-INS-94
chr7-58219994-INS-3236
chr7-58225342-INS-1408
chr7-58291618-INS-51
chr7-58306585-INS-170
chr7-58315029-INS-6425

chr7-58319356-INS-50
chr7-58325743-INS-96
chr7-58362157-INS-50
chr7-58373875-INS-52
chr7-58384916-INS-66
chr7-58518114-INS-305
chr7-58518543-INS-3244
chr7-58546678-INS-486
chr7-58572838-INS-129
chr7-58590357-INS-6427
chr7-58602988-INS-7335
chr7-58629119-INS-2213
chr7-58662110-INS-51
chr7-58662892-INS-50
chr7-58663018-INS-327
chr7-58686917-INS-71
chr7-58700544-INS-7097
chr7-58751740-INS-1461
chr7-58771662-INS-160
chr7-58813943-INS-1296
chr7-58843109-INS-171
chr7-58849754-INS-60
chr7-58964550-INS-151
chr7-58964849-INS-161
chr7-58973986-INS-130
chr7-59041968-INS-1870
chr7-59063792-INS-581
chr7-59064786-INS-370
chr7-59267933-INS-325
chr7-59271990-INS-180
chr7-59324369-INS-58
chr7-59555699-INS-77
chr7-59614892-INS-1492
chr7-59642455-INS-52
chr7-59647645-INS-50
chr7-59694306-INS-2195
chr7-59704272-INS-197
chr7-59707037-INS-246
chr7-59717266-INS-1297
chr7-59720032-INS-52
chr7-59784395-INS-6035
chr7-59831508-INS-1892
chr7-59831508-INS-400
chr7-59880543-INS-5635
chr7-59893891-INS-55
chr7-59939069-INS-700
chr7-60033087-INS-7248
chr7-60062850-INS-1462
chr7-60091983-INS-50
chr7-60109009-INS-65
chr7-60126004-INS-1514
chr7-60142472-INS-94
chr7-60149648-INS-361
chr7-60177664-INS-826
chr7-60191571-INS-72
chr7-60195927-INS-408
chr7-60210127-INS-172
chr7-60222684-INS-488
chr7-60224229-INS-71
chr7-60225470-INS-721
chr7-60341480-INS-2059
chr7-60343418-INS-499
chr7-60374822-INS-50
chr7-60383329-INS-118
chr7-60385681-INS-60
chr7-60386794-INS-1349
chr7-60388491-INS-167
chr7-60388743-INS-81
chr7-60457410-INS-230
chr7-60475548-INS-1100
chr7-60480071-INS-2390
chr7-60497288-INS-181
chr7-60497288-INS-56
chr7-60517171-INS-862
chr7-60536716-INS-62
chr7-60556931-INS-6515
chr7-60578627-INS-102

chr7-60579546-INS-83
chr7-60604258-INS-6401
chr7-60604916-INS-3714
chr7-60639285-INS-2639
chr7-60703969-INS-92
chr7-60729262-INS-99
chr7-60732639-INS-1022
chr7-60732669-INS-388
chr7-60763017-INS-63
chr7-60763383-INS-72
chr7-60779977-INS-109
chr7-60884327-INS-50
chr7-61028127-INS-183
chr7-61210972-INS-4776
chr7-61224867-INS-1785
chr7-61229059-INS-120
chr7-61273248-INS-217
chr7-61280358-INS-8852
chr7-61336793-INS-907
chr7-61384815-INS-415
chr7-61385630-INS-182
chr7-61390694-INS-851
chr7-61411672-INS-1672
chr7-61422928-INS-671
chr7-61469958-INS-109
chr7-61490234-INS-2400
chr7-61500590-INS-94
chr7-61501601-INS-54
chr7-61507209-INS-1223
chr7-61512609-INS-3033
chr7-61516644-INS-913
chr7-61532098-INS-1569
chr7-61537636-INS-1668
chr7-61558872-INS-57
chr7-61575288-INS-1015
chr7-61648735-INS-50
chr7-61668855-INS-1649
chr7-61688147-INS-73
chr7-61691529-INS-50
chr7-61720630-INS-927
chr7-61725541-INS-1635
chr7-61734415-INS-108
chr7-61770126-INS-64
chr7-61783015-INS-6178
chr7-61783163-INS-58
chr7-61798460-INS-55
chr7-61820052-INS-50
chr7-61879325-INS-57
chr7-61957835-INS-50
chr7-61961224-INS-93
chr7-61968070-INS-536
chr7-61985204-INS-213
chr7-62043350-INS-51
chr7-62143290-INS-11610
chr7-62152790-INS-4842
chr7-62181038-INS-56
chr7-62232947-INS-164
chr7-62261337-INS-100
chr7-62299607-INS-52
chr7-62320015-INS-402
chr7-62420765-INS-73
chr7-62464503-INS-7086
chr7-62521583-INS-102
chr7-62530466-INS-151
chr7-62556566-INS-6505
chr7-62577692-INS-6364
chr7-62581981-INS-1845
chr7-62591081-INS-76
chr7-62617344-INS-5821
chr7-62624065-INS-2445
chr7-62661118-INS-68
chr7-62695614-INS-72
chr7-62703046-INS-588
chr7-62727238-INS-1550
chr7-62727822-INS-130
chr7-62738019-INS-44097
chr7-62764383-INS-82

chr7-62800946-INS-464
chr7-62800946-INS-7569
chr7-62984923-INS-6690
chr7-62987880-INS-90
chr7-62991497-INS-2000
chr7-63019187-INS-50
chr7-63035699-INS-81
chr7-63166587-INS-51
chr7-63201201-INS-6226
chr7-63283119-INS-52
chr7-63289350-INS-163
chr7-63377599-INS-6715
chr7-63399145-INS-110
chr7-63425002-INS-1196
chr7-63425017-INS-313
chr7-63426956-INS-215
chr7-63464116-INS-50
chr7-63527030-INS-57
chr7-63527207-INS-177
chr7-63578469-INS-997
chr7-63578771-INS-1578
chr7-63604330-INS-50
chr7-63617225-INS-175
chr7-63624745-INS-50
chr7-63625016-INS-153
chr7-63625076-INS-318
chr7-63629389-INS-4091
chr7-63664866-INS-52
chr7-63681243-INS-219
chr7-63693985-INS-212
chr7-63694856-INS-263
chr7-63703922-INS-51
chr7-63710106-INS-51
chr7-63718021-INS-66
chr7-63747098-INS-1042
chr7-63747783-INS-3605
chr7-63765819-INS-496
chr7-63836921-INS-70
chr7-63865124-INS-194
chr7-63865124-INS-59
chr7-63866301-INS-78
chr7-63892861-INS-150
chr7-63991793-INS-63
chr7-64010169-INS-69
chr7-64055118-INS-144
chr7-64080910-INS-2754
chr7-64086413-INS-1510
chr7-64188821-INS-120
chr7-64193883-INS-277
chr7-64203896-INS-54
chr7-64217558-INS-3425
chr7-64287030-INS-6763
chr7-64302003-INS-6454
chr7-64366601-INS-3675
chr7-64368656-INS-63
chr7-64425639-INS-123
chr7-64461524-INS-54
chr7-64470384-INS-60
chr7-64500276-INS-498
chr7-64505491-INS-177
chr7-64543987-INS-56
chr7-64545095-INS-55
chr7-64548724-INS-6817
chr7-64563376-INS-50
chr7-64590824-INS-4749
chr7-64736744-INS-7093
chr7-64777294-INS-56
chr7-64799661-INS-548
chr7-64816267-INS-57
chr7-64817223-INS-158
chr7-64824660-INS-1888
chr7-64861711-INS-108
chr7-64875143-INS-205
chr7-64899769-INS-52
chr7-64932361-INS-55
chr7-64933196-INS-7184
chr7-64934866-INS-249

chr7-64975066-INS-52
chr7-65005544-INS-72
chr7-65008500-INS-76
chr7-65035916-INS-84
chr7-65053859-INS-121
chr7-65057601-INS-68
chr7-65085913-INS-319
chr7-65135923-INS-75
chr7-65184912-INS-54
chr7-65190647-INS-217
chr7-65192961-INS-57
chr7-65218208-INS-4839
chr7-65224611-INS-196
chr7-65225244-INS-392
chr7-65249284-INS-50
chr7-65272679-INS-63
chr7-65273353-INS-265
chr7-65308354-INS-1030
chr7-65316278-INS-51
chr7-65326473-INS-6403
chr7-65403366-INS-1642
chr7-65409246-INS-1054
chr7-65428281-INS-92
chr7-65441218-INS-120
chr7-65444133-INS-894
chr7-65471168-INS-194
chr7-65471308-INS-486
chr7-65476747-INS-55
chr7-65478695-INS-202
chr7-65482663-INS-179
chr7-65484766-INS-197
chr7-65489909-INS-198
chr7-65492560-INS-223
chr7-65493924-INS-196
chr7-65497437-INS-102
chr7-65499657-INS-58
chr7-65523593-INS-63
chr7-65561326-INS-544
chr7-65561364-INS-67
chr7-65607423-INS-50
chr7-65614924-INS-52
chr7-65663613-INS-52
chr7-65678457-INS-67
chr7-65713636-INS-98
chr7-65815834-INS-66
chr7-65866598-INS-51
chr7-65871473-INS-50
chr7-65881084-INS-206
chr7-65883729-INS-2465
chr7-65895456-INS-1640
chr7-65913511-INS-75
chr7-65927668-INS-78
chr7-65963789-INS-171
chr7-65969059-INS-80
chr7-65971621-INS-203
chr7-66003780-INS-55
chr7-66011487-INS-128
chr7-66012326-INS-652
chr7-66029940-INS-346
chr7-66030718-INS-2770
chr7-66160661-INS-672
chr7-66161210-INS-992
chr7-66210188-INS-342
chr7-66215396-INS-523
chr7-66245866-INS-166
chr7-66251400-INS-104
chr7-66377772-INS-52
chr7-66389607-INS-256
chr7-66409166-INS-111
chr7-66414715-INS-748
chr7-66440802-INS-72
chr7-66449808-INS-6524
chr7-66457094-INS-5299
chr7-66483380-INS-258
chr7-66506835-INS-130
chr7-66527131-INS-51
chr7-66560753-INS-54

chr7-66564283-INS-3056
chr7-66612252-INS-60
chr7-66620105-INS-265
chr7-66667756-INS-2777
chr7-66682885-INS-67
chr7-66691913-INS-635
chr7-66693599-INS-79
chr7-66695625-INS-115
chr7-66705245-INS-236
chr7-66709357-INS-192
chr7-66730027-INS-90
chr7-66735223-INS-5195
chr7-66795254-INS-110
chr7-66830146-INS-264
chr7-66830166-INS-105
chr7-66837638-INS-262
chr7-66837672-INS-85
chr7-66846720-INS-228
chr7-66898965-INS-546
chr7-67016741-INS-53
chr7-67077954-INS-225
chr7-67109137-INS-194
chr7-67123421-INS-401
chr7-67136299-INS-7283
chr7-67184104-INS-58
chr7-67192041-INS-155
chr7-67206516-INS-856
chr7-67312313-INS-197
chr7-67328232-INS-71
chr7-67427688-INS-50
chr7-67441867-INS-59
chr7-67531158-INS-60
chr7-67531203-INS-159
chr7-67660603-INS-156
chr7-67685661-INS-113
chr7-67704367-INS-79
chr7-67726590-INS-112
chr7-67734507-INS-93
chr7-67809028-INS-72
chr7-67821580-INS-238
chr7-67855350-INS-387
chr7-67858195-INS-56
chr7-67878907-INS-51
chr7-67892777-INS-67
chr7-67900525-INS-107
chr7-67903271-INS-1194
chr7-67903448-INS-101
chr7-67903448-INS-325
chr7-67916103-INS-205
chr7-67939359-INS-58
chr7-67954058-INS-50
chr7-67956222-INS-610
chr7-67959599-INS-62
chr7-67981410-INS-52
chr7-67986849-INS-188
chr7-67988453-INS-67
chr7-68013772-INS-56
chr7-68066042-INS-75
chr7-68077223-INS-60
chr7-68111928-INS-6174
chr7-68121473-INS-156
chr7-68148423-INS-148
chr7-68157451-INS-183
chr7-68164665-INS-826
chr7-68206770-INS-356
chr7-68215264-INS-54
chr7-68223771-INS-144
chr7-68231575-INS-98
chr7-68235700-INS-94
chr7-68266811-INS-121
chr7-68293687-INS-6268
chr7-68297615-INS-55
chr7-68334309-INS-1612
chr7-68352650-INS-50
chr7-68353263-INS-50
chr7-68353974-INS-91
chr7-68393571-INS-67

chr7-68431438-INS-2344
chr7-68451984-INS-1753
chr7-68559089-INS-1066
chr7-68604335-INS-131
chr7-68687746-INS-440
chr7-68707705-INS-2481
chr7-68760853-INS-12926
chr7-68773457-INS-50
chr7-68784429-INS-153
chr7-68816525-INS-4058
chr7-68828680-INS-95
chr7-68879744-INS-100
chr7-68881578-INS-2695
chr7-68891870-INS-105
chr7-68897390-INS-1940
chr7-68901299-INS-63
chr7-68905885-INS-50
chr7-68935383-INS-394
chr7-68941785-INS-136
chr7-68942387-INS-50
chr7-68943181-INS-6434
chr7-68963429-INS-161
chr7-68988129-INS-234
chr7-68993593-INS-371
chr7-69002772-INS-60
chr7-69033586-INS-896
chr7-69037676-INS-1018
chr7-69144167-INS-55
chr7-69213925-INS-103
chr7-69213925-INS-252
chr7-69221367-INS-61
chr7-69237195-INS-101
chr7-69240026-INS-80
chr7-69255467-INS-115
chr7-69256472-INS-4938
chr7-69272184-INS-6379
chr7-69280751-INS-1839
chr7-69283377-INS-2130
chr7-69326823-INS-327
chr7-69344552-INS-6152
chr7-69345672-INS-64
chr7-69369888-INS-247
chr7-69381565-INS-144
chr7-69392315-INS-2419
chr7-69455860-INS-70
chr7-69465160-INS-52
chr7-69475273-INS-58
chr7-69504506-INS-54
chr7-69514777-INS-70
chr7-69542482-INS-6614
chr7-69574228-INS-140
chr7-69667213-INS-60
chr7-69675151-INS-7325
chr7-69677599-INS-90
chr7-69732317-INS-68
chr7-69764323-INS-112
chr7-69774194-INS-163
chr7-69775921-INS-56
chr7-69793993-INS-52
chr7-69835860-INS-72
chr7-69850661-INS-51
chr7-69850880-INS-108
chr7-69855537-INS-362
chr7-69950144-INS-127
chr7-69974872-INS-66
chr7-70029491-INS-52
chr7-70056836-INS-52
chr7-70079397-INS-50
chr7-70118145-INS-1355
chr7-70140094-INS-6593
chr7-70212812-INS-104
chr7-70218070-INS-7988
chr7-70277622-INS-144
chr7-70292120-INS-199
chr7-70305736-INS-96
chr7-70370425-INS-177
chr7-70435545-INS-54

chr7-70461173-INS-50
chr7-70475577-INS-139
chr7-70593988-INS-467
chr7-70627526-INS-60
chr7-70638247-INS-1375
chr7-70646297-INS-498
chr7-70649693-INS-50
chr7-70650888-INS-50
chr7-70650889-INS-656
chr7-70666500-INS-531
chr7-70667830-INS-727
chr7-70678459-INS-167
chr7-70923113-INS-1040
chr7-70936504-INS-106
chr7-70936775-INS-214
chr7-70936775-INS-57
chr7-70976736-INS-50
chr7-70992630-INS-50
chr7-71052980-INS-232
chr7-71055461-INS-394
chr7-71055501-INS-156
chr7-71082341-INS-80
chr7-71159330-INS-380
chr7-71212388-INS-185
chr7-71244522-INS-107
chr7-71264217-INS-475
chr7-71264295-INS-100
chr7-71299700-INS-50
chr7-71328348-INS-6409
chr7-71361364-INS-145
chr7-71363162-INS-2615
chr7-71439442-INS-98
chr7-71443205-INS-253
chr7-71478200-INS-1626
chr7-71506484-INS-66
chr7-71507772-INS-118
chr7-71516299-INS-51
chr7-71561544-INS-6899
chr7-71615688-INS-50
chr7-71616072-INS-58
chr7-71625305-INS-718
chr7-71634604-INS-58
chr7-71647552-INS-104
chr7-71679702-INS-2231
chr7-71723268-INS-6261
chr7-71738182-INS-1908
chr7-71751761-INS-536
chr7-71751780-INS-51
chr7-71781162-INS-7520
chr7-71785672-INS-52
chr7-71791275-INS-6798
chr7-71812678-INS-1155
chr7-71844516-INS-2298
chr7-71886175-INS-231
chr7-71886266-INS-85
chr7-71888924-INS-80
chr7-71893141-INS-238
chr7-71936213-INS-213
chr7-71947771-INS-2205
chr7-72022727-INS-52
chr7-72052103-INS-64
chr7-72080883-INS-164
chr7-72092718-INS-6306
chr7-72095844-INS-59
chr7-72181878-INS-110
chr7-72187074-INS-50
chr7-72199383-INS-84
chr7-72200283-INS-294
chr7-72200516-INS-245
chr7-72203975-INS-2914
chr7-72215675-INS-2708
chr7-72241412-INS-1670
chr7-72253431-INS-528
chr7-72263550-INS-1497
chr7-72286909-INS-70
chr7-72320089-INS-80
chr7-72321305-INS-100

chr7-72321933-INS-215
chr7-72350590-INS-1696
chr7-72354856-INS-212
chr7-72382832-INS-195
chr7-72385936-INS-451
chr7-72406579-INS-866
chr7-72428342-INS-90
chr7-72447779-INS-6316
chr7-72459432-INS-1862
chr7-72463343-INS-65
chr7-72463753-INS-120
chr7-72465824-INS-6191
chr7-72505835-INS-84
chr7-72544572-INS-50
chr7-72556214-INS-50
chr7-72563789-INS-50
chr7-72564543-INS-4363
chr7-72573370-INS-2233
chr7-72585892-INS-52
chr7-72608123-INS-620
chr7-72619751-INS-6252
chr7-72626321-INS-1363
chr7-72631864-INS-218
chr7-72636324-INS-64
chr7-72669525-INS-1494
chr7-72777806-INS-58
chr7-72783385-INS-58
chr7-72796828-INS-67
chr7-72842123-INS-72
chr7-72855203-INS-60
chr7-72879342-INS-2246
chr7-72890664-INS-6401
chr7-72892837-INS-108
chr7-72905427-INS-2733
chr7-72929638-INS-86
chr7-72969469-INS-6818
chr7-72974458-INS-3328
chr7-72974458-INS-9123
chr7-72974465-INS-531
chr7-72981090-INS-58
chr7-72981105-INS-6856
chr7-72981179-INS-238
chr7-72989032-INS-197
chr7-72995689-INS-102
chr7-72998288-INS-56
chr7-73014058-INS-60
chr7-73051832-INS-82
chr7-73052238-INS-85
chr7-73053032-INS-781
chr7-73119683-INS-6367
chr7-73120063-INS-94
chr7-73120124-INS-343
chr7-73133453-INS-84
chr7-73135551-INS-2322
chr7-73160879-INS-214
chr7-73175983-INS-226
chr7-73181058-INS-206
chr7-73212122-INS-141
chr7-73212389-INS-53
chr7-73232250-INS-56
chr7-73248895-INS-208
chr7-73254626-INS-196
chr7-73254651-INS-465
chr7-73273268-INS-197
chr7-73278544-INS-493
chr7-73292173-INS-135
chr7-73308693-INS-3969
chr7-73313227-INS-50
chr7-73317917-INS-66
chr7-73328493-INS-200
chr7-73334417-INS-51
chr7-73360188-INS-4908
chr7-73362932-INS-181
chr7-73388064-INS-519
chr7-73497790-INS-100
chr7-73497790-INS-440
chr7-73498145-INS-574

chr7-73522634-INS-208
chr7-73547787-INS-6595
chr7-73580416-INS-4409
chr7-73692295-INS-298
chr7-73707982-INS-1663
chr7-73716997-INS-190
chr7-73716997-INS-53
chr7-73746960-INS-60
chr7-73754053-INS-4044
chr7-73756108-INS-233
chr7-73772639-INS-328
chr7-73772690-INS-54
chr7-73773287-INS-215
chr7-73782452-INS-1223
chr7-73783743-INS-399
chr7-73801749-INS-1428
chr7-73801795-INS-6636
chr7-73804653-INS-52
chr7-73832157-INS-64
chr7-73832588-INS-64
chr7-73832696-INS-56
chr7-73843105-INS-321
chr7-73843142-INS-897
chr7-73843401-INS-204
chr7-73843477-INS-62
chr7-73923086-INS-445
chr7-73970902-INS-60
chr7-73971828-INS-62
chr7-73993911-INS-50
chr7-73994077-INS-485
chr7-73995751-INS-74
chr7-74075356-INS-517
chr7-74221451-INS-88
chr7-74227747-INS-6415
chr7-74233796-INS-92
chr7-74302330-INS-375
chr7-74326250-INS-90
chr7-74331855-INS-66
chr7-74333096-INS-1104
chr7-74348323-INS-182
chr7-74359682-INS-60
chr7-74367963-INS-50
chr7-74385879-INS-52
chr7-74428204-INS-54
chr7-74474904-INS-67
chr7-74475952-INS-56
chr7-74498079-INS-59
chr7-74503865-INS-50
chr7-74576393-INS-50
chr7-74637922-INS-237
chr7-74661600-INS-6671
chr7-74670782-INS-7098
chr7-74710183-INS-918
chr7-74721655-INS-73
chr7-74744899-INS-185
chr7-74755032-INS-338
chr7-74757016-INS-526
chr7-74757207-INS-72
chr7-74767731-INS-166
chr7-74862940-INS-51
chr7-74862957-INS-180
chr7-74874264-INS-461
chr7-74874839-INS-54
chr7-74884908-INS-100
chr7-74907056-INS-51
chr7-74928050-INS-142
chr7-74971824-INS-57
chr7-74971898-INS-129
chr7-74982771-INS-157
chr7-74983330-INS-3478
chr7-74994350-INS-64
chr7-74997715-INS-52
chr7-75028460-INS-56
chr7-75044639-INS-50
chr7-75048552-INS-1147
chr7-75101915-INS-185
chr7-75123881-INS-79

chr7-75125321-INS-55
chr7-75162697-INS-206
chr7-75184404-INS-108
chr7-75325973-INS-230
chr7-75326184-INS-184
chr7-75363268-INS-80
chr7-75390352-INS-57
chr7-75391011-INS-56
chr7-75438024-INS-7347
chr7-75446968-INS-553
chr7-75470692-INS-202
chr7-75471161-INS-1414
chr7-75490193-INS-122
chr7-75518385-INS-119
chr7-75521989-INS-5548
chr7-75544474-INS-6240
chr7-75604783-INS-160
chr7-75716047-INS-207
chr7-75746144-INS-9726
chr7-75746155-INS-58
chr7-75788641-INS-71
chr7-75799246-INS-203
chr7-75813314-INS-52
chr7-75859092-INS-50
chr7-75864066-INS-517
chr7-75879957-INS-1540
chr7-75886825-INS-2132
chr7-75967473-INS-60
chr7-76081295-INS-57
chr7-76089635-INS-1252
chr7-76102093-INS-78
chr7-76114502-INS-56
chr7-76116066-INS-62
chr7-76134772-INS-1953
chr7-76134977-INS-96
chr7-76134994-INS-250
chr7-76154876-INS-170
chr7-76169420-INS-80
chr7-76174650-INS-6183
chr7-76229190-INS-200
chr7-76255889-INS-2733
chr7-76256290-INS-123
chr7-76265847-INS-63
chr7-76271115-INS-159
chr7-76277128-INS-185
chr7-76315196-INS-6817
chr7-76326885-INS-92
chr7-76327025-INS-240
chr7-76346398-INS-76
chr7-76367812-INS-50
chr7-76409110-INS-6119
chr7-76432374-INS-449
chr7-76508934-INS-2205
chr7-76529417-INS-769
chr7-76535445-INS-75
chr7-76576192-INS-6638
chr7-76624897-INS-6560
chr7-76626056-INS-689
chr7-76636914-INS-6287
chr7-76657582-INS-63
chr7-76657671-INS-266
chr7-76698987-INS-6617
chr7-76821528-INS-6510
chr7-76847919-INS-6226
chr7-76854003-INS-62
chr7-76914188-INS-140
chr7-76929971-INS-1892
chr7-76932719-INS-6644
chr7-76984543-INS-1142
chr7-77038041-INS-142
chr7-77088581-INS-6500
chr7-77203029-INS-84
chr7-77208028-INS-60
chr7-77210848-INS-74
chr7-77260503-INS-402
chr7-77279787-INS-416
chr7-77348394-INS-285

chr7-77353745-INS-6685
chr7-77358436-INS-1052
chr7-77385183-INS-124
chr7-77389532-INS-58
chr7-77406971-INS-1614
chr7-77409530-INS-52
chr7-77410719-INS-6277
chr7-77432347-INS-1494
chr7-77457914-INS-58
chr7-77474666-INS-50
chr7-77504044-INS-176
chr7-77534699-INS-703
chr7-77563145-INS-94
chr7-77586546-INS-183
chr7-77588917-INS-76
chr7-77626584-INS-65
chr7-77697893-INS-954
chr7-77714047-INS-2476
chr7-77741358-INS-370
chr7-77775131-INS-7321
chr7-77809704-INS-54
chr7-77809744-INS-481
chr7-77856222-INS-379
chr7-77900265-INS-350
chr7-77932409-INS-62
chr7-77947490-INS-58
chr7-78041028-INS-52
chr7-78090005-INS-66
chr7-78120452-INS-52
chr7-78151974-INS-166
chr7-78217670-INS-80
chr7-78313669-INS-113
chr7-78313715-INS-953
chr7-78336075-INS-52
chr7-78338646-INS-60
chr7-78338903-INS-216
chr7-78339547-INS-74
chr7-78394818-INS-168
chr7-78429779-INS-70
chr7-78429780-INS-224
chr7-78452173-INS-264
chr7-78452635-INS-172
chr7-78530258-INS-3534
chr7-78587743-INS-66
chr7-78593308-INS-125
chr7-78593698-INS-3352
chr7-78607789-INS-61
chr7-78607955-INS-205
chr7-78625763-INS-58
chr7-78633219-INS-93
chr7-78633422-INS-72
chr7-78634546-INS-63
chr7-78640062-INS-3268
chr7-78645352-INS-52
chr7-78647679-INS-151
chr7-78663796-INS-50
chr7-78685122-INS-2413
chr7-78696536-INS-280
chr7-78696536-INS-70
chr7-78696804-INS-101
chr7-78715686-INS-58
chr7-78717908-INS-132
chr7-78721025-INS-129
chr7-78728721-INS-179
chr7-78795222-INS-214
chr7-78807093-INS-54
chr7-78821507-INS-500
chr7-78835519-INS-109
chr7-78838494-INS-3010
chr7-78841140-INS-108
chr7-78865546-INS-104
chr7-78870960-INS-82
chr7-78907368-INS-344
chr7-78916465-INS-294
chr7-79024017-INS-214
chr7-79067448-INS-92
chr7-79089090-INS-138

chr7-79190511-INS-54
chr7-79195357-INS-84
chr7-79217761-INS-1000
chr7-79225885-INS-1618
chr7-79302246-INS-7468
chr7-79331161-INS-84
chr7-79364571-INS-196
chr7-79380298-INS-378
chr7-79418835-INS-67
chr7-79478665-INS-2538
chr7-79478815-INS-1242
chr7-79498354-INS-54
chr7-79567593-INS-232
chr7-79602578-INS-149
chr7-79602578-INS-71
chr7-79614353-INS-160
chr7-79634072-INS-75
chr7-79651219-INS-64
chr7-79684604-INS-102
chr7-79685164-INS-173
chr7-79751838-INS-54
chr7-79759836-INS-50
chr7-79770774-INS-64
chr7-79785227-INS-51
chr7-79811109-INS-120
chr7-79832149-INS-157
chr7-79832500-INS-451
chr7-79839829-INS-199
chr7-79862872-INS-50
chr7-79873310-INS-1002
chr7-79887684-INS-838
chr7-79895540-INS-253
chr7-79922161-INS-154
chr7-79922341-INS-180
chr7-79924206-INS-215
chr7-79937168-INS-196
chr7-79950413-INS-60
chr7-79950501-INS-240
chr7-79956710-INS-217
chr7-79956786-INS-98
chr7-79979843-INS-65
chr7-80035170-INS-116
chr7-80054334-INS-5371
chr7-80077652-INS-343
chr7-80078336-INS-76
chr7-80168727-INS-56
chr7-80181876-INS-68
chr7-80181914-INS-226
chr7-80223718-INS-62
chr7-80296141-INS-384
chr7-80304422-INS-216
chr7-80312801-INS-68
chr7-80344591-INS-50
chr7-80360663-INS-152
chr7-80387723-INS-63
chr7-80419659-INS-1038
chr7-80419764-INS-215
chr7-80420074-INS-478
chr7-80434722-INS-50
chr7-80443331-INS-451
chr7-80474968-INS-54
chr7-80478894-INS-196
chr7-80482501-INS-5043
chr7-80483821-INS-56
chr7-80497183-INS-8943
chr7-80502614-INS-502
chr7-80527235-INS-330
chr7-80527858-INS-52
chr7-80533691-INS-198
chr7-80613563-INS-290
chr7-80613942-INS-193
chr7-80615683-INS-249
chr7-80617100-INS-200
chr7-80620315-INS-179
chr7-80626052-INS-209
chr7-80626985-INS-58
chr7-80685058-INS-93

chr7-80689263-INS-59
chr7-80705166-INS-171
chr7-80737048-INS-521
chr7-80737103-INS-105
chr7-80737580-INS-384
chr7-80785647-INS-59
chr7-80786152-INS-58
chr7-80808238-INS-76
chr7-80814918-INS-88
chr7-80815180-INS-205
chr7-80822314-INS-78
chr7-80822472-INS-1286
chr7-80858033-INS-234
chr7-80952626-INS-52
chr7-80972986-INS-62
chr7-81009403-INS-200
chr7-81012553-INS-50
chr7-81012670-INS-56
chr7-81025635-INS-336
chr7-81040623-INS-65
chr7-81178539-INS-362
chr7-81242647-INS-99
chr7-81281879-INS-50
chr7-81305882-INS-59
chr7-81349220-INS-3171
chr7-81352361-INS-94
chr7-81373665-INS-400
chr7-81387076-INS-375
chr7-81407348-INS-4419
chr7-81441520-INS-208
chr7-81483060-INS-239
chr7-81483503-INS-573
chr7-81487882-INS-8672
chr7-81502537-INS-1741
chr7-81503798-INS-1093
chr7-81503887-INS-351
chr7-81528344-INS-64
chr7-81625981-INS-52
chr7-81708937-INS-216
chr7-81759082-INS-495
chr7-81789874-INS-69
chr7-81797009-INS-1902
chr7-81855032-INS-50
chr7-81857306-INS-1462
chr7-81884015-INS-99
chr7-81890406-INS-549
chr7-81900144-INS-134
chr7-81900951-INS-113
chr7-81914328-INS-148
chr7-81916563-INS-61
chr7-81917381-INS-139
chr7-81922355-INS-488
chr7-81922420-INS-52
chr7-81971384-INS-104
chr7-81979484-INS-55
chr7-82001226-INS-477
chr7-82001625-INS-886
chr7-82062060-INS-121
chr7-82069661-INS-112
chr7-82075802-INS-66
chr7-82087610-INS-53
chr7-82095237-INS-308
chr7-82113114-INS-450
chr7-82125330-INS-64
chr7-82131688-INS-2033
chr7-82144145-INS-75
chr7-82157775-INS-289
chr7-82167356-INS-55
chr7-82176830-INS-193
chr7-82210143-INS-6037
chr7-82216040-INS-71
chr7-82224094-INS-58
chr7-82224582-INS-165
chr7-82227631-INS-1138
chr7-82242106-INS-531
chr7-82242538-INS-747
chr7-82252316-INS-66

chr7-82272075-INS-405
chr7-82453053-INS-197
chr7-82463939-INS-304
chr7-82472082-INS-7167
chr7-82502790-INS-52
chr7-82521506-INS-642
chr7-82562439-INS-50
chr7-82580128-INS-50
chr7-82636153-INS-58
chr7-82657525-INS-54
chr7-82713974-INS-53
chr7-82747747-INS-200
chr7-82761780-INS-6697
chr7-82813981-INS-229
chr7-82940081-INS-5740
chr7-83067992-INS-818
chr7-83111869-INS-2332
chr7-83175982-INS-1887
chr7-83220008-INS-56
chr7-83231616-INS-76
chr7-83235816-INS-51
chr7-83240544-INS-1315
chr7-83247023-INS-69
chr7-83252429-INS-207
chr7-83345587-INS-1892
chr7-83345587-INS-396
chr7-83346162-INS-425
chr7-83353084-INS-204
chr7-83409355-INS-75
chr7-83415371-INS-4673
chr7-83416308-INS-157
chr7-83428839-INS-78
chr7-83432721-INS-428
chr7-83454729-INS-69
chr7-83454836-INS-78
chr7-83475479-INS-312
chr7-83480852-INS-127
chr7-83487280-INS-325
chr7-83487825-INS-68
chr7-83488901-INS-206
chr7-83498027-INS-50
chr7-83520268-INS-84
chr7-83521064-INS-54
chr7-83527070-INS-201
chr7-83528630-INS-70
chr7-83538230-INS-50
chr7-83561841-INS-54
chr7-83563773-INS-81
chr7-83564578-INS-1365
chr7-83587311-INS-338
chr7-83621018-INS-96
chr7-83667140-INS-120
chr7-83679247-INS-78
chr7-83684269-INS-70
chr7-83691143-INS-3126
chr7-83742208-INS-433
chr7-83755164-INS-872
chr7-83780436-INS-275
chr7-83809631-INS-4814
chr7-83812916-INS-2315
chr7-83838368-INS-7383
chr7-83851797-INS-229
chr7-83881104-INS-1991
chr7-83968070-INS-1904
chr7-84078250-INS-54
chr7-84078425-INS-84
chr7-84128180-INS-2211
chr7-84130300-INS-1737
chr7-84140416-INS-391
chr7-84142011-INS-97
chr7-84148421-INS-126
chr7-84160462-INS-4566
chr7-84170192-INS-1270
chr7-84193288-INS-499
chr7-84228956-INS-118
chr7-84229494-INS-341
chr7-84399624-INS-293

chr7-84401269-INS-440
chr7-84403515-INS-6207
chr7-84513590-INS-5705
chr7-84546139-INS-72
chr7-84649906-INS-1175
chr7-84696446-INS-1531
chr7-84721386-INS-6360
chr7-84765434-INS-4969
chr7-84775585-INS-2195
chr7-84779116-INS-54
chr7-84780204-INS-54
chr7-84797641-INS-54
chr7-84799541-INS-55
chr7-84824486-INS-205
chr7-84877959-INS-52
chr7-84889768-INS-106
chr7-84997299-INS-50
chr7-85076034-INS-90
chr7-85101489-INS-58
chr7-85149509-INS-66
chr7-85227991-INS-488
chr7-85238472-INS-213
chr7-85239515-INS-7193
chr7-85248409-INS-50
chr7-85332368-INS-71
chr7-85359973-INS-55
chr7-85520355-INS-421
chr7-85546242-INS-50
chr7-85566245-INS-739
chr7-85566685-INS-85
chr7-85599709-INS-54
chr7-85602963-INS-61
chr7-85603960-INS-114
chr7-85605975-INS-399
chr7-85621101-INS-1767
chr7-85635779-INS-7275
chr7-85643057-INS-498
chr7-85644015-INS-137
chr7-85650544-INS-6596
chr7-85716346-INS-2039
chr7-85826839-INS-5193
chr7-85866485-INS-2839
chr7-85910920-INS-7381
chr7-85911807-INS-135
chr7-85964592-INS-905
chr7-85982828-INS-50
chr7-86016939-INS-977
chr7-86024702-INS-116
chr7-86058944-INS-1775
chr7-86077784-INS-182
chr7-86096061-INS-6218
chr7-86101609-INS-113
chr7-86107887-INS-691
chr7-86108477-INS-1079
chr7-86110779-INS-1145
chr7-86139638-INS-81
chr7-86180473-INS-52
chr7-86183220-INS-3677
chr7-86226562-INS-52
chr7-86247027-INS-6208
chr7-86333905-INS-384
chr7-86334204-INS-543
chr7-86384038-INS-420
chr7-86387555-INS-67
chr7-86434693-INS-1072
chr7-86517250-INS-50
chr7-86526974-INS-70
chr7-86535405-INS-52
chr7-86566329-INS-7114
chr7-86570556-INS-3640
chr7-86573785-INS-52
chr7-86633554-INS-59
chr7-86655028-INS-97
chr7-86731008-INS-6222
chr7-86741030-INS-486
chr7-86765641-INS-2214
chr7-86786943-INS-14886

chr7-86804904-INS-54
chr7-86835248-INS-6236
chr7-86835497-INS-198
chr7-86848174-INS-6133
chr7-86858774-INS-399
chr7-86884648-INS-227
chr7-86897144-INS-692
chr7-86961627-INS-133
chr7-86965214-INS-585
chr7-87098453-INS-6698
chr7-87130019-INS-142
chr7-87132798-INS-205
chr7-87147716-INS-1483
chr7-87177874-INS-391
chr7-87179508-INS-68
chr7-87196329-INS-2243
chr7-87206695-INS-56
chr7-87207480-INS-607
chr7-87208158-INS-159
chr7-87225539-INS-118
chr7-87288141-INS-63
chr7-87339820-INS-498
chr7-87348664-INS-400
chr7-87374266-INS-6588
chr7-87507731-INS-80
chr7-87671245-INS-3143
chr7-87700127-INS-50
chr7-87812331-INS-353
chr7-87812331-INS-7095
chr7-87838092-INS-54
chr7-87883819-INS-190
chr7-87919467-INS-95
chr7-87974389-INS-58
chr7-87979060-INS-219
chr7-88005248-INS-6397
chr7-88019538-INS-474
chr7-88027778-INS-72
chr7-88047828-INS-443
chr7-88153889-INS-52
chr7-88166705-INS-1153
chr7-88166953-INS-51
chr7-88170110-INS-306
chr7-88170778-INS-60
chr7-88170887-INS-60
chr7-88251836-INS-471
chr7-88263484-INS-1941
chr7-88273655-INS-125
chr7-88386674-INS-56
chr7-88433545-INS-1035
chr7-88443728-INS-6430
chr7-88623248-INS-6403
chr7-88661006-INS-1384
chr7-88673008-INS-1110
chr7-88673008-INS-88
chr7-88675883-INS-3607
chr7-88679205-INS-131
chr7-88899864-INS-5148
chr7-88917076-INS-248
chr7-88917076-INS-80
chr7-88954815-INS-52
chr7-88981514-INS-1409
chr7-89004134-INS-58
chr7-89099832-INS-53
chr7-89101240-INS-59
chr7-89102466-INS-54
chr7-89110008-INS-202
chr7-89251671-INS-75
chr7-89267337-INS-114
chr7-89297782-INS-65
chr7-89321381-INS-1522
chr7-89370111-INS-6618
chr7-89375274-INS-313
chr7-89391277-INS-175
chr7-89421786-INS-98
chr7-89476328-INS-2628
chr7-89494692-INS-378
chr7-89524354-INS-452

chr7-89552345-INS-56
chr7-89553177-INS-50
chr7-89560504-INS-57
chr7-89561551-INS-205
chr7-89569890-INS-395
chr7-89569913-INS-96
chr7-89577134-INS-214
chr7-89581691-INS-384
chr7-89616392-INS-149
chr7-89616665-INS-65
chr7-89626333-INS-167
chr7-89631398-INS-55
chr7-89631597-INS-263
chr7-89637360-INS-183
chr7-89638065-INS-68
chr7-89638153-INS-143
chr7-89644339-INS-207
chr7-89648502-INS-111
chr7-89648637-INS-207
chr7-89649259-INS-3195
chr7-89652993-INS-87
chr7-89652997-INS-561
chr7-89653004-INS-2578
chr7-89666085-INS-182
chr7-89689837-INS-203
chr7-89715209-INS-210
chr7-89716001-INS-78
chr7-89729071-INS-80
chr7-89734673-INS-72
chr7-89754959-INS-207
chr7-89760134-INS-53
chr7-89761042-INS-60
chr7-89783258-INS-204
chr7-89812386-INS-7038
chr7-89846754-INS-84
chr7-89878511-INS-147
chr7-89935040-INS-1202
chr7-89937778-INS-66
chr7-89955019-INS-420
chr7-89970350-INS-153
chr7-89974286-INS-276
chr7-89974286-INS-54
chr7-90002275-INS-6221
chr7-90026328-INS-80
chr7-90087451-INS-152
chr7-90119523-INS-60
chr7-90129988-INS-182
chr7-90135250-INS-128
chr7-90136719-INS-413
chr7-90137096-INS-83
chr7-90154415-INS-174
chr7-90174055-INS-74
chr7-90195210-INS-72
chr7-90196490-INS-3240
chr7-90208125-INS-6320
chr7-90208827-INS-376
chr7-90228626-INS-2265
chr7-90249380-INS-866
chr7-90250385-INS-403
chr7-90317578-INS-6952
chr7-90350904-INS-6367
chr7-90356507-INS-1881
chr7-90394149-INS-717
chr7-90403067-INS-592
chr7-90408422-INS-59
chr7-90421054-INS-2940
chr7-90441521-INS-6712
chr7-90446464-INS-247
chr7-90466960-INS-50
chr7-90476149-INS-64
chr7-90484845-INS-50
chr7-90500591-INS-59
chr7-90527012-INS-210
chr7-90548744-INS-3496
chr7-90561447-INS-153
chr7-90564483-INS-6592
chr7-90595882-INS-7364

chr7-90604141-INS-155
chr7-90635848-INS-6173
chr7-90685948-INS-231
chr7-90778969-INS-75
chr7-90780298-INS-52
chr7-90783396-INS-4199
chr7-90812199-INS-124
chr7-90838419-INS-1402
chr7-90885313-INS-163
chr7-90943044-INS-120
chr7-90948445-INS-56
chr7-90967999-INS-87
chr7-90985481-INS-1078
chr7-91005704-INS-59
chr7-91042124-INS-877
chr7-91051695-INS-80
chr7-91067393-INS-448
chr7-91167395-INS-6916
chr7-91244480-INS-69
chr7-91267377-INS-7853
chr7-91282614-INS-57
chr7-91290538-INS-593
chr7-91305040-INS-7367
chr7-91310752-INS-235
chr7-91327762-INS-179
chr7-91336085-INS-50
chr7-91337331-INS-2582
chr7-91342795-INS-196
chr7-91350065-INS-64
chr7-91355302-INS-1397
chr7-91365298-INS-465
chr7-91365643-INS-74
chr7-91406252-INS-60
chr7-91444928-INS-1270
chr7-91448406-INS-154
chr7-91450070-INS-138
chr7-91468519-INS-1411
chr7-91485367-INS-569
chr7-91542009-INS-64
chr7-91583444-INS-245
chr7-91591000-INS-115
chr7-91634532-INS-428
chr7-91692471-INS-547
chr7-91697869-INS-81
chr7-91753431-INS-6730
chr7-91760594-INS-1434
chr7-91792666-INS-270
chr7-91795884-INS-209
chr7-91804391-INS-63
chr7-91841702-INS-50
chr7-91843977-INS-93
chr7-91943387-INS-3105
chr7-91950966-INS-6519
chr7-91967398-INS-62
chr7-91969767-INS-62
chr7-91993761-INS-52
chr7-92045329-INS-474
chr7-92061023-INS-2965
chr7-92084033-INS-205
chr7-92127162-INS-4601
chr7-92129725-INS-6417
chr7-92136394-INS-290
chr7-92136420-INS-116
chr7-92139598-INS-250
chr7-92151060-INS-7510
chr7-92180201-INS-2194
chr7-92181349-INS-50
chr7-92185018-INS-970
chr7-92193172-INS-6370
chr7-92194048-INS-110
chr7-92200664-INS-52
chr7-92212719-INS-250
chr7-92224923-INS-1906
chr7-92236617-INS-112
chr7-92352149-INS-230
chr7-92357900-INS-199
chr7-92364045-INS-64

chr7-92366690-INS-196
chr7-92369018-INS-117
chr7-92376711-INS-244
chr7-92398059-INS-272
chr7-92434098-INS-303
chr7-92434227-INS-58
chr7-92446504-INS-558
chr7-92534452-INS-50
chr7-92535970-INS-50
chr7-92591885-INS-106
chr7-92592493-INS-2206
chr7-92615274-INS-250
chr7-92615423-INS-76
chr7-92678413-INS-6912
chr7-92687703-INS-160
chr7-92687709-INS-78
chr7-92706230-INS-136
chr7-92707688-INS-2309
chr7-92707797-INS-943
chr7-92721297-INS-151
chr7-92735701-INS-50
chr7-92795215-INS-624
chr7-92911943-INS-5289
chr7-92927200-INS-199
chr7-92929449-INS-62
chr7-92936491-INS-1427
chr7-92980222-INS-94
chr7-92982606-INS-66
chr7-93039788-INS-53
chr7-93058935-INS-2991
chr7-93079307-INS-4273
chr7-93094715-INS-6448
chr7-93126426-INS-204
chr7-93127018-INS-546
chr7-93133215-INS-53
chr7-93252738-INS-90
chr7-93293335-INS-2227
chr7-93299781-INS-116
chr7-93372633-INS-199
chr7-93374183-INS-3997
chr7-93401534-INS-168
chr7-93401746-INS-153
chr7-93426344-INS-7227
chr7-93442122-INS-2158
chr7-93484202-INS-52
chr7-93486218-INS-200
chr7-93486220-INS-88
chr7-93500530-INS-140
chr7-93510472-INS-76
chr7-93611004-INS-136
chr7-93612826-INS-7731
chr7-93696045-INS-285
chr7-93708749-INS-62
chr7-93745745-INS-15044
chr7-93745745-INS-5017
chr7-93761610-INS-56
chr7-93816476-INS-1477
chr7-93857421-INS-74
chr7-93866066-INS-3209
chr7-93867613-INS-68
chr7-93891831-INS-6334
chr7-93919396-INS-7315
chr7-93941073-INS-60
chr7-93949774-INS-65
chr7-94052488-INS-96
chr7-94108596-INS-1478
chr7-94200055-INS-6544
chr7-94203021-INS-347
chr7-94207023-INS-2098
chr7-94221380-INS-1854
chr7-94231582-INS-6445
chr7-94372942-INS-4755
chr7-94410497-INS-204
chr7-94448196-INS-411
chr7-94450281-INS-1515
chr7-94452206-INS-2802
chr7-94460241-INS-2761

chr7-94553696-INS-52
chr7-94558426-INS-371
chr7-94585995-INS-956
chr7-94601871-INS-1422
chr7-94651803-INS-64
chr7-94694573-INS-106
chr7-94727336-INS-208
chr7-94747057-INS-50
chr7-94799546-INS-60
chr7-94799626-INS-60
chr7-94812962-INS-922
chr7-94846900-INS-4847
chr7-94849612-INS-114
chr7-94864320-INS-247
chr7-94864525-INS-114
chr7-94896328-INS-2768
chr7-94902657-INS-50
chr7-94972145-INS-59
chr7-94975746-INS-6395
chr7-95004270-INS-139
chr7-95004403-INS-486
chr7-95005399-INS-61
chr7-95071511-INS-2870
chr7-95099695-INS-6798
chr7-95102957-INS-68
chr7-95103087-INS-68
chr7-95111736-INS-5857
chr7-95121159-INS-2427
chr7-95122543-INS-2378
chr7-95138496-INS-109
chr7-95189251-INS-53
chr7-95189991-INS-85
chr7-95196875-INS-1399
chr7-95251028-INS-6798
chr7-95293978-INS-60
chr7-95295859-INS-1963
chr7-95300143-INS-68
chr7-95307007-INS-1912
chr7-95340340-INS-52
chr7-95424950-INS-105
chr7-95430802-INS-251
chr7-95460920-INS-5543
chr7-95461399-INS-4202
chr7-95495591-INS-65
chr7-95626994-INS-6178
chr7-95637375-INS-181
chr7-95649116-INS-102
chr7-95649201-INS-414
chr7-95700580-INS-50
chr7-95724402-INS-122
chr7-95748843-INS-2000
chr7-95856026-INS-60
chr7-95886499-INS-2876
chr7-95913806-INS-95
chr7-95916985-INS-52
chr7-96071500-INS-51
chr7-96110896-INS-1368
chr7-96232333-INS-64
chr7-96286528-INS-58
chr7-96341529-INS-80
chr7-96348402-INS-61
chr7-96393784-INS-56
chr7-96533149-INS-70
chr7-96576372-INS-61
chr7-96751783-INS-792
chr7-96778670-INS-51
chr7-96830007-INS-120
chr7-96910471-INS-64
chr7-97008079-INS-96
chr7-97008379-INS-201
chr7-97019675-INS-55
chr7-97041925-INS-72
chr7-97054185-INS-211
chr7-97055033-INS-545
chr7-97069576-INS-56
chr7-97074003-INS-64
chr7-97081744-INS-227

chr7-97082290-INS-55
chr7-97088602-INS-192
chr7-97150495-INS-149
chr7-97156614-INS-382
chr7-97171202-INS-123
chr7-97180740-INS-300
chr7-97187852-INS-59
chr7-97211631-INS-50
chr7-97214821-INS-7160
chr7-97256683-INS-96
chr7-97349031-INS-1238
chr7-97393693-INS-56
chr7-97401170-INS-194
chr7-97428426-INS-456
chr7-97446539-INS-208
chr7-97482366-INS-52
chr7-97519893-INS-108
chr7-97650877-INS-109
chr7-97693003-INS-219
chr7-97693026-INS-89
chr7-97741453-INS-99
chr7-97798204-INS-123
chr7-97804218-INS-288
chr7-97829405-INS-223
chr7-97837027-INS-88
chr7-97838181-INS-7119
chr7-97893801-INS-177
chr7-97919871-INS-59
chr7-97964374-INS-107
chr7-97964394-INS-50
chr7-97976613-INS-52
chr7-98010853-INS-50
chr7-98020794-INS-205
chr7-98059577-INS-60
chr7-98059940-INS-67
chr7-98062394-INS-159
chr7-98064237-INS-52
chr7-98069738-INS-60
chr7-98107127-INS-178
chr7-98107265-INS-137
chr7-98127117-INS-51
chr7-98186374-INS-5942
chr7-98210070-INS-206
chr7-98241597-INS-259
chr7-98378785-INS-126
chr7-98387608-INS-421
chr7-98412607-INS-114
chr7-98471938-INS-69
chr7-98622355-INS-4305
chr7-98640933-INS-4353
chr7-98646847-INS-1532
chr7-98646949-INS-143
chr7-98647037-INS-390
chr7-98683606-INS-87
chr7-98718123-INS-66
chr7-98718373-INS-186
chr7-98718373-INS-91
chr7-98724065-INS-172
chr7-98741760-INS-72
chr7-98741848-INS-360
chr7-98743680-INS-51
chr7-98768864-INS-591
chr7-98812500-INS-103
chr7-98818844-INS-94
chr7-98832672-INS-103
chr7-98858513-INS-74
chr7-98859164-INS-234
chr7-98859164-INS-507
chr7-98892650-INS-51
chr7-98906370-INS-4678
chr7-98913817-INS-56
chr7-98917972-INS-138
chr7-98953870-INS-184
chr7-98958246-INS-112
chr7-98958246-INS-568
chr7-98958850-INS-88
chr7-98958869-INS-192

chr7-98959281-INS-501
chr7-98982999-INS-63
chr7-99016859-INS-56
chr7-99054892-INS-125
chr7-99054892-INS-50
chr7-99061150-INS-2394
chr7-99061150-INS-7451
chr7-99155346-INS-124
chr7-99164743-INS-91
chr7-99172234-INS-56
chr7-99184574-INS-54
chr7-99199935-INS-423
chr7-99199935-INS-50
chr7-99211033-INS-270
chr7-99225584-INS-478
chr7-99257308-INS-183
chr7-99257984-INS-56
chr7-99295280-INS-91
chr7-99303069-INS-51
chr7-99325858-INS-608
chr7-99326702-INS-50
chr7-99363501-INS-80
chr7-99394216-INS-81
chr7-99394270-INS-163
chr7-99399002-INS-2112
chr7-99423725-INS-2869
chr7-99446617-INS-84
chr7-99518244-INS-104
chr7-99526014-INS-184
chr7-99526154-INS-358
chr7-99539159-INS-57
chr7-99540650-INS-88
chr7-99547542-INS-200
chr7-99547542-INS-50
chr7-99591774-INS-1726
chr7-99591884-INS-210
chr7-99609658-INS-1082
chr7-99636997-INS-196
chr7-99637177-INS-67
chr7-99719436-INS-113
chr7-99719771-INS-199
chr7-99782758-INS-571
chr7-99813195-INS-274
chr7-99825996-INS-54
chr7-99842630-INS-192
chr7-99857715-INS-50
chr7-99893303-INS-213
chr7-99922247-INS-52
chr7-99956075-INS-65
chr7-99957516-INS-51
chr7-99965113-INS-58
chr7-99991483-INS-182
chr7-100003295-INS-60
chr7-100022117-INS-86
chr7-100028035-INS-143
chr7-100035582-INS-85
chr7-100080854-INS-69
chr7-100106528-INS-214
chr7-100144614-INS-6487
chr7-100158365-INS-125
chr7-100159424-INS-339
chr7-100164465-INS-1619
chr7-100165721-INS-476
chr7-100171911-INS-3491
chr7-100174379-INS-125
chr7-100179204-INS-3407
chr7-100185460-INS-52
chr7-100200842-INS-206
chr7-100230113-INS-163
chr7-100281656-INS-68
chr7-100302015-INS-272
chr7-100348733-INS-66
chr7-100369654-INS-180
chr7-100391872-INS-219
chr7-100397050-INS-248
chr7-100397127-INS-725
chr7-100399392-INS-68

chr7-100400464-INS-62
chr7-100413570-INS-200
chr7-100414159-INS-218
chr7-100429569-INS-432
chr7-100456963-INS-461
chr7-100479014-INS-99
chr7-100540433-INS-209
chr7-100540596-INS-506
chr7-100540619-INS-76
chr7-100600455-INS-54
chr7-100600535-INS-4106
chr7-100600535-INS-785
chr7-100625459-INS-234
chr7-100631650-INS-463
chr7-100632010-INS-58
chr7-100674757-INS-223
chr7-100681182-INS-55
chr7-100708196-INS-148
chr7-100708807-INS-60
chr7-100731165-INS-87
chr7-100731240-INS-177
chr7-100944535-INS-112
chr7-101021132-INS-231
chr7-101043571-INS-3778
chr7-101058927-INS-80
chr7-101059209-INS-280
chr7-101078227-INS-64
chr7-101098452-INS-85
chr7-101126799-INS-88
chr7-101127532-INS-56
chr7-101150639-INS-69
chr7-101194546-INS-266
chr7-101194628-INS-63
chr7-101197912-INS-4562
chr7-101232156-INS-8758
chr7-101261253-INS-69
chr7-101287024-INS-60
chr7-101291402-INS-211
chr7-101310006-INS-56
chr7-101444899-INS-1805
chr7-101451545-INS-305
chr7-101478985-INS-208
chr7-101478992-INS-54
chr7-101496689-INS-212
chr7-101502948-INS-232
chr7-101521233-INS-286
chr7-101545177-INS-54
chr7-101563280-INS-237
chr7-101587418-INS-58
chr7-101587475-INS-285
chr7-101592024-INS-182
chr7-101627222-INS-268
chr7-101627454-INS-290
chr7-101706693-INS-208
chr7-101707329-INS-139
chr7-101712126-INS-9091
chr7-101758831-INS-51
chr7-101763728-INS-216
chr7-101798107-INS-2892
chr7-101809940-INS-50
chr7-101816407-INS-417
chr7-101842582-INS-92
chr7-101890781-INS-213
chr7-101910890-INS-79
chr7-101915116-INS-308
chr7-101929483-INS-50
chr7-101949020-INS-1248
chr7-101973607-INS-270
chr7-101974185-INS-72
chr7-102034390-INS-189
chr7-102034450-INS-87
chr7-102037811-INS-99
chr7-102042261-INS-55
chr7-102054016-INS-131
chr7-102090014-INS-50
chr7-102118757-INS-4792
chr7-102140418-INS-170

chr7-102140886-INS-200
chr7-102156618-INS-50
chr7-102169638-INS-51
chr7-102231370-INS-6256
chr7-102242389-INS-1936
chr7-102248900-INS-150
chr7-102253038-INS-56
chr7-102253251-INS-6423
chr7-102257920-INS-6792
chr7-102286194-INS-216
chr7-102315065-INS-72
chr7-102345742-INS-50
chr7-102372859-INS-7086
chr7-102374184-INS-1075
chr7-102379687-INS-7696
chr7-102384871-INS-548
chr7-102407865-INS-6452
chr7-102411299-INS-210
chr7-102430126-INS-272
chr7-102454653-INS-1427
chr7-102461988-INS-57
chr7-102462052-INS-130
chr7-102525258-INS-64
chr7-102531349-INS-1580
chr7-102534923-INS-7166
chr7-102562717-INS-106
chr7-102580169-INS-3434
chr7-102610349-INS-60
chr7-102610769-INS-121
chr7-102642115-INS-3770
chr7-102644874-INS-228
chr7-102669292-INS-443
chr7-102688078-INS-110
chr7-102719637-INS-102
chr7-102788682-INS-76
chr7-102788727-INS-346
chr7-102895351-INS-52
chr7-102931929-INS-832
chr7-103005750-INS-133
chr7-103022350-INS-1629
chr7-103064759-INS-12521
chr7-103078628-INS-101
chr7-103079908-INS-1289
chr7-103107909-INS-53
chr7-103112824-INS-696
chr7-103193673-INS-219
chr7-103202709-INS-50
chr7-103255334-INS-64
chr7-103281565-INS-81
chr7-103321968-INS-56
chr7-103367092-INS-375
chr7-103377520-INS-428
chr7-103377890-INS-1702
chr7-103387407-INS-52
chr7-103393210-INS-2909
chr7-103396930-INS-1848
chr7-103418113-INS-4438
chr7-103430911-INS-3039
chr7-103451670-INS-5138
chr7-103464705-INS-95
chr7-103481798-INS-2105
chr7-103487213-INS-56
chr7-103500090-INS-2001
chr7-103509306-INS-1815
chr7-103512750-INS-1520
chr7-103517279-INS-1576
chr7-103530046-INS-68
chr7-103594039-INS-125
chr7-103598805-INS-180
chr7-103600463-INS-1984
chr7-103622178-INS-1153
chr7-103638946-INS-6411
chr7-103640372-INS-6728
chr7-103648729-INS-66
chr7-103669351-INS-318
chr7-103721271-INS-6705
chr7-103724449-INS-54

chr7-103764590-INS-462
chr7-103780788-INS-131
chr7-103780822-INS-64
chr7-103781460-INS-210
chr7-103781613-INS-210
chr7-103787688-INS-186
chr7-103794367-INS-1272
chr7-103814791-INS-251
chr7-103815237-INS-52
chr7-103818136-INS-9298
chr7-103834668-INS-6481
chr7-103840656-INS-3445
chr7-103840656-INS-638
chr7-103847885-INS-187
chr7-103853411-INS-215
chr7-103862417-INS-3420
chr7-103863381-INS-112
chr7-103873402-INS-1076
chr7-103888101-INS-6256
chr7-103952842-INS-155
chr7-103971839-INS-50
chr7-103982817-INS-101
chr7-103983444-INS-84
chr7-103988456-INS-86
chr7-103995344-INS-53
chr7-104000980-INS-64
chr7-104002407-INS-1225
chr7-104013094-INS-1676
chr7-104038148-INS-3452
chr7-104055530-INS-4095
chr7-104231542-INS-104
chr7-104263865-INS-153
chr7-104270506-INS-4817
chr7-104330469-INS-1303
chr7-104330744-INS-9046
chr7-104379478-INS-2407
chr7-104415034-INS-8735
chr7-104417090-INS-50
chr7-104454835-INS-349
chr7-104497468-INS-50
chr7-104500810-INS-7292
chr7-104505622-INS-60
chr7-104511913-INS-100
chr7-104535117-INS-3110
chr7-104720740-INS-59
chr7-104759141-INS-662
chr7-104773509-INS-126
chr7-104811841-INS-52
chr7-104832463-INS-710
chr7-104835724-INS-710
chr7-104835734-INS-316
chr7-104842455-INS-2788
chr7-104858734-INS-193
chr7-104861102-INS-141
chr7-104906182-INS-52
chr7-104906389-INS-293
chr7-104910915-INS-518
chr7-104948611-INS-2352
chr7-104951442-INS-437
chr7-104951454-INS-98
chr7-104981313-INS-229
chr7-105042733-INS-148
chr7-105052434-INS-78
chr7-105074841-INS-56
chr7-105078788-INS-759
chr7-105097366-INS-382
chr7-105101424-INS-189
chr7-105108174-INS-351
chr7-105113985-INS-64
chr7-105158863-INS-1223
chr7-105179296-INS-82
chr7-105182318-INS-83
chr7-105192403-INS-136
chr7-105192577-INS-158
chr7-105202204-INS-6739
chr7-105221099-INS-57
chr7-105320449-INS-118

chr7-105320671-INS-56
chr7-105349697-INS-54
chr7-105377318-INS-188
chr7-105510441-INS-6415
chr7-105512181-INS-6439
chr7-105523976-INS-455
chr7-105526105-INS-400
chr7-105531505-INS-90
chr7-105531774-INS-864
chr7-105533639-INS-777
chr7-105541972-INS-1488
chr7-105543584-INS-6568
chr7-105553378-INS-54
chr7-105553746-INS-68
chr7-105567195-INS-54
chr7-105573332-INS-630
chr7-105945319-INS-2435
chr7-105991036-INS-1892
chr7-106004317-INS-2093
chr7-106037745-INS-1568
chr7-106052955-INS-2213
chr7-106084847-INS-50
chr7-106101918-INS-2090
chr7-106135929-INS-221
chr7-106211916-INS-178
chr7-106245962-INS-537
chr7-106249227-INS-524
chr7-106252444-INS-54
chr7-106282213-INS-150
chr7-106456219-INS-52
chr7-106463454-INS-112
chr7-106589885-INS-5243
chr7-106599690-INS-71
chr7-106668120-INS-3220
chr7-106678654-INS-7661
chr7-106678869-INS-1492
chr7-106758750-INS-75
chr7-106758860-INS-2965
chr7-106778872-INS-233
chr7-106778905-INS-1515
chr7-106794122-INS-200
chr7-106853706-INS-82
chr7-106888828-INS-4356
chr7-106898029-INS-75
chr7-106966675-INS-4902
chr7-107127382-INS-65
chr7-107150827-INS-96
chr7-107151564-INS-209
chr7-107179897-INS-50
chr7-107222063-INS-5195
chr7-107257041-INS-75
chr7-107365227-INS-57
chr7-107383475-INS-93
chr7-107407186-INS-209
chr7-107408667-INS-263
chr7-107415633-INS-70
chr7-107418288-INS-133
chr7-107440069-INS-20352
chr7-107487338-INS-916
chr7-107507788-INS-456
chr7-107535053-INS-63
chr7-107535614-INS-138
chr7-107580991-INS-383
chr7-107619177-INS-225
chr7-107667685-INS-95
chr7-107667807-INS-261
chr7-107684422-INS-88
chr7-107685162-INS-104
chr7-107686687-INS-56
chr7-107698484-INS-6196
chr7-107705893-INS-4355
chr7-107849050-INS-187
chr7-107868494-INS-6232
chr7-107872856-INS-52
chr7-107874568-INS-58
chr7-107905659-INS-135
chr7-107919595-INS-254

chr7-107963034-INS-2101
chr7-108006129-INS-1606
chr7-108008755-INS-61
chr7-108018753-INS-52
chr7-108033988-INS-70
chr7-108037377-INS-50
chr7-108045709-INS-51
chr7-108074372-INS-12782
chr7-108118966-INS-1657
chr7-108194220-INS-505
chr7-108205768-INS-1532
chr7-108238518-INS-75
chr7-108242097-INS-3340
chr7-108250237-INS-52
chr7-108274157-INS-6220
chr7-108289252-INS-1773
chr7-108293543-INS-56
chr7-108305339-INS-214
chr7-108358202-INS-60
chr7-108534674-INS-1812
chr7-108562482-INS-50
chr7-108569727-INS-181
chr7-108569786-INS-1061
chr7-108603639-INS-52
chr7-108607784-INS-50
chr7-108642372-INS-1881
chr7-108711015-INS-52
chr7-108730610-INS-752
chr7-108734045-INS-206
chr7-108748108-INS-172
chr7-108828150-INS-72
chr7-108830326-INS-92
chr7-108856341-INS-1648
chr7-108861789-INS-80
chr7-108862343-INS-50
chr7-108892504-INS-523
chr7-108950022-INS-56
chr7-108954616-INS-66
chr7-108956890-INS-250
chr7-108971413-INS-214
chr7-108997602-INS-1422
chr7-109061714-INS-240
chr7-109061714-INS-570
chr7-109065115-INS-98
chr7-109072015-INS-97
chr7-109103088-INS-448
chr7-109112021-INS-58
chr7-109137928-INS-52
chr7-109143148-INS-123
chr7-109143148-INS-54
chr7-109179471-INS-70
chr7-109182513-INS-60
chr7-109212579-INS-412
chr7-109212698-INS-119
chr7-109223860-INS-239
chr7-109285866-INS-200
chr7-109317777-INS-211
chr7-109329495-INS-69
chr7-109329606-INS-229
chr7-109333207-INS-50
chr7-109410165-INS-544
chr7-109444586-INS-90
chr7-109445303-INS-300
chr7-109461782-INS-6501
chr7-109465754-INS-50
chr7-109467036-INS-1460
chr7-109469375-INS-50
chr7-109469565-INS-60
chr7-109472598-INS-56
chr7-109473235-INS-209
chr7-109505920-INS-50
chr7-109538645-INS-114
chr7-109550324-INS-206
chr7-109553843-INS-64
chr7-109558078-INS-338
chr7-109566814-INS-52
chr7-109581144-INS-198

chr7-109597622-INS-203
chr7-109634389-INS-137
chr7-109666572-INS-208
chr7-109694710-INS-208
chr7-109699374-INS-749
chr7-109703113-INS-6410
chr7-109707842-INS-85
chr7-109717157-INS-241
chr7-109734062-INS-254
chr7-109762292-INS-104
chr7-109775523-INS-7111
chr7-109786761-INS-202
chr7-109787231-INS-52
chr7-109804112-INS-139
chr7-109804858-INS-88
chr7-109806388-INS-215
chr7-109816344-INS-231
chr7-109817544-INS-174
chr7-109817645-INS-50
chr7-109822658-INS-143
chr7-109840531-INS-59
chr7-109840968-INS-192
chr7-109892824-INS-117
chr7-109892903-INS-486
chr7-109893318-INS-197
chr7-109894283-INS-70
chr7-109941755-INS-414
chr7-109948683-INS-50
chr7-110032559-INS-6816
chr7-110045487-INS-1074
chr7-110048486-INS-212
chr7-110069124-INS-320
chr7-110069171-INS-117
chr7-110082231-INS-201
chr7-110083177-INS-6410
chr7-110111405-INS-1605
chr7-110148086-INS-372
chr7-110205202-INS-162
chr7-110205687-INS-162
chr7-110217621-INS-66
chr7-110334575-INS-56
chr7-110382062-INS-68
chr7-110484663-INS-148
chr7-110527509-INS-6669
chr7-110571283-INS-162
chr7-110730879-INS-1996
chr7-110784605-INS-57
chr7-110786235-INS-6434
chr7-110786776-INS-408
chr7-110792577-INS-11935
chr7-110820648-INS-124
chr7-110820789-INS-272
chr7-110839307-INS-6832
chr7-110873731-INS-6400
chr7-110880930-INS-69
chr7-110880931-INS-212
chr7-110881422-INS-498
chr7-110957182-INS-93
chr7-110957685-INS-413
chr7-111001783-INS-179
chr7-111029841-INS-7109
chr7-111051055-INS-208
chr7-111056520-INS-1659
chr7-111094809-INS-148
chr7-111113445-INS-53
chr7-111127095-INS-3048
chr7-111235145-INS-52
chr7-111278112-INS-52
chr7-111291594-INS-50
chr7-111299804-INS-2782
chr7-111318576-INS-151
chr7-111361638-INS-68
chr7-111405408-INS-56
chr7-111411597-INS-60
chr7-111458552-INS-50
chr7-111458864-INS-132
chr7-111462403-INS-5067

chr7-111499561-INS-138
chr7-111526839-INS-111
chr7-111526839-INS-306
chr7-111548142-INS-172
chr7-111572027-INS-52
chr7-111583330-INS-6557
chr7-111600811-INS-70
chr7-111659270-INS-80
chr7-111714228-INS-499
chr7-111759010-INS-145
chr7-111761484-INS-60
chr7-111767716-INS-7361
chr7-111781990-INS-54
chr7-111811592-INS-50
chr7-111817855-INS-50
chr7-111841602-INS-284
chr7-111864807-INS-118
chr7-111864807-INS-58
chr7-111894226-INS-151
chr7-111894268-INS-305
chr7-111926657-INS-118
chr7-111947534-INS-78
chr7-111950011-INS-148
chr7-111997832-INS-778
chr7-112104084-INS-50
chr7-112176355-INS-78
chr7-112177245-INS-129
chr7-112193697-INS-105
chr7-112193984-INS-365
chr7-112199429-INS-102
chr7-112237437-INS-3150
chr7-112237437-INS-870
chr7-112237786-INS-1410
chr7-112293103-INS-50
chr7-112301166-INS-295
chr7-112338050-INS-52
chr7-112372777-INS-50
chr7-112382838-INS-1732
chr7-112445965-INS-62
chr7-112447804-INS-69
chr7-112458581-INS-78
chr7-112494885-INS-82
chr7-112515871-INS-64
chr7-112541558-INS-148
chr7-112582212-INS-150
chr7-112597640-INS-84
chr7-112601713-INS-75
chr7-112601741-INS-228
chr7-112611642-INS-2634
chr7-112625874-INS-60
chr7-112651839-INS-64
chr7-112702243-INS-6450
chr7-112722290-INS-51
chr7-112774325-INS-498
chr7-112774325-INS-6475
chr7-112774581-INS-59
chr7-112830881-INS-134
chr7-112867425-INS-51
chr7-112923192-INS-59
chr7-112974544-INS-140
chr7-112979654-INS-190
chr7-112980720-INS-10206
chr7-112980720-INS-4900
chr7-113021712-INS-127
chr7-113044279-INS-130
chr7-113063691-INS-79
chr7-113156055-INS-209
chr7-113183454-INS-7268
chr7-113208586-INS-54
chr7-113232419-INS-486
chr7-113257281-INS-6165
chr7-113297865-INS-55
chr7-113305284-INS-3399
chr7-113337461-INS-156
chr7-113350311-INS-6365
chr7-113363398-INS-120
chr7-113376129-INS-101

chr7-113384300-INS-178
chr7-113412591-INS-68
chr7-113442794-INS-56
chr7-113480343-INS-6867
chr7-113517923-INS-104
chr7-113591276-INS-81
chr7-113619201-INS-58
chr7-113630871-INS-423
chr7-113659957-INS-166
chr7-113659989-INS-447
chr7-113660279-INS-202
chr7-113660817-INS-1022
chr7-113668962-INS-66
chr7-113676805-INS-5284
chr7-113747542-INS-69
chr7-113751805-INS-90
chr7-113764800-INS-52
chr7-113765302-INS-52
chr7-113799434-INS-154
chr7-113806095-INS-198
chr7-113806882-INS-213
chr7-113847983-INS-54
chr7-113851625-INS-208
chr7-113878934-INS-340
chr7-113920811-INS-196
chr7-113924749-INS-50
chr7-113944602-INS-111
chr7-114034109-INS-189
chr7-114050693-INS-3419
chr7-114123677-INS-54
chr7-114195732-INS-70
chr7-114335078-INS-59
chr7-114382497-INS-6113
chr7-114405526-INS-146
chr7-114413491-INS-4186
chr7-114447299-INS-128
chr7-114528583-INS-2037
chr7-114534278-INS-6212
chr7-114585149-INS-222
chr7-114620795-INS-258
chr7-114663277-INS-456
chr7-114699167-INS-60
chr7-114762714-INS-5616
chr7-114812241-INS-50
chr7-114835254-INS-3226
chr7-114840018-INS-50
chr7-114840540-INS-90
chr7-114868926-INS-59
chr7-114872485-INS-50
chr7-114895506-INS-244
chr7-114913714-INS-61
chr7-114913967-INS-58
chr7-114936917-INS-93
chr7-115023541-INS-370
chr7-115057378-INS-100
chr7-115102871-INS-415
chr7-115134037-INS-60
chr7-115188176-INS-206
chr7-115188382-INS-205
chr7-115238856-INS-509
chr7-115256570-INS-80
chr7-115270014-INS-88
chr7-115316876-INS-63
chr7-115317248-INS-217
chr7-115320274-INS-50
chr7-115339872-INS-57
chr7-115417183-INS-206
chr7-115439446-INS-50
chr7-115448599-INS-69
chr7-115449439-INS-6444
chr7-115468153-INS-76
chr7-115622854-INS-198
chr7-115632525-INS-54
chr7-115637041-INS-130
chr7-115648694-INS-168
chr7-115678961-INS-6509
chr7-115708343-INS-76

chr7-115741570-INS-50
chr7-115788234-INS-5496
chr7-115794921-INS-140
chr7-115828793-INS-377
chr7-115857388-INS-63
chr7-115893059-INS-83
chr7-115895877-INS-66
chr7-115913505-INS-67
chr7-115921550-INS-206
chr7-115921932-INS-72
chr7-115924036-INS-181
chr7-115924926-INS-151
chr7-115952244-INS-258
chr7-115996762-INS-50
chr7-116038461-INS-218
chr7-116085090-INS-147
chr7-116107486-INS-76
chr7-116201905-INS-6218
chr7-116204261-INS-62
chr7-116229990-INS-50
chr7-116250111-INS-6563
chr7-116272156-INS-68
chr7-116325462-INS-392
chr7-116327426-INS-52
chr7-116453514-INS-72
chr7-116467491-INS-2155
chr7-116517015-INS-60
chr7-116536623-INS-54
chr7-116542153-INS-58
chr7-116587419-INS-50
chr7-116596835-INS-78
chr7-116671170-INS-60
chr7-116699027-INS-483
chr7-116724232-INS-1056
chr7-116737635-INS-995
chr7-116756863-INS-64
chr7-116866280-INS-54
chr7-116905968-INS-216
chr7-116924437-INS-107
chr7-116933421-INS-6511
chr7-116970051-INS-53
chr7-116972866-INS-94
chr7-117073371-INS-269
chr7-117103282-INS-50
chr7-117138681-INS-50
chr7-117195920-INS-84
chr7-117329149-INS-6111
chr7-117351137-INS-510
chr7-117366024-INS-3131
chr7-117368573-INS-72
chr7-117372880-INS-105
chr7-117373485-INS-6452
chr7-117411533-INS-134
chr7-117482632-INS-2321
chr7-117491890-INS-193
chr7-117491957-INS-487
chr7-117491962-INS-190
chr7-117499747-INS-1887
chr7-117500661-INS-7101
chr7-117522152-INS-175
chr7-117530297-INS-58
chr7-117532641-INS-4859
chr7-117545407-INS-5129
chr7-117634687-INS-56
chr7-117640821-INS-1056
chr7-117640821-INS-8004
chr7-117643176-INS-70
chr7-117693287-INS-97
chr7-117786248-INS-469
chr7-117787105-INS-380
chr7-117787213-INS-95
chr7-117798717-INS-387
chr7-117798838-INS-94
chr7-117844161-INS-184
chr7-117851443-INS-79
chr7-117859732-INS-149
chr7-117897171-INS-72

chr7-117900956-INS-72
chr7-117901868-INS-401
chr7-117902325-INS-124
chr7-117903708-INS-58
chr7-117950645-INS-64
chr7-117962589-INS-93
chr7-117984752-INS-131
chr7-117994742-INS-63
chr7-118002553-INS-130
chr7-118002655-INS-872
chr7-118002790-INS-62
chr7-118011827-INS-690
chr7-118054573-INS-58
chr7-118054800-INS-162
chr7-118054807-INS-63
chr7-118066959-INS-1888
chr7-118067613-INS-52
chr7-118068969-INS-66
chr7-118072204-INS-3115
chr7-118073980-INS-154
chr7-118097709-INS-50
chr7-118115798-INS-168
chr7-118141648-INS-115
chr7-118153103-INS-6440
chr7-118157239-INS-102
chr7-118157996-INS-95
chr7-118158615-INS-50
chr7-118164084-INS-4970
chr7-118229710-INS-50
chr7-118259201-INS-447
chr7-118269775-INS-1761
chr7-118274433-INS-4438
chr7-118276066-INS-197
chr7-118285273-INS-126
chr7-118296866-INS-7135
chr7-118304005-INS-293
chr7-118313166-INS-225
chr7-118338039-INS-418
chr7-118339444-INS-72
chr7-118344863-INS-215
chr7-118369634-INS-52
chr7-118369973-INS-4140
chr7-118392128-INS-216
chr7-118412272-INS-210
chr7-118430685-INS-60
chr7-118453827-INS-1354
chr7-118472786-INS-53
chr7-118483910-INS-50
chr7-118494315-INS-6731
chr7-118495618-INS-1480
chr7-118499531-INS-131
chr7-118556475-INS-580
chr7-118561991-INS-51
chr7-118589825-INS-68
chr7-118599828-INS-6556
chr7-118615235-INS-1735
chr7-118615574-INS-1604
chr7-118624624-INS-235
chr7-118625022-INS-90
chr7-118626677-INS-885
chr7-118637293-INS-70
chr7-118658115-INS-1239
chr7-118663119-INS-193
chr7-118666720-INS-156
chr7-118704817-INS-65
chr7-118873386-INS-840
chr7-118892323-INS-7059
chr7-118909148-INS-52
chr7-118948486-INS-60
chr7-118960660-INS-2812
chr7-118961297-INS-57
chr7-119010594-INS-80
chr7-119105894-INS-3063
chr7-119116398-INS-59
chr7-119126182-INS-66
chr7-119183814-INS-302
chr7-119231327-INS-208

chr7-119263432-INS-1215
chr7-119265217-INS-563
chr7-119265338-INS-2425
chr7-119265816-INS-3870
chr7-119306946-INS-1558
chr7-119338527-INS-401
chr7-119485241-INS-50
chr7-119516988-INS-6404
chr7-119537744-INS-50
chr7-119552311-INS-56
chr7-119571355-INS-6512
chr7-119577013-INS-310
chr7-119585764-INS-68
chr7-119586650-INS-124
chr7-119586682-INS-280
chr7-119588265-INS-479
chr7-119652751-INS-55
chr7-119674620-INS-534
chr7-119736915-INS-51
chr7-119740179-INS-77
chr7-119792377-INS-194
chr7-119811111-INS-147
chr7-119813460-INS-90
chr7-119836216-INS-156
chr7-119854919-INS-363
chr7-119855017-INS-160
chr7-119870176-INS-1261
chr7-119884358-INS-7189
chr7-119923036-INS-2244
chr7-119954285-INS-210
chr7-119958505-INS-503
chr7-119959109-INS-196
chr7-119976554-INS-465
chr7-120041874-INS-1889
chr7-120043511-INS-2955
chr7-120070757-INS-1488
chr7-120159667-INS-68
chr7-120179949-INS-6947
chr7-120210075-INS-1349
chr7-120222799-INS-69
chr7-120223869-INS-60
chr7-120224264-INS-204
chr7-120226180-INS-7141
chr7-120230983-INS-2482
chr7-120303673-INS-62
chr7-120335026-INS-62
chr7-120361285-INS-189
chr7-120369298-INS-242
chr7-120380523-INS-337
chr7-120380627-INS-57
chr7-120387263-INS-1753
chr7-120396401-INS-80
chr7-120396628-INS-140
chr7-120419500-INS-366
chr7-120453827-INS-179
chr7-120475575-INS-96
chr7-120476593-INS-56
chr7-120491790-INS-65
chr7-120500342-INS-206
chr7-120509483-INS-306
chr7-120514241-INS-60
chr7-120545522-INS-102
chr7-120554451-INS-105
chr7-120554458-INS-247
chr7-120571290-INS-62
chr7-120597299-INS-6481
chr7-120610462-INS-73
chr7-120669065-INS-321
chr7-120689027-INS-6825
chr7-120706826-INS-6448
chr7-120723711-INS-92
chr7-120737458-INS-56
chr7-120769988-INS-202
chr7-120773768-INS-460
chr7-120792406-INS-7094
chr7-120807898-INS-50
chr7-120808643-INS-76

chr7-120829828-INS-158
chr7-120841992-INS-58
chr7-120853373-INS-235
chr7-120861393-INS-110
chr7-120874660-INS-50
chr7-120942841-INS-102
chr7-121008111-INS-53
chr7-121042075-INS-52
chr7-121071989-INS-98
chr7-121110292-INS-70
chr7-121115796-INS-253
chr7-121124856-INS-197
chr7-121133153-INS-50
chr7-121138263-INS-91
chr7-121163553-INS-5268
chr7-121166714-INS-88
chr7-121167367-INS-54
chr7-121174283-INS-84
chr7-121201254-INS-53
chr7-121213025-INS-245
chr7-121213298-INS-114
chr7-121223613-INS-62
chr7-121223704-INS-215
chr7-121224168-INS-379
chr7-121229511-INS-415
chr7-121229851-INS-83
chr7-121256270-INS-112
chr7-121328442-INS-231
chr7-121374695-INS-56
chr7-121387889-INS-68
chr7-121403243-INS-397
chr7-121424951-INS-63
chr7-121429264-INS-55
chr7-121449266-INS-61
chr7-121514584-INS-78
chr7-121514829-INS-498
chr7-121522461-INS-76
chr7-121544700-INS-2736
chr7-121565211-INS-632
chr7-121566833-INS-213
chr7-121598824-INS-68
chr7-121624219-INS-627
chr7-121624360-INS-130
chr7-121627431-INS-53
chr7-121630059-INS-224
chr7-121630777-INS-192
chr7-121641259-INS-50
chr7-121642932-INS-54
chr7-121648746-INS-1184
chr7-121686751-INS-52
chr7-121703022-INS-1686
chr7-121705706-INS-202
chr7-121719259-INS-214
chr7-121736886-INS-1666
chr7-121736968-INS-81
chr7-121741734-INS-172
chr7-121764251-INS-82
chr7-121777598-INS-166
chr7-121819090-INS-73
chr7-121857060-INS-258
chr7-121876420-INS-467
chr7-121885420-INS-80
chr7-121903825-INS-71
chr7-121934091-INS-52
chr7-121953728-INS-61
chr7-122022049-INS-52
chr7-122026304-INS-105
chr7-122047397-INS-944
chr7-122117889-INS-115
chr7-122153604-INS-90
chr7-122163415-INS-60
chr7-122205423-INS-54
chr7-122273889-INS-72
chr7-122298459-INS-2766
chr7-122313762-INS-231
chr7-122358872-INS-51
chr7-122366685-INS-114

chr7-122391202-INS-263
chr7-122436588-INS-50
chr7-122440770-INS-226
chr7-122440794-INS-76
chr7-122459551-INS-53
chr7-122469782-INS-189
chr7-122480668-INS-2915
chr7-122482358-INS-199
chr7-122483143-INS-106
chr7-122483425-INS-72
chr7-122483598-INS-234
chr7-122541928-INS-273
chr7-122616334-INS-127
chr7-122636178-INS-104
chr7-122663457-INS-4899
chr7-122677030-INS-219
chr7-122696122-INS-50
chr7-122700105-INS-129
chr7-122703199-INS-111
chr7-122739771-INS-72
chr7-122786160-INS-149
chr7-122818162-INS-756
chr7-122823745-INS-1699
chr7-122825340-INS-92
chr7-122829549-INS-50
chr7-122838314-INS-56
chr7-122855764-INS-56
chr7-122884453-INS-423
chr7-122884453-INS-94
chr7-122928249-INS-63
chr7-122931194-INS-66
chr7-123001027-INS-3248
chr7-123027983-INS-60
chr7-123043446-INS-724
chr7-123111661-INS-1160
chr7-123111816-INS-60
chr7-123169941-INS-64
chr7-123212583-INS-909
chr7-123232535-INS-2421
chr7-123252790-INS-122
chr7-123266109-INS-395
chr7-123269289-INS-1453
chr7-123290873-INS-9438
chr7-123336061-INS-4725
chr7-123368749-INS-2737
chr7-123386739-INS-52
chr7-123393540-INS-6477
chr7-123400260-INS-88
chr7-123421407-INS-58
chr7-123487823-INS-57
chr7-123521806-INS-6284
chr7-123533432-INS-84
chr7-123599230-INS-67
chr7-123625619-INS-56
chr7-123655774-INS-92
chr7-123716133-INS-58
chr7-123738046-INS-774
chr7-123743948-INS-694
chr7-123756276-INS-6179
chr7-123764320-INS-124
chr7-123773035-INS-58
chr7-123780045-INS-66
chr7-123783944-INS-70
chr7-123823803-INS-66
chr7-123845072-INS-51
chr7-123869815-INS-50
chr7-123871166-INS-73
chr7-123877550-INS-104
chr7-123877550-INS-501
chr7-123986741-INS-54
chr7-124004277-INS-6866
chr7-124008977-INS-10920
chr7-124036407-INS-131
chr7-124037180-INS-2180
chr7-124084884-INS-68
chr7-124111656-INS-70
chr7-124114163-INS-52

chr7-124191190-INS-52
chr7-124213905-INS-3338
chr7-124239930-INS-60
chr7-124249088-INS-133
chr7-124272145-INS-1673
chr7-124285882-INS-1220
chr7-124289706-INS-1043
chr7-124302410-INS-222
chr7-124302629-INS-257
chr7-124400462-INS-645
chr7-124401345-INS-2880
chr7-124407738-INS-4773
chr7-124420766-INS-82
chr7-124422526-INS-129
chr7-124442344-INS-52
chr7-124490072-INS-1179
chr7-124493534-INS-456
chr7-124511992-INS-148
chr7-124517362-INS-52
chr7-124536668-INS-182
chr7-124544398-INS-503
chr7-124566341-INS-158
chr7-124576224-INS-76
chr7-124592385-INS-9090
chr7-124625236-INS-120
chr7-124651288-INS-89
chr7-124703945-INS-1128
chr7-124716940-INS-80
chr7-124717540-INS-78
chr7-124771969-INS-152
chr7-124775145-INS-216
chr7-124830472-INS-53
chr7-124847022-INS-52
chr7-124853093-INS-2933
chr7-124880088-INS-62
chr7-124942359-INS-9053
chr7-124976713-INS-50
chr7-124981751-INS-112
chr7-124981892-INS-52
chr7-124981907-INS-248
chr7-124988105-INS-1026
chr7-125009934-INS-218
chr7-125146233-INS-201
chr7-125184448-INS-50
chr7-125317612-INS-172
chr7-125337413-INS-272
chr7-125361456-INS-65
chr7-125371691-INS-50
chr7-125389187-INS-3697
chr7-125462043-INS-96
chr7-125512730-INS-159
chr7-125552342-INS-61
chr7-125552379-INS-180
chr7-125569290-INS-52
chr7-125574947-INS-236
chr7-125603494-INS-240
chr7-125691681-INS-209
chr7-125691768-INS-112
chr7-125709740-INS-54
chr7-125739326-INS-155
chr7-125758970-INS-196
chr7-125766210-INS-228
chr7-125769280-INS-488
chr7-125800748-INS-130
chr7-125858877-INS-90
chr7-125860933-INS-1893
chr7-125920527-INS-87
chr7-125922167-INS-5223
chr7-125943362-INS-634
chr7-125959521-INS-54
chr7-126077759-INS-204
chr7-126109421-INS-204
chr7-126119911-INS-191
chr7-126138970-INS-1230
chr7-126170870-INS-305
chr7-126175316-INS-204
chr7-126191289-INS-205

chr7-126196232-INS-54
chr7-126406520-INS-50
chr7-126413613-INS-118
chr7-126413634-INS-56
chr7-126414048-INS-212
chr7-126476294-INS-58
chr7-126502879-INS-2032
chr7-126503423-INS-102
chr7-126507988-INS-168
chr7-126599721-INS-201
chr7-126659835-INS-103
chr7-126660125-INS-219
chr7-126684102-INS-52
chr7-126690048-INS-230
chr7-126724353-INS-187
chr7-126728001-INS-3344
chr7-126743023-INS-50
chr7-126759432-INS-133
chr7-126777911-INS-52
chr7-126781074-INS-50
chr7-126825850-INS-76
chr7-126833533-INS-55
chr7-126839749-INS-5759
chr7-126863247-INS-486
chr7-126865058-INS-137
chr7-126867302-INS-229
chr7-126883764-INS-7165
chr7-126888736-INS-173
chr7-126903129-INS-75
chr7-126915326-INS-207
chr7-126918971-INS-208
chr7-126925753-INS-60
chr7-126926162-INS-64
chr7-126935835-INS-3082
chr7-126937832-INS-64
chr7-126947946-INS-76
chr7-126985296-INS-65
chr7-126985349-INS-240
chr7-126989040-INS-177
chr7-127010993-INS-204
chr7-127092388-INS-54
chr7-127115026-INS-203
chr7-127170969-INS-50
chr7-127175459-INS-199
chr7-127204863-INS-173
chr7-127233822-INS-509
chr7-127234125-INS-215
chr7-127246754-INS-261
chr7-127278988-INS-436
chr7-127282562-INS-338
chr7-127282905-INS-187
chr7-127283185-INS-371
chr7-127305649-INS-242
chr7-127305649-INS-66
chr7-127318669-INS-2482
chr7-127320641-INS-220
chr7-127445902-INS-217
chr7-127463487-INS-52
chr7-127500339-INS-88
chr7-127538433-INS-152
chr7-127556466-INS-203
chr7-127642553-INS-505
chr7-127642722-INS-56
chr7-127654667-INS-129
chr7-127654667-INS-51
chr7-127655896-INS-54
chr7-127664284-INS-159
chr7-127677150-INS-57
chr7-127682209-INS-91
chr7-127683608-INS-279
chr7-127711052-INS-138
chr7-127748847-INS-62
chr7-127760454-INS-50
chr7-127764373-INS-73
chr7-127765178-INS-66
chr7-127795080-INS-224
chr7-127827984-INS-111

chr7-127842988-INS-346
chr7-127854986-INS-5469
chr7-127882641-INS-57
chr7-127889056-INS-2782
chr7-127907797-INS-172
chr7-127994747-INS-56
chr7-128067597-INS-107
chr7-128085774-INS-212
chr7-128102171-INS-434
chr7-128109860-INS-2575
chr7-128114165-INS-51
chr7-128131912-INS-81
chr7-128184460-INS-58
chr7-128196546-INS-54
chr7-128205961-INS-51
chr7-128206075-INS-113
chr7-128222866-INS-105
chr7-128272557-INS-90
chr7-128273011-INS-210
chr7-128323805-INS-125
chr7-128328007-INS-154
chr7-128414417-INS-50
chr7-128418396-INS-201
chr7-128418659-INS-151
chr7-128418723-INS-66
chr7-128436052-INS-303
chr7-128438853-INS-1230
chr7-128454584-INS-162
chr7-128454611-INS-67
chr7-128466273-INS-70
chr7-128471759-INS-56
chr7-128474243-INS-69
chr7-128488030-INS-6177
chr7-128501838-INS-457
chr7-128502580-INS-50
chr7-128507188-INS-607
chr7-128521899-INS-76
chr7-128539966-INS-1207
chr7-128551965-INS-6827
chr7-128560635-INS-92
chr7-128561470-INS-54
chr7-128606160-INS-60
chr7-128654398-INS-68
chr7-128661501-INS-7140
chr7-128662606-INS-1204
chr7-128678774-INS-6212
chr7-128701154-INS-3194
chr7-128711788-INS-192
chr7-128785147-INS-181
chr7-128786361-INS-198
chr7-128832086-INS-3154
chr7-128850533-INS-50
chr7-128869048-INS-7075
chr7-128898775-INS-50
chr7-128901622-INS-1765
chr7-128918505-INS-50
chr7-128948807-INS-3713
chr7-128953800-INS-5983
chr7-128967797-INS-51
chr7-129075126-INS-128
chr7-129106824-INS-7160
chr7-129200583-INS-75
chr7-129201526-INS-207
chr7-129209197-INS-4462
chr7-129248777-INS-159
chr7-129268890-INS-5983
chr7-129313028-INS-3483
chr7-129322087-INS-1891
chr7-129406735-INS-200
chr7-129419247-INS-87
chr7-129432131-INS-6199
chr7-129456377-INS-142
chr7-129463631-INS-306
chr7-129463670-INS-151
chr7-129465921-INS-312
chr7-129465923-INS-136
chr7-129467226-INS-53

chr7-129476809-INS-4206
chr7-129493243-INS-85
chr7-129501751-INS-78
chr7-129502002-INS-154
chr7-129558292-INS-108
chr7-129587419-INS-68
chr7-129600087-INS-6848
chr7-129677452-INS-999
chr7-129693036-INS-412
chr7-129704075-INS-68
chr7-129710554-INS-87
chr7-129719755-INS-184
chr7-129722997-INS-141
chr7-129791293-INS-54
chr7-129808146-INS-104
chr7-129822998-INS-201
chr7-129856299-INS-69
chr7-129934642-INS-98
chr7-129958635-INS-208
chr7-129973942-INS-50
chr7-129987654-INS-54
chr7-130004254-INS-50
chr7-130010699-INS-673
chr7-130010714-INS-171
chr7-130015239-INS-217
chr7-130059712-INS-2496
chr7-130059721-INS-129
chr7-130090622-INS-204
chr7-130127845-INS-83
chr7-130133592-INS-165
chr7-130147974-INS-81
chr7-130183710-INS-83
chr7-130193315-INS-115
chr7-130193316-INS-6435
chr7-130196390-INS-9572
chr7-130203044-INS-309
chr7-130207559-INS-56
chr7-130210679-INS-242
chr7-130241601-INS-74
chr7-130245057-INS-68
chr7-130262948-INS-410
chr7-130270413-INS-84
chr7-130274085-INS-69
chr7-130287175-INS-435
chr7-130354513-INS-68
chr7-130363421-INS-2778
chr7-130372957-INS-56
chr7-130376578-INS-62
chr7-130404802-INS-58
chr7-130423005-INS-217
chr7-130452963-INS-78
chr7-130453655-INS-57
chr7-130456288-INS-96
chr7-130464136-INS-62
chr7-130477330-INS-61
chr7-130508066-INS-209
chr7-130508857-INS-56
chr7-130524053-INS-123
chr7-130524624-INS-264
chr7-130530248-INS-64
chr7-130530254-INS-1393
chr7-130543639-INS-53
chr7-130558947-INS-70
chr7-130592798-INS-116
chr7-130611578-INS-50
chr7-130647673-INS-60
chr7-130672632-INS-564
chr7-130681371-INS-41737
chr7-130697748-INS-8795
chr7-130709205-INS-111
chr7-130733729-INS-92
chr7-130733845-INS-291
chr7-130773827-INS-316
chr7-130773913-INS-150
chr7-130792696-INS-261
chr7-130813596-INS-62
chr7-130821130-INS-263

chr7-130847572-INS-131
chr7-130848592-INS-765
chr7-130869593-INS-58
chr7-130872512-INS-178
chr7-130877393-INS-6259
chr7-130953920-INS-826
chr7-131028099-INS-4427
chr7-131028219-INS-790
chr7-131111559-INS-56
chr7-131112438-INS-65
chr7-131304460-INS-4140
chr7-131309578-INS-50
chr7-131316120-INS-240
chr7-131331047-INS-2396
chr7-131362225-INS-95
chr7-131374995-INS-6372
chr7-131383286-INS-303
chr7-131392433-INS-56
chr7-131445440-INS-64
chr7-131504042-INS-52
chr7-131603700-INS-261
chr7-131603736-INS-87
chr7-131611333-INS-7085
chr7-131618323-INS-95
chr7-131641254-INS-72
chr7-131669942-INS-7101
chr7-131677244-INS-481
chr7-131679184-INS-51
chr7-131687817-INS-160
chr7-131780679-INS-6367
chr7-131797831-INS-90
chr7-131838292-INS-70
chr7-131843645-INS-61
chr7-131844499-INS-58
chr7-131864347-INS-314
chr7-131899833-INS-131
chr7-131899833-INS-373
chr7-131913100-INS-66
chr7-131948072-INS-54
chr7-131966322-INS-6331
chr7-131979212-INS-381
chr7-131979359-INS-75
chr7-131982881-INS-7285
chr7-131985626-INS-54
chr7-131995801-INS-309
chr7-131995801-INS-80
chr7-132024786-INS-53
chr7-132032728-INS-435
chr7-132033128-INS-203
chr7-132035967-INS-66
chr7-132097917-INS-161
chr7-132108271-INS-116
chr7-132116574-INS-52
chr7-132127831-INS-51
chr7-132160596-INS-60
chr7-132172195-INS-54
chr7-132172195-INS-729
chr7-132200970-INS-67
chr7-132231725-INS-201
chr7-132245218-INS-169
chr7-132261772-INS-57
chr7-132263068-INS-112
chr7-132327625-INS-56
chr7-132328534-INS-207
chr7-132367798-INS-245
chr7-132367801-INS-532
chr7-132367803-INS-84
chr7-132419699-INS-84
chr7-132428002-INS-402
chr7-132447929-INS-99
chr7-132482659-INS-495
chr7-132482871-INS-66
chr7-132520431-INS-180
chr7-132523314-INS-52
chr7-132526898-INS-72
chr7-132546159-INS-852
chr7-132593679-INS-6189

chr7-132601321-INS-98
chr7-132707367-INS-52
chr7-132738108-INS-201
chr7-132749837-INS-363
chr7-132765127-INS-64
chr7-132770043-INS-65
chr7-132770328-INS-595
chr7-132799686-INS-62
chr7-132827758-INS-53
chr7-132849775-INS-65
chr7-132853867-INS-210
chr7-132867635-INS-54
chr7-132875466-INS-207
chr7-132909434-INS-92
chr7-132940859-INS-84
chr7-132970888-INS-140
chr7-132983657-INS-6245
chr7-133033992-INS-106
chr7-133040895-INS-385
chr7-133040907-INS-111
chr7-133091526-INS-75
chr7-133107916-INS-62
chr7-133129049-INS-52
chr7-133194152-INS-148
chr7-133218928-INS-50
chr7-133221806-INS-51
chr7-133250567-INS-350
chr7-133277898-INS-156
chr7-133277981-INS-51
chr7-133291977-INS-143
chr7-133306804-INS-199
chr7-133360906-INS-130
chr7-133467743-INS-100
chr7-133493870-INS-6217
chr7-133547574-INS-694
chr7-133623959-INS-148
chr7-133624462-INS-590
chr7-133630749-INS-68
chr7-133634126-INS-50
chr7-133647638-INS-56
chr7-133652091-INS-7206
chr7-133697480-INS-684
chr7-133722252-INS-6232
chr7-133744580-INS-3287
chr7-133756144-INS-91
chr7-133759352-INS-52
chr7-133762912-INS-409
chr7-133763233-INS-878
chr7-133779625-INS-55
chr7-133802461-INS-50
chr7-133810757-INS-50
chr7-133833217-INS-4214
chr7-133860678-INS-1078
chr7-133918198-INS-6478
chr7-133939414-INS-1438
chr7-133991688-INS-787
chr7-134042798-INS-2884
chr7-134075682-INS-59
chr7-134079008-INS-54
chr7-134096327-INS-6670
chr7-134118820-INS-7439
chr7-134141189-INS-1863
chr7-134147002-INS-6397
chr7-134171542-INS-50
chr7-134207284-INS-68
chr7-134243189-INS-58
chr7-134299098-INS-59
chr7-134312839-INS-1030
chr7-134330719-INS-213
chr7-134330781-INS-829
chr7-134357690-INS-7758
chr7-134414620-INS-4522
chr7-134417734-INS-101
chr7-134434727-INS-63
chr7-134446063-INS-512
chr7-134446144-INS-131
chr7-134460901-INS-142

chr7-134461405-INS-213
chr7-134461407-INS-703
chr7-134499972-INS-315
chr7-134581584-INS-71
chr7-134610303-INS-80
chr7-134632404-INS-56
chr7-134698455-INS-60
chr7-134712679-INS-64
chr7-134740809-INS-50
chr7-134743139-INS-54
chr7-134790773-INS-70
chr7-134795301-INS-73
chr7-134822774-INS-6276
chr7-134829019-INS-82
chr7-134857089-INS-64
chr7-134869361-INS-3478
chr7-134897804-INS-67
chr7-134916922-INS-5403
chr7-134951643-INS-94
chr7-134952903-INS-51
chr7-134960960-INS-3651
chr7-134971763-INS-66
chr7-134976994-INS-118
chr7-135005691-INS-1663
chr7-135015929-INS-53
chr7-135066965-INS-165
chr7-135066965-INS-55
chr7-135094364-INS-207
chr7-135132140-INS-503
chr7-135160675-INS-66
chr7-135168795-INS-84
chr7-135168907-INS-165
chr7-135176168-INS-60
chr7-135238910-INS-90
chr7-135270534-INS-452
chr7-135323814-INS-1892
chr7-135328936-INS-2252
chr7-135385350-INS-1602
chr7-135386740-INS-3155
chr7-135387787-INS-98
chr7-135387980-INS-149
chr7-135392340-INS-193
chr7-135401962-INS-290
chr7-135417001-INS-152
chr7-135425251-INS-172
chr7-135465895-INS-70
chr7-135466542-INS-132
chr7-135515535-INS-149
chr7-135519712-INS-52
chr7-135527470-INS-6581
chr7-135534262-INS-54
chr7-135576072-INS-115
chr7-135577746-INS-160
chr7-135586089-INS-996
chr7-135587713-INS-1407
chr7-135623927-INS-99
chr7-135713253-INS-401
chr7-135750940-INS-6622
chr7-135773459-INS-77
chr7-135780951-INS-119
chr7-135786829-INS-82
chr7-135787688-INS-101
chr7-135812562-INS-103
chr7-135822958-INS-181
chr7-135822978-INS-54
chr7-135852212-INS-54
chr7-135856164-INS-51
chr7-135966423-INS-53
chr7-135978556-INS-68
chr7-135978721-INS-272
chr7-135987116-INS-130
chr7-136002276-INS-94
chr7-136025545-INS-50
chr7-136028523-INS-147
chr7-136099381-INS-51
chr7-136162910-INS-50
chr7-136175203-INS-52

chr7-136175455-INS-559
chr7-136251492-INS-57
chr7-136263917-INS-67
chr7-136300464-INS-243
chr7-136300503-INS-81
chr7-136322556-INS-50
chr7-136326960-INS-140
chr7-136331450-INS-94
chr7-136372057-INS-573
chr7-136378123-INS-60
chr7-136404966-INS-199
chr7-136409381-INS-170
chr7-136409383-INS-346
chr7-136409517-INS-64
chr7-136438296-INS-64
chr7-136469795-INS-259
chr7-136513884-INS-2591
chr7-136518321-INS-74
chr7-136535387-INS-87
chr7-136611263-INS-72
chr7-136624086-INS-54
chr7-136624586-INS-67
chr7-136715024-INS-50
chr7-136716849-INS-420
chr7-136765652-INS-52
chr7-136777631-INS-1005
chr7-136842869-INS-56
chr7-136873911-INS-54
chr7-136883835-INS-60
chr7-136884074-INS-54
chr7-136927575-INS-60
chr7-136965577-INS-135
chr7-136974197-INS-118
chr7-136979404-INS-72
chr7-136996748-INS-208
chr7-137015449-INS-401
chr7-137023480-INS-328
chr7-137027795-INS-80
chr7-137032462-INS-58
chr7-137032676-INS-74
chr7-137044413-INS-485
chr7-137058011-INS-226
chr7-137076726-INS-79
chr7-137082180-INS-209
chr7-137112759-INS-122
chr7-137118391-INS-595
chr7-137134794-INS-120
chr7-137135205-INS-189
chr7-137167636-INS-1687
chr7-137198556-INS-61
chr7-137204642-INS-50
chr7-137211356-INS-50
chr7-137215726-INS-78
chr7-137260791-INS-54
chr7-137304356-INS-80
chr7-137312745-INS-52
chr7-137320940-INS-148
chr7-137321611-INS-349
chr7-137341223-INS-6843
chr7-137352894-INS-50
chr7-137364849-INS-876
chr7-137370923-INS-5264
chr7-137390365-INS-60
chr7-137400394-INS-56
chr7-137444848-INS-137
chr7-137449010-INS-80
chr7-137458091-INS-126
chr7-137488843-INS-84
chr7-137613983-INS-54
chr7-137627255-INS-140
chr7-137653794-INS-51
chr7-137675710-INS-848
chr7-137675904-INS-112
chr7-137696435-INS-3320
chr7-137706382-INS-90
chr7-137709958-INS-99
chr7-137716299-INS-50

chr7-137716312-INS-113
chr7-137718084-INS-6250
chr7-137745194-INS-51
chr7-137746769-INS-74
chr7-137747025-INS-66
chr7-137762795-INS-1895
chr7-137764365-INS-1000
chr7-137764419-INS-145
chr7-137764685-INS-58
chr7-137771044-INS-89
chr7-137776054-INS-578
chr7-137784384-INS-78
chr7-137840355-INS-2796
chr7-137924622-INS-50
chr7-137935920-INS-56
chr7-137942209-INS-6425
chr7-137980612-INS-70
chr7-137980838-INS-170
chr7-137988122-INS-63
chr7-138034125-INS-55
chr7-138083243-INS-60
chr7-138091777-INS-80
chr7-138100453-INS-6690
chr7-138111012-INS-74
chr7-138117625-INS-489
chr7-138124758-INS-67
chr7-138149097-INS-348
chr7-138166323-INS-52
chr7-138174013-INS-100
chr7-138201814-INS-512
chr7-138205734-INS-201
chr7-138230845-INS-50
chr7-138238586-INS-3423
chr7-138245178-INS-7268
chr7-138263777-INS-390
chr7-138263954-INS-65
chr7-138278356-INS-250
chr7-138278798-INS-62
chr7-138301699-INS-426
chr7-138314339-INS-1072
chr7-138315966-INS-58
chr7-138337732-INS-66
chr7-138343097-INS-96
chr7-138344798-INS-52
chr7-138352710-INS-50
chr7-138431070-INS-244
chr7-138445392-INS-153
chr7-138495139-INS-202
chr7-138531580-INS-189
chr7-138539055-INS-249
chr7-138557231-INS-201
chr7-138573615-INS-75
chr7-138574399-INS-64
chr7-138575565-INS-326
chr7-138576730-INS-1226
chr7-138584478-INS-76
chr7-138619838-INS-558
chr7-138634203-INS-56
chr7-138707556-INS-63
chr7-138762458-INS-252
chr7-138783840-INS-272
chr7-138785474-INS-145
chr7-138786248-INS-332
chr7-138812873-INS-152
chr7-138821515-INS-72
chr7-138824043-INS-140
chr7-138866754-INS-63
chr7-138867827-INS-1199
chr7-138895966-INS-1123
chr7-138906306-INS-366
chr7-138917081-INS-76
chr7-139035147-INS-50
chr7-139105066-INS-275
chr7-139119407-INS-374
chr7-139132990-INS-51
chr7-139149793-INS-615
chr7-139150341-INS-469

chr7-139204245-INS-182
chr7-139223197-INS-53
chr7-139237503-INS-181
chr7-139251055-INS-90
chr7-139280525-INS-250
chr7-139290523-INS-147
chr7-139293315-INS-60
chr7-139296756-INS-129
chr7-139299372-INS-2050
chr7-139321858-INS-52
chr7-139322170-INS-487
chr7-139326517-INS-80
chr7-139332614-INS-64
chr7-139425738-INS-164
chr7-139458363-INS-78
chr7-139505207-INS-105
chr7-139669459-INS-6371
chr7-139689212-INS-1035
chr7-139737660-INS-108
chr7-139749680-INS-3128
chr7-139762347-INS-197
chr7-139801159-INS-54
chr7-139829514-INS-3767
chr7-139844897-INS-196
chr7-139890374-INS-1667
chr7-139890836-INS-3433
chr7-139891665-INS-56
chr7-139894589-INS-50
chr7-139940358-INS-50
chr7-139968287-INS-66
chr7-140028829-INS-4053
chr7-140053834-INS-1587
chr7-140250215-INS-519
chr7-140399364-INS-55
chr7-140399460-INS-200
chr7-140399986-INS-393
chr7-140444040-INS-51
chr7-140451373-INS-96
chr7-140453027-INS-68
chr7-140455272-INS-220
chr7-140505893-INS-207
chr7-140505910-INS-82
chr7-140533518-INS-536
chr7-140533563-INS-56
chr7-140533567-INS-224
chr7-140544399-INS-202
chr7-140545455-INS-372
chr7-140552387-INS-121
chr7-140558240-INS-686
chr7-140559716-INS-52
chr7-140559806-INS-79
chr7-140580516-INS-121
chr7-140588160-INS-255
chr7-140608229-INS-60
chr7-140621277-INS-127
chr7-140621999-INS-188
chr7-140624331-INS-213
chr7-140666479-INS-219
chr7-140690942-INS-65
chr7-140760166-INS-156
chr7-140873547-INS-183
chr7-140899337-INS-197
chr7-140948738-INS-162
chr7-140956051-INS-65
chr7-140956608-INS-54
chr7-141059714-INS-60
chr7-141088639-INS-79
chr7-141111723-INS-50
chr7-141131248-INS-56
chr7-141152002-INS-192
chr7-141156047-INS-1544
chr7-141180564-INS-1911
chr7-141190521-INS-54
chr7-141195582-INS-353
chr7-141285305-INS-318
chr7-141291989-INS-318
chr7-141295064-INS-243

chr7-141303815-INS-84
chr7-141309812-INS-115
chr7-141314825-INS-119
chr7-141362385-INS-279
chr7-141396704-INS-120
chr7-141413545-INS-824
chr7-141586667-INS-80
chr7-141665675-INS-60
chr7-141666022-INS-90
chr7-141683973-INS-267
chr7-141691454-INS-165
chr7-141701154-INS-204
chr7-141701154-INS-610
chr7-141701614-INS-113
chr7-141702776-INS-63
chr7-141702785-INS-261
chr7-141708389-INS-96
chr7-141715851-INS-64
chr7-141743669-INS-78
chr7-141743887-INS-96
chr7-141743971-INS-204
chr7-141744852-INS-84
chr7-141746631-INS-55
chr7-141779929-INS-146
chr7-141783380-INS-60
chr7-141808874-INS-527
chr7-141809224-INS-214
chr7-141810134-INS-50
chr7-141810152-INS-136
chr7-141814805-INS-11034
chr7-141827333-INS-7023
chr7-141828045-INS-63
chr7-141831132-INS-207
chr7-141850353-INS-60
chr7-141850359-INS-144
chr7-141860350-INS-73
chr7-141860478-INS-206
chr7-141861948-INS-92
chr7-141872716-INS-1864
chr7-141886291-INS-1343
chr7-141897758-INS-62
chr7-141954159-INS-186
chr7-141959993-INS-110
chr7-142006095-INS-292
chr7-142007535-INS-95
chr7-142020070-INS-187
chr7-142073282-INS-60
chr7-142114472-INS-900
chr7-142114525-INS-291
chr7-142114674-INS-172
chr7-142324247-INS-1057
chr7-142360178-INS-7593
chr7-142469246-INS-403
chr7-142534939-INS-82
chr7-142543663-INS-150
chr7-142562739-INS-60
chr7-142563046-INS-130
chr7-142597475-INS-240
chr7-142613585-INS-88
chr7-142679339-INS-134
chr7-142679513-INS-398
chr7-142691498-INS-124
chr7-143134236-INS-136
chr7-143170707-INS-92
chr7-143181193-INS-187
chr7-143189411-INS-1753
chr7-143190801-INS-369
chr7-143223655-INS-266
chr7-143224894-INS-1001
chr7-143235762-INS-53
chr7-143241334-INS-1207
chr7-143267753-INS-50
chr7-143315675-INS-56
chr7-143322756-INS-98
chr7-143341054-INS-175
chr7-143362716-INS-107
chr7-143419351-INS-475

chr7-143439771-INS-105
chr7-143442757-INS-213
chr7-143452604-INS-3497
chr7-143452604-INS-413
chr7-143497912-INS-174
chr7-143498744-INS-6766
chr7-143528806-INS-50
chr7-143554223-INS-1738
chr7-143588382-INS-50
chr7-143644701-INS-126
chr7-143712196-INS-68
chr7-143717873-INS-199
chr7-143749627-INS-51
chr7-143801268-INS-217
chr7-143818360-INS-693
chr7-143874094-INS-85
chr7-144018746-INS-230
chr7-144025172-INS-267
chr7-144113093-INS-201
chr7-144126414-INS-51
chr7-144136541-INS-116
chr7-144137365-INS-4919
chr7-144152766-INS-475
chr7-144153296-INS-55
chr7-144157938-INS-135
chr7-144157938-INS-50
chr7-144228031-INS-54
chr7-144246383-INS-137
chr7-144310610-INS-50
chr7-144320126-INS-547
chr7-144364025-INS-200
chr7-144406330-INS-97
chr7-144448271-INS-62
chr7-144506595-INS-80
chr7-144506833-INS-192
chr7-144653979-INS-424
chr7-144765136-INS-52
chr8-3125596-INS-204
chr8-3156080-INS-118
chr8-3157953-INS-7164
chr8-3163896-INS-11994
chr8-3186330-INS-56
chr8-3205268-INS-73
chr8-3270562-INS-87
chr8-3270924-INS-105
chr8-3278517-INS-114
chr8-3287302-INS-50
chr8-3291077-INS-56
chr8-3317309-INS-72
chr8-3342316-INS-84
chr8-3350733-INS-52
chr8-3397593-INS-158
chr8-3433072-INS-203
chr8-3492391-INS-50
chr8-3536113-INS-75
chr8-3567716-INS-5916
chr8-3603976-INS-6474
chr8-3604182-INS-589
chr8-3604251-INS-280
chr8-3606235-INS-564
chr8-3606259-INS-243
chr8-3606362-INS-1734
chr8-3612444-INS-60
chr8-3626131-INS-55
chr8-3724896-INS-8422
chr8-3843162-INS-54
chr8-3961494-INS-2406
chr8-3970412-INS-54
chr8-3993149-INS-466
chr8-4162524-INS-3005
chr8-4166859-INS-66
chr8-4169232-INS-2111
chr8-4203504-INS-8323
chr8-4203770-INS-69
chr8-4231733-INS-7170
chr8-4240008-INS-77
chr8-4265945-INS-500

chr8-4324060-INS-53
chr8-4408473-INS-52
chr8-4434396-INS-210
chr8-4466974-INS-3210
chr8-4468281-INS-62
chr8-4485217-INS-574
chr8-4492945-INS-182
chr8-4499449-INS-2793
chr8-4563347-INS-288
chr8-4563347-INS-577
chr8-4577557-INS-144
chr8-4578174-INS-475
chr8-4645256-INS-1111
chr8-4703815-INS-134
chr8-4781026-INS-215
chr8-4781379-INS-1725
chr8-4860519-INS-8653
chr8-4900656-INS-113
chr8-4925289-INS-511
chr8-5040910-INS-50
chr8-5056409-INS-4989
chr8-5067854-INS-6382
chr8-5079781-INS-88
chr8-5126109-INS-1967
chr8-5133794-INS-64
chr8-5171468-INS-3019
chr8-5202605-INS-7230
chr8-5232725-INS-1961
chr8-5251256-INS-5542
chr8-5287944-INS-98
chr8-5314707-INS-54
chr8-5334369-INS-210
chr8-5394660-INS-1352
chr8-5396312-INS-6531
chr8-5397877-INS-1203
chr8-5415859-INS-537
chr8-5425529-INS-261
chr8-5432433-INS-1081
chr8-5435206-INS-6627
chr8-5448750-INS-8095
chr8-5492559-INS-992
chr8-5500687-INS-1001
chr8-5524946-INS-202
chr8-5576761-INS-117
chr8-5592432-INS-971
chr8-5655429-INS-206
chr8-5669623-INS-66
chr8-5699159-INS-66
chr8-5709941-INS-3197
chr8-5732407-INS-6589
chr8-5735930-INS-484
chr8-5803202-INS-52
chr8-5810182-INS-12358
chr8-5815031-INS-2030
chr8-5846183-INS-50
chr8-5867598-INS-7722
chr8-5884695-INS-65
chr8-5891499-INS-62
chr8-5891582-INS-195
chr8-5892392-INS-1284
chr8-5899465-INS-183
chr8-5912491-INS-156
chr8-5931814-INS-52
chr8-5936788-INS-7472
chr8-5949524-INS-50
chr8-5980646-INS-6229
chr8-5999898-INS-104
chr8-6011253-INS-56
chr8-6050498-INS-50
chr8-6064212-INS-408
chr8-6086396-INS-191
chr8-6093125-INS-60
chr8-6118421-INS-5275
chr8-6118945-INS-55
chr8-6187447-INS-188
chr8-6225869-INS-1674
chr8-6232114-INS-62

chr8-6302011-INS-52
chr8-6318259-INS-115
chr8-6328618-INS-52
chr8-6333300-INS-50
chr8-6346138-INS-50
chr8-6373193-INS-532
chr8-6385836-INS-6592
chr8-6400803-INS-56
chr8-6400909-INS-184
chr8-6401535-INS-77
chr8-6402403-INS-4337
chr8-6415921-INS-5016
chr8-6543284-INS-62
chr8-6602162-INS-72
chr8-6604893-INS-5411
chr8-6609188-INS-53
chr8-6638441-INS-115
chr8-6659568-INS-347
chr8-6695588-INS-53
chr8-6755522-INS-2532
chr8-6758497-INS-60
chr8-6766317-INS-137
chr8-6773090-INS-50
chr8-6783924-INS-7215
chr8-6859944-INS-52
chr8-6878320-INS-51
chr8-6916621-INS-2666
chr8-6936473-INS-6656
chr8-6961991-INS-4314
chr8-7108417-INS-51
chr8-7120403-INS-4258
chr8-7144958-INS-1394
chr8-7154525-INS-75
chr8-7266472-INS-3022
chr8-7331298-INS-60
chr8-7331715-INS-69
chr8-7335393-INS-3942
chr8-7428294-INS-84
chr8-7473547-INS-404
chr8-7493689-INS-1139
chr8-7533929-INS-54
chr8-7537474-INS-3344
chr8-7565805-INS-161
chr8-7601869-INS-57
chr8-7636396-INS-207
chr8-7638805-INS-184
chr8-7638805-INS-62
chr8-7639758-INS-54
chr8-7789128-INS-59
chr8-7791641-INS-56
chr8-7860643-INS-6205
chr8-7860729-INS-6259
chr8-7928833-INS-135
chr8-7999573-INS-56
chr8-7999735-INS-178
chr8-8053083-INS-1195
chr8-8070290-INS-55
chr8-8109658-INS-366
chr8-8109935-INS-949
chr8-8152390-INS-210
chr8-8192895-INS-6283
chr8-8217241-INS-50
chr8-8231450-INS-56
chr8-8253274-INS-1192
chr8-8266746-INS-461
chr8-8305849-INS-1884
chr8-8313229-INS-910
chr8-8411021-INS-50
chr8-8482749-INS-80
chr8-8488156-INS-106
chr8-8496901-INS-67
chr8-8496940-INS-60
chr8-8497119-INS-80
chr8-8504692-INS-195
chr8-8504692-INS-76
chr8-8509511-INS-51
chr8-8545134-INS-311

chr8-8545151-INS-1122
chr8-8555878-INS-123
chr8-8566999-INS-52
chr8-8592304-INS-152
chr8-8596101-INS-497
chr8-8611095-INS-86
chr8-8656832-INS-160
chr8-8795044-INS-2287
chr8-8799499-INS-375
chr8-8878224-INS-844
chr8-8910126-INS-52
chr8-8927944-INS-3586
chr8-8938946-INS-692
chr8-9016840-INS-567
chr8-9048537-INS-50
chr8-9060519-INS-6367
chr8-9080082-INS-52
chr8-9125419-INS-392
chr8-9126504-INS-162
chr8-9282358-INS-128
chr8-9304847-INS-2907
chr8-9357567-INS-194
chr8-9362587-INS-75
chr8-9362877-INS-240
chr8-9446275-INS-56
chr8-9454349-INS-50
chr8-9466382-INS-58
chr8-9471252-INS-1180
chr8-9477189-INS-75
chr8-9477306-INS-602
chr8-9493787-INS-448
chr8-9519591-INS-274
chr8-9542205-INS-67
chr8-9567500-INS-74
chr8-9614874-INS-51
chr8-9634073-INS-80
chr8-9654972-INS-56
chr8-9655709-INS-390
chr8-9731500-INS-52
chr8-9750347-INS-50
chr8-9753848-INS-51
chr8-9758721-INS-105
chr8-9799287-INS-205
chr8-9911733-INS-1104
chr8-9916442-INS-59
chr8-9924092-INS-495
chr8-9934355-INS-1069
chr8-9939726-INS-7630
chr8-9956956-INS-296
chr8-9961328-INS-6460
chr8-10003193-INS-60
chr8-10087720-INS-499
chr8-10158144-INS-78
chr8-10159684-INS-67
chr8-10161685-INS-420
chr8-10161705-INS-1320
chr8-10161730-INS-60
chr8-10163925-INS-59
chr8-10178147-INS-2182
chr8-10185722-INS-102
chr8-10238663-INS-300
chr8-10238665-INS-62
chr8-10240497-INS-94
chr8-10349633-INS-72
chr8-10355981-INS-1012
chr8-10374206-INS-96
chr8-10402549-INS-120
chr8-10402739-INS-339
chr8-10414773-INS-103
chr8-10473082-INS-112
chr8-10564923-INS-1891
chr8-10589190-INS-467
chr8-10655023-INS-50
chr8-10689372-INS-178
chr8-10723010-INS-60
chr8-10723194-INS-211
chr8-10732645-INS-64

chr8-10741251-INS-58
chr8-10746614-INS-124
chr8-10755084-INS-54
chr8-10775668-INS-247
chr8-10812855-INS-143
chr8-10830804-INS-189
chr8-10841617-INS-784
chr8-10847015-INS-76
chr8-10847366-INS-54
chr8-10853648-INS-54
chr8-10855624-INS-50
chr8-10869101-INS-111
chr8-10881334-INS-307
chr8-10903702-INS-666
chr8-10932127-INS-715
chr8-10932133-INS-3014
chr8-10950685-INS-119
chr8-10990624-INS-217
chr8-10994448-INS-50
chr8-10996920-INS-75
chr8-11045401-INS-650
chr8-11045430-INS-286
chr8-11047645-INS-599
chr8-11081008-INS-360
chr8-11081370-INS-109
chr8-11081370-INS-290
chr8-11100387-INS-221
chr8-11111317-INS-220
chr8-11114507-INS-374
chr8-11126326-INS-66
chr8-11131086-INS-7193
chr8-11165924-INS-50
chr8-11173109-INS-52
chr8-11290151-INS-426
chr8-11383485-INS-195
chr8-11474364-INS-55
chr8-11575738-INS-96
chr8-11578169-INS-58
chr8-11582737-INS-81
chr8-11595742-INS-50
chr8-11595962-INS-454
chr8-11617426-INS-76
chr8-11617560-INS-156
chr8-11738320-INS-1666
chr8-11748657-INS-1360
chr8-11764335-INS-130
chr8-11764839-INS-750
chr8-11835303-INS-108
chr8-11862071-INS-78
chr8-11900675-INS-395
chr8-11906226-INS-2643
chr8-11958143-INS-65
chr8-11958457-INS-98
chr8-11977302-INS-67
chr8-11978046-INS-121
chr8-12007114-INS-421
chr8-12017203-INS-86
chr8-12029987-INS-260
chr8-12055213-INS-240
chr8-12059362-INS-58
chr8-12069607-INS-2312
chr8-12071780-INS-58
chr8-12103330-INS-68
chr8-12135856-INS-56
chr8-12195635-INS-610
chr8-12208002-INS-53
chr8-12218672-INS-150
chr8-12237362-INS-509
chr8-12245035-INS-50
chr8-12270358-INS-122
chr8-12292797-INS-106
chr8-12292860-INS-466
chr8-12349719-INS-1140
chr8-12353323-INS-51
chr8-12381099-INS-6475
chr8-12406162-INS-611
chr8-12416770-INS-104

chr8-12428438-INS-179
chr8-12437839-INS-54
chr8-12459723-INS-52
chr8-12482171-INS-50
chr8-12487172-INS-50
chr8-12570020-INS-52
chr8-12571636-INS-115
chr8-12571648-INS-263
chr8-12588138-INS-308
chr8-12609597-INS-100
chr8-12623947-INS-67
chr8-12624050-INS-138
chr8-12670855-INS-52
chr8-12701829-INS-234
chr8-12704052-INS-1537
chr8-12722900-INS-91
chr8-12723848-INS-76
chr8-12746003-INS-131
chr8-12816455-INS-380
chr8-12845814-INS-384
chr8-12888350-INS-7242
chr8-12969106-INS-118
chr8-13021614-INS-127
chr8-13097022-INS-195
chr8-13121195-INS-172
chr8-13181795-INS-1278
chr8-13268145-INS-300
chr8-13293676-INS-65
chr8-13447277-INS-6447
chr8-13456662-INS-52
chr8-13478740-INS-90
chr8-13524159-INS-60
chr8-13569038-INS-507
chr8-13589735-INS-51
chr8-13662540-INS-97
chr8-13668931-INS-267
chr8-13671016-INS-56
chr8-13726954-INS-78
chr8-13834577-INS-357
chr8-13849065-INS-5984
chr8-13893649-INS-170
chr8-13893679-INS-72
chr8-13903021-INS-364
chr8-13965835-INS-78
chr8-13979558-INS-135
chr8-13990689-INS-75
chr8-14044159-INS-103
chr8-14061444-INS-6603
chr8-14088245-INS-53
chr8-14088607-INS-108
chr8-14169726-INS-194
chr8-14237833-INS-205
chr8-14283152-INS-69
chr8-14284235-INS-93
chr8-14343529-INS-122
chr8-14356702-INS-372
chr8-14356740-INS-10005
chr8-14357088-INS-497
chr8-14357113-INS-1611
chr8-14357843-INS-55
chr8-14404657-INS-60
chr8-14406230-INS-146
chr8-14416544-INS-83
chr8-14525975-INS-54
chr8-14590575-INS-68
chr8-14609436-INS-71
chr8-14612478-INS-50
chr8-14613167-INS-53
chr8-14654660-INS-192
chr8-14727929-INS-60
chr8-14783945-INS-140
chr8-14843681-INS-1106
chr8-14870921-INS-66
chr8-14930959-INS-68
chr8-14935261-INS-59
chr8-14986940-INS-720
chr8-14986982-INS-90

chr8-15085555-INS-54
chr8-15127196-INS-2116
chr8-15132098-INS-60
chr8-15151925-INS-115
chr8-15152290-INS-1000
chr8-15192204-INS-2304
chr8-15192204-INS-625
chr8-15192536-INS-978
chr8-15222309-INS-556
chr8-15239138-INS-84
chr8-15240679-INS-50
chr8-15278787-INS-375
chr8-15281382-INS-6396
chr8-15309470-INS-156
chr8-15309845-INS-57
chr8-15325367-INS-2704
chr8-15344624-INS-3107
chr8-15367040-INS-6451
chr8-15428321-INS-6753
chr8-15429531-INS-74
chr8-15429963-INS-60
chr8-15445493-INS-3327
chr8-15514954-INS-51
chr8-15526939-INS-401
chr8-15526982-INS-1846
chr8-15531999-INS-6188
chr8-15616796-INS-6318
chr8-15649493-INS-50
chr8-15697057-INS-2924
chr8-15710983-INS-81
chr8-15725670-INS-100
chr8-15729609-INS-68
chr8-15741349-INS-333
chr8-15758617-INS-2404
chr8-15873397-INS-54
chr8-15901342-INS-274
chr8-15901866-INS-72
chr8-15929311-INS-90
chr8-15939639-INS-76
chr8-15952265-INS-57
chr8-15952496-INS-260
chr8-15983357-INS-174
chr8-16007300-INS-51
chr8-16016366-INS-52
chr8-16065752-INS-64
chr8-16151826-INS-64
chr8-16152113-INS-54
chr8-16170519-INS-1034
chr8-16238052-INS-613
chr8-16238093-INS-126
chr8-16249793-INS-75
chr8-16295344-INS-998
chr8-16317725-INS-95
chr8-16334069-INS-55
chr8-16347184-INS-1211
chr8-16352883-INS-54
chr8-16458909-INS-3468
chr8-16462253-INS-50
chr8-16468310-INS-207
chr8-16478880-INS-3133
chr8-16528807-INS-50
chr8-16528872-INS-562
chr8-16533326-INS-1628
chr8-16605518-INS-500
chr8-16620933-INS-65
chr8-16648733-INS-112
chr8-16687589-INS-1349
chr8-16687879-INS-568
chr8-16717135-INS-75
chr8-16719827-INS-68
chr8-16733491-INS-3439
chr8-16766396-INS-20915
chr8-16816405-INS-345
chr8-16826130-INS-74
chr8-16847713-INS-66
chr8-16862165-INS-1499
chr8-16869637-INS-127

chr8-16869637-INS-743
chr8-16894853-INS-174
chr8-16916521-INS-84
chr8-16994641-INS-207
chr8-17019658-INS-123
chr8-17032972-INS-64
chr8-17105545-INS-217
chr8-17124883-INS-686
chr8-17125699-INS-1378
chr8-17151417-INS-308
chr8-17157136-INS-95
chr8-17187623-INS-392
chr8-17215782-INS-70
chr8-17235019-INS-647
chr8-17235671-INS-401
chr8-17252257-INS-53
chr8-17255486-INS-56
chr8-17264529-INS-171
chr8-17267665-INS-6487
chr8-17270950-INS-588
chr8-17271199-INS-2278
chr8-17271204-INS-1116
chr8-17309491-INS-2149
chr8-17315984-INS-322
chr8-17316022-INS-74
chr8-17324275-INS-1309
chr8-17324530-INS-217
chr8-17326017-INS-54
chr8-17399169-INS-258
chr8-17399169-INS-615
chr8-17527322-INS-270
chr8-17532730-INS-58
chr8-17538209-INS-264
chr8-17583921-INS-70
chr8-17613173-INS-74
chr8-17643670-INS-52
chr8-17647904-INS-56
chr8-17654721-INS-1122
chr8-17657610-INS-3071
chr8-17741155-INS-50
chr8-17741203-INS-3196
chr8-17746836-INS-648
chr8-17753192-INS-89
chr8-17794762-INS-1785
chr8-17810709-INS-54
chr8-17819065-INS-66
chr8-17835264-INS-3352
chr8-17841468-INS-50
chr8-17902106-INS-55
chr8-17962926-INS-6483
chr8-17977202-INS-4847
chr8-17982069-INS-6673
chr8-17989707-INS-62
chr8-18013527-INS-1148
chr8-18093286-INS-52
chr8-18185504-INS-74
chr8-18187238-INS-64
chr8-18222176-INS-3308
chr8-18248387-INS-198
chr8-18261748-INS-60
chr8-18305133-INS-80
chr8-18330492-INS-3138
chr8-18367582-INS-64
chr8-18395769-INS-68
chr8-18443981-INS-6395
chr8-18454899-INS-6196
chr8-18489722-INS-508
chr8-18496647-INS-164
chr8-18523858-INS-125
chr8-18549204-INS-80
chr8-18549252-INS-7178
chr8-18549858-INS-95
chr8-18575343-INS-605
chr8-18607731-INS-57
chr8-18617414-INS-50
chr8-18642332-INS-6801
chr8-18664072-INS-485

chr8-18713820-INS-221
chr8-18977601-INS-58
chr8-19042510-INS-470
chr8-19097068-INS-120
chr8-19100325-INS-467
chr8-19128036-INS-199
chr8-19130306-INS-137
chr8-19156289-INS-51
chr8-19186340-INS-7406
chr8-19194205-INS-175
chr8-19206487-INS-1332
chr8-19210154-INS-69
chr8-19272963-INS-11449
chr8-19290803-INS-7515
chr8-19303123-INS-52
chr8-19308892-INS-56
chr8-19333492-INS-64
chr8-19334718-INS-400
chr8-19357413-INS-401
chr8-19367395-INS-90
chr8-19384624-INS-76
chr8-19402734-INS-60
chr8-19461602-INS-78
chr8-19462465-INS-134
chr8-19509647-INS-3518
chr8-19531294-INS-2572
chr8-19570978-INS-74
chr8-19585113-INS-50
chr8-19594263-INS-270
chr8-19601698-INS-4868
chr8-19620953-INS-1603
chr8-19628018-INS-232
chr8-19628036-INS-67
chr8-19671416-INS-7182
chr8-19676494-INS-56
chr8-19711882-INS-76
chr8-19750853-INS-207
chr8-20050257-INS-136
chr8-20507697-INS-1508
chr8-21073389-INS-160
chr8-21394753-INS-81
chr8-21438729-INS-11200
chr8-21470653-INS-60
chr8-21491240-INS-344
chr8-21499736-INS-66
chr8-21499852-INS-56
chr8-21501554-INS-51
chr8-21548200-INS-498
chr8-21582711-INS-212
chr8-21681926-INS-552
chr8-22257994-INS-64
chr8-22302601-INS-2219
chr8-22314763-INS-52
chr8-22405289-INS-2770
chr8-22406055-INS-1278
chr8-22458532-INS-6488
chr8-22461328-INS-54
chr8-22467189-INS-55
chr8-22485864-INS-3000
chr8-22493555-INS-135
chr8-22580137-INS-74
chr8-22580258-INS-488
chr8-22580683-INS-134
chr8-22609096-INS-372
chr8-22686591-INS-185
chr8-22692811-INS-62
chr8-22695229-INS-476
chr8-22729972-INS-241
chr8-22756103-INS-78
chr8-22765273-INS-1090
chr8-22765629-INS-129
chr8-22765641-INS-60
chr8-22786987-INS-64
chr8-22820313-INS-51
chr8-22828915-INS-60
chr8-22842089-INS-114
chr8-22842860-INS-104

chr8-22845864-INS-85
chr8-22871081-INS-616
chr8-22876237-INS-64
chr8-22908907-INS-125
chr8-22909186-INS-451
chr8-22912867-INS-6907
chr8-22971690-INS-50
chr8-23039942-INS-52
chr8-23058292-INS-2000
chr8-23165264-INS-186
chr8-23198759-INS-3876
chr8-23231765-INS-7100
chr8-23269431-INS-150
chr8-23305324-INS-98
chr8-23326187-INS-274
chr8-23441851-INS-58
chr8-23449751-INS-53
chr8-23452455-INS-156
chr8-23456668-INS-200
chr8-23472798-INS-52
chr8-23514264-INS-52
chr8-23546146-INS-125
chr8-23550773-INS-190
chr8-23633359-INS-157
chr8-23674150-INS-62
chr8-23678213-INS-104
chr8-23700100-INS-4207
chr8-23761797-INS-1645
chr8-23767890-INS-52
chr8-23768034-INS-132
chr8-23768142-INS-62
chr8-23769416-INS-64
chr8-23770044-INS-65
chr8-23793012-INS-134
chr8-23812984-INS-211
chr8-23816645-INS-64
chr8-23842854-INS-52
chr8-23871040-INS-152
chr8-23887179-INS-171
chr8-23903896-INS-97
chr8-23951727-INS-5554
chr8-23971165-INS-6646
chr8-24016777-INS-162
chr8-24043495-INS-176
chr8-24043503-INS-80
chr8-24043605-INS-52
chr8-24082800-INS-71
chr8-24100049-INS-130
chr8-24114295-INS-125
chr8-24114307-INS-313
chr8-24223778-INS-52
chr8-24257163-INS-129
chr8-24285409-INS-142
chr8-24285409-INS-362
chr8-24381242-INS-77
chr8-24381689-INS-133
chr8-24387731-INS-54
chr8-24416206-INS-246
chr8-24424855-INS-152
chr8-24428701-INS-6434
chr8-24505258-INS-629
chr8-24510156-INS-50
chr8-24518168-INS-230
chr8-24549613-INS-54
chr8-24561947-INS-390
chr8-24568691-INS-405
chr8-24572697-INS-185
chr8-24613921-INS-64
chr8-24621390-INS-441
chr8-24621661-INS-318
chr8-24621899-INS-398
chr8-24654685-INS-62
chr8-24671534-INS-881
chr8-24737473-INS-81
chr8-24818226-INS-124
chr8-24834245-INS-1071
chr8-24870570-INS-135

chr8-24882319-INS-50
chr8-24895869-INS-68
chr8-24896594-INS-50
chr8-24900803-INS-68
chr8-24916389-INS-107
chr8-24940710-INS-202
chr8-24970331-INS-455
chr8-25033921-INS-60
chr8-25069430-INS-4349
chr8-25090553-INS-57
chr8-25154066-INS-2002
chr8-25160397-INS-1140
chr8-25160407-INS-348
chr8-25186700-INS-1422
chr8-25213219-INS-596
chr8-25217832-INS-52
chr8-25221509-INS-90
chr8-25238796-INS-111
chr8-25298416-INS-278
chr8-25363604-INS-433
chr8-25373418-INS-50
chr8-25414839-INS-55
chr8-25443324-INS-1373
chr8-25443416-INS-512
chr8-25459600-INS-1443
chr8-25478250-INS-117
chr8-25521011-INS-266
chr8-25531843-INS-100
chr8-25534549-INS-105
chr8-25538651-INS-92
chr8-25538827-INS-60
chr8-25556634-INS-80
chr8-25557105-INS-105
chr8-25589159-INS-106
chr8-25607751-INS-50
chr8-25612308-INS-56
chr8-25641331-INS-492
chr8-25677766-INS-151
chr8-25677775-INS-63
chr8-25702662-INS-57
chr8-25702734-INS-1092
chr8-25719361-INS-88
chr8-25724115-INS-166
chr8-25730450-INS-68
chr8-25731515-INS-4371
chr8-25732301-INS-243
chr8-25741387-INS-51
chr8-25745240-INS-93
chr8-25751305-INS-678
chr8-25770228-INS-50
chr8-25816623-INS-78
chr8-25827273-INS-74
chr8-25827401-INS-55
chr8-25827713-INS-180
chr8-25839444-INS-3153
chr8-25873741-INS-87
chr8-25877180-INS-52
chr8-25889660-INS-102
chr8-25918080-INS-115
chr8-25928969-INS-643
chr8-25964875-INS-120
chr8-25965736-INS-74
chr8-25970212-INS-72
chr8-25979890-INS-83
chr8-26009807-INS-108
chr8-26029576-INS-206
chr8-26051231-INS-146
chr8-26097136-INS-66
chr8-26147731-INS-539
chr8-26184085-INS-63
chr8-26325645-INS-88
chr8-26332954-INS-53
chr8-26361014-INS-231
chr8-26402533-INS-93
chr8-26425615-INS-162
chr8-26426343-INS-80
chr8-26441656-INS-56

chr8-26459568-INS-1696
chr8-26470166-INS-2266
chr8-26470166-INS-429
chr8-26481972-INS-68
chr8-26494868-INS-110
chr8-26523550-INS-593
chr8-26531484-INS-950
chr8-26540771-INS-129
chr8-26556631-INS-177
chr8-26566960-INS-6598
chr8-26583293-INS-280
chr8-26583433-INS-1540
chr8-26628820-INS-196
chr8-26636164-INS-1499
chr8-26659072-INS-1908
chr8-26660616-INS-117
chr8-26660729-INS-369
chr8-26660982-INS-995
chr8-26664290-INS-2700
chr8-26666056-INS-1868
chr8-26683849-INS-70
chr8-26726254-INS-79
chr8-26726983-INS-313
chr8-26735458-INS-278
chr8-26735520-INS-1276
chr8-26737508-INS-2345
chr8-26740462-INS-326
chr8-26740829-INS-4905
chr8-26775717-INS-183
chr8-26775978-INS-88
chr8-26791836-INS-101
chr8-26805301-INS-1455
chr8-26821437-INS-832
chr8-26855278-INS-69
chr8-26906573-INS-85
chr8-26912793-INS-52
chr8-26933521-INS-137
chr8-27007195-INS-6406
chr8-27043161-INS-193
chr8-27160109-INS-52
chr8-27190012-INS-3079
chr8-27192216-INS-52
chr8-27311173-INS-54
chr8-27420252-INS-3106
chr8-27426170-INS-262
chr8-27495626-INS-83
chr8-27507678-INS-52
chr8-27511492-INS-108
chr8-27517555-INS-68
chr8-27521044-INS-50
chr8-27567037-INS-216
chr8-27569081-INS-84
chr8-27569673-INS-125
chr8-27571202-INS-200
chr8-27590848-INS-210
chr8-27605102-INS-52
chr8-27654668-INS-50
chr8-27668728-INS-93
chr8-27676481-INS-114
chr8-27677497-INS-168
chr8-27682121-INS-87
chr8-27687499-INS-229
chr8-27770302-INS-244
chr8-27770763-INS-50
chr8-27788745-INS-191
chr8-27822641-INS-701
chr8-27822909-INS-2918
chr8-27823371-INS-70
chr8-27826892-INS-3082
chr8-27840559-INS-70
chr8-27841708-INS-84
chr8-27842084-INS-318
chr8-27848336-INS-6131
chr8-27850651-INS-68
chr8-27852181-INS-179
chr8-27852248-INS-78
chr8-27855604-INS-74

chr8-27856593-INS-1277
chr8-27862821-INS-118
chr8-27866673-INS-1029
chr8-27869237-INS-64
chr8-27878819-INS-366
chr8-27881453-INS-52
chr8-27914892-INS-3024
chr8-27928630-INS-4531
chr8-27928766-INS-63
chr8-28063515-INS-120
chr8-28085823-INS-56
chr8-28108978-INS-1020
chr8-28116865-INS-2293
chr8-28119055-INS-2727
chr8-28145190-INS-52
chr8-28167762-INS-60
chr8-28265654-INS-3104
chr8-28268379-INS-6392
chr8-28341591-INS-174
chr8-28391084-INS-6369
chr8-28501289-INS-1881
chr8-28534065-INS-54
chr8-28588715-INS-87
chr8-28599317-INS-62
chr8-28600242-INS-194
chr8-28644209-INS-660
chr8-28649643-INS-164
chr8-28677067-INS-58
chr8-28724476-INS-52
chr8-28735709-INS-52
chr8-28761911-INS-4170
chr8-28763684-INS-52
chr8-28825001-INS-52
chr8-28870657-INS-54
chr8-28908623-INS-6721
chr8-28975450-INS-66
chr8-28975919-INS-312
chr8-29014851-INS-72
chr8-29015965-INS-51
chr8-29017458-INS-173
chr8-29090297-INS-226
chr8-29097856-INS-125
chr8-29102834-INS-10581
chr8-29116865-INS-1157
chr8-29133240-INS-58
chr8-29148752-INS-62
chr8-29172297-INS-58
chr8-29175802-INS-195
chr8-29237682-INS-90
chr8-29252273-INS-215
chr8-29259673-INS-69
chr8-29428302-INS-1792
chr8-29435050-INS-52
chr8-29462907-INS-993
chr8-29489582-INS-475
chr8-29497282-INS-66
chr8-29514412-INS-198
chr8-29522038-INS-52
chr8-29559988-INS-60
chr8-29567494-INS-1124
chr8-29576236-INS-336
chr8-29598838-INS-52
chr8-29607023-INS-50
chr8-29614318-INS-54
chr8-29630439-INS-100
chr8-29720789-INS-386
chr8-29725352-INS-50
chr8-29759099-INS-64
chr8-29811410-INS-6265
chr8-29824493-INS-132
chr8-29835940-INS-2340
chr8-29839444-INS-72
chr8-29842579-INS-526
chr8-29854814-INS-52
chr8-29891116-INS-165
chr8-29926288-INS-3908
chr8-29946317-INS-6130

chr8-29954355-INS-130
chr8-29977628-INS-252
chr8-29979425-INS-53
chr8-30046512-INS-91
chr8-30098593-INS-7519
chr8-30161580-INS-3090
chr8-30174509-INS-52
chr8-30185812-INS-56
chr8-30208593-INS-688
chr8-30214230-INS-118
chr8-30221995-INS-6432
chr8-30239312-INS-1336
chr8-30246006-INS-333
chr8-30260841-INS-72
chr8-30308771-INS-3554
chr8-30366282-INS-408
chr8-30384607-INS-6237
chr8-30388891-INS-52
chr8-30407526-INS-7380
chr8-30457033-INS-1215
chr8-30457036-INS-2760
chr8-30503888-INS-92
chr8-30545092-INS-2869
chr8-30546029-INS-6499
chr8-30555389-INS-2855
chr8-30597199-INS-53
chr8-30621164-INS-7130
chr8-30658812-INS-214
chr8-30668518-INS-58
chr8-30677198-INS-7132
chr8-30700356-INS-79
chr8-30707334-INS-68
chr8-30733910-INS-260
chr8-30756937-INS-302
chr8-30797337-INS-221
chr8-30807158-INS-52
chr8-30903604-INS-63
chr8-30950015-INS-11076
chr8-31141827-INS-7170
chr8-31156107-INS-55
chr8-31164771-INS-1321
chr8-31169300-INS-1502
chr8-31236920-INS-7421
chr8-31260393-INS-609
chr8-31282319-INS-1864
chr8-31285101-INS-2736
chr8-31320632-INS-6088
chr8-31333969-INS-2228
chr8-31366227-INS-66
chr8-31438735-INS-1447
chr8-31453618-INS-98
chr8-31462727-INS-825
chr8-31469287-INS-111
chr8-31484822-INS-186
chr8-31488597-INS-55
chr8-31516908-INS-92
chr8-31552253-INS-3841
chr8-31552887-INS-124
chr8-31557616-INS-102
chr8-31558160-INS-54
chr8-31590633-INS-552
chr8-31634911-INS-60
chr8-31674461-INS-243
chr8-31699128-INS-882
chr8-31738385-INS-79
chr8-31777321-INS-72
chr8-31814401-INS-6401
chr8-31815124-INS-498
chr8-31815854-INS-2449
chr8-31825881-INS-6593
chr8-31827748-INS-55
chr8-31828160-INS-58
chr8-31831804-INS-445
chr8-31839720-INS-97
chr8-31844294-INS-1286
chr8-31853904-INS-6990
chr8-31855732-INS-2467

chr8-31873041-INS-95
chr8-31873149-INS-2205
chr8-31873261-INS-95
chr8-31887488-INS-6449
chr8-31887536-INS-50
chr8-31937271-INS-68
chr8-31939590-INS-6928
chr8-31950872-INS-4750
chr8-31957731-INS-1171
chr8-31958393-INS-4694
chr8-31961975-INS-358
chr8-31962294-INS-56
chr8-31973900-INS-223
chr8-31994032-INS-54
chr8-32017532-INS-7011
chr8-32054069-INS-3018
chr8-32079546-INS-6151
chr8-32088705-INS-836
chr8-32115162-INS-3117
chr8-32119476-INS-538
chr8-32119748-INS-75
chr8-32145623-INS-50
chr8-32151207-INS-1642
chr8-32186780-INS-88
chr8-32192148-INS-314
chr8-32197657-INS-108
chr8-32244453-INS-7524
chr8-32281867-INS-54
chr8-32297935-INS-50
chr8-32314644-INS-193
chr8-32416600-INS-121
chr8-32416620-INS-52
chr8-32422459-INS-3148
chr8-32445199-INS-77
chr8-32456406-INS-64
chr8-32535449-INS-64
chr8-32597057-INS-114
chr8-32606943-INS-2500
chr8-32633402-INS-7131
chr8-32651316-INS-52
chr8-32664042-INS-265
chr8-32730242-INS-68
chr8-32734337-INS-1665
chr8-32758330-INS-7010
chr8-32764678-INS-50
chr8-32792851-INS-50
chr8-32809154-INS-6358
chr8-32838847-INS-56
chr8-32870287-INS-84
chr8-32944684-INS-101
chr8-32953045-INS-2164
chr8-33010124-INS-2289
chr8-33036503-INS-52
chr8-33084329-INS-6239
chr8-33104068-INS-265
chr8-33131765-INS-50
chr8-33168122-INS-84
chr8-33172066-INS-7506
chr8-33175719-INS-234
chr8-33254276-INS-60
chr8-33254457-INS-64
chr8-33274777-INS-58
chr8-33274832-INS-200
chr8-33300856-INS-119
chr8-33312345-INS-326
chr8-33312445-INS-54
chr8-33312557-INS-112
chr8-33343939-INS-625
chr8-33379365-INS-1896
chr8-33394075-INS-150
chr8-33405164-INS-192
chr8-33414811-INS-63
chr8-33415523-INS-4525
chr8-33441982-INS-6326
chr8-33513669-INS-113
chr8-33530213-INS-1982
chr8-33532028-INS-6463

chr8-33555267-INS-72
chr8-33559405-INS-6707
chr8-33684577-INS-227
chr8-33705606-INS-63
chr8-33739174-INS-53
chr8-33750322-INS-101
chr8-33750370-INS-233
chr8-33781305-INS-108
chr8-33781682-INS-72
chr8-33852677-INS-601
chr8-33887454-INS-88
chr8-33910529-INS-72
chr8-33956131-INS-214
chr8-33987750-INS-3789
chr8-33999474-INS-111
chr8-34052366-INS-50
chr8-34059015-INS-424
chr8-34096250-INS-52
chr8-34096727-INS-130
chr8-34130307-INS-51
chr8-34146309-INS-50
chr8-34149421-INS-187
chr8-34150254-INS-220
chr8-34197828-INS-4241
chr8-34319401-INS-5593
chr8-34347027-INS-6402
chr8-34350392-INS-73
chr8-34361878-INS-149
chr8-34362758-INS-76
chr8-34382962-INS-1343
chr8-34433130-INS-1398
chr8-34444734-INS-2090
chr8-34445287-INS-235
chr8-34464393-INS-226
chr8-34489246-INS-161
chr8-34490502-INS-231
chr8-34491330-INS-216
chr8-34493137-INS-1273
chr8-34494477-INS-987
chr8-34498683-INS-340
chr8-34500483-INS-5218
chr8-34508444-INS-55
chr8-34538097-INS-59
chr8-34561341-INS-84
chr8-34562462-INS-263
chr8-34562533-INS-66
chr8-34573994-INS-3050
chr8-34590231-INS-215
chr8-34598572-INS-365
chr8-34598719-INS-205
chr8-34648009-INS-7110
chr8-34655335-INS-7086
chr8-34676828-INS-66
chr8-34696428-INS-50
chr8-34702796-INS-2578
chr8-34707902-INS-52
chr8-34726060-INS-68
chr8-34738031-INS-498
chr8-34751301-INS-230
chr8-34757587-INS-108
chr8-34773942-INS-131
chr8-34778295-INS-96
chr8-34806596-INS-80
chr8-34816784-INS-82
chr8-34821547-INS-86
chr8-34822989-INS-95
chr8-34828764-INS-4522
chr8-34832170-INS-72
chr8-34837514-INS-1919
chr8-34840335-INS-7084
chr8-34908678-INS-161
chr8-34910600-INS-58
chr8-34980304-INS-70
chr8-34986122-INS-54
chr8-35067023-INS-392
chr8-35067145-INS-993
chr8-35097937-INS-60

chr8-35104445-INS-157
chr8-35140027-INS-1057
chr8-35164568-INS-218
chr8-35190084-INS-164
chr8-35194516-INS-997
chr8-35195793-INS-134
chr8-35199106-INS-55
chr8-35211187-INS-198
chr8-35211369-INS-51
chr8-35257402-INS-4709
chr8-35306621-INS-57
chr8-35335876-INS-56
chr8-35459448-INS-202
chr8-35507608-INS-54
chr8-35512983-INS-66
chr8-35539475-INS-515
chr8-35611442-INS-68
chr8-35612092-INS-7082
chr8-35623301-INS-203
chr8-35623475-INS-78
chr8-35640159-INS-3211
chr8-35658058-INS-342
chr8-35674935-INS-5404
chr8-35676987-INS-56
chr8-35719060-INS-57
chr8-35724646-INS-50
chr8-35763118-INS-59
chr8-35771764-INS-67
chr8-35771881-INS-223
chr8-35798702-INS-88
chr8-35815087-INS-495
chr8-36045560-INS-54
chr8-36070943-INS-311
chr8-36089308-INS-63
chr8-36090768-INS-435
chr8-36123431-INS-181
chr8-36127413-INS-275
chr8-36148274-INS-52
chr8-36154898-INS-2740
chr8-36176700-INS-183
chr8-36183981-INS-1558
chr8-36201837-INS-2899
chr8-36217724-INS-602
chr8-36241332-INS-54
chr8-36288848-INS-175
chr8-36313384-INS-54
chr8-36314360-INS-164
chr8-36318005-INS-122
chr8-36322541-INS-50
chr8-36375849-INS-7121
chr8-36408572-INS-112
chr8-36408572-INS-54
chr8-36425159-INS-3150
chr8-36495272-INS-297
chr8-36527354-INS-123
chr8-36549447-INS-50
chr8-36549463-INS-150
chr8-36597664-INS-6679
chr8-36634223-INS-4819
chr8-36634708-INS-307
chr8-36644037-INS-2100
chr8-36656907-INS-102
chr8-36690417-INS-54
chr8-36788219-INS-6465
chr8-36794519-INS-50
chr8-36827548-INS-1912
chr8-36848792-INS-6401
chr8-36882181-INS-2174
chr8-36891925-INS-1541
chr8-36900965-INS-158
chr8-36903655-INS-56
chr8-36917243-INS-7918
chr8-36933041-INS-1480
chr8-36937807-INS-6466
chr8-36952159-INS-1031
chr8-36959393-INS-224
chr8-37005278-INS-279

chr8-37032333-INS-60
chr8-37062484-INS-51
chr8-37062780-INS-214
chr8-37102906-INS-6683
chr8-37207423-INS-58
chr8-37221743-INS-516
chr8-37245706-INS-55
chr8-37259685-INS-2562
chr8-37367282-INS-1169
chr8-37369208-INS-213
chr8-37377684-INS-196
chr8-37482917-INS-50
chr8-37495062-INS-7202
chr8-37502604-INS-1238
chr8-37511526-INS-52
chr8-37511733-INS-249
chr8-37538815-INS-204
chr8-37560215-INS-4158
chr8-37566822-INS-231
chr8-37572032-INS-10512
chr8-37572816-INS-32038
chr8-37577097-INS-1213
chr8-37600528-INS-189
chr8-37609112-INS-89
chr8-37623331-INS-174
chr8-37663220-INS-6387
chr8-37678985-INS-6499
chr8-37684186-INS-92
chr8-37709182-INS-1748
chr8-37799290-INS-1938
chr8-37820754-INS-361
chr8-37820815-INS-125
chr8-37826953-INS-4483
chr8-37975767-INS-6512
chr8-38021193-INS-205
chr8-38085807-INS-1838
chr8-38112125-INS-52
chr8-38126711-INS-50
chr8-38127190-INS-54
chr8-38131934-INS-208
chr8-38144532-INS-1041
chr8-38156700-INS-6297
chr8-38271161-INS-60
chr8-38322117-INS-215
chr8-38340027-INS-6761
chr8-38362063-INS-1898
chr8-38364775-INS-6147
chr8-38395127-INS-65
chr8-38400444-INS-5605
chr8-38410642-INS-50
chr8-38429860-INS-1616
chr8-38439926-INS-2542
chr8-38441316-INS-1281
chr8-38441677-INS-5413
chr8-38487246-INS-6682
chr8-38490062-INS-50
chr8-38493516-INS-1038
chr8-38493762-INS-2972
chr8-38540733-INS-1878
chr8-38587517-INS-4164
chr8-38607229-INS-1072
chr8-38608944-INS-1832
chr8-38619130-INS-2272
chr8-38625219-INS-6496
chr8-38629990-INS-66
chr8-38630280-INS-62
chr8-38638265-INS-1775
chr8-38646528-INS-58
chr8-38655579-INS-1678
chr8-38684436-INS-1503
chr8-38821157-INS-56
chr8-38821532-INS-1015
chr8-38826682-INS-56
chr8-38858994-INS-52
chr8-38895552-INS-208
chr8-38913070-INS-1368
chr8-38913498-INS-1207

chr8-38918049-INS-6403
chr8-38925219-INS-1722
chr8-39023865-INS-93
chr8-39024208-INS-3521
chr8-39045458-INS-138
chr8-39141718-INS-6174
chr8-39167220-INS-2762
chr8-39168637-INS-7102
chr8-39218107-INS-6004
chr8-39224552-INS-102
chr8-39234949-INS-657
chr8-39321285-INS-1607
chr8-39322344-INS-459
chr8-39322621-INS-128
chr8-39353084-INS-90
chr8-39442525-INS-2288
chr8-39449158-INS-6468
chr8-39495009-INS-1769
chr8-39512989-INS-78
chr8-39683791-INS-216
chr8-39729953-INS-1534
chr8-39742762-INS-70
chr8-39750925-INS-58
chr8-39753256-INS-6762
chr8-39815544-INS-220
chr8-39820216-INS-1510
chr8-39823968-INS-1821
chr8-39853757-INS-69
chr8-39890293-INS-5403
chr8-39905303-INS-1177
chr8-39931648-INS-55
chr8-39933654-INS-1175
chr8-39947837-INS-51
chr8-39959667-INS-50
chr8-39994436-INS-63
chr8-40028285-INS-1395
chr8-40035672-INS-6846
chr8-40048092-INS-110
chr8-40057120-INS-470
chr8-40112993-INS-80
chr8-40137105-INS-53
chr8-40153976-INS-7847
chr8-40189595-INS-89
chr8-40214158-INS-3080
chr8-40333007-INS-108
chr8-40385163-INS-6239
chr8-40391397-INS-966
chr8-40397878-INS-6090
chr8-40424446-INS-6394
chr8-40428940-INS-1764
chr8-40471022-INS-212
chr8-40471529-INS-552
chr8-40480412-INS-54
chr8-40494576-INS-383
chr8-40501385-INS-235
chr8-40528312-INS-1704
chr8-40533772-INS-332
chr8-40564307-INS-50
chr8-40641849-INS-54
chr8-40642098-INS-399
chr8-40727887-INS-64
chr8-40731425-INS-276
chr8-40743868-INS-454
chr8-40817265-INS-2035
chr8-40914853-INS-3104
chr8-41019251-INS-68
chr8-41033879-INS-67
chr8-41034114-INS-63
chr8-41069201-INS-82
chr8-41081300-INS-69
chr8-41082878-INS-58
chr8-41146921-INS-6655
chr8-41193773-INS-198
chr8-41301074-INS-87
chr8-41333181-INS-316
chr8-41366704-INS-1000
chr8-41378992-INS-2319

chr8-41393626-INS-50
chr8-41405727-INS-1903
chr8-41412829-INS-52
chr8-41413397-INS-175
chr8-41468391-INS-66
chr8-41473430-INS-56
chr8-41495115-INS-72
chr8-41514098-INS-508
chr8-41529908-INS-7131
chr8-41550842-INS-332
chr8-41607364-INS-190
chr8-41634957-INS-487
chr8-41675517-INS-50
chr8-41691160-INS-65
chr8-41691178-INS-251
chr8-41889057-INS-2571
chr8-41890757-INS-2034
chr8-41894959-INS-6487
chr8-41939810-INS-3682
chr8-41951017-INS-7004
chr8-41957031-INS-1913
chr8-41981369-INS-1762
chr8-42017340-INS-188
chr8-42035290-INS-104
chr8-42054477-INS-224
chr8-42076230-INS-409
chr8-42091971-INS-193
chr8-42214353-INS-6873
chr8-42239181-INS-78
chr8-42314401-INS-56
chr8-42336171-INS-6776
chr8-42398832-INS-1205
chr8-42423475-INS-56
chr8-42424706-INS-98
chr8-42431024-INS-5839
chr8-42446842-INS-6327
chr8-42447187-INS-99
chr8-42473375-INS-52
chr8-42473883-INS-61
chr8-42482861-INS-72
chr8-42492136-INS-162
chr8-42499357-INS-59
chr8-42570368-INS-72
chr8-42591471-INS-6415
chr8-42684747-INS-2657
chr8-42692332-INS-144
chr8-42745140-INS-1800
chr8-42767063-INS-78
chr8-42813762-INS-4992
chr8-42830062-INS-6059
chr8-42836228-INS-50
chr8-42861726-INS-3186
chr8-42869328-INS-6445
chr8-42878577-INS-84
chr8-42893627-INS-7538
chr8-42904142-INS-56
chr8-42906366-INS-627
chr8-42907707-INS-1013
chr8-42911977-INS-6270
chr8-42927687-INS-74
chr8-42937449-INS-52
chr8-42949548-INS-1547
chr8-42961066-INS-602
chr8-43065808-INS-4323
chr8-43087928-INS-321
chr8-43112794-INS-9659
chr8-43116085-INS-3253
chr8-43207145-INS-3216
chr8-43237080-INS-74
chr8-43310670-INS-974
chr8-43399859-INS-1721
chr8-43407472-INS-2593
chr8-43411339-INS-54
chr8-43420479-INS-1965
chr8-43436093-INS-6393
chr8-43436298-INS-778
chr8-43448083-INS-329

chr8-43511925-INS-4450
chr8-43526319-INS-6808
chr8-43536516-INS-341
chr8-43539198-INS-9959
chr8-43556996-INS-83
chr8-43560917-INS-6430
chr8-43593579-INS-64
chr8-43626595-INS-118
chr8-43627182-INS-541
chr8-43660407-INS-1329
chr8-43671487-INS-140
chr8-43683557-INS-1076
chr8-43691368-INS-710
chr8-43691655-INS-1567
chr8-43692198-INS-5406
chr8-43722414-INS-50
chr8-43732296-INS-233
chr8-43779760-INS-1266
chr8-43786450-INS-205
chr8-43791140-INS-65
chr8-43823809-INS-92
chr8-43837854-INS-65
chr8-43844893-INS-4721
chr8-43850258-INS-6727
chr8-43854212-INS-6851
chr8-43967801-INS-2448
chr8-44054298-INS-3850
chr8-44062461-INS-83
chr8-44066072-INS-255
chr8-44134212-INS-5513
chr8-44155603-INS-984
chr8-44187833-INS-2615
chr8-44222130-INS-54
chr8-44224808-INS-1377
chr8-44225137-INS-7061
chr8-44250864-INS-499
chr8-44308275-INS-3398
chr8-44316125-INS-184
chr8-44459034-INS-400
chr8-44465281-INS-50
chr8-44480220-INS-183
chr8-44517153-INS-327
chr8-44517467-INS-6800
chr8-44537878-INS-50
chr8-44642864-INS-218
chr8-44703260-INS-708
chr8-44713950-INS-132
chr8-44723402-INS-366
chr8-44772961-INS-72
chr8-44782267-INS-1435
chr8-44809032-INS-6327
chr8-44817693-INS-86
chr8-44818126-INS-280
chr8-44834798-INS-68
chr8-44874578-INS-146
chr8-44875199-INS-59
chr8-44892713-INS-52
chr8-44909195-INS-135
chr8-44909195-INS-379
chr8-44909195-INS-828
chr8-45017964-INS-117
chr8-45028685-INS-121
chr8-45078965-INS-6650
chr8-45079429-INS-182
chr8-45086752-INS-113
chr8-45100322-INS-112
chr8-45131629-INS-641
chr8-45269543-INS-373
chr8-45282618-INS-6398
chr8-45283322-INS-53
chr8-45357096-INS-67
chr8-45359446-INS-54
chr8-45380495-INS-120
chr8-45384212-INS-56
chr8-45453129-INS-52
chr8-45464183-INS-85
chr8-45514272-INS-50

chr8-45536109-INS-94
chr8-45553101-INS-152
chr8-45559612-INS-7099
chr8-45567395-INS-4901
chr8-45573162-INS-91
chr8-45574802-INS-6215
chr8-45629058-INS-421
chr8-45630075-INS-2435
chr8-45730181-INS-60
chr8-45740578-INS-7112
chr8-45821741-INS-3872
chr8-45835673-INS-252
chr8-45840341-INS-5554
chr8-45848389-INS-83
chr8-45858077-INS-216
chr8-45863113-INS-116
chr8-45882196-INS-400
chr8-45883716-INS-69
chr8-45891907-INS-175
chr8-45915262-INS-586
chr8-45915807-INS-72
chr8-45918296-INS-59
chr8-45956375-INS-381
chr8-45976988-INS-75
chr8-46001907-INS-239
chr8-46093637-INS-60
chr8-46101825-INS-130
chr8-46131706-INS-64
chr8-46132928-INS-112
chr8-46185080-INS-6801
chr8-46216758-INS-130
chr8-46222774-INS-60
chr8-46244911-INS-268
chr8-46261814-INS-55
chr8-46309277-INS-54
chr8-46321377-INS-286
chr8-46384454-INS-366
chr8-46407125-INS-484
chr8-46415762-INS-307
chr8-46459736-INS-193
chr8-46465196-INS-67
chr8-46498721-INS-55
chr8-46515354-INS-110
chr8-46547763-INS-6431
chr8-46559466-INS-61
chr8-46566832-INS-165
chr8-46567044-INS-73
chr8-46581047-INS-102
chr8-46645350-INS-199
chr8-46673475-INS-62
chr8-46728632-INS-54
chr8-46757530-INS-60
chr8-46759295-INS-78
chr8-46760436-INS-2133
chr8-46761232-INS-157
chr8-46771783-INS-51
chr8-46776460-INS-188
chr8-46789822-INS-1209
chr8-46858323-INS-151
chr8-46879635-INS-188
chr8-46880778-INS-71
chr8-46928931-INS-107
chr8-46968528-INS-306
chr8-46988407-INS-205
chr8-47002818-INS-69
chr8-47043477-INS-207
chr8-47043532-INS-2978
chr8-47043638-INS-359
chr8-47056839-INS-1039
chr8-47059035-INS-231
chr8-47148055-INS-1872
chr8-47157666-INS-59
chr8-47159624-INS-56
chr8-47162626-INS-120
chr8-47211482-INS-71
chr8-47309900-INS-3199
chr8-47320857-INS-1075

chr8-47348991-INS-124
chr8-47353375-INS-86
chr8-47353751-INS-1102
chr8-47356503-INS-70
chr8-47388517-INS-482
chr8-47420715-INS-52
chr8-47455505-INS-52
chr8-47459665-INS-254
chr8-47459667-INS-109
chr8-47464787-INS-50
chr8-47468931-INS-6211
chr8-47473276-INS-256
chr8-47473278-INS-88
chr8-47480500-INS-50
chr8-47499919-INS-69
chr8-47499979-INS-214
chr8-47504411-INS-1887
chr8-47508504-INS-186
chr8-47565785-INS-1175
chr8-47575104-INS-178
chr8-47579845-INS-95
chr8-47688050-INS-50
chr8-47700518-INS-53
chr8-47728103-INS-68
chr8-47744797-INS-3243
chr8-47782304-INS-60
chr8-47785231-INS-129
chr8-47807242-INS-368
chr8-47811100-INS-6805
chr8-47820604-INS-638
chr8-47824970-INS-426
chr8-47893278-INS-55
chr8-47918719-INS-13350
chr8-47938945-INS-160
chr8-47951646-INS-206
chr8-47952262-INS-81
chr8-48010764-INS-634
chr8-48035531-INS-6689
chr8-48094793-INS-984
chr8-48094835-INS-7086
chr8-48116931-INS-60
chr8-48141392-INS-198
chr8-48173368-INS-74
chr8-48179440-INS-72
chr8-48202278-INS-489
chr8-48202408-INS-66
chr8-48202753-INS-56
chr8-48205057-INS-58
chr8-48227063-INS-7097
chr8-48229692-INS-68
chr8-48255758-INS-58
chr8-48269288-INS-197
chr8-48270891-INS-121
chr8-48389814-INS-213
chr8-48394458-INS-75
chr8-48405540-INS-173
chr8-48411737-INS-3028
chr8-48417265-INS-59
chr8-48463442-INS-94
chr8-48464966-INS-3279
chr8-48468619-INS-54
chr8-48484200-INS-539
chr8-48486003-INS-52
chr8-48490449-INS-1053
chr8-48490704-INS-2101
chr8-48536109-INS-67
chr8-48546127-INS-3381
chr8-48549942-INS-96
chr8-48565408-INS-140
chr8-48640406-INS-477
chr8-48694973-INS-272
chr8-48786970-INS-94
chr8-48808948-INS-354
chr8-48853373-INS-73
chr8-48853936-INS-54
chr8-48935940-INS-73
chr8-48988929-INS-50

chr8-48992619-INS-180
chr8-48992627-INS-564
chr8-49017522-INS-52
chr8-49019202-INS-99
chr8-49027312-INS-1261
chr8-49071989-INS-618
chr8-49076244-INS-52
chr8-49087775-INS-52
chr8-49112464-INS-146
chr8-49180513-INS-62
chr8-49231559-INS-100
chr8-49264609-INS-52
chr8-49291242-INS-59
chr8-49365258-INS-60
chr8-49373839-INS-375
chr8-49413404-INS-52
chr8-49416904-INS-6677
chr8-49433559-INS-56
chr8-49474292-INS-714
chr8-49598111-INS-64
chr8-49643617-INS-62
chr8-49703447-INS-1845
chr8-49725805-INS-106
chr8-49750364-INS-292
chr8-49795956-INS-1556
chr8-49804360-INS-811
chr8-49821082-INS-1841
chr8-49842613-INS-4749
chr8-49855022-INS-245
chr8-49855222-INS-970
chr8-49906861-INS-61
chr8-49909564-INS-402
chr8-49912964-INS-100
chr8-49984873-INS-172
chr8-49985159-INS-655
chr8-50012726-INS-1293
chr8-50042617-INS-220
chr8-50068755-INS-50
chr8-50070992-INS-195
chr8-50084299-INS-213
chr8-50088044-INS-2831
chr8-50092576-INS-491
chr8-50124592-INS-50
chr8-50194744-INS-7219
chr8-50222409-INS-54
chr8-50224171-INS-1057
chr8-50229886-INS-190
chr8-50234199-INS-512
chr8-50276432-INS-124
chr8-50282664-INS-1328
chr8-50330002-INS-1007
chr8-50364411-INS-50
chr8-50367667-INS-452
chr8-50369915-INS-307
chr8-50452224-INS-68
chr8-50470047-INS-50
chr8-50491298-INS-55
chr8-50520157-INS-54
chr8-50544144-INS-73
chr8-50544149-INS-185
chr8-50640196-INS-56
chr8-50659729-INS-52
chr8-50660949-INS-336
chr8-50667105-INS-62
chr8-50688998-INS-572
chr8-50712054-INS-122
chr8-50722819-INS-210
chr8-50768846-INS-86
chr8-50786629-INS-660
chr8-50821502-INS-304
chr8-50838491-INS-94
chr8-50838878-INS-88
chr8-50887862-INS-70
chr8-50904158-INS-127
chr8-50916820-INS-304
chr8-51013527-INS-56
chr8-51014444-INS-116

chr8-51044933-INS-558
chr8-51044980-INS-194
chr8-51044980-INS-73
chr8-51048198-INS-1812
chr8-51066698-INS-201
chr8-51075608-INS-100
chr8-51075718-INS-278
chr8-51098954-INS-148
chr8-51116980-INS-50
chr8-51196322-INS-56
chr8-51207367-INS-52
chr8-51219107-INS-512
chr8-51235826-INS-52
chr8-51238573-INS-7186
chr8-51244647-INS-240
chr8-51251125-INS-54
chr8-51269518-INS-370
chr8-51292638-INS-99
chr8-51297276-INS-50
chr8-51302997-INS-50
chr8-51303103-INS-110
chr8-51325854-INS-694
chr8-51341612-INS-149
chr8-51347285-INS-738
chr8-51360314-INS-59
chr8-51360356-INS-236
chr8-51384454-INS-114
chr8-51414243-INS-147
chr8-51416084-INS-229
chr8-51435324-INS-213
chr8-51518785-INS-6291
chr8-51598352-INS-203
chr8-51610459-INS-7450
chr8-51620610-INS-59
chr8-51636434-INS-1679
chr8-51638733-INS-10384
chr8-51638733-INS-1575
chr8-51642335-INS-696
chr8-51665650-INS-6773
chr8-51671833-INS-79
chr8-51678074-INS-62
chr8-51716315-INS-54
chr8-51764753-INS-112
chr8-51787350-INS-264
chr8-51835850-INS-2129
chr8-51854742-INS-88
chr8-51898469-INS-1203
chr8-51905597-INS-80
chr8-51906019-INS-328
chr8-51929694-INS-109
chr8-51966928-INS-225
chr8-51972531-INS-6229
chr8-51981804-INS-3903
chr8-52043584-INS-114
chr8-52054459-INS-53
chr8-52084790-INS-58
chr8-52103941-INS-52
chr8-52124709-INS-92
chr8-52306513-INS-2068
chr8-52315927-INS-1516
chr8-52335669-INS-1252
chr8-52369959-INS-195
chr8-52372875-INS-60
chr8-52404008-INS-6357
chr8-52589049-INS-73
chr8-52598622-INS-2719
chr8-52670645-INS-1335
chr8-52677144-INS-2478
chr8-52691714-INS-189
chr8-52691893-INS-479
chr8-52692160-INS-54
chr8-52700257-INS-6570
chr8-52720979-INS-90
chr8-52786684-INS-52
chr8-52810899-INS-480
chr8-52820504-INS-90
chr8-52821298-INS-60

chr8-52834498-INS-142
chr8-52837783-INS-1560
chr8-52872546-INS-2876
chr8-52886378-INS-343
chr8-52908846-INS-2394
chr8-52920881-INS-279
chr8-52926797-INS-66
chr8-52935959-INS-54
chr8-52936883-INS-6425
chr8-52957305-INS-54
chr8-52987636-INS-51
chr8-53030133-INS-3308
chr8-53054225-INS-416
chr8-53154756-INS-56
chr8-53186146-INS-84
chr8-53186969-INS-6233
chr8-53196710-INS-62
chr8-53225939-INS-4818
chr8-53245984-INS-1937
chr8-53264088-INS-52
chr8-53277443-INS-51
chr8-53280219-INS-52
chr8-53332522-INS-51
chr8-53403100-INS-2349
chr8-53424936-INS-130
chr8-53485843-INS-2831
chr8-53516913-INS-304
chr8-53537853-INS-4837
chr8-53543572-INS-302
chr8-53545116-INS-1114
chr8-53596208-INS-6401
chr8-53616465-INS-1703
chr8-53638933-INS-54
chr8-53639294-INS-1405
chr8-53640920-INS-128
chr8-53672681-INS-124
chr8-53683780-INS-2904
chr8-53698718-INS-74
chr8-53698792-INS-639
chr8-53707720-INS-50
chr8-53748216-INS-1548
chr8-53778093-INS-800
chr8-53814829-INS-50
chr8-53890431-INS-1882
chr8-53910778-INS-54
chr8-53934159-INS-6382
chr8-54013623-INS-68
chr8-54020514-INS-50
chr8-54029493-INS-50
chr8-54048843-INS-158
chr8-54073838-INS-51
chr8-54102076-INS-429
chr8-54131597-INS-510
chr8-54158637-INS-1224
chr8-54211642-INS-123
chr8-54231657-INS-76
chr8-54239434-INS-768
chr8-54306621-INS-2279
chr8-54314063-INS-108
chr8-54319153-INS-67
chr8-54364237-INS-1499
chr8-54364731-INS-265
chr8-54364773-INS-282
chr8-54373628-INS-485
chr8-54376236-INS-803
chr8-54478903-INS-4590
chr8-54493522-INS-54
chr8-54493784-INS-78
chr8-54563573-INS-65
chr8-54580337-INS-6138
chr8-54640438-INS-74
chr8-54644233-INS-132
chr8-54672376-INS-68
chr8-54782481-INS-14181
chr8-54787742-INS-78
chr8-54792625-INS-4982
chr8-54793128-INS-7153

chr8-54817673-INS-3145
chr8-54906467-INS-397
chr8-54914758-INS-130
chr8-54931236-INS-408
chr8-54931982-INS-179
chr8-54933360-INS-6494
chr8-54956019-INS-111
chr8-54956293-INS-66
chr8-54956878-INS-50
chr8-54960267-INS-56
chr8-54969998-INS-90
chr8-55028514-INS-74
chr8-55037559-INS-147
chr8-55109741-INS-169
chr8-55134729-INS-2052
chr8-55134879-INS-440
chr8-55140045-INS-61
chr8-55186816-INS-81
chr8-55187179-INS-141
chr8-55202350-INS-374
chr8-55202463-INS-908
chr8-55205615-INS-72
chr8-55220468-INS-1718
chr8-55234998-INS-2341
chr8-55261967-INS-420
chr8-55397286-INS-3233
chr8-55470991-INS-60
chr8-55475029-INS-50
chr8-55486488-INS-144
chr8-55490200-INS-108
chr8-55542301-INS-50
chr8-55557809-INS-50
chr8-55564272-INS-1129
chr8-55861868-INS-4763
chr8-55862303-INS-91
chr8-55862783-INS-1086
chr8-55876527-INS-53
chr8-55901123-INS-474
chr8-55923218-INS-173
chr8-55924955-INS-248
chr8-55926329-INS-94
chr8-55959881-INS-499
chr8-56103403-INS-74
chr8-56121126-INS-56
chr8-56125052-INS-609
chr8-56140100-INS-198
chr8-56192772-INS-81
chr8-56226636-INS-4091
chr8-56259252-INS-214
chr8-56276938-INS-132
chr8-56327243-INS-502
chr8-56342689-INS-52
chr8-56345430-INS-2876
chr8-56348421-INS-1873
chr8-56392847-INS-70
chr8-56404317-INS-55
chr8-56465166-INS-1627
chr8-56528644-INS-6172
chr8-56555612-INS-200
chr8-56568729-INS-1215
chr8-56572582-INS-6385
chr8-56612654-INS-396
chr8-56614089-INS-195
chr8-56712982-INS-62
chr8-56759803-INS-6272
chr8-56786374-INS-3287
chr8-56788023-INS-309
chr8-56839697-INS-190
chr8-56879409-INS-4663
chr8-56895800-INS-62
chr8-56916589-INS-72
chr8-56917646-INS-166
chr8-57010502-INS-50
chr8-57034105-INS-451
chr8-57078317-INS-64
chr8-57115191-INS-119
chr8-57131657-INS-56

chr8-57149555-INS-144
chr8-57164668-INS-51
chr8-57177901-INS-55
chr8-57178770-INS-88
chr8-57196375-INS-317
chr8-57202316-INS-17916
chr8-57205808-INS-4100
chr8-57211367-INS-50
chr8-57224267-INS-530
chr8-57225884-INS-6182
chr8-57234140-INS-50
chr8-57284626-INS-55
chr8-57341225-INS-88
chr8-57342578-INS-1616
chr8-57357116-INS-83
chr8-57358701-INS-6569
chr8-57366747-INS-1186
chr8-57379031-INS-66
chr8-57385935-INS-153
chr8-57449425-INS-141
chr8-57449514-INS-115
chr8-57452639-INS-52
chr8-57507878-INS-325
chr8-57508170-INS-267
chr8-57508216-INS-822
chr8-57526412-INS-441
chr8-57600037-INS-133
chr8-57606736-INS-764
chr8-57681409-INS-417
chr8-57720069-INS-160
chr8-57736245-INS-316
chr8-57766470-INS-94
chr8-57791534-INS-50
chr8-57804325-INS-57
chr8-57811003-INS-312
chr8-57823830-INS-6224
chr8-57826617-INS-198
chr8-57838639-INS-1394
chr8-57861995-INS-64
chr8-57869080-INS-70
chr8-57869549-INS-81
chr8-57882967-INS-52
chr8-57883555-INS-56
chr8-57891200-INS-55
chr8-57895765-INS-250
chr8-57920477-INS-79
chr8-57928628-INS-533
chr8-57931775-INS-53
chr8-57932080-INS-58
chr8-57941895-INS-93
chr8-57953546-INS-8738
chr8-57969783-INS-192
chr8-57971449-INS-213
chr8-57997805-INS-702
chr8-57997820-INS-124
chr8-58020139-INS-241
chr8-58024566-INS-151
chr8-58024626-INS-397
chr8-58048527-INS-1177
chr8-58121603-INS-186
chr8-58170731-INS-183
chr8-58192625-INS-54
chr8-58193495-INS-84
chr8-58199231-INS-2696
chr8-58209761-INS-50
chr8-58258845-INS-500
chr8-58276545-INS-56
chr8-58281416-INS-2812
chr8-58285625-INS-52
chr8-58290961-INS-60
chr8-58321891-INS-3093
chr8-58322761-INS-4158
chr8-58322761-INS-8828
chr8-58341365-INS-7225
chr8-58341500-INS-63
chr8-58343496-INS-208
chr8-58484834-INS-994

chr8-58488135-INS-9091
chr8-58491847-INS-258
chr8-58492176-INS-1194
chr8-58497944-INS-60
chr8-58503762-INS-277
chr8-58510405-INS-125
chr8-58569254-INS-267
chr8-58595770-INS-83
chr8-58596091-INS-54
chr8-58651005-INS-1322
chr8-58656934-INS-303
chr8-58671274-INS-4477
chr8-58698503-INS-64
chr8-58702619-INS-58
chr8-58705069-INS-91
chr8-58705127-INS-491
chr8-58709798-INS-6430
chr8-58773458-INS-50
chr8-58820251-INS-5367
chr8-58932634-INS-8765
chr8-58961884-INS-1796
chr8-58973524-INS-330
chr8-58974849-INS-104
chr8-59031340-INS-1193
chr8-59053613-INS-392
chr8-59053726-INS-53
chr8-59053910-INS-464
chr8-59057741-INS-167
chr8-59101712-INS-6823
chr8-59102312-INS-56
chr8-59108460-INS-1678
chr8-59119510-INS-90
chr8-59156647-INS-50
chr8-59163271-INS-6449
chr8-59224497-INS-117
chr8-59224676-INS-704
chr8-59287835-INS-6628
chr8-59308774-INS-128
chr8-59308774-INS-380
chr8-59359619-INS-129
chr8-59415866-INS-721
chr8-59416333-INS-175
chr8-59435093-INS-2714
chr8-59436851-INS-61
chr8-59462595-INS-2228
chr8-59519191-INS-54
chr8-59527621-INS-142
chr8-59527647-INS-489
chr8-59544375-INS-52
chr8-59624442-INS-153
chr8-59624556-INS-483
chr8-59625165-INS-3161
chr8-59655059-INS-6781
chr8-59693545-INS-3359
chr8-59699922-INS-2642
chr8-59758365-INS-2765
chr8-59798412-INS-141
chr8-59798628-INS-77
chr8-59807904-INS-1693
chr8-59833600-INS-59
chr8-59890389-INS-500
chr8-59938030-INS-3129
chr8-59940743-INS-77
chr8-60011069-INS-94
chr8-60017237-INS-60
chr8-60026500-INS-4467
chr8-60031721-INS-697
chr8-60032268-INS-194
chr8-60032469-INS-129
chr8-60103172-INS-212
chr8-60131326-INS-1694
chr8-60162959-INS-2906
chr8-60175800-INS-796
chr8-60183708-INS-187
chr8-60210203-INS-399
chr8-60286427-INS-135
chr8-60304914-INS-1255

chr8-60341125-INS-4939
chr8-60347458-INS-79
chr8-60349121-INS-120
chr8-60370852-INS-1080
chr8-60392037-INS-52
chr8-60398029-INS-52
chr8-60409085-INS-225
chr8-60413968-INS-51
chr8-60414027-INS-6237
chr8-60424272-INS-2363
chr8-60434215-INS-4013
chr8-60453703-INS-4721
chr8-60474048-INS-5112
chr8-60514571-INS-433
chr8-60545274-INS-659
chr8-60547365-INS-200
chr8-60555818-INS-213
chr8-60556135-INS-616
chr8-60562646-INS-499
chr8-60580517-INS-6443
chr8-60582412-INS-52
chr8-60614523-INS-191
chr8-60614689-INS-100
chr8-60647247-INS-140
chr8-60663102-INS-412
chr8-60677206-INS-273
chr8-60677510-INS-66
chr8-60747719-INS-2757
chr8-60753056-INS-77
chr8-60797833-INS-6412
chr8-60819872-INS-1261
chr8-60847509-INS-1469
chr8-60918767-INS-1121
chr8-60945210-INS-1771
chr8-61016550-INS-8885
chr8-61081919-INS-1370
chr8-61134037-INS-505
chr8-61134128-INS-95
chr8-61149232-INS-224
chr8-61156109-INS-64
chr8-61168194-INS-58
chr8-61168211-INS-122
chr8-61213612-INS-1056
chr8-61224546-INS-68
chr8-61229316-INS-122
chr8-61313225-INS-82
chr8-61318466-INS-6422
chr8-61325608-INS-275
chr8-61342845-INS-56
chr8-61362779-INS-53
chr8-61596332-INS-108
chr8-61596906-INS-83
chr8-61613844-INS-52
chr8-61619527-INS-3760
chr8-61660791-INS-1053
chr8-61690756-INS-75
chr8-61717843-INS-72
chr8-61720571-INS-71
chr8-61740423-INS-4358
chr8-61759391-INS-6452
chr8-61767900-INS-1449
chr8-61781356-INS-539
chr8-61879978-INS-2798
chr8-61906520-INS-168
chr8-61926102-INS-263
chr8-61933987-INS-102
chr8-61946184-INS-62
chr8-61971755-INS-5284
chr8-62046044-INS-6656
chr8-62085325-INS-6988
chr8-62089742-INS-70
chr8-62112438-INS-60
chr8-62116403-INS-311
chr8-62136509-INS-59
chr8-62150844-INS-50
chr8-62175277-INS-82
chr8-62186620-INS-238

chr8-62215141-INS-164
chr8-62295863-INS-50
chr8-62369569-INS-350
chr8-62389119-INS-54
chr8-62398164-INS-361
chr8-62451096-INS-55
chr8-62479847-INS-723
chr8-62515074-INS-68
chr8-62518193-INS-72
chr8-62518729-INS-60
chr8-62582371-INS-50
chr8-62598746-INS-108
chr8-62690072-INS-3135
chr8-62740078-INS-438
chr8-62740568-INS-704
chr8-62741891-INS-84
chr8-62749412-INS-579
chr8-62768317-INS-355
chr8-62784965-INS-476
chr8-62786091-INS-138
chr8-62833327-INS-1312
chr8-62856231-INS-2910
chr8-62880287-INS-252
chr8-62884523-INS-494
chr8-62892319-INS-1324
chr8-62919835-INS-7111
chr8-62959088-INS-68
chr8-63065872-INS-352
chr8-63111225-INS-67
chr8-63111598-INS-252
chr8-63112407-INS-74
chr8-63113554-INS-164
chr8-63147681-INS-444
chr8-63150083-INS-75
chr8-63163628-INS-10011
chr8-63188320-INS-1148
chr8-63240640-INS-880
chr8-63273227-INS-250
chr8-63336346-INS-56
chr8-63347155-INS-5253
chr8-63395963-INS-6782
chr8-63400953-INS-514
chr8-63453747-INS-181
chr8-63460235-INS-672
chr8-63477476-INS-530
chr8-63478762-INS-92
chr8-63489462-INS-130
chr8-63489868-INS-70
chr8-63509991-INS-502
chr8-63553800-INS-52
chr8-63580779-INS-1270
chr8-63582280-INS-998
chr8-63592592-INS-3515
chr8-63596130-INS-772
chr8-63605496-INS-62
chr8-63634139-INS-214
chr8-63647542-INS-53
chr8-63656222-INS-52
chr8-63657189-INS-60
chr8-63664281-INS-106
chr8-63684123-INS-3423
chr8-63709623-INS-6190
chr8-63728811-INS-169
chr8-63740762-INS-78
chr8-63815907-INS-1222
chr8-63882062-INS-6849
chr8-63905338-INS-4871
chr8-63919756-INS-66
chr8-63922229-INS-436
chr8-63965423-INS-1324
chr8-63985891-INS-537
chr8-63989693-INS-6383
chr8-64002630-INS-146
chr8-64021395-INS-1798
chr8-64051950-INS-624
chr8-64063898-INS-4198
chr8-64064110-INS-339

chr8-64086702-INS-95
chr8-64125007-INS-1041
chr8-64181672-INS-159
chr8-64221167-INS-439
chr8-64232335-INS-68
chr8-64233445-INS-4487
chr8-64325519-INS-181
chr8-64326746-INS-6500
chr8-64341974-INS-3653
chr8-64348629-INS-308
chr8-64348629-INS-4353
chr8-64405632-INS-54
chr8-64492708-INS-378
chr8-64493678-INS-1048
chr8-64580503-INS-139
chr8-64580711-INS-4173
chr8-64585259-INS-60
chr8-64626267-INS-53
chr8-64632428-INS-60
chr8-64671314-INS-7289
chr8-64862631-INS-86
chr8-64873711-INS-52
chr8-64884693-INS-163
chr8-64898863-INS-636
chr8-64901852-INS-958
chr8-64952904-INS-2264
chr8-64961607-INS-53
chr8-64984948-INS-64
chr8-64989444-INS-54
chr8-64998662-INS-68
chr8-65006391-INS-50
chr8-65006769-INS-263
chr8-65006913-INS-50
chr8-65009219-INS-50
chr8-65016929-INS-108
chr8-65017404-INS-71
chr8-65033775-INS-56
chr8-65034554-INS-50
chr8-65036335-INS-60
chr8-65043218-INS-2606
chr8-65051193-INS-51
chr8-65054155-INS-162
chr8-65058058-INS-66
chr8-65058124-INS-363
chr8-65089334-INS-8099
chr8-65091723-INS-442
chr8-65229378-INS-4576
chr8-65259410-INS-3130
chr8-65269116-INS-1018
chr8-65387056-INS-494
chr8-65397998-INS-94
chr8-65413090-INS-2778
chr8-65456949-INS-1893
chr8-65462240-INS-50
chr8-65476078-INS-100
chr8-65483425-INS-74
chr8-65491130-INS-52
chr8-65535025-INS-206
chr8-65565320-INS-252
chr8-65572525-INS-195
chr8-65600666-INS-448
chr8-65609898-INS-458
chr8-65609955-INS-108
chr8-65610208-INS-323
chr8-65613709-INS-3070
chr8-65659303-INS-82
chr8-65677268-INS-1319
chr8-65679078-INS-156
chr8-65693671-INS-6587
chr8-65714486-INS-1153
chr8-65722967-INS-173
chr8-65722968-INS-68
chr8-65727432-INS-60
chr8-65743480-INS-2995
chr8-65785494-INS-3201
chr8-65794811-INS-3774
chr8-65808380-INS-654

chr8-65816702-INS-6584
chr8-65841398-INS-100
chr8-65842005-INS-418
chr8-65842532-INS-106
chr8-65862880-INS-464
chr8-65877398-INS-54
chr8-65878522-INS-955
chr8-65887966-INS-52
chr8-65898093-INS-128
chr8-65902846-INS-1798
chr8-65904654-INS-70
chr8-65953476-INS-4769
chr8-65963182-INS-6464
chr8-65988637-INS-1489
chr8-66013586-INS-223
chr8-66013647-INS-65
chr8-66123091-INS-522
chr8-66127378-INS-54
chr8-66134361-INS-6622
chr8-66135304-INS-50
chr8-66148024-INS-287
chr8-66148093-INS-780
chr8-66155412-INS-251
chr8-66182170-INS-58
chr8-66202514-INS-398
chr8-66212985-INS-52
chr8-66221483-INS-50
chr8-66245012-INS-7202
chr8-66254545-INS-280
chr8-66263048-INS-287
chr8-66264986-INS-53
chr8-66265595-INS-216
chr8-66302065-INS-72
chr8-66343873-INS-216
chr8-66389321-INS-1476
chr8-66435810-INS-1499
chr8-66435908-INS-6915
chr8-66448585-INS-451
chr8-66507942-INS-824
chr8-66507963-INS-389
chr8-66520062-INS-1491
chr8-66525291-INS-2552
chr8-66541100-INS-191
chr8-66565655-INS-995
chr8-66613735-INS-7199
chr8-66627800-INS-375
chr8-66637461-INS-50
chr8-66640357-INS-58
chr8-66646835-INS-889
chr8-66685132-INS-6360
chr8-66685151-INS-1700
chr8-66694605-INS-187
chr8-66733076-INS-57
chr8-66734082-INS-56
chr8-66774506-INS-71
chr8-66800247-INS-204
chr8-66805819-INS-130
chr8-66808485-INS-50
chr8-66852126-INS-60
chr8-66909339-INS-78
chr8-66909555-INS-127
chr8-66911584-INS-3261
chr8-66940201-INS-6647
chr8-66942726-INS-56
chr8-66945853-INS-112
chr8-67003325-INS-73
chr8-67008111-INS-52
chr8-67015850-INS-62
chr8-67028280-INS-5037
chr8-67091427-INS-175
chr8-67189147-INS-56
chr8-67197805-INS-7072
chr8-67227897-INS-6334
chr8-67242418-INS-1176
chr8-67254112-INS-5951
chr8-67304686-INS-57
chr8-67347657-INS-1239

chr8-67398187-INS-1262
chr8-67421014-INS-1534
chr8-67458861-INS-205
chr8-67464173-INS-6188
chr8-67495457-INS-229
chr8-67495650-INS-395
chr8-67535800-INS-6756
chr8-67537822-INS-3039
chr8-67570270-INS-200
chr8-67592380-INS-519
chr8-67605383-INS-3383
chr8-67679368-INS-2956
chr8-67682630-INS-274
chr8-67682680-INS-70
chr8-67691311-INS-1891
chr8-67726787-INS-8216
chr8-67772693-INS-64
chr8-67788284-INS-9200
chr8-67809913-INS-68
chr8-67810546-INS-241
chr8-67841293-INS-2294
chr8-67851909-INS-358
chr8-67912195-INS-60
chr8-67941739-INS-6113
chr8-67942494-INS-166
chr8-67942494-INS-66
chr8-67961858-INS-6528
chr8-68011627-INS-97
chr8-68031541-INS-6494
chr8-68070556-INS-537
chr8-68102465-INS-522
chr8-68130325-INS-434
chr8-68289031-INS-61
chr8-68305126-INS-130
chr8-68316128-INS-460
chr8-68316398-INS-464
chr8-68316891-INS-61
chr8-68348219-INS-9603
chr8-68365418-INS-6163
chr8-68410604-INS-50
chr8-68432180-INS-6961
chr8-68433234-INS-99
chr8-68448736-INS-58
chr8-68449181-INS-1082
chr8-68449827-INS-51
chr8-68507167-INS-82
chr8-68538644-INS-78
chr8-68566082-INS-4610
chr8-68583453-INS-56
chr8-68585535-INS-50
chr8-68592046-INS-80
chr8-68605034-INS-58
chr8-68612292-INS-1970
chr8-68614745-INS-6194
chr8-68628519-INS-1177
chr8-68734206-INS-77
chr8-68780201-INS-350
chr8-68780201-INS-60
chr8-68781047-INS-131
chr8-68794005-INS-458
chr8-68870472-INS-100
chr8-68888686-INS-52
chr8-68960276-INS-61
chr8-68960861-INS-437
chr8-68965526-INS-798
chr8-68978284-INS-54
chr8-69019280-INS-56
chr8-69039733-INS-105
chr8-69062325-INS-56
chr8-69105755-INS-110
chr8-69106657-INS-52
chr8-69159960-INS-1243
chr8-69167385-INS-1253
chr8-69194306-INS-1524
chr8-69206109-INS-168
chr8-69208118-INS-7000
chr8-69242978-INS-8809

chr8-69289738-INS-7140
chr8-69373946-INS-161
chr8-69380587-INS-63
chr8-69391529-INS-83
chr8-69397463-INS-61
chr8-69399446-INS-2173
chr8-69415529-INS-1199
chr8-69424015-INS-2003
chr8-69451195-INS-78
chr8-69460273-INS-79
chr8-69478458-INS-3125
chr8-69499760-INS-500
chr8-69516500-INS-57
chr8-69521643-INS-352
chr8-69525076-INS-144
chr8-69525096-INS-70
chr8-69549904-INS-174
chr8-69573141-INS-196
chr8-69583227-INS-1901
chr8-69689602-INS-96
chr8-69696001-INS-1498
chr8-69705129-INS-7251
chr8-69708656-INS-197
chr8-69709617-INS-498
chr8-69710982-INS-212
chr8-69713668-INS-54
chr8-69729111-INS-7915
chr8-69741661-INS-50
chr8-69785278-INS-5783
chr8-69791434-INS-6568
chr8-69909754-INS-187
chr8-69909811-INS-3177
chr8-70032616-INS-200
chr8-70151005-INS-230
chr8-70170407-INS-11512
chr8-70170407-INS-5012
chr8-70265345-INS-1232
chr8-70265348-INS-140
chr8-70290900-INS-291
chr8-70310503-INS-335
chr8-70310588-INS-480
chr8-70310795-INS-736
chr8-70316855-INS-194
chr8-70319801-INS-128
chr8-70320021-INS-421
chr8-70428131-INS-100
chr8-70435525-INS-289
chr8-70452121-INS-229
chr8-70460978-INS-2370
chr8-70462132-INS-8734
chr8-70567170-INS-8735
chr8-70577730-INS-203
chr8-70604929-INS-67
chr8-70619105-INS-205
chr8-70670945-INS-958
chr8-70671006-INS-200
chr8-70671006-INS-445
chr8-70698401-INS-52
chr8-70728133-INS-215
chr8-70730163-INS-84
chr8-70902582-INS-74
chr8-71090355-INS-77
chr8-71106597-INS-124
chr8-71106790-INS-221
chr8-71116793-INS-13287
chr8-71189366-INS-2237
chr8-71300105-INS-124
chr8-71329622-INS-182
chr8-71379481-INS-52
chr8-71453535-INS-152
chr8-71455655-INS-2871
chr8-71457400-INS-6873
chr8-71529944-INS-5536
chr8-71612961-INS-1432
chr8-71666878-INS-448
chr8-71677455-INS-77
chr8-71677561-INS-494

chr8-71715088-INS-337
chr8-71754360-INS-133
chr8-71842986-INS-59
chr8-71857831-INS-177
chr8-71862852-INS-51
chr8-71862958-INS-203
chr8-71866797-INS-200
chr8-71880166-INS-354
chr8-71882421-INS-216
chr8-71892558-INS-51
chr8-71908136-INS-55
chr8-71980336-INS-100
chr8-71992680-INS-2437
chr8-72027968-INS-564
chr8-72093181-INS-276
chr8-72093434-INS-484
chr8-72158482-INS-81
chr8-72210056-INS-443
chr8-72258682-INS-168
chr8-72703884-INS-8646
chr8-72725928-INS-89
chr8-72733395-INS-163
chr8-72755703-INS-11888
chr8-72811375-INS-105
chr8-72813315-INS-191
chr8-72817795-INS-314
chr8-72857396-INS-2407
chr8-72884161-INS-224
chr8-72943815-INS-2001
chr8-72946589-INS-5332
chr8-72958688-INS-203
chr8-73018897-INS-64
chr8-73021758-INS-5631
chr8-73039843-INS-170
chr8-73040658-INS-201
chr8-73049408-INS-203
chr8-73059529-INS-5150
chr8-73069682-INS-1681
chr8-73092955-INS-239
chr8-73211618-INS-493
chr8-73212246-INS-218
chr8-73346886-INS-60
chr8-73364940-INS-60
chr8-73367543-INS-802
chr8-73389362-INS-254
chr8-73398655-INS-66
chr8-73432471-INS-467
chr8-73440781-INS-72
chr8-73445737-INS-92
chr8-73481166-INS-1781
chr8-73486012-INS-62
chr8-73529061-INS-672
chr8-73566691-INS-329
chr8-73600471-INS-1211
chr8-73630344-INS-65
chr8-73635666-INS-58
chr8-73649583-INS-1116
chr8-73663105-INS-1280
chr8-73718911-INS-74
chr8-73719009-INS-89
chr8-73792280-INS-1880
chr8-73803080-INS-50
chr8-73831133-INS-216
chr8-73844721-INS-6577
chr8-73871149-INS-281
chr8-73889825-INS-1891
chr8-73892683-INS-77
chr8-73899232-INS-188
chr8-73901427-INS-50
chr8-73913650-INS-2899
chr8-73917757-INS-1196
chr8-73923209-INS-6172
chr8-73935859-INS-2064
chr8-74078267-INS-53
chr8-74092291-INS-4377
chr8-74165133-INS-221
chr8-74168544-INS-74

chr8-74200770-INS-1599
chr8-74212144-INS-6886
chr8-74235667-INS-196
chr8-74248498-INS-165
chr8-74248573-INS-63
chr8-74248910-INS-96
chr8-74270471-INS-199
chr8-74292776-INS-783
chr8-74301815-INS-485
chr8-74342903-INS-4698
chr8-74357440-INS-110
chr8-74403455-INS-6444
chr8-74403693-INS-99
chr8-74403862-INS-61
chr8-74404080-INS-65
chr8-74404119-INS-284
chr8-74429587-INS-1159
chr8-74440884-INS-56
chr8-74443018-INS-60
chr8-74443077-INS-4191
chr8-74462592-INS-281
chr8-74465545-INS-52
chr8-74492473-INS-73
chr8-74516630-INS-6405
chr8-74549713-INS-634
chr8-74550068-INS-3715
chr8-74550117-INS-853
chr8-74551699-INS-70
chr8-74562935-INS-1105
chr8-74564862-INS-6871
chr8-74569171-INS-116
chr8-74647285-INS-323
chr8-74649785-INS-50
chr8-74651573-INS-161
chr8-74668066-INS-65
chr8-74671252-INS-512
chr8-74672231-INS-74
chr8-74684581-INS-3500
chr8-74817229-INS-7243
chr8-74830291-INS-225
chr8-74832373-INS-78
chr8-74838871-INS-2912
chr8-74881133-INS-60
chr8-74901052-INS-2332
chr8-74934916-INS-2388
chr8-74979728-INS-66
chr8-74982190-INS-62
chr8-74996059-INS-107
chr8-74996954-INS-56
chr8-75000957-INS-99
chr8-75011414-INS-62
chr8-75025540-INS-124
chr8-75027891-INS-6239
chr8-75064794-INS-368
chr8-75067125-INS-152
chr8-75069053-INS-271
chr8-75071513-INS-208
chr8-75074123-INS-59
chr8-75074557-INS-123
chr8-75074557-INS-57
chr8-75091909-INS-2734
chr8-75229863-INS-104
chr8-75234074-INS-50
chr8-75288154-INS-52
chr8-75288311-INS-124
chr8-75308002-INS-156
chr8-75308609-INS-7224
chr8-75327016-INS-6485
chr8-75349173-INS-175
chr8-75355997-INS-322
chr8-75369880-INS-60
chr8-75394902-INS-6733
chr8-75409017-INS-376
chr8-75527044-INS-1672
chr8-75527080-INS-262
chr8-75527228-INS-604
chr8-75538617-INS-240

chr8-75560778-INS-332
chr8-75663672-INS-2522
chr8-75672395-INS-6254
chr8-75691665-INS-7301
chr8-75704481-INS-98
chr8-75752423-INS-50
chr8-75771577-INS-51
chr8-75771613-INS-186
chr8-75772263-INS-65
chr8-75793474-INS-7099
chr8-75798960-INS-144
chr8-75817730-INS-201
chr8-75819958-INS-130
chr8-75873783-INS-193
chr8-75933350-INS-7379
chr8-75946234-INS-92
chr8-75969588-INS-434
chr8-75974940-INS-217
chr8-76007710-INS-1856
chr8-76040326-INS-7128
chr8-76069692-INS-1052
chr8-76077605-INS-116
chr8-76105498-INS-4179
chr8-76142028-INS-10260
chr8-76142404-INS-551
chr8-76145513-INS-1068
chr8-76147825-INS-2770
chr8-76160073-INS-7678
chr8-76193413-INS-142
chr8-76197045-INS-72
chr8-76239845-INS-411
chr8-76241580-INS-132
chr8-76270068-INS-231
chr8-76272563-INS-6369
chr8-76280799-INS-50
chr8-76341930-INS-76
chr8-76342405-INS-50
chr8-76351909-INS-53
chr8-76360928-INS-50
chr8-76361667-INS-70
chr8-76387046-INS-60
chr8-76388052-INS-112
chr8-76494714-INS-494
chr8-76506909-INS-124
chr8-76558832-INS-96
chr8-76621550-INS-176
chr8-76651110-INS-7107
chr8-76653904-INS-445
chr8-76684333-INS-65
chr8-76906736-INS-54
chr8-76971919-INS-70
chr8-76981587-INS-96
chr8-76982094-INS-78
chr8-77056340-INS-94
chr8-77056874-INS-541
chr8-77057254-INS-70
chr8-77071345-INS-2406
chr8-77086838-INS-234
chr8-77091861-INS-467
chr8-77213688-INS-57
chr8-77226661-INS-6471
chr8-77241697-INS-230
chr8-77244214-INS-4791
chr8-77288161-INS-545
chr8-77321618-INS-75
chr8-77327879-INS-54
chr8-77333051-INS-50
chr8-77385586-INS-704
chr8-77391081-INS-1381
chr8-77392025-INS-416
chr8-77393275-INS-943
chr8-77426023-INS-148
chr8-77426027-INS-54
chr8-77500466-INS-7094
chr8-77520367-INS-6418
chr8-77595161-INS-63
chr8-77608190-INS-120

chr8-77608738-INS-1765
chr8-77612479-INS-84
chr8-77614967-INS-1122
chr8-77646462-INS-50
chr8-77725479-INS-50
chr8-77742077-INS-51
chr8-77781242-INS-94
chr8-77799434-INS-56
chr8-77809169-INS-50
chr8-77814409-INS-52
chr8-77824032-INS-122
chr8-77854508-INS-209
chr8-77854508-INS-510
chr8-77875408-INS-1412
chr8-77878440-INS-121
chr8-77981128-INS-184
chr8-78009816-INS-546
chr8-78022555-INS-141
chr8-78026743-INS-91
chr8-78036219-INS-62
chr8-78100059-INS-97
chr8-78109978-INS-210
chr8-78133958-INS-136
chr8-78149389-INS-562
chr8-78149389-INS-86
chr8-78150735-INS-4333
chr8-78152196-INS-159
chr8-78152340-INS-51
chr8-78158627-INS-433
chr8-78172217-INS-60
chr8-78175305-INS-364
chr8-78175433-INS-104
chr8-78176145-INS-261
chr8-78182251-INS-181
chr8-78184091-INS-1150
chr8-78219931-INS-1251
chr8-78246622-INS-198
chr8-78256978-INS-3245
chr8-78257338-INS-7080
chr8-78271561-INS-181
chr8-78298472-INS-82
chr8-78298476-INS-505
chr8-78301411-INS-185
chr8-78343558-INS-7329
chr8-78348383-INS-214
chr8-78355569-INS-52
chr8-78355791-INS-210
chr8-78356089-INS-458
chr8-78356546-INS-55
chr8-78358099-INS-74
chr8-78397481-INS-262
chr8-78406620-INS-478
chr8-78419104-INS-50
chr8-78424126-INS-50
chr8-78427146-INS-62
chr8-78435830-INS-273
chr8-78445660-INS-75
chr8-78456813-INS-1815
chr8-78469380-INS-56
chr8-78475265-INS-100
chr8-78477699-INS-133
chr8-78514826-INS-295
chr8-78534877-INS-1237
chr8-78547629-INS-54
chr8-78565126-INS-219
chr8-78565201-INS-58
chr8-78565492-INS-110
chr8-78603570-INS-147
chr8-78619251-INS-51
chr8-78619500-INS-362
chr8-78624480-INS-677
chr8-78668460-INS-4250
chr8-78726932-INS-172
chr8-78728637-INS-2841
chr8-78740269-INS-7145
chr8-78763654-INS-7115
chr8-78775619-INS-6158

chr8-78816471-INS-54
chr8-78854497-INS-56
chr8-78857304-INS-116
chr8-78857304-INS-310
chr8-78951766-INS-60
chr8-78982254-INS-56
chr8-78983117-INS-80
chr8-78984350-INS-50
chr8-78995266-INS-85
chr8-78995484-INS-64
chr8-79022294-INS-6497
chr8-79062237-INS-388
chr8-79121123-INS-1782
chr8-79136476-INS-382
chr8-79138446-INS-264
chr8-79140317-INS-597
chr8-79140617-INS-900
chr8-79216024-INS-205
chr8-79294074-INS-243
chr8-79348554-INS-8684
chr8-79361625-INS-217
chr8-79365852-INS-108
chr8-79457332-INS-76
chr8-79463085-INS-1952
chr8-79553788-INS-255
chr8-79600880-INS-1250
chr8-79602782-INS-56
chr8-79611031-INS-138
chr8-79642554-INS-4262
chr8-79672892-INS-7257
chr8-79678198-INS-2888
chr8-79711442-INS-63
chr8-79742247-INS-54
chr8-79744813-INS-149
chr8-79791976-INS-173
chr8-79791991-INS-80
chr8-79792183-INS-65
chr8-79848214-INS-52
chr8-79860828-INS-52
chr8-79866756-INS-54
chr8-79866907-INS-752
chr8-79960117-INS-95
chr8-80030942-INS-82
chr8-80081459-INS-176
chr8-80164172-INS-360
chr8-80164208-INS-78
chr8-80170846-INS-3543
chr8-80197899-INS-630
chr8-80253133-INS-58
chr8-80281563-INS-1968
chr8-80285073-INS-120
chr8-80285253-INS-360
chr8-80294891-INS-193
chr8-80319202-INS-107
chr8-80322842-INS-77
chr8-80331530-INS-1081
chr8-80348516-INS-54
chr8-80356621-INS-201
chr8-80357341-INS-50
chr8-80440167-INS-107
chr8-80481829-INS-2251
chr8-80484774-INS-2232
chr8-80517967-INS-5022
chr8-80518899-INS-146
chr8-80583366-INS-81
chr8-80597217-INS-67
chr8-80638260-INS-73
chr8-80638267-INS-154
chr8-80646921-INS-1140
chr8-80669634-INS-75
chr8-80674253-INS-5066
chr8-80687224-INS-161
chr8-80688579-INS-849
chr8-80740153-INS-82
chr8-80849136-INS-96
chr8-80914742-INS-430
chr8-81014762-INS-62

chr8-81108753-INS-102
chr8-81128009-INS-848
chr8-81162245-INS-6251
chr8-81164197-INS-3215
chr8-81200101-INS-3219
chr8-81279467-INS-1345
chr8-81280572-INS-230
chr8-81304698-INS-64
chr8-81310625-INS-354
chr8-81311698-INS-1153
chr8-81320591-INS-60
chr8-81326320-INS-249
chr8-81349621-INS-608
chr8-81379829-INS-170
chr8-81380035-INS-57
chr8-81396957-INS-64
chr8-81414384-INS-140
chr8-81453656-INS-477
chr8-81488742-INS-61
chr8-81489038-INS-169
chr8-81496570-INS-126
chr8-81500747-INS-541
chr8-81517905-INS-60
chr8-81543197-INS-92
chr8-81554907-INS-199
chr8-81579471-INS-93
chr8-81635625-INS-94
chr8-81658027-INS-569
chr8-81692830-INS-70
chr8-81712844-INS-304
chr8-81743410-INS-685
chr8-81770655-INS-88
chr8-81781010-INS-656
chr8-81822605-INS-2079
chr8-81852977-INS-54
chr8-81889013-INS-1060
chr8-81899680-INS-6818
chr8-81907313-INS-96
chr8-81933867-INS-125
chr8-81940600-INS-1895
chr8-81951071-INS-6401
chr8-81963839-INS-50
chr8-81981079-INS-6221
chr8-81997915-INS-316
chr8-81998673-INS-206
chr8-82022435-INS-120
chr8-82040183-INS-6855
chr8-82088897-INS-50
chr8-82089652-INS-50
chr8-82091168-INS-57
chr8-82095865-INS-108
chr8-82234649-INS-431
chr8-82281655-INS-495
chr8-82306776-INS-1336
chr8-82327891-INS-1362
chr8-82347449-INS-264
chr8-82347678-INS-51
chr8-82424188-INS-4419
chr8-82433107-INS-62
chr8-82455328-INS-83
chr8-82455442-INS-147
chr8-82498906-INS-50
chr8-82589145-INS-6401
chr8-82625901-INS-64
chr8-82626738-INS-68
chr8-82729702-INS-6583
chr8-82767327-INS-199
chr8-82841930-INS-6822
chr8-82874604-INS-7170
chr8-82895561-INS-5018
chr8-82898129-INS-4369
chr8-82945229-INS-2794
chr8-82963492-INS-7168
chr8-83003890-INS-55
chr8-83030192-INS-1639
chr8-83047875-INS-3126
chr8-83095255-INS-6560

chr8-83117134-INS-7548
chr8-83283920-INS-82
chr8-83346806-INS-6599
chr8-83347651-INS-52
chr8-83386421-INS-1372
chr8-83414853-INS-55
chr8-83471610-INS-7200
chr8-83475990-INS-7151
chr8-83476129-INS-235
chr8-83511048-INS-66
chr8-83511670-INS-84
chr8-83517456-INS-1635
chr8-83648842-INS-120
chr8-83664398-INS-66
chr8-83675618-INS-191
chr8-83682036-INS-299
chr8-83761821-INS-91
chr8-83839063-INS-191
chr8-83932624-INS-2573
chr8-83942945-INS-1358
chr8-83948275-INS-528
chr8-83949891-INS-65
chr8-83956502-INS-125
chr8-83969998-INS-120
chr8-84036463-INS-4525
chr8-84076878-INS-190
chr8-84093357-INS-671
chr8-84093468-INS-237
chr8-84093698-INS-302
chr8-84100671-INS-2877
chr8-84106543-INS-541
chr8-84119740-INS-80
chr8-84147489-INS-57
chr8-84157671-INS-213
chr8-84190236-INS-14488
chr8-84205887-INS-6553
chr8-84213490-INS-410
chr8-84214142-INS-57
chr8-84223100-INS-285
chr8-84245652-INS-68
chr8-84259496-INS-198
chr8-84296428-INS-2844
chr8-84296428-INS-444
chr8-84311517-INS-62
chr8-84327976-INS-140
chr8-84328342-INS-102
chr8-84329784-INS-478
chr8-84345866-INS-60
chr8-84373240-INS-202
chr8-84436453-INS-438
chr8-84484617-INS-112
chr8-84487503-INS-440
chr8-84487869-INS-84
chr8-84496411-INS-1888
chr8-84517442-INS-60
chr8-84517496-INS-260
chr8-84548380-INS-1061
chr8-84557755-INS-59
chr8-84561803-INS-50
chr8-84579359-INS-52
chr8-84580267-INS-93
chr8-84585862-INS-56
chr8-84585869-INS-3903
chr8-84761550-INS-225
chr8-84762357-INS-176
chr8-84839928-INS-1068
chr8-84839928-INS-498
chr8-84847133-INS-4055
chr8-84876215-INS-197
chr8-84936828-INS-105
chr8-85016971-INS-57
chr8-85033228-INS-172
chr8-85036249-INS-514
chr8-85075496-INS-502
chr8-85075690-INS-109
chr8-85075753-INS-238
chr8-85102865-INS-63

chr8-85125870-INS-419
chr8-85214290-INS-395
chr8-85214290-INS-79
chr8-85219616-INS-188
chr8-85356400-INS-157
chr8-85356400-INS-71
chr8-85357539-INS-52
chr8-85551104-INS-92
chr8-85603087-INS-449
chr8-85658334-INS-181
chr8-85660203-INS-64
chr8-85752835-INS-106
chr8-85842940-INS-89
chr8-85848604-INS-230
chr8-85848770-INS-1401
chr8-85848770-INS-600
chr8-85897083-INS-5620
chr8-85911304-INS-1457
chr8-85914772-INS-2055
chr8-85958846-INS-58
chr8-85978534-INS-2577
chr8-85991280-INS-199
chr8-86037570-INS-5550
chr8-86081373-INS-193
chr8-86126702-INS-7394
chr8-86171334-INS-215
chr8-86177756-INS-50
chr8-86206574-INS-123
chr8-86236558-INS-204
chr8-86236904-INS-216
chr8-86334082-INS-50
chr8-86342141-INS-148
chr8-86362886-INS-222
chr8-86387003-INS-6673
chr8-86387258-INS-295
chr8-86387878-INS-55
chr8-86437912-INS-6730
chr8-86446479-INS-378
chr8-86515464-INS-136
chr8-86527850-INS-2234
chr8-86527850-INS-468
chr8-86548769-INS-2996
chr8-86584300-INS-4106
chr8-86595578-INS-1164
chr8-86602920-INS-1555
chr8-86613467-INS-1290
chr8-86660404-INS-180
chr8-86670806-INS-58
chr8-86672116-INS-212
chr8-86774743-INS-128
chr8-86776923-INS-4960
chr8-86803831-INS-112
chr8-86815695-INS-52
chr8-86845364-INS-1676
chr8-86858771-INS-53
chr8-86858840-INS-917
chr8-86964528-INS-416
chr8-86999238-INS-52
chr8-86999650-INS-112
chr8-87001463-INS-50
chr8-87013347-INS-1364
chr8-87021654-INS-2572
chr8-87029469-INS-6392
chr8-87071740-INS-52
chr8-87094449-INS-54
chr8-87155540-INS-4059
chr8-87166279-INS-6430
chr8-87234215-INS-312
chr8-87291843-INS-438
chr8-87292832-INS-59
chr8-87334441-INS-288
chr8-87345920-INS-75
chr8-87345935-INS-690
chr8-87416567-INS-220
chr8-87468764-INS-3211
chr8-87488049-INS-83
chr8-87490252-INS-305

chr8-87500777-INS-54
chr8-87500836-INS-227
chr8-87505566-INS-6535
chr8-87508765-INS-502
chr8-87511355-INS-224
chr8-87530434-INS-361
chr8-87542184-INS-90
chr8-87545405-INS-64
chr8-87548304-INS-147
chr8-87549812-INS-89
chr8-87558507-INS-58
chr8-87569257-INS-62
chr8-87639923-INS-71
chr8-87640296-INS-155
chr8-87685198-INS-52
chr8-87686114-INS-156
chr8-87686204-INS-82
chr8-87777546-INS-380
chr8-87777574-INS-54
chr8-87779833-INS-228
chr8-87857989-INS-54
chr8-87892532-INS-64
chr8-87895736-INS-53
chr8-87928681-INS-54
chr8-88158634-INS-66
chr8-88193913-INS-158
chr8-88226502-INS-83
chr8-88247522-INS-52
chr8-88249611-INS-5239
chr8-88258625-INS-56
chr8-88265201-INS-97
chr8-88301392-INS-702
chr8-88361224-INS-98
chr8-88377226-INS-81
chr8-88377376-INS-96
chr8-88377547-INS-108
chr8-88436855-INS-52
chr8-88444020-INS-464
chr8-88500993-INS-82
chr8-88501116-INS-52
chr8-88517366-INS-2912
chr8-88522005-INS-147
chr8-88552135-INS-72
chr8-88552605-INS-213
chr8-88556628-INS-127
chr8-88556628-INS-285
chr8-88569046-INS-50
chr8-88609614-INS-327
chr8-88647837-INS-50
chr8-88683847-INS-1501
chr8-88692211-INS-227
chr8-88697483-INS-64
chr8-88697893-INS-204
chr8-88700179-INS-204
chr8-88710500-INS-88
chr8-88714762-INS-127
chr8-88739949-INS-214
chr8-88784219-INS-3139
chr8-88785543-INS-262
chr8-88800456-INS-240
chr8-88800469-INS-64
chr8-88806325-INS-4408
chr8-88886614-INS-92
chr8-88907784-INS-316
chr8-88907786-INS-82
chr8-88923109-INS-907
chr8-88929082-INS-3281
chr8-88929084-INS-486
chr8-88930554-INS-1416
chr8-88931023-INS-92
chr8-88931969-INS-999
chr8-88978042-INS-159
chr8-88990616-INS-209
chr8-88995490-INS-57
chr8-88995563-INS-279
chr8-89031880-INS-770
chr8-89032181-INS-1089

chr8-89047658-INS-66
chr8-89096308-INS-1480
chr8-89099185-INS-225
chr8-89099302-INS-461
chr8-89101247-INS-1060
chr8-89163127-INS-57
chr8-89203310-INS-2999
chr8-89304056-INS-78
chr8-89333598-INS-234
chr8-89339037-INS-62
chr8-89348148-INS-63
chr8-89348935-INS-263
chr8-89492708-INS-126
chr8-89497981-INS-2955
chr8-89530484-INS-382
chr8-89531088-INS-71
chr8-89550304-INS-165
chr8-89573769-INS-68
chr8-89574647-INS-70
chr8-89593580-INS-375
chr8-89594914-INS-117
chr8-89595098-INS-83
chr8-89648370-INS-199
chr8-89652655-INS-160
chr8-89652655-INS-68
chr8-89711971-INS-231
chr8-89793939-INS-2851
chr8-89815806-INS-130
chr8-89854717-INS-60
chr8-89877959-INS-7112
chr8-89885869-INS-1480
chr8-89885951-INS-104
chr8-89897670-INS-521
chr8-89899060-INS-316
chr8-89899060-INS-56
chr8-89915579-INS-58
chr8-89916901-INS-57
chr8-89940773-INS-214
chr8-89966607-INS-152
chr8-89966612-INS-53
chr8-90007098-INS-114
chr8-90208604-INS-192
chr8-90254849-INS-50
chr8-90265517-INS-60
chr8-90321238-INS-6346
chr8-90345360-INS-12853
chr8-90387754-INS-82
chr8-90402760-INS-90
chr8-90402995-INS-302
chr8-90418595-INS-226
chr8-90598110-INS-7382
chr8-90652410-INS-537
chr8-90717239-INS-193
chr8-90745541-INS-418
chr8-90745810-INS-191
chr8-90771246-INS-6211
chr8-90784934-INS-229
chr8-90785060-INS-64
chr8-90788507-INS-174
chr8-90846512-INS-54
chr8-90850675-INS-6594
chr8-90856058-INS-157
chr8-90880908-INS-1885
chr8-90924991-INS-10300
chr8-90989198-INS-130
chr8-91066347-INS-52
chr8-91069286-INS-129
chr8-91122211-INS-243
chr8-91132710-INS-98
chr8-91142830-INS-498
chr8-91147356-INS-6840
chr8-91183803-INS-59
chr8-91190242-INS-2290
chr8-91207578-INS-6167
chr8-91208059-INS-50
chr8-91228043-INS-50
chr8-91258336-INS-57

chr8-91258367-INS-2511
chr8-91258511-INS-264
chr8-91272066-INS-5532
chr8-91273154-INS-76
chr8-91291410-INS-692
chr8-91302232-INS-2576
chr8-91311958-INS-7300
chr8-91340142-INS-51
chr8-91348594-INS-7137
chr8-91360407-INS-9061
chr8-91420344-INS-164
chr8-91432918-INS-283
chr8-91438017-INS-66
chr8-91440102-INS-64
chr8-91459390-INS-63
chr8-91460691-INS-211
chr8-91469955-INS-67
chr8-91483057-INS-1891
chr8-91504689-INS-64
chr8-91505715-INS-52
chr8-91532357-INS-81
chr8-91537549-INS-170
chr8-91537619-INS-1174
chr8-91537634-INS-494
chr8-91550031-INS-222
chr8-91558693-INS-203
chr8-91559493-INS-260
chr8-91570275-INS-50
chr8-91575522-INS-51
chr8-91587870-INS-211
chr8-91593169-INS-266
chr8-91629941-INS-265
chr8-91676998-INS-188
chr8-91729182-INS-573
chr8-91734000-INS-254
chr8-91791152-INS-858
chr8-91817901-INS-58
chr8-91820783-INS-219
chr8-91830821-INS-1384
chr8-91840279-INS-249
chr8-91848589-INS-82
chr8-91879470-INS-6475
chr8-91914690-INS-156
chr8-91926951-INS-58
chr8-91962842-INS-184
chr8-91978279-INS-375
chr8-92078903-INS-153
chr8-92078903-INS-468
chr8-92094994-INS-1041
chr8-92114364-INS-4570
chr8-92139453-INS-486
chr8-92142249-INS-177
chr8-92182767-INS-52
chr8-92202110-INS-4728
chr8-92203594-INS-132
chr8-92217352-INS-70
chr8-92307869-INS-72
chr8-92347886-INS-130
chr8-92376362-INS-102
chr8-92408627-INS-1722
chr8-92408762-INS-72
chr8-92432185-INS-454
chr8-92513245-INS-133
chr8-92521850-INS-94
chr8-92594485-INS-306
chr8-92645761-INS-54
chr8-92737793-INS-66
chr8-92751361-INS-74
chr8-92777006-INS-7142
chr8-92800933-INS-846
chr8-92805172-INS-54
chr8-92852616-INS-225
chr8-92869872-INS-68
chr8-92872093-INS-1077
chr8-92876951-INS-79
chr8-92884440-INS-88
chr8-92942737-INS-54

chr8-92950418-INS-426
chr8-92998199-INS-52
chr8-93005859-INS-52
chr8-93057742-INS-88
chr8-93078815-INS-178
chr8-93102326-INS-351
chr8-93253585-INS-1876
chr8-93267460-INS-81
chr8-93291407-INS-7077
chr8-93292127-INS-238
chr8-93292127-INS-518
chr8-93292695-INS-200
chr8-93293362-INS-2914
chr8-93306084-INS-117
chr8-93306084-INS-241
chr8-93307318-INS-52
chr8-93318456-INS-1410
chr8-93320668-INS-59
chr8-93323580-INS-154
chr8-93323787-INS-178
chr8-93368734-INS-60
chr8-93370166-INS-203
chr8-93452625-INS-329
chr8-93452945-INS-50
chr8-93453308-INS-318
chr8-93459786-INS-156
chr8-93459806-INS-372
chr8-93490354-INS-5624
chr8-93493080-INS-58
chr8-93555944-INS-52
chr8-93556964-INS-58
chr8-93571342-INS-52
chr8-93657871-INS-177
chr8-93730878-INS-60
chr8-93808722-INS-188
chr8-93846460-INS-54
chr8-93866662-INS-1130
chr8-93880354-INS-1048
chr8-93884575-INS-1595
chr8-93889940-INS-815
chr8-93899475-INS-52
chr8-93926507-INS-96
chr8-93946839-INS-55
chr8-93982678-INS-12855
chr8-94029889-INS-56
chr8-94035971-INS-164
chr8-94035974-INS-66
chr8-94036539-INS-677
chr8-94039301-INS-3664
chr8-94047105-INS-172
chr8-94048009-INS-757
chr8-94048228-INS-934
chr8-94133465-INS-70
chr8-94134891-INS-72
chr8-94155362-INS-86
chr8-94156731-INS-7083
chr8-94157665-INS-528
chr8-94158309-INS-225
chr8-94169236-INS-103
chr8-94169637-INS-435
chr8-94172100-INS-225
chr8-94219329-INS-361
chr8-94221343-INS-486
chr8-94222473-INS-638
chr8-94222863-INS-174
chr8-94226608-INS-186
chr8-94226710-INS-256
chr8-94227228-INS-50
chr8-94270972-INS-2033
chr8-94281719-INS-63
chr8-94292237-INS-329
chr8-94315279-INS-165
chr8-94354818-INS-84
chr8-94398732-INS-852
chr8-94401945-INS-56
chr8-94413866-INS-171
chr8-94430217-INS-159

chr8-94440497-INS-214
chr8-94444652-INS-6268
chr8-94450248-INS-115
chr8-94460241-INS-1479
chr8-94519576-INS-91
chr8-94534818-INS-64
chr8-94534870-INS-132
chr8-94640073-INS-219
chr8-94641343-INS-75
chr8-94653132-INS-86
chr8-94653462-INS-296
chr8-94660542-INS-1488
chr8-94685617-INS-55
chr8-94729747-INS-56
chr8-94780178-INS-60
chr8-94795679-INS-243
chr8-94822462-INS-4774
chr8-94823490-INS-203
chr8-94833884-INS-112
chr8-94846875-INS-82
chr8-94848711-INS-1698
chr8-94858000-INS-52
chr8-94872042-INS-60
chr8-94884500-INS-56
chr8-94884840-INS-183
chr8-94884934-INS-878
chr8-94885991-INS-207
chr8-94890538-INS-53
chr8-94891092-INS-334
chr8-94896078-INS-59
chr8-94921910-INS-103
chr8-94926412-INS-53
chr8-94932894-INS-2828
chr8-94933845-INS-1056
chr8-94933845-INS-192
chr8-94933845-INS-60
chr8-94935514-INS-92
chr8-94947726-INS-192
chr8-94956864-INS-208
chr8-95000979-INS-8329
chr8-95005042-INS-91
chr8-95005332-INS-67
chr8-95018575-INS-70
chr8-95099339-INS-51
chr8-95137326-INS-315
chr8-95143287-INS-51
chr8-95154246-INS-6401
chr8-95187320-INS-60
chr8-95194362-INS-999
chr8-95240771-INS-7279
chr8-95243156-INS-50
chr8-95243292-INS-123
chr8-95271622-INS-50
chr8-95318240-INS-50
chr8-95320408-INS-74
chr8-95400385-INS-233
chr8-95428782-INS-59
chr8-95451255-INS-13460
chr8-95464364-INS-218
chr8-95485769-INS-6227
chr8-95496567-INS-120
chr8-95533839-INS-62
chr8-95556982-INS-1501
chr8-95604062-INS-57
chr8-95710449-INS-684
chr8-95743710-INS-150
chr8-95752926-INS-57
chr8-95779113-INS-50
chr8-95883069-INS-56
chr8-95894141-INS-340
chr8-95924227-INS-206
chr8-95934060-INS-183
chr8-95947044-INS-5274
chr8-95995865-INS-65
chr8-95996300-INS-50
chr8-96011864-INS-346
chr8-96012115-INS-347

chr8-96017839-INS-68
chr8-96105660-INS-212
chr8-96158027-INS-80
chr8-96221620-INS-53
chr8-96228301-INS-175
chr8-96242123-INS-52
chr8-96256470-INS-50
chr8-96276234-INS-2595
chr8-96276523-INS-164
chr8-96283626-INS-59
chr8-96295398-INS-528
chr8-96309892-INS-1489
chr8-96329931-INS-223
chr8-96348291-INS-191
chr8-96356739-INS-219
chr8-96485130-INS-116
chr8-96521242-INS-139
chr8-96535829-INS-50
chr8-96536529-INS-181
chr8-96539967-INS-67
chr8-96542554-INS-173
chr8-96544847-INS-209
chr8-96566484-INS-54
chr8-96577087-INS-80
chr8-96600074-INS-171
chr8-96618805-INS-50
chr8-96631579-INS-86
chr8-96632398-INS-208
chr8-96634175-INS-278
chr8-96642014-INS-214
chr8-96649544-INS-50
chr8-96652570-INS-66
chr8-96708106-INS-342
chr8-96712648-INS-56
chr8-96751179-INS-687
chr8-96760953-INS-1256
chr8-96768585-INS-138
chr8-96787131-INS-994
chr8-96794149-INS-558
chr8-96811943-INS-350
chr8-96823593-INS-7531
chr8-96824654-INS-152
chr8-96824683-INS-62
chr8-96830953-INS-3640
chr8-96832961-INS-6892
chr8-96851995-INS-136
chr8-96856155-INS-1102
chr8-96876615-INS-195
chr8-96946132-INS-1946
chr8-96946132-INS-3895
chr8-96994148-INS-498
chr8-97014426-INS-75
chr8-97019782-INS-62
chr8-97034090-INS-6554
chr8-97053778-INS-56
chr8-97066265-INS-612
chr8-97083309-INS-104
chr8-97084193-INS-224
chr8-97091252-INS-1024
chr8-97091941-INS-118
chr8-97095535-INS-123
chr8-97120787-INS-6806
chr8-97160899-INS-12678
chr8-97165088-INS-254
chr8-97168811-INS-6391
chr8-97179407-INS-52
chr8-97196425-INS-6534
chr8-97196511-INS-60
chr8-97270862-INS-112
chr8-97282998-INS-60
chr8-97285265-INS-374
chr8-97296331-INS-3031
chr8-97302944-INS-53
chr8-97345680-INS-54
chr8-97345707-INS-177
chr8-97386066-INS-115
chr8-97393286-INS-58

chr8-97393764-INS-282
chr8-97484948-INS-70
chr8-97485946-INS-66
chr8-97510672-INS-922
chr8-97512509-INS-62
chr8-97512845-INS-123
chr8-97513694-INS-8775
chr8-97517002-INS-56
chr8-97532180-INS-72
chr8-97534993-INS-1547
chr8-97546693-INS-6137
chr8-97563157-INS-52
chr8-97577678-INS-212
chr8-97582505-INS-60
chr8-97606065-INS-6169
chr8-97608088-INS-1011
chr8-97617043-INS-758
chr8-97643186-INS-5969
chr8-97649584-INS-63
chr8-97650070-INS-79
chr8-97656012-INS-60
chr8-97692287-INS-130
chr8-97730054-INS-1092
chr8-97736776-INS-85
chr8-97750749-INS-922
chr8-97751225-INS-508
chr8-97757340-INS-94
chr8-97776565-INS-7207
chr8-97781218-INS-6868
chr8-97798285-INS-59
chr8-97814583-INS-126
chr8-97849360-INS-460
chr8-97854914-INS-94
chr8-97861004-INS-50
chr8-97920278-INS-52
chr8-97924965-INS-1474
chr8-97945922-INS-19872
chr8-97945922-INS-7390
chr8-97953690-INS-2964
chr8-97987126-INS-12866
chr8-97999258-INS-240
chr8-98010085-INS-60
chr8-98029786-INS-8948
chr8-98044067-INS-85
chr8-98044679-INS-80
chr8-98075042-INS-62
chr8-98075654-INS-2421
chr8-98086316-INS-114
chr8-98106310-INS-180
chr8-98106310-INS-68
chr8-98205611-INS-80
chr8-98267873-INS-180
chr8-98285642-INS-1813
chr8-98300958-INS-7142
chr8-98347556-INS-1626
chr8-98373705-INS-54
chr8-98374990-INS-102
chr8-98406193-INS-6399
chr8-98414576-INS-70
chr8-98427489-INS-801
chr8-98431506-INS-52
chr8-98480585-INS-50
chr8-98510152-INS-199
chr8-98548823-INS-143
chr8-98573887-INS-593
chr8-98602103-INS-83
chr8-98622267-INS-62
chr8-98657558-INS-204
chr8-98659513-INS-1998
chr8-98664232-INS-77
chr8-98664934-INS-64
chr8-98689892-INS-1295
chr8-98745973-INS-60
chr8-98748048-INS-58
chr8-98758950-INS-287
chr8-98783220-INS-531
chr8-98801857-INS-54

chr8-98824934-INS-200
chr8-98825610-INS-74
chr8-98828136-INS-53
chr8-98924480-INS-56
chr8-98990353-INS-50
chr8-99082022-INS-52
chr8-99086784-INS-52
chr8-99101993-INS-937
chr8-99104771-INS-72
chr8-99121285-INS-1954
chr8-99179070-INS-86
chr8-99182111-INS-50
chr8-99208418-INS-52
chr8-99216344-INS-1728
chr8-99236596-INS-1779
chr8-99255249-INS-51
chr8-99260981-INS-76
chr8-99301166-INS-282
chr8-99301595-INS-53
chr8-99318622-INS-6253
chr8-99327975-INS-83
chr8-99334109-INS-52
chr8-99351490-INS-8696
chr8-99434921-INS-50
chr8-99443738-INS-2167
chr8-99457946-INS-104
chr8-99458782-INS-59
chr8-99473283-INS-4515
chr8-99485869-INS-59
chr8-99485896-INS-153
chr8-99487336-INS-6048
chr8-99494846-INS-829
chr8-99508123-INS-5956
chr8-99513436-INS-147
chr8-99571806-INS-1664
chr8-99607122-INS-50
chr8-99621519-INS-100
chr8-99621519-INS-284
chr8-99626134-INS-4112
chr8-99638775-INS-66
chr8-99646144-INS-8436
chr8-99652291-INS-56
chr8-99687790-INS-55
chr8-99703342-INS-52
chr8-99713439-INS-7073
chr8-99727553-INS-210
chr8-99790183-INS-52
chr8-99819089-INS-52
chr8-99869795-INS-56
chr8-99912010-INS-3234
chr8-99928234-INS-50
chr8-99934701-INS-1104
chr8-99949226-INS-271
chr8-99960427-INS-2869
chr8-99984645-INS-117
chr8-99992816-INS-54
chr8-100018643-INS-494
chr8-100022279-INS-52
chr8-100024553-INS-56
chr8-100051216-INS-369
chr8-100078382-INS-77
chr8-100146496-INS-268
chr8-100190834-INS-339
chr8-100263364-INS-3434
chr8-100263573-INS-84
chr8-100289858-INS-72
chr8-100308746-INS-7463
chr8-100310164-INS-412
chr8-100318721-INS-597
chr8-100325767-INS-50
chr8-100330562-INS-51
chr8-100332444-INS-56
chr8-100337914-INS-74
chr8-100338036-INS-1007
chr8-100356701-INS-6175
chr8-100393592-INS-1220
chr8-100466210-INS-118

chr8-100509875-INS-56
chr8-100512353-INS-202
chr8-100518247-INS-51
chr8-100551942-INS-55
chr8-100562495-INS-6845
chr8-100583261-INS-70
chr8-100583481-INS-50
chr8-100607863-INS-1819
chr8-100615693-INS-60
chr8-100617384-INS-64
chr8-100636103-INS-6568
chr8-100653183-INS-6616
chr8-100701387-INS-56
chr8-100708182-INS-170
chr8-100708182-INS-50
chr8-100709206-INS-70
chr8-100719035-INS-10983
chr8-100730988-INS-1751
chr8-100737843-INS-2963
chr8-100760460-INS-390
chr8-100793680-INS-84
chr8-100798512-INS-72
chr8-100800196-INS-80
chr8-100807181-INS-237
chr8-100811173-INS-50
chr8-100813438-INS-82
chr8-100858558-INS-194
chr8-100876148-INS-174
chr8-100911961-INS-125
chr8-100918329-INS-90
chr8-101033459-INS-50
chr8-101100006-INS-186
chr8-101108684-INS-72
chr8-101152173-INS-569
chr8-101156581-INS-56
chr8-101190339-INS-86
chr8-101225071-INS-64
chr8-101288961-INS-1681
chr8-101293371-INS-6148
chr8-101300912-INS-65
chr8-101303618-INS-397
chr8-101323498-INS-1553
chr8-101341098-INS-50
chr8-101350301-INS-66
chr8-101356352-INS-4108
chr8-101366566-INS-7081
chr8-101369439-INS-54
chr8-101418480-INS-6225
chr8-101433667-INS-2913
chr8-101440596-INS-3707
chr8-101444576-INS-52
chr8-101461247-INS-273
chr8-101466964-INS-88
chr8-101529317-INS-50
chr8-101532796-INS-70
chr8-101536113-INS-60
chr8-101552677-INS-6400
chr8-101554679-INS-1122
chr8-101555804-INS-260
chr8-101604292-INS-50
chr8-101604603-INS-175
chr8-101630504-INS-6046
chr8-101667885-INS-187
chr8-101678217-INS-52
chr8-101697432-INS-58
chr8-101702017-INS-71
chr8-101702846-INS-536
chr8-101725123-INS-50
chr8-101766370-INS-52
chr8-101801098-INS-264
chr8-101855603-INS-496
chr8-101876854-INS-341
chr8-101876857-INS-100
chr8-101893344-INS-126
chr8-102201027-INS-3484
chr8-102208550-INS-561
chr8-102214127-INS-52

chr8-102216215-INS-6201
chr8-102229711-INS-6642
chr8-102266152-INS-117
chr8-102283470-INS-1995
chr8-102283474-INS-4353
chr8-102285307-INS-1881
chr8-102356979-INS-135
chr8-102386408-INS-130
chr8-102390358-INS-2580
chr8-102418015-INS-50
chr8-102437182-INS-135
chr8-102448425-INS-1000
chr8-102472737-INS-4502
chr8-102475150-INS-81
chr8-102499145-INS-50
chr8-102503238-INS-62
chr8-102548532-INS-105
chr8-102548866-INS-591
chr8-102553326-INS-56
chr8-102566222-INS-70
chr8-102609896-INS-22792
chr8-102610891-INS-307
chr8-102624287-INS-4587
chr8-102632716-INS-60
chr8-102634113-INS-97
chr8-102657805-INS-70
chr8-102663550-INS-2476
chr8-102686774-INS-121
chr8-102755499-INS-6582
chr8-102765462-INS-6576
chr8-102782001-INS-255
chr8-102782221-INS-426
chr8-102800519-INS-6512
chr8-102836282-INS-59
chr8-102888936-INS-202
chr8-102898886-INS-70
chr8-102903471-INS-58
chr8-102904052-INS-77
chr8-102915506-INS-52
chr8-102923045-INS-106
chr8-102923873-INS-52
chr8-102944243-INS-888
chr8-103001114-INS-50
chr8-103026149-INS-387
chr8-103051689-INS-369
chr8-103081624-INS-211
chr8-103087902-INS-150
chr8-103089300-INS-52
chr8-103091245-INS-104
chr8-103091313-INS-554
chr8-103091317-INS-130
chr8-103099320-INS-436
chr8-103102434-INS-52
chr8-103109190-INS-61
chr8-103141299-INS-62
chr8-103196581-INS-1462
chr8-103211165-INS-106
chr8-103212694-INS-50
chr8-103222514-INS-87
chr8-103226773-INS-470
chr8-103231548-INS-138
chr8-103254770-INS-93
chr8-103267799-INS-55
chr8-103268583-INS-103
chr8-103273343-INS-236
chr8-103288893-INS-51
chr8-103288929-INS-338
chr8-103300490-INS-58
chr8-103380424-INS-368
chr8-103380708-INS-1107
chr8-103410497-INS-60
chr8-103411369-INS-437
chr8-103570347-INS-1526
chr8-103586099-INS-50
chr8-103595002-INS-7198
chr8-103611939-INS-125
chr8-103620708-INS-519

chr8-103628994-INS-163
chr8-103638646-INS-59
chr8-103649328-INS-6840
chr8-103680791-INS-54
chr8-103700648-INS-84
chr8-103723182-INS-68
chr8-103755674-INS-2326
chr8-103761499-INS-2224
chr8-103819710-INS-404
chr8-103819746-INS-115
chr8-103824090-INS-249
chr8-103824106-INS-111
chr8-103834674-INS-6807
chr8-103923993-INS-598
chr8-103931654-INS-1113
chr8-103940281-INS-50
chr8-103941871-INS-64
chr8-103944348-INS-56
chr8-103949809-INS-1173
chr8-103952591-INS-1876
chr8-104008875-INS-100
chr8-104023012-INS-207
chr8-104339345-INS-141
chr8-104341883-INS-68
chr8-104349882-INS-55
chr8-104354869-INS-231
chr8-104354869-INS-64
chr8-104366288-INS-1504
chr8-104392444-INS-53
chr8-104409712-INS-60
chr8-104447460-INS-2939
chr8-104463070-INS-6354
chr8-104502350-INS-52
chr8-104528058-INS-60
chr8-104537997-INS-53
chr8-104569154-INS-57
chr8-104570645-INS-100
chr8-104594195-INS-344
chr8-104607396-INS-528
chr8-104643282-INS-50
chr8-104654627-INS-68
chr8-104657847-INS-59
chr8-104676940-INS-133
chr8-104684929-INS-239
chr8-104684930-INS-76
chr8-104685042-INS-77
chr8-104690419-INS-7097
chr8-104819893-INS-261
chr8-104828610-INS-861
chr8-104874647-INS-73
chr8-104874721-INS-519
chr8-104879137-INS-1492
chr8-104886298-INS-128
chr8-104886606-INS-54
chr8-104945371-INS-298
chr8-104950026-INS-64
chr8-104984318-INS-268
chr8-104991418-INS-99
chr8-105007368-INS-67
chr8-105030482-INS-72
chr8-105042297-INS-56
chr8-105043818-INS-6453
chr8-105052884-INS-51
chr8-105105234-INS-1573
chr8-105113316-INS-182
chr8-105128989-INS-64
chr8-105170840-INS-6460
chr8-105176920-INS-982
chr8-105200857-INS-1894
chr8-105245133-INS-220
chr8-105248043-INS-63
chr8-105284557-INS-50
chr8-105296697-INS-54
chr8-105303749-INS-669
chr8-105315804-INS-113
chr8-105360516-INS-79
chr8-105370286-INS-84

chr8-105370804-INS-58
chr8-105422668-INS-50
chr8-105439820-INS-500
chr8-105444499-INS-110
chr8-105444687-INS-190
chr8-105463411-INS-60
chr8-105498599-INS-1339
chr8-105646290-INS-7479
chr8-105701465-INS-352
chr8-105702108-INS-192
chr8-105705728-INS-6957
chr8-105715168-INS-60
chr8-105726802-INS-62
chr8-105746433-INS-445
chr8-105751479-INS-1193
chr8-105791910-INS-2800
chr8-105821326-INS-50
chr8-105828029-INS-4607
chr8-105843447-INS-57
chr8-105846283-INS-347
chr8-105887053-INS-70
chr8-105889560-INS-1172
chr8-105892348-INS-241
chr8-105901035-INS-65
chr8-105910377-INS-599
chr8-105936981-INS-60
chr8-105948308-INS-6475
chr8-105970856-INS-215
chr8-106011022-INS-166
chr8-106037892-INS-54
chr8-106047480-INS-810
chr8-106086891-INS-214
chr8-106117793-INS-216
chr8-106131145-INS-62
chr8-106136373-INS-197
chr8-106152472-INS-304
chr8-106162437-INS-78
chr8-106173181-INS-467
chr8-106242304-INS-56
chr8-106243560-INS-57
chr8-106267677-INS-171
chr8-106268227-INS-143
chr8-106285310-INS-232
chr8-106329201-INS-54
chr8-106386150-INS-62
chr8-106445531-INS-68
chr8-106455903-INS-6357
chr8-106457041-INS-206
chr8-106459303-INS-106
chr8-106475104-INS-52
chr8-106475731-INS-211
chr8-106482538-INS-278
chr8-106483281-INS-56
chr8-106514012-INS-159
chr8-106533962-INS-209
chr8-106540708-INS-56
chr8-106541411-INS-68
chr8-106543269-INS-438
chr8-106543891-INS-152
chr8-106550311-INS-626
chr8-106551844-INS-106
chr8-106592368-INS-108
chr8-106626213-INS-172
chr8-106662089-INS-8747
chr8-106694520-INS-73
chr8-106751762-INS-5550
chr8-106753314-INS-198
chr8-106759801-INS-66
chr8-106760231-INS-183
chr8-106798897-INS-221
chr8-106842960-INS-203
chr8-106866396-INS-184
chr8-106917310-INS-1890
chr8-106948242-INS-80
chr8-106959288-INS-71
chr8-106966166-INS-3740
chr8-107073525-INS-422

chr8-107115823-INS-56
chr8-107122501-INS-133
chr8-107168827-INS-69
chr8-107197485-INS-118
chr8-107200295-INS-60
chr8-107222749-INS-2679
chr8-107257680-INS-400
chr8-107258309-INS-209
chr8-107294174-INS-77
chr8-107294601-INS-445
chr8-107320447-INS-199
chr8-107350060-INS-50
chr8-107398782-INS-4665
chr8-107411366-INS-199
chr8-107412189-INS-169
chr8-107425100-INS-779
chr8-107425229-INS-50
chr8-107428598-INS-52
chr8-107489379-INS-213
chr8-107492583-INS-494
chr8-107492937-INS-1145
chr8-107492995-INS-287
chr8-107493003-INS-103
chr8-107510077-INS-105
chr8-107513218-INS-1219
chr8-107514385-INS-58
chr8-107520311-INS-50
chr8-107520727-INS-537
chr8-107536648-INS-1024
chr8-107554764-INS-369
chr8-107556671-INS-187
chr8-107570322-INS-474
chr8-107590284-INS-308
chr8-107593098-INS-433
chr8-107645486-INS-123
chr8-107675040-INS-204
chr8-107693795-INS-92
chr8-107695933-INS-110
chr8-107720332-INS-209
chr8-107722902-INS-161
chr8-107725353-INS-101
chr8-107743815-INS-183
chr8-107749757-INS-200
chr8-107757542-INS-246
chr8-107765842-INS-172
chr8-107772536-INS-179
chr8-107789140-INS-96
chr8-107846863-INS-163
chr8-107861656-INS-62
chr8-107883262-INS-710
chr8-107889826-INS-87
chr8-107953381-INS-162
chr8-107962123-INS-86
chr8-107968557-INS-179
chr8-107980342-INS-228
chr8-107988420-INS-53
chr8-107993293-INS-50
chr8-107998265-INS-6468
chr8-108012163-INS-63
chr8-108123972-INS-66
chr8-108143861-INS-84
chr8-108143992-INS-226
chr8-108144310-INS-59
chr8-108156871-INS-3488
chr8-108188986-INS-55
chr8-108206682-INS-50
chr8-108209813-INS-1634
chr8-108252460-INS-185
chr8-108258747-INS-1257
chr8-108258768-INS-52
chr8-108268395-INS-76
chr8-108269335-INS-115
chr8-108318355-INS-408
chr8-108320926-INS-85
chr8-108355763-INS-133
chr8-108360160-INS-155
chr8-108360292-INS-626

chr8-108385953-INS-141
chr8-108397307-INS-208
chr8-108410160-INS-73
chr8-108447319-INS-60
chr8-108477986-INS-4725
chr8-108495282-INS-52
chr8-108495369-INS-222
chr8-108495386-INS-69
chr8-108502678-INS-53
chr8-108505401-INS-221
chr8-108536017-INS-375
chr8-108604939-INS-60
chr8-108616143-INS-460
chr8-108629937-INS-6793
chr8-108644051-INS-50
chr8-108692314-INS-647
chr8-108774951-INS-406
chr8-108774955-INS-83
chr8-108781182-INS-577
chr8-108810252-INS-50
chr8-108855608-INS-1556
chr8-108868724-INS-53
chr8-108868929-INS-129
chr8-108911568-INS-121
chr8-108938620-INS-114
chr8-108972908-INS-98
chr8-108977358-INS-3008
chr8-109088469-INS-50
chr8-109288584-INS-50
chr8-109298565-INS-2354
chr8-109336647-INS-174
chr8-109408580-INS-145
chr8-109411558-INS-52
chr8-109424849-INS-52
chr8-109445873-INS-64
chr8-109498973-INS-108
chr8-109509145-INS-102
chr8-109539987-INS-76
chr8-109540271-INS-170
chr8-109540271-INS-442
chr8-109546377-INS-54
chr8-109640251-INS-57
chr8-109738978-INS-58
chr8-109752267-INS-136
chr8-109760400-INS-270
chr8-109775052-INS-182
chr8-109775241-INS-774
chr8-109777794-INS-6264
chr8-109787690-INS-2078
chr8-109806318-INS-90
chr8-109834337-INS-50
chr8-109895032-INS-199
chr8-109923352-INS-112
chr8-109982489-INS-64
chr8-109986769-INS-199
chr8-109987268-INS-4259
chr8-110001620-INS-105
chr8-110006468-INS-346
chr8-110007338-INS-78
chr8-110039261-INS-82
chr8-110043490-INS-672
chr8-110077280-INS-527
chr8-110077292-INS-156
chr8-110133897-INS-55
chr8-110140906-INS-70
chr8-110157426-INS-101
chr8-110157572-INS-515
chr8-110173106-INS-60
chr8-110192529-INS-50
chr8-110253899-INS-64
chr8-110287708-INS-210
chr8-110363431-INS-109
chr8-110375964-INS-50
chr8-110389501-INS-56
chr8-110389695-INS-305
chr8-110389730-INS-110
chr8-110422184-INS-2103

chr8-110422367-INS-377
chr8-110476655-INS-2108
chr8-110509770-INS-219
chr8-110513104-INS-173
chr8-110526278-INS-50
chr8-110561554-INS-722
chr8-110576006-INS-84
chr8-110612748-INS-63
chr8-110613094-INS-105
chr8-110653178-INS-60
chr8-110732793-INS-74
chr8-110748727-INS-62
chr8-110751926-INS-448
chr8-110853406-INS-193
chr8-110863225-INS-3320
chr8-110873436-INS-64
chr8-110873633-INS-62
chr8-110884593-INS-54
chr8-110919632-INS-2621
chr8-110919632-INS-417
chr8-110932126-INS-55
chr8-110943261-INS-4927
chr8-110966398-INS-56
chr8-110979052-INS-122
chr8-110979052-INS-57
chr8-110983216-INS-541
chr8-110988795-INS-172
chr8-111000670-INS-1333
chr8-111024052-INS-52
chr8-111027934-INS-63
chr8-111030952-INS-96
chr8-111114058-INS-105
chr8-111128613-INS-85
chr8-111182673-INS-136
chr8-111213581-INS-498
chr8-111248176-INS-76
chr8-111256066-INS-193
chr8-111279573-INS-65
chr8-111298039-INS-111
chr8-111461366-INS-96
chr8-111474552-INS-190
chr8-111488558-INS-119
chr8-111499436-INS-202
chr8-111522219-INS-203
chr8-111522685-INS-64
chr8-111528591-INS-406
chr8-111553653-INS-50
chr8-111582929-INS-59
chr8-111583653-INS-204
chr8-111694487-INS-217
chr8-111717482-INS-62
chr8-111739457-INS-6245
chr8-111744766-INS-4167
chr8-111777663-INS-121
chr8-111829448-INS-154
chr8-111885864-INS-108
chr8-111913529-INS-223
chr8-111927290-INS-450
chr8-111980218-INS-274
chr8-111982696-INS-201
chr8-111983285-INS-494
chr8-112036760-INS-68
chr8-112040427-INS-80
chr8-112111955-INS-220
chr8-112114201-INS-1903
chr8-112122056-INS-54
chr8-112124128-INS-516
chr8-112151611-INS-68
chr8-112167147-INS-232
chr8-112178757-INS-1394
chr8-112191531-INS-72
chr8-112210181-INS-240
chr8-112213421-INS-180
chr8-112224948-INS-140
chr8-112289391-INS-135
chr8-112289501-INS-472
chr8-112291857-INS-52

chr8-112292535-INS-94
chr8-112321769-INS-182
chr8-112341419-INS-56
chr8-112341525-INS-144
chr8-112342349-INS-130
chr8-112358714-INS-120
chr8-112358811-INS-61
chr8-112368118-INS-131
chr8-112373858-INS-225
chr8-112373944-INS-1892
chr8-112376617-INS-8201
chr8-112405879-INS-196
chr8-112406568-INS-52
chr8-112414820-INS-126
chr8-112474156-INS-147
chr8-112477286-INS-617
chr8-112480094-INS-202
chr8-112484903-INS-244
chr8-112488369-INS-220
chr8-112491758-INS-141
chr8-112493320-INS-305
chr8-112510580-INS-195
chr8-112511231-INS-193
chr8-112524856-INS-52
chr8-112570306-INS-217
chr8-112573794-INS-127
chr8-112577902-INS-222
chr8-112586118-INS-161
chr8-112604273-INS-67
chr8-112613746-INS-185
chr8-112614516-INS-101
chr8-112626731-INS-52
chr8-112634159-INS-62
chr8-112635108-INS-88
chr8-112647063-INS-218
chr8-112663701-INS-124
chr8-112672667-INS-94
chr8-112677356-INS-56
chr8-112680533-INS-212
chr8-112681391-INS-122
chr8-112692197-INS-140
chr8-112728606-INS-77
chr8-112744454-INS-214
chr8-112745970-INS-139
chr8-112756935-INS-210
chr8-112757763-INS-78
chr8-112760740-INS-54
chr8-112765413-INS-205
chr8-112772097-INS-208
chr8-112772569-INS-50
chr8-112776941-INS-249
chr8-112778627-INS-176
chr8-112779224-INS-62
chr8-112780454-INS-344
chr8-112781140-INS-197
chr8-112781945-INS-172
chr8-112790402-INS-204
chr8-112814833-INS-57
chr8-112815523-INS-116
chr8-112822031-INS-56
chr8-112827636-INS-3024
chr8-112829314-INS-1108
chr8-112829322-INS-344
chr8-112842988-INS-76
chr8-112846430-INS-87
chr8-112849352-INS-122
chr8-112849886-INS-287
chr8-112862628-INS-52
chr8-112871480-INS-1284
chr8-112894648-INS-60
chr8-112933526-INS-365
chr8-112955621-INS-55
chr8-112955690-INS-416
chr8-112967865-INS-52
chr8-113009756-INS-50
chr8-113042202-INS-50
chr8-113046294-INS-52

chr8-113052708-INS-152
chr8-113052720-INS-351
chr8-113065835-INS-631
chr8-113066172-INS-1721
chr8-113089063-INS-342
chr8-113089063-INS-3583
chr8-113092887-INS-109
chr8-113115058-INS-6434
chr8-113142767-INS-67
chr8-113146970-INS-55
chr8-113147944-INS-50
chr8-113158116-INS-1081
chr8-113185685-INS-58
chr8-113235194-INS-54
chr8-113239308-INS-50
chr8-113258894-INS-88
chr8-113261316-INS-1864
chr8-113292921-INS-56
chr8-113294194-INS-100
chr8-113320117-INS-401
chr8-113339871-INS-499
chr8-113339904-INS-176
chr8-113379012-INS-537
chr8-113468724-INS-6823
chr8-113480943-INS-72
chr8-113512382-INS-52
chr8-113545685-INS-64
chr8-113545994-INS-436
chr8-113578257-INS-50
chr8-113588164-INS-211
chr8-113599167-INS-105
chr8-113661324-INS-53
chr8-113686923-INS-66
chr8-113687860-INS-5647
chr8-113696923-INS-2415
chr8-113713855-INS-78
chr8-113716841-INS-72
chr8-113725819-INS-147
chr8-113729364-INS-1175
chr8-113740686-INS-647
chr8-113774193-INS-114
chr8-113786063-INS-196
chr8-113788222-INS-68
chr8-113806167-INS-212
chr8-113807703-INS-6500
chr8-113821571-INS-455
chr8-113830911-INS-64
chr8-113835653-INS-6092
chr8-113846385-INS-171
chr8-113914390-INS-52
chr8-113937335-INS-100
chr8-114080758-INS-170
chr8-114083303-INS-337
chr8-114095180-INS-205
chr8-114100582-INS-61
chr8-114130794-INS-63
chr8-114140552-INS-243
chr8-114170027-INS-68
chr8-114172688-INS-174
chr8-114172688-INS-696
chr8-114174493-INS-6398
chr8-114177643-INS-208
chr8-114196863-INS-108
chr8-114212804-INS-103
chr8-114218396-INS-660
chr8-114245987-INS-7439
chr8-114250499-INS-1644
chr8-114260324-INS-166
chr8-114306275-INS-203
chr8-114320731-INS-76
chr8-114320766-INS-219
chr8-114327937-INS-277
chr8-114352588-INS-71
chr8-114353351-INS-50
chr8-114453348-INS-67
chr8-114475790-INS-64
chr8-114521053-INS-98

chr8-114525924-INS-358
chr8-114544942-INS-56
chr8-114569042-INS-1424
chr8-114586394-INS-57
chr8-114606039-INS-1380
chr8-114610265-INS-80
chr8-114658314-INS-64
chr8-114731629-INS-92
chr8-114748794-INS-2861
chr8-114756721-INS-307
chr8-114757184-INS-84
chr8-114768907-INS-56
chr8-114772497-INS-123
chr8-114789705-INS-56
chr8-114819030-INS-144
chr8-114872582-INS-51
chr8-114881658-INS-56
chr8-114881786-INS-147
chr8-114882903-INS-93
chr8-114885947-INS-219
chr8-114887284-INS-1202
chr8-114911290-INS-320
chr8-114925319-INS-102
chr8-114933090-INS-72
chr8-114974365-INS-56
chr8-114987338-INS-1892
chr8-114987653-INS-55
chr8-114988626-INS-929
chr8-115000668-INS-104
chr8-115015696-INS-195
chr8-115072533-INS-58
chr8-115083346-INS-55
chr8-115086720-INS-109
chr8-115126436-INS-538
chr8-115126804-INS-52
chr8-115142286-INS-4557
chr8-115150897-INS-106
chr8-115151212-INS-360
chr8-115164279-INS-112
chr8-115164577-INS-76
chr8-115184049-INS-442
chr8-115194383-INS-151
chr8-115243505-INS-163
chr8-115247699-INS-84
chr8-115330171-INS-52
chr8-115330475-INS-70
chr8-115348380-INS-1097
chr8-115354384-INS-50
chr8-115396569-INS-57
chr8-115462934-INS-77
chr8-115488525-INS-126
chr8-115488525-INS-302
chr8-115558012-INS-64
chr8-115568656-INS-206
chr8-115568669-INS-81
chr8-115570046-INS-52
chr8-115570581-INS-51
chr8-115590858-INS-182
chr8-115590858-INS-90
chr8-115637574-INS-52
chr8-115638615-INS-8748
chr8-115649634-INS-260
chr8-115658062-INS-557
chr8-115662729-INS-1299
chr8-115690589-INS-60
chr8-115730602-INS-224
chr8-115730680-INS-52
chr8-115730846-INS-50
chr8-115761004-INS-106
chr8-115774098-INS-482
chr8-115775685-INS-143
chr8-115783380-INS-63
chr8-115801889-INS-64
chr8-115802101-INS-60
chr8-115816470-INS-60
chr8-115817909-INS-70
chr8-115820818-INS-206

chr8-115835175-INS-1831
chr8-115898885-INS-508
chr8-115907400-INS-55
chr8-115917240-INS-74
chr8-115978011-INS-78
chr8-115978321-INS-210
chr8-115983882-INS-182
chr8-115986283-INS-76
chr8-115988653-INS-62
chr8-116028013-INS-140
chr8-116104591-INS-58
chr8-116113629-INS-74
chr8-116167595-INS-52
chr8-116243450-INS-88
chr8-116250555-INS-123
chr8-116274339-INS-50
chr8-116274494-INS-199
chr8-116280329-INS-254
chr8-116283488-INS-56
chr8-116283625-INS-195
chr8-116292018-INS-94
chr8-116476782-INS-67
chr8-116501054-INS-154
chr8-116501056-INS-60
chr8-116578250-INS-212
chr8-116620986-INS-185
chr8-116621907-INS-124
chr8-116642455-INS-1063
chr8-116674439-INS-114
chr8-116699569-INS-175
chr8-116717940-INS-72
chr8-116733933-INS-375
chr8-116734139-INS-368
chr8-116765600-INS-450
chr8-116774424-INS-1890
chr8-116784055-INS-183
chr8-116808390-INS-213
chr8-116843794-INS-282
chr8-116888807-INS-134
chr8-116892181-INS-61
chr8-117028279-INS-627
chr8-117034464-INS-452
chr8-117036306-INS-537
chr8-117052522-INS-50
chr8-117055355-INS-96
chr8-117055779-INS-68
chr8-117093075-INS-177
chr8-117112901-INS-7217
chr8-117171579-INS-387
chr8-117254218-INS-76
chr8-117261716-INS-590
chr8-117263201-INS-212
chr8-117264109-INS-322
chr8-117264165-INS-95
chr8-117268452-INS-211
chr8-117270974-INS-51
chr8-117273185-INS-51
chr8-117303199-INS-146
chr8-117317422-INS-515
chr8-117327399-INS-140
chr8-117327854-INS-71
chr8-117392690-INS-949
chr8-117394892-INS-58
chr8-117410531-INS-208
chr8-117416077-INS-56
chr8-117421591-INS-203
chr8-117465686-INS-58
chr8-117534877-INS-199
chr8-117540078-INS-315
chr8-117541853-INS-78
chr8-117559850-INS-160
chr8-117567917-INS-54
chr8-117599041-INS-5169
chr8-117609253-INS-60
chr8-117610466-INS-1134
chr8-117618810-INS-147
chr8-117635149-INS-178

chr8-117635190-INS-498
chr8-117720697-INS-112
chr8-117721431-INS-163
chr8-117728973-INS-284
chr8-117747510-INS-69
chr8-117748052-INS-997
chr8-117766722-INS-180
chr8-117778108-INS-5527
chr8-117783688-INS-84
chr8-117804871-INS-200
chr8-117814901-INS-50
chr8-117821450-INS-56
chr8-117857308-INS-83
chr8-117857465-INS-343
chr8-117946015-INS-64
chr8-117963401-INS-229
chr8-117965342-INS-53
chr8-117966952-INS-57
chr8-117971169-INS-216
chr8-118002014-INS-51
chr8-118021607-INS-54
chr8-118029700-INS-160
chr8-118032812-INS-80
chr8-118064939-INS-80
chr8-118070230-INS-1321
chr8-118076893-INS-1089
chr8-118128353-INS-105
chr8-118136848-INS-106
chr8-118152277-INS-50
chr8-118195774-INS-292
chr8-118196089-INS-60
chr8-118203811-INS-51
chr8-118203833-INS-53
chr8-118234523-INS-68
chr8-118245621-INS-78
chr8-118254972-INS-50
chr8-118257118-INS-62
chr8-118260748-INS-139
chr8-118262872-INS-224
chr8-118282633-INS-98
chr8-118283365-INS-2315
chr8-118299092-INS-90
chr8-118317441-INS-50
chr8-118336697-INS-155
chr8-118349207-INS-52
chr8-118435131-INS-400
chr8-118444431-INS-1361
chr8-118471136-INS-62
chr8-118550832-INS-50
chr8-118561185-INS-185
chr8-118664799-INS-2206
chr8-118665007-INS-195
chr8-118682046-INS-50
chr8-118686979-INS-50
chr8-118695955-INS-73
chr8-118740682-INS-1767
chr8-118770273-INS-102
chr8-118782180-INS-50
chr8-118786329-INS-55
chr8-118787166-INS-54
chr8-118799109-INS-92
chr8-118799784-INS-170
chr8-118830533-INS-1724
chr8-118920085-INS-117
chr8-118920085-INS-58
chr8-118923839-INS-80
chr8-118940066-INS-1329
chr8-118949597-INS-79
chr8-118949947-INS-50
chr8-118989002-INS-66
chr8-118993860-INS-6620
chr8-119076197-INS-701
chr8-119102548-INS-178
chr8-119108290-INS-84
chr8-119113831-INS-7253
chr8-119138971-INS-52
chr8-119139751-INS-62

chr8-119141380-INS-50
chr8-119157760-INS-76
chr8-119163024-INS-69
chr8-119184495-INS-52
chr8-119197751-INS-59
chr8-119256945-INS-59
chr8-119274631-INS-91
chr8-119340685-INS-900
chr8-119361344-INS-401
chr8-119364228-INS-7174
chr8-119405051-INS-89
chr8-119418243-INS-230
chr8-119454146-INS-362
chr8-119540984-INS-230
chr8-119541242-INS-70
chr8-119582337-INS-86
chr8-119599876-INS-520
chr8-119608683-INS-1551
chr8-119623525-INS-54
chr8-119648132-INS-167
chr8-119649879-INS-112
chr8-119651185-INS-80
chr8-119691105-INS-109
chr8-119723301-INS-68
chr8-119733093-INS-157
chr8-119733174-INS-100
chr8-119784919-INS-50
chr8-119785515-INS-1453
chr8-119789512-INS-67
chr8-119869354-INS-51
chr8-119879770-INS-90
chr8-119882849-INS-201
chr8-119897147-INS-60
chr8-119900359-INS-2170
chr8-119988006-INS-90
chr8-119991311-INS-134
chr8-120031770-INS-450
chr8-120039832-INS-167
chr8-120039912-INS-80
chr8-120093614-INS-149
chr8-120102524-INS-203
chr8-120111906-INS-60
chr8-120132134-INS-59
chr8-120143581-INS-55
chr8-120187925-INS-93
chr8-120216763-INS-99
chr8-120250051-INS-97
chr8-120251841-INS-236
chr8-120338375-INS-350
chr8-120346047-INS-385
chr8-120361751-INS-2978
chr8-120367370-INS-387
chr8-120377985-INS-107
chr8-120400570-INS-145
chr8-120411205-INS-324
chr8-120411785-INS-204
chr8-120422301-INS-52
chr8-120457679-INS-51
chr8-120491557-INS-69
chr8-120496798-INS-291
chr8-120526163-INS-50
chr8-120528286-INS-368
chr8-120528411-INS-943
chr8-120530498-INS-183
chr8-120532829-INS-429
chr8-120532831-INS-960
chr8-120568686-INS-64
chr8-120568786-INS-140
chr8-120569473-INS-153
chr8-120574854-INS-8297
chr8-120578391-INS-63
chr8-120578640-INS-237
chr8-120585569-INS-125
chr8-120624405-INS-84
chr8-120626800-INS-62
chr8-120634203-INS-54
chr8-120636162-INS-52

chr8-120644553-INS-72
chr8-120688125-INS-64
chr8-120713894-INS-312
chr8-120793049-INS-206
chr8-120793543-INS-212
chr8-120815064-INS-280
chr8-120822067-INS-168
chr8-120822067-INS-445
chr8-120847666-INS-53
chr8-120852373-INS-90
chr8-120876141-INS-5037
chr8-120900657-INS-629
chr8-120900735-INS-231
chr8-120909779-INS-136
chr8-120909995-INS-216
chr8-120927853-INS-86
chr8-120973081-INS-76
chr8-120982580-INS-184
chr8-120986871-INS-259
chr8-120989674-INS-79
chr8-121010362-INS-60
chr8-121012179-INS-323
chr8-121012191-INS-75
chr8-121038291-INS-84
chr8-121060266-INS-188
chr8-121063565-INS-302
chr8-121063635-INS-60
chr8-121089462-INS-59
chr8-121257779-INS-326
chr8-121317320-INS-1493
chr8-121320119-INS-249
chr8-121331108-INS-3537
chr8-121352315-INS-142
chr8-121377033-INS-103
chr8-121426703-INS-559
chr8-121457711-INS-100
chr8-121492842-INS-373
chr8-121498443-INS-104
chr8-121499034-INS-81
chr8-121581800-INS-275
chr8-121633310-INS-340
chr8-121671572-INS-64
chr8-121700788-INS-90
chr8-121746248-INS-95
chr8-121821645-INS-160
chr8-121837245-INS-103
chr8-121866021-INS-129
chr8-121866021-INS-51
chr8-121908786-INS-807
chr8-121933351-INS-140
chr8-121972575-INS-216
chr8-121980539-INS-129
chr8-121981382-INS-440
chr8-122000318-INS-64
chr8-122007994-INS-118
chr8-122035359-INS-68
chr8-122045493-INS-86
chr8-122045547-INS-464
chr8-122066256-INS-448
chr8-122079392-INS-132
chr8-122090032-INS-58
chr8-122190156-INS-80
chr8-122190480-INS-80
chr8-122198744-INS-1896
chr8-122199753-INS-156
chr8-122206330-INS-55
chr8-122237902-INS-52
chr8-122238232-INS-116
chr8-122241660-INS-123
chr8-122247754-INS-57
chr8-122251511-INS-72
chr8-122256416-INS-98
chr8-122262002-INS-110
chr8-122262706-INS-558
chr8-122262753-INS-232
chr8-122265049-INS-249
chr8-122265448-INS-557

chr8-122273650-INS-181
chr8-122274600-INS-62
chr8-122274890-INS-166
chr8-122275819-INS-213
chr8-122329921-INS-117
chr8-122362007-INS-626
chr8-122368596-INS-60
chr8-122382752-INS-72
chr8-122383035-INS-197
chr8-122404110-INS-66
chr8-122405414-INS-71
chr8-122406244-INS-307
chr8-122406294-INS-66
chr8-122406622-INS-234
chr8-122413600-INS-61
chr8-122414040-INS-84
chr8-122438699-INS-154
chr8-122530145-INS-52
chr8-122591111-INS-57
chr8-122632177-INS-1154
chr8-122648148-INS-182
chr8-122655619-INS-324
chr8-122699904-INS-1498
chr8-122706765-INS-120
chr8-122711886-INS-560
chr8-122711934-INS-105
chr8-122736882-INS-175
chr8-122745246-INS-225
chr8-122750186-INS-578
chr8-122787754-INS-192
chr8-122805041-INS-56
chr8-122809698-INS-139
chr8-122811153-INS-96
chr8-122847078-INS-64
chr8-122857096-INS-70
chr8-122865838-INS-146
chr8-122893124-INS-111
chr8-122893463-INS-322
chr8-122895205-INS-164
chr8-122901894-INS-68
chr8-122909879-INS-68
chr8-122912416-INS-63
chr8-122968673-INS-166
chr8-122983664-INS-128
chr8-123022516-INS-1370
chr8-123022545-INS-520
chr8-123044388-INS-64
chr8-123058660-INS-91
chr8-123080366-INS-212
chr8-123095338-INS-3689
chr8-123231781-INS-72
chr8-123281148-INS-62
chr8-123457880-INS-223
chr8-123510059-INS-50
chr8-123527802-INS-58
chr8-123529101-INS-175
chr8-123536579-INS-970
chr8-123557568-INS-111
chr8-123566444-INS-50
chr8-123602618-INS-86
chr8-123602834-INS-204
chr8-123639767-INS-90
chr8-123677210-INS-5499
chr8-123730934-INS-1160
chr8-123739594-INS-237
chr8-123741351-INS-870
chr8-123760796-INS-124
chr8-123760796-INS-558
chr8-123780027-INS-311
chr8-123780123-INS-168
chr8-123788696-INS-750
chr8-123813964-INS-55
chr8-123824847-INS-204
chr8-123841874-INS-156
chr8-123842248-INS-238
chr8-123851622-INS-86
chr8-123851671-INS-190

chr8-123868212-INS-163
chr8-123879364-INS-63
chr8-123902404-INS-205
chr8-123903211-INS-94
chr8-123943561-INS-232
chr8-123944253-INS-281
chr8-123944920-INS-197
chr8-123944959-INS-97
chr8-123996524-INS-91
chr8-124003040-INS-270
chr8-124004136-INS-275
chr8-124005927-INS-1135
chr8-124019103-INS-82
chr8-124023173-INS-205
chr8-124026476-INS-344
chr8-124026601-INS-61
chr8-124043446-INS-50
chr8-124046338-INS-184
chr8-124088617-INS-270
chr8-124127321-INS-58
chr8-124206426-INS-1447
chr8-124240690-INS-122
chr8-124247231-INS-200
chr8-124247231-INS-50
chr8-124253054-INS-55
chr8-124351781-INS-1897
chr8-124367201-INS-2800
chr8-124376186-INS-50
chr8-124399774-INS-3070
chr8-124401566-INS-57
chr8-124405588-INS-74
chr8-124410793-INS-68
chr8-124412092-INS-4835
chr8-124424708-INS-714
chr8-124485598-INS-66
chr8-124511010-INS-143
chr8-124519233-INS-70
chr8-124528453-INS-7195
chr8-124529825-INS-603
chr8-124565927-INS-50
chr8-124593560-INS-70
chr8-124594308-INS-83
chr8-124600414-INS-118
chr8-124603306-INS-52
chr8-124649743-INS-683
chr8-124674577-INS-205
chr8-124680653-INS-190
chr8-124681904-INS-55
chr8-124689281-INS-244
chr8-124690773-INS-429
chr8-124716059-INS-144
chr8-124717167-INS-245
chr8-124728616-INS-586
chr8-124728670-INS-181
chr8-124754408-INS-211
chr8-124762643-INS-250
chr8-124771498-INS-186
chr8-124830749-INS-51
chr8-124838057-INS-50
chr8-124878601-INS-51
chr8-124886132-INS-132
chr8-124887906-INS-66
chr8-124897761-INS-281
chr8-124904154-INS-928
chr8-124956495-INS-52
chr8-125007534-INS-240
chr8-125063685-INS-235
chr8-125123444-INS-335
chr8-125133411-INS-71
chr8-125136872-INS-60
chr8-125149518-INS-90
chr8-125149812-INS-150
chr8-125150063-INS-267
chr8-125153158-INS-182
chr8-125177592-INS-191
chr8-125198370-INS-320
chr8-125201865-INS-162

chr8-125251382-INS-99
chr8-125262977-INS-1064
chr8-125262977-INS-224
chr8-125299219-INS-61
chr8-125299581-INS-560
chr8-125323197-INS-50
chr8-125330188-INS-52
chr8-125384117-INS-711
chr8-125384218-INS-84
chr8-125384789-INS-199
chr8-125393754-INS-56
chr8-125402303-INS-5030
chr8-125417832-INS-59
chr8-125442111-INS-63
chr8-125449391-INS-132
chr8-125462297-INS-60
chr8-125463174-INS-64
chr8-125485046-INS-157
chr8-125486944-INS-3203
chr8-125528595-INS-372
chr8-125532717-INS-52
chr8-125545412-INS-70
chr8-125546644-INS-50
chr8-125561688-INS-83
chr8-125566344-INS-84
chr8-125601959-INS-6464
chr8-125709908-INS-54
chr8-125711705-INS-95
chr8-125712754-INS-283
chr8-125732948-INS-1889
chr8-125732948-INS-231
chr8-125732959-INS-112
chr8-125733320-INS-55
chr8-125738146-INS-202
chr8-125774602-INS-55
chr8-125804637-INS-98
chr8-125804849-INS-843
chr8-125811487-INS-221
chr8-125819968-INS-164
chr8-125868861-INS-3406
chr8-125889091-INS-76
chr8-125897312-INS-52
chr8-125897618-INS-84
chr8-125897662-INS-1652
chr8-125898127-INS-1568
chr8-125907745-INS-50
chr8-125909560-INS-50
chr8-125930936-INS-63
chr8-125974692-INS-348
chr8-125977119-INS-53
chr8-125992468-INS-8459
chr8-126028046-INS-96
chr8-126080588-INS-50
chr8-126088916-INS-65
chr8-126098977-INS-106
chr8-126098977-INS-402
chr8-126109098-INS-124
chr8-126113101-INS-50
chr8-126148270-INS-150
chr8-126156856-INS-94
chr8-126156928-INS-1080
chr8-126156938-INS-2857
chr8-126163418-INS-67
chr8-126163488-INS-198
chr8-126204149-INS-643
chr8-126204973-INS-590
chr8-126215326-INS-163
chr8-126256671-INS-95
chr8-126264487-INS-432
chr8-126264721-INS-660
chr8-126298673-INS-54
chr8-126326450-INS-86
chr8-126419992-INS-508
chr8-126447794-INS-97
chr8-126478712-INS-621
chr8-126491723-INS-50
chr8-126510556-INS-83

chr8-126536066-INS-426
chr8-126592041-INS-50
chr8-126599555-INS-52
chr8-126609985-INS-168
chr8-126626593-INS-1492
chr8-126629038-INS-1121
chr8-126630181-INS-160
chr8-126669613-INS-64
chr8-126747590-INS-98
chr8-126772746-INS-50
chr8-126809306-INS-406
chr8-126813540-INS-1063
chr8-126819268-INS-100
chr8-126847867-INS-51
chr8-126868713-INS-150
chr8-126889114-INS-110
chr8-126908879-INS-561
chr8-126995926-INS-53
chr8-127010188-INS-88
chr8-127022857-INS-50
chr8-127025914-INS-66
chr8-127112069-INS-62
chr8-127166720-INS-90
chr8-127186231-INS-9561
chr8-127206776-INS-62
chr8-127212306-INS-7158
chr8-127214103-INS-68
chr8-127247986-INS-249
chr8-127259072-INS-767
chr8-127302499-INS-137
chr8-127325977-INS-56
chr8-127369982-INS-140
chr8-127376885-INS-613
chr8-127517469-INS-86
chr8-127550643-INS-706
chr8-127583207-INS-1034
chr8-127583239-INS-448
chr8-127605209-INS-515
chr8-127632678-INS-87
chr8-127691559-INS-342
chr8-127696193-INS-197
chr8-127709247-INS-738
chr8-127719946-INS-747
chr8-127720240-INS-204
chr8-127720957-INS-233
chr8-127742934-INS-7129
chr8-127744354-INS-212
chr8-127748171-INS-2804
chr8-127752895-INS-214
chr8-127762471-INS-2803
chr8-127764281-INS-53
chr8-127773869-INS-198
chr8-127773874-INS-416
chr8-127842482-INS-58
chr8-127846981-INS-364
chr8-127859555-INS-390
chr8-127872916-INS-54
chr8-127943948-INS-6738
chr8-127945045-INS-51
chr8-127950641-INS-60
chr8-127961537-INS-97
chr8-127961544-INS-2210
chr8-127972956-INS-378
chr8-127983848-INS-460
chr8-127989646-INS-78
chr8-128084624-INS-50
chr8-128119945-INS-52
chr8-128240120-INS-104
chr8-128240159-INS-50
chr8-128260538-INS-50
chr8-128263741-INS-132
chr8-128293001-INS-52
chr8-128309973-INS-160
chr8-128316997-INS-54
chr8-128334392-INS-59
chr8-128370733-INS-64
chr8-128372525-INS-2329

chr8-128373340-INS-2139
chr8-128378495-INS-862
chr8-128378583-INS-64
chr8-128378919-INS-89
chr8-128378986-INS-273
chr8-128395103-INS-68
chr8-128417924-INS-1739
chr8-128431848-INS-6394
chr8-128445809-INS-82
chr8-128450241-INS-5382
chr8-128462727-INS-499
chr8-128462727-INS-5372
chr8-128523703-INS-70
chr8-128557654-INS-53
chr8-128561368-INS-162
chr8-128567093-INS-87
chr8-128574468-INS-153
chr8-128585282-INS-191
chr8-128588505-INS-100
chr8-128601934-INS-120
chr8-128627680-INS-3701
chr8-128659029-INS-56
chr8-128659825-INS-4189
chr8-128671119-INS-103
chr8-128687784-INS-52
chr8-128695521-INS-2735
chr8-128701733-INS-56
chr8-128735360-INS-116
chr8-128742740-INS-107
chr8-128742774-INS-266
chr8-128814058-INS-88
chr8-128832663-INS-58
chr8-128842484-INS-50
chr8-128856127-INS-50
chr8-128874061-INS-6605
chr8-128903266-INS-3323
chr8-128903266-INS-661
chr8-128937154-INS-59
chr8-128992546-INS-54
chr8-128996608-INS-50
chr8-129038006-INS-343
chr8-129040448-INS-50
chr8-129045264-INS-12870
chr8-129048124-INS-6214
chr8-129070203-INS-53
chr8-129116610-INS-398
chr8-129186714-INS-66
chr8-129222713-INS-58
chr8-129299042-INS-6182
chr8-129331931-INS-80
chr8-129393911-INS-181
chr8-129397658-INS-55
chr8-129407729-INS-1333
chr8-129430827-INS-1304
chr8-129502941-INS-2260
chr8-129517426-INS-734
chr8-129554939-INS-76
chr8-129620225-INS-3030
chr8-129631559-INS-152
chr8-129675537-INS-195
chr8-129684182-INS-2592
chr8-129690580-INS-1492
chr8-129701358-INS-155
chr8-129716621-INS-60
chr8-129784699-INS-6424
chr8-129802881-INS-1453
chr8-129924028-INS-121
chr8-129957788-INS-6364
chr8-129963559-INS-317
chr9-3003849-INS-60
chr9-3005905-INS-58
chr9-3010656-INS-60
chr9-3011300-INS-60
chr9-3015046-INS-60
chr9-3016800-INS-60
chr9-3017677-INS-60
chr9-3031091-INS-119

chr9-3033522-INS-60
chr9-3035948-INS-58
chr9-3053090-INS-213
chr9-3059434-INS-1088
chr9-3062273-INS-93
chr9-3093368-INS-4437
chr9-3100661-INS-69
chr9-3139622-INS-4133
chr9-3145934-INS-214
chr9-3241661-INS-565
chr9-3253938-INS-266
chr9-3283079-INS-573
chr9-3307500-INS-50
chr9-3308265-INS-516
chr9-3312707-INS-360
chr9-3366161-INS-57
chr9-3376263-INS-132
chr9-3437560-INS-56
chr9-3637434-INS-50
chr9-3672523-INS-378
chr9-3673017-INS-1059
chr9-3716247-INS-6404
chr9-3743182-INS-64
chr9-3823154-INS-79
chr9-3823472-INS-93
chr9-3823478-INS-219
chr9-4017904-INS-732
chr9-4060442-INS-55
chr9-4089355-INS-60
chr9-4098306-INS-60
chr9-4131379-INS-260
chr9-4245601-INS-233
chr9-4245678-INS-54
chr9-4262417-INS-3278
chr9-4296232-INS-180
chr9-4301841-INS-64
chr9-4394834-INS-490
chr9-4401202-INS-83
chr9-4405175-INS-555
chr9-4407330-INS-801
chr9-4408564-INS-144
chr9-4415937-INS-180
chr9-4483393-INS-1419
chr9-4510070-INS-67
chr9-4534547-INS-482
chr9-4534572-INS-190
chr9-4560191-INS-5286
chr9-4565567-INS-51
chr9-4572362-INS-1268
chr9-4645693-INS-253
chr9-4676582-INS-5535
chr9-4705161-INS-68
chr9-4708286-INS-69
chr9-4722846-INS-171
chr9-4804625-INS-50
chr9-4807615-INS-912
chr9-4812374-INS-85
chr9-4869161-INS-72
chr9-4873693-INS-65
chr9-4873713-INS-265
chr9-4896063-INS-56
chr9-4910778-INS-68
chr9-4933257-INS-108
chr9-4941802-INS-6419
chr9-4959067-INS-2853
chr9-4977056-INS-197
chr9-5007079-INS-2478
chr9-5010835-INS-1697
chr9-5013193-INS-7079
chr9-5067567-INS-642
chr9-5083678-INS-60
chr9-5104858-INS-50
chr9-5131201-INS-433
chr9-5275602-INS-794
chr9-5309384-INS-307
chr9-5309407-INS-56
chr9-5317003-INS-1012

chr9-5343195-INS-386
chr9-5363742-INS-440
chr9-5440152-INS-369
chr9-5468739-INS-81
chr9-5484723-INS-7375
chr9-5491013-INS-90
chr9-5491015-INS-237
chr9-5509180-INS-2818
chr9-5514377-INS-2749
chr9-5561996-INS-55
chr9-5563364-INS-67
chr9-5575844-INS-6548
chr9-5616451-INS-115
chr9-5693848-INS-2868
chr9-5717094-INS-488
chr9-5750803-INS-9780
chr9-5754826-INS-4600
chr9-5809016-INS-1921
chr9-5809016-INS-625
chr9-5849381-INS-80
chr9-5871316-INS-15939
chr9-5873156-INS-62
chr9-5873490-INS-279
chr9-5890827-INS-426
chr9-5907253-INS-62
chr9-5953763-INS-3173
chr9-5960467-INS-350
chr9-5963040-INS-3319
chr9-5980510-INS-116
chr9-5985011-INS-6002
chr9-5991920-INS-4801
chr9-5992278-INS-55
chr9-5992530-INS-268
chr9-6013093-INS-68
chr9-6017448-INS-68
chr9-6031279-INS-65
chr9-6035979-INS-243
chr9-6036391-INS-245
chr9-6046344-INS-790
chr9-6066588-INS-4761
chr9-6085177-INS-80
chr9-6140074-INS-1365
chr9-6179369-INS-56
chr9-6201585-INS-1670
chr9-6252265-INS-2551
chr9-6290139-INS-197
chr9-6318808-INS-6380
chr9-6342079-INS-52
chr9-6365594-INS-54
chr9-6369927-INS-5911
chr9-6404521-INS-87
chr9-6405012-INS-1117
chr9-6454971-INS-50
chr9-6469150-INS-605
chr9-6482749-INS-50
chr9-6487451-INS-405
chr9-6492928-INS-559
chr9-6518807-INS-2732
chr9-6523354-INS-263
chr9-6538398-INS-1701
chr9-6581864-INS-4438
chr9-6586591-INS-50
chr9-6613612-INS-6435
chr9-6712756-INS-723
chr9-6718617-INS-220
chr9-6727366-INS-65
chr9-6733081-INS-58
chr9-6752224-INS-6387
chr9-6801313-INS-59
chr9-6820133-INS-199
chr9-6828299-INS-1822
chr9-6835256-INS-50
chr9-6853122-INS-183
chr9-6853809-INS-297
chr9-6861692-INS-327
chr9-6870663-INS-251
chr9-6870663-INS-59

chr9-6902314-INS-201
chr9-6906672-INS-6868
chr9-6923407-INS-5377
chr9-6935035-INS-237
chr9-7012003-INS-4910
chr9-7024044-INS-274
chr9-7061532-INS-3000
chr9-7074492-INS-124
chr9-7123776-INS-78
chr9-7134905-INS-114
chr9-7143641-INS-178
chr9-7157353-INS-272
chr9-7163068-INS-66
chr9-7177641-INS-141
chr9-7204806-INS-54
chr9-7258036-INS-3595
chr9-7259903-INS-354
chr9-7279870-INS-199
chr9-7320594-INS-70
chr9-7320858-INS-93
chr9-7322270-INS-55
chr9-7331281-INS-199
chr9-7358980-INS-7508
chr9-7362070-INS-254
chr9-7436519-INS-498
chr9-7461009-INS-2291
chr9-7505178-INS-68
chr9-7520078-INS-398
chr9-7520078-INS-936
chr9-7528665-INS-1927
chr9-7542230-INS-175
chr9-7542256-INS-86
chr9-7554800-INS-102
chr9-7561830-INS-541
chr9-7596659-INS-51
chr9-7600168-INS-3803
chr9-7652334-INS-102
chr9-7655961-INS-298
chr9-7657750-INS-241
chr9-7685377-INS-111
chr9-7688582-INS-60
chr9-7716605-INS-68
chr9-7741672-INS-50
chr9-7749165-INS-186
chr9-7755207-INS-236
chr9-7796190-INS-505
chr9-7817040-INS-173
chr9-7827258-INS-1414
chr9-7840379-INS-173
chr9-7850271-INS-54
chr9-7854192-INS-85
chr9-7876747-INS-198
chr9-7877118-INS-437
chr9-7879232-INS-205
chr9-7889830-INS-60
chr9-7914571-INS-4449
chr9-7921320-INS-205
chr9-7934901-INS-702
chr9-7945015-INS-60
chr9-7970908-INS-192
chr9-7995408-INS-66
chr9-7996661-INS-116
chr9-8009006-INS-210
chr9-8015429-INS-202
chr9-8016326-INS-214
chr9-8018098-INS-65
chr9-8024686-INS-273
chr9-8024935-INS-202
chr9-8028861-INS-60
chr9-8028912-INS-148
chr9-8036205-INS-805
chr9-8040233-INS-142
chr9-8040233-INS-69
chr9-8062139-INS-50
chr9-8070629-INS-147
chr9-8070706-INS-53
chr9-8182494-INS-3590

chr9-8269882-INS-662
chr9-8311258-INS-102
chr9-8340369-INS-76
chr9-8343088-INS-416
chr9-8343089-INS-183
chr9-8348006-INS-3689
chr9-8400544-INS-89
chr9-8428618-INS-51
chr9-8428632-INS-159
chr9-8430120-INS-209
chr9-8449456-INS-570
chr9-8496216-INS-4656
chr9-8566394-INS-6326
chr9-8570192-INS-158
chr9-8575415-INS-234
chr9-8588819-INS-80
chr9-8591615-INS-291
chr9-8599050-INS-52
chr9-8652696-INS-68
chr9-8702143-INS-89
chr9-8708099-INS-563
chr9-8747733-INS-52
chr9-8805735-INS-117
chr9-8813273-INS-62
chr9-8826518-INS-7294
chr9-8841830-INS-68
chr9-8850387-INS-385
chr9-8850519-INS-962
chr9-8873194-INS-82
chr9-8905521-INS-693
chr9-8960644-INS-71
chr9-8976888-INS-101
chr9-8986143-INS-2120
chr9-9012770-INS-58
chr9-9022169-INS-50
chr9-9035684-INS-282
chr9-9038305-INS-2589
chr9-9042987-INS-173
chr9-9043054-INS-63
chr9-9054186-INS-51
chr9-9056316-INS-1181
chr9-9062715-INS-1880
chr9-9091074-INS-86
chr9-9096278-INS-67
chr9-9154972-INS-240
chr9-9157608-INS-220
chr9-9161550-INS-372
chr9-9173902-INS-158
chr9-9185969-INS-59
chr9-9226422-INS-116
chr9-9228695-INS-74
chr9-9229427-INS-199
chr9-9284935-INS-239
chr9-9287221-INS-3434
chr9-9290206-INS-116
chr9-9300778-INS-2344
chr9-9303419-INS-3329
chr9-9337965-INS-50
chr9-9338116-INS-309
chr9-9382905-INS-461
chr9-9430613-INS-144
chr9-9445041-INS-54
chr9-9490013-INS-2374
chr9-9510324-INS-2909
chr9-9512191-INS-647
chr9-9591142-INS-215
chr9-9600205-INS-495
chr9-9626259-INS-6591
chr9-9720849-INS-3363
chr9-9725973-INS-285
chr9-9771966-INS-60
chr9-9779557-INS-1122
chr9-9786412-INS-60
chr9-9804387-INS-3389
chr9-9879243-INS-2319
chr9-9905669-INS-172
chr9-9977061-INS-189

chr9-9979219-INS-1602
chr9-9991349-INS-108
chr9-9999141-INS-105
chr9-10005245-INS-6254
chr9-10008946-INS-83
chr9-10014614-INS-2093
chr9-10039769-INS-70
chr9-10040426-INS-319
chr9-10087186-INS-56
chr9-10202122-INS-129
chr9-10202122-INS-314
chr9-10208188-INS-1553
chr9-10225022-INS-268
chr9-10276539-INS-6680
chr9-10297420-INS-530
chr9-10298474-INS-62
chr9-10344961-INS-226
chr9-10356727-INS-84
chr9-10357046-INS-489
chr9-10425140-INS-6377
chr9-10427201-INS-160
chr9-10461126-INS-747
chr9-10474749-INS-51
chr9-10481206-INS-62
chr9-10483136-INS-878
chr9-10499738-INS-106
chr9-10508847-INS-90
chr9-10547136-INS-5544
chr9-10602685-INS-2818
chr9-10619179-INS-115
chr9-10639705-INS-73
chr9-10639975-INS-445
chr9-10674198-INS-5016
chr9-10881667-INS-6377
chr9-10916594-INS-3890
chr9-10917225-INS-498
chr9-10927175-INS-2875
chr9-10974004-INS-3713
chr9-11006023-INS-240
chr9-11006026-INS-73
chr9-11066025-INS-51
chr9-11078464-INS-8138
chr9-11103477-INS-498
chr9-11108354-INS-72
chr9-11115786-INS-229
chr9-11118926-INS-400
chr9-11199407-INS-61
chr9-11234328-INS-364
chr9-11261120-INS-6602
chr9-11294518-INS-5485
chr9-11329530-INS-53
chr9-11369606-INS-104
chr9-11388598-INS-1257
chr9-11408242-INS-1221
chr9-11420876-INS-198
chr9-11421634-INS-71
chr9-11428144-INS-156
chr9-11539170-INS-57
chr9-11586293-INS-1349
chr9-11595652-INS-3539
chr9-11649449-INS-66
chr9-11665687-INS-3332
chr9-11714960-INS-98
chr9-11751680-INS-72
chr9-11769002-INS-164
chr9-11769002-INS-54
chr9-11777296-INS-1880
chr9-11777720-INS-54
chr9-11808312-INS-1098
chr9-11870729-INS-51
chr9-11998698-INS-328
chr9-12001108-INS-70
chr9-12001511-INS-147
chr9-12051974-INS-940
chr9-12078687-INS-85
chr9-12106122-INS-7988
chr9-12155179-INS-54

chr9-12165609-INS-102
chr9-12206085-INS-118
chr9-12219079-INS-50
chr9-12393015-INS-74
chr9-12422567-INS-56
chr9-12426307-INS-6591
chr9-12458428-INS-247
chr9-12485063-INS-54
chr9-12546882-INS-209
chr9-12551052-INS-3125
chr9-12590128-INS-56
chr9-12593585-INS-213
chr9-12605800-INS-6317
chr9-12610088-INS-3103
chr9-12621473-INS-52
chr9-12633151-INS-326
chr9-12651709-INS-56
chr9-12657809-INS-152
chr9-12686962-INS-7124
chr9-12699454-INS-68
chr9-12719184-INS-6095
chr9-12820145-INS-605
chr9-12839033-INS-59
chr9-12860193-INS-484
chr9-12878346-INS-50
chr9-12879181-INS-3410
chr9-12890004-INS-2855
chr9-12897996-INS-3373
chr9-12911669-INS-76
chr9-12939142-INS-6391
chr9-12953332-INS-6239
chr9-12966110-INS-1855
chr9-12968293-INS-384
chr9-12973554-INS-6251
chr9-12973889-INS-6318
chr9-12974510-INS-168
chr9-12986689-INS-52
chr9-13013393-INS-64
chr9-13077894-INS-502
chr9-13078510-INS-7415
chr9-13079622-INS-259
chr9-13090475-INS-67
chr9-13132281-INS-293
chr9-13173584-INS-103
chr9-13194842-INS-154
chr9-13202714-INS-1467
chr9-13204512-INS-201
chr9-13220249-INS-4170
chr9-13234580-INS-252
chr9-13242564-INS-52
chr9-13325649-INS-103
chr9-13355789-INS-120
chr9-13389567-INS-533
chr9-13409856-INS-94
chr9-13415273-INS-1588
chr9-13415295-INS-70
chr9-13416810-INS-151
chr9-13446079-INS-612
chr9-13451655-INS-50
chr9-13473869-INS-56
chr9-13478090-INS-132
chr9-13478175-INS-352
chr9-13482271-INS-60
chr9-13529827-INS-58
chr9-13600802-INS-604
chr9-13689161-INS-54
chr9-13729233-INS-109
chr9-13735649-INS-75
chr9-13776954-INS-68
chr9-13817222-INS-82
chr9-13835847-INS-476
chr9-13861298-INS-74
chr9-13864617-INS-100
chr9-13884803-INS-50
chr9-13885192-INS-80
chr9-13902577-INS-5793
chr9-13904111-INS-2824

chr9-13904111-INS-361
chr9-13914937-INS-62
chr9-13943024-INS-50
chr9-13958613-INS-52
chr9-13958942-INS-999
chr9-13986229-INS-125
chr9-13986474-INS-100
chr9-13998722-INS-150
chr9-14029993-INS-170
chr9-14033393-INS-1965
chr9-14060313-INS-2813
chr9-14103233-INS-316
chr9-14104968-INS-126
chr9-14119958-INS-72
chr9-14129144-INS-50
chr9-14170110-INS-80
chr9-14170138-INS-193
chr9-14195083-INS-58
chr9-14206788-INS-56
chr9-14233079-INS-73
chr9-14258430-INS-6645
chr9-14288208-INS-1888
chr9-14317196-INS-68
chr9-14328055-INS-2752
chr9-14364209-INS-68
chr9-14426744-INS-294
chr9-14426768-INS-1885
chr9-14431808-INS-1564
chr9-14478195-INS-50
chr9-14497303-INS-76
chr9-14512105-INS-155
chr9-14524105-INS-77
chr9-14604250-INS-1176
chr9-14619721-INS-194
chr9-14658207-INS-64
chr9-14663809-INS-2858
chr9-14667354-INS-4001
chr9-14670609-INS-83
chr9-14676872-INS-912
chr9-14679382-INS-6301
chr9-14758406-INS-762
chr9-14788242-INS-5288
chr9-14833867-INS-677
chr9-14851653-INS-75
chr9-14852020-INS-210
chr9-14852072-INS-589
chr9-14872496-INS-8487
chr9-14875031-INS-107
chr9-14900213-INS-96
chr9-14915606-INS-177
chr9-14931098-INS-56
chr9-14932072-INS-168
chr9-14941591-INS-761
chr9-14967348-INS-124
chr9-14967348-INS-600
chr9-14974784-INS-173
chr9-14974810-INS-85
chr9-15007795-INS-440
chr9-15011256-INS-62
chr9-15029266-INS-6348
chr9-15032378-INS-1789
chr9-15056528-INS-7175
chr9-15098365-INS-215
chr9-15112507-INS-129
chr9-15113730-INS-195
chr9-15120057-INS-50
chr9-15130917-INS-219
chr9-15133451-INS-664
chr9-15133605-INS-1888
chr9-15145192-INS-99
chr9-15213175-INS-430
chr9-15213175-INS-86
chr9-15262713-INS-67
chr9-15300710-INS-713
chr9-15304956-INS-1718
chr9-15306174-INS-76
chr9-15309788-INS-401

chr9-15313424-INS-232
chr9-15337805-INS-7164
chr9-15355697-INS-97
chr9-15360577-INS-322
chr9-15422853-INS-226
chr9-15468775-INS-58
chr9-15477078-INS-123
chr9-15498370-INS-174
chr9-15502828-INS-79
chr9-15503027-INS-183
chr9-15551887-INS-55
chr9-15561056-INS-173
chr9-15562182-INS-58
chr9-15591623-INS-74
chr9-15598072-INS-1882
chr9-15627965-INS-1054
chr9-15637766-INS-168
chr9-15637934-INS-545
chr9-15644075-INS-626
chr9-15647408-INS-164
chr9-15676642-INS-56
chr9-15677405-INS-130
chr9-15782131-INS-7201
chr9-15857171-INS-295
chr9-15857314-INS-895
chr9-15884869-INS-52
chr9-15889411-INS-159
chr9-15915512-INS-50
chr9-15923628-INS-244
chr9-15923652-INS-687
chr9-15923842-INS-93
chr9-15971669-INS-344
chr9-16013576-INS-706
chr9-16034306-INS-50
chr9-16139225-INS-54
chr9-16161488-INS-380
chr9-16213022-INS-50
chr9-16240929-INS-50
chr9-16245684-INS-165
chr9-16259767-INS-52
chr9-16307835-INS-50
chr9-16313468-INS-56
chr9-16328050-INS-82
chr9-16329098-INS-87
chr9-16331683-INS-244
chr9-16344262-INS-72
chr9-16355533-INS-54
chr9-16399089-INS-2508
chr9-16421621-INS-6717
chr9-16444751-INS-60
chr9-16466014-INS-6902
chr9-16473877-INS-6208
chr9-16475481-INS-3048
chr9-16532177-INS-1274
chr9-16559160-INS-268
chr9-16569306-INS-59
chr9-16581375-INS-7617
chr9-16587187-INS-3783
chr9-16626229-INS-213
chr9-16653104-INS-6946
chr9-16659904-INS-5671
chr9-16675973-INS-1055
chr9-16680367-INS-50
chr9-16682935-INS-6561
chr9-16696628-INS-215
chr9-16727515-INS-2022
chr9-16764707-INS-70
chr9-16775031-INS-498
chr9-16779334-INS-4904
chr9-16796854-INS-84
chr9-16801117-INS-3241
chr9-16810767-INS-106
chr9-16820633-INS-56
chr9-16846331-INS-54
chr9-16886147-INS-6366
chr9-16955033-INS-61
chr9-16990237-INS-60

chr9-17014411-INS-159
chr9-17019300-INS-2039
chr9-17024023-INS-6593
chr9-17040261-INS-5103
chr9-17049600-INS-2905
chr9-17076393-INS-61
chr9-17082796-INS-193
chr9-17085882-INS-3169
chr9-17098477-INS-53
chr9-17102615-INS-112
chr9-17137641-INS-1175
chr9-17184374-INS-70
chr9-17198742-INS-74
chr9-17199944-INS-858
chr9-17201398-INS-79
chr9-17201504-INS-203
chr9-17220493-INS-2466
chr9-17227412-INS-7086
chr9-17231567-INS-58
chr9-17231622-INS-1853
chr9-17245871-INS-58
chr9-17283953-INS-52
chr9-17294057-INS-83
chr9-17306017-INS-1293
chr9-17361801-INS-3726
chr9-17370878-INS-53
chr9-17377661-INS-500
chr9-17383182-INS-5598
chr9-17384649-INS-59
chr9-17389858-INS-1085
chr9-17393275-INS-2352
chr9-17397965-INS-50
chr9-17428144-INS-6203
chr9-17438306-INS-3304
chr9-17451506-INS-892
chr9-17470731-INS-8754
chr9-17472486-INS-1456
chr9-17477926-INS-4481
chr9-17483815-INS-56
chr9-17492295-INS-562
chr9-17525520-INS-213
chr9-17540469-INS-1893
chr9-17549882-INS-398
chr9-17553406-INS-5501
chr9-17555347-INS-731
chr9-17555534-INS-53
chr9-17555817-INS-2334
chr9-17561408-INS-1325
chr9-17577784-INS-7114
chr9-17617547-INS-1623
chr9-17619076-INS-2456
chr9-17625555-INS-400
chr9-17688078-INS-506
chr9-17716959-INS-2650
chr9-17721535-INS-415
chr9-17722403-INS-3079
chr9-17735057-INS-1503
chr9-17783588-INS-400
chr9-17786765-INS-1424
chr9-17799819-INS-996
chr9-17815042-INS-83
chr9-17839782-INS-52
chr9-17852232-INS-60
chr9-17852697-INS-246
chr9-17861163-INS-242
chr9-17953818-INS-3132
chr9-17954241-INS-1254
chr9-17969830-INS-173
chr9-18012723-INS-206
chr9-18120838-INS-4247
chr9-18256593-INS-6669
chr9-18281077-INS-406
chr9-18281268-INS-139
chr9-18283157-INS-95
chr9-18295156-INS-1108
chr9-18329915-INS-51
chr9-18345592-INS-223

chr9-18351194-INS-3422
chr9-18452281-INS-1333
chr9-18535839-INS-6885
chr9-18551043-INS-1893
chr9-18551454-INS-3756
chr9-18573757-INS-6444
chr9-18627492-INS-50
chr9-18658047-INS-383
chr9-18660935-INS-6714
chr9-18664219-INS-57
chr9-18708358-INS-208
chr9-18709367-INS-963
chr9-18751938-INS-6458
chr9-18768471-INS-385
chr9-18772994-INS-116
chr9-18796113-INS-85
chr9-18796139-INS-175
chr9-18891865-INS-636
chr9-18894941-INS-498
chr9-18949812-INS-50
chr9-18966892-INS-10266
chr9-18971673-INS-1390
chr9-18996037-INS-663
chr9-18996787-INS-255
chr9-18997127-INS-439
chr9-19023143-INS-3721
chr9-19030941-INS-5068
chr9-19080269-INS-1896
chr9-19084370-INS-272
chr9-19158289-INS-54
chr9-19182107-INS-6803
chr9-19190860-INS-56
chr9-19190969-INS-57
chr9-19226172-INS-1101
chr9-19258342-INS-6933
chr9-19292204-INS-2201
chr9-19299578-INS-6591
chr9-19310477-INS-52
chr9-19314031-INS-55
chr9-19329573-INS-6371
chr9-19330831-INS-343
chr9-19340647-INS-415
chr9-19340881-INS-138
chr9-19380236-INS-440
chr9-19392792-INS-6357
chr9-19456570-INS-64
chr9-19499295-INS-1913
chr9-19539929-INS-54
chr9-19564610-INS-50
chr9-19596165-INS-6731
chr9-19604236-INS-56
chr9-19623562-INS-55
chr9-19692100-INS-12230
chr9-19700301-INS-445
chr9-19710757-INS-55
chr9-19718859-INS-75
chr9-19825621-INS-208
chr9-20029195-INS-222
chr9-20051031-INS-7502
chr9-20051819-INS-6262
chr9-20110414-INS-98
chr9-20111036-INS-53
chr9-20128010-INS-2102
chr9-20141702-INS-84
chr9-20178468-INS-54
chr9-20178581-INS-103
chr9-20203403-INS-4671
chr9-20232979-INS-208
chr9-20243017-INS-135
chr9-20254963-INS-293
chr9-20255309-INS-72
chr9-20326855-INS-226
chr9-20336236-INS-1871
chr9-20362689-INS-1409
chr9-20362689-INS-665
chr9-20404918-INS-64
chr9-20430871-INS-54

chr9-20493526-INS-60
chr9-20590445-INS-66
chr9-20610366-INS-95
chr9-20647404-INS-78
chr9-20647660-INS-54
chr9-20653637-INS-161
chr9-20669702-INS-268
chr9-20675769-INS-60
chr9-20680267-INS-135
chr9-20680677-INS-80
chr9-20700890-INS-50
chr9-20711621-INS-211
chr9-20717509-INS-214
chr9-20729155-INS-52
chr9-20730075-INS-110
chr9-20730396-INS-50
chr9-20732163-INS-50
chr9-20732708-INS-54
chr9-20736182-INS-5472
chr9-20774012-INS-125
chr9-20792185-INS-50
chr9-20866435-INS-56
chr9-20903052-INS-58
chr9-20927623-INS-50
chr9-20930868-INS-211
chr9-20941743-INS-61
chr9-20952524-INS-174
chr9-20954573-INS-72
chr9-20959998-INS-228
chr9-20960649-INS-345
chr9-20962573-INS-73
chr9-21012304-INS-92
chr9-21012798-INS-60
chr9-21088380-INS-169
chr9-21113797-INS-194
chr9-21130604-INS-65
chr9-21132209-INS-216
chr9-21133035-INS-256
chr9-21166304-INS-76
chr9-21168312-INS-71
chr9-21173804-INS-216
chr9-21204203-INS-219
chr9-21212641-INS-217
chr9-21220608-INS-172
chr9-21227959-INS-152
chr9-21236615-INS-58
chr9-21252307-INS-177
chr9-21275598-INS-4752
chr9-21332626-INS-184
chr9-21333800-INS-207
chr9-21408477-INS-50
chr9-21481294-INS-218
chr9-21537691-INS-181
chr9-21540950-INS-705
chr9-21601264-INS-278
chr9-21605580-INS-873
chr9-21605710-INS-624
chr9-21644195-INS-218
chr9-21652298-INS-217
chr9-21661697-INS-202
chr9-21677378-INS-132
chr9-21689858-INS-221
chr9-21693918-INS-144
chr9-21713186-INS-219
chr9-21719696-INS-72
chr9-21720380-INS-324
chr9-21720461-INS-84
chr9-21747721-INS-55
chr9-21749701-INS-90
chr9-21773334-INS-207
chr9-21806338-INS-144
chr9-21816665-INS-749
chr9-21833653-INS-2035
chr9-21833931-INS-777
chr9-21843325-INS-204
chr9-21906938-INS-199
chr9-22009802-INS-50

chr9-22044469-INS-301
chr9-22054017-INS-164
chr9-22054158-INS-4846
chr9-22057971-INS-3329
chr9-22062806-INS-64
chr9-22080548-INS-165
chr9-22100703-INS-1297
chr9-22107414-INS-54
chr9-22115136-INS-6819
chr9-22129515-INS-51
chr9-22132598-INS-71
chr9-22133651-INS-65
chr9-22136110-INS-53
chr9-22179862-INS-54
chr9-22182821-INS-1391
chr9-22183363-INS-8911
chr9-22211368-INS-828
chr9-22228263-INS-218
chr9-22228924-INS-514
chr9-22249155-INS-526
chr9-22284014-INS-85
chr9-22311478-INS-1130
chr9-22314092-INS-207
chr9-22316065-INS-8321
chr9-22318911-INS-2019
chr9-22352971-INS-7167
chr9-22356659-INS-55
chr9-22382084-INS-6385
chr9-22411551-INS-52
chr9-22426179-INS-126
chr9-22439680-INS-389
chr9-22441833-INS-64
chr9-22464752-INS-102
chr9-22471894-INS-1496
chr9-22481774-INS-141
chr9-22481777-INS-50
chr9-22486837-INS-157
chr9-22532728-INS-129
chr9-22537447-INS-55
chr9-22555272-INS-60
chr9-22577323-INS-3011
chr9-22588980-INS-208
chr9-22623797-INS-612
chr9-22624354-INS-2852
chr9-22654933-INS-295
chr9-22671495-INS-238
chr9-22699086-INS-70
chr9-22728084-INS-541
chr9-22759506-INS-58
chr9-22771006-INS-58
chr9-22788387-INS-61
chr9-22795320-INS-203
chr9-22805826-INS-228
chr9-22806487-INS-88
chr9-22820309-INS-220
chr9-22869960-INS-239
chr9-22899281-INS-466
chr9-22925575-INS-78
chr9-23002655-INS-6352
chr9-23031634-INS-59
chr9-23151617-INS-123
chr9-23169249-INS-150
chr9-23220615-INS-6513
chr9-23282618-INS-52
chr9-23286217-INS-58
chr9-23316129-INS-209
chr9-23316161-INS-52
chr9-23327860-INS-2117
chr9-23344470-INS-6411
chr9-23354797-INS-106
chr9-23423905-INS-50
chr9-23465036-INS-605
chr9-23468765-INS-174
chr9-23468796-INS-66
chr9-23476485-INS-230
chr9-23476485-INS-50
chr9-23481507-INS-1442

chr9-23501464-INS-3171
chr9-23525116-INS-6176
chr9-23526058-INS-3714
chr9-23527060-INS-1392
chr9-23611061-INS-602
chr9-23611062-INS-50
chr9-23623503-INS-54
chr9-23637536-INS-99
chr9-23680599-INS-5429
chr9-23688496-INS-1213
chr9-23736223-INS-6315
chr9-23761956-INS-7752
chr9-23766222-INS-5785
chr9-23771682-INS-2675
chr9-23780254-INS-1213
chr9-23783369-INS-7064
chr9-23783881-INS-112
chr9-23797935-INS-4828
chr9-23799247-INS-2103
chr9-23805639-INS-785
chr9-23842598-INS-171
chr9-23851181-INS-6002
chr9-23903056-INS-51
chr9-23924729-INS-50
chr9-23962083-INS-306
chr9-23966626-INS-1892
chr9-23982017-INS-6113
chr9-24021316-INS-1992
chr9-24027049-INS-1079
chr9-24046882-INS-52
chr9-24069609-INS-52
chr9-24088225-INS-50
chr9-24117239-INS-3036
chr9-24119721-INS-131
chr9-24120416-INS-1030
chr9-24122066-INS-1361
chr9-24126074-INS-200
chr9-24140370-INS-1210
chr9-24141067-INS-50
chr9-24159955-INS-60
chr9-24203356-INS-72
chr9-24211336-INS-53
chr9-24268219-INS-391
chr9-24268305-INS-106
chr9-24268579-INS-6396
chr9-24279649-INS-244
chr9-24279661-INS-85
chr9-24286939-INS-6319
chr9-24296861-INS-6181
chr9-24322158-INS-62
chr9-24359886-INS-50
chr9-24367390-INS-108
chr9-24392905-INS-484
chr9-24392927-INS-198
chr9-24392927-INS-67
chr9-24393182-INS-254
chr9-24416750-INS-306
chr9-24459909-INS-1472
chr9-24500671-INS-171
chr9-24511145-INS-317
chr9-24519052-INS-52
chr9-24579445-INS-135
chr9-24603055-INS-122
chr9-24725306-INS-136
chr9-24749484-INS-193
chr9-24760146-INS-94
chr9-24774448-INS-50
chr9-24793303-INS-551
chr9-24822502-INS-302
chr9-24896100-INS-466
chr9-24943654-INS-50
chr9-24950804-INS-201
chr9-25003462-INS-167
chr9-25097863-INS-4256
chr9-25099374-INS-56
chr9-25102226-INS-58
chr9-25102800-INS-102

chr9-25129000-INS-897
chr9-25229414-INS-100
chr9-25252005-INS-199
chr9-25252005-INS-99
chr9-25252574-INS-138
chr9-25276640-INS-5832
chr9-25280982-INS-54
chr9-25285987-INS-50
chr9-25286653-INS-326
chr9-25291569-INS-277
chr9-25355153-INS-1036
chr9-25355235-INS-408
chr9-25371939-INS-2000
chr9-25375917-INS-533
chr9-25376227-INS-78
chr9-25376253-INS-326
chr9-25376878-INS-312
chr9-25414344-INS-282
chr9-25441753-INS-108
chr9-25498444-INS-147
chr9-25504456-INS-96
chr9-25505224-INS-149
chr9-25505224-INS-330
chr9-25523797-INS-2092
chr9-25575515-INS-1228
chr9-25580434-INS-7519
chr9-25586463-INS-831
chr9-25591907-INS-140
chr9-25599060-INS-3794
chr9-25630843-INS-134
chr9-25638477-INS-467
chr9-25677621-INS-1350
chr9-25681486-INS-1187
chr9-25702105-INS-379
chr9-25719671-INS-58
chr9-25749091-INS-891
chr9-25793696-INS-1294
chr9-25794121-INS-15592
chr9-25810249-INS-2300
chr9-25810249-INS-5963
chr9-25819159-INS-6322
chr9-25820959-INS-56
chr9-25833796-INS-3013
chr9-25834592-INS-82
chr9-25861687-INS-2361
chr9-25871962-INS-9015
chr9-25875911-INS-58
chr9-25913157-INS-130
chr9-25915173-INS-50
chr9-25925578-INS-60
chr9-25962490-INS-58
chr9-25977974-INS-3428
chr9-26003271-INS-64
chr9-26024603-INS-106
chr9-26058652-INS-60
chr9-26060736-INS-313
chr9-26167284-INS-51
chr9-26167746-INS-62
chr9-26169549-INS-2682
chr9-26170769-INS-3377
chr9-26185590-INS-362
chr9-26241846-INS-1707
chr9-26280537-INS-128
chr9-26284110-INS-1476
chr9-26361382-INS-62
chr9-26371704-INS-113
chr9-26371756-INS-1278
chr9-26371948-INS-50
chr9-26506121-INS-2535
chr9-26523936-INS-212
chr9-26530767-INS-1771
chr9-26733357-INS-1965
chr9-26734229-INS-114
chr9-26740510-INS-2144
chr9-26757913-INS-860
chr9-26794112-INS-53
chr9-26844346-INS-1136

chr9-26853363-INS-723
chr9-26857183-INS-117
chr9-26858931-INS-6335
chr9-26898230-INS-88
chr9-26967787-INS-142
chr9-27047084-INS-73
chr9-27094540-INS-167
chr9-27192021-INS-151
chr9-27204150-INS-69
chr9-27306792-INS-54
chr9-27307488-INS-62
chr9-27315669-INS-62
chr9-27382648-INS-123
chr9-27385250-INS-749
chr9-27409729-INS-256
chr9-27431900-INS-72
chr9-27456655-INS-50
chr9-27458493-INS-73
chr9-27464581-INS-4116
chr9-27475230-INS-7054
chr9-27514119-INS-269
chr9-27557349-INS-73
chr9-27639581-INS-1332
chr9-27639680-INS-66
chr9-27727841-INS-646
chr9-27752025-INS-7144
chr9-27753785-INS-50
chr9-27770502-INS-89
chr9-27785476-INS-366
chr9-27785533-INS-149
chr9-27791452-INS-551
chr9-27805896-INS-466
chr9-27845279-INS-581
chr9-27908953-INS-63
chr9-27909158-INS-341
chr9-27914517-INS-6246
chr9-27939354-INS-141
chr9-28103393-INS-72
chr9-28173357-INS-58
chr9-28184714-INS-96
chr9-28246216-INS-60
chr9-28246263-INS-165
chr9-28246895-INS-50
chr9-28306466-INS-52
chr9-28362520-INS-152
chr9-28405959-INS-50
chr9-28410437-INS-64
chr9-28438582-INS-62
chr9-28516752-INS-88
chr9-28521700-INS-1265
chr9-28605684-INS-217
chr9-28608172-INS-60
chr9-28660102-INS-107
chr9-28684498-INS-2980
chr9-28699741-INS-258
chr9-28699851-INS-499
chr9-28738011-INS-357
chr9-28743539-INS-400
chr9-28743775-INS-51
chr9-28752622-INS-356
chr9-28791717-INS-477
chr9-28810083-INS-348
chr9-28846911-INS-941
chr9-28895040-INS-120
chr9-28929111-INS-141
chr9-28929735-INS-59
chr9-28949480-INS-55
chr9-29043167-INS-56
chr9-29058068-INS-514
chr9-29065659-INS-52
chr9-29157901-INS-56
chr9-29160527-INS-50
chr9-29268444-INS-561
chr9-29278265-INS-6804
chr9-29308505-INS-591
chr9-29331014-INS-58
chr9-29356211-INS-1924

chr9-29417426-INS-3107
chr9-29554682-INS-622
chr9-29559499-INS-62
chr9-29614635-INS-302
chr9-29671763-INS-102
chr9-29674122-INS-60
chr9-29743476-INS-52
chr9-29767759-INS-66
chr9-29815201-INS-52
chr9-29829183-INS-50
chr9-29921423-INS-52
chr9-29948420-INS-6385
chr9-29950933-INS-1923
chr9-29972524-INS-112
chr9-29995232-INS-59
chr9-30033361-INS-90
chr9-30041918-INS-57
chr9-30070131-INS-54
chr9-30074594-INS-1007
chr9-30107528-INS-58
chr9-30127319-INS-50
chr9-30246826-INS-54
chr9-30247494-INS-60
chr9-30260223-INS-965
chr9-30277959-INS-3088
chr9-30314353-INS-224
chr9-30317587-INS-1134
chr9-30439628-INS-2847
chr9-30462535-INS-60
chr9-30468385-INS-6476
chr9-30482141-INS-63
chr9-30618931-INS-74
chr9-30664620-INS-158
chr9-30689750-INS-200
chr9-30690220-INS-360
chr9-30690236-INS-107
chr9-30691178-INS-1611
chr9-30742349-INS-65
chr9-30744357-INS-75
chr9-30758916-INS-50
chr9-30770043-INS-1877
chr9-30771163-INS-372
chr9-30795775-INS-439
chr9-30796250-INS-312
chr9-30796341-INS-86
chr9-30825066-INS-55
chr9-30825179-INS-127
chr9-30843106-INS-468
chr9-30847794-INS-102
chr9-30849376-INS-431
chr9-30861239-INS-50
chr9-30892198-INS-56
chr9-30923286-INS-9101
chr9-30938669-INS-215
chr9-30993054-INS-62
chr9-31024914-INS-1364
chr9-31057937-INS-233
chr9-31064329-INS-235
chr9-31097088-INS-3006
chr9-31124460-INS-373
chr9-31131226-INS-192
chr9-31138330-INS-200
chr9-31148007-INS-50
chr9-31148310-INS-80
chr9-31148315-INS-320
chr9-31166836-INS-193
chr9-31167231-INS-66
chr9-31173102-INS-62
chr9-31173433-INS-111
chr9-31174149-INS-845
chr9-31218178-INS-80
chr9-31218525-INS-234
chr9-31218557-INS-78
chr9-31228554-INS-50
chr9-31248794-INS-52
chr9-31280835-INS-55
chr9-31319922-INS-108

chr9-31335165-INS-212
chr9-31342915-INS-72
chr9-31426898-INS-85
chr9-31429232-INS-607
chr9-31438567-INS-51
chr9-31456040-INS-54
chr9-31456499-INS-168
chr9-31537108-INS-1286
chr9-31569981-INS-2579
chr9-31587578-INS-6038
chr9-31679111-INS-101
chr9-31683883-INS-167
chr9-31691148-INS-5311
chr9-31692979-INS-84
chr9-31716839-INS-100
chr9-31736270-INS-50
chr9-31774453-INS-195
chr9-31774736-INS-56
chr9-31798267-INS-51
chr9-31866556-INS-1889
chr9-31867798-INS-65
chr9-31886453-INS-183
chr9-31887331-INS-58
chr9-31887955-INS-110
chr9-31892980-INS-932
chr9-31897960-INS-161
chr9-31910936-INS-317
chr9-31927943-INS-4369
chr9-31941877-INS-130
chr9-31961548-INS-172
chr9-32025864-INS-213
chr9-32026461-INS-80
chr9-32042811-INS-186
chr9-32045138-INS-1779
chr9-32053812-INS-50
chr9-32072379-INS-89
chr9-32090522-INS-1052
chr9-32096004-INS-56
chr9-32113876-INS-1058
chr9-32156163-INS-206
chr9-32164769-INS-797
chr9-32195627-INS-345
chr9-32198453-INS-290
chr9-32198783-INS-53
chr9-32199584-INS-86
chr9-32265816-INS-66
chr9-32279411-INS-236
chr9-32306501-INS-50
chr9-32406792-INS-52
chr9-32433751-INS-195
chr9-32481451-INS-201
chr9-32481647-INS-52
chr9-32486161-INS-369
chr9-32502642-INS-60
chr9-32504969-INS-380
chr9-32505200-INS-231
chr9-32566316-INS-64
chr9-32595130-INS-96
chr9-32595909-INS-54
chr9-32599580-INS-64
chr9-32599583-INS-298
chr9-32601157-INS-668
chr9-32609505-INS-148
chr9-32739786-INS-59
chr9-32792222-INS-86
chr9-32807261-INS-56
chr9-32840261-INS-86
chr9-32980088-INS-140
chr9-33001189-INS-1893
chr9-33005833-INS-1054
chr9-33091161-INS-2634
chr9-33124267-INS-2839
chr9-33143787-INS-130
chr9-33166292-INS-6751
chr9-33168645-INS-2872
chr9-33184249-INS-6289
chr9-33195378-INS-396

chr9-33195860-INS-53
chr9-33205335-INS-5062
chr9-33225849-INS-52
chr9-33277634-INS-471
chr9-33278308-INS-1706
chr9-33304627-INS-214
chr9-33320202-INS-56
chr9-33362914-INS-1891
chr9-33386716-INS-72
chr9-33392933-INS-4500
chr9-33415710-INS-1567
chr9-33427930-INS-6171
chr9-33440699-INS-3224
chr9-33469323-INS-89
chr9-33491870-INS-153
chr9-33492422-INS-85
chr9-33555187-INS-767
chr9-33555775-INS-329
chr9-33561791-INS-51
chr9-33562111-INS-6456
chr9-33590117-INS-969
chr9-33594588-INS-3129
chr9-33625763-INS-4053
chr9-33635012-INS-132
chr9-33681362-INS-55
chr9-33681387-INS-116
chr9-33695459-INS-50
chr9-33698148-INS-100
chr9-33716880-INS-54
chr9-33717396-INS-7098
chr9-33727867-INS-64
chr9-33743556-INS-52
chr9-33774970-INS-52
chr9-33797850-INS-496
chr9-33837387-INS-4790
chr9-33908012-INS-138
chr9-33908067-INS-361
chr9-33922549-INS-81
chr9-33931697-INS-1322
chr9-33943572-INS-124
chr9-33971043-INS-66
chr9-33998776-INS-74
chr9-34042404-INS-114
chr9-34206656-INS-2812
chr9-34211697-INS-6214
chr9-34264546-INS-52
chr9-34320979-INS-252
chr9-34336089-INS-6373
chr9-34361050-INS-3135
chr9-34424033-INS-50
chr9-34431200-INS-55
chr9-34438033-INS-114
chr9-34468190-INS-61
chr9-34477361-INS-65
chr9-34573909-INS-108
chr9-34603257-INS-51
chr9-34634020-INS-590
chr9-34661746-INS-54
chr9-34694402-INS-94
chr9-34719769-INS-215
chr9-34751801-INS-72
chr9-34795456-INS-278
chr9-34796949-INS-64
chr9-34797286-INS-69
chr9-34807630-INS-1656
chr9-34808071-INS-173
chr9-34815263-INS-69
chr9-34841145-INS-80
chr9-34888867-INS-199
chr9-34951174-INS-84
chr9-34971711-INS-4092
chr9-35040177-INS-3000
chr9-35074935-INS-50
chr9-35148225-INS-1162
chr9-35172555-INS-98
chr9-35225861-INS-60
chr9-35285643-INS-150

chr9-35338277-INS-53
chr9-35408871-INS-1078
chr9-35423207-INS-211
chr9-35453746-INS-52
chr9-35482298-INS-55
chr9-35612618-INS-8662
chr9-35613423-INS-107
chr9-35626062-INS-56
chr9-35631908-INS-6682
chr9-35650214-INS-56
chr9-35671067-INS-52
chr9-35675101-INS-56
chr9-35692191-INS-93
chr9-35725547-INS-9255
chr9-35857807-INS-403
chr9-35863814-INS-510
chr9-35874353-INS-91
chr9-35875244-INS-54
chr9-35909601-INS-2765
chr9-35911624-INS-226
chr9-35911624-INS-80
chr9-35911750-INS-4172
chr9-35940773-INS-85
chr9-36023878-INS-3212
chr9-36041599-INS-50
chr9-36119934-INS-4810
chr9-36130492-INS-87
chr9-36131300-INS-8131
chr9-36133419-INS-3150
chr9-36160399-INS-2040
chr9-36162356-INS-287
chr9-36174824-INS-14280
chr9-36186250-INS-3304
chr9-36223417-INS-4744
chr9-36237777-INS-27620
chr9-36328696-INS-240
chr9-36340572-INS-66
chr9-36422646-INS-143
chr9-36506751-INS-1184
chr9-36591415-INS-77
chr9-36623214-INS-182
chr9-36623450-INS-225
chr9-36653449-INS-172
chr9-36658796-INS-1418
chr9-36667465-INS-220
chr9-36668518-INS-201
chr9-36669783-INS-212
chr9-36720771-INS-54
chr9-36724002-INS-401
chr9-36724102-INS-284
chr9-36726279-INS-226
chr9-36748192-INS-314
chr9-36749101-INS-181
chr9-36758694-INS-64
chr9-36763759-INS-142
chr9-36775560-INS-54
chr9-36790343-INS-212
chr9-36798923-INS-58
chr9-36838432-INS-58
chr9-36910115-INS-113
chr9-37021252-INS-64
chr9-37041421-INS-225
chr9-37042389-INS-1889
chr9-37066329-INS-183
chr9-37094045-INS-6007
chr9-37104980-INS-280
chr9-37104980-INS-80
chr9-37106902-INS-54
chr9-37135932-INS-60
chr9-37201339-INS-93
chr9-37228451-INS-218
chr9-37349415-INS-50
chr9-37378583-INS-162
chr9-37391004-INS-56
chr9-37393710-INS-107
chr9-37407482-INS-225
chr9-37462522-INS-150

chr9-37465766-INS-135
chr9-37470214-INS-152
chr9-37539369-INS-368
chr9-37573439-INS-204
chr9-37579887-INS-74
chr9-37607729-INS-66
chr9-37608538-INS-3072
chr9-37609484-INS-6309
chr9-37632903-INS-713
chr9-37634856-INS-212
chr9-37646071-INS-6537
chr9-37651011-INS-95
chr9-37669075-INS-138
chr9-37728677-INS-6167
chr9-37741564-INS-204
chr9-37774094-INS-62
chr9-37836639-INS-51
chr9-37837443-INS-58
chr9-37841350-INS-805
chr9-37841690-INS-1035
chr9-37863650-INS-377
chr9-37864029-INS-195
chr9-37864972-INS-4597
chr9-37900451-INS-6858
chr9-37911028-INS-56
chr9-37981017-INS-52
chr9-38098201-INS-7447
chr9-38109899-INS-184
chr9-38116616-INS-1856
chr9-38126820-INS-348
chr9-38134196-INS-6341
chr9-38136756-INS-55
chr9-38148028-INS-4813
chr9-38148442-INS-88
chr9-38165700-INS-218
chr9-38251160-INS-1472
chr9-38264131-INS-6413
chr9-38276052-INS-3995
chr9-38298439-INS-738
chr9-38303472-INS-2956
chr9-38305326-INS-77
chr9-38311188-INS-522
chr9-38348641-INS-50
chr9-38361672-INS-411
chr9-38378751-INS-50
chr9-38509664-INS-1112
chr9-38510147-INS-800
chr9-38551538-INS-739
chr9-38562400-INS-1074
chr9-38574378-INS-6265
chr9-38587403-INS-253
chr9-38600551-INS-6605
chr9-38600830-INS-51
chr9-38612286-INS-316
chr9-38683020-INS-69
chr9-38685125-INS-168
chr9-38686232-INS-52
chr9-38692511-INS-53
chr9-38744774-INS-550
chr9-38767608-INS-587
chr9-38783336-INS-50
chr9-38808009-INS-2020
chr9-38834229-INS-61
chr9-38900552-INS-6472
chr9-38903048-INS-221
chr9-38925761-INS-612
chr9-38932874-INS-18510
chr9-38938432-INS-69
chr9-38942777-INS-141
chr9-38944686-INS-1777
chr9-38945762-INS-195
chr9-38969924-INS-76
chr9-38995989-INS-87
chr9-39019509-INS-8769
chr9-39025236-INS-52
chr9-39042587-INS-7212
chr9-39112610-INS-1287

chr9-39112748-INS-195
chr9-39119289-INS-50
chr9-39131670-INS-56
chr9-39185357-INS-8301
chr9-39196088-INS-2273
chr9-39302817-INS-107
chr9-39362084-INS-114
chr9-39383012-INS-5235
chr9-39383913-INS-868
chr9-39454306-INS-12763
chr9-39456907-INS-1900
chr9-39573506-INS-2283
chr9-39612646-INS-6836
chr9-39615067-INS-79
chr9-39636305-INS-52
chr9-39639326-INS-4444
chr9-39656257-INS-76
chr9-39658297-INS-7193
chr9-39701569-INS-139
chr9-39705544-INS-68
chr9-39705584-INS-306
chr9-39762817-INS-420
chr9-39816031-INS-134
chr9-39860196-INS-55
chr9-39860488-INS-2419
chr9-39864825-INS-63
chr9-39909490-INS-4199
chr9-39920495-INS-54
chr9-40029799-INS-1668
chr9-40033783-INS-257
chr9-40043849-INS-54
chr9-40049670-INS-54
chr9-40065180-INS-1221
chr9-40068400-INS-50
chr9-40071026-INS-75
chr9-40075639-INS-204
chr9-40106241-INS-222
chr9-40134168-INS-258
chr9-40148216-INS-82
chr9-40155645-INS-112
chr9-40159089-INS-3197
chr9-40178558-INS-196
chr9-40196371-INS-70
chr9-40307934-INS-72
chr9-40318969-INS-54
chr9-40382807-INS-55
chr9-40424849-INS-54
chr9-40428018-INS-438
chr9-40479030-INS-205
chr9-40491590-INS-591
chr9-40491654-INS-235
chr9-40500420-INS-7488
chr9-40501005-INS-272
chr9-40501069-INS-110
chr9-40517238-INS-61
chr9-40520682-INS-50
chr9-40534733-INS-60
chr9-40542560-INS-3142
chr9-40558227-INS-77
chr9-40582370-INS-277
chr9-40633016-INS-1892
chr9-40635574-INS-6401
chr9-40648152-INS-64
chr9-40648349-INS-164
chr9-40652772-INS-213
chr9-40676264-INS-2896
chr9-40681919-INS-59
chr9-40684073-INS-178
chr9-40708594-INS-58
chr9-40730421-INS-1979
chr9-40736627-INS-1451
chr9-40742897-INS-1092
chr9-40747044-INS-187
chr9-40836531-INS-51
chr9-40847505-INS-83
chr9-40859253-INS-50
chr9-40966319-INS-75

chr9-40966463-INS-169
chr9-41078583-INS-91
chr9-41090176-INS-67
chr9-41090344-INS-118
chr9-41116138-INS-56
chr9-41141992-INS-52
chr9-41174321-INS-1127
chr9-41174599-INS-78
chr9-41174609-INS-307
chr9-41192686-INS-288
chr9-41223254-INS-1371
chr9-41292038-INS-66
chr9-41297885-INS-54
chr9-41310274-INS-131
chr9-41376374-INS-125
chr9-41513190-INS-61
chr9-41534123-INS-116
chr9-41543530-INS-1701
chr9-41547328-INS-2546
chr9-41558889-INS-6393
chr9-41589233-INS-1862
chr9-41639639-INS-167
chr9-41639642-INS-62
chr9-41653572-INS-1897
chr9-41676418-INS-644
chr9-41686213-INS-153
chr9-41698911-INS-50
chr9-41739894-INS-68
chr9-41907029-INS-70
chr9-41912618-INS-243
chr9-41932001-INS-1825
chr9-41932735-INS-306
chr9-41962953-INS-52
chr9-41977729-INS-170
chr9-41987646-INS-2057
chr9-42003508-INS-181
chr9-42020360-INS-50
chr9-42043474-INS-52
chr9-42063833-INS-63
chr9-42065935-INS-63
chr9-42066036-INS-154
chr9-42078574-INS-7156
chr9-42097237-INS-305
chr9-42110266-INS-93
chr9-42110423-INS-221
chr9-42111258-INS-243
chr9-42117792-INS-55
chr9-42171486-INS-220
chr9-42281338-INS-118
chr9-42327994-INS-51
chr9-42347192-INS-166
chr9-42363796-INS-52
chr9-42385526-INS-6383
chr9-42387467-INS-95
chr9-42444104-INS-135
chr9-42465752-INS-59
chr9-42521900-INS-51
chr9-42544649-INS-6680
chr9-42548145-INS-99
chr9-42555454-INS-309
chr9-42561006-INS-59
chr9-42576340-INS-50
chr9-42590771-INS-50
chr9-42610070-INS-2889
chr9-42738329-INS-52
chr9-42741540-INS-74
chr9-42778439-INS-50
chr9-42792133-INS-217
chr9-42863730-INS-60
chr9-42930534-INS-878
chr9-42953378-INS-66
chr9-42963397-INS-205
chr9-43007976-INS-1866
chr9-43072723-INS-56
chr9-43075569-INS-252
chr9-43091286-INS-212
chr9-43124629-INS-50

chr9-43149887-INS-81
chr9-43156590-INS-212
chr9-43159356-INS-50
chr9-43186466-INS-68
chr9-43282012-INS-51
chr9-43297677-INS-214
chr9-43357494-INS-112
chr9-43364145-INS-559
chr9-43369697-INS-62
chr9-43430490-INS-206
chr9-43444403-INS-462
chr9-43492738-INS-92
chr9-43520941-INS-217
chr9-43584384-INS-92
chr9-43595737-INS-136
chr9-43595872-INS-487
chr9-43732091-INS-54
chr9-43766354-INS-77
chr9-43784578-INS-50
chr9-43801639-INS-192
chr9-43851179-INS-249
chr9-43861766-INS-65
chr9-43889485-INS-53
chr9-43956917-INS-136
chr9-43965964-INS-98
chr9-43971615-INS-2402
chr9-43972688-INS-73
chr9-44012482-INS-224
chr9-44057485-INS-195
chr9-44057737-INS-211
chr9-44072111-INS-609
chr9-44098580-INS-206
chr9-44117784-INS-225
chr9-44140748-INS-194
chr9-44141424-INS-81
chr9-44181104-INS-94
chr9-44223722-INS-105
chr9-44306195-INS-76
chr9-44336477-INS-75
chr9-44344507-INS-50
chr9-44355461-INS-57
chr9-44355479-INS-194
chr9-44431748-INS-217
chr9-44441098-INS-101
chr9-44441396-INS-383
chr9-44455104-INS-1600
chr9-44466250-INS-129
chr9-44491144-INS-55
chr9-44494190-INS-51
chr9-44525313-INS-221
chr9-44540221-INS-76
chr9-44552765-INS-195
chr9-44571077-INS-198
chr9-44577679-INS-112
chr9-44588984-INS-733
chr9-44606495-INS-199
chr9-44661809-INS-229
chr9-44814168-INS-628
chr9-44822265-INS-240
chr9-44886837-INS-62
chr9-44888064-INS-485
chr9-44929414-INS-9498
chr9-44937492-INS-458
chr9-44942762-INS-116
chr9-45023673-INS-777
chr9-45028434-INS-72
chr9-45167961-INS-160
chr9-45175303-INS-121
chr9-45175303-INS-245
chr9-45175367-INS-51
chr9-45222464-INS-429
chr9-45235758-INS-52
chr9-45239763-INS-66
chr9-45249325-INS-81
chr9-45249334-INS-217
chr9-45277671-INS-77
chr9-45284506-INS-229

chr9-45339415-INS-123
chr9-45356271-INS-75
chr9-45377773-INS-206
chr9-45437836-INS-54
chr9-45443102-INS-58
chr9-45479300-INS-262
chr9-45542149-INS-84
chr9-45570186-INS-68
chr9-45584338-INS-70
chr9-45629584-INS-210
chr9-45641234-INS-98
chr9-45643744-INS-55
chr9-45687714-INS-50
chr9-45706046-INS-76
chr9-45707434-INS-137
chr9-45722315-INS-167
chr9-45735410-INS-849
chr9-45810650-INS-110
chr9-45855265-INS-65
chr9-45865125-INS-3060
chr9-45888286-INS-52
chr9-45903001-INS-1280
chr9-45914319-INS-224
chr9-45936059-INS-50
chr9-45945182-INS-56
chr9-45945185-INS-122
chr9-45945619-INS-198
chr9-45956816-INS-65
chr9-45957990-INS-6442
chr9-45967893-INS-148
chr9-45968035-INS-1940
chr9-45968082-INS-700
chr9-45989113-INS-432
chr9-45989616-INS-116
chr9-46078147-INS-89
chr9-46079273-INS-104
chr9-46084100-INS-60
chr9-46109285-INS-60
chr9-46121085-INS-61
chr9-46160470-INS-181
chr9-46162495-INS-52
chr9-46164381-INS-120
chr9-46164689-INS-435
chr9-46183340-INS-206
chr9-46190573-INS-51
chr9-46194837-INS-552
chr9-46207509-INS-78
chr9-46245055-INS-65
chr9-46252207-INS-210
chr9-46311542-INS-419
chr9-46311550-INS-73
chr9-46355448-INS-232
chr9-46365633-INS-1195
chr9-46371990-INS-64
chr9-46400000-INS-125
chr9-46515302-INS-177
chr9-46560267-INS-54
chr9-46563557-INS-1574
chr9-46569852-INS-79
chr9-46633164-INS-1089
chr9-46687222-INS-56
chr9-46703503-INS-52
chr9-47017638-INS-177
chr9-47056460-INS-95
chr9-47061217-INS-98
chr9-47088620-INS-60
chr9-47171140-INS-233
chr9-47209807-INS-162
chr9-47210284-INS-234
chr9-47254559-INS-52
chr9-47338714-INS-228
chr9-47358093-INS-50
chr9-47366835-INS-158
chr9-47447557-INS-52
chr9-47456948-INS-7143
chr9-47461657-INS-19941
chr9-47670243-INS-217

chr9-47883850-INS-75
chr9-47883900-INS-165
chr9-47967493-INS-876
chr9-47967508-INS-388
chr9-48118930-INS-116
chr9-48119106-INS-230
chr9-48168062-INS-263
chr9-48215704-INS-661
chr9-48226568-INS-480
chr9-48226805-INS-1044
chr9-48227128-INS-95
chr9-48244919-INS-56
chr9-48260708-INS-6270
chr9-48280004-INS-56
chr9-48289016-INS-492
chr9-48296075-INS-50
chr9-48300900-INS-6384
chr9-48302299-INS-72
chr9-48332891-INS-310
chr9-48333290-INS-112
chr9-48352246-INS-294
chr9-48366627-INS-70
chr9-48469100-INS-63
chr9-48486938-INS-1893
chr9-48509784-INS-159
chr9-48638099-INS-52
chr9-48645124-INS-58
chr9-48718647-INS-178
chr9-48801927-INS-80
chr9-48802617-INS-213
chr9-48811797-INS-50
chr9-48900802-INS-203
chr9-48979402-INS-260
chr9-49012106-INS-1131
chr9-49088026-INS-124
chr9-49088165-INS-369
chr9-49095936-INS-56
chr9-49131906-INS-71
chr9-49132357-INS-108
chr9-49198822-INS-600
chr9-49199058-INS-677
chr9-49199067-INS-2099
chr9-49210135-INS-2740
chr9-49214791-INS-2500
chr9-49328907-INS-75
chr9-49329006-INS-176
chr9-49345385-INS-1492
chr9-49345751-INS-51
chr9-49379643-INS-912
chr9-49407817-INS-108
chr9-49446657-INS-54
chr9-49455448-INS-3036
chr9-49500895-INS-622
chr9-49965188-INS-6366
chr9-49990493-INS-129
chr9-50065057-INS-88
chr9-50206407-INS-52
chr9-50251935-INS-60
chr9-50281210-INS-191
chr9-50327413-INS-400
chr9-50327413-INS-89
chr9-50327681-INS-651
chr9-50401418-INS-225
chr9-50448524-INS-68
chr9-50487140-INS-214
chr9-50672526-INS-51
chr9-50730379-INS-673
chr9-50744397-INS-408
chr9-50906714-INS-1141
chr9-50923340-INS-58
chr9-50926834-INS-6798
chr9-50957257-INS-6758
chr9-51001249-INS-162
chr9-51011969-INS-160
chr9-51019834-INS-69
chr9-51039472-INS-352
chr9-51039504-INS-96

chr9-51064270-INS-51
chr9-51086711-INS-112
chr9-51106312-INS-50
chr9-51138935-INS-125
chr9-51234788-INS-2166
chr9-51252823-INS-62
chr9-51291975-INS-74
chr9-51303498-INS-142
chr9-51303498-INS-345
chr9-51340340-INS-204
chr9-51340511-INS-67
chr9-51349855-INS-1885
chr9-51404097-INS-100
chr9-51458332-INS-66
chr9-51474827-INS-1037
chr9-51490659-INS-70
chr9-51515082-INS-367
chr9-51515089-INS-140
chr9-51515757-INS-52
chr9-51587443-INS-4054
chr9-51644788-INS-68
chr9-51646168-INS-3739
chr9-51649564-INS-50
chr9-51672476-INS-100
chr9-51770934-INS-2013
chr9-51815327-INS-164
chr9-51823436-INS-274
chr9-51823497-INS-4895
chr9-51838488-INS-82
chr9-51865649-INS-6589
chr9-51882750-INS-64
chr9-51889755-INS-218
chr9-51889957-INS-259
chr9-51946413-INS-318
chr9-52085539-INS-135
chr9-52098183-INS-507
chr9-52145376-INS-574
chr9-52156085-INS-79
chr9-52156567-INS-269
chr9-52166443-INS-2683
chr9-52174633-INS-77
chr9-52178791-INS-86
chr9-52216405-INS-53
chr9-52228025-INS-128
chr9-52259612-INS-498
chr9-52316424-INS-61
chr9-52381916-INS-50
chr9-52409587-INS-120
chr9-52423679-INS-105
chr9-52497244-INS-149
chr9-52497244-INS-621
chr9-52499717-INS-6208
chr9-52521813-INS-422
chr9-52521832-INS-162
chr9-52541418-INS-4049
chr9-52544558-INS-52
chr9-52545012-INS-3519
chr9-52603024-INS-5415
chr9-52606813-INS-5642
chr9-52660228-INS-168
chr9-52660423-INS-284
chr9-52675024-INS-51
chr9-52710592-INS-50
chr9-52753341-INS-625
chr9-52833215-INS-75
chr9-52844334-INS-549
chr9-52853524-INS-58
chr9-52889757-INS-56
chr9-52919608-INS-3873
chr9-53009318-INS-65
chr9-53040329-INS-83
chr9-53041578-INS-52
chr9-53046775-INS-312
chr9-53057934-INS-4053
chr9-53065750-INS-2075
chr9-53117105-INS-64
chr9-53117874-INS-120

chr9-53151118-INS-50
chr9-53156058-INS-205
chr9-53226009-INS-176
chr9-53243936-INS-208
chr9-53294890-INS-54
chr9-53372497-INS-86
chr9-53380046-INS-221
chr9-53388042-INS-285
chr9-53414208-INS-84
chr9-53418183-INS-54
chr9-53458449-INS-310
chr9-53538098-INS-50
chr9-53545639-INS-205
chr9-53550529-INS-117
chr9-53561707-INS-264
chr9-53566841-INS-186
chr9-53588651-INS-50
chr9-53591342-INS-216
chr9-53627182-INS-88
chr9-53665030-INS-52
chr9-53683474-INS-99
chr9-53725550-INS-54
chr9-53743957-INS-55
chr9-53743987-INS-946
chr9-53744699-INS-52
chr9-53767934-INS-2639
chr9-53785305-INS-250
chr9-53822545-INS-6390
chr9-53852031-INS-58
chr9-53901974-INS-6382
chr9-53914960-INS-52
chr9-53919675-INS-72
chr9-53952518-INS-79
chr9-53954301-INS-6730
chr9-53958903-INS-1848
chr9-53999671-INS-218
chr9-54009398-INS-52
chr9-54221474-INS-60
chr9-54287876-INS-2539
chr9-54335171-INS-52
chr9-54336981-INS-54
chr9-54340814-INS-238
chr9-54354795-INS-173
chr9-54377336-INS-7091
chr9-54389515-INS-353
chr9-54391603-INS-115
chr9-54441073-INS-189
chr9-54475314-INS-95
chr9-54594121-INS-2319
chr9-54629876-INS-178
chr9-54718413-INS-61
chr9-54733866-INS-136
chr9-54767279-INS-58
chr9-54769108-INS-167
chr9-54796397-INS-218
chr9-54850059-INS-215
chr9-54859675-INS-223
chr9-54865688-INS-64
chr9-54868842-INS-3864
chr9-54870741-INS-67
chr9-54999618-INS-246
chr9-55025995-INS-219
chr9-55136174-INS-50
chr9-55145217-INS-283
chr9-55166442-INS-50
chr9-55170312-INS-130
chr9-55175280-INS-162
chr9-55251341-INS-207
chr9-55282712-INS-50
chr9-55309130-INS-152
chr9-55310718-INS-50
chr9-55321368-INS-6224
chr9-55362481-INS-132
chr9-55366963-INS-236
chr9-55382800-INS-207
chr9-55425722-INS-6361
chr9-55486333-INS-982

chr9-55570462-INS-57
chr9-55631868-INS-56
chr9-55632097-INS-72
chr9-55669808-INS-80
chr9-55739302-INS-1598
chr9-55742427-INS-104
chr9-55762695-INS-2659
chr9-55833972-INS-100
chr9-55899970-INS-58
chr9-55926501-INS-793
chr9-55959026-INS-177
chr9-55980086-INS-1000
chr9-55988208-INS-5519
chr9-56019746-INS-68
chr9-56024707-INS-204
chr9-56057911-INS-247
chr9-56157188-INS-116
chr9-56224398-INS-53
chr9-56227968-INS-2853
chr9-56262200-INS-56
chr9-56298430-INS-1011
chr9-56359927-INS-214
chr9-56395708-INS-201
chr9-56495561-INS-201
chr9-56511503-INS-252
chr9-56624909-INS-66
chr9-56628494-INS-118
chr9-56716262-INS-135
chr9-56841192-INS-89
chr9-56846425-INS-95
chr9-56885660-INS-239
chr9-56948475-INS-456
chr9-56950362-INS-211
chr9-56954751-INS-107
chr9-56955350-INS-182
chr9-57070848-INS-205
chr9-57071529-INS-200
chr9-57072440-INS-2702
chr9-57072472-INS-235
chr9-57078741-INS-52
chr9-57120690-INS-203
chr9-57172224-INS-126
chr9-57192348-INS-498
chr9-57248624-INS-202
chr9-57280733-INS-3884
chr9-57312116-INS-50
chr9-57317408-INS-192
chr9-57347571-INS-167
chr9-57379724-INS-4842
chr9-57504341-INS-217
chr9-57572187-INS-2058
chr9-57572191-INS-2382
chr9-57579878-INS-68
chr9-57602908-INS-466
chr9-57632125-INS-51
chr9-57636126-INS-1027
chr9-57636225-INS-153
chr9-57753300-INS-51
chr9-57757912-INS-304
chr9-57757912-INS-85
chr9-57771308-INS-3885
chr9-57772052-INS-142
chr9-57787592-INS-130
chr9-57790972-INS-89
chr9-57806978-INS-62
chr9-57831400-INS-215
chr9-57843698-INS-210
chr9-57878703-INS-56
chr9-57880008-INS-3744
chr9-57899906-INS-85
chr9-57910674-INS-513
chr9-57929671-INS-399
chr9-57936370-INS-5797
chr9-58003038-INS-51
chr9-58148049-INS-168
chr9-58168438-INS-285
chr9-58173439-INS-202

chr9-58181935-INS-225
chr9-58189851-INS-183
chr9-58191157-INS-128
chr9-58191254-INS-221
chr9-58191540-INS-64
chr9-58227086-INS-209
chr9-58238355-INS-211
chr9-58239314-INS-704
chr9-58284475-INS-168
chr9-58288802-INS-198
chr9-58296254-INS-57
chr9-58319638-INS-50
chr9-58356411-INS-156
chr9-58390084-INS-167
chr9-58477845-INS-50
chr9-58479670-INS-408
chr9-58480580-INS-452
chr9-58483272-INS-249
chr9-58483350-INS-78
chr9-58499952-INS-113
chr9-58518153-INS-60
chr9-58518153-INS-615
chr9-58602480-INS-124
chr9-58625692-INS-4174
chr9-58645685-INS-52
chr9-58654398-INS-68
chr9-58663474-INS-62
chr9-58674761-INS-51
chr9-58674774-INS-139
chr9-58674821-INS-2344
chr9-58674874-INS-69
chr9-58696473-INS-74
chr9-58706127-INS-245
chr9-58706139-INS-931
chr9-58713733-INS-5628
chr9-58717109-INS-1063
chr9-58785057-INS-50
chr9-58786128-INS-84
chr9-58800773-INS-179
chr9-58837093-INS-56
chr9-58846302-INS-1984
chr9-58910997-INS-1196
chr9-59013798-INS-105
chr9-59022784-INS-50
chr9-59042818-INS-60
chr9-59059116-INS-126
chr9-59067994-INS-114
chr9-59069542-INS-6508
chr9-59086383-INS-646
chr9-59086794-INS-902
chr9-59087675-INS-249
chr9-59091405-INS-106
chr9-59160976-INS-84
chr9-59163562-INS-500
chr9-59189624-INS-222
chr9-59203633-INS-60
chr9-59211719-INS-6442
chr9-59212328-INS-452
chr9-59213007-INS-76
chr9-59322829-INS-51
chr9-59351231-INS-57
chr9-59377951-INS-90
chr9-59407247-INS-498
chr9-59416423-INS-92
chr9-59420445-INS-6147
chr9-59533730-INS-180
chr9-59546458-INS-219
chr9-59546497-INS-60
chr9-59546829-INS-207
chr9-59584713-INS-53
chr9-59625329-INS-378
chr9-59625731-INS-1296
chr9-59638365-INS-52
chr9-59652188-INS-197
chr9-59724495-INS-68
chr9-59913165-INS-4396
chr9-59931092-INS-50

chr9-60110162-INS-50
chr9-60161416-INS-51
chr9-60166489-INS-70
chr9-60216014-INS-128
chr9-60238563-INS-85
chr9-60241896-INS-58
chr9-60264857-INS-219
chr9-60293821-INS-50
chr9-60357907-INS-6786
chr9-60413911-INS-73
chr9-60415174-INS-372
chr9-60446878-INS-334
chr9-60469554-INS-50
chr9-60474528-INS-1475
chr9-60498355-INS-50
chr9-60510832-INS-62
chr9-60548944-INS-71
chr9-60564202-INS-165
chr9-60606445-INS-140
chr9-60694641-INS-157
chr9-60706472-INS-70
chr9-60715986-INS-50
chr9-60792880-INS-149
chr9-60805348-INS-100
chr9-60810996-INS-240
chr9-60913706-INS-65
chr9-60935176-INS-107
chr9-60935405-INS-119
chr9-60947320-INS-126
chr9-60965888-INS-3435
chr9-61049543-INS-50
chr9-61113093-INS-50
chr9-61126664-INS-54
chr9-61168523-INS-52
chr9-61332265-INS-714
chr9-61337731-INS-52
chr9-61361918-INS-80
chr9-61369659-INS-54
chr9-61369961-INS-68
chr9-61371676-INS-468
chr9-61379523-INS-64
chr9-61380281-INS-205
chr9-61381051-INS-345
chr9-61388461-INS-100
chr9-61402085-INS-1460
chr9-61415537-INS-507
chr9-61428325-INS-1859
chr9-61492689-INS-774
chr9-61492701-INS-136
chr9-61494560-INS-3813
chr9-61514380-INS-172
chr9-61541111-INS-62
chr9-61566001-INS-56
chr9-61571300-INS-108
chr9-61674905-INS-1733
chr9-61694460-INS-62
chr9-61732968-INS-168
chr9-61802730-INS-52
chr9-61818282-INS-188
chr9-61830689-INS-193
chr9-61846168-INS-90
chr9-61847394-INS-96
chr9-61862410-INS-206
chr9-61887510-INS-138
chr9-61943116-INS-180
chr9-61943116-INS-384
chr9-62109100-INS-57
chr9-62198696-INS-12791
chr9-62217841-INS-78
chr9-62217939-INS-83
chr9-62222202-INS-55
chr9-62222610-INS-252
chr9-62227501-INS-166
chr9-62227739-INS-361
chr9-62238419-INS-2722
chr9-62239356-INS-7165
chr9-62299817-INS-50

chr9-62304007-INS-97
chr9-62356571-INS-2599
chr9-62375265-INS-176
chr9-62395355-INS-116
chr9-62395659-INS-55
chr9-62396005-INS-262
chr9-62419812-INS-2796
chr9-62488514-INS-7273
chr9-62511357-INS-240
chr9-62512258-INS-3494
chr9-62516418-INS-81
chr9-62531740-INS-65
chr9-62541855-INS-77
chr9-62544155-INS-172
chr9-62554658-INS-156
chr9-62556890-INS-256
chr9-62561251-INS-210
chr9-62575873-INS-323
chr9-62597595-INS-138
chr9-62598237-INS-89
chr9-62623656-INS-87
chr9-62640253-INS-340
chr9-62653847-INS-111
chr9-62667312-INS-50
chr9-62749417-INS-174
chr9-62781914-INS-120
chr9-62898266-INS-352
chr9-62933429-INS-306
chr9-62951558-INS-232
chr9-62955193-INS-88
chr9-62992274-INS-55
chr9-63063632-INS-118
chr9-63078273-INS-82
chr9-63147011-INS-170
chr9-63186511-INS-6513
chr9-63215478-INS-504
chr9-63374728-INS-1088
chr9-63382540-INS-259
chr9-63474158-INS-330
chr9-63485224-INS-51
chr9-63492293-INS-56
chr9-63493248-INS-8790
chr9-63522912-INS-85
chr9-63588158-INS-62
chr9-63591337-INS-86
chr9-63597361-INS-473
chr9-63597420-INS-1118
chr9-63607750-INS-180
chr9-63631427-INS-98
chr9-63655992-INS-143
chr9-63728446-INS-122
chr9-63739145-INS-281
chr9-63755800-INS-421
chr9-63769634-INS-50
chr9-63781712-INS-400
chr9-63781833-INS-86
chr9-63858407-INS-210
chr9-63892812-INS-7275
chr9-63972342-INS-216
chr9-64012493-INS-140
chr9-64013344-INS-57
chr9-64069517-INS-226
chr9-64077645-INS-124
chr9-64106863-INS-252
chr9-64107710-INS-226
chr9-64114374-INS-207
chr9-64146114-INS-201
chr9-64146308-INS-174
chr9-64147898-INS-58
chr9-64167167-INS-252
chr9-64180580-INS-215
chr9-64185283-INS-216
chr9-64233507-INS-127
chr9-64233520-INS-54
chr9-64258523-INS-240
chr9-64268182-INS-157
chr9-64268574-INS-50

chr9-64280180-INS-185
chr9-64283386-INS-52
chr9-64342817-INS-56
chr9-64373457-INS-74
chr9-64374452-INS-94
chr9-64404812-INS-296
chr9-64464116-INS-232
chr9-64474005-INS-56
chr9-64496594-INS-64
chr9-64498563-INS-192
chr9-64510846-INS-141
chr9-64526438-INS-50
chr9-64661122-INS-4683
chr9-64667313-INS-126
chr9-64668927-INS-50
chr9-64673393-INS-51
chr9-64742463-INS-217
chr9-64747861-INS-204
chr9-64776975-INS-282
chr9-64792825-INS-56
chr9-64883338-INS-120
chr9-64914586-INS-54
chr9-64926840-INS-186
chr9-64978590-INS-56
chr9-65007689-INS-60
chr9-65021527-INS-104
chr9-65058791-INS-123
chr9-65082331-INS-240
chr9-65108010-INS-203
chr9-65108442-INS-261
chr9-65157079-INS-214
chr9-65200771-INS-64
chr9-65220143-INS-2612
chr9-65239181-INS-84
chr9-65251320-INS-70
chr9-65284668-INS-227
chr9-65321643-INS-51
chr9-65331427-INS-52
chr9-65379881-INS-64
chr9-65395554-INS-383
chr9-65431302-INS-217
chr9-65452259-INS-78
chr9-65510184-INS-60
chr9-65512207-INS-205
chr9-65526141-INS-103
chr9-65551041-INS-446
chr9-65714929-INS-50
chr9-65728414-INS-50
chr9-65728835-INS-201
chr9-65744765-INS-103
chr9-65808297-INS-5130
chr9-65844914-INS-214
chr9-65873724-INS-56
chr9-65893238-INS-52
chr9-65942248-INS-54
chr9-66019433-INS-55
chr9-66019987-INS-62
chr9-66060465-INS-50
chr9-66084303-INS-50
chr9-66102312-INS-92
chr9-66123822-INS-52
chr9-66130658-INS-108
chr9-66155850-INS-186
chr9-66170762-INS-309
chr9-66184991-INS-56
chr9-66189832-INS-4717
chr9-66190343-INS-746
chr9-66203394-INS-2625
chr9-66241381-INS-200
chr9-66253058-INS-64
chr9-66365037-INS-86
chr9-66410425-INS-480
chr9-66475104-INS-150
chr9-66527149-INS-499
chr9-66540472-INS-1138
chr9-66551403-INS-176
chr9-66570966-INS-452

chr9-66570972-INS-124
chr9-66571701-INS-100
chr9-66582203-INS-79
chr9-66582337-INS-90
chr9-66591465-INS-2832
chr9-66597606-INS-1845
chr9-66602690-INS-290
chr9-66612546-INS-338
chr9-66612593-INS-124
chr9-66616101-INS-131
chr9-66616101-INS-56
chr9-66618758-INS-123
chr9-66689230-INS-102
chr9-66721635-INS-2686
chr9-66739955-INS-60
chr9-66866250-INS-50
chr9-66885124-INS-66
chr9-66885444-INS-60
chr9-66888254-INS-134
chr9-66968652-INS-60
chr9-67002634-INS-6460
chr9-67030102-INS-62
chr9-67034497-INS-232
chr9-67035712-INS-7127
chr9-67068264-INS-6626
chr9-67167476-INS-372
chr9-67169018-INS-146
chr9-67182188-INS-191
chr9-67207575-INS-82
chr9-67315896-INS-52
chr9-67329124-INS-50
chr9-67329188-INS-125
chr9-67348428-INS-739
chr9-67357062-INS-64
chr9-67369814-INS-64
chr9-67374074-INS-2937
chr9-67400765-INS-52
chr9-67405573-INS-117
chr9-67407641-INS-137
chr9-67407724-INS-542
chr9-67425412-INS-50
chr9-67488015-INS-6403
chr9-67489675-INS-65
chr9-67494899-INS-70
chr9-67498305-INS-94
chr9-67498495-INS-243
chr9-67527457-INS-68
chr9-67580761-INS-1868
chr9-67586604-INS-6323
chr9-67587417-INS-70
chr9-67663386-INS-196
chr9-67678269-INS-434
chr9-67684937-INS-50
chr9-67700205-INS-59
chr9-67713023-INS-494
chr9-67713147-INS-102
chr9-67730223-INS-52
chr9-67747485-INS-162
chr9-67750056-INS-53
chr9-67756373-INS-188
chr9-67763687-INS-210
chr9-67774029-INS-52
chr9-67796515-INS-52
chr9-67798756-INS-136
chr9-68004844-INS-54
chr9-68020224-INS-57
chr9-68024929-INS-50
chr9-68073232-INS-66
chr9-68073268-INS-159
chr9-68089379-INS-1640
chr9-68142945-INS-98
chr9-68156623-INS-66
chr9-68190099-INS-148
chr9-68206987-INS-64
chr9-68216088-INS-56
chr9-68235469-INS-55
chr9-68263219-INS-15748

chr9-68339293-INS-482
chr9-68365208-INS-2698
chr9-68391152-INS-3641
chr9-68401135-INS-183
chr9-68442531-INS-52
chr9-68460156-INS-2663
chr9-68470883-INS-2708
chr9-68547146-INS-54
chr9-68578965-INS-69
chr9-68587872-INS-71
chr9-68624043-INS-149
chr9-68635916-INS-52
chr9-68681608-INS-67
chr9-68779714-INS-63
chr9-68793925-INS-148
chr9-68804111-INS-60
chr9-68810272-INS-155
chr9-68841904-INS-113
chr9-68863764-INS-1862
chr9-68890288-INS-978
chr9-68900130-INS-6413
chr9-68912771-INS-131
chr9-68917480-INS-100
chr9-68925280-INS-54
chr9-68926091-INS-51
chr9-68974318-INS-60
chr9-68982725-INS-75
chr9-69007232-INS-75
chr9-69007432-INS-850
chr9-69024238-INS-189
chr9-69035589-INS-243
chr9-69041540-INS-105
chr9-69064662-INS-58
chr9-69090991-INS-72
chr9-69091106-INS-114
chr9-69091106-INS-55
chr9-69144993-INS-50
chr9-69148571-INS-68
chr9-69157438-INS-50
chr9-69158063-INS-263
chr9-69163401-INS-89
chr9-69189638-INS-78
chr9-69189771-INS-239
chr9-69209916-INS-71
chr9-69262373-INS-56
chr9-69283154-INS-56
chr9-69310682-INS-207
chr9-69315011-INS-498
chr9-69329838-INS-54
chr9-69372719-INS-1854
chr9-69381726-INS-56
chr9-69390022-INS-50
chr9-69412462-INS-60
chr9-69429895-INS-66
chr9-69478682-INS-1574
chr9-69508964-INS-1972
chr9-69509921-INS-170
chr9-69522356-INS-1069
chr9-69572363-INS-81
chr9-69573339-INS-60
chr9-69625038-INS-59
chr9-69644559-INS-60
chr9-69644625-INS-72
chr9-69649556-INS-236
chr9-69712404-INS-309
chr9-69784961-INS-7127
chr9-69799208-INS-62
chr9-69807088-INS-54
chr9-69825689-INS-5903
chr9-69842691-INS-121
chr9-69846354-INS-98
chr9-69846654-INS-75
chr9-69857527-INS-262
chr9-70024273-INS-56
chr9-70133982-INS-50
chr9-70140855-INS-191
chr9-70153750-INS-60

chr9-70202726-INS-56
chr9-70243284-INS-182
chr9-70252531-INS-174
chr9-70257547-INS-114
chr9-70269264-INS-57
chr9-70288519-INS-100
chr9-70300611-INS-76
chr9-70305495-INS-50
chr9-70396379-INS-60
chr9-70428787-INS-73
chr9-70439003-INS-89
chr9-70444969-INS-191
chr9-70448436-INS-201
chr9-70484800-INS-200
chr9-70530365-INS-387
chr9-70530843-INS-315
chr9-70566075-INS-203
chr9-70609459-INS-52
chr9-70646119-INS-52
chr9-70689492-INS-6591
chr9-70700733-INS-54
chr9-70703133-INS-6285
chr9-70718077-INS-298
chr9-70721286-INS-251
chr9-70721286-INS-81
chr9-70768001-INS-50
chr9-70820217-INS-176
chr9-70917604-INS-54
chr9-70933662-INS-225
chr9-70936223-INS-72
chr9-70938281-INS-448
chr9-70938334-INS-2864
chr9-70938398-INS-74
chr9-71003746-INS-1354
chr9-71009334-INS-120
chr9-71050149-INS-1536
chr9-71089650-INS-1875
chr9-71091209-INS-142
chr9-71112067-INS-202
chr9-71141484-INS-82
chr9-71143302-INS-50
chr9-71151058-INS-204
chr9-71184077-INS-191
chr9-71234506-INS-58
chr9-71240019-INS-60
chr9-71250509-INS-272
chr9-71273492-INS-65
chr9-71274662-INS-88
chr9-71313150-INS-177
chr9-71323997-INS-58
chr9-71356074-INS-282
chr9-71356102-INS-104
chr9-71357375-INS-50
chr9-71374499-INS-72
chr9-71379654-INS-53
chr9-71399037-INS-160
chr9-71402008-INS-60
chr9-71410494-INS-51
chr9-71460571-INS-114
chr9-71460619-INS-462
chr9-71469969-INS-209
chr9-71477142-INS-159
chr9-71479212-INS-1034
chr9-71488288-INS-2485
chr9-71493628-INS-202
chr9-71518515-INS-151
chr9-71552676-INS-90
chr9-71575610-INS-2994
chr9-71575610-INS-374
chr9-71593690-INS-84
chr9-71652610-INS-66
chr9-71695206-INS-73
chr9-71699006-INS-57
chr9-71700825-INS-247
chr9-71707765-INS-127
chr9-71711449-INS-58
chr9-71722480-INS-1492

chr9-71782657-INS-220
chr9-71811714-INS-62
chr9-71927502-INS-81
chr9-72078302-INS-69
chr9-72117955-INS-237
chr9-72125530-INS-247
chr9-72125530-INS-78
chr9-72129522-INS-64
chr9-72147899-INS-6120
chr9-72153074-INS-6228
chr9-72172042-INS-54
chr9-72178302-INS-4751
chr9-72228645-INS-208
chr9-72249242-INS-345
chr9-72281977-INS-75
chr9-72288698-INS-320
chr9-72290410-INS-730
chr9-72303728-INS-393
chr9-72320305-INS-78
chr9-72322130-INS-343
chr9-72365950-INS-1941
chr9-72378648-INS-260
chr9-72382224-INS-727
chr9-72385829-INS-95
chr9-72393870-INS-2587
chr9-72416525-INS-56
chr9-72433942-INS-259
chr9-72474854-INS-209
chr9-72493631-INS-208
chr9-72494243-INS-143
chr9-72502225-INS-185
chr9-72533030-INS-207
chr9-72536200-INS-214
chr9-72538810-INS-133
chr9-72538871-INS-307
chr9-72539583-INS-236
chr9-72540521-INS-7215
chr9-72542044-INS-268
chr9-72557643-INS-63
chr9-72571283-INS-57
chr9-72585733-INS-199
chr9-72595955-INS-177
chr9-72608396-INS-222
chr9-72616332-INS-374
chr9-72622997-INS-1671
chr9-72658652-INS-584
chr9-72660820-INS-52
chr9-72660969-INS-50
chr9-72669949-INS-2775
chr9-72691774-INS-298
chr9-72704612-INS-352
chr9-72712517-INS-52
chr9-72723051-INS-200
chr9-72725166-INS-222
chr9-72728350-INS-129
chr9-72743410-INS-488
chr9-72771154-INS-125
chr9-72784983-INS-580
chr9-72791771-INS-1885
chr9-72826475-INS-303
chr9-72840814-INS-192
chr9-72841773-INS-218
chr9-72856494-INS-74
chr9-72864532-INS-200
chr9-72898640-INS-142
chr9-72913845-INS-128
chr9-72929105-INS-50
chr9-72939300-INS-207
chr9-72940183-INS-185
chr9-72960670-INS-56
chr9-72970591-INS-246
chr9-73007107-INS-88
chr9-73010723-INS-291
chr9-73010797-INS-78
chr9-73021678-INS-588
chr9-73025307-INS-197
chr9-73026161-INS-179

chr9-73057074-INS-322
chr9-73058952-INS-76
chr9-73066775-INS-50
chr9-73075028-INS-4667
chr9-73095356-INS-72
chr9-73107294-INS-106
chr9-73109058-INS-6451
chr9-73128419-INS-540
chr9-73130305-INS-1130
chr9-73152407-INS-151
chr9-73153181-INS-104
chr9-73157244-INS-199
chr9-73163465-INS-1885
chr9-73172025-INS-1336
chr9-73182733-INS-54
chr9-73191345-INS-203
chr9-73216847-INS-2578
chr9-73230436-INS-52
chr9-73238335-INS-1080
chr9-73250174-INS-6250
chr9-73286787-INS-108
chr9-73323760-INS-1034
chr9-73349299-INS-72
chr9-73359963-INS-212
chr9-73359963-INS-92
chr9-73368377-INS-199
chr9-73444232-INS-1967
chr9-73450923-INS-105
chr9-73451341-INS-165
chr9-73451374-INS-1494
chr9-73456827-INS-5142
chr9-73468940-INS-60
chr9-73469101-INS-124
chr9-73469738-INS-194
chr9-73475278-INS-68
chr9-73522510-INS-1974
chr9-73549949-INS-232
chr9-73552703-INS-877
chr9-73555838-INS-95
chr9-73567045-INS-60
chr9-73615287-INS-126
chr9-73660347-INS-146
chr9-73662588-INS-156
chr9-73672963-INS-51
chr9-73825207-INS-74
chr9-73852725-INS-1775
chr9-73860156-INS-76
chr9-73860960-INS-1245
chr9-73868964-INS-66
chr9-73883539-INS-6643
chr9-73891659-INS-60
chr9-73904274-INS-6441
chr9-73910790-INS-55
chr9-73935267-INS-527
chr9-73963592-INS-387
chr9-73966752-INS-71
chr9-73978255-INS-2138
chr9-73984257-INS-128
chr9-73997376-INS-3144
chr9-74000308-INS-104
chr9-74005707-INS-142
chr9-74005707-INS-326
chr9-74005707-INS-52
chr9-74024720-INS-50
chr9-74038383-INS-1108
chr9-74116937-INS-56
chr9-74117167-INS-2557
chr9-74224821-INS-60
chr9-74228406-INS-164
chr9-74231067-INS-51
chr9-74279281-INS-54
chr9-74311333-INS-60
chr9-74314326-INS-57
chr9-74314746-INS-223
chr9-74317043-INS-86
chr9-74337524-INS-108
chr9-74347692-INS-72

chr9-74370926-INS-52
chr9-74403577-INS-7524
chr9-74435813-INS-75
chr9-74455375-INS-54
chr9-74465794-INS-50
chr9-74476896-INS-50
chr9-74480394-INS-71
chr9-74520546-INS-50
chr9-74534937-INS-213
chr9-74566783-INS-152
chr9-74610902-INS-52
chr9-74618253-INS-110
chr9-74627143-INS-58
chr9-74635125-INS-197
chr9-74656701-INS-54
chr9-74675335-INS-178
chr9-74675535-INS-246
chr9-74722103-INS-6254
chr9-74804833-INS-5109
chr9-74849566-INS-66
chr9-74873806-INS-209
chr9-74909563-INS-65
chr9-74925710-INS-107
chr9-74942417-INS-211
chr9-74946502-INS-59
chr9-74954574-INS-216
chr9-74955049-INS-147
chr9-74989461-INS-2357
chr9-74995591-INS-1591
chr9-75001582-INS-50
chr9-75010471-INS-2498
chr9-75156605-INS-52
chr9-75172145-INS-54
chr9-75206191-INS-495
chr9-75206309-INS-231
chr9-75216276-INS-54
chr9-75231932-INS-52
chr9-75259943-INS-222
chr9-75261209-INS-8930
chr9-75346377-INS-139
chr9-75356814-INS-122
chr9-75363151-INS-287
chr9-75366519-INS-248
chr9-75381899-INS-291
chr9-75382045-INS-121
chr9-75395471-INS-50
chr9-75420672-INS-204
chr9-75421190-INS-200
chr9-75426354-INS-3458
chr9-75433755-INS-1845
chr9-75434651-INS-763
chr9-75443474-INS-191
chr9-75484224-INS-50
chr9-75515860-INS-224
chr9-75527355-INS-102
chr9-75546315-INS-52
chr9-75563786-INS-51
chr9-75603340-INS-255
chr9-75645478-INS-7005
chr9-75715358-INS-64
chr9-75751147-INS-50
chr9-75767393-INS-7103
chr9-75782038-INS-52
chr9-75783554-INS-92
chr9-75787947-INS-99
chr9-75798561-INS-865
chr9-75799248-INS-50
chr9-75807112-INS-50
chr9-75852711-INS-84
chr9-75859352-INS-102
chr9-75863426-INS-51
chr9-75888131-INS-83
chr9-75901451-INS-72
chr9-75912448-INS-2822
chr9-75913031-INS-7218
chr9-75928691-INS-85
chr9-75944075-INS-110

chr9-75970048-INS-54
chr9-75990464-INS-729
chr9-76022422-INS-302
chr9-76078224-INS-104
chr9-76205425-INS-774
chr9-76205437-INS-316
chr9-76217679-INS-191
chr9-76217699-INS-64
chr9-76265251-INS-120
chr9-76265251-INS-52
chr9-76288978-INS-137
chr9-76288986-INS-65
chr9-76311833-INS-143
chr9-76323959-INS-167
chr9-76357024-INS-60
chr9-76384555-INS-50
chr9-76385314-INS-6767
chr9-76404806-INS-68
chr9-76415180-INS-58
chr9-76418156-INS-57
chr9-76471627-INS-334
chr9-76521093-INS-2810
chr9-76582975-INS-62
chr9-76587894-INS-105
chr9-76608492-INS-291
chr9-76674917-INS-6413
chr9-76731339-INS-50
chr9-76741656-INS-1226
chr9-76771880-INS-389
chr9-76779745-INS-156
chr9-76865375-INS-1027
chr9-76866487-INS-142
chr9-76867221-INS-120
chr9-76896709-INS-104
chr9-76990077-INS-5300
chr9-76990186-INS-494
chr9-77022686-INS-1347
chr9-77058968-INS-1458
chr9-77095609-INS-1073
chr9-77099138-INS-200
chr9-77115981-INS-3216
chr9-77116694-INS-1023
chr9-77205291-INS-123
chr9-77237699-INS-380
chr9-77287456-INS-119
chr9-77287967-INS-361
chr9-77289341-INS-229
chr9-77293164-INS-205
chr9-77302867-INS-282
chr9-77303064-INS-58
chr9-77328473-INS-79
chr9-77331746-INS-408
chr9-77353872-INS-103
chr9-77368881-INS-159
chr9-77388348-INS-7494
chr9-77396764-INS-155
chr9-77438508-INS-317
chr9-77439433-INS-334
chr9-77463248-INS-66
chr9-77501214-INS-1178
chr9-77514900-INS-70
chr9-77542236-INS-498
chr9-77558574-INS-54
chr9-77560089-INS-1088
chr9-77575385-INS-138
chr9-77578820-INS-934
chr9-77591821-INS-63
chr9-77600929-INS-7154
chr9-77629324-INS-73
chr9-77643358-INS-168
chr9-77650264-INS-238
chr9-77718161-INS-158
chr9-77718852-INS-74
chr9-77719316-INS-82
chr9-77724944-INS-68
chr9-77772058-INS-3541
chr9-77780941-INS-839

chr9-77790928-INS-1444
chr9-77803387-INS-377
chr9-77803405-INS-124
chr9-77803803-INS-62
chr9-77807531-INS-91
chr9-77807834-INS-68
chr9-77813851-INS-203
chr9-77815271-INS-311
chr9-77820668-INS-51
chr9-77828701-INS-52
chr9-77840501-INS-490
chr9-77841087-INS-66
chr9-77848772-INS-50
chr9-77850194-INS-109
chr9-77884998-INS-57
chr9-77910145-INS-56
chr9-77940755-INS-56
chr9-77947719-INS-420
chr9-77967311-INS-435
chr9-77967657-INS-100
chr9-77985403-INS-50
chr9-77995564-INS-112
chr9-78012110-INS-71
chr9-78029285-INS-203
chr9-78044273-INS-50
chr9-78062178-INS-79
chr9-78072608-INS-234
chr9-78079431-INS-133
chr9-78079656-INS-469
chr9-78082091-INS-126
chr9-78120507-INS-110
chr9-78134056-INS-216
chr9-78145832-INS-241
chr9-78146571-INS-207
chr9-78217970-INS-248
chr9-78226580-INS-129
chr9-78227118-INS-67
chr9-78227163-INS-4663
chr9-78228473-INS-7156
chr9-78230100-INS-310
chr9-78238127-INS-128
chr9-78252206-INS-315
chr9-78252495-INS-209
chr9-78271582-INS-70
chr9-78279365-INS-53
chr9-78280005-INS-174
chr9-78300331-INS-3795
chr9-78324197-INS-782
chr9-78324347-INS-299
chr9-78352580-INS-552
chr9-78362141-INS-258
chr9-78373696-INS-116
chr9-78375203-INS-204
chr9-78377496-INS-203
chr9-78402756-INS-61
chr9-78412722-INS-236
chr9-78414855-INS-186
chr9-78442839-INS-199
chr9-78456835-INS-98
chr9-78460958-INS-68
chr9-78460966-INS-142
chr9-78502596-INS-8458
chr9-78503032-INS-172
chr9-78508318-INS-60
chr9-78515813-INS-234
chr9-78527470-INS-50
chr9-78531475-INS-120
chr9-78581608-INS-318
chr9-78585395-INS-1022
chr9-78599535-INS-70
chr9-78599569-INS-145
chr9-78639807-INS-1547
chr9-78723243-INS-147
chr9-78759647-INS-3447
chr9-78773876-INS-6708
chr9-78801361-INS-147
chr9-78803113-INS-135

chr9-78814244-INS-6940
chr9-78819092-INS-50
chr9-78841082-INS-1877
chr9-78847325-INS-6288
chr9-78889497-INS-64
chr9-78892607-INS-70
chr9-78893227-INS-311
chr9-78902927-INS-68
chr9-78904704-INS-4463
chr9-78962325-INS-76
chr9-78966373-INS-343
chr9-78976403-INS-60
chr9-78976737-INS-78
chr9-78982671-INS-402
chr9-79010183-INS-50
chr9-79012423-INS-449
chr9-79013411-INS-2034
chr9-79043601-INS-1286
chr9-79093421-INS-198
chr9-79111216-INS-58
chr9-79186114-INS-4903
chr9-79187788-INS-7103
chr9-79202426-INS-68
chr9-79206932-INS-7107
chr9-79231453-INS-2068
chr9-79270481-INS-69
chr9-79295119-INS-13289
chr9-79306677-INS-191
chr9-79321547-INS-50
chr9-79335338-INS-1488
chr9-79362906-INS-60
chr9-79368442-INS-6782
chr9-79426698-INS-230
chr9-79426698-INS-690
chr9-79444709-INS-646
chr9-79464546-INS-68
chr9-79464590-INS-228
chr9-79484931-INS-6460
chr9-79510155-INS-99
chr9-79518359-INS-54
chr9-79525715-INS-59
chr9-79545100-INS-128
chr9-79555349-INS-1437
chr9-79643614-INS-389
chr9-79679632-INS-211
chr9-79695972-INS-216
chr9-79696642-INS-164
chr9-79718804-INS-54
chr9-79723856-INS-186
chr9-79731111-INS-52
chr9-79779183-INS-64
chr9-79785776-INS-60
chr9-79794928-INS-74
chr9-79837181-INS-65
chr9-79839780-INS-325
chr9-79839942-INS-461
chr9-79852429-INS-1282
chr9-79864654-INS-1568
chr9-79869099-INS-50
chr9-79877090-INS-4828
chr9-79950374-INS-68
chr9-79958581-INS-1011
chr9-79973393-INS-104
chr9-79995625-INS-194
chr9-79996967-INS-74
chr9-80027324-INS-368
chr9-80061655-INS-232
chr9-80061830-INS-166
chr9-80087938-INS-70
chr9-80103056-INS-76
chr9-80112522-INS-500
chr9-80135026-INS-50
chr9-80256358-INS-6345
chr9-80265673-INS-3772
chr9-80268103-INS-55
chr9-80268684-INS-52
chr9-80278711-INS-315

chr9-80294818-INS-94
chr9-80316001-INS-1168
chr9-80318858-INS-5217
chr9-80326735-INS-6356
chr9-80344284-INS-76
chr9-80378821-INS-781
chr9-80378895-INS-286
chr9-80414308-INS-1682
chr9-80438833-INS-178
chr9-80445490-INS-51
chr9-80449964-INS-1777
chr9-80467030-INS-278
chr9-80503952-INS-6295
chr9-80508016-INS-979
chr9-80626070-INS-50
chr9-80650281-INS-170
chr9-80668066-INS-6333
chr9-80699120-INS-51
chr9-80712178-INS-6790
chr9-80720342-INS-4516
chr9-80763321-INS-272
chr9-80778695-INS-60
chr9-80796645-INS-55
chr9-80808339-INS-64
chr9-80838383-INS-424
chr9-80847833-INS-50
chr9-80891242-INS-2355
chr9-80896184-INS-78
chr9-80900501-INS-50
chr9-80917363-INS-5898
chr9-80951450-INS-1679
chr9-80961441-INS-50
chr9-81014012-INS-479
chr9-81038426-INS-431
chr9-81106446-INS-76
chr9-81110874-INS-1799
chr9-81127684-INS-1057
chr9-81157238-INS-160
chr9-81207101-INS-6479
chr9-81241196-INS-4627
chr9-81258272-INS-50
chr9-81305648-INS-2932
chr9-81312675-INS-117
chr9-81443981-INS-4156
chr9-81450568-INS-703
chr9-81491576-INS-215
chr9-81593708-INS-50
chr9-81607368-INS-162
chr9-81607384-INS-350
chr9-81629638-INS-60
chr9-81660408-INS-50
chr9-81703887-INS-4815
chr9-81716636-INS-1656
chr9-81726737-INS-10640
chr9-81770694-INS-2156
chr9-81775086-INS-162
chr9-81795325-INS-647
chr9-81827610-INS-1272
chr9-81881555-INS-2302
chr9-81887159-INS-1461
chr9-81899947-INS-2414
chr9-81935176-INS-111
chr9-81937663-INS-73
chr9-81957872-INS-551
chr9-82009458-INS-1869
chr9-82061488-INS-90
chr9-82089923-INS-59
chr9-82090370-INS-491
chr9-82105235-INS-109
chr9-82108877-INS-62
chr9-82146774-INS-72
chr9-82146812-INS-173
chr9-82190510-INS-76
chr9-82203057-INS-7104
chr9-82217649-INS-2580
chr9-82236871-INS-811
chr9-82296854-INS-50

chr9-82306558-INS-50
chr9-82309626-INS-1863
chr9-82334542-INS-203
chr9-82366305-INS-1508
chr9-82524167-INS-1386
chr9-82557113-INS-105
chr9-82564001-INS-6373
chr9-82565290-INS-1372
chr9-82567350-INS-50
chr9-82616550-INS-83
chr9-82641577-INS-53
chr9-82646869-INS-108
chr9-82672208-INS-53
chr9-82689638-INS-2283
chr9-82725170-INS-162
chr9-82785503-INS-6189
chr9-82878836-INS-93
chr9-82878911-INS-492
chr9-82879459-INS-200
chr9-82881995-INS-182
chr9-82926569-INS-4390
chr9-83012194-INS-57
chr9-83034910-INS-50
chr9-83038192-INS-90
chr9-83099057-INS-50
chr9-83137867-INS-93
chr9-83137912-INS-647
chr9-83137952-INS-1363
chr9-83158918-INS-51
chr9-83243387-INS-2084
chr9-83258895-INS-549
chr9-83303655-INS-180
chr9-83336230-INS-66
chr9-83354633-INS-1519
chr9-83396546-INS-146
chr9-83400358-INS-399
chr9-83417169-INS-210
chr9-83467459-INS-1352
chr9-83472763-INS-603
chr9-83488730-INS-493
chr9-83500711-INS-1964
chr9-83537147-INS-449
chr9-83563071-INS-1568
chr9-83570237-INS-569
chr9-83572820-INS-6514
chr9-83581477-INS-2622
chr9-83588438-INS-51
chr9-83685145-INS-301
chr9-83697296-INS-52
chr9-83699054-INS-70
chr9-83742954-INS-61
chr9-83777810-INS-6444
chr9-83803183-INS-51
chr9-83821319-INS-176
chr9-83851322-INS-59
chr9-83852944-INS-62
chr9-83864501-INS-50
chr9-83895366-INS-63
chr9-83906497-INS-6332
chr9-83917136-INS-71
chr9-83988598-INS-1734
chr9-83997408-INS-55
chr9-84000640-INS-64
chr9-84011918-INS-54
chr9-84036088-INS-52
chr9-84056176-INS-52
chr9-84092663-INS-3477
chr9-84167143-INS-6596
chr9-84195989-INS-163
chr9-84196018-INS-332
chr9-84240704-INS-3516
chr9-84243647-INS-6467
chr9-84337523-INS-56
chr9-84337976-INS-50
chr9-84381678-INS-678
chr9-84414965-INS-66
chr9-84429105-INS-2133

chr9-84482225-INS-58
chr9-84490679-INS-52
chr9-84547301-INS-6427
chr9-84551562-INS-53
chr9-84554412-INS-78
chr9-84563148-INS-4994
chr9-84634485-INS-139
chr9-84634485-INS-401
chr9-84659876-INS-243
chr9-84691659-INS-202
chr9-84724149-INS-707
chr9-84724159-INS-310
chr9-84740310-INS-2945
chr9-84745386-INS-247
chr9-84746951-INS-168
chr9-84746951-INS-350
chr9-84852638-INS-77
chr9-84852786-INS-122
chr9-84899224-INS-294
chr9-84997027-INS-84
chr9-84998442-INS-92
chr9-85019826-INS-84
chr9-85041800-INS-241
chr9-85041983-INS-90
chr9-85053485-INS-350
chr9-85053819-INS-1532
chr9-85065358-INS-642
chr9-85084906-INS-187
chr9-85087588-INS-615
chr9-85114625-INS-176
chr9-85139559-INS-477
chr9-85142860-INS-62
chr9-85143283-INS-134
chr9-85164146-INS-1020
chr9-85193469-INS-233
chr9-85195426-INS-150
chr9-85255058-INS-168
chr9-85271284-INS-1693
chr9-85305189-INS-53
chr9-85305348-INS-181
chr9-85342059-INS-7086
chr9-85367788-INS-6590
chr9-85378452-INS-6175
chr9-85408830-INS-237
chr9-85419770-INS-51
chr9-85422869-INS-4410
chr9-85425698-INS-72
chr9-85426267-INS-70
chr9-85438009-INS-7270
chr9-85446595-INS-123
chr9-85466722-INS-92
chr9-85473598-INS-50
chr9-85475788-INS-6455
chr9-85488809-INS-5879
chr9-85493328-INS-202
chr9-85530003-INS-77
chr9-85539448-INS-85
chr9-85545236-INS-313
chr9-85549888-INS-191
chr9-85550144-INS-272
chr9-85553286-INS-52
chr9-85572176-INS-6961
chr9-85620051-INS-248
chr9-85641355-INS-59
chr9-85641891-INS-126
chr9-85642339-INS-825
chr9-85655848-INS-3915
chr9-85659873-INS-201
chr9-85666288-INS-81
chr9-85666762-INS-559
chr9-85768085-INS-638
chr9-85772379-INS-1939
chr9-85781310-INS-212
chr9-85790112-INS-208
chr9-85805214-INS-141
chr9-85883665-INS-1809
chr9-85918955-INS-50

chr9-86109658-INS-56
chr9-86117184-INS-6712
chr9-86131591-INS-395
chr9-86132159-INS-1595
chr9-86132729-INS-81
chr9-86159539-INS-94
chr9-86160367-INS-64
chr9-86216019-INS-109
chr9-86218196-INS-52
chr9-86240245-INS-498
chr9-86254815-INS-288
chr9-86271698-INS-50
chr9-86274598-INS-50
chr9-86302645-INS-733
chr9-86326812-INS-77
chr9-86426304-INS-149
chr9-86435946-INS-1194
chr9-86470856-INS-485
chr9-86475493-INS-60
chr9-86479377-INS-60
chr9-86482359-INS-190
chr9-86490287-INS-7088
chr9-86494363-INS-224
chr9-86539382-INS-140
chr9-86541233-INS-929
chr9-86542797-INS-618
chr9-86547073-INS-54
chr9-86553966-INS-270
chr9-86570761-INS-61
chr9-86651126-INS-52
chr9-86690131-INS-59
chr9-86726206-INS-106
chr9-86752866-INS-605
chr9-86805993-INS-220
chr9-86807395-INS-6473
chr9-86812318-INS-51
chr9-86876103-INS-64
chr9-86887359-INS-86
chr9-86937401-INS-6610
chr9-86950868-INS-194
chr9-86957731-INS-200
chr9-86958487-INS-58
chr9-86966065-INS-327
chr9-86977983-INS-403
chr9-87006323-INS-50
chr9-87029242-INS-171
chr9-87034848-INS-167
chr9-87034896-INS-83
chr9-87035441-INS-72
chr9-87058539-INS-6918
chr9-87059107-INS-328
chr9-87075194-INS-1268
chr9-87087477-INS-202
chr9-87094750-INS-243
chr9-87125167-INS-108
chr9-87126892-INS-427
chr9-87129958-INS-1150
chr9-87150405-INS-168
chr9-87161224-INS-1376
chr9-87173110-INS-117
chr9-87218120-INS-78
chr9-87218154-INS-337
chr9-87270214-INS-1145
chr9-87272116-INS-151
chr9-87294421-INS-555
chr9-87294497-INS-1945
chr9-87298357-INS-50
chr9-87306242-INS-91
chr9-87307338-INS-52
chr9-87307400-INS-115
chr9-87356723-INS-156
chr9-87369708-INS-201
chr9-87417823-INS-50
chr9-87447834-INS-103
chr9-87498644-INS-335
chr9-87502226-INS-6862
chr9-87567305-INS-128

chr9-87635572-INS-84
chr9-87638156-INS-2220
chr9-87654685-INS-6485
chr9-87661474-INS-238
chr9-87802320-INS-1030
chr9-87821364-INS-56
chr9-87842728-INS-2783
chr9-87845587-INS-1214
chr9-87845613-INS-86
chr9-87856894-INS-84
chr9-87863795-INS-125
chr9-87884715-INS-72
chr9-87915858-INS-116
chr9-87933087-INS-348
chr9-87941773-INS-52
chr9-87948990-INS-52
chr9-87953599-INS-51
chr9-87960214-INS-140
chr9-87978453-INS-196
chr9-87978453-INS-404
chr9-87978453-INS-6393
chr9-87978685-INS-272
chr9-87979393-INS-416
chr9-88018550-INS-71
chr9-88058913-INS-399
chr9-88072622-INS-217
chr9-88105727-INS-152
chr9-88113097-INS-255
chr9-88118636-INS-62
chr9-88135253-INS-170
chr9-88148077-INS-6253
chr9-88167816-INS-1152
chr9-88171662-INS-84
chr9-88173279-INS-120
chr9-88177048-INS-56
chr9-88194154-INS-55
chr9-88194295-INS-117
chr9-88194425-INS-153
chr9-88255548-INS-1524
chr9-88259099-INS-60
chr9-88265277-INS-159
chr9-88285953-INS-193
chr9-88297221-INS-610
chr9-88297308-INS-220
chr9-88297911-INS-207
chr9-88306735-INS-405
chr9-88306740-INS-1057
chr9-88315235-INS-182
chr9-88341168-INS-209
chr9-88383028-INS-589
chr9-88384104-INS-336
chr9-88392696-INS-56
chr9-88398981-INS-269
chr9-88420734-INS-53
chr9-88433903-INS-574
chr9-88456729-INS-69
chr9-88491194-INS-978
chr9-88507660-INS-418
chr9-88603664-INS-53
chr9-88875442-INS-204
chr9-88910650-INS-255
chr9-89002334-INS-152
chr9-89078707-INS-56
chr9-89084041-INS-62
chr9-89088538-INS-341
chr9-89103192-INS-750
chr9-89117574-INS-175
chr9-89117574-INS-70
chr9-89127277-INS-54
chr9-89129106-INS-91
chr9-89138112-INS-7102
chr9-89172647-INS-400
chr9-89238875-INS-114
chr9-89254280-INS-6794
chr9-89261946-INS-51
chr9-89272454-INS-80
chr9-89279777-INS-105

chr9-89297028-INS-60
chr9-89298127-INS-129
chr9-89311217-INS-121
chr9-89323181-INS-6665
chr9-89348090-INS-66
chr9-89351162-INS-209
chr9-89361445-INS-2581
chr9-89388027-INS-62
chr9-89470322-INS-170
chr9-89473764-INS-108
chr9-89498263-INS-68
chr9-89515005-INS-106
chr9-89515134-INS-112
chr9-89516260-INS-54
chr9-89520626-INS-72
chr9-89531891-INS-70
chr9-89536251-INS-2881
chr9-89547275-INS-1290
chr9-89557209-INS-1530
chr9-89566524-INS-50
chr9-89576147-INS-784
chr9-89580209-INS-8325
chr9-89605303-INS-718
chr9-89627957-INS-6494
chr9-89630468-INS-547
chr9-89645898-INS-100
chr9-89713088-INS-5330
chr9-89719421-INS-723
chr9-89762853-INS-200
chr9-89763476-INS-246
chr9-89772534-INS-52
chr9-89779048-INS-103
chr9-89788340-INS-76
chr9-89820356-INS-183
chr9-89842176-INS-289
chr9-89845260-INS-88
chr9-89861825-INS-72
chr9-89874891-INS-671
chr9-89881731-INS-290
chr9-89907954-INS-173
chr9-89940584-INS-6180
chr9-89954505-INS-52
chr9-89955297-INS-113
chr9-89965464-INS-55
chr9-89990833-INS-203
chr9-90007445-INS-70
chr9-90019953-INS-3121
chr9-90072217-INS-496
chr9-90089044-INS-211
chr9-90114012-INS-50
chr9-90136886-INS-475
chr9-90138366-INS-50
chr9-90149837-INS-64
chr9-90163563-INS-7084
chr9-90212520-INS-5347
chr9-90216137-INS-104
chr9-90230329-INS-495
chr9-90231384-INS-83
chr9-90231433-INS-715
chr9-90238547-INS-54
chr9-90255813-INS-86
chr9-90260875-INS-54
chr9-90327567-INS-6884
chr9-90330410-INS-496
chr9-90360433-INS-50
chr9-90370178-INS-1282
chr9-90408387-INS-421
chr9-90441359-INS-6404
chr9-90458362-INS-54
chr9-90458852-INS-88
chr9-90481962-INS-150
chr9-90488415-INS-6648
chr9-90514905-INS-66
chr9-90516959-INS-766
chr9-90545188-INS-50
chr9-90618184-INS-72
chr9-90625029-INS-60

chr9-90628679-INS-1082
chr9-90649659-INS-54
chr9-90659124-INS-70
chr9-90682009-INS-62
chr9-90688143-INS-259
chr9-90791467-INS-6215
chr9-90829873-INS-524
chr9-90829935-INS-254
chr9-90855343-INS-52
chr9-90947118-INS-4958
chr9-90992291-INS-119
chr9-91042731-INS-317
chr9-91043322-INS-2693
chr9-91056594-INS-56
chr9-91061681-INS-50
chr9-91067461-INS-64
chr9-91079987-INS-88
chr9-91082510-INS-111
chr9-91179330-INS-495
chr9-91209473-INS-6171
chr9-91217164-INS-54
chr9-91248222-INS-57
chr9-91260230-INS-57
chr9-91263047-INS-69
chr9-91304048-INS-52
chr9-91334812-INS-954
chr9-91361172-INS-2245
chr9-91362139-INS-68
chr9-91366190-INS-526
chr9-91379157-INS-98
chr9-91406162-INS-384
chr9-91431392-INS-67
chr9-91449603-INS-51
chr9-91456625-INS-2106
chr9-91460956-INS-70
chr9-91485169-INS-51
chr9-91518085-INS-1395
chr9-91530831-INS-663
chr9-91554387-INS-6307
chr9-91585223-INS-80
chr9-91589707-INS-57
chr9-91629803-INS-1358
chr9-91659762-INS-75
chr9-91659765-INS-154
chr9-91705832-INS-7516
chr9-91729347-INS-1484
chr9-91734265-INS-90
chr9-91736423-INS-7078
chr9-91757363-INS-257
chr9-91798515-INS-223
chr9-91831117-INS-58
chr9-91898980-INS-52
chr9-91899129-INS-54
chr9-91914621-INS-3771
chr9-91916312-INS-5969
chr9-91988861-INS-58
chr9-92010086-INS-647
chr9-92034954-INS-60
chr9-92038418-INS-5012
chr9-92044217-INS-50
chr9-92067770-INS-2999
chr9-92082671-INS-60
chr9-92112372-INS-89
chr9-92116597-INS-94
chr9-92116974-INS-386
chr9-92138269-INS-80
chr9-92196609-INS-199
chr9-92225136-INS-90
chr9-92227875-INS-101
chr9-92270742-INS-1742
chr9-92279792-INS-2683
chr9-92320615-INS-3543
chr9-92391478-INS-479
chr9-92418668-INS-70
chr9-92443040-INS-7027
chr9-92457459-INS-140
chr9-92564504-INS-52

chr9-92635694-INS-595
chr9-92658881-INS-9552
chr9-92686997-INS-66
chr9-92689233-INS-2832
chr9-92728482-INS-1888
chr9-92728769-INS-99
chr9-92731507-INS-486
chr9-92745696-INS-3970
chr9-92753578-INS-4021
chr9-92763449-INS-7089
chr9-92769696-INS-2095
chr9-92769726-INS-224
chr9-92955150-INS-4916
chr9-92967412-INS-2837
chr9-92971000-INS-56
chr9-92984386-INS-14989
chr9-93058452-INS-51
chr9-93113920-INS-51
chr9-93127184-INS-1450
chr9-93133892-INS-50
chr9-93136491-INS-6461
chr9-93174027-INS-838
chr9-93198538-INS-56
chr9-93205578-INS-3782
chr9-93227816-INS-1244
chr9-93334446-INS-50
chr9-93347322-INS-66
chr9-93347885-INS-151
chr9-93370644-INS-1777
chr9-93371422-INS-54
chr9-93372996-INS-4867
chr9-93387223-INS-6393
chr9-93392722-INS-50
chr9-93399440-INS-2538
chr9-93411366-INS-1453
chr9-93430692-INS-51
chr9-93493275-INS-409
chr9-93551775-INS-50
chr9-93566036-INS-57
chr9-93566577-INS-1323
chr9-93570256-INS-7142
chr9-93603670-INS-269
chr9-93652445-INS-295
chr9-93652545-INS-211
chr9-93655925-INS-54
chr9-93682113-INS-6308
chr9-93693139-INS-51
chr9-93704560-INS-6100
chr9-93716422-INS-1054
chr9-93754237-INS-7219
chr9-93765199-INS-55
chr9-93849734-INS-56
chr9-93878124-INS-6634
chr9-93885392-INS-222
chr9-93897396-INS-212
chr9-93989802-INS-50
chr9-93990843-INS-395
chr9-93992112-INS-5850
chr9-94022419-INS-50
chr9-94095253-INS-50
chr9-94159680-INS-7094
chr9-94176811-INS-6588
chr9-94181019-INS-56
chr9-94227720-INS-50
chr9-94292606-INS-54
chr9-94309644-INS-52
chr9-94312811-INS-52
chr9-94323414-INS-50
chr9-94335979-INS-1411
chr9-94403222-INS-1137
chr9-94411875-INS-52
chr9-94456470-INS-1377
chr9-94479852-INS-1059
chr9-94500863-INS-52
chr9-94505850-INS-201
chr9-94533667-INS-268
chr9-94533905-INS-2301

chr9-94538224-INS-50
chr9-94539855-INS-6465
chr9-94541630-INS-72
chr9-94580940-INS-6572
chr9-94591641-INS-1041
chr9-94592358-INS-80
chr9-94610517-INS-50
chr9-94622910-INS-521
chr9-94625671-INS-72
chr9-94632004-INS-122
chr9-94771556-INS-151
chr9-94794976-INS-62
chr9-94816714-INS-88
chr9-94819476-INS-999
chr9-94826754-INS-126
chr9-94860641-INS-324
chr9-94861959-INS-62
chr9-94883749-INS-125
chr9-94900425-INS-50
chr9-94963936-INS-59
chr9-94964276-INS-330
chr9-94964502-INS-504
chr9-94975922-INS-314
chr9-94979636-INS-52
chr9-94991696-INS-83
chr9-95006617-INS-50
chr9-95099731-INS-202
chr9-95101604-INS-51
chr9-95109929-INS-55
chr9-95132945-INS-1287
chr9-95139043-INS-8233
chr9-95147611-INS-192
chr9-95220757-INS-14299
chr9-95227004-INS-73
chr9-95235092-INS-6432
chr9-95240611-INS-6387
chr9-95246531-INS-407
chr9-95252470-INS-4960
chr9-95260098-INS-2574
chr9-95261002-INS-5059
chr9-95261002-INS-525
chr9-95265564-INS-610
chr9-95282592-INS-131
chr9-95296963-INS-7405
chr9-95317297-INS-60
chr9-95324006-INS-134
chr9-95333160-INS-237
chr9-95362799-INS-200
chr9-95415112-INS-2104
chr9-95426720-INS-449
chr9-95428124-INS-57
chr9-95434394-INS-95
chr9-95471103-INS-434
chr9-95474877-INS-53
chr9-95487971-INS-210
chr9-95495841-INS-52
chr9-95500705-INS-71
chr9-95502558-INS-1621
chr9-95543317-INS-50
chr9-95544800-INS-327
chr9-95594728-INS-148
chr9-95594728-INS-65
chr9-95648078-INS-468
chr9-95659555-INS-105
chr9-95672120-INS-68
chr9-95672254-INS-501
chr9-95690831-INS-222
chr9-95702536-INS-50
chr9-95776350-INS-3377
chr9-95843446-INS-200
chr9-95851772-INS-108
chr9-95852266-INS-380
chr9-95859238-INS-240
chr9-95871597-INS-450
chr9-95909574-INS-105
chr9-95957395-INS-432
chr9-95958031-INS-77

chr9-95959665-INS-156
chr9-95960890-INS-57
chr9-95969370-INS-52
chr9-95975641-INS-52
chr9-96004366-INS-138
chr9-96035855-INS-222
chr9-96039725-INS-999
chr9-96077121-INS-108
chr9-96077541-INS-208
chr9-96084450-INS-187
chr9-96085075-INS-243
chr9-96085495-INS-52
chr9-96090689-INS-195
chr9-96121614-INS-64
chr9-96134767-INS-51
chr9-96162083-INS-113
chr9-96164914-INS-1005
chr9-96174257-INS-55
chr9-96198176-INS-218
chr9-96253871-INS-235
chr9-96276049-INS-50
chr9-96277983-INS-554
chr9-96296597-INS-200
chr9-96300023-INS-566
chr9-96300741-INS-822
chr9-96329742-INS-4050
chr9-96335282-INS-3020
chr9-96336879-INS-512
chr9-96336879-INS-6456
chr9-96345058-INS-187
chr9-96345323-INS-207
chr9-96363087-INS-52
chr9-96391580-INS-55
chr9-96391953-INS-492
chr9-96411729-INS-64
chr9-96429383-INS-61
chr9-96501614-INS-72
chr9-96510735-INS-50
chr9-96551659-INS-55
chr9-96562140-INS-1108
chr9-96640076-INS-84
chr9-96640159-INS-230
chr9-96641781-INS-156
chr9-96642059-INS-317
chr9-96648893-INS-54
chr9-96649979-INS-4181
chr9-96675777-INS-108
chr9-96682894-INS-56
chr9-96684291-INS-216
chr9-96688132-INS-5546
chr9-96716666-INS-114
chr9-96717060-INS-224
chr9-96720918-INS-7142
chr9-96734456-INS-137
chr9-96735170-INS-282
chr9-96808534-INS-998
chr9-96828840-INS-239
chr9-96916447-INS-56
chr9-96917223-INS-98
chr9-96930075-INS-7143
chr9-96965659-INS-264
chr9-96965659-INS-726
chr9-96965921-INS-66
chr9-96965987-INS-924
chr9-97000664-INS-236
chr9-97006026-INS-73
chr9-97010290-INS-2913
chr9-97044247-INS-76
chr9-97051881-INS-52
chr9-97152110-INS-1892
chr9-97172583-INS-7916
chr9-97173352-INS-1586
chr9-97189420-INS-56
chr9-97205543-INS-53
chr9-97208149-INS-98
chr9-97229509-INS-129
chr9-97252938-INS-134

chr9-97252977-INS-766
chr9-97261238-INS-4393
chr9-97276340-INS-1231
chr9-97351966-INS-55
chr9-97358714-INS-63
chr9-97377478-INS-62
chr9-97381401-INS-50
chr9-97395722-INS-1008
chr9-97409872-INS-6304
chr9-97530071-INS-243
chr9-97547627-INS-6932
chr9-97583092-INS-145
chr9-97592686-INS-53
chr9-97621761-INS-362
chr9-97649265-INS-3093
chr9-97772156-INS-369
chr9-97790948-INS-12997
chr9-97846430-INS-173
chr9-97894969-INS-50
chr9-98030577-INS-2980
chr9-98064270-INS-120
chr9-98113010-INS-6858
chr9-98149449-INS-1277
chr9-98164116-INS-57
chr9-98192685-INS-51
chr9-98246512-INS-458
chr9-98296328-INS-57
chr9-98341218-INS-132
chr9-98348704-INS-212
chr9-98400827-INS-100
chr9-98403300-INS-72
chr9-98430108-INS-6230
chr9-98505255-INS-55
chr9-98575077-INS-119
chr9-98595308-INS-135
chr9-98615289-INS-186
chr9-98625463-INS-51
chr9-98666502-INS-274
chr9-98737459-INS-50
chr9-98769756-INS-52
chr9-98770012-INS-9396
chr9-98771436-INS-18808
chr9-98811136-INS-985
chr9-98850292-INS-66
chr9-98868286-INS-270
chr9-98899576-INS-50
chr9-98945833-INS-677
chr9-98951190-INS-190
chr9-98956251-INS-78
chr9-98991024-INS-131
chr9-98992621-INS-5991
chr9-98999998-INS-105
chr9-99050945-INS-56
chr9-99083067-INS-373
chr9-99083563-INS-50
chr9-99088287-INS-132
chr9-99101098-INS-520
chr9-99105351-INS-132
chr9-99105351-INS-330
chr9-99133064-INS-5366
chr9-99141430-INS-64
chr9-99144163-INS-54
chr9-99192839-INS-56
chr9-99201322-INS-7474
chr9-99221142-INS-246
chr9-99295326-INS-52
chr9-99322788-INS-108
chr9-99322788-INS-648
chr9-99349165-INS-92
chr9-99359700-INS-81
chr9-99376547-INS-111
chr9-99389899-INS-81
chr9-99427711-INS-175
chr9-99440230-INS-288
chr9-99445652-INS-209
chr9-99476737-INS-207
chr9-99586193-INS-52

chr9-99615804-INS-269
chr9-99657609-INS-88
chr9-99668899-INS-406
chr9-99669516-INS-1528
chr9-99697321-INS-4585
chr9-99703017-INS-67
chr9-99705642-INS-1299
chr9-99843668-INS-3132
chr9-99853283-INS-52
chr9-99858215-INS-65
chr9-99866967-INS-2960
chr9-99871276-INS-216
chr9-99922898-INS-65
chr9-99939598-INS-342
chr9-99983129-INS-6752
chr9-100004233-INS-50
chr9-100005121-INS-74
chr9-100104299-INS-1733
chr9-100125247-INS-2536
chr9-100146259-INS-6459
chr9-100218579-INS-3188
chr9-100242904-INS-128
chr9-100244025-INS-99
chr9-100255426-INS-114
chr9-100268275-INS-274
chr9-100268275-INS-583
chr9-100277315-INS-1029
chr9-100281864-INS-50
chr9-100302876-INS-65
chr9-100328999-INS-7402
chr9-100346675-INS-138
chr9-100418991-INS-138
chr9-100418991-INS-1791
chr9-100439083-INS-109
chr9-100460158-INS-54
chr9-100464679-INS-54
chr9-100476083-INS-58
chr9-100477482-INS-98
chr9-100496313-INS-52
chr9-100497536-INS-128
chr9-100499934-INS-326
chr9-100504368-INS-50
chr9-100522238-INS-76
chr9-100533050-INS-176
chr9-100533070-INS-60
chr9-100585734-INS-194
chr9-100617412-INS-595
chr9-100623272-INS-3123
chr9-100623272-INS-480
chr9-100679857-INS-380
chr9-100795366-INS-121
chr9-100800511-INS-323
chr9-100825854-INS-135
chr9-100844622-INS-203
chr9-100845412-INS-376
chr9-100859467-INS-296
chr9-100860435-INS-156
chr9-100860435-INS-63
chr9-100860830-INS-50
chr9-100868446-INS-76
chr9-100883133-INS-50
chr9-100926177-INS-52
chr9-100927506-INS-6433
chr9-100940062-INS-230
chr9-100950584-INS-1011
chr9-100967716-INS-51
chr9-100967802-INS-112
chr9-100974984-INS-2898
chr9-100974984-INS-789
chr9-101051229-INS-51
chr9-101068801-INS-54
chr9-101106576-INS-459
chr9-101126131-INS-440
chr9-101163378-INS-657
chr9-101190012-INS-87
chr9-101193783-INS-1491
chr9-101205011-INS-6460

chr9-101221839-INS-62
chr9-101221878-INS-159
chr9-101222104-INS-215
chr9-101245590-INS-132
chr9-101282752-INS-56
chr9-101292978-INS-60
chr9-101293119-INS-301
chr9-101348819-INS-54
chr9-101367244-INS-60
chr9-101367959-INS-88
chr9-101368766-INS-129
chr9-101370446-INS-62
chr9-101375318-INS-1216
chr9-101416081-INS-1992
chr9-101427411-INS-180
chr9-101428710-INS-51
chr9-101431700-INS-51
chr9-101433832-INS-80
chr9-101434596-INS-80
chr9-101466746-INS-6546
chr9-101482309-INS-292
chr9-101507611-INS-60
chr9-101541910-INS-84
chr9-101647341-INS-6551
chr9-101683953-INS-56
chr9-101803313-INS-216
chr9-101956958-INS-156
chr9-101965610-INS-142
chr9-101983563-INS-141
chr9-101988070-INS-74
chr9-101999432-INS-375
chr9-102006271-INS-215
chr9-102053435-INS-114
chr9-102116669-INS-121
chr9-102159813-INS-428
chr9-102180756-INS-697
chr9-102325533-INS-117
chr9-102334990-INS-298
chr9-102337343-INS-50
chr9-102344127-INS-289
chr9-102418565-INS-276
chr9-102449418-INS-50
chr9-102495569-INS-65
chr9-102547441-INS-72
chr9-102558316-INS-70
chr9-102575178-INS-52
chr9-102578839-INS-190
chr9-102584516-INS-1000
chr9-102634722-INS-56
chr9-102678719-INS-230
chr9-102678839-INS-213
chr9-102690988-INS-594
chr9-102693389-INS-71
chr9-102693507-INS-52
chr9-102729396-INS-58
chr9-102813461-INS-1843
chr9-102826458-INS-128
chr9-102830615-INS-92
chr9-102830621-INS-191
chr9-102849456-INS-130
chr9-102860035-INS-3405
chr9-102875700-INS-69
chr9-102893012-INS-52
chr9-102908325-INS-213
chr9-102912962-INS-1729
chr9-102915904-INS-52
chr9-102920751-INS-999
chr9-102964317-INS-7185
chr9-102966600-INS-140
chr9-102981900-INS-637
chr9-102996373-INS-6236
chr9-103057866-INS-160
chr9-103132230-INS-206
chr9-103135074-INS-146
chr9-103144658-INS-7110
chr9-103174981-INS-1489
chr9-103190640-INS-60

chr9-103212768-INS-350
chr9-103212771-INS-64
chr9-103250298-INS-56
chr9-103302404-INS-72
chr9-103330167-INS-568
chr9-103350318-INS-50
chr9-103491765-INS-56
chr9-103508020-INS-62
chr9-103509075-INS-4671
chr9-103509455-INS-352
chr9-103510384-INS-1036
chr9-103526482-INS-800
chr9-103529566-INS-50
chr9-103535826-INS-7428
chr9-103669241-INS-93
chr9-103727814-INS-103
chr9-103764895-INS-111
chr9-103768484-INS-2680
chr9-103791359-INS-4092
chr9-103794054-INS-1176
chr9-103829510-INS-1508
chr9-103831967-INS-50
chr9-103842496-INS-62
chr9-103863035-INS-231
chr9-103877107-INS-79
chr9-103883157-INS-70
chr9-103883775-INS-146
chr9-103942448-INS-115
chr9-103955507-INS-7198
chr9-103957979-INS-187
chr9-104022346-INS-102
chr9-104043495-INS-137
chr9-104054856-INS-258
chr9-104063448-INS-54
chr9-104187355-INS-62
chr9-104190310-INS-4141
chr9-104226158-INS-100
chr9-104226864-INS-80
chr9-104227242-INS-1159
chr9-104268745-INS-156
chr9-104297369-INS-81
chr9-104300359-INS-155
chr9-104301892-INS-145
chr9-104319317-INS-3123
chr9-104372471-INS-5610
chr9-104399202-INS-76
chr9-104406134-INS-2419
chr9-104408996-INS-205
chr9-104425805-INS-672
chr9-104461166-INS-623
chr9-104480806-INS-72
chr9-104497504-INS-216
chr9-104498270-INS-105
chr9-104504658-INS-57
chr9-104572528-INS-1469
chr9-104572595-INS-101
chr9-104576619-INS-128
chr9-104576630-INS-51
chr9-104681662-INS-6482
chr9-104747728-INS-349
chr9-104762337-INS-1776
chr9-104763000-INS-530
chr9-104801137-INS-50
chr9-104806800-INS-6428
chr9-104816890-INS-1123
chr9-104827887-INS-4758
chr9-104832273-INS-52
chr9-104834989-INS-50
chr9-104845081-INS-2536
chr9-104861849-INS-80
chr9-104865493-INS-80
chr9-104865508-INS-203
chr9-104880338-INS-1797
chr9-104897434-INS-60
chr9-104967373-INS-1662
chr9-104986092-INS-191
chr9-105050624-INS-53

chr9-105059288-INS-54
chr9-105070872-INS-52
chr9-105071757-INS-136
chr9-105076700-INS-92
chr9-105080678-INS-132
chr9-105080809-INS-630
chr9-105085289-INS-58
chr9-105113282-INS-7238
chr9-105165760-INS-53
chr9-105199757-INS-50
chr9-105234058-INS-83
chr9-105252547-INS-93
chr9-105257748-INS-4788
chr9-105258828-INS-1445
chr9-105315095-INS-73
chr9-105318961-INS-68
chr9-105326396-INS-136
chr9-105326427-INS-150
chr9-105332841-INS-108
chr9-105410216-INS-294
chr9-105464113-INS-2563
chr9-105506744-INS-208
chr9-105513221-INS-424
chr9-105514677-INS-498
chr9-105537903-INS-596
chr9-105554841-INS-382
chr9-105554902-INS-1881
chr9-105584178-INS-153
chr9-105587460-INS-68
chr9-105591337-INS-6386
chr9-105607411-INS-80
chr9-105627444-INS-72
chr9-105665329-INS-50
chr9-105668453-INS-372
chr9-105677645-INS-7089
chr9-105678221-INS-1681
chr9-105691162-INS-626
chr9-105692667-INS-6450
chr9-105702091-INS-100
chr9-105706800-INS-1631
chr9-105735772-INS-52
chr9-105778649-INS-66
chr9-105778970-INS-352
chr9-105820331-INS-129
chr9-105824271-INS-52
chr9-105828838-INS-5412
chr9-105839771-INS-5327
chr9-105850086-INS-56
chr9-105860705-INS-283
chr9-105879072-INS-104
chr9-105903437-INS-144
chr9-105907725-INS-428
chr9-105949054-INS-234
chr9-105950545-INS-604
chr9-105958016-INS-6247
chr9-105986047-INS-6230
chr9-105989194-INS-1016
chr9-106038327-INS-81
chr9-106119815-INS-72
chr9-106135941-INS-71
chr9-106162917-INS-54
chr9-106163723-INS-53
chr9-106301616-INS-502
chr9-106301800-INS-905
chr9-106355932-INS-54
chr9-106414865-INS-186
chr9-106430570-INS-52
chr9-106475063-INS-1040
chr9-106505889-INS-76
chr9-106548150-INS-55
chr9-106607769-INS-66
chr9-106609016-INS-136
chr9-106622167-INS-214
chr9-106651988-INS-106
chr9-106680937-INS-98
chr9-106719162-INS-229
chr9-106728764-INS-140

chr9-106755198-INS-556
chr9-106994436-INS-447
chr9-106995137-INS-1888
chr9-107001868-INS-471
chr9-107117247-INS-99
chr9-107155099-INS-186
chr9-107172413-INS-959
chr9-107183916-INS-58
chr9-107241240-INS-153
chr9-107241414-INS-50
chr9-107272804-INS-64
chr9-107309621-INS-215
chr9-107343593-INS-52
chr9-107407052-INS-1766
chr9-107423243-INS-207
chr9-107434604-INS-272
chr9-107530417-INS-62
chr9-107538934-INS-214
chr9-107631556-INS-301
chr9-107638108-INS-278
chr9-107643774-INS-168
chr9-107655914-INS-50
chr9-107685293-INS-201
chr9-107688316-INS-210
chr9-107689787-INS-169
chr9-107694344-INS-205
chr9-107727615-INS-210
chr9-107739238-INS-204
chr9-107742073-INS-68
chr9-107752632-INS-181
chr9-107752632-INS-52
chr9-107799918-INS-60
chr9-107829988-INS-201
chr9-107866729-INS-50
chr9-107889125-INS-226
chr9-107892787-INS-215
chr9-107977650-INS-105
chr9-107977755-INS-544
chr9-108001056-INS-500
chr9-108023452-INS-58
chr9-108026807-INS-127
chr9-108037072-INS-52
chr9-108142706-INS-210
chr9-108149145-INS-80
chr9-108158942-INS-211
chr9-108160017-INS-98
chr9-108169057-INS-202
chr9-108169267-INS-376
chr9-108169655-INS-433
chr9-108169935-INS-78
chr9-108204083-INS-106
chr9-108223995-INS-57
chr9-108252697-INS-72
chr9-108291961-INS-79
chr9-108292573-INS-197
chr9-108308624-INS-58
chr9-108310511-INS-94
chr9-108311274-INS-228
chr9-108320506-INS-142
chr9-108344480-INS-205
chr9-108483648-INS-72
chr9-108499096-INS-631
chr9-108499687-INS-892
chr9-108527416-INS-7635
chr9-108545427-INS-217
chr9-108597474-INS-156
chr9-108597527-INS-52
chr9-108613996-INS-72
chr9-108615528-INS-541
chr9-108629498-INS-7293
chr9-108647116-INS-220
chr9-108725280-INS-55
chr9-108749569-INS-50
chr9-108752429-INS-60
chr9-108848956-INS-150
chr9-108849509-INS-176
chr9-108913779-INS-192

chr9-108914302-INS-68
chr9-109029076-INS-60
chr9-109044686-INS-50
chr9-109129740-INS-107
chr9-109144296-INS-204
chr9-109200112-INS-142
chr9-109239615-INS-5977
chr9-109353775-INS-52
chr9-109389483-INS-62
chr9-109498158-INS-7113
chr9-109514585-INS-258
chr9-109531313-INS-175
chr9-109531448-INS-71
chr9-109591787-INS-52
chr9-109603987-INS-309
chr9-109627676-INS-460
chr9-109639659-INS-1890
chr9-109644682-INS-127
chr9-109644876-INS-60
chr9-109685770-INS-208
chr9-109686767-INS-154
chr9-109697685-INS-138
chr9-109737685-INS-58
chr9-109739955-INS-77
chr9-109742066-INS-195
chr9-109744703-INS-52
chr9-109748013-INS-50
chr9-109748715-INS-254
chr9-109752400-INS-144
chr9-109752914-INS-457
chr9-109770107-INS-1085
chr9-109776193-INS-54
chr9-109776680-INS-58
chr9-109836524-INS-59
chr9-109836533-INS-1418
chr9-109851814-INS-184
chr9-109852520-INS-142
chr9-109971569-INS-50
chr9-109990583-INS-101
chr9-110063524-INS-211
chr9-110107617-INS-811
chr9-110118828-INS-165
chr9-110130041-INS-231
chr9-110141849-INS-77
chr9-110151498-INS-210
chr9-110188314-INS-453
chr9-110188318-INS-80
chr9-110197835-INS-94
chr9-110198186-INS-161
chr9-110239889-INS-71
chr9-110252606-INS-746
chr9-110259451-INS-151
chr9-110279176-INS-923
chr9-110279393-INS-72
chr9-110287035-INS-231
chr9-110310759-INS-180
chr9-110311119-INS-81
chr9-110328380-INS-72
chr9-110331824-INS-62
chr9-110340077-INS-206
chr9-110342490-INS-450
chr9-110347254-INS-86
chr9-110347809-INS-61
chr9-110381255-INS-87
chr9-110405375-INS-191
chr9-110445736-INS-488
chr9-110492199-INS-62
chr9-110503242-INS-176
chr9-110503298-INS-52
chr9-110503482-INS-344
chr9-110519189-INS-2040
chr9-110519303-INS-900
chr9-110537343-INS-52
chr9-110552348-INS-183
chr9-110594025-INS-119
chr9-110623046-INS-55
chr9-110632433-INS-245

chr9-110637637-INS-859
chr9-110649949-INS-56
chr9-110661995-INS-308
chr9-110689275-INS-75
chr9-110742356-INS-68
chr9-110786254-INS-82
chr9-110786659-INS-150
chr9-110808408-INS-52
chr9-110814366-INS-54
chr9-110824671-INS-464
chr9-110834466-INS-90
chr9-110840097-INS-60
chr9-110844307-INS-115
chr9-110845221-INS-74
chr9-110848553-INS-639
chr9-110881847-INS-57
chr9-110898886-INS-96
chr9-110906103-INS-247
chr9-110912401-INS-102
chr9-110914032-INS-50
chr9-110923138-INS-358
chr9-110963093-INS-432
chr9-110974804-INS-204
chr9-110983310-INS-2000
chr9-111005511-INS-943
chr9-111005829-INS-246
chr9-111037815-INS-406
chr9-111052171-INS-111
chr9-111075034-INS-426
chr9-111075198-INS-1162
chr9-111110762-INS-3100
chr9-111138981-INS-64
chr9-111145109-INS-236
chr9-111167306-INS-1612
chr9-111186849-INS-181
chr9-111187504-INS-50
chr9-111224099-INS-55
chr9-111238881-INS-4830
chr9-111283969-INS-60
chr9-111290044-INS-100
chr9-111325123-INS-823
chr9-111334135-INS-148
chr9-111445752-INS-230
chr9-111446246-INS-70
chr9-111457777-INS-50
chr9-111464711-INS-2701
chr9-111527285-INS-7490
chr9-111551287-INS-86
chr9-111577212-INS-1042
chr9-111585867-INS-133
chr9-111590841-INS-50
chr9-111612437-INS-58
chr9-111672528-INS-93
chr9-111685408-INS-1553
chr9-111699449-INS-54
chr9-111719170-INS-114
chr9-111741496-INS-6499
chr9-111742419-INS-53
chr9-111744486-INS-50
chr9-111744817-INS-5412
chr9-111752781-INS-64
chr9-111756499-INS-153
chr9-111777515-INS-90
chr9-111790447-INS-144
chr9-111812048-INS-52
chr9-111819333-INS-3173
chr9-111831331-INS-76
chr9-111842712-INS-144
chr9-111887513-INS-247
chr9-111920302-INS-52
chr9-111921046-INS-1146
chr9-111935407-INS-81
chr9-111977865-INS-80
chr9-112012549-INS-1378
chr9-112014458-INS-99
chr9-112041118-INS-1000
chr9-112118657-INS-54

chr9-112126598-INS-90
chr9-112126990-INS-358
chr9-112130921-INS-6147
chr9-112148946-INS-66
chr9-112150586-INS-449
chr9-112151193-INS-114
chr9-112178589-INS-439
chr9-112178919-INS-50
chr9-112180384-INS-52
chr9-112206033-INS-6857
chr9-112222399-INS-278
chr9-112225987-INS-82
chr9-112264001-INS-1420
chr9-112264277-INS-103
chr9-112267109-INS-66
chr9-112267555-INS-254
chr9-112282545-INS-1183
chr9-112293328-INS-52
chr9-112304735-INS-55
chr9-112308725-INS-50
chr9-112324493-INS-220
chr9-112331209-INS-8539
chr9-112347860-INS-1463
chr9-112350748-INS-4177
chr9-112358827-INS-71
chr9-112358895-INS-250
chr9-112379591-INS-6518
chr9-112391565-INS-160
chr9-112467647-INS-274
chr9-112468253-INS-500
chr9-112617577-INS-54
chr9-112637587-INS-64
chr9-112650457-INS-3186
chr9-112666636-INS-50
chr9-112678621-INS-467
chr9-112685908-INS-860
chr9-112732104-INS-132
chr9-112753037-INS-756
chr9-112781915-INS-144
chr9-112846571-INS-208
chr9-112846737-INS-582
chr9-112888771-INS-50
chr9-112937707-INS-80
chr9-112937845-INS-120
chr9-112950486-INS-68
chr9-112966339-INS-924
chr9-112974680-INS-72
chr9-112998516-INS-56
chr9-113022810-INS-128
chr9-113023179-INS-65
chr9-113034324-INS-6392
chr9-113044364-INS-50
chr9-113044399-INS-969
chr9-113062200-INS-70
chr9-113064523-INS-6793
chr9-113074947-INS-65
chr9-113141620-INS-2769
chr9-113176632-INS-60
chr9-113186250-INS-50
chr9-113209137-INS-83
chr9-113210287-INS-348
chr9-113227925-INS-3907
chr9-113227936-INS-889
chr9-113227977-INS-57
chr9-113239939-INS-770
chr9-113248993-INS-142
chr9-113250013-INS-251
chr9-113311717-INS-50
chr9-113322447-INS-156
chr9-113322451-INS-72
chr9-113325457-INS-2775
chr9-113342322-INS-408
chr9-113342718-INS-54
chr9-113373124-INS-6485
chr9-113398952-INS-370
chr9-113421928-INS-7139
chr9-113438605-INS-1017

chr9-113438700-INS-172
chr9-113451968-INS-72
chr9-113459334-INS-160
chr9-113478703-INS-523
chr9-113490783-INS-615
chr9-113551903-INS-119
chr9-113564266-INS-69
chr9-113723543-INS-208
chr9-113741387-INS-284
chr9-113763343-INS-59
chr9-113789789-INS-52
chr9-113817761-INS-308
chr9-113830673-INS-113
chr9-113858409-INS-50
chr9-113878191-INS-148
chr9-113879749-INS-218
chr9-113882076-INS-226
chr9-113899791-INS-251
chr9-113900143-INS-60
chr9-113914174-INS-60
chr9-113938964-INS-220
chr9-113957822-INS-496
chr9-113968059-INS-62
chr9-113994749-INS-13404
chr9-113999531-INS-636
chr9-114012458-INS-350
chr9-114041724-INS-99
chr9-114076008-INS-186
chr9-114076020-INS-70
chr9-114094718-INS-194
chr9-114097983-INS-1259
chr9-114124453-INS-130
chr9-114134760-INS-3390
chr9-114161321-INS-178
chr9-114198435-INS-160
chr9-114282703-INS-269
chr9-114282725-INS-65
chr9-114320192-INS-63
chr9-114349623-INS-234
chr9-114350782-INS-1132
chr9-114374412-INS-65
chr9-114397639-INS-90
chr9-114397808-INS-213
chr9-114398371-INS-128
chr9-114398662-INS-54
chr9-114400192-INS-209
chr9-114404393-INS-67
chr9-114409153-INS-134
chr9-114411783-INS-189
chr9-114412451-INS-209
chr9-114418605-INS-205
chr9-114438702-INS-74
chr9-114446183-INS-1261
chr9-114452257-INS-222
chr9-114480761-INS-90
chr9-114480927-INS-2953
chr9-114495679-INS-54
chr9-114502912-INS-185
chr9-114529057-INS-2260
chr9-114539517-INS-412
chr9-114550116-INS-52
chr9-114572675-INS-1083
chr9-114572703-INS-456
chr9-114575390-INS-219
chr9-114594761-INS-232
chr9-114598647-INS-103
chr9-114618133-INS-50
chr9-114618621-INS-148
chr9-114630812-INS-80
chr9-114631277-INS-4745
chr9-114638927-INS-195
chr9-114661962-INS-54
chr9-114662465-INS-55
chr9-114670984-INS-847
chr9-114685749-INS-52
chr9-114686119-INS-65
chr9-114694989-INS-80

chr9-114724030-INS-7129
chr9-114727141-INS-103
chr9-114741577-INS-84
chr9-114757394-INS-66
chr9-114767611-INS-69
chr9-114768733-INS-498
chr9-114775246-INS-91
chr9-114784821-INS-557
chr9-114784873-INS-65
chr9-114785262-INS-142
chr9-114794449-INS-379
chr9-114794953-INS-189
chr9-114795593-INS-120
chr9-114813283-INS-52
chr9-114820013-INS-214
chr9-114820038-INS-106
chr9-114868298-INS-80
chr9-114868576-INS-222
chr9-114877047-INS-63
chr9-114887052-INS-56
chr9-114889218-INS-50
chr9-114898781-INS-100
chr9-114906414-INS-60
chr9-114914869-INS-205
chr9-114957578-INS-185
chr9-114957583-INS-84
chr9-114965722-INS-339
chr9-114970525-INS-72
chr9-115016565-INS-94
chr9-115028329-INS-1512
chr9-115028354-INS-59
chr9-115035126-INS-1500
chr9-115091939-INS-219
chr9-115112253-INS-238
chr9-115128574-INS-528
chr9-115154821-INS-53
chr9-115155622-INS-7242
chr9-115168975-INS-1829
chr9-115171265-INS-7214
chr9-115181257-INS-162
chr9-115196774-INS-209
chr9-115218882-INS-84
chr9-115242869-INS-233
chr9-115243742-INS-208
chr9-115260202-INS-3968
chr9-115272546-INS-167
chr9-115279440-INS-772
chr9-115315053-INS-140
chr9-115329027-INS-2079
chr9-115389380-INS-65
chr9-115535926-INS-66
chr9-115544660-INS-416
chr9-115551026-INS-52
chr9-115565139-INS-50
chr9-115596682-INS-56
chr9-115616212-INS-70
chr9-115693224-INS-78
chr9-115693385-INS-596
chr9-115694192-INS-371
chr9-115718066-INS-391
chr9-115718990-INS-224
chr9-115759009-INS-84
chr9-115764550-INS-335
chr9-115767476-INS-1632
chr9-115770931-INS-3030
chr9-115782994-INS-52
chr9-115844659-INS-6481
chr9-115851346-INS-258
chr9-115863951-INS-155
chr9-115863951-INS-341
chr9-115864105-INS-55
chr9-115871548-INS-426
chr9-115888613-INS-1184
chr9-115908837-INS-66
chr9-115911635-INS-80
chr9-115915966-INS-75
chr9-115924997-INS-144

chr9-115947037-INS-5548
chr9-116005462-INS-50
chr9-116024249-INS-50
chr9-116026600-INS-52
chr9-116080603-INS-54
chr9-116080863-INS-190
chr9-116091878-INS-94
chr9-116172988-INS-220
chr9-116184527-INS-117
chr9-116191744-INS-65
chr9-116214279-INS-56
chr9-116221764-INS-73
chr9-116227156-INS-557
chr9-116238646-INS-90
chr9-116285323-INS-57
chr9-116290842-INS-2160
chr9-116294115-INS-1049
chr9-116296082-INS-108
chr9-116315786-INS-1180
chr9-116324769-INS-222
chr9-116329722-INS-5662
chr9-116354003-INS-104
chr9-116366901-INS-1569
chr9-116478090-INS-234
chr9-116501993-INS-136
chr9-116534833-INS-53
chr9-116610700-INS-200
chr9-116655080-INS-68
chr9-116707411-INS-282
chr9-116707941-INS-62
chr9-116719126-INS-534
chr9-116766427-INS-55
chr9-116766953-INS-60
chr9-116834660-INS-50
chr9-116870488-INS-246
chr9-116924982-INS-67
chr9-116929039-INS-6440
chr9-116935987-INS-306
chr9-116937961-INS-418
chr9-116938018-INS-133
chr9-116943427-INS-106
chr9-116954402-INS-52
chr9-116958645-INS-60
chr9-117015656-INS-77
chr9-117021651-INS-54
chr9-117061108-INS-1325
chr9-117090801-INS-50
chr9-117116643-INS-82
chr9-117161947-INS-7257
chr9-117162491-INS-154
chr9-117173068-INS-122
chr9-117175180-INS-68
chr9-117200061-INS-207
chr9-117218735-INS-84
chr9-117246385-INS-6210
chr9-117268547-INS-55
chr9-117282308-INS-140
chr9-117328859-INS-129
chr9-117331080-INS-57
chr9-117376451-INS-6591
chr9-117386827-INS-84
chr9-117398591-INS-120
chr9-117432077-INS-58
chr9-117434014-INS-50
chr9-117438813-INS-50
chr9-117464887-INS-82
chr9-117467090-INS-56
chr9-117476149-INS-68
chr9-117526642-INS-420
chr9-117546052-INS-124
chr9-117547485-INS-138
chr9-117594977-INS-50
chr9-117623235-INS-62
chr9-117630108-INS-500
chr9-117633990-INS-50
chr9-117651299-INS-411
chr9-117672351-INS-60

chr9-117678900-INS-2634
chr9-117693375-INS-66
chr9-117693428-INS-245
chr9-117695120-INS-74
chr9-117746575-INS-1110
chr9-117746639-INS-510
chr9-117754178-INS-1470
chr9-117754402-INS-511
chr9-117798786-INS-60
chr9-117801937-INS-250
chr9-117804154-INS-3456
chr9-117810504-INS-3225
chr9-117820705-INS-50
chr9-117839909-INS-632
chr9-117879454-INS-135
chr9-117897293-INS-7162
chr9-117929689-INS-151
chr9-117929689-INS-56
chr9-117933782-INS-62
chr9-117953745-INS-1007
chr9-118009682-INS-54
chr9-118013846-INS-2055
chr9-118021607-INS-52
chr9-118021710-INS-362
chr9-118022134-INS-52
chr9-118022265-INS-520
chr9-118049966-INS-65
chr9-118052332-INS-150
chr9-118062215-INS-356
chr9-118109518-INS-57
chr9-118143796-INS-240
chr9-118150630-INS-1264
chr9-118151976-INS-78
chr9-118154524-INS-3220
chr9-118163198-INS-56
chr9-118171914-INS-215
chr9-118182036-INS-242
chr9-118187264-INS-128
chr9-118187612-INS-77
chr9-118215353-INS-77
chr9-118239152-INS-152
chr9-118239221-INS-506
chr9-118239264-INS-138
chr9-118244050-INS-77
chr9-118244234-INS-275
chr9-118244358-INS-51
chr9-118249306-INS-133
chr9-118279656-INS-102
chr9-118291548-INS-212
chr9-118294132-INS-56
chr9-118318405-INS-96
chr9-118319919-INS-1457
chr9-118329910-INS-122
chr9-118364128-INS-204
chr9-118399901-INS-157
chr9-118403428-INS-111
chr9-118454860-INS-67
chr9-118460408-INS-55
chr9-118460471-INS-175
chr9-118485005-INS-392
chr9-118485005-INS-56
chr9-118502766-INS-58
chr9-118541293-INS-57
chr9-118587284-INS-341
chr9-118598512-INS-82
chr9-118632336-INS-114
chr9-118636969-INS-174
chr9-118653209-INS-226
chr9-118684365-INS-51
chr9-118719451-INS-2522
chr9-118738350-INS-63
chr9-118746708-INS-713
chr9-118750281-INS-84
chr9-118785195-INS-687
chr9-118817210-INS-1115
chr9-118831912-INS-192
chr9-118832300-INS-680

chr9-118918085-INS-54
chr9-118946609-INS-73
chr9-118951449-INS-406
chr9-119016708-INS-199
chr9-119016888-INS-187
chr9-119037518-INS-58
chr9-119049856-INS-921
chr9-119054385-INS-278
chr9-119077741-INS-50
chr9-119100444-INS-359
chr9-119119395-INS-84
chr9-119175968-INS-62
chr9-119182174-INS-158
chr9-119182967-INS-498
chr9-119201900-INS-60
chr9-119203632-INS-54
chr9-119305671-INS-610
chr9-119333848-INS-379
chr9-119356685-INS-831
chr9-119357263-INS-343
chr9-119357319-INS-711
chr9-119357440-INS-68
chr9-119442455-INS-1158
chr9-119455018-INS-1495
chr9-119455753-INS-234
chr9-119462427-INS-232
chr9-119477720-INS-54
chr9-119540997-INS-126
chr9-119541401-INS-58
chr9-119586578-INS-76
chr9-119629014-INS-108
chr9-119664985-INS-61
chr9-119715825-INS-513
chr9-119741692-INS-202
chr9-119768575-INS-88
chr9-119775459-INS-96
chr9-119776531-INS-193
chr9-119815560-INS-50
chr9-119822499-INS-264
chr9-119822499-INS-561
chr9-119832592-INS-486
chr9-119832650-INS-183
chr9-119836134-INS-216
chr9-119858527-INS-301
chr9-119865525-INS-108
chr9-119873663-INS-53
chr9-119885731-INS-156
chr9-119910863-INS-135
chr9-119914470-INS-143
chr9-119932486-INS-201
chr9-119960985-INS-156
chr9-119971637-INS-295
chr9-119971827-INS-118
chr9-119980085-INS-194
chr9-120004236-INS-58
chr9-120024438-INS-70
chr9-120024787-INS-327
chr9-120041285-INS-113
chr9-120116336-INS-132
chr9-120183010-INS-5524
chr9-120208078-INS-50
chr9-120225237-INS-52
chr9-120236212-INS-136
chr9-120236722-INS-180
chr9-120241847-INS-90
chr9-120262545-INS-66
chr9-120269343-INS-60
chr9-120272758-INS-52
chr9-120274099-INS-50
chr9-120279540-INS-52
chr9-120287459-INS-64
chr9-120298774-INS-52
chr9-120305387-INS-176
chr9-120329904-INS-113
chr9-120335368-INS-115
chr9-120416654-INS-470
chr9-120417800-INS-386

chr9-120436086-INS-120
chr9-120436304-INS-218
chr9-120439942-INS-78
chr9-120443048-INS-66
chr9-120443400-INS-122
chr9-120502901-INS-342
chr9-120515500-INS-7066
chr9-120516113-INS-62
chr9-120516879-INS-220
chr9-120516879-INS-85
chr9-120517666-INS-244
chr9-120536346-INS-60
chr9-120537156-INS-208
chr9-120549629-INS-384
chr9-120554125-INS-92
chr9-120558372-INS-63
chr9-120571117-INS-184
chr9-120571965-INS-94
chr9-120575506-INS-109
chr9-120595892-INS-2487
chr9-120597053-INS-434
chr9-120598948-INS-67
chr9-120606473-INS-414
chr9-120606473-INS-9018
chr9-120632175-INS-84
chr9-120633229-INS-111
chr9-120657866-INS-96
chr9-120671302-INS-207
chr9-120707885-INS-56
chr9-120708837-INS-60
chr9-120732651-INS-117
chr9-120735888-INS-58
chr9-120740354-INS-114
chr9-120796610-INS-51
chr9-120799499-INS-837
chr9-120871135-INS-336
chr9-120876083-INS-1548
chr9-120883146-INS-97
chr9-120883336-INS-137
chr9-120894148-INS-70
chr9-120903324-INS-60
chr9-120904244-INS-207
chr9-120904345-INS-898
chr9-120918897-INS-75
chr9-120933707-INS-55
chr9-120933726-INS-243
chr9-120948684-INS-58
chr9-120949749-INS-150
chr9-120950110-INS-6168
chr9-120961014-INS-80
chr9-120963151-INS-52
chr9-120963166-INS-174
chr9-120976415-INS-334
chr9-120976530-INS-75
chr9-120982301-INS-1959
chr9-121013977-INS-270
chr9-121014300-INS-150
chr9-121017583-INS-356
chr9-121026672-INS-331
chr9-121044323-INS-106
chr9-121079708-INS-154
chr9-121080977-INS-242
chr9-121087417-INS-320
chr9-121164016-INS-110
chr9-121194145-INS-650
chr9-121194167-INS-298
chr9-121194167-INS-78
chr9-121194696-INS-95
chr9-121198342-INS-1159
chr9-121204801-INS-106
chr9-121218205-INS-60
chr9-121219689-INS-298
chr9-121243984-INS-211
chr9-121248211-INS-183
chr9-121260023-INS-458
chr9-121266315-INS-60
chr9-121276958-INS-139

chr9-121288983-INS-209
chr9-121327067-INS-494
chr9-121333407-INS-561
chr9-121342390-INS-219
chr9-121352605-INS-6389
chr9-121374715-INS-275
chr9-121383038-INS-387
chr9-121390705-INS-788
chr9-121466271-INS-56
chr9-121506065-INS-315
chr9-121515493-INS-56
chr9-121518196-INS-72
chr9-121538120-INS-507
chr9-121632458-INS-4335
chr9-121752810-INS-50
chr9-121775567-INS-153
chr9-121788196-INS-610
chr9-121789796-INS-338
chr9-121855932-INS-50
chr9-121865536-INS-62
chr9-121873006-INS-83
chr9-121897682-INS-66
chr9-121915519-INS-96
chr9-121934770-INS-945
chr9-122001608-INS-2751
chr9-122060104-INS-76
chr9-122062563-INS-64
chr9-122080923-INS-1514
chr9-122089926-INS-52
chr9-122102113-INS-111
chr9-122103748-INS-1786
chr9-122114544-INS-104
chr9-122125898-INS-198
chr9-122169792-INS-74
chr9-122183590-INS-203
chr9-122213758-INS-380
chr9-122219760-INS-76
chr9-122242710-INS-72
chr9-122243040-INS-146
chr9-122274078-INS-610
chr9-122286664-INS-2495
chr9-122315031-INS-91
chr9-122327399-INS-51
chr9-122336145-INS-152
chr9-122347717-INS-483
chr9-122359648-INS-379
chr9-122367352-INS-2058
chr9-122375021-INS-635
chr9-122405883-INS-55
chr9-122435844-INS-92
chr9-122462581-INS-70
chr9-122477444-INS-594
chr9-122477445-INS-93
chr9-122498106-INS-112
chr9-122515048-INS-273
chr9-122583532-INS-224
chr9-122590256-INS-91
chr9-122590618-INS-59
chr9-122591052-INS-92
chr9-122591516-INS-425
chr9-122606701-INS-704
chr9-122606725-INS-337
chr9-122609220-INS-92
chr9-122609522-INS-105
chr9-122693241-INS-196
chr9-122702165-INS-458
chr9-122703371-INS-4807
chr9-122727128-INS-305
chr9-122735540-INS-58
chr9-122749439-INS-62
chr9-122769464-INS-74
chr9-122802497-INS-287
chr9-122823355-INS-968
chr9-122830878-INS-66
chr9-122859320-INS-97
chr9-122860069-INS-52
chr9-122872232-INS-111

chr9-122895029-INS-189
chr9-122922836-INS-1579
chr9-122969096-INS-50
chr9-122982628-INS-82
chr9-123004701-INS-528
chr9-123010520-INS-1088
chr9-123032634-INS-187
chr9-123040033-INS-325
chr9-123056715-INS-5613
chr9-123070929-INS-192
chr9-123077486-INS-134
chr9-123077749-INS-162
chr9-123080227-INS-168
chr9-123141084-INS-314
chr9-123142083-INS-102
chr9-123153752-INS-186
chr9-123159214-INS-2556
chr9-123159216-INS-7987
chr9-123179911-INS-2738
chr9-123261928-INS-750
chr9-123274562-INS-60
chr9-123279732-INS-7264
chr9-123307678-INS-50
chr9-123327579-INS-1462
chr9-123327751-INS-136
chr9-123327752-INS-606
chr9-123338077-INS-72
chr9-123338681-INS-129
chr9-123339191-INS-176
chr9-123343078-INS-63
chr9-123368696-INS-205
chr9-123383442-INS-1891
chr9-123421992-INS-441
chr9-123453968-INS-343
chr9-123481690-INS-2202
chr9-123546538-INS-210
chr9-123563788-INS-128
chr9-123604440-INS-578
chr9-123625028-INS-96
chr9-123631208-INS-190
chr9-123653820-INS-54
chr9-123722031-INS-382
chr9-123739678-INS-58
chr9-123760044-INS-277
chr9-123783809-INS-7089
chr9-123792139-INS-171
chr9-123792458-INS-293
chr9-123796335-INS-66
chr9-123840201-INS-64
chr9-123864633-INS-6625
chr9-123892327-INS-87
chr9-123913592-INS-5705
chr9-124031852-INS-84
chr9-124113608-INS-186
chr9-124114644-INS-498
chr9-124115608-INS-1421
chr9-124129559-INS-399
chr9-124176804-INS-1325
chrX-4530040-INS-366
chrX-5413413-INS-64
chrX-5551059-INS-98
chrX-5624612-INS-417
chrX-5625168-INS-294
chrX-5625211-INS-126
chrX-5626475-INS-372
chrX-5653745-INS-3003
chrX-5672505-INS-76
chrX-5841033-INS-523
chrX-5846765-INS-1174
chrX-5957066-INS-610
chrX-6121685-INS-50
chrX-6121754-INS-112
chrX-6154385-INS-64
chrX-6316052-INS-276
chrX-6338546-INS-412
chrX-6339492-INS-224
chrX-6353106-INS-384

chrX-6385395-INS-1344
chrX-6385795-INS-54
chrX-6481531-INS-1000
chrX-6501503-INS-74
chrX-6516401-INS-53
chrX-6538995-INS-6680
chrX-6597537-INS-198
chrX-6616643-INS-80
chrX-6629977-INS-2949
chrX-6690893-INS-72
chrX-6695296-INS-372
chrX-6775232-INS-174
chrX-6775232-INS-78
chrX-6782639-INS-216
chrX-6817905-INS-90
chrX-6886977-INS-54
chrX-6899236-INS-65
chrX-6935516-INS-132
chrX-7040503-INS-79
chrX-7073283-INS-72
chrX-7097302-INS-6376
chrX-7108492-INS-2998
chrX-7119903-INS-2918
chrX-7136195-INS-430
chrX-7137152-INS-1908
chrX-7147043-INS-1705
chrX-7319691-INS-60
chrX-7319761-INS-50
chrX-7326209-INS-1412
chrX-7342654-INS-324
chrX-7396505-INS-58
chrX-7418623-INS-65
chrX-7462929-INS-54
chrX-7466773-INS-99
chrX-7468897-INS-148
chrX-7615246-INS-145
chrX-7618471-INS-205
chrX-7692620-INS-145
chrX-7699866-INS-52
chrX-7844438-INS-340
chrX-7896358-INS-1478
chrX-7896458-INS-218
chrX-7909266-INS-148
chrX-7914117-INS-323
chrX-7965097-INS-792
chrX-7965629-INS-814
chrX-7973995-INS-208
chrX-8038871-INS-58
chrX-8063557-INS-60
chrX-8158379-INS-3656
chrX-8159016-INS-6726
chrX-8180149-INS-156
chrX-8216386-INS-168
chrX-8221029-INS-50
chrX-8232381-INS-434
chrX-8368401-INS-297
chrX-8369295-INS-201
chrX-8446639-INS-1498
chrX-8448136-INS-998
chrX-8458974-INS-106
chrX-8490466-INS-2317
chrX-8557167-INS-6866
chrX-8574922-INS-16180
chrX-8703358-INS-539
chrX-8740353-INS-2904
chrX-8829545-INS-364
chrX-8836018-INS-215
chrX-8843196-INS-71
chrX-8849833-INS-54
chrX-8899247-INS-355
chrX-8899389-INS-498
chrX-8913588-INS-91
chrX-8983551-INS-192
chrX-8989406-INS-56
chrX-9005905-INS-6452
chrX-9007688-INS-158
chrX-9062916-INS-180

chrX-9063361-INS-479
chrX-9067274-INS-100
chrX-9075142-INS-39197
chrX-9084493-INS-56
chrX-9169359-INS-526
chrX-9169390-INS-242
chrX-9211519-INS-63
chrX-9270137-INS-700
chrX-9273127-INS-858
chrX-9311695-INS-56
chrX-9328791-INS-86
chrX-9339175-INS-370
chrX-9356838-INS-60
chrX-9394752-INS-8074
chrX-9397826-INS-502
chrX-9419924-INS-1467
chrX-9487258-INS-57
chrX-9500748-INS-104
chrX-9524696-INS-50
chrX-9546667-INS-6600
chrX-9554304-INS-6520
chrX-9559062-INS-746
chrX-9559136-INS-1523
chrX-9559949-INS-134
chrX-9623030-INS-6270
chrX-9632490-INS-4153
chrX-9654905-INS-60
chrX-9740745-INS-1188
chrX-9783637-INS-142
chrX-9790686-INS-6801
chrX-9798537-INS-54
chrX-9822086-INS-2705
chrX-9908021-INS-56
chrX-9944868-INS-492
chrX-9945217-INS-712
chrX-10072838-INS-62
chrX-10077155-INS-212
chrX-10088546-INS-4356
chrX-10097595-INS-91
chrX-10098158-INS-163
chrX-10114389-INS-51
chrX-10204961-INS-7388
chrX-10219192-INS-53
chrX-10224822-INS-57
chrX-10331168-INS-50
chrX-10391117-INS-212
chrX-10398595-INS-123
chrX-10485484-INS-6462
chrX-10505504-INS-69
chrX-10512264-INS-80
chrX-10570484-INS-77
chrX-10588115-INS-214
chrX-10595254-INS-50
chrX-10615941-INS-7271
chrX-10678023-INS-50
chrX-10714714-INS-72
chrX-10760782-INS-150
chrX-10807464-INS-6894
chrX-10826429-INS-56
chrX-10869181-INS-423
chrX-10889712-INS-52
chrX-10889958-INS-1162
chrX-10931216-INS-120
chrX-10969170-INS-152
chrX-10979381-INS-474
chrX-11028503-INS-108
chrX-11061544-INS-3061
chrX-11066733-INS-6342
chrX-11092823-INS-424
chrX-11130575-INS-219
chrX-11164600-INS-3172
chrX-11254431-INS-157
chrX-11254510-INS-509
chrX-11318385-INS-593
chrX-11424067-INS-76
chrX-11436515-INS-52
chrX-11453809-INS-54

chrX-11571151-INS-227
chrX-11577452-INS-51
chrX-11597635-INS-50
chrX-11618075-INS-59
chrX-11658385-INS-131
chrX-11685887-INS-4494
chrX-11754398-INS-52
chrX-11806363-INS-130
chrX-11876397-INS-70
chrX-11880650-INS-121
chrX-11889836-INS-197
chrX-11890568-INS-132
chrX-11894270-INS-52
chrX-11897564-INS-52
chrX-11898930-INS-90
chrX-11974885-INS-116
chrX-11986760-INS-206
chrX-12032964-INS-120
chrX-12038503-INS-218
chrX-12039453-INS-63
chrX-12045057-INS-93
chrX-12045192-INS-210
chrX-12045490-INS-220
chrX-12058259-INS-284
chrX-12119876-INS-64
chrX-12131170-INS-52
chrX-12158867-INS-218
chrX-12172423-INS-330
chrX-12230825-INS-929
chrX-12271446-INS-523
chrX-12295612-INS-50
chrX-12303216-INS-60
chrX-12343323-INS-507
chrX-12408568-INS-100
chrX-12449971-INS-1488
chrX-12481251-INS-211
chrX-12497848-INS-6502
chrX-12576817-INS-207
chrX-12577028-INS-74
chrX-12577397-INS-201
chrX-12641686-INS-220
chrX-12655520-INS-145
chrX-12656217-INS-88
chrX-12656272-INS-149
chrX-12747045-INS-56
chrX-12797910-INS-6499
chrX-12840135-INS-514
chrX-12896768-INS-69
chrX-12896816-INS-341
chrX-12932649-INS-1572
chrX-12957923-INS-120
chrX-13081222-INS-65
chrX-13083860-INS-57
chrX-13090399-INS-265
chrX-13110175-INS-60
chrX-13130301-INS-187
chrX-13174155-INS-50
chrX-13206438-INS-444
chrX-13215290-INS-213
chrX-13270852-INS-156
chrX-13270855-INS-349
chrX-13307418-INS-407
chrX-13321586-INS-168
chrX-13323917-INS-87
chrX-13360984-INS-225
chrX-13376369-INS-270
chrX-13429245-INS-6554
chrX-13456787-INS-56
chrX-13469806-INS-369
chrX-13524259-INS-56
chrX-13644353-INS-2625
chrX-13748298-INS-68
chrX-13766080-INS-50
chrX-13828422-INS-134
chrX-13834496-INS-149
chrX-13856380-INS-2760
chrX-13888864-INS-500

chrX-13920142-INS-4956
chrX-13980352-INS-240
chrX-13988223-INS-110
chrX-13989735-INS-255
chrX-14077208-INS-3164
chrX-14087855-INS-2195
chrX-14111298-INS-51
chrX-14111784-INS-3319
chrX-14179651-INS-50
chrX-14188893-INS-186
chrX-14235874-INS-102
chrX-14250741-INS-2993
chrX-14478127-INS-625
chrX-14478132-INS-1526
chrX-14528194-INS-7078
chrX-14574177-INS-6530
chrX-14618025-INS-6714
chrX-14641379-INS-2314
chrX-14717891-INS-2853
chrX-14765312-INS-2491
chrX-14795653-INS-75
chrX-14814858-INS-1070
chrX-14958857-INS-4942
chrX-14993788-INS-7294
chrX-15009336-INS-7268
chrX-15038113-INS-57
chrX-15053941-INS-701
chrX-15072293-INS-252
chrX-15083288-INS-52
chrX-15104313-INS-55
chrX-15121294-INS-74
chrX-15128322-INS-54
chrX-15128577-INS-2631
chrX-15151011-INS-138
chrX-15155910-INS-55
chrX-15185751-INS-1371
chrX-15187395-INS-2528
chrX-15205032-INS-66
chrX-15215283-INS-212
chrX-15223371-INS-11073
chrX-15239987-INS-129
chrX-15246581-INS-241
chrX-15385106-INS-426
chrX-15411179-INS-50
chrX-15421424-INS-6446
chrX-15432544-INS-53
chrX-15445605-INS-79
chrX-15446526-INS-8734
chrX-15476397-INS-60
chrX-15507004-INS-211
chrX-15509087-INS-62
chrX-15531903-INS-8740
chrX-15601520-INS-6763
chrX-15655145-INS-83
chrX-15662286-INS-381
chrX-15721028-INS-65
chrX-15743824-INS-54
chrX-15809714-INS-156
chrX-15848318-INS-103
chrX-15848427-INS-50
chrX-15852457-INS-107
chrX-15857679-INS-82
chrX-15912502-INS-101
chrX-15916483-INS-138
chrX-15963766-INS-140
chrX-16009913-INS-7291
chrX-16078842-INS-485
chrX-16083153-INS-64
chrX-16100255-INS-2743
chrX-16165478-INS-201
chrX-16268923-INS-201
chrX-16303968-INS-717
chrX-16337470-INS-72
chrX-16364509-INS-2847
chrX-16377858-INS-80
chrX-16410013-INS-50
chrX-16416699-INS-7157

chrX-16419438-INS-1600
chrX-16474778-INS-52
chrX-16480474-INS-7104
chrX-16577893-INS-796
chrX-16578275-INS-84
chrX-16623490-INS-6805
chrX-16659038-INS-3074
chrX-16687854-INS-52
chrX-16750839-INS-2182
chrX-16878301-INS-65
chrX-16954002-INS-15046
chrX-16989197-INS-1223
chrX-17189599-INS-56
chrX-17193981-INS-6783
chrX-17206524-INS-6572
chrX-17242434-INS-99
chrX-17261547-INS-88
chrX-17346096-INS-208
chrX-17346096-INS-85
chrX-17372920-INS-3459
chrX-17379556-INS-85
chrX-17413303-INS-104
chrX-17421550-INS-4706
chrX-17487860-INS-84
chrX-17540298-INS-144
chrX-17554535-INS-516
chrX-17557812-INS-1945
chrX-17719370-INS-1053
chrX-17731494-INS-64
chrX-17752333-INS-567
chrX-17754780-INS-225
chrX-17784288-INS-1945
chrX-17826843-INS-55
chrX-17831598-INS-103
chrX-17912477-INS-6487
chrX-17923593-INS-3254
chrX-17924197-INS-244
chrX-17970059-INS-3976
chrX-17985289-INS-828
chrX-18064555-INS-1162
chrX-18078535-INS-7204
chrX-18098054-INS-59
chrX-18140218-INS-85
chrX-18180292-INS-99
chrX-18187095-INS-50
chrX-18238771-INS-252
chrX-18245844-INS-55
chrX-18268586-INS-6958
chrX-18296242-INS-2585
chrX-18303069-INS-123
chrX-18326687-INS-73
chrX-18369564-INS-1888
chrX-18428536-INS-416
chrX-18472509-INS-2779
chrX-18478726-INS-1854
chrX-18499009-INS-437
chrX-18525290-INS-66
chrX-18573823-INS-2493
chrX-18626375-INS-1500
chrX-18690497-INS-1500
chrX-18748302-INS-145
chrX-18754762-INS-7137
chrX-18865824-INS-52
chrX-18895231-INS-142
chrX-18910370-INS-58
chrX-18924435-INS-12538
chrX-18999654-INS-348
chrX-19007487-INS-574
chrX-19079824-INS-302
chrX-19114316-INS-1155
chrX-19136776-INS-1259
chrX-19184424-INS-51
chrX-19207373-INS-16900
chrX-19207373-INS-5005
chrX-19214129-INS-53
chrX-19215293-INS-934
chrX-19224101-INS-88

chrX-19228773-INS-212
chrX-19249806-INS-55
chrX-19264287-INS-1357
chrX-19302670-INS-1006
chrX-19313091-INS-1755
chrX-19321003-INS-191
chrX-19322818-INS-5060
chrX-19426723-INS-2943
chrX-19506498-INS-387
chrX-19551928-INS-4189
chrX-19571739-INS-71
chrX-19574042-INS-351
chrX-19584151-INS-6219
chrX-19625414-INS-6172
chrX-19641554-INS-278
chrX-19733572-INS-56
chrX-19876836-INS-1732
chrX-19941963-INS-55
chrX-19983636-INS-80
chrX-19987855-INS-7157
chrX-20037832-INS-6828
chrX-20038602-INS-118
chrX-20074812-INS-4777
chrX-20081600-INS-270
chrX-20110109-INS-54
chrX-20142568-INS-1059
chrX-20170667-INS-56
chrX-20198452-INS-172
chrX-20224583-INS-203
chrX-20327838-INS-350
chrX-20390050-INS-51
chrX-20420123-INS-60
chrX-20451213-INS-74
chrX-20526607-INS-678
chrX-20594827-INS-88
chrX-20599700-INS-51
chrX-20611301-INS-120
chrX-20611301-INS-300
chrX-20621416-INS-239
chrX-20651715-INS-196
chrX-20661186-INS-294
chrX-20669508-INS-3081
chrX-20669590-INS-374
chrX-20679312-INS-195
chrX-20681009-INS-195
chrX-20692086-INS-362
chrX-20693053-INS-201
chrX-20773797-INS-889
chrX-20830911-INS-50
chrX-20841492-INS-1194
chrX-20872929-INS-2657
chrX-20882404-INS-542
chrX-20900233-INS-6810
chrX-20912521-INS-50
chrX-20961990-INS-128
chrX-20991195-INS-7497
chrX-21049145-INS-165
chrX-21049733-INS-274
chrX-21054368-INS-306
chrX-21208414-INS-88
chrX-21222863-INS-62
chrX-21224858-INS-59
chrX-21268476-INS-223
chrX-21280656-INS-50
chrX-21377181-INS-2136
chrX-21386618-INS-55
chrX-21390128-INS-591
chrX-21390978-INS-50
chrX-21660717-INS-349
chrX-21686517-INS-1603
chrX-21809313-INS-50
chrX-21962498-INS-6428
chrX-21975180-INS-3318
chrX-22129130-INS-1279
chrX-22144365-INS-236
chrX-22148526-INS-6319
chrX-22229494-INS-8694

chrX-22460400-INS-50
chrX-22473818-INS-177
chrX-22523556-INS-104
chrX-22578310-INS-84
chrX-22638083-INS-6082
chrX-22683657-INS-56
chrX-22697349-INS-7090
chrX-22702165-INS-86
chrX-22728838-INS-3161
chrX-22774646-INS-5481
chrX-22829568-INS-123
chrX-22867615-INS-13043
chrX-22879591-INS-6761
chrX-22901870-INS-7198
chrX-22966991-INS-3102
chrX-22975814-INS-2983
chrX-22978041-INS-360
chrX-23044550-INS-84
chrX-23046762-INS-6432
chrX-23049098-INS-56
chrX-23083998-INS-133
chrX-23140259-INS-159
chrX-23167256-INS-52
chrX-23270456-INS-3617
chrX-23281018-INS-352
chrX-23284964-INS-231
chrX-23305084-INS-197
chrX-23322592-INS-1301
chrX-23325259-INS-148
chrX-23325475-INS-696
chrX-23331348-INS-54
chrX-23345732-INS-6701
chrX-23433003-INS-54
chrX-23461699-INS-76
chrX-23471473-INS-5075
chrX-23560171-INS-58
chrX-23604379-INS-148
chrX-23716802-INS-6724
chrX-23822500-INS-96
chrX-23900620-INS-1493
chrX-23915820-INS-60
chrX-23956386-INS-6833
chrX-24108128-INS-565
chrX-24521432-INS-217
chrX-24589714-INS-148
chrX-25758649-INS-509
chrX-25888123-INS-428
chrX-25914380-INS-257
chrX-26039018-INS-52
chrX-26076033-INS-158
chrX-26206536-INS-417
chrX-26581159-INS-514
chrX-29377956-INS-443
chrX-30883268-INS-295
chrX-31499263-INS-55
chrX-32450952-INS-220
chrX-33156116-INS-66
chrX-33238314-INS-56
chrX-33732445-INS-1355
chrX-35028054-INS-872
chrX-35043779-INS-2388
chrX-35045817-INS-72
chrX-35058034-INS-5042
chrX-35075167-INS-58
chrX-35349708-INS-91
chrX-35367938-INS-222
chrX-35390052-INS-204
chrX-35402787-INS-505
chrX-35456648-INS-430
chrX-35463324-INS-280
chrX-35493613-INS-331
chrX-35598617-INS-702
chrX-35656991-INS-1430
chrX-35684915-INS-137
chrX-35690538-INS-96
chrX-35758673-INS-58
chrX-35964614-INS-106

chrX-36014180-INS-644
chrX-36033058-INS-200
chrX-36043910-INS-186
chrX-36053960-INS-54
chrX-36067199-INS-192
chrX-36081935-INS-54
chrX-36082341-INS-219
chrX-36103548-INS-58
chrX-36119135-INS-178
chrX-36134624-INS-193
chrX-36168484-INS-200
chrX-36172413-INS-10523
chrX-36299544-INS-200
chrX-36299584-INS-80
chrX-36386804-INS-5190
chrX-36394138-INS-4428
chrX-36430519-INS-72
chrX-36456124-INS-64
chrX-36465332-INS-158
chrX-36499801-INS-196
chrX-36524767-INS-126
chrX-36759411-INS-608
chrX-36761956-INS-58
chrX-36765066-INS-211
chrX-36812659-INS-62
chrX-36821516-INS-56
chrX-36837219-INS-128
chrX-36841372-INS-202
chrX-36861251-INS-90
chrX-36861940-INS-360
chrX-36928562-INS-350
chrX-37039627-INS-7024
chrX-37042641-INS-3008
chrX-37069917-INS-2059
chrX-37130433-INS-50
chrX-37162033-INS-13698
chrX-37183718-INS-1191
chrX-37209861-INS-341
chrX-37231028-INS-90
chrX-37263823-INS-202
chrX-37305897-INS-52
chrX-37308833-INS-207
chrX-37379353-INS-1095
chrX-37395786-INS-6653
chrX-37495284-INS-80
chrX-37522577-INS-64
chrX-37534154-INS-120
chrX-37552966-INS-348
chrX-37608531-INS-1645
chrX-37712164-INS-99
chrX-37721519-INS-61
chrX-37763229-INS-72
chrX-37766905-INS-397
chrX-37767481-INS-506
chrX-37768085-INS-57
chrX-37777215-INS-58
chrX-37880726-INS-636
chrX-37900175-INS-776
chrX-37947699-INS-1071
chrX-38082495-INS-160
chrX-38120138-INS-52
chrX-38280369-INS-78
chrX-38348456-INS-6383
chrX-38357339-INS-1398
chrX-38476700-INS-6503
chrX-38484252-INS-2287
chrX-38487080-INS-54
chrX-38637920-INS-1801
chrX-38652855-INS-2904
chrX-38661427-INS-50
chrX-38741417-INS-84
chrX-38796157-INS-6760
chrX-38888863-INS-6393
chrX-39077549-INS-1804
chrX-39137145-INS-4839
chrX-39186052-INS-215
chrX-39208533-INS-3480

chrX-39225787-INS-78
chrX-39232718-INS-74
chrX-39248565-INS-198
chrX-39249710-INS-1309
chrX-39256813-INS-944
chrX-39263088-INS-60
chrX-39371301-INS-2223
chrX-39442399-INS-432
chrX-39489287-INS-3195
chrX-39611140-INS-6592
chrX-39629237-INS-5175
chrX-39670044-INS-52
chrX-39702812-INS-52
chrX-39743492-INS-7051
chrX-39915395-INS-60
chrX-39953706-INS-199
chrX-39959209-INS-1143
chrX-40062724-INS-8949
chrX-40095418-INS-198
chrX-40097635-INS-50
chrX-40118595-INS-104
chrX-40144120-INS-918
chrX-40196010-INS-3535
chrX-40224211-INS-548
chrX-40228741-INS-247
chrX-40242331-INS-4592
chrX-40246496-INS-6555
chrX-40339135-INS-2071
chrX-40402867-INS-1422
chrX-40419948-INS-6564
chrX-40468372-INS-6446
chrX-40561149-INS-1543
chrX-40648907-INS-498
chrX-40843177-INS-184
chrX-40905993-INS-56
chrX-41054203-INS-398
chrX-41054539-INS-87
chrX-41069980-INS-54
chrX-41082830-INS-1808
chrX-41089141-INS-167
chrX-41124203-INS-2472
chrX-41131434-INS-50
chrX-41151720-INS-60
chrX-41216411-INS-5360
chrX-41246776-INS-52
chrX-41267157-INS-1339
chrX-41304862-INS-57
chrX-41312433-INS-191
chrX-41344995-INS-60
chrX-41375618-INS-1564
chrX-41410694-INS-635
chrX-41410759-INS-100
chrX-41434593-INS-214
chrX-41439309-INS-62
chrX-41463520-INS-7299
chrX-41481634-INS-6249
chrX-41488634-INS-54
chrX-41551773-INS-58
chrX-41602336-INS-1620
chrX-41695885-INS-66
chrX-41780888-INS-320
chrX-41796604-INS-591
chrX-41850799-INS-1492
chrX-41890884-INS-6382
chrX-41946365-INS-7227
chrX-42012068-INS-1090
chrX-42101608-INS-3245
chrX-42116139-INS-198
chrX-42120227-INS-844
chrX-42163701-INS-53
chrX-42241656-INS-51
chrX-42286620-INS-173
chrX-42300352-INS-1151
chrX-42315919-INS-3346
chrX-42424048-INS-50
chrX-42424172-INS-201
chrX-42605491-INS-54

chrX-42712540-INS-371
chrX-42714849-INS-7182
chrX-42724275-INS-2819
chrX-42740904-INS-83
chrX-42863000-INS-53
chrX-42918326-INS-75
chrX-42939682-INS-56
chrX-43037679-INS-140
chrX-43133739-INS-7840
chrX-43244545-INS-50
chrX-43270763-INS-6567
chrX-43324290-INS-155
chrX-43324915-INS-199
chrX-43358215-INS-624
chrX-43413501-INS-50
chrX-43425328-INS-6219
chrX-43622556-INS-201
chrX-43628923-INS-120
chrX-43667106-INS-6503
chrX-43701994-INS-56
chrX-43727686-INS-51
chrX-43727848-INS-60
chrX-43760171-INS-52
chrX-43804194-INS-82
chrX-43928859-INS-6496
chrX-44018110-INS-80
chrX-44034510-INS-56
chrX-44036926-INS-370
chrX-44128539-INS-500
chrX-44158359-INS-62
chrX-44169889-INS-82
chrX-44391216-INS-539
chrX-44440086-INS-2848
chrX-44495681-INS-6384
chrX-44552206-INS-52
chrX-44617649-INS-68
chrX-44617651-INS-137
chrX-44621408-INS-201
chrX-44666516-INS-2379
chrX-44850976-INS-6368
chrX-44856219-INS-62
chrX-44884884-INS-3562
chrX-44902544-INS-3368
chrX-44952722-INS-50
chrX-44959286-INS-6937
chrX-44994449-INS-1765
chrX-45121386-INS-58
chrX-45178777-INS-1550
chrX-45217707-INS-6681
chrX-45318568-INS-20006
chrX-45331391-INS-2851
chrX-45383435-INS-6143
chrX-45414107-INS-6489
chrX-45432119-INS-60
chrX-45537053-INS-1004
chrX-45552516-INS-7519
chrX-45587322-INS-6661
chrX-45639921-INS-56
chrX-45649665-INS-7727
chrX-45664682-INS-81
chrX-45694386-INS-6603
chrX-45703385-INS-50
chrX-45729741-INS-96
chrX-45791164-INS-7249
chrX-45890525-INS-1000
chrX-46033069-INS-52
chrX-46067406-INS-1539
chrX-46079134-INS-498
chrX-46113549-INS-226
chrX-46136927-INS-6225
chrX-46148899-INS-54
chrX-46156554-INS-1731
chrX-46174117-INS-6211
chrX-46220234-INS-1150
chrX-46249110-INS-1573
chrX-46267273-INS-2008
chrX-46303752-INS-51

chrX-46306052-INS-72
chrX-46393936-INS-93
chrX-46401972-INS-170
chrX-46558794-INS-55
chrX-46609120-INS-6659
chrX-46625825-INS-6443
chrX-46650676-INS-1067
chrX-46664295-INS-124
chrX-46763715-INS-186
chrX-46773821-INS-4746
chrX-46815197-INS-3714
chrX-46834738-INS-4000
chrX-46892242-INS-120
chrX-46920833-INS-1000
chrX-46937494-INS-50
chrX-47014235-INS-56
chrX-47034396-INS-64
chrX-47037660-INS-960
chrX-47050820-INS-205
chrX-47091587-INS-241
chrX-47091587-INS-55
chrX-47096754-INS-50
chrX-47197608-INS-77
chrX-47203114-INS-119
chrX-47283642-INS-60
chrX-47305019-INS-69
chrX-47335323-INS-58
chrX-47390163-INS-66
chrX-47390973-INS-54
chrX-47393444-INS-196
chrX-47421227-INS-111
chrX-47515518-INS-182
chrX-47544749-INS-50
chrX-47585309-INS-86
chrX-47590013-INS-124
chrX-47640283-INS-8030
chrX-47684693-INS-84
chrX-47796872-INS-121
chrX-47797193-INS-120
chrX-47929043-INS-6374
chrX-47956404-INS-50
chrX-48026667-INS-3868
chrX-48026991-INS-1030
chrX-48028299-INS-58
chrX-48031454-INS-183
chrX-48031908-INS-6911
chrX-48105044-INS-191
chrX-48114846-INS-1534
chrX-48117206-INS-1861
chrX-48120621-INS-218
chrX-48139471-INS-6796
chrX-48143138-INS-110
chrX-48160940-INS-6666
chrX-48256121-INS-2828
chrX-48259787-INS-6580
chrX-48280318-INS-52
chrX-48392671-INS-25740
chrX-48421201-INS-53
chrX-48457388-INS-248
chrX-48560917-INS-109
chrX-48581385-INS-53
chrX-48731941-INS-2846
chrX-48738427-INS-2112
chrX-48747960-INS-995
chrX-48830239-INS-66
chrX-48945155-INS-2877
chrX-48962817-INS-52
chrX-48987064-INS-644
chrX-48995302-INS-159
chrX-49043410-INS-416
chrX-49045173-INS-2600
chrX-49148704-INS-743
chrX-49224168-INS-312
chrX-49253956-INS-15606
chrX-49348268-INS-4075
chrX-49379267-INS-2134
chrX-49381348-INS-50

chrX-49395972-INS-5559
chrX-49397149-INS-1043
chrX-49441693-INS-1454
chrX-49467220-INS-124
chrX-49510332-INS-56
chrX-49552233-INS-4474
chrX-49604860-INS-324
chrX-49620802-INS-698
chrX-49635652-INS-2473
chrX-49702701-INS-1358
chrX-49702993-INS-7906
chrX-49762379-INS-52
chrX-49774915-INS-6442
chrX-49798737-INS-87
chrX-49814326-INS-51
chrX-49816953-INS-51
chrX-49825360-INS-456
chrX-49825468-INS-170
chrX-49834693-INS-379
chrX-49872903-INS-3425
chrX-49911274-INS-316
chrX-49926797-INS-52
chrX-49996892-INS-131
chrX-50036031-INS-1808
chrX-50058221-INS-204
chrX-50086626-INS-706
chrX-50125929-INS-50
chrX-50196069-INS-6678
chrX-50269700-INS-110
chrX-50318385-INS-50
chrX-50336632-INS-68
chrX-50460699-INS-50
chrX-50547219-INS-105
chrX-50580609-INS-56
chrX-50638117-INS-154
chrX-50671631-INS-50
chrX-50682715-INS-66
chrX-50691685-INS-175
chrX-50708909-INS-181
chrX-50719424-INS-68
chrX-50758328-INS-64
chrX-50784974-INS-60
chrX-50805729-INS-58
chrX-50830024-INS-56
chrX-50857927-INS-811
chrX-50880311-INS-60
chrX-50891157-INS-64
chrX-51027054-INS-54
chrX-51098584-INS-68
chrX-51112554-INS-85
chrX-51156163-INS-8340
chrX-51165045-INS-52
chrX-51220738-INS-56
chrX-51220802-INS-148
chrX-51272123-INS-155
chrX-51272123-INS-539
chrX-51272393-INS-56
chrX-51272443-INS-774
chrX-51294838-INS-4964
chrX-51294932-INS-11770
chrX-51348884-INS-167
chrX-51348888-INS-79
chrX-51379786-INS-140
chrX-51389173-INS-163
chrX-51564850-INS-200
chrX-51623117-INS-221
chrX-51686653-INS-52
chrX-51742763-INS-51
chrX-51761046-INS-119
chrX-51804626-INS-174
chrX-51824041-INS-8272
chrX-51857887-INS-126
chrX-51859136-INS-212
chrX-51859512-INS-2472
chrX-51939715-INS-54
chrX-51941463-INS-1891
chrX-51944967-INS-50

chrX-51949680-INS-75
chrX-51956575-INS-149
chrX-51973681-INS-998
chrX-51974172-INS-51
chrX-51974809-INS-12561
chrX-52039108-INS-66
chrX-52059972-INS-86
chrX-52081090-INS-52
chrX-52083119-INS-181
chrX-52092758-INS-81
chrX-52095756-INS-203
chrX-52106086-INS-216
chrX-52120365-INS-209
chrX-52129548-INS-55
chrX-52233613-INS-58
chrX-52244816-INS-88
chrX-52245162-INS-54
chrX-52462563-INS-62
chrX-52472544-INS-78
chrX-52485797-INS-76
chrX-52519272-INS-72
chrX-52530895-INS-270
chrX-52535783-INS-1473
chrX-52566402-INS-50
chrX-52595992-INS-977
chrX-52596671-INS-1154
chrX-52601579-INS-61
chrX-52654899-INS-62
chrX-52682790-INS-10881
chrX-52727826-INS-212
chrX-52839940-INS-5986
chrX-52883448-INS-72
chrX-52901207-INS-2310
chrX-52911132-INS-147
chrX-53074194-INS-3478
chrX-53087354-INS-53
chrX-53165849-INS-154
chrX-53195774-INS-2579
chrX-54556659-INS-290
chrX-54602155-INS-59
chrX-54697053-INS-6364
chrX-54725036-INS-2672
chrX-54878663-INS-175
chrX-54906168-INS-120
chrX-54937680-INS-92
chrX-55002273-INS-52
chrX-55161069-INS-199
chrX-55171332-INS-7175
chrX-55215287-INS-62
chrX-55312719-INS-50
chrX-55351824-INS-100
chrX-55428460-INS-58
chrX-55433146-INS-3036
chrX-55438317-INS-3220
chrX-55441465-INS-7086
chrX-55587197-INS-135
chrX-55596731-INS-52
chrX-55604155-INS-1455
chrX-55608502-INS-1091
chrX-55612735-INS-50
chrX-55612826-INS-684
chrX-55687996-INS-3437
chrX-55730347-INS-3190
chrX-55733789-INS-258
chrX-55777331-INS-56
chrX-55814029-INS-55
chrX-55852990-INS-120
chrX-55857186-INS-1832
chrX-55874758-INS-144
chrX-55877812-INS-67
chrX-55884859-INS-6865
chrX-55921457-INS-1739
chrX-56004615-INS-66
chrX-56120579-INS-65
chrX-56125306-INS-56
chrX-56154677-INS-186
chrX-56160309-INS-1825

chrX-56167967-INS-86
chrX-56169249-INS-107
chrX-56172152-INS-54
chrX-56177841-INS-6469
chrX-56239739-INS-60
chrX-56298162-INS-64
chrX-56379377-INS-136
chrX-56389322-INS-217
chrX-56409630-INS-58
chrX-56421288-INS-210
chrX-56440157-INS-64
chrX-56447498-INS-50
chrX-56455925-INS-136
chrX-56466159-INS-56
chrX-56492551-INS-6836
chrX-56504048-INS-896
chrX-56507611-INS-211
chrX-56577075-INS-68
chrX-56630689-INS-829
chrX-56632979-INS-2760
chrX-56637125-INS-2369
chrX-56678175-INS-776
chrX-56768526-INS-4811
chrX-56855696-INS-88
chrX-56870024-INS-60
chrX-56912307-INS-50
chrX-56965720-INS-50
chrX-56976898-INS-6596
chrX-57038961-INS-57
chrX-57039181-INS-80
chrX-57044093-INS-83
chrX-57057958-INS-193
chrX-57057959-INS-1040
chrX-57088832-INS-1041
chrX-57123225-INS-52
chrX-57145394-INS-149
chrX-57145394-INS-359
chrX-57150018-INS-69
chrX-57174779-INS-120
chrX-57217278-INS-194
chrX-57230397-INS-296
chrX-57245692-INS-1317
chrX-57382317-INS-3890
chrX-57390648-INS-176
chrX-57395759-INS-658
chrX-57522004-INS-1439
chrX-57542206-INS-50
chrX-57554303-INS-821
chrX-57644322-INS-198
chrX-57702857-INS-6570
chrX-57780119-INS-68
chrX-57803171-INS-3130
chrX-57883548-INS-50
chrX-57904413-INS-96
chrX-57932060-INS-80
chrX-57957730-INS-84
chrX-57991640-INS-163
chrX-58016699-INS-2988
chrX-58055831-INS-206
chrX-58063265-INS-54
chrX-58069180-INS-4901
chrX-58069608-INS-52
chrX-58092024-INS-1729
chrX-58093609-INS-192
chrX-58219247-INS-420
chrX-58253133-INS-52
chrX-58278492-INS-51
chrX-58312700-INS-352
chrX-58411345-INS-663
chrX-58435502-INS-53
chrX-58451922-INS-69
chrX-58530342-INS-344
chrX-58550022-INS-340
chrX-58615551-INS-52
chrX-58681918-INS-1996
chrX-58807040-INS-159
chrX-58826729-INS-381

chrX-58862360-INS-6466
chrX-58903564-INS-261
chrX-58956718-INS-78
chrX-58959217-INS-6415
chrX-58970298-INS-52
chrX-58978910-INS-2152
chrX-58984901-INS-6672
chrX-58988145-INS-3185
chrX-59008535-INS-287
chrX-59009173-INS-58
chrX-59025630-INS-4096
chrX-59135986-INS-6593
chrX-59159860-INS-53
chrX-59178609-INS-86
chrX-59178924-INS-344
chrX-59196914-INS-63
chrX-59204959-INS-6221
chrX-59254602-INS-450
chrX-59266654-INS-2870
chrX-59394762-INS-54
chrX-59413967-INS-664
chrX-59414713-INS-636
chrX-59523950-INS-1453
chrX-59573173-INS-3000
chrX-59613453-INS-52
chrX-59630249-INS-1484
chrX-59681158-INS-548
chrX-59769901-INS-51
chrX-59774364-INS-56
chrX-59777209-INS-443
chrX-59799577-INS-104
chrX-59800525-INS-66
chrX-59823889-INS-6500
chrX-59829150-INS-64
chrX-59845434-INS-52
chrX-59846290-INS-122
chrX-59883815-INS-79
chrX-59898583-INS-105
chrX-59984359-INS-54
chrX-60011247-INS-411
chrX-60026368-INS-344
chrX-60032306-INS-71
chrX-60102871-INS-369
chrX-60125767-INS-6410
chrX-60229118-INS-163
chrX-60287438-INS-52
chrX-60300829-INS-1831
chrX-60345591-INS-72
chrX-60389189-INS-212
chrX-60405143-INS-104
chrX-60428535-INS-182
chrX-60511774-INS-6455
chrX-60545216-INS-6417
chrX-60554581-INS-1752
chrX-60582984-INS-82
chrX-60602896-INS-199
chrX-60632761-INS-174
chrX-60704337-INS-588
chrX-60787502-INS-1489
chrX-60831523-INS-6395
chrX-60874363-INS-6386
chrX-60920794-INS-93
chrX-60922167-INS-125
chrX-60942760-INS-6530
chrX-60997978-INS-2500
chrX-61087141-INS-129
chrX-61102626-INS-94
chrX-61186710-INS-4041
chrX-61240752-INS-1464
chrX-61241612-INS-1051
chrX-61315405-INS-75
chrX-61364547-INS-648
chrX-61448941-INS-5027
chrX-61476404-INS-1108
chrX-61488261-INS-1961
chrX-61561907-INS-898
chrX-61564889-INS-783

chrX-61570629-INS-1528
chrX-61587229-INS-746
chrX-61641578-INS-5542
chrX-61647897-INS-748
chrX-61714168-INS-124
chrX-61756370-INS-1489
chrX-61823740-INS-387
chrX-61977983-INS-1500
chrX-62071127-INS-99
chrX-62072619-INS-257
chrX-62165663-INS-5362
chrX-62195231-INS-56
chrX-62213204-INS-52
chrX-62229683-INS-58
chrX-62332603-INS-1650
chrX-62523711-INS-2796
chrX-62715638-INS-57
chrX-62738868-INS-126
chrX-62749930-INS-7126
chrX-62799965-INS-117
chrX-62803644-INS-4774
chrX-62838698-INS-2421
chrX-62895921-INS-60
chrX-62912145-INS-174
chrX-62933130-INS-541
chrX-63069978-INS-884
chrX-63152828-INS-1280
chrX-63174865-INS-1003
chrX-63190505-INS-64
chrX-63238486-INS-60
chrX-63366049-INS-353
chrX-63418508-INS-2912
chrX-63477858-INS-6672
chrX-63514792-INS-129
chrX-63568731-INS-6689
chrX-63607864-INS-50
chrX-63695025-INS-52
chrX-63708477-INS-148
chrX-63743086-INS-7333
chrX-63928069-INS-2630
chrX-63964848-INS-52
chrX-64221521-INS-74
chrX-64252504-INS-50
chrX-64270118-INS-6078
chrX-64372652-INS-52
chrX-64399756-INS-250
chrX-64400127-INS-52
chrX-64462027-INS-2920
chrX-64529988-INS-8726
chrX-64599656-INS-383
chrX-64650060-INS-167
chrX-64712950-INS-1620
chrX-64769426-INS-148
chrX-64903816-INS-9618
chrX-65018640-INS-416
chrX-65086511-INS-6309
chrX-65121258-INS-618
chrX-65205431-INS-54
chrX-65227978-INS-6287
chrX-65344216-INS-6457
chrX-65410476-INS-5806
chrX-65421335-INS-63
chrX-65426021-INS-11544
chrX-65456483-INS-55
chrX-65493617-INS-159
chrX-65510289-INS-50
chrX-65539479-INS-604
chrX-65582378-INS-3085
chrX-65608849-INS-215
chrX-65676025-INS-58
chrX-65923904-INS-1113
chrX-65986076-INS-168
chrX-66108438-INS-526
chrX-66108438-INS-74
chrX-66119864-INS-2136
chrX-66144481-INS-1182
chrX-66145195-INS-1459

chrX-66167234-INS-348
chrX-66229576-INS-2702
chrX-66249235-INS-52
chrX-66261699-INS-7569
chrX-66514710-INS-428
chrX-66515083-INS-428
chrX-66575652-INS-200
chrX-66593738-INS-50
chrX-66648180-INS-4338
chrX-66722452-INS-54
chrX-66764665-INS-1369
chrX-66783347-INS-351
chrX-66794506-INS-70
chrX-66844214-INS-7651
chrX-66935260-INS-760
chrX-67004095-INS-2313
chrX-67044682-INS-6799
chrX-67056386-INS-6799
chrX-67116973-INS-3472
chrX-67178919-INS-6665
chrX-67247284-INS-80
chrX-67292906-INS-56
chrX-67360370-INS-1599
chrX-67367182-INS-194
chrX-67413183-INS-1432
chrX-67441959-INS-2426
chrX-67483773-INS-5998
chrX-67576186-INS-261
chrX-67581679-INS-3953
chrX-67643541-INS-92
chrX-67650427-INS-151
chrX-67871450-INS-2963
chrX-67950947-INS-14235
chrX-67975002-INS-2151
chrX-68016041-INS-110
chrX-68016158-INS-85
chrX-68041316-INS-2318
chrX-68046741-INS-658
chrX-68079157-INS-114
chrX-68083877-INS-4083
chrX-68188251-INS-4767
chrX-68188667-INS-147
chrX-68196266-INS-1303
chrX-68229658-INS-4013
chrX-68265164-INS-7256
chrX-68302625-INS-2608
chrX-68433891-INS-92
chrX-68439114-INS-57
chrX-68471944-INS-50
chrX-68525293-INS-60
chrX-68538588-INS-50
chrX-68579363-INS-1514
chrX-68579689-INS-481
chrX-68649933-INS-52
chrX-68685786-INS-57
chrX-68715073-INS-206
chrX-68748918-INS-362
chrX-68751930-INS-50
chrX-68753194-INS-50
chrX-68761512-INS-106
chrX-68761589-INS-52
chrX-68789356-INS-150
chrX-68924044-INS-59
chrX-68924610-INS-216
chrX-68924665-INS-57
chrX-68932043-INS-371
chrX-68960725-INS-11292
chrX-68961489-INS-208
chrX-68981962-INS-848
chrX-69039980-INS-216
chrX-69040489-INS-204
chrX-69040489-INS-524
chrX-69078530-INS-63
chrX-69078558-INS-400
chrX-69124673-INS-1325
chrX-69198320-INS-58
chrX-69235370-INS-199

chrX-69255882-INS-126
chrX-69319308-INS-58
chrX-69341472-INS-5730
chrX-69437753-INS-60
chrX-69474375-INS-644
chrX-69500628-INS-54
chrX-69552718-INS-1293
chrX-69552810-INS-430
chrX-69607269-INS-53
chrX-69617915-INS-962
chrX-69618528-INS-1127
chrX-69795115-INS-2832
chrX-69822923-INS-181
chrX-69886002-INS-268
chrX-70005504-INS-64
chrX-70013628-INS-107
chrX-70045015-INS-204
chrX-70058516-INS-76
chrX-70079029-INS-63
chrX-70079160-INS-166
chrX-70092216-INS-50
chrX-70136212-INS-56
chrX-70220236-INS-2366
chrX-70287269-INS-222
chrX-70316686-INS-50
chrX-70335835-INS-54
chrX-70341235-INS-2910
chrX-70344200-INS-1919
chrX-70390905-INS-12960
chrX-70418486-INS-187
chrX-70418563-INS-570
chrX-70506171-INS-95
chrX-70544343-INS-84
chrX-70566139-INS-179
chrX-70566363-INS-59
chrX-70566835-INS-53
chrX-70591980-INS-2768
chrX-70621895-INS-577
chrX-70641670-INS-205
chrX-70719259-INS-613
chrX-70735728-INS-70
chrX-70768023-INS-60
chrX-70812494-INS-78
chrX-70854533-INS-54
chrX-70871284-INS-54
chrX-70874255-INS-69
chrX-70874875-INS-199
chrX-70877567-INS-6423
chrX-70893490-INS-328
chrX-70955318-INS-6171
chrX-70960191-INS-52
chrX-70960784-INS-350
chrX-70961363-INS-1085
chrX-70990156-INS-102
chrX-71014294-INS-50
chrX-71063193-INS-6896
chrX-71088326-INS-1275
chrX-71101666-INS-2013
chrX-71110787-INS-3103
chrX-71186054-INS-52
chrX-71231214-INS-6390
chrX-71241894-INS-84
chrX-71253987-INS-94
chrX-71268908-INS-703
chrX-71321631-INS-6655
chrX-71350696-INS-12803
chrX-71359587-INS-7684
chrX-71531664-INS-52
chrX-71563049-INS-150
chrX-71601841-INS-50
chrX-71706618-INS-3587
chrX-71737582-INS-2730
chrX-71765551-INS-608
chrX-71770451-INS-2095
chrX-71775068-INS-108
chrX-71802700-INS-4861
chrX-71811726-INS-4833

chrX-71847928-INS-188
chrX-71865217-INS-273
chrX-71865226-INS-55
chrX-71865909-INS-66
chrX-71906086-INS-356
chrX-71927106-INS-1034
chrX-71948525-INS-86
chrX-72011999-INS-138
chrX-72018596-INS-69
chrX-72037586-INS-64
chrX-72126937-INS-76
chrX-72238811-INS-123
chrX-72302907-INS-748
chrX-72318220-INS-848
chrX-72338213-INS-79
chrX-72377757-INS-115
chrX-72381787-INS-60
chrX-72404358-INS-4316
chrX-72411622-INS-5072
chrX-72562312-INS-70
chrX-72574439-INS-172
chrX-72581224-INS-286
chrX-72613288-INS-72
chrX-72676176-INS-206
chrX-72785779-INS-251
chrX-72791737-INS-77
chrX-72872311-INS-54
chrX-72875422-INS-176
chrX-72907348-INS-110
chrX-72929856-INS-58
chrX-72951871-INS-205
chrX-72966821-INS-211
chrX-72968369-INS-223
chrX-72977628-INS-195
chrX-72981554-INS-184
chrX-73063469-INS-52
chrX-73086355-INS-172
chrX-73106880-INS-148
chrX-73228203-INS-56
chrX-73259579-INS-593
chrX-73260259-INS-622
chrX-73451387-INS-204
chrX-73521106-INS-77
chrX-73606148-INS-135
chrX-73606222-INS-505
chrX-73626422-INS-52
chrX-73740254-INS-98
chrX-73749231-INS-110
chrX-73759977-INS-82
chrX-73769104-INS-50
chrX-73801924-INS-487
chrX-73802015-INS-6377
chrX-73812468-INS-61
chrX-73814322-INS-60
chrX-73824828-INS-4803
chrX-73830170-INS-497
chrX-73857641-INS-68
chrX-73877868-INS-53
chrX-73880462-INS-442
chrX-73888488-INS-502
chrX-73906864-INS-56
chrX-73957101-INS-78
chrX-74058844-INS-7025
chrX-74181179-INS-169
chrX-74190326-INS-60
chrX-74237123-INS-165
chrX-74258895-INS-3059
chrX-74374388-INS-7183
chrX-74429047-INS-50
chrX-74528277-INS-2042
chrX-74541540-INS-53
chrX-74663690-INS-2387
chrX-74687566-INS-60
chrX-74706529-INS-3203
chrX-74709429-INS-999
chrX-74723538-INS-66
chrX-74751844-INS-6576

chrX-74782666-INS-103
chrX-74783646-INS-64
chrX-74788465-INS-74
chrX-74792607-INS-961
chrX-74797515-INS-989
chrX-74812772-INS-63
chrX-74818132-INS-62
chrX-74880447-INS-54
chrX-75051271-INS-7684
chrX-75108781-INS-7908
chrX-75193097-INS-216
chrX-75193841-INS-76
chrX-75204173-INS-50
chrX-75207647-INS-52
chrX-75208120-INS-5668
chrX-75273552-INS-54
chrX-75367558-INS-50
chrX-75406745-INS-424
chrX-75413204-INS-391
chrX-75516921-INS-50
chrX-75581202-INS-4837
chrX-75586162-INS-1053
chrX-75608271-INS-204
chrX-75608432-INS-857
chrX-75616142-INS-50
chrX-75639512-INS-18221
chrX-75725747-INS-7971
chrX-75846522-INS-51
chrX-75855503-INS-6401
chrX-75866063-INS-624
chrX-75893223-INS-177
chrX-75908336-INS-7831
chrX-75912261-INS-2738
chrX-75924102-INS-6589
chrX-75963891-INS-2852
chrX-76023766-INS-1688
chrX-76077372-INS-50
chrX-76080248-INS-624
chrX-76089158-INS-64
chrX-76112596-INS-188
chrX-76146833-INS-6586
chrX-76185357-INS-3124
chrX-76185357-INS-445
chrX-76218754-INS-2958
chrX-76320195-INS-2322
chrX-76349501-INS-92
chrX-76366206-INS-52
chrX-76369404-INS-1278
chrX-76408276-INS-6711
chrX-76420530-INS-160
chrX-76430323-INS-94
chrX-76441366-INS-432
chrX-76441962-INS-424
chrX-76464130-INS-61
chrX-76499429-INS-50
chrX-76576304-INS-80
chrX-76721250-INS-1377
chrX-76722745-INS-55
chrX-76819989-INS-231
chrX-76839796-INS-50
chrX-76877080-INS-2741
chrX-76982836-INS-2321
chrX-77050787-INS-70
chrX-77063097-INS-1483
chrX-77066857-INS-510
chrX-77102569-INS-60
chrX-77138815-INS-51
chrX-77139244-INS-4035
chrX-77155170-INS-5757
chrX-77192269-INS-53
chrX-77235266-INS-1489
chrX-77242398-INS-8744
chrX-77246749-INS-117
chrX-77288031-INS-60
chrX-77290261-INS-1706
chrX-77291861-INS-58
chrX-77302763-INS-83

chrX-77303022-INS-80
chrX-77407570-INS-5627
chrX-77415969-INS-1287
chrX-77450971-INS-8859
chrX-77542202-INS-6819
chrX-77650306-INS-433
chrX-77656333-INS-309
chrX-77713956-INS-275
chrX-77773769-INS-105
chrX-77949362-INS-50
chrX-78141470-INS-6369
chrX-78172192-INS-3194
chrX-78241312-INS-50
chrX-78251741-INS-114
chrX-78336982-INS-144
chrX-78342438-INS-1455
chrX-78342444-INS-380
chrX-78604093-INS-188
chrX-78610255-INS-55
chrX-78620054-INS-52
chrX-78702397-INS-568
chrX-78794309-INS-522
chrX-78821961-INS-1410
chrX-78823543-INS-66
chrX-78824076-INS-54
chrX-78919044-INS-4750
chrX-78922811-INS-76
chrX-78965194-INS-201
chrX-78965420-INS-203
chrX-78969714-INS-4729
chrX-79017341-INS-8723
chrX-79139810-INS-6895
chrX-79168868-INS-51
chrX-79196761-INS-70
chrX-79307728-INS-11904
chrX-79323341-INS-347
chrX-79334392-INS-4815
chrX-79374424-INS-52
chrX-79381595-INS-574
chrX-79422540-INS-201
chrX-79569664-INS-6369
chrX-79654119-INS-2326
chrX-79661854-INS-116
chrX-79681682-INS-6483
chrX-79768688-INS-3151
chrX-79788730-INS-110
chrX-79835948-INS-6790
chrX-79928684-INS-211
chrX-79966618-INS-6642
chrX-80089438-INS-50
chrX-80111393-INS-206
chrX-80177102-INS-82
chrX-80191747-INS-62
chrX-80219573-INS-6147
chrX-80227214-INS-204
chrX-80256544-INS-910
chrX-80314878-INS-305
chrX-80314880-INS-125
chrX-80336816-INS-90
chrX-80456716-INS-208
chrX-80487172-INS-103
chrX-80618727-INS-1487
chrX-80822728-INS-79
chrX-80826839-INS-56
chrX-80862239-INS-6520
chrX-80934917-INS-584
chrX-80951717-INS-50
chrX-80976451-INS-533
chrX-80989344-INS-1830
chrX-81065081-INS-3152
chrX-81068218-INS-118
chrX-81068701-INS-320
chrX-81073648-INS-73
chrX-81159513-INS-50
chrX-81174893-INS-50
chrX-81178254-INS-3048
chrX-81180805-INS-54

chrX-81229684-INS-2381
chrX-81280698-INS-6560
chrX-81370565-INS-475
chrX-81392300-INS-458
chrX-81393633-INS-406
chrX-81438761-INS-200
chrX-81443359-INS-418
chrX-81465928-INS-67
chrX-81479855-INS-2836
chrX-81612857-INS-6854
chrX-81625228-INS-54
chrX-81637428-INS-96
chrX-81691321-INS-52
chrX-81719463-INS-52
chrX-81796000-INS-7045
chrX-81911999-INS-140
chrX-81938639-INS-170
chrX-81996709-INS-1440
chrX-82030853-INS-179
chrX-82030853-INS-59
chrX-82039082-INS-465
chrX-82103136-INS-54
chrX-82121816-INS-57
chrX-82136496-INS-56
chrX-82136500-INS-160
chrX-82150822-INS-68
chrX-82169344-INS-90
chrX-82204936-INS-1094
chrX-82220701-INS-56
chrX-82231262-INS-59
chrX-82233961-INS-6691
chrX-82242588-INS-50
chrX-82269177-INS-992
chrX-82278540-INS-6710
chrX-82318592-INS-62
chrX-82342538-INS-506
chrX-82351847-INS-753
chrX-82390077-INS-94
chrX-82421491-INS-2020
chrX-82464243-INS-100
chrX-82530664-INS-171
chrX-82557685-INS-3417
chrX-82647475-INS-56
chrX-82652904-INS-72
chrX-82690709-INS-203
chrX-82698118-INS-656
chrX-82707322-INS-282
chrX-82709251-INS-201
chrX-82725538-INS-810
chrX-82747477-INS-6718
chrX-82776618-INS-1887
chrX-82794697-INS-132
chrX-82797960-INS-2970
chrX-82842741-INS-2684
chrX-82880051-INS-178
chrX-82892137-INS-53
chrX-82892217-INS-3321
chrX-82934910-INS-3288
chrX-82987585-INS-476
chrX-83030952-INS-261
chrX-83180627-INS-70
chrX-83270647-INS-68
chrX-83279024-INS-2998
chrX-83279024-INS-382
chrX-83297745-INS-114
chrX-83314072-INS-56
chrX-83317613-INS-575
chrX-83356963-INS-178
chrX-83362840-INS-6256
chrX-83392013-INS-9821
chrX-83395981-INS-50
chrX-83444637-INS-60
chrX-83558324-INS-52
chrX-83569078-INS-164
chrX-83573126-INS-9618
chrX-83605535-INS-4811
chrX-83651346-INS-624

chrX-83724666-INS-51
chrX-83750464-INS-1069
chrX-83842543-INS-172
chrX-83881157-INS-63
chrX-83889071-INS-521
chrX-83940259-INS-240
chrX-83940312-INS-55
chrX-83951972-INS-106
chrX-84031355-INS-54
chrX-84063601-INS-226
chrX-84075645-INS-89
chrX-84076518-INS-489
chrX-84102176-INS-106
chrX-84172174-INS-68
chrX-84172317-INS-58
chrX-84229918-INS-6757
chrX-84236722-INS-303
chrX-84257101-INS-6696
chrX-84302430-INS-500
chrX-84344287-INS-2322
chrX-84389458-INS-319
chrX-84401318-INS-6928
chrX-84477407-INS-1726
chrX-84483693-INS-6891
chrX-84560244-INS-6473
chrX-84569203-INS-57
chrX-84684578-INS-90
chrX-84685077-INS-70
chrX-84704640-INS-164
chrX-84722031-INS-1775
chrX-84728424-INS-6402
chrX-84728677-INS-2705
chrX-84737644-INS-52
chrX-84798305-INS-202
chrX-84805615-INS-118
chrX-84808953-INS-6914
chrX-84817779-INS-412
chrX-84821524-INS-213
chrX-84821541-INS-61
chrX-84830291-INS-90
chrX-84830417-INS-572
chrX-84844017-INS-449
chrX-84882449-INS-4648
chrX-84900104-INS-50
chrX-84900305-INS-189
chrX-84964289-INS-2658
chrX-84983994-INS-843
chrX-85015261-INS-1614
chrX-85033723-INS-2048
chrX-85033723-INS-443
chrX-85088961-INS-54
chrX-85123950-INS-2652
chrX-85161160-INS-138
chrX-85161826-INS-2435
chrX-85165806-INS-3812
chrX-85187129-INS-293
chrX-85268315-INS-669
chrX-85304377-INS-999
chrX-85307792-INS-4701
chrX-85333637-INS-6298
chrX-85349911-INS-2999
chrX-85602715-INS-64
chrX-85610673-INS-64
chrX-85736290-INS-52
chrX-85759718-INS-104
chrX-85907855-INS-63
chrX-86036322-INS-51
chrX-86045667-INS-50
chrX-86050411-INS-744
chrX-86117316-INS-234
chrX-86152821-INS-264
chrX-86152894-INS-66
chrX-86174416-INS-53
chrX-86178734-INS-52
chrX-86210766-INS-57
chrX-86444963-INS-6490
chrX-86471781-INS-651

chrX-86549085-INS-54
chrX-86563391-INS-50
chrX-86587828-INS-498
chrX-86656603-INS-57
chrX-86710019-INS-240
chrX-86749142-INS-55
chrX-87063980-INS-80
chrX-87141808-INS-329
chrX-87200194-INS-6613
chrX-87210679-INS-224
chrX-87214454-INS-6395
chrX-87237279-INS-6596
chrX-87320342-INS-137
chrX-87332410-INS-6559
chrX-87347894-INS-549
chrX-87370226-INS-6372
chrX-87377209-INS-52
chrX-87421370-INS-7968
chrX-87487127-INS-6237
chrX-87506336-INS-256
chrX-87507136-INS-411
chrX-87541082-INS-68
chrX-87559713-INS-7567
chrX-87583479-INS-114
chrX-87629063-INS-172
chrX-87734496-INS-994
chrX-87751078-INS-50
chrX-87762119-INS-197
chrX-87806519-INS-6712
chrX-87817545-INS-124
chrX-87826510-INS-7691
chrX-87835086-INS-340
chrX-87853021-INS-1578
chrX-87866444-INS-6389
chrX-87883860-INS-58
chrX-87895861-INS-7098
chrX-87936086-INS-412
chrX-87956800-INS-1461
chrX-88083934-INS-55
chrX-88127541-INS-1756
chrX-88141341-INS-5212
chrX-88230610-INS-66
chrX-88271682-INS-9892
chrX-88417029-INS-2974
chrX-88436254-INS-52
chrX-88440603-INS-52
chrX-88506149-INS-50
chrX-88524981-INS-1608
chrX-88529640-INS-55
chrX-88530134-INS-428
chrX-88549806-INS-78
chrX-88612014-INS-73
chrX-88705470-INS-1468
chrX-88781262-INS-5304
chrX-88820944-INS-73
chrX-88820945-INS-415
chrX-88837109-INS-722
chrX-88905757-INS-192
chrX-88940505-INS-52
chrX-89025989-INS-118
chrX-89040919-INS-6692
chrX-89044394-INS-7002
chrX-89048293-INS-440
chrX-89157990-INS-131
chrX-89180841-INS-71
chrX-89202296-INS-6405
chrX-89223617-INS-839
chrX-89223851-INS-91
chrX-89240224-INS-6532
chrX-89256633-INS-5015
chrX-89284387-INS-710
chrX-89329047-INS-6275
chrX-89370003-INS-1479
chrX-89429436-INS-5219
chrX-89437791-INS-3997
chrX-89506655-INS-3062
chrX-89608921-INS-8332

chrX-89656683-INS-52
chrX-89707710-INS-1022
chrX-89796579-INS-20826
chrX-89819008-INS-848
chrX-89886071-INS-11997
chrX-89886684-INS-997
chrX-89918714-INS-4018
chrX-90085058-INS-325
chrX-90127276-INS-6716
chrX-90128338-INS-1226
chrX-90148256-INS-50
chrX-90215510-INS-62
chrX-90238507-INS-3346
chrX-90244232-INS-1762
chrX-90251280-INS-11775
chrX-90251280-INS-4650
chrX-90295260-INS-106
chrX-90303674-INS-1610
chrX-90330821-INS-296
chrX-90357735-INS-57
chrX-90362072-INS-84
chrX-90367017-INS-57
chrX-90367640-INS-57
chrX-90369969-INS-628
chrX-90370175-INS-3538
chrX-90370992-INS-5597
chrX-90435810-INS-414
chrX-90452278-INS-53
chrX-90458306-INS-341
chrX-90463816-INS-71
chrX-90482492-INS-77
chrX-90555572-INS-160
chrX-90583141-INS-644
chrX-90602232-INS-6413
chrX-90623844-INS-335
chrX-90623955-INS-142
chrX-90691840-INS-54
chrX-90696426-INS-67
chrX-90702292-INS-1800
chrX-90948246-INS-84
chrX-90961707-INS-3873
chrX-91057683-INS-1000
chrX-91063125-INS-81
chrX-91078866-INS-1439
chrX-91107539-INS-1882
chrX-91150971-INS-164
chrX-91151267-INS-132
chrX-91157581-INS-578
chrX-91195888-INS-54
chrX-91362444-INS-60
chrX-91400156-INS-146
chrX-91486084-INS-2886
chrX-91552919-INS-132
chrX-91654137-INS-58
chrX-91865810-INS-74
chrX-91866135-INS-68
chrX-91928566-INS-604
chrX-91942497-INS-1500
chrX-92026838-INS-51
chrX-92063774-INS-139
chrX-92075139-INS-649
chrX-92137238-INS-63
chrX-92137262-INS-288
chrX-92159124-INS-1772
chrX-92159565-INS-6254
chrX-92198310-INS-52
chrX-92222636-INS-587
chrX-92235232-INS-78
chrX-92312641-INS-117
chrX-92387417-INS-1981
chrX-92394035-INS-76
chrX-92493397-INS-64
chrX-92545080-INS-122
chrX-92690853-INS-362
chrX-92695414-INS-173
chrX-92714125-INS-208
chrX-92714720-INS-163

chrX-92725865-INS-116
chrX-92782387-INS-51
chrX-92790470-INS-53
chrX-92830509-INS-107
chrX-92882675-INS-163
chrX-92892861-INS-197
chrX-92937029-INS-70
chrX-92965973-INS-319
chrX-92981922-INS-91
chrX-93021067-INS-196
chrX-93021647-INS-71
chrX-93040141-INS-54
chrX-93040935-INS-88
chrX-93041776-INS-50
chrX-93088401-INS-90
chrX-93097159-INS-201
chrX-93105720-INS-61
chrX-93133766-INS-144
chrX-93134165-INS-197
chrX-93228840-INS-196
chrX-93236233-INS-63
chrX-93238335-INS-108
chrX-93263360-INS-89
chrX-93302928-INS-70
chrX-93436663-INS-68
chrX-93437439-INS-104
chrX-93449990-INS-2167
chrX-93507886-INS-164
chrX-93592893-INS-6502
chrX-93834891-INS-68
chrX-93886342-INS-116
chrX-93888353-INS-2870
chrX-93902503-INS-193
chrX-94014254-INS-696
chrX-94033208-INS-59
chrX-94064847-INS-68
chrX-94129857-INS-2851
chrX-94195803-INS-56
chrX-94287294-INS-6479
chrX-94302496-INS-1544
chrX-94342920-INS-171
chrX-94361262-INS-74
chrX-94362117-INS-50
chrX-94391704-INS-51
chrX-94493278-INS-77
chrX-94522935-INS-6403
chrX-94526506-INS-7759
chrX-94531330-INS-212
chrX-94554497-INS-259
chrX-94559299-INS-6585
chrX-94581684-INS-2807
chrX-94673669-INS-65
chrX-94674553-INS-83
chrX-94739567-INS-702
chrX-94829766-INS-357
chrX-94830077-INS-1269
chrX-94850995-INS-360
chrX-94919336-INS-1460
chrX-94976367-INS-171
chrX-95120043-INS-56
chrX-95122451-INS-551
chrX-95131381-INS-124
chrX-95207875-INS-160
chrX-95233223-INS-2269
chrX-95236818-INS-203
chrX-95242714-INS-547
chrX-95299094-INS-6753
chrX-95330707-INS-6659
chrX-95341395-INS-490
chrX-95389548-INS-100
chrX-95411035-INS-1132
chrX-95438410-INS-1487
chrX-95507605-INS-80
chrX-95565803-INS-50
chrX-95576568-INS-54
chrX-95577690-INS-4720
chrX-95651041-INS-67

chrX-95737612-INS-321
chrX-95737667-INS-1828
chrX-95746913-INS-995
chrX-95774634-INS-60
chrX-95797996-INS-771
chrX-95822492-INS-358
chrX-95921786-INS-6957
chrX-96134927-INS-1381
chrX-96166739-INS-52
chrX-96185348-INS-4522
chrX-96269666-INS-1261
chrX-96272813-INS-213
chrX-96315243-INS-100
chrX-96535309-INS-597
chrX-96584871-INS-2257
chrX-96624939-INS-7006
chrX-96651569-INS-489
chrX-96651572-INS-3594
chrX-96687892-INS-164
chrX-96740552-INS-362
chrX-96771901-INS-212
chrX-96838392-INS-50
chrX-96850925-INS-50
chrX-96971284-INS-1270
chrX-96984531-INS-402
chrX-96992366-INS-8274
chrX-97120955-INS-98
chrX-97147247-INS-212
chrX-97226103-INS-711
chrX-97232242-INS-112
chrX-97258080-INS-136
chrX-97374595-INS-7233
chrX-97390557-INS-2240
chrX-97463681-INS-916
chrX-97533319-INS-3271
chrX-97578832-INS-475
chrX-97583180-INS-4167
chrX-97638992-INS-1039
chrX-97681002-INS-74
chrX-97818551-INS-2749
chrX-97824882-INS-75
chrX-97826859-INS-1561
chrX-97848628-INS-1735
chrX-97906868-INS-193
chrX-97913967-INS-853
chrX-97930588-INS-185
chrX-97931089-INS-269
chrX-98012216-INS-6389
chrX-98019072-INS-589
chrX-98065575-INS-55
chrX-98066177-INS-328
chrX-98146275-INS-327
chrX-98217780-INS-163
chrX-98311706-INS-105
chrX-98335224-INS-128
chrX-98385613-INS-55
chrX-98385686-INS-111
chrX-98387130-INS-82
chrX-98411068-INS-50
chrX-98491242-INS-95
chrX-98493928-INS-52
chrX-98540615-INS-56
chrX-98601737-INS-64
chrX-98617110-INS-52
chrX-98644277-INS-58
chrX-98659734-INS-4119
chrX-98701143-INS-181
chrX-98720307-INS-154
chrX-98760959-INS-55
chrX-98785044-INS-54
chrX-98786647-INS-214
chrX-98816601-INS-1107
chrX-98847864-INS-50
chrX-98856486-INS-50
chrX-98943520-INS-4316
chrX-98963476-INS-58
chrX-98986438-INS-64

chrX-98986681-INS-63
chrX-98986793-INS-117
chrX-99059845-INS-2387
chrX-99080463-INS-1533
chrX-99092208-INS-1640
chrX-99175833-INS-63
chrX-99191433-INS-216
chrX-99225226-INS-17158
chrX-99237106-INS-80
chrX-99283513-INS-60
chrX-99425830-INS-109
chrX-99617701-INS-202
chrX-99676235-INS-191
chrX-99676930-INS-162
chrX-99699470-INS-1111
chrX-99706628-INS-198
chrX-99718420-INS-54
chrX-99783993-INS-153
chrX-99844661-INS-238
chrX-99905816-INS-1367
chrX-99915792-INS-198
chrX-99996750-INS-2917
chrX-100043262-INS-333
chrX-100049387-INS-61
chrX-100076880-INS-85
chrX-100112902-INS-50
chrX-100117531-INS-161
chrX-100142128-INS-52
chrX-100142207-INS-273
chrX-100144218-INS-52
chrX-100153085-INS-53
chrX-100153635-INS-209
chrX-100164029-INS-61
chrX-100242105-INS-210
chrX-100262125-INS-56
chrX-100281343-INS-50
chrX-100287459-INS-1244
chrX-100294700-INS-305
chrX-100295202-INS-225
chrX-100323279-INS-177
chrX-100330488-INS-218
chrX-100354889-INS-210
chrX-100400409-INS-241
chrX-100406001-INS-54
chrX-100407490-INS-112
chrX-100509082-INS-220
chrX-100551344-INS-200
chrX-100609603-INS-306
chrX-100658419-INS-220
chrX-100670805-INS-273
chrX-100723958-INS-139
chrX-100724223-INS-180
chrX-100739759-INS-328
chrX-100788442-INS-461
chrX-100847989-INS-4774
chrX-100882639-INS-7234
chrX-100892848-INS-53
chrX-100896517-INS-52
chrX-100969464-INS-300
chrX-100981996-INS-198
chrX-100982395-INS-148
chrX-101036634-INS-72
chrX-101085124-INS-55
chrX-101138218-INS-50
chrX-101142827-INS-52
chrX-101184456-INS-7233
chrX-101215924-INS-557
chrX-101255441-INS-56
chrX-101262447-INS-136
chrX-101308691-INS-51
chrX-101313683-INS-1174
chrX-101313855-INS-56
chrX-101313978-INS-2245
chrX-101315055-INS-80
chrX-101323620-INS-63
chrX-101339101-INS-52
chrX-101343497-INS-538

chrX-101374557-INS-112
chrX-101439259-INS-52
chrX-101439608-INS-98
chrX-101440045-INS-69
chrX-101443959-INS-58
chrX-101545538-INS-2904
chrX-101550184-INS-116
chrX-101555860-INS-1890
chrX-101555959-INS-750
chrX-101556375-INS-660
chrX-101620185-INS-62
chrX-101625763-INS-1142
chrX-101671177-INS-182
chrX-101721094-INS-6610
chrX-101741286-INS-21153
chrX-101743348-INS-184
chrX-101745585-INS-1501
chrX-101760589-INS-289
chrX-101773096-INS-4803
chrX-101780932-INS-19341
chrX-101795269-INS-55
chrX-101799844-INS-84
chrX-101875765-INS-176
chrX-101971569-INS-504
chrX-102016012-INS-1281
chrX-102022979-INS-53
chrX-102044066-INS-50
chrX-102050407-INS-1375
chrX-102061622-INS-2843
chrX-102117741-INS-1375
chrX-102121371-INS-195
chrX-102158025-INS-109
chrX-102241637-INS-70
chrX-102275848-INS-1020
chrX-102295267-INS-107
chrX-102296018-INS-56
chrX-102321214-INS-240
chrX-102329082-INS-1197
chrX-102379764-INS-214
chrX-102391063-INS-51
chrX-102396056-INS-225
chrX-102438848-INS-54
chrX-102477655-INS-59
chrX-102498832-INS-85
chrX-102500496-INS-612
chrX-102511056-INS-55
chrX-102547132-INS-207
chrX-102588862-INS-117
chrX-102606360-INS-52
chrX-102612384-INS-57
chrX-102673519-INS-278
chrX-102674069-INS-51
chrX-102689269-INS-135
chrX-102717798-INS-55
chrX-102827244-INS-289
chrX-102827326-INS-69
chrX-102831814-INS-6601
chrX-102955653-INS-1749
chrX-102994986-INS-52
chrX-103042518-INS-52
chrX-103064363-INS-57
chrX-103079389-INS-60
chrX-103116086-INS-492
chrX-103141717-INS-245
chrX-103187533-INS-1901
chrX-103258127-INS-52
chrX-103270568-INS-3070
chrX-103273717-INS-3103
chrX-103279266-INS-105
chrX-103290693-INS-196
chrX-103300936-INS-8681
chrX-103360620-INS-2263
chrX-103379831-INS-1502
chrX-103441228-INS-1002
chrX-103472473-INS-54
chrX-103496929-INS-90
chrX-103500727-INS-10218

chrX-103527710-INS-110
chrX-103578097-INS-59
chrX-103609915-INS-68
chrX-103627163-INS-6563
chrX-103869339-INS-110
chrX-103990279-INS-2542
chrX-104003253-INS-212
chrX-104057960-INS-50
chrX-104094346-INS-455
chrX-104130543-INS-371
chrX-104168935-INS-50
chrX-104188434-INS-6421
chrX-104244749-INS-3030
chrX-104247326-INS-6323
chrX-104257635-INS-56
chrX-104288551-INS-1825
chrX-104291857-INS-111
chrX-104313294-INS-68
chrX-104372475-INS-1795
chrX-104401311-INS-53
chrX-104422842-INS-637
chrX-104502848-INS-9597
chrX-104514567-INS-1685
chrX-104644068-INS-188
chrX-104679230-INS-500
chrX-104775958-INS-4469
chrX-104777061-INS-51
chrX-104809546-INS-72
chrX-104872097-INS-200
chrX-104877923-INS-186
chrX-104901666-INS-108
chrX-104961848-INS-280
chrX-104965354-INS-255
chrX-104987579-INS-80
chrX-105003040-INS-2146
chrX-105037797-INS-162
chrX-105060518-INS-199
chrX-105081186-INS-504
chrX-105083672-INS-158
chrX-105089557-INS-80
chrX-105115854-INS-1792
chrX-105125859-INS-406
chrX-105146692-INS-1723
chrX-105173172-INS-2121
chrX-105181842-INS-150
chrX-105182030-INS-75
chrX-105197089-INS-206
chrX-105234233-INS-59
chrX-105276606-INS-50
chrX-105285070-INS-72
chrX-105285254-INS-64
chrX-105415524-INS-56
chrX-105462562-INS-7003
chrX-105742422-INS-72
chrX-105744913-INS-72
chrX-105745193-INS-165
chrX-105770350-INS-391
chrX-105830673-INS-1515
chrX-105896993-INS-58
chrX-105987855-INS-207
chrX-106042459-INS-52
chrX-106077037-INS-500
chrX-106121206-INS-2449
chrX-106137566-INS-232
chrX-106163010-INS-1923
chrX-106164432-INS-196
chrX-106164534-INS-81
chrX-106169277-INS-932
chrX-106198519-INS-59
chrX-106223316-INS-408
chrX-106250001-INS-2842
chrX-106270045-INS-135
chrX-106285706-INS-392
chrX-106360252-INS-262
chrX-106425614-INS-699
chrX-106430263-INS-50
chrX-106449301-INS-50

chrX-106597609-INS-51
chrX-106661755-INS-1657
chrX-106750327-INS-6579
chrX-106790283-INS-536
chrX-106794684-INS-70
chrX-106800087-INS-419
chrX-106800355-INS-4673
chrX-106952284-INS-60
chrX-106963488-INS-71
chrX-107115143-INS-208
chrX-107216281-INS-4831
chrX-107231412-INS-17001
chrX-107280293-INS-65
chrX-107310655-INS-3495
chrX-107385263-INS-3901
chrX-107448614-INS-4467
chrX-107449145-INS-50
chrX-107539168-INS-52
chrX-107546807-INS-50
chrX-107605149-INS-6365
chrX-107614790-INS-2895
chrX-107623299-INS-99
chrX-107745786-INS-174
chrX-107791382-INS-78
chrX-107850586-INS-50
chrX-107864309-INS-211
chrX-107893947-INS-54
chrX-107942038-INS-60
chrX-107950232-INS-60
chrX-108059706-INS-155
chrX-108139659-INS-55
chrX-108147980-INS-52
chrX-108175021-INS-187
chrX-108250266-INS-420
chrX-108254071-INS-5548
chrX-108255338-INS-10537
chrX-108274264-INS-198
chrX-108279986-INS-1482
chrX-108290454-INS-266
chrX-108304167-INS-1665
chrX-108334453-INS-2683
chrX-108374796-INS-3373
chrX-108571082-INS-50
chrX-108574625-INS-138
chrX-108645689-INS-7988
chrX-108701282-INS-52
chrX-108782831-INS-6524
chrX-108787801-INS-71
chrX-109193168-INS-205
chrX-109203138-INS-50
chrX-109227511-INS-7923
chrX-109251309-INS-4908
chrX-109259450-INS-306
chrX-109335292-INS-210
chrX-109346126-INS-72
chrX-109375147-INS-11819
chrX-109585722-INS-56
chrX-109613267-INS-55
chrX-109706510-INS-52
chrX-109732007-INS-50
chrX-109808842-INS-6934
chrX-109976246-INS-75
chrX-110087281-INS-20499
chrX-110114255-INS-6333
chrX-110136365-INS-437
chrX-110142520-INS-208
chrX-110180726-INS-6692
chrX-110200809-INS-4791
chrX-110208090-INS-2984
chrX-110220996-INS-9692
chrX-110221009-INS-1428
chrX-110228044-INS-3958
chrX-110286038-INS-694
chrX-110416140-INS-294
chrX-110848416-INS-484
chrX-111044028-INS-315
chrX-111044706-INS-363

chrX-111060873-INS-408
chrX-111202669-INS-1839
chrX-111205521-INS-6191
chrX-111372289-INS-3214
chrX-111449221-INS-538
chrX-111458860-INS-52
chrX-111485172-INS-84
chrX-111539028-INS-556
chrX-111594912-INS-150
chrX-111594991-INS-89
chrX-111623488-INS-52
chrX-111868392-INS-636
chrX-111908448-INS-50
chrX-111977859-INS-57
chrX-112054687-INS-1596
chrX-112068477-INS-81
chrX-112073306-INS-60
chrX-112128294-INS-56
chrX-112203100-INS-52
chrX-112207142-INS-50
chrX-112269071-INS-120
chrX-112275311-INS-164
chrX-112342949-INS-876
chrX-112372358-INS-425
chrX-112473307-INS-209
chrX-112488498-INS-54
chrX-112606903-INS-252
chrX-112639946-INS-5470
chrX-112763227-INS-8623
chrX-112802115-INS-131
chrX-112802406-INS-121
chrX-112900894-INS-4419
chrX-112920895-INS-7477
chrX-112944524-INS-134
chrX-113378445-INS-1889
chrX-113406419-INS-101
chrX-113412031-INS-1465
chrX-113452661-INS-52
chrX-113674487-INS-214
chrX-113710553-INS-2000
chrX-113751855-INS-14006
chrX-113751855-INS-7197
chrX-113814908-INS-476
chrX-113854831-INS-342
chrX-113879362-INS-52
chrX-113888854-INS-68
chrX-113953940-INS-1489
chrX-113953952-INS-4951
chrX-113981441-INS-702
chrX-114013016-INS-61
chrX-114018642-INS-2748
chrX-114031293-INS-54
chrX-114091460-INS-50
chrX-114143821-INS-2839
chrX-114242714-INS-221
chrX-114367045-INS-3099
chrX-114370676-INS-52
chrX-114413572-INS-9340
chrX-114424758-INS-1496
chrX-114467381-INS-446
chrX-114473680-INS-1131
chrX-114568304-INS-1033
chrX-114660757-INS-3842
chrX-114661379-INS-7691
chrX-114699216-INS-1183
chrX-114740588-INS-85
chrX-114749437-INS-3132
chrX-114805980-INS-70
chrX-114829298-INS-116
chrX-114847228-INS-50
chrX-114906996-INS-54
chrX-114920154-INS-54
chrX-114940188-INS-252
chrX-115006468-INS-114
chrX-115031528-INS-826
chrX-115032819-INS-56
chrX-115047255-INS-4998

chrX-115075416-INS-56
chrX-115099362-INS-4500
chrX-115245183-INS-166
chrX-115281748-INS-171
chrX-115314118-INS-69
chrX-115468425-INS-1531
chrX-115536149-INS-248
chrX-115546994-INS-3305
chrX-115609607-INS-52
chrX-115798618-INS-104
chrX-115798618-INS-50
chrX-115803246-INS-68
chrX-115932204-INS-216
chrX-115989547-INS-70
chrX-116028500-INS-6516
chrX-116094713-INS-2748
chrX-116098626-INS-194
chrX-116166318-INS-98
chrX-116215637-INS-327
chrX-116215693-INS-106
chrX-116232198-INS-557
chrX-116287510-INS-6801
chrX-116411785-INS-6199
chrX-116560921-INS-123
chrX-116598854-INS-5325
chrX-116607055-INS-2146
chrX-116794598-INS-76
chrX-116846968-INS-2948
chrX-116901233-INS-70
chrX-116959408-INS-2490
chrX-117026314-INS-7086
chrX-117076891-INS-6376
chrX-117086463-INS-209
chrX-117091184-INS-315
chrX-117248940-INS-52
chrX-117515442-INS-52
chrX-117650697-INS-56
chrX-117742856-INS-205
chrX-117758637-INS-1181
chrX-117774367-INS-109
chrX-117866544-INS-110
chrX-117903084-INS-90
chrX-118008783-INS-62
chrX-118052144-INS-3395
chrX-118095354-INS-5452
chrX-118098601-INS-50
chrX-118100754-INS-7437
chrX-118180620-INS-52
chrX-118226024-INS-67
chrX-118238291-INS-376
chrX-118238291-INS-54
chrX-118316261-INS-58
chrX-118392990-INS-50
chrX-118742749-INS-50
chrX-118744220-INS-6386
chrX-118777049-INS-205
chrX-118930347-INS-2846
chrX-118941818-INS-1252
chrX-119018835-INS-3204
chrX-119038610-INS-75
chrX-119064879-INS-4049
chrX-119147603-INS-477
chrX-119173160-INS-6831
chrX-119278991-INS-306
chrX-119433403-INS-6392
chrX-119434204-INS-204
chrX-119629182-INS-7086
chrX-119680274-INS-535
chrX-119756728-INS-2823
chrX-119978617-INS-3016
chrX-119986267-INS-2989
chrX-120063675-INS-570
chrX-120070464-INS-1130
chrX-120174362-INS-73
chrX-120212013-INS-915
chrX-120416395-INS-101
chrX-120707829-INS-66

chrX-120746765-INS-1548
chrX-120809089-INS-51
chrX-120810862-INS-7099
chrX-120925825-INS-135
chrX-120925825-INS-295
chrX-120992200-INS-50
chrX-121028590-INS-50
chrX-121061471-INS-3108
chrX-121078504-INS-1092
chrX-121078759-INS-458
chrX-121187794-INS-252
chrX-121282642-INS-56
chrX-121283267-INS-115
chrX-121288261-INS-1439
chrX-121302172-INS-264
chrX-121380141-INS-1465
chrX-121387302-INS-50
chrX-121713619-INS-54
chrX-121859557-INS-50
chrX-121875412-INS-71
chrX-121907161-INS-40319
chrX-121950543-INS-59
chrX-121961614-INS-76
chrX-121963969-INS-652
chrX-121966375-INS-56
chrX-121971195-INS-84
chrX-121994540-INS-1237
chrX-121998404-INS-6702
chrX-122284534-INS-208
chrX-122398743-INS-92
chrX-122565420-INS-459
chrX-122577645-INS-1167
chrX-123093301-INS-4652
chrX-123096707-INS-52
chrX-123096815-INS-126
chrX-123111489-INS-1147
chrX-123122594-INS-56
chrX-123123819-INS-52
chrX-123263817-INS-6744
chrX-124989962-INS-259
chrX-125026737-INS-7079
chrX-125066710-INS-2881
chrX-125089238-INS-2449
chrX-125201081-INS-6647
chrX-125309090-INS-855
chrX-125381972-INS-500
chrX-125400094-INS-416
chrX-125557823-INS-7225
chrX-125582844-INS-52
chrX-125679836-INS-327
chrX-125729098-INS-416
chrX-125775634-INS-1397
chrX-125904587-INS-50
chrX-125972288-INS-1841
chrX-125987459-INS-58
chrX-126019188-INS-501
chrX-126130649-INS-60
chrX-126143586-INS-6385
chrX-126157701-INS-1300
chrX-126191353-INS-391
chrX-126350787-INS-195
chrX-126395009-INS-67
chrX-126409605-INS-141
chrX-126535208-INS-6478
chrX-126662541-INS-853
chrX-126667474-INS-1078
chrX-126682572-INS-52
chrX-126731179-INS-51
chrX-126765685-INS-471
chrX-126799193-INS-50
chrX-126909328-INS-74
chrX-126923800-INS-55
chrX-126936103-INS-81
chrX-126951149-INS-52
chrX-127018341-INS-76
chrX-127088845-INS-122
chrX-127090446-INS-216

chrX-127097160-INS-227
chrX-127278933-INS-7082
chrX-127303996-INS-414
chrX-127330368-INS-53
chrX-127359525-INS-50
chrX-127417527-INS-52
chrX-127433631-INS-980
chrX-127436722-INS-2960
chrX-127659218-INS-1181
chrX-127829567-INS-125
chrX-127829712-INS-75
chrX-127829766-INS-711
chrX-127871928-INS-2909
chrX-127994473-INS-467
chrX-128053907-INS-5428
chrX-128057644-INS-55
chrX-128069687-INS-226
chrX-128099666-INS-719
chrX-128117712-INS-6887
chrX-128147469-INS-205
chrX-128179382-INS-115
chrX-128199112-INS-50
chrX-128221232-INS-500
chrX-128233237-INS-54
chrX-128374488-INS-507
chrX-128401363-INS-327
chrX-128440408-INS-8665
chrX-128452102-INS-68
chrX-128465026-INS-898
chrX-128470231-INS-125
chrX-128490224-INS-7259
chrX-128493640-INS-274
chrX-128493850-INS-260
chrX-128496087-INS-1207
chrX-128547101-INS-8670
chrX-128584298-INS-3186
chrX-128586380-INS-660
chrX-128691679-INS-7013
chrX-128734273-INS-76
chrX-128735604-INS-107
chrX-128954590-INS-6379
chrX-129041486-INS-244
chrX-129101214-INS-508
chrX-129184279-INS-66
chrX-129342111-INS-348
chrX-129388793-INS-1489
chrX-129412083-INS-6763
chrX-129432854-INS-6538
chrX-129470159-INS-4172
chrX-129592768-INS-61
chrX-129624293-INS-178
chrX-129627553-INS-339
chrX-129693802-INS-66
chrX-129754051-INS-7250
chrX-129782163-INS-157
chrX-129944123-INS-50
chrX-129984393-INS-213
chrX-130051295-INS-2344
chrX-130163523-INS-120
chrX-130201614-INS-7416
chrX-130296386-INS-450
chrX-130391018-INS-52
chrX-130409247-INS-6388
chrX-130441972-INS-61
chrX-130457683-INS-203
chrX-130474452-INS-52
chrX-130490519-INS-54
chrX-130491180-INS-78
chrX-130493378-INS-8690
chrX-130550613-INS-2131
chrX-130558214-INS-6543
chrX-130583828-INS-203
chrX-130606423-INS-82
chrX-130638026-INS-62
chrX-130689719-INS-85
chrX-130691910-INS-55
chrX-130707269-INS-51

chrX-130846016-INS-6804
chrX-130862953-INS-204
chrX-130883649-INS-11435
chrX-130890870-INS-3019
chrX-130901596-INS-66
chrX-130908642-INS-67
chrX-130950738-INS-78
chrX-131032858-INS-9395
chrX-131052272-INS-6484
chrX-131075058-INS-550
chrX-131130498-INS-55
chrX-131385840-INS-6811
chrX-131430392-INS-1622
chrX-131637093-INS-513
chrX-131646315-INS-198
chrX-131670610-INS-4597
chrX-131712827-INS-743
chrX-131715385-INS-52
chrX-131725036-INS-614
chrX-131732087-INS-72
chrX-131860670-INS-72
chrX-131866799-INS-2860
chrX-131956810-INS-3025
chrX-131961342-INS-87
chrX-132110645-INS-9949
chrX-132123494-INS-256
chrX-132125703-INS-339
chrX-132171542-INS-2217
chrX-132178480-INS-190
chrX-132198117-INS-88
chrX-132209794-INS-1128
chrX-132285222-INS-6565
chrX-132411762-INS-1056
chrX-132415852-INS-6544
chrX-132432998-INS-6553
chrX-132470637-INS-6711
chrX-132475599-INS-70
chrX-132543898-INS-62
chrX-132557654-INS-50
chrX-132590461-INS-67
chrX-132641834-INS-60
chrX-132745319-INS-1324
chrX-132750184-INS-58
chrX-132840115-INS-54
chrX-132883421-INS-133
chrX-132934506-INS-52
chrX-132935246-INS-7143
chrX-132950733-INS-17034
chrX-133005827-INS-59
chrX-133069682-INS-203
chrX-133114477-INS-252
chrX-133175842-INS-108
chrX-133176296-INS-60
chrX-133184318-INS-112
chrX-133230443-INS-63
chrX-133230755-INS-196
chrX-133237706-INS-166
chrX-133242252-INS-242
chrX-133272342-INS-120
chrX-133272702-INS-200
chrX-133277849-INS-211
chrX-133281851-INS-3538
chrX-133312460-INS-7148
chrX-133385063-INS-721
chrX-133415571-INS-1055
chrX-133417608-INS-56
chrX-133420590-INS-405
chrX-133477949-INS-70
chrX-133482919-INS-375
chrX-133525587-INS-56
chrX-133560574-INS-60
chrX-133721454-INS-86
chrX-133750428-INS-7096
chrX-133790787-INS-50
chrX-133864114-INS-55
chrX-133864251-INS-82
chrX-133885416-INS-76

chrX-133909454-INS-5031
chrX-133949350-INS-18817
chrX-133968166-INS-592
chrX-133976251-INS-62
chrX-134000943-INS-117
chrX-134014354-INS-129
chrX-134092409-INS-86
chrX-134103497-INS-6588
chrX-134221361-INS-95
chrX-134268938-INS-3182
chrX-134277781-INS-57
chrX-134328257-INS-52
chrX-134467497-INS-92
chrX-134467797-INS-68
chrX-134535470-INS-1999
chrX-134572478-INS-138
chrX-134574936-INS-84
chrX-134576523-INS-79
chrX-134586297-INS-55
chrX-134586637-INS-156
chrX-134599903-INS-6154
chrX-134623542-INS-56
chrX-134635396-INS-6295
chrX-134698289-INS-226
chrX-134741500-INS-700
chrX-134775378-INS-2011
chrX-134777180-INS-6540
chrX-134807421-INS-78
chrX-134879752-INS-300
chrX-134892331-INS-328
chrX-134912133-INS-2002
chrX-134922475-INS-400
chrX-134930298-INS-50
chrX-135064119-INS-369
chrX-135156776-INS-55
chrX-135370925-INS-3713
chrX-135383769-INS-1187
chrX-135432350-INS-1588
chrX-135467535-INS-491
chrX-135505640-INS-357
chrX-135506105-INS-241
chrX-135506408-INS-50
chrX-135509519-INS-50
chrX-135518896-INS-52
chrX-135622125-INS-136
chrX-135706250-INS-66
chrX-135735451-INS-57
chrX-135840336-INS-102
chrX-135846249-INS-4238
chrX-135847823-INS-542
chrX-135851122-INS-329
chrX-135856463-INS-78
chrX-135858256-INS-19656
chrX-135872834-INS-1393
chrX-135940726-INS-7110
chrX-135988104-INS-79
chrX-136010292-INS-4126
chrX-136054606-INS-55
chrX-136057447-INS-70
chrX-136071833-INS-515
chrX-136127398-INS-82
chrX-136145896-INS-6212
chrX-136154573-INS-145
chrX-136184173-INS-695
chrX-136211842-INS-1500
chrX-136232183-INS-55
chrX-136243547-INS-170
chrX-136314662-INS-680
chrX-136322096-INS-54
chrX-136378907-INS-67
chrX-136427096-INS-103
chrX-136505015-INS-1253
chrX-136513269-INS-144
chrX-136560255-INS-56
chrX-136565512-INS-937
chrX-136576865-INS-491
chrX-136577290-INS-2227

chrX-136589378-INS-60
chrX-136589628-INS-120
chrX-136623559-INS-155
chrX-136625054-INS-1886
chrX-136644670-INS-5496
chrX-136662575-INS-50
chrX-136767451-INS-933
chrX-136767658-INS-217
chrX-136796144-INS-59
chrX-136814760-INS-56
chrX-136833387-INS-6985
chrX-136930238-INS-2498
chrX-136976968-INS-180
chrX-137012189-INS-52
chrX-137014394-INS-50
chrX-137023676-INS-342
chrX-137032547-INS-201
chrX-137045587-INS-7982
chrX-137051264-INS-720
chrX-137065516-INS-51
chrX-137096663-INS-997
chrX-137099696-INS-6874
chrX-137140508-INS-52
chrX-137215327-INS-3793
chrX-137269760-INS-130
chrX-137270084-INS-6732
chrX-137358130-INS-60
chrX-137449657-INS-1413
chrX-137465406-INS-4841
chrX-137500676-INS-205
chrX-137595275-INS-4636
chrX-137636985-INS-90
chrX-137666861-INS-180
chrX-137684758-INS-50
chrX-137747815-INS-107
chrX-137852516-INS-74
chrX-137863622-INS-1058
chrX-137919564-INS-342
chrX-137978924-INS-70
chrX-137981518-INS-347
chrX-138001688-INS-404
chrX-138016579-INS-6935
chrX-138060996-INS-4034
chrX-138061024-INS-1468
chrX-138099060-INS-116
chrX-138099142-INS-314
chrX-138151487-INS-56
chrX-138157914-INS-148
chrX-138291135-INS-50
chrX-138328569-INS-67
chrX-138349365-INS-50
chrX-138366487-INS-132
chrX-138397257-INS-68
chrX-138432962-INS-6594
chrX-138449288-INS-54
chrX-138485401-INS-50
chrX-138533223-INS-57
chrX-138576401-INS-87
chrX-138590013-INS-60
chrX-138618927-INS-526
chrX-138665602-INS-90
chrX-138716686-INS-56
chrX-138788620-INS-120
chrX-138788620-INS-52
chrX-138788927-INS-195
chrX-138866440-INS-157
chrX-138951180-INS-394
chrX-139007236-INS-3345
chrX-139032345-INS-158
chrX-139082495-INS-198
chrX-139129533-INS-204
chrX-139170293-INS-72
chrX-139312756-INS-129
chrX-139342619-INS-4746
chrX-139397763-INS-56
chrX-139447602-INS-85
chrX-139510249-INS-215

chrX-139541548-INS-1467
chrX-139551235-INS-3554
chrX-139633264-INS-7844
chrX-139640218-INS-81
chrX-139655383-INS-51
chrX-139655838-INS-69
chrX-139716172-INS-2359
chrX-139872795-INS-2925
chrX-139898097-INS-75
chrX-139915366-INS-328
chrX-139918281-INS-80
chrX-139923015-INS-1312
chrX-139980864-INS-81
chrX-140036253-INS-363
chrX-140073235-INS-66
chrX-140073428-INS-174
chrX-140098662-INS-52
chrX-140122250-INS-977
chrX-140147577-INS-85
chrX-140202030-INS-1389
chrX-140295424-INS-52
chrX-140318421-INS-721
chrX-140320905-INS-201
chrX-140336301-INS-90
chrX-140336369-INS-80
chrX-140401711-INS-1501
chrX-140452472-INS-55
chrX-140459474-INS-172
chrX-140459925-INS-681
chrX-140459938-INS-134
chrX-140481840-INS-110
chrX-140485607-INS-50
chrX-140535861-INS-50
chrX-140537573-INS-62
chrX-140558148-INS-161
chrX-140574444-INS-68
chrX-140633581-INS-54
chrX-140660773-INS-6599
chrX-140666231-INS-451
chrX-140704000-INS-68
chrX-140718698-INS-158
chrX-140767339-INS-6905
chrX-140855014-INS-252
chrX-140882520-INS-151
chrX-140931585-INS-2220
chrX-140984384-INS-3358
chrX-141110475-INS-2038
chrX-141124798-INS-55
chrX-141148071-INS-1364
chrX-141170543-INS-198
chrX-141272572-INS-204
chrX-141301721-INS-144
chrX-141309918-INS-72
chrX-141326038-INS-128
chrX-141351837-INS-190
chrX-141374399-INS-120
chrX-141383904-INS-1071
chrX-141406356-INS-82
chrX-141409926-INS-67
chrX-141410688-INS-167
chrX-141447377-INS-2806
chrX-141495655-INS-1017
chrX-141547078-INS-53
chrX-141557210-INS-195
chrX-141564563-INS-195
chrX-141565284-INS-430
chrX-141594433-INS-284
chrX-141603758-INS-61
chrX-141684356-INS-52
chrX-141723389-INS-56
chrX-141790094-INS-56
chrX-141798853-INS-71
chrX-141856078-INS-362
chrX-141921539-INS-10499
chrX-141921618-INS-2767
chrX-141937354-INS-53
chrX-141970782-INS-77

chrX-142034080-INS-60
chrX-142152530-INS-51
chrX-142155527-INS-135
chrX-142203953-INS-502
chrX-142227329-INS-210
chrX-142228148-INS-384
chrX-142332703-INS-1253
chrX-142389720-INS-6550
chrX-142475465-INS-405
chrX-142492420-INS-66
chrX-142544552-INS-98
chrX-142659717-INS-7227
chrX-142683971-INS-459
chrX-142721028-INS-6166
chrX-142846754-INS-2451
chrX-142849206-INS-2447
chrX-142886429-INS-63
chrX-142899865-INS-208
chrX-142933972-INS-77
chrX-142961498-INS-110
chrX-142967460-INS-50
chrX-142975172-INS-7145
chrX-142978816-INS-1245
chrX-142980244-INS-6174
chrX-143028477-INS-127
chrX-143039221-INS-58
chrX-143047960-INS-52
chrX-143082986-INS-52
chrX-143090259-INS-115
chrX-143092662-INS-341
chrX-143092869-INS-407
chrX-143166200-INS-52
chrX-143175327-INS-77
chrX-143193625-INS-153
chrX-143213886-INS-1813
chrX-143215254-INS-75
chrX-143233398-INS-120
chrX-143233398-INS-348
chrX-143365596-INS-57
chrX-143400441-INS-4175
chrX-143506808-INS-432
chrX-143555283-INS-465
chrX-143565648-INS-148
chrX-143583486-INS-2715
chrX-143638193-INS-50
chrX-143647355-INS-6407
chrX-143657294-INS-1824
chrX-143686575-INS-159
chrX-143811802-INS-2772
chrX-143856627-INS-3225
chrX-143862881-INS-6439
chrX-143884375-INS-110
chrX-143895724-INS-2742
chrX-143926902-INS-487
chrX-144000445-INS-5505
chrX-144050044-INS-8699
chrX-144132146-INS-175
chrX-144177467-INS-2882
chrX-144243321-INS-60
chrX-144301143-INS-7098
chrX-144316231-INS-88
chrX-144321708-INS-6439
chrX-144379989-INS-215
chrX-144379989-INS-83
chrX-144387844-INS-6892
chrX-144397204-INS-6169
chrX-144407511-INS-135
chrX-144410669-INS-1622
chrX-144415218-INS-1193
chrX-144417080-INS-114
chrX-144596518-INS-70
chrX-144622338-INS-485
chrX-144629297-INS-3444
chrX-144639141-INS-70
chrX-144653130-INS-1329
chrX-144653154-INS-6212
chrX-144658700-INS-232

chrX-144677957-INS-199
chrX-144713285-INS-1273
chrX-144713509-INS-60
chrX-144720281-INS-60
chrX-144750023-INS-304
chrX-144798765-INS-52
chrX-144842557-INS-54
chrX-144878148-INS-58
chrX-144969463-INS-849
chrX-145017034-INS-6481
chrX-145041705-INS-56
chrX-145041914-INS-127
chrX-145063705-INS-61
chrX-145084166-INS-59
chrX-145173760-INS-52
chrX-145182402-INS-58
chrX-145182482-INS-273
chrX-145185748-INS-64
chrX-145220586-INS-3100
chrX-145237458-INS-60
chrX-145265747-INS-50
chrX-145306163-INS-159
chrX-145414042-INS-50
chrX-145458198-INS-2398
chrX-145465371-INS-7017
chrX-145567868-INS-6984
chrX-145583782-INS-258
chrX-145630532-INS-6423
chrX-145666570-INS-1563
chrX-145689957-INS-199
chrX-145721775-INS-1788
chrX-145738275-INS-2226
chrX-145790030-INS-602
chrX-145811746-INS-51
chrX-145811775-INS-139
chrX-145852288-INS-6606
chrX-145938559-INS-9154
chrX-145986339-INS-52
chrX-146006117-INS-509
chrX-146011978-INS-4447
chrX-146018295-INS-3122
chrX-146067620-INS-119
chrX-146130301-INS-3305
chrX-146155588-INS-50
chrX-146155622-INS-172
chrX-146237181-INS-6417
chrX-146262499-INS-62
chrX-146314527-INS-7177
chrX-146316510-INS-426
chrX-146321439-INS-90
chrX-146323008-INS-397
chrX-146487214-INS-69
chrX-146595338-INS-1982
chrX-146617877-INS-115
chrX-146742461-INS-50
chrX-148172909-INS-106
chrX-148421423-INS-57
chrX-149007976-INS-6676
chrX-149144740-INS-478
chrX-149162656-INS-214
chrX-149175872-INS-64
chrX-149215449-INS-98
chrX-149262570-INS-428
chrX-149377815-INS-1012
chrX-149460943-INS-50
chrX-149503691-INS-7001
chrX-149517690-INS-948
chrX-149557832-INS-6358
chrX-149600611-INS-68
chrX-149606485-INS-755
chrX-149773927-INS-50
chrX-149791823-INS-134
chrX-149802654-INS-196
chrX-149823757-INS-210
chrX-149826756-INS-51
chrX-149844039-INS-457
chrX-149915017-INS-262

chrX-149947203-INS-51
chrX-150001892-INS-186
chrX-150010137-INS-500
chrX-150080767-INS-56
chrX-150271145-INS-204
chrX-150276350-INS-61
chrX-150435130-INS-3630
chrX-150543614-INS-566
chrX-150655686-INS-203
chrX-150775885-INS-55
chrX-150815024-INS-52
chrX-150853941-INS-181
chrX-150856055-INS-239
chrX-150891690-INS-445
chrX-150901607-INS-516
chrX-150976179-INS-59
chrX-151040482-INS-66
chrX-151067183-INS-52
chrX-151067740-INS-89
chrX-151131192-INS-55
chrX-151141965-INS-80
chrX-151215035-INS-51
chrX-151253026-INS-289
chrX-151253095-INS-52
chrX-151370036-INS-352
chrX-151373726-INS-561
chrX-151401045-INS-857
chrX-151483101-INS-202
chrX-151511169-INS-729
chrX-151516116-INS-60
chrX-151529878-INS-195
chrX-151572334-INS-255
chrX-151588425-INS-137
chrX-151588425-INS-54
chrX-151630567-INS-225
chrX-151631907-INS-204
chrX-151666578-INS-557
chrX-151672543-INS-419
chrX-151777072-INS-50
chrX-151881522-INS-135
chrX-151884525-INS-50
chrX-151896637-INS-7286
chrX-151918759-INS-68
chrX-151921948-INS-3137
chrX-151958706-INS-101
chrX-151959666-INS-203
chrX-151982079-INS-229
chrX-151989941-INS-183
chrX-151999493-INS-202
chrX-152082785-INS-216
chrX-152094348-INS-2935
chrX-152139141-INS-70
chrX-152139231-INS-335
chrX-152262409-INS-449
chrX-152331464-INS-50
chrX-152356288-INS-6599
chrX-152362425-INS-70
chrX-152401649-INS-205
chrX-152441387-INS-52
chrX-152464983-INS-4973
chrX-152479364-INS-105
chrX-152544682-INS-54
chrX-152549466-INS-7216
chrX-152626316-INS-6343
chrX-152675437-INS-184
chrX-152711059-INS-3123
chrX-152867990-INS-52
chrX-152869950-INS-5018
chrX-152929071-INS-6476
chrX-152987159-INS-3275
chrX-153051750-INS-161
chrX-153377257-INS-68
chrX-153404850-INS-5055
chrX-153417333-INS-54
chrX-153427732-INS-144
chrX-153461273-INS-123
chrX-153721438-INS-20595

chrX-153722639-INS-111
chrX-153724418-INS-1511
chrX-153735491-INS-2572
chrX-153800216-INS-114
chrX-153884046-INS-51
chrX-153886436-INS-3506
chrX-153891672-INS-62
chrX-153899106-INS-1883
chrX-153900301-INS-6220
chrX-153953925-INS-52
chrX-153963036-INS-120
chrX-153971935-INS-1853
chrX-154054697-INS-225
chrX-154088808-INS-145
chrX-154088970-INS-336
chrX-154126890-INS-72
chrX-154129878-INS-498
chrX-154160432-INS-8438
chrX-154179482-INS-6632
chrX-154208033-INS-3185
chrX-154226416-INS-51
chrX-154233938-INS-1348
chrX-154338400-INS-51
chrX-154347269-INS-6915
chrX-154436966-INS-51
chrX-154446231-INS-2577
chrX-154464630-INS-1021
chrX-154472017-INS-10413
chrX-154529613-INS-213
chrX-154547988-INS-6174
chrX-154592017-INS-193
chrX-154593860-INS-7585
chrX-154609205-INS-154
chrX-154646356-INS-66
chrX-154718144-INS-193
chrX-154727764-INS-1115
chrX-154773569-INS-141
chrX-154879020-INS-822
chrX-154907095-INS-62
chrX-154921267-INS-445
chrX-154960079-INS-266
chrX-154975936-INS-758
chrX-154976476-INS-321
chrX-155003707-INS-6620
chrX-155016273-INS-6539
chrX-155072331-INS-621
chrX-155112853-INS-6717
chrX-155170683-INS-238
chrX-155177557-INS-50
chrX-155210974-INS-56
chrX-155222858-INS-4495
chrX-155284392-INS-84
chrX-155372624-INS-69
chrX-155505563-INS-62
chrX-155536671-INS-54
chrX-155670653-INS-56
chrX-155709104-INS-231
chrX-155760585-INS-224
chrX-155818934-INS-50
chrX-155827983-INS-818
chrX-155864211-INS-7205
chrX-155864336-INS-1977
chrX-155893724-INS-6401
chrX-155902754-INS-3496
chrX-155911459-INS-5313
chrX-155948289-INS-95
chrX-155981498-INS-154
chrX-155990498-INS-920
chrX-155990630-INS-64
chrX-156018258-INS-7849
chrX-156040960-INS-199
chrX-156104797-INS-1552
chrX-156156597-INS-472
chrX-156177829-INS-671
chrX-156209539-INS-2823
chrX-156387700-INS-343
chrX-156396810-INS-299

chrX-156397433-INS-391
chrX-156427447-INS-50
chrX-156430325-INS-1108
chrX-156430538-INS-1912
chrX-156433117-INS-152
chrX-156433233-INS-6152
chrX-156433556-INS-60
chrX-156436676-INS-6301
chrX-156436911-INS-138
chrX-156436911-INS-68
chrX-156576828-INS-116
chrX-156664671-INS-6250
chrX-156680867-INS-6493
chrX-156746744-INS-1882
chrX-156843724-INS-404
chrX-156875795-INS-2706
chrX-156907623-INS-52
chrX-156920455-INS-70
chrX-157044210-INS-1140
chrX-157044273-INS-161
chrX-157066680-INS-566
chrX-157077730-INS-2624
chrX-157089560-INS-50
chrX-157229541-INS-204
chrX-157290665-INS-6584
chrX-157308858-INS-8506
chrX-157327596-INS-56
chrX-157328191-INS-75
chrX-157373506-INS-53
chrX-157382506-INS-1401
chrX-157473244-INS-1221
chrX-157510021-INS-65
chrX-157536541-INS-9986
chrX-157538834-INS-61
chrX-157557046-INS-100
chrX-157563088-INS-108
chrX-157690241-INS-579
chrX-157902971-INS-2150
chrX-158019718-INS-2872
chrX-158022917-INS-79
chrX-158053216-INS-168
chrX-158173534-INS-488
chrX-158174323-INS-546
chrX-158179388-INS-53
chrX-158213293-INS-1688
chrX-158222196-INS-4725
chrX-158251752-INS-198
chrX-158300031-INS-94
chrX-158359868-INS-202
chrX-158445130-INS-211
chrX-158453217-INS-1223
chrX-158455203-INS-63
chrX-158507782-INS-78
chrX-158508147-INS-136
chrX-158537458-INS-84
chrX-158618634-INS-52
chrX-158646546-INS-162
chrX-158668038-INS-110
chrX-158675256-INS-2809
chrX-158834992-INS-222
chrX-158835545-INS-81
chrX-158840955-INS-117
chrX-158841382-INS-77
chrX-158962140-INS-177
chrX-158965484-INS-52
chrX-159022129-INS-412
chrX-159022889-INS-5832
chrX-159102919-INS-50
chrX-159108059-INS-53
chrX-159160404-INS-116
chrX-159215033-INS-7187
chrX-159219899-INS-60
chrX-159251112-INS-58
chrX-159251115-INS-438
chrX-159251500-INS-78
chrX-159277668-INS-228
chrX-159284065-INS-7033

chrX-159378024-INS-471
chrX-159439792-INS-2810
chrX-159463200-INS-204
chrX-159692070-INS-537
chrX-159718172-INS-264
chrX-159791288-INS-54
chrX-159824339-INS-72
chrX-159828030-INS-71
chrX-159893931-INS-221
chrX-159905865-INS-106
chrX-159908480-INS-149
chrX-159934121-INS-50
chrX-159995996-INS-6036
chrX-160011626-INS-570
chrX-160045420-INS-7731
chrX-160069858-INS-665
chrX-160069980-INS-70
chrX-160158069-INS-371
chrX-160180015-INS-54
chrX-160181567-INS-1141
chrX-160191856-INS-75
chrX-160370810-INS-131
chrX-160373072-INS-126
chrX-160418716-INS-52
chrX-160495428-INS-208
chrX-160632050-INS-51
chrX-160826395-INS-98
chrX-160858778-INS-366
chrX-160858778-INS-78
chrX-160867842-INS-4063
chrX-161080746-INS-59
chrX-161110305-INS-6834
chrX-161165053-INS-809
chrX-161320001-INS-108
chrX-161342951-INS-244
chrX-161380706-INS-110
chrX-161412899-INS-8546
chrX-161486518-INS-50
chrX-161503080-INS-158
chrX-161537100-INS-273
chrX-161580405-INS-93
chrX-161580993-INS-131
chrX-161614738-INS-810
chrX-161624017-INS-97
chrX-161646801-INS-59
chrX-161691131-INS-173
chrX-161694559-INS-50
chrX-161700337-INS-194
chrX-161720525-INS-172
chrX-161720536-INS-467
chrX-161754167-INS-66
chrX-161776802-INS-50
chrX-161779233-INS-218
chrX-161787213-INS-185
chrX-161841247-INS-594
chrX-161872703-INS-6247
chrX-161876688-INS-52
chrX-161885866-INS-6819
chrX-161902789-INS-201
chrX-161913774-INS-167
chrX-161937239-INS-107
chrX-161998076-INS-52
chrX-162021138-INS-192
chrX-162059429-INS-150
chrX-162119644-INS-78
chrX-162152096-INS-2782
chrX-162159270-INS-492
chrX-162195754-INS-7056
chrX-162213976-INS-68
chrX-162216809-INS-50
chrX-162321975-INS-89
chrX-162326540-INS-6261
chrX-162361358-INS-76
chrX-162421135-INS-140
chrX-162422059-INS-192
chrX-162494162-INS-7229
chrX-162519810-INS-164

chrX-162610904-INS-207
chrX-162737810-INS-50
chrX-162748926-INS-999
chrX-162749179-INS-60
chrX-162751322-INS-454
chrX-162774085-INS-677
chrX-162774705-INS-500
chrX-162786798-INS-51
chrX-162787634-INS-70
chrX-162871156-INS-170
chrX-162944967-INS-52
chrX-162962715-INS-948
chrX-162989241-INS-205
chrX-163016856-INS-72
chrX-163056018-INS-69
chrX-163147894-INS-1500
chrX-163172742-INS-59
chrX-163245492-INS-1993
chrX-163307686-INS-186
chrX-163337662-INS-3436
chrX-163360512-INS-1328
chrX-163453661-INS-198
chrX-163489610-INS-2040
chrX-163573456-INS-6867
chrX-163692415-INS-60
chrX-163695737-INS-1343
chrX-163696808-INS-4832
chrX-163708545-INS-66
chrX-163797902-INS-5143
chrX-163842395-INS-2758
chrX-163867979-INS-150
chrX-163919809-INS-6336
chrX-163932493-INS-7188
chrX-163969081-INS-67
chrX-164052815-INS-314
chrX-164098294-INS-6675
chrX-164126207-INS-50
chrX-164135022-INS-50
chrX-164220755-INS-126
chrX-164387031-INS-74
chrX-164764495-INS-76
chrX-164898993-INS-10491
chrX-164926044-INS-52
chrX-164934458-INS-133
chrX-165100146-INS-58
chrX-165269890-INS-179
chrX-165277994-INS-3190
chrX-165282104-INS-54
chrX-165339537-INS-1476
chrX-165340936-INS-53
chrX-165443014-INS-151
chrX-165494125-INS-7071
chrX-165498765-INS-3430
chrX-165516506-INS-198
chrX-165516976-INS-50
chrX-165525383-INS-291
chrX-165539154-INS-50
chrX-165618150-INS-464
chrX-165624592-INS-2847
chrX-165677264-INS-50
chrX-165691676-INS-452
chrX-165730968-INS-1764
chrX-165790134-INS-78
chrX-165802383-INS-1317
chrX-165807431-INS-50
chrX-165819604-INS-108
chrX-165819978-INS-52
chrX-165842833-INS-53
chrX-165855213-INS-1633
chrX-165864284-INS-573
chrX-165865644-INS-1750
chrX-165966068-INS-2065
chrX-166029209-INS-214
chrX-166067721-INS-87
chrX-166107943-INS-59
chrX-166179060-INS-76
chrX-166189903-INS-75

chrX-166190020-INS-133
chrX-166236094-INS-4107
chrX-166244375-INS-129
chrX-166380910-INS-100
chrX-166382130-INS-1190
chrX-166481528-INS-56
chrX-166581202-INS-6993
chrX-166598298-INS-1338
chrX-166615502-INS-56
chrX-166622243-INS-75
chrX-166624674-INS-914
chrX-166641378-INS-52
chrX-166644326-INS-7228
chrX-166658511-INS-85
chrX-166672331-INS-56
chrX-166709615-INS-672
chrX-166734740-INS-6143
chrX-166756480-INS-124
chrX-166756530-INS-59
chrX-166783619-INS-55
chrX-166784134-INS-1757
chrX-166792526-INS-66
chrX-166857986-INS-51
chrX-166891187-INS-75
chrX-166891226-INS-154
chrX-166980312-INS-52
chrX-167072491-INS-70
chrX-167112700-INS-251
chrX-167194689-INS-6207
chrX-167237335-INS-6804
chrX-167391497-INS-7106
chrX-167543322-INS-2752
chrX-167571043-INS-2848
chrX-167584425-INS-55
chrX-167600312-INS-5050
chrX-167613247-INS-2759
chrX-167652705-INS-107
chrX-167664118-INS-1449
chrX-167862304-INS-525
chrX-167933713-INS-66
chrX-168191433-INS-2015
chrX-168206973-INS-460
chrX-168366061-INS-2844
chrX-168372641-INS-51
chrX-168600581-INS-52
chrX-168669110-INS-1326
chrX-168670751-INS-723
chrX-168750263-INS-204
chrX-168756122-INS-247
chrX-168771206-INS-113
chrX-168772511-INS-63
chrX-168772772-INS-247
chrX-168773893-INS-405
chrX-168773893-INS-932
chrX-168777324-INS-62
chrX-168779902-INS-306
chrX-168781409-INS-198
chrX-168791880-INS-996
chrX-168793019-INS-243
chrX-169072094-INS-73
chrX-169077103-INS-158
chrX-169077289-INS-636
chrX-169077412-INS-164
chrX-169077901-INS-93
chrX-169078180-INS-89
chrX-169078243-INS-186
chrX-169079722-INS-54
chrX-169080422-INS-177
chrX-169082420-INS-341
chrX-169083534-INS-279
chrX-169086549-INS-63
chrX-169086962-INS-62
chrX-169093444-INS-52
chrX-169095008-INS-62
chrX-169096148-INS-19649
chrX-169107279-INS-744
chrX-169107592-INS-214

chrX-169108790-INS-63
chrX-169110908-INS-95
chrX-169111356-INS-79
chrX-169116835-INS-219
chrX-169295496-INS-578
chrX-169307217-INS-93

| Identified_By_Us | Identified_By_JAX |
|---|---|
| chr1-55525702-INV-320 | chr1-55525701-INV-328 |
| chr1-73270460-INV-954 | chr1-73270462-INV-977 |
| chr1-110137904-INV-346 | chr1-110137885-INV-363 |
| chr1-129554314-INV-303 | chr1-129554291-INV-346 |
| chr1-167394713-INV-1078 | chr1-167394724-INV-1088 |
| chr1-175205685-INV-629 | chr1-175205643-INV-660 |
| chr1-182330754-INV-2128 | chr1-182330756-INV-2127 |
| chr10-42755478-INV-4679 | chr10-42755495-INV-4659 |
| chr11-8257235-INV-509 | chr11-8257238-INV-513 |
| chr11-18130963-INV-514 | chr11-18130961-INV-515 |
| chr11-65546990-INV-2679 | chr11-65546988-INV-2686 |
| chr12-60910841-INV-1069 | chr12-60910838-INV-1074 |
| chr12-73702591-INV-925 | chr12-73702582-INV-922 |
| chr12-76109065-INV-1664 | chr12-76109061-INV-1673 |
| chr12-81865341-INV-810 | chr12-81865323-INV-828 |
| chr13-23187383-INV-1701 | chr13-23187368-INV-1705 |
| chr13-53593987-INV-2238 | chr13-53593993-INV-2232 |
| chr13-54468753-INV-698 | chr13-54468755-INV-697 |
| chr13-108324811-INV-4841 | chr13-108324805-INV-4849 |
| chr13-112999776-INV-2187 | chr13-112999761-INV-2189 |
| chr14-6099128-INV-747 | chr14-6099130-INV-751 |
| chr14-21317153-INV-925 | chr14-21317150-INV-927 |
| chr15-91931041-INV-1973 | chr15-91931055-INV-1949 |
| chr15-94705924-INV-337 | chr15-94705920-INV-334 |
| chr16-40570447-INV-1231 | chr16-40570452-INV-1270 |
| chr16-47357811-INV-567 | chr16-47357790-INV-605 |
| chr16-52468246-INV-567 | chr16-52468230-INV-597 |
| chr17-69068525-INV-930 | chr17-69068536-INV-921 |
| chr17-82956817-INV-2269 | chr17-82956857-INV-2232 |
| chr17-85856064-INV-422 | chr17-85856061-INV-461 |
| chr18-7052921-INV-538 | chr18-7052922-INV-568 |
| chr18-13789443-INV-359 | chr18-13789456-INV-342 |
| chr18-55470533-INV-398 | chr18-55470524-INV-400 |
| chr19-52932484-INV-1069 | chr19-52932482-INV-1065 |
| chr2-75548031-INV-1594 | chr2-75548028-INV-1612 |
| chr2-77251976-INV-388 | chr2-77251983-INV-399 |
| chr2-92057174-INV-386 | chr2-92057153-INV-416 |
| chr2-154451791-INV-2243 | chr2-154451793-INV-2279 |
| chr3-27527909-INV-2460 | chr3-27527924-INV-2442 |
| chr3-96145778-INV-8904 | chr3-96145803-INV-8871 |
| chr3-117227878-INV-450 | chr3-117227877-INV-445 |
| chr3-126580362-INV-1361 | chr3-126580358-INV-1407 |
| chr3-135283838-INV-2805 | chr3-135283850-INV-2798 |
| chr4-100586751-INV-839 | chr4-100586749-INV-846 |
| chr4-108635034-INV-320 | chr4-108635043-INV-301 |
| chr4-129557150-INV-1257 | chr4-129557151-INV-1264 |

| | |
|---|---|
| chr5-37561810-INV-921 | chr5-37561813-INV-918 |
| chr5-72391577-INV-1499 | chr5-72391556-INV-1517 |
| chr7-85453808-INV-1643 | chr7-85453811-INV-1658 |
| chr7-105108322-INV-1175 | chr7-105108320-INV-1188 |
| chr7-109368734-INV-306 | chr7-109368731-INV-306 |
| chr7-127827009-INV-1306 | chr7-127827010-INV-1304 |
| chr8-70305999-INV-1278 | chr8-70305996-INV-1327 |
| chr9-29477078-INV-2936 | chr9-29477074-INV-2941 |
| chrX-69357636-INV-433 | chrX-69357636-INV-431 |
| chrX-102260883-INV-326 | chrX-102260906-INV-347 |

**Identified_By_Us**
chr1-3192284-INV-1569
chr1-4498582-INV-3382
chr1-7910818-INV-1290
chr1-8577511-INV-359
chr1-9101845-INV-286
chr1-9558025-INV-8583
chr1-9716534-INV-518
chr1-9946199-INV-253
chr1-11384470-INV-1499
chr1-14410529-INV-1469
chr1-14411390-INV-1350
chr1-14546450-INV-6631
chr1-16388574-INV-14041
chr1-16389346-INV-13268
chr1-16394170-INV-9316
chr1-17358782-INV-3533
chr1-19418764-INV-555
chr1-22301046-INV-6209
chr1-24468568-INV-3349
chr1-26034302-INV-2141
chr1-27353869-INV-4577
chr1-27794175-INV-6231
chr1-27795574-INV-4864
chr1-29179906-INV-3015
chr1-30134538-INV-1111
chr1-30135020-INV-632
chr1-31512524-INV-2945
chr1-31514711-INV-1378
chr1-31611064-INV-1228
chr1-31755028-INV-802
chr1-33093914-INV-1850
chr1-33338680-INV-6877
chr1-34941022-INV-774
chr1-35624414-INV-2008
chr1-35865216-INV-1513
chr1-35865616-INV-2540
chr1-38027683-INV-627
chr1-40188818-INV-939
chr1-40189016-INV-2015
chr1-40606335-INV-685
chr1-41833374-INV-4104
chr1-41833399-INV-2485
chr1-43679870-INV-1975
chr1-44311450-INV-1058
chr1-44311670-INV-2006
chr1-44439467-INV-1807
chr1-44440400-INV-1246
chr1-45308398-INV-1948
chr1-45437972-INV-410
chr1-47013665-INV-341
chr1-47161561-INV-4006
chr1-47291023-INV-443
chr1-47579784-INV-1207
chr1-47626851-INV-1529
chr1-47866555-INV-1132
chr1-48357392-INV-2016
chr1-48811258-INV-529
chr1-49822900-INV-1001
chr1-50717802-INV-226
chr1-50720379-INV-1084
chr1-50894398-INV-303
chr1-53683073-INV-229
chr1-54698971-INV-447
chr1-57495558-INV-2611
chr1-58073081-INV-2142
chr1-59332577-INV-2457
chr1-62267242-INV-1446
chr1-62441195-INV-727
chr1-63260644-INV-1440
chr1-63261313-INV-771
chr1-63878921-INV-1787
chr1-64510712-INV-2715
chr1-65475776-INV-870
chr1-66054893-INV-1876
chr1-66831537-INV-248
chr1-66831537-INV-4231

chr1-69284325-INV-2409
chr1-69286163-INV-584
chr1-74155678-INV-2408
chr1-74360600-INV-2991
chr1-75945121-INV-2283
chr1-79010149-INV-1911
chr1-79049228-INV-2093
chr1-79225828-INV-9427
chr1-79227348-INV-5545
chr1-79227348-INV-6386
chr1-79227349-INV-7897
chr1-79229464-INV-4270
chr1-80102274-INV-3200
chr1-83117877-INV-5030
chr1-83208566-INV-4150
chr1-88158026-INV-1084
chr1-90327974-INV-271
chr1-91663953-INV-3273
chr1-94043886-INV-586
chr1-94952669-INV-2386
chr1-95777026-INV-1248
chr1-96423208-INV-1217
chr1-98923482-INV-1723
chr1-99583884-INV-1985
chr1-99584332-INV-4348
chr1-99628600-INV-1424
chr1-99837338-INV-714
chr1-99837346-INV-1994
chr1-101287217-INV-4100
chr1-101287301-INV-6189
chr1-102148878-INV-639
chr1-102441331-INV-2643
chr1-102446190-INV-10713
chr1-103182078-INV-4342
chr1-103819873-INV-1421
chr1-106119816-INV-628
chr1-110290644-INV-3876
chr1-111068753-INV-4620
chr1-111576416-INV-2651
chr1-111643496-INV-382
chr1-113652278-INV-4582
chr1-113822909-INV-2868
chr1-114692240-INV-6863
chr1-114730090-INV-7355
chr1-119623617-INV-3227
chr1-122065818-INV-1128
chr1-122283100-INV-253
chr1-122851020-INV-2208
chr1-123516894-INV-5135
chr1-123518424-INV-4508
chr1-123518458-INV-5026
chr1-124449685-INV-524
chr1-125533371-INV-1032
chr1-129294033-INV-1069
chr1-129294240-INV-862
chr1-130518233-INV-278
chr1-132031302-INV-6632
chr1-133078132-INV-587
chr1-133395416-INV-590
chr1-133461806-INV-946
chr1-134557055-INV-348
chr1-136827825-INV-552
chr1-139273370-INV-556
chr1-139727825-INV-694
chr1-139752079-INV-1057
chr1-139762728-INV-5200
chr1-139765230-INV-7491
chr1-141183299-INV-838
chr1-143014748-INV-968
chr1-143016519-INV-271
chr1-147687495-INV-2197
chr1-147708987-INV-3223
chr1-149359784-INV-8476
chr1-150008728-INV-1108
chr1-150074280-INV-955
chr1-150173354-INV-350
chr1-150615700-INV-1735

chr1-150615702-INV-827
chr1-152952979-INV-6928
chr1-156757035-INV-1389
chr1-161229553-INV-1091
chr1-162091608-INV-7012
chr1-162647273-INV-6598
chr1-162647781-INV-6090
chr1-162648831-INV-5073
chr1-164971369-INV-322
chr1-168386809-INV-1305
chr1-170442569-INV-799
chr1-171089657-INV-450
chr1-174694278-INV-845
chr1-175107988-INV-5476
chr1-177534659-INV-279
chr1-177534675-INV-2043
chr1-181056751-INV-1172
chr1-181364448-INV-1092
chr1-181365228-INV-360
chr1-181587270-INV-706
chr1-183802323-INV-2338
chr1-186587760-INV-385
chr1-187805483-INV-2399
chr1-187806776-INV-2591
chr1-188671832-INV-747
chr1-188672351-INV-231
chr1-190244156-INV-2971
chr1-191319489-INV-327
chr1-191382994-INV-3287
chr1-191383081-INV-4807
chr1-193946703-INV-1175
chr1-194871917-INV-545
chr10-3436046-INV-757
chr10-6066074-INV-951
chr10-6223173-INV-15797
chr10-6224968-INV-4895
chr10-6227799-INV-2172
chr10-7439694-INV-593
chr10-14392626-INV-3900
chr10-14947910-INV-1212
chr10-15002521-INV-443
chr10-16647127-INV-8361
chr10-17904207-INV-2413
chr10-18098448-INV-282
chr10-23464703-INV-1733
chr10-23721483-INV-464
chr10-24823314-INV-2904
chr10-26484875-INV-2060
chr10-26524922-INV-3527
chr10-27209713-INV-790
chr10-27209718-INV-1327
chr10-27229937-INV-5323
chr10-27229937-INV-6668
chr10-28102399-INV-3251
chr10-28104912-INV-737
chr10-28292106-INV-4248
chr10-28899715-INV-634
chr10-29069753-INV-739
chr10-29732943-INV-1195
chr10-29732943-INV-735
chr10-30118674-INV-650
chr10-30325801-INV-1631
chr10-30925793-INV-691
chr10-31069992-INV-357
chr10-32333134-INV-739
chr10-35312293-INV-437
chr10-36480006-INV-1593
chr10-36481035-INV-746
chr10-36807624-INV-356
chr10-37931265-INV-781
chr10-43643646-INV-219
chr10-45851423-INV-1748
chr10-46182329-INV-4914
chr10-46186057-INV-1318
chr10-46471994-INV-551
chr10-47270793-INV-412
chr10-48647679-INV-335

chr10-49464020-INV-378
chr10-51512731-INV-2827
chr10-51515201-INV-447
chr10-52761951-INV-615
chr10-54831974-INV-6087
chr10-59311531-INV-531
chr10-59790660-INV-1324
chr10-64741379-INV-1699
chr10-64998139-INV-1466
chr10-65219129-INV-7762
chr10-65451892-INV-427
chr10-70138398-INV-533
chr10-72661504-INV-1057
chr10-74932002-INV-1276
chr10-77435955-INV-3427
chr10-77450145-INV-672
chr10-77450147-INV-1392
chr10-79034286-INV-4651
chr10-79035252-INV-3686
chr10-82106419-INV-1574
chr10-82188009-INV-1919
chr10-83623157-INV-5123
chr10-83637861-INV-5974
chr10-83638155-INV-4794
chr10-84455748-INV-1150
chr10-86483827-INV-2795
chr10-87470264-INV-1385
chr10-88660071-INV-381
chr10-93389643-INV-635
chr10-95001302-INV-674
chr10-95297336-INV-621
chr10-95297338-INV-1077
chr10-100074100-INV-1418
chr10-100074939-INV-918
chr10-100114815-INV-304
chr10-100666965-INV-2169
chr10-100666965-INV-244
chr10-101813294-INV-392
chr10-102159443-INV-6429
chr10-102788171-INV-491
chr10-103176433-INV-1704
chr10-103780096-INV-3149
chr10-104626693-INV-1868
chr10-104626702-INV-797
chr10-105868876-INV-1886
chr10-105870133-INV-727
chr10-106496096-INV-3663
chr10-106497875-INV-460
chr10-110112347-INV-472
chr10-110949978-INV-364
chr10-111881224-INV-3671
chr10-112061798-INV-293
chr10-112448580-INV-4308
chr10-112959251-INV-1116
chr10-112959419-INV-946
chr10-115637350-INV-781
chr10-115637558-INV-3286
chr10-117161539-INV-690
chr10-117161545-INV-1131
chr10-120643604-INV-247
chr10-122467476-INV-6597
chr10-122557213-INV-1622
chr10-124065390-INV-5536
chr10-124069702-INV-1213
chr10-125880282-INV-613
chr10-126340151-INV-600
chr10-129423174-INV-7092
chr11-4419801-INV-990
chr11-5515369-INV-1763
chr11-7349554-INV-1761
chr11-8951691-INV-751
chr11-9885191-INV-605
chr11-11321836-INV-394
chr11-11420893-INV-530
chr11-13201662-INV-1493
chr11-13500625-INV-3116
chr11-14381766-INV-529

chr11-15753256-INV-2091
chr11-16698481-INV-370
chr11-17346227-INV-895
chr11-30703150-INV-1724
chr11-30983233-INV-961
chr11-31997466-INV-897
chr11-33128799-INV-806
chr11-35798170-INV-1815
chr11-37643088-INV-7268
chr11-39093459-INV-858
chr11-43828351-INV-861
chr11-46489321-INV-6573
chr11-46848246-INV-573
chr11-49089014-INV-895
chr11-49250825-INV-1144
chr11-52296688-INV-267
chr11-56609432-INV-288
chr11-58158302-INV-2620
chr11-59443188-INV-381
chr11-59844941-INV-723
chr11-61222339-INV-1499
chr11-61311196-INV-7210
chr11-63726137-INV-2210
chr11-65438162-INV-321
chr11-68577950-INV-341
chr11-72988202-INV-785
chr11-72988226-INV-1007
chr11-74656199-INV-308
chr11-74666164-INV-696
chr11-76049730-INV-1712
chr11-79375229-INV-3675
chr11-85959835-INV-564
chr11-92017998-INV-254
chr11-92343272-INV-472
chr11-92382582-INV-2791
chr11-92471956-INV-3420
chr11-92808159-INV-2565
chr11-95799329-INV-401
chr11-96277388-INV-1076
chr11-97786811-INV-1266
chr11-99638798-INV-4762
chr11-99638870-INV-733
chr11-101533239-INV-2851
chr11-103805032-INV-266
chr11-106757732-INV-488
chr11-108411681-INV-1751
chr11-108804510-INV-1433
chr11-115632447-INV-851
chr12-3554091-INV-1113
chr12-3755463-INV-909
chr12-4822860-INV-357
chr12-5671474-INV-5497
chr12-6327015-INV-1592
chr12-6692416-INV-4761
chr12-6693787-INV-2805
chr12-7691528-INV-1049
chr12-7692313-INV-796
chr12-10657122-INV-1299
chr12-10897961-INV-2456
chr12-12115437-INV-3605
chr12-13540513-INV-2775
chr12-13542254-INV-1107
chr12-14894381-INV-6112
chr12-14894382-INV-7277
chr12-14898409-INV-4641
chr12-15241483-INV-7187
chr12-17034833-INV-1234
chr12-24314131-INV-2359
chr12-24314895-INV-2685
chr12-24915882-INV-1226
chr12-25652919-INV-360
chr12-26938081-INV-1660
chr12-31449901-INV-512
chr12-35412504-INV-358
chr12-36300480-INV-1574
chr12-39703063-INV-916
chr12-39835337-INV-1266

chr12-40929655-INV-2931
chr12-42270436-INV-3522
chr12-42866445-INV-3066
chr12-43262569-INV-1705
chr12-43264028-INV-762
chr12-43390098-INV-7450
chr12-43746510-INV-589
chr12-43949158-INV-605
chr12-44995009-INV-242
chr12-45343015-INV-6885
chr12-45384479-INV-1123
chr12-45808090-INV-675
chr12-45836212-INV-214
chr12-47058966-INV-588
chr12-47849592-INV-2257
chr12-48361848-INV-1998
chr12-48361851-INV-3119
chr12-48889824-INV-6877
chr12-49394675-INV-2288
chr12-49394676-INV-3632
chr12-49395461-INV-1502
chr12-51349353-INV-7155
chr12-51575931-INV-3603
chr12-52080810-INV-872
chr12-53520850-INV-261
chr12-55447707-INV-5121
chr12-55447717-INV-1090
chr12-55922817-INV-3294
chr12-56396522-INV-485
chr12-57984794-INV-4738
chr12-58212613-INV-2564
chr12-59041213-INV-1188
chr12-59422935-INV-1691
chr12-59999686-INV-5107
chr12-60873574-INV-2163
chr12-60888626-INV-1461
chr12-61417904-INV-240
chr12-62494803-INV-309
chr12-62726260-INV-1367
chr12-63154815-INV-2563
chr12-63155837-INV-1558
chr12-64033675-INV-902
chr12-69988886-INV-1617
chr12-70112476-INV-3617
chr12-77363806-INV-1150
chr12-79062007-INV-339
chr12-82609624-INV-2561
chr12-86697875-INV-4968
chr12-91084153-INV-4783
chr12-92590991-INV-1383
chr12-92592046-INV-360
chr12-92691971-INV-759
chr12-92692014-INV-1843
chr12-93342922-INV-1793
chr12-93343893-INV-2734
chr12-95248728-INV-327
chr12-95293494-INV-1749
chr12-96388849-INV-601
chr12-97156975-INV-433
chr12-97901156-INV-389
chr12-98037001-INV-364
chr12-98495849-INV-788
chr12-100479785-INV-700
chr12-101432473-INV-1771
chr12-101458016-INV-2323
chr12-104385147-INV-218
chr12-104385149-INV-1748
chr12-107284140-INV-3350
chr12-107286406-INV-2487
chr12-109087134-INV-1835
chr12-110152713-INV-3059
chr12-111196316-INV-600
chr12-111220148-INV-423
chr12-111448964-INV-500
chr12-111786743-INV-1194
chr12-114561058-INV-4675
chr12-116025469-INV-648

chr13-3663633-INV-5276
chr13-6009238-INV-767
chr13-10351676-INV-6450
chr13-10860650-INV-11702
chr13-11261735-INV-1069
chr13-11261912-INV-885
chr13-11555482-INV-568
chr13-15605812-INV-7269
chr13-16559807-INV-753
chr13-17640543-INV-868
chr13-17964286-INV-446
chr13-18138971-INV-1024
chr13-18158296-INV-288
chr13-18642029-INV-492
chr13-18721547-INV-816
chr13-18721548-INV-1271
chr13-19865276-INV-3458
chr13-22251586-INV-15036
chr13-22253937-INV-10336
chr13-22792216-INV-709
chr13-33392550-INV-6204
chr13-34725691-INV-484
chr13-35273126-INV-366
chr13-36842421-INV-552
chr13-42022996-INV-687
chr13-42022997-INV-1025
chr13-45493929-INV-1486
chr13-53593730-INV-967
chr13-58062240-INV-630
chr13-58062382-INV-1019
chr13-60393869-INV-245
chr13-62298840-INV-2535
chr13-62477404-INV-10938
chr13-62833917-INV-1037
chr13-62839048-INV-5365
chr13-64872089-INV-2029
chr13-64873182-INV-938
chr13-77290511-INV-2455
chr13-78419449-INV-1631
chr13-78633859-INV-7942
chr13-79481834-INV-1431
chr13-84534090-INV-2346
chr13-85874778-INV-6311
chr13-86375129-INV-1958
chr13-86426954-INV-1935
chr13-86428557-INV-1519
chr13-86614870-INV-2604
chr13-88635716-INV-1819
chr13-89495911-INV-1414
chr13-89593365-INV-1083
chr13-89594142-INV-1920
chr13-90752181-INV-6723
chr13-92595229-INV-1201
chr13-97627458-INV-910
chr13-97627480-INV-5444
chr13-98478960-INV-665
chr13-99092092-INV-1208
chr13-100643186-INV-3767
chr13-100710260-INV-694
chr13-102375038-INV-557
chr13-103122040-INV-896
chr13-103804798-INV-1181
chr13-105591747-INV-623
chr13-105853367-INV-1358
chr13-106035168-INV-789
chr13-106263466-INV-3000
chr13-106263468-INV-1818
chr13-106448509-INV-728
chr13-106815968-INV-6622
chr13-106821684-INV-2153
chr13-110362227-INV-1229
chr13-110362227-INV-1974
chr13-111491923-INV-569
chr13-111657762-INV-494
chr13-111699760-INV-247
chr13-113480947-INV-543
chr13-116227229-INV-7016

chr13-116973729-INV-6212
chr13-117189657-INV-259
chr13-118396227-INV-3064
chr13-118799481-INV-1598
chr13-119093853-INV-357
chr14-9333848-INV-7307
chr14-11741807-INV-371
chr14-13969259-INV-947
chr14-17644594-INV-2196
chr14-18170523-INV-2086
chr14-21630753-INV-2639
chr14-23187885-INV-337
chr14-26054547-INV-2571
chr14-27286233-INV-751
chr14-27479951-INV-353
chr14-28403641-INV-2853
chr14-28405400-INV-2067
chr14-31844835-INV-1234
chr14-31845558-INV-525
chr14-35799619-INV-529
chr14-35811315-INV-7341
chr14-36216813-INV-295
chr14-36565708-INV-998
chr14-38032280-INV-847
chr14-38107794-INV-1990
chr14-40448555-INV-5053
chr14-40479404-INV-6444
chr14-40483243-INV-3563
chr14-42003455-INV-2390
chr14-45079707-INV-2250
chr14-49447816-INV-1343
chr14-50175914-INV-7352
chr14-51665970-INV-2643
chr14-51667721-INV-1470
chr14-52205724-INV-5207
chr14-55772228-INV-706
chr14-56432657-INV-205
chr14-58895915-INV-1975
chr14-58896977-INV-1132
chr14-62675471-INV-4853
chr14-62677509-INV-3589
chr14-62720587-INV-902
chr14-67173441-INV-3996
chr14-67176576-INV-890
chr14-68620912-INV-262
chr14-74501862-INV-1526
chr14-76207154-INV-515
chr14-82974971-INV-1290
chr14-83634414-INV-856
chr14-84337208-INV-5340
chr14-84453217-INV-7249
chr14-84454423-INV-4855
chr14-86521552-INV-557
chr14-86846008-INV-294
chr14-87820672-INV-543
chr14-89555111-INV-682
chr14-89555123-INV-5698
chr14-89990467-INV-876
chr14-90577037-INV-2011
chr14-90577058-INV-7556
chr14-94269136-INV-1742
chr14-95367152-INV-2731
chr14-96646843-INV-2226
chr14-97813395-INV-396
chr14-98679331-INV-1353
chr14-98679573-INV-2433
chr14-101639151-INV-1144
chr14-101639151-INV-697
chr14-102389939-INV-931
chr14-102423410-INV-1210
chr14-102658976-INV-3622
chr14-102659828-INV-3409
chr14-103106690-INV-3280
chr14-103108787-INV-1765
chr14-103676076-INV-7602
chr14-103682334-INV-1731
chr14-103682715-INV-967

chr14-104583957-INV-320
chr14-104744106-INV-524
chr14-105509650-INV-1012
chr14-105981480-INV-2591
chr14-106056533-INV-1648
chr14-107366800-INV-1000
chr14-108381876-INV-5059
chr14-110280758-INV-7107
chr14-110639907-INV-3599
chr14-110911564-INV-299
chr14-111848070-INV-2379
chr14-113228280-INV-1232
chr14-113823583-INV-1067
chr14-114657746-INV-4161
chr14-116440842-INV-560
chr14-117276158-INV-2175
chr14-117896199-INV-237
chr14-118313260-INV-3615
chr14-119329349-INV-1901
chr14-119864737-INV-932
chr14-120170590-INV-6977
chr14-120846339-INV-1913
chr14-122300638-INV-289
chr14-124004412-INV-293
chr14-124167458-INV-2934
chr15-6011820-INV-3144
chr15-6011848-INV-1876
chr15-10146040-INV-255
chr15-10853388-INV-4387
chr15-13787376-INV-291
chr15-13841338-INV-411
chr15-14804237-INV-275
chr15-16787611-INV-517
chr15-18028077-INV-957
chr15-18318454-INV-2157
chr15-19676713-INV-1088
chr15-20042932-INV-7978
chr15-20217449-INV-785
chr15-20867589-INV-1734
chr15-20868128-INV-2888
chr15-21067678-INV-1598
chr15-21982193-INV-1373
chr15-22930339-INV-1389
chr15-24481816-INV-949
chr15-25952428-INV-776
chr15-27384221-INV-6061
chr15-27834888-INV-1683
chr15-29340847-INV-811
chr15-29350540-INV-529
chr15-31124479-INV-660
chr15-32148049-INV-1311
chr15-32833543-INV-6314
chr15-32835802-INV-5218
chr15-33399073-INV-1290
chr15-34824153-INV-1254
chr15-36093590-INV-672
chr15-38808400-INV-440
chr15-42844205-INV-6787
chr15-42847421-INV-3065
chr15-42871261-INV-918
chr15-45866017-INV-12027
chr15-47403380-INV-556
chr15-47991941-INV-722
chr15-50004415-INV-1402
chr15-52516993-INV-2047
chr15-53608505-INV-366
chr15-57184319-INV-3426
chr15-57507225-INV-348
chr15-58104487-INV-317
chr15-59988065-INV-947
chr15-60907784-INV-1706
chr15-60909212-INV-512
chr15-65260754-INV-1905
chr15-67010100-INV-511
chr15-68416042-INV-1201
chr15-69083140-INV-306
chr15-71097098-INV-3839

chr15-74297001-INV-985
chr15-74839049-INV-2495
chr15-86603902-INV-1092
chr15-94303520-INV-475
chr15-95995388-INV-2355
chr15-98653193-INV-727
chr15-98653195-INV-2162
chr15-99795448-INV-2136
chr15-103598024-INV-1162
chr16-3331671-INV-327
chr16-7506356-INV-546
chr16-8789692-INV-299
chr16-10192645-INV-1196
chr16-10192645-INV-693
chr16-10926091-INV-420
chr16-13773476-INV-1201
chr16-19638342-INV-553
chr16-21673452-INV-319
chr16-23096050-INV-1344
chr16-24843978-INV-271
chr16-26120097-INV-3272
chr16-26121321-INV-2210
chr16-26627223-INV-7037
chr16-27951988-INV-1644
chr16-27953409-INV-245
chr16-30720799-INV-630
chr16-33127464-INV-729
chr16-38090836-INV-2130
chr16-42252370-INV-3281
chr16-42830689-INV-1629
chr16-47114985-INV-10388
chr16-47121493-INV-3835
chr16-48118428-INV-992
chr16-53029061-INV-317
chr16-53407810-INV-6011
chr16-53460065-INV-312
chr16-53469811-INV-1293
chr16-54484487-INV-4032
chr16-54484491-INV-1552
chr16-54584534-INV-1077
chr16-54870271-INV-651
chr16-55412097-INV-1362
chr16-56022950-INV-1288
chr16-57406443-INV-1907
chr16-57406473-INV-7209
chr16-58584286-INV-2073
chr16-59582043-INV-7594
chr16-60592823-INV-7167
chr16-61554281-INV-1748
chr16-61679299-INV-1415
chr16-61911949-INV-939
chr16-62152828-INV-924
chr16-63505768-INV-2319
chr16-63507468-INV-648
chr16-66314919-INV-693
chr16-66677672-INV-315
chr16-67730954-INV-6579
chr16-67735556-INV-2462
chr16-68621308-INV-471
chr16-70737736-INV-7898
chr16-71168699-INV-3493
chr16-73527086-INV-6217
chr16-73527117-INV-6860
chr16-73952188-INV-1014
chr16-74537603-INV-2752
chr16-74537829-INV-1465
chr16-76585789-INV-6145
chr16-76588073-INV-4173
chr16-78419888-INV-2164
chr16-82623561-INV-6315
chr16-82627105-INV-2775
chr16-82763754-INV-256
chr16-83785324-INV-843
chr16-84565406-INV-312
chr16-86127998-INV-6784
chr16-86528571-INV-3769
chr16-86598553-INV-2468

chr16-87340509-INV-1301
chr16-87341176-INV-3024
chr16-93453383-INV-1103
chr16-95546143-INV-1044
chr16-97147593-INV-6198
chr17-3486982-INV-238
chr17-4798113-INV-234
chr17-4798115-INV-1389
chr17-8824942-INV-332
chr17-10106363-INV-3634
chr17-10369561-INV-295
chr17-11998543-INV-1397
chr17-12589498-INV-748
chr17-15702459-INV-2333
chr17-15703826-INV-966
chr17-18598238-INV-2225
chr17-19000106-INV-817
chr17-19340899-INV-345
chr17-19818121-INV-1479
chr17-19845665-INV-1829
chr17-19999841-INV-2698
chr17-21294553-INV-1823
chr17-21295621-INV-4129
chr17-21826144-INV-1890
chr17-21826808-INV-4505
chr17-22373080-INV-938
chr17-23642650-INV-7428
chr17-23753073-INV-1169
chr17-24099297-INV-4877
chr17-24166988-INV-2758
chr17-27961024-INV-763
chr17-28087897-INV-2369
chr17-28089383-INV-876
chr17-29001138-INV-978
chr17-29323106-INV-777
chr17-29323126-INV-1120
chr17-29485202-INV-827
chr17-29529828-INV-1450
chr17-29664735-INV-1000
chr17-29664735-INV-1001
chr17-29799264-INV-884
chr17-31004249-INV-1440
chr17-31004250-INV-634
chr17-32317448-INV-420
chr17-32930561-INV-785
chr17-36792193-INV-4675
chr17-36876837-INV-6511
chr17-37018173-INV-6921
chr17-37093114-INV-3974
chr17-38804383-INV-1092
chr17-38805027-INV-1924
chr17-38898261-INV-3625
chr17-39225832-INV-1123
chr17-40197297-INV-1412
chr17-41836147-INV-748
chr17-41998169-INV-6204
chr17-42858507-INV-461
chr17-50958935-INV-447
chr17-51727670-INV-1560
chr17-51856632-INV-442
chr17-52534072-INV-368
chr17-52681840-INV-1542
chr17-53581050-INV-6527
chr17-54013822-INV-1600
chr17-54350920-INV-6912
chr17-54574468-INV-861
chr17-54676189-INV-785
chr17-55323217-INV-1001
chr17-55323221-INV-995
chr17-56226866-INV-312
chr17-56572767-INV-1372
chr17-59103433-INV-3269
chr17-60250033-INV-2661
chr17-60251771-INV-976
chr17-64062206-INV-739
chr17-66295362-INV-835
chr17-66295581-INV-1755

chr17-69005665-INV-2679
chr17-70009556-INV-1892
chr17-70009557-INV-6784
chr17-70527821-INV-1322
chr17-70568547-INV-728
chr17-76231229-INV-1035
chr17-76386364-INV-2306
chr17-76387971-INV-1024
chr17-78040288-INV-502
chr17-80763897-INV-598
chr17-82157691-INV-1366
chr17-83947816-INV-516
chr17-89302197-INV-4995
chr17-89744065-INV-1326
chr17-93437577-INV-2150
chr17-93700276-INV-1784
chr17-94155912-INV-2081
chr18-3972158-INV-1146
chr18-6695113-INV-1931
chr18-6716346-INV-682
chr18-8340170-INV-6457
chr18-8341245-INV-5391
chr18-8391036-INV-5360
chr18-9150192-INV-4779
chr18-9439761-INV-943
chr18-10384409-INV-1725
chr18-10492134-INV-2228
chr18-10493475-INV-890
chr18-12496564-INV-893
chr18-13341437-INV-2416
chr18-13342076-INV-5224
chr18-14257141-INV-1364
chr18-14540792-INV-1131
chr18-16182537-INV-1205
chr18-16270502-INV-1055
chr18-16270613-INV-945
chr18-17526926-INV-251
chr18-23151259-INV-1132
chr18-23151280-INV-363
chr18-24598038-INV-888
chr18-25253870-INV-426
chr18-28067180-INV-769
chr18-28142650-INV-975
chr18-28797951-INV-402
chr18-29023301-INV-905
chr18-29064738-INV-2934
chr18-29201844-INV-1856
chr18-29845493-INV-6861
chr18-30821235-INV-2495
chr18-31054182-INV-3162
chr18-31175897-INV-375
chr18-31379302-INV-2274
chr18-31380342-INV-1352
chr18-31487792-INV-6875
chr18-31516392-INV-1629
chr18-31826533-INV-1704
chr18-32107341-INV-368
chr18-33219876-INV-836
chr18-35305029-INV-394
chr18-35899483-INV-2241
chr18-36232694-INV-239
chr18-36831907-INV-497
chr18-41716673-INV-925
chr18-42886825-INV-314
chr18-44920930-INV-1136
chr18-51941902-INV-6605
chr18-53356002-INV-1633
chr18-58515821-INV-2184
chr18-58998405-INV-847
chr18-60338394-INV-2744
chr18-64156869-INV-1721
chr18-64182620-INV-6746
chr18-64960074-INV-1252
chr18-68507487-INV-6834
chr18-69168042-INV-4561
chr18-70446828-INV-4237
chr18-73205984-INV-1572

chr18-73207142-INV-739
chr18-75115443-INV-848
chr18-80885752-INV-238
chr18-81820166-INV-306
chr18-84204694-INV-1081
chr18-85156849-INV-2814
chr18-85279972-INV-1820
chr18-85811997-INV-1336
chr18-86925872-INV-286
chr18-87274519-INV-1187
chr18-88365904-INV-4666
chr18-88591605-INV-213
chr18-89363768-INV-736
chr18-90513761-INV-2047
chr19-6048217-INV-1788
chr19-9165708-INV-6477
chr19-9166232-INV-5953
chr19-9166951-INV-5397
chr19-11692787-INV-1353
chr19-13997596-INV-3122
chr19-14000115-INV-603
chr19-15994182-INV-3208
chr19-16351102-INV-1175
chr19-17280986-INV-6562
chr19-19269938-INV-4323
chr19-21168163-INV-609
chr19-22390640-INV-435
chr19-25049891-INV-1944
chr19-26948491-INV-846
chr19-28804903-INV-4224
chr19-28807938-INV-1217
chr19-30734697-INV-7615
chr19-33963779-INV-3772
chr19-34464820-INV-1053
chr19-36699478-INV-1503
chr19-38631392-INV-3543
chr19-38634292-INV-656
chr19-38962018-INV-990
chr19-41064356-INV-344
chr19-41207064-INV-2400
chr19-42737369-INV-266
chr19-42869844-INV-953
chr19-43275345-INV-1666
chr19-48019063-INV-350
chr19-52443372-INV-1019
chr19-52443372-INV-988
chr19-54224704-INV-436
chr19-55674914-INV-1744
chr19-55676111-INV-558
chr19-57007632-INV-596
chr19-59913368-INV-2411
chr19-59948040-INV-294
chr19-60659320-INV-5015
chr19-60660714-INV-3626
chr19-60894574-INV-893
chr19-60894962-INV-1950
chr2-4686601-INV-695
chr2-4686612-INV-2176
chr2-5269621-INV-1704
chr2-7123268-INV-670
chr2-8910385-INV-993
chr2-11950150-INV-3972
chr2-11990376-INV-7011
chr2-11991747-INV-6770
chr2-13919992-INV-4160
chr2-14203836-INV-1737
chr2-14495814-INV-1976
chr2-15697055-INV-1122
chr2-19967329-INV-1031
chr2-20878372-INV-2343
chr2-21720254-INV-3599
chr2-21723632-INV-222
chr2-22090297-INV-6687
chr2-35667696-INV-575
chr2-37020841-INV-1269
chr2-39708928-INV-291
chr2-39749349-INV-1083

chr2-39749440-INV-908
chr2-39791865-INV-368
chr2-40653375-INV-6595
chr2-41820484-INV-573
chr2-41934323-INV-1338
chr2-42030914-INV-515
chr2-42155884-INV-2792
chr2-42438092-INV-313
chr2-43298615-INV-6458
chr2-43503164-INV-2225
chr2-46603637-INV-2063
chr2-46716171-INV-226
chr2-47612504-INV-10626
chr2-49556524-INV-896
chr2-50654204-INV-6470
chr2-51514985-INV-1045
chr2-53488689-INV-1456
chr2-53489383-INV-1598
chr2-55087471-INV-16193
chr2-55088965-INV-14685
chr2-55093985-INV-12522
chr2-56051101-INV-566
chr2-56310941-INV-808
chr2-56505120-INV-3215
chr2-56505697-INV-2385
chr2-56923442-INV-4485
chr2-56924216-INV-3551
chr2-56925476-INV-2875
chr2-57462012-INV-2353
chr2-58115411-INV-280
chr2-63265291-INV-6296
chr2-63436805-INV-2935
chr2-63478323-INV-2020
chr2-63548530-INV-7653
chr2-64843534-INV-1421
chr2-64844309-INV-674
chr2-65233902-INV-1164
chr2-66703996-INV-8301
chr2-68016234-INV-1174
chr2-69547764-INV-259
chr2-71838941-INV-352
chr2-71838982-INV-944
chr2-72543355-INV-4303
chr2-76009387-INV-1846
chr2-78163651-INV-6204
chr2-79682207-INV-1204
chr2-80031178-INV-1549
chr2-80052591-INV-605
chr2-80052616-INV-1505
chr2-81919369-INV-738
chr2-82014520-INV-309
chr2-85396312-INV-1733
chr2-87845428-INV-3925
chr2-88420094-INV-664
chr2-94547966-INV-398
chr2-95599599-INV-505
chr2-96617030-INV-574
chr2-96698436-INV-2256
chr2-96698437-INV-644
chr2-97966251-INV-5415
chr2-98636694-INV-751
chr2-99356054-INV-3675
chr2-99359500-INV-1721
chr2-100030794-INV-1115
chr2-101155731-INV-4402
chr2-101606511-INV-804
chr2-103897178-INV-1214
chr2-106252287-INV-1709
chr2-106918052-INV-3629
chr2-107587312-INV-499
chr2-109113541-INV-950
chr2-109281729-INV-3116
chr2-109655355-INV-491
chr2-110675899-INV-862
chr2-111202660-INV-2893
chr2-111204253-INV-1299
chr2-111429858-INV-1547

chr2-111627536-INV-3552
chr2-111627560-INV-1193
chr2-113684273-INV-1030
chr2-117308436-INV-1633
chr2-117519284-INV-360
chr2-118394410-INV-1928
chr2-118394539-INV-582
chr2-119438960-INV-374
chr2-119664301-INV-846
chr2-121144878-INV-972
chr2-122038350-INV-3018
chr2-122038350-INV-552
chr2-123421760-INV-2201
chr2-125947759-INV-4448
chr2-128196848-INV-2220
chr2-131097998-INV-589
chr2-131203843-INV-945
chr2-131203849-INV-1339
chr2-132734540-INV-533
chr2-137266406-INV-1675
chr2-137267792-INV-289
chr2-139058641-INV-1390
chr2-141186911-INV-723
chr2-141186929-INV-1437
chr2-141431981-INV-655
chr2-141883974-INV-420
chr2-143064418-INV-1002
chr2-145051160-INV-1710
chr2-145052007-INV-872
chr2-146997876-INV-2747
chr2-147717729-INV-2866
chr2-149081802-INV-383
chr2-149316751-INV-1664
chr2-149568175-INV-599
chr2-150362640-INV-3951
chr2-153689126-INV-3355
chr2-153690014-INV-2962
chr2-155899201-INV-309
chr2-157543141-INV-1203
chr2-160765011-INV-880
chr2-163004043-INV-1231
chr2-163750977-INV-290
chr2-164043007-INV-2368
chr2-165745263-INV-482
chr2-165760523-INV-318
chr2-166908238-INV-313
chr2-171354331-INV-530
chr2-173335565-INV-2270
chr2-173983253-INV-1170
chr2-181342035-INV-506
chr3-4140392-INV-2373
chr3-4217153-INV-3227
chr3-6274194-INV-2513
chr3-6424084-INV-7450
chr3-9044377-INV-300
chr3-10654629-INV-1689
chr3-10854917-INV-6703
chr3-10988140-INV-3473
chr3-11505584-INV-1073
chr3-11505584-INV-845
chr3-12234164-INV-858
chr3-13013156-INV-768
chr3-15269098-INV-1193
chr3-15297826-INV-2191
chr3-15299245-INV-821
chr3-18402343-INV-2096
chr3-19020321-INV-817
chr3-20350851-INV-4190
chr3-22951995-INV-2668
chr3-23544498-INV-390
chr3-25016629-INV-7014
chr3-26468513-INV-1277
chr3-26658970-INV-12108
chr3-26667003-INV-6055
chr3-27262040-INV-2264
chr3-27525928-INV-3393
chr3-28805774-INV-549

chr3-29508123-INV-2795
chr3-29800550-INV-1593
chr3-33052867-INV-1007
chr3-33053080-INV-811
chr3-33653812-INV-3672
chr3-33750729-INV-6833
chr3-33778952-INV-1191
chr3-34646234-INV-566
chr3-35042058-INV-4127
chr3-35827056-INV-3780
chr3-35830572-INV-622
chr3-37703361-INV-428
chr3-39702554-INV-171
chr3-40467275-INV-8534
chr3-42040451-INV-1102
chr3-44757645-INV-4125
chr3-44978398-INV-497
chr3-45730310-INV-2137
chr3-45975560-INV-805
chr3-47530760-INV-1674
chr3-47577000-INV-4041
chr3-48654870-INV-2333
chr3-48655969-INV-1308
chr3-48837656-INV-300
chr3-48934230-INV-309
chr3-49069485-INV-252
chr3-49273803-INV-8798
chr3-49274489-INV-8118
chr3-49308972-INV-6055
chr3-54481975-INV-3297
chr3-54483215-INV-2226
chr3-57021414-INV-257
chr3-57107557-INV-3843
chr3-57107561-INV-1739
chr3-57257044-INV-1603
chr3-58688005-INV-171
chr3-59794822-INV-2561
chr3-60775631-INV-2985
chr3-62393877-INV-445
chr3-62780416-INV-6997
chr3-67442623-INV-877
chr3-67442649-INV-6515
chr3-67629647-INV-3160
chr3-69477523-INV-1692
chr3-69477523-INV-283
chr3-69821873-INV-5706
chr3-70527623-INV-987
chr3-71179551-INV-919
chr3-71179575-INV-2050
chr3-71362763-INV-4843
chr3-71932412-INV-5731
chr3-72225318-INV-1278
chr3-73877923-INV-1602
chr3-74467170-INV-2264
chr3-75677497-INV-554
chr3-75835005-INV-5142
chr3-76280921-INV-579
chr3-77182267-INV-1920
chr3-77573997-INV-2345
chr3-77574639-INV-7707
chr3-78400887-INV-2800
chr3-80197196-INV-6274
chr3-81827725-INV-357
chr3-83236047-INV-4239
chr3-87483699-INV-292
chr3-88298265-INV-383
chr3-90284759-INV-585
chr3-91725467-INV-2700
chr3-91909234-INV-412
chr3-92431401-INV-2553
chr3-92431402-INV-4201
chr3-93658104-INV-1416
chr3-96147881-INV-4689
chr3-96147882-INV-6802
chr3-96361845-INV-1573
chr3-96362964-INV-530
chr3-96394526-INV-1029

chr3-96807201-INV-575
chr3-99296164-INV-3356
chr3-100290630-INV-1648
chr3-104058766-INV-509
chr3-106704233-INV-2117
chr3-106705344-INV-5543
chr3-108851201-INV-2465
chr3-108851344-INV-3489
chr3-109832361-INV-9751
chr3-110742646-INV-1655
chr3-112188470-INV-6503
chr3-112497920-INV-1507
chr3-113142245-INV-1355
chr3-114147157-INV-1850
chr3-114605192-INV-2283
chr3-115859915-INV-237
chr3-118664387-INV-920
chr3-118664586-INV-1523
chr3-121396114-INV-399
chr3-121923426-INV-224
chr3-122584760-INV-1115
chr3-124917732-INV-1427
chr3-124917732-INV-228
chr3-129148426-INV-7941
chr3-130819897-INV-401
chr3-133917164-INV-771
chr3-133935516-INV-2707
chr3-133936163-INV-2107
chr3-134236670-INV-6771
chr3-134597004-INV-2436
chr3-134597015-INV-5993
chr3-135180404-INV-227
chr3-135661204-INV-3178
chr3-135811411-INV-1284
chr3-135811825-INV-2040
chr3-137234880-INV-4020
chr3-139092383-INV-1398
chr3-140021631-INV-1127
chr3-140390215-INV-2308
chr3-140396631-INV-395
chr3-144066482-INV-829
chr3-144651875-INV-2180
chr3-144743485-INV-7454
chr3-149685766-INV-850
chr3-150900607-INV-2356
chr3-151627377-INV-952
chr3-151663788-INV-1858
chr3-151717636-INV-6671
chr3-151723184-INV-1132
chr3-151745052-INV-214
chr3-151754327-INV-4539
chr3-154339887-INV-298
chr3-154961240-INV-618
chr3-157117706-INV-2640
chr4-4152381-INV-277
chr4-4310289-INV-9295
chr4-8121396-INV-1328
chr4-8479535-INV-6364
chr4-11738701-INV-453
chr4-12431211-INV-6764
chr4-15993614-INV-773
chr4-16798245-INV-2034
chr4-17383804-INV-581
chr4-20198839-INV-596
chr4-21452313-INV-2364
chr4-22601210-INV-425
chr4-22749880-INV-1583
chr4-24025452-INV-1025
chr4-25192402-INV-981
chr4-26522509-INV-288
chr4-27588386-INV-982
chr4-28408135-INV-3813
chr4-30252025-INV-1642
chr4-30370047-INV-300
chr4-30532738-INV-3685
chr4-32902341-INV-279
chr4-35616316-INV-3109

chr4-36583293-INV-624
chr4-36652396-INV-452
chr4-36756676-INV-591
chr4-37912484-INV-591
chr4-38582216-INV-1818
chr4-38795676-INV-559
chr4-50316351-INV-310
chr4-50684881-INV-351
chr4-52791831-INV-402
chr4-53608064-INV-6485
chr4-53767590-INV-243
chr4-54148030-INV-335
chr4-59139859-INV-6410
chr4-59142636-INV-3634
chr4-64530685-INV-2596
chr4-64532068-INV-1823
chr4-67029762-INV-7964
chr4-67052557-INV-1754
chr4-67053400-INV-1214
chr4-68052438-INV-1240
chr4-68444331-INV-3137
chr4-69716847-INV-2171
chr4-69993376-INV-5185
chr4-71833760-INV-4481
chr4-73161771-INV-2887
chr4-73972725-INV-5944
chr4-75310629-INV-576
chr4-77601804-INV-5111
chr4-77664169-INV-730
chr4-78894866-INV-1530
chr4-78998963-INV-844
chr4-79742400-INV-1724
chr4-81413530-INV-2446
chr4-81413546-INV-4690
chr4-81523402-INV-1596
chr4-81928730-INV-1127
chr4-81928733-INV-2684
chr4-81929192-INV-8384
chr4-82844417-INV-1831
chr4-83086287-INV-2478
chr4-85390046-INV-3589
chr4-86731785-INV-903
chr4-88818568-INV-877
chr4-90066706-INV-2890
chr4-91622375-INV-2181
chr4-91782677-INV-7326
chr4-93274454-INV-341
chr4-94224929-INV-3574
chr4-94647929-INV-3538
chr4-96190967-INV-6779
chr4-96197214-INV-993
chr4-97274443-INV-4548
chr4-97277171-INV-1817
chr4-98867052-INV-641
chr4-98867062-INV-1004
chr4-101587861-INV-436
chr4-105491777-INV-2502
chr4-107101938-INV-255
chr4-110984534-INV-1567
chr4-112460259-INV-3107
chr4-112547807-INV-582
chr4-112974468-INV-4385
chr4-112978183-INV-2492
chr4-115124837-INV-7013
chr4-116523053-INV-935
chr4-116526335-INV-807
chr4-117792527-INV-1283
chr4-117792530-INV-622
chr4-118064106-INV-342
chr4-118327768-INV-537
chr4-123788679-INV-2331
chr4-124281876-INV-3984
chr4-126829024-INV-491
chr4-127044638-INV-966
chr4-128963100-INV-238
chr4-129433012-INV-230
chr4-130132391-INV-616

chr4-135547033-INV-957
chr4-135787829-INV-4741
chr4-136377563-INV-994
chr4-137066185-INV-1246
chr4-137066717-INV-714
chr4-141730445-INV-222
chr4-144478474-INV-8527
chr4-144498569-INV-1074
chr4-152333926-INV-1061
chr4-152824280-INV-902
chr4-156296404-INV-981
chr4-156406558-INV-500
chr5-3282177-INV-444
chr5-3365861-INV-2910
chr5-5219334-INV-481
chr5-5867926-INV-2071
chr5-5869480-INV-1262
chr5-6262520-INV-3219
chr5-6760044-INV-909
chr5-7644912-INV-221
chr5-8013397-INV-225
chr5-8013404-INV-1957
chr5-8335754-INV-1311
chr5-9338913-INV-4608
chr5-10109143-INV-910
chr5-12864046-INV-3554
chr5-15014075-INV-6479
chr5-15663097-INV-1459
chr5-15726265-INV-1023
chr5-15829599-INV-2813
chr5-18249791-INV-1970
chr5-19029112-INV-6795
chr5-19185974-INV-2655
chr5-19314278-INV-2484
chr5-21913140-INV-1012
chr5-23223380-INV-1313
chr5-24155800-INV-1013
chr5-25673806-INV-363
chr5-28155131-INV-929
chr5-28251143-INV-692
chr5-28251160-INV-1562
chr5-30417348-INV-478
chr5-30698219-INV-331
chr5-31597150-INV-539
chr5-36585801-INV-1465
chr5-37602282-INV-384
chr5-38286734-INV-5458
chr5-38286735-INV-953
chr5-39502943-INV-783
chr5-42380239-INV-400
chr5-43049304-INV-620
chr5-43049306-INV-2200
chr5-43312890-INV-296
chr5-44037288-INV-456
chr5-44411353-INV-6379
chr5-47438629-INV-7889
chr5-49541896-INV-502
chr5-49902527-INV-3522
chr5-50050444-INV-596
chr5-51112162-INV-867
chr5-51112189-INV-3569
chr5-51270921-INV-3606
chr5-54489325-INV-1054
chr5-54489519-INV-864
chr5-55538755-INV-2613
chr5-56262629-INV-989
chr5-56439299-INV-6311
chr5-57030181-INV-1169
chr5-57907257-INV-1984
chr5-58557564-INV-6936
chr5-59123699-INV-6592
chr5-59727573-INV-872
chr5-60961199-INV-5185
chr5-61140576-INV-3629
chr5-61169684-INV-704
chr5-61257422-INV-1502
chr5-62482446-INV-1116

chr5-66186420-INV-497
chr5-66863982-INV-1073
chr5-66972360-INV-337
chr5-68487489-INV-669
chr5-69691173-INV-2019
chr5-71403344-INV-919
chr5-73094629-INV-708
chr5-74338690-INV-915
chr5-74470416-INV-305
chr5-74660593-INV-4106
chr5-74661433-INV-6568
chr5-77115494-INV-910
chr5-77379648-INV-336
chr5-77915142-INV-1275
chr5-79798289-INV-5505
chr5-79957976-INV-5085
chr5-80664778-INV-855
chr5-80876635-INV-2747
chr5-80876638-INV-1170
chr5-82704992-INV-1234
chr5-83309858-INV-1507
chr5-84889909-INV-2083
chr5-86846824-INV-2220
chr5-88145352-INV-631
chr5-88308994-INV-494
chr5-88513790-INV-819
chr5-91057574-INV-6451
chr5-98466498-INV-3722
chr5-99258201-INV-503
chr5-99465770-INV-703
chr5-100593790-INV-257
chr5-102497453-INV-1412
chr5-103385392-INV-720
chr5-103819475-INV-1853
chr5-104736052-INV-219956
chr5-104752780-INV-240797
chr5-104922219-INV-83226
chr5-105252845-INV-96991
chr5-107393133-INV-3464
chr5-107393835-INV-2798
chr5-110188303-INV-344
chr5-113568044-INV-361
chr5-116733726-INV-4422
chr5-118926434-INV-235
chr5-121241226-INV-621
chr5-121692042-INV-6483
chr5-121696946-INV-1578
chr5-124574781-INV-735
chr5-125443646-INV-788
chr5-130436391-INV-1772
chr5-135652140-INV-5698
chr5-137445799-INV-394
chr5-137741414-INV-960
chr5-138543038-INV-5198
chr5-142348379-INV-259
chr5-146510985-INV-606
chr5-148050415-INV-393
chr5-148050480-INV-975
chr5-148897740-INV-1970
chr5-149284241-INV-846
chr5-150577170-INV-283
chr6-3057218-INV-3298
chr6-6734331-INV-2281
chr6-6734498-INV-1021
chr6-7390548-INV-652
chr6-7390550-INV-1750
chr6-7498542-INV-6870
chr6-10047463-INV-307
chr6-11013784-INV-822
chr6-11962991-INV-6939
chr6-13566499-INV-499
chr6-14440710-INV-6248
chr6-15459619-INV-310
chr6-16657317-INV-926
chr6-19973397-INV-1415
chr6-21749708-INV-4079
chr6-23668693-INV-6410

chr6-24289418-INV-6778
chr6-26231787-INV-722
chr6-26612486-INV-2922
chr6-31584582-INV-430
chr6-33129575-INV-518
chr6-34959070-INV-372
chr6-40958638-INV-853
chr6-41202361-INV-351
chr6-41299874-INV-421
chr6-41474414-INV-1456
chr6-43673801-INV-360
chr6-44077392-INV-929
chr6-44077405-INV-1980
chr6-45510748-INV-2663
chr6-46840800-INV-1622
chr6-48017054-INV-583
chr6-48123499-INV-5976
chr6-48254274-INV-955
chr6-56394390-INV-1181
chr6-57008619-INV-1454
chr6-58130494-INV-795
chr6-58650586-INV-323
chr6-59244126-INV-1787
chr6-59244774-INV-1097
chr6-60411944-INV-4690
chr6-60412577-INV-3404
chr6-61265685-INV-1100
chr6-62135952-INV-368
chr6-62547014-INV-226
chr6-62843551-INV-1288
chr6-64507761-INV-644
chr6-65192192-INV-6999
chr6-65444756-INV-7093
chr6-65991376-INV-2394
chr6-66214446-INV-1598
chr6-66704751-INV-7803
chr6-66765067-INV-606
chr6-68911189-INV-1915
chr6-69272376-INV-740
chr6-69430420-INV-6680
chr6-69474845-INV-652
chr6-69531060-INV-3570
chr6-69533170-INV-6954
chr6-70879430-INV-440
chr6-73337697-INV-268
chr6-73983969-INV-3216
chr6-74215683-INV-1416
chr6-74313657-INV-5051
chr6-74316227-INV-2487
chr6-74353659-INV-2080
chr6-74586368-INV-455
chr6-74895470-INV-1177
chr6-76011325-INV-259
chr6-76222119-INV-5433
chr6-76775272-INV-2563
chr6-78494361-INV-1445
chr6-78825404-INV-636
chr6-79284022-INV-6249
chr6-79289360-INV-929
chr6-79738992-INV-390
chr6-80616300-INV-1433
chr6-86332761-INV-606
chr6-88009542-INV-450
chr6-88438872-INV-842
chr6-88438946-INV-1745
chr6-90953713-INV-1253
chr6-93246633-INV-2121
chr6-95388800-INV-13230
chr6-95389653-INV-12377
chr6-95389745-INV-9421
chr6-96738420-INV-6816
chr6-98497599-INV-256
chr6-98862268-INV-410
chr6-101567362-INV-9911
chr6-101569547-INV-5065
chr6-103454245-INV-4804
chr6-103466663-INV-2498

chr6-104047473-INV-1446
chr6-105158253-INV-441
chr6-105190962-INV-1915
chr6-105473249-INV-355
chr6-105921660-INV-1550
chr6-107116490-INV-6930
chr6-107149074-INV-1166
chr6-107249103-INV-810
chr6-108816190-INV-1044
chr6-109005495-INV-654
chr6-110386966-INV-13771
chr6-110667166-INV-1450
chr6-116008530-INV-3462
chr6-121563750-INV-7487
chr6-121674145-INV-6868
chr6-122520256-INV-1290
chr6-123788349-INV-2150
chr6-123948562-INV-259
chr6-123948582-INV-3362
chr6-124539172-INV-1748
chr6-126432422-INV-7288
chr6-128862405-INV-573
chr6-130096234-INV-2579
chr6-130353430-INV-634
chr6-130726785-INV-6616
chr6-131217016-INV-1194
chr6-132245958-INV-1847
chr6-132246016-INV-6869
chr6-132246036-INV-4254
chr6-133516636-INV-265
chr6-133857156-INV-8812
chr6-136395041-INV-920
chr6-139396096-INV-611
chr6-140516528-INV-927
chr6-143040099-INV-757
chr6-149273876-INV-734
chr6-149273884-INV-1493
chr7-3146706-INV-265
chr7-4042770-INV-1968
chr7-4053950-INV-975
chr7-4912221-INV-325
chr7-6033982-INV-1300
chr7-7219420-INV-6716
chr7-10125392-INV-283
chr7-10190603-INV-814
chr7-10439866-INV-1841
chr7-11422409-INV-1605
chr7-15010191-INV-6981
chr7-15557134-INV-262
chr7-25281833-INV-1985
chr7-28333588-INV-1057
chr7-29464241-INV-627
chr7-31064904-INV-6817
chr7-31093455-INV-439
chr7-35194979-INV-1048
chr7-39264150-INV-1897
chr7-40655095-INV-3758
chr7-40657738-INV-2451
chr7-41396821-INV-3364
chr7-41696855-INV-427
chr7-42013287-INV-5225
chr7-42491400-INV-7876
chr7-42493721-INV-3243
chr7-42883207-INV-994
chr7-42969199-INV-409
chr7-43549729-INV-13131
chr7-43551188-INV-12942
chr7-43553200-INV-8918
chr7-44582922-INV-347
chr7-44591717-INV-310
chr7-46844210-INV-1522
chr7-47456469-INV-4346
chr7-47456589-INV-6112
chr7-48796518-INV-1190
chr7-49581195-INV-521
chr7-50128161-INV-663
chr7-50292081-INV-1631

chr7-50809685-INV-711
chr7-52418974-INV-6652
chr7-52499454-INV-6431
chr7-52654009-INV-1396
chr7-52703647-INV-1781
chr7-52952454-INV-958
chr7-54520511-INV-2068
chr7-54573208-INV-6716
chr7-56209546-INV-2826
chr7-56211710-INV-1111
chr7-58409510-INV-394
chr7-61038869-INV-2134
chr7-61091250-INV-1386
chr7-61423546-INV-314
chr7-62170643-INV-11215
chr7-62170644-INV-9501
chr7-62172694-INV-7112
chr7-62357533-INV-2480
chr7-62427581-INV-271
chr7-62528034-INV-1564
chr7-62528034-INV-2095
chr7-63311788-INV-724
chr7-70645034-INV-643
chr7-70703806-INV-4831
chr7-72532336-INV-457
chr7-72806354-INV-350
chr7-78071906-INV-702
chr7-78889244-INV-1633
chr7-78919023-INV-372
chr7-83398998-INV-1015
chr7-83399794-INV-250
chr7-84217241-INV-2271
chr7-85009842-INV-8472
chr7-85015785-INV-2920
chr7-85843847-INV-6623
chr7-85849401-INV-669
chr7-86711376-INV-899
chr7-87325600-INV-552
chr7-87568283-INV-330
chr7-92837854-INV-6826
chr7-92946885-INV-2856
chr7-93393172-INV-3299
chr7-93936284-INV-273
chr7-94631413-INV-390
chr7-96818878-INV-566
chr7-102088450-INV-486
chr7-102088700-INV-1315
chr7-102384272-INV-601
chr7-102448218-INV-3666
chr7-103172128-INV-1575
chr7-103546639-INV-587
chr7-103611140-INV-414
chr7-104124590-INV-611
chr7-104386571-INV-1257
chr7-108272067-INV-1279
chr7-108684201-INV-723
chr7-108684248-INV-1896
chr7-111549136-INV-1212
chr7-113637287-INV-1260
chr7-113779804-INV-269
chr7-120317018-INV-2564
chr7-121418566-INV-364
chr7-125574260-INV-1000
chr7-139444998-INV-1491
chr7-140737759-INV-1348
chr7-142356731-INV-3451
chr7-143182928-INV-299
chr8-4062270-INV-10545
chr8-4112663-INV-408
chr8-5060036-INV-5913
chr8-5741853-INV-778
chr8-5757490-INV-6080
chr8-10766303-INV-370
chr8-12316761-INV-2414
chr8-13657917-INV-792
chr8-14238314-INV-282
chr8-14840490-INV-2321

chr8-14842407-INV-404
chr8-17928718-INV-335
chr8-18037398-INV-230
chr8-18356857-INV-411
chr8-22457958-INV-1533
chr8-22458663-INV-832
chr8-26386015-INV-473
chr8-27357198-INV-331
chr8-27693995-INV-377
chr8-27852617-INV-2040
chr8-28689049-INV-498
chr8-31717299-INV-4731
chr8-31717827-INV-2424
chr8-31718762-INV-2613
chr8-31842247-INV-2047
chr8-33634523-INV-692
chr8-34001855-INV-2285
chr8-34229229-INV-1036
chr8-36223018-INV-1145
chr8-37200393-INV-796
chr8-38689887-INV-449
chr8-42691009-INV-690
chr8-42985202-INV-1020
chr8-45559151-INV-4114
chr8-45560739-INV-3320
chr8-45659053-INV-4047
chr8-46408321-INV-509
chr8-47825560-INV-4713
chr8-50306061-INV-6727
chr8-56446375-INV-969
chr8-56477864-INV-4043
chr8-58695969-INV-262
chr8-58837386-INV-482
chr8-60552222-INV-1198
chr8-60552905-INV-516
chr8-60717879-INV-498
chr8-60777498-INV-1054
chr8-63750890-INV-3379
chr8-63824277-INV-4630
chr8-66558267-INV-379
chr8-72857181-INV-4987
chr8-74611075-INV-1794
chr8-74765149-INV-1797
chr8-74914508-INV-7928
chr8-74918166-INV-6198
chr8-75339940-INV-513
chr8-75751557-INV-2216
chr8-76782073-INV-4482
chr8-79502973-INV-10500
chr8-81866485-INV-1591
chr8-86323117-INV-4314
chr8-88881717-INV-340
chr8-88961779-INV-674
chr8-89703762-INV-915
chr8-90300459-INV-588
chr8-92415770-INV-973
chr8-94208568-INV-561
chr8-102757027-INV-1862
chr8-102758291-INV-630
chr8-105954219-INV-879
chr8-106640828-INV-347
chr8-109992956-INV-4523
chr8-109995028-INV-3446
chr8-111873705-INV-4295
chr8-113108924-INV-1196
chr8-118553760-INV-315
chr8-119583699-INV-874
chr8-129792443-INV-709
chr8-129986989-INV-1206
chr8-129987565-INV-642
chr9-3652205-INV-372
chr9-4182475-INV-6500
chr9-5170075-INV-2463
chr9-5219529-INV-1228
chr9-5220539-INV-219
chr9-6585857-INV-385
chr9-6742772-INV-6311

chr9-7954596-INV-873
chr9-8723189-INV-859
chr9-8725706-INV-642
chr9-10343171-INV-1744
chr9-10344356-INV-558
chr9-11995388-INV-276
chr9-12373256-INV-6501
chr9-12983761-INV-500
chr9-13068133-INV-965
chr9-14036535-INV-708
chr9-15161264-INV-477
chr9-16744355-INV-7253
chr9-17721479-INV-275
chr9-18981126-INV-3210
chr9-21628225-INV-587
chr9-22750874-INV-5017
chr9-24448106-INV-2429
chr9-25571467-INV-1878
chr9-26263949-INV-819
chr9-26880719-INV-2414
chr9-27743739-INV-695
chr9-36087757-INV-1721
chr9-36284050-INV-4313
chr9-37743456-INV-2768
chr9-38175868-INV-1001
chr9-41193411-INV-527
chr9-41394654-INV-1092
chr9-50838195-INV-3635
chr9-51882147-INV-248
chr9-52067530-INV-321
chr9-53777543-INV-589
chr9-54759362-INV-362
chr9-56125616-INV-558
chr9-56183564-INV-634
chr9-57134250-INV-1233
chr9-60459251-INV-671
chr9-60459259-INV-1400
chr9-62371628-INV-304
chr9-62544943-INV-1281
chr9-63343627-INV-4230
chr9-63345195-INV-3323
chr9-65753564-INV-1601
chr9-66201246-INV-5733
chr9-68398173-INV-1727
chr9-68398180-INV-3061
chr9-72554701-INV-2735
chr9-75915505-INV-508
chr9-76514367-INV-635
chr9-78223497-INV-8549
chr9-78223955-INV-7634
chr9-78225554-INV-4214
chr9-79063211-INV-258
chr9-79747426-INV-1419
chr9-80211692-INV-560
chr9-80225149-INV-958
chr9-82704989-INV-3795
chr9-82704989-INV-4534
chr9-93173123-INV-904
chr9-93176693-INV-1958
chr9-93487793-INV-1559
chr9-94886939-INV-450
chr9-96234706-INV-1742
chr9-96445462-INV-1075
chr9-97291281-INV-2391
chr9-97779605-INV-322
chr9-99937370-INV-1110
chr9-100694239-INV-1354
chr9-103558721-INV-2890
chr9-105377767-INV-1430
chr9-105580183-INV-738
chr9-107002195-INV-363
chr9-112797554-INV-570
chr9-113069170-INV-642
chr9-114061527-INV-697
chr9-114685083-INV-298
chr9-118507743-INV-526
chr9-119030479-INV-1158

chr9-119319278-INV-577
chr9-122966093-INV-1680
chrX-5389200-INV-1727
chrX-5430352-INV-305
chrX-5776993-INV-1353
chrX-5870161-INV-246
chrX-5904766-INV-402
chrX-9625897-INV-427
chrX-10045261-INV-323
chrX-14017605-INV-738
chrX-15111212-INV-5405
chrX-15546613-INV-1947
chrX-15761377-INV-2731
chrX-15762483-INV-528
chrX-17907364-INV-1228
chrX-18048161-INV-921
chrX-18048494-INV-4666
chrX-21977119-INV-4441
chrX-21980175-INV-365
chrX-23795973-INV-6176
chrX-32098874-INV-3061
chrX-36667556-INV-5416
chrX-37164419-INV-15599
chrX-37878738-INV-891
chrX-41850018-INV-5460
chrX-41850267-INV-7918
chrX-42484284-INV-834
chrX-44361236-INV-348
chrX-44449900-INV-349
chrX-45394537-INV-1830
chrX-45997881-INV-358
chrX-49300724-INV-6317
chrX-49326255-INV-2242
chrX-49326874-INV-2884
chrX-49477875-INV-3268
chrX-49792793-INV-2079
chrX-52660376-INV-6595
chrX-53273610-INV-1976
chrX-60383059-INV-6400
chrX-60510545-INV-5355
chrX-60511250-INV-4650
chrX-60991508-INV-7824
chrX-62371462-INV-373
chrX-63566842-INV-2035
chrX-63927322-INV-7034
chrX-66074977-INV-6302
chrX-66293937-INV-7530
chrX-66295963-INV-5534
chrX-68874752-INV-370
chrX-69158921-INV-2065
chrX-69163571-INV-3676
chrX-70365641-INV-6965
chrX-71411368-INV-4156
chrX-71412542-INV-3028
chrX-74699377-INV-535
chrX-74871492-INV-1344
chrX-74871492-INV-234
chrX-74989139-INV-739
chrX-77145043-INV-1869
chrX-77467042-INV-1569
chrX-77711687-INV-701
chrX-77711691-INV-1665
chrX-78123760-INV-360
chrX-78781471-INV-7477
chrX-79077783-INV-2189
chrX-79466032-INV-1463
chrX-80192500-INV-2007
chrX-81153490-INV-1268
chrX-81632739-INV-1696
chrX-83280848-INV-796
chrX-84856307-INV-1380
chrX-85810889-INV-2005
chrX-85935275-INV-6961
chrX-88432207-INV-955
chrX-88970816-INV-1125
chrX-89257273-INV-1389
chrX-89257932-INV-2016

chrX-90068246-INV-2119
chrX-90291811-INV-651
chrX-91086901-INV-668
chrX-91583584-INV-1343
chrX-95122257-INV-214
chrX-97026840-INV-1954
chrX-97259589-INV-631
chrX-100517022-INV-1431
chrX-100518233-INV-371
chrX-100630751-INV-416
chrX-101379852-INV-638
chrX-101675176-INV-461
chrX-101821339-INV-2739
chrX-103646755-INV-312
chrX-104380261-INV-771
chrX-108161940-INV-651
chrX-108756910-INV-558
chrX-108979458-INV-254
chrX-111650672-INV-265
chrX-112254733-INV-1246
chrX-112598642-INV-463
chrX-112687331-INV-1250
chrX-113900762-INV-6816
chrX-115436722-INV-467
chrX-116358513-INV-938
chrX-116358549-INV-1098
chrX-116416311-INV-4833
chrX-117072618-INV-2585
chrX-117073645-INV-2387
chrX-118651675-INV-1980
chrX-120272550-INV-215
chrX-120780046-INV-453
chrX-120830933-INV-8019
chrX-126710022-INV-863
chrX-129571105-INV-238
chrX-130115201-INV-708
chrX-130602799-INV-678
chrX-131980919-INV-803
chrX-133073592-INV-1139
chrX-134878192-INV-1491
chrX-137171207-INV-1153
chrX-138483419-INV-4103
chrX-140375135-INV-1029
chrX-140375619-INV-545
chrX-140988506-INV-217
chrX-141240408-INV-1330
chrX-145678269-INV-2153
chrX-145679192-INV-1253
chrX-146219745-INV-423
chrX-146251290-INV-1727
chrX-154011370-INV-1301
chrX-155632928-INV-815
chrX-155757905-INV-247
chrX-156845317-INV-1513
chrX-161632242-INV-996
chrX-162239200-INV-610
chrX-168856708-INV-2321

**Identified_By_JAX**
chr1-3192115-INV-1750
chr1-8576742-INV-1881
chr1-9716444-INV-584
chr1-31755041-INV-514
chr1-48811313-INV-498
chr1-48911119-INV-478839
chr1-49992237-INV-1100
chr1-61531315-INV-29879
chr1-79226513-INV-7729
chr1-98811352-INV-17746
chr1-114565317-INV-2137
chr1-132031270-INV-6601
chr1-141262162-INV-51250
chr1-142942778-INV-19294
chr1-147760005-INV-16956
chr1-156756886-INV-1791
chr1-167438810-INV-709
chr1-170442338-INV-1163
chr1-191853081-INV-16227
chr10-6226667-INV-4359
chr10-61469084-INV-390
chr10-103780635-INV-5267
chr11-7701330-INV-24043
chr11-13609979-INV-19906
chr11-17345865-INV-1554
chr11-28941704-INV-14838
chr11-45492246-INV-18101
chr11-48093703-INV-36416
chr11-48220952-INV-16437
chr11-48503363-INV-20405
chr11-58763391-INV-1405
chr11-63726235-INV-2118
chr11-101532092-INV-5235
chr11-108804515-INV-1381
chr12-6692373-INV-5284
chr12-14894225-INV-6413
chr12-21211013-INV-410
chr12-54750776-INV-40915
chr12-63694889-INV-4544
chr12-93080849-INV-27523
chr12-100479786-INV-764
chr12-111094926-INV-755
chr12-113198213-INV-15776
chr13-8991329-INV-18489
chr13-21978201-INV-55632
chr13-22252591-INV-13026

chr13-23091505-INV-76591
chr13-86375038-INV-2273
chr14-3955924-INV-23504
chr14-10202691-INV-4532
chr14-12577191-INV-12648
chr14-40448358-INV-5314
chr14-56426138-INV-13261
chr14-84453443-INV-6906
chr15-10145842-INV-688
chr15-85921028-INV-500
chr15-103666863-INV-23960
chr16-24843839-INV-543
chr16-26146535-INV-24567
chr16-52967857-INV-47407
chr16-54584462-INV-1312
chr16-84565354-INV-415
chr17-19811556-INV-14614
chr17-23751408-INV-4654
chr17-24163669-INV-9880
chr17-28089091-INV-1441
chr17-29529849-INV-1506
chr17-29799100-INV-1189
chr17-33456470-INV-66889
chr17-56572704-INV-1440
chr17-82157592-INV-1503
chr17-89302141-INV-5191
chr17-94285118-INV-9992
chr18-6716104-INV-1179
chr18-8703368-INV-29066
chr18-11013189-INV-30914
chr18-29064740-INV-3030
chr18-48575103-INV-44400
chr18-80885589-INV-590
chr18-84204659-INV-1158
chr18-85808238-INV-107298
chr18-90433218-INV-24296
chr19-57007528-INV-913
chr2-7123261-INV-732
chr2-10940849-INV-2310
chr2-14203057-INV-3283
chr2-21720190-INV-6931
chr2-22095508-INV-1930
chr2-35667693-INV-763
chr2-43503330-INV-2060
chr2-56505345-INV-2054
chr2-72543204-INV-4705
chr2-81107177-INV-36136

chr2-85394721-INV-3322
chr2-94547911-INV-426
chr2-100542833-INV-53386
chr2-103896974-INV-1622
chr2-117308236-INV-2074
chr2-150926364-INV-21671
chr2-154773797-INV-22627
chr2-173981926-INV-4002
chr3-11505682-INV-740
chr3-20834692-INV-27311
chr3-20889662-INV-16029
chr3-28804161-INV-2176
chr3-33651854-INV-7691
chr3-34646203-INV-664
chr3-40662192-INV-24131
chr3-75085481-INV-16819
chr3-82064187-INV-125601
chr3-90722739-INV-120367
chr3-96354231-INV-16822
chr3-111358441-INV-20318
chr3-115263860-INV-27409
chr3-151656670-INV-16090
chr3-151744847-INV-627
chr3-154961334-INV-627
chr4-4310255-INV-9082
chr4-20805044-INV-28375
chr4-24318648-INV-2617
chr4-53958899-INV-15867
chr4-67417831-INV-13044
chr4-81923115-INV-14013
chr4-115006011-INV-15517
chr4-144331300-INV-42044
chr5-12530018-INV-2725
chr5-57649017-INV-12359
chr5-66137396-INV-14022
chr5-80664747-INV-924
chr5-100593656-INV-537
chr5-105345071-INV-43191
chr6-41299750-INV-619
chr6-66764748-INV-1171
chr6-70878652-INV-2005
chr6-105921710-INV-2527
chr6-107642504-INV-23683
chr6-108815834-INV-1726
chr6-110386916-INV-13955
chr6-132243279-INV-9598
chr7-42491965-INV-6792

chr7-42883113-INV-1090
chr7-43551424-INV-12467
chr7-50807472-INV-5415
chr7-53036846-INV-35746
chr7-59704465-INV-16686
chr7-61423444-INV-447
chr7-62172199-INV-7982
chr7-66905044-INV-342
chr7-84557319-INV-109584
chr7-102670207-INV-14269
chr8-34001740-INV-2535
chr8-52472589-INV-162792
chr8-53780855-INV-38904
chr8-88961648-INV-927
chr8-97259772-INV-13908
chr8-111873540-INV-4330
chr8-129789001-INV-6734
chr9-5169507-INV-3029
chr9-99937000-INV-1773
chr9-105580051-INV-921
chrX-5386830-INV-6332
chrX-5770609-INV-14112
chrX-8178316-INV-44966
chrX-8825242-INV-89699
chrX-36922628-INV-47637
chrX-40833291-INV-32557
chrX-41094153-INV-22511
chrX-49838397-INV-21242
chrX-55224741-INV-49654
chrX-68178742-INV-31817
chrX-69416088-INV-175848
chrX-73742314-INV-153178
chrX-75879411-INV-114642
chrX-76907116-INV-64214
chrX-79622822-INV-24847
chrX-93488372-INV-23940
chrX-93719144-INV-179596
chrX-97070933-INV-87925
chrX-100630601-INV-797
chrX-100830923-INV-14393
chrX-101840091-INV-68700
chrX-102030528-INV-106521
chrX-103645966-INV-1905
chrX-133073362-INV-1661
chrX-135270657-INV-40430
chrX-135370515-INV-48826
chrX-138482698-INV-5459

chrX-142012758-INV-33073
chrX-151197417-INV-62349
chrX-155714533-INV-24250
chrX-156842759-INV-6435

| Identified_By_Us | Identified_By_JAX |
|---|---|
| chr1-16151971-DUP-237 | chr1-16151971-DUP-237 |
| chr1-28106534-DUP-1432 | chr1-28106534-DUP-1438 |
| chr1-66227553-DUP-376 | chr1-66227553-DUP-378 |
| chr1-98743395-DUP-286 | chr1-98743395-DUP-296 |
| chr1-149902315-DUP-319 | chr1-149902315-DUP-321 |
| chr1-161036363-DUP-256 | chr1-161036360-DUP-253 |
| chr10-70885864-DUP-362 | chr10-70885864-DUP-366 |
| chr10-107930140-DUP-2748 | chr10-107930141-DUP-2748 |
| chr11-23164256-DUP-322 | chr11-23164256-DUP-324 |
| chr11-91118437-DUP-405 | chr11-91118437-DUP-407 |
| chr11-101597224-DUP-256 | chr11-101597236-DUP-251 |
| chr12-71041499-DUP-494 | chr12-71041498-DUP-496 |
| chr12-76284548-DUP-282 | chr12-76284548-DUP-286 |
| chr13-96043904-DUP-237 | chr13-96043904-DUP-239 |
| chr13-110884164-DUP-206 | chr13-110884164-DUP-208 |
| chr14-23468674-DUP-2633 | chr14-23468674-DUP-2633 |
| chr14-81691116-DUP-216 | chr14-81691116-DUP-218 |
| chr14-86977382-DUP-210 | chr14-86977382-DUP-212 |
| chr14-114822067-DUP-366 | chr14-114822067-DUP-368 |
| chr14-122148495-DUP-260 | chr14-122148495-DUP-262 |
| chr15-66955835-DUP-456 | chr15-66955835-DUP-461 |
| chr15-76034637-DUP-200 | chr15-76034637-DUP-202 |
| chr15-80503720-DUP-234 | chr15-80503720-DUP-239 |
| chr15-84982496-DUP-320 | chr15-84982496-DUP-359 |
| chr16-17420418-DUP-925 | chr16-17420417-DUP-932 |
| chr16-31241520-DUP-3887 | chr16-31241521-DUP-3902 |
| chr16-56658694-DUP-2641 | chr16-56658691-DUP-2653 |
| chr17-20650685-DUP-461 | chr17-20650685-DUP-463 |
| chr17-32139961-DUP-300 | chr17-32139968-DUP-326 |
| chr17-33683862-DUP-244 | chr17-33683861-DUP-281 |
| chr18-25505934-DUP-5632 | chr18-25505932-DUP-5633 |
| chr18-38431334-DUP-411 | chr18-38431334-DUP-413 |
| chr19-51879797-DUP-5783 | chr19-51879797-DUP-5768 |
| chr2-3426224-DUP-203 | chr2-3426224-DUP-205 |
| chr2-21067661-DUP-544 | chr2-21067661-DUP-546 |
| chr2-76257056-DUP-232 | chr2-76257056-DUP-234 |
| chr2-82177223-DUP-774 | chr2-82177223-DUP-770 |
| chr2-99499211-DUP-263 | chr2-99499211-DUP-287 |
| chr2-123881426-DUP-359 | chr2-123881426-DUP-361 |
| chr2-126197088-DUP-2231 | chr2-126197083-DUP-2246 |
| chr2-157679074-DUP-1299 | chr2-157679074-DUP-1301 |
| chr3-48329611-DUP-984 | chr3-48329611-DUP-996 |
| chr4-154211763-DUP-271 | chr4-154211763-DUP-302 |
| chr5-14648139-DUP-267 | chr5-14648139-DUP-269 |
| chr5-79742143-DUP-215 | chr5-79742143-DUP-217 |
| chr5-115855664-DUP-257 | chr5-115855664-DUP-259 |
| chr5-140357484-DUP-606 | chr5-140357484-DUP-608 |
| chr5-146119602-DUP-2635 | chr5-146119603-DUP-2635 |
| chr6-20368976-DUP-869 | chr6-20368976-DUP-871 |
| chr6-86993296-DUP-897 | chr6-86993274-DUP-897 |
| chr6-132075459-DUP-2992 | chr6-132075455-DUP-2988 |
| chr7-25473079-DUP-268 | chr7-25473078-DUP-268 |
| chr7-70344691-DUP-235 | chr7-70344691-DUP-237 |
| chr7-94992751-DUP-289 | chr7-94992751-DUP-291 |
| chr7-114528583-DUP-2032 | chr7-114528583-DUP-2037 |
| chr8-9284996-DUP-304 | chr8-9284996-DUP-306 |
| chr8-10327481-DUP-291 | chr8-10327481-DUP-293 |
| chr9-31124460-DUP-371 | chr9-31124460-DUP-373 |
| chr9-41192688-DUP-288 | chr9-41192686-DUP-288 |
| chr9-48841852-DUP-286 | chr9-48841852-DUP-288 |
| chr9-100950584-DUP-1010 | chr9-100950584-DUP-1011 |
| chrX-12449971-DUP-1486 | chrX-12449971-DUP-1488 |

**Identified_By_Us**
chr1-3600450-DUP-139
chr1-4421459-DUP-77
chr1-4904188-DUP-100
chr1-4905015-DUP-177
chr1-4905107-DUP-675
chr1-4905735-DUP-105
chr1-5142660-DUP-388
chr1-5163646-DUP-60
chr1-9020433-DUP-98
chr1-10325078-DUP-103
chr1-10325119-DUP-130
chr1-10356639-DUP-76
chr1-10376322-DUP-60
chr1-10829488-DUP-405
chr1-11130907-DUP-62
chr1-11163462-DUP-192
chr1-11726206-DUP-61
chr1-11874097-DUP-79
chr1-12727751-DUP-199
chr1-12727865-DUP-187
chr1-12911215-DUP-386
chr1-13022222-DUP-585
chr1-13032699-DUP-56
chr1-13193382-DUP-60
chr1-13273598-DUP-138
chr1-13328311-DUP-78
chr1-13511872-DUP-73
chr1-13721164-DUP-70
chr1-13841465-DUP-190
chr1-13847297-DUP-27407
chr1-14122244-DUP-87
chr1-14336242-DUP-113
chr1-14506623-DUP-187
chr1-14554932-DUP-93
chr1-15758001-DUP-90
chr1-16142474-DUP-372
chr1-16143267-DUP-1100
chr1-16158661-DUP-748
chr1-16203214-DUP-137
chr1-16203288-DUP-244
chr1-17119693-DUP-367
chr1-17419827-DUP-52
chr1-18506568-DUP-139
chr1-19226543-DUP-58
chr1-20046273-DUP-131
chr1-20735346-DUP-122
chr1-20890135-DUP-90
chr1-20890172-DUP-126
chr1-21094998-DUP-72
chr1-21095027-DUP-52
chr1-21301283-DUP-101
chr1-21301357-DUP-61
chr1-21988483-DUP-150
chr1-21988515-DUP-76
chr1-22431761-DUP-172
chr1-22814649-DUP-113
chr1-22814720-DUP-115
chr1-22839112-DUP-85
chr1-22962782-DUP-214
chr1-23099955-DUP-284
chr1-24128459-DUP-193
chr1-24129313-DUP-122
chr1-24224609-DUP-213
chr1-24247612-DUP-72
chr1-24323381-DUP-83
chr1-24445762-DUP-110
chr1-24723146-DUP-138
chr1-24723187-DUP-67
chr1-24738563-DUP-68
chr1-24738594-DUP-70
chr1-25210767-DUP-64
chr1-25245567-DUP-139
chr1-25668640-DUP-119
chr1-25668721-DUP-71
chr1-26177016-DUP-190
chr1-26313696-DUP-284

chr1-26313740-DUP-126
chr1-27612460-DUP-218
chr1-28106555-DUP-1388
chr1-29343825-DUP-26759
chr1-29346843-DUP-68
chr1-30253618-DUP-81
chr1-31018312-DUP-621
chr1-31996035-DUP-78
chr1-32320239-DUP-9893
chr1-32485073-DUP-88
chr1-32485098-DUP-117
chr1-32712842-DUP-57
chr1-33606710-DUP-154
chr1-33607161-DUP-232
chr1-33608097-DUP-154
chr1-33608643-DUP-388
chr1-33720652-DUP-121
chr1-33720735-DUP-164
chr1-33822868-DUP-83
chr1-34641939-DUP-2560
chr1-34647327-DUP-71
chr1-34837060-DUP-94
chr1-34919487-DUP-113
chr1-35271291-DUP-70
chr1-35691521-DUP-177
chr1-35744443-DUP-299
chr1-35953541-DUP-126
chr1-36001084-DUP-62
chr1-36035568-DUP-96
chr1-36198885-DUP-92
chr1-36302314-DUP-93
chr1-36302340-DUP-72
chr1-36658751-DUP-52
chr1-36658938-DUP-425
chr1-37170503-DUP-81
chr1-37253237-DUP-67
chr1-37352516-DUP-128
chr1-37435463-DUP-98
chr1-37435486-DUP-166
chr1-37473728-DUP-289
chr1-37883267-DUP-138
chr1-38426006-DUP-178
chr1-38469487-DUP-60
chr1-38605824-DUP-96
chr1-38625551-DUP-87
chr1-38848322-DUP-328
chr1-38887114-DUP-66
chr1-38887136-DUP-68
chr1-38912228-DUP-55
chr1-39092004-DUP-67
chr1-39284035-DUP-189
chr1-40360823-DUP-62
chr1-40424230-DUP-94
chr1-40716844-DUP-66
chr1-40724098-DUP-121
chr1-40831515-DUP-78
chr1-40851439-DUP-61
chr1-41158974-DUP-113
chr1-41372300-DUP-54
chr1-41923995-DUP-163
chr1-42774309-DUP-53
chr1-42982287-DUP-102
chr1-43338978-DUP-59
chr1-43816778-DUP-1327
chr1-43817119-DUP-58
chr1-43817766-DUP-119
chr1-45416670-DUP-80
chr1-45732518-DUP-82
chr1-45921287-DUP-166
chr1-46151396-DUP-1318
chr1-46570174-DUP-181
chr1-46855216-DUP-70
chr1-49059443-DUP-125
chr1-49059469-DUP-77
chr1-50011013-DUP-1682
chr1-50340824-DUP-93
chr1-50600815-DUP-51

chr1-51161682-DUP-152
chr1-51161753-DUP-104
chr1-51318756-DUP-147
chr1-51520977-DUP-145
chr1-52114925-DUP-55
chr1-52223506-DUP-81
chr1-52318926-DUP-81
chr1-52522382-DUP-62
chr1-52892104-DUP-262
chr1-53234769-DUP-183
chr1-53468179-DUP-154
chr1-53491853-DUP-55
chr1-53611647-DUP-63
chr1-53800760-DUP-65
chr1-53872091-DUP-145
chr1-54531616-DUP-58
chr1-54652804-DUP-53
chr1-54799239-DUP-60
chr1-54801370-DUP-84
chr1-54813504-DUP-118
chr1-54921462-DUP-122
chr1-54996443-DUP-54
chr1-55248346-DUP-95
chr1-55533436-DUP-63
chr1-55731401-DUP-60
chr1-55772402-DUP-58
chr1-56605908-DUP-190
chr1-56605970-DUP-110
chr1-56717221-DUP-71
chr1-57022225-DUP-126
chr1-57164683-DUP-134
chr1-57184635-DUP-62
chr1-57386866-DUP-55
chr1-57455110-DUP-68
chr1-57605962-DUP-198
chr1-57635813-DUP-94
chr1-57744654-DUP-84
chr1-57909591-DUP-75
chr1-58125514-DUP-83
chr1-58129195-DUP-89
chr1-58134460-DUP-76
chr1-58254649-DUP-106
chr1-58254697-DUP-65
chr1-58414060-DUP-205
chr1-58690090-DUP-60251
chr1-58998441-DUP-67
chr1-59045982-DUP-99
chr1-59046167-DUP-53
chr1-59137176-DUP-157
chr1-59178772-DUP-66
chr1-59217189-DUP-58
chr1-59439800-DUP-87
chr1-59519770-DUP-79
chr1-60048068-DUP-57
chr1-60109927-DUP-64
chr1-60182354-DUP-110
chr1-60227965-DUP-147
chr1-61040963-DUP-89
chr1-61085644-DUP-394
chr1-61085665-DUP-346
chr1-61138785-DUP-58
chr1-61335067-DUP-591
chr1-61409186-DUP-55
chr1-61561742-DUP-2408
chr1-61605933-DUP-98
chr1-61928167-DUP-62
chr1-62437189-DUP-83
chr1-63025447-DUP-93
chr1-63278857-DUP-88
chr1-63322017-DUP-59
chr1-63322043-DUP-61
chr1-63585330-DUP-111
chr1-63668892-DUP-1241
chr1-63669211-DUP-67
chr1-63669692-DUP-136
chr1-63963225-DUP-163
chr1-63963298-DUP-60

chr1-64158928-DUP-119
chr1-64583022-DUP-343
chr1-64680908-DUP-72
chr1-64837748-DUP-358
chr1-64839399-DUP-238
chr1-64887543-DUP-162
chr1-64887597-DUP-78
chr1-65601900-DUP-63
chr1-66227591-DUP-277
chr1-66535783-DUP-385
chr1-66535895-DUP-250
chr1-66567335-DUP-69
chr1-66755378-DUP-72
chr1-66881344-DUP-65
chr1-67024337-DUP-57
chr1-67551598-DUP-136
chr1-67798182-DUP-93
chr1-67879616-DUP-77
chr1-68141875-DUP-39983
chr1-68592576-DUP-77
chr1-69709912-DUP-77
chr1-69803305-DUP-168
chr1-70199753-DUP-67
chr1-70285468-DUP-60
chr1-71756844-DUP-77
chr1-71960553-DUP-76
chr1-71977705-DUP-120
chr1-72217788-DUP-54
chr1-72753957-DUP-99
chr1-73025688-DUP-115
chr1-73108954-DUP-240
chr1-73109002-DUP-202
chr1-73524044-DUP-126
chr1-73870602-DUP-56
chr1-73877327-DUP-143
chr1-74463222-DUP-196
chr1-74463246-DUP-198
chr1-74536336-DUP-60
chr1-75015548-DUP-81
chr1-75084942-DUP-333
chr1-75084998-DUP-239
chr1-75171595-DUP-52
chr1-75203954-DUP-71
chr1-75367139-DUP-147
chr1-75568213-DUP-57
chr1-75568290-DUP-60
chr1-76547092-DUP-394
chr1-76807147-DUP-82
chr1-76901164-DUP-67
chr1-77952598-DUP-92
chr1-78550330-DUP-969
chr1-78550365-DUP-931
chr1-78646556-DUP-97
chr1-78855051-DUP-59
chr1-79450771-DUP-74
chr1-79570205-DUP-95
chr1-79817072-DUP-358
chr1-79817144-DUP-283
chr1-80633947-DUP-64
chr1-80901861-DUP-75865
chr1-81103907-DUP-64
chr1-81410154-DUP-57
chr1-81528353-DUP-131
chr1-81765027-DUP-78
chr1-82052458-DUP-222
chr1-82430647-DUP-61
chr1-82545324-DUP-53
chr1-83101624-DUP-72
chr1-84278524-DUP-284
chr1-84581495-DUP-275
chr1-84655487-DUP-88
chr1-84836893-DUP-167
chr1-85011393-DUP-64
chr1-85079028-DUP-12962
chr1-85286537-DUP-64
chr1-85472573-DUP-64
chr1-85595696-DUP-251

chr1-85595772-DUP-207
chr1-85742142-DUP-77
chr1-86082783-DUP-95
chr1-86118498-DUP-68
chr1-86419448-DUP-71
chr1-86752165-DUP-56
chr1-86812948-DUP-246
chr1-87034011-DUP-126
chr1-87382588-DUP-148
chr1-87382633-DUP-72
chr1-87968099-DUP-4316
chr1-87974775-DUP-4342
chr1-88162322-DUP-144
chr1-88259084-DUP-192
chr1-88313712-DUP-118
chr1-88397459-DUP-53
chr1-88460364-DUP-78
chr1-88515543-DUP-74
chr1-88552478-DUP-60
chr1-88741224-DUP-10836
chr1-89047796-DUP-56
chr1-89062557-DUP-103
chr1-89429377-DUP-254
chr1-89430252-DUP-446
chr1-89498867-DUP-69
chr1-89579188-DUP-68
chr1-89607326-DUP-154
chr1-89637812-DUP-475
chr1-89637845-DUP-427
chr1-89904052-DUP-61
chr1-89974745-DUP-122
chr1-90111832-DUP-58
chr1-90375325-DUP-80
chr1-90378592-DUP-69
chr1-90466085-DUP-129
chr1-90550731-DUP-5623
chr1-90551275-DUP-5077
chr1-90684145-DUP-139
chr1-90941019-DUP-85
chr1-91066045-DUP-80
chr1-91108640-DUP-57
chr1-91248754-DUP-93
chr1-91349843-DUP-456
chr1-91365270-DUP-74
chr1-91370008-DUP-84
chr1-91443071-DUP-53
chr1-91595354-DUP-54
chr1-91666908-DUP-62
chr1-91714723-DUP-53
chr1-91909440-DUP-79
chr1-92063506-DUP-54
chr1-92290634-DUP-87
chr1-92417258-DUP-17834
chr1-92444673-DUP-210
chr1-92448954-DUP-55
chr1-92666432-DUP-53
chr1-92819191-DUP-189
chr1-92909451-DUP-137
chr1-93057555-DUP-4432
chr1-93375193-DUP-104
chr1-93745033-DUP-115
chr1-94066715-DUP-56
chr1-94367690-DUP-427
chr1-94595628-DUP-57
chr1-94616674-DUP-1241
chr1-97238166-DUP-352
chr1-97274740-DUP-96
chr1-97714758-DUP-137
chr1-97847326-DUP-91
chr1-99080841-DUP-73
chr1-99314830-DUP-2979
chr1-99552034-DUP-128
chr1-99789256-DUP-165
chr1-99857505-DUP-52
chr1-99892985-DUP-257
chr1-101898194-DUP-68
chr1-102279690-DUP-66

chr1-102635050-DUP-83
chr1-102976828-DUP-128
chr1-103497996-DUP-674
chr1-104190494-DUP-92
chr1-104210129-DUP-193
chr1-104457211-DUP-187
chr1-104761317-DUP-1571
chr1-105107557-DUP-220
chr1-105152434-DUP-454
chr1-106271868-DUP-108
chr1-106289985-DUP-53
chr1-106549749-DUP-56
chr1-106645583-DUP-201
chr1-107246281-DUP-58
chr1-107691132-DUP-107
chr1-107871529-DUP-2142
chr1-107920684-DUP-59
chr1-108382984-DUP-69
chr1-108731380-DUP-662
chr1-109010009-DUP-57
chr1-110698390-DUP-71
chr1-110816417-DUP-15502
chr1-111591653-DUP-67
chr1-112321746-DUP-116
chr1-112844249-DUP-50
chr1-113308869-DUP-355
chr1-113308891-DUP-293
chr1-113600294-DUP-92
chr1-113648691-DUP-1632
chr1-114354339-DUP-175
chr1-116262158-DUP-67
chr1-116316136-DUP-2726
chr1-116838685-DUP-98
chr1-117075667-DUP-77
chr1-117709473-DUP-315081
chr1-118387705-DUP-97
chr1-118899545-DUP-51
chr1-119458438-DUP-88
chr1-119712010-DUP-98
chr1-120138724-DUP-89
chr1-120849174-DUP-93
chr1-120906028-DUP-62
chr1-120977580-DUP-53
chr1-121369714-DUP-71
chr1-121459488-DUP-66
chr1-121700465-DUP-1262
chr1-121741421-DUP-67
chr1-121814738-DUP-83
chr1-122462288-DUP-8575
chr1-122913260-DUP-58
chr1-123088124-DUP-5481
chr1-123274755-DUP-53
chr1-123369594-DUP-815
chr1-123369659-DUP-773
chr1-123751155-DUP-113
chr1-124823077-DUP-72
chr1-125317987-DUP-55
chr1-125656404-DUP-84
chr1-125809902-DUP-121
chr1-126282641-DUP-54
chr1-126824807-DUP-103
chr1-126824838-DUP-59
chr1-126964915-DUP-81
chr1-126968798-DUP-54
chr1-127036906-DUP-370
chr1-127043157-DUP-105
chr1-127183264-DUP-86
chr1-127254249-DUP-75
chr1-127421839-DUP-270
chr1-127422557-DUP-58
chr1-127992793-DUP-1808
chr1-127993103-DUP-512
chr1-128768586-DUP-97
chr1-129049574-DUP-521
chr1-129082908-DUP-85
chr1-129196709-DUP-146
chr1-129372229-DUP-308

chr1-129448001-DUP-128
chr1-129689186-DUP-51
chr1-129783408-DUP-382
chr1-130376642-DUP-194
chr1-130376668-DUP-193
chr1-130639412-DUP-57
chr1-130881212-DUP-51
chr1-131016466-DUP-78
chr1-131040678-DUP-574
chr1-131045502-DUP-191
chr1-131087746-DUP-200
chr1-131153843-DUP-67
chr1-131754759-DUP-82
chr1-131971361-DUP-92
chr1-132201960-DUP-80
chr1-132606969-DUP-92
chr1-132649954-DUP-52
chr1-133272290-DUP-95
chr1-133522182-DUP-878
chr1-133522185-DUP-1705
chr1-133522223-DUP-1537
chr1-133522347-DUP-100
chr1-133522389-DUP-1253
chr1-133523262-DUP-283
chr1-133523580-DUP-51
chr1-133786501-DUP-55
chr1-133894563-DUP-168
chr1-134231204-DUP-84
chr1-134231273-DUP-66
chr1-134241532-DUP-77
chr1-134282171-DUP-85
chr1-134284065-DUP-530
chr1-134318868-DUP-253
chr1-134663674-DUP-281
chr1-134663799-DUP-189
chr1-135201184-DUP-72
chr1-135233433-DUP-129
chr1-135293644-DUP-66
chr1-135363018-DUP-135
chr1-135422224-DUP-64
chr1-135493287-DUP-64
chr1-135849965-DUP-119
chr1-136006537-DUP-277
chr1-136014138-DUP-117
chr1-136606992-DUP-136
chr1-136799444-DUP-230
chr1-137031775-DUP-57
chr1-137251505-DUP-59
chr1-137323151-DUP-344
chr1-137537292-DUP-58
chr1-137881727-DUP-197
chr1-137956584-DUP-78
chr1-138193098-DUP-162
chr1-138218138-DUP-64
chr1-138487107-DUP-56
chr1-139250900-DUP-54
chr1-139325079-DUP-583
chr1-139549434-DUP-292365
chr1-140198006-DUP-172
chr1-140255444-DUP-51
chr1-141220393-DUP-446
chr1-141672748-DUP-72
chr1-142621280-DUP-70
chr1-143001368-DUP-120
chr1-144205482-DUP-86
chr1-144753719-DUP-87
chr1-144915483-DUP-1308
chr1-145443540-DUP-86
chr1-145830817-DUP-3413
chr1-146021172-DUP-72
chr1-146038977-DUP-25927
chr1-148740434-DUP-122
chr1-148787476-DUP-170
chr1-148787537-DUP-79
chr1-149418315-DUP-253
chr1-149964568-DUP-70
chr1-150622112-DUP-243

chr1-150795205-DUP-93
chr1-150846842-DUP-78
chr1-151011510-DUP-62
chr1-151028572-DUP-61
chr1-151060176-DUP-27555
chr1-151112009-DUP-75
chr1-151381429-DUP-121
chr1-151450481-DUP-360
chr1-151526925-DUP-66
chr1-151563762-DUP-128
chr1-151672690-DUP-52
chr1-151677344-DUP-85
chr1-152154708-DUP-65
chr1-152387167-DUP-96
chr1-152396171-DUP-216
chr1-152481364-DUP-180
chr1-152500244-DUP-54
chr1-152535910-DUP-59
chr1-152549358-DUP-54
chr1-152555815-DUP-158
chr1-152555865-DUP-110
chr1-153090649-DUP-258
chr1-153100456-DUP-91
chr1-153169575-DUP-80
chr1-153175548-DUP-65
chr1-153286329-DUP-4828
chr1-153289589-DUP-4667
chr1-153429838-DUP-161
chr1-153497219-DUP-83
chr1-153794961-DUP-64
chr1-153881469-DUP-72
chr1-154031685-DUP-359
chr1-154031706-DUP-311
chr1-154083374-DUP-168
chr1-154381680-DUP-7551
chr1-154657238-DUP-56
chr1-154795211-DUP-94
chr1-155150728-DUP-117
chr1-155900142-DUP-127
chr1-156383707-DUP-104
chr1-156689392-DUP-127
chr1-156962362-DUP-233
chr1-156962426-DUP-203
chr1-157021353-DUP-100
chr1-157386300-DUP-493
chr1-157974597-DUP-123
chr1-158075175-DUP-198
chr1-158337855-DUP-100
chr1-158665236-DUP-54
chr1-158670643-DUP-67
chr1-158890589-DUP-81
chr1-159901795-DUP-61159
chr1-161060583-DUP-182
chr1-161129012-DUP-92
chr1-161290520-DUP-52
chr1-161315813-DUP-303
chr1-161700159-DUP-62
chr1-161803182-DUP-150
chr1-162089911-DUP-60
chr1-162199314-DUP-52
chr1-163133021-DUP-83
chr1-163582331-DUP-61
chr1-163669430-DUP-109
chr1-163700677-DUP-100
chr1-163805127-DUP-51
chr1-164006864-DUP-62
chr1-164021231-DUP-54
chr1-164464853-DUP-111
chr1-164592469-DUP-100
chr1-164725160-DUP-51
chr1-164872111-DUP-111
chr1-164872136-DUP-81
chr1-164937701-DUP-185
chr1-164937734-DUP-51
chr1-165156624-DUP-61
chr1-165429553-DUP-71492
chr1-165438637-DUP-61

chr1-165507609-DUP-144
chr1-165690938-DUP-917
chr1-166577277-DUP-83
chr1-166827102-DUP-57
chr1-166839187-DUP-178
chr1-167047392-DUP-114
chr1-167094744-DUP-90
chr1-167329248-DUP-225
chr1-167591121-DUP-201
chr1-167591147-DUP-237
chr1-167907917-DUP-116
chr1-168121035-DUP-160
chr1-168134337-DUP-75
chr1-168236200-DUP-93
chr1-169345172-DUP-675
chr1-169735131-DUP-62
chr1-170558469-DUP-82
chr1-170601567-DUP-187
chr1-170631782-DUP-213
chr1-170671162-DUP-107
chr1-170708348-DUP-155
chr1-171305545-DUP-98
chr1-171390151-DUP-1647
chr1-171907684-DUP-207
chr1-171907715-DUP-159
chr1-171943670-DUP-52
chr1-171949155-DUP-55
chr1-172296229-DUP-58
chr1-173556688-DUP-82
chr1-173569235-DUP-5422
chr1-173587714-DUP-59
chr1-173591669-DUP-214386
chr1-173611739-DUP-82308
chr1-173733716-DUP-59
chr1-173737663-DUP-68391
chr1-173927390-DUP-11519
chr1-174041842-DUP-122
chr1-174424033-DUP-122
chr1-174643484-DUP-316
chr1-175034771-DUP-4946
chr1-175430578-DUP-62
chr1-175556504-DUP-113
chr1-175897032-DUP-85
chr1-176269100-DUP-22030
chr1-176549674-DUP-214
chr1-176549737-DUP-221
chr1-176850818-DUP-192
chr1-176868037-DUP-216
chr1-176868123-DUP-256
chr1-177498279-DUP-87
chr1-177498326-DUP-59
chr1-177643152-DUP-325
chr1-177685585-DUP-147
chr1-177685620-DUP-151
chr1-177826683-DUP-158
chr1-177924667-DUP-136
chr1-178376597-DUP-246
chr1-178376728-DUP-62
chr1-178867278-DUP-54
chr1-178975903-DUP-54
chr1-179064041-DUP-150
chr1-179125336-DUP-54
chr1-179167363-DUP-56
chr1-179201369-DUP-80
chr1-179289073-DUP-75
chr1-179421609-DUP-62
chr1-179667321-DUP-214
chr1-179885279-DUP-92
chr1-179948601-DUP-539
chr1-179948673-DUP-432
chr1-180056382-DUP-68
chr1-180071223-DUP-122
chr1-180072142-DUP-172
chr1-180296901-DUP-59
chr1-180363238-DUP-62
chr1-180618341-DUP-501
chr1-180797691-DUP-109

chr1-180958218-DUP-66
chr1-180966021-DUP-61
chr1-181032735-DUP-139
chr1-181467089-DUP-68
chr1-181570552-DUP-1080
chr1-181632622-DUP-123
chr1-181654089-DUP-108
chr1-181759378-DUP-111
chr1-181963409-DUP-53
chr1-182332695-DUP-218
chr1-182332727-DUP-170
chr1-182403133-DUP-111
chr1-182712014-DUP-177
chr1-182712087-DUP-60
chr1-182733177-DUP-52
chr1-182763366-DUP-125
chr1-182765366-DUP-460
chr1-182765396-DUP-412
chr1-182884899-DUP-674
chr1-182887648-DUP-111
chr1-182943324-DUP-410
chr1-182962449-DUP-69
chr1-183267859-DUP-80
chr1-183267887-DUP-148
chr1-183305553-DUP-50
chr1-183316710-DUP-142
chr1-183350116-DUP-72
chr1-183600933-DUP-301
chr1-183877631-DUP-85
chr1-183877669-DUP-87
chr1-184179176-DUP-63
chr1-184206051-DUP-255
chr1-184223957-DUP-136
chr1-184495510-DUP-118
chr1-184547225-DUP-358
chr1-184658921-DUP-176
chr1-184708040-DUP-112
chr1-184708093-DUP-56
chr1-185216914-DUP-96
chr1-185481803-DUP-900
chr1-185529208-DUP-75
chr1-185827370-DUP-131
chr1-186462119-DUP-755
chr1-186462837-DUP-895
chr1-186463369-DUP-159
chr1-186464781-DUP-62
chr1-186767571-DUP-68
chr1-186888964-DUP-228
chr1-187085374-DUP-122
chr1-187351958-DUP-73
chr1-187357703-DUP-83
chr1-187933485-DUP-166
chr1-187933511-DUP-168
chr1-188505263-DUP-57
chr1-188714537-DUP-74
chr1-188765063-DUP-70
chr1-188789354-DUP-53
chr1-188846077-DUP-144
chr1-188846124-DUP-57
chr1-189239388-DUP-118
chr1-189244630-DUP-100
chr1-189254317-DUP-283
chr1-189269424-DUP-176
chr1-189330964-DUP-70
chr1-189356610-DUP-150
chr1-189365933-DUP-147
chr1-189468134-DUP-114
chr1-189478526-DUP-287
chr1-189478611-DUP-166
chr1-189674172-DUP-86
chr1-189674194-DUP-83
chr1-189712877-DUP-57
chr1-189723430-DUP-84
chr1-189935183-DUP-256
chr1-190010016-DUP-72
chr1-190032203-DUP-4325
chr1-190032567-DUP-322

chr1-190033512-DUP-1144
chr1-190033648-DUP-368
chr1-190034112-DUP-97
chr1-190034436-DUP-52
chr1-190035404-DUP-1121
chr1-190036261-DUP-94
chr1-190136850-DUP-80
chr1-190150280-DUP-58
chr1-190406909-DUP-107
chr1-190512513-DUP-69
chr1-190513379-DUP-885
chr1-190564686-DUP-66
chr1-190583458-DUP-237
chr1-190604386-DUP-186
chr1-190634968-DUP-82
chr1-190689073-DUP-134
chr1-190783021-DUP-151
chr1-190783925-DUP-291
chr1-190954904-DUP-81
chr1-191132309-DUP-75
chr1-191183452-DUP-328
chr1-191224282-DUP-60
chr1-191224406-DUP-58
chr1-191240280-DUP-228
chr1-191287395-DUP-603
chr1-191490918-DUP-51
chr1-191693144-DUP-55
chr1-191799655-DUP-170
chr1-191880103-DUP-6832
chr1-191884842-DUP-119
chr1-191885020-DUP-119
chr1-191912582-DUP-307
chr1-191913273-DUP-341
chr1-191934091-DUP-66
chr1-192047570-DUP-118
chr1-192047603-DUP-92
chr1-192275621-DUP-54
chr1-192320772-DUP-98
chr1-192475753-DUP-59
chr1-192527892-DUP-106
chr1-193007613-DUP-64
chr1-193057215-DUP-65
chr1-193109853-DUP-158
chr1-193159703-DUP-65
chr1-193180171-DUP-345
chr1-193557706-DUP-57
chr1-193603834-DUP-52
chr1-194331585-DUP-79
chr1-194805605-DUP-53542
chr10-3061650-DUP-152
chr10-3062533-DUP-118
chr10-3063768-DUP-237
chr10-3064511-DUP-52
chr10-3066959-DUP-89
chr10-3092902-DUP-63
chr10-3536941-DUP-67
chr10-3658951-DUP-119
chr10-3703426-DUP-99
chr10-3719916-DUP-63
chr10-3816567-DUP-82
chr10-3952415-DUP-135
chr10-4031747-DUP-73
chr10-4255929-DUP-65
chr10-4500977-DUP-256
chr10-4501155-DUP-80
chr10-4668643-DUP-113
chr10-4843602-DUP-70
chr10-5019330-DUP-112
chr10-5019366-DUP-76
chr10-5489517-DUP-108
chr10-5489580-DUP-51
chr10-6464744-DUP-130
chr10-6661277-DUP-54
chr10-6743404-DUP-321
chr10-6830443-DUP-61
chr10-6985617-DUP-84
chr10-7098826-DUP-83

chr10-7099222-DUP-295
chr10-7123499-DUP-94
chr10-7207531-DUP-64
chr10-7224269-DUP-139
chr10-7224379-DUP-72
chr10-7312991-DUP-84
chr10-7321445-DUP-510
chr10-7405164-DUP-1994
chr10-7484379-DUP-72
chr10-7496518-DUP-99
chr10-7511552-DUP-538
chr10-7535041-DUP-60
chr10-7580786-DUP-133
chr10-7609045-DUP-226
chr10-7690552-DUP-65
chr10-7783802-DUP-62
chr10-8224549-DUP-145
chr10-8224579-DUP-74
chr10-8600277-DUP-86
chr10-8894000-DUP-106
chr10-9470347-DUP-112
chr10-9982970-DUP-57
chr10-10333849-DUP-124
chr10-10333908-DUP-55
chr10-10347118-DUP-58
chr10-10423598-DUP-159
chr10-10661900-DUP-52
chr10-10899250-DUP-72
chr10-10899981-DUP-104
chr10-10900086-DUP-538
chr10-10900622-DUP-141
chr10-10914228-DUP-97
chr10-11149736-DUP-325
chr10-11263817-DUP-85
chr10-11417633-DUP-139
chr10-11644744-DUP-63
chr10-12216683-DUP-355
chr10-12250894-DUP-68
chr10-12414539-DUP-75
chr10-12424155-DUP-192
chr10-12513208-DUP-72
chr10-12555254-DUP-138
chr10-13083978-DUP-94
chr10-13607421-DUP-59
chr10-13886159-DUP-89
chr10-13945299-DUP-50
chr10-14525799-DUP-64
chr10-14908816-DUP-66
chr10-15417464-DUP-55
chr10-15455100-DUP-3426
chr10-15535470-DUP-215
chr10-15535496-DUP-186
chr10-15915921-DUP-267
chr10-16216548-DUP-148
chr10-16403468-DUP-77
chr10-17590395-DUP-84
chr10-17952820-DUP-103
chr10-17976807-DUP-79
chr10-18167141-DUP-62
chr10-18370702-DUP-118
chr10-18737950-DUP-60
chr10-18793613-DUP-250
chr10-19580870-DUP-376
chr10-19637810-DUP-231
chr10-19905483-DUP-78
chr10-20062978-DUP-107
chr10-20112112-DUP-70
chr10-20304410-DUP-79
chr10-20314636-DUP-61
chr10-20353180-DUP-67
chr10-20396463-DUP-86
chr10-20596779-DUP-70
chr10-20790301-DUP-156
chr10-20836336-DUP-155
chr10-20890480-DUP-100
chr10-20928845-DUP-70
chr10-20952783-DUP-59

chr10-21330808-DUP-56
chr10-21334916-DUP-355
chr10-21891371-DUP-210
chr10-22059301-DUP-189212
chr10-22133975-DUP-55
chr10-22161621-DUP-28474
chr10-22346731-DUP-155
chr10-22693869-DUP-54
chr10-22791953-DUP-79
chr10-23219016-DUP-99
chr10-23460521-DUP-65
chr10-23728313-DUP-73
chr10-23824389-DUP-55
chr10-23840943-DUP-79
chr10-24614233-DUP-129
chr10-24753557-DUP-80
chr10-24755016-DUP-263
chr10-24845415-DUP-60
chr10-24883942-DUP-154
chr10-24968138-DUP-549
chr10-25134253-DUP-60
chr10-25322351-DUP-68
chr10-25334521-DUP-423
chr10-25538696-DUP-55
chr10-26308475-DUP-157593
chr10-26432660-DUP-96
chr10-26432689-DUP-176
chr10-26755495-DUP-56
chr10-27354433-DUP-236
chr10-27354531-DUP-172
chr10-28310188-DUP-227
chr10-28429583-DUP-61
chr10-28848832-DUP-56494
chr10-29163189-DUP-87
chr10-29424646-DUP-58
chr10-29640910-DUP-259
chr10-30066822-DUP-278
chr10-30463182-DUP-146
chr10-30596073-DUP-76
chr10-31714010-DUP-51
chr10-33318349-DUP-68
chr10-33340355-DUP-3738
chr10-33340488-DUP-1863
chr10-33342164-DUP-1928
chr10-34192175-DUP-171
chr10-34381377-DUP-5898
chr10-35454151-DUP-52
chr10-36856165-DUP-82
chr10-38656317-DUP-91
chr10-39015890-DUP-316
chr10-39281266-DUP-109
chr10-39281302-DUP-199
chr10-39519694-DUP-90
chr10-40712710-DUP-66
chr10-40973556-DUP-160
chr10-41068834-DUP-75
chr10-41100821-DUP-297
chr10-41101406-DUP-102
chr10-41101925-DUP-324
chr10-41298597-DUP-53
chr10-41605820-DUP-113
chr10-42052119-DUP-60
chr10-42102872-DUP-88
chr10-42199987-DUP-87
chr10-42302710-DUP-53
chr10-42305509-DUP-71
chr10-42451054-DUP-53
chr10-42531946-DUP-58
chr10-43493872-DUP-166
chr10-43652749-DUP-237
chr10-44505232-DUP-67
chr10-44677916-DUP-85
chr10-44986947-DUP-138
chr10-45167441-DUP-53
chr10-45318939-DUP-1814
chr10-45322575-DUP-110
chr10-45581551-DUP-125

chr10-50938549-DUP-3440
chr10-51941916-DUP-247
chr10-53048276-DUP-147187
chr10-53147450-DUP-281
chr10-53328052-DUP-145
chr10-53649019-DUP-51
chr10-53858906-DUP-65
chr10-54227596-DUP-105
chr10-54900439-DUP-3525
chr10-55802014-DUP-57
chr10-55978913-DUP-58
chr10-57551854-DUP-182
chr10-57551921-DUP-111
chr10-57576991-DUP-433
chr10-57977509-DUP-9818
chr10-57982430-DUP-4899
chr10-58056666-DUP-4881
chr10-58057567-DUP-4884
chr10-58209131-DUP-1448
chr10-58726893-DUP-73
chr10-58842045-DUP-54
chr10-59295548-DUP-63
chr10-59300115-DUP-374
chr10-59347647-DUP-68
chr10-59636863-DUP-67
chr10-59672735-DUP-153
chr10-60202607-DUP-83
chr10-60919635-DUP-439
chr10-60982942-DUP-66
chr10-61130486-DUP-84
chr10-61287342-DUP-87
chr10-61424860-DUP-145
chr10-61431672-DUP-55
chr10-61664059-DUP-59
chr10-61665045-DUP-97
chr10-61859265-DUP-92
chr10-61979206-DUP-104
chr10-61984424-DUP-169
chr10-61984448-DUP-69
chr10-62133219-DUP-246
chr10-62239123-DUP-58
chr10-62293814-DUP-104
chr10-62409539-DUP-110
chr10-63035214-DUP-1638
chr10-63176176-DUP-80
chr10-63428937-DUP-350
chr10-64242215-DUP-132
chr10-64242265-DUP-85
chr10-65109948-DUP-54
chr10-66135503-DUP-61
chr10-66347548-DUP-232
chr10-66347578-DUP-234
chr10-66412620-DUP-88
chr10-67372681-DUP-116
chr10-67605969-DUP-54
chr10-67626375-DUP-133
chr10-67869441-DUP-314
chr10-67869496-DUP-267
chr10-68204198-DUP-91
chr10-68451206-DUP-55
chr10-68549143-DUP-61
chr10-68697478-DUP-204
chr10-68740755-DUP-59
chr10-68799639-DUP-230
chr10-69690485-DUP-76
chr10-69720249-DUP-99
chr10-70072440-DUP-77
chr10-70147544-DUP-317
chr10-70261900-DUP-118
chr10-70782101-DUP-50
chr10-71035925-DUP-187
chr10-71048277-DUP-1711
chr10-71061229-DUP-97
chr10-71076853-DUP-58
chr10-71341127-DUP-90
chr10-73331411-DUP-71
chr10-73394245-DUP-127

chr10-74844027-DUP-76
chr10-74844107-DUP-74
chr10-74903599-DUP-163
chr10-75184669-DUP-69
chr10-75313199-DUP-59
chr10-75421654-DUP-53
chr10-75518272-DUP-97
chr10-75572861-DUP-54
chr10-75627406-DUP-53
chr10-75774846-DUP-75
chr10-76061555-DUP-96
chr10-76147007-DUP-1413
chr10-76166677-DUP-208
chr10-76272034-DUP-76
chr10-76320439-DUP-54
chr10-76387018-DUP-101
chr10-76519806-DUP-80
chr10-76834873-DUP-110
chr10-77018840-DUP-69
chr10-77037434-DUP-78
chr10-77039888-DUP-84
chr10-77352260-DUP-51
chr10-77444836-DUP-197
chr10-77828459-DUP-163
chr10-77986352-DUP-150
chr10-78288984-DUP-51
chr10-78348188-DUP-73392
chr10-79383888-DUP-145
chr10-79384080-DUP-68
chr10-79434449-DUP-224
chr10-79566186-DUP-91
chr10-80438452-DUP-59
chr10-80638307-DUP-140
chr10-80747899-DUP-59
chr10-80910356-DUP-121
chr10-81048108-DUP-134
chr10-81132812-DUP-95
chr10-81816592-DUP-59
chr10-82542457-DUP-78
chr10-82669534-DUP-53
chr10-82729667-DUP-52
chr10-82756597-DUP-154
chr10-82756665-DUP-52
chr10-82999637-DUP-54
chr10-83016772-DUP-65
chr10-83080064-DUP-56
chr10-83082700-DUP-102
chr10-83096128-DUP-122
chr10-83120963-DUP-86
chr10-83572224-DUP-72
chr10-83709041-DUP-334
chr10-84214353-DUP-51
chr10-84351896-DUP-50
chr10-84426487-DUP-58
chr10-84553384-DUP-132
chr10-84574349-DUP-236
chr10-84959906-DUP-55
chr10-85250671-DUP-219
chr10-85252124-DUP-98
chr10-85253590-DUP-219
chr10-85265019-DUP-68
chr10-85265046-DUP-70
chr10-85472525-DUP-73
chr10-85793889-DUP-344
chr10-85887153-DUP-84
chr10-86203236-DUP-88
chr10-86623866-DUP-68
chr10-86681266-DUP-334
chr10-86764742-DUP-397
chr10-87202697-DUP-51
chr10-88046331-DUP-328
chr10-88124116-DUP-211
chr10-88195508-DUP-83
chr10-88283843-DUP-57
chr10-88561635-DUP-4175
chr10-88610115-DUP-142
chr10-88617032-DUP-119

chr10-88617454-DUP-474
chr10-89092237-DUP-393
chr10-89453888-DUP-50
chr10-89506194-DUP-426
chr10-90140788-DUP-126
chr10-90260083-DUP-117
chr10-91339499-DUP-143
chr10-91434668-DUP-1086
chr10-91497295-DUP-3098
chr10-91956406-DUP-53
chr10-91962215-DUP-96
chr10-92481516-DUP-137
chr10-92705903-DUP-130
chr10-92805000-DUP-137
chr10-92861196-DUP-61
chr10-92887672-DUP-148
chr10-93909361-DUP-75
chr10-93909386-DUP-157
chr10-93995634-DUP-22611
chr10-94119490-DUP-76
chr10-94243464-DUP-91
chr10-94669898-DUP-62
chr10-94744727-DUP-61
chr10-94832074-DUP-298
chr10-95122156-DUP-67
chr10-95174523-DUP-121
chr10-95387064-DUP-60
chr10-96376467-DUP-264
chr10-96376622-DUP-128
chr10-97383084-DUP-280
chr10-97383230-DUP-109
chr10-99006837-DUP-150
chr10-99859480-DUP-111
chr10-100139210-DUP-803
chr10-100139775-DUP-110
chr10-100956667-DUP-52
chr10-101301162-DUP-57
chr10-101961494-DUP-244
chr10-102556495-DUP-99
chr10-105290115-DUP-58
chr10-105354760-DUP-78
chr10-105610652-DUP-57
chr10-106119853-DUP-60
chr10-106284231-DUP-7596
chr10-106552571-DUP-4871
chr10-106624285-DUP-102
chr10-106704645-DUP-121
chr10-107239936-DUP-60
chr10-108067590-DUP-111
chr10-109113422-DUP-139
chr10-110433604-DUP-83
chr10-110572389-DUP-74
chr10-111814720-DUP-157
chr10-112654975-DUP-81
chr10-113322117-DUP-66
chr10-113476432-DUP-402
chr10-113577123-DUP-61
chr10-113582133-DUP-2472
chr10-113662180-DUP-77
chr10-114729290-DUP-7781
chr10-115039621-DUP-54
chr10-115178772-DUP-61
chr10-115368365-DUP-95
chr10-115491177-DUP-112
chr10-116067194-DUP-147
chr10-116548058-DUP-104
chr10-116623631-DUP-72
chr10-116714301-DUP-74
chr10-116715092-DUP-138
chr10-116846105-DUP-116
chr10-116846186-DUP-59
chr10-116853382-DUP-163
chr10-116853408-DUP-165
chr10-116892589-DUP-90
chr10-117472498-DUP-296
chr10-117563656-DUP-91
chr10-117604979-DUP-300

chr10-117850546-DUP-91
chr10-117852546-DUP-127
chr10-117866295-DUP-2426
chr10-117884744-DUP-60
chr10-117894577-DUP-172
chr10-117989092-DUP-99
chr10-117989126-DUP-68
chr10-118567450-DUP-108
chr10-119378104-DUP-53
chr10-119583736-DUP-88
chr10-119583760-DUP-77
chr10-119695914-DUP-166
chr10-119749014-DUP-54
chr10-119856798-DUP-135
chr10-120052701-DUP-58
chr10-120058963-DUP-52
chr10-120109682-DUP-217
chr10-120202232-DUP-62
chr10-120417806-DUP-135
chr10-120417829-DUP-112
chr10-120427717-DUP-53
chr10-120438355-DUP-382
chr10-120654262-DUP-59
chr10-120656252-DUP-66
chr10-120737403-DUP-79
chr10-121020144-DUP-69
chr10-121414686-DUP-258
chr10-121415499-DUP-361
chr10-121435213-DUP-52
chr10-121567337-DUP-79
chr10-121733244-DUP-67
chr10-121802816-DUP-58
chr10-122295214-DUP-62
chr10-122295304-DUP-65
chr10-122408325-DUP-57
chr10-122533429-DUP-82
chr10-122679962-DUP-141
chr10-122882978-DUP-1246
chr10-122954798-DUP-70
chr10-122974208-DUP-92
chr10-122974230-DUP-94
chr10-123137484-DUP-58
chr10-123262186-DUP-275
chr10-123262238-DUP-218
chr10-123601442-DUP-69
chr10-123628292-DUP-204
chr10-123628335-DUP-70
chr10-124096852-DUP-70
chr10-124801027-DUP-78
chr10-124801073-DUP-156
chr10-124953729-DUP-117
chr10-125434621-DUP-362
chr10-125471678-DUP-516
chr10-125537104-DUP-72
chr10-125537125-DUP-61
chr10-125754955-DUP-1355
chr10-125754965-DUP-55
chr10-125755342-DUP-358
chr10-125755916-DUP-93
chr10-125975850-DUP-184
chr10-125975873-DUP-86
chr10-126313969-DUP-53
chr10-126333486-DUP-117
chr10-126498188-DUP-151
chr10-126586387-DUP-125
chr10-126722990-DUP-57
chr10-126819646-DUP-54
chr10-126976855-DUP-56
chr10-127070212-DUP-309
chr10-127070788-DUP-178
chr10-127071288-DUP-51
chr10-127135644-DUP-102
chr10-127283933-DUP-70
chr10-127329261-DUP-74
chr10-127329302-DUP-218
chr10-127395729-DUP-78
chr10-127503841-DUP-60

chr10-127552371-DUP-91
chr10-127578252-DUP-51
chr10-127753200-DUP-156
chr10-127753296-DUP-70
chr10-127774532-DUP-177
chr10-127878231-DUP-135
chr10-127932015-DUP-97
chr10-128069501-DUP-70
chr10-128072956-DUP-70
chr10-128217109-DUP-335
chr10-128877908-DUP-1417
chr10-128886484-DUP-59
chr11-3101763-DUP-248
chr11-3299678-DUP-242
chr11-3299754-DUP-88
chr11-3473857-DUP-295
chr11-3518454-DUP-105
chr11-3711230-DUP-60
chr11-3737287-DUP-103
chr11-3737330-DUP-105
chr11-3737897-DUP-120
chr11-3737922-DUP-97
chr11-4015478-DUP-89
chr11-4373003-DUP-62
chr11-4523401-DUP-660
chr11-4790779-DUP-114
chr11-4971088-DUP-558
chr11-5118237-DUP-2254
chr11-5118251-DUP-970
chr11-5118489-DUP-1023
chr11-5119082-DUP-140
chr11-5119493-DUP-224
chr11-5138947-DUP-55
chr11-5172191-DUP-78
chr11-5239476-DUP-66
chr11-5420910-DUP-243
chr11-5532136-DUP-70
chr11-5673029-DUP-97
chr11-5854541-DUP-55
chr11-5855906-DUP-68
chr11-5890217-DUP-115
chr11-5907344-DUP-90
chr11-5962595-DUP-175
chr11-6720159-DUP-53
chr11-6780720-DUP-58
chr11-6820042-DUP-107
chr11-6867688-DUP-74
chr11-7427259-DUP-78
chr11-7427314-DUP-80
chr11-7590888-DUP-88
chr11-8490275-DUP-90
chr11-8490298-DUP-92
chr11-8902831-DUP-56
chr11-9164941-DUP-69
chr11-9451681-DUP-93
chr11-9568513-DUP-60
chr11-10146890-DUP-232
chr11-10152478-DUP-70
chr11-10342452-DUP-55
chr11-10478129-DUP-52
chr11-11495295-DUP-63
chr11-11753711-DUP-901
chr11-12263492-DUP-348
chr11-12263619-DUP-232
chr11-12325023-DUP-189
chr11-12806313-DUP-7283
chr11-12807246-DUP-6348
chr11-13075339-DUP-54
chr11-13415789-DUP-51
chr11-17164569-DUP-138
chr11-17309570-DUP-51
chr11-17319444-DUP-71
chr11-17436234-DUP-547
chr11-17436281-DUP-454
chr11-17950114-DUP-118
chr11-17950236-DUP-55
chr11-18261238-DUP-2345

chr11-18514161-DUP-67
chr11-19129523-DUP-82
chr11-20315237-DUP-537
chr11-20380454-DUP-51
chr11-20414907-DUP-86
chr11-20624557-DUP-96
chr11-20784458-DUP-69
chr11-20859682-DUP-98
chr11-20879116-DUP-124
chr11-20883290-DUP-152
chr11-20980867-DUP-57
chr11-21220906-DUP-55
chr11-21768275-DUP-54
chr11-22177094-DUP-79
chr11-22625564-DUP-332
chr11-22796708-DUP-67
chr11-22907488-DUP-68
chr11-23521135-DUP-72
chr11-24107425-DUP-66
chr11-24926426-DUP-104
chr11-25235917-DUP-65
chr11-25330669-DUP-92
chr11-25583221-DUP-226
chr11-25691660-DUP-68
chr11-26019602-DUP-53
chr11-26157440-DUP-70
chr11-26289885-DUP-102
chr11-26363100-DUP-102
chr11-26363156-DUP-54
chr11-26538293-DUP-186
chr11-26668655-DUP-172
chr11-26904758-DUP-76
chr11-26924415-DUP-94
chr11-27814139-DUP-151
chr11-28440096-DUP-52
chr11-29104848-DUP-90
chr11-29732330-DUP-86
chr11-29878820-DUP-68
chr11-31899635-DUP-233
chr11-31986207-DUP-112
chr11-32145245-DUP-65
chr11-32145266-DUP-59
chr11-32222282-DUP-338
chr11-32534420-DUP-77
chr11-32577910-DUP-76
chr11-32647256-DUP-128
chr11-32647312-DUP-74
chr11-32716721-DUP-106
chr11-32792869-DUP-79
chr11-32863492-DUP-66
chr11-32986118-DUP-169
chr11-33105546-DUP-113
chr11-33233530-DUP-128
chr11-33451218-DUP-77
chr11-33511837-DUP-118
chr11-33555726-DUP-1378
chr11-33617068-DUP-159
chr11-33618640-DUP-100
chr11-33652109-DUP-74
chr11-33658902-DUP-123
chr11-33682648-DUP-52
chr11-33688315-DUP-79
chr11-33805521-DUP-54
chr11-33980094-DUP-501
chr11-34332713-DUP-251
chr11-34619976-DUP-93
chr11-34658156-DUP-76
chr11-34701628-DUP-182
chr11-34701650-DUP-172
chr11-34703320-DUP-156
chr11-34703491-DUP-488
chr11-35083709-DUP-109
chr11-35272567-DUP-51
chr11-35586156-DUP-83
chr11-35586291-DUP-115
chr11-35622674-DUP-542
chr11-35801927-DUP-53

chr11-35896399-DUP-133
chr11-36885723-DUP-117
chr11-37411372-DUP-63
chr11-37550378-DUP-56
chr11-37610454-DUP-145
chr11-38201532-DUP-58
chr11-38394927-DUP-55
chr11-39555834-DUP-74
chr11-39901615-DUP-192
chr11-40600783-DUP-123
chr11-40600938-DUP-63
chr11-41435421-DUP-130
chr11-41958431-DUP-164
chr11-42318523-DUP-100
chr11-43088068-DUP-66
chr11-43594316-DUP-70
chr11-43668747-DUP-145
chr11-44335297-DUP-205
chr11-44353501-DUP-74
chr11-45107718-DUP-73
chr11-45400056-DUP-157
chr11-45400086-DUP-159
chr11-45540469-DUP-72
chr11-45747418-DUP-58
chr11-45934343-DUP-300
chr11-46230295-DUP-114
chr11-46518295-DUP-22827
chr11-46672628-DUP-102
chr11-48309251-DUP-51
chr11-48355385-DUP-181
chr11-49000217-DUP-55
chr11-49164560-DUP-147
chr11-49319754-DUP-51
chr11-49414269-DUP-106171
chr11-49552814-DUP-180
chr11-49655965-DUP-52
chr11-49819521-DUP-75
chr11-49880349-DUP-109
chr11-49880373-DUP-56
chr11-49955406-DUP-60
chr11-50046182-DUP-94
chr11-50103316-DUP-55
chr11-50374296-DUP-221
chr11-50588672-DUP-51
chr11-50669762-DUP-105
chr11-51313043-DUP-52
chr11-51401470-DUP-62
chr11-51465341-DUP-107
chr11-51625183-DUP-127
chr11-51832853-DUP-64
chr11-51920921-DUP-198
chr11-52554175-DUP-107
chr11-53144670-DUP-58
chr11-53317452-DUP-54
chr11-53652595-DUP-57
chr11-53677514-DUP-107
chr11-53737444-DUP-227
chr11-54211450-DUP-62
chr11-54217304-DUP-466
chr11-54573423-DUP-748
chr11-54784160-DUP-249
chr11-55100479-DUP-116
chr11-55341551-DUP-67
chr11-55408575-DUP-52
chr11-55698584-DUP-264
chr11-55892285-DUP-167
chr11-55892307-DUP-118
chr11-56326120-DUP-56
chr11-56607147-DUP-66
chr11-56758211-DUP-54
chr11-57344711-DUP-174
chr11-57384299-DUP-72
chr11-57559113-DUP-51
chr11-57902755-DUP-67
chr11-58029546-DUP-121
chr11-58241782-DUP-53
chr11-58319654-DUP-102

chr11-58519496-DUP-111
chr11-58706891-DUP-65
chr11-58957406-DUP-53
chr11-59003366-DUP-3406
chr11-59259318-DUP-62
chr11-59264576-DUP-75
chr11-59359039-DUP-92
chr11-59835923-DUP-11444
chr11-59836025-DUP-11300
chr11-60247368-DUP-73
chr11-60267491-DUP-55
chr11-60360582-DUP-108
chr11-60479413-DUP-9108
chr11-60480242-DUP-9154
chr11-60516102-DUP-13380
chr11-60549015-DUP-22231
chr11-60692697-DUP-107
chr11-60725535-DUP-69
chr11-61299356-DUP-60
chr11-61548633-DUP-177
chr11-61657476-DUP-14353
chr11-61657518-DUP-14244
chr11-61721687-DUP-52
chr11-61773319-DUP-60
chr11-61809563-DUP-165
chr11-61911582-DUP-212
chr11-62051123-DUP-191
chr11-62390453-DUP-55
chr11-62480887-DUP-52
chr11-62706428-DUP-61
chr11-62733908-DUP-144
chr11-62872408-DUP-64
chr11-62899773-DUP-3266
chr11-62912199-DUP-113
chr11-63298011-DUP-106
chr11-63665451-DUP-199
chr11-63665473-DUP-123
chr11-63835981-DUP-65
chr11-63845182-DUP-81
chr11-63866713-DUP-149
chr11-64054338-DUP-104
chr11-64664946-DUP-63
chr11-64705291-DUP-72
chr11-64892697-DUP-73
chr11-64901518-DUP-55
chr11-65049921-DUP-93
chr11-65177765-DUP-52
chr11-65319829-DUP-79
chr11-65601499-DUP-54
chr11-65639502-DUP-58
chr11-65838063-DUP-128
chr11-66133021-DUP-59
chr11-66762822-DUP-114
chr11-66762862-DUP-74
chr11-66983592-DUP-267
chr11-67756169-DUP-56
chr11-67873510-DUP-69
chr11-68007754-DUP-60
chr11-68149110-DUP-527
chr11-68149153-DUP-479
chr11-68457325-DUP-63
chr11-68643209-DUP-1331
chr11-68678025-DUP-61
chr11-68876105-DUP-141
chr11-69049693-DUP-145
chr11-69049724-DUP-148
chr11-69104255-DUP-73
chr11-69209956-DUP-56
chr11-69210405-DUP-118
chr11-69340439-DUP-60
chr11-69353938-DUP-63
chr11-69756661-DUP-276
chr11-70243875-DUP-31191
chr11-70500650-DUP-77
chr11-70500673-DUP-67
chr11-70664453-DUP-1045
chr11-70682750-DUP-76

chr11-70806897-DUP-56
chr11-71455289-DUP-121
chr11-71580871-DUP-88
chr11-71600894-DUP-67
chr11-71736384-DUP-165
chr11-72140814-DUP-87
chr11-72140873-DUP-137
chr11-72449040-DUP-73
chr11-72550424-DUP-151
chr11-72655850-DUP-1533
chr11-72656955-DUP-561
chr11-72724872-DUP-53
chr11-72956691-DUP-207
chr11-73350524-DUP-326
chr11-73523283-DUP-57
chr11-73523314-DUP-121
chr11-74430331-DUP-132
chr11-74491665-DUP-70
chr11-75554064-DUP-1710
chr11-75554108-DUP-1812
chr11-75575420-DUP-155
chr11-75650598-DUP-191
chr11-75756354-DUP-147
chr11-75896459-DUP-211
chr11-75896495-DUP-163
chr11-76008173-DUP-62
chr11-76085199-DUP-261
chr11-76187276-DUP-464
chr11-76219824-DUP-159
chr11-76310813-DUP-62
chr11-76311626-DUP-82
chr11-76353441-DUP-109
chr11-76460699-DUP-58
chr11-76521027-DUP-92
chr11-76579744-DUP-76
chr11-77028569-DUP-117
chr11-77028630-DUP-60
chr11-77503283-DUP-161
chr11-77637082-DUP-52
chr11-77669540-DUP-52
chr11-77779357-DUP-1309
chr11-78591041-DUP-102
chr11-78591086-DUP-63
chr11-78817864-DUP-134
chr11-78899348-DUP-108
chr11-78933121-DUP-54
chr11-78988642-DUP-100
chr11-79310456-DUP-77
chr11-79607172-DUP-65
chr11-79682167-DUP-427
chr11-79689103-DUP-292
chr11-79842778-DUP-385
chr11-79842802-DUP-337
chr11-80176175-DUP-168
chr11-81427226-DUP-74
chr11-81452190-DUP-290
chr11-81452229-DUP-97
chr11-81458484-DUP-211
chr11-81590493-DUP-54
chr11-82821098-DUP-82
chr11-82822115-DUP-221
chr11-82823087-DUP-101
chr11-83193308-DUP-289
chr11-83272801-DUP-51
chr11-83298108-DUP-98
chr11-83489239-DUP-56
chr11-83577929-DUP-16535
chr11-83591836-DUP-3505
chr11-83633861-DUP-72
chr11-83976645-DUP-52
chr11-84302106-DUP-90
chr11-84927024-DUP-53
chr11-85409817-DUP-88
chr11-85459966-DUP-59
chr11-85484035-DUP-69
chr11-85525786-DUP-151
chr11-85525878-DUP-122

chr11-85879617-DUP-92
chr11-86125144-DUP-62
chr11-86861507-DUP-116
chr11-87353406-DUP-9083
chr11-87413380-DUP-72
chr11-87693069-DUP-181
chr11-87717049-DUP-166
chr11-87935018-DUP-180
chr11-88197304-DUP-250
chr11-88741184-DUP-94
chr11-88889467-DUP-86
chr11-88920761-DUP-3516
chr11-88920785-DUP-3426
chr11-88934251-DUP-63
chr11-88940228-DUP-83
chr11-88940251-DUP-64
chr11-88970745-DUP-63
chr11-89442646-DUP-144
chr11-89697186-DUP-54
chr11-90225819-DUP-66
chr11-90225851-DUP-79
chr11-90316394-DUP-102
chr11-90426481-DUP-56
chr11-90928966-DUP-104
chr11-91695086-DUP-392
chr11-91695246-DUP-362
chr11-93283243-DUP-59
chr11-93452354-DUP-83
chr11-93507252-DUP-70
chr11-93545497-DUP-56
chr11-93774924-DUP-25749
chr11-93917661-DUP-547
chr11-93994581-DUP-138
chr11-93994619-DUP-84
chr11-94177123-DUP-71
chr11-94426930-DUP-62
chr11-94430679-DUP-57
chr11-94638180-DUP-97
chr11-95097970-DUP-52
chr11-95452000-DUP-153
chr11-95863068-DUP-248
chr11-95990842-DUP-103
chr11-96007738-DUP-1887
chr11-96007819-DUP-1719
chr11-96126013-DUP-54
chr11-96400219-DUP-59
chr11-96404626-DUP-80
chr11-96831542-DUP-65
chr11-97225110-DUP-187
chr11-97266992-DUP-60
chr11-97345667-DUP-74
chr11-97481689-DUP-53
chr11-97510499-DUP-76
chr11-98192672-DUP-90
chr11-98226696-DUP-104
chr11-98480871-DUP-3199
chr11-98486756-DUP-70
chr11-98496326-DUP-35301
chr11-98679104-DUP-84
chr11-98934252-DUP-93
chr11-99157039-DUP-117
chr11-99166981-DUP-174
chr11-99276823-DUP-58
chr11-99500214-DUP-438
chr11-99549563-DUP-75
chr11-99564024-DUP-26201
chr11-99575616-DUP-40587
chr11-99683648-DUP-73
chr11-99705987-DUP-103
chr11-99839625-DUP-199
chr11-99890226-DUP-95
chr11-100051954-DUP-129
chr11-100145725-DUP-53
chr11-100293868-DUP-66
chr11-100450115-DUP-249
chr11-100715225-DUP-61
chr11-100748281-DUP-101

chr11-100748342-DUP-103
chr11-100809710-DUP-60
chr11-101016757-DUP-58
chr11-101290784-DUP-2586
chr11-101290941-DUP-2591
chr11-102738469-DUP-55
chr11-102738510-DUP-54
chr11-102763533-DUP-292
chr11-102999335-DUP-60
chr11-103277427-DUP-65
chr11-103610023-DUP-53
chr11-103656945-DUP-182
chr11-103734897-DUP-70
chr11-104012221-DUP-52
chr11-104095433-DUP-60
chr11-104261638-DUP-154
chr11-104261681-DUP-119
chr11-104621307-DUP-130
chr11-104705572-DUP-408
chr11-104978627-DUP-122
chr11-104978698-DUP-64
chr11-105061431-DUP-166
chr11-105225493-DUP-102
chr11-105316031-DUP-54
chr11-105508093-DUP-127
chr11-106398634-DUP-53
chr11-106697403-DUP-250
chr11-106724892-DUP-75
chr11-106859040-DUP-55
chr11-106939513-DUP-103
chr11-106939662-DUP-576
chr11-107027807-DUP-3396
chr11-107030422-DUP-7064
chr11-107030491-DUP-6945
chr11-107390501-DUP-56
chr11-107451259-DUP-57
chr11-107656840-DUP-61
chr11-107751339-DUP-109633
chr11-107797683-DUP-160
chr11-107819915-DUP-291
chr11-107983242-DUP-114
chr11-108210223-DUP-68
chr11-108333419-DUP-51
chr11-108416459-DUP-97
chr11-108497738-DUP-76
chr11-108621796-DUP-91
chr11-108824531-DUP-63
chr11-109248385-DUP-72
chr11-109409164-DUP-181
chr11-109464588-DUP-54
chr11-109496651-DUP-53
chr11-109923348-DUP-245
chr11-110092842-DUP-67
chr11-110195362-DUP-87
chr11-110308153-DUP-92
chr11-110344543-DUP-70
chr11-110392191-DUP-78395
chr11-110861695-DUP-152
chr11-110937352-DUP-97
chr11-111509845-DUP-66
chr11-112589374-DUP-138
chr11-113060461-DUP-92
chr11-113064437-DUP-271
chr11-113312943-DUP-55
chr11-113391966-DUP-111
chr11-113434687-DUP-59
chr11-113551867-DUP-51
chr11-113582221-DUP-77
chr11-113582272-DUP-79
chr11-114018141-DUP-63
chr11-114109871-DUP-98
chr11-114318866-DUP-89
chr11-114421901-DUP-157
chr11-114593538-DUP-199
chr11-114593569-DUP-193
chr11-114645822-DUP-78
chr11-115210540-DUP-89

chr11-115368625-DUP-211
chr11-115368665-DUP-213
chr11-115431732-DUP-204
chr11-115448313-DUP-2998
chr11-115448384-DUP-2904
chr11-115725408-DUP-63
chr11-116053627-DUP-69
chr11-116053653-DUP-72
chr11-116178010-DUP-123
chr11-116190372-DUP-150
chr11-116348178-DUP-118
chr11-116416344-DUP-159
chr11-116460607-DUP-59
chr11-116665799-DUP-60
chr11-116859881-DUP-110
chr11-116957915-DUP-1242
chr11-116958163-DUP-181
chr11-116958813-DUP-227
chr11-117707062-DUP-130
chr11-117771350-DUP-61
chr11-117791677-DUP-102
chr11-117812883-DUP-65
chr11-117970708-DUP-75
chr11-118082039-DUP-277
chr11-118165019-DUP-54
chr11-118226206-DUP-72
chr11-118236216-DUP-58
chr11-118267304-DUP-139
chr11-118451539-DUP-81
chr11-119044272-DUP-113
chr11-119173405-DUP-264
chr11-119238302-DUP-108
chr11-119251669-DUP-111
chr11-119281696-DUP-104
chr11-119281717-DUP-106
chr11-119319746-DUP-59
chr11-119473850-DUP-300
chr11-119606762-DUP-51
chr11-119654288-DUP-120
chr11-119654359-DUP-172
chr11-119669900-DUP-152
chr11-119961527-DUP-74
chr11-120075848-DUP-68
chr11-120578792-DUP-76
chr11-120598446-DUP-59
chr11-120745903-DUP-303
chr11-120746685-DUP-138
chr11-121010547-DUP-3338
chr11-121072630-DUP-87
chr11-121072656-DUP-63
chr11-121085418-DUP-92
chr11-121123539-DUP-12453
chr11-121260518-DUP-71852
chr11-121551524-DUP-78
chr11-121596252-DUP-66
chr12-3099638-DUP-92
chr12-3142862-DUP-58
chr12-3284491-DUP-172
chr12-3446213-DUP-364
chr12-3543825-DUP-91
chr12-3993201-DUP-68
chr12-4023336-DUP-129
chr12-4057302-DUP-80
chr12-4057326-DUP-82
chr12-4159420-DUP-72
chr12-4160946-DUP-1388
chr12-4161018-DUP-462
chr12-4161402-DUP-114
chr12-4161593-DUP-764
chr12-4212380-DUP-226
chr12-4235180-DUP-112
chr12-4237302-DUP-240
chr12-4254475-DUP-78
chr12-4269320-DUP-1662
chr12-4269548-DUP-1265
chr12-4269694-DUP-2010
chr12-4269959-DUP-231

chr12-4270216-DUP-1856
chr12-4271216-DUP-830
chr12-4398618-DUP-93
chr12-4398653-DUP-51
chr12-4415213-DUP-15130
chr12-4442027-DUP-112
chr12-4444064-DUP-56
chr12-4487466-DUP-142
chr12-4495713-DUP-71
chr12-4500343-DUP-152
chr12-4516653-DUP-168
chr12-4517095-DUP-780
chr12-4517323-DUP-167
chr12-4660323-DUP-98
chr12-5098359-DUP-68
chr12-5105821-DUP-495
chr12-5106973-DUP-990
chr12-5401345-DUP-109
chr12-5441440-DUP-78
chr12-5462639-DUP-52
chr12-5556485-DUP-98
chr12-5741891-DUP-64
chr12-7715035-DUP-166
chr12-7715098-DUP-56
chr12-7971608-DUP-62
chr12-8275246-DUP-104
chr12-8891044-DUP-92
chr12-8990445-DUP-97
chr12-9018016-DUP-639
chr12-9280322-DUP-85
chr12-9476201-DUP-140
chr12-9959255-DUP-266
chr12-10421293-DUP-57
chr12-10607290-DUP-200
chr12-10662572-DUP-107
chr12-11272269-DUP-1639
chr12-12061993-DUP-187
chr12-12081507-DUP-264
chr12-12081529-DUP-258
chr12-12432854-DUP-56
chr12-12558856-DUP-85
chr12-13323468-DUP-72
chr12-13613377-DUP-74
chr12-13617573-DUP-78
chr12-13677045-DUP-4019
chr12-14387881-DUP-57
chr12-14454400-DUP-110
chr12-15272917-DUP-71
chr12-15688230-DUP-65
chr12-16349397-DUP-148
chr12-16624262-DUP-60
chr12-16686879-DUP-203
chr12-16687027-DUP-104
chr12-16707383-DUP-77
chr12-17724794-DUP-84
chr12-17936534-DUP-57
chr12-17936566-DUP-227
chr12-18625637-DUP-74
chr12-18625666-DUP-152
chr12-19908215-DUP-945
chr12-20251288-DUP-112
chr12-20251327-DUP-97
chr12-20471192-DUP-112
chr12-20669772-DUP-105
chr12-21045519-DUP-164
chr12-21265147-DUP-132
chr12-21265175-DUP-135
chr12-21455420-DUP-121
chr12-21823190-DUP-58
chr12-21943772-DUP-60
chr12-22076012-DUP-227
chr12-22790823-DUP-56
chr12-23234905-DUP-169
chr12-23846067-DUP-110
chr12-24014223-DUP-108
chr12-24142287-DUP-87
chr12-24405652-DUP-169

chr12-24769955-DUP-135
chr12-24810128-DUP-93
chr12-24943642-DUP-54
chr12-24947923-DUP-153
chr12-25254823-DUP-72
chr12-25387442-DUP-304
chr12-25464610-DUP-66
chr12-25860144-DUP-75
chr12-26489214-DUP-62
chr12-26585667-DUP-61
chr12-26587347-DUP-69
chr12-26844613-DUP-57
chr12-26858270-DUP-250
chr12-27069544-DUP-66
chr12-27183019-DUP-112
chr12-27309618-DUP-84
chr12-28604840-DUP-174
chr12-28604894-DUP-149
chr12-28744382-DUP-78
chr12-28891916-DUP-55
chr12-28917614-DUP-240
chr12-28924268-DUP-105
chr12-29098207-DUP-68
chr12-29287949-DUP-57
chr12-29364462-DUP-85
chr12-29671776-DUP-64
chr12-29737920-DUP-61
chr12-29829768-DUP-53
chr12-29899533-DUP-69
chr12-30015309-DUP-176
chr12-30296660-DUP-51
chr12-30523993-DUP-61
chr12-30631431-DUP-64
chr12-31121279-DUP-2648
chr12-31352598-DUP-10946
chr12-31426257-DUP-70
chr12-31439033-DUP-120
chr12-31453184-DUP-71
chr12-31875955-DUP-179
chr12-32014224-DUP-56
chr12-32059899-DUP-239
chr12-32206581-DUP-119
chr12-32293251-DUP-416
chr12-32636027-DUP-332
chr12-33110815-DUP-103
chr12-33303544-DUP-71
chr12-33431078-DUP-200
chr12-33431107-DUP-77
chr12-33607937-DUP-396
chr12-34680452-DUP-192
chr12-34729870-DUP-94
chr12-34948601-DUP-214
chr12-35053712-DUP-61
chr12-35598928-DUP-64
chr12-36340311-DUP-69
chr12-36410428-DUP-75
chr12-36561637-DUP-88
chr12-36649832-DUP-95
chr12-37030204-DUP-4126
chr12-37221052-DUP-154780
chr12-38306412-DUP-796
chr12-38880709-DUP-57
chr12-39982007-DUP-73
chr12-40505940-DUP-80
chr12-41394591-DUP-94
chr12-41801743-DUP-112
chr12-41801775-DUP-64
chr12-41914954-DUP-2115
chr12-42098184-DUP-397
chr12-43700752-DUP-71
chr12-44263841-DUP-66
chr12-45449210-DUP-515
chr12-45646847-DUP-51
chr12-45654212-DUP-151
chr12-48807926-DUP-114
chr12-48819412-DUP-67
chr12-49503114-DUP-55

chr12-49766504-DUP-83
chr12-50059967-DUP-162
chr12-50582352-DUP-229
chr12-50781128-DUP-86
chr12-52189318-DUP-318
chr12-52380537-DUP-64
chr12-52380656-DUP-124
chr12-52438398-DUP-75
chr12-52770057-DUP-53
chr12-52907770-DUP-99
chr12-52910299-DUP-202
chr12-52953572-DUP-119
chr12-52973664-DUP-112
chr12-52974377-DUP-82
chr12-52975927-DUP-207
chr12-52977272-DUP-54
chr12-52977559-DUP-594
chr12-52977673-DUP-1024
chr12-52978324-DUP-539
chr12-52978869-DUP-83
chr12-53077483-DUP-231
chr12-53273854-DUP-107
chr12-53273879-DUP-143
chr12-53306492-DUP-56
chr12-53847845-DUP-115
chr12-54220400-DUP-319
chr12-54361840-DUP-53
chr12-54373501-DUP-91
chr12-54728056-DUP-2060
chr12-54728119-DUP-753
chr12-54728959-DUP-417
chr12-54729463-DUP-628
chr12-54807103-DUP-57
chr12-55049505-DUP-2793
chr12-55125488-DUP-226677
chr12-55585207-DUP-404
chr12-55585244-DUP-381
chr12-55713270-DUP-65
chr12-55912369-DUP-92
chr12-55955758-DUP-338
chr12-56468406-DUP-61
chr12-56469730-DUP-137
chr12-56469761-DUP-63
chr12-57140804-DUP-51
chr12-57244574-DUP-139
chr12-57608577-DUP-1875
chr12-57998241-DUP-52
chr12-58080933-DUP-1927
chr12-58266028-DUP-54
chr12-59159686-DUP-1230
chr12-59197757-DUP-276
chr12-59247014-DUP-321
chr12-59955744-DUP-75
chr12-60177313-DUP-2768
chr12-60978138-DUP-277
chr12-61614943-DUP-128
chr12-61632490-DUP-65
chr12-63186299-DUP-70
chr12-64072991-DUP-55
chr12-64073023-DUP-51
chr12-64433959-DUP-74
chr12-65034431-DUP-51
chr12-66287687-DUP-194
chr12-66297596-DUP-225
chr12-66638613-DUP-5194
chr12-66957402-DUP-683
chr12-67064519-DUP-55
chr12-67322008-DUP-197
chr12-67322039-DUP-99
chr12-67542413-DUP-56
chr12-67542439-DUP-54
chr12-67884726-DUP-54
chr12-68145875-DUP-59
chr12-69015468-DUP-120
chr12-69015563-DUP-58
chr12-69489974-DUP-320
chr12-69633926-DUP-62

chr12-69659415-DUP-106
chr12-69671018-DUP-902
chr12-69830388-DUP-102
chr12-69916306-DUP-58
chr12-69960118-DUP-3204
chr12-69979229-DUP-77
chr12-70003197-DUP-184
chr12-70211760-DUP-55
chr12-70318491-DUP-72
chr12-70693628-DUP-223
chr12-70787125-DUP-80
chr12-71138419-DUP-99
chr12-71138948-DUP-315
chr12-71472294-DUP-221
chr12-71689289-DUP-63
chr12-71718032-DUP-80
chr12-72060816-DUP-72
chr12-72380390-DUP-67
chr12-72380418-DUP-69
chr12-72682653-DUP-113
chr12-72751149-DUP-80
chr12-72751197-DUP-78
chr12-72802942-DUP-110
chr12-73192737-DUP-70
chr12-73338479-DUP-59
chr12-73575012-DUP-168
chr12-73601408-DUP-67
chr12-73771718-DUP-52
chr12-74274605-DUP-274
chr12-74772949-DUP-158
chr12-75186910-DUP-55
chr12-75403216-DUP-80
chr12-75896134-DUP-198
chr12-76040402-DUP-71
chr12-76211407-DUP-114
chr12-77138193-DUP-72
chr12-77218664-DUP-62
chr12-77243827-DUP-72
chr12-77379717-DUP-70
chr12-77700158-DUP-76
chr12-78736802-DUP-108
chr12-78979782-DUP-57
chr12-79106290-DUP-53
chr12-79161080-DUP-2073
chr12-79181089-DUP-142
chr12-79289937-DUP-56
chr12-79310284-DUP-90
chr12-79481127-DUP-111
chr12-80137248-DUP-65
chr12-81118693-DUP-120
chr12-81118741-DUP-78
chr12-81530575-DUP-58
chr12-81855749-DUP-178
chr12-81855843-DUP-54
chr12-81903059-DUP-76
chr12-81932339-DUP-175
chr12-82032349-DUP-180
chr12-82032443-DUP-238
chr12-82241055-DUP-76
chr12-82241089-DUP-78
chr12-82328090-DUP-1858
chr12-82331901-DUP-132
chr12-82364736-DUP-433
chr12-82365461-DUP-349
chr12-82606425-DUP-410
chr12-82691202-DUP-62
chr12-82694291-DUP-111
chr12-82894096-DUP-140
chr12-82894123-DUP-138
chr12-82974824-DUP-77
chr12-82974921-DUP-74
chr12-83223771-DUP-194
chr12-83244913-DUP-60
chr12-83774403-DUP-100
chr12-84027318-DUP-62
chr12-85263533-DUP-154
chr12-86268147-DUP-66

chr12-86491087-DUP-195
chr12-86549787-DUP-56
chr12-86812179-DUP-212
chr12-86812316-DUP-168
chr12-87147352-DUP-73
chr12-87502275-DUP-56
chr12-87531359-DUP-102
chr12-87832207-DUP-101
chr12-87994165-DUP-62
chr12-88249802-DUP-50
chr12-89471242-DUP-119
chr12-90203847-DUP-217
chr12-91114573-DUP-215
chr12-91201942-DUP-358
chr12-91260028-DUP-238
chr12-91550324-DUP-56
chr12-91683878-DUP-105
chr12-91931729-DUP-180
chr12-91979713-DUP-61
chr12-93381203-DUP-6372
chr12-94431127-DUP-111
chr12-95533875-DUP-5425
chr12-95725137-DUP-82
chr12-96591467-DUP-57
chr12-96896783-DUP-10939
chr12-98379610-DUP-75
chr12-98993997-DUP-72
chr12-98994794-DUP-64
chr12-99471382-DUP-435
chr12-99474220-DUP-210
chr12-99474720-DUP-119
chr12-99671566-DUP-293
chr12-99770359-DUP-66
chr12-99773566-DUP-128
chr12-99975667-DUP-124
chr12-100449476-DUP-60
chr12-100456967-DUP-293
chr12-100457126-DUP-756
chr12-100457659-DUP-325
chr12-100812515-DUP-55
chr12-100933776-DUP-55
chr12-101449315-DUP-77
chr12-101806244-DUP-345
chr12-101806933-DUP-267
chr12-101807577-DUP-175
chr12-101843377-DUP-283
chr12-101843452-DUP-157
chr12-101858764-DUP-111
chr12-102158509-DUP-75
chr12-102188050-DUP-66
chr12-102533558-DUP-112
chr12-102625430-DUP-155
chr12-102976739-DUP-215
chr12-104086348-DUP-108
chr12-104188013-DUP-321
chr12-104281509-DUP-57
chr12-104371641-DUP-135
chr12-104371670-DUP-167
chr12-104478331-DUP-85
chr12-104501009-DUP-56
chr12-104527097-DUP-62
chr12-104611102-DUP-142
chr12-104634262-DUP-51
chr12-104698303-DUP-83
chr12-104720417-DUP-61
chr12-104786494-DUP-114
chr12-104786520-DUP-68
chr12-104877470-DUP-166
chr12-105043435-DUP-115
chr12-105290960-DUP-51
chr12-105391494-DUP-52
chr12-105516264-DUP-59
chr12-105621848-DUP-196
chr12-105638502-DUP-145
chr12-105920024-DUP-333
chr12-105925705-DUP-232
chr12-105925743-DUP-198

chr12-105979368-DUP-56
chr12-105984139-DUP-80
chr12-106036251-DUP-56
chr12-106269804-DUP-163
chr12-106350194-DUP-54
chr12-106897131-DUP-69
chr12-107745556-DUP-64
chr12-107758572-DUP-521
chr12-107764081-DUP-53
chr12-107764176-DUP-95
chr12-108258781-DUP-114
chr12-108304062-DUP-59
chr12-108315584-DUP-412
chr12-108648628-DUP-55
chr12-108664288-DUP-94
chr12-108810263-DUP-13316
chr12-108875444-DUP-94
chr12-108898990-DUP-213
chr12-108932227-DUP-392
chr12-109415752-DUP-140
chr12-109540677-DUP-139
chr12-109540723-DUP-284
chr12-109730448-DUP-221
chr12-109730478-DUP-96
chr12-109977263-DUP-96
chr12-109979878-DUP-57
chr12-110020974-DUP-60
chr12-110116954-DUP-60
chr12-110335142-DUP-543
chr12-110335270-DUP-405
chr12-110335455-DUP-1187
chr12-110336268-DUP-269
chr12-110336986-DUP-266
chr12-110468254-DUP-97
chr12-110468292-DUP-100
chr12-110784679-DUP-154
chr12-110784745-DUP-156
chr12-111107056-DUP-55
chr12-111165485-DUP-109
chr12-111165535-DUP-87
chr12-111258159-DUP-65
chr12-111278987-DUP-110
chr12-111295831-DUP-173
chr12-111314680-DUP-141
chr12-111377174-DUP-271
chr12-111377287-DUP-213
chr12-111759424-DUP-19506
chr12-111816409-DUP-286
chr12-111956060-DUP-77
chr12-111990121-DUP-64
chr12-112002119-DUP-146
chr12-112002265-DUP-102
chr12-112014209-DUP-257
chr12-112128600-DUP-59
chr12-112146600-DUP-72
chr12-112260819-DUP-60
chr12-112263771-DUP-404
chr12-112292208-DUP-239
chr12-112292245-DUP-162
chr12-112347216-DUP-67
chr12-112374194-DUP-158
chr12-112374229-DUP-160
chr12-112411319-DUP-74
chr12-112513558-DUP-347
chr12-112514432-DUP-274
chr12-112573168-DUP-91
chr12-112679875-DUP-59
chr12-112710711-DUP-78
chr12-112736955-DUP-68
chr12-112742279-DUP-2076
chr12-112742280-DUP-6147
chr12-112742722-DUP-516
chr12-112743715-DUP-4665
chr12-112744476-DUP-516
chr12-112744798-DUP-3630
chr12-112745836-DUP-2592
chr12-112747707-DUP-563

chr12-112757204-DUP-80
chr12-112819377-DUP-116
chr12-112949667-DUP-212
chr12-113159063-DUP-63
chr12-113197548-DUP-6243
chr12-113257968-DUP-280
chr12-113274116-DUP-341
chr12-113298601-DUP-1074
chr12-113298642-DUP-271
chr12-113299706-DUP-143
chr12-113327174-DUP-1221
chr12-113427675-DUP-9675
chr12-113660690-DUP-85
chr12-114410125-DUP-170
chr12-114436771-DUP-43415
chr12-114703896-DUP-109076
chr12-114704710-DUP-5866
chr12-114807139-DUP-5833
chr12-115026693-DUP-396296
chr12-115137882-DUP-557
chr12-115139000-DUP-110
chr12-115422164-DUP-315
chr12-115565277-DUP-188219
chr12-115677359-DUP-361
chr12-115719123-DUP-315
chr12-115753232-DUP-5861
chr12-116167143-DUP-77
chr12-116177464-DUP-63
chr12-116205682-DUP-615
chr12-116212654-DUP-110
chr12-116213510-DUP-72
chr12-116270221-DUP-74
chr12-116338091-DUP-95
chr12-116622574-DUP-227
chr12-116641861-DUP-214
chr12-116970922-DUP-78
chr12-116971382-DUP-147
chr12-116984031-DUP-97
chr12-117059181-DUP-132
chr12-117059218-DUP-100
chr12-117103696-DUP-52
chr12-117403664-DUP-98
chr12-117414548-DUP-51
chr12-117551994-DUP-318
chr12-118072252-DUP-118
chr12-118462043-DUP-105
chr12-119010651-DUP-64
chr12-119153279-DUP-88
chr12-119220555-DUP-171
chr13-3089980-DUP-115
chr13-3117664-DUP-79
chr13-3237246-DUP-94
chr13-3931857-DUP-70
chr13-3951276-DUP-1175
chr13-3951812-DUP-204
chr13-3952331-DUP-232
chr13-4828405-DUP-170
chr13-5693498-DUP-2183
chr13-5844217-DUP-104
chr13-6099217-DUP-71
chr13-6185343-DUP-248
chr13-6310482-DUP-54
chr13-6446858-DUP-60
chr13-6550338-DUP-163
chr13-6658497-DUP-117
chr13-6660518-DUP-118
chr13-6661827-DUP-957
chr13-6662141-DUP-117
chr13-7031510-DUP-146
chr13-7031663-DUP-62
chr13-7154218-DUP-226
chr13-7362871-DUP-71
chr13-8375573-DUP-53
chr13-8879538-DUP-68
chr13-8910783-DUP-67
chr13-9164059-DUP-212
chr13-9201832-DUP-150

chr13-9377213-DUP-243
chr13-9403699-DUP-3970
chr13-9703054-DUP-75
chr13-10431767-DUP-55
chr13-10892751-DUP-117
chr13-11039692-DUP-55
chr13-11459748-DUP-135
chr13-11748411-DUP-309
chr13-11748451-DUP-60
chr13-11785831-DUP-53
chr13-11974529-DUP-54
chr13-11974558-DUP-51
chr13-12220500-DUP-55
chr13-12370414-DUP-58
chr13-12417811-DUP-97
chr13-12528634-DUP-60
chr13-12575641-DUP-143
chr13-12612368-DUP-107
chr13-12639786-DUP-64244
chr13-12652811-DUP-198
chr13-13927522-DUP-2620
chr13-13927631-DUP-2483
chr13-13927871-DUP-66
chr13-13928983-DUP-439
chr13-14031908-DUP-62
chr13-14332511-DUP-72
chr13-15913398-DUP-1129
chr13-16150776-DUP-62
chr13-16981342-DUP-58
chr13-16982068-DUP-178
chr13-17170872-DUP-333
chr13-17524537-DUP-55
chr13-17928517-DUP-61
chr13-18185998-DUP-78
chr13-18186057-DUP-75
chr13-18338903-DUP-132
chr13-18762146-DUP-60
chr13-18953760-DUP-341
chr13-19216009-DUP-146
chr13-20064953-DUP-148
chr13-20073720-DUP-71
chr13-20101600-DUP-110
chr13-20102135-DUP-73
chr13-20329019-DUP-202
chr13-20420675-DUP-302
chr13-20661901-DUP-258
chr13-21413400-DUP-171
chr13-22280294-DUP-284
chr13-23225244-DUP-154
chr13-23225835-DUP-348
chr13-23407945-DUP-127
chr13-23408379-DUP-288
chr13-23584992-DUP-12175
chr13-24071599-DUP-97
chr13-24288722-DUP-288
chr13-24339894-DUP-129
chr13-24579156-DUP-82
chr13-24641866-DUP-53
chr13-25075116-DUP-193
chr13-25249914-DUP-102
chr13-25275267-DUP-52
chr13-25352540-DUP-82
chr13-25362440-DUP-82
chr13-25364699-DUP-91
chr13-25380529-DUP-385
chr13-25385722-DUP-61
chr13-26251284-DUP-85
chr13-26447658-DUP-4321
chr13-27387374-DUP-63
chr13-28977418-DUP-63
chr13-29678833-DUP-111
chr13-29760237-DUP-150
chr13-29797392-DUP-53
chr13-29934534-DUP-309
chr13-30434004-DUP-231
chr13-30434102-DUP-68
chr13-30518942-DUP-54

chr13-30524444-DUP-70
chr13-30680526-DUP-445
chr13-30680565-DUP-397
chr13-30971689-DUP-90
chr13-30997643-DUP-108
chr13-30997675-DUP-56
chr13-31069293-DUP-78
chr13-31069319-DUP-133
chr13-31175786-DUP-57
chr13-31252976-DUP-68
chr13-31438561-DUP-70
chr13-31576829-DUP-78
chr13-31608659-DUP-134
chr13-31803504-DUP-67
chr13-32250866-DUP-61
chr13-34286928-DUP-1098
chr13-34287469-DUP-558
chr13-35125309-DUP-508
chr13-35156836-DUP-50893
chr13-35379465-DUP-82
chr13-35393902-DUP-80
chr13-36217370-DUP-54
chr13-36318826-DUP-64
chr13-36757355-DUP-53
chr13-37149095-DUP-58
chr13-37332821-DUP-83
chr13-37666466-DUP-140
chr13-37697709-DUP-131
chr13-37698446-DUP-53
chr13-38043269-DUP-116
chr13-38148453-DUP-62
chr13-38173719-DUP-60
chr13-38241280-DUP-87
chr13-38802016-DUP-202
chr13-39373507-DUP-86
chr13-40327048-DUP-136
chr13-40359359-DUP-180
chr13-40520669-DUP-69
chr13-40539629-DUP-126
chr13-40888751-DUP-51
chr13-41346596-DUP-238
chr13-41346617-DUP-212
chr13-41758000-DUP-55
chr13-41868412-DUP-60
chr13-42105171-DUP-69
chr13-42293497-DUP-262
chr13-42383855-DUP-73
chr13-43189515-DUP-64
chr13-44467369-DUP-94
chr13-44608164-DUP-70
chr13-44710213-DUP-79
chr13-45085908-DUP-98
chr13-45177597-DUP-55
chr13-45215890-DUP-88
chr13-46051464-DUP-191
chr13-46672408-DUP-153
chr13-46967878-DUP-53
chr13-47563918-DUP-121
chr13-47693581-DUP-572
chr13-48899076-DUP-56
chr13-49276758-DUP-69
chr13-49408411-DUP-725
chr13-49423416-DUP-88
chr13-49524413-DUP-124
chr13-49553008-DUP-140
chr13-50128237-DUP-116
chr13-50129094-DUP-55
chr13-50436011-DUP-125116
chr13-51096715-DUP-141
chr13-51527240-DUP-277
chr13-51527282-DUP-262
chr13-51743064-DUP-76
chr13-51817994-DUP-325
chr13-51832177-DUP-212
chr13-53021649-DUP-52
chr13-53085179-DUP-78
chr13-53213888-DUP-80

chr13-53500554-DUP-116
chr13-53515790-DUP-116
chr13-53940091-DUP-83
chr13-54027723-DUP-246
chr13-54327263-DUP-106
chr13-54409735-DUP-53
chr13-54580257-DUP-59
chr13-54731133-DUP-108
chr13-56039800-DUP-61
chr13-56068857-DUP-64
chr13-56169359-DUP-52
chr13-56190213-DUP-50
chr13-56246386-DUP-152
chr13-56246772-DUP-350
chr13-56624344-DUP-122
chr13-56838030-DUP-205
chr13-56838077-DUP-182
chr13-56878482-DUP-52
chr13-57152832-DUP-165
chr13-57524970-DUP-64
chr13-57635071-DUP-80
chr13-57806257-DUP-225
chr13-58044442-DUP-74
chr13-58210004-DUP-86
chr13-58432814-DUP-121
chr13-58432911-DUP-74
chr13-58802466-DUP-160
chr13-58802582-DUP-58
chr13-58872435-DUP-117
chr13-59287255-DUP-358
chr13-59344514-DUP-1596
chr13-59391836-DUP-214
chr13-59400522-DUP-222
chr13-59490523-DUP-52
chr13-59652678-DUP-54
chr13-59673626-DUP-333
chr13-59767510-DUP-195
chr13-59953137-DUP-112
chr13-60084305-DUP-145
chr13-60157490-DUP-58
chr13-60218179-DUP-525
chr13-60343382-DUP-154
chr13-60383543-DUP-5986
chr13-60586584-DUP-97
chr13-60829317-DUP-198
chr13-60900300-DUP-85
chr13-61090356-DUP-282
chr13-61228689-DUP-5578
chr13-61420121-DUP-162
chr13-61622306-DUP-70
chr13-61786641-DUP-306385
chr13-62804884-DUP-136
chr13-62966847-DUP-60
chr13-62986454-DUP-130
chr13-63311776-DUP-180
chr13-63312332-DUP-81
chr13-63312823-DUP-139
chr13-63313330-DUP-206
chr13-63495730-DUP-1227
chr13-63527218-DUP-111
chr13-63527438-DUP-684
chr13-63548181-DUP-66
chr13-63614216-DUP-170
chr13-63668971-DUP-57
chr13-63726764-DUP-88
chr13-63883087-DUP-126
chr13-63961281-DUP-118
chr13-64000304-DUP-51
chr13-64339265-DUP-114
chr13-64408749-DUP-4899
chr13-64408812-DUP-4853
chr13-64457058-DUP-54
chr13-65218088-DUP-92
chr13-65289205-DUP-65
chr13-65494357-DUP-69
chr13-65766852-DUP-89
chr13-66215801-DUP-380763

chr13-67563133-DUP-347
chr13-67563184-DUP-199
chr13-68016680-DUP-80
chr13-68054176-DUP-158
chr13-68232745-DUP-188
chr13-68584945-DUP-76
chr13-68894455-DUP-65
chr13-68996367-DUP-104
chr13-69485042-DUP-184
chr13-70133350-DUP-61
chr13-70247064-DUP-61
chr13-70532444-DUP-1090
chr13-70678088-DUP-200
chr13-71168363-DUP-181
chr13-71227065-DUP-93
chr13-71575172-DUP-7859
chr13-71621270-DUP-69
chr13-72149407-DUP-54
chr13-72257467-DUP-59
chr13-72658511-DUP-180
chr13-72758380-DUP-52
chr13-72884898-DUP-53
chr13-73122142-DUP-65
chr13-73271011-DUP-110
chr13-73313885-DUP-53
chr13-73585964-DUP-54
chr13-73853081-DUP-114
chr13-73925093-DUP-73
chr13-74141337-DUP-143
chr13-74195460-DUP-78
chr13-74504358-DUP-85
chr13-76591608-DUP-157
chr13-77088411-DUP-183
chr13-77257644-DUP-206
chr13-77710502-DUP-54
chr13-78604166-DUP-100
chr13-78643078-DUP-220
chr13-78643132-DUP-205
chr13-79834413-DUP-212
chr13-79941334-DUP-5533
chr13-80856503-DUP-127
chr13-80895442-DUP-82
chr13-81092815-DUP-78
chr13-81551938-DUP-170
chr13-81562037-DUP-54
chr13-81622022-DUP-155
chr13-81622066-DUP-74
chr13-83361742-DUP-559
chr13-84839375-DUP-53
chr13-84850211-DUP-52
chr13-84906936-DUP-59
chr13-85674804-DUP-58
chr13-86164667-DUP-54
chr13-88339802-DUP-131
chr13-88982232-DUP-81
chr13-89898576-DUP-227
chr13-89898649-DUP-93
chr13-91172240-DUP-106
chr13-91322682-DUP-393
chr13-91323341-DUP-75
chr13-91783089-DUP-197
chr13-91783120-DUP-149
chr13-91786693-DUP-104
chr13-91788869-DUP-61
chr13-91895918-DUP-221
chr13-92366187-DUP-502
chr13-92366302-DUP-432
chr13-92385308-DUP-109
chr13-92396534-DUP-52
chr13-92448487-DUP-151
chr13-92452475-DUP-490
chr13-92874382-DUP-59
chr13-92906794-DUP-164
chr13-93183671-DUP-1349
chr13-93218957-DUP-97
chr13-93233240-DUP-154
chr13-93576851-DUP-128

chr13-93576895-DUP-138
chr13-93838614-DUP-99
chr13-93864631-DUP-601
chr13-93864655-DUP-553
chr13-93894084-DUP-81
chr13-94022901-DUP-58
chr13-94278790-DUP-53
chr13-94327083-DUP-112
chr13-94365343-DUP-80
chr13-94434102-DUP-141
chr13-94747987-DUP-75
chr13-94792854-DUP-99
chr13-95004155-DUP-473
chr13-95148993-DUP-67
chr13-95240906-DUP-66
chr13-95528920-DUP-62
chr13-95731391-DUP-426
chr13-95957867-DUP-134
chr13-96257965-DUP-64
chr13-96269790-DUP-51
chr13-96473735-DUP-60
chr13-96708756-DUP-257
chr13-97322373-DUP-223
chr13-98104534-DUP-58
chr13-98395498-DUP-101
chr13-98614025-DUP-113
chr13-98697327-DUP-120
chr13-98697373-DUP-92
chr13-98785244-DUP-172
chr13-98963531-DUP-75
chr13-99354813-DUP-59
chr13-99492017-DUP-315
chr13-99508838-DUP-101
chr13-99921800-DUP-112
chr13-100764908-DUP-63
chr13-100808854-DUP-1520
chr13-101127312-DUP-61
chr13-101192216-DUP-297
chr13-103031715-DUP-56
chr13-103254128-DUP-138
chr13-104355664-DUP-103
chr13-104355691-DUP-56
chr13-104880101-DUP-64
chr13-105102832-DUP-178
chr13-105418210-DUP-356
chr13-105665249-DUP-219
chr13-106532628-DUP-50
chr13-106682933-DUP-63
chr13-106989644-DUP-147
chr13-107197513-DUP-56
chr13-107324658-DUP-148
chr13-107528816-DUP-72
chr13-107603836-DUP-79
chr13-107994771-DUP-1797
chr13-107994893-DUP-699
chr13-107995256-DUP-1313
chr13-108312611-DUP-129
chr13-108783196-DUP-254
chr13-108989275-DUP-170
chr13-109335346-DUP-100
chr13-109435864-DUP-51
chr13-109754378-DUP-9537
chr13-110165333-DUP-83
chr13-110605426-DUP-54
chr13-110862770-DUP-81
chr13-111398027-DUP-182
chr13-111442789-DUP-84
chr13-111628891-DUP-92
chr13-111911122-DUP-85
chr13-112287154-DUP-304
chr13-112287198-DUP-286
chr13-112728990-DUP-75
chr13-112783332-DUP-51
chr13-112854206-DUP-53
chr13-112860964-DUP-52
chr13-112861523-DUP-216
chr13-112862074-DUP-58

chr13-112866098-DUP-183
chr13-113228363-DUP-9772
chr13-113354966-DUP-83
chr13-113523262-DUP-57
chr13-113666207-DUP-55
chr13-113941166-DUP-261
chr13-113943010-DUP-634
chr13-114035273-DUP-56
chr13-114220776-DUP-99
chr13-114422103-DUP-51
chr13-114500958-DUP-242
chr13-114517690-DUP-91
chr13-114567827-DUP-68
chr13-114581277-DUP-184
chr13-114581325-DUP-112
chr13-116422229-DUP-75
chr13-117220618-DUP-230
chr13-117246294-DUP-65
chr13-119012556-DUP-8075
chr13-119566279-DUP-100
chr13-120058380-DUP-6678
chr13-120058659-DUP-60
chr13-120060385-DUP-207
chr13-120062095-DUP-119
chr13-120063659-DUP-90
chr13-120064338-DUP-91
chr13-120064415-DUP-124
chr13-120072411-DUP-46754
chr13-120366998-DUP-6346
chr13-120368879-DUP-6251
chr13-120368962-DUP-6136
chr13-120486361-DUP-12655
chr13-120489484-DUP-1013
chr13-120493007-DUP-6032
chr13-120495040-DUP-3999
chr13-120497758-DUP-336
chr13-120497784-DUP-1264
chr13-120498531-DUP-494
chr14-3432618-DUP-58
chr14-3690222-DUP-186
chr14-4082640-DUP-519
chr14-4082662-DUP-471
chr14-4095294-DUP-54
chr14-4558848-DUP-163
chr14-4632064-DUP-72
chr14-4650079-DUP-59
chr14-5213808-DUP-76
chr14-5316179-DUP-54
chr14-5528524-DUP-2687
chr14-5978852-DUP-132
chr14-5978878-DUP-141
chr14-6058341-DUP-95
chr14-6058362-DUP-79
chr14-6072386-DUP-92
chr14-6124103-DUP-126
chr14-6283230-DUP-54
chr14-7176877-DUP-159
chr14-7485481-DUP-406
chr14-8445181-DUP-60
chr14-8445693-DUP-313
chr14-8446480-DUP-60
chr14-8513578-DUP-63
chr14-8890173-DUP-66
chr14-9390820-DUP-66
chr14-9531408-DUP-51
chr14-10534598-DUP-63
chr14-11358845-DUP-62
chr14-11420016-DUP-502
chr14-11420466-DUP-3060
chr14-11421159-DUP-136
chr14-11422176-DUP-101
chr14-11423080-DUP-424
chr14-11492389-DUP-282
chr14-11672722-DUP-145
chr14-11802648-DUP-53
chr14-12138093-DUP-56
chr14-12509454-DUP-76

chr14-12688843-DUP-69
chr14-12783977-DUP-327
chr14-12798326-DUP-70
chr14-13649686-DUP-54
chr14-13755824-DUP-57
chr14-13781828-DUP-70
chr14-13820355-DUP-96
chr14-13855001-DUP-268
chr14-14089123-DUP-51
chr14-14295457-DUP-158
chr14-14534228-DUP-95
chr14-14598046-DUP-1226
chr14-14598930-DUP-245
chr14-15095943-DUP-79
chr14-15832044-DUP-445
chr14-15923561-DUP-486934
chr14-16156822-DUP-133
chr14-16848597-DUP-131
chr14-17010519-DUP-107
chr14-17500399-DUP-267
chr14-18170214-DUP-2876
chr14-18546009-DUP-266
chr14-18546214-DUP-760
chr14-18547009-DUP-176
chr14-18923692-DUP-132
chr14-18923831-DUP-622
chr14-19162642-DUP-681
chr14-19660427-DUP-78
chr14-19853343-DUP-61
chr14-19897462-DUP-74
chr14-19962259-DUP-178
chr14-20007373-DUP-52
chr14-20469411-DUP-106
chr14-20471026-DUP-2812
chr14-20524981-DUP-518
chr14-20525136-DUP-471
chr14-20618855-DUP-121
chr14-20762578-DUP-96
chr14-20762599-DUP-93
chr14-21130074-DUP-58
chr14-21361862-DUP-1655
chr14-21377228-DUP-363
chr14-21538149-DUP-617
chr14-21806886-DUP-357
chr14-21827817-DUP-149
chr14-21827846-DUP-141
chr14-21949574-DUP-142
chr14-22078253-DUP-231
chr14-22103559-DUP-71
chr14-22474537-DUP-85
chr14-22818945-DUP-101
chr14-22970092-DUP-88
chr14-23672552-DUP-81
chr14-23922869-DUP-127
chr14-23922892-DUP-112
chr14-24246174-DUP-65
chr14-24713392-DUP-75
chr14-24989329-DUP-133
chr14-25421213-DUP-71
chr14-25532074-DUP-56
chr14-25912223-DUP-1760
chr14-25912792-DUP-1734
chr14-25913817-DUP-1734
chr14-26371566-DUP-17289
chr14-26467516-DUP-374
chr14-26467631-DUP-374
chr14-26478061-DUP-102
chr14-26585705-DUP-73
chr14-26600548-DUP-76
chr14-26885450-DUP-115
chr14-27753492-DUP-161
chr14-28522365-DUP-88
chr14-28522980-DUP-324
chr14-29070686-DUP-56
chr14-29414901-DUP-66
chr14-29806547-DUP-150
chr14-30252531-DUP-95

chr14-30419144-DUP-52
chr14-30628646-DUP-62
chr14-30990586-DUP-95
chr14-31161575-DUP-51
chr14-31651473-DUP-74
chr14-32085206-DUP-148
chr14-32130114-DUP-77
chr14-32296574-DUP-57
chr14-33009258-DUP-115
chr14-33086150-DUP-96
chr14-33289610-DUP-52
chr14-33683048-DUP-127
chr14-33754144-DUP-63
chr14-33921447-DUP-213
chr14-34273148-DUP-151
chr14-34634928-DUP-79
chr14-34634970-DUP-78
chr14-34697626-DUP-163
chr14-34954861-DUP-66
chr14-36603354-DUP-150
chr14-36603387-DUP-51
chr14-36624427-DUP-70
chr14-36628361-DUP-110
chr14-36632458-DUP-3293
chr14-36641168-DUP-86
chr14-36696539-DUP-348
chr14-37494642-DUP-51
chr14-37906878-DUP-51
chr14-38091647-DUP-94
chr14-38101355-DUP-54
chr14-39132036-DUP-713
chr14-39133067-DUP-60
chr14-39550444-DUP-66
chr14-39680303-DUP-82
chr14-40634585-DUP-185
chr14-40872810-DUP-54
chr14-40893675-DUP-76
chr14-40899473-DUP-108
chr14-40904343-DUP-147
chr14-41244349-DUP-56
chr14-42366445-DUP-159640
chr14-43765978-DUP-90007
chr14-44738255-DUP-117
chr14-45232481-DUP-60
chr14-45240799-DUP-67
chr14-45552249-DUP-76
chr14-45562261-DUP-384
chr14-45562288-DUP-336
chr14-46024153-DUP-183
chr14-46162195-DUP-70
chr14-46522186-DUP-70
chr14-46926937-DUP-149
chr14-47608402-DUP-319
chr14-47725469-DUP-82
chr14-47791663-DUP-94
chr14-47890205-DUP-127
chr14-48126430-DUP-128
chr14-48281933-DUP-174
chr14-48301315-DUP-65
chr14-48380745-DUP-85
chr14-48424387-DUP-60
chr14-48474586-DUP-330
chr14-48586213-DUP-153
chr14-48713134-DUP-122
chr14-48740215-DUP-78
chr14-48754891-DUP-76
chr14-48817728-DUP-83
chr14-49456523-DUP-67
chr14-49457120-DUP-101
chr14-49457149-DUP-69
chr14-49860093-DUP-127
chr14-49860128-DUP-64
chr14-51322698-DUP-131
chr14-51524484-DUP-126
chr14-51759277-DUP-428
chr14-52074876-DUP-25126
chr14-52076260-DUP-25149

chr14-52522974-DUP-57
chr14-53450954-DUP-389278
chr14-54105008-DUP-7376
chr14-54152462-DUP-75
chr14-54163653-DUP-131
chr14-54293437-DUP-141
chr14-54452286-DUP-76
chr14-54543529-DUP-1374
chr14-54647088-DUP-156
chr14-54647119-DUP-120
chr14-55606259-DUP-20112
chr14-55817708-DUP-92
chr14-56183147-DUP-107991
chr14-56711447-DUP-97
chr14-56797768-DUP-164
chr14-56871597-DUP-110
chr14-57073497-DUP-154
chr14-57252548-DUP-7234
chr14-57254041-DUP-7278
chr14-58165198-DUP-221
chr14-58224850-DUP-58
chr14-58266819-DUP-1064
chr14-58267260-DUP-76
chr14-58550116-DUP-54
chr14-58795628-DUP-70
chr14-58855142-DUP-73
chr14-59290527-DUP-72
chr14-59683188-DUP-285
chr14-59849194-DUP-1912
chr14-60148985-DUP-273
chr14-60190355-DUP-69
chr14-60230535-DUP-162
chr14-60232274-DUP-139
chr14-60261397-DUP-68
chr14-60261426-DUP-67
chr14-60431335-DUP-59
chr14-61335441-DUP-58
chr14-61459442-DUP-82
chr14-61674657-DUP-67
chr14-62040200-DUP-78
chr14-62040226-DUP-56
chr14-62043454-DUP-146
chr14-62301656-DUP-69
chr14-62318175-DUP-71
chr14-62417988-DUP-114
chr14-62424610-DUP-120
chr14-62642475-DUP-87
chr14-62787968-DUP-60
chr14-62798167-DUP-310
chr14-62817747-DUP-200
chr14-62817804-DUP-85
chr14-62877235-DUP-69
chr14-62939929-DUP-127
chr14-63214264-DUP-92
chr14-63639268-DUP-143
chr14-64455395-DUP-198
chr14-64455432-DUP-180
chr14-64461567-DUP-413
chr14-64461625-DUP-351
chr14-64524249-DUP-68
chr14-64869324-DUP-798
chr14-65304637-DUP-81
chr14-65772357-DUP-56
chr14-65863315-DUP-90
chr14-65865258-DUP-334
chr14-65867397-DUP-94
chr14-65920999-DUP-201
chr14-66467958-DUP-68
chr14-66767782-DUP-62
chr14-67266165-DUP-121
chr14-67266205-DUP-63
chr14-67741817-DUP-54
chr14-67777178-DUP-188
chr14-67913127-DUP-80
chr14-67999405-DUP-614
chr14-68174836-DUP-72
chr14-68440951-DUP-77

chr14-68581399-DUP-71
chr14-68581421-DUP-73
chr14-68925015-DUP-500
chr14-68960057-DUP-110
chr14-69941703-DUP-140
chr14-70101815-DUP-179
chr14-70255428-DUP-75
chr14-70708465-DUP-75
chr14-71205530-DUP-52
chr14-71684496-DUP-56
chr14-71893115-DUP-76
chr14-72008642-DUP-64
chr14-72094054-DUP-60
chr14-72560418-DUP-130
chr14-72681392-DUP-77
chr14-73165188-DUP-62
chr14-73389617-DUP-63
chr14-73392945-DUP-187
chr14-73412434-DUP-312
chr14-73412491-DUP-333
chr14-74825461-DUP-10060
chr14-74855398-DUP-344
chr14-74974086-DUP-54
chr14-74992400-DUP-257
chr14-75028133-DUP-81
chr14-75095778-DUP-58
chr14-75095810-DUP-85
chr14-75179275-DUP-57
chr14-75442969-DUP-163
chr14-75515459-DUP-63
chr14-75547509-DUP-107
chr14-75911403-DUP-204
chr14-75948188-DUP-264
chr14-75955266-DUP-70
chr14-76003665-DUP-170
chr14-76026997-DUP-33323
chr14-76041547-DUP-72
chr14-76147125-DUP-148
chr14-76620015-DUP-114
chr14-77527869-DUP-2623
chr14-78210328-DUP-57
chr14-78743371-DUP-50
chr14-78947730-DUP-216
chr14-79222870-DUP-142
chr14-79223400-DUP-96
chr14-79232918-DUP-261
chr14-79808647-DUP-54
chr14-81517704-DUP-347
chr14-81933702-DUP-4150
chr14-82067360-DUP-88
chr14-83382445-DUP-84
chr14-83383721-DUP-12829
chr14-84994591-DUP-170
chr14-86287159-DUP-58
chr14-86347545-DUP-1354
chr14-86868531-DUP-73
chr14-87169390-DUP-62
chr14-87319242-DUP-79
chr14-87358787-DUP-2554
chr14-87444983-DUP-5330
chr14-87700276-DUP-176
chr14-87721626-DUP-61
chr14-88592464-DUP-53
chr14-90642737-DUP-87
chr14-90913639-DUP-53
chr14-90979582-DUP-1458
chr14-91491326-DUP-516
chr14-91644169-DUP-989
chr14-91644197-DUP-1051
chr14-91813253-DUP-355
chr14-93686487-DUP-317
chr14-96221241-DUP-155
chr14-97736228-DUP-294
chr14-97798304-DUP-57
chr14-99297733-DUP-141
chr14-100318852-DUP-71
chr14-100482529-DUP-88

chr14-100654698-DUP-88
chr14-100781245-DUP-61
chr14-100990512-DUP-79
chr14-101897636-DUP-115
chr14-101991635-DUP-173
chr14-102149001-DUP-274
chr14-102149038-DUP-237
chr14-102879528-DUP-85
chr14-103061277-DUP-56
chr14-103211234-DUP-57
chr14-103492345-DUP-124
chr14-103701833-DUP-356
chr14-103914482-DUP-421
chr14-103914535-DUP-1062
chr14-104044578-DUP-115
chr14-104135563-DUP-87
chr14-104370250-DUP-58
chr14-105192084-DUP-61
chr14-105552835-DUP-84
chr14-105612926-DUP-114
chr14-105612961-DUP-86
chr14-105670857-DUP-61
chr14-106140892-DUP-5304
chr14-106140914-DUP-5185
chr14-106769194-DUP-120
chr14-106769219-DUP-72
chr14-107418672-DUP-57
chr14-107799823-DUP-169
chr14-107817136-DUP-1268
chr14-108884309-DUP-173
chr14-111525998-DUP-208
chr14-111526086-DUP-160
chr14-112356221-DUP-366
chr14-112356261-DUP-329
chr14-113762192-DUP-426
chr14-114154910-DUP-89
chr14-114718415-DUP-57
chr14-114753279-DUP-55
chr14-114760419-DUP-211
chr14-114822119-DUP-318
chr14-115253966-DUP-70
chr14-115643036-DUP-298
chr14-115805365-DUP-295
chr14-116425639-DUP-60
chr14-116500948-DUP-51
chr14-116513166-DUP-84
chr14-116799953-DUP-84
chr14-117397223-DUP-81
chr14-117958873-DUP-58
chr14-117959720-DUP-65
chr14-118069495-DUP-186
chr14-118110105-DUP-99
chr14-118199435-DUP-74
chr14-118248971-DUP-219
chr14-118382717-DUP-175
chr14-118393233-DUP-369
chr14-118436545-DUP-146
chr14-118450975-DUP-74
chr14-118466575-DUP-77
chr14-118573348-DUP-141
chr14-118784987-DUP-135
chr14-118878333-DUP-166
chr14-119118881-DUP-444
chr14-119132625-DUP-64
chr14-119272771-DUP-117
chr14-119272803-DUP-114
chr14-119318206-DUP-134
chr14-119406742-DUP-244
chr14-119596608-DUP-192
chr14-119669117-DUP-194
chr14-119905672-DUP-66
chr14-119905753-DUP-68
chr14-120062098-DUP-69
chr14-120064573-DUP-10730
chr14-120187191-DUP-121
chr14-120366174-DUP-81
chr14-120588148-DUP-132

chr14-120740136-DUP-357
chr14-121067953-DUP-206
chr14-121197146-DUP-52
chr14-121322348-DUP-51
chr14-121344262-DUP-53
chr14-121369101-DUP-155
chr14-121369197-DUP-59
chr14-121439383-DUP-96
chr14-121471444-DUP-144
chr14-121503449-DUP-55
chr14-121818203-DUP-217
chr14-121894030-DUP-94
chr14-121894073-DUP-123
chr14-121969358-DUP-51
chr14-122245567-DUP-72
chr14-122423837-DUP-127
chr14-122540490-DUP-80
chr14-122669955-DUP-70
chr14-122939942-DUP-131
chr14-122940000-DUP-192
chr14-123286451-DUP-253
chr14-123402826-DUP-95
chr14-124377891-DUP-63
chr14-124691779-DUP-2288
chr15-3298353-DUP-37732
chr15-3433947-DUP-79
chr15-4164058-DUP-103
chr15-4492885-DUP-271
chr15-5128216-DUP-58
chr15-5190562-DUP-351
chr15-5191947-DUP-191
chr15-5239928-DUP-180
chr15-5283341-DUP-142
chr15-5283406-DUP-83
chr15-5360852-DUP-104
chr15-5616767-DUP-60
chr15-6101934-DUP-64
chr15-6176232-DUP-408
chr15-6348645-DUP-75
chr15-6381188-DUP-418
chr15-6426818-DUP-154
chr15-6557182-DUP-65
chr15-6637348-DUP-77
chr15-6890100-DUP-80
chr15-6962296-DUP-151
chr15-7362472-DUP-51
chr15-7382031-DUP-58
chr15-7418647-DUP-88
chr15-7452273-DUP-151
chr15-7452836-DUP-235
chr15-7830647-DUP-1468
chr15-7878156-DUP-60
chr15-7932631-DUP-52
chr15-7933163-DUP-1459
chr15-8021257-DUP-444
chr15-8075775-DUP-59
chr15-8116516-DUP-56
chr15-8205901-DUP-359
chr15-8333780-DUP-64
chr15-8768799-DUP-108
chr15-8770426-DUP-79
chr15-8790732-DUP-61
chr15-8829706-DUP-6713
chr15-8830965-DUP-6705
chr15-8917015-DUP-97
chr15-9048216-DUP-66
chr15-9085369-DUP-93
chr15-9085430-DUP-133
chr15-9103168-DUP-154
chr15-9863276-DUP-59
chr15-10312507-DUP-151
chr15-10538256-DUP-124
chr15-10583617-DUP-57
chr15-10661546-DUP-9452
chr15-10676092-DUP-68
chr15-10922232-DUP-54
chr15-11314943-DUP-67

chr15-11413706-DUP-66
chr15-11611070-DUP-365
chr15-11629992-DUP-93
chr15-12049791-DUP-53
chr15-12256428-DUP-203
chr15-12562773-DUP-65
chr15-12585396-DUP-54
chr15-12872770-DUP-71
chr15-12893901-DUP-53
chr15-13870225-DUP-65
chr15-14100592-DUP-1001
chr15-14140621-DUP-67
chr15-14238728-DUP-78
chr15-14445783-DUP-178
chr15-15364535-DUP-70
chr15-16069282-DUP-54
chr15-17700793-DUP-1286
chr15-17700951-DUP-116
chr15-18222596-DUP-9189
chr15-18917455-DUP-2430
chr15-19355692-DUP-58
chr15-19852947-DUP-110
chr15-19868059-DUP-74
chr15-20611770-DUP-82
chr15-20891321-DUP-1308
chr15-20891359-DUP-1145
chr15-20942142-DUP-246
chr15-20942171-DUP-148
chr15-21101967-DUP-134
chr15-21188473-DUP-74
chr15-21188495-DUP-126
chr15-22578503-DUP-80
chr15-23124605-DUP-77
chr15-25093900-DUP-847
chr15-25146098-DUP-57
chr15-25436010-DUP-1289
chr15-25436171-DUP-590
chr15-25436228-DUP-404
chr15-25436691-DUP-146
chr15-25570235-DUP-120
chr15-25746880-DUP-51
chr15-25903468-DUP-64
chr15-25903490-DUP-68
chr15-26071228-DUP-59
chr15-26421483-DUP-1432
chr15-26709421-DUP-272
chr15-26900979-DUP-60
chr15-26923918-DUP-73
chr15-27551269-DUP-65
chr15-27825572-DUP-69
chr15-27855491-DUP-61
chr15-28011491-DUP-160
chr15-30244848-DUP-67
chr15-30334157-DUP-66
chr15-30483620-DUP-365
chr15-30623914-DUP-51
chr15-30649061-DUP-323
chr15-30759060-DUP-204
chr15-30780229-DUP-120
chr15-30798902-DUP-68
chr15-30850242-DUP-208
chr15-30850332-DUP-56
chr15-30947518-DUP-82
chr15-31171887-DUP-141
chr15-31570550-DUP-51
chr15-32419636-DUP-187
chr15-32728727-DUP-71
chr15-32735619-DUP-126
chr15-33342600-DUP-348
chr15-34130766-DUP-81
chr15-34630241-DUP-238
chr15-35015449-DUP-85
chr15-35075939-DUP-158
chr15-35286929-DUP-4560
chr15-35515709-DUP-58
chr15-35863865-DUP-58
chr15-35909151-DUP-309

chr15-36056363-DUP-51
chr15-36169272-DUP-90
chr15-36189882-DUP-171
chr15-36237230-DUP-136
chr15-36392217-DUP-123
chr15-36392270-DUP-105
chr15-36453585-DUP-84
chr15-36457976-DUP-171
chr15-36537101-DUP-75
chr15-36624480-DUP-72
chr15-36842206-DUP-53
chr15-36909190-DUP-218
chr15-36929608-DUP-144
chr15-36977032-DUP-247
chr15-36977073-DUP-129
chr15-36995825-DUP-1407
chr15-37309643-DUP-165
chr15-37446364-DUP-182
chr15-37537971-DUP-272
chr15-37715728-DUP-72
chr15-37716752-DUP-116
chr15-37908981-DUP-72
chr15-37950889-DUP-299
chr15-38119799-DUP-360
chr15-38172784-DUP-209
chr15-38206865-DUP-51
chr15-38250539-DUP-85
chr15-38492530-DUP-924
chr15-38709070-DUP-114
chr15-39105240-DUP-5134
chr15-39307738-DUP-51
chr15-39684025-DUP-70
chr15-39870056-DUP-73
chr15-39921564-DUP-51
chr15-40015220-DUP-56
chr15-40265036-DUP-128
chr15-40797926-DUP-1653
chr15-40811637-DUP-85
chr15-41056628-DUP-85
chr15-42352694-DUP-235
chr15-42352861-DUP-135
chr15-42478893-DUP-75
chr15-43910030-DUP-87
chr15-44735558-DUP-88
chr15-45974883-DUP-103
chr15-48751973-DUP-75
chr15-49945686-DUP-73
chr15-50352595-DUP-3193
chr15-50353140-DUP-2645
chr15-51401140-DUP-56
chr15-51445123-DUP-87
chr15-51749594-DUP-126
chr15-52229349-DUP-51
chr15-53463726-DUP-87
chr15-54812685-DUP-148
chr15-54812760-DUP-145
chr15-55561313-DUP-121
chr15-55892283-DUP-364
chr15-56194728-DUP-1967
chr15-56473812-DUP-50
chr15-56558201-DUP-587
chr15-57707028-DUP-240
chr15-57707122-DUP-105
chr15-57753209-DUP-54
chr15-57848574-DUP-54
chr15-57926787-DUP-152
chr15-58178715-DUP-84
chr15-58309636-DUP-2975
chr15-58309796-DUP-286
chr15-58309951-DUP-1821
chr15-58310113-DUP-764
chr15-58310685-DUP-125
chr15-58310753-DUP-924
chr15-58311674-DUP-221
chr15-58312310-DUP-102
chr15-58376050-DUP-86
chr15-58400766-DUP-126

chr15-58422163-DUP-66
chr15-58693353-DUP-92
chr15-58988926-DUP-80
chr15-59052489-DUP-906
chr15-59219420-DUP-328
chr15-59219492-DUP-330
chr15-59346227-DUP-91
chr15-59681496-DUP-164
chr15-59729009-DUP-81
chr15-60197359-DUP-60
chr15-60209375-DUP-613
chr15-60667877-DUP-324
chr15-60828103-DUP-86
chr15-61215028-DUP-96
chr15-61420891-DUP-114
chr15-61685564-DUP-149
chr15-61946780-DUP-51
chr15-61997416-DUP-78
chr15-62157239-DUP-411147
chr15-63692681-DUP-37977
chr15-63726447-DUP-4212
chr15-64080503-DUP-65
chr15-64621797-DUP-82
chr15-64777662-DUP-72
chr15-65560432-DUP-165
chr15-66591912-DUP-184
chr15-66792799-DUP-79
chr15-66992059-DUP-54
chr15-67090223-DUP-135
chr15-67272910-DUP-59
chr15-67934853-DUP-161
chr15-67934918-DUP-81
chr15-67961021-DUP-52
chr15-68026941-DUP-309
chr15-68086383-DUP-166
chr15-68331347-DUP-107
chr15-68772972-DUP-51
chr15-69034538-DUP-1486
chr15-70428411-DUP-78
chr15-70629260-DUP-68
chr15-71425016-DUP-72
chr15-71586559-DUP-116
chr15-71723850-DUP-267
chr15-71984773-DUP-1989
chr15-71984795-DUP-2137
chr15-72722463-DUP-66
chr15-72735234-DUP-32997
chr15-73099693-DUP-64
chr15-73656045-DUP-297
chr15-74211339-DUP-1198
chr15-74211411-DUP-64
chr15-74212384-DUP-82
chr15-74367260-DUP-84
chr15-74424285-DUP-539
chr15-74695567-DUP-137
chr15-75034247-DUP-52
chr15-75336929-DUP-72386
chr15-75353236-DUP-22205
chr15-75519874-DUP-57
chr15-75794854-DUP-53
chr15-76193588-DUP-70
chr15-76271487-DUP-155
chr15-76271524-DUP-74
chr15-76309151-DUP-821
chr15-76633675-DUP-118
chr15-76643317-DUP-148
chr15-76841046-DUP-72
chr15-76860409-DUP-52
chr15-76886355-DUP-349
chr15-76912505-DUP-72
chr15-76922487-DUP-23589
chr15-77638097-DUP-106
chr15-77865980-DUP-177
chr15-78315389-DUP-106
chr15-78593752-DUP-77
chr15-78660539-DUP-66
chr15-78954272-DUP-150

chr15-79066711-DUP-66
chr15-79168509-DUP-52
chr15-79345749-DUP-1894
chr15-79521577-DUP-54
chr15-79606502-DUP-79
chr15-79920105-DUP-431
chr15-80072369-DUP-119
chr15-81068868-DUP-333
chr15-81331480-DUP-166
chr15-81475187-DUP-137
chr15-81807074-DUP-6030
chr15-81807124-DUP-5936
chr15-81846748-DUP-177
chr15-82525231-DUP-201
chr15-82781631-DUP-162
chr15-82932925-DUP-54
chr15-83203428-DUP-189
chr15-83297989-DUP-170
chr15-83298067-DUP-72
chr15-83804413-DUP-70
chr15-83854693-DUP-151
chr15-83924069-DUP-245
chr15-84069907-DUP-168
chr15-84474448-DUP-157
chr15-84524500-DUP-64
chr15-85070206-DUP-805
chr15-85184298-DUP-108
chr15-85578285-DUP-149
chr15-85611471-DUP-150
chr15-85800266-DUP-163
chr15-85817969-DUP-51
chr15-86135527-DUP-177
chr15-86151701-DUP-70
chr15-86183423-DUP-67
chr15-86288356-DUP-218
chr15-86304387-DUP-117
chr15-86344867-DUP-243
chr15-86515208-DUP-58
chr15-86898274-DUP-56
chr15-87065815-DUP-2880
chr15-87240369-DUP-117
chr15-87845569-DUP-1539
chr15-87845608-DUP-1437
chr15-87957977-DUP-59
chr15-88460128-DUP-445
chr15-88491132-DUP-125
chr15-88641697-DUP-64
chr15-88796453-DUP-147
chr15-88859783-DUP-375
chr15-89103416-DUP-108
chr15-89103660-DUP-418
chr15-89103702-DUP-420
chr15-89182948-DUP-263
chr15-89424178-DUP-95
chr15-89424200-DUP-55
chr15-89886873-DUP-238
chr15-89929649-DUP-65
chr15-90094919-DUP-58
chr15-90450657-DUP-61
chr15-90453606-DUP-90
chr15-91221349-DUP-73
chr15-91303686-DUP-2323
chr15-91670124-DUP-342
chr15-91670267-DUP-221
chr15-91728970-DUP-3569
chr15-91730541-DUP-3613
chr15-92045988-DUP-776
chr15-92046073-DUP-688
chr15-92065367-DUP-69
chr15-92528857-DUP-154
chr15-92528923-DUP-308
chr15-93039613-DUP-309
chr15-93045332-DUP-56
chr15-93319757-DUP-66
chr15-93365754-DUP-897
chr15-93365755-DUP-1387
chr15-93366225-DUP-918

chr15-93366376-DUP-592
chr15-93465926-DUP-86
chr15-93605436-DUP-72
chr15-93694952-DUP-70
chr15-93745199-DUP-75
chr15-93829118-DUP-228
chr15-93874089-DUP-91
chr15-93899948-DUP-112
chr15-93996108-DUP-2230
chr15-94171712-DUP-60
chr15-94393692-DUP-90
chr15-94674281-DUP-63
chr15-94837891-DUP-81
chr15-94852551-DUP-55
chr15-94928731-DUP-524
chr15-95131571-DUP-75
chr15-95716123-DUP-6284
chr15-95789815-DUP-134
chr15-95839272-DUP-70
chr15-95839310-DUP-77
chr15-96306034-DUP-1439
chr15-96330854-DUP-56
chr15-96385414-DUP-77
chr15-96478518-DUP-2421
chr15-96555965-DUP-55
chr15-96656130-DUP-89
chr15-96692682-DUP-407
chr15-96721612-DUP-51
chr15-96952360-DUP-72
chr15-97226984-DUP-57
chr15-97233251-DUP-132
chr15-97311694-DUP-64
chr15-97311791-DUP-58
chr15-97545042-DUP-127
chr15-97845095-DUP-484
chr15-97953179-DUP-63
chr15-97977596-DUP-57
chr15-98018995-DUP-57
chr15-98022146-DUP-100
chr15-98022176-DUP-134
chr15-98027425-DUP-123
chr15-98084722-DUP-20590
chr15-98145836-DUP-58
chr15-98314711-DUP-248
chr15-98641963-DUP-55
chr15-98748903-DUP-85
chr15-99030790-DUP-105
chr15-99030843-DUP-54
chr15-99328358-DUP-57
chr15-99382106-DUP-52
chr15-99516258-DUP-264
chr15-99516423-DUP-128
chr15-99578898-DUP-57
chr15-99611896-DUP-58
chr15-99671779-DUP-699
chr15-99671807-DUP-697
chr15-99840402-DUP-306
chr15-99925690-DUP-320
chr15-100002893-DUP-163
chr15-100096365-DUP-141
chr15-100096387-DUP-143
chr15-100189645-DUP-120
chr15-100385717-DUP-80
chr15-100412455-DUP-71
chr15-100412480-DUP-73
chr15-100482723-DUP-121
chr15-100513001-DUP-53
chr15-100543952-DUP-170
chr15-100544995-DUP-1464
chr15-100545259-DUP-197
chr15-100570491-DUP-130
chr15-100611709-DUP-252
chr15-100666552-DUP-122
chr15-100729721-DUP-231
chr15-100814874-DUP-874
chr15-100818505-DUP-90
chr15-100884408-DUP-2319

chr15-101325330-DUP-146
chr15-101354423-DUP-145
chr15-101381120-DUP-131
chr15-101582329-DUP-224
chr15-101607797-DUP-68
chr15-101791763-DUP-55
chr15-101855780-DUP-264
chr15-101856286-DUP-22159
chr15-102082923-DUP-90
chr15-103083656-DUP-101
chr15-103720618-DUP-64814
chr16-3054680-DUP-323
chr16-3321746-DUP-77
chr16-3469247-DUP-230
chr16-3469274-DUP-68
chr16-3479182-DUP-58
chr16-3480064-DUP-218
chr16-3779147-DUP-119
chr16-3779212-DUP-102
chr16-3792452-DUP-97
chr16-4057794-DUP-53
chr16-4092563-DUP-64
chr16-4118932-DUP-195
chr16-4614264-DUP-577
chr16-4680036-DUP-100
chr16-4737836-DUP-69
chr16-5122496-DUP-73
chr16-5152506-DUP-72
chr16-5298782-DUP-370
chr16-5302367-DUP-60
chr16-5347123-DUP-101
chr16-5625946-DUP-240
chr16-6348900-DUP-126
chr16-6614690-DUP-70
chr16-6872297-DUP-91
chr16-7050180-DUP-100
chr16-7440613-DUP-102
chr16-7544008-DUP-52
chr16-7755960-DUP-160
chr16-7803723-DUP-194
chr16-8210320-DUP-84
chr16-8334199-DUP-52
chr16-8637964-DUP-1946
chr16-8638510-DUP-541
chr16-8844611-DUP-57
chr16-8920358-DUP-167
chr16-8957431-DUP-58
chr16-8981882-DUP-458
chr16-8981908-DUP-320
chr16-9167074-DUP-68
chr16-9170239-DUP-56
chr16-9278110-DUP-92
chr16-9465442-DUP-51
chr16-9541004-DUP-93
chr16-9638597-DUP-130
chr16-9638625-DUP-132
chr16-9660692-DUP-109
chr16-9894250-DUP-52
chr16-10246703-DUP-240
chr16-10381906-DUP-422
chr16-10704245-DUP-392
chr16-11066475-DUP-910
chr16-11379506-DUP-55
chr16-11744358-DUP-61
chr16-11875641-DUP-178
chr16-12646660-DUP-2181
chr16-12647749-DUP-307
chr16-12782418-DUP-54
chr16-12913656-DUP-282
chr16-13130416-DUP-249
chr16-13267706-DUP-69
chr16-13359213-DUP-124
chr16-13510712-DUP-110
chr16-13822225-DUP-76
chr16-13822248-DUP-51
chr16-14856470-DUP-53887
chr16-15504114-DUP-75

chr16-15535991-DUP-58
chr16-15578378-DUP-311
chr16-15732149-DUP-85
chr16-16376917-DUP-74
chr16-16929557-DUP-301
chr16-17243886-DUP-208
chr16-17296699-DUP-200
chr16-17420250-DUP-1059
chr16-17723007-DUP-100
chr16-17767779-DUP-13067
chr16-18386613-DUP-53
chr16-18449374-DUP-379
chr16-18449395-DUP-331
chr16-18504158-DUP-51
chr16-19539939-DUP-164
chr16-19816853-DUP-10897
chr16-20217290-DUP-110
chr16-20538016-DUP-196
chr16-20899258-DUP-77
chr16-21160687-DUP-56
chr16-21161158-DUP-85
chr16-21370209-DUP-356
chr16-21370235-DUP-350
chr16-21615585-DUP-67
chr16-21616169-DUP-176
chr16-21616275-DUP-52
chr16-21622229-DUP-59
chr16-21912817-DUP-130
chr16-21912881-DUP-139
chr16-21973634-DUP-205
chr16-22649500-DUP-71
chr16-22805837-DUP-69
chr16-23182827-DUP-109
chr16-23255312-DUP-149
chr16-23324139-DUP-52
chr16-24414257-DUP-4308
chr16-25382523-DUP-126
chr16-26789808-DUP-10794
chr16-27074037-DUP-155
chr16-28573830-DUP-57
chr16-28580044-DUP-90
chr16-29013786-DUP-86
chr16-29236792-DUP-252
chr16-29236823-DUP-116
chr16-29731660-DUP-209
chr16-29963260-DUP-218
chr16-30052054-DUP-64
chr16-30098375-DUP-229
chr16-30098436-DUP-70
chr16-30109260-DUP-60
chr16-30800873-DUP-54
chr16-30974181-DUP-142
chr16-30975221-DUP-172
chr16-30977175-DUP-52
chr16-30980001-DUP-61
chr16-30981355-DUP-450
chr16-30981504-DUP-430
chr16-31363162-DUP-95
chr16-31462611-DUP-819
chr16-31462704-DUP-946
chr16-31487386-DUP-51
chr16-31703645-DUP-78
chr16-32284135-DUP-23274
chr16-32284178-DUP-23147
chr16-32309752-DUP-81
chr16-32428677-DUP-63
chr16-32553938-DUP-80
chr16-32573705-DUP-742
chr16-33290670-DUP-76
chr16-33372782-DUP-170
chr16-33483093-DUP-135
chr16-33509610-DUP-70
chr16-34211263-DUP-63
chr16-34796943-DUP-65
chr16-34998081-DUP-104
chr16-35001481-DUP-138
chr16-35001506-DUP-92

chr16-35092781-DUP-232
chr16-35440174-DUP-66
chr16-35704262-DUP-186
chr16-35704315-DUP-194
chr16-36298869-DUP-138
chr16-36310813-DUP-723
chr16-36348824-DUP-66
chr16-38980242-DUP-69
chr16-39399475-DUP-97
chr16-39821941-DUP-6421
chr16-40449564-DUP-75
chr16-41546983-DUP-231
chr16-41547009-DUP-207
chr16-41847435-DUP-532
chr16-41939907-DUP-54
chr16-42116914-DUP-52
chr16-42953195-DUP-99
chr16-43084768-DUP-203
chr16-43161358-DUP-54
chr16-43376036-DUP-53
chr16-43732690-DUP-472
chr16-43733414-DUP-1601
chr16-43734053-DUP-1097
chr16-43734357-DUP-411
chr16-43988466-DUP-77
chr16-44034909-DUP-102
chr16-44087607-DUP-100
chr16-44091367-DUP-97
chr16-44223954-DUP-62
chr16-44522799-DUP-222
chr16-44537629-DUP-669
chr16-44622168-DUP-279
chr16-44921824-DUP-90
chr16-45291719-DUP-129
chr16-45508410-DUP-53
chr16-46349959-DUP-243
chr16-46854938-DUP-117
chr16-48401382-DUP-143
chr16-49318264-DUP-85
chr16-50556191-DUP-54
chr16-50586589-DUP-54
chr16-51903787-DUP-169
chr16-52060025-DUP-56
chr16-53158989-DUP-2793
chr16-54396960-DUP-3557
chr16-55137565-DUP-73
chr16-55729719-DUP-60
chr16-56025916-DUP-2551
chr16-56197723-DUP-103
chr16-56296338-DUP-69
chr16-56606305-DUP-286
chr16-56606342-DUP-251
chr16-56610258-DUP-82
chr16-56657721-DUP-2631
chr16-56658644-DUP-2589
chr16-56659318-DUP-54
chr16-56865887-DUP-109
chr16-57360394-DUP-96
chr16-57360418-DUP-68
chr16-58093213-DUP-2079
chr16-58242845-DUP-122
chr16-58242887-DUP-93
chr16-58264884-DUP-70
chr16-58647253-DUP-144
chr16-59429263-DUP-302
chr16-59610934-DUP-92
chr16-59796085-DUP-85
chr16-60338490-DUP-121
chr16-60455535-DUP-14375
chr16-61867076-DUP-85
chr16-62535504-DUP-192
chr16-63048479-DUP-88
chr16-63104331-DUP-128
chr16-63149571-DUP-51
chr16-63177926-DUP-57
chr16-63287874-DUP-105
chr16-65549552-DUP-465

chr16-68424063-DUP-116
chr16-68442008-DUP-89
chr16-69446823-DUP-73
chr16-71301594-DUP-61
chr16-71633632-DUP-234
chr16-72413606-DUP-101
chr16-72898838-DUP-155
chr16-73150745-DUP-63
chr16-73320290-DUP-156
chr16-73628530-DUP-86
chr16-74331870-DUP-118
chr16-74486960-DUP-2623
chr16-74538048-DUP-56
chr16-74657096-DUP-481
chr16-74986314-DUP-162
chr16-74994032-DUP-736
chr16-75102111-DUP-111
chr16-76142871-DUP-55
chr16-76388423-DUP-62
chr16-76852476-DUP-65
chr16-77767767-DUP-62
chr16-78122315-DUP-65
chr16-78247987-DUP-174
chr16-78248108-DUP-68
chr16-78265447-DUP-51
chr16-78440131-DUP-62
chr16-78620561-DUP-58
chr16-78930387-DUP-52
chr16-79859384-DUP-77
chr16-79886048-DUP-78
chr16-80323221-DUP-128
chr16-81046732-DUP-104
chr16-81793000-DUP-65
chr16-83129144-DUP-1475
chr16-83641077-DUP-75
chr16-84323068-DUP-70
chr16-84355370-DUP-104
chr16-84619250-DUP-51
chr16-84861779-DUP-56
chr16-85254983-DUP-145
chr16-85475994-DUP-58
chr16-85493915-DUP-101
chr16-85787509-DUP-77
chr16-85868677-DUP-51
chr16-86189004-DUP-26146
chr16-86326797-DUP-57
chr16-86541125-DUP-4831
chr16-86602217-DUP-117
chr16-86617719-DUP-104
chr16-86963054-DUP-98
chr16-87787229-DUP-51
chr16-87844601-DUP-58
chr16-88293446-DUP-270
chr16-88293496-DUP-86
chr16-88395098-DUP-234
chr16-88666970-DUP-4854
chr16-88671053-DUP-3730
chr16-89316422-DUP-158
chr16-89986142-DUP-65
chr16-90201165-DUP-170
chr16-90224987-DUP-128
chr16-90425241-DUP-54
chr16-90699336-DUP-171
chr16-90737571-DUP-130
chr16-90738181-DUP-73
chr16-90771960-DUP-71
chr16-90782391-DUP-230
chr16-90782451-DUP-308
chr16-90926533-DUP-77
chr16-91031874-DUP-73
chr16-91054269-DUP-51
chr16-91139142-DUP-1211
chr16-91139249-DUP-316
chr16-91139798-DUP-374
chr16-91200068-DUP-100
chr16-91364743-DUP-252
chr16-91377301-DUP-70

chr16-91518359-DUP-138
chr16-91546630-DUP-335
chr16-91546653-DUP-380
chr16-91672774-DUP-74
chr16-91776197-DUP-100
chr16-91787325-DUP-94
chr16-91795612-DUP-61
chr16-91808127-DUP-189
chr16-92014733-DUP-178
chr16-92045637-DUP-76
chr16-92210648-DUP-1433
chr16-92210770-DUP-1309
chr16-92211122-DUP-202
chr16-92255331-DUP-51
chr16-92327944-DUP-144
chr16-92448251-DUP-72
chr16-92457569-DUP-444
chr16-92511239-DUP-105
chr16-92584447-DUP-86
chr16-92584482-DUP-69
chr16-92744422-DUP-68
chr16-93199592-DUP-87
chr16-93205737-DUP-323
chr16-93474898-DUP-77
chr16-93575189-DUP-238
chr16-93586095-DUP-188
chr16-93599952-DUP-149
chr16-93811612-DUP-52
chr16-93885469-DUP-72
chr16-94038461-DUP-74
chr16-94057412-DUP-113
chr16-94059958-DUP-112
chr16-94429659-DUP-64
chr16-94430845-DUP-52
chr16-94468084-DUP-69
chr16-94476755-DUP-307
chr16-94477522-DUP-519
chr16-94581302-DUP-192
chr16-94581326-DUP-190
chr16-94599289-DUP-107
chr16-94657759-DUP-171
chr16-94680336-DUP-75
chr16-94706306-DUP-143
chr16-94833410-DUP-101
chr16-94833436-DUP-182
chr16-94834865-DUP-51
chr16-94995454-DUP-105
chr16-95009734-DUP-64
chr16-95118520-DUP-78
chr16-95156863-DUP-77
chr16-95475999-DUP-73
chr16-95669886-DUP-226
chr16-96165368-DUP-81
chr16-96452641-DUP-57
chr16-96565804-DUP-58
chr16-96584879-DUP-166
chr16-96757820-DUP-110
chr16-96764463-DUP-115
chr16-96938024-DUP-88
chr16-97588369-DUP-60
chr17-3060923-DUP-163
chr17-3062188-DUP-185
chr17-3062690-DUP-8283
chr17-3070929-DUP-61
chr17-3072061-DUP-1858
chr17-3072167-DUP-1916
chr17-3076772-DUP-373
chr17-3077006-DUP-1459
chr17-3078163-DUP-4917
chr17-3214932-DUP-85
chr17-3300373-DUP-112
chr17-3411554-DUP-118
chr17-3450438-DUP-91
chr17-3516608-DUP-54
chr17-3944267-DUP-101
chr17-4275621-DUP-850
chr17-4290104-DUP-1464

chr17-4404718-DUP-55
chr17-4549474-DUP-93
chr17-4676452-DUP-343
chr17-4676474-DUP-320
chr17-4846004-DUP-192
chr17-4911807-DUP-195
chr17-5054330-DUP-69
chr17-5061928-DUP-108
chr17-5088573-DUP-102
chr17-5097938-DUP-51
chr17-5133290-DUP-117
chr17-5206892-DUP-93
chr17-5291285-DUP-295
chr17-5406447-DUP-110
chr17-5471226-DUP-52
chr17-5617921-DUP-52
chr17-5791772-DUP-102
chr17-5807375-DUP-66
chr17-5815500-DUP-136
chr17-5901936-DUP-83
chr17-5930064-DUP-54
chr17-5930811-DUP-79
chr17-5932121-DUP-323
chr17-5934019-DUP-162
chr17-5938386-DUP-138
chr17-6090030-DUP-62
chr17-6111033-DUP-90
chr17-6111122-DUP-133
chr17-6174181-DUP-82
chr17-6255379-DUP-85
chr17-6269871-DUP-171
chr17-6440627-DUP-104
chr17-6658101-DUP-191
chr17-6824221-DUP-502
chr17-6825021-DUP-151
chr17-6961385-DUP-536
chr17-7012646-DUP-93
chr17-7016590-DUP-87
chr17-7031190-DUP-63
chr17-7065263-DUP-72
chr17-7181978-DUP-62
chr17-7200674-DUP-547
chr17-7201922-DUP-98
chr17-7202919-DUP-151
chr17-7257801-DUP-68
chr17-7262500-DUP-77
chr17-7320801-DUP-119
chr17-7407987-DUP-70
chr17-7409599-DUP-128
chr17-7454489-DUP-62
chr17-7596967-DUP-1159
chr17-7610359-DUP-73
chr17-7757012-DUP-61
chr17-7782605-DUP-93
chr17-7934817-DUP-132
chr17-7962362-DUP-90
chr17-7985894-DUP-72
chr17-8004072-DUP-356
chr17-8053983-DUP-63
chr17-8175061-DUP-986
chr17-8205807-DUP-96
chr17-8205870-DUP-61
chr17-8252723-DUP-77
chr17-8291661-DUP-116
chr17-8291698-DUP-118
chr17-8470386-DUP-222
chr17-8566026-DUP-222
chr17-8642071-DUP-148
chr17-8656748-DUP-91
chr17-8656777-DUP-87
chr17-8847257-DUP-82
chr17-8847781-DUP-82
chr17-8889782-DUP-393
chr17-8923135-DUP-213
chr17-8956807-DUP-141
chr17-9012535-DUP-62
chr17-9052729-DUP-182299

chr17-9067808-DUP-278
chr17-9071617-DUP-82
chr17-9681130-DUP-124
chr17-9789105-DUP-51
chr17-9963669-DUP-363
chr17-10218830-DUP-80
chr17-10663895-DUP-196
chr17-10663923-DUP-264
chr17-10700020-DUP-76
chr17-10743985-DUP-129
chr17-10744009-DUP-72
chr17-11126447-DUP-185
chr17-11165979-DUP-143
chr17-11268055-DUP-114
chr17-11300116-DUP-73
chr17-11326136-DUP-105
chr17-11749220-DUP-124
chr17-11847168-DUP-92
chr17-11872415-DUP-125
chr17-11874031-DUP-58
chr17-12113997-DUP-70
chr17-12377278-DUP-188
chr17-12399235-DUP-54
chr17-12508589-DUP-306
chr17-12584001-DUP-126
chr17-12674470-DUP-146
chr17-12690119-DUP-799
chr17-12693587-DUP-181
chr17-12855151-DUP-582
chr17-12856274-DUP-59
chr17-13330352-DUP-66
chr17-13391050-DUP-70
chr17-13445529-DUP-9357
chr17-13447013-DUP-9375
chr17-13536063-DUP-138
chr17-13688235-DUP-75151
chr17-13807842-DUP-112
chr17-13808625-DUP-131
chr17-13809370-DUP-160
chr17-13875322-DUP-184
chr17-13994916-DUP-82
chr17-14148015-DUP-220
chr17-14156821-DUP-51
chr17-14441549-DUP-127
chr17-14469172-DUP-54
chr17-14511711-DUP-138
chr17-14601881-DUP-53
chr17-14807815-DUP-186
chr17-14862584-DUP-59
chr17-15081991-DUP-185
chr17-15096087-DUP-123
chr17-15545295-DUP-85
chr17-15630043-DUP-83
chr17-15647278-DUP-133
chr17-16145064-DUP-148
chr17-16211620-DUP-474
chr17-16211654-DUP-437
chr17-16273439-DUP-205
chr17-16378531-DUP-100
chr17-16704763-DUP-52
chr17-18075990-DUP-51
chr17-18653141-DUP-147549
chr17-20650725-DUP-463
chr17-21807254-DUP-96
chr17-22314477-DUP-93093
chr17-23086388-DUP-191
chr17-23870965-DUP-54
chr17-24213401-DUP-55
chr17-24459302-DUP-260
chr17-24650878-DUP-95
chr17-24844118-DUP-85
chr17-24896602-DUP-95
chr17-24988439-DUP-62
chr17-25290968-DUP-61
chr17-25310429-DUP-201
chr17-25314802-DUP-51
chr17-25877939-DUP-371

chr17-25918814-DUP-630
chr17-26210830-DUP-64
chr17-26315699-DUP-5157
chr17-26328781-DUP-199
chr17-26443988-DUP-143
chr17-26479033-DUP-75651
chr17-26663394-DUP-60
chr17-26767068-DUP-370
chr17-26852062-DUP-201
chr17-26852101-DUP-58
chr17-26898124-DUP-188
chr17-26983429-DUP-518
chr17-26986548-DUP-70
chr17-27216040-DUP-118
chr17-27324813-DUP-121
chr17-27351974-DUP-117
chr17-27416345-DUP-154
chr17-27425601-DUP-128056
chr17-27455285-DUP-183262
chr17-27689345-DUP-78
chr17-27818253-DUP-52
chr17-27918812-DUP-65
chr17-28115621-DUP-180
chr17-28138474-DUP-124
chr17-28218930-DUP-96
chr17-28240439-DUP-332
chr17-28245497-DUP-288
chr17-28330218-DUP-70
chr17-28339694-DUP-1558
chr17-28503910-DUP-282
chr17-28577742-DUP-109
chr17-28646950-DUP-82
chr17-28721043-DUP-94
chr17-29128004-DUP-66
chr17-29212543-DUP-320
chr17-29282059-DUP-62
chr17-29321223-DUP-6992
chr17-29321396-DUP-113
chr17-29322233-DUP-170
chr17-29325206-DUP-3009
chr17-29326278-DUP-1929
chr17-29327745-DUP-169
chr17-29504839-DUP-503
chr17-29504954-DUP-622
chr17-29505522-DUP-116
chr17-29505602-DUP-54
chr17-29595220-DUP-113
chr17-29789875-DUP-205
chr17-29857236-DUP-55
chr17-29930920-DUP-54
chr17-30302585-DUP-157
chr17-30440508-DUP-56
chr17-30556383-DUP-1543
chr17-30556515-DUP-1706
chr17-30666348-DUP-476837
chr17-30702447-DUP-57
chr17-30725784-DUP-385500
chr17-30737935-DUP-56
chr17-30936999-DUP-173
chr17-30994182-DUP-1544
chr17-31016966-DUP-213
chr17-31115045-DUP-76
chr17-31136903-DUP-65
chr17-31217564-DUP-32473
chr17-31252886-DUP-79
chr17-31345652-DUP-52
chr17-31376194-DUP-63
chr17-31437907-DUP-550
chr17-31441857-DUP-57999
chr17-31472596-DUP-56150
chr17-31473829-DUP-88449
chr17-31474505-DUP-324
chr17-31524719-DUP-55
chr17-31614699-DUP-54
chr17-31682496-DUP-87
chr17-31867576-DUP-69
chr17-31982442-DUP-2122

chr17-32232812-DUP-96
chr17-32320368-DUP-54
chr17-32434316-DUP-60
chr17-32742711-DUP-55
chr17-32742761-DUP-64
chr17-33047446-DUP-286
chr17-33117523-DUP-82
chr17-33578345-DUP-96
chr17-33655013-DUP-106
chr17-33744468-DUP-128
chr17-33744489-DUP-106
chr17-34057356-DUP-298
chr17-34163412-DUP-64
chr17-34326489-DUP-82
chr17-34326515-DUP-147
chr17-34412341-DUP-91
chr17-34421177-DUP-148
chr17-34421909-DUP-318
chr17-34472957-DUP-244
chr17-34473001-DUP-188
chr17-34572658-DUP-137
chr17-34577287-DUP-78
chr17-34653958-DUP-77
chr17-34669256-DUP-182
chr17-34669782-DUP-148
chr17-34669854-DUP-67
chr17-34752905-DUP-103
chr17-34856434-DUP-54
chr17-35194336-DUP-62
chr17-35285660-DUP-51
chr17-35474575-DUP-2030
chr17-35474609-DUP-1912
chr17-35826490-DUP-70
chr17-35931530-DUP-313
chr17-36301269-DUP-272
chr17-36341360-DUP-112132
chr17-36387349-DUP-62369
chr17-36552391-DUP-80
chr17-37342382-DUP-4455
chr17-37342417-DUP-4359
chr17-37384168-DUP-8700
chr17-37868707-DUP-4293
chr17-38519624-DUP-1394
chr17-39429376-DUP-7784
chr17-39610960-DUP-179
chr17-40534750-DUP-159
chr17-41218581-DUP-72
chr17-41526134-DUP-1089
chr17-41526284-DUP-935
chr17-42880594-DUP-65
chr17-43005307-DUP-53
chr17-43054349-DUP-709
chr17-43054963-DUP-264
chr17-43067392-DUP-128
chr17-43181394-DUP-51
chr17-43908009-DUP-63
chr17-43961822-DUP-95
chr17-44799998-DUP-96
chr17-44966332-DUP-120
chr17-45037724-DUP-113
chr17-45132973-DUP-62
chr17-45453782-DUP-78
chr17-45469902-DUP-58
chr17-45548737-DUP-69
chr17-45770093-DUP-64
chr17-45831665-DUP-69
chr17-45870791-DUP-51
chr17-46025998-DUP-96
chr17-46300484-DUP-60
chr17-46380429-DUP-54
chr17-46444146-DUP-82
chr17-46570366-DUP-115
chr17-46572882-DUP-57
chr17-46623053-DUP-91
chr17-46758854-DUP-87
chr17-47126595-DUP-337
chr17-47130877-DUP-55

chr17-47247240-DUP-85
chr17-47596991-DUP-74
chr17-47602706-DUP-58
chr17-47681622-DUP-79
chr17-47967218-DUP-135
chr17-47967255-DUP-87
chr17-48023903-DUP-1914
chr17-48100062-DUP-114
chr17-48112840-DUP-57
chr17-48503348-DUP-89
chr17-48505740-DUP-75
chr17-48525938-DUP-259
chr17-48525993-DUP-256
chr17-48647902-DUP-58
chr17-48737864-DUP-68
chr17-48753379-DUP-280
chr17-48772944-DUP-132
chr17-48822454-DUP-75
chr17-49044075-DUP-65
chr17-49190958-DUP-61
chr17-49260731-DUP-54
chr17-49851517-DUP-647
chr17-49851544-DUP-595
chr17-49986424-DUP-98
chr17-50423398-DUP-109
chr17-50451264-DUP-521
chr17-50621769-DUP-57
chr17-50701873-DUP-120
chr17-50892468-DUP-53
chr17-51375625-DUP-322
chr17-51696946-DUP-84
chr17-51737941-DUP-116
chr17-51859833-DUP-60
chr17-52809919-DUP-92
chr17-53117256-DUP-92
chr17-53152747-DUP-319
chr17-53553855-DUP-752
chr17-54130157-DUP-90
chr17-54233832-DUP-443
chr17-54336086-DUP-104
chr17-55140780-DUP-94
chr17-55979720-DUP-54
chr17-56411925-DUP-196
chr17-56592083-DUP-275
chr17-56727055-DUP-142
chr17-56729823-DUP-56
chr17-56962210-DUP-56
chr17-57101818-DUP-59
chr17-57148620-DUP-90
chr17-57158297-DUP-87
chr17-57355648-DUP-77
chr17-57527786-DUP-72
chr17-58787979-DUP-118
chr17-59176981-DUP-504
chr17-59383773-DUP-614
chr17-60219135-DUP-4788
chr17-61702831-DUP-59
chr17-61842816-DUP-162
chr17-63357754-DUP-83
chr17-63426907-DUP-85
chr17-63587512-DUP-105
chr17-63641362-DUP-82
chr17-63647014-DUP-74
chr17-63703314-DUP-286
chr17-63730200-DUP-285
chr17-63999577-DUP-357
chr17-64033977-DUP-182
chr17-64034132-DUP-590
chr17-64256827-DUP-61
chr17-64302327-DUP-52
chr17-64321994-DUP-55
chr17-64438528-DUP-348
chr17-64741901-DUP-51
chr17-64923635-DUP-346
chr17-65614522-DUP-126
chr17-65725264-DUP-106
chr17-66227822-DUP-531

chr17-66268615-DUP-144
chr17-67556537-DUP-65
chr17-67570204-DUP-92
chr17-67694791-DUP-56
chr17-68027222-DUP-119
chr17-68194312-DUP-56
chr17-68521351-DUP-267
chr17-68551041-DUP-51
chr17-68642096-DUP-79
chr17-68747081-DUP-925
chr17-69122499-DUP-105
chr17-69162097-DUP-62
chr17-69498324-DUP-54
chr17-69542031-DUP-101
chr17-69605835-DUP-91
chr17-69741266-DUP-94
chr17-69741296-DUP-96
chr17-69802379-DUP-154
chr17-70090030-DUP-63
chr17-70147343-DUP-101
chr17-70330211-DUP-68
chr17-70468794-DUP-53
chr17-70726527-DUP-201
chr17-70807035-DUP-109
chr17-71235755-DUP-166
chr17-71341857-DUP-133
chr17-71480063-DUP-60
chr17-71535182-DUP-229
chr17-71552338-DUP-52
chr17-71687490-DUP-54
chr17-71703088-DUP-1869
chr17-71745500-DUP-66
chr17-71745752-DUP-354
chr17-72108438-DUP-144
chr17-72211831-DUP-162
chr17-72369213-DUP-138
chr17-72690348-DUP-277
chr17-73122611-DUP-529
chr17-73761056-DUP-2909
chr17-73855534-DUP-118
chr17-74340233-DUP-104
chr17-74441503-DUP-75
chr17-74516587-DUP-231
chr17-74516633-DUP-338
chr17-74519440-DUP-60
chr17-74610046-DUP-83
chr17-75148530-DUP-63
chr17-75148554-DUP-56
chr17-75371864-DUP-77
chr17-75520777-DUP-166
chr17-75770287-DUP-196
chr17-75786656-DUP-90
chr17-76775062-DUP-121
chr17-77309687-DUP-5385
chr17-78367569-DUP-91
chr17-78367651-DUP-96
chr17-78607752-DUP-77
chr17-78926606-DUP-99
chr17-79021329-DUP-56
chr17-79021374-DUP-50
chr17-79058293-DUP-138
chr17-79068741-DUP-130
chr17-79198964-DUP-273
chr17-79198996-DUP-227
chr17-79202563-DUP-101
chr17-79641144-DUP-68
chr17-79646900-DUP-70
chr17-79659600-DUP-59
chr17-80189633-DUP-58
chr17-80347088-DUP-776
chr17-80709173-DUP-84
chr17-80728999-DUP-491
chr17-80906336-DUP-411
chr17-80906391-DUP-380
chr17-81157660-DUP-63
chr17-81329985-DUP-114
chr17-81354853-DUP-145

chr17-81743498-DUP-51
chr17-81814828-DUP-103
chr17-81879279-DUP-282
chr17-82361613-DUP-93
chr17-82778184-DUP-13236
chr17-83142885-DUP-103
chr17-83241131-DUP-80
chr17-83363294-DUP-63
chr17-83561733-DUP-114
chr17-83696707-DUP-123
chr17-83754852-DUP-72
chr17-83898335-DUP-60
chr17-83910065-DUP-66
chr17-83997112-DUP-7908
chr17-84043861-DUP-87
chr17-84080043-DUP-87
chr17-84143651-DUP-54
chr17-84434610-DUP-63
chr17-84473104-DUP-94
chr17-84576819-DUP-82
chr17-84630718-DUP-55
chr17-84635079-DUP-75
chr17-84769246-DUP-166
chr17-84771269-DUP-273
chr17-84813937-DUP-64
chr17-84854957-DUP-234
chr17-84855001-DUP-190
chr17-85005546-DUP-72
chr17-85077185-DUP-100
chr17-85204953-DUP-56
chr17-85308857-DUP-179
chr17-85448240-DUP-158
chr17-85448328-DUP-83
chr17-85560693-DUP-175
chr17-85560789-DUP-88
chr17-85692399-DUP-74
chr17-86084451-DUP-149
chr17-86211333-DUP-281
chr17-86262339-DUP-89
chr17-86351952-DUP-58
chr17-86458415-DUP-66
chr17-86498285-DUP-283
chr17-86536195-DUP-81
chr17-86559649-DUP-58
chr17-86565115-DUP-54
chr17-86775424-DUP-59
chr17-86797957-DUP-100
chr17-86845694-DUP-59
chr17-86913509-DUP-143
chr17-87278261-DUP-54
chr17-87320057-DUP-262
chr17-87321476-DUP-195
chr17-87355579-DUP-83
chr17-87370317-DUP-62
chr17-87425692-DUP-193
chr17-87604571-DUP-103
chr17-87681833-DUP-180
chr17-87681901-DUP-121
chr17-87875715-DUP-86
chr17-87984560-DUP-54
chr17-88002201-DUP-60
chr17-88021020-DUP-216
chr17-88152733-DUP-111
chr17-88153895-DUP-55
chr17-88227454-DUP-86
chr17-88238101-DUP-141
chr17-88617623-DUP-71
chr17-88881028-DUP-132
chr17-88932306-DUP-96
chr17-88964212-DUP-53
chr17-88965736-DUP-172
chr17-88980292-DUP-99
chr17-89008484-DUP-69
chr17-89025380-DUP-78
chr17-89085920-DUP-71
chr17-89136937-DUP-53
chr17-89153301-DUP-320

chr17-89320319-DUP-1729
chr17-89483918-DUP-154
chr17-89617647-DUP-52
chr17-89629283-DUP-78
chr17-89840206-DUP-88
chr17-90099253-DUP-72
chr17-90519256-DUP-86
chr17-90870683-DUP-85
chr17-90900051-DUP-658
chr17-90900250-DUP-1131
chr17-90900631-DUP-641
chr17-91204277-DUP-151
chr17-91475272-DUP-111
chr17-91527469-DUP-250
chr17-91657766-DUP-63
chr17-92178179-DUP-502
chr17-92407822-DUP-132
chr17-92553388-DUP-201
chr17-93732999-DUP-198
chr17-93867028-DUP-179
chr18-3115903-DUP-3976
chr18-3116877-DUP-58
chr18-3117648-DUP-108
chr18-3266980-DUP-54
chr18-3501608-DUP-73
chr18-3514499-DUP-53
chr18-3934512-DUP-115
chr18-3934538-DUP-206
chr18-4276742-DUP-950
chr18-4290746-DUP-82
chr18-4367421-DUP-198
chr18-4367473-DUP-177
chr18-4550686-DUP-75
chr18-5164924-DUP-62
chr18-5402462-DUP-91
chr18-5440320-DUP-101
chr18-5495538-DUP-75
chr18-5681372-DUP-173
chr18-5778181-DUP-97
chr18-5909594-DUP-68
chr18-6033658-DUP-310
chr18-6046894-DUP-79
chr18-6148739-DUP-57
chr18-6444818-DUP-62
chr18-6787724-DUP-244
chr18-6882270-DUP-56
chr18-7437117-DUP-173
chr18-7547489-DUP-86
chr18-7770233-DUP-6368
chr18-8288422-DUP-70
chr18-9373688-DUP-72
chr18-10361096-DUP-51
chr18-10508422-DUP-120
chr18-10595058-DUP-55
chr18-10612915-DUP-60
chr18-10653286-DUP-255
chr18-10653354-DUP-174
chr18-10663930-DUP-113
chr18-10664006-DUP-230
chr18-10939031-DUP-96
chr18-10939062-DUP-68
chr18-10984793-DUP-136
chr18-11193527-DUP-116
chr18-11415481-DUP-94
chr18-11517587-DUP-129
chr18-11544592-DUP-90
chr18-11556645-DUP-64
chr18-11721379-DUP-71
chr18-12014532-DUP-195
chr18-12018854-DUP-51
chr18-12033334-DUP-62
chr18-12056880-DUP-178
chr18-12056907-DUP-180
chr18-12120679-DUP-136
chr18-12166766-DUP-183
chr18-12310644-DUP-74
chr18-12484152-DUP-70

chr18-12507059-DUP-59
chr18-12585449-DUP-137
chr18-12603462-DUP-127
chr18-12604722-DUP-62
chr18-12651316-DUP-128
chr18-12668658-DUP-147
chr18-12684042-DUP-59
chr18-12823690-DUP-56
chr18-12847216-DUP-90
chr18-12853395-DUP-278
chr18-12939587-DUP-237
chr18-12963751-DUP-220
chr18-12963778-DUP-1351
chr18-12987980-DUP-85
chr18-13058909-DUP-133
chr18-13239271-DUP-204
chr18-13279158-DUP-173
chr18-13279248-DUP-180
chr18-13383890-DUP-68
chr18-13429518-DUP-355
chr18-13430467-DUP-714
chr18-13430692-DUP-2089
chr18-14090263-DUP-113
chr18-14149342-DUP-436
chr18-14149416-DUP-340
chr18-14266137-DUP-157
chr18-14493711-DUP-15996
chr18-14798522-DUP-66
chr18-15051404-DUP-53
chr18-15172605-DUP-304
chr18-15580518-DUP-88
chr18-15791738-DUP-139
chr18-16070543-DUP-273
chr18-17578823-DUP-125
chr18-17687223-DUP-68
chr18-17877046-DUP-62
chr18-18119031-DUP-52
chr18-18495687-DUP-67
chr18-20541369-DUP-298
chr18-20567286-DUP-74
chr18-20761204-DUP-68
chr18-20762396-DUP-64
chr18-20913866-DUP-129
chr18-21178699-DUP-59
chr18-21804929-DUP-71
chr18-23237074-DUP-92
chr18-23727092-DUP-117
chr18-23796020-DUP-89
chr18-24392300-DUP-108
chr18-24582207-DUP-101
chr18-24755840-DUP-88
chr18-24804843-DUP-670
chr18-24856453-DUP-83
chr18-24947640-DUP-78
chr18-25320455-DUP-77
chr18-25434524-DUP-82
chr18-26370259-DUP-74
chr18-26567657-DUP-104
chr18-26567689-DUP-100
chr18-26769741-DUP-111
chr18-27027108-DUP-59
chr18-28409365-DUP-84
chr18-28793150-DUP-2615
chr18-29245548-DUP-976
chr18-30139908-DUP-124
chr18-30460041-DUP-413
chr18-30492276-DUP-87
chr18-30521908-DUP-51
chr18-31748109-DUP-23251
chr18-31993935-DUP-212
chr18-31994101-DUP-64
chr18-32285988-DUP-87
chr18-32514161-DUP-170
chr18-32750239-DUP-113
chr18-32994729-DUP-58
chr18-33943926-DUP-64
chr18-34402282-DUP-173

chr18-34440308-DUP-63
chr18-34701830-DUP-97
chr18-35009142-DUP-241
chr18-35217186-DUP-2733
chr18-35416236-DUP-180
chr18-36145364-DUP-152
chr18-36245503-DUP-71
chr18-36407884-DUP-135
chr18-36407974-DUP-88
chr18-36692158-DUP-166
chr18-36806425-DUP-92
chr18-36806943-DUP-99
chr18-36903685-DUP-103
chr18-37548178-DUP-116
chr18-38032865-DUP-238
chr18-38032951-DUP-168
chr18-38369371-DUP-98
chr18-38509856-DUP-57
chr18-38609450-DUP-935
chr18-38609537-DUP-894
chr18-38610201-DUP-157
chr18-38957682-DUP-69
chr18-39049151-DUP-160
chr18-39470693-DUP-55
chr18-39763666-DUP-105
chr18-39855439-DUP-62
chr18-39949855-DUP-77
chr18-41775619-DUP-89
chr18-42016212-DUP-174
chr18-42016252-DUP-92
chr18-42225821-DUP-53
chr18-42363869-DUP-82
chr18-43109006-DUP-56
chr18-43112110-DUP-99
chr18-43339586-DUP-54
chr18-43484821-DUP-316
chr18-43485021-DUP-318
chr18-44544411-DUP-306
chr18-44544439-DUP-488
chr18-44666589-DUP-80
chr18-45311753-DUP-77
chr18-45819929-DUP-114
chr18-45820081-DUP-116
chr18-46432126-DUP-166
chr18-46559590-DUP-52
chr18-46685266-DUP-102
chr18-47038430-DUP-54
chr18-47421178-DUP-626
chr18-47467022-DUP-61
chr18-47855230-DUP-64
chr18-48086059-DUP-135
chr18-48291990-DUP-80
chr18-49618489-DUP-214
chr18-49867687-DUP-262
chr18-49985507-DUP-439
chr18-50088393-DUP-266
chr18-50088467-DUP-120
chr18-50149647-DUP-147
chr18-50293705-DUP-377
chr18-50591240-DUP-107
chr18-51152260-DUP-74
chr18-51732796-DUP-154
chr18-51732989-DUP-110
chr18-51808470-DUP-160
chr18-52282904-DUP-53
chr18-52607060-DUP-62
chr18-53488411-DUP-243
chr18-53559042-DUP-99
chr18-53732661-DUP-103
chr18-53849545-DUP-76
chr18-53856027-DUP-90
chr18-55364760-DUP-97
chr18-55364781-DUP-61
chr18-55393265-DUP-56
chr18-55404004-DUP-8916
chr18-56159495-DUP-113
chr18-56503154-DUP-57

chr18-56660670-DUP-80
chr18-57066982-DUP-136
chr18-57496873-DUP-52
chr18-57611364-DUP-55
chr18-57611432-DUP-64
chr18-57791270-DUP-83298
chr18-57912289-DUP-91
chr18-57916046-DUP-167
chr18-58366449-DUP-93
chr18-58808918-DUP-65
chr18-59262155-DUP-120
chr18-59498780-DUP-108
chr18-59498804-DUP-69
chr18-60770826-DUP-55
chr18-60923005-DUP-218
chr18-60923623-DUP-414
chr18-60966164-DUP-150
chr18-60966191-DUP-60
chr18-61167107-DUP-170
chr18-61328158-DUP-57
chr18-61592541-DUP-107422
chr18-61737046-DUP-211
chr18-61833872-DUP-197
chr18-62092315-DUP-615
chr18-62092346-DUP-567
chr18-62122039-DUP-75
chr18-62193592-DUP-126
chr18-62362381-DUP-76
chr18-62886580-DUP-822
chr18-62895128-DUP-70
chr18-63119704-DUP-164
chr18-63204538-DUP-377
chr18-63204576-DUP-219
chr18-63240667-DUP-344
chr18-63512311-DUP-229
chr18-63512336-DUP-258
chr18-63530426-DUP-102
chr18-64602013-DUP-55
chr18-64607349-DUP-326
chr18-64844013-DUP-130
chr18-65131891-DUP-117
chr18-65148660-DUP-63
chr18-65170246-DUP-67
chr18-65200244-DUP-127
chr18-65250009-DUP-55
chr18-65260166-DUP-61
chr18-65510668-DUP-72
chr18-65844942-DUP-111
chr18-66369562-DUP-62
chr18-66665650-DUP-59
chr18-66837358-DUP-128
chr18-66837397-DUP-92
chr18-67252677-DUP-69
chr18-67354969-DUP-70
chr18-67511469-DUP-144
chr18-67702277-DUP-86
chr18-67844830-DUP-68
chr18-67926738-DUP-85
chr18-68410399-DUP-61
chr18-68605612-DUP-158
chr18-68891638-DUP-76
chr18-69602547-DUP-78
chr18-69618279-DUP-115
chr18-70162864-DUP-74
chr18-70400571-DUP-242
chr18-70400597-DUP-123
chr18-70407804-DUP-52
chr18-70907506-DUP-121
chr18-71424777-DUP-303
chr18-71633190-DUP-170
chr18-72179959-DUP-57
chr18-72639010-DUP-77
chr18-72661719-DUP-54
chr18-72822343-DUP-65
chr18-73056500-DUP-59
chr18-73428997-DUP-65
chr18-73448896-DUP-103

chr18-73553087-DUP-111
chr18-73557638-DUP-293
chr18-73667802-DUP-51
chr18-73691077-DUP-53
chr18-73774275-DUP-51
chr18-73823262-DUP-63
chr18-73908926-DUP-52
chr18-74232982-DUP-85
chr18-74334262-DUP-98
chr18-74434101-DUP-70
chr18-74596994-DUP-98
chr18-74623065-DUP-154
chr18-74623138-DUP-78
chr18-74714086-DUP-65
chr18-74720623-DUP-120
chr18-74743864-DUP-64
chr18-74819370-DUP-242
chr18-74847096-DUP-82
chr18-74895964-DUP-151
chr18-74968620-DUP-107
chr18-74980090-DUP-89
chr18-75218780-DUP-174
chr18-75433649-DUP-56
chr18-75748707-DUP-109
chr18-75748811-DUP-196
chr18-75946887-DUP-213
chr18-75949487-DUP-73
chr18-75949517-DUP-54
chr18-75991333-DUP-52
chr18-76474282-DUP-51
chr18-76614852-DUP-325
chr18-76681115-DUP-128
chr18-76684946-DUP-971
chr18-76684947-DUP-1036
chr18-76689862-DUP-166
chr18-76745592-DUP-84
chr18-76745656-DUP-58
chr18-76776076-DUP-56
chr18-77012074-DUP-55
chr18-77270130-DUP-318
chr18-77270159-DUP-388
chr18-77430384-DUP-61
chr18-77599003-DUP-58
chr18-77795532-DUP-72
chr18-77859032-DUP-103
chr18-77994072-DUP-82
chr18-77994128-DUP-118
chr18-78019880-DUP-114
chr18-78044007-DUP-205
chr18-78044095-DUP-92
chr18-78059703-DUP-688
chr18-78060491-DUP-141
chr18-78185672-DUP-70
chr18-78208746-DUP-89
chr18-78542490-DUP-118
chr18-78628532-DUP-154
chr18-78629486-DUP-287
chr18-78984943-DUP-172
chr18-79113357-DUP-88
chr18-79485096-DUP-532
chr18-79823285-DUP-76
chr18-80062010-DUP-386
chr18-80068508-DUP-387
chr18-80071817-DUP-384
chr18-80299133-DUP-53
chr18-80349769-DUP-103
chr18-80479641-DUP-58
chr18-80479669-DUP-57
chr18-80497649-DUP-147
chr18-80568917-DUP-169
chr18-80655815-DUP-103
chr18-80793730-DUP-84
chr18-80844353-DUP-182
chr18-80844398-DUP-186
chr18-80850764-DUP-157
chr18-80850875-DUP-262
chr18-80938617-DUP-129

chr18-80941448-DUP-90
chr18-80941478-DUP-70
chr18-81139449-DUP-127
chr18-81140492-DUP-230
chr18-81242836-DUP-135
chr18-81557797-DUP-99
chr18-81712276-DUP-171
chr18-81712313-DUP-133
chr18-81740259-DUP-102
chr18-81742179-DUP-389
chr18-81838494-DUP-675
chr18-81916051-DUP-178
chr18-81984452-DUP-68
chr18-82011441-DUP-72
chr18-82036093-DUP-273
chr18-82051472-DUP-477
chr18-82204318-DUP-79
chr18-82205856-DUP-161
chr18-82206958-DUP-116
chr18-82207386-DUP-845
chr18-82207693-DUP-194
chr18-82207908-DUP-724
chr18-82209180-DUP-286
chr18-82209742-DUP-77
chr18-82210807-DUP-77
chr18-82212634-DUP-80
chr18-82232991-DUP-118
chr18-82235792-DUP-77
chr18-82238047-DUP-159
chr18-82694986-DUP-302
chr18-82695722-DUP-132
chr18-82695772-DUP-84
chr18-82942047-DUP-164
chr18-83080273-DUP-71
chr18-83449753-DUP-117
chr18-83463444-DUP-128
chr18-83503208-DUP-290
chr18-83572855-DUP-86
chr18-83901040-DUP-188
chr18-84048550-DUP-73
chr18-84083979-DUP-106
chr18-84221220-DUP-85
chr18-84230901-DUP-70
chr18-84247762-DUP-60
chr18-84294445-DUP-75
chr18-84315973-DUP-230
chr18-84326281-DUP-122
chr18-84471365-DUP-83
chr18-84536142-DUP-65
chr18-84571126-DUP-44416
chr18-84581865-DUP-136
chr18-84582707-DUP-127
chr18-84583057-DUP-361
chr18-84751928-DUP-111
chr18-84881345-DUP-63
chr18-86510637-DUP-73
chr18-87088604-DUP-51
chr18-87696182-DUP-188
chr18-88576512-DUP-2479
chr18-88989654-DUP-89
chr18-89178273-DUP-121
chr18-90179356-DUP-823
chr18-90180271-DUP-76
chr19-3174710-DUP-75
chr19-3255655-DUP-147
chr19-3348275-DUP-97
chr19-3422422-DUP-134
chr19-3579131-DUP-157
chr19-3656537-DUP-114
chr19-3656561-DUP-116
chr19-4280902-DUP-160
chr19-4352906-DUP-60
chr19-4636903-DUP-233
chr19-4696074-DUP-58
chr19-4707579-DUP-66
chr19-5033238-DUP-104
chr19-5033278-DUP-80

chr19-5561991-DUP-81
chr19-5623673-DUP-63
chr19-5650643-DUP-136
chr19-5825673-DUP-85
chr19-5911934-DUP-92
chr19-5971299-DUP-74
chr19-5971323-DUP-114
chr19-6061448-DUP-62
chr19-6071860-DUP-80
chr19-6171474-DUP-97
chr19-6243949-DUP-125
chr19-6298566-DUP-434
chr19-6397387-DUP-54
chr19-6531920-DUP-52
chr19-6558827-DUP-216
chr19-6649299-DUP-100
chr19-6685918-DUP-51
chr19-6941423-DUP-120
chr19-6980953-DUP-149
chr19-7050828-DUP-213
chr19-7050857-DUP-101
chr19-7423235-DUP-53
chr19-7903341-DUP-87
chr19-8041286-DUP-73
chr19-9064390-DUP-221
chr19-9905466-DUP-86
chr19-10204323-DUP-87
chr19-10581490-DUP-98
chr19-10676772-DUP-82
chr19-11280599-DUP-75
chr19-11781241-DUP-52
chr19-11806412-DUP-26927
chr19-11938820-DUP-87
chr19-11980635-DUP-125
chr19-11990460-DUP-55
chr19-12089382-DUP-93
chr19-14159691-DUP-78
chr19-14343460-DUP-68
chr19-14343590-DUP-99
chr19-15445781-DUP-115
chr19-15530845-DUP-56
chr19-16014169-DUP-96
chr19-16250955-DUP-110
chr19-16313166-DUP-84
chr19-16677429-DUP-52
chr19-17228349-DUP-64
chr19-17354789-DUP-52
chr19-17354865-DUP-156
chr19-17708523-DUP-54
chr19-17729334-DUP-106
chr19-18277190-DUP-68
chr19-18497976-DUP-57
chr19-18683752-DUP-55
chr19-19043268-DUP-73
chr19-19160745-DUP-58
chr19-19419849-DUP-60
chr19-19546891-DUP-145
chr19-20181480-DUP-249
chr19-20181688-DUP-214
chr19-20605725-DUP-68
chr19-20705121-DUP-72
chr19-20924470-DUP-83
chr19-20927799-DUP-26889
chr19-21440409-DUP-73
chr19-21526757-DUP-73
chr19-21580719-DUP-291
chr19-21583746-DUP-84
chr19-21918536-DUP-78
chr19-22617945-DUP-100
chr19-23250385-DUP-128
chr19-23413186-DUP-116
chr19-23413214-DUP-62
chr19-24020081-DUP-186
chr19-24274159-DUP-145
chr19-24323213-DUP-201
chr19-24327043-DUP-58
chr19-24444815-DUP-80

chr19-24516938-DUP-118
chr19-24640508-DUP-203
chr19-24640578-DUP-60
chr19-24776072-DUP-63
chr19-25102148-DUP-56
chr19-25102295-DUP-78
chr19-25225655-DUP-150
chr19-25274080-DUP-53
chr19-25301764-DUP-167
chr19-25310791-DUP-64
chr19-25340338-DUP-78
chr19-25687577-DUP-621
chr19-26615020-DUP-109
chr19-26631446-DUP-52
chr19-26697171-DUP-73
chr19-26748079-DUP-86
chr19-26748100-DUP-109
chr19-27119409-DUP-50
chr19-27178171-DUP-147
chr19-27256726-DUP-80
chr19-27644970-DUP-52
chr19-27747013-DUP-309
chr19-28819434-DUP-55
chr19-28980477-DUP-159
chr19-29298023-DUP-184
chr19-29674352-DUP-99
chr19-30153879-DUP-128
chr19-30174976-DUP-32555
chr19-30193272-DUP-119
chr19-30831967-DUP-57
chr19-30832009-DUP-53
chr19-30835698-DUP-239
chr19-31438524-DUP-79
chr19-32354873-DUP-63
chr19-32479885-DUP-458
chr19-32479939-DUP-448
chr19-32663902-DUP-128
chr19-33765200-DUP-234
chr19-33866252-DUP-57
chr19-34057134-DUP-70
chr19-34426797-DUP-90
chr19-34547445-DUP-87
chr19-35239082-DUP-226
chr19-35579690-DUP-75
chr19-35964592-DUP-181
chr19-36096560-DUP-67
chr19-36713955-DUP-59
chr19-36908878-DUP-113
chr19-37078218-DUP-120
chr19-37205254-DUP-153
chr19-37276802-DUP-142
chr19-37277846-DUP-84
chr19-37371461-DUP-83
chr19-37371492-DUP-62
chr19-37389744-DUP-75
chr19-37954768-DUP-55
chr19-37991643-DUP-81
chr19-38655883-DUP-90
chr19-38739537-DUP-94
chr19-38804598-DUP-219
chr19-38804629-DUP-221
chr19-39664393-DUP-253232
chr19-39754395-DUP-127
chr19-39897529-DUP-2222
chr19-39897530-DUP-4394
chr19-40317686-DUP-300
chr19-40496956-DUP-306
chr19-40553981-DUP-121
chr19-40558476-DUP-52
chr19-40626876-DUP-86
chr19-40658065-DUP-107
chr19-40702099-DUP-117
chr19-40706917-DUP-52
chr19-40761751-DUP-84
chr19-40802557-DUP-81
chr19-40830979-DUP-262
chr19-40846551-DUP-105

chr19-41236981-DUP-113
chr19-41298565-DUP-190
chr19-41490052-DUP-3301
chr19-41571696-DUP-123
chr19-41643195-DUP-128
chr19-41670022-DUP-68
chr19-41749320-DUP-110
chr19-41758964-DUP-89
chr19-41861538-DUP-70
chr19-41861562-DUP-142
chr19-41872250-DUP-77
chr19-41872306-DUP-74
chr19-41983194-DUP-52
chr19-42135929-DUP-72
chr19-42263932-DUP-109
chr19-42519997-DUP-63
chr19-42703935-DUP-126
chr19-42703972-DUP-187
chr19-42717656-DUP-69
chr19-42762020-DUP-126
chr19-42875646-DUP-232
chr19-43191462-DUP-62
chr19-43471862-DUP-57
chr19-43704970-DUP-183
chr19-43752477-DUP-78
chr19-43879403-DUP-53
chr19-43896683-DUP-80
chr19-44077805-DUP-55
chr19-44114218-DUP-165
chr19-44174669-DUP-127
chr19-44175205-DUP-136
chr19-44363759-DUP-233
chr19-44532449-DUP-76
chr19-45006177-DUP-178
chr19-45006238-DUP-145
chr19-45389979-DUP-51
chr19-45600080-DUP-64
chr19-45842359-DUP-66
chr19-46218015-DUP-182
chr19-46775649-DUP-275
chr19-46775673-DUP-220
chr19-46804254-DUP-79
chr19-46821317-DUP-69
chr19-46908536-DUP-52
chr19-47004977-DUP-55
chr19-47090130-DUP-81
chr19-47466788-DUP-102
chr19-47534089-DUP-64
chr19-47623057-DUP-105
chr19-47649666-DUP-50
chr19-47849667-DUP-274
chr19-48021907-DUP-114
chr19-48546153-DUP-111
chr19-48618855-DUP-54
chr19-48722804-DUP-67
chr19-49079883-DUP-72
chr19-49637373-DUP-159
chr19-49637405-DUP-84
chr19-49919520-DUP-792
chr19-50072269-DUP-60
chr19-50335465-DUP-57
chr19-50725116-DUP-123
chr19-50831197-DUP-88
chr19-50981588-DUP-103
chr19-51137377-DUP-75
chr19-51409470-DUP-56
chr19-51761323-DUP-106
chr19-52909932-DUP-57
chr19-53203711-DUP-51
chr19-53214355-DUP-74
chr19-53513830-DUP-337
chr19-53656981-DUP-67
chr19-53687858-DUP-87
chr19-53737253-DUP-61
chr19-53765323-DUP-424
chr19-53910629-DUP-91
chr19-53928525-DUP-219

chr19-54139178-DUP-521
chr19-54139871-DUP-216
chr19-54406111-DUP-93
chr19-54406134-DUP-64
chr19-54965547-DUP-60
chr19-54965571-DUP-57
chr19-55083184-DUP-104
chr19-55135292-DUP-57
chr19-55335496-DUP-120
chr19-55357396-DUP-194
chr19-55516892-DUP-54
chr19-55582024-DUP-53
chr19-55759009-DUP-430
chr19-55976489-DUP-58
chr19-56094562-DUP-118
chr19-56372652-DUP-76
chr19-56477768-DUP-71
chr19-56496189-DUP-83
chr19-56561285-DUP-134
chr19-56805103-DUP-84
chr19-56805137-DUP-86
chr19-56835979-DUP-856
chr19-56836012-DUP-772
chr19-56836119-DUP-142
chr19-56921493-DUP-52
chr19-57148076-DUP-56
chr19-57220497-DUP-150
chr19-57233206-DUP-80
chr19-57446842-DUP-143
chr19-57447021-DUP-147
chr19-57570905-DUP-91
chr19-57622644-DUP-233
chr19-57623139-DUP-153
chr19-57667955-DUP-62
chr19-57696250-DUP-86
chr19-57696286-DUP-88
chr19-57783859-DUP-264
chr19-57783944-DUP-190
chr19-58251137-DUP-103
chr19-58506936-DUP-219
chr19-58541105-DUP-68
chr19-58828798-DUP-124
chr19-58930452-DUP-80
chr19-58990061-DUP-56
chr19-59076920-DUP-142
chr19-59076962-DUP-73
chr19-59268605-DUP-166
chr19-59313320-DUP-72
chr19-59328515-DUP-115
chr19-59337774-DUP-2143
chr19-59433682-DUP-2249
chr19-59724438-DUP-336
chr19-59858527-DUP-61
chr19-59989486-DUP-52
chr19-59995656-DUP-81
chr19-60180239-DUP-61
chr2-3174123-DUP-147
chr2-3218731-DUP-119
chr2-3434216-DUP-243
chr2-3435027-DUP-80
chr2-3704221-DUP-145
chr2-3779689-DUP-51
chr2-3786267-DUP-51
chr2-3909831-DUP-106
chr2-3942696-DUP-300
chr2-3965525-DUP-2759
chr2-3965526-DUP-3905
chr2-3965608-DUP-1676
chr2-3965986-DUP-291
chr2-3966712-DUP-2715
chr2-3967484-DUP-1948
chr2-3967512-DUP-165
chr2-3968034-DUP-1397
chr2-3968418-DUP-374
chr2-3969133-DUP-205
chr2-4050887-DUP-267
chr2-4387279-DUP-102

chr2-4783005-DUP-145
chr2-4783036-DUP-105
chr2-5014585-DUP-454
chr2-5163509-DUP-139
chr2-5323586-DUP-342
chr2-5407064-DUP-144
chr2-5467316-DUP-53
chr2-5587173-DUP-385
chr2-5587355-DUP-258
chr2-5606765-DUP-66
chr2-5650863-DUP-74
chr2-5672446-DUP-99
chr2-5713655-DUP-60
chr2-5803230-DUP-193
chr2-6332018-DUP-307
chr2-6332294-DUP-711
chr2-6349739-DUP-72
chr2-6477631-DUP-82
chr2-6637588-DUP-112
chr2-6695478-DUP-58
chr2-6783380-DUP-54
chr2-6831794-DUP-643
chr2-7209015-DUP-137
chr2-7731696-DUP-76
chr2-7859745-DUP-90
chr2-8509177-DUP-978
chr2-8713663-DUP-54
chr2-8951475-DUP-215
chr2-9532734-DUP-53
chr2-9935564-DUP-96
chr2-10176645-DUP-93
chr2-10190783-DUP-135
chr2-10462942-DUP-72
chr2-10511595-DUP-88
chr2-10755690-DUP-162
chr2-10860747-DUP-97
chr2-11065219-DUP-296
chr2-11067128-DUP-87
chr2-11284057-DUP-132
chr2-11284172-DUP-55
chr2-11454888-DUP-53
chr2-11731442-DUP-80
chr2-13348724-DUP-392
chr2-13348746-DUP-516
chr2-13495356-DUP-51
chr2-14602972-DUP-52
chr2-14834711-DUP-68
chr2-14873903-DUP-57
chr2-14880922-DUP-110
chr2-15047173-DUP-116
chr2-15528702-DUP-90
chr2-15571542-DUP-386
chr2-16061158-DUP-187
chr2-17034973-DUP-81
chr2-17778059-DUP-110
chr2-17778095-DUP-145
chr2-17862471-DUP-92
chr2-18783618-DUP-81
chr2-19274581-DUP-73
chr2-19781790-DUP-112
chr2-20037852-DUP-71
chr2-20280138-DUP-173
chr2-20412405-DUP-80
chr2-20582506-DUP-181
chr2-20630968-DUP-66
chr2-20845606-DUP-66
chr2-21022131-DUP-153
chr2-21640960-DUP-109
chr2-21640995-DUP-61
chr2-22575990-DUP-144
chr2-22576012-DUP-146
chr2-22658911-DUP-54
chr2-23962891-DUP-64
chr2-24350915-DUP-81
chr2-24353899-DUP-58
chr2-24435794-DUP-60
chr2-24452557-DUP-60

chr2-24519912-DUP-102
chr2-24557665-DUP-54
chr2-24624140-DUP-869
chr2-24624336-DUP-416
chr2-24624781-DUP-113
chr2-24930799-DUP-62
chr2-24934286-DUP-1618
chr2-24999336-DUP-126
chr2-25334717-DUP-71
chr2-26248348-DUP-66
chr2-26909695-DUP-131
chr2-26954496-DUP-443
chr2-26954574-DUP-510
chr2-27019546-DUP-174
chr2-27165828-DUP-341
chr2-27765428-DUP-6715
chr2-27802428-DUP-59
chr2-27831399-DUP-72
chr2-27851394-DUP-122
chr2-28056111-DUP-52
chr2-28092445-DUP-215
chr2-28656379-DUP-131
chr2-29024857-DUP-89
chr2-29051533-DUP-102
chr2-29325005-DUP-142
chr2-29417904-DUP-58
chr2-29439671-DUP-92
chr2-29439693-DUP-64
chr2-29885603-DUP-58
chr2-30317871-DUP-104
chr2-30382949-DUP-158
chr2-30445414-DUP-86
chr2-30461080-DUP-159
chr2-30655486-DUP-165
chr2-30737542-DUP-1616
chr2-30987486-DUP-277
chr2-30987520-DUP-233
chr2-31086627-DUP-54
chr2-31326502-DUP-137
chr2-31648352-DUP-124
chr2-31759881-DUP-61
chr2-31823130-DUP-179
chr2-32135487-DUP-47758
chr2-32650865-DUP-112
chr2-32804205-DUP-61
chr2-33416114-DUP-121
chr2-33594584-DUP-125
chr2-33919303-DUP-78
chr2-33941389-DUP-64
chr2-34049667-DUP-4247
chr2-34376271-DUP-53
chr2-34558894-DUP-73
chr2-34611344-DUP-54
chr2-35134038-DUP-196
chr2-35440959-DUP-102
chr2-37676881-DUP-146
chr2-38301848-DUP-162
chr2-38490851-DUP-125
chr2-38498550-DUP-89
chr2-41393343-DUP-59
chr2-41943847-DUP-58
chr2-42246390-DUP-70
chr2-43668877-DUP-98
chr2-43720030-DUP-135
chr2-43788827-DUP-68
chr2-44248182-DUP-74
chr2-46171832-DUP-463
chr2-46661142-DUP-209
chr2-47716541-DUP-55
chr2-47896789-DUP-88
chr2-48453888-DUP-56
chr2-48501632-DUP-101
chr2-48825344-DUP-149
chr2-49607857-DUP-146
chr2-49607889-DUP-75
chr2-50631422-DUP-170
chr2-50631459-DUP-69

chr2-51706918-DUP-81
chr2-51730084-DUP-195
chr2-51730105-DUP-100
chr2-51851095-DUP-126
chr2-52070082-DUP-218
chr2-52297086-DUP-95
chr2-52305964-DUP-122
chr2-52305996-DUP-69
chr2-52562153-DUP-112
chr2-52562212-DUP-76
chr2-52586416-DUP-1153
chr2-52586417-DUP-158
chr2-52587343-DUP-1709
chr2-52747345-DUP-119
chr2-54162354-DUP-218
chr2-56666644-DUP-130
chr2-57147313-DUP-185
chr2-57284626-DUP-136
chr2-57461776-DUP-129
chr2-58170042-DUP-168
chr2-58170077-DUP-88
chr2-58366019-DUP-55
chr2-58550309-DUP-53
chr2-59083258-DUP-228
chr2-59083280-DUP-233
chr2-59115981-DUP-158
chr2-59545756-DUP-104
chr2-59641315-DUP-86
chr2-59801836-DUP-951
chr2-59802032-DUP-858
chr2-60020229-DUP-376
chr2-60193676-DUP-85
chr2-60231420-DUP-173
chr2-60299744-DUP-132
chr2-60821825-DUP-106
chr2-60921737-DUP-68
chr2-60947598-DUP-68
chr2-61014553-DUP-51
chr2-61031511-DUP-75
chr2-61057363-DUP-53
chr2-61358598-DUP-53
chr2-61626884-DUP-52
chr2-63183841-DUP-56
chr2-64446146-DUP-76
chr2-65613980-DUP-129
chr2-66689818-DUP-19669
chr2-66752257-DUP-84
chr2-67347980-DUP-106
chr2-67518825-DUP-67
chr2-67649688-DUP-53
chr2-67827348-DUP-148
chr2-67827388-DUP-80
chr2-68279404-DUP-159
chr2-68279435-DUP-110
chr2-68718520-DUP-62
chr2-69151583-DUP-51
chr2-69301864-DUP-78
chr2-69502896-DUP-65507
chr2-69570075-DUP-188
chr2-69570116-DUP-99
chr2-69896548-DUP-119
chr2-69985179-DUP-51
chr2-70227820-DUP-88
chr2-70368629-DUP-174
chr2-70833028-DUP-133
chr2-71065219-DUP-58
chr2-71094982-DUP-108
chr2-71095044-DUP-59
chr2-71299347-DUP-644
chr2-71329702-DUP-71
chr2-71358305-DUP-70
chr2-72008407-DUP-51
chr2-72130568-DUP-68
chr2-72325944-DUP-68
chr2-72389468-DUP-102
chr2-72389509-DUP-100
chr2-72578177-DUP-51

chr2-72731955-DUP-81
chr2-72931517-DUP-91
chr2-73055569-DUP-55
chr2-73194434-DUP-70
chr2-73615029-DUP-94
chr2-73770826-DUP-61
chr2-73992417-DUP-76
chr2-74295133-DUP-175
chr2-74899913-DUP-70
chr2-74908803-DUP-60
chr2-74940002-DUP-60
chr2-74949639-DUP-278
chr2-75960273-DUP-172
chr2-76038253-DUP-1737
chr2-77001378-DUP-54
chr2-77030886-DUP-124
chr2-77033121-DUP-116
chr2-77082251-DUP-98
chr2-77428941-DUP-162
chr2-77428987-DUP-79
chr2-77700000-DUP-157702
chr2-77801540-DUP-112
chr2-78061241-DUP-164
chr2-78192134-DUP-125
chr2-78852527-DUP-100
chr2-78934773-DUP-200932
chr2-79163316-DUP-128
chr2-80094618-DUP-184
chr2-80359011-DUP-71
chr2-80780406-DUP-168
chr2-82208041-DUP-187
chr2-83531861-DUP-23108
chr2-83532041-DUP-352
chr2-84229915-DUP-576
chr2-84301261-DUP-150
chr2-84460171-DUP-198
chr2-85030185-DUP-107
chr2-85671181-DUP-170
chr2-86614586-DUP-1898
chr2-86643135-DUP-117
chr2-87225360-DUP-98
chr2-87901689-DUP-185
chr2-87901728-DUP-137
chr2-88157366-DUP-200
chr2-88218843-DUP-2262
chr2-88483738-DUP-859
chr2-88483764-DUP-943
chr2-88699357-DUP-89
chr2-89894491-DUP-3935
chr2-90211114-DUP-131
chr2-90523629-DUP-295
chr2-90564981-DUP-186
chr2-90584441-DUP-31096
chr2-90584495-DUP-30963
chr2-90793382-DUP-67
chr2-90813945-DUP-80
chr2-90925838-DUP-101
chr2-91631811-DUP-92
chr2-91798950-DUP-57
chr2-91971808-DUP-242
chr2-92248084-DUP-51
chr2-92459185-DUP-119
chr2-92621202-DUP-158
chr2-92714426-DUP-122
chr2-92714474-DUP-155
chr2-92916471-DUP-297
chr2-92940248-DUP-58
chr2-93080409-DUP-92
chr2-93499516-DUP-54
chr2-93603776-DUP-172
chr2-93606623-DUP-91
chr2-93675956-DUP-134
chr2-93735531-DUP-78
chr2-93816710-DUP-971
chr2-94155228-DUP-57
chr2-94279576-DUP-65
chr2-94324616-DUP-74

chr2-94549369-DUP-60
chr2-95308167-DUP-2684
chr2-96030859-DUP-615
chr2-97313786-DUP-1693
chr2-97367058-DUP-79
chr2-97427619-DUP-57
chr2-97935338-DUP-5870
chr2-98487974-DUP-3311
chr2-99349972-DUP-346
chr2-100702227-DUP-104
chr2-101196330-DUP-142
chr2-101357040-DUP-184
chr2-101780163-DUP-113
chr2-101842133-DUP-56
chr2-101956789-DUP-87
chr2-102198994-DUP-258
chr2-102200960-DUP-67
chr2-103261030-DUP-111
chr2-103420148-DUP-112
chr2-103877182-DUP-53
chr2-104126461-DUP-130
chr2-104297981-DUP-66
chr2-104501723-DUP-54
chr2-104851556-DUP-86
chr2-104984271-DUP-74
chr2-105089026-DUP-267
chr2-105445286-DUP-200
chr2-105492413-DUP-69
chr2-105576865-DUP-152
chr2-105760044-DUP-133
chr2-106390098-DUP-103
chr2-106607555-DUP-80
chr2-106618948-DUP-64
chr2-106956030-DUP-832
chr2-106956136-DUP-102
chr2-107655518-DUP-99
chr2-107778874-DUP-62
chr2-107976063-DUP-52
chr2-107990451-DUP-284
chr2-108433895-DUP-96
chr2-108433930-DUP-72
chr2-108690649-DUP-109
chr2-109070923-DUP-1227
chr2-109410242-DUP-267
chr2-109432192-DUP-677
chr2-109585484-DUP-121
chr2-109735859-DUP-53
chr2-110160177-DUP-106
chr2-110310762-DUP-3001
chr2-112420928-DUP-325
chr2-112434678-DUP-109
chr2-112548247-DUP-74
chr2-112824716-DUP-1258
chr2-112847156-DUP-457
chr2-113119021-DUP-104
chr2-113152987-DUP-303
chr2-113649730-DUP-55
chr2-113659749-DUP-56
chr2-113674867-DUP-116
chr2-113696753-DUP-106
chr2-113866920-DUP-141
chr2-113866943-DUP-71
chr2-113880489-DUP-9677
chr2-114066206-DUP-85
chr2-114066228-DUP-87
chr2-114324177-DUP-60
chr2-115381875-DUP-197
chr2-115972666-DUP-53
chr2-116306883-DUP-90
chr2-116391327-DUP-1176
chr2-116399414-DUP-1704
chr2-116891518-DUP-93
chr2-117063981-DUP-1206
chr2-117064038-DUP-416
chr2-117132051-DUP-57
chr2-117247118-DUP-55
chr2-117289247-DUP-51

chr2-117508719-DUP-76
chr2-117605818-DUP-76
chr2-117610223-DUP-245
chr2-117610322-DUP-58
chr2-117943224-DUP-51
chr2-118275107-DUP-662
chr2-118533766-DUP-168587
chr2-118730228-DUP-103
chr2-119149498-DUP-107
chr2-119209132-DUP-56
chr2-119316764-DUP-2876
chr2-119324939-DUP-66
chr2-119432339-DUP-82
chr2-119618550-DUP-306
chr2-119652663-DUP-114
chr2-119652713-DUP-135
chr2-119849813-DUP-74
chr2-120037855-DUP-80
chr2-120127796-DUP-93
chr2-120127822-DUP-244
chr2-120173538-DUP-52
chr2-120451349-DUP-79
chr2-120875188-DUP-70
chr2-120899999-DUP-100
chr2-121257395-DUP-193
chr2-121384630-DUP-95
chr2-121590924-DUP-108
chr2-121925655-DUP-60
chr2-122181029-DUP-24761
chr2-122946132-DUP-80
chr2-123294184-DUP-71
chr2-123468402-DUP-226
chr2-124108507-DUP-133
chr2-124902675-DUP-70
chr2-125336764-DUP-66
chr2-125349847-DUP-592
chr2-125429459-DUP-81
chr2-125443776-DUP-216
chr2-125586224-DUP-53
chr2-125608265-DUP-457
chr2-125608848-DUP-99
chr2-125621102-DUP-63
chr2-125659793-DUP-298
chr2-125740797-DUP-103
chr2-125893469-DUP-71
chr2-126123963-DUP-80
chr2-126212361-DUP-59
chr2-126361945-DUP-60
chr2-126754724-DUP-74
chr2-126798107-DUP-81
chr2-126879382-DUP-83
chr2-126965219-DUP-52
chr2-127067151-DUP-52
chr2-128060616-DUP-195
chr2-128251636-DUP-294
chr2-128251661-DUP-248
chr2-128296087-DUP-76
chr2-128403928-DUP-50
chr2-128404255-DUP-335
chr2-128839916-DUP-315
chr2-128902276-DUP-60
chr2-129212772-DUP-208
chr2-129218704-DUP-51
chr2-129262222-DUP-60
chr2-129271872-DUP-101
chr2-129284429-DUP-100
chr2-129500098-DUP-200
chr2-129500166-DUP-91
chr2-129782824-DUP-158
chr2-130653328-DUP-311
chr2-130653354-DUP-271
chr2-130675937-DUP-68
chr2-130795218-DUP-55
chr2-130976169-DUP-166
chr2-131043147-DUP-64
chr2-131146757-DUP-371
chr2-131247262-DUP-290

chr2-131491180-DUP-72
chr2-131511413-DUP-5064
chr2-131562510-DUP-61
chr2-131584176-DUP-137
chr2-131618643-DUP-52
chr2-131693474-DUP-60
chr2-131701592-DUP-289
chr2-131709026-DUP-246
chr2-131880354-DUP-60
chr2-131909119-DUP-97
chr2-132323684-DUP-94
chr2-132353540-DUP-54
chr2-133011808-DUP-52
chr2-133567087-DUP-52
chr2-135504669-DUP-7039
chr2-135520186-DUP-90
chr2-135520324-DUP-457
chr2-135600662-DUP-389
chr2-135838084-DUP-73
chr2-135994288-DUP-62
chr2-136413381-DUP-157
chr2-137105408-DUP-123
chr2-138117800-DUP-60
chr2-138209654-DUP-66
chr2-138483799-DUP-550
chr2-139171525-DUP-271
chr2-139526895-DUP-258
chr2-139863914-DUP-115
chr2-139996894-DUP-58
chr2-140325604-DUP-36814
chr2-140487225-DUP-3159
chr2-140516640-DUP-83
chr2-140587711-DUP-1889
chr2-140824150-DUP-91
chr2-141543680-DUP-148
chr2-141774921-DUP-72
chr2-142181784-DUP-62
chr2-142735324-DUP-85
chr2-143355932-DUP-90
chr2-143481856-DUP-61
chr2-143556260-DUP-90
chr2-143659996-DUP-52
chr2-143835835-DUP-899
chr2-143835938-DUP-700
chr2-143923652-DUP-231
chr2-143923693-DUP-299
chr2-144206687-DUP-58
chr2-144755031-DUP-121
chr2-144755058-DUP-73
chr2-145540892-DUP-177
chr2-145876061-DUP-109
chr2-145893639-DUP-59
chr2-146424055-DUP-144
chr2-146497976-DUP-77
chr2-146557759-DUP-100
chr2-147403534-DUP-127
chr2-147439754-DUP-64
chr2-147776476-DUP-55
chr2-147914026-DUP-160
chr2-148621283-DUP-175
chr2-148927714-DUP-108673
chr2-149156623-DUP-1970
chr2-150035793-DUP-118
chr2-150663819-DUP-273
chr2-151085977-DUP-154936
chr2-151556595-DUP-102825
chr2-151685704-DUP-91
chr2-151904623-DUP-327
chr2-151904666-DUP-171
chr2-152235052-DUP-103
chr2-152236668-DUP-119
chr2-152674272-DUP-55
chr2-152744674-DUP-51
chr2-152880956-DUP-62
chr2-152936393-DUP-626
chr2-152937693-DUP-1033
chr2-152938354-DUP-269

chr2-152938751-DUP-626
chr2-152940073-DUP-756
chr2-152940142-DUP-181
chr2-152941162-DUP-216
chr2-152964881-DUP-54
chr2-153123895-DUP-292
chr2-153395660-DUP-186
chr2-153593262-DUP-73
chr2-154003242-DUP-53
chr2-154162684-DUP-147
chr2-154266758-DUP-324
chr2-154266866-DUP-348
chr2-154359700-DUP-415
chr2-154444200-DUP-4160
chr2-154444258-DUP-4049
chr2-154506867-DUP-58
chr2-154881050-DUP-268
chr2-155085102-DUP-344
chr2-155508660-DUP-52
chr2-155743593-DUP-128
chr2-155983989-DUP-84
chr2-156195820-DUP-123
chr2-156252002-DUP-126
chr2-156299927-DUP-243
chr2-156351267-DUP-72
chr2-156551955-DUP-489
chr2-156619881-DUP-52
chr2-156887759-DUP-4583
chr2-157323379-DUP-124
chr2-157571730-DUP-265
chr2-157591849-DUP-68
chr2-157709186-DUP-148
chr2-157807813-DUP-72
chr2-157892562-DUP-600
chr2-158122002-DUP-68
chr2-158146982-DUP-2913
chr2-158486167-DUP-410
chr2-158723385-DUP-261
chr2-158742440-DUP-62
chr2-160238782-DUP-500
chr2-160238821-DUP-452
chr2-160815713-DUP-68
chr2-161030641-DUP-53
chr2-161299476-DUP-104
chr2-161404035-DUP-586
chr2-161523314-DUP-64
chr2-162200151-DUP-117
chr2-162285720-DUP-55
chr2-162398223-DUP-5076
chr2-162797432-DUP-287
chr2-162845659-DUP-71
chr2-162879664-DUP-810
chr2-163200887-DUP-86
chr2-163335921-DUP-51
chr2-163396591-DUP-127
chr2-163396624-DUP-80
chr2-163543800-DUP-242
chr2-163699291-DUP-82
chr2-163852210-DUP-660
chr2-163877308-DUP-58
chr2-163984651-DUP-69
chr2-164070898-DUP-56
chr2-164341590-DUP-92
chr2-164402368-DUP-55
chr2-164457544-DUP-82
chr2-164501335-DUP-136
chr2-164501369-DUP-105
chr2-164742655-DUP-53
chr2-165132751-DUP-110
chr2-165133300-DUP-112
chr2-165134652-DUP-143
chr2-165265499-DUP-131
chr2-165277146-DUP-175
chr2-165281498-DUP-60
chr2-165353357-DUP-85
chr2-165522209-DUP-61
chr2-165577717-DUP-137

chr2-165626713-DUP-80
chr2-165638317-DUP-96
chr2-165644027-DUP-207
chr2-165644053-DUP-215
chr2-165883669-DUP-67
chr2-166002643-DUP-78
chr2-166003144-DUP-68
chr2-166044851-DUP-117
chr2-166289595-DUP-265
chr2-166446207-DUP-215
chr2-166544645-DUP-178
chr2-166785818-DUP-72
chr2-166887307-DUP-159
chr2-166908357-DUP-1133
chr2-166908452-DUP-1011
chr2-167105146-DUP-96
chr2-167123228-DUP-51
chr2-167124398-DUP-53
chr2-167132695-DUP-137
chr2-167549325-DUP-295
chr2-167655319-DUP-75
chr2-167718681-DUP-226
chr2-167820390-DUP-194
chr2-168061314-DUP-277
chr2-168068524-DUP-71
chr2-168165535-DUP-118
chr2-168244806-DUP-54
chr2-168438263-DUP-64
chr2-168480418-DUP-173
chr2-168590700-DUP-126
chr2-168638536-DUP-58
chr2-169317055-DUP-77
chr2-169379506-DUP-6115
chr2-169409049-DUP-136
chr2-169459814-DUP-93
chr2-169485726-DUP-63
chr2-169485747-DUP-65
chr2-169487011-DUP-234
chr2-169658562-DUP-176
chr2-169680757-DUP-74
chr2-169715897-DUP-55
chr2-169847679-DUP-67
chr2-170191757-DUP-57
chr2-170346539-DUP-97
chr2-170732624-DUP-107
chr2-170745407-DUP-59
chr2-171005125-DUP-860
chr2-171182108-DUP-191
chr2-171235931-DUP-751
chr2-171236771-DUP-205
chr2-171347907-DUP-163
chr2-171480961-DUP-75
chr2-172204807-DUP-100
chr2-172206049-DUP-62
chr2-172484229-DUP-58
chr2-172835917-DUP-72
chr2-172920599-DUP-5270
chr2-172945045-DUP-96
chr2-173029649-DUP-59
chr2-173112734-DUP-98
chr2-173173770-DUP-79
chr2-173173818-DUP-78
chr2-173279924-DUP-73
chr2-173306697-DUP-54
chr2-173384192-DUP-78
chr2-173475381-DUP-304
chr2-173478604-DUP-143
chr2-173541944-DUP-137
chr2-173595435-DUP-99
chr2-173804000-DUP-163
chr2-173935059-DUP-126
chr2-174157458-DUP-126
chr2-174189097-DUP-1295
chr2-174189514-DUP-199
chr2-174190449-DUP-77
chr2-174314229-DUP-66
chr2-174326408-DUP-83

chr2-174363010-DUP-106
chr2-174417793-DUP-93
chr2-174750388-DUP-65
chr2-176402896-DUP-366512
chr2-176493556-DUP-1519
chr2-177802969-DUP-91
chr2-177803003-DUP-95
chr2-177871293-DUP-52
chr2-178104006-DUP-140
chr2-178112559-DUP-76
chr2-178170027-DUP-433
chr2-178170647-DUP-56
chr2-178175242-DUP-103
chr2-178225544-DUP-78
chr2-178283526-DUP-207
chr2-178343078-DUP-60
chr2-178645024-DUP-108
chr2-178951568-DUP-361
chr2-178954921-DUP-56
chr2-178992692-DUP-78
chr2-179134182-DUP-70
chr2-179155494-DUP-150
chr2-179280498-DUP-65
chr2-179341370-DUP-176
chr2-179358319-DUP-106
chr2-179361211-DUP-64
chr2-179368996-DUP-99
chr2-179406506-DUP-67
chr2-179406645-DUP-156
chr2-179472696-DUP-65
chr2-179483114-DUP-66
chr2-179708424-DUP-413
chr2-179708514-DUP-429
chr2-179765396-DUP-94
chr2-180918040-DUP-56
chr2-181085563-DUP-58
chr3-3034855-DUP-1637
chr3-3043268-DUP-604
chr3-3075768-DUP-3564
chr3-3080756-DUP-61
chr3-3852073-DUP-94
chr3-4374627-DUP-538
chr3-7424240-DUP-77
chr3-7657178-DUP-232
chr3-8617510-DUP-63
chr3-8766658-DUP-82
chr3-8830772-DUP-62
chr3-8835793-DUP-252
chr3-8835876-DUP-64
chr3-9017813-DUP-149
chr3-9017835-DUP-95
chr3-9018638-DUP-110
chr3-9237574-DUP-136
chr3-9264324-DUP-84
chr3-9396813-DUP-93
chr3-9516232-DUP-57
chr3-9566799-DUP-142
chr3-9605596-DUP-119
chr3-9667297-DUP-73
chr3-9682418-DUP-88
chr3-9682448-DUP-85
chr3-9748125-DUP-108
chr3-9792532-DUP-99
chr3-9830157-DUP-71
chr3-9864011-DUP-126
chr3-9991320-DUP-55
chr3-9994565-DUP-81
chr3-10049173-DUP-59
chr3-10100709-DUP-195
chr3-10270008-DUP-113
chr3-10471681-DUP-113
chr3-10779932-DUP-277
chr3-10795108-DUP-113
chr3-11240321-DUP-91
chr3-12661253-DUP-112
chr3-13297753-DUP-78
chr3-13617985-DUP-8024

chr3-14749349-DUP-318
chr3-14907083-DUP-201
chr3-14940362-DUP-84
chr3-14964667-DUP-100
chr3-15064020-DUP-88
chr3-15244384-DUP-62
chr3-15363162-DUP-590
chr3-16321680-DUP-168
chr3-16985451-DUP-457
chr3-17143799-DUP-65
chr3-18245406-DUP-108
chr3-18277180-DUP-310
chr3-18547258-DUP-51
chr3-18831090-DUP-86
chr3-19307555-DUP-200
chr3-19491996-DUP-53
chr3-19720850-DUP-235
chr3-19737629-DUP-56
chr3-20383522-DUP-115
chr3-20431471-DUP-62
chr3-20552251-DUP-802
chr3-20552284-DUP-557
chr3-20835912-DUP-68
chr3-20860904-DUP-55
chr3-22171170-DUP-63
chr3-22262056-DUP-75
chr3-22563027-DUP-81
chr3-24419140-DUP-138
chr3-24419196-DUP-92
chr3-26515265-DUP-53
chr3-26655820-DUP-331
chr3-27051607-DUP-915
chr3-27052268-DUP-249
chr3-27333889-DUP-138
chr3-27341653-DUP-55
chr3-27358461-DUP-4481
chr3-27360303-DUP-4512
chr3-27700985-DUP-54
chr3-27806579-DUP-77
chr3-28065578-DUP-146
chr3-28463469-DUP-116
chr3-29932858-DUP-190
chr3-30041588-DUP-151
chr3-30239434-DUP-376
chr3-30787684-DUP-270
chr3-30984654-DUP-118
chr3-31075783-DUP-55
chr3-31126729-DUP-72
chr3-31244789-DUP-2858
chr3-31267495-DUP-63
chr3-31279071-DUP-127
chr3-31456449-DUP-75
chr3-31490940-DUP-129
chr3-31491097-DUP-801
chr3-31741179-DUP-51
chr3-31856722-DUP-63
chr3-32106041-DUP-88
chr3-32544096-DUP-82
chr3-32799863-DUP-161
chr3-32941726-DUP-64
chr3-33508378-DUP-53
chr3-34241313-DUP-108
chr3-34274434-DUP-64
chr3-34290070-DUP-193
chr3-34447688-DUP-227020
chr3-34493083-DUP-170
chr3-34493105-DUP-132
chr3-34508349-DUP-68
chr3-34782948-DUP-112
chr3-34857847-DUP-73
chr3-34858805-DUP-68
chr3-34948748-DUP-218
chr3-34952966-DUP-54
chr3-34969468-DUP-87
chr3-35035012-DUP-67
chr3-35164800-DUP-444168
chr3-35548379-DUP-83

chr3-35887025-DUP-125
chr3-35963887-DUP-96
chr3-36128569-DUP-469
chr3-36350498-DUP-94
chr3-36515623-DUP-56
chr3-36815335-DUP-203
chr3-36815411-DUP-123
chr3-37159634-DUP-61
chr3-37488120-DUP-153
chr3-37504057-DUP-68
chr3-37665153-DUP-834
chr3-37699886-DUP-51
chr3-37735292-DUP-58
chr3-37827124-DUP-73
chr3-37853719-DUP-151
chr3-37887678-DUP-137
chr3-38110390-DUP-345
chr3-38552137-DUP-66
chr3-38722658-DUP-52
chr3-39867436-DUP-170
chr3-39902033-DUP-63
chr3-39917986-DUP-8216
chr3-39918134-DUP-8216
chr3-40604298-DUP-1386
chr3-40604507-DUP-400
chr3-40818698-DUP-46959
chr3-41080331-DUP-117
chr3-41368912-DUP-95
chr3-41514448-DUP-2166
chr3-41532442-DUP-159
chr3-42057011-DUP-817
chr3-42057067-DUP-759
chr3-45413043-DUP-525
chr3-45428756-DUP-191
chr3-47009767-DUP-117
chr3-47034118-DUP-60
chr3-47074513-DUP-111
chr3-47428025-DUP-4611
chr3-47715031-DUP-586
chr3-48154373-DUP-119
chr3-49083122-DUP-181
chr3-49299096-DUP-89
chr3-49456313-DUP-107
chr3-49971412-DUP-105
chr3-50593968-DUP-181
chr3-50893155-DUP-125
chr3-51179917-DUP-177
chr3-51179963-DUP-100
chr3-51209339-DUP-140
chr3-51263783-DUP-113
chr3-51271725-DUP-74
chr3-52235514-DUP-471
chr3-52261289-DUP-222
chr3-52730940-DUP-77
chr3-52738661-DUP-84
chr3-53141063-DUP-69
chr3-53158044-DUP-75
chr3-53419147-DUP-61
chr3-53438281-DUP-312
chr3-53438346-DUP-963
chr3-53442312-DUP-57
chr3-53590836-DUP-375
chr3-53590928-DUP-294
chr3-54305854-DUP-109
chr3-54585992-DUP-373
chr3-54618758-DUP-79
chr3-55039380-DUP-115
chr3-55323661-DUP-223
chr3-55354211-DUP-74
chr3-55594408-DUP-134
chr3-55594468-DUP-68
chr3-55902027-DUP-120
chr3-55902052-DUP-61
chr3-55973162-DUP-196
chr3-56483911-DUP-946
chr3-56540123-DUP-1626
chr3-57667610-DUP-98

chr3-61853932-DUP-55
chr3-62179629-DUP-152
chr3-63756700-DUP-176
chr3-63756727-DUP-128
chr3-64700539-DUP-549
chr3-65124383-DUP-356
chr3-65249460-DUP-390
chr3-65437507-DUP-80
chr3-66156575-DUP-54
chr3-66351177-DUP-222
chr3-66458324-DUP-85
chr3-66556824-DUP-76
chr3-66601276-DUP-580
chr3-66630254-DUP-226
chr3-67027456-DUP-70
chr3-67075981-DUP-82
chr3-67135236-DUP-174
chr3-67489489-DUP-53
chr3-68647260-DUP-85
chr3-68647320-DUP-58
chr3-68704630-DUP-57046
chr3-68901744-DUP-60
chr3-68951947-DUP-52
chr3-69682792-DUP-570
chr3-69865504-DUP-77
chr3-69865874-DUP-291
chr3-69873024-DUP-123
chr3-70221192-DUP-63
chr3-71767829-DUP-86
chr3-73623712-DUP-4532
chr3-73623734-DUP-4477
chr3-75854094-DUP-56
chr3-76343071-DUP-961
chr3-76943798-DUP-99
chr3-77197747-DUP-399
chr3-77847359-DUP-72
chr3-78385103-DUP-44988
chr3-78633452-DUP-60897
chr3-79150074-DUP-146
chr3-79273155-DUP-158
chr3-79273192-DUP-123
chr3-79347819-DUP-286
chr3-79378056-DUP-78
chr3-79404180-DUP-62
chr3-79806626-DUP-177
chr3-79902674-DUP-111
chr3-80774175-DUP-318
chr3-80799473-DUP-90
chr3-81238380-DUP-69
chr3-81715126-DUP-384
chr3-81983791-DUP-62
chr3-82254209-DUP-75
chr3-82287657-DUP-222
chr3-82562388-DUP-53
chr3-82894959-DUP-95
chr3-82971997-DUP-681
chr3-82972068-DUP-633
chr3-83121643-DUP-151
chr3-83215321-DUP-67
chr3-83215352-DUP-69
chr3-83753952-DUP-171
chr3-83755737-DUP-58
chr3-83848838-DUP-57
chr3-83861550-DUP-62
chr3-83870201-DUP-88
chr3-84022001-DUP-222
chr3-84295477-DUP-145
chr3-84731655-DUP-6019
chr3-84752140-DUP-581
chr3-85241904-DUP-172
chr3-85327110-DUP-64
chr3-85413669-DUP-116
chr3-85514841-DUP-112
chr3-85865241-DUP-91
chr3-86544014-DUP-524
chr3-86564780-DUP-70
chr3-86732532-DUP-149

chr3-86817629-DUP-79
chr3-87530457-DUP-152
chr3-87668066-DUP-260
chr3-88003740-DUP-61
chr3-88020123-DUP-228
chr3-88503963-DUP-274
chr3-88635374-DUP-174
chr3-89025275-DUP-60
chr3-89050855-DUP-61
chr3-89088325-DUP-70
chr3-89271903-DUP-190
chr3-89305049-DUP-192
chr3-89596333-DUP-54
chr3-89673433-DUP-211
chr3-89751149-DUP-206
chr3-89751212-DUP-104
chr3-90174216-DUP-59
chr3-90275408-DUP-569
chr3-91020896-DUP-109037
chr3-92479161-DUP-79
chr3-92591826-DUP-200
chr3-93110593-DUP-72
chr3-93113206-DUP-238
chr3-93189922-DUP-735
chr3-93198590-DUP-762
chr3-93231839-DUP-529
chr3-93237575-DUP-537
chr3-93304211-DUP-106
chr3-94234734-DUP-142
chr3-94376048-DUP-53
chr3-94612788-DUP-7669
chr3-94613330-DUP-7676
chr3-94619237-DUP-7622
chr3-94781010-DUP-68
chr3-95118873-DUP-2603
chr3-95374578-DUP-55
chr3-95597293-DUP-82
chr3-95772987-DUP-110
chr3-95820491-DUP-76
chr3-95923965-DUP-58
chr3-96117835-DUP-66
chr3-96117856-DUP-68
chr3-96136643-DUP-748
chr3-96177631-DUP-2976
chr3-96748562-DUP-380
chr3-96749307-DUP-315
chr3-96750139-DUP-209
chr3-96751594-DUP-293
chr3-96751801-DUP-727
chr3-96752014-DUP-362
chr3-96753964-DUP-51
chr3-96797345-DUP-79
chr3-96979143-DUP-58
chr3-97067871-DUP-81
chr3-97110110-DUP-70
chr3-97168330-DUP-126
chr3-97478533-DUP-53
chr3-97651365-DUP-409
chr3-97665884-DUP-67
chr3-97837964-DUP-54
chr3-97885660-DUP-60
chr3-98039143-DUP-310
chr3-98102497-DUP-110
chr3-98126026-DUP-64
chr3-98359665-DUP-65
chr3-98722724-DUP-174
chr3-98917221-DUP-4628
chr3-99070401-DUP-238
chr3-99502010-DUP-257
chr3-99502049-DUP-167
chr3-99938742-DUP-303
chr3-99971887-DUP-65
chr3-99993754-DUP-350
chr3-100067409-DUP-96
chr3-100381251-DUP-74
chr3-100448153-DUP-152
chr3-100448204-DUP-112

chr3-101227016-DUP-104
chr3-101347293-DUP-54
chr3-101584634-DUP-118
chr3-101584691-DUP-57
chr3-101778491-DUP-112
chr3-102318933-DUP-243
chr3-102404882-DUP-84
chr3-102553374-DUP-123
chr3-102644641-DUP-79
chr3-102821931-DUP-60
chr3-103029267-DUP-136
chr3-103210120-DUP-80
chr3-103570758-DUP-82
chr3-103668339-DUP-359
chr3-103670917-DUP-55
chr3-103808815-DUP-73
chr3-103883667-DUP-51
chr3-104208083-DUP-62
chr3-104450005-DUP-75
chr3-104638794-DUP-77
chr3-104962523-DUP-70
chr3-105213277-DUP-51
chr3-105390256-DUP-71
chr3-105531785-DUP-56
chr3-105622161-DUP-54
chr3-105667026-DUP-67
chr3-105732172-DUP-59
chr3-105893684-DUP-205
chr3-106696015-DUP-366
chr3-107017328-DUP-94
chr3-107045957-DUP-51
chr3-107092905-DUP-69
chr3-107180867-DUP-62
chr3-107302277-DUP-3487
chr3-107306292-DUP-83
chr3-107386327-DUP-1036
chr3-107386420-DUP-179
chr3-107387130-DUP-432
chr3-108125893-DUP-118
chr3-108201396-DUP-133
chr3-108297762-DUP-164
chr3-108612376-DUP-198
chr3-108715225-DUP-117
chr3-108723975-DUP-145
chr3-108724789-DUP-295
chr3-108923418-DUP-53
chr3-109265957-DUP-144
chr3-109333163-DUP-167
chr3-109776538-DUP-96
chr3-110629695-DUP-197
chr3-112732186-DUP-70
chr3-113570339-DUP-82
chr3-114843552-DUP-51
chr3-115581931-DUP-76
chr3-115700113-DUP-1082
chr3-115700680-DUP-153
chr3-115837433-DUP-63
chr3-116227326-DUP-58
chr3-116553476-DUP-211
chr3-116660040-DUP-157
chr3-116685225-DUP-60
chr3-116844204-DUP-171
chr3-117046382-DUP-58
chr3-117707693-DUP-119
chr3-118315021-DUP-86
chr3-118888311-DUP-54
chr3-120758003-DUP-814
chr3-121156215-DUP-127
chr3-121523308-DUP-473
chr3-121523336-DUP-479
chr3-121715088-DUP-52
chr3-121902274-DUP-51
chr3-121987149-DUP-1485
chr3-121987266-DUP-1335
chr3-122124849-DUP-78
chr3-122133648-DUP-64
chr3-122171402-DUP-67

chr3-122273461-DUP-202
chr3-122318301-DUP-118
chr3-122398397-DUP-52
chr3-122598821-DUP-627
chr3-122856862-DUP-216
chr3-123156813-DUP-87
chr3-123317156-DUP-52
chr3-123338899-DUP-54
chr3-123377517-DUP-160
chr3-123470346-DUP-257
chr3-123471354-DUP-127
chr3-123471743-DUP-128
chr3-123472699-DUP-1907
chr3-123472851-DUP-1753
chr3-123473016-DUP-295
chr3-123474176-DUP-324
chr3-123579830-DUP-809
chr3-124134400-DUP-53
chr3-124381846-DUP-213
chr3-126066989-DUP-92
chr3-126067030-DUP-89
chr3-126334734-DUP-110
chr3-126474954-DUP-55
chr3-126626161-DUP-57
chr3-126631037-DUP-52
chr3-127491922-DUP-348
chr3-127602730-DUP-78
chr3-127669050-DUP-56
chr3-127799530-DUP-84
chr3-127799574-DUP-86
chr3-128092839-DUP-52
chr3-128660426-DUP-141
chr3-128777649-DUP-108
chr3-129084923-DUP-80
chr3-129118877-DUP-233
chr3-129378285-DUP-53
chr3-129672810-DUP-69
chr3-129725558-DUP-132
chr3-129867298-DUP-73
chr3-129869470-DUP-76
chr3-130153353-DUP-64
chr3-130296357-DUP-61
chr3-130335851-DUP-51
chr3-130338190-DUP-57
chr3-130402723-DUP-109
chr3-130417994-DUP-64
chr3-130425212-DUP-57
chr3-130964766-DUP-77
chr3-131994091-DUP-340
chr3-132067766-DUP-53
chr3-132327620-DUP-273
chr3-132598257-DUP-102
chr3-132598281-DUP-73
chr3-132601119-DUP-64
chr3-132646556-DUP-69
chr3-133025896-DUP-200
chr3-133058717-DUP-558
chr3-133062848-DUP-110
chr3-133241580-DUP-92
chr3-133250239-DUP-123
chr3-133373224-DUP-57
chr3-133473454-DUP-67
chr3-134093976-DUP-1631
chr3-134216968-DUP-58
chr3-134829850-DUP-91
chr3-134956393-DUP-220
chr3-135078674-DUP-63
chr3-135102894-DUP-73
chr3-135120554-DUP-66
chr3-135125954-DUP-175
chr3-135192064-DUP-81
chr3-135354387-DUP-62
chr3-135558874-DUP-92
chr3-135978300-DUP-218
chr3-137230776-DUP-13247
chr3-137558703-DUP-219
chr3-137558817-DUP-110

chr3-137812089-DUP-163
chr3-137812117-DUP-132
chr3-137816855-DUP-57
chr3-137820023-DUP-78
chr3-137836923-DUP-51
chr3-138048930-DUP-86
chr3-138060264-DUP-51
chr3-138464262-DUP-204
chr3-139067136-DUP-97
chr3-139605675-DUP-254
chr3-141824032-DUP-51
chr3-141897313-DUP-212
chr3-141932831-DUP-65
chr3-141990844-DUP-147
chr3-141991035-DUP-95
chr3-142202266-DUP-195
chr3-142203262-DUP-56
chr3-142475366-DUP-60
chr3-142709970-DUP-331
chr3-143015264-DUP-95
chr3-143015339-DUP-60
chr3-143016721-DUP-228
chr3-143862062-DUP-110
chr3-143866382-DUP-130
chr3-143986962-DUP-66
chr3-144195796-DUP-154
chr3-144195874-DUP-621
chr3-144232760-DUP-142
chr3-144252908-DUP-834
chr3-144279033-DUP-86
chr3-144304464-DUP-435
chr3-144420916-DUP-54
chr3-144426256-DUP-133
chr3-145401818-DUP-58
chr3-145409906-DUP-225
chr3-145556437-DUP-87
chr3-145659943-DUP-60
chr3-145660352-DUP-124
chr3-145660394-DUP-123
chr3-145728759-DUP-85
chr3-145834758-DUP-51
chr3-145858817-DUP-54
chr3-146182922-DUP-1414
chr3-146183111-DUP-182
chr3-146184048-DUP-287
chr3-146265841-DUP-333
chr3-146301572-DUP-63
chr3-146434817-DUP-80
chr3-146565704-DUP-89
chr3-146893091-DUP-315
chr3-146893146-DUP-296
chr3-147050052-DUP-87
chr3-147225827-DUP-55
chr3-147515035-DUP-145
chr3-147515531-DUP-584
chr3-148556263-DUP-77
chr3-148817160-DUP-4225
chr3-148817186-DUP-4176
chr3-148934435-DUP-52
chr3-149103974-DUP-187
chr3-149152129-DUP-88
chr3-149488103-DUP-258
chr3-149557288-DUP-219
chr3-149557449-DUP-55
chr3-149960589-DUP-4387
chr3-150270374-DUP-93
chr3-150270449-DUP-57
chr3-150282617-DUP-96
chr3-150457815-DUP-53
chr3-151286438-DUP-325
chr3-151406659-DUP-190
chr3-151406692-DUP-192
chr3-151558073-DUP-60
chr3-151672917-DUP-69
chr3-151673728-DUP-116
chr3-151673775-DUP-120
chr3-151793594-DUP-81

chr3-152191849-DUP-70
chr3-152355451-DUP-75
chr3-152431550-DUP-100
chr3-152447786-DUP-133
chr3-152447812-DUP-135
chr3-152461062-DUP-139
chr3-152531682-DUP-52
chr3-152560534-DUP-100
chr3-152569190-DUP-156
chr3-152709219-DUP-89
chr3-152726439-DUP-59
chr3-152726476-DUP-56
chr3-152771050-DUP-99
chr3-152813836-DUP-213
chr3-152879426-DUP-57
chr3-152927952-DUP-149
chr3-153060094-DUP-57
chr3-153112605-DUP-186
chr3-153126192-DUP-65
chr3-153139874-DUP-60
chr3-153327602-DUP-53
chr3-153439080-DUP-122
chr3-153459143-DUP-120
chr3-153499889-DUP-200
chr3-153500079-DUP-50
chr3-153557353-DUP-243
chr3-153583201-DUP-53
chr3-153627523-DUP-220
chr3-153627567-DUP-147
chr3-153902343-DUP-971
chr3-153924005-DUP-443
chr3-153925256-DUP-56
chr3-154013929-DUP-63
chr3-154158991-DUP-110
chr3-154159026-DUP-141
chr3-154178774-DUP-127
chr3-154232106-DUP-77
chr3-154283843-DUP-525
chr3-154287688-DUP-52
chr3-154323560-DUP-84
chr3-154459114-DUP-83
chr3-154468185-DUP-62
chr3-154552094-DUP-394
chr3-154651262-DUP-53
chr3-154663034-DUP-105
chr3-154733755-DUP-44767
chr3-154739195-DUP-110
chr3-154798565-DUP-96
chr3-154835716-DUP-71
chr3-154851599-DUP-6993
chr3-154964391-DUP-70
chr3-155386994-DUP-134
chr3-155522670-DUP-78
chr3-156139014-DUP-115
chr3-156444948-DUP-106
chr3-156745112-DUP-143
chr3-156857114-DUP-122
chr3-156939491-DUP-126
chr3-156948362-DUP-69
chr3-157146759-DUP-1361
chr3-159193916-DUP-54
chr4-3062230-DUP-88
chr4-3175224-DUP-1218
chr4-3175355-DUP-612
chr4-3271135-DUP-70
chr4-3422900-DUP-74
chr4-3926934-DUP-72
chr4-4023961-DUP-134
chr4-4545346-DUP-118
chr4-4628721-DUP-61
chr4-4723196-DUP-66
chr4-4832158-DUP-448
chr4-5535384-DUP-69
chr4-5611595-DUP-80
chr4-5611630-DUP-83
chr4-5660881-DUP-223
chr4-5748386-DUP-376

chr4-5783902-DUP-58
chr4-6230734-DUP-166
chr4-6431904-DUP-477
chr4-6465802-DUP-86
chr4-6685410-DUP-52
chr4-7222168-DUP-80
chr4-7856838-DUP-54
chr4-8152070-DUP-255
chr4-8152248-DUP-71
chr4-8212841-DUP-56
chr4-8604636-DUP-69
chr4-9025740-DUP-138
chr4-9025779-DUP-90
chr4-9072703-DUP-359
chr4-9085494-DUP-79
chr4-9378947-DUP-70
chr4-9459342-DUP-63
chr4-9502697-DUP-167
chr4-9502721-DUP-132
chr4-9580066-DUP-92
chr4-9620229-DUP-101
chr4-9665151-DUP-53
chr4-9748983-DUP-51
chr4-9824294-DUP-104
chr4-10137220-DUP-434
chr4-10989522-DUP-82
chr4-11171028-DUP-52
chr4-11532029-DUP-152
chr4-11590019-DUP-51
chr4-11687166-DUP-17526
chr4-12233564-DUP-112
chr4-12233589-DUP-87
chr4-12459777-DUP-199
chr4-12764418-DUP-291
chr4-12873965-DUP-60
chr4-12913944-DUP-80
chr4-13087672-DUP-96
chr4-13262411-DUP-58
chr4-15024013-DUP-54
chr4-15169131-DUP-103
chr4-15415624-DUP-56
chr4-15981104-DUP-59
chr4-16870999-DUP-286
chr4-17492379-DUP-57
chr4-18919598-DUP-51
chr4-19134754-DUP-62
chr4-19202450-DUP-127
chr4-19368590-DUP-65
chr4-19694265-DUP-133
chr4-20191897-DUP-62
chr4-20449351-DUP-256
chr4-20608620-DUP-92
chr4-20814336-DUP-87
chr4-21576772-DUP-8678
chr4-21885672-DUP-170
chr4-23775685-DUP-178
chr4-23898752-DUP-489
chr4-24624867-DUP-183
chr4-25275514-DUP-249
chr4-26592395-DUP-550
chr4-26897176-DUP-162
chr4-27601456-DUP-75
chr4-28470576-DUP-52
chr4-28688956-DUP-1479
chr4-28688998-DUP-1481
chr4-29306032-DUP-62
chr4-29788625-DUP-104
chr4-29864031-DUP-4059
chr4-30162262-DUP-109
chr4-30666701-DUP-7633
chr4-31303097-DUP-588
chr4-31986955-DUP-271
chr4-32303596-DUP-52
chr4-32610354-DUP-317
chr4-32862568-DUP-73
chr4-33073500-DUP-72
chr4-33358493-DUP-278

chr4-34544870-DUP-56
chr4-34856914-DUP-58
chr4-34992765-DUP-101
chr4-36708464-DUP-205
chr4-37768063-DUP-197
chr4-37832006-DUP-67726
chr4-37925181-DUP-62
chr4-40446137-DUP-123
chr4-40505359-DUP-63
chr4-41592884-DUP-102
chr4-41775803-DUP-256
chr4-41883740-DUP-363606
chr4-41886613-DUP-360733
chr4-41909576-DUP-327
chr4-42233286-DUP-2552
chr4-42241596-DUP-5750
chr4-43969127-DUP-111
chr4-44046045-DUP-98
chr4-44627684-DUP-114
chr4-44966580-DUP-208
chr4-44966660-DUP-132
chr4-44974925-DUP-405
chr4-44974953-DUP-357
chr4-44986396-DUP-71
chr4-45190784-DUP-100
chr4-45404312-DUP-78
chr4-45417864-DUP-69
chr4-45883646-DUP-150
chr4-46024038-DUP-359
chr4-46104332-DUP-89
chr4-46483788-DUP-4076
chr4-46483836-DUP-3972
chr4-46745947-DUP-88
chr4-46777799-DUP-339
chr4-46777876-DUP-218
chr4-47296252-DUP-53
chr4-47319976-DUP-80
chr4-48312230-DUP-60
chr4-48391497-DUP-53
chr4-48524766-DUP-54
chr4-48685033-DUP-61
chr4-49240425-DUP-57
chr4-49533358-DUP-64
chr4-52937325-DUP-5241
chr4-53336919-DUP-316
chr4-53759524-DUP-69
chr4-53840332-DUP-91
chr4-53929193-DUP-53
chr4-53933734-DUP-260
chr4-53933757-DUP-252
chr4-53986611-DUP-132
chr4-53986634-DUP-90
chr4-54880098-DUP-2804
chr4-54961059-DUP-4749
chr4-55204850-DUP-60
chr4-55370519-DUP-2226
chr4-55552680-DUP-20424
chr4-55584747-DUP-123
chr4-56776973-DUP-64
chr4-57199004-DUP-149
chr4-57283271-DUP-135
chr4-57283352-DUP-57
chr4-57463652-DUP-259
chr4-57463677-DUP-247
chr4-57501138-DUP-3278
chr4-57811150-DUP-289
chr4-57811186-DUP-434
chr4-58382818-DUP-56
chr4-58443167-DUP-54
chr4-58505017-DUP-118
chr4-58916213-DUP-114
chr4-59020858-DUP-158
chr4-59556491-DUP-328
chr4-59679705-DUP-240
chr4-62007380-DUP-70477
chr4-62232159-DUP-54
chr4-62413147-DUP-28919

chr4-62562617-DUP-56
chr4-62921538-DUP-143
chr4-62949995-DUP-60
chr4-63194715-DUP-72
chr4-63304274-DUP-143
chr4-63581844-DUP-184
chr4-63582503-DUP-300
chr4-63602712-DUP-70
chr4-63619894-DUP-77
chr4-63619936-DUP-110
chr4-63927928-DUP-247
chr4-64174908-DUP-1452
chr4-64346607-DUP-105
chr4-64981793-DUP-55
chr4-65848287-DUP-112
chr4-66503927-DUP-349
chr4-66770667-DUP-219
chr4-67021440-DUP-95
chr4-67627160-DUP-907
chr4-68030357-DUP-81
chr4-68445792-DUP-56
chr4-68660183-DUP-66
chr4-68675691-DUP-100
chr4-68999855-DUP-58
chr4-70139206-DUP-184
chr4-71197304-DUP-200
chr4-71197330-DUP-170
chr4-72612295-DUP-313
chr4-76037491-DUP-52
chr4-76154320-DUP-79
chr4-76763262-DUP-458
chr4-76998317-DUP-62
chr4-77177584-DUP-56
chr4-77190793-DUP-178
chr4-77721225-DUP-128
chr4-77969921-DUP-59
chr4-78524722-DUP-105
chr4-79794207-DUP-200
chr4-79823128-DUP-370
chr4-79823154-DUP-322
chr4-80849706-DUP-123
chr4-81424346-DUP-160
chr4-82171255-DUP-65
chr4-82564315-DUP-64
chr4-82630165-DUP-173
chr4-82654414-DUP-119
chr4-83760537-DUP-238
chr4-83910401-DUP-62
chr4-84080074-DUP-76
chr4-84251814-DUP-86
chr4-84698610-DUP-76
chr4-85451634-DUP-229
chr4-85837169-DUP-86
chr4-85956299-DUP-156
chr4-85956326-DUP-114
chr4-85966290-DUP-111
chr4-86398299-DUP-55
chr4-86417515-DUP-60
chr4-87121110-DUP-2272
chr4-87161095-DUP-71
chr4-87734884-DUP-159
chr4-87889852-DUP-77
chr4-88035325-DUP-119
chr4-88308683-DUP-206
chr4-88308752-DUP-203
chr4-88549949-DUP-75
chr4-88674350-DUP-2973
chr4-88734801-DUP-3731
chr4-88735104-DUP-3581
chr4-88745704-DUP-58
chr4-88812860-DUP-121
chr4-88991580-DUP-51
chr4-89203191-DUP-563
chr4-89790615-DUP-56
chr4-89898526-DUP-1204
chr4-91167561-DUP-111
chr4-91280915-DUP-53

chr4-91450733-DUP-3510
chr4-91599664-DUP-206
chr4-93558584-DUP-91
chr4-94571223-DUP-93
chr4-95047677-DUP-60
chr4-95447109-DUP-60
chr4-95567895-DUP-161
chr4-95668198-DUP-273
chr4-95706755-DUP-315
chr4-95807983-DUP-65
chr4-96135645-DUP-142
chr4-96396207-DUP-6347
chr4-97518361-DUP-133
chr4-98366743-DUP-71
chr4-98584099-DUP-169
chr4-99136684-DUP-482
chr4-99203886-DUP-217
chr4-99203912-DUP-178
chr4-99629292-DUP-63
chr4-99629423-DUP-218
chr4-99633871-DUP-51
chr4-99883289-DUP-160
chr4-99892664-DUP-280
chr4-99993346-DUP-90
chr4-100573932-DUP-132
chr4-100932147-DUP-68
chr4-101010497-DUP-66
chr4-101169823-DUP-2020
chr4-101170363-DUP-1162
chr4-101170751-DUP-253
chr4-101609598-DUP-120
chr4-101626571-DUP-910
chr4-101627091-DUP-456
chr4-102053976-DUP-51
chr4-102252094-DUP-61
chr4-102658723-DUP-348
chr4-103064558-DUP-20207
chr4-103303475-DUP-166
chr4-103303540-DUP-82
chr4-103378383-DUP-71
chr4-103662761-DUP-58214
chr4-104506004-DUP-163
chr4-104552880-DUP-1210
chr4-104994970-DUP-62
chr4-105230434-DUP-90
chr4-105544896-DUP-67
chr4-105904024-DUP-81
chr4-106059027-DUP-106
chr4-106059059-DUP-66
chr4-106235582-DUP-390
chr4-106235636-DUP-237
chr4-106236752-DUP-104
chr4-106277316-DUP-72
chr4-106317172-DUP-266
chr4-106483585-DUP-57
chr4-106542451-DUP-260
chr4-106683220-DUP-140
chr4-106899978-DUP-92
chr4-106971158-DUP-55
chr4-107028855-DUP-243
chr4-107253221-DUP-177
chr4-107419025-DUP-90
chr4-107580016-DUP-202
chr4-107834887-DUP-150
chr4-108450893-DUP-57
chr4-108834973-DUP-57
chr4-109144559-DUP-80
chr4-109286445-DUP-74
chr4-109290008-DUP-106
chr4-109651070-DUP-88
chr4-110080850-DUP-55
chr4-110155157-DUP-96
chr4-110646591-DUP-8495
chr4-110988755-DUP-76
chr4-111280571-DUP-163
chr4-111328208-DUP-86
chr4-111336714-DUP-82

chr4-111728405-DUP-69
chr4-111745471-DUP-146
chr4-114165343-DUP-56
chr4-114501005-DUP-80
chr4-115228053-DUP-83
chr4-115558184-DUP-171
chr4-115760607-DUP-69
chr4-115776030-DUP-56407
chr4-115973105-DUP-59
chr4-116155604-DUP-92
chr4-116375398-DUP-186
chr4-116924942-DUP-65
chr4-117071025-DUP-136
chr4-117071066-DUP-79
chr4-117239974-DUP-78
chr4-117317913-DUP-76
chr4-117366150-DUP-57
chr4-117396624-DUP-122
chr4-117564483-DUP-872
chr4-117654319-DUP-770
chr4-117654906-DUP-318
chr4-117875942-DUP-1360
chr4-118277038-DUP-166
chr4-118277148-DUP-85
chr4-118404254-DUP-859
chr4-118404258-DUP-1669
chr4-118405529-DUP-261
chr4-118413418-DUP-458
chr4-118490799-DUP-127
chr4-118490861-DUP-52
chr4-118832645-DUP-123
chr4-119715683-DUP-70
chr4-119766327-DUP-133
chr4-119766404-DUP-74
chr4-119936571-DUP-64
chr4-119936594-DUP-136
chr4-120035025-DUP-98
chr4-120072769-DUP-71
chr4-120380440-DUP-68
chr4-120623843-DUP-115
chr4-120773375-DUP-85
chr4-120804557-DUP-153
chr4-120834766-DUP-53
chr4-122836735-DUP-66
chr4-122844775-DUP-72
chr4-123024601-DUP-125
chr4-123159122-DUP-278
chr4-123548249-DUP-51
chr4-123559092-DUP-54
chr4-123572163-DUP-76
chr4-123647266-DUP-70
chr4-123647769-DUP-118
chr4-123989900-DUP-79
chr4-124031613-DUP-81
chr4-124378862-DUP-133
chr4-124724353-DUP-91
chr4-124746849-DUP-152219
chr4-124764068-DUP-72
chr4-124768375-DUP-57
chr4-124829123-DUP-851
chr4-124829146-DUP-803
chr4-125228130-DUP-311
chr4-125282343-DUP-92
chr4-125839808-DUP-57
chr4-125953083-DUP-150
chr4-126309764-DUP-56
chr4-126345368-DUP-68
chr4-126396856-DUP-117
chr4-126483161-DUP-442
chr4-126485138-DUP-122
chr4-126492362-DUP-80
chr4-126740652-DUP-69
chr4-126891147-DUP-4389
chr4-127159706-DUP-122
chr4-127170858-DUP-91
chr4-127295115-DUP-78
chr4-127304083-DUP-121

chr4-127304116-DUP-82
chr4-127778116-DUP-220
chr4-127784331-DUP-210
chr4-128195464-DUP-79
chr4-128317474-DUP-139
chr4-128398429-DUP-62
chr4-128410829-DUP-84
chr4-128475644-DUP-76
chr4-128591447-DUP-107
chr4-128596870-DUP-67
chr4-128646891-DUP-92
chr4-128753710-DUP-990
chr4-128759323-DUP-95
chr4-128822255-DUP-358
chr4-128903790-DUP-95
chr4-128903817-DUP-63
chr4-128922663-DUP-274
chr4-129813651-DUP-63
chr4-129836473-DUP-87
chr4-130006888-DUP-80
chr4-130569863-DUP-141
chr4-130613944-DUP-117
chr4-130691096-DUP-297
chr4-130708995-DUP-54
chr4-130722757-DUP-139
chr4-130786590-DUP-119
chr4-130823264-DUP-73
chr4-130826545-DUP-117
chr4-131180098-DUP-75
chr4-131237633-DUP-152
chr4-131472913-DUP-173
chr4-131504889-DUP-93
chr4-131872298-DUP-104
chr4-131950649-DUP-54
chr4-132016395-DUP-230
chr4-133312963-DUP-129
chr4-133692302-DUP-177
chr4-133991724-DUP-86
chr4-134312411-DUP-99
chr4-134408966-DUP-72
chr4-134458181-DUP-6893
chr4-134678880-DUP-76
chr4-134881353-DUP-61
chr4-135166668-DUP-51
chr4-135406470-DUP-103
chr4-135406526-DUP-126
chr4-135418052-DUP-94
chr4-135460129-DUP-60
chr4-135471823-DUP-86
chr4-135996021-DUP-105
chr4-136059975-DUP-105
chr4-136081060-DUP-57
chr4-136136616-DUP-218
chr4-136136703-DUP-91
chr4-136149219-DUP-94
chr4-136433145-DUP-98
chr4-136688659-DUP-59
chr4-136868133-DUP-2115
chr4-136868619-DUP-55
chr4-136929325-DUP-60
chr4-137334923-DUP-197
chr4-137334986-DUP-205
chr4-137379137-DUP-79
chr4-137436125-DUP-175
chr4-137632955-DUP-109
chr4-137759750-DUP-69
chr4-137784627-DUP-73
chr4-137861176-DUP-85
chr4-137924818-DUP-76
chr4-138011491-DUP-144
chr4-138016587-DUP-61
chr4-138265871-DUP-55
chr4-138274981-DUP-121
chr4-138558729-DUP-91
chr4-138624533-DUP-137
chr4-138627164-DUP-93
chr4-138635466-DUP-133

chr4-138648270-DUP-63
chr4-138716427-DUP-102
chr4-138753308-DUP-787
chr4-138758424-DUP-63
chr4-138789616-DUP-67
chr4-138934150-DUP-85
chr4-138983867-DUP-208
chr4-138983910-DUP-168
chr4-139051344-DUP-60
chr4-139082523-DUP-92
chr4-139082556-DUP-94
chr4-139090005-DUP-83
chr4-139144762-DUP-173
chr4-139260707-DUP-64
chr4-139263037-DUP-116
chr4-139416125-DUP-155
chr4-139416222-DUP-57
chr4-139578188-DUP-63
chr4-139768732-DUP-308
chr4-139828471-DUP-109
chr4-139856470-DUP-355
chr4-139898781-DUP-100
chr4-140042183-DUP-262
chr4-140175278-DUP-85
chr4-140236900-DUP-188
chr4-140302420-DUP-101
chr4-140338281-DUP-64
chr4-140439204-DUP-86
chr4-140450559-DUP-310
chr4-140494252-DUP-57
chr4-140596546-DUP-67
chr4-140912259-DUP-66
chr4-141121556-DUP-98
chr4-141289493-DUP-51
chr4-141395163-DUP-105
chr4-141477878-DUP-208
chr4-141674396-DUP-100
chr4-141674491-DUP-69
chr4-141827065-DUP-52
chr4-141827127-DUP-50
chr4-141999867-DUP-660
chr4-142317053-DUP-406
chr4-142317118-DUP-374
chr4-142353492-DUP-297
chr4-142419599-DUP-57
chr4-142508343-DUP-57
chr4-142930078-DUP-308
chr4-142936582-DUP-84
chr4-143184457-DUP-298
chr4-143660682-DUP-32675
chr4-143783909-DUP-84276
chr4-143820689-DUP-31748
chr4-144248481-DUP-69
chr4-144869652-DUP-87
chr4-145045649-DUP-70
chr4-145158084-DUP-152752
chr4-145159161-DUP-152753
chr4-145311899-DUP-227
chr4-145437168-DUP-459
chr4-145563573-DUP-70
chr4-146355038-DUP-90
chr4-146559570-DUP-70
chr4-147412718-DUP-79
chr4-147570012-DUP-5069
chr4-147958725-DUP-62
chr4-147958746-DUP-65
chr4-148063491-DUP-147
chr4-148129805-DUP-55
chr4-148301434-DUP-241
chr4-148538848-DUP-51
chr4-148789511-DUP-333
chr4-149087066-DUP-80
chr4-149126574-DUP-93
chr4-149179868-DUP-100
chr4-149432844-DUP-108
chr4-149432925-DUP-83
chr4-149551090-DUP-117

chr4-149612915-DUP-417
chr4-149703835-DUP-60
chr4-149894963-DUP-180
chr4-149951125-DUP-58
chr4-150051212-DUP-80
chr4-150051237-DUP-58
chr4-150143849-DUP-202
chr4-150268461-DUP-98
chr4-150409215-DUP-146
chr4-150570228-DUP-377
chr4-150570256-DUP-359
chr4-150598207-DUP-492
chr4-150598309-DUP-488
chr4-150598847-DUP-74
chr4-150665546-DUP-68
chr4-150756654-DUP-134
chr4-150778729-DUP-75
chr4-150824002-DUP-191
chr4-150926257-DUP-58
chr4-150932571-DUP-60
chr4-151072419-DUP-459
chr4-151361120-DUP-131
chr4-151393589-DUP-118
chr4-151414238-DUP-85
chr4-151460044-DUP-56
chr4-151569396-DUP-148
chr4-151616184-DUP-834
chr4-151616869-DUP-173
chr4-151706360-DUP-111
chr4-151762409-DUP-56
chr4-151895814-DUP-52
chr4-151929912-DUP-209
chr4-151957226-DUP-124
chr4-151957281-DUP-168
chr4-152328575-DUP-58
chr4-152348375-DUP-55
chr4-152432567-DUP-117
chr4-152451966-DUP-110
chr4-152559935-DUP-80
chr4-152651607-DUP-151
chr4-152693824-DUP-84
chr4-152760690-DUP-135
chr4-152801147-DUP-85
chr4-152978588-DUP-53
chr4-153109822-DUP-137
chr4-153224801-DUP-57
chr4-153446585-DUP-94
chr4-153446613-DUP-52
chr4-153481769-DUP-172
chr4-153525818-DUP-200
chr4-153525978-DUP-138
chr4-153537686-DUP-74
chr4-153611905-DUP-65
chr4-153652394-DUP-98
chr4-153663842-DUP-60
chr4-153707033-DUP-4040
chr4-153754828-DUP-64
chr4-153826816-DUP-63
chr4-153844454-DUP-56
chr4-153888942-DUP-57
chr4-153889972-DUP-230
chr4-153890032-DUP-190
chr4-154070536-DUP-94
chr4-154085727-DUP-238
chr4-154087326-DUP-133
chr4-154099988-DUP-100
chr4-154144366-DUP-183
chr4-154176291-DUP-54
chr4-154177296-DUP-68
chr4-154178143-DUP-109
chr4-154233402-DUP-318
chr4-154374091-DUP-117
chr4-154462575-DUP-115
chr4-154462602-DUP-67
chr4-154595382-DUP-109
chr4-154623347-DUP-62
chr4-154930495-DUP-74

chr4-154931481-DUP-149
chr4-155111090-DUP-128
chr4-155139793-DUP-190
chr4-155139915-DUP-74
chr4-155273976-DUP-91
chr4-155358256-DUP-275
chr4-155358449-DUP-260
chr4-155377784-DUP-79
chr4-155526452-DUP-51
chr4-155548796-DUP-217
chr4-155548834-DUP-114
chr4-155608075-DUP-141
chr4-155981958-DUP-333
chr4-156077150-DUP-172
chr4-156077191-DUP-211
chr4-156085936-DUP-410
chr4-156302483-DUP-66
chr4-156302516-DUP-97
chr4-156370136-DUP-62
chr5-3663728-DUP-62
chr5-3738829-DUP-57
chr5-3745952-DUP-78
chr5-3802089-DUP-100
chr5-3886096-DUP-201
chr5-3917256-DUP-92
chr5-4012617-DUP-412
chr5-4012654-DUP-356
chr5-4102302-DUP-193
chr5-4103030-DUP-1496
chr5-4103215-DUP-299
chr5-4104060-DUP-51
chr5-4322177-DUP-82
chr5-4597817-DUP-370
chr5-4910497-DUP-56
chr5-4962679-DUP-147
chr5-5037769-DUP-103
chr5-5198745-DUP-81
chr5-5201661-DUP-70
chr5-5278251-DUP-376
chr5-5432030-DUP-202
chr5-5981587-DUP-57
chr5-6011264-DUP-71
chr5-6120795-DUP-69
chr5-6869430-DUP-51
chr5-8146815-DUP-303
chr5-8230960-DUP-97
chr5-8230988-DUP-99
chr5-8645907-DUP-3730
chr5-8647104-DUP-3713
chr5-8751924-DUP-178
chr5-8753559-DUP-105
chr5-9502239-DUP-214
chr5-9502374-DUP-325
chr5-9789847-DUP-79
chr5-11448768-DUP-498971
chr5-13317897-DUP-52
chr5-14192352-DUP-93
chr5-14378356-DUP-160
chr5-15068849-DUP-130
chr5-16110225-DUP-66
chr5-16417103-DUP-66
chr5-16446417-DUP-117
chr5-16446457-DUP-112
chr5-16501603-DUP-172
chr5-16808416-DUP-238
chr5-18521918-DUP-68
chr5-19438620-DUP-72
chr5-19828074-DUP-124
chr5-19830900-DUP-100
chr5-20818627-DUP-54
chr5-20877219-DUP-184576
chr5-20991714-DUP-313
chr5-21715104-DUP-106
chr5-21822778-DUP-136
chr5-21985643-DUP-157
chr5-22028002-DUP-52
chr5-22114221-DUP-193

chr5-22133729-DUP-66
chr5-22174756-DUP-88
chr5-22306355-DUP-76
chr5-22355690-DUP-58
chr5-22467623-DUP-62
chr5-22652339-DUP-898
chr5-22697132-DUP-1266
chr5-22698824-DUP-151
chr5-22854141-DUP-54
chr5-22947114-DUP-51
chr5-23016997-DUP-51
chr5-23017033-DUP-66
chr5-23222681-DUP-57
chr5-23242810-DUP-134
chr5-23386086-DUP-214
chr5-23530524-DUP-4840
chr5-23589207-DUP-618
chr5-23754874-DUP-147
chr5-23844329-DUP-250
chr5-23984691-DUP-57
chr5-24019934-DUP-152
chr5-24020041-DUP-110
chr5-24094450-DUP-308
chr5-24130959-DUP-62
chr5-24446752-DUP-59
chr5-24487403-DUP-56
chr5-24752582-DUP-58
chr5-24818008-DUP-317
chr5-25110526-DUP-56
chr5-25178091-DUP-75
chr5-25220384-DUP-332
chr5-25251682-DUP-86
chr5-25282354-DUP-148
chr5-25642152-DUP-80
chr5-25939735-DUP-147
chr5-26020197-DUP-115
chr5-26020229-DUP-93
chr5-26038254-DUP-107
chr5-26112974-DUP-83
chr5-26113009-DUP-90
chr5-26563117-DUP-55
chr5-27046363-DUP-104
chr5-27046395-DUP-66
chr5-27127025-DUP-57
chr5-27127067-DUP-75
chr5-27451038-DUP-217
chr5-27451085-DUP-196
chr5-27526720-DUP-53
chr5-27670780-DUP-80
chr5-27881939-DUP-84
chr5-28077708-DUP-143
chr5-28462009-DUP-280
chr5-28462172-DUP-114
chr5-28904629-DUP-70
chr5-29539046-DUP-225
chr5-29901773-DUP-62716
chr5-30196416-DUP-95
chr5-30583007-DUP-56
chr5-30794538-DUP-56
chr5-30886159-DUP-72
chr5-31114330-DUP-200
chr5-31238735-DUP-88
chr5-31560454-DUP-118
chr5-31560490-DUP-111
chr5-32202956-DUP-132
chr5-32302077-DUP-62
chr5-32399952-DUP-62
chr5-32411792-DUP-208
chr5-32411864-DUP-95
chr5-32618643-DUP-216
chr5-33304756-DUP-91
chr5-33378969-DUP-81
chr5-33821832-DUP-284
chr5-33838073-DUP-95
chr5-33914870-DUP-67
chr5-33992544-DUP-70
chr5-35161480-DUP-52

chr5-35473427-DUP-126
chr5-35532564-DUP-53
chr5-35590583-DUP-52
chr5-36200700-DUP-57
chr5-36296299-DUP-75
chr5-36614532-DUP-103
chr5-36614731-DUP-77
chr5-36943840-DUP-54
chr5-37261375-DUP-90
chr5-37292650-DUP-77
chr5-37753645-DUP-141
chr5-37937311-DUP-83
chr5-38266889-DUP-71
chr5-38272546-DUP-66
chr5-38643673-DUP-79
chr5-39224159-DUP-312
chr5-40847489-DUP-81
chr5-41341093-DUP-63
chr5-41886191-DUP-82
chr5-42628656-DUP-98
chr5-43182894-DUP-141
chr5-43182932-DUP-235
chr5-43777732-DUP-51
chr5-43850998-DUP-246
chr5-43966039-DUP-67
chr5-44016517-DUP-64
chr5-44178312-DUP-51
chr5-44329515-DUP-188
chr5-44592246-DUP-98
chr5-44631920-DUP-86
chr5-45517603-DUP-51
chr5-46268688-DUP-65
chr5-47175070-DUP-58
chr5-47254686-DUP-1741
chr5-48575356-DUP-138
chr5-48659574-DUP-223
chr5-49020468-DUP-56
chr5-49790447-DUP-89
chr5-50189934-DUP-134
chr5-50362763-DUP-1873
chr5-50960760-DUP-54
chr5-51358762-DUP-634
chr5-51563662-DUP-53
chr5-51614733-DUP-59
chr5-52495433-DUP-55
chr5-52594057-DUP-89
chr5-52679061-DUP-79
chr5-52835030-DUP-2965
chr5-53106154-DUP-60
chr5-53171639-DUP-77
chr5-53171669-DUP-79
chr5-53191677-DUP-237
chr5-54493318-DUP-267
chr5-54805785-DUP-197
chr5-54805843-DUP-108
chr5-54959678-DUP-136
chr5-55595942-DUP-68
chr5-56344735-DUP-122
chr5-56763436-DUP-53
chr5-57237623-DUP-74
chr5-57892687-DUP-55
chr5-58048555-DUP-70
chr5-59188156-DUP-1651
chr5-61049175-DUP-473
chr5-61815799-DUP-72
chr5-62124859-DUP-77
chr5-63085538-DUP-401
chr5-63085565-DUP-353
chr5-64030958-DUP-77
chr5-64123983-DUP-78
chr5-65202028-DUP-227
chr5-65407038-DUP-57
chr5-66258407-DUP-311
chr5-66501893-DUP-232
chr5-66501963-DUP-183
chr5-66714272-DUP-156
chr5-66838063-DUP-158

chr5-66922289-DUP-63
chr5-67052009-DUP-97
chr5-67481530-DUP-162
chr5-67579254-DUP-206
chr5-67672390-DUP-56
chr5-67788168-DUP-58
chr5-67815473-DUP-61
chr5-67818659-DUP-55
chr5-67956151-DUP-64
chr5-68177178-DUP-192
chr5-68847780-DUP-405
chr5-68847810-DUP-357
chr5-70101604-DUP-260
chr5-70166727-DUP-777
chr5-71333153-DUP-109
chr5-72086447-DUP-27279
chr5-72998592-DUP-123
chr5-73556578-DUP-275
chr5-73556612-DUP-93
chr5-73613195-DUP-60
chr5-73709614-DUP-223
chr5-74426051-DUP-168
chr5-74510870-DUP-164
chr5-74545584-DUP-133
chr5-74965193-DUP-78
chr5-74981153-DUP-127
chr5-75337888-DUP-58
chr5-75622698-DUP-148
chr5-75622812-DUP-262
chr5-75860642-DUP-86
chr5-76314228-DUP-51
chr5-76495418-DUP-105
chr5-76632060-DUP-76
chr5-76690959-DUP-51
chr5-77023019-DUP-51
chr5-77150427-DUP-57
chr5-77260536-DUP-52
chr5-77355863-DUP-222
chr5-77925994-DUP-139
chr5-80067942-DUP-123
chr5-80651436-DUP-1134
chr5-80772519-DUP-76
chr5-80783279-DUP-94
chr5-82250930-DUP-62
chr5-84612442-DUP-95270
chr5-85099482-DUP-62
chr5-86113768-DUP-279
chr5-86330486-DUP-106
chr5-86492393-DUP-64
chr5-86499946-DUP-150
chr5-87289732-DUP-86
chr5-87726730-DUP-6466
chr5-87823685-DUP-249
chr5-88649969-DUP-96
chr5-88819291-DUP-174
chr5-88869296-DUP-62
chr5-89670155-DUP-76
chr5-89767921-DUP-1132
chr5-89768844-DUP-81
chr5-89769607-DUP-207
chr5-90051477-DUP-70
chr5-90060177-DUP-95
chr5-90077434-DUP-53
chr5-90213287-DUP-56
chr5-90368827-DUP-192
chr5-91942313-DUP-194
chr5-92043578-DUP-555
chr5-92485073-DUP-78
chr5-92777725-DUP-172
chr5-93091418-DUP-129
chr5-93290949-DUP-51
chr5-93325918-DUP-114
chr5-93391565-DUP-67
chr5-96363448-DUP-61
chr5-96514946-DUP-66
chr5-96584072-DUP-118
chr5-97161346-DUP-143

chr5-97161379-DUP-58
chr5-97167022-DUP-66
chr5-97167050-DUP-159
chr5-97544916-DUP-869
chr5-97691499-DUP-61
chr5-98132659-DUP-51
chr5-98221048-DUP-78
chr5-98338395-DUP-59
chr5-98350553-DUP-110
chr5-98350579-DUP-58
chr5-98518648-DUP-301
chr5-98970874-DUP-72
chr5-99396193-DUP-52
chr5-99509727-DUP-63
chr5-99686124-DUP-65
chr5-99883018-DUP-113
chr5-99979647-DUP-133
chr5-100091752-DUP-581
chr5-100216368-DUP-67
chr5-100330470-DUP-932
chr5-100330620-DUP-277
chr5-100331254-DUP-96
chr5-100332019-DUP-336
chr5-100370564-DUP-1108
chr5-100370692-DUP-112
chr5-100392447-DUP-67
chr5-100425063-DUP-56
chr5-100700292-DUP-548
chr5-100844922-DUP-100
chr5-100844975-DUP-54
chr5-100909141-DUP-404
chr5-101002245-DUP-65
chr5-101039750-DUP-100
chr5-101484823-DUP-59
chr5-101605885-DUP-51
chr5-101783438-DUP-52
chr5-101864079-DUP-74
chr5-102028272-DUP-53
chr5-102056728-DUP-99
chr5-102087467-DUP-136
chr5-102093293-DUP-52
chr5-102349446-DUP-62
chr5-102529213-DUP-111
chr5-102623256-DUP-231
chr5-102843478-DUP-255
chr5-102843508-DUP-132
chr5-103686695-DUP-72
chr5-103717819-DUP-56
chr5-103718221-DUP-212
chr5-103727742-DUP-51
chr5-104006347-DUP-359
chr5-104040391-DUP-57
chr5-104236883-DUP-86
chr5-104251081-DUP-81
chr5-104396931-DUP-84
chr5-104496801-DUP-105
chr5-104707006-DUP-66
chr5-104979954-DUP-65
chr5-104991134-DUP-258
chr5-105006672-DUP-60728
chr5-105154100-DUP-61579
chr5-105201501-DUP-5297
chr5-105574624-DUP-52
chr5-105686944-DUP-173
chr5-105687017-DUP-98
chr5-105780719-DUP-52
chr5-105807121-DUP-117
chr5-105916688-DUP-70
chr5-106006134-DUP-154
chr5-106077198-DUP-418
chr5-106122000-DUP-76
chr5-106358709-DUP-66
chr5-106393605-DUP-124
chr5-106516482-DUP-193
chr5-106516520-DUP-91
chr5-106884376-DUP-91
chr5-106974273-DUP-215

chr5-107135926-DUP-72
chr5-107151631-DUP-208
chr5-107354969-DUP-158
chr5-107355028-DUP-68
chr5-107534399-DUP-78
chr5-107992506-DUP-54
chr5-108207657-DUP-2622
chr5-108574751-DUP-67
chr5-108830535-DUP-92
chr5-108874590-DUP-64
chr5-109832833-DUP-55
chr5-109911257-DUP-1395
chr5-109911334-DUP-1314
chr5-110399866-DUP-74
chr5-110553588-DUP-157
chr5-110743095-DUP-57
chr5-111042437-DUP-55
chr5-111109986-DUP-208
chr5-111111997-DUP-52
chr5-111164144-DUP-129
chr5-111266460-DUP-162
chr5-111282732-DUP-215
chr5-111282769-DUP-186
chr5-111340713-DUP-62
chr5-111394962-DUP-177
chr5-111399732-DUP-58
chr5-111420419-DUP-93
chr5-111592901-DUP-215
chr5-111709765-DUP-92
chr5-111737921-DUP-98
chr5-111886305-DUP-146
chr5-111886337-DUP-148
chr5-111900799-DUP-274
chr5-111948999-DUP-139
chr5-112443768-DUP-91
chr5-112677943-DUP-71
chr5-112770041-DUP-98
chr5-112854994-DUP-94
chr5-113358566-DUP-115
chr5-113365655-DUP-85
chr5-113416269-DUP-120
chr5-113675775-DUP-53
chr5-113742298-DUP-358
chr5-113742340-DUP-356
chr5-114013381-DUP-243
chr5-114013431-DUP-155
chr5-114199226-DUP-172
chr5-114234182-DUP-877
chr5-114346576-DUP-174
chr5-114346604-DUP-165
chr5-114602835-DUP-68
chr5-114809544-DUP-204
chr5-115017898-DUP-2322
chr5-115793879-DUP-122
chr5-116028863-DUP-138
chr5-116130962-DUP-77
chr5-116225826-DUP-56
chr5-116230287-DUP-235
chr5-116912231-DUP-51
chr5-117074008-DUP-83
chr5-117074061-DUP-202
chr5-117236914-DUP-70
chr5-117236939-DUP-72
chr5-117286745-DUP-461
chr5-117369742-DUP-263
chr5-118122255-DUP-75
chr5-118159718-DUP-59
chr5-118330475-DUP-58
chr5-118385316-DUP-57
chr5-118424399-DUP-184
chr5-118462095-DUP-168
chr5-118583104-DUP-189
chr5-118733534-DUP-92
chr5-118764387-DUP-58
chr5-118852049-DUP-209
chr5-118886188-DUP-85
chr5-118982915-DUP-170

chr5-119658021-DUP-61
chr5-119873895-DUP-64
chr5-119904012-DUP-52
chr5-119904035-DUP-54
chr5-119988972-DUP-61
chr5-120032771-DUP-297
chr5-120032983-DUP-120
chr5-120158824-DUP-8985
chr5-120388963-DUP-69
chr5-120480287-DUP-421
chr5-120687830-DUP-109
chr5-120866776-DUP-70
chr5-121310251-DUP-60
chr5-121352196-DUP-461
chr5-121352389-DUP-363
chr5-121506672-DUP-144
chr5-121506740-DUP-104
chr5-121672831-DUP-271
chr5-121692830-DUP-63
chr5-122682424-DUP-137
chr5-122700039-DUP-174
chr5-122700087-DUP-114
chr5-122964750-DUP-2511
chr5-122964771-DUP-2637
chr5-123028712-DUP-59
chr5-123258222-DUP-90
chr5-123384714-DUP-182
chr5-123710135-DUP-855
chr5-123921643-DUP-51
chr5-124191727-DUP-140
chr5-124315544-DUP-168
chr5-124701814-DUP-101
chr5-124702971-DUP-74
chr5-124774181-DUP-57
chr5-124786563-DUP-465
chr5-124855401-DUP-52
chr5-125243434-DUP-167
chr5-125369558-DUP-96
chr5-125463413-DUP-227
chr5-125574202-DUP-308
chr5-125788916-DUP-51
chr5-125800900-DUP-180
chr5-125801018-DUP-60
chr5-126286805-DUP-76
chr5-126417818-DUP-187
chr5-126616700-DUP-132
chr5-126827979-DUP-66
chr5-126997462-DUP-161
chr5-127160813-DUP-82
chr5-127396840-DUP-80
chr5-127712745-DUP-83
chr5-127893718-DUP-130
chr5-128250615-DUP-120
chr5-128660922-DUP-54
chr5-128743550-DUP-79
chr5-128989271-DUP-77
chr5-129062020-DUP-120
chr5-129169344-DUP-110
chr5-129307059-DUP-68
chr5-129331274-DUP-140
chr5-129331793-DUP-123
chr5-129392747-DUP-82
chr5-129489028-DUP-158
chr5-129645556-DUP-60
chr5-130060948-DUP-2641
chr5-130125743-DUP-59
chr5-130179323-DUP-112
chr5-130423680-DUP-83
chr5-131398117-DUP-89
chr5-131826609-DUP-118
chr5-131856826-DUP-159
chr5-132281709-DUP-77
chr5-132463853-DUP-113
chr5-132463931-DUP-58
chr5-132566799-DUP-74
chr5-132850884-DUP-121
chr5-134812644-DUP-154

chr5-134836464-DUP-452
chr5-134836946-DUP-86
chr5-135153139-DUP-157
chr5-135296108-DUP-245
chr5-135417359-DUP-119
chr5-135460672-DUP-175
chr5-135540872-DUP-276
chr5-135541064-DUP-80
chr5-135587298-DUP-82
chr5-135613074-DUP-105
chr5-135860344-DUP-54
chr5-136033110-DUP-228
chr5-136278641-DUP-260
chr5-136374594-DUP-54
chr5-136858485-DUP-103
chr5-137115760-DUP-97
chr5-137152811-DUP-490
chr5-137182292-DUP-88
chr5-137259311-DUP-174
chr5-137461985-DUP-99
chr5-137462091-DUP-96
chr5-137486942-DUP-261
chr5-137596972-DUP-145
chr5-137629325-DUP-112
chr5-137754829-DUP-119
chr5-137809373-DUP-161
chr5-137852738-DUP-94
chr5-138147774-DUP-518
chr5-138471917-DUP-93149
chr5-138606433-DUP-81
chr5-138640217-DUP-51
chr5-138673976-DUP-300
chr5-138674010-DUP-304
chr5-138748492-DUP-79
chr5-139169544-DUP-62
chr5-139264893-DUP-109
chr5-139264935-DUP-58
chr5-139358295-DUP-176
chr5-139739536-DUP-54
chr5-140033059-DUP-94
chr5-140259150-DUP-419
chr5-140423479-DUP-59
chr5-140424608-DUP-196
chr5-140439736-DUP-59
chr5-140540328-DUP-104
chr5-140982957-DUP-102
chr5-140982988-DUP-55
chr5-141039195-DUP-56
chr5-141526984-DUP-85
chr5-141723352-DUP-74
chr5-142023344-DUP-222
chr5-142322922-DUP-100
chr5-142428963-DUP-206
chr5-142429025-DUP-74
chr5-142676642-DUP-134
chr5-142794557-DUP-312
chr5-143462359-DUP-229
chr5-143462553-DUP-90
chr5-143692192-DUP-85
chr5-143747127-DUP-88
chr5-143747153-DUP-60
chr5-143751585-DUP-56
chr5-143904701-DUP-114
chr5-144104498-DUP-209
chr5-144104626-DUP-138
chr5-144228769-DUP-59
chr5-144376924-DUP-214
chr5-144382912-DUP-103
chr5-144447444-DUP-57
chr5-144659795-DUP-365
chr5-144724912-DUP-115
chr5-144863657-DUP-57
chr5-144864532-DUP-74
chr5-144906948-DUP-85
chr5-145054217-DUP-103
chr5-145223832-DUP-67
chr5-145535468-DUP-553

chr5-146226349-DUP-270
chr5-146872578-DUP-112
chr5-146918765-DUP-79
chr5-146923313-DUP-99
chr5-146931528-DUP-57
chr5-147126502-DUP-56
chr5-147167246-DUP-58
chr5-147351069-DUP-99
chr5-147351118-DUP-68
chr5-147357957-DUP-113
chr5-147596566-DUP-98
chr5-147637045-DUP-53
chr5-147689000-DUP-262
chr5-147815835-DUP-104
chr5-147960987-DUP-57
chr5-148159413-DUP-117
chr5-148674324-DUP-82
chr5-148739440-DUP-811
chr5-148830070-DUP-108
chr5-148857042-DUP-438
chr5-148857092-DUP-717
chr5-149108544-DUP-515
chr5-150132730-DUP-107
chr5-150554329-DUP-61
chr5-151044962-DUP-2396
chr5-151298906-DUP-81
chr5-151326786-DUP-110
chr6-3135254-DUP-3693
chr6-3193828-DUP-88
chr6-3290963-DUP-155
chr6-3384101-DUP-39811
chr6-3539767-DUP-71
chr6-3560076-DUP-76
chr6-3573917-DUP-87
chr6-3894633-DUP-77
chr6-4083837-DUP-126
chr6-4465628-DUP-53
chr6-4762384-DUP-55
chr6-4762433-DUP-93
chr6-4789274-DUP-80
chr6-4959292-DUP-74
chr6-5008455-DUP-153
chr6-5056870-DUP-56
chr6-5106309-DUP-179
chr6-5106421-DUP-78
chr6-5113869-DUP-57
chr6-5263789-DUP-60
chr6-5346405-DUP-55
chr6-5356809-DUP-203
chr6-5356920-DUP-103
chr6-5704908-DUP-490
chr6-5704930-DUP-1184
chr6-5705650-DUP-416
chr6-5911468-DUP-82
chr6-6032524-DUP-320
chr6-6192430-DUP-56
chr6-6283543-DUP-508
chr6-6284563-DUP-1213
chr6-6285463-DUP-92
chr6-6293122-DUP-102
chr6-6543545-DUP-68
chr6-7554205-DUP-87
chr6-7566791-DUP-70
chr6-7681572-DUP-100
chr6-7757861-DUP-407
chr6-7962895-DUP-121
chr6-8004406-DUP-92
chr6-8459385-DUP-130
chr6-9245654-DUP-155
chr6-9245725-DUP-94
chr6-11911201-DUP-1117
chr6-11922923-DUP-123
chr6-12604667-DUP-85
chr6-12648861-DUP-68
chr6-13600409-DUP-57
chr6-13861325-DUP-7732
chr6-16219408-DUP-118

chr6-16949865-DUP-240
chr6-16989285-DUP-244
chr6-17117343-DUP-73
chr6-17147057-DUP-56
chr6-17233371-DUP-161
chr6-17711445-DUP-76
chr6-17738436-DUP-54
chr6-17766174-DUP-202
chr6-17935874-DUP-187
chr6-18131274-DUP-78
chr6-18254793-DUP-86
chr6-18254863-DUP-84
chr6-18401999-DUP-55
chr6-18717343-DUP-95
chr6-19332406-DUP-85
chr6-19559051-DUP-53
chr6-19688616-DUP-53859
chr6-20398202-DUP-821
chr6-21860414-DUP-73
chr6-21888088-DUP-99
chr6-21888134-DUP-51
chr6-22046967-DUP-69
chr6-22118731-DUP-63
chr6-22263229-DUP-67
chr6-22527405-DUP-159
chr6-23042037-DUP-52
chr6-23121747-DUP-266
chr6-23155938-DUP-7476
chr6-23161811-DUP-7340
chr6-23243557-DUP-62
chr6-23246307-DUP-120
chr6-23945909-DUP-77
chr6-24373119-DUP-85
chr6-24889232-DUP-97
chr6-25548254-DUP-1866
chr6-25776515-DUP-64
chr6-27963769-DUP-60
chr6-28094006-DUP-68
chr6-28189789-DUP-61
chr6-28300931-DUP-78
chr6-28431422-DUP-61
chr6-29037274-DUP-54
chr6-29283756-DUP-1296
chr6-29497130-DUP-52
chr6-29830584-DUP-248
chr6-30862848-DUP-346
chr6-31060013-DUP-70
chr6-31811387-DUP-68
chr6-32724096-DUP-196
chr6-32730881-DUP-60
chr6-32784293-DUP-88
chr6-33595641-DUP-185
chr6-33595703-DUP-71
chr6-33661406-DUP-57
chr6-33797135-DUP-115
chr6-33820044-DUP-70
chr6-33989203-DUP-122
chr6-34330785-DUP-58
chr6-34411941-DUP-58
chr6-34841237-DUP-92
chr6-34898476-DUP-152
chr6-35280456-DUP-64
chr6-35741482-DUP-65
chr6-35980459-DUP-72
chr6-36502834-DUP-66
chr6-36808211-DUP-54
chr6-37919605-DUP-97
chr6-38048036-DUP-57
chr6-38135216-DUP-92
chr6-38189530-DUP-79
chr6-38474949-DUP-68
chr6-38662912-DUP-142
chr6-38691806-DUP-100
chr6-39960143-DUP-91
chr6-40105588-DUP-53
chr6-40354676-DUP-58
chr6-40409009-DUP-82

chr6-40844009-DUP-118
chr6-40844545-DUP-118
chr6-41316423-DUP-89
chr6-41316464-DUP-91
chr6-41670448-DUP-69
chr6-42092855-DUP-58
chr6-42190349-DUP-86
chr6-42224220-DUP-110
chr6-42845081-DUP-111
chr6-43093074-DUP-187
chr6-47623554-DUP-5413
chr6-47961180-DUP-58
chr6-48131095-DUP-5463
chr6-48131792-DUP-5466
chr6-48725612-DUP-134
chr6-48769428-DUP-138
chr6-49393034-DUP-114
chr6-49448339-DUP-79
chr6-49591864-DUP-77
chr6-49723885-DUP-180
chr6-49723962-DUP-106
chr6-50173863-DUP-387
chr6-50334642-DUP-9698
chr6-50478315-DUP-114
chr6-50597468-DUP-7840
chr6-50612901-DUP-346
chr6-50615983-DUP-303
chr6-50781420-DUP-82
chr6-51118073-DUP-101
chr6-51619835-DUP-75
chr6-51681744-DUP-165
chr6-51787286-DUP-51
chr6-52046984-DUP-193
chr6-52074721-DUP-186
chr6-52074833-DUP-64
chr6-52770247-DUP-78
chr6-52801189-DUP-240
chr6-52865589-DUP-92
chr6-52882744-DUP-80
chr6-52897777-DUP-89
chr6-52908841-DUP-69
chr6-52947943-DUP-92
chr6-53109008-DUP-103
chr6-53249724-DUP-81
chr6-53688944-DUP-66
chr6-53723306-DUP-253
chr6-54094856-DUP-58
chr6-54143293-DUP-254
chr6-54191250-DUP-77
chr6-54337179-DUP-168
chr6-54445827-DUP-150
chr6-54565441-DUP-253
chr6-54596707-DUP-51
chr6-54600269-DUP-52
chr6-54752267-DUP-54
chr6-55007857-DUP-52
chr6-55687989-DUP-135
chr6-57232325-DUP-434
chr6-58167951-DUP-287
chr6-61325643-DUP-113
chr6-61399249-DUP-67
chr6-61447726-DUP-37379
chr6-61849282-DUP-73
chr6-62147279-DUP-108
chr6-62578413-DUP-147
chr6-63802023-DUP-233
chr6-64795852-DUP-62
chr6-64849835-DUP-237
chr6-65856968-DUP-52
chr6-67393650-DUP-135
chr6-67520391-DUP-145
chr6-68235142-DUP-60
chr6-68747424-DUP-524
chr6-68747448-DUP-504
chr6-70193207-DUP-156037
chr6-71778665-DUP-53
chr6-72254995-DUP-198

chr6-72565457-DUP-70
chr6-72734941-DUP-221
chr6-72734989-DUP-92
chr6-72979576-DUP-75
chr6-73166902-DUP-52
chr6-73181026-DUP-65
chr6-73214566-DUP-142
chr6-73214620-DUP-93
chr6-73338984-DUP-191
chr6-73340505-DUP-86
chr6-74050236-DUP-224
chr6-74686252-DUP-60
chr6-75406846-DUP-96
chr6-76918869-DUP-58
chr6-77664545-DUP-55
chr6-78737995-DUP-68
chr6-79623411-DUP-94
chr6-80363501-DUP-517
chr6-80440839-DUP-59
chr6-80697728-DUP-58
chr6-80739585-DUP-302
chr6-81323694-DUP-129
chr6-81347241-DUP-54
chr6-81614928-DUP-51
chr6-81783099-DUP-109
chr6-81783125-DUP-74
chr6-81993568-DUP-65
chr6-82111605-DUP-227
chr6-82686245-DUP-166
chr6-82718384-DUP-155
chr6-82891604-DUP-112
chr6-82959662-DUP-51
chr6-82964762-DUP-105
chr6-83127300-DUP-61
chr6-83193119-DUP-126
chr6-83194706-DUP-352
chr6-83362892-DUP-58
chr6-83378255-DUP-246
chr6-83430466-DUP-59
chr6-83661983-DUP-324
chr6-84081653-DUP-195
chr6-84166818-DUP-87
chr6-84248438-DUP-254
chr6-84992451-DUP-458
chr6-84992473-DUP-411
chr6-85162516-DUP-80
chr6-85251520-DUP-59
chr6-85537918-DUP-53
chr6-85994108-DUP-59
chr6-86091238-DUP-113
chr6-86369661-DUP-232
chr6-86369689-DUP-240
chr6-86768557-DUP-91
chr6-86769464-DUP-87
chr6-86993274-DUP-963
chr6-87080361-DUP-64
chr6-87080382-DUP-66
chr6-87141936-DUP-70
chr6-87193819-DUP-79
chr6-87434008-DUP-1084
chr6-87434869-DUP-87
chr6-87498791-DUP-68
chr6-87560872-DUP-62
chr6-87607283-DUP-56
chr6-87643825-DUP-104
chr6-87774287-DUP-122
chr6-87774309-DUP-70
chr6-88040272-DUP-125
chr6-88094815-DUP-554
chr6-88161758-DUP-105
chr6-88220402-DUP-90
chr6-88220428-DUP-62
chr6-88364535-DUP-56
chr6-88471498-DUP-158
chr6-88471720-DUP-476
chr6-88498478-DUP-172
chr6-89058398-DUP-74

chr6-89181994-DUP-7929
chr6-89297052-DUP-165
chr6-89297076-DUP-201
chr6-89388879-DUP-122
chr6-89630184-DUP-53
chr6-89657878-DUP-86
chr6-89688256-DUP-61
chr6-89902662-DUP-232623
chr6-90521435-DUP-57
chr6-90522564-DUP-341
chr6-90550938-DUP-57
chr6-90635724-DUP-96
chr6-91121756-DUP-120
chr6-92002551-DUP-452
chr6-92151525-DUP-199
chr6-92863075-DUP-60
chr6-93444147-DUP-56
chr6-93684478-DUP-63
chr6-93696704-DUP-64
chr6-93900894-DUP-59
chr6-93911305-DUP-96
chr6-93958902-DUP-54
chr6-93982912-DUP-50
chr6-94027450-DUP-58
chr6-94152488-DUP-90
chr6-94495281-DUP-57
chr6-94561285-DUP-129
chr6-94568162-DUP-53
chr6-94685576-DUP-165
chr6-94851567-DUP-55
chr6-94851596-DUP-53
chr6-94865599-DUP-2111
chr6-94918345-DUP-81
chr6-95147158-DUP-63
chr6-95630775-DUP-52
chr6-96208255-DUP-104
chr6-96968930-DUP-174
chr6-97151711-DUP-72
chr6-97213509-DUP-53
chr6-97986248-DUP-86
chr6-98039829-DUP-93
chr6-98041220-DUP-717
chr6-98380821-DUP-116
chr6-98464269-DUP-52
chr6-99314000-DUP-98
chr6-99331280-DUP-65
chr6-99423005-DUP-116
chr6-99574297-DUP-148
chr6-99628795-DUP-98
chr6-99843531-DUP-64
chr6-99974251-DUP-57
chr6-100043774-DUP-74
chr6-100441906-DUP-93
chr6-100474080-DUP-95
chr6-100602258-DUP-64
chr6-100635047-DUP-60
chr6-100690873-DUP-145
chr6-101753171-DUP-8017
chr6-101831763-DUP-126
chr6-102116356-DUP-120
chr6-102600886-DUP-73
chr6-103386384-DUP-3671
chr6-105449941-DUP-69
chr6-106488674-DUP-88
chr6-107439804-DUP-88
chr6-108217514-DUP-66
chr6-108336287-DUP-303
chr6-108671838-DUP-51
chr6-108893095-DUP-94
chr6-108901377-DUP-61
chr6-108978217-DUP-362
chr6-109633116-DUP-4486
chr6-110215693-DUP-227
chr6-111780700-DUP-211
chr6-112345209-DUP-138
chr6-112457432-DUP-59
chr6-112477772-DUP-63

chr6-113058796-DUP-58
chr6-113226641-DUP-135
chr6-113226723-DUP-51
chr6-113746620-DUP-109
chr6-114423360-DUP-79
chr6-114656200-DUP-114
chr6-115458333-DUP-91
chr6-115550503-DUP-166
chr6-115550531-DUP-168
chr6-115843148-DUP-8300
chr6-115862798-DUP-131
chr6-115898061-DUP-130
chr6-115972179-DUP-134
chr6-116207958-DUP-177
chr6-116208038-DUP-73
chr6-116215750-DUP-190
chr6-116228100-DUP-797
chr6-116228171-DUP-81
chr6-116228686-DUP-91
chr6-116275357-DUP-58
chr6-117639638-DUP-243
chr6-117639817-DUP-107
chr6-117684927-DUP-143
chr6-117716691-DUP-55
chr6-117776576-DUP-152
chr6-117821717-DUP-120
chr6-117939047-DUP-107
chr6-118057759-DUP-78
chr6-118494566-DUP-74
chr6-118494647-DUP-70
chr6-118627974-DUP-52
chr6-118689679-DUP-104
chr6-118689716-DUP-57
chr6-119081952-DUP-54
chr6-119220491-DUP-231
chr6-119488972-DUP-95
chr6-120778204-DUP-275
chr6-121025081-DUP-239
chr6-121026330-DUP-67
chr6-122358901-DUP-63
chr6-122746227-DUP-75
chr6-123719737-DUP-74028
chr6-124419476-DUP-238
chr6-124506719-DUP-213
chr6-124725776-DUP-246
chr6-125049091-DUP-51
chr6-125082346-DUP-497
chr6-125082444-DUP-352
chr6-125317768-DUP-54
chr6-125923661-DUP-69
chr6-126066221-DUP-152
chr6-126411857-DUP-259
chr6-126813003-DUP-66
chr6-126937657-DUP-62
chr6-126965384-DUP-144
chr6-126965460-DUP-253
chr6-127286988-DUP-144
chr6-127419345-DUP-86
chr6-127445339-DUP-516
chr6-127548975-DUP-71
chr6-128096346-DUP-76
chr6-128183859-DUP-68
chr6-128562480-DUP-69
chr6-128574152-DUP-88
chr6-128602471-DUP-107
chr6-128611559-DUP-159
chr6-128820365-DUP-10976
chr6-128834436-DUP-85
chr6-128864454-DUP-83140
chr6-129023307-DUP-248
chr6-129158774-DUP-65
chr6-130020546-DUP-218618
chr6-130022077-DUP-280
chr6-131285746-DUP-245
chr6-131285800-DUP-196
chr6-132091680-DUP-52
chr6-132415581-DUP-52

chr6-132549054-DUP-121
chr6-133083404-DUP-53
chr6-133740042-DUP-24201
chr6-134547074-DUP-522
chr6-134644183-DUP-173
chr6-134644289-DUP-88
chr6-134651050-DUP-73
chr6-134999161-DUP-204
chr6-134999233-DUP-121
chr6-135159519-DUP-59
chr6-135294829-DUP-64
chr6-135914928-DUP-54
chr6-136032270-DUP-158
chr6-136032313-DUP-120
chr6-136140843-DUP-236
chr6-136299594-DUP-2571
chr6-136799377-DUP-101
chr6-136828097-DUP-149
chr6-136828645-DUP-88
chr6-136903537-DUP-70
chr6-137225957-DUP-64
chr6-137286204-DUP-379
chr6-137317395-DUP-214
chr6-137598444-DUP-145
chr6-137687628-DUP-100
chr6-137718607-DUP-62
chr6-137766123-DUP-62
chr6-137788642-DUP-122
chr6-137900137-DUP-89
chr6-138279260-DUP-186
chr6-138407110-DUP-382
chr6-138809441-DUP-58
chr6-139249918-DUP-177
chr6-140213780-DUP-68
chr6-140286500-DUP-87
chr6-140379148-DUP-1693
chr6-140979438-DUP-368
chr6-141270896-DUP-78
chr6-141366778-DUP-1890
chr6-141383042-DUP-58
chr6-141946120-DUP-84
chr6-142322077-DUP-57
chr6-142345224-DUP-260
chr6-142414684-DUP-51
chr6-142471496-DUP-62
chr6-142884940-DUP-84
chr6-142959452-DUP-78
chr6-143003065-DUP-357
chr6-143113069-DUP-114
chr6-143306753-DUP-63
chr6-143334543-DUP-106
chr6-143373452-DUP-84
chr6-143470065-DUP-58
chr6-143788419-DUP-67
chr6-144165049-DUP-57
chr6-144275191-DUP-186
chr6-144316590-DUP-72
chr6-144464837-DUP-399
chr6-144502649-DUP-80
chr6-144592562-DUP-166
chr6-144593402-DUP-56
chr6-144593431-DUP-109
chr6-144605298-DUP-67
chr6-144615229-DUP-62
chr6-144628997-DUP-2833
chr6-144698249-DUP-63
chr6-144771355-DUP-93
chr6-144979645-DUP-133
chr6-145021166-DUP-87
chr6-145040888-DUP-58
chr6-145084478-DUP-503
chr6-145114573-DUP-262
chr6-145208779-DUP-109
chr6-145208844-DUP-134
chr6-145283462-DUP-318
chr6-145385848-DUP-92
chr6-145400103-DUP-406

chr6-145400147-DUP-371
chr6-145439734-DUP-80
chr6-145454176-DUP-193
chr6-145562173-DUP-66
chr6-145599230-DUP-68
chr6-145624794-DUP-346
chr6-145719107-DUP-87
chr6-146153285-DUP-119
chr6-146210171-DUP-186
chr6-146222737-DUP-94
chr6-146303631-DUP-168
chr6-146341516-DUP-106
chr6-146432795-DUP-52
chr6-146616306-DUP-81
chr6-146877544-DUP-57
chr6-146977952-DUP-190
chr6-147020626-DUP-558
chr6-147066677-DUP-54
chr6-147410493-DUP-178
chr6-147518202-DUP-298
chr6-147527978-DUP-162
chr6-147528092-DUP-80
chr6-148195917-DUP-66
chr6-148418184-DUP-274
chr6-148418215-DUP-226
chr6-148701448-DUP-58
chr6-148840098-DUP-61868
chr6-148957640-DUP-57
chr6-149182392-DUP-530
chr6-149182413-DUP-500
chr7-3158316-DUP-92
chr7-3223417-DUP-4527
chr7-3235277-DUP-5653
chr7-3242104-DUP-3054
chr7-3242175-DUP-2915
chr7-3393433-DUP-82
chr7-4128941-DUP-113
chr7-4317206-DUP-62
chr7-4343554-DUP-71
chr7-4345049-DUP-88
chr7-4464409-DUP-68
chr7-4551970-DUP-52
chr7-4575378-DUP-1460
chr7-4652284-DUP-538
chr7-4652928-DUP-153
chr7-4653476-DUP-113
chr7-4653938-DUP-343
chr7-4672475-DUP-274
chr7-4727700-DUP-66
chr7-4853380-DUP-4062
chr7-5030770-DUP-56
chr7-5057462-DUP-2180
chr7-5318854-DUP-52
chr7-5414494-DUP-52
chr7-5494876-DUP-112
chr7-6077954-DUP-108
chr7-6198579-DUP-105
chr7-6252372-DUP-3094
chr7-6312434-DUP-85
chr7-6321588-DUP-153
chr7-6368791-DUP-130
chr7-6735898-DUP-72
chr7-6792463-DUP-64
chr7-6886488-DUP-53
chr7-7106319-DUP-160
chr7-7130029-DUP-3600
chr7-7281704-DUP-6164
chr7-7282693-DUP-6156
chr7-7303508-DUP-93
chr7-9455229-DUP-362672
chr7-11421408-DUP-1868
chr7-11889440-DUP-51
chr7-12508560-DUP-156
chr7-12508637-DUP-56
chr7-14279054-DUP-167
chr7-14316410-DUP-163
chr7-14717813-DUP-7215

chr7-15628082-DUP-11499
chr7-15628146-DUP-11404
chr7-15634426-DUP-2440
chr7-15666556-DUP-99
chr7-15783608-DUP-65
chr7-16376884-DUP-25376
chr7-16377603-DUP-25463
chr7-18270292-DUP-184
chr7-18270342-DUP-136
chr7-18700574-DUP-96
chr7-19030571-DUP-55
chr7-19464280-DUP-136
chr7-19583340-DUP-51
chr7-19584441-DUP-88
chr7-19685937-DUP-93
chr7-19685960-DUP-63
chr7-19704755-DUP-321
chr7-19704892-DUP-226
chr7-23037463-DUP-86
chr7-23632906-DUP-62
chr7-23786630-DUP-108
chr7-23989827-DUP-115
chr7-23990339-DUP-77
chr7-24048950-DUP-280
chr7-24062301-DUP-123
chr7-24074570-DUP-421
chr7-24138993-DUP-118
chr7-24147283-DUP-210
chr7-24389409-DUP-1729
chr7-24409953-DUP-88
chr7-24419004-DUP-17177
chr7-24434281-DUP-6381
chr7-24492897-DUP-68
chr7-24539979-DUP-65
chr7-24697344-DUP-63
chr7-24824697-DUP-138
chr7-24891566-DUP-182
chr7-25052473-DUP-57
chr7-26137071-DUP-96
chr7-26528264-DUP-280
chr7-26921930-DUP-104
chr7-27043148-DUP-96
chr7-27049417-DUP-112
chr7-27334649-DUP-162
chr7-27388475-DUP-160
chr7-27397388-DUP-232
chr7-27623145-DUP-78
chr7-27630869-DUP-134
chr7-27909228-DUP-1054
chr7-27929761-DUP-6219
chr7-27946478-DUP-261
chr7-27989647-DUP-59
chr7-28248845-DUP-50
chr7-28251957-DUP-201
chr7-28529034-DUP-89
chr7-28540406-DUP-62
chr7-28604114-DUP-150
chr7-28912167-DUP-214
chr7-29113770-DUP-149
chr7-29113801-DUP-114
chr7-29159911-DUP-53
chr7-29211423-DUP-70
chr7-29275407-DUP-243
chr7-29275438-DUP-232
chr7-29516608-DUP-132
chr7-29516639-DUP-86
chr7-29781001-DUP-113
chr7-29781126-DUP-72
chr7-30111355-DUP-105
chr7-30142372-DUP-51
chr7-30150721-DUP-200
chr7-30150810-DUP-162
chr7-30177391-DUP-75
chr7-30437971-DUP-4291
chr7-30451470-DUP-106
chr7-30451584-DUP-166
chr7-30452385-DUP-52

chr7-30452432-DUP-81
chr7-30518388-DUP-66
chr7-30618519-DUP-19256
chr7-31102728-DUP-143631
chr7-31170086-DUP-104
chr7-32516946-DUP-352876
chr7-33273692-DUP-88
chr7-33438167-DUP-352
chr7-33438201-DUP-354
chr7-33808447-DUP-84
chr7-34046622-DUP-154
chr7-34241702-DUP-25568
chr7-34322779-DUP-62
chr7-34726089-DUP-76
chr7-34859210-DUP-1113
chr7-34921337-DUP-62
chr7-35006799-DUP-51
chr7-35310341-DUP-363
chr7-35335235-DUP-113
chr7-35404474-DUP-194
chr7-37115115-DUP-170
chr7-37115169-DUP-58
chr7-37576030-DUP-229
chr7-37576041-DUP-987
chr7-37863903-DUP-51
chr7-38097867-DUP-222
chr7-38103549-DUP-62
chr7-38412584-DUP-53
chr7-39049776-DUP-53
chr7-39403503-DUP-735
chr7-39403652-DUP-58
chr7-40767319-DUP-421
chr7-40839285-DUP-57
chr7-41685103-DUP-402577
chr7-42114211-DUP-56
chr7-42647877-DUP-75
chr7-42648511-DUP-77
chr7-42722528-DUP-82
chr7-42938961-DUP-76
chr7-43258033-DUP-124
chr7-43258058-DUP-77
chr7-43326426-DUP-112
chr7-43868848-DUP-73
chr7-43890320-DUP-134
chr7-43898363-DUP-111
chr7-44048693-DUP-58
chr7-44054936-DUP-142
chr7-44068919-DUP-339
chr7-44086321-DUP-85
chr7-44133447-DUP-137
chr7-44264613-DUP-86
chr7-44417690-DUP-60
chr7-44482961-DUP-266
chr7-44659359-DUP-53
chr7-45243180-DUP-19916
chr7-45265326-DUP-65
chr7-45335159-DUP-79
chr7-45669698-DUP-139
chr7-45720369-DUP-147
chr7-46181558-DUP-83
chr7-46181581-DUP-54
chr7-46585773-DUP-72
chr7-46676769-DUP-129
chr7-46676794-DUP-101
chr7-47276891-DUP-1542
chr7-48777807-DUP-53
chr7-48780390-DUP-569
chr7-48780450-DUP-571
chr7-49000098-DUP-64
chr7-49016330-DUP-217
chr7-49565277-DUP-55
chr7-49615570-DUP-261
chr7-49893931-DUP-54
chr7-49905541-DUP-91
chr7-50087190-DUP-64
chr7-50135767-DUP-160
chr7-50163454-DUP-94

chr7-50359068-DUP-112
chr7-50784849-DUP-156
chr7-51327696-DUP-84
chr7-51547300-DUP-243
chr7-51547348-DUP-171
chr7-51711736-DUP-613
chr7-51722885-DUP-1462
chr7-52374940-DUP-197
chr7-52710684-DUP-112
chr7-53023197-DUP-130
chr7-55777768-DUP-58
chr7-55811298-DUP-1214
chr7-55811792-DUP-453
chr7-55811944-DUP-867
chr7-55812628-DUP-111
chr7-56259710-DUP-64
chr7-56633539-DUP-91
chr7-57244254-DUP-129
chr7-57775931-DUP-70
chr7-57900200-DUP-54
chr7-59040099-DUP-1866
chr7-59040895-DUP-1869
chr7-59055079-DUP-1867
chr7-59252607-DUP-1870
chr7-59322688-DUP-2526
chr7-60440676-DUP-5193
chr7-60441286-DUP-5127
chr7-60820359-DUP-148
chr7-61383877-DUP-6834
chr7-61387756-DUP-3014
chr7-61388656-DUP-3015
chr7-61389844-DUP-3015
chr7-61452975-DUP-17829
chr7-61921782-DUP-54
chr7-62903776-DUP-66
chr7-63866217-DUP-72
chr7-64134152-DUP-91
chr7-64135605-DUP-180
chr7-64144102-DUP-135
chr7-64425652-DUP-78
chr7-64642113-DUP-891
chr7-64666763-DUP-51
chr7-65313017-DUP-85
chr7-65523604-DUP-60
chr7-65561337-DUP-54
chr7-65627407-DUP-84
chr7-65713663-DUP-75
chr7-65795397-DUP-760
chr7-66011415-DUP-997
chr7-66160669-DUP-286
chr7-66210206-DUP-141
chr7-66283747-DUP-76
chr7-66389618-DUP-158
chr7-66695648-DUP-90
chr7-67063273-DUP-137
chr7-67660603-DUP-100
chr7-68051874-DUP-72
chr7-68067732-DUP-266
chr7-68231064-DUP-89
chr7-68363765-DUP-984
chr7-68493989-DUP-2145
chr7-69071307-DUP-145
chr7-69748679-DUP-68
chr7-69868431-DUP-66
chr7-69880311-DUP-128
chr7-70032831-DUP-70
chr7-70437623-DUP-75
chr7-70475528-DUP-60
chr7-70622708-DUP-103
chr7-70808215-DUP-103
chr7-70945681-DUP-62
chr7-71509810-DUP-115
chr7-71516267-DUP-57
chr7-71561041-DUP-1596
chr7-71647244-DUP-229
chr7-71967259-DUP-829
chr7-72165932-DUP-68

chr7-72165972-DUP-68
chr7-72316281-DUP-285
chr7-72339304-DUP-75
chr7-72793520-DUP-52
chr7-72869550-DUP-158
chr7-73062166-DUP-61
chr7-73489254-DUP-131
chr7-74071093-DUP-120
chr7-74139959-DUP-60
chr7-74736434-DUP-212
chr7-75184404-DUP-69
chr7-75344220-DUP-69861
chr7-75568962-DUP-154
chr7-75745530-DUP-109
chr7-76308214-DUP-91
chr7-76326838-DUP-232
chr7-76732696-DUP-99
chr7-76738404-DUP-123
chr7-77602234-DUP-6028
chr7-77604378-DUP-6026
chr7-77703303-DUP-425
chr7-78215050-DUP-160
chr7-78728612-DUP-232
chr7-78871698-DUP-307
chr7-79067448-DUP-90
chr7-79602555-DUP-111
chr7-79626033-DUP-111
chr7-80097753-DUP-132
chr7-80358198-DUP-1350
chr7-80359497-DUP-127
chr7-80374307-DUP-57
chr7-80443275-DUP-90
chr7-80443927-DUP-110
chr7-80463941-DUP-121
chr7-80562855-DUP-51
chr7-80780007-DUP-492
chr7-80783054-DUP-87
chr7-80808276-DUP-75
chr7-80884432-DUP-66
chr7-81017439-DUP-60
chr7-81164603-DUP-85
chr7-81239645-DUP-191
chr7-81352361-DUP-92
chr7-81528342-DUP-64
chr7-81759070-DUP-129
chr7-81759121-DUP-57
chr7-82155853-DUP-202
chr7-82489214-DUP-72
chr7-82747658-DUP-204
chr7-82905562-DUP-143
chr7-83147327-DUP-202
chr7-83147464-DUP-65
chr7-83346250-DUP-131
chr7-83547623-DUP-156
chr7-83742134-DUP-454
chr7-84062549-DUP-68
chr7-85983550-DUP-53
chr7-86961627-DUP-131
chr7-87391091-DUP-80
chr7-87915761-DUP-358
chr7-87916380-DUP-2118
chr7-87917181-DUP-112
chr7-87918354-DUP-219
chr7-87947473-DUP-672
chr7-88027778-DUP-357
chr7-89194640-DUP-126
chr7-89846765-DUP-121
chr7-89938255-DUP-57
chr7-89964319-DUP-51
chr7-90135251-DUP-64
chr7-90195138-DUP-214
chr7-90195180-DUP-96
chr7-91041963-DUP-307
chr7-91197629-DUP-106
chr7-92136394-DUP-288
chr7-92136902-DUP-56
chr7-92224982-DUP-66

chr7-92828690-DUP-119
chr7-93695996-DUP-225
chr7-93744639-DUP-5022
chr7-93745180-DUP-5014
chr7-94503606-DUP-5842
chr7-95004273-DUP-94
chr7-95444108-DUP-450
chr7-95637363-DUP-50
chr7-95951193-DUP-54
chr7-96542924-DUP-142
chr7-97169055-DUP-390
chr7-97360413-DUP-86
chr7-97445001-DUP-59
chr7-97446050-DUP-188
chr7-97780612-DUP-76
chr7-97798178-DUP-79
chr7-98403223-DUP-94
chr7-98415837-DUP-56
chr7-98711757-DUP-96
chr7-98741796-DUP-70
chr7-98917972-DUP-135
chr7-98947330-DUP-275
chr7-98948364-DUP-346
chr7-98969814-DUP-66
chr7-99054951-DUP-51
chr7-99199880-DUP-401
chr7-99284153-DUP-72
chr7-99394222-DUP-74
chr7-99586116-DUP-55
chr7-99707808-DUP-190
chr7-100301822-DUP-306
chr7-100301947-DUP-304
chr7-100359969-DUP-194
chr7-100584738-DUP-96
chr7-100584775-DUP-98
chr7-100600239-DUP-78
chr7-100600263-DUP-162
chr7-101020527-DUP-130
chr7-101041149-DUP-61
chr7-101206784-DUP-25372
chr7-101321944-DUP-87
chr7-101397998-DUP-68
chr7-101435635-DUP-64
chr7-101796730-DUP-3506
chr7-102426377-DUP-51
chr7-103882738-DUP-83
chr7-103950073-DUP-86
chr7-104419177-DUP-2269
chr7-104419767-DUP-4562
chr7-104420720-DUP-2296
chr7-104699404-DUP-62
chr7-105090822-DUP-56
chr7-105144686-DUP-2982
chr7-106463454-DUP-110
chr7-106994051-DUP-176
chr7-107095398-DUP-98
chr7-108266621-DUP-28283
chr7-108430955-DUP-166
chr7-108525739-DUP-78
chr7-108764619-DUP-77
chr7-108764641-DUP-59
chr7-108817726-DUP-56
chr7-108861789-DUP-78
chr7-109352598-DUP-94
chr7-109605069-DUP-60
chr7-109816209-DUP-207
chr7-109816296-DUP-88
chr7-110526698-DUP-57
chr7-110552107-DUP-56
chr7-110552140-DUP-89
chr7-110784611-DUP-6311
chr7-110913308-DUP-88
chr7-111261052-DUP-69
chr7-111304625-DUP-100
chr7-111325687-DUP-92
chr7-111325717-DUP-63
chr7-111411548-DUP-87

chr7-111411592-DUP-60
chr7-111567085-DUP-55
chr7-111596260-DUP-192
chr7-112236466-DUP-1405
chr7-112236496-DUP-298
chr7-112236663-DUP-1749
chr7-112237297-DUP-1117
chr7-112256906-DUP-58
chr7-112352958-DUP-60
chr7-112403571-DUP-79
chr7-112597974-DUP-51
chr7-112735667-DUP-86
chr7-112802042-DUP-57
chr7-113297439-DUP-282
chr7-113337432-DUP-68
chr7-113660321-DUP-448
chr7-113660700-DUP-195
chr7-113800137-DUP-159
chr7-113806093-DUP-56
chr7-113815959-DUP-208
chr7-113815988-DUP-162
chr7-113821369-DUP-51
chr7-113847983-DUP-51
chr7-114034164-DUP-149
chr7-114198433-DUP-128
chr7-114204922-DUP-89
chr7-114620772-DUP-225
chr7-115169303-DUP-157
chr7-115238851-DUP-52
chr7-115515021-DUP-91
chr7-115893052-DUP-56
chr7-116724264-DUP-254
chr7-117056632-DUP-61
chr7-117727618-DUP-91
chr7-117808521-DUP-119
chr7-118259230-DUP-65
chr7-118386147-DUP-131
chr7-118386183-DUP-130
chr7-118410077-DUP-91
chr7-118410124-DUP-102
chr7-118511969-DUP-61
chr7-118540079-DUP-51
chr7-118873386-DUP-67
chr7-119257732-DUP-399
chr7-119315944-DUP-52
chr7-119631086-DUP-67
chr7-119837170-DUP-110
chr7-119837200-DUP-56
chr7-120424703-DUP-95
chr7-120449183-DUP-116
chr7-120514205-DUP-52
chr7-120545515-DUP-817
chr7-120861350-DUP-53
chr7-121104568-DUP-115
chr7-121507384-DUP-54
chr7-121514585-DUP-78
chr7-121514677-DUP-63
chr7-121544343-DUP-168
chr7-121564803-DUP-116
chr7-121564860-DUP-272
chr7-121596769-DUP-59
chr7-122636178-DUP-102
chr7-122636232-DUP-54
chr7-122884453-DUP-92
chr7-122992136-DUP-63
chr7-122992162-DUP-99
chr7-123014941-DUP-286
chr7-123015102-DUP-384
chr7-123598941-DUP-187
chr7-124024363-DUP-52
chr7-124302428-DUP-223
chr7-124566308-DUP-157
chr7-124587963-DUP-114
chr7-124703851-DUP-1125
chr7-124719623-DUP-163
chr7-125074781-DUP-53
chr7-125242902-DUP-57

chr7-125512271-DUP-155
chr7-125757773-DUP-159
chr7-125757844-DUP-219
chr7-126413613-DUP-110
chr7-126499343-DUP-126
chr7-127306835-DUP-71
chr7-127317786-DUP-2521
chr7-127317821-DUP-2473
chr7-127718300-DUP-219
chr7-127786082-DUP-110
chr7-127786750-DUP-417
chr7-127787461-DUP-222
chr7-127795080-DUP-56
chr7-127842758-DUP-343
chr7-127868874-DUP-182
chr7-128297801-DUP-51
chr7-128810056-DUP-81
chr7-129007286-DUP-95
chr7-129244762-DUP-58
chr7-129469931-DUP-2045
chr7-129642491-DUP-74
chr7-129943622-DUP-77
chr7-129970408-DUP-75
chr7-129992786-DUP-193
chr7-130193303-DUP-3082
chr7-130194306-DUP-75656
chr7-130195247-DUP-3057
chr7-130286959-DUP-184
chr7-130475691-DUP-62
chr7-130535959-DUP-188
chr7-130695013-DUP-4420
chr7-130733717-DUP-274
chr7-130885662-DUP-3606
chr7-131603734-DUP-141
chr7-131615881-DUP-501
chr7-131843645-DUP-59
chr7-131844499-DUP-56
chr7-131995801-DUP-63
chr7-132160609-DUP-87
chr7-132326055-DUP-59
chr7-132482799-DUP-67
chr7-132545655-DUP-855
chr7-133469964-DUP-64
chr7-133547582-DUP-64
chr7-133633523-DUP-106
chr7-134243010-DUP-82
chr7-134429787-DUP-51
chr7-134460754-DUP-74
chr7-134548056-DUP-56
chr7-134795108-DUP-79
chr7-134911181-DUP-56
chr7-135240126-DUP-56
chr7-135300768-DUP-813
chr7-135465895-DUP-68
chr7-136028595-DUP-94
chr7-136398266-DUP-58
chr7-136471603-DUP-371
chr7-136619222-DUP-75
chr7-136619254-DUP-57
chr7-136737256-DUP-109
chr7-136740767-DUP-251
chr7-137060618-DUP-97
chr7-137071557-DUP-199
chr7-137393002-DUP-93
chr7-137775872-DUP-537
chr7-137798217-DUP-110
chr7-137877594-DUP-68
chr7-137888713-DUP-80
chr7-138165899-DUP-131
chr7-138174025-DUP-82
chr7-138263814-DUP-117
chr7-138263959-DUP-63
chr7-138344798-DUP-66
chr7-138421204-DUP-57
chr7-138431874-DUP-235
chr7-138465075-DUP-81
chr7-138622719-DUP-91

chr7-138640257-DUP-189
chr7-138640361-DUP-168
chr7-138646534-DUP-86
chr7-138738213-DUP-79
chr7-138775275-DUP-104
chr7-138876529-DUP-73
chr7-139015137-DUP-222
chr7-139027776-DUP-551
chr7-139090866-DUP-123
chr7-139132990-DUP-66
chr7-139251055-DUP-83
chr7-139296638-DUP-115
chr7-139580718-DUP-187
chr7-139794636-DUP-72
chr7-139998710-DUP-30114
chr7-140285691-DUP-97
chr7-140318114-DUP-2862
chr7-140443990-DUP-54
chr7-140507818-DUP-54
chr7-140533565-DUP-56
chr7-140649249-DUP-119
chr7-140681232-DUP-122
chr7-140681257-DUP-62
chr7-140690917-DUP-84
chr7-140854254-DUP-61
chr7-141088669-DUP-63
chr7-141224990-DUP-509
chr7-141296808-DUP-289
chr7-141363058-DUP-189
chr7-141412183-DUP-886
chr7-141665869-DUP-88
chr7-141743689-DUP-201
chr7-141755477-DUP-96
chr7-141783413-DUP-55
chr7-141827607-DUP-111
chr7-141834048-DUP-177
chr7-141834085-DUP-90
chr7-141850232-DUP-114
chr7-142403749-DUP-895
chr7-142548788-DUP-88
chr7-143190378-DUP-109
chr7-143644669-DUP-58
chr7-144322616-DUP-181
chr7-144575658-DUP-52
chr7-144757807-DUP-136
chr8-3156020-DUP-117
chr8-3205261-DUP-69
chr8-3272588-DUP-67
chr8-4138794-DUP-108
chr8-4463137-DUP-57
chr8-4581991-DUP-311
chr8-4729024-DUP-100
chr8-4915738-DUP-88
chr8-5134465-DUP-157
chr8-5548227-DUP-127
chr8-5699159-DUP-66
chr8-6605010-DUP-116
chr8-6700281-DUP-59
chr8-6783581-DUP-11503
chr8-7649625-DUP-103
chr8-8360074-DUP-129
chr8-8432567-DUP-90
chr8-8509440-DUP-55
chr8-8555902-DUP-121
chr8-8555949-DUP-82
chr8-8794417-DUP-609
chr8-8799486-DUP-192
chr8-9282297-DUP-80
chr8-9781201-DUP-63
chr8-9916391-DUP-54
chr8-9975556-DUP-77
chr8-10242348-DUP-73
chr8-10325427-DUP-65
chr8-10374155-DUP-142
chr8-10374193-DUP-96
chr8-10402672-DUP-114
chr8-10476417-DUP-62

chr8-10573374-DUP-122
chr8-10620230-DUP-61
chr8-10852626-DUP-52
chr8-10853648-DUP-51
chr8-10868895-DUP-201
chr8-11196617-DUP-92
chr8-11384543-DUP-63
chr8-11449181-DUP-114
chr8-11449210-DUP-273
chr8-11459270-DUP-102
chr8-11552276-DUP-142
chr8-11552346-DUP-73
chr8-11563430-DUP-76
chr8-11885584-DUP-66
chr8-11988076-DUP-58
chr8-12017287-DUP-89
chr8-12049766-DUP-125
chr8-12054658-DUP-189
chr8-12054703-DUP-312
chr8-12121549-DUP-62
chr8-12295724-DUP-64
chr8-12582852-DUP-249
chr8-12596975-DUP-104
chr8-12637268-DUP-173
chr8-12701740-DUP-186
chr8-12703994-DUP-113
chr8-12816440-DUP-106
chr8-12845714-DUP-131
chr8-12917049-DUP-89
chr8-13068280-DUP-96
chr8-13553345-DUP-61
chr8-13658543-DUP-143
chr8-13662540-DUP-95
chr8-13693243-DUP-256
chr8-13699592-DUP-86
chr8-13806359-DUP-56
chr8-13841315-DUP-371
chr8-13990665-DUP-71
chr8-14555712-DUP-71
chr8-14908896-DUP-406
chr8-14908949-DUP-350
chr8-15044791-DUP-81
chr8-15076779-DUP-55
chr8-15110851-DUP-54
chr8-15176518-DUP-63
chr8-15682307-DUP-61
chr8-16072213-DUP-91
chr8-16989959-DUP-20660
chr8-17307217-DUP-51
chr8-17396016-DUP-67
chr8-17613173-DUP-72
chr8-17849432-DUP-60
chr8-17927048-DUP-54
chr8-18167642-DUP-75
chr8-18464560-DUP-784
chr8-18563267-DUP-99
chr8-18772367-DUP-146
chr8-19084738-DUP-52
chr8-20112308-DUP-102
chr8-20507541-DUP-102
chr8-20507921-DUP-256
chr8-20645571-DUP-199
chr8-20647139-DUP-256
chr8-20660346-DUP-137
chr8-20660739-DUP-291
chr8-20672126-DUP-512
chr8-21053020-DUP-826
chr8-21053536-DUP-153
chr8-21064571-DUP-1029
chr8-21064685-DUP-357
chr8-21065227-DUP-393
chr8-21187880-DUP-1963
chr8-21188041-DUP-195
chr8-21188886-DUP-361
chr8-21189675-DUP-153
chr8-21217819-DUP-195
chr8-21222398-DUP-594

chr8-21222992-DUP-669
chr8-21224074-DUP-150
chr8-21236644-DUP-153
chr8-21237278-DUP-151
chr8-21238048-DUP-354
chr8-21239543-DUP-273
chr8-21240622-DUP-154
chr8-21241668-DUP-366
chr8-21242356-DUP-368
chr8-21244063-DUP-509
chr8-21244555-DUP-317
chr8-22580684-DUP-121
chr8-22653711-DUP-106
chr8-22841948-DUP-126
chr8-22871042-DUP-56
chr8-22876142-DUP-93
chr8-22908964-DUP-408
chr8-22951839-DUP-58
chr8-23269413-DUP-148
chr8-23575979-DUP-109
chr8-23576003-DUP-86
chr8-23605591-DUP-57
chr8-23734849-DUP-51
chr8-23903896-DUP-95
chr8-23916533-DUP-55
chr8-24247016-DUP-90
chr8-24424620-DUP-228
chr8-24424741-DUP-74
chr8-24621757-DUP-314
chr8-24621803-DUP-223
chr8-24623923-DUP-122
chr8-25492223-DUP-172
chr8-25534451-DUP-158
chr8-25759051-DUP-54
chr8-26183959-DUP-104
chr8-26425753-DUP-81
chr8-26426343-DUP-78
chr8-26493475-DUP-165
chr8-26523550-DUP-72
chr8-26735458-DUP-162
chr8-26736575-DUP-2345
chr8-26872594-DUP-84
chr8-27011545-DUP-124
chr8-27198387-DUP-51
chr8-27768282-DUP-39579
chr8-28032045-DUP-59
chr8-30091927-DUP-134
chr8-30519332-DUP-110
chr8-31003997-DUP-58
chr8-31484841-DUP-200
chr8-31552887-DUP-112
chr8-32197581-DUP-104
chr8-32197629-DUP-114
chr8-32389343-DUP-54
chr8-32568957-DUP-239
chr8-32838674-DUP-171
chr8-32838706-DUP-93
chr8-33910481-DUP-64
chr8-34110054-DUP-79
chr8-34402990-DUP-95
chr8-34507913-DUP-152
chr8-34562484-DUP-108
chr8-34598622-DUP-63
chr8-34729559-DUP-106
chr8-34729623-DUP-72
chr8-34908809-DUP-103
chr8-35209861-DUP-105
chr8-35209909-DUP-84
chr8-35409674-DUP-57
chr8-35581951-DUP-59
chr8-35662730-DUP-236
chr8-35888123-DUP-60
chr8-36234906-DUP-57
chr8-36647652-DUP-89
chr8-36911676-DUP-8004
chr8-38263067-DUP-115
chr8-38547456-DUP-60

chr8-39231416-DUP-52
chr8-40545217-DUP-105
chr8-41055888-DUP-90
chr8-41851586-DUP-153
chr8-41980475-DUP-11137
chr8-42372368-DUP-73
chr8-44764286-DUP-77
chr8-45017901-DUP-67
chr8-45269306-DUP-116
chr8-45269339-DUP-139
chr8-45867842-DUP-70
chr8-45927967-DUP-163
chr8-46407316-DUP-62
chr8-46481335-DUP-53
chr8-46514972-DUP-157
chr8-46566850-DUP-182
chr8-46690942-DUP-251
chr8-47143976-DUP-76
chr8-47456642-DUP-170
chr8-47821470-DUP-118
chr8-48271277-DUP-91
chr8-48394454-DUP-78
chr8-48440845-DUP-145
chr8-48651909-DUP-89
chr8-48770380-DUP-6357
chr8-49328920-DUP-291
chr8-49328952-DUP-243
chr8-49643561-DUP-59
chr8-49691410-DUP-241
chr8-49795113-DUP-96
chr8-49803334-DUP-201
chr8-50635785-DUP-73
chr8-50820855-DUP-55
chr8-50859693-DUP-174
chr8-52692141-DUP-52
chr8-53091592-DUP-405
chr8-53972037-DUP-90
chr8-54321949-DUP-72
chr8-54672281-DUP-99
chr8-54938915-DUP-70
chr8-55037559-DUP-145
chr8-56348065-DUP-1936
chr8-56348207-DUP-1936
chr8-56981309-DUP-60
chr8-57009985-DUP-67
chr8-57090204-DUP-60
chr8-57178255-DUP-133
chr8-57758187-DUP-52
chr8-58024652-DUP-413
chr8-58024684-DUP-404
chr8-58099365-DUP-74
chr8-58717598-DUP-55
chr8-58839224-DUP-115
chr8-60920446-DUP-47810
chr8-61155172-DUP-151
chr8-61829598-DUP-138
chr8-62443862-DUP-64
chr8-63240803-DUP-34908
chr8-64044988-DUP-56
chr8-64176710-DUP-116
chr8-65057926-DUP-293
chr8-65194683-DUP-51
chr8-65499282-DUP-104
chr8-66478180-DUP-72
chr8-66478214-DUP-66
chr8-66733076-DUP-55
chr8-66909920-DUP-69
chr8-67091434-DUP-134
chr8-67091497-DUP-100
chr8-68100882-DUP-61
chr8-68201720-DUP-134
chr8-68296449-DUP-119
chr8-68371675-DUP-115
chr8-68371698-DUP-190
chr8-68414757-DUP-110
chr8-68583453-DUP-53
chr8-68643327-DUP-79

chr8-68992758-DUP-51
chr8-69516500-DUP-55
chr8-69549904-DUP-53
chr8-69874430-DUP-84
chr8-70165526-DUP-328
chr8-70319539-DUP-286
chr8-70353472-DUP-221
chr8-70360084-DUP-59
chr8-70586623-DUP-51
chr8-70604863-DUP-57
chr8-70619102-DUP-82
chr8-70741098-DUP-628
chr8-70741185-DUP-630
chr8-70826743-DUP-58
chr8-70986118-DUP-180
chr8-70986179-DUP-177
chr8-71090283-DUP-87
chr8-71280821-DUP-2026
chr8-71280873-DUP-1953
chr8-71448252-DUP-10040
chr8-71451313-DUP-75
chr8-71453535-DUP-149
chr8-72028078-DUP-134
chr8-72028153-DUP-64
chr8-72050200-DUP-247
chr8-72050223-DUP-199
chr8-72116411-DUP-64
chr8-72209450-DUP-355
chr8-72755702-DUP-98210
chr8-72934873-DUP-115
chr8-72934906-DUP-62
chr8-73005431-DUP-178
chr8-73018647-DUP-116
chr8-73356999-DUP-9982
chr8-73357169-DUP-383
chr8-73364940-DUP-58
chr8-73365051-DUP-110
chr8-73848228-DUP-73
chr8-74569139-DUP-58
chr8-74647285-DUP-318
chr8-75694458-DUP-100
chr8-75937308-DUP-103
chr8-75993107-DUP-133
chr8-76755046-DUP-67
chr8-77034259-DUP-156
chr8-77071457-DUP-53
chr8-77630157-DUP-56
chr8-77647300-DUP-189
chr8-77854418-DUP-207
chr8-77854484-DUP-60
chr8-78100025-DUP-71
chr8-78175322-DUP-456
chr8-78175399-DUP-309
chr8-78397433-DUP-89
chr8-78414729-DUP-98
chr8-78510862-DUP-84
chr8-78550787-DUP-82
chr8-78550813-DUP-79
chr8-78559181-DUP-57
chr8-78567681-DUP-393
chr8-78704223-DUP-64
chr8-78781191-DUP-93
chr8-78783454-DUP-51
chr8-78979537-DUP-115
chr8-79062337-DUP-134
chr8-79138350-DUP-132
chr8-79318342-DUP-148
chr8-79318980-DUP-71
chr8-79319488-DUP-52
chr8-79404911-DUP-84
chr8-79437980-DUP-50
chr8-79520785-DUP-445
chr8-79532286-DUP-52
chr8-79553434-DUP-251
chr8-79773758-DUP-57
chr8-79773816-DUP-93
chr8-79822307-DUP-128

chr8-79866902-DUP-243
chr8-79870841-DUP-68
chr8-80081620-DUP-99
chr8-80263008-DUP-97
chr8-80366576-DUP-109
chr8-80646889-DUP-358
chr8-80688440-DUP-308
chr8-80774019-DUP-88
chr8-80915270-DUP-157
chr8-81002178-DUP-233
chr8-81048843-DUP-60
chr8-81320591-DUP-59
chr8-81564205-DUP-54
chr8-82234598-DUP-181
chr8-82325477-DUP-268
chr8-82625839-DUP-56
chr8-83609596-DUP-51
chr8-83618641-DUP-60
chr8-83802463-DUP-171
chr8-83955651-DUP-235
chr8-84076886-DUP-80
chr8-84093245-DUP-298
chr8-84297671-DUP-118
chr8-84538380-DUP-62
chr8-84839932-DUP-374
chr8-85016639-DUP-92
chr8-85103237-DUP-61
chr8-85152449-DUP-146
chr8-85166205-DUP-196
chr8-85200712-DUP-119
chr8-85600067-DUP-1324
chr8-86194117-DUP-86
chr8-86236905-DUP-108
chr8-86387876-DUP-54
chr8-86446223-DUP-141
chr8-86446402-DUP-334
chr8-86446475-DUP-456
chr8-86672030-DUP-73
chr8-86738967-DUP-63
chr8-86739042-DUP-59
chr8-87040740-DUP-54
chr8-87119358-DUP-107
chr8-87568775-DUP-58
chr8-87649468-DUP-84
chr8-88394257-DUP-53
chr8-88936839-DUP-184
chr8-89031866-DUP-268
chr8-89195725-DUP-51
chr8-89333368-DUP-52
chr8-89613116-DUP-107
chr8-89643069-DUP-54
chr8-89652699-DUP-116
chr8-89806382-DUP-6809
chr8-90049510-DUP-77
chr8-90467792-DUP-51
chr8-90792177-DUP-66
chr8-90911678-DUP-150
chr8-91695025-DUP-77
chr8-91746519-DUP-98
chr8-91820332-DUP-505
chr8-91830545-DUP-55
chr8-91831189-DUP-52
chr8-91863263-DUP-140
chr8-91962233-DUP-150
chr8-91962265-DUP-90
chr8-92071283-DUP-53
chr8-92155780-DUP-93
chr8-92166282-DUP-52
chr8-92233162-DUP-60
chr8-92253515-DUP-51
chr8-92736752-DUP-51
chr8-92987943-DUP-71
chr8-93078824-DUP-70
chr8-93197520-DUP-238
chr8-93429888-DUP-163
chr8-93586835-DUP-66
chr8-93898036-DUP-210

chr8-94379691-DUP-59
chr8-94450167-DUP-73
chr8-94522994-DUP-55
chr8-94858000-DUP-108
chr8-94872050-DUP-100
chr8-94884794-DUP-90
chr8-94884822-DUP-89
chr8-94933834-DUP-214
chr8-94943798-DUP-189
chr8-95470424-DUP-73
chr8-95534645-DUP-116
chr8-95603193-DUP-100
chr8-95763837-DUP-93
chr8-95763878-DUP-60
chr8-96002562-DUP-76
chr8-96158003-DUP-384
chr8-96320677-DUP-64
chr8-96326799-DUP-75
chr8-97009751-DUP-864
chr8-98044699-DUP-78
chr8-98044746-DUP-140
chr8-98470420-DUP-77
chr8-98826956-DUP-285
chr8-100460889-DUP-1137
chr8-101893579-DUP-88
chr8-102382064-DUP-83
chr8-102587567-DUP-169
chr8-103758927-DUP-52
chr8-103923871-DUP-149
chr8-104003017-DUP-6316
chr8-104139936-DUP-96
chr8-104991331-DUP-97
chr8-105076826-DUP-66
chr8-105245177-DUP-130
chr8-106047420-DUP-403
chr8-106137584-DUP-217
chr8-106347322-DUP-93
chr8-106647628-DUP-150
chr8-106920635-DUP-51
chr8-106949515-DUP-193
chr8-107129354-DUP-60
chr8-107142830-DUP-4550
chr8-107197360-DUP-1865
chr8-107197383-DUP-1833
chr8-107510016-DUP-70
chr8-107636107-DUP-992
chr8-107636264-DUP-1019
chr8-107637168-DUP-80
chr8-107640719-DUP-166
chr8-107953502-DUP-70
chr8-108232343-DUP-114
chr8-108232381-DUP-78
chr8-108273716-DUP-136
chr8-110304740-DUP-55
chr8-110327084-DUP-54
chr8-110389518-DUP-243
chr8-110814541-DUP-153
chr8-111670668-DUP-106
chr8-111820177-DUP-57506
chr8-112341352-DUP-56
chr8-112728567-DUP-75
chr8-112884792-DUP-65
chr8-113040914-DUP-58
chr8-113092887-DUP-107
chr8-114934001-DUP-3947
chr8-115073142-DUP-132
chr8-115311140-DUP-64
chr8-115590610-DUP-274
chr8-116280329-DUP-213
chr8-117261172-DUP-588
chr8-117430662-DUP-406
chr8-117630545-DUP-101
chr8-117630593-DUP-56
chr8-118022281-DUP-61
chr8-118287403-DUP-66
chr8-118561181-DUP-195
chr8-119395298-DUP-64

chr8-120131053-DUP-114
chr8-120301190-DUP-209
chr8-120301213-DUP-111
chr8-120433993-DUP-92
chr8-120552459-DUP-421
chr8-120645224-DUP-194
chr8-120672837-DUP-75
chr8-120815350-DUP-356
chr8-120900346-DUP-425
chr8-120900380-DUP-254
chr8-120986918-DUP-168
chr8-121650513-DUP-136
chr8-121700479-DUP-146
chr8-121981326-DUP-60
chr8-122092631-DUP-230
chr8-122092689-DUP-67
chr8-122199760-DUP-151
chr8-122205504-DUP-60
chr8-122361876-DUP-51
chr8-122441728-DUP-57
chr8-122548819-DUP-1015
chr8-122548919-DUP-354
chr8-122549578-DUP-1936
chr8-122553052-DUP-150
chr8-122596273-DUP-92
chr8-122629046-DUP-100
chr8-122736822-DUP-63
chr8-122736909-DUP-87
chr8-122909769-DUP-73
chr8-123022493-DUP-208
chr8-123022602-DUP-152
chr8-123038838-DUP-61
chr8-123535688-DUP-976
chr8-123536214-DUP-483
chr8-123684821-DUP-206
chr8-123751858-DUP-93
chr8-123756175-DUP-63
chr8-123788525-DUP-142
chr8-123791023-DUP-81
chr8-124529724-DUP-269
chr8-124689290-DUP-69
chr8-124716059-DUP-142
chr8-124988947-DUP-184
chr8-125062845-DUP-74
chr8-125063453-DUP-279
chr8-125063653-DUP-184
chr8-125088825-DUP-61
chr8-125136938-DUP-78
chr8-125141904-DUP-80
chr8-125149888-DUP-249
chr8-125153158-DUP-180
chr8-125177385-DUP-73
chr8-125209266-DUP-164
chr8-125225217-DUP-55
chr8-125414497-DUP-74
chr8-125417687-DUP-115
chr8-125417743-DUP-57
chr8-125444746-DUP-51
chr8-125456524-DUP-72
chr8-125555435-DUP-145
chr8-125746737-DUP-62
chr8-125751730-DUP-66
chr8-125756560-DUP-138
chr8-125756597-DUP-112
chr8-125849340-DUP-72
chr8-125897848-DUP-280
chr8-125918678-DUP-342
chr8-125930936-DUP-61
chr8-126058878-DUP-74
chr8-126111075-DUP-104
chr8-126156807-DUP-142
chr8-126163418-DUP-65
chr8-126203984-DUP-132
chr8-126264403-DUP-55
chr8-126264487-DUP-97
chr8-126326453-DUP-127
chr8-126326476-DUP-87

chr8-126532419-DUP-143
chr8-127025930-DUP-67
chr8-127121098-DUP-51
chr8-127302475-DUP-55
chr8-127404636-DUP-856
chr8-127583389-DUP-234
chr8-127638789-DUP-57
chr8-128260121-DUP-54
chr8-128574468-DUP-151
chr8-128700627-DUP-85
chr8-129717984-DUP-64
chr9-3057569-DUP-56
chr9-3076289-DUP-542
chr9-3077108-DUP-180
chr9-3596782-DUP-149
chr9-4446194-DUP-492
chr9-5062675-DUP-5064
chr9-6268475-DUP-2319
chr9-6496604-DUP-109
chr9-7157252-DUP-137
chr9-7157301-DUP-84
chr9-7657756-DUP-80
chr9-7934775-DUP-486
chr9-7977811-DUP-54
chr9-8040087-DUP-217
chr9-8183818-DUP-97
chr9-8575220-DUP-232
chr9-8976683-DUP-75
chr9-9196393-DUP-360026
chr9-9201261-DUP-66
chr9-9766586-DUP-72
chr9-9964397-DUP-268
chr9-10442781-DUP-54
chr9-10755443-DUP-51
chr9-10825336-DUP-2886
chr9-11269997-DUP-87
chr9-11283980-DUP-58
chr9-11348967-DUP-87
chr9-12277399-DUP-1545
chr9-13445575-DUP-684
chr9-13472565-DUP-97
chr9-13749119-DUP-51
chr9-13803440-DUP-2052
chr9-13906770-DUP-969
chr9-13967910-DUP-59
chr9-14134149-DUP-89
chr9-14511750-DUP-245
chr9-14833834-DUP-468
chr9-14916731-DUP-115
chr9-15112446-DUP-99
chr9-15113489-DUP-243
chr9-15398745-DUP-155
chr9-15502828-DUP-187
chr9-15502999-DUP-318
chr9-15519509-DUP-474
chr9-15520053-DUP-201
chr9-15627687-DUP-102
chr9-15637432-DUP-501
chr9-15810036-DUP-177
chr9-18549616-DUP-1303
chr9-18549684-DUP-1893
chr9-18549811-DUP-1890
chr9-19031907-DUP-51
chr9-19539930-DUP-54
chr9-20264255-DUP-71
chr9-20404918-DUP-61
chr9-20593303-DUP-153
chr9-20680838-DUP-76
chr9-20866345-DUP-88
chr9-21411715-DUP-443
chr9-21411772-DUP-1018
chr9-21412292-DUP-96
chr9-21412748-DUP-173
chr9-21713186-DUP-65
chr9-21833118-DUP-68
chr9-21833395-DUP-590
chr9-22101030-DUP-182

chr9-22101636-DUP-465
chr9-22104313-DUP-53
chr9-22133881-DUP-4105
chr9-22133918-DUP-3950
chr9-22179864-DUP-58
chr9-22411465-DUP-102
chr9-22438849-DUP-115
chr9-22438937-DUP-111
chr9-22588929-DUP-59
chr9-22623193-DUP-588
chr9-22671496-DUP-68
chr9-22862755-DUP-71
chr9-22935785-DUP-69
chr9-23491926-DUP-255
chr9-23667714-DUP-309
chr9-24169656-DUP-71
chr9-24659011-DUP-58
chr9-24924062-DUP-70
chr9-25102700-DUP-96
chr9-25188910-DUP-118
chr9-25325542-DUP-63
chr9-26534776-DUP-94
chr9-26704580-DUP-60
chr9-26794089-DUP-104
chr9-26794112-DUP-53
chr9-27271599-DUP-117
chr9-27640262-DUP-100
chr9-27640296-DUP-51
chr9-27727468-DUP-294
chr9-28869595-DUP-61
chr9-29646984-DUP-52
chr9-29647009-DUP-102
chr9-29757225-DUP-3483
chr9-29943208-DUP-95
chr9-31066255-DUP-96
chr9-31124485-DUP-323
chr9-31167231-DUP-64
chr9-31354040-DUP-112
chr9-31774439-DUP-57
chr9-32042811-DUP-184
chr9-32220048-DUP-141
chr9-32453339-DUP-94
chr9-32739786-DUP-56
chr9-33007622-DUP-73
chr9-34633942-DUP-57
chr9-34836717-DUP-70
chr9-35018786-DUP-66
chr9-35147593-DUP-873
chr9-35147953-DUP-289
chr9-35201553-DUP-366
chr9-35201583-DUP-318
chr9-35254445-DUP-104
chr9-35874357-DUP-118
chr9-35874385-DUP-100
chr9-36697077-DUP-68
chr9-36726130-DUP-3751
chr9-36726171-DUP-3674
chr9-37393677-DUP-71
chr9-37393701-DUP-67
chr9-37451997-DUP-56
chr9-37462522-DUP-64
chr9-37465772-DUP-87
chr9-37678584-DUP-135
chr9-37959421-DUP-14098
chr9-39237584-DUP-78
chr9-39339307-DUP-43705
chr9-40269543-DUP-56
chr9-40666563-DUP-90
chr9-40666604-DUP-51
chr9-40966441-DUP-166
chr9-41086691-DUP-2027
chr9-41274229-DUP-74
chr9-41932615-DUP-132
chr9-42110429-DUP-181
chr9-42110891-DUP-236
chr9-42347193-DUP-225
chr9-42375656-DUP-194

chr9-42375685-DUP-153
chr9-42626254-DUP-110
chr9-42684356-DUP-63
chr9-42959287-DUP-80
chr9-43301406-DUP-154
chr9-43766299-DUP-55
chr9-44007256-DUP-76
chr9-44356002-DUP-3355
chr9-44496679-DUP-110
chr9-44727151-DUP-554
chr9-45023481-DUP-193
chr9-45239778-DUP-64
chr9-45245129-DUP-80
chr9-45293237-DUP-62
chr9-45349777-DUP-98
chr9-45997808-DUP-51
chr9-46044152-DUP-206
chr9-46055846-DUP-328
chr9-46077494-DUP-51
chr9-46782333-DUP-288367
chr9-46865245-DUP-56
chr9-48219096-DUP-50
chr9-48354422-DUP-51
chr9-48694347-DUP-61
chr9-48718604-DUP-100
chr9-49087916-DUP-95
chr9-49141541-DUP-185
chr9-49141702-DUP-105
chr9-49407818-DUP-54
chr9-49731741-DUP-304
chr9-50376636-DUP-579
chr9-50462072-DUP-109
chr9-50744131-DUP-528
chr9-50744286-DUP-386
chr9-51011866-DUP-187
chr9-51011967-DUP-96
chr9-51252624-DUP-76
chr9-51744971-DUP-56
chr9-51889733-DUP-165
chr9-52174620-DUP-77
chr9-52710614-DUP-97
chr9-53167157-DUP-64
chr9-53380569-DUP-105
chr9-54068825-DUP-92
chr9-54489966-DUP-10331
chr9-54596506-DUP-90
chr9-54629887-DUP-127
chr9-54963063-DUP-177
chr9-55175348-DUP-136
chr9-55309130-DUP-75
chr9-55481373-DUP-215
chr9-55499280-DUP-111
chr9-56181380-DUP-87
chr9-56224228-DUP-68
chr9-56521877-DUP-88
chr9-56607833-DUP-44303
chr9-57806967-DUP-57
chr9-57830109-DUP-64
chr9-57920916-DUP-104
chr9-57920968-DUP-56
chr9-58085488-DUP-158
chr9-59066761-DUP-142
chr9-59255096-DUP-115
chr9-59608105-DUP-244
chr9-59906025-DUP-73
chr9-60166418-DUP-70
chr9-60238563-DUP-168
chr9-60510777-DUP-60
chr9-60564233-DUP-167
chr9-60753990-DUP-69
chr9-60769809-DUP-237
chr9-60825499-DUP-58
chr9-60937181-DUP-57
chr9-60947305-DUP-120
chr9-60947346-DUP-60
chr9-61906111-DUP-59
chr9-62437074-DUP-57

chr9-62575843-DUP-317
chr9-62575873-DUP-323
chr9-62781854-DUP-147
chr9-62781897-DUP-70
chr9-62933385-DUP-367
chr9-62983307-DUP-150
chr9-63265544-DUP-468
chr9-63351392-DUP-52
chr9-63463010-DUP-113
chr9-63463094-DUP-174
chr9-63522524-DUP-253
chr9-63607217-DUP-56
chr9-63679632-DUP-111
chr9-63958825-DUP-73
chr9-64077585-DUP-84
chr9-64256697-DUP-65
chr9-64388734-DUP-62
chr9-65239181-DUP-76
chr9-65270129-DUP-53
chr9-65335878-DUP-6667
chr9-65335879-DUP-3604
chr9-65395313-DUP-269
chr9-65499888-DUP-64
chr9-66371553-DUP-80
chr9-66781697-DUP-85
chr9-67167518-DUP-84
chr9-67239459-DUP-58
chr9-67347782-DUP-760
chr9-67348362-DUP-53
chr9-67506036-DUP-57
chr9-67798733-DUP-93
chr9-68061427-DUP-53
chr9-68073172-DUP-100
chr9-69158063-DUP-261
chr9-69189562-DUP-73
chr9-70252375-DUP-53
chr9-70323709-DUP-54
chr9-70501409-DUP-68
chr9-70553593-DUP-91
chr9-70553631-DUP-155
chr9-71552645-DUP-53
chr9-71925989-DUP-65
chr9-72288712-DUP-54
chr9-72743410-DUP-63
chr9-72822674-DUP-94
chr9-73010744-DUP-137
chr9-73346356-DUP-4687
chr9-73963380-DUP-402
chr9-74883608-DUP-83
chr9-75433640-DUP-979
chr9-75798205-DUP-66
chr9-77371814-DUP-65
chr9-77389527-DUP-72
chr9-77438479-DUP-132
chr9-77438501-DUP-179
chr9-77439166-DUP-196
chr9-77440153-DUP-117
chr9-77850178-DUP-58
chr9-78324324-DUP-159
chr9-78570186-DUP-86
chr9-78806140-DUP-212
chr9-80019090-DUP-281
chr9-80444661-DUP-57
chr9-80490753-DUP-109
chr9-80649269-DUP-50
chr9-80778698-DUP-51
chr9-80859756-DUP-72
chr9-81831208-DUP-138
chr9-82601368-DUP-54
chr9-82725245-DUP-160
chr9-83038192-DUP-58
chr9-83107579-DUP-348
chr9-83107619-DUP-338
chr9-83492046-DUP-94
chr9-83852947-DUP-57
chr9-83855685-DUP-83
chr9-83906713-DUP-191

chr9-83964248-DUP-262
chr9-85303308-DUP-5805
chr9-86151348-DUP-168
chr9-87018902-DUP-80
chr9-87034939-DUP-81
chr9-87173110-DUP-63
chr9-87270199-DUP-1155
chr9-87305906-DUP-193
chr9-87661476-DUP-60
chr9-88297172-DUP-251
chr9-88424788-DUP-60
chr9-88676877-DUP-52
chr9-89762346-DUP-259
chr9-89762917-DUP-77
chr9-90072659-DUP-51
chr9-90093235-DUP-60
chr9-92338839-DUP-58
chr9-92424818-DUP-64
chr9-92499302-DUP-196
chr9-92851772-DUP-62
chr9-93077588-DUP-241
chr9-93555236-DUP-186
chr9-93652294-DUP-348
chr9-93713452-DUP-68
chr9-94379009-DUP-56
chr9-94485274-DUP-60
chr9-94500128-DUP-192
chr9-94505850-DUP-199
chr9-94533718-DUP-422
chr9-94534347-DUP-79
chr9-94541624-DUP-65
chr9-94875715-DUP-111
chr9-95420063-DUP-319
chr9-95500290-DUP-56
chr9-95648048-DUP-465
chr9-95648110-DUP-503
chr9-95657007-DUP-88
chr9-95659450-DUP-102
chr9-95677606-DUP-121
chr9-96452139-DUP-97
chr9-96734924-DUP-249
chr9-96734995-DUP-210
chr9-96962198-DUP-75
chr9-97805155-DUP-60
chr9-98037949-DUP-82
chr9-98614917-DUP-184
chr9-98730942-DUP-208
chr9-99359025-DUP-1271
chr9-99359237-DUP-1059
chr9-99360050-DUP-78
chr9-99376501-DUP-220
chr9-99407587-DUP-214
chr9-99427711-DUP-173
chr9-99550276-DUP-72
chr9-99690094-DUP-426
chr9-99731121-DUP-182
chr9-99911143-DUP-56
chr9-99913513-DUP-103
chr9-99937017-DUP-66
chr9-100100995-DUP-654
chr9-100129881-DUP-68
chr9-100171679-DUP-52
chr9-100257501-DUP-59
chr9-100719644-DUP-127
chr9-101079140-DUP-113
chr9-101285678-DUP-625
chr9-102690926-DUP-267
chr9-102690992-DUP-139
chr9-102732466-DUP-127
chr9-103173596-DUP-525
chr9-103174946-DUP-525
chr9-103175658-DUP-285
chr9-103176325-DUP-1101
chr9-103176583-DUP-51
chr9-103176918-DUP-526
chr9-103401873-DUP-78
chr9-103461202-DUP-52

chr9-103537934-DUP-85
chr9-103548002-DUP-134
chr9-103571455-DUP-87
chr9-103617911-DUP-52
chr9-103994977-DUP-84
chr9-104022245-DUP-100
chr9-104043495-DUP-60
chr9-104167499-DUP-59
chr9-104304153-DUP-58
chr9-104823347-DUP-94
chr9-105059340-DUP-50
chr9-105174736-DUP-81
chr9-105318975-DUP-235
chr9-105332877-DUP-159
chr9-105448750-DUP-280
chr9-105448842-DUP-286
chr9-105625258-DUP-92
chr9-106044993-DUP-79
chr9-106078615-DUP-3783
chr9-106093108-DUP-1713
chr9-106093141-DUP-1830
chr9-106559737-DUP-78
chr9-106607630-DUP-194
chr9-107841363-DUP-63
chr9-108002291-DUP-189
chr9-108132092-DUP-66
chr9-108169655-DUP-323
chr9-108371872-DUP-20895
chr9-109643612-DUP-50
chr9-109644730-DUP-125
chr9-109878762-DUP-91
chr9-109975946-DUP-57
chr9-110215209-DUP-68
chr9-110290806-DUP-73
chr9-110347809-DUP-115
chr9-110492216-DUP-62
chr9-110503182-DUP-109
chr9-110519017-DUP-204
chr9-110552376-DUP-159
chr9-110552417-DUP-88
chr9-110623048-DUP-55
chr9-110694117-DUP-130
chr9-110815523-DUP-369
chr9-110844251-DUP-62
chr9-111005466-DUP-133
chr9-111283811-DUP-62
chr9-111310465-DUP-58
chr9-111699399-DUP-98
chr9-112497232-DUP-51
chr9-112650811-DUP-67
chr9-112706433-DUP-144
chr9-112931711-DUP-64
chr9-112931735-DUP-155
chr9-113043747-DUP-59
chr9-113115609-DUP-87
chr9-113342092-DUP-126
chr9-113809113-DUP-40796
chr9-113817772-DUP-350
chr9-114001567-DUP-93
chr9-114004392-DUP-386
chr9-114004443-DUP-282
chr9-114007482-DUP-454
chr9-114447829-DUP-67
chr9-114572659-DUP-114
chr9-114784834-DUP-76
chr9-114785288-DUP-140
chr9-114849455-DUP-54
chr9-115112158-DUP-236
chr9-115112252-DUP-144
chr9-115175530-DUP-67
chr9-115326508-DUP-75
chr9-115362665-DUP-61
chr9-115472886-DUP-169
chr9-115724432-DUP-54
chr9-115871211-DUP-359
chr9-116354003-DUP-100
chr9-116354050-DUP-52

chr9-116645580-DUP-69
chr9-116844831-DUP-109
chr9-116937999-DUP-55
chr9-117283794-DUP-57
chr9-117546045-DUP-60
chr9-117546068-DUP-74
chr9-117619168-DUP-166
chr9-117619313-DUP-246
chr9-117629880-DUP-64
chr9-117754117-DUP-126
chr9-117879342-DUP-409
chr9-117920618-DUP-153
chr9-118021724-DUP-463
chr9-118061906-DUP-403
chr9-118181877-DUP-212
chr9-118438830-DUP-57
chr9-118622486-DUP-96
chr9-118622509-DUP-132
chr9-118632103-DUP-112
chr9-119100291-DUP-51
chr9-119100318-DUP-100
chr9-119218407-DUP-63
chr9-119255932-DUP-86
chr9-119338899-DUP-115
chr9-119462352-DUP-117
chr9-119526838-DUP-59
chr9-119607815-DUP-164
chr9-119607853-DUP-91
chr9-119971482-DUP-245
chr9-120236795-DUP-90
chr9-120383527-DUP-53
chr9-120517849-DUP-241
chr9-120715099-DUP-172
chr9-120715255-DUP-113
chr9-120801940-DUP-177
chr9-120862096-DUP-52
chr9-120871471-DUP-82
chr9-120884719-DUP-122
chr9-120884751-DUP-63
chr9-120904244-DUP-238
chr9-120949713-DUP-60
chr9-120991493-DUP-61
chr9-121013988-DUP-150
chr9-121014030-DUP-177
chr9-121017520-DUP-50
chr9-121032275-DUP-92
chr9-121044296-DUP-56
chr9-121084521-DUP-56
chr9-121390592-DUP-185
chr9-121390615-DUP-167
chr9-121403462-DUP-59
chr9-121466384-DUP-76
chr9-121469334-DUP-98
chr9-121602317-DUP-232
chr9-121662586-DUP-142
chr9-121782807-DUP-217
chr9-121789927-DUP-205
chr9-121797944-DUP-518
chr9-121934607-DUP-76
chr9-122079352-DUP-125
chr9-122371819-DUP-413
chr9-122372862-DUP-97
chr9-122374127-DUP-180
chr9-122590130-DUP-124
chr9-122591234-DUP-90
chr9-122609284-DUP-88
chr9-122734345-DUP-75
chr9-122735229-DUP-88
chr9-122988487-DUP-64
chr9-123032555-DUP-106
chr9-123274564-DUP-55
chr9-123722742-DUP-89
chr9-123817233-DUP-57
chr9-123892530-DUP-54
chrX-4439283-DUP-233
chrX-5079362-DUP-66
chrX-5304086-DUP-66

chrX-5531415-DUP-67
chrX-6096192-DUP-54
chrX-6227264-DUP-80
chrX-6353106-DUP-85
chrX-6501454-DUP-111
chrX-6501486-DUP-72
chrX-6690672-DUP-51
chrX-6736419-DUP-54
chrX-6775139-DUP-229
chrX-6935455-DUP-130
chrX-6937156-DUP-100
chrX-7074632-DUP-59
chrX-7523246-DUP-106
chrX-7623820-DUP-21075
chrX-8038493-DUP-57
chrX-8399025-DUP-191
chrX-8507982-DUP-35444
chrX-8829612-DUP-214
chrX-8976450-DUP-77
chrX-8989294-DUP-54
chrX-9029359-DUP-317
chrX-9029389-DUP-269
chrX-9244517-DUP-106
chrX-9489016-DUP-104
chrX-9515578-DUP-102
chrX-9558850-DUP-1097
chrX-10077106-DUP-205
chrX-10077180-DUP-126
chrX-10538903-DUP-68
chrX-10952837-DUP-52
chrX-11028467-DUP-70
chrX-12932641-DUP-685
chrX-12971153-DUP-162
chrX-14582060-DUP-7745
chrX-17118517-DUP-52
chrX-20607107-DUP-58
chrX-21386620-DUP-130
chrX-21573409-DUP-62
chrX-22078012-DUP-494
chrX-22498239-DUP-62
chrX-23000569-DUP-480
chrX-23083391-DUP-66
chrX-23432970-DUP-89
chrX-25624489-DUP-221
chrX-31364693-DUP-57
chrX-35169284-DUP-17136
chrX-35300763-DUP-93
chrX-35367848-DUP-119
chrX-35534905-DUP-80
chrX-36812659-DUP-60
chrX-36861228-DUP-87
chrX-36861940-DUP-358
chrX-37158246-DUP-6733
chrX-37238102-DUP-136
chrX-39117240-DUP-543
chrX-39163755-DUP-54
chrX-40228553-DUP-186
chrX-41509102-DUP-117
chrX-41989802-DUP-97
chrX-43141869-DUP-126
chrX-43143471-DUP-99
chrX-43233415-DUP-60
chrX-44782924-DUP-83
chrX-45142162-DUP-85
chrX-47141642-DUP-93
chrX-47152486-DUP-52
chrX-48778403-DUP-54
chrX-50890333-DUP-403
chrX-52556793-DUP-142
chrX-52589954-DUP-92
chrX-52595708-DUP-148
chrX-52595749-DUP-150
chrX-55257114-DUP-133
chrX-55257162-DUP-263
chrX-55729589-DUP-56
chrX-56881958-DUP-134
chrX-64787761-DUP-55

chrX-66026814-DUP-91
chrX-66430058-DUP-51
chrX-66998423-DUP-2210
chrX-67000669-DUP-2292
chrX-67001800-DUP-2289
chrX-67004353-DUP-2304
chrX-67925047-DUP-71
chrX-68324485-DUP-1307
chrX-68324541-DUP-1137
chrX-68381773-DUP-100
chrX-69552718-DUP-429
chrX-69757840-DUP-248
chrX-70774875-DUP-128
chrX-70774896-DUP-94
chrX-70961480-DUP-354
chrX-71531652-DUP-103
chrX-72161433-DUP-135
chrX-72192063-DUP-135
chrX-72192111-DUP-137
chrX-72220603-DUP-60
chrX-72336638-DUP-133
chrX-72799392-DUP-85
chrX-73370302-DUP-74
chrX-73521106-DUP-76
chrX-73636423-DUP-125
chrX-73814274-DUP-81
chrX-73830151-DUP-155
chrX-73847496-DUP-55
chrX-74107760-DUP-2059
chrX-74423514-DUP-57
chrX-74434085-DUP-68
chrX-74787864-DUP-993
chrX-74801363-DUP-951
chrX-75400674-DUP-155
chrX-75754945-DUP-127
chrX-77466781-DUP-1805
chrX-78176010-DUP-84
chrX-80119696-DUP-596
chrX-80976284-DUP-411
chrX-81996715-DUP-1141
chrX-82351205-DUP-697
chrX-82485345-DUP-181
chrX-82530610-DUP-187
chrX-83713909-DUP-70
chrX-83908527-DUP-52
chrX-85343694-DUP-250
chrX-86477365-DUP-132
chrX-86477388-DUP-137
chrX-86871009-DUP-56
chrX-87064827-DUP-60
chrX-87762119-DUP-195
chrX-88313062-DUP-104
chrX-88640451-DUP-194
chrX-89778934-DUP-56
chrX-90356144-DUP-270
chrX-90362805-DUP-1245
chrX-90591503-DUP-139
chrX-90810183-DUP-67
chrX-91068464-DUP-622
chrX-91072970-DUP-1249
chrX-91370762-DUP-118
chrX-91372975-DUP-1418
chrX-92063702-DUP-134
chrX-92830509-DUP-105
chrX-92848596-DUP-464
chrX-92848619-DUP-466
chrX-92965862-DUP-210
chrX-94342913-DUP-69
chrX-96114473-DUP-282
chrX-96453031-DUP-52
chrX-97257844-DUP-232
chrX-98598529-DUP-185
chrX-98705125-DUP-77
chrX-98953441-DUP-781
chrX-100739840-DUP-326
chrX-100866551-DUP-79
chrX-101440036-DUP-75

chrX-101534001-DUP-156
chrX-101555959-DUP-447
chrX-101834857-DUP-557
chrX-101834884-DUP-559
chrX-102295379-DUP-104
chrX-102498916-DUP-83
chrX-102499748-DUP-621
chrX-102680271-DUP-150
chrX-105140548-DUP-55
chrX-105492162-DUP-63
chrX-105986450-DUP-76
chrX-107330016-DUP-59
chrX-107433181-DUP-80
chrX-108503671-DUP-104
chrX-109290846-DUP-116
chrX-110425042-DUP-231
chrX-112112759-DUP-1331
chrX-112334328-DUP-65
chrX-112734949-DUP-69
chrX-113153349-DUP-115
chrX-113720225-DUP-750
chrX-114266321-DUP-51
chrX-114851444-DUP-92914
chrX-115855438-DUP-456
chrX-118159022-DUP-139
chrX-119733936-DUP-29372
chrX-119844804-DUP-274
chrX-119941198-DUP-58
chrX-121963943-DUP-620
chrX-121993331-DUP-1254
chrX-121993833-DUP-1256
chrX-122147348-DUP-1149
chrX-122151372-DUP-378
chrX-122176131-DUP-488
chrX-122375045-DUP-1147
chrX-122376552-DUP-1159
chrX-122377466-DUP-1138
chrX-122378109-DUP-1146
chrX-122383686-DUP-1147
chrX-122388116-DUP-378
chrX-122609047-DUP-379
chrX-122614955-DUP-85
chrX-122819165-DUP-2296
chrX-124794800-DUP-85
chrX-125955009-DUP-79
chrX-126270740-DUP-50
chrX-129342031-DUP-116
chrX-129921543-DUP-2613
chrX-130394188-DUP-65
chrX-132032858-DUP-70
chrX-132539363-DUP-128043
chrX-133237706-DUP-164
chrX-133272460-DUP-122
chrX-133462513-DUP-78
chrX-133671671-DUP-79
chrX-133710308-DUP-108
chrX-133852723-DUP-76
chrX-133946809-DUP-528
chrX-133948157-DUP-720
chrX-133949943-DUP-74
chrX-133950341-DUP-682
chrX-133952390-DUP-74
chrX-133953642-DUP-200
chrX-133955974-DUP-130
chrX-133956505-DUP-224
chrX-133987774-DUP-155
chrX-134000514-DUP-234
chrX-134381442-DUP-170
chrX-134572134-DUP-84
chrX-134593573-DUP-519
chrX-134698290-DUP-135
chrX-134698468-DUP-215
chrX-134741200-DUP-223
chrX-135249267-DUP-90006
chrX-135369979-DUP-1854
chrX-135674842-DUP-175
chrX-135674876-DUP-77

chrX-135705236-DUP-69
chrX-135780871-DUP-8808
chrX-136007510-DUP-2727
chrX-136008871-DUP-4092
chrX-136010238-DUP-2725
chrX-136513255-DUP-134
chrX-136991216-DUP-107
chrX-137269703-DUP-125
chrX-139307493-DUP-74
chrX-139823996-DUP-127
chrX-139872489-DUP-2931
chrX-140078297-DUP-101
chrX-140237188-DUP-57
chrX-140243298-DUP-56
chrX-141148060-DUP-676
chrX-141415912-DUP-52
chrX-141723471-DUP-86
chrX-142707100-DUP-2635
chrX-142734787-DUP-54
chrX-142820733-DUP-105
chrX-143129710-DUP-141
chrX-143362029-DUP-570
chrX-144871548-DUP-62
chrX-145408559-DUP-29049
chrX-145506957-DUP-235
chrX-145929347-DUP-9170
chrX-145930614-DUP-9178
chrX-145931652-DUP-5939
chrX-145932525-DUP-5936
chrX-146563905-DUP-4759
chrX-147017857-DUP-54
chrX-147506790-DUP-66
chrX-147804756-DUP-54
chrX-148587855-DUP-72
chrX-148680979-DUP-52
chrX-148766876-DUP-121
chrX-149175873-DUP-96
chrX-149175904-DUP-62
chrX-149602267-DUP-1490
chrX-150157459-DUP-64
chrX-150437190-DUP-93
chrX-150994477-DUP-120
chrX-150994547-DUP-56
chrX-152039902-DUP-81
chrX-152177710-DUP-75
chrX-152734344-DUP-335
chrX-154088970-DUP-110
chrX-156040814-DUP-1601
chrX-156040928-DUP-2905
chrX-156042189-DUP-92
chrX-156042317-DUP-1396
chrX-156482756-DUP-53
chrX-156786991-DUP-72
chrX-157563296-DUP-91
chrX-159239190-DUP-63
chrX-159633031-DUP-156
chrX-159751967-DUP-73
chrX-159759885-DUP-81
chrX-161075050-DUP-67
chrX-161358118-DUP-63
chrX-161623941-DUP-93
chrX-162052175-DUP-68
chrX-162442556-DUP-222
chrX-162678440-DUP-54
chrX-163338352-DUP-3399
chrX-164836353-DUP-57
chrX-165250373-DUP-66
chrX-165819590-DUP-118
chrX-166740568-DUP-97
chrX-167862638-DUP-102
chrX-167887235-DUP-956
chrX-168151963-DUP-118
chrX-168609362-DUP-286
chrX-168769136-DUP-89
chrX-168772477-DUP-89
chrX-168774190-DUP-6854
chrX-168774260-DUP-150

chrX-168776822-DUP-172
chrX-168777591-DUP-432
chrX-168779618-DUP-188
chrX-168780993-DUP-147
chrX-168783331-DUP-337
chrX-168790055-DUP-60
chrX-168794499-DUP-222
chrX-168796644-DUP-88
chrX-168797508-DUP-175
chrX-169072177-DUP-71
chrX-169077950-DUP-260
chrX-169078091-DUP-180
chrX-169081128-DUP-180
chrX-169082141-DUP-60
chrX-169088655-DUP-66
chrX-169088677-DUP-170
chrX-169090519-DUP-107
chrX-169091723-DUP-213
chrX-169091749-DUP-215
chrX-169095996-DUP-139
chrX-169097199-DUP-53
chrX-169104311-DUP-68
chrX-169107091-DUP-181
chrX-169107178-DUP-93
chrX-169107592-DUP-220
chrX-169118561-DUP-87
chrX-169119383-DUP-2626
chrX-169260409-DUP-58
chrX-169263695-DUP-60
chrX-169266391-DUP-104
chrX-169270303-DUP-1412
chrX-169270653-DUP-1775
chrX-169288811-DUP-1976
chrX-169304604-DUP-950

**Identified_By_JAX**
chr1-5555045-DUP-1811
chr1-6623341-DUP-252
chr1-9567046-DUP-300
chr1-16158644-DUP-839
chr1-26313900-DUP-3168
chr1-26725896-DUP-17946
chr1-26725969-DUP-25663
chr1-28341813-DUP-1260
chr1-29324559-DUP-59135
chr1-34641782-DUP-5254
chr1-34644503-DUP-5075
chr1-45135001-DUP-3008
chr1-46622552-DUP-212
chr1-47797988-DUP-798
chr1-50276138-DUP-4547
chr1-53712901-DUP-2224
chr1-54819586-DUP-2348
chr1-57347504-DUP-2616
chr1-61163897-DUP-358
chr1-63328316-DUP-255
chr1-65738594-DUP-1274
chr1-65738594-DUP-3813
chr1-79817071-DUP-292
chr1-87974104-DUP-4473
chr1-90550728-DUP-5077
chr1-92434908-DUP-18491
chr1-95340475-DUP-16655
chr1-104270910-DUP-3096
chr1-104664664-DUP-2779
chr1-107872347-DUP-2140
chr1-110810468-DUP-3621
chr1-114492133-DUP-7141
chr1-124017190-DUP-8175
chr1-133274793-DUP-31118
chr1-136006598-DUP-287
chr1-137810458-DUP-228
chr1-138790916-DUP-14538
chr1-138794145-DUP-33926
chr1-143326762-DUP-6583
chr1-145832879-DUP-3408
chr1-153272293-DUP-7674
chr1-166706035-DUP-255
chr1-170601556-DUP-244
chr1-185482708-DUP-987
chr10-7405534-DUP-4001
chr10-10449299-DUP-5495
chr10-16053924-DUP-1152
chr10-40501403-DUP-288
chr10-48883789-DUP-3541
chr10-54906950-DUP-3475
chr10-54967393-DUP-2192
chr10-58209131-DUP-2900
chr10-66261180-DUP-14165
chr10-88498515-DUP-2886
chr10-95769211-DUP-399
chr10-101961717-DUP-248
chr10-117868993-DUP-2456
chr10-130321710-DUP-2491
chr11-8280557-DUP-929
chr11-11394169-DUP-496
chr11-21473030-DUP-6023
chr11-32222716-DUP-345
chr11-33998818-DUP-413
chr11-43728093-DUP-3017
chr11-49591027-DUP-264
chr11-59191615-DUP-2136
chr11-69757087-DUP-278
chr11-70252929-DUP-42845
chr11-74256640-DUP-13279
chr11-83936205-DUP-626
chr11-92703415-DUP-1680
chr11-99195141-DUP-4441
chr11-107029876-DUP-2936
chr11-108569863-DUP-826
chr11-115442521-DUP-8776
chr11-119292599-DUP-4473

chr12-5102046-DUP-2492
chr12-8671029-DUP-6678
chr12-41915391-DUP-4230
chr12-57609220-DUP-1873
chr12-58081240-DUP-1883
chr12-71041824-DUP-497
chr12-79163040-DUP-2271
chr12-109931905-DUP-477
chr12-111515265-DUP-311
chr12-115677241-DUP-345
chr12-118942432-DUP-208
chr12-119805844-DUP-601
chr13-10224814-DUP-11673
chr13-14900724-DUP-2031
chr13-16006246-DUP-4925
chr13-23595355-DUP-12100
chr13-26361890-DUP-3297
chr13-51527559-DUP-277
chr13-60218400-DUP-515
chr13-68478086-DUP-3537
chr13-73291096-DUP-3066
chr13-78373453-DUP-2921
chr13-88747257-DUP-6898
chr13-99471636-DUP-1608
chr13-100810392-DUP-1466
chr13-108123572-DUP-477
chr13-111732665-DUP-867
chr13-113230099-DUP-29035
chr13-113230261-DUP-9974
chr13-119018487-DUP-8114
chr14-21361841-DUP-3294
chr14-21363515-DUP-3300
chr14-21538261-DUP-567
chr14-36632505-DUP-15343
chr14-36632924-DUP-7380
chr14-55625472-DUP-20153
chr14-57254558-DUP-7240
chr14-57254850-DUP-14611
chr14-74855462-DUP-349
chr14-84592966-DUP-536
chr14-86348825-DUP-3856
chr14-87361180-DUP-2597
chr14-90978439-DUP-2664
chr14-109579325-DUP-928
chr14-116623386-DUP-1954
chr14-120066649-DUP-10695
chr14-123242362-DUP-229
chr14-124693885-DUP-2316
chr14-124693885-DUP-4668
chr15-8834943-DUP-13419
chr15-9970470-DUP-14801
chr15-18231723-DUP-18572
chr15-20686575-DUP-224
chr15-34226754-DUP-2850
chr15-50332341-DUP-2410
chr15-52916126-DUP-662
chr15-67741470-DUP-364
chr15-76309457-DUP-2794
chr15-76309457-DUP-694
chr15-84982819-DUP-359
chr15-86681675-DUP-395
chr15-95716938-DUP-6333
chr15-96480853-DUP-2484
chr15-98097015-DUP-20457
chr15-100886752-DUP-2218
chr15-101878401-DUP-22035
chr16-3080193-DUP-2657
chr16-3168158-DUP-2122
chr16-5178422-DUP-2133
chr16-27486116-DUP-544
chr16-31243732-DUP-3917
chr16-36144721-DUP-6765
chr16-40101208-DUP-228
chr16-53122179-DUP-468
chr16-54397287-DUP-3586
chr16-56659151-DUP-5316
chr16-58095314-DUP-4220

chr16-59158154-DUP-6359
chr16-59187204-DUP-268
chr16-60469035-DUP-28769
chr16-61177684-DUP-371
chr16-61391227-DUP-334
chr16-62417126-DUP-2136
chr16-65549998-DUP-467
chr16-69241555-DUP-619
chr16-74160167-DUP-201
chr16-74657374-DUP-549
chr16-87183269-DUP-870
chr16-87183719-DUP-504
chr16-97286645-DUP-339
chr17-3670108-DUP-319
chr17-21473096-DUP-15652
chr17-21503704-DUP-25978
chr17-25261516-DUP-696
chr17-26318312-DUP-10309
chr17-35474643-DUP-5813
chr17-41526280-DUP-1081
chr17-55068114-DUP-9876
chr17-74610178-DUP-443
chr17-76517267-DUP-3947
chr18-9177787-DUP-10380
chr18-11288664-DUP-19809
chr18-11895930-DUP-1633
chr18-18108095-DUP-5720
chr18-29154932-DUP-22361
chr18-52585134-DUP-8958
chr18-60463152-DUP-215
chr18-60593659-DUP-1651
chr18-74443672-DUP-4300
chr18-89739982-DUP-993
chr19-22911849-DUP-1871
chr19-29335239-DUP-631
chr19-41490933-DUP-3300
chr19-47830393-DUP-7786
chr19-47836275-DUP-4987
chr19-56168385-DUP-356
chr19-57016257-DUP-1891
chr2-8011661-DUP-8368
chr2-23636640-DUP-2953
chr2-24934739-DUP-1650
chr2-36493977-DUP-317
chr2-39320700-DUP-341
chr2-76038957-DUP-1736
chr2-76038957-DUP-3476
chr2-86137133-DUP-669
chr2-96031337-DUP-625
chr2-104780241-DUP-393
chr2-113501533-DUP-2229
chr2-127851315-DUP-317
chr2-135504669-DUP-14067
chr2-139005792-DUP-365
chr2-162400427-DUP-5092
chr2-162687854-DUP-6196
chr2-168884628-DUP-1495
chr3-5808721-DUP-3584
chr3-10517885-DUP-4942
chr3-11175553-DUP-219
chr3-13621977-DUP-8022
chr3-16681553-DUP-2410
chr3-16683857-DUP-12056
chr3-16683857-DUP-4823
chr3-21422846-DUP-11945
chr3-27333784-DUP-14384
chr3-33751501-DUP-18995
chr3-41515213-DUP-2170
chr3-44517832-DUP-36663
chr3-45751659-DUP-1499
chr3-57621193-DUP-250
chr3-64672821-DUP-901
chr3-64776295-DUP-288
chr3-73395668-DUP-1489
chr3-73626376-DUP-17879
chr3-78180126-DUP-575
chr3-80521073-DUP-236

chr3-84734317-DUP-6327
chr3-86184684-DUP-477
chr3-90400601-DUP-1413
chr3-114774541-DUP-2059
chr3-133359787-DUP-478
chr3-134879152-DUP-403
chr3-142307427-DUP-31458
chr3-153562424-DUP-203
chr3-155083157-DUP-3416
chr4-3732870-DUP-6327
chr4-10247123-DUP-552
chr4-12133153-DUP-4434
chr4-12189513-DUP-4082
chr4-14758877-DUP-211
chr4-22141203-DUP-354
chr4-22627721-DUP-11478
chr4-29933269-DUP-579
chr4-30405131-DUP-2943
chr4-43758064-DUP-4454
chr4-46486390-DUP-3947
chr4-51325228-DUP-2638
chr4-54962855-DUP-4741
chr4-57463914-DUP-297
chr4-57858994-DUP-210
chr4-62413148-DUP-16989
chr4-62432834-DUP-24495
chr4-64176041-DUP-1420
chr4-67628089-DUP-924
chr4-71974899-DUP-4196
chr4-97277585-DUP-18944
chr4-99296552-DUP-234
chr4-109553331-DUP-9208
chr4-115525521-DUP-6395
chr4-115541088-DUP-6109
chr4-117876421-DUP-1368
chr4-144482090-DUP-14071
chr4-145278143-DUP-9310
chr5-8951597-DUP-1214
chr5-16543020-DUP-520
chr5-23972915-DUP-504
chr5-40912429-DUP-566
chr5-47255805-DUP-1764
chr5-59189185-DUP-1656
chr5-61434229-DUP-6325
chr5-77526622-DUP-327
chr5-79640745-DUP-513
chr5-108972502-DUP-4710
chr5-110978095-DUP-1047
chr5-129916971-DUP-3132
chr5-138674067-DUP-312
chr5-148303167-DUP-2148
chr5-148739833-DUP-811
chr5-151046474-DUP-4778
chr6-20143026-DUP-3345
chr6-20787380-DUP-299
chr6-25244439-DUP-209
chr6-31756400-DUP-5262
chr6-51204757-DUP-315
chr6-70253753-DUP-3554
chr6-75318748-DUP-431
chr6-86154875-DUP-2093
chr6-89709488-DUP-3153
chr6-94865599-DUP-2002
chr6-101759053-DUP-16026
chr6-101777862-DUP-504
chr6-118905986-DUP-33309
chr6-118911843-DUP-8329
chr6-120637243-DUP-784
chr6-128820366-DUP-6031
chr6-128831306-DUP-21371
chr6-128945139-DUP-4171
chr6-136302330-DUP-25788
chr6-148200497-DUP-243
chr7-4857432-DUP-4192
chr7-5058496-DUP-2186
chr7-13857554-DUP-493
chr7-15638201-DUP-11405

chr7-16395247-DUP-25585
chr7-19781349-DUP-1956
chr7-24435860-DUP-12797
chr7-26359340-DUP-2194
chr7-27528556-DUP-909
chr7-30443000-DUP-4278
chr7-31082046-DUP-4002
chr7-31815325-DUP-309
chr7-40436594-DUP-1492
chr7-46394133-DUP-12960
chr7-48053599-DUP-3356
chr7-54079626-DUP-4484
chr7-61422928-DUP-671
chr7-62624065-DUP-2445
chr7-63629389-DUP-4091
chr7-93745745-DUP-15044
chr7-93745745-DUP-5017
chr7-99199935-DUP-423
chr7-101798107-DUP-2892
chr7-109473235-DUP-209
chr7-110786235-DUP-6434
chr7-114663277-DUP-456
chr7-120509483-DUP-306
chr7-124407738-DUP-4773
chr7-127318669-DUP-2482
chr7-130193316-DUP-6435
chr7-130196390-DUP-9572
chr7-131028099-DUP-4427
chr7-132546159-DUP-852
chr7-141413545-DUP-824
chr7-141827333-DUP-7023
chr8-21438729-DUP-11200
chr8-26737508-DUP-2345
chr8-32569196-DUP-243
chr8-34059015-DUP-424
chr8-36917243-DUP-7918
chr8-44187833-DUP-2615
chr8-56345430-DUP-2876
chr8-57202316-DUP-17916
chr8-58569254-DUP-267
chr8-61781356-DUP-539
chr8-65387056-DUP-494
chr8-66491468-DUP-4050
chr8-71457400-DUP-6873
chr8-81246122-DUP-232
chr8-86548769-DUP-2996
chr8-87013347-DUP-1364
chr8-96011864-DUP-346
chr8-96012115-DUP-347
chr9-12278944-DUP-2023
chr9-18971673-DUP-1390
chr9-19112219-DUP-362
chr9-19292204-DUP-2201
chr9-24240342-DUP-348
chr9-36506751-DUP-1184
chr9-64661122-DUP-4683
chr9-93205578-DUP-3782
chr9-101286301-DUP-220
chr9-105410216-DUP-294
chr9-110661995-DUP-308
chr9-113044399-DUP-969
chrX-12932649-DUP-1572
chrX-22728838-DUP-3161
chrX-22867615-DUP-13043
chrX-37162033-DUP-13698
chrX-42120227-DUP-844
chrX-57088832-DUP-1041
chrX-65410476-DUP-5806
chrX-66648180-DUP-4338
chrX-67360370-DUP-1599
chrX-67441959-DUP-2426
chrX-71350696-DUP-12803
chrX-92690853-DUP-362
chrX-100287459-DUP-1244
chrX-113751855-DUP-14006
chrX-116607055-DUP-2146
chrX-126667474-DUP-1078

chrX-135370925-DUP-3713
chrX-135858256-DUP-19656
chrX-136010292-DUP-4126
chrX-137096663-DUP-997
chrX-139872795-DUP-2925
chrX-142846754-DUP-2451
chrX-142849206-DUP-2447
chrX-145938559-DUP-9154
chrX-150543025-DUP-566
chrX-150543614-DUP-566
chrX-156104797-DUP-1552
chrX-157902971-DUP-2150
chrX-167543322-DUP-2752

**Table S7. Lists of high impact DELs present in all 39 or in the 35 classical inbred strains.** The chromosome, starting and ending position, size, gene symbol, strains with the variant allele, and the predicted effect for each DEL are shown.

| CHR | START | END | SIZE | SYMBOL | STRAINS with VARIANT SV ALLELE | CONSEQUENCE |
|---|---|---|---|---|---|---|
| 1 | 4851851 | 4851919 | 68 | Mrpl15 | C3H,CBA,MRL,RF | feature_truncation,3_prime_UTR_variant |
| 1 | 13721098 | 13721166 | 68 | Lactb2 | AJ,AKR,BALB,C3H,CBA,DBA1J,DBA2J,FVB,ILNJ2,KK,LP,MRL,NOD,NZB,RF,RH2,SEA,SJL,SMJ | feature_truncation,coding_sequence_variant,intron_variant |
| 1 | 34484669 | 34484738 | 69 | Imp4 | RF | feature_truncation,3_prime_UTR_variant |
| 1 | 37664302 | 37664375 | 73 | Cracdl | AJ,B10J,C3H,CE,MA/My,MRL,NOD,SJL | frameshift_variant,feature_truncation |
| 1 | 38833129 | 38833189 | 60 | Lonrf2 | 129S1,AJ,AKR,BTBR,BUB,C3H,C57L,CBA,CE,DBA1J,DBA2J,ILNJ2,KK,MRL,NOD,NOR,NZB,NZW,PJ,PL,RH2,SJL,SMJ,TH | feature_truncation,3_prime_UTR_variant |
| 1 | 52697296 | 52697354 | 58 | Mfsd6 | C58J | feature_truncation,3_prime_UTR_variant |
| 1 | 59045610 | 59045661 | 51 | C2cd6 | MA/My | feature_truncation,inframe_deletion |
| 1 | 59756069 | 59756169 | 100 | Nop58 | CE,LP | feature_truncation,3_prime_UTR_variant |
| 1 | 60023658 | 60023710 | 52 | Fam117b | CE,LP | feature_truncation,3_prime_UTR_variant |
| 1 | 64597703 | 64598084 | 381 | Creb1 | 129S1,AJ,AKR,BTBR,C3H,C57L,CBA,CE,DBA1J,DBA2J,FVB,ILNJ2,KK,LP,MA/My,MRL,NOD,NOR,NZB,NZO,NZW,PJ,PL,RF,SJL,SMJ,SWR,TH | feature_truncation,coding_sequence_variant,intron_variant |
| 1 | 74199029 | 74199222 | 193 | Cxcr2 | 129S1,AJ,BTBR,C3H,C57L,CBA,MRL,NOD,NZB,NZO,PJ,RH2,SEA,SMJ,SWR | feature_truncation,3_prime_UTR_variant |
| 1 | 78648418 | 78648480 | 62 | Utp14b | LP | feature_truncation,3_prime_UTR_variant |
| 1 | 79584844 | 79585161 | 317 | Ap1s3 | 129S1,AJ,BALB,BTBR,BUB,C3H,C57L,C58J,CBA,CE,DBA1J,DBA2J,FVB,ILNJ2,LP,MRL,NOD,NZB,NZO,PJ,RF,RH2,SEA,SJL,SMJ,SWR,TH | feature_truncation,3_prime_UTR_variant |
| 1 | 82938077 | 82938140 | 63 | Gm47955 | PJ,RF,SMJ | feature_truncation,inframe_deletion |
| 1 | 82978485 | 82978553 | 68 | Gm47959 | AKR,BUB,CBA,MRL,NOD,PJ,PL,RF,SMJ | frameshift_variant,feature_truncation |
| 1 | 83036905 | 83036959 | 54 | A030005K14Rik | RF | feature_truncation,inframe_deletion |
| 1 | 83038865 | 83038922 | 57 | Krtap28-13 | PJ,SMJ | feature_truncation,inframe_deletion |
| 1 | 85191260 | 85191510 | 250 | Gm7609 | AKR,BUB,CBA,MRL,NOD,PJ,PL,SMJ | stop_lost,feature_truncation,coding_sequence_variant,3_prime_UTR_variant |
| 1 | 86205774 | 86205847 | 73 | Armc9 | BUB,NZB,RF | feature_truncation,3_prime_UTR_variant |
| 1 | 86354309 | 86354385 | 76 | Tex44 | 129S1,AJ,BTBR,C3H,C57L,C58J,CE,DBA1J,DBA2J,FVB,ILNJ2,KK,NZB,NZO,NZW,RH2,SJL,SMJ,SWR,TH | frameshift_variant,feature_truncation |
| 1 | 86535314 | 86535380 | 66 | Cops7b | BUB | feature_truncation,3_prime_UTR_variant |
| 1 | 90703524 | 90703587 | 63 | Col6a3 | 129S1,AKR,BUB,C3H,C57L,CBA,FVB,KK,LP,MA/My,MRL,NZB,NZO,NZW,PJ,PL,RF,SJL,SMJ,SWR,TH | feature_truncation,inframe_deletion |
| 1 | 97552793 | 97552927 | 134 | Macir | MRL,PL | feature_truncation,3_prime_UTR_variant |
| 1 | 97570705 | 97570768 | 63 | Macir | MRL,PL | feature_truncation,3_prime_UTR_variant |
| 1 | 121432868 | 121433033 | 165 | Ccdc93 | BALB,BTBR,PL,RF,SEA,SMJ | feature_truncation,3_prime_UTR_variant |
| 1 | 131106721 | 131106797 | 76 | Eif2d | 129S1,BALB,C57L,DBA1J,DBA2J,FVB,ILNJ2,KK,MA/My,NZO,NZW,SEA,SJL,SWR,TH | feature_truncation,3_prime_UTR_variant |
| 1 | 131697925 | 131697984 | 59 | Slc26a9 | PL | feature_truncation,3_prime_UTR_variant |
| 1 | 134436505 | 134436706 | 201 | Rabif | 129S1,AJ,AKR,BTBR,BUB,C3H,CBA,CE,DBA1J,DBA2J,FVB,ILNJ2,KK,MA/My,MRL,NOD,NOR,NZB,NZO,PJ,PL,RF,RH2,SJL,SMJ,SWR,TH | feature_truncation,3_prime_UTR_variant |
| 1 | 134818444 | 134818499 | 55 | Ppp1r12b | 129S1,AJ,AKR,BTBR,BUB,C3H,CBA,CE,DBA1J,DBA2J,FVB,ILNJ2,KK,MA/My,MRL,NOD,NOR,NZB,NZO,PJ,PL,RF,RH2,SJL,SMJ,SWR,TH | feature_truncation,coding_sequence_variant,intron_variant |
| 1 | 150330660 | 150330876 | 216 | Prg4 | AJ,AKR,BTBR,BUB,C3H,C57L,CBA,CE,DBA1J,DBA2J,ILNJ2,MRL,PL,RF,RH2,SMJ | feature_truncation,inframe_deletion |
| 1 | 153752072 | 153752275 | 203 | Teddm1b | BUB,C57L,DBA1J,DBA2J,FVB,MA/My,RF,SJL,SWR,TH | feature_truncation,3_prime_UTR_variant |
| 1 | 156193594 | 156194101 | 507 | Rpl35rt | 129S1,AJ,AKR,BALB,BTBR,BUB,C3H,C57L,C58J,CBA,CE,DBA1J,DBA2J,FVB,ILNJ2,KK,LP,MA/My,MRL,NOD,NOR,NZB,NZO,NZW,PJ,PL,RF,RH2,SEA,SWR,TH | transcript_ablation |
| 1 | 159753853 | 159753920 | 67 | Tnr | BALB,C58J,DBA1J,DBA2J,LP | feature_truncation,3_prime_UTR_variant |
| 1 | 160995589 | 160995711 | 122 | Ankrd45 | 129S1,AJ,BALB,BUB,C3H,C57L,C58J,CBA,CE,DBA1J,DBA2J,ILNJ2,LP,NZW,PJ,RH2,SEA,SMJ | feature_truncation,3_prime_UTR_variant |
| 1 | 163976790 | 163976866 | 76 | Selp | DBA1J,DBA2J,ILNJ2 | feature_truncation,3_prime_UTR_variant |
| 1 | 164175683 | 164175880 | 197 | Nme7 | BUB,FVB,NOD,NOR,PJ,RF,SEA,SJL,SMJ,SWR | stop_lost,feature_truncation,coding_sequence_variant,3_prime_UTR_variant,intron_variant |
| 1 | 165004861 | 165004921 | 60 | Sft2d2 | DBA1J,DBA2J,ILNJ2 | feature_truncation,3_prime_UTR_variant |
| 1 | 167426126 | 167426300 | 174 | Rxrg | BUB,SWR,TH | feature_truncation,5_prime_UTR_variant,feature_truncation,5_prime_UTR_variant,intron_variant |
| 1 | 171831327 | 171831453 | 126 | Vangl2 | 129S1,AJ,AKR,BALB,BUB,C3H,C57L,CBA,CE,DBA1J,DBA2J,FVB,KK,MRL,NOD,NOR,NZB,NZO,NZW,PJ,PL,RF,RH2,SEA,SJL,SMJ,SWR,TH | feature_truncation,3_prime_UTR_variant |
| 1 | 172101601 | 172101653 | 52 | Atp1a2 | PL | feature_truncation,3_prime_UTR_variant |
| 1 | 172198493 | 172198547 | 54 | Kcnj10 | 129S1,AJ,AKR,BALB,BUB,C3H,C57L,C58J,CBA,CE,DBA1J,DBA2J,FVB,KK,MRL,NOD,NOR,NZB,NZO,NZW,PJ,PL,RF,RH2,SEA,SJL,SMJ,SWR,TH | feature_truncation,3_prime_UTR_variant |
| 1 | 172206793 | 172207134 | 341 | Pigm | AKR,RF | feature_truncation,3_prime_UTR_variant |
| 1 | 172207321 | 172207790 | 469 | Pigm | CE,NZB,PL,SMJ,TH | feature_truncation,3_prime_UTR_variant |
| 1 | 172527106 | 172527166 | 60 | Crp | 129S1,AJ,AKR,BALB,BTBR,BUB,C3H,C57L,C58J,CBA,CE,DBA1J,DBA2J,FVB,ILNJ2,KK,MRL,NOD,NOR,NZB,NZO,NZW,PJ,PL,RF,RH2,SEA,SJL,SMJ,SWR,TH | feature_truncation,3_prime_UTR_variant |
| 1 | 173562682 | 173562745 | 63 | Ifi207 | AJ,FVB,PL,RH2,TH | feature_truncation,inframe_deletion |
| 1 | 173870799 | 173870877 | 78 | Or10aa1 | AKR,BTBR,BUB,C58J,CBA,CE,DBA1J,DBA2J,ILNJ2,KK,NZB,NZO,PJ,PL,RF,SEA,SMJ,SWR,TH | feature_truncation,3_prime_UTR_variant |
| 1 | 173988263 | 173988555 | 292 | Or6k2 | 129S1,AJ,AKR,BALB,BTBR,BUB,C3H,C57L,C58J,CBA,CE,DBA1J,DBA2J,FVB,ILNJ2,KK,MRL,NOD,NOR,NZB,NZO,PJ,PL,RF,RH2,SEA,SJL,SMJ,SWR,TH | feature_truncation,3_prime_UTR_variant |
| 1 | 174041803 | 174041864 | 61 | Spta1 | 129S1,AJ,AKR,BALB,BTBR,BUB,C3H,C57L,C58J,CBA,CE,DBA1J,DBA2J,FVB,ILNJ2,KK,MRL,NOD,NOR,NZB,NZO,NZW,PJ,PL,RF,RH2,SEA,SJL,SMJ,SWR,TH | feature_truncation,coding_sequence_variant,intron_variant |

| | | | | | | |
|---|---|---|---|---|---|---|
| 1 | 175488632 | 175488700 | 68 | Kmo | 129S1,BALB,BTBR,C3H,C57L,MRL,NOD,NOR,NZO,NZW,PL,SMJ,TH | feature_truncation,3_prime_UTR_variant |
| 1 | 179335900 | 179335960 | 60 | Smyd3 | AKR | feature_truncation,3_prime_UTR_variant |
| 1 | 181337202 | 181337293 | 91 | Ccdc121rt1 | TH | feature_truncation,3_prime_UTR_variant |
| 1 | 181729876 | 181729942 | 66 | Enah | NZO | feature_truncation,3_prime_UTR_variant |
| 1 | 190815834 | 190815887 | 53 | Nsl1 | 129S1,BUB,CBA,DBA1J,DBA2J,ILNJ2,KK,MA/My,MRL,NOD,NOR,NZO,NZW,PL,SWR | feature_truncation,3_prime_UTR_variant |
| 1 | 194720240 | 194720307 | 67 | Cd46 | NOD,NOR,SWR | feature_truncation,3_prime_UTR_variant |
| 1 | 194821674 | 194821725 | 51 | Cr2 | NZO,NZW,RF | feature_truncation,3_prime_UTR_variant |
| 10 | 13006405 | 13006463 | 58 | Plagl1 | DBA1J,DBA2J | feature_truncation,3_prime_UTR_variant |
| 10 | 33470537 | 33470617 | 80 | Clvs2 | KK | feature_truncation,3_prime_UTR_variant |
| 10 | 43400069 | 43400639 | 570 | Pam16l | 129S1,BTBR,C57L,ILNJ2,KK | stop_lost,feature_truncation,coding_sequence_variant,5_prime_UTR_variant,3_prime_UTR_variant,stop_lost,feature_truncation,coding_sequence_variant,5_prime_UTR_variant,3_prime_UTR_variant,intron_variant |
| 10 | 45043240 | 45043735 | 495 | Prep | KK | feature_truncation,3_prime_UTR_variant |
| 10 | 53152674 | 53152728 | 54 | Cep85l | CE | feature_truncation,3_prime_UTR_variant |
| 10 | 62849148 | 62849214 | 66 | Rufy2 | RF | feature_truncation,3_prime_UTR_variant |
| 10 | 62913117 | 62913323 | 206 | Pbld1 | CE,RF | feature_truncation,3_prime_UTR_variant |
| 10 | 66848254 | 66848309 | 55 | Reep3 | ILNJ2,SMJ | feature_truncation,3_prime_UTR_variant |
| 10 | 75386588 | 75386684 | 96 | Lrrc75b | NZW | feature_truncation,3_prime_UTR_variant |
| 10 | 76881787 | 76882153 | 366 | Slc19a1 | BUB | feature_truncation,3_prime_UTR_variant |
| 10 | 78716400 | 78716878 | 478 | Or7a39 | TH | feature_truncation,3_prime_UTR_variant |
| 10 | 81171660 | 81171714 | 54 | Gipc3 | RF | feature_truncation,3_prime_UTR_variant |
| 10 | 83227179 | 83227256 | 77 | Nopchap1 | C58J,LP | feature_truncation,3_prime_UTR_variant |
| 10 | 83254239 | 83254314 | 75 | Nopchap1 | BUB,SMJ | feature_truncation,3_prime_UTR_variant |
| 10 | 83295351 | 83295453 | 102 | Nopchap1 | BUB,SMJ | feature_truncation,3_prime_UTR_variant |
| 10 | 83322300 | 83322355 | 55 | Nopchap1 | BUB,SMJ | feature_truncation,3_prime_UTR_variant |
| 10 | 83431733 | 83431936 | 203 | Washc4 | BUB | feature_truncation,3_prime_UTR_variant |
| 10 | 89245617 | 89245824 | 207 | Gas2l3 | AJ,AKR,BALB,BTBR,BUB,C3H,C58J,CBA,CE,FVB,ILNJ2,KK,LP,MRL,NOD,NOR,NZB,NZO,NZW,PJ,PL,RF,RH2,SEA,SWR,TH | feature_truncation,3_prime_UTR_variant |
| 10 | 89475032 | 89475092 | 60 | Scyl2 | AJ,AKR,BALB,CE,FVB,ILNJ2,MRL,NOD,NOR,NZO,PJ,PL,RF,RH2,SEA | feature_truncation,3_prime_UTR_variant |
| 10 | 93353273 | 93353366 | 93 | Hal | 129S1,AJ,AKR,BALB,BTBR,BUB,C3H,C57L,C58J,CBA,CE,FVB,ILNJ2,KK,LP,MA/My,NOD,NOR,PJ,RF,RH2,SEA,SJL,SMJ,SWR,TH | feature_truncation,3_prime_UTR_variant |
| 10 | 93688439 | 93688583 | 144 | Usp44 | 129S1,AJ,AKR,BALB,BTBR,BUB,C3H,C57L,C58J,CBA,CE,DBA1J,DBA2J,FVB,ILNJ2,KK,LP,MA/My,MRL,NOD,NOR,NZO,NZW,PJ,PL,RF,RH2,SEA,SJL,SMJ,SWR,TH | feature_truncation,coding_sequence_variant,intron_variant |
| 10 | 117569715 | 117569769 | 54 | Slc35e3 | SMJ | feature_truncation,3_prime_UTR_variant |
| 10 | 122779223 | 122779788 | 565 | Ppm1h | AKR,C3H,CBA,DBA2J,RF,SJL,SWR | feature_truncation,3_prime_UTR_variant |
| 10 | 122828302 | 122828409 | 107 | Mon2 | BTBR,C57L,KK,NZW,TH | feature_truncation,3_prime_UTR_variant |
| 10 | 126846496 | 126846579 | 83 | Avil | CE,FVB,NOD,NOR | feature_truncation,coding_sequence_variant,intron_variant |
| 10 | 127582641 | 127582766 | 125 | Zbtb39 | CE | feature_truncation,3_prime_UTR_variant |
| 10 | 128945525 | 128945924 | 399 | Or6c2b | BUB,C3H,CBA,CE,KK,RF,SMJ | feature_truncation,3_prime_UTR_variant |
| 10 | 130217036 | 130217090 | 54 | Vmn2r84 | AKR,MRL,SJL,SWR | feature_truncation,3_prime_UTR_variant |
| 11 | 4172015 | 4172068 | 53 | Castor1 | NZB | feature_truncation,3_prime_UTR_variant |
| 11 | 4890448 | 4890520 | 72 | Nefh | AJ,BUB,NOD,PL | feature_truncation,inframe_deletion |
| 11 | 5040420 | 5040704 | 284 | Ewsr1 | BALB,BTBR,ILNJ2,MA/My,PJ,SEA,TH | feature_truncation,3_prime_UTR_variant |
| 11 | 9199731 | 9199838 | 107 | Abca13 | BUB,FVB,KK,NZB,NZO,SJL,SWR | splice_donor_variant,splice_donor_5th_base_variant,coding_sequence_variant,intron_variant |
| 11 | 17889094 | 17889185 | 91 | Etaa1 | 129S1,AJ,AKR,BALB,BTBR,BUB,C3H,C57L,C58J,CE,DBA1J,DBA2J,FVB,ILNJ2,KK,MA/My,MRL,NOD,NOR,NZB,NZO,NZW,PJ,PL,RF,SEA,SJL,SWR,TH | feature_truncation,3_prime_UTR_variant |
| 11 | 22469179 | 22470091 | 912 | Tmem17 | BALB,C57L,NOR,RH2,SEA | feature_truncation,3_prime_UTR_variant |
| 11 | 30904145 | 30904217 | 72 | Erlec1 | AJ,AKR,BUB,CBA,CE,DBA1J,DBA2J,FVB,ILNJ2,NOD,PJ,PL,RF,SEA,SJL,SMJ,SWR | feature_truncation,5_prime_UTR_variant |
| 11 | 34039920 | 34039978 | 58 | Lcp2 | ILNJ2,KK,MA/My,NOD,NZW,PL,RF | feature_truncation,3_prime_UTR_variant |
| 11 | 40592461 | 40592805 | 344 | Hmmr | B10D2,B10J,BTBR,C58J,LP,MA/My | feature_truncation,3_prime_UTR_variant |
| 11 | 46297545 | 46297608 | 63 | Garin3 | B10D2,B10J,C58J,NZO | feature_truncation,inframe_deletion |
| 11 | 46669383 | 46669515 | 132 | Havcr1 | NZB | stop_lost,feature_truncation,coding_sequence_variant,3_prime_UTR_variant |
| 11 | 50918795 | 50919250 | 455 | Or1x2 | B10D2,B10J,DBA2J,NOR,PJ,SEA | feature_truncation,3_prime_UTR_variant |
| 11 | 50919433 | 50919513 | 80 | Or1x2 | B10D2,B10J,NOR,TH | feature_truncation,3_prime_UTR_variant |
| 11 | 53492677 | 53492822 | 145 | Kif3a | B10D2,B10J,BUB,C58J,CE,KK,MA/My,NOD,NOR,NZW,PL,SJL,SWR,TH | feature_truncation,3_prime_UTR_variant |
| 11 | 58352928 | 58352984 | 56 | Or2z2 | 129S1,AJ,AKR,BALB,BTBR,C3H,C57L,CBA,CE,DBA1J,DBA2J,FVB,ILNJ2,KK,LP,MA/My,MRL,NOD,NOR,NZB,NZO,NZW,PJ,PL,RF,RH2,SJL,SMJ,SWR,TH | feature_truncation,5_prime_UTR_variant |
| 11 | 60418338 | 60418537 | 199 | Myo15a | AJ,C3H,CBA,MRL,NZB,NZO,PJ,SEA | feature_truncation,3_prime_UTR_variant |
| 11 | 62110476 | 62110555 | 79 | Specc1 | FVB,NOD,SWR | feature_truncation,3_prime_UTR_variant |
| 11 | 66818848 | 66819424 | 576 | Pirt | KK | feature_truncation,3_prime_UTR_variant |
| 11 | 68097177 | 68097568 | 391 | Stx8 | BALB,C3H,CBA,DBA1J,DBA2J,KK,MA/My,MRL,NOR,NZW,PJ,RH2,SEA,SJL,SMJ,SWR,TH | feature_truncation,3_prime_UTR_variant |
| 11 | 70345504 | 70345584 | 80 | Cxcl16 | CE | feature_truncation,3_prime_UTR_variant |
| 11 | 72202807 | 72203055 | 248 | Xaf1 | 129S1,BTBR,C57L,CE,DBA1J,DBA2J,NOR,PJ,SEA | stop_lost,feature_truncation,coding_sequence_variant,3_prime_UTR_variant,frameshift_variant,stop_lost,feature_truncation,3_prime_UTR_variant |
| 11 | 72662701 | 72662902 | 201 | Ankfy1 | DBA1J,DBA2J,NOR,NZB,PJ,SEA,TH | feature_truncation,3_prime_UTR_variant |
| 11 | 72734165 | 72734460 | 295 | Zzef1 | DBA1J,DBA2J,NOR,NZB,PJ,SEA,TH | feature_truncation,coding_sequence_variant,intron_variant |
| 11 | 72974200 | 72974427 | 227 | Ncbp3 | C58J,CE,DBA1J,DBA2J,NOR,NZB,PJ,SEA,TH | feature_truncation,3_prime_UTR_variant |
| 11 | 77678268 | 77678781 | 513 | Gm10277 | PJ | feature_truncation,5_prime_UTR_variant |
| 11 | 82854589 | 82854823 | 234 | Slfn5 | 129S1,AJ,BTBR,C3H,C57L,C58J,CBA,MA/My,MRL,NZO | feature_truncation,3_prime_UTR_variant |
| 11 | 84913691 | 84913886 | 195 | Usp32 | AKR,DBA1J,DBA2J,NZB,NZO,RF | feature_truncation,coding_sequence_variant,intron_variant |
| 11 | 93983798 | 93983901 | 103 | Spag9 | AJ,AKR,C3H,CBA,CE,MRL,RF | feature_truncation,coding_sequence_variant,intron_variant |
| 11 | 97884678 | 97884797 | 119 | Arl5c | NZW,SWR | feature_truncation,3_prime_UTR_variant |
| 11 | 97978623 | 97978765 | 142 | Fbxl20 | 129S1,BTBR,C3H,C57L,CE,DBA1J,DBA2J,FVB,ILNJ2,NZW,SWR | feature_truncation,3_prime_UTR_variant |
| 11 | 99676258 | 99676390 | 132 | Krtap4-9 | AJ,SMJ | feature_truncation,inframe_deletion |

| | | | | | | | |
|---|---|---|---|---|---|---|---|
| 11 | 99839973 | 99840045 | 72 | Krtap9-5 | NZB | feature_truncation,inframe_deletion | |
| 11 | 102781403 | 102781761 | 358 | Gfap | 129S1,AJ,BTBR,C3H,C57L,CBA,DBA1J,DBA2J,ILNJ2 | feature_truncation,3_prime_UTR_variant | |
| 11 | 107624631 | 107624690 | 59 | Cacng4 | ILNJ2,RF | feature_truncation,3_prime_UTR_variant | |
| 11 | 109287786 | 109287870 | 84 | Gna13 | BALB,ILNJ2,RF,SEA | feature_truncation,3_prime_UTR_variant | |
| 11 | 109823924 | 109823992 | 68 | Abca8b | BUB | feature_truncation,3_prime_UTR_variant | |
| 11 | 114847215 | 114847279 | 64 | Cd300c | 129S1,BALB,C58J,CE,ILNJ2,KK,MRL,NOD,NOR,NZB,NZO,NZW,RF,SEA,SMJ,TH | feature_truncation,coding_sequence_variant,intron_variant | |
| 11 | 115135638 | 115135760 | 122 | Tmem104 | 129S1,BALB,C58J,KK,NOD,NOR,NZB,NZO,NZW,RF,SEA,SMJ,TH | feature_truncation,3_prime_UTR_variant | |
| 11 | 116766283 | 116766498 | 215 | Mfsd11 | 129S1,AJ,AKR,BALB,BTBR,C3H,C57L,C58J,CBA,CE,DBA1J,DBA2J,FVB,ILNJ2,KK,MA/My,NZB,NZO,NZW,PL,RF,RH2,SEA,SJL,SMJ,SWR,TH | feature_truncation,3_prime_UTR_variant | |
| 11 | 120672103 | 120672202 | 99 | Rfng | SJL | feature_truncation,3_prime_UTR_variant | |
| 11 | 121105563 | 121105746 | 183 | Hexd | 129S1,AJ,AKR,B10D2,B10J,BALB,BTBR,BUB,C3H,C57L,C58J,CBA,CE,DBA1J,DBA2J,FVB,ILNJ2,KK,LP,MA/My,MRL,NOD,NOR,NZB,NZO,NZW,PJ,PL,RF,RH2,SEA,SJL,SMJ,SWR,TH | feature_truncation,coding_sequence_variant,intron_variant | |
| 12 | 4159495 | 4159571 | 76 | Dnajc27 | MRL | feature_truncation,3_prime_UTR_variant | |
| 12 | 4957632 | 4957685 | 53 | Ubxn2a | C3H,CBA,CE,TH | feature_truncation,5_prime_UTR_variant | |
| 12 | 28520377 | 28520437 | 60 | Dcdc2c | 129S1,AKR,CE,ILNJ2,RF | feature_truncation,inframe_deletion,feature_truncation,3_prime_UTR_variant | |
| 12 | 30640423 | 30640798 | 375 | Tmem18 | 129S1,BUB,MRL | feature_truncation,3_prime_UTR_variant | |
| 12 | 31124581 | 31124721 | 140 | Fam110c | AJ,AKR,BALB,BTBR,BUB,C3H,CBA,PJ,PL,RF,RH2,SEA | frameshift_variant,feature_truncation | |
| 12 | 33029543 | 33029719 | 176 | Sypl1 | 129S1,AJ,AKR,BALB,BTBR,BUB,C3H,CBA,PJ,PL,RF,RH2,SEA | feature_truncation,3_prime_UTR_variant | |
| 12 | 34098620 | 34098686 | 66 | Hdac9 | NZW | feature_truncation,3_prime_UTR_variant | |
| 12 | 40110311 | 40110364 | 53 | Scin | CE | feature_truncation,3_prime_UTR_variant | |
| 12 | 52357820 | 52358088 | 268 | Nubpl | 129S1,C57L,NZB | feature_truncation,3_prime_UTR_variant | |
| 12 | 53198896 | 53198983 | 87 | Akap6 | 129S1,C57L,NZB | feature_truncation,3_prime_UTR_variant | |
| 12 | 54718634 | 54718850 | 216 | Eapp | AJ,AKR,BALB,C3H,CBA,DBA1J,DBA2J,ILNJ2,KK,NOD,NOR,PJ,RF,RH2,SEA,SJL,SWR,TH | feature_truncation,3_prime_UTR_variant | |
| 12 | 71009568 | 71009792 | 224 | Actr10 | FVB,NZW,SJL,SWR | feature_truncation,3_prime_UTR_variant | |
| 12 | 75677656 | 75678346 | 690 | Wdr89,Rplp2-ps1 | FVB,PJ,SEA,SMJ,SWR | feature_truncation,3_prime_UTR_variant,transcript_ablation | |
| 12 | 76376197 | 76376269 | 72 | Akap5 | AKR,C58J,CE,DBA1J,DBA2J,ILNJ2,PJ,RF,SEA,SMJ | feature_truncation,inframe_deletion | |
| 12 | 78887751 | 78887809 | 58 | Atp6v1d | AKR,BALB,C58J,CE,KK,LP,MRL,NOD,NOR,NZO,PJ,PL,RF,SEA,SJL,SMJ,SWR,TH | feature_truncation,3_prime_UTR_variant | |
| 12 | 86562228 | 86562470 | 242 | Esrrb | CE,NOD,NOR,SWR | feature_truncation,3_prime_UTR_variant | |
| 12 | 87353190 | 87353322 | 132 | Sptlc2 | 129S1,AJ,AKR,BTBR,BUB,C3H,C57L,CBA,CE,DBA1J,DBA2J,FVB,ILNJ2,MRL,NOD,NOR,PJ,PL,RF,SEA,SJL,SWR,TH | feature_truncation,3_prime_UTR_variant | |
| 12 | 87994245 | 87994426 | 181 | Eif1ad11 | NOD,SWR | feature_truncation,3_prime_UTR_variant | |
| 12 | 102719709 | 102719911 | 202 | Gon7 | CE,DBA1J,DBA2J,PJ,SEA | feature_truncation,3_prime_UTR_variant | |
| 12 | 103582618 | 103582670 | 52 | Serpina10 | NZO,PJ,SEA | feature_truncation,3_prime_UTR_variant | |
| 12 | 103961819 | 103961876 | 57 | Serpina9 | 129S1,AKR,BALB,BTBR,BUB,C57L,CBA,DBA1J,DBA2J,FVB,ILNJ2,KK,LP,MRL,NOD,NOR,NZB,PJ,PL,RF,SEA,SJL,SWR | feature_truncation,3_prime_UTR_variant | |
| 12 | 103979533 | 103980223 | 690 | Serpina9 | FVB | feature_truncation,5_prime_UTR_variant,intron_variant | |
| 12 | 104078583 | 104079100 | 517 | Serpina3a | SJL,SWR | feature_truncation,5_prime_UTR_variant | FALSE |
| 12 | 104181329 | 104181724 | 395 | Serpina3f | AKR,BALB,BUB,CBA,CE,DBA1J,DBA2J,FVB,ILNJ2,KK,LP,MRL,NOD,NOR,PJ,PL,RF,SEA,SJL,SWR | feature_truncation,5_prime_UTR_variant,intron_variant | |
| 12 | 105668802 | 105668862 | 60 | Gskip | BALB,CE,FVB,KK,LP,MA/My,NZB,NZO,NZW,PJ,SEA,SJL,TH | feature_truncation,3_prime_UTR_variant | |
| 12 | 111770465 | 111770531 | 66 | Xrcc3 | AJ,AKR,C3H,CBA,CE,DBA1J,DBA2J,FVB,ILNJ2,MRL,NZB,NZO,NZW,PL,SMJ,SWR,TH | feature_truncation,3_prime_UTR_variant | |
| 12 | 113036095 | 113036185 | 90 | Pacs2 | AJ,C3H,CBA,CE,FVB,KK,MRL,NZB,NZO,NZW,PJ,PL,SEA,SMJ,SWR,TH | feature_truncation,3_prime_UTR_variant | |
| 13 | 3903966 | 3904316 | 350 | Calm5 | KK,LP,NZB,NZW | feature_truncation,5_prime_UTR_variant,feature_truncation,coding_sequence_variant,5_prime_UTR_variant,intron_variant | |
| 13 | 8854303 | 8854382 | 79 | Wdr37 | AKR,BTBR,C57L,KK,LP,NOD,NOR,NZB,NZO,NZW,RF,TH | feature_truncation,3_prime_UTR_variant | |
| 13 | 12487978 | 12488032 | 54 | Edaradd | AKR,ILNJ2,LP,RF | feature_truncation,3_prime_UTR_variant | |
| 13 | 21301498 | 21301622 | 124 | Or2w1b | AJ,AKR,B10D2,B10J,BALB,BTBR,BUB,C3H,C57L,C58J,CBA,DBA1J,DBA2J,FVB,ILNJ2,MA/My,MRL,NOD,NOR,PL,RF,RH2,SJL,SMJ,SWR,TH | feature_truncation,3_prime_UTR_variant | |
| 13 | 21303102 | 21303339 | 237 | Or2w1b | AJ,AKR,B10D2,B10J,BALB,BTBR,BUB,C3H,C57L,C58J,CBA,CE,DBA1J,DBA2J,FVB,ILNJ2,KK,LP,MA/My,MRL,NOD,NOR,NZB,PJ,PL,RF,RH2,SEA,SJL,SMJ,SWR,TH | feature_truncation,3_prime_UTR_variant | |
| 13 | 21627333 | 21627392 | 59 | Zscan26 | LP,PJ,SEA | feature_truncation,3_prime_UTR_variant | |
| 13 | 21685082 | 21685621 | 539 | Cox5b-ps | LP,PJ,SEA | transcript_ablation | |
| 13 | 22909627 | 22909721 | 94 | Vmn1r207 | BTBR,C57L,KK,LP,NZB,NZO,NZW,PJ,SEA | feature_truncation,3_prime_UTR_variant | |
| 13 | 52802452 | 52802554 | 102 | Syk | C58J,LP | feature_truncation,3_prime_UTR_variant | |
| 13 | 55848023 | 55848198 | 175 | B230219D22Rik | AJ,BALB,C3H,CBA,CE,SWR | feature_truncation,3_prime_UTR_variant | |
| 13 | 56620247 | 56620359 | 112 | Slc25a48 | 129S1,AJ,BALB,BTBR,BUB,C3H,C57L,CBA,CE,DBA1J,DBA2J,ILNJ2,MRL,NOD,NOR,NZB,NZW,PJ,PL,SEA,SMJ,SWR | feature_truncation,3_prime_UTR_variant | |
| 13 | 56621552 | 56622058 | 506 | Slc25a48 | AJ,BALB,C3H,CBA,CE | feature_truncation,3_prime_UTR_variant | |
| 13 | 58701552 | 58701724 | 172 | Slc28a3 | FVB,SWR | feature_truncation,3_prime_UTR_variant | |
| 13 | 61049014 | 61049120 | 106 | Tpbpb | PJ,SEA | feature_truncation,3_prime_UTR_variant | |
| 13 | 62922242 | 62922326 | 84 | Gm5141 | BALB | feature_truncation,inframe_deletion | |
| 13 | 64509628 | 64509855 | 227 | Ctsl | 129S1,AJ,AKR,BALB,BTBR,BUB,C3H,C57L,C58J,CBA,CE,DBA1J,DBA2J,FVB,ILNJ2,KK,MRL,NOD,NOR,NZB,NZO,NZW,PJ,PL,RF,SEA,SJL,SMJ,SWR,TH | feature_truncation,3_prime_UTR_variant | |
| 13 | 64510883 | 64510969 | 86 | Ctsl | 129S1,AJ,AKR,BALB,BTBR,BUB,C3H,C57L,C58J,CBA,CE,DBA1J,DBA2J,ILNJ2,KK,MRL,NOD,NOR,NZB,NZO,NZW,PL,RF,SJL,SMJ,TH | feature_truncation,3_prime_UTR_variant | |
| 13 | 65302570 | 65302645 | 75 | Or9s18 | 129S1,AJ,AKR,BALB,BTBR,BUB,C3H,C57L,C58J,CBA,CE,DBA1J,DBA2J,FVB,ILNJ2,KK,MRL,NOD,NOR,NZB,NZO,NZW,PJ,PL,RF,SEA,SJL,SMJ,SWR,TH | feature_truncation,3_prime_UTR_variant | |

| | | | | | | |
|---|---|---|---|---|---|---|
| 13 | 67286835 | 67286919 | 84 | Zfp759 | AJ,BALB,BUB,C3H,C58J,CBA,CE,DBA1J,DBA2J,FVB,ILNJ2,KK,NOD,NOR,NZB,NZO,PL,SJL,SMJ,SWR,TH | feature_truncation,inframe_deletion |
| 13 | 67329899 | 67330319 | 420 | Rsl1 | SWR | feature_truncation,inframe_deletion |
| 13 | 67439814 | 67440687 | 873 | Zfp457 | 129S1,AJ,AKR,BALB,BTBR,BUB,C3H,C57L,C58J,CBA,CE,DBA1J,DBA2J,FVB,ILNJ2,KK,MRL,NOD,NOR,NZB,NZO,NZW,PJ,PL,RF,SEA,SJL,SMJ,SWR,TH | stop_lost,feature_truncation,coding_sequence_variant,3_prime_UTR_variant |
| 13 | 67441674 | 67441758 | 84 | Zfp457 | 129S1,AJ,AKR,BALB,BTBR,BUB,C3H,C57L,C58J,CBA,CE,DBA1J,DBA2J,FVB,ILNJ2,KK,MRL,NOD,NOR,NZB,NZO,NZW,PJ,PL,RF,SEA,SJL,SMJ,SWR,TH | feature_truncation,inframe_deletion |
| 13 | 67537503 | 67537818 | 315 | Zfp429 | 129S1,AKR,MRL,NZW,PL,RF | feature_truncation,3_prime_UTR_variant |
| 13 | 67579561 | 67579683 | 122 | Zfp874a | 129S1,AJ,AKR,BALB,BTBR,BUB,C3H,C57L,C58J,CBA,CE,DBA1J,DBA2J,FVB,ILNJ2,KK,MRL,NOD,NOR,NZB,NZO,NZW,PJ,PL,RF,SEA,SJL,SMJ,SWR,TH | splice_acceptor_variant,splice_donor_variant,splice_donor_5th_base_variant,coding_sequence_variant,intron_variant |
| 13 | 67812299 | 67813239 | 940 | Zfp738 | 129S1,AJ,AKR,BALB,BTBR,BUB,C3H,C57L,C58J,CBA,CE,DBA1J,DBA2J,FVB,ILNJ2,KK,MRL,NOD,NOR,NZB,NZO,NZW,PJ,PL,RF,SEA,SJL,SMJ,SWR,TH | feature_truncation,3_prime_UTR_variant |
| 13 | 67814255 | 67814585 | 330 | Zfp738 | 129S1,AJ,AKR,BALB,BTBR,BUB,C3H,C57L,C58J,CBA,CE,DBA1J,DBA2J,FVB,ILNJ2,KK,MRL,NOD,NOR,NZB,NZO,NZW,PJ,PL,RF,SEA,SJL,SMJ,SWR,TH | feature_truncation,3_prime_UTR_variant |
| 13 | 67940346 | 67940431 | 85 | Zfp493 | 129S1,AJ,AKR,BALB,BTBR,BUB,C3H,C57L,C58J,CBA,CE,DBA1J,DBA2J,FVB,ILNJ2,MA/My,MRL,NOD,NOR,NZB,NZO,NZW,PJ,PL,RF,SEA,SJL,SMJ,SWR,TH | feature_truncation,3_prime_UTR_variant |
| 13 | 74554501 | 74555477 | 976 | Ftl1-ps1 | ILNJ2,SWR | transcript_ablation |
| 13 | 76150376 | 76150486 | 110 | Rfesd | 129S1,AJ,AKR,BALB,BTBR,BUB,C3H,C57L,CBA,CE,DBA1J,DBA2J,FVB,ILNJ2,KK,MA/My,MRL,NOD,NOR,NZB,NZO,NZW,PJ,PL,RF,SEA,SJL,SMJ,SWR,TH | feature_truncation,3_prime_UTR_variant |
| 13 | 95149011 | 95149659 | 648 | Wdr41 | C58J,DBA1J,KK,MA/My,RF | feature_truncation,coding_sequence_variant,intron_variant |
| 13 | 96097243 | 96097323 | 80 | Sv2c | 129S1,AJ,AKR,BALB,BTBR,BUB,C3H,C57L,CBA,DBA1J,DBA2J,FVB,KK,MRL,NZB,NZO,NZW,PJ,PL,SEA,SJL,SMJ,SWR,TH | feature_truncation,3_prime_UTR_variant |
| 13 | 103877384 | 103877511 | 127 | Srek1 | C58J,CE,MA/My,NZB,NZO | feature_truncation,3_prime_UTR_variant |
| 13 | 108189182 | 108189235 | 53 | Ndufaf2 | CE,MA/My,NZO,TH | feature_truncation,3_prime_UTR_variant |
| 13 | 114291977 | 114292041 | 64 | Arl15 | AJ,BUB,CE,DBA1J,FVB,KK,MA/My,NZB,NZO,NZW,PL,RH2,SJL,SWR | feature_truncation,3_prime_UTR_variant |
| 13 | 120249500 | 120249664 | 164 | Zfp131 | 129S1,BTBR,C3H,C57L,CBA,CE,DBA2J,KK,MA/My,NZO,SMJ | frameshift_variant,stop_lost,feature_truncation,3_prime_UTR_variant |
| 14 | 7666136 | 7666457 | 321 | Nek10 | BALB,SEA | feature_truncation,5_prime_UTR_variant |
| 14 | 10229143 | 10229364 | 221 | Ptprg | 129S1,C3H,C57L,CBA,DBA1J,DBA2J,KK,MRL,NZW,PJ,PL | feature_truncation,3_prime_UTR_variant |
| 14 | 42353052 | 42353649 | 597 | Gm8024 | C57L,NZO,PJ,PL | feature_truncation,coding_sequence_variant,5_prime_UTR_variant,intron_variant,feature_truncation,5_prime_UTR_variant,intron_variant |
| 14 | 45370863 | 45371079 | 216 | Txndc16 | C57L,KK,NZO,PJ,PL,SMJ | feature_truncation,3_prime_UTR_variant |
| 14 | 47005526 | 47005812 | 286 | Cdkn3 | 129S1,AJ,AKR,BALB,BUB,C3H,CBA,CE,DBA1J,DBA2J,FVB,ILNJ2,MA/My,MRL,NOD,NOR,NZB,NZO,NZW,PJ,RF,RH2,SEA,SJL,SMJ,SWR,TH | feature_truncation,3_prime_UTR_variant |
| 14 | 51201913 | 51202326 | 413 | Pnp2 | 129S1,AJ,AKR,BALB,BUB,C3H,C57L,CBA,CE,DBA1J,DBA2J,FVB,ILNJ2,KK,MA/My,MRL,NOD,NOR,NZB,NZO,NZW,PJ,PL,RF,RH2,SEA,SJL,SMJ,SWR,TH | stop_lost,feature_truncation,coding_sequence_variant,3_prime_UTR_variant |
| 14 | 54781224 | 54781286 | 62 | Haus4 | AJ,AKR,BALB,BTBR,BUB,C3H,C57L,CBA,CE,DBA1J,DBA2J,FVB,ILNJ2,KK,MA/My,MRL,NOD,NOR,NZB,NZO,NZW,PJ,PL,RF,RH2,SEA,SJL,SWR,TH | splice_donor_variant,splice_donor_5th_base_variant,coding_sequence_variant,intron_variant |
| 14 | 54865097 | 54865157 | 60 | Psmb11 | 129S1,AJ,AKR,BALB,BTBR,BUB,C3H,C57L,CBA,CE,DBA1J,DBA2J,FVB,ILNJ2,KK,MA/My,MRL,NOD,NOR,NZB,NZO,NZW,PJ,PL,RF,RH2,SEA,SJL,SMJ,SWR,TH | feature_truncation,3_prime_UTR_variant |
| 14 | 55260795 | 55260890 | 95 | Ngdn | BUB,SMJ | feature_truncation,coding_sequence_variant,intron_variant |
| 14 | 56885180 | 56885468 | 288 | Parp4 | BTBR,BUB,DBA2J,KK,NZB,SJL | feature_truncation,inframe_deletion |
| 14 | 57930795 | 57931002 | 207 | Lats2 | BTBR,BUB,DBA2J,KK | feature_truncation,3_prime_UTR_variant |
| 14 | 58348378 | 58348433 | 55 | Fgf9 | BTBR,BUB,DBA2J | feature_truncation,3_prime_UTR_variant |
| 14 | 69932944 | 69933149 | 205 | Loxl2 | 129S1,C57L,KK | feature_truncation,3_prime_UTR_variant |
| 14 | 70821114 | 70821242 | 128 | Fhip2b | 129S1,AJ,BALB,C57L,KK,PL,SEA | feature_truncation,3_prime_UTR_variant |
| 14 | 76372870 | 76372990 | 120 | Nufip1 | NZB,NZO | feature_truncation,3_prime_UTR_variant |
| 14 | 78799970 | 78800030 | 60 | Dgkh | ILNJ2 | feature_truncation,3_prime_UTR_variant |
| 14 | 78969919 | 78969976 | 57 | Dgkh | 129S1,AJ,AKR,BALB,BTBR,BUB,C3H,C57L,CBA,CE,DBA1J,DBA2J,FVB,ILNJ2,KK,LP,NZB,NZO,NZW,PJ,PL,RF,SEA,SJL,SMJ,SWR,TH | feature_truncation,5_prime_UTR_variant |
| 14 | 79563601 | 79563676 | 75 | Naa16 | CE,DBA1J,DBA2J,PJ | feature_truncation,3_prime_UTR_variant |
| 14 | 87694359 | 87694451 | 92 | Tdrd3 | AKR,DBA1J,DBA2J,NZO,PJ | feature_truncation,coding_sequence_variant,5_prime_UTR_variant,intron_variant |
| 14 | 96341930 | 96341987 | 57 | Klhl1 | AJ,AKR,BALB,BTBR,BUB,C3H,CBA,CE,DBA1J,DBA2J,FVB,MRL,NOD,PJ,PL,RF,RH2,SEA,SJL,SMJ,SWR,TH | feature_truncation,3_prime_UTR_variant |
| 14 | 103480616 | 103480715 | 99 | Mycbp2 | 129S1,C57L,ILNJ2,NZB,NZO,PJ | feature_truncation,coding_sequence_variant,intron_variant |
| 14 | 118504615 | 118505195 | 580 | Gm9376 | 129S1,KK | stop_lost,feature_truncation,coding_sequence_variant,3_prime_UTR_variant |
| 15 | 7222191 | 7222257 | 66 | Lifr | AKR,LP,RF,TH | feature_truncation,3_prime_UTR_variant |
| 15 | 7223231 | 7223295 | 64 | Lifr | LP,TH | feature_truncation,3_prime_UTR_variant |
| 15 | 10498390 | 10498748 | 358 | Rad1 | TH | feature_truncation,3_prime_UTR_variant |
| 15 | 35178886 | 35179287 | 401 | Stk3 | 129S1,BUB,C57L,KK,LP,NOD,NOR,PJ,SWR | feature_truncation,coding_sequence_variant,5_prime_UTR_variant,intron_variant |
| 15 | 54616278 | 54616346 | 68 | Ccn3 | 129S1,C57L,NZB,TH | feature_truncation,3_prime_UTR_variant |
| 15 | 58422145 | 58422343 | 198 | Fer1l6 | ILNJ2,NZB,NZW,SMJ,TH | feature_truncation,coding_sequence_variant,intron_variant |
| 15 | 75088229 | 75088412 | 183 | Ly6g2 | ILNJ2 | feature_truncation,5_prime_UTR_variant,intron_variant |
| 15 | 76935187 | 76935339 | 152 | Apol6 | MA/My | frameshift_variant,feature_truncation |

| | | | | | | |
|---|---|---|---|---|---|---|
| 15 | 77373859 | 77374389 | 530 | Apol10a | MA/My | feature_truncation,3_prime_UTR_variant |
| 15 | 78660360 | 78660426 | 66 | Card10 | 129S1,AJ,AKR,BALB,BUB,C3H,C57L,C58J,CBA,CE,DBA1J,DBA2J,FVB,ILNJ2,KK,LP,MA/My,MRL,NOD,NOR,NZW,PJ,PL,SEA,SMJ,TH | feature_truncation,3_prime_UTR_variant |
| 15 | 78851187 | 78851259 | 72 | Triobp | MA/My | feature_truncation,inframe_deletion |
| 15 | 88896186 | 88896285 | 99 | Mov10l1 | 129S1,C57L,CE,FVB,ILNJ2,PL | feature_truncation,inframe_deletion |
| 15 | 98395993 | 98396311 | 318 | Or11m3 | AJ,C3H,CE,RH2 | stop_lost,3_prime_UTR_variant |
| 15 | 98399500 | 98399560 | 60 | Or11m3 | BUB,MA/My,SWR | feature_truncation,3_prime_UTR_variant |
| 15 | 98748887 | 98748947 | 60 | Kmt2d | C3H | feature_truncation,inframe_deletion |
| 15 | 99842052 | 99842166 | 114 | Fam186a | BALB,CBA | feature_truncation,inframe_deletion |
| 15 | 100286368 | 100286487 | 119 | Slc11a2 | 129S1,BALB,BTBR,BUB,C57L,C58J,CBA,ILNJ2,KK,MA/My,NOD,NOR,NZB,NZO,NZW,PJ,SEA,SMJ,SWR,TH | feature_truncation,3_prime_UTR_variant |
| 15 | 100513326 | 100513380 | 54 | Dazap2 | 129S1,BALB,BTBR,BUB,C57L,C58J,CBA,ILNJ2,KK,MA/My,NOD,NOR,NZB,NZO,NZW,PJ,SEA,SWR,TH | feature_truncation,5_prime_UTR_variant |
| 15 | 102441494 | 102441763 | 269 | Atf7 | AKR,DBA1J,DBA2J,FVB,ILNJ2,LP,MRL,NOD,NOR,PJ,PL,SEA,SJL,SMJ | feature_truncation,3_prime_UTR_variant |
| 16 | 3764271 | 3764559 | 288 | Nlrc3 | AJ,BTBR,BUB,CE,FVB,MRL,PL,SJL,SMJ,SWR,TH | feature_truncation,3_prime_UTR_variant |
| 16 | 3764832 | 3764927 | 95 | Nlrc3 | SMJ | feature_truncation,3_prime_UTR_variant |
| 16 | 11479218 | 11479723 | 505 | Snx29 | KK | feature_truncation,3_prime_UTR_variant |
| 16 | 13573589 | 13573763 | 174 | A930007A09Rik | BUB,KK | feature_truncation,inframe_deletion |
| 16 | 13767014 | 13767241 | 227 | Mpv17l | C3H,CBA,DBA1J,DBA2J,KK | feature_truncation,3_prime_UTR_variant |
| 16 | 18226049 | 18226279 | 230 | Comt | C3H,C57L,C58J,CBA,DBA1J,DBA2J,FVB,ILNJ2,KK,LP,MA/My,NZB,NZO,NZW,PJ,RF,SJL,SWR,TH | feature_truncation,3_prime_UTR_variant |
| 16 | 23298178 | 23298268 | 90 | Masp1 | PL,SMJ,TH | feature_truncation,3_prime_UTR_variant |
| 16 | 27270344 | 27270402 | 58 | Ccdc50 | BALB | feature_truncation,3_prime_UTR_variant |
| 16 | 31114801 | 31115056 | 255 | Apod | 129S1,AJ,BALB,FVB,ILNJ2,KK,LP,MRL,NOD,NOR,NZB,NZO,NZW,PL,RH2,SEA,SJL,TH | feature_truncation,3_prime_UTR_variant |
| 16 | 48139206 | 48139371 | 165 | Dppa2 | C3H,CBA | feature_truncation,3_prime_UTR_variant |
| 16 | 55853913 | 55854007 | 94 | Trmt10c | CBA,NZW,SMJ | feature_truncation,3_prime_UTR_variant |
| 16 | 56435066 | 56435990 | 924 | Abi3bp | C58J,LP | feature_truncation,coding_sequence_variant,intron_variant |
| 16 | 84659881 | 84659932 | 51 | Gabpa | BALB,BUB,C58J,CBA,LP,MRL,NZO,RH2 | feature_truncation,3_prime_UTR_variant |
| 16 | 88570804 | 88570864 | 60 | 2310057N15Rik | 129S1,C57L,C58J,KK,NZB,NZO,NZW,PL | feature_truncation,inframe_deletion |
| 16 | 88670829 | 88670991 | 162 | Krtap19-2 | 129S1,AKR,C57L,C58J,CBA,CE,DBA1J,DBA2J,ILNJ2,KK,MRL,NZB,NZO,NZW,PJ,PL,RF,SEA,SMJ,TH | feature_truncation,inframe_deletion |
| 16 | 91627395 | 91627476 | 81 | Itsn1 | CBA,CE,DBA1J,MA/My,PL | feature_truncation,3_prime_UTR_variant |
| 16 | 92144628 | 92144820 | 192 | Kcne1 | BUB,CE,DBA1J,FVB,NZB,PL,SJL,SWR,TH | feature_truncation,3_prime_UTR_variant |
| 16 | 94549489 | 94549543 | 54 | Kcnj6 | NZW | feature_truncation,3_prime_UTR_variant |
| 16 | 95058697 | 95058802 | 105 | Kcnj15 | 129S1,AJ,AKR,B10D2,B10J,BALB,BTBR,BUB,C3H,C57L,C58J,CBA,CE,DBA1J,DBA2J,FVB,ILNJ2,KK,LP,MA/My,MRL,NOD,NOR,NZB,NZO,NZW,PJ,PL,RF,RH2,SEA,SJL,SMJ,SWR,TH | feature_truncation,5_prime_UTR_variant |
| 17 | 6973005 | 6973229 | 224 | Sytl3 | 129S1,AJ,AKR,BALB,C3H,C57L,CBA,CE,DBA1J,DBA2J,ILNJ2,MRL,NOD,NOR,NZO,PJ,PL,RF,RH2,SEA,SMJ,TH | feature_truncation,5_prime_UTR_variant |
| 17 | 15764864 | 15764948 | 84 | Prdm9 | ILNJ2 | feature_truncation,inframe_deletion |
| 17 | 17307825 | 17307980 | 155 | Zfp960 | C58J,PL | frameshift_variant,feature_truncation |
| 17 | 17366495 | 17366747 | 252 | Zfp97 | LP | feature_truncation,3_prime_UTR_variant |
| 17 | 21262502 | 21262553 | 51 | Zfp160 | AKR,C3H,CBA,FVB,MA/My,MRL,PJ | feature_truncation,3_prime_UTR_variant |
| 17 | 21807273 | 21807372 | 99 | Zfp948 | 129S1,AJ,BALB,C57L,CE,DBA1J,DBA2J,NZO,RH2,SEA | feature_truncation,inframe_deletion |
| 17 | 23847162 | 23847450 | 288 | Mmp25 | BTBR,BUB,C58J,KK,LP,NOD,NOR,NZW,PL,SWR | feature_truncation,3_prime_UTR_variant |
| 17 | 24397374 | 24397425 | 51 | Tbc1d24 | AKR | feature_truncation,3_prime_UTR_variant |
| 17 | 24399026 | 24399152 | 126 | Tbc1d24 | 129S1,AJ,AKR,B10D2,BALB,BTBR,BUB,C3H,C57L,CBA,CE,DBA1J,DBA2J,FVB,ILNJ2,KK,MA/My,MRL,NOD,NOR,NZB,NZO,NZW,PJ,RF,RH2,SEA,SJL,SMJ,SWR,TH | feature_truncation,3_prime_UTR_variant |
| 17 | 24912587 | 24913136 | 549 | Gfer | 129S1,AJ,AKR,B10D2,BALB,BTBR,BUB,C3H,C57L,CBA,CE,DBA1J,DBA2J,FVB,ILNJ2,KK,MA/My,MRL,NOD,NOR,NZB,NZO,NZW,PJ,RF,RH2,SEA,SJL,SMJ,SWR,TH | feature_truncation,3_prime_UTR_variant |
| 17 | 26663356 | 26663419 | 63 | Neurl1b | 129S1,AJ,AKR,B10D2,BALB,C3H,C57L,CBA,CE,DBA1J,DBA2J,ILNJ2,MRL,NOD,NOR,NZO,PJ,RF,RH2,SEA,TH | feature_truncation,3_prime_UTR_variant |
| 17 | 28056819 | 28057312 | 493 | Snrpc | KK,NZB,RF | feature_truncation,5_prime_UTR_variant,intron_variant |
| 17 | 29108791 | 29109372 | 581 | Pnpla1 | KK,RF | feature_truncation,3_prime_UTR_variant |
| 17 | 29863485 | 29864099 | 614 | Rnf8 | NOD,NOR,NZB | feature_truncation,3_prime_UTR_variant |
| 17 | 29923672 | 29923786 | 114 | Cmtr1 | KK,RF | feature_truncation,3_prime_UTR_variant |
| 17 | 30863677 | 30863913 | 236 | Dnah8 | AJ,AKR,BALB,BTBR,BUB,C3H,C58J,CBA,CE,FVB,ILNJ2,KK,MA/My,MRL,NOD,NOR,NZB,NZO,NZW,PJ,PL,RF,SJL,SMJ,SWR,TH | feature_truncation,coding_sequence_variant,intron_variant |
| 17 | 31095907 | 31096007 | 100 | Dnah8 | 129S1,AJ,AKR,B10D2,BALB,BTBR,BUB,C3H,C57L,C58J,CBA,CE,DBA1J,DBA2J,ILNJ2,KK,MA/My,MRL,NOD,NOR,NZO,NZW,PJ,PL,RF,RH2,SEA,SMJ,SWR | feature_truncation,3_prime_UTR_variant |
| 17 | 31463761 | 31463959 | 198 | Ubash3a | C58J,KK,NOD,NOR,NZB,NZO,NZW | feature_truncation,3_prime_UTR_variant |
| 17 | 32402972 | 32403078 | 106 | Ephx3 | PL | feature_truncation,3_prime_UTR_variant |
| 17 | 33461224 | 33461320 | 96 | Zfp955a | TH | feature_truncation,inframe_deletion |
| 17 | 34662424 | 34662516 | 92 | Tsbp1 | ILNJ2 | feature_truncation,coding_sequence_variant,intron_variant |
| 17 | 35912503 | 35912595 | 92 | Sfta2 | DBA1J,ILNJ2,PJ,SJL,TH | feature_truncation,5_prime_UTR_variant |
| 17 | 35930919 | 35931009 | 90 | Muc21 | 129S1,AKR,BTBR,C3H,C57L,CBA,CE,LP,MA/My,MRL,NZO,NZW,RF,SMJ,TH | feature_truncation,inframe_deletion |
| 17 | 35931846 | 35932115 | 269 | Muc21 | ILNJ2 | frameshift_variant,feature_truncation |
| 17 | 35932248 | 35932788 | 540 | Muc21 | 129S1,AKR,B10D2,BALB,BTBR,BUB,C3H,C57L,C58J,CBA,CE,DBA1J,DBA2J,FVB,ILNJ2,KK,LP,MA/My,MRL,NOD,NOR,NZB,NZO,NZW,PJ,PL,RF,SMJ,SWR | feature_truncation,inframe_deletion |
| 17 | 35933391 | 35933548 | 157 | Muc21 | BUB,CE,MA/My | frameshift_variant,feature_truncation |
| 17 | 36241478 | 36241609 | 131 | Ppp1r10 | NZW | frameshift_variant,feature_truncation |
| 17 | 36268176 | 36268236 | 60 | Abcf1 | B10D2,BALB,DBA2J,PJ,RH2,SEA | feature_truncation,3_prime_UTR_variant |

| | | | | | | |
|---|---|---|---|---|---|---|
| 17 | 37394629 | 37395252 | 623 | Or2h2 | 129S1,AJ,AKR,B10D2,BALB,BTBR,BUB,C3H,C57L,C58J,CBA,CE,DBA1J,DBA2J,FVB,ILNJ2,KK,LP,MA/My,MRL,NOD,NOR,NZB,NZO,NZW,PJ,PL,RF,RH2,SEA,SJL,SWR,TH | feature_truncation,3_prime_UTR_variant |
| 17 | 37461066 | 37461124 | 58 | Or2h1b | AJ,B10D2,BALB,BUB,C58J,DBA1J,DBA2J,KK,NOD,NOR,NZB,NZO,NZW,PL,RH2,SEA,SJL,SWR | feature_truncation,3_prime_UTR_variant |
| 17 | 37690906 | 37691241 | 335 | Or12d15 | B10D2,BALB,DBA1J,DBA2J,NZO,NZW,RH2,SEA | feature_truncation,coding_sequence_variant,intron_variant |
| 17 | 38059742 | 38059828 | 86 | Or10al5 | B10D2,BALB,DBA1J,DBA2J,FVB,ILNJ2,NZO,NZW,RH2,SEA,TH | feature_truncation,5_prime_UTR_variant |
| 17 | 44446111 | 44446195 | 84 | Clic5 | AKR,BALB,C3H,CBA,CE,DBA2J,ILNJ2,MA/My,PJ,RF,RH2,SEA | feature_truncation,inframe_deletion |
| 17 | 46563358 | 46563463 | 105 | Yipf3 | 129S1,BUB,FVB,NOD,NOR,NZW,SJL,SWR | feature_truncation,3_prime_UTR_variant |
| 17 | 47084813 | 47084958 | 145 | Rpl7l1 | NOD,NOR,NZW,SJL,SMJ | feature_truncation,3_prime_UTR_variant |
| 17 | 53981999 | 53982235 | 236 | Sgo1 | BTBR,MA/My,TH | feature_truncation,3_prime_UTR_variant |
| 17 | 56882590 | 56882671 | 81 | Safb2 | SEA | feature_truncation,inframe_deletion |
| 17 | 74732730 | 74732913 | 183 | Nlrc4 | 129S1,BTBR,BUB,C57L,ILNJ2,KK,NOD,NOR,NZW,SWR | feature_truncation,3_prime_UTR_variant |
| 17 | 79153607 | 79153816 | 209 | Gpatch11 | CE,ILNJ2,NZO,NZW | feature_truncation,3_prime_UTR_variant |
| 17 | 87139069 | 87139125 | 56 | Epas1 | PL | feature_truncation,3_prime_UTR_variant |
| 18 | 6202202 | 6202371 | 169 | Kif5b | 129S1,AKR,BALB,C3H,C57L,CBA,ILNJ2,PJ,RF,RH2,SEA,SMJ | feature_truncation,3_prime_UTR_variant |
| 18 | 10725780 | 10725837 | 57 | Mib1 | 129S1,C57L | feature_truncation,5_prime_UTR_variant |
| 18 | 34732619 | 34732834 | 215 | Brd8 | 129S1,AJ,BALB,BUB,C3H,C57L,C58J,CBA,CE,DBA1J,DBA2J,ILNJ2,MRL,NOD,NOR,PL,RF,RH2,SEA,SJL,TH | feature_truncation,3_prime_UTR_variant |
| 18 | 37437165 | 37437271 | 106 | Pcdhb3 | 129S1,AJ,AKR,BALB,BUB,C3H,C57L,C58J,CBA,CE,DBA1J,DBA2J,FVB,ILNJ2,KK,MA/My,NOD,NOR,NZB,NZO,NZW,PL,RF,RH2,SEA,SJL,SMJ,SWR,TH | feature_truncation,3_prime_UTR_variant |
| 18 | 37636474 | 37636527 | 53 | Pcdhb19 | KK | feature_truncation,3_prime_UTR_variant |
| 18 | 37828371 | 37828516 | 145 | Pcdhga5 | BUB,C3H,CE,DBA1J,MA/My,NOD,NOR,PL,RF | frameshift_variant,feature_truncation |
| 18 | 44559174 | 44559226 | 52 | Mcc | BALB,C3H,CBA,DBA1J,DBA2J,KK,SEA | feature_truncation,3_prime_UTR_variant |
| 18 | 47245193 | 47245746 | 553 | Hspe1-rs1 | BALB,C3H,C58J,CBA,DBA1J,DBA2J,ILNJ2,KK,LP,MA/My,NZO,NZW,SEA | transcript_ablation |
| 18 | 52624796 | 52624857 | 61 | Srfbp1 | 129S1,AJ,AKR,BALB,BUB,C3H,C57L,C58J,CBA,CE,DBA1J,DBA2J,FVB,ILNJ2,LP,MA/My,MRL,NOD,NOR,NZB,NZO,NZW,PL,RF,RH2,SEA,SJL,SWR | feature_truncation,3_prime_UTR_variant |
| 18 | 56720630 | 56720750 | 120 | Phax | C3H,CE,DBA1J,DBA2J,PJ | feature_truncation,3_prime_UTR_variant |
| 18 | 60402190 | 60402373 | 183 | Gm4841 | KK,NOD,NOR | feature_truncation,3_prime_UTR_variant |
| 18 | 60403399 | 60404196 | 797 | Gm4841 | NZO,NZW | feature_truncation,coding_sequence_variant,5_prime_UTR_variant,intron_variant |
| 18 | 61327045 | 61327097 | 52 | Slc26a2 | NZW | feature_truncation,3_prime_UTR_variant |
| 18 | 62800804 | 62801103 | 299 | Spink10 | AJ,BALB,CBA,ILNJ2,PL,RH2,SEA | feature_truncation,3_prime_UTR_variant |
| 18 | 75219140 | 75219201 | 61 | Dym | 129S1,AKR,BTBR,BUB,C3H,C57L,CBA,DBA1J,DBA2J,FVB,NZO,PJ,PL,SEA,TH | feature_truncation,3_prime_UTR_variant |
| 18 | 75219642 | 75219824 | 182 | Dym | ILNJ2,NZW,RF | feature_truncation,3_prime_UTR_variant |
| 18 | 77545284 | 77545414 | 130 | Ark2c | KK,NZO,NZW,TH | feature_truncation,3_prime_UTR_variant |
| 18 | 80266867 | 80267036 | 169 | Txnl4a | 129S1,AJ,AKR,BALB,BTBR,BUB,C57L,FVB,ILNJ2,MA/My,MRL,NOD,NOR,NZO,RH2,SJL,TH | feature_truncation,3_prime_UTR_variant |
| 18 | 80317493 | 80317553 | 60 | Slc66a2 | AKR,NZO | feature_truncation,3_prime_UTR_variant |
| 18 | 84963612 | 84964422 | 810 | Timm21 | CE | feature_truncation,3_prime_UTR_variant |
| 19 | 3439428 | 3439510 | 82 | Tesmin | FVB,SJL | feature_truncation,5_prime_UTR_variant |
| 19 | 4438398 | 4438633 | 235 | Kdm2a | AJ,AKR,BALB,BUB,C3H,C58J,CBA,CE,DBA1J,DBA2J,FVB,ILNJ2,KK,LP,MA/My,MRL,NOD,NOR,NZW,PJ,PL,RF,RH2,SEA,SJL,SMJ,SWR,TH | feature_truncation,3_prime_UTR_variant,intron_variant |
| 19 | 5545018 | 5545244 | 226 | Cfl1 | FVB,LP,SJL | feature_truncation,3_prime_UTR_variant |
| 19 | 5991093 | 5991275 | 182 | Pola2 | BUB,C58J,FVB,LP,MRL,SJL,TH | feature_truncation,3_prime_UTR_variant |
| 19 | 6113093 | 6113149 | 56 | Tm7sf2 | BUB | feature_truncation,coding_sequence_variant,intron_variant |
| 19 | 10695081 | 10695297 | 216 | Cd5 | BUB,C3H,CBA,DBA1J,DBA2J,FVB,ILNJ2,LP,MA/My,SWR | feature_truncation,3_prime_UTR_variant |
| 19 | 11439876 | 11440757 | 881 | Ms4a4b | FVB,SWR | stop_lost,feature_truncation,coding_sequence_variant,3_prime_UTR_variant,intron_variant |
| 19 | 12010978 | 12011174 | 196 | Or4d11 | FVB,SWR | feature_truncation,3_prime_UTR_variant |
| 19 | 12261512 | 12261614 | 102 | Or5an1 | AJ,BALB,CE,MA/My,NOD,NOR,PL,RH2,SEA | feature_truncation,3_prime_UTR_variant |
| 19 | 12442401 | 12442479 | 78 | Mpeg1 | AJ,BALB,CE,FVB,LP,MA/My,NOD,NOR,PL,RH2,SEA,SWR | feature_truncation,3_prime_UTR_variant |
| 19 | 12660755 | 12661212 | 457 | Or5b95 | ILNJ2 | feature_truncation,3_prime_UTR_variant |
| 19 | 13071041 | 13071260 | 219 | Or5b104 | LP | feature_truncation,3_prime_UTR_variant |
| 19 | 17410102 | 17410168 | 66 | Pcsk5 | CBA,DBA1J,DBA2J | feature_truncation,3_prime_UTR_variant |
| 19 | 21369844 | 21370026 | 182 | Gda | BUB,MRL,NZB,NZO,NZW | feature_truncation,3_prime_UTR_variant |
| 19 | 30544002 | 30544116 | 114 | Prkg1 | 129S1,BTBR,C57L,C58J,NZW | feature_truncation,3_prime_UTR_variant |
| 19 | 40804758 | 40805018 | 260 | Cc2d2b | C3H,CBA,DBA1J,DBA2J,FVB,ILNJ2,MA/My,NZB,NZO | feature_truncation,coding_sequence_variant,intron_variant |
| 19 | 41757716 | 41757848 | 132 | Arhgap19 | AJ,BALB,CE,RH2 | feature_truncation,3_prime_UTR_variant |
| 19 | 44333627 | 44334087 | 460 | Scd4 | 129S1,BUB,C57L,FVB,KK,SMJ,SWR | feature_truncation,3_prime_UTR_variant |
| 19 | 53576561 | 53577493 | 932 | Nutf2-ps1 | 129S1,AJ,AKR,BALB,BTBR,C3H,C57L,CBA,CE,DBA1J,DBA2J,FVB,ILNJ2,KK,MA/My,MRL,NOD,NOR,NZB,NZO,NZW,PJ,PL,RF,RH2,SEA,SJL,SWR,TH | stop_lost,feature_truncation,coding_sequence_variant,5_prime_UTR_variant,3_prime_UTR_variant |
| 19 | 60745538 | 60745670 | 132 | Nanos1 | TH | feature_truncation,3_prime_UTR_variant |
| 19 | 61107079 | 61107148 | 69 | Zfp950 | 129S1,AJ,AKR,BALB,BTBR,C3H,C57L,CBA,DBA1J,DBA2J,FVB,KK,MA/My,MRL,NOD,NOR,NZB,NZW,PJ,PL,RF,RH2,SEA,SMJ,SWR,TH | feature_truncation,3_prime_UTR_variant |
| 2 | 13513769 | 13514234 | 465 | Trdmt1 | SJL | feature_truncation,3_prime_UTR_variant |
| 2 | 24980606 | 24980661 | 55 | Noxa1 | FVB | feature_truncation,coding_sequence_variant,intron_variant |
| 2 | 27001749 | 27001839 | 90 | Fam163b | 129S1,C57L | feature_truncation,3_prime_UTR_variant |
| 2 | 28976913 | 28977126 | 213 | Ttf1 | 129S1,C57L | feature_truncation,3_prime_UTR_variant |
| 2 | 32000584 | 32000641 | 57 | Plpp7 | SWR | feature_truncation,3_prime_UTR_variant |
| 2 | 37112090 | 37112150 | 60 | Or1af1 | BUB | feature_truncation,3_prime_UTR_variant |
| 2 | 62529547 | 62529651 | 104 | Kcnh7 | AJ,AKR,BTBR,BUB,C3H,C58J,CBA,CE,DBA1J,DBA2J,FVB,ILNJ2,KK,LP,MA/My,MRL,NOD,NOR,NZB,NZO,NZW,PJ,PL,RF,RH2,SEA,SMJ,SWR,TH | feature_truncation,3_prime_UTR_variant |

| | | | | | | |
|---|---|---|---|---|---|---|
| 2 | 65183600 | 65184411 | 811 | Slc38a11 | BTBR,BUB,NOD,NOR,SJL,SWR | feature_truncation,coding_sequence_variant,intron_variant |
| 2 | 68943464 | 68943516 | 52 | Cers6 | 129S1,AJ,AKR,BUB,FVB,ILNJ2,KK,MA/My,MRL,NOD,NOR,NZB,NZO,NZW,RH2 | feature_truncation,3_prime_UTR_variant |
| 2 | 69514147 | 69514321 | 174 | Klhl41 | BALB,BTBR,BUB,C3H,C57L,C58J,CBA,CE,DBA1J,DBA2J,MA/My,NZB,NZO,NZW,PJ,PL,RF,SEA,SJL,SMJ,SWR,TH | feature_truncation,3_prime_UTR_variant |
| 2 | 71103323 | 71103969 | 646 | Slc25a12 | AJ,AKR,BUB,C3H,FVB,ILNJ2,MA/My,MRL,NOD,NOR,NZW,PL,RH2,SMJ | feature_truncation,3_prime_UTR_variant |
| 2 | 73394670 | 73394856 | 186 | Chrna1 | NZB | feature_truncation,3_prime_UTR_variant |
| 2 | 85587643 | 85587996 | 353 | Or9g3 | AJ,AKR,BALB,BUB,C3H,C57L,C58J,CBA,CE,DBA1J,DBA2J,ILNJ2,MA/My,MRL,NOD,NOR,NZB,NZO,NZW,PJ,PL,RF,RH2,SEA,SJL,SMJ,SWR | feature_truncation,3_prime_UTR_variant |
| 2 | 85823816 | 85824755 | 939 | Or5m8 | AJ,AKR,BALB,BUB,C3H,C57L,C58J,CBA,CE,DBA1J,DBA2J,FVB,ILNJ2,KK,MA/My,MRL,NOD,NOR,NZB,NZO,NZW,PJ,PL,RF,RH2,SEA,SJL,SMJ,SWR | feature_truncation,3_prime_UTR_variant |
| 2 | 85959961 | 85960018 | 57 | Or5al5 | FVB,KK | feature_truncation,3_prime_UTR_variant |
| 2 | 86286204 | 86286263 | 59 | Or8k28 | AJ,AKR,BALB,BUB,C3H,C57L,CBA,CE,DBA1J,DBA2J,FVB,KK,MA/My,MRL,NOD,NOR,NZB,NZO,NZW,PJ,PL,RF,RH2,SEA,SMJ | frameshift_variant,feature_truncation |
| 2 | 86936253 | 86936445 | 192 | Or5aq7 | AJ,AKR,BALB,BUB,C3H,C57L,CBA,CE,DBA1J,DBA2J,FVB,KK,MA/My,MRL,NOD,NOR,NZB,NZO,NZW,PJ,PL,RF,RH2,SEA,SMJ | feature_truncation,3_prime_UTR_variant |
| 2 | 87221154 | 87221338 | 184 | Or10ag57 | AJ,AKR,BALB,C3H,C57L,C58J,CBA,CE,DBA1J,DBA2J,FVB,ILNJ2,KK,MA/My,MRL,NOD,NOR,NZB,NZO,NZW,PJ,RF,RH2,SEA,SJL,SMJ,SWR | feature_truncation,3_prime_UTR_variant |
| 2 | 87253131 | 87253191 | 60 | Or10ag2 | C3H,CBA,CE,ILNJ2,MA/My,NZW,RF,SJL,SWR | feature_truncation,3_prime_UTR_variant |
| 2 | 87696672 | 87697062 | 390 | Or5w19 | C58J,SJL,SWR | feature_truncation,5_prime_UTR_variant,intron_variant |
| 2 | 89356549 | 89356604 | 55 | Or4a71 | C3H,CBA,CE,MA/My,NZW | feature_truncation,3_prime_UTR_variant |
| 2 | 89850335 | 89850433 | 98 | Or4c3 | C58J,ILNJ2,KK,SJL,SWR | feature_truncation,3_prime_UTR_variant |
| 2 | 89978745 | 89979321 | 576 | Or4b1 | BUB,C58J,ILNJ2,KK,PL,SJL,SWR | feature_truncation,3_prime_UTR_variant,feature_truncation,3_prime_UTR_variant,intron_variant |
| 2 | 104661840 | 104661934 | 94 | Prrg4 | 129S1,AJ,AKR,BALB,BTBR,BUB,C3H,C57L,CBA,CE,DBA1J,DBA2J,ILNJ2,KK,MRL,NOD,NOR,PJ,PL,RH2,SEA,SJL | feature_truncation,3_prime_UTR_variant |
| 2 | 110467927 | 110468654 | 727 | Slc5a12 | C3H,CE,PL | feature_truncation,coding_sequence_variant,intron_variant |
| 2 | 113662677 | 113663049 | 372 | Arhgap11a | AJ,AKR,BALB,BTBR,C3H,CBA,CE,DBA1J,DBA2J,FVB,ILNJ2,KK,MA/My,MRL,NOD,NOR,NZB,NZO,NZW,PJ,PL,RF,RH2,SEA,SJL,SMJ,SWR,TH | feature_truncation,3_prime_UTR_variant |
| 2 | 115894816 | 115894968 | 152 | Meis2 | 129S1,AJ,AKR,BALB,BTBR,BUB,C3H,C57L,C58J,CBA,CE,DBA1J,DBA2J,FVB,ILNJ2,KK,MA/My,MRL,NOD,NOR,NZB,NZO,NZW,PJ,PL,RF,RH2,SEA,SJL,SMJ,SWR,TH | feature_truncation,5_prime_UTR_variant |
| 2 | 118608388 | 118608451 | 63 | Phgr1 | 129S1,AJ,AKR,BALB,BTBR,BUB,C3H,C57L,C58J,CBA,CE,DBA1J,DBA2J,FVB,ILNJ2,MA/My,NOD,NOR,NZB,NZO,NZW,PJ,PL,RF,RH2,SEA,SJL,SMJ,SWR,TH | feature_truncation,inframe_deletion |
| 2 | 119878007 | 119878070 | 63 | Sptbn5 | NZW,PL | feature_truncation,inframe_deletion |
| 2 | 121766495 | 121766698 | 203 | Golm2 | 129S1,AJ,AKR,BALB,BTBR,BUB,C3H,C57L,C58J,CBA,CE,DBA1J,DBA2J,FVB,ILNJ2,KK,MA/My,NOD,NZB,NZO,NZW,PJ,PL,RF,RH2,SEA,SJL,SMJ,SWR,TH | feature_truncation,3_prime_UTR_variant |
| 2 | 122316312 | 122316789 | 477 | Slc28a2b | 129S1,AJ,AKR,BALB,BTBR,BUB,C3H,C57L,C58J,CBA,CE,DBA1J,DBA2J,FVB,ILNJ2,MA/My,NOD,NZB,NZO,NZW,PJ,PL,RF,RH2,SEA,SJL,SMJ,SWR,TH | feature_truncation,5_prime_UTR_variant,intron_variant |
| 2 | 130409296 | 130409512 | 216 | Mrps26 | BALB,C3H,CBA,DBA1J,FVB,NOD,PL,SEA | feature_truncation,3_prime_UTR_variant |
| 2 | 131833380 | 131833436 | 56 | Rassf2 | 129S1,AKR,BTBR,C3H,C57L,CBA,CE,DBA1J,DBA2J,ILNJ2,KK,MRL,NZO,NZW,PJ,PL,RF,SJL,SWR | feature_truncation,3_prime_UTR_variant |
| 2 | 132147684 | 132147785 | 101 | Cds2 | 129S1,BTBR,C57L,ILNJ2,MA/My,NOD,PJ | feature_truncation,3_prime_UTR_variant |
| 2 | 134449205 | 134449805 | 600 | Tmx4 | 129S1,BTBR,C57L,CE,FVB,MRL,NZB,SMJ,SWR | feature_truncation,3_prime_UTR_variant |
| 2 | 150080901 | 150080985 | 84 | Zfp937 | NOD,NZB,NZW,RF | feature_truncation,inframe_deletion |
| 2 | 153460232 | 153460336 | 104 | Commd7 | 129S1,AJ,BALB,BTBR,C3H,C57L,C58J,CBA,DBA1J,DBA2J,ILNJ2,NOD,PL,RF,SEA,TH | feature_truncation,3_prime_UTR_variant |
| 2 | 154402269 | 154402323 | 54 | E2f1 | AJ,BTBR,C58J,DBA1J,DBA2J,KK,NZB,NZO,NZW | feature_truncation,3_prime_UTR_variant |
| 2 | 154714089 | 154714158 | 69 | Eif2s2 | 129S1,BALB,C3H,C57L,CBA,CE,FVB,MA/My,NOD,NZO,NZW,PL,SEA,SJL,SMJ,SWR,TH | feature_truncation,3_prime_UTR_variant |
| 2 | 155812023 | 155812086 | 63 | Cep250 | 129S1,BALB,C3H,C57L,CBA,CE,FVB,MA/My,NOD,NOR,NZO,NZW,PL,SEA,SJL,SMJ,SWR,TH | feature_truncation,inframe_deletion,feature_truncation,coding_sequence_variant,intron_variant |
| 2 | 156778497 | 156778697 | 200 | Ndrg3 | AKR,BTBR,BUB,DBA1J,DBA2J,ILNJ2,KK,MRL,NZB,PJ,RF,SJL | feature_truncation,3_prime_UTR_variant |
| 2 | 158226536 | 158226627 | 91 | Snhg11 | DBA1J,DBA2J,ILNJ2,MA/My,NZB,SMJ | feature_truncation,3_prime_UTR_variant |
| 2 | 158606731 | 158606960 | 229 | Ppp1r16b | AJ,CE,DBA1J,DBA2J,ILNJ2,KK,MA/My,NZB,SJL,SMJ | feature_truncation,3_prime_UTR_variant |
| 2 | 163165522 | 163165672 | 150 | Tox2 | KK | feature_truncation,3_prime_UTR_variant |
| 2 | 164349424 | 164349616 | 192 | Pigt | 129S1,AJ,BALB,BTBR,BUB,C3H,C57L,C58J,CBA,CE,DBA1J,DBA2J,FVB,KK,MA/My,NOD,NOR,NZO,NZW,PJ,PL,SEA,SMJ,SWR,TH | stop_lost,feature_truncation,coding_sequence_variant,3_prime_UTR_variant |
| 2 | 165218319 | 165218373 | 54 | Zfp334 | 129S1,AJ,AKR,BALB,BTBR,C57L,CBA,CE,ILNJ2,KK,MA/My,MRL,NOD,NOR,NZB,PL,RF,SEA,SJL,SMJ,TH | feature_truncation,3_prime_UTR_variant |
| 2 | 165669633 | 165669847 | 214 | Zmynd8 | 129S1,AJ,AKR,BALB,BTBR,BUB,C3H,C57L,C58J,CBA,CE,DBA1J,DBA2J,FVB,ILNJ2,KK,MA/My,NOD,NOR,NZB,NZO,PJ,PL,RF,SEA,SJL,SMJ,SWR,TH | feature_truncation,3_prime_UTR_variant |
| 2 | 172710979 | 172711085 | 106 | Bmp7 | 129S1,AJ,AKR,BUB,C57L,CE,MA/My,MRL,NZO,NZW,PL,RF,RH2,SMJ | feature_truncation,3_prime_UTR_variant |
| 2 | 175010857 | 175011445 | 588 | Gm14443 | CE,FVB,NZB,SJL,SWR | feature_truncation,inframe_deletion |
| 2 | 176372294 | 176372546 | 252 | Zfp973 | BALB,NZB,PJ | stop_lost,inframe_deletion |
| 2 | 177167503 | 177167587 | 84 | Zfp970 | SEA | feature_truncation,inframe_deletion |
| 2 | 177598805 | 177599620 | 815 | Gm14326 | BUB,KK,NOD,NOR | feature_truncation,5_prime_UTR_variant,intron_variant |

| | | | | | | |
|---|---|---|---|---|---|---|
| 2 | 177674926 | 177675010 | 84 | Zfp971 | CE,KK,PJ | feature_truncation,inframe_deletion |
| 2 | 177709457 | 177709876 | 419 | Zfp931 | PJ | stop_lost,feature_truncation,coding_sequence_variant,3_prime_UTR_variant,feature_truncation,3_prime_UTR_variant |
| 3 | 27396549 | 27396624 | 75 | Tnfsf10 | 129S1,AJ,AKR,BTBR,BUB,C3H,C57L,C58J,CBA,CE,DBA1J,DBA2J,FVB,ILNJ2,MRL,NOD,NOR,NZW,PJ,PL,RF,SEA,SJL,SWR | feature_truncation,3_prime_UTR_variant |
| 3 | 31260361 | 31260563 | 202 | Slc7a14 | CE,SWR | feature_truncation,3_prime_UTR_variant |
| 3 | 36220607 | 36220801 | 194 | Zfp267 | BUB,CE,DBA1J,FVB,NZB,NZO,NZW,SWR | feature_truncation,3_prime_UTR_variant |
| 3 | 37365937 | 37366028 | 91 | Cetn4 | 129S1,AJ,AKR,BTBR,C3H,C57L,CBA,CE,DBA1J,DBA2J,ILNJ2,MA/My,MRL,NOD,NOR,NZB,NZO,NZW,PL,RF,SJL,SMJ,TH | feature_truncation,5_prime_UTR_variant |
| 3 | 37768844 | 37769376 | 532 | Gm5148 | 129S1,AJ,AKR,BTBR,C3H,C57L,CBA,CE,DBA1J,DBA2J,ILNJ2,MA/My,MRL,NOD,NOR,NZB,NZO,NZW,PL,RF,SJL,SMJ,SWR,TH | feature_truncation,coding_sequence_variant,5_prime_UTR_variant,intron_variant |
| 3 | 40657076 | 40657475 | 399 | Intu | AJ,C3H,CBA,ILNJ2,KK,NZB,NZW,PL | feature_truncation,3_prime_UTR_variant |
| 3 | 40994230 | 40994295 | 65 | Larp1b | KK,NZW | feature_truncation,3_prime_UTR_variant |
| 3 | 51403438 | 51403650 | 212 | Rab33b | CE,NZW,SJL,SWR | feature_truncation,3_prime_UTR_variant |
| 3 | 72955110 | 72955174 | 64 | Slitrk3 | C58J,LP,NZW | feature_truncation,3_prime_UTR_variant |
| 3 | 79365318 | 79365428 | 110 | Fnip2 | RF | feature_truncation,3_prime_UTR_variant |
| 3 | 87348471 | 87349333 | 862 | Fcrl5 | FVB,MA/My,NOD,NOR,SJL,SWR,TH | feature_truncation,coding_sequence_variant,intron_variant |
| 3 | 89764484 | 89764603 | 119 | She | KK | feature_truncation,3_prime_UTR_variant |
| 3 | 89775195 | 89775631 | 436 | Il6ra | KK | feature_truncation,3_prime_UTR_variant |
| 3 | 92479212 | 92479299 | 87 | Ivl | CBA,KK | feature_truncation,inframe_deletion |
| 3 | 93116829 | 93117770 | 941 | Flg2 | AJ,C3H,CBA,CE | frameshift_variant,feature_truncation |
| 3 | 93117682 | 93118151 | 469 | Flg2 | BALB,MA/My,NZB,RH2 | frameshift_variant,feature_truncation |
| 3 | 93119292 | 93119649 | 357 | Flg2 | BALB,RH2 | feature_truncation,inframe_deletion |
| 3 | 93120694 | 93120922 | 228 | Flg2 | AJ,BALB,C3H,CBA,CE,MA/My,NZB,NZO,RF,RH2 | stop_lost,inframe_deletion |
| 3 | 93122837 | 93123064 | 227 | Flg2 | 129S1,AJ,BTBR,C3H,C57L,CBA,CE,ILNJ2,PJ,PL,SEA,TH | frameshift_variant,feature_truncation |
| 3 | 93124476 | 93124710 | 234 | Flg2 | AKR,B10D2,B10J,BUB,C58J,DBA1J,DBA2J,FVB,LP,MRL,NZW,RF,SJL,SMJ,SWR | feature_truncation,inframe_deletion |
| 3 | 93125285 | 93125642 | 357 | Flg2 | AKR,B10D2,B10J,BUB,C58J,CE,DBA1J,DBA2J,FVB,LP,MRL,NZW,SJL | feature_truncation,inframe_deletion |
| 3 | 93126118 | 93126339 | 221 | Flg2 | 129S1,BTBR,CE,NOD,RF,SMJ,SWR | frameshift_variant |
| 3 | 93126139 | 93126834 | 695 | Flg2 | NOR | frameshift_variant,feature_truncation |
| 3 | 93235829 | 93236342 | 513 | Hrnr | 129S1,AKR,BTBR,BUB,C57L,FVB,KK,MRL,NZW,SJL,SWR | feature_truncation,inframe_deletion |
| 3 | 93353127 | 93353247 | 120 | Tchh | BALB,NZB,NZO,RH2 | feature_truncation,inframe_deletion |
| 3 | 107141635 | 107142334 | 699 | Prok1 | SMJ | feature_truncation,3_prime_UTR_variant |
| 3 | 115736194 | 115736345 | 151 | Slc30a7 | 129S1,AJ,AKR,BALB,BUB,C3H,C57L,CBA,CE,DBA1J,DBA2J,FVB,ILNJ2,LP,MA/My,MRL,NZB,NZO,NZW,PL,RF,RH2,SJL,SWR,TH | feature_truncation,3_prime_UTR_variant |
| 3 | 116387842 | 116387895 | 53 | Trmt13 | SMJ | feature_truncation,3_prime_UTR_variant |
| 3 | 123166915 | 123167120 | 205 | Mettl14 | AJ,BALB,C3H,CBA,CE,DBA1J,DBA2J,KK,MA/My,NZB,PJ,RF,RH2,SEA,SMJ | feature_truncation,3_prime_UTR_variant |
| 3 | 124203263 | 124203403 | 140 | 1700006A11Rik | 129S1,BTBR,C57L,CE,PJ,SEA,TH | feature_truncation,coding_sequence_variant |
| 3 | 131082645 | 131082983 | 338 | Cyp2u1 | NZB,NZO | feature_truncation,3_prime_UTR_variant |
| 3 | 135048620 | 135048868 | 248 | Slc9b2 | 129S1,AKR,BTBR,BUB,C57L,C58J,LP,NZB,NZO,NZW,PJ,RF,SEA | feature_truncation,3_prime_UTR_variant |
| 3 | 138612058 | 138612152 | 94 | Tspan5 | 129S1,AJ,BALB,BTBR,C3H,C57L,CBA,CE,DBA1J,ILNJ2,KK,MA/My,MRL,NOD,NOR,NZB,NZO,NZW,PJ,PL,RH2,SEA,SJL,SMJ,SWR,TH | feature_truncation,3_prime_UTR_variant |
| 3 | 144394421 | 144394477 | 56 | Sh3glb1 | BUB,C58J,DBA2J,LP,MRL,RF | feature_truncation,3_prime_UTR_variant |
| 3 | 144528604 | 144528766 | 162 | Clca3b | C58J,LP,SJL | feature_truncation,inframe_deletion |
| 3 | 144533602 | 144533798 | 196 | Clca3b | SJL | feature_truncation,coding_sequence_variant,intron_variant |
| 3 | 152491894 | 152492044 | 150 | Pigk | 129S1,AJ,AKR,BALB,BTBR,C3H,C57L,CBA,CE,DBA1J,ILNJ2,PL,RH2,SEA,SMJ,TH | feature_truncation,3_prime_UTR_variant |
| 3 | 152524988 | 152525339 | 351 | St6galnac5 | TH | feature_truncation,3_prime_UTR_variant |
| 3 | 154284010 | 154284158 | 148 | Tyw3 | KK | feature_truncation,3_prime_UTR_variant |
| 3 | 154468054 | 154468123 | 69 | Erich3 | AJ,BALB,BUB,C3H,DBA2J,FVB,ILNJ2,KK,MA/My,NOD,NOR,NZB,NZO,NZW,RH2,SEA,SJL,SWR,TH | feature_truncation,inframe_deletion |
| 4 | 3973082 | 3973596 | 514 | Uba52rt | KK | transcript_ablation |
| 4 | 11262864 | 11262919 | 55 | Dpy19l4 | 129S1,AKR,BALB,BTBR,BUB,C3H,C57L,CBA,CE,DBA1J,DBA2J,FVB,ILNJ2,KK,MA/My,MRL,NZB,NZO,PJ,PL,RF,RH2,SEA,SJL,SMJ,SWR | feature_truncation,3_prime_UTR_variant |
| 4 | 12981638 | 12981694 | 56 | Triqk | AJ,AKR,BUB,NOD,NOR,NZB,NZO,PJ,SEA | feature_truncation,3_prime_UTR_variant |
| 4 | 20243831 | 20243994 | 163 | Nkain3 | KK,NZB,NZO | feature_truncation,3_prime_UTR_variant |
| 4 | 24503268 | 24503367 | 99 | Mms22l | NZB,NZO,NZW,RF | feature_truncation,3_prime_UTR_variant |
| 4 | 24613669 | 24613722 | 53 | Klhl32 | ILNJ2 | feature_truncation,3_prime_UTR_variant |
| 4 | 24903265 | 24903430 | 165 | Ndufaf4 | C57L | feature_truncation,3_prime_UTR_variant |
| 4 | 34950694 | 34950760 | 66 | Mob3b | AJ,AKR,BTBR,BUB,C3H,CBA,DBA1J,DBA2J,FVB,ILNJ2,LP,MRL,NOD,NOR,RF,SJL,SMJ,SWR | feature_truncation,3_prime_UTR_variant |
| 4 | 51228991 | 51229072 | 81 | Cylc2 | 129S1,C57L,NZW | feature_truncation,inframe_deletion |
| 4 | 52852930 | 52852981 | 51 | Or13c3 | MA/My | feature_truncation,3_prime_UTR_variant |
| 4 | 56875664 | 56875876 | 212 | Tmem245 | 129S1,AJ,AKR,BALB,BTBR,BUB,C3H,C57L,CBA,CE,DBA1J,DBA2J,ILNJ2,LP,MRL,NZB,NZO,NZW,PJ,PL,RF,RH2,SEA,SMJ | feature_truncation,3_prime_UTR_variant |
| 4 | 57854020 | 57854217 | 197 | Pakap | KK | feature_truncation,5_prime_UTR_variant |
| 4 | 59138810 | 59139313 | 503 | Shoc1 | 129S1,AJ,AKR,BALB,BTBR,BUB,C3H,C57L,CBA,CE,DBA1J,DBA2J,FVB,ILNJ2,KK,LP,MA/My,MRL,NOD,NOR,NZB,NZO,NZW,PJ,RF,RH2,SEA,SJL,SWR,TH | feature_truncation,5_prime_UTR_variant,intron_variant |
| 4 | 59753707 | 59753802 | 95 | E130308A19Rik | BTBR | feature_truncation,3_prime_UTR_variant |
| 4 | 98342641 | 98342999 | 358 | Patj | SMJ | stop_lost,feature_truncation,coding_sequence_variant,3_prime_UTR_variant |
| 4 | 99147442 | 99147510 | 68 | Atg4c | AJ,BALB,BTBR,C3H,CBA,MA/My,SEA | feature_truncation,3_prime_UTR_variant |
| 4 | 107834868 | 107834967 | 99 | Slc1a7 | 129S1,AJ,BALB,BUB,C3H,C57L,CBA,CE,FVB,NOD,PL,RF,SEA,SJL | feature_truncation,coding_sequence_variant,intron_variant |
| 4 | 114470634 | 114470746 | 112 | Trabd2b | 129S1,AJ,BTBR,C57L,DBA1J,DBA2J,FVB,ILNJ2,MA/My,SWR | feature_truncation,3_prime_UTR_variant |
| 4 | 115625960 | 115626678 | 718 | Efcab14 | BALB,BUB,C3H,CBA,FVB,KK,NOD,NOR,RF,SJL,SWR,TH | feature_truncation,3_prime_UTR_variant |
| 4 | 116557362 | 116557424 | 62 | Prdx1 | AKR,CE,KK,MRL,NZB,NZO,NZW,PJ,SEA,SMJ | feature_truncation,3_prime_UTR_variant |

| | | | | | | |
|---|---|---|---|---|---|---|
| 4 | 119077411 | 119077591 | 180 | AU022252 | AJ,B10D2,B10J,BALB,BUB,C3H,C58J,CBA,CE,DBA1J,DBA2J,FVB,ILNJ2,MA/My,NOD,NOR,NZB,PJ,PL,RF,SEA,SJL,SWR,TH | feature_truncation,3_prime_UTR_variant |
| 4 | 119079093 | 119079296 | 203 | AU022252 | ILNJ2 | feature_truncation,3_prime_UTR_variant |
| 4 | 122761262 | 122761433 | 171 | Cap1 | AJ,B10D2,B10J,BUB,C58J,CBA,CE,DBA1J,DBA2J,FVB,ILNJ2,MA/My,NOD,NOR,NZB,NZW,PL,RF,SJL,SMJ,SWR,TH | splice_acceptor_variant,splice_donor_variant,splice_donor_5th_base_variant,coding_sequence_variant,intron_variant |
| 4 | 123018893 | 123018951 | 58 | Bmp8b | B10D2,B10J,C58J | feature_truncation,3_prime_UTR_variant |
| 4 | 125604595 | 125604647 | 52 | Grik3 | ILNJ2 | feature_truncation,3_prime_UTR_variant |
| 4 | 129200886 | 129201173 | 287 | Rbbp4 | BUB,FVB,SJL,SWR | feature_truncation,3_prime_UTR_variant |
| 4 | 135177430 | 135177546 | 116 | Nipal3 | B10D2,B10J,DBA1J,KK,NZO,PJ,SEA | feature_truncation,3_prime_UTR_variant |
| 4 | 137455081 | 137455174 | 93 | Rap1gap | SMJ | feature_truncation,coding_sequence_variant,intron_variant |
| 4 | 138167370 | 138167433 | 63 | Mul1 | AJ,BALB,BUB,C3H,CBA,CE,DBA1J,DBA2J,FVB,ILNJ2,KK,MA/My,MRL,NZB,NZW,RH2,SJL,SMJ,SWR,TH | feature_truncation,3_prime_UTR_variant |
| 4 | 139104558 | 139104628 | 70 | Emc1 | AKR,BALB,BUB,C3H,CBA,DBA1J,DBA2J,ILNJ2,KK,MA/My,MRL,NOR,NZB,PL,RF,SJL,SMJ,TH | feature_truncation,3_prime_UTR_variant |
| 4 | 140785315 | 140785388 | 73 | Crocc | 129S1,BTBR,C57L,CE,DBA1J,DBA2J,FVB,ILNJ2,KK,NOD,NOR,NZB,NZO,PL,RF,SJL,SWR,TH | feature_truncation,5_prime_UTR_variant,intron_variant |
| 4 | 145349581 | 145349902 | 321 | Zfp268 | C58J,DBA2J,PJ,SEA | stop_lost,feature_truncation,coding_sequence_variant,3_prime_UTR_variant,feature_truncation,inframe_deletion |
| 4 | 145349853 | 145350525 | 672 | Zfp268 | DBA2J,KK,LP,TH | feature_truncation,inframe_deletion,feature_truncation,coding_sequence_variant,intron_variant |
| 4 | 146061591 | 146061674 | 83 | Zfp987 | DBA1J,MRL,NOD,NZB,NZW,SJL | frameshift_variant,feature_truncation |
| 4 | 146132071 | 146132154 | 83 | Zfp600 | BTBR,DBA1J,NOD,SWR | frameshift_variant,feature_truncation |
| 4 | 146133114 | 146133198 | 84 | Zfp600 | CE,KK,MA/My,SMJ | stop_lost,inframe_deletion |
| 4 | 146623297 | 146623557 | 260 | Zfp981 | PJ,TH | feature_truncation,3_prime_UTR_variant |
| 4 | 147074248 | 147074499 | 251 | Zfp989 | PL | frameshift_variant,feature_truncation |
| 4 | 147075026 | 147075109 | 83 | Zfp989 | FVB,LP,SEA | frameshift_variant,feature_truncation |
| 4 | 147142351 | 147142435 | 84 | Rex2 | C58J,LP,PJ,SEA | feature_truncation,inframe_deletion |
| 4 | 147416497 | 147416580 | 83 | Zfp988 | SMJ | frameshift_variant,feature_truncation |
| 4 | 147597252 | 147597419 | 167 | Zfp982 | 129S1,BTBR,C57L,C58J,LP,SEA,TH | frameshift_variant,feature_truncation |
| 4 | 147668773 | 147668846 | 73 | Zfp985 | 129S1,BTBR,KK | feature_truncation,3_prime_UTR_variant |
| 4 | 147759462 | 147759881 | 419 | Zfp534 | NZO | frameshift_variant,feature_truncation |
| 4 | 147909370 | 147909545 | 175 | Zfp933 | 129S1,AKR,BTBR,BUB,C3H,C57L,C58J,CBA,CE,DBA1J,DBA2J,KK,LP,MA/My,MRL,NOD,NZB,NZO,NZW,PJ,PL,RF,SEA,SJL,SMJ,SWR,TH | feature_truncation,3_prime_UTR_variant |
| 4 | 149212801 | 149212883 | 82 | Cenps | 129S1,AJ,AKR,BALB,BTBR,BUB,C3H,C57L,C58J,CBA,CE,DBA1J,DBA2J,KK,LP,MA/My,MRL,NOD,NZB,NZO,NZW,PJ,PL,RF,SEA,SJL,SMJ,SWR,TH | feature_truncation,3_prime_UTR_variant |
| 4 | 149464780 | 149464876 | 96 | Ube4b | 129S1,AKR,BTBR,C3H,C57L,CBA,DBA1J,DBA2J,MA/My,MRL,NOD,PL,SJL,TH | feature_truncation,coding_sequence_variant,intron_variant |
| 4 | 155506133 | 155506506 | 373 | Cfap74 | CE,FVB,ILNJ2,NOD,NOR,SJL,SMJ,SWR,TH | feature_truncation,3_prime_UTR_variant |
| 5 | 4131076 | 4131238 | 162 | Cyp51,Akap9 | BUB,NOD,NOR | feature_truncation,3_prime_UTR_variant |
| 5 | 14571967 | 14572057 | 90 | Pclo | 129S1,BALB,BTBR,C57L,C58J,CE,FVB,MA/My,PJ,RF,RH2,SEA,SJL,SWR | feature_truncation,inframe_deletion |
| 5 | 43999503 | 43999567 | 64 | Bst1 | PJ,PL | feature_truncation,3_prime_UTR_variant |
| 5 | 44151234 | 44151310 | 76 | Prom1 | PJ,PL | feature_truncation,3_prime_UTR_variant |
| 5 | 65604667 | 65604802 | 135 | Smim14 | C3H,CBA,DBA1J,DBA2J,ILNJ2,KK,NZB,PJ | feature_truncation,3_prime_UTR_variant |
| 5 | 88476262 | 88476316 | 54 | Prol1 | MA/My,NZB,NZO | feature_truncation,inframe_deletion |
| 5 | 103174854 | 103174933 | 79 | Mapk10 | 129S1,AKR,BUB,C3H,C57L,CBA,CE,DBA1J,DBA2J,MRL,NOD,NOR,NZO,NZW,PJ,PL,RF,SMJ,TH | feature_truncation,5_prime_UTR_variant |
| 5 | 104962344 | 104962430 | 86 | Zfp951 | AJ,BALB,BUB,C3H,CBA,DBA1J,DBA2J,FVB,ILNJ2,KK,NZB,NZO,RH2,SEA,SJL,SWR,TH | frameshift_variant,stop_lost |
| 5 | 105007539 | 105007811 | 272 | Zfp951 | AJ,BALB,BUB,C3H,CBA,DBA1J,DBA2J,FVB,ILNJ2,KK,NZB,NZO,RH2,SEA,SJL,SWR,TH | frameshift_variant,start_lost,feature_truncation,start_retained_variant,5_prime_UTR_variant |
| 5 | 108266887 | 108267090 | 203 | Tmed5 | 129S1,AJ,AKR,BALB,BTBR,BUB,C3H,C57L,CBA,CE,DBA1J,DBA2J,FVB,ILNJ2,KK,LP,MA/My,MRL,NOD,NOR,NZB,NZO,NZW,PJ,PL,RF,RH2,SEA,SJL,SMJ,SWR,TH | feature_truncation,3_prime_UTR_variant |
| 5 | 109363045 | 109363591 | 546 | Vmn2r14 | 129S1,AKR,BUB,C3H,C57L,CBA,FVB,KK,MA/My,NOD,NOR,NZB,NZO,NZW,PJ,PL,RF,SJL,SWR,TH | stop_lost,feature_truncation,coding_sequence_variant,3_prime_UTR_variant |
| 5 | 109823483 | 109823567 | 84 | Zfp1007 | BUB,C3H,CBA,FVB,PJ,PL,SJL,SWR,TH | feature_truncation,inframe_deletion |
| 5 | 110226068 | 110226152 | 84 | Gm35315 | 129S1,AJ,AKR,BALB,BTBR,BUB,C3H,C57L,CBA,FVB,ILNJ2,KK,MA/My,MRL,NOD,NOR,NZB,NZO,NZW,PJ,PL,RF,RH2,SEA,SJL,SMJ,SWR,TH | feature_truncation,inframe_deletion |
| 5 | 113957727 | 113957787 | 60 | Selplg | 129S1,BALB,BUB,C3H,C57L,CBA,FVB,ILNJ2,MRL,PJ,PL,RH2,SEA,SJL,SMJ,SWR | feature_truncation,inframe_deletion |
| 5 | 118095046 | 118095108 | 62 | Nos1 | BUB,C3H,CBA,ILNJ2,KK,NZB,NZO,PL,SJL,TH | feature_truncation,3_prime_UTR_variant |
| 5 | 118399986 | 118400127 | 141 | Spring1 | RF | feature_truncation,3_prime_UTR_variant |
| 5 | 123153130 | 123153191 | 61 | Orai1 | B10D2 | feature_truncation,5_prime_UTR_variant |
| 5 | 125464210 | 125464438 | 228 | Ubc | FVB,MA/My,SWR,TH | feature_truncation,inframe_deletion |
| 5 | 134736365 | 134736425 | 60 | Eln | KK | feature_truncation,inframe_deletion |
| 5 | 137140644 | 137140697 | 53 | Trim56 | LP | feature_truncation,3_prime_UTR_variant |
| 5 | 137178167 | 137178311 | 144 | Muc17 | LP | feature_truncation,inframe_deletion |
| 5 | 137181310 | 137181399 | 89 | Muc17 | 129S1,AJ,BUB,C57L,CBA,FVB,MA/My,NOR | frameshift_variant,feature_truncation |
| 5 | 137182489 | 137182632 | 143 | Muc17 | LP | frameshift_variant,feature_truncation |
| 5 | 137186295 | 137186402 | 107 | Muc17 | LP | frameshift_variant,feature_truncation |
| 5 | 137189044 | 137189178 | 134 | Muc17 | B10D2,B10J,C58J,LP,PL,SMJ,TH | frameshift_variant,feature_truncation |
| 5 | 137462596 | 137462833 | 237 | Zan | 129S1,BUB,LP,RF,SWR | feature_truncation,inframe_deletion |
| 5 | 137594987 | 137595080 | 93 | Mospd3 | 129S1,AKR,BUB,KK,LP,MA/My,MRL,NZB,NZO,NZW,RF,SWR | feature_truncation,3_prime_UTR_variant |
| 5 | 138250795 | 138250865 | 70 | Nxpe5 | BUB,LP | feature_truncation,3_prime_UTR_variant |
| 5 | 138646892 | 138646958 | 66 | A430033K04Rik | 129S1,C3H,CBA,CE,DBA1J,DBA2J,KK,LP,MA/My,NZB,NZO,NZW | feature_truncation,3_prime_UTR_variant |
| 5 | 139789876 | 139789968 | 92 | Tmem184a | ILNJ2 | feature_truncation,3_prime_UTR_variant |
| 5 | 144793877 | 144794033 | 156 | Trrap | 129S1,AKR,BTBR,C57L,CBA,PL,SJL,SMJ,TH | feature_truncation,coding_sequence_variant,intron_variant |
| 6 | 7040524 | 7040706 | 182 | Sdhaf3 | PJ | feature_truncation,3_prime_UTR_variant |
| 6 | 23003608 | 23003713 | 105 | Ptprz1 | TH | feature_truncation,3_prime_UTR_variant |
| 6 | 23344469 | 23345112 | 643 | Cadps2 | C3H,CBA,CE,FVB,NOD,NOR | feature_truncation,coding_sequence_variant,intron_variant |

| | | | | | | |
|---|---|---|---|---|---|---|
| 6 | 38481156 | 38481653 | 497 | Ubn2 | KK | feature_truncation,3_prime_UTR_variant |
| 6 | 48478165 | 48478779 | 614 | Sspo | AJ,AKR,BALB,BUB,C3H,CBA,DBA1J,DBA2J,FVB,MRL,NOD,NOR,PJ,PL,RF,RH2,SEA,SJL,SMJ,SWR,TH | feature_truncation,3_prime_UTR_variant |
| 6 | 48599337 | 48599429 | 92 | Zfp775 | AKR,BALB,C3H,C58J,CBA,DBA1J,DBA2J,FVB,MA/My,MRL,PJ,PL,RF,RH2,SEA,SJL,SMJ,SWR | feature_truncation,3_prime_UTR_variant |
| 6 | 56057449 | 56057507 | 58 | Pde1c | 129S1,C57L,KK,NZB,NZO,NZW | feature_truncation,3_prime_UTR_variant |
| 6 | 57217815 | 57217872 | 57 | Vmn1r14 | 129S1,C57L,CE,ILNJ2,KK,NZB,NZO,NZW,PJ | feature_truncation,3_prime_UTR_variant |
| 6 | 57874406 | 57875178 | 772 | Vmn1r22 | CE,ILNJ2,PJ | feature_truncation,3_prime_UTR_variant |
| 6 | 67271531 | 67271584 | 53 | Serbp1 | C58J,KK,NOD,NOR,NZB,NZO | feature_truncation,3_prime_UTR_variant |
| 6 | 71790241 | 71790839 | 598 | Mrpl35 | C3H,CBA,DBA1J,DBA2J,MA/My,SMJ | feature_truncation,3_prime_UTR_variant |
| 6 | 72291034 | 72291085 | 51 | Sftpb | AKR,C58J,KK,MRL,NOD,NOR,PL,RF | feature_truncation,3_prime_UTR_variant |
| 6 | 82535044 | 82535143 | 99 | Tacr1 | KK,NZB,NZO,TH | feature_truncation,3_prime_UTR_variant |
| 6 | 83281554 | 83281633 | 79 | Slc4a5 | KK,NZB,NZO,TH | feature_truncation,3_prime_UTR_variant |
| 6 | 84166799 | 84166940 | 141 | Dysf | SJL | feature_truncation,coding_sequence_variant,intron_variant |
| 6 | 87605886 | 87606146 | 260 | Aplf | PJ,SEA | feature_truncation,3_prime_UTR_variant |
| 6 | 87768641 | 87768740 | 99 | Rab43 | AKR,BALB,BUB,CBA,FVB,KK,LP,MA/My,MRL,NOD,NOR,PJ,PL,RH2,SEA,SJL,SWR | feature_truncation,3_prime_UTR_variant |
| 6 | 91439106 | 91439757 | 651 | Chchd4 | CE,NZB,NZW,PJ,SEA | feature_truncation,3_prime_UTR_variant |
| 6 | 100547923 | 100548581 | 658 | Shq1 | BUB,NZB,NZO,NZW,SWR | feature_truncation,3_prime_UTR_variant |
| 6 | 100783038 | 100783090 | 52 | Gxylt2 | SWR | feature_truncation,3_prime_UTR_variant |
| 6 | 115782071 | 115782245 | 174 | Cand2 | AKR,C3H,CBA,ILNJ2,MRL,PJ,PL,RF | feature_truncation,3_prime_UTR_variant |
| 6 | 123781229 | 123782005 | 776 | Vmn2r24 | FVB,NZO,SWR | feature_truncation,coding_sequence_variant,intron_variant |
| 6 | 128586551 | 128586683 | 132 | Klrb1a | NOD,NOR,SWR | stop_lost,feature_truncation,coding_sequence_variant,3_prime_UTR_variant |
| 6 | 128947891 | 128948266 | 375 | Clec2g | 129S1,AJ,AKR,BALB,BUB,C3H,C57L,CBA,DBA1J,DBA2J,ILNJ2,MRL,PL,RF,SEA,SMJ | feature_truncation,5_prime_UTR_variant,feature_truncation,coding_sequence_variant,5_prime_UTR_variant,intron_variant |
| 6 | 129842165 | 129842870 | 705 | Klra17 | 129S1,C58J,CE,FVB,LP,MA/My,SJL,TH | feature_truncation,coding_sequence_variant,intron_variant |
| 6 | 132577154 | 132577277 | 123 | Prp2 | 129S1,AJ,AKR,BALB,BUB,C3H,C57L,CBA,DBA1J,DBA2J,ILNJ2,MRL,NZB,NZO,PL,RF,RH2,SEA,SMJ | feature_truncation,inframe_deletion |
| 6 | 134424796 | 134424859 | 63 | Lrp6 | FVB,NZO,SWR | feature_truncation,3_prime_UTR_variant |
| 6 | 135701084 | 135701375 | 291 | Grin2b | 129S1,AJ,AKR,BALB,BTBR,BUB,C3H,C57L,CBA,CE,DBA1J,DBA2J,FVB,KK,MA/My,MRL,NOD,NOR,NZB,NZW,PJ,PL,RF,RH2,SEA,SJL,SWR,TH | feature_truncation,3_prime_UTR_variant |
| 6 | 136584359 | 136584431 | 72 | Atf7ip | 129S1,AJ,AKR,BALB,BTBR,BUB,C3H,C57L,CBA,CE,DBA1J,DBA2J,FVB,KK,MA/My,MRL,NOD,NOR,NZO,NZW,PJ,PL,RF,RH2,SEA,SJL,SWR,TH | feature_truncation,3_prime_UTR_variant |
| 6 | 136778675 | 136779043 | 368 | H4c16 | 129S1,AJ,AKR,BALB,BTBR,BUB,C3H,C57L,CBA,CE,DBA1J,DBA2J,FVB,KK,MA/My,MRL,NOD,NOR,NZO,NZW,PJ,PL,RF,RH2,SEA,SJL,SWR,TH | feature_truncation,3_prime_UTR_variant |
| 6 | 136805813 | 136806284 | 471 | BC049715 | 129S1,BALB,BTBR,C57L,CE,DBA1J,FVB,MRL,NZW,SEA | feature_truncation,5_prime_UTR_variant,intron_variant |
| 6 | 136941033 | 136941125 | 92 | Pde6h | BALB,FVB,KK,MA/My,MRL,NZO,NZW,RH2,SEA | feature_truncation,3_prime_UTR_variant |
| 6 | 136944910 | 136945030 | 120 | Pde6h | 129S1,AJ,AKR,BALB,BTBR,BUB,C3H,C57L,CBA,CE,DBA1J,DBA2J,FVB,KK,MA/My,MRL,NOD,NOR,NZO,NZW,PJ,PL,RF,RH2,SEA,SJL,SWR,TH | feature_truncation,3_prime_UTR_variant |
| 6 | 140627277 | 140627450 | 173 | Aebp2 | BTBR,C57L,C58J,FVB,PJ,SMJ,SWR | feature_truncation,3_prime_UTR_variant |
| 6 | 145396408 | 145396459 | 51 | Lmntd1 | AKR,BUB,C3H,CE,DBA1J,DBA2J,MRL,NZB,NZO,SJL | feature_truncation,5_prime_UTR_variant |
| 6 | 145910552 | 145910624 | 72 | Sspn | AKR,BUB,C3H,CE,DBA1J,DBA2J,FVB,MRL,NOD,NOR,NZB,NZO,NZW,SJL,SWR | feature_truncation,3_prime_UTR_variant |
| 6 | 148891544 | 148891764 | 220 | Dennd5b | KK,PL | feature_truncation,3_prime_UTR_variant |
| 7 | 3268668 | 3268721 | 53 | Nlrp12 | 129S1,BALB,BTBR,C3H,C57L,CBA,CE,DBA1J,DBA2J,ILNJ2,MRL,NZB,NZO,PJ,SEA,SMJ | feature_truncation,3_prime_UTR_variant |
| 7 | 4551997 | 4552072 | 75 | Ptprh | CE,SWR,TH | feature_truncation,3_prime_UTR_variant |
| 7 | 5612759 | 5612840 | 81 | Vmn1r61 | BUB,CE,NZO,SWR,TH | feature_truncation,3_prime_UTR_variant |
| 7 | 6477604 | 6477733 | 129 | Or6z5 | MRL,SMJ | feature_truncation,inframe_deletion |
| 7 | 11586237 | 11586330 | 93 | Vmn1r74 | AKR,FVB,MRL,NOD,NOR,RH2 | feature_truncation,3_prime_UTR_variant |
| 7 | 11889405 | 11889507 | 102 | Vmn1r78 | AKR,FVB,MRL,NOD,NOR,RH2 | feature_truncation,3_prime_UTR_variant |
| 7 | 12739245 | 12739461 | 216 | Zbtb45 | C58J | feature_truncation,3_prime_UTR_variant |
| 7 | 16718848 | 16719052 | 204 | Pnma8c | 129S1,AJ,AKR,BALB,BTBR,BUB,C3H,C57L,C58J,CBA,CE,ILNJ2,LP,MRL,NOD,NOR,NZB,NZO,NZW,PL,RF,RH2,SJL,SWR,TH | feature_truncation,3_prime_UTR_variant |
| 7 | 18340201 | 18340688 | 487 | Psg23 | 129S1,BTBR,C3H,C57L,CBA,CE,DBA1J,DBA2J,FVB,MA/My,NZB,NZW,PJ,PL,RF,SEA,SMJ | feature_truncation,3_prime_UTR_variant |
| 7 | 23246223 | 23246292 | 69 | Vmn1r168 | AJ,BALB | feature_truncation,3_prime_UTR_variant |
| 7 | 23674231 | 23674350 | 119 | Vmn1r181 | 129S1,AJ,BALB,BTBR,BUB,C3H,C57L,CBA,CE,NOD,NOR,NZB,NZW,SJL | feature_truncation,5_prime_UTR_variant,intron_variant |
| 7 | 23824648 | 23825176 | 528 | Zfp112 | LP,MA/My,PJ,SEA,SMJ | feature_truncation,inframe_deletion |
| 7 | 27355493 | 27355583 | 90 | Ttc9b | BUB,NZB,SJL | feature_truncation,3_prime_UTR_variant |
| 7 | 30578325 | 30578393 | 68 | Cd22 | DBA1J,DBA2J,NZB,NZO,NZW,SMJ | feature_truncation,5_prime_UTR_variant |
| 7 | 35279910 | 35279966 | 56 | Rgs9bp | AJ,BALB,CE,ILNJ2,SEA | feature_truncation,3_prime_UTR_variant |
| 7 | 35416095 | 35416189 | 94 | Dpy19l3 | BTBR,C57L,NZB,NZO,NZW | feature_truncation,coding_sequence_variant,intron_variant |
| 7 | 39187834 | 39188222 | 388 | Zfp619 | 129S1,BTBR,BUB,C3H,C57L,CBA,KK,MRL,NZW,PL,SJL,SMJ | feature_truncation,3_prime_UTR_variant |
| 7 | 40693731 | 40693818 | 87 | Gm4884 | DBA1J,DBA2J,ILNJ2,NZB,NZO,NZW,PJ,SMJ | feature_truncation,inframe_deletion |
| 7 | 41300718 | 41301089 | 371 | Zfp788 | DBA1J,DBA2J,ILNJ2,NZW,PJ | feature_truncation,3_prime_UTR_variant |
| 7 | 42466167 | 42466687 | 520 | Gm2381 | SMJ | feature_truncation,3_prime_UTR_variant |
| 7 | 42552292 | 42552488 | 196 | Vmn2r63 | DBA1J,DBA2J,ILNJ2,NZB,NZO,NZW,PJ,SJL,SMJ | feature_truncation,3_prime_UTR_variant |
| 7 | 42672702 | 42672912 | 210 | Vmn2r-ps158 | NZB,SJL | feature_truncation,coding_sequence_variant,intron_variant |
| 7 | 42838929 | 42838983 | 54 | Zfp936 | DBA1J,DBA2J,ILNJ2,NZW,PJ | feature_truncation,inframe_deletion |
| 7 | 43326434 | 43326491 | 57 | Ctu1 | C57L,SMJ | feature_truncation,inframe_deletion |
| 7 | 44917465 | 44918139 | 674 | Slc6a16 | AKR | feature_truncation,coding_sequence_variant,intron_variant |
| 7 | 45122617 | 45122827 | 210 | Dhdh | CBA,ILNJ2,NZW,PJ,SJL | feature_truncation,3_prime_UTR_variant |

| | | | | | | | |
|---|---|---|---|---|---|---|---|
| 7 | 45944696 | 45945037 | 341 | Otog | ILNJ2,NZW,PJ,SJL | feature_truncation,5_prime_UTR_variant,intron_variant | |
| 7 | 46377942 | 46377998 | 56 | Saa4 | CE | feature_truncation,3_prime_UTR_variant | |
| 7 | 46573083 | 46573231 | 148 | Uevld | ILNJ2,NZW,PJ,SJL | feature_truncation,3_prime_UTR_variant | |
| 7 | 47631732 | 47631808 | 76 | Mrgpra4 | CE | feature_truncation,5_prime_UTR_variant | |
| 7 | 48095305 | 48095372 | 67 | Mrgprb1 | ILNJ2,NZW,PJ,SJL | feature_truncation,3_prime_UTR_variant | |
| 7 | 66058248 | 66058447 | 199 | Aldh1a3 | BALB,BUB,C57L,CE,DBA1J,DBA2J,ILNJ2,KK,LP,NZB,NZO,PJ,SEA,SMJ,TH | feature_truncation,3_prime_UTR_variant | |
| 7 | 77828543 | 77828609 | 66 | Ntrk3 | 129S1,AJ,AKR,BALB,BTBR,BUB,C3H,C57L,C58J,CBA,CE,DBA1J,DBA2J,FVB,KK,MA/My,MRL,NOD,NOR,NZB,NZO,NZW,PJ,PL,RF,RH2,SEA,SJL,SMJ,SWR,TH | feature_truncation,3_prime_UTR_variant | |
| 7 | 79059320 | 79059563 | 243 | Fanci | 129S1,C3H,CBA,DBA1J,DBA2J,KK,PJ,SEA | feature_truncation,3_prime_UTR_variant | |
| 7 | 79966293 | 79966438 | 145 | Rccd1 | 129S1,AJ,AKR,BALB,BTBR,BUB,C3H,C57L,CBA,CE,DBA1J,DBA2J,FVB,ILNJ2,KK,MA/My,MRL,NOD,NOR,NZB,NZO,NZW,PJ,PL,RF,RH2,SEA,SJL,SMJ,SWR,TH | feature_truncation,3_prime_UTR_variant | |
| 7 | 81542164 | 81542648 | 484 | Hdgfl3 | 129S1,AJ,AKR,BALB,BTBR,BUB,C3H,C57L,CBA,DBA1J,DBA2J,FVB,KK,NOD,NOR,NZW,PJ,PL,RF,RH2,SEA,SJL,SWR | feature_truncation,3_prime_UTR_variant | |
| 7 | 85852601 | 85852657 | 56 | Vmn2r76 | 129S1,BTBR,C3H,C57L,CBA,DBA1J,DBA2J,ILNJ2,KK,NZB,NZO,NZW,PJ,SEA,SMJ | feature_truncation,3_prime_UTR_variant | |
| 7 | 86061957 | 86062047 | 90 | Or14c44 | 129S1,BTBR,C3H,C57L,CBA,DBA1J,DBA2J,ILNJ2,KK,NZB,NZO,NZW,PJ,SEA | feature_truncation,inframe_deletion | |
| 7 | 97108055 | 97108627 | 572 | Kctd14 | C3H,CBA,DBA1J,DBA2J,ILNJ2,NZO,PJ,SEA | feature_truncation,3_prime_UTR_variant | |
| 7 | 99809663 | 99809726 | 63 | Lipt2 | 129S1,AJ,AKR,BALB,BTBR,BUB,C3H,C57L,CBA,CE,DBA1J,DBA2J,FVB,ILNJ2,KK,MA/My,MRL,NZB,NZW,PJ,PL,RF,RH2,SEA,SJL,TH | feature_truncation,3_prime_UTR_variant | |
| 7 | 101794147 | 101794351 | 204 | Nup98 | SMJ | feature_truncation,3_prime_UTR_variant | |
| 7 | 102140779 | 102140833 | 54 | Or55b10 | KK,MA/My,NOD,SJL,SMJ,SWR | feature_truncation,3_prime_UTR_variant | |
| 7 | 102362252 | 102362309 | 57 | Or51a39 | KK,NOD,SJL,SWR | feature_truncation,3_prime_UTR_variant | |
| 7 | 102452456 | 102453159 | 703 | Or51f23 | KK,NOD,SJL,SWR | feature_truncation,coding_sequence_variant,5_prime_UTR_variant | |
| 7 | 102692549 | 102692607 | 58 | Or52r1b | TH | feature_truncation,3_prime_UTR_variant | |
| 7 | 102767700 | 102768254 | 554 | Or51a5 | KK,NOD,SJL,SWR | feature_truncation,3_prime_UTR_variant | |
| 7 | 102970764 | 102970871 | 107 | Or52ab2 | KK,NOD,SJL,SWR | feature_truncation,3_prime_UTR_variant | |
| 7 | 102993876 | 102993936 | 60 | Or52ad1 | KK,NOD,SJL,SWR | feature_truncation,3_prime_UTR_variant | |
| 7 | 103107759 | 103108411 | 652 | Or52d13 | KK,NOD,SJL,SWR | feature_truncation,3_prime_UTR_variant | |
| 7 | 103247632 | 103248575 | 943 | Or52z13 | SMJ | feature_truncation,3_prime_UTR_variant | |
| 7 | 103368491 | 103368555 | 64 | Or52a20 | 129S1,AJ,AKR,BALB,BTBR,BUB,C3H,C57L,CBA,DBA1J,DBA2J,FVB,ILNJ2,KK,MA/My,MRL,NOD,NZB,NZO,NZW,PJ,PL,RF,RH2,SEA,SJL,SMJ,SWR,TH | feature_truncation,3_prime_UTR_variant | |
| 7 | 103382175 | 103382235 | 60 | Or52a24 | 129S1,AJ,AKR,BALB,BTBR,BUB,C3H,C57L,CBA,DBA1J,DBA2J,FVB,ILNJ2,MRL,NZB,NZW,PJ,PL,RF,RH2,SEA,TH | feature_truncation,3_prime_UTR_variant | |
| 7 | 103529539 | 103529609 | 70 | Or51b4 | TH | feature_truncation,3_prime_UTR_variant | |
| 7 | 103539225 | 103539302 | 77 | Or51b17 | 129S1,AJ,AKR,BALB,BTBR,BUB,C3H,C57L,CBA,DBA1J,DBA2J,FVB,ILNJ2,MRL,NZB,NZW,PJ,PL,RF,RH2,SEA,TH | feature_truncation,3_prime_UTR_variant | |
| 7 | 103758007 | 103758548 | 541 | Or52d1 | KK,NOD,SJL,SMJ,SWR,TH | feature_truncation,3_prime_UTR_variant | |
| 7 | 103826704 | 103827642 | 938 | Or52h1 | KK,NOD,SJL,SMJ,SWR,TH | feature_truncation,3_prime_UTR_variant | |
| 7 | 103832235 | 103832912 | 677 | Or52h1 | KK | feature_truncation,5_prime_UTR_variant,intron_variant | |
| 7 | 103882658 | 103882748 | 90 | Trim6 | CE,NOD,NZO,SJL,SMJ,SWR | feature_truncation,3_prime_UTR_variant | |
| 7 | 103988993 | 103989055 | 62 | Trim12c | 129S1,AJ,BALB,BTBR,BUB,C3H,C57L,CBA,DBA1J,DBA2J,FVB,ILNJ2,NOD,NZB,NZW,PJ,PL,RH2,SEA,SJL,SMJ,SWR | feature_truncation,3_prime_UTR_variant | |
| 7 | 104005701 | 104005882 | 181 | Trim30b | 129S1,AJ,AKR,BALB,BTBR,BUB,C3H,C57L,CBA,CE,DBA1J,DBA2J,FVB,ILNJ2,MRL,NZB,NZO,NZW,PJ,PL,RF,RH2,SEA | feature_truncation,3_prime_UTR_variant | |
| 7 | 104232791 | 104232974 | 183 | Or52d3 | NOD,SJL,SWR | feature_truncation,3_prime_UTR_variant | |
| 7 | 104736640 | 104736890 | 250 | Or56a3 | NOD,NZW,SWR | feature_truncation,3_prime_UTR_variant | |
| 7 | 106500427 | 106500520 | 93 | Or2d3 | KK,NOD,NZO,SMJ,SWR | feature_truncation,5_prime_UTR_variant | FALSE |
| 7 | 106599543 | 106599674 | 131 | Or6a2 | B10D2,B10J,BUB,C58J,CE,FVB,KK,LP,MA/My,NOD,NOR,SJL,SMJ,SWR,TH | feature_truncation,3_prime_UTR_variant | |
| 7 | 106749425 | 106749845 | 420 | Or2d36 | 129S1,AJ,AKR,BALB,BTBR,C3H,C57L,CBA,CE,DBA1J,DBA2J,ILNJ2,KK,MRL,NOD,NZB,NZO,NZW,PJ,PL,RF,RH2,SEA,SMJ,SWR | feature_truncation,3_prime_UTR_variant | |
| 7 | 107629728 | 107629779 | 51 | Or5p6 | NOD,SMJ,SWR | feature_truncation,3_prime_UTR_variant | |
| 7 | 107655793 | 107655863 | 70 | Or10ab4 | NOD,NZO,SMJ,SWR | feature_truncation,3_prime_UTR_variant | |
| 7 | 107681812 | 107681901 | 89 | Or5p4 | NOD,NZO,SMJ,SWR | feature_truncation,3_prime_UTR_variant | |
| 7 | 108144669 | 108145160 | 491 | Or52n4b | NOD,SMJ,SWR | stop_lost,feature_truncation,coding_sequence_variant,3_prime_UTR_variant | |
| 7 | 108214189 | 108214940 | 751 | Or5p78 | NOD,SMJ,SWR | feature_truncation,3_prime_UTR_variant | |
| 7 | 108269414 | 108269548 | 134 | Or5p81 | NOD,SMJ,SWR | feature_truncation,3_prime_UTR_variant | |
| 7 | 118221483 | 118221992 | 509 | Tmc5 | 129S1,C3H,C57L,CBA,CE,DBA1J,DBA2J,NOR | feature_truncation,5_prime_UTR_variant,intron_variant | |
| 7 | 119314574 | 119314923 | 349 | Thumpd1 | AJ,BALB,C3H,CBA,DBA1J,DBA2J,ILNJ2,KK,NOR,NZB,NZO,PL,RH2 | feature_truncation,3_prime_UTR_variant | |
| 7 | 120545683 | 120546159 | 476 | Polr3e | 129S1,AJ,AKR,B10D2,B10J,BTBR,BUB,C3H,C58J,CBA,CE,DBA1J,DBA2J,ILNJ2,LP,MA/My,MRL,NOD,NZB,NZO,NZW,PJ,RF,RH2,SEA,SJL,SMJ,SWR,TH | feature_truncation,3_prime_UTR_variant | FALSE |
| 7 | 120545853 | 120546033 | 180 | Polr3e | AJ,BUB,FVB,NOR,PL | feature_truncation,3_prime_UTR_variant | FALSE |
| 7 | 126289624 | 126289835 | 211 | Slx1b | 129S1,AJ,AKR,BALB,BTBR,BUB,C57L,C58J,FVB,ILNJ2,KK,LP,MA/My,MRL,NOD,NZB,NZO,NZW,PJ,PL,RF,RH2,SEA,SJL,SMJ,SWR,TH | feature_truncation,3_prime_UTR_variant | |
| 7 | 127477030 | 127477087 | 57 | Zfp646 | TH | feature_truncation,5_prime_UTR_variant | |
| 7 | 128385885 | 128386096 | 211 | Sec23ip | 129S1,BTBR,C57L,ILNJ2,LP,PJ | feature_truncation,3_prime_UTR_variant | |
| 7 | 140443989 | 140444042 | 53 | Sirt3 | CE | feature_truncation,coding_sequence_variant,intron_variant | |
| 7 | 141031310 | 141031390 | 80 | Rplp2 | ILNJ2 | feature_truncation,3_prime_UTR_variant | |
| 7 | 141219584 | 141219790 | 206 | Muc6 | NZW | frameshift_variant,feature_truncation | |
| 7 | 141219965 | 141220483 | 518 | Muc6 | NZW | frameshift_variant,feature_truncation | |
| 7 | 141292749 | 141292882 | 133 | Muc2 | ILNJ2,NZW | frameshift_variant,feature_truncation | |
| 7 | 141295011 | 141295140 | 129 | Muc2 | NZW | feature_truncation,inframe_deletion | |
| 7 | 141363194 | 141363482 | 288 | Muc5ac | NZO,PJ | feature_truncation,inframe_deletion | |
| 7 | 141412901 | 141413842 | 941 | Muc5b | NZW | frameshift_variant,feature_truncation | |

| | | | | | | | |
|---|---|---|---|---|---|---|---|
| 7 | 141413519 | 141413573 | 54 | Muc5b | 129S1,AJ,AKR,B10D2,B10J,BALB,BTBR,BUB,C3H,C57L,C58J,CBA,CE,DBA1J,DBA2J,FVB,ILNJ2,KK,LP,MA/My,MRL,NOD,NOR,NZB,NZO,PJ,PL,RF,RH2,SMJ,SWR,TH | feature_truncation,inframe_deletion | |
| 7 | 141413523 | 141414020 | 497 | Muc5b | NZW,SJL | frameshift_variant,feature_truncation | FALSE |
| 7 | 141729057 | 141729192 | 135 | Krtap5-2 | BALB,MRL,RF,RH2,SEA | feature_truncation,inframe_deletion | |
| 7 | 141755267 | 141755555 | 288 | Krtap5-3 | BALB,MRL,RF,RH2,SEA | stop_lost,inframe_deletion | |
| 7 | 141782953 | 141783043 | 90 | Krtap5-5 | BALB,MRL,RF,RH2,SEA | feature_truncation,inframe_deletion | |
| 7 | 143333719 | 143333783 | 64 | Mrgpre | C58J,LP,MA/My,PJ,SMJ | feature_truncation,3_prime_UTR_variant | |
| 8 | 3440010 | 3440353 | 343 | Arhgef18 | 129S1,NZB | feature_truncation,3_prime_UTR_variant | |
| 8 | 3633737 | 3634248 | 511 | Rps23rg1 | 129S1,NZB,NZO,NZW | feature_truncation,coding_sequence_variant,intron_variant,feature_truncation,coding_sequence_variant,5_prime_UTR_variant,intron_variant | |
| 8 | 3727602 | 3727689 | 87 | Trappc5 | 129S1,NZB | feature_truncation,5_prime_UTR_variant | |
| 8 | 5135425 | 5135479 | 54 | Slc10a2 | 129S1,NZB | feature_truncation,3_prime_UTR_variant | |
| 8 | 13125567 | 13125732 | 165 | Proz | C57L,ILNJ2,NZB,NZW,PJ,PL,SEA,SMJ,TH | feature_truncation,3_prime_UTR_variant | |
| 8 | 13466907 | 13467274 | 367 | Grk1 | 129S1,C57L,CE,ILNJ2,LP,NZB,NZO,NZW,PJ,PL,SEA,SMJ,TH | feature_truncation,3_prime_UTR_variant | |
| 8 | 21161992 | 21162217 | 225 | Potefam3b | 129S1,AJ,AKR,BTBR,BUB,C3H,CBA,KK,MA/My,MRL,NOD,NOR,PL,RF,RH2,SMJ,SWR,TH | splice_donor_variant,splice_donor_5th_base_variant,coding_sequence_variant,intron_variant | |
| 8 | 25452880 | 25452934 | 54 | Adam9 | 129S1,AJ,AKR,BALB,BTBR,BUB,C3H,C57L,CBA,DBA1J,DBA2J,FVB,ILNJ2,KK,MA/My,MRL,NOD,NOR,NZB,NZO,NZW,PL,RF,RH2,SJL,SMJ,SWR,TH | feature_truncation,inframe_deletion | |
| 8 | 26516856 | 26516910 | 54 | Hook3 | 129S1,AJ,AKR,BALB,BTBR,BUB,C3H,C57L,CBA,DBA1J,DBA2J,FVB,ILNJ2,KK,MA/My,MRL,NOD,NOR,NZB,NZO,NZW,RF,RH2,SJL,SMJ,SWR,TH | feature_truncation,3_prime_UTR_variant | |
| 8 | 27765814 | 27765877 | 63 | Eif4ebp1 | AKR,C3H,CE,DBA1J,DBA2J,MA/My,PL,RF | feature_truncation,3_prime_UTR_variant | |
| 8 | 34578103 | 34578892 | 789 | Erhrt-ps | 129S1,AJ,AKR,BALB,BUB,C3H,C57L,CBA,CE,DBA1J,DBA2J,MA/My,MRL,NOD,NOR,PL,RF,RH2,SJL,SMJ,SWR,TH | transcript_ablation | |
| 8 | 47964092 | 47964152 | 60 | Trappc11 | C58J,LP | feature_truncation,5_prime_UTR_variant | |
| 8 | 60996624 | 60996763 | 139 | Aadat | AJ,AKR,BALB,BTBR,BUB,C3H,C57L,CBA,CE,DBA1J,DBA2J,FVB,ILNJ2,MA/My,NOD,NOR,NZB,NZO,NZW,PJ,PL,RF,RH2,SEA,SMJ | feature_truncation,coding_sequence_variant,intron_variant | |
| 8 | 69358096 | 69358147 | 51 | Lpl | 129S1,AJ,AKR,BALB,BTBR,BUB,C3H,C57L,C58J,CBA,CE,DBA1J,DBA2J,FVB,ILNJ2,KK,MA/My,MRL,NOD,NOR,NZB,NZO,NZW,PJ,PL,RF,RH2,SEA,SJL,SMJ,SWR,TH | feature_truncation,3_prime_UTR_variant | |
| 8 | 70637878 | 70637942 | 64 | Slc25a42 | 129S1,AKR,BALB,BUB,C3H,C57L,C58J,CBA,CE,DBA1J,DBA2J,FVB,ILNJ2,KK,MA/My,MRL,NZB,NZO,NZW,PJ,PL,RF,SEA,SJL,SMJ,SWR,TH | feature_truncation,3_prime_UTR_variant | |
| 8 | 72855851 | 72856077 | 226 | Or2z9 | 129S1,AKR,BALB,BUB,C3H,C57L,C58J,CBA,CE,DBA1J,DBA2J,FVB,ILNJ2,KK,MA/My,MRL,NZB,NZO,NZW,PJ,PL,RF,SEA,SJL,SMJ,SWR,TH | feature_truncation,3_prime_UTR_variant | |
| 8 | 72865267 | 72865879 | 612 | Or1ab2 | 129S1,AKR,BALB,BUB,C3H,C57L,C58J,CBA,CE,DBA1J,DBA2J,FVB,ILNJ2,KK,MA/My,MRL,NZB,NZO,NZW,PJ,PL,RF,SEA,SJL,SMJ,SWR,TH | feature_truncation,3_prime_UTR_variant | |
| 8 | 72867481 | 72867545 | 64 | Or1ab2 | 129S1,AKR,BALB,BUB,C3H,C57L,C58J,CBA,CE,DBA1J,DBA2J,FVB,ILNJ2,KK,MA/My,MRL,NZB,NZO,NZW,PJ,PL,RF,SEA,SJL,SMJ,SWR,TH | feature_truncation,3_prime_UTR_variant | |
| 8 | 94685489 | 94685616 | 127 | Gnao1 | 129S1,AKR,BUB,C57L,MA/My,PL,SJL | feature_truncation,3_prime_UTR_variant | |
| 8 | 96551968 | 96552785 | 817 | Sap18b | SWR | transcript_ablation | |
| 8 | 105076806 | 105076865 | 59 | Cmtm4 | PJ | feature_truncation,3_prime_UTR_variant | |
| 8 | 123209487 | 123209554 | 67 | Ctu2 | ILNJ2 | feature_truncation,3_prime_UTR_variant | |
| 8 | 123354870 | 123354931 | 61 | Cbfa2t3 | AJ,AKR,BALB,C3H,C58J,CE,ILNJ2,LP,MRL,PL,RF,RH2,SEA,SMJ | feature_truncation,3_prime_UTR_variant | |
| 8 | 125565614 | 125565705 | 91 | 2810004N23Rik | AJ,BALB,ILNJ2,RH2,SEA | feature_truncation,3_prime_UTR_variant | |
| 8 | 125756560 | 125756672 | 112 | Tsnax | AJ,BALB,ILNJ2,RH2,SEA | feature_truncation,3_prime_UTR_variant | |
| 8 | 127659877 | 127659972 | 95 | Tomm20 | KK | feature_truncation,3_prime_UTR_variant | |
| 9 | 20340121 | 20340178 | 57 | Zfp26 | 129S1 | feature_truncation,3_prime_UTR_variant | |
| 9 | 20509869 | 20509960 | 91 | Zfp846 | 129S1,BUB,C58J,FVB,KK,LP,MA/My,NZB,NZO,NZW,RF,SJL,SWR | feature_truncation,3_prime_UTR_variant | |
| 9 | 21048531 | 21048590 | 59 | Cdc37 | 129S1,BUB,C58J,CBA,CE,FVB,ILNJ2,KK,LP,MA/My,MRL,NOD,NOR,NZB,NZO,NZW,PJ,PL,RF,SEA,SJL,SMJ,SWR,TH | feature_truncation,3_prime_UTR_variant | |
| 9 | 21666898 | 21667094 | 196 | Spc24 | BTBR,C57L,C58J,FVB,MA/My,NOD,NOR,NZB,NZW,RF,SJL,SWR | feature_truncation,3_prime_UTR_variant | |
| 9 | 22111677 | 22111845 | 168 | Zfp872 | BTBR,C57L,FVB,MA/My,NOD,NOR,NZB,NZW,RF,SJL,SWR | feature_truncation,inframe_deletion | |
| 9 | 28907226 | 28907291 | 65 | Ntm | NZO | feature_truncation,3_prime_UTR_variant | |
| 9 | 38204881 | 38204947 | 66 | Or8c18 | 129S1,AJ,AKR,BALB,BTBR,BUB,C3H,CBA,DBA2J,NOD,NOR,PL,RF,RH2,SJL,SMJ,SWR,TH | feature_truncation,3_prime_UTR_variant | |
| 9 | 38205751 | 38206293 | 542 | Or8c18 | 129S1,AJ,AKR,BALB,BTBR,BUB,C3H,CBA,DBA2J,NOD,NOR,PL,RF,RH2,SJL,SMJ,SWR,TH | feature_truncation,3_prime_UTR_variant | |
| 9 | 38380165 | 38381160 | 995 | Or8b1b | BTBR,CBA,DBA1J,ILNJ2,MA/My,NOD,NOR,PL,SJL,TH | feature_truncation,3_prime_UTR_variant | |
| 9 | 38506298 | 38507207 | 909 | Or8b49 | 129S1,AJ,AKR,BALB,BTBR,BUB,C3H,DBA2J,FVB,MA/My,NOD,NOR,RF,RH2,SMJ,SWR,TH | stop_lost,feature_truncation,coding_sequence_variant,3_prime_UTR_variant | |
| 9 | 38566705 | 38567280 | 575 | Or8b51 | CBA,MA/My,PL,SJL | feature_truncation,3_prime_UTR_variant | |
| 9 | 39149428 | 39149513 | 85 | Or8g18 | PL,SJL | frameshift_variant,feature_truncation | |
| 9 | 39860337 | 39860968 | 631 | Or10d5 | CBA,CE,KK,MRL,NZB,NZW,PJ,PL,SEA,SJL | feature_truncation,3_prime_UTR_variant | |
| 9 | 39934770 | 39934822 | 52 | Or10g6 | CE,MRL,PL,SJL | feature_truncation,3_prime_UTR_variant | |
| 9 | 40100912 | 40101610 | 698 | Or6x1 | CBA,KK,MA/My,MRL,NZB,NZW,PJ,PL,SEA,SJL | feature_truncation,3_prime_UTR_variant | |
| 9 | 44055627 | 44055679 | 52 | Cbl | SMJ | feature_truncation,3_prime_UTR_variant | |
| 9 | 45040035 | 45040903 | 868 | Scn2b | AKR,BUB,FVB,MA/My,NZB,PJ,PL,RF,SEA | feature_truncation,3_prime_UTR_variant | |
| 9 | 49428618 | 49428698 | 80 | Ncam1 | KK | feature_truncation,3_prime_UTR_variant | |
| 9 | 50255488 | 50256284 | 796 | Rpl10-ps3 | FVB,SWR | transcript_ablation | |
| 9 | 53372416 | 53372474 | 58 | Atm | 129S1,C57L,PJ,SEA | feature_truncation,coding_sequence_variant,intron_variant | |

| | | | | | | |
|---|---|---|---|---|---|---|
| 9 | 59625223 | 59625433 | 210 | Gramd2a | 129S1,BTBR,C57L,CBA,KK,MA/My,NZB,NZO,NZW,PL,SMJ | feature_truncation,3_prime_UTR_variant |
| 9 | 65462881 | 65463095 | 214 | Plekho2 | B10D2,B10J,LP,MA/My | feature_truncation,3_prime_UTR_variant |
| 9 | 67838029 | 67838085 | 56 | Vps13c | CE | feature_truncation,coding_sequence_variant,intron_variant |
| 9 | 69292063 | 69292119 | 56 | Rora | NOD,NOR | feature_truncation,3_prime_UTR_variant |
| 9 | 75477846 | 75477947 | 101 | Tmod2 | 129S1,AJ,AKR,BALB,BTBR,BUB,C3H,C57L,C58J,CBA,CE,DBA1J,DBA2J,FVB,ILNJ2,KK,LP,MA/My,MRL,NOD,NOR,NZB,NZO,NZW,PJ,PL,RF,RH2,SEA,SJL,SMJ,SWR,TH | feature_truncation,3_prime_UTR_variant |
| 9 | 86406393 | 86406514 | 121 | Dop1a | DBA1J,DBA2J,FVB,KK,MA/My,NZO,NZW,PL,RF,SJL,SWR | feature_truncation,coding_sequence_variant,intron_variant |
| 9 | 95282592 | 95282704 | 112 | Chst2 | CE,MRL,NZO,NZW | feature_truncation,3_prime_UTR_variant |
| 9 | 96742621 | 96742890 | 269 | Pxylp1 | AKR,BUB,RF,SMJ,TH | feature_truncation,5_prime_UTR_variant,intron_variant |
| 9 | 103066684 | 103066782 | 98 | Srprb | 129S1,AJ,AKR,BALB,BTBR,BUB,C3H,C57L,CBA,FVB,ILNJ2,MA/My,NOD,NOR,NZB,NZW,PJ,PL,RF,RH2,SEA,SJL,SMJ,SWR,TH | feature_truncation,3_prime_UTR_variant |
| 9 | 103227574 | 103227625 | 51 | Topbp1 | CE,SMJ | feature_truncation,3_prime_UTR_variant |
| 9 | 103924354 | 103924418 | 64 | Uba5 | MA/My | feature_truncation,3_prime_UTR_variant |
| 9 | 106322858 | 106322973 | 115 | Abhd14a | 129S1,AJ,AKR,BALB,BUB,C3H,C57L,CE,DBA1J,DBA2J,FVB,ILNJ2,MA/My,MRL,NOD,NOR,NZB,NZW,PL,RH2,SJL,SMJ,SWR,TH | feature_truncation,5_prime_UTR_variant,feature_truncation,5_prime_UTR_variant,intron_variant |
| 9 | 106757415 | 106757467 | 52 | Dcaf1 | 129S1,AJ,AKR,BALB,BUB,C3H,C57L,CE,FVB,ILNJ2,MA/My,MRL,NOD,NOR,NZB,NZO,NZW,PL,RF,RH2,SJL,SMJ,SWR,TH | feature_truncation,3_prime_UTR_variant |
| 9 | 108280803 | 108280917 | 114 | Iho1 | CBA | feature_truncation,3_prime_UTR_variant |
| 9 | 110138005 | 110138171 | 166 | Elp6 | BTBR,CBA,FVB,NZW,SJL,SWR | feature_truncation,5_prime_UTR_variant,intron_variant |
| 9 | 110815512 | 110815698 | 186 | Rtp3 | 129S1,AJ,AKR,BALB,BTBR,BUB,C3H,C57L,CBA,CE,DBA1J,DBA2J,FVB,ILNJ2,KK,MA/My,MRL,NOD,NOR,NZB,NZO,PJ,PL,RF,RH2,SEA,TH | feature_truncation,inframe_deletion,feature_truncation,coding_sequence_variant,intron_variant |
| 9 | 114811420 | 114811808 | 388 | Rps27rt | 129S1,AJ,AKR,BALB,BTBR,C3H,C57L,CE,FVB,ILNJ2,KK,MRL,NOD,NOR,NZB,NZO,NZW,PJ,PL,RF,RH2,SEA,SJL,SMJ,SWR,TH | transcript_ablation |
| 9 | 115869588 | 115870198 | 610 | Gadl1 | 129S1,C57L,CE,NZB,NZO | stop_lost,feature_truncation,coding_sequence_variant,3_prime_UTR_variant,intron_variant |
| 9 | 121510106 | 121510157 | 51 | Sec22c | 129S1,AJ,AKR,B10D2,B10J,BALB,BTBR,BUB,C3H,C57L,C58J,CBA,CE,DBA1J,DBA2J,FVB,ILNJ2,KK,LP,MA/My,MRL,NOD,NOR,NZB,NZO,NZW,PJ,PL,RF,RH2,SEA,SJL,SMJ,SWR,TH | feature_truncation,3_prime_UTR_variant |
| 9 | 122986486 | 122986707 | 221 | Clec3b | 129S1,AJ,BALB,C3H,C57L,C58J,CE,FVB,MA/My,PJ,PL,SEA,SMJ | feature_truncation,3_prime_UTR_variant |
| 9 | 123620960 | 123621095 | 135 | Fyco1 | 129S1,C57L,C58J,CE,MA/My,SMJ,TH | feature_truncation,3_prime_UTR_variant |
| 9 | 123680961 | 123681081 | 120 | Fyco1 | C58J,MA/My,SMJ,TH | feature_truncation,5_prime_UTR_variant |
| 9 | 123932413 | 123932996 | 583 | Ccr5 | 129S1,C57L,C58J,CBA,CE,DBA1J,DBA2J,FVB,MA/My,NOD,NOR,SJL,SMJ,SWR | feature_truncation,3_prime_UTR_variant |
| X | 88452357 | 88452616 | 259 | Dcaf8l | CE | feature_truncation,5_prime_UTR_variant,intron_variant |
| X | 99486056 | 99486199 | 143 | Awat2 | NZB,NZO | feature_truncation,5_prime_UTR_variant |
| X | 149431903 | 149431963 | 60 | Tro | AJ,BALB,FVB,MA/My,MRL,NOD,NOR,NZB,NZW,RH2,SEA,SJL,SMJ,SWR | feature_truncation,inframe_deletion |
| X | 159258543 | 159258619 | 76 | Adgrg2 | BALB,MRL,NOD,NOR,SEA | feature_truncation,coding_sequence_variant,intron_variant |
| X | 162761874 | 162761942 | 68 | Car5b | TH | feature_truncation,3_prime_UTR_variant |
| X | 164959996 | 164960217 | 221 | Gemin8 | 129S1,AJ,AKR,BALB,BTBR,BUB,C3H,C57L,C58J,CBA,CE,DBA1J,DBA2J,FVB,ILNJ2,KK,MA/My,MRL,NOD,NOR,NZB,NZO,NZW,PJ,PL,RF,RH2,SEA,SJL,SMJ,SWR,TH | splice_donor_variant,splice_donor_5th_base_variant,5_prime_UTR_variant,intron_variant |
| X | 168772538 | 168772707 | 169 | Mid1 | 129S1,BTBR,C3H,CE,DBA2J,FVB,ILNJ2,KK,MA/My,MRL,NZB,NZO,NZW,PJ,PL,RF,SEA,SMJ,SWR | feature_truncation,3_prime_UTR_variant |

| CHR | START | END | SIZE | SYMBOL | STRAINS with VARIANT SV ALLELE | CONSEQUENCE | |
|---|---|---|---|---|---|---|---|
| 1 | 31304335 | 31326680 | 22345 | Gm6489 | RF | feature_truncation,coding_sequence_variant,5_prime_UTR_variant | |
| 1 | 58679924 | 58689654 | 9730 | Flacc1 | CE,SWR | stop_lost,feature_truncation,coding_sequence_variant,3_prime_UTR_variant,intron_variant | |
| 1 | 58685740 | 58687490 | 1750 | Flacc1 | CBA,DBA1J,DBA2J,ILNJ2,MA/My,NZB,NZO,NZW | stop_lost,feature_truncation,coding_sequence_variant,3_prime_UTR_variant,intron_variant | |
| 1 | 79693270 | 79700696 | 7426 | Wdfy1 | DBA2J | stop_lost,feature_truncation,coding_sequence_variant,3_prime_UTR_variant,intron_variant,feature_truncation,coding_sequence_variant,intron_variant | |
| 1 | 79696043 | 79703491 | 7448 | Wdfy1 | C57L,PJ | stop_lost,feature_truncation,coding_sequence_variant,3_prime_UTR_variant,intron_variant,feature_truncation,coding_sequence_variant,intron_variant | |
| 1 | 79698356 | 79705791 | 7435 | Wdfy1 | CBA | stop_lost,feature_truncation,coding_sequence_variant,3_prime_UTR_variant,intron_variant,feature_truncation,coding_sequence_variant,intron_variant | |
| 1 | 79698614 | 79700632 | 2018 | Wdfy1 | FVB | feature_truncation,coding_sequence_variant,intron_variant | |
| 1 | 85224521 | 85234100 | 9579 | Sp140l2 | MRL,PJ,PL | stop_lost,feature_truncation,coding_sequence_variant,3_prime_UTR_variant,intron_variant | |
| 1 | 85500108 | 85508544 | 8436 | Sp110 | NOD | stop_lost,feature_truncation,coding_sequence_variant,3_prime_UTR_variant,intron_variant | |
| 1 | 85516811 | 85517859 | 1048 | Sp110 | RF | feature_truncation,coding_sequence_variant,intron_variant | |
| 1 | 85527137 | 85531478 | 4341 | Sp140 | CBA,MRL,SMJ | feature_truncation,coding_sequence_variant,5_prime_UTR_variant,intron_variant | |
| 1 | 85651674 | 85653397 | 1723 | A630001G21Rik | RF | feature_truncation,coding_sequence_variant,intron_variant | |
| 1 | 88156516 | 88158359 | 1843 | Mroh2a | DBA1J,DBA2J | feature_truncation,coding_sequence_variant,intron_variant | |
| 1 | 90892310 | 90895296 | 2986 | Rab17 | AJ,BUB,C3H,CBA,CE,MA/My,MRL,PJ,RH2 | feature_truncation,5_prime_UTR_variant,intron_variant | |
| 1 | 92414945 | 92417170 | 2225 | Or6b2b | PL | feature_truncation,3_prime_UTR_variant | |
| 1 | 130740894 | 130747469 | 6575 | Fcamr | SEA | stop_lost,feature_truncation,coding_sequence_variant,3_prime_UTR_variant,intron_variant | |
| 1 | 151058084 | 151067891 | 9807 | Gm47985 | NZO,TH | transcript_ablation | |
| 1 | 156292059 | 156298875 | 6816 | Soat1 | AKR | feature_truncation,coding_sequence_variant,5_prime_UTR_variant,intron_variant | |
| 1 | 160841518 | 160843318 | 1800 | Zbtb37 | BALB,C58J,DBA1J,DBA2J,LP | feature_truncation,3_prime_UTR_variant | |
| 1 | 162668101 | 162675212 | 7111 | Fmo1 | 129S1,AJ,BALB,BTBR,BUB,C3H,C57L,C58J,CBA,CE,DBA1J,DBA2J,ILNJ2,KK,LP,MA/My,MRL,NZB,NZO,NZW,PJ,PL,RH2,SEA,SMJ,TH | stop_lost,feature_truncation,coding_sequence_variant,3_prime_UTR_variant,intron_variant | |
| 1 | 170510268 | 170516315 | 6047 | Olfml2b | ILNJ2 | feature_truncation,3_prime_UTR_variant | |
| 1 | 173745676 | 173748075 | 2399 | Ifi203 | AKR,BALB,BUB,C58J,CBA,CE,DBA1J,DBA2J,ILNJ2,NZB,NZO,PJ,RF,SEA,SJL,SWR | feature_truncation,3_prime_UTR_variant | |
| 1 | 181983461 | 181999423 | 15962 | Vmn1r1 | AKR,CBA,DBA1J,DBA2J,FVB,KK,PL,RF,SWR | transcript_ablation,stop_lost,feature_truncation,coding_sequence_variant,5_prime_UTR_variant,3_prime_UTR_variant,intron_variant | |
| 10 | 39849656 | 39853245 | 3589 | Mfsd4b5 | SJL | feature_truncation,coding_sequence_variant,intron_variant | |
| 10 | 77606404 | 77617620 | 11216 | Gm7138,Gm3238 | TH | transcript_ablation | |
| 10 | 77620397 | 77627052 | 6655 | Gm7137 | BUB | transcript_ablation | |
| 10 | 103976218 | 103979898 | 3680 | Gm21293 | PL | feature_truncation,5_prime_UTR_variant,intron_variant | |
| 10 | 118074185 | 118082776 | 8591 | Cdc5lrt1 | 129S1,AJ,AKR,BALB,BTBR,C3H,C57L,C58J,CBA,CE,DBA1J,DBA2J,KK,MA/My,MRL,NOD,NOR,NZB,NZO,NZW,PJ,PL,RF,RH2,SEA,SJL,TH | transcript_ablation | |
| 10 | 118164126 | 118187278 | 23152 | Cdc5lrt9,Cdc5lrt6,Cdc5lrt8,Cdc5lrt7 | NZB,SJL | transcript_ablation | FALSE |
| 10 | 118169550 | 118181852 | 12302 | Cdc5lrt8,Cdc5lrt7 | 129S1,DBA1J | transcript_ablation | |
| 10 | 129074977 | 129077579 | 2602 | Or6c5 | BUB,C3H,CBA,CE,KK,RF,SMJ | feature_truncation,3_prime_UTR_variant | |
| 10 | 129224680 | 129231985 | 7305 | Or6c208 | BUB,C3H,CBA,CE,KK,LP,NOD,NOR,RF,SMJ | feature_truncation,3_prime_UTR_variant | |
| 10 | 129559439 | 129564211 | 4772 | Or6c212 | CE,RF | feature_truncation,5_prime_UTR_variant,intron_variant | |
| 10 | 129798887 | 129801272 | 2385 | Or10u4 | KK,SMJ | feature_truncation,3_prime_UTR_variant | |
| 10 | 130248103 | 130250013 | 1910 | Vmn2r85 | AKR,CE,MRL,RF,SJL,SMJ,SWR | feature_truncation,3_prime_UTR_variant | |
| 11 | 34041269 | 34042413 | 1144 | Lcp2,4930469K13Rik | ILNJ2,KK,MA/My,NOD,NZW,PL,RF | feature_truncation,3_prime_UTR_variant,stop_lost,feature_truncation,coding_sequence_variant,3_prime_UTR_variant,intron_variant | |
| 11 | 46557281 | 46560322 | 3041 | Timd2 | 129S1,AJ,BALB,BTBR,C3H,C57L,CBA,DBA1J,ILNJ2,KK,MRL,NZB,NZW,PL,RH2,SJL,SMJ | feature_truncation,3_prime_UTR_variant | |
| 11 | 48716160 | 48717571 | 1411 | Cdk2ap1rt,Trim7 | CE,NZO,NZW,PJ,SEA | transcript_ablation,feature_truncation,5_prime_UTR_variant,intron_variant | |
| 11 | 49336004 | 49343486 | 7482 | Or2y16 | AKR,B10D2,B10J,BUB,C58J,CE,KK,MA/My,NOR,NZB,NZO,NZW,PJ,RF,SEA,SWR,TH | feature_truncation,3_prime_UTR_variant | |
| 11 | 58649769 | 58651709 | 1940 | Or2ak4 | AJ,BTBR,BUB,C3H,C57L,CBA,DBA1J,DBA2J,FVB,ILNJ2,KK,MA/My,MRL,NOR,NZB,NZO,NZW,PJ,SEA,SJL,SWR,TH | feature_truncation,3_prime_UTR_variant | |
| 11 | 71034946 | 71037667 | 2721 | Nlrp1a | 129S1,BALB,BTBR,C3H,C57L,C58J,CBA,FVB,LP,MA/My,MRL,NZB,NZO,RH2,SWR | feature_truncation,5_prime_UTR_variant,intron_variant | |
| 11 | 71058273 | 71086157 | 27884 | Nlrp1b | KK,NZW | feature_truncation,coding_sequence_variant,intron_variant | |
| 11 | 71091867 | 71118755 | 26888 | Nlrp1b | C3H,PL | feature_truncation,coding_sequence_variant,intron_variant,feature_truncation,coding_sequence_variant,5_prime_UTR_variant,intron_variant | |
| 11 | 71093037 | 71174694 | 81657 | Nlrp1b | CBA | feature_truncation,coding_sequence_variant,5_prime_UTR_variant,intron_variant | |
| 11 | 73675791 | 73679175 | 3384 | Or1e30 | AJ,NZO,NZW | transcript_ablation,stop_lost,feature_truncation,coding_sequence_variant,5_prime_UTR_variant,3_prime_UTR_variant,intron_variant | |
| 11 | 95957709 | 95960790 | 3081 | Atp5mc1 | KK | feature_truncation,3_prime_UTR_variant | |
| 11 | 99688712 | 99694195 | 5483 | Gm11569 | BALB,CBA,CE,MA/My,MRL,NZO,SEA | transcript_ablation | |
| 11 | 99696077 | 99701435 | 5358 | Krtap4-13 | AKR,RF | transcript_ablation | |
| 11 | 100222817 | 100230307 | 7490 | Gast | MA/My | transcript_ablation | |
| 11 | 108571632 | 108574270 | 2638 | Cep112 | BUB | feature_truncation,coding_sequence_variant,5_prime_UTR_variant,intron_variant | |
| 11 | 114908101 | 114932941 | 24840 | Cd300ld5,Cd300ld4 | SJL | transcript_ablation | |
| 11 | 116640381 | 116656483 | 16102 | St6galnac1 | AJ,BALB,C3H,C57L,CBA,DBA1J,DBA2J,MRL,SEA,TH | feature_truncation,3_prime_UTR_variant | |
| 11 | 120653604 | 120655245 | 1641 | Hmga1b | SJL | transcript_ablation | |

| | | | | | | |
|---|---|---|---|---|---|---|
| 12 | 44278085 | 44284684 | 6599 | Pnpla8 | AJ,AKR,BALB,BTBR,C3H,CBA,KK,PJ,PL,RF,RH2,SEA,SMJ,SWR | feature_truncation,5_prime_UTR_variant,intron_variant |
| 12 | 55287457 | 55300491 | 13034 | Srp54c | CE | feature_truncation,coding_sequence_variant,intron_variant |
| 12 | 69790997 | 69793449 | 2452 | Dmac2l | SWR | feature_truncation,3_prime_UTR_variant |
| 12 | 73357761 | 73362635 | 4874 | Slc38a6 | 129S1,AJ,BALB,BTBR,BUB,C3H,C57L,C58J,CBA,CE,FVB,KK,MA/My,NOD,NOR,NZW,PJ,RH2,SEA,SJL,SMJ,SWR,TH | stop_lost,feature_truncation,coding_sequence_variant,3_prime_UTR_variant |
| 12 | 84689815 | 84691609 | 1794 | Vrtn | PL,SJL | feature_truncation,5_prime_UTR_variant,intron_variant |
| 12 | 87669306 | 87671605 | 2299 | Eif1ad6 | NOD | feature_truncation,3_prime_UTR_variant |
| 12 | 87787027 | 87789326 | 2299 | Eif1ad2 | C57L | feature_truncation,3_prime_UTR_variant |
| 12 | 87880685 | 87885626 | 4941 | Eif1ad14 | CE,NOD,SWR | feature_truncation,3_prime_UTR_variant |
| 12 | 103407138 | 103415137 | 7999 | Ifi27l2a | PJ,SEA | transcript_ablation |
| 12 | 103562465 | 103563694 | 1229 | Ppp4r4 | NZO | feature_truncation,coding_sequence_variant,intron_variant |
| 13 | 21679379 | 21681317 | 1938 | AK157302 | LP,PJ,SEA | transcript_ablation |
| 13 | 21698190 | 21699475 | 1285 | Zkscan8 | 129S1,ILNJ2 | feature_truncation,3_prime_UTR_variant |
| 13 | 21738733 | 21742278 | 3545 | Or2b7 | LP,PJ,SEA | transcript_ablation,stop_lost,feature_truncation,coding_sequence_variant,5_prime_UTR_variant,3_prime_UTR_variant,intron_variant |
| 13 | 21813982 | 21821080 | 7098 | Or2b6 | 129S1,AJ,AKR,B10D2,B10J,BALB,BTBR,BUB,C3H,C57L,C58J,CBA,CE,DBA1J,DBA2J,FVB,ILNJ2,KK,LP,MA/My,MRL,NOD,NOR,NZB,PJ,PL,RF,RH2,SEA,SJL,SMJ,SWR,TH | feature_truncation,3_prime_UTR_variant |
| 13 | 21919800 | 21921069 | 1269 | H4c11 | LP,PJ,SEA | feature_truncation,3_prime_UTR_variant |
| 13 | 22270041 | 22273284 | 3243 | Vmn1r188 | 129S1,AJ,B10D2,B10J,BALB,BUB,C3H,C58J,CBA,DBA1J,DBA2J,FVB,MA/My,MRL,NOD,NOR,PL,RH2,SJL,SMJ,SWR,TH | transcript_ablation,stop_lost,feature_truncation,coding_sequence_variant,5_prime_UTR_variant,3_prime_UTR_variant,intron_variant |
| 13 | 22516447 | 22519483 | 3036 | Vmn1r197 | BTBR,C57L,KK,LP,NZB,NZO,NZW | feature_truncation,3_prime_UTR_variant |
| 13 | 22808311 | 22809745 | 1434 | Vmn1r206 | BTBR,C57L,KK,LP,NZB,NZO,NZW,PJ,SEA | feature_truncation,5_prime_UTR_variant |
| 13 | 43540263 | 43547352 | 7089 | Sirt5 | MRL,PJ,SEA | stop_lost,feature_truncation,coding_sequence_variant,3_prime_UTR_variant,intron_variant |
| 13 | 62849226 | 62850473 | 1247 | Platr25 | FVB,PJ,SEA,SWR | stop_lost,feature_truncation,coding_sequence_variant,3_prime_UTR_variant,intron_variant |
| 13 | 62874305 | 62880780 | 6475 | Platr25 | 129S1,AJ,BALB,BTBR,BUB,C57L,FVB,KK,MA/My,MRL,NOD,NOR,NZB,NZW,PJ,SEA,SMJ,SWR | feature_truncation,5_prime_UTR_variant,intron_variant |
| 13 | 67054010 | 67055581 | 1571 | Ramacl | PJ,SEA | feature_truncation,5_prime_UTR_variant,intron_variant |
| 13 | 67419469 | 67427196 | 7727 | Zfp458 | SWR | feature_truncation,5_prime_UTR_variant,intron_variant |
| 13 | 67541178 | 67543435 | 2257 | Zfp429 | 129S1,AKR,MRL,NZW,PL,RF | feature_truncation,coding_sequence_variant,intron_variant |
| 13 | 67588657 | 67589780 | 1123 | Zfp874a | 129S1,AJ,AKR,BALB,BTBR,BUB,C3H,C57L,C58J,CBA,CE,DBA1J,DBA2J,FVB,ILNJ2,KK,MRL,NOD,NOR,NZB,NZO,NZW,PJ,PL,RF,SEA,SJL,SMJ,SWR,TH | feature_truncation,3_prime_UTR_variant |
| 13 | 96215717 | 96217884 | 2167 | Btf3l4b | 129S1,AJ,BUB,CBA,DBA1J,DBA2J,SMJ | transcript_ablation |
| 13 | 97234707 | 97235850 | 1143 | Prp2rt | 129S1,AKR,C3H,C57L,CE,FVB,ILNJ2,KK,MRL,NZB,NZO,PL,SJL,SWR | transcript_ablation |
| 13 | 100486340 | 100557660 | 71320 | Naip1 | BTBR | stop_lost,feature_truncation,coding_sequence_variant,3_prime_UTR_variant,intron_variant |
| 13 | 113452593 | 113456924 | 4331 | Cspg4b | MRL | feature_truncation,coding_sequence_variant,intron_variant |
| 13 | 120411491 | 120415738 | 4247 | B020031M17Rik | NZW | feature_truncation,coding_sequence_variant,5_prime_UTR_variant,feature_truncation,coding_sequence_variant,5_prime_UTR_variant,intron_variant |
| 13 | 120635651 | 120748120 | 112469 | Gm21762,AF067061,Gm21731 | 129S1 | transcript_ablation |
| 14 | 11187752 | 11189046 | 1294 | Gapdhrt | 129S1,C3H,C57L,CBA,DBA1J,DBA2J,KK,MRL,NZW,PJ,PL | stop_lost,feature_truncation,coding_sequence_variant,5_prime_UTR_variant,3_prime_UTR_variant,intron_variant |
| 14 | 16736141 | 16750175 | 14034 | Gm3373 | 129S1 | feature_truncation,coding_sequence_variant,5_prime_UTR_variant,intron_variant |
| 14 | 18462888 | 18465895 | 3007 | Gm3099 | FVB,KK,MA/My,NOD,NOR,NZB,NZO,NZW,SJL,SMJ,SWR | stop_lost,feature_truncation,coding_sequence_variant,3_prime_UTR_variant,intron_variant |
| 14 | 18503785 | 18506828 | 3043 | Gm3095 | FVB,MA/My,NOD,NOR,SJL,SWR | feature_truncation,coding_sequence_variant,5_prime_UTR_variant,intron_variant |
| 14 | 19582823 | 19584297 | 1474 | Gm2244 | KK | stop_lost,feature_truncation,coding_sequence_variant,3_prime_UTR_variant,intron_variant |
| 14 | 40535381 | 40536584 | 1203 | Sh2d4b | CBA,DBA1J,DBA2J,ILNJ2 | feature_truncation,3_prime_UTR_variant |
| 14 | 41073567 | 41078154 | 4587 | Gm5798 | NZB | stop_lost,feature_truncation,coding_sequence_variant,3_prime_UTR_variant,intron_variant |
| 14 | 41881269 | 42209050 | 327781 | Gm17026,Gm3573,Gm7995 | MA/My | transcript_ablation |
| 14 | 42354691 | 42362885 | 8194 | Gm8024 | C57L,NZO,PJ,PL | feature_truncation,5_prime_UTR_variant,intron_variant |
| 14 | 42866512 | 42867966 | 1454 | Gm10376 | ILNJ2 | stop_lost,feature_truncation,coding_sequence_variant,3_prime_UTR_variant |
| 14 | 96337827 | 96340615 | 2788 | Klhl1 | AJ,AKR,BALB,BTBR,BUB,C3H,CBA,CE,DBA1J,DBA2J,FVB,MRL,NOD,PJ,PL,RF,SEA,SJL,SMJ,SWR,TH | feature_truncation,3_prime_UTR_variant |
| 14 | 113552025 | 113554169 | 2144 | Tpm3-rs7 | FVB,SJL | stop_lost,feature_truncation,coding_sequence_variant,5_prime_UTR_variant,3_prime_UTR_variant |
| 15 | 9314182 | 9321393 | 7211 | Ugt3a2 | RF,SJL,TH | stop_lost,feature_truncation,coding_sequence_variant,3_prime_UTR_variant,intron_variant |
| 15 | 9315071 | 9322280 | 7209 | Ugt3a2 | BTBR,C57L,CBA,KK,LP,NOR,NZW,PJ,PL,SEA | feature_truncation,3_prime_UTR_variant |
| 15 | 36995803 | 36997602 | 1799 | Zfp706 | 129S1,AJ,AKR,BUB,C57L,CE,FVB,LP,MA/My,MRL,NOD,NOR,PJ,RF,RH2,SJL,SMJ,SWR,TH | feature_truncation,3_prime_UTR_variant |
| 15 | 75981188 | 75988903 | 7715 | Eppk1 | BTBR | feature_truncation,coding_sequence_variant,intron_variant |
| 15 | 77464042 | 77486566 | 22524 | Apol10b,Gm36245 | MA/My | transcript_ablation,feature_truncation,coding_sequence_variant,5_prime_UTR_variant,intron_variant |
| 15 | 90520665 | 90527825 | 7160 | Cpne8 | AKR,BALB,BUB,DBA1J,DBA2J,MRL,NZW,PL,SJL | feature_truncation,3_prime_UTR_variant |
| 15 | 94419528 | 94421141 | 1613 | Pus7l | BTBR | feature_truncation,3_prime_UTR_variant |
| 15 | 100226497 | 100245831 | 19334 | Tmt1a3,Tmt1a | BUB,MA/My,SWR | transcript_ablation,feature_truncation,3_prime_UTR_variant |
| 15 | 100228159 | 100264668 | 36509 | Tmt1a3,Tmt1a2,Higd1c | NOD | transcript_ablation,feature_truncation,coding_sequence_variant,5_prime_UTR_variant,intron_variant |
| 15 | 100232589 | 100275332 | 42743 | Tmt1a3,Tmt1a2,Higd1c | C58J,KK,NOR,NZB,NZO,NZW,TH | transcript_ablation,feature_truncation,coding_sequence_variant,5_prime_UTR_variant,intron_variant |

| | | | | | | | |
|---|---|---|---|---|---|---|---|
| 15 | 100266044 | 100275331 | 9287 | Higd1c | NOD,NOR,NZO | feature_truncation,coding_sequence_variant,intron_variant | |
| 15 | 101535571 | 101554698 | 19127 | Gm5414,Gm5478 | KK,NZB,NZO,RF,TH | splice_acceptor_variant,splice_donor_variant,splice_donor_5th_base_variant,coding_sequence_variant,5_prime_UTR_variant,intron_variant,splice_acceptor_variant,splice_donor_variant,splice_donor_5th_base_variant,coding_sequence_variant,5_prime_UTR_variant,3_prime_UTR_variant,intron_variant,splice_acceptor_variant,splice_donor_variant,splice_donor_5th_base_variant,coding_sequence_variant,3_prime_UTR_variant,intron_variant | |
| 16 | 4573788 | 4574963 | 1175 | Hmox2 | AJ,BTBR,CE,FVB,NOD,NOR,SMJ,TH | feature_truncation,5_prime_UTR_variant | |
| 16 | 4675678 | 4678510 | 2832 | 4930562C15Rik | AJ,BTBR,CE,FVB,NOD,NOR,SMJ,TH | feature_truncation,coding_sequence_variant,5_prime_UTR_variant,intron_variant | |
| 16 | 45973324 | 45975448 | 2124 | Ahcyl | 129S1,C3H,C58J,CBA,CE,FVB,SJL,SWR,TH | transcript_ablation | |
| 16 | 89862422 | 89866040 | 3618 | Tiam1 | DBA1J,DBA2J | feature_truncation,5_prime_UTR_variant,intron_variant | |
| 17 | 6347859 | 6349361 | 1502 | Tmem181a | SEA | feature_truncation,coding_sequence_variant,intron_variant | |
| 17 | 13580338 | 13581449 | 1111 | Tcp10c | 129S1,AJ,AKR,B10D2,BALB,C3H,C57L,C58J,CBA,DBA1J,DBA2J,LP,NZO,RH2,SEA | feature_truncation,coding_sequence_variant,intron_variant | |
| 17 | 15037908 | 15041020 | 3112 | Wdr27 | PL | feature_truncation,3_prime_UTR_variant,intron_variant | |
| 17 | 18821788 | 18828341 | 6553 | Vmn2r96 | 129S1,AJ,AKR,BALB,C57L,CE,DBA1J,DBA2J,FVB,ILNJ2,KK,MA/My,NZO,RH2,SEA | feature_truncation,3_prime_UTR_variant | |
| 17 | 19611375 | 19612625 | 1250 | Vmn2r99 | 129S1,AJ,AKR,BALB,C57L,CE,DBA1J,DBA2J,FVB,ILNJ2,KK,MA/My,NZO,RH2,SEA | feature_truncation,coding_sequence_variant,intron_variant | |
| 17 | 19924820 | 19930804 | 5984 | Vmn2r102 | AKR,FVB,MA/My | feature_truncation,3_prime_UTR_variant | |
| 17 | 24505657 | 24508795 | 3138 | Abca17 | BTBR | feature_truncation,coding_sequence_variant,intron_variant | |
| 17 | 24509137 | 24514054 | 4917 | Abca17 | BTBR | feature_truncation,coding_sequence_variant,intron_variant | |
| 17 | 32588390 | 32595491 | 7101 | Wiz | 129S1,AJ,AKR,B10D2,BALB,BUB,C3H,C57L,CBA,CE,DBA1J,DBA2J,FVB,ILNJ2,MA/My,MRL,NOD,NOR,NZB,NZO,NZW,PJ,PL,RF,RH2,SEA,SJL,SMJ,SWR,TH | feature_truncation,coding_sequence_variant,intron_variant | |
| 17 | 33198118 | 33199861 | 1743 | Zfp472 | BUB,DBA1J,FVB,NOD,NOR,PL,SMJ,SWR | feature_truncation,3_prime_UTR_variant | |
| 17 | 35536985 | 35565596 | 28611 | H2-Q1,H2-Q2 | TH | transcript_ablation | FALSE |
| 17 | 36359906 | 36367019 | 7113 | H2-T27 | 129S1,AJ,AKR,B10D2,BALB,BTBR,BUB,C3H,C57L,C58J,CBA,CE,DBA1J,DBA2J,FVB,ILNJ2,KK,LP,MA/My,MRL,NOD,NOR,NZB,NZO,NZW,PJ,PL,RF,RH2,SEA,SJL,SWR,TH | stop_lost,feature_truncation,coding_sequence_variant,3_prime_UTR_variant | |
| 17 | 36967898 | 36995302 | 27404 | H2-M1 | AKR,C3H,CBA,CE,MA/My,MRL,RF | transcript_ablation | |
| 17 | 39387032 | 39388986 | 1954 | Esp23 | PJ | feature_truncation,5_prime_UTR_variant,intron_variant | |
| 17 | 40231761 | 40478211 | 246450 | Esp38,Gm49835 | 129S1 | transcript_ablation | |
| 17 | 44445972 | 44454889 | 8917 | Clic5 | FVB,SWR | feature_truncation,coding_sequence_variant,intron_variant | |
| 17 | 56814103 | 56824003 | 9900 | Znrf4 | SJL | transcript_ablation | |
| 17 | 70274708 | 70276324 | 1616 | Dlgap1 | C3H,MRL | feature_truncation,5_prime_UTR_variant | |
| 18 | 43605703 | 43610924 | 5221 | Eif3j2 | 129S1,AJ,AKR,BTBR,BUB,C57L,CE,FVB,ILNJ2,KK,MA/My,MRL,NOD,NOR,NZB,PJ,PL,RF,RH2,SJL,SWR,TH | transcript_ablation | |
| 18 | 48179700 | 48181479 | 1779 | Eno1b | 129S1,AJ,AKR,BUB,C57L,C58J,CE,FVB,ILNJ2,KK,LP,MA/My,MRL,NOD,NOR,NZB,NZO,NZW,PL,RF,RH2,SJL,SMJ,SWR,TH | stop_lost,feature_truncation,coding_sequence_variant,5_prime_UTR_variant,3_prime_UTR_variant,intron_variant,transcript_ablation | |
| 18 | 49822380 | 49829499 | 7119 | Dtwd2 | C58J,KK,LP | feature_truncation,3_prime_UTR_variant | |
| 18 | 87773570 | 87777505 | 3935 | Bhmt1b | KK,TH | transcript_ablation | |
| 19 | 11433992 | 11436262 | 2270 | Ms4a4b | KK,LP,TH | feature_truncation,coding_sequence_variant,intron_variant | |
| 19 | 12156202 | 12205503 | 49301 | Or5an9,Or5bb10 | AJ,PJ | transcript_ablation,feature_truncation,3_prime_UTR_variant | |
| 19 | 20593514 | 20602695 | 9181 | Aldh1a1 | DBA1J | feature_truncation,coding_sequence_variant,intron_variant | |
| 19 | 24852973 | 24854244 | 1271 | Gm10053 | AJ,AKR,BALB,BUB,C3H,C58J,CBA,CE,DBA1J,DBA2J,FVB,ILNJ2,KK,LP,MA/My,MRL,PJ,PL,RF,RH2,SEA,SJL,SWR,TH | transcript_ablation | |
| 19 | 33489300 | 33492348 | 3048 | Lipo4 | FVB,NZW | feature_truncation,coding_sequence_variant,intron_variant | |
| 19 | 52973976 | 52979378 | 5402 | Xpnpep1 | 129S1,AJ,AKR,BALB,BTBR,BUB,C3H,C57L,CBA,CE,DBA1J,DBA2J,ILNJ2,KK,MA/My,MRL,NOD,NOR,NZB,NZO,NZW,PJ,PL,RF,RH2,SEA,SMJ,TH | feature_truncation,3_prime_UTR_variant | |
| 19 | 61157208 | 61163676 | 6468 | Mplkipl1 | 129S1,AJ,AKR,BALB,BTBR,C3H,C57L,CBA,DBA1J,DBA2J,FVB,KK,MA/My,MRL,NOD,NOR,NZB,NZW,PJ,PL,RF,SEA,SJL,SMJ,SWR,TH | feature_truncation,3_prime_UTR_variant | |
| 2 | 3397298 | 3401214 | 3916 | Olah | MA/My,NZO,NZW | feature_truncation,5_prime_UTR_variant,intron_variant | |
| 2 | 11652968 | 11654570 | 1602 | Il2ra | SJL | feature_truncation,3_prime_UTR_variant | |
| 2 | 24083290 | 24089025 | 5735 | Il36g | AJ,AKR,DBA2J,FVB,PL,RH2,SJL | feature_truncation,3_prime_UTR_variant | |
| 2 | 26888304 | 26894330 | 6026 | Adamts13 | 129S1,BUB,C57L,FVB,MA/My,NOD,NOR,RF,SMJ | stop_lost,feature_truncation,coding_sequence_variant,3_prime_UTR_variant | |
| 2 | 49819282 | 49823487 | 4205 | Lypd6b | C57L | feature_truncation,5_prime_UTR_variant,intron_variant | |
| 2 | 83789414 | 83818979 | 29565 | Cwc22rt7,Cwc22rt3,Cwc22rt5,Cwc22rt2,Cwc22rt4 | NZO | transcript_ablation | |
| 2 | 85183702 | 85201516 | 17814 | Or5ak20 | C58J,NZB,SJL,SWR | feature_truncation,coding_sequence_variant,5_prime_UTR_variant,feature_truncation,coding_sequence_variant,5_prime_UTR_variant,intron_variant | |
| 2 | 85507681 | 85508744 | 1063 | Or9g4 | SJL,SWR | feature_truncation,5_prime_UTR_variant,intron_variant | |
| 2 | 85848706 | 85853929 | 5223 | Or5m3b | AJ,AKR,BALB,BUB,C3H,C57L,CBA,CE,DBA1J,DBA2J,ILNJ2,KK,MA/My,MRL,NOD,NOR,NZB,NZO,NZW,PJ,PL,RF,RH2,SEA,SMJ | feature_truncation,5_prime_UTR_variant,intron_variant | |
| 2 | 86170085 | 86177170 | 7085 | Or8k53 | AJ,AKR,BALB,BUB,C3H,C57L,CBA,CE,DBA1J,DBA2J,FVB,KK,MA/My,MRL,NOD,NOR,NZB,NZO,NZW,PJ,PL,RF,RH2,SEA,SMJ | stop_lost,feature_truncation,coding_sequence_variant,3_prime_UTR_variant | |
| 2 | 86217770 | 86245506 | 27736 | Or8k25,Or8k24 | FVB | transcript_ablation,feature_truncation,5_prime_UTR_variant,intron_variant | |
| 2 | 87702904 | 87705569 | 2665 | Or5w19 | C58J,SWR | feature_truncation,3_prime_UTR_variant | |
| 2 | 89997115 | 90079547 | 82432 | Or4x15 | BUB,C58J,ILNJ2,KK,PL,SJL,SWR | transcript_ablation | |
| 2 | 90003559 | 90073065 | 69506 | Or4x15 | 129S1,AJ,AKR,BALB,BTBR,C3H,C57L,CBA,CE,DBA1J,DBA2J,FVB,NOD,NOR,NZO,PJ,RH2 | transcript_ablation | |

| | | | | | | |
|---|---|---|---|---|---|---|
| 2 | 122254711 | 122258831 | 4120 | Slc28a2 | 129S1,AJ,AKR,BALB,BTBR,BUB,C3H,C57L,C58J,CBA,CE,DBA1J,DBA2J,FVB,ILNJ2,MA/My,NOD,NZB,NZO,NZW,PJ,PL,RF,RH2,SEA,SJL,SMJ,SWR,TH | feature_truncation,5_prime_UTR_variant,intron_variant |
| 2 | 122350748 | 122352139 | 1391 | Slc28a2b | 129S1,AJ,AKR,BALB,BTBR,BUB,C3H,C57L,C58J,CBA,CE,DBA1J,DBA2J,FVB,ILNJ2,MA/My,NOD,NZB,NZO,NZW,PJ,PL,RF,RH2,SEA,SJL,SMJ,SWR,TH | feature_truncation,coding_sequence_variant,intron_variant |
| 2 | 130249325 | 130250574 | 1249 | Tmem239 | BALB,DBA1J,PL,SEA | feature_truncation,3_prime_UTR_variant |
| 2 | 150936711 | 150944533 | 7822 | Gm14147 | NOD,NZB,NZW,RF | feature_truncation,coding_sequence_variant,5_prime_UTR_variant,intron_variant |
| 2 | 156198472 | 156205526 | 7054 | Cnbd2 | 129S1,BALB,C3H,C57L,CBA,CE,FVB,MA/My,NOD,NZO,NZW,PL,SEA,SJL,SMJ,SWR,TH | stop_lost,feature_truncation,coding_sequence_variant,3_prime_UTR_variant |
| 2 | 181645559 | 181649985 | 4426 | Gm14496 | BUB,CE,PJ,RF | feature_truncation,3_prime_UTR_variant |
| 3 | 10151563 | 10154671 | 3108 | Myef2l | 129S1,AJ,AKR,BALB,BTBR,C3H,C57L,C58J,CBA,DBA1J,FVB,ILNJ2,KK,LP,MRL,NZB,NZO,NZW,PJ,PL,RF,SEA,SJL,SWR | feature_truncation,coding_sequence_variant,5_prime_UTR_variant |
| 3 | 15567466 | 15570916 | 3450 | Sirpb1b | AJ,BALB,BUB,C3H,CBA,FVB,PL,SJL,SWR,TH | stop_lost,feature_truncation,coding_sequence_variant,3_prime_UTR_variant,intron_variant |
| 3 | 15882313 | 15885531 | 3218 | Sirpb1c | CE,DBA2J,LP | feature_truncation,3_prime_UTR_variant |
| 3 | 51147383 | 51152723 | 5340 | Noct | 129S1,BTBR,BUB,C3H,C57L,C58J,CBA,CE,FVB,LP,NZB,NZW,SJL,SMJ,SWR,TH | feature_truncation,5_prime_UTR_variant |
| 3 | 88280795 | 88286515 | 5720 | Bglap2 | RF | transcript_ablation |
| 3 | 91935247 | 91938771 | 3524 | Pglyrp3 | NOD,NOR | feature_truncation,coding_sequence_variant,intron_variant |
| 3 | 93111394 | 93113038 | 1644 | Flg2 | AJ,BALB,C3H,CBA,MA/My,NZB,NZO,RH2 | feature_truncation,inframe_deletion |
| 3 | 93123742 | 93125146 | 1404 | Flg2 | BALB,MA/My,NZB,NZO,RH2 | feature_truncation,inframe_deletion |
| 3 | 93124241 | 93125872 | 1631 | Flg2 | KK | frameshift_variant,feature_truncation |
| 3 | 93187612 | 93189850 | 2238 | Flg | C58J,LP,RF | transcript_ablation,stop_lost,feature_truncation,coding_sequence_variant,3_prime_UTR_variant |
| 3 | 93189614 | 93191090 | 1476 | Flg | SMJ | transcript_ablation |
| 3 | 93196332 | 93199355 | 3023 | Flg | CBA | transcript_ablation |
| 3 | 93234278 | 93235321 | 1043 | Hrnr | ILNJ2,PJ,PL,SEA,TH | frameshift_variant,feature_truncation |
| 3 | 103551238 | 103553386 | 2148 | Atg4a-ps,Syt6 | 129S1,AKR,BUB,C57L,C58J,CE,FVB,ILNJ2,KK,LP,PJ,RF,SEA,SJL,SWR,TH | transcript_ablation,stop_lost,feature_truncation,coding_sequence_variant,3_prime_UTR_variant,intron_variant |
| 3 | 131273078 | 131274456 | 1378 | Papss1 | AKR,BUB,C3H,C58J,CE,DBA1J,DBA2J,FVB,KK,LP,MA/My,NOD,NOR,NZB,NZO,NZW,PJ,PL,RF,SEA,SJL,SMJ,SWR,TH | feature_truncation,5_prime_UTR_variant,intron_variant |
| 3 | 152028773 | 152037820 | 9047 | Miga1 | 129S1,AJ,BALB,BTBR,C3H,C57L,CBA,CE,DBA1J,ILNJ2,KK,MRL,PL,RH2,SEA,SMJ | feature_truncation,3_prime_UTR_variant |
| 3 | 157351195 | 157353621 | 2426 | Ptger3 | RF | feature_truncation,3_prime_UTR_variant |
| 4 | 3994609 | 3997118 | 2509 | Sdr16c5 | 129S1,BALB,BTBR,C3H,C57L,CBA,CE,DBA1J,DBA2J,MRL,NZB,NZO,PL,RH2,SEA | stop_lost,feature_truncation,coding_sequence_variant,3_prime_UTR_variant,intron_variant |
| 4 | 60218186 | 60221626 | 3440 | Mup8 | 129S1,AJ,AKR,BALB,C3H,C57L,CBA,CE,DBA1J,DBA2J,FVB,ILNJ2,MA/My,MRL,NOD,NOR,NZB,NZO,NZW,RF,RH2,SEA,SJL,SWR | stop_lost,feature_truncation,coding_sequence_variant,3_prime_UTR_variant,intron_variant |
| 4 | 61433395 | 61436814 | 3419 | Mup16 | 129S1,BALB,C3H,C57L,CBA,DBA1J,DBA2J,FVB,MA/My,NOD,NOR,NZO,NZW,RF,RH2,SEA,SJL,SWR | stop_lost,feature_truncation,coding_sequence_variant,3_prime_UTR_variant,intron_variant |
| 4 | 88708973 | 88711909 | 2936 | Gm13278 | AJ | transcript_ablation |
| 4 | 101832107 | 101838549 | 6442 | Gm12790 | AJ,AKR,BALB,BUB,C3H,C58J,CBA,CE,KK,LP,MA/My,MRL,NZB,NZO,NZW,PJ,PL,RF,SEA,SJL,SMJ | feature_truncation,5_prime_UTR_variant,intron_variant |
| 4 | 106588722 | 106599282 | 10560 | Fam151a | PL | feature_truncation,coding_sequence_variant,5_prime_UTR_variant,intron_variant |
| 4 | 112401341 | 112468388 | 67047 | Skint2 | AKR,SMJ,SWR | feature_truncation,5_prime_UTR_variant,intron_variant |
| 4 | 112998167 | 113069759 | 71592 | Skint6 | LP | feature_truncation,coding_sequence_variant,intron_variant |
| 4 | 113016893 | 113031363 | 14470 | Skint6 | BALB,BUB,NOR,SEA,SJL,TH | feature_truncation,coding_sequence_variant,intron_variant |
| 4 | 113364578 | 113392236 | 27658 | Skint5 | AJ | feature_truncation,coding_sequence_variant,intron_variant |
| 4 | 113403579 | 113410500 | 6921 | Skint5 | AKR,C57L,DBA1J,DBA2J,LP,MRL,TH | feature_truncation,coding_sequence_variant,intron_variant |
| 4 | 113704823 | 113711685 | 6862 | Skint5 | 129S1,AJ,AKR,BTBR,C57L,DBA1J,DBA2J,FVB,LP,MRL,SWR | feature_truncation,coding_sequence_variant,intron_variant |
| 4 | 141161788 | 141163788 | 2000 | Srarp | TH | feature_truncation,coding_sequence_variant,5_prime_UTR_variant,intron_variant |
| 4 | 143574797 | 143588157 | 13360 | Pramel27 | BUB,CE,DBA2J,KK,MA/My,NZB,NZW,PL,RF,SMJ,SWR,TH | stop_lost,feature_truncation,coding_sequence_variant,3_prime_UTR_variant,intron_variant |
| 4 | 143668501 | 143683902 | 15401 | Pramel16 | CE,SWR | transcript_ablation |
| 4 | 143881566 | 143898718 | 17152 | Oog3 | AKR,C3H,CBA | transcript_ablation |
| 4 | 145265575 | 145268487 | 2912 | Zfp990 | PJ,SEA | feature_truncation,3_prime_UTR_variant |
| 4 | 146926154 | 146976189 | 50035 | Gm21411 | C57L | stop_lost,feature_truncation,coding_sequence_variant,3_prime_UTR_variant |
| 4 | 147954746 | 147956088 | 1342 | Fv1 | 129S1,AKR,BTBR,BUB,C3H,C57L,C58J,CBA,CE,DBA1J,DBA2J,KK,LP,MA/My,MRL,NOD,NZB,NZO,NZW,PJ,PL,RF,SEA,SJL,SWR,TH | stop_lost,feature_truncation,coding_sequence_variant,3_prime_UTR_variant |
| 5 | 15857413 | 15862680 | 5267 | 4930572O03Rik | MA/My | stop_lost,feature_truncation,coding_sequence_variant,5_prime_UTR_variant,3_prime_UTR_variant,intron_variant |
| 5 | 26141225 | 26153250 | 12025 | Gm21663 | AJ,AKR,B10D2,B10J,BALB,C3H,CBA,DBA1J,DBA2J,FVB,ILNJ2,MA/My,MRL,NZB,NZO,PJ,RF,SJL,TH | stop_lost,feature_truncation,coding_sequence_variant,5_prime_UTR_variant,3_prime_UTR_variant,intron_variant |
| 5 | 26146974 | 26159011 | 12037 | Gm21671,Gm21663 | BUB,C58J,LP,NOR | stop_lost,feature_truncation,coding_sequence_variant,3_prime_UTR_variant,intron_variant,feature_truncation,coding_sequence_variant,5_prime_UTR_variant,intron_variant |
| 5 | 26148326 | 26153283 | 4957 | Gm21671 | 129S1,BTBR,CE,NZW | feature_truncation,3_prime_UTR_variant |
| 5 | 26153438 | 26154461 | 1023 | Gm21671 | BTBR | feature_truncation,3_prime_UTR_variant |
| 5 | 26166046 | 26182146 | 16100 | Gm21680 | 129S1,BTBR,C57L,CE,NZW | transcript_ablation |
| 5 | 26311481 | 26327539 | 16058 | Gm10220 | BTBR,NZW | transcript_ablation |
| 5 | 26320634 | 26326624 | 5990 | Gm10220 | KK | stop_lost,feature_truncation,coding_sequence_variant,5_prime_UTR_variant,3_prime_UTR_variant,intron_variant |
| 5 | 44069651 | 44071575 | 1924 | Cd38,Ppihl | PJ,PL | feature_truncation,3_prime_UTR_variant,transcript_ablation |
| 5 | 87569388 | 87573834 | 4446 | Ugt2b38 | 129S1,C57L,SMJ | feature_truncation,coding_sequence_variant,5_prime_UTR_variant,intron_variant |
| 5 | 92707354 | 92711152 | 3798 | Fam47e | DBA2J | feature_truncation,coding_sequence_variant,intron_variant |

| | | | | | | | |
|---|---|---|---|---|---|---|---|
| 5 | 109293847 | 109300849 | 7002 | Vmn2r13 | 129S1,AKR,BUB,C3H,C57L,CBA,FVB,KK,MA/My,NOD,NOR,NZB,NZO,NZW,PJ,PL,RF,SJL,SWR,TH | feature_truncation,3_prime_UTR_variant | |
| 5 | 137836131 | 137869140 | 33009 | Pilrb1,Pilrb2 | BUB,RF | transcript_ablation,stop_lost,feature_truncation,coding_sequence_variant,3_prime_UTR_variant,intron_variant | |
| 5 | 145280983 | 145285149 | 4166 | Cyp3a57 | TH | feature_truncation,coding_sequence_variant,5_prime_UTR_variant,intron_variant | |
| 5 | 146418233 | 146427634 | 9401 | Gm6408 | TH | transcript_ablation | |
| 6 | 3388053 | 3420796 | 32743 | Samd9l | SEA | feature_truncation,5_prime_UTR_variant,intron_variant | |
| 6 | 65093219 | 65094389 | 1170 | Hpgds | KK | feature_truncation,3_prime_UTR_variant | |
| 6 | 71287151 | 71289038 | 1887 | Smyd1 | AKR,C3H,C58J,CBA,DBA1J,DBA2J,KK,MA/My,MRL,NOD,NOR,PL,RF,SMJ | feature_truncation,3_prime_UTR_variant | |
| 6 | 76469330 | 76476656 | 7326 | Rnf26rt | KK,PJ | transcript_ablation | |
| 6 | 102549922 | 102557101 | 7179 | Cntn3 | BUB,C58J,LP,NOD,NOR,NZB,NZO,NZW,SWR | feature_truncation,5_prime_UTR_variant,intron_variant | |
| 6 | 112335959 | 112337523 | 1564 | Ssu2 | SWR | stop_lost,feature_truncation,coding_sequence_variant,3_prime_UTR_variant,intron_variant | |
| 6 | 123044870 | 123046268 | 1398 | Clec4b1 | PJ,SMJ,SWR | feature_truncation,coding_sequence_variant,intron_variant | |
| 6 | 129574505 | 129576889 | 2384 | Klrd1 | DBA2J | stop_lost,feature_truncation,coding_sequence_variant,3_prime_UTR_variant,intron_variant | |
| 6 | 129716471 | 129718157 | 1686 | Klri2 | 129S1,SMJ | feature_truncation,coding_sequence_variant,5_prime_UTR_variant,intron_variant | |
| 6 | 129738259 | 129744140 | 5881 | Klrh1 | 129S1,C58J,KK,NOD,NOR,NZB,NZW,SMJ,SWR | stop_lost,feature_truncation,coding_sequence_variant,3_prime_UTR_variant,intron_variant,feature_truncation,3_prime_UTR_variant | |
| 6 | 132221716 | 132368953 | 147237 | Prb1b,Prb1c | NZO | transcript_ablation | |
| 6 | 142103592 | 142107519 | 3927 | Slco1a6 | PJ | feature_truncation,coding_sequence_variant,5_prime_UTR_variant,intron_variant | |
| 7 | 5224666 | 5230251 | 5585 | Vmn1r57 | BUB,CE,MRL,NZO,SMJ,SWR,TH | feature_truncation,3_prime_UTR_variant | |
| 7 | 5523289 | 5552569 | 29280 | Vmn1r60 | BUB,NZO,SWR,TH | transcript_ablation | |
| 7 | 9028486 | 9033280 | 4794 | Vmn2r39 | 129S1,BTBR | feature_truncation,coding_sequence_variant,intron_variant | |
| 7 | 11611459 | 11612691 | 1232 | Vmn1r75 | AKR,FVB,MRL,NOD,NOR,RH2 | feature_truncation,5_prime_UTR_variant,intron_variant | |
| 7 | 11663538 | 11667578 | 4040 | Vmn1r76 | SWR | transcript_ablation,stop_lost,feature_truncation,coding_sequence_variant,5_prime_UTR_variant,3_prime_UTR_variant,intron_variant | |
| 7 | 12837099 | 12838577 | 1478 | Vmn1r86 | NZO | feature_truncation,5_prime_UTR_variant,intron_variant | |
| 7 | 21086069 | 21098918 | 12849 | Vmn1r129 | ILNJ2,MRL,PL | transcript_ablation | |
| 7 | 23194111 | 23203075 | 8964 | Vmn1r167 | 129S1,BTBR,BUB,C3H,C57L,CBA,NOD,NOR,NZB,NZW,SJL | feature_truncation,3_prime_UTR_variant | |
| 7 | 24277585 | 24283556 | 5971 | Zfp575 | BUB,C3H,CBA,CE,NOD,NOR,SJL | feature_truncation,3_prime_UTR_variant | |
| 7 | 25688012 | 25897736 | 209724 | Cyp2b13,Cyp2b9 | 129S1,BTBR,C57L | transcript_ablation,feature_truncation,coding_sequence_variant,5_prime_UTR_variant,intron_variant | |
| 7 | 27387653 | 27389195 | 1542 | Zfp607b | KK,NZO | feature_truncation,5_prime_UTR_variant,intron_variant | |
| 7 | 42537169 | 42547477 | 10308 | Vmn2r63 | DBA1J,DBA2J,ILNJ2,NZW,PJ | feature_truncation,3_prime_UTR_variant | |
| 7 | 46835720 | 46838404 | 2684 | Mrgpra6 | C57L,CE,ILNJ2,NZW,PJ,RF,SJL | feature_truncation,coding_sequence_variant,intron_variant | |
| 7 | 47063585 | 47103412 | 39827 | Mrgpra2a | C57L,FVB,NOR | transcript_ablation | |
| 7 | 47215078 | 47243884 | 28806 | Mrgpra3 | CE,ILNJ2,NZW,PJ | stop_lost,feature_truncation,coding_sequence_variant,3_prime_UTR_variant,intron_variant | |
| 7 | 79993836 | 79995220 | 1384 | Hddc3 | 129S1,AJ,AKR,BALB,BTBR,BUB,C3H,C57L,CBA,CE,DBA1J,DBA2J,FVB,ILNJ2,KK,MA/My,MRL,NOD,NOR,NZB,NZO,NZW,PJ,PL,RF,RH2,SEA,SJL,SMJ,SWR,TH | stop_lost,feature_truncation,coding_sequence_variant,3_prime_UTR_variant,intron_variant | |
| 7 | 85036649 | 85043084 | 6435 | Vmn2r69 | 129S1,AJ,AKR,BALB,BTBR,BUB,C3H,C57L,CBA,DBA1J,DBA2J,FVB,ILNJ2,KK,MA/My,MRL,NOD,NOR,NZB,NZO,NZW,PJ,PL,RF,RH2,SEA,SJL,SMJ,SWR,TH | feature_truncation,3_prime_UTR_variant | |
| 7 | 97858997 | 97860488 | 1491 | Acer3 | C3H,CBA,DBA1J,DBA2J,ILNJ2,PJ,SEA | feature_truncation,3_prime_UTR_variant | |
| 7 | 101206786 | 101229154 | 22368 | Art2b | BTBR,C57L,KK,MA/My,NZW | stop_lost,feature_truncation,coding_sequence_variant,3_prime_UTR_variant,intron_variant | |
| 7 | 103198912 | 103200102 | 1190 | Or51ab3 | 129S1,AJ,AKR,BALB,BTBR,BUB,C3H,C57L,CBA,DBA1J,DBA2J,FVB,ILNJ2,MA/My,MRL,NZB,NZO,NZW,PJ,PL,RF,RH2,SEA,SMJ,TH | feature_truncation,5_prime_UTR_variant | |
| 7 | 103314791 | 103316670 | 1879 | Or51v8 | 129S1,AJ,AKR,BALB,BTBR,BUB,C3H,C57L,CBA,DBA1J,DBA2J,FVB,ILNJ2,MRL,NZB,NZW,PJ,PL,RF,RH2,SEA,TH | feature_truncation,3_prime_UTR_variant | |
| 7 | 103657752 | 103660760 | 3008 | Or51k1 | KK,NOD,SJL,SMJ,SWR,TH | feature_truncation,3_prime_UTR_variant | |
| 7 | 103705421 | 103706676 | 1255 | Or51a42 | SJL,SWR,TH | feature_truncation,3_prime_UTR_variant | |
| 7 | 103916051 | 103949854 | 33803 | Trim5,Trim12a | NZW,SJL | feature_truncation,coding_sequence_variant,5_prime_UTR_variant,intron_variant,feature_truncation,3_prime_UTR_variant,intron_variant | |
| 7 | 104105961 | 104153641 | 47680 | Trim30d,Trim30a | CE | stop_lost,feature_truncation,coding_sequence_variant,5_prime_UTR_variant,3_prime_UTR_variant,intron_variant,feature_truncation,5_prime_UTR_variant,intron_variant | FALSE |
| 7 | 105589360 | 105805589 | 216229 | Gvin1,Gvin2 | SMJ,SWR | stop_lost,feature_truncation,coding_sequence_variant,5_prime_UTR_variant,3_prime_UTR_variant,intron_variant,feature_truncation,5_prime_UTR_variant,intron_variant | |
| 7 | 106519056 | 106522231 | 3175 | Or2d3c | 129S1,AJ,AKR,BALB,BTBR,C3H,C57L,CBA,CE,DBA1J,DBA2J,ILNJ2,KK,MRL,NOD,NZB,NZO,NZW,PJ,PL,RF,RH2,SEA,SMJ,SWR | feature_truncation,3_prime_UTR_variant | |
| 7 | 106717743 | 106726092 | 8349 | Or2d2 | BALB,NZB,NZO | feature_truncation,3_prime_UTR_variant | |
| 7 | 107357231 | 107358831 | 1600 | Cyb5r2 | KK | feature_truncation,5_prime_UTR_variant | |
| 7 | 107436245 | 107458637 | 22392 | Or5p51 | CE | transcript_ablation | |
| 7 | 107765252 | 107775239 | 9987 | Or5p62 | NOD,SMJ,SWR | transcript_ablation | |
| 7 | 127331135 | 127337587 | 6452 | Ctf2 | AJ,TH | feature_truncation,5_prime_UTR_variant,intron_variant | |
| 7 | 135193862 | 135196479 | 2617 | Ptpre | BALB,SEA | feature_truncation,coding_sequence_variant,5_prime_UTR_variant,intron_variant | |
| 7 | 140194139 | 140200732 | 6593 | Or13a18 | NZW,PJ | feature_truncation,3_prime_UTR_variant | |
| 7 | 140273044 | 140276112 | 3068 | Or13a17 | NZW,PJ | feature_truncation,3_prime_UTR_variant | |
| 7 | 141224174 | 141225499 | 1325 | Muc6 | NZW | frameshift_variant,feature_truncation | |
| 7 | 141224929 | 141226464 | 1535 | Muc6 | ILNJ2 | frameshift_variant,feature_truncation | |
| 8 | 3995958 | 4002932 | 6974 | Cd209c | 129S1,NZB,NZO,NZW | feature_truncation,5_prime_UTR_variant | |
| 8 | 21171157 | 21179109 | 7952 | Potefam3b | MA/My | feature_truncation,coding_sequence_variant,intron_variant | FALSE |

| | | | | | | |
|---|---|---|---|---|---|---|
| 8 | 21748107 | 22122931 | 374824 | Defa26,Defa37,Defa32,Defa20,Defa36,Defa28,Defa29,Defa5,Defa3,Defa43,Defa42,Defa33,Defa27,Defa2 | BTBR,NOR,NZB,NZW | transcript_ablation |
| 8 | 21796427 | 22037852 | 241425 | Defa32,Defa20,Defa36,Defa29,Defa42,Defa33,Defa27,Defa2 | NZB | transcript_ablation |
| 8 | 21888729 | 21921018 | 32289 | Defa36 | BUB,CBA,DBA1J,DBA2J,FVB,MRL,NOR,PL,RH2 | transcript_ablation |
| 8 | 21925976 | 22037852 | 111876 | Defa32,Defa20,Defa42 | DBA1J,NOR,RH2,SWR,TH | transcript_ablation |
| 8 | 22003546 | 22023356 | 19810 | Defa32 | C3H | transcript_ablation |
| 8 | 22071558 | 22077184 | 5626 | Defa28 | C57L,PJ | transcript_ablation |
| 8 | 43981634 | 44021497 | 39863 | Adam26a | NZB,NZO,NZW | stop_lost,feature_truncation,coding_sequence_variant,3_prime_UTR_variant |
| 8 | 72870306 | 72871835 | 1529 | Or1ab2 | C58J,PJ,SEA | feature_truncation,3_prime_UTR_variant |
| 8 | 73057910 | 73059159 | 1249 | Hmgn2-ps | 129S1,AKR,BALB,BUB,C3H,C57L,C58J,CBA,CE,DBA1J,DBA2J,FVB,ILNJ2,KK,MA/My,MRL,NZB,NZO,NZW,PJ,PL,RF,SEA,SJL,SMJ,SWR,TH | transcript_ablation |
| 8 | 105579697 | 105613844 | 34147 | Ces2c | KK | stop_lost,feature_truncation,coding_sequence_variant,3_prime_UTR_variant,intron_variant |
| 8 | 117747892 | 117749935 | 2043 | Pkd1l2 | SWR | feature_truncation,coding_sequence_variant,intron_variant |
| 9 | 8628262 | 8633605 | 5343 | Trpc6 | 129S1,AKR,BTBR,C57L,MRL,NZB,NZW,PL,RF | feature_truncation,3_prime_UTR_variant |
| 9 | 14540233 | 14559307 | 19074 | Amotl1 | NZO | feature_truncation,coding_sequence_variant,5_prime_UTR_variant,intron_variant |
| 9 | 37764761 | 37766052 | 1291 | Or8b9 | CBA,CE,MRL | feature_truncation,5_prime_UTR_variant,intron_variant |
| 9 | 38024168 | 38054263 | 30095 | Or8b40,Or8b41 | KK,NZB,NZW,PJ,SEA | transcript_ablation,feature_truncation,5_prime_UTR_variant,intron_variant |
| 9 | 38090878 | 38106922 | 16044 | Or8c13 | MA/My | transcript_ablation |
| 9 | 38411089 | 38417577 | 6488 | Or8b44 | 129S1,AJ,AKR,BALB,BTBR,BUB,C3H,CBA,CE,DBA1J,DBA2J,FVB,ILNJ2,KK,MA/My,MRL,NOD,NOR,NZB,NZW,PJ,PL,RF,RH2,SEA,SJL,SMJ,SWR,TH | feature_truncation,3_prime_UTR_variant |
| 9 | 38582877 | 38585657 | 2780 | Or8b3b | CBA,PL,SJL | transcript_ablation,stop_lost,feature_truncation,coding_sequence_variant,5_prime_UTR_variant,3_prime_UTR_variant,intron_variant |
| 9 | 39295852 | 39361330 | 65478 | Or8g33,Or8g32 | SJL | transcript_ablation |
| 9 | 39300641 | 39365213 | 64572 | Or8g33,Or8g32 | MRL | transcript_ablation,stop_lost,feature_truncation,coding_sequence_variant,5_prime_UTR_variant,3_prime_UTR_variant,intron_variant |
| 9 | 50963962 | 50966735 | 2773 | Layn | 129S1,C57L | stop_lost,feature_truncation,coding_sequence_variant,3_prime_UTR_variant,intron_variant |
| 9 | 57315270 | 57316753 | 1483 | Ppcdc | 129S1,C57L,CBA | stop_lost,feature_truncation,coding_sequence_variant,3_prime_UTR_variant,intron_variant |
| 9 | 83824595 | 83831922 | 7327 | Bckdhb | 129S1,AJ,AKR,BALB,BTBR,BUB,C3H,C57L,C58J,CBA,CE,DBA1J,DBA2J,FVB,ILNJ2,KK,LP,MRL,NOD,NOR,NZB,NZO,NZW,PJ,PL,RF,RH2,SEA,SJL,SMJ,SWR,TH | feature_truncation,5_prime_UTR_variant,intron_variant |
| 9 | 103330317 | 103336614 | 6297 | Bfsp2 | 129S1,CBA,FVB,NZB,NZW | feature_truncation,coding_sequence_variant,intron_variant |
| 9 | 105286553 | 105288170 | 1617 | Atp2c1 | 129S1,AJ,AKR,BALB,BTBR,BUB,C3H,C57L,CBA,CE,DBA1J,DBA2J,FVB,ILNJ2,KK,MRL,NOD,NOR,NZB,NZO,NZW,PJ,PL,RF,RH2,SEA,SJL,SMJ,SWR,TH | feature_truncation,3_prime_UTR_variant |
| X | 67864689 | 67868938 | 4249 | Styx-ps | AJ,BALB,BUB,C3H,C58J,ILNJ2,LP,MRL,NOD,NOR,NZB,NZO,NZW,PJ,PL,RH2,SEA,SJL,TH | transcript_ablation |
| X | 146235555 | 146236864 | 1309 | Vmn1r239-ps | NZB | transcript_ablation |
| X | 146955392 | 146965040 | 9648 | Gm15107 | FVB,MRL,NOD | feature_truncation,coding_sequence_variant,5_prime_UTR_variant,intron_variant |
| X | 147636439 | 147640761 | 4322 | Luzp4 | NZB | feature_truncation,coding_sequence_variant,5_prime_UTR_variant,intron_variant |
| X | 160006956 | 160011172 | 4216 | Scml2 | MRL | feature_truncation,coding_sequence_variant,intron_variant |

| CHR | START | END | SIZE | SYMBOL | STRAINS with VARIANT SV ALLELE | CONSEQUENCE |
|---|---|---|---|---|---|---|
| 1 | 4851851 | 4851919 | 68 | Mrpl15 | C3H,CAST,CBA,MRL,RF,SPRET,WSB | feature_truncation,3_prime_UTR_variant |
| 1 | 4980086 | 4980142 | 56 | Rgs20 | SPRET | feature_truncation,3_prime_UTR_variant |
| 1 | 9990991 | 9991192 | 201 | Mcmdc2 | SPRET | stop_lost,feature_truncation,coding_sequence_variant,3_prime_UTR_variant,intron_variant,feature_truncation,3_prime_UTR_variant |
| 1 | 10011298 | 10011481 | 183 | Mcmdc2 | SPRET | feature_truncation,3_prime_UTR_variant |
| 1 | 11369858 | 11369980 | 122 | Prex2 | SPRET | feature_truncation,3_prime_UTR_variant |
| 1 | 11669125 | 11669192 | 67 | A830018L16Rik | SPRET | feature_truncation,3_prime_UTR_variant |
| 1 | 13721098 | 13721166 | 68 | Lactb2 | AJ,AKR,BALB,C3H,CAST,CBA,DBA1J,DBA2J,FVB,ILNJ2,KK,LP,MRL,NOD,NZB,RF,RH2,SEA,SJL,SMJ,SPRET,WSB | feature_truncation,coding_sequence_variant,intron_variant |
| 1 | 15788129 | 15788181 | 52 | Kcnb2 | MOLF | feature_truncation,3_prime_UTR_variant |
| 1 | 24247155 | 24247255 | 100 | Col9a1 | SPRET | splice_acceptor_variant,coding_sequence_variant,intron_variant |
| 1 | 32284559 | 32284611 | 52 | Khdrbs2 | SPRET | feature_truncation,3_prime_UTR_variant |
| 1 | 32583596 | 32583769 | 173 | Semp2l1 | SPRET | feature_truncation,3_prime_UTR_variant |
| 1 | 34175229 | 34175547 | 318 | Dst | CAST | feature_truncation,3_prime_UTR_variant |
| 1 | 34484669 | 34484738 | 69 | Imp4 | RF | feature_truncation,3_prime_UTR_variant |
| 1 | 34487468 | 34487532 | 64 | Imp4 | CAST | feature_truncation,3_prime_UTR_variant |
| 1 | 37458653 | 37458747 | 94 | Coa5 | SPRET | feature_truncation,3_prime_UTR_variant |
| 1 | 37664302 | 37664375 | 73 | Cracdl | AJ,B10J,C3H,CE,MA/My,MOLF,MRL,NOD,SJL,SPRET,WSB | frameshift_variant,feature_truncation |
| 1 | 38833129 | 38833189 | 60 | Lonrf2 | 129S1,AJ,AKR,BTBR,BUB,C3H,C57L,CBA,CE,DBA1J,DBA2J,ILNJ2,KK,MRL,NOD,NOR,NZB,NZW,PJ,PL,RH2,SJL,SMJ,TH | feature_truncation,3_prime_UTR_variant |
| 1 | 38906385 | 38906580 | 195 | Chst10 | MOLF | feature_truncation,3_prime_UTR_variant |
| 1 | 39591037 | 39591249 | 212 | Rnf149 | SPRET | feature_truncation,3_prime_UTR_variant |
| 1 | 52697296 | 52697354 | 58 | Mfsd6 | C58J | feature_truncation,3_prime_UTR_variant |
| 1 | 52828264 | 52828422 | 158 | Inpp1 | SPRET | feature_truncation,3_prime_UTR_variant |
| 1 | 54435782 | 54436128 | 346 | Gtf3c3 | CAST,MOLF,SPRET | feature_truncation,3_prime_UTR_variant |
| 1 | 58484558 | 58484613 | 55 | Ppil3,Nif3l1 | MOLF,SPRET | feature_truncation,5_prime_UTR_variant |
| 1 | 58796296 | 58796495 | 199 | Cflar | SPRET | feature_truncation,3_prime_UTR_variant |
| 1 | 59045610 | 59045661 | 51 | C2cd6 | MA/My | feature_truncation,inframe_deletion |
| 1 | 59391564 | 59391775 | 211 | Cdk15 | CAST,SPRET | feature_truncation,3_prime_UTR_variant |
| 1 | 59754392 | 59754594 | 202 | Nop58 | SPRET | feature_truncation,3_prime_UTR_variant |
| 1 | 59756069 | 59756169 | 100 | Nop58 | CE,LP | feature_truncation,3_prime_UTR_variant |
| 1 | 60023658 | 60023710 | 52 | Fam117b | CE,LP,MOLF | feature_truncation,3_prime_UTR_variant |
| 1 | 60113780 | 60113977 | 197 | Wdr12 | CAST,SPRET | feature_truncation,3_prime_UTR_variant |
| 1 | 66692401 | 66692487 | 86 | Unc80 | SPRET | feature_truncation,coding_sequence_variant,intron_variant |
| 1 | 71068087 | 71068370 | 283 | Bard1 | SPRET | feature_truncation,3_prime_UTR_variant |
| 1 | 74199029 | 74199222 | 193 | Cxcr2 | 129S1,AJ,BTBR,C3H,C57L,CAST,CBA,MOLF,MRL,NOD,NZB,NZO,PJ,RH2,SEA,SMJ,SPRET,SWR,WSB | feature_truncation,3_prime_UTR_variant |
| 1 | 75285758 | 75285839 | 81 | Dnpep | SPRET | feature_truncation,3_prime_UTR_variant |
| 1 | 78648418 | 78648480 | 62 | Utp14b | CAST,LP | feature_truncation,3_prime_UTR_variant |
| 1 | 79584844 | 79585161 | 317 | Ap1s3 | 129S1,AJ,BALB,BTBR,BUB,C3H,C57L,C58J,CAST,CBA,CE,DBA1J,DBA2J,FVB,ILNJ2,LP,MOLF,MRL,NOD,NZB,NZO,PJ,RF,RH2,SEA,SJL,SMJ,SPRET,SWR,TH,WSB | feature_truncation,3_prime_UTR_variant |
| 1 | 79772438 | 79772518 | 80 | Serpine2 | SPRET | feature_truncation,3_prime_UTR_variant |
| 1 | 82938077 | 82938140 | 63 | Gm47955 | CAST,MOLF,PJ,RF,SMJ,SPRET | feature_truncation,inframe_deletion |
| 1 | 82978485 | 82978553 | 68 | Gm47959 | AKR,BUB,CAST,CBA,MOLF,MRL,NOD,PJ,PL,RF,SMJ,SPRET,WSB | frameshift_variant,feature_truncation |
| 1 | 83036905 | 83036959 | 54 | A030005K14Rik | MOLF,RF | feature_truncation,inframe_deletion |
| 1 | 83038865 | 83038922 | 57 | Krtap28-13 | PJ,SMJ | feature_truncation,inframe_deletion |
| 1 | 85191260 | 85191510 | 250 | Gm7609 | AKR,BUB,CBA,MRL,NOD,PJ,PL,SMJ | stop_lost,feature_truncation,coding_sequence_variant,3_prime_UTR_variant |
| 1 | 86205774 | 86205847 | 73 | Armc9 | BUB,CAST,MOLF,NZB,RF,WSB | feature_truncation,3_prime_UTR_variant |
| 1 | 86354309 | 86354385 | 76 | Tex44 | 129S1,AJ,BTBR,C3H,C57L,C58J,CAST,CE,DBA1J,DBA2J,FVB,ILNJ2,KK,NZB,NZO,NZW,RH2,SJL,SMJ,SWR,TH,WSB | frameshift_variant,feature_truncation |
| 1 | 86535314 | 86535380 | 66 | Cops7b | BUB | feature_truncation,3_prime_UTR_variant |
| 1 | 88430869 | 88431127 | 258 | Glrp1 | CAST,MOLF,SPRET,WSB | feature_truncation,inframe_deletion |
| 1 | 89878533 | 89879016 | 483 | Asb18 | CAST,MOLF | feature_truncation,3_prime_UTR_variant |
| 1 | 90703524 | 90703587 | 63 | Col6a3 | 129S1,AKR,BUB,C3H,C57L,CBA,FVB,KK,LP,MA/My,MOLF,MRL,NZB,NZO,NZW,PJ,PL,RF,SJL,SMJ,SPRET,SWR,TH,WSB | feature_truncation,inframe_deletion |
| 1 | 91214464 | 91214573 | 109 | Ube2f | SPRET | feature_truncation,3_prime_UTR_variant |
| 1 | 92404191 | 92404269 | 78 | Or6b2 | SPRET | feature_truncation,3_prime_UTR_variant |
| 1 | 92903095 | 92903538 | 443 | Gpr35 | SPRET | feature_truncation,5_prime_UTR_variant,intron_variant |
| 1 | 93717642 | 93717948 | 306 | Atg4b | SPRET | feature_truncation,3_prime_UTR_variant |
| 1 | 93917816 | 93918286 | 470 | Gal3st2c | CAST,SPRET | feature_truncation,5_prime_UTR_variant |
| 1 | 95549060 | 95549347 | 287 | St8sia4 | SPRET | feature_truncation,3_prime_UTR_variant |
| 1 | 97552793 | 97552927 | 134 | Macir | CAST,MRL,PL,SPRET | feature_truncation,3_prime_UTR_variant |
| 1 | 97560516 | 97560702 | 186 | Macir | MOLF,SPRET | feature_truncation,3_prime_UTR_variant |
| 1 | 97570705 | 97570768 | 63 | Macir | CAST,MOLF,MRL,PL,SPRET | feature_truncation,3_prime_UTR_variant |
| 1 | 106650934 | 106650997 | 63 | Kdsr | MOLF | feature_truncation,3_prime_UTR_variant |
| 1 | 117728949 | 117729117 | 168 | B020011L13Rik | MOLF | feature_truncation,inframe_deletion |
| 1 | 118580078 | 118580249 | 171 | Tfcp2l1 | CAST,MOLF,SPRET | splice_donor_variant,splice_donor_5th_base_variant,coding_sequence_variant,intron_variant |
| 1 | 120153189 | 120153279 | 90 | Steap3 | SPRET | feature_truncation,3_prime_UTR_variant |
| 1 | 121432868 | 121433033 | 165 | Ccdc93 | BALB,BTBR,PL,RF,SEA,SMJ | feature_truncation,3_prime_UTR_variant |
| 1 | 127415512 | 127415564 | 52 | Mgat5 | CAST | feature_truncation,3_prime_UTR_variant |
| 1 | 130625118 | 130625188 | 70 | Pfkfb2 | SPRET | feature_truncation,3_prime_UTR_variant |
| 1 | 131106829 | 131106902 | 73 | Eif2d | 129S1,BALB,C57L,CAST,DBA1J,DBA2J,FVB,ILNJ2,KK,MA/My,MOLF,NZO,NZW,SEA,SJL,SPRET,SWR,TH | feature_truncation,3_prime_UTR_variant |
| 1 | 131697925 | 131697984 | 59 | Slc26a9 | PL | feature_truncation,3_prime_UTR_variant |
| 1 | 131955146 | 131955286 | 140 | Mfsd4a | SPRET | feature_truncation,3_prime_UTR_variant,intron_variant,feature_truncation,3_prime_UTR_variant |

| | | | | | | |
|---|---|---|---|---|---|---|
| 1 | 134436505 | 134436706 | 201 | Rabif | 129S1,AJ,AKR,BTBR,BUB,C3H,CAST,CBA,CE,DBA1J,DBA2J,FVB,ILNJ2,KK,MA/My,MRL,NOD,NOR,NZB,NZO,PJ,PL,RF,RH2,SJL,SMJ,SPRET,SWR,TH | feature_truncation,3_prime_UTR_variant |
| 1 | 135224344 | 135224458 | 114 | Timm17a | SPRET | feature_truncation,3_prime_UTR_variant |
| 1 | 150330660 | 150330876 | 216 | Prg4 | AJ,AKR,BTBR,BUB,C3H,C57L,CBA,CE,DBA1J,DBA2J,ILNJ2,MOLF,MRL,PL,RF,RH2,SMJ | feature_truncation,inframe_deletion |
| 1 | 150331112 | 150331256 | 144 | Prg4 | SPRET | feature_truncation,inframe_deletion |
| 1 | 153752072 | 153752275 | 203 | Teddm1b | BUB,C57L,CAST,DBA1J,DBA2J,FVB,MA/My,RF,SJL,SPRET,SWR,TH,WSB | feature_truncation,3_prime_UTR_variant |
| 1 | 154739345 | 154740314 | 969 | Cacna1e | SPRET | feature_truncation,coding_sequence_variant,5_prime_UTR_variant,intron_variant |
| 1 | 156193594 | 156194101 | 507 | Rpl35rt | 129S1,AJ,AKR,BALB,BTBR,BUB,C3H,C57L,C58J,CAST,CBA,CE,DBA1J,DBA2J,FVB,ILNJ2,KK,LP,MA/My,MOLF,MRL,NOD,NOR,NZB,NZO,NZW,PJ,PL,RF,RH2,SEA,SPRET,SWR,TH,WSB | transcript_ablation |
| 1 | 156476172 | 156476230 | 58 | Abl2 | SPRET | feature_truncation,3_prime_UTR_variant |
| 1 | 156966252 | 156966353 | 101 | Rasal2 | CAST | feature_truncation,3_prime_UTR_variant |
| 1 | 159753853 | 159753920 | 67 | Tnr | BALB,C58J,DBA1J,DBA2J,LP,MOLF,SPRET | feature_truncation,3_prime_UTR_variant |
| 1 | 160995589 | 160995711 | 122 | Ankrd45 | 129S1,AJ,BALB,BUB,C3H,C57L,C58J,CBA,CE,DBA1J,DBA2J,ILNJ2,LP,MOLF,NZW,PJ,RH2,SEA,SMJ,SPRET | feature_truncation,3_prime_UTR_variant |
| 1 | 161245883 | 161246212 | 329 | Tnfsf4 | SPRET | feature_truncation,3_prime_UTR_variant |
| 1 | 163213419 | 163213514 | 95 | Gorab | SPRET | feature_truncation,3_prime_UTR_variant |
| 1 | 163976790 | 163976866 | 76 | Selp | DBA1J,DBA2J,ILNJ2,MOLF | feature_truncation,3_prime_UTR_variant |
| 1 | 164175683 | 164175880 | 197 | Nme7 | BUB,FVB,NOD,NOR,PJ,RF,SEA,SJL,SMJ,SPRET,SWR,WSB | stop_lost,feature_truncation,coding_sequence_variant,3_prime_UTR_variant,intron_variant |
| 1 | 164965771 | 164965839 | 68 | Tbx19 | SPRET | feature_truncation,3_prime_UTR_variant |
| 1 | 165004861 | 165004921 | 60 | Sft2d2 | CAST,DBA1J,DBA2J,ILNJ2,MOLF,SPRET | feature_truncation,3_prime_UTR_variant |
| 1 | 165156378 | 165156682 | 304 | Dcaf6 | CAST,SPRET,WSB | feature_truncation,3_prime_UTR_variant |
| 1 | 166206771 | 166206851 | 80 | Tada1 | SPRET | feature_truncation,5_prime_UTR_variant |
| 1 | 166223125 | 166223190 | 65 | Pogk | CAST,SPRET | feature_truncation,3_prime_UTR_variant |
| 1 | 167426126 | 167426300 | 174 | Rxrg | BUB,SWR,TH | feature_truncation,5_prime_UTR_variant,feature_truncation,5_prime_UTR_variant,intron_variant |
| 1 | 171043632 | 171043696 | 64 | Tomm40l | SPRET | feature_truncation,3_prime_UTR_variant |
| 1 | 171776994 | 171777059 | 65 | Slamf6 | SPRET | feature_truncation,3_prime_UTR_variant |
| 1 | 171831327 | 171831453 | 126 | Vangl2 | 129S1,AJ,AKR,BALB,BUB,C3H,C57L,CBA,CE,DBA1J,DBA2J,FVB,KK,MRL,NOD,NOR,NZB,NZO,NZW,PJ,PL,RF,RH2,SEA,SJL,SMJ,SPRET,SWR,TH,WSB | feature_truncation,3_prime_UTR_variant |
| 1 | 172101601 | 172101653 | 52 | Atp1a2 | PL | feature_truncation,3_prime_UTR_variant |
| 1 | 172198491 | 172198547 | 56 | Kcnj10 | 129S1,AJ,AKR,BALB,BUB,C3H,C57L,C58J,CBA,CE,DBA1J,DBA2J,FVB,KK,MRL,NOD,NOR,NZB,NZO,NZW,PJ,PL,RF,RH2,SEA,SJL,SMJ,SWR,TH,WSB | feature_truncation,3_prime_UTR_variant |
| 1 | 172206793 | 172207134 | 341 | Pigm | AKR,RF | feature_truncation,3_prime_UTR_variant |
| 1 | 172207321 | 172207790 | 469 | Pigm | CE,NZB,PL,SMJ,TH | feature_truncation,3_prime_UTR_variant |
| 1 | 172527106 | 172527166 | 60 | Crp | 129S1,AJ,AKR,BALB,BTBR,BUB,C3H,C57L,C58J,CAST,CBA,CE,DBA1J,DBA2J,FVB,ILNJ2,KK,MRL,NOD,NOR,NZB,NZO,NZW,PJ,PL,RF,RH2,SEA,SJL,SMJ,SPRET,SWR,TH,WSB | feature_truncation,3_prime_UTR_variant |
| 1 | 173308092 | 173308518 | 426 | Ifi206 | SPRET | feature_truncation,inframe_deletion |
| 1 | 173562682 | 173562745 | 63 | Ifi207 | AJ,FVB,PL,RH2,TH | feature_truncation,inframe_deletion |
| 1 | 173870781 | 173870868 | 87 | Or10aa1 | AKR,BTBR,BUB,C58J,CBA,CE,DBA1J,DBA2J,ILNJ2,KK,NZB,NZO,PJ,PL,RF,SEA,SMJ,SPRET,SWR,TH | feature_truncation,3_prime_UTR_variant |
| 1 | 173940951 | 173941185 | 234 | Or6k6 | SPRET | feature_truncation,3_prime_UTR_variant |
| 1 | 173988263 | 173988555 | 292 | Or6k2 | 129S1,AJ,AKR,BALB,BTBR,BUB,C3H,C57L,C58J,CAST,CBA,CE,DBA1J,DBA2J,FVB,ILNJ2,KK,MOLF,MRL,NOD,NOR,NZB,NZO,PJ,PL,RF,RH2,SEA,SJL,SMJ,SPRET,SWR,TH,WSB | feature_truncation,3_prime_UTR_variant |
| 1 | 174041803 | 174041864 | 61 | Spta1 | 129S1,AJ,AKR,BALB,BTBR,BUB,C3H,C57L,C58J,CBA,CE,DBA1J,DBA2J,FVB,ILNJ2,KK,MRL,NOD,NOR,NZB,NZO,NZW,PJ,PL,RF,RH2,SEA,SJL,SMJ,SPRET,SWR,TH,WSB | feature_truncation,coding_sequence_variant,intron_variant |
| 1 | 175488632 | 175488700 | 68 | Kmo | 129S1,BALB,BTBR,C3H,C57L,CAST,MOLF,MRL,NOD,NOR,NZO,NZW,PL,SMJ,SPRET,TH | feature_truncation,3_prime_UTR_variant |
| 1 | 175739009 | 175739590 | 581 | Exo1 | SPRET | feature_truncation,3_prime_UTR_variant |
| 1 | 178081238 | 178081975 | 737 | Desi2 | SPRET | feature_truncation,3_prime_UTR_variant |
| 1 | 179335900 | 179335960 | 60 | Smyd3 | AKR,CAST,MOLF | feature_truncation,3_prime_UTR_variant |
| 1 | 179420411 | 179420666 | 255 | Cnst | SPRET | feature_truncation,coding_sequence_variant,intron_variant |
| 1 | 181337202 | 181337293 | 91 | Ccdc121rt1 | MOLF,TH | feature_truncation,3_prime_UTR_variant |
| 1 | 181729866 | 181729950 | 84 | Enah | NZO,SPRET | feature_truncation,3_prime_UTR_variant |
| 1 | 185089442 | 185089594 | 152 | Bpnt1 | SPRET | feature_truncation,3_prime_UTR_variant |
| 1 | 189606610 | 189606683 | 73 | Ptpn14 | MOLF | feature_truncation,3_prime_UTR_variant |
| 1 | 190815834 | 190815887 | 53 | Nsl1 | 129S1,BUB,CBA,DBA1J,DBA2J,ILNJ2,KK,MA/My,MOLF,MRL,NOD,NOR,NZO,NZW,PL,SWR | feature_truncation,3_prime_UTR_variant |
| 1 | 192195949 | 192196013 | 64 | Hhat | CAST,MOLF | feature_truncation,3_prime_UTR_variant |
| 1 | 194720240 | 194720307 | 67 | Cd46 | NOD,NOR,SWR | feature_truncation,3_prime_UTR_variant |
| 1 | 194821674 | 194821725 | 51 | Cr2 | MOLF,NZO,NZW,RF | feature_truncation,3_prime_UTR_variant |
| 10 | 4402353 | 4402421 | 68 | Armt1 | CAST,MOLF,SPRET | feature_truncation,3_prime_UTR_variant |
| 10 | 6838588 | 6838642 | 54 | Ipcef1 | SPRET | feature_truncation,3_prime_UTR_variant |
| 10 | 7396513 | 7396579 | 66 | Ulbp1 | MOLF | feature_truncation,coding_sequence_variant,intron_variant |
| 10 | 11084704 | 11084867 | 163 | Shprh | SPRET | feature_truncation,3_prime_UTR_variant |
| 10 | 11333299 | 11333355 | 56 | Epm2a | SPRET | feature_truncation,3_prime_UTR_variant |
| 10 | 12815033 | 12815109 | 76 | Stx11 | SPRET | feature_truncation,3_prime_UTR_variant |
| 10 | 13006405 | 13006463 | 58 | Plagl1 | CAST,DBA1J,DBA2J,MOLF,SPRET | feature_truncation,3_prime_UTR_variant |
| 10 | 24065206 | 24065308 | 102 | Stx7 | SPRET | feature_truncation,3_prime_UTR_variant |
| 10 | 29106242 | 29106299 | 57 | 9330159F19Rik | CAST,MOLF | feature_truncation,3_prime_UTR_variant |
| 10 | 33470537 | 33470617 | 80 | Clvs2 | KK,WSB | feature_truncation,3_prime_UTR_variant |

| | | | | | | |
|---|---|---|---|---|---|---|
| 10 | 33655256 | 33655642 | 386 | Sult3a2 | SPRET | feature_truncation,coding_sequence_variant,intron_variant |
| 10 | 40130352 | 40130560 | 208 | Gtf3c6 | CAST,MOLF,SPRET | splice_acceptor_variant,splice_donor_variant,splice_donor_5th_base_variant,coding_sequence_variant,intron_variant |
| 10 | 40708502 | 40708722 | 220 | Cdc40 | SPRET | feature_truncation,3_prime_UTR_variant |
| 10 | 41310040 | 41310259 | 219 | Ak9 | MOLF,SPRET | feature_truncation,3_prime_UTR_variant |
| 10 | 43400069 | 43400639 | 570 | Pam16l | 129S1,BTBR,C57L,CAST,ILNJ2,KK,MOLF,SPRET,WSB | stop_lost,feature_truncation,coding_sequence_variant,5_prime_UTR_variant,3_prime_UTR_variant,stop_lost,feature_truncation,coding_sequence_variant,5_prime_UTR_variant,3_prime_UTR variant,intron variant |
| 10 | 45043240 | 45043735 | 495 | Prep | KK,WSB | feature_truncation,3_prime_UTR_variant |
| 10 | 49454256 | 49454473 | 217 | Grik2 | BALB,BTBR,CAST,KK,MOLF,NZW,PJ,SEA,SPRET | feature_truncation,coding_sequence_variant,intron_variant |
| 10 | 49454256 | 49454468 | 212 | Grik2 | BALB,BTBR,CAST,KK,MOLF,NZW,PJ,SEA,SPRET | splice_acceptor_variant,coding_sequence_variant,intron_variant |
| 10 | 50494545 | 50494604 | 59 | Ascc3 | SPRET | feature_truncation,3_prime_UTR_variant |
| 10 | 53152663 | 53152750 | 87 | Cep85l | CE,SPRET,WSB | feature_truncation,3_prime_UTR_variant |
| 10 | 58972891 | 58972967 | 76 | Sh3rf3 | CAST | feature_truncation,3_prime_UTR_variant |
| 10 | 60548867 | 60548925 | 58 | Slc29a3 | SPRET | feature_truncation,3_prime_UTR_variant |
| 10 | 62849148 | 62849214 | 66 | Rufy2 | MOLF,RF | feature_truncation,3_prime_UTR_variant |
| 10 | 62913117 | 62913323 | 206 | Pbld1 | CAST,CE,MOLF,RF,SPRET | feature_truncation,3_prime_UTR_variant |
| 10 | 66848254 | 66848309 | 55 | Reep3 | ILNJ2,SMJ,SPRET | feature_truncation,3_prime_UTR_variant |
| 10 | 67849562 | 67849776 | 214 | Rtkn2 | SPRET | splice_acceptor_variant,splice_donor_variant,splice_donor_5th_base_variant,coding_sequence_variant,intron_variant |
| 10 | 71218545 | 71218643 | 98 | Ipmk | MOLF | feature_truncation,3_prime_UTR_variant |
| 10 | 75386588 | 75386684 | 96 | Lrrc75b | CAST,NZW,SPRET | feature_truncation,3_prime_UTR_variant |
| 10 | 75667476 | 75667663 | 187 | Gstt2 | SPRET | feature_truncation,3_prime_UTR_variant |
| 10 | 76295602 | 76295828 | 226 | Ybey | SPRET | feature_truncation,3_prime_UTR_variant |
| 10 | 76297020 | 76297093 | 73 | Ybey | MOLF,SPRET | feature_truncation,3_prime_UTR_variant |
| 10 | 76881787 | 76882153 | 366 | Slc19a1 | BUB | feature_truncation,3_prime_UTR_variant |
| 10 | 77650634 | 77650694 | 60 | Gm49918 | SPRET | feature_truncation,inframe_deletion |
| 10 | 78548574 | 78548629 | 55 | Casp14 | SPRET | feature_truncation,3_prime_UTR_variant |
| 10 | 78716400 | 78716878 | 478 | Or7a39 | TH | feature_truncation,3_prime_UTR_variant |
| 10 | 78855227 | 78855283 | 56 | Or7a35 | SPRET | feature_truncation,3_prime_UTR_variant |
| 10 | 79730628 | 79730728 | 100 | Med16 | MOLF,SPRET | feature_truncation,3_prime_UTR_variant |
| 10 | 80737014 | 80737160 | 146 | Lmnb2 | SPRET | feature_truncation,3_prime_UTR_variant |
| 10 | 81171660 | 81171744 | 84 | Gipc3 | RF,SPRET | feature_truncation,3_prime_UTR_variant |
| 10 | 81500343 | 81500544 | 201 | Gm10778 | CAST,MOLF,SPRET | feature_truncation,3_prime_UTR_variant |
| 10 | 82124464 | 82124590 | 126 | 4932415D10Rik | SPRET | feature_truncation,inframe_deletion |
| 10 | 82213337 | 82213421 | 84 | Gm4924 | MOLF,SPRET | feature_truncation,inframe_deletion |
| 10 | 83227179 | 83227256 | 77 | Nopchap1 | C58J,LP | feature_truncation,3_prime_UTR_variant |
| 10 | 83254239 | 83254314 | 75 | Nopchap1 | BUB,SMJ | feature_truncation,3_prime_UTR_variant |
| 10 | 83295351 | 83295453 | 102 | Nopchap1 | BUB,SMJ | feature_truncation,3_prime_UTR_variant |
| 10 | 83322269 | 83322345 | 76 | Nopchap1 | BUB,CAST,MOLF,SMJ,SPRET | feature_truncation,3_prime_UTR_variant |
| 10 | 83382759 | 83382967 | 208 | Washc4 | SPRET | feature_truncation,coding_sequence_variant,intron_variant |
| 10 | 83431733 | 83431936 | 203 | Washc4 | BUB,CAST,MOLF,SPRET,WSB | feature_truncation,3_prime_UTR_variant |
| 10 | 85794925 | 85795427 | 502 | Bpifc | SPRET | feature_truncation,3_prime_UTR_variant |
| 10 | 85887211 | 85887292 | 81 | Fbxo7 | SPRET | feature_truncation,3_prime_UTR_variant |
| 10 | 88322891 | 88323108 | 217 | Chpt1 | SPRET | stop_lost,feature_truncation,coding_sequence_variant,3_prime_UTR_variant |
| 10 | 89245617 | 89245824 | 207 | Gas2l3 | AJ,AKR,BALB,BTBR,BUB,C3H,C58J,CAST,CBA,CE,FVB,ILNJ2,KK,LP,MOLF,MRL,NOD,NOR,NZB,NZO,NZW,PJ,PL,RF,RH2,SEA,SPRET,SWR,TH,WSB | feature_truncation,3_prime_UTR_variant |
| 10 | 89475032 | 89475092 | 60 | Scyl2 | AJ,AKR,BALB,CE,FVB,ILNJ2,MRL,NOD,NOR,NZO,PJ,PL,RF,RH2,SEA | feature_truncation,3_prime_UTR_variant |
| 10 | 93353273 | 93353366 | 93 | Hal | 129S1,AJ,AKR,BALB,BTBR,BUB,C3H,C57L,C58J,CBA,CE,FVB,ILNJ2,KK,LP,MA/My,NOD,NOR,PJ,RF,RH2,SEA,SJL,SMJ,SPRET,SWR,TH | feature_truncation,3_prime_UTR_variant |
| 10 | 93419675 | 93419739 | 64 | Ccdc38 | SPRET,WSB | feature_truncation,3_prime_UTR_variant |
| 10 | 93688439 | 93688583 | 144 | Usp44 | 129S1,AJ,AKR,BALB,BTBR,BUB,C3H,C57L,C58J,CAST,CBA,CE,DBA1J,DBA2J,FVB,ILNJ2,KK,LP,MA/My,MRL,NOD,NOR,NZO,NZW,PJ,PL,RF,RH2,SEA,SJL,SMJ,SPRET,SWR,TH,WSB | feature_truncation,coding_sequence_variant,intron_variant |
| 10 | 93803099 | 93803159 | 60 | Vezt | CAST,SPRET | feature_truncation,3_prime_UTR_variant |
| 10 | 116119512 | 116119617 | 105 | Ptprb | SPRET | feature_truncation,inframe_deletion |
| 10 | 117569715 | 117569769 | 54 | Slc35e3 | MOLF,SMJ,SPRET | feature_truncation,3_prime_UTR_variant |
| 10 | 120617715 | 120617779 | 64 | Msrb3 | CAST,MOLF,WSB | feature_truncation,3_prime_UTR_variant |
| 10 | 121916392 | 121916593 | 201 | Rxylt1 | SPRET | feature_truncation,3_prime_UTR_variant |
| 10 | 122779223 | 122779788 | 565 | Ppm1h | AKR,C3H,CBA,DBA2J,RF,SJL,SWR | feature_truncation,3_prime_UTR_variant |
| 10 | 122828302 | 122828409 | 107 | Mon2 | BTBR,C57L,CAST,KK,MOLF,NZW,SPRET,TH,WSB | feature_truncation,3_prime_UTR_variant |
| 10 | 126835212 | 126835353 | 141 | Ctdsp2 | CAST,MOLF,SPRET | feature_truncation,3_prime_UTR_variant |
| 10 | 126846496 | 126846579 | 83 | Avil | CAST,CE,FVB,MOLF,NOD,NOR,SPRET | feature_truncation,coding_sequence_variant,intron_variant |
| 10 | 126868014 | 126868210 | 196 | Eef1akmt3 | CAST,SPRET,WSB | feature_truncation,3_prime_UTR_variant |
| 10 | 127494015 | 127494136 | 121 | Stat6 | CAST,SPRET | feature_truncation,coding_sequence_variant,intron_variant |
| 10 | 127582641 | 127582766 | 125 | Zbtb39 | CE,WSB | feature_truncation,3_prime_UTR_variant |
| 10 | 127651695 | 127651754 | 59 | Rdh16 | SPRET | feature_truncation,3_prime_UTR_variant |
| 10 | 127719977 | 127720361 | 384 | Rdh7 | SPRET | feature_truncation,3_prime_UTR_variant |
| 10 | 128126583 | 128126799 | 216 | Stat2 | CAST,SPRET | feature_truncation,inframe_deletion |
| 10 | 128635079 | 128635218 | 139 | Mmp19 | SPRET | feature_truncation,3_prime_UTR_variant |
| 10 | 128945525 | 128945924 | 399 | Or6c2b | BUB,C3H,CAST,CBA,CE,KK,MOLF,RF,SMJ,SPRET,WSB | feature_truncation,3_prime_UTR_variant |
| 10 | 129220072 | 129220232 | 160 | Or6c208 | SPRET | feature_truncation,5_prime_UTR_variant |
| 10 | 129311798 | 129311858 | 60 | Or6c3 | BUB,KK,MOLF,SMJ | feature_truncation,3_prime_UTR_variant |
| 10 | 130217036 | 130217090 | 54 | Vmn2r84 | AKR,MRL,SJL,SPRET,SWR | feature_truncation,3_prime_UTR_variant |
| 11 | 3929665 | 3930007 | 342 | Pes1 | CAST,SPRET | feature_truncation,3_prime_UTR_variant |

| | | | | | | |
|---|---|---|---|---|---|---|
| 11 | 4172015 | 4172068 | 53 | Castor1 | CAST,NZB | feature_truncation,3_prime_UTR_variant |
| 11 | 4890448 | 4890520 | 72 | Nefh | AJ,BUB,NOD,PL | feature_truncation,inframe_deletion |
| 11 | 5040420 | 5040704 | 284 | Ewsr1 | BALB,BTBR,ILNJ2,MA/My,PJ,SEA,TH | feature_truncation,3_prime_UTR_variant |
| 11 | 6221315 | 6221382 | 67 | Tmed4 | SPRET | feature_truncation,3_prime_UTR_variant |
| 11 | 7120088 | 7120154 | 66 | Adcy1 | SPRET | feature_truncation,3_prime_UTR_variant |
| 11 | 9199731 | 9199838 | 107 | Abca13 | BUB,FVB,KK,NZB,NZO,SJL,SPRET,SWR | splice_donor_variant,splice_donor_5th_base_variant,coding_sequence_variant,intron_variant |
| 11 | 11848112 | 11848178 | 66 | Ddc | WSB | feature_truncation,5_prime_UTR_variant |
| 11 | 17889094 | 17889185 | 91 | Etaa1 | 129S1,AJ,AKR,BALB,BTBR,BUB,C3H,C57L,C58J,CE,DBA1J,DBA2J,FVB,ILNJ2,KK,MA/My,MRL,NOD,NOR,NZB,NZO,NZW,PJ,PL,RF,SEA,SJL,SWR,TH,WSB | feature_truncation,3_prime_UTR_variant |
| 11 | 22469179 | 22470091 | 912 | Tmem17 | BALB,C57L,NOR,RH2,SEA | feature_truncation,3_prime_UTR_variant |
| 11 | 29692939 | 29693017 | 78 | Rtn4 | SPRET | feature_truncation,3_prime_UTR_variant |
| 11 | 30904145 | 30904202 | 57 | Erlec1 | AJ,AKR,BUB,CBA,CE,DBA1J,DBA2J,FVB,ILNJ2,NOD,PJ,PL,RF,SEA,SJL,SMJ,SWR,WSB | feature_truncation,5_prime_UTR_variant |
| 11 | 33422676 | 33422733 | 57 | Ranbp17 | CAST | feature_truncation,3_prime_UTR_variant |
| 11 | 34039920 | 34039978 | 58 | Lcp2 | CAST,ILNJ2,KK,MA/My,MOLF,NOD,NZW,PL,RF,SPRET | feature_truncation,3_prime_UTR_variant |
| 11 | 34176803 | 34177041 | 238 | Dock2 | SPRET | feature_truncation,3_prime_UTR_variant |
| 11 | 40592461 | 40592805 | 344 | Hmmr | B10D2,B10J,BTBR,C58J,CAST,LP,MA/My,MOLF | feature_truncation,3_prime_UTR_variant |
| 11 | 44444646 | 44444702 | 56 | Rnf145 | SPRET | feature_truncation,3_prime_UTR_variant |
| 11 | 45942179 | 45943177 | 998 | Sft2d1rt | 129S1,AJ,BALB,BTBR,C3H,C57L,CAST,CBA,CE,DBA1J,DBA2J,ILNJ2,KK,MOLF,MRL,NOD,NZB,NZW,PJ,PL,RH2,SEA,SJL,SMJ,SPRET | transcript_ablation |
| 11 | 46297545 | 46297608 | 63 | Garin3 | B10D2,B10J,C58J,NZO,WSB | feature_truncation,inframe_deletion |
| 11 | 46669383 | 46669515 | 132 | Havcr1 | NZB,WSB | stop_lost,feature_truncation,coding_sequence_variant,3_prime_UTR_variant |
| 11 | 49353572 | 49353642 | 70 | Or2y15 | CAST,SPRET | feature_truncation,3_prime_UTR_variant |
| 11 | 49456578 | 49457042 | 464 | Or2y10 | MOLF,SPRET | feature_truncation,3_prime_UTR_variant |
| 11 | 50918795 | 50919250 | 455 | Or1x2 | B10D2,B10J,DBA2J,NOR,PJ,SEA | feature_truncation,3_prime_UTR_variant |
| 11 | 50919433 | 50919513 | 80 | Or1x2 | B10D2,B10J,NOR,TH | feature_truncation,3_prime_UTR_variant |
| 11 | 53492677 | 53492822 | 145 | Kif3a | B10D2,B10J,BUB,C58J,CAST,CE,KK,MA/My,MOLF,NOD,NOR,NZW,PL,SJL,SPRET,SWR,TH | feature_truncation,3_prime_UTR_variant |
| 11 | 58352954 | 58353030 | 76 | Or2z2 | 129S1,AJ,AKR,BALB,BTBR,C3H,C57L,CBA,CE,DBA1J,DBA2J,FVB,ILNJ2,KK,LP,MA/My,MOLF,MRL,NOD,NOR,NZB,NZO,NZW,PJ,PL,RF,RH2,SJL,SMJ,SWR,TH,WSB | feature_truncation,5_prime_UTR_variant |
| 11 | 58443519 | 58444018 | 499 | Or2t47 | SPRET | feature_truncation,5_prime_UTR_variant |
| 11 | 58543014 | 58543082 | 68 | Trim58 | MOLF | feature_truncation,3_prime_UTR_variant |
| 11 | 58815346 | 58815437 | 91 | Btnl10 | SPRET | feature_truncation,3_prime_UTR_variant |
| 11 | 60418338 | 60418537 | 199 | Myo15a | AJ,C3H,CAST,CBA,MOLF,MRL,NZB,NZO,PJ,SEA,SPRET | feature_truncation,3_prime_UTR_variant |
| 11 | 62110476 | 62110555 | 79 | Specc1 | FVB,MOLF,NOD,SPRET,SWR | feature_truncation,3_prime_UTR_variant |
| 11 | 62491161 | 62491337 | 176 | Trpv2 | MOLF | feature_truncation,3_prime_UTR_variant |
| 11 | 66818848 | 66819424 | 576 | Pirt | KK | feature_truncation,3_prime_UTR_variant |
| 11 | 68097177 | 68097568 | 391 | Stx8 | BALB,C3H,CAST,CBA,DBA1J,DBA2J,KK,MA/My,MOLF,MRL,NOR,NZW,PJ,RH2,SEA,SJL,SMJ,SPRET,SWR,TH | feature_truncation,3_prime_UTR_variant |
| 11 | 69014498 | 69014594 | 96 | Hes7 | SPRET | feature_truncation,3_prime_UTR_variant |
| 11 | 69925000 | 69925183 | 183 | Dlg4 | SPRET | feature_truncation,3_prime_UTR_variant |
| 11 | 70028145 | 70028486 | 341 | Mgl2 | SPRET | feature_truncation,3_prime_UTR_variant |
| 11 | 70345504 | 70345584 | 80 | Cxcl16 | CE | feature_truncation,3_prime_UTR_variant |
| 11 | 72202807 | 72203055 | 248 | Xaf1 | 129S1,BTBR,C57L,CAST,CE,DBA1J,DBA2J,MOLF,NOR,PJ,SEA,SPRET | stop_lost,feature_truncation,coding_sequence_variant,3_prime_UTR_variant,frameshift_variant,stop_lost,feature_truncation,3_prime_UTR_variant |
| 11 | 72339788 | 72339851 | 63 | Mybbp1a | SPRET | feature_truncation,coding_sequence_variant,intron_variant |
| 11 | 72662701 | 72662902 | 201 | Ankfy1 | DBA1J,DBA2J,MOLF,NOR,NZB,PJ,SEA,SPRET,TH | feature_truncation,3_prime_UTR_variant |
| 11 | 72734165 | 72734460 | 295 | Zzef1 | DBA1J,DBA2J,MOLF,NOR,NZB,PJ,SEA,SPRET,TH | feature_truncation,coding_sequence_variant,intron_variant |
| 11 | 72974200 | 72974427 | 227 | Ncbp3 | C58J,CE,DBA1J,DBA2J,MOLF,NOR,NZB,PJ,SEA,SPRET,TH,WSB | feature_truncation,3_prime_UTR_variant |
| 11 | 73796117 | 73796169 | 52 | Or1e35 | WSB | feature_truncation,3_prime_UTR_variant |
| 11 | 73873651 | 73874198 | 547 | Or1r1 | SPRET | feature_truncation,3_prime_UTR_variant |
| 11 | 74714282 | 74714354 | 72 | Mettl16 | SPRET | feature_truncation,3_prime_UTR_variant |
| 11 | 74718635 | 74718697 | 62 | Mettl16 | SPRET | feature_truncation,3_prime_UTR_variant |
| 11 | 75360926 | 75361326 | 400 | Tlcd2 | CAST,MOLF,SPRET | feature_truncation,coding_sequence_variant,intron_variant,stop_lost,feature_truncation,coding_sequence_variant,3_prime_UTR_variant,intron_variant |
| 11 | 75467166 | 75467217 | 51 | Slc43a2 | SPRET | feature_truncation,3_prime_UTR_variant |
| 11 | 76305433 | 76305532 | 99 | Timm22 | SPRET | feature_truncation,3_prime_UTR_variant |
| 11 | 77678268 | 77678781 | 513 | Gm10277 | PJ | feature_truncation,5_prime_UTR_variant |
| 11 | 80158041 | 80158393 | 352 | Rhot1 | SPRET | feature_truncation,3_prime_UTR_variant |
| 11 | 82845570 | 82845627 | 57 | Slfn5 | MOLF | feature_truncation,5_prime_UTR_variant,intron_variant |
| 11 | 82854589 | 82854823 | 234 | Slfn5 | 129S1,AJ,BTBR,C3H,C57L,C58J,CAST,CBA,MA/My,MOLF,MRL,NZO,SPRET,WSB | feature_truncation,3_prime_UTR_variant |
| 11 | 83932711 | 83932914 | 203 | Synrg | SPRET | feature_truncation,3_prime_UTR_variant |
| 11 | 84913691 | 84913886 | 195 | Usp32 | AKR,CAST,DBA1J,DBA2J,MOLF,NZB,NZO,RF,SPRET,WSB | feature_truncation,coding_sequence_variant,intron_variant |
| 11 | 87268328 | 87268437 | 109 | Rad51c | SPRET | feature_truncation,3_prime_UTR_variant |
| 11 | 87780987 | 87781039 | 52 | Or4d2b | CAST | feature_truncation,5_prime_UTR_variant |
| 11 | 87870886 | 87870960 | 74 | Dynll2 | SPRET | feature_truncation,3_prime_UTR_variant |
| 11 | 88864229 | 88864570 | 341 | Coil | MOLF | feature_truncation,5_prime_UTR_variant,intron_variant |
| 11 | 88928298 | 88928462 | 164 | Dgke | SPRET | feature_truncation,3_prime_UTR_variant |
| 11 | 93983798 | 93983901 | 103 | Spag9 | AJ,AKR,C3H,CAST,CBA,CE,MOLF,MRL,RF,SPRET | feature_truncation,coding_sequence_variant,intron_variant |
| 11 | 94380652 | 94380790 | 138 | Epn3 | SPRET | feature_truncation,3_prime_UTR_variant |
| 11 | 96822144 | 96822262 | 118 | Prr15l | SPRET | feature_truncation,5_prime_UTR_variant |
| 11 | 97884678 | 97884797 | 119 | Arl5c | MOLF,NZW,SPRET,SWR | feature_truncation,3_prime_UTR_variant |

| | | | | | | |
|---|---|---|---|---|---|---|
| 11 | 97978623 | 97978765 | 142 | Fbxl20 | 129S1,BTBR,C3H,C57L,CE,DBA1J,DBA2J,FVB,ILNJ2,MOLF,NZW,SPRET,SWR,WSB | feature_truncation,3_prime_UTR_variant |
| 11 | 99525474 | 99525534 | 60 | Krtap4-2 | SPRET | feature_truncation,inframe_deletion |
| 11 | 99663094 | 99663236 | 142 | Gm11595 | CAST,MOLF | frameshift_variant,feature_truncation |
| 11 | 99676258 | 99676390 | 132 | Krtap4-9 | AJ,SMJ | feature_truncation,inframe_deletion |
| 11 | 99728031 | 99728208 | 177 | 2300003K06Rik | SPRET | stop_lost,feature_truncation,coding_sequence_variant,3_prime_UTR_variant |
| 11 | 99741968 | 99742025 | 57 | Krtap4-16 | CAST,SPRET | feature_truncation,inframe_deletion |
| 11 | 99839973 | 99840045 | 72 | Krtap9-5 | NZB | feature_truncation,inframe_deletion |
| 11 | 100079682 | 100079772 | 90 | Krt9 | SPRET | feature_truncation,inframe_deletion |
| 11 | 100212722 | 100212799 | 77 | Eif1 | WSB | feature_truncation,3_prime_UTR_variant |
| 11 | 100657087 | 100657151 | 64 | Ghdc | SPRET | feature_truncation,3_prime_UTR_variant |
| 11 | 100953305 | 100953357 | 52 | Atp6v0a1 | SPRET | feature_truncation,3_prime_UTR_variant |
| 11 | 101349357 | 101349439 | 82 | Ifi35 | SPRET | feature_truncation,3_prime_UTR_variant |
| 11 | 102781403 | 102781761 | 358 | Gfap | 129S1,AJ,BTBR,C3H,C57L,CAST,CBA,DBA1J,DBA2J,ILNJ2,SPRET,WSB | feature_truncation,3_prime_UTR_variant |
| 11 | 104558968 | 104559034 | 66 | Itgb3 | SPRET | feature_truncation,3_prime_UTR_variant |
| 11 | 104611782 | 104611980 | 198 | Gm11639 | SPRET | feature_truncation,inframe_deletion |
| 11 | 106659339 | 106660338 | 999 | Milr1 | SPRET | feature_truncation,3_prime_UTR_variant |
| 11 | 107561116 | 107561179 | 63 | Helz | SPRET | feature_truncation,inframe_deletion |
| 11 | 107624631 | 107624690 | 59 | Cacng4 | ILNJ2,RF | feature_truncation,3_prime_UTR_variant |
| 11 | 109116208 | 109116262 | 54 | Rgs9 | SPRET | feature_truncation,3_prime_UTR_variant |
| 11 | 109146766 | 109146848 | 82 | Rgs9 | CAST | feature_truncation,3_prime_UTR_variant |
| 11 | 109148571 | 109148677 | 106 | Rgs9 | CAST | feature_truncation,3_prime_UTR_variant |
| 11 | 109287786 | 109287870 | 84 | Gna13 | BALB,ILNJ2,RF,SEA,SPRET | feature_truncation,3_prime_UTR_variant |
| 11 | 109823924 | 109823992 | 68 | Abca8b | BUB,CAST,SPRET | feature_truncation,3_prime_UTR_variant |
| 11 | 114792961 | 114793431 | 470 | Cd300a | CAST,SPRET,WSB | feature_truncation,3_prime_UTR_variant |
| 11 | 114847215 | 114847279 | 64 | Cd300c | 129S1,BALB,C58J,CE,ILNJ2,KK,MRL,NOD,NOR,NZB,NZO,NZW,RF,SEA,SMJ,TH,WSB | feature_truncation,coding_sequence_variant,intron_variant |
| 11 | 115135638 | 115135760 | 122 | Tmem104 | 129S1,BALB,C58J,KK,NOD,NOR,NZB,NZO,NZW,RF,SEA,SMJ,TH,WSB | feature_truncation,3_prime_UTR_variant |
| 11 | 115137160 | 115137215 | 55 | Tmem104 | SPRET | feature_truncation,3_prime_UTR_variant |
| 11 | 116014900 | 116014964 | 64 | Trim65 | MOLF,SPRET | feature_truncation,3_prime_UTR_variant |
| 11 | 116033581 | 116033643 | 62 | Fbf1 | SPRET | feature_truncation,3_prime_UTR_variant |
| 11 | 116694928 | 116695082 | 154 | Mxra7 | CAST,MOLF | feature_truncation,3_prime_UTR_variant |
| 11 | 116766283 | 116766498 | 215 | Mfsd11 | 129S1,AJ,AKR,BALB,BTBR,C3H,C57L,C58J,CAST,CBA,CE,DBA1J,DBA2J,FVB,ILNJ2,KK,MA/My,MOLF,NZB,NZO,NZW,PL,RF,RH2,SEA,SJL,SMJ,SPRET,SWR,TH,WSB | feature_truncation,3_prime_UTR_variant |
| 11 | 116869300 | 116869373 | 73 | Mgat5b | SPRET | feature_truncation,coding_sequence_variant,intron_variant |
| 11 | 117048532 | 117048605 | 73 | Sec14l1 | SPRET | feature_truncation,3_prime_UTR_variant |
| 11 | 117745616 | 117745798 | 182 | Birc5 | CAST,SPRET | feature_truncation,3_prime_UTR_variant |
| 11 | 119401531 | 119401657 | 126 | Endov | SPRET | feature_truncation,3_prime_UTR_variant |
| 11 | 120672103 | 120672202 | 99 | Rfng | CAST,MOLF,SJL,SPRET | feature_truncation,3_prime_UTR_variant |
| 11 | 121105563 | 121105746 | 183 | Hexd | 129S1,AJ,AKR,B10D2,B10J,BALB,BTBR,BUB,C3H,C57L,C58J,CAST,CBA,CE,DBA1J,DBA2J,FVB,ILNJ2,KK,LP,MA/My,MOLF,MRL,NOD,NOR,NZB,NZO,NZW,PJ,PL,RF,RH2,SEA,SJL,SMJ,SPRET,SWR,TH,WSB | feature_truncation,coding_sequence_variant,intron_variant |
| 12 | 3547187 | 3547462 | 275 | Asxl2 | SPRET | feature_truncation,3_prime_UTR_variant |
| 12 | 4159495 | 4159571 | 76 | Dnajc27 | MRL | feature_truncation,3_prime_UTR_variant |
| 12 | 4909060 | 4909265 | 205 | Wdcp | SPRET | feature_truncation,3_prime_UTR_variant |
| 12 | 4957569 | 4957622 | 53 | Ubxn2a | C3H,CBA,CE,MOLF,SPRET,TH | feature_truncation,5_prime_UTR_variant |
| 12 | 8601579 | 8602398 | 819 | Slc7a15 | MOLF | feature_truncation,5_prime_UTR_variant,intron_variant |
| 12 | 12427778 | 12428116 | 338 | Cyria | SPRET | feature_truncation,3_prime_UTR_variant |
| 12 | 28520377 | 28520437 | 60 | Dcdc2c | 129S1,AKR,CAST,CE,ILNJ2,RF | feature_truncation,inframe_deletion,feature_truncation,3_prime_UTR_variant |
| 12 | 28917531 | 28917715 | 184 | Eipr1 | CAST | feature_truncation,3_prime_UTR_variant |
| 12 | 30066277 | 30066330 | 53 | Pxdn | CAST | feature_truncation,3_prime_UTR_variant |
| 12 | 30640423 | 30640798 | 375 | Tmem18 | 129S1,BUB,MRL,WSB | feature_truncation,3_prime_UTR_variant |
| 12 | 31124618 | 31124759 | 141 | Fam110c | AJ,AKR,BALB,BTBR,BUB,C3H,CBA,MOLF,PJ,PL,RF,RH2,SEA,WSB | feature_truncation,inframe_deletion |
| 12 | 33029543 | 33029719 | 176 | Sypl1 | 129S1,AJ,AKR,BALB,BTBR,BUB,C3H,CAST,CBA,PJ,PL,RF,RH2,SEA,WSB | feature_truncation,3_prime_UTR_variant |
| 12 | 33378522 | 33379005 | 483 | Atxn7l1 | SPRET | feature_truncation,3_prime_UTR_variant |
| 12 | 34098620 | 34098686 | 66 | Hdac9 | MOLF,NZW | feature_truncation,3_prime_UTR_variant |
| 12 | 38918319 | 38918371 | 52 | Etv1 | CAST,MOLF,WSB | feature_truncation,3_prime_UTR_variant |
| 12 | 40110255 | 40110338 | 83 | Scin | CAST,CE,MOLF,SPRET | feature_truncation,3_prime_UTR_variant |
| 12 | 52357820 | 52358088 | 268 | Nubpl | 129S1,C57L,CAST,MOLF,NZB | feature_truncation,3_prime_UTR_variant |
| 12 | 53198896 | 53198983 | 87 | Akap6 | 129S1,C57L,CAST,MOLF,NZB,SPRET | feature_truncation,3_prime_UTR_variant |
| 12 | 54718634 | 54718850 | 216 | Eapp | AJ,AKR,BALB,C3H,CAST,CBA,DBA1J,DBA2J,ILNJ2,KK,MOLF,NOD,NOR,PJ,RF,RH2,SEA,SJL,SPRET,SWR,TH | feature_truncation,3_prime_UTR_variant |
| 12 | 55349987 | 55350506 | 519 | Prorp | SPRET | feature_truncation,5_prime_UTR_variant,feature_truncation,5_prime_UTR_variant,intron_variant |
| 12 | 59237537 | 59237767 | 230 | Mia2 | CAST,MOLF,SPRET | feature_truncation,3_prime_UTR_variant |
| 12 | 71009568 | 71009792 | 224 | Actr10 | CAST,FVB,MOLF,NZW,SJL,SPRET,SWR,WSB | feature_truncation,3_prime_UTR_variant |
| 12 | 75677656 | 75678346 | 690 | Wdr89,Rplp2-ps1 | CAST,FVB,MOLF,PJ,SEA,SMJ,SPRET,SWR | feature_truncation,3_prime_UTR_variant,transcript_ablation |
| 12 | 76376364 | 76376436 | 72 | Akap5 | AKR,C58J,CE,DBA1J,DBA2J,ILNJ2,MOLF,PJ,RF,SEA,SMJ,WSB | feature_truncation,inframe_deletion |
| 12 | 76829724 | 76829921 | 197 | Churc1 | CAST,SPRET | feature_truncation,3_prime_UTR_variant |
| 12 | 78245676 | 78246500 | 824 | Ccdc196 | MOLF | feature_truncation,coding_sequence_variant,intron_variant |
| 12 | 78887751 | 78887809 | 58 | Atp6v1d | AKR,BALB,C58J,CE,KK,LP,MRL,NOD,NOR,NZO,PJ,PL,RF,SEA,SJL,SMJ,SWR,TH | feature_truncation,3_prime_UTR_variant |
| 12 | 84040418 | 84040484 | 66 | Acot2 | SPRET | feature_truncation,3_prime_UTR_variant |
| 12 | 84093154 | 84093247 | 93 | Acot4 | SPRET | feature_truncation,3_prime_UTR_variant |
| 12 | 84696964 | 84697018 | 54 | Vrtn | SPRET | feature_truncation,inframe_deletion |
| 12 | 86562228 | 86562470 | 242 | Esrrb | CAST,CE,MOLF,NOD,NOR,SPRET,SWR,WSB | feature_truncation,3_prime_UTR_variant |

| | | | | | | |
|---|---|---|---|---|---|---|
| 12 | 87353190 | 87353322 | 132 | Sptlc2 | 129S1,AJ,AKR,BTBR,BUB,C3H,C57L,CAST,CBA,CE,DBA1J,DBA2J,FVB,ILNJ2,MOLF,MRL,NOD,NOR,PJ,PL,RF,SEA,SJL,SPRET,SWR,TH,WSB | feature_truncation,3_prime_UTR_variant |
| 12 | 87994245 | 87994426 | 181 | Eif1ad11 | NOD,SWR | feature_truncation,3_prime_UTR_variant |
| 12 | 91602959 | 91603019 | 60 | Ston2 | SPRET | feature_truncation,3_prime_UTR_variant |
| 12 | 101613684 | 101613886 | 202 | Tc2n | SPRET | feature_truncation,3_prime_UTR_variant |
| 12 | 102719709 | 102719911 | 202 | Gon7 | CAST,CE,DBA1J,DBA2J,MOLF,PJ,SEA,SPRET,WSB | feature_truncation,3_prime_UTR_variant |
| 12 | 103582618 | 103582670 | 52 | Serpina10 | NZO,PJ,SEA,SPRET | feature_truncation,3_prime_UTR_variant |
| 12 | 103961819 | 103961876 | 57 | Serpina9 | 129S1,AKR,BALB,BTBR,BUB,C57L,CAST,CBA,DBA1J,DBA2J,FVB,ILNJ2,KK,LP,MRL,NOD,NOR,NZB,PJ,PL,RF,SEA,SJL,SWR,WSB | feature_truncation,3_prime_UTR_variant |
| 12 | 103979533 | 103980223 | 690 | Serpina9 | FVB | feature_truncation,5_prime_UTR_variant,intron_variant |
| 12 | 104078583 | 104079100 | 517 | Serpina3a | SJL,SWR | feature_truncation,5_prime_UTR_variant |
| 12 | 104181329 | 104181724 | 395 | Serpina3f | AKR,BALB,BUB,CAST,CBA,CE,DBA1J,DBA2J,FVB,ILNJ2,KK,LP,MRL,NOD,NOR,PJ,PL,RF,SEA,SJL,SPRET,SWR,WSB | feature_truncation,5_prime_UTR_variant,intron_variant |
| 12 | 105008707 | 105008780 | 73 | Glrx5 | SPRET | feature_truncation,3_prime_UTR_variant |
| 12 | 105455538 | 105455590 | 52 | D430019H16Rik | CAST | feature_truncation,3_prime_UTR_variant |
| 12 | 105668802 | 105668862 | 60 | Gskip | BALB,CAST,CE,FVB,KK,LP,MA/My,MOLF,NZB,NZO,NZW,PJ,SEA,SJL,SPRET,TH,WSB | feature_truncation,3_prime_UTR_variant |
| 12 | 108993869 | 108993923 | 54 | Wdr25 | SPRET | feature_truncation,3_prime_UTR_variant |
| 12 | 109561378 | 109561444 | 66 | Rtl1 | SPRET | feature_truncation,inframe_deletion |
| 12 | 111770485 | 111770553 | 68 | Xrcc3 | AJ,AKR,C3H,CBA,CE,DBA1J,DBA2J,FVB,ILNJ2,MOLF,MRL,NZB,NZO,NZW,PL,SMJ,SWR,TH,WSB | feature_truncation,3_prime_UTR_variant |
| 12 | 113036095 | 113036185 | 90 | Pacs2 | AJ,C3H,CBA,CE,FVB,KK,MRL,NZB,NZO,NZW,PJ,PL,SEA,SMJ,SPRET,SWR,TH,WSB | feature_truncation,3_prime_UTR_variant |
| 12 | 119433122 | 119433274 | 152 | Macc1 | SPRET,WSB | feature_truncation,3_prime_UTR_variant |
| 13 | 3903966 | 3904316 | 350 | Calm5 | KK,LP,NZB,NZW | feature_truncation,5_prime_UTR_variant,feature_truncation,coding_sequence_variant,5_prime_UTR_variant,intron_variant |
| 13 | 8854303 | 8854382 | 79 | Wdr37 | AKR,BTBR,C57L,CAST,KK,LP,MOLF,NOD,NOR,NZB,NZO,NZW,RF,SPRET,TH,WSB | feature_truncation,3_prime_UTR_variant |
| 13 | 9706573 | 9706678 | 105 | Dip2c | CAST | feature_truncation,coding_sequence_variant,intron_variant |
| 13 | 12487978 | 12488032 | 54 | Edaradd | AKR,CAST,ILNJ2,LP,MOLF,RF | feature_truncation,3_prime_UTR_variant |
| 13 | 12489988 | 12490090 | 102 | Edaradd | CAST | feature_truncation,3_prime_UTR_variant |
| 13 | 14226422 | 14226473 | 51 | Ggps1 | SPRET | feature_truncation,3_prime_UTR_variant |
| 13 | 14228072 | 14228140 | 68 | Ggps1 | SPRET | feature_truncation,3_prime_UTR_variant |
| 13 | 16202266 | 16202336 | 70 | Inhba | CAST,SPRET | feature_truncation,3_prime_UTR_variant |
| 13 | 21301498 | 21301622 | 124 | Or2w1b | AJ,AKR,B10D2,B10J,BALB,BTBR,BUB,C3H,C57L,C58J,CAST,CBA,DBA1J,DBA2J,FVB,ILNJ2,MA/My,MOLF,MRL,NOD,NOR,PL,RF,RH2,SJL,SMJ,SPRET,SWR,TH | feature_truncation,3_prime_UTR_variant |
| 13 | 21303102 | 21303339 | 237 | Or2w1b | AJ,AKR,B10D2,B10J,BALB,BTBR,BUB,C3H,C57L,C58J,CAST,CBA,CE,DBA1J,DBA2J,FVB,ILNJ2,KK,LP,MA/My,MOLF,MRL,NOD,NOR,NZB,PJ,PL,RF,RH2,SEA,SJL,SMJ,SPRET,SWR,TH,WSB | feature_truncation,3_prime_UTR_variant |
| 13 | 21471096 | 21471157 | 61 | Gpx5 | SPRET | feature_truncation,3_prime_UTR_variant |
| 13 | 21554669 | 21555381 | 712 | Zscan12 | SPRET | feature_truncation,3_prime_UTR_variant |
| 13 | 21575893 | 21576065 | 172 | Zkscan3 | CAST,SPRET | feature_truncation,3_prime_UTR_variant |
| 13 | 21627333 | 21627392 | 59 | Zscan26 | LP,MOLF,PJ,SEA,SPRET | feature_truncation,3_prime_UTR_variant |
| 13 | 21685082 | 21685621 | 539 | Cox5b-ps | CAST,LP,MOLF,PJ,SEA,SPRET | transcript_ablation |
| 13 | 22224769 | 22224855 | 86 | H4c9 | SPRET | feature_truncation,3_prime_UTR_variant |
| 13 | 22357979 | 22358291 | 312 | Vmn1r191 | SPRET | feature_truncation,3_prime_UTR_variant |
| 13 | 22682811 | 22683243 | 432 | Vmn1r202 | SPRET | feature_truncation,3_prime_UTR_variant |
| 13 | 22909627 | 22909721 | 94 | Vmn1r207 | BTBR,C57L,KK,LP,MOLF,NZB,NZO,NZW,PJ,SEA,SPRET | feature_truncation,3_prime_UTR_variant |
| 13 | 23295571 | 23296551 | 980 | Vmn1r217 | SPRET | feature_truncation,3_prime_UTR_variant |
| 13 | 23886827 | 23887227 | 400 | Hfe | MOLF,SPRET | feature_truncation,3_prime_UTR_variant |
| 13 | 25000071 | 25000273 | 202 | BC005537 | CAST | feature_truncation,3_prime_UTR_variant |
| 13 | 25084847 | 25084904 | 57 | D130043K22Rik | CAST | feature_truncation,3_prime_UTR_variant |
| 13 | 25172956 | 25173232 | 276 | Mrs2 | SPRET | feature_truncation,3_prime_UTR_variant |
| 13 | 32888557 | 32888611 | 54 | Mylk4 | SPRET | feature_truncation,3_prime_UTR_variant |
| 13 | 33149330 | 33149400 | 70 | Serpinb6b | SPRET | feature_truncation,5_prime_UTR_variant |
| 13 | 36079265 | 36079389 | 124 | Rpp40 | SPRET | feature_truncation,3_prime_UTR_variant |
| 13 | 38157942 | 38158158 | 216 | Ssr1 | SPRET | feature_truncation,3_prime_UTR_variant |
| 13 | 41175785 | 41175848 | 63 | Tmem14c | MOLF | feature_truncation,3_prime_UTR_variant |
| 13 | 46830946 | 46831046 | 100 | Fam8a1 | WSB | feature_truncation,3_prime_UTR_variant |
| 13 | 47399686 | 47399737 | 51 | Rnf144b | SPRET | feature_truncation,3_prime_UTR_variant |
| 13 | 49741092 | 49741325 | 233 | Omd | SPRET | feature_truncation,5_prime_UTR_variant,intron_variant |
| 13 | 52802452 | 52802554 | 102 | Syk | C58J,LP,MOLF | feature_truncation,3_prime_UTR_variant |
| 13 | 54527796 | 54527895 | 99 | Cplx2 | SPRET | feature_truncation,3_prime_UTR_variant |
| 13 | 54673081 | 54673144 | 63 | Simc1 | MOLF | feature_truncation,inframe_deletion |
| 13 | 55848023 | 55848198 | 175 | B230219D22Rik | AJ,BALB,C3H,CAST,CBA,CE,MOLF,SPRET,SWR | feature_truncation,3_prime_UTR_variant |
| 13 | 55849613 | 55849666 | 53 | B230219D22Rik | SPRET | feature_truncation,3_prime_UTR_variant |
| 13 | 56322790 | 56322874 | 84 | Tifab | CAST | feature_truncation,3_prime_UTR_variant |
| 13 | 56620235 | 56620362 | 127 | Slc25a48 | 129S1,AJ,BALB,BTBR,BUB,C3H,C57L,CAST,CBA,CE,DBA1J,DBA2J,ILNJ2,MOLF,MRL,NOD,NOR,NZB,NZW,PJ,PL,SEA,SMJ,SPRET,SWR,WSB | feature_truncation,3_prime_UTR_variant |
| 13 | 56621552 | 56622058 | 506 | Slc25a48 | AJ,BALB,C3H,CBA,CE | feature_truncation,3_prime_UTR_variant |
| 13 | 58149676 | 58149767 | 91 | Klhl3 | WSB | feature_truncation,3_prime_UTR_variant |
| 13 | 58701925 | 58702052 | 127 | Slc28a3 | CAST,FVB,SPRET,SWR | feature_truncation,3_prime_UTR_variant |
| 13 | 59083472 | 59083548 | 76 | Ntrk2 | SPRET | feature_truncation,3_prime_UTR_variant |
| 13 | 59847526 | 59847741 | 215 | Spata31d1a | SPRET | feature_truncation,3_prime_UTR_variant |
| 13 | 61049014 | 61049120 | 106 | Tpbpb | PJ,SEA | feature_truncation,3_prime_UTR_variant |
| 13 | 62319236 | 62319319 | 83 | Zfp808 | WSB | frameshift_variant,feature_truncation,frameshift_variant,stop_lost,feature_truncation,3_prime_UTR_variant |
| 13 | 62320193 | 62320444 | 251 | Zfp808 | CAST,MOLF,WSB | frameshift_variant,feature_truncation |

| | | | | | | |
|---|---|---|---|---|---|---|
| 13 | 62600852 | 62601018 | 166 | Zfp935 | SPRET | feature_truncation,3_prime_UTR_variant |
| 13 | 62922242 | 62922326 | 84 | Gm5141 | BALB | feature_truncation,inframe_deletion |
| 13 | 64509628 | 64509855 | 227 | Ctsl | 129S1,AJ,AKR,BALB,BTBR,BUB,C3H,C57L,C58J,CAST,CBA,CE,DBA1J,DBA2J,FVB,ILNJ2,KK,MRL,NOD,NOR,NZB,NZO,NZW,PJ,PL,RF,SEA,SJL,SMJ,SPRET,SWR,TH,WSB | feature_truncation,3_prime_UTR_variant |
| 13 | 64510883 | 64510969 | 86 | Ctsl | 129S1,AJ,AKR,BALB,BTBR,BUB,C3H,C57L,C58J,CBA,CE,DBA1J,DBA2J,ILNJ2,KK,MRL,NOD,NOR,NZB,NZO,NZW,PL,RF,SJL,SMJ,TH,WSB | feature_truncation,3_prime_UTR_variant |
| 13 | 65302548 | 65302612 | 64 | Or9s18 | 129S1,AJ,AKR,BALB,BTBR,BUB,C3H,C57L,C58J,CAST,CBA,CE,DBA1J,DBA2J,FVB,ILNJ2,KK,MOLF,MRL,NOD,NOR,NZB,NZO,NZW,PJ,PL,RF,SEA,SJL,SMJ,SPRET,SWR,TH,WSB | feature_truncation,3_prime_UTR_variant |
| 13 | 66375101 | 66375226 | 125 | Zfp999 | B10D2,B10J,C3H,C58J,CBA,CE,NZO,PL,SMJ,SPRET | stop_lost,feature_truncation,coding_sequence_variant,3_prime_UTR_variant |
| 13 | 67246856 | 67246940 | 84 | Rslcan18 | SPRET | feature_truncation,inframe_deletion |
| 13 | 67286835 | 67286919 | 84 | Zfp759 | AJ,BALB,BUB,C3H,C58J,CBA,CE,DBA1J,DBA2J,FVB,ILNJ2,KK,NOD,NOR,NZB,NZO,PL,SJL,SMJ,SWR,TH,WSB | feature_truncation,inframe_deletion |
| 13 | 67329899 | 67330319 | 420 | Rsl1 | SWR | feature_truncation,inframe_deletion |
| 13 | 67439814 | 67440687 | 873 | Zfp457 | 129S1,AJ,AKR,BALB,BTBR,BUB,C3H,C57L,C58J,CAST,CBA,CE,DBA1J,DBA2J,FVB,ILNJ2,KK,MOLF,MRL,NOD,NOR,NZB,NZO,NZW,PJ,PL,RF,SEA,SJL,SMJ,SPRET,SWR,TH,WSB | stop_lost,feature_truncation,coding_sequence_variant,3_prime_UTR_variant |
| 13 | 67441674 | 67441758 | 84 | Zfp457 | 129S1,AJ,AKR,BALB,BTBR,BUB,C3H,C57L,C58J,CAST,CBA,CE,DBA1J,DBA2J,FVB,ILNJ2,KK,MOLF,MRL,NOD,NOR,NZB,NZO,NZW,PJ,PL,RF,SEA,SJL,SMJ,SPRET,SWR,TH,WSB | feature_truncation,inframe_deletion |
| 13 | 67490890 | 67490989 | 99 | Zfp953 | SPRET | feature_truncation,3_prime_UTR_variant |
| 13 | 67537503 | 67537818 | 315 | Zfp429 | 129S1,AKR,MRL,NZW,PL,RF,WSB | feature_truncation,3_prime_UTR_variant |
| 13 | 67812299 | 67813239 | 940 | Zfp738 | 129S1,AJ,AKR,BALB,BTBR,BUB,C3H,C57L,C58J,CAST,CBA,CE,DBA1J,DBA2J,FVB,ILNJ2,KK,MRL,NOD,NOR,NZB,NZO,NZW,PJ,PL,RF,SEA,SJL,SMJ,SWR,TH | feature_truncation,3_prime_UTR_variant |
| 13 | 67814255 | 67814585 | 330 | Zfp738 | 129S1,AJ,AKR,BALB,BTBR,BUB,C3H,C57L,C58J,CAST,CBA,CE,DBA1J,DBA2J,FVB,ILNJ2,KK,MOLF,MRL,NOD,NOR,NZB,NZO,NZW,PJ,PL,RF,SEA,SJL,SMJ,SPRET,SWR,TH | feature_truncation,3_prime_UTR_variant |
| 13 | 67940346 | 67940431 | 85 | Zfp493 | 129S1,AJ,AKR,BALB,BTBR,BUB,C3H,C57L,C58J,CAST,CBA,CE,DBA1J,DBA2J,FVB,ILNJ2,MA/My,MOLF,MRL,NOD,NOR,NZB,NZO,NZW,PJ,PL,RF,SEA,SJL,SMJ,SPRET,SWR,TH,WSB | feature_truncation,3_prime_UTR_variant |
| 13 | 74554515 | 74555477 | 962 | Ftl1-ps1 | CAST,ILNJ2,MOLF,SPRET,SWR,WSB | transcript_ablation |
| 13 | 76019674 | 76019725 | 51 | Rhobtb3 | SPRET | feature_truncation,3_prime_UTR_variant |
| 13 | 76150376 | 76150486 | 110 | Rfesd | 129S1,AJ,AKR,BALB,BTBR,BUB,C3H,C57L,CBA,CE,DBA1J,DBA2J,FVB,ILNJ2,KK,MA/My,MRL,NOD,NOR,NZB,NZO,NZW,PJ,PL,RF,SEA,SJL,SMJ,SPRET,SWR,TH,WSB | feature_truncation,3_prime_UTR_variant |
| 13 | 86194079 | 86194285 | 206 | Cox7c | MOLF | feature_truncation,5_prime_UTR_variant |
| 13 | 92845597 | 92845815 | 218 | Serinc5 | SPRET | feature_truncation,3_prime_UTR_variant |
| 13 | 93791839 | 93792619 | 780 | Bhmt2 | SPRET | feature_truncation,3_prime_UTR_variant |
| 13 | 95149234 | 95149666 | 432 | Wdr41 | C58J,CAST,DBA1J,KK,MA/My,MOLF,RF,SPRET | feature_truncation,coding_sequence_variant,intron_variant,feature_truncation,inframe_deletion |
| 13 | 96095994 | 96096049 | 55 | Sv2c | SPRET | feature_truncation,3_prime_UTR_variant |
| 13 | 96097251 | 96097313 | 62 | Sv2c | 129S1,AJ,AKR,BALB,BTBR,BUB,C3H,C57L,CBA,DBA1J,DBA2J,FVB,KK,MOLF,MRL,NZB,NZO,NZW,PJ,PL,SEA,SJL,SMJ,SWR,TH,WSB | feature_truncation,3_prime_UTR_variant |
| 13 | 97896574 | 97897141 | 567 | Rps18-ps6 | CAST,SPRET | transcript_ablation |
| 13 | 99384421 | 99384651 | 230 | Zfp366 | CAST,SPRET | feature_truncation,3_prime_UTR_variant |
| 13 | 99385316 | 99385448 | 132 | Zfp366 | CAST | feature_truncation,3_prime_UTR_variant |
| 13 | 99386354 | 99386477 | 123 | Zfp366 | SPRET | feature_truncation,3_prime_UTR_variant |
| 13 | 103877384 | 103877511 | 127 | Srek1 | C58J,CE,MA/My,NZB,NZO | feature_truncation,3_prime_UTR_variant |
| 13 | 105285984 | 105286086 | 102 | Rnf180 | SPRET | feature_truncation,coding_sequence_variant,intron_variant |
| 13 | 108189182 | 108189235 | 53 | Ndufaf2 | CE,MA/My,NZO,TH | feature_truncation,3_prime_UTR_variant |
| 13 | 110196729 | 110196786 | 57 | Rab3c | SPRET | feature_truncation,3_prime_UTR_variant |
| 13 | 112870587 | 112870807 | 220 | Slc38a9 | SPRET | feature_truncation,3_prime_UTR_variant |
| 13 | 114291977 | 114292041 | 64 | Arl15 | AJ,BUB,CAST,CE,DBA1J,FVB,KK,MA/My,MOLF,NZB,NZO,NZW,PL,RH2,SJL,SPRET,SWR | feature_truncation,3_prime_UTR_variant |
| 13 | 120249500 | 120249664 | 164 | Zfp131 | 129S1,BTBR,C3H,C57L,CAST,CBA,CE,DBA2J,KK,MA/My,NZO,SMJ,SPRET | frameshift_variant,stop_lost,feature_truncation,3_prime_UTR_variant |
| 14 | 7666136 | 7666457 | 321 | Nek10 | BALB,SEA,SPRET | feature_truncation,5_prime_UTR_variant |
| 14 | 10229143 | 10229364 | 221 | Ptprg | 129S1,C3H,C57L,CBA,DBA1J,DBA2J,KK,MOLF,MRL,NZW,PJ,PL | feature_truncation,3_prime_UTR_variant |
| 14 | 15463569 | 15463655 | 86 | Gm10406 | SPRET | feature_truncation,3_prime_UTR_variant |
| 14 | 17685235 | 17685333 | 98 | Gm9602 | SPRET | feature_truncation,3_prime_UTR_variant |
| 14 | 19137320 | 19137593 | 273 | Gm2956 | MOLF | feature_truncation,5_prime_UTR_variant |
| 14 | 20421211 | 20421398 | 187 | Fam149b | CAST,MOLF,SPRET | feature_truncation,3_prime_UTR_variant |
| 14 | 20709125 | 20709210 | 85 | Synpo2l | SPRET | feature_truncation,3_prime_UTR_variant |
| 14 | 24510130 | 24510220 | 90 | Polr3a | CAST,MOLF,SPRET | feature_truncation,coding_sequence_variant,intron_variant |
| 14 | 31199422 | 31199822 | 400 | Mettl6 | SPRET | feature_truncation,3_prime_UTR_variant |
| 14 | 32017605 | 32017661 | 56 | Parg | CAST | feature_truncation,3_prime_UTR_variant |
| 14 | 32019318 | 32019519 | 201 | Parg | CAST | feature_truncation,3_prime_UTR_variant |
| 14 | 32069705 | 32069928 | 223 | Ogdhl | SPRET | feature_truncation,3_prime_UTR_variant |
| 14 | 33296525 | 33296797 | 272 | Frmpd2 | SPRET | feature_truncation,3_prime_UTR_variant |
| 14 | 33689306 | 33690270 | 964 | Zfp488 | SPRET | feature_truncation,3_prime_UTR_variant |
| 14 | 42353052 | 42353649 | 597 | Gm8024 | C57L,CAST,MOLF,NZO,PJ,PL | feature_truncation,coding_sequence_variant,5_prime_UTR_variant,intron_variant,feature_truncation,5_prime_UTR_variant,intron_variant |
| 14 | 45370863 | 45371079 | 216 | Txndc16 | C57L,CAST,KK,MOLF,NZO,PJ,PL,SMJ,SPRET | feature_truncation,3_prime_UTR_variant |
| 14 | 45405129 | 45405238 | 109 | Txndc16 | SPRET | feature_truncation,3_prime_UTR_variant |

| | | | | | | |
|---|---|---|---|---|---|---|
| 14 | 45620092 | 45620175 | 83 | Styx | SPRET | feature_truncation,3_prime_UTR_variant |
| 14 | 45827391 | 45828163 | 772 | Ddhd1 | SPRET | feature_truncation,3_prime_UTR_variant |
| 14 | 47005526 | 47005812 | 286 | Cdkn3 | 129S1,AJ,AKR,BALB,BUB,C3H,CBA,CE,DBA1J,DBA2J,FVB,ILNJ2,MA/My,MRL,NOD,NOR,NZB,NZO,NZW,PJ,RF,RH2,SEA,SJL,SMJ,SWR,TH,WSB | feature_truncation,3_prime_UTR_variant |
| 14 | 47495333 | 47495577 | 244 | Wdhd1 | SPRET | feature_truncation,3_prime_UTR_variant |
| 14 | 48759613 | 48760254 | 641 | Tmem260 | SPRET | feature_truncation,3_prime_UTR_variant |
| 14 | 49325254 | 49325324 | 70 | Ap5m1 | SPRET | feature_truncation,3_prime_UTR_variant |
| 14 | 49456542 | 49457241 | 699 | Ccdc198 | CAST,SPRET | feature_truncation,3_prime_UTR_variant |
| 14 | 50664454 | 50665245 | 791 | Or11g24 | 129S1,AJ,AKR,BALB,BUB,C3H,C57L,CBA,CE,DBA1J,DBA2J,FVB,ILNJ2,KK,MA/My,MOLF,MRL,NOD,NOR,NZB,NZO,NZW,PJ,PL,RF,RH2,SEA,SJL,SPRET,SWR,TH,WSB | feature_truncation,3_prime_UTR_variant |
| 14 | 51151296 | 51151666 | 370 | Osgep | SPRET | feature_truncation,3_prime_UTR_variant |
| 14 | 51201913 | 51202326 | 413 | Pnp2 | 129S1,AJ,AKR,BALB,BUB,C3H,C57L,CBA,CE,DBA1J,DBA2J,FVB,ILNJ2,KK,MA/My,MRL,NOD,NOR,NZB,NZO,NZW,PJ,PL,RF,RH2,SEA,SJL,SMJ,SPRET,SWR,TH,WSB | stop_lost,feature_truncation,coding_sequence_variant,3_prime_UTR_variant |
| 14 | 54781224 | 54781286 | 62 | Haus4 | AJ,AKR,BALB,BTBR,BUB,C3H,C57L,CBA,CE,DBA1J,DBA2J,FVB,ILNJ2,KK,MA/My,MRL,NOD,NOR,NZB,NZO,NZW,PJ,PL,RF,RH2,SEA,SJL,SWR,TH,WSB | splice_donor_variant,splice_donor_5th_base_variant,coding_sequence_variant,intron_variant |
| 14 | 54865097 | 54865157 | 60 | Psmb11 | 129S1,AJ,AKR,BALB,BTBR,BUB,C3H,C57L,CAST,CBA,CE,DBA1J,DBA2J,FVB,ILNJ2,KK,MA/My,MRL,NOD,NOR,NZB,NZO,NZW,PJ,PL,RF,RH2,SEA,SJL,SMJ,SPRET,SWR,TH,WSB | feature_truncation,3_prime_UTR_variant |
| 14 | 55095072 | 55095138 | 66 | Homez | SPRET | feature_truncation,inframe_deletion |
| 14 | 55461227 | 55461282 | 55 | Dhrs2 | MOLF | feature_truncation,5_prime_UTR_variant |
| 14 | 56265413 | 56265536 | 123 | Mcpt9 | WSB | feature_truncation,coding_sequence_variant,intron_variant |
| 14 | 56885180 | 56885468 | 288 | Parp4 | BTBR,BUB,CAST,DBA2J,KK,MOLF,NZB,SJL,SPRET,WSB | feature_truncation,inframe_deletion |
| 14 | 57930796 | 57931002 | 206 | Lats2 | BTBR,BUB,CAST,DBA2J,KK,MOLF,SPRET,WSB | feature_truncation,3_prime_UTR_variant |
| 14 | 58040729 | 58040794 | 65 | Sap18 | CAST,MOLF | feature_truncation,3_prime_UTR_variant |
| 14 | 58348370 | 58348429 | 59 | Fgf9 | BTBR,BUB,CAST,DBA2J,MOLF,SPRET | feature_truncation,3_prime_UTR_variant |
| 14 | 62642069 | 62642156 | 87 | Gucy1b2 | MOLF,SPRET | feature_truncation,coding_sequence_variant,intron_variant |
| 14 | 63701081 | 63701481 | 400 | Fam167a | SPRET | feature_truncation,3_prime_UTR_variant |
| 14 | 65634325 | 65634377 | 52 | Zfp395 | MOLF | feature_truncation,3_prime_UTR_variant |
| 14 | 67937689 | 67937749 | 60 | Cdca2 | SPRET | feature_truncation,inframe_deletion |
| 14 | 67991616 | 67991806 | 190 | Dock5 | SPRET | feature_truncation,3_prime_UTR_variant |
| 14 | 67992672 | 67992728 | 56 | Dock5 | CAST,MOLF,SPRET | feature_truncation,3_prime_UTR_variant |
| 14 | 69932944 | 69933149 | 205 | Loxl2 | 129S1,C57L,CAST,KK,MOLF,WSB | feature_truncation,3_prime_UTR_variant |
| 14 | 70819011 | 70819085 | 74 | Nudt18 | SPRET | feature_truncation,3_prime_UTR_variant |
| 14 | 70821114 | 70821242 | 128 | Fhip2b | 129S1,AJ,BALB,C57L,CAST,KK,MOLF,PL,SEA,SPRET | feature_truncation,3_prime_UTR_variant |
| 14 | 70838994 | 70839072 | 78 | Dmtn | SPRET | feature_truncation,3_prime_UTR_variant |
| 14 | 73478285 | 73478337 | 52 | Lpar6 | SPRET | feature_truncation,3_prime_UTR_variant |
| 14 | 73758526 | 73758771 | 245 | Nudt15 | SPRET | feature_truncation,3_prime_UTR_variant |
| 14 | 76372870 | 76372990 | 120 | Nufip1 | CAST,MOLF,NZB,NZO | feature_truncation,3_prime_UTR_variant |
| 14 | 77273380 | 77273504 | 124 | Lacc1 | SPRET | feature_truncation,5_prime_UTR_variant,intron_variant |
| 14 | 78798216 | 78798278 | 62 | Dgkh | SPRET | feature_truncation,3_prime_UTR_variant |
| 14 | 78799970 | 78800030 | 60 | Dgkh | ILNJ2 | feature_truncation,3_prime_UTR_variant |
| 14 | 78969919 | 78969976 | 57 | Dgkh | 129S1,AJ,AKR,BALB,BTBR,BUB,C3H,C57L,CAST,CBA,CE,DBA1J,DBA2J,FVB,ILNJ2,KK,LP,MOLF,NZB,NZO,NZW,PJ,PL,RF,SEA,SJL,SMJ,SPRET,SWR,TH,WSB | feature_truncation,5_prime_UTR_variant |
| 14 | 79450820 | 79450949 | 129 | Zfp957 | SPRET | feature_truncation,inframe_deletion |
| 14 | 79563601 | 79563676 | 75 | Naa16 | CAST,CE,DBA1J,DBA2J,MOLF,PJ,SPRET | feature_truncation,3_prime_UTR_variant |
| 14 | 79868338 | 79868414 | 76 | Sugt1 | MOLF,WSB | feature_truncation,3_prime_UTR_variant |
| 14 | 79870477 | 79870625 | 148 | Sugt1 | SPRET | feature_truncation,3_prime_UTR_variant |
| 14 | 96341930 | 96341987 | 57 | Klhl1 | AJ,AKR,BALB,BTBR,BUB,C3H,CBA,CE,DBA1J,DBA2J,FVB,MRL,NOD,PJ,PL,RF,RH2,SEA,SJL,SMJ,SPRET,SWR,TH,WSB | feature_truncation,3_prime_UTR_variant |
| 14 | 99312966 | 99313606 | 640 | Dis3 | SPRET | feature_truncation,3_prime_UTR_variant |
| 14 | 103480616 | 103480715 | 99 | Mycbp2 | 129S1,C57L,CAST,ILNJ2,MOLF,NZB,NZO,PJ,SPRET,WSB | feature_truncation,coding_sequence_variant,intron_variant |
| 14 | 118504615 | 118505195 | 580 | Gm9376 | 129S1,CAST,KK,MOLF,SPRET | stop_lost,feature_truncation,coding_sequence_variant,3_prime_UTR_variant |
| 15 | 5128064 | 5128154 | 90 | Card6 | SPRET | feature_truncation,inframe_deletion |
| 15 | 5241596 | 5241660 | 64 | Ttc33 | SPRET | feature_truncation,coding_sequence_variant,intron_variant,feature_truncation,3_prime_UTR_variant,intron_variant |
| 15 | 7222163 | 7222229 | 66 | Lifr | AKR,CAST,LP,MOLF,RF,SPRET,TH | feature_truncation,3_prime_UTR_variant |
| 15 | 7223231 | 7223295 | 64 | Lifr | LP,MOLF,TH | feature_truncation,3_prime_UTR_variant |
| 15 | 10493479 | 10493560 | 81 | Rad1 | CAST,MOLF | feature_truncation,3_prime_UTR_variant |
| 15 | 10978881 | 10978985 | 104 | C1qtnf3 | MOLF,SPRET | feature_truncation,3_prime_UTR_variant |
| 15 | 35178886 | 35179287 | 401 | Stk3 | 129S1,BUB,C57L,KK,LP,MOLF,NOD,NOR,PJ,SPRET,SWR | feature_truncation,coding_sequence_variant,5_prime_UTR_variant,intron_variant |
| 15 | 38291894 | 38292054 | 160 | Klf10 | MOLF,SPRET | feature_truncation,3_prime_UTR_variant |
| 15 | 43728515 | 43728601 | 86 | Tmem74 | MOLF,SPRET | feature_truncation,3_prime_UTR_variant |
| 15 | 44236852 | 44237640 | 788 | Nudcd1 | MOLF,SPRET | feature_truncation,3_prime_UTR_variant |
| 15 | 51747029 | 51747088 | 59 | Utp23 | SPRET | feature_truncation,3_prime_UTR_variant |
| 15 | 53317763 | 53317823 | 60 | Samd12 | CAST | feature_truncation,3_prime_UTR_variant |
| 15 | 54616278 | 54616346 | 68 | Ccn3 | 129S1,C57L,MOLF,NZB,SPRET,TH | feature_truncation,3_prime_UTR_variant |
| 15 | 58422145 | 58422343 | 198 | Fer1l6 | ILNJ2,MOLF,NZB,NZW,SMJ,TH,WSB | feature_truncation,coding_sequence_variant,intron_variant |
| 15 | 66976624 | 66976785 | 161 | St3gal1 | SPRET | feature_truncation,3_prime_UTR_variant |
| 15 | 71303389 | 71303476 | 87 | Fam135b | CAST,MOLF,SPRET | feature_truncation,3_prime_UTR_variant |
| 15 | 71305991 | 71306126 | 135 | Fam135b | MOLF,SPRET | feature_truncation,3_prime_UTR_variant |
| 15 | 71668570 | 71668745 | 175 | Col22a1 | SPRET | feature_truncation,3_prime_UTR_variant |
| 15 | 75088229 | 75088412 | 183 | Ly6g2 | ILNJ2 | feature_truncation,5_prime_UTR_variant,intron_variant |
| 15 | 75874613 | 75874679 | 66 | Fam83h | SPRET | feature_truncation,inframe_deletion |

| | | | | | | |
|---|---|---|---|---|---|---|
| 15 | 76322212 | 76322298 | 86 | Mroh1 | MOLF | stop_lost,feature_truncation,coding_sequence_variant,3_prime_UTR_variant,intron_variant |
| 15 | 76599400 | 76599457 | 57 | Lrrc14 | SPRET | feature_truncation,3_prime_UTR_variant |
| 15 | 76785420 | 76785737 | 317 | Commd5 | MOLF | feature_truncation,3_prime_UTR_variant |
| 15 | 76935187 | 76935339 | 152 | Apol6 | MA/My | frameshift_variant,feature_truncation |
| 15 | 77373859 | 77374389 | 530 | Apol10a | MA/My,MOLF | feature_truncation,3_prime_UTR_variant |
| 15 | 77800010 | 77800362 | 352 | Txn2 | SPRET | feature_truncation,coding_sequence_variant,intron_variant |
| 15 | 78168155 | 78168231 | 76 | Csf2rb2 | SPRET | feature_truncation,3_prime_UTR_variant |
| 15 | 78364407 | 78364477 | 70 | Il2rb | SPRET | feature_truncation,3_prime_UTR_variant |
| 15 | 78660165 | 78660253 | 88 | Card10 | CAST,CBA,CE,DBA2J,FVB,ILNJ2,KK,LP,MA/My,MOLF,NOD,NOR,NZW,PL,SEA,SMJ,SPRET,TH,WSB | feature_truncation,3_prime_UTR_variant |
| 15 | 78660378 | 78660433 | 55 | Card10 | 129S1,AJ,AKR,BALB,BUB,C3H,C57L,C58J,DBA1J,MRL,PJ | feature_truncation,3_prime_UTR_variant |
| 15 | 78851187 | 78851259 | 72 | Triobp | MA/My,MOLF | feature_truncation,inframe_deletion |
| 15 | 80895939 | 80895995 | 56 | Sgsm3 | SPRET | feature_truncation,3_prime_UTR_variant |
| 15 | 81901135 | 81901272 | 137 | Xrcc6 | CAST | feature_truncation,5_prime_UTR_variant |
| 15 | 82255053 | 82255105 | 52 | Cyp2d22 | MOLF | feature_truncation,3_prime_UTR_variant |
| 15 | 83011472 | 83011580 | 108 | Poldip3 | MOLF,SPRET | feature_truncation,3_prime_UTR_variant |
| 15 | 83376047 | 83376134 | 87 | Ttll1 | CAST,SPRET | feature_truncation,3_prime_UTR_variant |
| 15 | 83459657 | 83459798 | 141 | Ttll12 | CAST | feature_truncation,3_prime_UTR_variant |
| 15 | 84226040 | 84226269 | 229 | Parvg | MOLF | feature_truncation,3_prime_UTR_variant |
| 15 | 88896186 | 88896285 | 99 | Mov10l1 | 129S1,C57L,CAST,CE,FVB,ILNJ2,PL,WSB | feature_truncation,inframe_deletion |
| 15 | 88987343 | 88987424 | 81 | Tubgcp6 | SPRET | feature_truncation,inframe_deletion |
| 15 | 89305602 | 89305652 | 50 | Cpt1b | MOLF,SPRET | frameshift_variant,feature_truncation |
| 15 | 93140727 | 93140896 | 169 | Gxylt1 | SPRET | feature_truncation,3_prime_UTR_variant |
| 15 | 98207421 | 98207731 | 310 | Or8s16 | CAST | feature_truncation,3_prime_UTR_variant |
| 15 | 98395993 | 98396311 | 318 | Or11m3 | AJ,C3H,CE,RH2 | stop_lost,3_prime_UTR_variant |
| 15 | 98399500 | 98399560 | 60 | Or11m3 | BUB,CAST,MA/My,SWR | feature_truncation,3_prime_UTR_variant |
| 15 | 98636348 | 98636508 | 160 | Arf3 | SPRET | feature_truncation,3_prime_UTR_variant |
| 15 | 98748887 | 98748947 | 60 | Kmt2d | C3H | feature_truncation,inframe_deletion |
| 15 | 99842052 | 99842166 | 114 | Fam186a | BALB,CAST,CBA,SPRET | feature_truncation,inframe_deletion |
| 15 | 99914036 | 99914127 | 91 | Larp4 | SPRET | feature_truncation,3_prime_UTR_variant |
| 15 | 100286368 | 100286487 | 119 | Slc11a2 | 129S1,BALB,BTBR,BUB,C57L,C58J,CAST,CBA,ILNJ2,KK,MA/My,MOLF,NOD,NOR,NZB,NZO,NZW,PJ,SEA,SMJ,SPRET,SWR,TH,WSB | feature_truncation,3_prime_UTR_variant |
| 15 | 100513326 | 100513380 | 54 | Dazap2 | 129S1,BALB,BTBR,BUB,C57L,C58J,CBA,ILNJ2,KK,MA/My,NOD,NOR,NZB,NZO,NZW,PJ,SEA,SWR,TH,WSB | feature_truncation,5_prime_UTR_variant |
| 15 | 100539142 | 100539205 | 63 | Bin2 | MOLF,SPRET | feature_truncation,3_prime_UTR_variant |
| 15 | 100720330 | 100720502 | 172 | Slc4a8 | MOLF,SPRET | feature_truncation,3_prime_UTR_variant |
| 15 | 101855641 | 101855740 | 99 | Krt78 | CAST,WSB | feature_truncation,inframe_deletion |
| 15 | 101998724 | 101998776 | 52 | Eif4b | CAST | feature_truncation,3_prime_UTR_variant |
| 15 | 102441494 | 102441763 | 269 | Atf7 | AKR,DBA1J,DBA2J,FVB,ILNJ2,LP,MRL,NOD,NOR,PJ,PL,SEA,SJL,SMJ | feature_truncation,3_prime_UTR_variant |
| 15 | 103783595 | 103783757 | 162 | Mucl1 | SPRET | stop_lost,feature_truncation,coding_sequence_variant,3_prime_UTR_variant,intron_variant |
| 16 | 3763468 | 3763655 | 187 | Nlrc3 | SPRET | feature_truncation,3_prime_UTR_variant |
| 16 | 3764271 | 3764559 | 288 | Nlrc3 | AJ,BTBR,BUB,CAST,CE,FVB,MOLF,MRL,PL,SJL,SMJ,SPRET,SWR,TH,WSB | feature_truncation,3_prime_UTR_variant |
| 16 | 3764832 | 3764927 | 95 | Nlrc3 | SMJ | feature_truncation,3_prime_UTR_variant |
| 16 | 10102097 | 10102148 | 51 | Emp2 | SPRET | feature_truncation,3_prime_UTR_variant |
| 16 | 10309100 | 10309178 | 78 | Ciita | CAST,MOLF,SPRET | feature_truncation,3_prime_UTR_variant |
| 16 | 11479218 | 11479723 | 505 | Snx29 | CAST,KK,MOLF,SPRET | feature_truncation,3_prime_UTR_variant |
| 16 | 13573589 | 13573763 | 174 | A930007A09Rik | BUB,CAST,KK,MOLF | feature_truncation,inframe_deletion |
| 16 | 13767014 | 13767241 | 227 | Mpv17l | C3H,CAST,CBA,DBA1J,DBA2J,KK,MOLF,SPRET,WSB | feature_truncation,3_prime_UTR_variant |
| 16 | 16126721 | 16126778 | 57 | Yars2 | CAST | feature_truncation,3_prime_UTR_variant |
| 16 | 17889821 | 17889879 | 58 | Prodh | SPRET | feature_truncation,3_prime_UTR_variant |
| 16 | 18226049 | 18226279 | 230 | Comt | C3H,C57L,C58J,CAST,CBA,DBA1J,DBA2J,FVB,ILNJ2,KK,LP,MA/My,MOLF,NZB,NZO,NZW,PJ,RF,SJL,SPRET,SWR,TH,WSB | feature_truncation,3_prime_UTR_variant |
| 16 | 19225205 | 19225636 | 431 | Or2m13 | MOLF | feature_truncation,3_prime_UTR_variant |
| 16 | 19472681 | 19472766 | 85 | Lamp3 | CAST,MOLF,SPRET | feature_truncation,3_prime_UTR_variant |
| 16 | 19473775 | 19473847 | 72 | Lamp3 | SPRET | feature_truncation,3_prime_UTR_variant |
| 16 | 19945092 | 19945272 | 180 | Klhl24 | CAST,SPRET,WSB | feature_truncation,3_prime_UTR_variant |
| 16 | 19946738 | 19946935 | 197 | Klhl24 | CAST,SPRET | feature_truncation,3_prime_UTR_variant |
| 16 | 20351292 | 20351394 | 102 | Dvl3 | CAST,MOLF,SPRET | feature_truncation,3_prime_UTR_variant |
| 16 | 21727519 | 21727753 | 234 | Map3k13 | CAST,MOLF,SPRET,WSB | feature_truncation,coding_sequence_variant,intron_variant |
| 16 | 23298178 | 23298268 | 90 | Masp1 | CAST,MOLF,PL,SMJ,SPRET,TH,WSB | feature_truncation,3_prime_UTR_variant |
| 16 | 27269139 | 27269209 | 70 | Ccdc50 | SPRET | feature_truncation,3_prime_UTR_variant |
| 16 | 27270344 | 27270402 | 58 | Ccdc50 | BALB | feature_truncation,3_prime_UTR_variant |
| 16 | 31114801 | 31115056 | 255 | Apod | 129S1,AJ,BALB,CAST,FVB,ILNJ2,KK,LP,MOLF,MRL,NOD,NOR,NZB,NZO,NZW,PL,RH2,SEA,SJL,TH,WSB | feature_truncation,3_prime_UTR_variant |
| 16 | 32569539 | 32569880 | 341 | Muc4 | CAST | frameshift_variant,feature_truncation |
| 16 | 32571430 | 32571801 | 371 | Muc4 | SPRET | frameshift_variant,feature_truncation |
| 16 | 32614682 | 32614745 | 63 | Muc20 | SPRET | feature_truncation,inframe_deletion |
| 16 | 38063414 | 38063480 | 66 | Gsk3b | MOLF,SPRET | feature_truncation,3_prime_UTR_variant |
| 16 | 38198175 | 38198353 | 178 | Popdc2 | MOLF | feature_truncation,3_prime_UTR_variant |
| 16 | 38265010 | 38265062 | 52 | Adprh | SPRET | feature_truncation,3_prime_UTR_variant |
| 16 | 43469230 | 43469290 | 60 | Tigit | CAST | feature_truncation,3_prime_UTR_variant |
| 16 | 44780332 | 44780648 | 316 | Cd200r3 | SPRET | feature_truncation,3_prime_UTR_variant |
| 16 | 45312779 | 45313590 | 811 | Cd200l1 | SPRET | feature_truncation,coding_sequence_variant,5_prime_UTR_variant,intron_variant |
| 16 | 48139206 | 48139371 | 165 | Dppa2 | C3H,CAST,CBA,MOLF,SPRET | feature_truncation,3_prime_UTR_variant |
| 16 | 55853913 | 55854007 | 94 | Trmt10c | CBA,MOLF,NZW,SMJ | feature_truncation,3_prime_UTR_variant |
| 16 | 56435066 | 56435990 | 924 | Abi3bp | C58J,LP | feature_truncation,coding_sequence_variant,intron_variant |
| 16 | 58579779 | 58579989 | 210 | Or5k1b | SPRET | feature_truncation,3_prime_UTR_variant |
| 16 | 58709159 | 58709213 | 54 | Or5k15 | SPRET | feature_truncation,3_prime_UTR_variant |

| | | | | | | |
|---|---|---|---|---|---|---|
| 16 | 59124597 | 59124685 | 88 | Or5ac21 | SPRET | feature_truncation,3_prime_UTR_variant |
| 16 | 65350819 | 65351120 | 301 | Chmp2b | CAST | feature_truncation,coding_sequence_variant,5_prime_UTR_variant,intron_variant |
| 16 | 81394611 | 81394841 | 230 | Ncam2 | CAST,MOLF,SPRET | feature_truncation,3_prime_UTR_variant |
| 16 | 84659881 | 84659932 | 51 | Gabpa | BALB,BUB,C58J,CBA,LP,MRL,NZO,RH2,SPRET | feature_truncation,3_prime_UTR_variant |
| 16 | 85253511 | 85253574 | 63 | Cyyr1 | SPRET | feature_truncation,3_prime_UTR_variant |
| 16 | 87162106 | 87162320 | 214 | N6amt1 | CAST,MOLF,SPRET | feature_truncation,3_prime_UTR_variant |
| 16 | 88570804 | 88570864 | 60 | 2310057N15Rik | 129S1,C57L,C58J,KK,NZB,NZO,NZW,PL | feature_truncation,inframe_deletion |
| 16 | 88670780 | 88670933 | 153 | Krtap19-2 | 129S1,AKR,C57L,C58J,CAST,CBA,CE,DBA1J,DBA2J,ILNJ2,KK,MOLF,MRL,NZB,NZO,NZW,PJ,PL,RF,SEA,SMJ,SPRET,TH | feature_truncation,inframe_deletion |
| 16 | 89216724 | 89216844 | 120 | Krtap6-2 | CAST,MOLF,SPRET | feature_truncation,inframe_deletion |
| 16 | 91411899 | 91412052 | 153 | Dnajc28 | SPRET | feature_truncation,3_prime_UTR_variant |
| 16 | 91627395 | 91627476 | 81 | Itsn1 | CBA,CE,DBA1J,MA/My,PL | feature_truncation,3_prime_UTR_variant |
| 16 | 91714191 | 91714253 | 62 | Itsn1 | SPRET | feature_truncation,3_prime_UTR_variant |
| 16 | 92108278 | 92108354 | 76 | Smim11 | SPRET | feature_truncation,3_prime_UTR_variant |
| 16 | 92144628 | 92144820 | 192 | Kcne1 | BUB,CAST,CE,DBA1J,FVB,MOLF,NZB,PL,SJL,SPRET,SWR,TH | feature_truncation,3_prime_UTR_variant |
| 16 | 92255346 | 92255397 | 51 | Rcan1 | CE,DBA1J,MOLF,NZO,NZW,PL,TH | feature_truncation,5_prime_UTR_variant |
| 16 | 94549489 | 94549543 | 54 | Kcnj6 | MOLF,NZW,SPRET | feature_truncation,3_prime_UTR_variant |
| 16 | 95058705 | 95058802 | 97 | Kcnj15 | 129S1,AJ,AKR,B10D2,B10J,BALB,BTBR,BUB,C3H,C57L,C58J,CAST,CBA,CE,DBA1J,DBA2J,FVB,ILNJ2,KK,LP,MA/My,MOLF,MRL,NOD,NOR,NZB,NZO,NZW,PJ,PL,RF,RH2,SEA,SJL,SMJ,SPRET,SWR,TH,WSB | feature_truncation,5_prime_UTR_variant |
| 16 | 97314701 | 97314834 | 133 | Fam3b | SPRET | feature_truncation,5_prime_UTR_variant,intron_variant |
| 17 | 5482495 | 5483354 | 859 | Tmem242 | SPRET | feature_truncation,3_prime_UTR_variant,stop_lost,feature_truncation,coding_sequence_variant,3_prime_UTR_variant,intron_variant |
| 17 | 5978501 | 5978657 | 156 | Snx9 | SPRET | feature_truncation,coding_sequence_variant,intron_variant |
| 17 | 6973005 | 6973229 | 224 | Sytl3 | 129S1,AJ,AKR,BALB,C3H,C57L,CAST,CBA,CE,DBA1J,DBA2J,ILNJ2,MOLF,MRL,NOD,NOR,NZO,PJ,PL,RF,RH2,SEA,SMJ,SPRET,TH,WSB | feature_truncation,5_prime_UTR_variant |
| 17 | 13449114 | 13449178 | 64 | Smok2b | SPRET | feature_truncation,5_prime_UTR_variant |
| 17 | 15764864 | 15764948 | 84 | Prdm9 | CAST,ILNJ2,SPRET | feature_truncation,inframe_deletion |
| 17 | 17307825 | 17307980 | 155 | Zfp960 | C58J,PL | frameshift_variant,feature_truncation |
| 17 | 19619005 | 19619121 | 116 | Vmn2r99 | SPRET | feature_truncation,3_prime_UTR_variant |
| 17 | 19927994 | 19928048 | 54 | Vmn2r102 | SPRET | feature_truncation,3_prime_UTR_variant |
| 17 | 20426023 | 20426410 | 387 | Vmn2r105 | SPRET | feature_truncation,3_prime_UTR_variant |
| 17 | 20676884 | 20676986 | 102 | Vmn2r108 | SPRET | feature_truncation,3_prime_UTR_variant |
| 17 | 20958875 | 20959232 | 357 | Vmn1r227 | SPRET | feature_truncation,3_prime_UTR_variant |
| 17 | 21213494 | 21213562 | 68 | Vmn1r233 | SPRET | feature_truncation,3_prime_UTR_variant |
| 17 | 21262502 | 21262553 | 51 | Zfp160 | AKR,C3H,CBA,FVB,MA/My,MRL,PJ,WSB | feature_truncation,3_prime_UTR_variant |
| 17 | 21618789 | 21618957 | 168 | Zfp677 | CAST,MOLF,WSB | stop_lost,feature_truncation,coding_sequence_variant,3_prime_UTR_variant |
| 17 | 21807273 | 21807372 | 99 | Zfp948 | 129S1,AJ,BALB,C57L,CE,DBA1J,DBA2J,NZO,RH2,SEA,SPRET | feature_truncation,inframe_deletion |
| 17 | 22037627 | 22037711 | 84 | Zfp820 | SPRET | stop_lost,feature_truncation,coding_sequence_variant,3_prime_UTR_variant |
| 17 | 22766131 | 22766239 | 108 | Vmn2r111 | CAST,MOLF,SPRET | feature_truncation,3_prime_UTR_variant |
| 17 | 23847162 | 23847450 | 288 | Mmp25 | BTBR,BUB,C58J,KK,LP,NOD,NOR,NZW,PL,SWR | feature_truncation,3_prime_UTR_variant |
| 17 | 24397374 | 24397425 | 51 | Tbc1d24 | AKR | feature_truncation,3_prime_UTR_variant |
| 17 | 24399026 | 24399173 | 147 | Tbc1d24 | 129S1,AJ,AKR,B10D2,BALB,BTBR,BUB,C3H,C57L,CAST,CBA,CE,DBA1J,DBA2J,FVB,ILNJ2,KK,MA/My,MRL,NOD,NOR,NZB,NZO,NZW,PJ,RF,RH2,SEA,SJL,SMJ,SPRET,SWR,TH,WSB | feature_truncation,3_prime_UTR_variant |
| 17 | 24912587 | 24913136 | 549 | Gfer | 129S1,AJ,AKR,B10D2,BALB,BTBR,BUB,C3H,C57L,CAST,CBA,CE,DBA1J,DBA2J,FVB,ILNJ2,KK,MA/My,MRL,NOD,NOR,NZB,NZO,NZW,PJ,RF,RH2,SEA,SJL,SMJ,SPRET,SWR,TH,WSB | feature_truncation,3_prime_UTR_variant |
| 17 | 24940687 | 24940775 | 88 | Rps2 | CAST | feature_truncation,coding_sequence_variant,intron_variant |
| 17 | 25314823 | 25314878 | 55 | Ift140 | SPRET | feature_truncation,3_prime_UTR_variant |
| 17 | 26287760 | 26287829 | 69 | Rab11fip3 | SPRET | feature_truncation,inframe_deletion |
| 17 | 26663356 | 26663419 | 63 | Neurl1b | 129S1,AJ,AKR,B10D2,BALB,C3H,C57L,CBA,CE,DBA1J,DBA2J,ILNJ2,MRL,NOD,NOR,NZO,PJ,RF,RH2,SEA,TH | feature_truncation,3_prime_UTR_variant |
| 17 | 28056819 | 28057312 | 493 | Snrpc | KK,NZB,RF,SPRET | feature_truncation,5_prime_UTR_variant,intron_variant |
| 17 | 28115605 | 28115713 | 108 | Bltp3a | SPRET | feature_truncation,3_prime_UTR_variant |
| 17 | 28519321 | 28519379 | 58 | Ppard | SPRET | feature_truncation,3_prime_UTR_variant |
| 17 | 28656388 | 28656916 | 528 | Fkbp5 | SPRET | feature_truncation,3_prime_UTR_variant |
| 17 | 28808323 | 28808395 | 72 | Srpk1 | CAST,SPRET,WSB | feature_truncation,3_prime_UTR_variant |
| 17 | 29058682 | 29059118 | 436 | Brpf3 | SPRET | feature_truncation,3_prime_UTR_variant |
| 17 | 29108791 | 29109372 | 581 | Pnpla1 | CAST,KK,MOLF,RF | feature_truncation,3_prime_UTR_variant |
| 17 | 29531134 | 29531282 | 148 | BC004004 | SPRET | feature_truncation,3_prime_UTR_variant |
| 17 | 29536476 | 29536748 | 272 | Pi16 | SPRET | feature_truncation,5_prime_UTR_variant |
| 17 | 29863485 | 29864099 | 614 | Rnf8 | CAST,NOD,NOR,NZB | feature_truncation,3_prime_UTR_variant |
| 17 | 29923672 | 29923786 | 114 | Cmtr1 | KK,MOLF,RF | feature_truncation,3_prime_UTR_variant |
| 17 | 30810681 | 30810882 | 201 | Glo1 | SPRET | feature_truncation,3_prime_UTR_variant |
| 17 | 30863677 | 30863913 | 236 | Dnah8 | AJ,AKR,BALB,BTBR,BUB,C3H,C58J,CAST,CBA,CE,FVB,ILNJ2,KK,MA/My,MOLF,MRL,NOD,NOR,NZB,NZO,NZW,PJ,PL,RF,SJL,SMJ,SPRET,SWR,TH,WSB | feature_truncation,coding_sequence_variant,intron_variant |
| 17 | 30936864 | 30937172 | 308 | Dnah8 | SPRET | feature_truncation,coding_sequence_variant,intron_variant |

| | | | | | | |
|---|---|---|---|---|---|---|
| 17 | 31095907 | 31096007 | 100 | Dnah8 | 129S1,AJ,AKR,B10D2,BALB,BTBR,BUB,C3H,C57L,C58J,CAST,CBA,CE,DBA1J,DBA2J,ILNJ2,KK,MA/My,MOLF,MRL,NOD,NOR,NZO,NZW,PJ,PL,RF,RH2,SEA,SMJ,SPRET,SWR,WSB | feature_truncation,3_prime_UTR_variant |
| 17 | 31463761 | 31463959 | 198 | Ubash3a | C58J,CAST,KK,MOLF,NOD,NOR,NZB,NZO,NZW,SPRET | feature_truncation,3_prime_UTR_variant |
| 17 | 31828624 | 31828688 | 64 | Cbs | SPRET | feature_truncation,3_prime_UTR_variant |
| 17 | 32402972 | 32403078 | 106 | Ephx3 | MOLF,PL,SPRET | feature_truncation,3_prime_UTR_variant |
| 17 | 33035551 | 33035610 | 59 | Zfp799 | SPRET | feature_truncation,3_prime_UTR_variant |
| 17 | 33098389 | 33098611 | 222 | Zfp870 | MOLF | feature_truncation,3_prime_UTR_variant |
| 17 | 33461224 | 33461320 | 96 | Zfp955a | MOLF,TH | feature_truncation,inframe_deletion |
| 17 | 33492125 | 33492337 | 212 | Or10h28 | SPRET | feature_truncation,3_prime_UTR_variant |
| 17 | 33524255 | 33524310 | 55 | Zfp955b | SPRET | feature_truncation,3_prime_UTR_variant |
| 17 | 34171285 | 34171375 | 90 | Rps18 | SPRET | feature_truncation,coding_sequence_variant,intron_variant |
| 17 | 34181759 | 34181824 | 65 | Vps52 | SPRET | feature_truncation,coding_sequence_variant,intron_variant |
| 17 | 35912503 | 35912595 | 92 | Sfta2 | CAST,DBA1J,ILNJ2,PJ,SJL,TH,WSB | feature_truncation,5_prime_UTR_variant |
| 17 | 35930158 | 35930270 | 112 | Muc21 | CAST,SPRET | frameshift_variant,feature_truncation |
| 17 | 35930919 | 35931009 | 90 | Muc21 | 129S1,AKR,BTBR,C3H,C57L,CBA,CE,LP,MA/My,MRL,NZO,NZW,RF,SMJ,TH | feature_truncation,inframe_deletion |
| 17 | 35931846 | 35932115 | 269 | Muc21 | ILNJ2 | frameshift_variant,feature_truncation |
| 17 | 35932248 | 35932788 | 540 | Muc21 | 129S1,AKR,B10D2,BALB,BTBR,BUB,C3H,C57L,C58J,CBA,CE,DBA1J,DBA2J,FVB,ILNJ2,KK,LP,MA/My,MRL,NOD,NOR,NZB,NZO,NZW,PJ,PL,RF,SMJ,SWR | feature_truncation,inframe_deletion |
| 17 | 35933391 | 35933548 | 157 | Muc21 | BUB,CE,MA/My | frameshift_variant,feature_truncation |
| 17 | 35934039 | 35934292 | 253 | Muc21 | CAST,MOLF,SPRET,TH | feature_truncation,coding_sequence_variant,intron_variant |
| 17 | 36241478 | 36241609 | 131 | Ppp1r10 | NZW | frameshift_variant,feature_truncation |
| 17 | 36268176 | 36268236 | 60 | Abcf1 | B10D2,BALB,DBA2J,PJ,RH2,SEA,SPRET | feature_truncation,3_prime_UTR_variant |
| 17 | 37394629 | 37395252 | 623 | Or2h2 | 129S1,AJ,AKR,B10D2,BALB,BTBR,BUB,C3H,C57L,C58J,CAST,CBA,CE,DBA1J,DBA2J,FVB,ILNJ2,KK,LP,MA/My,MRL,NOD,NOR,NZB,NZO,NZW,PJ,PL,RF,RH2,SEA,SJL,SPRET,SWR,TH | feature_truncation,3_prime_UTR_variant |
| 17 | 37461066 | 37461124 | 58 | Or2h1b | AJ,B10D2,BALB,BUB,C58J,DBA1J,DBA2J,KK,NOD,NOR,NZB,NZO,NZW,PL,RH2,SEA,SJL,SWR | feature_truncation,3_prime_UTR_variant |
| 17 | 37690906 | 37691241 | 335 | Or12d15 | B10D2,BALB,CAST,DBA1J,DBA2J,NZO,NZW,RH2,SEA,SPRET | feature_truncation,coding_sequence_variant,intron_variant |
| 17 | 37978562 | 37979057 | 495 | Or10al2 | SPRET | feature_truncation,5_prime_UTR_variant,intron_variant |
| 17 | 38059742 | 38059828 | 86 | Or10al5 | B10D2,BALB,DBA1J,DBA2J,FVB,ILNJ2,NZO,NZW,RH2,SEA,TH | feature_truncation,5_prime_UTR_variant |
| 17 | 38364192 | 38364442 | 250 | Or10al7 | MOLF | feature_truncation,3_prime_UTR_variant |
| 17 | 38381597 | 38382496 | 899 | Or2g7 | MOLF | feature_truncation,3_prime_UTR_variant |
| 17 | 41105387 | 41105440 | 53 | Crisp2 | MOLF | feature_truncation,5_prime_UTR_variant |
| 17 | 44446111 | 44446195 | 84 | Clic5 | AKR,BALB,C3H,CBA,CE,DBA2J,ILNJ2,MA/My,PJ,RF,RH2,SEA,SPRET | feature_truncation,inframe_deletion |
| 17 | 46563358 | 46563463 | 105 | Yipf3 | 129S1,BUB,FVB,NOD,NOR,NZW,SJL,SPRET,SWR | feature_truncation,3_prime_UTR_variant |
| 17 | 47084813 | 47084958 | 145 | Rpl7l1 | CAST,MOLF,NOD,NOR,NZW,SJL,SMJ,SPRET,WSB | feature_truncation,3_prime_UTR_variant |
| 17 | 53981999 | 53982235 | 236 | Sgo1 | BTBR,MA/My,TH | feature_truncation,3_prime_UTR_variant |
| 17 | 56411965 | 56412955 | 990 | Plin4 | CAST,PL,SPRET | feature_truncation,coding_sequence_variant,intron_variant,feature_truncation,inframe_deletion |
| 17 | 56483127 | 56483183 | 56 | Mydgf | CAST | feature_truncation,3_prime_UTR_variant |
| 17 | 56882590 | 56882671 | 81 | Safb2 | SEA | feature_truncation,inframe_deletion |
| 17 | 65945416 | 65945506 | 90 | Txndc2 | SPRET | feature_truncation,inframe_deletion |
| 17 | 74324144 | 74324202 | 58 | Srd5a2 | CAST,MOLF | feature_truncation,3_prime_UTR_variant |
| 17 | 74732730 | 74732913 | 183 | Nlrc4 | 129S1,BTBR,BUB,C57L,ILNJ2,KK,NOD,NOR,NZW,SPRET,SWR | feature_truncation,3_prime_UTR_variant |
| 17 | 79153607 | 79153816 | 209 | Gpatch11 | CAST,CE,ILNJ2,NZO,NZW,SPRET,WSB | feature_truncation,3_prime_UTR_variant |
| 17 | 87139069 | 87139125 | 56 | Epas1 | MOLF,PL,SPRET | feature_truncation,3_prime_UTR_variant |
| 18 | 3515915 | 3515987 | 72 | Bambi | CAST | feature_truncation,3_prime_UTR_variant |
| 18 | 6202202 | 6202371 | 169 | Kif5b | 129S1,AKR,BALB,C3H,C57L,CBA,ILNJ2,PJ,RF,RH2,SEA,SMJ,SPRET | feature_truncation,3_prime_UTR_variant |
| 18 | 10639639 | 10639709 | 70 | Snrpd1 | CAST,MOLF,SPRET | frameshift_variant |
| 18 | 10725780 | 10725837 | 57 | Mib1 | 129S1,C57L,MOLF | feature_truncation,5_prime_UTR_variant |
| 18 | 20736592 | 20736647 | 55 | Dsg2 | CAST | feature_truncation,3_prime_UTR_variant |
| 18 | 25269296 | 25269355 | 59 | Tpgs2 | MOLF | feature_truncation,3_prime_UTR_variant |
| 18 | 25886998 | 25887058 | 60 | Celf4 | SPRET | feature_truncation,5_prime_UTR_variant |
| 18 | 34402304 | 34402410 | 106 | Apc | SPRET | feature_truncation,coding_sequence_variant,intron_variant |
| 18 | 34479272 | 34479373 | 101 | Reep5 | SPRET | feature_truncation,3_prime_UTR_variant |
| 18 | 34732619 | 34732834 | 215 | Brd8 | 129S1,AJ,BALB,BUB,C3H,C57L,C58J,CAST,CBA,CE,DBA1J,DBA2J,ILNJ2,MOLF,MRL,NOD,NOR,PL,RF,RH2,SEA,SJL,SPRET,TH,WSB | feature_truncation,3_prime_UTR_variant |
| 18 | 34870406 | 34871105 | 699 | Rpl27rt | CAST,SPRET | stop_lost,feature_truncation,coding_sequence_variant,5_prime_UTR_variant,3_prime_UTR_variant |
| 18 | 35726068 | 35726293 | 225 | Matr3 | CAST,MOLF,SPRET | feature_truncation,3_prime_UTR_variant |
| 18 | 37437165 | 37437271 | 106 | Pcdhb3 | 129S1,AJ,AKR,BALB,BUB,C3H,C57L,C58J,CAST,CBA,CE,DBA1J,DBA2J,FVB,ILNJ2,KK,MA/My,MOLF,NOD,NOR,NZB,NZO,NZW,PL,RF,RH2,SEA,SJL,SMJ,SWR,TH,WSB | feature_truncation,3_prime_UTR_variant |
| 18 | 37636474 | 37636527 | 53 | Pcdhb19 | CAST,KK,MOLF,SPRET | feature_truncation,3_prime_UTR_variant |
| 18 | 37654957 | 37655039 | 82 | Pcdhb22 | CAST,MOLF,SPRET | feature_truncation,3_prime_UTR_variant |
| 18 | 37828371 | 37828516 | 145 | Pcdhga5 | BUB,C3H,CE,DBA1J,MA/My,NOD,NOR,PL,RF | frameshift_variant,feature_truncation |
| 18 | 40336899 | 40337227 | 328 | Yipf5 | SPRET | feature_truncation,3_prime_UTR_variant |
| 18 | 44559174 | 44559226 | 52 | Mcc | BALB,C3H,CAST,CBA,DBA1J,DBA2J,KK,MOLF,SEA,SPRET | feature_truncation,3_prime_UTR_variant |
| 18 | 47245193 | 47245746 | 553 | Hspe1-rs1 | BALB,C3H,C58J,CAST,CBA,DBA1J,DBA2J,ILNJ2,KK,LP,MA/My,MOLF,NZO,NZW,SEA,SPRET | transcript_ablation |

| | | | | | | |
|---|---|---|---|---|---|---|
| 18 | 50279640 | 50279755 | 115 | Hsd17b4 | SPRET | feature_truncation,coding_sequence_variant,intron_variant |
| 18 | 52624796 | 52624857 | 61 | Srfbp1 | 129S1,AJ,AKR,BALB,BUB,C3H,C57L,C58J,CAST,CBA,CE,DBA1J,DBA2J,FVB,ILNJ2,LP,MA/My,MRL,NOD,NOR,NZB,NZO,NZW,PL,RF,RH2,SEA,SJL,SPRET,SWR | feature_truncation,3_prime_UTR_variant |
| 18 | 56720630 | 56720750 | 120 | Phax | C3H,CE,DBA1J,DBA2J,PJ,SPRET | feature_truncation,3_prime_UTR_variant |
| 18 | 56866168 | 56866317 | 149 | Lmnb1 | SPRET | feature_truncation,coding_sequence_variant,intron_variant |
| 18 | 60402190 | 60402373 | 183 | Gm4841 | KK,MOLF,NOD,NOR | feature_truncation,3_prime_UTR_variant |
| 18 | 60403399 | 60404196 | 797 | Gm4841 | NZO,NZW | feature_truncation,coding_sequence_variant,5_prime_UTR_variant,intron_variant |
| 18 | 60709194 | 60709252 | 58 | Myoz3 | CAST | feature_truncation,3_prime_UTR_variant |
| 18 | 61327045 | 61327097 | 52 | Slc26a2 | NZW | feature_truncation,3_prime_UTR_variant |
| 18 | 61421432 | 61421487 | 55 | Pde6a | CAST | feature_truncation,3_prime_UTR_variant |
| 18 | 62156526 | 62156819 | 293 | Sh3tc2 | SPRET | feature_truncation,3_prime_UTR_variant |
| 18 | 62800804 | 62801103 | 299 | Spink10 | AJ,BALB,CBA,ILNJ2,PL,RH2,SEA | feature_truncation,3_prime_UTR_variant |
| 18 | 63092781 | 63092863 | 82 | Apcdd1 | CAST,MOLF,SPRET | feature_truncation,3_prime_UTR_variant |
| 18 | 65610665 | 65610801 | 136 | Malt1 | SPRET | feature_truncation,3_prime_UTR_variant |
| 18 | 75219533 | 75219757 | 224 | Dym | 129S1,AKR,BTBR,BUB,C3H,C57L,CBA,DBA1J,DBA2J,FVB,NZO,PJ,PL,SEA,SPRET,TH,WSB | feature_truncation,3_prime_UTR_variant |
| 18 | 75219780 | 75219892 | 112 | Dym | CAST,ILNJ2,MOLF,NZW,RF | feature_truncation,3_prime_UTR_variant |
| 18 | 75565905 | 75565967 | 62 | Ctif | SPRET | feature_truncation,3_prime_UTR_variant |
| 18 | 76440876 | 76441477 | 601 | Smad2 | CAST | feature_truncation,3_prime_UTR_variant |
| 18 | 76443116 | 76443177 | 61 | Smad2 | SPRET | feature_truncation,3_prime_UTR_variant |
| 18 | 77343636 | 77343693 | 57 | St8sia5 | CAST | feature_truncation,3_prime_UTR_variant |
| 18 | 77545200 | 77545368 | 168 | Ark2c | CAST,KK,MOLF,NZO,NZW,TH | feature_truncation,3_prime_UTR_variant |
| 18 | 78795972 | 78796032 | 60 | Setbp1 | CAST,MOLF,SPRET | feature_truncation,3_prime_UTR_variant |
| 18 | 80266867 | 80267036 | 169 | Txnl4a | 129S1,AJ,AKR,BALB,BTBR,BUB,C57L,CAST,FVB,ILNJ2,MA/My,MOLF,MRL,NOD,NOR,NZO,RH2,SJL,SPRET,TH,WSB | feature_truncation,3_prime_UTR_variant |
| 18 | 80268726 | 80268815 | 89 | Txnl4a | SPRET | feature_truncation,3_prime_UTR_variant |
| 18 | 80317493 | 80317553 | 60 | Slc66a2 | AKR,CAST,MOLF,NZO,SPRET | feature_truncation,3_prime_UTR_variant |
| 18 | 84963685 | 84964495 | 810 | Timm21 | CE,MOLF | feature_truncation,3_prime_UTR_variant |
| 19 | 3439428 | 3439510 | 82 | Tesmin | FVB,SJL | feature_truncation,5_prime_UTR_variant |
| 19 | 3857463 | 3857529 | 66 | Kmt5b | SPRET | feature_truncation,3_prime_UTR_variant |
| 19 | 4438398 | 4438633 | 235 | Kdm2a | AJ,AKR,BALB,BUB,C3H,C58J,CAST,CBA,CE,DBA1J,DBA2J,FVB,ILNJ2,KK,LP,MA/My,MOLF,MRL,NOD,NOR,NZW,PJ,PL,RF,RH2,SEA,SJL,SMJ,SPRET,SWR,TH,WSB | feature_truncation,3_prime_UTR_variant,intron_variant |
| 19 | 4918611 | 4918804 | 193 | Actn3 | CAST,MOLF,SPRET | splice_donor_variant,splice_donor_5th_base_variant,coding_sequence_variant,intron_variant |
| 19 | 5015000 | 5015751 | 751 | Mrpl11 | SPRET | feature_truncation,3_prime_UTR_variant |
| 19 | 5512637 | 5512711 | 74 | Fibp | SPRET | feature_truncation,coding_sequence_variant,intron_variant |
| 19 | 5545018 | 5545244 | 226 | Cfl1 | CAST,FVB,LP,MOLF,SJL,SPRET | feature_truncation,3_prime_UTR_variant |
| 19 | 5991093 | 5991275 | 182 | Pola2 | BUB,C58J,CAST,FVB,LP,MOLF,MRL,SJL,SPRET,TH | feature_truncation,3_prime_UTR_variant |
| 19 | 6113093 | 6113149 | 56 | Tm7sf2 | BUB,CAST | feature_truncation,coding_sequence_variant,intron_variant |
| 19 | 6441070 | 6441174 | 104 | Pygm | CAST,MOLF | feature_truncation,coding_sequence_variant,intron_variant |
| 19 | 8684627 | 8684814 | 187 | Slc3a2 | CAST,SPRET | feature_truncation,3_prime_UTR_variant |
| 19 | 8830584 | 8830660 | 76 | Lrrn4cl | SPRET | feature_truncation,3_prime_UTR_variant |
| 19 | 10695081 | 10695297 | 216 | Cd5 | BUB,C3H,CAST,CBA,DBA1J,DBA2J,FVB,ILNJ2,LP,MA/My,MOLF,SPRET,SWR | feature_truncation,3_prime_UTR_variant |
| 19 | 10885770 | 10885829 | 59 | Prpf19 | SPRET | feature_truncation,3_prime_UTR_variant |
| 19 | 10926167 | 10926347 | 180 | Ccdc86 | CAST,MOLF,SPRET | feature_truncation,inframe_deletion |
| 19 | 11439876 | 11440757 | 881 | Ms4a4b | FVB,SWR | stop_lost,feature_truncation,coding_sequence_variant,3_prime_UTR_variant,intron_variant |
| 19 | 11441000 | 11441052 | 52 | Ms4a4b | SPRET | feature_truncation,3_prime_UTR_variant |
| 19 | 11508483 | 11509455 | 972 | Ms4a6b | SPRET | feature_truncation,3_prime_UTR_variant |
| 19 | 11863707 | 11864212 | 505 | Or10v1 | MOLF | feature_truncation,5_prime_UTR_variant |
| 19 | 12010978 | 12011174 | 196 | Or4d11 | FVB,SPRET,SWR | feature_truncation,3_prime_UTR_variant |
| 19 | 12261512 | 12261614 | 102 | Or5an1 | AJ,BALB,CE,MA/My,NOD,NOR,PL,RH2,SEA,WSB | feature_truncation,3_prime_UTR_variant |
| 19 | 12307186 | 12307329 | 143 | Or5a21 | MOLF,SPRET | feature_truncation,3_prime_UTR_variant |
| 19 | 12660755 | 12661212 | 457 | Or5b95 | ILNJ2,SPRET,WSB | feature_truncation,3_prime_UTR_variant |
| 19 | 13071041 | 13071260 | 219 | Or5b104 | CAST,LP,WSB | feature_truncation,3_prime_UTR_variant |
| 19 | 13682735 | 13682801 | 66 | Or10q1b | MOLF | feature_truncation,inframe_deletion |
| 19 | 15881904 | 15882075 | 171 | Psat1 | CAST,MOLF,SPRET | feature_truncation,3_prime_UTR_variant |
| 19 | 17410102 | 17410168 | 66 | Pcsk5 | CAST,CBA,DBA1J,DBA2J,MOLF | feature_truncation,3_prime_UTR_variant |
| 19 | 21369844 | 21370026 | 182 | Gda | BUB,MOLF,MRL,NZB,NZO,NZW | feature_truncation,3_prime_UTR_variant |
| 19 | 27313243 | 27313323 | 80 | Kcnv2 | CAST,MOLF | feature_truncation,3_prime_UTR_variant |
| 19 | 29364414 | 29364466 | 52 | Cd274 | SPRET | feature_truncation,3_prime_UTR_variant |
| 19 | 30544002 | 30544116 | 114 | Prkg1 | 129S1,BTBR,C57L,C58J,MOLF,NZW,SPRET | feature_truncation,3_prime_UTR_variant |
| 19 | 31062515 | 31062727 | 212 | Cstf2t | SPRET | feature_truncation,3_prime_UTR_variant |
| 19 | 32443255 | 32443977 | 722 | Rpl9-ps6 | CAST,MOLF,SPRET | stop_lost,feature_truncation,coding_sequence_variant,5_prime_UTR_variant,3_prime_UTR_variant |
| 19 | 32676036 | 32676104 | 68 | Atad1 | SPRET | feature_truncation,coding_sequence_variant,intron_variant |
| 19 | 38957417 | 38957503 | 86 | Hells | SPRET | feature_truncation,3_prime_UTR_variant |
| 19 | 40728338 | 40728412 | 74 | Entpd1 | CAST | feature_truncation,3_prime_UTR_variant |
| 19 | 40743189 | 40743928 | 739 | Cc2d2b | CAST | feature_truncation,coding_sequence_variant,intron_variant |
| 19 | 40804758 | 40805018 | 260 | Cc2d2b | C3H,CAST,CBA,DBA1J,DBA2J,FVB,ILNJ2,MA/My,MOLF,NZB,NZO,SPRET | feature_truncation,coding_sequence_variant,intron_variant |
| 19 | 41757716 | 41757782 | 66 | Arhgap19 | AJ,BALB,CE,MOLF,RH2,SPRET,WSB | feature_truncation,3_prime_UTR_variant |
| 19 | 41910492 | 41911098 | 606 | Exosc1 | MOLF | feature_truncation,3_prime_UTR_variant |
| 19 | 42579432 | 42579500 | 68 | R3hcc1l | SPRET | feature_truncation,3_prime_UTR_variant |
| 19 | 44333627 | 44334087 | 460 | Scd4 | 129S1,BUB,C57L,CAST,FVB,KK,SMJ,SWR | feature_truncation,3_prime_UTR_variant |
| 19 | 44993345 | 44993562 | 217 | Mrpl43 | CAST | feature_truncation,3_prime_UTR_variant |

| | | | | | | |
|---|---|---|---|---|---|---|
| 19 | 45425911 | 45426081 | 170 | Btrc | MOLF | frameshift_variant,stop_lost,feature_truncation,3_prime_UTR_variant |
| 19 | 46713428 | 46713523 | 95 | As3mt | CAST,MOLF,SPRET | splice_donor_variant,splice_donor_5th_base_variant,coding_sequence_variant,intron_variant |
| 19 | 47073659 | 47073830 | 171 | Atp5mk | CAST,SPRET | feature_truncation,3_prime_UTR_variant |
| 19 | 47721141 | 47721237 | 96 | Sfr1 | CAST,SPRET | feature_truncation,inframe_deletion |
| 19 | 53028184 | 53028250 | 66 | Xpnpep1 | SPRET | feature_truncation,5_prime_UTR_variant |
| 19 | 53576561 | 53577493 | 932 | Nutf2-ps1 | 129S1,AJ,AKR,BALB,BTBR,C3H,C57L,CAST,CBA,CE,DBA1J,DBA2J,FVB,ILNJ2,KK,MA/My,MOLF,MRL,NOD,NOR,NZB,NZO,NZW,PJ,PL,RF,RH2,SEA,SJL,SPRET,SWR,TH,WSB | stop_lost,feature_truncation,coding_sequence_variant,5_prime_UTR_variant,3_prime_UTR_variant |
| 19 | 56588125 | 56588179 | 54 | Nhlrc2 | CAST | feature_truncation,3_prime_UTR_variant |
| 19 | 56590328 | 56590434 | 106 | Nhlrc2 | SPRET | feature_truncation,3_prime_UTR_variant |
| 19 | 59224418 | 59224497 | 79 | Kcnk18 | SPRET | feature_truncation,3_prime_UTR_variant |
| 19 | 60745538 | 60745670 | 132 | Nanos1 | TH | feature_truncation,3_prime_UTR_variant |
| 19 | 61107079 | 61107148 | 69 | Zfp950 | 129S1,AJ,AKR,BALB,BTBR,C3H,C57L,CAST,CBA,DBA1J,DBA2J,FVB,KK,MA/My,MRL,NOD,NOR,NZB,NZW,PJ,PL,RF,RH2,SEA,SMJ,SWR,TH,WSB | feature_truncation,3_prime_UTR_variant |
| 2 | 3398217 | 3398624 | 407 | Olah | SPRET | feature_truncation,5_prime_UTR_variant |
| 2 | 3525292 | 3525450 | 158 | Cdnf | CAST,SPRET | feature_truncation,3_prime_UTR_variant |
| 2 | 4627845 | 4627936 | 91 | Prpf18 | SPRET | feature_truncation,3_prime_UTR_variant |
| 2 | 5900929 | 5901018 | 89 | Dhtkd1 | SPRET | feature_truncation,3_prime_UTR_variant |
| 2 | 11477684 | 11477735 | 51 | Pfkfb3 | CAST,MOLF,SPRET | feature_truncation,3_prime_UTR_variant |
| 2 | 13513811 | 13513929 | 118 | Trdmt1 | SJL,SPRET | feature_truncation,3_prime_UTR_variant |
| 2 | 13516180 | 13516249 | 69 | Trdmt1 | CAST | feature_truncation,3_prime_UTR_variant |
| 2 | 24849930 | 24850001 | 71 | Zmynd19 | MOLF | feature_truncation,3_prime_UTR_variant |
| 2 | 24980606 | 24980661 | 55 | Noxa1 | CAST,FVB,MOLF,SPRET | feature_truncation,coding_sequence_variant,intron_variant |
| 2 | 25250436 | 25250593 | 157 | Uap1l1 | SPRET | feature_truncation,3_prime_UTR_variant |
| 2 | 25408519 | 25408613 | 94 | Traf2 | SPRET | feature_truncation,3_prime_UTR_variant |
| 2 | 26909743 | 26909870 | 127 | Cacfd1 | SPRET | feature_truncation,3_prime_UTR_variant |
| 2 | 27001749 | 27001839 | 90 | Fam163b | 129S1,C57L,SPRET | feature_truncation,3_prime_UTR_variant |
| 2 | 28976913 | 28977126 | 213 | Ttf1 | 129S1,C57L,CAST,MOLF,SPRET | feature_truncation,3_prime_UTR_variant |
| 2 | 29666262 | 29666691 | 429 | Trub2 | CAST,SPRET | feature_truncation,3_prime_UTR_variant |
| 2 | 30896780 | 30897378 | 598 | Usp20 | SPRET | stop_lost,feature_truncation,coding_sequence_variant,3_prime_UTR_variant,intron_variant |
| 2 | 31106902 | 31106962 | 60 | Gpr107 | SPRET | feature_truncation,3_prime_UTR_variant |
| 2 | 31571131 | 31571338 | 207 | Exosc2 | MOLF,SPRET | feature_truncation,3_prime_UTR_variant |
| 2 | 31684637 | 31684763 | 126 | Abl1 | MOLF,SPRET | feature_truncation,coding_sequence_variant,intron_variant |
| 2 | 32000584 | 32000641 | 57 | Plpp7 | SWR | feature_truncation,3_prime_UTR_variant |
| 2 | 32304895 | 32305001 | 106 | Slc25a25 | MOLF | feature_truncation,3_prime_UTR_variant |
| 2 | 32652031 | 32652105 | 74 | Tor2a | SPRET | feature_truncation,3_prime_UTR_variant |
| 2 | 33298087 | 33298213 | 126 | Zbtb34 | SPRET | feature_truncation,3_prime_UTR_variant |
| 2 | 36270998 | 36271164 | 166 | Or1j10 | SPRET | feature_truncation,3_prime_UTR_variant |
| 2 | 36463903 | 36464436 | 533 | Or1j15 | SPRET | feature_truncation,3_prime_UTR_variant |
| 2 | 36465889 | 36466013 | 124 | Or1j15 | SPRET | feature_truncation,3_prime_UTR_variant |
| 2 | 37094643 | 37094731 | 88 | Or1l4 | SPRET | feature_truncation,3_prime_UTR_variant |
| 2 | 37112090 | 37112150 | 60 | Or1af1 | BUB | feature_truncation,3_prime_UTR_variant |
| 2 | 40741582 | 40741809 | 227 | Lrp1b | BUB,SPRET,WSB | splice_donor_variant,splice_donor_5th_base_variant,coding_sequence_variant,intron_variant |
| 2 | 58456869 | 58456975 | 106 | Acvr1 | MOLF | feature_truncation,5_prime_UTR_variant |
| 2 | 62529547 | 62529651 | 104 | Kcnh7 | AJ,AKR,BTBR,BUB,C3H,C58J,CBA,CE,DBA1J,DBA2J,FVB,ILNJ2,KK,LP,MA/My,MRL,NOD,NOR,NZB,NZO,NZW,PJ,PL,RF,RH2,SEA,SMJ,SWR,TH | feature_truncation,3_prime_UTR_variant |
| 2 | 65183600 | 65184411 | 811 | Slc38a11 | BTBR,BUB,NOD,NOR,SJL,SWR | feature_truncation,coding_sequence_variant,intron_variant |
| 2 | 68943721 | 68943847 | 126 | Cers6 | 129S1,AJ,AKR,BUB,CAST,FVB,ILNJ2,KK,MA/My,MOLF,MRL,NOD,NOR,NZB,NZO,NZW,RH2,SPRET | feature_truncation,3_prime_UTR_variant |
| 2 | 69514147 | 69514321 | 174 | Klhl41 | BALB,BTBR,BUB,C3H,C57L,C58J,CAST,CBA,CE,DBA1J,DBA2J,MA/My,MOLF,NZB,NZO,NZW,PJ,PL,RF,SEA,SJL,SMJ,SPRET,SWR,TH,WSB | feature_truncation,3_prime_UTR_variant |
| 2 | 70258616 | 70258765 | 149 | Myo3b | MOLF | feature_truncation,3_prime_UTR_variant |
| 2 | 70924538 | 70924736 | 198 | Dcaf17 | SPRET | feature_truncation,3_prime_UTR_variant |
| 2 | 71103323 | 71103969 | 646 | Slc25a12 | AJ,AKR,BUB,C3H,FVB,ILNJ2,MA/My,MRL,NOD,NOR,NZW,PL,RH2,SMJ | feature_truncation,3_prime_UTR_variant |
| 2 | 73394670 | 73394856 | 186 | Chrna1 | CAST,MOLF,NZB,SPRET | feature_truncation,3_prime_UTR_variant |
| 2 | 76525840 | 76525927 | 87 | Plekha3 | SPRET | stop_lost,feature_truncation,coding_sequence_variant,3_prime_UTR_variant |
| 2 | 77358739 | 77358806 | 67 | Zfp385b | CAST | feature_truncation,3_prime_UTR_variant |
| 2 | 82820868 | 82820976 | 108 | Fsip2 | SPRET | feature_truncation,inframe_deletion |
| 2 | 84639919 | 84640429 | 510 | Ube2l6 | SPRET | feature_truncation,3_prime_UTR_variant |
| 2 | 85192622 | 85193267 | 645 | Or5ak20 | SPRET | feature_truncation,5_prime_UTR_variant,intron_variant |
| 2 | 85437998 | 85438060 | 62 | Or5g23 | CAST | feature_truncation,3_prime_UTR_variant |
| 2 | 85587643 | 85587996 | 353 | Or9g3 | AJ,AKR,BALB,BUB,C3H,C57L,C58J,CBA,CE,DBA1J,DBA2J,ILNJ2,MA/My,MOLF,MRL,NOD,NOR,NZB,NZO,NZW,PJ,PL,RF,RH2,SEA,SJL,SMJ,SPRET,SWR | feature_truncation,3_prime_UTR_variant |
| 2 | 85592112 | 85592602 | 490 | Or9g3 | MOLF | feature_truncation,5_prime_UTR_variant,intron_variant |
| 2 | 85823816 | 85824755 | 939 | Or5m8 | AJ,AKR,BALB,BUB,C3H,C57L,C58J,CBA,CE,DBA1J,DBA2J,FVB,ILNJ2,KK,MA/My,MOLF,MRL,NOD,NOR,NZB,NZO,NZW,PJ,PL,RF,RH2,SEA,SJL,SMJ,SPRET,SWR,WSB | feature_truncation,3_prime_UTR_variant |
| 2 | 85873592 | 85873762 | 170 | Or5m3b | SPRET | feature_truncation,3_prime_UTR_variant |
| 2 | 85960030 | 85960176 | 146 | Or5al5 | FVB,KK,SPRET,WSB | feature_truncation,3_prime_UTR_variant |
| 2 | 85999317 | 85999495 | 178 | Or8u9 | SPRET | feature_truncation,3_prime_UTR_variant |
| 2 | 86286204 | 86286263 | 59 | Or8k28 | AJ,AKR,BALB,BUB,C3H,C57L,CBA,CE,DBA1J,DBA2J,FVB,KK,MA/My,MOLF,MRL,NOD,NOR,NZB,NZO,NZW,PJ,PL,RF,RH2,SEA,SMJ,WSB | frameshift_variant,feature_truncation |

| | | | | | | |
|---|---|---|---|---|---|---|
| 2 | 86486959 | 86487010 | 51 | Or8k38 | SPRET | feature_truncation,3_prime_UTR_variant |
| 2 | 86807883 | 86807976 | 93 | Or8h10 | SPRET | feature_truncation,3_prime_UTR_variant |
| 2 | 86863133 | 86863258 | 125 | Or5be3 | SPRET | feature_truncation,3_prime_UTR_variant |
| 2 | 86936249 | 86936441 | 192 | Or5aq7 | AJ,AKR,BALB,BUB,C3H,C57L,CAST,CBA,CE,DBA1J,DBA2J,FVB,KK,MA/My,MOLF,MRL,NOD,NOR,NZB,NZO,NZW,PJ,PL,RF,RH2,SEA,SMJ,WSB | feature_truncation,3_prime_UTR_variant |
| 2 | 87140880 | 87140950 | 70 | Or10ag56 | SPRET | feature_truncation,3_prime_UTR_variant |
| 2 | 87221154 | 87221338 | 184 | Or10ag57 | AJ,AKR,BALB,C3H,C57L,C58J,CBA,CE,DBA1J,DBA2J,FVB,ILNJ2,KK,MA/My,MOLF,MRL,NOD,NOR,NZB,NZO,NZW,PJ,RF,RH2,SEA,SJL,SMJ,SWR | feature_truncation,3_prime_UTR_variant |
| 2 | 87253131 | 87253191 | 60 | Or10ag2 | C3H,CBA,CE,ILNJ2,MA/My,NZW,RF,SJL,SPRET,SWR | feature_truncation,3_prime_UTR_variant |
| 2 | 87319617 | 87319704 | 87 | Pramel7 | MOLF | feature_truncation,3_prime_UTR_variant |
| 2 | 87696672 | 87697062 | 390 | Or5w19 | C58J,SJL,SWR,WSB | feature_truncation,5_prime_UTR_variant,intron_variant |
| 2 | 88226128 | 88226588 | 460 | Or4p7 | SPRET | feature_truncation,3_prime_UTR_variant |
| 2 | 88252492 | 88252705 | 213 | Or4p20 | SPRET | feature_truncation,3_prime_UTR_variant |
| 2 | 88472034 | 88472089 | 55 | Or4s2 | SPRET | feature_truncation,5_prime_UTR_variant |
| 2 | 88511732 | 88511799 | 67 | Or4c103 | SPRET | feature_truncation,3_prime_UTR_variant |
| 2 | 88574697 | 88575097 | 400 | Or4p23 | SPRET | feature_truncation,3_prime_UTR_variant |
| 2 | 88687724 | 88687796 | 72 | Or4c106 | WSB | feature_truncation,3_prime_UTR_variant |
| 2 | 88857545 | 88857896 | 351 | Or4c112 | SPRET | feature_truncation,5_prime_UTR_variant,intron_variant |
| 2 | 88909910 | 88910906 | 996 | Or4c114 | SPRET | feature_truncation,5_prime_UTR_variant,intron_variant |
| 2 | 89111142 | 89111636 | 494 | Or4c15b | MOLF | feature_truncation,3_prime_UTR_variant |
| 2 | 89356549 | 89356604 | 55 | Or4a71 | C3H,CBA,CE,MA/My,NZW | feature_truncation,3_prime_UTR_variant |
| 2 | 89618152 | 89618275 | 123 | Or4a81 | SPRET | feature_truncation,3_prime_UTR_variant |
| 2 | 89850335 | 89850433 | 98 | Or4c3 | C58J,ILNJ2,KK,MOLF,SJL,SPRET,SWR,WSB | feature_truncation,3_prime_UTR_variant |
| 2 | 89978745 | 89979321 | 576 | Or4b1 | BUB,C58J,ILNJ2,KK,MOLF,PL,SJL,SPRET,SWR,WSB | feature_truncation,3_prime_UTR_variant,feature_truncation,3_prime_UTR_variant,intron_variant |
| 2 | 90155776 | 90156275 | 499 | Or4x18 | MOLF | feature_truncation,3_prime_UTR_variant |
| 2 | 93033490 | 93033554 | 64 | Tspan18 | CAST,MOLF,SPRET | feature_truncation,3_prime_UTR_variant |
| 2 | 93509992 | 93510217 | 225 | Alx4 | CAST | feature_truncation,3_prime_UTR_variant |
| 2 | 93664700 | 93665651 | 951 | Accs | CAST | feature_truncation,3_prime_UTR_variant |
| 2 | 102612545 | 102612601 | 56 | Slc1a2 | CAST | feature_truncation,3_prime_UTR_variant |
| 2 | 104661840 | 104661934 | 94 | Prrg4 | 129S1,AJ,AKR,BALB,BTBR,BUB,C3H,C57L,CAST,CBA,CE,DBA1J,DBA2J,ILNJ2,KK,MOLF,MRL,NOD,NOR,PJ,PL,RH2,SEA,SJL | feature_truncation,3_prime_UTR_variant |
| 2 | 110467927 | 110468654 | 727 | Slc5a12 | C3H,CE,MOLF,PL,SPRET | feature_truncation,coding_sequence_variant,intron_variant |
| 2 | 111091610 | 111091703 | 93 | Or4f53 | SPRET | feature_truncation,3_prime_UTR_variant |
| 2 | 111318658 | 111318942 | 284 | Or4k42 | CAST | feature_truncation,3_prime_UTR_variant |
| 2 | 111333164 | 111333485 | 321 | Or4f4-ps1 | MOLF | feature_truncation,3_prime_UTR_variant |
| 2 | 111763700 | 111764340 | 640 | Or4f14b | SPRET | feature_truncation,3_prime_UTR_variant |
| 2 | 111770121 | 111770358 | 237 | Or4f14b | MOLF | feature_truncation,3_prime_UTR_variant |
| 2 | 112298819 | 112298898 | 79 | Emc7 | SPRET | feature_truncation,3_prime_UTR_variant |
| 2 | 113578657 | 113578722 | 65 | Grem1 | SPRET | feature_truncation,3_prime_UTR_variant |
| 2 | 113662677 | 113663049 | 372 | Arhgap11a | AJ,AKR,BALB,BTBR,C3H,CAST,CBA,CE,DBA1J,DBA2J,FVB,ILNJ2,KK,MA/My,MOLF,MRL,NOD,NOR,NZB,NZO,NZW,PJ,PL,RF,RH2,SEA,SJL,SMJ,SPRET,SWR,TH,WSB | feature_truncation,3_prime_UTR_variant |
| 2 | 115894828 | 115894983 | 155 | Meis2 | 129S1,AJ,AKR,BALB,BTBR,BUB,C3H,C57L,C58J,CAST,CBA,CE,DBA1J,DBA2J,FVB,ILNJ2,KK,MA/My,MOLF,MRL,NOD,NOR,NZB,NZO,NZW,PJ,PL,RF,RH2,SEA,SJL,SMJ,SPRET,SWR,TH,WSB | feature_truncation,5_prime_UTR_variant |
| 2 | 118608388 | 118608451 | 63 | Phgr1 | 129S1,AJ,AKR,BALB,BTBR,BUB,C3H,C57L,C58J,CAST,CBA,CE,DBA1J,DBA2J,FVB,ILNJ2,MA/My,NOD,NOR,NZB,NZO,NZW,PJ,PL,RF,RH2,SEA,SJL,SMJ,SPRET,SWR,TH,WSB | feature_truncation,inframe_deletion |
| 2 | 119878007 | 119878070 | 63 | Sptbn5 | CAST,NZW,PL,SPRET | feature_truncation,inframe_deletion |
| 2 | 120281375 | 120281434 | 59 | Ganc | SPRET | feature_truncation,coding_sequence_variant,intron_variant |
| 2 | 121766495 | 121766698 | 203 | Golm2 | 129S1,AJ,AKR,BALB,BTBR,BUB,C3H,C57L,C58J,CAST,CBA,CE,DBA1J,DBA2J,FVB,ILNJ2,KK,MA/My,NOD,NZB,NZO,NZW,PJ,PL,RF,RH2,SEA,SJL,SMJ,SPRET,SWR,TH,WSB | feature_truncation,3_prime_UTR_variant |
| 2 | 122316312 | 122316789 | 477 | Slc28a2b | 129S1,AJ,AKR,BALB,BTBR,BUB,C3H,C57L,C58J,CBA,CE,DBA1J,DBA2J,FVB,ILNJ2,MA/My,NOD,NZB,NZO,NZW,PJ,PL,RF,RH2,SEA,SJL,SMJ,SWR,TH,WSB | feature_truncation,5_prime_UTR_variant,intron_variant |
| 2 | 122425382 | 122425458 | 76 | Gatm | CAST,SPRET | feature_truncation,3_prime_UTR_variant |
| 2 | 130409296 | 130409512 | 216 | Mrps26 | BALB,C3H,CAST,CBA,DBA1J,FVB,MOLF,NOD,PL,SEA,SPRET,WSB | feature_truncation,3_prime_UTR_variant |
| 2 | 130913952 | 130914153 | 201 | Siglec1 | SPRET | feature_truncation,3_prime_UTR_variant |
| 2 | 131797244 | 131797539 | 295 | Prnd | WSB | feature_truncation,3_prime_UTR_variant |
| 2 | 131833375 | 131833441 | 66 | Rassf2 | 129S1,AKR,BTBR,C3H,C57L,CBA,CE,DBA1J,DBA2J,ILNJ2,KK,MOLF,MRL,NZO,NZW,PJ,PL,RF,SJL,SWR | feature_truncation,3_prime_UTR_variant |
| 2 | 132147674 | 132147808 | 134 | Cds2 | 129S1,BTBR,C57L,CAST,ILNJ2,MA/My,MOLF,NOD,PJ,SPRET | feature_truncation,3_prime_UTR_variant |
| 2 | 132746666 | 132746948 | 282 | Fermt1 | CAST | feature_truncation,3_prime_UTR_variant |
| 2 | 134449205 | 134449805 | 600 | Tmx4 | 129S1,BTBR,C57L,CE,FVB,MRL,NZB,SMJ,SPRET,SWR,WSB | feature_truncation,3_prime_UTR_variant |
| 2 | 146084938 | 146085015 | 77 | Ralgapa2 | SPRET | feature_truncation,3_prime_UTR_variant |
| 2 | 150080901 | 150080985 | 84 | Zfp937 | CAST,MOLF,NOD,NZB,NZW,RF,SPRET | feature_truncation,inframe_deletion |
| 2 | 151312874 | 151312935 | 61 | 4921509C19Rik | SPRET | feature_truncation,3_prime_UTR_variant |
| 2 | 152327767 | 152327857 | 90 | Defb22 | SPRET | feature_truncation,inframe_deletion |
| 2 | 152349821 | 152349905 | 84 | Defb26 | SPRET | feature_truncation,inframe_deletion |
| 2 | 153459323 | 153459584 | 261 | Commd7 | SPRET | feature_truncation,3_prime_UTR_variant |
| 2 | 153460232 | 153460336 | 104 | Commd7 | 129S1,AJ,BALB,BTBR,C3H,C57L,C58J,CAST,CBA,DBA1J,DBA2J,ILNJ2,MOLF,NOD,PL,RF,SEA,SPRET,TH,WSB | feature_truncation,3_prime_UTR_variant |

| | | | | | | |
|---|---|---|---|---|---|---|
| 2 | 154402269 | 154402323 | 54 | E2f1 | AJ,BTBR,C58J,DBA1J,DBA2J,KK,MOLF,NZB,NZO,NZW | feature_truncation,3_prime_UTR_variant |
| 2 | 154714089 | 154714158 | 69 | Eif2s2 | 129S1,BALB,C3H,C57L,CBA,CE,FVB,MA/My,NOD,NZO,NZW,PL,SEA,SJL,SMJ,SWR,TH,WSB | feature_truncation,3_prime_UTR_variant |
| 2 | 155812023 | 155812086 | 63 | Cep250 | 129S1,BALB,C3H,C57L,CBA,CE,FVB,MA/My,NOD,NOR,NZO,NZW,PL,SEA,SJL,SMJ,SPRET,SWR,TH,WSB | feature_truncation,inframe_deletion,feature_truncation,coding_sequence_variant,intron_variant |
| 2 | 156778497 | 156778697 | 200 | Ndrg3 | AKR,BTBR,BUB,CAST,DBA1J,DBA2J,ILNJ2,KK,MOLF,MRL,NZB,PJ,RF,SJL | feature_truncation,3_prime_UTR_variant |
| 2 | 158226536 | 158226627 | 91 | Snhg11 | DBA1J,DBA2J,ILNJ2,MA/My,NZB,SMJ,WSB | feature_truncation,3_prime_UTR_variant |
| 2 | 158606727 | 158606953 | 226 | Ppp1r16b | AJ,CAST,CE,DBA1J,DBA2J,ILNJ2,KK,MA/My,MOLF,NZB,SJL,SMJ,SPRET,WSB | feature_truncation,3_prime_UTR_variant |
| 2 | 163165520 | 163165769 | 249 | Tox2 | KK,SPRET | feature_truncation,3_prime_UTR_variant |
| 2 | 164079460 | 164079529 | 69 | Semg1 | SPRET | feature_truncation,inframe_deletion |
| 2 | 164333153 | 164333213 | 60 | Dbndd2 | CAST | feature_truncation,3_prime_UTR_variant |
| 2 | 164349585 | 164349777 | 192 | Pigt | 129S1,AJ,BALB,BTBR,BUB,C3H,C57L,C58J,CAST,CBA,CE,DBA1J,DBA2J,FVB,KK,MA/My,NOD,NOR,NZO,NZW,PJ,PL,SEA,SMJ,SPRET,SWR,TH,WSB | feature_truncation,3_prime_UTR_variant |
| 2 | 165218320 | 165218373 | 53 | Zfp334 | 129S1,AJ,AKR,BALB,BTBR,C57L,CAST,CBA,CE,ILNJ2,KK,MA/My,MRL,NOD,NOR,NZB,PL,RF,SEA,SJL,SMJ,SPRET,TH,WSB | feature_truncation,3_prime_UTR_variant |
| 2 | 165669633 | 165669847 | 214 | Zmynd8 | 129S1,AJ,AKR,BALB,BTBR,BUB,C3H,C57L,C58J,CAST,CBA,CE,DBA1J,DBA2J,FVB,ILNJ2,KK,MA/My,NOD,NOR,NZB,NZO,PJ,PL,RF,SEA,SJL,SMJ,SPRET,SWR,TH,WSB | feature_truncation,3_prime_UTR_variant |
| 2 | 172710968 | 172711196 | 228 | Bmp7 | 129S1,AJ,AKR,BUB,C57L,CAST,CE,MA/My,MOLF,MRL,NZO,NZW,PL,RF,RH2,SMJ,SPRET | feature_truncation,3_prime_UTR_variant |
| 2 | 174972834 | 174973210 | 376 | Gm14399 | SPRET | frameshift_variant,feature_truncation,frameshift_variant,stop_lost,feature_truncation,3_prime_UTR_variant |
| 2 | 175010853 | 175011189 | 336 | Gm14443 | CE,FVB,NZB,SJL,SPRET,SWR | feature_truncation,inframe_deletion |
| 2 | 176372294 | 176372546 | 252 | Zfp973 | BALB,MOLF,NZB,PJ,WSB | stop_lost,inframe_deletion |
| 2 | 176502163 | 176502582 | 419 | Gm14295 | SPRET | frameshift_variant,feature_truncation |
| 2 | 177167206 | 177167332 | 126 | Zfp970 | DBA2J,MOLF,SEA | feature_truncation,inframe_deletion |
| 2 | 177201034 | 177201118 | 84 | Gm14403 | SPRET | stop_lost,inframe_deletion |
| 2 | 177587649 | 177587816 | 167 | Gm14326 | SPRET | frameshift_variant,feature_truncation |
| 2 | 177598805 | 177599620 | 815 | Gm14326 | BUB,KK,NOD,NOR | feature_truncation,5_prime_UTR_variant,intron_variant |
| 2 | 177675218 | 177675428 | 210 | Zfp971 | CE,KK,PJ,SPRET | feature_truncation,inframe_deletion |
| 2 | 177709457 | 177709876 | 419 | Zfp931 | PJ | stop_lost,feature_truncation,coding_sequence_variant,3_prime_UTR_variant,feature_truncation,3_prime_UTR_variant |
| 3 | 3723510 | 3723566 | 56 | Hnf4g | CAST,MOLF | feature_truncation,3_prime_UTR_variant |
| 3 | 5815850 | 5816351 | 501 | Gm8797 | SPRET | stop_lost,feature_truncation,coding_sequence_variant,5_prime_UTR_variant,3_prime_UTR_variant |
| 3 | 7637475 | 7637576 | 101 | Il7 | MOLF | feature_truncation,3_prime_UTR_variant |
| 3 | 9501714 | 9501880 | 166 | Zfp704 | SPRET | feature_truncation,3_prime_UTR_variant |
| 3 | 9753507 | 9753560 | 53 | Pag1 | SPRET,WSB | feature_truncation,3_prime_UTR_variant |
| 3 | 10380640 | 10380717 | 77 | Impa1 | CAST | feature_truncation,3_prime_UTR_variant |
| 3 | 20050012 | 20050245 | 233 | Hps3 | CAST,MOLF | feature_truncation,3_prime_UTR_variant |
| 3 | 27333413 | 27333645 | 232 | Nceh1 | SPRET | feature_truncation,coding_sequence_variant,intron_variant |
| 3 | 27391852 | 27392127 | 275 | Tnfsf10 | SPRET | feature_truncation,3_prime_UTR_variant |
| 3 | 27396549 | 27396624 | 75 | Tnfsf10 | 129S1,AJ,AKR,BTBR,BUB,C3H,C57L,C58J,CBA,CE,DBA1J,DBA2J,FVB,ILNJ2,MRL,NOD,NOR,NZW,PJ,PL,RF,SEA,SJL,SPRET,SWR | feature_truncation,3_prime_UTR_variant |
| 3 | 30956909 | 30956963 | 54 | Phc3 | SPRET | feature_truncation,3_prime_UTR_variant |
| 3 | 31107103 | 31107223 | 120 | Prkci | SPRET | feature_truncation,3_prime_UTR_variant |
| 3 | 31260304 | 31260597 | 293 | Slc7a14 | CAST,CE,MOLF,SPRET,SWR,WSB | feature_truncation,3_prime_UTR_variant |
| 3 | 32585822 | 32586012 | 190 | Mfn1 | CAST,MOLF,SPRET | feature_truncation,5_prime_UTR_variant |
| 3 | 32635811 | 32636199 | 388 | Gnb4 | MOLF | feature_truncation,3_prime_UTR_variant |
| 3 | 32990626 | 32990722 | 96 | Usp13 | SPRET | feature_truncation,3_prime_UTR_variant |
| 3 | 34110673 | 34110985 | 312 | Dnajc19 | SPRET | feature_truncation,3_prime_UTR_variant |
| 3 | 36220607 | 36220801 | 194 | Zfp267 | BUB,CAST,CE,DBA1J,FVB,MOLF,NZB,NZO,NZW,SPRET,SWR,WSB | feature_truncation,3_prime_UTR_variant |
| 3 | 37277246 | 37277299 | 53 | Il21 | AKR,BTBR,DBA1J,ILNJ2,MA/My,MOLF,MRL,NOR,PL,RF | feature_truncation,3_prime_UTR_variant |
| 3 | 37365937 | 37366028 | 91 | Cetn4 | 129S1,AJ,AKR,BTBR,C3H,C57L,CBA,CE,DBA1J,DBA2J,ILNJ2,MA/My,MRL,NOD,NOR,NZB,NZO,NZW,PL,RF,SJL,SMJ,TH,WSB | feature_truncation,5_prime_UTR_variant |
| 3 | 37768844 | 37769376 | 532 | Gm5148 | 129S1,AJ,AKR,BTBR,C3H,C57L,CAST,CBA,CE,DBA1J,DBA2J,ILNJ2,MA/My,MRL,NOD,NOR,NZB,NZO,NZW,PL,RF,SJL,SMJ,SPRET,SWR,TH,WSB | feature_truncation,coding_sequence_variant,5_prime_UTR_variant,intron_variant |
| 3 | 40657076 | 40657475 | 399 | Intu | AJ,C3H,CAST,CBA,ILNJ2,KK,MOLF,NZB,NZW,PL,SPRET | feature_truncation,3_prime_UTR_variant |
| 3 | 40994230 | 40994295 | 65 | Larp1b | CAST,KK,MOLF,NZW,SPRET | feature_truncation,3_prime_UTR_variant |
| 3 | 51403642 | 51403790 | 148 | Rab33b | CAST,CE,MOLF,NZW,SJL,SPRET,SWR | feature_truncation,3_prime_UTR_variant |
| 3 | 53358004 | 53358446 | 442 | Nhlrc3 | SPRET | feature_truncation,3_prime_UTR_variant |
| 3 | 53422162 | 53422389 | 227 | Frem2 | CAST,SPRET | feature_truncation,3_prime_UTR_variant |
| 3 | 54642531 | 54642632 | 101 | Exosc8 | SPRET | feature_truncation,5_prime_UTR_variant |
| 3 | 57573890 | 57573949 | 59 | Ankub1 | MOLF | feature_truncation,3_prime_UTR_variant |
| 3 | 62376635 | 62376857 | 222 | Dhx36 | MOLF | feature_truncation,3_prime_UTR_variant |
| 3 | 64016140 | 64017023 | 883 | Vmn2r1 | SPRET | feature_truncation,3_prime_UTR_variant |
| 3 | 64438406 | 64438793 | 387 | Vmn2r6 | CAST,SPRET | feature_truncation,3_prime_UTR_variant |
| 3 | 72955110 | 72955174 | 64 | Slitrk3 | C58J,LP,MOLF,NZW | feature_truncation,3_prime_UTR_variant |
| 3 | 79365318 | 79365428 | 110 | Fnip2 | RF | feature_truncation,3_prime_UTR_variant |
| 3 | 85580066 | 85580129 | 63 | Fhip1a | SPRET | feature_truncation,inframe_deletion |
| 3 | 87348471 | 87349333 | 862 | Fcrl5 | FVB,MA/My,NOD,NOR,SJL,SWR,TH | feature_truncation,coding_sequence_variant,intron_variant |
| 3 | 87760112 | 87760265 | 153 | Sh2d2a | SPRET | feature_truncation,3_prime_UTR_variant |
| 3 | 87884050 | 87884182 | 132 | Nes | MOLF | feature_truncation,inframe_deletion |
| 3 | 88653811 | 88654006 | 195 | Syt11 | SPRET | feature_truncation,3_prime_UTR_variant |
| 3 | 89137754 | 89137880 | 126 | Muc1 | CAST | feature_truncation,inframe_deletion |

| | | | | | | |
|---|---|---|---|---|---|---|
| 3 | 89764484 | 89764603 | 119 | She | CAST,KK,MOLF | feature_truncation,3_prime_UTR_variant |
| 3 | 89775195 | 89775631 | 436 | Il6ra | CAST,KK,MOLF,SPRET | feature_truncation,3_prime_UTR_variant |
| 3 | 90565423 | 90565586 | 163 | S100a7a | SPRET | feature_truncation,3_prime_UTR_variant |
| 3 | 92224827 | 92224881 | 54 | Sprr2b | CAST | feature_truncation,inframe_deletion |
| 3 | 92479212 | 92479299 | 87 | Ivl | CBA,KK,WSB | feature_truncation,inframe_deletion |
| 3 | 92626240 | 92626300 | 60 | Lce1f | SPRET | feature_truncation,inframe_deletion |
| 3 | 92645180 | 92645240 | 60 | Gm38119 | MOLF | feature_truncation,inframe_deletion |
| 3 | 92684750 | 92684810 | 60 | Lce1i | CAST,SPRET | feature_truncation,inframe_deletion |
| 3 | 92925574 | 92925625 | 51 | Lce1m | CAST | feature_truncation,inframe_deletion |
| 3 | 93111349 | 93111570 | 221 | Flg2 | CAST,MOLF,SPRET | frameshift_variant,feature_truncation |
| 3 | 93114716 | 93114955 | 239 | Flg2 | MOLF,SPRET | frameshift_variant,feature_truncation |
| 3 | 93116829 | 93117770 | 941 | Flg2 | AJ,C3H,CBA,CE,MOLF,SPRET,WSB | frameshift_variant,feature_truncation |
| 3 | 93117682 | 93118151 | 469 | Flg2 | BALB,MA/My,NZB,RH2 | frameshift_variant,feature_truncation |
| 3 | 93119292 | 93119649 | 357 | Flg2 | BALB,RH2 | feature_truncation,inframe_deletion |
| 3 | 93120807 | 93121515 | 708 | Flg2 | AJ,BALB,C3H,CBA,CE,MA/My,NZO,RH2,WSB | feature_truncation,inframe_deletion |
| 3 | 93120979 | 93121444 | 465 | Flg2 | CAST,MA/My,NZB,RF | feature_truncation,inframe_deletion |
| 3 | 93121913 | 93122848 | 935 | Flg2 | SPRET | frameshift_variant,feature_truncation |
| 3 | 93122837 | 93123064 | 227 | Flg2 | 129S1,AJ,BTBR,C3H,C57L,CBA,CE,ILNJ2,PJ,PL,SEA,TH,WSB | frameshift_variant,feature_truncation |
| 3 | 93124476 | 93124710 | 234 | Flg2 | AKR,B10D2,B10J,BUB,C58J,DBA1J,DBA2J,FVB,LP,MRL,NZW,RF,SJL,SMJ,SWR | feature_truncation,inframe_deletion |
| 3 | 93125385 | 93125859 | 474 | Flg2 | AKR,B10D2,B10J,BUB,C58J,CE,DBA1J,DBA2J,FVB,LP,MOLF,MRL,NZW,SJL,SPRET | feature_truncation,inframe_deletion |
| 3 | 93126115 | 93126342 | 227 | Flg2 | NOR,PJ,SEA,SPRET | frameshift_variant,feature_truncation |
| 3 | 93126118 | 93126339 | 221 | Flg2 | 129S1,BTBR,CAST,CE,NOD,RF,SMJ,SWR | frameshift_variant |
| 3 | 93126545 | 93126779 | 234 | Flg2 | MOLF | feature_truncation,inframe_deletion |
| 3 | 93232012 | 93232542 | 530 | Hrnr | MOLF | frameshift_variant,feature_truncation |
| 3 | 93235829 | 93236342 | 513 | Hrnr | 129S1,AKR,BTBR,BUB,C57L,CAST,FVB,KK,MRL,NZW,SJL,SWR | feature_truncation,inframe_deletion |
| 3 | 93303697 | 93303769 | 72 | Rptn | MOLF | feature_truncation,inframe_deletion |
| 3 | 93304757 | 93304829 | 72 | Rptn | CAST | feature_truncation,inframe_deletion |
| 3 | 93353127 | 93353247 | 120 | Tchh | BALB,NZB,NZO,RH2 | feature_truncation,inframe_deletion |
| 3 | 94341230 | 94341297 | 67 | Tdrkh | SPRET | feature_truncation,3_prime_UTR_variant |
| 3 | 94979313 | 94979598 | 285 | Pip5k1a | SPRET | feature_truncation,coding_sequence_variant,intron_variant |
| 3 | 95737286 | 95737475 | 189 | Prpf3 | MOLF,SPRET | feature_truncation,3_prime_UTR_variant |
| 3 | 96105790 | 96106118 | 328 | Bola1 | MOLF,SPRET | feature_truncation,5_prime_UTR_variant |
| 3 | 96169919 | 96170137 | 218 | H4c14 | MOLF,SPRET | feature_truncation,3_prime_UTR_variant |
| 3 | 97315739 | 97316031 | 292 | Or13l2 | SPRET | feature_truncation,3_prime_UTR_variant |
| 3 | 97829563 | 97829684 | 121 | Sec22b | SPRET | feature_truncation,3_prime_UTR_variant |
| 3 | 99126566 | 99126647 | 81 | Wars2 | MOLF | feature_truncation,3_prime_UTR_variant |
| 3 | 100839237 | 100839973 | 736 | Trim45 | SPRET | feature_truncation,3_prime_UTR_variant,stop_lost,feature_truncation,coding_sequence_variant,3_prime_UTR_variant |
| 3 | 102061633 | 102061714 | 81 | Vangl1 | SPRET | feature_truncation,3_prime_UTR_variant |
| 3 | 102062489 | 102062721 | 232 | Vangl1 | SPRET | feature_truncation,3_prime_UTR_variant |
| 3 | 103086279 | 103086478 | 199 | Bcas2 | CAST,MOLF | feature_truncation,3_prime_UTR_variant |
| 3 | 107141635 | 107142334 | 699 | Prok1 | CAST,SMJ | feature_truncation,3_prime_UTR_variant |
| 3 | 108762045 | 108762208 | 163 | Fndc7 | SPRET | feature_truncation,3_prime_UTR_variant |
| 3 | 108880836 | 108881520 | 684 | Eeig2 | CAST | feature_truncation,3_prime_UTR_variant |
| 3 | 115722404 | 115722479 | 75 | Dph5 | CAST | feature_truncation,3_prime_UTR_variant |
| 3 | 115736194 | 115736345 | 151 | Slc30a7 | 129S1,AJ,AKR,BALB,BUB,C3H,C57L,CBA,CE,DBA1J,DBA2J,FVB,ILNJ2,LP,MA/My,MRL,NZB,NZO,NZW,PL,RF,RH2,SJL,SWR,TH | feature_truncation,3_prime_UTR_variant |
| 3 | 116342790 | 116342873 | 83 | Dbt | MOLF | feature_truncation,3_prime_UTR_variant |
| 3 | 116387842 | 116387895 | 53 | Trmt13 | MOLF,SMJ,WSB | feature_truncation,3_prime_UTR_variant |
| 3 | 116534227 | 116534349 | 122 | Agl | MOLF,SPRET | feature_truncation,3_prime_UTR_variant |
| 3 | 116698134 | 116698204 | 70 | Frrs1 | CAST,MOLF,SPRET | feature_truncation,3_prime_UTR_variant |
| 3 | 119225366 | 119225418 | 52 | Dpyd | CAST,MOLF,SPRET | feature_truncation,3_prime_UTR_variant |
| 3 | 122905195 | 122905264 | 69 | Synpo2 | CAST,MOLF,SPRET | feature_truncation,3_prime_UTR_variant |
| 3 | 123166915 | 123167120 | 205 | Mettl14 | AJ,BALB,C3H,CAST,CBA,CE,DBA1J,DBA2J,KK,MA/My,MOLF,NZB,PJ,RF,RH2,SEA,SMJ,SPRET | feature_truncation,3_prime_UTR_variant |
| 3 | 124203263 | 124203403 | 140 | 1700006A11Rik | 129S1,BTBR,C57L,CE,PJ,SEA,TH | feature_truncation,coding_sequence_variant |
| 3 | 127356830 | 127356911 | 81 | Zgrf1 | SPRET | feature_truncation,coding_sequence_variant,intron_variant,feature_truncation,inframe_deletion |
| 3 | 127663170 | 127663306 | 136 | Fam241a | SPRET | feature_truncation,3_prime_UTR_variant |
| 3 | 129598133 | 129598749 | 616 | Rrh | SPRET | feature_truncation,3_prime_UTR_variant,intron_variant |
| 3 | 131082645 | 131082983 | 338 | Cyp2u1 | CAST,MOLF,NZB,NZO,SPRET,WSB | feature_truncation,3_prime_UTR_variant |
| 3 | 134955481 | 134955592 | 111 | Cenpe | CAST,MOLF | splice_donor_variant,splice_donor_5th_base_variant,coding_sequence_variant,intron_variant |
| 3 | 135048618 | 135048898 | 280 | Slc9b2 | 129S1,AKR,BTBR,BUB,C57L,C58J,CAST,LP,MOLF,NZB,NZO,NZW,PJ,RF,SEA,SPRET | feature_truncation,3_prime_UTR_variant |
| 3 | 135253395 | 135254085 | 690 | Manba | CAST | feature_truncation,coding_sequence_variant,intron_variant |
| 3 | 137615435 | 137615513 | 78 | Dnajb14 | SPRET | feature_truncation,3_prime_UTR_variant |
| 3 | 137870052 | 137870118 | 66 | 4930579F01Rik | SPRET | feature_truncation,3_prime_UTR_variant |
| 3 | 138612058 | 138612152 | 94 | Tspan5 | 129S1,AJ,BALB,BTBR,C3H,C57L,CAST,CBA,CE,DBA1J,ILNJ2,KK,MA/My,MOLF,MRL,NOD,NOR,NZB,NZO,NZW,PJ,PL,RH2,SEA,SJL,SMJ,SPRET,SWR,TH | feature_truncation,3_prime_UTR_variant |
| 3 | 144394421 | 144394477 | 56 | Sh3glb1 | BUB,C58J,DBA2J,LP,MRL,RF | feature_truncation,3_prime_UTR_variant |
| 3 | 144528604 | 144528766 | 162 | Clca3b | C58J,CAST,LP,MOLF,SJL,SPRET,WSB | feature_truncation,inframe_deletion |
| 3 | 144533602 | 144533798 | 196 | Clca3b | SJL | feature_truncation,coding_sequence_variant,intron_variant |
| 3 | 144616439 | 144616720 | 281 | Clca4b | MOLF | feature_truncation,3_prime_UTR_variant |
| 3 | 152038192 | 152038291 | 99 | Miga1 | SPRET | feature_truncation,3_prime_UTR_variant |
| 3 | 152492085 | 152492325 | 240 | Pigk | 129S1,AJ,AKR,BALB,BTBR,C3H,C57L,CAST,CBA,CE,DBA1J,ILNJ2,MOLF,PL,RH2,SEA,SMJ,SPRET,TH,WSB | feature_truncation,3_prime_UTR_variant |
| 3 | 152524988 | 152525339 | 351 | St6galnac5 | CAST,TH | feature_truncation,3_prime_UTR_variant |
| 3 | 154283947 | 154284453 | 506 | Tyw3 | KK,MOLF,SPRET | feature_truncation,3_prime_UTR_variant |
| 3 | 154328550 | 154328602 | 52 | Cryz | MOLF,SPRET | feature_truncation,3_prime_UTR_variant |

| | | | | | | |
|---|---|---|---|---|---|---|
| 3 | 154468054 | 154468123 | 69 | Erich3 | AJ,BALB,BUB,C3H,DBA2J,FVB,ILNJ2,KK,MA/My,MOLF,NOD,NOR,NZB,NZO,NZW,RH2,SEA,SJL,SWR,TH | feature_truncation,inframe_deletion |
| 4 | 3973082 | 3973596 | 514 | Uba52rt | CAST,KK,SPRET | transcript_ablation |
| 4 | 8144070 | 8144144 | 74 | Car8 | SPRET | feature_truncation,3_prime_UTR_variant |
| 4 | 11171008 | 11171086 | 78 | Trp53inp1 | SPRET | feature_truncation,3_prime_UTR_variant |
| 4 | 11262864 | 11262919 | 55 | Dpy19l4 | 129S1,AKR,BALB,BTBR,BUB,C3H,C57L,CBA,CE,DBA1J,DBA2J,FVB,ILNJ2,KK,MA/My,MRL,NZB,NZO,PJ,PL,RF,RH2,SEA,SJL,SMJ,SWR | feature_truncation,3_prime_UTR_variant |
| 4 | 12981638 | 12981694 | 56 | Triqk | AJ,AKR,BUB,NOD,NOR,NZB,NZO,PJ,SEA,WSB | feature_truncation,3_prime_UTR_variant |
| 4 | 20243831 | 20243994 | 163 | Nkain3 | CAST,KK,NZB,NZO,SPRET,WSB | feature_truncation,3_prime_UTR_variant |
| 4 | 24503268 | 24503367 | 99 | Mms22l | NZB,NZO,NZW,RF | feature_truncation,3_prime_UTR_variant |
| 4 | 24613669 | 24613722 | 53 | Klhl32 | ILNJ2 | feature_truncation,3_prime_UTR_variant |
| 4 | 24903265 | 24903430 | 165 | Ndufaf4 | C57L | feature_truncation,3_prime_UTR_variant |
| 4 | 24904811 | 24904955 | 144 | Ndufaf4 | CAST | feature_truncation,3_prime_UTR_variant |
| 4 | 25618783 | 25619104 | 321 | Fut9 | SPRET | feature_truncation,3_prime_UTR_variant |
| 4 | 33174177 | 33174237 | 60 | Pm20d2 | CAST,MOLF,SPRET | feature_truncation,3_prime_UTR_variant |
| 4 | 34950694 | 34950760 | 66 | Mob3b | AJ,AKR,BTBR,BUB,C3H,CBA,DBA1J,DBA2J,FVB,ILNJ2,LP,MOLF,MRL,NOD,NOR,RF,SJL,SMJ,SWR | feature_truncation,3_prime_UTR_variant |
| 4 | 40685609 | 40685930 | 321 | Aptx | CAST,SPRET | feature_truncation,3_prime_UTR_variant |
| 4 | 41033339 | 41033393 | 54 | Aqp7 | SPRET | feature_truncation,3_prime_UTR_variant |
| 4 | 43666002 | 43666071 | 69 | Fam221b | CAST,MOLF | feature_truncation,inframe_deletion |
| 4 | 43734217 | 43734273 | 56 | Spaar | SPRET | feature_truncation,3_prime_UTR_variant |
| 4 | 51228991 | 51229072 | 81 | Cylc2 | 129S1,C57L,NZW | feature_truncation,inframe_deletion |
| 4 | 52852930 | 52852981 | 51 | Or13c3 | MA/My | feature_truncation,3_prime_UTR_variant |
| 4 | 53549982 | 53550179 | 197 | Slc44a1 | SPRET | stop_lost,feature_truncation,coding_sequence_variant,3_prime_UTR_variant,intron_variant |
| 4 | 56875664 | 56875876 | 212 | Tmem245 | 129S1,AJ,AKR,BALB,BTBR,BUB,C3H,C57L,CAST,CBA,CE,DBA1J,DBA2J,ILNJ2,LP,MOLF,MRL,NZB,NZO,NZW,PJ,PL,RF,RH2,SEA,SMJ,SPRET | feature_truncation,3_prime_UTR_variant |
| 4 | 57854017 | 57854221 | 204 | Pakap | CAST,KK,MOLF,SPRET | feature_truncation,5_prime_UTR_variant |
| 4 | 59138810 | 59139313 | 503 | Shoc1 | 129S1,AJ,AKR,BALB,BTBR,BUB,C3H,C57L,CAST,CBA,CE,DBA1J,DBA2J,FVB,ILNJ2,KK,LP,MA/My,MOLF,MRL,NOD,NOR,NZB,NZO,NZW,PJ,RF,RH2,SEA,SJL,SPRET,SWR,TH,WSB | feature_truncation,5_prime_UTR_variant,intron_variant |
| 4 | 59610573 | 59610627 | 54 | Hsdl2 | SPRET | feature_truncation,inframe_deletion |
| 4 | 59753707 | 59753802 | 95 | E130308A19Rik | BTBR | feature_truncation,3_prime_UTR_variant |
| 4 | 62607859 | 62607913 | 54 | Rgs3 | SPRET | feature_truncation,inframe_deletion |
| 4 | 86117863 | 86117975 | 112 | Adamtsl1 | CAST,SPRET | feature_truncation,3_prime_UTR_variant |
| 4 | 86837691 | 86837761 | 70 | Acer2 | SPRET | feature_truncation,3_prime_UTR_variant |
| 4 | 95931805 | 95931878 | 73 | Cyp2j13 | SPRET | feature_truncation,3_prime_UTR_variant |
| 4 | 98342641 | 98342999 | 358 | Patj | CAST,MOLF,SMJ,SPRET | stop_lost,feature_truncation,coding_sequence_variant,3_prime_UTR_variant |
| 4 | 99147442 | 99147510 | 68 | Atg4c | AJ,BALB,BTBR,C3H,CBA,MA/My,SEA,SPRET | feature_truncation,3_prime_UTR_variant |
| 4 | 101701019 | 101701497 | 478 | Pramel17 | SPRET | feature_truncation,5_prime_UTR_variant,intron_variant |
| 4 | 106602217 | 106602302 | 85 | Acot11 | CAST | feature_truncation,3_prime_UTR_variant |
| 4 | 107834868 | 107834967 | 99 | Slc1a7 | 129S1,AJ,BALB,BUB,C3H,C57L,CBA,CE,FVB,NOD,PL,RF,SEA,SJL,SPRET,SWR | feature_truncation,coding_sequence_variant,intron_variant |
| 4 | 114470634 | 114470746 | 112 | Trabd2b | 129S1,AJ,BTBR,C57L,DBA1J,DBA2J,FVB,ILNJ2,MA/My,SWR | feature_truncation,3_prime_UTR_variant |
| 4 | 114963839 | 114964149 | 310 | Cyp4x1 | CAST,SPRET | feature_truncation,3_prime_UTR_variant |
| 4 | 114965352 | 114965495 | 143 | Cyp4x1 | SPRET | feature_truncation,3_prime_UTR_variant |
| 4 | 115625960 | 115626678 | 718 | Efcab14 | BALB,BUB,C3H,CBA,FVB,KK,NOD,NOR,RF,SJL,SWR,TH,WSB | feature_truncation,3_prime_UTR_variant |
| 4 | 116557362 | 116557424 | 62 | Prdx1 | AKR,CE,KK,MRL,NZB,NZO,NZW,PJ,SEA,SMJ | feature_truncation,3_prime_UTR_variant |
| 4 | 118546185 | 118546452 | 267 | Or13p4 | SPRET | feature_truncation,3_prime_UTR_variant |
| 4 | 118790200 | 118790691 | 491 | Or10ak7 | WSB | feature_truncation,3_prime_UTR_variant |
| 4 | 119077411 | 119077591 | 180 | AU022252 | AJ,B10D2,B10J,BALB,BUB,C3H,C58J,CAST,CBA,CE,DBA1J,DBA2J,FVB,ILNJ2,MA/My,MOLF,NOD,NOR,NZB,PJ,PL,RF,SEA,SJL,SPRET,SWR,TH | feature_truncation,3_prime_UTR_variant |
| 4 | 119079093 | 119079296 | 203 | AU022252 | CAST,ILNJ2,MOLF,SPRET | feature_truncation,3_prime_UTR_variant |
| 4 | 119249867 | 119250100 | 233 | Ccdc30 | SPRET | feature_truncation,3_prime_UTR_variant |
| 4 | 120804506 | 120804584 | 78 | Zfp69 | SPRET | feature_truncation,inframe_deletion |
| 4 | 122761262 | 122761433 | 171 | Cap1 | AJ,B10D2,B10J,BUB,C58J,CAST,CBA,CE,DBA1J,DBA2J,FVB,ILNJ2,MA/My,MOLF,NOD,NOR,NZB,NZW,PL,RF,SJL,SMJ,SPRET,SWR,TH,WSB | splice_acceptor_variant,splice_donor_variant,splice_donor_5th_base_variant,coding_sequence_variant,intron_variant |
| 4 | 122895414 | 122895550 | 136 | Mycl | SPRET | feature_truncation,3_prime_UTR_variant |
| 4 | 123018882 | 123018954 | 72 | Bmp8b | B10D2,B10J,C58J,CAST,MOLF,SPRET,WSB | feature_truncation,3_prime_UTR_variant |
| 4 | 125604595 | 125604647 | 52 | Grik3 | ILNJ2 | feature_truncation,3_prime_UTR_variant |
| 4 | 129200886 | 129201173 | 287 | Rbbp4 | BUB,FVB,SJL,SWR | feature_truncation,3_prime_UTR_variant |
| 4 | 129242088 | 129242277 | 189 | Zbtb8os | SPRET | feature_truncation,3_prime_UTR_variant |
| 4 | 130662441 | 130662765 | 324 | Laptm5 | SPRET | feature_truncation,3_prime_UTR_variant |
| 4 | 131949573 | 131949777 | 204 | Gmeb1 | SPRET | feature_truncation,3_prime_UTR_variant |
| 4 | 132449982 | 132450044 | 62 | Eya3 | SPRET | feature_truncation,3_prime_UTR_variant |
| 4 | 133929523 | 133929772 | 249 | Zpld2 | SPRET | feature_truncation,inframe_deletion |
| 4 | 135177430 | 135177546 | 116 | Nipal3 | B10D2,B10J,DBA1J,KK,NZO,PJ,SEA | feature_truncation,3_prime_UTR_variant |
| 4 | 136077883 | 136078750 | 867 | Hnrnpr | SPRET | feature_truncation,3_prime_UTR_variant |
| 4 | 136078853 | 136078984 | 131 | Hnrnpr | SPRET | feature_truncation,3_prime_UTR_variant |
| 4 | 138019655 | 138020619 | 964 | Kif17 | SPRET | feature_truncation,coding_sequence_variant,intron_variant |
| 4 | 138167370 | 138167433 | 63 | Mul1 | AJ,BALB,BUB,C3H,CBA,CE,DBA1J,DBA2J,FVB,ILNJ2,KK,MA/My,MRL,NZB,NZW,RH2,SJL,SMJ,SWR,TH,WSB | feature_truncation,3_prime_UTR_variant |
| 4 | 139104558 | 139104628 | 70 | Emc1 | AKR,BALB,BUB,C3H,CBA,DBA1J,DBA2J,ILNJ2,KK,MA/My,MRL,NOR,NZB,PL,RF,SJL,SMJ,TH | feature_truncation,3_prime_UTR_variant |
| 4 | 139688685 | 139689123 | 438 | Klhdc7a | CAST,SPRET | feature_truncation,3_prime_UTR_variant |

| | | | | | | |
|---|---|---|---|---|---|---|
| 4 | 140785315 | 140785388 | 73 | Crocc | 129S1,BTBR,C57L,CE,DBA1J,DBA2J,FVB,ILNJ2,KK,NOD,NOR,NZB,NZO,PL,RF,SJL,SPRET,SWR,TH,WSB | feature_truncation,5_prime_UTR_variant,intron_variant |
| 4 | 145348866 | 145349525 | 659 | Zfp268 | WSB | frameshift_variant,feature_truncation,feature_truncation,coding_sequence_variant,intron_variant |
| 4 | 145349581 | 145349902 | 321 | Zfp268 | C58J,DBA2J,PJ,SEA,TH | stop_lost,feature_truncation,coding_sequence_variant,3_prime_UTR_variant,feature_truncation,inframe_deletion |
| 4 | 145350209 | 145350293 | 84 | Zfp268 | DBA2J,KK,LP | feature_truncation,inframe_deletion |
| 4 | 146061591 | 146061674 | 83 | Zfp987 | DBA1J,MRL,NOD,NZB,NZW,SJL | frameshift_variant,feature_truncation |
| 4 | 146132071 | 146132154 | 83 | Zfp600 | BTBR,DBA1J,NOD,SWR,WSB | frameshift_variant,feature_truncation |
| 4 | 146133114 | 146133198 | 84 | Zfp600 | CE,KK,MA/My,SMJ | stop_lost,inframe_deletion |
| 4 | 146623297 | 146623557 | 260 | Zfp981 | PJ,TH | feature_truncation,3_prime_UTR_variant |
| 4 | 147074248 | 147074499 | 251 | Zfp989 | PL | frameshift_variant,feature_truncation |
| 4 | 147075026 | 147075109 | 83 | Zfp989 | FVB,LP,SEA | frameshift_variant,feature_truncation |
| 4 | 147142351 | 147142435 | 84 | Rex2 | C58J,LP,PJ,SEA,SPRET | feature_truncation,inframe_deletion |
| 4 | 147416497 | 147416580 | 83 | Zfp988 | SMJ | frameshift_variant,feature_truncation |
| 4 | 147597252 | 147597421 | 169 | Zfp982 | 129S1,BTBR,C57L,C58J,CAST,LP,MOLF,SEA,SPRET,TH | frameshift_variant,feature_truncation |
| 4 | 147668773 | 147668846 | 73 | Zfp985 | 129S1,BTBR,KK | feature_truncation,3_prime_UTR_variant |
| 4 | 147759462 | 147759881 | 419 | Zfp534 | NZO | frameshift_variant,feature_truncation |
| 4 | 147909370 | 147909545 | 175 | Zfp933 | 129S1,AKR,BTBR,BUB,C3H,C57L,C58J,CAST,CBA,CE,DBA1J,DBA2J,KK,LP,MA/My,MRL,NOD,NZB,NZO,NZW,PJ,PL,RF,SEA,SJL,SMJ,SPRET,SWR,TH,WSB | feature_truncation,3_prime_UTR_variant |
| 4 | 147958725 | 147958789 | 64 | Mfn2 | SPRET | feature_truncation,3_prime_UTR_variant |
| 4 | 148140073 | 148140125 | 52 | Mthfr | CAST | feature_truncation,3_prime_UTR_variant |
| 4 | 148519573 | 148519705 | 132 | Ubiad1 | SPRET | feature_truncation,3_prime_UTR_variant |
| 4 | 149212801 | 149212883 | 82 | Cenps | 129S1,AJ,AKR,BALB,BTBR,BUB,C3H,C57L,C58J,CBA,CE,DBA1J,DBA2J,KK,LP,MA/My,MRL,NOD,NZB,NZO,NZW,PJ,PL,RF,SEA,SJL,SMJ,SPRET,SWR,TH,WSB | feature_truncation,3_prime_UTR_variant |
| 4 | 149464780 | 149464876 | 96 | Ube4b | 129S1,AKR,BTBR,C3H,C57L,CAST,CBA,DBA1J,DBA2J,MA/My,MOLF,MRL,NOD,PL,SJL,SPRET,TH | feature_truncation,coding_sequence_variant,intron_variant |
| 4 | 149552720 | 149552924 | 204 | Nmnat1 | SPRET | feature_truncation,3_prime_UTR_variant |
| 4 | 152101423 | 152101526 | 103 | Klhl21 | CAST,MOLF,SPRET | feature_truncation,3_prime_UTR_variant |
| 4 | 154104690 | 154104814 | 124 | Lrrc47 | MOLF | feature_truncation,3_prime_UTR_variant |
| 4 | 154106247 | 154106456 | 209 | Smim1 | CAST | feature_truncation,3_prime_UTR_variant |
| 4 | 155506133 | 155506506 | 373 | Cfap74 | CAST,CE,FVB,ILNJ2,MOLF,NOD,NOR,SJL,SMJ,SPRET,SWR,TH,WSB | feature_truncation,3_prime_UTR_variant |
| 4 | 155551095 | 155551151 | 56 | Cfap74 | SPRET | feature_truncation,3_prime_UTR_variant |
| 4 | 155705802 | 155705855 | 53 | Slc35e2 | CAST | feature_truncation,3_prime_UTR_variant |
| 4 | 155853115 | 155853176 | 61 | Vwa1 | CAST | feature_truncation,3_prime_UTR_variant |
| 4 | 156077194 | 156077382 | 188 | Sdf4 | SPRET | feature_truncation,5_prime_UTR_variant |
| 5 | 3575955 | 3576781 | 826 | Cdk6 | SPRET | feature_truncation,3_prime_UTR_variant |
| 5 | 3943592 | 3943645 | 53 | Mterf1a | SPRET | feature_truncation,5_prime_UTR_variant |
| 5 | 4131127 | 4131289 | 162 | Cyp51,Akap9 | BUB,CAST,MOLF,NOD,NOR | feature_truncation,3_prime_UTR_variant |
| 5 | 5721320 | 5721379 | 59 | Steap2 | SPRET | feature_truncation,3_prime_UTR_variant |
| 5 | 10298041 | 10298853 | 812 | Gm5152 | CAST | feature_truncation,coding_sequence_variant,5_prime_UTR_variant,intron_variant |
| 5 | 14571967 | 14572057 | 90 | Pclo | 129S1,BALB,BTBR,C57L,C58J,CE,FVB,MA/My,PJ,RF,RH2,SEA,SJL,SWR,WSB | feature_truncation,inframe_deletion |
| 5 | 17685392 | 17685482 | 90 | Speer4f1 | CAST,SPRET | stop_lost,feature_truncation,inframe_deletion |
| 5 | 21160843 | 21161617 | 774 | Rsbn1l | CAST | feature_truncation,5_prime_UTR_variant |
| 5 | 22115798 | 22115890 | 92 | Reln | SPRET | splice_donor_variant,splice_donor_5th_base_variant,coding_sequence_variant,intron_variant |
| 5 | 24645751 | 24645808 | 57 | Slc4a2 | SPRET | feature_truncation,3_prime_UTR_variant |
| 5 | 25895640 | 25895824 | 184 | Xrcc2 | SPRET | feature_truncation,3_prime_UTR_variant |
| 5 | 29657217 | 29658027 | 810 | Nom1 | SPRET | feature_truncation,3_prime_UTR_variant |
| 5 | 31034275 | 31034354 | 79 | Tmem214 | SPRET | feature_truncation,3_prime_UTR_variant |
| 5 | 31035827 | 31035927 | 100 | Tmem214 | SPRET | feature_truncation,3_prime_UTR_variant |
| 5 | 31654848 | 31655028 | 180 | Gpn1 | SPRET | feature_truncation,3_prime_UTR_variant |
| 5 | 31675093 | 31675300 | 207 | Supt7l | CAST,MOLF,SPRET | feature_truncation,3_prime_UTR_variant |
| 5 | 31711480 | 31711597 | 117 | Slc4a1ap | SPRET | feature_truncation,3_prime_UTR_variant |
| 5 | 33574942 | 33574999 | 57 | Uvssa | MOLF | feature_truncation,3_prime_UTR_variant |
| 5 | 33935838 | 33935892 | 54 | Letm1 | CAST,MOLF,SPRET | feature_truncation,3_prime_UTR_variant |
| 5 | 35772116 | 35772268 | 152 | Acox3 | CAST | feature_truncation,3_prime_UTR_variant |
| 5 | 37604833 | 37604896 | 63 | Stk32b | WSB | feature_truncation,3_prime_UTR_variant |
| 5 | 43999503 | 43999567 | 64 | Bst1 | PJ,PL | feature_truncation,3_prime_UTR_variant |
| 5 | 44001094 | 44001276 | 182 | Bst1 | MOLF,SPRET | feature_truncation,3_prime_UTR_variant |
| 5 | 44070476 | 44071278 | 802 | Ppihl | SPRET | transcript_ablation |
| 5 | 44151234 | 44151310 | 76 | Prom1 | CAST,MOLF,PJ,PL | feature_truncation,3_prime_UTR_variant |
| 5 | 45684420 | 45684718 | 298 | Med28 | MOLF | feature_truncation,3_prime_UTR_variant |
| 5 | 48464242 | 48464311 | 69 | Slit2 | SPRET | feature_truncation,3_prime_UTR_variant |
| 5 | 52311190 | 52311322 | 132 | Dhx15 | SPRET | feature_truncation,3_prime_UTR_variant |
| 5 | 53758093 | 53758305 | 212 | Rbpj | SPRET | feature_truncation,coding_sequence_variant,5_prime_UTR_variant,intron_variant |
| 5 | 65199049 | 65199192 | 143 | Fam114a1 | SPRET | feature_truncation,3_prime_UTR_variant |
| 5 | 65547016 | 65547123 | 107 | Rpl9 | CAST,MOLF | feature_truncation,3_prime_UTR_variant |
| 5 | 65604667 | 65604802 | 135 | Smim14 | C3H,CAST,CBA,DBA1J,DBA2J,ILNJ2,KK,NZB,PJ,SPRET,WSB | feature_truncation,3_prime_UTR_variant |
| 5 | 65605886 | 65606181 | 295 | Smim14 | SPRET | feature_truncation,3_prime_UTR_variant |
| 5 | 72314508 | 72314850 | 342 | Commd8 | CAST | feature_truncation,3_prime_UTR_variant |
| 5 | 73164998 | 73165303 | 305 | Slc10a4 | SPRET | feature_truncation,coding_sequence_variant,intron_variant |
| 5 | 77429676 | 77429730 | 54 | Rest | SPRET | feature_truncation,inframe_deletion |
| 5 | 77499256 | 77500136 | 880 | Igfbp7 | SPRET | feature_truncation,coding_sequence_variant,intron_variant |
| 5 | 86252645 | 86252776 | 131 | Stap1 | SPRET | feature_truncation,3_prime_UTR_variant |
| 5 | 87159503 | 87159573 | 70 | Ugt2b35 | SPRET | feature_truncation,3_prime_UTR_variant |
| 5 | 88476262 | 88476316 | 54 | Prol1 | MA/My,NZB,NZO,SPRET | feature_truncation,inframe_deletion |
| 5 | 90908395 | 90908461 | 66 | Cxcl5 | BALB,BTBR,BUB,C3H,CBA,CE,DBA1J,DBA2J,FVB,ILNJ2,KK,MOLF,MRL,NOD,NOR,PJ,PL,RH2,SEA,SJL,SMJ,SPRET | feature_truncation,3_prime_UTR_variant |

| | | | | | | |
|---|---|---|---|---|---|---|
| 5 | 91052523 | 91052697 | 174 | Cxcl2 | CAST | feature_truncation,3_prime_UTR_variant |
| 5 | 91176036 | 91176146 | 110 | Epgn | SPRET | splice_acceptor_variant,5_prime_UTR_variant,intron_variant |
| 5 | 100105965 | 100106067 | 102 | Hnrnpd | CAST | feature_truncation,3_prime_UTR_variant |
| 5 | 103174854 | 103174933 | 79 | Mapk10 | 129S1,AKR,BUB,C3H,C57L,CAST,CBA,CE,DBA1J,DBA2J,MOLF,MRL,NOD,NOR,NZO,NZW,PJ,PL,RF,SMJ,SPRET,TH,WSB | feature_truncation,5_prime_UTR_variant |
| 5 | 104010755 | 104010853 | 98 | Klhl8 | WSB | feature_truncation,3_prime_UTR_variant |
| 5 | 104962344 | 104962430 | 86 | Zfp951 | AJ,BALB,BUB,C3H,CBA,DBA1J,DBA2J,FVB,ILNJ2,KK,MOLF,NZB,NZO,RH2,SEA,SJL,SWR,TH,WSB | frameshift_variant,stop_lost |
| 5 | 105007539 | 105007811 | 272 | Zfp951 | AJ,BALB,BUB,C3H,CBA,DBA1J,DBA2J,FVB,ILNJ2,KK,MOLF,NZB,NZO,RH2,SEA,SJL,SWR,TH,WSB | frameshift_variant,start_lost,feature_truncation,start_retained_variant,5_prime_UTR_variant |
| 5 | 107534367 | 107534583 | 216 | Brdt | SPRET | feature_truncation,3_prime_UTR_variant |
| 5 | 107697541 | 107697898 | 357 | 1700028K03Rik | SPRET | feature_truncation,3_prime_UTR_variant |
| 5 | 108266887 | 108267090 | 203 | Tmed5 | 129S1,AJ,AKR,BALB,BTBR,BUB,C3H,C57L,CAST,CBA,CE,DBA1J,DBA2J,FVB,ILNJ2,KK,LP,MA/My,MOLF,MRL,NOD,NOR,NZB,NZO,NZW,PJ,PL,RF,RH2,SEA,SJL,SMJ,SPRET,SWR,TH,WSB | feature_truncation,3_prime_UTR_variant |
| 5 | 108791781 | 108791949 | 168 | Tmem175 | SPRET | feature_truncation,3_prime_UTR_variant |
| 5 | 109233298 | 109233528 | 230 | Vmn2r12 | SPRET | feature_truncation,3_prime_UTR_variant |
| 5 | 109363045 | 109363591 | 546 | Vmn2r14 | 129S1,AKR,BUB,C3H,C57L,CAST,CBA,FVB,KK,MA/My,MOLF,NOD,NOR,NZB,NZO,NZW,PJ,PL,RF,SJL,SPRET,SWR,TH,WSB | stop_lost,feature_truncation,coding_sequence_variant,3_prime_UTR_variant |
| 5 | 109602149 | 109602218 | 69 | Vmn2r17 | SPRET | feature_truncation,3_prime_UTR_variant |
| 5 | 109603277 | 109603427 | 150 | Vmn2r17 | SPRET | feature_truncation,3_prime_UTR_variant |
| 5 | 109823483 | 109823567 | 84 | Zfp1007 | BUB,C3H,CBA,FVB,MOLF,PJ,PL,SJL,SWR,TH | feature_truncation,inframe_deletion |
| 5 | 110090065 | 110090149 | 84 | Gm15446 | CAST,SPRET | feature_truncation,inframe_deletion |
| 5 | 113915459 | 113915670 | 211 | Iscu | CAST,MOLF,SPRET | feature_truncation,3_prime_UTR_variant |
| 5 | 113957727 | 113957787 | 60 | Selplg | 129S1,BALB,BUB,C3H,C57L,CBA,FVB,ILNJ2,MOLF,MRL,PJ,PL,RH2,SEA,SJL,SMJ,SWR,WSB | feature_truncation,inframe_deletion |
| 5 | 114336159 | 114336435 | 276 | Acacb | MOLF,SPRET | splice_acceptor_variant,splice_donor_variant,splice_donor_5th_base_variant,coding_sequence_variant,intron_variant |
| 5 | 114959633 | 114960589 | 956 | 1500011B03Rik | SPRET | stop_lost,feature_truncation,coding_sequence_variant,3_prime_UTR_variant |
| 5 | 117912335 | 117912397 | 62 | Ksr2 | SPRET | feature_truncation,3_prime_UTR_variant |
| 5 | 118095046 | 118095108 | 62 | Nos1 | BUB,C3H,CBA,ILNJ2,KK,NZB,NZO,PL,SJL,TH,WSB | feature_truncation,3_prime_UTR_variant |
| 5 | 118399986 | 118400127 | 141 | Spring1 | CAST,MOLF,RF | feature_truncation,3_prime_UTR_variant |
| 5 | 120578819 | 120578885 | 66 | Lhx5 | SPRET | feature_truncation,3_prime_UTR_variant |
| 5 | 120883856 | 120883917 | 61 | Oas2 | PL,SPRET | frameshift_variant,feature_truncation |
| 5 | 120892059 | 120892135 | 76 | Oas3 | SPRET | feature_truncation,3_prime_UTR_variant |
| 5 | 121564209 | 121564270 | 61 | Naa25 | SPRET | feature_truncation,coding_sequence_variant,intron_variant |
| 5 | 121579261 | 121579446 | 185 | Naa25,Erp29 | SPRET | feature_truncation,3_prime_UTR_variant |
| 5 | 121583045 | 121583098 | 53 | Erp29 | SPRET | feature_truncation,3_prime_UTR_variant |
| 5 | 121704696 | 121704836 | 140 | Aldh2 | CAST | feature_truncation,3_prime_UTR_variant |
| 5 | 121736567 | 121736702 | 135 | Acad12 | CAST,MOLF | feature_truncation,3_prime_UTR_variant |
| 5 | 122375909 | 122376097 | 188 | Tctn1 | SPRET | feature_truncation,3_prime_UTR_variant |
| 5 | 122821404 | 122821474 | 70 | P2rx7 | MOLF,SPRET | feature_truncation,3_prime_UTR_variant |
| 5 | 123153130 | 123153191 | 61 | Orai1 | B10D2 | feature_truncation,5_prime_UTR_variant |
| 5 | 125464210 | 125464438 | 228 | Ubc | FVB,MA/My,SWR,TH | feature_truncation,inframe_deletion |
| 5 | 125535326 | 125535409 | 83 | Bri3bp | SPRET | feature_truncation,3_prime_UTR_variant |
| 5 | 129062030 | 129062093 | 63 | Stx2 | SPRET | feature_truncation,3_prime_UTR_variant |
| 5 | 129871527 | 129871643 | 116 | Cct6a | MOLF,SPRET | feature_truncation,coding_sequence_variant,intron_variant |
| 5 | 134736365 | 134736425 | 60 | Eln | KK | feature_truncation,inframe_deletion |
| 5 | 135991116 | 135991171 | 55 | Ssc4d | SPRET | feature_truncation,5_prime_UTR_variant |
| 5 | 136944065 | 136944132 | 67 | Ift22 | CAST,MOLF,SPRET | feature_truncation,3_prime_UTR_variant |
| 5 | 137140644 | 137140697 | 53 | Trim56 | CAST,LP,SPRET | feature_truncation,3_prime_UTR_variant |
| 5 | 137178194 | 137178428 | 234 | Muc17 | LP,SPRET | feature_truncation,inframe_deletion |
| 5 | 137178731 | 137178803 | 72 | Muc17 | CAST | feature_truncation,inframe_deletion |
| 5 | 137181487 | 137181712 | 225 | Muc17 | CAST,CBA,CE,DBA2J,LP,NZW,PJ,SJL | feature_truncation,inframe_deletion |
| 5 | 137181933 | 137182051 | 118 | Muc17 | 129S1,AJ,BALB,BTBR,BUB,C57L,MA/My,NOR,RF,SEA,SPRET,SWR | frameshift_variant,feature_truncation |
| 5 | 137184544 | 137184625 | 81 | Muc17 | CAST,LP | feature_truncation,inframe_deletion |
| 5 | 137186467 | 137186521 | 54 | Muc17 | CAST,MOLF | feature_truncation,inframe_deletion |
| 5 | 137186823 | 137187633 | 810 | Muc17 | CAST,MOLF,SPRET | feature_truncation,inframe_deletion |
| 5 | 137188130 | 137188831 | 701 | Muc17 | CAST | frameshift_variant,feature_truncation |
| 5 | 137189347 | 137189401 | 54 | Muc17 | B10D2,B10J,C58J,LP,MOLF,PL,SMJ,SPRET,TH | feature_truncation,inframe_deletion |
| 5 | 137189539 | 137189970 | 431 | Muc17 | CAST | frameshift_variant,feature_truncation |
| 5 | 137462698 | 137462868 | 170 | Zan | 129S1,BUB,CAST,LP,MOLF,RF,SPRET,SWR,WSB | frameshift_variant,feature_truncation |
| 5 | 137594987 | 137595142 | 155 | Mospd3 | 129S1,AKR,BUB,CAST,KK,LP,MA/My,MOLF,MRL,NZB,NZO,NZW,RF,SPRET,SWR | feature_truncation,3_prime_UTR_variant |
| 5 | 137821257 | 137821486 | 229 | Pilra | SPRET | feature_truncation,3_prime_UTR_variant |
| 5 | 138139712 | 138139825 | 113 | Zfp113 | SPRET | feature_truncation,3_prime_UTR_variant |
| 5 | 138162563 | 138162614 | 51 | Cops6 | MOLF | feature_truncation,3_prime_UTR_variant |
| 5 | 138250815 | 138250985 | 170 | Nxpe5 | MOLF,SPRET | feature_truncation,3_prime_UTR_variant |
| 5 | 138646892 | 138646976 | 84 | A430033K04Rik | 129S1,C3H,CBA,CE,DBA1J,DBA2J,KK,LP,MA/My,NZB,NZO,NZW,SPRET,WSB | feature_truncation,3_prime_UTR_variant |
| 5 | 139787733 | 139787838 | 105 | Mafk | CAST,MOLF | feature_truncation,3_prime_UTR_variant |
| 5 | 139789876 | 139789968 | 92 | Tmem184a | ILNJ2 | feature_truncation,3_prime_UTR_variant |
| 5 | 143119878 | 143120258 | 380 | Or10ah1-ps1 | CAST,MOLF,SPRET,WSB | feature_truncation,3_prime_UTR_variant |
| 5 | 143120167 | 143120378 | 211 | Or10ah1-ps1 | SPRET | feature_truncation,3_prime_UTR_variant |
| 5 | 143122013 | 143122080 | 67 | Or10ah1-ps1 | SPRET | feature_truncation,3_prime_UTR_variant |
| 5 | 143869099 | 143869152 | 53 | Pms2 | MOLF,SPRET | feature_truncation,3_prime_UTR_variant |
| 5 | 144673098 | 144673207 | 109 | Tmem130 | SPRET | feature_truncation,3_prime_UTR_variant |

| | | | | | | |
|---|---|---|---|---|---|---|
| 5 | 144793877 | 144793981 | 104 | Trrap | 129S1,AKR,BTBR,C57L,CAST,CBA,MOLF,PL,SJL,SMJ,SPRET,TH,WSB | feature_truncation,coding_sequence_variant,intron_variant |
| 5 | 147806521 | 147806597 | 76 | Pomp | MOLF,SPRET | feature_truncation,3_prime_UTR_variant |
| 6 | 7040524 | 7040706 | 182 | Sdhaf3 | CAST,MOLF,PJ | feature_truncation,3_prime_UTR_variant |
| 6 | 23003608 | 23003713 | 105 | Ptprz1 | TH | feature_truncation,3_prime_UTR_variant |
| 6 | 23344469 | 23345112 | 643 | Cadps2 | C3H,CAST,CBA,CE,FVB,MOLF,NOD,NOR,SPRET | feature_truncation,coding_sequence_variant,intron_variant |
| 6 | 25758353 | 25758535 | 182 | Pot1a | CAST,MOLF,SPRET | feature_truncation,3_prime_UTR_variant |
| 6 | 31493642 | 31493702 | 60 | Mkln1 | SPRET | feature_truncation,3_prime_UTR_variant |
| 6 | 34079026 | 34079086 | 60 | Lrguk | CAST,MOLF,SPRET | feature_truncation,inframe_deletion |
| 6 | 34085773 | 34086464 | 691 | Lrguk | CAST,MOLF | feature_truncation,3_prime_UTR_variant |
| 6 | 34482142 | 34482194 | 52 | Bpgm | SPRET | feature_truncation,3_prime_UTR_variant |
| 6 | 37544619 | 37545433 | 814 | Akr1d1 | SPRET | feature_truncation,3_prime_UTR_variant |
| 6 | 38481156 | 38481653 | 497 | Ubn2 | CAST,KK,MOLF,SPRET | feature_truncation,3_prime_UTR_variant |
| 6 | 38583066 | 38583298 | 232 | Luc7l2 | SPRET | feature_truncation,3_prime_UTR_variant |
| 6 | 39116719 | 39116893 | 174 | Kdm7a | SPRET | feature_truncation,3_prime_UTR_variant |
| 6 | 41747969 | 41748139 | 170 | Or9a2 | SPRET | feature_truncation,3_prime_UTR_variant |
| 6 | 43132250 | 43132609 | 359 | Or2a14 | SPRET | feature_truncation,3_prime_UTR_variant |
| 6 | 47848947 | 47849040 | 93 | Zfp398 | SPRET | feature_truncation,3_prime_UTR_variant |
| 6 | 48478165 | 48478779 | 614 | Sspo | AJ,AKR,BALB,BUB,C3H,CBA,DBA1J,DBA2J,FVB,MRL,NOD,NOR,PJ,PL,RF,RH2,SEA,SJL,SMJ,SWR,TH,WSB | feature_truncation,3_prime_UTR_variant |
| 6 | 48599337 | 48599429 | 92 | Zfp775 | AKR,BALB,C3H,C58J,CBA,DBA1J,DBA2J,FVB,MA/My,MOLF,MRL,PJ,PL,RF,RH2,SEA,SJL,SMJ,SPRET,SWR | feature_truncation,3_prime_UTR_variant |
| 6 | 48637433 | 48637799 | 366 | Gimap8 | SPRET | feature_truncation,3_prime_UTR_variant |
| 6 | 50173491 | 50173841 | 350 | Pals2 | CAST,SPRET | splice_acceptor_variant,splice_donor_variant,splice_donor_5th_base_variant,coding_sequence_variant,intron_variant |
| 6 | 56048851 | 56048908 | 57 | Pde1c | SPRET | feature_truncation,3_prime_UTR_variant |
| 6 | 56057449 | 56057507 | 58 | Pde1c | 129S1,C57L,KK,MOLF,NZB,NZO,NZW,SPRET | feature_truncation,3_prime_UTR_variant |
| 6 | 57217815 | 57217872 | 57 | Vmn1r14 | 129S1,C57L,CE,ILNJ2,KK,MOLF,NZB,NZO,NZW,PJ,WSB | feature_truncation,3_prime_UTR_variant |
| 6 | 57336745 | 57336808 | 63 | Vmn1r17 | SPRET | feature_truncation,3_prime_UTR_variant |
| 6 | 57737449 | 57737529 | 80 | Vopp1 | SPRET | feature_truncation,3_prime_UTR_variant |
| 6 | 57874406 | 57875178 | 772 | Vmn1r22 | CAST,CE,ILNJ2,MOLF,PJ,SPRET,WSB | feature_truncation,3_prime_UTR_variant |
| 6 | 57875776 | 57876069 | 293 | Vmn1r22 | CAST | feature_truncation,3_prime_UTR_variant |
| 6 | 57877937 | 57878487 | 550 | Vmn1r22 | SPRET | feature_truncation,coding_sequence_variant,5_prime_UTR_variant,intron_variant,feature_truncation,coding_sequence_variant,5_prime_UTR_variant |
| 6 | 58238919 | 58239035 | 116 | Vmn1r28 | SPRET | feature_truncation,5_prime_UTR_variant |
| 6 | 58415937 | 58416884 | 947 | Vmn1r30 | CAST,MOLF | feature_truncation,5_prime_UTR_variant,intron_variant |
| 6 | 58448231 | 58448973 | 742 | Vmn1r31 | CAST | frameshift_variant,stop_lost,feature_truncation,3_prime_UTR_variant,stop_lost,feature_truncation,coding_sequence_variant,3_prime_UTR_variant |
| 6 | 59186872 | 59186983 | 111 | Tigd2 | SPRET | feature_truncation,5_prime_UTR_variant |
| 6 | 67268914 | 67268970 | 56 | Serbp1,Il12rb2 | MOLF | feature_truncation,3_prime_UTR_variant |
| 6 | 67271531 | 67271584 | 53 | Serbp1 | C58J,KK,NOD,NOR,NZB,NZO | feature_truncation,3_prime_UTR_variant |
| 6 | 71314315 | 71314516 | 201 | Cd8b1 | SPRET | feature_truncation,3_prime_UTR_variant |
| 6 | 71790241 | 71790839 | 598 | Mrpl35 | C3H,CBA,DBA1J,DBA2J,MA/My,SMJ | feature_truncation,3_prime_UTR_variant |
| 6 | 72291034 | 72291085 | 51 | Sftpb | AKR,C58J,KK,MRL,NOD,NOR,PL,RF,SPRET | feature_truncation,3_prime_UTR_variant |
| 6 | 81938557 | 81938737 | 180 | Mrpl19 | CAST,SPRET | feature_truncation,3_prime_UTR_variant |
| 6 | 82535044 | 82535143 | 99 | Tacr1 | KK,MOLF,NZB,NZO,SPRET,TH | feature_truncation,3_prime_UTR_variant |
| 6 | 83029357 | 83029425 | 68 | Loxl3 | CAST | feature_truncation,3_prime_UTR_variant |
| 6 | 83281554 | 83281633 | 79 | Slc4a5 | CAST,KK,MOLF,NZB,NZO,SPRET,TH | feature_truncation,3_prime_UTR_variant |
| 6 | 84166799 | 84166940 | 141 | Dysf | CAST,SJL | feature_truncation,coding_sequence_variant,intron_variant |
| 6 | 85479719 | 85479775 | 56 | Fbxo41 | SPRET | feature_truncation,5_prime_UTR_variant |
| 6 | 85920302 | 85921165 | 863 | Dusp11 | SPRET | feature_truncation,3_prime_UTR_variant |
| 6 | 87605572 | 87605981 | 409 | Aplf | CAST,MOLF,PJ,SEA,SPRET | feature_truncation,3_prime_UTR_variant |
| 6 | 87606199 | 87606279 | 80 | Aplf | MA/My,MOLF,NZB,NZO,SEA,TH | feature_truncation,3_prime_UTR_variant |
| 6 | 87768641 | 87768740 | 99 | Rab43 | AKR,BALB,BUB,CAST,CBA,FVB,KK,LP,MA/My,MOLF,MRL,NOD,NOR,PJ,PL,RH2,SEA,SJL,SPRET,SWR,WSB | feature_truncation,3_prime_UTR_variant |
| 6 | 87792249 | 87792488 | 239 | Isy1 | SPRET | feature_truncation,3_prime_UTR_variant |
| 6 | 89842289 | 89843112 | 823 | Vmn1r43 | SPRET | feature_truncation,3_prime_UTR_variant |
| 6 | 91439106 | 91439757 | 651 | Chchd4 | CE,MOLF,NZB,NZW,PJ,SEA | feature_truncation,3_prime_UTR_variant |
| 6 | 92164712 | 92164842 | 130 | Rbsn | SPRET | feature_truncation,3_prime_UTR_variant |
| 6 | 100547923 | 100548581 | 658 | Shq1 | BUB,CAST,MOLF,NZB,NZO,NZW,SPRET,SWR,WSB | feature_truncation,3_prime_UTR_variant |
| 6 | 100783038 | 100783090 | 52 | Gxylt2 | SPRET,SWR | feature_truncation,3_prime_UTR_variant |
| 6 | 112626567 | 112626639 | 72 | Rad18 | MOLF,SPRET | feature_truncation,inframe_deletion |
| 6 | 113146197 | 113146286 | 89 | Lhfpl4 | CAST,MOLF | feature_truncation,3_prime_UTR_variant |
| 6 | 113267465 | 113267549 | 84 | Cpne9 | SPRET | feature_truncation,coding_sequence_variant,intron_variant |
| 6 | 113342874 | 113343060 | 186 | Tada3 | CAST,MOLF,SPRET | feature_truncation,3_prime_UTR_variant |
| 6 | 115782071 | 115782245 | 174 | Cand2 | AKR,C3H,CBA,ILNJ2,MOLF,MRL,PJ,PL,RF,SPRET,WSB | feature_truncation,3_prime_UTR_variant |
| 6 | 116483186 | 116484002 | 816 | Or6d12 | SPRET | feature_truncation,5_prime_UTR_variant,intron_variant |
| 6 | 116657145 | 116657203 | 58 | Tmem72 | SPRET | feature_truncation,3_prime_UTR_variant |
| 6 | 120467712 | 120467776 | 64 | Tmem121b | MOLF | feature_truncation,3_prime_UTR_variant |
| 6 | 120747364 | 120747553 | 189 | Cecr2 | SPRET | feature_truncation,3_prime_UTR_variant |
| 6 | 122582737 | 122582845 | 108 | Gdf3 | SPRET | feature_truncation,3_prime_UTR_variant |
| 6 | 122825993 | 122826057 | 64 | C3ar1 | SPRET | feature_truncation,3_prime_UTR_variant |
| 6 | 123781229 | 123782005 | 776 | Vmn2r24 | FVB,NZO,SWR | feature_truncation,coding_sequence_variant,intron_variant |
| 6 | 124167291 | 124167724 | 433 | Vmn2r27 | SPRET | feature_truncation,3_prime_UTR_variant,frameshift_variant,stop_lost,feature_truncation,3_prime_UTR_variant |
| 6 | 126952675 | 126952876 | 201 | D6Wsu163e | SPRET | feature_truncation,3_prime_UTR_variant |
| 6 | 128299898 | 128299952 | 54 | Tulp3 | CAST,MOLF | feature_truncation,3_prime_UTR_variant |
| 6 | 128384603 | 128384883 | 280 | Nrip2 | CAST | feature_truncation,coding_sequence_variant,intron_variant |

| | | | | | | |
|---|---|---|---|---|---|---|
| 6 | 128586551 | 128586683 | 132 | Klrb1a | NOD,NOR,SPRET,SWR | stop_lost,feature_truncation,coding_sequence_variant,3_prime_UTR_variant |
| 6 | 128947891 | 128948266 | 375 | Clec2g | 129S1,AJ,AKR,BALB,BUB,C3H,C57L,CBA,DBA1J,DBA2J,ILNJ2,MRL,PL,RF,SEA,SMJ,WSB | feature_truncation,5_prime_UTR_variant,feature_truncation,coding_sequence_variant,5_prime_UTR_variant,intron_variant |
| 6 | 129463671 | 129463732 | 61 | Olr1 | MOLF | feature_truncation,3_prime_UTR_variant |
| 6 | 129706780 | 129706972 | 192 | Klri2 | MOLF | feature_truncation,3_prime_UTR_variant |
| 6 | 129842165 | 129842870 | 705 | Klra17 | 129S1,C58J,CE,FVB,LP,MA/My,SJL,TH,WSB | feature_truncation,coding_sequence_variant,intron_variant |
| 6 | 132577154 | 132577277 | 123 | Prp2 | 129S1,AJ,AKR,BALB,BUB,C3H,C57L,CBA,DBA1J,DBA2J,ILNJ2,MRL,NZB,NZO,PL,RF,RH2,SEA,SMJ,SPRET | feature_truncation,inframe_deletion |
| 6 | 134424796 | 134424859 | 63 | Lrp6 | FVB,NZO,SWR | feature_truncation,3_prime_UTR_variant |
| 6 | 134835596 | 134835663 | 67 | Crebl2 | SPRET | feature_truncation,3_prime_UTR_variant |
| 6 | 135701084 | 135701375 | 291 | Grin2b | 129S1,AJ,AKR,BALB,BTBR,BUB,C3H,C57L,CBA,CE,DBA1J,DBA2J,FVB,KK,MA/My,MRL,NOD,NOR,NZB,NZW,PJ,PL,RF,RH2,SEA,SJL,SWR,TH,WSB | feature_truncation,3_prime_UTR_variant |
| 6 | 136584359 | 136584419 | 60 | Atf7ip | 129S1,AJ,AKR,BALB,BTBR,BUB,C3H,C57L,CAST,CBA,CE,DBA1J,DBA2J,FVB,KK,MA/My,MOLF,MRL,NOD,NOR,NZO,NZW,PJ,PL,RF,RH2,SEA,SJL,SPRET,SWR,TH,WSB | feature_truncation,3_prime_UTR_variant |
| 6 | 136778675 | 136779043 | 368 | H4c16 | 129S1,AJ,AKR,BALB,BTBR,BUB,C3H,C57L,CAST,CBA,CE,DBA1J,DBA2J,FVB,KK,MA/My,MRL,NOD,NOR,NZO,NZW,PJ,PL,RF,RH2,SEA,SJL,SPRET,SWR,TH,WSB | feature_truncation,3_prime_UTR_variant |
| 6 | 136785953 | 136786039 | 86 | H2aj | SPRET | feature_truncation,3_prime_UTR_variant |
| 6 | 136805813 | 136806284 | 471 | BC049715 | 129S1,BALB,BTBR,C57L,CE,DBA1J,FVB,MRL,NZW,SEA,WSB | feature_truncation,5_prime_UTR_variant,intron_variant |
| 6 | 136941033 | 136941125 | 92 | Pde6h | BALB,FVB,KK,MA/My,MRL,NZO,NZW,RH2,SEA | feature_truncation,3_prime_UTR_variant |
| 6 | 136944910 | 136945030 | 120 | Pde6h | 129S1,AJ,AKR,BALB,BTBR,BUB,C3H,C57L,CAST,CBA,CE,DBA1J,DBA2J,FVB,KK,MA/My,MRL,NOD,NOR,NZO,NZW,PJ,PL,RF,RH2,SEA,SJL,SPRET,SWR,TH,WSB | feature_truncation,3_prime_UTR_variant |
| 6 | 140540430 | 140540646 | 216 | Plekha5 | SPRET | feature_truncation,3_prime_UTR_variant |
| 6 | 140627277 | 140627450 | 173 | Aebp2 | BTBR,C57L,C58J,FVB,MOLF,PJ,SMJ,SWR,WSB | feature_truncation,3_prime_UTR_variant |
| 6 | 142767576 | 142767627 | 51 | St8sia1 | MOLF | feature_truncation,3_prime_UTR_variant |
| 6 | 143776898 | 143776958 | 60 | Sox5 | SPRET | feature_truncation,3_prime_UTR_variant |
| 6 | 145396408 | 145396459 | 51 | Lmntd1 | AKR,BUB,C3H,CE,DBA1J,DBA2J,MRL,NZB,NZO,SJL | feature_truncation,5_prime_UTR_variant |
| 6 | 145910552 | 145910624 | 72 | Sspn | AKR,BUB,C3H,CAST,CE,DBA1J,DBA2J,FVB,MOLF,MRL,NOD,NOR,NZB,NZO,NZW,SJL,SPRET,SWR | feature_truncation,3_prime_UTR_variant |
| 6 | 146935539 | 146935742 | 203 | Rep15 | SPRET | feature_truncation,3_prime_UTR_variant |
| 6 | 148891544 | 148891764 | 220 | Dennd5b | CAST,KK,MOLF,PL,SPRET,WSB | feature_truncation,3_prime_UTR_variant |
| 6 | 149226210 | 149226268 | 58 | Resf1 | SPRET | feature_truncation,5_prime_UTR_variant,intron_variant |
| 7 | 3268668 | 3268721 | 53 | Nlrp12 | 129S1,BALB,BTBR,C3H,C57L,CBA,CE,DBA1J,DBA2J,ILNJ2,MRL,NZB,NZO,PJ,SEA,SMJ,WSB | feature_truncation,3_prime_UTR_variant |
| 7 | 4153034 | 4153085 | 51 | Leng9 | SPRET | feature_truncation,5_prime_UTR_variant |
| 7 | 4369438 | 4369882 | 444 | Gp6 | SPRET | feature_truncation,3_prime_UTR_variant |
| 7 | 4551890 | 4552032 | 142 | Ptprh | CE,MOLF,SPRET,SWR,TH | feature_truncation,3_prime_UTR_variant,intron_variant,feature_truncation,3_prime_UTR_variant |
| 7 | 4898526 | 4898580 | 54 | Isoc2a | SPRET | feature_truncation,3_prime_UTR_variant |
| 7 | 4969974 | 4970029 | 55 | Sbk3 | SPRET | feature_truncation,3_prime_UTR_variant |
| 7 | 5612759 | 5612840 | 81 | Vmn1r61 | BUB,CE,NZO,SWR,TH,WSB | feature_truncation,3_prime_UTR_variant |
| 7 | 6358330 | 6358417 | 87 | Zfp582 | CAST,MOLF | feature_truncation,3_prime_UTR_variant |
| 7 | 6469397 | 6469566 | 169 | Or6z3 | CAST,MOLF | feature_truncation,3_prime_UTR_variant |
| 7 | 6477604 | 6477733 | 129 | Or6z5 | MRL,SMJ | feature_truncation,inframe_deletion |
| 7 | 6676604 | 6676678 | 74 | Zim1 | CAST | feature_truncation,3_prime_UTR_variant |
| 7 | 10366632 | 10366684 | 52 | Vmn1r70 | SPRET | feature_truncation,5_prime_UTR_variant |
| 7 | 10486171 | 10486389 | 218 | Vmn1r71 | SPRET | feature_truncation,5_prime_UTR_variant,intron_variant |
| 7 | 11583133 | 11583209 | 76 | Vmn1r74 | SPRET | feature_truncation,3_prime_UTR_variant |
| 7 | 11586237 | 11586326 | 89 | Vmn1r74 | AKR,CAST,FVB,MRL,NOD,NOR,RH2 | feature_truncation,3_prime_UTR_variant |
| 7 | 11615774 | 11615960 | 186 | Vmn1r75 | CAST | feature_truncation,3_prime_UTR_variant |
| 7 | 11889405 | 11889507 | 102 | Vmn1r78 | AKR,FVB,MRL,NOD,NOR,RH2,SPRET | feature_truncation,3_prime_UTR_variant |
| 7 | 12053095 | 12053926 | 831 | Vmn1r83 | CAST,SPRET | feature_truncation,3_prime_UTR_variant |
| 7 | 12449323 | 12449640 | 317 | Vmn2r56 | SPRET | frameshift_variant,feature_truncation |
| 7 | 12508676 | 12508739 | 63 | Zscan18 | SPRET | feature_truncation,inframe_deletion |
| 7 | 12739245 | 12739461 | 216 | Zbtb45 | C58J,SPRET | feature_truncation,3_prime_UTR_variant |
| 7 | 12822162 | 12822231 | 69 | Vmn1r85 | SPRET | feature_truncation,5_prime_UTR_variant |
| 7 | 14294211 | 14294398 | 187 | Vmn1r90 | CAST | feature_truncation,3_prime_UTR_variant |
| 7 | 15705070 | 15705128 | 58 | Bicra | BTBR,BUB,C58J,MOLF,NOD,NOR | feature_truncation,3_prime_UTR_variant |
| 7 | 16718848 | 16719052 | 204 | Pnma8c | 129S1,AJ,AKR,BALB,BTBR,BUB,C3H,C57L,C58J,CAST,CBA,CE,ILNJ2,LP,MOLF,MRL,NOD,NOR,NZB,NZO,NZW,PL,RF,RH2,SJL,SPRET,SWR,TH | feature_truncation,3_prime_UTR_variant |
| 7 | 16949385 | 16949732 | 347 | Psg29 | SPRET | feature_truncation,3_prime_UTR_variant |
| 7 | 18340201 | 18340688 | 487 | Psg23 | 129S1,BTBR,C3H,C57L,CAST,CBA,CE,DBA1J,DBA2J,FVB,MA/My,MOLF,NZB,NZW,PJ,PL,RF,SEA,SMJ,SPRET,WSB | feature_truncation,3_prime_UTR_variant |
| 7 | 19464567 | 19464823 | 256 | Nectin2 | SPRET | splice_acceptor_variant,coding_sequence_variant,intron_variant |
| 7 | 23246202 | 23246286 | 84 | Vmn1r168 | AJ,BALB,CAST,SPRET | feature_truncation,3_prime_UTR_variant |
| 7 | 23674231 | 23674350 | 119 | Vmn1r181 | 129S1,AJ,BALB,BTBR,BUB,C3H,C57L,CAST,CBA,CE,MOLF,NOD,NOR,NZB,NZW,SJL,SPRET,WSB | feature_truncation,5_prime_UTR_variant,intron_variant |
| 7 | 23687438 | 23688061 | 623 | Vmn1r181 | SPRET | feature_truncation,3_prime_UTR_variant |
| 7 | 23824648 | 23825176 | 528 | Zfp112 | LP,MA/My,PJ,SEA,SMJ | feature_truncation,inframe_deletion |
| 7 | 24236240 | 24236293 | 53 | Irgq | CAST,MOLF,SPRET | feature_truncation,3_prime_UTR_variant |
| 7 | 24561614 | 24562121 | 507 | Tmsb10b | CAST,SPRET | stop_lost,feature_truncation,coding_sequence_variant,5_prime_UTR_variant,3_prime_UTR_variant |
| 7 | 25095581 | 25095639 | 58 | Lipe | MOLF | feature_truncation,5_prime_UTR_variant |

| | | | | | | |
|---|---|---|---|---|---|---|
| 7 | 25219000 | 25219599 | 599 | Ceacam2 | SPRET | feature_truncation,coding_sequence_variant,intron_variant |
| 7 | 27355493 | 27355583 | 90 | Ttc9b | BUB,NZB,SJL | feature_truncation,3_prime_UTR_variant |
| 7 | 27405244 | 27405549 | 305 | Zfp607b | SPRET | feature_truncation,3_prime_UTR_variant |
| 7 | 27449594 | 27449676 | 82 | Zfp60 | CAST,MOLF | feature_truncation,3_prime_UTR_variant |
| 7 | 27451504 | 27451703 | 199 | Zfp60 | SPRET | feature_truncation,3_prime_UTR_variant |
| 7 | 27521269 | 27521491 | 222 | Zfp626 | CAST,SPRET | feature_truncation,3_prime_UTR_variant |
| 7 | 27606070 | 27606941 | 871 | Zfp974 | MOLF | feature_truncation,3_prime_UTR_variant |
| 7 | 27659732 | 27660068 | 336 | Zfp780b | SPRET | stop_lost,feature_truncation,coding_sequence_variant,3_prime_UTR_variant,intron_variant |
| 7 | 30578325 | 30578393 | 68 | Cd22 | CAST,DBA1J,DBA2J,MOLF,NZB,NZO,NZW,SMJ,SPRET | feature_truncation,5_prime_UTR_variant |
| 7 | 35279910 | 35279966 | 56 | Rgs9bp | AJ,BALB,CE,ILNJ2,SEA,SPRET | feature_truncation,3_prime_UTR_variant |
| 7 | 35416095 | 35416189 | 94 | Dpy19l3 | BTBR,C57L,MOLF,NZB,NZO,NZW,SPRET | feature_truncation,coding_sequence_variant,intron_variant |
| 7 | 39187834 | 39188222 | 388 | Zfp619 | 129S1,BTBR,BUB,C3H,C57L,CBA,KK,MRL,NZW,PL,SJL,SMJ,WSB | feature_truncation,3_prime_UTR_variant |
| 7 | 40693731 | 40693818 | 87 | Gm4884 | DBA1J,DBA2J,ILNJ2,NZB,NZO,NZW,PJ,SMJ | feature_truncation,inframe_deletion |
| 7 | 41195866 | 41195950 | 84 | Gm6871 | MOLF | feature_truncation,inframe_deletion |
| 7 | 41300718 | 41301089 | 371 | Zfp788 | DBA1J,DBA2J,ILNJ2,NZW,PJ | feature_truncation,3_prime_UTR_variant |
| 7 | 42356424 | 42356675 | 251 | Gm17067 | SPRET | feature_truncation,3_prime_UTR_variant |
| 7 | 42466167 | 42466687 | 520 | Gm2381 | SMJ | feature_truncation,3_prime_UTR_variant |
| 7 | 42552292 | 42552488 | 196 | Vmn2r63 | CAST,DBA1J,DBA2J,ILNJ2,MOLF,NZB,NZO,NZW,PJ,SJL,SMJ,SPRET | feature_truncation,3_prime_UTR_variant |
| 7 | 42672702 | 42672912 | 210 | Vmn2r-ps158 | NZB,SJL,SPRET | feature_truncation,coding_sequence_variant,intron_variant |
| 7 | 42838929 | 42838983 | 54 | Zfp936 | CAST,DBA1J,DBA2J,ILNJ2,MOLF,NZW,PJ,SPRET | feature_truncation,inframe_deletion |
| 7 | 43120553 | 43120625 | 72 | Vsig10l | MOLF | feature_truncation,3_prime_UTR_variant |
| 7 | 43326426 | 43326540 | 114 | Ctu1 | C57L,MOLF,SMJ | feature_truncation,inframe_deletion |
| 7 | 44008185 | 44008270 | 85 | Shank1 | CAST,SPRET | feature_truncation,3_prime_UTR_variant |
| 7 | 44917465 | 44918139 | 674 | Slc6a16 | AKR | feature_truncation,coding_sequence_variant,intron_variant |
| 7 | 45122617 | 45122827 | 210 | Dhdh | CAST,CBA,ILNJ2,MOLF,NZW,PJ,SJL,SPRET | feature_truncation,3_prime_UTR_variant |
| 7 | 45944696 | 45945037 | 341 | Otog | ILNJ2,MOLF,NZW,PJ,SJL | feature_truncation,5_prime_UTR_variant,intron_variant |
| 7 | 46363541 | 46363603 | 62 | Saa3 | SPRET | feature_truncation,3_prime_UTR_variant |
| 7 | 46377942 | 46377998 | 56 | Saa4 | CE,WSB | feature_truncation,3_prime_UTR_variant |
| 7 | 46573083 | 46573231 | 148 | Uevld | ILNJ2,MOLF,NZW,PJ,SJL,SPRET | feature_truncation,3_prime_UTR_variant |
| 7 | 47631732 | 47631808 | 76 | Mrgpra4 | CE | feature_truncation,5_prime_UTR_variant |
| 7 | 48095305 | 48095372 | 67 | Mrgprb1 | ILNJ2,MOLF,NZW,PJ,SJL | feature_truncation,3_prime_UTR_variant |
| 7 | 48131262 | 48131389 | 127 | Mrgprx2 | SPRET | feature_truncation,3_prime_UTR_variant |
| 7 | 66058248 | 66058447 | 199 | Aldh1a3 | BALB,BUB,C57L,CAST,CE,DBA1J,DBA2J,ILNJ2,KK,LP,MOLF,NZB,NZO,PJ,SEA,SMJ,SPRET,TH | feature_truncation,3_prime_UTR_variant |
| 7 | 77828566 | 77828687 | 121 | Ntrk3 | 129S1,AJ,AKR,BALB,BTBR,BUB,C3H,C57L,C58J,CAST,CBA,CE,DBA1J,DBA2J,FVB,KK,MA/My,MOLF,MRL,NOD,NOR,NZB,NZO,NZW,PJ,PL,RF,RH2,SEA,SJL,SMJ,SPRET,SWR,TH | feature_truncation,3_prime_UTR_variant |
| 7 | 77832387 | 77832673 | 286 | Ntrk3 | SPRET | feature_truncation,3_prime_UTR_variant |
| 7 | 79059320 | 79059563 | 243 | Fanci | 129S1,C3H,CAST,CBA,DBA1J,DBA2J,KK,MOLF,PJ,SEA,SPRET | feature_truncation,3_prime_UTR_variant |
| 7 | 79527735 | 79528457 | 722 | Ap3s2 | CAST | feature_truncation,3_prime_UTR_variant |
| 7 | 79876202 | 79876312 | 110 | Sema4b | MOLF,SPRET | feature_truncation,3_prime_UTR_variant |
| 7 | 79889674 | 79889790 | 116 | Gdpgp1 | SPRET | feature_truncation,3_prime_UTR_variant |
| 7 | 79941060 | 79941132 | 72 | Vps33b | SPRET | feature_truncation,3_prime_UTR_variant |
| 7 | 79966283 | 79966440 | 157 | Rccd1 | 129S1,AJ,AKR,BALB,BTBR,BUB,C3H,C57L,CAST,CBA,CE,DBA1J,DBA2J,FVB,ILNJ2,KK,MA/My,MOLF,MRL,NOD,NOR,NZB,NZO,NZW,PJ,PL,RF,RH2,SEA,SJL,SMJ,SPRET,SWR,TH,WSB | feature_truncation,3_prime_UTR_variant |
| 7 | 80967185 | 80967296 | 111 | Pde8a | CAST,MOLF | feature_truncation,coding_sequence_variant,intron_variant |
| 7 | 81253273 | 81253324 | 51 | Homer2 | CAST,MOLF | feature_truncation,3_prime_UTR_variant |
| 7 | 81542164 | 81542648 | 484 | Hdgfl3 | 129S1,AJ,AKR,BALB,BTBR,BUB,C3H,C57L,CBA,DBA1J,DBA2J,FVB,KK,NOD,NOR,NZW,PJ,PL,RF,RH2,SEA,SJL,SWR | feature_truncation,3_prime_UTR_variant |
| 7 | 81616816 | 81616963 | 147 | Bnc1 | SPRET | feature_truncation,3_prime_UTR_variant |
| 7 | 82298224 | 82298438 | 214 | Efl1 | CAST | feature_truncation,5_prime_UTR_variant |
| 7 | 83547623 | 83547749 | 126 | Mesd | SPRET | feature_truncation,3_prime_UTR_variant |
| 7 | 83583359 | 83583425 | 66 | Cemip | SPRET | feature_truncation,3_prime_UTR_variant |
| 7 | 84228764 | 84228887 | 123 | Ctxnd1 | SPRET | feature_truncation,3_prime_UTR_variant |
| 7 | 84231175 | 84231469 | 294 | Ctxnd1 | CAST,MOLF,SPRET | feature_truncation,3_prime_UTR_variant |
| 7 | 85064600 | 85064705 | 105 | Vmn2r69 | CAST | feature_truncation,coding_sequence_variant,intron_variant |
| 7 | 85272753 | 85272855 | 102 | Vmn2r71 | WSB | feature_truncation,coding_sequence_variant,intron_variant |
| 7 | 85852601 | 85852657 | 56 | Vmn2r76 | 129S1,BTBR,C3H,C57L,CBA,DBA1J,DBA2J,ILNJ2,KK,NZB,NZO,NZW,PJ,SEA,SMJ | feature_truncation,3_prime_UTR_variant |
| 7 | 85968282 | 85968448 | 166 | Or6f1 | SPRET | feature_truncation,3_prime_UTR_variant |
| 7 | 86061957 | 86062047 | 90 | Or14c44 | 129S1,BTBR,C3H,C57L,CBA,DBA1J,DBA2J,ILNJ2,KK,NZB,NZO,NZW,PJ,SEA | feature_truncation,inframe_deletion |
| 7 | 86314268 | 86314748 | 480 | Or14c40 | CAST,MOLF | feature_truncation,3_prime_UTR_variant |
| 7 | 86345930 | 86346165 | 235 | Or14c39 | SPRET | feature_truncation,3_prime_UTR_variant |
| 7 | 86654846 | 86654944 | 98 | Vmn2r79 | SPRET | feature_truncation,3_prime_UTR_variant |
| 7 | 87075781 | 87075986 | 205 | Tyr | SPRET | feature_truncation,3_prime_UTR_variant |
| 7 | 89613765 | 89613945 | 180 | Eed | SPRET | splice_donor_variant,splice_donor_5th_base_variant,coding_sequence_variant,intron_variant |
| 7 | 96956208 | 96956287 | 79 | Gab2 | CAST | feature_truncation,3_prime_UTR_variant |
| 7 | 97108055 | 97108627 | 572 | Kctd14 | C3H,CAST,CBA,DBA1J,DBA2J,ILNJ2,MOLF,NZO,PJ,SEA,SPRET | feature_truncation,3_prime_UTR_variant |
| 7 | 99809663 | 99809726 | 63 | Lipt2 | 129S1,AJ,AKR,BALB,BTBR,BUB,C3H,C57L,CAST,CBA,CE,DBA1J,DBA2J,FVB,ILNJ2,KK,MA/My,MOLF,MRL,NZB,NZW,PJ,PL,RF,RH2,SEA,SJL,SPRET,TH | feature_truncation,3_prime_UTR_variant |

| | | | | | | |
|---|---|---|---|---|---|---|
| 7 | 100196724 | 100196876 | 152 | Mrpl48 | SPRET | feature_truncation,5_prime_UTR_variant,feature_truncation,3_prime_UTR_variant |
| 7 | 101443712 | 101444109 | 397 | Clpb | CAST | feature_truncation,3_prime_UTR_variant |
| 7 | 101794147 | 101794351 | 204 | Nup98 | SMJ,SPRET | feature_truncation,3_prime_UTR_variant |
| 7 | 102140779 | 102140833 | 54 | Or55b10 | KK,MA/My,NOD,SJL,SMJ,SWR | feature_truncation,3_prime_UTR_variant |
| 7 | 102362252 | 102362309 | 57 | Or51a39 | CAST,KK,MOLF,NOD,SJL,SPRET,SWR | feature_truncation,3_prime_UTR_variant |
| 7 | 102452456 | 102453159 | 703 | Or51f23 | KK,NOD,SJL,SWR | feature_truncation,coding_sequence_variant,5_prime_UTR_variant |
| 7 | 102692549 | 102692607 | 58 | Or52r1b | CAST,MOLF,TH,WSB | feature_truncation,3_prime_UTR_variant |
| 7 | 102767700 | 102768254 | 554 | Or51a5 | KK,NOD,SJL,SPRET,SWR | feature_truncation,3_prime_UTR_variant |
| 7 | 102768204 | 102769043 | 839 | Or51a5 | MOLF | feature_truncation,3_prime_UTR_variant |
| 7 | 102769776 | 102770285 | 509 | Or51a5 | CAST,MOLF | feature_truncation,3_prime_UTR_variant |
| 7 | 102970764 | 102970871 | 107 | Or52ab2 | KK,NOD,SJL,SPRET,SWR | feature_truncation,3_prime_UTR_variant |
| 7 | 102978367 | 102978439 | 72 | Or52ab7 | MOLF | feature_truncation,inframe_deletion |
| 7 | 102993876 | 102993936 | 60 | Or52ad1 | KK,NOD,SJL,SPRET,SWR | feature_truncation,3_prime_UTR_variant |
| 7 | 103107759 | 103108411 | 652 | Or52d13 | KK,NOD,SJL,SPRET,SWR | feature_truncation,3_prime_UTR_variant |
| 7 | 103247632 | 103248575 | 943 | Or52z13 | SMJ | feature_truncation,3_prime_UTR_variant |
| 7 | 103382175 | 103382235 | 60 | Or52a24 | 129S1,AJ,AKR,BALB,BTBR,BUB,C3H,C57L,CAST,CBA,DBA1J,DBA2J,FVB,ILNJ2,MOLF,MRL,NZB,NZW,PJ,PL,RF,RH2,SEA,TH,WSB | feature_truncation,3_prime_UTR_variant |
| 7 | 103529539 | 103529609 | 70 | Or51b4 | CAST,TH,WSB | feature_truncation,3_prime_UTR_variant |
| 7 | 103539225 | 103539302 | 77 | Or51b17 | 129S1,AJ,AKR,BALB,BTBR,BUB,C3H,C57L,CBA,DBA1J,DBA2J,FVB,ILNJ2,MRL,NZB,NZW,PJ,PL,RF,RH2,SEA,SPRET,TH,WSB | feature_truncation,3_prime_UTR_variant |
| 7 | 103655517 | 103656193 | 676 | Or51k1 | CAST,KK,MOLF,SJL,SMJ,SPRET,SWR | feature_truncation,3_prime_UTR_variant |
| 7 | 103656461 | 103656525 | 64 | Or51k1 | SPRET | feature_truncation,3_prime_UTR_variant |
| 7 | 103704833 | 103704884 | 51 | Or51a42 | MOLF | feature_truncation,3_prime_UTR_variant |
| 7 | 103758007 | 103758548 | 541 | Or52d1 | CAST,KK,MOLF,NOD,SJL,SMJ,SWR,TH | feature_truncation,3_prime_UTR_variant |
| 7 | 103826704 | 103827642 | 938 | Or52h1 | CAST,KK,MOLF,NOD,SJL,SMJ,SWR,TH | feature_truncation,3_prime_UTR_variant |
| 7 | 103832235 | 103832912 | 677 | Or52h1 | KK | feature_truncation,5_prime_UTR_variant,intron_variant |
| 7 | 103882658 | 103882748 | 90 | Trim6 | CE,MOLF,NOD,NZO,SJL,SMJ,SPRET,SWR,WSB | feature_truncation,3_prime_UTR_variant |
| 7 | 103988993 | 103989055 | 62 | Trim12c | 129S1,AJ,BALB,BTBR,BUB,C3H,C57L,CBA,DBA1J,DBA2J,FVB,ILNJ2,MOLF,NOD,NZB,NZW,PJ,PL,RH2,SEA,SJL,SMJ,SWR | feature_truncation,3_prime_UTR_variant |
| 7 | 104005701 | 104005882 | 181 | Trim30b | 129S1,AJ,AKR,BALB,BTBR,BUB,C3H,C57L,CBA,CE,DBA1J,DBA2J,FVB,ILNJ2,MRL,NZB,NZO,NZW,PJ,PL,RF,RH2,SEA | feature_truncation,3_prime_UTR_variant |
| 7 | 104232791 | 104232974 | 183 | Or52d3 | MOLF,NOD,SJL,SWR | feature_truncation,3_prime_UTR_variant |
| 7 | 104736640 | 104736890 | 250 | Or56a3 | CAST,NOD,NZW,SWR | feature_truncation,3_prime_UTR_variant |
| 7 | 104737817 | 104737871 | 54 | Or56a3 | WSB | feature_truncation,3_prime_UTR_variant |
| 7 | 104773094 | 104773955 | 861 | Or56a3b | MOLF | feature_truncation,3_prime_UTR_variant |
| 7 | 104965401 | 104965827 | 426 | Or56b35 | SPRET | feature_truncation,3_prime_UTR_variant |
| 7 | 105023593 | 105023647 | 54 | Fhip1b | SPRET | feature_truncation,3_prime_UTR_variant |
| 7 | 105291780 | 105291947 | 167 | Timm10b | SPRET | feature_truncation,3_prime_UTR_variant |
| 7 | 106500427 | 106500520 | 93 | Or2d3 | CAST,KK,NOD,NZO,SMJ,SWR,WSB | feature_truncation,5_prime_UTR_variant |
| 7 | 106599543 | 106599666 | 123 | Or6a2 | B10D2,B10J,BUB,C58J,CAST,CE,FVB,KK,LP,MA/My,MOLF,NOD,NOR,SJL,SMJ,SPRET,SWR,TH,WSB | feature_truncation,3_prime_UTR_variant |
| 7 | 106749425 | 106749845 | 420 | Or2d36 | 129S1,AJ,AKR,BALB,BTBR,C3H,C57L,CAST,CBA,CE,DBA1J,DBA2J,ILNJ2,KK,MOLF,MRL,NOD,NZB,NZO,NZW,PJ,PL,RF,RH2,SEA,SMJ,SPRET,SWR,WSB | feature_truncation,3_prime_UTR_variant |
| 7 | 107415514 | 107415572 | 58 | Or5p5 | SPRET | feature_truncation,3_prime_UTR_variant |
| 7 | 107629728 | 107629779 | 51 | Or5p6 | CAST,NOD,SMJ,SWR | feature_truncation,3_prime_UTR_variant |
| 7 | 107655793 | 107655863 | 70 | Or10ab4 | CAST,NOD,NZO,SMJ,SPRET,SWR | feature_truncation,3_prime_UTR_variant |
| 7 | 107681814 | 107681896 | 82 | Or5p4 | CAST,NOD,NZO,SMJ,SWR | feature_truncation,3_prime_UTR_variant |
| 7 | 108144669 | 108145160 | 491 | Or52n4b | CAST,NOD,SMJ,SWR | stop_lost,feature_truncation,coding_sequence_variant,3_prime_UTR_variant |
| 7 | 108214189 | 108214940 | 751 | Or5p78 | CAST,MOLF,NOD,SMJ,SPRET,SWR,WSB | feature_truncation,3_prime_UTR_variant |
| 7 | 108269414 | 108269548 | 134 | Or5p81 | CAST,NOD,SMJ,SWR | feature_truncation,3_prime_UTR_variant |
| 7 | 108522397 | 108522519 | 122 | Nlrp10 | SPRET | feature_truncation,3_prime_UTR_variant |
| 7 | 110650116 | 110650214 | 98 | Ctr9 | SPRET | feature_truncation,coding_sequence_variant,intron_variant |
| 7 | 112180098 | 112180184 | 86 | Parva | CAST,ILNJ2,MOLF,NZO,SMJ | feature_truncation,coding_sequence_variant,intron_variant |
| 7 | 118221483 | 118221992 | 509 | Tmc5 | 129S1,C3H,C57L,CAST,CBA,CE,DBA1J,DBA2J,MOLF,NOR,SPRET | feature_truncation,5_prime_UTR_variant,intron_variant |
| 7 | 119198339 | 119198562 | 223 | Acsm2 | MOLF | feature_truncation,3_prime_UTR_variant |
| 7 | 119314574 | 119314923 | 349 | Thumpd1 | AJ,BALB,C3H,CBA,DBA1J,DBA2J,ILNJ2,KK,NOR,NZB,NZO,PL,RH2 | feature_truncation,3_prime_UTR_variant |
| 7 | 119552275 | 119552375 | 100 | Lyrm1 | SPRET | feature_truncation,3_prime_UTR_variant |
| 7 | 119553746 | 119554157 | 411 | Lyrm1 | SPRET | feature_truncation,3_prime_UTR_variant |
| 7 | 120545660 | 120546033 | 373 | Polr3e | 129S1,AJ,AKR,B10D2,B10J,BTBR,BUB,C3H,C58J,CAST,CBA,CE,DBA1J,DBA2J,ILNJ2,LP,MA/My,MOLF,MRL,NOD,NZB,NZO,NZW,PJ,RF,RH2,SEA,SJL,SMJ,SPRET,SWR,TH,WSB | feature_truncation,3_prime_UTR_variant |
| 7 | 120545853 | 120546033 | 180 | Polr3e | AJ,BUB,FVB,NOR,PL | feature_truncation,3_prime_UTR_variant |
| 7 | 126289624 | 126289835 | 211 | Slx1b | 129S1,AJ,AKR,BALB,BTBR,BUB,C57L,C58J,CAST,FVB,ILNJ2,KK,LP,MA/My,MOLF,MRL,NOD,NZB,NZO,NZW,PJ,PL,RF,RH2,SEA,SJL,SMJ,SPRET,SWR,TH,WSB | feature_truncation,3_prime_UTR_variant |
| 7 | 126462973 | 126463027 | 54 | Hirip3 | SPRET | feature_truncation,inframe_deletion |
| 7 | 126984371 | 126984538 | 167 | 9130019O22Rik | CAST | frameshift_variant,feature_truncation,feature_truncation,3_prime_UTR_variant |
| 7 | 127402519 | 127402570 | 51 | Hsd3b7 | SPRET | feature_truncation,3_prime_UTR_variant |
| 7 | 127477030 | 127477087 | 57 | Zfp646 | MOLF,TH | feature_truncation,5_prime_UTR_variant |
| 7 | 127716685 | 127716737 | 52 | Itgam | SPRET | feature_truncation,3_prime_UTR_variant |
| 7 | 128385885 | 128386096 | 211 | Sec23ip | 129S1,BTBR,C57L,CAST,ILNJ2,LP,MOLF,PJ,SPRET | feature_truncation,3_prime_UTR_variant |
| 7 | 130943663 | 130943729 | 66 | 2310057M21Rik | CAST | feature_truncation,3_prime_UTR_variant |
| 7 | 131592138 | 131592349 | 211 | Gpr26 | CAST,SPRET | feature_truncation,3_prime_UTR_variant |
| 7 | 132198218 | 132198292 | 74 | Nkx1-2 | CAST,MOLF | feature_truncation,3_prime_UTR_variant |
| 7 | 132315080 | 132315140 | 60 | Fam53b | SPRET | feature_truncation,3_prime_UTR_variant |

| | | | | | | |
|---|---|---|---|---|---|---|
| 7 | 134860183 | 134860515 | 332 | Nps | SPRET | feature_truncation,5_prime_UTR_variant |
| 7 | 135286048 | 135286123 | 75 | Ptpre | MOLF | feature_truncation,3_prime_UTR_variant |
| 7 | 139218718 | 139218781 | 63 | Cfap46 | SPRET | feature_truncation,inframe_deletion |
| 7 | 139917745 | 139917827 | 82 | Or12j2 | CAST,SPRET | feature_truncation,3_prime_UTR_variant |
| 7 | 140194381 | 140194595 | 214 | Or13a18 | WSB | feature_truncation,3_prime_UTR_variant |
| 7 | 140252169 | 140252918 | 749 | Or13a25 | SPRET | feature_truncation,3_prime_UTR_variant |
| 7 | 140275206 | 140275418 | 212 | Or13a17 | CAST,MOLF,SPRET,WSB | feature_truncation,3_prime_UTR_variant |
| 7 | 140443989 | 140444042 | 53 | Sirt3 | CAST,CE | feature_truncation,coding_sequence_variant,intron_variant |
| 7 | 140681271 | 140681331 | 60 | Ano9 | SPRET | feature_truncation,3_prime_UTR_variant |
| 7 | 141031310 | 141031390 | 80 | Rplp2 | ILNJ2 | feature_truncation,3_prime_UTR_variant |
| 7 | 141051747 | 141051942 | 195 | Polr2l | CAST,SPRET | feature_truncation,3_prime_UTR_variant |
| 7 | 141218119 | 141218638 | 519 | Muc6 | CAST | feature_truncation,inframe_deletion |
| 7 | 141218343 | 141219161 | 818 | Muc6 | WSB | frameshift_variant,feature_truncation |
| 7 | 141219584 | 141219790 | 206 | Muc6 | NZW | frameshift_variant,feature_truncation |
| 7 | 141219965 | 141220483 | 518 | Muc6 | MOLF,NZW | frameshift_variant,feature_truncation |
| 7 | 141222246 | 141222454 | 208 | Muc6 | CAST,ILNJ2 | frameshift_variant |
| 7 | 141223274 | 141223480 | 206 | Muc6 | MOLF,WSB | frameshift_variant,feature_truncation |
| 7 | 141292749 | 141292882 | 133 | Muc2 | ILNJ2,NZW | frameshift_variant,feature_truncation |
| 7 | 141295011 | 141295140 | 129 | Muc2 | NZW | feature_truncation,inframe_deletion |
| 7 | 141296256 | 141296389 | 133 | Muc2 | CAST,MOLF,WSB | frameshift_variant,feature_truncation |
| 7 | 141297283 | 141297427 | 144 | Muc2 | CAST,MOLF,WSB | feature_truncation,inframe_deletion |
| 7 | 141363194 | 141363482 | 288 | Muc5ac | NZO,PJ | feature_truncation,inframe_deletion |
| 7 | 141412901 | 141413842 | 941 | Muc5b | NZW | frameshift_variant,feature_truncation |
| 7 | 141413519 | 141413573 | 54 | Muc5b | 129S1,AJ,AKR,B10D2,B10J,BALB,BTBR,BUB,C3H,C57L,C58J,CAST,CBA,CE,DBA1J,DBA2J,FVB,ILNJ2,KK,LP,MA/My,MOLF,MRL,NOD,NOR,NZB,NZO,PJ,PL,RF,RH2,SMJ,SWR,TH,WSB | feature_truncation,inframe_deletion |
| 7 | 141413523 | 141414020 | 497 | Muc5b | NZW,SJL | frameshift_variant,feature_truncation |
| 7 | 141416004 | 141416946 | 942 | Muc5b | SPRET | feature_truncation,inframe_deletion |
| 7 | 141674180 | 141674240 | 60 | Gm45618 | CAST | feature_truncation,inframe_deletion |
| 7 | 141729057 | 141729192 | 135 | Krtap5-2 | BALB,CAST,MOLF,MRL,RF,RH2,SEA,SPRET | feature_truncation,inframe_deletion |
| 7 | 141755588 | 141755741 | 153 | Krtap5-3 | BALB,CAST,MOLF,MRL,RF,RH2,SEA,SPRET | feature_truncation,inframe_deletion |
| 7 | 141782953 | 141783043 | 90 | Krtap5-5 | BALB,MRL,RF,RH2,SEA | feature_truncation,inframe_deletion |
| 7 | 141794525 | 141794585 | 60 | Gm40460 | CAST | feature_truncation,inframe_deletion |
| 7 | 143190465 | 143190516 | 51 | Tnfrsf22 | CAST,MOLF | feature_truncation,3_prime_UTR_variant |
| 7 | 143333528 | 143333600 | 72 | Mrgpre | C58J,CAST,LP,MA/My,MOLF,PJ,SMJ,SPRET | feature_truncation,3_prime_UTR_variant |
| 7 | 144133039 | 144133117 | 78 | Fadd | SPRET | feature_truncation,3_prime_UTR_variant |
| 8 | 3440010 | 3440353 | 343 | Arhgef18 | 129S1,CAST,NZB,SPRET | feature_truncation,3_prime_UTR_variant |
| 8 | 3633737 | 3634248 | 511 | Rps23rg1 | 129S1,CAST,NZB,NZO,NZW,SPRET,WSB | feature_truncation,coding_sequence_variant,intron_variant,feature_truncation,coding_sequence_variant,5_prime_UTR_variant,intron_variant |
| 8 | 3727602 | 3727689 | 87 | Trappc5 | 129S1,NZB | feature_truncation,5_prime_UTR_variant |
| 8 | 5135425 | 5135479 | 54 | Slc10a2 | 129S1,NZB,SPRET | feature_truncation,3_prime_UTR_variant |
| 8 | 10620251 | 10620314 | 63 | Myo16 | SPRET | feature_truncation,coding_sequence_variant,intron_variant,feature_truncation,inframe_deletion |
| 8 | 11663477 | 11663637 | 160 | Ankrd10 | SPRET | feature_truncation,3_prime_UTR_variant |
| 8 | 11875668 | 11875739 | 71 | Arhgef7 | SPRET | feature_truncation,3_prime_UTR_variant |
| 8 | 13068378 | 13068432 | 54 | Mcf2l | CAST,SPRET | feature_truncation,3_prime_UTR_variant |
| 8 | 13106216 | 13106288 | 72 | F10 | SPRET | feature_truncation,3_prime_UTR_variant |
| 8 | 13125567 | 13125732 | 165 | Proz | C57L,CAST,ILNJ2,MOLF,NZB,NZW,PJ,PL,SEA,SMJ,TH | feature_truncation,3_prime_UTR_variant |
| 8 | 13466907 | 13467274 | 367 | Grk1 | 129S1,C57L,CAST,CE,ILNJ2,LP,MOLF,NZB,NZO,NZW,PJ,PL,SEA,SMJ,SPRET,TH,WSB | feature_truncation,3_prime_UTR_variant |
| 8 | 15091975 | 15092056 | 81 | BB014433 | SPRET | feature_truncation,inframe_deletion |
| 8 | 21161992 | 21162217 | 225 | Potefam3b | 129S1,AJ,AKR,BTBR,BUB,C3H,CBA,KK,MA/My,MRL,NOD,NOR,PL,RF,RH2,SMJ,SPRET,SWR,TH,WSB | splice_donor_variant,splice_donor_5th_base_variant,coding_sequence_variant,intron_variant |
| 8 | 25023426 | 25023541 | 115 | Ido2 | SPRET | feature_truncation,3_prime_UTR_variant |
| 8 | 25171509 | 25171569 | 60 | Adam3 | MOLF | feature_truncation,inframe_deletion |
| 8 | 25452880 | 25452934 | 54 | Adam9 | 129S1,AJ,AKR,BALB,BTBR,BUB,C3H,C57L,CBA,DBA1J,DBA2J,FVB,ILNJ2,KK,MA/My,MRL,NOD,NOR,NZB,NZO,NZW,PL,RF,RH2,SJL,SMJ,SWR,TH | feature_truncation,inframe_deletion |
| 8 | 26516856 | 26516910 | 54 | Hook3 | 129S1,AJ,AKR,BALB,BTBR,BUB,C3H,C57L,CBA,DBA1J,DBA2J,FVB,ILNJ2,KK,MA/My,MRL,NOD,NOR,NZB,NZO,NZW,RF,RH2,SJL,SMJ,SPRET,SWR,TH | feature_truncation,3_prime_UTR_variant |
| 8 | 27765814 | 27765877 | 63 | Eif4ebp1 | AKR,C3H,CE,DBA1J,DBA2J,MA/My,PL,RF,WSB | feature_truncation,3_prime_UTR_variant |
| 8 | 34578103 | 34578892 | 789 | Erhrt-ps | 129S1,AJ,AKR,BALB,BUB,C3H,C57L,CAST,CBA,CE,DBA1J,DBA2J,MA/My,MRL,NOD,NOR,PL,RF,RH2,SJL,SMJ,SPRET,SWR,TH,WSB | transcript_ablation |
| 8 | 40999686 | 40999775 | 89 | Vps37a | SPRET | feature_truncation,3_prime_UTR_variant |
| 8 | 46230856 | 46230981 | 125 | Sorbs2 | CAST,MOLF,SPRET | feature_truncation,3_prime_UTR_variant |
| 8 | 47964092 | 47964152 | 60 | Trappc11 | C58J,CAST,LP,MOLF,SPRET | feature_truncation,5_prime_UTR_variant |
| 8 | 60996624 | 60996763 | 139 | Aadat | AJ,AKR,BALB,BTBR,BUB,C3H,C57L,CBA,CE,DBA1J,DBA2J,FVB,ILNJ2,MA/My,NOD,NOR,NZB,NZO,NZW,PJ,PL,RF,RH2,SEA,SMJ,SPRET | feature_truncation,coding_sequence_variant,intron_variant |
| 8 | 62510285 | 62510352 | 67 | Anxa10 | SPRET | feature_truncation,3_prime_UTR_variant |
| 8 | 67313463 | 67313592 | 129 | Naf1 | CAST,MOLF,SPRET | feature_truncation,inframe_deletion |
| 8 | 69280808 | 69281010 | 202 | Ints10 | CAST | feature_truncation,3_prime_UTR_variant |
| 8 | 69358096 | 69358147 | 51 | Lpl | 129S1,AJ,AKR,BALB,BTBR,BUB,C3H,C57L,C58J,CBA,CE,DBA1J,DBA2J,FVB,ILNJ2,KK,MA/My,MRL,NOD,NOR,NZB,NZO,NZW,PJ,PL,RF,RH2,SEA,SJL,SMJ,SWR,TH,WSB | feature_truncation,3_prime_UTR_variant |
| 8 | 70072014 | 70072074 | 60 | Zfp868 | SPRET | feature_truncation,5_prime_UTR_variant |
| 8 | 70637884 | 70637942 | 58 | Slc25a42 | 129S1,AKR,BALB,BUB,C3H,C57L,C58J,CAST,CBA,CE,DBA1J,DBA2J,FVB,ILNJ2,KK,MA/My,MRL,NZB,NZO,NZW,PJ,PL,RF,SEA,SJL,SMJ,SPRET,SWR,TH,WSB | feature_truncation,3_prime_UTR_variant |
| 8 | 70963349 | 70963653 | 304 | Uba52 | SPRET | feature_truncation,5_prime_UTR_variant |

| | | | | | | |
|---|---|---|---|---|---|---|
| 8 | 71827642 | 71827987 | 345 | Ocel1 | MOLF | feature_truncation,3_prime_UTR_variant |
| 8 | 72687694 | 72687862 | 168 | Zfp617 | SPRET | feature_truncation,3_prime_UTR_variant |
| 8 | 72855851 | 72856077 | 226 | Or2z9 | 129S1,AKR,BALB,BUB,C3H,C57L,C58J,CAST,CBA,CE,DBA1J,DBA2J,FVB,ILNJ2,KK,MA/My,MRL,NZB,NZO,NZW,PJ,PL,RF,SEA,SJL,SMJ,SPRET,SWR,TH,WSB | feature_truncation,3_prime_UTR_variant |
| 8 | 72865267 | 72865879 | 612 | Or1ab2 | 129S1,AKR,BALB,BUB,C3H,C57L,C58J,CAST,CBA,CE,DBA1J,DBA2J,FVB,ILNJ2,KK,MA/My,MRL,NZB,NZO,NZW,PJ,PL,RF,SEA,SJL,SMJ,SPRET,SWR,TH,WSB | feature_truncation,3_prime_UTR_variant |
| 8 | 72867488 | 72867558 | 70 | Or1ab2 | 129S1,AKR,BALB,BUB,C3H,C57L,C58J,CAST,CBA,CE,DBA1J,DBA2J,FVB,ILNJ2,KK,MA/My,MRL,NZB,NZO,NZW,PJ,PL,RF,SEA,SJL,SMJ,SPRET,SWR,TH,WSB | feature_truncation,3_prime_UTR_variant |
| 8 | 75949363 | 75949422 | 59 | Rasd2 | CAST,MOLF | feature_truncation,3_prime_UTR_variant |
| 8 | 84957708 | 84957787 | 79 | Zswim4 | SPRET | feature_truncation,coding_sequence_variant,intron_variant |
| 8 | 94685579 | 94685707 | 128 | Gnao1 | 129S1,AKR,BUB,C57L,CAST,MA/My,PL,SJL,SPRET,WSB | feature_truncation,3_prime_UTR_variant |
| 8 | 95247594 | 95247981 | 387 | Nlrc5 | SPRET | feature_truncation,coding_sequence_variant,intron_variant |
| 8 | 95351026 | 95351175 | 149 | Rspry1 | SPRET | feature_truncation,3_prime_UTR_variant |
| 8 | 96551968 | 96552785 | 817 | Sap18b | CAST,MOLF,SPRET,SWR,WSB | transcript_ablation |
| 8 | 96572897 | 96573345 | 448 | Slc38a7 | CAST | feature_truncation,coding_sequence_variant,intron_variant |
| 8 | 104985050 | 104985408 | 358 | Cklf | SPRET | feature_truncation,3_prime_UTR_variant |
| 8 | 105076806 | 105076865 | 59 | Cmtm4 | CAST,MOLF,PJ,SPRET | feature_truncation,3_prime_UTR_variant |
| 8 | 106664850 | 106664930 | 80 | Psmb10 | SPRET | feature_truncation,5_prime_UTR_variant |
| 8 | 107741304 | 107741447 | 143 | Sntb2 | SPRET | feature_truncation,3_prime_UTR_variant |
| 8 | 110349864 | 110349942 | 78 | Pkd1l3 | CAST,MOLF | feature_truncation,inframe_deletion |
| 8 | 123209487 | 123209554 | 67 | Ctu2 | ILNJ2,MOLF | feature_truncation,3_prime_UTR_variant |
| 8 | 123354870 | 123354931 | 61 | Cbfa2t3 | AJ,AKR,BALB,C3H,C58J,CE,ILNJ2,LP,MOLF,MRL,PL,RF,RH2,SEA,SMJ | feature_truncation,3_prime_UTR_variant |
| 8 | 125262071 | 125262141 | 70 | Cog2 | SPRET | feature_truncation,coding_sequence_variant,intron_variant |
| 8 | 125565614 | 125565705 | 91 | 2810004N23Rik | AJ,BALB,CAST,ILNJ2,MOLF,RH2,SEA | feature_truncation,3_prime_UTR_variant |
| 8 | 125756560 | 125756672 | 112 | Tsnax | AJ,BALB,CAST,ILNJ2,MOLF,RH2,SEA | feature_truncation,3_prime_UTR_variant |
| 8 | 127659877 | 127659972 | 95 | Tomm20 | CAST,KK,MOLF,SPRET | feature_truncation,3_prime_UTR_variant |
| 8 | 129772122 | 129772176 | 54 | Ccdc7a | SPRET | feature_truncation,3_prime_UTR_variant |
| 9 | 14454711 | 14454762 | 51 | Amotl1 | SPRET | feature_truncation,3_prime_UTR_variant |
| 9 | 14690833 | 14690967 | 134 | Ankrd49 | CAST | feature_truncation,3_prime_UTR_variant |
| 9 | 14913979 | 14914047 | 68 | Panx1 | MOLF | feature_truncation,3_prime_UTR_variant |
| 9 | 14916731 | 14916824 | 93 | Panx1 | SPRET | feature_truncation,3_prime_UTR_variant |
| 9 | 15009270 | 15009497 | 227 | Hephl1 | CAST,MOLF,SPRET | feature_truncation,coding_sequence_variant,intron_variant |
| 9 | 15653786 | 15653871 | 85 | Slc36a4 | SPRET | feature_truncation,3_prime_UTR_variant |
| 9 | 18323861 | 18323921 | 60 | Ubtfl1 | CAST,MOLF | feature_truncation,3_prime_UTR_variant |
| 9 | 18789554 | 18789823 | 269 | Or7g18 | SPRET | feature_truncation,3_prime_UTR_variant |
| 9 | 19543329 | 19543813 | 484 | Zfp317 | CAST | feature_truncation,5_prime_UTR_variant,intron_variant |
| 9 | 19839096 | 19839620 | 524 | Or7d10 | SPRET | feature_truncation,3_prime_UTR_variant |
| 9 | 19964703 | 19964802 | 99 | Or7d11 | MOLF,SPRET | feature_truncation,3_prime_UTR_variant |
| 9 | 20339973 | 20340282 | 309 | Zfp26 | 129S1,CAST,MOLF,SPRET | feature_truncation,3_prime_UTR_variant |
| 9 | 20342386 | 20342452 | 66 | Zfp26 | CAST | feature_truncation,3_prime_UTR_variant |
| 9 | 20509869 | 20509960 | 91 | Zfp846 | 129S1,BUB,C58J,FVB,KK,LP,MA/My,NZB,NZO,NZW,RF,SJL,SPRET,SWR,WSB | feature_truncation,3_prime_UTR_variant |
| 9 | 21047047 | 21048029 | 982 | Cdc37 | WSB | feature_truncation,3_prime_UTR_variant |
| 9 | 21048531 | 21048590 | 59 | Cdc37 | 129S1,BUB,C58J,CBA,CE,FVB,ILNJ2,KK,LP,MA/My,MOLF,MRL,NOD,NOR,NZB,NZO,NZW,PJ,PL,RF,SEA,SJL,SMJ,SPRET,SWR,TH | feature_truncation,3_prime_UTR_variant |
| 9 | 21666898 | 21667094 | 196 | Spc24 | BTBR,C57L,C58J,CAST,FVB,MA/My,NOD,NOR,NZB,NZW,RF,SJL,SWR,WSB | feature_truncation,3_prime_UTR_variant |
| 9 | 22111677 | 22111845 | 168 | Zfp872 | BTBR,C57L,CAST,FVB,MA/My,NOD,NOR,NZB,NZW,RF,SJL,SPRET,SWR | feature_truncation,inframe_deletion |
| 9 | 24225936 | 24225988 | 52 | Npsr1 | SPRET | feature_truncation,3_prime_UTR_variant |
| 9 | 24556419 | 24556581 | 162 | Dpy19l2 | CAST,SPRET,WSB | feature_truncation,coding_sequence_variant,intron_variant |
| 9 | 28907226 | 28907291 | 65 | Ntm | MOLF,NZO,SPRET | feature_truncation,3_prime_UTR_variant |
| 9 | 34941443 | 34941512 | 69 | Kirrel3 | SPRET | feature_truncation,3_prime_UTR_variant |
| 9 | 37622275 | 37622332 | 57 | Or8a1b | SPRET | feature_truncation,3_prime_UTR_variant |
| 9 | 38028910 | 38029123 | 213 | Or8b40 | SPRET | feature_truncation,3_prime_UTR_variant |
| 9 | 38204881 | 38204947 | 66 | Or8c18 | 129S1,AJ,AKR,BALB,BTBR,BUB,C3H,CAST,CBA,DBA2J,MOLF,NOD,NOR,PL,RF,RH2,SJL,SMJ,SPRET,SWR,TH,WSB | feature_truncation,3_prime_UTR_variant |
| 9 | 38205751 | 38206293 | 542 | Or8c18 | 129S1,AJ,AKR,BALB,BTBR,BUB,C3H,CBA,DBA2J,NOD,NOR,PL,RF,RH2,SJL,SMJ,SPRET,SWR,TH,WSB | feature_truncation,3_prime_UTR_variant |
| 9 | 38261752 | 38261936 | 184 | Or8c20 | CAST,MOLF,SPRET | feature_truncation,3_prime_UTR_variant |
| 9 | 38262726 | 38263100 | 374 | Or8c20 | CAST,MOLF,SPRET | feature_truncation,3_prime_UTR_variant |
| 9 | 38380165 | 38381160 | 995 | Or8b1b | BTBR,CBA,DBA1J,ILNJ2,MA/My,MOLF,NOD,NOR,PL,SJL,TH,WSB | feature_truncation,3_prime_UTR_variant |
| 9 | 38506298 | 38507207 | 909 | Or8b49 | 129S1,AJ,AKR,BALB,BTBR,BUB,C3H,DBA2J,FVB,MA/My,NOD,NOR,RF,RH2,SMJ,SWR,TH | stop_lost,feature_truncation,coding_sequence_variant,3_prime_UTR_variant |
| 9 | 38566705 | 38567280 | 575 | Or8b51 | CBA,MA/My,PL,SJL,WSB | feature_truncation,3_prime_UTR_variant |
| 9 | 38567255 | 38567366 | 111 | Or8b51 | MOLF | feature_truncation,3_prime_UTR_variant |
| 9 | 39149428 | 39149513 | 85 | Or8g18 | PL,SJL | frameshift_variant,feature_truncation |
| 9 | 39612641 | 39612989 | 348 | Or10d1b | CAST,MOLF,SPRET | feature_truncation,3_prime_UTR_variant |
| 9 | 39662615 | 39662943 | 328 | Or8g4 | SPRET | stop_lost,feature_truncation,coding_sequence_variant,3_prime_UTR_variant |
| 9 | 39823207 | 39823405 | 198 | Or8g2 | SPRET | feature_truncation,3_prime_UTR_variant |
| 9 | 39860337 | 39860968 | 631 | Or10d5 | CAST,CBA,CE,KK,MOLF,MRL,NZB,NZW,PJ,PL,SEA,SJL,SPRET,WSB | feature_truncation,3_prime_UTR_variant |
| 9 | 39871982 | 39872690 | 708 | Or10d5j | MOLF | feature_truncation,5_prime_UTR_variant,intron_variant |
| 9 | 39934770 | 39934822 | 52 | Or10g6 | CAST,CE,MRL,PL,SJL,WSB | feature_truncation,3_prime_UTR_variant |
| 9 | 40008818 | 40009257 | 439 | Or4d5 | SPRET | feature_truncation,3_prime_UTR_variant |

| | | | | | | |
|---|---|---|---|---|---|---|
| 9 | 40100912 | 40101610 | 698 | Or6x1 | CBA,KK,MA/My,MOLF,MRL,NZB,NZW,PJ,PL,SEA,SJL,SPRET,WSB | feature_truncation,3_prime_UTR_variant |
| 9 | 40207519 | 40207590 | 71 | Gramd1b | SPRET | feature_truncation,3_prime_UTR_variant |
| 9 | 44055627 | 44055679 | 52 | Cbl | SMJ | feature_truncation,3_prime_UTR_variant |
| 9 | 45040035 | 45040903 | 868 | Scn2b | AKR,BUB,FVB,MA/My,NZB,PJ,PL,RF,SEA | feature_truncation,3_prime_UTR_variant |
| 9 | 49348204 | 49348540 | 336 | Ttc12 | SPRET | feature_truncation,3_prime_UTR_variant |
| 9 | 50255488 | 50256284 | 796 | Rpl10-ps3 | CAST,FVB,MOLF,SPRET,SWR,WSB | transcript_ablation |
| 9 | 50517131 | 50517189 | 58 | Nkapd1 | SPRET | feature_truncation,3_prime_UTR_variant |
| 9 | 50519500 | 50519565 | 65 | Nkapd1 | SPRET | feature_truncation,3_prime_UTR_variant |
| 9 | 51762794 | 51762905 | 111 | Arhgap20 | MOLF | feature_truncation,3_prime_UTR_variant |
| 9 | 53372416 | 53372474 | 58 | Atm | 129S1,C57L,PJ,SEA | feature_truncation,coding_sequence_variant,intron_variant |
| 9 | 53757407 | 53757459 | 52 | Sln | SPRET | feature_truncation,5_prime_UTR_variant |
| 9 | 53932288 | 53932340 | 52 | Tnfaip8l3 | SPRET | feature_truncation,3_prime_UTR_variant |
| 9 | 54073489 | 54073593 | 104 | Cyp19a1 | CAST | feature_truncation,3_prime_UTR_variant |
| 9 | 55234133 | 55234233 | 100 | Tmem266 | MOLF | feature_truncation,5_prime_UTR_variant |
| 9 | 59625223 | 59625433 | 210 | Gramd2a | 129S1,BTBR,C57L,CAST,CBA,KK,MA/My,MOLF,NZB,NZO,NZW,PL,SMJ,SPRET,WSB | feature_truncation,3_prime_UTR_variant |
| 9 | 61861059 | 61861508 | 449 | Paqr5 | SPRET | feature_truncation,3_prime_UTR_variant |
| 9 | 62326935 | 62327037 | 102 | Coro2b | SPRET | feature_truncation,3_prime_UTR_variant |
| 9 | 64764972 | 64765160 | 188 | Dennd4a | SPRET | splice_donor_variant,splice_donor_5th_base_variant,coding_sequence_variant,intron_variant |
| 9 | 65462881 | 65463095 | 214 | Plekho2 | B10D2,B10J,LP,MA/My | feature_truncation,3_prime_UTR_variant |
| 9 | 67838029 | 67838085 | 56 | Vps13c | CE | feature_truncation,coding_sequence_variant,intron_variant |
| 9 | 69292063 | 69292119 | 56 | Rora | NOD,NOR | feature_truncation,3_prime_UTR_variant |
| 9 | 72366458 | 72366609 | 151 | Mns1,Tex9 | SPRET | feature_truncation,3_prime_UTR_variant |
| 9 | 72575945 | 72576150 | 205 | Nedd4 | CAST,MOLF,SPRET | feature_truncation,5_prime_UTR_variant |
| 9 | 75428006 | 75428183 | 177 | Tmod3 | MOLF,SPRET | feature_truncation,3_prime_UTR_variant |
| 9 | 75477846 | 75477947 | 101 | Tmod2 | 129S1,AJ,AKR,BALB,BTBR,BUB,C3H,C57L,C58J,CBA,CE,DBA1J,DBA2J,FVB,ILNJ2,KK,LP,MA/My,MRL,NOD,NOR,NZB,NZO,NZW,PJ,PL,RF,RH2,SEA,SJL,SMJ,SWR,TH | feature_truncation,3_prime_UTR_variant |
| 9 | 75480383 | 75480445 | 62 | Tmod2 | CAST | feature_truncation,3_prime_UTR_variant |
| 9 | 86406393 | 86406514 | 121 | Dop1a | CAST,DBA1J,DBA2J,FVB,KK,MA/My,MOLF,NZO,NZW,PL,RF,SJL,SPRET,SWR | feature_truncation,coding_sequence_variant,intron_variant |
| 9 | 88965807 | 88966788 | 981 | Trim43b | SPRET | feature_truncation,3_prime_UTR_variant |
| 9 | 90086446 | 90086503 | 57 | Tbc1d2b | CAST | feature_truncation,3_prime_UTR_variant |
| 9 | 95282592 | 95282704 | 112 | Chst2 | CE,MRL,NZO,NZW,SPRET | feature_truncation,3_prime_UTR_variant |
| 9 | 96742621 | 96742890 | 269 | Pxylp1 | AKR,BUB,RF,SMJ,TH | feature_truncation,5_prime_UTR_variant,intron_variant |
| 9 | 103066684 | 103066782 | 98 | Srprb | 129S1,AJ,AKR,BALB,BTBR,BUB,C3H,C57L,CAST,CBA,FVB,ILNJ2,MA/My,MOLF,NOD,NOR,NZB,NZW,PJ,PL,RF,RH2,SEA,SJL,SMJ,SPRET,SWR,TH,WSB | feature_truncation,3_prime_UTR_variant |
| 9 | 103227574 | 103227625 | 51 | Topbp1 | CE,SMJ | feature_truncation,3_prime_UTR_variant |
| 9 | 103924354 | 103924418 | 64 | Uba5 | CAST,MA/My,SPRET | feature_truncation,3_prime_UTR_variant |
| 9 | 106322858 | 106322973 | 115 | Abhd14a | 129S1,AJ,AKR,BALB,BUB,C3H,C57L,CAST,CE,DBA1J,DBA2J,FVB,ILNJ2,MA/My,MRL,NOD,NOR,NZB,NZW,PL,RH2,SJL,SMJ,SWR,TH | feature_truncation,5_prime_UTR_variant,feature_truncation,5_prime_UTR_variant,intron_variant |
| 9 | 106757415 | 106757467 | 52 | Dcaf1 | 129S1,AJ,AKR,BALB,BUB,C3H,C57L,CE,FVB,ILNJ2,MA/My,MRL,NOD,NOR,NZB,NZO,NZW,PL,RF,RH2,SJL,SMJ,SWR,TH,WSB | feature_truncation,3_prime_UTR_variant |
| 9 | 108280803 | 108280917 | 114 | Iho1 | CAST,CBA,MOLF,SPRET,WSB | feature_truncation,3_prime_UTR_variant |
| 9 | 110137713 | 110138206 | 493 | Elp6 | BTBR,CAST,CBA,FVB,MOLF,NZW,SJL,SPRET,SWR | splice_acceptor_variant,5_prime_UTR_variant,intron_variant |
| 9 | 110812424 | 110812639 | 215 | Lrrc2 | CAST | feature_truncation,3_prime_UTR_variant |
| 9 | 110815750 | 110815936 | 186 | Rtp3 | 129S1,AJ,AKR,BALB,BTBR,BUB,C3H,C57L,CAST,CBA,CE,DBA1J,DBA2J,FVB,ILNJ2,KK,MA/My,MOLF,MRL,NOD,NOR,NZB,NZO,PJ,PL,RF,RH2,SEA,SPRET,TH,WSB | feature_truncation,coding_sequence_variant,intron_variant |
| 9 | 114811420 | 114811808 | 388 | Rps27rt | 129S1,AJ,AKR,BALB,BTBR,C3H,C57L,CAST,CE,FVB,ILNJ2,KK,MOLF,MRL,NOD,NOR,NZB,NZO,NZW,PJ,PL,RF,RH2,SEA,SJL,SMJ,SPRET,SWR,TH,WSB | transcript_ablation |
| 9 | 115869588 | 115870198 | 610 | Gadl1 | 129S1,C57L,CAST,CE,MOLF,NZB,NZO,SPRET,WSB | stop_lost,feature_truncation,coding_sequence_variant,3_prime_UTR_variant,intron_variant |
| 9 | 119897638 | 119897700 | 62 | Cx3cr1 | SPRET | feature_truncation,5_prime_UTR_variant |
| 9 | 121500697 | 121500817 | 120 | Vipr1 | CAST,MOLF | feature_truncation,3_prime_UTR_variant |
| 9 | 121510106 | 121510157 | 51 | Sec22c | 129S1,AJ,AKR,B10D2,B10J,BALB,BTBR,BUB,C3H,C57L,C58J,CAST,CBA,CE,DBA1J,DBA2J,FVB,ILNJ2,KK,LP,MA/My,MOLF,MRL,NOD,NOR,NZB,NZO,NZW,PJ,PL,RF,RH2,SEA,SJL,SMJ,SPRET,SWR,TH,WSB | feature_truncation,3_prime_UTR_variant |
| 9 | 122986486 | 122986707 | 221 | Clec3b | 129S1,AJ,BALB,C3H,C57L,C58J,CAST,CE,FVB,MA/My,MOLF,PJ,PL,SEA,SMJ,SPRET,WSB | feature_truncation,3_prime_UTR_variant |
| 9 | 123620986 | 123621099 | 113 | Fyco1 | 129S1,C57L,C58J,CAST,CE,MA/My,MOLF,SMJ,SPRET,WSB | feature_truncation,3_prime_UTR_variant |
| 9 | 123680961 | 123681081 | 120 | Fyco1 | C58J,CAST,MA/My,SMJ,SPRET,TH | feature_truncation,5_prime_UTR_variant |
| 9 | 123932423 | 123932996 | 573 | Ccr5 | 129S1,C57L,C58J,CBA,CE,DBA1J,DBA2J,FVB,MA/My,MOLF,NOD,NOR,SJL,SMJ,SPRET,SWR,WSB | feature_truncation,3_prime_UTR_variant |
| X | 7518728 | 7518806 | 78 | Syp | SPRET | feature_truncation,3_prime_UTR_variant |
| X | 7991318 | 7991426 | 108 | Wdr13 | CAST,SPRET | feature_truncation,3_prime_UTR_variant |
| X | 36696044 | 36696127 | 83 | Rhox4d | MOLF | feature_truncation,5_prime_UTR_variant |
| X | 47500587 | 47500692 | 105 | Elf4 | SPRET | feature_truncation,3_prime_UTR_variant |
| X | 52776594 | 52776666 | 72 | Zfp36l3 | SPRET | feature_truncation,inframe_deletion |
| X | 63312436 | 63312499 | 63 | Slitrk4 | MOLF | feature_truncation,3_prime_UTR_variant |
| X | 71313875 | 71313959 | 84 | Gabre | CAST | feature_truncation,inframe_deletion |
| X | 72220572 | 72220702 | 130 | Spin2d | MOLF,SPRET | feature_truncation,5_prime_UTR_variant |
| X | 72378544 | 72378971 | 427 | Gm45015 | CAST,MOLF,SPRET | feature_truncation,3_prime_UTR_variant |
| X | 72467674 | 72467732 | 58 | Zfp92 | MOLF | feature_truncation,3_prime_UTR_variant |
| X | 73482661 | 73482745 | 84 | Ikbkg | CAST,MOLF,SPRET | feature_truncation,coding_sequence_variant,intron_variant |
| X | 73496138 | 73496343 | 205 | Ikbkg | CAST,MOLF,SPRET | feature_truncation,3_prime_UTR_variant |

| | | | | | | |
|---|---|---|---|---|---|---|
| X | 76815478 | 76815558 | 80 | Prkx | CAST,MOLF | feature_truncation,3_prime_UTR_variant |
| X | 84249533 | 84249744 | 211 | Dmd | CAST,MOLF | feature_truncation,3_prime_UTR_variant |
| X | 85294549 | 85294858 | 309 | Mageb4 | CAST,SPRET | feature_truncation,inframe_deletion |
| X | 88452357 | 88452616 | 259 | Dcaf8l | CE | feature_truncation,5_prime_UTR_variant,intron_variant |
| X | 99486056 | 99486199 | 143 | Awat2 | CAST,MOLF,NZB,NZO,SPRET | feature_truncation,5_prime_UTR_variant |
| X | 101134302 | 101134365 | 63 | Nhsl2 | CAST,SPRET | feature_truncation,3_prime_UTR_variant |
| X | 101185264 | 101185650 | 386 | Ercc6l | CAST,SPRET | feature_truncation,3_prime_UTR_variant |
| X | 102438713 | 102438818 | 105 | Chic1 | SPRET | feature_truncation,3_prime_UTR_variant |
| X | 133050866 | 133050980 | 114 | Xkrx | SPRET | feature_truncation,3_prime_UTR_variant |
| X | 135738438 | 135738504 | 66 | Plp1 | CAST,SPRET | feature_truncation,3_prime_UTR_variant |
| X | 135828926 | 135829021 | 95 | H2bw2 | MOLF | feature_truncation,3_prime_UTR_variant |
| X | 135874181 | 135875089 | 908 | Tmsb15b1 | MOLF | stop_lost,feature_truncation,coding_sequence_variant,3_prime_UTR_variant,intron_variant |
| X | 135882654 | 135882706 | 52 | Slc25a53 | SPRET | feature_truncation,3_prime_UTR_variant |
| X | 140471379 | 140471520 | 141 | Col4a5 | SPRET | feature_truncation,3_prime_UTR_variant |
| X | 149431903 | 149431963 | 60 | Tro | AJ,BALB,CAST,FVB,MA/My,MOLF,MRL,NOD,NOR,NZB,NZW,RH2,SEA,SJL,SMJ,SWR,WSB | feature_truncation,inframe_deletion |
| X | 149766735 | 149766894 | 159 | Gnl3l | CAST,MOLF,SPRET | feature_truncation,3_prime_UTR_variant |
| X | 155126147 | 155126206 | 59 | Gm15155 | SPRET | feature_truncation,coding_sequence_variant,intron_variant |
| X | 158269284 | 158269463 | 179 | Map7d2 | SPRET | splice_acceptor_variant,coding_sequence_variant,3_prime_UTR_variant,intron_variant |
| X | 159258543 | 159258619 | 76 | Adgrg2 | BALB,CAST,MRL,NOD,NOR,SEA,SPRET,WSB | feature_truncation,coding_sequence_variant,intron_variant |
| X | 159579337 | 159579389 | 52 | Rs1 | SPRET | feature_truncation,3_prime_UTR_variant |
| X | 161198224 | 161198288 | 64 | Reps2 | SPRET | feature_truncation,3_prime_UTR_variant |
| X | 162761874 | 162761942 | 68 | Car5b | MOLF,TH | feature_truncation,3_prime_UTR_variant |
| X | 164960017 | 164960490 | 473 | Gemin8 | 129S1,AJ,AKR,BALB,BTBR,BUB,C3H,C57L,C58J,CAST,CBA,CE,DBA1J,DBA2J,FVB,ILNJ2,KK,MA/My,MOLF,MRL,NOD,NOR,NZB,NZO,NZW,PJ,PL,RF,RH2,SEA,SJL,SMJ,SPRET,SWR,TH,WSB | feature_truncation,5_prime_UTR_variant,intron_variant |
| X | 168772538 | 168772707 | 169 | Mid1 | 129S1,BTBR,C3H,CE,DBA2J,FVB,ILNJ2,KK,MA/My,MRL,NZB,NZO,NZW,PJ,PL,RF,SEA,SMJ,SWR | feature_truncation,3_prime_UTR_variant |

| CHR | START | END | SIZE | SYMBOL | STRAINS with VARIANT SV ALLELE | CONSEQUENCE |
|---|---|---|---|---|---|---|
| 1 | 16581616 | 16585868 | 4252 | Stau2 | WSB | feature_truncation,5_prime_UTR_variant,intron_variant |
| 1 | 31304335 | 31326680 | 22345 | Gm6489 | RF | feature_truncation,coding_sequence_variant,5_prime_UTR_variant |
| 1 | 31315348 | 31330883 | 15535 | Gm6489 | SPRET | transcript_ablation |
| 1 | 37089787 | 37123263 | 33476 | Vwa3b | SPRET | stop_lost,feature_truncation,coding_sequence_variant,3_prime_UTR_variant,intron_variant,feature_truncation,coding_sequence_variant,intron_variant |
| 1 | 39035925 | 39037147 | 1222 | Pdcl3 | SPRET | feature_truncation,3_prime_UTR_variant |
| 1 | 40355828 | 40356966 | 1138 | Il1r1 | SPRET | feature_truncation,3_prime_UTR_variant |
| 1 | 40486213 | 40488665 | 2452 | Il1rl1 | CAST | feature_truncation,3_prime_UTR_variant |
| 1 | 50947871 | 50950171 | 2300 | Tmeff2 | SPRET | feature_truncation,5_prime_UTR_variant,intron_variant |
| 1 | 54494118 | 54496167 | 2049 | Gm34066 | MOLF | feature_truncation,5_prime_UTR_variant,intron_variant |
| 1 | 58679920 | 58690084 | 10164 | Flacc1 | CE,SPRET,SWR | stop_lost,feature_truncation,coding_sequence_variant,3_prime_UTR_variant,intron_variant |
| 1 | 58685740 | 58687490 | 1750 | Flacc1 | CAST,CBA,DBA1J,DBA2J,ILNJ2,MA/My,NZB,NZO,NZW | stop_lost,feature_truncation,coding_sequence_variant,3_prime_UTR_variant,intron_variant |
| 1 | 79696043 | 79703491 | 7448 | Wdfy1 | C57L,PJ | stop_lost,feature_truncation,coding_sequence_variant,3_prime_UTR_variant,intron_variant,feature_truncation,coding_sequence_variant,intron_variant |
| 1 | 79698356 | 79705791 | 7435 | Wdfy1 | CBA | stop_lost,feature_truncation,coding_sequence_variant,3_prime_UTR_variant,intron_variant,feature_truncation,coding_sequence_variant,intron_variant |
| 1 | 79698614 | 79700632 | 2018 | Wdfy1 | FVB | feature_truncation,coding_sequence_variant,intron_variant |
| 1 | 84313765 | 84315332 | 1567 | Pid1 | SPRET | feature_truncation,5_prime_UTR_variant |
| 1 | 85224521 | 85234100 | 9579 | Sp140l2 | MRL,PJ,PL | stop_lost,feature_truncation,coding_sequence_variant,3_prime_UTR_variant,intron_variant |
| 1 | 85500108 | 85508544 | 8436 | Sp110 | CAST,NOD | stop_lost,feature_truncation,coding_sequence_variant,3_prime_UTR_variant,intron_variant |
| 1 | 85516811 | 85517859 | 1048 | Sp110 | CAST,RF | feature_truncation,coding_sequence_variant,intron_variant |
| 1 | 85527137 | 85531478 | 4341 | Sp140 | CBA,MRL,SMJ | feature_truncation,coding_sequence_variant,5_prime_UTR_variant,intron_variant |
| 1 | 85651674 | 85653397 | 1723 | A630001G21Rik | MOLF,RF | feature_truncation,coding_sequence_variant,intron_variant |
| 1 | 88156516 | 88158359 | 1843 | Mroh2a | DBA1J,DBA2J | feature_truncation,coding_sequence_variant,intron_variant |
| 1 | 90892310 | 90895296 | 2986 | Rab17 | AJ,BUB,C3H,CBA,CE,MA/My,MRL,PJ,RH2,WSB | feature_truncation,5_prime_UTR_variant,intron_variant |
| 1 | 92414945 | 92417170 | 2225 | Or6b2b | MOLF,PL,SPRET | feature_truncation,3_prime_UTR_variant |
| 1 | 105482811 | 105484591 | 1780 | Pign | SPRET | feature_truncation,3_prime_UTR_variant |
| 1 | 107034970 | 107137292 | 102322 | Serpinb3b | SPRET | transcript_ablation,stop_lost,feature_truncation,coding_sequence_variant,5_prime_UTR_variant,3_prime_UTR_variant,intron_variant |
| 1 | 130740894 | 130747469 | 6575 | Fcamr | SEA | stop_lost,feature_truncation,coding_sequence_variant,3_prime_UTR_variant,intron_variant |
| 1 | 135226312 | 135227591 | 1279 | Timm17a | CAST | feature_truncation,3_prime_UTR_variant |
| 1 | 151058084 | 151067891 | 9807 | Gm47985 | NZO,TH | transcript_ablation |
| 1 | 156292059 | 156298875 | 6816 | Soat1 | AKR | feature_truncation,coding_sequence_variant,5_prime_UTR_variant,intron_variant |
| 1 | 160841518 | 160843318 | 1800 | Zbtb37 | BALB,C58J,DBA1J,DBA2J,LP,MOLF | feature_truncation,3_prime_UTR_variant |
| 1 | 162668101 | 162675212 | 7111 | Fmo1 | 129S1,AJ,BALB,BTBR,BUB,C3H,C57L,C58J,CAST,CBA,CE,DBA1J,DBA2J,ILNJ2,KK,LP,MA/My,MOLF,MRL,NZB,NZO,NZW,PJ,PL,RH2,SEA,SMJ,SPRET,TH,WSB | stop_lost,feature_truncation,coding_sequence_variant,3_prime_UTR_variant,intron_variant |
| 1 | 170510268 | 170516315 | 6047 | Olfml2b | ILNJ2 | feature_truncation,3_prime_UTR_variant |
| 1 | 173745676 | 173748075 | 2399 | Ifi203 | AKR,BALB,BUB,C58J,CBA,CE,DBA1J,DBA2J,ILNJ2,NZB,NZO,PJ,RF,SEA,SJL,SWR,WSB | feature_truncation,3_prime_UTR_variant |
| 1 | 181975324 | 181984273 | 8949 | Vmn1r1 | CAST | feature_truncation,3_prime_UTR_variant |
| 1 | 181983461 | 181999423 | 15962 | Vmn1r1 | AKR,CBA,DBA1J,DBA2J,FVB,KK,PL,RF,SWR | transcript_ablation,stop_lost,feature_truncation,coding_sequence_variant,5_prime_UTR_variant,3_prime_UTR_variant,intron_variant |
| 1 | 189372045 | 189373424 | 1379 | Cenpf | SPRET | feature_truncation,3_prime_UTR_variant |
| 10 | 22160867 | 22165223 | 4356 | H60b | MOLF | stop_lost,feature_truncation,coding_sequence_variant,3_prime_UTR_variant,intron_variant |
| 10 | 24692123 | 24693339 | 1216 | Enpp3 | CAST,MOLF | feature_truncation,3_prime_UTR_variant |
| 10 | 35584604 | 35588977 | 4373 | Amd2 | CAST,MOLF,SPRET | transcript_ablation |
| 10 | 35584631 | 35587889 | 3258 | Amd2 | WSB | transcript_ablation |
| 10 | 39849656 | 39853245 | 3589 | Mfsd4b5 | SJL | feature_truncation,coding_sequence_variant,intron_variant |
| 10 | 77596418 | 77616787 | 20369 | Gm7138,Gm3238 | SPRET | transcript_ablation |
| 10 | 77606404 | 77617620 | 11216 | Gm7138,Gm3238 | MOLF,TH | transcript_ablation |
| 10 | 77609794 | 77615386 | 5592 | Gm7138 | CAST | transcript_ablation |
| 10 | 77620397 | 77627052 | 6655 | Gm7137 | BUB | transcript_ablation |
| 10 | 78724172 | 78790489 | 66317 | Or7a38,Or7a42 | MOLF | transcript_ablation,feature_truncation,5_prime_UTR_variant,intron_variant |
| 10 | 79049962 | 79130285 | 80323 | Vmn2r81 | SPRET | stop_lost,feature_truncation,coding_sequence_variant,5_prime_UTR_variant,3_prime_UTR_variant,intron_variant |
| 10 | 83246227 | 83261974 | 15747 | Nopchap1 | CAST,MOLF | feature_truncation,3_prime_UTR_variant |
| 10 | 83274173 | 83276904 | 2731 | Nopchap1 | CAST,MOLF | feature_truncation,3_prime_UTR_variant |
| 10 | 83295819 | 83310363 | 14544 | Nopchap1 | SPRET | feature_truncation,3_prime_UTR_variant |
| 10 | 83317068 | 83318814 | 1746 | Nopchap1 | CAST,MOLF,SPRET | feature_truncation,3_prime_UTR_variant |
| 10 | 103976218 | 103979898 | 3680 | Gm21293 | PL | feature_truncation,5_prime_UTR_variant,intron_variant |
| 10 | 112209191 | 112220482 | 11291 | Kcnc2 | CAST | feature_truncation,5_prime_UTR_variant,intron_variant |
| 10 | 118074185 | 118082776 | 8591 | Cdc5lrt1 | 129S1,AJ,AKR,BALB,BTBR,C3H,C57L,C58J,CAST,CBA,CE,DBA1J,DBA2J,KK,MA/My,MOLF,MRL,NOD,NOR,NZB,NZO,NZW,PJ,PL,RF,RH2,SEA,SJL,SPRET,TH,WSB | transcript_ablation |
| 10 | 118093852 | 118111417 | 17565 | Cdc5lrt4 | MRL,NZO,NZW,WSB | transcript_ablation |
| 10 | 118164126 | 118187278 | 23152 | Cdc5lrt9,Cdc5lrt6,Cdc5lrt8,Cdc5lrt7 | MOLF,NZB,SJL | transcript_ablation |
| 10 | 118169550 | 118181852 | 12302 | Cdc5lrt8,Cdc5lrt7 | 129S1,DBA1J | transcript_ablation |
| 10 | 129074977 | 129077579 | 2602 | Or6c5 | BUB,C3H,CAST,CBA,CE,KK,MOLF,RF,SMJ,SPRET,WSB | feature_truncation,3_prime_UTR_variant |

| | | | | | | |
|---|---|---|---|---|---|---|
| 10 | 129161870 | 129168897 | 7027 | Or6c35 | CAST | feature_truncation,coding_sequence_variant,5_prime_UTR_variant,feature_truncation,coding_sequence_variant,5_prime_UTR_variant,intron_variant |
| 10 | 129224680 | 129231985 | 7305 | Or6c208 | BUB,C3H,CAST,CBA,CE,KK,LP,MOLF,NOD,NOR,RF,SMJ,SPRET,WSB | feature_truncation,3_prime_UTR_variant |
| 10 | 129559439 | 129564211 | 4772 | Or6c212 | CE,RF | feature_truncation,5_prime_UTR_variant,intron_variant |
| 10 | 129713548 | 129715537 | 1989 | Or6c70 | SPRET | feature_truncation,5_prime_UTR_variant,intron_variant |
| 10 | 129726275 | 129737308 | 11033 | Or6c217 | SPRET | feature_truncation,3_prime_UTR_variant |
| 10 | 129798887 | 129801272 | 2385 | Or10u4 | CAST,KK,SMJ | feature_truncation,3_prime_UTR_variant |
| 10 | 129804354 | 129816701 | 12347 | Or10u3,Gm10310 | SPRET | transcript_ablation,feature_truncation,3_prime_UTR_variant |
| 10 | 129809690 | 129818564 | 8874 | Or10u3,Gm10310 | WSB | transcript_ablation,feature_truncation,3_prime_UTR_variant |
| 10 | 130247005 | 130250104 | 3099 | Vmn2r85 | CAST | feature_truncation,3_prime_UTR_variant |
| 10 | 130248103 | 130250013 | 1910 | Vmn2r85 | AKR,CE,MOLF,MRL,RF,SJL,SMJ,SPRET,SWR | feature_truncation,3_prime_UTR_variant |
| 10 | 130252452 | 130273905 | 21453 | Vmn2r85 | SPRET | stop_lost,feature_truncation,coding_sequence_variant,5_prime_UTR_variant,3_prime_UTR_variant,intron_variant |
| 11 | 11576688 | 11583590 | 6902 | 4930512M02Rik | CAST | feature_truncation,coding_sequence_variant,5_prime_UTR_variant,intron_variant |
| 11 | 17887419 | 17889833 | 2414 | Etaa1 | SPRET | feature_truncation,3_prime_UTR_variant |
| 11 | 34041269 | 34042413 | 1144 | Lcp2,4930469K13Rik | ILNJ2,KK,MA/My,MOLF,NOD,NZW,PL,RF | feature_truncation,3_prime_UTR_variant,stop_lost,feature_truncation,coding_sequence_variant,3_prime_UTR_variant,intron_variant |
| 11 | 46557281 | 46560322 | 3041 | Timd2 | 129S1,AJ,BALB,BTBR,C3H,C57L,CBA,DBA1J,ILNJ2,KK,MRL,NZB,NZW,PL,RH2,SJL,SMJ,WSB | feature_truncation,3_prime_UTR_variant |
| 11 | 48716160 | 48717571 | 1411 | Cdk2ap1rt,Trim7 | CAST,CE,MOLF,NZO,NZW,PJ,SEA,SPRET | transcript_ablation,feature_truncation,5_prime_UTR_variant,intron_variant |
| 11 | 49228597 | 49269989 | 41392 | Or2y17 | CAST | transcript_ablation,stop_lost,feature_truncation,coding_sequence_variant,5_prime_UTR_variant,3_prime_UTR_variant,intron_variant |
| 11 | 49336004 | 49343486 | 7482 | Or2y16 | AKR,B10D2,B10J,BUB,C58J,CAST,CE,KK,MA/My,MOLF,NOR,NZB,NZO,NZW,PJ,RF,SEA,SPRET,SWR,TH,WSB | feature_truncation,3_prime_UTR_variant |
| 11 | 49387223 | 49388381 | 1158 | Or2y1 | MOLF,SPRET | feature_truncation,3_prime_UTR_variant |
| 11 | 58649769 | 58651709 | 1940 | Or2ak4 | AJ,BTBR,BUB,C3H,C57L,CBA,DBA1J,DBA2J,FVB,ILNJ2,KK,MA/My,MRL,NOR,NZB,NZO,NZW,PJ,SEA,SJL,SWR,TH | feature_truncation,3_prime_UTR_variant |
| 11 | 71034946 | 71037667 | 2721 | Nlrp1a | 129S1,BALB,BTBR,C3H,C57L,C58J,CBA,FVB,LP,MA/My,MRL,NZB,NZO,RH2,SWR | feature_truncation,5_prime_UTR_variant,intron_variant |
| 11 | 71058273 | 71086157 | 27884 | Nlrp1b | KK,NZW | feature_truncation,coding_sequence_variant,intron_variant |
| 11 | 71091867 | 71118755 | 26888 | Nlrp1b | C3H,PL | feature_truncation,coding_sequence_variant,intron_variant,feature_truncation,coding_sequence_variant,5_prime_UTR_variant,intron_variant |
| 11 | 71093037 | 71174694 | 81657 | Nlrp1b | CBA | feature_truncation,coding_sequence_variant,5_prime_UTR_variant,intron_variant |
| 11 | 73675791 | 73679175 | 3384 | Or1e30 | AJ,NZO,NZW,WSB | transcript_ablation,stop_lost,feature_truncation,coding_sequence_variant,5_prime_UTR_variant,3_prime_UTR_variant,intron_variant |
| 11 | 74011762 | 74013112 | 1350 | Or3a1b | CAST | transcript_ablation,stop_lost,feature_truncation,coding_sequence_variant,5_prime_UTR_variant,3_prime_UTR_variant,intron_variant |
| 11 | 74038719 | 74146019 | 107300 | Or3a1c,Or1a1,Or1a1b | MOLF | transcript_ablation |
| 11 | 95957709 | 95960790 | 3081 | Atp5mc1 | KK | feature_truncation,3_prime_UTR_variant |
| 11 | 99686280 | 99692048 | 5768 | Gm11569 | MOLF | transcript_ablation |
| 11 | 99688712 | 99694195 | 5483 | Gm11569 | BALB,CBA,CE,MA/My,MRL,NZO,SEA | transcript_ablation |
| 11 | 99696077 | 99701435 | 5358 | Krtap4-13 | AKR,RF | transcript_ablation |
| 11 | 100222817 | 100230307 | 7490 | Gast | MA/My | transcript_ablation |
| 11 | 108571632 | 108574270 | 2638 | Cep112 | BUB | feature_truncation,coding_sequence_variant,5_prime_UTR_variant,intron_variant |
| 11 | 114908101 | 114932941 | 24840 | Cd300ld5,Cd300ld4 | SJL | transcript_ablation |
| 11 | 116640381 | 116656483 | 16102 | St6galnac1 | AJ,BALB,C3H,C57L,CBA,DBA1J,DBA2J,MRL,SEA,TH | feature_truncation,3_prime_UTR_variant |
| 11 | 120653604 | 120655245 | 1641 | Hmga1b | CAST,MOLF,SJL,SPRET | transcript_ablation |
| 12 | 44278084 | 44284683 | 6599 | Pnpla8 | AJ,AKR,BALB,BTBR,C3H,CAST,CBA,KK,MOLF,PJ,PL,RF,RH2,SEA,SMJ,SPRET,SWR | feature_truncation,5_prime_UTR_variant,intron_variant |
| 12 | 55287457 | 55300491 | 13034 | Srp54c | CE | feature_truncation,coding_sequence_variant,intron_variant |
| 12 | 69790997 | 69793449 | 2452 | Dmac2l | SWR | feature_truncation,3_prime_UTR_variant |
| 12 | 70230408 | 70236599 | 6191 | Abhd12b | CAST,SPRET | feature_truncation,3_prime_UTR_variant |
| 12 | 73357761 | 73362635 | 4874 | Slc38a6 | 129S1,AJ,BALB,BTBR,BUB,C3H,C57L,C58J,CAST,CBA,CE,FVB,KK,MA/My,MOLF,NOD,NOR,NZW,PJ,RH2,SEA,SJL,SMJ,SPRET,SWR,TH,WSB | stop_lost,feature_truncation,coding_sequence_variant,3_prime_UTR_variant |
| 12 | 84099355 | 84100562 | 1207 | Acot3 | MOLF | feature_truncation,coding_sequence_variant,5_prime_UTR_variant,intron_variant,feature_truncation,coding_sequence_variant,5_prime_UTR_variant |
| 12 | 84381847 | 84382905 | 1058 | Zfp410 | MOLF | stop_lost,feature_truncation,coding_sequence_variant,3_prime_UTR_variant |
| 12 | 84689815 | 84691609 | 1794 | Vrtn | PL,SJL | feature_truncation,5_prime_UTR_variant,intron_variant |
| 12 | 87531926 | 87543245 | 11319 | Eif1ad12 | CAST | transcript_ablation |
| 12 | 87538209 | 87543259 | 5050 | Eif1ad12 | SPRET | stop_lost,feature_truncation,coding_sequence_variant,5_prime_UTR_variant,3_prime_UTR_variant,intron_variant |
| 12 | 87669306 | 87671605 | 2299 | Eif1ad6 | NOD | feature_truncation,3_prime_UTR_variant |
| 12 | 87787027 | 87789326 | 2299 | Eif1ad2 | C57L | feature_truncation,3_prime_UTR_variant |
| 12 | 87880685 | 87885626 | 4941 | Eif1ad14 | CAST,CE,MOLF,NOD,SWR | feature_truncation,3_prime_UTR_variant |
| 12 | 103407138 | 103415137 | 7999 | Ifi27l2a | PJ,SEA | transcript_ablation |
| 12 | 103562465 | 103563694 | 1229 | Ppp4r4 | NZO | feature_truncation,coding_sequence_variant,intron_variant |
| 12 | 103731323 | 103739180 | 7857 | Serpina1d | CAST | feature_truncation,coding_sequence_variant,5_prime_UTR_variant,intron_variant |

| | | | | | | |
|---|---|---|---|---|---|---|
| 12 | 104079540 | 104082379 | 2839 | Serpina3a | SPRET | feature_truncation,coding_sequence_variant,5_prime_UTR_variant,feature_truncation,coding_sequence_variant,5_prime_UTR_variant,intron_variant |
| 13 | 21679379 | 21681317 | 1938 | AK157302 | CAST,LP,MOLF,PJ,SEA,SPRET | transcript_ablation |
| 13 | 21698190 | 21699475 | 1285 | Zkscan8 | 129S1,ILNJ2 | feature_truncation,3_prime_UTR_variant |
| 13 | 21738733 | 21742278 | 3545 | Or2b7 | LP,PJ,SEA | transcript_ablation,stop_lost,feature_truncation,coding_sequence_variant,5_prime_UTR_variant,3_prime_UTR_variant,intron_variant |
| 13 | 21813982 | 21821080 | 7098 | Or2b6 | 129S1,AJ,AKR,B10D2,B10J,BALB,BTBR,BUB,C3H,C57L,C58J,CAST,CBA,CE,DBA1J,DBA2J,FVB,ILNJ2,KK,LP,MA/My,MOLF,MRL,NOD,NOR,NZB,PJ,PL,RF,RH2,SEA,SJL,SMJ,SPRET,SWR,TH,WSB | feature_truncation,3_prime_UTR_variant |
| 13 | 21919800 | 21921069 | 1269 | H4c11 | CAST,LP,MOLF,PJ,SEA | feature_truncation,3_prime_UTR_variant |
| 13 | 22270041 | 22273284 | 3243 | Vmn1r188 | 129S1,AJ,B10D2,B10J,BALB,BUB,C3H,C58J,CBA,DBA1J,DBA2J,FVB,MA/My,MRL,NOD,NOR,PL,RH2,SJL,SMJ,SWR,TH | transcript_ablation,stop_lost,feature_truncation,coding_sequence_variant,5_prime_UTR_variant,3_prime_UTR_variant,intron_variant |
| 13 | 22478283 | 22479678 | 1395 | Vmn1r196 | SPRET | feature_truncation,3_prime_UTR_variant |
| 13 | 22516447 | 22519483 | 3036 | Vmn1r197 | BTBR,C57L,KK,LP,MOLF,NZB,NZO,NZW | feature_truncation,3_prime_UTR_variant |
| 13 | 22808311 | 22809745 | 1434 | Vmn1r206 | BTBR,C57L,KK,LP,MOLF,NZB,NZO,NZW,PJ,SEA | feature_truncation,5_prime_UTR_variant |
| 13 | 23262120 | 23264486 | 2366 | Vmn1r215 | SPRET | feature_truncation,3_prime_UTR_variant |
| 13 | 23350941 | 23361414 | 10473 | Vmn1r219 | SPRET | feature_truncation,3_prime_UTR_variant |
| 13 | 23362258 | 23363437 | 1179 | Vmn1r220 | SPRET | feature_truncation,3_prime_UTR_variant |
| 13 | 24655279 | 24660484 | 5205 | Cmah | SPRET | feature_truncation,3_prime_UTR_variant |
| 13 | 27758270 | 27759478 | 1208 | Prl8a1 | MOLF | stop_lost,feature_truncation,coding_sequence_variant,3_prime_UTR_variant |
| 13 | 27889905 | 27892284 | 2379 | Prl7d1 | MOLF | feature_truncation,3_prime_UTR_variant |
| 13 | 34922623 | 34925735 | 3112 | Fam50b | SPRET | feature_truncation,5_prime_UTR_variant,intron_variant |
| 13 | 43540263 | 43547352 | 7089 | Sirt5 | CAST,MOLF,MRL,PJ,SEA,SPRET,WSB | stop_lost,feature_truncation,coding_sequence_variant,3_prime_UTR_variant,intron_variant |
| 13 | 43540271 | 43543640 | 3369 | Sirt5 | MOLF | stop_lost,feature_truncation,coding_sequence_variant,3_prime_UTR_variant,intron_variant |
| 13 | 49647062 | 49649874 | 2812 | Tes3-ps | SPRET | frameshift_variant,stop_lost,start_lost,feature_truncation,start_retained_variant,5_prime_UTR_variant,3_prime_UTR_variant |
| 13 | 53061093 | 53062506 | 1413 | Auh | MOLF | feature_truncation,3_prime_UTR_variant |
| 13 | 53062842 | 53064192 | 1350 | Auh | MOLF | stop_lost,feature_truncation,coding_sequence_variant,3_prime_UTR_variant,intron_variant |
| 13 | 55155638 | 55166514 | 10876 | Hk3 | MOLF | feature_truncation,coding_sequence_variant,intron_variant,feature_truncation,coding_sequence_variant,5_prime_UTR_variant,intron_variant |
| 13 | 62849226 | 62850473 | 1247 | Platr25 | FVB,PJ,SEA,SWR | stop_lost,feature_truncation,coding_sequence_variant,3_prime_UTR_variant,intron_variant |
| 13 | 62874305 | 62880780 | 6475 | Platr25 | 129S1,AJ,BALB,BTBR,BUB,C57L,CAST,FVB,KK,MA/My,MOLF,MRL,NOD,NOR,NZB,NZW,PJ,SEA,SMJ,SPRET,SWR,WSB | feature_truncation,5_prime_UTR_variant,intron_variant |
| 13 | 67054010 | 67055581 | 1571 | Ramacl | PJ,SEA | feature_truncation,5_prime_UTR_variant,intron_variant |
| 13 | 67419469 | 67427196 | 7727 | Zfp458 | SPRET,SWR | feature_truncation,5_prime_UTR_variant,intron_variant |
| 13 | 67541178 | 67543435 | 2257 | Zfp429 | 129S1,AKR,MRL,NZW,PL,RF | feature_truncation,coding_sequence_variant,intron_variant |
| 13 | 67544276 | 67546224 | 1948 | Zfp429 | WSB | feature_truncation,coding_sequence_variant,intron_variant |
| 13 | 67553705 | 67559411 | 5706 | Zfp459 | SPRET | stop_lost,feature_truncation,coding_sequence_variant,3_prime_UTR_variant,intron_variant |
| 13 | 67588657 | 67589780 | 1123 | Zfp874a | 129S1,AJ,AKR,BALB,BTBR,BUB,C3H,C57L,C58J,CAST,CBA,CE,DBA1J,DBA2J,FVB,ILNJ2,KK,MRL,NOD,NOR,NZB,NZO,NZW,PJ,PL,RF,SEA,SJL,SMJ,SWR,TH,WSB | feature_truncation,3_prime_UTR_variant |
| 13 | 67804073 | 67840964 | 36891 | Zfp738 | WSB | transcript_ablation |
| 13 | 67988653 | 67993920 | 5267 | Krbox5 | MOLF | stop_lost,feature_truncation,coding_sequence_variant,3_prime_UTR_variant,intron_variant |
| 13 | 96215717 | 96217884 | 2167 | Btf3l4b | 129S1,AJ,BUB,CAST,CBA,DBA1J,DBA2J,MOLF,SMJ,SPRET,WSB | transcript_ablation |
| 13 | 97234707 | 97235850 | 1143 | Prp2rt | 129S1,AKR,C3H,C57L,CAST,CE,FVB,ILNJ2,KK,MOLF,MRL,NZB,NZO,PL,SJL,SPRET,SWR | transcript_ablation |
| 13 | 100486340 | 100557660 | 71320 | Naip1 | BTBR | stop_lost,feature_truncation,coding_sequence_variant,3_prime_UTR_variant,intron_variant |
| 13 | 113452593 | 113456924 | 4331 | Cspg4b | MRL | feature_truncation,coding_sequence_variant,intron_variant |
| 13 | 120353175 | 120358454 | 5279 | Tcstv1 | SPRET | transcript_ablation |
| 13 | 120411491 | 120415738 | 4247 | B020031M17Rik | NZW | feature_truncation,coding_sequence_variant,5_prime_UTR_variant,feature_truncation,coding_sequence_variant,5_prime_UTR_variant,intron_variant |
| 13 | 120492531 | 120497885 | 5354 | Anxa2r1 | WSB | stop_lost,feature_truncation,coding_sequence_variant,5_prime_UTR_variant,3_prime_UTR_variant,intron_variant |
| 13 | 120635651 | 120748120 | 112469 | Gm21762,AF067061,Gm21731 | 129S1 | transcript_ablation |
| 14 | 8532720 | 8537144 | 4424 | Gm11100 | CAST | feature_truncation,5_prime_UTR_variant,intron_variant |
| 14 | 11187752 | 11189046 | 1294 | Gapdhrt | 129S1,C3H,C57L,CAST,CBA,DBA1J,DBA2J,KK,MOLF,MRL,NZW,PJ,PL,SPRET | stop_lost,feature_truncation,coding_sequence_variant,5_prime_UTR_variant,3_prime_UTR_variant,intron_variant |
| 14 | 11739408 | 11740633 | 1225 | Fhit | WSB | feature_truncation,coding_sequence_variant,intron_variant,feature_truncation,5_prime_UTR_variant,intron_variant |
| 14 | 15578288 | 15582383 | 4095 | Gm3667 | SPRET | feature_truncation,5_prime_UTR_variant,intron_variant |
| 14 | 16736141 | 16750175 | 14034 | Gm3373 | 129S1 | feature_truncation,coding_sequence_variant,5_prime_UTR_variant,intron_variant |
| 14 | 17614413 | 17615944 | 1531 | Gm3264 | MOLF | feature_truncation,5_prime_UTR_variant,intron_variant |
| 14 | 18462888 | 18465895 | 3007 | Gm3099 | FVB,KK,MA/My,NOD,NOR,NZB,NZO,NZW,SJL,SMJ,SWR | stop_lost,feature_truncation,coding_sequence_variant,3_prime_UTR_variant,intron_variant |
| 14 | 18503785 | 18506828 | 3043 | Gm3095 | FVB,MA/My,NOD,NOR,SJL,SWR | feature_truncation,coding_sequence_variant,5_prime_UTR_variant,intron_variant |
| 14 | 19336932 | 19339016 | 2084 | Gm10340 | SPRET | feature_truncation,5_prime_UTR_variant |
| 14 | 19582823 | 19584297 | 1474 | Gm2244 | KK | stop_lost,feature_truncation,coding_sequence_variant,3_prime_UTR_variant,intron_variant |
| 14 | 40535381 | 40536584 | 1203 | Sh2d4b | CBA,DBA1J,DBA2J,ILNJ2 | feature_truncation,3_prime_UTR_variant |

| | | | | | | |
|---|---|---|---|---|---|---|
| 14 | 41073567 | 41078154 | 4587 | Gm5798 | NZB | stop_lost,feature_truncation,coding_sequence_variant,3_prime_UTR_variant,intron_variant |
| 14 | 41747740 | 41748804 | 1064 | 1700049E17Rik1 | BTBR,FVB,KK,NOD,NOR,NZB,NZW,WSB | feature_truncation,coding_sequence_variant,intron_variant |
| 14 | 41881269 | 42209050 | 327781 | Gm17026,Gm3573,Gm7995 | MA/My | transcript_ablation |
| 14 | 42354691 | 42362885 | 8194 | Gm8024 | C57L,CAST,MOLF,NZO,PJ,PL | feature_truncation,5_prime_UTR_variant,intron_variant |
| 14 | 42866512 | 42867966 | 1454 | Gm10376 | ILNJ2 | stop_lost,feature_truncation,coding_sequence_variant,3_prime_UTR_variant |
| 14 | 44575046 | 44581602 | 6556 | Gm5930 | SPRET | feature_truncation,coding_sequence_variant,5_prime_UTR_variant,intron_variant |
| 14 | 44893819 | 44898644 | 4825 | Gm8257 | SPRET | feature_truncation,coding_sequence_variant,intron_variant,feature_truncation,coding_sequence_variant,5_prime_UTR_variant,intron_variant |
| 14 | 44986759 | 45000946 | 14187 | Gm8267 | SPRET | feature_truncation,5_prime_UTR_variant,intron_variant |
| 14 | 45405247 | 45406260 | 1013 | Txndc16 | CAST | feature_truncation,3_prime_UTR_variant |
| 14 | 50380818 | 50382382 | 1564 | Or4k5 | SPRET | feature_truncation,3_prime_UTR_variant |
| 14 | 57026582 | 57028191 | 1609 | Zmym5 | SPRET | feature_truncation,3_prime_UTR_variant |
| 14 | 96337827 | 96340615 | 2788 | Klhl1 | AJ,AKR,BALB,BTBR,BUB,C3H,CBA,CE,DBA1J,DBA2J,FVB,MRL,NOD,PJ,PL,RF,SEA,SJL,SMJ,SWR,TH,WSB | feature_truncation,3_prime_UTR_variant |
| 14 | 113552025 | 113554169 | 2144 | Tpm3-rs7 | FVB,MOLF,SJL,SPRET | stop_lost,feature_truncation,coding_sequence_variant,5_prime_UTR_variant,3_prime_UTR_variant |
| 15 | 4831614 | 4833559 | 1945 | C6 | SPRET | stop_lost,feature_truncation,coding_sequence_variant,3_prime_UTR_variant,intron_variant |
| 15 | 9313785 | 9320993 | 7208 | Ugt3a2 | CAST,MOLF,RF,SJL,TH | stop_lost,feature_truncation,coding_sequence_variant,3_prime_UTR_variant,intron_variant |
| 15 | 9315071 | 9322280 | 7209 | Ugt3a2 | BTBR,C57L,CBA,KK,LP,NOR,NZW,PJ,PL,SEA | feature_truncation,3_prime_UTR_variant |
| 15 | 36995803 | 36997602 | 1799 | Zfp706 | 129S1,AJ,AKR,BUB,C57L,CAST,CE,FVB,LP,MA/My,MRL,NOD,NOR,PJ,RF,RH2,SJL,SMJ,SPRET,SWR,TH | feature_truncation,3_prime_UTR_variant |
| 15 | 75981188 | 75988903 | 7715 | Eppk1 | BTBR | feature_truncation,coding_sequence_variant,intron_variant |
| 15 | 75983266 | 75989437 | 6171 | Eppk1 | SPRET | feature_truncation,coding_sequence_variant,intron_variant |
| 15 | 77464042 | 77486566 | 22524 | Apol10b,Gm36245 | MA/My,MOLF | transcript_ablation,feature_truncation,coding_sequence_variant,5_prime_UTR_variant,intron_variant |
| 15 | 84226995 | 84229893 | 2898 | Parvg | SPRET | feature_truncation,3_prime_UTR_variant |
| 15 | 90520665 | 90527825 | 7160 | Cpne8 | AKR,BALB,BUB,CAST,DBA1J,DBA2J,MOLF,MRL,NZW,PL,SJL,SPRET,WSB | feature_truncation,3_prime_UTR_variant |
| 15 | 94419528 | 94421141 | 1613 | Pus7l | BTBR | feature_truncation,3_prime_UTR_variant |
| 15 | 100226497 | 100245831 | 19334 | Tmt1a3,Tmt1a | BUB,MA/My,SWR | transcript_ablation,feature_truncation,3_prime_UTR_variant |
| 15 | 100228159 | 100264668 | 36509 | Tmt1a3,Tmt1a2,Higd1c | NOD | transcript_ablation,feature_truncation,coding_sequence_variant,5_prime_UTR_variant,intron_variant |
| 15 | 100232589 | 100275332 | 42743 | Tmt1a3,Tmt1a2,Higd1c | C58J,KK,NOR,NZB,NZO,NZW,TH | transcript_ablation,feature_truncation,coding_sequence_variant,5_prime_UTR_variant,intron_variant |
| 15 | 100266044 | 100275331 | 9287 | Higd1c | NOD,NOR,NZO | feature_truncation,coding_sequence_variant,intron_variant |
| 15 | 101535571 | 101554698 | 19127 | Gm5414,Gm5478 | KK,NZB,NZO,RF,TH | splice_acceptor_variant,splice_donor_variant,splice_donor_5th_base_variant,coding_sequence_variant,5_prime_UTR_variant,intron_variant,splice_acceptor_variant,splice_donor_variant,splice_donor_5th_base_variant,coding_sequence_variant,5_prime_UTR_variant,3_prime_UTR_variant,intron_variant,splice_acceptor_variant,splice_donor_variant,splice_donor_5th_base_variant,coding_sequence_variant,3_prime_UTR_variant,intron_variant |
| 15 | 101550520 | 101580496 | 29976 | Gm5478 | SPRET | transcript_ablation |
| 16 | 4573788 | 4574963 | 1175 | Hmox2 | AJ,BTBR,CAST,CE,FVB,MOLF,NOD,NOR,SMJ,SPRET,TH,WSB | feature_truncation,5_prime_UTR_variant |
| 16 | 4675678 | 4678510 | 2832 | 4930562C15Rik | AJ,BTBR,CAST,CE,FVB,MOLF,NOD,NOR,SMJ,SPRET,TH | feature_truncation,coding_sequence_variant,5_prime_UTR_variant,intron_variant |
| 16 | 10040291 | 10045295 | 5004 | Atf7ip2 | SPRET | stop_lost,feature_truncation,coding_sequence_variant,3_prime_UTR_variant,intron_variant |
| 16 | 16973109 | 16975706 | 2597 | Ube2l3 | SPRET | feature_truncation,3_prime_UTR_variant |
| 16 | 17019918 | 17022822 | 2904 | Ube2l3 | CAST | feature_truncation,5_prime_UTR_variant,intron_variant |
| 16 | 19385793 | 19442637 | 56844 | Or2aj5,Or2aj4 | SPRET | transcript_ablation,feature_truncation,5_prime_UTR_variant,intron_variant |
| 16 | 30378591 | 30379802 | 1211 | Lsg1 | SPRET | feature_truncation,3_prime_UTR_variant |
| 16 | 36081298 | 36108348 | 27050 | Stfa1 | MOLF,SPRET | transcript_ablation |
| 16 | 44729296 | 44731799 | 2503 | Cd200r2 | CAST | feature_truncation,coding_sequence_variant,intron_variant |
| 16 | 45973316 | 45975441 | 2125 | Ahcyl | 129S1,C3H,C58J,CAST,CBA,CE,FVB,MOLF,SJL,SPRET,SWR,TH,WSB | transcript_ablation |
| 16 | 88929874 | 88937911 | 8037 | Gm6358 | CAST | feature_truncation,coding_sequence_variant,5_prime_UTR_variant |
| 16 | 89862422 | 89866040 | 3618 | Tiam1 | DBA1J,DBA2J | feature_truncation,5_prime_UTR_variant,intron_variant |
| 16 | 91625377 | 91626519 | 1142 | Itsn1 | SPRET | feature_truncation,3_prime_UTR_variant |
| 16 | 97817755 | 97819076 | 1321 | B230307C23Rik | SPRET | feature_truncation,3_prime_UTR_variant |
| 17 | 6277281 | 6289462 | 12181 | Tulp4 | CAST | stop_lost,feature_truncation,coding_sequence_variant,3_prime_UTR_variant,intron_variant |
| 17 | 6347859 | 6349361 | 1502 | Tmem181a | SEA | feature_truncation,coding_sequence_variant,intron_variant |
| 17 | 7602259 | 7605118 | 2859 | Tcp10a | SPRET | feature_truncation,coding_sequence_variant,intron_variant |
| 17 | 13190585 | 13191645 | 1060 | Wtap | SPRET | feature_truncation,3_prime_UTR_variant |
| 17 | 13580338 | 13581449 | 1111 | Tcp10c | 129S1,AJ,AKR,B10D2,BALB,C3H,C57L,C58J,CBA,DBA1J,DBA2J,LP,NZO,RH2,SEA | feature_truncation,coding_sequence_variant,intron_variant |
| 17 | 15037908 | 15041020 | 3112 | Wdr27 | MOLF,PL | feature_truncation,3_prime_UTR_variant,intron_variant |
| 17 | 15200268 | 15203593 | 3325 | Ermardl2 | SPRET | feature_truncation,coding_sequence_variant,intron_variant |
| 17 | 15263015 | 15266340 | 3325 | Ermard | SPRET | feature_truncation,coding_sequence_variant,intron_variant,stop_lost,feature_truncation,coding_sequence_variant,3_prime_UTR_variant,intron_variant |
| 17 | 18296289 | 18297348 | 1059 | Vmn2r124 | SPRET | feature_truncation,3_prime_UTR_variant |

| | | | | | | |
|---|---|---|---|---|---|---|
| 17 | 18821788 | 18828341 | 6553 | Vmn2r96 | 129S1,AJ,AKR,BALB,C57L,CAST,CE,DBA1J,DBA2J,FVB,ILNJ2,KK,MA/My,MOLF,NZO,RH2,SEA,SPRET,WSB | feature_truncation,3_prime_UTR_variant |
| 17 | 19611375 | 19612625 | 1250 | Vmn2r99 | 129S1,AJ,AKR,BALB,C57L,CAST,CE,DBA1J,DBA2J,FVB,ILNJ2,KK,MA/My,MOLF,NZO,RH2,SEA,SPRET,WSB | feature_truncation,coding_sequence_variant,intron_variant |
| 17 | 19734130 | 19750560 | 16430 | Vmn2r100 | SPRET | feature_truncation,coding_sequence_variant,intron_variant |
| 17 | 19924820 | 19930804 | 5984 | Vmn2r102 | AKR,FVB,MA/My | feature_truncation,3_prime_UTR_variant |
| 17 | 19954193 | 20029813 | 75620 | Vmn2r103 | MOLF | feature_truncation,coding_sequence_variant,5_prime_UTR_variant,intron_variant |
| 17 | 20071048 | 20250726 | 179678 | Vmn2r104 | SPRET | stop_lost,feature_truncation,coding_sequence_variant,3_prime_UTR_variant,intron_variant |
| 17 | 20242560 | 20248732 | 6172 | Vmn2r104 | CAST,MOLF | feature_truncation,3_prime_UTR_variant |
| 17 | 20486580 | 20500319 | 13739 | Vmn2r106 | SPRET | stop_lost,feature_truncation,coding_sequence_variant,3_prime_UTR_variant,intron_variant |
| 17 | 21204583 | 21210070 | 5487 | Vmn1r233 | CAST,MOLF,SPRET | feature_truncation,3_prime_UTR_variant |
| 17 | 21255485 | 21261029 | 5544 | Zfp160 | SPRET | stop_lost,feature_truncation,coding_sequence_variant,3_prime_UTR_variant |
| 17 | 21454813 | 21457704 | 2891 | Vmn1r234 | SPRET | feature_truncation,3_prime_UTR_variant |
| 17 | 21512656 | 21514361 | 1705 | Vmn1r236 | CAST,MOLF | feature_truncation,3_prime_UTR_variant |
| 17 | 21532696 | 21535010 | 2314 | Vmn1r237 | CAST | feature_truncation,coding_sequence_variant,5_prime_UTR_variant,feature_truncation,coding_sequence_variant,5_prime_UTR_variant,intron_variant |
| 17 | 22159816 | 22160910 | 1094 | Zfp942 | MOLF | feature_truncation,5_prime_UTR_variant,intron_variant |
| 17 | 22528494 | 22531620 | 3126 | Gm7072 | SPRET | feature_truncation,5_prime_UTR_variant,intron_variant |
| 17 | 24505657 | 24508795 | 3138 | Abca17 | BTBR | feature_truncation,coding_sequence_variant,intron_variant |
| 17 | 24509137 | 24514054 | 4917 | Abca17 | BTBR | feature_truncation,coding_sequence_variant,intron_variant |
| 17 | 30842603 | 30845160 | 2557 | Dnah8 | SPRET | feature_truncation,5_prime_UTR_variant,intron_variant |
| 17 | 30844034 | 30846501 | 2467 | Dnah8,Glo1 | KK,MOLF,NOD,NOR,SPRET | feature_truncation,5_prime_UTR_variant,intron_variant |
| 17 | 32588390 | 32595491 | 7101 | Wiz | 129S1,AJ,AKR,B10D2,BALB,BUB,C3H,C57L,CAST,CBA,CE,DBA1J,DBA2J,FVB,ILNJ2,MA/My,MOLF,MRL,NOD,NOR,NZB,NZO,NZW,PJ,PL,RF,RH2,SEA,SJL,SMJ,SPRET,SWR,TH,WSB | feature_truncation,coding_sequence_variant,intron_variant |
| 17 | 32921992 | 32927504 | 5512 | Cyp4f15 | SPRET | feature_truncation,3_prime_UTR_variant |
| 17 | 33198118 | 33199861 | 1743 | Zfp472 | BUB,DBA1J,FVB,MOLF,NOD,NOR,PL,SMJ,SWR | feature_truncation,3_prime_UTR_variant |
| 17 | 35536985 | 35565596 | 28611 | H2-Q1,H2-Q2 | TH | transcript_ablation |
| 17 | 36359906 | 36367019 | 7113 | H2-T27 | 129S1,AJ,AKR,B10D2,BALB,BTBR,BUB,C3H,C57L,C58J,CAST,CBA,CE,DBA1J,DBA2J,FVB,ILNJ2,KK,LP,MA/My,MOLF,MRL,NOD,NOR,NZB,NZO,NZW,PJ,PL,RF,RH2,SEA,SJL,SPRET,SWR,TH,WSB | stop_lost,feature_truncation,coding_sequence_variant,3_prime_UTR_variant |
| 17 | 36967898 | 36995302 | 27404 | H2-M1 | AKR,C3H,CBA,CE,MA/My,MRL,RF | transcript_ablation |
| 17 | 38865681 | 38874259 | 8578 | Esp34 | MOLF | stop_lost,feature_truncation,coding_sequence_variant,3_prime_UTR_variant,intron_variant,stop_lost,feature_truncation,coding_sequence_variant,5_prime_UTR_variant,3_prime_UTR_variant,intron_variant |
| 17 | 39387032 | 39388986 | 1954 | Esp23 | PJ | feature_truncation,5_prime_UTR_variant,intron_variant |
| 17 | 40231761 | 40478211 | 246450 | Esp38,Gm49835 | 129S1 | transcript_ablation |
| 17 | 40889325 | 40900179 | 10854 | Esp5 | MOLF | stop_lost,feature_truncation,coding_sequence_variant,3_prime_UTR_variant,intron_variant |
| 17 | 44445972 | 44454889 | 8917 | Clic5 | FVB,SWR | feature_truncation,coding_sequence_variant,intron_variant |
| 17 | 56814103 | 56824003 | 9900 | Znrf4 | SJL | transcript_ablation |
| 17 | 70274708 | 70276324 | 1616 | Dlgap1 | C3H,MRL | feature_truncation,5_prime_UTR_variant |
| 17 | 90215869 | 90217871 | 2002 | Kpna2rt | CAST,MOLF,SPRET | stop_lost,feature_truncation,coding_sequence_variant,5_prime_UTR_variant,3_prime_UTR_variant |
| 18 | 3121554 | 3126392 | 4838 | Vmn1r238 | SPRET | transcript_ablation |
| 18 | 43605703 | 43610924 | 5221 | Eif3j2 | 129S1,AJ,AKR,BTBR,BUB,C57L,CAST,CE,FVB,ILNJ2,KK,MA/My,MOLF,MRL,NOD,NOR,NZB,PJ,PL,RF,RH2,SJL,SPRET,SWR,TH | transcript_ablation |
| 18 | 48179700 | 48181479 | 1779 | Eno1b | 129S1,AJ,AKR,BUB,C57L,C58J,CAST,CE,FVB,ILNJ2,KK,LP,MA/My,MOLF,MRL,NOD,NOR,NZB,NZO,NZW,PL,RF,RH2,SJL,SMJ,SPRET,SWR,TH,WSB | stop_lost,feature_truncation,coding_sequence_variant,5_prime_UTR_variant,3_prime_UTR_variant,intron_variant,transcript_ablation |
| 18 | 49822387 | 49829499 | 7112 | Dtwd2 | C58J,CAST,KK,LP,MOLF,SPRET | feature_truncation,3_prime_UTR_variant |
| 18 | 60388134 | 60525341 | 137207 | Iigp1,Gm4841,F830016B08Rik | CAST | stop_lost,feature_truncation,coding_sequence_variant,5_prime_UTR_variant,3_prime_UTR_variant,intron_variant,transcript_ablation |
| 18 | 60493893 | 60510765 | 16872 | Iigp1 | MOLF,SPRET | feature_truncation,5_prime_UTR_variant,intron_variant |
| 18 | 62794471 | 62796063 | 1592 | Spink10 | SPRET | feature_truncation,3_prime_UTR_variant,transcript_ablation |
| 18 | 87773570 | 87777505 | 3935 | Bhmt1b | KK,TH | transcript_ablation |
| 18 | 87774401 | 87776293 | 1892 | Bhmt1b | CAST,MOLF,SPRET | transcript_ablation |
| 19 | 7496480 | 7497499 | 1019 | Atl3 | CAST,MOLF | stop_lost,feature_truncation,coding_sequence_variant,3_prime_UTR_variant,intron_variant |
| 19 | 8175803 | 8188911 | 13108 | Slc22a29 | SPRET | feature_truncation,coding_sequence_variant,intron_variant |
| 19 | 10490140 | 10492139 | 1999 | Sdhaf2 | SPRET | feature_truncation,3_prime_UTR_variant |
| 19 | 11387108 | 11472656 | 85548 | Ms4a4b,Ms4a4c,Gm8369,Ms4a6c | MOLF | transcript_ablation,stop_lost,feature_truncation,coding_sequence_variant,5_prime_UTR_variant,3_prime_UTR_variant,intron_variant,feature_truncation,5_prime_UTR_variant,intron_variant |
| 19 | 11399525 | 11432010 | 32485 | Ms4a4b,Ms4a4c | WSB | feature_truncation,coding_sequence_variant,5_prime_UTR_variant,intron_variant,stop_lost,feature_truncation,coding_sequence_variant,3_prime_UTR_variant,intron_variant |
| 19 | 11433992 | 11436262 | 2270 | Ms4a4b | KK,LP,TH | feature_truncation,coding_sequence_variant,intron_variant |
| 19 | 11484001 | 11488023 | 4022 | Gm8369 | SPRET | feature_truncation,coding_sequence_variant,intron_variant |

| | | | | | | |
|---|---|---|---|---|---|---|
| 19 | 11570392 | 11577199 | 6807 | Ms4a6d | AJ,BALB,BUB,CAST,CE,FVB,LP,MOLF,NOD,NOR,PJ,PL,RH2,SEA,SPRET,SWR,WSB | feature_truncation,5_prime_UTR_variant,intron_variant |
| 19 | 12033188 | 12048122 | 14934 | Or4d10b | MOLF | transcript_ablation |
| 19 | 12156202 | 12205503 | 49301 | Or5an9,Or5bb10 | AJ,PJ | transcript_ablation,feature_truncation,3_prime_UTR_variant |
| 19 | 12894960 | 12896542 | 1582 | Or5b12 | CAST | feature_truncation,3_prime_UTR_variant |
| 19 | 13115263 | 13122711 | 7448 | Or5b106 | CAST,SPRET | feature_truncation,3_prime_UTR_variant |
| 19 | 20593514 | 20602695 | 9181 | Aldh1a1 | DBA1J | feature_truncation,coding_sequence_variant,intron_variant |
| 19 | 24852973 | 24854244 | 1271 | Gm10053 | AJ,AKR,BALB,BUB,C3H,C58J,CAST,CBA,CE,DBA1J,DBA2J,FVB,ILNJ2,KK,LP,MA/My,MOLF,MRL,PJ,PL,RF,RH2,SEA,SJL,SPRET,SWR,TH,WSB | transcript_ablation |
| 19 | 33489300 | 33492348 | 3048 | Lipo4 | FVB,NZW | feature_truncation,coding_sequence_variant,intron_variant |
| 19 | 33638580 | 33788972 | 150392 | Lipo3,Lipo2,Lipo1 | WSB | feature_truncation,5_prime_UTR_variant,intron_variant,transcript_ablation |
| 19 | 41044790 | 41047470 | 2680 | Dntt | CAST | feature_truncation,coding_sequence_variant,intron_variant |
| 19 | 52973976 | 52979378 | 5402 | Xpnpep1 | 129S1,AJ,AKR,BALB,BTBR,BUB,C3H,C57L,CAST,CBA,CE,DBA1J,DBA2J,ILNJ2,KK,MA/My,MOLF,MRL,NOD,NOR,NZB,NZO,NZW,PJ,PL,RF,RH2,SEA,SMJ,SPRET,TH,WSB | feature_truncation,3_prime_UTR_variant |
| 19 | 61157208 | 61163676 | 6468 | Mplkipl1 | 129S1,AJ,AKR,BALB,BTBR,C3H,C57L,CAST,CBA,DBA1J,DBA2J,FVB,KK,MA/My,MRL,NOD,NOR,NZB,NZW,PJ,PL,RF,SEA,SJL,SMJ,SPRET,SWR,TH,WSB | feature_truncation,3_prime_UTR_variant |
| 2 | 3397298 | 3401214 | 3916 | Olah | MA/My,NZO,NZW | feature_truncation,5_prime_UTR_variant,intron_variant |
| 2 | 11652968 | 11654570 | 1602 | Il2ra | SJL,WSB | feature_truncation,3_prime_UTR_variant |
| 2 | 22011295 | 22235429 | 224134 | Myo3a | WSB | feature_truncation,5_prime_UTR_variant,intron_variant |
| 2 | 24083290 | 24089025 | 5735 | Il36g | AJ,AKR,CAST,DBA2J,FVB,PL,RH2,SJL,SPRET,WSB | feature_truncation,3_prime_UTR_variant |
| 2 | 26888304 | 26894330 | 6026 | Adamts13 | 129S1,BUB,C57L,CAST,FVB,MA/My,MOLF,NOD,NOR,RF,SMJ,SPRET,WSB | stop_lost,feature_truncation,coding_sequence_variant,3_prime_UTR_variant |
| 2 | 36561265 | 36577970 | 16705 | Or1j17 | SPRET | feature_truncation,5_prime_UTR_variant,intron_variant |
| 2 | 36884034 | 36885693 | 1659 | Or1q1 | CAST | feature_truncation,5_prime_UTR_variant,intron_variant |
| 2 | 49819282 | 49823487 | 4205 | Lypd6b | C57L | feature_truncation,5_prime_UTR_variant,intron_variant |
| 2 | 58014395 | 58019662 | 5267 | Cytip | SPRET | feature_truncation,3_prime_UTR_variant |
| 2 | 83789414 | 83818463 | 29049 | Cwc22rt7,Cwc22rt3,Cwc22rt5,Cwc22rt2,Cwc22rt4 | MOLF | transcript_ablation |
| 2 | 83789414 | 83818979 | 29565 | Cwc22rt7,Cwc22rt3,Cwc22rt5,Cwc22rt2,Cwc22rt4 | MOLF | transcript_ablation |
| 2 | 85183702 | 85201516 | 17814 | Or5ak20 | C58J,NZB,SJL,SWR | feature_truncation,coding_sequence_variant,5_prime_UTR_variant,feature_truncation,coding_sequence_variant,5_prime_UTR_variant,intron_variant |
| 2 | 85507681 | 85508744 | 1063 | Or9g4 | SJL,SWR | feature_truncation,5_prime_UTR_variant,intron_variant |
| 2 | 85848706 | 85853929 | 5223 | Or5m3b | AJ,AKR,BALB,BUB,C3H,C57L,CAST,CBA,CE,DBA1J,DBA2J,ILNJ2,KK,MA/My,MOLF,MRL,NOD,NOR,NZB,NZO,NZW,PJ,PL,RF,RH2,SEA,SMJ,SPRET | feature_truncation,5_prime_UTR_variant,intron_variant |
| 2 | 85880318 | 85881607 | 1289 | Or5m9 | CAST | feature_truncation,3_prime_UTR_variant |
| 2 | 86085857 | 86103647 | 17790 | Or8k18 | CAST | feature_truncation,coding_sequence_variant,5_prime_UTR_variant,feature_truncation,coding_sequence_variant,5_prime_UTR_variant,intron_variant |
| 2 | 86170085 | 86177170 | 7085 | Or8k53 | AJ,AKR,BALB,BUB,C3H,C57L,CBA,CE,DBA1J,DBA2J,FVB,KK,MA/My,MOLF,MRL,NOD,NOR,NZB,NZO,NZW,PJ,PL,RF,RH2,SEA,SMJ,SPRET,WSB | stop_lost,feature_truncation,coding_sequence_variant,3_prime_UTR_variant |
| 2 | 86217778 | 86245506 | 27728 | Or8k25,Or8k24 | FVB,WSB | transcript_ablation,feature_truncation,5_prime_UTR_variant,intron_variant |
| 2 | 86424224 | 86446693 | 22469 | Or8k35 | MOLF | stop_lost,feature_truncation,coding_sequence_variant,5_prime_UTR_variant,3_prime_UTR_variant,intron_variant |
| 2 | 86527800 | 86529205 | 1405 | Or8k3b | CAST | feature_truncation,5_prime_UTR_variant,intron_variant |
| 2 | 86561921 | 86563941 | 2020 | Or8k39 | SPRET | stop_lost,feature_truncation,coding_sequence_variant,3_prime_UTR_variant |
| 2 | 87199813 | 87201708 | 1895 | Or12e9 | CAST | feature_truncation,5_prime_UTR_variant,intron_variant |
| 2 | 87678884 | 87680577 | 1693 | Or12e14 | CAST | feature_truncation,3_prime_UTR_variant |
| 2 | 87702900 | 87705569 | 2669 | Or5w19 | MOLF,SPRET | feature_truncation,3_prime_UTR_variant |
| 2 | 88054907 | 88057040 | 2133 | Or5d41 | MOLF | feature_truncation,coding_sequence_variant,5_prime_UTR_variant,feature_truncation,coding_sequence_variant,5_prime_UTR_variant,intron_variant |
| 2 | 88147629 | 88152624 | 4995 | Or5d45 | SPRET | feature_truncation,3_prime_UTR_variant |
| 2 | 88533662 | 88534890 | 1228 | Or4a66 | SPRET | feature_truncation,5_prime_UTR_variant,intron_variant |
| 2 | 89484610 | 89485625 | 1015 | Or4a77 | SPRET | feature_truncation,3_prime_UTR_variant |
| 2 | 89997115 | 90079547 | 82432 | Or4x15 | BUB,C58J,ILNJ2,KK,MOLF,PL,SJL,SWR,WSB | transcript_ablation |
| 2 | 90003559 | 90073065 | 69506 | Or4x15 | 129S1,AJ,AKR,BALB,BTBR,C3H,C57L,CBA,CE,DBA1J,DBA2J,FVB,NOD,NOR,NZO,PJ,RH2 | transcript_ablation |
| 2 | 110553226 | 110554358 | 1132 | Muc15 | SPRET | feature_truncation,5_prime_UTR_variant,intron_variant |
| 2 | 111718985 | 111758442 | 39457 | Or4f14 | MOLF | transcript_ablation |
| 2 | 111946032 | 111958803 | 12771 | Or4f14d | SPRET | feature_truncation,3_prime_UTR_variant |
| 2 | 122254711 | 122258831 | 4120 | Slc28a2 | 129S1,AJ,AKR,BALB,BTBR,BUB,C3H,C57L,C58J,CBA,CE,DBA1J,DBA2J,FVB,ILNJ2,MA/My,NOD,NZB,NZO,NZW,PJ,PL,RF,RH2,SEA,SJL,SMJ,SWR,TH,WSB | feature_truncation,5_prime_UTR_variant,intron_variant |
| 2 | 122294058 | 122347241 | 53183 | Slc28a2b | SPRET | feature_truncation,coding_sequence_variant,5_prime_UTR_variant,intron_variant |
| 2 | 122350748 | 122352139 | 1391 | Slc28a2b | 129S1,AJ,AKR,BALB,BTBR,BUB,C3H,C57L,C58J,CAST,CBA,CE,DBA1J,DBA2J,FVB,ILNJ2,MA/My,NOD,NZB,NZO,NZW,PJ,PL,RF,RH2,SEA,SJL,SMJ,SWR,TH,WSB | feature_truncation,coding_sequence_variant,intron_variant |
| 2 | 130249325 | 130250574 | 1249 | Tmem239 | BALB,CAST,DBA1J,MOLF,PL,SEA | feature_truncation,3_prime_UTR_variant |

| | | | | | | |
|---|---|---|---|---|---|---|
| 2 | 148844283 | 148845359 | 1076 | Gm1330 | MOLF | feature_truncation,coding_sequence_variant,5_prime_UTR_variant,intron_variant |
| 2 | 150926160 | 150938776 | 12616 | Gm14151 | CAST | transcript_ablation |
| 2 | 150936711 | 150944533 | 7822 | Gm14147 | NOD,NZB,NZW,RF | feature_truncation,coding_sequence_variant,5_prime_UTR_variant,intron_variant |
| 2 | 156198472 | 156205526 | 7054 | Cnbd2 | 129S1,BALB,C3H,C57L,CAST,CBA,CE,FVB,MA/My,NOD,NZO,NZW,PL,SEA,SJL,SMJ,SPRET,SWR,TH,WSB | stop_lost,feature_truncation,coding_sequence_variant,3_prime_UTR_variant |
| 2 | 164592726 | 164595428 | 2702 | Dnttip1 | SPRET | stop_lost,feature_truncation,coding_sequence_variant,3_prime_UTR_variant,intron_variant |
| 2 | 170194922 | 170205925 | 11003 | Bcas1 | WSB | feature_truncation,coding_sequence_variant,intron_variant |
| 2 | 181645559 | 181649985 | 4426 | Gm14496 | BUB,CAST,CE,MOLF,PJ,RF,SPRET,WSB | feature_truncation,3_prime_UTR_variant |
| 3 | 10151563 | 10154671 | 3108 | Myef2l | 129S1,AJ,AKR,BALB,BTBR,C3H,C57L,C58J,CBA,DBA1J,FVB,ILNJ2,KK,LP,MRL,NZB,NZO,NZW,PJ,PL,RF,SEA,SJL,SWR | feature_truncation,coding_sequence_variant,5_prime_UTR_variant |
| 3 | 10153155 | 10155286 | 2131 | Myef2l | CAST,MOLF,SPRET | feature_truncation,coding_sequence_variant,5_prime_UTR_variant,intron_variant |
| 3 | 15567466 | 15570916 | 3450 | Sirpb1b | AJ,BALB,BUB,C3H,CBA,FVB,PL,SJL,SWR,TH | stop_lost,feature_truncation,coding_sequence_variant,3_prime_UTR_variant,intron_variant |
| 3 | 15882313 | 15885531 | 3218 | Sirpb1c | CE,DBA2J,LP | feature_truncation,3_prime_UTR_variant |
| 3 | 51147383 | 51152723 | 5340 | Noct | 129S1,BTBR,BUB,C3H,C57L,C58J,CAST,CBA,CE,FVB,LP,MOLF,NZB,NZW,SJL,SMJ,SPRET,SWR,TH,WSB | feature_truncation,5_prime_UTR_variant |
| 3 | 59001528 | 59002913 | 1385 | Med12l | SPRET | feature_truncation,3_prime_UTR_variant |
| 3 | 87374725 | 87386123 | 11398 | Fcrl5 | SPRET | feature_truncation,3_prime_UTR_variant |
| 3 | 88280618 | 88286324 | 5706 | Bglap2 | RF,SPRET | transcript_ablation |
| 3 | 90683058 | 90684925 | 1867 | Gm5849 | SPRET | stop_lost,feature_truncation,coding_sequence_variant,3_prime_UTR_variant,intron_variant |
| 3 | 91935247 | 91938771 | 3524 | Pglyrp3 | NOD,NOR | feature_truncation,coding_sequence_variant,intron_variant |
| 3 | 92114662 | 92131332 | 16670 | Sprr2a1,Sprr2a2 | MOLF | transcript_ablation,feature_truncation,5_prime_UTR_variant,intron_variant |
| 3 | 92335325 | 92336731 | 1406 | Adrm1b | CAST,MOLF | transcript_ablation |
| 3 | 92644976 | 92647207 | 2231 | Gm38119 | SPRET | stop_lost,feature_truncation,coding_sequence_variant,5_prime_UTR_variant,3_prime_UTR_variant,intron_variant |
| 3 | 93111394 | 93113038 | 1644 | Flg2 | AJ,BALB,C3H,CBA,MA/My,NZB,NZO,RH2 | feature_truncation,inframe_deletion |
| 3 | 93123742 | 93125146 | 1404 | Flg2 | BALB,MA/My,NZB,NZO,RH2 | feature_truncation,inframe_deletion |
| 3 | 93124241 | 93125872 | 1631 | Flg2 | KK | frameshift_variant,feature_truncation |
| 3 | 93187612 | 93189850 | 2238 | Flg | C58J,LP,RF | transcript_ablation,stop_lost,feature_truncation,coding_sequence_variant,3_prime_UTR_variant |
| 3 | 93196332 | 93199355 | 3023 | Flg | CBA | transcript_ablation |
| 3 | 93234278 | 93235321 | 1043 | Hrnr | ILNJ2,PJ,PL,SEA,TH | frameshift_variant,feature_truncation |
| 3 | 98616561 | 98617634 | 1073 | Hsd3b2 | SPRET | feature_truncation,3_prime_UTR_variant |
| 3 | 98766517 | 98767986 | 1469 | Hsd3b1 | SPRET | feature_truncation,5_prime_UTR_variant,intron_variant |
| 3 | 103551238 | 103553386 | 2148 | Atg4a-ps,Syt6 | 129S1,AKR,BUB,C57L,C58J,CAST,CE,FVB,ILNJ2,KK,LP,MOLF,PJ,RF,SEA,SJL,SPRET,SWR,TH | transcript_ablation,stop_lost,feature_truncation,coding_sequence_variant,3_prime_UTR_variant,intron_variant |
| 3 | 106722594 | 106728874 | 6280 | Or11i1 | MOLF | feature_truncation,3_prime_UTR_variant |
| 3 | 116488402 | 116489699 | 1297 | Slc35a3 | CAST,MOLF | feature_truncation,5_prime_UTR_variant,intron_variant |
| 3 | 122064075 | 122066511 | 2436 | Gclm | MOLF | feature_truncation,3_prime_UTR_variant |
| 3 | 131273078 | 131274456 | 1378 | Papss1 | AKR,BUB,C3H,C58J,CAST,CE,DBA1J,DBA2J,FVB,KK,LP,MA/My,MOLF,NOD,NOR,NZB,NZO,NZW,PJ,PL,RF,SEA,SJL,SMJ,SPRET,SWR,TH,WSB | feature_truncation,5_prime_UTR_variant,intron_variant |
| 3 | 152028773 | 152037820 | 9047 | Miga1 | 129S1,AJ,BALB,BTBR,C3H,C57L,CAST,CBA,CE,DBA1J,ILNJ2,KK,MOLF,MRL,PL,RH2,SEA,SMJ,SPRET,WSB | feature_truncation,3_prime_UTR_variant |
| 3 | 157351195 | 157353621 | 2426 | Ptger3 | RF | feature_truncation,3_prime_UTR_variant |
| 4 | 3994609 | 3997118 | 2509 | Sdr16c5 | 129S1,BALB,BTBR,C3H,C57L,CBA,CE,DBA1J,DBA2J,MRL,NZB,NZO,PL,RH2,SEA | stop_lost,feature_truncation,coding_sequence_variant,3_prime_UTR_variant,intron_variant |
| 4 | 44033284 | 44034459 | 1175 | Gne | MOLF | feature_truncation,3_prime_UTR_variant |
| 4 | 46631855 | 46632926 | 1071 | Tbc1d2 | SPRET | feature_truncation,3_prime_UTR_variant |
| 4 | 60218186 | 60221626 | 3440 | Mup8 | 129S1,AJ,AKR,BALB,C3H,C57L,CBA,CE,DBA1J,DBA2J,FVB,ILNJ2,MA/My,MRL,NOD,NOR,NZB,NZO,NZW,RF,RH2,SEA,SJL,SWR | stop_lost,feature_truncation,coding_sequence_variant,3_prime_UTR_variant,intron_variant |
| 4 | 61433395 | 61436814 | 3419 | Mup16 | 129S1,BALB,C3H,C57L,CBA,DBA1J,DBA2J,FVB,MA/My,NOD,NOR,NZO,NZW,RF,RH2,SEA,SJL,SWR | stop_lost,feature_truncation,coding_sequence_variant,3_prime_UTR_variant,intron_variant |
| 4 | 61999946 | 62061498 | 61552 | Mup3 | SPRET | transcript_ablation |
| 4 | 62002432 | 62005284 | 2852 | Mup3 | MOLF | stop_lost,feature_truncation,coding_sequence_variant,3_prime_UTR_variant,intron_variant |
| 4 | 62065813 | 62068682 | 2869 | Mup21 | SPRET | stop_lost,feature_truncation,coding_sequence_variant,3_prime_UTR_variant,intron_variant |
| 4 | 73522675 | 73527750 | 5075 | Gm38423 | CAST | feature_truncation,coding_sequence_variant,5_prime_UTR_variant,intron_variant |
| 4 | 73791545 | 73793389 | 1844 | Msantd5f7 | MOLF | feature_truncation,coding_sequence_variant,intron_variant |
| 4 | 88708973 | 88711909 | 2936 | Gm13278 | AJ | transcript_ablation |
| 4 | 92078941 | 92080082 | 1141 | Larp7-ps | CAST,SPRET | transcript_ablation |
| 4 | 99717713 | 99719244 | 1531 | Efcab7 | MOLF | feature_truncation,5_prime_UTR_variant,intron_variant |
| 4 | 101832107 | 101838549 | 6442 | Gm12790 | AJ,AKR,BALB,BUB,C3H,C58J,CAST,CBA,CE,KK,LP,MA/My,MOLF,MRL,NZB,NZO,NZW,PJ,PL,RF,SEA,SJL,SMJ,SPRET,WSB | feature_truncation,5_prime_UTR_variant,intron_variant |
| 4 | 106588722 | 106599282 | 10560 | Fam151a | PL | feature_truncation,coding_sequence_variant,5_prime_UTR_variant,intron_variant |
| 4 | 111730790 | 111734325 | 3535 | Slc5a9 | SPRET | feature_truncation,3_prime_UTR_variant |
| 4 | 112401341 | 112468388 | 67047 | Skint2 | AKR,SMJ,SWR | feature_truncation,5_prime_UTR_variant,intron_variant |
| 4 | 112998167 | 113069759 | 71592 | Skint6 | LP | feature_truncation,coding_sequence_variant,intron_variant |
| 4 | 113016893 | 113031363 | 14470 | Skint6 | BALB,BUB,NOR,SEA,SJL,TH,WSB | feature_truncation,coding_sequence_variant,intron_variant |
| 4 | 113364578 | 113392236 | 27658 | Skint5 | AJ | feature_truncation,coding_sequence_variant,intron_variant |

| | | | | | | |
|---|---|---|---|---|---|---|
| 4 | 113403578 | 113410796 | 7218 | Skint5 | AKR,C57L,CAST,DBA1J,DBA2J,LP,MOLF,MRL,TH,WSB | feature_truncation,coding_sequence_variant,intron_variant |
| 4 | 113704823 | 113711685 | 6862 | Skint5 | 129S1,AJ,AKR,BTBR,C57L,DBA1J,DBA2J,FVB,LP,MOLF,MRL,SPRET,SWR,WSB | feature_truncation,coding_sequence_variant,intron_variant |
| 4 | 121164950 | 121217435 | 52485 | Gm12888 | MOLF | transcript_ablation |
| 4 | 121476181 | 121481378 | 5197 | Gm12887 | MOLF | feature_truncation,coding_sequence_variant,5_prime_UTR_variant,intron_variant |
| 4 | 138163332 | 138164802 | 1470 | Mul1 | SPRET | feature_truncation,coding_sequence_variant,5_prime_UTR_variant,intron_variant |
| 4 | 138600623 | 138602103 | 1480 | Gm13030 | CAST | feature_truncation,coding_sequence_variant,5_prime_UTR_variant,intron_variant |
| 4 | 141161788 | 141163788 | 2000 | Srarp | TH | feature_truncation,coding_sequence_variant,5_prime_UTR_variant,intron_variant |
| 4 | 143253176 | 143255472 | 2296 | Gm13040 | CAST | feature_truncation,5_prime_UTR_variant,intron_variant |
| 4 | 143293580 | 143295504 | 1924 | Pramel20 | CAST | feature_truncation,5_prime_UTR_variant,intron_variant |
| 4 | 143574797 | 143588157 | 13360 | Pramel27 | BUB,CE,DBA2J,KK,MA/My,NZB,NZW,PL,RF,SMJ,SWR,TH | stop_lost,feature_truncation,coding_sequence_variant,3_prime_UTR_variant,intron_variant |
| 4 | 143668501 | 143683902 | 15401 | Pramel16 | CE,SWR | transcript_ablation |
| 4 | 143881566 | 143898718 | 17152 | Oog3 | AKR,C3H,CBA | transcript_ablation |
| 4 | 146926154 | 146976189 | 50035 | Gm21411 | C57L | stop_lost,feature_truncation,coding_sequence_variant,3_prime_UTR_variant |
| 4 | 147954746 | 147956088 | 1342 | Fv1 | 129S1,AKR,BTBR,BUB,C3H,C57L,C58J,CBA,CE,DBA1J,DBA2J,KK,LP,MA/My,MRL,NOD,NZB,NZO,NZW,PJ,PL,RF,SEA,SJL,SWR,TH,WSB | stop_lost,feature_truncation,coding_sequence_variant,3_prime_UTR_variant |
| 4 | 154982644 | 154984700 | 2056 | Prxl2b | CAST | feature_truncation,coding_sequence_variant,5_prime_UTR_variant,intron_variant |
| 5 | 7358132 | 7370164 | 12032 | Tex47 | SPRET | feature_truncation,3_prime_UTR_variant,intron_variant |
| 5 | 15857413 | 15862680 | 5267 | 4930572O03Rik | MA/My | stop_lost,feature_truncation,coding_sequence_variant,5_prime_UTR_variant,3_prime_UTR_variant,intron_variant |
| 5 | 26141225 | 26153250 | 12025 | Gm21663 | AJ,AKR,B10D2,B10J,BALB,C3H,CBA,DBA1J,DBA2J,FVB,ILNJ2,MA/My,MRL,NZB,NZO,PJ,RF,SJL,TH | stop_lost,feature_truncation,coding_sequence_variant,5_prime_UTR_variant,3_prime_UTR_variant,intron_variant |
| 5 | 26146974 | 26159011 | 12037 | Gm21671,Gm21663 | BUB,C58J,LP,NOR | stop_lost,feature_truncation,coding_sequence_variant,3_prime_UTR_variant,intron_variant,feature_truncation,coding_sequence_variant,5_prime_UTR_variant,intron_variant |
| 5 | 26148326 | 26153283 | 4957 | Gm21671 | 129S1,BTBR,CE,NZW | feature_truncation,3_prime_UTR_variant |
| 5 | 26153438 | 26154461 | 1023 | Gm21671 | BTBR | feature_truncation,3_prime_UTR_variant |
| 5 | 26166507 | 26182606 | 16099 | Gm21680 | MOLF | transcript_ablation |
| 5 | 26311481 | 26327539 | 16058 | Gm10220 | BTBR,NZW | transcript_ablation |
| 5 | 30831838 | 30832870 | 1032 | Cenpa | CAST | feature_truncation,3_prime_UTR_variant |
| 5 | 44069651 | 44071575 | 1924 | Cd38,Ppihl | CAST,MOLF,PJ,PL | feature_truncation,3_prime_UTR_variant,transcript_ablation |
| 5 | 82268850 | 82270264 | 1414 | 1700031L13Rik | SPRET | feature_truncation,3_prime_UTR_variant |
| 5 | 87569388 | 87573834 | 4446 | Ugt2b38 | 129S1,C57L,SMJ | feature_truncation,coding_sequence_variant,5_prime_UTR_variant,intron_variant |
| 5 | 92707354 | 92711152 | 3798 | Fam47e | DBA2J | feature_truncation,coding_sequence_variant,intron_variant |
| 5 | 93955360 | 94004439 | 49079 | Pramel35 | MOLF | transcript_ablation |
| 5 | 104725128 | 104736051 | 10923 | Zfp33b | MOLF | stop_lost,feature_truncation,coding_sequence_variant,3_prime_UTR_variant,intron_variant,stop_lost,feature_truncation,coding_sequence_variant,5_prime_UTR_variant,3_prime_UTR_variant,intron_variant |
| 5 | 108953013 | 108960905 | 7892 | Vmn2r8 | SPRET | feature_truncation,coding_sequence_variant,5_prime_UTR_variant,intron_variant |
| 5 | 109293847 | 109300849 | 7002 | Vmn2r13 | 129S1,AKR,BUB,C3H,C57L,CAST,CBA,FVB,KK,MA/My,MOLF,NOD,NOR,NZB,NZO,NZW,PJ,PL,RF,SJL,SPRET,SWR,TH,WSB | feature_truncation,3_prime_UTR_variant |
| 5 | 109320509 | 109334167 | 13658 | Vmn2r13 | CAST | feature_truncation,coding_sequence_variant,intron_variant |
| 5 | 109427385 | 109433600 | 6215 | Vmn2r15 | SPRET | feature_truncation,3_prime_UTR_variant |
| 5 | 109516405 | 109518566 | 2161 | Vmn2r16 | CAST | feature_truncation,3_prime_UTR_variant |
| 5 | 110081353 | 110085226 | 3873 | Gm15446 | SPRET | feature_truncation,5_prime_UTR_variant,intron_variant |
| 5 | 110100371 | 110171878 | 71507 | Zfp932 | SPRET | transcript_ablation |
| 5 | 120661799 | 120665366 | 3567 | Slc8b1 | CAST | feature_truncation,coding_sequence_variant,intron_variant |
| 5 | 120915788 | 120977701 | 61913 | Oas1e,Oas1c | SPRET | transcript_ablation |
| 5 | 121347219 | 121350324 | 3105 | Rpl6 | SPRET | feature_truncation,3_prime_UTR_variant |
| 5 | 137179893 | 137181004 | 1111 | Muc17 | SPRET | frameshift_variant,feature_truncation |
| 5 | 137184350 | 137186063 | 1713 | Muc17 | MOLF | feature_truncation,inframe_deletion |
| 5 | 137836131 | 137869140 | 33009 | Pilrb1,Pilrb2 | BUB,RF | transcript_ablation,stop_lost,feature_truncation,coding_sequence_variant,3_prime_UTR_variant,intron_variant |
| 5 | 145280983 | 145285149 | 4166 | Cyp3a57 | MOLF,TH | feature_truncation,coding_sequence_variant,5_prime_UTR_variant,intron_variant |
| 5 | 146101171 | 146118623 | 17452 | Gm6309 | SPRET | transcript_ablation |
| 5 | 146418233 | 146427634 | 9401 | Gm6408 | CAST,TH | transcript_ablation |
| 5 | 146424095 | 146432919 | 8824 | Gm6370 | MOLF | transcript_ablation |
| 5 | 146430450 | 146441487 | 11037 | 4930449I24Rik,Gm6370 | WSB | feature_truncation,coding_sequence_variant,5_prime_UTR_variant,intron_variant,stop_lost,feature_truncation,coding_sequence_variant,3_prime_UTR_variant |
| 5 | 146773218 | 146774398 | 1180 | Rpl21 | SPRET | feature_truncation,3_prime_UTR_variant |
| 6 | 3388053 | 3420796 | 32743 | Samd9l | SEA | feature_truncation,5_prime_UTR_variant,intron_variant |
| 6 | 8401090 | 8405284 | 4194 | Umad1 | WSB | feature_truncation,coding_sequence_variant,intron_variant |
| 6 | 35239775 | 35241223 | 1448 | Stmp1 | SPRET | feature_truncation,3_prime_UTR_variant |
| 6 | 40957543 | 40977611 | 20068 | Gm2663 | SPRET | transcript_ablation |
| 6 | 42875807 | 42877290 | 1483 | Or2a5 | CAST | feature_truncation,3_prime_UTR_variant |
| 6 | 43079902 | 43083978 | 4076 | Or2a54 | MOLF | feature_truncation,5_prime_UTR_variant,intron_variant |
| 6 | 48740281 | 48741701 | 1420 | Gimap3 | SPRET | feature_truncation,3_prime_UTR_variant |
| 6 | 57735110 | 57736650 | 1540 | Vopp1 | SPRET | feature_truncation,3_prime_UTR_variant |
| 6 | 65093219 | 65094389 | 1170 | Hpgds | KK | feature_truncation,3_prime_UTR_variant |
| 6 | 66659059 | 66660208 | 1149 | Vmn1r35 | SPRET | feature_truncation,5_prime_UTR_variant,intron_variant |

| | | | | | | |
|---|---|---|---|---|---|---|
| 6 | 71287151 | 71289038 | 1887 | Smyd1 | AKR,C3H,C58J,CAST,CBA,DBA1J,DBA2J,KK,MA/My,MOLF,MRL,NOD,NOR,PL,RF,SMJ,SPRET,WSB | feature_truncation,3_prime_UTR_variant |
| 6 | 76469330 | 76476656 | 7326 | Rnf26rt | KK,MOLF,PJ | transcript_ablation |
| 6 | 76472403 | 76475091 | 2688 | Rnf26rt | CAST,SPRET | transcript_ablation |
| 6 | 78397181 | 78399717 | 2536 | Reg1 | SPRET | feature_truncation,5_prime_UTR_variant |
| 6 | 87992471 | 87994463 | 1992 | Rab7 | SPRET | feature_truncation,5_prime_UTR_variant,intron_variant |
| 6 | 102549922 | 102557101 | 7179 | Cntn3 | BUB,C58J,CAST,LP,MOLF,NOD,NOR,NZB,NZO,NZW,SPRET,SWR,WSB | feature_truncation,5_prime_UTR_variant,intron_variant |
| 6 | 112335959 | 112337523 | 1564 | Ssu2 | SWR | stop_lost,feature_truncation,coding_sequence_variant,3_prime_UTR_variant,intron_variant |
| 6 | 121761447 | 121762502 | 1055 | Gm7298 | SPRET | feature_truncation,coding_sequence_variant,intron_variant |
| 6 | 123044870 | 123046268 | 1398 | Clec4b1 | PJ,SMJ,SWR | feature_truncation,coding_sequence_variant,intron_variant |
| 6 | 123487697 | 123489253 | 1556 | Vmn2r21 | SPRET,WSB | feature_truncation,coding_sequence_variant,intron_variant |
| 6 | 128855784 | 128857791 | 2007 | BC035044 | CAST | stop_lost,feature_truncation,coding_sequence_variant,3_prime_UTR_variant,intron_variant |
| 6 | 129574505 | 129576889 | 2384 | Klrd1 | DBA2J | stop_lost,feature_truncation,coding_sequence_variant,3_prime_UTR_variant,intron_variant |
| 6 | 129716471 | 129718157 | 1686 | Klri2 | 129S1,SMJ | feature_truncation,coding_sequence_variant,5_prime_UTR_variant,intron_variant |
| 6 | 129738259 | 129744140 | 5881 | Klrh1 | 129S1,C58J,KK,NOD,NOR,NZB,NZW,SMJ,SWR | stop_lost,feature_truncation,coding_sequence_variant,3_prime_UTR_variant,intron_variant,feature_truncation,3_prime_UTR_variant |
| 6 | 132221716 | 132368953 | 147237 | Prb1b,Prb1c | NZO | transcript_ablation |
| 6 | 142103592 | 142107519 | 3927 | Slco1a6 | PJ | feature_truncation,coding_sequence_variant,5_prime_UTR_variant,intron_variant |
| 7 | 5223245 | 5231238 | 7993 | Vmn1r57 | SPRET | transcript_ablation,stop_lost,feature_truncation,coding_sequence_variant,5_prime_UTR_variant,3_prime_UTR_variant |
| 7 | 5224666 | 5230251 | 5585 | Vmn1r57 | BUB,CAST,CE,MOLF,MRL,NZO,SMJ,SWR,TH,WSB | feature_truncation,3_prime_UTR_variant |
| 7 | 5523289 | 5552569 | 29280 | Vmn1r60 | BUB,NZO,SWR,TH,WSB | transcript_ablation |
| 7 | 6358780 | 6359953 | 1173 | Zfp582 | CAST,MOLF | feature_truncation,3_prime_UTR_variant |
| 7 | 6480823 | 6482863 | 2040 | Or6z5,Or6z7 | SPRET | feature_truncation,3_prime_UTR_variant |
| 7 | 9028486 | 9033280 | 4794 | Vmn2r39 | 129S1,BTBR | feature_truncation,coding_sequence_variant,intron_variant |
| 7 | 9884418 | 9891406 | 6988 | Vmn2r52 | CAST,MOLF | feature_truncation,3_prime_UTR_variant |
| 7 | 10368463 | 10369798 | 1335 | Vmn1r70 | SPRET | feature_truncation,3_prime_UTR_variant |
| 7 | 10788201 | 10886187 | 97986 | Zscan4-ps1 | FVB,MOLF,SPRET | transcript_ablation |
| 7 | 10800159 | 10801882 | 1723 | Zscan4-ps1 | MOLF | feature_truncation,coding_sequence_variant,intron_variant |
| 7 | 11611459 | 11612691 | 1232 | Vmn1r75 | AKR,FVB,MRL,NOD,NOR,RH2,WSB | feature_truncation,5_prime_UTR_variant,intron_variant |
| 7 | 11663538 | 11667578 | 4040 | Vmn1r76 | SWR | transcript_ablation,stop_lost,feature_truncation,coding_sequence_variant,5_prime_UTR_variant,3_prime_UTR_variant,intron_variant |
| 7 | 11776411 | 11778634 | 2223 | Vmn1r77 | SPRET | feature_truncation,3_prime_UTR_variant |
| 7 | 11887755 | 11892686 | 4931 | Vmn1r78 | MOLF | feature_truncation,3_prime_UTR_variant |
| 7 | 12719071 | 12720074 | 1003 | Zfp446 | SPRET | feature_truncation,3_prime_UTR_variant |
| 7 | 12837099 | 12838577 | 1478 | Vmn1r86 | NZO | feature_truncation,5_prime_UTR_variant,intron_variant |
| 7 | 13628298 | 13658591 | 30293 | Sult2a4 | MOLF | stop_lost,feature_truncation,coding_sequence_variant,3_prime_UTR_variant,intron_variant |
| 7 | 14289437 | 14290939 | 1502 | Vmn1r90 | SPRET | feature_truncation,3_prime_UTR_variant |
| 7 | 18191252 | 18230102 | 38850 | Psg26 | SPRET | transcript_ablation |
| 7 | 18323150 | 18369273 | 46123 | Psg23 | SPRET | transcript_ablation |
| 7 | 19156123 | 19157256 | 1133 | Ckm | SPRET | feature_truncation,3_prime_UTR_variant |
| 7 | 21086262 | 21098918 | 12656 | Vmn1r129 | ILNJ2,MRL,PL,WSB | transcript_ablation |
| 7 | 23194111 | 23203075 | 8964 | Vmn1r167 | 129S1,BTBR,BUB,C3H,C57L,CAST,CBA,MOLF,NOD,NOR,NZB,NZW,SJL,SPRET | feature_truncation,3_prime_UTR_variant |
| 7 | 24277585 | 24283556 | 5971 | Zfp575 | BUB,C3H,CAST,CBA,CE,MOLF,NOD,NOR,SJL,SPRET | feature_truncation,3_prime_UTR_variant |
| 7 | 25622438 | 25624849 | 2411 | Cyp2b10 | SPRET | feature_truncation,coding_sequence_variant,intron_variant |
| 7 | 25688012 | 25897736 | 209724 | Cyp2b13,Cyp2b9 | 129S1,BTBR,C57L | transcript_ablation,feature_truncation,coding_sequence_variant,5_prime_UTR_variant,intron_variant |
| 7 | 25908859 | 25911425 | 2566 | Cyp2b9 | MOLF | stop_lost,feature_truncation,coding_sequence_variant,3_prime_UTR_variant,intron_variant |
| 7 | 25967641 | 25968694 | 1053 | Vmn1r184 | MOLF | feature_truncation,3_prime_UTR_variant |
| 7 | 27387653 | 27389195 | 1542 | Zfp607b | KK,NZO | feature_truncation,5_prime_UTR_variant,intron_variant |
| 7 | 29532474 | 29535082 | 2608 | Zfp940 | SPRET | stop_lost,feature_truncation,coding_sequence_variant,3_prime_UTR_variant |
| 7 | 42375688 | 42379517 | 3829 | Gm17067 | SPRET | feature_truncation,coding_sequence_variant,5_prime_UTR_variant,intron_variant |
| 7 | 42537169 | 42547477 | 10308 | Vmn2r63 | DBA1J,DBA2J,ILNJ2,NZW,PJ | feature_truncation,3_prime_UTR_variant |
| 7 | 46835720 | 46838404 | 2684 | Mrgpra6 | C57L,CAST,CE,ILNJ2,MOLF,NZW,PJ,RF,SJL,SPRET,WSB | feature_truncation,coding_sequence_variant,intron_variant |
| 7 | 47053331 | 47098567 | 45236 | Mrgpra2a | SPRET | stop_lost,feature_truncation,coding_sequence_variant,3_prime_UTR_variant,intron_variant |
| 7 | 47063585 | 47103412 | 39827 | Mrgpra2a | C57L,FVB,NOR | transcript_ablation |
| 7 | 47178066 | 47179399 | 1333 | Mrgpra2b | SPRET | feature_truncation,5_prime_UTR_variant,intron_variant |
| 7 | 47215078 | 47243884 | 28806 | Mrgpra3 | CE,ILNJ2,NZW,PJ | stop_lost,feature_truncation,coding_sequence_variant,3_prime_UTR_variant,intron_variant |
| 7 | 47215085 | 47337115 | 122030 | Mrgpra3 | WSB | transcript_ablation |
| 7 | 64515547 | 64517288 | 1741 | Nsmce3 | SPRET | feature_truncation,3_prime_UTR_variant |
| 7 | 79993836 | 79995220 | 1384 | Hddc3 | 129S1,AJ,AKR,BALB,BTBR,BUB,C3H,C57L,CAST,CBA,CE,DBA1J,DBA2J,FVB,ILNJ2,KK,MA/My,MOLF,MRL,NOD,NOR,NZB,NZO,NZW,PJ,PL,RF,RH2,SEA,SJL,SMJ,SPRET,SWR,TH,WSB | stop_lost,feature_truncation,coding_sequence_variant,3_prime_UTR_variant,intron_variant |
| 7 | 85036649 | 85043084 | 6435 | Vmn2r69 | 129S1,AJ,AKR,BALB,BTBR,BUB,C3H,C57L,CAST,CBA,DBA1J,DBA2J,FVB,ILNJ2,KK,MA/My,MOLF,MRL,NOD,NOR,NZB,NZO,NZW,PJ,PL,RF,RH2,SEA,SJL,SMJ,SWR,TH | feature_truncation,3_prime_UTR_variant |

| | | | | | | |
|---|---|---|---|---|---|---|
| 7 | 85193998 | 85200880 | 6882 | Vmn2r70 | SPRET | feature_truncation,3_prime_UTR_variant |
| 7 | 86041401 | 86043029 | 1628 | Or6aa1 | CAST,MOLF | feature_truncation,3_prime_UTR_variant |
| 7 | 97858997 | 97860488 | 1491 | Acer3 | C3H,CBA,DBA1J,DBA2J,ILNJ2,MOLF,PJ,SEA | feature_truncation,3_prime_UTR_variant |
| 7 | 101206786 | 101229154 | 22368 | Art2b | BTBR,C57L,KK,MA/My,NZW,WSB | stop_lost,feature_truncation,coding_sequence_variant,3_prime_UTR_variant,intron_variant |
| 7 | 102320769 | 102322093 | 1324 | Or52i2 | MOLF | feature_truncation,3_prime_UTR_variant |
| 7 | 102451562 | 102452987 | 1425 | Or51f23 | SPRET | feature_truncation,coding_sequence_variant,5_prime_UTR_variant |
| 7 | 102974327 | 102977616 | 3289 | Or52ab7 | MOLF | feature_truncation,5_prime_UTR_variant,intron_variant |
| 7 | 103063822 | 103065352 | 1530 | Usp17lc | SPRET | feature_truncation,coding_sequence_variant,5_prime_UTR_variant,intron_variant |
| 7 | 103198912 | 103200102 | 1190 | Or51ab3 | 129S1,AJ,AKR,BALB,BTBR,BUB,C3H,C57L,CAST,CBA,DBA1J,DBA2J,FVB,ILNJ2,MA/My,MOLF,MRL,NZB,NZO,NZW,PJ,PL,RF,RH2,SEA,SMJ,TH,WSB | feature_truncation,5_prime_UTR_variant |
| 7 | 103314791 | 103316670 | 1879 | Or51v8 | 129S1,AJ,AKR,BALB,BTBR,BUB,C3H,C57L,CAST,CBA,DBA1J,DBA2J,FVB,ILNJ2,MOLF,MRL,NZB,NZW,PJ,PL,RF,RH2,SEA,TH,WSB | feature_truncation,3_prime_UTR_variant |
| 7 | 103629508 | 103652288 | 22780 | Or51q1,Or51q1c | SPRET | feature_truncation,3_prime_UTR_variant,feature_truncation,5_prime_UTR_variant,intron_variant |
| 7 | 103630195 | 103636596 | 6401 | Or51q1 | MOLF | feature_truncation,3_prime_UTR_variant |
| 7 | 103657752 | 103660760 | 3008 | Or51k1 | CAST,KK,MOLF,NOD,SJL,SMJ,SPRET,SWR,TH | feature_truncation,3_prime_UTR_variant |
| 7 | 103705421 | 103706676 | 1255 | Or51a42 | CAST,MOLF,SJL,SWR,TH | feature_truncation,3_prime_UTR_variant |
| 7 | 103916051 | 103949854 | 33803 | Trim5,Trim12a | NZW,SJL | feature_truncation,coding_sequence_variant,5_prime_UTR_variant,intron_variant,feature_truncation,3_prime_UTR_variant,intron_variant |
| 7 | 104105961 | 104153641 | 47680 | Trim30d,Trim30a | CE | stop_lost,feature_truncation,coding_sequence_variant,5_prime_UTR_variant,3_prime_UTR_variant,intron_variant,feature_truncation,5_prime_UTR_variant,intron_variant |
| 7 | 105589360 | 105805589 | 216229 | Gvin1,Gvin2 | SMJ,SWR | stop_lost,feature_truncation,coding_sequence_variant,5_prime_UTR_variant,3_prime_UTR_variant,intron_variant,feature_truncation,5_prime_UTR_variant,intron_variant |
| 7 | 106519056 | 106522231 | 3175 | Or2d3c | 129S1,AJ,AKR,BALB,BTBR,C3H,C57L,CAST,CBA,CE,DBA1J,DBA2J,ILNJ2,KK,MOLF,MRL,NOD,NZB,NZO,NZW,PJ,PL,RF,RH2,SEA,SMJ,SPRET,SWR,WSB | feature_truncation,3_prime_UTR_variant |
| 7 | 106717743 | 106726092 | 8349 | Or2d2 | BALB,NZB,NZO | feature_truncation,3_prime_UTR_variant |
| 7 | 107357231 | 107358831 | 1600 | Cyb5r2 | KK,WSB | feature_truncation,5_prime_UTR_variant |
| 7 | 107436245 | 107458637 | 22392 | Or5p51 | CE | transcript_ablation |
| 7 | 107692380 | 107693788 | 1408 | Or5p58 | CAST,NOD,NZO,SMJ,SPRET,SWR | feature_truncation,3_prime_UTR_variant |
| 7 | 107765252 | 107775239 | 9987 | Or5p62 | CAST,NOD,SMJ,SWR | transcript_ablation |
| 7 | 107765253 | 107777657 | 12404 | Or5p62 | SPRET | transcript_ablation |
| 7 | 108221358 | 108222498 | 1140 | Or5p79 | MOLF | stop_lost,feature_truncation,coding_sequence_variant,3_prime_UTR_variant |
| 7 | 108417748 | 108424465 | 6717 | Or10a48 | MOLF | stop_lost,feature_truncation,coding_sequence_variant,3_prime_UTR_variant |
| 7 | 110428404 | 110430368 | 1964 | Rnf141 | CAST | stop_lost,feature_truncation,coding_sequence_variant,3_prime_UTR_variant,intron_variant |
| 7 | 127331135 | 127337587 | 6452 | Ctf2 | AJ,CAST,MOLF,SPRET,TH | feature_truncation,5_prime_UTR_variant,intron_variant |
| 7 | 130803523 | 130804567 | 1044 | Cdcp3 | CAST,SPRET | feature_truncation,coding_sequence_variant,intron_variant |
| 7 | 130879728 | 130881029 | 1301 | Cdcp3 | SPRET | feature_truncation,coding_sequence_variant,intron_variant |
| 7 | 133929468 | 133946539 | 17071 | D7Ertd443e | CAST | feature_truncation,coding_sequence_variant,5_prime_UTR_variant,intron_variant |
| 7 | 135193862 | 135196479 | 2617 | Ptpre | BALB,SEA | feature_truncation,coding_sequence_variant,5_prime_UTR_variant,intron_variant |
| 7 | 135298938 | 135300336 | 1398 | Mki67 | SPRET | feature_truncation,inframe_deletion |
| 7 | 139841570 | 139848073 | 6503 | Scart2 | CAST | feature_truncation,coding_sequence_variant,intron_variant |
| 7 | 140194139 | 140200732 | 6593 | Or13a18 | CAST,MOLF,NZW,PJ,SPRET | feature_truncation,3_prime_UTR_variant |
| 7 | 140250445 | 140253346 | 2901 | Or13a25 | CAST,MOLF | feature_truncation,3_prime_UTR_variant |
| 7 | 140257190 | 140258599 | 1409 | Or13a25 | SPRET | feature_truncation,3_prime_UTR_variant |
| 7 | 140273044 | 140276112 | 3068 | Or13a17 | NZW,PJ | feature_truncation,3_prime_UTR_variant |
| 7 | 141224174 | 141225499 | 1325 | Muc6 | NZW | frameshift_variant,feature_truncation |
| 7 | 141224929 | 141226464 | 1535 | Muc6 | ILNJ2,WSB | frameshift_variant,feature_truncation |
| 8 | 3995958 | 4002932 | 6974 | Cd209c | 129S1,CAST,NZB,NZO,NZW,SPRET,WSB | feature_truncation,5_prime_UTR_variant |
| 8 | 15059979 | 15061126 | 1147 | Kbtbd11 | SPRET | feature_truncation,5_prime_UTR_variant |
| 8 | 21171157 | 21179109 | 7952 | Potefam3b | MA/My | feature_truncation,coding_sequence_variant,intron_variant |
| 8 | 21748114 | 22122932 | 374818 | Defa26,Defa37,Defa32,Defa20,Defa36,Defa28,Defa29,Defa5,Defa3,Defa43,Defa42,Defa33,Defa27,Defa2 | BTBR,CAST,MOLF,NOR,NZB,NZW | transcript_ablation |
| 8 | 21796427 | 22037852 | 241425 | Defa32,Defa20,Defa36,Defa29,Defa42,Defa33,Defa27,Defa2 | NZB | transcript_ablation |
| 8 | 21888729 | 21921018 | 32289 | Defa36 | BUB,CBA,DBA1J,DBA2J,FVB,MRL,NOR,PL,RH2 | transcript_ablation |
| 8 | 21925976 | 22037852 | 111876 | Defa32,Defa20,Defa42 | DBA1J,NOR,RH2,SWR,TH | transcript_ablation |
| 8 | 22003546 | 22023356 | 19810 | Defa32 | C3H | transcript_ablation |
| 8 | 22071558 | 22077184 | 5626 | Defa28 | C57L,PJ | transcript_ablation |
| 8 | 27544055 | 27545086 | 1031 | Plpbp | CAST | feature_truncation,3_prime_UTR_variant |
| 8 | 43981634 | 44021497 | 39863 | Adam26a | NZB,NZO,NZW,WSB | stop_lost,feature_truncation,coding_sequence_variant,3_prime_UTR_variant |
| 8 | 44073189 | 44098987 | 25798 | Adam34l | MOLF | transcript_ablation |
| 8 | 44161171 | 44163551 | 2380 | Adam34 | MOLF | feature_truncation,5_prime_UTR_variant,intron_variant |
| 8 | 69945916 | 69960121 | 14205 | Potefam3d | MOLF | feature_truncation,coding_sequence_variant,intron_variant |
| 8 | 72870306 | 72871835 | 1529 | Or1ab2 | C58J,PJ,SEA | feature_truncation,3_prime_UTR_variant |

| | | | | | | |
|---|---|---|---|---|---|---|
| 8 | 73057910 | 73059159 | 1249 | Hmgn2-ps | 129S1,AKR,BALB,BUB,C3H,C57L,C58J,CBA,CE,DBA1J,DBA2J,FVB,ILNJ2,KK,MA/My,MRL,NZB,NZO,NZW,PJ,PL,RF,SEA,SJL,SMJ,SPRET,SWR,TH,WSB | transcript_ablation |
| 8 | 76131920 | 76133217 | 1297 | Gapdhrt2 | MOLF,SPRET | transcript_ablation |
| 8 | 105579697 | 105613844 | 34147 | Ces2c | KK | stop_lost,feature_truncation,coding_sequence_variant,3_prime_UTR_variant,intron_variant |
| 8 | 106738396 | 106741278 | 2882 | Ddx28 | SPRET | feature_truncation,5_prime_UTR_variant |
| 8 | 107740089 | 107742883 | 2794 | Sntb2 | MOLF | feature_truncation,3_prime_UTR_variant |
| 8 | 107785997 | 107787089 | 1092 | Nip7 | CAST,SPRET | feature_truncation,3_prime_UTR_variant |
| 8 | 117747892 | 117749935 | 2043 | Pkd1l2 | SWR | feature_truncation,coding_sequence_variant,intron_variant |
| 9 | 8628262 | 8633605 | 5343 | Trpc6 | 129S1,AKR,BTBR,C57L,CAST,MOLF,MRL,NZB,NZW,PL,RF,WSB | feature_truncation,3_prime_UTR_variant |
| 9 | 14540233 | 14559307 | 19074 | Amotl1 | NZO | feature_truncation,coding_sequence_variant,5_prime_UTR_variant,intron_variant |
| 9 | 18856994 | 18865515 | 8521 | Or7g19 | CAST | feature_truncation,3_prime_UTR_variant |
| 9 | 20198629 | 20200980 | 2351 | Or7d9 | CAST,SPRET | feature_truncation,3_prime_UTR_variant |
| 9 | 37764761 | 37766052 | 1291 | Or8b9 | CBA,CE,MRL | feature_truncation,5_prime_UTR_variant,intron_variant |
| 9 | 38024168 | 38054263 | 30095 | Or8b40,Or8b41 | KK,NZB,NZW,PJ,SEA | transcript_ablation,feature_truncation,5_prime_UTR_variant,intron_variant |
| 9 | 38029280 | 38030924 | 1644 | Or8b40 | SPRET | feature_truncation,3_prime_UTR_variant |
| 9 | 38090878 | 38106922 | 16044 | Or8c13 | MA/My | transcript_ablation |
| 9 | 38411089 | 38417577 | 6488 | Or8b44 | 129S1,AJ,AKR,BALB,BTBR,BUB,C3H,CAST,CBA,CE,DBA1J,DBA2J,FVB,ILNJ2,KK,MA/My,MOLF,MRL,NOD,NOR,NZB,NZW,PJ,PL,RF,RH2,SEA,SJL,SMJ,SPRET,SWR,TH,WSB | feature_truncation,3_prime_UTR_variant |
| 9 | 38582877 | 38585657 | 2780 | Or8b3b | CBA,PL,SJL,WSB | transcript_ablation,stop_lost,feature_truncation,coding_sequence_variant,5_prime_UTR_variant,3_prime_UTR_variant,intron_variant |
| 9 | 39098326 | 39101525 | 3199 | Or8g26 | SPRET | feature_truncation,3_prime_UTR_variant |
| 9 | 39295852 | 39361330 | 65478 | Or8g33,Or8g32 | SJL,WSB | transcript_ablation |
| 9 | 39300641 | 39365213 | 64572 | Or8g33,Or8g32 | MRL | transcript_ablation,stop_lost,feature_truncation,coding_sequence_variant,5_prime_UTR_variant,3_prime_UTR_variant,intron_variant |
| 9 | 40015693 | 40018734 | 3041 | Or4d5 | MOLF | feature_truncation,5_prime_UTR_variant |
| 9 | 42876076 | 42877740 | 1664 | Arhgef12 | SPRET | feature_truncation,3_prime_UTR_variant |
| 9 | 48264338 | 48268082 | 3744 | Nxpe4 | SPRET,WSB | feature_truncation,5_prime_UTR_variant,intron_variant |
| 9 | 50963962 | 50966735 | 2773 | Layn | 129S1,C57L,MOLF | stop_lost,feature_truncation,coding_sequence_variant,3_prime_UTR_variant,intron_variant |
| 9 | 56344696 | 56345735 | 1039 | Ube2srt | CAST,MOLF,SPRET | transcript_ablation |
| 9 | 57315270 | 57316753 | 1483 | Ppcdc | 129S1,C57L,CAST,CBA,SPRET | stop_lost,feature_truncation,coding_sequence_variant,3_prime_UTR_variant,intron_variant |
| 9 | 57594101 | 57602719 | 8618 | Cyp1a1 | SPRET | feature_truncation,5_prime_UTR_variant,intron_variant |
| 9 | 83824595 | 83831922 | 7327 | Bckdhb | 129S1,AJ,AKR,BALB,BTBR,BUB,C3H,C57L,C58J,CAST,CBA,CE,DBA1J,DBA2J,FVB,ILNJ2,KK,LP,MOLF,MRL,NOD,NOR,NZB,NZO,NZW,PJ,PL,RF,RH2,SEA,SJL,SMJ,SPRET,SWR,TH,WSB | feature_truncation,5_prime_UTR_variant,intron_variant |
| 9 | 89608712 | 89609916 | 1204 | Ankrd34c | SPRET | feature_truncation,3_prime_UTR_variant |
| 9 | 103330317 | 103336614 | 6297 | Bfsp2 | 129S1,CBA,FVB,NZB,NZW | feature_truncation,coding_sequence_variant,intron_variant |
| 9 | 105286553 | 105288170 | 1617 | Atp2c1 | 129S1,AJ,AKR,BALB,BTBR,BUB,C3H,C57L,CBA,CE,DBA1J,DBA2J,FVB,ILNJ2,KK,MRL,NOD,NOR,NZB,NZO,NZW,PJ,PL,RF,RH2,SEA,SJL,SMJ,SWR,TH,WSB | feature_truncation,3_prime_UTR_variant |
| 9 | 106109614 | 106110728 | 1114 | Alas1 | SPRET | feature_truncation,3_prime_UTR_variant |
| 9 | 106431184 | 106434532 | 3348 | Iqcf3 | SPRET | feature_truncation,coding_sequence_variant,intron_variant |
| X | 9202102 | 9239969 | 37867 | H2al1k | MOLF | transcript_ablation |
| X | 11169029 | 11189905 | 20876 | H2al1h,H2al1d,H2al1c,H2al1e,H2al1f,H2al1g,H2al1b | SPRET | transcript_ablation,stop_lost,feature_truncation,coding_sequence_variant,3_prime_UTR_variant |
| X | 13354065 | 13356592 | 2527 | Nyx | CAST | feature_truncation,3_prime_UTR_variant |
| X | 26264591 | 26267098 | 2507 | Gm2030 | MOLF | feature_truncation,5_prime_UTR_variant,intron_variant |
| X | 34688045 | 34698608 | 10563 | Gm14819 | MOLF | stop_lost,feature_truncation,coding_sequence_variant,3_prime_UTR_variant,intron_variant |
| X | 36827797 | 36829726 | 1929 | Rhox4g | SPRET | stop_lost,feature_truncation,coding_sequence_variant,3_prime_UTR_variant,intron_variant |
| X | 67864689 | 67868938 | 4249 | Styx-ps | AJ,BALB,BUB,C3H,C58J,CAST,ILNJ2,LP,MOLF,MRL,NOD,NOR,NZB,NZO,NZW,PJ,PL,RH2,SEA,SJL,SPRET,TH | transcript_ablation |
| X | 73210403 | 73211416 | 1013 | Tex28 | CAST,MOLF,SPRET | feature_truncation,5_prime_UTR_variant,intron_variant |
| X | 112746047 | 112747744 | 1697 | Dach2 | SPRET | feature_truncation,3_prime_UTR_variant |
| X | 117337791 | 117339513 | 1722 | Tgif2lx2 | SPRET | feature_truncation,3_prime_UTR_variant |
| X | 122151905 | 122387321 | 235416 | Cldn34c6,- | MOLF | transcript_ablation,stop_lost,feature_truncation,coding_sequence_variant,3_prime_UTR_variant |
| X | 146235555 | 146236864 | 1309 | Vmn1r239-ps | CAST,MOLF,NZB | transcript_ablation |
| X | 146571391 | 146576281 | 4890 | Gm15128 | WSB | feature_truncation,coding_sequence_variant,5_prime_UTR_variant,intron_variant |
| X | 146955392 | 146965040 | 9648 | Gm15107 | FVB,MRL,NOD | feature_truncation,coding_sequence_variant,5_prime_UTR_variant,intron_variant |
| X | 147636439 | 147640761 | 4322 | Luzp4 | NZB | feature_truncation,coding_sequence_variant,5_prime_UTR_variant,intron_variant |
| X | 147856586 | 147860892 | 4306 | Ott | MOLF | feature_truncation,coding_sequence_variant,5_prime_UTR_variant,intron_variant |
| X | 148365435 | 148379303 | 13868 | Gm10439 | MOLF | feature_truncation,coding_sequence_variant,5_prime_UTR_variant,intron_variant |
| X | 160006956 | 160011172 | 4216 | Scml2 | MRL | feature_truncation,coding_sequence_variant,intron_variant |
| X | 160008011 | 160009537 | 1526 | Scml2 | CAST,CE,MOLF,WSB | splice_acceptor_variant,splice_donor_variant,splice_donor_5th_base_variant,coding_sequence_variant,intron_variant |

**Table S8. Lists of high impact INS present in all 39 or in the 35 classical inbred strains.** The chromosome, starting and ending position, size, gene symbol, strains with the variant allele, and the predicted effect for each INS are shown.

| CHR | START | END | SIZE | SYMBOL | STRAINS with VARIANT SV ALLELE | CONSEQUENCE |
|---|---|---|---|---|---|---|
| 1 | 4846259 | 4846472 | 213 | Mrpl15 | DBA1J,DBA2J | feature_elongation,3_prime_UTR_variant |
| 1 | 4851852 | 4851988 | 136 | Mrpl15 | DBA1J,DBA2J | feature_elongation,3_prime_UTR_variant |
| 1 | 11670519 | 11670931 | 412 | A830018L16Rik | 129S1,AJ,AKR,BALB,BTBR,BUB,C3H,C57L,CBA,CE,FVB,KK,MRL,NOD,NZB,NZO,NZW,PL,RF,RH2,SEA,SJL,SWR,TH | feature_elongation,3_prime_UTR_variant |
| 1 | 13721098 | 13721166 | 68 | Lactb2 | TH | feature_elongation,coding_sequence_variant |
| 1 | 16203253 | 16203497 | 244 | Rdh10 | LP | feature_elongation,3_prime_UTR_variant |
| 1 | 30980183 | 30980237 | 54 | Ptp4a1 | SWR | feature_elongation,3_prime_UTR_variant |
| 1 | 33493000 | 33493058 | 58 | Prim2 | 129S1,AKR,BALB,BTBR,C57L,CBA,CE,LP,NOD,RF,SEA,SWR | feature_elongation,3_prime_UTR_variant |
| 1 | 34174632 | 34174702 | 70 | Dst | RF | feature_elongation,3_prime_UTR_variant |
| 1 | 34175238 | 34175443 | 205 | Dst | CE | feature_elongation,3_prime_UTR_variant |
| 1 | 34303931 | 34304071 | 140 | Dst | NZW | feature_elongation,coding_sequence_variant |
| 1 | 34484167 | 34484226 | 59 | Imp4 | 129S1,BTBR,C57L,ILNJ2,KK | feature_elongation,3_prime_UTR_variant |
| 1 | 37457066 | 37457126 | 60 | Coa5 | MA/My | feature_elongation,3_prime_UTR_variant |
| 1 | 38906630 | 38906708 | 78 | Chst10 | BALB,BTBR,C57L,DBA1J,DBA2J,FVB,ILNJ2,LP,MA/My,NOR,NZB,NZO,PL,SEA,SMJ,TH | feature_elongation,3_prime_UTR_variant |
| 1 | 39585134 | 39585339 | 205 | Cnot11 | RF | feature_elongation,3_prime_UTR_variant |
| 1 | 44183845 | 44183929 | 84 | Bivm | AKR,FVB,MA/My,NOD,NOR,RF,RH2 | feature_elongation,3_prime_UTR_variant |
| 1 | 44244716 | 44244820 | 104 | Mettl21e | 129S1,AJ,AKR,BUB,C3H,CBA,CE,DBA1J,DBA2J,FVB,ILNJ2,KK,MA/My,MRL,NOD,NOR,PJ,PL,RF,RH2,SJL,SMJ,SWR,TH | feature_elongation,3_prime_UTR_variant |
| 1 | 57455110 | 57455180 | 70 | Maip1 | SMJ,SWR | feature_elongation,3_prime_UTR_variant |
| 1 | 58566540 | 58566769 | 229 | Hycc2 | CBA,DBA1J,DBA2J,ILNJ2,NZB,NZO,NZW,RH2 | feature_elongation,3_prime_UTR_variant |
| 1 | 59752225 | 59752423 | 198 | Nop58 | ILNJ2,MA/My,NZB,NZO,NZW | feature_elongation,3_prime_UTR_variant |
| 1 | 59755468 | 59755688 | 220 | Nop58 | ILNJ2,MA/My,NZB,NZO,NZW,SMJ | feature_elongation,3_prime_UTR_variant |
| 1 | 59911861 | 59912068 | 207 | Bmpr2 | CE | feature_elongation,3_prime_UTR_variant |
| 1 | 60109960 | 60110026 | 66 | Wdr12 | CE,LP | feature_elongation,3_prime_UTR_variant |
| 1 | 60111845 | 60112397 | 552 | Wdr12 | ILNJ2,NZB,NZO,NZW | feature_elongation,3_prime_UTR_variant |
| 1 | 63350797 | 63350995 | 198 | Zdbf2 | C58J,NZB | feature_elongation,3_prime_UTR_variant |
| 1 | 67041321 | 67041376 | 55 | Lancl1 | PJ | feature_elongation,3_prime_UTR_variant |
| 1 | 82748860 | 82748935 | 75 | Gm47791 | PJ | feature_elongation,coding_sequence_variant |
| 1 | 82933542 | 82933603 | 61 | Gm7544 | RF | feature_elongation,coding_sequence_variant |
| 1 | 82938140 | 82938302 | 162 | Gm47955 | AKR,CBA,LP,MRL,NOD | feature_elongation,coding_sequence_variant |
| 1 | 84563271 | 84563325 | 54 | Dner | AKR,CBA,MRL,NOD,PJ,PL | feature_elongation,coding_sequence_variant |
| 1 | 86205808 | 86205882 | 74 | Armc9 | SMJ | feature_elongation,3_prime_UTR_variant |
| 1 | 88431142 | 88431199 | 57 | Glrp1 | SMJ | feature_elongation,coding_sequence_variant |
| 1 | 89818413 | 89818633 | 220 | Agap1 | C58J | feature_elongation,3_prime_UTR_variant |
| 1 | 90877734 | 90877823 | 89 | Mlph | B10D2,B10J,BALB,BTBR,BUB,C3H,C58J,CBA,CE,DBA1J,DBA2J,ILNJ2,MA/My,MRL,NOD,NOR,PJ,PL,RH2,SEA,SMJ | feature_elongation,3_prime_UTR_variant |
| 1 | 91483493 | 91483709 | 216 | Asb1 | 129S1,AKR,C57L,FVB,KK,LP,NZB,NZO,NZW,RF,SJL,SWR,TH | feature_elongation,3_prime_UTR_variant |
| 1 | 91486902 | 91486972 | 70 | Asb1 | 129S1,AKR,BALB,BTBR,C57L,DBA1J,DBA2J,FVB,ILNJ2,KK,LP,NOD,NOR,NZB,NZO,NZW,PJ,PL,RF,SEA,SJL,SMJ,SWR,TH | feature_elongation,3_prime_UTR_variant |
| 1 | 92404220 | 92404292 | 72 | Or6b2 | PL | feature_elongation,3_prime_UTR_variant |
| 1 | 92405288 | 92405458 | 170 | Or6b2 | BUB,C3H,CBA,DBA1J,DBA2J,PJ,PL,SEA | feature_elongation,3_prime_UTR_variant |
| 1 | 92540499 | 92540612 | 113 | Or9s14 | BUB,C3H,CBA | feature_elongation,3_prime_UTR_variant |
| 1 | 95253264 | 95253794 | 530 | Fam174a | PL | feature_elongation,3_prime_UTR_variant |
| 1 | 97545428 | 97545988 | 560 | Macir | MRL,PL | feature_elongation,3_prime_UTR_variant |
| 1 | 97551636 | 97551720 | 84 | Macir | MRL,PL | feature_elongation,3_prime_UTR_variant |
| 1 | 105773115 | 105773498 | 383 | Tnfrsf11a | 129S1,BUB,C57L,DBA1J,DBA2J,ILNJ2,NOD,NOR,SJL,TH | feature_elongation,3_prime_UTR_variant |
| 1 | 107536119 | 107536318 | 199 | Serpinb8 | 129S1,C57L,DBA1J,DBA2J,SJL,SWR,TH | feature_elongation,3_prime_UTR_variant |
| 1 | 118128515 | 118128683 | 168 | Gm29106 | SWR | feature_elongation,3_prime_UTR_variant |
| 1 | 130779619 | 130779676 | 57 | Pigr | 129S1,BALB,C57L,DBA1J,DBA2J,FVB,ILNJ2,KK,MA/My,NZO,NZW,SEA,SJL,SWR,TH | feature_elongation,3_prime_UTR_variant |
| 1 | 131640794 | 131641204 | 410 | Rab7b | 129S1,AKR,BALB,BTBR,BUB,CBA,DBA1J,DBA2J,FVB,ILNJ2,KK,MA/My,MRL,NZO,NZW,PJ,PL,RF,SEA,SJL,SMJ,SWR,TH | feature_elongation,3_prime_UTR_variant |
| 1 | 135363073 | 135363189 | 116 | Nav1 | 129S1,AJ,AKR,BTBR,C3H,CE,DBA1J,DBA2J,ILNJ2,KK,MA/My,NZB,NZO,NZW,PJ,RF,RH2,SJL,SWR,TH | feature_elongation,3_prime_UTR_variant |
| 1 | 143625218 | 143625436 | 218 | Glrx2 | AJ,C3H,CBA,CE,FVB,ILNJ2,PL,RH2 | feature_elongation,3_prime_UTR_variant |
| 1 | 143630205 | 143630404 | 199 | Ro60 | 129S1,NZO,NZW,SJL,SWR,TH | feature_elongation,3_prime_UTR_variant |
| 1 | 143682922 | 143683140 | 218 | Uchl5 | AJ,C3H,CBA,CE,FVB,ILNJ2,MRL,PL,RH2 | feature_elongation,3_prime_UTR_variant |
| 1 | 144395319 | 144395381 | 62 | Rgs21 | AJ,C3H,CBA,CE,FVB,ILNJ2,NZO,PL,RH2,SJL,SWR | feature_elongation,3_prime_UTR_variant |
| 1 | 150331351 | 150331441 | 90 | Prg4 | FVB,SJL,SWR | feature_elongation,coding_sequence_variant |
| 1 | 155003932 | 155004152 | 220 | Mr1 | C57L | feature_elongation,3_prime_UTR_variant |
| 1 | 155574699 | 155574795 | 96 | Lhx4 | C57L | feature_elongation,3_prime_UTR_variant |
| 1 | 159753917 | 159754025 | 108 | Tnr | BUB,C3H,CBA,CE,PJ,SEA | feature_elongation,3_prime_UTR_variant |
| 1 | 160835404 | 160835615 | 211 | Zbtb37 | BALB,C58J,DBA1J,DBA2J,LP | feature_elongation,3_prime_UTR_variant |
| 1 | 160917163 | 160917243 | 80 | Klhl20 | BALB,C58J,DBA1J,DBA2J,ILNJ2,LP | feature_elongation,3_prime_UTR_variant |
| 1 | 164021151 | 164021310 | 159 | F5 | DBA1J,DBA2J,ILNJ2 | feature_elongation,coding_sequence_variant |
| 1 | 165152131 | 165152187 | 56 | Gpr161 | AKR,BUB,C3H,C58J,CBA,LP,MRL,RH2 | feature_elongation,3_prime_UTR_variant |
| 1 | 165156494 | 165156629 | 135 | Dcaf6 | DBA1J,DBA2J,ILNJ2,PJ,RF,SEA,SJL | feature_elongation,3_prime_UTR_variant |
| 1 | 165178573 | 165178648 | 75 | Dcaf6 | DBA1J,DBA2J,ILNJ2 | feature_elongation,coding_sequence_variant |
| 1 | 167426225 | 167426281 | 56 | Rxrg | SMJ | feature_elongation,5_prime_UTR_variant |
| 1 | 169800493 | 169800660 | 167 | Ddr2 | 129S1,AJ,BTBR,BUB,C3H,C57L,CE,FVB,KK,MRL,NOD,NOR,NZB,NZO,NZW,PJ,PL,RF,RH2,SEA,SJL,SMJ,SWR,TH | feature_elongation,3_prime_UTR_variant |
| 1 | 171460903 | 171460968 | 65 | Slamf7 | 129S1,AJ,AKR,BALB,BUB,C3H,C57L,CBA,CE,DBA1J,DBA2J,FVB,KK,MRL,NOD,NOR,NZB,NZO,NZW,PJ,PL,RF,RH2,SEA,SJL,SMJ,SWR,TH | feature_elongation,3_prime_UTR_variant |
| 1 | 171780686 | 171780894 | 208 | Slamf6 | 129S1,AJ,AKR,BALB,C3H,C57L,CBA,CE,DBA1J,DBA2J,KK,MRL,NOD,NOR,NZO,NZW,PJ,PL,RF,RH2,SEA,SJL,SMJ,TH | feature_elongation,3_prime_UTR_variant |
| 1 | 172526846 | 172526900 | 54 | Crp | NOD,NOR | feature_elongation,3_prime_UTR_variant |
| 1 | 173099816 | 173100376 | 560 | Or10j2 | 129S1,AJ,AKR,BALB,BTBR,BUB,C3H,C57L,C58J,CBA,CE,DBA1J,DBA2J,FVB,ILNJ2,KK,MRL,NOD,NOR,NZB,NZO,NZW,PJ,PL,RF,RH2,SEA,SJL,SMJ,SWR,TH | feature_elongation,3_prime_UTR_variant |

| | | | | | | |
|---|---|---|---|---|---|---|
| 1 | 173556584 | 173556668 | 84 | Ifi207 | 129S1,AJ,AKR,BALB,BTBR,BUB,C3H,C57L,C58J,CBA,CE,DBA1J,DBA2J,FVB,ILNJ2,KK,MRL,NOD,NOR,NZB,NZO,PJ,PL,RF,RH2,SEA,SJL,SMJ,SWR,TH | stop_gained,protein_altering_variant |
| 1 | 173587714 | 173587777 | 63 | Ifi204 | 129S1,AKR,BALB,BUB,C3H,C57L,C58J,CBA,CE,DBA1J,DBA2J,ILNJ2,MRL,NOD,NOR,NZB,NZO,PJ,RF,SEA,SJL,SMJ,SWR | feature_elongation,coding_sequence_variant |
| 1 | 173733734 | 173733796 | 62 | Ifi211 | 129S1,AKR,BALB,BUB,C3H,C57L,C58J,CBA,CE,DBA1J,DBA2J,ILNJ2,MRL,NOD,NOR,NZB,NZO,PJ,RF,SEA,SJL,SMJ | frameshift_variant |
| 1 | 173896677 | 173896743 | 66 | Or6n2 | AJ,AKR,BTBR,BUB,C58J,CBA,CE,DBA1J,DBA2J,FVB,ILNJ2,KK,NZB,NZO,PJ,PL,RF,RH2,SEA,SMJ,SWR,TH | feature_elongation,5_prime_UTR_variant |
| 1 | 175805702 | 175805914 | 212 | Rbm8a2 | NZO | feature_elongation,3_prime_UTR_variant |
| 1 | 176850818 | 176851012 | 194 | Akt3 | SMJ | feature_elongation,3_prime_UTR_variant |
| 1 | 180994765 | 180994985 | 220 | Cnih4 | CBA,DBA1J,DBA2J | feature_elongation,3_prime_UTR_variant |
| 1 | 194723637 | 194723868 | 231 | Cd46 | NOD,NOR,NZO,NZW,RF,SWR | feature_elongation,3_prime_UTR_variant |
| 2 | 3458321 | 3458493 | 172 | Suv39h2 | MA/My,NZO,NZW | feature_elongation,3_prime_UTR_variant |
| 2 | 5799150 | 5799259 | 109 | Cdc123 | SMJ | feature_elongation,3_prime_UTR_variant |
| 2 | 11478428 | 11478502 | 74 | Pfkfb3 | MA/My | feature_elongation,3_prime_UTR_variant |
| 2 | 18221937 | 18222139 | 202 | Dnajc1 | MA/My,RF | feature_elongation,3_prime_UTR_variant |
| 2 | 20749272 | 20749471 | 199 | Etl4 | CE | feature_elongation,3_prime_UTR_variant |
| 2 | 24862157 | 24862362 | 205 | Dph7 | FVB | feature_elongation,3_prime_UTR_variant |
| 2 | 26909673 | 26909906 | 233 | Cacfd1 | 129S1,AJ,AKR,B10D2,B10J,BALB,BTBR,BUB,C3H,C57L,C58J,CBA,CE,DBA1J,DBA2J,FVB,ILNJ2,KK,LP,MA/My,MRL,NOD,NOR,NZB,NZO,NZW,PJ,PL,RF,RH2,SEA,SJL,SMJ,SWR,TH | feature_elongation,3_prime_UTR_variant |
| 2 | 26998572 | 26998630 | 58 | Adamtsl2 | 129S1,BUB,C57L,FVB,MA/My,RF,SMJ | feature_elongation,3_prime_UTR_variant |
| 2 | 32489325 | 32490150 | 825 | St6galnac4 | AJ,AKR,BTBR,BUB,C58J,CE,DBA2J,FVB,ILNJ2,KK,MRL,NOD,NOR,NZB,NZW,PL,RH2,SMJ,TH | feature_elongation,3_prime_UTR_variant |
| 2 | 36956125 | 36956185 | 60 | Or12k7 | BUB | feature_elongation,5_prime_UTR_variant |
| 2 | 37037656 | 37037811 | 155 | Or1l4b | BUB,TH | feature_elongation,3_prime_UTR_variant |
| 2 | 37325068 | 37325462 | 394 | Zbtb26 | DBA1J | feature_elongation,3_prime_UTR_variant |
| 2 | 43525678 | 43525878 | 200 | Kynu | 129S1,AJ,AKR,B10D2,B10J,BALB,BTBR,BUB,C3H,C57L,C58J,CBA,CE,DBA1J,DBA2J,FVB,ILNJ2,KK,LP,MA/My,MRL,NOD,NOR,NZB,NZO,NZW,PJ,PL,RF,RH2,SEA,SJL,SMJ,SWR,TH | feature_elongation,3_prime_UTR_variant |
| 2 | 51024040 | 51024096 | 56 | Rnd3 | 129S1,AJ,AKR,BUB,C3H,C58J,CBA,CE,DBA1J,DBA2J,FVB,ILNJ2,KK,MRL,NOD,NOR,NZW,PJ,RF,SEA,SJL,SMJ,SWR,TH | feature_elongation,coding_sequence_variant |
| 2 | 52306089 | 52306227 | 138 | Arl5a | 129S1,AJ,AKR,BUB,C3H,CBA,CE,DBA1J,DBA2J,ILNJ2,MRL,NOD,NOR,NZW,PJ,PL,RF,SEA,SJL,SMJ,SWR,TH | feature_elongation,coding_sequence_variant |
| 2 | 58020800 | 58020860 | 60 | Cytip | 129S1,AJ,AKR,BUB,C3H,C58J,CBA,DBA1J,DBA2J,FVB,NOD,NOR,PJ,PL,RF,SEA,SJL,SWR,TH | feature_elongation,3_prime_UTR_variant |
| 2 | 58022231 | 58022300 | 69 | Cytip | 129S1,AJ,AKR,BUB,C3H,C58J,CBA,DBA1J,DBA2J,FVB,NOD,NOR,PJ,PL,RF,SEA,SJL,SWR,TH | feature_elongation,3_prime_UTR_variant |
| 2 | 70969337 | 70969397 | 60 | Cybrd1 | AJ,AKR,BUB,C3H,FVB,ILNJ2,MA/My,MRL,NOD,NOR,NZW,PL,RH2,SMJ | feature_elongation,3_prime_UTR_variant |
| 2 | 73262517 | 73262593 | 76 | Wipf1 | KK,MA/My,NZB | feature_elongation,3_prime_UTR_variant |
| 2 | 80359011 | 80359084 | 73 | Nckap1 | AJ,AKR,BALB,BUB,C3H,C57L,C58J,CBA,CE,DBA1J,DBA2J,FVB,KK,MA/My,MRL,NOD,NOR,PJ,PL,RF,RH2,SEA,SJL,SMJ,SWR,TH | feature_elongation,coding_sequence_variant |
| 2 | 83816023 | 83816261 | 238 | Cwc22rt7 | ILNJ2 | feature_elongation,coding_sequence_variant |
| 2 | 85617082 | 85617297 | 215 | Or9g4b | AJ,AKR,BALB,BUB,C3H,C57L,C58J,CBA,CE,DBA1J,DBA2J,ILNJ2,MA/My,MRL,NOD,NOR,NZB,NZO,NZW,PJ,PL,RF,RH2,SEA,SMJ | feature_elongation,3_prime_UTR_variant |
| 2 | 85671181 | 85671353 | 172 | Or5ar1 | KK | feature_elongation,3_prime_UTR_variant |
| 2 | 87189139 | 87189203 | 64 | Or12e8 | SJL,SWR | feature_elongation,3_prime_UTR_variant |
| 2 | 87251332 | 87251558 | 226 | Or10ag2 | AJ,AKR,BALB,C57L,DBA1J,DBA2J,ILNJ2,MRL,NOD,NOR,NZB,NZO,PJ,RH2,SEA,SJL,SMJ,SWR | feature_elongation,3_prime_UTR_variant |
| 2 | 87540998 | 87541052 | 54 | Or5w14 | C3H,CBA,CE,KK,MA/My,NZW,RF | feature_elongation,3_prime_UTR_variant |
| 2 | 87614117 | 87614191 | 74 | Or5i1 | C3H,C58J,CBA,CE,KK,MA/My,NZW,RF,SJL,SWR | feature_elongation,3_prime_UTR_variant |
| 2 | 87789889 | 87790049 | 160 | Or5l14 | AJ,AKR,BALB,C57L,C58J,DBA1J,DBA2J,FVB,ILNJ2,KK,MRL,NOD,NOR,NZB,NZO,PJ,RH2,SEA,SJL,SMJ,SWR | feature_elongation,3_prime_UTR_variant |
| 2 | 88738457 | 88739070 | 613 | Or4c29 | C3H,CBA,CE,MA/My,NZW | feature_elongation,3_prime_UTR_variant |
| 2 | 88757583 | 88758085 | 502 | Or4c15 | AJ,AKR,BALB,C57L,DBA1J,DBA2J,FVB,KK,MRL,NOD,NOR,NZB,NZO,PJ,RF,RH2,SEA,SMJ | feature_elongation,3_prime_UTR_variant |
| 2 | 89771112 | 89771574 | 462 | Or4c12 | ILNJ2 | feature_elongation,3_prime_UTR_variant |
| 2 | 104495673 | 104495773 | 100 | Cstf3 | SMJ | feature_elongation,3_prime_UTR_variant |
| 2 | 105003264 | 105003325 | 61 | Wt1 | PL | feature_elongation,3_prime_UTR_variant |
| 2 | 111248366 | 111248462 | 96 | Or4k40 | C3H,CE,PL | feature_elongation,3_prime_UTR_variant |
| 2 | 111250384 | 111250526 | 142 | Or4k40 | C3H,CE,PL | feature_elongation,coding_sequence_variant |
| 2 | 111332178 | 111332311 | 133 | Or4f4-ps1 | C3H,CE,PL | feature_elongation,3_prime_UTR_variant |
| 2 | 111447524 | 111447592 | 68 | Or4k47 | C3H,CE,PL | feature_elongation,3_prime_UTR_variant |
| 2 | 111836264 | 111836316 | 52 | Or4f6 | 129S1,AKR,BALB,BUB,C57L,KK,MA/My,MRL,NOD,NOR,NZO,NZW,PJ,SEA,SWR,TH | feature_elongation,3_prime_UTR_variant |
| 2 | 112186126 | 112186463 | 337 | Slc12a6 | C3H,CE,PL | feature_elongation,3_prime_UTR_variant |
| 2 | 113578622 | 113578687 | 65 | Grem1 | C3H,CE,PL | feature_elongation,3_prime_UTR_variant |
| 2 | 118712816 | 118712867 | 51 | Ivd | 129S1,AJ,AKR,BALB,BTBR,BUB,C3H,C57L,C58J,CBA,CE,DBA1J,DBA2J,FVB,ILNJ2,MA/My,NOD,NOR,NZB,NZO,NZW,PJ,PL,RF,RH2,SEA,SJL,SMJ,SWR,TH | feature_elongation,3_prime_UTR_variant |
| 2 | 119048863 | 119049089 | 226 | Ppp1r14d | AKR | feature_elongation,coding_sequence_variant |
| 2 | 119233190 | 119233653 | 463 | Ino80 | 129S1,AKR,BTBR,BUB,C3H,C57L,C58J,CBA,CE,FVB,NOD,NZB,NZO,SJL,SWR | feature_elongation,3_prime_UTR_variant |
| 2 | 119481204 | 119481399 | 195 | Nusap1 | 129S1,AJ,AKR,BALB,BTBR,BUB,C3H,C57L,C58J,CBA,CE,DBA1J,DBA2J,FVB,ILNJ2,KK,MA/My,NOD,NZB,NZO,NZW,PJ,PL,RF,RH2,SEA,SJL,SMJ,SWR,TH | feature_elongation,3_prime_UTR_variant |
| 2 | 119798755 | 119798976 | 221 | Mga | AJ,BALB,DBA1J,DBA2J,ILNJ2,PJ,PL,RF,RH2,SEA,SMJ,TH | feature_elongation,3_prime_UTR_variant |

| | | | | | | |
|---|---|---|---|---|---|---|
| 2 | 119849812 | 119849889 | 77 | Mapkbp1 | 129S1,AJ,AKR,BALB,BTBR,BUB,C3H,C57L,C58J,CBA,CE,DBA1J,DBA2J,FVB,ILNJ2,KK,MA/My,NOD,NZB,NZO,PJ,PL,RF,RH2,SEA,SJL,SMJ,SWR,TH | feature_elongation,coding_sequence_variant |
| 2 | 121136770 | 121136830 | 60 | Map1a | 129S1,AJ,AKR,BALB,BTBR,BUB,C3H,C57L,C58J,CBA,CE,DBA1J,FVB,ILNJ2,KK,MA/My,NZB,NZO,NZW,PJ,PL,RF,RH2,SEA,SJL,SMJ,SWR,TH | feature_elongation,coding_sequence_variant |
| 2 | 121285306 | 121285362 | 56 | Hypk | MA/My | feature_elongation,5_prime_UTR_variant |
| 2 | 121291564 | 121291773 | 209 | Mfap1b | 129S1,AJ,AKR,BALB,BTBR,BUB,C3H,C57L,C58J,CBA,CE,DBA1J,DBA2J,FVB,ILNJ2,KK,MA/My,NZB,NZO,NZW,PJ,PL,RF,RH2,SEA,SJL,SMJ,SWR,TH | feature_elongation,3_prime_UTR_variant |
| 2 | 121764678 | 121764814 | 136 | Golm2 | 129S1,BALB,BTBR,BUB,C57L,CE,NZW | feature_elongation,3_prime_UTR_variant |
| 2 | 122095395 | 122095712 | 317 | Sord | AJ,BALB,C3H,C58J,CBA,DBA1J,DBA2J,ILNJ2,NZB,NZO,PJ,PL,RF,RH2,SEA,SMJ,TH | feature_elongation,3_prime_UTR_variant |
| 2 | 122590418 | 122590823 | 405 | Bloc1s6 | MA/My | feature_elongation,3_prime_UTR_variant |
| 2 | 126471182 | 126471879 | 697 | Gabpb1 | SMJ | feature_elongation,3_prime_UTR_variant |
| 2 | 130259695 | 130259907 | 212 | Pced1a | NOD | feature_elongation,3_prime_UTR_variant |
| 2 | 131701592 | 131701883 | 291 | Erv3 | MA/My | feature_elongation,5_prime_UTR_variant |
| 2 | 132091098 | 132091288 | 190 | Pcna | ILNJ2,PJ | feature_elongation,3_prime_UTR_variant |
| 2 | 150110249 | 150110418 | 169 | Zfp1005 | NOD,NZB,NZW,RF | feature_elongation,coding_sequence_variant |
| 2 | 152236730 | 152236854 | 124 | Sox12 | AJ,C58J,NOD,NZB,NZW,RF | feature_elongation,3_prime_UTR_variant |
| 2 | 154359700 | 154360117 | 417 | Cbfa2t2 | FVB,SWR | feature_elongation,coding_sequence_variant |
| 2 | 156940944 | 156941020 | 76 | Samhd1 | CE | feature_elongation,3_prime_UTR_variant |
| 2 | 158225136 | 158225265 | 129 | Snhg11 | DBA1J,DBA2J,ILNJ2,MA/My,SMJ | feature_elongation,3_prime_UTR_variant |
| 2 | 164032497 | 164032704 | 207 | Wfdc12 | KK | feature_elongation,5_prime_UTR_variant |
| 2 | 165626756 | 165626838 | 82 | Zmynd8 | AJ,AKR,BALB,BTBR,BUB,C3H,C58J,CBA,CE,DBA1J,DBA2J,FVB,ILNJ2,KK,MA/My,NOD,NOR,NZB,NZO,PJ,PL,RF,SEA,SJL,SMJ,SWR,TH | feature_elongation,3_prime_UTR_variant |
| 2 | 167820390 | 167820586 | 196 | Ptpn1 | 129S1,C57L,NZB | feature_elongation,3_prime_UTR_variant |
| 2 | 168590700 | 168590828 | 128 | Sall4 | B10D2 | feature_elongation,3_prime_UTR_variant |
| 2 | 172204807 | 172204909 | 102 | Aurka | FVB,SJL | feature_elongation,coding_sequence_variant |
| 2 | 175037635 | 175037719 | 84 | Gm14391 | BUB,CE,FVB,NZB,SJL,SWR | feature_elongation,coding_sequence_variant |
| 2 | 175672079 | 175672163 | 84 | Gm6710 | BUB,CE,FVB,KK,NOD,NOR,NZB,PJ,SJL,SWR | feature_elongation,coding_sequence_variant |
| 2 | 175772093 | 175772301 | 208 | Gm2026 | CE,FVB,NZB,SJL,SWR | feature_elongation,3_prime_UTR_variant |
| 2 | 178064237 | 178064305 | 68 | Fam217b | PJ | feature_elongation,3_prime_UTR_variant |
| 3 | 10377772 | 10378270 | 498 | Impa1 | SMJ | feature_elongation,3_prime_UTR_variant |
| 3 | 10378403 | 10378474 | 71 | Impa1 | BUB,CE,DBA2J,SMJ | feature_elongation,3_prime_UTR_variant |
| 3 | 10455705 | 10455768 | 63 | Chmp4c | BUB,CE,DBA2J | feature_elongation,3_prime_UTR_variant |
| 3 | 19186037 | 19186101 | 64 | Armc1 | AJ,C3H,C58J,CBA,CE,DBA2J,LP,MRL,NZB,NZO,NZW,PL | feature_elongation,3_prime_UTR_variant |
| 3 | 32394611 | 32394802 | 191 | Zmat3 | SJL | feature_elongation,3_prime_UTR_variant |
| 3 | 32520798 | 32520880 | 82 | Pik3ca | CE | feature_elongation,3_prime_UTR_variant |
| 3 | 32991174 | 32991602 | 428 | Usp13 | 129S1,AJ,BTBR,BUB,C3H,C57L,CBA,FVB,MRL,NOD,NOR,NZB,NZO,NZW,PJ,SEA,SWR | feature_elongation,3_prime_UTR_variant |
| 3 | 36145884 | 36146200 | 316 | Acad9 | BUB,FVB | feature_elongation,3_prime_UTR_variant |
| 3 | 51403622 | 51403780 | 158 | Rab33b | NZW | feature_elongation,3_prime_UTR_variant |
| 3 | 67386628 | 67386756 | 128 | Rarres1 | AKR,BALB,BTBR,CBA,CE,DBA1J,DBA2J,FVB,NOD,NOR,NZO,RF,SJL,SWR,TH | feature_elongation,3_prime_UTR_variant |
| 3 | 68951947 | 68952001 | 54 | Trim59 | 129S1,BTBR,C57L,CE,FVB,SJL,SWR,TH | feature_elongation,5_prime_UTR_variant |
| 3 | 69658344 | 69658491 | 147 | Nmd3 | AJ,AKR,BUB,C3H,C58J,CBA,DBA1J,DBA2J,KK,LP,MRL,NOD,NOR,NZO,NZW,RF | feature_elongation,3_prime_UTR_variant |
| 3 | 81864031 | 81864101 | 70 | Ctso | TH | feature_elongation,3_prime_UTR_variant |
| 3 | 88020123 | 88020353 | 230 | Iqgap3 | NZB,NZO | feature_elongation,coding_sequence_variant |
| 3 | 89655278 | 89655332 | 54 | Chrnb2 | KK | feature_elongation,coding_sequence_variant |
| 3 | 89773373 | 89773562 | 189 | Il6ra | KK | feature_elongation,3_prime_UTR_variant |
| 3 | 92479165 | 92479597 | 432 | Ivl | NOD,NOR,RF | feature_elongation,coding_sequence_variant |
| 3 | 93110593 | 93110671 | 78 | Flg2 | 129S1,AJ,BTBR,C3H,C57L,CBA,CE,ILNJ2,PJ,PL,SEA,TH | feature_elongation,coding_sequence_variant |
| 3 | 93304211 | 93304319 | 108 | Rptn | BALB,ILNJ2,KK,NOD,NOR,NZB,NZO,PJ,PL,RH2,SEA,TH | feature_elongation,coding_sequence_variant |
| 3 | 96541096 | 96541303 | 207 | Rbm8a | 129S1,AJ,BALB,C3H,C57L,CBA,CE,FVB,MA/My,NOD,NOR,NZB,NZO,NZW,PJ,RH2,SEA,SJL,SMJ,SWR | feature_elongation,3_prime_UTR_variant |
| 3 | 97561481 | 97561705 | 224 | Fmo5 | NOD,NOR | feature_elongation,3_prime_UTR_variant |
| 3 | 98056712 | 98056768 | 56 | Notch2 | 129S1,C57L,FVB,PJ,RF,SEA,SJL,SWR | feature_elongation,3_prime_UTR_variant |
| 3 | 101721087 | 101721139 | 52 | Mab21l3 | MA/My | feature_elongation,3_prime_UTR_variant |
| 3 | 103783078 | 103783132 | 54 | Ptpn22 | AJ,AKR,C3H,CBA,FVB,ILNJ2,RF,SMJ | feature_elongation,coding_sequence_variant |
| 3 | 105626786 | 105627088 | 302 | Inka2 | MA/My,NZB,NZO,NZW | feature_elongation,3_prime_UTR_variant |
| 3 | 105893744 | 105893954 | 210 | Ovgp1 | AJ,C3H,CBA | feature_elongation,coding_sequence_variant |
| 3 | 107017328 | 107017430 | 102 | Kcna2 | AJ,AKR,BALB,C3H,CBA,DBA1J,DBA2J,ILNJ2,MA/My,MRL,NOD,NOR,NZB,NZO,NZW,PL,RF,RH2,TH | feature_elongation,3_prime_UTR_variant |
| 3 | 108464895 | 108465292 | 397 | Tmem167b | BALB,MRL,PJ,RH2,SEA | feature_elongation,3_prime_UTR_variant |
| 3 | 110156115 | 110157084 | 969 | Prmt6 | MA/My | feature_elongation,3_prime_UTR_variant |
| 3 | 122719386 | 122719476 | 90 | 1810037I17Rik | 129S1,BTBR,C57L,CE,DBA1J,DBA2J,KK,MA/My,NZB,PJ,RF,SEA,SMJ,TH | feature_elongation,3_prime_UTR_variant |
| 3 | 122749802 | 122750779 | 977 | Usp53 | 129S1,BTBR,C57L,CE,DBA1J,DBA2J,KK,MA/My,NZB,PJ,RF,SEA,SMJ,TH | feature_elongation,3_prime_UTR_variant |
| 3 | 123165690 | 123166034 | 344 | Mettl14 | AJ,BALB,C3H,CBA,DBA1J,DBA2J,NZB,RH2,SMJ | feature_elongation,3_prime_UTR_variant |
| 3 | 137615461 | 137615513 | 52 | Dnajb14 | AJ,AKR,BALB,C3H,CBA,CE,DBA1J,FVB,ILNJ2,KK,MA/My,MRL,NOD,NOR,PJ,RH2,SEA,SJL,SWR,TH | feature_elongation,3_prime_UTR_variant |
| 3 | 138035995 | 138036178 | 183 | Adh6a | 129S1,AJ,AKR,BALB,BTBR,BUB,C3H,C57L,C58J,CBA,CE,DBA1J,DBA2J,FVB,ILNJ2,KK,LP,MA/My,MRL,NOD,NOR,NZB,NZO,NZW,PJ,PL,RF,RH2,SEA,SJL,SMJ,SWR,TH | feature_elongation,3_prime_UTR_variant |
| 3 | 138610731 | 138610947 | 216 | Tspan5 | AKR,FVB | feature_elongation,3_prime_UTR_variant |
| 3 | 142210254 | 142211029 | 775 | Gbp5 | CBA,DBA1J,MRL,NZW | feature_elongation,coding_sequence_variant |
| 3 | 152038258 | 152038324 | 66 | Miga1 | 129S1,BTBR,C57L,CBA,DBA1J,SMJ | feature_elongation,3_prime_UTR_variant |
| 3 | 154283843 | 154284435 | 592 | Tyw3 | CE,RF | feature_elongation,3_prime_UTR_variant |
| 4 | 43832556 | 43833528 | 972 | Or13c7c | 129S1,BUB,PJ,PL,SEA,SJL | feature_elongation,3_prime_UTR_variant |
| 4 | 43856121 | 43856173 | 52 | Or13c7 | AJ,AKR,BALB,C3H,CE,KK,MA/My,MRL,NOD,NOR,NZB,NZO,PL,RF,RH2 | feature_elongation,3_prime_UTR_variant |
| 4 | 46633093 | 46633265 | 172 | Tbc1d2 | 129S1,AJ,BALB,C3H,C57L,PJ,RH2,SEA,SJL | feature_elongation,3_prime_UTR_variant |

| | | | | | | | |
|---|---|---|---|---|---|---|---|
| 4 | 51228991 | 51229072 | 81 | Cylc2 | AKR,CBA,CE,DBA1J,DBA2J,MRL,RF,SMJ | feature_elongation,coding_sequence_variant | |
| 4 | 52853755 | 52853820 | 65 | Or13c3 | MA/My | feature_elongation,3_prime_UTR_variant | |
| 4 | 59757794 | 59757902 | 108 | E130308A19Rik | CE | feature_elongation,3_prime_UTR_variant | |
| 4 | 62607859 | 62607913 | 54 | Rgs3 | CE | feature_elongation,coding_sequence_variant | |
| 4 | 65274381 | 65274458 | 77 | Pappa | PJ | feature_elongation,3_prime_UTR_variant | |
| 4 | 70139317 | 70139480 | 163 | Cdk5rap2 | 129S1,C57L,ILNJ2,KK,PJ | feature_elongation,3_prime_UTR_variant | |
| 4 | 86902280 | 86902345 | 65 | Slc24a2 | CE | feature_elongation,3_prime_UTR_variant | |
| 4 | 88745704 | 88745764 | 60 | Ifna6 | CE | feature_elongation,coding_sequence_variant | |
| 4 | 89192816 | 89192872 | 56 | Cdkn2a | MA/My | feature_elongation,3_prime_UTR_variant | |
| 4 | 99147955 | 99148173 | 218 | Atg4c | 129S1,AJ,AKR,BALB,BTBR,BUB,C3H,C57L,CBA,CE,FVB,KK,MA/My,MRL,NOD,NOR,NZB,PJ,PL,RF,RH2,SEA,SJL,SMJ,SWR,TH | feature_elongation,3_prime_UTR_variant | |
| 4 | 101747714 | 101747805 | 91 | C130073F10Rik | NZB,NZO | feature_elongation,coding_sequence_variant | |
| 4 | 103030463 | 103030515 | 52 | Slc35d1 | CE,MA/My,NZW,SJL | feature_elongation,3_prime_UTR_variant | |
| 4 | 115669326 | 115669560 | 234 | Atpaf1 | AJ,CE,DBA1J,DBA2J | feature_elongation,3_prime_UTR_variant | |
| 4 | 115990157 | 115990361 | 204 | Lurap1 | B10D2,B10J,C58J,KK,PL,TH | feature_elongation,3_prime_UTR_variant | |
| 4 | 117746600 | 117746652 | 52 | Dph2 | ILNJ2 | feature_elongation,3_prime_UTR_variant | |
| 4 | 119117520 | 119117735 | 215 | Cldn19 | NZO,NZW | feature_elongation,3_prime_UTR_variant | |
| 4 | 120804674 | 120804830 | 156 | Zfp69 | ILNJ2 | feature_elongation,coding_sequence_variant | |
| 4 | 124836904 | 124837109 | 205 | Airim | BUB,FVB,NOD,SJL,SWR | feature_elongation,3_prime_UTR_variant | |
| 4 | 125006469 | 125006526 | 57 | Meaf6 | AJ | feature_elongation,3_prime_UTR_variant | |
| 4 | 126229814 | 126229906 | 92 | Ago3 | B10D2,B10J,NZW | feature_elongation,3_prime_UTR_variant | |
| 4 | 126231620 | 126232012 | 392 | Ago3 | AJ,B10D2,B10J,NZW,RH2 | feature_elongation,3_prime_UTR_variant | |
| 4 | 128662818 | 128662983 | 165 | A3galt2 | B10D2,B10J,DBA1J,KK,NOR,PJ,SEA,SMJ | feature_elongation,3_prime_UTR_variant | |
| 4 | 130662673 | 130662780 | 107 | Laptm5 | 129S1,B10D2,B10J,BTBR,BUB,C57L,DBA1J,FVB,ILNJ2,KK,MA/My,NOR,NZB,PJ,PL,SEA,SJL,SMJ,SWR,TH | feature_elongation,3_prime_UTR_variant | |
| 4 | 131950649 | 131950705 | 56 | Gmeb1 | DBA1J,KK,NZB,SMJ | feature_elongation,3_prime_UTR_variant | |
| 4 | 132083758 | 132083811 | 53 | Phactr4 | DBA1J,KK,SMJ | feature_elongation,3_prime_UTR_variant | |
| 4 | 132452249 | 132452447 | 198 | Xkr8 | 129S1,AJ,AKR,BALB,BTBR,C3H,C57L,CBA,CE,DBA2J,ILNJ2,MRL,NOD,NZO,NZW,RF,RH2,SJL,SWR,TH | feature_elongation,3_prime_UTR_variant | |
| 4 | 132729192 | 132729370 | 178 | Fgr | DBA1J,KK,NZB,NZO,SMJ | feature_elongation,3_prime_UTR_variant | |
| 4 | 133389205 | 133389409 | 204 | Pigv | FVB | feature_elongation,3_prime_UTR_variant | |
| 4 | 133692302 | 133692481 | 179 | Hmgn2 | 129S1,BALB,BTBR,C57L,DBA1J,DBA2J,ILNJ2,NOD,PL,SMJ | feature_elongation,3_prime_UTR_variant | |
| 4 | 136019007 | 136019077 | 70 | Zfp46 | 129S1,B10D2,B10J,C57L,DBA1J,KK,NZO,PJ,SEA | feature_elongation,3_prime_UTR_variant | |
| 4 | 136081060 | 136081119 | 59 | Hnrnpr | 129S1,B10D2,B10J,C57L,DBA1J,KK,NZO,PJ,SEA | feature_elongation,3_prime_UTR_variant | |
| 4 | 136086167 | 136086640 | 473 | Hnrnpr | 129S1,B10D2,B10J,C57L,DBA1J,KK,NZO,PJ,SEA | feature_elongation,3_prime_UTR_variant | |
| 4 | 136365877 | 136366085 | 208 | Lactbl1 | AKR,BUB,C3H,CBA,CE,ILNJ2,MRL,NZB,NZW,RF | feature_elongation,3_prime_UTR_variant | |
| 4 | 137408315 | 137408378 | 63 | Rap1gap | AJ,AKR,BALB,BUB,C3H,CBA,CE,DBA2J,FVB,KK,MRL,NZB,NZW,PL,RF,RH2,SJL,SWR,TH | feature_elongation,5_prime_UTR_variant | |
| 4 | 138043683 | 138043866 | 183 | Pink1 | BALB,DBA2J,FVB,KK,SJL,SWR | feature_elongation,5_prime_UTR_variant | |
| 4 | 138184295 | 138184349 | 54 | Camk2n1 | BALB,BUB,C3H,CBA,CE,DBA1J,DBA2J,ILNJ2,KK,MA/My,MRL,NZB,NZW,SJL,SMJ,TH | feature_elongation,3_prime_UTR_variant | |
| 4 | 139465648 | 139465704 | 56 | Pax7 | AKR,BALB,BUB,C3H,CBA,DBA1J,DBA2J,MRL,NOR,PL,RF,SMJ | feature_elongation,3_prime_UTR_variant | |
| 4 | 146061338 | 146061422 | 84 | Zfp987 | 129S1,BTBR,C57L,C58J,LP,PJ,SEA | feature_elongation,coding_sequence_variant | |
| 4 | 147759389 | 147759556 | 167 | Zfp534 | AKR,BUB,C3H,CBA,CE,DBA1J,KK,MA/My,MRL,NOD,NZB,NZO,NZW,PL,RF,SJL,SMJ | feature_elongation,coding_sequence_variant | FALSE |
| 4 | 147958725 | 147958789 | 64 | Mfn2 | 129S1,AKR,BTBR,BUB,C3H,C57L,C58J,CBA,CE,DBA1J,DBA2J,KK,LP,MA/My,MRL,NOD,NZB,NZO,NZW,PJ,PL,RF,SEA,SJL,SMJ,SWR,TH | feature_elongation,3_prime_UTR_variant | |
| 4 | 148163867 | 148164002 | 135 | Agtrap | DBA2J,TH | feature_elongation,3_prime_UTR_variant | |
| 4 | 148184523 | 148184746 | 223 | Draxin | BUB,C58J,KK,LP,MA/My,NOD,NZB,NZW,PJ,RF,SEA,SJL,SMJ,SWR | feature_elongation,3_prime_UTR_variant | |
| 4 | 149557829 | 149558010 | 181 | Nmnat1 | SWR | feature_elongation,coding_sequence_variant | |
| 4 | 154950251 | 154950595 | 344 | Ttc34 | 129S1,AJ,AKR,BALB,BTBR,BUB,C3H,C57L,CBA,CE,DBA1J,DBA2J,FVB,ILNJ2,KK,MA/My,MRL,NOD,NOR,NZB,NZO,NZW,PJ,PL,RF,RH2,SEA,SJL,SMJ,SWR,TH | feature_elongation,3_prime_UTR_variant | |
| 4 | 155867086 | 155867289 | 203 | Tmem88b | C57L,CE,ILNJ2,SMJ | feature_elongation,3_prime_UTR_variant | |
| 4 | 155917576 | 155917785 | 209 | Aurkaip1 | C57L,CE,FVB,NOD,NOR,SJL,SWR,TH | feature_elongation,3_prime_UTR_variant | |
| 4 | 155973321 | 155973531 | 210 | Ints11,Pusl1 | 129S1,AJ,AKR,BALB,BUB,C3H,C57L,CBA,DBA1J,DBA2J,ILNJ2,NZO,PL,RF,RH2,SMJ | feature_elongation,3_prime_UTR_variant | |
| 4 | 156043815 | 156044270 | 455 | Ube2j2 | 129S1,AJ,AKR,BALB,BUB,C3H,CBA,CE,DBA1J,DBA2J,FVB,ILNJ2,NOD,NOR,NZO,PL,RF,RH2,SJL,SMJ,SWR,TH | feature_elongation,3_prime_UTR_variant | |
| 4 | 156302483 | 156302582 | 99 | Perm1 | 129S1,AJ,AKR,BALB,C3H,C57L,C58J,DBA1J,DBA2J,FVB,ILNJ2,MA/My,MRL,NOD,NOR,NZO,PL,RF,RH2,SJL,SMJ,SWR | feature_elongation,coding_sequence_variant | |
| 4 | 156332083 | 156332270 | 187 | Samd11 | NZO | feature_elongation,3_prime_UTR_variant | |
| 5 | 3576001 | 3576092 | 91 | Cdk6 | BUB,NOD,NOR | feature_elongation,3_prime_UTR_variant | |
| 5 | 3577768 | 3577838 | 70 | Cdk6 | BUB,NOD,NOR | feature_elongation,3_prime_UTR_variant | |
| 5 | 8751924 | 8752104 | 180 | Abcb1a | BUB,NOD,NOR | feature_elongation,coding_sequence_variant | |
| 5 | 21962539 | 21962756 | 217 | Dnajc2 | NZO | feature_elongation,3_prime_UTR_variant | |
| 5 | 31301623 | 31301839 | 216 | Mpv17 | NZO,NZW | feature_elongation,3_prime_UTR_variant | |
| 5 | 43775222 | 43775278 | 56 | C1qtnf7 | MA/My,PJ,PL | feature_elongation,3_prime_UTR_variant | |
| 5 | 44068957 | 44069475 | 518 | Cd38 | BALB,BUB,DBA1J,DBA2J,FVB,ILNJ2,KK,LP,MA/My,RH2,SEA,SWR | feature_elongation,3_prime_UTR_variant | |
| 5 | 52925614 | 52925823 | 209 | Pi4k2b | AJ,AKR,BALB,BTBR,BUB,C3H,CBA,CE,DBA1J,DBA2J,FVB,ILNJ2,KK,MA/My,MRL,NOD,NZB,NZO,NZW,PJ,PL,RF,RH2,SEA,SJL,SMJ,SWR,TH | feature_elongation,3_prime_UTR_variant | |
| 5 | 67514499 | 67514996 | 497 | Slc30a9 | BUB,CE,FVB,PJ | feature_elongation,3_prime_UTR_variant | |
| 5 | 74161669 | 74161721 | 52 | Usp46 | C3H,CBA,DBA1J,DBA2J,ILNJ2,NZB | feature_elongation,3_prime_UTR_variant | |
| 5 | 76632096 | 76632182 | 86 | Exoc1l | AJ,AKR,BUB,CE,FVB,KK,MRL,PJ,PL,SJL,SMJ | feature_elongation,5_prime_UTR_variant | |
| 5 | 84214579 | 84214656 | 77 | Epha5 | 129S1,AJ,AKR,B10D2,B10J,BALB,BTBR,BUB,C3H,C57L,C58J,CBA,CE,DBA1J,DBA2J,FVB,ILNJ2,KK,LP,MA/My,MRL,NOD,NOR,NZB,NZO,NZW,PJ,PL,RF,RH2,SEA,SJL,SMJ,SWR,TH | feature_elongation,3_prime_UTR_variant | |
| 5 | 87965877 | 87965931 | 54 | Csn1s2b | SMJ | feature_elongation,3_prime_UTR_variant | |
| 5 | 93189123 | 93189326 | 203 | Sowahb | AKR,BALB,BTBR,BUB,C3H,CBA,CE,DBA1J,DBA2J,FVB,MA/My,MRL,NOD,NOR,NZW,PJ,RH2,SEA,SJL,SWR,TH | feature_elongation,3_prime_UTR_variant | |

| | | | | | | | |
|---|---|---|---|---|---|---|---|
| 5 | 97230339 | 97230438 | 99 | Paqr3 | 129S1,AJ,BTBR,BUB,C57L,KK,MA/My,NOD,NOR,NZB,NZO,SWR,TH | feature_elongation,3_prime_UTR_variant | |
| 5 | 104325526 | 104325796 | 270 | Dspp | NZB | feature_elongation,coding_sequence_variant | |
| 5 | 104699955 | 104700168 | 213 | Thoc2l | AJ,BALB,BUB,C3H,CBA,DBA1J,DBA2J,FVB,ILNJ2,KK,NZB,NZO,RH2,SEA,SJL,SWR,TH | feature_elongation,3_prime_UTR_variant | |
| 5 | 104730384 | 104730636 | 252 | Zfp33b | 129S1,AJ,AKR,B10D2,B10J,BALB,BTBR,C57L,C58J,CE,DBA1J,DBA2J,ILNJ2,KK,LP,MA/My,MRL,NOD,NOR,NZB,NZO,NZW,PJ,PL,RF,RH2,SEA,SMJ,SWR | feature_elongation,coding_sequence_variant | |
| 5 | 107697640 | 107697691 | 51 | 1700028K03Rik | BUB,C3H,CBA,FVB,PJ,PL,SJL,SWR,TH | feature_elongation,3_prime_UTR_variant | |
| 5 | 109886154 | 109886453 | 299 | 4930522L14Rik | CE,DBA1J,DBA2J,LP | feature_elongation,3_prime_UTR_variant | |
| 5 | 110224606 | 110224816 | 210 | Gm35315 | 129S1,AJ,AKR,BALB,BTBR,C57L,ILNJ2,KK,MA/My,MRL,NOD,NOR,NZB,NZO,NZW,RF,RH2,SEA,SMJ | feature_elongation,3_prime_UTR_variant | |
| 5 | 110275243 | 110275580 | 337 | Zfp605 | BUB,C3H,CBA,FVB,PJ,PL,SJL,SWR,TH | feature_elongation,coding_sequence_variant | |
| 5 | 117492869 | 117493050 | 181 | Vsig10 | RF | feature_elongation,3_prime_UTR_variant | |
| 5 | 117885493 | 117885863 | 370 | Ksr2 | 129S1,AJ,AKR,BALB,BUB,C57L,CE,DBA1J,DBA2J,PJ,PL,RF,RH2,SEA,SJL,SMJ | feature_elongation,coding_sequence_variant | |
| 5 | 118330475 | 118330535 | 60 | Rnft2 | 129S1,AJ,AKR,BALB,BUB,C3H,C57L,CBA,CE,DBA1J,DBA2J,ILNJ2,KK,NOR,NZB,NZO,PJ,PL,RH2,SEA,SJL,SMJ,TH | feature_elongation,3_prime_UTR_variant | |
| 5 | 121735584 | 121735639 | 55 | Acad12 | AKR,BUB,ILNJ2,KK,MA/My,PL,RF,SJL,TH | feature_elongation,3_prime_UTR_variant | |
| 5 | 122867592 | 122868247 | 655 | P2rx4 | RF | feature_elongation,3_prime_UTR_variant | |
| 5 | 123584316 | 123584547 | 231 | Mlxip | 129S1,AJ,AKR,BALB,BUB,C3H,C57L,CBA,CE,DBA1J,DBA2J,KK,MA/My,MRL,NOD,NOR,NZW,PJ,PL,RH2,SEA,SJL,SMJ | feature_elongation,3_prime_UTR_variant | |
| 5 | 124734957 | 124735144 | 187 | Gtf2h3 | 129S1,AJ,BALB,C57L,MRL,PL,RH2,SEA,SJL | feature_elongation,3_prime_UTR_variant | |
| 5 | 129062083 | 129062209 | 126 | Stx2 | AJ,BALB,C58J,FVB,LP,MRL,NOD,NOR,NZB,NZO,NZW,RH2,SEA,SMJ,SWR,TH | feature_elongation,3_prime_UTR_variant | |
| 5 | 130028471 | 130028677 | 206 | Gusb | AJ,BALB,MRL,RH2,SEA,SMJ | feature_elongation,3_prime_UTR_variant | |
| 5 | 130903567 | 130903641 | 74 | Galnt17 | NOD,NOR | feature_elongation,3_prime_UTR_variant | |
| 5 | 137178802 | 137178892 | 90 | Muc17 | LP | feature_elongation,coding_sequence_variant | |
| 5 | 137461977 | 137462076 | 99 | Zan | 129S1,BUB,LP,RF,SWR | feature_elongation,coding_sequence_variant | |
| 5 | 137641877 | 137642090 | 213 | Irs3 | ILNJ2,MA/My,NZW | feature_elongation,3_prime_UTR_variant | |
| 5 | 138192168 | 138192369 | 201 | Cnpy4 | 129S1,AJ,BALB,BTBR,C3H,C57L,CBA,CE,DBA1J,DBA2J,FVB,LP,MA/My,NOD,NOR,NZW,PJ,RH2,SEA,SJL | feature_elongation,3_prime_UTR_variant | |
| 5 | 138250128 | 138250187 | 59 | Nxpe5 | BUB,KK,LP,NZB,NZO,NZW,RF | feature_elongation,3_prime_UTR_variant | |
| 5 | 139323258 | 139323432 | 174 | Cox19 | NZW,SMJ | feature_elongation,3_prime_UTR_variant | |
| 5 | 139381580 | 139381784 | 204 | Gpr146 | ILNJ2,NZW,SMJ | feature_elongation,3_prime_UTR_variant | |
| 5 | 144493796 | 144493858 | 62 | Nptx2 | CE | feature_elongation,3_prime_UTR_variant | |
| 5 | 146409596 | 146409657 | 61 | Wasf3 | TH | feature_elongation,3_prime_UTR_variant | |
| 5 | 147498981 | 147499129 | 148 | Flt1 | TH | feature_elongation,3_prime_UTR_variant | |
| 5 | 147815847 | 147816008 | 161 | Slc46a3 | CE | feature_elongation,3_prime_UTR_variant | |
| 5 | 148252153 | 148252211 | 58 | Mtus2 | AKR,BUB,MA/My,MRL,NOD,NOR | feature_elongation,3_prime_UTR_variant | |
| 5 | 149215909 | 149216151 | 242 | Alox5ap | AKR,BUB,CE,MRL,SJL | feature_elongation,5_prime_UTR_variant | |
| 5 | 150565406 | 150565579 | 173 | N4bp2l2 | CE,ILNJ2,SEA,SWR | feature_elongation,3_prime_UTR_variant | |
| 6 | 4756453 | 4756534 | 81 | Peg10 | 129S1,AJ,AKR,B10D2,B10J,BALB,BTBR,BUB,C3H,C57L,C58J,CBA,CE,DBA1J,DBA2J,FVB,ILNJ2,KK,LP,MA/My,MRL,NOD,NOR,NZB,NZO,NZW,PL,RF,RH2,SEA,SJL,SMJ,SWR,TH | feature_elongation,3_prime_UTR_variant,feature_elongation,coding_sequence_variant | |
| 6 | 5008455 | 5008610 | 155 | Ppp1r9a | PJ | feature_elongation,3_prime_UTR_variant | |
| 6 | 15770578 | 15770629 | 51 | Mdfic | KK | feature_elongation,coding_sequence_variant | |
| 6 | 40844545 | 40844665 | 120 | Mgam2-ps | BUB,FVB,LP,SJL,SWR,TH | feature_elongation,coding_sequence_variant | |
| 6 | 47941607 | 47941665 | 58 | Zfp956 | AJ,AKR,BALB,BUB,C3H,CBA,CE,DBA1J,DBA2J,FVB,ILNJ2,MRL,NOD,NOR,RF,RH2,SJL,SMJ,SWR | feature_elongation,3_prime_UTR_variant | |
| 6 | 48964956 | 48965010 | 54 | Aoc1l3 | CE,ILNJ2 | feature_elongation,coding_sequence_variant | |
| 6 | 57046064 | 57046120 | 56 | Vmn1r9 | NZW | feature_elongation,5_prime_UTR_variant | |
| 6 | 57364776 | 57364976 | 200 | Vmn1r18 | 129S1,C57L,CE,ILNJ2,KK,NZB,NZO,NZW,PJ | feature_elongation,3_prime_UTR_variant | |
| 6 | 57382791 | 57383243 | 452 | Vmn1r19 | 129S1,C57L,KK,NZB,NZO,NZW | feature_elongation,3_prime_UTR_variant | |
| 6 | 65433857 | 65434003 | 146 | Qrfprl | KK | feature_elongation,3_prime_UTR_variant | |
| 6 | 67263640 | 67263736 | 96 | Serbp1 | MA/My | feature_elongation,3_prime_UTR_variant | |
| 6 | 72329092 | 72329153 | 61 | 0610030E20Rik | AKR,C58J,KK,NOD,NOR,PL,RF | feature_elongation,3_prime_UTR_variant | |
| 6 | 72407385 | 72407451 | 66 | Ggcx | AKR,C3H,C58J,CBA,DBA1J,DBA2J,KK,MA/My,MRL,NOD,NOR,PL,RF,SMJ | feature_elongation,3_prime_UTR_variant | |
| 6 | 72584962 | 72585190 | 228 | Retsat | C58J,KK,NOD,NOR | feature_elongation,3_prime_UTR_variant | |
| 6 | 82535170 | 82535224 | 54 | Tacr1 | AJ,AKR,BALB,BTBR,BUB,CE,FVB,ILNJ2,LP,MA/My,MRL,NOD,NOR,NZW,PL,RH2,SEA,SJL,SWR | feature_elongation,3_prime_UTR_variant | |
| 6 | 83127309 | 83127433 | 124 | Wdr54 | KK,NZB,NZO,TH | feature_elongation,3_prime_UTR_variant | |
| 6 | 86992589 | 86992720 | 131 | Nfu1 | AKR,BALB,KK,MA/My,MRL,NZB,NZO,PL,RH2,TH | feature_elongation,coding_sequence_variant | FALSE |
| 6 | 87413046 | 87413130 | 84 | Bmp10 | AJ,AKR,B10D2,B10J,BALB,BTBR,BUB,C57L,C58J,FVB,ILNJ2,KK,LP,MRL,NOD,NOR,NZO,PL,RF,RH2,SJL,SMJ,TH | feature_elongation,3_prime_UTR_variant | |
| 6 | 87791450 | 87791648 | 198 | Isy1 | SWR | feature_elongation,3_prime_UTR_variant | |
| 6 | 89691868 | 89692283 | 415 | Vmn1r40 | NZO | feature_elongation,coding_sequence_variant | |
| 6 | 90247433 | 90247497 | 64 | Vmn1r54 | CE,PJ,SEA | feature_elongation,3_prime_UTR_variant | |
| 6 | 90285570 | 90285657 | 87 | Chst13 | CE,PJ,SEA | feature_elongation,3_prime_UTR_variant | |
| 6 | 90635724 | 90635822 | 98 | Iqsec1 | CE,PJ,SEA | feature_elongation,3_prime_UTR_variant | |
| 6 | 91441264 | 91441479 | 215 | Chchd4 | AJ,BALB,BTBR,BUB,RF,RH2,SJL | feature_elongation,3_prime_UTR_variant | |
| 6 | 92151624 | 92151824 | 200 | Mrps25 | CE,PJ,SEA | feature_elongation,3_prime_UTR_variant | |
| 6 | 93653633 | 93653800 | 167 | Magi1 | C3H,CBA,CE,ILNJ2,NOD,NOR,SMJ | feature_elongation,3_prime_UTR_variant | |
| 6 | 100784144 | 100784202 | 58 | Gxylt2 | SJL | feature_elongation,3_prime_UTR_variant | |
| 6 | 115599045 | 115599735 | 690 | Mkrn2 | BALB | feature_elongation,3_prime_UTR_variant | |
| 6 | 116557792 | 116558112 | 320 | Or6d15 | BUB,CE,FVB,KK,NZB,NZW,PJ,SJL,SMJ,SWR | feature_elongation,3_prime_UTR_variant | |
| 6 | 116659873 | 116659924 | 51 | Tmem72 | 129S1,AKR,BTBR,BUB,C3H,C57L,C58J,CBA,DBA1J,DBA2J,FVB,KK,LP,MRL,NOD,NOR,NZO,PJ,PL,RF,RH2,SJL,SMJ,SWR | feature_elongation,3_prime_UTR_variant | |
| 6 | 116668790 | 116668878 | 88 | Tmem72 | C3H,CBA,PL | feature_elongation,3_prime_UTR_variant | |
| 6 | 117154173 | 117154235 | 62 | Cxcl12 | SWR | feature_elongation,3_prime_UTR_variant | |
| 6 | 123119769 | 123119823 | 54 | Clec4a2 | SWR | feature_elongation,3_prime_UTR_variant | |
| 6 | 123361261 | 123361746 | 485 | Vmn2r20 | SMJ | feature_elongation,3_prime_UTR_variant | |
| 6 | 125523986 | 125524168 | 182 | Vwf | KK | feature_elongation,coding_sequence_variant | |
| 6 | 126509828 | 126509883 | 55 | Kcna5 | NOD,NOR | feature_elongation,3_prime_UTR_variant | |
| 6 | 128791418 | 128791623 | 205 | Klrb1b | NZO | feature_elongation,3_prime_UTR_variant | |

| | | | | | | |
|---|---|---|---|---|---|---|
| 6 | 129463734 | 129463912 | 178 | Olr1 | AJ,AKR,BALB,BUB,C3H,C57L,CBA,DBA1J,DBA2J,ILNJ2,LP,MRL,PL,RF,RH2,SEA | feature_elongation,3_prime_UTR_variant |
| 6 | 129625027 | 129625297 | 270 | Klrc2 | 129S1,AJ,AKR,BALB,BUB,C3H,C57L,C58J,CBA,CE,DBA1J,DBA2J,FVB,ILNJ2,LP,MA/My,MRL,NOD,NOR,NZO,PL,RF,RH2,SEA,SJL,SMJ,SWR,TH | feature_elongation,3_prime_UTR_variant |
| 6 | 129744568 | 129744622 | 54 | Klrh1 | AKR,C3H,NZO,RF | feature_elongation,3_prime_UTR_variant |
| 6 | 132289137 | 132289344 | 207 | Prb1b | AJ,AKR,BALB,BUB,C3H,C57L,CBA,DBA1J,DBA2J,ILNJ2,MRL,NOD,NOR,NZB,PL,RF,RH2,SEA,SWR | feature_elongation,coding_sequence_variant |
| 6 | 132549261 | 132549384 | 123 | Prh1 | AJ,AKR,BALB,BUB,C3H,C57L,CBA,DBA1J,DBA2J,ILNJ2,MRL,NZB,NZO,PL,RF,RH2,SEA | feature_elongation,coding_sequence_variant |
| 6 | 133083395 | 133083498 | 103 | Smim10l1 | 129S1,C58J,FVB,KK,LP,NOD,NOR,NZW,SJL,SMJ,SWR,TH | feature_elongation,3_prime_UTR_variant |
| 6 | 134425852 | 134426096 | 244 | Lrp6 | AJ,AKR,BUB,C3H,C58J,CBA,CE,DBA1J,DBA2J,KK,MRL,NOD,NOR,NZB,NZW,PL,RF,SJL,TH | feature_elongation,3_prime_UTR_variant |
| 6 | 135705656 | 135705717 | 61 | Grin2b | 129S1,AJ,AKR,BALB,BTBR,BUB,C3H,C57L,CBA,CE,DBA1J,DBA2J,FVB,KK,MA/My,MRL,NOD,NOR,NZB,NZW,PJ,PL,RF,RH2,SEA,SJL,SWR,TH | feature_elongation,3_prime_UTR_variant |
| 6 | 140623945 | 140624171 | 226 | Aebp2 | PJ,SWR | feature_elongation,3_prime_UTR_variant |
| 6 | 140626101 | 140626153 | 52 | Aebp2 | FVB | feature_elongation,3_prime_UTR_variant |
| 6 | 144155263 | 144155316 | 53 | Sox5 | AKR,BUB,C3H,CBA,CE,DBA2J,ILNJ2,MRL,NZO,RF,SJL | feature_elongation,5_prime_UTR_variant |
| 6 | 146734471 | 146734596 | 125 | Bmal2 | 129S1,AJ,BALB,BTBR,C57L,CBA,FVB,ILNJ2,MA/My,NOD,NOR,PJ,RF,SEA,SMJ,SWR,TH | feature_elongation,3_prime_UTR_variant |
| 7 | 3537506 | 3538199 | 693 | Tarm1 | 129S1,AJ,AKR,BALB,BTBR,BUB,C57L,FVB,KK,LP,MA/My,MRL,NOD,NOR,NZB,NZO,NZW,PL,RF,RH2,SJL,SMJ,SWR,TH | feature_elongation,3_prime_UTR_variant |
| 7 | 3824543 | 3824763 | 220 | Pira13 | MRL | feature_elongation,coding_sequence_variant |
| 7 | 4371488 | 4371768 | 280 | Gp6 | 129S1,AJ,AKR,BALB,BTBR,BUB,C3H,C57L,CBA,CE,DBA1J,DBA2J,FVB,ILNJ2,KK,LP,MA/My,MRL,NOD,NOR,NZB,NZO,NZW,PJ,PL,RF,RH2,SEA,SJL,SMJ,SWR,TH | feature_elongation,3_prime_UTR_variant |
| 7 | 4482739 | 4483014 | 275 | Eps8l1 | CE,MRL,SWR,TH | feature_elongation,3_prime_UTR_variant |
| 7 | 4695285 | 4695400 | 115 | Brsk1 | BUB,CE,MRL,NZO,SMJ | feature_elongation,5_prime_UTR_variant |
| 7 | 4968267 | 4968441 | 174 | Sbk3 | 129S1,AJ,AKR,BALB,BTBR,BUB,C3H,C57L,CBA,DBA1J,DBA2J,FVB,ILNJ2,KK,LP,MA/My,NOD,NOR,NZW,PJ,PL,RF,RH2,SEA,SJL | feature_elongation,3_prime_UTR_variant |
| 7 | 5543514 | 5543604 | 90 | Vmn1r60 | MRL | feature_elongation,3_prime_UTR_variant |
| 7 | 10007619 | 10007673 | 54 | Vmn1r66 | SWR,TH | feature_elongation,3_prime_UTR_variant |
| 7 | 11583302 | 11583392 | 90 | Vmn1r74 | AKR,FVB,MRL,NOD,NOR,RH2 | feature_elongation,3_prime_UTR_variant |
| 7 | 11782418 | 11783162 | 744 | Vmn1r77 | AKR,FVB,MRL,NOD,NOR,RH2 | feature_elongation,3_prime_UTR_variant |
| 7 | 11887617 | 11887679 | 62 | Vmn1r78 | AKR,FVB,MRL,NOD,NOR | feature_elongation,3_prime_UTR_variant |
| 7 | 12911808 | 12912362 | 554 | Vmn1r88 | C58J | feature_elongation,coding_sequence_variant |
| 7 | 12951497 | 12951688 | 191 | Vmn1r89 | C58J | feature_elongation,5_prime_UTR_variant |
| 7 | 13093681 | 13093853 | 172 | Pla2g4c | C58J | feature_elongation,3_prime_UTR_variant |
| 7 | 15671212 | 15671380 | 168 | Nop53 | BTBR,BUB,C58J,CE,FVB,KK,NOD,NOR,NZO,PL | feature_elongation,3_prime_UTR_variant |
| 7 | 15968971 | 15969031 | 60 | C5ar2 | 129S1,AJ,AKR,BALB,BTBR,C3H,C57L,C58J,CBA,DBA1J,DBA2J,ILNJ2,KK,LP,MA/My,MRL,NZB,NZO,NZW,PJ,RF,RH2,SEA,SJL,SMJ,TH | feature_elongation,3_prime_UTR_variant |
| 7 | 15970294 | 15970376 | 82 | C5ar2 | 129S1,AJ,AKR,BALB,BTBR,C3H,C57L,CBA,DBA1J,DBA2J,ILNJ2,KK,LP,MA/My,MRL,NZB,NZO,NZW,PJ,RF,RH2,SEA,SJL,SMJ,TH | feature_elongation,3_prime_UTR_variant |
| 7 | 16054274 | 16054361 | 87 | Sae1 | 129S1,BTBR,C3H,C57L,C58J,CBA,DBA1J,DBA2J,MA/My,NZB,PJ,RF,SEA,SMJ | feature_elongation,3_prime_UTR_variant |
| 7 | 16459558 | 16459628 | 70 | Ceacam9 | NOD,NOR,SJL | feature_elongation,3_prime_UTR_variant |
| 7 | 18958097 | 18958311 | 214 | Gpr4 | 129S1,BTBR,C3H,C57L,CBA,NOD,NOR,NZB,NZW,RF,SJL | feature_elongation,3_prime_UTR_variant |
| 7 | 19078720 | 19078784 | 64 | Ercc1 | AKR,BUB,C58J,CE,MRL,NZO,TH | feature_elongation,5_prime_UTR_variant |
| 7 | 19724368 | 19724661 | 293 | Ceacam20 | 129S1,BTBR,C3H,C57L,CBA,NOD,NOR,NZB,RF,SJL | feature_elongation,3_prime_UTR_variant |
| 7 | 23203747 | 23203841 | 94 | Vmn1r167 | 129S1,BTBR,BUB,C3H,C57L,CBA,NOD,NOR,NZB,NZW,SJL | feature_elongation,3_prime_UTR_variant |
| 7 | 23536610 | 23537179 | 569 | Vmn1r176 | 129S1,BTBR,BUB,C3H,C57L,CBA,NOD,NOR,NZB,NZW,SJL | feature_elongation,5_prime_UTR_variant |
| 7 | 23880563 | 23880638 | 75 | Zfp114 | 129S1,BTBR,BUB,C3H,C57L,CBA,NOD,NOR,NZB,NZW,SJL | feature_elongation,coding_sequence_variant |
| 7 | 27450569 | 27450782 | 213 | Zfp60 | TH | feature_elongation,3_prime_UTR_variant |
| 7 | 30451828 | 30451882 | 54 | Sbsn | 129S1,AJ,AKR,BALB,BTBR,C57L,C58J,CE,DBA1J,DBA2J,ILNJ2,LP,MA/My,MRL,NOD,NOR,NZB,NZO,NZW,PJ,PL,RF,SEA,SJL,SMJ,TH | feature_elongation,coding_sequence_variant |
| 7 | 30577837 | 30578631 | 794 | Cd22 | AKR,BTBR,C57L,DBA1J,DBA2J,NZB,NZO,NZW,PJ,PL,SEA,SMJ,TH | feature_elongation,5_prime_UTR_variant |
| 7 | 35246858 | 35247033 | 175 | Nudt19 | BTBR,C57L,C58J,CE,DBA1J,DBA2J,FVB,ILNJ2,LP,MA/My,MRL,NOD,NOR,PJ,RF,SJL,SMJ,SWR | feature_elongation,3_prime_UTR_variant |
| 7 | 41300737 | 41300915 | 178 | Zfp788 | AJ,BALB,C57L,CE,FVB,MA/My,NOD,NOR,NZB,NZO,PL,RH2,SEA,SMJ,SWR,TH | feature_elongation,3_prime_UTR_variant |
| 7 | 43142044 | 43142244 | 200 | 4931406B18Rik | DBA1J,DBA2J,ILNJ2,NZW,PJ | feature_elongation,3_prime_UTR_variant |
| 7 | 43959688 | 43960022 | 334 | Shank1 | 129S1,AJ,AKR,B10D2,B10J,BALB,BTBR,BUB,C3H,C57L,C58J,CBA,CE,DBA1J,DBA2J,FVB,ILNJ2,KK,LP,MA/My,MRL,NOD,NOR,NZB,NZO,NZW,PJ,PL,RF,RH2,SEA,SJL,SMJ,SWR,TH | feature_elongation,5_prime_UTR_variant |
| 7 | 44385782 | 44386152 | 370 | Zfp473 | ILNJ2,NZW,PJ,SJL | feature_elongation,3_prime_UTR_variant |
| 7 | 44387256 | 44387457 | 201 | Zfp473 | ILNJ2,NZW,PJ,SJL | feature_elongation,3_prime_UTR_variant |
| 7 | 44906960 | 44907025 | 65 | Slc6a16 | ILNJ2,NZW,PJ,SJL | frameshift_variant,splice_region_variant |
| 7 | 46079660 | 46079740 | 80 | Kcnc1 | C58J | feature_elongation,3_prime_UTR_variant |
| 7 | 49258159 | 49258210 | 51 | Nav2 | ILNJ2,SJL | feature_elongation,3_prime_UTR_variant |
| 7 | 49609827 | 49609883 | 56 | Slc6a5 | ILNJ2,NZW,PJ,SJL | feature_elongation,3_prime_UTR_variant |
| 7 | 50249178 | 50249503 | 325 | 4933405O20Rik | KK | feature_elongation,coding_sequence_variant |
| 7 | 55428213 | 55428375 | 162 | Siglech | ILNJ2,KK,NZW,PJ,SJL | feature_elongation,3_prime_UTR_variant |
| 7 | 84587416 | 84588411 | 995 | Vmn2r65 | SMJ | feature_elongation,3_prime_UTR_variant |
| 7 | 85597826 | 85598755 | 929 | Vmn2r74 | 129S1,AJ,AKR,BALB,BTBR,BUB,C3H,C57L,CBA,DBA1J,DBA2J,FVB,ILNJ2,KK,MA/My,MRL,NOD,NOR,NZB,NZO,NZW,PJ,PL,RF,RH2,SEA,SJL,SWR,TH | feature_elongation,3_prime_UTR_variant |

| | | | | | | |
|---|---|---|---|---|---|---|
| 7 | 85983563 | 85983618 | 55 | Or14a260 | 129S1,BTBR,C3H,C57L,CBA,DBA1J,DBA2J,ILNJ2,KK,NZB,NZO,NZW,PJ,SEA | feature_elongation,3_prime_UTR_variant |
| 7 | 89176234 | 89176316 | 82 | Prss23 | SMJ | feature_elongation,5_prime_UTR_variant |
| 7 | 89502170 | 89502411 | 241 | Me3 | SMJ | feature_elongation,3_prime_UTR_variant |
| 7 | 97108129 | 97108312 | 183 | Kctd14 | 129S1,AJ,AKR,BALB,BTBR,BUB,C57L,CE,FVB,KK,MA/My,MRL,NOD,NOR,NZB,NZW,PL,RF,RH2,SJL,SMJ,SWR,TH | feature_elongation,3_prime_UTR_variant |
| 7 | 97856120 | 97856647 | 527 | Acer3 | CE | feature_elongation,3_prime_UTR_variant |
| 7 | 98918041 | 98918179 | 138 | Map6 | C3H,CBA,DBA1J,DBA2J,ILNJ2,PJ,SEA | feature_elongation,coding_sequence_variant |
| 7 | 101530539 | 101530894 | 355 | Anapc15 | C3H,CBA,DBA1J,DBA2J,ILNJ2,PJ,SEA | feature_elongation,5_prime_UTR_variant |
| 7 | 102141979 | 102142159 | 180 | Or55b10 | KK,NOD,SJL,SMJ,SWR | feature_elongation,3_prime_UTR_variant |
| 7 | 102204962 | 102205166 | 204 | Or52b3 | 129S1,AJ,AKR,BALB,BTBR,BUB,C3H,C57L,CBA,DBA1J,DBA2J,FVB,ILNJ2,MRL,NZB,NZO,NZW,PJ,PL,RF,RH2,SEA,SMJ,TH | feature_elongation,3_prime_UTR_variant |
| 7 | 102255617 | 102255900 | 283 | Or52k2 | KK,NOD,SJL,SWR | feature_elongation,3_prime_UTR_variant |
| 7 | 102389211 | 102389353 | 142 | Or51e2 | MA/My,NZO,SMJ,TH | feature_elongation,3_prime_UTR_variant |
| 7 | 102955782 | 102955978 | 196 | Or52e19 | KK,NOD,SJL,SWR | feature_elongation,5_prime_UTR_variant |
| 7 | 103101635 | 103102174 | 539 | Or51l14 | NZO,SMJ,TH | feature_elongation,3_prime_UTR_variant |
| 7 | 103367005 | 103367226 | 221 | Or52a20 | KK,MA/My,NOD,NZO,SJL,SMJ,SWR | feature_elongation,3_prime_UTR_variant |
| 7 | 103595519 | 103595607 | 88 | Or51k2 | KK,NOD,SJL,SMJ,SWR,TH | feature_elongation,5_prime_UTR_variant |
| 7 | 103696558 | 103696634 | 76 | Or51a10 | KK,NOD,SJL,SMJ,SWR,TH | feature_elongation,3_prime_UTR_variant |
| 7 | 103732677 | 103733108 | 431 | Or51a24 | KK,NOD,SJL,SMJ,SWR,TH | feature_elongation,3_prime_UTR_variant |
| 7 | 103911127 | 103911331 | 204 | Trim34a | 129S1,AJ,AKR,BALB,BTBR,BUB,C3H,C57L,CBA,DBA1J,DBA2J,FVB,ILNJ2,MRL,NZB,NZW,PJ,RF,SEA | feature_elongation,3_prime_UTR_variant |
| 7 | 104012324 | 104012705 | 381 | Trim30b | 129S1,AJ,AKR,BALB,BTBR,BUB,C3H,C57L,CBA,CE,DBA1J,DBA2J,FVB,ILNJ2,MRL,NZB,NZO,NZW,PJ,PL,RF,RH2,SEA | feature_elongation,3_prime_UTR_variant |
| 7 | 104231542 | 104231639 | 97 | Or52d3 | KK,NZO,SMJ | feature_elongation,3_prime_UTR_variant |
| 7 | 105259731 | 105259950 | 219 | Trim3 | CE,KK | feature_elongation,3_prime_UTR_variant |
| 7 | 105292481 | 105292541 | 60 | Timm10b | CE | feature_elongation,3_prime_UTR_variant |
| 7 | 106572863 | 106573200 | 337 | Or6b6 | KK | feature_elongation,5_prime_UTR_variant |
| 7 | 106599825 | 106599898 | 73 | Or6a2 | 129S1,AJ,AKR,BALB,BTBR,C3H,C57L,CBA,DBA1J,DBA2J,ILNJ2,MRL,NZB,NZO,NZW,PJ,PL,RF,RH2,SEA | feature_elongation,3_prime_UTR_variant |
| 7 | 106636676 | 106636747 | 71 | Or10a5 | NZO | feature_elongation,3_prime_UTR_variant |
| 7 | 107630450 | 107630530 | 80 | Or5p6 | NOD,SMJ,SWR | feature_elongation,3_prime_UTR_variant |
| 7 | 107681963 | 107682063 | 100 | Or5p4 | NOD,NZO,SMJ,SWR | feature_elongation,3_prime_UTR_variant |
| 7 | 107721160 | 107721947 | 787 | Or5p60 | NOD,SMJ,SWR | feature_elongation,3_prime_UTR_variant |
| 7 | 107918020 | 107918108 | 88 | Or5p1 | NOD,SMJ,SWR | feature_elongation,3_prime_UTR_variant |
| 7 | 109907897 | 109907973 | 76 | Sbf2 | NOD,SWR | feature_elongation,3_prime_UTR_variant |
| 7 | 110655290 | 110655345 | 55 | Ctr9 | SMJ | feature_elongation,3_prime_UTR_variant |
| 7 | 119552375 | 119552431 | 56 | Lyrm1 | AJ,BALB,BTBR,C3H,CBA,DBA1J,DBA2J,ILNJ2,KK,NOR,NZB,NZO,PL,RH2,TH | feature_elongation,3_prime_UTR_variant |
| 7 | 119783295 | 119783346 | 51 | Anks4b | 129S1,AKR,B10D2,B10J,C57L,C58J,FVB,LP,MA/My,MRL,NOD,NZW,PJ,RF,SEA,SJL,SMJ,SWR | feature_elongation,3_prime_UTR_variant |
| 7 | 120996884 | 120996936 | 52 | Hs3st2 | BTBR | feature_elongation,3_prime_UTR_variant |
| 7 | 126186855 | 126186921 | 66 | Apobr | AJ,NZB,NZO,TH | feature_elongation,coding_sequence_variant |
| 7 | 126989123 | 126989303 | 180 | E430018J23Rik | AJ | feature_elongation,3_prime_UTR_variant |
| 7 | 127463487 | 127463539 | 52 | Zfp668 | TH | feature_elongation,3_prime_UTR_variant |
| 7 | 135286041 | 135286241 | 200 | Ptpre | ILNJ2 | feature_elongation,3_prime_UTR_variant |
| 7 | 139740901 | 139741790 | 889 | Or6ae1 | PJ | feature_elongation,3_prime_UTR_variant |
| 7 | 139917699 | 139917823 | 124 | Or12j2 | NZW,PJ | feature_elongation,3_prime_UTR_variant |
| 7 | 139924124 | 139924219 | 95 | Or13a27 | CE | feature_elongation,3_prime_UTR_variant |
| 7 | 139952248 | 139952443 | 195 | Or12j3 | NZW,PJ | feature_elongation,3_prime_UTR_variant |
| 7 | 140507818 | 140507874 | 56 | Nlrp6 | ILNJ2 | feature_elongation,3_prime_UTR_variant |
| 7 | 140681232 | 140681356 | 124 | Ano9 | NZW | feature_elongation,3_prime_UTR_variant |
| 7 | 141297148 | 141297295 | 147 | Muc2 | NZW | feature_elongation,coding_sequence_variant |
| 7 | 141743604 | 141743807 | 203 | Gm10153 | BALB,MRL,RF,RH2,SEA | stop_gained,frameshift_variant |
| 7 | 141755630 | 141755707 | 77 | Krtap5-3 | NZW | feature_elongation,coding_sequence_variant |
| 7 | 143190695 | 143190906 | 211 | Tnfrsf22 | C58J,LP,MA/My,PJ,SMJ | feature_elongation,3_prime_UTR_variant |
| 8 | 4577557 | 4577701 | 144 | Cers4 | 129S1,NZB | feature_elongation,3_prime_UTR_variant |
| 8 | 11035636 | 11035696 | 60 | Irs2 | BTBR | feature_elongation,3_prime_UTR_variant |
| 8 | 11662311 | 11662381 | 70 | Ankrd10 | NZB,SMJ,TH | feature_elongation,3_prime_UTR_variant |
| 8 | 12917049 | 12917140 | 91 | Atp11a | C57L,ILNJ2,NZB,NZW,PJ,PL,SEA,SMJ,TH | feature_elongation,3_prime_UTR_variant |
| 8 | 13068432 | 13068530 | 98 | Mcf2l | 129S1,C57L,CE,ILNJ2,LP,NZB,NZO,NZW,PJ,PL,SEA,SMJ,TH | feature_elongation,3_prime_UTR_variant |
| 8 | 15091988 | 15092072 | 84 | BB014433 | AJ,AKR,BALB,BTBR,BUB,C3H,CBA,CE,DBA1J,DBA2J,FVB,KK,LP,MRL,NOD,NOR,PL,RF,RH2,SJL,SMJ,SWR,TH | feature_elongation,coding_sequence_variant |
| 8 | 25531894 | 25532001 | 107 | Plekha2 | PL | feature_elongation,3_prime_UTR_variant |
| 8 | 27538594 | 27538668 | 74 | Plpbp | PL | feature_elongation,3_prime_UTR_variant |
| 8 | 31750969 | 31751053 | 84 | Fut10 | 129S1,AJ,BALB,BUB,C57L,CBA,FVB,MRL,NOD,NOR,PL,RH2,SJL,SMJ,SWR,TH | feature_elongation,3_prime_UTR_variant |
| 8 | 34110054 | 34110135 | 81 | Ubxn8 | 129S1,AJ,AKR,BALB,BUB,C3H,C57L,CBA,CE,DBA1J,DBA2J,FVB,MA/My,MRL,NOD,NOR,PL,RF,RH2,SJL,SMJ,SWR,TH | feature_elongation,3_prime_UTR_variant |
| 8 | 34637775 | 34637903 | 128 | Saraf | 129S1,AJ,AKR,BALB,C3H,C57L,CBA,CE,DBA1J,DBA2J,MA/My,MRL,PL,RF,RH2,SMJ | feature_elongation,3_prime_UTR_variant |
| 8 | 41370766 | 41370828 | 62 | Slc7a2 | 129S1,AJ,AKR,BALB,BTBR,BUB,C3H,C57L,CBA,CE,DBA1J,DBA2J,FVB,ILNJ2,KK,MA/My,MRL,NOD,NOR,NZB,NZO,NZW,PJ,PL,RF,RH2,SEA,SJL,SMJ,SWR,TH | feature_elongation,3_prime_UTR_variant |
| 8 | 65194683 | 65194736 | 53 | Klhl2 | PJ,SEA | feature_elongation,3_prime_UTR_variant |
| 8 | 65465819 | 65465883 | 64 | Trim61 | AKR,BALB,C3H,C57L,CBA,CE,DBA1J,DBA2J,MA/My,MRL,NZB,NZO,PJ,PL,RF,SEA,SMJ | feature_elongation,3_prime_UTR_variant |
| 8 | 69823455 | 69823517 | 62 | Gm10033 | AKR,BUB,C3H,C58J,CE,DBA1J,DBA2J,MA/My,MRL,NZB,PL,SMJ | feature_elongation,3_prime_UTR_variant |
| 8 | 69823978 | 69824631 | 653 | Gm10033 | NZB,NZO | feature_elongation,3_prime_UTR_variant |
| 8 | 70157767 | 70157971 | 204 | Zfp869 | PJ,SEA | feature_elongation,3_prime_UTR_variant |
| 8 | 70280950 | 70281152 | 202 | Lpar2 | 129S1,AKR,BALB,BUB,C3H,C57L,C58J,CBA,CE,DBA1J,DBA2J,FVB,ILNJ2,KK,MA/My,MRL,NZB,NZO,NZW,PJ,PL,RF,SEA,SJL,SMJ,SWR,TH | feature_elongation,3_prime_UTR_variant |
| 8 | 70360078 | 70360187 | 109 | Gatad2a | 129S1,BALB,C3H,C57L,CBA,DBA1J,DBA2J,FVB,ILNJ2,KK,NZB,NZO,RF,SJL,SMJ,SWR,TH | feature_elongation,3_prime_UTR_variant |
| 8 | 71295045 | 71295253 | 208 | Kcnn1 | C58J,NZW | feature_elongation,3_prime_UTR_variant |

| | | | | | | |
|---|---|---|---|---|---|---|
| 8 | 71347378 | 71347436 | 58 | Rpl18a | 129S1,AKR,BALB,BUB,C3H,C57L,C58J,CBA,CE,DBA1J,DBA2J,FVB,ILNJ2,KK,MA/My,MRL,NZB,NZO,NZW,PJ,PL,RF,SEA,SJL,SMJ,SWR,TH | feature_elongation,3_prime_UTR_variant |
| 8 | 72144752 | 72144957 | 205 | B3gnt3 | AKR,BUB,CE,MA/My,MRL,PL | feature_elongation,3_prime_UTR_variant |
| 8 | 72955236 | 72955295 | 59 | Hsh2d | NZO,PJ,SEA,SMJ | feature_elongation,3_prime_UTR_variant |
| 8 | 80068725 | 80068884 | 159 | Smad1 | C58J,FVB,KK,NOD,NOR,NZB,NZO,PL,RF,SWR,TH | feature_elongation,3_prime_UTR_variant |
| 8 | 81339751 | 81339856 | 105 | Frem3 | TH | feature_elongation,coding_sequence_variant |
| 8 | 84270472 | 84270524 | 52 | Or10k2 | AJ,C3H,CBA,CE,DBA1J,DBA2J,ILNJ2,KK,MA/My,RH2 | feature_elongation,3_prime_UTR_variant |
| 8 | 85898990 | 85899229 | 239 | Cks1brt | NOD,NOR,NZW,SJL | feature_elongation,3_prime_UTR_variant |
| 8 | 88898585 | 88898652 | 67 | Heatr3 | BUB,C57L,C58J,SMJ,SWR | feature_elongation,3_prime_UTR_variant |
| 8 | 92171248 | 92171443 | 195 | Fto | C58J | feature_elongation,3_prime_UTR_variant |
| 8 | 104991418 | 104991517 | 99 | Cklf | KK,PL | feature_elongation,3_prime_UTR_variant |
| 8 | 105076889 | 105076958 | 69 | Cmtm4 | KK,PL | feature_elongation,3_prime_UTR_variant |
| 8 | 107741831 | 107742019 | 188 | Sntb2 | RH2 | feature_elongation,3_prime_UTR_variant |
| 8 | 111732359 | 111732539 | 180 | Ddx19b | BALB,SEA | feature_elongation,3_prime_UTR_variant |
| 8 | 121821645 | 121821779 | 134 | Mthfsd | ILNJ2 | feature_elongation,3_prime_UTR_variant |
| 8 | 121824908 | 121824978 | 70 | Mthfsd | ILNJ2 | feature_elongation,3_prime_UTR_variant |
| 8 | 122318669 | 122319001 | 332 | Map1lc3b | ILNJ2 | feature_elongation,3_prime_UTR_variant |
| 8 | 123868239 | 123868293 | 54 | Sult5a1 | PJ | feature_elongation,3_prime_UTR_variant |
| 8 | 125631096 | 125631152 | 56 | Sprtn | AJ,BALB,ILNJ2,RH2,SEA | feature_elongation,3_prime_UTR_variant |
| 8 | 125756560 | 125756700 | 140 | Tsnax | NZB | feature_elongation,3_prime_UTR_variant |
| 9 | 7476708 | 7476891 | 183 | Mmp1a | MA/My | feature_elongation,3_prime_UTR_variant |
| 9 | 7834925 | 7835039 | 114 | Birc2 | AKR,MRL,RF | feature_elongation,5_prime_UTR_variant |
| 9 | 7857174 | 7857230 | 56 | Birc3 | AKR,MRL,RF | feature_elongation,3_prime_UTR_variant |
| 9 | 14661930 | 14662107 | 177 | Fut4 | C58J,CE,ILNJ2,PJ,SEA,SMJ,TH | feature_elongation,3_prime_UTR_variant |
| 9 | 14780837 | 14780922 | 85 | Gpr83 | C58J,CE,ILNJ2,PJ,SEA,SMJ,TH | feature_elongation,3_prime_UTR_variant |
| 9 | 20343416 | 20343769 | 353 | Zfp26 | 129S1 | feature_elongation,3_prime_UTR_variant |
| 9 | 32177234 | 32177298 | 64 | Arhgap32 | BTBR,C57L | feature_elongation,3_prime_UTR_variant |
| 9 | 36640891 | 36640943 | 52 | Stt3a | 129S1,AJ,C3H,C57L,CBA,FVB,MA/My,NZO,RH2,SJL,SMJ,SWR,TH | feature_elongation,3_prime_UTR_variant |
| 9 | 37560397 | 37560755 | 358 | Tbrg1,Siae | PL,SJL | feature_elongation,3_prime_UTR_variant |
| 9 | 37768016 | 37768482 | 466 | Or8b9 | CBA,CE,MRL | feature_elongation,3_prime_UTR_variant |
| 9 | 37904690 | 37905085 | 395 | Or8b35 | CE | feature_elongation,coding_sequence_variant |
| 9 | 38244247 | 38245104 | 857 | Or8c9 | DBA1J,ILNJ2,MA/My | feature_elongation,3_prime_UTR_variant |
| 9 | 38385024 | 38385748 | 724 | Or8b1c | 129S1,AJ,AKR,BALB,BTBR,BUB,C3H,CBA,DBA1J,DBA2J,ILNJ2,MA/My,NOD,NOR,PL,RF,RH2,SJL,SMJ,SWR,TH | feature_elongation,3_prime_UTR_variant |
| 9 | 39482517 | 39482619 | 102 | Or10d1 | MRL | feature_elongation,3_prime_UTR_variant |
| 9 | 39860196 | 39860325 | 129 | Or10d5 | KK,NZB,NZW,PJ,SEA | feature_elongation,3_prime_UTR_variant |
| 9 | 40884141 | 40884354 | 213 | Crtam | CE,NOD,NOR | feature_elongation,3_prime_UTR_variant |
| 9 | 44057485 | 44057575 | 90 | Cbl | 129S1,C57L,ILNJ2,SMJ | feature_elongation,3_prime_UTR_variant |
| 9 | 44323576 | 44323771 | 195 | Cenatac | DBA1J,NZO | feature_elongation,3_prime_UTR_variant |
| 9 | 45039966 | 45040060 | 94 | Scn2b | 129S1,BTBR,C57L,KK | feature_elongation,3_prime_UTR_variant |
| 9 | 51857752 | 51857981 | 229 | Fdx1 | FVB,SWR | feature_elongation,3_prime_UTR_variant |
| 9 | 64214093 | 64214302 | 209 | Dis3l | AJ,BUB,C3H,CBA,DBA1J,DBA2J,FVB,ILNJ2,KK,MRL,NZB,NZO,NZW,PJ,RH2,SEA,SJL,SWR | feature_elongation,3_prime_UTR_variant |
| 9 | 65502748 | 65503008 | 260 | Pif1 | AJ | feature_elongation,3_prime_UTR_variant |
| 9 | 86476658 | 86477026 | 368 | Me1 | 129S1,BTBR,C57L,CE,MRL | feature_elongation,coding_sequence_variant |
| 9 | 95939144 | 95939200 | 56 | Xrn1 | MRL,NZO,NZW | feature_elongation,3_prime_UTR_variant |
| 9 | 99550276 | 99550350 | 74 | Dzip1l | NOD,NOR | feature_elongation,3_prime_UTR_variant |
| 9 | 102966548 | 102966634 | 86 | Slco2a1 | CE,MRL,SMJ | feature_elongation,3_prime_UTR_variant |
| 9 | 103231359 | 103231417 | 58 | Cdv3 | SMJ | feature_elongation,3_prime_UTR_variant |
| 9 | 105039659 | 105039946 | 287 | Nek11 | 129S1,AJ,AKR,BALB,BTBR,BUB,C3H,C57L,CBA,CE,DBA1J,DBA2J,FVB,ILNJ2,KK,MRL,NOD,NOR,NZB,NZO,NZW,PJ,PL,RF,RH2,SEA,SJL,SMJ,SWR,TH | feature_elongation,3_prime_UTR_variant |
| 9 | 106125767 | 106125977 | 210 | Alas1 | 129S1,AJ,AKR,BALB,BTBR,BUB,C3H,C57L,CE,DBA1J,DBA2J,FVB,ILNJ2,MA/My,MRL,NOD,NOR,NZB,NZO,NZW,PJ,PL,RF,RH2,SEA,SJL,SMJ,SWR,TH | feature_elongation,5_prime_UTR_variant |
| 9 | 106227284 | 106227366 | 82 | Poc1a | 129S1,AJ,AKR,BALB,BUB,C3H,C57L,DBA1J,DBA2J,FVB,ILNJ2,MA/My,MRL,NOD,NOR,NZB,NZW,PL,RH2,SJL,SMJ,SWR,TH | feature_elongation,3_prime_UTR_variant |
| 9 | 110090077 | 110090557 | 480 | Cspg5 | NZW | feature_elongation,3_prime_UTR_variant |
| 9 | 110215211 | 110215281 | 70 | Ptpn23 | CBA | feature_elongation,coding_sequence_variant |
| 9 | 110815636 | 110815824 | 188 | Rtp3 | NZW,SJL,SWR | feature_elongation,coding_sequence_variant |
| 9 | 119749896 | 119750103 | 207 | Wdr48 | BTBR,MRL | feature_elongation,3_prime_UTR_variant |
| 9 | 119878210 | 119878294 | 84 | Cx3cr1 | BTBR,MRL | feature_elongation,3_prime_UTR_variant |
| 9 | 119960737 | 119961102 | 365 | Rpsa | AKR,BTBR,BUB,C58J,CBA,DBA1J,DBA2J,MRL,PL,RF,RH2 | feature_elongation,3_prime_UTR_variant |
| 9 | 123421992 | 123422484 | 492 | Slc6a20b | TH | feature_elongation,3_prime_UTR_variant |
| 9 | 123423249 | 123423321 | 72 | Slc6a20b | 129S1,C57L | feature_elongation,3_prime_UTR_variant |
| 9 | 123680958 | 123681078 | 120 | Fyco1 | 129S1,C57L,CE | feature_elongation,5_prime_UTR_variant |
| 10 | 4352193 | 4352414 | 221 | Rmnd1 | 129S1,AJ,AKR,BALB,BTBR,BUB,C57L,ILNJ2,PL,RF,RH2,SJL,SMJ | feature_elongation,3_prime_UTR_variant |
| 10 | 4353384 | 4353619 | 235 | Rmnd1 | NOD,NOR | feature_elongation,3_prime_UTR_variant |
| 10 | 6837057 | 6837121 | 64 | Ipcef1 | BUB | feature_elongation,3_prime_UTR_variant |
| 10 | 7393517 | 7393574 | 57 | Ulbp1 | C3H,CBA,DBA1J,DBA2J,FVB,LP,MA/My,MRL,NZW,SWR | feature_elongation,3_prime_UTR_variant |
| 10 | 9503431 | 9503584 | 153 | Samd5 | 129S1,BUB,CE,DBA1J,DBA2J,ILNJ2,KK,MA/My,NOD,NOR,NZB,NZO,NZW,PJ,PL,SEA,TH | feature_elongation,3_prime_UTR_variant |
| 10 | 11334909 | 11334967 | 58 | Epm2a | AJ,AKR,BALB,BTBR,C3H,C57L,CBA,MRL,NOD,NOR,PL,RF,RH2,SJL,SMJ | feature_elongation,3_prime_UTR_variant |
| 10 | 12423895 | 12424036 | 141 | Utrn | CE,DBA1J,DBA2J,LP,MA/My,TH | feature_elongation,5_prime_UTR_variant |
| 10 | 13005992 | 13006050 | 58 | Plagl1 | DBA1J,DBA2J | feature_elongation,3_prime_UTR_variant |
| 10 | 29221207 | 29221275 | 68 | Echdc1,Rnf146 | FVB,PJ,SEA | feature_elongation,3_prime_UTR_variant |
| 10 | 55892150 | 55892266 | 116 | Tbc1d32 | KK | feature_elongation,3_prime_UTR_variant |
| 10 | 55893463 | 55893625 | 162 | Tbc1d32 | KK | feature_elongation,3_prime_UTR_variant |
| 10 | 62936780 | 62936968 | 188 | Atoh7 | RF | feature_elongation,3_prime_UTR_variant |
| 10 | 66848321 | 66848402 | 81 | Reep3 | C57L,MRL | feature_elongation,3_prime_UTR_variant |
| 10 | 80527313 | 80527606 | 293 | Mob3a | BUB | feature_elongation,coding_sequence_variant |
| 10 | 81200565 | 81200617 | 52 | Mfsd12 | BUB | feature_elongation,3_prime_UTR_variant |

| | | | | | | |
|---|---|---|---|---|---|---|
| 10 | 82214042 | 82214293 | 251 | Gm4924 | 129S1,AJ,AKR,BALB,BTBR,C3H,C57L,CBA,CE,DBA1J,DBA2J,FVB,ILNJ2,KK,MA/My,MRL,NOD,NOR,NZB,NZO,NZW,PJ,PL,RF,RH2,SEA,SJL,SMJ,SWR,TH | feature_elongation,coding_sequence_variant |
| 10 | 82577481 | 82577543 | 62 | Hcfc2 | 129S1,AJ,AKR,BALB,BTBR,BUB,C3H,C57L,CBA,CE,DBA1J,DBA2J,FVB,ILNJ2,KK,MA/My,MRL,NOD,NOR,NZB,NZO,NZW,PJ,PL,RF,RH2,SEA,SJL,SMJ,SWR,TH | feature_elongation,3_prime_UTR_variant |
| 10 | 83260968 | 83261378 | 410 | Nopchap1 | BUB,SMJ | feature_elongation,3_prime_UTR_variant |
| 10 | 85887193 | 85887487 | 294 | Fbxo7 | C58J,LP,NZB | feature_elongation,3_prime_UTR_variant |
| 10 | 86128971 | 86129079 | 108 | Syn3 | AJ,AKR,BALB,C3H,CBA,CE,ILNJ2,MA/My,MRL,NOD,NOR,NZB,NZO,PJ,PL,RF,RH2,SEA,SWR,TH | feature_elongation,3_prime_UTR_variant |
| 10 | 93076134 | 93076197 | 63 | Cdk17 | MRL,NZW | feature_elongation,3_prime_UTR_variant |
| 10 | 93803102 | 93803162 | 60 | Vezt | AJ,BALB,BUB,PJ,RH2,SEA,SJL | feature_elongation,3_prime_UTR_variant |
| 10 | 94056874 | 94056927 | 53 | Ndufa12 | AKR,MA/My,MRL,NZO,PL,RF | feature_elongation,3_prime_UTR_variant |
| 10 | 111821118 | 111821282 | 164 | Krr1 | CE,LP,MRL,NZB,PL,SJL | feature_elongation,3_prime_UTR_variant |
| 10 | 117068411 | 117068483 | 72 | Lyz3 | SMJ | feature_elongation,3_prime_UTR_variant |
| 10 | 120618611 | 120618687 | 76 | Msrb3 | TH | feature_elongation,3_prime_UTR_variant |
| 10 | 121212151 | 121212295 | 144 | Gns | DBA1J,KK,MA/My,SMJ | feature_elongation,coding_sequence_variant |
| 10 | 127578251 | 127578302 | 51 | Zbtb39 | AJ,BALB,FVB,NOD,NOR,PL,RH2 | feature_elongation,coding_sequence_variant |
| 10 | 127581270 | 127581384 | 114 | Zbtb39 | CE | feature_elongation,3_prime_UTR_variant |
| 10 | 127602382 | 127602593 | 211 | Rdh1 | CE | feature_elongation,3_prime_UTR_variant |
| 10 | 127702392 | 127702444 | 52 | Rdh16f2 | SMJ | feature_elongation,5_prime_UTR_variant |
| 10 | 128104228 | 128104426 | 198 | Apof | CE | feature_elongation,5_prime_UTR_variant |
| 10 | 128378812 | 128378985 | 173 | Zc3h10 | BUB,KK,SMJ | feature_elongation,3_prime_UTR_variant |
| 10 | 128635147 | 128635427 | 280 | Mmp19 | BUB,KK,SMJ | feature_elongation,3_prime_UTR_variant |
| 10 | 128912802 | 128912976 | 174 | Or6c8 | C3H,CBA,CE,RF | feature_elongation,3_prime_UTR_variant |
| 10 | 129303015 | 129303078 | 63 | Or6c5c | LP,NOD,NOR | feature_elongation,3_prime_UTR_variant |
| 10 | 130043117 | 130043442 | 325 | Or9k7 | SMJ | feature_elongation,3_prime_UTR_variant |
| 11 | 3836543 | 3836651 | 108 | 4921536K21Rik | B10D2,B10J,C58J,KK | feature_elongation,3_prime_UTR_variant |
| 11 | 3849068 | 3849235 | 167 | Dusp18 | B10D2,B10J,C58J,KK | feature_elongation,3_prime_UTR_variant |
| 11 | 4172107 | 4172160 | 53 | Castor1 | B10D2,B10J,C58J,KK | feature_elongation,3_prime_UTR_variant |
| 11 | 5144115 | 5144285 | 170 | Kremen1 | B10D2,B10J,C58J,KK,NOR | feature_elongation,3_prime_UTR_variant |
| 11 | 5854541 | 5854598 | 57 | Gck | AJ,AKR,B10D2,B10J,BALB,BTBR,BUB,C3H,C57L,C58J,CE,FVB,KK,MA/My,MRL,NOD,NOR,NZB,NZO,PJ,PL,RF,SEA,SJL,SWR,TH | feature_elongation,coding_sequence_variant |
| 11 | 9068596 | 9068653 | 57 | Upp1 | C58J | feature_elongation,5_prime_UTR_variant |
| 11 | 29104848 | 29104940 | 92 | Pnpt1 | AJ,C3H,DBA1J,DBA2J,ILNJ2,MA/My,MRL,NZB,NZO,PL | feature_elongation,coding_sequence_variant |
| 11 | 29171710 | 29171912 | 202 | Cfap36 | AKR,BALB,BUB,C57L,C58J,CE,FVB,KK,NOD,NZW,RF,RH2 | feature_elongation,3_prime_UTR_variant |
| 11 | 40601035 | 40601161 | 126 | Hmmr | B10D2,B10J,BTBR,C58J,LP,MA/My | feature_elongation,coding_sequence_variant |
| 11 | 45821507 | 45821688 | 181 | Lsm11 | CE,PL,SMJ | feature_elongation,3_prime_UTR_variant |
| 11 | 48796639 | 48796835 | 196 | Gm12185 | NZW | feature_elongation,3_prime_UTR_variant |
| 11 | 54731188 | 54731244 | 56 | Lyrm7 | B10D2,B10J,C58J,NOR,NZB,NZO,NZW | feature_elongation,3_prime_UTR_variant |
| 11 | 58490275 | 58490337 | 62 | Or2ab1 | 129S1,AJ,AKR,BALB,BTBR,BUB,C3H,C57L,CBA,DBA1J,DBA2J,FVB,ILNJ2,KK,LP,MA/My,MRL,NOD,NOR,NZB,NZO,NZW,PJ,PL,RF,RH2,SEA,SJL,SWR,TH | feature_elongation,3_prime_UTR_variant |
| 11 | 58853329 | 58853428 | 99 | H3f4 | 129S1,AKR,BALB,BTBR,C57L,LP,RF,RH2 | feature_elongation,3_prime_UTR_variant |
| 11 | 58951824 | 58952453 | 629 | Obscn | AJ,BTBR,BUB,C3H,C57L,CBA,CE,DBA1J,DBA2J,FVB,ILNJ2,KK,MA/My,MRL,NOD,NOR,NZW,PJ,PL,RF,SEA,SJL,SMJ,SWR,TH | feature_elongation,coding_sequence_variant |
| 11 | 59047601 | 59048311 | 710 | Iba57 | ILNJ2 | feature_elongation,3_prime_UTR_variant |
| 11 | 60267491 | 60267776 | 285 | Drc3 | AJ,C3H,CBA,MRL,NZB,NZO,PJ,SEA | feature_elongation,3_prime_UTR_variant |
| 11 | 60679177 | 60679406 | 229 | Shmt1 | 129S1,AKR,BTBR,C57L,LP,MA/My,NOD,PL,RF | feature_elongation,3_prime_UTR_variant |
| 11 | 60750131 | 60750267 | 136 | Dhrs7b | AJ,C3H,CBA,MRL,NZB,NZO,PJ,SEA | feature_elongation,3_prime_UTR_variant |
| 11 | 64892697 | 64892772 | 75 | Elac2 | CE,ILNJ2 | feature_elongation,3_prime_UTR_variant |
| 11 | 68838660 | 68838900 | 240 | Arhgef15 | DBA1J,DBA2J,NOR,PJ,SEA,SMJ | stop_gained,inframe_insertion |
| 11 | 69111732 | 69111795 | 63 | Gucy2e | DBA1J,DBA2J,NOR,PJ,SEA,SMJ | feature_elongation,3_prime_UTR_variant |
| 11 | 69514502 | 69514669 | 167 | Sat2 | DBA1J,DBA2J,ILNJ2,NOR,PJ,SEA,SMJ | feature_elongation,3_prime_UTR_variant |
| 11 | 70028268 | 70028489 | 221 | Mgl2 | CE,DBA1J,DBA2J,FVB,ILNJ2,KK,NOR,PJ,SEA,SMJ | feature_elongation,3_prime_UTR_variant |
| 11 | 70235307 | 70235375 | 68 | Alox15 | CE,FVB,ILNJ2,NZW | feature_elongation,3_prime_UTR_variant |
| 11 | 73703017 | 73703273 | 256 | Or1e32 | AJ,NZO,NZW | feature_elongation,3_prime_UTR_variant |
| 11 | 73975983 | 73976235 | 252 | Zfp616 | AJ,CE,NZO,NZW | feature_elongation,3_prime_UTR_variant,feature_elongation,coding_sequence_variant |
| 11 | 77028564 | 77028624 | 60 | Efcab5 | TH | feature_elongation,coding_sequence_variant |
| 11 | 79392356 | 79392570 | 214 | Omg | AKR,BALB,C3H,CBA,CE,ILNJ2,MRL,PL,RF,RH2,SMJ | feature_elongation,3_prime_UTR_variant |
| 11 | 80371768 | 80372207 | 439 | Cdk5r1 | C3H | feature_elongation,3_prime_UTR_variant |
| 11 | 83166239 | 83166644 | 405 | Slfn14 | 129S1,AJ,AKR,B10D2,B10J,BALB,BTBR,BUB,C3H,C57L,C58J,CBA,CE,DBA1J,DBA2J,FVB,ILNJ2,KK,LP,MA/My,MRL,NOD,NOR,NZB,NZO,NZW,PJ,PL,RF,RH2,SEA,SJL,SMJ,SWR,TH | feature_elongation,3_prime_UTR_variant |
| 11 | 83193427 | 83193718 | 291 | Ap2b1 | KK | feature_elongation,5_prime_UTR_variant |
| 11 | 86389501 | 86389703 | 202 | Rnft1 | 129S1,AJ,AKR,BTBR,C3H,C57L,C58J,CBA,DBA1J,DBA2J,KK,MRL,NZB,NZO,PL,RF,SJL,SWR | feature_elongation,3_prime_UTR_variant |
| 11 | 88930820 | 88930996 | 176 | Dgke | BALB,CE,KK,MRL,RH2,SJL | feature_elongation,3_prime_UTR_variant |
| 11 | 94180656 | 94180720 | 64 | Luc7l3 | AJ,AKR,C3H,CBA,CE,MRL,RF | feature_elongation,3_prime_UTR_variant |
| 11 | 94430679 | 94430735 | 56 | Rsad1 | 129S1,BALB,BTBR,C57L,MA/My,NOD,NOR,PL,SJL,SMJ | feature_elongation,3_prime_UTR_variant |
| 11 | 94448825 | 94448960 | 135 | Acsf2 | AJ,AKR,C3H,CBA,CE,MRL,RF | feature_elongation,3_prime_UTR_variant |
| 11 | 95660269 | 95660483 | 214 | Zfp652 | DBA1J,DBA2J,ILNJ2,KK,NZB,NZO,PJ,SEA | feature_elongation,3_prime_UTR_variant |
| 11 | 95990891 | 95991017 | 126 | Calcoco2 | AJ,AKR,BUB,C3H,CBA,DBA1J,DBA2J,ILNJ2,MRL,NZB,NZO,PJ,RH2,SEA | feature_elongation,coding_sequence_variant |
| 11 | 97606053 | 97606275 | 222 | Pip4k2b | SEA | feature_elongation,3_prime_UTR_variant |
| 11 | 97884768 | 97884845 | 77 | Arl5c | 129S1,BTBR,C3H,C57L,CE,DBA1J,DBA2J,FVB,ILNJ2 | feature_elongation,3_prime_UTR_variant |
| 11 | 97979155 | 97979207 | 52 | Fbxl20 | NZW,SWR | feature_elongation,3_prime_UTR_variant |
| 11 | 98723431 | 98723653 | 222 | Casc3 | CE | feature_elongation,3_prime_UTR_variant |
| 11 | 98754599 | 98754689 | 90 | Wipf2 | SMJ | feature_elongation,5_prime_UTR_variant |
| 11 | 99276823 | 99276883 | 60 | Krt10 | AKR,RF,RH2,SJL | feature_elongation,coding_sequence_variant |
| 11 | 101862617 | 101862838 | 221 | Dusp3 | AJ,TH | feature_elongation,3_prime_UTR_variant |
| 11 | 103255689 | 103255840 | 151 | Plekhm1 | NZW | feature_elongation,3_prime_UTR_variant |

| | | | | | | |
|---|---|---|---|---|---|---|
| 11 | 109992711 | 109992770 | 59 | Abca9 | BALB,BUB,RF,SEA | feature_elongation,3_prime_UTR_variant |
| 11 | 114935004 | 114935184 | 180 | Cd300ld3 | FVB,SJL | feature_elongation,coding_sequence_variant |
| 11 | 116566129 | 116566411 | 282 | St6galnac2 | AJ,AKR,BTBR,C3H,C57L,CBA,CE,FVB,ILNJ2,MA/My,RH2,SJL,SMJ,SWR | feature_elongation,3_prime_UTR_variant |
| 11 | 119301052 | 119301106 | 54 | Rnf213 | CE | feature_elongation,coding_sequence_variant |
| 11 | 120014944 | 120015176 | 232 | Slc38a10 | SJL | feature_elongation,3_prime_UTR_variant |
| 11 | 120356316 | 120356527 | 211 | Arl16 | SJL | feature_elongation,3_prime_UTR_variant |
| 11 | 120501231 | 120501285 | 54 | Pcyt2 | SJL | feature_elongation,3_prime_UTR_variant |
| 12 | 4013159 | 4013359 | 200 | Efr3b | MRL | feature_elongation,3_prime_UTR_variant |
| 12 | 4819152 | 4819254 | 102 | Fam228b,Pfn4 | MRL | feature_elongation,5_prime_UTR_variant |
| 12 | 18584005 | 18584172 | 167 | 5730507C01Rik | MRL | feature_elongation,coding_sequence_variant |
| 12 | 24716919 | 24716977 | 58 | Cys1 | AKR,BTBR,BUB,C57L,CE,DBA2J,FVB,KK,NZO,RF,SJL,SMJ,SWR,TH | feature_elongation,3_prime_UTR_variant |
| 12 | 52035390 | 52035596 | 206 | Dtd2 | 129S1,AJ,AKR,BALB,BUB,C3H,C57L,CBA,CE,MRL,NOD,NOR,NZB,PJ,PL,RF,RH2,SEA,SJL,SMJ,SWR | feature_elongation,3_prime_UTR_variant |
| 12 | 52044760 | 52045130 | 370 | Dtd2 | AJ,AKR,BALB,BUB,C3H,CBA,NOD,NOR,PJ,PL,RF,RH2,SEA,SJL,SWR | feature_elongation,3_prime_UTR_variant |
| 12 | 54719914 | 54720158 | 244 | Eapp | AJ,AKR,BALB,C3H,CBA,DBA1J,DBA2J,ILNJ2,KK,NOD,NOR,PJ,RF,RH2,SEA | feature_elongation,3_prime_UTR_variant |
| 12 | 55327273 | 55327509 | 236 | Ppp2r3c | FVB,KK,NZW,SMJ,TH | feature_elongation,3_prime_UTR_variant |
| 12 | 64986678 | 64987010 | 332 | Klhl28 | BUB,SJL,SWR | feature_elongation,3_prime_UTR_variant |
| 12 | 69358598 | 69358783 | 185 | Nemf | MRL | feature_elongation,3_prime_UTR_variant |
| 12 | 71041697 | 71042192 | 495 | Psma3 | NOD,NOR | feature_elongation,3_prime_UTR_variant |
| 12 | 72808314 | 72808366 | 52 | Ppm1a | NZW | feature_elongation,5_prime_UTR_variant |
| 12 | 75677401 | 75677624 | 223 | Wdr89 | CE | feature_elongation,3_prime_UTR_variant |
| 12 | 87310996 | 87311200 | 204 | Vipas39 | SWR | feature_elongation,5_prime_UTR_variant |
| 12 | 87994165 | 87994229 | 64 | Eif1ad11 | 129S1,AJ,AKR,BTBR,BUB,C3H,C57L,CBA,DBA1J,DBA2J,FVB,ILNJ2,MRL,NOR,PJ,RF,SEA,SJL,TH | feature_elongation,coding_sequence_variant |
| 12 | 91604862 | 91604920 | 58 | Ston2 | LP,SWR | feature_elongation,3_prime_UTR_variant |
| 12 | 98946123 | 98946190 | 67 | Ttc8 | PJ,SEA | feature_elongation,coding_sequence_variant |
| 12 | 98949223 | 98949411 | 188 | Ttc8 | 129S1,BTBR,BUB,C57L,C58J,CBA,FVB,ILNJ2,KK,LP,MRL,NZB,NZO,NZW,RF,SJL,SMJ,SWR | feature_elongation,3_prime_UTR_variant |
| 12 | 103962610 | 103962747 | 137 | Serpina9 | 129S1,AKR,BALB,BTBR,BUB,C57L,CBA,CE,DBA1J,DBA2J,FVB,ILNJ2,KK,LP,MRL,NOD,NOR,NZB,PJ,PL,RF,SEA,SJL,SWR | feature_elongation,3_prime_UTR_variant |
| 12 | 104204055 | 104204486 | 431 | Serpina3g | AKR,BALB,BUB,CBA,CE,DBA1J,DBA2J,FVB,ILNJ2,KK,LP,MRL,NOD,NOR,PJ,PL,RF,SEA,SJL,SWR | feature_elongation,5_prime_UTR_variant |
| 12 | 107877986 | 107878042 | 56 | Bcl11b | AJ,BUB,CE,FVB,MA/My,NZB,NZO,NZW,PJ,PL,RF,SEA,SJL,SMJ,SWR,TH | feature_elongation,3_prime_UTR_variant |
| 13 | 3703272 | 3703347 | 75 | Asb13 | 129S1,AJ,AKR,B10D2,B10J,BALB,BTBR,BUB,C3H,C57L,C58J,CBA,CE,DBA1J,DBA2J,FVB,ILNJ2,KK,LP,MA/My,MRL,NOD,NOR,NZB,NZO,NZW,PJ,PL,RF,RH2,SEA,SJL,SMJ,SWR,TH | feature_elongation,3_prime_UTR_variant |
| 13 | 6630709 | 6630794 | 85 | Pfkp | AKR,BUB,C57L,KK,LP,MA/My,NOD,NOR,NZB,NZO,NZW,RF,TH | feature_elongation,3_prime_UTR_variant |
| 13 | 16202344 | 16202576 | 232 | Inhba | AKR,ILNJ2,LP,PJ,RF,SEA | feature_elongation,3_prime_UTR_variant |
| 13 | 19815987 | 19816206 | 219 | Sfrp4 | LP,PJ,SEA | feature_elongation,3_prime_UTR_variant |
| 13 | 21254785 | 21254990 | 205 | Or2p2 | LP,PJ,SEA | feature_elongation,3_prime_UTR_variant |
| 13 | 21841863 | 21841991 | 128 | Or2w6 | LP,PJ,SEA | feature_elongation,3_prime_UTR_variant |
| 13 | 22277102 | 22277162 | 60 | Vmn1r188 | ILNJ2 | feature_elongation,3_prime_UTR_variant |
| 13 | 23323506 | 23323597 | 91 | Vmn1r218 | BTBR,C57L,KK,LP,NOD,NOR,NZB,NZO,NZW,PJ,SEA | feature_elongation,3_prime_UTR_variant |
| 13 | 27343095 | 27343265 | 170 | Prl3d3 | KK,PJ,SEA | feature_elongation,coding_sequence_variant |
| 13 | 36163130 | 36163331 | 201 | Lyrm4 | CE,KK,LP | feature_elongation,3_prime_UTR_variant |
| 13 | 54527805 | 54527878 | 73 | Cplx2 | AJ,BALB,C3H,CBA,CE | feature_elongation,3_prime_UTR_variant |
| 13 | 55565773 | 55565996 | 223 | F12 | AJ,BALB,C3H,CBA,CE | feature_elongation,3_prime_UTR_variant |
| 13 | 58308759 | 58308847 | 88 | Idnk | BTBR,C57L,C58J,DBA1J,DBA2J,ILNJ2,KK,MA/My,MRL,SWR | feature_elongation,3_prime_UTR_variant |
| 13 | 62848408 | 62848498 | 90 | Platr25 | FVB,PJ,SEA,SWR | feature_elongation,3_prime_UTR_variant,feature_elongation,coding_sequence_variant |
| 13 | 64508650 | 64508870 | 220 | Ctsl | FVB,PJ,SEA,SWR | feature_elongation,3_prime_UTR_variant |
| 13 | 65303139 | 65303339 | 200 | Or9s18 | 129S1,AJ,AKR,BALB,BTBR,BUB,C3H,C57L,C58J,CBA,CE,DBA1J,DBA2J,FVB,ILNJ2,KK,MRL,NOD,NOR,NZB,NZO,NZW,PJ,PL,RF,SEA,SJL,SMJ,SWR,TH | feature_elongation,3_prime_UTR_variant |
| 13 | 67056673 | 67056864 | 191 | Ramacl | PJ,SEA,SWR | feature_elongation,3_prime_UTR_variant |
| 13 | 67246775 | 67246859 | 84 | Rslcan18 | 129S1,AKR,BTBR,BUB,C3H,C57L,C58J,CBA,CE,DBA1J,DBA2J,ILNJ2,MRL,NOD,NOR,NZW,PL,RF,SMJ,SWR,TH | feature_elongation,coding_sequence_variant |
| 13 | 67441178 | 67441262 | 84 | Zfp457 | 129S1,AJ,AKR,BALB,BTBR,BUB,C3H,C57L,C58J,CBA,CE,DBA1J,DBA2J,FVB,ILNJ2,KK,MRL,NOD,NOR,NZB,NZO,NZW,PJ,PL,RF,SEA,SJL,SMJ,SWR,TH | feature_elongation,coding_sequence_variant |
| 13 | 67590027 | 67590298 | 271 | Zfp874a | 129S1,AJ,AKR,BALB,BTBR,BUB,C3H,C57L,C58J,CBA,CE,DBA1J,DBA2J,FVB,ILNJ2,KK,MRL,NOD,NOR,NZB,NZO,NZW,PJ,PL,RF,SEA,SJL,SMJ,SWR,TH | feature_elongation,3_prime_UTR_variant |
| 13 | 67622508 | 67622750 | 242 | Zfp874b | MA/My | feature_elongation,coding_sequence_variant |
| 13 | 67853736 | 67853925 | 189 | Zfp65 | KK,NZB,NZO,SWR | feature_elongation,3_prime_UTR_variant |
| 13 | 70935189 | 70935544 | 355 | Adamts16 | MRL | feature_elongation,coding_sequence_variant |
| 13 | 74519204 | 74519400 | 196 | Zfp72 | ILNJ2,NZW | feature_elongation,3_prime_UTR_variant |
| 13 | 75919518 | 75919575 | 57 | Ell2 | BALB,NZW | feature_elongation,3_prime_UTR_variant |
| 13 | 93233253 | 93233398 | 145 | Cmya5 | 129S1,AJ,AKR,BALB,BTBR,BUB,C3H,C57L,C58J,CBA,CE,DBA1J,DBA2J,FVB,ILNJ2,KK,MA/My,MRL,NZB,NZO,NZW,PJ,PL,RF,SEA,SJL,SMJ,SWR | feature_elongation,coding_sequence_variant |
| 13 | 97265288 | 97265543 | 255 | Fam169a | AJ,BUB | feature_elongation,3_prime_UTR_variant |
| 13 | 105087715 | 105087779 | 64 | Rgs7bp | NZB | feature_elongation,3_prime_UTR_variant |
| 13 | 108310797 | 108310862 | 65 | Ercc8 | CE,MA/My,NZO,TH | feature_elongation,3_prime_UTR_variant |
| 13 | 110489382 | 110489446 | 64 | Gapt | AKR,BALB,DBA2J,ILNJ2,RF,SEA | feature_elongation,3_prime_UTR_variant |
| 13 | 112871130 | 112871196 | 66 | Slc38a9 | AJ,DBA1J,FVB,KK,MA/My,NZO,PL,RF,RH2,SJL,SWR | feature_elongation,3_prime_UTR_variant |
| 13 | 116435508 | 116435568 | 60 | Isl1 | AJ,DBA1J,MA/My,RH2 | feature_elongation,3_prime_UTR_variant |
| 13 | 120114279 | 120114547 | 268 | Ccl28 | 129S1,BTBR,C3H,C57L,CBA,CE,DBA2J,KK,MA/My,NZO,SMJ | feature_elongation,3_prime_UTR_variant |

| | | | | | | |
|---|---|---|---|---|---|---|
| 14 | 8296045 | 8296097 | 52 | Or2t6 | 129S1,C3H,C57L,DBA1J,DBA2J,KK,MRL,PJ,PL | feature_elongation,3_prime_UTR_variant |
| 14 | 8520344 | 8520520 | 176 | Gm11100 | 129S1,AJ,AKR,BALB,BTBR,BUB,C3H,C57L,C58J,CBA,CE,DBA1J,DBA2J,ILNJ2,KK,MRL,NZB,NZO,NZW,PJ,PL,RF,SEA,SMJ | feature_elongation,3_prime_UTR_variant |
| 14 | 33914887 | 33915721 | 834 | Gprin2 | PJ | feature_elongation,3_prime_UTR_variant |
| 14 | 45240799 | 45240868 | 69 | Ptger2 | C57L,NZO,PJ,PL | feature_elongation,3_prime_UTR_variant |
| 14 | 47533031 | 47533294 | 263 | Socs4 | SWR | feature_elongation,3_prime_UTR_variant |
| 14 | 49247026 | 49247234 | 208 | Exoc5 | NZW | feature_elongation,3_prime_UTR_variant |
| 14 | 50382781 | 50383701 | 920 | Or4k5 | 129S1,AJ,AKR,BALB,BUB,C3H,C57L,CBA,CE,DBA1J,DBA2J,FVB,ILNJ2,KK,MA/My,MRL,NOD,NOR,NZB,NZO,NZW,PJ,PL,RF,RH2,SEA,SJL,SMJ,SWR,TH | feature_elongation,3_prime_UTR_variant |
| 14 | 50471627 | 50471678 | 51 | Or4k6 | AJ,AKR,C3H,C57L,CBA,CE,MRL,NOD,NZB,PJ,PL,RF,RH2,SEA | feature_elongation,3_prime_UTR_variant |
| 14 | 50555329 | 50555380 | 51 | Or4m1 | DBA1J,DBA2J | feature_elongation,3_prime_UTR_variant |
| 14 | 50692959 | 50693155 | 196 | Or11g7 | 129S1,AJ,AKR,BALB,BUB,C3H,C57L,CBA,CE,DBA1J,DBA2J,FVB,ILNJ2,KK,MA/My,MRL,NOD,NOR,NZB,NZO,NZW,PJ,PL,RF,RH2,SEA,SJL,SMJ,SWR,TH | feature_elongation,3_prime_UTR_variant |
| 14 | 50771843 | 50771958 | 115 | Or11g27 | 129S1,AJ,AKR,BALB,BUB,C3H,C57L,CBA,CE,DBA1J,DBA2J,FVB,ILNJ2,KK,MA/My,MRL,NOD,NOR,NZB,NZO,NZW,PJ,PL,RF,RH2,SEA,SJL,SMJ,SWR,TH | feature_elongation,3_prime_UTR_variant |
| 14 | 51128258 | 51128347 | 89 | Klhl33 | 129S1,AJ,AKR,BALB,BUB,C3H,C57L,CBA,CE,DBA1J,DBA2J,FVB,ILNJ2,KK,MA/My,MRL,NOD,NOR,NZB,NZO,NZW,PJ,PL,RF,RH2,SEA,SJL,SMJ,SWR,TH | feature_elongation,3_prime_UTR_variant |
| 14 | 52549564 | 52549800 | 236 | Sall2 | FVB | feature_elongation,3_prime_UTR_variant |
| 14 | 54517221 | 54517404 | 183 | Or6e1 | 129S1,AJ,AKR,BALB,BTBR,BUB,C3H,C57L,CBA,CE,DBA1J,DBA2J,FVB,ILNJ2,KK,MA/My,MRL,NOD,NOR,NZB,NZO,NZW,PJ,PL,RF,RH2,SEA,SJL,SMJ,SWR,TH | feature_elongation,3_prime_UTR_variant |
| 14 | 54960250 | 54960472 | 222 | Slc7a8 | 129S1,AJ,AKR,BALB,BTBR,C3H,C57L,CBA,CE,DBA1J,KK,MRL,NZO,PJ,RF,RH2,SEA,TH | feature_elongation,3_prime_UTR_variant |
| 14 | 57029250 | 57029454 | 204 | Zmym5 | KK | feature_elongation,3_prime_UTR_variant |
| 14 | 58072440 | 58072643 | 203 | Zdhhc20 | BTBR,BUB | feature_elongation,3_prime_UTR_variant |
| 14 | 59785333 | 59785763 | 430 | Cab39l | C3H,CE,DBA2J | feature_elongation,3_prime_UTR_variant |
| 14 | 69913127 | 69913387 | 260 | Loxl2 | 129S1,BTBR,KK,LP,SJL,TH | feature_elongation,coding_sequence_variant |
| 14 | 69930749 | 69930831 | 82 | Loxl2 | BTBR,LP,TH | feature_elongation,3_prime_UTR_variant |
| 14 | 70821124 | 70821361 | 237 | Fhip2b | BTBR,BUB,LP | feature_elongation,3_prime_UTR_variant |
| 14 | 71216059 | 71216135 | 76 | Gfra2 | FVB,RH2,SJL | feature_elongation,3_prime_UTR_variant |
| 14 | 73477904 | 73478124 | 220 | Lpar6 | SJL | feature_elongation,3_prime_UTR_variant |
| 14 | 74992492 | 74992747 | 255 | Lrch1 | BUB,KK | feature_elongation,3_prime_UTR_variant |
| 14 | 75044978 | 75045966 | 988 | Lrch1 | BUB,KK,TH | feature_elongation,coding_sequence_variant |
| 14 | 78318844 | 78319069 | 225 | Fam216b | FVB,MA/My,MRL,NOD,NOR | feature_elongation,3_prime_UTR_variant |
| 14 | 121780426 | 121780574 | 148 | Dock9 | RF | feature_elongation,3_prime_UTR_variant |
| 15 | 8189758 | 8189810 | 52 | Nup155 | AKR,RF | feature_elongation,3_prime_UTR_variant |
| 15 | 9115886 | 9116147 | 261 | Skp2 | AJ,BUB,FVB,NOD,NOR,RH2 | feature_elongation,3_prime_UTR_variant |
| 15 | 9199833 | 9200219 | 386 | Lmbrd2 | LP,SJL | feature_elongation,3_prime_UTR_variant |
| 15 | 10570618 | 10570827 | 209 | Rai14 | NZB | feature_elongation,3_prime_UTR_variant |
| 15 | 31294865 | 31294918 | 53 | Ankrd33b | 129S1,BTBR,C57L,KK,LP,SJL | feature_elongation,3_prime_UTR_variant |
| 15 | 38492530 | 38493370 | 840 | Azin1 | NZB,NZW | feature_elongation,coding_sequence_variant |
| 15 | 55119103 | 55119265 | 162 | Deptor | 129S1,AJ,AKR,C57L,CE,DBA1J,DBA2J,FVB,ILNJ2,MRL,NZB,PJ,RH2,SJL,SMJ,TH | feature_elongation,3_prime_UTR_variant |
| 15 | 59528651 | 59528929 | 278 | Trib1 | 129S1 | feature_elongation,3_prime_UTR_variant |
| 15 | 60693155 | 60693508 | 353 | Lratd2 | ILNJ2 | feature_elongation,3_prime_UTR_variant |
| 15 | 65740942 | 65741005 | 63 | Efr3a | BALB,C3H,CBA,CE,SEA | feature_elongation,3_prime_UTR_variant |
| 15 | 72835468 | 72835755 | 287 | Trappc9 | AJ,BALB,C3H,CBA,CE,DBA1J,DBA2J,ILNJ2,NOD,NOR,NZB,NZO,SEA | stop_gained,frameshift_variant |
| 15 | 74597301 | 74597366 | 65 | Slurp1 | 129S1,ILNJ2,NZW | feature_elongation,3_prime_UTR_variant |
| 15 | 76835096 | 76835208 | 112 | 1110038F14Rik | MA/My | feature_elongation,3_prime_UTR_variant |
| 15 | 84198149 | 84198200 | 51 | Parvb | BUB,DBA1J,DBA2J,ILNJ2 | feature_elongation,3_prime_UTR_variant |
| 15 | 85170191 | 85171170 | 979 | Fbln1 | 129S1,AJ,AKR,BALB,BTBR,BUB,C3H,C57L,CBA,CE,DBA1J,DBA2J,FVB,ILNJ2,KK,MA/My,MRL,NOD,NOR,NZB,NZO,NZW,PJ,PL,RF,SEA,SJL,SMJ,SWR,TH | feature_elongation,3_prime_UTR_variant |
| 15 | 85377219 | 85377563 | 344 | 7530416G11Rik | SJL,SMJ,SWR | feature_elongation,3_prime_UTR_variant |
| 15 | 85698981 | 85699136 | 155 | Pkdrej | BUB,DBA1J,DBA2J,NZW,PJ,SEA | feature_elongation,3_prime_UTR_variant |
| 15 | 94460947 | 94461037 | 90 | Irak4 | 129S1,AJ,AKR,BTBR,BUB,C3H,C57L,CBA,CE,DBA1J,DBA2J,FVB,ILNJ2,KK,MA/My,NOD,NOR,NZB,NZO,NZW,PJ,PL,RF,RH2,SEA,SJL,SMJ,SWR,TH | feature_elongation,coding_sequence_variant |
| 15 | 97937995 | 97938049 | 54 | Senp1 | C57L,DBA2J,ILNJ2,KK,NOD,NOR,NZB,NZO,PJ,SEA,SMJ,TH | feature_elongation,3_prime_UTR_variant |
| 15 | 98358221 | 98358662 | 441 | Or8s8 | BUB,MA/My,SWR | feature_elongation,3_prime_UTR_variant |
| 15 | 98748903 | 98748990 | 87 | Kmt2d | ILNJ2,NOD,NOR | feature_elongation,coding_sequence_variant |
| 15 | 99626507 | 99626737 | 230 | Cox14 | 129S1,C57L | feature_elongation,3_prime_UTR_variant |
| 15 | 102246337 | 102246392 | 55 | Myg1 | 129S1,AJ,AKR,BTBR,C3H,C57L,C58J,CBA,CE,DBA1J,DBA2J,FVB,ILNJ2,KK,LP,MRL,NOD,NOR,NZB,NZO,PJ,PL,RF,RH2,SEA,SJL,SMJ,TH | feature_elongation,3_prime_UTR_variant |
| 16 | 10658800 | 10659040 | 240 | Rmi2 | BUB | feature_elongation,3_prime_UTR_variant |
| 16 | 22976702 | 22976902 | 200 | Adipoq | TH | feature_elongation,3_prime_UTR_variant |
| 16 | 30379362 | 30379570 | 208 | Lsg1 | SMJ | feature_elongation,3_prime_UTR_variant |
| 16 | 32572401 | 32572773 | 372 | Muc4 | AJ,BALB,BTBR,BUB,C3H,C57L,CBA,CE,DBA1J,DBA2J,FVB,ILNJ2,KK,MRL,PL,SEA,SJL,SMJ,TH | feature_elongation,coding_sequence_variant |
| 16 | 32574492 | 32575236 | 744 | Muc4 | C57L,CE,DBA1J,DBA2J,KK,SMJ | feature_elongation,coding_sequence_variant |
| 16 | 32576214 | 32576586 | 372 | Muc4 | AJ,BALB,BTBR,BUB,C3H,C57L,CBA,CE,DBA1J,DBA2J,FVB,ILNJ2,KK,MRL,PL,RH2,SEA,SJL,SMJ,TH | feature_elongation,coding_sequence_variant |
| 16 | 33619651 | 33619725 | 74 | Muc13 | SMJ | feature_elongation,coding_sequence_variant |
| 16 | 37656311 | 37656411 | 100 | Fstl1 | AKR,MRL,RF,SMJ | feature_elongation,3_prime_UTR_variant |
| 16 | 45202855 | 45202909 | 54 | Cd200 | AKR,PL,RF | feature_elongation,3_prime_UTR_variant |
| 16 | 48412913 | 48412982 | 69 | Morc1 | C3H,CBA,CE | feature_elongation,coding_sequence_variant |
| 16 | 59134137 | 59134188 | 51 | Or5ac22 | SMJ | feature_elongation,3_prime_UTR_variant |
| 16 | 84514601 | 84514665 | 64 | Mrpl39 | AJ,C3H | feature_elongation,3_prime_UTR_variant |

| | | | | | | |
|---|---|---|---|---|---|---|
| 16 | 84622328 | 84622793 | 465 | Jam2 | C58J | feature_elongation,3_prime_UTR_variant |
| 16 | 84659872 | 84659940 | 68 | Gabpa | AJ,C3H,DBA2J,FVB,NOD,NOR,NZB,NZW,PL,SJL,SWR | feature_elongation,3_prime_UTR_variant |
| 16 | 88574947 | 88575007 | 60 | Gm5965 | 129S1,C57L,C58J,KK,NZB,NZO,NZW,PL | feature_elongation,coding_sequence_variant |
| 16 | 88666138 | 88666228 | 90 | Krtap19-1 | 129S1,C57L,C58J,ILNJ2,KK,NZB,NZO,NZW,PL | feature_elongation,coding_sequence_variant |
| 16 | 88971754 | 88971832 | 78 | 1110057P08Rik | 129S1,C57L | feature_elongation,coding_sequence_variant |
| 16 | 91714256 | 91714394 | 138 | Itsn1 | AKR,BUB,CBA,CE,DBA1J,FVB,ILNJ2,MA/My,NZB,NZO,PL,SJL,SWR,TH | feature_elongation,3_prime_UTR_variant |
| 17 | 7172289 | 7172343 | 54 | Rsph3b | 129S1,AJ,AKR,BALB,BUB,C3H,C57L,CBA,DBA1J,DBA2J,FVB,ILNJ2,KK,MA/My,MRL,NOD,NOR,NZO,PJ,PL,RF,SEA,SMJ,TH | feature_elongation,3_prime_UTR_variant |
| 17 | 8164849 | 8164903 | 54 | Rsph3a | AKR,C57L,ILNJ2,NZO,PJ,PL,RF | feature_elongation,coding_sequence_variant |
| 17 | 10538307 | 10538412 | 105 | Qki | 129S1,AJ,AKR,B10D2,B10J,BALB,BTBR,BUB,C3H,C57L,C58J,CBA,CE,DBA1J,DBA2J,FVB,ILNJ2,KK,LP,MA/My,MRL,NOD,NOR,NZB,NZO,NZW,PJ,PL,RF,RH2,SEA,SJL,SMJ,SWR,TH | feature_elongation,5_prime_UTR_variant |
| 17 | 15165800 | 15165864 | 64 | 1600012H06Rik | PL | feature_elongation,3_prime_UTR_variant |
| 17 | 15695567 | 15695876 | 309 | Psmb1 | BTBR,BUB,C3H,CBA,MRL,NOD,NOR,PJ,RF,SJL,SMJ,SWR | feature_elongation,3_prime_UTR_variant |
| 17 | 15719822 | 15720371 | 549 | Psmb1 | PL | feature_elongation,5_prime_UTR_variant |
| 17 | 15738956 | 15739402 | 446 | Tbp | BTBR,BUB,C3H,CBA,CE,ILNJ2,MRL,NOD,NOR,PJ,PL,RF,SJL,SMJ,SWR,TH | feature_elongation,3_prime_UTR_variant |
| 17 | 15764370 | 15764454 | 84 | Prdm9 | B10J,BTBR,BUB,C3H,CBA,CE,MRL,NOD,NOR,PJ,PL,RF,SJL,SMJ,TH | feature_elongation,coding_sequence_variant |
| 17 | 19752938 | 19752993 | 55 | Vmn2r100 | AJ,BALB,CE,DBA1J,DBA2J,KK,RH2 | feature_elongation,3_prime_UTR_variant |
| 17 | 20789725 | 20790181 | 456 | Vmn2r110 | 129S1,AJ,AKR,BALB,C3H,C57L,CBA,CE,DBA1J,DBA2J,FVB,KK,MA/My,MRL,NZO,PJ,RH2,SEA | feature_elongation,3_prime_UTR_variant |
| 17 | 21904218 | 21904454 | 236 | Gm10509 | 129S1,AJ,BALB,C57L,CE,DBA1J,DBA2J,NZO,RH2,SEA | feature_elongation,5_prime_UTR_variant |
| 17 | 24394457 | 24394664 | 207 | Tbc1d24 | 129S1,AJ,AKR,B10D2,BALB,BTBR,BUB,C3H,C57L,CBA,CE,DBA1J,DBA2J,FVB,ILNJ2,KK,MA/My,MRL,NOD,NOR,NZB,NZO,NZW,PJ,RF,RH2,SEA,SJL,SMJ,SWR,TH | feature_elongation,3_prime_UTR_variant |
| 17 | 24659265 | 24659473 | 208 | Dnase1l2 | 129S1,AJ,AKR,B10D2,BALB,BTBR,BUB,C3H,C57L,CBA,CE,DBA1J,DBA2J,FVB,ILNJ2,KK,MA/My,MRL,NOD,NOR,NZB,NZO,NZW,PJ,RF,RH2,SEA,SJL,SMJ,SWR,TH | feature_elongation,3_prime_UTR_variant |
| 17 | 25314766 | 25314820 | 54 | Ift140 | 129S1,AJ,AKR,B10D2,BALB,BTBR,BUB,C3H,C57L,CBA,CE,DBA1J,DBA2J,FVB,ILNJ2,KK,MA/My,MRL,NOD,NOR,NZB,NZO,RF,RH2,SEA,SJL,SWR,TH | feature_elongation,3_prime_UTR_variant |
| 17 | 28115605 | 28115659 | 54 | Bltp3a | 129S1,AJ,AKR,B10D2,BALB,BTBR,C3H,C57L,CBA,CE,DBA1J,DBA2J,FVB,ILNJ2,KK,MA/My,MRL,NZB,NZW,RF,RH2,SEA,SJL,SMJ,TH | feature_elongation,3_prime_UTR_variant |
| 17 | 29537235 | 29537456 | 221 | Pi16 | AJ,AKR,BTBR,C3H,CBA,CE,MRL | feature_elongation,5_prime_UTR_variant |
| 17 | 30047234 | 30047300 | 66 | Mdga1 | KK,RF,TH | feature_elongation,3_prime_UTR_variant |
| 17 | 30810875 | 30811111 | 236 | Glo1 | KK,NOD,NOR,RF | feature_elongation,3_prime_UTR_variant |
| 17 | 30936864 | 30936974 | 110 | Dnah8 | 129S1,B10D2,BALB,BUB,C57L,DBA1J,DBA2J,MA/My,PL,RH2,SEA,SWR | feature_elongation,coding_sequence_variant |
| 17 | 31465694 | 31465897 | 203 | Ubash3a | KK,NOD,NOR | feature_elongation,3_prime_UTR_variant |
| 17 | 32911607 | 32911769 | 162 | Cyp4f15 | AJ,AKR,C3H,CBA,CE,MRL,NOD,NOR,PJ,RF,TH | feature_elongation,coding_sequence_variant |
| 17 | 33034992 | 33035097 | 105 | Zfp799 | MA/My,PL,SJL,TH | feature_elongation,3_prime_UTR_variant |
| 17 | 33433730 | 33433795 | 65 | Or10h5 | AJ,AKR,B10J,BTBR,BUB,C3H,C58J,CBA,CE,FVB,ILNJ2,KK,LP,MA/My,MRL,NOD,NOR,PJ,PL,RF,SJL,SWR,TH | feature_elongation,3_prime_UTR_variant |
| 17 | 34068697 | 34068970 | 273 | Cd320 | B10D2,BALB,BUB,DBA1J,DBA2J,FVB,ILNJ2,NOD,NOR,NZB,NZO,NZW,PJ,PL,RH2,SEA,SJL,SMJ,SWR,TH | feature_elongation,3_prime_UTR_variant |
| 17 | 34463785 | 34463861 | 76 | H2-Ob | AJ,AKR,C3H,C58J,CBA,CE,MA/My,MRL,PJ,RF,SMJ | feature_elongation,3_prime_UTR_variant |
| 17 | 34559744 | 34559871 | 127 | H2-Eb2 | PJ | feature_elongation,3_prime_UTR_variant |
| 17 | 35930978 | 35931180 | 202 | Muc21 | AJ,BALB,DBA2J,NZB | feature_elongation,coding_sequence_variant |
| 17 | 35931729 | 35931864 | 135 | Muc21 | PJ | feature_elongation,coding_sequence_variant |
| 17 | 35933305 | 35933395 | 90 | Muc21 | AJ,B10D2,BALB,DBA2J,NZB,PL,RH2,SEA,SJL | feature_elongation,coding_sequence_variant |
| 17 | 35947124 | 35947194 | 70 | Mucl3 | AJ,B10D2,BALB,C58J,DBA2J,ILNJ2,NZB,PL,RH2,SEA,SJL,TH | feature_elongation,3_prime_UTR_variant |
| 17 | 43399052 | 43399108 | 56 | Tnfrsf21 | C58J,ILNJ2,TH | feature_elongation,3_prime_UTR_variant |
| 17 | 44349956 | 44350009 | 53 | Rcan2 | C3H | feature_elongation,3_prime_UTR_variant |
| 17 | 44916090 | 44916171 | 81 | Runx2 | 129S1,AJ,BTBR,BUB,C57L,C58J,DBA1J,KK,LP,MRL,NOD,NOR,PL,TH | feature_elongation,3_prime_UTR_variant |
| 17 | 45831404 | 45831615 | 211 | Aars2 | AKR,BALB,C3H,CBA,CE,MRL,PJ,PL,RH2,SEA | feature_elongation,3_prime_UTR_variant |
| 17 | 46570366 | 46570483 | 117 | Tjap1 | BTBR,C57L,DBA1J,KK,LP,MA/My,NZO,RF | feature_elongation,coding_sequence_variant |
| 17 | 46633183 | 46633282 | 99 | Abcc10 | NOD,NOR,NZW,SJL,SWR | feature_elongation,coding_sequence_variant |
| 17 | 46723441 | 46723492 | 51 | Zfp318 | NOD,NOR,NZW,SJL,SWR | feature_elongation,coding_sequence_variant |
| 17 | 46728315 | 46728405 | 90 | Zfp318 | NOD,NOR,NZW,SJL,SWR | feature_elongation,3_prime_UTR_variant |
| 17 | 48694945 | 48694999 | 54 | Nfya | SWR | feature_elongation,3_prime_UTR_variant |
| 17 | 63806660 | 63806751 | 91 | Fbxl17 | SMJ | feature_elongation,coding_sequence_variant |
| 17 | 73800373 | 73800435 | 62 | Galnt14 | C58J,MA/My,NZO,PL | feature_elongation,3_prime_UTR_variant |
| 17 | 84498780 | 84498991 | 211 | Thada | DBA1J,DBA2J,ILNJ2 | feature_elongation,3_prime_UTR_variant |
| 17 | 84928028 | 84928201 | 173 | Plekhh2 | DBA1J,DBA2J,ILNJ2 | feature_elongation,3_prime_UTR_variant |
| 17 | 87139125 | 87139211 | 86 | Epas1 | AJ,BALB,BTBR,C3H,C57L,CBA,DBA1J,DBA2J,ILNJ2,LP,RH2,SEA | feature_elongation,3_prime_UTR_variant |
| 18 | 9881189 | 9881646 | 457 | Colec12 | 129S1,C57L | feature_elongation,3_prime_UTR_variant |
| 18 | 10595058 | 10595115 | 57 | Esco1 | 129S1,C57L | feature_elongation,coding_sequence_variant |
| 18 | 15102333 | 15102387 | 54 | Kctd1 | BALB,C3H,CBA,NOD,RH2 | feature_elongation,3_prime_UTR_variant |
| 18 | 21264277 | 21264501 | 224 | Garem1 | CBA,CE,DBA1J,DBA2J,FVB,KK,NOD,NOR,RF,SMJ,SWR,TH | feature_elongation,3_prime_UTR_variant |
| 18 | 30463861 | 30464535 | 674 | Pik3c3 | BALB,C3H,C58J,CE,MRL,PL,RF,RH2,SEA | feature_elongation,3_prime_UTR_variant |
| 18 | 32236955 | 32237362 | 407 | Iws1 | 129S1,C57L,ILNJ2 | feature_elongation,3_prime_UTR_variant |
| 18 | 37626589 | 37626786 | 197 | Pcdhb18 | KK | feature_elongation,3_prime_UTR_variant |
| 18 | 37850906 | 37850963 | 57 | Pcdhga7 | BUB,C3H,CE,DBA1J,MA/My,NOD,NOR,PL,RF,SMJ | feature_elongation,3_prime_UTR_variant |
| 18 | 43449949 | 43450008 | 59 | Stk32a | MA/My | feature_elongation,3_prime_UTR_variant |

| | | | | | | |
|---|---|---|---|---|---|---|
| 18 | 43910273 | 43910743 | 470 | Gm94 | 129S1,AJ,AKR,BALB,BTBR,C3H,C57L,CBA,CE,DBA1J,DBA2J,FVB,ILNJ2,KK,MA/My,MRL,NZB,NZW,PJ,PL,RF,RH2,SEA,SJL,SWR,TH | feature_elongation,3_prime_UTR_variant |
| 18 | 46370482 | 46370545 | 63 | Pggt1b | KK | feature_elongation,3_prime_UTR_variant |
| 18 | 46742644 | 46742702 | 58 | Eif1a | KK | feature_elongation,3_prime_UTR_variant |
| 18 | 53522829 | 53523225 | 396 | Snx24 | KK | feature_elongation,3_prime_UTR_variant |
| 18 | 60709282 | 60709390 | 108 | Myoz3 | AKR,B10D2,B10J,BTBR,BUB,C3H,C58J,CE,DBA1J,DBA2J,FVB,LP,MA/My,MRL,NZB,NZO,NZW,PJ,PL,RF,SJL | feature_elongation,3_prime_UTR_variant |
| 18 | 61328154 | 61328318 | 164 | Slc26a2 | KK,NZO,NZW | feature_elongation,3_prime_UTR_variant |
| 18 | 64409590 | 64410234 | 644 | St8sia3 | AJ,AKR,BALB,BUB,C3H,CBA,FVB,ILNJ2,KK,MRL,NOD,NOR,NZB,NZW,PL,RF,RH2,SJL | feature_elongation,3_prime_UTR_variant |
| 18 | 67926767 | 67926825 | 58 | Seh1l | AJ,BALB,FVB,ILNJ2,KK,MRL,NOD,NOR,NZB,NZO,NZW,PL,RF,RH2,SJL,SMJ,SWR,TH | feature_elongation,3_prime_UTR_variant |
| 18 | 70663594 | 70663648 | 54 | Poli | BUB | feature_elongation,5_prime_UTR_variant |
| 18 | 74012017 | 74012067 | 50 | Mro | BUB,NZO | feature_elongation,3_prime_UTR_variant |
| 18 | 75218775 | 75218841 | 66 | Dym | AJ,AKR,BALB,ILNJ2,KK,MA/My,MRL,NOD,NOR,NZB,RF,RH2,SJL,SWR | feature_elongation,coding_sequence_variant |
| 18 | 75219500 | 75219640 | 140 | Dym | C3H,CBA,DBA1J,DBA2J,PJ,PL,SEA | feature_elongation,3_prime_UTR_variant |
| 18 | 77345665 | 77345820 | 155 | St8sia5 | 129S1,AJ,AKR,B10D2,B10J,BALB,BTBR,BUB,C3H,C57L,C58J,CBA,CE,DBA1J,DBA2J,FVB,ILNJ2,KK,LP,MA/My,MRL,NOD,NOR,NZB,NZO,NZW,PJ,PL,RF,RH2,SEA,SJL,SMJ,SWR,TH | feature_elongation,3_prime_UTR_variant |
| 18 | 78076990 | 78077190 | 200 | Epg5 | KK,NZB,TH | feature_elongation,3_prime_UTR_variant |
| 18 | 80335070 | 80335127 | 57 | Slc66a2 | AKR,NZO | feature_elongation,3_prime_UTR_variant |
| 18 | 90610533 | 90610747 | 214 | Gm45871 | KK,NZB,NZO,RF,TH | feature_elongation,3_prime_UTR_variant |
| 19 | 4234544 | 4234645 | 101 | Tbc1d10c | C58J,LP,TH | feature_elongation,3_prime_UTR_variant |
| 19 | 4938502 | 4938662 | 160 | Bbs1 | BUB,NZW | feature_elongation,3_prime_UTR_variant |
| 19 | 6171474 | 6171573 | 99 | Snx15 | TH | feature_elongation,coding_sequence_variant |
| 19 | 6586075 | 6586162 | 87 | Slc22a12 | BUB,C58J,CBA,DBA1J,DBA2J,FVB,LP,MA/My | feature_elongation,3_prime_UTR_variant |
| 19 | 9970358 | 9970589 | 231 | Best1 | AJ,BALB,C3H,C58J,CE,LP,MRL,RH2,SEA | feature_elongation,coding_sequence_variant |
| 19 | 11752806 | 11752909 | 103 | Stx3 | AJ,BALB,BUB,CE,FVB,ILNJ2,KK,LP,NOD,NOR,PJ,PL,RH2,SEA,SWR,TH | feature_elongation,3_prime_UTR_variant |
| 19 | 12307200 | 12307297 | 97 | Or5a21 | AJ,BALB,CE,MA/My,NOD,NOR,PL,RH2,SEA | feature_elongation,3_prime_UTR_variant |
| 19 | 12577399 | 12577525 | 126 | Gm4952 | AJ,BALB,CE,MA/My,NOD,NOR,PL,RH2,SEA | feature_elongation,5_prime_UTR_variant |
| 19 | 18683752 | 18683809 | 57 | Carnmt1 | NZB,NZO,NZW,PJ | feature_elongation,3_prime_UTR_variant |
| 19 | 38776697 | 38776905 | 208 | Noc3l | 129S1,AJ,AKR,BALB,BUB,C3H,C57L,CBA,CE,DBA1J,DBA2J,FVB,ILNJ2,KK,MRL,NOD,NOR,NZB,NZO,PJ,PL,RF,RH2,SEA,SJL,SMJ,SWR,TH | feature_elongation,3_prime_UTR_variant |
| 19 | 43723763 | 43723819 | 56 | Cox15 | PL,SMJ | feature_elongation,3_prime_UTR_variant |
| 19 | 46874692 | 46875183 | 491 | Nt5c2 | NZB | feature_elongation,3_prime_UTR_variant |
| 19 | 47230093 | 47230926 | 833 | Neurl1a | CE | feature_elongation,3_prime_UTR_variant |
| 19 | 57474127 | 57474194 | 67 | Trub1 | BALB,MA/My,PL,RF | feature_elongation,3_prime_UTR_variant |
| 19 | 61113578 | 61113816 | 238 | Zfp950 | 129S1,AJ,AKR,BALB,BTBR,C3H,C57L,CBA,DBA1J,DBA2J,FVB,KK,MA/My,MRL,NOD,NOR,NZB,NZW,PJ,PL,RF,RH2,SEA,SMJ,SWR,TH | feature_elongation,3_prime_UTR_variant |
| X | 70977785 | 70977863 | 78 | Pasd1 | ILNJ2 | feature_elongation,5_prime_UTR_variant,feature_elongation,coding_sequence_variant |
| X | 74423514 | 74423573 | 59 | F8 | 129S1,C57L,NZB | feature_elongation,5_prime_UTR_variant |
| X | 74439103 | 74439860 | 757 | Fundc2 | 129S1,C57L,NZB | feature_elongation,3_prime_UTR_variant |
| X | 82530664 | 82530835 | 171 | Tsga8 | AJ,AKR,BALB,BUB,C3H,CBA,CE,DBA1J,DBA2J,ILNJ2,KK,MA/My,MRL,NZB,PJ,PL,RF,RH2,SEA,SMJ | feature_elongation,coding_sequence_variant |
| X | 125721041 | 125721112 | 71 | Dmtf1l | AJ,BALB,C3H,CBA,MRL,RH2 | feature_elongation,3_prime_UTR_variant |

| CHR | START | END | SIZE | SYMBOL | STRAINS with VARIANT SV ALLELE | CONSEQUENCE |
|---|---|---|---|---|---|---|
| 1 | 4159116 | 4166266 | 7150 | Rp1 | 129S1,AKR,BTBR,BUB,C57L,CE,FVB,ILNJ2,LP,MA/My,NOD,NOR,NZB,PJ,PL,SJL,SWR | feature_elongation,coding_sequence_variant,intron_variant |
| 1 | 6454042 | 6460962 | 6920 | Alkal1 | KK | feature_elongation,coding_sequence_variant,intron_variant |
| 1 | 8532733 | 8539247 | 6514 | Sntg1 | 129S1,AJ,AKR,BALB,BTBR,C57L,FVB,ILNJ2,KK,LP,NOD,NOR,NZO,PJ,PL,RH2,SEA,SJL,SMJ,SWR,TH | feature_elongation,coding_sequence_variant,intron_variant |
| 1 | 9987587 | 9994690 | 7103 | Mcmdc2 | C3H,CBA,MRL,RF | feature_elongation,coding_sequence_variant,intron_variant |
| 1 | 14951181 | 14952762 | 1581 | Trpa1 | 129S1,BTBR,C57L,CE,DBA1J,DBA2J,MA/My,NOR | feature_elongation,coding_sequence_variant,intron_variant |
| 1 | 20672026 | 20678856 | 6830 | Pkhd1 | SMJ | feature_elongation,coding_sequence_variant,intron_variant |
| 1 | 21572030 | 21578724 | 6694 | Kcnq5 | BALB,CE,KK,MA/My,NOD,SEA | feature_elongation,coding_sequence_variant,intron_variant |
| 1 | 22506002 | 22512707 | 6705 | Rims1 | AJ,AKR,BALB,BTBR,BUB,C3H,C57L,CBA,DBA1J,DBA2J,FVB,KK,LP,NOD,RF,RH2,SEA,SWR | feature_elongation,coding_sequence_variant,intron_variant |
| 1 | 34899533 | 34906230 | 6697 | Plekhb2 | 129S1,AKR,BTBR,C57L,CBA,KK,MA/My,PJ,PL,SJL,TH | feature_elongation,coding_sequence_variant,5_prime_UTR_variant,intron_variant |
| 1 | 36533774 | 36540226 | 6452 | Cnnm4 | AJ,BUB,C3H,CE,MRL,NZW,SMJ | feature_elongation,coding_sequence_variant,intron_variant |
| 1 | 46387602 | 46392840 | 5238 | Dnah7b | 129S1,AJ,BUB,C3H,CBA,CE,DBA1J,DBA2J,KK,MRL,PJ,PL,SJL,SMJ | feature_elongation,coding_sequence_variant,intron_variant |
| 1 | 53531846 | 53536404 | 4558 | Dnah7a | SMJ | feature_elongation,coding_sequence_variant,intron_variant |
| 1 | 53606388 | 53611930 | 5542 | Dnah7a | KK | stop_gained,frameshift_variant |
| 1 | 55209800 | 55213959 | 4159 | Rftn2 | SMJ | feature_elongation,coding_sequence_variant,3_prime_UTR_variant,intron_variant |
| 1 | 57998655 | 58005591 | 6936 | Kctd18 | NZB | feature_elongation,coding_sequence_variant,intron_variant |
| 1 | 60557982 | 60561238 | 3256 | Raph1 | SWR | feature_elongation,coding_sequence_variant,intron_variant |
| 1 | 65298180 | 65304433 | 6253 | Pikfyve | CBA,DBA1J,DBA2J,LP,PJ,SMJ | feature_elongation,coding_sequence_variant,intron_variant |
| 1 | 69933467 | 69939870 | 6403 | Spag16 | C58J,SWR | feature_elongation,coding_sequence_variant,intron_variant |
| 1 | 72633305 | 72639759 | 6454 | Smarcal1 | SEA | feature_elongation,coding_sequence_variant,intron_variant |
| 1 | 86122837 | 86124727 | 1890 | Armc9 | NZB | feature_elongation,coding_sequence_variant,intron_variant |
| 1 | 86512642 | 86519352 | 6710 | Cops7b | 129S1,AJ,BTBR,C3H,C57L,CE,FVB,ILNJ2,KK,NZO,NZW,RH2,SWR,TH | feature_elongation,coding_sequence_variant,5_prime_UTR_variant,intron_variant |
| 1 | 86714327 | 86721036 | 6709 | Dis3l2 | CE | feature_elongation,coding_sequence_variant,intron_variant |
| 1 | 91232035 | 91233910 | 1875 | Scly | PJ | feature_elongation,coding_sequence_variant,intron_variant |
| 1 | 92417891 | 92422948 | 5057 | Or6b2b | BUB,C3H,CBA | feature_elongation,coding_sequence_variant,5_prime_UTR_variant,3_prime_UTR_variant,intron_variant |
| 1 | 97041495 | 97048031 | 6536 | Slco6c1 | AKR,BUB,SJL | feature_elongation,coding_sequence_variant,intron_variant |
| 1 | 97540138 | 97546620 | 6482 | Macir | PL | feature_elongation,3_prime_UTR_variant |
| 1 | 97548040 | 97554632 | 6592 | Macir | MRL,PL | feature_elongation,3_prime_UTR_variant |
| 1 | 98422756 | 98424925 | 2169 | Slco6d1 | MRL,PL | feature_elongation,coding_sequence_variant,intron_variant |
| 1 | 105489808 | 105496042 | 6234 | Pign | AJ,AKR,BALB,BTBR,BUB,C3H,CBA,CE,FVB,ILNJ2,KK,LP,MA/My,MRL,NZB,NZO,NZW,PJ,PL,RF,RH2,SEA,SJL,SMJ,SWR | feature_elongation,coding_sequence_variant,intron_variant |
| 1 | 106212050 | 106216540 | 4490 | Phlpp1 | ILNJ2 | feature_elongation,coding_sequence_variant,intron_variant |
| 1 | 106715108 | 106719949 | 4841 | Vps4b | BUB,NZO,NZW | feature_elongation,coding_sequence_variant,intron_variant |
| 1 | 127229449 | 127236563 | 7114 | Mgat5 | CBA,NOD,NOR,SMJ | feature_elongation,coding_sequence_variant,5_prime_UTR_variant,intron_variant |
| 1 | 128194794 | 128199991 | 5197 | Ubxn4 | MRL,NZB,SWR | feature_elongation,coding_sequence_variant,intron_variant |
| 1 | 134231291 | 134232523 | 1232 | Ppfia4 | AJ,C3H,CE,KK,NZB,NZO,PJ,RH2,SJL,SWR,TH | feature_elongation,coding_sequence_variant,intron_variant |
| 1 | 139482282 | 139484882 | 2600 | Cfhr1 | BTBR | feature_elongation,coding_sequence_variant,intron_variant |
| 1 | 140494719 | 140501872 | 7153 | Kcnt2 | KK,SWR | feature_elongation,coding_sequence_variant,intron_variant |
| 1 | 150604322 | 150610933 | 6611 | Hmcn1 | BUB,DBA1J,DBA2J,RF | feature_elongation,coding_sequence_variant,intron_variant |
| 1 | 152682288 | 152684181 | 1893 | Ncf2 | C57L | feature_elongation,coding_sequence_variant,5_prime_UTR_variant,intron_variant |
| 1 | 152698928 | 152700918 | 1990 | Ncf2 | AKR,BTBR,KK,NZB,NZO,NZW,SMJ | feature_elongation,coding_sequence_variant,intron_variant |
| 1 | 156448172 | 156449794 | 1622 | Abl2 | C58J,FVB,ILNJ2,RH2 | feature_elongation,coding_sequence_variant,intron_variant |
| 1 | 159970355 | 159976791 | 6436 | Tnn | ILNJ2,MA/My,NZW,PL | feature_elongation,coding_sequence_variant,5_prime_UTR_variant,intron_variant |
| 1 | 160925218 | 160926861 | 1643 | Klhl20 | 129S1,AJ,BALB,BUB,C3H,C57L,C58J,CBA,CE,DBA1J,DBA2J,ILNJ2,LP,NZW,PJ,RH2,SEA,SMJ | feature_elongation,coding_sequence_variant,intron_variant |
| 1 | 162869470 | 162875286 | 5816 | Mroh9 | 129S1,AJ,BALB,BUB,C3H,C57L,C58J,CBA,CE,ILNJ2,LP,MA/My,MRL,PL,RH2,TH | feature_elongation,coding_sequence_variant,intron_variant |
| 1 | 173408768 | 173414009 | 5241 | Ifi213 | 129S1,BALB,C3H,C57L,MRL,NOD,NOR | feature_elongation,coding_sequence_variant,intron_variant |
| 1 | 174517445 | 174523806 | 6361 | Fmn2 | PL,TH | feature_elongation,coding_sequence_variant,intron_variant |
| 1 | 174947470 | 174951990 | 4520 | Rgs7 | C58J,CBA,CE,DBA1J,DBA2J,ILNJ2,NZB,NZW,PJ,SEA,SMJ | feature_elongation,coding_sequence_variant,intron_variant |
| 1 | 175586380 | 175593247 | 6867 | Wdr64 | 129S1,BALB,C3H,C57L,MRL,NOD,NOR,NZW,PL,SMJ | feature_elongation,coding_sequence_variant,intron_variant |
| 1 | 177708269 | 177714708 | 6439 | Catspere1 | AJ,BUB,RH2 | feature_elongation,coding_sequence_variant,intron_variant |
| 1 | 177843026 | 177850124 | 7098 | Catspere2 | CBA,DBA1J,DBA2J,KK | feature_elongation,coding_sequence_variant,intron_variant |

| | | | | | | |
|---|---|---|---|---|---|---|
| 1 | 178023051 | 178029491 | 6440 | Desi2 | AJ,BUB,RH2 | feature_elongation,5_prime_UTR_variant,intron_variant |
| 1 | 179928080 | 179931075 | 2995 | Cdc42bpa | C58J,LP | feature_elongation,coding_sequence_variant,intron_variant |
| 1 | 182428980 | 182435381 | 6401 | Capn8 | 129S1,BALB,C3H,C57L,CBA,DBA1J,DBA2J,KK,MA/My,PJ,PL,SEA | feature_elongation,coding_sequence_variant,intron_variant |
| 1 | 189529611 | 189531623 | 2012 | Ptpn14 | AKR,NZO,NZW | feature_elongation,coding_sequence_variant,intron_variant |
| 2 | 5835759 | 5842232 | 6473 | Cdc123 | FVB,NZB | feature_elongation,coding_sequence_variant,intron_variant |
| 2 | 13218105 | 13223361 | 5256 | Rsu1 | SMJ | feature_elongation,coding_sequence_variant,intron_variant |
| 2 | 14036516 | 14041745 | 5229 | Hacd1 | C3H | feature_elongation,coding_sequence_variant,3_prime_UTR_variant,intron_variant,feature_elongation,coding_sequence_variant,intron_variant |
| 2 | 14408055 | 14414458 | 6403 | Slc39a12 | SJL | feature_elongation,coding_sequence_variant,intron_variant |
| 2 | 30971780 | 30976980 | 5200 | Fnbp1 | MA/My | feature_elongation,coding_sequence_variant,intron_variant |
| 2 | 31663079 | 31670199 | 7120 | Abl1 | NZO | feature_elongation,coding_sequence_variant,intron_variant |
| 2 | 35272195 | 35278411 | 6216 | 4930402F06Rik | BUB,C58J,MA/My,NZO | feature_elongation,coding_sequence_variant,intron_variant |
| 2 | 37910710 | 37917185 | 6475 | Dennd1a | BUB | feature_elongation,coding_sequence_variant,intron_variant |
| 2 | 38845661 | 38852810 | 7149 | Olfml2a | 129S1,AJ,AKR,BTBR,C3H,C57L,C58J,CBA,CE,DBA1J,DBA2J,FVB,ILNJ2,KK,MA/My,MRL,NOD,NOR,NZB,NZO,NZW,PJ,PL,RF,SEA,SMJ,SWR | feature_elongation,coding_sequence_variant,3_prime_UTR_variant,intron_variant |
| 2 | 39014823 | 39020372 | 5549 | Scai | NOD | feature_elongation,coding_sequence_variant,intron_variant |
| 2 | 40789273 | 40795948 | 6675 | Lrp1b | BUB | feature_elongation,coding_sequence_variant,intron_variant |
| 2 | 41196725 | 41203526 | 6801 | Lrp1b | 129S1,AJ,AKR,BTBR,C3H,C57L,C58J,CBA,DBA1J,DBA2J,FVB,ILNJ2,KK,MA/My,MRL,NOD,NOR,NZB,NZO,NZW,PJ,PL,RF,SEA,SWR | feature_elongation,coding_sequence_variant,intron_variant |
| 2 | 48813144 | 48820224 | 7080 | Orc4 | FVB,SJL,TH | feature_elongation,coding_sequence_variant,intron_variant |
| 2 | 62108530 | 62114972 | 6442 | Slc4a10 | NOD,NOR,NZB,NZO,NZW,SJL | feature_elongation,coding_sequence_variant,intron_variant |
| 2 | 62562405 | 62572147 | 9742 | Kcnh7 | NZO | feature_elongation,coding_sequence_variant,intron_variant |
| 2 | 65527192 | 65534481 | 7289 | Scn2a | C3H,C58J,CBA,MRL,PL,SMJ,TH | feature_elongation,coding_sequence_variant,intron_variant |
| 2 | 69086415 | 69092671 | 6256 | Abcb11 | 129S1,C3H,C58J,CBA,NZW,SJL | feature_elongation,coding_sequence_variant,intron_variant |
| 2 | 70822284 | 70827832 | 5548 | Mettl8 | MA/My | feature_elongation,coding_sequence_variant,intron_variant |
| 2 | 83582299 | 83588614 | 6315 | Itgav | AJ,AKR,BALB,C3H,C57L,C58J,CBA,CE,DBA1J,DBA2J,MA/My,MRL,NZW,PJ,RF,RH2,SEA,SMJ | feature_elongation,coding_sequence_variant,intron_variant |
| 2 | 88150536 | 88156976 | 6440 | Or5d45 | C58J,ILNJ2,KK | feature_elongation,coding_sequence_variant,3_prime_UTR_variant,intron_variant |
| 2 | 88574843 | 88576333 | 1490 | Or4p23 | AJ,AKR,BALB,C57L,DBA1J,DBA2J,FVB,KK,MRL,NOD,NOR,NZB,NZO,PJ,RF,RH2,SEA,SMJ | feature_elongation,coding_sequence_variant,3_prime_UTR_variant |
| 2 | 102407053 | 102413573 | 6520 | Pamr1 | ILNJ2,NZB,RF,SMJ | feature_elongation,coding_sequence_variant,intron_variant |
| 2 | 113391854 | 113398105 | 6251 | Fmn1 | C3H,CE,PL | feature_elongation,coding_sequence_variant,intron_variant |
| 2 | 118465340 | 118468847 | 3507 | Bub1b | 129S1,AJ,AKR,BALB,BUB,C57L,CE,DBA1J,DBA2J,FVB,ILNJ2,MA/My,NZW,PJ,PL,RF,RH2,SEA,SJL,SMJ,SWR | feature_elongation,coding_sequence_variant,intron_variant |
| 2 | 120325114 | 120331556 | 6442 | Capn3 | PL | feature_elongation,coding_sequence_variant,intron_variant |
| 2 | 125727666 | 125736484 | 8818 | Galk2 | C3H,CBA,DBA2J,ILNJ2,PJ | feature_elongation,coding_sequence_variant,intron_variant |
| 2 | 128922836 | 128929455 | 6619 | Ttl | BALB,C3H,C58J,CBA,NZB,SEA,SMJ | feature_elongation,coding_sequence_variant,intron_variant |
| 2 | 152822451 | 152828893 | 6442 | Ttll9 | AJ,ILNJ2,NOD,NZW,RF | feature_elongation,coding_sequence_variant,intron_variant |
| 2 | 153699769 | 153701377 | 1608 | Sun5 | AKR,B10D2,B10J,BALB,BUB,C3H,CBA,DBA1J,DBA2J,FVB,KK,LP,NOR,NZO,PL,RH2,SEA,SJL,SWR | feature_elongation,coding_sequence_variant,intron_variant |
| 2 | 154480584 | 154485696 | 5112 | Zfp341 | CE | feature_elongation,coding_sequence_variant,intron_variant |
| 2 | 166704183 | 166709472 | 5289 | Arfgef2 | SMJ | feature_elongation,coding_sequence_variant,intron_variant |
| 2 | 167274094 | 167281308 | 7214 | Slc9a8 | CE | feature_elongation,coding_sequence_variant,intron_variant |
| 2 | 177006961 | 177009397 | 2436 | Gm14412 | DBA2J,SEA | feature_elongation,coding_sequence_variant,intron_variant |
| 2 | 178014598 | 178018743 | 4145 | Sycp2 | PJ | feature_elongation,coding_sequence_variant,intron_variant |
| 3 | 14717811 | 14722126 | 4315 | Car13 | NZO | feature_elongation,coding_sequence_variant,intron_variant |
| 3 | 19595838 | 19602237 | 6399 | Dnajc5b | DBA2J,LP,MA/My,NOD,NOR,NZB,NZO,NZW | feature_elongation,coding_sequence_variant,5_prime_UTR_variant,intron_variant |
| 3 | 41670269 | 41673642 | 3373 | Sclt1 | SJL,SWR | feature_elongation,coding_sequence_variant,intron_variant |
| 3 | 41685568 | 41692192 | 6624 | Sclt1 | 129S1 | feature_elongation,coding_sequence_variant,intron_variant |
| 3 | 58530722 | 58536279 | 5557 | Erich6 | BUB,CE,FVB,SJL,SWR | feature_elongation,coding_sequence_variant,intron_variant |
| 3 | 69923383 | 69930088 | 6705 | Otol1 | 129S1,BTBR,C57L,FVB,SWR,TH | feature_elongation,coding_sequence_variant,5_prime_UTR_variant,intron_variant |
| 3 | 72832576 | 72834170 | 1594 | Sis | 129S1,AJ,AKR,BALB,BTBR,BUB,C57L,FVB,KK,MA/My,MRL,NOD,NOR,NZB,NZO,PJ,RF,RH2,SEA,SJL,SWR,TH | feature_elongation,coding_sequence_variant,intron_variant |
| 3 | 76439469 | 76447058 | 7589 | Fstl5 | AKR,DBA1J,DBA2J,MRL,PL,RF | feature_elongation,coding_sequence_variant,intron_variant |

| | | | | | | |
|---|---|---|---|---|---|---|
| 3 | 86206998 | 86214144 | 7146 | Lrba | NZB,NZO | feature_elongation,coding_sequence_variant,intron_variant |
| 3 | 90039969 | 90047084 | 7115 | Nup210l | NOD,NOR | feature_elongation,coding_sequence_variant,intron_variant |
| 3 | 90047557 | 90053075 | 5518 | Nup210l | RF | feature_elongation,coding_sequence_variant,intron_variant |
| 3 | 92865525 | 92881535 | 16010 | Lce3d | DBA1J,DBA2J | stop_gained,inframe_insertion |
| 3 | 97629422 | 97636323 | 6901 | Pde4dip | NOD,NOR,RF | feature_elongation,coding_sequence_variant,intron_variant |
| 3 | 99954703 | 99957016 | 2313 | Spag17 | 129S1,B10D2,B10J,BTBR,C57L,C58J,DBA1J,DBA2J,LP,MA/My,NOD,RH2,TH | feature_elongation,coding_sequence_variant,intron_variant |
| 3 | 100021433 | 100028546 | 7113 | Spag17 | BTBR,DBA1J,DBA2J,NOD,NOR | feature_elongation,coding_sequence_variant,intron_variant |
| 3 | 100043841 | 100050157 | 6316 | Spag17 | KK | feature_elongation,3_prime_UTR_variant,intron_variant |
| 3 | 102835780 | 102840386 | 4606 | Sycp1 | 129S1,AKR,BALB,BUB,C57L,C58J,FVB,ILNJ2,KK,LP,MA/My,MRL,NOD,NOR,NZB,NZO,NZW,PL,RF,RH2,SWR | feature_elongation,coding_sequence_variant,intron_variant |
| 3 | 104369175 | 104370923 | 1748 | Lrig2 | 129S1,AKR,BUB,C57L,C58J,FVB,ILNJ2,KK,PL,SJL,SWR,TH | feature_elongation,coding_sequence_variant,intron_variant |
| 3 | 104745458 | 104751918 | 6460 | Capza1 | CE,NZB,PJ,SEA | feature_elongation,coding_sequence_variant,intron_variant |
| 3 | 108276191 | 108281443 | 5252 | Mybphl | MRL | feature_elongation,coding_sequence_variant,intron_variant |
| 3 | 113361945 | 113363972 | 2027 | Amy1 | CE,NOD,NOR | feature_elongation,coding_sequence_variant,5_prime_UTR_variant,intron_variant |
| 3 | 118403388 | 118409612 | 6224 | Dpyd | BUB,LP | feature_elongation,coding_sequence_variant,intron_variant |
| 3 | 123334910 | 123341435 | 6525 | Ndst3 | PJ,SEA | feature_elongation,coding_sequence_variant,intron_variant |
| 3 | 129514326 | 129519521 | 5195 | Egf | TH | feature_elongation,coding_sequence_variant,intron_variant |
| 3 | 135217939 | 135224412 | 6473 | Manba | KK | feature_elongation,coding_sequence_variant,intron_variant |
| 3 | 135551040 | 135555571 | 4531 | Slc39a8 | CBA,CE,DBA1J,DBA2J,FVB,KK,MA/My,NOD,NOR,SJL,SMJ,SWR,TH | feature_elongation,coding_sequence_variant,intron_variant |
| 3 | 138127419 | 138128907 | 1488 | Adh4 | 129S1,AJ,BALB,BTBR,BUB,C3H,C57L,C58J,CBA,CE,DBA1J,DBA2J,ILNJ2,KK,LP,MRL,NZB,NZO,NZW,RF,RH2 | feature_elongation,coding_sequence_variant,intron_variant |
| 3 | 139225878 | 139228933 | 3055 | Stpg2 | CE | feature_elongation,coding_sequence_variant,intron_variant |
| 3 | 153596052 | 153603829 | 7777 | Msh4 | 129S1,BTBR,C57L | feature_elongation,coding_sequence_variant,intron_variant,feature_elongation,coding_sequence_variant,5_prime_UTR_variant,intron_variant |
| 3 | 154465114 | 154469660 | 4546 | Erich3 | FVB,SJL,SWR | feature_elongation,coding_sequence_variant,intron_variant |
| 3 | 154660368 | 154667563 | 7195 | Tnni3k | AJ,BALB,BUB,C3H,DBA2J,ILNJ2,KK,MA/My,NOD,NOR,NZB,NZO,NZW,RH2,SEA | feature_elongation,coding_sequence_variant,intron_variant |
| 4 | 19833785 | 19840259 | 6474 | Slc7a13 | ILNJ2 | feature_elongation,coding_sequence_variant,intron_variant |
| 4 | 43319825 | 43325374 | 5549 | Atp8b5 | C58J,LP | feature_elongation,coding_sequence_variant,3_prime_UTR_variant,intron_variant,feature_elongation,coding_sequence_variant,intron_variant |
| 4 | 59483832 | 59490259 | 6427 | Ptbp3 | CE | feature_elongation,coding_sequence_variant,intron_variant |
| 4 | 59604688 | 59611432 | 6744 | Hsdl2 | BUB | feature_elongation,coding_sequence_variant,intron_variant |
| 4 | 83494262 | 83499810 | 5548 | Ccdc171 | PL | feature_elongation,coding_sequence_variant,intron_variant |
| 4 | 96352448 | 96359866 | 7418 | Cyp2j8 | ILNJ2 | feature_elongation,coding_sequence_variant,intron_variant |
| 4 | 99108714 | 99115160 | 6446 | Atg4c | CE | feature_elongation,coding_sequence_variant,intron_variant |
| 4 | 100717005 | 100723565 | 6560 | Cachd1 | 129S1,AJ,BALB,C3H,C57L,CBA,CE,NOD,NOR,RH2,SEA,SWR,TH | feature_elongation,coding_sequence_variant,intron_variant |
| 4 | 107021998 | 107025544 | 3546 | Tceanc2 | 129S1,AJ,C57L,FVB,TH | feature_elongation,coding_sequence_variant,intron_variant |
| 4 | 108778770 | 108784207 | 5437 | Rab3b | BALB,BUB,C3H,CBA,NOD,PL,RF,SEA,SJL | feature_elongation,coding_sequence_variant,intron_variant |
| 4 | 108866720 | 108872269 | 5549 | Nrdc | NOD | feature_elongation,coding_sequence_variant,intron_variant |
| 4 | 109608115 | 109615273 | 7158 | Faf1 | CE | feature_elongation,coding_sequence_variant,intron_variant |
| 4 | 111467612 | 111475125 | 7513 | Agbl4 | CE,SWR | feature_elongation,coding_sequence_variant,intron_variant |
| 4 | 115667258 | 115668862 | 1604 | Atpaf1 | AJ,BALB,BUB,C3H,CBA,CE,DBA1J,DBA2J,FVB,KK,NOD,NOR,RF,SJL,SWR,TH | feature_elongation,coding_sequence_variant,3_prime_UTR_variant,intron_variant |
| 4 | 116387118 | 116389413 | 2295 | Ipp | AKR,B10D2,B10J,C58J,KK,MRL,NZB,NZO,NZW,PJ,PL,SEA,SMJ,TH | feature_elongation,coding_sequence_variant,intron_variant |
| 4 | 116835780 | 116837481 | 1701 | Zswim5 | ILNJ2,TH | feature_elongation,coding_sequence_variant,intron_variant |
| 4 | 119334594 | 119337427 | 2833 | Rimkla | 129S1,BALB,C3H,C57L,CBA,CE,SJL | feature_elongation,coding_sequence_variant,intron_variant |
| 4 | 120361057 | 120364675 | 3618 | Scmh1 | FVB,SWR | feature_elongation,coding_sequence_variant,intron_variant |
| 4 | 126808396 | 126809434 | 1038 | Zmym4 | B10D2,B10J,ILNJ2,NOD,PJ,SEA | feature_elongation,coding_sequence_variant,intron_variant |
| 4 | 132642063 | 132646120 | 4057 | Fam76a | SJL,SWR | feature_elongation,coding_sequence_variant,intron_variant |
| 4 | 134627035 | 134628943 | 1908 | Tmem50a | AJ,AKR,C3H,CBA,CE,MRL,RF,RH2 | feature_elongation,coding_sequence_variant,intron_variant |
| 4 | 135991461 | 135995831 | 4370 | Tcea3 | ILNJ2,PL,SMJ,TH | feature_elongation,coding_sequence_variant,intron_variant |
| 4 | 148523644 | 148529192 | 5548 | Ubiad1 | AKR,C3H,CBA,MRL,PL | feature_elongation,coding_sequence_variant,5_prime_UTR_variant,intron_variant |
| 4 | 155074247 | 155079792 | 5545 | Plch2 | AJ | stop_gained,frameshift_variant |
| 4 | 155085820 | 155091254 | 5434 | Plch2 | LP | feature_elongation,coding_sequence_variant,intron_variant |
| 4 | 155730736 | 155733282 | 2546 | Cdk11b | FVB,NOD,NOR,SJL,SWR | feature_elongation,coding_sequence_variant,intron_variant |

| | | | | | | |
|---|---|---|---|---|---|---|
| 5 | 8487519 | 8493980 | 6461 | Slc25a40 | BUB,NOD,NOR,SMJ | feature_elongation,coding_sequence_variant,intron_variant |
| 5 | 8739787 | 8744956 | 5169 | Abcb1a | SWR | feature_elongation,coding_sequence_variant,intron_variant |
| 5 | 8900711 | 8905213 | 4502 | Abcb1b | BUB,NOD,NOR | feature_elongation,coding_sequence_variant,intron_variant |
| 5 | 12544406 | 12550954 | 6548 | Sema3d | AKR,BUB,DBA1J,DBA2J,FVB,ILNJ2,KK,MRL,NOD,NOR,NZB,NZO,NZW,SJL,SWR | feature_elongation,coding_sequence_variant,intron_variant |
| 5 | 13625539 | 13632451 | 6912 | Sema3a | AJ,AKR,BUB,C3H,CBA,DBA1J,DBA2J,FVB,ILNJ2,KK,LP,MRL,NOD,NOR,SJL,SMJ,SWR | feature_elongation,coding_sequence_variant,intron_variant |
| 5 | 24004981 | 24011926 | 6945 | Rint1 | AKR,BUB,NOD,NOR,NZW,PJ,RF | feature_elongation,coding_sequence_variant,intron_variant |
| 5 | 25395815 | 25403182 | 7367 | Galntl5 | BUB | feature_elongation,coding_sequence_variant,intron_variant |
| 5 | 35675727 | 35682185 | 6458 | Cpz | AKR | feature_elongation,coding_sequence_variant,intron_variant |
| 5 | 54000357 | 54004334 | 3977 | Tbc1d19 | PL | feature_elongation,coding_sequence_variant,intron_variant |
| 5 | 66446372 | 66447667 | 1295 | Nsun7 | AKR,FVB,RF | feature_elongation,coding_sequence_variant,intron_variant |
| 5 | 76986101 | 76993231 | 7130 | Cracd | SMJ | feature_elongation,coding_sequence_variant,5_prime_UTR_variant,intron_variant,feature_elongation,coding_sequence_variant,intron_variant |
| 5 | 86169634 | 86171498 | 1864 | Cenpc1 | 129S1,C57L | feature_elongation,coding_sequence_variant,intron_variant |
| 5 | 88422444 | 88429154 | 6710 | Smr2l | NZO | feature_elongation,coding_sequence_variant,5_prime_UTR_variant,intron variant |
| 5 | 92169112 | 92170976 | 1864 | Cdkl2 | AJ,NOD,NOR,NZO | feature_elongation,coding_sequence_variant,intron_variant |
| 5 | 96782852 | 96784350 | 1498 | Fras1 | 129S1,AJ,AKR,BALB,BTBR,BUB,C3H,C57L,CBA,CE,DBA1J,DBA2J,FVB,ILNJ2,KK,MA/My,MRL,NOD,NOR,NZB,NZO,NZW,PJ,RF,RH2,SEA,SJL,SMJ,SWR,TH | feature_elongation,coding_sequence_variant,intron_variant |
| 5 | 100673917 | 100679362 | 5445 | Cops4 | 129S1,C57L | feature_elongation,coding_sequence_variant,intron_variant |
| 5 | 100956115 | 100960637 | 4522 | Abraxas1 | NZB | feature_elongation,coding_sequence_variant,intron_variant |
| 5 | 104480092 | 104485412 | 5320 | Mepe | 129S1,AKR,B10D2,B10J,BTBR,C57L,C58J,CE,LP,MA/My,MRL,NOD,NOR,NZW,PJ,PL,RF,SMJ | feature_elongation,coding_sequence_variant,intron_variant |
| 5 | 104644958 | 104652045 | 7087 | Pkd2 | AJ,BALB,DBA1J,DBA2J,ILNJ2,KK,NZB,NZO,RH2,SEA,SWR | feature_elongation,coding_sequence_variant,3_prime_UTR_variant,intron_variant |
| 5 | 104673296 | 104675622 | 2326 | Thoc2l | AJ,BALB,BUB,C3H,CBA,DBA1J,DBA2J,FVB,ILNJ2,KK,NZB,NZO,RH2,SEA,SJL,SWR,TH | feature_elongation,3_prime_UTR_variant,intron_variant |
| 5 | 107631378 | 107638487 | 7109 | Btbd8 | 129S1,C57L,NZB,NZO,NZW | feature_elongation,coding_sequence_variant,intron_variant |
| 5 | 107935033 | 107941685 | 6652 | Evi5 | DBA1J,DBA2J | feature_elongation,coding_sequence_variant,intron_variant |
| 5 | 109484995 | 109487947 | 2952 | Vmn2r16 | CE,DBA1J,DBA2J,LP | feature_elongation,coding_sequence_variant,intron_variant |
| 5 | 110393829 | 110402386 | 8557 | Ankle2 | AJ,BALB,BTBR,CE,DBA1J,DBA2J,ILNJ2,LP,MRL,RH2,SEA,SMJ | feature_elongation,coding_sequence_variant,intron_variant |
| 5 | 112869082 | 112876169 | 7087 | Myo18b | AKR,CE,DBA1J,DBA2J,NZW,RF | feature_elongation,coding_sequence_variant,intron_variant |
| 5 | 118038045 | 118040303 | 2258 | Nos1 | NOD | feature_elongation,coding_sequence_variant,intron_variant |
| 5 | 118038137 | 118039289 | 1152 | Nos1 | NOD | feature_elongation,coding_sequence_variant,intron_variant |
| 5 | 124224113 | 124229625 | 5512 | Abcb9 | BUB | feature_elongation,coding_sequence_variant,5_prime_UTR_variant,intron_variant |
| 5 | 129182490 | 129188817 | 6327 | Adgrd1 | C58J,LP | feature_elongation,coding_sequence_variant,intron_variant |
| 5 | 129234247 | 129245106 | 10859 | Adgrd1 | 129S1,C57L,ILNJ2,KK,NZB,NZO,NZW,SWR | feature_elongation,coding_sequence_variant,intron_variant |
| 5 | 131111542 | 131117098 | 5556 | Galnt17 | NOR | feature_elongation,coding_sequence_variant,intron_variant |
| 5 | 143681535 | 143688103 | 6568 | Cyth3 | CBA | feature_elongation,coding_sequence_variant,intron_variant |
| 5 | 144486739 | 144491912 | 5173 | Nptx2 | AJ,BALB,C3H,CBA,DBA1J,DBA2J,PJ,RH2,SEA | feature_elongation,coding_sequence_variant,intron_variant |
| 5 | 146044158 | 146049933 | 5775 | Cyp3a59 | AJ,DBA2J,SEA | feature_elongation,coding_sequence_variant,3_prime_UTR_variant,intron_variant |
| 5 | 147030871 | 147038099 | 7228 | Polr1d | NZB | feature_elongation,coding_sequence_variant,intron_variant |
| 5 | 148222820 | 148230151 | 7331 | Mtus2 | CE,FVB | feature_elongation,coding_sequence_variant,intron_variant |
| 5 | 149526731 | 149533696 | 6965 | Wdr95 | NZB | feature_elongation,coding_sequence_variant,intron_variant |
| 5 | 150232003 | 150234359 | 2356 | Fry | AKR,BUB,FVB,MRL,NOD,NOR,PJ,SJL | feature_elongation,coding_sequence_variant,intron_variant |
| 5 | 150398620 | 150405035 | 6415 | Fry | NZO | feature_elongation,coding_sequence_variant,intron_variant |
| 6 | 4078847 | 4085777 | 6930 | Bet1 | MA/My,NZB,NZO,NZW | feature_elongation,coding_sequence_variant,intron_variant |
| 6 | 38268735 | 38274054 | 5319 | Zc3hav1l | C3H | feature_elongation,coding_sequence_variant,3_prime_UTR_variant,intron_variant |
| 6 | 40785680 | 40792158 | 6478 | Mgam2-ps | BUB,FVB,LP,SJL,SWR,TH | feature_elongation,coding_sequence_variant,intron_variant |
| 6 | 47838550 | 47844992 | 6442 | Zfp398 | C58J,MA/My | feature_elongation,coding_sequence_variant,3_prime_UTR_variant,intron_variant |
| 6 | 49377580 | 49383124 | 5544 | Stk31 | AJ,BUB | feature_elongation,coding_sequence_variant,intron_variant |
| 6 | 49440591 | 49443178 | 2587 | Stk31 | AJ,AKR,BALB,BUB,C3H,CBA,DBA1J,DBA2J,FVB,MRL,NOD,NOR,PJ,PL,RF,RH2,SEA,SJL,SMJ,SWR | feature_elongation,coding_sequence_variant,intron_variant |
| 6 | 57708248 | 57709925 | 1677 | Lancl2 | 129S1,C57L,KK,NZB,NZO,NZW | feature_elongation,coding_sequence_variant,intron_variant |
| 6 | 70746700 | 70753153 | 6453 | Rpia | MA/My,SJL | feature_elongation,coding_sequence_variant,intron_variant |

| | | | | | | |
|---|---|---|---|---|---|---|
| 6 | 73098133 | 73101265 | 3132 | Dnah6 | AKR,BALB,MRL,PL,RF,RH2,SEA | feature_elongation,coding_sequence_variant,intron_variant |
| 6 | 93917940 | 93920912 | 2972 | Magi1 | AKR,MRL,PL | feature_elongation,coding_sequence_variant,intron_variant |
| 6 | 100725694 | 100732117 | 6423 | Gxylt2 | FVB,SJL | feature_elongation,coding_sequence_variant,intron_variant |
| 6 | 113360269 | 113363084 | 2815 | Arpc4 | AKR,C3H,CBA,FVB,ILNJ2,MRL,NOD,NOR,NZB,PJ,PL,RF,SJL | feature_elongation,coding_sequence_variant,intron_variant,feature_elongation,coding_sequence_variant,5_prime_UTR_variant,intron_variant |
| 6 | 118135292 | 118142893 | 7601 | Ret | SJL,SWR | feature_elongation,coding_sequence_variant,intron_variant |
| 6 | 119606680 | 119613093 | 6413 | Erc1 | TH | feature_elongation,coding_sequence_variant,intron_variant |
| 6 | 119709461 | 119715892 | 6431 | Erc1 | SJL,SWR | feature_elongation,coding_sequence_variant,intron_variant |
| 6 | 122032947 | 122039779 | 6832 | Mug2 | SWR | feature_elongation,coding_sequence_variant,intron_variant |
| 6 | 123153694 | 123160557 | 6863 | Clec4b2 | SWR | feature_elongation,coding_sequence_variant,intron_variant |
| 6 | 123368079 | 123375620 | 7541 | Vmn2r20 | BALB | feature_elongation,coding_sequence_variant,intron_variant |
| 6 | 123602269 | 123604044 | 1775 | Vmn2r22 | SWR | feature_elongation,coding_sequence_variant,intron_variant |
| 6 | 126910486 | 126915226 | 4740 | Rad51ap1 | AJ,AKR,BALB,BUB,C3H,C57L,C58J,CBA,CE,DBA1J,DBA2J,FVB,ILNJ2,KK,LP,MA/My,MRL,NOD,NOR,NZO,PJ,PL,RF,SEA,SJL,TH | feature_elongation,coding_sequence_variant,intron_variant |
| 6 | 140517588 | 140519907 | 2319 | Plekha5 | BTBR,C57L,C58J,FVB,PJ,SMJ,SWR | feature_elongation,coding_sequence_variant,intron_variant |
| 6 | 141953235 | 141955886 | 2651 | Slco1a8 | BALB | feature_elongation,coding_sequence_variant,intron_variant,feature_elongation,coding_sequence_variant,5_prime_UTR_variant,intron_variant |
| 7 | 3275616 | 3283094 | 7478 | Nlrp12 | MA/My | feature_elongation,coding_sequence_variant,intron_variant |
| 7 | 4007749 | 4013999 | 6250 | Lair1 | MRL | feature_elongation,coding_sequence_variant,3_prime_UTR_variant,intron_variant |
| 7 | 6475563 | 6477433 | 1870 | Or6z5 | MRL | feature_elongation,coding_sequence_variant,5_prime_UTR_variant,intron_variant |
| 7 | 7133836 | 7137428 | 3592 | Zfp773 | C58J | feature_elongation,coding_sequence_variant,3_prime_UTR_variant,intron_variant,feature_elongation,coding_sequence_variant,intron_variant |
| 7 | 9772768 | 9779182 | 6414 | Vmn2r50 | AKR,MRL,RH2 | feature_elongation,coding_sequence_variant,intron_variant |
| 7 | 11985137 | 11991353 | 6216 | Vmn1r81 | AKR,MRL,RH2 | feature_elongation,3_prime_UTR_variant |
| 7 | 12900283 | 12907709 | 7426 | Vmn1r88 | PL | feature_elongation,5_prime_UTR_variant,intron_variant |
| 7 | 15712441 | 15721175 | 8734 | Bicra | AKR,RH2 | feature_elongation,coding_sequence_variant,intron_variant |
| 7 | 17616160 | 17623028 | 6868 | Ceacam23 | CE,NOD,NOR,SJL | feature_elongation,coding_sequence_variant,intron_variant |
| 7 | 23534278 | 23541885 | 7607 | Vmn1r176 | CE | stop_gained,frameshift_variant |
| 7 | 25783833 | 25790418 | 6585 | Cyp2b13 | AJ | feature_elongation,coding_sequence_variant,intron_variant |
| 7 | 34980617 | 34987875 | 7258 | Gpatch1 | AJ,BALB,SEA | feature_elongation,coding_sequence_variant,intron_variant |
| 7 | 34987750 | 34989647 | 1897 | Gpatch1 | NZW | feature_elongation,coding_sequence_variant,intron_variant |
| 7 | 35414261 | 35420703 | 6442 | Dpy19l3 | ILNJ2 | feature_elongation,coding_sequence_variant,intron_variant |
| 7 | 44270299 | 44271587 | 1288 | Myh14 | CE | feature_elongation,coding_sequence_variant,intron_variant |
| 7 | 49926932 | 49932060 | 5128 | Nell1 | ILNJ2,KK,NZW,PJ,SJL | feature_elongation,coding_sequence_variant,intron_variant |
| 7 | 55774972 | 55781611 | 6639 | Herc2 | 129S1,AJ,AKR,BALB,BTBR,BUB,C3H,C57L,CBA,DBA1J,DBA2J,FVB,MA/My,MRL,NOD,NOR,NZB,NZO,PL,RF,SEA,SWR,TH | feature_elongation,coding_sequence_variant,intron_variant |
| 7 | 56381566 | 56388934 | 7368 | Gabrg3 | 129S1,AJ,AKR,BTBR,BUB,C3H,C57L,CBA,MA/My,MRL,NOD,NOR,PL,TH | feature_elongation,coding_sequence_variant,intron_variant |
| 7 | 57064504 | 57071933 | 7429 | Gabra5 | AJ,BUB,FVB,MA/My,MRL,NZB,NZO,SMJ,SWR | feature_elongation,coding_sequence_variant,intron_variant |
| 7 | 66065765 | 66072930 | 7165 | Aldh1a3 | 129S1,AJ,AKR,BTBR,C3H,CBA,FVB,MA/My,MRL,NOD,NOR,NZW,PL,RF,SJL,SWR | feature_elongation,coding_sequence_variant,intron_variant,feature_elongation,5_prime_UTR_variant,intron_variant |
| 7 | 66554812 | 66561462 | 6650 | Adamts17 | BALB,C57L,CE,FVB,SEA,SWR | feature_elongation,coding_sequence_variant,intron_variant |
| 7 | 68002086 | 68009177 | 7091 | Fam169b | LP,SMJ | feature_elongation,coding_sequence_variant,intron_variant |
| 7 | 74850255 | 74859045 | 8790 | Sv2b | FVB,SWR,TH | feature_elongation,coding_sequence_variant,5_prime_UTR_variant,intron_variant |
| 7 | 77853132 | 77854852 | 1720 | Ntrk3 | TH | feature_elongation,coding_sequence_variant,intron_variant |
| 7 | 78122462 | 78129777 | 7315 | Ntrk3 | 129S1,C3H,CBA,DBA1J,DBA2J,KK,PJ,SEA | feature_elongation,coding_sequence_variant,intron_variant |
| 7 | 82026250 | 82032524 | 6274 | Adamtsl3 | CE,SMJ | feature_elongation,coding_sequence_variant,intron_variant |
| 7 | 86303732 | 86309080 | 5348 | Or14c40 | 129S1,BTBR,C3H,C57L,CBA,DBA1J,DBA2J,ILNJ2,KK,NZB,NZO,NZW,PJ,SEA | feature_elongation,5_prime_UTR_variant,intron_variant |
| 7 | 87126761 | 87133441 | 6680 | Tyr | 129S1,BTBR,C3H,C57L,CBA,DBA1J,DBA2J,ILNJ2,KK,NZB,NZO,PJ,SEA | feature_elongation,coding_sequence_variant,intron_variant |
| 7 | 89812385 | 89818999 | 6614 | Picalm | FVB,MA/My,NZW,SJL,SWR | feature_elongation,coding_sequence_variant,intron_variant |
| 7 | 91714292 | 91719624 | 5332 | Dlg2 | AJ | feature_elongation,coding_sequence_variant,5_prime_UTR_variant,intron_variant |
| 7 | 92019060 | 92026090 | 7030 | Dlg2 | SMJ | feature_elongation,coding_sequence_variant,intron_variant |
| 7 | 97377402 | 97382833 | 5431 | Aqp11 | NOD,NOR,SWR | feature_elongation,coding_sequence_variant,intron_variant |
| 7 | 99900118 | 99907320 | 7202 | Pgm2l1 | NOD,NOR,SWR | feature_elongation,coding_sequence_variant,intron_variant |
| 7 | 100022730 | 100023809 | 1079 | C2cd3 | NOD,NOR,SWR | feature_elongation,coding_sequence_variant,intron_variant |

| | | | | | | |
|---|---|---|---|---|---|---|
| 7 | 100930140 | 100932001 | 1861 | Fchsd2 | 129S1,AJ,AKR,BALB,BTBR,BUB,C3H,C57L,CBA,DBA1J,DBA2J,FVB,ILNJ2,KK,MA/My,MRL,NOD,NOR,NZB,NZO,NZW,PJ,PL,RF,RH2,SEA,SJL,SMJ,SWR,TH | feature_elongation,coding_sequence_variant,3_prime_UTR_variant,intron_variant |
| 7 | 108645500 | 108651814 | 6314 | Ric3 | KK | feature_elongation,coding_sequence_variant,intron_variant |
| 7 | 119587555 | 119594728 | 7173 | Dnah3 | SJL,SWR | feature_elongation,coding_sequence_variant,intron_variant |
| 7 | 130198187 | 130204494 | 6307 | Tacc2 | RF | feature_elongation,coding_sequence_variant,intron_variant |
| 7 | 139103466 | 139109871 | 6405 | Inpp5a | 129S1,AJ,BALB,BTBR,C3H,C57L,C58J,CE,DBA1J,DBA2J,FVB,ILNJ2,LP,MA/My,NOD,NOR,PJ,PL,RF,SEA,SJL,SMJ,SWR,TH | feature_elongation,coding_sequence_variant,intron_variant |
| 8 | 3474812 | 3481211 | 6399 | Arhgef18 | NZO,NZW | feature_elongation,coding_sequence_variant,5_prime_UTR_variant,intron_variant,feature_elongation,coding_sequence_variant,intron_variant |
| 8 | 15951457 | 15954172 | 2715 | Csmd1 | 129S1,NZB,NZO,NZW | feature_elongation,coding_sequence_variant,intron_variant |
| 8 | 23595074 | 23597826 | 2752 | Ank1 | 129S1,AJ,AKR,B10D2,B10J,BALB,BTBR,BUB,C3H,CBA,CE,DBA1J,DBA2J,FVB,KK,LP,MA/My,MRL,NOD,NOR,NZB,PL,RF,RH2,SJL,SMJ,SWR,TH | feature_elongation,coding_sequence_variant,intron_variant |
| 8 | 34630260 | 34637376 | 7116 | Saraf | NOD,NOR | feature_elongation,coding_sequence_variant,3_prime_UTR_variant,intron_variant |
| 8 | 40813982 | 40820268 | 6286 | Micu3 | SMJ | feature_elongation,coding_sequence_variant,intron_variant |
| 8 | 46551412 | 46558770 | 7358 | Snx25 | C58J,LP | feature_elongation,coding_sequence_variant,intron_variant |
| 8 | 58406818 | 58414053 | 7235 | Galntl6 | 129S1,AJ,CBA,KK,MA/My,NOD,NOR,NZW,RH2,SMJ | feature_elongation,coding_sequence_variant,intron_variant |
| 8 | 61461051 | 61468261 | 7210 | Nek1 | C58J,LP | feature_elongation,coding_sequence_variant,intron_variant |
| 8 | 61843531 | 61848749 | 5218 | Sh3rf1 | NZW | feature_elongation,coding_sequence_variant,3_prime_UTR_variant,intron_variant |
| 8 | 62549471 | 62556501 | 7030 | Anxa10 | CE | feature_elongation,coding_sequence_variant,intron_variant |
| 8 | 68571683 | 68577720 | 6037 | Psd3 | 129S1,AJ,AKR,BALB,BUB,C3H,C57L,CBA,CE,DBA1J,DBA2J,FVB,ILNJ2,KK,MA/My,MRL,NOD,NOR,NZW,PL,RF,RH2,SJL,SMJ,SWR,TH | feature_elongation,coding_sequence_variant,5_prime_UTR_variant,intron_variant |
| 8 | 71765081 | 71772517 | 7436 | Myo9b | 129S1,AKR,BALB,BUB,C3H,C57L,C58J,CBA,CE,DBA1J,DBA2J,FVB,ILNJ2,KK,MA/My,MRL,NZB,NZO,NZW,PJ,PL,RF,SEA,SJL,SMJ,SWR,TH | feature_elongation,coding_sequence_variant,intron_variant |
| 8 | 73211625 | 73213148 | 1523 | 1700030K09Rik | AJ,B10D2,B10J,BTBR,LP,NOD,NOR,RH2 | feature_elongation,coding_sequence_variant,intron_variant |
| 8 | 73235894 | 73240723 | 4829 | Slc35e1 | PJ | feature_elongation,coding_sequence_variant,3_prime_UTR_variant,intron_variant |
| 8 | 82841929 | 82848543 | 6614 | Inpp4b | 129S1,AJ,AKR,BALB,BTBR,BUB,C3H,C57L,CBA,CE,DBA1J,DBA2J,ILNJ2,KK,MA/My,MRL,NOD,NOR,NZB,NZO,NZW,PJ,PL,RF,RH2,SEA,SJL,SWR | feature_elongation,coding_sequence_variant,3_prime_UTR_variant,intron_variant |
| 8 | 84946144 | 84951502 | 5358 | Zswim4 | C3H | feature_elongation,coding_sequence_variant,intron_variant |
| 8 | 92167246 | 92173659 | 6413 | Fto | 129S1,AKR,BALB,BUB,C57L,CE,FVB,ILNJ2,MRL,NZB,NZO,NZW,PJ,RF,SEA,TH | feature_elongation,coding_sequence_variant,intron_variant |
| 8 | 107423550 | 107426678 | 3128 | Tango6 | PL | feature_elongation,coding_sequence_variant,intron_variant |
| 8 | 107425099 | 107426240 | 1141 | Tango6 | RF | feature_elongation,coding_sequence_variant,intron_variant |
| 8 | 123526757 | 123534042 | 7285 | Acsf3 | AJ,AKR,BALB,C3H,C58J,CE,LP,MRL,PL,RF,RH2,SEA,SMJ | feature_elongation,coding_sequence_variant,intron_variant |
| 8 | 123866983 | 123871810 | 4827 | Sult5a1 | 129S1,BTBR,C57L,MA/My,NZB,PJ,TH | feature_elongation,coding_sequence_variant,3_prime_UTR_variant,intron_variant |
| 8 | 126399158 | 126406641 | 7483 | Map10 | TH | stop_gained,frameshift_variant |
| 8 | 128186231 | 128187496 | 1265 | Pard3 | CBA,DBA1J,DBA2J,NZW | feature_elongation,coding_sequence_variant,intron_variant |
| 8 | 129579892 | 129586561 | 6669 | Ccdc7a | 129S1,BTBR,C57L,NZB | feature_elongation,coding_sequence_variant,intron_variant |
| 8 | 129690408 | 129691901 | 1493 | Ccdc7a | KK | feature_elongation,coding_sequence_variant,intron_variant |
| 8 | 129831960 | 129836887 | 4927 | Ccdc7b | 129S1,BTBR,C57L,NZB | feature_elongation,coding_sequence_variant,intron_variant |
| 9 | 25458611 | 25465262 | 6651 | Eepd1 | BUB,C58J,FVB,LP,NOD,NOR,NZO,RF,SJL,SWR | feature_elongation,coding_sequence_variant,intron_variant |
| 9 | 32177263 | 32178797 | 1534 | Arhgap32 | KK,MRL | feature_elongation,3_prime_UTR_variant |
| 9 | 34934094 | 34940535 | 6441 | Kirrel3 | CBA | feature_elongation,coding_sequence_variant,intron_variant |
| 9 | 44461954 | 44469031 | 7077 | Cxcr5 | BTBR,NZO | feature_elongation,5_prime_UTR_variant,intron_variant |
| 9 | 49350769 | 49356022 | 5253 | Ttc12 | BTBR,ILNJ2 | feature_elongation,coding_sequence_variant,intron_variant |
| 9 | 58885449 | 58887024 | 1575 | Neo1 | MA/My,NZB,PL | feature_elongation,coding_sequence_variant,intron_variant |
| 9 | 65416148 | 65421137 | 4989 | Ankdd1a | B10D2,B10J,LP,MA/My,NZW,RF | feature_elongation,coding_sequence_variant,intron_variant |
| 9 | 65763893 | 65765757 | 1864 | Trip4 | 129S1,B10D2,B10J,BTBR,BUB,C57L,KK,LP,MA/My,MRL,NOD,NOR,NZO,PJ,SEA,SMJ,SWR,TH | feature_elongation,coding_sequence_variant,intron_variant |
| 9 | 67195756 | 67201335 | 5579 | Tln2 | AJ,AKR,BALB,C3H,MRL,PL,RF,RH2 | feature_elongation,coding_sequence_variant,intron_variant |
| 9 | 67209184 | 67213618 | 4434 | Tln2 | B10D2,B10J,LP,MA/My | feature_elongation,coding_sequence_variant,intron_variant |
| 9 | 70280778 | 70287221 | 6443 | Myo1e | 129S1,BTBR,C57L | feature_elongation,coding_sequence_variant,intron_variant |
| 9 | 71426701 | 71433784 | 7083 | Myzap | AJ,BALB,BUB,C3H,C58J,CBA,CE,DBA1J,DBA2J,FVB,ILNJ2,KK,MA/My,NOD,NOR,NZB,NZO,NZW,PJ,RH2,SEA,SJL,SMJ,SWR,TH | feature_elongation,coding_sequence_variant,intron_variant |
| 9 | 71626673 | 71629073 | 2400 | Cgnl1 | 129S1,AJ,BALB,BTBR,BUB,C3H,C57L,C58J,CBA,CE,DBA1J,DBA2J,FVB,ILNJ2,KK,LP,MA/My,NOD,NOR,NZB,NZO,NZW,PJ,RH2,SEA,SJL,SMJ,SWR,TH | feature_elongation,coding_sequence_variant,intron_variant |

| | | | | | | | |
|---|---|---|---|---|---|---|---|
| 9 | 75555251 | 75562386 | 7135 | Scg3 | AJ,AKR,BALB,BTBR,BUB,C3H,C58J,CBA,CE,DBA1J,DBA2J,FVB,ILNJ2,KK,LP,MA/My,MRL,NOD,NOR,NZB,NZO,NZW,PJ,PL,RF,RH2,SEA,SJL,SMJ,SWR,TH | feature_elongation,coding_sequence_variant,intron_variant | |
| 9 | 78003607 | 78008243 | 4636 | Fbxo9 | KK | feature_elongation,coding_sequence_variant,intron_variant | |
| 9 | 80016316 | 80022756 | 6440 | Senp6 | AJ,AKR,BALB,BUB,C3H,CBA,CE,DBA1J,DBA2J,FVB,KK,MA/My,MRL,NOD,NOR,NZB,NZO,PL,RF,RH2,SEA,SJL,SWR,TH | feature_elongation,coding_sequence_variant,intron_variant | |
| 9 | 86923585 | 86930290 | 6705 | Cyb5r4 | 129S1,BTBR,C57L,CE | feature_elongation,coding_sequence_variant,intron_variant | |
| 9 | 87098664 | 87105354 | 6690 | Cep162 | FVB,SJL,SWR | feature_elongation,coding_sequence_variant,intron_variant | |
| 9 | 87101420 | 87103193 | 1773 | Cep162 | PJ,SEA | feature_elongation,coding_sequence_variant,intron_variant | |
| 9 | 99208834 | 99210698 | 1864 | Esyt3 | AJ,BALB,C3H,CBA,FVB,NZB,PJ,RH2,SEA,SJL,SWR | feature_elongation,coding_sequence_variant,intron_variant | |
| 9 | 99377180 | 99383835 | 6655 | Armc8 | CE | feature_elongation,coding_sequence_variant,intron_variant | |
| 9 | 100868551 | 100870247 | 1696 | Pccb | AKR,BUB,CBA,CE,FVB,ILNJ2,NOD,NOR,PJ,PL,RF,SEA,SJL,SMJ,SWR,TH | feature_elongation,coding_sequence_variant,intron_variant | |
| 9 | 101837481 | 101843536 | 6055 | Ephb1 | 129S1,AJ,AKR,BALB,BUB,C3H,CBA,DBA1J,DBA2J,ILNJ2,NOD,NOR,PL,RH2 | feature_elongation,coding_sequence_variant,intron_variant | |
| 9 | 103037310 | 103038370 | 1060 | Rab6b | 129S1,AJ,AKR,BALB,BTBR,BUB,C3H,C57L,CBA,CE,FVB,ILNJ2,MA/My,NOD,NOR,NZB,NZO,NZW,PJ,PL,RF,RH2,SEA,SJL,SMJ,SWR,TH | feature_elongation,coding_sequence_variant,intron_variant,feature_elongation,coding_sequence_variant,5_prime_UTR_variant,intron_variant | |
| 9 | 103305563 | 103310343 | 4780 | Bfsp2 | MA/My,NZO | feature_elongation,coding_sequence_variant,intron_variant | |
| 9 | 105222423 | 105228915 | 6492 | Nek11 | CE,FVB,SJL,SWR | feature_elongation,coding_sequence_variant,intron_variant | |
| 9 | 106860606 | 106869247 | 8641 | Dock3 | 129S1,AJ,AKR,BALB,BTBR,BUB,C3H,C57L,FVB,ILNJ2,MA/My,MRL,NOD,NOR,NZB,NZO,NZW,PJ,PL,RF,RH2,SEA,SJL,SMJ,SWR,TH | feature_elongation,coding_sequence_variant,intron_variant | |
| 9 | 108254919 | 108256783 | 1864 | Usp4 | NZB | feature_elongation,coding_sequence_variant,intron_variant | |
| 9 | 111091866 | 111098320 | 6454 | Mlh1 | CE,PJ,SEA | feature_elongation,coding_sequence_variant,intron_variant | |
| 9 | 116934080 | 116941198 | 7118 | Rbms3 | CE | feature_elongation,coding_sequence_variant,intron_variant | |
| 9 | 119111977 | 119118066 | 6089 | Oxsr1 | CE | feature_elongation,coding_sequence_variant,intron_variant | |
| 9 | 119623861 | 119625725 | 1864 | Scn11a | 129S1,C57L | feature_elongation,coding_sequence_variant,intron_variant | |
| 10 | 4805115 | 4811579 | 6464 | Esr1 | ILNJ2 | feature_elongation,coding_sequence_variant,intron_variant | |
| 10 | 4885531 | 4891973 | 6442 | Esr1 | KK | feature_elongation,coding_sequence_variant,intron_variant | |
| 10 | 4912221 | 4916604 | 4383 | Esr1 | C3H,CBA,CE,DBA1J,DBA2J,FVB,ILNJ2,LP,MA/My,MRL,NZW,PJ,SEA,SWR | feature_elongation,coding_sequence_variant,3_prime_UTR_variant,intron_variant | |
| 10 | 5271250 | 5273138 | 1888 | Syne1 | NZB,NZO | feature_elongation,coding_sequence_variant,intron_variant | |
| 10 | 10285780 | 10292266 | 6486 | Adgb | 129S1,DBA1J,DBA2J,FVB,MA/My,NZB,NZO,NZW,SWR,TH | feature_elongation,coding_sequence_variant,intron_variant | |
| 10 | 13105639 | 13110997 | 5358 | Phactr2 | AKR,RF | feature_elongation,coding_sequence_variant,intron_variant | |
| 10 | 19630696 | 19632587 | 1891 | Il20ra | AKR,ILNJ2,RF | feature_elongation,coding_sequence_variant,intron_variant | |
| 10 | 28652858 | 28659320 | 6462 | Themis | AKR,C3H,C58J,CBA,DBA1J,DBA2J,FVB,MRL,PJ,SEA | feature_elongation,coding_sequence_variant,intron_variant | |
| 10 | 50855108 | 50862406 | 7298 | Sim1 | TH | feature_elongation,coding_sequence_variant,3_prime_UTR_variant,intron_variant | |
| 10 | 53790112 | 53796525 | 6413 | Man1a | KK | feature_elongation,coding_sequence_variant,intron_variant | |
| 10 | 60220120 | 60225686 | 5566 | Cdh23 | AJ | feature_elongation,coding_sequence_variant,intron_variant | |
| 10 | 71206366 | 71211717 | 5351 | Ipmk | MRL | feature_elongation,coding_sequence_variant,intron_variant | |
| 10 | 75432601 | 75439000 | 6399 | Ggt5 | NZW | feature_elongation,coding_sequence_variant,intron_variant | |
| 10 | 79014882 | 79017847 | 2965 | Vmn2r80 | TH | feature_elongation,coding_sequence_variant,intron_variant | |
| 10 | 83234636 | 83240227 | 5591 | Nopchap1 | BUB,SMJ | feature_elongation,3_prime_UTR_variant | |
| 10 | 83320368 | 83322200 | 1832 | Nopchap1 | BUB | feature_elongation,3_prime_UTR_variant | |
| 10 | 85873092 | 85879526 | 6434 | Fbxo7 | BUB | feature_elongation,coding_sequence_variant,intron_variant | |
| 10 | 86647925 | 86651434 | 3509 | Nt5dc3 | BUB,FVB | feature_elongation,coding_sequence_variant,intron_variant | |
| 10 | 86741080 | 86742203 | 1123 | Stab2 | BUB,FVB | feature_elongation,coding_sequence_variant,intron_variant | |
| 10 | 90882456 | 90884900 | 2444 | Apaf1 | AJ,AKR,BALB,C3H,C57L,CBA,CE,FVB,ILNJ2,KK,MRL,NOD,NOR,NZB,NZO,NZW,PJ,PL,RF,RH2,SEA,SMJ,SWR,TH | feature_elongation,coding_sequence_variant,intron_variant | |
| 10 | 106391638 | 106396058 | 4420 | Ppfia2 | BTBR,BUB,FVB,NZB | feature_elongation,coding_sequence_variant,intron_variant | |
| 10 | 121403056 | 121411382 | 8326 | Tbk1 | DBA1J,KK,SMJ | feature_elongation,coding_sequence_variant,intron_variant | FALSE |
| 10 | 121427758 | 121430249 | 2491 | Xpot | AJ,BALB,DBA2J,MA/My,RH2 | feature_elongation,coding_sequence_variant,intron_variant | |
| 10 | 128911048 | 128913069 | 2021 | Or6c8 | BUB,C3H,CBA,CE,KK,RF,SMJ | feature_elongation,3_prime_UTR_variant | |
| 11 | 4828982 | 4833474 | 4492 | Nipsnap1 | NZW,PJ | feature_elongation,coding_sequence_variant,intron_variant | |
| 11 | 6291396 | 6292540 | 1144 | Ogdh | 129S1,AJ,AKR,BALB,BTBR,C57L,MA/My,MRL,NOD,NZW,PL,RF,SEA,TH | feature_elongation,coding_sequence_variant,intron_variant | |
| 11 | 29796158 | 29803157 | 6999 | Eml6 | FVB,SJL,SWR | feature_elongation,coding_sequence_variant,intron_variant | |
| 11 | 31964491 | 31968858 | 4367 | Nsg2 | BALB,BTBR,C57L,C58J,RH2 | feature_elongation,coding_sequence_variant,intron_variant | |
| 11 | 32687011 | 32693451 | 6440 | Fbxw11 | NOD,SWR,TH | feature_elongation,coding_sequence_variant,3_prime_UTR_variant,intron_variant | |

| | | | | | | |
|---|---|---|---|---|---|---|
| 11 | 43120151 | 43121375 | 1224 | Atp10b | B10D2,B10J,BTBR,BUB,C58J,CE,DBA2J,ILNJ2,KK,MA/My,NZB,NZO,NZW,PJ,SEA,SJL,SMJ | feature_elongation,coding_sequence_variant,intron_variant |
| 11 | 45989071 | 45990964 | 1893 | Adam19 | 129S1,AJ,BALB,BTBR,BUB,C3H,C57L,CBA,DBA1J,DBA2J,FVB,MA/My,MRL,PJ,PL,RH2,SEA,SWR,TH | feature_elongation,coding_sequence_variant,intron_variant |
| 11 | 49014463 | 49025581 | 11118 | Or2v1 | SJL | feature_elongation,coding_sequence_variant,5_prime_UTR_variant,intron_variant |
| 11 | 50084006 | 50090444 | 6438 | Mrnip | NZW | feature_elongation,coding_sequence_variant,intron_variant |
| 11 | 53857131 | 53862835 | 5704 | Slc22a21 | B10D2,B10J,C58J,DBA2J,KK,NOR,NZB,NZO,NZW,PL | feature_elongation,coding_sequence_variant,intron_variant |
| 11 | 53887533 | 53892557 | 5024 | Slc22a4 | KK,NZB,NZO | feature_elongation,coding_sequence_variant,intron_variant |
| 11 | 59315024 | 59327404 | 12380 | Snap47 | NZB,NZO,NZW | feature_elongation,coding_sequence_variant,intron_variant |
| 11 | 62301149 | 62302493 | 1344 | Ncor1 | AJ,B10D2,B10J,C3H,CBA,MA/My,MRL,PJ,RH2 | feature_elongation,coding_sequence_variant,intron_variant |
| 11 | 64940776 | 64947176 | 6400 | Arhgap44 | ILNJ2 | feature_elongation,coding_sequence_variant,intron_variant |
| 11 | 65071969 | 65078388 | 6419 | Myocd | FVB,ILNJ2,NOD,NZB,NZO,SWR | feature_elongation,coding_sequence_variant,intron_variant |
| 11 | 68089194 | 68095641 | 6447 | Stx8 | ILNJ2 | feature_elongation,coding_sequence_variant,3_prime_UTR_variant,intron_variant |
| 11 | 71067330 | 71074830 | 7500 | Nlrp1b | CE | feature_elongation,coding_sequence_variant,intron_variant |
| 11 | 73829158 | 73833468 | 4310 | Or1e17 | AJ,NZO,NZW | feature_elongation,coding_sequence_variant,5_prime_UTR_variant,3_prime_UTR_variant,intron_variant |
| 11 | 104172201 | 104173575 | 1374 | Mapt | 129S1,BTBR,C3H,C57L,DBA1J,DBA2J | feature_elongation,coding_sequence_variant,5_prime_UTR_variant,intron_variant |
| 11 | 104809410 | 104810595 | 1185 | Gm11639 | AJ,CBA | feature_elongation,coding_sequence_variant,intron_variant |
| 11 | 104815836 | 104822440 | 6604 | Gm11639 | 129S1,AJ,AKR,BTBR,C3H,C57L,CBA,DBA1J,DBA2J,ILNJ2,KK,MRL,PL,RF,RH2 | feature_elongation,coding_sequence_variant,intron_variant |
| 11 | 105699939 | 105706816 | 6877 | Tanc2 | NZO | feature_elongation,coding_sequence_variant,intron_variant |
| 11 | 117696191 | 117697259 | 1068 | 6030468B19Rik | 129S1,AJ,AKR,BTBR,BUB,C3H,C57L,C58J,CBA,CE,DBA1J,DBA2J,ILNJ2,KK,MA/My,MRL,NZB,NZO,NZW,PL,RH2,SJL,SMJ | feature_elongation,coding_sequence_variant,intron_variant |
| 11 | 120339835 | 120341723 | 1888 | Pde6g | PL | feature_elongation,coding_sequence_variant,5_prime_UTR_variant,intron_variant |
| 12 | 4389333 | 4396483 | 7150 | Ncoa1 | KK | feature_elongation,coding_sequence_variant,5_prime_UTR_variant,intron_variant |
| 12 | 4720804 | 4727301 | 6497 | Itsn2 | CE | feature_elongation,coding_sequence_variant,intron_variant |
| 12 | 5033196 | 5040045 | 6849 | Atad2b | MRL | feature_elongation,coding_sequence_variant,intron_variant |
| 12 | 9056598 | 9061563 | 4965 | Wdr35 | ILNJ2 | feature_elongation,coding_sequence_variant,intron_variant |
| 12 | 11352411 | 11357722 | 5311 | Smc6 | CBA | feature_elongation,coding_sequence_variant,intron_variant |
| 12 | 13440349 | 13443952 | 3603 | Nbas | MRL | feature_elongation,coding_sequence_variant,intron_variant |
| 12 | 24571840 | 24576760 | 4920 | Taf1b | AKR,RF | feature_elongation,coding_sequence_variant,intron_variant |
| 12 | 51668314 | 51669597 | 1283 | Strn3 | BUB,SJL,SWR | feature_elongation,coding_sequence_variant,intron_variant |
| 12 | 53055852 | 53061396 | 5544 | Akap6 | AJ,AKR,BALB,C3H,CBA,NOD,NOR,PJ,RF,RH2,SEA | feature_elongation,coding_sequence_variant,intron_variant |
| 12 | 59147077 | 59154165 | 7088 | Mia2 | AJ,BALB,C3H,CBA,DBA1J,DBA2J,ILNJ2,KK,MA/My,RH2,SEA | feature_elongation,coding_sequence_variant,intron_variant |
| 12 | 59259283 | 59266376 | 7093 | Fbxo33 | BUB,FVB | feature_elongation,coding_sequence_variant,5_prime_UTR_variant,intron_variant |
| 12 | 65020078 | 65026628 | 6550 | Togaram1 | MRL | feature_elongation,coding_sequence_variant,intron_variant |
| 12 | 69779239 | 69784759 | 5520 | Dmac2l | 129S1,AJ,AKR,BALB,BTBR,BUB,C3H,C57L,CBA,CE,DBA1J,DBA2J,FVB,ILNJ2,KK,MA/My,MRL,NOD,NOR,NZB,NZO,NZW,PL,RF,RH2,SEA,SJL,SMJ,SWR,TH | feature_elongation,coding_sequence_variant,5_prime_UTR_variant,intron_variant |
| 12 | 69920999 | 69928121 | 7122 | Map4k5 | AJ,AKR,BALB,BTBR,BUB,C3H,C57L,CBA,CE,DBA1J,DBA2J,FVB,ILNJ2,KK,MA/My,NOD,NOR,NZW,PL,RF,RH2,SEA,SJL,SMJ,TH | feature_elongation,coding_sequence_variant,intron_variant |
| 12 | 72586554 | 72588199 | 1645 | Pcnx4 | BUB,PJ,SEA,SMJ | feature_elongation,5_prime_UTR_variant,intron_variant |
| 12 | 89494564 | 89501750 | 7186 | Nrxn3 | NOD | feature_elongation,coding_sequence_variant,intron_variant |
| 12 | 101486055 | 101489068 | 3013 | Catsperb | 129S1,AKR,BALB,BTBR,BUB,C57L,C58J,CBA,CE,DBA1J,DBA2J,FVB,ILNJ2,KK,LP,MA/My,NOD,NOR,NZB,NZO,NZW,PJ,PL,RF,SEA,SJL,SMJ,SWR,TH | feature_elongation,coding_sequence_variant,intron_variant |
| 12 | 101506107 | 101512559 | 6452 | Catsperb | 129S1,AKR,BALB,BTBR,BUB,C57L,C58J,CBA,DBA1J,DBA2J,FVB,ILNJ2,KK,LP,MA/My,NOD,NOR,NZB,NZO,NZW,PL,RF,SJL,SMJ,SWR,TH | feature_elongation,coding_sequence_variant,intron_variant |
| 12 | 105453718 | 105459031 | 5313 | D430019H16Rik | C3H | feature_elongation,coding_sequence_variant,3_prime_UTR_variant,intron_variant |
| 12 | 108149835 | 108156929 | 7094 | Ccnk | DBA1J,DBA2J,ILNJ2 | feature_elongation,coding_sequence_variant,5_prime_UTR_variant,intron_variant |
| 12 | 108279409 | 108285848 | 6439 | Hhipl1 | DBA1J,DBA2J,ILNJ2 | feature_elongation,coding_sequence_variant,intron_variant |
| 12 | 110861265 | 110863480 | 2215 | Tecpr2 | C3H,CBA,CE | feature_elongation,coding_sequence_variant,5_prime_UTR_variant,intron_variant |
| 12 | 116055986 | 116057402 | 1416 | Vipr2 | AKR,FVB,NZB,NZO,NZW,SWR,TH | feature_elongation,coding_sequence_variant,intron_variant |
| 12 | 118138657 | 118145510 | 6853 | Dnah11 | AJ | feature_elongation,coding_sequence_variant,intron_variant |
| 13 | 14540168 | 14546977 | 6809 | Hecw1 | BUB,FVB,NOD,NOR,SJL,SWR | feature_elongation,coding_sequence_variant,intron_variant |
| 13 | 17835846 | 17842329 | 6483 | Sugct | AJ,B10D2,B10J,BALB,BTBR,C3H,C57L,C58J,CBA,CE,DBA1J,DBA2J,KK,MA/My,MRL,NZB,PL,RH2,SMJ | feature_elongation,coding_sequence_variant,intron_variant |

| | | | | | | |
|---|---|---|---|---|---|---|
| 13 | 23295677 | 23300988 | 5311 | Vmn1r217 | CE | feature_elongation,coding_sequence_variant,5_prime_UTR_variant,3_prime_UTR_variant,intron_variant |
| 13 | 23886935 | 23888410 | 1475 | Hfe | ILNJ2,PJ,SEA | feature_elongation,coding_sequence_variant,3_prime_UTR_variant |
| 13 | 33150681 | 33155965 | 5284 | Serpinb6b | CE | feature_elongation,coding_sequence_variant,5_prime_UTR_variant,intron_variant,feature_elongation,5_prime_UTR_variant,intron_variant |
| 13 | 40324587 | 40330135 | 5548 | Ofcc1 | MRL | feature_elongation,coding_sequence_variant,intron_variant |
| 13 | 55933261 | 55940422 | 7161 | Catsper3 | BTBR,C57L | feature_elongation,coding_sequence_variant,intron_variant |
| 13 | 58170236 | 58178694 | 8458 | Klhl3 | AKR,BALB,C3H,CBA,CE,RF | feature_elongation,coding_sequence_variant,intron_variant |
| 13 | 59768965 | 59774347 | 5382 | Naa35 | BTBR,BUB,C57L,SMJ | feature_elongation,coding_sequence_variant,intron_variant |
| 13 | 63331803 | 63338929 | 7126 | Aopep | BALB | feature_elongation,coding_sequence_variant,intron_variant |
| 13 | 63500488 | 63509474 | 8986 | Fancc | BUB,DBA1J,DBA2J,ILNJ2,MA/My,NOD,NOR,SMJ | feature_elongation,coding_sequence_variant,intron_variant |
| 13 | 64925805 | 64932200 | 6395 | Cntnap3 | FVB,SWR | feature_elongation,coding_sequence_variant,intron_variant |
| 13 | 65241394 | 65243811 | 2417 | Mfsd14b | 129S1,AKR,BUB,C3H,C58J,CBA,CE,DBA1J,DBA2J,ILNJ2,KK,MRL,NOD,NOR,NZB,NZO,NZW,PJ,PL,RF,SEA,SMJ,SWR,TH | feature_elongation,coding_sequence_variant,intron_variant |
| 13 | 65431122 | 65433014 | 1892 | Zfp369 | BTBR,C57L | feature_elongation,coding_sequence_variant,intron_variant |
| 13 | 67512180 | 67514116 | 1936 | Zfp456 | 129S1,AJ,AKR,BALB,BTBR,BUB,C3H,C57L,C58J,CBA,CE,DBA1J,DBA2J,FVB,ILNJ2,KK,MRL,NOD,NOR,NZB,NZO,NZW,PJ,PL,RF,SEA,SJL,SMJ,SWR,TH | feature_elongation,3_prime_UTR_variant |
| 13 | 67538581 | 67541868 | 3287 | Zfp429 | KK,NZB,NZO,PJ,SEA | feature_elongation,coding_sequence_variant,intron_variant,feature_elongation,coding_sequence_variant,3_prime_UTR_variant,intron_variant |
| 13 | 74345563 | 74348302 | 2739 | Exoc3 | ILNJ2 | feature_elongation,coding_sequence_variant,intron_variant |
| 13 | 85402195 | 85408637 | 6442 | Rasa1 | AJ,AKR,BALB,BTBR,BUB,C3H,C57L,CBA,CE,DBA1J,DBA2J,FVB,ILNJ2,KK,MRL,NOD,NOR,NZB,NZO,NZW,PJ,PL,RF,SEA,SJL,TH | feature_elongation,coding_sequence_variant,intron_variant |
| 13 | 93184906 | 93187672 | 2766 | Cmya5 | BUB | feature_elongation,coding_sequence_variant,intron_variant |
| 13 | 96928951 | 96934560 | 5609 | Ankrd31 | BTBR,C58J,CBA,DBA1J,DBA2J,MA/My,SMJ | feature_elongation,coding_sequence_variant,intron_variant |
| 13 | 107114965 | 107120340 | 5375 | Kif2a | C3H | feature_elongation,coding_sequence_variant,intron_variant |
| 13 | 114100533 | 114105355 | 4822 | Arl15 | AJ,BUB,CE,DBA1J,FVB,KK,MA/My,NZO,PL,RF,RH2,SJL,SWR | feature_elongation,coding_sequence_variant,intron_variant |
| 14 | 20728880 | 20734125 | 5245 | Sec24c | SEA | feature_elongation,coding_sequence_variant,intron_variant |
| 14 | 51601148 | 51608919 | 7771 | Gm7247 | FVB,ILNJ2,MA/My,NOD,NOR,SJL,SMJ,SWR | feature_elongation,coding_sequence_variant,5_prime_UTR_variant,intron_variant |
| 14 | 70471225 | 70476493 | 5268 | Ppp3cc | LP | feature_elongation,coding_sequence_variant,intron_variant |
| 14 | 73268545 | 73275221 | 6676 | Cysltr2 | 129S1,C57L | feature_elongation,5_prime_UTR_variant,intron_variant |
| 14 | 73451490 | 73458577 | 7087 | Rb1 | SJL | feature_elongation,coding_sequence_variant,intron_variant |
| 14 | 75504764 | 75506336 | 1572 | Cpb2 | SJL | feature_elongation,coding_sequence_variant,intron_variant |
| 14 | 78853423 | 78854793 | 1370 | Dgkh | 129S1,AJ,AKR,B10D2,B10J,BALB,BTBR,BUB,C3H,C57L,C58J,CBA,FVB,ILNJ2,KK,LP,MA/My,MRL,NOD,NOR,NZB,NZO,NZW,PL,RF,RH2,SEA,SJL,SMJ,SWR,TH | feature_elongation,coding_sequence_variant,intron_variant |
| 14 | 87007888 | 87010695 | 2807 | Diaph3 | NZO | feature_elongation,coding_sequence_variant,intron_variant |
| 14 | 103558971 | 103565370 | 6399 | Mycbp2 | C57L,ILNJ2 | feature_elongation,coding_sequence_variant,intron_variant |
| 14 | 115365919 | 115372381 | 6462 | Gpc5 | FVB,SJL | feature_elongation,coding_sequence_variant,intron_variant |
| 14 | 118106971 | 118108464 | 1493 | Gpc6 | 129S1,KK | feature_elongation,coding_sequence_variant,intron_variant |
| 14 | 118252902 | 118259336 | 6434 | Dct | BTBR,C57L,C58J | feature_elongation,coding_sequence_variant,intron_variant |
| 14 | 118903115 | 118908368 | 5253 | Abcc4 | C58J | feature_elongation,coding_sequence_variant,intron_variant |
| 14 | 119304641 | 119311845 | 7204 | Uggt2 | AJ,AKR,B10D2,B10J,BALB,BUB,C3H,CBA,CE,DBA1J,DBA2J,FVB,ILNJ2,MRL,NOD,NOR,PL,RF,RH2,SEA,SJL,SMJ,SWR | feature_elongation,coding_sequence_variant,intron_variant |
| 14 | 122545427 | 122552593 | 7166 | Clybl | C3H | feature_elongation,coding_sequence_variant,intron_variant |
| 15 | 8262124 | 8265441 | 3317 | Cplane1 | LP | feature_elongation,coding_sequence_variant,intron_variant |
| 15 | 12831626 | 12838086 | 6460 | Drosha | BUB,CE,ILNJ2 | feature_elongation,coding_sequence_variant,intron_variant |
| 15 | 16776577 | 16780953 | 4376 | Cdh9 | 129S1,BTBR,C57L | feature_elongation,coding_sequence_variant,5_prime_UTR_variant,intron_variant |
| 15 | 18982766 | 18989843 | 7077 | Cdh10 | CE,ILNJ2 | feature_elongation,coding_sequence_variant,intron_variant |
| 15 | 35469070 | 35476166 | 7096 | Vps13b | CE | feature_elongation,coding_sequence_variant,intron_variant |
| 15 | 39337809 | 39344527 | 6718 | Rims2 | C3H,PL | feature_elongation,coding_sequence_variant,intron_variant |
| 15 | 42356705 | 42362972 | 6267 | Angpt1 | BTBR,ILNJ2,LP | feature_elongation,coding_sequence_variant,intron_variant |
| 15 | 44253825 | 44260923 | 7098 | Nudcd1 | 129S1,C57L,CBA,DBA1J,DBA2J,MA/My,NOD,NOR,PJ,SMJ | feature_elongation,coding_sequence_variant,intron_variant |
| 15 | 58413379 | 58415037 | 1658 | Fer1l6 | BALB,NOD,NOR,NZB,NZW,SEA | feature_elongation,coding_sequence_variant,5_prime_UTR_variant,intron_variant |
| 15 | 65812756 | 65818067 | 5311 | Hhla1 | C3H | feature_elongation,coding_sequence_variant,intron_variant |
| 15 | 67975993 | 67982395 | 6402 | Zfat | AKR,NOD,NOR,SMJ,SWR | feature_elongation,coding_sequence_variant,intron_variant |

| | | | | | | |
|---|---|---|---|---|---|---|
| 15 | 71323866 | 71326564 | 2698 | Fam135b | BTBR,BUB,ILNJ2,KK,MA/My,TH | feature_elongation,coding_sequence_variant,intron_variant |
| 15 | 78650535 | 78656213 | 5678 | Mfng | NZB | feature_elongation,coding_sequence_variant,intron_variant |
| 15 | 79468038 | 79475148 | 7110 | Dmc1 | 129S1,AJ,AKR,BALB,C3H,C57L,C58J,CBA,CE,DBA1J,DBA2J,FVB,ILNJ2,KK,LP,MA/My,MRL,NOD,NOR,NZW,PJ,PL,SEA,SJL,SMJ,SWR,TH | feature_elongation,coding_sequence_variant,intron_variant |
| 15 | 81978546 | 81980427 | 1881 | Mei1 | FVB,MRL | feature_elongation,coding_sequence_variant,intron_variant |
| 15 | 82001267 | 82007709 | 6442 | Mei1 | MA/My | feature_elongation,coding_sequence_variant,intron_variant |
| 15 | 82062776 | 82064357 | 1581 | Srebf2 | FVB,MRL | feature_elongation,coding_sequence_variant,intron_variant |
| 15 | 82907639 | 82914112 | 6473 | Nfam1 | 129S1,AJ,AKR,BALB,BTBR,BUB,C3H,C57L,CBA,CE,DBA1J,DBA2J,ILNJ2,KK,LP,MA/My,NOD,NOR,NZB,NZO,NZW,PJ,RF,SEA,SJL,SMJ,SWR | feature_elongation,5_prime_UTR_variant,intron_variant |
| 15 | 84839130 | 84843942 | 4812 | 5031439G07Rik | NZO | feature_elongation,coding_sequence_variant,intron_variant |
| 15 | 84971640 | 84978124 | 6484 | Smc1b | AKR,BTBR,FVB,MRL,NOD,NOR,PL,TH | feature_elongation,coding_sequence_variant,intron_variant |
| 15 | 87564968 | 87571419 | 6451 | Tafa5 | PJ,SEA | feature_elongation,coding_sequence_variant,intron_variant |
| 15 | 100089979 | 100091869 | 1890 | Dip2b | AJ,C3H,CE,MRL,RH2 | feature_elongation,coding_sequence_variant,intron_variant |
| 16 | 14253080 | 14259522 | 6442 | Abcc1 | BALB,SEA | feature_elongation,coding_sequence_variant,intron_variant |
| 16 | 15489060 | 15490842 | 1782 | Prkdc | B10D2,B10J,BTBR,C57L,CBA,FVB,KK,PJ,RH2,SJL | feature_elongation,coding_sequence_variant,intron_variant |
| 16 | 25798325 | 25803592 | 5267 | P3h2 | BUB,CE,LP | feature_elongation,coding_sequence_variant,intron_variant |
| 16 | 29061846 | 29068081 | 6235 | Atp13a5 | SMJ | feature_elongation,coding_sequence_variant,intron_variant |
| 16 | 29250403 | 29257301 | 6898 | Atp13a4 | 129S1,AJ,AKR,BALB,BUB,C57L,CE,FVB,ILNJ2,KK,LP,MA/My,MRL,NZB,RF,RH2,SEA,SJL,SWR,TH | feature_elongation,coding_sequence_variant,intron_variant |
| 16 | 31905012 | 31910527 | 5515 | Pigx | NZB | feature_elongation,coding_sequence_variant,intron_variant,feature_elongation,coding_sequence_variant,5_prime_UTR_variant,intron_variant |
| 16 | 34865144 | 34872286 | 7142 | Hacd2 | C3H,CBA,DBA1J,DBA2J | feature_elongation,coding_sequence_variant,intron_variant |
| 16 | 36908419 | 36914774 | 6355 | Polq | SMJ | feature_elongation,coding_sequence_variant,intron_variant |
| 16 | 38154924 | 38156158 | 1234 | Cfap91 | AJ,BALB,CE,DBA1J,FVB,MA/My,RH2,SEA,SWR | feature_elongation,coding_sequence_variant,intron_variant |
| 16 | 43721249 | 43727473 | 6224 | Ccdc191 | PL | feature_elongation,coding_sequence_variant,intron_variant |
| 16 | 48952196 | 48959342 | 7146 | Myh15 | C3H,CBA | feature_elongation,coding_sequence_variant,intron_variant |
| 16 | 56050360 | 56055240 | 4880 | Impg2 | 129S1,AJ,BALB,DBA1J,DBA2J,MRL,RH2,SEA,SJL | feature_elongation,coding_sequence_variant,intron_variant |
| 16 | 56740684 | 56748205 | 7521 | Lnp1 | AKR,C3H,MA/My,NOD,NOR,RF,TH | feature_elongation,coding_sequence_variant,intron_variant |
| 16 | 90264555 | 90271700 | 7145 | Hunk | SEA | feature_elongation,coding_sequence_variant,intron_variant |
| 16 | 90769671 | 90772524 | 2853 | Synj1 | 129S1,AKR,C57L,ILNJ2,SMJ | feature_elongation,coding_sequence_variant,intron_variant |
| 16 | 91706700 | 91713103 | 6403 | Itsn1 | BUB,FVB,NZB,NZO,SJL,SWR | feature_elongation,coding_sequence_variant,3_prime_UTR_variant,intron_variant |
| 16 | 93937343 | 93942384 | 5041 | Hlcs | NZB | feature_elongation,coding_sequence_variant,intron_variant |
| 17 | 12681775 | 12685093 | 3318 | Slc22a3 | PL | feature_elongation,coding_sequence_variant,intron_variant |
| 17 | 13199051 | 13200501 | 1450 | Wtap | 129S1,AJ,AKR,B10D2,BALB,C57L,C58J,DBA1J,DBA2J,LP,NZO,RH2,SEA | feature_elongation,coding_sequence_variant,intron_variant |
| 17 | 14594307 | 14600986 | 6679 | Smoc2 | NZW | feature_elongation,coding_sequence_variant,intron_variant |
| 17 | 16039026 | 16042866 | 3840 | Rgmb | MA/My,NZB | feature_elongation,coding_sequence_variant,intron_variant |
| 17 | 19910745 | 19917707 | 6962 | Vmn2r102 | 129S1,AJ,BALB,C57L,CE,DBA1J,DBA2J,KK,NZO,RH2,SEA | feature_elongation,coding_sequence_variant,3_prime_UTR_variant,intron_variant |
| 17 | 20490592 | 20498918 | 8326 | Vmn2r106 | AKR,C3H,CBA,FVB,MA/My,MRL,NZB,PJ,RF,SJL,SMJ,TH | feature_elongation,coding_sequence_variant,intron_variant |
| 17 | 20588484 | 20595679 | 7195 | Vmn2r107 | ILNJ2 | feature_elongation,coding_sequence_variant,intron_variant |
| 17 | 22785707 | 22790868 | 5161 | Vmn2r111 | RF | feature_elongation,coding_sequence_variant,intron_variant |
| 17 | 24949232 | 24951625 | 2393 | Rpl3l | FVB,SJL | feature_elongation,coding_sequence_variant,intron_variant |
| 17 | 30806457 | 30812006 | 5549 | Glo1 | BTBR | feature_elongation,3_prime_UTR_variant |
| 17 | 30993144 | 30996142 | 2998 | Dnah8 | SMJ | feature_elongation,coding_sequence_variant,intron_variant |
| 17 | 31745172 | 31746422 | 1250 | Ndufv3 | FVB,SJL | feature_elongation,coding_sequence_variant,intron_variant |
| 17 | 32548706 | 32554254 | 5548 | Akap8l | NZB | feature_elongation,coding_sequence_variant,intron_variant |
| 17 | 32550639 | 32551950 | 1311 | Akap8l | MA/My,SJL | feature_elongation,coding_sequence_variant,intron_variant |
| 17 | 35120933 | 35122077 | 1144 | Ehmt2 | ILNJ2 | feature_elongation,coding_sequence_variant,intron_variant |
| 17 | 37542613 | 37549499 | 6886 | Or1o2 | PJ | stop_gained,frameshift_variant |
| 17 | 41221508 | 41223462 | 1954 | Glyatl3 | BALB,DBA2J,RH2,SEA | feature_elongation,coding_sequence_variant,intron_variant |
| 17 | 42902282 | 42909505 | 7223 | Opn5 | BALB,BUB,DBA1J,DBA2J,PJ,RH2,SEA,SJL,SWR | feature_elongation,coding_sequence_variant,intron_variant |
| 17 | 44011717 | 44017930 | 6213 | Cyp39a1 | 129S1,AJ,BALB,BTBR,C57L,C58J,DBA2J,ILNJ2,KK,LP,MRL,NOD,NOR,NZB,NZO,NZW,PJ,PL,RH2,SEA,SJL,TH | feature_elongation,coding_sequence_variant,intron_variant |

| | | | | | | |
|---|---|---|---|---|---|---|
| 17 | 46433248 | 46439649 | 6401 | Mrps18a | BTBR,C57L,LP | feature_elongation,coding_sequence_variant,3_prime_UTR_variant,intron_variant |
| 17 | 66229189 | 66236300 | 7111 | Twsg1 | AJ,BALB,BTBR,BUB,C58J,CE,DBA1J,DBA2J,FVB,KK,NOD,NOR,NZB,NZW,PJ,PL,RH2,SEA,SJL,SMJ,TH | feature_elongation,coding_sequence_variant,3_prime_UTR_variant,intron_variant |
| 17 | 68727558 | 68734842 | 7284 | L3mbtl4 | C3H,CBA,MRL | feature_elongation,coding_sequence_variant,intron_variant |
| 17 | 71704897 | 71706791 | 1894 | Smchd1 | MA/My,NZB,NZO | feature_elongation,coding_sequence_variant,intron_variant |
| 17 | 72253489 | 72259892 | 6403 | Alk | C3H,CBA,CE,DBA1J,DBA2J,MA/My,MRL,NZB,NZO | feature_elongation,coding_sequence_variant,intron_variant |
| 17 | 75133846 | 75140289 | 6443 | Ttc27 | AKR,MA/My,NZB,NZO | feature_elongation,coding_sequence_variant,intron_variant |
| 17 | 88870919 | 88876224 | 5305 | Ppp1r21 | C3H | feature_elongation,coding_sequence_variant,intron_variant |
| 18 | 14888731 | 14895173 | 6442 | Psma8 | MA/My | feature_elongation,coding_sequence_variant,3_prime_UTR_variant,intron_variant |
| 18 | 22234362 | 22241286 | 6924 | Ccdc178 | SJL | feature_elongation,coding_sequence_variant,intron_variant |
| 18 | 22236981 | 22242498 | 5517 | Ccdc178 | CE | feature_elongation,coding_sequence_variant,intron_variant |
| 18 | 42679207 | 42681855 | 2648 | Tcerg1 | 129S1,AJ,AKR,BALB,BTBR,BUB,C3H,C57L,C58J,CBA,CE,DBA1J,DBA2J,FVB,KK,MA/My,MRL,NOD,NOR,NZW,PJ,PL,RF,RH2,SEA,SJL,SMJ,SWR,TH | feature_elongation,coding_sequence_variant,intron_variant |
| 18 | 43441860 | 43448263 | 6403 | Stk32a | KK | feature_elongation,coding_sequence_variant,3_prime_UTR_variant,intron_variant |
| 18 | 44971844 | 44978510 | 6666 | Ythdc2 | 129S1,BTBR,C57L,FVB,ILNJ2,MA/My,NZB,NZW,PJ,SWR | feature_elongation,coding_sequence_variant,intron_variant |
| 18 | 46716685 | 46723084 | 6399 | Tmed7 | C58J,LP,NZO,NZW | feature_elongation,coding_sequence_variant,intron_variant |
| 18 | 54044604 | 54051073 | 6469 | Csnk1g3 | 129S1,C57L | feature_elongation,coding_sequence_variant,intron_variant |
| 18 | 60940102 | 60941340 | 1238 | Cd74 | AJ,BALB,CBA,ILNJ2,RH2,SEA,SMJ | feature_elongation,coding_sequence_variant,intron_variant |
| 18 | 61656199 | 61658087 | 1888 | Arhgef37 | AJ,AKR,BALB,BUB,CBA,ILNJ2,MA/My,MRL,NOD,NOR,PL,RF,RH2,SEA | feature_elongation,coding_sequence_variant,5_prime_UTR_variant,intron_variant |
| 18 | 62559001 | 62562031 | 3030 | Htr4 | SMJ,SWR | feature_elongation,coding_sequence_variant,intron_variant |
| 18 | 63147047 | 63153489 | 6442 | Piezo2 | AJ,AKR,BALB,BUB,CBA,ILNJ2,MA/My,MRL,NOD,NOR,PL,RF,RH2,SEA | feature_elongation,coding_sequence_variant,intron_variant |
| 18 | 63955699 | 63958500 | 2801 | Wdr7 | AJ,BALB,C3H,CBA,ILNJ2,RH2 | feature_elongation,coding_sequence_variant,intron_variant |
| 18 | 80886501 | 80887835 | 1334 | Atp9b | NOD,NOR,SJL | feature_elongation,coding_sequence_variant,intron_variant |
| 18 | 89211224 | 89217704 | 6480 | Cd226 | FVB,KK,TH | feature_elongation,5_prime_UTR_variant,intron_variant |
| 19 | 9973915 | 9975034 | 1119 | Best1 | 129S1,AKR,B10D2,B10J,BTBR,C57L,DBA1J,DBA2J,FVB,MA/My,NOD,NOR,NZB,NZO,NZW,PJ,PL,RF,SJL,SMJ,SWR | feature_elongation,coding_sequence_variant,5_prime_UTR_variant,intron_variant |
| 19 | 11160491 | 11162381 | 1890 | Ms4a13 | TH | feature_elongation,coding_sequence_variant,intron_variant |
| 19 | 11569669 | 11571181 | 1512 | Ms4a6d | AJ,BALB,CE,LP,NOD,NOR,PJ,PL,RH2,SEA | feature_elongation,coding_sequence_variant,intron_variant |
| 19 | 29004271 | 29009829 | 5558 | Ak3 | AKR,BUB,MRL | feature_elongation,coding_sequence_variant,intron_variant |
| 19 | 30560616 | 30562598 | 1982 | Prkg1 | 129S1,BTBR,C57L,C58J,NZW | feature_elongation,coding_sequence_variant,intron_variant |
| 19 | 37560109 | 37561529 | 1420 | Exoc6 | B10D2,B10J,BTBR,C58J,LP,MA/My | feature_elongation,coding_sequence_variant,intron_variant |
| 19 | 37936898 | 37943339 | 6441 | Myof | NZW | feature_elongation,coding_sequence_variant,intron_variant |
| 19 | 42973009 | 42979700 | 6691 | Hpse2 | C3H,CBA,ILNJ2,NZB,NZO,NZW | feature_elongation,coding_sequence_variant,intron_variant |
| 19 | 50237087 | 50243693 | 6606 | Sorcs1 | NZB,NZO,NZW,RF | feature_elongation,coding_sequence_variant,intron_variant |
| 19 | 52920575 | 52921890 | 1315 | Xpnpep1 | NZB,NZW | feature_elongation,coding_sequence_variant,3_prime_UTR_variant,intron_variant |
| 19 | 57442463 | 57449600 | 7137 | Trub1 | AJ,RH2 | feature_elongation,coding_sequence_variant,intron_variant |
| X | 6432543 | 6437856 | 5313 | Shroom4 | C3H | feature_elongation,coding_sequence_variant,intron_variant |
| X | 7770740 | 7774174 | 3434 | Timm17b | PL | feature_elongation,coding_sequence_variant,3_prime_UTR_variant,intron_variant |
| X | 73202453 | 73205783 | 3330 | Tex28 | CE | feature_elongation,coding_sequence_variant,intron_variant |
| X | 76812714 | 76815188 | 2474 | Prkx | CE | feature_elongation,coding_sequence_variant,intron_variant |
| X | 76829470 | 76836581 | 7111 | Prkx | CE | feature_elongation,coding_sequence_variant,intron_variant |
| X | 82918179 | 82924682 | 6503 | Dmd | C58J | feature_elongation,coding_sequence_variant,intron_variant |
| X | 84229917 | 84236441 | 6524 | Dmd | 129S1,BUB,C57L,DBA1J,DBA2J,PJ | feature_elongation,coding_sequence_variant,intron_variant |
| X | 134695760 | 134697648 | 1888 | Gprasp1 | 129S1,AKR,BTBR,BUB,C3H,C57L,C58J,CBA,CE,DBA1J,DBA2J,LP,PJ,RF,SEA,SMJ | feature_elongation,5_prime_UTR_variant,intron_variant |
| X | 135412343 | 135419013 | 6670 | Kir3dl2 | KK | feature_elongation,5_prime_UTR_variant,intron_variant |
| X | 140409210 | 140415591 | 6381 | Col4a5 | CE,FVB,NOD,NOR,TH | feature_elongation,coding_sequence_variant,intron_variant |
| X | 147911202 | 147914137 | 2935 | Ott | MRL,SJL | feature_elongation,coding_sequence_variant,intron_variant |
| X | 150644747 | 150647160 | 2413 | Huwe1 | AJ,BALB,FVB,KK,MA/My,MRL,NOD,NOR,NZB,RH2,SEA,SJL,SMJ,SWR,TH | feature_elongation,coding_sequence_variant,intron_variant |
| X | 155084689 | 155089187 | 4498 | Gm15155 | AJ,BALB,MA/My,NOD,NOR,NZB,NZW,RH2,SEA,TH | feature_elongation,coding_sequence_variant,intron_variant |
| X | 159215032 | 159220771 | 5739 | Adgrg2 | BALB,CE,MRL,NOD,NOR,SEA | feature_elongation,5_prime_UTR_variant,intron_variant |
| X | 161700620 | 161705839 | 5219 | Ctps2 | AKR,BALB,RF,SEA | feature_elongation,coding_sequence_variant,intron_variant |
| X | 165227230 | 165232778 | 5548 | Trappc2 | TH | feature_elongation,coding_sequence_variant,5_prime_UTR_variant,intron_variant |

| CHR | START | END | SIZE | SYMBOL | STRAINS with VARIANT SV ALLELE | CONSEQUENCE |
|---|---|---|---|---|---|---|
| 1 | 4846259 | 4846472 | 213 | Mrpl15 | DBA1J,DBA2J | feature_elongation,3_prime_UTR_variant |
| 1 | 4851852 | 4851988 | 136 | Mrpl15 | DBA1J,DBA2J | feature_elongation,3_prime_UTR_variant |
| 1 | 9951878 | 9952390 | 512 | Sgk3 | SPRET | feature_elongation,coding_sequence_variant |
| 1 | 10010784 | 10010921 | 137 | Mcmdc2 | SPRET | feature_elongation,3_prime_UTR_variant |
| 1 | 10033327 | 10033526 | 199 | Tcf24 | CAST | feature_elongation,3_prime_UTR_variant |
| 1 | 10259804 | 10259881 | 77 | Arfgef1 | SPRET | feature_elongation,coding_sequence_variant |
| 1 | 11670519 | 11670931 | 412 | A830018L16Rik | 129S1,AJ,AKR,BALB,BTBR,BUB,C3H,C57L,CBA,CE,FVB,KK,MOLF,MRL,NOD,NZB,NZO,NZW,PL,RF,RH2,SEA,SJL,SPRET,SWR,TH | feature_elongation,3_prime_UTR_variant |
| 1 | 13721098 | 13721166 | 68 | Lactb2 | MOLF,TH | feature_elongation,coding_sequence_variant |
| 1 | 15793503 | 15793740 | 237 | Kcnb2 | SPRET | feature_elongation,3_prime_UTR_variant |
| 1 | 15914133 | 15914326 | 193 | Terf1 | SPRET | feature_elongation,3_prime_UTR_variant |
| 1 | 16151971 | 16152207 | 236 | 4930444P10Rik | WSB | feature_elongation,5_prime_UTR_variant |
| 1 | 16203422 | 16203560 | 138 | Rdh10 | CAST,LP,MOLF,SPRET | feature_elongation,3_prime_UTR_variant |
| 1 | 16640141 | 16640203 | 62 | Ube2w | CAST | feature_elongation,3_prime_UTR_variant |
| 1 | 16747739 | 16747923 | 184 | Tmem70 | CAST,MOLF | feature_elongation,3_prime_UTR_variant |
| 1 | 17819835 | 17819926 | 91 | Crispld1 | 129S1,BTBR,C57L,CE,MOLF,SPRET | feature_elongation,coding_sequence_variant |
| 1 | 30980183 | 30980237 | 54 | Ptp4a1 | SWR | feature_elongation,3_prime_UTR_variant |
| 1 | 31262931 | 31263139 | 208 | Dnaaf6rt | SPRET | feature_elongation,3_prime_UTR_variant |
| 1 | 33493000 | 33493058 | 58 | Prim2 | 129S1,AKR,BALB,BTBR,C57L,CAST,CBA,CE,LP,MOLF,NOD,RF,SEA,SPRET,SWR,WSB | feature_elongation,3_prime_UTR_variant |
| 1 | 34174632 | 34174702 | 70 | Dst | CAST,RF,SPRET | feature_elongation,3_prime_UTR_variant |
| 1 | 34175238 | 34175444 | 206 | Dst | CE,MOLF | feature_elongation,3_prime_UTR_variant |
| 1 | 34303931 | 34304071 | 140 | Dst | NZW | feature_elongation,coding_sequence_variant |
| 1 | 34484167 | 34484226 | 59 | Imp4 | 129S1,BTBR,C57L,ILNJ2,KK | feature_elongation,3_prime_UTR_variant |
| 1 | 36181589 | 36181685 | 96 | Uggt1 | MOLF | feature_elongation,3_prime_UTR_variant |
| 1 | 36732507 | 36732562 | 55 | Cox5b | SPRET | feature_elongation,3_prime_UTR_variant |
| 1 | 37457066 | 37457130 | 64 | Coa5 | MA/My,MOLF | feature_elongation,3_prime_UTR_variant |
| 1 | 38833629 | 38833804 | 175 | Lonrf2 | SPRET | feature_elongation,3_prime_UTR_variant |
| 1 | 38906630 | 38906708 | 78 | Chst10 | BALB,BTBR,C57L,DBA1J,DBA2J,FVB,ILNJ2,LP,MA/My,NOR,NZB,NZO,PL,SEA,SMJ,TH | feature_elongation,3_prime_UTR_variant |
| 1 | 39585134 | 39585339 | 205 | Cnot11 | RF | feature_elongation,3_prime_UTR_variant |
| 1 | 44183845 | 44183916 | 71 | Bivm | AKR,CAST,FVB,MA/My,MOLF,NOD,NOR,RF,RH2 | feature_elongation,3_prime_UTR_variant |
| 1 | 44244716 | 44244820 | 104 | Mettl21e | 129S1,AJ,AKR,BUB,C3H,CAST,CBA,CE,DBA1J,DBA2J,FVB,ILNJ2,KK,MA/My,MOLF,MRL,NOD,NOR,PJ,PL,RF,RH2,SJL,SMJ,SWR,TH | feature_elongation,3_prime_UTR_variant |
| 1 | 52223569 | 52223654 | 85 | Gls | SPRET | feature_elongation,3_prime_UTR_variant |
| 1 | 52825479 | 52825617 | 138 | Inpp1 | SPRET | feature_elongation,3_prime_UTR_variant |
| 1 | 53795206 | 53795416 | 210 | Stk17b | CAST,MOLF | feature_elongation,3_prime_UTR_variant |
| 1 | 57455110 | 57455180 | 70 | Maip1 | SMJ,SWR | feature_elongation,3_prime_UTR_variant |
| 1 | 58566540 | 58566769 | 229 | Hycc2 | CBA,DBA1J,DBA2J,ILNJ2,NZB,NZO,NZW,RH2 | feature_elongation,3_prime_UTR_variant |
| 1 | 59046000 | 59046895 | 895 | C2cd6 | 129S1,BTBR,BUB,C3H,CBA,DBA2J,ILNJ2,KK,LP,NZO,NZW,SMJ,SPRET,SWR | feature_elongation,coding_sequence_variant |
| 1 | 59391679 | 59391753 | 74 | Cdk15 | CE,MOLF | feature_elongation,3_prime_UTR_variant |
| 1 | 59752225 | 59752424 | 199 | Nop58 | ILNJ2,MA/My,NZB,NZO,NZW,WSB | feature_elongation,3_prime_UTR_variant |
| 1 | 59755468 | 59755688 | 220 | Nop58 | ILNJ2,MA/My,NZB,NZO,NZW,SMJ,WSB | feature_elongation,3_prime_UTR_variant |
| 1 | 59757121 | 59757180 | 59 | Nop58 | SPRET | feature_elongation,3_prime_UTR_variant |
| 1 | 59911861 | 59912068 | 207 | Bmpr2 | CE | feature_elongation,3_prime_UTR_variant |
| 1 | 60109960 | 60110026 | 66 | Wdr12 | CE,LP,MOLF,SPRET | feature_elongation,3_prime_UTR_variant |
| 1 | 60111845 | 60112397 | 552 | Wdr12 | ILNJ2,NZB,NZO,NZW | feature_elongation,3_prime_UTR_variant |
| 1 | 60375404 | 60375597 | 193 | Nbeal1 | SPRET | feature_elongation,3_prime_UTR_variant |
| 1 | 63218917 | 63219120 | 203 | Eef1b2 | CAST | feature_elongation,3_prime_UTR_variant |
| 1 | 63350797 | 63350994 | 197 | Zdbf2 | C58J,MOLF,NZB | feature_elongation,3_prime_UTR_variant |
| 1 | 65317295 | 65317347 | 52 | Pikfyve | SPRET | feature_elongation,3_prime_UTR_variant |
| 1 | 67041321 | 67041381 | 60 | Lancl1 | PJ,WSB | feature_elongation,3_prime_UTR_variant |
| 1 | 72467364 | 72467444 | 80 | Marchf4 | SPRET | feature_elongation,3_prime_UTR_variant |
| 1 | 74778567 | 74778619 | 52 | Prkag3 | SPRET | feature_elongation,3_prime_UTR_variant |
| 1 | 78646556 | 78646656 | 100 | Utp14b | CAST | feature_elongation,3_prime_UTR_variant |
| 1 | 82430647 | 82430713 | 66 | Col4a4 | CAST,MOLF | feature_elongation,3_prime_UTR_variant |
| 1 | 82748860 | 82748935 | 75 | Gm47791 | PJ | feature_elongation,coding_sequence_variant |
| 1 | 82933542 | 82933603 | 61 | Gm7544 | RF | feature_elongation,coding_sequence_variant |
| 1 | 82938140 | 82938293 | 153 | Gm47955 | AKR,CBA,LP,MRL,NOD,WSB | feature_elongation,coding_sequence_variant |
| 1 | 83233045 | 83233162 | 117 | Sphkap | CAST | feature_elongation,3_prime_UTR_variant |
| 1 | 84563271 | 84563325 | 54 | Dner | AKR,CBA,MRL,NOD,PJ,PL | feature_elongation,coding_sequence_variant |
| 1 | 85637380 | 85637529 | 149 | Sp100 | SPRET | feature_elongation,3_prime_UTR_variant |
| 1 | 86204807 | 86204883 | 76 | Armc9 | SPRET | feature_elongation,3_prime_UTR_variant |
| 1 | 86205808 | 86205882 | 74 | Armc9 | SMJ | feature_elongation,3_prime_UTR_variant |
| 1 | 86535261 | 86535319 | 58 | Cops7b | WSB | feature_elongation,3_prime_UTR_variant |
| 1 | 88431142 | 88431199 | 57 | Glrp1 | SMJ | feature_elongation,coding_sequence_variant |
| 1 | 89818413 | 89818633 | 220 | Agap1 | C58J | feature_elongation,3_prime_UTR_variant |
| 1 | 89880000 | 89880190 | 190 | Asb18 | SPRET | feature_elongation,3_prime_UTR_variant |
| 1 | 90877734 | 90877823 | 89 | Mlph | B10D2,B10J,BALB,BTBR,BUB,C3H,C58J,CBA,CE,DBA1J,DBA2J,ILNJ2,MA/My,MRL,NOD,NOR,PJ,PL,RH2,SEA,SMJ,WSB | feature_elongation,3_prime_UTR_variant |
| 1 | 91483493 | 91483709 | 216 | Asb1 | 129S1,AKR,C57L,FVB,KK,LP,NZB,NZO,NZW,RF,SJL,SWR,TH | feature_elongation,3_prime_UTR_variant |
| 1 | 91486902 | 91486972 | 70 | Asb1 | 129S1,AKR,BALB,BTBR,C57L,CAST,DBA1J,DBA2J,FVB,ILNJ2,KK,LP,MOLF,NOD,NOR,NZB,NZO,NZW,PJ,PL,RF,SEA,SJL,SMJ,SPRET,SWR,TH,WSB | feature_elongation,3_prime_UTR_variant |
| 1 | 92404232 | 92404302 | 70 | Or6b2 | CAST,MOLF,PL | feature_elongation,3_prime_UTR_variant |
| 1 | 92405287 | 92405457 | 170 | Or6b2 | BUB,C3H,CAST,CBA,DBA1J,DBA2J,MOLF,PJ,PL,SEA,SPRET,WSB | feature_elongation,3_prime_UTR_variant |
| 1 | 92436382 | 92436582 | 200 | Or6b3 | SPRET | feature_elongation,3_prime_UTR_variant |
| 1 | 92540499 | 92540612 | 113 | Or9s14 | BUB,C3H,CBA | feature_elongation,3_prime_UTR_variant |
| 1 | 92907020 | 92907179 | 159 | Gpr35 | SPRET | feature_elongation,5_prime_UTR_variant |
| 1 | 93630119 | 93630176 | 57 | Thap4 | SPRET | feature_elongation,3_prime_UTR_variant |
| 1 | 95253264 | 95253795 | 531 | Fam174a | PL,SPRET | feature_elongation,3_prime_UTR_variant |
| 1 | 95254107 | 95254187 | 80 | Fam174a | SPRET | feature_elongation,3_prime_UTR_variant |
| 1 | 97545428 | 97545988 | 560 | Macir | MOLF,MRL,PL | feature_elongation,3_prime_UTR_variant |
| 1 | 97551636 | 97551720 | 84 | Macir | MRL,PL | feature_elongation,3_prime_UTR_variant |

| | | | | | | |
|---|---|---|---|---|---|---|
| 1 | 97565164 | 97565313 | 149 | Macir | SPRET | feature_elongation,3_prime_UTR_variant |
| 1 | 105772969 | 105773134 | 165 | Tnfrsf11a | 129S1,BUB,C57L,CAST,DBA1J,DBA2J,ILNJ2,MOLF,NOD,NOR,SJL,SPRET,TH,WSB | feature_elongation,3_prime_UTR_variant |
| 1 | 107536119 | 107536318 | 199 | Serpinb8 | 129S1,C57L,CAST,DBA1J,DBA2J,MOLF,SJL,SWR,TH | feature_elongation,3_prime_UTR_variant |
| 1 | 117914188 | 117914524 | 336 | Gm7145 | MOLF,SPRET | feature_elongation,coding_sequence_variant |
| 1 | 118128515 | 118128683 | 168 | Gm29106 | MOLF,SWR | feature_elongation,3_prime_UTR_variant |
| 1 | 118222543 | 118222739 | 196 | Tsn | MOLF | feature_elongation,3_prime_UTR_variant |
| 1 | 118608036 | 118608106 | 70 | Tfcp2l1 | CAST | feature_elongation,3_prime_UTR_variant |
| 1 | 123254401 | 123255352 | 951 | Dpp10 | SPRET | feature_elongation,3_prime_UTR_variant |
| 1 | 127415541 | 127415597 | 56 | Mgat5 | MOLF,RF | feature_elongation,3_prime_UTR_variant |
| 1 | 127695241 | 127695709 | 468 | Acmsd | SPRET | feature_elongation,3_prime_UTR_variant |
| 1 | 130671871 | 130672018 | 147 | AA986860 | CAST | feature_elongation,3_prime_UTR_variant |
| 1 | 130779619 | 130779676 | 57 | Pigr | 129S1,BALB,C57L,DBA1J,DBA2J,FVB,ILNJ2,KK,MA/My,NZO,NZW,SEA,SJL,SWR,TH | feature_elongation,3_prime_UTR_variant |
| 1 | 131106911 | 131107063 | 152 | Eif2d | WSB | feature_elongation,3_prime_UTR_variant |
| 1 | 131213773 | 131214005 | 232 | Srgap2 | CAST,MOLF,SPRET | feature_elongation,3_prime_UTR_variant |
| 1 | 131640794 | 131641204 | 410 | Rab7b | 129S1,AKR,BALB,BTBR,BUB,CBA,DBA1J,DBA2J,FVB,ILNJ2,KK,MA/My,MRL,NZO,NZW,PJ,PL,RF,SEA,SJL,SMJ,SWR,TH,WSB | feature_elongation,3_prime_UTR_variant |
| 1 | 132889916 | 132890054 | 138 | Mdm4 | CAST,MOLF | feature_elongation,3_prime_UTR_variant |
| 1 | 133065684 | 133065874 | 190 | Ppp1r15b | WSB | feature_elongation,3_prime_UTR_variant |
| 1 | 133627625 | 133627712 | 87 | Atp2b4 | SPRET | feature_elongation,3_prime_UTR_variant |
| 1 | 133629537 | 133629607 | 70 | Atp2b4 | CAST,MOLF,MRL,SJL,SWR | frameshift_variant |
| 1 | 134339327 | 134339397 | 70 | Cyb5r1 | SPRET | feature_elongation,3_prime_UTR_variant |
| 1 | 135226013 | 135226111 | 98 | Timm17a | CAST | feature_elongation,3_prime_UTR_variant |
| 1 | 135363118 | 135363253 | 135 | Nav1 | 129S1,AJ,AKR,BTBR,C3H,CAST,CE,DBA1J,DBA2J,ILNJ2,KK,MA/My,MOLF,NZB,NZO,NZW,PJ,RF,RH2,SJL,SPRET,SWR,TH,WSB | feature_elongation,3_prime_UTR_variant |
| 1 | 138754651 | 138754735 | 84 | Lhx9 | SPRET | feature_elongation,3_prime_UTR_variant |
| 1 | 143625218 | 143625436 | 218 | Glrx2 | AJ,C3H,CBA,CE,FVB,ILNJ2,PL,RH2 | feature_elongation,3_prime_UTR_variant |
| 1 | 143630205 | 143630404 | 199 | Ro60 | 129S1,NZO,NZW,SJL,SWR,TH | feature_elongation,3_prime_UTR_variant |
| 1 | 143682922 | 143683140 | 218 | Uchl5 | AJ,C3H,CBA,CE,FVB,ILNJ2,MRL,PL,RH2 | feature_elongation,3_prime_UTR_variant |
| 1 | 144395319 | 144395373 | 54 | Rgs21 | AJ,C3H,CBA,CE,FVB,ILNJ2,NZO,PL,RH2,SJL,SWR,WSB | feature_elongation,3_prime_UTR_variant |
| 1 | 150331351 | 150331441 | 90 | Prg4 | FVB,SJL,SWR | feature_elongation,coding_sequence_variant |
| 1 | 152993614 | 152993709 | 95 | Nmnat2 | CAST | feature_elongation,3_prime_UTR_variant |
| 1 | 153638463 | 153638537 | 74 | Rnasel | CAST,SPRET,WSB | feature_elongation,3_prime_UTR_variant |
| 1 | 153776450 | 153776501 | 51 | Glul | CAST | feature_elongation,5_prime_UTR_variant |
| 1 | 155003932 | 155004152 | 220 | Mr1 | C57L | feature_elongation,3_prime_UTR_variant |
| 1 | 155574699 | 155574795 | 96 | Lhx4 | C57L,CAST,MOLF | feature_elongation,3_prime_UTR_variant |
| 1 | 155577550 | 155577673 | 123 | Lhx4 | CAST | feature_elongation,3_prime_UTR_variant |
| 1 | 156254652 | 156254851 | 199 | Soat1 | SPRET | feature_elongation,3_prime_UTR_variant |
| 1 | 156666175 | 156666236 | 61 | Angptl1 | CAST | feature_elongation,5_prime_UTR_variant |
| 1 | 156965251 | 156965388 | 137 | Rasal2 | SPRET | feature_elongation,3_prime_UTR_variant |
| 1 | 159174439 | 159175221 | 782 | Cop1 | SPRET | feature_elongation,3_prime_UTR_variant |
| 1 | 159753917 | 159754025 | 108 | Tnr | BUB,C3H,CBA,CE,PJ,SEA | feature_elongation,3_prime_UTR_variant |
| 1 | 160028902 | 160029275 | 373 | Mrps14 | SPRET | feature_elongation,3_prime_UTR_variant |
| 1 | 160835404 | 160835615 | 211 | Zbtb37 | BALB,C58J,DBA1J,DBA2J,LP,MOLF | feature_elongation,3_prime_UTR_variant |
| 1 | 160841070 | 160841240 | 170 | Zbtb37 | SPRET | feature_elongation,3_prime_UTR_variant |
| 1 | 160917163 | 160917243 | 80 | Klhl20 | BALB,C58J,DBA1J,DBA2J,ILNJ2,LP,MOLF | feature_elongation,3_prime_UTR_variant |
| 1 | 162851898 | 162852101 | 203 | Mroh9 | SPRET | feature_elongation,3_prime_UTR_variant |
| 1 | 163084650 | 163084719 | 69 | Prrx1 | SPRET | feature_elongation,3_prime_UTR_variant |
| 1 | 164021215 | 164021375 | 160 | F5 | CAST,DBA1J,DBA2J,ILNJ2,MOLF,SPRET | feature_elongation,coding_sequence_variant |
| 1 | 164965805 | 164965873 | 68 | Tbx19 | CAST | feature_elongation,3_prime_UTR_variant |
| 1 | 165039901 | 165039957 | 56 | Tiprl | SPRET | feature_elongation,3_prime_UTR_variant |
| 1 | 165152131 | 165152187 | 56 | Gpr161 | AKR,BUB,C3H,C58J,CBA,LP,MRL,RH2 | feature_elongation,3_prime_UTR_variant |
| 1 | 165156494 | 165156629 | 135 | Dcaf6 | FVB,PJ,RF,SEA,SJL | feature_elongation,3_prime_UTR_variant |
| 1 | 165156607 | 165157053 | 446 | Dcaf6 | DBA1J,DBA2J,ILNJ2,MOLF | feature_elongation,3_prime_UTR_variant |
| 1 | 165178573 | 165178648 | 75 | Dcaf6 | CAST,DBA1J,DBA2J,ILNJ2,MOLF | feature_elongation,coding_sequence_variant |
| 1 | 167426225 | 167426281 | 56 | Rxrg | SMJ | feature_elongation,5_prime_UTR_variant |
| 1 | 169800493 | 169800660 | 167 | Ddr2 | 129S1,AJ,BTBR,BUB,C3H,C57L,CE,FVB,KK,MRL,NOD,NOR,NZB,NZO,NZW,PJ,PL,RF,RH2,SEA,SJL,SMJ,SWR,TH,WSB | feature_elongation,3_prime_UTR_variant |
| 1 | 171460903 | 171460968 | 65 | Slamf7 | 129S1,AJ,AKR,BALB,BUB,C3H,C57L,CBA,CE,DBA1J,DBA2J,FVB,KK,MRL,NOD,NOR,NZB,NZO,NZW,PJ,PL,RF,RH2,SEA,SJL,SMJ,SWR,TH,WSB | feature_elongation,3_prime_UTR_variant |
| 1 | 171628587 | 171628969 | 382 | Slamf1 | SPRET | feature_elongation,3_prime_UTR_variant |
| 1 | 171776737 | 171776797 | 60 | Slamf6 | SPRET | feature_elongation,3_prime_UTR_variant |
| 1 | 171780686 | 171780894 | 208 | Slamf6 | 129S1,AJ,AKR,BALB,C3H,C57L,CBA,CE,DBA1J,DBA2J,KK,MRL,NOD,NOR,NZO,NZW,PJ,PL,RF,RH2,SEA,SJL,SMJ,TH | feature_elongation,3_prime_UTR_variant |
| 1 | 172526846 | 172526900 | 54 | Crp | NOD,NOR | feature_elongation,3_prime_UTR_variant |
| 1 | 173099728 | 173100329 | 601 | Or10j2 | 129S1,AJ,AKR,BALB,BTBR,BUB,C3H,C57L,C58J,CAST,CBA,CE,DBA1J,DBA2J,FVB,ILNJ2,KK,MRL,NOD,NOR,NZB,NZO,NZW,PJ,PL,RF,RH2,SEA,SJL,SMJ,SPRET,SWR,TH,WSB | feature_elongation,3_prime_UTR_variant |
| 1 | 173556779 | 173557110 | 331 | Ifi207 | 129S1,AJ,AKR,BALB,BTBR,BUB,C3H,C57L,C58J,CBA,CE,DBA1J,DBA2J,FVB,ILNJ2,KK,MRL,NOD,NOR,NZB,NZO,PJ,PL,RF,RH2,SEA,SJL,SMJ,SPRET,SWR,TH,WSB | feature_elongation,coding_sequence_variant |
| 1 | 173587714 | 173587777 | 63 | Ifi204 | 129S1,AKR,BALB,BUB,C3H,C57L,C58J,CBA,CE,DBA1J,DBA2J,ILNJ2,MRL,NOD,NOR,NZB,NZO,PJ,RF,SEA,SJL,SMJ,SWR,WSB | feature_elongation,coding_sequence_variant |
| 1 | 173733734 | 173733796 | 62 | Ifi211 | 129S1,AKR,BALB,BUB,C3H,C57L,C58J,CBA,CE,DBA1J,DBA2J,ILNJ2,MRL,NOD,NOR,NZB,NZO,PJ,RF,SEA,SJL,SMJ,WSB | frameshift_variant |
| 1 | 173896677 | 173896743 | 66 | Or6n2 | AJ,AKR,BTBR,BUB,C58J,CBA,CE,DBA1J,DBA2J,FVB,ILNJ2,KK,NZB,NZO,PJ,PL,RF,RH2,SEA,SMJ,SPRET,SWR,TH | feature_elongation,5_prime_UTR_variant |
| 1 | 175556503 | 175556666 | 163 | Wdr64 | SPRET | feature_elongation,coding_sequence_variant |
| 1 | 175805702 | 175805914 | 212 | Rbm8a2 | MOLF,NZO | feature_elongation,3_prime_UTR_variant |
| 1 | 176850818 | 176851012 | 194 | Akt3 | SMJ | feature_elongation,3_prime_UTR_variant |

| | | | | | | |
|---|---|---|---|---|---|---|
| 1 | 177875630 | 177875713 | 83 | Catspere2 | MOLF | feature_elongation,3_prime_UTR_variant |
| 1 | 180521856 | 180522394 | 538 | Mixl1 | CAST | feature_elongation,3_prime_UTR_variant |
| 1 | 180916439 | 180916505 | 66 | Nvl | SPRET | feature_elongation,3_prime_UTR_variant |
| 1 | 180994765 | 180994985 | 220 | Cnih4 | CBA,DBA1J,DBA2J | feature_elongation,3_prime_UTR_variant |
| 1 | 189605316 | 189605469 | 153 | Ptpn14 | CAST,MOLF,SPRET | feature_elongation,3_prime_UTR_variant |
| 1 | 190738816 | 190738992 | 176 | Flvcr1 | SPRET | feature_elongation,3_prime_UTR_variant |
| 1 | 192196105 | 192196201 | 96 | Hhat | SPRET | feature_elongation,3_prime_UTR_variant |
| 1 | 192527962 | 192528080 | 118 | Sertad4 | SPRET | feature_elongation,3_prime_UTR_variant |
| 1 | 194723637 | 194723868 | 231 | Cd46 | CAST,MOLF,NOD,NOR,NZO,NZW,RF,SPRET,SWR,WSB | feature_elongation,3_prime_UTR_variant |
| 2 | 3458321 | 3458493 | 172 | Suv39h2 | MA/My,NZO,NZW,SPRET | feature_elongation,3_prime_UTR_variant |
| 2 | 3462645 | 3462835 | 190 | Dclre1c | SPRET | feature_elongation,3_prime_UTR_variant |
| 2 | 3614893 | 3614972 | 79 | Fam107b | SPRET | feature_elongation,5_prime_UTR_variant |
| 2 | 5799150 | 5799259 | 109 | Cdc123 | SMJ | feature_elongation,3_prime_UTR_variant |
| 2 | 5803295 | 5803584 | 289 | Cdc123 | SPRET | feature_elongation,coding_sequence_variant |
| 2 | 5874425 | 5874502 | 77 | Nudt5 | CAST | feature_elongation,3_prime_UTR_variant |
| 2 | 6548271 | 6548338 | 67 | Celf2 | SPRET | feature_elongation,3_prime_UTR_variant |
| 2 | 10258849 | 10259061 | 212 | Itih5 | SPRET | feature_elongation,3_prime_UTR_variant |
| 2 | 11478428 | 11478502 | 74 | Pfkfb3 | MA/My | feature_elongation,3_prime_UTR_variant |
| 2 | 13516249 | 13516317 | 68 | Trdmt1 | SPRET | feature_elongation,3_prime_UTR_variant |
| 2 | 14224520 | 14224724 | 204 | Tmem236 | MOLF | feature_elongation,3_prime_UTR_variant |
| 2 | 18221937 | 18222139 | 202 | Dnajc1 | MA/My,MOLF,RF,SPRET | feature_elongation,3_prime_UTR_variant |
| 2 | 20749272 | 20749471 | 199 | Etl4 | CE | feature_elongation,3_prime_UTR_variant |
| 2 | 24076660 | 24076756 | 96 | Il36g | CAST | feature_elongation,5_prime_UTR_variant |
| 2 | 24240782 | 24240970 | 188 | Il1rn | SPRET | feature_elongation,3_prime_UTR_variant |
| 2 | 24862157 | 24862362 | 205 | Dph7 | FVB,WSB | feature_elongation,3_prime_UTR_variant |
| 2 | 25465701 | 25465789 | 88 | Ajm1 | CAST,MOLF,SPRET | feature_elongation,3_prime_UTR_variant |
| 2 | 26100029 | 26100082 | 53 | Qsox2 | SPRET | feature_elongation,3_prime_UTR_variant |
| 2 | 26909673 | 26909906 | 233 | Cacfd1 | 129S1,AJ,AKR,B10D2,B10J,BALB,BTBR,BUB,C3H,C57L,C58J,CAST,CBA,CE,DBA1J,DBA2J,FVB,ILNJ2,KK,LP,MA/My,MRL,NOD,NOR,NZB,NZO,NZW,PJ,PL,RF,RH2,SEA,SJL,SMJ,SWR,TH,WSB | feature_elongation,3_prime_UTR_variant |
| 2 | 26998572 | 26998630 | 58 | Adamtsl2 | 129S1,BUB,C57L,CAST,FVB,MA/My,MOLF,RF,SMJ,SPRET | feature_elongation,3_prime_UTR_variant |
| 2 | 28548588 | 28548790 | 202 | Tsc1 | SPRET | feature_elongation,5_prime_UTR_variant |
| 2 | 29051533 | 29051637 | 104 | Setx | MOLF | feature_elongation,coding_sequence_variant |
| 2 | 30719321 | 30719523 | 202 | Asb6 | SPRET | feature_elongation,5_prime_UTR_variant |
| 2 | 30920068 | 30920157 | 89 | Fnbp1 | CAST,MOLF | feature_elongation,3_prime_UTR_variant |
| 2 | 31693592 | 31693702 | 110 | Abl1 | SPRET | feature_elongation,3_prime_UTR_variant |
| 2 | 32489325 | 32490150 | 825 | St6galnac4 | AJ,AKR,BTBR,BUB,C58J,CE,DBA2J,FVB,ILNJ2,KK,MRL,NOD,NOR,NZB,NZW,PL,RH2,SMJ,TH | feature_elongation,3_prime_UTR_variant |
| 2 | 33621717 | 33621768 | 51 | Mvb12b | SPRET | feature_elongation,3_prime_UTR_variant |
| 2 | 34666682 | 34666874 | 192 | Hspa5 | CAST | feature_elongation,3_prime_UTR_variant |
| 2 | 34832954 | 34833009 | 55 | Traf1 | SPRET | feature_elongation,3_prime_UTR_variant |
| 2 | 34851672 | 34851743 | 71 | Traf1 | SPRET | feature_elongation,5_prime_UTR_variant |
| 2 | 36466038 | 36466090 | 52 | Or1j15 | MOLF | feature_elongation,3_prime_UTR_variant |
| 2 | 36748377 | 36748428 | 51 | Or1n1 | SPRET | feature_elongation,3_prime_UTR_variant |
| 2 | 36956125 | 36956185 | 60 | Or12k7 | BUB | feature_elongation,5_prime_UTR_variant |
| 2 | 37037656 | 37037811 | 155 | Or1l4b | BUB,CAST,MOLF,SPRET,TH | feature_elongation,3_prime_UTR_variant |
| 2 | 37325068 | 37325462 | 394 | Zbtb26 | DBA1J | feature_elongation,3_prime_UTR_variant |
| 2 | 37409688 | 37409786 | 98 | Gpr21 | CAST,SPRET | feature_elongation,3_prime_UTR_variant |
| 2 | 37465079 | 37465137 | 58 | Strbp | SPRET | feature_elongation,3_prime_UTR_variant |
| 2 | 43525676 | 43525866 | 190 | Kynu | 129S1,AJ,AKR,B10D2,B10J,BALB,BTBR,BUB,C3H,C57L,C58J,CAST,CBA,CE,DBA1J,DBA2J,FVB,ILNJ2,KK,LP,MA/My,MOLF,MRL,NOD,NOR,NZB,NZO,NZW,PJ,PL,RF,RH2,SEA,SJL,SMJ,SPRET,SWR,TH,WSB | feature_elongation,3_prime_UTR_variant |
| 2 | 44876189 | 44876304 | 115 | Zeb2 | SPRET | feature_elongation,3_prime_UTR_variant |
| 2 | 51024040 | 51024096 | 56 | Rnd3 | 129S1,AJ,AKR,BUB,C3H,C58J,CAST,CBA,CE,DBA1J,DBA2J,FVB,ILNJ2,KK,MOLF,MRL,NOD,NOR,NZW,PJ,RF,SEA,SJL,SMJ,SWR,TH | feature_elongation,coding_sequence_variant |
| 2 | 52306089 | 52306227 | 138 | Arl5a | 129S1,AJ,AKR,BUB,C3H,CBA,CE,DBA1J,DBA2J,ILNJ2,MRL,NOD,NOR,NZW,PJ,PL,RF,SEA,SJL,SMJ,SWR,TH,WSB | feature_elongation,coding_sequence_variant |
| 2 | 52581930 | 52582135 | 205 | Stam2 | MOLF | feature_elongation,3_prime_UTR_variant |
| 2 | 55005401 | 55005501 | 100 | Galnt13 | SPRET | feature_elongation,3_prime_UTR_variant |
| 2 | 58020800 | 58020860 | 60 | Cytip | 129S1,AJ,AKR,BUB,C3H,C58J,CAST,CBA,DBA1J,DBA2J,FVB,MOLF,NOD,NOR,PJ,PL,RF,SEA,SJL,SPRET,SWR,TH | feature_elongation,3_prime_UTR_variant |
| 2 | 58022231 | 58022300 | 69 | Cytip | 129S1,AJ,AKR,BUB,C3H,C58J,CAST,CBA,DBA1J,DBA2J,FVB,MOLF,NOD,NOR,PJ,PL,RF,SEA,SJL,SWR,TH | feature_elongation,3_prime_UTR_variant |
| 2 | 59135482 | 59136025 | 543 | Pkp4 | SPRET | feature_elongation,coding_sequence_variant,feature_elongation,5_prime_UTR_variant |
| 2 | 60430033 | 60430405 | 372 | Itgb6 | SPRET | feature_elongation,3_prime_UTR_variant |
| 2 | 70969337 | 70969397 | 60 | Cybrd1 | AJ,AKR,BUB,C3H,CAST,FVB,ILNJ2,MA/My,MOLF,MRL,NOD,NOR,NZW,PL,RH2,SMJ | feature_elongation,3_prime_UTR_variant |
| 2 | 73150116 | 73150270 | 154 | Scrn3 | SPRET | feature_elongation,3_prime_UTR_variant |
| 2 | 73262517 | 73262593 | 76 | Wipf1 | CAST,KK,MA/My,MOLF,NZB,SPRET | feature_elongation,3_prime_UTR_variant |
| 2 | 73393585 | 73394030 | 445 | Chrna1 | SPRET | feature_elongation,3_prime_UTR_variant |
| 2 | 76842952 | 76843043 | 91 | Ccdc141 | WSB | feature_elongation,3_prime_UTR_variant |
| 2 | 77726734 | 77726885 | 151 | Cwc22 | SPRET,WSB | feature_elongation,coding_sequence_variant |
| 2 | 79163316 | 79163446 | 130 | Itga4 | MOLF | feature_elongation,3_prime_UTR_variant |
| 2 | 80359011 | 80359084 | 73 | Nckap1 | AJ,AKR,BALB,BUB,C3H,C57L,C58J,CAST,CBA,CE,DBA1J,DBA2J,FVB,KK,MA/My,MRL,NOD,NOR,PJ,PL,RF,RH2,SEA,SJL,SMJ,SWR,TH,WSB | feature_elongation,coding_sequence_variant |
| 2 | 83816023 | 83816261 | 238 | Cwc22rt7 | ILNJ2 | feature_elongation,coding_sequence_variant |
| 2 | 85437422 | 85437476 | 54 | Or5g23 | SPRET,WSB | feature_elongation,3_prime_UTR_variant |
| 2 | 85617082 | 85617297 | 215 | Or9g4b | AJ,AKR,BALB,BUB,C3H,C57L,C58J,CBA,CE,DBA1J,DBA2J,ILNJ2,MA/My,MRL,NOD,NOR,NZB,NZO,NZW,PJ,PL,RF,RH2,SEA,SMJ | feature_elongation,3_prime_UTR_variant |

| | | | | | | |
|---|---|---|---|---|---|---|
| 2 | 85628337 | 85628401 | 64 | Or9g20 | SPRET | feature_elongation,3_prime_UTR_variant |
| 2 | 85671181 | 85671353 | 172 | Or5ar1 | KK | feature_elongation,3_prime_UTR_variant |
| 2 | 86180436 | 86181103 | 667 | Or8k53 | MOLF | feature_elongation,5_prime_UTR_variant |
| 2 | 86863115 | 86863239 | 124 | Or5be3 | MOLF | feature_elongation,3_prime_UTR_variant |
| 2 | 86936591 | 86937364 | 773 | Or5aq7 | CAST | feature_elongation,3_prime_UTR_variant |
| 2 | 87189135 | 87189266 | 131 | Or12e8 | MOLF,SJL,SPRET,SWR | feature_elongation,3_prime_UTR_variant |
| 2 | 87251332 | 87251558 | 226 | Or10ag2 | AJ,AKR,BALB,C57L,DBA1J,DBA2J,ILNJ2,MOLF,MRL,NOD,NOR,NZB,NZO,PJ,RH2,SEA,SJL,SMJ,SPRET,SWR | feature_elongation,3_prime_UTR_variant |
| 2 | 87540998 | 87541052 | 54 | Or5w14 | C3H,CBA,CE,KK,MA/My,MOLF,NZW,RF | feature_elongation,3_prime_UTR_variant |
| 2 | 87614057 | 87614126 | 69 | Or5i1 | C3H,C58J,CBA,CE,KK,MA/My,MOLF,NZW,RF,SJL,SPRET,SWR,WSB | feature_elongation,3_prime_UTR_variant |
| 2 | 87633743 | 87634049 | 306 | Or5w18 | MOLF | feature_elongation,3_prime_UTR_variant |
| 2 | 87789889 | 87790049 | 160 | Or5l14 | AJ,AKR,BALB,C57L,C58J,DBA1J,DBA2J,FVB,ILNJ2,KK,MOLF,MRL,NOD,NOR,NZB,NZO,PJ,RH2,SEA,SJL,SMJ,SPRET,SWR,WSB | feature_elongation,3_prime_UTR_variant |
| 2 | 87878498 | 87878566 | 68 | Or5d14 | SPRET | feature_elongation,3_prime_UTR_variant |
| 2 | 87901689 | 87901876 | 187 | Or5d36 | MOLF | feature_elongation,coding_sequence_variant |
| 2 | 88224627 | 88224682 | 55 | Or4p7 | MOLF | feature_elongation,3_prime_UTR_variant |
| 2 | 88227534 | 88227618 | 84 | Or4p7 | MOLF | feature_elongation,3_prime_UTR_variant |
| 2 | 88512171 | 88512225 | 54 | Or4c103 | MOLF,SPRET | feature_elongation,3_prime_UTR_variant |
| 2 | 88558710 | 88558768 | 58 | Or4a27 | SPRET | feature_elongation,3_prime_UTR_variant |
| 2 | 88738457 | 88739070 | 613 | Or4c29 | C3H,CBA,CE,MA/My,NZW | feature_elongation,3_prime_UTR_variant |
| 2 | 88757583 | 88758085 | 502 | Or4c15 | AJ,AKR,BALB,C57L,DBA1J,DBA2J,FVB,KK,MOLF,MRL,NOD,NOR,NZB,NZO,PJ,RF,RH2,SEA,SMJ,WSB | feature_elongation,3_prime_UTR_variant |
| 2 | 88941244 | 88941334 | 90 | Or4c116 | SPRET | feature_elongation,3_prime_UTR_variant |
| 2 | 88974252 | 88974437 | 185 | Or4c118 | SPRET | feature_elongation,3_prime_UTR_variant |
| 2 | 89025504 | 89025800 | 296 | Or4c121 | WSB | feature_elongation,5_prime_UTR_variant |
| 2 | 89026363 | 89026545 | 182 | Or4c121 | SPRET | feature_elongation,5_prime_UTR_variant |
| 2 | 89080475 | 89080676 | 201 | Or4c122 | MOLF | feature_elongation,5_prime_UTR_variant |
| 2 | 89191671 | 89191736 | 65 | Or4a15 | SPRET | feature_elongation,3_prime_UTR_variant |
| 2 | 89235377 | 89235507 | 130 | Or4a39 | SPRET | feature_elongation,3_prime_UTR_variant |
| 2 | 89268659 | 89268715 | 56 | Or4a68 | SPRET | feature_elongation,3_prime_UTR_variant |
| 2 | 89435933 | 89436390 | 457 | Or4a74 | SPRET | feature_elongation,3_prime_UTR_variant |
| 2 | 89771112 | 89771574 | 462 | Or4c12 | ILNJ2,WSB | feature_elongation,3_prime_UTR_variant |
| 2 | 89835610 | 89836075 | 465 | Or4c127 | C58J,ILNJ2,KK,SJL,SWR,WSB | feature_elongation,3_prime_UTR_variant |
| 2 | 89838307 | 89838359 | 52 | Or4c127 | SPRET | feature_elongation,3_prime_UTR_variant |
| 2 | 91474547 | 91474603 | 56 | Zfp408 | CAST | feature_elongation,3_prime_UTR_variant |
| 2 | 91545552 | 91545771 | 219 | Harbi1 | SPRET | feature_elongation,3_prime_UTR_variant |
| 2 | 92191726 | 92191794 | 68 | Phf21a | CAST,MOLF | feature_elongation,3_prime_UTR_variant |
| 2 | 103915009 | 103915063 | 54 | Cd59b | SPRET | feature_elongation,3_prime_UTR_variant |
| 2 | 104495673 | 104495773 | 100 | Cstf3 | SMJ | feature_elongation,3_prime_UTR_variant |
| 2 | 105003264 | 105003325 | 61 | Wt1 | PL | feature_elongation,3_prime_UTR_variant |
| 2 | 105503294 | 105503348 | 54 | Pax6 | SPRET | feature_elongation,5_prime_UTR_variant |
| 2 | 111091390 | 111091584 | 194 | Or4f53 | SPRET | feature_elongation,3_prime_UTR_variant |
| 2 | 111150483 | 111150548 | 65 | Or4k36 | MOLF | feature_elongation,3_prime_UTR_variant |
| 2 | 111160723 | 111160877 | 154 | Or4k37 | SPRET | feature_elongation,3_prime_UTR_variant |
| 2 | 111248366 | 111248462 | 96 | Or4k40 | C3H,CE,PL | feature_elongation,3_prime_UTR_variant |
| 2 | 111248951 | 111249019 | 68 | Or4k40 | MOLF | feature_elongation,3_prime_UTR_variant |
| 2 | 111250381 | 111250522 | 141 | Or4k40 | C3H,CE,MOLF,PL,SPRET | feature_elongation,coding_sequence_variant |
| 2 | 111332178 | 111332311 | 133 | Or4f4-ps1 | C3H,CE,PL | feature_elongation,3_prime_UTR_variant |
| 2 | 111447524 | 111447600 | 76 | Or4k47 | C3H,CE,MOLF,PL | feature_elongation,3_prime_UTR_variant |
| 2 | 111612043 | 111612243 | 200 | Or4k52 | CAST | feature_elongation,3_prime_UTR_variant |
| 2 | 111836264 | 111836316 | 52 | Or4f6 | 129S1,AKR,BALB,BUB,C57L,KK,MA/My,MOLF,MRL,NOD,NOR,NZO,NZW,PJ,SEA,SPRET,SWR,TH | feature_elongation,3_prime_UTR_variant |
| 2 | 111837346 | 111837406 | 60 | Or4f6 | SPRET | feature_elongation,3_prime_UTR_variant |
| 2 | 112093283 | 112094263 | 980 | Nop10 | SPRET | feature_elongation,3_prime_UTR_variant |
| 2 | 112186126 | 112186463 | 337 | Slc12a6 | C3H,CE,MOLF,PL,SPRET | feature_elongation,3_prime_UTR_variant |
| 2 | 113578622 | 113578687 | 65 | Grem1 | C3H,CE,MOLF,PL | feature_elongation,3_prime_UTR_variant |
| 2 | 114348410 | 114348476 | 66 | Dph6 | AKR,C3H,CAST,CBA,CE,FVB,MOLF,MRL,NOD,NOR,SJL,SPRET,SWR | feature_elongation,3_prime_UTR_variant |
| 2 | 118269381 | 118269457 | 76 | Eif2ak4 | SPRET | feature_elongation,coding_sequence_variant |
| 2 | 118712816 | 118712867 | 51 | Ivd | 129S1,AJ,AKR,BALB,BTBR,BUB,C3H,C57L,C58J,CAST,CBA,CE,DBA1J,DBA2J,FVB,ILNJ2,MA/My,NOD,NOR,NZB,NZO,NZW,PJ,PL,RF,RH2,SEA,SJL,SMJ,SPRET,SWR,TH,WSB | feature_elongation,3_prime_UTR_variant |
| 2 | 118966994 | 118967066 | 72 | Rad51 | SPRET | feature_elongation,3_prime_UTR_variant |
| 2 | 119048863 | 119049089 | 226 | Ppp1r14d | AKR | feature_elongation,coding_sequence_variant |
| 2 | 119233190 | 119233653 | 463 | Ino80 | 129S1,AKR,BTBR,BUB,C3H,C57L,C58J,CBA,CE,FVB,NOD,NZB,NZO,SJL,SWR,WSB | feature_elongation,3_prime_UTR_variant |
| 2 | 119481204 | 119481399 | 195 | Nusap1 | 129S1,AJ,AKR,BALB,BTBR,BUB,C3H,C57L,C58J,CAST,CBA,CE,DBA1J,DBA2J,FVB,ILNJ2,KK,MA/My,NOD,NZB,NZO,NZW,PJ,PL,RF,RH2,SEA,SJL,SMJ,SWR,TH,WSB | feature_elongation,3_prime_UTR_variant |
| 2 | 119798755 | 119798976 | 221 | Mga | AJ,BALB,DBA1J,DBA2J,ILNJ2,PJ,PL,RF,RH2,SEA,SMJ,TH | feature_elongation,3_prime_UTR_variant |
| 2 | 119849812 | 119849966 | 154 | Mapkbp1 | 129S1,AJ,AKR,BALB,BTBR,BUB,C3H,C57L,C58J,CBA,CE,DBA1J,DBA2J,FVB,ILNJ2,KK,MA/My,NOD,NZB,NZO,PJ,PL,RF,RH2,SEA,SJL,SMJ,SWR,TH,WSB | feature_elongation,coding_sequence_variant |
| 2 | 119997909 | 119998192 | 283 | Pla2g4e | SPRET | feature_elongation,3_prime_UTR_variant |
| 2 | 120451360 | 120451441 | 81 | Haus2 | SPRET | feature_elongation,3_prime_UTR_variant |
| 2 | 121136770 | 121136830 | 60 | Map1a | 129S1,AJ,AKR,BALB,BTBR,BUB,C3H,C57L,C58J,CAST,CBA,CE,DBA1J,FVB,ILNJ2,KK,MA/My,NZB,NZO,NZW,PJ,PL,RF,RH2,SEA,SJL,SMJ,SWR,TH,WSB | feature_elongation,coding_sequence_variant |
| 2 | 121285306 | 121285362 | 56 | Hypk | MA/My | feature_elongation,5_prime_UTR_variant |
| 2 | 121291564 | 121291773 | 209 | Mfap1b | 129S1,AJ,AKR,BALB,BTBR,BUB,C3H,C57L,C58J,CBA,CE,DBA1J,DBA2J,FVB,ILNJ2,KK,MA/My,NZB,NZO,NZW,PJ,PL,RF,RH2,SEA,SJL,SMJ,SWR,TH,WSB | feature_elongation,3_prime_UTR_variant |
| 2 | 121323404 | 121323468 | 64 | Mfap1a | SPRET | feature_elongation,3_prime_UTR_variant |

| | | | | | | |
|---|---|---|---|---|---|---|
| 2 | 121764678 | 121764814 | 136 | Golm2 | 129S1,BALB,BTBR,BUB,C57L,CE,NZW | feature_elongation,3_prime_UTR_variant |
| 2 | 122095395 | 122095712 | 317 | Sord | AJ,BALB,C3H,C58J,CBA,DBA1J,DBA2J,ILNJ2,NZB,NZO,PJ,PL,RF,RH2,SEA,SMJ,TH | feature_elongation,3_prime_UTR_variant |
| 2 | 122482733 | 122482806 | 73 | AA467197 | SPRET | feature_elongation,3_prime_UTR_variant |
| 2 | 122590418 | 122590823 | 405 | Bloc1s6 | CAST,MA/My | feature_elongation,3_prime_UTR_variant |
| 2 | 126471182 | 126471879 | 697 | Gabpb1 | SMJ | feature_elongation,3_prime_UTR_variant |
| 2 | 126490886 | 126491103 | 217 | Gabpb1 | SPRET | feature_elongation,3_prime_UTR_variant |
| 2 | 127083040 | 127083245 | 205 | Ciao1 | SPRET | feature_elongation,3_prime_UTR_variant |
| 2 | 127420623 | 127420691 | 68 | Zfp661 | SPRET | feature_elongation,coding_sequence_variant |
| 2 | 130259695 | 130259908 | 213 | Pced1a | NOD,WSB | feature_elongation,3_prime_UTR_variant |
| 2 | 130409207 | 130409478 | 271 | Mrps26 | SPRET | feature_elongation,3_prime_UTR_variant |
| 2 | 130418923 | 130418981 | 58 | Oxt | WSB | feature_elongation,3_prime_UTR_variant |
| 2 | 131696881 | 131696956 | 75 | Erv3 | CAST | feature_elongation,3_prime_UTR_variant |
| 2 | 131701592 | 131701883 | 291 | Erv3 | MA/My | feature_elongation,5_prime_UTR_variant |
| 2 | 132091098 | 132091286 | 188 | Pcna | ILNJ2,MOLF,PJ | feature_elongation,3_prime_UTR_variant |
| 2 | 132180245 | 132181106 | 861 | Prokr2 | SPRET | feature_elongation,3_prime_UTR_variant |
| 2 | 132747632 | 132747764 | 132 | Fermt1 | SPRET | feature_elongation,3_prime_UTR_variant |
| 2 | 136923947 | 136924005 | 58 | Jag1 | SPRET | feature_elongation,3_prime_UTR_variant |
| 2 | 142459720 | 142459818 | 98 | Kif16b | SPRET | feature_elongation,3_prime_UTR_variant |
| 2 | 146085071 | 146085135 | 64 | Ralgapa2 | SPRET | feature_elongation,3_prime_UTR_variant |
| 2 | 150034492 | 150034744 | 252 | Zfp1004 | CAST | feature_elongation,coding_sequence_variant |
| 2 | 150110249 | 150110417 | 168 | Zfp1005 | CAST,MOLF,NOD,NZB,NZW,RF | feature_elongation,coding_sequence_variant |
| 2 | 152236730 | 152236854 | 124 | Sox12 | AJ,C58J,MOLF,NOD,NZB,NZW,RF | feature_elongation,3_prime_UTR_variant |
| 2 | 154359700 | 154360117 | 417 | Cbfa2t2 | FVB,SWR | feature_elongation,coding_sequence_variant |
| 2 | 154429139 | 154429342 | 203 | Pxmp4 | CAST | feature_elongation,3_prime_UTR_variant |
| 2 | 155876941 | 155877221 | 280 | Fer1l4 | SPRET | stop_gained,frameshift_variant |
| 2 | 155983989 | 155984077 | 88 | Nfs1 | WSB | feature_elongation,coding_sequence_variant |
| 2 | 156940944 | 156941020 | 76 | Samhd1 | CE | feature_elongation,3_prime_UTR_variant |
| 2 | 158225523 | 158225652 | 129 | Snhg11 | DBA1J,DBA2J,ILNJ2,SMJ | feature_elongation,3_prime_UTR_variant |
| 2 | 164032497 | 164032704 | 207 | Wfdc12 | KK | feature_elongation,5_prime_UTR_variant |
| 2 | 164455495 | 164455796 | 301 | Wfdc6b,Wfdc8 | SPRET | feature_elongation,5_prime_UTR_variant |
| 2 | 164677635 | 164677723 | 88 | Ctsa | SPRET | feature_elongation,coding_sequence_variant |
| 2 | 165133922 | 165134114 | 192 | Elmo2 | CAST | feature_elongation,coding_sequence_variant |
| 2 | 165218530 | 165218584 | 54 | Zfp334 | CAST | feature_elongation,3_prime_UTR_variant |
| 2 | 165626756 | 165626838 | 82 | Zmynd8 | AJ,AKR,BALB,BTBR,BUB,C3H,C58J,CAST,CBA,CE,DBA1J,DBA2J,FVB,ILNJ2,KK,MA/My,NOD,NOR,NZB,NZO,PJ,PL,RF,SEA,SJL,SMJ,SWR,TH,WSB | feature_elongation,3_prime_UTR_variant |
| 2 | 166635482 | 166635546 | 64 | Trp53rkb | CAST | feature_elongation,5_prime_UTR_variant |
| 2 | 167820390 | 167820586 | 196 | Ptpn1 | 129S1,C57L,NZB | feature_elongation,3_prime_UTR_variant |
| 2 | 168321826 | 168321944 | 118 | Nfatc2 | SPRET | feature_elongation,3_prime_UTR_variant |
| 2 | 168349249 | 168349322 | 73 | Nfatc2 | CAST,SPRET | feature_elongation,coding_sequence_variant |
| 2 | 168590700 | 168590828 | 128 | Sall4 | B10D2 | feature_elongation,3_prime_UTR_variant |
| 2 | 170721356 | 170721424 | 68 | Dok5 | CAST | feature_elongation,3_prime_UTR_variant |
| 2 | 172204807 | 172204909 | 102 | Aurka | FVB,SJL | feature_elongation,coding_sequence_variant |
| 2 | 175037635 | 175037719 | 84 | Gm14391 | BUB,CE,FVB,NZB,SJL,SWR | feature_elongation,coding_sequence_variant |
| 2 | 175672079 | 175672163 | 84 | Gm6710 | BUB,CAST,CE,FVB,KK,MOLF,NOD,NOR,NZB,PJ,SJL,SWR,WSB | feature_elongation,coding_sequence_variant |
| 2 | 175772093 | 175772301 | 208 | Gm2026 | CE,FVB,MOLF,NZB,SJL,SWR | feature_elongation,3_prime_UTR_variant |
| 2 | 176412779 | 176412946 | 167 | Gm14305 | CAST,WSB | frameshift_variant |
| 2 | 181000451 | 181000550 | 99 | Arfrp1 | SPRET | feature_elongation,3_prime_UTR_variant |
| 3 | 7638112 | 7638192 | 80 | Il7 | SPRET | feature_elongation,3_prime_UTR_variant |
| 3 | 8991916 | 8992012 | 96 | Tpd52 | MOLF,SPRET | feature_elongation,3_prime_UTR_variant |
| 3 | 8993538 | 8993594 | 56 | Tpd52 | MOLF | feature_elongation,3_prime_UTR_variant |
| 3 | 9756624 | 9757356 | 732 | Pag1 | SPRET | feature_elongation,3_prime_UTR_variant |
| 3 | 10377772 | 10378270 | 498 | Impa1 | SMJ | feature_elongation,3_prime_UTR_variant |
| 3 | 10378403 | 10378474 | 71 | Impa1 | BUB,CAST,CE,DBA2J,SMJ,SPRET | feature_elongation,3_prime_UTR_variant |
| 3 | 10455705 | 10455768 | 63 | Chmp4c | BUB,CAST,CE,DBA2J,MOLF,SPRET | feature_elongation,3_prime_UTR_variant |
| 3 | 19186037 | 19186101 | 64 | Armc1 | AJ,C3H,C58J,CBA,CE,DBA2J,LP,MRL,NZB,NZO,NZW,PL,SPRET | feature_elongation,3_prime_UTR_variant |
| 3 | 19584884 | 19585312 | 428 | Dnajc5b | CAST,MOLF | feature_elongation,5_prime_UTR_variant |
| 3 | 19746585 | 19746649 | 64 | Trim55 | MOLF,SPRET | feature_elongation,3_prime_UTR_variant |
| 3 | 20303251 | 20303516 | 265 | Cpb1 | SPRET | feature_elongation,3_prime_UTR_variant |
| 3 | 23854063 | 23854231 | 168 | Naaladl2 | CAST,MOLF,SPRET,WSB | feature_elongation,3_prime_UTR_variant |
| 3 | 23856230 | 23856426 | 196 | Naaladl2 | CAST,MOLF,WSB | feature_elongation,3_prime_UTR_variant |
| 3 | 25484846 | 25485038 | 192 | Nlgn1 | MOLF | feature_elongation,3_prime_UTR_variant |
| 3 | 27393036 | 27393444 | 408 | Tnfsf10 | MOLF,SPRET | feature_elongation,3_prime_UTR_variant |
| 3 | 30951105 | 30951221 | 116 | Gpr160 | SPRET | feature_elongation,3_prime_UTR_variant |
| 3 | 32394572 | 32394911 | 339 | Zmat3 | SJL,SPRET | feature_elongation,3_prime_UTR_variant |
| 3 | 32520798 | 32520880 | 82 | Pik3ca | CAST,CE | feature_elongation,3_prime_UTR_variant |
| 3 | 32781320 | 32781642 | 322 | Mrpl47 | SPRET | feature_elongation,3_prime_UTR_variant |
| 3 | 32991174 | 32991602 | 428 | Usp13 | 129S1,AJ,BTBR,BUB,C3H,C57L,CBA,FVB,MRL,NOD,NOR,NZB,NZO,NZW,PJ,SEA,SWR | feature_elongation,3_prime_UTR_variant |
| 3 | 33865114 | 33865494 | 380 | Ttc14 | MOLF | feature_elongation,3_prime_UTR_variant |
| 3 | 34111197 | 34111839 | 642 | Dnajc19 | CAST | feature_elongation,3_prime_UTR_variant |
| 3 | 36145884 | 36146200 | 316 | Acad9 | BUB,FVB,MOLF | feature_elongation,3_prime_UTR_variant |
| 3 | 36619691 | 36619745 | 54 | Exosc9,Ccna2 | CAST | feature_elongation,3_prime_UTR_variant |
| 3 | 40878112 | 40878428 | 316 | Abhd18 | MOLF | feature_elongation,3_prime_UTR_variant |
| 3 | 51403454 | 51403590 | 136 | Rab33b | CAST,MOLF,NZW | feature_elongation,3_prime_UTR_variant |
| 3 | 53356740 | 53357219 | 479 | Nhlrc3 | SPRET | feature_elongation,3_prime_UTR_variant |
| 3 | 54635070 | 54635124 | 54 | Supt20 | SPRET | feature_elongation,coding_sequence_variant |
| 3 | 58752558 | 58752634 | 76 | Clrn1 | SPRET | feature_elongation,3_prime_UTR_variant |
| 3 | 62512450 | 62512507 | 57 | Gpr149 | MOLF | feature_elongation,5_prime_UTR_variant |
| 3 | 64048477 | 64049317 | 840 | Vmn2r2 | SPRET | feature_elongation,5_prime_UTR_variant |
| 3 | 67386628 | 67386756 | 128 | Rarres1 | AKR,BALB,BTBR,CAST,CBA,CE,DBA1J,DBA2J,FVB,MOLF,NOD,NOR,NZO,RF,SJL,SPRET,SWR,TH,WSB | feature_elongation,3_prime_UTR_variant |
| 3 | 68951947 | 68952001 | 54 | Trim59 | 129S1,BTBR,C57L,CAST,CE,FVB,MOLF,SJL,SWR,TH | feature_elongation,5_prime_UTR_variant |
| 3 | 69658344 | 69658491 | 147 | Nmd3 | AJ,AKR,BUB,C3H,C58J,CBA,DBA1J,DBA2J,KK,LP,MRL,NOD,NOR,NZO,NZW,RF,SPRET | feature_elongation,3_prime_UTR_variant |
| 3 | 75549913 | 75549977 | 64 | Serpini1 | SPRET | feature_elongation,3_prime_UTR_variant |

| | | | | | | |
|---|---|---|---|---|---|---|
| 3 | 79550684 | 79550878 | 194 | Rxfp1 | CAST,MOLF | feature_elongation,3_prime_UTR_variant |
| 3 | 79551698 | 79551764 | 66 | Rxfp1 | CAST,MOLF,SPRET | feature_elongation,3_prime_UTR_variant |
| 3 | 80589804 | 80589864 | 60 | Gria2 | CAST,SPRET | feature_elongation,3_prime_UTR_variant |
| 3 | 81864031 | 81864101 | 70 | Ctso | CAST,TH | feature_elongation,3_prime_UTR_variant |
| 3 | 82802722 | 82802773 | 51 | Lrat | SPRET | feature_elongation,3_prime_UTR_variant |
| 3 | 84081950 | 84082148 | 198 | Gm6525 | SPRET | feature_elongation,5_prime_UTR_variant |
| 3 | 88020123 | 88020353 | 230 | Iqgap3 | NZB,NZO | feature_elongation,coding_sequence_variant |
| 3 | 89655278 | 89655332 | 54 | Chrnb2 | CAST,KK,MOLF,SPRET | feature_elongation,coding_sequence_variant |
| 3 | 89773373 | 89773562 | 189 | Il6ra | KK,MOLF | feature_elongation,3_prime_UTR_variant |
| 3 | 92273313 | 92273502 | 189 | Sprr2f | SPRET | feature_elongation,coding_sequence_variant |
| 3 | 92479165 | 92479597 | 432 | Ivl | CAST,MOLF,NOD,NOR,RF,SPRET | feature_elongation,coding_sequence_variant |
| 3 | 92593092 | 92593152 | 60 | Lce1d | MOLF | feature_elongation,coding_sequence_variant |
| 3 | 92657469 | 92657667 | 198 | Lce1g | SPRET | feature_elongation,3_prime_UTR_variant |
| 3 | 92731645 | 92731729 | 84 | Kprp | SPRET | feature_elongation,coding_sequence_variant |
| 3 | 92865688 | 92866623 | 935 | Lce3d | SPRET | feature_elongation,3_prime_UTR_variant |
| 3 | 93110593 | 93110671 | 78 | Flg2 | 129S1,AJ,BTBR,C3H,C57L,CAST,CBA,CE,ILNJ2,MOLF,PJ,PL,SEA,TH,WSB | feature_elongation,coding_sequence_variant |
| 3 | 93304211 | 93304319 | 108 | Rptn | BALB,ILNJ2,KK,NOD,NOR,NZB,NZO,PJ,PL,RH2,SEA,SPRET,TH,WSB | feature_elongation,coding_sequence_variant |
| 3 | 94311307 | 94311367 | 60 | Lingo4 | MOLF | feature_elongation,3_prime_UTR_variant |
| 3 | 94340312 | 94340421 | 109 | Tdrkh | SPRET | feature_elongation,3_prime_UTR_variant |
| 3 | 95110706 | 95110805 | 99 | Gabpb2 | SPRET | feature_elongation,3_prime_UTR_variant |
| 3 | 95670833 | 95670917 | 84 | Rprd2 | SPRET | feature_elongation,3_prime_UTR_variant |
| 3 | 96541096 | 96541303 | 207 | Rbm8a | 129S1,AJ,BALB,C3H,C57L,CAST,CBA,CE,FVB,MA/My,NOD,NOR,NZB,NZO,NZW,PJ,RH2,SEA,SJL,SMJ,SWR,WSB | feature_elongation,3_prime_UTR_variant |
| 3 | 97560524 | 97560680 | 156 | Fmo5 | SPRET | feature_elongation,3_prime_UTR_variant |
| 3 | 97561481 | 97561705 | 224 | Fmo5 | CAST,NOD,NOR | feature_elongation,3_prime_UTR_variant |
| 3 | 98056712 | 98056768 | 56 | Notch2 | 129S1,C57L,CAST,FVB,MOLF,PJ,RF,SEA,SJL,SPRET,SWR,WSB | feature_elongation,3_prime_UTR_variant |
| 3 | 98071382 | 98071435 | 53 | Adam30 | SPRET | feature_elongation,3_prime_UTR_variant |
| 3 | 101721087 | 101721139 | 52 | Mab21l3 | CAST,MA/My,MOLF | feature_elongation,3_prime_UTR_variant |
| 3 | 101765094 | 101765258 | 164 | Slc22a15 | MOLF,SPRET | feature_elongation,3_prime_UTR_variant |
| 3 | 103640211 | 103640267 | 56 | Olfml3 | SPRET | feature_elongation,3_prime_UTR_variant |
| 3 | 103783078 | 103783132 | 54 | Ptpn22 | AJ,AKR,C3H,CBA,FVB,ILNJ2,RF,SMJ | feature_elongation,coding_sequence_variant |
| 3 | 105626786 | 105627088 | 302 | Inka2 | MA/My,MOLF,NZB,NZO,NZW,SPRET | feature_elongation,3_prime_UTR_variant |
| 3 | 105893744 | 105893891 | 147 | Ovgp1 | AJ,C3H,CBA,SPRET | feature_elongation,coding_sequence_variant |
| 3 | 107015182 | 107015256 | 74 | Kcna2 | MOLF | feature_elongation,3_prime_UTR_variant |
| 3 | 107017328 | 107017430 | 102 | Kcna2 | AJ,AKR,BALB,C3H,CAST,CBA,DBA1J,DBA2J,ILNJ2,MA/My,MRL,NOD,NOR,NZB,NZO,NZW,PL,RF,RH2,TH,WSB | feature_elongation,3_prime_UTR_variant |
| 3 | 108097271 | 108097331 | 60 | Amigo1 | SPRET | feature_elongation,3_prime_UTR_variant |
| 3 | 108464895 | 108465292 | 397 | Tmem167b | BALB,MRL,PJ,RH2,SEA | feature_elongation,3_prime_UTR_variant |
| 3 | 108490235 | 108490437 | 202 | Taf13 | CAST | feature_elongation,3_prime_UTR_variant |
| 3 | 110156115 | 110157084 | 969 | Prmt6 | MA/My | feature_elongation,3_prime_UTR_variant |
| 3 | 113155555 | 113155945 | 390 | Amy2a5 | SPRET | feature_elongation,3_prime_UTR_variant |
| 3 | 115732931 | 115733204 | 273 | Slc30a7 | SPRET | feature_elongation,3_prime_UTR_variant |
| 3 | 116343070 | 116343130 | 60 | Dbt | MOLF | feature_elongation,3_prime_UTR_variant |
| 3 | 116466633 | 116466687 | 54 | Slc35a3 | MOLF,SPRET | feature_elongation,3_prime_UTR_variant |
| 3 | 116536621 | 116536672 | 51 | Agl | SPRET | feature_elongation,3_prime_UTR_variant |
| 3 | 116777958 | 116778280 | 322 | 4930455H04Rik | SPRET | feature_elongation,3_prime_UTR_variant |
| 3 | 117154446 | 117154548 | 102 | Plppr4 | MOLF | feature_elongation,5_prime_UTR_variant |
| 3 | 121156371 | 121156501 | 130 | Alg14 | SPRET | feature_elongation,3_prime_UTR_variant |
| 3 | 122719386 | 122719476 | 90 | 1810037I17Rik | 129S1,BTBR,C57L,CAST,CE,DBA1J,DBA2J,KK,MA/My,MOLF,NZB,PJ,RF,SEA,SMJ,SPRET,TH | feature_elongation,3_prime_UTR_variant |
| 3 | 122749802 | 122750779 | 977 | Usp53 | 129S1,BTBR,C57L,CAST,CE,DBA1J,DBA2J,KK,MA/My,MOLF,NZB,PJ,RF,SEA,SMJ,SPRET,TH | feature_elongation,3_prime_UTR_variant |
| 3 | 122903772 | 122903848 | 76 | Synpo2 | MOLF | feature_elongation,3_prime_UTR_variant |
| 3 | 123165690 | 123166034 | 344 | Mettl14 | AJ,BALB,C3H,CBA,DBA1J,DBA2J,MOLF,NZB,RH2,SMJ,SPRET | feature_elongation,3_prime_UTR_variant |
| 3 | 127601624 | 127601682 | 58 | Ap1ar | SPRET | feature_elongation,3_prime_UTR_variant |
| 3 | 129582455 | 129582518 | 63 | Lrit3 | SPRET | feature_elongation,coding_sequence_variant |
| 3 | 131017292 | 131017359 | 67 | Lef1 | CAST | feature_elongation,3_prime_UTR_variant |
| 3 | 134638689 | 134638841 | 152 | Tacr3 | SPRET | feature_elongation,3_prime_UTR_variant |
| 3 | 135531445 | 135531507 | 62 | Slc39a8 | CAST,MOLF | feature_elongation,5_prime_UTR_variant |
| 3 | 137615461 | 137615513 | 52 | Dnajb14 | AJ,AKR,BALB,C3H,CAST,CBA,CE,DBA1J,FVB,ILNJ2,KK,MA/My,MOLF,MRL,NOD,NOR,PJ,RH2,SEA,SJL,SWR,TH | feature_elongation,3_prime_UTR_variant |
| 3 | 137864328 | 137864426 | 98 | Trmt10a | SPRET | feature_elongation,3_prime_UTR_variant |
| 3 | 138035995 | 138036166 | 171 | Adh6a | 129S1,AJ,AKR,BALB,BTBR,BUB,C3H,C57L,C58J,CAST,CBA,CE,DBA1J,DBA2J,FVB,ILNJ2,KK,LP,MA/My,MOLF,MRL,NOD,NOR,NZB,NZO,NZW,PJ,PL,RF,RH2,SEA,SJL,SMJ,SPRET,SWR,TH,WSB | feature_elongation,3_prime_UTR_variant |
| 3 | 138610731 | 138610947 | 216 | Tspan5 | AKR,FVB | feature_elongation,3_prime_UTR_variant |
| 3 | 142210254 | 142211031 | 777 | Gbp5 | CBA,DBA1J,MOLF,MRL,NZW | feature_elongation,coding_sequence_variant |
| 3 | 142228031 | 142228289 | 258 | Gbp5 | CAST,SPRET | feature_elongation,3_prime_UTR_variant |
| 3 | 142266031 | 142266424 | 393 | Gbp3 | WSB | feature_elongation,5_prime_UTR_variant |
| 3 | 144135490 | 144135677 | 187 | Hs2st1 | SPRET | feature_elongation,3_prime_UTR_variant |
| 3 | 146123385 | 146123437 | 52 | Ssx2ip | CAST,MOLF,SPRET | feature_elongation,5_prime_UTR_variant |
| 3 | 146170415 | 146170478 | 63 | Ctbs | SPRET | feature_elongation,3_prime_UTR_variant |
| 3 | 146351619 | 146351671 | 52 | 4930503B20Rik,Samd13 | SPRET | feature_elongation,3_prime_UTR_variant |
| 3 | 151941632 | 151941708 | 76 | Fubp1 | CAST,SPRET,WSB | feature_elongation,3_prime_UTR_variant |
| 3 | 152038258 | 152038324 | 66 | Miga1 | 129S1,BTBR,C57L,CBA,DBA1J,SMJ | feature_elongation,3_prime_UTR_variant |
| 3 | 154283843 | 154284435 | 592 | Tyw3 | CAST,RF | feature_elongation,3_prime_UTR_variant |
| 3 | 154284099 | 154284839 | 740 | Tyw3 | CE | feature_elongation,3_prime_UTR_variant |
| 3 | 154468181 | 154468247 | 66 | Erich3 | SPRET | feature_elongation,coding_sequence_variant |
| 4 | 8144991 | 8145044 | 53 | Car8 | SPRET | feature_elongation,3_prime_UTR_variant |
| 4 | 10989522 | 10989606 | 84 | Plekhf2 | CAST | feature_elongation,3_prime_UTR_variant |
| 4 | 11171049 | 11171101 | 52 | Trp53inp1 | CAST | feature_elongation,3_prime_UTR_variant |

| | | | | | | |
|---|---|---|---|---|---|---|
| 4 | 11584491 | 11584557 | 66 | Rad54b,Fsbp | SPRET | feature_elongation,3_prime_UTR_variant |
| 4 | 11810464 | 11810534 | 70 | Cdh17 | SPRET | feature_elongation,coding_sequence_variant |
| 4 | 25618531 | 25618586 | 55 | Fut9 | SPRET | feature_elongation,3_prime_UTR_variant |
| 4 | 33232541 | 33232603 | 62 | Srsf12 | SPRET | feature_elongation,3_prime_UTR_variant |
| 4 | 41033324 | 41033382 | 58 | Aqp7 | CAST | feature_elongation,3_prime_UTR_variant |
| 4 | 41389867 | 41390069 | 202 | Ubap1 | WSB | feature_elongation,3_prime_UTR_variant |
| 4 | 41633372 | 41633575 | 203 | Dnai1 | SPRET | feature_elongation,5_prime_UTR_variant |
| 4 | 43513460 | 43513556 | 96 | Car9 | CAST | feature_elongation,3_prime_UTR_variant |
| 4 | 43832556 | 43833528 | 972 | Or13c7c | 129S1,BUB,PJ,PL,SEA,SJL | feature_elongation,3_prime_UTR_variant |
| 4 | 43856121 | 43856173 | 52 | Or13c7 | AJ,AKR,BALB,C3H,CE,KK,MA/My,MRL,NOD,NOR,NZB,NZO,PL,RF,RH2,WSB | feature_elongation,3_prime_UTR_variant |
| 4 | 44035984 | 44036100 | 116 | Gne | CAST,MOLF | feature_elongation,3_prime_UTR_variant |
| 4 | 44036487 | 44036706 | 219 | Gne | WSB | feature_elongation,3_prime_UTR_variant |
| 4 | 45397739 | 45397855 | 116 | Slc25a51 | WSB | feature_elongation,3_prime_UTR_variant |
| 4 | 46633094 | 46633303 | 209 | Tbc1d2 | 129S1,AJ,BALB,C3H,C57L,MOLF,PJ,RH2,SEA,SJL,WSB | feature_elongation,3_prime_UTR_variant |
| 4 | 49059422 | 49059482 | 60 | Plppr1 | MOLF | feature_elongation,5_prime_UTR_variant |
| 4 | 51228991 | 51229072 | 81 | Cylc2 | AKR,CAST,CBA,CE,DBA1J,DBA2J,MOLF,MRL,RF,SMJ,SPRET | feature_elongation,coding_sequence_variant |
| 4 | 52853755 | 52853820 | 65 | Or13c3 | CAST,MA/My,MOLF | feature_elongation,3_prime_UTR_variant |
| 4 | 53703738 | 53703910 | 172 | Fsd1l | CAST | feature_elongation,3_prime_UTR_variant |
| 4 | 56877599 | 56877755 | 156 | Tmem245 | CAST | feature_elongation,3_prime_UTR_variant |
| 4 | 56879115 | 56879166 | 51 | Tmem245 | SPRET | feature_elongation,3_prime_UTR_variant |
| 4 | 58783517 | 58784502 | 985 | Or2k2 | SPRET | feature_elongation,3_prime_UTR_variant |
| 4 | 59757794 | 59757902 | 108 | E130308A19Rik | CE | feature_elongation,3_prime_UTR_variant |
| 4 | 62607859 | 62607913 | 54 | Rgs3 | CAST,CE,MOLF | feature_elongation,coding_sequence_variant |
| 4 | 65274381 | 65274457 | 76 | Pappa | CAST,MOLF,PJ | feature_elongation,3_prime_UTR_variant |
| 4 | 70137888 | 70138669 | 781 | Cdk5rap2 | MOLF | feature_elongation,3_prime_UTR_variant |
| 4 | 70138495 | 70138616 | 121 | Cdk5rap2 | MOLF | feature_elongation,3_prime_UTR_variant |
| 4 | 70139317 | 70139480 | 163 | Cdk5rap2 | 129S1,C57L,CAST,ILNJ2,KK,PJ | feature_elongation,3_prime_UTR_variant |
| 4 | 73635246 | 73635424 | 178 | Rasef | SPRET | feature_elongation,3_prime_UTR_variant |
| 4 | 83139996 | 83140050 | 54 | Ttc39b | MOLF,SPRET | feature_elongation,3_prime_UTR_variant |
| 4 | 83141892 | 83142047 | 155 | Ttc39b | MOLF | feature_elongation,3_prime_UTR_variant |
| 4 | 85306687 | 85306741 | 54 | Sh3gl2 | SPRET | feature_elongation,3_prime_UTR_variant |
| 4 | 86902280 | 86902345 | 65 | Slc24a2 | CE | feature_elongation,3_prime_UTR_variant |
| 4 | 88745704 | 88745764 | 60 | Ifna6 | CE | feature_elongation,coding_sequence_variant |
| 4 | 89192816 | 89192872 | 56 | Cdkn2a | MA/My | feature_elongation,3_prime_UTR_variant |
| 4 | 90113937 | 90113989 | 52 | Zfp352 | CAST | feature_elongation,3_prime_UTR_variant |
| 4 | 94525409 | 94525592 | 183 | Lrrc19 | CAST,MOLF,SPRET | feature_elongation,3_prime_UTR_variant |
| 4 | 99147435 | 99147529 | 94 | Atg4c | CAST,MOLF | feature_elongation,3_prime_UTR_variant |
| 4 | 99147955 | 99148173 | 218 | Atg4c | 129S1,AJ,AKR,BALB,BTBR,BUB,C3H,C57L,CBA,CE,FVB,KK,MA/My,MRL,NOD,NOR,NZB,PJ,PL,RF,RH2,SEA,SJL,SMJ,SWR,TH,WSB | feature_elongation,3_prime_UTR_variant |
| 4 | 100860939 | 100861002 | 63 | Cachd1 | SPRET | feature_elongation,3_prime_UTR_variant |
| 4 | 101010497 | 101010565 | 68 | Jak1 | SPRET | feature_elongation,3_prime_UTR_variant |
| 4 | 101747714 | 101747805 | 91 | C130073F10Rik | NZB,NZO | feature_elongation,coding_sequence_variant |
| 4 | 102895672 | 102895754 | 82 | Dnai4 | CAST | feature_elongation,3_prime_UTR_variant |
| 4 | 103030463 | 103030515 | 52 | Slc35d1 | CE,MA/My,NZW,SJL | feature_elongation,3_prime_UTR_variant |
| 4 | 107752922 | 107753091 | 169 | Czib | SPRET | feature_elongation,3_prime_UTR_variant |
| 4 | 109280760 | 109280812 | 52 | Ttc39a | SPRET | feature_elongation,3_prime_UTR_variant |
| 4 | 115669326 | 115669560 | 234 | Atpaf1 | AJ,CE,DBA1J,DBA2J | feature_elongation,3_prime_UTR_variant |
| 4 | 115774095 | 115774213 | 118 | Dmbx1 | SPRET | feature_elongation,3_prime_UTR_variant |
| 4 | 115774672 | 115774726 | 54 | Dmbx1 | MOLF | feature_elongation,3_prime_UTR_variant |
| 4 | 115990157 | 115990361 | 204 | Lurap1 | B10D2,B10J,C58J,KK,PL,TH | feature_elongation,3_prime_UTR_variant |
| 4 | 117746600 | 117746657 | 57 | Dph2 | ILNJ2,SPRET | feature_elongation,3_prime_UTR_variant |
| 4 | 119117520 | 119117735 | 215 | Cldn19 | NZO,NZW,WSB | feature_elongation,3_prime_UTR_variant |
| 4 | 119250231 | 119250335 | 104 | Ccdc30 | SPRET | feature_elongation,3_prime_UTR_variant |
| 4 | 119993915 | 119994011 | 96 | Hivep3 | MOLF | feature_elongation,3_prime_UTR_variant |
| 4 | 120804582 | 120804777 | 195 | Zfp69 | ILNJ2,MOLF | feature_elongation,coding_sequence_variant |
| 4 | 122845118 | 122845178 | 60 | Mfsd2a | MOLF,SPRET | feature_elongation,coding_sequence_variant |
| 4 | 123019762 | 123019829 | 67 | Bmp8b | CAST | feature_elongation,3_prime_UTR_variant |
| 4 | 124572355 | 124572413 | 58 | Utp11 | CAST | feature_elongation,3_prime_UTR_variant |
| 4 | 124836904 | 124837109 | 205 | Airim | BUB,FVB,NOD,SJL,SWR | feature_elongation,3_prime_UTR_variant |
| 4 | 125006469 | 125006526 | 57 | Meaf6 | AJ | feature_elongation,3_prime_UTR_variant |
| 4 | 125983342 | 125983396 | 54 | Oscp1 | CAST | feature_elongation,3_prime_UTR_variant |
| 4 | 125983928 | 125984137 | 209 | Oscp1 | SPRET | feature_elongation,3_prime_UTR_variant |
| 4 | 126229814 | 126229906 | 92 | Ago3 | B10D2,B10J,MOLF,NZW | feature_elongation,3_prime_UTR_variant |
| 4 | 126231620 | 126232012 | 392 | Ago3 | AJ,B10D2,B10J,MOLF,NZW,RH2 | feature_elongation,3_prime_UTR_variant |
| 4 | 126997431 | 126997544 | 113 | Zmym6 | SPRET | feature_elongation,coding_sequence_variant |
| 4 | 128662818 | 128663021 | 203 | A3galt2 | B10D2,B10J,DBA1J,KK,MOLF,NOR,PJ,SEA,SMJ | feature_elongation,3_prime_UTR_variant |
| 4 | 129501477 | 129501694 | 217 | Tmem234 | SPRET | feature_elongation,3_prime_UTR_variant |
| 4 | 130662744 | 130662852 | 108 | Laptm5 | 129S1,B10D2,B10J,BTBR,BUB,C57L,DBA1J,FVB,ILNJ2,KK,MA/My,MOLF,NOR,NZB,PJ,PL,SEA,SJL,SMJ,SWR,TH,WSB | feature_elongation,3_prime_UTR_variant |
| 4 | 130663308 | 130663378 | 70 | Laptm5 | SPRET | feature_elongation,3_prime_UTR_variant |
| 4 | 131949085 | 131949238 | 153 | Gmeb1 | SPRET | feature_elongation,3_prime_UTR_variant |
| 4 | 131950649 | 131950705 | 56 | Gmeb1 | DBA1J,KK,MOLF,NZB,SMJ | feature_elongation,3_prime_UTR_variant |
| 4 | 131951379 | 131951524 | 145 | Gmeb1 | SPRET | feature_elongation,3_prime_UTR_variant |
| 4 | 132083758 | 132083854 | 96 | Phactr4 | DBA1J,KK,MOLF,SMJ | feature_elongation,3_prime_UTR_variant |
| 4 | 132452249 | 132452447 | 198 | Xkr8 | 129S1,AJ,AKR,BALB,BTBR,C3H,C57L,CBA,CE,DBA2J,ILNJ2,MRL,NOD,NZO,NZW,RF,RH2,SJL,SWR,TH | feature_elongation,3_prime_UTR_variant |
| 4 | 132729192 | 132729370 | 178 | Fgr | DBA1J,KK,NZB,NZO,SMJ | feature_elongation,3_prime_UTR_variant |
| 4 | 133389205 | 133389409 | 204 | Pigv | FVB | feature_elongation,3_prime_UTR_variant |
| 4 | 133692302 | 133692481 | 179 | Hmgn2 | 129S1,BALB,BTBR,C57L,DBA1J,DBA2J,ILNJ2,NOD,PL,SMJ | feature_elongation,3_prime_UTR_variant |
| 4 | 134080419 | 134080623 | 204 | Slc30a2 | SPRET | feature_elongation,3_prime_UTR_variant |
| 4 | 134290526 | 134291070 | 544 | Man1c1 | SPRET | feature_elongation,3_prime_UTR_variant |
| 4 | 136019007 | 136019077 | 70 | Zfp46 | 129S1,B10D2,B10J,C57L,DBA1J,KK,MOLF,NZO,PJ,SEA | feature_elongation,3_prime_UTR_variant |
| 4 | 136076419 | 136076473 | 54 | Hnrnpr | 129S1,B10D2,B10J,C57L,DBA1J,KK,MOLF,NZO,PJ,SEA | feature_elongation,3_prime_UTR_variant |

| | | | | | | |
|---|---|---|---|---|---|---|
| 4 | 136081060 | 136081119 | 59 | Hnrnpr | 129S1,B10D2,B10J,C57L,DBA1J,KK,MOLF,NZO,PJ,SEA | feature_elongation,3_prime_UTR_variant |
| 4 | 136086167 | 136086640 | 473 | Hnrnpr | 129S1,B10D2,B10J,C57L,DBA1J,KK,MOLF,NZO,PJ,SEA | feature_elongation,3_prime_UTR_variant |
| 4 | 136365877 | 136366085 | 208 | Lactbl1 | AKR,BUB,C3H,CBA,CE,ILNJ2,MRL,NZB,NZW,RF,WSB | feature_elongation,3_prime_UTR_variant |
| 4 | 136381170 | 136381266 | 96 | Ephb2 | SPRET | feature_elongation,3_prime_UTR_variant |
| 4 | 137408315 | 137408385 | 70 | Rap1gap | AJ,AKR,BALB,BUB,C3H,CBA,CE,DBA2J,FVB,KK,MRL,NZB,NZW,PL,RF,RH2,SJL,SWR,TH,WSB | feature_elongation,5_prime_UTR_variant |
| 4 | 138016587 | 138016644 | 57 | Kif17 | CAST,SPRET | feature_elongation,coding_sequence_variant |
| 4 | 138043683 | 138043866 | 183 | Pink1 | BALB,DBA2J,FVB,KK,SJL,SWR | feature_elongation,5_prime_UTR_variant |
| 4 | 138184295 | 138184349 | 54 | Camk2n1 | BALB,BUB,C3H,CBA,CE,DBA1J,DBA2J,ILNJ2,KK,MA/My,MRL,NZB,NZW,SJL,SMJ,SPRET,TH | feature_elongation,3_prime_UTR_variant |
| 4 | 139465648 | 139465704 | 56 | Pax7 | AKR,BALB,BUB,C3H,CBA,DBA1J,DBA2J,MRL,NOR,PL,RF,SMJ | feature_elongation,3_prime_UTR_variant |
| 4 | 141312026 | 141312093 | 67 | Fblim1 | CAST | feature_elongation,coding_sequence_variant |
| 4 | 141346603 | 141346655 | 52 | Slc25a34 | CAST | feature_elongation,3_prime_UTR_variant |
| 4 | 145428989 | 145429198 | 209 | Zfp980 | SWR,WSB | feature_elongation,coding_sequence_variant |
| 4 | 146061338 | 146061422 | 84 | Zfp987 | 129S1,BTBR,C57L,C58J,LP,PJ,SEA | feature_elongation,coding_sequence_variant |
| 4 | 147759389 | 147759556 | 167 | Zfp534 | AKR,BUB,C3H,CAST,CBA,CE,DBA1J,KK,MA/My,MRL,NOD,NZB,NZO,NZW,PL,RF,SJL,SMJ,WSB | feature_elongation,coding_sequence_variant |
| 4 | 147839024 | 147839150 | 126 | Zfp984 | DBA2J,SPRET,TH | feature_elongation,3_prime_UTR_variant |
| 4 | 147958725 | 147958789 | 64 | Mfn2 | 129S1,AKR,BTBR,BUB,C3H,C57L,C58J,CBA,CE,DBA1J,DBA2J,KK,LP,MA/My,MRL,NOD,NZB,NZO,NZW,PJ,PL,RF,SEA,SJL,SMJ,SWR,TH,WSB | feature_elongation,3_prime_UTR_variant |
| 4 | 148164115 | 148164340 | 225 | Agtrap | DBA2J | feature_elongation,3_prime_UTR_variant |
| 4 | 148184523 | 148184746 | 223 | Draxin | BUB,C58J,KK,LP,MA/My,NOD,NZB,NZW,PJ,RF,SEA,SJL,SMJ,SWR,WSB | feature_elongation,3_prime_UTR_variant |
| 4 | 148325647 | 148325714 | 67 | Disp3 | SPRET | feature_elongation,3_prime_UTR_variant |
| 4 | 149038110 | 149038164 | 54 | Casz1 | SPRET | feature_elongation,3_prime_UTR_variant |
| 4 | 149209632 | 149209840 | 208 | Cort | CAST | feature_elongation,3_prime_UTR_variant |
| 4 | 149557829 | 149558010 | 181 | Nmnat1 | CAST,SWR | feature_elongation,coding_sequence_variant |
| 4 | 152101404 | 152101458 | 54 | Klhl21 | 129S1,WSB | feature_elongation,3_prime_UTR_variant |
| 4 | 152144208 | 152144468 | 260 | Nol9 | SPRET | feature_elongation,3_prime_UTR_variant |
| 4 | 154104501 | 154104553 | 52 | Lrrc47 | WSB | feature_elongation,3_prime_UTR_variant |
| 4 | 154359259 | 154359388 | 129 | Megf6 | SPRET | feature_elongation,3_prime_UTR_variant |
| 4 | 154950251 | 154950595 | 344 | Ttc34 | 129S1,AJ,AKR,BALB,BTBR,BUB,C3H,C57L,CAST,CBA,CE,DBA1J,DBA2J,FVB,ILNJ2,KK,MA/My,MRL,NOD,NOR,NZB,NZO,NZW,PJ,PL,RF,RH2,SEA,SJL,SMJ,SPRET,SWR,TH,WSB | feature_elongation,3_prime_UTR_variant |
| 4 | 155703487 | 155703550 | 63 | Slc35e2 | CAST | feature_elongation,3_prime_UTR_variant |
| 4 | 155707022 | 155707232 | 210 | Slc35e2 | CAST,MOLF,SPRET | feature_elongation,3_prime_UTR_variant |
| 4 | 155867086 | 155867289 | 203 | Tmem88b | C57L,CE,ILNJ2,SMJ,WSB | feature_elongation,3_prime_UTR_variant |
| 4 | 155917576 | 155917785 | 209 | Aurkaip1 | C57L,CE,FVB,NOD,NOR,SJL,SWR,TH,WSB | feature_elongation,3_prime_UTR_variant |
| 4 | 155944152 | 155944715 | 563 | Tas1r3 | SPRET | feature_elongation,3_prime_UTR_variant |
| 4 | 155973321 | 155973531 | 210 | Ints11,Pusl1 | 129S1,AJ,AKR,BALB,BUB,C3H,C57L,CBA,DBA1J,DBA2J,ILNJ2,NZO,PL,RF,RH2,SMJ,WSB | feature_elongation,3_prime_UTR_variant |
| 4 | 156043815 | 156044270 | 455 | Ube2j2 | 129S1,AJ,AKR,BALB,BUB,C3H,CBA,CE,DBA1J,DBA2J,FVB,ILNJ2,NOD,NOR,NZO,PL,RF,RH2,SJL,SMJ,SWR,TH,WSB | feature_elongation,3_prime_UTR_variant |
| 4 | 156095351 | 156095441 | 90 | Sdf4 | SPRET | feature_elongation,3_prime_UTR_variant |
| 4 | 156119673 | 156119727 | 54 | Ttll10 | CAST,MOLF | feature_elongation,coding_sequence_variant |
| 4 | 156302625 | 156303021 | 396 | Perm1 | 129S1,AJ,AKR,BALB,C3H,C57L,C58J,DBA1J,DBA2J,FVB,ILNJ2,MA/My,MOLF,MRL,NOD,NOR,NZO,PL,RF,RH2,SJL,SMJ,SWR,WSB | feature_elongation,coding_sequence_variant |
| 4 | 156332083 | 156332270 | 187 | Samd11 | MOLF,NZO | feature_elongation,3_prime_UTR_variant |
| 5 | 3576016 | 3576080 | 64 | Cdk6 | BUB,CAST,MOLF,NOD,NOR | feature_elongation,3_prime_UTR_variant |
| 5 | 3577768 | 3577838 | 70 | Cdk6 | BUB,NOD,NOR,SPRET | feature_elongation,3_prime_UTR_variant |
| 5 | 8751924 | 8752104 | 180 | Abcb1a | BUB,NOD,NOR | feature_elongation,coding_sequence_variant |
| 5 | 14305239 | 14305309 | 70 | Sema3e | SPRET | feature_elongation,3_prime_UTR_variant |
| 5 | 21159637 | 21159709 | 72 | Rsbn1l | SPRET | feature_elongation,5_prime_UTR_variant |
| 5 | 21775365 | 21775481 | 116 | Lrrc17 | CAST,MOLF | feature_elongation,coding_sequence_variant |
| 5 | 21865663 | 21865877 | 214 | Armc10 | CAST,MOLF | feature_elongation,coding_sequence_variant |
| 5 | 21962539 | 21962756 | 217 | Dnajc2 | NZO | feature_elongation,3_prime_UTR_variant |
| 5 | 24025077 | 24025294 | 217 | Rint1 | MOLF | feature_elongation,3_prime_UTR_variant |
| 5 | 29435337 | 29435714 | 377 | Lmbr1 | SPRET | feature_elongation,3_prime_UTR_variant |
| 5 | 29436756 | 29436934 | 178 | Lmbr1 | SPRET | feature_elongation,3_prime_UTR_variant |
| 5 | 29660025 | 29660894 | 869 | Nom1 | SPRET | feature_elongation,3_prime_UTR_variant |
| 5 | 30263959 | 30264450 | 491 | Tyms | SPRET | feature_elongation,3_prime_UTR_variant |
| 5 | 30322625 | 30322684 | 59 | Garem2 | SPRET | feature_elongation,3_prime_UTR_variant |
| 5 | 31108227 | 31108648 | 421 | Preb | MOLF | feature_elongation,3_prime_UTR_variant |
| 5 | 31133869 | 31134299 | 430 | Tcf23 | CAST,MOLF | feature_elongation,3_prime_UTR_variant |
| 5 | 31301623 | 31301839 | 216 | Mpv17 | NZO,NZW | feature_elongation,3_prime_UTR_variant |
| 5 | 31668616 | 31668667 | 51 | Gpn1 | CAST,MOLF,SPRET | feature_elongation,3_prime_UTR_variant |
| 5 | 33574922 | 33575042 | 120 | Uvssa | CAST | feature_elongation,3_prime_UTR_variant |
| 5 | 33576487 | 33577014 | 527 | Uvssa | SPRET | feature_elongation,3_prime_UTR_variant |
| 5 | 33821797 | 33821977 | 180 | Tacc3 | CAST,SPRET | feature_elongation,coding_sequence_variant |
| 5 | 33934128 | 33934214 | 86 | Letm1 | MOLF | feature_elongation,3_prime_UTR_variant |
| 5 | 43775222 | 43775278 | 56 | C1qtnf7 | MA/My,PJ,PL | feature_elongation,3_prime_UTR_variant |
| 5 | 44068370 | 44068436 | 66 | Cd38 | MOLF | feature_elongation,3_prime_UTR_variant |
| 5 | 44068957 | 44069475 | 518 | Cd38 | BALB,BUB,DBA1J,DBA2J,FVB,ILNJ2,KK,LP,MA/My,RH2,SEA,SWR | feature_elongation,3_prime_UTR_variant |
| 5 | 45684925 | 45684986 | 61 | Med28 | CAST | feature_elongation,3_prime_UTR_variant |
| 5 | 51614796 | 51614910 | 114 | Ppargc1a | MOLF,SPRET | feature_elongation,3_prime_UTR_variant |
| 5 | 52925614 | 52925823 | 209 | Pi4k2b | AJ,AKR,BALB,BTBR,BUB,C3H,CBA,CE,DBA1J,DBA2J,FVB,ILNJ2,KK,MA/My,MRL,NOD,NZB,NZO,NZW,PJ,PL,RF,RH2,SEA,SJL,SMJ,SWR,TH | feature_elongation,3_prime_UTR_variant |
| 5 | 66052883 | 66052976 | 93 | Rhoh | SPRET | feature_elongation,3_prime_UTR_variant |
| 5 | 67514499 | 67514996 | 497 | Slc30a9 | BUB,CE,FVB,PJ | feature_elongation,3_prime_UTR_variant |

| | | | | | | |
|---|---|---|---|---|---|---|
| 5 | 67552082 | 67552140 | 58 | Bend4 | CAST | feature_elongation,3_prime_UTR_variant |
| 5 | 71118148 | 71118369 | 221 | Gabra2 | SPRET | feature_elongation,3_prime_UTR_variant |
| 5 | 74161669 | 74161721 | 52 | Usp46 | C3H,CAST,CBA,DBA1J,DBA2J,ILNJ2,NZB,SPRET | feature_elongation,3_prime_UTR_variant |
| 5 | 74365618 | 74365676 | 58 | Scfd2 | SPRET | feature_elongation,3_prime_UTR_variant |
| 5 | 76358799 | 76358852 | 53 | Clock | SPRET | feature_elongation,3_prime_UTR_variant |
| 5 | 76632096 | 76632182 | 86 | Exoc1l | AJ,AKR,BUB,CE,FVB,KK,MOLF,MRL,PJ,PL,SJL,SMJ | feature_elongation,5_prime_UTR_variant |
| 5 | 77235071 | 77235131 | 60 | Hopx | SPRET | feature_elongation,3_prime_UTR_variant |
| 5 | 84214579 | 84214647 | 68 | Epha5 | 129S1,AJ,AKR,B10D2,B10J,BALB,BTBR,BUB,C3H,C57L,C58J,CAST,CBA,CE,DBA1J,DBA2J,FVB,ILNJ2,KK,LP,MA/My,MOLF,MRL,NOD,NOR,NZB,NZO,NZW,PJ,PL,RF,RH2,SEA,SJL,SMJ,SWR,TH,WSB | feature_elongation,3_prime_UTR_variant |
| 5 | 87039782 | 87040277 | 495 | Ugt2b34 | CAST,SPRET | feature_elongation,3_prime_UTR_variant |
| 5 | 87159512 | 87159574 | 62 | Ugt2b35 | CAST | feature_elongation,3_prime_UTR_variant |
| 5 | 87662205 | 87662657 | 452 | Sult1b1 | CAST | feature_elongation,3_prime_UTR_variant |
| 5 | 87965877 | 87965931 | 54 | Csn1s2b | SMJ | feature_elongation,3_prime_UTR_variant |
| 5 | 88649969 | 88650068 | 99 | Enam | CAST | feature_elongation,coding_sequence_variant |
| 5 | 88929556 | 88929638 | 82 | Dck | CAST | feature_elongation,3_prime_UTR_variant |
| 5 | 89189933 | 89190013 | 80 | Slc4a4 | SPRET | feature_elongation,3_prime_UTR_variant |
| 5 | 90908769 | 90909043 | 274 | Cxcl5 | CAST | feature_elongation,3_prime_UTR_variant |
| 5 | 92097131 | 92097205 | 74 | Rchy1 | SPRET | feature_elongation,3_prime_UTR_variant |
| 5 | 93189123 | 93189326 | 203 | Sowahb | AKR,BALB,BTBR,BUB,C3H,CBA,CE,DBA1J,DBA2J,FVB,MA/My,MRL,NOD,NOR,NZW,PJ,RH2,SEA,SJL,SWR,TH | feature_elongation,3_prime_UTR_variant |
| 5 | 97230339 | 97230438 | 99 | Paqr3 | 129S1,AJ,BTBR,BUB,C57L,KK,MA/My,NOD,NOR,NZB,NZO,SPRET,SWR,TH | feature_elongation,3_prime_UTR_variant |
| 5 | 99445711 | 99445769 | 58 | Rasgef1b | CAST | feature_elongation,3_prime_UTR_variant |
| 5 | 100216368 | 100216439 | 71 | Enoph1 | CAST,SPRET | feature_elongation,3_prime_UTR_variant |
| 5 | 101954473 | 101954745 | 272 | Cds1 | CAST | feature_elongation,coding_sequence_variant |
| 5 | 101970994 | 101971134 | 140 | Cds1 | SPRET | feature_elongation,3_prime_UTR_variant |
| 5 | 104325526 | 104325796 | 270 | Dspp | NZB | feature_elongation,coding_sequence_variant |
| 5 | 104699955 | 104700168 | 213 | Thoc2l | AJ,BALB,BUB,C3H,CBA,DBA1J,DBA2J,FVB,ILNJ2,KK,MOLF,NZB,NZO,RH2,SEA,SJL,SWR,TH,WSB | feature_elongation,3_prime_UTR_variant |
| 5 | 104730384 | 104730636 | 252 | Zfp33b | 129S1,AJ,AKR,B10D2,B10J,BALB,BTBR,C57L,C58J,CAST,CE,DBA1J,DBA2J,ILNJ2,KK,LP,MA/My,MOLF,MRL,NOD,NOR,NZB,NZO,NZW,PJ,PL,RF,RH2,SEA,SMJ,SPRET,SWR,WSB | feature_elongation,coding_sequence_variant |
| 5 | 107534561 | 107534642 | 81 | Brdt | CAST | feature_elongation,3_prime_UTR_variant |
| 5 | 107697640 | 107697691 | 51 | 1700028K03Rik | BUB,C3H,CBA,FVB,MOLF,PJ,PL,SJL,SWR,TH | feature_elongation,3_prime_UTR_variant |
| 5 | 108574751 | 108574820 | 69 | Pde6b | CAST | feature_elongation,coding_sequence_variant |
| 5 | 109514834 | 109515687 | 853 | Vmn2r16 | SPRET | feature_elongation,3_prime_UTR_variant |
| 5 | 109886154 | 109886453 | 299 | 4930522L14Rik | CE,DBA1J,DBA2J,LP | feature_elongation,3_prime_UTR_variant |
| 5 | 110224606 | 110224816 | 210 | Gm35315 | 129S1,AJ,AKR,BALB,BTBR,C57L,ILNJ2,KK,MA/My,MRL,NOD,NOR,NZB,NZO,NZW,RF,RH2,SEA,SMJ | feature_elongation,3_prime_UTR_variant |
| 5 | 110225458 | 110225586 | 128 | Gm35315 | SPRET | feature_elongation,3_prime_UTR_variant |
| 5 | 110275243 | 110275580 | 337 | Zfp605 | BUB,C3H,CBA,FVB,MOLF,PJ,PL,SJL,SWR,TH | feature_elongation,coding_sequence_variant |
| 5 | 110403780 | 110404000 | 220 | Ankle2 | SPRET | feature_elongation,3_prime_UTR_variant |
| 5 | 111437201 | 111437257 | 56 | Ttc28 | CAST | feature_elongation,3_prime_UTR_variant |
| 5 | 113061325 | 113061382 | 57 | Grk3 | SPRET | feature_elongation,3_prime_UTR_variant |
| 5 | 113108412 | 113108579 | 167 | Grk3 | SPRET | feature_elongation,3_prime_UTR_variant |
| 5 | 114569745 | 114569952 | 207 | Mmab | MOLF | feature_elongation,3_prime_UTR_variant |
| 5 | 114946515 | 114946568 | 53 | 1500011B03Rik | CAST | feature_elongation,3_prime_UTR_variant |
| 5 | 115049680 | 115049749 | 69 | Oasl2 | MOLF,SPRET | feature_elongation,3_prime_UTR_variant |
| 5 | 116177512 | 116177683 | 171 | Tmem233 | CAST,MOLF | feature_elongation,3_prime_UTR_variant |
| 5 | 117492869 | 117493050 | 181 | Vsig10 | MOLF,RF,SPRET | feature_elongation,3_prime_UTR_variant |
| 5 | 117912382 | 117912438 | 56 | Ksr2 | CAST | feature_elongation,3_prime_UTR_variant |
| 5 | 118330475 | 118330535 | 60 | Rnft2 | 129S1,AJ,AKR,BALB,BUB,C3H,C57L,CBA,CE,DBA1J,DBA2J,ILNJ2,KK,NOR,NZB,NZO,PJ,PL,RH2,SEA,SJL,SMJ,TH,WSB | feature_elongation,3_prime_UTR_variant |
| 5 | 118401413 | 118401871 | 458 | Spring1 | SPRET | feature_elongation,3_prime_UTR_variant |
| 5 | 120578757 | 120578826 | 69 | Lhx5 | MOLF | feature_elongation,3_prime_UTR_variant |
| 5 | 121024042 | 121024175 | 133 | Oas1g | MOLF,SPRET | feature_elongation,coding_sequence_variant |
| 5 | 121580004 | 121580070 | 66 | Naa25,Erp29 | MOLF | feature_elongation,3_prime_UTR_variant |
| 5 | 121580741 | 121581210 | 469 | Erp29 | SPRET | feature_elongation,3_prime_UTR_variant |
| 5 | 121683726 | 121683777 | 51 | Mapkapk5,- | MOLF | feature_elongation,5_prime_UTR_variant,feature_elongation,coding_sequence_variant |
| 5 | 121735584 | 121735645 | 61 | Acad12 | AKR,BUB,CAST,ILNJ2,KK,MA/My,MOLF,PL,RF,SJL,SPRET,TH | feature_elongation,3_prime_UTR_variant |
| 5 | 122377814 | 122377931 | 117 | Tctn1 | SPRET | feature_elongation,3_prime_UTR_variant |
| 5 | 122502342 | 122502538 | 196 | Vps29 | SPRET | feature_elongation,3_prime_UTR_variant |
| 5 | 122867592 | 122868247 | 655 | P2rx4 | RF | feature_elongation,3_prime_UTR_variant |
| 5 | 123008495 | 123008680 | 185 | Rnf34 | CAST | feature_elongation,3_prime_UTR_variant |
| 5 | 123465542 | 123465753 | 211 | Cfap251 | SPRET | feature_elongation,3_prime_UTR_variant |
| 5 | 123584316 | 123584548 | 232 | Mlxip | 129S1,AJ,AKR,BALB,BUB,C3H,C57L,CBA,CE,DBA1J,DBA2J,KK,MA/My,MRL,NOD,NOR,NZW,PJ,PL,RH2,SEA,SJL,SMJ,WSB | feature_elongation,3_prime_UTR_variant |
| 5 | 123836758 | 123836896 | 138 | Zcchc8 | SPRET | feature_elongation,3_prime_UTR_variant |
| 5 | 124066631 | 124066802 | 171 | Denr | SPRET | feature_elongation,3_prime_UTR_variant |
| 5 | 124479112 | 124479260 | 148 | Mtrfr | SPRET | feature_elongation,3_prime_UTR_variant |
| 5 | 124734957 | 124735144 | 187 | Gtf2h3 | 129S1,AJ,BALB,C57L,MRL,PL,RH2,SEA,SJL | feature_elongation,3_prime_UTR_variant |
| 5 | 125464955 | 125465183 | 228 | Ubc | PL,SPRET,WSB | feature_elongation,coding_sequence_variant |
| 5 | 125533912 | 125534114 | 202 | Bri3bp | MOLF | feature_elongation,3_prime_UTR_variant |
| 5 | 125868893 | 125868949 | 56 | Tmem132b | CAST,SPRET | feature_elongation,3_prime_UTR_variant |
| 5 | 128831697 | 128831770 | 73 | Piwil1 | SPRET | feature_elongation,3_prime_UTR_variant |
| 5 | 129062083 | 129062209 | 126 | Stx2 | AJ,BALB,C58J,CAST,FVB,LP,MOLF,MRL,NOD,NOR,NZB,NZO,NZW,RH2,SEA,SMJ,SWR,TH | feature_elongation,3_prime_UTR_variant |
| 5 | 130028471 | 130028677 | 206 | Gusb | AJ,BALB,MRL,RH2,SEA,SMJ | feature_elongation,3_prime_UTR_variant |

| | | | | | | |
|---|---|---|---|---|---|---|
| 5 | 130903567 | 130903641 | 74 | Galnt17 | CAST,NOD,NOR | feature_elongation,3_prime_UTR_variant |
| 5 | 135939421 | 135939482 | 61 | Ywhag | MOLF | feature_elongation,3_prime_UTR_variant |
| 5 | 136177363 | 136177470 | 107 | Orai2 | SPRET | feature_elongation,3_prime_UTR_variant |
| 5 | 136207651 | 136207820 | 169 | Prkrip1 | CAST,MOLF,SPRET | feature_elongation,3_prime_UTR_variant |
| 5 | 136943024 | 136943243 | 219 | Ift22 | SPRET | feature_elongation,3_prime_UTR_variant |
| 5 | 137178802 | 137178892 | 90 | Muc17 | LP | feature_elongation,coding_sequence_variant |
| 5 | 137180234 | 137180288 | 54 | Muc17 | CAST,LP,MOLF | feature_elongation,coding_sequence_variant |
| 5 | 137462190 | 137462289 | 99 | Zan | 129S1,BUB,CAST,LP,MOLF,RF,SWR,WSB | feature_elongation,coding_sequence_variant |
| 5 | 137641877 | 137642090 | 213 | Irs3 | ILNJ2,MA/My,NZW | feature_elongation,3_prime_UTR_variant |
| 5 | 138192168 | 138192369 | 201 | Cnpy4 | 129S1,AJ,BALB,BTBR,C3H,C57L,CBA,CE,DBA1J,DBA2J,FVB,LP,MA/My,NOD,NOR,NZW,PJ,RH2,SEA,SJL | feature_elongation,3_prime_UTR_variant |
| 5 | 138250128 | 138250187 | 59 | Nxpe5 | BUB,CAST,KK,LP,NZB,NZO,NZW,RF,SPRET | feature_elongation,3_prime_UTR_variant |
| 5 | 138646471 | 138646547 | 76 | A430033K04Rik | SPRET | feature_elongation,3_prime_UTR_variant |
| 5 | 138646714 | 138647195 | 481 | A430033K04Rik | CAST | feature_elongation,3_prime_UTR_variant |
| 5 | 139171554 | 139171615 | 61 | Dnaaf5 | CAST | feature_elongation,3_prime_UTR_variant |
| 5 | 139323258 | 139323432 | 174 | Cox19 | NZW,SMJ | feature_elongation,3_prime_UTR_variant |
| 5 | 139345696 | 139345847 | 151 | 3110082I17Rik | ILNJ2,SPRET | feature_elongation,3_prime_UTR_variant |
| 5 | 139364158 | 139364735 | 577 | Gpr146 | MOLF | feature_elongation,5_prime_UTR_variant |
| 5 | 139381580 | 139381784 | 204 | Gpr146 | CAST,ILNJ2,MOLF,NZW,SMJ,SPRET | feature_elongation,3_prime_UTR_variant |
| 5 | 139457720 | 139457774 | 54 | Zfand2a | SPRET | feature_elongation,3_prime_UTR_variant |
| 5 | 142654018 | 142654070 | 52 | Wipi2 | SPRET | feature_elongation,3_prime_UTR_variant |
| 5 | 143233789 | 143233872 | 83 | Zfp12 | SPRET | feature_elongation,3_prime_UTR_variant |
| 5 | 143694717 | 143694769 | 52 | Cyth3 | MOLF | feature_elongation,3_prime_UTR_variant |
| 5 | 144129236 | 144129302 | 66 | Bhlha15 | MOLF | feature_elongation,3_prime_UTR_variant |
| 5 | 144493796 | 144493858 | 62 | Nptx2 | CE | feature_elongation,3_prime_UTR_variant |
| 5 | 144783274 | 144783351 | 77 | Trrap | SPRET | feature_elongation,coding_sequence_variant |
| 5 | 146409596 | 146409657 | 61 | Wasf3 | MOLF,TH | feature_elongation,3_prime_UTR_variant |
| 5 | 147498981 | 147499129 | 148 | Flt1 | MOLF,TH | feature_elongation,3_prime_UTR_variant |
| 5 | 147815752 | 147815859 | 107 | Slc46a3 | CAST,CE,MOLF,SPRET,WSB | feature_elongation,3_prime_UTR_variant |
| 5 | 148252153 | 148252211 | 58 | Mtus2 | AKR,BUB,MA/My,MRL,NOD,NOR | feature_elongation,3_prime_UTR_variant |
| 5 | 149502597 | 149502673 | 76 | Wdr95 | SPRET | feature_elongation,coding_sequence_variant |
| 5 | 150565406 | 150565579 | 173 | N4bp2l2 | CE,ILNJ2,SEA,SWR | feature_elongation,3_prime_UTR_variant |
| 5 | 150962420 | 150962559 | 139 | Stard13 | CAST,MOLF,SPRET | feature_elongation,3_prime_UTR_variant |
| 6 | 4756453 | 4756534 | 81 | Peg10 | 129S1,AJ,AKR,B10D2,B10J,BALB,BTBR,BUB,C3H,C57L,C58J,CAST,CBA,CE,DBA1J,DBA2J,FVB,ILNJ2,KK,LP,MA/My,MOLF,MRL,NOD,NOR,NZB,NZO,NZW,PL,RF,RH2,SEA,SJL,SMJ,SPRET,SWR,TH,WSB | feature_elongation,3_prime_UTR_variant,feature_elongation,coding_sequence_variant |
| 6 | 5008455 | 5008610 | 155 | Ppp1r9a | PJ | feature_elongation,3_prime_UTR_variant |
| 6 | 5164175 | 5164253 | 78 | Ppp1r9a | SPRET | feature_elongation,3_prime_UTR_variant |
| 6 | 8459385 | 8459516 | 131 | Umad1 | SPRET | feature_elongation,3_prime_UTR_variant |
| 6 | 15770578 | 15770629 | 51 | Mdfic | KK | feature_elongation,coding_sequence_variant |
| 6 | 17665665 | 17665760 | 95 | Capza2 | SPRET | feature_elongation,3_prime_UTR_variant |
| 6 | 23097157 | 23097270 | 113 | Aass | SPRET | feature_elongation,coding_sequence_variant |
| 6 | 24952618 | 24952713 | 95 | Tmem229a | SPRET | feature_elongation,3_prime_UTR_variant |
| 6 | 25666442 | 25666537 | 95 | Gpr37 | CAST | feature_elongation,3_prime_UTR_variant |
| 6 | 28934267 | 28934413 | 146 | Snd1 | MOLF | feature_elongation,3_prime_UTR_variant |
| 6 | 29959675 | 29959889 | 214 | Strip2 | MOLF | feature_elongation,3_prime_UTR_variant |
| 6 | 31485167 | 31485735 | 568 | Mkln1 | SPRET | feature_elongation,3_prime_UTR_variant |
| 6 | 34083063 | 34083136 | 73 | Lrguk | CAST | feature_elongation,3_prime_UTR_variant |
| 6 | 34849404 | 34849896 | 492 | Cyren | SPRET | feature_elongation,3_prime_UTR_variant |
| 6 | 35238807 | 35239642 | 835 | Stmp1 | MOLF | feature_elongation,3_prime_UTR_variant |
| 6 | 35238889 | 35239104 | 215 | Stmp1 | SPRET | feature_elongation,3_prime_UTR_variant |
| 6 | 36502833 | 36502899 | 66 | Chrm2 | SPRET | feature_elongation,3_prime_UTR_variant |
| 6 | 36717738 | 36717831 | 93 | Ptn | SPRET | feature_elongation,3_prime_UTR_variant |
| 6 | 39113576 | 39113814 | 238 | Kdm7a | SPRET | feature_elongation,3_prime_UTR_variant |
| 6 | 39578527 | 39578580 | 53 | Ndufb2 | SPRET | feature_elongation,3_prime_UTR_variant |
| 6 | 40519625 | 40519684 | 59 | Or9a7 | MOLF | feature_elongation,3_prime_UTR_variant |
| 6 | 40549878 | 40549942 | 64 | Or9a4 | CAST | feature_elongation,3_prime_UTR_variant |
| 6 | 40844545 | 40844665 | 120 | Mgam2-ps | BUB,FVB,LP,MOLF,SJL,SWR,TH,WSB | feature_elongation,coding_sequence_variant |
| 6 | 40844849 | 40845206 | 357 | Mgam2-ps | CAST | feature_elongation,coding_sequence_variant |
| 6 | 41747084 | 41747180 | 96 | Or9a2 | CAST | feature_elongation,3_prime_UTR_variant |
| 6 | 42890356 | 42890416 | 60 | Or2a25 | MOLF | feature_elongation,3_prime_UTR_variant |
| 6 | 42906828 | 42907089 | 261 | Or2a12 | CAST,MOLF | feature_elongation,3_prime_UTR_variant |
| 6 | 43125095 | 43125174 | 79 | Or2a14 | CAST,SPRET | feature_elongation,5_prime_UTR_variant |
| 6 | 43213539 | 43213901 | 362 | Or2a57 | SPRET | feature_elongation,3_prime_UTR_variant |
| 6 | 47941607 | 47941665 | 58 | Zfp956 | AJ,AKR,BALB,BUB,C3H,CBA,CE,DBA1J,DBA2J,FVB,ILNJ2,MRL,NOD,NOR,RF,RH2,SJL,SMJ,SWR | feature_elongation,3_prime_UTR_variant |
| 6 | 48964956 | 48965010 | 54 | Aoc1l3 | CAST,CE,ILNJ2,MOLF | feature_elongation,coding_sequence_variant |
| 6 | 54554861 | 54555046 | 185 | Fkbp14 | CAST | feature_elongation,3_prime_UTR_variant |
| 6 | 54619376 | 54619586 | 210 | Plekha8 | MOLF | feature_elongation,3_prime_UTR_variant |
| 6 | 56983797 | 56983876 | 79 | Vmn1r6 | CAST | feature_elongation,3_prime_UTR_variant |
| 6 | 57046064 | 57046120 | 56 | Vmn1r9 | MOLF,NZW | feature_elongation,5_prime_UTR_variant |
| 6 | 57138718 | 57139246 | 528 | Vmn1r12 | SPRET | feature_elongation,3_prime_UTR_variant |
| 6 | 57364776 | 57364976 | 200 | Vmn1r18 | 129S1,C57L,CAST,CE,ILNJ2,KK,MOLF,NZB,NZO,NZW,PJ,WSB | feature_elongation,3_prime_UTR_variant |
| 6 | 57382791 | 57383243 | 452 | Vmn1r19 | 129S1,C57L,KK,MOLF,NZB,NZO,NZW | feature_elongation,3_prime_UTR_variant |
| 6 | 57410872 | 57411227 | 355 | Vmn1r20 | SPRET | feature_elongation,3_prime_UTR_variant |
| 6 | 57738303 | 57738499 | 196 | Vopp1 | SPRET | feature_elongation,3_prime_UTR_variant |
| 6 | 58191583 | 58191635 | 52 | Vmn1r27 | SPRET | feature_elongation,3_prime_UTR_variant |
| 6 | 58672239 | 58672597 | 358 | Abcg2 | SPRET | feature_elongation,3_prime_UTR_variant |
| 6 | 65094723 | 65095124 | 401 | Hpgds | SPRET | feature_elongation,3_prime_UTR_variant |
| 6 | 65433857 | 65434003 | 146 | Qrfprl | CAST,KK,SPRET | feature_elongation,3_prime_UTR_variant |
| 6 | 66519241 | 66519924 | 683 | Mad2l1 | SPRET | feature_elongation,3_prime_UTR_variant |
| 6 | 66520391 | 66520483 | 92 | Mad2l1 | SPRET | feature_elongation,3_prime_UTR_variant |
| 6 | 67263640 | 67263736 | 96 | Serbp1 | MA/My | feature_elongation,3_prime_UTR_variant |
| 6 | 71951860 | 71951950 | 90 | Polr1a | SPRET | feature_elongation,coding_sequence_variant |
| 6 | 72329092 | 72329153 | 61 | 0610030E20Rik | AKR,C58J,KK,MOLF,NOD,NOR,PL,RF | feature_elongation,3_prime_UTR_variant |

| | | | | | | |
|---|---|---|---|---|---|---|
| 6 | 72407385 | 72407444 | 59 | Ggcx | AKR,C3H,C58J,CAST,CBA,DBA1J,DBA2J,KK,MA/My,MOLF,MRL,NOD,NOR,PL,RF,SMJ,SPRET,WSB | feature_elongation,3_prime_UTR_variant |
| 6 | 72584962 | 72585189 | 227 | Retsat | C58J,KK,MOLF,NOD,NOR | feature_elongation,3_prime_UTR_variant |
| 6 | 82535164 | 82535226 | 62 | Tacr1 | AJ,AKR,BALB,BTBR,BUB,CE,FVB,ILNJ2,LP,MA/My,MRL,NOD,NOR,NZW,PL,RH2,SEA,SJL,SWR,WSB | feature_elongation,3_prime_UTR_variant |
| 6 | 83127309 | 83127433 | 124 | Wdr54 | CAST,KK,NZB,NZO,TH | feature_elongation,3_prime_UTR_variant |
| 6 | 83965422 | 83965477 | 55 | Zfp638 | SPRET | feature_elongation,3_prime_UTR_variant |
| 6 | 85807449 | 85807647 | 198 | Nat8 | MOLF | feature_elongation,coding_sequence_variant |
| 6 | 86992589 | 86992720 | 131 | Nfu1 | AKR,BALB,KK,MA/My,MRL,NZB,NZO,PL,RH2,TH,WSB | feature_elongation,coding_sequence_variant |
| 6 | 87067470 | 87067534 | 64 | Gfpt1 | SPRET | feature_elongation,3_prime_UTR_variant |
| 6 | 87413046 | 87413130 | 84 | Bmp10 | AJ,AKR,B10D2,B10J,BALB,BTBR,BUB,C57L,C58J,FVB,ILNJ2,KK,LP,MRL,NOD,NOR,NZO,PL,RF,RH2,SJL,SMJ,TH | feature_elongation,3_prime_UTR_variant |
| 6 | 87791450 | 87791648 | 198 | Isy1 | SWR | feature_elongation,3_prime_UTR_variant |
| 6 | 89691868 | 89692283 | 415 | Vmn1r40 | NZO | feature_elongation,coding_sequence_variant |
| 6 | 90247433 | 90247499 | 66 | Vmn1r54 | CAST,CE,MOLF,PJ,SEA | feature_elongation,3_prime_UTR_variant |
| 6 | 90285570 | 90285657 | 87 | Chst13 | CAST,CE,MOLF,PJ,SEA,SPRET | feature_elongation,3_prime_UTR_variant |
| 6 | 90635724 | 90635920 | 196 | Iqsec1 | CAST,CE,MOLF,PJ,SEA | feature_elongation,3_prime_UTR_variant |
| 6 | 91441264 | 91441479 | 215 | Chchd4 | AJ,BALB,BTBR,BUB,RF,RH2,SJL | feature_elongation,3_prime_UTR_variant |
| 6 | 92151624 | 92151824 | 200 | Mrps25 | CE,PJ,SEA | feature_elongation,3_prime_UTR_variant |
| 6 | 93653633 | 93653800 | 167 | Magi1 | C3H,CBA,CE,ILNJ2,NOD,NOR,SMJ | feature_elongation,3_prime_UTR_variant |
| 6 | 100784144 | 100784202 | 58 | Gxylt2 | CAST,SJL,SPRET | feature_elongation,3_prime_UTR_variant |
| 6 | 103707086 | 103707138 | 52 | Chl1 | CAST | feature_elongation,3_prime_UTR_variant |
| 6 | 106687367 | 106687528 | 161 | Il5ra | SPRET | feature_elongation,3_prime_UTR_variant |
| 6 | 112452787 | 112452882 | 95 | Oxtr | SPRET | feature_elongation,3_prime_UTR_variant |
| 6 | 113147543 | 113147597 | 54 | Lhfpl4 | CAST | feature_elongation,3_prime_UTR_variant |
| 6 | 113366881 | 113366965 | 84 | Arpc4 | SPRET | feature_elongation,3_prime_UTR_variant |
| 6 | 115599045 | 115599735 | 690 | Mkrn2 | BALB | feature_elongation,3_prime_UTR_variant |
| 6 | 115819180 | 115819390 | 210 | Mbd4 | SPRET | feature_elongation,3_prime_UTR_variant |
| 6 | 116557792 | 116558112 | 320 | Or6d15 | BUB,CE,FVB,KK,NZB,NZW,PJ,SJL,SMJ,SWR | feature_elongation,3_prime_UTR_variant |
| 6 | 116659873 | 116659924 | 51 | Tmem72 | 129S1,AKR,BTBR,BUB,C3H,C57L,C58J,CAST,CBA,DBA1J,DBA2J,FVB,KK,LP,MOLF,MRL,NOD,NOR,NZO,PJ,PL,RF,RH2,SJL,SMJ,SPRET,SWR | feature_elongation,3_prime_UTR_variant |
| 6 | 116668790 | 116668878 | 88 | Tmem72 | C3H,CBA,PL,SPRET | feature_elongation,3_prime_UTR_variant |
| 6 | 116669456 | 116669560 | 104 | Tmem72 | SPRET | feature_elongation,3_prime_UTR_variant |
| 6 | 117154173 | 117154235 | 62 | Cxcl12 | SWR | feature_elongation,3_prime_UTR_variant |
| 6 | 119749794 | 119749920 | 126 | Erc1 | CAST,MOLF,SPRET | feature_elongation,3_prime_UTR_variant |
| 6 | 122946761 | 122946813 | 52 | Clec4a3 | WSB | feature_elongation,3_prime_UTR_variant |
| 6 | 123119769 | 123119823 | 54 | Clec4a2 | SWR | feature_elongation,3_prime_UTR_variant |
| 6 | 123352787 | 123352843 | 56 | Vmn2r20 | MOLF | feature_elongation,3_prime_UTR_variant |
| 6 | 123361261 | 123361746 | 485 | Vmn2r20 | SMJ | feature_elongation,3_prime_UTR_variant |
| 6 | 123794634 | 123794694 | 60 | Vmn2r24 | MOLF | feature_elongation,3_prime_UTR_variant |
| 6 | 123800922 | 123801365 | 443 | Vmn2r24 | SPRET | feature_elongation,3_prime_UTR_variant |
| 6 | 125145148 | 125145205 | 57 | Ncapd2 | SPRET | feature_elongation,3_prime_UTR_variant |
| 6 | 125217444 | 125217747 | 303 | Vamp1 | MOLF,SPRET | feature_elongation,coding_sequence_variant |
| 6 | 125523986 | 125524168 | 182 | Vwf | KK | feature_elongation,coding_sequence_variant |
| 6 | 126509828 | 126509883 | 55 | Kcna5 | NOD,NOR | feature_elongation,3_prime_UTR_variant |
| 6 | 128791418 | 128791623 | 205 | Klrb1b | CAST,NZO | feature_elongation,3_prime_UTR_variant |
| 6 | 128948169 | 128948221 | 52 | Clec2g | CAST,SPRET | feature_elongation,5_prime_UTR_variant |
| 6 | 129163185 | 129163295 | 110 | Clec2d | SPRET | feature_elongation,3_prime_UTR_variant |
| 6 | 129463734 | 129463912 | 178 | Olr1 | AJ,AKR,BALB,BUB,C3H,C57L,CAST,CBA,DBA1J,DBA2J,ILNJ2,LP,MRL,PL,RF,RH2,SEA,SPRET | feature_elongation,3_prime_UTR_variant |
| 6 | 129589514 | 129589916 | 402 | Klrk1 | SPRET | feature_elongation,3_prime_UTR_variant |
| 6 | 129625027 | 129625297 | 270 | Klrc2 | 129S1,AJ,AKR,BALB,BUB,C3H,C57L,C58J,CAST,CBA,CE,DBA1J,DBA2J,FVB,ILNJ2,LP,MA/My,MOLF,MRL,NOD,NOR,NZO,PL,RF,RH2,SEA,SJL,SMJ,SPRET,SWR,TH,WSB | feature_elongation,3_prime_UTR_variant |
| 6 | 129744568 | 129744622 | 54 | Klrh1 | AKR,C3H,MOLF,NZO,RF,WSB | feature_elongation,3_prime_UTR_variant |
| 6 | 131261388 | 131261589 | 201 | Magohb | CAST | feature_elongation,3_prime_UTR_variant |
| 6 | 131276974 | 131277039 | 65 | Styk1 | SPRET | feature_elongation,3_prime_UTR_variant |
| 6 | 132289152 | 132289304 | 152 | Prb1b | AJ,AKR,BALB,BUB,C3H,C57L,CBA,DBA1J,DBA2J,ILNJ2,MOLF,MRL,NOD,NOR,NZB,PL,RF,RH2,SEA,SWR,WSB | frameshift_variant |
| 6 | 132338666 | 132339022 | 356 | Prb1c | CAST,MOLF | feature_elongation,coding_sequence_variant |
| 6 | 132549261 | 132549384 | 123 | Prh1 | AJ,AKR,BALB,BUB,C3H,C57L,CBA,DBA1J,DBA2J,ILNJ2,MRL,NZB,NZO,PL,RF,RH2,SEA | feature_elongation,coding_sequence_variant |
| 6 | 133083395 | 133083498 | 103 | Smim10l1 | 129S1,C58J,FVB,KK,LP,NOD,NOR,NZW,SJL,SMJ,SPRET,SWR,TH,WSB | feature_elongation,3_prime_UTR_variant |
| 6 | 134425852 | 134426096 | 244 | Lrp6 | AJ,AKR,BUB,C3H,C58J,CBA,CE,DBA1J,DBA2J,KK,MRL,NOD,NOR,NZB,NZW,PL,RF,SJL,TH | feature_elongation,3_prime_UTR_variant |
| 6 | 135286151 | 135286344 | 193 | Pbp2 | SPRET | feature_elongation,3_prime_UTR_variant |
| 6 | 135698056 | 135698246 | 190 | Grin2b | CAST,MOLF,SPRET | feature_elongation,3_prime_UTR_variant |
| 6 | 135705656 | 135705717 | 61 | Grin2b | 129S1,AJ,AKR,BALB,BTBR,BUB,C3H,C57L,CBA,CE,DBA1J,DBA2J,FVB,KK,MA/My,MRL,NOD,NOR,NZB,NZW,PJ,PL,RF,RH2,SEA,SJL,SWR,TH,WSB | feature_elongation,3_prime_UTR_variant |
| 6 | 136943892 | 136943944 | 52 | Pde6h | CAST | feature_elongation,3_prime_UTR_variant |
| 6 | 140623945 | 140624171 | 226 | Aebp2 | PJ,SWR | feature_elongation,3_prime_UTR_variant |
| 6 | 140626136 | 140626190 | 54 | Aebp2 | CAST,FVB,SPRET,WSB | feature_elongation,3_prime_UTR_variant |
| 6 | 141853442 | 141853497 | 55 | Slco1a1 | SPRET | feature_elongation,3_prime_UTR_variant |
| 6 | 142131844 | 142132403 | 559 | Slco1a6 | SPRET | feature_elongation,5_prime_UTR_variant |
| 6 | 142308005 | 142308297 | 292 | Pyroxd1,Recql | SPRET | feature_elongation,3_prime_UTR_variant |
| 6 | 142322561 | 142322683 | 122 | Recql | SPRET | frameshift_variant |
| 6 | 142365203 | 142365413 | 210 | Spx | SPRET | feature_elongation,3_prime_UTR_variant |
| 6 | 142510838 | 142510912 | 74 | Kcnj8 | SPRET | feature_elongation,3_prime_UTR_variant |
| 6 | 143113069 | 143113185 | 116 | Etnk1 | WSB | feature_elongation,5_prime_UTR_variant |
| 6 | 144155263 | 144155321 | 58 | Sox5 | AKR,BUB,C3H,CBA,CE,DBA2J,ILNJ2,MOLF,MRL,NZO,RF,SJL | feature_elongation,5_prime_UTR_variant |

| | | | | | | |
|---|---|---|---|---|---|---|
| 6 | 146734471 | 146734600 | 129 | Bmal2 | 129S1,AJ,BALB,BTBR,C57L,CAST,CBA,FVB,ILNJ2,MA/My,MOLF,NOD,NOR,PJ,RF,SEA,SMJ,SWR,TH,WSB | feature_elongation,3_prime_UTR_variant |
| 6 | 146975278 | 146975363 | 85 | Mrps35 | SPRET | feature_elongation,3_prime_UTR_variant |
| 6 | 148135941 | 148136317 | 376 | Tmtc1 | CAST | feature_elongation,3_prime_UTR_variant |
| 6 | 149046056 | 149046269 | 213 | Etfbkmt | CAST | feature_elongation,coding_sequence_variant |
| 7 | 3537506 | 3538199 | 693 | Tarm1 | 129S1,AJ,AKR,BALB,BTBR,BUB,C57L,CAST,FVB,KK,LP,MA/My,MOLF,MRL,NOD,NOR,NZB,NZO,NZW,PL,RF,RH2,SJL,SMJ,SPRET,SWR,TH | feature_elongation,3_prime_UTR_variant |
| 7 | 3824543 | 3824763 | 220 | Pira13 | MOLF,MRL | feature_elongation,coding_sequence_variant |
| 7 | 4371488 | 4371768 | 280 | Gp6 | 129S1,AJ,AKR,BALB,BTBR,BUB,C3H,C57L,CBA,CE,DBA1J,DBA2J,FVB,ILNJ2,KK,LP,MA/My,MRL,NOD,NOR,NZB,NZO,NZW,PJ,PL,RF,RH2,SEA,SJL,SMJ,SWR,TH,WSB | feature_elongation,3_prime_UTR_variant |
| 7 | 4482739 | 4483014 | 275 | Eps8l1 | CE,MRL,SWR,TH | feature_elongation,3_prime_UTR_variant |
| 7 | 4695285 | 4695400 | 115 | Brsk1 | BUB,CAST,CE,MRL,NZO,SMJ,SPRET | feature_elongation,5_prime_UTR_variant |
| 7 | 4968267 | 4968441 | 174 | Sbk3 | 129S1,AJ,AKR,BALB,BTBR,BUB,C3H,C57L,CBA,DBA1J,DBA2J,FVB,ILNJ2,KK,LP,MA/My,NOD,NOR,NZW,PJ,PL,RF,RH2,SEA,SJL | feature_elongation,3_prime_UTR_variant |
| 7 | 5480399 | 5480579 | 180 | Vmn2r28 | CAST | feature_elongation,3_prime_UTR_variant |
| 7 | 6358672 | 6358730 | 58 | Zfp582 | SPRET | feature_elongation,3_prime_UTR_variant |
| 7 | 6674784 | 6674848 | 64 | Zim1 | CAST | feature_elongation,3_prime_UTR_variant |
| 7 | 10007619 | 10007673 | 54 | Vmn1r66 | SWR,TH | feature_elongation,3_prime_UTR_variant |
| 7 | 11495664 | 11495727 | 63 | Vmn1r73 | CAST,SPRET | feature_elongation,3_prime_UTR_variant |
| 7 | 11583302 | 11583392 | 90 | Vmn1r74 | AKR,CAST,FVB,MRL,NOD,NOR,RH2,WSB | feature_elongation,3_prime_UTR_variant |
| 7 | 11583843 | 11584172 | 329 | Vmn1r74 | SPRET | feature_elongation,3_prime_UTR_variant |
| 7 | 11586296 | 11586493 | 197 | Vmn1r74 | SPRET | feature_elongation,3_prime_UTR_variant |
| 7 | 11782418 | 11783162 | 744 | Vmn1r77 | AKR,FVB,MRL,NOD,NOR,RH2,WSB | feature_elongation,3_prime_UTR_variant |
| 7 | 11887617 | 11887679 | 62 | Vmn1r78 | AKR,FVB,MOLF,MRL,NOD,NOR,SPRET | feature_elongation,3_prime_UTR_variant |
| 7 | 11889432 | 11889483 | 51 | Vmn1r78 | CAST,WSB | feature_elongation,3_prime_UTR_variant |
| 7 | 11912839 | 11913167 | 328 | Vmn1r79 | MOLF | feature_elongation,3_prime_UTR_variant |
| 7 | 11932668 | 11932735 | 67 | Vmn1r80 | MOLF | feature_elongation,3_prime_UTR_variant |
| 7 | 11985708 | 11986217 | 509 | Vmn1r81 | CAST,MOLF | feature_elongation,3_prime_UTR_variant |
| 7 | 11992270 | 11992442 | 172 | Vmn1r81 | CAST | feature_elongation,3_prime_UTR_variant |
| 7 | 12042016 | 12042213 | 197 | Vmn1r82 | SPRET | feature_elongation,3_prime_UTR_variant |
| 7 | 12508438 | 12508510 | 72 | Zscan18 | CAST,MOLF,SPRET | feature_elongation,coding_sequence_variant |
| 7 | 12706566 | 12706790 | 224 | Zfp324 | SPRET | feature_elongation,3_prime_UTR_variant |
| 7 | 12821540 | 12821869 | 329 | Vmn1r85 | SPRET | feature_elongation,5_prime_UTR_variant |
| 7 | 12822179 | 12822238 | 59 | Vmn1r85 | CAST,MOLF | feature_elongation,5_prime_UTR_variant |
| 7 | 12869164 | 12869218 | 54 | Vmn1r87 | CAST | feature_elongation,5_prime_UTR_variant |
| 7 | 12911808 | 12912362 | 554 | Vmn1r88 | C58J | feature_elongation,coding_sequence_variant |
| 7 | 12951497 | 12951688 | 191 | Vmn1r89 | C58J | feature_elongation,5_prime_UTR_variant |
| 7 | 13093681 | 13093853 | 172 | Pla2g4c | C58J,CAST,MOLF,SPRET | feature_elongation,3_prime_UTR_variant |
| 7 | 13142498 | 13142584 | 86 | Cabp5 | SPRET | feature_elongation,3_prime_UTR_variant |
| 7 | 15671590 | 15671749 | 159 | Nop53 | BTBR,BUB,C58J,CE,FVB,KK,MOLF,NOD,NOR,NZO,PL,WSB | feature_elongation,3_prime_UTR_variant |
| 7 | 15704946 | 15705004 | 58 | Bicra | SPRET | feature_elongation,3_prime_UTR_variant |
| 7 | 15968971 | 15969031 | 60 | C5ar2 | 129S1,AJ,AKR,BALB,BTBR,C3H,C57L,C58J,CBA,DBA1J,DBA2J,ILNJ2,KK,LP,MA/My,MRL,NZB,NZO,NZW,PJ,RF,RH2,SEA,SJL,SMJ,TH,WSB | feature_elongation,3_prime_UTR_variant |
| 7 | 15970294 | 15970376 | 82 | C5ar2 | 129S1,AJ,AKR,BALB,BTBR,C3H,C57L,CBA,DBA1J,DBA2J,ILNJ2,KK,LP,MA/My,MRL,NZB,NZO,NZW,PJ,RF,RH2,SEA,SJL,SMJ,TH,WSB | feature_elongation,3_prime_UTR_variant |
| 7 | 16054663 | 16054731 | 68 | Sae1 | 129S1,BTBR,C3H,C57L,C58J,CBA,DBA1J,DBA2J,MA/My,MOLF,NZB,PJ,RF,SEA,SMJ,SPRET | feature_elongation,3_prime_UTR_variant |
| 7 | 16459558 | 16459628 | 70 | Ceacam9 | CAST,MOLF,NOD,NOR,SJL,SPRET | feature_elongation,3_prime_UTR_variant |
| 7 | 16729484 | 16729580 | 96 | Ccdc8 | SPRET | feature_elongation,coding_sequence_variant |
| 7 | 16774896 | 16775053 | 157 | Hif3a | 129S1,AJ,BALB,BTBR,BUB,C3H,C57L,CAST,CBA,CE,FVB,ILNJ2,LP,MOLF,MRL,NZB,NZW,PL,RF,SPRET,WSB | stop_gained,frameshift_variant |
| 7 | 18958097 | 18958311 | 214 | Gpr4 | 129S1,BTBR,C3H,C57L,CBA,MOLF,NOD,NOR,NZB,NZW,RF,SJL | feature_elongation,3_prime_UTR_variant |
| 7 | 19078720 | 19078784 | 64 | Ercc1 | AKR,BUB,C58J,CE,MRL,NZO,TH | feature_elongation,5_prime_UTR_variant |
| 7 | 19724354 | 19724654 | 300 | Ceacam20 | 129S1,BTBR,C3H,C57L,CAST,CBA,MOLF,NOD,NOR,NZB,RF,SJL | feature_elongation,3_prime_UTR_variant |
| 7 | 23203747 | 23203841 | 94 | Vmn1r167 | 129S1,BTBR,BUB,C3H,C57L,CAST,CBA,MOLF,NOD,NOR,NZB,NZW,SJL | feature_elongation,3_prime_UTR_variant |
| 7 | 23277823 | 23277877 | 54 | Vmn1r169 | CAST | feature_elongation,3_prime_UTR_variant |
| 7 | 23461753 | 23461962 | 209 | Vmn1r174 | SPRET | feature_elongation,3_prime_UTR_variant |
| 7 | 23536610 | 23537179 | 569 | Vmn1r176 | 129S1,BTBR,BUB,C3H,C57L,CBA,MOLF,NOD,NOR,NZB,NZW,SJL | feature_elongation,5_prime_UTR_variant |
| 7 | 23758811 | 23758865 | 54 | Vmn1r183 | CAST | feature_elongation,3_prime_UTR_variant |
| 7 | 23880563 | 23880638 | 75 | Zfp114 | 129S1,BTBR,BUB,C3H,C57L,CBA,MOLF,NOD,NOR,NZB,NZW,SJL,SPRET,WSB | feature_elongation,coding_sequence_variant |
| 7 | 24588613 | 24588782 | 169 | Rps19 | MOLF | feature_elongation,coding_sequence_variant |
| 7 | 25967615 | 25967774 | 159 | Vmn1r184 | WSB | feature_elongation,3_prime_UTR_variant |
| 7 | 26906579 | 26906789 | 210 | Actmap | SPRET | feature_elongation,3_prime_UTR_variant |
| 7 | 27367754 | 27367811 | 57 | Map3k10 | MOLF | feature_elongation,3_prime_UTR_variant |
| 7 | 27450569 | 27450782 | 213 | Zfp60 | TH | feature_elongation,3_prime_UTR_variant |
| 7 | 27556978 | 27557327 | 349 | Zfp607a | CAST | feature_elongation,5_prime_UTR_variant |
| 7 | 28922343 | 28922476 | 133 | Kcnk6 | MOLF | feature_elongation,3_prime_UTR_variant |
| 7 | 29484201 | 29484326 | 125 | Zfp30 | CAST,MOLF | feature_elongation,5_prime_UTR_variant |
| 7 | 29494037 | 29494221 | 184 | Zfp30 | SPRET | feature_elongation,3_prime_UTR_variant |
| 7 | 29759642 | 29759817 | 175 | Zfp82 | SPRET | feature_elongation,3_prime_UTR_variant |
| 7 | 30451812 | 30451905 | 93 | Sbsn | 129S1,AJ,AKR,BALB,BTBR,C57L,C58J,CAST,CE,DBA1J,DBA2J,ILNJ2,LP,MA/My,MOLF,MRL,NOD,NOR,NZB,NZO,NZW,PJ,PL,RF,SEA,SJL,SMJ,SPRET,TH,WSB | feature_elongation,coding_sequence_variant |
| 7 | 30518388 | 30518465 | 77 | Ffar2 | SPRET | feature_elongation,3_prime_UTR_variant |
| 7 | 30557530 | 30557584 | 54 | Ffar1 | SPRET | feature_elongation,3_prime_UTR_variant |

| | | | | | | |
|---|---|---|---|---|---|---|
| 7 | 30577837 | 30578631 | 794 | Cd22 | AKR,BTBR,C57L,CAST,DBA1J,DBA2J,MOLF,NZB,NZO,NZW,PJ,PL,SEA,SMJ,SPRET,TH,WSB | feature_elongation,5_prime_UTR_variant |
| 7 | 35246858 | 35247033 | 175 | Nudt19 | BTBR,C57L,C58J,CE,DBA1J,DBA2J,FVB,ILNJ2,LP,MA/My,MRL,NOD,NOR,PJ,RF,SJL,SMJ,SWR | feature_elongation,3_prime_UTR_variant |
| 7 | 35387258 | 35387396 | 138 | Dpy19l3 | CAST,SPRET | feature_elongation,3_prime_UTR_variant |
| 7 | 41300737 | 41300915 | 178 | Zfp788 | AJ,BALB,C57L,CAST,CE,FVB,MA/My,MOLF,NOD,NOR,NZB,NZO,PL,RH2,SEA,SMJ,SPRET,SWR,TH,WSB | feature_elongation,3_prime_UTR_variant |
| 7 | 42123693 | 42124209 | 516 | Zfp141 | CAST | feature_elongation,3_prime_UTR_variant |
| 7 | 42841210 | 42841301 | 91 | Zfp936 | SPRET | feature_elongation,3_prime_UTR_variant |
| 7 | 43142044 | 43142244 | 200 | 4931406B18Rik | DBA1J,DBA2J,ILNJ2,NZW,PJ | feature_elongation,3_prime_UTR_variant |
| 7 | 43175234 | 43175447 | 213 | Cd33 | SPRET | feature_elongation,3_prime_UTR_variant |
| 7 | 43326426 | 43326540 | 114 | Ctu1 | SPRET | feature_elongation,coding_sequence_variant |
| 7 | 43344596 | 43344720 | 124 | Klk14 | 129S1,AJ,AKR,B10D2,B10J,BALB,BTBR,BUB,C3H,C57L,C58J,CAST,CBA,CE,DBA1J,DBA2J,FVB,ILNJ2,KK,LP,MA/My,MOLF,MRL,NOD,NOR,NZB,NZO,NZW,PJ,PL,RF,RH2,SEA,SJL,SMJ,SWR,TH,WSB | feature_elongation,3_prime_UTR_variant |
| 7 | 43898363 | 43898476 | 113 | 2410002F23Rik | CAST | feature_elongation,5_prime_UTR_variant |
| 7 | 43952237 | 43952308 | 71 | Clec11a | MOLF | feature_elongation,3_prime_UTR_variant |
| 7 | 43959688 | 43960021 | 333 | Shank1 | 129S1,AJ,AKR,B10D2,B10J,BALB,BTBR,BUB,C3H,C57L,C58J,CAST,CBA,CE,DBA1J,DBA2J,FVB,ILNJ2,KK,LP,MA/My,MOLF,MRL,NOD,NOR,NZB,NZO,NZW,PJ,PL,RF,RH2,SEA,SJL,SMJ,SPRET,SWR,TH,WSB | feature_elongation,5_prime_UTR_variant |
| 7 | 44385782 | 44386153 | 371 | Zfp473 | ILNJ2,MOLF,NZW,PJ,SJL | feature_elongation,3_prime_UTR_variant |
| 7 | 44727578 | 44727787 | 209 | Nosip | CAST,MOLF | feature_elongation,3_prime_UTR_variant |
| 7 | 44906960 | 44907025 | 65 | Slc6a16 | ILNJ2,NZW,PJ,SJL,SPRET | frameshift_variant,splice_region_variant |
| 7 | 45122165 | 45122382 | 217 | Dhdh | CAST,MOLF | feature_elongation,3_prime_UTR_variant |
| 7 | 45173863 | 45174334 | 471 | Ppp1r15a | SPRET | feature_elongation,coding_sequence_variant |
| 7 | 45456389 | 45456461 | 72 | Cyth2 | SPRET | feature_elongation,3_prime_UTR_variant |
| 7 | 46079660 | 46079740 | 80 | Kcnc1 | C58J | feature_elongation,3_prime_UTR_variant |
| 7 | 46574276 | 46574702 | 426 | Uevld | CAST | feature_elongation,3_prime_UTR_variant |
| 7 | 46575252 | 46575325 | 73 | Uevld | MOLF | feature_elongation,3_prime_UTR_variant |
| 7 | 49258159 | 49258214 | 55 | Nav2 | ILNJ2,MOLF,SJL | feature_elongation,3_prime_UTR_variant |
| 7 | 49609827 | 49609883 | 56 | Slc6a5 | ILNJ2,MOLF,NZW,PJ,SJL | feature_elongation,3_prime_UTR_variant |
| 7 | 50249178 | 50249503 | 325 | 4933405O20Rik | KK | feature_elongation,coding_sequence_variant |
| 7 | 55428213 | 55428375 | 162 | Siglech | CAST,ILNJ2,KK,MOLF,NZW,PJ,SJL,SPRET | feature_elongation,3_prime_UTR_variant |
| 7 | 56185816 | 56185876 | 60 | Oca2 | CAST | feature_elongation,3_prime_UTR_variant |
| 7 | 66058853 | 66058913 | 60 | Aldh1a3 | CAST | feature_elongation,coding_sequence_variant |
| 7 | 66297327 | 66297383 | 56 | Asb7 | SPRET | feature_elongation,3_prime_UTR_variant |
| 7 | 77830704 | 77830906 | 202 | Ntrk3 | SPRET | feature_elongation,3_prime_UTR_variant |
| 7 | 79059137 | 79059354 | 217 | Fanci | SPRET | feature_elongation,3_prime_UTR_variant |
| 7 | 79096138 | 79096348 | 210 | Polg | SPRET | feature_elongation,3_prime_UTR_variant |
| 7 | 79527986 | 79528065 | 79 | Ap3s2 | SPRET | feature_elongation,3_prime_UTR_variant |
| 7 | 79575868 | 79576007 | 139 | Arpin | SPRET | feature_elongation,3_prime_UTR_variant |
| 7 | 79585003 | 79585057 | 54 | Arpin | SPRET | feature_elongation,5_prime_UTR_variant |
| 7 | 83547698 | 83547866 | 168 | Mesd | CAST | feature_elongation,3_prime_UTR_variant |
| 7 | 84587416 | 84588411 | 995 | Vmn2r65 | SMJ | feature_elongation,3_prime_UTR_variant |
| 7 | 85597826 | 85598755 | 929 | Vmn2r74 | 129S1,AJ,AKR,BALB,BTBR,BUB,C3H,C57L,CBA,DBA1J,DBA2J,FVB,ILNJ2,KK,MA/My,MRL,NOD,NOR,NZB,NZO,NZW,PJ,PL,RF,RH2,SEA,SJL,SPRET,SWR,TH | feature_elongation,3_prime_UTR_variant |
| 7 | 85599024 | 85599088 | 64 | Vmn2r74 | SPRET | feature_elongation,3_prime_UTR_variant |
| 7 | 85796144 | 85797032 | 888 | Vmn2r75 | SPRET | feature_elongation,3_prime_UTR_variant |
| 7 | 85851863 | 85851939 | 76 | Vmn2r76 | CAST,MOLF | feature_elongation,3_prime_UTR_variant |
| 7 | 85874046 | 85874106 | 60 | Vmn2r76 | SPRET | feature_elongation,3_prime_UTR_variant |
| 7 | 85983563 | 85983618 | 55 | Or14a260 | 129S1,BTBR,C3H,C57L,CBA,DBA1J,DBA2J,ILNJ2,KK,NZB,NZO,NZW,PJ,SEA,SPRET | feature_elongation,3_prime_UTR_variant |
| 7 | 86010456 | 86011085 | 629 | Or14a259 | SPRET | feature_elongation,3_prime_UTR_variant |
| 7 | 86655159 | 86655211 | 52 | Vmn2r79 | CAST,MOLF | feature_elongation,3_prime_UTR_variant |
| 7 | 89176234 | 89176316 | 82 | Prss23 | SMJ | feature_elongation,5_prime_UTR_variant |
| 7 | 89502170 | 89502411 | 241 | Me3 | SMJ | feature_elongation,3_prime_UTR_variant |
| 7 | 97100578 | 97100757 | 179 | Kctd14 | CAST | feature_elongation,5_prime_UTR_variant |
| 7 | 97108129 | 97108312 | 183 | Kctd14 | 129S1,AJ,AKR,BALB,BTBR,BUB,C57L,CE,FVB,KK,MA/My,MRL,NOD,NOR,NZB,NZW,PL,RF,RH2,SJL,SMJ,SPRET,SWR,TH | feature_elongation,3_prime_UTR_variant |
| 7 | 97856120 | 97856647 | 527 | Acer3 | CE | feature_elongation,3_prime_UTR_variant |
| 7 | 98534443 | 98534595 | 152 | Uvrag | SPRET | feature_elongation,3_prime_UTR_variant |
| 7 | 98918049 | 98918187 | 138 | Map6 | C3H,CBA,DBA1J,DBA2J,ILNJ2,MOLF,PJ,SEA | feature_elongation,coding_sequence_variant |
| 7 | 101471993 | 101472080 | 87 | Inppl1 | CAST,SPRET | feature_elongation,3_prime_UTR_variant |
| 7 | 101530539 | 101530902 | 363 | Anapc15 | C3H,CAST,CBA,DBA1J,DBA2J,ILNJ2,MOLF,PJ,SEA,SPRET | feature_elongation,5_prime_UTR_variant |
| 7 | 102141979 | 102142159 | 180 | Or55b10 | KK,NOD,SJL,SMJ,SPRET,SWR | feature_elongation,3_prime_UTR_variant |
| 7 | 102204962 | 102205166 | 204 | Or52b3 | 129S1,AJ,AKR,BALB,BTBR,BUB,C3H,C57L,CAST,CBA,DBA1J,DBA2J,FVB,ILNJ2,MOLF,MRL,NZB,NZO,NZW,PJ,PL,RF,RH2,SEA,SMJ,TH,WSB | feature_elongation,3_prime_UTR_variant |
| 7 | 102255617 | 102255900 | 283 | Or52k2 | KK,NOD,SJL,SPRET,SWR | feature_elongation,3_prime_UTR_variant |
| 7 | 102320663 | 102320716 | 53 | Or52i2 | MOLF | feature_elongation,3_prime_UTR_variant |
| 7 | 102389211 | 102389353 | 142 | Or51e2 | MA/My,NZO,SMJ,TH,WSB | feature_elongation,3_prime_UTR_variant |
| 7 | 102390313 | 102390830 | 517 | Or51e2 | SPRET | feature_elongation,3_prime_UTR_variant |
| 7 | 102504826 | 102504885 | 59 | Or51f1 | SPRET | feature_elongation,3_prime_UTR_variant |
| 7 | 102557424 | 102558227 | 803 | Or51s1 | SPRET | feature_elongation,3_prime_UTR_variant |
| 7 | 102747915 | 102748357 | 442 | Or51f1e | CAST,MOLF | feature_elongation,3_prime_UTR_variant |
| 7 | 102796672 | 102797077 | 405 | Or51p1 | SPRET | feature_elongation,3_prime_UTR_variant |
| 7 | 102955782 | 102955978 | 196 | Or52e19 | CAST,KK,NOD,SJL,SPRET,SWR | feature_elongation,5_prime_UTR_variant |
| 7 | 103101635 | 103102174 | 539 | Or51l14 | NZO,SMJ,TH,WSB | feature_elongation,3_prime_UTR_variant |
| 7 | 103186506 | 103186558 | 52 | Or51aa2 | CAST | feature_elongation,3_prime_UTR_variant |
| 7 | 103367005 | 103367226 | 221 | Or52a20 | KK,MA/My,NOD,NZO,SJL,SMJ,SWR | feature_elongation,3_prime_UTR_variant |
| 7 | 103595519 | 103595607 | 88 | Or51k2 | KK,NOD,SJL,SMJ,SWR,TH | feature_elongation,5_prime_UTR_variant |

| | | | | | | |
|---|---|---|---|---|---|---|
| 7 | 103696576 | 103696698 | 122 | Or51a10 | KK,MOLF,NOD,SJL,SMJ,SWR,TH | feature_elongation,3_prime_UTR_variant |
| 7 | 103732677 | 103733110 | 433 | Or51a24 | CAST,KK,MOLF,NOD,SJL,SMJ,SWR,TH | feature_elongation,3_prime_UTR_variant |
| 7 | 103882736 | 103882823 | 87 | Trim6 | CAST | feature_elongation,3_prime_UTR_variant |
| 7 | 103911127 | 103911331 | 204 | Trim34a | 129S1,AJ,AKR,BALB,BTBR,BUB,C3H,C57L,CBA,DBA1J,DBA2J,FVB,ILNJ2,MRL,NZB,NZW,PJ,RF,SEA | feature_elongation,3_prime_UTR_variant |
| 7 | 103913718 | 103913796 | 78 | Trim5 | CAST,MOLF,SPRET | feature_elongation,3_prime_UTR_variant |
| 7 | 104012324 | 104012705 | 381 | Trim30b | 129S1,AJ,AKR,BALB,BTBR,BUB,C3H,C57L,CBA,CE,DBA1J,DBA2J,FVB,ILNJ2,MRL,NZB,NZO,NZW,PJ,PL,RF,RH2,SEA,SPRET,WSB | feature_elongation,3_prime_UTR_variant |
| 7 | 104060223 | 104060327 | 104 | Trim30a | CAST,SPRET | feature_elongation,3_prime_UTR_variant |
| 7 | 104231490 | 104231560 | 70 | Or52d3 | KK,MOLF,NZO,SMJ,SPRET,WSB | feature_elongation,3_prime_UTR_variant |
| 7 | 104671777 | 104671833 | 56 | Or52e8b | MOLF | feature_elongation,3_prime_UTR_variant |
| 7 | 104967496 | 104967594 | 98 | Or56b35 | SPRET | feature_elongation,3_prime_UTR_variant |
| 7 | 104978342 | 104978404 | 62 | Or52b1 | SPRET | feature_elongation,3_prime_UTR_variant |
| 7 | 105259731 | 105259950 | 219 | Trim3 | CAST,CE,KK,WSB | feature_elongation,3_prime_UTR_variant |
| 7 | 105292476 | 105292536 | 60 | Timm10b | CE,WSB | feature_elongation,3_prime_UTR_variant |
| 7 | 106525049 | 106525105 | 56 | Or2d3c | MOLF | feature_elongation,3_prime_UTR_variant |
| 7 | 106572863 | 106573200 | 337 | Or6b6 | KK,WSB | feature_elongation,5_prime_UTR_variant |
| 7 | 106599825 | 106599898 | 73 | Or6a2 | 129S1,AJ,AKR,BALB,BTBR,C3H,C57L,CBA,DBA1J,DBA2J,ILNJ2,MOLF,MRL,NZB,NZO,NZW,PJ,PL,RF,RH2,SEA | feature_elongation,3_prime_UTR_variant |
| 7 | 106636676 | 106636747 | 71 | Or10a5 | CAST,NZO | feature_elongation,3_prime_UTR_variant |
| 7 | 107630450 | 107630530 | 80 | Or5p6 | CAST,NOD,SMJ,SWR | feature_elongation,3_prime_UTR_variant |
| 7 | 107681963 | 107682063 | 100 | Or5p4 | CAST,NOD,NZO,SMJ,SWR | feature_elongation,3_prime_UTR_variant |
| 7 | 107721160 | 107721947 | 787 | Or5p60 | CAST,MOLF,NOD,SMJ,SPRET,SWR | feature_elongation,3_prime_UTR_variant |
| 7 | 107723055 | 107723405 | 350 | Or5p60 | MOLF,SPRET | feature_elongation,3_prime_UTR_variant |
| 7 | 107851327 | 107851522 | 195 | Or5p64 | WSB | feature_elongation,3_prime_UTR_variant |
| 7 | 107918020 | 107918108 | 88 | Or5p1 | CAST,NOD,SMJ,SWR | feature_elongation,3_prime_UTR_variant |
| 7 | 109358293 | 109358394 | 101 | Nrip3 | SPRET | feature_elongation,3_prime_UTR_variant |
| 7 | 109907897 | 109907973 | 76 | Sbf2 | MOLF,NOD,SWR | feature_elongation,3_prime_UTR_variant |
| 7 | 110415273 | 110415351 | 78 | Rnf141 | SPRET | feature_elongation,3_prime_UTR_variant |
| 7 | 110655290 | 110655345 | 55 | Ctr9 | CAST,SMJ | feature_elongation,3_prime_UTR_variant |
| 7 | 118339516 | 118339590 | 74 | Vps35l | WSB | feature_elongation,5_prime_UTR_variant |
| 7 | 119315944 | 119315998 | 54 | Thumpd1 | CAST | feature_elongation,coding_sequence_variant |
| 7 | 119552375 | 119552431 | 56 | Lyrm1 | AJ,BALB,BTBR,C3H,CBA,DBA1J,DBA2J,ILNJ2,KK,MOLF,NOR,NZB,NZO,PL,RH2,TH,WSB | feature_elongation,3_prime_UTR_variant |
| 7 | 119783295 | 119783346 | 51 | Anks4b | 129S1,AKR,B10D2,B10J,C57L,C58J,FVB,LP,MA/My,MRL,NOD,NZW,PJ,RF,SEA,SJL,SMJ,SWR | feature_elongation,3_prime_UTR_variant |
| 7 | 119784151 | 119784247 | 96 | Anks4b | SPRET | feature_elongation,3_prime_UTR_variant |
| 7 | 119924543 | 119924910 | 367 | Abca14 | SPRET | feature_elongation,3_prime_UTR_variant |
| 7 | 120996884 | 120996936 | 52 | Hs3st2 | BTBR | feature_elongation,3_prime_UTR_variant |
| 7 | 122583034 | 122583217 | 183 | Rbbp6 | MOLF | feature_elongation,3_prime_UTR_variant |
| 7 | 123076797 | 123077155 | 358 | Zkscan2 | SPRET | feature_elongation,3_prime_UTR_variant |
| 7 | 126170690 | 126171104 | 414 | Cln3 | CAST,SPRET | feature_elongation,3_prime_UTR_variant |
| 7 | 126186855 | 126186921 | 66 | Apobr | AJ,CAST,MOLF,NZB,NZO,SPRET,TH | feature_elongation,coding_sequence_variant |
| 7 | 126989120 | 126989300 | 180 | E430018J23Rik | AJ,WSB | feature_elongation,3_prime_UTR_variant |
| 7 | 127463663 | 127463818 | 155 | Zfp668 | MOLF,SPRET,TH | feature_elongation,3_prime_UTR_variant |
| 7 | 127870808 | 127870920 | 112 | Rusf1 | MOLF | feature_elongation,3_prime_UTR_variant |
| 7 | 128297801 | 128297854 | 53 | Inpp5f | SPRET | feature_elongation,3_prime_UTR_variant |
| 7 | 130940306 | 130940377 | 71 | 2310057M21Rik | MOLF,SPRET | feature_elongation,3_prime_UTR_variant |
| 7 | 131049260 | 131049376 | 116 | Acadsb | SPRET | feature_elongation,3_prime_UTR_variant |
| 7 | 135286041 | 135286159 | 118 | Ptpre | ILNJ2,WSB | feature_elongation,3_prime_UTR_variant |
| 7 | 136988930 | 136989096 | 166 | 9430038I01Rik | CAST | feature_elongation,3_prime_UTR_variant |
| 7 | 139740901 | 139741790 | 889 | Or6ae1 | MOLF,PJ | feature_elongation,3_prime_UTR_variant |
| 7 | 139917699 | 139917823 | 124 | Or12j2 | MOLF,NZW,PJ | feature_elongation,3_prime_UTR_variant |
| 7 | 139924440 | 139924653 | 213 | Or13a27 | CE,SPRET | feature_elongation,3_prime_UTR_variant |
| 7 | 139952248 | 139952443 | 195 | Or12j3 | MOLF,NZW,PJ | feature_elongation,3_prime_UTR_variant |
| 7 | 140253094 | 140253158 | 64 | Or13a25 | SPRET | feature_elongation,3_prime_UTR_variant |
| 7 | 140507818 | 140507874 | 56 | Nlrp6 | ILNJ2,WSB | feature_elongation,3_prime_UTR_variant |
| 7 | 140681232 | 140681356 | 124 | Ano9 | CAST,MOLF,NZW | feature_elongation,3_prime_UTR_variant |
| 7 | 141219084 | 141219335 | 251 | Muc6 | NZW,SPRET,WSB | feature_elongation,coding_sequence_variant |
| 7 | 141221227 | 141221527 | 300 | Muc6 | ILNJ2,SPRET | feature_elongation,coding_sequence_variant |
| 7 | 141225447 | 141226248 | 801 | Muc6 | SPRET | feature_elongation,coding_sequence_variant |
| 7 | 141292037 | 141292122 | 85 | Muc2 | CAST,MOLF,SPRET | frameshift_variant |
| 7 | 141362384 | 141362527 | 143 | Muc5ac | CAST,MOLF,SPRET | feature_elongation,coding_sequence_variant |
| 7 | 141363453 | 141363645 | 192 | Muc5ac | 129S1,AKR,B10D2,B10J,BALB,BTBR,BUB,C3H,C57L,C58J,CAST,CBA,CE,DBA1J,DBA2J,FVB,ILNJ2,LP,MA/My,MOLF,MRL,NOD,NOR,NZB,PL,RF,RH2,SEA,SJL,SMJ,SPRET,SWR,TH,WSB | feature_elongation,coding_sequence_variant |
| 7 | 141412631 | 141413156 | 525 | Muc5b | SPRET | feature_elongation,coding_sequence_variant |
| 7 | 141689104 | 141689164 | 60 | Gm39115 | SPRET | feature_elongation,coding_sequence_variant |
| 7 | 141697858 | 141697912 | 54 | Gm45337 | MOLF | feature_elongation,coding_sequence_variant |
| 7 | 141710654 | 141710714 | 60 | Gm29735 | SPRET | feature_elongation,coding_sequence_variant |
| 7 | 141718768 | 141718852 | 84 | Gm4553 | SPRET | feature_elongation,coding_sequence_variant |
| 7 | 141743604 | 141743807 | 203 | Gm10153 | BALB,CAST,MRL,RF,RH2,SEA,SPRET | stop_gained,frameshift_variant |
| 7 | 141755630 | 141755707 | 77 | Krtap5-3 | NZW | feature_elongation,coding_sequence_variant |
| 7 | 141783349 | 141783532 | 183 | Krtap5-5 | CAST,SPRET | feature_elongation,coding_sequence_variant |
| 7 | 141827637 | 141827751 | 114 | Gm4559 | CAST,MOLF,WSB | feature_elongation,coding_sequence_variant |
| 7 | 141850268 | 141850322 | 54 | Krtap5-1 | MOLF,SPRET,WSB | feature_elongation,coding_sequence_variant |
| 7 | 143190695 | 143190906 | 211 | Tnfrsf22 | C58J,CAST,LP,MA/My,MOLF,PJ,SMJ | feature_elongation,3_prime_UTR_variant |
| 8 | 3205325 | 3205449 | 124 | Insr | CAST,SPRET | feature_elongation,3_prime_UTR_variant |
| 8 | 4577865 | 4578156 | 291 | Cers4 | 129S1,CAST,NZB | feature_elongation,3_prime_UTR_variant |
| 8 | 5134129 | 5134233 | 104 | Slc10a2 | CAST,SPRET | feature_elongation,3_prime_UTR_variant |
| 8 | 10027595 | 10027863 | 268 | Lig4 | SPRET | feature_elongation,5_prime_UTR_variant |
| 8 | 10620272 | 10620335 | 63 | Myo16 | CAST,MOLF | feature_elongation,coding_sequence_variant |
| 8 | 11035636 | 11035696 | 60 | Irs2 | BTBR | feature_elongation,3_prime_UTR_variant |
| 8 | 11453015 | 11453135 | 120 | Col4a2 | CAST | feature_elongation,coding_sequence_variant |
| 8 | 11662311 | 11662381 | 70 | Ankrd10 | MOLF,NZB,SMJ,TH | feature_elongation,3_prime_UTR_variant |
| 8 | 12917049 | 12917140 | 91 | Atp11a | C57L,ILNJ2,NZB,NZW,PJ,PL,SEA,SMJ,TH | feature_elongation,3_prime_UTR_variant |

| | | | | | | |
|---|---|---|---|---|---|---|
| 8 | 13068432 | 13068530 | 98 | Mcf2l | 129S1,C57L,CE,ILNJ2,LP,MOLF,NZB,NZO,NZW,PJ,PL,SEA,SMJ,TH | feature_elongation,3_prime_UTR_variant |
| 8 | 13106216 | 13106276 | 60 | F10 | WSB | feature_elongation,3_prime_UTR_variant |
| 8 | 13125751 | 13125871 | 120 | Proz | SPRET | feature_elongation,3_prime_UTR_variant |
| 8 | 15076836 | 15076894 | 58 | Kbtbd11 | SPRET | feature_elongation,5_prime_UTR_variant |
| 8 | 15091988 | 15092072 | 84 | BB014433 | AJ,AKR,BALB,BTBR,BUB,C3H,CAST,CBA,CE,DBA1J,DBA2J,FVB,KK,LP,MRL,NOD,NOR,PL,RF,RH2,SJL,SMJ,SWR,TH | feature_elongation,coding_sequence_variant |
| 8 | 25171513 | 25171573 | 60 | Adam3 | SPRET | feature_elongation,coding_sequence_variant |
| 8 | 25531894 | 25531995 | 101 | Plekha2 | CAST,MOLF,PL | feature_elongation,3_prime_UTR_variant |
| 8 | 26471208 | 26471337 | 129 | Pomk | SPRET | feature_elongation,3_prime_UTR_variant |
| 8 | 26515715 | 26515887 | 172 | Hook3 | SPRET | feature_elongation,3_prime_UTR_variant |
| 8 | 26516864 | 26516918 | 54 | Hook3 | CAST | feature_elongation,3_prime_UTR_variant |
| 8 | 26517819 | 26517905 | 86 | Hook3 | SPRET | feature_elongation,3_prime_UTR_variant |
| 8 | 26518839 | 26519008 | 169 | Hook3 | CAST | feature_elongation,3_prime_UTR_variant |
| 8 | 27538594 | 27538668 | 74 | Plpbp | CAST,MOLF,PL | feature_elongation,3_prime_UTR_variant |
| 8 | 27605370 | 27605423 | 53 | Adgra2 | WSB | feature_elongation,coding_sequence_variant |
| 8 | 31750969 | 31751053 | 84 | Fut10 | 129S1,AJ,BALB,BUB,C57L,CBA,FVB,MRL,NOD,NOR,PL,RH2,SJL,SMJ,SWR,TH,WSB | feature_elongation,3_prime_UTR_variant |
| 8 | 34110054 | 34110135 | 81 | Ubxn8 | 129S1,AJ,AKR,BALB,BUB,C3H,C57L,CBA,CE,DBA1J,DBA2J,FVB,MA/My,MRL,NOD,NOR,PL,RF,RH2,SJL,SMJ,SWR,TH,WSB | feature_elongation,3_prime_UTR_variant |
| 8 | 34637775 | 34637903 | 128 | Saraf | 129S1,AJ,AKR,BALB,C3H,C57L,CBA,CE,DBA1J,DBA2J,MA/My,MRL,PL,RF,RH2,SMJ | feature_elongation,3_prime_UTR_variant |
| 8 | 36145482 | 36145542 | 60 | Mfhas1 | CAST,WSB | feature_elongation,3_prime_UTR_variant |
| 8 | 39618525 | 39618656 | 131 | Tusc3 | SPRET | feature_elongation,3_prime_UTR_variant |
| 8 | 40963716 | 40963929 | 213 | Cnot7 | SPRET | feature_elongation,5_prime_UTR_variant |
| 8 | 41370766 | 41370828 | 62 | Slc7a2 | 129S1,AJ,AKR,BALB,BTBR,BUB,C3H,C57L,CBA,CE,DBA1J,DBA2J,FVB,ILNJ2,KK,MA/My,MRL,NOD,NOR,NZB,NZO,NZW,PJ,PL,RF,RH2,SEA,SJL,SMJ,SWR,TH,WSB | feature_elongation,3_prime_UTR_variant |
| 8 | 45757987 | 45758338 | 351 | Cyp4v3 | WSB | feature_elongation,3_prime_UTR_variant |
| 8 | 46427220 | 46427289 | 69 | Cfap96 | SPRET | feature_elongation,5_prime_UTR_variant |
| 8 | 46481334 | 46481497 | 163 | Lrp2bp | SPRET | feature_elongation,3_prime_UTR_variant |
| 8 | 47635353 | 47635409 | 56 | Stox2 | CAST | feature_elongation,3_prime_UTR_variant |
| 8 | 47964107 | 47964347 | 240 | Trappc11 | WSB | feature_elongation,5_prime_UTR_variant |
| 8 | 61957362 | 61957429 | 67 | Cbr4 | SPRET | feature_elongation,3_prime_UTR_variant |
| 8 | 61959421 | 61959496 | 75 | Cbr4 | SPRET | feature_elongation,3_prime_UTR_variant |
| 8 | 65194683 | 65194736 | 53 | Klhl2 | PJ,SEA,WSB | feature_elongation,3_prime_UTR_variant |
| 8 | 65434887 | 65435017 | 130 | Trim75 | SPRET | feature_elongation,3_prime_UTR_variant |
| 8 | 65465819 | 65465883 | 64 | Trim61 | AKR,BALB,C3H,C57L,CBA,CE,DBA1J,DBA2J,MA/My,MRL,NZB,NZO,PJ,PL,RF,SEA,SMJ,SPRET,WSB | feature_elongation,3_prime_UTR_variant |
| 8 | 69281315 | 69281481 | 166 | Ints10 | SPRET | feature_elongation,3_prime_UTR_variant |
| 8 | 69681772 | 69681982 | 210 | Zfp930 | CAST,MOLF | feature_elongation,3_prime_UTR_variant |
| 8 | 69755842 | 69756413 | 571 | D130040H23Rik | SPRET | feature_elongation,3_prime_UTR_variant |
| 8 | 69820870 | 69821295 | 425 | Gm10033 | CAST,MOLF | feature_elongation,3_prime_UTR_variant |
| 8 | 69823455 | 69823517 | 62 | Gm10033 | AKR,BUB,C3H,C58J,CE,DBA1J,DBA2J,MA/My,MRL,NZB,PL,SMJ | feature_elongation,3_prime_UTR_variant |
| 8 | 69823978 | 69824631 | 653 | Gm10033 | NZB,NZO | feature_elongation,3_prime_UTR_variant |
| 8 | 70117125 | 70117525 | 400 | Zfp964 | SPRET | feature_elongation,3_prime_UTR_variant |
| 8 | 70157767 | 70157971 | 204 | Zfp869 | PJ,SEA | feature_elongation,3_prime_UTR_variant |
| 8 | 70280950 | 70281152 | 202 | Lpar2 | 129S1,AKR,BALB,BUB,C3H,C57L,C58J,CBA,CE,DBA1J,DBA2J,FVB,ILNJ2,KK,MA/My,MRL,NZB,NZO,NZW,PJ,PL,RF,SEA,SJL,SMJ,SWR,TH,WSB | feature_elongation,3_prime_UTR_variant |
| 8 | 70281367 | 70281751 | 384 | Lpar2 | SPRET | feature_elongation,3_prime_UTR_variant |
| 8 | 70360078 | 70360187 | 109 | Gatad2a | 129S1,BALB,C3H,C57L,CBA,DBA1J,DBA2J,FVB,ILNJ2,KK,NZB,NZO,RF,SJL,SMJ,SWR,TH | feature_elongation,3_prime_UTR_variant |
| 8 | 70541798 | 70541892 | 94 | Hapln4 | SPRET | feature_elongation,3_prime_UTR_variant |
| 8 | 70745517 | 70745750 | 233 | Ddx49 | SPRET | feature_elongation,3_prime_UTR_variant |
| 8 | 71295045 | 71295253 | 208 | Kcnn1 | C58J,NZW | feature_elongation,3_prime_UTR_variant |
| 8 | 71347378 | 71347436 | 58 | Rpl18a | 129S1,AKR,BALB,BUB,C3H,C57L,C58J,CAST,CBA,CE,DBA1J,DBA2J,FVB,ILNJ2,KK,MA/My,MRL,NZB,NZO,NZW,PJ,PL,RF,SEA,SJL,SMJ,SPRET,SWR,TH,WSB | feature_elongation,3_prime_UTR_variant |
| 8 | 72144752 | 72144957 | 205 | B3gnt3 | AKR,BUB,CE,MA/My,MRL,PL | feature_elongation,3_prime_UTR_variant |
| 8 | 72811377 | 72811581 | 204 | Or2z8 | SPRET | feature_elongation,5_prime_UTR_variant |
| 8 | 72934873 | 72934991 | 118 | Rab8a | SPRET | feature_elongation,3_prime_UTR_variant |
| 8 | 72955236 | 72955295 | 59 | Hsh2d | NZO,PJ,SEA,SMJ | feature_elongation,3_prime_UTR_variant |
| 8 | 73199314 | 73199542 | 228 | 1700030K09Rik | SPRET | feature_elongation,coding_sequence_variant |
| 8 | 73442020 | 73442075 | 55 | Nwd1 | SPRET | feature_elongation,3_prime_UTR_variant |
| 8 | 78322614 | 78322833 | 219 | Tmem184c | SPRET | feature_elongation,3_prime_UTR_variant |
| 8 | 79532283 | 79532339 | 56 | Lsm6 | SPRET | feature_elongation,3_prime_UTR_variant |
| 8 | 80068725 | 80068884 | 159 | Smad1 | C58J,FVB,KK,MOLF,NOD,NOR,NZB,NZO,PL,RF,SWR,TH,WSB | feature_elongation,3_prime_UTR_variant |
| 8 | 81237045 | 81237258 | 213 | Gypa | CAST | feature_elongation,3_prime_UTR_variant |
| 8 | 81339751 | 81339856 | 105 | Frem3 | TH | feature_elongation,coding_sequence_variant |
| 8 | 83811828 | 83811884 | 56 | Rnf150 | SPRET | feature_elongation,3_prime_UTR_variant |
| 8 | 84270472 | 84270524 | 52 | Or10k2 | AJ,C3H,CBA,CE,DBA1J,DBA2J,ILNJ2,KK,MA/My,RH2 | feature_elongation,3_prime_UTR_variant |
| 8 | 85898990 | 85899229 | 239 | Cks1brt | NOD,NOR,NZW,SJL | feature_elongation,3_prime_UTR_variant |
| 8 | 87568772 | 87568830 | 58 | N4bp1 | CAST,SPRET | feature_elongation,3_prime_UTR_variant |
| 8 | 88898585 | 88898652 | 67 | Heatr3 | BUB,C57L,C58J,SMJ,SWR | feature_elongation,3_prime_UTR_variant |
| 8 | 92171248 | 92171447 | 199 | Fto | C58J,WSB | feature_elongation,3_prime_UTR_variant |
| 8 | 94685722 | 94685780 | 58 | Gnao1 | MOLF | feature_elongation,3_prime_UTR_variant |
| 8 | 94880503 | 94880559 | 56 | Mt3 | MOLF | feature_elongation,3_prime_UTR_variant |
| 8 | 96148203 | 96148257 | 54 | Cfap20 | SPRET | feature_elongation,3_prime_UTR_variant |
| 8 | 104984701 | 104984985 | 284 | Cklf | MOLF | feature_elongation,3_prime_UTR_variant |
| 8 | 104991331 | 104991430 | 99 | Cklf | KK,PL,SPRET,WSB | feature_elongation,3_prime_UTR_variant |
| 8 | 105076889 | 105076958 | 69 | Cmtm4 | KK,PL,WSB | feature_elongation,3_prime_UTR_variant |
| 8 | 106441069 | 106441271 | 202 | Gfod2 | SPRET | feature_elongation,3_prime_UTR_variant |
| 8 | 107282983 | 107283060 | 77 | Cdh3 | CAST,MOLF | feature_elongation,3_prime_UTR_variant |

| | | | | | | |
|---|---|---|---|---|---|---|
| 8 | 107741831 | 107742019 | 188 | Sntb2 | RH2 | feature_elongation,3_prime_UTR_variant |
| 8 | 107743364 | 107743570 | 206 | Sntb2 | MOLF,RH2 | feature_elongation,3_prime_UTR_variant |
| 8 | 107773249 | 107773437 | 188 | Cog8,Pdf | SPRET | feature_elongation,3_prime_UTR_variant |
| 8 | 108103104 | 108103255 | 151 | Nfat5 | MOLF | feature_elongation,3_prime_UTR_variant |
| 8 | 108103767 | 108103979 | 212 | Nfat5 | SPRET | feature_elongation,3_prime_UTR_variant |
| 8 | 110350673 | 110350907 | 234 | Pkd1l3 | SPRET | feature_elongation,coding_sequence_variant |
| 8 | 110668614 | 110668677 | 63 | Phlpp2 | SPRET | feature_elongation,3_prime_UTR_variant |
| 8 | 110725750 | 110725847 | 97 | Tat | SPRET | feature_elongation,3_prime_UTR_variant |
| 8 | 111698804 | 111698985 | 181 | St3gal2 | CAST | feature_elongation,3_prime_UTR_variant |
| 8 | 111732359 | 111732539 | 180 | Ddx19b | BALB,SEA | feature_elongation,3_prime_UTR_variant |
| 8 | 112082035 | 112082286 | 251 | Fa2h | SPRET | feature_elongation,coding_sequence_variant |
| 8 | 112745970 | 112746161 | 191 | Terf2ip | CAST,MOLF | feature_elongation,3_prime_UTR_variant |
| 8 | 120414764 | 120414845 | 81 | Wfdc1 | CAST,MOLF,SPRET | feature_elongation,3_prime_UTR_variant |
| 8 | 121821645 | 121821753 | 108 | Mthfsd | ILNJ2,MOLF | feature_elongation,3_prime_UTR_variant |
| 8 | 121824908 | 121824978 | 70 | Mthfsd | ILNJ2,MOLF | feature_elongation,3_prime_UTR_variant |
| 8 | 122318669 | 122319001 | 332 | Map1lc3b | CAST,ILNJ2,MOLF,SPRET | feature_elongation,3_prime_UTR_variant |
| 8 | 122643026 | 122643077 | 51 | Car5a | SPRET | feature_elongation,3_prime_UTR_variant |
| 8 | 123209338 | 123209402 | 64 | Ctu2 | CAST | feature_elongation,3_prime_UTR_variant |
| 8 | 123354402 | 123354470 | 68 | Cbfa2t3 | CAST,SPRET | feature_elongation,3_prime_UTR_variant |
| 8 | 123868239 | 123868293 | 54 | Sult5a1 | CAST,PJ | feature_elongation,3_prime_UTR_variant |
| 8 | 125283287 | 125283797 | 510 | Agt | SPRET | feature_elongation,3_prime_UTR_variant |
| 8 | 125558153 | 125558245 | 92 | Trim67 | SPRET | feature_elongation,3_prime_UTR_variant |
| 8 | 125631096 | 125631152 | 56 | Sprtn | AJ,BALB,ILNJ2,RH2,SEA | feature_elongation,3_prime_UTR_variant |
| 8 | 125756560 | 125756700 | 140 | Tsnax | NZB | feature_elongation,3_prime_UTR_variant |
| 8 | 126564532 | 126564636 | 104 | Pcnx2 | SPRET | feature_elongation,coding_sequence_variant |
| 8 | 126757252 | 126757366 | 114 | Kcnk1 | MOLF | feature_elongation,3_prime_UTR_variant |
| 9 | 6263498 | 6263582 | 84 | Ddi1 | MOLF | feature_elongation,3_prime_UTR_variant |
| 9 | 7476708 | 7476891 | 183 | Mmp1a | MA/My | feature_elongation,3_prime_UTR_variant |
| 9 | 7834925 | 7835039 | 114 | Birc2 | AKR,MRL,RF | feature_elongation,5_prime_UTR_variant |
| 9 | 7857174 | 7857230 | 56 | Birc3 | AKR,CAST,MOLF,MRL,RF,SPRET | feature_elongation,3_prime_UTR_variant |
| 9 | 13719637 | 13719862 | 225 | Cep57 | CAST | feature_elongation,3_prime_UTR_variant |
| 9 | 14661930 | 14662107 | 177 | Fut4 | C58J,CE,ILNJ2,PJ,SEA,SMJ,SPRET,TH,WSB | feature_elongation,3_prime_UTR_variant |
| 9 | 14780837 | 14780937 | 100 | Gpr83 | C58J,CAST,CE,ILNJ2,MOLF,PJ,SEA,SMJ,TH | feature_elongation,3_prime_UTR_variant |
| 9 | 14915606 | 14915675 | 69 | Panx1 | MOLF | feature_elongation,3_prime_UTR_variant |
| 9 | 14916712 | 14916774 | 62 | Panx1 | MOLF,WSB | feature_elongation,3_prime_UTR_variant |
| 9 | 15223329 | 15223493 | 164 | Taf1d | SPRET | feature_elongation,3_prime_UTR_variant |
| 9 | 18233630 | 18234333 | 703 | Naalad2 | SPRET | feature_elongation,3_prime_UTR_variant |
| 9 | 19031899 | 19031956 | 57 | Or7g21 | MOLF | feature_elongation,5_prime_UTR_variant |
| 9 | 19033550 | 19033767 | 217 | Or7g21 | CAST | feature_elongation,3_prime_UTR_variant |
| 9 | 19159635 | 19159687 | 52 | Or7g25 | CAST | feature_elongation,3_prime_UTR_variant |
| 9 | 19268742 | 19268797 | 55 | Or7g28 | MOLF,SPRET | feature_elongation,3_prime_UTR_variant |
| 9 | 19355883 | 19356273 | 390 | Or7g30 | MOLF | feature_elongation,3_prime_UTR_variant |
| 9 | 19410579 | 19410771 | 192 | Or7g32 | MOLF | feature_elongation,3_prime_UTR_variant |
| 9 | 19447563 | 19447959 | 396 | Or7g33 | MOLF | feature_elongation,3_prime_UTR_variant |
| 9 | 19796066 | 19796166 | 100 | Or7e170 | CAST | feature_elongation,5_prime_UTR_variant |
| 9 | 20343416 | 20343760 | 344 | Zfp26 | 129S1,CAST,MOLF,SPRET | feature_elongation,3_prime_UTR_variant |
| 9 | 21045769 | 21045877 | 108 | Cdc37 | MOLF | feature_elongation,3_prime_UTR_variant |
| 9 | 21185625 | 21185862 | 237 | Atg4d | CAST,SPRET | feature_elongation,3_prime_UTR_variant |
| 9 | 21615062 | 21615272 | 210 | Smarca4 | SPRET | feature_elongation,3_prime_UTR_variant |
| 9 | 22110992 | 22111076 | 84 | Zfp872 | MOLF | feature_elongation,coding_sequence_variant |
| 9 | 22160092 | 22160156 | 64 | Zfp599 | SPRET | feature_elongation,3_prime_UTR_variant |
| 9 | 25188910 | 25189030 | 120 | Septin7 | MOLF,SPRET,WSB | feature_elongation,coding_sequence_variant |
| 9 | 26967362 | 26967523 | 161 | Ncapd3 | MOLF,SPRET | feature_elongation,coding_sequence_variant |
| 9 | 32177234 | 32177310 | 76 | Arhgap32 | BTBR,C57L,MOLF,SPRET | feature_elongation,3_prime_UTR_variant |
| 9 | 32296571 | 32296831 | 260 | Kcnj1 | SPRET | feature_elongation,5_prime_UTR_variant |
| 9 | 36640891 | 36640943 | 52 | Stt3a | 129S1,AJ,C3H,C57L,CAST,CBA,FVB,MA/My,MOLF,NZO,RH2,SJL,SMJ,SWR,TH | feature_elongation,3_prime_UTR_variant |
| 9 | 37138912 | 37139014 | 102 | Slc37a2 | SPRET | feature_elongation,3_prime_UTR_variant |
| 9 | 37560397 | 37560755 | 358 | Tbrg1,Siae | PL,SJL,WSB | feature_elongation,3_prime_UTR_variant |
| 9 | 37622244 | 37622452 | 208 | Or8a1b | CAST,MOLF | feature_elongation,3_prime_UTR_variant |
| 9 | 37659517 | 37659960 | 443 | Or8b12 | SPRET | feature_elongation,3_prime_UTR_variant |
| 9 | 37716993 | 37717491 | 498 | Or8b12c | CAST | feature_elongation,3_prime_UTR_variant |
| 9 | 37768016 | 37768482 | 466 | Or8b9 | CBA,CE,MRL | feature_elongation,3_prime_UTR_variant |
| 9 | 37904690 | 37905085 | 395 | Or8b35 | CE | feature_elongation,coding_sequence_variant |
| 9 | 37997577 | 37997651 | 74 | Or8b39 | SPRET | feature_elongation,3_prime_UTR_variant |
| 9 | 38031902 | 38031985 | 83 | Or8b40 | SPRET | feature_elongation,3_prime_UTR_variant |
| 9 | 38165866 | 38165981 | 115 | Or8c8 | CAST,MOLF,SPRET | feature_elongation,3_prime_UTR_variant |
| 9 | 38244247 | 38245104 | 857 | Or8c9 | DBA1J,ILNJ2,MA/My | feature_elongation,3_prime_UTR_variant |
| 9 | 38385024 | 38385748 | 724 | Or8b1c | 129S1,AJ,AKR,BALB,BTBR,BUB,C3H,CAST,CBA,DBA1J,DBA2J,ILNJ2,MA/My,MOLF,NOD,NOR,PL,RF,RH2,SJL,SMJ,SPRET,SWR,TH,WSB | feature_elongation,3_prime_UTR_variant |
| 9 | 38729469 | 38729533 | 64 | Or8b55 | SPRET | feature_elongation,3_prime_UTR_variant |
| 9 | 38761418 | 38761807 | 389 | Or8d2 | SPRET | feature_elongation,3_prime_UTR_variant |
| 9 | 39482517 | 39482619 | 102 | Or10d1 | MRL | feature_elongation,3_prime_UTR_variant |
| 9 | 39860196 | 39860325 | 129 | Or10d5 | CAST,KK,MOLF,NZB,NZW,PJ,SEA,WSB | feature_elongation,3_prime_UTR_variant |
| 9 | 39934749 | 39934915 | 166 | Or10g6 | SPRET | feature_elongation,3_prime_UTR_variant |
| 9 | 39989289 | 39989341 | 52 | Or10s1 | MOLF,SPRET | feature_elongation,3_prime_UTR_variant |
| 9 | 40202577 | 40202629 | 52 | Scn3b | SPRET | feature_elongation,3_prime_UTR_variant |
| 9 | 40884141 | 40884354 | 213 | Crtam | CE,NOD,NOR | feature_elongation,3_prime_UTR_variant |
| 9 | 43021323 | 43021399 | 76 | Tlcd5 | MOLF,SPRET | feature_elongation,3_prime_UTR_variant |
| 9 | 43110358 | 43110484 | 126 | Pou2f3 | SPRET | feature_elongation,coding_sequence_variant |
| 9 | 44057485 | 44057575 | 90 | Cbl | 129S1,C57L,CAST,ILNJ2,MOLF,SMJ | feature_elongation,3_prime_UTR_variant |
| 9 | 44205786 | 44205993 | 207 | Hinfp | SPRET | feature_elongation,3_prime_UTR_variant |
| 9 | 44315199 | 44315283 | 84 | Trappc4 | SPRET | feature_elongation,3_prime_UTR_variant |
| 9 | 44323576 | 44323771 | 195 | Cenatac | DBA1J,NZO | feature_elongation,3_prime_UTR_variant |
| 9 | 44898148 | 44898254 | 106 | Cd3d | CAST,SPRET | feature_elongation,3_prime_UTR_variant |
| 9 | 45039966 | 45040060 | 94 | Scn2b | 129S1,BTBR,C57L,KK,MOLF | feature_elongation,3_prime_UTR_variant |
| 9 | 45064224 | 45064318 | 94 | Scn4b | SPRET | feature_elongation,3_prime_UTR_variant |
| 9 | 45239816 | 45239897 | 81 | Tmprss13 | CAST,SPRET | feature_elongation,coding_sequence_variant |
| 9 | 51857752 | 51857981 | 229 | Fdx1 | FVB,SWR | feature_elongation,3_prime_UTR_variant |
| 9 | 53345304 | 53345436 | 132 | 4930550C14Rik | SPRET | feature_elongation,3_prime_UTR_variant |

| | | | | | | |
|---|---|---|---|---|---|---|
| 9 | 53414397 | 53414606 | 209 | Atm | CAST,SPRET | stop_gained,frameshift_variant |
| 9 | 56396585 | 56396645 | 60 | Hmg20a | SPRET | feature_elongation,3_prime_UTR_variant |
| 9 | 57508067 | 57508271 | 204 | Cplx3 | SPRET | feature_elongation,3_prime_UTR_variant |
| 9 | 59874853 | 59874943 | 90 | Thsd4 | SPRET | feature_elongation,3_prime_UTR_variant |
| 9 | 64214093 | 64214302 | 209 | Dis3l | AJ,BUB,C3H,CBA,DBA1J,DBA2J,FVB,ILNJ2,KK,MRL,NZB,NZO,NZW,PJ,RH2,SEA,SJL,SWR | feature_elongation,3_prime_UTR_variant |
| 9 | 65395554 | 65395828 | 274 | Spg21 | CAST | feature_elongation,3_prime_UTR_variant |
| 9 | 65502748 | 65503008 | 260 | Pif1 | AJ | feature_elongation,3_prime_UTR_variant |
| 9 | 66931471 | 66931523 | 52 | Tpm1 | SPRET | feature_elongation,3_prime_UTR_variant |
| 9 | 70508246 | 70508322 | 76 | Mindy2 | CAST,MOLF | feature_elongation,3_prime_UTR_variant |
| 9 | 72822673 | 72822770 | 97 | Prtg | SPRET | feature_elongation,3_prime_UTR_variant |
| 9 | 73010723 | 73010801 | 78 | Khdc3 | CAST,MOLF,SPRET | feature_elongation,coding_sequence_variant |
| 9 | 78283945 | 78284147 | 202 | Ooep | SPRET | feature_elongation,3_prime_UTR_variant |
| 9 | 86476658 | 86477026 | 368 | Me1 | 129S1,BTBR,C57L,CE,MRL | feature_elongation,coding_sequence_variant |
| 9 | 87586253 | 87586332 | 79 | Tbx18 | CAST | feature_elongation,3_prime_UTR_variant |
| 9 | 89470342 | 89470566 | 224 | Minar1 | SPRET | feature_elongation,3_prime_UTR_variant |
| 9 | 89795860 | 89795919 | 59 | Rasgrf1 | SPRET | feature_elongation,3_prime_UTR_variant |
| 9 | 95939144 | 95939200 | 56 | Xrn1 | MRL,NZO,NZW | feature_elongation,3_prime_UTR_variant |
| 9 | 99550276 | 99550350 | 74 | Dzip1l | NOD,NOR | feature_elongation,3_prime_UTR_variant |
| 9 | 100375611 | 100376018 | 407 | Nck1 | CAST | feature_elongation,3_prime_UTR_variant |
| 9 | 102965388 | 102965472 | 84 | Slco2a1 | SPRET | feature_elongation,3_prime_UTR_variant |
| 9 | 102966548 | 102966634 | 86 | Slco2a1 | CAST,CE,MRL,SMJ,WSB | feature_elongation,3_prime_UTR_variant |
| 9 | 103231359 | 103231417 | 58 | Cdv3 | SMJ | feature_elongation,3_prime_UTR_variant |
| 9 | 104167499 | 104167560 | 61 | Acp3 | SPRET | feature_elongation,3_prime_UTR_variant |
| 9 | 105039659 | 105039946 | 287 | Nek11 | 129S1,AJ,AKR,BALB,BTBR,BUB,C3H,C57L,CBA,CE,DBA1J,DBA2J,FVB,ILNJ2,KK,MRL,NOD,NOR,NZB,NZO,NZW,PJ,PL,RF,RH2,SEA,SJL,SMJ,SWR,TH,WSB | feature_elongation,3_prime_UTR_variant |
| 9 | 106125767 | 106125977 | 210 | Alas1 | 129S1,AJ,AKR,BALB,BTBR,BUB,C3H,C57L,CE,DBA1J,DBA2J,FVB,ILNJ2,MA/My,MRL,NOD,NOR,NZB,NZO,NZW,PJ,PL,RF,RH2,SEA,SJL,SMJ,SWR,TH,WSB | feature_elongation,5_prime_UTR_variant |
| 9 | 106227284 | 106227366 | 82 | Poc1a | 129S1,AJ,AKR,BALB,BUB,C3H,C57L,DBA1J,DBA2J,FVB,ILNJ2,MA/My,MRL,NOD,NOR,NZB,NZW,PL,RH2,SJL,SMJ,SWR,TH | feature_elongation,3_prime_UTR_variant |
| 9 | 107910574 | 107910699 | 125 | Ip6k1 | SPRET | feature_elongation,3_prime_UTR_variant |
| 9 | 110085847 | 110085908 | 61 | Cspg5 | CAST | feature_elongation,3_prime_UTR_variant |
| 9 | 110090077 | 110090557 | 480 | Cspg5 | NZW | feature_elongation,3_prime_UTR_variant |
| 9 | 110215209 | 110215279 | 70 | Ptpn23 | CBA,MOLF | feature_elongation,coding_sequence_variant |
| 9 | 110769543 | 110769723 | 180 | Cripto | SPRET | feature_elongation,3_prime_UTR_variant |
| 9 | 110814665 | 110814858 | 193 | Rtp3 | SPRET | feature_elongation,3_prime_UTR_variant |
| 9 | 110815636 | 110815824 | 188 | Rtp3 | NZW,SJL,SWR | feature_elongation,coding_sequence_variant |
| 9 | 114320110 | 114320289 | 179 | Ccr4 | CAST,SPRET | feature_elongation,3_prime_UTR_variant |
| 9 | 114728811 | 114728940 | 129 | Gpd1l | MOLF | feature_elongation,3_prime_UTR_variant |
| 9 | 114732701 | 114732781 | 80 | Gpd1l | SPRET | feature_elongation,3_prime_UTR_variant |
| 9 | 115915157 | 115915209 | 52 | Tgfbr2 | SPRET | feature_elongation,3_prime_UTR_variant |
| 9 | 119437617 | 119437674 | 57 | Scn10a | CAST,MOLF | feature_elongation,3_prime_UTR_variant |
| 9 | 119607984 | 119608348 | 364 | Scn11a | BTBR,CAST,MOLF,MRL | feature_elongation,coding_sequence_variant |
| 9 | 119749896 | 119750101 | 205 | Wdr48 | BTBR,MOLF,MRL | feature_elongation,3_prime_UTR_variant |
| 9 | 119878210 | 119878294 | 84 | Cx3cr1 | BTBR,MRL | feature_elongation,3_prime_UTR_variant |
| 9 | 119960737 | 119961102 | 365 | Rpsa | AKR,BTBR,BUB,C58J,CBA,DBA1J,DBA2J,MOLF,MRL,PL,RF,RH2 | feature_elongation,3_prime_UTR_variant |
| 9 | 121291103 | 121291516 | 413 | Trak1 | CAST,MOLF,SPRET | feature_elongation,3_prime_UTR_variant |
| 9 | 121805529 | 121805641 | 112 | Gask1a | SPRET | feature_elongation,coding_sequence_variant |
| 9 | 122691254 | 122691358 | 104 | Zfp445 | CAST,MOLF | feature_elongation,5_prime_UTR_variant |
| 9 | 122723588 | 122723796 | 208 | Zkscan7 | MOLF | feature_elongation,3_prime_UTR_variant |
| 9 | 122727128 | 122727318 | 190 | Zkscan7 | CAST | feature_elongation,3_prime_UTR_variant |
| 9 | 122903408 | 122903476 | 68 | Zdhhc3 | MOLF,SPRET | feature_elongation,3_prime_UTR_variant |
| 9 | 122904409 | 122904719 | 310 | Zdhhc3 | MOLF | feature_elongation,3_prime_UTR_variant |
| 9 | 123421992 | 123422484 | 492 | Slc6a20b | TH | feature_elongation,3_prime_UTR_variant |
| 9 | 123423249 | 123423321 | 72 | Slc6a20b | 129S1,C57L,MOLF | feature_elongation,3_prime_UTR_variant |
| 9 | 123609993 | 123610081 | 88 | Ccr9 | SPRET | feature_elongation,3_prime_UTR_variant |
| 9 | 123680961 | 123681081 | 120 | Fyco1 | 129S1,C57L,CE,MOLF | feature_elongation,5_prime_UTR_variant |
| 10 | 3504512 | 3504800 | 288 | Iyd | SPRET | feature_elongation,3_prime_UTR_variant |
| 10 | 4352193 | 4352414 | 221 | Rmnd1 | 129S1,AJ,AKR,BALB,BTBR,BUB,C57L,ILNJ2,PL,RF,RH2,SJL,SMJ | feature_elongation,3_prime_UTR_variant |
| 10 | 4353384 | 4353619 | 235 | Rmnd1 | NOD,NOR | feature_elongation,3_prime_UTR_variant |
| 10 | 4401750 | 4402205 | 455 | Armt1 | CAST,MOLF | feature_elongation,3_prime_UTR_variant |
| 10 | 4512211 | 4512295 | 84 | Ccdc170 | CAST | feature_elongation,3_prime_UTR_variant |
| 10 | 5782225 | 5782297 | 72 | Rgs17 | CAST,MOLF | feature_elongation,3_prime_UTR_variant |
| 10 | 6837057 | 6837121 | 64 | Ipcef1 | BUB | feature_elongation,3_prime_UTR_variant |
| 10 | 6838622 | 6838750 | 128 | Ipcef1 | CAST,MOLF | feature_elongation,3_prime_UTR_variant |
| 10 | 7393517 | 7393574 | 57 | Ulbp1 | C3H,CBA,DBA1J,DBA2J,FVB,LP,MA/My,MRL,NZW,SPRET,SWR | feature_elongation,3_prime_UTR_variant |
| 10 | 7745520 | 7745594 | 74 | Zc3h12d | SPRET,WSB | feature_elongation,3_prime_UTR_variant |
| 10 | 8600307 | 8600397 | 90 | Sash1 | CAST,MOLF | feature_elongation,3_prime_UTR_variant |
| 10 | 9503431 | 9503584 | 153 | Samd5 | 129S1,BUB,CE,DBA1J,DBA2J,ILNJ2,KK,MA/My,NOD,NOR,NZB,NZO,NZW,PJ,PL,SEA,TH | feature_elongation,3_prime_UTR_variant |
| 10 | 10306889 | 10306952 | 63 | Adgb | SPRET | feature_elongation,coding_sequence_variant |
| 10 | 10564500 | 10564599 | 99 | Grm1 | SPRET | feature_elongation,3_prime_UTR_variant |
| 10 | 11334909 | 11334967 | 58 | Epm2a | AJ,AKR,BALB,BTBR,C3H,C57L,CAST,CBA,MRL,NOD,NOR,PL,RF,RH2,SJL,SMJ | feature_elongation,3_prime_UTR_variant |
| 10 | 12423895 | 12424036 | 141 | Utrn | CE,DBA1J,DBA2J,LP,MA/My,TH | feature_elongation,5_prime_UTR_variant |
| 10 | 12883625 | 12883688 | 63 | Sf3b5 | SPRET | feature_elongation,5_prime_UTR_variant |
| 10 | 13005992 | 13006050 | 58 | Plagl1 | DBA1J,DBA2J,MOLF | feature_elongation,3_prime_UTR_variant |
| 10 | 13083976 | 13084072 | 96 | Phactr2 | WSB | feature_elongation,3_prime_UTR_variant |
| 10 | 13086458 | 13086518 | 60 | Phactr2 | SPRET | feature_elongation,3_prime_UTR_variant |
| 10 | 18458599 | 18458663 | 64 | Arfgef3 | SPRET | feature_elongation,3_prime_UTR_variant |
| 10 | 19733982 | 19734039 | 57 | Pex7 | SPRET | feature_elongation,3_prime_UTR_variant |
| 10 | 22979363 | 22979760 | 397 | Eya4 | SPRET | feature_elongation,3_prime_UTR_variant |
| 10 | 29221207 | 29221275 | 68 | Echdc1,Rnf146 | CAST,FVB,MOLF,PJ,SEA,WSB | feature_elongation,3_prime_UTR_variant |
| 10 | 33965445 | 33965577 | 132 | Calhm5 | MOLF,SPRET | feature_elongation,3_prime_UTR_variant |
| 10 | 40124320 | 40124939 | 619 | Gtf3c6 | CAST,MOLF,SPRET | feature_elongation,3_prime_UTR_variant |

| | | | | | | |
|---|---|---|---|---|---|---|
| 10 | 40125002 | 40125110 | 108 | Gtf3c6 | CAST,MOLF,SPRET | feature_elongation,3_prime_UTR_variant |
| 10 | 40358788 | 40358849 | 61 | Cdk19 | SPRET | feature_elongation,3_prime_UTR_variant |
| 10 | 40479848 | 40479920 | 72 | Slc22a16 | SPRET | feature_elongation,coding_sequence_variant |
| 10 | 40708505 | 40708635 | 130 | Cdc40 | CAST,MOLF | feature_elongation,3_prime_UTR_variant |
| 10 | 41310473 | 41310689 | 216 | Ak9 | CAST | feature_elongation,3_prime_UTR_variant |
| 10 | 41341303 | 41341373 | 70 | Zbtb24 | CAST,MOLF,SPRET | feature_elongation,3_prime_UTR_variant |
| 10 | 41358676 | 41358762 | 86 | Mical1 | CAST,MOLF | frameshift_variant |
| 10 | 43389955 | 43390009 | 54 | Bend3 | SPRET | feature_elongation,3_prime_UTR_variant |
| 10 | 45035402 | 45035463 | 61 | Prep | SPRET | feature_elongation,3_prime_UTR_variant |
| 10 | 45041543 | 45042399 | 856 | Prep | CAST,MOLF,SPRET | feature_elongation,3_prime_UTR_variant |
| 10 | 52987478 | 52987601 | 123 | Slc35f1 | SPRET | feature_elongation,3_prime_UTR_variant |
| 10 | 53413249 | 53413309 | 60 | Mcm9 | SPRET | feature_elongation,3_prime_UTR_variant |
| 10 | 55892160 | 55892232 | 72 | Tbc1d32 | KK,MOLF | feature_elongation,3_prime_UTR_variant |
| 10 | 55893463 | 55893625 | 162 | Tbc1d32 | KK | feature_elongation,3_prime_UTR_variant |
| 10 | 58116513 | 58116680 | 167 | Gcc2 | MOLF | feature_elongation,coding_sequence_variant |
| 10 | 58972893 | 58973375 | 482 | Sh3rf3 | MOLF | feature_elongation,3_prime_UTR_variant |
| 10 | 60548742 | 60548806 | 64 | Slc29a3 | SPRET | feature_elongation,3_prime_UTR_variant |
| 10 | 61943520 | 61943583 | 63 | Fam241b | SPRET | feature_elongation,3_prime_UTR_variant |
| 10 | 62645032 | 62645084 | 52 | Tet1 | CAST | feature_elongation,3_prime_UTR_variant |
| 10 | 62936780 | 62936968 | 188 | Atoh7 | MOLF,RF | feature_elongation,3_prime_UTR_variant |
| 10 | 66848321 | 66848402 | 81 | Reep3 | C57L,MRL,WSB | feature_elongation,3_prime_UTR_variant |
| 10 | 68616690 | 68616743 | 53 | Tmem26 | CAST,MOLF,SPRET | feature_elongation,3_prime_UTR_variant |
| 10 | 69171773 | 69171864 | 91 | Cdk1 | SPRET | feature_elongation,3_prime_UTR_variant |
| 10 | 71062118 | 71062426 | 308 | Tfam | SPRET | feature_elongation,3_prime_UTR_variant |
| 10 | 75409430 | 75409505 | 75 | Ggt1 | MOLF | feature_elongation,5_prime_UTR_variant |
| 10 | 75451077 | 75451479 | 402 | Ggt5 | CAST,MOLF,WSB | feature_elongation,3_prime_UTR_variant |
| 10 | 75800319 | 75800514 | 195 | Gm5134 | BUB,RF,SPRET,WSB | stop_gained,protein_altering_variant |
| 10 | 75893634 | 75893844 | 210 | Slc5a4b | SPRET | feature_elongation,3_prime_UTR_variant |
| 10 | 76434963 | 76435090 | 127 | Col6a2 | SPRET | feature_elongation,3_prime_UTR_variant |
| 10 | 76880957 | 76881181 | 224 | Slc19a1 | SPRET | feature_elongation,3_prime_UTR_variant |
| 10 | 77551738 | 77551798 | 60 | Gm10142 | SPRET | feature_elongation,coding_sequence_variant |
| 10 | 77578270 | 77578345 | 75 | Gm18596 | CAST | feature_elongation,coding_sequence_variant |
| 10 | 77697982 | 77698043 | 61 | Gm3285 | SPRET | frameshift_variant |
| 10 | 77743840 | 77743896 | 56 | Trpm2 | SPRET | feature_elongation,3_prime_UTR_variant |
| 10 | 78274594 | 78275242 | 648 | Pdxk | SPRET | feature_elongation,3_prime_UTR_variant |
| 10 | 78681181 | 78681474 | 293 | Or7c70 | MOLF | feature_elongation,3_prime_UTR_variant |
| 10 | 78753852 | 78754041 | 189 | Or7a38 | WSB | feature_elongation,3_prime_UTR_variant |
| 10 | 80527313 | 80527606 | 293 | Mob3a | BUB | feature_elongation,coding_sequence_variant |
| 10 | 80786478 | 80786529 | 51 | Gng7 | SPRET | feature_elongation,3_prime_UTR_variant |
| 10 | 81034466 | 81034659 | 193 | Nmrk2 | CAST | feature_elongation,3_prime_UTR_variant |
| 10 | 81200565 | 81200617 | 52 | Mfsd12 | BUB | feature_elongation,3_prime_UTR_variant |
| 10 | 82214042 | 82214293 | 251 | Gm4924 | 129S1,AJ,AKR,BALB,BTBR,C3H,C57L,CBA,CE,DBA1J,DBA2J,FVB,ILNJ2,KK,MA/My,MRL,NOD,NOR,NZB,NZO,NZW,PJ,PL,RF,RH2,SEA,SJL,SMJ,SWR,TH | feature_elongation,coding_sequence_variant |
| 10 | 82577481 | 82577543 | 62 | Hcfc2 | 129S1,AJ,AKR,BALB,BTBR,BUB,C3H,C57L,CBA,CE,DBA1J,DBA2J,FVB,ILNJ2,KK,MA/My,MRL,NOD,NOR,NZB,NZO,NZW,PJ,PL,RF,RH2,SEA,SJL,SMJ,SWR,TH,WSB | feature_elongation,3_prime_UTR_variant |
| 10 | 82733216 | 82733300 | 84 | Txnrd1 | CAST | feature_elongation,3_prime_UTR_variant |
| 10 | 83213067 | 83213554 | 487 | Nopchap1 | SPRET | feature_elongation,3_prime_UTR_variant |
| 10 | 83260968 | 83261378 | 410 | Nopchap1 | BUB,SMJ | feature_elongation,3_prime_UTR_variant |
| 10 | 83316129 | 83316500 | 371 | Nopchap1 | SPRET | feature_elongation,3_prime_UTR_variant |
| 10 | 85887193 | 85887487 | 294 | Fbxo7 | C58J,LP,NZB | feature_elongation,3_prime_UTR_variant |
| 10 | 86128971 | 86129079 | 108 | Syn3 | AJ,AKR,BALB,C3H,CBA,CE,ILNJ2,MA/My,MRL,NOD,NOR,NZB,NZO,PJ,PL,RF,RH2,SEA,SWR,TH | feature_elongation,3_prime_UTR_variant |
| 10 | 87327557 | 87327657 | 100 | Ascl1 | CAST | feature_elongation,3_prime_UTR_variant |
| 10 | 89411558 | 89411610 | 52 | Slc17a8 | SPRET | feature_elongation,3_prime_UTR_variant |
| 10 | 93076134 | 93076197 | 63 | Cdk17 | MRL,NZW | feature_elongation,3_prime_UTR_variant |
| 10 | 93727012 | 93727068 | 56 | Metap2 | SPRET | feature_elongation,5_prime_UTR_variant |
| 10 | 93803102 | 93803162 | 60 | Vezt | AJ,BALB,BUB,PJ,RH2,SEA,SJL | feature_elongation,3_prime_UTR_variant |
| 10 | 93979486 | 93980310 | 824 | Fgd6 | SPRET | feature_elongation,3_prime_UTR_variant |
| 10 | 94056874 | 94056927 | 53 | Ndufa12 | AKR,MA/My,MRL,NZO,PL,RF,WSB | feature_elongation,3_prime_UTR_variant |
| 10 | 94424087 | 94424179 | 92 | Tmcc3 | SPRET | feature_elongation,3_prime_UTR_variant |
| 10 | 102383797 | 102383875 | 78 | Rassf9 | SPRET | feature_elongation,3_prime_UTR_variant |
| 10 | 111131285 | 111131347 | 62 | Osbpl8 | SPRET | feature_elongation,3_prime_UTR_variant |
| 10 | 111821118 | 111821282 | 164 | Krr1 | CAST,CE,LP,MOLF,MRL,NZB,PL,SJL,WSB | feature_elongation,3_prime_UTR_variant |
| 10 | 115123744 | 115123800 | 56 | Rab21 | MOLF | feature_elongation,3_prime_UTR_variant |
| 10 | 117068378 | 117068666 | 288 | Lyz3 | CAST,MOLF,SMJ | feature_elongation,3_prime_UTR_variant |
| 10 | 119978598 | 119978659 | 61 | Irak3 | SPRET | feature_elongation,3_prime_UTR_variant |
| 10 | 120053590 | 120053661 | 71 | Tmbim4 | SPRET | feature_elongation,3_prime_UTR_variant |
| 10 | 120618611 | 120618687 | 76 | Msrb3 | MOLF,SPRET,TH,WSB | feature_elongation,3_prime_UTR_variant |
| 10 | 120619799 | 120619916 | 117 | Msrb3 | MOLF | feature_elongation,3_prime_UTR_variant |
| 10 | 121212151 | 121212295 | 144 | Gns | DBA1J,KK,MA/My,MOLF,SMJ | feature_elongation,coding_sequence_variant |
| 10 | 121916389 | 121916463 | 74 | Rxylt1 | CAST | feature_elongation,3_prime_UTR_variant |
| 10 | 122757661 | 122757862 | 201 | Ppm1h | MOLF | feature_elongation,3_prime_UTR_variant |
| 10 | 122828785 | 122829001 | 216 | Mon2 | SPRET | feature_elongation,3_prime_UTR_variant |
| 10 | 125062208 | 125062274 | 66 | Slc16a7 | CAST,SPRET | feature_elongation,3_prime_UTR_variant |
| 10 | 126846496 | 126846579 | 83 | Avil | WSB | feature_elongation,coding_sequence_variant |
| 10 | 127533459 | 127533513 | 54 | Nemp1 | CAST | feature_elongation,3_prime_UTR_variant |
| 10 | 127578251 | 127578302 | 51 | Zbtb39 | AJ,BALB,FVB,NOD,NOR,PL,RH2 | feature_elongation,coding_sequence_variant |
| 10 | 127581270 | 127581322 | 52 | Zbtb39 | CAST,CE,MOLF,WSB | feature_elongation,3_prime_UTR_variant |
| 10 | 127602382 | 127602593 | 211 | Rdh1 | CE | feature_elongation,3_prime_UTR_variant |
| 10 | 127602899 | 127603122 | 223 | Rdh1 | WSB | feature_elongation,3_prime_UTR_variant |
| 10 | 127649579 | 127650048 | 469 | Rdh16 | SPRET | feature_elongation,3_prime_UTR_variant |
| 10 | 127702392 | 127702444 | 52 | Rdh16f2 | MOLF,SMJ | feature_elongation,5_prime_UTR_variant |
| 10 | 128104228 | 128104426 | 198 | Apof | CE | feature_elongation,5_prime_UTR_variant |
| 10 | 128378812 | 128378986 | 174 | Zc3h10 | BUB,KK,MOLF,SMJ | feature_elongation,3_prime_UTR_variant |
| 10 | 128635147 | 128635427 | 280 | Mmp19 | BUB,KK,MOLF,SMJ | feature_elongation,3_prime_UTR_variant |
| 10 | 128825545 | 128825638 | 93 | Or10p22 | CAST | feature_elongation,5_prime_UTR_variant |
| 10 | 128912802 | 128912976 | 174 | Or6c8 | C3H,CBA,CE,RF | feature_elongation,3_prime_UTR_variant |

| | | | | | | |
|---|---|---|---|---|---|---|
| 10 | 129274265 | 129274339 | 74 | Or6c1b | CAST | feature_elongation,3_prime_UTR_variant |
| 10 | 129303015 | 129303078 | 63 | Or6c5c | LP,NOD,NOR | feature_elongation,3_prime_UTR_variant |
| 10 | 129515126 | 129515626 | 500 | Or6c1 | SPRET | feature_elongation,3_prime_UTR_variant |
| 10 | 129539364 | 129539706 | 342 | Or6c6c | SPRET | feature_elongation,5_prime_UTR_variant |
| 10 | 129625345 | 129625411 | 66 | Or6c69b | SPRET | feature_elongation,3_prime_UTR_variant |
| 10 | 130043117 | 130043442 | 325 | Or9k7 | MOLF,SMJ | feature_elongation,3_prime_UTR_variant |
| 11 | 3292245 | 3292307 | 62 | Pik3ip1 | SPRET | feature_elongation,3_prime_UTR_variant |
| 11 | 3836537 | 3836669 | 132 | 4921536K21Rik | B10D2,B10J,C58J,CAST,KK,SPRET | feature_elongation,3_prime_UTR_variant |
| 11 | 3849068 | 3849235 | 167 | Dusp18 | B10D2,B10J,C58J,KK | feature_elongation,3_prime_UTR_variant |
| 11 | 4027718 | 4027845 | 127 | Sec14l3 | SPRET | feature_elongation,3_prime_UTR_variant |
| 11 | 4047704 | 4047768 | 64 | Sec14l2 | SPRET | feature_elongation,3_prime_UTR_variant |
| 11 | 4172107 | 4172160 | 53 | Castor1 | B10D2,B10J,C58J,KK,SPRET | feature_elongation,3_prime_UTR_variant |
| 11 | 4717995 | 4718047 | 52 | Nf2 | SPRET | feature_elongation,3_prime_UTR_variant |
| 11 | 5144064 | 5144311 | 247 | Kremen1 | B10D2,B10J,C58J,KK,NOR,SPRET | feature_elongation,3_prime_UTR_variant |
| 11 | 5854541 | 5854598 | 57 | Gck | AJ,AKR,B10D2,B10J,BALB,BTBR,BUB,C3H,C57L,C58J,CE,FVB,KK,MA/My,MOLF,MRL,NOD,NOR,NZB,NZO,PJ,PL,RF,SEA,SJL,SPRET,SWR,TH,WSB | feature_elongation,coding_sequence_variant |
| 11 | 6220547 | 6220752 | 205 | Tmed4 | SPRET | feature_elongation,3_prime_UTR_variant |
| 11 | 9068596 | 9068653 | 57 | Upp1 | C58J | feature_elongation,5_prime_UTR_variant |
| 11 | 11577641 | 11577707 | 66 | 4930512M02Rik | SPRET | feature_elongation,coding_sequence_variant |
| 11 | 16867412 | 16867506 | 94 | Egfr | CAST | feature_elongation,3_prime_UTR_variant |
| 11 | 23596671 | 23596896 | 225 | Pex13 | CAST | feature_elongation,3_prime_UTR_variant |
| 11 | 29104848 | 29104940 | 92 | Pnpt1 | AJ,C3H,CAST,DBA1J,DBA2J,ILNJ2,MA/My,MOLF,MRL,NZB,NZO,PL | feature_elongation,coding_sequence_variant |
| 11 | 29171710 | 29171912 | 202 | Cfap36 | AKR,BALB,BUB,C57L,C58J,CE,FVB,KK,NOD,NZW,RF,RH2,WSB | feature_elongation,3_prime_UTR_variant |
| 11 | 30057849 | 30057966 | 117 | Sptbn1 | SPRET | feature_elongation,3_prime_UTR_variant |
| 11 | 33923106 | 33923547 | 441 | Kcnmb1 | SPRET | feature_elongation,3_prime_UTR_variant |
| 11 | 33997038 | 33997171 | 133 | Lcp2 | CAST | feature_elongation,5_prime_UTR_variant |
| 11 | 40601035 | 40601161 | 126 | Hmmr | B10D2,B10J,BTBR,C58J,CAST,LP,MA/My,MOLF,SPRET | feature_elongation,coding_sequence_variant |
| 11 | 45821507 | 45821681 | 174 | Lsm11 | CE,PL,SMJ,WSB | feature_elongation,3_prime_UTR_variant |
| 11 | 45838219 | 45838271 | 52 | Thg1l | CAST | feature_elongation,3_prime_UTR_variant |
| 11 | 46039757 | 46039819 | 62 | Nipal4 | SPRET | feature_elongation,3_prime_UTR_variant |
| 11 | 46203651 | 46203973 | 322 | Cyfip2 | SPRET | feature_elongation,5_prime_UTR_variant |
| 11 | 46560312 | 46560511 | 199 | Timd2 | CAST | feature_elongation,3_prime_UTR_variant |
| 11 | 48796639 | 48796835 | 196 | Gm12185 | MOLF,NZW | feature_elongation,3_prime_UTR_variant |
| 11 | 48909649 | 48909849 | 200 | 9930111J21Rik2 | SPRET | feature_elongation,3_prime_UTR_variant |
| 11 | 54730080 | 54730271 | 191 | Lyrm7 | MOLF,SPRET | feature_elongation,3_prime_UTR_variant |
| 11 | 54731188 | 54731242 | 54 | Lyrm7 | B10D2,B10J,C58J,NOR,NZB,NZO,NZW,SPRET | feature_elongation,3_prime_UTR_variant |
| 11 | 57219741 | 57219794 | 53 | Gria1 | SPRET | feature_elongation,3_prime_UTR_variant |
| 11 | 58490275 | 58490337 | 62 | Or2ab1 | 129S1,AJ,AKR,BALB,BTBR,BUB,C3H,C57L,CBA,DBA1J,DBA2J,FVB,ILNJ2,KK,LP,MA/My,MRL,NOD,NOR,NZB,NZO,NZW,PJ,PL,RF,RH2,SEA,SJL,SPRET,SWR,TH,WSB | feature_elongation,3_prime_UTR_variant |
| 11 | 58780350 | 58780498 | 148 | Zfp39 | SPRET | feature_elongation,3_prime_UTR_variant |
| 11 | 58853329 | 58853428 | 99 | H3f4 | 129S1,AKR,BALB,BTBR,C57L,LP,RF,RH2,WSB | feature_elongation,3_prime_UTR_variant |
| 11 | 58951870 | 58952489 | 619 | Obscn | AJ,BTBR,BUB,C3H,C57L,CAST,CBA,CE,DBA1J,DBA2J,FVB,ILNJ2,KK,MA/My,MOLF,MRL,NOD,NOR,NZW,PJ,PL,RF,SEA,SJL,SMJ,SWR,TH,WSB | stop_gained,frameshift_variant |
| 11 | 59047601 | 59048311 | 710 | Iba57 | ILNJ2 | feature_elongation,3_prime_UTR_variant |
| 11 | 60267491 | 60267776 | 285 | Drc3 | AJ,C3H,CAST,CBA,MOLF,MRL,NZB,NZO,PJ,SEA,SPRET | feature_elongation,3_prime_UTR_variant |
| 11 | 60679177 | 60679406 | 229 | Shmt1 | 129S1,AKR,BTBR,C57L,LP,MA/My,NOD,PL,RF | feature_elongation,3_prime_UTR_variant |
| 11 | 60750131 | 60750269 | 138 | Dhrs7b | AJ,C3H,CAST,CBA,MRL,NZB,NZO,PJ,SEA | feature_elongation,3_prime_UTR_variant |
| 11 | 60793081 | 60793286 | 205 | Natd1 | MOLF | feature_elongation,3_prime_UTR_variant |
| 11 | 62404327 | 62404387 | 60 | Pigl | CAST,MOLF | feature_elongation,3_prime_UTR_variant |
| 11 | 62404965 | 62405170 | 205 | Pigl | SPRET | feature_elongation,3_prime_UTR_variant |
| 11 | 64892697 | 64892772 | 75 | Elac2 | CE,ILNJ2 | feature_elongation,3_prime_UTR_variant |
| 11 | 68095918 | 68095974 | 56 | Stx8 | MOLF | feature_elongation,3_prime_UTR_variant |
| 11 | 68387617 | 68387799 | 182 | Pik3r5 | CAST | feature_elongation,3_prime_UTR_variant |
| 11 | 68834800 | 68834853 | 53 | Arhgef15 | CAST | feature_elongation,3_prime_UTR_variant |
| 11 | 68838660 | 68838900 | 240 | Arhgef15 | DBA1J,DBA2J,MOLF,NOR,PJ,SEA,SMJ,SPRET | stop_gained,inframe_insertion |
| 11 | 68877637 | 68877770 | 133 | Pfas | SPRET | feature_elongation,3_prime_UTR_variant |
| 11 | 68963909 | 68964024 | 115 | Tmem107 | CAST | feature_elongation,3_prime_UTR_variant |
| 11 | 69111732 | 69111795 | 63 | Gucy2e | DBA1J,DBA2J,NOR,PJ,SEA,SMJ | feature_elongation,3_prime_UTR_variant |
| 11 | 69514502 | 69514669 | 167 | Sat2 | DBA1J,DBA2J,ILNJ2,MOLF,NOR,PJ,SEA,SMJ | feature_elongation,3_prime_UTR_variant |
| 11 | 69687918 | 69687986 | 68 | Fgf11 | CAST,MOLF | feature_elongation,3_prime_UTR_variant |
| 11 | 70028268 | 70028489 | 221 | Mgl2 | CAST,CE,DBA1J,DBA2J,FVB,ILNJ2,KK,MOLF,NOR,PJ,SEA,SMJ,WSB | feature_elongation,3_prime_UTR_variant |
| 11 | 70235307 | 70235375 | 68 | Alox15 | CAST,CE,FVB,ILNJ2,NZW,SPRET,WSB | feature_elongation,3_prime_UTR_variant |
| 11 | 72204137 | 72204320 | 183 | Xaf1 | SPRET | feature_elongation,3_prime_UTR_variant |
| 11 | 72972930 | 72972982 | 52 | Ncbp3 | SPRET | feature_elongation,3_prime_UTR_variant |
| 11 | 73375727 | 73375783 | 56 | Or1e22 | SPRET | feature_elongation,3_prime_UTR_variant |
| 11 | 73679985 | 73680284 | 299 | Or1e30 | SPRET | feature_elongation,3_prime_UTR_variant |
| 11 | 73703017 | 73703273 | 256 | Or1e32 | AJ,CAST,NZO,NZW,WSB | feature_elongation,3_prime_UTR_variant |
| 11 | 73738005 | 73738586 | 581 | Or1e33 | SPRET | feature_elongation,3_prime_UTR_variant |
| 11 | 73832246 | 73832316 | 70 | Or1e17 | SPRET | feature_elongation,3_prime_UTR_variant |
| 11 | 73975983 | 73976235 | 252 | Zfp616 | AJ,CAST,CE,NZO,NZW,SPRET,WSB | feature_elongation,3_prime_UTR_variant,feature_elongation,coding_sequence_variant |
| 11 | 74099565 | 74099623 | 58 | Or1a1b | CAST | feature_elongation,5_prime_UTR_variant |
| 11 | 76669478 | 76669575 | 97 | Cpd | SPRET | feature_elongation,3_prime_UTR_variant |
| 11 | 76936391 | 76936502 | 111 | Nsrp1 | MOLF | feature_elongation,3_prime_UTR_variant |
| 11 | 77028661 | 77028780 | 119 | Efcab5 | CAST,MOLF,SPRET,TH | feature_elongation,coding_sequence_variant |
| 11 | 77964371 | 77964540 | 169 | Eral1 | CAST | feature_elongation,3_prime_UTR_variant |
| 11 | 79130574 | 79130777 | 203 | Wsb1 | CAST | feature_elongation,3_prime_UTR_variant |

| | | | | | | |
|---|---|---|---|---|---|---|
| 11 | 79392356 | 79392570 | 214 | Omg | AKR,BALB,C3H,CBA,CE,ILNJ2,MRL,PL,RF,RH2,SMJ | feature_elongation,3_prime_UTR_variant |
| 11 | 80371768 | 80372207 | 439 | Cdk5r1 | C3H | feature_elongation,3_prime_UTR_variant |
| 11 | 83166239 | 83166614 | 375 | Slfn14 | 129S1,AJ,AKR,B10D2,B10J,BALB,BTBR,BUB,C3H,C57L,C58J,CAST,CBA,CE,DBA1J,DBA2J,FVB,ILNJ2,KK,LP,MA/My,MOLF,MRL,NOD,NOR,NZB,NZO,NZW,PJ,PL,RF,RH2,SEA,SJL,SMJ,SPRET,SWR,TH,WSB | feature_elongation,3_prime_UTR_variant |
| 11 | 83185853 | 83186060 | 207 | Pex12 | CAST | feature_elongation,3_prime_UTR_variant |
| 11 | 83193427 | 83193718 | 291 | Ap2b1 | KK | feature_elongation,5_prime_UTR_variant |
| 11 | 86388627 | 86388909 | 282 | Rnft1 | SPRET | feature_elongation,3_prime_UTR_variant |
| 11 | 86389501 | 86389703 | 202 | Rnft1 | 129S1,AJ,AKR,BTBR,C3H,C57L,C58J,CBA,DBA1J,DBA2J,KK,MOLF,MRL,NZB,NZO,PL,RF,SJL,SWR | feature_elongation,3_prime_UTR_variant |
| 11 | 86980703 | 86980945 | 242 | Prr11 | CAST,MOLF,SPRET | feature_elongation,3_prime_UTR_variant |
| 11 | 87413380 | 87413452 | 72 | Tex14 | SPRET | feature_elongation,coding_sequence_variant |
| 11 | 87780987 | 87781093 | 106 | Or4d2b | SPRET | feature_elongation,5_prime_UTR_variant |
| 11 | 88930820 | 88930996 | 176 | Dgke | BALB,CE,KK,MRL,RH2,SJL,WSB | feature_elongation,3_prime_UTR_variant |
| 11 | 93491476 | 93491568 | 92 | Car10 | SPRET | feature_elongation,3_prime_UTR_variant |
| 11 | 94180656 | 94180720 | 64 | Luc7l3 | AJ,AKR,C3H,CAST,CBA,CE,MRL,RF | feature_elongation,3_prime_UTR_variant |
| 11 | 94390670 | 94390730 | 60 | Epn3 | SPRET | feature_elongation,5_prime_UTR_variant |
| 11 | 94430679 | 94430735 | 56 | Rsad1 | 129S1,BALB,BTBR,C57L,MA/My,NOD,NOR,PL,SJL,SMJ | feature_elongation,3_prime_UTR_variant |
| 11 | 94448825 | 94448960 | 135 | Acsf2 | AJ,AKR,C3H,CAST,CBA,CE,MOLF,MRL,RF | feature_elongation,3_prime_UTR_variant |
| 11 | 95660269 | 95660483 | 214 | Zfp652 | DBA1J,DBA2J,ILNJ2,KK,NZB,NZO,PJ,SEA | feature_elongation,3_prime_UTR_variant |
| 11 | 95960534 | 95960586 | 52 | Atp5mc1 | CAST | feature_elongation,3_prime_UTR_variant |
| 11 | 95990891 | 95991017 | 126 | Calcoco2 | AJ,AKR,BUB,C3H,CAST,CBA,DBA1J,DBA2J,ILNJ2,MOLF,MRL,NZB,NZO,PJ,RH2,SEA,SPRET | feature_elongation,coding_sequence_variant |
| 11 | 96025593 | 96025662 | 69 | Ttll6 | SPRET | feature_elongation,5_prime_UTR_variant |
| 11 | 97225126 | 97225359 | 233 | Gpr179 | MOLF,NZW,SPRET,SWR | feature_elongation,3_prime_UTR_variant |
| 11 | 97408859 | 97409073 | 214 | Srcin1 | SPRET | feature_elongation,3_prime_UTR_variant |
| 11 | 97606053 | 97606275 | 222 | Pip4k2b | SEA | feature_elongation,3_prime_UTR_variant |
| 11 | 97884768 | 97884845 | 77 | Arl5c | 129S1,BTBR,C3H,C57L,CE,DBA1J,DBA2J,FVB,ILNJ2,WSB | feature_elongation,3_prime_UTR_variant |
| 11 | 97975264 | 97975318 | 54 | Fbxl20 | CAST | feature_elongation,3_prime_UTR_variant |
| 11 | 97979155 | 97979207 | 52 | Fbxl20 | CAST,MOLF,NZW,SWR | feature_elongation,3_prime_UTR_variant |
| 11 | 98109981 | 98110162 | 181 | Cdk12 | MOLF,NZW,SWR | feature_elongation,coding_sequence_variant |
| 11 | 98567803 | 98568301 | 498 | Gsdma | CAST,MOLF,SPRET | feature_elongation,3_prime_UTR_variant |
| 11 | 98723431 | 98723653 | 222 | Casc3 | CE | feature_elongation,3_prime_UTR_variant |
| 11 | 98754599 | 98754689 | 90 | Wipf2 | SMJ | feature_elongation,5_prime_UTR_variant |
| 11 | 99276823 | 99276883 | 60 | Krt10 | AKR,RF,RH2,SJL | feature_elongation,coding_sequence_variant |
| 11 | 99549559 | 99549709 | 150 | Gm11563 | SPRET | feature_elongation,coding_sequence_variant |
| 11 | 99683648 | 99683723 | 75 | Gm11596 | CAST,MOLF | feature_elongation,coding_sequence_variant |
| 11 | 99700282 | 99700372 | 90 | Krtap4-13 | CAST,SPRET,WSB | feature_elongation,coding_sequence_variant |
| 11 | 99705987 | 99706092 | 105 | Gm11564 | SPRET | feature_elongation,coding_sequence_variant |
| 11 | 99839625 | 99839829 | 204 | Krtap9-5 | SPRET | feature_elongation,coding_sequence_variant |
| 11 | 100657125 | 100657187 | 62 | Ghdc | CAST | feature_elongation,3_prime_UTR_variant |
| 11 | 101862617 | 101862838 | 221 | Dusp3 | AJ,TH | feature_elongation,3_prime_UTR_variant |
| 11 | 102170036 | 102170108 | 72 | Asb16 | SPRET | feature_elongation,3_prime_UTR_variant |
| 11 | 102781900 | 102781981 | 81 | Gfap | MOLF | feature_elongation,3_prime_UTR_variant |
| 11 | 103255689 | 103255840 | 151 | Plekhm1 | MOLF,NZW | feature_elongation,3_prime_UTR_variant |
| 11 | 103619021 | 103619109 | 88 | Wnt9b | CAST,SPRET | feature_elongation,3_prime_UTR_variant |
| 11 | 104611644 | 104611743 | 99 | Gm11639 | CAST | feature_elongation,coding_sequence_variant |
| 11 | 106074870 | 106074930 | 60 | Strada | CAST | feature_elongation,3_prime_UTR_variant |
| 11 | 106075885 | 106076081 | 196 | Strada | SPRET | feature_elongation,3_prime_UTR_variant |
| 11 | 106088956 | 106089462 | 506 | Ccdc47 | SPRET | feature_elongation,3_prime_UTR_variant |
| 11 | 106304869 | 106305090 | 221 | Ern1 | SPRET | feature_elongation,3_prime_UTR_variant |
| 11 | 107394759 | 107394899 | 140 | Psmd12 | SPRET | feature_elongation,3_prime_UTR_variant |
| 11 | 109147074 | 109147144 | 70 | Rgs9 | CAST,SPRET | feature_elongation,3_prime_UTR_variant |
| 11 | 109289650 | 109289848 | 198 | Gna13 | SPRET | feature_elongation,3_prime_UTR_variant |
| 11 | 109464601 | 109464655 | 54 | Wipi1 | SPRET | feature_elongation,3_prime_UTR_variant |
| 11 | 109992711 | 109992770 | 59 | Abca9 | BALB,BUB,CAST,RF,SEA,SPRET | feature_elongation,3_prime_UTR_variant |
| 11 | 110142597 | 110142657 | 60 | Abca6 | CAST | feature_elongation,5_prime_UTR_variant |
| 11 | 110964534 | 110964590 | 56 | Kcnj2 | SPRET | feature_elongation,3_prime_UTR_variant |
| 11 | 114935004 | 114935184 | 180 | Cd300ld3 | FVB,SJL | feature_elongation,coding_sequence_variant |
| 11 | 115320364 | 115320578 | 214 | Kctd2 | SPRET | feature_elongation,coding_sequence_variant |
| 11 | 116013202 | 116013401 | 199 | Trim65 | MOLF | feature_elongation,3_prime_UTR_variant |
| 11 | 116348263 | 116348399 | 136 | Qrich2 | CAST,MOLF | feature_elongation,coding_sequence_variant |
| 11 | 116566129 | 116566411 | 282 | St6galnac2 | AJ,AKR,BTBR,C3H,C57L,CBA,CE,FVB,ILNJ2,MA/My,RH2,SJL,SMJ,SWR,WSB | feature_elongation,3_prime_UTR_variant |
| 11 | 119301052 | 119301124 | 72 | Rnf213 | CAST,CE,SPRET,WSB | feature_elongation,coding_sequence_variant |
| 11 | 120014944 | 120015176 | 232 | Slc38a10 | CAST,MOLF,SJL | feature_elongation,3_prime_UTR_variant |
| 11 | 120356316 | 120356527 | 211 | Arl16 | MOLF,SJL | feature_elongation,3_prime_UTR_variant |
| 11 | 120501231 | 120501285 | 54 | Pcyt2 | MOLF,SJL | feature_elongation,3_prime_UTR_variant |
| 11 | 121321701 | 121321772 | 71 | Fn3krp | MOLF | feature_elongation,3_prime_UTR_variant |
| 12 | 4013159 | 4013359 | 200 | Efr3b | MRL | feature_elongation,3_prime_UTR_variant |
| 12 | 4159420 | 4159496 | 76 | Dnajc27 | CAST,MOLF,SPRET | feature_elongation,3_prime_UTR_variant |
| 12 | 4819152 | 4819254 | 102 | Fam228b,Pfn4 | MOLF,MRL | feature_elongation,5_prime_UTR_variant |
| 12 | 4877441 | 4877612 | 171 | Sf3b6 | CAST,MOLF,SPRET | feature_elongation,3_prime_UTR_variant |
| 12 | 4902860 | 4902982 | 122 | Wdcp | MOLF,SPRET | feature_elongation,3_prime_UTR_variant |
| 12 | 18584005 | 18584172 | 167 | 5730507C01Rik | MRL | feature_elongation,coding_sequence_variant |
| 12 | 24716919 | 24716977 | 58 | Cys1 | AKR,BTBR,BUB,C57L,CE,DBA2J,FVB,KK,NZO,RF,SJL,SMJ,SWR,TH | feature_elongation,3_prime_UTR_variant |
| 12 | 28917555 | 28917806 | 251 | Eipr1 | SPRET | feature_elongation,3_prime_UTR_variant |
| 12 | 28919861 | 28919916 | 55 | Eipr1 | CAST,SPRET | feature_elongation,3_prime_UTR_variant |
| 12 | 31641043 | 31641514 | 471 | Bcap29 | SPRET | feature_elongation,3_prime_UTR_variant |
| 12 | 33377260 | 33378020 | 760 | Atxn7l1 | SPRET | feature_elongation,3_prime_UTR_variant |
| 12 | 44361112 | 44361206 | 94 | Pnpla8 | CAST | feature_elongation,3_prime_UTR_variant |
| 12 | 52035390 | 52035596 | 206 | Dtd2 | 129S1,AJ,AKR,BALB,BUB,C3H,C57L,CAST,CBA,CE,MOLF,MRL,NOD,NOR,NZB,PJ,PL,RF,RH2,SEA,SJL,SMJ,SPRET,SWR,WSB | feature_elongation,3_prime_UTR_variant |

| | | | | | | |
|---|---|---|---|---|---|---|
| 12 | 52044760 | 52045128 | 368 | Dtd2 | AJ,AKR,BALB,BUB,C3H,CBA,NOD,NOR,PJ,PL,RF,RH2,SEA,SJL,SWR,WSB | feature_elongation,3_prime_UTR_variant |
| 12 | 54692546 | 54692658 | 112 | Sptssa | MOLF | feature_elongation,3_prime_UTR_variant |
| 12 | 54719914 | 54720158 | 244 | Eapp | AJ,AKR,BALB,C3H,CBA,DBA1J,DBA2J,ILNJ2,KK,NOD,NOR,PJ,RF,RH2,SEA | feature_elongation,3_prime_UTR_variant |
| 12 | 55309193 | 55309400 | 207 | Srp54c | SPRET | feature_elongation,3_prime_UTR_variant |
| 12 | 55327273 | 55327509 | 236 | Ppp2r3c | FVB,KK,NZW,SMJ,TH | feature_elongation,3_prime_UTR_variant |
| 12 | 56579572 | 56579672 | 100 | Nkx2-1 | CAST,SPRET | feature_elongation,3_prime_UTR_variant |
| 12 | 64519538 | 64519682 | 144 | Fscb | SPRET | feature_elongation,coding_sequence_variant |
| 12 | 64986678 | 64987010 | 332 | Klhl28 | BUB,SJL,SWR | feature_elongation,3_prime_UTR_variant |
| 12 | 65178769 | 65178865 | 96 | Fancm | SPRET | feature_elongation,3_prime_UTR_variant |
| 12 | 69358598 | 69358783 | 185 | Nemf | MRL | feature_elongation,3_prime_UTR_variant |
| 12 | 69790853 | 69790907 | 54 | Dmac2l | SPRET | feature_elongation,3_prime_UTR_variant |
| 12 | 71041697 | 71042192 | 495 | Psma3 | NOD,NOR,SPRET | feature_elongation,3_prime_UTR_variant |
| 12 | 71179525 | 71180121 | 596 | Timm9 | SPRET | feature_elongation,5_prime_UTR_variant |
| 12 | 72808314 | 72808366 | 52 | Ppm1a | NZW | feature_elongation,5_prime_UTR_variant |
| 12 | 73149707 | 73149803 | 96 | Six4 | SPRET | feature_elongation,3_prime_UTR_variant |
| 12 | 73154413 | 73154597 | 184 | Six4 | SPRET | feature_elongation,3_prime_UTR_variant |
| 12 | 75447810 | 75448312 | 502 | Rhoj | SPRET | feature_elongation,3_prime_UTR_variant |
| 12 | 75677401 | 75677624 | 223 | Wdr89 | CE | feature_elongation,3_prime_UTR_variant |
| 12 | 76438766 | 76438964 | 198 | Zbtb1 | MOLF | feature_elongation,3_prime_UTR_variant |
| 12 | 76486248 | 76486463 | 215 | Ppp1r36 | SPRET | feature_elongation,3_prime_UTR_variant |
| 12 | 79222324 | 79223198 | 874 | Rdh11 | MOLF | feature_elongation,3_prime_UTR_variant |
| 12 | 82496141 | 82496193 | 52 | Sipa1l1 | SPRET | feature_elongation,coding_sequence_variant |
| 12 | 84040013 | 84040067 | 54 | Acot2 | MOLF | feature_elongation,3_prime_UTR_variant |
| 12 | 84091656 | 84091724 | 68 | Acot4 | SPRET | feature_elongation,3_prime_UTR_variant |
| 12 | 84382057 | 84382961 | 904 | Zfp410 | SPRET | feature_elongation,3_prime_UTR_variant |
| 12 | 87310996 | 87311200 | 204 | Vipas39 | SWR | feature_elongation,5_prime_UTR_variant |
| 12 | 87994165 | 87994229 | 64 | Eif1ad11 | 129S1,AJ,AKR,BTBR,BUB,C3H,C57L,CBA,DBA1J,DBA2J,FVB,ILNJ2,MRL,NOR,PJ,RF,SEA,SJL,TH,WSB | feature_elongation,coding_sequence_variant |
| 12 | 88051105 | 88051290 | 185 | Eif1ad18 | CAST,MOLF | feature_elongation,3_prime_UTR_variant |
| 12 | 91505427 | 91505482 | 55 | Tshr | SPRET | feature_elongation,3_prime_UTR_variant |
| 12 | 91604862 | 91604920 | 58 | Ston2 | LP,SWR | feature_elongation,3_prime_UTR_variant |
| 12 | 98946123 | 98946190 | 67 | Ttc8 | PJ,SEA | feature_elongation,coding_sequence_variant |
| 12 | 98949223 | 98949411 | 188 | Ttc8 | 129S1,BTBR,BUB,C57L,C58J,CBA,FVB,ILNJ2,KK,LP,MOLF,MRL,NZB,NZO,NZW,RF,SJL,SMJ,SPRET,SWR,WSB | feature_elongation,3_prime_UTR_variant |
| 12 | 101631837 | 101632209 | 372 | Tc2n | SPRET | stop_gained,protein_altering_variant |
| 12 | 101800769 | 101800907 | 138 | Trip11 | MOLF | feature_elongation,3_prime_UTR_variant |
| 12 | 101887119 | 101887977 | 858 | Atxn3 | MOLF | feature_elongation,3_prime_UTR_variant |
| 12 | 101887123 | 101887330 | 207 | Atxn3 | CAST | feature_elongation,3_prime_UTR_variant |
| 12 | 102536101 | 102536163 | 62 | Itpk1 | MOLF | feature_elongation,3_prime_UTR_variant |
| 12 | 103417476 | 103417557 | 81 | Ifi27l2b | CAST | feature_elongation,coding_sequence_variant |
| 12 | 103962610 | 103962747 | 137 | Serpina9 | 129S1,AKR,BALB,BTBR,BUB,C57L,CAST,CBA,CE,DBA1J,DBA2J,FVB,ILNJ2,KK,LP,MRL,NOD,NOR,NZB,PJ,PL,RF,SEA,SJL,SWR,WSB | feature_elongation,3_prime_UTR_variant |
| 12 | 104204054 | 104204474 | 420 | Serpina3g | AKR,BALB,BUB,CAST,CBA,CE,DBA1J,DBA2J,FVB,ILNJ2,KK,LP,MRL,NOD,NOR,PJ,PL,RF,SEA,SJL | feature_elongation,5_prime_UTR_variant |
| 12 | 104204055 | 104204486 | 431 | Serpina3g | SPRET,SWR,WSB | feature_elongation,5_prime_UTR_variant |
| 12 | 104735101 | 104735375 | 274 | Clmn | MOLF,SPRET | feature_elongation,3_prime_UTR_variant |
| 12 | 107877986 | 107878042 | 56 | Bcl11b | AJ,BUB,CE,FVB,MA/My,NZB,NZO,NZW,PJ,PL,RF,SEA,SJL,SMJ,SWR,TH | feature_elongation,3_prime_UTR_variant |
| 12 | 109561092 | 109561158 | 66 | Rtl1 | WSB | feature_elongation,coding_sequence_variant |
| 12 | 111768933 | 111769057 | 124 | Klc1 | SPRET | feature_elongation,3_prime_UTR_variant |
| 12 | 111778592 | 111778665 | 73 | Xrcc3 | CAST | feature_elongation,5_prime_UTR_variant |
| 12 | 116354584 | 116354763 | 179 | Esyt2 | CAST,SPRET | feature_elongation,3_prime_UTR_variant |
| 13 | 3701161 | 3701220 | 59 | Asb13 | SPRET | feature_elongation,3_prime_UTR_variant |
| 13 | 3703272 | 3703347 | 75 | Asb13 | 129S1,AJ,AKR,B10D2,B10J,BALB,BTBR,BUB,C3H,C57L,C58J,CAST,CBA,CE,DBA1J,DBA2J,FVB,ILNJ2,KK,LP,MA/My,MOLF,MRL,NOD,NOR,NZB,NZO,NZW,PJ,PL,RF,RH2,SEA,SJL,SMJ,SPRET,SWR,TH,WSB | feature_elongation,3_prime_UTR_variant |
| 13 | 4298723 | 4299055 | 332 | Akr1c19 | SPRET | feature_elongation,3_prime_UTR_variant |
| 13 | 4299323 | 4300005 | 682 | Akr1c19 | CAST | feature_elongation,3_prime_UTR_variant |
| 13 | 6630709 | 6630794 | 85 | Pfkp | AKR,BUB,C57L,KK,LP,MA/My,MOLF,NOD,NOR,NZB,NZO,NZW,RF,TH,WSB | feature_elongation,3_prime_UTR_variant |
| 13 | 8901609 | 8901911 | 302 | Wdr37 | SPRET | feature_elongation,3_prime_UTR_variant |
| 13 | 13952677 | 13952730 | 53 | Lyst | SPRET | feature_elongation,3_prime_UTR_variant |
| 13 | 14173341 | 14173397 | 56 | B3galnt2 | SPRET | feature_elongation,3_prime_UTR_variant |
| 13 | 14228184 | 14228238 | 54 | Ggps1 | SPRET | feature_elongation,3_prime_UTR_variant |
| 13 | 16202288 | 16202530 | 242 | Inhba | AKR,ILNJ2,LP,MOLF,PJ,RF,SEA | feature_elongation,3_prime_UTR_variant |
| 13 | 19815987 | 19816206 | 219 | Sfrp4 | LP,MOLF,PJ,SEA,SPRET | feature_elongation,3_prime_UTR_variant |
| 13 | 21254785 | 21254990 | 205 | Or2p2 | LP,PJ,SEA | feature_elongation,3_prime_UTR_variant |
| 13 | 21739708 | 21739899 | 191 | Or2b7 | MOLF | feature_elongation,coding_sequence_variant |
| 13 | 21841863 | 21841991 | 128 | Or2w6 | CAST,LP,MOLF,PJ,SEA,SPRET | feature_elongation,3_prime_UTR_variant |
| 13 | 22277102 | 22277162 | 60 | Vmn1r188 | ILNJ2 | feature_elongation,3_prime_UTR_variant |
| 13 | 22278750 | 22278865 | 115 | Vmn1r188 | CAST | feature_elongation,3_prime_UTR_variant |
| 13 | 22368063 | 22368129 | 66 | Vmn1r191 | SPRET | feature_elongation,5_prime_UTR_variant |
| 13 | 22803420 | 22803609 | 189 | Vmn1r206 | SPRET | feature_elongation,3_prime_UTR_variant |
| 13 | 23323506 | 23323597 | 91 | Vmn1r218 | BTBR,C57L,KK,LP,NOD,NOR,NZB,NZO,NZW,PJ,SEA | feature_elongation,3_prime_UTR_variant |
| 13 | 23765971 | 23766753 | 782 | H4c4 | SPRET | feature_elongation,3_prime_UTR_variant |
| 13 | 25394525 | 25394579 | 54 | Dcdc2a | SPRET | feature_elongation,3_prime_UTR_variant |
| 13 | 26953497 | 26953599 | 102 | Hdgfl1 | WSB | feature_elongation,coding_sequence_variant |
| 13 | 27343095 | 27343265 | 170 | Prl3d3 | CAST,KK,PJ,SEA | feature_elongation,coding_sequence_variant |
| 13 | 29986484 | 29986587 | 103 | Cdkal1 | SPRET | feature_elongation,3_prime_UTR_variant |
| 13 | 30891142 | 30891195 | 53 | Dusp22 | MOLF | feature_elongation,3_prime_UTR_variant |
| 13 | 33200855 | 33201213 | 358 | Serpinb9 | SPRET | feature_elongation,3_prime_UTR_variant |
| 13 | 34220869 | 34221197 | 328 | Ripk1 | SPRET | feature_elongation,3_prime_UTR_variant |
| 13 | 36079965 | 36080178 | 213 | Rpp40 | SPRET | feature_elongation,3_prime_UTR_variant |
| 13 | 36163130 | 36163331 | 201 | Lyrm4 | CE,KK,LP | feature_elongation,3_prime_UTR_variant |

| | | | | | | |
|---|---|---|---|---|---|---|
| 13 | 37602923 | 37602978 | 55 | Ly86 | MOLF | feature_elongation,3_prime_UTR_variant |
| 13 | 38117453 | 38117524 | 71 | Rreb1 | SPRET | feature_elongation,3_prime_UTR_variant |
| 13 | 38157733 | 38157800 | 67 | Ssr1 | CAST | feature_elongation,3_prime_UTR_variant |
| 13 | 40869382 | 40869452 | 70 | Tfap2a | SPRET | feature_elongation,3_prime_UTR_variant |
| 13 | 43288845 | 43288897 | 52 | Phactr1 | SPRET | feature_elongation,3_prime_UTR_variant |
| 13 | 43289862 | 43289937 | 75 | Phactr1 | MOLF | feature_elongation,3_prime_UTR_variant |
| 13 | 43539727 | 43540166 | 439 | Sirt5 | SPRET | feature_elongation,3_prime_UTR_variant |
| 13 | 46583232 | 46583298 | 66 | Rbm24 | CAST | feature_elongation,3_prime_UTR_variant |
| 13 | 49415975 | 49416459 | 484 | Fgd3 | SPRET | feature_elongation,3_prime_UTR_variant |
| 13 | 51574313 | 51574373 | 60 | S1pr3 | CAST | feature_elongation,3_prime_UTR_variant |
| 13 | 51819651 | 51819824 | 173 | Secisbp2 | CAST | feature_elongation,coding_sequence_variant |
| 13 | 52797282 | 52797348 | 66 | Syk | CAST | feature_elongation,3_prime_UTR_variant |
| 13 | 54527805 | 54527878 | 73 | Cplx2 | AJ,BALB,C3H,CBA,CE | feature_elongation,3_prime_UTR_variant |
| 13 | 54530818 | 54530876 | 58 | Cplx2 | SPRET | feature_elongation,3_prime_UTR_variant |
| 13 | 54935993 | 54936074 | 81 | Eif4e1b | MOLF,SPRET | feature_elongation,3_prime_UTR_variant |
| 13 | 55565773 | 55565996 | 223 | F12 | AJ,BALB,C3H,CBA,CE | feature_elongation,3_prime_UTR_variant |
| 13 | 55848635 | 55848731 | 96 | B230219D22Rik | MOLF,SPRET | feature_elongation,3_prime_UTR_variant |
| 13 | 58308759 | 58308847 | 88 | Idnk | BTBR,C57L,C58J,CAST,DBA1J,DBA2J,ILNJ2,KK,MA/My,MOLF,MRL,SPRET,SWR,WSB | feature_elongation,3_prime_UTR_variant |
| 13 | 58703031 | 58703089 | 58 | Slc28a3 | SPRET | feature_elongation,3_prime_UTR_variant |
| 13 | 59035623 | 59035786 | 163 | Ntrk2 | MOLF | feature_elongation,3_prime_UTR_variant |
| 13 | 59080500 | 59080564 | 64 | Ntrk2 | MOLF | feature_elongation,3_prime_UTR_variant |
| 13 | 59788052 | 59788248 | 196 | Golm1 | SPRET | stop_gained,frameshift_variant |
| 13 | 59904415 | 59904490 | 75 | Isca1 | CAST | feature_elongation,3_prime_UTR_variant |
| 13 | 59919667 | 59919896 | 229 | Tut7 | SPRET | feature_elongation,3_prime_UTR_variant |
| 13 | 59953137 | 59953260 | 123 | Tut7 | SPRET | feature_elongation,coding_sequence_variant |
| 13 | 62517890 | 62518058 | 168 | Gm3604 | CAST,MOLF | feature_elongation,coding_sequence_variant,feature_elongation,3_prime_UTR_variant |
| 13 | 62601933 | 62602017 | 84 | Zfp935 | SPRET | feature_elongation,3_prime_UTR_variant |
| 13 | 62752055 | 62752139 | 84 | Zfp1008 | CAST,MOLF,SPRET | feature_elongation,3_prime_UTR_variant |
| 13 | 62848408 | 62848498 | 90 | Platr25 | FVB,PJ,SEA,SWR | feature_elongation,3_prime_UTR_variant,feature_elongation,coding_sequence_variant |
| 13 | 63657237 | 63657309 | 72 | Ptch1 | SPRET | feature_elongation,3_prime_UTR_variant |
| 13 | 64289079 | 64289131 | 52 | Zfp367 | SPRET | feature_elongation,3_prime_UTR_variant |
| 13 | 64341493 | 64341712 | 219 | Cdc14b | CAST | feature_elongation,3_prime_UTR_variant |
| 13 | 64508650 | 64508870 | 220 | Ctsl | FVB,PJ,SEA,SWR | feature_elongation,3_prime_UTR_variant |
| 13 | 65303139 | 65303339 | 200 | Or9s18 | 129S1,AJ,AKR,BALB,BTBR,BUB,C3H,C57L,C58J,CBA,CE,DBA1J,DBA2J,FVB,ILNJ2,KK,MRL,NOD,NOR,NZB,NZO,NZW,PJ,PL,RF,SEA,SJL,SMJ,SWR,TH,WSB | feature_elongation,3_prime_UTR_variant |
| 13 | 67056673 | 67056864 | 191 | Ramacl | PJ,SEA,SPRET,SWR | feature_elongation,3_prime_UTR_variant |
| 13 | 67246775 | 67246859 | 84 | Rslcan18 | 129S1,AKR,BTBR,BUB,C3H,C57L,C58J,CBA,CE,DBA1J,DBA2J,ILNJ2,MRL,NOD,NOR,NZW,PL,RF,SMJ,SWR,TH | feature_elongation,coding_sequence_variant |
| 13 | 67441178 | 67441262 | 84 | Zfp457 | 129S1,AJ,AKR,BALB,BTBR,BUB,C3H,C57L,C58J,CAST,CBA,CE,DBA1J,DBA2J,FVB,ILNJ2,KK,MOLF,MRL,NOD,NOR,NZB,NZO,NZW,PJ,PL,RF,SEA,SJL,SMJ,SPRET,SWR,TH,WSB | feature_elongation,coding_sequence_variant |
| 13 | 67574468 | 67575274 | 806 | Zfp874a | MOLF,SPRET | feature_elongation,coding_sequence_variant |
| 13 | 67588647 | 67588816 | 169 | Zfp874a | MOLF,SPRET | feature_elongation,3_prime_UTR_variant |
| 13 | 67590027 | 67590299 | 272 | Zfp874a | 129S1,AJ,AKR,BALB,BTBR,BUB,C3H,C57L,C58J,CAST,CBA,CE,DBA1J,DBA2J,FVB,ILNJ2,KK,MOLF,MRL,NOD,NOR,NZB,NZO,NZW,PJ,PL,RF,SEA,SJL,SMJ,SPRET,SWR,TH,WSB | feature_elongation,3_prime_UTR_variant |
| 13 | 67622508 | 67622750 | 242 | Zfp874b | MA/My | feature_elongation,coding_sequence_variant |
| 13 | 67764195 | 67764881 | 686 | Zfp729a | SPRET | feature_elongation,3_prime_UTR_variant |
| 13 | 67814761 | 67814815 | 54 | Zfp738 | MOLF | feature_elongation,3_prime_UTR_variant |
| 13 | 67853736 | 67853925 | 189 | Zfp65 | KK,NZB,NZO,SWR | feature_elongation,3_prime_UTR_variant |
| 13 | 70935189 | 70935544 | 355 | Adamts16 | MRL | feature_elongation,coding_sequence_variant |
| 13 | 73810256 | 73810325 | 69 | Slc6a18 | MOLF,SPRET | feature_elongation,3_prime_UTR_variant |
| 13 | 74519204 | 74519400 | 196 | Zfp72 | ILNJ2,NZW,WSB | feature_elongation,3_prime_UTR_variant |
| 13 | 75919518 | 75919575 | 57 | Ell2 | BALB,NZW,WSB | feature_elongation,3_prime_UTR_variant |
| 13 | 76020334 | 76020517 | 183 | Rhobtb3 | ILNJ2,WSB | feature_elongation,3_prime_UTR_variant |
| 13 | 90256752 | 90256925 | 173 | Tmem167 | SPRET | feature_elongation,3_prime_UTR_variant |
| 13 | 91072951 | 91073158 | 207 | Rps23 | SPRET | feature_elongation,3_prime_UTR_variant |
| 13 | 92847135 | 92847187 | 52 | Serinc5 | SPRET | feature_elongation,3_prime_UTR_variant |
| 13 | 93233253 | 93233398 | 145 | Cmya5 | 129S1,AJ,AKR,BALB,BTBR,BUB,C3H,C57L,C58J,CAST,CBA,CE,DBA1J,DBA2J,FVB,ILNJ2,KK,MA/My,MOLF,MRL,NZB,NZO,NZW,PJ,PL,RF,SEA,SJL,SMJ,SPRET,SWR | feature_elongation,coding_sequence_variant |
| 13 | 97170738 | 97170822 | 84 | 1700029F12Rik | SPRET | feature_elongation,coding_sequence_variant |
| 13 | 97265288 | 97265543 | 255 | Fam169a | AJ,BUB,WSB | feature_elongation,3_prime_UTR_variant |
| 13 | 97266755 | 97266961 | 206 | Nsa2,Fam169a | CAST | feature_elongation,3_prime_UTR_variant |
| 13 | 100155479 | 100155781 | 302 | Bdp1 | CAST,SPRET | feature_elongation,3_prime_UTR_variant |
| 13 | 100247419 | 100247949 | 530 | Serf1 | SPRET | feature_elongation,3_prime_UTR_variant |
| 13 | 100349053 | 100349258 | 205 | Naip5 | SPRET | feature_elongation,3_prime_UTR_variant |
| 13 | 101821104 | 101821165 | 61 | Pik3r1 | MOLF | feature_elongation,3_prime_UTR_variant |
| 13 | 105087715 | 105087779 | 64 | Rgs7bp | NZB | feature_elongation,3_prime_UTR_variant |
| 13 | 108310797 | 108310862 | 65 | Ercc8 | CAST,CE,MA/My,MOLF,NZO,SPRET,TH | feature_elongation,3_prime_UTR_variant |
| 13 | 110489382 | 110489446 | 64 | Gapt | AKR,BALB,DBA2J,ILNJ2,RF,SEA,WSB | feature_elongation,3_prime_UTR_variant |
| 13 | 112871130 | 112871196 | 66 | Slc38a9 | AJ,CAST,DBA1J,FVB,KK,MA/My,MOLF,NZO,PL,RF,RH2,SJL,SPRET,SWR | feature_elongation,3_prime_UTR_variant |
| 13 | 114425008 | 114425288 | 280 | Ndufs4 | SPRET | feature_elongation,3_prime_UTR_variant |
| 13 | 114965308 | 114965373 | 65 | Mocs2 | SPRET | feature_elongation,3_prime_UTR_variant |
| 13 | 115092561 | 115092633 | 72 | Itga1 | SPRET | feature_elongation,3_prime_UTR_variant |
| 13 | 116435508 | 116435568 | 60 | Isl1 | AJ,DBA1J,MA/My,RH2 | feature_elongation,3_prime_UTR_variant |
| 13 | 120114279 | 120114547 | 268 | Ccl28 | 129S1,BTBR,C3H,C57L,CAST,CBA,CE,DBA2J,KK,MA/My,MOLF,NZO,SMJ | feature_elongation,3_prime_UTR_variant |
| 14 | 6036784 | 6036939 | 155 | Top2b | SPRET | feature_elongation,3_prime_UTR_variant |
| 14 | 6230449 | 6230530 | 81 | Oxsm | SPRET | feature_elongation,3_prime_UTR_variant |
| 14 | 8296045 | 8296105 | 60 | Or2t6 | 129S1,C3H,C57L,DBA1J,DBA2J,KK,MOLF,MRL,PJ,PL,SPRET | feature_elongation,3_prime_UTR_variant |
| 14 | 8349776 | 8350049 | 273 | Psmd6 | CAST | feature_elongation,coding_sequence_variant |

| | | | | | | |
|---|---|---|---|---|---|---|
| 14 | 8365514 | 8365584 | 70 | Atxn7 | SPRET | feature_elongation,3_prime_UTR_variant |
| 14 | 8520344 | 8520520 | 176 | Gm11100 | 129S1,AJ,AKR,BALB,BTBR,BUB,C3H,C57L,C58J,CAST,CBA,CE,DBA1J,DBA2J,ILNJ2,KK,MOLF,MRL,NZB,NZO,NZW,PJ,PL,RF,SEA,SMJ,SPRET,WSB | feature_elongation,3_prime_UTR_variant |
| 14 | 20393410 | 20393637 | 227 | Ecd | SPRET | feature_elongation,coding_sequence_variant |
| 14 | 22766361 | 22766416 | 55 | Lrmda | CAST,SPRET | feature_elongation,3_prime_UTR_variant |
| 14 | 25988362 | 25988659 | 297 | Duxbl1 | SPRET | feature_elongation,3_prime_UTR_variant |
| 14 | 26825568 | 26825624 | 56 | Il17rd | SPRET | feature_elongation,3_prime_UTR_variant |
| 14 | 28753176 | 28753292 | 116 | Lrtm1 | SPRET | feature_elongation,3_prime_UTR_variant |
| 14 | 29781833 | 29781973 | 140 | Cacna1d | CAST,SPRET | feature_elongation,3_prime_UTR_variant |
| 14 | 30296584 | 30296793 | 209 | Tkt | SPRET | feature_elongation,3_prime_UTR_variant |
| 14 | 31245136 | 31245364 | 228 | Colq | WSB | feature_elongation,3_prime_UTR_variant |
| 14 | 31899196 | 31899457 | 261 | Ncoa4 | MOLF | feature_elongation,coding_sequence_variant |
| 14 | 32130165 | 32130245 | 80 | Chat | CAST | feature_elongation,3_prime_UTR_variant |
| 14 | 33656466 | 33656530 | 64 | Gdf10 | SPRET | feature_elongation,coding_sequence_variant |
| 14 | 33659319 | 33659595 | 276 | Gdf10 | SPRET | feature_elongation,3_prime_UTR_variant |
| 14 | 33914887 | 33915721 | 834 | Gprin2 | PJ,SPRET | feature_elongation,3_prime_UTR_variant |
| 14 | 38091720 | 38091826 | 106 | Nrg3 | MOLF | feature_elongation,3_prime_UTR_variant |
| 14 | 40858294 | 40858505 | 211 | Sftpa1 | SPRET | feature_elongation,3_prime_UTR_variant |
| 14 | 45240799 | 45240868 | 69 | Ptger2 | C57L,CAST,MOLF,NZO,PJ,PL | feature_elongation,3_prime_UTR_variant |
| 14 | 47533031 | 47533294 | 263 | Socs4 | SWR | feature_elongation,3_prime_UTR_variant |
| 14 | 47974917 | 47975131 | 214 | Ktn1 | SPRET | feature_elongation,3_prime_UTR_variant |
| 14 | 49245870 | 49246032 | 162 | Exoc5 | SPRET | feature_elongation,3_prime_UTR_variant |
| 14 | 49247026 | 49247234 | 208 | Exoc5 | NZW | feature_elongation,3_prime_UTR_variant |
| 14 | 50382781 | 50383702 | 921 | Or4k5 | 129S1,AJ,AKR,BALB,BUB,C3H,C57L,CAST,CBA,CE,DBA1J,DBA2J,FVB,ILNJ2,KK,MA/My,MRL,NOD,NOR,NZB,NZO,NZW,PJ,PL,RF,RH2,SEA,SJL,SMJ,SWR,TH | feature_elongation,3_prime_UTR_variant |
| 14 | 50421901 | 50422364 | 463 | Or4k2 | SPRET | feature_elongation,3_prime_UTR_variant |
| 14 | 50471627 | 50471678 | 51 | Or4k6 | AJ,AKR,C3H,C57L,CBA,CE,MRL,NOD,NZB,PJ,PL,RF,RH2,SEA | feature_elongation,3_prime_UTR_variant |
| 14 | 50555329 | 50555380 | 51 | Or4m1 | DBA1J,DBA2J | feature_elongation,3_prime_UTR_variant |
| 14 | 50600974 | 50601408 | 434 | Tlr11 | SPRET | feature_elongation,3_prime_UTR_variant |
| 14 | 50663298 | 50663364 | 66 | Or11g24 | MOLF | feature_elongation,3_prime_UTR_variant |
| 14 | 50692959 | 50693155 | 196 | Or11g7 | 129S1,AJ,AKR,BALB,BUB,C3H,C57L,CAST,CBA,CE,DBA1J,DBA2J,FVB,ILNJ2,KK,MA/My,MOLF,MRL,NOD,NOR,NZB,NZO,NZW,PJ,PL,RF,RH2,SEA,SJL,SMJ,SPRET,SWR,TH,WSB | feature_elongation,3_prime_UTR_variant |
| 14 | 50771843 | 50771958 | 115 | Or11g27 | 129S1,AJ,AKR,BALB,BUB,C3H,C57L,CBA,CE,DBA1J,DBA2J,FVB,ILNJ2,KK,MA/My,MOLF,MRL,NOD,NOR,NZB,NZO,NZW,PJ,PL,RF,RH2,SEA,SJL,SMJ,SPRET,SWR,TH,WSB | feature_elongation,3_prime_UTR_variant |
| 14 | 50880883 | 50881471 | 588 | Or11h6 | CAST | feature_elongation,3_prime_UTR_variant |
| 14 | 51128283 | 51128343 | 60 | Klhl33 | 129S1,AJ,AKR,BALB,BUB,C3H,C57L,CAST,CBA,CE,DBA1J,DBA2J,FVB,ILNJ2,KK,MA/My,MOLF,MRL,NOD,NOR,NZB,NZO,NZW,PJ,PL,RF,RH2,SEA,SJL,SMJ,SPRET,SWR,TH | feature_elongation,3_prime_UTR_variant |
| 14 | 51759277 | 51759705 | 428 | Gm7247 | SPRET | feature_elongation,coding_sequence_variant |
| 14 | 52549564 | 52549800 | 236 | Sall2 | FVB | feature_elongation,3_prime_UTR_variant |
| 14 | 52697396 | 52697595 | 199 | Or4e1 | SPRET | feature_elongation,3_prime_UTR_variant |
| 14 | 52727084 | 52727210 | 126 | Or4e5 | SPRET | feature_elongation,3_prime_UTR_variant |
| 14 | 54517221 | 54517404 | 183 | Or6e1 | 129S1,AJ,AKR,BALB,BTBR,BUB,C3H,C57L,CAST,CBA,CE,DBA1J,DBA2J,FVB,ILNJ2,KK,MA/My,MRL,NOD,NOR,NZB,NZO,NZW,PJ,PL,RF,RH2,SEA,SJL,SMJ,SWR,TH,WSB | feature_elongation,3_prime_UTR_variant |
| 14 | 54866850 | 54867343 | 493 | Psmb11 | SPRET | feature_elongation,3_prime_UTR_variant |
| 14 | 54960250 | 54960472 | 222 | Slc7a8 | 129S1,AJ,AKR,BALB,BTBR,C3H,C57L,CBA,CE,DBA1J,KK,MRL,NZO,PJ,RF,RH2,SEA,TH | feature_elongation,3_prime_UTR_variant |
| 14 | 56496344 | 56496510 | 166 | Gzmb | MOLF | feature_elongation,3_prime_UTR_variant |
| 14 | 57029250 | 57029457 | 207 | Zmym5 | KK,WSB | feature_elongation,3_prime_UTR_variant |
| 14 | 58072440 | 58072643 | 203 | Zdhhc20 | BTBR,BUB,CAST | feature_elongation,3_prime_UTR_variant |
| 14 | 59473853 | 59473943 | 90 | Rcbtb1 | SPRET | feature_elongation,3_prime_UTR_variant |
| 14 | 59785333 | 59785763 | 430 | Cab39l | C3H,CE,DBA2J | feature_elongation,3_prime_UTR_variant |
| 14 | 60622162 | 60622289 | 127 | Amer2 | SPRET | feature_elongation,3_prime_UTR_variant |
| 14 | 61456878 | 61457196 | 318 | Sgcg | MOLF | feature_elongation,3_prime_UTR_variant |
| 14 | 61477931 | 61478049 | 118 | Sacs | MOLF | feature_elongation,3_prime_UTR_variant |
| 14 | 61576057 | 61576109 | 52 | Ebpl | CAST | feature_elongation,3_prime_UTR_variant |
| 14 | 62939949 | 62940076 | 127 | Ints6 | MOLF | feature_elongation,coding_sequence_variant |
| 14 | 65634248 | 65634302 | 54 | Zfp395 | CAST | feature_elongation,3_prime_UTR_variant |
| 14 | 67251877 | 67252102 | 225 | Ppp2r2a | CAST | feature_elongation,3_prime_UTR_variant |
| 14 | 67991625 | 67991853 | 228 | Dock5 | CAST | feature_elongation,3_prime_UTR_variant |
| 14 | 67992901 | 67992989 | 88 | Dock5 | MOLF | feature_elongation,3_prime_UTR_variant |
| 14 | 69930749 | 69930827 | 78 | Loxl2 | BTBR,LP,TH,WSB | feature_elongation,3_prime_UTR_variant |
| 14 | 70820980 | 70821055 | 75 | Fhip2b | BTBR,BUB,LP,WSB | feature_elongation,3_prime_UTR_variant |
| 14 | 70892733 | 70892992 | 259 | Xpo7 | CAST | feature_elongation,3_prime_UTR_variant |
| 14 | 70893780 | 70893979 | 199 | Xpo7 | SPRET | feature_elongation,3_prime_UTR_variant |
| 14 | 71216059 | 71216135 | 76 | Gfra2 | FVB,RH2,SJL | feature_elongation,3_prime_UTR_variant |
| 14 | 73477904 | 73478125 | 221 | Lpar6 | SJL,WSB | feature_elongation,3_prime_UTR_variant |
| 14 | 74992492 | 74992747 | 255 | Lrch1 | BUB,KK,MOLF | feature_elongation,3_prime_UTR_variant |
| 14 | 75044978 | 75045966 | 988 | Lrch1 | BUB,KK,TH | feature_elongation,coding_sequence_variant |
| 14 | 78318844 | 78319069 | 225 | Fam216b | FVB,MA/My,MRL,NOD,NOR | feature_elongation,3_prime_UTR_variant |
| 14 | 78731272 | 78731370 | 98 | Akap11 | CAST | feature_elongation,3_prime_UTR_variant |
| 14 | 79871495 | 79871554 | 59 | Sugt1 | CAST | feature_elongation,3_prime_UTR_variant |
| 14 | 106343868 | 106343947 | 79 | Trim52 | CAST | feature_elongation,5_prime_UTR_variant |
| 14 | 121780426 | 121780574 | 148 | Dock9 | CAST,MOLF,RF,SPRET | feature_elongation,3_prime_UTR_variant |
| 15 | 5128246 | 5128306 | 60 | Card6 | MOLF | feature_elongation,coding_sequence_variant |
| 15 | 8189758 | 8189810 | 52 | Nup155 | AKR,RF | feature_elongation,3_prime_UTR_variant |
| 15 | 9115886 | 9116147 | 261 | Skp2 | AJ,BUB,FVB,NOD,NOR,RH2 | feature_elongation,3_prime_UTR_variant |
| 15 | 9199833 | 9200219 | 386 | Lmbrd2 | LP,SJL | feature_elongation,3_prime_UTR_variant |
| 15 | 10570618 | 10570827 | 209 | Rai14 | NZB | feature_elongation,3_prime_UTR_variant |
| 15 | 30847536 | 30847646 | 110 | Ctnnd2 | SPRET | feature_elongation,coding_sequence_variant |
| 15 | 31294865 | 31294918 | 53 | Ankrd33b | 129S1,BTBR,C57L,CAST,KK,LP,SJL | feature_elongation,3_prime_UTR_variant |

| | | | | | | |
|---|---|---|---|---|---|---|
| 15 | 38492530 | 38493370 | 840 | Azin1 | NZB,NZW | feature_elongation,coding_sequence_variant |
| 15 | 38692037 | 38692091 | 54 | Atp6v1c1 | SPRET | feature_elongation,3_prime_UTR_variant |
| 15 | 51746533 | 51746617 | 84 | Utp23 | MOLF,SPRET,WSB | feature_elongation,3_prime_UTR_variant |
| 15 | 55119103 | 55119269 | 166 | Deptor | 129S1,AJ,AKR,C57L,CE,DBA1J,DBA2J,FVB,ILNJ2,MOLF,MRL,NZB,PJ,RH2,SJL,SMJ,TH,WSB | feature_elongation,3_prime_UTR_variant |
| 15 | 59528651 | 59528929 | 278 | Trib1 | 129S1 | feature_elongation,3_prime_UTR_variant |
| 15 | 60693155 | 60693508 | 353 | Lratd2 | ILNJ2 | feature_elongation,3_prime_UTR_variant |
| 15 | 64569697 | 64569766 | 69 | Adcy8 | MOLF | feature_elongation,3_prime_UTR_variant |
| 15 | 65740942 | 65741005 | 63 | Efr3a | BALB,C3H,CAST,CBA,CE,SEA | feature_elongation,3_prime_UTR_variant |
| 15 | 66792799 | 66792884 | 85 | Ccn4 | CAST | feature_elongation,3_prime_UTR_variant |
| 15 | 66975686 | 66975806 | 120 | St3gal1 | CAST,SPRET | feature_elongation,3_prime_UTR_variant |
| 15 | 71310029 | 71310115 | 86 | Fam135b | SPRET | feature_elongation,3_prime_UTR_variant |
| 15 | 71313280 | 71313680 | 400 | Fam135b | SPRET | feature_elongation,3_prime_UTR_variant |
| 15 | 71668527 | 71668643 | 116 | Col22a1 | CAST | feature_elongation,3_prime_UTR_variant |
| 15 | 72835468 | 72835759 | 291 | Trappc9 | AJ,BALB,C3H,CAST,CBA,CE,DBA1J,DBA2J,ILNJ2,MOLF,NOD,NOR,NZB,NZO,SEA,SPRET,WSB | stop_gained,protein_altering_variant |
| 15 | 72865945 | 72866001 | 56 | Trappc9 | MOLF | feature_elongation,3_prime_UTR_variant |
| 15 | 74597301 | 74597366 | 65 | Slurp1 | 129S1,CAST,ILNJ2,MOLF,NZW | feature_elongation,3_prime_UTR_variant |
| 15 | 74706276 | 74706405 | 129 | Cyp11b1 | CAST | feature_elongation,3_prime_UTR_variant |
| 15 | 76426155 | 76426369 | 214 | Slc52a2 | CAST | feature_elongation,3_prime_UTR_variant |
| 15 | 76499062 | 76499116 | 54 | Slc39a4 | CAST | feature_elongation,coding_sequence_variant |
| 15 | 76835096 | 76835208 | 112 | 1110038F14Rik | MA/My | feature_elongation,3_prime_UTR_variant |
| 15 | 79009676 | 79009811 | 135 | Micall1 | SPRET | feature_elongation,coding_sequence_variant |
| 15 | 80000503 | 80000559 | 56 | Syngr1 | SPRET | feature_elongation,3_prime_UTR_variant |
| 15 | 80535258 | 80535339 | 81 | Grap2 | SPRET | feature_elongation,3_prime_UTR_variant |
| 15 | 80815026 | 80815081 | 55 | Tnrc6b | SPRET | feature_elongation,3_prime_UTR_variant |
| 15 | 80854321 | 80854730 | 409 | Adsl | MOLF,SPRET | feature_elongation,3_prime_UTR_variant |
| 15 | 81708389 | 81708449 | 60 | Tef | SPRET | feature_elongation,3_prime_UTR_variant |
| 15 | 81733700 | 81733767 | 67 | Tob2 | CAST,SPRET | feature_elongation,3_prime_UTR_variant |
| 15 | 81748736 | 81748901 | 165 | Phf5a | CAST | feature_elongation,3_prime_UTR_variant |
| 15 | 81799567 | 81799756 | 189 | Polr3h | WSB | feature_elongation,3_prime_UTR_variant |
| 15 | 81924977 | 81925192 | 215 | Snu13 | SPRET,WSB | feature_elongation,3_prime_UTR_variant |
| 15 | 84198149 | 84198200 | 51 | Parvb | BUB,DBA1J,DBA2J,ILNJ2 | feature_elongation,3_prime_UTR_variant |
| 15 | 85170191 | 85171170 | 979 | Fbln1 | 129S1,AJ,AKR,BALB,BTBR,BUB,C3H,C57L,CAST,CBA,CE,DBA1J,DBA2J,FVB,ILNJ2,KK,MA/My,MRL,NOD,NOR,NZB,NZO,NZW,PJ,PL,RF,SEA,SJL,SMJ,SWR,TH,WSB | feature_elongation,3_prime_UTR_variant |
| 15 | 85377219 | 85377563 | 344 | 7530416G11Rik | MOLF,SJL,SMJ,SPRET,SWR | feature_elongation,3_prime_UTR_variant |
| 15 | 85698981 | 85699136 | 155 | Pkdrej | BUB,DBA1J,DBA2J,NZW,PJ,SEA,SPRET | feature_elongation,3_prime_UTR_variant |
| 15 | 94438560 | 94438620 | 60 | Pus7l | SPRET | feature_elongation,coding_sequence_variant |
| 15 | 94460947 | 94461037 | 90 | Irak4 | 129S1,AJ,AKR,BTBR,BUB,C3H,C57L,CAST,CBA,CE,DBA1J,DBA2J,FVB,ILNJ2,KK,MA/My,MOLF,NOD,NOR,NZB,NZO,NZW,PJ,PL,RF,RH2,SEA,SJL,SMJ,SWR,TH | feature_elongation,coding_sequence_variant |
| 15 | 95553600 | 95553774 | 174 | Dbx2 | SPRET | feature_elongation,5_prime_UTR_variant |
| 15 | 96317779 | 96317875 | 96 | Scaf11 | SPRET | feature_elongation,coding_sequence_variant |
| 15 | 96512847 | 96513396 | 549 | Slc38a1 | MOLF | feature_elongation,3_prime_UTR_variant |
| 15 | 96513926 | 96513989 | 63 | Slc38a1 | MOLF | feature_elongation,3_prime_UTR_variant |
| 15 | 97937995 | 97938049 | 54 | Senp1 | C57L,DBA2J,ILNJ2,KK,NOD,NOR,NZB,NZO,PJ,SEA,SMJ,TH | feature_elongation,3_prime_UTR_variant |
| 15 | 98132205 | 98132277 | 72 | Or10ad1b | SPRET | feature_elongation,5_prime_UTR_variant |
| 15 | 98355643 | 98356180 | 537 | Or8s8 | MOLF,SPRET | feature_elongation,3_prime_UTR_variant |
| 15 | 98358221 | 98358662 | 441 | Or8s8 | BUB,MA/My,SWR | feature_elongation,3_prime_UTR_variant |
| 15 | 98397761 | 98397821 | 60 | Or11m3 | MOLF | feature_elongation,3_prime_UTR_variant |
| 15 | 98398774 | 98398844 | 70 | Or11m3 | CAST,MOLF,SPRET | feature_elongation,3_prime_UTR_variant |
| 15 | 98748903 | 98748960 | 57 | Kmt2d | ILNJ2 | feature_elongation,coding_sequence_variant |
| 15 | 99502346 | 99502486 | 140 | Aqp6 | CAST,MOLF | feature_elongation,3_prime_UTR_variant |
| 15 | 99626507 | 99626737 | 230 | Cox14 | 129S1,C57L | feature_elongation,3_prime_UTR_variant |
| 15 | 99841182 | 99841405 | 223 | Fam186a | C57L,CAST,MOLF | feature_elongation,coding_sequence_variant |
| 15 | 100378530 | 100378585 | 55 | Csrnp2 | CAST | feature_elongation,3_prime_UTR_variant |
| 15 | 100385887 | 100386697 | 810 | Csrnp2 | MOLF | stop_gained,protein_altering_variant |
| 15 | 101351722 | 101351774 | 52 | Krt88 | MOLF | feature_elongation,3_prime_UTR_variant |
| 15 | 102246337 | 102246392 | 55 | Myg1 | 129S1,AJ,AKR,BTBR,C3H,C57L,C58J,CAST,CBA,CE,DBA1J,DBA2J,FVB,ILNJ2,KK,LP,MOLF,MRL,NOD,NOR,NZB,NZO,PJ,PL,RF,RH2,SEA,SJL,SMJ,SPRET,TH,WSB | feature_elongation,3_prime_UTR_variant |
| 15 | 102406585 | 102406766 | 181 | Map3k12 | WSB | feature_elongation,3_prime_UTR_variant |
| 15 | 102615418 | 102615635 | 217 | Calcoco1 | CAST,MOLF,SPRET | feature_elongation,3_prime_UTR_variant |
| 16 | 8453280 | 8453812 | 532 | Tmem186 | SPRET | feature_elongation,3_prime_UTR_variant |
| 16 | 8501584 | 8501785 | 201 | Litafd | SPRET | feature_elongation,3_prime_UTR_variant |
| 16 | 10658800 | 10659040 | 240 | Rmi2 | BUB | feature_elongation,3_prime_UTR_variant |
| 16 | 13632243 | 13632463 | 220 | Rrn3 | SPRET | feature_elongation,3_prime_UTR_variant |
| 16 | 13651545 | 13651673 | 128 | Pdxdc1 | CAST,SPRET | feature_elongation,3_prime_UTR_variant |
| 16 | 15680818 | 15680872 | 54 | Mzt2 | CAST,MOLF,SPRET | feature_elongation,5_prime_UTR_variant |
| 16 | 15964695 | 15964769 | 74 | Spidr | CAST | feature_elongation,5_prime_UTR_variant |
| 16 | 16958680 | 16959474 | 794 | Ccdc116 | SPRET | feature_elongation,3_prime_UTR_variant |
| 16 | 17363250 | 17363307 | 57 | Lrrc74b | SPRET | feature_elongation,3_prime_UTR_variant |
| 16 | 18226056 | 18226257 | 201 | Comt | SPRET | feature_elongation,3_prime_UTR_variant |
| 16 | 19328704 | 19329177 | 473 | Or2l5 | CAST | feature_elongation,3_prime_UTR_variant |
| 16 | 20127711 | 20127852 | 141 | Cyp2ab1 | SPRET | feature_elongation,3_prime_UTR_variant |
| 16 | 21031036 | 21031187 | 151 | Ephb3 | CAST,SPRET | feature_elongation,5_prime_UTR_variant |
| 16 | 21750227 | 21750426 | 199 | Map3k13 | CAST | feature_elongation,3_prime_UTR_variant |
| 16 | 22976702 | 22976902 | 200 | Adipoq | MOLF,TH | feature_elongation,3_prime_UTR_variant |
| 16 | 25778718 | 25778825 | 107 | P3h2 | SPRET | feature_elongation,3_prime_UTR_variant |
| 16 | 26520888 | 26521217 | 329 | Il1rap | SPRET | feature_elongation,3_prime_UTR_variant |
| 16 | 27207686 | 27207809 | 123 | Ccdc50 | SPRET | feature_elongation,5_prime_UTR_variant |
| 16 | 30379362 | 30379570 | 208 | Lsg1 | CAST,SMJ,WSB | feature_elongation,3_prime_UTR_variant |
| 16 | 30419162 | 30419290 | 128 | Fam43a | MOLF | feature_elongation,5_prime_UTR_variant |
| 16 | 31692479 | 31692674 | 195 | Dlg1 | SPRET | feature_elongation,3_prime_UTR_variant |

| | | | | | | |
|---|---|---|---|---|---|---|
| 16 | 32572401 | 32572773 | 372 | Muc4 | AJ,BALB,BTBR,BUB,C3H,C57L,CBA,CE,DBA1J,DBA2J,FVB,ILNJ2,KK,MRL,PL,SEA,SJL,SMJ,TH,WSB | feature_elongation,coding_sequence_variant |
| 16 | 32574492 | 32575236 | 744 | Muc4 | C57L,CE,DBA1J,DBA2J,KK,SMJ,WSB | feature_elongation,coding_sequence_variant |
| 16 | 32576214 | 32576586 | 372 | Muc4 | AJ,BALB,BTBR,BUB,C3H,C57L,CBA,CE,DBA1J,DBA2J,FVB,ILNJ2,KK,MRL,PL,RH2,SEA,SJL,SMJ,TH,WSB | feature_elongation,coding_sequence_variant |
| 16 | 33587537 | 33587674 | 137 | Heg1 | SPRET | feature_elongation,3_prime_UTR_variant |
| 16 | 33619609 | 33619681 | 72 | Muc13 | SMJ,SPRET | feature_elongation,coding_sequence_variant |
| 16 | 37656311 | 37656411 | 100 | Fstl1 | AKR,CAST,MOLF,MRL,RF,SMJ,SPRET,WSB | feature_elongation,3_prime_UTR_variant |
| 16 | 38589353 | 38589469 | 116 | B4galt4 | SPRET | feature_elongation,3_prime_UTR_variant |
| 16 | 43906509 | 43906665 | 156 | Atp6v1a | SPRET | feature_elongation,3_prime_UTR_variant |
| 16 | 45202855 | 45202907 | 52 | Cd200 | AKR,PL,RF,SPRET | feature_elongation,3_prime_UTR_variant |
| 16 | 48412913 | 48412982 | 69 | Morc1 | C3H,CAST,CBA,CE,MOLF | feature_elongation,coding_sequence_variant |
| 16 | 49734156 | 49734285 | 129 | Cd47 | SPRET | feature_elongation,3_prime_UTR_variant |
| 16 | 52070044 | 52070100 | 56 | Alcam | CAST,MOLF | feature_elongation,3_prime_UTR_variant |
| 16 | 55724259 | 55724572 | 313 | Cep97 | SPRET | feature_elongation,3_prime_UTR_variant |
| 16 | 56858850 | 56859846 | 996 | Tmem45a2 | SPRET | feature_elongation,3_prime_UTR_variant |
| 16 | 58709104 | 58709305 | 201 | Or5k15 | SPRET | feature_elongation,3_prime_UTR_variant |
| 16 | 59134137 | 59134188 | 51 | Or5ac22 | MOLF,SMJ | feature_elongation,3_prime_UTR_variant |
| 16 | 59296459 | 59296583 | 124 | Riox2 | SPRET | feature_elongation,3_prime_UTR_variant |
| 16 | 78337552 | 78337609 | 57 | D16Ertd472e | CAST | feature_elongation,3_prime_UTR_variant |
| 16 | 81393413 | 81393478 | 65 | Ncam2 | SPRET | feature_elongation,3_prime_UTR_variant |
| 16 | 84514601 | 84514665 | 64 | Mrpl39 | AJ,C3H | feature_elongation,3_prime_UTR_variant |
| 16 | 84622328 | 84622793 | 465 | Jam2 | C58J | feature_elongation,3_prime_UTR_variant |
| 16 | 84659872 | 84659940 | 68 | Gabpa | AJ,C3H,DBA2J,FVB,NOD,NOR,NZB,NZW,PL,SJL,SWR | feature_elongation,3_prime_UTR_variant |
| 16 | 87391566 | 87391622 | 56 | Map3k7cl | MOLF | feature_elongation,3_prime_UTR_variant |
| 16 | 88574947 | 88575007 | 60 | Gm5965 | 129S1,C57L,C58J,KK,NZB,NZO,NZW,PL | feature_elongation,coding_sequence_variant |
| 16 | 88666138 | 88666228 | 90 | Krtap19-1 | 129S1,C57L,C58J,CAST,ILNJ2,KK,MOLF,NZB,NZO,NZW,PL | feature_elongation,coding_sequence_variant |
| 16 | 88971754 | 88971832 | 78 | 1110057P08Rik | 129S1,C57L,WSB | feature_elongation,coding_sequence_variant |
| 16 | 91222297 | 91222351 | 54 | Il10rb | MOLF | feature_elongation,3_prime_UTR_variant |
| 16 | 91710188 | 91710242 | 54 | Itsn1 | SPRET | feature_elongation,3_prime_UTR_variant |
| 16 | 91714256 | 91714394 | 138 | Itsn1 | AKR,BUB,CAST,CBA,CE,DBA1J,FVB,ILNJ2,MA/My,MOLF,NZB,NZO,PL,SJL,SWR,TH | feature_elongation,3_prime_UTR_variant |
| 16 | 94495066 | 94495123 | 57 | Dyrk1a | SPRET | feature_elongation,3_prime_UTR_variant |
| 16 | 95099854 | 95099910 | 56 | Kcnj15 | SPRET | feature_elongation,3_prime_UTR_variant |
| 16 | 97659126 | 97659262 | 136 | C2cd2 | SPRET | feature_elongation,3_prime_UTR_variant |
| 16 | 97817267 | 97817444 | 177 | B230307C23Rik | SPRET | feature_elongation,3_prime_UTR_variant |
| 17 | 8164849 | 8164903 | 54 | Rsph3a | AKR,C57L,ILNJ2,NZO,PJ,PL,RF | feature_elongation,coding_sequence_variant |
| 17 | 8198180 | 8198234 | 54 | Rsph3a | SPRET | feature_elongation,3_prime_UTR_variant |
| 17 | 8414745 | 8414833 | 88 | Cep43 | SPRET | feature_elongation,3_prime_UTR_variant |
| 17 | 8642071 | 8642221 | 150 | T2 | SPRET | feature_elongation,3_prime_UTR_variant |
| 17 | 10538307 | 10538412 | 105 | Qki | 129S1,AJ,AKR,B10D2,B10J,BALB,BTBR,BUB,C3H,C57L,C58J,CAST,CBA,CE,DBA1J,DBA2J,FVB,ILNJ2,KK,LP,MA/My,MOLF,MRL,NOD,NOR,NZB,NZO,NZW,PJ,PL,RF,RH2,SEA,SJL,SMJ,SPRET,SWR,TH,WSB | feature_elongation,5_prime_UTR_variant |
| 17 | 15165800 | 15165864 | 64 | 1600012H06Rik | MOLF,PL,SPRET | feature_elongation,3_prime_UTR_variant |
| 17 | 15695567 | 15695876 | 309 | Psmb1 | BTBR,BUB,C3H,CBA,MRL,NOD,NOR,PJ,RF,SJL,SMJ,SWR,WSB | feature_elongation,3_prime_UTR_variant |
| 17 | 15719822 | 15720371 | 549 | Psmb1 | MOLF,PL | feature_elongation,5_prime_UTR_variant |
| 17 | 15738956 | 15739402 | 446 | Tbp | BTBR,BUB,C3H,CAST,CBA,CE,ILNJ2,MOLF,MRL,NOD,NOR,PJ,PL,RF,SJL,SMJ,SPRET,SWR,TH,WSB | feature_elongation,3_prime_UTR_variant |
| 17 | 15764370 | 15764454 | 84 | Prdm9 | B10J,BTBR,BUB,C3H,CBA,CE,MOLF,MRL,NOD,NOR,PJ,PL,RF,SJL,SMJ,TH,WSB | feature_elongation,coding_sequence_variant |
| 17 | 17679274 | 17679345 | 71 | Lix1 | SPRET | feature_elongation,3_prime_UTR_variant |
| 17 | 17746778 | 17746832 | 54 | Lnpep | SPRET | feature_elongation,3_prime_UTR_variant |
| 17 | 19619184 | 19619678 | 494 | Vmn2r99 | SPRET | feature_elongation,3_prime_UTR_variant |
| 17 | 19752938 | 19752993 | 55 | Vmn2r100 | AJ,BALB,CE,DBA1J,DBA2J,KK,RH2,SPRET | feature_elongation,3_prime_UTR_variant |
| 17 | 19922243 | 19922742 | 499 | Vmn2r102 | SPRET | feature_elongation,3_prime_UTR_variant |
| 17 | 19923711 | 19923851 | 140 | Vmn2r102 | SPRET | feature_elongation,3_prime_UTR_variant |
| 17 | 20424319 | 20424419 | 100 | Vmn2r105 | SPRET | feature_elongation,3_prime_UTR_variant |
| 17 | 20644763 | 20644873 | 110 | Vmn1r224 | MOLF | feature_elongation,3_prime_UTR_variant |
| 17 | 20789725 | 20790181 | 456 | Vmn2r110 | 129S1,AJ,AKR,BALB,C3H,C57L,CBA,CE,DBA1J,DBA2J,FVB,KK,MA/My,MRL,NZO,PJ,RH2,SEA,WSB | feature_elongation,3_prime_UTR_variant |
| 17 | 20792837 | 20793421 | 584 | Vmn2r110 | SPRET | feature_elongation,3_prime_UTR_variant |
| 17 | 20995804 | 20996261 | 457 | Vmn1r228 | MOLF,SPRET | feature_elongation,3_prime_UTR_variant |
| 17 | 21072360 | 21072861 | 501 | Vmn1r230 | MOLF | feature_elongation,3_prime_UTR_variant |
| 17 | 21213543 | 21213873 | 330 | Vmn1r233 | CAST,MOLF | feature_elongation,3_prime_UTR_variant |
| 17 | 21730514 | 21731012 | 498 | Zfp53 | SPRET | feature_elongation,3_prime_UTR_variant |
| 17 | 21807353 | 21807452 | 99 | Zfp948 | MOLF | feature_elongation,coding_sequence_variant |
| 17 | 21904218 | 21904454 | 236 | Gm10509 | 129S1,AJ,BALB,C57L,CE,DBA1J,DBA2J,NZO,RH2,SEA | feature_elongation,5_prime_UTR_variant |
| 17 | 21910182 | 21910266 | 84 | Gm10509 | SPRET | feature_elongation,coding_sequence_variant |
| 17 | 21967126 | 21967201 | 75 | Zfp229 | SPRET | feature_elongation,3_prime_UTR_variant |
| 17 | 22418930 | 22419092 | 162 | Zfp994 | SPRET | feature_elongation,3_prime_UTR_variant |
| 17 | 23183467 | 23183582 | 115 | Vmn2r113 | CAST,MOLF | feature_elongation,3_prime_UTR_variant |
| 17 | 23622023 | 23622208 | 185 | Vmn2r116 | MOLF | feature_elongation,3_prime_UTR_variant |
| 17 | 23776149 | 23776261 | 112 | Zfp213 | MOLF | feature_elongation,3_prime_UTR_variant |
| 17 | 24394457 | 24394666 | 209 | Tbc1d24 | 129S1,AJ,AKR,B10D2,BALB,BTBR,BUB,C3H,C57L,CBA,CE,DBA1J,DBA2J,FVB,ILNJ2,KK,MA/My,MRL,NOD,NOR,NZB,NZO,NZW,PJ,RF,RH2,SEA,SJL,SMJ,SWR,TH,WSB | feature_elongation,3_prime_UTR_variant |
| 17 | 24659265 | 24659474 | 209 | Dnase1l2 | 129S1,AJ,AKR,B10D2,BALB,BTBR,BUB,C3H,C57L,CBA,CE,DBA1J,DBA2J,FVB,ILNJ2,KK,MA/My,MRL,NOD,NOR,NZB,NZO,NZW,PJ,RF,RH2,SEA,SJL,SMJ,SPRET,SWR,TH,WSB | feature_elongation,3_prime_UTR_variant |

| | | | | | | |
|---|---|---|---|---|---|---|
| 17 | 25314802 | 25314856 | 54 | Ift140 | 129S1,AJ,AKR,B10D2,BALB,BTBR,BUB,C3H,C57L,CBA,CE,DBA1J,DBA2J,FVB,ILNJ2,KK,MA/My,MOLF,MRL,NOD,NOR,NZB,NZO,RF,RH2,SEA,SJL,SWR,TH,WSB | feature_elongation,3_prime_UTR_variant |
| 17 | 25452109 | 25452349 | 240 | Unkl | CAST | feature_elongation,3_prime_UTR_variant |
| 17 | 25488860 | 25488953 | 93 | Ube2i | SPRET | feature_elongation,5_prime_UTR_variant |
| 17 | 26039831 | 26039882 | 51 | Fbxl16 | SPRET | feature_elongation,3_prime_UTR_variant |
| 17 | 26663457 | 26663520 | 63 | Neurl1b | SPRET | feature_elongation,3_prime_UTR_variant |
| 17 | 26915178 | 26915393 | 215 | Atp6v0e | SPRET | feature_elongation,3_prime_UTR_variant |
| 17 | 27970648 | 27970712 | 64 | Ilrun | SPRET | feature_elongation,3_prime_UTR_variant |
| 17 | 27977874 | 27978008 | 134 | Ilrun | SPRET | feature_elongation,3_prime_UTR_variant |
| 17 | 28115605 | 28115677 | 72 | Bltp3a | 129S1,AJ,AKR,B10D2,BALB,BTBR,C3H,C57L,CAST,CBA,CE,DBA1J,DBA2J,FVB,ILNJ2,KK,MA/My,MOLF,MRL,NZB,NZW,RF,RH2,SEA,SJL,SMJ,TH,WSB | feature_elongation,3_prime_UTR_variant |
| 17 | 28619187 | 28619240 | 53 | Fkbp5 | MOLF,SPRET | feature_elongation,3_prime_UTR_variant |
| 17 | 28656128 | 28656200 | 72 | Fkbp5 | SPRET | feature_elongation,3_prime_UTR_variant |
| 17 | 29058890 | 29058956 | 66 | Brpf3 | CAST | feature_elongation,3_prime_UTR_variant |
| 17 | 29488280 | 29488486 | 206 | BC004004 | SPRET | feature_elongation,5_prime_UTR_variant |
| 17 | 29525994 | 29526068 | 74 | BC004004 | CAST,SPRET | feature_elongation,3_prime_UTR_variant |
| 17 | 29537235 | 29537456 | 221 | Pi16 | AJ,AKR,BTBR,C3H,CBA,CE,MRL | feature_elongation,5_prime_UTR_variant |
| 17 | 29547263 | 29547377 | 114 | Pi16 | CAST | feature_elongation,3_prime_UTR_variant |
| 17 | 29923321 | 29923372 | 51 | Cmtr1,Ccdc167 | SPRET | feature_elongation,3_prime_UTR_variant |
| 17 | 30810875 | 30811111 | 236 | Glo1 | KK,MOLF,NOD,NOR,RF,SPRET | feature_elongation,3_prime_UTR_variant |
| 17 | 30936864 | 30936974 | 110 | Dnah8 | 129S1,B10D2,BALB,BUB,C57L,DBA1J,DBA2J,MA/My,PL,RH2,SEA,SWR | feature_elongation,coding_sequence_variant |
| 17 | 31464509 | 31464568 | 59 | Ubash3a | SPRET | feature_elongation,3_prime_UTR_variant |
| 17 | 31465694 | 31465897 | 203 | Ubash3a | CAST,KK,NOD,NOR | feature_elongation,3_prime_UTR_variant |
| 17 | 31830886 | 31831075 | 189 | Cbs | SPRET | feature_elongation,3_prime_UTR_variant |
| 17 | 32855093 | 32855296 | 203 | Cyp4f37 | SPRET | feature_elongation,3_prime_UTR_variant |
| 17 | 32911607 | 32911769 | 162 | Cyp4f15 | AJ,AKR,C3H,CBA,CE,MRL,NOD,NOR,PJ,RF,TH | feature_elongation,coding_sequence_variant |
| 17 | 33015840 | 33015900 | 60 | Zfp811 | SPRET | feature_elongation,3_prime_UTR_variant |
| 17 | 33034992 | 33035097 | 105 | Zfp799 | CAST,MA/My,MOLF,PL,SJL,SPRET,TH,WSB | feature_elongation,3_prime_UTR_variant |
| 17 | 33433730 | 33433795 | 65 | Or10h5 | AJ,AKR,B10J,BTBR,BUB,C3H,C58J,CAST,CBA,CE,FVB,ILNJ2,KK,LP,MA/My,MOLF,MRL,NOD,NOR,PJ,PL,RF,SJL,SWR,TH | feature_elongation,3_prime_UTR_variant |
| 17 | 33552511 | 33552609 | 98 | Zfp81 | SPRET | feature_elongation,3_prime_UTR_variant |
| 17 | 33963305 | 33963522 | 217 | Rab11b | SPRET | feature_elongation,3_prime_UTR_variant |
| 17 | 34068697 | 34068970 | 273 | Cd320 | B10D2,BALB,BUB,DBA1J,DBA2J,FVB,ILNJ2,MOLF,NOD,NOR,NZB,NZO,NZW,PJ,PL,RH2,SEA,SJL,SMJ,SWR,TH,WSB | feature_elongation,3_prime_UTR_variant |
| 17 | 34463785 | 34463861 | 76 | H2-Ob | AJ,AKR,C3H,C58J,CBA,CE,MA/My,MOLF,MRL,PJ,RF,SMJ,WSB | feature_elongation,3_prime_UTR_variant |
| 17 | 34559749 | 34559876 | 127 | H2-Eb2 | PJ,WSB | feature_elongation,3_prime_UTR_variant |
| 17 | 35930978 | 35931180 | 202 | Muc21 | AJ,BALB,DBA2J,NZB | feature_elongation,coding_sequence_variant |
| 17 | 35931729 | 35931864 | 135 | Muc21 | MOLF,PJ,WSB | feature_elongation,coding_sequence_variant |
| 17 | 35933305 | 35933395 | 90 | Muc21 | AJ,B10D2,BALB,DBA2J,NZB,PL,RH2,SEA,SJL | feature_elongation,coding_sequence_variant |
| 17 | 35933865 | 35934043 | 178 | Muc21 | CAST,MOLF,TH | feature_elongation,coding_sequence_variant |
| 17 | 35947124 | 35947194 | 70 | Mucl3 | AJ,B10D2,BALB,C58J,DBA2J,ILNJ2,NZB,PL,RH2,SEA,SJL,TH,WSB | feature_elongation,3_prime_UTR_variant |
| 17 | 36207068 | 36207249 | 181 | 2310061I04Rik | SPRET | feature_elongation,5_prime_UTR_variant |
| 17 | 37461721 | 37461893 | 172 | Or2h1b | B10D2,BALB,DBA1J,DBA2J,MOLF,NZO,NZW,RH2,SEA | feature_elongation,3_prime_UTR_variant |
| 17 | 37779511 | 37779695 | 184 | Or12d17 | MOLF | feature_elongation,3_prime_UTR_variant |
| 17 | 38012894 | 38013004 | 110 | Or10al3 | MOLF | feature_elongation,3_prime_UTR_variant |
| 17 | 38083865 | 38083969 | 104 | Or10al6 | MOLF | feature_elongation,3_prime_UTR_variant |
| 17 | 38363451 | 38364077 | 626 | Or10al7 | MOLF,SPRET | feature_elongation,3_prime_UTR_variant |
| 17 | 38520407 | 38520675 | 268 | Or2n1c | MOLF | feature_elongation,3_prime_UTR_variant |
| 17 | 43399052 | 43399108 | 56 | Tnfrsf21 | C58J,CAST,ILNJ2,TH | feature_elongation,3_prime_UTR_variant |
| 17 | 43978322 | 43978420 | 98 | Cyp39a1 | SPRET | feature_elongation,5_prime_UTR_variant |
| 17 | 44349909 | 44349980 | 71 | Rcan2 | C3H,SPRET | feature_elongation,3_prime_UTR_variant |
| 17 | 44916090 | 44916171 | 81 | Runx2 | 129S1,AJ,BTBR,BUB,C57L,C58J,DBA1J,KK,LP,MRL,NOD,NOR,PL,SPRET,TH | feature_elongation,3_prime_UTR_variant |
| 17 | 45831404 | 45831615 | 211 | Aars2 | AKR,BALB,C3H,CBA,CE,MRL,PJ,PL,RH2,SEA | feature_elongation,3_prime_UTR_variant |
| 17 | 46570366 | 46570483 | 117 | Tjap1 | BTBR,C57L,DBA1J,KK,LP,MA/My,NZO,RF | feature_elongation,coding_sequence_variant |
| 17 | 46633183 | 46633282 | 99 | Abcc10 | CAST,MOLF,NOD,NOR,NZW,SJL,SWR | feature_elongation,coding_sequence_variant |
| 17 | 46723441 | 46723492 | 51 | Zfp318 | CAST,NOD,NOR,NZW,SJL,SWR | feature_elongation,coding_sequence_variant |
| 17 | 46728315 | 46728405 | 90 | Zfp318 | MOLF,NOD,NOR,NZW,SJL,SPRET,SWR | feature_elongation,3_prime_UTR_variant |
| 17 | 48556642 | 48556695 | 53 | Trem3 | MOLF | feature_elongation,coding_sequence_variant |
| 17 | 48618530 | 48618655 | 125 | Treml2 | SPRET | feature_elongation,3_prime_UTR_variant |
| 17 | 48694945 | 48694999 | 54 | Nfya | SWR | feature_elongation,3_prime_UTR_variant |
| 17 | 48769376 | 48769435 | 59 | Unc5cl | SPRET | feature_elongation,coding_sequence_variant |
| 17 | 55959962 | 55960174 | 212 | Pot1b | SPRET | feature_elongation,3_prime_UTR_variant |
| 17 | 56411826 | 56412024 | 198 | Plin4 | SPRET | feature_elongation,coding_sequence_variant |
| 17 | 56585671 | 56585853 | 182 | Plin3 | CAST | feature_elongation,3_prime_UTR_variant |
| 17 | 59306934 | 59307032 | 98 | Nudt12 | CAST | feature_elongation,3_prime_UTR_variant |
| 17 | 63806660 | 63806751 | 91 | Fbxl17 | SMJ | feature_elongation,coding_sequence_variant |
| 17 | 66229294 | 66229484 | 190 | Twsg1 | MOLF | feature_elongation,3_prime_UTR_variant |
| 17 | 69086673 | 69086779 | 106 | L3mbtl4 | CAST,MOLF,SPRET | feature_elongation,3_prime_UTR_variant |
| 17 | 73800373 | 73800435 | 62 | Galnt14 | C58J,MA/My,MOLF,NZO,PL,SPRET | feature_elongation,3_prime_UTR_variant |
| 17 | 84498780 | 84498991 | 211 | Thada | DBA1J,DBA2J,ILNJ2,MOLF | feature_elongation,3_prime_UTR_variant |
| 17 | 84928028 | 84928201 | 173 | Plekhh2 | DBA1J,DBA2J,ILNJ2 | feature_elongation,3_prime_UTR_variant |
| 17 | 86962873 | 86962948 | 75 | Prkce | CAST | feature_elongation,3_prime_UTR_variant |
| 17 | 86964409 | 86964475 | 66 | Prkce | SPRET | feature_elongation,3_prime_UTR_variant |
| 17 | 87139125 | 87139211 | 86 | Epas1 | AJ,BALB,BTBR,C3H,C57L,CBA,DBA1J,DBA2J,ILNJ2,LP,RH2,SEA | feature_elongation,3_prime_UTR_variant |
| 18 | 3266988 | 3267092 | 104 | Crem | SPRET | feature_elongation,3_prime_UTR_variant |
| 18 | 3514498 | 3514551 | 53 | Bambi | SPRET | feature_elongation,3_prime_UTR_variant |
| 18 | 9878344 | 9878457 | 113 | Colec12 | MOLF | feature_elongation,3_prime_UTR_variant |
| 18 | 9881189 | 9881646 | 457 | Colec12 | 129S1,C57L,CAST,MOLF,SPRET | feature_elongation,3_prime_UTR_variant |
| 18 | 10595058 | 10595115 | 57 | Esco1 | 129S1,C57L,CAST,MOLF | feature_elongation,coding_sequence_variant |

| | | | | | | |
|---|---|---|---|---|---|---|
| 18 | 12711187 | 12711509 | 322 | Lama3 | CAST | feature_elongation,coding_sequence_variant |
| 18 | 15102333 | 15102387 | 54 | Kctd1 | BALB,C3H,CBA,NOD,RH2 | feature_elongation,3_prime_UTR_variant |
| 18 | 20217837 | 20217888 | 51 | Dsc1 | MOLF | feature_elongation,3_prime_UTR_variant |
| 18 | 20475735 | 20475787 | 52 | Dsg1a | CAST | feature_elongation,3_prime_UTR_variant |
| 18 | 20735919 | 20735989 | 70 | Dsg2 | MOLF,WSB | feature_elongation,3_prime_UTR_variant |
| 18 | 20818354 | 20818551 | 197 | B4galt6 | CAST | feature_elongation,3_prime_UTR_variant |
| 18 | 21264277 | 21264501 | 224 | Garem1 | CBA,CE,DBA1J,DBA2J,FVB,KK,NOD,NOR,RF,SMJ,SWR,TH,WSB | feature_elongation,3_prime_UTR_variant |
| 18 | 24239332 | 24239541 | 209 | Ino80c | SPRET | feature_elongation,3_prime_UTR_variant |
| 18 | 30463861 | 30464535 | 674 | Pik3c3 | BALB,C3H,C58J,CE,MRL,PL,RF,RH2,SEA | feature_elongation,3_prime_UTR_variant |
| 18 | 32236955 | 32237362 | 407 | Iws1 | 129S1,C57L,CAST,ILNJ2,MOLF,SPRET | feature_elongation,3_prime_UTR_variant |
| 18 | 33596726 | 33596792 | 66 | Nrep | MOLF | feature_elongation,5_prime_UTR_variant |
| 18 | 35804943 | 35805013 | 70 | Dnajc18 | MOLF | feature_elongation,3_prime_UTR_variant |
| 18 | 37626589 | 37626786 | 197 | Pcdhb18 | KK | feature_elongation,3_prime_UTR_variant |
| 18 | 37850906 | 37850963 | 57 | Pcdhga7 | BUB,C3H,CAST,CE,DBA1J,MA/My,MOLF,NOD,NOR,PL,RF,SMJ,SPRET | feature_elongation,3_prime_UTR_variant |
| 18 | 43449949 | 43450008 | 59 | Stk32a | MA/My | feature_elongation,3_prime_UTR_variant |
| 18 | 43910273 | 43910747 | 474 | Gm94 | 129S1,AJ,AKR,BALB,BTBR,C3H,C57L,CAST,CBA,CE,DBA1J,DBA2J,FVB,ILNJ2,KK,MA/My,MOLF,MRL,NZB,NZW,PJ,PL,RF,RH2,SEA,SJL,SPRET,SWR,TH,WSB | feature_elongation,3_prime_UTR_variant |
| 18 | 44559399 | 44559460 | 61 | Mcc | SPRET | feature_elongation,3_prime_UTR_variant |
| 18 | 46370482 | 46370545 | 63 | Pggt1b | CAST,KK,SPRET | feature_elongation,3_prime_UTR_variant |
| 18 | 46690624 | 46690812 | 188 | Ticam2 | CAST | feature_elongation,3_prime_UTR_variant |
| 18 | 46742644 | 46742698 | 54 | Eif1a | KK,MOLF | feature_elongation,3_prime_UTR_variant |
| 18 | 46866041 | 46866157 | 116 | Atg12 | SPRET | feature_elongation,3_prime_UTR_variant |
| 18 | 47038430 | 47038486 | 56 | Lvrn | CAST | feature_elongation,3_prime_UTR_variant |
| 18 | 53522829 | 53523225 | 396 | Snx24 | KK,MOLF | feature_elongation,3_prime_UTR_variant |
| 18 | 57089769 | 57090502 | 733 | C330018D20Rik | SPRET | feature_elongation,3_prime_UTR_variant |
| 18 | 60709286 | 60709382 | 96 | Myoz3 | AKR,B10D2,B10J,BTBR,BUB,C3H,C58J,CE,DBA1J,DBA2J,FVB,LP,MA/My,MRL,NZB,NZO,NZW,PJ,PL,RF,SJL,SPRET,WSB | feature_elongation,3_prime_UTR_variant |
| 18 | 61328154 | 61328318 | 164 | Slc26a2 | CAST,KK,MOLF,NZO,NZW,SPRET | feature_elongation,3_prime_UTR_variant |
| 18 | 62800928 | 62801600 | 672 | Spink10 | SPRET | feature_elongation,3_prime_UTR_variant |
| 18 | 63091134 | 63091207 | 73 | Apcdd1 | CAST | feature_elongation,3_prime_UTR_variant |
| 18 | 64409590 | 64410234 | 644 | St8sia3 | AJ,AKR,BALB,BUB,C3H,CAST,CBA,FVB,ILNJ2,KK,MOLF,MRL,NOD,NOR,NZB,NZW,PL,RF,RH2,SJL,SPRET,WSB | feature_elongation,3_prime_UTR_variant |
| 18 | 64524615 | 64524693 | 78 | Onecut2 | SPRET | feature_elongation,3_prime_UTR_variant |
| 18 | 64528851 | 64529251 | 400 | Onecut2 | MOLF | feature_elongation,3_prime_UTR_variant |
| 18 | 67926767 | 67926825 | 58 | Seh1l | AJ,BALB,FVB,ILNJ2,KK,MOLF,MRL,NOD,NOR,NZB,NZO,NZW,PL,RF,RH2,SJL,SMJ,SPRET,SWR,TH | feature_elongation,3_prime_UTR_variant |
| 18 | 68472874 | 68473405 | 531 | Mc5r | SPRET | feature_elongation,3_prime_UTR_variant |
| 18 | 70663594 | 70663648 | 54 | Poli | BUB | feature_elongation,5_prime_UTR_variant |
| 18 | 74012017 | 74012067 | 50 | Mro | BUB,NZO | feature_elongation,3_prime_UTR_variant |
| 18 | 75218775 | 75218841 | 66 | Dym | AJ,AKR,BALB,ILNJ2,KK,MA/My,MOLF,MRL,NOD,NOR,NZB,RF,RH2,SJL,SWR | feature_elongation,coding_sequence_variant |
| 18 | 75219500 | 75219640 | 140 | Dym | C3H,CBA,DBA1J,DBA2J,PJ,PL,SEA | feature_elongation,3_prime_UTR_variant |
| 18 | 77345665 | 77345820 | 155 | St8sia5 | 129S1,AJ,AKR,B10D2,B10J,BALB,BTBR,BUB,C3H,C57L,C58J,CAST,CBA,CE,DBA1J,DBA2J,FVB,ILNJ2,KK,LP,MA/My,MOLF,MRL,NOD,NOR,NZB,NZO,NZW,PJ,PL,RF,RH2,SEA,SJL,SMJ,SWR,TH,WSB | feature_elongation,3_prime_UTR_variant |
| 18 | 78076990 | 78077190 | 200 | Epg5 | KK,MOLF,NZB,TH | feature_elongation,3_prime_UTR_variant |
| 18 | 80318192 | 80318406 | 214 | Slc66a2 | CAST | feature_elongation,3_prime_UTR_variant |
| 18 | 80335070 | 80335127 | 57 | Slc66a2 | AKR,NZO | feature_elongation,3_prime_UTR_variant |
| 18 | 82411112 | 82411206 | 94 | Galr1 | CAST,MOLF,SPRET | feature_elongation,3_prime_UTR_variant |
| 18 | 82575080 | 82575240 | 160 | Mbp | MOLF | feature_elongation,3_prime_UTR_variant |
| 18 | 90610533 | 90610765 | 232 | Gm45871 | KK,NZB,NZO,RF,TH,WSB | feature_elongation,3_prime_UTR_variant |
| 19 | 3939054 | 3939126 | 72 | Chka | SPRET | feature_elongation,coding_sequence_variant |
| 19 | 4193537 | 4193681 | 144 | Gpr152 | MOLF,SPRET | feature_elongation,coding_sequence_variant |
| 19 | 4234544 | 4234645 | 101 | Tbc1d10c | C58J,LP,TH | feature_elongation,3_prime_UTR_variant |
| 19 | 4938502 | 4938662 | 160 | Bbs1 | BUB,NZW | feature_elongation,3_prime_UTR_variant |
| 19 | 5016755 | 5016875 | 120 | Mrpl11 | CAST,MOLF,SPRET | feature_elongation,3_prime_UTR_variant |
| 19 | 5358425 | 5358485 | 60 | Gal3st3 | MOLF | feature_elongation,3_prime_UTR_variant |
| 19 | 5899943 | 5900116 | 173 | Frmd8 | CAST | feature_elongation,3_prime_UTR_variant |
| 19 | 6113093 | 6113233 | 140 | Tm7sf2 | SPRET | feature_elongation,coding_sequence_variant |
| 19 | 6148029 | 6148155 | 126 | Gm550 | CAST,MOLF | feature_elongation,coding_sequence_variant |
| 19 | 6171474 | 6171573 | 99 | Snx15 | TH | feature_elongation,coding_sequence_variant |
| 19 | 6582687 | 6582817 | 130 | Nrxn2 | SPRET | feature_elongation,3_prime_UTR_variant |
| 19 | 6586075 | 6586138 | 63 | Slc22a12 | BUB,C58J,CBA,DBA1J,DBA2J,FVB,LP,MA/My,SPRET,WSB | feature_elongation,3_prime_UTR_variant |
| 19 | 7515247 | 7515307 | 60 | Atl3 | SPRET | feature_elongation,3_prime_UTR_variant |
| 19 | 8814306 | 8814508 | 202 | Gng3 | SPRET | feature_elongation,3_prime_UTR_variant |
| 19 | 9970358 | 9970588 | 230 | Best1 | AJ,BALB,C3H,C58J,CE,LP,MRL,RH2,SEA,WSB | feature_elongation,coding_sequence_variant |
| 19 | 10433404 | 10433460 | 56 | Lrrc10b | SPRET | feature_elongation,3_prime_UTR_variant |
| 19 | 10569921 | 10569989 | 68 | Tkfc | MOLF | feature_elongation,3_prime_UTR_variant |
| 19 | 11459183 | 11459268 | 85 | Ms4a6c | SPRET | feature_elongation,3_prime_UTR_variant |
| 19 | 11489276 | 11489695 | 419 | Gm8369 | CAST,MOLF | feature_elongation,3_prime_UTR_variant |
| 19 | 11752806 | 11752909 | 103 | Stx3 | AJ,BALB,BUB,CE,FVB,ILNJ2,KK,LP,NOD,NOR,PJ,PL,RH2,SEA,SWR,TH,WSB | feature_elongation,3_prime_UTR_variant |
| 19 | 12010378 | 12010496 | 118 | Or4d11 | CAST,FVB,SWR | feature_elongation,3_prime_UTR_variant |
| 19 | 12307200 | 12307297 | 97 | Or5a21 | AJ,BALB,CE,MA/My,NOD,NOR,PL,RH2,SEA,WSB | feature_elongation,3_prime_UTR_variant |
| 19 | 12577399 | 12577525 | 126 | Gm4952 | AJ,BALB,CE,MA/My,NOD,NOR,PL,RH2,SEA | feature_elongation,5_prime_UTR_variant |
| 19 | 12630365 | 12630425 | 60 | Glyat | CAST,MOLF,SPRET | feature_elongation,3_prime_UTR_variant |
| 19 | 12894110 | 12894168 | 58 | Or5b12 | SPRET | feature_elongation,3_prime_UTR_variant |
| 19 | 12908254 | 12908405 | 151 | Or5b24 | CAST | feature_elongation,5_prime_UTR_variant |
| 19 | 12932204 | 12932282 | 78 | Or5b98 | CAST,SPRET | feature_elongation,3_prime_UTR_variant |
| 19 | 13041142 | 13041811 | 669 | Or5b102 | MOLF | feature_elongation,coding_sequence_variant |
| 19 | 13813100 | 13813298 | 198 | Or9i2 | CAST,MOLF | feature_elongation,3_prime_UTR_variant |

| | | | | | | |
|---|---|---|---|---|---|---|
| 19 | 17377364 | 17377440 | 76 | Rfk | SPRET | feature_elongation,3_prime_UTR_variant |
| 19 | 18683752 | 18683809 | 57 | Carnmt1 | MOLF,NZB,NZO,NZW,PJ | feature_elongation,3_prime_UTR_variant |
| 19 | 21370072 | 21370235 | 163 | Gda | SPRET | feature_elongation,3_prime_UTR_variant |
| 19 | 21575570 | 21575638 | 68 | 1110059E24Rik | SPRET | feature_elongation,coding_sequence_variant |
| 19 | 22967785 | 22967838 | 53 | Trpm3 | SPRET | feature_elongation,3_prime_UTR_variant |
| 19 | 22968860 | 22968920 | 60 | Trpm3 | SPRET | feature_elongation,3_prime_UTR_variant |
| 19 | 23703166 | 23703257 | 91 | Ptar1 | SPRET | feature_elongation,3_prime_UTR_variant |
| 19 | 24661489 | 24661571 | 82 | Pgm5 | SPRET | feature_elongation,3_prime_UTR_variant |
| 19 | 27312233 | 27312294 | 61 | Kcnv2 | MOLF | feature_elongation,3_prime_UTR_variant |
| 19 | 29364518 | 29364596 | 78 | Cd274 | CAST,MOLF | feature_elongation,3_prime_UTR_variant |
| 19 | 30544041 | 30544167 | 126 | Prkg1 | CAST | feature_elongation,3_prime_UTR_variant |
| 19 | 31962363 | 31962415 | 52 | Asah2 | MOLF | feature_elongation,3_prime_UTR_variant |
| 19 | 32078991 | 32079057 | 66 | Asah2 | SPRET | feature_elongation,5_prime_UTR_variant |
| 19 | 36713955 | 36714016 | 61 | Ppp1r3c | CAST | feature_elongation,coding_sequence_variant |
| 19 | 37927812 | 37927934 | 122 | Myof | SPRET | feature_elongation,coding_sequence_variant |
| 19 | 38776697 | 38776905 | 208 | Noc3l | 129S1,AJ,AKR,BALB,BUB,C3H,C57L,CBA,CE,DBA1J,DBA2J,FVB,ILNJ2,KK,MRL,NOD,NOR,NZB,NZO,PJ,PL,RF,RH2,SEA,SJL,SMJ,SWR,TH,WSB | feature_elongation,3_prime_UTR_variant |
| 19 | 40595626 | 40595686 | 60 | Tctn3 | SPRET,WSB | feature_elongation,coding_sequence_variant |
| 19 | 40835883 | 40836248 | 365 | Ccnj | SPRET | feature_elongation,3_prime_UTR_variant |
| 19 | 41538078 | 41538141 | 63 | Lcor | MOLF,SPRET | feature_elongation,5_prime_UTR_variant |
| 19 | 41571696 | 41571822 | 126 | Lcor | SPRET | feature_elongation,coding_sequence_variant |
| 19 | 41930615 | 41930701 | 86 | Mms19 | CAST | feature_elongation,3_prime_UTR_variant |
| 19 | 41931852 | 41932048 | 196 | Mms19 | MOLF | feature_elongation,3_prime_UTR_variant |
| 19 | 42135929 | 42136014 | 85 | Marveld1 | MOLF | feature_elongation,5_prime_UTR_variant |
| 19 | 43723763 | 43723819 | 56 | Cox15 | PL,SMJ | feature_elongation,3_prime_UTR_variant |
| 19 | 44231699 | 44231844 | 145 | Scd3 | SPRET | feature_elongation,3_prime_UTR_variant |
| 19 | 44517328 | 44517457 | 129 | Sec31b | MOLF | feature_elongation,coding_sequence_variant |
| 19 | 46583742 | 46583803 | 61 | Sfxn2 | SPRET | feature_elongation,3_prime_UTR_variant |
| 19 | 46874692 | 46875183 | 491 | Nt5c2 | NZB | feature_elongation,3_prime_UTR_variant |
| 19 | 47095234 | 47095446 | 212 | Calhm2 | SPRET | feature_elongation,3_prime_UTR_variant |
| 19 | 47230093 | 47230926 | 833 | Neurl1a | CE | feature_elongation,3_prime_UTR_variant |
| 19 | 53028509 | 53028623 | 114 | Xpnpep1 | SPRET | feature_elongation,5_prime_UTR_variant |
| 19 | 53880788 | 53880848 | 60 | Pdcd4 | SPRET | feature_elongation,5_prime_UTR_variant |
| 19 | 53919158 | 53919260 | 102 | Bbip1 | SPRET | feature_elongation,3_prime_UTR_variant |
| 19 | 55186569 | 55186994 | 425 | Gucy2g | SPRET | feature_elongation,3_prime_UTR_variant |
| 19 | 56778374 | 56778530 | 156 | Ccdc186 | SPRET | feature_elongation,3_prime_UTR_variant |
| 19 | 57474127 | 57474194 | 67 | Trub1 | BALB,MA/My,PL,RF | feature_elongation,3_prime_UTR_variant |
| 19 | 61040023 | 61040534 | 511 | Grk5 | SPRET | feature_elongation,coding_sequence_variant |
| 19 | 61113313 | 61113440 | 127 | Zfp950 | SPRET | feature_elongation,3_prime_UTR_variant |
| 19 | 61113578 | 61113816 | 238 | Zfp950 | 129S1,AJ,AKR,BALB,BTBR,C3H,C57L,CAST,CBA,DBA1J,DBA2J,FVB,KK,MA/My,MRL,NOD,NOR,NZB,NZW,PJ,PL,RF,RH2,SEA,SMJ,SWR,TH,WSB | feature_elongation,3_prime_UTR_variant |
| X | 5994379 | 5994457 | 78 | Ezhip | MOLF | feature_elongation,coding_sequence_variant |
| X | 6227278 | 6227359 | 81 | Bmp15 | CAST | feature_elongation,3_prime_UTR_variant |
| X | 6548630 | 6548686 | 56 | Shroom4 | MOLF | feature_elongation,3_prime_UTR_variant |
| X | 7760832 | 7760995 | 163 | Pqbp1 | SPRET | feature_elongation,3_prime_UTR_variant |
| X | 7764955 | 7765119 | 164 | Pqbp1 | CAST | feature_elongation,5_prime_UTR_variant |
| X | 7773420 | 7773635 | 215 | Timm17b | SPRET | feature_elongation,3_prime_UTR_variant |
| X | 10181215 | 10181778 | 563 | Otc | WSB | feature_elongation,coding_sequence_variant |
| X | 10456869 | 10456927 | 58 | Tspan7 | SPRET | feature_elongation,3_prime_UTR_variant |
| X | 10458978 | 10459045 | 67 | Tspan7 | SPRET | feature_elongation,3_prime_UTR_variant |
| X | 35433928 | 35433997 | 69 | Il13ra1 | MOLF | feature_elongation,3_prime_UTR_variant |
| X | 36967277 | 36967358 | 81 | Rhox8 | SPRET | feature_elongation,coding_sequence_variant |
| X | 41198010 | 41198061 | 51 | Xiap | MOLF,SPRET | feature_elongation,3_prime_UTR_variant |
| X | 46964017 | 46964176 | 159 | Smarca1 | SPRET | feature_elongation,coding_sequence_variant |
| X | 59355129 | 59355292 | 163 | Atp11c | CAST | feature_elongation,coding_sequence_variant |
| X | 70977785 | 70977863 | 78 | Pasd1 | CAST,ILNJ2,MOLF,SPRET,WSB | feature_elongation,5_prime_UTR_variant,feature_elongation,coding_sequence_variant |
| X | 71300593 | 71300645 | 52 | Gabre | CAST | feature_elongation,3_prime_UTR_variant |
| X | 72002095 | 72002172 | 77 | Nsdhl | CAST | feature_elongation,3_prime_UTR_variant |
| X | 72220578 | 72220694 | 116 | Spin2d | CAST | feature_elongation,5_prime_UTR_variant |
| X | 72326721 | 72327665 | 944 | Xlr5c | MOLF | feature_elongation,3_prime_UTR_variant |
| X | 73194601 | 73194975 | 374 | Tex28 | SPRET | feature_elongation,3_prime_UTR_variant |
| X | 73497246 | 73497412 | 166 | Ikbkg | SPRET | feature_elongation,3_prime_UTR_variant |
| X | 74423514 | 74423573 | 59 | F8 | 129S1,C57L,NZB | feature_elongation,5_prime_UTR_variant |
| X | 74439103 | 74439860 | 757 | Fundc2 | 129S1,C57L,NZB | feature_elongation,3_prime_UTR_variant |
| X | 82530664 | 82530781 | 117 | Tsga8 | AJ,AKR,BALB,BUB,C3H,CAST,CBA,CE,DBA1J,DBA2J,ILNJ2,KK,MA/My,MOLF,MRL,NZB,PJ,PL,RF,RH2,SEA,SMJ,SPRET,WSB | feature_elongation,coding_sequence_variant |
| X | 84933109 | 84933606 | 497 | Tasl | SPRET | feature_elongation,3_prime_UTR_variant |
| X | 88450669 | 88450930 | 261 | Dcaf8l | SPRET | feature_elongation,5_prime_UTR_variant |
| X | 93232416 | 93232616 | 200 | Eif2s3x | SPRET | feature_elongation,3_prime_UTR_variant |
| X | 94470890 | 94470950 | 60 | Amer1 | CAST | feature_elongation,coding_sequence_variant |
| X | 97992403 | 97992973 | 570 | Yipf6 | SPRET | feature_elongation,3_prime_UTR_variant |
| X | 104128288 | 104128490 | 202 | Pbdc1 | CAST | feature_elongation,3_prime_UTR_variant |
| X | 105012665 | 105012795 | 130 | Magt1 | CAST,MOLF | feature_elongation,3_prime_UTR_variant |
| X | 111951008 | 111951772 | 764 | Chm | SPRET | feature_elongation,3_prime_UTR_variant |
| X | 123034727 | 123035084 | 357 | Vmn2r121 | SPRET | feature_elongation,3_prime_UTR_variant |
| X | 125721041 | 125721112 | 71 | Dmtf1l | AJ,BALB,C3H,CBA,MOLF,MRL,RH2 | feature_elongation,3_prime_UTR_variant |
| X | 134741426 | 134741661 | 235 | Gprasp2 | SPRET | feature_elongation,5_prime_UTR_variant |
| X | 135070290 | 135070494 | 204 | Tceal8 | MOLF | feature_elongation,3_prime_UTR_variant |
| X | 143129710 | 143129853 | 143 | Alg13 | SPRET,WSB | feature_elongation,coding_sequence_variant |
| X | 152177707 | 152177832 | 125 | Klf8 | MOLF | feature_elongation,3_prime_UTR_variant |
| X | 152479354 | 152479519 | 165 | Kctd12b | MOLF | feature_elongation,5_prime_UTR_variant |
| X | 155948322 | 155948816 | 494 | Phex | MOLF,SPRET | feature_elongation,3_prime_UTR_variant |
| X | 158172162 | 158172275 | 113 | Eif1ax | CAST,MOLF,SPRET | feature_elongation,3_prime_UTR_variant |
| X | 159685942 | 159686425 | 483 | Gja6 | SPRET | feature_elongation,3_prime_UTR_variant |
| X | 160147085 | 160147308 | 223 | Bend2 | SPRET | feature_elongation,3_prime_UTR_variant |
| X | 161580671 | 161580767 | 96 | Txlng | CAST,MOLF,SPRET,TH | feature_elongation,coding_sequence_variant |

| | | | | | | |
|---|---|---|---|---|---|---|
| X | 168087229 | 168087289 | 60 | Arhgap6 | MOLF | feature_elongation,3_prime_UTR_variant |

| CHR | START | END | SIZE | SYMBOL | STRAINS with VARIANT SV ALLELE | CONSEQUENCE |
|---|---|---|---|---|---|---|
| 1 | 4159116 | 4166266 | 7150 | Rp1 | 129S1,AKR,BTBR,BUB,C57L,CE,FVB,ILNJ2,LP,MA/My,NOD,NOR,NZB,PJ,PL,SJL,SWR | feature_elongation,coding_sequence_variant,intron_variant |
| 1 | 6454042 | 6460963 | 6921 | Alkal1 | KK,MOLF | feature_elongation,coding_sequence_variant,intron_variant |
| 1 | 8532733 | 8539247 | 6514 | Sntg1 | 129S1,AJ,AKR,BALB,BTBR,C57L,FVB,ILNJ2,KK,LP,MOLF,NOD,NOR,NZO,PJ,PL,RH2,SEA,SJL,SMJ,SWR,TH | feature_elongation,coding_sequence_variant,intron_variant |
| 1 | 9987587 | 9994690 | 7103 | Mcmdc2 | C3H,CBA,MRL,RF | feature_elongation,coding_sequence_variant,intron_variant |
| 1 | 14951181 | 14952762 | 1581 | Trpa1 | 129S1,BTBR,C57L,CE,DBA1J,DBA2J,MA/My,NOR | feature_elongation,coding_sequence_variant,intron_variant |
| 1 | 20185953 | 20187275 | 1322 | Pkhd1 | MOLF | feature_elongation,coding_sequence_variant,intron_variant |
| 1 | 20672026 | 20678856 | 6830 | Pkhd1 | SMJ | feature_elongation,coding_sequence_variant,intron_variant |
| 1 | 21572030 | 21578724 | 6694 | Kcnq5 | BALB,CE,KK,MA/My,NOD,SEA | feature_elongation,coding_sequence_variant,intron_variant |
| 1 | 22506002 | 22512707 | 6705 | Rims1 | AJ,AKR,BALB,BTBR,BUB,C3H,C57L,CBA,DBA1J,DBA2J,FVB,KK,LP,NOD,RF,RH2,SEA,SWR | feature_elongation,coding_sequence_variant,intron_variant |
| 1 | 34899533 | 34906230 | 6697 | Plekhb2 | 129S1,AKR,BTBR,C57L,CBA,KK,MA/My,PJ,PL,SJL,TH | feature_elongation,coding_sequence_variant,5_prime_UTR_variant,intron_variant |
| 1 | 36354084 | 36363060 | 8976 | Arid5a | CAST | feature_elongation,coding_sequence_variant,3_prime_UTR_variant,intron_variant |
| 1 | 36533774 | 36540226 | 6452 | Cnnm4 | AJ,BUB,C3H,CE,MRL,NZW,SMJ | feature_elongation,coding_sequence_variant,intron_variant |
| 1 | 39725057 | 39731487 | 6430 | Rfx8 | MOLF | feature_elongation,coding_sequence_variant,intron_variant |
| 1 | 44006353 | 44010006 | 3653 | Tpp2 | SPRET | feature_elongation,coding_sequence_variant,intron_variant |
| 1 | 46387602 | 46392840 | 5238 | Dnah7b | 129S1,AJ,BUB,C3H,CBA,CE,DBA1J,DBA2J,KK,MRL,PJ,PL,SJL,SMJ | feature_elongation,coding_sequence_variant,intron_variant |
| 1 | 46674161 | 46675651 | 1490 | Dnah7c | CAST,MOLF | feature_elongation,coding_sequence_variant,intron_variant |
| 1 | 46691054 | 46697423 | 6369 | Dnah7c | MOLF | feature_elongation,coding_sequence_variant,intron_variant |
| 1 | 46784020 | 46791374 | 7354 | Dnah7c | WSB | feature_elongation,coding_sequence_variant,intron_variant |
| 1 | 53531846 | 53536404 | 4558 | Dnah7a | SMJ | feature_elongation,coding_sequence_variant,intron_variant |
| 1 | 53567066 | 53573350 | 6284 | Dnah7a | SPRET | feature_elongation,coding_sequence_variant,intron_variant |
| 1 | 53606388 | 53611930 | 5542 | Dnah7a | KK | stop_gained,frameshift_variant |
| 1 | 53988425 | 53994867 | 6442 | Hecw2 | CAST | feature_elongation,coding_sequence_variant,intron_variant |
| 1 | 54581038 | 54588347 | 7309 | Pgap1 | SPRET | feature_elongation,coding_sequence_variant,intron_variant |
| 1 | 55209800 | 55213959 | 4159 | Rftn2 | SMJ | feature_elongation,coding_sequence_variant,3_prime_UTR_variant,intron_variant |
| 1 | 57998655 | 58005591 | 6936 | Kctd18 | NZB | feature_elongation,coding_sequence_variant,intron_variant |
| 1 | 60474347 | 60477562 | 3215 | Abi2 | SPRET | feature_elongation,coding_sequence_variant,intron_variant |
| 1 | 60557982 | 60561238 | 3256 | Raph1 | SWR | feature_elongation,coding_sequence_variant,intron_variant |
| 1 | 62195723 | 62202464 | 6741 | Pard3b | CAST | feature_elongation,coding_sequence_variant,intron_variant |
| 1 | 62202361 | 62208674 | 6313 | Pard3b | CAST | feature_elongation,coding_sequence_variant,intron_variant |
| 1 | 63746593 | 63753537 | 6944 | Mdh1b | CAST | feature_elongation,coding_sequence_variant,intron_variant |
| 1 | 65298180 | 65304433 | 6253 | Pikfyve | CBA,DBA1J,DBA2J,LP,PJ,SMJ | feature_elongation,coding_sequence_variant,intron_variant |
| 1 | 66536961 | 66544167 | 7206 | Unc80 | MOLF | feature_elongation,coding_sequence_variant,intron_variant |
| 1 | 69933467 | 69939870 | 6403 | Spag16 | C58J,SWR | feature_elongation,coding_sequence_variant,intron_variant |
| 1 | 72633305 | 72639759 | 6454 | Smarcal1 | SEA | feature_elongation,coding_sequence_variant,intron_variant |
| 1 | 72683907 | 72691036 | 7129 | Ankar | WSB | feature_elongation,coding_sequence_variant,intron_variant |
| 1 | 86122837 | 86124727 | 1890 | Armc9 | NZB | feature_elongation,coding_sequence_variant,intron_variant |
| 1 | 86129389 | 86131281 | 1892 | Armc9 | SPRET | feature_elongation,coding_sequence_variant,intron_variant |
| 1 | 86512642 | 86519352 | 6710 | Cops7b | 129S1,AJ,BTBR,C3H,C57L,CE,FVB,ILNJ2,KK,NZO,NZW,RH2,SWR,TH | feature_elongation,coding_sequence_variant,5_prime_UTR_variant,intron_variant |
| 1 | 86714327 | 86721036 | 6709 | Dis3l2 | CE | feature_elongation,coding_sequence_variant,intron_variant |
| 1 | 91208275 | 91214714 | 6439 | Ube2f | WSB | feature_elongation,coding_sequence_variant,3_prime_UTR_variant,intron_variant |
| 1 | 91232035 | 91233910 | 1875 | Scly | PJ | feature_elongation,coding_sequence_variant,intron_variant |
| 1 | 92417891 | 92422948 | 5057 | Or6b2b | BUB,C3H,CBA | feature_elongation,coding_sequence_variant,5_prime_UTR_variant,3_prime_UTR_variant,intron_variant |
| 1 | 93220124 | 93226559 | 6435 | Sned1 | SPRET | feature_elongation,coding_sequence_variant,3_prime_UTR_variant,intron_variant |
| 1 | 93243221 | 93245105 | 1884 | Pask | SPRET | feature_elongation,coding_sequence_variant,intron_variant |
| 1 | 97041495 | 97048031 | 6536 | Slco6c1 | AKR,BUB,SJL | feature_elongation,coding_sequence_variant,intron_variant |
| 1 | 97540138 | 97546620 | 6482 | Macir | PL | feature_elongation,3_prime_UTR_variant |
| 1 | 97548040 | 97554632 | 6592 | Macir | MRL,PL | feature_elongation,3_prime_UTR_variant |
| 1 | 98422756 | 98424925 | 2169 | Slco6d1 | MOLF,MRL,PL | feature_elongation,coding_sequence_variant,intron_variant |

| | | | | | | |
|---|---|---|---|---|---|---|
| 1 | 105489808 | 105496042 | 6234 | Pign | AJ,AKR,BALB,BTBR,BUB,C3H,CBA,CE,FVB,ILNJ2,KK,LP,MA/My,MRL,NZB,NZO,NZW,PJ,PL,RF,RH2,SEA,SJL,SMJ,SWR,WSB | feature_elongation,coding_sequence_variant,intron_variant |
| 1 | 106212050 | 106216540 | 4490 | Phlpp1 | ILNJ2 | feature_elongation,coding_sequence_variant,intron_variant |
| 1 | 106715108 | 106719949 | 4841 | Vps4b | BUB,NZO,NZW | feature_elongation,coding_sequence_variant,intron_variant |
| 1 | 117911094 | 117913563 | 2469 | Gm7145 | CAST | feature_elongation,coding_sequence_variant,intron_variant |
| 1 | 127229449 | 127236563 | 7114 | Mgat5 | CBA,NOD,NOR,SMJ | feature_elongation,coding_sequence_variant,5_prime_UTR_variant,intron_variant |
| 1 | 128194794 | 128199991 | 5197 | Ubxn4 | CAST,MRL,NZB,SWR,WSB | feature_elongation,coding_sequence_variant,intron_variant |
| 1 | 134231291 | 134232523 | 1232 | Ppfia4 | AJ,C3H,CE,KK,NZB,NZO,PJ,RH2,SJL,SWR,TH | feature_elongation,coding_sequence_variant,intron_variant |
| 1 | 139166673 | 139172384 | 5711 | Crb1 | SPRET | feature_elongation,coding_sequence_variant,intron_variant |
| 1 | 139482282 | 139484882 | 2600 | Cfhr1 | BTBR | feature_elongation,coding_sequence_variant,intron_variant |
| 1 | 140494719 | 140501872 | 7153 | Kcnt2 | KK,SWR | feature_elongation,coding_sequence_variant,intron_variant |
| 1 | 150604322 | 150610933 | 6611 | Hmcn1 | BUB,DBA1J,DBA2J,RF,WSB | feature_elongation,coding_sequence_variant,intron_variant |
| 1 | 151588619 | 151591816 | 3197 | Niban1 | CAST | feature_elongation,coding_sequence_variant,intron_variant |
| 1 | 152682288 | 152684181 | 1893 | Ncf2 | C57L | feature_elongation,coding_sequence_variant,5_prime_UTR_variant,intron_variant |
| 1 | 152698928 | 152700918 | 1990 | Ncf2 | AKR,BTBR,KK,MOLF,NZB,NZO,NZW,SMJ | feature_elongation,coding_sequence_variant,intron_variant |
| 1 | 153704017 | 153705075 | 1058 | Rgsl1 | WSB | feature_elongation,coding_sequence_variant,intron_variant |
| 1 | 155254314 | 155260707 | 6393 | Xpr1 | WSB | feature_elongation,coding_sequence_variant,intron_variant |
| 1 | 156448172 | 156449794 | 1622 | Abl2 | C58J,FVB,ILNJ2,RH2 | feature_elongation,coding_sequence_variant,intron_variant |
| 1 | 159970355 | 159976791 | 6436 | Tnn | ILNJ2,MA/My,NZW,PL | feature_elongation,coding_sequence_variant,5_prime_UTR_variant,intron_variant |
| 1 | 160925218 | 160926861 | 1643 | Klhl20 | 129S1,AJ,BALB,BUB,C3H,C57L,C58J,CAST,CBA,CE,DBA1J,DBA2J,ILNJ2,LP,MOLF,NZW,PJ,RH2,SEA,SMJ | feature_elongation,coding_sequence_variant,intron_variant |
| 1 | 162869470 | 162875286 | 5816 | Mroh9 | 129S1,AJ,BALB,BUB,C3H,C57L,C58J,CBA,CE,ILNJ2,LP,MA/My,MRL,PL,RH2,TH | feature_elongation,coding_sequence_variant,intron_variant |
| 1 | 162869727 | 162874062 | 4335 | Mroh9 | CAST | feature_elongation,coding_sequence_variant,intron_variant |
| 1 | 172054186 | 172055309 | 1123 | Atp1a4 | WSB | feature_elongation,coding_sequence_variant,intron_variant |
| 1 | 173408768 | 173414009 | 5241 | Ifi213 | 129S1,BALB,C3H,C57L,MRL,NOD,NOR | feature_elongation,coding_sequence_variant,intron_variant |
| 1 | 174517445 | 174523806 | 6361 | Fmn2 | PL,TH | feature_elongation,coding_sequence_variant,intron_variant |
| 1 | 174947470 | 174951990 | 4520 | Rgs7 | C58J,CBA,CE,DBA1J,DBA2J,ILNJ2,NZB,NZW,PJ,SEA,SMJ | feature_elongation,coding_sequence_variant,intron_variant |
| 1 | 175586380 | 175593247 | 6867 | Wdr64 | 129S1,BALB,C3H,C57L,MRL,NOD,NOR,NZW,PL,SMJ | feature_elongation,coding_sequence_variant,intron_variant |
| 1 | 176565838 | 176568758 | 2920 | Cep170 | CAST,MOLF,SPRET | feature_elongation,coding_sequence_variant,intron_variant |
| 1 | 176919103 | 176925514 | 6411 | Akt3 | SPRET | feature_elongation,coding_sequence_variant,intron_variant |
| 1 | 177708269 | 177714708 | 6439 | Catspere1 | AJ,BUB,RH2,WSB | feature_elongation,coding_sequence_variant,intron_variant |
| 1 | 177843026 | 177850124 | 7098 | Catspere2 | CBA,DBA1J,DBA2J,KK | feature_elongation,coding_sequence_variant,intron_variant |
| 1 | 178023051 | 178029491 | 6440 | Desi2 | AJ,BUB,RH2,WSB | feature_elongation,5_prime_UTR_variant,intron_variant |
| 1 | 179865697 | 179872798 | 7101 | Cdc42bpa | WSB | feature_elongation,coding_sequence_variant,intron_variant |
| 1 | 179928080 | 179931075 | 2995 | Cdc42bpa | C58J,LP | feature_elongation,coding_sequence_variant,intron_variant |
| 1 | 182428980 | 182435381 | 6401 | Capn8 | 129S1,BALB,C3H,C57L,CBA,DBA1J,DBA2J,KK,MA/My,PJ,PL,SEA | feature_elongation,coding_sequence_variant,intron_variant |
| 1 | 189529611 | 189531623 | 2012 | Ptpn14 | AKR,NZO,NZW | feature_elongation,coding_sequence_variant,intron_variant |
| 1 | 194821520 | 194829506 | 7986 | Cr2 | CAST,WSB | feature_elongation,coding_sequence_variant,3_prime_UTR_variant,intron_variant |
| 2 | 5835759 | 5842232 | 6473 | Cdc123 | FVB,NZB,WSB | feature_elongation,coding_sequence_variant,intron_variant |
| 2 | 6408932 | 6413882 | 4950 | Usp6nl | CAST | feature_elongation,coding_sequence_variant,intron_variant |
| 2 | 13218105 | 13223361 | 5256 | Rsu1 | SMJ | feature_elongation,coding_sequence_variant,intron_variant |
| 2 | 14036516 | 14041745 | 5229 | Hacd1 | C3H | feature_elongation,coding_sequence_variant,3_prime_UTR_variant,intron_variant,feature_elongation,coding_sequence_variant,intron_variant |
| 2 | 14408055 | 14414458 | 6403 | Slc39a12 | SJL | feature_elongation,coding_sequence_variant,intron_variant |
| 2 | 15755617 | 15761876 | 6259 | Malrd1 | CAST | feature_elongation,coding_sequence_variant,intron_variant |
| 2 | 15836518 | 15842967 | 6449 | Malrd1 | MOLF | feature_elongation,coding_sequence_variant,intron_variant |
| 2 | 20121233 | 20124229 | 2996 | Etl4 | WSB | feature_elongation,5_prime_UTR_variant,intron_variant |
| 2 | 24504892 | 24511679 | 6787 | Cacna1b | MOLF | feature_elongation,coding_sequence_variant,intron_variant |
| 2 | 25822206 | 25824607 | 2401 | Camsap1 | SPRET | feature_elongation,coding_sequence_variant,intron_variant |
| 2 | 26112834 | 26118964 | 6130 | Qsox2 | CAST | feature_elongation,coding_sequence_variant,intron_variant |
| 2 | 27407927 | 27410935 | 3008 | Wdr5 | MOLF | feature_elongation,coding_sequence_variant,5_prime_UTR_variant,intron_variant |

| | | | | | | |
|---|---|---|---|---|---|---|
| 2 | 28390006 | 28394539 | 4533 | Gbgt1 | CAST | feature_elongation,coding_sequence_variant,intron_variant |
| 2 | 30971780 | 30976980 | 5200 | Fnbp1 | MA/My | feature_elongation,coding_sequence_variant,intron_variant |
| 2 | 31368289 | 31369779 | 1490 | Ass1 | CAST | feature_elongation,coding_sequence_variant,intron_variant |
| 2 | 31663079 | 31670199 | 7120 | Abl1 | NZO | feature_elongation,coding_sequence_variant,intron_variant |
| 2 | 35272195 | 35278411 | 6216 | 4930402F06Rik | BUB,C58J,MA/My,NZO | feature_elongation,coding_sequence_variant,intron_variant |
| 2 | 36461901 | 36464385 | 2484 | Or1j15 | CAST | feature_elongation,3_prime_UTR_variant |
| 2 | 37910710 | 37917186 | 6476 | Dennd1a | BUB,WSB | feature_elongation,coding_sequence_variant,intron_variant |
| 2 | 38845661 | 38852810 | 7149 | Olfml2a | 129S1,AJ,AKR,BTBR,C3H,C57L,C58J,CBA,CE,DBA1J,DBA2J,FVB,ILNJ2,KK,MA/My,MRL,NOD,NOR,NZB,NZO,NZW,PJ,PL,RF,SEA,SMJ,SWR | feature_elongation,coding_sequence_variant,3_prime_UTR_variant,intron_variant |
| 2 | 39014823 | 39020372 | 5549 | Scai | NOD | feature_elongation,coding_sequence_variant,intron_variant |
| 2 | 40498603 | 40503449 | 4846 | Lrp1b | CAST | feature_elongation,coding_sequence_variant,intron_variant |
| 2 | 40789273 | 40795948 | 6675 | Lrp1b | BUB | feature_elongation,coding_sequence_variant,intron_variant |
| 2 | 41196725 | 41203526 | 6801 | Lrp1b | 129S1,AJ,AKR,BTBR,C3H,C57L,C58J,CBA,DBA1J,DBA2J,FVB,ILNJ2,KK,MA/My,MRL,NOD,NOR,NZB,NZO,NZW,PJ,PL,RF,SEA,SWR | feature_elongation,coding_sequence_variant,intron_variant |
| 2 | 48813144 | 48820224 | 7080 | Orc4 | FVB,SJL,TH | feature_elongation,coding_sequence_variant,intron_variant |
| 2 | 59761062 | 59767526 | 6464 | Baz2b | MOLF | feature_elongation,coding_sequence_variant,intron_variant |
| 2 | 62108530 | 62114972 | 6442 | Slc4a10 | NOD,NOR,NZB,NZO,NZW,SJL | feature_elongation,coding_sequence_variant,intron_variant |
| 2 | 62562405 | 62572147 | 9742 | Kcnh7 | NZO | feature_elongation,coding_sequence_variant,intron_variant |
| 2 | 65527192 | 65534481 | 7289 | Scn2a | C3H,C58J,CBA,MRL,PL,SMJ,TH | feature_elongation,coding_sequence_variant,intron_variant |
| 2 | 65828294 | 65835847 | 7553 | Csrnp3 | MOLF | feature_elongation,coding_sequence_variant,intron_variant |
| 2 | 68437584 | 68445413 | 7829 | 4933409G03Rik | MOLF | feature_elongation,coding_sequence_variant,intron_variant |
| 2 | 68565718 | 68571943 | 6225 | 4932414N04Rik | CAST | feature_elongation,coding_sequence_variant,intron_variant |
| 2 | 69086415 | 69092671 | 6256 | Abcb11 | 129S1,C3H,C58J,CBA,NZW,SJL | feature_elongation,coding_sequence_variant,intron_variant |
| 2 | 70822284 | 70827832 | 5548 | Mettl8 | MA/My | feature_elongation,coding_sequence_variant,intron_variant |
| 2 | 77750054 | 77756292 | 6238 | Cwc22 | CAST | feature_elongation,coding_sequence_variant,intron_variant |
| 2 | 83582299 | 83588614 | 6315 | Itgav | AJ,AKR,BALB,C3H,C57L,C58J,CBA,CE,DBA1J,DBA2J,MA/My,MRL,NZW,PJ,RF,RH2,SEA,SMJ | feature_elongation,coding_sequence_variant,intron_variant |
| 2 | 85905403 | 85910600 | 5197 | Or5m9b | MOLF | stop_gained,frameshift_variant |
| 2 | 88150536 | 88156976 | 6440 | Or5d45 | C58J,CAST,ILNJ2,KK,WSB | feature_elongation,coding_sequence_variant,3_prime_UTR_variant,intron_variant |
| 2 | 88574843 | 88576333 | 1490 | Or4p23 | AJ,AKR,BALB,C57L,DBA1J,DBA2J,FVB,KK,MRL,NOD,NOR,NZB,NZO,PJ,RF,RH2,SEA,SMJ,WSB | feature_elongation,coding_sequence_variant,3_prime_UTR_variant |
| 2 | 91062224 | 91065214 | 2990 | Ddb2 | WSB | feature_elongation,coding_sequence_variant,intron_variant,feature_elongation,coding_sequence_variant,5_prime_UTR_variant,intron_variant |
| 2 | 101436210 | 101442778 | 6568 | Iftap | MOLF | feature_elongation,coding_sequence_variant,5_prime_UTR_variant,intron_variant |
| 2 | 102407053 | 102413573 | 6520 | Pamr1 | ILNJ2,NZB,RF,SMJ,WSB | feature_elongation,coding_sequence_variant,intron_variant |
| 2 | 104073912 | 104079287 | 5375 | D430041D05Rik | CAST | feature_elongation,coding_sequence_variant,intron_variant |
| 2 | 111868509 | 111874132 | 5623 | Or4f59 | MOLF | feature_elongation,coding_sequence_variant,5_prime_UTR_variant,3_prime_UTR_variant,intron_variant |
| 2 | 113391854 | 113398105 | 6251 | Fmn1 | C3H,CE,MOLF,PL | feature_elongation,coding_sequence_variant,intron_variant |
| 2 | 117097132 | 117098925 | 1793 | Fam98b | SPRET | feature_elongation,coding_sequence_variant,intron_variant |
| 2 | 118465340 | 118468847 | 3507 | Bub1b | 129S1,AJ,AKR,BALB,BUB,C57L,CAST,CE,DBA1J,DBA2J,FVB,ILNJ2,MA/My,NZW,PJ,PL,RF,RH2,SEA,SJL,SMJ,SWR | feature_elongation,coding_sequence_variant,intron_variant |
| 2 | 120325114 | 120331556 | 6442 | Capn3 | PL | feature_elongation,coding_sequence_variant,intron_variant |
| 2 | 125727666 | 125736484 | 8818 | Galk2 | C3H,CBA,DBA2J,ILNJ2,PJ | feature_elongation,coding_sequence_variant,intron_variant |
| 2 | 125770136 | 125783247 | 13111 | Galk2 | SPRET | feature_elongation,coding_sequence_variant,intron_variant |
| 2 | 128922836 | 128929455 | 6619 | Ttl | BALB,C3H,C58J,CBA,NZB,SEA,SMJ | feature_elongation,coding_sequence_variant,intron_variant |
| 2 | 152822451 | 152828893 | 6442 | Ttll9 | AJ,ILNJ2,MOLF,NOD,NZW,RF | feature_elongation,coding_sequence_variant,intron_variant |
| 2 | 153699754 | 153701566 | 1812 | Sun5 | AKR,B10D2,B10J,BALB,BUB,C3H,CBA,DBA1J,DBA2J,FVB,KK,LP,NOR,NZO,PL,RH2,SEA,SJL,SWR,WSB | feature_elongation,coding_sequence_variant,intron_variant |
| 2 | 154480584 | 154485696 | 5112 | Zfp341 | CE | feature_elongation,coding_sequence_variant,intron_variant |
| 2 | 160880413 | 160887773 | 7360 | Chd6 | CAST | feature_elongation,coding_sequence_variant,intron_variant |
| 2 | 166704183 | 166709472 | 5289 | Arfgef2 | SMJ | feature_elongation,coding_sequence_variant,intron_variant |
| 2 | 167274094 | 167281308 | 7214 | Slc9a8 | CE | feature_elongation,coding_sequence_variant,intron_variant |

| | | | | | | |
|---|---|---|---|---|---|---|
| 2 | 177006961 | 177009397 | 2436 | Gm14412 | DBA2J,SEA | feature_elongation,coding_sequence_variant,intron_variant |
| 2 | 178014598 | 178018743 | 4145 | Sycp2 | CAST,MOLF,PJ | feature_elongation,coding_sequence_variant,intron_variant |
| 3 | 14717811 | 14722126 | 4315 | Car13 | NZO | feature_elongation,coding_sequence_variant,intron_variant |
| 3 | 19595838 | 19602237 | 6399 | Dnajc5b | DBA2J,LP,MA/My,NOD,NOR,NZB,NZO,NZW | feature_elongation,coding_sequence_variant,5_prime_UTR_variant,intron_variant |
| 3 | 32532777 | 32537268 | 4491 | Kcnmb3 | CAST | feature_elongation,coding_sequence_variant,intron_variant |
| 3 | 37039339 | 37045897 | 6558 | Bltp1 | CAST | feature_elongation,coding_sequence_variant,intron_variant |
| 3 | 41670269 | 41673642 | 3373 | Sclt1 | SJL,SWR | feature_elongation,coding_sequence_variant,intron_variant |
| 3 | 41685568 | 41692192 | 6624 | Sclt1 | 129S1 | feature_elongation,coding_sequence_variant,intron_variant |
| 3 | 58530722 | 58536279 | 5557 | Erich6 | BUB,CE,FVB,SJL,SWR | feature_elongation,coding_sequence_variant,intron_variant |
| 3 | 64026998 | 64035415 | 8417 | Vmn2r2 | CAST | feature_elongation,coding_sequence_variant,intron_variant |
| 3 | 69923383 | 69930088 | 6705 | Otol1 | 129S1,BTBR,C57L,FVB,SWR,TH | feature_elongation,coding_sequence_variant,5_prime_UTR_variant,intron_variant |
| 3 | 72832576 | 72834170 | 1594 | Sis | 129S1,AJ,AKR,BALB,BTBR,BUB,C57L,FVB,KK,MA/My,MRL,NOD,NOR,NZB,NZO,PJ,RF,RH2,SEA,SJL,SWR,TH | feature_elongation,coding_sequence_variant,intron_variant |
| 3 | 74974583 | 74982298 | 7715 | Zbbx | CAST,MOLF | feature_elongation,coding_sequence_variant,intron_variant |
| 3 | 74997023 | 75002190 | 5167 | Zbbx | WSB | feature_elongation,coding_sequence_variant,intron_variant |
| 3 | 76439469 | 76447058 | 7589 | Fstl5 | AKR,DBA1J,DBA2J,MRL,PL,RF | feature_elongation,coding_sequence_variant,intron_variant |
| 3 | 79838148 | 79844368 | 6220 | Gask1b | MOLF | feature_elongation,coding_sequence_variant,intron_variant |
| 3 | 86206998 | 86214144 | 7146 | Lrba | NZB,NZO | feature_elongation,coding_sequence_variant,intron_variant |
| 3 | 90039969 | 90047084 | 7115 | Nup210l | NOD,NOR | feature_elongation,coding_sequence_variant,intron_variant |
| 3 | 90047557 | 90053075 | 5518 | Nup210l | RF | feature_elongation,coding_sequence_variant,intron_variant |
| 3 | 91932359 | 91933543 | 1184 | Pglyrp3 | CAST | feature_elongation,coding_sequence_variant,intron_variant |
| 3 | 92865525 | 92881535 | 16010 | Lce3d | DBA1J,DBA2J | stop_gained,inframe_insertion |
| 3 | 96059074 | 96060960 | 1886 | Otud7b | SPRET | feature_elongation,coding_sequence_variant,intron_variant |
| 3 | 97629422 | 97636321 | 6899 | Pde4dip | NOD,NOR,RF,WSB | feature_elongation,coding_sequence_variant,intron_variant |
| 3 | 99923377 | 99929630 | 6253 | Spag17 | CAST | feature_elongation,coding_sequence_variant,intron_variant |
| 3 | 99954703 | 99957016 | 2313 | Spag17 | 129S1,B10D2,B10J,BTBR,C57L,C58J,DBA1J,DBA2J,LP,MA/My,NOD,RH2,TH | feature_elongation,coding_sequence_variant,intron_variant |
| 3 | 100016143 | 100019411 | 3268 | Spag17 | WSB | feature_elongation,coding_sequence_variant,intron_variant |
| 3 | 100021433 | 100028546 | 7113 | Spag17 | BTBR,DBA1J,DBA2J,NOD,NOR | feature_elongation,coding_sequence_variant,intron_variant |
| 3 | 100043841 | 100050157 | 6316 | Spag17 | KK,MOLF | feature_elongation,3_prime_UTR_variant,intron_variant |
| 3 | 102835780 | 102840386 | 4606 | Sycp1 | 129S1,AKR,BALB,BUB,C57L,C58J,FVB,ILNJ2,KK,LP,MA/My,MRL,NOD,NOR,NZB,NZO,NZW,PL,RF,RH2,SWR | feature_elongation,coding_sequence_variant,intron_variant |
| 3 | 104369175 | 104370923 | 1748 | Lrig2 | 129S1,AKR,BUB,C57L,C58J,FVB,ILNJ2,KK,PL,SJL,SWR,TH | feature_elongation,coding_sequence_variant,intron_variant |
| 3 | 104745458 | 104751918 | 6460 | Capza1 | CE,NZB,PJ,SEA | feature_elongation,coding_sequence_variant,intron_variant |
| 3 | 106062555 | 106064917 | 2362 | Chil3 | MOLF | feature_elongation,coding_sequence_variant,intron_variant |
| 3 | 108244175 | 108246066 | 1891 | Sort1 | WSB | feature_elongation,coding_sequence_variant,intron_variant |
| 3 | 108276191 | 108281443 | 5252 | Mybphl | MRL | feature_elongation,coding_sequence_variant,intron_variant |
| 3 | 113361945 | 113363972 | 2027 | Amy1 | CE,NOD,NOR | feature_elongation,coding_sequence_variant,5_prime_UTR_variant,intron_variant |
| 3 | 118403388 | 118409612 | 6224 | Dpyd | BUB,LP | feature_elongation,coding_sequence_variant,intron_variant |
| 3 | 123334910 | 123341435 | 6525 | Ndst3 | PJ,SEA | feature_elongation,coding_sequence_variant,intron_variant |
| 3 | 129514326 | 129519521 | 5195 | Egf | TH | feature_elongation,coding_sequence_variant,intron_variant |
| 3 | 133070040 | 133076525 | 6485 | Ppa2 | CAST | feature_elongation,coding_sequence_variant,intron_variant |
| 3 | 135217939 | 135224412 | 6473 | Manba | KK | feature_elongation,coding_sequence_variant,intron_variant |
| 3 | 135551040 | 135555571 | 4531 | Slc39a8 | CBA,CE,DBA1J,DBA2J,FVB,KK,MA/My,NOD,NOR,SJL,SMJ,SWR,TH | feature_elongation,coding_sequence_variant,intron_variant |
| 3 | 138127419 | 138128907 | 1488 | Adh4 | 129S1,AJ,BALB,BTBR,BUB,C3H,C57L,C58J,CBA,CE,DBA1J,DBA2J,ILNJ2,KK,LP,MRL,NZB,NZO,NZW,RF,RH2 | feature_elongation,coding_sequence_variant,intron_variant |
| 3 | 138660637 | 138662497 | 1860 | Rap1gds1 | CAST | feature_elongation,coding_sequence_variant,intron_variant |
| 3 | 139225878 | 139228933 | 3055 | Stpg2 | CE | feature_elongation,coding_sequence_variant,intron_variant |
| 3 | 141561465 | 141562707 | 1242 | Bmpr1b | CAST,MOLF | feature_elongation,coding_sequence_variant,intron_variant |
| 3 | 145152504 | 145159192 | 6688 | Col24a1 | CAST | feature_elongation,coding_sequence_variant,intron_variant |
| 3 | 145162513 | 145169116 | 6603 | Col24a1 | CAST | feature_elongation,coding_sequence_variant,intron_variant |
| 3 | 146178954 | 146186268 | 7314 | Spata1 | WSB | feature_elongation,coding_sequence_variant,intron_variant |

| | | | | | | |
|---|---|---|---|---|---|---|
| 3 | 153596052 | 153603829 | 7777 | Msh4 | 129S1,BTBR,C57L | feature_elongation,coding_sequence_variant,intron_variant,feature_elongation,coding_sequence_variant,5_prime_UTR_variant,intron_variant |
| 3 | 154465114 | 154469660 | 4546 | Erich3 | FVB,SJL,SWR | feature_elongation,coding_sequence_variant,intron_variant |
| 3 | 154660368 | 154667563 | 7195 | Tnni3k | AJ,BALB,BUB,C3H,DBA2J,ILNJ2,KK,MA/My,NOD,NOR,NZB,NZO,NZW,RH2,SEA | feature_elongation,coding_sequence_variant,intron_variant |
| 4 | 14971827 | 14978007 | 6180 | Necab1 | CAST | feature_elongation,coding_sequence_variant,intron_variant |
| 4 | 19833785 | 19840259 | 6474 | Slc7a13 | ILNJ2 | feature_elongation,coding_sequence_variant,intron_variant |
| 4 | 36135172 | 36141074 | 5902 | Lingo2 | WSB | feature_elongation,5_prime_UTR_variant,intron_variant |
| 4 | 43319825 | 43325374 | 5549 | Atp8b5 | C58J,LP | feature_elongation,coding_sequence_variant,3_prime_UTR_variant,intron_variant,feature_elongation,coding_sequence_variant,intron_variant |
| 4 | 43353242 | 43354470 | 1228 | Atp8b5 | SPRET,WSB | feature_elongation,coding_sequence_variant,intron_variant |
| 4 | 43359052 | 43362673 | 3621 | Atp8b5 | SPRET | feature_elongation,coding_sequence_variant,intron_variant |
| 4 | 49298178 | 49304792 | 6614 | Plppr1 | CAST | feature_elongation,coding_sequence_variant,intron_variant |
| 4 | 59483832 | 59490259 | 6427 | Ptbp3 | CE | feature_elongation,coding_sequence_variant,intron_variant |
| 4 | 59604688 | 59611432 | 6744 | Hsdl2 | BUB | feature_elongation,coding_sequence_variant,intron_variant |
| 4 | 73988436 | 73994596 | 6160 | Frmd3 | CAST | feature_elongation,coding_sequence_variant,intron_variant |
| 4 | 83494262 | 83499810 | 5548 | Ccdc171 | PL | feature_elongation,coding_sequence_variant,intron_variant |
| 4 | 86702179 | 86705012 | 2833 | Dennd4c | MOLF | feature_elongation,coding_sequence_variant,intron_variant |
| 4 | 88679502 | 88686847 | 7345 | Gm13283 | WSB | stop_gained,frameshift_variant |
| 4 | 96352448 | 96359866 | 7418 | Cyp2j8 | ILNJ2 | feature_elongation,coding_sequence_variant,intron_variant |
| 4 | 99108714 | 99115160 | 6446 | Atg4c | CE,WSB | feature_elongation,coding_sequence_variant,intron_variant |
| 4 | 100717005 | 100723565 | 6560 | Cachd1 | 129S1,AJ,BALB,C3H,C57L,CBA,CE,NOD,NOR,RH2,SEA,SWR,TH | feature_elongation,coding_sequence_variant,intron_variant |
| 4 | 104904755 | 104911199 | 6444 | Prkaa2 | MOLF | feature_elongation,coding_sequence_variant,intron_variant |
| 4 | 107021998 | 107025544 | 3546 | Tceanc2 | 129S1,AJ,C57L,FVB,TH | feature_elongation,coding_sequence_variant,intron_variant |
| 4 | 107763105 | 107770221 | 7116 | Cpt2 | CAST | feature_elongation,coding_sequence_variant,intron_variant |
| 4 | 108778770 | 108784207 | 5437 | Rab3b | BALB,BUB,C3H,CBA,NOD,PL,RF,SEA,SJL | feature_elongation,coding_sequence_variant,intron_variant |
| 4 | 108866720 | 108872269 | 5549 | Nrdc | NOD | feature_elongation,coding_sequence_variant,intron_variant |
| 4 | 109608115 | 109615273 | 7158 | Faf1 | CE | feature_elongation,coding_sequence_variant,intron_variant |
| 4 | 111467612 | 111475125 | 7513 | Agbl4 | CE,SWR,WSB | feature_elongation,coding_sequence_variant,intron_variant |
| 4 | 115495252 | 115503009 | 7757 | Cyp4b1 | WSB | feature_elongation,coding_sequence_variant,intron_variant |
| 4 | 115667258 | 115668862 | 1604 | Atpaf1 | AJ,BALB,BUB,C3H,CAST,CBA,CE,DBA1J,DBA2J,FVB,KK,MOLF,NOD,NOR,RF,SJL,SWR,TH,WSB | feature_elongation,coding_sequence_variant,3_prime_UTR_variant,intron_variant |
| 4 | 116387118 | 116389413 | 2295 | Ipp | AKR,B10D2,B10J,C58J,KK,MRL,NZB,NZO,NZW,PJ,PL,SEA,SMJ,TH | feature_elongation,coding_sequence_variant,intron_variant |
| 4 | 116835780 | 116837481 | 1701 | Zswim5 | ILNJ2,MOLF,TH | feature_elongation,coding_sequence_variant,intron_variant |
| 4 | 118449769 | 118456761 | 6992 | Cfap57 | WSB | feature_elongation,coding_sequence_variant,intron_variant |
| 4 | 119334594 | 119337427 | 2833 | Rimkla | 129S1,BALB,C3H,C57L,CBA,CE,SJL | feature_elongation,coding_sequence_variant,intron_variant |
| 4 | 120361057 | 120364675 | 3618 | Scmh1 | FVB,SWR | feature_elongation,coding_sequence_variant,intron_variant |
| 4 | 124683414 | 124685636 | 2222 | Inpp5b | MOLF | feature_elongation,coding_sequence_variant,intron_variant |
| 4 | 126256110 | 126264841 | 8731 | Ago3 | MOLF | feature_elongation,coding_sequence_variant,intron_variant |
| 4 | 126808396 | 126809447 | 1051 | Zmym4 | B10D2,B10J,CAST,ILNJ2,MOLF,NOD,PJ,SEA | feature_elongation,coding_sequence_variant,intron_variant |
| 4 | 132642063 | 132646120 | 4057 | Fam76a | SJL,SWR | feature_elongation,coding_sequence_variant,intron_variant |
| 4 | 134627035 | 134628943 | 1908 | Tmem50a | AJ,AKR,C3H,CBA,CE,MRL,RF,RH2 | feature_elongation,coding_sequence_variant,intron_variant |
| 4 | 135991461 | 135995831 | 4370 | Tcea3 | ILNJ2,PL,SMJ,TH | feature_elongation,coding_sequence_variant,intron_variant |
| 4 | 137921672 | 137926147 | 4475 | Eif4g3 | WSB | feature_elongation,coding_sequence_variant,intron_variant |
| 4 | 137993833 | 137997242 | 3409 | Kif17 | MOLF | feature_elongation,coding_sequence_variant,intron_variant |
| 4 | 147878952 | 147885992 | 7040 | Zfp984 | MOLF | feature_elongation,5_prime_UTR_variant,intron_variant |
| 4 | 148523644 | 148529192 | 5548 | Ubiad1 | AKR,C3H,CBA,MRL,PL | feature_elongation,coding_sequence_variant,5_prime_UTR_variant,intron_variant |
| 4 | 149805277 | 149810276 | 4999 | Tmem201 | CAST | feature_elongation,coding_sequence_variant,intron_variant |
| 4 | 155074247 | 155079792 | 5545 | Plch2 | AJ | stop_gained,frameshift_variant |
| 4 | 155085820 | 155091254 | 5434 | Plch2 | LP | feature_elongation,coding_sequence_variant,intron_variant |
| 4 | 155730736 | 155733282 | 2546 | Cdk11b | FVB,MOLF,NOD,NOR,SJL,SWR | feature_elongation,coding_sequence_variant,intron_variant |
| 5 | 5672370 | 5679717 | 7347 | Cfap69 | WSB | feature_elongation,coding_sequence_variant,intron_variant |
| 5 | 8487519 | 8493980 | 6461 | Slc25a40 | BUB,NOD,NOR,SMJ | feature_elongation,coding_sequence_variant,intron_variant |

| | | | | | | |
|---|---|---|---|---|---|---|
| 5 | 8650450 | 8657915 | 7465 | Rundc3b | CAST | feature_elongation,coding_sequence_variant,intron_variant |
| 5 | 8739787 | 8744956 | 5169 | Abcb1a | SWR | feature_elongation,coding_sequence_variant,intron_variant |
| 5 | 8900711 | 8905213 | 4502 | Abcb1b | BUB,MOLF,NOD,NOR | feature_elongation,coding_sequence_variant,intron_variant |
| 5 | 12544406 | 12550954 | 6548 | Sema3d | AKR,BUB,DBA1J,DBA2J,FVB,ILNJ2,KK,MRL,NOD,NOR,NZB,NZO,NZW,SJL,SWR,WSB | feature_elongation,coding_sequence_variant,intron_variant |
| 5 | 13625539 | 13632451 | 6912 | Sema3a | AJ,AKR,BUB,C3H,CBA,DBA1J,DBA2J,FVB,ILNJ2,KK,LP,MRL,NOD,NOR,SJL,SMJ,SWR | feature_elongation,coding_sequence_variant,intron_variant |
| 5 | 14271156 | 14277736 | 6580 | Sema3e | WSB | feature_elongation,coding_sequence_variant,intron_variant |
| 5 | 24004981 | 24011926 | 6945 | Rint1 | AKR,BUB,NOD,NOR,NZW,PJ,RF | feature_elongation,coding_sequence_variant,intron_variant |
| 5 | 25395815 | 25403182 | 7367 | Galntl5 | BUB | feature_elongation,coding_sequence_variant,intron_variant |
| 5 | 30264281 | 30271408 | 7127 | Tyms | CAST | feature_elongation,coding_sequence_variant,3_prime_UTR_variant,intron_variant |
| 5 | 32937467 | 32942966 | 5499 | Pisd | WSB | feature_elongation,coding_sequence_variant,5_prime_UTR_variant,intron_variant |
| 5 | 35675727 | 35682185 | 6458 | Cpz | AKR | feature_elongation,coding_sequence_variant,intron_variant |
| 5 | 35748551 | 35749808 | 1257 | Acox3 | MOLF | feature_elongation,coding_sequence_variant,intron_variant |
| 5 | 53279804 | 53281692 | 1888 | Sel1l3 | WSB | feature_elongation,coding_sequence_variant,intron_variant |
| 5 | 54000357 | 54004330 | 3973 | Tbc1d19 | CAST,MOLF,PL | feature_elongation,coding_sequence_variant,intron_variant |
| 5 | 66446372 | 66447667 | 1295 | Nsun7 | AKR,FVB,RF | feature_elongation,coding_sequence_variant,intron_variant |
| 5 | 76986101 | 76993231 | 7130 | Cracd | SMJ | feature_elongation,coding_sequence_variant,5_prime_UTR_variant,intron_variant,feature_elongation,coding_sequence_variant,intron_variant |
| 5 | 77043136 | 77044292 | 1156 | Aasdh | CAST | feature_elongation,coding_sequence_variant,intron_variant |
| 5 | 86169634 | 86171498 | 1864 | Cenpc1 | 129S1,C57L | feature_elongation,coding_sequence_variant,intron_variant |
| 5 | 88422444 | 88429154 | 6710 | Smr2l | NZO | feature_elongation,coding_sequence_variant,5_prime_UTR_variant,intron_variant |
| 5 | 89360565 | 89367855 | 7290 | Slc4a4 | WSB | feature_elongation,coding_sequence_variant,intron_variant |
| 5 | 92169112 | 92170976 | 1864 | Cdkl2 | AJ,NOD,NOR,NZO | feature_elongation,coding_sequence_variant,intron_variant |
| 5 | 96782852 | 96784350 | 1498 | Fras1 | 129S1,AJ,AKR,BALB,BTBR,BUB,C3H,C57L,CAST,CBA,CE,DBA1J,DBA2J,FVB,ILNJ2,KK,MA/My,MRL,NOD,NOR,NZB,NZO,NZW,PJ,RF,RH2,SEA,SJL,SMJ,SWR,TH,WSB | feature_elongation,coding_sequence_variant,intron_variant |
| 5 | 100673917 | 100679362 | 5445 | Cops4 | 129S1,C57L | feature_elongation,coding_sequence_variant,intron_variant |
| 5 | 100956115 | 100960637 | 4522 | Abraxas1 | NZB | feature_elongation,coding_sequence_variant,intron_variant |
| 5 | 104480092 | 104485412 | 5320 | Mepe | 129S1,AKR,B10D2,B10J,BTBR,C57L,C58J,CE,LP,MA/My,MOLF,MRL,NOD,NOR,NZW,PJ,PL,RF,SMJ | feature_elongation,coding_sequence_variant,intron_variant |
| 5 | 104644958 | 104652045 | 7087 | Pkd2 | AJ,BALB,DBA1J,DBA2J,ILNJ2,KK,MOLF,NZB,NZO,RH2,SEA,SWR,WSB | feature_elongation,coding_sequence_variant,3_prime_UTR_variant,intron_variant |
| 5 | 104673296 | 104675622 | 2326 | Thoc2l | AJ,BALB,BUB,C3H,CBA,DBA1J,DBA2J,FVB,ILNJ2,KK,MOLF,NZB,NZO,RH2,SEA,SJL,SWR,TH,WSB | feature_elongation,3_prime_UTR_variant,intron_variant |
| 5 | 107631378 | 107638487 | 7109 | Btbd8 | 129S1,C57L,NZB,NZO,NZW | feature_elongation,coding_sequence_variant,intron_variant |
| 5 | 107935033 | 107941685 | 6652 | Evi5 | DBA1J,DBA2J | feature_elongation,coding_sequence_variant,intron_variant |
| 5 | 109484995 | 109487947 | 2952 | Vmn2r16 | CE,DBA1J,DBA2J,LP | feature_elongation,coding_sequence_variant,intron_variant |
| 5 | 110393829 | 110402386 | 8557 | Ankle2 | AJ,BALB,BTBR,CE,DBA1J,DBA2J,ILNJ2,LP,MRL,RH2,SEA,SMJ | feature_elongation,coding_sequence_variant,intron_variant |
| 5 | 112869082 | 112876169 | 7087 | Myo18b | AKR,CE,DBA1J,DBA2J,NZW,RF | feature_elongation,coding_sequence_variant,intron_variant |
| 5 | 115743858 | 115745016 | 1158 | Gcn1 | CAST,MOLF | feature_elongation,coding_sequence_variant,intron_variant |
| 5 | 118038045 | 118040303 | 2258 | Nos1 | NOD | feature_elongation,coding_sequence_variant,intron_variant |
| 5 | 118038137 | 118039289 | 1152 | Nos1 | NOD | feature_elongation,coding_sequence_variant,intron_variant |
| 5 | 121593520 | 121602309 | 8789 | Tmem116 | MOLF | feature_elongation,5_prime_UTR_variant,intron_variant |
| 5 | 124224113 | 124229625 | 5512 | Abcb9 | BUB | feature_elongation,coding_sequence_variant,5_prime_UTR_variant,intron_variant |
| 5 | 124439912 | 124445277 | 5365 | Mphosph9 | MOLF | feature_elongation,coding_sequence_variant,intron_variant |
| 5 | 129182490 | 129188817 | 6327 | Adgrd1 | C58J,LP | feature_elongation,coding_sequence_variant,intron_variant |
| 5 | 129234247 | 129245106 | 10859 | Adgrd1 | 129S1,C57L,ILNJ2,KK,NZB,NZO,NZW,SWR | feature_elongation,coding_sequence_variant,intron_variant |
| 5 | 131111542 | 131117098 | 5556 | Galnt17 | NOR | feature_elongation,coding_sequence_variant,intron_variant |
| 5 | 143681535 | 143688103 | 6568 | Cyth3 | CBA,WSB | feature_elongation,coding_sequence_variant,intron_variant |
| 5 | 144486739 | 144491912 | 5173 | Nptx2 | AJ,BALB,C3H,CBA,DBA1J,DBA2J,PJ,RH2,SEA | feature_elongation,coding_sequence_variant,intron_variant |
| 5 | 146044158 | 146049933 | 5775 | Cyp3a59 | AJ,DBA2J,SEA,WSB | feature_elongation,coding_sequence_variant,3_prime_UTR_variant,intron_variant |
| 5 | 147030871 | 147038099 | 7228 | Polr1d | NZB | feature_elongation,coding_sequence_variant,intron_variant |
| 5 | 148222820 | 148230151 | 7331 | Mtus2 | CE,FVB | feature_elongation,coding_sequence_variant,intron_variant |

| | | | | | | |
|---|---|---|---|---|---|---|
| 5 | 149526731 | 149533696 | 6965 | Wdr95 | NZB | feature_elongation,coding_sequence_variant,intron_variant |
| 5 | 150232003 | 150234359 | 2356 | Fry | AKR,BUB,FVB,MRL,NOD,NOR,PJ,SJL | feature_elongation,coding_sequence_variant,intron_variant |
| 5 | 150398620 | 150405035 | 6415 | Fry | NZO | feature_elongation,coding_sequence_variant,intron_variant |
| 5 | 150645295 | 150647172 | 1877 | Pds5b | CAST,MOLF,SPRET | feature_elongation,coding_sequence_variant,intron_variant |
| 6 | 4078847 | 4085777 | 6930 | Bet1 | MA/My,NZB,NZO,NZW,WSB | feature_elongation,coding_sequence_variant,intron_variant |
| 6 | 5273775 | 5278351 | 4576 | Pon2 | WSB | feature_elongation,coding_sequence_variant,intron_variant |
| 6 | 8079237 | 8081442 | 2205 | Col28a1 | MOLF | feature_elongation,coding_sequence_variant,intron_variant |
| 6 | 25776462 | 25779744 | 3282 | Pot1a | CAST | feature_elongation,coding_sequence_variant,intron_variant |
| 6 | 25793879 | 25800639 | 6760 | Pot1a | CAST | feature_elongation,coding_sequence_variant,5_prime_UTR_variant,intron_variant |
| 6 | 38268735 | 38274054 | 5319 | Zc3hav1l | C3H | feature_elongation,coding_sequence_variant,3_prime_UTR_variant,intron_variant |
| 6 | 40548384 | 40555486 | 7102 | Or9a4 | WSB | stop_gained,frameshift_variant |
| 6 | 40785680 | 40792158 | 6478 | Mgam2-ps | BUB,FVB,LP,SJL,SWR,TH,WSB | feature_elongation,coding_sequence_variant,intron_variant |
| 6 | 46996972 | 46998876 | 1904 | Cntnap2 | CAST | feature_elongation,coding_sequence_variant,intron_variant |
| 6 | 47838550 | 47844992 | 6442 | Zfp398 | C58J,MA/My | feature_elongation,coding_sequence_variant,3_prime_UTR_variant,intron_variant |
| 6 | 49377580 | 49383124 | 5544 | Stk31 | AJ,BUB | feature_elongation,coding_sequence_variant,intron_variant |
| 6 | 49440591 | 49443178 | 2587 | Stk31 | AJ,AKR,BALB,BUB,C3H,CBA,DBA1J,DBA2J,FVB,MOLF,MRL,NOD,NOR,PJ,PL,RF,RH2,SEA,SJL,SMJ,SWR,WSB | feature_elongation,coding_sequence_variant,intron_variant |
| 6 | 51538467 | 51543502 | 5035 | Snx10 | WSB | feature_elongation,coding_sequence_variant,5_prime_UTR_variant,intron_variant |
| 6 | 52728915 | 52730491 | 1576 | Tax1bp1 | CAST | feature_elongation,coding_sequence_variant,intron_variant |
| 6 | 57708248 | 57709925 | 1677 | Lancl2 | 129S1,C57L,KK,NZB,NZO,NZW | feature_elongation,coding_sequence_variant,intron_variant |
| 6 | 61350008 | 61351799 | 1791 | Ccser1 | CAST,SPRET | feature_elongation,coding_sequence_variant,intron_variant |
| 6 | 61543743 | 61550153 | 6410 | Ccser1 | WSB | feature_elongation,coding_sequence_variant,intron_variant |
| 6 | 70746700 | 70753153 | 6453 | Rpia | MA/My,SJL | feature_elongation,coding_sequence_variant,intron_variant |
| 6 | 73098133 | 73101265 | 3132 | Dnah6 | AKR,BALB,MRL,PL,RF,RH2,SEA | feature_elongation,coding_sequence_variant,intron_variant |
| 6 | 89956581 | 89958076 | 1495 | Vmn1r46 | WSB | feature_elongation,3_prime_UTR_variant |
| 6 | 92542987 | 92544305 | 1318 | Prickle2 | CAST | feature_elongation,5_prime_UTR_variant,intron_variant |
| 6 | 93917940 | 93920912 | 2972 | Magi1 | AKR,MRL,PL | feature_elongation,coding_sequence_variant,intron_variant |
| 6 | 100725694 | 100732117 | 6423 | Gxylt2 | FVB,SJL | feature_elongation,coding_sequence_variant,intron_variant |
| 6 | 113360269 | 113363084 | 2815 | Arpc4 | AKR,C3H,CBA,FVB,ILNJ2,MRL,NOD,NOR,NZB,PJ,PL,RF,SJL | feature_elongation,coding_sequence_variant,intron_variant,feature_elongation,coding_sequence_variant,5_prime_UTR_variant,intron_variant |
| 6 | 118135292 | 118142893 | 7601 | Ret | SJL,SWR | feature_elongation,coding_sequence_variant,intron_variant |
| 6 | 119606680 | 119613093 | 6413 | Erc1 | TH | feature_elongation,coding_sequence_variant,intron_variant |
| 6 | 119709461 | 119715892 | 6431 | Erc1 | SJL,SWR | feature_elongation,coding_sequence_variant,intron_variant |
| 6 | 121402673 | 121405349 | 2676 | Iqsec3 | SPRET | feature_elongation,coding_sequence_variant,intron_variant |
| 6 | 122032947 | 122039779 | 6832 | Mug2 | SWR | feature_elongation,coding_sequence_variant,intron_variant |
| 6 | 123153694 | 123160557 | 6863 | Clec4b2 | SWR | feature_elongation,coding_sequence_variant,intron_variant |
| 6 | 123368079 | 123375620 | 7541 | Vmn2r20 | BALB | feature_elongation,coding_sequence_variant,intron_variant |
| 6 | 123602269 | 123604044 | 1775 | Vmn2r22 | SWR | feature_elongation,coding_sequence_variant,intron_variant |
| 6 | 124196211 | 124203065 | 6854 | Vmn2r27 | MOLF | feature_elongation,coding_sequence_variant,intron_variant |
| 6 | 126910486 | 126915226 | 4740 | Rad51ap1 | AJ,AKR,BALB,BUB,C3H,C57L,C58J,CBA,CE,DBA1J,DBA2J,FVB,ILNJ2,KK,LP,MA/My,MRL,NOD,NOR,NZO,PJ,PL,RF,SEA,SJL,TH | feature_elongation,coding_sequence_variant,intron_variant |
| 6 | 140517588 | 140520026 | 2438 | Plekha5 | BTBR,C57L,C58J,CAST,FVB,MOLF,PJ,SMJ,SWR,WSB | feature_elongation,coding_sequence_variant,intron_variant |
| 6 | 140537001 | 140538387 | 1386 | Plekha5 | WSB | feature_elongation,coding_sequence_variant,intron_variant |
| 6 | 141953235 | 141955886 | 2651 | Slco1a8 | BALB | feature_elongation,coding_sequence_variant,intron_variant,feature_elongation,coding_sequence_variant,5_prime_UTR_variant,intron_variant |
| 6 | 146272171 | 146273682 | 1511 | Itpr2 | CAST | feature_elongation,coding_sequence_variant,intron_variant |
| 6 | 146889377 | 146896053 | 6676 | Ppfibp1 | WSB | feature_elongation,coding_sequence_variant,intron_variant |
| 7 | 3275616 | 3283094 | 7478 | Nlrp12 | MA/My | feature_elongation,coding_sequence_variant,intron_variant |
| 7 | 4007749 | 4013999 | 6250 | Lair1 | MRL | feature_elongation,coding_sequence_variant,3_prime_UTR_variant,intron_variant |
| 7 | 6475563 | 6477433 | 1870 | Or6z5 | MRL | feature_elongation,coding_sequence_variant,5_prime_UTR_variant,intron_variant |
| 7 | 7133836 | 7137428 | 3592 | Zfp773 | C58J | feature_elongation,coding_sequence_variant,3_prime_UTR_variant,intron_variant,feature_elongation,coding_sequence_variant,intron_variant |

| | | | | | | |
|---|---|---|---|---|---|---|
| 7 | 9772768 | 9779182 | 6414 | Vmn2r50 | AKR,MRL,RH2 | feature_elongation,coding_sequence_variant,intron_variant |
| 7 | 11985137 | 11991353 | 6216 | Vmn1r81 | AKR,MRL,RH2 | feature_elongation,3_prime_UTR_variant |
| 7 | 12313921 | 12318765 | 4844 | Vmn2r53 | WSB | feature_elongation,coding_sequence_variant,3_prime_UTR_variant,intron_variant |
| 7 | 12355077 | 12361789 | 6712 | Vmn2r54 | CAST | feature_elongation,coding_sequence_variant,intron_variant |
| 7 | 12363238 | 12371453 | 8215 | Vmn2r54 | CAST | feature_elongation,coding_sequence_variant,intron_variant |
| 7 | 12389632 | 12396948 | 7316 | Vmn2r55 | WSB | feature_elongation,coding_sequence_variant,intron_variant |
| 7 | 12900283 | 12907709 | 7426 | Vmn1r88 | PL | feature_elongation,5_prime_UTR_variant,intron_variant |
| 7 | 13393997 | 13397194 | 3197 | Sult2a5 | CAST | feature_elongation,coding_sequence_variant,intron_variant |
| 7 | 13723471 | 13728227 | 4756 | Sult2a4 | WSB | stop_gained,frameshift_variant |
| 7 | 15712441 | 15721175 | 8734 | Bicra | AKR,RH2 | feature_elongation,coding_sequence_variant,intron_variant |
| 7 | 17616160 | 17623028 | 6868 | Ceacam23 | CE,NOD,NOR,SJL | feature_elongation,coding_sequence_variant,intron_variant |
| 7 | 23534278 | 23541885 | 7607 | Vmn1r176 | CE | stop_gained,frameshift_variant |
| 7 | 25783833 | 25790418 | 6585 | Cyp2b13 | AJ | feature_elongation,coding_sequence_variant,intron_variant |
| 7 | 29407721 | 29413712 | 5991 | Catsperg2 | SPRET | feature_elongation,coding_sequence_variant,intron_variant |
| 7 | 34980617 | 34987875 | 7258 | Gpatch1 | AJ,BALB,SEA | feature_elongation,coding_sequence_variant,intron_variant |
| 7 | 34987750 | 34989647 | 1897 | Gpatch1 | NZW | feature_elongation,coding_sequence_variant,intron_variant |
| 7 | 35414261 | 35420703 | 6442 | Dpy19l3 | ILNJ2 | feature_elongation,coding_sequence_variant,intron_variant |
| 7 | 44270299 | 44271587 | 1288 | Myh14 | CE | feature_elongation,coding_sequence_variant,intron_variant |
| 7 | 49926932 | 49932060 | 5128 | Nell1 | ILNJ2,KK,NZW,PJ,SJL | feature_elongation,coding_sequence_variant,intron_variant |
| 7 | 50496858 | 50505433 | 8575 | Nell1 | WSB | feature_elongation,coding_sequence_variant,intron_variant |
| 7 | 54703098 | 54709484 | 6386 | Luzp2 | MOLF | feature_elongation,coding_sequence_variant,intron_variant |
| 7 | 55774972 | 55781611 | 6639 | Herc2 | 129S1,AJ,AKR,BALB,BTBR,BUB,C3H,C57L,CBA,DBA1J,DBA2J,FVB,MA/My,MRL,NOD,NOR,NZB,NZO,PL,RF,SEA,SWR,TH | feature_elongation,coding_sequence_variant,intron_variant |
| 7 | 56381566 | 56388934 | 7368 | Gabrg3 | 129S1,AJ,AKR,BTBR,BUB,C3H,C57L,CBA,MA/My,MRL,NOD,NOR,PL,TH | feature_elongation,coding_sequence_variant,intron_variant |
| 7 | 57064504 | 57071933 | 7429 | Gabra5 | AJ,BUB,FVB,MA/My,MRL,NZB,NZO,SMJ,SWR | feature_elongation,coding_sequence_variant,intron_variant |
| 7 | 57436969 | 57439582 | 2613 | Gabrb3 | CAST | feature_elongation,coding_sequence_variant,intron_variant |
| 7 | 66065765 | 66072930 | 7165 | Aldh1a3 | 129S1,AJ,AKR,BTBR,C3H,CBA,FVB,MA/My,MRL,NOD,NOR,NZW,PL,RF,SJL,SWR | feature_elongation,coding_sequence_variant,intron_variant,feature_elongation,5_prime_UTR_variant,intron_variant |
| 7 | 66554812 | 66561462 | 6650 | Adamts17 | BALB,C57L,CE,FVB,SEA,SWR | feature_elongation,coding_sequence_variant,intron_variant |
| 7 | 68002086 | 68009177 | 7091 | Fam169b | LP,SMJ | feature_elongation,coding_sequence_variant,intron_variant |
| 7 | 74007117 | 74013551 | 6434 | Slco3a1 | WSB | feature_elongation,coding_sequence_variant,intron_variant |
| 7 | 74850255 | 74859045 | 8790 | Sv2b | FVB,SWR,TH | feature_elongation,coding_sequence_variant,5_prime_UTR_variant,intron_variant |
| 7 | 77853132 | 77854852 | 1720 | Ntrk3 | TH | feature_elongation,coding_sequence_variant,intron_variant |
| 7 | 78122462 | 78129777 | 7315 | Ntrk3 | 129S1,C3H,CBA,DBA1J,DBA2J,KK,PJ,SEA | feature_elongation,coding_sequence_variant,intron_variant |
| 7 | 81085402 | 81091636 | 6234 | Cpeb1 | SPRET | feature_elongation,coding_sequence_variant,intron_variant |
| 7 | 82026250 | 82032521 | 6271 | Adamtsl3 | CE,MOLF,SMJ | feature_elongation,coding_sequence_variant,intron_variant |
| 7 | 86303732 | 86309080 | 5348 | Or14c40 | 129S1,BTBR,C3H,C57L,CBA,DBA1J,DBA2J,ILNJ2,KK,NZB,NZO,NZW,PJ,SEA | feature_elongation,5_prime_UTR_variant,intron_variant |
| 7 | 87126761 | 87133441 | 6680 | Tyr | 129S1,BTBR,C3H,C57L,CBA,DBA1J,DBA2J,ILNJ2,KK,NZB,NZO,PJ,SEA | feature_elongation,coding_sequence_variant,intron_variant |
| 7 | 89812385 | 89818999 | 6614 | Picalm | FVB,MA/My,NZW,SJL,SWR | feature_elongation,coding_sequence_variant,intron_variant |
| 7 | 91714292 | 91719624 | 5332 | Dlg2 | AJ | feature_elongation,coding_sequence_variant,5_prime_UTR_variant,intron_variant |
| 7 | 92019060 | 92026090 | 7030 | Dlg2 | SMJ | feature_elongation,coding_sequence_variant,intron_variant |
| 7 | 97377402 | 97382833 | 5431 | Aqp11 | NOD,NOR,SWR | feature_elongation,coding_sequence_variant,intron_variant |
| 7 | 97856119 | 97861196 | 5077 | Acer3 | CAST,WSB | feature_elongation,3_prime_UTR_variant |
| 7 | 99826704 | 99833806 | 7102 | Kcne3 | MOLF | feature_elongation,coding_sequence_variant,5_prime_UTR_variant,3_prime_UTR_variant,intron_variant |
| 7 | 99900118 | 99907320 | 7202 | Pgm2l1 | NOD,NOR,SWR | feature_elongation,coding_sequence_variant,intron_variant |
| 7 | 100022730 | 100023809 | 1079 | C2cd3 | NOD,NOR,SWR | feature_elongation,coding_sequence_variant,intron_variant |
| 7 | 100930140 | 100932015 | 1875 | Fchsd2 | 129S1,AJ,AKR,BALB,BTBR,BUB,C3H,C57L,CAST,CBA,DBA1J,DBA2J,FVB,ILNJ2,KK,MA/My,MOLF,MRL,NOD,NOR,NZB,NZO,NZW,PJ,PL,RF,RH2,SEA,SJL,SMJ,SWR,TH,WSB | feature_elongation,coding_sequence_variant,3_prime_UTR_variant,intron_variant |
| 7 | 103997139 | 104004204 | 7065 | Trim12c | CAST | stop_gained |
| 7 | 107500300 | 107506773 | 6473 | Or5p52 | SPRET | feature_elongation,coding_sequence_variant,5_prime_UTR_variant,3_prime_UTR_variant,intron_variant |
| 7 | 108645500 | 108651814 | 6314 | Ric3 | KK | feature_elongation,coding_sequence_variant,intron_variant |
| 7 | 119587555 | 119594728 | 7173 | Dnah3 | SJL,SWR | feature_elongation,coding_sequence_variant,intron_variant |

| | | | | | | |
|---|---|---|---|---|---|---|
| 7 | 121493895 | 121499290 | 5395 | Scnn1b | WSB | feature_elongation,coding_sequence_variant,5_prime_UTR_variant,intron_variant |
| 7 | 121654676 | 121660101 | 5425 | Ears2 | WSB | feature_elongation,coding_sequence_variant,intron_variant |
| 7 | 130198187 | 130204494 | 6307 | Tacc2 | RF | feature_elongation,coding_sequence_variant,intron_variant |
| 7 | 133506933 | 133513538 | 6605 | Adam12 | SPRET | feature_elongation,coding_sequence_variant,intron_variant |
| 7 | 139103466 | 139109871 | 6405 | Inpp5a | 129S1,AJ,BALB,BTBR,C3H,C57L,C58J,CE,DBA1J,DBA2J,FVB,ILNJ2,LP,MA/My,NOD,NOR,PJ,PL,RF,SEA,SJL,SMJ,SWR,TH | feature_elongation,coding_sequence_variant,intron_variant |
| 8 | 3474812 | 3481211 | 6399 | Arhgef18 | NZO,NZW | feature_elongation,coding_sequence_variant,5_prime_UTR_variant,intron_variant,feature_elongation,coding_sequence_variant,intron_variant |
| 8 | 10544479 | 10551065 | 6586 | Myo16 | MOLF | feature_elongation,coding_sequence_variant,intron_variant |
| 8 | 15951457 | 15954172 | 2715 | Csmd1 | 129S1,NZB,NZO,NZW | feature_elongation,coding_sequence_variant,intron_variant |
| 8 | 23595074 | 23597826 | 2752 | Ank1 | 129S1,AJ,AKR,B10D2,B10J,BALB,BTBR,BUB,C3H,CBA,CE,DBA1J,DBA2J,FVB,KK,LP,MA/My,MRL,NOD,NOR,NZB,PL,RF,RH2,SJL,SMJ,SWR,TH,WSB | feature_elongation,coding_sequence_variant,intron_variant |
| 8 | 34352557 | 34353701 | 1144 | Rbpms | CAST | feature_elongation,coding_sequence_variant,intron_variant |
| 8 | 34630260 | 34637376 | 7116 | Saraf | NOD,NOR | feature_elongation,coding_sequence_variant,3_prime_UTR_variant,intron_variant |
| 8 | 40813982 | 40820268 | 6286 | Micu3 | SMJ | feature_elongation,coding_sequence_variant,intron_variant |
| 8 | 46551412 | 46558770 | 7358 | Snx25 | C58J,LP | feature_elongation,coding_sequence_variant,intron_variant |
| 8 | 47025158 | 47027048 | 1890 | Cenpu | WSB | feature_elongation,coding_sequence_variant,intron_variant |
| 8 | 55496909 | 55504095 | 7186 | Gpm6a | CAST | feature_elongation,coding_sequence_variant,intron_variant |
| 8 | 58406818 | 58414053 | 7235 | Galntl6 | 129S1,AJ,CBA,KK,MA/My,NOD,NOR,NZW,RH2,SMJ | feature_elongation,coding_sequence_variant,intron_variant |
| 8 | 60964879 | 60971965 | 7086 | Aadat | CAST | feature_elongation,coding_sequence_variant,intron_variant |
| 8 | 61461051 | 61468261 | 7210 | Nek1 | C58J,LP | feature_elongation,coding_sequence_variant,intron_variant |
| 8 | 61843531 | 61848749 | 5218 | Sh3rf1 | NZW | feature_elongation,coding_sequence_variant,3_prime_UTR_variant,intron_variant |
| 8 | 62549471 | 62556501 | 7030 | Anxa10 | CE | feature_elongation,coding_sequence_variant,intron_variant |
| 8 | 68571683 | 68577720 | 6037 | Psd3 | 129S1,AJ,AKR,BALB,BUB,C3H,C57L,CBA,CE,DBA1J,DBA2J,FVB,ILNJ2,KK,MA/My,MRL,NOD,NOR,NZW,PL,RF,RH2,SJL,SMJ,SWR,TH,WSB | feature_elongation,coding_sequence_variant,5_prime_UTR_variant,intron_variant |
| 8 | 68743111 | 68749364 | 6253 | Sh2d4a | MOLF | feature_elongation,coding_sequence_variant,intron_variant |
| 8 | 71765081 | 71772517 | 7436 | Myo9b | 129S1,AKR,BALB,BUB,C3H,C57L,C58J,CBA,CE,DBA1J,DBA2J,FVB,ILNJ2,KK,MA/My,MRL,NZB,NZO,NZW,PJ,PL,RF,SEA,SJL,SMJ,SWR,TH,WSB | feature_elongation,coding_sequence_variant,intron_variant |
| 8 | 73211626 | 73213131 | 1505 | 1700030K09Rik | AJ,B10D2,B10J,BTBR,CAST,LP,NOD,NOR,RH2 | feature_elongation,coding_sequence_variant,intron_variant |
| 8 | 73235894 | 73240723 | 4829 | Slc35e1 | PJ | feature_elongation,coding_sequence_variant,3_prime_UTR_variant,intron_variant |
| 8 | 82841929 | 82848543 | 6614 | Inpp4b | 129S1,AJ,AKR,BALB,BTBR,BUB,C3H,C57L,CBA,CE,DBA1J,DBA2J,ILNJ2,KK,MA/My,MRL,NOD,NOR,NZB,NZO,NZW,PJ,PL,RF,RH2,SEA,SJL,SWR,WSB | feature_elongation,coding_sequence_variant,3_prime_UTR_variant,intron_variant |
| 8 | 84946144 | 84951502 | 5358 | Zswim4 | C3H | feature_elongation,coding_sequence_variant,intron_variant |
| 8 | 86731689 | 86738161 | 6472 | Phkb | WSB | feature_elongation,coding_sequence_variant,intron_variant |
| 8 | 86766399 | 86771115 | 4716 | Phkb | CAST | feature_elongation,coding_sequence_variant,intron_variant |
| 8 | 91975055 | 91976631 | 1576 | Rpgrip1l | SPRET | feature_elongation,coding_sequence_variant,intron_variant |
| 8 | 92167246 | 92173659 | 6413 | Fto | 129S1,AKR,BALB,BUB,C57L,CE,FVB,ILNJ2,MRL,NZB,NZO,NZW,PJ,RF,SEA,TH | feature_elongation,coding_sequence_variant,intron_variant |
| 8 | 93795963 | 93797317 | 1354 | Ces1b | SPRET | stop_gained,frameshift_variant |
| 8 | 95190121 | 95197850 | 7729 | Nlrc5 | CAST | feature_elongation,5_prime_UTR_variant,intron_variant |
| 8 | 107423550 | 107426678 | 3128 | Tango6 | PL | feature_elongation,coding_sequence_variant,intron_variant |
| 8 | 107425099 | 107426240 | 1141 | Tango6 | RF | feature_elongation,coding_sequence_variant,intron_variant |
| 8 | 123526757 | 123534042 | 7285 | Acsf3 | AJ,AKR,BALB,C3H,C58J,CE,LP,MRL,PL,RF,RH2,SEA,SMJ | feature_elongation,coding_sequence_variant,intron_variant |
| 8 | 123866983 | 123871810 | 4827 | Sult5a1 | 129S1,BTBR,C57L,MA/My,NZB,PJ,TH | feature_elongation,coding_sequence_variant,3_prime_UTR_variant,intron_variant |
| 8 | 126399158 | 126406641 | 7483 | Map10 | TH | stop_gained,frameshift_variant |
| 8 | 128186231 | 128187496 | 1265 | Pard3 | CBA,DBA1J,DBA2J,NZW | feature_elongation,coding_sequence_variant,intron_variant |
| 8 | 129158095 | 129159167 | 1072 | Nrp1 | SPRET | feature_elongation,coding_sequence_variant,intron_variant |
| 8 | 129579892 | 129586561 | 6669 | Ccdc7a | 129S1,BTBR,C57L,NZB | feature_elongation,coding_sequence_variant,intron_variant |
| 8 | 129690643 | 129692136 | 1493 | Ccdc7a | KK,MOLF | feature_elongation,coding_sequence_variant,intron_variant |
| 8 | 129831960 | 129836887 | 4927 | Ccdc7b | 129S1,BTBR,C57L,NZB | feature_elongation,coding_sequence_variant,intron_variant |
| 9 | 4378439 | 4384912 | 6473 | Msantd4 | SPRET | feature_elongation,coding_sequence_variant,5_prime_UTR_variant,intron_variant |
| 9 | 7356607 | 7357910 | 1303 | Mmp12 | CAST | feature_elongation,coding_sequence_variant,intron_variant |

| | | | | | | |
|---|---|---|---|---|---|---|
| 9 | 8620555 | 8627041 | 6486 | Trpc6 | MOLF | feature_elongation,coding_sequence_variant,intron_variant,feature_elongation,coding_sequence_variant,3_prime_UTR_variant,intron_variant |
| 9 | 24331415 | 24333308 | 1893 | Dpy19l1 | WSB | feature_elongation,coding_sequence_variant,intron_variant |
| 9 | 25458611 | 25465262 | 6651 | Eepd1 | BUB,C58J,FVB,LP,NOD,NOR,NZO,RF,SJL,SWR | feature_elongation,coding_sequence_variant,intron_variant |
| 9 | 26945299 | 26951730 | 6431 | Ncapd3 | SPRET | feature_elongation,coding_sequence_variant,intron_variant |
| 9 | 28922752 | 28929998 | 7246 | Ntm | SPRET | feature_elongation,coding_sequence_variant,intron_variant |
| 9 | 32177263 | 32178797 | 1534 | Arhgap32 | KK,MRL | feature_elongation,3_prime_UTR_variant |
| 9 | 34934094 | 34940535 | 6441 | Kirrel3 | CBA | feature_elongation,coding_sequence_variant,intron_variant |
| 9 | 35516866 | 35521241 | 4375 | Pate2 | MOLF | feature_elongation,5_prime_UTR_variant,intron_variant |
| 9 | 35589981 | 35597079 | 7098 | Pate2 | WSB | feature_elongation,coding_sequence_variant,intron_variant |
| 9 | 36344193 | 36346060 | 1867 | Pate12 | SPRET | stop_gained,frameshift_variant,splice_region_variant |
| 9 | 44461954 | 44469031 | 7077 | Cxcr5 | BTBR,NZO | feature_elongation,5_prime_UTR_variant,intron_variant |
| 9 | 49350769 | 49356022 | 5253 | Ttc12 | BTBR,ILNJ2 | feature_elongation,coding_sequence_variant,intron_variant |
| 9 | 58885449 | 58887024 | 1575 | Neo1 | MA/My,MOLF,NZB,PL | feature_elongation,coding_sequence_variant,intron_variant |
| 9 | 63232497 | 63237565 | 5068 | Map2k5 | SPRET | feature_elongation,coding_sequence_variant,intron_variant |
| 9 | 65416148 | 65421137 | 4989 | Ankdd1a | B10D2,B10J,LP,MA/My,NZW,RF | feature_elongation,coding_sequence_variant,intron_variant |
| 9 | 65763893 | 65765757 | 1864 | Trip4 | 129S1,B10D2,B10J,BTBR,BUB,C57L,KK,LP,MA/My,MRL,NOD,NOR,NZO,PJ,SEA,SMJ,SWR,TH | feature_elongation,coding_sequence_variant,intron_variant |
| 9 | 67195756 | 67201335 | 5579 | Tln2 | AJ,AKR,BALB,C3H,MRL,PL,RF,RH2 | feature_elongation,coding_sequence_variant,intron_variant |
| 9 | 67209184 | 67213618 | 4434 | Tln2 | B10D2,B10J,LP,MA/My | feature_elongation,coding_sequence_variant,intron_variant |
| 9 | 70280778 | 70287221 | 6443 | Myo1e | 129S1,BTBR,C57L,WSB | feature_elongation,coding_sequence_variant,intron_variant |
| 9 | 71426701 | 71433784 | 7083 | Myzap | AJ,BALB,BUB,C3H,C58J,CBA,CE,DBA1J,DBA2J,FVB,ILNJ2,KK,MA/My,NOD,NOR,NZB,NZO,NZW,PJ,RH2,SEA,SJL,SMJ,SWR,TH | feature_elongation,coding_sequence_variant,intron_variant |
| 9 | 71626673 | 71629099 | 2426 | Cgnl1 | 129S1,AJ,BALB,BTBR,BUB,C3H,C57L,C58J,CBA,CE,DBA1J,DBA2J,FVB,ILNJ2,KK,LP,MA/My,MOLF,NOD,NOR,NZB,NZO,NZW,PJ,RH2,SEA,SJL,SMJ,SWR,TH | feature_elongation,coding_sequence_variant,intron_variant |
| 9 | 71791407 | 71798588 | 7181 | Tcf12 | CAST | feature_elongation,coding_sequence_variant,intron_variant |
| 9 | 74039543 | 74046727 | 7184 | Wdr72 | WSB | feature_elongation,coding_sequence_variant,5_prime_UTR_variant,intron_variant |
| 9 | 75555251 | 75562386 | 7135 | Scg3 | AJ,AKR,BALB,BTBR,BUB,C3H,C58J,CBA,CE,DBA1J,DBA2J,FVB,ILNJ2,KK,LP,MA/My,MRL,NOD,NOR,NZB,NZO,NZW,PJ,PL,RF,RH2,SEA,SJL,SMJ,SWR,TH,WSB | feature_elongation,coding_sequence_variant,intron_variant |
| 9 | 76867257 | 76873416 | 6159 | Tinag | CAST | feature_elongation,coding_sequence_variant,intron_variant |
| 9 | 78003607 | 78008243 | 4636 | Fbxo9 | KK | feature_elongation,coding_sequence_variant,intron_variant |
| 9 | 80016316 | 80022756 | 6440 | Senp6 | AJ,AKR,BALB,BUB,C3H,CBA,CE,DBA1J,DBA2J,FVB,KK,MA/My,MRL,NOD,NOR,NZB,NZO,PL,RF,RH2,SEA,SJL,SWR,TH | feature_elongation,coding_sequence_variant,intron_variant |
| 9 | 86923585 | 86930290 | 6705 | Cyb5r4 | 129S1,BTBR,C57L,CE | feature_elongation,coding_sequence_variant,intron_variant |
| 9 | 87098664 | 87105354 | 6690 | Cep162 | FVB,SJL,SWR | feature_elongation,coding_sequence_variant,intron_variant |
| 9 | 87101420 | 87103193 | 1773 | Cep162 | PJ,SEA | feature_elongation,coding_sequence_variant,intron_variant |
| 9 | 99208834 | 99210698 | 1864 | Esyt3 | AJ,BALB,C3H,CBA,FVB,MOLF,NZB,PJ,RH2,SEA,SJL,SWR,WSB | feature_elongation,coding_sequence_variant,intron_variant |
| 9 | 99377180 | 99383835 | 6655 | Armc8 | CE | feature_elongation,coding_sequence_variant,intron_variant |
| 9 | 100868551 | 100870247 | 1696 | Pccb | AKR,BUB,CBA,CE,FVB,ILNJ2,NOD,NOR,PJ,PL,RF,SEA,SJL,SMJ,SWR,TH | feature_elongation,coding_sequence_variant,intron_variant |
| 9 | 101837481 | 101843536 | 6055 | Ephb1 | 129S1,AJ,AKR,BALB,BUB,C3H,CBA,DBA1J,DBA2J,ILNJ2,NOD,NOR,PL,RH2 | feature_elongation,coding_sequence_variant,intron_variant |
| 9 | 103037310 | 103038370 | 1060 | Rab6b | 129S1,AJ,AKR,BALB,BTBR,BUB,C3H,C57L,CAST,CBA,CE,FVB,ILNJ2,MA/My,MOLF,NOD,NOR,NZB,NZO,NZW,PJ,PL,RF,RH2,SEA,SJL,SMJ,SWR,TH,WSB | feature_elongation,coding_sequence_variant,intron_variant,feature_elongation,coding_sequence_variant,5_prime_UTR_variant,intron_variant |
| 9 | 103305563 | 103310343 | 4780 | Bfsp2 | MA/My,NZO | feature_elongation,coding_sequence_variant,intron_variant |
| 9 | 105198524 | 105201292 | 2768 | Nek11 | WSB | feature_elongation,coding_sequence_variant,intron_variant |
| 9 | 105222423 | 105228915 | 6492 | Nek11 | CE,FVB,SJL,SWR | feature_elongation,coding_sequence_variant,intron_variant |
| 9 | 106860606 | 106869247 | 8641 | Dock3 | 129S1,AJ,AKR,BALB,BTBR,BUB,C3H,C57L,FVB,ILNJ2,MA/My,MRL,NOD,NOR,NZB,NZO,NZW,PJ,PL,RF,RH2,SEA,SJL,SMJ,SWR,TH,WSB | feature_elongation,coding_sequence_variant,intron_variant |
| 9 | 108254919 | 108256783 | 1864 | Usp4 | NZB | feature_elongation,coding_sequence_variant,intron_variant |
| 9 | 111091866 | 111098320 | 6454 | Mlh1 | CE,PJ,SEA,WSB | feature_elongation,coding_sequence_variant,intron_variant |
| 9 | 116934080 | 116941194 | 7114 | Rbms3 | CE,WSB | feature_elongation,coding_sequence_variant,intron_variant |
| 9 | 119111977 | 119118066 | 6089 | Oxsr1 | CAST,CE | feature_elongation,coding_sequence_variant,intron_variant |
| 9 | 119623861 | 119625725 | 1864 | Scn11a | 129S1,C57L | feature_elongation,coding_sequence_variant,intron_variant |

| | | | | | | |
|---|---|---|---|---|---|---|
| 10 | 4805115 | 4811579 | 6464 | Esr1 | ILNJ2 | feature_elongation,coding_sequence_variant,intron_variant |
| 10 | 4885531 | 4891973 | 6442 | Esr1 | KK | feature_elongation,coding_sequence_variant,intron_variant |
| 10 | 4912221 | 4916604 | 4383 | Esr1 | C3H,CBA,CE,DBA1J,DBA2J,FVB,ILNJ2,LP,MA/My,MRL,NZW,PJ,SEA,SWR | feature_elongation,coding_sequence_variant,3_prime_UTR_variant,intron_variant |
| 10 | 5271250 | 5273138 | 1888 | Syne1 | NZB,NZO | feature_elongation,coding_sequence_variant,intron_variant |
| 10 | 10285780 | 10292266 | 6486 | Adgb | 129S1,DBA1J,DBA2J,FVB,MA/My,NZB,NZO,NZW,SWR,TH | feature_elongation,coding_sequence_variant,intron_variant |
| 10 | 13105639 | 13110997 | 5358 | Phactr2 | AKR,RF | feature_elongation,coding_sequence_variant,intron_variant |
| 10 | 13434194 | 13437377 | 3183 | Adat2 | MOLF | feature_elongation,coding_sequence_variant,intron_variant |
| 10 | 17687938 | 17691354 | 3416 | Txlnb | MOLF | feature_elongation,coding_sequence_variant,intron_variant |
| 10 | 19630696 | 19632587 | 1891 | Il20ra | AKR,ILNJ2,RF,WSB | feature_elongation,coding_sequence_variant,intron_variant |
| 10 | 22567249 | 22573785 | 6536 | Slc2a12 | MOLF | feature_elongation,coding_sequence_variant,intron_variant |
| 10 | 28652858 | 28659320 | 6462 | Themis | AKR,C3H,C58J,CBA,DBA1J,DBA2J,FVB,MRL,PJ,SEA,WSB | feature_elongation,coding_sequence_variant,intron_variant |
| 10 | 29054008 | 29060596 | 6588 | Mtcl3 | CAST | feature_elongation,coding_sequence_variant,intron_variant |
| 10 | 40522997 | 40524770 | 1773 | Ddo | WSB | feature_elongation,coding_sequence_variant,3_prime_UTR_variant,intron_variant |
| 10 | 50698151 | 50709310 | 11159 | Ascc3 | SPRET | feature_elongation,coding_sequence_variant,intron_variant |
| 10 | 50855108 | 50862406 | 7298 | Sim1 | TH | feature_elongation,coding_sequence_variant,3_prime_UTR_variant,intron_variant |
| 10 | 53185912 | 53192817 | 6905 | Cep85l | MOLF | feature_elongation,coding_sequence_variant,intron_variant |
| 10 | 53186360 | 53192662 | 6302 | Cep85l | SPRET | feature_elongation,coding_sequence_variant,intron_variant |
| 10 | 53190510 | 53197925 | 7415 | Cep85l | SPRET | feature_elongation,coding_sequence_variant,intron_variant |
| 10 | 53437731 | 53444632 | 6901 | Mcm9 | MOLF | feature_elongation,coding_sequence_variant,intron_variant,feature_elongation,5_prime_UTR_variant,intron_variant |
| 10 | 53790112 | 53796525 | 6413 | Man1a | KK | feature_elongation,coding_sequence_variant,intron_variant |
| 10 | 58392198 | 58399477 | 7279 | Ccdc138 | WSB | feature_elongation,coding_sequence_variant,intron_variant |
| 10 | 59182218 | 59184056 | 1838 | P4ha1 | SPRET | feature_elongation,coding_sequence_variant,intron_variant |
| 10 | 60220120 | 60225686 | 5566 | Cdh23 | AJ | feature_elongation,coding_sequence_variant,intron_variant |
| 10 | 63176997 | 63181512 | 4515 | Sirt1 | WSB | feature_elongation,5_prime_UTR_variant,intron_variant |
| 10 | 71206366 | 71211717 | 5351 | Ipmk | MRL | feature_elongation,coding_sequence_variant,intron_variant |
| 10 | 75432601 | 75439000 | 6399 | Ggt5 | NZW | feature_elongation,coding_sequence_variant,intron_variant |
| 10 | 78624154 | 78625327 | 1173 | Slc1a6 | CAST | feature_elongation,coding_sequence_variant,intron_variant |
| 10 | 79014882 | 79017847 | 2965 | Vmn2r80 | MOLF,TH | feature_elongation,coding_sequence_variant,intron_variant |
| 10 | 83234636 | 83240227 | 5591 | Nopchap1 | BUB,SMJ | feature_elongation,3_prime_UTR_variant |
| 10 | 83320368 | 83322200 | 1832 | Nopchap1 | BUB | feature_elongation,3_prime_UTR_variant |
| 10 | 85873092 | 85879523 | 6431 | Fbxo7 | BUB,CAST | feature_elongation,coding_sequence_variant,intron_variant |
| 10 | 86647925 | 86651434 | 3509 | Nt5dc3 | BUB,FVB | feature_elongation,coding_sequence_variant,intron_variant |
| 10 | 86741080 | 86742203 | 1123 | Stab2 | BUB,FVB | feature_elongation,coding_sequence_variant,intron_variant |
| 10 | 90882456 | 90884900 | 2444 | Apaf1 | AJ,AKR,BALB,C3H,C57L,CAST,CBA,CE,FVB,ILNJ2,KK,MRL,NOD,NOR,NZB,NZO,NZW,PJ,PL,RF,RH2,SEA,SMJ,SWR,TH,WSB | feature_elongation,coding_sequence_variant,intron_variant |
| 10 | 92851390 | 92852506 | 1116 | Cfap54 | WSB | feature_elongation,coding_sequence_variant,intron_variant |
| 10 | 106391638 | 106396058 | 4420 | Ppfia2 | BTBR,BUB,FVB,NZB,WSB | feature_elongation,coding_sequence_variant,intron_variant |
| 10 | 121403056 | 121411382 | 8326 | Tbk1 | DBA1J,KK,SMJ | feature_elongation,coding_sequence_variant,intron_variant |
| 10 | 121427758 | 121430249 | 2491 | Xpot | AJ,BALB,DBA2J,MA/My,RH2 | feature_elongation,coding_sequence_variant,intron_variant |
| 10 | 128911048 | 128913069 | 2021 | Or6c8 | BUB,C3H,CAST,CBA,CE,KK,MOLF,RF,SMJ | feature_elongation,3_prime_UTR_variant |
| 10 | 130326556 | 130333261 | 6705 | Vmn2r87 | CAST | feature_elongation,coding_sequence_variant,5_prime_UTR_variant,intron_variant |
| 11 | 3380781 | 3384767 | 3986 | Rnf185 | CAST | feature_elongation,coding_sequence_variant,intron_variant,feature_elongation,coding_sequence_variant,5_prime_UTR_variant,intron_variant |
| 11 | 4828982 | 4833474 | 4492 | Nipsnap1 | NZW,PJ,WSB | feature_elongation,coding_sequence_variant,intron_variant |
| 11 | 6291396 | 6292540 | 1144 | Ogdh | 129S1,AJ,AKR,BALB,BTBR,C57L,CAST,MA/My,MOLF,MRL,NOD,NZW,PL,RF,SEA,TH | feature_elongation,coding_sequence_variant,intron_variant |
| 11 | 9302337 | 9308806 | 6469 | Abca13 | CAST | feature_elongation,coding_sequence_variant,intron_variant |
| 11 | 9343024 | 9350409 | 7385 | Abca13 | CAST | feature_elongation,coding_sequence_variant,intron_variant |
| 11 | 11817131 | 11824233 | 7102 | Ddc | CAST | feature_elongation,coding_sequence_variant,intron_variant |
| 11 | 29750180 | 29755516 | 5336 | Eml6 | SPRET | feature_elongation,coding_sequence_variant,intron_variant |
| 11 | 29796158 | 29803157 | 6999 | Eml6 | FVB,SJL,SWR | feature_elongation,coding_sequence_variant,intron_variant |
| 11 | 31964491 | 31968858 | 4367 | Nsg2 | BALB,BTBR,C57L,C58J,RH2 | feature_elongation,coding_sequence_variant,intron_variant |

| | | | | | | |
|---|---|---|---|---|---|---|
| 11 | 32687011 | 32693451 | 6440 | Fbxw11 | NOD,SWR,TH | feature_elongation,coding_sequence_variant,3_prime_UTR_variant,intron_variant |
| 11 | 43120151 | 43121375 | 1224 | Atp10b | B10D2,B10J,BTBR,BUB,C58J,CE,DBA2J,ILNJ2,KK,MA/My,NZB,NZO,NZW,PJ,SEA,SJL,SMJ | feature_elongation,coding_sequence_variant,intron_variant |
| 11 | 45989071 | 45990964 | 1893 | Adam19 | 129S1,AJ,BALB,BTBR,BUB,C3H,C57L,CBA,DBA1J,DBA2J,FVB,MA/My,MRL,PJ,PL,RH2,SEA,SWR,TH | feature_elongation,coding_sequence_variant,intron_variant |
| 11 | 49014463 | 49025581 | 11118 | Or2v1 | SJL | feature_elongation,coding_sequence_variant,5_prime_UTR_variant,intron_variant |
| 11 | 49148389 | 49153849 | 5460 | Mgat1 | WSB | feature_elongation,coding_sequence_variant,5_prime_UTR_variant,3_prime_UTR_variant,intron_variant |
| 11 | 50084006 | 50090444 | 6438 | Mrnip | NZW | feature_elongation,coding_sequence_variant,intron_variant |
| 11 | 53857215 | 53862927 | 5712 | Slc22a21 | B10D2,B10J,C58J,CAST,DBA2J,KK,MOLF,NOR,NZB,NZO,NZW,PL,WSB | feature_elongation,coding_sequence_variant,intron_variant |
| 11 | 53887533 | 53892557 | 5024 | Slc22a4 | KK,MOLF,NZB,NZO | feature_elongation,coding_sequence_variant,intron_variant |
| 11 | 59315024 | 59327404 | 12380 | Snap47 | NZB,NZO,NZW | feature_elongation,coding_sequence_variant,intron_variant |
| 11 | 62301149 | 62302493 | 1344 | Ncor1 | AJ,B10D2,B10J,C3H,CBA,MA/My,MRL,PJ,RH2 | feature_elongation,coding_sequence_variant,intron_variant |
| 11 | 64940776 | 64947176 | 6400 | Arhgap44 | ILNJ2 | feature_elongation,coding_sequence_variant,intron_variant |
| 11 | 65071969 | 65078388 | 6419 | Myocd | FVB,ILNJ2,NOD,NZB,NZO,SWR | feature_elongation,coding_sequence_variant,intron_variant |
| 11 | 68089194 | 68095641 | 6447 | Stx8 | ILNJ2 | feature_elongation,coding_sequence_variant,3_prime_UTR_variant,intron_variant |
| 11 | 71067330 | 71074831 | 7501 | Nlrp1b | CE,WSB | feature_elongation,coding_sequence_variant,intron_variant |
| 11 | 73829158 | 73833468 | 4310 | Or1e17 | AJ,NZO,NZW,WSB | feature_elongation,coding_sequence_variant,5_prime_UTR_variant,3_prime_UTR_variant,intron_variant |
| 11 | 74805634 | 74812903 | 7269 | Srr | CAST | feature_elongation,5_prime_UTR_variant,intron_variant |
| 11 | 80088311 | 80089688 | 1377 | Rnf135 | WSB | feature_elongation,coding_sequence_variant,intron_variant |
| 11 | 94240465 | 94245667 | 5202 | Abcc3 | CAST | feature_elongation,coding_sequence_variant,intron_variant |
| 11 | 96562699 | 96569098 | 6399 | Skap1 | WSB | feature_elongation,coding_sequence_variant,3_prime_UTR_variant,intron_variant |
| 11 | 98518324 | 98523427 | 5103 | Gsdma3 | CAST | feature_elongation,coding_sequence_variant,5_prime_UTR_variant,intron_variant |
| 11 | 104172201 | 104173575 | 1374 | Mapt | 129S1,BTBR,C3H,C57L,DBA1J,DBA2J | feature_elongation,coding_sequence_variant,5_prime_UTR_variant,intron_variant |
| 11 | 104809410 | 104810595 | 1185 | Gm11639 | AJ,CBA | feature_elongation,coding_sequence_variant,intron_variant |
| 11 | 104815836 | 104822440 | 6604 | Gm11639 | 129S1,AJ,AKR,BTBR,C3H,C57L,CBA,DBA1J,DBA2J,ILNJ2,KK,MRL,PL,RF,RH2 | feature_elongation,coding_sequence_variant,intron_variant |
| 11 | 105743116 | 105752085 | 8969 | Tanc2 | CAST | feature_elongation,coding_sequence_variant,intron_variant |
| 11 | 117696191 | 117697259 | 1068 | 6030468B19Rik | 129S1,AJ,AKR,BTBR,BUB,C3H,C57L,C58J,CBA,CE,DBA1J,DBA2J,ILNJ2,KK,MA/My,MRL,NZB,NZO,NZW,PL,RH2,SJL,SMJ | feature_elongation,coding_sequence_variant,intron_variant |
| 11 | 120339835 | 120341723 | 1888 | Pde6g | PL | feature_elongation,coding_sequence_variant,5_prime_UTR_variant,intron_variant |
| 12 | 4389333 | 4396483 | 7150 | Ncoa1 | KK | feature_elongation,coding_sequence_variant,5_prime_UTR_variant,intron_variant |
| 12 | 4403149 | 4404403 | 1254 | Ncoa1 | CAST,MOLF | feature_elongation,5_prime_UTR_variant,intron_variant |
| 12 | 4720804 | 4727301 | 6497 | Itsn2 | CE | feature_elongation,coding_sequence_variant,intron_variant |
| 12 | 4949632 | 4955013 | 5381 | Ubxn2a | WSB | feature_elongation,coding_sequence_variant,5_prime_UTR_variant,intron_variant |
| 12 | 5033196 | 5040045 | 6849 | Atad2b | MRL | feature_elongation,coding_sequence_variant,intron_variant |
| 12 | 9056598 | 9061563 | 4965 | Wdr35 | ILNJ2 | feature_elongation,coding_sequence_variant,intron_variant |
| 12 | 11352411 | 11357722 | 5311 | Smc6 | CBA | feature_elongation,coding_sequence_variant,intron_variant |
| 12 | 13440349 | 13443952 | 3603 | Nbas | MRL | feature_elongation,coding_sequence_variant,intron_variant |
| 12 | 24571840 | 24576760 | 4920 | Taf1b | AKR,CAST,RF | feature_elongation,coding_sequence_variant,intron_variant |
| 12 | 36228248 | 36235350 | 7102 | Ankmy2 | WSB | feature_elongation,coding_sequence_variant,intron_variant |
| 12 | 51668314 | 51669597 | 1283 | Strn3 | BUB,SJL,SWR | feature_elongation,coding_sequence_variant,intron_variant |
| 12 | 53055852 | 53061396 | 5544 | Akap6 | AJ,AKR,BALB,C3H,CBA,NOD,NOR,PJ,RF,RH2,SEA | feature_elongation,coding_sequence_variant,intron_variant |
| 12 | 57694173 | 57700950 | 6777 | Ttc6 | MOLF | feature_elongation,coding_sequence_variant,intron_variant |
| 12 | 59147077 | 59154165 | 7088 | Mia2 | AJ,BALB,C3H,CBA,DBA1J,DBA2J,ILNJ2,KK,MA/My,RH2,SEA | feature_elongation,coding_sequence_variant,intron_variant |
| 12 | 59259283 | 59266376 | 7093 | Fbxo33 | BUB,FVB | feature_elongation,coding_sequence_variant,5_prime_UTR_variant,intron_variant |
| 12 | 65020078 | 65026628 | 6550 | Togaram1 | MRL | feature_elongation,coding_sequence_variant,intron_variant |
| 12 | 65134196 | 65136286 | 2090 | Fancm | CAST | feature_elongation,coding_sequence_variant,intron_variant |
| 12 | 69779239 | 69784759 | 5520 | Dmac2l | 129S1,AJ,AKR,BALB,BTBR,BUB,C3H,C57L,CAST,CBA,CE,DBA1J,DBA2J,FVB,ILNJ2,KK,MA/My,MOLF,MRL,NOD,NOR,NZB,NZO,NZW,PL,RF,RH2,SEA,SJL,SMJ,SWR,TH,WSB | feature_elongation,coding_sequence_variant,5_prime_UTR_variant,intron_variant |
| 12 | 69920999 | 69928121 | 7122 | Map4k5 | AJ,AKR,BALB,BTBR,BUB,C3H,C57L,CBA,CE,DBA1J,DBA2J,FVB,ILNJ2,KK,MA/My,NOD,NOR,NZW,PL,RF,RH2,SEA,SJL,SMJ,TH | feature_elongation,coding_sequence_variant,intron_variant |
| 12 | 72224909 | 72226963 | 2054 | Ccdc175 | CAST | feature_elongation,coding_sequence_variant,intron_variant |
| 12 | 72586554 | 72588199 | 1645 | Pcnx4 | BUB,CAST,MOLF,PJ,SEA,SMJ | feature_elongation,5_prime_UTR_variant,intron_variant |

| | | | | | | |
|---|---|---|---|---|---|---|
| 12 | 76199975 | 76206428 | 6453 | Esr2 | WSB | feature_elongation,coding_sequence_variant,intron_variant |
| 12 | 77407974 | 77414544 | 6570 | Fut8 | CAST,MOLF | feature_elongation,coding_sequence_variant,intron_variant |
| 12 | 89494564 | 89501750 | 7186 | Nrxn3 | NOD | feature_elongation,coding_sequence_variant,intron_variant |
| 12 | 101486055 | 101489068 | 3013 | Catsperb | 129S1,AKR,BALB,BTBR,BUB,C57L,C58J,CAST,CBA,CE,DBA1J,DBA2J,FVB,ILNJ2,KK,LP,MA/My,NOD,NOR,NZB,NZO,NZW,PJ,PL,RF,SEA,SJL,SMJ,SWR,TH,WSB | feature_elongation,coding_sequence_variant,intron_variant |
| 12 | 101506107 | 101512559 | 6452 | Catsperb | 129S1,AKR,BALB,BTBR,BUB,C57L,C58J,CBA,DBA1J,DBA2J,FVB,ILNJ2,KK,LP,MA/My,MOLF,NOD,NOR,NZB,NZO,NZW,PL,RF,SJL,SMJ,SWR,TH | feature_elongation,coding_sequence_variant,intron_variant |
| 12 | 105453718 | 105459031 | 5313 | D430019H16Rik | C3H | feature_elongation,coding_sequence_variant,3_prime_UTR_variant,intron_variant |
| 12 | 108149835 | 108156929 | 7094 | Ccnk | DBA1J,DBA2J,ILNJ2 | feature_elongation,coding_sequence_variant,5_prime_UTR_variant,intron_variant |
| 12 | 108279409 | 108285848 | 6439 | Hhipl1 | DBA1J,DBA2J,ILNJ2 | feature_elongation,coding_sequence_variant,intron_variant |
| 12 | 110861265 | 110863480 | 2215 | Tecpr2 | C3H,CBA,CE,WSB | feature_elongation,coding_sequence_variant,5_prime_UTR_variant,intron_variant |
| 12 | 116055986 | 116057402 | 1416 | Vipr2 | AKR,FVB,NZB,NZO,NZW,SWR,TH,WSB | feature_elongation,coding_sequence_variant,intron_variant |
| 12 | 118138657 | 118145510 | 6853 | Dnah11 | AJ | feature_elongation,coding_sequence_variant,intron_variant |
| 12 | 119295793 | 119302028 | 6235 | Macc1 | SPRET | feature_elongation,5_prime_UTR_variant,intron_variant |
| 12 | 119401311 | 119407751 | 6440 | Macc1 | MOLF | feature_elongation,coding_sequence_variant,5_prime_UTR_variant,intron_variant |
| 13 | 4103053 | 4109456 | 6403 | Akr1c14 | WSB | feature_elongation,5_prime_UTR_variant,intron_variant |
| 13 | 4556978 | 4561315 | 4337 | Akr1c20 | SPRET | feature_elongation,coding_sequence_variant,intron_variant |
| 13 | 10077777 | 10084482 | 6705 | Chrm3 | WSB | feature_elongation,5_prime_UTR_variant,intron_variant |
| 13 | 14540168 | 14546977 | 6809 | Hecw1 | BUB,FVB,NOD,NOR,SJL,SWR | feature_elongation,coding_sequence_variant,intron_variant |
| 13 | 17835846 | 17842329 | 6483 | Sugct | AJ,B10D2,B10J,BALB,BTBR,C3H,C57L,C58J,CBA,CE,DBA1J,DBA2J,KK,MA/My,MRL,NZB,PL,RH2,SMJ | feature_elongation,coding_sequence_variant,intron_variant |
| 13 | 21610900 | 21618013 | 7113 | Pgbd1 | CAST | feature_elongation,coding_sequence_variant,intron_variant |
| 13 | 23295677 | 23300988 | 5311 | Vmn1r217 | CE | feature_elongation,coding_sequence_variant,5_prime_UTR_variant,3_prime_UTR_variant,intron_variant |
| 13 | 23886935 | 23888410 | 1475 | Hfe | CAST,ILNJ2,PJ,SEA | feature_elongation,coding_sequence_variant,3_prime_UTR_variant |
| 13 | 25068868 | 25072786 | 3918 | D130043K22Rik | CAST | feature_elongation,coding_sequence_variant,intron_variant |
| 13 | 31065599 | 31072694 | 7095 | Exoc2 | WSB | feature_elongation,coding_sequence_variant,intron_variant |
| 13 | 33150681 | 33155965 | 5284 | Serpinb6b | CE | feature_elongation,coding_sequence_variant,5_prime_UTR_variant,intron_variant,feature_elongation,5_prime_UTR_variant,intron_variant |
| 13 | 37559596 | 37561484 | 1888 | Ly86 | WSB | feature_elongation,coding_sequence_variant,intron_variant |
| 13 | 40247202 | 40251885 | 4683 | Ofcc1 | WSB | feature_elongation,coding_sequence_variant,intron_variant |
| 13 | 40324587 | 40330135 | 5548 | Ofcc1 | MRL | feature_elongation,coding_sequence_variant,intron_variant |
| 13 | 41314884 | 41322442 | 7558 | Sycp2l | WSB | feature_elongation,coding_sequence_variant,intron_variant |
| 13 | 55933261 | 55940422 | 7161 | Catsper3 | BTBR,C57L | feature_elongation,coding_sequence_variant,intron_variant |
| 13 | 58170236 | 58178694 | 8458 | Klhl3 | AKR,BALB,C3H,CBA,CE,RF | feature_elongation,coding_sequence_variant,intron_variant |
| 13 | 59768965 | 59774347 | 5382 | Naa35 | BTBR,BUB,C57L,SMJ | feature_elongation,coding_sequence_variant,intron_variant |
| 13 | 63331803 | 63338929 | 7126 | Aopep | BALB | feature_elongation,coding_sequence_variant,intron_variant |
| 13 | 63500488 | 63509474 | 8986 | Fancc | BUB,DBA1J,DBA2J,ILNJ2,MA/My,NOD,NOR,SMJ | feature_elongation,coding_sequence_variant,intron_variant |
| 13 | 64925805 | 64932200 | 6395 | Cntnap3 | FVB,SWR | feature_elongation,coding_sequence_variant,intron_variant |
| 13 | 65241394 | 65243811 | 2417 | Mfsd14b | 129S1,AKR,BUB,C3H,C58J,CBA,CE,DBA1J,DBA2J,ILNJ2,KK,MOLF,MRL,NOD,NOR,NZB,NZO,NZW,PJ,PL,RF,SEA,SMJ,SWR,TH | feature_elongation,coding_sequence_variant,intron_variant |
| 13 | 65431122 | 65433014 | 1892 | Zfp369 | BTBR,C57L | feature_elongation,coding_sequence_variant,intron_variant |
| 13 | 67512180 | 67514116 | 1936 | Zfp456 | 129S1,AJ,AKR,BALB,BTBR,BUB,C3H,C57L,C58J,CBA,CE,DBA1J,DBA2J,FVB,ILNJ2,KK,MRL,NOD,NOR,NZB,NZO,NZW,PJ,PL,RF,SEA,SJL,SMJ,SWR,TH | feature_elongation,3_prime_UTR_variant |
| 13 | 67538581 | 67541868 | 3287 | Zfp429 | KK,NZB,NZO,PJ,SEA | feature_elongation,coding_sequence_variant,intron_variant,feature_elongation,coding_sequence_variant,3_prime_UTR_variant,intron_variant |
| 13 | 70933680 | 70935154 | 1474 | Adamts16 | MOLF,WSB | feature_elongation,coding_sequence_variant,intron_variant |
| 13 | 74345563 | 74348302 | 2739 | Exoc3 | ILNJ2 | feature_elongation,coding_sequence_variant,intron_variant |
| 13 | 74492034 | 74494920 | 2886 | Sdha | MOLF | feature_elongation,coding_sequence_variant,intron_variant |
| 13 | 85402195 | 85408637 | 6442 | Rasa1 | AJ,AKR,BALB,BTBR,BUB,C3H,C57L,CBA,CE,DBA1J,DBA2J,FVB,ILNJ2,KK,MRL,NOD,NOR,NZB,NZO,NZW,PJ,PL,RF,SEA,SJL,TH | feature_elongation,coding_sequence_variant,intron_variant |
| 13 | 93184906 | 93187672 | 2766 | Cmya5 | BUB | feature_elongation,coding_sequence_variant,intron_variant |
| 13 | 96928951 | 96934560 | 5609 | Ankrd31 | BTBR,C58J,CBA,DBA1J,DBA2J,MA/My,SMJ | feature_elongation,coding_sequence_variant,intron_variant |

| | | | | | | |
|---|---|---|---|---|---|---|
| 13 | 104005097 | 104006222 | 1125 | Erbin | SPRET | feature_elongation,coding_sequence_variant,intron_variant |
| 13 | 104373526 | 104380142 | 6616 | Cenpk | CAST | feature_elongation,coding_sequence_variant,intron_variant |
| 13 | 107114965 | 107120340 | 5375 | Kif2a | C3H | feature_elongation,coding_sequence_variant,intron_variant |
| 13 | 114100533 | 114105355 | 4822 | Arl15 | AJ,BUB,CAST,CE,DBA1J,FVB,KK,MA/My,NZO,PL,RF,RH2,SJL,SPRET,SWR | feature_elongation,coding_sequence_variant,intron_variant |
| 14 | 20728880 | 20734125 | 5245 | Sec24c | SEA | feature_elongation,coding_sequence_variant,intron_variant |
| 14 | 27169295 | 27170530 | 1235 | Tasor | CAST,MOLF | feature_elongation,coding_sequence_variant,intron_variant |
| 14 | 32291209 | 32292936 | 1727 | Ercc6 | CAST | feature_elongation,coding_sequence_variant,intron_variant |
| 14 | 51009114 | 51015524 | 6410 | Ttc5 | MOLF | feature_elongation,coding_sequence_variant,intron_variant |
| 14 | 51601148 | 51608919 | 7771 | Gm7247 | FVB,ILNJ2,MA/My,NOD,NOR,SJL,SMJ,SWR | feature_elongation,coding_sequence_variant,5_prime_UTR_variant,intron_variant |
| 14 | 51890143 | 51896581 | 6438 | Gm5622 | CAST | feature_elongation,coding_sequence_variant,intron_variant |
| 14 | 56889309 | 56896078 | 6769 | Parp4 | CAST | feature_elongation,coding_sequence_variant,intron_variant |
| 14 | 60262640 | 60263856 | 1216 | Atp8a2 | CAST | feature_elongation,coding_sequence_variant,intron_variant |
| 14 | 60942003 | 60949423 | 7420 | Spata13 | MOLF | feature_elongation,coding_sequence_variant,intron_variant |
| 14 | 63253809 | 63255302 | 1493 | Defb43 | CAST | feature_elongation,coding_sequence_variant,intron_variant |
| 14 | 70471225 | 70476493 | 5268 | Ppp3cc | LP | feature_elongation,coding_sequence_variant,intron_variant |
| 14 | 73268545 | 73275221 | 6676 | Cysltr2 | 129S1,C57L | feature_elongation,5_prime_UTR_variant,intron_variant |
| 14 | 73451490 | 73458577 | 7087 | Rb1 | SJL,WSB | feature_elongation,coding_sequence_variant,intron_variant |
| 14 | 75044977 | 75046121 | 1144 | Lrch1 | MOLF | stop_gained,frameshift_variant |
| 14 | 75504764 | 75506336 | 1572 | Cpb2 | SJL | feature_elongation,coding_sequence_variant,intron_variant |
| 14 | 78853423 | 78854770 | 1347 | Dgkh | 129S1,AJ,AKR,B10D2,B10J,BALB,BTBR,BUB,C3H,C57L,C58J,CBA,FVB,ILNJ2,KK,LP,MA/My,MRL,NOD,NOR,NZB,NZO,NZW,PL,RF,RH2,SEA,SJL,SMJ,SWR,TH,WSB | feature_elongation,coding_sequence_variant,intron_variant |
| 14 | 78853448 | 78855070 | 1622 | Dgkh | MOLF | feature_elongation,coding_sequence_variant,intron_variant |
| 14 | 86901279 | 86907905 | 6626 | Diaph3 | MOLF | feature_elongation,5_prime_UTR_variant,intron_variant |
| 14 | 87007888 | 87010695 | 2807 | Diaph3 | NZO | feature_elongation,coding_sequence_variant,intron_variant |
| 14 | 101697342 | 101706289 | 8947 | Tbc1d4 | WSB | feature_elongation,coding_sequence_variant,intron_variant |
| 14 | 103558971 | 103565370 | 6399 | Mycbp2 | C57L,CAST,ILNJ2,MOLF | feature_elongation,coding_sequence_variant,intron_variant |
| 14 | 115365919 | 115372381 | 6462 | Gpc5 | FVB,SJL | feature_elongation,coding_sequence_variant,intron_variant |
| 14 | 118106971 | 118108464 | 1493 | Gpc6 | 129S1,KK | feature_elongation,coding_sequence_variant,intron_variant |
| 14 | 118252902 | 118259336 | 6434 | Dct | BTBR,C57L,C58J | feature_elongation,coding_sequence_variant,intron_variant |
| 14 | 118903115 | 118908368 | 5253 | Abcc4 | C58J | feature_elongation,coding_sequence_variant,intron_variant |
| 14 | 119241895 | 119249147 | 7252 | Uggt2 | CAST | feature_elongation,coding_sequence_variant,intron_variant |
| 14 | 119304641 | 119311845 | 7204 | Uggt2 | AJ,AKR,B10D2,B10J,BALB,BUB,C3H,CBA,CE,DBA1J,DBA2J,FVB,ILNJ2,MRL,NOD,NOR,PL,RF,RH2,SEA,SJL,SMJ,SWR,WSB | feature_elongation,coding_sequence_variant,intron_variant |
| 14 | 122545427 | 122552593 | 7166 | Clybl | C3H | feature_elongation,coding_sequence_variant,intron_variant |
| 15 | 4958539 | 4960368 | 1829 | Mroh2b | SPRET | feature_elongation,coding_sequence_variant,intron_variant |
| 15 | 5057511 | 5064782 | 7271 | C7 | CAST | feature_elongation,coding_sequence_variant,intron_variant |
| 15 | 8262124 | 8265441 | 3317 | Cplane1 | LP,MOLF | feature_elongation,coding_sequence_variant,intron_variant |
| 15 | 9665856 | 9672226 | 6370 | Spef2 | MOLF | feature_elongation,coding_sequence_variant,intron_variant |
| 15 | 10290627 | 10295640 | 5013 | Prlr | MOLF | feature_elongation,5_prime_UTR_variant,intron_variant |
| 15 | 12386565 | 12392968 | 6403 | Pdzd2 | WSB | feature_elongation,coding_sequence_variant,intron_variant |
| 15 | 12831626 | 12838086 | 6460 | Drosha | BUB,CE,ILNJ2,WSB | feature_elongation,coding_sequence_variant,intron_variant |
| 15 | 16776563 | 16782663 | 6100 | Cdh9 | WSB | feature_elongation,coding_sequence_variant,5_prime_UTR_variant,intron_variant |
| 15 | 16776577 | 16780953 | 4376 | Cdh9 | 129S1,BTBR,C57L | feature_elongation,coding_sequence_variant,5_prime_UTR_variant,intron_variant |
| 15 | 18982766 | 18989843 | 7077 | Cdh10 | CE,ILNJ2 | feature_elongation,coding_sequence_variant,intron_variant |
| 15 | 18982889 | 18986383 | 3494 | Cdh10 | MOLF | feature_elongation,coding_sequence_variant,intron_variant |
| 15 | 35469070 | 35476166 | 7096 | Vps13b | CE | feature_elongation,coding_sequence_variant,intron_variant |
| 15 | 35623421 | 35626737 | 3316 | Vps13b | CAST | feature_elongation,coding_sequence_variant,intron_variant |
| 15 | 39337809 | 39344527 | 6718 | Rims2 | C3H,PL | feature_elongation,coding_sequence_variant,intron_variant |
| 15 | 42356705 | 42362972 | 6267 | Angpt1 | BTBR,ILNJ2,LP | feature_elongation,coding_sequence_variant,intron_variant |
| 15 | 44253825 | 44260923 | 7098 | Nudcd1 | 129S1,C57L,CBA,DBA1J,DBA2J,MA/My,NOD,NOR,PJ,SMJ | feature_elongation,coding_sequence_variant,intron_variant |
| 15 | 58413379 | 58415037 | 1658 | Fer1l6 | BALB,NOD,NOR,NZB,NZW,SEA | feature_elongation,coding_sequence_variant,5_prime_UTR_variant,intron_variant |

| | | | | | | |
|---|---|---|---|---|---|---|
| 15 | 59287544 | 59288567 | 1023 | Nsmce2 | CAST,MOLF | feature_elongation,coding_sequence_variant,intron_variant |
| 15 | 65812756 | 65818067 | 5311 | Hhla1 | C3H | feature_elongation,coding_sequence_variant,intron_variant |
| 15 | 66320664 | 66322023 | 1359 | Dnaaf11 | MOLF | feature_elongation,coding_sequence_variant,intron_variant |
| 15 | 67975993 | 67982395 | 6402 | Zfat | AKR,NOD,NOR,SMJ,SWR | feature_elongation,coding_sequence_variant,intron_variant |
| 15 | 71323866 | 71326564 | 2698 | Fam135b | BTBR,BUB,ILNJ2,KK,MA/My,TH | feature_elongation,coding_sequence_variant,intron_variant |
| 15 | 71728469 | 71735760 | 7291 | Col22a1 | WSB | feature_elongation,coding_sequence_variant,intron_variant |
| 15 | 76299598 | 76306037 | 6439 | Mroh1 | WSB | feature_elongation,coding_sequence_variant,intron_variant |
| 15 | 78650535 | 78656213 | 5678 | Mfng | NZB | feature_elongation,coding_sequence_variant,intron_variant |
| 15 | 79468038 | 79475148 | 7110 | Dmc1 | 129S1,AJ,AKR,BALB,C3H,C57L,C58J,CBA,CE,DBA1J,DBA2J,FVB,ILNJ2,KK,LP,MA/My,MRL,NOD,NOR,NZW,PJ,PL,SEA,SJL,SMJ,SWR,TH | feature_elongation,coding_sequence_variant,intron_variant |
| 15 | 81978546 | 81980427 | 1881 | Mei1 | FVB,MRL | feature_elongation,coding_sequence_variant,intron_variant |
| 15 | 82001267 | 82007709 | 6442 | Mei1 | MA/My | feature_elongation,coding_sequence_variant,intron_variant |
| 15 | 82062776 | 82064345 | 1569 | Srebf2 | CAST,FVB,MOLF,MRL,WSB | feature_elongation,coding_sequence_variant,intron_variant |
| 15 | 82907639 | 82914112 | 6473 | Nfam1 | 129S1,AJ,AKR,BALB,BTBR,BUB,C3H,C57L,CBA,CE,DBA1J,DBA2J,ILNJ2,KK,LP,MA/My,NOD,NOR,NZB,NZO,NZW,PJ,RF,SEA,SJL,SMJ,SWR | feature_elongation,5_prime_UTR_variant,intron_variant |
| 15 | 84839130 | 84843942 | 4812 | 5031439G07Rik | NZO | feature_elongation,coding_sequence_variant,intron_variant |
| 15 | 84971640 | 84978124 | 6484 | Smc1b | AKR,BTBR,FVB,MRL,NOD,NOR,PL,TH | feature_elongation,coding_sequence_variant,intron_variant |
| 15 | 85022609 | 85026746 | 4137 | Ribc2 | CAST | feature_elongation,coding_sequence_variant,intron_variant |
| 15 | 85149040 | 85155480 | 6440 | Fbln1 | WSB | feature_elongation,coding_sequence_variant,intron_variant |
| 15 | 87564968 | 87571419 | 6451 | Tafa5 | PJ,SEA | feature_elongation,coding_sequence_variant,intron_variant |
| 15 | 100089979 | 100091869 | 1890 | Dip2b | AJ,C3H,CE,MRL,RH2,WSB | feature_elongation,coding_sequence_variant,intron_variant |
| 15 | 103429377 | 103435417 | 6040 | Pde1b | CAST | feature_elongation,coding_sequence_variant,intron_variant |
| 16 | 13992505 | 13999826 | 7321 | Nde1 | CAST | feature_elongation,coding_sequence_variant,intron_variant |
| 16 | 14253080 | 14259522 | 6442 | Abcc1 | BALB,SEA,WSB | feature_elongation,coding_sequence_variant,intron_variant |
| 16 | 15489060 | 15490842 | 1782 | Prkdc | B10D2,B10J,BTBR,C57L,CBA,FVB,KK,PJ,RH2,SJL | feature_elongation,coding_sequence_variant,intron_variant |
| 16 | 15935948 | 15943435 | 7487 | Spidr | MOLF | feature_elongation,coding_sequence_variant,intron_variant |
| 16 | 23135326 | 23136542 | 1216 | St6gal1 | CAST | feature_elongation,5_prime_UTR_variant,intron_variant |
| 16 | 25798325 | 25803592 | 5267 | P3h2 | BUB,CE,LP | feature_elongation,coding_sequence_variant,intron_variant |
| 16 | 29061846 | 29068081 | 6235 | Atp13a5 | CAST,SMJ,WSB | feature_elongation,coding_sequence_variant,intron_variant |
| 16 | 29065824 | 29072186 | 6362 | Atp13a5 | MOLF | feature_elongation,coding_sequence_variant,intron_variant |
| 16 | 29250403 | 29257301 | 6898 | Atp13a4 | 129S1,AJ,AKR,BALB,BUB,C57L,CE,FVB,ILNJ2,KK,LP,MA/My,MRL,NZB,RF,RH2,SEA,SJL,SWR,TH | feature_elongation,coding_sequence_variant,intron_variant |
| 16 | 31905012 | 31910527 | 5515 | Pigx | NZB | feature_elongation,coding_sequence_variant,intron_variant,feature_elongation,coding_sequence_variant,5_prime_UTR_variant,intron_variant |
| 16 | 32863549 | 32870630 | 7081 | Iqcg | WSB | feature_elongation,coding_sequence_variant,intron_variant |
| 16 | 32868700 | 32871969 | 3269 | Iqcg | MOLF | feature_elongation,coding_sequence_variant,intron_variant |
| 16 | 33769116 | 33776334 | 7218 | Itgb5 | CAST | stop_gained,inframe_insertion |
| 16 | 34865144 | 34872286 | 7142 | Hacd2 | C3H,CBA,DBA1J,DBA2J | feature_elongation,coding_sequence_variant,intron_variant |
| 16 | 36431683 | 36436490 | 4807 | Cd86 | MOLF | feature_elongation,coding_sequence_variant,intron_variant |
| 16 | 36908419 | 36914774 | 6355 | Polq | CAST,SMJ,WSB | feature_elongation,coding_sequence_variant,intron_variant |
| 16 | 37104507 | 37110237 | 5730 | Stxbp5l | SPRET | feature_elongation,coding_sequence_variant,intron_variant |
| 16 | 38154924 | 38156158 | 1234 | Cfap91 | AJ,BALB,CE,DBA1J,FVB,MA/My,RH2,SEA,SWR | feature_elongation,coding_sequence_variant,intron_variant |
| 16 | 43721249 | 43727473 | 6224 | Ccdc191 | PL | feature_elongation,coding_sequence_variant,intron_variant |
| 16 | 48761176 | 48763458 | 2282 | Dzip3 | CAST | feature_elongation,coding_sequence_variant,intron_variant |
| 16 | 48952196 | 48959342 | 7146 | Myh15 | C3H,CBA | feature_elongation,coding_sequence_variant,intron_variant |
| 16 | 48994078 | 48997556 | 3478 | Myh15 | SPRET | feature_elongation,coding_sequence_variant,intron_variant |
| 16 | 49005182 | 49011435 | 6253 | Myh15 | SPRET | feature_elongation,coding_sequence_variant,intron_variant |
| 16 | 56050360 | 56055240 | 4880 | Impg2 | 129S1,AJ,BALB,DBA1J,DBA2J,MRL,RH2,SEA,SJL | feature_elongation,coding_sequence_variant,intron_variant |
| 16 | 56740684 | 56748205 | 7521 | Lnp1 | AKR,C3H,MA/My,NOD,NOR,RF,TH | feature_elongation,coding_sequence_variant,intron_variant |
| 16 | 58930404 | 58936992 | 6588 | Or5h25 | MOLF | stop_gained,inframe_insertion |
| 16 | 75679802 | 75681026 | 1224 | Samsn1 | MOLF | feature_elongation,coding_sequence_variant,intron_variant |

| | | | | | | |
|---|---|---|---|---|---|---|
| 16 | 90264555 | 90271700 | 7145 | Hunk | SEA | feature_elongation,coding_sequence_variant,intron_variant |
| 16 | 90769671 | 90772524 | 2853 | Synj1 | 129S1,AKR,C57L,ILNJ2,SMJ | feature_elongation,coding_sequence_variant,intron_variant |
| 16 | 91706700 | 91713103 | 6403 | Itsn1 | BUB,FVB,NZB,NZO,SJL,SWR | feature_elongation,coding_sequence_variant,3_prime_UTR_variant,intron_variant |
| 16 | 93937343 | 93942384 | 5041 | Hlcs | NZB | feature_elongation,coding_sequence_variant,intron_variant |
| 16 | 96171905 | 96178152 | 6247 | Igsf5 | WSB | feature_elongation,coding_sequence_variant,intron_variant |
| 17 | 12681775 | 12685093 | 3318 | Slc22a3 | PL | feature_elongation,coding_sequence_variant,intron_variant |
| 17 | 13199051 | 13200501 | 1450 | Wtap | 129S1,AJ,AKR,B10D2,BALB,C57L,C58J,DBA1J,DBA2J,LP,NZO,RH2,SEA | feature_elongation,coding_sequence_variant,intron_variant |
| 17 | 14594307 | 14600986 | 6679 | Smoc2 | NZW | feature_elongation,coding_sequence_variant,intron_variant |
| 17 | 16039026 | 16042866 | 3840 | Rgmb | MA/My,NZB | feature_elongation,coding_sequence_variant,intron_variant |
| 17 | 18292398 | 18293622 | 1224 | Vmn2r124 | SPRET | feature_elongation,coding_sequence_variant,intron_variant |
| 17 | 19910745 | 19917707 | 6962 | Vmn2r102 | 129S1,AJ,BALB,C57L,CE,DBA1J,DBA2J,KK,NZO,RH2,SEA,WSB | feature_elongation,coding_sequence_variant,3_prime_UTR_variant,intron_variant |
| 17 | 20490592 | 20498918 | 8326 | Vmn2r106 | AKR,C3H,CBA,FVB,MA/My,MRL,NZB,PJ,RF,SJL,SMJ,TH | feature_elongation,coding_sequence_variant,intron_variant |
| 17 | 20588484 | 20595679 | 7195 | Vmn2r107 | ILNJ2 | feature_elongation,coding_sequence_variant,intron_variant |
| 17 | 20903169 | 20904432 | 1263 | Vmn1r226 | SPRET | feature_elongation,5_prime_UTR_variant,intron_variant |
| 17 | 22773766 | 22780567 | 6801 | Vmn2r111 | SPRET | feature_elongation,coding_sequence_variant,intron_variant |
| 17 | 22785707 | 22790868 | 5161 | Vmn2r111 | RF | feature_elongation,coding_sequence_variant,intron_variant |
| 17 | 24949232 | 24951625 | 2393 | Rpl3l | FVB,SJL | feature_elongation,coding_sequence_variant,intron_variant |
| 17 | 30806457 | 30812006 | 5549 | Glo1 | BTBR | feature_elongation,3_prime_UTR_variant |
| 17 | 30993144 | 30996142 | 2998 | Dnah8 | SMJ | feature_elongation,coding_sequence_variant,intron_variant |
| 17 | 31745172 | 31746422 | 1250 | Ndufv3 | FVB,SJL | feature_elongation,coding_sequence_variant,intron_variant |
| 17 | 32548706 | 32554254 | 5548 | Akap8l | NZB | feature_elongation,coding_sequence_variant,intron_variant |
| 17 | 32550639 | 32551950 | 1311 | Akap8l | MA/My,SJL | feature_elongation,coding_sequence_variant,intron_variant |
| 17 | 34930571 | 34932804 | 2233 | Tnxb | SPRET | stop_gained,frameshift_variant |
| 17 | 35120933 | 35122077 | 1144 | Ehmt2 | ILNJ2 | feature_elongation,coding_sequence_variant,intron_variant |
| 17 | 37542613 | 37549499 | 6886 | Or1o2 | PJ | stop_gained,frameshift_variant |
| 17 | 41221508 | 41223462 | 1954 | Glyatl3 | BALB,DBA2J,RH2,SEA | feature_elongation,coding_sequence_variant,intron_variant |
| 17 | 42902282 | 42909505 | 7223 | Opn5 | BALB,BUB,DBA1J,DBA2J,PJ,RH2,SEA,SJL,SWR,WSB | feature_elongation,coding_sequence_variant,intron_variant |
| 17 | 44011717 | 44017930 | 6213 | Cyp39a1 | 129S1,AJ,BALB,BTBR,C57L,C58J,DBA2J,ILNJ2,KK,LP,MRL,NOD,NOR,NZB,NZO,NZW,PJ,PL,RH2,SEA,SJL,TH | feature_elongation,coding_sequence_variant,intron_variant |
| 17 | 46433248 | 46439649 | 6401 | Mrps18a | BTBR,C57L,LP | feature_elongation,coding_sequence_variant,3_prime_UTR_variant,intron_variant |
| 17 | 49768357 | 49771135 | 2778 | Daam2 | MOLF | feature_elongation,coding_sequence_variant,intron_variant |
| 17 | 57713452 | 57715253 | 1801 | Adgre1 | CAST,MOLF | feature_elongation,coding_sequence_variant,intron_variant |
| 17 | 66229189 | 66236300 | 7111 | Twsg1 | AJ,BALB,BTBR,BUB,C58J,CE,DBA1J,DBA2J,FVB,KK,NOD,NOR,NZB,NZW,PJ,PL,RH2,SEA,SJL,SMJ,TH | feature_elongation,coding_sequence_variant,3_prime_UTR_variant,intron_variant |
| 17 | 68727558 | 68734842 | 7284 | L3mbtl4 | C3H,CBA,MRL | feature_elongation,coding_sequence_variant,intron_variant |
| 17 | 71704897 | 71706791 | 1894 | Smchd1 | MA/My,NZB,NZO | feature_elongation,coding_sequence_variant,intron_variant |
| 17 | 72253489 | 72259892 | 6403 | Alk | C3H,CBA,CE,DBA1J,DBA2J,MA/My,MRL,NZB,NZO | feature_elongation,coding_sequence_variant,intron_variant |
| 17 | 75133846 | 75140289 | 6443 | Ttc27 | AKR,MA/My,NZB,NZO | feature_elongation,coding_sequence_variant,intron_variant |
| 17 | 88870919 | 88876224 | 5305 | Ppp1r21 | C3H | feature_elongation,coding_sequence_variant,intron_variant |
| 18 | 11866036 | 11868620 | 2584 | Rbbp8 | SPRET | feature_elongation,coding_sequence_variant,intron_variant |
| 18 | 14888731 | 14895173 | 6442 | Psma8 | MA/My | feature_elongation,coding_sequence_variant,3_prime_UTR_variant,intron_variant |
| 18 | 22234362 | 22241286 | 6924 | Ccdc178 | SJL | feature_elongation,coding_sequence_variant,intron_variant |
| 18 | 22236981 | 22242498 | 5517 | Ccdc178 | CAST,CE,MOLF | feature_elongation,coding_sequence_variant,intron_variant |
| 18 | 34053147 | 34055032 | 1885 | Epb41l4a | WSB | feature_elongation,coding_sequence_variant,intron_variant |
| 18 | 42679198 | 42681846 | 2648 | Tcerg1 | 129S1,AJ,AKR,BALB,BTBR,BUB,C3H,C57L,C58J,CBA,CE,DBA1J,DBA2J,FVB,KK,MA/My,MRL,NOD,NOR,NZW,PJ,PL,RF,RH2,SEA,SJL,SMJ,SWR,TH,WSB | feature_elongation,coding_sequence_variant,intron_variant |
| 18 | 43441860 | 43448263 | 6403 | Stk32a | KK | feature_elongation,coding_sequence_variant,3_prime_UTR_variant,intron_variant |
| 18 | 44971844 | 44978510 | 6666 | Ythdc2 | 129S1,BTBR,C57L,FVB,ILNJ2,MA/My,NZB,NZW,PJ,SWR | feature_elongation,coding_sequence_variant,intron_variant |
| 18 | 46716685 | 46723084 | 6399 | Tmed7 | C58J,LP,NZO,NZW | feature_elongation,coding_sequence_variant,intron_variant |
| 18 | 54044604 | 54051073 | 6469 | Csnk1g3 | 129S1,C57L,WSB | feature_elongation,coding_sequence_variant,intron_variant |
| 18 | 60940102 | 60941340 | 1238 | Cd74 | AJ,BALB,CBA,ILNJ2,RH2,SEA,SMJ | feature_elongation,coding_sequence_variant,intron_variant |

| | | | | | | |
|---|---|---|---|---|---|---|
| 18 | 61656199 | 61658087 | 1888 | Arhgef37 | AJ,AKR,BALB,BUB,CBA,ILNJ2,MA/My,MRL,NOD,NOR,PL,RF,RH2,SEA | feature_elongation,coding_sequence_variant,5_prime_UTR_variant,intron_variant |
| 18 | 62559001 | 62562031 | 3030 | Htr4 | SMJ,SWR | feature_elongation,coding_sequence_variant,intron_variant |
| 18 | 63147047 | 63153489 | 6442 | Piezo2 | AJ,AKR,BALB,BUB,CBA,ILNJ2,MA/My,MRL,NOD,NOR,PL,RF,RH2,SEA,WSB | feature_elongation,coding_sequence_variant,intron_variant |
| 18 | 63955699 | 63958500 | 2801 | Wdr7 | AJ,BALB,C3H,CBA,ILNJ2,RH2 | feature_elongation,coding_sequence_variant,intron_variant |
| 18 | 80886501 | 80887835 | 1334 | Atp9b | NOD,NOR,SJL | feature_elongation,coding_sequence_variant,intron_variant |
| 18 | 89211224 | 89217704 | 6480 | Cd226 | FVB,KK,TH | feature_elongation,5_prime_UTR_variant,intron_variant |
| 19 | 3560265 | 3562212 | 1947 | Ppp6r3 | WSB | feature_elongation,coding_sequence_variant,intron_variant |
| 19 | 9973915 | 9975034 | 1119 | Best1 | 129S1,AKR,B10D2,B10J,BTBR,C57L,DBA1J,DBA2J,FVB,MA/My,NOD,NOR,NZB,NZO,NZW,PJ,PL,RF,SJL,SMJ,SWR | feature_elongation,coding_sequence_variant,5_prime_UTR_variant,intron_variant |
| 19 | 11160491 | 11162381 | 1890 | Ms4a13 | TH | feature_elongation,coding_sequence_variant,intron_variant |
| 19 | 11569669 | 11571181 | 1512 | Ms4a6d | AJ,BALB,CE,LP,NOD,NOR,PJ,PL,RH2,SEA,WSB | feature_elongation,coding_sequence_variant,intron_variant |
| 19 | 11830868 | 11838058 | 7190 | Or10v9 | WSB | start_lost,feature_elongation,start_retained_variant,coding_sequence_variant,5_prime_UTR_variant,3_prime_UTR_variant,intron_variant |
| 19 | 12182729 | 12189229 | 6500 | Or5an9 | MOLF | feature_elongation,coding_sequence_variant,5_prime_UTR_variant,3_prime_UTR_variant,intron_variant |
| 19 | 12621225 | 12625245 | 4020 | Glyat | CAST | feature_elongation,coding_sequence_variant,intron_variant,feature_elongation,coding_sequence_variant,5_prime_UTR_variant,intron_variant |
| 19 | 12894781 | 12900798 | 6017 | Or5b12 | MOLF | feature_elongation,coding_sequence_variant,5_prime_UTR_variant,3_prime_UTR_variant,intron_variant |
| 19 | 29004271 | 29009829 | 5558 | Ak3 | AKR,BUB,MRL | feature_elongation,coding_sequence_variant,intron_variant |
| 19 | 29013709 | 29016566 | 2857 | Ak3 | SPRET | feature_elongation,coding_sequence_variant,intron_variant |
| 19 | 29928466 | 29935063 | 6597 | Il33 | CAST | feature_elongation,coding_sequence_variant,intron_variant |
| 19 | 30560616 | 30562598 | 1982 | Prkg1 | 129S1,BTBR,C57L,C58J,NZW | feature_elongation,coding_sequence_variant,intron_variant |
| 19 | 37560109 | 37561529 | 1420 | Exoc6 | B10D2,B10J,BTBR,C58J,LP,MA/My,WSB | feature_elongation,coding_sequence_variant,intron_variant |
| 19 | 37936898 | 37943339 | 6441 | Myof | NZW | feature_elongation,coding_sequence_variant,intron_variant |
| 19 | 42973009 | 42979700 | 6691 | Hpse2 | C3H,CBA,ILNJ2,NZB,NZO,NZW,WSB | feature_elongation,coding_sequence_variant,intron_variant |
| 19 | 50237087 | 50243693 | 6606 | Sorcs1 | NZB,NZO,NZW,RF | feature_elongation,coding_sequence_variant,intron_variant |
| 19 | 52920575 | 52921890 | 1315 | Xpnpep1 | NZB,NZW,WSB | feature_elongation,coding_sequence_variant,3_prime_UTR_variant,intron_variant |
| 19 | 57442463 | 57449600 | 7137 | Trub1 | AJ,RH2 | feature_elongation,coding_sequence_variant,intron_variant |
| X | 6432543 | 6437856 | 5313 | Shroom4 | C3H | feature_elongation,coding_sequence_variant,intron_variant |
| X | 7770740 | 7774174 | 3434 | Timm17b | PL | feature_elongation,coding_sequence_variant,3_prime_UTR_variant,intron_variant |
| X | 13005400 | 13007404 | 2004 | Usp9x | CAST | feature_elongation,coding_sequence_variant,intron_variant |
| X | 23597124 | 23600612 | 3488 | Wdr44 | MOLF | feature_elongation,coding_sequence_variant,intron_variant |
| X | 52791843 | 52798193 | 6350 | Xlr | CAST | feature_elongation,coding_sequence_variant,5_prime_UTR_variant,intron_variant |
| X | 59317284 | 59323264 | 5980 | Atp11c | SPRET | feature_elongation,coding_sequence_variant,intron_variant |
| X | 70333844 | 70337164 | 3320 | Mtm1 | WSB | feature_elongation,coding_sequence_variant,intron_variant |
| X | 73202453 | 73205783 | 3330 | Tex28 | CE | feature_elongation,coding_sequence_variant,intron_variant |
| X | 74335118 | 74341310 | 6192 | F8 | MOLF | feature_elongation,coding_sequence_variant,intron_variant |
| X | 74374387 | 74381057 | 6670 | F8 | CAST | feature_elongation,coding_sequence_variant,intron_variant |
| X | 74392277 | 74399063 | 6786 | F8 | SPRET | feature_elongation,coding_sequence_variant,intron_variant |
| X | 76812714 | 76815188 | 2474 | Prkx | CE | feature_elongation,coding_sequence_variant,intron_variant |
| X | 76829470 | 76836581 | 7111 | Prkx | CE | feature_elongation,coding_sequence_variant,intron_variant |
| X | 78499748 | 78506212 | 6464 | Cfap47 | MOLF | feature_elongation,coding_sequence_variant,intron_variant |
| X | 82667019 | 82672196 | 5177 | Dmd | CAST | feature_elongation,coding_sequence_variant,intron_variant |
| X | 82918179 | 82924682 | 6503 | Dmd | C58J | feature_elongation,coding_sequence_variant,intron_variant |
| X | 82948980 | 82951502 | 2522 | Dmd | MOLF | feature_elongation,coding_sequence_variant,intron_variant |
| X | 83344454 | 83355356 | 10902 | Dmd | CAST,MOLF | feature_elongation,coding_sequence_variant,intron_variant |
| X | 84229917 | 84236441 | 6524 | Dmd | 129S1,BUB,C57L,DBA1J,DBA2J,PJ | feature_elongation,coding_sequence_variant,intron_variant |
| X | 95536795 | 95544131 | 7336 | Heph | SPRET | feature_elongation,coding_sequence_variant,intron_variant |
| X | 99467527 | 99474370 | 6843 | Awat2 | MOLF | feature_elongation,5_prime_UTR_variant,intron_variant |
| X | 99972202 | 99978707 | 6505 | Tex11 | SPRET | feature_elongation,coding_sequence_variant,intron_variant |
| X | 101125174 | 101127016 | 1842 | Nhsl2 | SPRET | feature_elongation,coding_sequence_variant,3_prime_UTR_variant,intron_variant |
| X | 103587601 | 103589485 | 1884 | Zdhhc15 | SPRET | feature_elongation,coding_sequence_variant,3_prime_UTR_variant,intron_variant |

| | | | | | | |
|---|---|---|---|---|---|---|
| X | 105509044 | 105516566 | 7522 | Fndc3c1 | WSB | feature_elongation,coding_sequence_variant,intron_variant |
| X | 105715147 | 105721461 | 6314 | Gm5127 | MOLF | feature_elongation,5_prime_UTR_variant,intron_variant |
| X | 128720198 | 128727131 | 6933 | Diaph2 | CAST | feature_elongation,coding_sequence_variant,intron_variant |
| X | 134695760 | 134697648 | 1888 | Gprasp1 | 129S1,AKR,BTBR,BUB,C3H,C57L,C58J,CBA,CE,DBA1J,DBA2J,LP,PJ,RF,SEA,SMJ | feature_elongation,5_prime_UTR_variant,intron_variant |
| X | 135366553 | 135373218 | 6665 | Kir3dl2 | WSB | feature_elongation,coding_sequence_variant,5_prime_UTR_variant,intron_variant |
| X | 135412343 | 135419013 | 6670 | Kir3dl2 | KK | feature_elongation,5_prime_UTR_variant,intron_variant |
| X | 136552053 | 136556105 | 4052 | Il1rapl2 | MOLF | feature_elongation,coding_sequence_variant,5_prime_UTR_variant,intron_variant |
| X | 140409210 | 140415591 | 6381 | Col4a5 | CE,FVB,NOD,NOR,TH | feature_elongation,coding_sequence_variant,intron_variant |
| X | 147911202 | 147914137 | 2935 | Ott | MOLF,MRL,SJL | feature_elongation,coding_sequence_variant,intron_variant |
| X | 149424673 | 149425838 | 1165 | Pfkfb1 | CAST,MOLF | feature_elongation,coding_sequence_variant,intron_variant |
| X | 150644747 | 150647160 | 2413 | Huwe1 | AJ,BALB,CAST,FVB,KK,MA/My,MOLF,MRL,NOD,NOR,NZB,RH2,SEA,SJL,SMJ,SWR,TH,WSB | feature_elongation,coding_sequence_variant,intron_variant |
| X | 155084689 | 155089187 | 4498 | Gm15155 | AJ,BALB,MA/My,NOD,NOR,NZB,NZW,RH2,SEA,TH | feature_elongation,coding_sequence_variant,intron_variant |
| X | 158123758 | 158127414 | 3656 | Rps6ka3 | MOLF | feature_elongation,coding_sequence_variant,intron_variant |
| X | 158130962 | 158134072 | 3110 | Rps6ka3 | CAST | feature_elongation,coding_sequence_variant,intron_variant |
| X | 159215032 | 159220771 | 5739 | Adgrg2 | BALB,CE,MRL,NOD,NOR,SEA | feature_elongation,5_prime_UTR_variant,intron_variant |
| X | 159452603 | 159459190 | 6587 | Ppef1 | MOLF | feature_elongation,coding_sequence_variant,intron_variant |
| X | 161578643 | 161583648 | 5005 | Txlng | WSB | feature_elongation,coding_sequence_variant,intron_variant |
| X | 161700620 | 161705839 | 5219 | Ctps2 | AKR,BALB,RF,SEA | feature_elongation,coding_sequence_variant,intron_variant |
| X | 165191983 | 165198365 | 6382 | Ofd1 | MOLF | feature_elongation,coding_sequence_variant,intron_variant |
| X | 165227230 | 165232778 | 5548 | Trappc2 | TH | feature_elongation,coding_sequence_variant,5_prime_UTR_variant,intron_variant |
| X | 165341103 | 165347580 | 6477 | Egfl6 | WSB | feature_elongation,coding_sequence_variant,intron_variant |

**Table S9. Lists of INVs present in all 39 or in the 35 classical inbred strains.** The chromosome, starting and ending position, size, gene symbol, strains with the variant allele, and the predicted effect for each INV are shown.

| CHR | START | END | SIZE | SYMBOL | STRAINS with VARIANT SV ALLELE | CONSEQUENCE |
|---|---|---|---|---|---|---|
| 1 | 8577513 | 8577872 | 359 | Sntg1 | 129S1,AJ,AKR,BALB,BTBR,C3H,C57L,CBA,DBA1J,DBA2J,FVB,ILNJ2,KK,LP,MRL,NOD,NOR,NZO,PJ,PL,RF,RH2,SEA,SJL,SMJ,SWR,TH | intron_variant |
| 1 | 9101845 | 9102131 | 286 | Sntg1 | 129S1,AKR,BTBR,C57L,ILNJ2,LP,NOD,NOR,PJ,PL,SMJ,SWR,TH | intron_variant |
| 1 | 19418764 | 19419319 | 555 | - | PL | intergenic_variant |
| 1 | 31755029 | 31755830 | 801 | - | CE | intergenic_variant |
| 1 | 40188818 | 40189757 | 939 | - | 129S1,BTBR,C57L,NZB,NZO | intergenic_variant |
| 1 | 40606335 | 40607020 | 685 | - | C3H,KK,NZB,PJ,TH | intergenic_variant |
| 1 | 45437972 | 45438382 | 410 | Col5a2 | FVB,MA/My,NOD,NOR,RH2 | intron_variant |
| 1 | 47013665 | 47014006 | 341 | - | AKR,BUB,C58J,RF | intergenic_variant |
| 1 | 48811258 | 48811787 | 529 | - | AKR,BALB,BUB,C58J,MA/My,RF,RH2,SEA | intergenic_variant |
| 1 | 54698971 | 54699418 | 447 | Ankrd44 | NZO,NZW | intron_variant |
| 1 | 55525702 | 55526022 | 320 | Plcl1 | AKR,BALB,BUB,MRL,RH2,SEA,SJL | intron_variant |
| 1 | 66831537 | 66831785 | 248 | Kansl1l | AJ,CBA,DBA1J,NZW,SWR | intron_variant |
| 1 | 69286163 | 69286747 | 584 | - | 129S1,AJ,BTBR,C3H,C57L,DBA2J,FVB,KK,LP,MA/My,MRL,NOD,NOR,NZB,NZO,NZW,SMJ,TH | regulatory_region_variant,intergenic_variant |
| 1 | 94043886 | 94044472 | 586 | - | NZO,PL | intergenic_variant |
| 1 | 99837338 | 99838052 | 714 | Cntnap5b | 129S1,AKR,BUB,C57L,DBA1J,DBA2J,LP,MA/My,NOD,NOR,NZO,PJ,SEA,SJL,TH | intron_variant |
| 1 | 150074280 | 150075235 | 955 | - | BTBR,SMJ | intergenic_variant |
| 1 | 170442569 | 170443368 | 799 | - | 129S1,AJ,BTBR,BUB,C3H,C57L,CE,FVB,KK,MRL,NOD,NOR,NZB,NZO,NZW,PJ,PL,RF,RH2,SEA,SJL,SMJ,SWR,TH | intergenic_variant |
| 1 | 171089657 | 171090107 | 450 | - | CBA,DBA1J,DBA2J,KK,NZW,TH | regulatory_region_variant,intergenic_variant |
| 1 | 175205685 | 175206314 | 629 | Rgs7,5730408A14Rik | 129S1,AJ,AKR,BALB,BUB,C3H,C57L,C58J,CBA,CE,DBA1J,DBA2J,FVB,ILNJ2,KK,MRL,NOD,NOR,NZB,NZW,PJ,RF,RH2,SEA,SJL,SMJ,SWR | intron_variant,downstream_gene_variant |
| 1 | 177534659 | 177534938 | 279 | - | FVB,NZO | intergenic_variant |
| 1 | 186587760 | 186588145 | 385 | - | 129S1,AKR,BALB,BTBR,C3H,C57L,CBA,CE,DBA1J,DBA2J,ILNJ2,KK,MRL,NZW,PJ,PL,RH2,SEA,SMJ | regulatory_region_variant,intergenic_variant |
| 1 | 191319489 | 191319816 | 327 | Ints7 | TH | intron_variant,regulatory_region_variant |
| 1 | 194871917 | 194872462 | 545 | - | NZO,NZW,RF | intergenic_variant |
| 10 | 18098448 | 18098730 | 282 | Ccdc28a | AJ,BALB,BTBR,C3H,C57L,CBA,CE,DBA1J,DBA2J,MA/My,MRL,PL,RH2,SJL,SMJ | intron_variant |
| 10 | 30182071 | 30182918 | 847 | - | MA/My | intergenic_variant |
| 10 | 48647679 | 48648014 | 335 | - | BALB,BTBR,KK,NZW,PJ,SEA,TH | intergenic_variant |
| 10 | 88660071 | 88660452 | 381 | Utp20 | BUB | intron_variant |
| 10 | 93389643 | 93390278 | 635 | Ccdc38 | 129S1,AJ,AKR,BALB,BTBR,BUB,C3H,C57L,C58J,CBA,CE,DBA1J,DBA2J,FVB,ILNJ2,KK,LP,MA/My,MRL,NOD,NOR,NZO,NZW,PJ,PL,RF,RH2,SEA,SJL,SMJ,SWR,TH | intron_variant,upstream_gene_variant |
| 10 | 95297336 | 95297957 | 621 | Gm47599 | PL | upstream_gene_variant |
| 10 | 100074939 | 100075857 | 918 | - | AKR,BUB,C3H,CBA,FVB,MA/My,NOD,NOR,PL,SMJ | intergenic_variant |
| 10 | 100666965 | 100667209 | 244 | - | CBA,NZB | intergenic_variant |
| 10 | 104626702 | 104627499 | 797 | - | AJ,BALB,BTBR,CE,DBA1J,DBA2J,MRL,NZB,PJ,RH2,SEA,SJL | intergenic_variant |
| 10 | 106497875 | 106498335 | 460 | Ppfia2 | BTBR,NZB | intron_variant |
| 10 | 110112347 | 110112819 | 472 | Nav3 | SMJ | intron_variant |
| 10 | 112959440 | 112960375 | 935 | - | MA/My | intergenic_variant |
| 10 | 126340151 | 126340751 | 600 | - | AJ,BALB,ILNJ2,KK,LP,MA/My,NZW,PJ,RF,RH2,SEA | intergenic_variant |
| 11 | 4419801 | 4420791 | 990 | - | AKR,B10D2,B10J,C58J,CE,FVB,KK,MRL,RF,SJL,SWR | intergenic_variant |
| 11 | 8257235 | 8257744 | 509 | - | B10D2,B10J,BUB,C3H,C58J,CE,NOR,NZB,PJ,SJL,SWR | intergenic_variant |
| 11 | 16698481 | 16698851 | 370 | - | SEA | intergenic_variant |
| 11 | 18130963 | 18131477 | 514 | - | 129S1,AJ,AKR,BALB,BTBR,BUB,C3H,C57L,C58J,CBA,CE,DBA1J,DBA2J,FVB,ILNJ2,KK,MA/My,MRL,NOD,NOR,NZB,NZO,NZW,PJ,PL,RF,SEA,SJL,SMJ,SWR,TH | intergenic_variant |
| 11 | 31997466 | 31998363 | 897 | Nsg2 | AKR,BALB,BTBR,BUB,C3H,C57L,C58J,CE,FVB,ILNJ2,KK,MRL,NOD,NZO,PJ,PL,RH2,SJL,SMJ,SWR | intron_variant |
| 11 | 46848246 | 46848819 | 573 | Sgcd | B10J,C58J,NZB,PJ,SEA | intron_variant |
| 11 | 49089014 | 49089909 | 895 | - | AKR,B10D2,B10J,BUB,C58J,CE,KK,MA/My,NOR,NZO,NZW,RF,SJL,SWR,TH | intergenic_variant |
| 11 | 56609432 | 56609720 | 288 | - | CE | intergenic_variant |
| 11 | 59844941 | 59845664 | 723 | Med9 | AJ,C3H,CBA,MRL,PJ,SEA | intron_variant |
| 11 | 65438162 | 65438483 | 321 | - | CE,LP,NOD,SWR | intergenic_variant |
| 11 | 68577950 | 68578291 | 341 | - | SEA | intergenic_variant |
| 11 | 72988202 | 72988987 | 785 | Itgae | TH | intron_variant |
| 11 | 92017998 | 92018252 | 254 | - | BUB | intergenic_variant |
| 11 | 95799329 | 95799730 | 401 | B4galnt2 | AJ,AKR,BUB,C3H,CBA,KK,MA/My,MRL,RF,RH2,SWR,TH | intron_variant |
| 11 | 103805034 | 103805300 | 266 | Nsf | AJ,CBA | intron_variant |
| 11 | 115632448 | 115633298 | 850 | - | AKR,MRL,RH2,SMJ | intergenic_variant |
| 12 | 7692313 | 7693109 | 796 | Gm32828 | CE,ILNJ2,MRL,NZW | intron_variant,non_coding_transcript_variant |
| 12 | 25652919 | 25653279 | 360 | - | C57L,NZW | intergenic_variant |
| 12 | 44995009 | 44995251 | 242 | Stxbp6 | BALB | intron_variant,regulatory_region_variant |
| 12 | 45836212 | 45836426 | 214 | - | KK,NZO,NZW | intergenic_variant |
| 12 | 76740992 | 76741176 | 184 | Sptb | 129S1 | intron_variant |
| 12 | 92592046 | 92592406 | 360 | - | SMJ | intergenic_variant |
| 12 | 116025469 | 116026117 | 648 | Zfp386 | DBA1J,DBA2J,RF | 3_prime_UTR_variant |
| 13 | 6009238 | 6010005 | 767 | Gm46401 | AKR,BUB,MA/My,NOD,NOR,RF | intron_variant,non_coding_transcript_variant |
| 13 | 16559807 | 16560560 | 753 | Gm48493 | AKR,PJ,RF,SEA | non_coding_transcript_exon_variant |
| 13 | 18721547 | 18722363 | 816 | Pou6f2 | AKR,ILNJ2,LP,PJ,RF,SEA | intron_variant |

| | | | | | | |
|---|---|---|---|---|---|---|
| 13 | 22792216 | 22792925 | 709 | - | CE | intergenic_variant |
| 13 | 35273126 | 35273492 | 366 | - | AJ,AKR,B10D2,B10J,BALB,BTBR,C3H,C57L,C58J,CBA,DBA1J,DBA2J,ILNJ2,KK,MA/My,MRL,NZB,NZO,NZW,PL,RF,RH2,SMJ,TH | intergenic_variant |
| 13 | 58062240 | 58062870 | 630 | - | BUB,SMJ,SWR | intergenic_variant |
| 13 | 97627458 | 97628368 | 910 | Lncenc1 | AKR | intron_variant,non_coding_transcript_variant,non_coding_transcript_exon_variant |
| 13 | 98478960 | 98479625 | 665 | - | AKR,C58J,CBA,DBA1J,DBA2J,MA/My,PL,SMJ | intergenic_variant |
| 13 | 105591747 | 105592370 | 623 | - | CE,MA/My,NZO | intergenic_variant |
| 13 | 106035168 | 106035957 | 789 | Gm10739 | CE,MA/My,NZO | intron_variant,non_coding_transcript_variant |
| 13 | 106448509 | 106449237 | 728 | - | NZB | intergenic_variant |
| 13 | 111491923 | 111492492 | 569 | - | KK | intergenic_variant |
| 13 | 111699760 | 111700007 | 247 | Gm15288 | MA/My,NZB | splice_polypyrimidine_tract_variant,intron_variant,non_coding_transcript_variant |
| 13 | 113480947 | 113481490 | 543 | Cspg4b | MRL | intron_variant |
| 14 | 6099128 | 6099875 | 747 | Top2b | AJ,AKR,BTBR,BUB,C58J,CE,ILNJ2,NOD,NOR,RF,SMJ | intron_variant |
| 14 | 36216813 | 36217108 | 295 | - | 129S1,C57L | intergenic_variant |
| 14 | 56432657 | 56432862 | 205 | Gm4022 | 129S1,AJ,AKR,B10D2,B10J,BALB,C3H,C57L,C58J,CBA,CE,DBA1J,FVB,KK,LP,MA/My,MRL,NOD,NOR,NZW,PJ,RF,SEA,SWR | intron_variant,non_coding_transcript_variant |
| 14 | 83634414 | 83635270 | 856 | - | 129S1,BUB,C57L,C58J,ILNJ2,KK,NOD,NOR,NZW,SJL | intergenic_variant |
| 14 | 86041295 | 86041712 | 417 | - | C57L,C58J | intergenic_variant |
| 14 | 86521552 | 86522109 | 557 | 4930529K09Rik,Gm32815 | AJ,AKR,BALB,BTBR,BUB,C3H,CBA,DBA1J,DBA2J,FVB,NOD,NZO,PJ,RF,SEA,SJL,SMJ,SWR,TH | downstream_gene_variant,intron_variant,non_coding_transcript_variant |
| 14 | 89990467 | 89991343 | 876 | - | PJ | intergenic_variant |
| 14 | 101639151 | 101639848 | 697 | - | 129S1,AKR,BALB,BTBR,C3H,C57L,C58J,DBA1J,DBA2J,FVB,ILNJ2,KK,MA/My,NZB,NZO,PJ,RF,SEA,SJL,SMJ,SWR,TH | intergenic_variant |
| 14 | 104744106 | 104744630 | 524 | Obi1 | FVB | intron_variant |
| 14 | 116440842 | 116441402 | 560 | Gpc5 | KK | intron_variant,regulatory_region_variant |
| 14 | 117896199 | 117896436 | 237 | Gpc6 | PJ | intron_variant |
| 15 | 10146041 | 10146295 | 254 | - | LP,SJL,SWR,TH | intergenic_variant |
| 15 | 14804237 | 14804512 | 275 | - | BUB,CE,ILNJ2,MRL,PL | intergenic_variant |
| 15 | 24481816 | 24482765 | 949 | - | NZB,NZO,NZW | intergenic_variant |
| 15 | 31124471 | 31125139 | 668 | - | C3H,CBA,CE,DBA1J,DBA2J,MRL,NOD,NOR,SWR | intergenic_variant |
| 15 | 36093591 | 36094262 | 671 | Rgs22 | 129S1,C57L,KK,LP,NOD,NOR,SWR | intron_variant |
| 15 | 57507225 | 57507573 | 348 | - | ILNJ2 | intergenic_variant |
| 15 | 59988065 | 59989012 | 947 | - | ILNJ2 | intergenic_variant |
| 15 | 94705924 | 94706261 | 337 | Tmem117 | 129S1,AJ,AKR,BUB,C3H,C57L,CBA,CE,DBA1J,DBA2J,FVB,ILNJ2,KK,MA/My,NOD,NOR,NZB,NZO,NZW,PJ,PL,RF,RH2,SEA,SJL,SMJ,SWR,TH | intron_variant |
| 16 | 3331671 | 3332000 | 329 | - | BUB,MRL,PL,SMJ | intergenic_variant |
| 16 | 21673452 | 21673771 | 319 | Map3k13 | PL,SMJ | intron_variant |
| 16 | 24843978 | 24844249 | 271 | - | BUB,CE,NOD,NOR | regulatory_region_variant,intergenic_variant |
| 16 | 27953409 | 27953654 | 245 | - | CE,KK | intergenic_variant |
| 16 | 47357811 | 47358378 | 567 | - | C3H,C58J,CBA | intergenic_variant |
| 16 | 48118428 | 48119420 | 992 | - | C3H,CBA | intergenic_variant |
| 16 | 54870271 | 54870922 | 651 | - | CBA,NZW | intergenic_variant |
| 16 | 66677672 | 66677987 | 315 | Cadm2 | SMJ | intron_variant |
| 16 | 82763754 | 82764010 | 256 | - | C58J,KK,LP,SMJ | intergenic_variant |
| 17 | 3486982 | 3487220 | 238 | Tiam2 | 129S1,CE | intron_variant |
| 17 | 8824942 | 8825274 | 332 | Pde10a | BTBR,PL | intron_variant |
| 17 | 15703826 | 15704792 | 966 | Psmb1 | BTBR,BUB,C3H,CBA,MRL,NOD,NOR,PJ,PL,RF,SJL,SMJ,SWR | intron_variant |
| 17 | 19000106 | 19000923 | 817 | - | 129S1,AJ,BALB,C57L,CE,DBA1J,DBA2J,NZO,RH2,SEA | intergenic_variant |
| 17 | 22373080 | 22374018 | 938 | Zfp947 | NZB | intron_variant |
| 17 | 28089383 | 28090259 | 876 | Bltp3a | KK,NZB,RF | intron_variant |
| 17 | 29323106 | 29323883 | 777 | - | TH | intergenic_variant |
| 17 | 29485202 | 29486029 | 827 | - | KK,RF | intergenic_variant |
| 17 | 29664735 | 29665735 | 1000 | Gm36199 | KK | intron_variant,non_coding_transcript_variant |
| 17 | 32317448 | 32317868 | 420 | Pdxk-ps | PJ | intron_variant,non_coding_transcript_variant |
| 17 | 32930561 | 32931346 | 785 | - | CE | intergenic_variant |
| 17 | 64062206 | 64062945 | 739 | - | C58J,KK,LP,NZO | intergenic_variant |
| 17 | 85856064 | 85856486 | 422 | - | 129S1,PJ,RF | intergenic_variant |
| 18 | 7052921 | 7053459 | 538 | - | BALB,C3H,CBA,ILNJ2,RH2 | intergenic_variant |
| 18 | 17526926 | 17527177 | 251 | - | SMJ | intergenic_variant |
| 18 | 28067180 | 28067949 | 769 | - | AKR,KK,MA/My,NZB,NZO,SJL | intergenic_variant |
| 18 | 55470523 | 55470909 | 386 | - | KK | intergenic_variant |
| 18 | 73207142 | 73207881 | 739 | - | BUB,DBA1J,DBA2J,PJ,SEA | intergenic_variant |
| 18 | 88591605 | 88591818 | 213 | - | NZB,NZO | intergenic_variant |
| 18 | 89363769 | 89364504 | 735 | Dok6 | 129S1,CE,DBA1J,DBA2J,FVB,RF,SMJ | intron_variant |
| 19 | 14000115 | 14000718 | 603 | - | FVB,ILNJ2,LP,SJL,SWR | intergenic_variant |
| 19 | 42737369 | 42737635 | 266 | Pyroxd2,Gm19437 | DBA1J | intron_variant,non_coding_transcript_variant |
| 19 | 59948040 | 59948334 | 294 | - | BTBR | intergenic_variant |
| 19 | 60894574 | 60895467 | 893 | Grk5 | TH | intron_variant |
| 2 | 21723632 | 21723854 | 222 | Gpr158 | AJ,AKR,BTBR,C3H,DBA2J,PL,RH2 | intron_variant |
| 2 | 39708928 | 39709219 | 291 | - | SMJ | intergenic_variant |
| 2 | 39791865 | 39792233 | 368 | - | SMJ | intergenic_variant |
| 2 | 42438092 | 42438405 | 313 | Lrp1b | TH | intron_variant |
| 2 | 46716171 | 46716397 | 226 | - | CE,KK,NZB,SMJ,TH | intergenic_variant |
| 2 | 56310941 | 56311749 | 808 | - | KK,NZO | intergenic_variant |
| 2 | 69547764 | 69548023 | 259 | - | 129S1,AJ,AKR,B10D2,B10J,BALB,BTBR,BUB,C3H,C57L,C58J,CBA,CE,DBA1J,DBA2J,FVB,ILNJ2,KK,LP,MA/My,MRL,NOD,NOR,NZB,NZO,NZW,PJ,PL,RF,RH2,SEA,SJL,SMJ,SWR,TH | intergenic_variant |
| 2 | 71838941 | 71839293 | 352 | Rapgef4 | AJ,AKR,BUB,C57L,FVB,ILNJ2,MA/My,NOD,NOR,PJ,PL,RH2 | intron_variant |

| | | | | | | |
|---|---|---|---|---|---|---|
| 2 | 71838982 | 71839926 | 944 | Rapgef4 | BALB,BTBR,MRL,SEA | intron_variant,regulatory_region_variant |
| 2 | 77251976 | 77252364 | 388 | Zfp385b | NZW | intron_variant |
| 2 | 109113542 | 109114493 | 951 | Kif18a | C3H,CE,PL,RF | intron_variant |
| 2 | 109655356 | 109655846 | 490 | - | C3H,CE,PL | intergenic_variant |
| 2 | 118394539 | 118395121 | 582 | - | BTBR,C3H,C58J,CBA,NOD,NOR,NZB,NZO,TH | intergenic_variant |
| 2 | 119438960 | 119439334 | 374 | Oip5os1 | RF | downstream_gene_variant |
| 2 | 119664301 | 119665147 | 846 | - | AJ,BALB,DBA1J,DBA2J,ILNJ2,PJ,PL,RF,RH2,SEA,SMJ,TH | intergenic_variant |
| 2 | 122038350 | 122038902 | 552 | Gm14050 | AJ,BALB,C3H,C58J,CBA,DBA1J,DBA2J,ILNJ2,NZB,NZO,PJ,PL,RF,RH2,SEA,SMJ,TH | non_coding_transcript_variant |
| 2 | 131203843 | 131204788 | 945 | - | DBA1J,DBA2J,ILNJ2,PJ,PL | intergenic_variant |
| 2 | 141431981 | 141432636 | 655 | Macrod2 | C58J,DBA1J,DBA2J,NZO,NZW | intron_variant |
| 2 | 141883974 | 141884394 | 420 | Macrod2 | AJ | intron_variant |
| 2 | 149081802 | 149082185 | 383 | - | NOD | intergenic_variant |
| 2 | 155899201 | 155899510 | 309 | Gm14240 | 129S1,BALB,C3H,C57L,CBA,CE,FVB,MA/My,NOD,NZO,NZW,PL,SEA,SJL,SMJ,SWR,TH | downstream_gene_variant |
| 2 | 163750977 | 163751267 | 290 | Rims4 | AJ,B10D2,B10J,BALB,BTBR,C3H,CBA,KK,SEA,SMJ | intron_variant |
| 2 | 165760523 | 165760841 | 318 | - | KK | intergenic_variant |
| 2 | 166908238 | 166908551 | 313 | Snord12,Zfas1 | KK | downstream_gene_variant |
| 2 | 171354331 | 171354861 | 530 | - | CE | intergenic_variant |
| 3 | 9044377 | 9044677 | 300 | Tpd52 | AKR,BUB,C58J,DBA1J,FVB,KK,MRL,NOD | intron_variant,regulatory_region_variant |
| 3 | 12234164 | 12235022 | 858 | - | SMJ | intergenic_variant |
| 3 | 15299245 | 15300066 | 821 | - | CE,DBA2J,LP | intergenic_variant |
| 3 | 34646234 | 34646800 | 566 | Sox2ot | BUB,FVB | intron_variant,non_coding_transcript_variant |
| 3 | 35830572 | 35831194 | 622 | Atp11b | DBA1J,RF | intron_variant |
| 3 | 39702554 | 39702725 | 171 | - | KK,NZW | intergenic_variant |
| 3 | 45975560 | 45976365 | 805 | - | KK | intergenic_variant |
| 3 | 67442623 | 67443500 | 877 | - | SJL | intergenic_variant |
| 3 | 69477523 | 69477806 | 283 | - | BUB,C58J,SEA | intergenic_variant |
| 3 | 76280935 | 76281484 | 549 | Fstl5 | 129S1,BTBR,C57L,NZB,NZO,SWR,TH | intron_variant |
| 3 | 87483699 | 87483991 | 292 | - | 129S1,C57L,FVB,MA/My,NOD,NOR,SJL,SWR,TH | intergenic_variant |
| 3 | 115859915 | 115860152 | 237 | Gm26344 | AJ,AKR,BALB,BUB,C3H,CBA,CE,DBA1J,DBA2J,FVB,ILNJ2,LP,MRL,NOD,NOR,NZB,NZO,NZW,PL,RF,RH2,SJL,SWR,TH | upstream_gene_variant |
| 3 | 117227878 | 117228328 | 450 | - | AJ,AKR,BALB,BUB,C3H,C58J,CBA,DBA1J,DBA2J,FVB,ILNJ2,MRL,NZO,NZW,PL,RF,RH2,SJL,SMJ,SWR,TH | intergenic_variant |
| 3 | 121923426 | 121923650 | 224 | Abca4,Gm43158 | BTBR,DBA1J,DBA2J,TH | intron_variant,non_coding_transcript_variant,regulatory_region_variant |
| 3 | 124917732 | 124917960 | 228 | - | AJ,BALB,C3H,CBA,NZB,RH2,SEA,SMJ | intergenic_variant |
| 3 | 140396631 | 140397026 | 395 | Gm43049 | BALB,BUB,C3H,ILNJ2,MA/My,MRL,SEA,SMJ,SWR | downstream_gene_variant |
| 3 | 151745052 | 151745266 | 214 | - | B10D2,BUB,NOD,NOR,NZB,NZO,NZW,TH | intergenic_variant |
| 4 | 15993614 | 15994387 | 773 | - | NZW | intergenic_variant |
| 4 | 17383804 | 17384385 | 581 | - | NZW | intergenic_variant |
| 4 | 22601210 | 22601635 | 425 | - | RF | intergenic_variant |
| 4 | 26522509 | 26522797 | 288 | - | 129S1,AJ,AKR,BALB,BTBR,BUB,C57L,CE,FVB,ILNJ2,MA/My,MRL,NOD,NOR,NZB,NZO,NZW,PJ,PL,RF,RH2,SEA,SJL,SWR | intergenic_variant |
| 4 | 27588386 | 27589368 | 982 | - | C57L | intergenic_variant |
| 4 | 30370047 | 30370347 | 300 | - | KK | intergenic_variant |
| 4 | 36652396 | 36652848 | 452 | Lingo2 | TH | intron_variant |
| 4 | 37912484 | 37913075 | 591 | - | TH | intergenic_variant |
| 4 | 54148030 | 54148365 | 335 | - | MA/My | intergenic_variant |
| 4 | 86731785 | 86732688 | 903 | Dennd4c | SEA | coding_sequence_variant,intron_variant |
| 4 | 93274454 | 93274795 | 341 | - | 129S1,C57L,RH2 | intergenic_variant |
| 4 | 96197214 | 96198207 | 993 | Cyp2j11 | ILNJ2 | intron_variant |
| 4 | 98867052 | 98867693 | 641 | Dock7 | MRL,NOD,RH2,SWR | intron_variant |
| 4 | 100586751 | 100587590 | 839 | - | AJ,AKR,B10D2,B10J,BALB,BTBR,BUB,C3H,C58J,CBA,CE,FVB,ILNJ2,MA/My,MRL,NZW,PJ,PL,RF,SEA,SJL | intergenic_variant |
| 4 | 101587861 | 101588297 | 436 | Lepr | NZB,NZO | intron_variant |
| 4 | 107101938 | 107102193 | 255 | Lrrc42 | MA/My | intron_variant |
| 4 | 112547807 | 112548389 | 582 | - | SEA | intergenic_variant |
| 4 | 116526335 | 116527142 | 807 | - | CE | regulatory_region_variant,intergenic_variant |
| 4 | 117792530 | 117793152 | 622 | St3gal3 | AKR,ILNJ2,KK,MRL | intron_variant |
| 4 | 118327768 | 118328305 | 537 | Tie1 | ILNJ2 | downstream_gene_variant,regulatory_region_variant |
| 4 | 128963100 | 128963338 | 238 | Rnf19b | FVB | intron_variant |
| 4 | 137066717 | 137067431 | 714 | Cdc42 | BUB,MRL,NZW | intron_variant,regulatory_region_variant |
| 4 | 141730445 | 141730667 | 222 | Fhad1 | SWR | intron_variant |
| 5 | 3282177 | 3282621 | 444 | - | FVB,SJL,SWR | intergenic_variant |
| 5 | 6760044 | 6760953 | 909 | - | AJ,BTBR,C3H,C57L,DBA1J,DBA2J,MA/My,MRL,RF,SJL,SWR | intergenic_variant |
| 5 | 7644919 | 7645133 | 214 | - | 129S1,AKR,BALB,BTBR,C57L,C58J,CE,FVB,LP,MA/My,NZB,NZO,NZW,PJ,RF,RH2,SEA,SJL,SWR | intergenic_variant |
| 5 | 8013397 | 8013622 | 225 | Steap4 | 129S1,BALB,BTBR,C57L,CE,NZB,NZO,NZW,PJ,PL,RF,SEA,SJL,SWR | intron_variant |
| 5 | 10109143 | 10110053 | 910 | - | AKR,NZB,NZO,NZW,SMJ | intergenic_variant |
| 5 | 42380239 | 42380639 | 400 | - | NZB,NZO | intergenic_variant |
| 5 | 44037288 | 44037744 | 456 | Cd38 | PJ,PL | intron_variant |
| 5 | 50050444 | 50051040 | 596 | - | KK,NOD,NZB,NZO,NZW,PJ,TH | intergenic_variant |
| 5 | 51112162 | 51113029 | 867 | - | KK,MA/My,NOD,NZB,NZO,NZW,PJ,SJL,SWR,TH | intergenic_variant |
| 5 | 59727573 | 59728445 | 872 | Gm43042 | C3H,CBA,DBA1J,DBA2J,ILNJ2 | intron_variant,non_coding_transcript_variant |
| 5 | 66186420 | 66186917 | 497 | Rbm47 | AJ,AKR,BUB,C58J,CE,FVB,MA/My,NOD,NOR,PL,RF,SJL,SMJ,SWR | intron_variant |
| 5 | 80664780 | 80665633 | 853 | - | SWR | intergenic_variant |
| 5 | 88513790 | 88514609 | 819 | - | MA/My | intergenic_variant |
| 5 | 118926434 | 118926669 | 235 | - | RF | intergenic_variant |
| 5 | 137445799 | 137446193 | 394 | Zan | BUB,RF | intron_variant |

| | | | | | | |
|---|---|---|---|---|---|---|
| 5 | 142348379 | 142348638 | 259 | - | NOD | regulatory_region_variant,intergenic_variant |
| 5 | 146510985 | 146511591 | 606 | Gm8494 | TH | upstream_gene_variant |
| 5 | 148050415 | 148050808 | 393 | Mtus2 | AJ,AKR,BALB,BUB,C3H,ILNJ2,MA/My,NOD,NOR,NZB,SEA | intron_variant |
| 5 | 148050480 | 148051455 | 975 | Mtus2 | MRL | intron_variant,regulatory_region_variant |
| 6 | 10047463 | 10047770 | 307 | - | 129S1,AJ,BUB,C57L,CE,DBA1J,DBA2J,ILNJ2,KK,LP,NOD,NOR,NZW,SMJ,TH | intergenic_variant |
| 6 | 13566499 | 13566998 | 499 | - | BALB,C3H,CBA,SEA,SMJ,SWR | intergenic_variant |
| 6 | 40958638 | 40959491 | 853 | - | FVB,SJL,TH | intergenic_variant |
| 6 | 48254274 | 48255229 | 955 | - | MA/My | intergenic_variant |
| 6 | 66765067 | 66765673 | 606 | - | KK | intergenic_variant |
| 6 | 69474845 | 69475497 | 652 | - | NOR | intergenic_variant |
| 6 | 70879409 | 70879859 | 450 | Eif2ak3 | 129S1,AJ,AKR,B10D2,B10J,BALB,BTBR,BUB,C3H,C57L,C58J,CBA,CE,DBA1J,DBA2J,FVB,ILNJ2,KK,LP,MA/My,MRL,NOD,NOR,NZB,NZO,NZW,PJ,PL,RF,RH2,SEA,SJL,SMJ,SWR,TH | intron_variant |
| 6 | 73337697 | 73337965 | 268 | - | AKR,C58J,NOD | intergenic_variant |
| 6 | 76011318 | 76011584 | 266 | - | KK,NOD,NOR | intergenic_variant |
| 6 | 78825404 | 78826040 | 636 | - | MRL | intergenic_variant |
| 6 | 79289360 | 79290289 | 929 | - | KK,NZB,NZO,NZW,TH | intergenic_variant |
| 6 | 98497599 | 98497855 | 256 | Gm44196 | SEA | intron_variant,non_coding_transcript_variant |
| 6 | 123948562 | 123948821 | 259 | A330044P14Rik | CE | intron_variant,non_coding_transcript_variant,regulatory_region_variant |
| 6 | 128862405 | 128862978 | 573 | BC035044,Gm8774 | NZO | intron_variant,downstream_gene_variant |
| 6 | 130353430 | 130354064 | 634 | Klra1 | KK,NOD,NOR,NZB,NZW,SWR | intron_variant |
| 6 | 133516637 | 133516903 | 266 | - | AJ,AKR,BALB,BUB,C3H,C57L,CBA,CE,DBA1J,DBA2J,ILNJ2,MRL,NZB,NZO,PL,RF,RH2,SEA | intergenic_variant |
| 6 | 139396096 | 139396707 | 611 | - | C3H | intergenic_variant |
| 7 | 10125392 | 10125675 | 283 | - | NZW | intergenic_variant |
| 7 | 10190603 | 10191417 | 814 | - | C58J,ILNJ2,NOD,SJL,TH | intergenic_variant |
| 7 | 15557134 | 15557396 | 262 | Obox4-ps35 | C58J | intron_variant,non_coding_transcript_variant |
| 7 | 29464241 | 29464868 | 627 | Gm44954 | 129S1,LP,NZW,RF,SJL | upstream_gene_variant |
| 7 | 31093455 | 31093894 | 439 | - | DBA1J,DBA2J,NZW | intergenic_variant |
| 7 | 42883207 | 42884088 | 881 | - | DBA1J,DBA2J,ILNJ2,NZW,PJ | intergenic_variant |
| 7 | 44582922 | 44583269 | 347 | - | 129S1,AJ,AKR,BALB,C3H,CBA,DBA1J,DBA2J,KK,MA/My,PL,RH2,SEA,SMJ | intergenic_variant |
| 7 | 44591717 | 44592027 | 310 | Tsks | 129S1,AJ,AKR,BALB,C3H,CBA,CE,DBA1J,DBA2J,KK,MA/My,PL,RH2,SEA,SMJ | upstream_gene_variant |
| 7 | 50128157 | 50128823 | 666 | Nell1 | ILNJ2,KK,NZW,PJ | intron_variant |
| 7 | 50809685 | 50810396 | 711 | - | 129S1,C3H,DBA1J,DBA2J | intergenic_variant |
| 7 | 52952454 | 52953412 | 958 | - | DBA2J | intergenic_variant |
| 7 | 72532336 | 72532793 | 457 | - | CBA,DBA1J,DBA2J,ILNJ2,NZB,NZO,PJ,SEA | intergenic_variant |
| 7 | 94631413 | 94631803 | 390 | - | NZB,NZO | intergenic_variant |
| 7 | 96818878 | 96819444 | 566 | Gab2 | 129S1,AJ,AKR,BALB,BTBR,BUB,C57L,FVB,KK,MA/My,MRL,NOD,NOR,NZB,NZW,PL,RF,RH2,SJL,SMJ,SWR,TH | intron_variant |
| 7 | 103546639 | 103547226 | 587 | - | 129S1,AJ,AKR,BALB,BTBR,BUB,C3H,C57L,CBA,DBA1J,DBA2J,FVB,ILNJ2,MRL,NZB,NZW,PJ,PL,RF,RH2,SEA,TH | intergenic_variant |
| 7 | 103611140 | 103611554 | 414 | - | KK,NOD,SJL,SWR,TH | intergenic_variant |
| 7 | 108684201 | 108684924 | 723 | - | NOD,SMJ,SWR | intergenic_variant |
| 7 | 109368734 | 109369040 | 306 | Nrip3 | NOD,SWR | intron_variant,regulatory_region_variant |
| 7 | 113779804 | 113780073 | 269 | Gm57208 | 129S1,AJ,AKR,BALB,BUB,C57L,MRL,NOD,NZB,NZW,PL,RH2,SJL | upstream_gene_variant,regulatory_region_variant |
| 8 | 14238314 | 14238596 | 282 | Dlgap2 | AKR,BALB,C57L,CE,ILNJ2,LP,MRL,NZB,NZW,PJ,PL,RF,RH2,SEA,SMJ,TH | intron_variant |
| 8 | 14842407 | 14842811 | 404 | Dlgap2 | SEA | intron_variant |
| 8 | 27357198 | 27357529 | 331 | 2310008N11Rik | PL | intron_variant,non_coding_transcript_variant |
| 8 | 38689887 | 38690336 | 449 | Sgcz | 129S1,AJ,AKR,BALB,BTBR,BUB,C3H,C57L,CBA,CE,DBA1J,DBA2J,FVB,ILNJ2,KK,MA/My,MRL,NOD,NOR,NZB,NZO,NZW,PJ,PL,RF,RH2,SEA,SJL,SMJ,SWR,TH | intron_variant |
| 8 | 60552905 | 60553421 | 516 | - | PL | intergenic_variant |
| 8 | 60717879 | 60718377 | 498 | Gm34623 | KK | intron_variant,non_coding_transcript_variant |
| 8 | 75339941 | 75340455 | 514 | Gm11033 | 129S1,BALB,BUB,C3H,CBA,DBA1J,DBA2J,FVB,ILNJ2,NZB,NZO,RF,SJL,SMJ,SWR | intron_variant,non_coding_transcript_variant |
| 8 | 88961779 | 88962453 | 674 | Tent4b | AJ,RH2 | intron_variant |
| 8 | 89703762 | 89704677 | 915 | - | 129S1,AKR,BALB,BTBR,BUB,C57L,C58J,CBA,CE,DBA1J,DBA2J,FVB,ILNJ2,KK,MA/My,MRL,NOD,NOR,NZB,NZO,NZW,PJ,PL,RF,SEA,SJL,SMJ,SWR,TH | intergenic_variant |
| 8 | 92415770 | 92416743 | 973 | Gm2875,Gm9619 | 129S1,AKR,BALB,BUB,C57L,C58J,CBA,CE,DBA1J,DBA2J,FVB,ILNJ2,MA/My,MRL,NOD,NOR,NZB,NZO,NZW,PJ,PL,RF,SEA,SJL,SMJ,SWR,TH | non_coding_transcript_variant,upstream_gene_variant |
| 8 | 106640828 | 106641175 | 347 | Pskh1 | PJ | intron_variant |
| 8 | 129792443 | 129793152 | 709 | Ccdc7b,Ccdc7a | CBA,DBA1J,DBA2J,MA/My,NOD,NOR,PJ | upstream_gene_variant |
| 8 | 129987565 | 129988207 | 642 | - | KK | intergenic_variant |
| 9 | 40669708 | 40670132 | 424 | Clmp,Gm48293 | FVB | intron_variant,non_coding_transcript_variant |
| 9 | 41193411 | 41193938 | 527 | - | 129S1,AJ,AKR,BALB,BTBR,BUB,C57L,CBA,CE,DBA1J,DBA2J,FVB,ILNJ2,KK,MRL,NOD,NOR,NZB,NZO,PJ,PL,RF,RH2,SEA,SWR,TH | intergenic_variant |
| 9 | 56183564 | 56184198 | 634 | Peak1 | KK | intron_variant |
| 9 | 62371628 | 62371932 | 304 | Coro2b | AKR,NZB,RF | intron_variant,regulatory_region_variant |
| 9 | 80225149 | 80226107 | 958 | Impg1 | 129S1,AJ,BALB,BTBR,C3H,C57L,CBA,CE,DBA1J,DBA2J,KK,MA/My,NZB,NZO,PJ,RH2,SEA,SMJ,SWR,TH | intron_variant,regulatory_region_variant |
| 9 | 94886939 | 94887389 | 450 | Slc9a9 | MRL,NZO,NZW | intron_variant |
| 9 | 105580183 | 105580921 | 738 | Col6a6 | 129S1,AJ,AKR,BALB,BTBR,BUB,C3H,C57L,CBA,CE,DBA1J,DBA2J,FVB,ILNJ2,MA/My,MRL,NOD,NOR,NZB,NZO,NZW,PJ,PL,RF,RH2,SEA,SJL,SMJ,SWR,TH | intron_variant |
| 9 | 112797554 | 112798124 | 570 | - | ILNJ2,NZB,NZO,NZW | intergenic_variant |

| | | | | | | |
|---|---|---|---|---|---|---|
| 9 | 114685094 | 114685353 | 259 | - | AJ,AKR,BALB,C3H,CE,FVB,MRL,PJ,PL,RF,RH2,SEA,SMJ | intergenic_variant |
| 9 | 118507743 | 118508269 | 526 | Itga9 | MRL,PJ | intron_variant |
| X | 66738128 | 66738349 | 221 | Gm14692,Gm1140 | AJ,BALB,C3H,ILNJ2,MRL,NZB,NZO,NZW,PL,SEA,SJL | upstream_gene_variant |
| X | 74699377 | 74699912 | 535 | - | B10D2,B10J,BTBR,MA/My,NZB | intergenic_variant |
| X | 74989139 | 74989878 | 739 | - | CE | intergenic_variant |
| X | 108161940 | 108162591 | 651 | Sh3bgrl | CE,NZB,NZO,NZW | intron_variant |
| X | 111650672 | 111650937 | 265 | - | AJ,BALB,C3H,CBA,MRL,RH2 | intergenic_variant |
| X | 116358513 | 116359451 | 938 | - | AJ,BALB,C3H,CBA,MRL,RH2 | intergenic_variant |
| X | 146219745 | 146220168 | 423 | - | NZB | intergenic_variant |

| CHR | START | END | SIZE | SYMBOL | STRAINS with VARIANT SV ALLELE | CONSEQUENCE |
|---|---|---|---|---|---|---|
| 1 | 3192284 | 3193853 | 1569 | - | C3H,CBA,MRL,RF | regulatory_region_variant,intergenic_variant |
| 1 | 4498582 | 4501964 | 3382 | - | MRL | intergenic_variant |
| 1 | 7910818 | 7912108 | 1290 | - | C3H,CBA,MRL,RF | intergenic_variant |
| 1 | 14410529 | 14411998 | 1469 | - | 129S1,AJ,AKR,BALB,BTBR,C3H,C57L,CBA,CE,DBA1J,DBA2J,FVB,ILNJ2,KK,LP,MRL,NOD,NOR,NZB,NZO,NZW,PJ,PL,RF,RH2,SEA,SJL,SMJ,SWR,TH | intergenic_variant |
| 1 | 17358782 | 17362315 | 3533 | - | BTBR,BUB,LP | intergenic_variant |
| 1 | 22301046 | 22307255 | 6209 | Gm28239 | MRL | upstream_gene_variant |
| 1 | 26034302 | 26036443 | 2141 | Gm18443 | 129S1,BTBR,C57L,NZW | non_coding_transcript_variant |
| 1 | 29179906 | 29182921 | 3015 | - | RF | intergenic_variant |
| 1 | 31512524 | 31515469 | 2945 | - | BUB,FVB,NZB,NZO,PJ,SJL,SWR | intergenic_variant |
| 1 | 31514711 | 31516089 | 1378 | - | BUB,FVB,NZB,NZO,PJ,SJL,SWR | intergenic_variant |
| 1 | 31611064 | 31612292 | 1228 | - | CE | intergenic_variant |
| 1 | 33338680 | 33345557 | 6877 | - | AJ,BALB,BUB,C3H,DBA1J,DBA2J,FVB,ILNJ2,KK,LP,MRL,NOR,NZB,NZO,PJ,PL,RF,RH2,SEA,SMJ | intergenic_variant |
| 1 | 35865216 | 35866729 | 1513 | - | AJ,AKR,BUB,C3H,CBA,CE,FVB,KK,MRL,NZB,NZO,NZW,RF,RH2,SMJ,SWR | regulatory_region_variant,intergenic_variant |
| 1 | 35865616 | 35868156 | 2540 | - | AJ,AKR,BUB,C3H,CBA,CE,FVB,KK,MRL,NZB,NZO,NZW,RF,RH2,SMJ,SWR | regulatory_region_variant,intergenic_variant |
| 1 | 40189016 | 40191031 | 2015 | - | 129S1,BTBR,C57L,NZB,NZO | regulatory_region_variant,intergenic_variant |
| 1 | 47626851 | 47628380 | 1529 | - | SWR | intergenic_variant |
| 1 | 50720379 | 50721461 | 1082 | - | AKR,BALB,BUB,C58J,MA/My,RF,RH2,SEA,SWR | intergenic_variant |
| 1 | 59332577 | 59335034 | 2457 | Cdk15 | CE | intron_variant |
| 1 | 66054893 | 66056769 | 1876 | - | 129S1,AJ,AKR,BTBR,BUB,C3H,C57L,C58J,CBA,CE,DBA1J,DBA2J,FVB,ILNJ2,KK,LP,MA/My,MRL,NOD,NOR,NZB,PJ,PL,RF,SJL,SMJ,SWR,TH | regulatory_region_variant,intergenic_variant |
| 1 | 66831537 | 66835768 | 4231 | Kansl1l,6820402A03Rik | AJ,CBA,DBA1J,NZW,SWR | intron_variant,non_coding_transcript_variant,regulatory_region_variant |
| 1 | 69284325 | 69286734 | 2409 | - | 129S1,AJ,BTBR,C3H,C57L,DBA2J,FVB,KK,LP,MA/My,MRL,NOD,NOR,NZB,NZO,NZW,SMJ,TH | regulatory_region_variant,intergenic_variant |
| 1 | 79010149 | 79012060 | 1911 | Gm29536 | AJ,BALB,C3H,ILNJ2,NZB,PJ,RH2,SEA,SJL,SWR | intron_variant,non_coding_transcript_variant |
| 1 | 79225828 | 79235255 | 9427 | - | AKR,B10J,BUB,C3H,C57L,NOR,PL,RH2,SEA | intergenic_variant |
| 1 | 79227348 | 79232893 | 5545 | - | PJ,SWR | intergenic_variant |
| 1 | 79227348 | 79233734 | 6386 | - | 129S1,AJ,AKR,B10D2,B10J,BALB,BTBR,BUB,C3H,C57L,CBA,CE,DBA1J,DBA2J,ILNJ2,KK,MA/My,MRL,NOD,NOR,NZO,NZW,PL,RF,RH2,SEA,SJL,TH | intergenic_variant |
| 1 | 79227349 | 79235246 | 7897 | - | PL,RH2,SEA | intergenic_variant |
| 1 | 79229464 | 79233734 | 4270 | - | CBA,CE,DBA1J,DBA2J,MRL,NOD | intergenic_variant |
| 1 | 83208566 | 83212716 | 4150 | - | LP | intergenic_variant |
| 1 | 88158034 | 88159149 | 1115 | Mroh2a | 129S1,AJ,AKR,B10D2,B10J,BALB,BTBR,BUB,C3H,C57L,C58J,CBA,CE,DBA1J,DBA2J,FVB,ILNJ2,KK,LP,MA/My,MRL,NOD,NOR,NZB,NZO,NZW,PJ,PL,RH2,SEA,SJL,SMJ,SWR,TH | coding_sequence_variant,intron_variant,regulatory_region_variant |
| 1 | 96423208 | 96424425 | 1217 | - | AKR,C57L,DBA2J,ILNJ2,NOD,SMJ | intergenic_variant |
| 1 | 99837346 | 99839340 | 1994 | Cntnap5b | 129S1,AKR,BUB,C57L,DBA1J,DBA2J,LP,MA/My,NOD,NOR,NZO,PJ,SEA,SJL,TH | intron_variant |
| 1 | 101287217 | 101291317 | 4100 | - | 129S1,AKR,C57L,DBA1J,DBA2J,LP,MA/My,NOD,NOR,PJ,SEA,TH | intergenic_variant |
| 1 | 101287301 | 101293490 | 6189 | - | 129S1,AKR,C57L,DBA1J,DBA2J,LP,MA/My,NOD,NOR,PJ,SEA,TH | intergenic_variant |
| 1 | 122065818 | 122066946 | 1128 | - | CE,DBA2J,SMJ | intergenic_variant |
| 1 | 122851020 | 122853228 | 2208 | - | BUB | intergenic_variant |
| 1 | 123516894 | 123522029 | 5135 | Dpp10 | AJ,BUB,C3H,CBA,CE,KK,NZO,RH2,SMJ,SWR | intron_variant |
| 1 | 123518424 | 123522932 | 4508 | Dpp10 | NZW | intron_variant |
| 1 | 123518458 | 123523484 | 5026 | Dpp10 | 129S1,AJ,BALB,BTBR,BUB,C3H,C57L,CBA,CE,ILNJ2,KK,NZO,NZW,PL,RF,RH2,SEA,SMJ,SWR | intron_variant |
| 1 | 132031302 | 132037934 | 6632 | Gm37120 | KK,MA/My,NZO | non_coding_transcript_exon_variant,regulatory_region_variant |
| 1 | 139752079 | 139753136 | 1057 | Cfhr2 | MRL | intron_variant |
| 1 | 147076762 | 147080366 | 3604 | - | BTBR,SMJ | intergenic_variant |
| 1 | 149359784 | 149368260 | 8476 | - | BTBR,SMJ | intergenic_variant |
| 1 | 167394713 | 167395791 | 1078 | - | BTBR,BUB,FVB,ILNJ2,KK,NZB,NZO,NZW,RF,SMJ,SWR,TH | regulatory_region_variant,intergenic_variant |
| 1 | 175107988 | 175113464 | 5476 | Rgs7 | 129S1,AJ,AKR,BALB,BTBR,BUB,C3H,C57L,C58J,CBA,CE,DBA1J,DBA2J,FVB,ILNJ2,KK,MRL,NOD,NOR,NZB,NZO,NZW,PJ,PL,RF,RH2,SEA,SJL,SMJ,SWR,TH | intron_variant |
| 1 | 177534675 | 177536718 | 2043 | - | FVB,NZO | regulatory_region_variant,intergenic_variant |
| 1 | 181056751 | 181057923 | 1172 | A430110L20Rik,Gm57146 | C57L | downstream_gene_variant,intron_variant,non_coding_transcript_variant |
| 1 | 182330754 | 182332882 | 2128 | Capn2 | CE,NZB | intron_variant,regulatory_region_variant |
| 10 | 6223173 | 6238970 | 15797 | - | BUB | intergenic_variant |
| 10 | 6224968 | 6229966 | 4998 | - | BUB | intergenic_variant |
| 10 | 6227799 | 6229971 | 2172 | - | BUB,KK,NOD,NOR,TH | intergenic_variant |
| 10 | 16647127 | 16655488 | 8361 | - | BUB,LP,SWR | intergenic_variant |
| 10 | 45851423 | 45853171 | 1748 | - | TH | regulatory_region_variant,intergenic_variant |
| 10 | 51512731 | 51515558 | 2827 | - | KK,NZW | intergenic_variant |
| 10 | 77435955 | 77439382 | 3427 | - | TH | regulatory_region_variant,intergenic_variant |
| 10 | 84455748 | 84456898 | 1150 | Gm17249 | BTBR,C58J,LP,NZB,NZO,SMJ | non_coding_transcript_exon_variant |
| 10 | 87470264 | 87471649 | 1385 | - | BUB,FVB | intergenic_variant |
| 10 | 95297338 | 95298415 | 1077 | Gm47599 | PL | upstream_gene_variant |
| 10 | 98112002 | 98115458 | 3456 | - | NZB | intergenic_variant |
| 10 | 100074100 | 100075518 | 1418 | - | AKR,BUB,C3H,CBA,FVB,MA/My,NOD,NOR,PL,SMJ | intergenic_variant |
| 10 | 100666965 | 100669134 | 2169 | - | CBA,NZB | intergenic_variant |

| | | | | | | |
|---|---|---|---|---|---|---|
| 10 | 103780096 | 103783245 | 3149 | - | AJ,BALB,CE,DBA1J,DBA2J,ILNJ2,MRL,PJ,RH2,SEA,SJL | intergenic_variant |
| 10 | 104626693 | 104628561 | 1868 | - | AJ,BALB,BTBR,CE,DBA1J,DBA2J,MRL,NZB,PJ,RH2,SEA,SJL | intergenic_variant |
| 10 | 112448580 | 112452888 | 4308 | - | PL | intergenic_variant |
| 10 | 122557213 | 122558835 | 1622 | Ppm1h | KK | intron_variant |
| 10 | 124065390 | 124070926 | 5536 | - | PL | intergenic_variant |
| 10 | 124069702 | 124070915 | 1213 | - | PL | intergenic_variant |
| 10 | 129423174 | 129430266 | 7092 | - | C3H,CBA | regulatory_region_variant,intergenic_variant |
| 11 | 7349554 | 7351315 | 1761 | - | KK | intergenic_variant |
| 11 | 30703150 | 30704874 | 1724 | Gm12101,Gm12100 | BALB,BTBR,C57L,RH2 | non_coding_transcript_exon_variant,non_coding_transcript_exon_variant,intron_variant,regulatory_region_variant |
| 11 | 38970120 | 38971140 | 1020 | - | CE | intergenic_variant |
| 11 | 61311196 | 61318406 | 7210 | - | FVB,NOD,SWR | intergenic_variant |
| 11 | 63726137 | 63728347 | 2210 | - | FVB,NOD,SWR | intergenic_variant |
| 11 | 65546990 | 65549669 | 2679 | - | 129S1,BTBR,C57L,NZW | intergenic_variant |
| 11 | 76049730 | 76051442 | 1712 | Vps53 | 129S1,AJ,AKR,BALB,BTBR,BUB,C3H,C57L,C58J,CBA,DBA1J,DBA2J,FVB,ILNJ2,KK,MA/My,MRL,NOD,NOR,NZB,PJ,PL,RF,RH2,SEA,SJL,SMJ,SWR,TH | intron_variant,regulatory_region_variant |
| 11 | 92808159 | 92810724 | 2565 | - | 129S1,BTBR,C57L,CE | intergenic_variant |
| 11 | 97786811 | 97788077 | 1266 | - | NZW,SWR | regulatory_region_variant,intergenic_variant |
| 11 | 101533239 | 101536090 | 2851 | Gm24950,Gm23971 | 129S1,AJ,BTBR,BUB,C3H,C57L,CE,DBA1J,DBA2J,ILNJ2,KK,NOD,NOR,NZB,NZO | downstream_gene_variant |
| 12 | 3554091 | 3555204 | 1113 | Asxl2 | C58J,LP | 3_prime_UTR_variant |
| 12 | 7691528 | 7692577 | 1049 | Gm32828 | MRL,NZW | intron_variant,non_coding_transcript_variant |
| 12 | 14894381 | 14900493 | 6112 | - | AJ,BALB,CE,ILNJ2,MRL | intergenic_variant |
| 12 | 14898409 | 14903050 | 4641 | - | C58J,DBA1J,DBA2J,NOD,NOR,NZB,NZW,SMJ | intergenic_variant |
| 12 | 36300480 | 36302054 | 1574 | Lrrc72 | CE | intron_variant |
| 12 | 40929655 | 40932586 | 2931 | - | BUB,DBA1J,DBA2J,FVB,ILNJ2,MRL,NOD,NOR,NZB,NZW,SJL | regulatory_region_variant,intergenic_variant |
| 12 | 42270436 | 42273957 | 3521 | - | BUB,DBA1J,DBA2J,FVB,ILNJ2,MRL,NOD,NOR,NZB,SJL | intergenic_variant |
| 12 | 42866445 | 42869511 | 3066 | - | KK,SWR | intergenic_variant |
| 12 | 45384479 | 45385602 | 1123 | - | KK,NZO,NZW | intergenic_variant |
| 12 | 51349354 | 51356510 | 7156 | - | 129S1,C57L,NZB | regulatory_region_variant,intergenic_variant |
| 12 | 60910841 | 60911910 | 1069 | - | AJ,BALB,BTBR,C3H,C57L,CBA,DBA1J,DBA2J,ILNJ2,KK,MA/My,NZB,NZO,NZW,PJ,RH2,SEA,SMJ,TH | intergenic_variant |
| 12 | 82609624 | 82612185 | 2561 | - | PJ,SEA | intergenic_variant |
| 12 | 92590991 | 92592374 | 1383 | - | AJ,BUB,C3H,C58J,DBA1J,DBA2J,ILNJ2,KK,NZO,PJ,PL,SEA | intergenic_variant |
| 12 | 93342922 | 93344715 | 1793 | - | PJ,SEA | intergenic_variant |
| 12 | 93343893 | 93346627 | 2734 | - | PJ,SEA | intergenic_variant |
| 12 | 101458016 | 101460339 | 2323 | Catsperb | C58J,SMJ | intron_variant |
| 12 | 107284140 | 107287490 | 3350 | - | BALB,BUB,CE,MA/My,NZW,PJ,SEA,SJL | intergenic_variant |
| 12 | 107286406 | 107288893 | 2487 | - | BALB,BUB,CE,MA/My,NZW,PJ,SEA,SJL | intergenic_variant |
| 12 | 109087134 | 109088969 | 1835 | - | KK,NOD,NOR | intergenic_variant |
| 12 | 110152713 | 110155772 | 3059 | Gm34785 | DBA2J | non_coding_transcript_exon_variant,intron_variant,regulatory_region_variant |
| 12 | 114561058 | 114565733 | 4675 | Ighv1-10 | 129S1,BALB,BUB,C58J,KK,LP,MA/My,PJ,SEA,SMJ,TH | non_coding_transcript_variant |
| 13 | 3663633 | 3668872 | 5239 | - | C57L,KK,LP,NZB,NZO,NZW | intergenic_variant |
| 13 | 15605812 | 15613081 | 7269 | - | NZO,NZW | intergenic_variant |
| 13 | 18721548 | 18722819 | 1271 | Pou6f2 | AKR,ILNJ2,LP,PJ,RF,SEA | intron_variant |
| 13 | 22251586 | 22266622 | 15036 | Vmn1r-ps93 | 129S1,AJ,B10D2,B10J,BALB,BTBR,BUB,C3H,C57L,C58J,CBA,DBA1J,DBA2J,FVB,ILNJ2,KK,LP,MA/My,MRL,NOD,NOR,NZB,PJ,PL,RH2,SEA,SJL,SMJ,SWR,TH | non_coding_transcript_variant |
| 13 | 22253937 | 22264273 | 10336 | Vmn1r-ps93 | 129S1,AJ,B10D2,B10J,BALB,BTBR,BUB,C3H,C57L,C58J,CBA,DBA1J,DBA2J,FVB,ILNJ2,KK,LP,MA/My,MRL,NOD,NOR,NZB,PJ,PL,RH2,SEA,SJL,SMJ,SWR,TH | non_coding_transcript_variant |
| 13 | 23187383 | 23189084 | 1701 | - | BTBR,C57L,KK,LP,NOD,NOR,NZB,NZO,NZW,PJ,SEA | intergenic_variant |
| 13 | 53593986 | 53596209 | 2223 | Gm2762 | 129S1,AJ,AKR,BALB,BTBR,BUB,C3H,C57L,CBA,CE,DBA1J,DBA2J,FVB,ILNJ2,KK,MA/My,MRL,NOD,NOR,NZB,NZO,NZW,PJ,PL,RF,RH2,SEA,SJL,SMJ,SWR,TH | intron_variant,non_coding_transcript_variant,non_coding_transcript_exon_variant,regulatory_region_variant |
| 13 | 58062382 | 58063401 | 1019 | - | BUB | intergenic_variant |
| 13 | 62833917 | 62834954 | 1037 | Platr25 | BUB | intron_variant |
| 13 | 78419448 | 78421080 | 1632 | A830082K12Rik,Gm5042 | BUB,C3H,CBA,CE,DBA1J,DBA2J,NOD,NOR,TH | intron_variant,non_coding_transcript_variant,downstream_gene_variant |
| 13 | 79481834 | 79483265 | 1431 | - | SWR | intergenic_variant |
| 13 | 84534090 | 84536436 | 2346 | - | SWR | intergenic_variant |
| 13 | 86375129 | 86377087 | 1958 | - | 129S1,AKR,BALB,BTBR,BUB,C3H,C57L,C58J,CBA,CE,DBA1J,DBA2J,FVB,ILNJ2,KK,MA/My,MRL,NOD,NOR,NZB,NZO,NZW,PJ,PL,RF,SEA,SJL,SMJ,TH | intergenic_variant |
| 13 | 89593365 | 89594448 | 1083 | - | BALB,DBA1J,FVB,MRL,SEA | intergenic_variant |
| 13 | 89594142 | 89596062 | 1920 | - | BALB,DBA1J,FVB,MRL,SEA | intergenic_variant |
| 13 | 97627480 | 97632924 | 5444 | Lncenc1 | AKR | non_coding_transcript_exon_variant,intron_variant,non_coding_transcript_exon_variant,regulatory_region_variant |
| 13 | 106815968 | 106822590 | 6622 | - | CE,MA/My | intergenic_variant |
| 13 | 106821684 | 106823837 | 2153 | - | CE,MA/My | intergenic_variant |
| 13 | 108324811 | 108329652 | 4841 | Ercc8 | 129S1,BTBR,C57L,KK,NZB,NZW,PL | intron_variant,regulatory_region_variant |
| 13 | 110362227 | 110363456 | 1229 | Rab3c | CE,MA/My,NOD,NOR,NZO | intron_variant,regulatory_region_variant |
| 13 | 112999776 | 113001963 | 2187 | Plpp1 | AJ,DBA1J,FVB,KK,MA/My,NZO,PL,RF,RH2,SJL,SWR | intron_variant |
| 13 | 116973729 | 116979941 | 6212 | 4930435F18Rik | 129S1,BTBR,BUB,C3H,C57L,CE,DBA2J,KK,NZB,NZO,NZW | non_coding_transcript_exon_variant,intron_variant,regulatory_region_variant |
| 14 | 9333848 | 9341155 | 7307 | - | PL | intergenic_variant |
| 14 | 17644594 | 17646790 | 2196 | - | 129S1,C57L,KK,PL | intergenic_variant |

| | | | | | | |
|---|---|---|---|---|---|---|
| 14 | 18170523 | 18172609 | 2086 | - | C57L,KK | intergenic_variant |
| 14 | 35811315 | 35818656 | 7341 | Gm7853,4930474N05Rik | 129S1,C57L,NZW | non_coding_transcript_variant,coding_sequence_variant,5_prime_UTR_variant,intron_variant |
| 14 | 42003455 | 42005845 | 2390 | - | SMJ | intergenic_variant |
| 14 | 45079707 | 45081965 | 2258 | - | C57L,NZO,PJ,PL | intergenic_variant |
| 14 | 51665970 | 51668613 | 2643 | Gm7247,Vmn2r-ps111 | 129S1,AJ,AKR,BALB,BUB,C3H,C57L,CBA,CE,KK,MRL,NZB,NZO,NZW,PJ,PL,RF,RH2,SEA,TH | non_coding_transcript_exon_variant,intron_variant |
| 14 | 51667721 | 51669191 | 1470 | Gm7247,Vmn2r-ps111 | 129S1,AJ,AKR,BALB,BUB,C3H,C57L,CBA,CE,KK,MRL,NZB,NZO,NZW,PJ,PL,RF,RH2,SEA,TH | non_coding_transcript_exon_variant,intron_variant |
| 14 | 52205724 | 52210931 | 5207 | - | 129S1,AJ,AKR,BALB,BUB,C3H,C57L,CBA,CE,DBA2J,KK,MRL,NZB,NZO,NZW,PJ,PL,RF,RH2,SEA,TH | intergenic_variant |
| 14 | 74501864 | 74503388 | 1524 | Gm49171 | BUB,KK,NZO,TH | intron_variant,non_coding_transcript_variant |
| 14 | 84453219 | 84460464 | 7245 | - | BUB,C57L,C58J | intergenic_variant |
| 14 | 84454423 | 84459278 | 4855 | - | 129S1,BUB,C57L,C58J,ILNJ2,KK,NOD,NOR,NZW,SJL | intergenic_variant |
| 14 | 90577042 | 90579048 | 2006 | - | AJ,AKR,BALB,BUB,C3H,CBA,CE,DBA1J,DBA2J,FVB,NOD,NZO,PJ,RF,SEA,SJL,SMJ,SWR,TH | intergenic_variant |
| 14 | 90577058 | 90584539 | 7481 | - | NOD,NZO | intergenic_variant |
| 14 | 96646843 | 96649069 | 2226 | Klhl1 | AJ,AKR,BALB,BTBR,BUB,C3H,CBA,CE,DBA1J,DBA2J,FVB,MRL,PJ,PL,RF,SEA,SJL,SMJ,SWR | intron_variant |
| 14 | 101639151 | 101640296 | 1145 | - | C57L,C58J,ILNJ2,NZB,NZO,PJ | intergenic_variant |
| 14 | 105981480 | 105984071 | 2591 | - | 129S1,BTBR,BUB,C57L,C58J,FVB,ILNJ2,LP,MA/My,NOD,NOR,NZW,SWR | regulatory_region_variant,intergenic_variant |
| 14 | 111848070 | 111850449 | 2379 | Gm30439 | 129S1,AJ,AKR,BUB,C57L,C58J,FVB,ILNJ2,LP,MA/My,NOD,NOR,NZB,NZO,NZW,PL,RF,SJL,SMJ,SWR,TH | intron_variant,non_coding_transcript_variant |
| 14 | 114657746 | 114661907 | 4161 | - | SWR | intergenic_variant |
| 14 | 119329349 | 119331250 | 1901 | Uggt2 | 129S1,KK | intron_variant |
| 15 | 15908554 | 15914624 | 6070 | - | LP | intergenic_variant |
| 15 | 19676713 | 19677801 | 1088 | Gm49166 | AJ,BALB,BUB,C3H,CBA,CE,DBA1J,DBA2J,FVB,ILNJ2,KK,LP,MA/My,MRL,NZB,NZO,NZW,PL,RH2,SJL,SWR,TH | upstream_gene_variant |
| 15 | 20042932 | 20050910 | 7978 | - | 129S1,BTBR,C57L | intergenic_variant |
| 15 | 20867589 | 20869323 | 1734 | Gm49685 | 129S1,BALB,BTBR,BUB,C3H,C57L,DBA2J,KK,LP,MA/My,NZB,NZO,NZW,PJ,SEA,SJL,SMJ | intron_variant,non_coding_transcript_variant |
| 15 | 20868128 | 20871016 | 2888 | Gm49685 | 129S1,BALB,BTBR,BUB,C3H,C57L,DBA2J,KK,LP,MA/My,NZB,NZO,NZW,PJ,SEA,SJL,SMJ | intron_variant,non_coding_transcript_variant |
| 15 | 21067678 | 21069276 | 1598 | - | CE,ILNJ2,MRL | intergenic_variant |
| 15 | 27384221 | 27390282 | 6061 | - | MRL | intergenic_variant |
| 15 | 32833543 | 32839857 | 6314 | - | C3H,CBA,CE,DBA1J | intergenic_variant |
| 15 | 32835802 | 32841020 | 5218 | - | C3H,CBA,DBA1J,DBA2J | intergenic_variant |
| 15 | 42844205 | 42850992 | 6787 | - | MRL | regulatory_region_variant,intergenic_variant |
| 15 | 42847421 | 42850486 | 3065 | - | MRL | regulatory_region_variant,intergenic_variant |
| 15 | 52516993 | 52519040 | 2047 | - | KK | intergenic_variant |
| 16 | 23096050 | 23097394 | 1344 | St6gal1 | TH | intron_variant |
| 16 | 26627223 | 26634260 | 7037 | - | ILNJ2,NZB,NZO,NZW | intergenic_variant |
| 16 | 27951988 | 27953632 | 1644 | - | CE,KK | intergenic_variant |
| 16 | 40570447 | 40571678 | 1231 | Lsamp | 129S1,BTBR,BUB,C3H,C57L,CBA,CE,ILNJ2,KK,LP,MRL,NZB,NZO,NZW,PL,SMJ,TH | intron_variant |
| 16 | 47114993 | 47125317 | 10324 | - | 129S1,C3H,C58J,CBA,CE,SJL,TH | intergenic_variant |
| 16 | 47121660 | 47125360 | 3700 | - | AKR,FVB,LP,MRL,PJ,RF,SWR | intergenic_variant |
| 16 | 53407810 | 53413821 | 6011 | - | C3H,C58J,CBA,FVB,KK,LP,MRL,NZW,SWR,TH | intergenic_variant |
| 16 | 53469811 | 53471104 | 1293 | - | SMJ | intergenic_variant |
| 16 | 54484489 | 54488519 | 4030 | - | CBA,NZW | intergenic_variant |
| 16 | 54484491 | 54486043 | 1552 | - | CBA,NZW | intergenic_variant |
| 16 | 54584534 | 54585611 | 1077 | - | CBA,NZW | intergenic_variant |
| 16 | 59582043 | 59589637 | 7594 | Epha6 | PL | intron_variant |
| 16 | 67730954 | 67737533 | 6579 | - | AJ,C3H,C58J,CBA,DBA2J,ILNJ2,KK,LP,MA/My,MRL,NZB,NZO,PL,RH2,SMJ | intergenic_variant |
| 16 | 67735556 | 67738018 | 2462 | - | AJ,BALB,C3H,C58J,CBA,DBA1J,DBA2J,ILNJ2,KK,LP,MA/My,MRL,NZB,NZO,NZW,PL,RH2,SMJ | intergenic_variant |
| 16 | 73952188 | 73953202 | 1014 | Robo2 | KK | intron_variant |
| 16 | 76585789 | 76591934 | 6145 | - | C58J,NZB,NZO,NZW | intergenic_variant |
| 16 | 76588073 | 76592246 | 4173 | - | C58J,NZB,NZO,NZW | intergenic_variant |
| 16 | 87340509 | 87341810 | 1301 | - | 129S1,C57L,ILNJ2 | intergenic_variant |
| 16 | 87341176 | 87344200 | 3024 | - | 129S1,C57L,ILNJ2 | intergenic_variant |
| 16 | 97147593 | 97153791 | 6198 | - | NZO | intergenic_variant |
| 17 | 15702459 | 15704792 | 2333 | Psmb1 | BTBR,BUB,C3H,CBA,MRL,NOD,NOR,PJ,PL,RF,SJL,SMJ,SWR | intron_variant |
| 17 | 18598238 | 18600463 | 2225 | - | AKR,FVB,ILNJ2,KK,MA/My | intergenic_variant |
| 17 | 19818121 | 19819600 | 1479 | Vmn2r101 | B10D2,B10J,BTBR,BUB,C3H,C58J,CBA,LP,MRL,NOD,NOR,NZB,NZW,PJ,PL,RF,SJL,SWR,TH | intron_variant |
| 17 | 23753073 | 23754242 | 1169 | - | PJ,PL | intergenic_variant |
| 17 | 28087897 | 28090266 | 2369 | Bltp3a,Gm49803 | KK,NZB | intron_variant,non_coding_transcript_exon_variant,intron_variant |
| 17 | 29323126 | 29324246 | 1120 | - | B10J,BUB,C58J,FVB,LP,MA/My,NZO,NZW,PL,SJL,SMJ,TH | intergenic_variant |
| 17 | 29529828 | 29531278 | 1450 | BC004004 | C58J,KK,LP,NZO,NZW,RF,SMJ,TH | 3_prime_UTR_variant |
| 17 | 29664735 | 29665736 | 1001 | Gm36199 | RF | intron_variant,non_coding_transcript_variant |
| 17 | 36792193 | 36796868 | 4675 | - | PJ | regulatory_region_variant,intergenic_variant |
| 17 | 37018173 | 37025094 | 6921 | Gm20523 | AKR,B10D2,BALB,C3H,CBA,CE,DBA1J,DBA2J,MA/My,MRL,NZO,NZW,RF,RH2,SEA,TH | upstream_gene_variant,regulatory_region_variant |
| 17 | 38804383 | 38805475 | 1092 | - | DBA1J,NZW,RH2,SEA | intergenic_variant |
| 17 | 38805027 | 38806951 | 1924 | - | AKR,DBA1J,NZW,RH2,SEA | intergenic_variant |
| 17 | 40197297 | 40198709 | 1412 | - | AKR,C3H,CBA,CE,MA/My,MRL,RF | intergenic_variant |

| | | | | | | |
|---|---|---|---|---|---|---|
| 17 | 41998169 | 42004373 | 6204 | - | 129S1,AJ,AKR,BALB,BTBR,BUB,C3H,C57L,C58J,CBA,CE,DBA1J,DBA2J,FVB,ILNJ2,KK,LP,MA/My,MRL,NOD,NOR,NZB,NZO,NZW,PJ,PL,RF,RH2,SEA,SJL,SWR,TH | intergenic_variant |
| 17 | 52681840 | 52683382 | 1542 | - | AJ,AKR,BALB,C3H,CBA,DBA1J,DBA2J,ILNJ2,KK,LP,MA/My,MRL,NOD,NOR,NZB,PJ,PL,RF,RH2,SEA,SMJ,SWR | intergenic_variant |
| 17 | 70009556 | 70011448 | 1892 | - | AKR,NOR,PL,SMJ | intergenic_variant |
| 17 | 70009557 | 70016341 | 6784 | - | AKR,NOD,NOR,PL,SMJ | intergenic_variant |
| 17 | 76231229 | 76232264 | 1035 | - | AJ,AKR,BALB,C3H,CBA,DBA1J,DBA2J,PJ,RF,SEA,SMJ | intergenic_variant |
| 17 | 82157690 | 82159058 | 1368 | - | MA/My,NZW,PL | intergenic_variant |
| 17 | 82956817 | 82959086 | 2269 | - | BUB,CE | intergenic_variant |
| 17 | 89302197 | 89307192 | 4995 | Fshr | KK,NOD,NOR,PJ,RF,SMJ | intron_variant |
| 18 | 13341437 | 13343853 | 2416 | Gm22251 | 129S1,AJ,AKR,BALB,BTBR,BUB,C3H,C57L,CBA,FVB,ILNJ2,KK,RF,RH2,SJL,SMJ,SWR,TH | non_coding_transcript_variant |
| 18 | 13342076 | 13347300 | 5224 | Gm22251 | 129S1,AJ,AKR,BALB,BTBR,BUB,C3H,C57L,CBA,FVB,ILNJ2,KK,RF,RH2,SJL,SMJ,SWR,TH | non_coding_transcript_variant |
| 18 | 30821235 | 30823730 | 2495 | - | 129S1,C57L | intergenic_variant |
| 18 | 69168042 | 69172603 | 4561 | - | KK | intergenic_variant |
| 18 | 70446828 | 70451065 | 4237 | - | BALB,KK,NZB,RF,RH2 | regulatory_region_variant,intergenic_variant |
| 18 | 73205984 | 73207556 | 1572 | - | BUB,CBA,DBA1J,DBA2J,MA/My,PJ,SEA | intergenic_variant |
| 18 | 84204694 | 84205775 | 1081 | - | 129S1,AJ,AKR,B10D2,B10J,BALB,BTBR,BUB,C3H,C57L,C58J,CBA,DBA1J,DBA2J,FVB,ILNJ2,KK,LP,MA/My,MRL,NOR,NZW,PJ,PL,RF,RH2,SEA,SJL,SMJ,SWR,TH | regulatory_region_variant,intergenic_variant |
| 18 | 85156849 | 85159663 | 2814 | - | 129S1,BTBR,C57L | intergenic_variant |
| 18 | 90513761 | 90515808 | 2047 | - | CE | intergenic_variant |
| 19 | 13997596 | 14000718 | 3122 | - | FVB,ILNJ2,LP,SJL,SWR | intergenic_variant |
| 19 | 17280986 | 17287548 | 6562 | - | BALB,BTBR,C57L,CBA,DBA1J,DBA2J,ILNJ2,SEA | intergenic_variant |
| 19 | 19269938 | 19274257 | 4319 | Gm41815 | NZB,NZO,NZW,PJ | intron_variant,non_coding_transcript_variant |
| 19 | 52932484 | 52933553 | 1069 | Xpnpep1 | 129S1,AJ,BALB,BTBR,BUB,C3H,C57L,CBA,CE,DBA1J,DBA2J,ILNJ2,KK,MA/My,MRL,NOD,NOR,NZO,PJ,RF,RH2,SEA,SMJ | intron_variant |
| 19 | 59913368 | 59915779 | 2411 | Rab11fip2 | BTBR,FVB,NZB,PJ,SEA,SMJ,TH | intron_variant |
| 19 | 60894962 | 60896912 | 1950 | Grk5 | TH | intron_variant |
| 2 | 20878372 | 20880715 | 2343 | Arhgap21 | AJ,AKR,DBA2J,PL,RH2 | intron_variant |
| 2 | 21720254 | 21723853 | 3599 | Gpr158 | AJ,AKR,DBA2J,PL,RH2 | intron_variant |
| 2 | 22090297 | 22096984 | 6687 | - | AJ,AKR,CE,DBA2J,FVB,PL,RH2,SJL | intergenic_variant |
| 2 | 42155884 | 42158676 | 2792 | Lrp1b | 129S1,AJ,AKR,BUB,C57L,C58J,DBA1J,ILNJ2,MA/My,MRL,NOD,NOR,NZB,NZO,NZW,PJ,RF,SEA,SWR,TH | intron_variant |
| 2 | 43298615 | 43305073 | 6458 | - | SMJ,TH | intergenic_variant |
| 2 | 43503164 | 43505388 | 2224 | Kynu | BUB | intron_variant,regulatory_region_variant |
| 2 | 50654204 | 50660674 | 6470 | - | AJ,AKR,BUB,C3H,CBA,CE,DBA1J,DBA2J,ILNJ2,MRL,NOD,NOR,PJ,PL,RF,SWR | intergenic_variant |
| 2 | 65233901 | 65235066 | 1165 | - | 129S1,BTBR,BUB,CE,DBA1J,DBA2J,NOD,NOR,RF,SJL,SWR | intergenic_variant |
| 2 | 72543355 | 72547658 | 4303 | 8430437L04Rik | MA/My,NZB | non_coding_transcript_exon_variant,intron_variant |
| 2 | 100030794 | 100031909 | 1115 | - | 129S1,AJ,AKR,BALB,BTBR,BUB,C3H,C57L,CBA,CE,DBA1J,DBA2J,FVB,ILNJ2,KK,MRL,NOD,NOR,NZB,NZO,NZW,PJ,RF,RH2,SEA,SJL,SMJ,SWR | intergenic_variant |
| 2 | 106252287 | 106253996 | 1709 | - | C3H,CE | intergenic_variant |
| 2 | 111429858 | 111431405 | 1547 | - | AKR,BTBR,MRL | intergenic_variant |
| 2 | 111627534 | 111631088 | 3554 | - | C3H,CE,PL | intergenic_variant |
| 2 | 111627579 | 111628753 | 1174 | - | C3H,CE,PL | intergenic_variant |
| 2 | 117308436 | 117310069 | 1633 | - | KK,NZB,TH | regulatory_region_variant,intergenic_variant |
| 2 | 118394410 | 118396338 | 1928 | - | CBA | intergenic_variant |
| 2 | 122038350 | 122041368 | 3018 | Gm14050 | AJ,BALB,C3H,C58J,CBA,DBA1J,DBA2J,ILNJ2,NZB,NZO,PJ,PL,RF,RH2,SEA,SMJ,TH | non_coding_transcript_variant |
| 2 | 128196849 | 128199166 | 2317 | Morrbid | C3H,CBA,DBA1J,DBA2J,ILNJ2,PJ,PL | intron_variant,non_coding_transcript_variant,non_coding_transcript_exon_variant,intron_variant |
| 2 | 131203849 | 131205188 | 1339 | - | DBA1J,DBA2J,ILNJ2,PJ,PL | intergenic_variant |
| 2 | 147717729 | 147720595 | 2866 | Gm25516 | FVB | upstream_gene_variant |
| 2 | 150362640 | 150366591 | 3951 | Gm14128 | BUB,MA/My | non_coding_transcript_variant |
| 2 | 153689126 | 153692481 | 3355 | - | AJ,BTBR,C58J,ILNJ2,MRL,NOD,NZB,NZW,PJ,RF | regulatory_region_variant,intergenic_variant |
| 2 | 153690014 | 153692969 | 2955 | - | AJ,BTBR,C58J,ILNJ2,MRL,NOD,NZB,NZW,PJ,RF | regulatory_region_variant,intergenic_variant |
| 2 | 154451791 | 154454034 | 2243 | 4930519P11Rik | AJ,BTBR,C58J,DBA1J,DBA2J,KK,NZB,NZO,NZW | non_coding_transcript_exon_variant |
| 2 | 157543144 | 157544344 | 1200 | - | 129S1,AJ,AKR,BALB,BUB,C3H,C57L,C58J,CBA,CE,DBA1J,DBA2J,FVB,ILNJ2,MA/My,MRL,NOD,NZB,NZO,NZW,PJ,PL,RF,SEA,SMJ,SWR | regulatory_region_variant,intergenic_variant |
| 2 | 163004043 | 163005274 | 1231 | - | KK | intergenic_variant |
| 2 | 173983253 | 173984423 | 1170 | Gm10714 | PJ | intron_variant,non_coding_transcript_variant |
| 3 | 6274194 | 6276707 | 2513 | - | C58J,CE,DBA1J,DBA2J,FVB,MA/My,MRL,NOD,NOR,PL,SJL,SWR | intergenic_variant |
| 3 | 15297826 | 15300017 | 2191 | - | CE,DBA2J,LP | intergenic_variant |
| 3 | 22951995 | 22954663 | 2668 | - | MA/My,NOD,NOR,PL,SMJ | regulatory_region_variant,intergenic_variant |
| 3 | 27525928 | 27529321 | 3393 | Fndc3b | AJ,AKR,C3H,CBA,CE,PL,RF | intron_variant,regulatory_region_variant |
| 3 | 27527909 | 27530369 | 2460 | Fndc3b | AJ,AKR,C3H,CBA,CE,PL,RF | intron_variant,regulatory_region_variant |
| 3 | 29170943 | 29186408 | 15465 | Egfem1 | PJ | intron_variant |
| 3 | 35827056 | 35830836 | 3780 | Atp11b | DBA1J,RF | intron_variant,regulatory_region_variant |
| 3 | 40467275 | 40475809 | 8534 | - | CE | regulatory_region_variant,intergenic_variant |
| 3 | 48654870 | 48657203 | 2333 | - | CE,SJL,SWR | intergenic_variant |
| 3 | 48655969 | 48657277 | 1308 | - | CE,SJL,SWR | intergenic_variant |
| 3 | 49308972 | 49315027 | 6055 | Gm20557 | AJ,AKR,BALB,BUB,C3H,CBA,CE,DBA1J,DBA2J,FVB,ILNJ2,MRL,NOD,NOR,NZB,PL,RF,RH2,SMJ,TH | non_coding_transcript_exon_variant,intron_variant |
| 3 | 57107557 | 57111400 | 3843 | - | CE,FVB,SJL,SMJ,SWR | regulatory_region_variant,intergenic_variant |

| | | | | | | |
|---|---|---|---|---|---|---|
| 3 | 57107561 | 57109300 | 1739 | - | CE,FVB,SJL,SMJ,SWR | intergenic_variant |
| 3 | 60775631 | 60778616 | 2985 | - | AKR | intergenic_variant |
| 3 | 67442649 | 67449164 | 6515 | - | SJL | intergenic_variant |
| 3 | 69477523 | 69479215 | 1692 | - | BUB,SEA | intergenic_variant |
| 3 | 73877923 | 73879525 | 1602 | - | AKR,FVB,RF,SJL | intergenic_variant |
| 3 | 77573997 | 77576342 | 2345 | - | 129S1,AKR,BALB,BTBR,BUB,C57L,CE,DBA1J,DBA2J,FVB,ILNJ2,KK,MA/My,MRL,NOD,NOR,NZB,NZO,NZW,PJ,PL,RF,RH2,SEA,SMJ,SWR,TH | intergenic_variant |
| 3 | 77574640 | 77582346 | 7706 | - | 129S1,AKR,BALB,BTBR,BUB,C57L,CE,DBA1J,DBA2J,FVB,ILNJ2,KK,MA/My,MRL,NOD,NOR,NZB,NZO,NZW,PJ,PL,RF,RH2,SEA,SMJ,SWR,TH | intergenic_variant |
| 3 | 78400887 | 78403687 | 2800 | - | B10D2,B10J,C58J,LP | regulatory_region_variant,intergenic_variant |
| 3 | 83236047 | 83240286 | 4239 | Dchs2 | NZB,NZO,NZW | intron_variant |
| 3 | 92431401 | 92433954 | 2553 | - | NOD,NOR | regulatory_region_variant,intergenic_variant |
| 3 | 92431402 | 92435603 | 4201 | - | NOD,NOR | regulatory_region_variant,intergenic_variant |
| 3 | 96145778 | 96154682 | 8904 | H2ac19,H2ac18,H3c15,H3c14,Gm20633,Gm20632,Gm54987,Gm54988 | 129S1,AJ,BALB,C3H,C57L,CBA,CE,FVB,KK,MA/My,NOR,NZB,NZO,NZW,PJ,RH2,SEA,SJL,SMJ,SWR | coding_transcript_variant,coding_sequence_variant,5_prime_UTR_variant,3_prime_UTR_variant,non_coding_transcript_variant,regulatory_region_variant |
| 3 | 96147881 | 96152570 | 4689 | H2ac19,H2ac18,Gm20633,Gm20632,Gm54987,Gm54988 | 129S1,AJ,BALB,C3H,C57L,CBA,CE,FVB,KK,MA/My,NOD,NOR,NZB,NZO,NZW,PJ,RH2,SEA,SJL,SMJ,SWR | upstream_gene_variant,non_coding_transcript_exon_variant,downstream_gene_variant,regulatory_region_variant |
| 3 | 96147882 | 96154684 | 6802 | H2ac19,H2ac18,H3c14,Gm20633,Gm20632,Gm54987,Gm54988 | AJ,BALB,C57L,CBA,CE,FVB,KK,NZO,NZW,SJL,SMJ,SWR | upstream_gene_variant,coding_transcript_variant,coding_sequence_variant,5_prime_UTR_variant,3_prime_UTR_variant,non_coding_transcript_variant,non_coding_transcript_exon_variant,downstream_gene_variant,regulatory_region_variant |
| 3 | 96361845 | 96363418 | 1573 | Gm9207 | 129S1,B10J,C57L,CBA,DBA2J,FVB,KK,NOR,NZO,NZW,SJL | non_coding_transcript_variant |
| 3 | 96394526 | 96395555 | 1029 | Gm25890,Gm22614 | 129S1,AJ,B10D2,BALB,C3H,C57L,CBA,CE,FVB,KK,MA/My,NOD,NOR,NZB,NZO,NZW,PJ,RH2,SEA,SJL,SMJ,SWR | upstream_gene_variant,regulatory_region_variant |
| 3 | 109832361 | 109842112 | 9751 | Ntng1 | DBA2J | coding_sequence_variant,intron_variant,regulatory_region_variant |
| 3 | 114147157 | 114149007 | 1850 | - | KK | intergenic_variant |
| 3 | 124917732 | 124919159 | 1427 | - | AJ,BALB,C3H,CBA,NZB,RH2,SEA,SMJ | intergenic_variant |
| 3 | 126580362 | 126581723 | 1361 | Camk2d | 129S1,AJ,BALB,C3H,C57L,CBA,NZB,RH2,SMJ | intron_variant |
| 3 | 129148426 | 129156367 | 7941 | Gm42629 | PJ,SEA | non_coding_transcript_exon_variant,regulatory_region_variant |
| 3 | 137234880 | 137238900 | 4020 | - | BUB,C58J,DBA2J,NZB,NZO,NZW,PL,RF,SMJ | regulatory_region_variant,intergenic_variant |
| 3 | 151663788 | 151665646 | 1858 | Gm2939 | AJ,AKR,B10D2,B10J,BALB,BUB,C3H,C58J,DBA2J,FVB,ILNJ2,LP,MA/My,NOD,NOR,NZB,NZO,NZW,PJ,PL,RF,RH2,SEA,SJL,SWR,TH | non_coding_transcript_variant |
| 3 | 157117706 | 157120346 | 2640 | - | RF | intergenic_variant |
| 4 | 4310289 | 4319584 | 9295 | - | AJ | regulatory_region_variant,intergenic_variant |
| 4 | 8121396 | 8122724 | 1328 | - | 129S1,C58J,LP,MA/My,MRL,RF,SJL | regulatory_region_variant,intergenic_variant |
| 4 | 8479535 | 8485899 | 6364 | - | AJ,KK,NOD,NOR | intergenic_variant |
| 4 | 12431211 | 12437975 | 6764 | - | BUB,FVB,NZB,NZO,NZW,PJ,SEA,SJL,SWR,TH | intergenic_variant |
| 4 | 16798245 | 16800279 | 2034 | - | NOD,NOR | intergenic_variant |
| 4 | 22749880 | 22751463 | 1583 | - | TH | intergenic_variant |
| 4 | 24025454 | 24026479 | 1025 | - | 129S1,AJ,AKR,BALB,BTBR,BUB,CE,FVB,MRL,NOD,NOR,NZO,PJ,PL,RH2,SJL,SWR,TH | intergenic_variant |
| 4 | 67029762 | 67037726 | 7964 | - | 129S1,C57L,PJ | intergenic_variant |
| 4 | 67053400 | 67054616 | 1216 | - | 129S1,C57L,PJ | intergenic_variant |
| 4 | 69993376 | 69998561 | 5185 | - | 129S1,C57L,DBA1J,DBA2J,ILNJ2,KK,NZB,NZO,NZW,PJ,SMJ | intergenic_variant |
| 4 | 71833760 | 71838241 | 4481 | - | ILNJ2,PJ | intergenic_variant |
| 4 | 81413530 | 81415976 | 2446 | - | 129S1,BTBR,C57L | intergenic_variant |
| 4 | 81413546 | 81418236 | 4690 | - | 129S1,BTBR,C57L | regulatory_region_variant,intergenic_variant |
| 4 | 81928733 | 81931417 | 2684 | - | AKR,B10D2,B10J,BUB,C3H,CBA,CE,DBA1J,DBA2J,FVB,KK,NOD,NOR,NZB,NZO,RF,SJL,TH | intergenic_variant |
| 4 | 81929192 | 81937576 | 8384 | - | 129S1,AJ,BALB,BTBR,C57L,RH2,SEA,SWR | regulatory_region_variant,intergenic_variant |
| 4 | 85390046 | 85393635 | 3589 | Sh3gl2 | B10D2,B10J,C58J,DBA1J,DBA2J,ILNJ2,PJ,RF | intron_variant |
| 4 | 91782677 | 91790003 | 7326 | - | CE,MA/My,NZB | intergenic_variant |
| 4 | 98867062 | 98868066 | 1004 | Dock7 | 129S1,BTBR,C57L,MA/My,NOR,SJL,SWR,TH | intron_variant |
| 4 | 110984534 | 110986101 | 1567 | Agbl4 | AKR,KK,MRL,SWR | intron_variant |
| 4 | 112460259 | 112463366 | 3107 | Skint2 | AJ,BUB,C3H,CE,PL,SEA,SMJ,SWR | intron_variant |
| 4 | 112974468 | 112978853 | 4385 | Skint6 | AJ | intron_variant |
| 4 | 112978183 | 112980675 | 2492 | Skint6 | AJ,DBA1J,FVB | intron_variant |
| 4 | 117792527 | 117793810 | 1283 | St3gal3 | AKR,ILNJ2,KK,MRL | intron_variant |
| 4 | 129557150 | 129558407 | 1257 | Kpna6 | 129S1,AJ,AKR,BALB,BTBR,BUB,C3H,C57L,CBA,CE,DBA2J,FVB,ILNJ2,MA/My,MRL,NOD,NZO,NZW,RF,RH2,SJL,SWR,TH | upstream_gene_variant,intron_variant |
| 4 | 137066185 | 137067431 | 1246 | Cdc42 | C3H,CBA,NZW | intron_variant,regulatory_region_variant |
| 4 | 144478474 | 144487001 | 8527 | - | BTBR | regulatory_region_variant,intergenic_variant |
| 4 | 144498569 | 144499643 | 1074 | - | 129S1,BTBR,C57L | intergenic_variant |
| 5 | 3365861 | 3368771 | 2910 | - | BUB,NOD,NOR | intergenic_variant |
| 5 | 6262520 | 6265739 | 3219 | - | C58J,MA/My,MRL | intergenic_variant |
| 5 | 8013404 | 8015361 | 1957 | Steap4 | 129S1,BALB,BTBR,C57L,CE,NZB,NZO,NZW,PJ,PL,RF,SEA,SJL,SWR | intron_variant,regulatory_region_variant |
| 5 | 9338913 | 9343521 | 4608 | Elapor2 | NOD,NOR | intron_variant |
| 5 | 15663097 | 15664556 | 1459 | - | TH | regulatory_region_variant,intergenic_variant |
| 5 | 15726265 | 15727288 | 1023 | Gm21847 | NZO | intron_variant,non_coding_transcript_variant,upstream_gene_variant,regulatory_region_variant |
| 5 | 15829599 | 15832412 | 2813 | Speer4d,Gm21847 | FVB,MA/My,RF,SJL,SWR | downstream_gene_variant,intron_variant,non_coding_transcript_variant,regulatory_region_variant |
| 5 | 18249791 | 18251423 | 1632 | - | AKR,TH | intergenic_variant |
| 5 | 21913140 | 21914152 | 1012 | - | 129S1,BTBR,C57L,CE,DBA1J,DBA2J,NOD,NOR,NZO | intergenic_variant |

| | | | | | | |
|---|---|---|---|---|---|---|
| 5 | 23223380 | 23224676 | 1296 | Lhfpl3 | 129S1,AKR,BALB,BTBR,BUB,C57L,C58J,CE,DBA1J,DBA2J,FVB,KK,MA/My,MRL,NOD,NOR,NZO,NZW,PJ,PL,RF,RH2,SEA,SJL,SMJ,SWR,TH | intron_variant |
| 5 | 47438629 | 47446518 | 7889 | - | FVB,SJL,SWR | intergenic_variant |
| 5 | 49902527 | 49906049 | 3522 | - | NOR | intergenic_variant |
| 5 | 51112189 | 51115758 | 3569 | - | KK,MA/My,NOD,NZB,NZO,NZW,PJ,SJL,SWR,TH | intergenic_variant |
| 5 | 57907257 | 57909241 | 1984 | Pcdh7 | NZO | intron_variant |
| 5 | 62482446 | 62483562 | 1116 | - | TH | intergenic_variant |
| 5 | 66863982 | 66865055 | 1073 | - | BALB,RH2,SEA | intergenic_variant |
| 5 | 69691173 | 69693192 | 2019 | Yipf7 | BTBR,PL | intron_variant |
| 5 | 77915142 | 77916417 | 1275 | - | SWR | intergenic_variant |
| 5 | 79798289 | 79803794 | 5505 | - | 129S1,BALB,C57L,C58J,LP,NZO,NZW,RH2,SEA | intergenic_variant |
| 5 | 83309858 | 83311365 | 1507 | - | KK,NZB,NZO,NZW | intergenic_variant |
| 5 | 102497453 | 102498865 | 1412 | - | AKR,CE,ILNJ2,MRL,NOD,NOR,NZO,NZW,PJ,PL,RF,SMJ,TH | intergenic_variant |
| 5 | 104736052 | 104956008 | 219956 | Gm8258 | AJ,BALB,BUB,C3H,CBA,DBA1J,DBA2J,FVB,ILNJ2,NZB,NZO,RH2,SEA,SJL,SWR,TH | non_coding_transcript_variant,regulatory_region_variant |
| 5 | 104752780 | 104993577 | 240797 | Gm8258,Zfp951 | SWR | non_coding_transcript_variant,coding_transcript_variant,coding_sequence_variant,3_prime_UTR_variant,intron_variant,regulatory_region_variant |
| 5 | 104922219 | 105005445 | 83226 | Gm8258,Zfp951 | BUB,NZO,RH2 | non_coding_transcript_variant,coding_transcript_variant,coding_sequence_variant,3_prime_UTR_variant,intron_variant,regulatory_region_variant |
| 5 | 107393133 | 107396597 | 3464 | Tgfbr3 | B10J | intron_variant,regulatory_region_variant |
| 5 | 107393835 | 107396633 | 2798 | Tgfbr3 | B10J | intron_variant,regulatory_region_variant |
| 5 | 135652140 | 135657838 | 5698 | - | AKR,BUB,ILNJ2,MRL | regulatory_region_variant,intergenic_variant |
| 6 | 14440710 | 14446958 | 6248 | - | BALB,C3H,CBA,SWR | intergenic_variant |
| 6 | 19973397 | 19974812 | 1415 | - | SMJ | intergenic_variant |
| 6 | 57008619 | 57010073 | 1454 | Vmn1r8 | 129S1,KK,NZO,NZW,PJ | 5_prime_UTR_variant,intron_variant |
| 6 | 60411944 | 60416634 | 4690 | - | CE,ILNJ2,PJ | intergenic_variant |
| 6 | 60411945 | 60415983 | 4038 | - | CE | intergenic_variant |
| 6 | 60412577 | 60415981 | 3404 | - | CE,ILNJ2,PJ | intergenic_variant |
| 6 | 65444756 | 65451849 | 7093 | - | KK | intergenic_variant |
| 6 | 68911192 | 68913106 | 1914 | - | AKR,C58J,KK,MRL,NOD,NOR,PL,RF | intergenic_variant |
| 6 | 73983966 | 73987187 | 3221 | - | C58J,KK,NOD,NOR | intergenic_variant |
| 6 | 74313657 | 74318708 | 5051 | Gm26179 | AKR,BALB,MRL,PL,RF,RH2,SEA | non_coding_transcript_variant |
| 6 | 74316227 | 74318714 | 2487 | Gm26179 | AKR,BALB,MRL,PL,RF,RH2,SEA | non_coding_transcript_variant |
| 6 | 74895470 | 74896651 | 1181 | - | C58J,KK,NOD,NOR | intergenic_variant |
| 6 | 79284022 | 79290271 | 6249 | - | KK,NZB,NZO,NZW,TH | intergenic_variant |
| 6 | 90953713 | 90954966 | 1253 | Iqsec1 | CE,NZB,NZW,PJ,SEA | intron_variant |
| 6 | 95388800 | 95402030 | 13230 | - | CE | intergenic_variant |
| 6 | 95389653 | 95402030 | 12377 | - | CE | intergenic_variant |
| 6 | 95389745 | 95399166 | 9421 | - | CE | intergenic_variant |
| 6 | 103454245 | 103459049 | 4804 | - | NOD | intergenic_variant |
| 6 | 103466663 | 103469161 | 2498 | - | BUB,NOD,NOR | intergenic_variant |
| 6 | 105921660 | 105923210 | 1550 | Cntn4 | AJ,AKR,BALB,BUB,C3H,CBA,CE,DBA1J,ILNJ2,KK,MRL,NZB,NZO,NZW,PJ,PL,RF,RH2,SEA | intron_variant |
| 6 | 110386966 | 110400737 | 13771 | - | 129S1,AJ,BALB,BTBR,BUB,C57L,CE,DBA1J,DBA2J,FVB,NZB,NZO,NZW,PJ,RH2,SEA,SJL,SMJ,SWR,TH | intergenic_variant |
| 6 | 116008530 | 116011993 | 3463 | Tmcc1 | AKR,C3H,CBA,ILNJ2,MRL,PL,RF | intron_variant,regulatory_region_variant |
| 6 | 121674145 | 121681013 | 6868 | - | CE | regulatory_region_variant,intergenic_variant |
| 6 | 123948582 | 123951944 | 3362 | A330044P14Rik | CE | intron_variant,non_coding_transcript_variant,regulatory_region_variant |
| 6 | 124539172 | 124540920 | 1748 | Gm18840 | SWR | non_coding_transcript_exon_variant,intron_variant |
| 6 | 130726785 | 130733401 | 6616 | - | NZW | intergenic_variant |
| 6 | 131217016 | 131218210 | 1194 | Klra2 | AJ,AKR,BALB,C3H,C57L,CBA,CE,DBA1J,DBA2J,FVB,ILNJ2,KK,LP,MA/My,MRL,NZB,NZO,NZW,PL,RF,RH2,SEA,SJL,TH | intron_variant |
| 6 | 132245958 | 132247805 | 1847 | - | 129S1,AJ,AKR,BALB,BUB,C3H,C57L,CBA,DBA1J,DBA2J,ILNJ2,MRL,NZB,PL,RF,RH2,SEA,SMJ | intergenic_variant |
| 6 | 132246016 | 132252885 | 6869 | - | 129S1,AJ,AKR,BALB,BUB,C3H,C57L,CBA,DBA1J,DBA2J,ILNJ2,MRL,NZB,PL,RF,RH2,SEA | intergenic_variant |
| 6 | 132246036 | 132250290 | 4254 | - | 129S1,AJ,AKR,BALB,BUB,C3H,C57L,CBA,DBA1J,DBA2J,ILNJ2,MRL,NZB,PL,RF,RH2,SEA,SMJ | intergenic_variant |
| 7 | 7219420 | 7226136 | 6716 | - | 129S1,AJ,BALB,BTBR,BUB,C57L,C58J,CBA,CE,ILNJ2,KK,LP,NOD,NOR,NZB,NZO,NZW,PL,RF,SJL,SWR,TH | regulatory_region_variant,intergenic_variant |
| 7 | 15010191 | 15017172 | 6981 | - | AJ,AKR,BTBR,ILNJ2,LP,MRL,RH2,SJL | intergenic_variant |
| 7 | 31064904 | 31071721 | 6817 | - | 129S1,C58J,LP,MA/My,MRL,RF,SJL | intergenic_variant |
| 7 | 40655095 | 40658853 | 3758 | - | NZB,NZO,SMJ | intergenic_variant |
| 7 | 40657738 | 40660189 | 2451 | - | NZB,NZO,SMJ | intergenic_variant |
| 7 | 43551188 | 43564130 | 12942 | - | AJ,BALB,C57L,CE,NZB,NZO,NZW,RH2,SEA,SJL,SMJ,TH | intergenic_variant |
| 7 | 43553200 | 43562118 | 8918 | - | AJ,BALB,C57L,CE,DBA1J,DBA2J,ILNJ2,NZB,NZO,NZW,PJ,RH2,SEA,SJL,SMJ,TH | intergenic_variant |
| 7 | 46844210 | 46845732 | 1522 | Mrgpra6 | ILNJ2,NZW,PJ,SJL | intron_variant |
| 7 | 47456469 | 47460815 | 4346 | Mrgprc7-ps | C57L | non_coding_transcript_variant |
| 7 | 48796515 | 48797705 | 1190 | Nav2,9130015G15Rik | ILNJ2,NZW,PJ,SJL | intron_variant,non_coding_transcript_variant,regulatory_region_variant |
| 7 | 52418974 | 52425626 | 6652 | - | MA/My,SJL,TH | intergenic_variant |
| 7 | 54520508 | 54522578 | 2070 | Luzp2 | CE,ILNJ2,KK,MA/My,NZB,NZO,NZW,PJ,SJL,TH | intron_variant |
| 7 | 56209546 | 56212372 | 2826 | - | ILNJ2,KK,NZW,PJ,SJL | intergenic_variant |
| 7 | 56211710 | 56212821 | 1111 | - | ILNJ2,KK,NZW,PJ,SJL | intergenic_variant |
| 7 | 62528034 | 62529598 | 1564 | - | KK | intergenic_variant |
| 7 | 62528034 | 62529598 | 1564 | - | KK | intergenic_variant |
| 7 | 70703806 | 70708637 | 4831 | Gm42397 | NZB,NZO | intron_variant,non_coding_transcript_variant |

| | | | | | | |
|---|---|---|---|---|---|---|
| 7 | 85453808 | 85455451 | 1643 | - | 129S1,BTBR,C3H,C57L,CBA,DBA1J,DBA2J,ILNJ2,KK,NZB,NZO,NZW,PJ,SEA,SMJ | intergenic_variant |
| 7 | 102448218 | 102451884 | 3666 | Or51f23 | KK,NOD,SJL | upstream_gene_variant |
| 7 | 105108322 | 105109497 | 1175 | Cckbr | BUB,FVB,NZO,SJL,TH | intron_variant |
| 7 | 108684248 | 108686144 | 1896 | - | NOD,SMJ,SWR | intergenic_variant |
| 7 | 111549136 | 111550348 | 1212 | - | NZO | intergenic_variant |
| 7 | 113637287 | 113638547 | 1260 | Spon1 | CE | intron_variant |
| 7 | 127827009 | 127828315 | 1306 | - | 129S1,BTBR,C57L,ILNJ2,KK,PJ,TH | intergenic_variant |
| 7 | 140737759 | 140739107 | 1348 | Ptdss2 | NZW | downstream_gene_variant |
| 8 | 14840490 | 14842811 | 2321 | Dlgap2 | NZB | intron_variant |
| 8 | 27852617 | 27854657 | 2040 | - | PL | regulatory_region_variant,intergenic_variant |
| 8 | 31717827 | 31720241 | 2414 | Fut10 | 129S1,AJ,AKR,BALB,BUB,C3H,C57L,CBA,DBA1J,DBA2J,FVB,MA/My,MRL,NOD,NOR,NZW,PL,RF,RH2,SJL,SMJ,SWR,TH | intron_variant |
| 8 | 31718757 | 31721377 | 2620 | Fut10 | 129S1,AJ,AKR,BALB,BUB,C3H,C57L,CBA,DBA1J,DBA2J,FVB,MA/My,MRL,NOD,NOR,PL,RF,RH2,SJL,SMJ,SWR,TH | intron_variant |
| 8 | 31842247 | 31844294 | 2047 | - | 129S1,AJ,AKR,BALB,BUB,C3H,C57L,CBA,DBA1J,DBA2J,FVB,ILNJ2,MA/My,MRL,NOD,NOR,NZW,PL,RF,RH2,SJL,SMJ,SWR,TH | intergenic_variant |
| 8 | 60552222 | 60553420 | 1198 | - | AKR,BALB,BTBR,BUB,C57L,CBA,CE,DBA1J,DBA2J,ILNJ2,NZO,PJ,RF,SEA | intergenic_variant |
| 8 | 60777498 | 60778552 | 1054 | - | 129S1,BALB,C57L,CE,DBA2J,KK,NZO,SMJ | intergenic_variant |
| 8 | 63750890 | 63754269 | 3379 | Spock3 | KK | intron_variant,regulatory_region_variant |
| 8 | 74611075 | 74612869 | 1794 | - | AKR,C57L,CE,MA/My,MRL,NOD,NOR,PL | intergenic_variant |
| 8 | 74765149 | 74766945 | 1796 | - | 129S1,BALB,BUB,C3H,CBA,DBA1J,DBA2J,FVB,ILNJ2,NZB,NZO,RF,SJL,SMJ,SWR | intergenic_variant |
| 8 | 81866485 | 81868076 | 1591 | - | 129S1,AKR,BALB,BTBR,BUB,C57L,MRL,NOD,NOR,NZW,PJ,PL,RF,SEA,SJL | intergenic_variant |
| 8 | 86323117 | 86327431 | 4314 | Gm45424 | 129S1,AJ,AKR,BALB,BTBR,C3H,C57L,CBA,CE,DBA1J,DBA2J,FVB,ILNJ2,KK,MA/My,MRL,PJ,PL,RF,RH2,SEA,SMJ | non_coding_transcript_exon_variant,regulatory_region_variant |
| 8 | 113108924 | 113110120 | 1196 | - | BALB,NZW,RH2,SEA | intergenic_variant |
| 8 | 129986988 | 129988195 | 1207 | - | KK | intergenic_variant |
| 9 | 5170075 | 5172538 | 2463 | - | 129S1,AKR,CBA,RF | intergenic_variant |
| 9 | 12373256 | 12379757 | 6501 | - | AKR,MRL,RF | intergenic_variant |
| 9 | 36087762 | 36089469 | 1707 | - | C57L,NZO | intergenic_variant |
| 9 | 36284050 | 36288364 | 4314 | Gm48401 | C57L,NZO | non_coding_transcript_exon_variant |
| 9 | 41394654 | 41395746 | 1092 | Mir100hg,Gm48742 | CE | intron_variant,non_coding_transcript_variant,regulatory_region_variant |
| 9 | 57134250 | 57135483 | 1233 | - | MA/My | regulatory_region_variant,intergenic_variant |
| 9 | 65753564 | 65755165 | 1601 | Trip4 | 129S1,AJ,AKR,B10D2,B10J,BALB,BTBR,BUB,C3H,C57L,CE,DBA1J,DBA2J,FVB,ILNJ2,KK,LP,MA/My,MRL,NOD,NOR,NZB,NZO,PJ,PL,RF,RH2,SEA,SJL,SMJ,SWR,TH | intron_variant,regulatory_region_variant |
| 9 | 68398173 | 68399900 | 1727 | 4930502A04Rik | AJ,BALB,BUB,C3H,CBA,DBA1J,DBA2J,FVB,ILNJ2,MA/My,NOD,NOR,NZO,PJ,RH2,SEA,SJL,SWR | intron_variant,non_coding_transcript_variant |
| 9 | 68398180 | 68401241 | 3061 | 4930502A04Rik | AJ,BALB,BUB,C3H,CBA,DBA1J,DBA2J,FVB,ILNJ2,MA/My,NOD,NOR,NZO,PJ,RH2,SEA,SJL,SWR | intron_variant,non_coding_transcript_variant |
| 9 | 78223497 | 78232046 | 8549 | Gm8074 | LP,NZW | non_coding_transcript_variant,regulatory_region_variant |
| 9 | 78225547 | 78229776 | 4229 | Gm8074 | C58J,ILNJ2,LP,NZW | non_coding_transcript_variant,regulatory_region_variant |
| 9 | 79747426 | 79748845 | 1419 | Filip1 | 129S1,BTBR,C57L,C58J,ILNJ2,LP,NZW | intron_variant |
| 9 | 96234706 | 96236448 | 1742 | Atp1b3 | MRL,NZO,NZW | intron_variant,regulatory_region_variant |
| 9 | 99937370 | 99938480 | 1110 | - | NOD,NOR,RF | intergenic_variant |
| 9 | 105377767 | 105379197 | 1430 | Atp2c1 | 129S1,AJ,AKR,BALB,BTBR,BUB,C3H,C57L,CBA,CE,DBA1J,DBA2J,FVB,ILNJ2,KK,MRL,NOD,NOR,NZB,NZO,NZW,PJ,PL,RF,RH2,SEA,SJL,SMJ,SWR,TH | intron_variant |
| X | 5776993 | 5778346 | 1353 | - | PL | intergenic_variant |
| X | 32098857 | 32101935 | 3078 | - | C57L,KK,NZB,NZO | intergenic_variant |
| X | 79466032 | 79467495 | 1463 | - | AJ,AKR,BALB,BUB,C3H,CBA,DBA1J,DBA2J,KK,MRL,NZO,NZW,PJ,PL,RF,RH2,SEA,SJL,SMJ | intergenic_variant |
| X | 113900762 | 113907578 | 6816 | - | AJ,BALB,C3H,CBA,MRL | regulatory_region_variant,intergenic_variant |
| X | 116358549 | 116359647 | 1098 | - | AJ,BALB,C3H,CBA,MRL,RH2 | intergenic_variant |
| X | 117072618 | 117075203 | 2585 | - | AJ,BALB,C3H,CBA,MRL | intergenic_variant |
| X | 117073645 | 117076032 | 2387 | - | RH2 | intergenic_variant |
| X | 145678270 | 145680428 | 2158 | - | NZB | intergenic_variant |
| X | 145679192 | 145680445 | 1253 | - | NZB | intergenic_variant |
| X | 151482254 | 151487885 | 5631 | Shroom2 | AKR,BUB,RF | intron_variant,regulatory_region_variant |
| X | 168856708 | 168859029 | 2321 | - | C58J,MA/My,PL | coding_transcript_variant |

| CHR | START | END | SIZE | SYMBOL | STRAINS with VARIANT SV ALLELE | CONSEQUENCE |
|---|---|---|---|---|---|---|
| 1 | 8577511 | 8577870 | 359 | Sntg1 | MOLF | intron_variant |
| 1 | 9101845 | 9102131 | 286 | Sntg1 | 129S1,AKR,BTBR,C57L,ILNJ2,LP,NOD,NOR,PJ,PL,SMJ,SWR,TH | intron_variant |
| 1 | 9716534 | 9717052 | 518 | - | CAST | intergenic_variant |
| 1 | 9946199 | 9946452 | 253 | Sgk3,Snhg6 | CAST | intron_variant,non_coding_transcript_variant |
| 1 | 19418764 | 19419319 | 555 | - | PL | intergenic_variant |
| 1 | 30135020 | 30135652 | 632 | - | SPRET | intergenic_variant |
| 1 | 31755028 | 31755830 | 802 | - | CE,MOLF | intergenic_variant |
| 1 | 34941022 | 34941796 | 774 | - | SPRET | intergenic_variant |
| 1 | 38027683 | 38028310 | 627 | Txndc9 | SPRET | intron_variant,3_prime_UTR_variant |
| 1 | 40188818 | 40189757 | 939 | - | 129S1,BTBR,C57L,NZB,NZO | intergenic_variant |
| 1 | 40606335 | 40607020 | 685 | - | C3H,KK,NZB,PJ,TH | intergenic_variant |
| 1 | 45437972 | 45438382 | 410 | Col5a2 | CAST,FVB,MA/My,MOLF,NOD,NOR,RH2,WSB | intron_variant |
| 1 | 47013665 | 47014006 | 341 | - | AKR,BUB,C58J,RF | intergenic_variant |
| 1 | 47291023 | 47291466 | 443 | - | SPRET | intergenic_variant |
| 1 | 48811258 | 48811787 | 529 | - | AKR,BALB,BUB,C58J,MA/My,RF,RH2,SEA | intergenic_variant |
| 1 | 50717802 | 50718028 | 226 | - | MOLF | intergenic_variant |
| 1 | 50894398 | 50894701 | 303 | - | CAST,SPRET | intergenic_variant |
| 1 | 53683073 | 53683302 | 229 | Dnah7a | SPRET | intron_variant |
| 1 | 54698971 | 54699418 | 447 | Ankrd44 | CAST,MOLF,NZO,NZW | intron_variant |
| 1 | 55525702 | 55526022 | 320 | Plcl1 | AKR,BALB,BUB,CAST,MRL,RH2,SEA,SJL | intron_variant |
| 1 | 62441195 | 62441922 | 727 | Pard3b | SPRET | intron_variant |
| 1 | 63261313 | 63262084 | 771 | - | SPRET | intergenic_variant |
| 1 | 65475776 | 65476646 | 870 | - | SPRET | intergenic_variant |
| 1 | 66831537 | 66831785 | 248 | Kansl1l | AJ,CBA,DBA1J,NZW,SWR | intron_variant |
| 1 | 69286163 | 69286747 | 584 | - | 129S1,AJ,BTBR,C3H,C57L,DBA2J,FVB,KK,LP,MA/My,MRL,NOD,NOR,NZB,NZO,NZW,SMJ,TH | regulatory_region_variant,intergenic_variant |
| 1 | 73270460 | 73271414 | 954 | - | CAST | intergenic_variant |
| 1 | 90327974 | 90328245 | 271 | - | CAST | intergenic_variant |
| 1 | 94043886 | 94044472 | 586 | - | MOLF,NZO,PL | intergenic_variant |
| 1 | 99837338 | 99838052 | 714 | Cntnap5b | 129S1,AKR,BUB,C57L,DBA1J,DBA2J,LP,MA/My,NOD,NOR,NZO,PJ,SEA,SJL,TH,WSB | intron_variant |
| 1 | 102148878 | 102149517 | 639 | - | SPRET | intergenic_variant |
| 1 | 106119816 | 106120444 | 628 | Phlpp1 | SPRET | intron_variant,regulatory_region_variant |
| 1 | 110137904 | 110138250 | 346 | - | CAST | intergenic_variant |
| 1 | 111643496 | 111643878 | 382 | - | SPRET | intergenic_variant |
| 1 | 122283100 | 122283353 | 253 | - | MOLF,SPRET | intergenic_variant |
| 1 | 124449685 | 124450209 | 524 | Dpp10 | SPRET | intron_variant |
| 1 | 129294240 | 129295102 | 862 | Thsd7b | MOLF | intron_variant |
| 1 | 129554314 | 129554617 | 303 | Thsd7b | SPRET | intron_variant |
| 1 | 130518233 | 130518511 | 278 | Zp3r | MOLF | intron_variant |
| 1 | 133078132 | 133078719 | 587 | - | SPRET | regulatory_region_variant,intergenic_variant |
| 1 | 133395416 | 133396006 | 590 | - | SPRET | regulatory_region_variant,intergenic_variant |
| 1 | 133461806 | 133462752 | 946 | - | SPRET | intergenic_variant |
| 1 | 134557055 | 134557403 | 348 | Kdm5b | SPRET | intron_variant |
| 1 | 136827825 | 136828377 | 552 | Nr5a2 | CAST | intron_variant |
| 1 | 139273370 | 139273926 | 556 | Crb1 | CAST,MOLF | intron_variant,regulatory_region_variant |
| 1 | 139727825 | 139728519 | 694 | - | SPRET | intergenic_variant |
| 1 | 141183299 | 141184137 | 838 | Gm4845 | SPRET | downstream_gene_variant |
| 1 | 143014748 | 143015716 | 968 | - | MOLF | intergenic_variant |
| 1 | 143016519 | 143016790 | 271 | - | SPRET | intergenic_variant |
| 1 | 150074280 | 150075235 | 955 | - | BTBR,CAST,MOLF,SMJ,SPRET | intergenic_variant |
| 1 | 150173354 | 150173704 | 350 | - | CAST | intergenic_variant |
| 1 | 150615702 | 150616529 | 827 | Hmcn1 | SPRET | intron_variant |
| 1 | 164971369 | 164971691 | 322 | Tbx19 | CAST | intron_variant |
| 1 | 170442569 | 170443368 | 799 | - | 129S1,AJ,BTBR,BUB,C3H,C57L,CAST,CE,FVB,KK,MRL,NOD,NOR,NZB,NZO,NZW,PJ,PL,RF,RH2,SEA,SJL,SMJ,SPRET,SWR,TH,WSB | intergenic_variant |
| 1 | 171089657 | 171090107 | 450 | - | CBA,DBA1J,DBA2J,KK,NZW,TH | regulatory_region_variant,intergenic_variant |
| 1 | 174694278 | 174695123 | 845 | Grem2 | CAST | intron_variant |
| 1 | 175205685 | 175206314 | 629 | Rgs7,5730408A14Rik | 129S1,AJ,AKR,BALB,BUB,C3H,C57L,C58J,CBA,CE,DBA1J,DBA2J,FVB,ILNJ2,KK,MRL,NOD,NOR,NZB,NZW,PJ,RF,RH2,SEA,SJL,SMJ,SWR,WSB | intron_variant,downstream_gene_variant |
| 1 | 177534659 | 177534938 | 279 | - | FVB,NZO | intergenic_variant |
| 1 | 181365228 | 181365588 | 360 | - | SPRET | intergenic_variant |
| 1 | 181587270 | 181587976 | 706 | Dnah14 | SPRET | intron_variant |
| 1 | 186587760 | 186588145 | 385 | - | 129S1,AKR,BALB,BTBR,C3H,C57L,CAST,CBA,CE,DBA1J,DBA2J,ILNJ2,KK,MRL,NZW,PJ,PL,RH2,SEA,SMJ | regulatory_region_variant,intergenic_variant |
| 1 | 188671832 | 188672579 | 747 | Ush2a | SPRET | intron_variant |
| 1 | 188672351 | 188672582 | 231 | Ush2a | SPRET | intron_variant |
| 1 | 191319489 | 191319816 | 327 | Ints7 | TH | intron_variant,regulatory_region_variant |
| 1 | 194871917 | 194872462 | 545 | - | NZO,NZW,RF | intergenic_variant |
| 10 | 3436046 | 3436803 | 757 | - | MOLF | intergenic_variant |
| 10 | 6066074 | 6067025 | 951 | - | SPRET | intergenic_variant |
| 10 | 7439694 | 7440287 | 593 | - | SPRET | intergenic_variant |
| 10 | 15002521 | 15002964 | 443 | - | CAST | intergenic_variant |
| 10 | 18098448 | 18098730 | 282 | Ccdc28a | AJ,BALB,BTBR,C3H,C57L,CAST,CBA,CE,DBA1J,DBA2J,MA/My,MRL,PL,RH2,SJL,SMJ,SPRET | intron_variant |
| 10 | 23721483 | 23721947 | 464 | - | MOLF | intergenic_variant |
| 10 | 27209713 | 27210503 | 790 | Lama2 | MOLF | intron_variant |
| 10 | 28104912 | 28105649 | 737 | Ptprk | SPRET | intron_variant,regulatory_region_variant |
| 10 | 28899715 | 28900349 | 634 | - | SPRET | intergenic_variant |
| 10 | 29069753 | 29070492 | 739 | Mtcl3 | SPRET | intron_variant |
| 10 | 29732943 | 29733678 | 735 | - | SPRET | intergenic_variant |
| 10 | 30118674 | 30119324 | 650 | - | SPRET | intergenic_variant |
| 10 | 30925793 | 30926484 | 691 | Gm30676 | SPRET | intron_variant,non_coding_transcript_variant,regulatory_region_variant |

| | | | | | | |
|---|---|---|---|---|---|---|
| 10 | 31069992 | 31070349 | 357 | - | SPRET | intergenic_variant |
| 10 | 32333134 | 32333873 | 739 | Nkain2 | MOLF | intron_variant |
| 10 | 35312293 | 35312730 | 437 | - | MOLF | intergenic_variant |
| 10 | 36481035 | 36481781 | 746 | Hs3st5 | MOLF | intron_variant |
| 10 | 36807624 | 36807980 | 356 | - | CAST,MOLF,SPRET | intergenic_variant |
| 10 | 37931265 | 37932046 | 781 | - | CAST | intergenic_variant |
| 10 | 43643646 | 43643865 | 219 | - | CAST,MOLF | regulatory_region_variant,intergenic_variant |
| 10 | 46471994 | 46472545 | 551 | - | CAST | intergenic_variant |
| 10 | 47270793 | 47271205 | 412 | - | SPRET | intergenic_variant |
| 10 | 48647679 | 48648014 | 335 | - | BALB,BTBR,CAST,KK,MOLF,NZW,PJ,SEA,SPRET,TH,WSB | intergenic_variant |
| 10 | 49464020 | 49464398 | 378 | Grik2 | SPRET | intron_variant |
| 10 | 51515201 | 51515648 | 447 | - | MOLF | intergenic_variant |
| 10 | 52761951 | 52762566 | 615 | Slc35f1 | SPRET | intron_variant |
| 10 | 59311531 | 59312062 | 531 | Mcu | SPRET | intron_variant,regulatory_region_variant |
| 10 | 65451892 | 65452319 | 427 | - | SPRET | intergenic_variant |
| 10 | 70138398 | 70138931 | 533 | - | SPRET | regulatory_region_variant,intergenic_variant |
| 10 | 77450145 | 77450817 | 672 | Sumo3 | SPRET | intron_variant,regulatory_region_variant |
| 10 | 88660071 | 88660452 | 381 | Utp20 | BUB,MOLF | intron_variant |
| 10 | 93389643 | 93390278 | 635 | Ccdc38 | 129S1,AJ,AKR,BALB,BTBR,BUB,C3H,C57L,C58J,CBA,CE,DBA1J,DBA2J,FVB,ILNJ2,KK,LP,MA/My,MRL,NOD,NOR,NZO,NZW,PJ,PL,RF,RH2,SEA,SJL,SMJ,SWR,TH,WSB | intron_variant,upstream_gene_variant |
| 10 | 95001302 | 95001976 | 674 | Gm48874 | CAST | non_coding_transcript_exon_variant |
| 10 | 95297336 | 95297957 | 621 | Gm47599 | PL | upstream_gene_variant |
| 10 | 100074939 | 100075857 | 918 | - | AKR,BUB,C3H,CBA,FVB,MA/My,NOD,NOR,PL,SMJ | intergenic_variant |
| 10 | 100114815 | 100115119 | 304 | - | CAST | intergenic_variant |
| 10 | 100666965 | 100667209 | 244 | - | CBA,NZB | intergenic_variant |
| 10 | 101813294 | 101813686 | 392 | - | CAST | intergenic_variant |
| 10 | 102788171 | 102788662 | 491 | - | SPRET | intergenic_variant |
| 10 | 104626702 | 104627499 | 797 | - | AJ,BALB,BTBR,CE,DBA1J,DBA2J,MRL,NZB,PJ,RH2,SEA,SJL | intergenic_variant |
| 10 | 105870133 | 105870860 | 727 | - | WSB | intergenic_variant |
| 10 | 106497875 | 106498335 | 460 | Ppfia2 | BTBR,CAST,NZB | intron_variant |
| 10 | 110112347 | 110112819 | 472 | Nav3 | MOLF,SMJ | intron_variant |
| 10 | 110949978 | 110950342 | 364 | - | SPRET | intergenic_variant |
| 10 | 112061798 | 112062091 | 293 | Gm47419 | MOLF,SPRET | intron_variant,non_coding_transcript_variant |
| 10 | 112959419 | 112960365 | 946 | - | MA/My,MOLF,WSB | intergenic_variant |
| 10 | 115637350 | 115638131 | 781 | 4930422I22Rik | SPRET | downstream_gene_variant |
| 10 | 117161539 | 117162229 | 690 | - | SPRET | intergenic_variant |
| 10 | 120643604 | 120643851 | 247 | Msrb3 | SPRET | intron_variant |
| 10 | 125880282 | 125880895 | 613 | - | SPRET | intergenic_variant |
| 10 | 126340151 | 126340751 | 600 | - | AJ,BALB,CAST,ILNJ2,KK,LP,MA/My,MOLF,NZW,PJ,RF,RH2,SEA | intergenic_variant |
| 11 | 4419801 | 4420791 | 990 | - | AKR,B10D2,B10J,C58J,CAST,CE,FVB,KK,MOLF,MRL,RF,SJL,SWR | intergenic_variant |
| 11 | 8257235 | 8257744 | 509 | - | B10D2,B10J,BUB,C3H,C58J,CAST,CE,NOR,NZB,PJ,SJL,SPRET,SWR,WSB | intergenic_variant |
| 11 | 8951691 | 8952442 | 751 | Hus1 | SPRET | intron_variant |
| 11 | 9885191 | 9885796 | 605 | - | SPRET | intergenic_variant |
| 11 | 11321836 | 11322230 | 394 | Zpbp | SPRET | intron_variant |
| 11 | 11420893 | 11421423 | 530 | Spmip7 | SPRET | intron_variant |
| 11 | 14381766 | 14382295 | 529 | - | SPRET | intergenic_variant |
| 11 | 16698481 | 16698851 | 370 | - | SEA | intergenic_variant |
| 11 | 17346227 | 17347122 | 895 | - | SPRET | intergenic_variant |
| 11 | 18130963 | 18131477 | 514 | - | 129S1,AJ,AKR,BALB,BTBR,BUB,C3H,C57L,C58J,CAST,CBA,CE,DBA1J,DBA2J,FVB,ILNJ2,KK,MA/My,MOLF,MRL,NOD,NOR,NZB,NZO,NZW,PJ,PL,RF,SEA,SJL,SMJ,SWR,TH,WSB | intergenic_variant |
| 11 | 30983233 | 30984194 | 961 | Asb3 | SPRET | intron_variant |
| 11 | 31997466 | 31998363 | 897 | Nsg2 | AKR,BALB,BTBR,BUB,C3H,C57L,C58J,CE,FVB,ILNJ2,KK,MRL,NOD,NZO,PJ,PL,RH2,SJL,SMJ,SWR | intron_variant |
| 11 | 33128799 | 33129605 | 806 | - | SPRET | intergenic_variant |
| 11 | 39093459 | 39094317 | 858 | - | SPRET | intergenic_variant |
| 11 | 43828351 | 43829212 | 861 | - | CAST | intergenic_variant |
| 11 | 46848246 | 46848819 | 573 | Sgcd | B10J,C58J,NZB,PJ,SEA | intron_variant |
| 11 | 49089014 | 49089909 | 895 | - | AKR,B10D2,B10J,BUB,C58J,CE,KK,MA/My,NOR,NZO,NZW,RF,SJL,SWR,TH | intergenic_variant |
| 11 | 52296688 | 52296955 | 267 | 9530068E07Rik,Gm12208 | CAST | intron_variant,downstream_gene_variant |
| 11 | 56609432 | 56609720 | 288 | - | CE | intergenic_variant |
| 11 | 59443188 | 59443569 | 381 | Nlrp3 | SPRET | intron_variant |
| 11 | 59844941 | 59845664 | 723 | Med9 | AJ,C3H,CAST,CBA,MOLF,MRL,PJ,SEA | intron_variant |
| 11 | 65438162 | 65438483 | 321 | - | CE,LP,NOD,SWR,WSB | intergenic_variant |
| 11 | 68577950 | 68578291 | 341 | - | SEA | intergenic_variant |
| 11 | 72988202 | 72988987 | 785 | Itgae | TH | intron_variant |
| 11 | 74656199 | 74656507 | 308 | - | MOLF | regulatory_region_variant,intergenic_variant |
| 11 | 74666164 | 74666860 | 696 | Mettl16 | SPRET | intron_variant |
| 11 | 85959835 | 85960399 | 564 | Brip1 | MOLF | intron_variant,regulatory_region_variant |
| 11 | 92017998 | 92018252 | 254 | - | BUB | intergenic_variant |
| 11 | 92343272 | 92343744 | 472 | - | MOLF | intergenic_variant |
| 11 | 95799329 | 95799730 | 401 | B4galnt2 | AJ,AKR,BUB,C3H,CBA,KK,MA/My,MOLF,MRL,RF,RH2,SWR,TH | intron_variant |
| 11 | 99638870 | 99639603 | 733 | Gm52963 | SPRET | upstream_gene_variant |
| 11 | 103805032 | 103805298 | 266 | Nsf | SPRET | intron_variant |
| 11 | 106757732 | 106758220 | 488 | Smurf2 | CAST | intron_variant |
| 11 | 115632447 | 115633298 | 851 | - | AKR,CAST,MOLF,MRL,RH2,SMJ | intergenic_variant |
| 12 | 3755463 | 3756372 | 909 | Dtnb | SPRET | intron_variant |
| 12 | 4822860 | 4823217 | 357 | Pfn4 | SPRET | intron_variant |
| 12 | 7692313 | 7693109 | 796 | Gm32828 | CE,ILNJ2,MRL,NZW,WSB | intron_variant,non_coding_transcript_variant |
| 12 | 25652919 | 25653279 | 360 | - | C57L,NZW | intergenic_variant |

| | | | | | | |
|---|---|---|---|---|---|---|
| 12 | 31449901 | 31450413 | 512 | - | SPRET | intergenic_variant |
| 12 | 35412504 | 35412862 | 358 | - | SPRET | intergenic_variant |
| 12 | 39703063 | 39703979 | 916 | - | SPRET | intergenic_variant |
| 12 | 43264028 | 43264790 | 762 | - | CAST,MOLF | intergenic_variant |
| 12 | 43746510 | 43747099 | 589 | - | SPRET | intergenic_variant |
| 12 | 43949158 | 43949763 | 605 | - | SPRET | intergenic_variant |
| 12 | 44995009 | 44995251 | 242 | Stxbp6 | BALB | intron_variant,regulatory_region_variant |
| 12 | 45808090 | 45808765 | 675 | - | SPRET | intergenic_variant |
| 12 | 45836212 | 45836426 | 214 | - | KK,NZO,NZW | intergenic_variant |
| 12 | 47058966 | 47059554 | 588 | - | MOLF | intergenic_variant |
| 12 | 52080810 | 52081682 | 872 | - | SPRET | intergenic_variant |
| 12 | 53520850 | 53521111 | 261 | Npas3 | CAST | intron_variant |
| 12 | 56396522 | 56397007 | 485 | - | MOLF | intergenic_variant |
| 12 | 61417904 | 61418144 | 240 | - | WSB | intergenic_variant |
| 12 | 62494803 | 62495112 | 309 | - | SPRET | intergenic_variant |
| 12 | 64033675 | 64034577 | 902 | - | MOLF | intergenic_variant |
| 12 | 73702591 | 73703516 | 925 | Prkch | CAST,MOLF,SPRET | intron_variant,regulatory_region_variant |
| 12 | 79062007 | 79062346 | 339 | - | MOLF,SPRET | intergenic_variant |
| 12 | 81865341 | 81866151 | 810 | - | CAST,MOLF | regulatory_region_variant,intergenic_variant |
| 12 | 92592046 | 92592406 | 360 | - | SMJ | intergenic_variant |
| 12 | 92691971 | 92692730 | 759 | - | CAST,MOLF | intergenic_variant |
| 12 | 95248728 | 95249055 | 327 | - | CAST,MOLF | intergenic_variant |
| 12 | 96388849 | 96389450 | 601 | - | SPRET | intergenic_variant |
| 12 | 97156975 | 97157408 | 433 | - | SPRET | intergenic_variant |
| 12 | 97901156 | 97901545 | 389 | - | CAST | intergenic_variant |
| 12 | 98037001 | 98037365 | 364 | - | SPRET | intergenic_variant |
| 12 | 98495849 | 98496637 | 788 | Kcnk10 | SPRET | intron_variant |
| 12 | 100479785 | 100480485 | 700 | Ttc7b | CAST,MOLF | intron_variant |
| 12 | 104385147 | 104385365 | 218 | Gm24334 | CAST | downstream_gene_variant,regulatory_region_variant |
| 12 | 111196316 | 111196916 | 600 | Traf3 | CAST,SPRET | intron_variant |
| 12 | 111220148 | 111220571 | 423 | Traf3 | SPRET | intron_variant |
| 12 | 111448964 | 111449464 | 500 | Gm40578 | SPRET | intron_variant,non_coding_transcript_variant |
| 12 | 116025469 | 116026117 | 648 | Zfp386 | DBA1J,DBA2J,RF,WSB | 3_prime_UTR_variant |
| 13 | 6009238 | 6010005 | 767 | Gm46401 | AKR,BUB,MA/My,NOD,NOR,RF | intron_variant,non_coding_transcript_variant |
| 13 | 11261912 | 11262797 | 885 | - | SPRET | regulatory_region_variant,intergenic_variant |
| 13 | 11555482 | 11556050 | 568 | - | SPRET | regulatory_region_variant,intergenic_variant |
| 13 | 16559807 | 16560560 | 753 | Gm48493 | AKR,PJ,RF,SEA | non_coding_transcript_exon_variant |
| 13 | 17640543 | 17641411 | 868 | Sugct | CAST,MOLF | intron_variant |
| 13 | 17964286 | 17964732 | 446 | Cdk13 | SPRET | intron_variant |
| 13 | 18158296 | 18158584 | 288 | Yae1d1 | SPRET | 3_prime_UTR_variant |
| 13 | 18642029 | 18642521 | 492 | Pou6f2 | SPRET | intron_variant |
| 13 | 18721547 | 18722363 | 816 | Pou6f2 | AKR,ILNJ2,LP,PJ,RF,SEA | intron_variant |
| 13 | 22792216 | 22792925 | 709 | - | CE | intergenic_variant |
| 13 | 34725691 | 34726175 | 484 | - | CAST | intergenic_variant |
| 13 | 35273126 | 35273492 | 366 | - | AJ,AKR,B10D2,B10J,BALB,BTBR,C3H,C57L,C58J,CBA,DBA1J,DBA2J,ILNJ2,KK,MA/My,MRL,NZB,NZO,NZW,PL,RF,RH2,SMJ,SPRET,TH | intergenic_variant |
| 13 | 36842421 | 36842973 | 552 | - | CAST | intergenic_variant |
| 13 | 42022996 | 42023683 | 687 | Gm10790 | SPRET | intron_variant,non_coding_transcript_variant |
| 13 | 53593730 | 53594697 | 967 | Gm2762 | CAST,MOLF | intron_variant,non_coding_transcript_variant,non_coding_transcript_exon_variant |
| 13 | 54468753 | 54469451 | 698 | Gm48622 | CAST,MOLF | intron_variant,non_coding_transcript_variant |
| 13 | 58062240 | 58062870 | 630 | - | BUB,SMJ,SWR | intergenic_variant |
| 13 | 60393869 | 60394114 | 245 | - | SPRET | intergenic_variant |
| 13 | 64873182 | 64874120 | 938 | - | SPRET | intergenic_variant |
| 13 | 97627458 | 97628368 | 910 | Lncenc1 | AKR | intron_variant,non_coding_transcript_variant,non_coding_transcript_exon_variant |
| 13 | 98478960 | 98479625 | 665 | - | AKR,C58J,CAST,CBA,DBA1J,DBA2J,MA/My,PL,SMJ,WSB | intergenic_variant |
| 13 | 100710260 | 100710954 | 694 | - | SPRET | intergenic_variant |
| 13 | 102375038 | 102375595 | 557 | - | SPRET | intergenic_variant |
| 13 | 103122040 | 103122936 | 896 | Mast4 | SPRET | intron_variant |
| 13 | 105591747 | 105592370 | 623 | - | CE,MA/My,MOLF,NZO | intergenic_variant |
| 13 | 106035168 | 106035957 | 789 | Gm10739 | CE,MA/My,MOLF,NZO | intron_variant,non_coding_transcript_variant |
| 13 | 106448509 | 106449237 | 728 | - | CAST,NZB,SPRET,WSB | intergenic_variant |
| 13 | 111491923 | 111492492 | 569 | - | KK | intergenic_variant |
| 13 | 111657762 | 111658256 | 494 | Gm15290 | CAST | downstream_gene_variant |
| 13 | 111699760 | 111700007 | 247 | Gm15288 | MA/My,MOLF,NZB | splice_polypyrimidine_tract_variant,intron_variant,non_coding_transcript_variant |
| 13 | 113480947 | 113481490 | 543 | Cspg4b | MRL | intron_variant |
| 13 | 117189657 | 117189916 | 259 | - | SPRET | regulatory_region_variant,intergenic_variant |
| 13 | 119093853 | 119094210 | 357 | - | SPRET | intergenic_variant |
| 14 | 6099128 | 6099875 | 747 | Top2b | AJ,AKR,BTBR,BUB,C58J,CE,ILNJ2,NOD,NOR,RF,SMJ | intron_variant |
| 14 | 11741807 | 11742178 | 371 | Fhit | SPRET | intron_variant |
| 14 | 13969259 | 13970206 | 947 | Cfap20dc | SPRET | intron_variant |
| 14 | 21317153 | 21318078 | 925 | Adk | WSB | intron_variant |
| 14 | 23187885 | 23188222 | 337 | - | SPRET | intergenic_variant |
| 14 | 27286233 | 27286984 | 751 | - | CAST,MOLF | intergenic_variant |
| 14 | 27479951 | 27480304 | 353 | Erc2 | SPRET | intron_variant,regulatory_region_variant |
| 14 | 31845558 | 31846083 | 525 | - | SPRET | intergenic_variant |
| 14 | 35799619 | 35800148 | 529 | - | SPRET | intergenic_variant |
| 14 | 36216813 | 36217108 | 295 | - | 129S1,C57L,MOLF | intergenic_variant |
| 14 | 36565708 | 36566706 | 998 | - | SPRET | intergenic_variant |
| 14 | 38032280 | 38033127 | 847 | - | MOLF | intergenic_variant |
| 14 | 55772228 | 55772934 | 706 | Gm10876 | MOLF | intron_variant,non_coding_transcript_variant |
| 14 | 56432657 | 56432862 | 205 | Gm4022 | 129S1,AJ,AKR,B10D2,B10J,BALB,C3H,C57L,C58J,CBA,CE,DBA1J,FVB,KK,LP,MA/My,MRL,NOD,NOR,NZW,PJ,RF,SEA,SWR | intron_variant,non_coding_transcript_variant |
| 14 | 62720587 | 62721489 | 902 | - | SPRET | intergenic_variant |
| 14 | 67176576 | 67177466 | 890 | 4930578I07Rik | SPRET | downstream_gene_variant |
| 14 | 68620912 | 68621174 | 262 | - | SPRET | regulatory_region_variant,intergenic_variant |

| | | | | | | |
|---|---|---|---|---|---|---|
| 14 | 76207154 | 76207669 | 515 | Gtf2f2 | SPRET | intron_variant |
| 14 | 83634414 | 83635270 | 856 | - | 129S1,BUB,C57L,C58J,ILNJ2,KK,MOLF,NOD,NOR,NZW,SJL | intergenic_variant |
| 14 | 86521552 | 86522109 | 557 | 4930529K09Rik,Gm32815 | AJ,AKR,BALB,BTBR,BUB,C3H,CBA,DBA1J,DBA2J,FVB,MOLF,NOD,NZO,PJ,RF,SEA,SJL,SMJ,SWR,TH,WSB | downstream_gene_variant,intron_variant,non_coding_transcript_variant |
| 14 | 86846008 | 86846302 | 294 | - | CAST | intergenic_variant |
| 14 | 87820672 | 87821215 | 543 | - | CAST | intergenic_variant |
| 14 | 89555111 | 89555793 | 682 | - | SPRET | intergenic_variant |
| 14 | 89990467 | 89991343 | 876 | - | PJ | intergenic_variant |
| 14 | 97813395 | 97813791 | 396 | - | SPRET | intergenic_variant |
| 14 | 101639151 | 101639848 | 697 | - | 129S1,AKR,BALB,BTBR,C3H,C57L,C58J,CAST,DBA1J,DBA2J,FVB,ILNJ2,KK,MA/My,MOLF,NZB,NZO,PJ,RF,SEA,SJL,SMJ,SWR,TH | intergenic_variant |
| 14 | 102389939 | 102390870 | 931 | - | CAST,SPRET | intergenic_variant |
| 14 | 103682715 | 103683682 | 967 | 1700087I21Rik | CAST,MOLF | intron_variant,non_coding_transcript_variant |
| 14 | 104583957 | 104584277 | 320 | D130079A08Rik | SPRET | intron_variant,non_coding_transcript_variant |
| 14 | 104744106 | 104744630 | 524 | Obi1 | FVB | intron_variant |
| 14 | 107366800 | 107367800 | 1000 | - | SPRET | intergenic_variant |
| 14 | 110911564 | 110911863 | 299 | - | SPRET | intergenic_variant |
| 14 | 116440842 | 116441402 | 560 | Gpc5 | KK | intron_variant,regulatory_region_variant |
| 14 | 117896199 | 117896436 | 237 | Gpc6 | PJ | intron_variant |
| 14 | 119864737 | 119865669 | 932 | Hs6st3 | SPRET | intron_variant |
| 14 | 122300638 | 122300927 | 289 | Gm49236 | SPRET | downstream_gene_variant,regulatory_region_variant |
| 14 | 124004412 | 124004705 | 293 | Itgbl1 | SPRET | intron_variant |
| 15 | 10146040 | 10146295 | 255 | - | CAST,LP,MOLF,SJL,SPRET,SWR,TH | intergenic_variant |
| 15 | 13787376 | 13787667 | 291 | - | CAST | intergenic_variant |
| 15 | 13841338 | 13841749 | 411 | Gm49114 | CAST | downstream_gene_variant |
| 15 | 14804237 | 14804512 | 275 | - | BUB,CAST,CE,ILNJ2,MRL,PL,WSB | intergenic_variant |
| 15 | 16787611 | 16788128 | 517 | Cdh9 | SPRET | intron_variant |
| 15 | 18028077 | 18029034 | 957 | - | SPRET | intergenic_variant |
| 15 | 20217449 | 20218234 | 785 | - | SPRET | intergenic_variant |
| 15 | 24481816 | 24482765 | 949 | - | MOLF,NZB,NZO,NZW,SPRET | intergenic_variant |
| 15 | 25952428 | 25953204 | 776 | Retreg1 | CAST | intron_variant |
| 15 | 29340847 | 29341658 | 811 | - | CAST | intergenic_variant |
| 15 | 29350540 | 29351069 | 529 | - | CAST | intergenic_variant |
| 15 | 31124479 | 31125139 | 660 | - | C3H,CBA,CE,DBA1J,DBA2J,MRL,NOD,NOR,SWR,WSB | intergenic_variant |
| 15 | 36093590 | 36094262 | 672 | Rgs22 | 129S1,C57L,CAST,KK,LP,MOLF,NOD,NOR,SPRET,SWR | intron_variant |
| 15 | 38808400 | 38808840 | 440 | Baalc | SPRET | intron_variant |
| 15 | 42871261 | 42872179 | 918 | - | SPRET | intergenic_variant |
| 15 | 47403380 | 47403936 | 556 | - | SPRET | regulatory_region_variant,intergenic_variant |
| 15 | 47991941 | 47992663 | 722 | Csmd3 | SPRET | intron_variant |
| 15 | 53608505 | 53608871 | 366 | Samd12 | CAST,MOLF,SPRET | intron_variant |
| 15 | 57507225 | 57507573 | 348 | - | CAST,ILNJ2,MOLF,SPRET | intergenic_variant |
| 15 | 58104487 | 58104804 | 317 | Gm15943 | CAST | intron_variant,non_coding_transcript_variant |
| 15 | 59988065 | 59989012 | 947 | - | ILNJ2 | intergenic_variant |
| 15 | 60909212 | 60909724 | 512 | - | SPRET | intergenic_variant |
| 15 | 67010100 | 67010611 | 511 | St3gal1 | MOLF,SPRET | intron_variant |
| 15 | 69083140 | 69083446 | 306 | - | SPRET | intergenic_variant |
| 15 | 74297001 | 74297986 | 985 | Gm53029 | MOLF | intron_variant,non_coding_transcript_variant |
| 15 | 94303520 | 94303995 | 475 | - | SPRET | intergenic_variant |
| 15 | 94705924 | 94706261 | 337 | Tmem117 | 129S1,AJ,AKR,BUB,C3H,C57L,CAST,CBA,CE,DBA1J,DBA2J,FVB,ILNJ2,KK,MA/My,MOLF,NOD,NOR,NZB,NZO,NZW,PJ,PL,RF,RH2,SEA,SJL,SMJ,SPRET,SWR,TH,WSB | intron_variant |
| 15 | 98653193 | 98653920 | 727 | Arf3 | MOLF | intron_variant,regulatory_region_variant |
| 16 | 3331671 | 3331998 | 327 | - | BUB,CAST,MOLF,MRL,PL,SMJ,WSB | intergenic_variant |
| 16 | 7506356 | 7506902 | 546 | - | CAST | intergenic_variant |
| 16 | 8789692 | 8789991 | 299 | - | CAST,SPRET | intergenic_variant |
| 16 | 10192645 | 10193338 | 693 | Tekt5 | MOLF | intron_variant |
| 16 | 10926091 | 10926511 | 420 | Txndc11 | SPRET | intron_variant |
| 16 | 19638342 | 19638895 | 553 | - | SPRET | intergenic_variant |
| 16 | 21673452 | 21673771 | 319 | Map3k13 | PL,SMJ | intron_variant |
| 16 | 24843978 | 24844249 | 271 | - | BUB,CE,NOD,NOR,WSB | regulatory_region_variant,intergenic_variant |
| 16 | 27953409 | 27953654 | 245 | - | CE,KK | intergenic_variant |
| 16 | 30720799 | 30721429 | 630 | - | SPRET | regulatory_region_variant,intergenic_variant |
| 16 | 33127464 | 33128193 | 729 | - | CAST | regulatory_region_variant,intergenic_variant |
| 16 | 47357811 | 47358378 | 567 | - | C3H,C58J,CBA,MOLF | intergenic_variant |
| 16 | 48118428 | 48119420 | 992 | - | C3H,CBA,MOLF | intergenic_variant |
| 16 | 52468246 | 52468813 | 567 | - | SPRET | intergenic_variant |
| 16 | 53029061 | 53029378 | 317 | - | SPRET | intergenic_variant |
| 16 | 53460065 | 53460377 | 312 | - | SPRET | intergenic_variant |
| 16 | 54870271 | 54870922 | 651 | - | CAST,CBA,MOLF,NZW,SPRET | intergenic_variant |
| 16 | 61911949 | 61912888 | 939 | - | SPRET | intergenic_variant |
| 16 | 62152828 | 62153752 | 924 | - | SPRET | intergenic_variant |
| 16 | 63507468 | 63508116 | 648 | Epha3 | SPRET | intron_variant,regulatory_region_variant |
| 16 | 66314919 | 66315612 | 693 | - | CAST | intergenic_variant |
| 16 | 66677672 | 66677987 | 315 | Cadm2 | SMJ | intron_variant |
| 16 | 68621308 | 68621779 | 471 | - | MOLF | intergenic_variant |
| 16 | 82763754 | 82764010 | 256 | - | C58J,KK,LP,SMJ | intergenic_variant |
| 16 | 83785324 | 83786167 | 843 | - | CAST | intergenic_variant |
| 16 | 84565406 | 84565718 | 312 | - | WSB | intergenic_variant |
| 17 | 3486982 | 3487220 | 238 | Tiam2 | 129S1,CE | intron_variant |
| 17 | 4798113 | 4798347 | 234 | - | CAST | intergenic_variant |
| 17 | 8824942 | 8825274 | 332 | Pde10a | BTBR,MOLF,PL,SPRET | intron_variant |
| 17 | 10369561 | 10369856 | 295 | - | SPRET | regulatory_region_variant,intergenic_variant |
| 17 | 12589498 | 12590246 | 748 | - | SPRET | regulatory_region_variant,intergenic_variant |
| 17 | 15703826 | 15704792 | 966 | Psmb1 | BTBR,BUB,C3H,CAST,CBA,MOLF,MRL,NOD,NOR,PJ,PL,RF,SJL,SMJ,SWR,WSB | intron_variant |

| | | | | | | |
|---|---|---|---|---|---|---|
| 17 | 19000106 | 19000923 | 817 | - | 129S1,AJ,BALB,C57L,CAST,CE,DBA1J,DBA2J,MOLF,NZO,RH2,SEA | intergenic_variant |
| 17 | 19340899 | 19341244 | 345 | - | SPRET | intergenic_variant |
| 17 | 22373080 | 22374018 | 938 | Zfp947 | NZB | intron_variant |
| 17 | 27961024 | 27961787 | 763 | Gm15458 | SPRET | downstream_gene_variant |
| 17 | 28089383 | 28090259 | 876 | Bltp3a | CAST,KK,MOLF,NZB,RF,SPRET | intron_variant |
| 17 | 29001138 | 29002116 | 978 | - | CAST | intergenic_variant |
| 17 | 29323106 | 29323883 | 777 | - | TH | intergenic_variant |
| 17 | 29485202 | 29486029 | 827 | - | KK,MOLF,RF | intergenic_variant |
| 17 | 29664735 | 29665735 | 1000 | Gm36199 | KK,MOLF,SPRET | intron_variant,non_coding_transcript_variant |
| 17 | 29799264 | 29800148 | 884 | Tbc1d22b,Tbc1d22bos | CAST | intron_variant,intron_variant,non_coding_transcript_variant,regulatory_region_variant |
| 17 | 31004250 | 31004884 | 634 | Dnah8 | SPRET | intron_variant,regulatory_region_variant |
| 17 | 32317448 | 32317868 | 420 | Pdxk-ps | PJ | intron_variant,non_coding_transcript_variant |
| 17 | 32930561 | 32931346 | 785 | - | CE | intergenic_variant |
| 17 | 41836147 | 41836895 | 748 | - | SPRET | intergenic_variant |
| 17 | 42858507 | 42858968 | 461 | - | SPRET | intergenic_variant |
| 17 | 50958935 | 50959382 | 447 | Plcl2 | MOLF | intron_variant |
| 17 | 51856632 | 51857074 | 442 | - | SPRET | intergenic_variant |
| 17 | 52534072 | 52534440 | 368 | - | SPRET | intergenic_variant |
| 17 | 54574468 | 54575329 | 861 | - | SPRET | intergenic_variant |
| 17 | 54676189 | 54676974 | 785 | 1700025K24Rik | SPRET | non_coding_transcript_exon_variant,intron_variant |
| 17 | 55323221 | 55324216 | 995 | - | SPRET | intergenic_variant |
| 17 | 56226866 | 56227178 | 312 | - | CAST,SPRET | intergenic_variant |
| 17 | 60251771 | 60252747 | 976 | Gm49853 | CAST | intron_variant,non_coding_transcript_variant |
| 17 | 64062206 | 64062945 | 739 | - | C58J,CAST,KK,LP,MOLF,NZO,SPRET,WSB | intergenic_variant |
| 17 | 66295362 | 66296197 | 835 | Ankrd12 | SPRET | intron_variant,regulatory_region_variant |
| 17 | 69068525 | 69069455 | 930 | L3mbtl4 | CAST | intron_variant |
| 17 | 70568547 | 70569275 | 728 | Dlgap1 | SPRET | intron_variant |
| 17 | 78040288 | 78040790 | 502 | - | SPRET | intergenic_variant |
| 17 | 80763897 | 80764495 | 598 | Sos1 | SPRET | intron_variant |
| 17 | 83947816 | 83948332 | 516 | - | SPRET | regulatory_region_variant,intergenic_variant |
| 17 | 85856064 | 85856486 | 422 | - | 129S1,PJ,RF | intergenic_variant |
| 18 | 6716346 | 6717028 | 682 | - | CAST,MOLF,SPRET | intergenic_variant |
| 18 | 7052921 | 7053459 | 538 | - | BALB,C3H,CBA,ILNJ2,RH2 | intergenic_variant |
| 18 | 9439761 | 9440704 | 943 | Ccny | SPRET | intron_variant |
| 18 | 10493475 | 10494365 | 890 | Greb1l | SPRET | intron_variant |
| 18 | 12496564 | 12497457 | 893 | Lama3 | SPRET | intron_variant |
| 18 | 13789443 | 13789802 | 359 | - | MOLF | intergenic_variant |
| 18 | 16270613 | 16271558 | 945 | Gm50076 | CAST,MOLF,SPRET,WSB | intron_variant,non_coding_transcript_variant |
| 18 | 17526926 | 17527177 | 251 | - | SMJ | intergenic_variant |
| 18 | 23151280 | 23151643 | 363 | Nol4 | CAST | intron_variant |
| 18 | 24598038 | 24598926 | 888 | - | SPRET | intergenic_variant |
| 18 | 25253870 | 25254296 | 426 | Fhod3 | SPRET | intron_variant |
| 18 | 28067180 | 28067949 | 769 | - | AKR,KK,MA/My,NZB,NZO,SJL,WSB | intergenic_variant |
| 18 | 28142650 | 28143625 | 975 | - | SPRET | regulatory_region_variant,intergenic_variant |
| 18 | 28797951 | 28798353 | 402 | - | SPRET | intergenic_variant |
| 18 | 29023301 | 29024206 | 905 | Gm33948 | SPRET | intron_variant,non_coding_transcript_variant |
| 18 | 31175897 | 31176272 | 375 | Rit2 | SPRET | intron_variant |
| 18 | 32107341 | 32107709 | 368 | Myo7b | MOLF | intron_variant |
| 18 | 33219876 | 33220712 | 836 | Camk4 | WSB | intron_variant |
| 18 | 35305029 | 35305423 | 394 | Ctnna1 | SPRET | intron_variant |
| 18 | 36232694 | 36232933 | 239 | Nrg2 | CAST,MOLF,SPRET | intron_variant,regulatory_region_variant |
| 18 | 36831907 | 36832404 | 497 | - | CAST | intergenic_variant |
| 18 | 41716673 | 41717598 | 925 | Gm50409 | SPRET | intron_variant,non_coding_transcript_variant |
| 18 | 42886825 | 42887139 | 314 | Ppp2r2b | SPRET | intron_variant |
| 18 | 55470533 | 55470931 | 398 | - | CAST | intergenic_variant |
| 18 | 58998405 | 58999252 | 847 | Adamts19 | SPRET | intron_variant |
| 18 | 73207142 | 73207881 | 739 | - | BUB,DBA1J,DBA2J,PJ,SEA | intergenic_variant |
| 18 | 75115443 | 75116291 | 848 | 9030625G05Rik | SPRET | upstream_gene_variant |
| 18 | 80885752 | 80885990 | 238 | Atp9b | CAST | intron_variant |
| 18 | 81820166 | 81820472 | 306 | Gm46614 | SPRET | intron_variant,non_coding_transcript_variant |
| 18 | 86925872 | 86926158 | 286 | - | CAST,MOLF,SPRET,WSB | intergenic_variant |
| 18 | 88591605 | 88591818 | 213 | - | NZB,NZO | intergenic_variant |
| 18 | 89363768 | 89364504 | 736 | Dok6 | 129S1,CE,DBA1J,DBA2J,FVB,RF,SMJ,WSB | intron_variant |
| 19 | 14000115 | 14000718 | 603 | - | FVB,ILNJ2,LP,SJL,SWR,WSB | intergenic_variant |
| 19 | 21168163 | 21168772 | 609 | Gm52981 | SPRET | intron_variant,non_coding_transcript_variant |
| 19 | 22390640 | 22391075 | 435 | Trpm3 | SPRET | intron_variant |
| 19 | 26948491 | 26949337 | 846 | - | SPRET | intergenic_variant |
| 19 | 38634292 | 38634948 | 656 | Plce1 | CAST | intron_variant,regulatory_region_variant |
| 19 | 38962018 | 38963008 | 990 | - | MOLF | intergenic_variant |
| 19 | 41064356 | 41064700 | 344 | Opalin | CAST | intron_variant |
| 19 | 42737369 | 42737635 | 266 | Pyroxd2,Gm19437 | DBA1J | intron_variant,non_coding_transcript_variant |
| 19 | 42869844 | 42870797 | 953 | Hpse2 | SPRET | intron_variant |
| 19 | 48019063 | 48019413 | 350 | Cfap58 | SPRET | intron_variant |
| 19 | 52443372 | 52444360 | 988 | - | CAST | intergenic_variant |
| 19 | 54224704 | 54225140 | 436 | - | CAST,MOLF,SPRET,WSB | intergenic_variant |
| 19 | 55676111 | 55676669 | 558 | - | MOLF | regulatory_region_variant,intergenic_variant |
| 19 | 57007632 | 57008228 | 596 | - | CAST | intergenic_variant |
| 19 | 59948040 | 59948334 | 294 | - | BTBR,WSB | intergenic_variant |
| 19 | 60894574 | 60895467 | 893 | Grk5 | TH | intron_variant |
| 2 | 4686601 | 4687296 | 695 | - | SPRET | intergenic_variant |
| 2 | 7123268 | 7123938 | 670 | Celf2 | SPRET | intron_variant |
| 2 | 8910385 | 8911378 | 993 | - | MOLF | intergenic_variant |
| 2 | 21723632 | 21723854 | 222 | Gpr158 | AJ,AKR,BTBR,C3H,DBA2J,PL,RH2 | intron_variant |
| 2 | 35667696 | 35668271 | 575 | Ttll11 | CAST | intron_variant |
| 2 | 39708928 | 39709219 | 291 | - | SMJ | intergenic_variant |
| 2 | 39749440 | 39750348 | 908 | - | SPRET | intergenic_variant |
| 2 | 39791865 | 39792233 | 368 | - | SMJ | intergenic_variant |
| 2 | 41820484 | 41821057 | 573 | Lrp1b | SPRET | intron_variant |
| 2 | 42030914 | 42031429 | 515 | Lrp1b | SPRET | intron_variant |
| 2 | 42438092 | 42438405 | 313 | Lrp1b | TH | intron_variant |

| | | | | | | |
|---|---|---|---|---|---|---|
| 2 | 46716171 | 46716397 | 226 | - | CAST,CE,KK,NZB,SMJ,TH,WSB | intergenic_variant |
| 2 | 49556524 | 49557420 | 896 | Kif5c | SPRET | intron_variant |
| 2 | 56051101 | 56051667 | 566 | - | SPRET | intergenic_variant |
| 2 | 56310941 | 56311749 | 808 | - | KK,NZO | intergenic_variant |
| 2 | 58115411 | 58115691 | 280 | - | SPRET | intergenic_variant |
| 2 | 64844309 | 64844983 | 674 | Grb14 | SPRET | intron_variant,regulatory_region_variant |
| 2 | 69547764 | 69548023 | 259 | - | 129S1,AJ,AKR,B10D2,B10J,BALB,BTBR,BUB,C3H,C57L,C58J,CAST,CBA,CE,DBA1J,DBA2J,FVB,ILNJ2,KK,LP,MA/My,MOLF,MRL,NOD,NOR,NZB,NZO,NZW,PJ,PL,RF,RH2,SEA,SJL,SMJ,SPRET,SWR,TH,WSB | intergenic_variant |
| 2 | 71838941 | 71839293 | 352 | Rapgef4 | AJ,AKR,BUB,C57L,FVB,ILNJ2,MA/My,MOLF,NOD,NOR,PJ,PL,RH2 | intron_variant |
| 2 | 71838982 | 71839926 | 944 | Rapgef4 | BALB,BTBR,CAST,MRL,SEA | intron_variant,regulatory_region_variant |
| 2 | 77251976 | 77252364 | 388 | Zfp385b | MOLF,NZW | intron_variant |
| 2 | 80052591 | 80053196 | 605 | - | SPRET | intergenic_variant |
| 2 | 81919369 | 81920107 | 738 | Zfp804a | SPRET | intron_variant |
| 2 | 82014520 | 82014829 | 309 | Zfp804a | SPRET | intron_variant |
| 2 | 88420094 | 88420758 | 664 | Or4c102 | SPRET | intron_variant |
| 2 | 92057174 | 92057560 | 386 | Phf21a | CAST,SPRET | intron_variant |
| 2 | 94547966 | 94548364 | 398 | - | CAST | intergenic_variant |
| 2 | 95599599 | 95600104 | 505 | - | SPRET | intergenic_variant |
| 2 | 96617030 | 96617604 | 574 | Lrrc4c | CAST,MOLF | intron_variant |
| 2 | 96698437 | 96699081 | 644 | Lrrc4c | CAST,MOLF | intron_variant |
| 2 | 98636694 | 98637445 | 751 | - | CAST,MOLF | intergenic_variant |
| 2 | 101606511 | 101607315 | 804 | Prr5l | SPRET | intron_variant |
| 2 | 107587312 | 107587811 | 499 | - | SPRET | intergenic_variant |
| 2 | 109113541 | 109114491 | 950 | Kif18a | C3H,CAST,CE,MOLF,PL,RF,SPRET | intron_variant |
| 2 | 109655355 | 109655846 | 491 | - | C3H,CE,MOLF,PL | intergenic_variant |
| 2 | 110675899 | 110676761 | 862 | Ano3 | SPRET | intron_variant |
| 2 | 117519284 | 117519644 | 360 | - | SPRET | intergenic_variant |
| 2 | 118394539 | 118395121 | 582 | - | BTBR,C3H,C58J,CBA,NOD,NOR,NZB,NZO,TH,WSB | intergenic_variant |
| 2 | 119438960 | 119439334 | 374 | Oip5os1 | RF | downstream_gene_variant |
| 2 | 119664301 | 119665147 | 846 | - | AJ,BALB,DBA1J,DBA2J,ILNJ2,PJ,PL,RF,RH2,SEA,SMJ,TH | intergenic_variant |
| 2 | 121144878 | 121145850 | 972 | Ppip5k1 | MOLF | intron_variant |
| 2 | 122038350 | 122038902 | 552 | Gm14050 | AJ,BALB,C3H,C58J,CBA,DBA1J,DBA2J,ILNJ2,NZB,NZO,PJ,PL,RF,RH2,SEA,SMJ,TH | non_coding_transcript_variant |
| 2 | 131097998 | 131098587 | 589 | - | SPRET | regulatory_region_variant,intergenic_variant |
| 2 | 131203843 | 131204788 | 945 | - | CAST,DBA1J,DBA2J,ILNJ2,MOLF,PJ,PL | intergenic_variant |
| 2 | 132734540 | 132735073 | 533 | - | SPRET | intergenic_variant |
| 2 | 137267792 | 137268081 | 289 | - | SPRET | intergenic_variant |
| 2 | 141186911 | 141187634 | 723 | Macrod2 | CAST | intron_variant |
| 2 | 141431981 | 141432636 | 655 | Macrod2 | C58J,CAST,DBA1J,DBA2J,MOLF,NZO,NZW | intron_variant |
| 2 | 141883974 | 141884394 | 420 | Macrod2 | AJ | intron_variant |
| 2 | 145052007 | 145052879 | 872 | Slc24a3 | SPRET | intron_variant |
| 2 | 149081802 | 149082185 | 383 | - | NOD | intergenic_variant |
| 2 | 149568175 | 149568774 | 599 | - | SPRET | intergenic_variant |
| 2 | 155899201 | 155899510 | 309 | Gm14240 | 129S1,BALB,C3H,C57L,CBA,CE,FVB,MA/My,NOD,NZO,NZW,PL,SEA,SJL,SMJ,SWR,TH,WSB | downstream_gene_variant |
| 2 | 160765011 | 160765891 | 880 | - | SPRET | intergenic_variant |
| 2 | 163750977 | 163751267 | 290 | Rims4 | AJ,B10D2,B10J,BALB,BTBR,C3H,CBA,KK,SEA,SMJ | intron_variant |
| 2 | 165745263 | 165745745 | 482 | - | CAST | intergenic_variant |
| 2 | 165760523 | 165760841 | 318 | - | KK | intergenic_variant |
| 2 | 166908238 | 166908551 | 313 | Snord12,Zfas1 | CAST,KK | downstream_gene_variant |
| 2 | 171354331 | 171354861 | 530 | - | CAST,CE | intergenic_variant |
| 2 | 181342035 | 181342541 | 506 | - | SPRET | intergenic_variant |
| 3 | 9044377 | 9044677 | 300 | Tpd52 | AKR,BUB,C58J,DBA1J,FVB,KK,MRL,NOD | intron_variant,regulatory_region_variant |
| 3 | 11505584 | 11506429 | 845 | - | MOLF | intergenic_variant |
| 3 | 12234164 | 12235022 | 858 | - | SMJ | intergenic_variant |
| 3 | 13013156 | 13013924 | 768 | - | SPRET | intergenic_variant |
| 3 | 15299245 | 15300066 | 821 | - | CE,DBA2J,LP | intergenic_variant |
| 3 | 19020321 | 19021138 | 817 | - | MOLF | intergenic_variant |
| 3 | 23544498 | 23544888 | 390 | 4930412E21Rik | SPRET | intron_variant,non_coding_transcript_variant |
| 3 | 28805774 | 28806323 | 549 | 1700112D23Rik | CAST,MOLF,SPRET | intron_variant,non_coding_transcript_variant |
| 3 | 33053080 | 33053891 | 811 | Pex5l | SPRET | intron_variant |
| 3 | 34646234 | 34646800 | 566 | Sox2ot | BUB,FVB,MOLF | intron_variant,non_coding_transcript_variant |
| 3 | 35830572 | 35831194 | 622 | Atp11b | DBA1J,RF | intron_variant |
| 3 | 37703361 | 37703789 | 428 | - | SPRET | intergenic_variant |
| 3 | 39702554 | 39702725 | 171 | - | KK,NZW | intergenic_variant |
| 3 | 44978398 | 44978895 | 497 | - | CAST | intergenic_variant |
| 3 | 45975560 | 45976365 | 805 | - | KK | intergenic_variant |
| 3 | 48837656 | 48837956 | 300 | - | SPRET | intergenic_variant |
| 3 | 48934230 | 48934539 | 309 | - | SPRET | intergenic_variant |
| 3 | 49069485 | 49069737 | 252 | - | SPRET | intergenic_variant |
| 3 | 57021414 | 57021671 | 257 | - | SPRET | intergenic_variant |
| 3 | 58688005 | 58688176 | 171 | 4930593A02Rik | MOLF | intron_variant,non_coding_transcript_variant |
| 3 | 62393877 | 62394322 | 445 | Dhx36 | SPRET | intron_variant |
| 3 | 67442623 | 67443500 | 877 | - | SJL | intergenic_variant |
| 3 | 69477523 | 69477806 | 283 | - | BUB,C58J,SEA | intergenic_variant |
| 3 | 70527623 | 70528610 | 987 | - | SPRET | intergenic_variant |
| 3 | 71179551 | 71180470 | 919 | - | SPRET | intergenic_variant |
| 3 | 75677497 | 75678051 | 554 | Lrrc77 | CAST,MOLF | intron_variant |
| 3 | 76280921 | 76281500 | 579 | Fstl5 | CAST | intron_variant |
| 3 | 81827725 | 81828082 | 357 | Gm9989 | SPRET | intron_variant,non_coding_transcript_variant |
| 3 | 87483699 | 87483991 | 292 | - | 129S1,C57L,FVB,MA/My,NOD,NOR,SJL,SWR,TH | intergenic_variant |
| 3 | 88298265 | 88298648 | 383 | - | SPRET | intergenic_variant |
| 3 | 90284759 | 90285344 | 585 | - | SPRET | intergenic_variant |

| | | | | | | |
|---|---|---|---|---|---|---|
| 3 | 91909234 | 91909646 | 412 | - | CAST | intergenic_variant |
| 3 | 96362964 | 96363494 | 530 | Gm9207 | MOLF | non_coding_transcript_exon_variant |
| 3 | 96807201 | 96807776 | 575 | Gpr89 | MOLF | intron_variant |
| 3 | 104058766 | 104059275 | 509 | Magi3,Gm43258 | SPRET | intron_variant,non_coding_transcript_exon_variant |
| 3 | 115859915 | 115860152 | 237 | Gm26344 | AJ,AKR,BALB,BUB,C3H,CBA,CE,DBA1J,DBA2J,FVB,ILNJ2,LP,MRL,NOD,NOR,NZB,NZO,NZW,PL,RF,RH2,SJL,SWR,TH | upstream_gene_variant |
| 3 | 117227878 | 117228328 | 450 | - | AJ,AKR,BALB,BUB,C3H,C58J,CBA,DBA1J,DBA2J,FVB,ILNJ2,MRL,NZO,NZW,PL,RF,RH2,SJL,SMJ,SWR,TH,WSB | intergenic_variant |
| 3 | 118664387 | 118665307 | 920 | Dpyd,Gm43410 | CAST | intron_variant,downstream_gene_variant |
| 3 | 121396114 | 121396513 | 399 | - | MOLF | regulatory_region_variant,intergenic_variant |
| 3 | 121923426 | 121923650 | 224 | Abca4,Gm43158 | BTBR,DBA1J,DBA2J,TH | intron_variant,non_coding_transcript_variant,regulatory_region_variant |
| 3 | 124917732 | 124917960 | 228 | - | AJ,BALB,C3H,CAST,CBA,MOLF,NZB,RH2,SEA,SMJ,WSB | intergenic_variant |
| 3 | 130819897 | 130820298 | 401 | A430072C10Rik | CAST,SPRET | non_coding_transcript_exon_variant |
| 3 | 133917164 | 133917935 | 771 | Gm26691,Gm30648 | CAST,MOLF | intron_variant,non_coding_transcript_variant,non_coding_transcript_exon_variant,intron_variant |
| 3 | 135180404 | 135180631 | 227 | - | CAST | intergenic_variant |
| 3 | 140396631 | 140397026 | 395 | Gm43049 | BALB,BUB,C3H,ILNJ2,MA/My,MRL,SEA,SMJ,SWR | downstream_gene_variant |
| 3 | 144066482 | 144067311 | 829 | - | CAST,MOLF,SPRET | intergenic_variant |
| 3 | 149685766 | 149686616 | 850 | - | CAST | intergenic_variant |
| 3 | 151627377 | 151628329 | 952 | - | CAST | intergenic_variant |
| 3 | 151745052 | 151745266 | 214 | - | B10D2,BUB,NOD,NOR,NZB,NZO,NZW,TH | intergenic_variant |
| 3 | 154339887 | 154340185 | 298 | - | SPRET | intergenic_variant |
| 3 | 154961240 | 154961858 | 618 | - | CAST,MOLF,SPRET | intergenic_variant |
| 4 | 4152381 | 4152658 | 277 | A830012C17Rik | CAST | intron_variant,non_coding_transcript_variant |
| 4 | 11738701 | 11739154 | 453 | - | SPRET | intergenic_variant |
| 4 | 15993614 | 15994387 | 773 | - | CAST,NZW | intergenic_variant |
| 4 | 17383804 | 17384385 | 581 | - | NZW | intergenic_variant |
| 4 | 20198839 | 20199435 | 596 | Nkain3 | SPRET | intron_variant |
| 4 | 22601210 | 22601635 | 425 | - | RF | intergenic_variant |
| 4 | 25192402 | 25193383 | 981 | Gm11906 | CAST | intron_variant,non_coding_transcript_variant,regulatory_region variant |
| 4 | 26522509 | 26522797 | 288 | - | 129S1,AJ,AKR,BALB,BTBR,BUB,C57L,CE,FVB,ILNJ2,MA/My,MOLF,MRL,NOD,NOR,NZB,NZO,NZW,PJ,PL,RF,RH2,SEA,SJL,SPRET,SWR | intergenic_variant |
| 4 | 27588386 | 27589368 | 982 | - | C57L,WSB | intergenic_variant |
| 4 | 30370047 | 30370347 | 300 | - | KK | intergenic_variant |
| 4 | 32902341 | 32902620 | 279 | Ankrd6 | MOLF | intron_variant |
| 4 | 36583293 | 36583917 | 624 | Lingo2 | SPRET | intron_variant |
| 4 | 36652396 | 36652848 | 452 | Lingo2 | SPRET,TH | intron_variant |
| 4 | 36756676 | 36757267 | 591 | Lingo2,Gm56187 | SPRET | intron_variant,downstream_gene_variant |
| 4 | 37912484 | 37913075 | 591 | - | TH | intergenic_variant |
| 4 | 38795676 | 38796235 | 559 | Gm12381 | WSB | intron_variant,non_coding_transcript_variant |
| 4 | 50316351 | 50316661 | 310 | - | WSB | intergenic_variant |
| 4 | 50684881 | 50685232 | 351 | - | SPRET | intergenic_variant |
| 4 | 52791831 | 52792233 | 402 | - | SPRET | intergenic_variant |
| 4 | 53767590 | 53767833 | 243 | - | CAST | intergenic_variant |
| 4 | 54148030 | 54148365 | 335 | - | MA/My,MOLF | intergenic_variant |
| 4 | 75310629 | 75311205 | 576 | - | SPRET | intergenic_variant |
| 4 | 77664169 | 77664899 | 730 | - | SPRET | intergenic_variant |
| 4 | 78998963 | 78999807 | 844 | - | SPRET | intergenic_variant |
| 4 | 86731785 | 86732688 | 903 | Dennd4c | SEA | coding_sequence_variant,intron_variant |
| 4 | 88818568 | 88819445 | 877 | - | SPRET | intergenic_variant |
| 4 | 93274454 | 93274795 | 341 | - | 129S1,C57L,RH2 | intergenic_variant |
| 4 | 96197214 | 96198207 | 993 | Cyp2j11 | CAST,ILNJ2,MOLF | intron_variant |
| 4 | 98867052 | 98867693 | 641 | Dock7 | MRL,NOD,RH2,SWR | intron_variant |
| 4 | 100586751 | 100587590 | 839 | - | AJ,AKR,B10D2,B10J,BALB,BTBR,BUB,C3H,C58J,CBA,CE,FVB,ILNJ2,MA/My,MRL,NZW,PJ,PL,RF,SEA,SJL | intergenic_variant |
| 4 | 101587861 | 101588297 | 436 | Lepr | CAST,MOLF,NZB,NZO | intron_variant |
| 4 | 107101938 | 107102193 | 255 | Lrrc42 | CAST,MA/My,MOLF,SPRET | intron_variant |
| 4 | 108635034 | 108635354 | 320 | Zfyve9 | CAST | intron_variant |
| 4 | 112547807 | 112548389 | 582 | - | SEA | intergenic_variant |
| 4 | 116523053 | 116523988 | 935 | - | CAST,MOLF | regulatory_region_variant,intergenic_variant |
| 4 | 116526335 | 116527142 | 807 | - | CE | regulatory_region_variant,intergenic_variant |
| 4 | 117792530 | 117793152 | 622 | St3gal3 | AKR,ILNJ2,KK,MRL | intron_variant |
| 4 | 118064106 | 118064448 | 342 | Ptprf | SPRET | downstream_gene_variant,regulatory_region_variant |
| 4 | 118327768 | 118328305 | 537 | Tie1 | CAST,ILNJ2,MOLF,SPRET | downstream_gene_variant,regulatory_region_variant |
| 4 | 126829024 | 126829515 | 491 | Zmym4 | SPRET | intron_variant |
| 4 | 127044638 | 127045604 | 966 | - | SPRET | intergenic_variant |
| 4 | 128963100 | 128963338 | 238 | Rnf19b | FVB | intron_variant |
| 4 | 129433012 | 129433242 | 230 | Hdac1 | CAST | intron_variant |
| 4 | 130132391 | 130133007 | 616 | - | SPRET | intergenic_variant |
| 4 | 135547033 | 135547990 | 957 | - | SPRET | intergenic_variant |
| 4 | 136377563 | 136378557 | 994 | Ephb2 | SPRET | 3_prime_UTR_variant |
| 4 | 137066717 | 137067431 | 714 | Cdc42 | BUB,MRL,NZW | intron_variant,regulatory_region_variant |
| 4 | 141730445 | 141730667 | 222 | Fhad1 | SWR | intron_variant |
| 4 | 152824280 | 152825182 | 902 | - | SPRET | intergenic_variant |
| 4 | 156296404 | 156297385 | 981 | - | SPRET | intergenic_variant |
| 4 | 156406558 | 156407058 | 500 | - | SPRET | intergenic_variant |
| 5 | 3282177 | 3282621 | 444 | - | FVB,SJL,SWR | intergenic_variant |
| 5 | 5219334 | 5219815 | 481 | Cdk14,Gm10484 | SPRET | intron_variant,downstream_gene_variant |
| 5 | 6760044 | 6760953 | 909 | - | AJ,BTBR,C3H,C57L,DBA1J,DBA2J,MA/My,MRL,RF,SJL,SWR | intergenic_variant |
| 5 | 7644912 | 7645133 | 221 | - | 129S1,AKR,BALB,BTBR,C57L,C58J,CE,FVB,LP,MA/My,NZB,NZO,NZW,PJ,RF,RH2,SEA,SJL,SWR,WSB | intergenic_variant |
| 5 | 8013397 | 8013622 | 225 | Steap4 | 129S1,BALB,BTBR,C57L,CE,NZB,NZO,NZW,PJ,PL,RF,SEA,SJL,SWR,WSB | intron_variant |

| | | | | | | |
|---|---|---|---|---|---|---|
| 5 | 10109143 | 10110053 | 910 | - | AKR,CAST,NZB,NZO,NZW,SMJ,SPRET | intergenic_variant |
| 5 | 25673806 | 25674169 | 363 | Kmt2c | SPRET | intron_variant |
| 5 | 28155131 | 28156060 | 929 | - | MOLF | intergenic_variant |
| 5 | 28251143 | 28251835 | 692 | Gm26608 | MOLF | intron_variant,non_coding_transcript_variant |
| 5 | 30417348 | 30417826 | 478 | - | SPRET | intergenic_variant |
| 5 | 30698219 | 30698550 | 331 | Cib4,Gm42765 | WSB | intron_variant,non_coding_transcript_exon_variant,intron_variant |
| 5 | 31597150 | 31597689 | 539 | - | CAST | intergenic_variant |
| 5 | 37561810 | 37562731 | 921 | Evc2,Gm15732 | MOLF | intron_variant,non_coding_transcript_exon_variant |
| 5 | 37602282 | 37602666 | 384 | - | SPRET | intergenic_variant |
| 5 | 38286735 | 38287688 | 953 | Stx18 | CAST,SPRET | intron_variant |
| 5 | 39502943 | 39503726 | 783 | Gm55843 | SPRET | upstream_gene_variant |
| 5 | 42380239 | 42380639 | 400 | - | NZB,NZO | intergenic_variant |
| 5 | 43049304 | 43049924 | 620 | - | MOLF | regulatory_region_variant,intergenic_variant |
| 5 | 43312890 | 43313186 | 296 | Gm7854 | CAST,MOLF,SPRET | intron_variant,non_coding_transcript_variant |
| 5 | 44037288 | 44037744 | 456 | Cd38 | PJ,PL | intron_variant |
| 5 | 49541896 | 49542398 | 502 | Kcnip4 | SPRET | intron_variant |
| 5 | 50050444 | 50051040 | 596 | - | KK,MOLF,NOD,NZB,NZO,NZW,PJ,TH | intergenic_variant |
| 5 | 51112162 | 51113029 | 867 | - | KK,MA/My,MOLF,NOD,NZB,NZO,NZW,PJ,SJL,SWR,TH,WSB | intergenic_variant |
| 5 | 54489519 | 54490383 | 864 | - | CAST | intergenic_variant |
| 5 | 56262629 | 56263618 | 989 | - | SPRET | intergenic_variant |
| 5 | 59727573 | 59728445 | 872 | Gm43042 | C3H,CBA,DBA1J,DBA2J,ILNJ2,WSB | intron_variant,non_coding_transcript_variant |
| 5 | 61169684 | 61170388 | 704 | - | SPRET | intergenic_variant |
| 5 | 66186420 | 66186917 | 497 | Rbm47 | AJ,AKR,BUB,C58J,CE,FVB,MA/My,NOD,NOR,PL,RF,SJL,SMJ,SPRET,SWR | intron_variant |
| 5 | 66972360 | 66972697 | 337 | Limch1 | CAST | intron_variant |
| 5 | 68487489 | 68488158 | 669 | - | SPRET | intergenic_variant |
| 5 | 71403344 | 71404263 | 919 | - | SPRET | intergenic_variant |
| 5 | 73094629 | 73095337 | 708 | Slain2 | CAST,SPRET,WSB | intron_variant |
| 5 | 74338690 | 74339605 | 915 | - | SPRET | intergenic_variant |
| 5 | 74470416 | 74470721 | 305 | Scfd2 | SPRET | intron_variant |
| 5 | 77115494 | 77116404 | 910 | Paics | SPRET | downstream_gene_variant |
| 5 | 77379648 | 77379984 | 336 | - | CAST | intergenic_variant |
| 5 | 80664778 | 80665633 | 855 | - | SWR,WSB | intergenic_variant |
| 5 | 88145352 | 88145983 | 631 | - | SPRET | intergenic_variant |
| 5 | 88308994 | 88309488 | 494 | - | SPRET | intergenic_variant |
| 5 | 88513790 | 88514609 | 819 | - | MA/My | intergenic_variant |
| 5 | 99258201 | 99258704 | 503 | Gm43251 | CAST | intron_variant,non_coding_transcript_variant |
| 5 | 99465770 | 99466473 | 703 | Rasgef1b | SPRET | intron_variant |
| 5 | 100593790 | 100594047 | 257 | Lin54 | CAST | intron_variant |
| 5 | 103385392 | 103386112 | 720 | 4930429D17Rik | SPRET | intron_variant,non_coding_transcript_variant |
| 5 | 110188303 | 110188647 | 344 | - | SPRET | intergenic_variant |
| 5 | 113568044 | 113568405 | 361 | - | SPRET | regulatory_region_variant,intergenic_variant |
| 5 | 118926434 | 118926669 | 235 | - | MOLF,RF,SPRET | intergenic_variant |
| 5 | 121241226 | 121241847 | 621 | - | SPRET | intergenic_variant |
| 5 | 124574781 | 124575516 | 735 | - | CAST | intergenic_variant |
| 5 | 125443646 | 125444434 | 788 | Gm40323 | SPRET | intron_variant,non_coding_transcript_variant |
| 5 | 137445799 | 137446193 | 394 | Zan | BUB,MOLF,RF | intron_variant |
| 5 | 137741414 | 137742374 | 960 | - | SPRET | intergenic_variant |
| 5 | 142348379 | 142348638 | 259 | - | NOD,SPRET | regulatory_region_variant,intergenic_variant |
| 5 | 146510985 | 146511591 | 606 | Gm8494 | CAST,MOLF,TH | upstream_gene_variant |
| 5 | 148050415 | 148050808 | 393 | Mtus2 | AJ,AKR,BALB,BUB,C3H,ILNJ2,MA/My,NOD,NOR,NZB,SEA,WSB | intron_variant |
| 5 | 148050480 | 148051455 | 975 | Mtus2 | MRL | intron_variant,regulatory_region_variant |
| 5 | 149284241 | 149285087 | 846 | - | CAST,MOLF,SPRET | intergenic_variant |
| 5 | 150577170 | 150577453 | 283 | N4bp2l2 | SPRET | intron_variant |
| 6 | 7390548 | 7391200 | 652 | - | CAST | intergenic_variant |
| 6 | 10047463 | 10047770 | 307 | - | 129S1,AJ,BUB,C57L,CE,DBA1J,DBA2J,ILNJ2,KK,LP,NOD,NOR,NZW,SMJ,TH | intergenic_variant |
| 6 | 11013784 | 11014606 | 822 | - | SPRET | intergenic_variant |
| 6 | 13566499 | 13566998 | 499 | - | BALB,C3H,CBA,SEA,SMJ,SWR | intergenic_variant |
| 6 | 15459619 | 15459929 | 310 | - | CAST | intergenic_variant |
| 6 | 16657317 | 16658243 | 926 | - | WSB | intergenic_variant |
| 6 | 26231787 | 26232509 | 722 | - | SPRET | intergenic_variant |
| 6 | 31584582 | 31585012 | 430 | Gm43154 | SPRET | intron_variant,non_coding_transcript_variant |
| 6 | 33129575 | 33130093 | 518 | - | SPRET | intergenic_variant |
| 6 | 34959070 | 34959442 | 372 | Slc23a4 | SPRET | intron_variant |
| 6 | 40958638 | 40959491 | 853 | - | CAST,FVB,SJL,TH | intergenic_variant |
| 6 | 41202361 | 41202712 | 351 | - | MOLF,SPRET | intergenic_variant |
| 6 | 41299874 | 41300295 | 421 | - | MOLF | intergenic_variant |
| 6 | 43673801 | 43674161 | 360 | - | MOLF | intergenic_variant |
| 6 | 44077392 | 44078321 | 929 | - | SPRET | intergenic_variant |
| 6 | 48017054 | 48017637 | 583 | Zfp777 | CAST,MOLF | intron_variant |
| 6 | 48254274 | 48255229 | 955 | - | MA/My | intergenic_variant |
| 6 | 58130494 | 58131289 | 795 | - | SPRET | intergenic_variant |
| 6 | 58650586 | 58650909 | 323 | Abcg2 | SPRET | intron_variant |
| 6 | 62135952 | 62136320 | 368 | Ccser1 | SPRET | intron_variant |
| 6 | 62547014 | 62547240 | 226 | - | SPRET | intergenic_variant |
| 6 | 64507761 | 64508405 | 644 | Grid2 | CAST | intron_variant |
| 6 | 66765067 | 66765673 | 606 | - | KK | intergenic_variant |
| 6 | 69272376 | 69273116 | 740 | - | WSB | intergenic_variant |
| 6 | 69474845 | 69475497 | 652 | - | NOR | intergenic_variant |
| 6 | 70879430 | 70879870 | 440 | Eif2ak3 | 129S1,AJ,AKR,B10D2,B10J,BALB,BTBR,BUB,C3H,C57L,C58J,CAST,CBA,CE,DBA1J,DBA2J,FVB,ILNJ2,KK,LP,MA/My,MOLF,MRL,NOD,NOR,NZB,NZO,NZW,PJ,PL,RF,RH2,SEA,SJL,SMJ,SPRET,SWR,TH,WSB | intron_variant |
| 6 | 73337697 | 73337965 | 268 | - | AKR,C58J,CAST,NOD | intergenic_variant |
| 6 | 74586368 | 74586823 | 455 | - | SPRET | intergenic_variant |
| 6 | 76011325 | 76011584 | 259 | - | CAST,KK,MOLF,NOD,NOR | intergenic_variant |
| 6 | 78825404 | 78826040 | 636 | - | MRL | intergenic_variant |

| | | | | | | |
|---|---|---|---|---|---|---|
| 6 | 79289360 | 79290289 | 929 | - | KK,MOLF,NZB,NZO,NZW,TH | intergenic_variant |
| 6 | 79738992 | 79739382 | 390 | - | SPRET | intergenic_variant |
| 6 | 86332761 | 86333367 | 606 | - | SPRET | intergenic_variant |
| 6 | 88009542 | 88009992 | 450 | Rab7 | SPRET | intron_variant,regulatory_region_variant |
| 6 | 88438872 | 88439714 | 842 | - | CAST,MOLF | intergenic_variant |
| 6 | 98497599 | 98497855 | 256 | Gm44196 | SEA | intron_variant,non_coding_transcript_variant |
| 6 | 98862268 | 98862678 | 410 | - | CAST | intergenic_variant |
| 6 | 105158253 | 105158694 | 441 | - | SPRET | intergenic_variant |
| 6 | 105473249 | 105473604 | 355 | - | CAST | intergenic_variant |
| 6 | 107249103 | 107249913 | 810 | Gm34933 | MOLF | intron_variant,non_coding_transcript_variant |
| 6 | 109005495 | 109006149 | 654 | - | SPRET | intergenic_variant |
| 6 | 123948562 | 123948821 | 259 | A330044P14Rik | CE | intron_variant,non_coding_transcript_variant,regulatory_region_variant |
| 6 | 128862405 | 128862978 | 573 | BC035044,Gm8774 | NZO | intron_variant,downstream_gene_variant |
| 6 | 130353430 | 130354064 | 634 | Klra1 | KK,NOD,NOR,NZB,NZW,SWR,WSB | intron_variant |
| 6 | 133516636 | 133516901 | 265 | - | SPRET | intergenic_variant |
| 6 | 136395041 | 136395961 | 920 | Gm55605 | MOLF | downstream_gene_variant,regulatory_region_variant |
| 6 | 139396096 | 139396707 | 611 | - | C3H | intergenic_variant |
| 6 | 140516528 | 140517455 | 927 | Plekha5 | SPRET | intron_variant |
| 6 | 143040099 | 143040856 | 757 | C2cd5 | SPRET | intron_variant,regulatory_region_variant |
| 6 | 149273876 | 149274610 | 734 | - | SPRET | intergenic_variant |
| 7 | 3146706 | 3146971 | 265 | Gm18198 | SPRET | upstream_gene_variant |
| 7 | 4053950 | 4054925 | 975 | Lair1 | MOLF,SPRET | intron_variant |
| 7 | 4912221 | 4912546 | 325 | Gm5320,Gm44983 | MOLF | non_coding_transcript_exon_variant,non_coding_transcript_exon_variant,intron_variant |
| 7 | 10125392 | 10125675 | 283 | - | CAST,NZW,SPRET | intergenic_variant |
| 7 | 10190603 | 10191417 | 814 | - | C58J,ILNJ2,NOD,SJL,TH | intergenic_variant |
| 7 | 15557134 | 15557396 | 262 | Obox4-ps35 | C58J,MOLF | intron_variant,non_coding_transcript_variant |
| 7 | 29464241 | 29464868 | 627 | Gm44954 | 129S1,CAST,LP,NZW,RF,SJL | upstream_gene_variant |
| 7 | 31093455 | 31093894 | 439 | - | DBA1J,DBA2J,MOLF,NZW | intergenic_variant |
| 7 | 41696855 | 41697282 | 427 | Gm7224,Vmn2r59 | SPRET | downstream_gene_variant,intron_variant |
| 7 | 42883207 | 42884201 | 994 | - | DBA1J,DBA2J,ILNJ2,MOLF,NZW,PJ | intergenic_variant |
| 7 | 42969199 | 42969608 | 409 | - | MOLF | regulatory_region_variant,intergenic_variant |
| 7 | 44582922 | 44583269 | 347 | - | 129S1,AJ,AKR,BALB,C3H,CBA,DBA1J,DBA2J,KK,MA/My,PL,RH2,SEA,SMJ,WSB | intergenic_variant |
| 7 | 44591717 | 44592027 | 310 | Tsks | 129S1,AJ,AKR,BALB,C3H,CBA,CE,DBA1J,DBA2J,KK,MA/My,PL,RH2,SEA,SMJ,WSB | upstream_gene_variant |
| 7 | 49581195 | 49581716 | 521 | Slc6a5 | CAST | intron_variant |
| 7 | 50128161 | 50128824 | 663 | Nell1 | ILNJ2,KK,MOLF,NZW,PJ | intron_variant |
| 7 | 50809685 | 50810396 | 711 | - | 129S1,C3H,DBA1J,DBA2J | intergenic_variant |
| 7 | 52952454 | 52953412 | 958 | - | DBA2J | intergenic_variant |
| 7 | 58409510 | 58409904 | 394 | Atp10a | SPRET | intron_variant |
| 7 | 61423546 | 61423860 | 314 | A230057D06Rik | CAST | intron_variant,non_coding_transcript_variant |
| 7 | 62427581 | 62427852 | 271 | - | SPRET | intergenic_variant |
| 7 | 63311788 | 63312512 | 724 | Otud7a | SPRET | intron_variant |
| 7 | 70645034 | 70645677 | 643 | Gm42397 | SPRET | intron_variant,non_coding_transcript_variant |
| 7 | 72532336 | 72532793 | 457 | - | CBA,DBA1J,DBA2J,ILNJ2,MOLF,NZB,NZO,PJ,SEA | intergenic_variant |
| 7 | 72806354 | 72806704 | 350 | - | CAST | intergenic_variant |
| 7 | 78071906 | 78072608 | 702 | Ntrk3 | MOLF | intron_variant,regulatory_region_variant |
| 7 | 78919023 | 78919395 | 372 | - | SPRET | intergenic_variant |
| 7 | 83399794 | 83400044 | 250 | - | CAST | intergenic_variant |
| 7 | 85849401 | 85850070 | 669 | - | CAST | intergenic_variant |
| 7 | 86711376 | 86712275 | 899 | - | SPRET | intergenic_variant |
| 7 | 87325600 | 87326152 | 552 | Grm5 | SPRET | intron_variant |
| 7 | 87568283 | 87568613 | 330 | Grm5 | SPRET | intron_variant |
| 7 | 93936284 | 93936557 | 273 | Gm32647 | MOLF | intron_variant,non_coding_transcript_variant |
| 7 | 94631413 | 94631803 | 390 | - | CAST,MOLF,NZB,NZO,SPRET | intergenic_variant |
| 7 | 96818878 | 96819444 | 566 | Gab2 | 129S1,AJ,AKR,BALB,BTBR,BUB,C57L,FVB,KK,MA/My,MRL,NOD,NOR,NZB,NZW,PL,RF,RH2,SJL,SMJ,SWR,TH | intron_variant |
| 7 | 102088450 | 102088936 | 486 | - | CAST | intergenic_variant |
| 7 | 102384272 | 102384873 | 601 | Gm24210 | CAST,MOLF | upstream_gene_variant |
| 7 | 103546639 | 103547226 | 587 | - | 129S1,AJ,AKR,BALB,BTBR,BUB,C3H,C57L,CBA,DBA1J,DBA2J,FVB,ILNJ2,MRL,NZB,NZW,PJ,PL,RF,RH2,SEA,TH,WSB | intergenic_variant |
| 7 | 103611140 | 103611554 | 414 | - | KK,NOD,SJL,SWR,TH | intergenic_variant |
| 7 | 104124590 | 104125201 | 611 | Trim30d | CAST | intron_variant |
| 7 | 108684201 | 108684924 | 723 | - | NOD,SMJ,SWR | intergenic_variant |
| 7 | 109368734 | 109369040 | 306 | Nrip3 | CAST,MOLF,NOD,SPRET,SWR | intron_variant,regulatory_region_variant |
| 7 | 113779804 | 113780073 | 269 | Gm57208 | 129S1,AJ,AKR,BALB,BUB,C57L,MRL,NOD,NZB,NZW,PL,RH2,SJL,WSB | upstream_gene_variant,regulatory_region_variant |
| 7 | 121418566 | 121418930 | 364 | - | CAST,MOLF,SPRET | intergenic_variant |
| 7 | 125574260 | 125575260 | 1000 | Gsg1l | WSB | intron_variant,regulatory_region_variant |
| 7 | 143182928 | 143183227 | 299 | E230032D23Rik,Tnfrsf26 | CAST,MOLF | intron_variant,non_coding_transcript_variant,non_coding_transcript_exon_variant,intron_variant,upstream_gene_variant,downstream_gene_variant |
| 8 | 4112663 | 4113071 | 408 | - | SPRET | intergenic_variant |
| 8 | 5741853 | 5742631 | 778 | - | CAST,SPRET | intergenic_variant |
| 8 | 10766303 | 10766673 | 370 | - | SPRET | intergenic_variant |
| 8 | 13657917 | 13658709 | 792 | Rasa3 | SPRET | intron_variant,regulatory_region_variant |
| 8 | 14238314 | 14238596 | 282 | Dlgap2 | AKR,BALB,C57L,CAST,CE,ILNJ2,LP,MOLF,MRL,NZB,NZW,PJ,PL,RF,RH2,SEA,SMJ,SPRET,TH,WSB | intron_variant |
| 8 | 14842407 | 14842811 | 404 | Dlgap2 | SEA | intron_variant |
| 8 | 17928718 | 17929053 | 335 | - | SPRET | intergenic_variant |
| 8 | 18037398 | 18037628 | 230 | - | WSB | intergenic_variant |
| 8 | 18356857 | 18357268 | 411 | - | SPRET | intergenic_variant |
| 8 | 22458663 | 22459495 | 832 | Ccdc70 | SPRET | upstream_gene_variant |
| 8 | 26386015 | 26386488 | 473 | Kcnu1 | SPRET | intron_variant |
| 8 | 27357198 | 27357529 | 331 | 2310008N11Rik | PL | intron_variant,non_coding_transcript_variant |
| 8 | 27693995 | 27694372 | 377 | - | SPRET | regulatory_region_variant,intergenic_variant |
| 8 | 28689049 | 28689547 | 498 | - | SPRET | intergenic_variant |

| | | | | | | |
|---|---|---|---|---|---|---|
| 8 | 33634523 | 33635215 | 692 | - | SPRET | intergenic_variant |
| 8 | 37200393 | 37201189 | 796 | Dlc1 | CAST | upstream_gene_variant,intron_variant,regulatory_region_variant |
| 8 | 38689887 | 38690336 | 449 | Sgcz | 129S1,AJ,AKR,BALB,BTBR,BUB,C3H,C57L,CBA,CE,DBA1J,DBA2J,FVB,ILNJ2,KK,MA/My,MRL,NOD,NOR,NZB,NZO,NZW,PJ,PL,RF,RH2,SEA,SJL,SMJ,SWR,TH,WSB | intron_variant |
| 8 | 42691009 | 42691699 | 690 | - | SPRET | intergenic_variant |
| 8 | 46408321 | 46408830 | 509 | Cfap96 | SPRET | intron_variant |
| 8 | 56446375 | 56447344 | 969 | Glra3 | CAST | intron_variant |
| 8 | 58695969 | 58696231 | 262 | Galntl6 | CAST,SPRET | intron_variant |
| 8 | 58837386 | 58837868 | 482 | Galntl6 | SPRET | intron_variant |
| 8 | 60552905 | 60553421 | 516 | - | PL | intergenic_variant |
| 8 | 60717879 | 60718377 | 498 | Gm34623 | KK | intron_variant,non_coding_transcript_variant |
| 8 | 66558267 | 66558646 | 379 | Marchf1 | SPRET | intron_variant |
| 8 | 75339940 | 75340453 | 513 | Gm11033 | CAST,SPRET | intron_variant,non_coding_transcript_variant |
| 8 | 88881717 | 88882057 | 340 | Heatr3 | SPRET | intron_variant |
| 8 | 88961779 | 88962453 | 674 | Tent4b | AJ,RH2 | intron_variant |
| 8 | 89703762 | 89704677 | 915 | - | 129S1,AKR,BALB,BTBR,BUB,C57L,C58J,CAST,CBA,CE,DBA1J,DBA2J,FVB,ILNJ2,KK,MA/My,MRL,NOD,NOR,NZB,NZO,NZW,PJ,PL,RF,SEA,SJL,SMJ,SPRET,SWR,TH,WSB | intergenic_variant |
| 8 | 90300459 | 90301047 | 588 | - | SPRET | intergenic_variant |
| 8 | 92415770 | 92416743 | 973 | Gm2875,Gm9619 | 129S1,AKR,BALB,BUB,C57L,C58J,CAST,CBA,CE,DBA1J,DBA2J,FVB,ILNJ2,MA/My,MOLF,MRL,NOD,NOR,NZB,NZO,NZW,PJ,PL,RF,SEA,SJL,SMJ,SPRET,SWR,TH,WSB | non_coding_transcript_variant,upstream_gene_variant |
| 8 | 94208568 | 94209129 | 561 | Gm53022 | CAST,MOLF | intron_variant,non_coding_transcript_variant |
| 8 | 102758291 | 102758921 | 630 | - | MOLF | intergenic_variant |
| 8 | 105954219 | 105955098 | 879 | Phaf1 | SPRET | intron_variant |
| 8 | 106640828 | 106641175 | 347 | Pskh1 | PJ | intron_variant |
| 8 | 118553760 | 118554075 | 315 | - | CAST | intergenic_variant |
| 8 | 119583699 | 119584573 | 874 | Cdh13 | MOLF | intron_variant |
| 8 | 129792443 | 129793152 | 709 | Ccdc7b,Ccdc7a | CAST,CBA,DBA1J,DBA2J,MA/My,NOD,NOR,PJ,SPRET | upstream_gene_variant |
| 8 | 129987565 | 129988207 | 642 | - | KK,MOLF | intergenic_variant |
| 9 | 3652205 | 3652577 | 372 | Gucy1a2 | SPRET | intron_variant |
| 9 | 5220539 | 5220758 | 219 | - | CAST | intergenic_variant |
| 9 | 6585857 | 6586242 | 385 | - | MOLF | intergenic_variant |
| 9 | 7954596 | 7955469 | 873 | Yap1 | SPRET | intron_variant,regulatory_region_variant |
| 9 | 8723189 | 8724048 | 859 | Gm32014 | CAST,SPRET | non_coding_transcript_exon_variant,intron_variant,non_coding_transcript_variant |
| 9 | 8725706 | 8726348 | 642 | Gm32014 | CAST,SPRET | intron_variant,non_coding_transcript_variant |
| 9 | 10344356 | 10344914 | 558 | Cntn5 | CAST,WSB | intron_variant |
| 9 | 11995388 | 11995664 | 276 | - | CAST,SPRET,WSB | intergenic_variant |
| 9 | 12983761 | 12984261 | 500 | - | SPRET | intergenic_variant |
| 9 | 13068133 | 13069098 | 965 | - | SPRET | intergenic_variant |
| 9 | 14036535 | 14037243 | 708 | - | MOLF,SPRET | intergenic_variant |
| 9 | 15161264 | 15161741 | 477 | Vstm5 | MOLF | intron_variant,regulatory_region_variant |
| 9 | 17721479 | 17721754 | 275 | - | MOLF | intergenic_variant |
| 9 | 21628225 | 21628812 | 587 | Gm26511 | SPRET | downstream_gene_variant |
| 9 | 26263949 | 26264768 | 819 | - | MOLF,SPRET | intergenic_variant |
| 9 | 27743739 | 27744434 | 695 | Opcml | SPRET | intron_variant |
| 9 | 41193411 | 41193938 | 527 | - | 129S1,AJ,AKR,BALB,BTBR,BUB,C57L,CAST,CBA,CE,DBA1J,DBA2J,FVB,ILNJ2,KK,MRL,NOD,NOR,NZB,NZO,PJ,PL,RF,RH2,SEA,SWR,TH,WSB | intergenic_variant |
| 9 | 51882147 | 51882395 | 248 | - | SPRET | intergenic_variant |
| 9 | 52067530 | 52067851 | 321 | Zc3h12c | MOLF | intron_variant |
| 9 | 53777543 | 53778132 | 589 | - | CAST,MOLF | intergenic_variant |
| 9 | 54759362 | 54759724 | 362 | - | SPRET | intergenic_variant |
| 9 | 56125616 | 56126174 | 558 | Peak1 | SPRET | intron_variant |
| 9 | 56183564 | 56184198 | 634 | Peak1 | KK | intron_variant |
| 9 | 60459251 | 60459922 | 671 | - | SPRET | intergenic_variant |
| 9 | 62371628 | 62371932 | 304 | Coro2b | AKR,MOLF,NZB,RF,SPRET | intron_variant,regulatory_region_variant |
| 9 | 75915505 | 75916013 | 508 | Gm26903 | MOLF | intron_variant,non_coding_transcript_variant,regulatory_region_variant |
| 9 | 76514367 | 76515002 | 635 | - | MOLF | intergenic_variant |
| 9 | 79063211 | 79063469 | 258 | - | SPRET | intergenic_variant |
| 9 | 80211692 | 80212252 | 560 | Myo6 | MOLF | intron_variant |
| 9 | 80225149 | 80226107 | 958 | Impg1 | 129S1,AJ,BALB,BTBR,C3H,C57L,CAST,CBA,CE,DBA1J,DBA2J,KK,MA/My,MOLF,NZB,NZO,PJ,RH2,SEA,SMJ,SWR,TH | intron_variant,regulatory_region_variant |
| 9 | 93173123 | 93174027 | 904 | - | CAST | intergenic_variant |
| 9 | 94886939 | 94887389 | 450 | Slc9a9 | MRL,NZO,NZW,SPRET | intron_variant |
| 9 | 97779605 | 97779927 | 322 | Clstn2 | SPRET | intron_variant |
| 9 | 105580183 | 105580921 | 738 | Col6a6 | 129S1,AJ,AKR,BALB,BTBR,BUB,C3H,C57L,CAST,CBA,CE,DBA1J,DBA2J,FVB,ILNJ2,MA/My,MRL,NOD,NOR,NZB,NZO,NZW,PJ,PL,RF,RH2,SEA,SJL,SMJ,SPRET,SWR,TH,WSB | intron_variant |
| 9 | 107002195 | 107002558 | 363 | Dock3 | SPRET | intron_variant |
| 9 | 112797554 | 112798124 | 570 | - | ILNJ2,NZB,NZO,NZW | intergenic_variant |
| 9 | 113069170 | 113069812 | 642 | - | SPRET | intergenic_variant |
| 9 | 114061527 | 114062224 | 697 | - | SPRET | intergenic_variant |
| 9 | 114685083 | 114685381 | 298 | - | AJ,AKR,BALB,C3H,CE,FVB,MRL,PJ,PL,RF,RH2,SEA,SMJ,WSB | intergenic_variant |
| 9 | 118507743 | 118508269 | 526 | Itga9 | MOLF,MRL,PJ,SPRET | intron_variant |
| 9 | 119319278 | 119319855 | 577 | Scn5a | SPRET | intron_variant |
| X | 5430352 | 5430657 | 305 | - | MOLF | intergenic_variant |
| X | 5870161 | 5870407 | 246 | - | SPRET | intergenic_variant |
| X | 5904766 | 5905168 | 402 | - | SPRET | intergenic_variant |
| X | 9625897 | 9626324 | 427 | - | SPRET | intergenic_variant |
| X | 10045261 | 10045584 | 323 | Rpgr | SPRET | intron_variant |

| | | | | | | |
|---|---|---|---|---|---|---|
| X | 14017605 | 14018343 | 738 | - | SPRET | intergenic_variant |
| X | 15762483 | 15763011 | 528 | Gm23516 | SPRET | non_coding_transcript_variant |
| X | 18048161 | 18049082 | 921 | Kdm6a | SPRET | intron_variant |
| X | 21980175 | 21980540 | 365 | - | SPRET | intergenic_variant |
| X | 37878738 | 37879629 | 891 | 6030498E09Rik | WSB | intron_variant |
| X | 42484284 | 42485118 | 834 | - | MOLF,SPRET | intergenic_variant |
| X | 44361236 | 44361584 | 348 | - | SPRET | intergenic_variant |
| X | 44449900 | 44450249 | 349 | - | SPRET | intergenic_variant |
| X | 45997881 | 45998239 | 358 | - | SPRET | intergenic_variant |
| X | 62371462 | 62371835 | 373 | - | CAST | intergenic_variant |
| X | 68874752 | 68875122 | 370 | Aff2 | SPRET | intron_variant |
| X | 69357636 | 69358069 | 433 | - | CAST,MOLF | intergenic_variant |
| X | 74699377 | 74699912 | 535 | - | B10D2,B10J,BTBR,MA/My,NZB | intergenic_variant |
| X | 74871492 | 74871726 | 234 | Pls3 | SPRET | intron_variant,upstream_gene_variant |
| X | 74989139 | 74989878 | 739 | - | CE,WSB | intergenic_variant |
| X | 77711687 | 77712388 | 701 | - | CAST | intergenic_variant |
| X | 78123760 | 78124120 | 360 | - | SPRET | intergenic_variant |
| X | 83280848 | 83281644 | 796 | Dmd | WSB | intron_variant |
| X | 88432207 | 88433162 | 955 | - | SPRET | intergenic_variant |
| X | 90291811 | 90292462 | 651 | - | SPRET | intergenic_variant |
| X | 91086901 | 91087569 | 668 | - | SPRET | intergenic_variant |
| X | 95122257 | 95122471 | 214 | - | MOLF | intergenic_variant |
| X | 97259589 | 97260220 | 631 | Ar | SPRET | intron_variant |
| X | 100518233 | 100518604 | 371 | - | MOLF | intergenic_variant |
| X | 100630751 | 100631167 | 416 | Taf1 | MOLF | intron_variant |
| X | 101379852 | 101380490 | 638 | Hdac8,Gm14858 | SPRET | intron_variant,non_coding_transcript_variant |
| X | 101675176 | 101675637 | 461 | Phka1 | SPRET | intron_variant |
| X | 102260883 | 102261209 | 326 | - | MOLF | intergenic_variant |
| X | 103646755 | 103647067 | 312 | Zdhhc15 | CAST,MOLF | intron_variant |
| X | 104380261 | 104381032 | 771 | - | CAST,MOLF | intergenic_variant |
| X | 108161940 | 108162591 | 651 | Sh3bgrl | CAST,CE,MOLF,NZB,NZO,NZW,SPRET | intron_variant |
| X | 108756910 | 108757468 | 558 | - | SPRET | intergenic_variant |
| X | 108979458 | 108979712 | 254 | - | SPRET | intergenic_variant |
| X | 111650672 | 111650937 | 265 | - | AJ,BALB,C3H,CBA,MOLF,MRL,RH2,SPRET | intergenic_variant |
| X | 112598642 | 112599105 | 463 | Dach2 | MOLF | intron_variant |
| X | 115436722 | 115437189 | 467 | - | MOLF | intergenic_variant |
| X | 116358513 | 116359451 | 938 | - | AJ,BALB,C3H,CBA,MOLF,MRL,RH2 | intergenic_variant |
| X | 120272550 | 120272765 | 215 | - | WSB | intergenic_variant |
| X | 120780046 | 120780499 | 453 | - | SPRET | intergenic_variant |
| X | 126710022 | 126710885 | 863 | - | SPRET | intergenic_variant |
| X | 129571105 | 129571343 | 238 | - | SPRET | intergenic_variant |
| X | 130115201 | 130115909 | 708 | - | SPRET | intergenic_variant |
| X | 130602799 | 130603477 | 678 | - | MOLF,SPRET | intergenic_variant |
| X | 131980919 | 131981722 | 803 | - | SPRET | intergenic_variant |
| X | 140375619 | 140376164 | 545 | Col4a5 | SPRET | intron_variant |
| X | 140988506 | 140988723 | 217 | - | CAST | intergenic_variant |
| X | 146219745 | 146220168 | 423 | - | CAST,MOLF,NZB | intergenic_variant |
| X | 155632928 | 155633743 | 815 | - | SPRET | intergenic_variant |
| X | 155757905 | 155758152 | 247 | - | CAST | intergenic_variant |
| X | 161632242 | 161633238 | 996 | Gm23901 | SPRET | non_coding_transcript_exon_variant |
| X | 162239200 | 162239810 | 610 | - | SPRET | intergenic_variant |

| CHR | START | END | SIZE | SYMBOL | STRAINS with VARIANT SV ALLELE | CONSEQUENCE |
|---|---|---|---|---|---|---|
| 1 | 3192284 | 3193853 | 1569 | - | C3H,CBA,MRL,RF | regulatory_region_variant,intergenic_variant |
| 1 | 4498582 | 4501964 | 3382 | - | MRL | intergenic_variant |
| 1 | 7910818 | 7912108 | 1290 | - | C3H,CBA,MRL,RF | intergenic_variant |
| 1 | 9558025 | 9566608 | 8583 | - | SPRET | intergenic_variant |
| 1 | 11384470 | 11385969 | 1499 | - | SPRET | intergenic_variant |
| 1 | 14410529 | 14411998 | 1469 | - | 129S1,AJ,AKR,BALB,BTBR,C3H,C57L,CBA,CE,DBA1J,DBA2J,FVB,ILNJ2,KK,LP,MRL,NOD,NOR,NZB,NZO,NZW,PJ,PL,RF,RH2,SEA,SJL,SMJ,SWR,TH | intergenic_variant |
| 1 | 14411390 | 14412740 | 1350 | - | CAST | intergenic_variant |
| 1 | 14546450 | 14553081 | 6631 | - | SPRET | intergenic_variant |
| 1 | 16388574 | 16402615 | 14041 | Stau2 | CAST,MOLF | intron_variant |
| 1 | 16389346 | 16402614 | 13268 | Stau2 | CAST,MOLF | intron_variant |
| 1 | 16394170 | 16403486 | 9316 | Stau2 | CAST,MOLF | intron_variant |
| 1 | 17358782 | 17362315 | 3533 | - | BTBR,BUB,LP | intergenic_variant |
| 1 | 22301046 | 22307255 | 6209 | Gm28239 | MRL | upstream_gene_variant |
| 1 | 24468568 | 24471917 | 3349 | Col19a1 | SPRET | intron_variant |
| 1 | 26034302 | 26036443 | 2141 | Gm18443 | 129S1,BTBR,C57L,NZW | non_coding_transcript_variant |
| 1 | 27353869 | 27358446 | 4577 | - | SPRET | intergenic_variant |
| 1 | 27794175 | 27800406 | 6231 | - | SPRET | intergenic_variant |
| 1 | 27795574 | 27800438 | 4864 | - | SPRET | intergenic_variant |
| 1 | 29179906 | 29182921 | 3015 | - | CAST,RF | intergenic_variant |
| 1 | 30134538 | 30135649 | 1111 | - | SPRET | intergenic_variant |
| 1 | 31512524 | 31515469 | 2945 | - | BUB,CAST,FVB,NZB,NZO,PJ,SJL,SWR | intergenic_variant |
| 1 | 31514711 | 31516089 | 1378 | - | BUB,CAST,FVB,NZB,NZO,PJ,SJL,SWR | intergenic_variant |
| 1 | 31611064 | 31612292 | 1228 | - | CE,MOLF | intergenic_variant |
| 1 | 33093914 | 33095764 | 1850 | - | SPRET | intergenic_variant |
| 1 | 33338680 | 33345557 | 6877 | - | AJ,BALB,BUB,C3H,DBA1J,DBA2J,FVB,ILNJ2,KK,LP,MRL,NOR,NZB,NZO,PJ,PL,RF,RH2,SEA,SMJ | intergenic_variant |
| 1 | 35624414 | 35626422 | 2008 | - | MOLF | regulatory_region_variant,intergenic_variant |
| 1 | 35865216 | 35866729 | 1513 | - | AJ,AKR,BUB,C3H,CBA,CE,FVB,KK,MRL,NZB,NZO,NZW,RF,RH2,SMJ,SWR,WSB | regulatory_region_variant,intergenic_variant |
| 1 | 35865616 | 35868156 | 2540 | - | AJ,AKR,BUB,C3H,CBA,CE,FVB,KK,MRL,NZB,NZO,NZW,RF,RH2,SMJ,SWR,WSB | regulatory_region_variant,intergenic_variant |
| 1 | 40189016 | 40191031 | 2015 | - | 129S1,BTBR,C57L,CAST,MOLF,NZB,NZO | regulatory_region_variant,intergenic_variant |
| 1 | 41833374 | 41837478 | 4104 | Gm29260 | CAST,MOLF | intron_variant,non_coding_transcript_variant,regulatory_region_variant |
| 1 | 41833399 | 41835884 | 2485 | Gm29260 | CAST,MOLF | intron_variant,non_coding_transcript_variant |
| 1 | 43679870 | 43681845 | 1975 | - | CAST | intergenic_variant |
| 1 | 44311450 | 44312508 | 1058 | - | MOLF | intergenic_variant |
| 1 | 44311670 | 44313676 | 2006 | - | MOLF | intergenic_variant |
| 1 | 44439467 | 44441274 | 1807 | - | SPRET | intergenic_variant |
| 1 | 44440400 | 44441646 | 1246 | - | SPRET | intergenic_variant |
| 1 | 45308398 | 45310346 | 1948 | - | MOLF,SPRET | intergenic_variant |
| 1 | 47161561 | 47165567 | 4006 | - | SPRET | intergenic_variant |
| 1 | 47579784 | 47580991 | 1207 | - | SPRET | intergenic_variant |
| 1 | 47626851 | 47628380 | 1529 | - | SWR | intergenic_variant |
| 1 | 47866555 | 47867687 | 1132 | - | SPRET | intergenic_variant |
| 1 | 48357392 | 48359408 | 2016 | - | SPRET | intergenic_variant |
| 1 | 49822900 | 49823901 | 1001 | - | CAST,SPRET | intergenic_variant |
| 1 | 50720379 | 50721463 | 1084 | - | AKR,BALB,BUB,C58J,MA/My,RF,RH2,SEA,SWR,WSB | intergenic_variant |
| 1 | 57495558 | 57498169 | 2611 | - | CAST | intergenic_variant |
| 1 | 58073081 | 58075223 | 2142 | Aox1 | CAST | intron_variant |
| 1 | 59332577 | 59335034 | 2457 | Cdk15 | CE,MOLF | intron_variant |
| 1 | 62267242 | 62268688 | 1446 | Pard3b | SPRET | intron_variant |
| 1 | 63260644 | 63262084 | 1440 | - | SPRET | intergenic_variant |
| 1 | 63878921 | 63880708 | 1787 | - | MOLF | intergenic_variant |
| 1 | 64510712 | 64513427 | 2715 | - | CAST,MOLF | intergenic_variant |
| 1 | 66054893 | 66056769 | 1876 | - | 129S1,AJ,AKR,BTBR,BUB,C3H,C57L,C58J,CBA,CE,DBA1J,DBA2J,FVB,ILNJ2,KK,LP,MA/My,MRL,NOD,NOR,NZB,PJ,PL,RF,SJL,SMJ,SWR,TH,WSB | regulatory_region_variant,intergenic_variant |
| 1 | 66831537 | 66835768 | 4231 | Kansl1l,6820402A03Ril | AJ,CBA,DBA1J,NZW,SWR | intron_variant,non_coding_transcript_variant,regulatory_region_variant |
| 1 | 69284325 | 69286734 | 2409 | - | 129S1,AJ,BTBR,C3H,C57L,DBA2J,FVB,KK,LP,MA/My,MRL,NOD,NOR,NZB,NZO,NZW,SMJ,TH | regulatory_region_variant,intergenic_variant |
| 1 | 74155678 | 74158086 | 2408 | Tns1 | SPRET | intron_variant,regulatory_region_variant |
| 1 | 74360600 | 74363591 | 2991 | Pnkd | CAST,SPRET | intron_variant,regulatory_region_variant |
| 1 | 75945121 | 75947404 | 2283 | - | SPRET | intergenic_variant |
| 1 | 79010149 | 79014077 | 3928 | Gm29536 | CAST | intron_variant,non_coding_transcript_variant |
| 1 | 79010149 | 79012060 | 1911 | Gm29536 | AJ,BALB,C3H,ILNJ2,NZB,PJ,RH2,SEA,SJL,SWR,WSB | intron_variant,non_coding_transcript_variant |
| 1 | 79049228 | 79051321 | 2093 | Gm29536 | SPRET | intron_variant,non_coding_transcript_variant |
| 1 | 79225828 | 79235255 | 9427 | - | AKR,B10J,BUB,C3H,C57L,NOR,PL,RH2,SEA | intergenic_variant |
| 1 | 79227348 | 79232893 | 5545 | - | PJ,SWR | intergenic_variant |
| 1 | 79227348 | 79233734 | 6386 | - | 129S1,AJ,AKR,B10D2,B10J,BALB,BTBR,BUB,C3H,C57L,CBA,CE,DBA1J,DBA2J,ILNJ2,KK,MA/My,MRL,NOD,NOR,NZO,NZW,PL,RF,RH2,SEA,SJL,TH | intergenic_variant |
| 1 | 79227349 | 79235246 | 7897 | - | PL,RH2,SEA | intergenic_variant |
| 1 | 79229464 | 79233734 | 4270 | - | CBA,CE,DBA1J,DBA2J,MRL,NOD | intergenic_variant |
| 1 | 80102274 | 80105474 | 3200 | - | CAST | intergenic_variant |
| 1 | 83117877 | 83122907 | 5030 | - | SPRET | regulatory_region_variant,intergenic_variant |
| 1 | 83208566 | 83212716 | 4150 | - | LP | intergenic_variant |
| 1 | 88158026 | 88159110 | 1084 | Mroh2a | 129S1,AJ,AKR,B10D2,B10J,BALB,BTBR,BUB,C3H,C57L,C58J,CBA,CE,DBA1J,DBA2J,FVB,ILNJ2,KK,LP,MA/My,MOLF,MRL,NOD,NOR,NZB,NZO,NZW,PJ,PL,RH2,SEA,SJL,SMJ,SWR,TH,WSB | coding_sequence_variant,intron_variant,regulatory_region_variant |

| | | | | | | |
|---|---|---|---|---|---|---|
| 1 | 91663953 | 91667226 | 3273 | - | SPRET | intergenic_variant |
| 1 | 94952669 | 94955055 | 2386 | - | MOLF | regulatory_region_variant,intergenic_variant |
| 1 | 95777026 | 95778274 | 1248 | - | CAST | intergenic_variant |
| 1 | 96423208 | 96424425 | 1217 | - | AKR,C57L,DBA2J,ILNJ2,NOD,SMJ | intergenic_variant |
| 1 | 98923482 | 98925205 | 1723 | - | CAST | intergenic_variant |
| 1 | 99583884 | 99585869 | 1985 | - | WSB | intergenic_variant |
| 1 | 99584332 | 99588680 | 4348 | - | WSB | intergenic_variant |
| 1 | 99628600 | 99630024 | 1424 | - | MOLF | intergenic_variant |
| 1 | 99837346 | 99839340 | 1994 | Cntnap5b | 129S1,AKR,BUB,C57L,DBA1J,DBA2J,LP,MA/My,NOD,NOR,NZO,PJ,SEA,SJL,TH,WSB | intron_variant |
| 1 | 101287217 | 101291317 | 4100 | - | 129S1,AKR,C57L,DBA1J,DBA2J,LP,MA/My,NOD,NOR,PJ,SEA,SPRET,TH | intergenic_variant |
| 1 | 101287301 | 101293490 | 6189 | - | 129S1,AKR,C57L,DBA1J,DBA2J,LP,MA/My,NOD,NOR,PJ,SEA,SPRET,TH | intergenic_variant |
| 1 | 102441331 | 102443974 | 2643 | Gm53054 | CAST | intron_variant,non_coding_transcript_variant |
| 1 | 102446190 | 102456903 | 10713 | Gm53054 | WSB | non_coding_transcript_exon_variant,intron_variant |
| 1 | 103182078 | 103186420 | 4342 | Gm18728 | SPRET | intron_variant,non_coding_transcript_variant |
| 1 | 103819873 | 103821294 | 1421 | - | MOLF,SPRET | intergenic_variant |
| 1 | 110290644 | 110294520 | 3876 | - | SPRET | intergenic_variant |
| 1 | 111068753 | 111073373 | 4620 | - | SPRET | intergenic_variant |
| 1 | 111576416 | 111579067 | 2651 | - | SPRET | intergenic_variant |
| 1 | 113652278 | 113656860 | 4582 | - | CAST | intergenic_variant |
| 1 | 113822909 | 113825777 | 2868 | 9330185C12Rik | MOLF | intron_variant,non_coding_transcript_variant |
| 1 | 114692240 | 114699103 | 6863 | - | WSB | intergenic_variant |
| 1 | 114730090 | 114737445 | 7355 | - | SPRET | intergenic_variant |
| 1 | 119623617 | 119626844 | 3227 | Ptpn4 | MOLF | intron_variant |
| 1 | 122065818 | 122066946 | 1128 | - | CE,DBA2J,SMJ | intergenic_variant |
| 1 | 122851020 | 122853228 | 2208 | - | BUB | intergenic_variant |
| 1 | 123516894 | 123522029 | 5135 | Dpp10 | AJ,BUB,C3H,CAST,CBA,CE,KK,MOLF,NZO,RH2,SMJ,SPRET,SWR,WSB | intron_variant |
| 1 | 123518424 | 123522932 | 4508 | Dpp10 | NZW | intron_variant |
| 1 | 123518458 | 123523484 | 5026 | Dpp10 | 129S1,AJ,BALB,BTBR,BUB,C3H,C57L,CAST,CBA,CE,ILNJ2,KK,MOLF,NZO,NZW,PL,RF,RH2,SEA,SMJ,SPRET,SWR,WSB | intron_variant |
| 1 | 125533371 | 125534403 | 1032 | - | SPRET | intergenic_variant |
| 1 | 129294033 | 129295102 | 1069 | Thsd7b | MOLF | intron_variant |
| 1 | 132031302 | 132037934 | 6632 | Gm37120 | KK,MA/My,NZO | non_coding_transcript_exon_variant,regulatory_region_variant |
| 1 | 139752079 | 139753136 | 1057 | Cfhr2 | MRL | intron_variant |
| 1 | 139762728 | 139767928 | 5200 | Cfhr2 | CAST,MOLF | intron_variant |
| 1 | 139765230 | 139772721 | 7491 | Cfhr2 | CAST,MOLF | intron_variant |
| 1 | 147687495 | 147689692 | 2197 | - | SPRET | intergenic_variant |
| 1 | 147708987 | 147712210 | 3223 | - | BTBR,CAST,MOLF,SMJ | intergenic_variant |
| 1 | 149359784 | 149368260 | 8476 | - | BTBR,CAST,MOLF,SMJ | intergenic_variant |
| 1 | 150008728 | 150009836 | 1108 | - | SPRET | intergenic_variant |
| 1 | 150615700 | 150617435 | 1735 | Hmcn1 | SPRET | intron_variant |
| 1 | 152952979 | 152959907 | 6928 | Nmnat2,Gm28960 | CAST | intron_variant,intron_variant,non_coding_transcript_variant,regulatory_region_variant |
| 1 | 156757035 | 156758424 | 1389 | Ralgps2 | CAST | intron_variant |
| 1 | 161229553 | 161230644 | 1091 | Tnfsf4 | SPRET | intron_variant |
| 1 | 162091608 | 162098620 | 7012 | Dnm3 | SPRET | intron_variant |
| 1 | 162647273 | 162653871 | 6598 | - | SPRET | intergenic_variant |
| 1 | 162647781 | 162653871 | 6090 | - | SPRET | intergenic_variant |
| 1 | 162648831 | 162653904 | 5073 | - | SPRET | intergenic_variant |
| 1 | 167394713 | 167395791 | 1078 | - | BTBR,BUB,CAST,FVB,ILNJ2,KK,NZB,NZO,NZW,RF,SMJ,SWR,TH | regulatory_region_variant,intergenic_variant |
| 1 | 168386809 | 168388114 | 1305 | - | SPRET | intergenic_variant |
| 1 | 175107988 | 175113464 | 5476 | Rgs7 | 129S1,AJ,AKR,BALB,BTBR,BUB,C3H,C57L,C58J,CBA,CE,DBA1J,DBA2J,FVB,ILNJ2,KK,MRL,NOD,NOR,NZB,NZO,NZW,PJ,PL,RF,RH2,SEA,SJL,SMJ,SWR,TH,WSB | intron_variant |
| 1 | 177534675 | 177536718 | 2043 | - | FVB,NZO | regulatory_region_variant,intergenic_variant |
| 1 | 181056751 | 181057923 | 1172 | A430110L20Rik,Gm57 | C57L | downstream_gene_variant,intron_variant,non_coding_transcript_variant |
| 1 | 181364448 | 181365540 | 1092 | - | SPRET | regulatory_region_variant,intergenic_variant |
| 1 | 182330754 | 182332882 | 2128 | Capn2 | CE,NZB | intron_variant,regulatory_region_variant |
| 1 | 183802323 | 183804661 | 2338 | Dusp10 | SPRET | intron_variant,regulatory_region_variant |
| 1 | 187805483 | 187807882 | 2399 | Esrrg | SPRET | intron_variant |
| 1 | 187806776 | 187809367 | 2591 | Esrrg | SPRET | intron_variant |
| 1 | 190244156 | 190247127 | 2971 | - | SPRET | intergenic_variant |
| 1 | 191382994 | 191386281 | 3287 | - | CAST | intergenic_variant |
| 1 | 191383081 | 191387888 | 4807 | - | CAST | intergenic_variant |
| 1 | 193946703 | 193947878 | 1175 | - | SPRET | intergenic_variant |
| 10 | 6223173 | 6238970 | 15797 | - | BUB | intergenic_variant |
| 10 | 6224968 | 6229863 | 4895 | - | BUB,CAST,MOLF,SPRET | intergenic_variant |
| 10 | 6227799 | 6229971 | 2172 | - | BUB,CAST,KK,MOLF,NOD,NOR,SPRET,TH | intergenic_variant |
| 10 | 14392626 | 14396526 | 3900 | Adgrg6 | SPRET | intron_variant |
| 10 | 14947910 | 14949122 | 1212 | - | SPRET | intergenic_variant |
| 10 | 16647127 | 16655488 | 8361 | - | BUB,CAST,LP,MOLF,SWR | intergenic_variant |
| 10 | 17904207 | 17906620 | 2413 | - | SPRET | intergenic_variant |
| 10 | 23464703 | 23466436 | 1733 | Gm47843 | CAST,MOLF | intron_variant,non_coding_transcript_variant |
| 10 | 24823314 | 24826218 | 2904 | - | WSB | regulatory_region_variant,intergenic_variant |
| 10 | 26484875 | 26486935 | 2060 | Gm48864 | CAST | upstream_gene_variant |
| 10 | 26524922 | 26528449 | 3527 | - | SPRET | intergenic_variant |
| 10 | 27209718 | 27211045 | 1327 | Lama2 | MOLF | intron_variant |
| 10 | 27229937 | 27235260 | 5323 | Lama2 | SPRET | intron_variant,regulatory_region_variant |
| 10 | 27229937 | 27235260 | 5323 | Lama2 | SPRET | intron_variant |
| 10 | 28102399 | 28105650 | 3251 | Ptprk | SPRET | intron_variant,regulatory_region_variant |
| 10 | 28292106 | 28296354 | 4248 | Ptprk | SPRET | intron_variant |
| 10 | 29732943 | 29734138 | 1195 | - | SPRET | intergenic_variant |
| 10 | 30325801 | 30327432 | 1631 | - | SPRET | intergenic_variant |
| 10 | 36480006 | 36481599 | 1593 | Hs3st5 | MOLF | intron_variant |
| 10 | 42755478 | 42760157 | 4679 | Scml4 | CAST,MOLF,SPRET | intron_variant,regulatory_region_variant |

| | | | | | | |
|---|---|---|---|---|---|---|
| 10 | 45851423 | 45853171 | 1748 | - | TH | regulatory_region_variant,intergenic_variant |
| 10 | 46182329 | 46187243 | 4914 | Gm35430 | SPRET | intron_variant,non_coding_transcript_variant |
| 10 | 46186057 | 46187375 | 1318 | Gm35430 | SPRET | intron_variant,non_coding_transcript_variant |
| 10 | 51512731 | 51515558 | 2827 | - | KK,NZW,WSB | intergenic_variant |
| 10 | 54831974 | 54838061 | 6087 | - | SPRET | intergenic_variant |
| 10 | 59790660 | 59791984 | 1324 | - | SPRET | intergenic_variant |
| 10 | 64741379 | 64743078 | 1699 | Ctnna3 | SPRET | intron_variant |
| 10 | 64998139 | 64999605 | 1466 | - | CAST | intergenic_variant |
| 10 | 65219129 | 65226891 | 7762 | - | SPRET | intergenic_variant |
| 10 | 72661504 | 72662561 | 1057 | - | MOLF | intergenic_variant |
| 10 | 74932002 | 74933278 | 1276 | Bcr | SPRET | intron_variant,regulatory_region_variant |
| 10 | 77435955 | 77439382 | 3427 | - | TH | regulatory_region_variant,intergenic_variant |
| 10 | 77450147 | 77451539 | 1392 | Sumo3 | SPRET | intron_variant,regulatory_region_variant |
| 10 | 79034286 | 79038937 | 4651 | - | CAST | intergenic_variant |
| 10 | 79035252 | 79038938 | 3686 | - | CAST | intergenic_variant |
| 10 | 82106419 | 82107993 | 1574 | - | SPRET | intergenic_variant |
| 10 | 82188009 | 82189928 | 1919 | Gm4924 | CAST,SPRET | upstream_gene_variant,regulatory_region_variant |
| 10 | 83623157 | 83628280 | 5123 | - | MOLF | regulatory_region_variant,intergenic_variant |
| 10 | 83637861 | 83643835 | 5974 | - | SPRET | regulatory_region_variant,intergenic_variant |
| 10 | 83638155 | 83642949 | 4794 | - | SPRET | regulatory_region_variant,intergenic_variant |
| 10 | 84455748 | 84456898 | 1150 | Gm17249 | BTBR,C58J,LP,NZB,NZO,SMJ | non_coding_transcript_exon_variant |
| 10 | 86483827 | 86486622 | 2795 | - | CAST | intergenic_variant |
| 10 | 87470264 | 87471649 | 1385 | - | BUB,CAST,FVB | intergenic_variant |
| 10 | 95297338 | 95298415 | 1077 | Gm47599 | PL | upstream_gene_variant |
| 10 | 100074100 | 100075518 | 1418 | - | AKR,BUB,C3H,CBA,FVB,MA/My,NOD,NOR,PL,SMJ | intergenic_variant |
| 10 | 100666965 | 100669134 | 2169 | - | CBA,NZB | intergenic_variant |
| 10 | 102159443 | 102165872 | 6429 | Mgat4c | CAST | intron_variant |
| 10 | 103176433 | 103178137 | 1704 | - | SPRET | intergenic_variant |
| 10 | 103780096 | 103783245 | 3149 | - | AJ,BALB,CE,DBA1J,DBA2J,ILNJ2,MRL,PJ,RH2,SEA,SJL | intergenic_variant |
| 10 | 104626693 | 104628561 | 1868 | - | AJ,BALB,BTBR,CE,DBA1J,DBA2J,MRL,NZB,PJ,RH2,SEA,SJL | intergenic_variant |
| 10 | 105868876 | 105870762 | 1886 | - | WSB | intergenic_variant |
| 10 | 106496096 | 106499759 | 3663 | Ppfia2 | WSB | intron_variant |
| 10 | 111881224 | 111884895 | 3671 | Gm7269 | SPRET | non_coding_transcript_exon_variant,intron_variant,regulatory_region_variant |
| 10 | 112448580 | 112452888 | 4308 | - | PL,WSB | intergenic_variant |
| 10 | 112959251 | 112960367 | 1116 | - | CAST,MOLF | intergenic_variant |
| 10 | 115637558 | 115640844 | 3286 | 4930422I22Rik | SPRET | non_coding_transcript_exon_variant,intron_variant,regulatory_region_variant |
| 10 | 117161545 | 117162676 | 1131 | - | SPRET | intergenic_variant |
| 10 | 122467476 | 122474073 | 6597 | - | SPRET | intergenic_variant |
| 10 | 122557213 | 122558835 | 1622 | Ppm1h | KK,MOLF | intron_variant |
| 10 | 124065390 | 124070926 | 5536 | - | PL | intergenic_variant |
| 10 | 124069702 | 124070915 | 1213 | - | PL | intergenic_variant |
| 10 | 129423174 | 129430266 | 7092 | - | C3H,CBA | regulatory_region_variant,intergenic_variant |
| 11 | 5515369 | 5517132 | 1763 | - | SPRET | intergenic_variant |
| 11 | 7349554 | 7351315 | 1761 | - | KK | intergenic_variant |
| 11 | 13201662 | 13203155 | 1493 | - | SPRET | intergenic_variant |
| 11 | 13500625 | 13503741 | 3116 | - | MOLF | intergenic_variant |
| 11 | 15753256 | 15755347 | 2091 | - | SPRET | intergenic_variant |
| 11 | 30703150 | 30704874 | 1724 | Gm12101,Gm12100 | BALB,BTBR,C57L,RH2,WSB | non_coding_transcript_exon_variant,non_coding_transcript_exon_variant,intron_variant,regulatory_region_variant |
| 11 | 35798170 | 35799985 | 1815 | Wwc1 | SPRET | intron_variant |
| 11 | 37643088 | 37650356 | 7268 | - | SPRET | regulatory_region_variant,intergenic_variant |
| 11 | 46489321 | 46495894 | 6573 | - | MOLF | intergenic_variant |
| 11 | 49250825 | 49251969 | 1144 | - | MOLF | regulatory_region_variant,intergenic_variant |
| 11 | 58158302 | 58160922 | 2620 | - | SPRET | regulatory_region_variant,intergenic_variant |
| 11 | 61222339 | 61223838 | 1499 | Slc47a2 | SPRET | intron_variant,regulatory_region_variant |
| 11 | 61311196 | 61318406 | 7210 | - | FVB,NOD,SWR | intergenic_variant |
| 11 | 63726137 | 63728347 | 2210 | - | FVB,NOD,SWR,WSB | intergenic_variant |
| 11 | 65546990 | 65549669 | 2679 | - | 129S1,BTBR,C57L,NZW | intergenic_variant |
| 11 | 72988226 | 72989233 | 1007 | Itgae | DBA1J,DBA2J,MOLF,NOR,NZB,PJ,SEA | intron_variant |
| 11 | 76049730 | 76051442 | 1712 | Vps53 | 129S1,AJ,AKR,BALB,BTBR,BUB,C3H,C57L,C58J,CBA,DBA1J,DBA2J,FVB,ILNJ2,KK,MA/My,MRL,NOD,NOR,NZB,PJ,PL,RF,RH2,SEA,SJL,SMJ,SWR,TH | intron_variant,regulatory_region_variant |
| 11 | 79375229 | 79378904 | 3675 | Nf1,AU040972 | SPRET | intron_variant,upstream_gene_variant |
| 11 | 92382582 | 92385373 | 2791 | - | SPRET | intergenic_variant |
| 11 | 92471956 | 92475376 | 3420 | - | SPRET | intergenic_variant |
| 11 | 92808159 | 92810724 | 2565 | - | 129S1,BTBR,C57L,CE | intergenic_variant |
| 11 | 96277388 | 96278464 | 1076 | - | SPRET | intergenic_variant |
| 11 | 97786811 | 97788077 | 1266 | - | MOLF,NZW,SWR | regulatory_region_variant,intergenic_variant |
| 11 | 99638798 | 99643560 | 4762 | Gm52963 | SPRET | upstream_gene_variant |
| 11 | 101533239 | 101536090 | 2851 | Gm24950,Gm23971 | 129S1,AJ,BTBR,BUB,C3H,C57L,CE,DBA1J,DBA2J,ILNJ2,KK,NOD,NOR,NZB,NZO | downstream_gene_variant |
| 11 | 108411681 | 108413432 | 1751 | Cep112 | CAST | intron_variant |
| 11 | 108804510 | 108805943 | 1433 | - | CAST,SPRET | intergenic_variant |
| 12 | 3554091 | 3555204 | 1113 | Asxl2 | C58J,LP,MOLF | 3_prime_UTR_variant |
| 12 | 5671474 | 5676971 | 5497 | - | BALB,ILNJ2,MRL,SEA,SPRET,WSB | regulatory_region_variant,intergenic_variant |
| 12 | 6327015 | 6328607 | 1592 | - | MOLF | intergenic_variant |
| 12 | 6692416 | 6697177 | 4761 | - | CAST | intergenic_variant |
| 12 | 6693787 | 6696592 | 2805 | - | CAST | intergenic_variant |
| 12 | 7691528 | 7692577 | 1049 | Gm32828 | MRL,NZW,WSB | intron_variant,non_coding_transcript_variant |
| 12 | 10657122 | 10658421 | 1299 | - | SPRET | intergenic_variant |
| 12 | 10897961 | 10900417 | 2456 | - | CAST,MOLF | intergenic_variant |
| 12 | 12115437 | 12119042 | 3605 | - | MOLF | intergenic_variant |
| 12 | 13540513 | 13543288 | 2775 | Nbas,Gm24142 | SPRET | intron_variant,non_coding_transcript_exon_variant |
| 12 | 13542254 | 13543361 | 1107 | Nbas | SPRET | intron_variant |
| 12 | 14894381 | 14900493 | 6112 | - | AJ,BALB,CE,ILNJ2,MRL | intergenic_variant |
| 12 | 14894382 | 14901659 | 7277 | - | MOLF | intergenic_variant |

| | | | | | | |
|---|---|---|---|---|---|---|
| 12 | 14898409 | 14903050 | 4641 | - | C58J,CAST,DBA1J,DBA2J,MOLF,NOD,NOR,NZB,NZW,SMJ,SPRET | intergenic_variant |
| 12 | 15241483 | 15248670 | 7187 | - | CAST | intergenic_variant |
| 12 | 17034833 | 17036067 | 1234 | Rock2 | SPRET | 3_prime_UTR_variant,regulatory_region_variant |
| 12 | 24314131 | 24316490 | 2359 | - | SPRET,WSB | intergenic_variant |
| 12 | 24314895 | 24317580 | 2685 | - | SPRET | intergenic_variant |
| 12 | 24915882 | 24917108 | 1226 | Mboat2 | SPRET | intron_variant,regulatory_region_variant |
| 12 | 26938081 | 26939741 | 1660 | - | SPRET | regulatory_region_variant,intergenic_variant |
| 12 | 36300480 | 36302054 | 1574 | Lrrc72 | CE | intron_variant |
| 12 | 39835337 | 39836603 | 1266 | - | MOLF | intergenic_variant |
| 12 | 40929655 | 40932586 | 2931 | - | BUB,DBA1J,DBA2J,FVB,ILNJ2,MRL,NOD,NOR,NZB,NZW,SJL | regulatory_region_variant,intergenic_variant |
| 12 | 42270436 | 42273958 | 3522 | - | BUB,CAST,DBA1J,DBA2J,FVB,ILNJ2,MOLF,MRL,NOD,NOR,NZB,SJL,SPRET | intergenic_variant |
| 12 | 42866445 | 42869511 | 3066 | - | CAST,KK,SWR | intergenic_variant |
| 12 | 43262569 | 43264274 | 1705 | - | CAST,MOLF | intergenic_variant |
| 12 | 43390098 | 43397548 | 7450 | Gm47030 | SPRET | intron_variant,non_coding_transcript_variant |
| 12 | 45343015 | 45349900 | 6885 | - | CAST,MOLF | intergenic_variant |
| 12 | 45384479 | 45385602 | 1123 | - | KK,NZO,NZW | intergenic_variant |
| 12 | 47849592 | 47851849 | 2257 | - | CAST,MOLF,SPRET | intergenic_variant |
| 12 | 48361848 | 48363846 | 1998 | - | CAST,MOLF | intergenic_variant |
| 12 | 48361851 | 48364970 | 3119 | - | CAST,MOLF | intergenic_variant |
| 12 | 48889824 | 48896701 | 6877 | - | SPRET | intergenic_variant |
| 12 | 49394675 | 49396963 | 2288 | - | MOLF | intergenic_variant |
| 12 | 49394676 | 49398308 | 3632 | - | CAST,MOLF,SPRET | intergenic_variant |
| 12 | 49395461 | 49396963 | 1502 | - | CAST,MOLF | intergenic_variant |
| 12 | 51349353 | 51356508 | 7155 | - | 129S1,C57L,CAST,MOLF,NZB,SPRET | regulatory_region_variant,intergenic_variant |
| 12 | 51575931 | 51579534 | 3603 | - | SPRET | intergenic_variant |
| 12 | 55447707 | 55452828 | 5121 | Psma6 | MOLF | intron_variant |
| 12 | 55447717 | 55448807 | 1090 | Psma6 | MOLF | intron_variant |
| 12 | 55922817 | 55926111 | 3294 | - | SPRET | intergenic_variant |
| 12 | 57984794 | 57989532 | 4738 | - | MOLF | regulatory_region_variant,intergenic_variant |
| 12 | 58212613 | 58215177 | 2564 | - | SPRET | intergenic_variant |
| 12 | 59041213 | 59042401 | 1188 | Sec23a | SPRET | intron_variant,regulatory_region_variant |
| 12 | 59422935 | 59424626 | 1691 | - | SPRET | intergenic_variant |
| 12 | 59999686 | 60004793 | 5107 | - | CAST | intergenic_variant |
| 12 | 60873574 | 60875737 | 2163 | - | CAST,MOLF | intergenic_variant |
| 12 | 60888626 | 60890087 | 1461 | - | MOLF | intergenic_variant |
| 12 | 60910841 | 60911910 | 1069 | - | AJ,BALB,BTBR,C3H,C57L,CBA,DBA1J,DBA2J,ILNJ2,KK,MA/My,NZB,NZO,NZW,PJ,RH2,SEA,SMJ,TH,WSB | intergenic_variant |
| 12 | 62726260 | 62727627 | 1367 | - | MOLF | intergenic_variant |
| 12 | 63154815 | 63157378 | 2563 | - | CAST,MOLF | intergenic_variant |
| 12 | 63155837 | 63157395 | 1558 | - | SPRET | intergenic_variant |
| 12 | 69988886 | 69990503 | 1617 | Atl1 | SPRET | intron_variant,regulatory_region_variant |
| 12 | 70112476 | 70116093 | 3617 | Nin | SPRET | intron_variant,regulatory_region_variant |
| 12 | 76109065 | 76110729 | 1664 | Syne2 | CAST | intron_variant |
| 12 | 77363806 | 77364956 | 1150 | Fut8 | SPRET | intron_variant |
| 12 | 82609624 | 82612185 | 2561 | - | PJ,SEA | intergenic_variant |
| 12 | 86697875 | 86702843 | 4968 | Gm22004,Gm8504,Gm | MOLF | non_coding_transcript_variant,non_coding_transcript_exon_variant,regulatory_region_variant |
| 12 | 91084153 | 91088936 | 4783 | Cep128 | CAST,MOLF,SPRET | intron_variant,regulatory_region_variant |
| 12 | 92590991 | 92592374 | 1383 | - | AJ,BUB,C3H,C58J,DBA1J,DBA2J,ILNJ2,KK,NZO,PJ,PL,SEA | intergenic_variant |
| 12 | 92692014 | 92693857 | 1843 | - | CAST,MOLF,SPRET | intergenic_variant |
| 12 | 93342922 | 93344715 | 1793 | - | PJ,SEA | intergenic_variant |
| 12 | 93343893 | 93346627 | 2734 | - | PJ,SEA | intergenic_variant |
| 12 | 95293494 | 95295243 | 1749 | - | CAST,MOLF | intergenic_variant |
| 12 | 101432473 | 101434244 | 1771 | Catsperb | SPRET | intron_variant |
| 12 | 101458016 | 101460339 | 2323 | Catsperb | C58J,SMJ | intron_variant |
| 12 | 104385149 | 104386897 | 1748 | Gm24334 | CAST | non_coding_transcript_variant,regulatory_region_variant |
| 12 | 107284140 | 107287490 | 3350 | - | BALB,BUB,CE,MA/My,NZW,PJ,SEA,SJL | intergenic_variant |
| 12 | 107286406 | 107288893 | 2487 | - | BALB,BUB,CE,MA/My,NZW,PJ,SEA,SJL | intergenic_variant |
| 12 | 109087134 | 109088969 | 1835 | - | KK,MOLF,NOD,NOR,SPRET | intergenic_variant |
| 12 | 110152713 | 110155772 | 3059 | Gm34785 | DBA2J | non_coding_transcript_exon_variant,intron_variant,regulatory_region_variant |
| 12 | 111786743 | 111787937 | 1194 | Zfyve21 | MOLF | intron_variant |
| 12 | 114561058 | 114565733 | 4675 | Ighv1-10 | 129S1,BALB,BUB,C58J,KK,LP,MA/My,PJ,SEA,SMJ,TH | non_coding_transcript_variant |
| 13 | 3663633 | 3668909 | 5276 | - | C57L,KK,LP,MOLF,NZB,NZO,NZW | intergenic_variant |
| 13 | 10351676 | 10358126 | 6450 | Chrm3 | CAST | intron_variant |
| 13 | 10860650 | 10872352 | 11702 | Gm47408 | SPRET | intron_variant,non_coding_transcript_variant |
| 13 | 11261735 | 11262804 | 1069 | - | SPRET | regulatory_region_variant,intergenic_variant |
| 13 | 15605812 | 15613081 | 7269 | - | NZO,NZW | intergenic_variant |
| 13 | 18138971 | 18139995 | 1024 | - | SPRET | intergenic_variant |
| 13 | 18721548 | 18722819 | 1271 | Pou6f2 | AKR,ILNJ2,LP,PJ,RF,SEA | intron_variant |
| 13 | 19865276 | 19868734 | 3458 | Nme8 | SPRET | intron_variant,regulatory_region_variant |
| 13 | 22251586 | 22266622 | 15036 | Vmn1r-ps93 | 129S1,AJ,B10D2,B10J,BALB,BTBR,BUB,C3H,C57L,C58J,CAST,CBA,DBA1J,DBA2J,FVB,ILNJ2,KK,LP,MA/My,MOLF,MRL,NOD,NOR,NZB,PJ,PL,RH2,SEA,SJL,SMJ,SPRET,SWR,TH | non_coding_transcript_variant |
| 13 | 22253937 | 22264273 | 10336 | Vmn1r-ps93 | 129S1,AJ,B10D2,B10J,BALB,BTBR,BUB,C3H,C57L,C58J,CAST,CBA,DBA1J,DBA2J,FVB,ILNJ2,KK,LP,MA/My,MOLF,MRL,NOD,NOR,NZB,PJ,PL,RH2,SEA,SJL,SMJ,SPRET,SWR,TH | non_coding_transcript_variant |
| 13 | 23187383 | 23189084 | 1701 | - | BTBR,C57L,KK,LP,MOLF,NOD,NOR,NZB,NZO,NZW,PJ,SEA | intergenic_variant |
| 13 | 33392550 | 33398754 | 6204 | - | SPRET | intergenic_variant |
| 13 | 42022997 | 42024022 | 1025 | Gm10790 | SPRET | intron_variant,non_coding_transcript_variant |
| 13 | 45493929 | 45495415 | 1486 | - | SPRET | intergenic_variant |

| | | | | | | |
|---|---|---|---|---|---|---|
| 13 | 53593987 | 53596225 | 2238 | Gm2762 | 129S1,AJ,AKR,BALB,BTBR,BUB,C3H,C57L,CAST,CBA,CE,DBA1J,DBA2J,FVB,ILNJ2,KK,MA/My,MOLF,MRL,NOD,NOR,NZB,NZO,NZW,PJ,PL,RF,RH2,SEA,SJL,SMJ,SPRET,SWR,TH,WSB | intron_variant,non_coding_transcript_variant,non_coding_transcript_exon_variant,regulatory_region_variant |
| 13 | 58062382 | 58063401 | 1019 | - | BUB | intergenic_variant |
| 13 | 62298840 | 62301375 | 2535 | Zfp808,Gm36501 | CAST,MOLF | intron_variant,non_coding_transcript_variant,regulatory_region_variant |
| 13 | 62477404 | 62488342 | 10938 | Gm19792 | CAST,MOLF | non_coding_transcript_exon_variant,intron_variant,regulatory_region_variant |
| 13 | 62833917 | 62834954 | 1037 | Platr25 | BUB | intron_variant |
| 13 | 62839048 | 62844413 | 5365 | Platr25 | SPRET | intron_variant |
| 13 | 64872089 | 64874118 | 2029 | - | SPRET | intergenic_variant |
| 13 | 77290511 | 77292966 | 2455 | 2210408I21Rik | SPRET | intron_variant |
| 13 | 78419449 | 78421080 | 1631 | A830082K12Rik,Gm50 | BUB,C3H,CBA,CE,DBA1J,DBA2J,NOD,NOR,TH,WSB | intron_variant,non_coding_transcript_variant,downstream_gene_variant |
| 13 | 78633859 | 78641801 | 7942 | Gm32067 | CAST | intron_variant,non_coding_transcript_variant,regulatory_region_variant |
| 13 | 79481834 | 79483265 | 1431 | - | SWR | intergenic_variant |
| 13 | 84534090 | 84536436 | 2346 | - | CAST,MOLF,SWR | intergenic_variant |
| 13 | 85874778 | 85881089 | 6311 | - | SPRET | intergenic_variant |
| 13 | 86375129 | 86377087 | 1958 | - | 129S1,AKR,BALB,BTBR,BUB,C3H,C57L,C58J,CBA,CE,DBA1J,DBA2J,FVB,ILNJ2,KK,MA/My,MRL,NOD,NOR,NZB,NZO,NZW,PJ,PL,RF,SEA,SJL,SMJ,TH | intergenic_variant |
| 13 | 86426954 | 86428889 | 1935 | - | CAST,MOLF | intergenic_variant |
| 13 | 86428557 | 86430076 | 1519 | - | SPRET | intergenic_variant |
| 13 | 86614870 | 86617474 | 2604 | - | CAST | intergenic_variant |
| 13 | 88635716 | 88637535 | 1819 | - | SPRET | intergenic_variant |
| 13 | 89495911 | 89497325 | 1414 | - | SPRET | intergenic_variant |
| 13 | 89593365 | 89594448 | 1083 | - | BALB,DBA1J,FVB,MRL,SEA | intergenic_variant |
| 13 | 89594142 | 89596062 | 1920 | - | BALB,DBA1J,FVB,MRL,SEA | intergenic_variant |
| 13 | 90752181 | 90758904 | 6723 | - | SPRET | regulatory_region_variant,intergenic_variant |
| 13 | 92595229 | 92596430 | 1201 | Fam151b | SPRET | intron_variant |
| 13 | 97627480 | 97632924 | 5444 | Lncenc1 | AKR | non_coding_transcript_exon_variant,intron_variant,non_coding_transcript_exon_variant,regulatory_region_variant |
| 13 | 99092092 | 99093300 | 1208 | - | CAST,SPRET | regulatory_region_variant,intergenic_variant |
| 13 | 100643186 | 100646953 | 3767 | Ocln | MOLF | intron_variant |
| 13 | 103804798 | 103805979 | 1181 | Gm30551 | CAST,MOLF | intron_variant,non_coding_transcript_variant |
| 13 | 105853367 | 105854725 | 1358 | - | SPRET | intergenic_variant |
| 13 | 106263466 | 106266466 | 3000 | - | SPRET | intergenic_variant |
| 13 | 106263468 | 106265286 | 1818 | - | SPRET | intergenic_variant |
| 13 | 106815968 | 106822590 | 6622 | - | CAST,CE,MA/My,MOLF | intergenic_variant |
| 13 | 106821684 | 106823837 | 2153 | - | CAST,CE,MA/My,MOLF | intergenic_variant |
| 13 | 108324811 | 108329652 | 4841 | Ercc8 | 129S1,BTBR,C57L,KK,NZB,NZW,PL | intron_variant,regulatory_region_variant |
| 13 | 110362227 | 110363456 | 1229 | Rab3c | CAST,CE,MA/My,MOLF,NOD,NOR,NZO | intron_variant,regulatory_region_variant |
| 13 | 110362227 | 110363456 | 1229 | Rab3c | CAST,CE,MA/My,MOLF,NOD,NOR,NZO | intron_variant,regulatory_region_variant |
| 13 | 112999776 | 113001963 | 2187 | Plpp1 | AJ,DBA1J,FVB,KK,MA/My,MOLF,NZO,PL,RF,RH2,SJL,SWR | intron_variant |
| 13 | 116227229 | 116234245 | 7016 | Gm47893 | SPRET | non_coding_transcript_variant,regulatory_region_variant |
| 13 | 116973729 | 116979941 | 6212 | 4930435F18Rik | 129S1,BTBR,BUB,C3H,C57L,CE,DBA2J,KK,NZB,NZO,NZW,WSB | non_coding_transcript_exon_variant,intron_variant,regulatory_region_variant |
| 13 | 118396227 | 118399291 | 3064 | - | SPRET | intergenic_variant |
| 13 | 118799481 | 118801079 | 1598 | - | SPRET | intergenic_variant |
| 14 | 9333848 | 9341155 | 7307 | - | PL | intergenic_variant |
| 14 | 17644594 | 17646790 | 2196 | - | 129S1,C57L,KK,MOLF,PL | intergenic_variant |
| 14 | 18170523 | 18172609 | 2086 | - | C57L,KK,MOLF | intergenic_variant |
| 14 | 21630753 | 21633392 | 2639 | Kat6b | MOLF | intron_variant,regulatory_region_variant |
| 14 | 26054547 | 26057118 | 2571 | - | CAST,MOLF | intergenic_variant |
| 14 | 28403641 | 28406494 | 2853 | Gm35164 | SPRET | intron_variant,non_coding_transcript_variant |
| 14 | 28405400 | 28407467 | 2067 | Gm35164 | SPRET | intron_variant,non_coding_transcript_variant |
| 14 | 31844835 | 31846069 | 1234 | - | SPRET | intergenic_variant |
| 14 | 35811315 | 35818656 | 7341 | Gm7853,4930474N05R | 129S1,C57L,NZW | non_coding_transcript_variant,coding_sequence_variant,5_prime_UTR_variant,intron_variant |
| 14 | 38107794 | 38109784 | 1990 | Nrg3 | SPRET | intron_variant |
| 14 | 40448555 | 40453608 | 5053 | - | CAST | intergenic_variant |
| 14 | 40479404 | 40485848 | 6444 | - | SPRET | intergenic_variant |
| 14 | 40483243 | 40486806 | 3563 | - | SPRET | intergenic_variant |
| 14 | 42003455 | 42005845 | 2390 | - | SMJ | intergenic_variant |
| 14 | 45079707 | 45081957 | 2250 | - | C57L,CAST,MOLF,NZO,PJ,PL | intergenic_variant |
| 14 | 49447816 | 49449159 | 1343 | - | SPRET | intergenic_variant |
| 14 | 50175914 | 50183266 | 7352 | - | SPRET | intergenic_variant |
| 14 | 51665970 | 51668613 | 2643 | Gm7247,Vmn2r-ps111 | 129S1,AJ,AKR,BALB,BUB,C3H,C57L,CBA,CE,KK,MRL,NZB,NZO,NZW,PJ,PL,RF,RH2,SEA,TH | non_coding_transcript_exon_variant,intron_variant,intron_variant |
| 14 | 51667721 | 51669191 | 1470 | Gm7247,Vmn2r-ps111 | 129S1,AJ,AKR,BALB,BUB,C3H,C57L,CBA,CE,KK,MRL,NZB,NZO,NZW,PJ,PL,RF,RH2,SEA,TH | non_coding_transcript_exon_variant,intron_variant,intron_variant |
| 14 | 52205724 | 52210931 | 5207 | - | 129S1,AJ,AKR,BALB,BUB,C3H,C57L,CBA,CE,DBA2J,KK,MRL,NZB,NZO,NZW,PJ,PL,RF,RH2,SEA,TH | intergenic_variant |
| 14 | 58895915 | 58897890 | 1975 | - | SPRET | intergenic_variant |
| 14 | 58896977 | 58898109 | 1132 | - | SPRET | intergenic_variant |
| 14 | 62675471 | 62680324 | 4853 | Gucy1b2 | SPRET | intron_variant,regulatory_region_variant |
| 14 | 62677509 | 62681098 | 3589 | Gucy1b2 | SPRET | intron_variant,regulatory_region_variant |
| 14 | 67173441 | 67177437 | 3996 | 4930578I07Rik | SPRET | non_coding_transcript_variant |
| 14 | 74501862 | 74503388 | 1526 | Gm49171 | BUB,CAST,KK,MOLF,NZO,SPRET,TH | intron_variant,non_coding_transcript_variant |
| 14 | 82974971 | 82976261 | 1290 | - | SPRET | intergenic_variant |
| 14 | 84337208 | 84342548 | 5340 | - | SPRET | intergenic_variant |
| 14 | 84453217 | 84460466 | 7249 | - | BUB,C57L,C58J,CAST,SPRET | intergenic_variant |
| 14 | 84454423 | 84459278 | 4855 | - | 129S1,BUB,C57L,C58J,CAST,ILNJ2,KK,NOD,NOR,NZW,SJL,SPRET | intergenic_variant |
| 14 | 89555123 | 89560821 | 5698 | - | SPRET | intergenic_variant |

| | | | | | | |
|---|---|---|---|---|---|---|
| 14 | 90577037 | 90579048 | 2011 | - | AJ,AKR,BALB,BUB,C3H,CBA,CE,DBA1J,DBA2J,FVB,NOD,NZO,PJ,RF,SEA,SJL,SMJ,SPRET,SWR,TH,WSB | intergenic_variant |
| 14 | 90577058 | 90584614 | 7556 | - | NOD,NZO,SPRET,WSB | intergenic_variant |
| 14 | 94269136 | 94270878 | 1742 | - | CAST | intergenic_variant |
| 14 | 95367152 | 95369883 | 2731 | - | CAST | intergenic_variant |
| 14 | 96646843 | 96649069 | 2226 | Klhl1 | AJ,AKR,BALB,BTBR,BUB,C3H,CBA,CE,DBA1J,DBA2J,FVB,MRL,PJ,PL,RF,SEA,SJL,SMJ,SWR | intron_variant |
| 14 | 98679331 | 98680684 | 1353 | - | SPRET | intergenic_variant |
| 14 | 98679573 | 98682006 | 2433 | - | SPRET | intergenic_variant |
| 14 | 101639151 | 101640295 | 1144 | - | C57L,C58J,ILNJ2,MOLF,NZB,NZO,PJ | intergenic_variant |
| 14 | 102423410 | 102424620 | 1210 | - | SPRET | intergenic_variant |
| 14 | 102658976 | 102662598 | 3622 | - | CAST | intergenic_variant |
| 14 | 102659828 | 102663237 | 3409 | - | CAST | intergenic_variant |
| 14 | 103106690 | 103109970 | 3280 | - | CAST | intergenic_variant |
| 14 | 103108787 | 103110552 | 1765 | - | CAST | intergenic_variant |
| 14 | 103676076 | 103683678 | 7602 | 1700087I21Rik | CAST,MOLF | non_coding_transcript_exon_variant,intron_variant |
| 14 | 103682334 | 103684065 | 1731 | 1700087I21Rik | CAST | intron_variant,non_coding_transcript_variant |
| 14 | 105509650 | 105510662 | 1012 | Ndfip2 | SPRET | intron_variant,regulatory_region_variant |
| 14 | 105981480 | 105984071 | 2591 | - | 129S1,BTBR,BUB,C57L,C58J,CAST,FVB,ILNJ2,LP,MA/My,MOLF,NOD,NOR,NZW,SPRET,SWR,WSB | regulatory_region_variant,intergenic_variant |
| 14 | 106056533 | 106058181 | 1648 | - | MOLF | intergenic_variant |
| 14 | 108381876 | 108386935 | 5059 | - | SPRET | intergenic_variant |
| 14 | 110280758 | 110287865 | 7107 | - | SPRET | intergenic_variant |
| 14 | 110639907 | 110643506 | 3599 | - | SPRET | intergenic_variant |
| 14 | 111848070 | 111850449 | 2379 | Gm30439 | 129S1,AJ,AKR,BUB,C57L,C58J,CAST,FVB,ILNJ2,LP,MA/My,MOLF,NOD,NOR,NZB,NZO,NZW,PL,RF,SJL,SMJ,SPRET,SWR,TH,WSB | intron_variant,non_coding_transcript_variant |
| 14 | 113228280 | 113229512 | 1232 | - | SPRET | intergenic_variant |
| 14 | 113823583 | 113824650 | 1067 | - | MOLF | intergenic_variant |
| 14 | 114657746 | 114661907 | 4161 | - | SWR | intergenic_variant |
| 14 | 117276158 | 117278333 | 2175 | Gpc6 | MOLF | intron_variant |
| 14 | 118313260 | 118316875 | 3615 | - | SPRET | intergenic_variant |
| 14 | 119329349 | 119331250 | 1901 | Uggt2 | 129S1,KK,MOLF | intron_variant |
| 14 | 120170590 | 120177567 | 6977 | - | SPRET | intergenic_variant |
| 14 | 120846339 | 120848252 | 1913 | - | SPRET | intergenic_variant |
| 14 | 124167458 | 124170392 | 2934 | Itgbl1,Gm25272 | SPRET | intron_variant,upstream_gene_variant |
| 15 | 6011820 | 6014964 | 3144 | - | CAST | intergenic_variant |
| 15 | 6011848 | 6013724 | 1876 | - | CAST | intergenic_variant |
| 15 | 10853388 | 10857775 | 4387 | - | CAST | regulatory_region_variant,intergenic_variant |
| 15 | 18318454 | 18320611 | 2157 | 4921515E04Rik | WSB | intron_variant,non_coding_transcript_variant |
| 15 | 19676713 | 19677801 | 1088 | Gm49166 | AJ,BALB,BUB,C3H,CBA,CE,DBA1J,DBA2J,FVB,ILNJ2,KK,LP,MA/My,MRL,NZB,NZO,NZW,PL,RH2,SJL,SWR,TH,WSB | upstream_gene_variant |
| 15 | 20042932 | 20050910 | 7978 | - | 129S1,BTBR,C57L | intergenic_variant |
| 15 | 20867589 | 20869323 | 1734 | Gm49685 | 129S1,BALB,BTBR,BUB,C3H,C57L,DBA2J,KK,LP,MA/My,NZB,NZO,NZW,PJ,SEA,SJL,SMJ | intron_variant,non_coding_transcript_variant |
| 15 | 20868128 | 20871016 | 2888 | Gm49685 | 129S1,BALB,BTBR,BUB,C3H,C57L,DBA2J,KK,LP,MA/My,NZB,NZO,NZW,PJ,SEA,SJL,SMJ | intron_variant,non_coding_transcript_variant |
| 15 | 21067678 | 21069276 | 1598 | - | CE,ILNJ2,MOLF,MRL,SPRET,WSB | intergenic_variant |
| 15 | 21982193 | 21983566 | 1373 | - | WSB | intergenic_variant |
| 15 | 22930339 | 22931728 | 1389 | - | SPRET | intergenic_variant |
| 15 | 27384221 | 27390282 | 6061 | - | MRL | intergenic_variant |
| 15 | 27834888 | 27836571 | 1683 | Trio | CAST | intron_variant |
| 15 | 32148049 | 32149360 | 1311 | - | SPRET | intergenic_variant |
| 15 | 32833543 | 32839857 | 6314 | - | C3H,CBA,CE,DBA1J,WSB | intergenic_variant |
| 15 | 32835802 | 32841020 | 5218 | - | C3H,CBA,DBA1J,DBA2J | intergenic_variant |
| 15 | 33399073 | 33400363 | 1290 | Cpq | SPRET | intron_variant |
| 15 | 34824153 | 34825407 | 1254 | Gm48932 | CAST | non_coding_transcript_exon_variant,regulatory_region_variant |
| 15 | 42844205 | 42850992 | 6787 | - | MRL | regulatory_region_variant,intergenic_variant |
| 15 | 42847421 | 42850486 | 3065 | - | MRL | regulatory_region_variant,intergenic_variant |
| 15 | 45866017 | 45878044 | 12027 | - | CAST | intergenic_variant |
| 15 | 50004415 | 50005817 | 1402 | - | SPRET | intergenic_variant |
| 15 | 52516993 | 52519040 | 2047 | - | KK | intergenic_variant |
| 15 | 57184319 | 57187745 | 3426 | Slc22a22 | CAST,MOLF | intron_variant |
| 15 | 60907784 | 60909490 | 1706 | - | SPRET | intergenic_variant |
| 15 | 65260754 | 65262659 | 1905 | - | SPRET | intergenic_variant |
| 15 | 68416042 | 68417243 | 1201 | - | CAST | intergenic_variant |
| 15 | 71097098 | 71100937 | 3839 | - | SPRET | intergenic_variant |
| 15 | 74839049 | 74841544 | 2495 | - | CAST,SPRET | regulatory_region_variant,intergenic_variant |
| 15 | 86603902 | 86604994 | 1092 | - | SPRET | intergenic_variant |
| 15 | 91931041 | 91933014 | 1973 | - | MOLF | intergenic_variant |
| 15 | 95995388 | 95997743 | 2355 | - | SPRET | intergenic_variant |
| 15 | 98653195 | 98655357 | 2162 | Arf3 | MOLF | intron_variant,regulatory_region_variant |
| 15 | 99795448 | 99797584 | 2136 | Uqcrh-ps1 | CAST | non_coding_transcript_variant,regulatory_region_variant |
| 15 | 103598024 | 103599186 | 1162 | - | SPRET | intergenic_variant |
| 16 | 10192645 | 10193841 | 1196 | Tekt5 | MOLF | intron_variant |
| 16 | 13773476 | 13774677 | 1201 | - | SPRET | intergenic_variant |
| 16 | 23096050 | 23097394 | 1344 | St6gal1 | CAST,MOLF,SPRET,TH | intron_variant |
| 16 | 26120097 | 26123369 | 3272 | - | CAST,MOLF | intergenic_variant |
| 16 | 26121321 | 26123531 | 2210 | - | CAST,MOLF | intergenic_variant |
| 16 | 26627223 | 26634260 | 7037 | - | ILNJ2,NZB,NZO,NZW | intergenic_variant |
| 16 | 27951988 | 27953632 | 1644 | - | CE,KK | intergenic_variant |
| 16 | 38090836 | 38092966 | 2130 | Nr1i2 | SPRET | intron_variant,regulatory_region_variant |
| 16 | 40570447 | 40571678 | 1231 | Lsamp | 129S1,BTBR,BUB,C3H,C57L,CAST,CBA,CE,ILNJ2,KK,LP,MOLF,MRL,NZB,NZO,NZW,PL,SMJ,SPRET,TH,WSB | intron_variant |
| 16 | 42252370 | 42255651 | 3281 | - | MOLF | regulatory_region_variant,intergenic_variant |
| 16 | 42830689 | 42832318 | 1629 | Zbtb20 | SPRET | intron_variant |

| | | | | | | |
|---|---|---|---|---|---|---|
| 16 | 47114985 | 47125373 | 10388 | - | 129S1,C3H,C58J,CAST,CBA,CE,MOLF,SJL,SPRET,TH | intergenic_variant |
| 16 | 47121493 | 47125328 | 3835 | - | AKR,FVB,LP,MRL,PJ,RF,SWR,WSB | intergenic_variant |
| 16 | 53407810 | 53413821 | 6011 | - | C3H,C58J,CAST,CBA,FVB,KK,LP,MOLF,MRL,NZW,SPRET,SWR,TH,WSB | intergenic_variant |
| 16 | 53469811 | 53471104 | 1293 | - | SMJ | intergenic_variant |
| 16 | 54484487 | 54488519 | 4032 | - | CBA,MOLF,NZW | intergenic_variant |
| 16 | 54484491 | 54486043 | 1552 | - | CBA,MOLF,NZW | intergenic_variant |
| 16 | 54584534 | 54585611 | 1077 | - | CAST,CBA,MOLF,NZW,SPRET | intergenic_variant |
| 16 | 55412097 | 55413459 | 1362 | - | SPRET | intergenic_variant |
| 16 | 56022950 | 56024238 | 1288 | Impg2 | CAST | upstream_gene_variant |
| 16 | 57406443 | 57408350 | 1907 | Cmss1 | SPRET | intron_variant |
| 16 | 57406473 | 57413682 | 7209 | Cmss1 | SPRET | intron_variant,regulatory_region_variant |
| 16 | 58584286 | 58586359 | 2073 | Or5k1b | SPRET | upstream_gene_variant,regulatory_region_variant |
| 16 | 59582043 | 59589637 | 7594 | Epha6 | PL | intron_variant |
| 16 | 60592823 | 60599990 | 7167 | - | CAST | intergenic_variant |
| 16 | 61554281 | 61556029 | 1748 | - | CAST,MOLF,SPRET | intergenic_variant |
| 16 | 61679299 | 61680714 | 1415 | - | SPRET | intergenic_variant |
| 16 | 63505768 | 63508087 | 2319 | Epha3 | SPRET | intron_variant,regulatory_region_variant |
| 16 | 67730954 | 67737533 | 6579 | - | AJ,C3H,C58J,CBA,DBA2J,ILNJ2,KK,LP,MA/My,MRL,NZB,NZO,PL,RH2,SMJ | intergenic_variant |
| 16 | 67735556 | 67738018 | 2462 | - | AJ,BALB,C3H,C58J,CBA,DBA1J,DBA2J,ILNJ2,KK,LP,MA/My,MRL,NZB,NZO,NZW,PL,RH2,SMJ | intergenic_variant |
| 16 | 70737736 | 70745634 | 7898 | - | SPRET | intergenic_variant |
| 16 | 71168699 | 71172192 | 3493 | - | SPRET | intergenic_variant |
| 16 | 73527086 | 73533303 | 6217 | - | CAST | intergenic_variant |
| 16 | 73527117 | 73533977 | 6860 | - | CAST,MOLF | intergenic_variant |
| 16 | 73952188 | 73953202 | 1014 | Robo2 | KK | intron_variant |
| 16 | 74537603 | 74540355 | 2752 | Robo2 | SPRET | intron_variant |
| 16 | 74537829 | 74539294 | 1465 | Robo2 | SPRET | intron_variant |
| 16 | 76585789 | 76591934 | 6145 | - | C58J,NZB,NZO,NZW | intergenic_variant |
| 16 | 76588073 | 76592246 | 4173 | - | C58J,NZB,NZO,NZW | intergenic_variant |
| 16 | 78419888 | 78422052 | 2164 | Gm24861 | MOLF | non_coding_transcript_variant |
| 16 | 82623561 | 82629876 | 6315 | - | SPRET | intergenic_variant |
| 16 | 82627105 | 82629880 | 2775 | - | SPRET | intergenic_variant |
| 16 | 86127998 | 86134782 | 6784 | - | SPRET | intergenic_variant |
| 16 | 86528571 | 86532340 | 3769 | - | SPRET | intergenic_variant |
| 16 | 86598553 | 86601021 | 2468 | Gm32461 | SPRET | non_coding_transcript_exon_variant,intron_variant |
| 16 | 87340509 | 87341810 | 1301 | - | 129S1,C57L,ILNJ2 | intergenic_variant |
| 16 | 87341176 | 87344200 | 3024 | - | 129S1,C57L,ILNJ2 | intergenic_variant |
| 16 | 93453383 | 93454486 | 1103 | Gm52975 | SPRET | non_coding_transcript_variant |
| 16 | 95546143 | 95547187 | 1044 | - | CAST | regulatory_region_variant,intergenic_variant |
| 16 | 97147593 | 97153791 | 6198 | - | MOLF,NZO | intergenic_variant |
| 17 | 4798115 | 4799504 | 1389 | - | CAST | intergenic_variant |
| 17 | 10106363 | 10109997 | 3634 | - | SPRET | regulatory_region_variant,intergenic_variant |
| 17 | 11998543 | 11999940 | 1397 | Prkn | WSB | intron_variant,regulatory_region_variant |
| 17 | 15702459 | 15704792 | 2333 | Psmb1 | BTBR,BUB,C3H,CAST,CBA,MOLF,MRL,NOD,NOR,PJ,PL,RF,SJL,SMJ,SWR,WSB | intron_variant |
| 17 | 18598238 | 18600463 | 2225 | - | AKR,FVB,ILNJ2,KK,MA/My,WSB | intergenic_variant |
| 17 | 19818121 | 19819600 | 1479 | Vmn2r101 | B10D2,B10J,BTBR,BUB,C3H,C58J,CBA,LP,MRL,NOD,NOR,NZB,NZW,PJ,PL,RF,SJL,SWR,TH | intron_variant |
| 17 | 19845665 | 19847494 | 1829 | - | CAST | intergenic_variant |
| 17 | 19999841 | 20002539 | 2698 | Vmn2r103 | SPRET | intron_variant |
| 17 | 21294553 | 21296376 | 1823 | - | SPRET | intergenic_variant |
| 17 | 21295621 | 21299750 | 4129 | Vmn1r-ps146 | SPRET | downstream_gene_variant |
| 17 | 21826144 | 21828034 | 1890 | - | CAST,MOLF | intergenic_variant |
| 17 | 21826808 | 21831313 | 4505 | - | CAST,MOLF | intergenic_variant |
| 17 | 23642650 | 23650078 | 7428 | - | SPRET | intergenic_variant |
| 17 | 23753073 | 23754242 | 1169 | - | PJ,PL,SPRET | intergenic_variant |
| 17 | 24099297 | 24104174 | 4877 | Dcpp1,Dcpp2 | SPRET | coding_transcript_variant,5_prime_UTR_variant,intron_variant |
| 17 | 24166988 | 24169746 | 2758 | - | WSB | intergenic_variant |
| 17 | 28087897 | 28090266 | 2369 | Bltp3a,Gm49803 | KK,NZB | intron_variant,non_coding_transcript_exon_variant,intron_variant |
| 17 | 29323126 | 29324246 | 1120 | - | B10J,BUB,C58J,FVB,LP,MA/My,NZO,NZW,PL,SJL,SMJ,TH,WSB | intergenic_variant |
| 17 | 29529828 | 29531278 | 1450 | BC004004 | C58J,CAST,KK,LP,MOLF,NZO,NZW,RF,SMJ,TH,WSB | 3_prime_UTR_variant |
| 17 | 29664735 | 29665736 | 1001 | Gm36199 | MOLF,RF,SPRET | intron_variant,non_coding_transcript_variant |
| 17 | 31004249 | 31005689 | 1440 | Dnah8 | SPRET | intron_variant,regulatory_region_variant |
| 17 | 36792193 | 36796868 | 4675 | - | MOLF,PJ,SPRET | regulatory_region_variant,intergenic_variant |
| 17 | 36876837 | 36883348 | 6511 | - | SPRET | intergenic_variant |
| 17 | 37018173 | 37025094 | 6921 | Gm20523 | AKR,B10D2,BALB,C3H,CBA,CE,DBA1J,DBA2J,MA/My,MRL,NZO,NZW,RF,RH2,SEA,TH | upstream_gene_variant,regulatory_region_variant |
| 17 | 37093114 | 37097088 | 3974 | - | SPRET | regulatory_region_variant,intergenic_variant |
| 17 | 38804383 | 38805475 | 1092 | - | DBA1J,MOLF,NZW,RH2,SEA | intergenic_variant |
| 17 | 38805027 | 38806951 | 1924 | - | AKR,DBA1J,NZW,RH2,SEA | intergenic_variant |
| 17 | 38898261 | 38901886 | 3625 | Gm49813 | SPRET | upstream_gene_variant |
| 17 | 39225832 | 39226955 | 1123 | - | WSB | intergenic_variant |
| 17 | 40197297 | 40198709 | 1412 | - | AKR,C3H,CBA,CE,MA/My,MRL,RF | intergenic_variant |
| 17 | 41998169 | 42004373 | 6204 | - | 129S1,AJ,AKR,BALB,BTBR,BUB,C3H,C57L,C58J,CAST,CBA,CE,DBA1J,DBA2J,FVB,ILNJ2,KK,LP,MA/My,MRL,NOD,NOR,NZB,NZO,NZW,PJ,PL,RF,RH2,SEA,SJL,SWR,TH,WSB | intergenic_variant |
| 17 | 51727670 | 51729230 | 1560 | - | SPRET | intergenic_variant |
| 17 | 52681840 | 52683382 | 1542 | - | AJ,AKR,BALB,C3H,CBA,DBA1J,DBA2J,ILNJ2,KK,LP,MA/My,MRL,NOD,NOR,NZB,PJ,PL,RF,RH2,SEA,SMJ,SWR | intergenic_variant |
| 17 | 53581050 | 53587577 | 6527 | - | MOLF | intergenic_variant |
| 17 | 54013822 | 54015422 | 1600 | - | SPRET | intergenic_variant |
| 17 | 54350920 | 54357832 | 6912 | - | CAST,MOLF | intergenic_variant |

| | | | | | | |
|---|---|---|---|---|---|---|
| 17 | 55323217 | 55324218 | 1001 | - | SPRET | intergenic_variant |
| 17 | 56572767 | 56574139 | 1372 | - | CAST,MOLF | intergenic_variant |
| 17 | 59103433 | 59106702 | 3269 | A330072L02Rik | CAST,SPRET | intron_variant,non_coding_transcript_variant |
| 17 | 60250033 | 60252694 | 2661 | Gm49853 | CAST | intron_variant,non_coding_transcript_variant |
| 17 | 66295581 | 66297336 | 1755 | Ankrd12 | SPRET | intron_variant,regulatory_region_variant |
| 17 | 69005665 | 69008344 | 2679 | L3mbtl4,Gm17921 | SPRET | intron_variant,non_coding_transcript_variant |
| 17 | 70009556 | 70011448 | 1892 | - | AKR,NOR,PL,SMJ | intergenic_variant |
| 17 | 70009557 | 70016341 | 6784 | - | AKR,NOD,NOR,PL,SMJ | intergenic_variant |
| 17 | 70527821 | 70529143 | 1322 | Dlgap1 | MOLF | intron_variant |
| 17 | 76231229 | 76232264 | 1035 | - | AJ,AKR,BALB,C3H,CBA,DBA1J,DBA2J,PJ,RF,SEA,SMJ,WSB | intergenic_variant |
| 17 | 76386364 | 76388670 | 2306 | - | SPRET | intergenic_variant |
| 17 | 76387971 | 76388995 | 1024 | - | SPRET | intergenic_variant |
| 17 | 82157691 | 82159057 | 1366 | - | MA/My,NZW,PL,WSB | intergenic_variant |
| 17 | 82956817 | 82959086 | 2269 | - | BUB,CE,WSB | intergenic_variant |
| 17 | 89302197 | 89307192 | 4995 | Fshr | KK,NOD,NOR,PJ,RF,SMJ | intron_variant |
| 17 | 89744065 | 89745391 | 1326 | - | SPRET | intergenic_variant |
| 17 | 93437577 | 93439727 | 2150 | - | SPRET | intergenic_variant |
| 17 | 93700276 | 93702060 | 1784 | - | SPRET | intergenic_variant |
| 17 | 94155912 | 94157993 | 2081 | - | SPRET | intergenic_variant |
| 18 | 3972158 | 3973304 | 1146 | - | SPRET | intergenic_variant |
| 18 | 6695113 | 6697044 | 1931 | - | CAST,MOLF | intergenic_variant |
| 18 | 8340170 | 8346627 | 6457 | - | SPRET | intergenic_variant |
| 18 | 8341245 | 8346636 | 5391 | - | SPRET | intergenic_variant |
| 18 | 8391036 | 8396396 | 5360 | - | SPRET | intergenic_variant |
| 18 | 9150192 | 9154971 | 4779 | - | SPRET | intergenic_variant |
| 18 | 10384409 | 10386134 | 1725 | Greb1l | SPRET | intron_variant |
| 18 | 10492134 | 10494362 | 2228 | Greb1l | SPRET | intron_variant |
| 18 | 13341437 | 13343853 | 2416 | Gm22251 | 129S1,AJ,AKR,BALB,BTBR,BUB,C3H,C57L,CBA,FVB,ILNJ2,KK,RF,RH2,SJL,SMJ,SWR,TH,WSB | non_coding_transcript_variant |
| 18 | 13342076 | 13347300 | 5224 | Gm22251 | 129S1,AJ,AKR,BALB,BTBR,BUB,C3H,C57L,CBA,FVB,ILNJ2,KK,RF,RH2,SJL,SMJ,SWR,TH,WSB | non_coding_transcript_variant |
| 18 | 14257141 | 14258505 | 1364 | - | MOLF | intergenic_variant |
| 18 | 14540792 | 14541923 | 1131 | - | SPRET | intergenic_variant |
| 18 | 16182537 | 16183742 | 1205 | Gm41673 | SPRET | intron_variant,non_coding_transcript_variant |
| 18 | 16270502 | 16271557 | 1055 | Gm50076 | CAST,WSB | intron_variant,non_coding_transcript_variant |
| 18 | 23151259 | 23152391 | 1132 | Nol4 | CAST | intron_variant |
| 18 | 29064738 | 29067672 | 2934 | Gm33948 | CAST,MOLF | intron_variant,non_coding_transcript_variant |
| 18 | 29201844 | 29203700 | 1856 | - | WSB | intergenic_variant |
| 18 | 29845493 | 29852354 | 6861 | - | SPRET | regulatory_region_variant,intergenic_variant |
| 18 | 30821235 | 30823730 | 2495 | - | 129S1,C57L | intergenic_variant |
| 18 | 31054182 | 31057344 | 3162 | - | MOLF | intergenic_variant |
| 18 | 31379302 | 31381576 | 2274 | Rit2 | MOLF | intron_variant |
| 18 | 31380342 | 31381694 | 1352 | Rit2 | MOLF | intron_variant |
| 18 | 31487792 | 31494667 | 6875 | - | SPRET | intergenic_variant |
| 18 | 31516392 | 31518021 | 1629 | - | SPRET | intergenic_variant |
| 18 | 31826533 | 31828237 | 1704 | Sap130 | MOLF | intron_variant |
| 18 | 35899483 | 35901724 | 2241 | - | SPRET | intergenic_variant |
| 18 | 44920930 | 44922066 | 1136 | Mcc | CAST | intron_variant |
| 18 | 51941902 | 51948507 | 6605 | - | CAST | regulatory_region_variant,intergenic_variant |
| 18 | 53356002 | 53357635 | 1633 | - | SPRET | intergenic_variant |
| 18 | 58515821 | 58518005 | 2184 | - | CAST,MOLF | regulatory_region_variant,intergenic_variant |
| 18 | 60338394 | 60341138 | 2744 | - | SPRET | regulatory_region_variant,intergenic_variant |
| 18 | 64156869 | 64158590 | 1721 | - | MOLF | intergenic_variant |
| 18 | 64182620 | 64189366 | 6746 | - | MOLF | intergenic_variant |
| 18 | 64960074 | 64961326 | 1252 | - | SPRET | intergenic_variant |
| 18 | 68507487 | 68514321 | 6834 | - | SPRET | intergenic_variant |
| 18 | 69168042 | 69172603 | 4561 | - | KK | intergenic_variant |
| 18 | 70446828 | 70451065 | 4237 | - | BALB,KK,MOLF,NZB,RF,RH2 | regulatory_region_variant,intergenic_variant |
| 18 | 73205984 | 73207556 | 1572 | - | BUB,CBA,DBA1J,DBA2J,MA/My,PJ,SEA | intergenic_variant |
| 18 | 84204694 | 84205775 | 1081 | - | 129S1,AJ,AKR,B10D2,B10J,BALB,BTBR,BUB,C3H,C57L,C58J,CAST,CBA,DBA1J,DBA2J,FVB,ILNJ2,KK,LP,MA/My,MOLF,MRL,NOR,NZW,PJ,PL,RF,RH2,SEA,SJL,SMJ,SPRET,SWR,TH | regulatory_region_variant,intergenic_variant |
| 18 | 85156849 | 85159663 | 2814 | - | 129S1,BTBR,C57L | intergenic_variant |
| 18 | 85279972 | 85281792 | 1820 | - | SPRET | intergenic_variant |
| 18 | 85811997 | 85813333 | 1336 | - | CAST,MOLF,SPRET | intergenic_variant |
| 18 | 87274519 | 87275706 | 1187 | - | SPRET | regulatory_region_variant,intergenic_variant |
| 18 | 88365904 | 88370570 | 4666 | - | SPRET | intergenic_variant |
| 18 | 90513761 | 90515808 | 2047 | - | CE | intergenic_variant |
| 19 | 6048217 | 6050005 | 1788 | Capn1 | MOLF | intron_variant,regulatory_region_variant |
| 19 | 9165708 | 9172185 | 6477 | Gm9750 | CAST | intron_variant,non_coding_transcript_variant |
| 19 | 9166232 | 9172185 | 5953 | Gm9750 | CAST | intron_variant,non_coding_transcript_variant |
| 19 | 9166951 | 9172348 | 5397 | Gm9750 | CAST | intron_variant,non_coding_transcript_variant |
| 19 | 11692787 | 11694140 | 1353 | - | CAST | intergenic_variant |
| 19 | 13997596 | 14000718 | 3122 | - | FVB,ILNJ2,LP,SJL,SWR,WSB | intergenic_variant |
| 19 | 15994182 | 15997390 | 3208 | - | MOLF | regulatory_region_variant,intergenic_variant |
| 19 | 16351102 | 16352277 | 1175 | Gnaq | MOLF,SPRET | intron_variant,regulatory_region_variant |
| 19 | 17280986 | 17287548 | 6562 | - | BALB,BTBR,C57L,CBA,DBA1J,DBA2J,ILNJ2,MOLF,SEA,WSB | intergenic_variant |
| 19 | 19269938 | 19274261 | 4323 | Gm41815 | MOLF,NZB,NZO,NZW,PJ | intron_variant,non_coding_transcript_variant |
| 19 | 25049891 | 25051835 | 1944 | Dock8 | SPRET | intron_variant,regulatory_region_variant |
| 19 | 28804903 | 28809127 | 4224 | - | CAST | intergenic_variant |
| 19 | 28807938 | 28809155 | 1217 | - | CAST | intergenic_variant |
| 19 | 30734697 | 30742312 | 7615 | Prkg1 | MOLF | intron_variant |
| 19 | 33963779 | 33967551 | 3772 | - | CAST,SPRET | intergenic_variant |
| 19 | 34464820 | 34465873 | 1053 | - | CAST,MOLF | intergenic_variant |
| 19 | 36699478 | 36700981 | 1503 | - | SPRET | intergenic_variant |
| 19 | 38631392 | 38634935 | 3543 | Plce1 | CAST | intron_variant,regulatory_region_variant |

| | | | | | | |
|---|---|---|---|---|---|---|
| 19 | 41207064 | 41209464 | 2400 | Tm9sf3 | SPRET | intron_variant |
| 19 | 43275345 | 43277011 | 1666 | Hpse2 | SPRET | intron_variant |
| 19 | 52443372 | 52444391 | 1019 | - | SPRET | intergenic_variant |
| 19 | 52932484 | 52933553 | 1069 | Xpnpep1 | 129S1,AJ,BALB,BTBR,BUB,C3H,C57L,CBA,CE,DBA1J,DBA2J,ILNJ2,KK,MA/My,MRL,NOD,NOR,NZO,PJ,RF,RH2,SEA,SMJ | intron_variant |
| 19 | 55674914 | 55676658 | 1744 | - | MOLF | regulatory_region_variant,intergenic_variant |
| 19 | 59913368 | 59915779 | 2411 | Rab11fip2 | BTBR,CAST,FVB,NZB,PJ,SEA,SMJ,TH,WSB | intron_variant |
| 19 | 60659320 | 60664335 | 5015 | - | SPRET | regulatory_region_variant,intergenic_variant |
| 19 | 60660714 | 60664340 | 3626 | - | SPRET | regulatory_region_variant,intergenic_variant |
| 19 | 60894962 | 60896912 | 1950 | Grk5 | TH | intron_variant |
| 2 | 4686612 | 4688788 | 2176 | - | SPRET | intergenic_variant |
| 2 | 5269621 | 5271325 | 1704 | - | SPRET | intergenic_variant |
| 2 | 11950150 | 11954122 | 3972 | - | CAST,SPRET | intergenic_variant |
| 2 | 11990376 | 11997387 | 7011 | - | CAST | intergenic_variant |
| 2 | 11991747 | 11998517 | 6770 | - | CAST | intergenic_variant |
| 2 | 13919992 | 13924152 | 4160 | - | CAST,MOLF | intergenic_variant |
| 2 | 14203836 | 14205573 | 1737 | Tmem236 | SPRET,WSB | intron_variant,regulatory_region_variant |
| 2 | 14495814 | 14497790 | 1976 | Slc39a12 | CAST | intron_variant |
| 2 | 15697055 | 15698177 | 1122 | Malrd1 | SPRET | intron_variant |
| 2 | 19967329 | 19968360 | 1031 | Etl4 | CAST | intron_variant |
| 2 | 20878372 | 20880715 | 2343 | Arhgap21 | AJ,AKR,DBA2J,PL,RH2 | intron_variant |
| 2 | 21720254 | 21723853 | 3599 | Gpr158 | AJ,AKR,DBA2J,PL,RH2 | intron_variant |
| 2 | 22090297 | 22096984 | 6687 | - | AJ,AKR,CE,DBA2J,FVB,PL,RH2,SJL,SPRET | intergenic_variant |
| 2 | 37020841 | 37022110 | 1269 | - | SPRET | intergenic_variant |
| 2 | 39749349 | 39750432 | 1083 | - | SPRET | intergenic_variant |
| 2 | 40653375 | 40659970 | 6595 | Lrp1b | SPRET | intron_variant,regulatory_region_variant |
| 2 | 41934323 | 41935661 | 1338 | Lrp1b | MOLF | intron_variant |
| 2 | 42155884 | 42158676 | 2792 | Lrp1b | 129S1,AJ,AKR,BUB,C57L,C58J,DBA1J,ILNJ2,MA/My,MOLF,MRL,NOD,NOR,NZB,NZO,NZW,PJ,RF,SEA,SPRET,SWR,TH,WSB | intron_variant |
| 2 | 43298615 | 43305073 | 6458 | - | SMJ,TH | intergenic_variant |
| 2 | 43503164 | 43505389 | 2225 | Kynu | BUB,CAST,WSB | intron_variant,regulatory_region_variant |
| 2 | 46603637 | 46605700 | 2063 | - | SPRET | intergenic_variant |
| 2 | 47612504 | 47623130 | 10626 | - | SPRET | intergenic_variant |
| 2 | 50654204 | 50660674 | 6470 | - | AJ,AKR,BUB,C3H,CBA,CE,DBA1J,DBA2J,ILNJ2,MRL,NOD,NOR,PJ,PL,RF,SWR | intergenic_variant |
| 2 | 51514985 | 51516030 | 1045 | - | CAST | intergenic_variant |
| 2 | 53488689 | 53490145 | 1456 | - | MOLF | intergenic_variant |
| 2 | 53489383 | 53490981 | 1598 | - | CAST | intergenic_variant |
| 2 | 55087471 | 55103664 | 16193 | - | MOLF,SPRET | regulatory_region_variant,intergenic_variant |
| 2 | 55088965 | 55103650 | 14685 | - | MOLF,SPRET | regulatory_region_variant,intergenic_variant |
| 2 | 55093985 | 55106507 | 12522 | - | CAST,MOLF,SPRET | intergenic_variant |
| 2 | 56505120 | 56508335 | 3215 | - | CAST,SPRET | intergenic_variant |
| 2 | 56505697 | 56508082 | 2385 | - | SPRET | intergenic_variant |
| 2 | 56923442 | 56927927 | 4485 | - | CAST | intergenic_variant |
| 2 | 56924216 | 56927767 | 3551 | - | MOLF | intergenic_variant |
| 2 | 56925476 | 56928351 | 2875 | - | CAST,MOLF | intergenic_variant |
| 2 | 57462012 | 57464365 | 2353 | - | SPRET | intergenic_variant |
| 2 | 63265291 | 63271587 | 6296 | - | SPRET | intergenic_variant |
| 2 | 63436805 | 63439740 | 2935 | - | MOLF | intergenic_variant |
| 2 | 63478323 | 63480343 | 2020 | - | CAST,MOLF,SPRET | intergenic_variant |
| 2 | 63548530 | 63556183 | 7653 | - | SPRET | intergenic_variant |
| 2 | 64843534 | 64844955 | 1421 | Grb14 | SPRET | intron_variant,regulatory_region_variant |
| 2 | 65233902 | 65235066 | 1164 | - | 129S1,BTBR,BUB,CAST,CE,DBA1J,DBA2J,MOLF,NOD,NOR,RF,SJL,SPRET,SWR,WSB | intergenic_variant |
| 2 | 66703996 | 66712297 | 8301 | - | SPRET | intergenic_variant |
| 2 | 68016234 | 68017408 | 1174 | - | WSB | intergenic_variant |
| 2 | 72543355 | 72547658 | 4303 | 8430437L04Rik | MA/My,MOLF,NZB,SPRET | non_coding_transcript_exon_variant,intron_variant |
| 2 | 75548031 | 75549625 | 1594 | - | CAST | regulatory_region_variant,intergenic_variant |
| 2 | 76009387 | 76011233 | 1846 | Pde11a | SPRET | intron_variant |
| 2 | 78163651 | 78169855 | 6204 | - | MOLF | intergenic_variant |
| 2 | 79682207 | 79683411 | 1204 | Pde1a | MOLF | intron_variant |
| 2 | 80031178 | 80032727 | 1549 | - | SPRET | intergenic_variant |
| 2 | 80052616 | 80054121 | 1505 | - | SPRET | intergenic_variant |
| 2 | 85396312 | 85398045 | 1733 | - | MOLF,SPRET,WSB | intergenic_variant |
| 2 | 87845428 | 87849353 | 3925 | - | SPRET | regulatory_region_variant,intergenic_variant |
| 2 | 96698436 | 96700692 | 2256 | Lrrc4c | CAST,MOLF | intron_variant |
| 2 | 97966251 | 97971666 | 5415 | - | SPRET | intergenic_variant |
| 2 | 99356054 | 99359729 | 3675 | - | SPRET | intergenic_variant |
| 2 | 99359500 | 99361221 | 1721 | - | SPRET | intergenic_variant |
| 2 | 100030794 | 100031909 | 1115 | - | 129S1,AJ,AKR,BALB,BTBR,BUB,C3H,C57L,CAST,CBA,CE,DBA1J,DBA2J,FVB,ILNJ2,KK,MOLF,MRL,NOD,NOR,NZB,NZO,NZW,PJ,RF,RH2,SEA,SJL,SMJ,SPRET,SWR,WSB | intergenic_variant |
| 2 | 101155731 | 101160133 | 4402 | - | CAST | intergenic_variant |
| 2 | 103897178 | 103898392 | 1214 | Cd59b | MOLF,SPRET | 5_prime_UTR_variant,intron_variant |
| 2 | 106252287 | 106253996 | 1709 | - | C3H,CE,MOLF | intergenic_variant |
| 2 | 106918052 | 106921681 | 3629 | - | SPRET | intergenic_variant |
| 2 | 109281729 | 109284845 | 3116 | - | SPRET | intergenic_variant |
| 2 | 111202660 | 111205553 | 2893 | Or4g17 | MOLF | upstream_gene_variant |
| 2 | 111204253 | 111205552 | 1299 | Or4g17 | MOLF | upstream_gene_variant |
| 2 | 111429858 | 111431405 | 1547 | - | AKR,BTBR,MRL | intergenic_variant |
| 2 | 111627536 | 111631088 | 3552 | - | C3H,CE,MOLF,PL | intergenic_variant |
| 2 | 111627560 | 111628753 | 1193 | - | C3H,CE,MOLF,PL | intergenic_variant |
| 2 | 113684273 | 113685303 | 1030 | - | SPRET | intergenic_variant |
| 2 | 117308436 | 117310069 | 1633 | - | CAST,KK,NZB,TH | regulatory_region_variant,intergenic_variant |
| 2 | 118394410 | 118396338 | 1928 | - | CBA | intergenic_variant |
| 2 | 122038350 | 122041368 | 3018 | Gm14050 | AJ,BALB,C3H,C58J,CBA,DBA1J,DBA2J,ILNJ2,NZB,NZO,PJ,PL,RF,RH2,SEA,SMJ,TH | non_coding_transcript_variant |
| 2 | 123421760 | 123423961 | 2201 | - | SPRET | intergenic_variant |
| 2 | 125947759 | 125952207 | 4448 | Fam227b,Gm26496 | SPRET | intron_variant,non_coding_transcript_variant |

| | | | | | | |
|---|---|---|---|---|---|---|
| 2 | 128196848 | 128199068 | 2220 | Morrbid | C3H,CBA,DBA1J,DBA2J,ILNJ2,MOLF,PJ,PL | intron_variant,non_coding_transcript_variant,non_coding_transcript_exon_variant,intron_variant |
| 2 | 131203849 | 131205188 | 1339 | - | CAST,DBA1J,DBA2J,ILNJ2,MOLF,PJ,PL | intergenic_variant |
| 2 | 137266406 | 137268081 | 1675 | - | SPRET | intergenic_variant |
| 2 | 139058641 | 139060031 | 1390 | - | SPRET | intergenic_variant |
| 2 | 141186929 | 141188366 | 1437 | Macrod2 | CAST | intron_variant |
| 2 | 143064418 | 143065420 | 1002 | - | SPRET | intergenic_variant |
| 2 | 145051160 | 145052870 | 1710 | Slc24a3 | SPRET | intron_variant |
| 2 | 146997876 | 147000623 | 2747 | - | SPRET | intergenic_variant |
| 2 | 147717729 | 147720595 | 2866 | Gm25516 | FVB,SPRET | upstream_gene_variant |
| 2 | 149316751 | 149318415 | 1664 | - | SPRET | intergenic_variant |
| 2 | 150362640 | 150366591 | 3951 | Gm14128 | BUB,MA/My,SPRET | non_coding_transcript_variant |
| 2 | 153689126 | 153692481 | 3355 | - | AJ,BTBR,C58J,ILNJ2,MOLF,MRL,NOD,NZB,NZW,PJ,RF | regulatory_region_variant,intergenic_variant |
| 2 | 153690014 | 153692976 | 2962 | - | AJ,BTBR,C58J,ILNJ2,MOLF,MRL,NOD,NZB,NZW,PJ,RF | regulatory_region_variant,intergenic_variant |
| 2 | 154451791 | 154454034 | 2243 | 4930519P11Rik | AJ,BTBR,C58J,CAST,DBA1J,DBA2J,KK,MOLF,NZB,NZO,NZW,SPRET | non_coding_transcript_exon_variant |
| 2 | 157543141 | 157544344 | 1203 | - | 129S1,AJ,AKR,BALB,BUB,C3H,C57L,C58J,CAST,CBA,CE,DBA1J,DBA2J,FVB,ILNJ2,MA/My,MOLF,MRL,NOD,NZB,NZO,NZW,PJ,PL,RF,SEA,SMJ,SPRET,SWR,WSB | regulatory_region_variant,intergenic_variant |
| 2 | 163004043 | 163005274 | 1231 | - | KK | intergenic_variant |
| 2 | 164043007 | 164045375 | 2368 | Wfdc15a | SPRET | upstream_gene_variant |
| 2 | 173335565 | 173337835 | 2270 | - | CAST,SPRET | intergenic_variant |
| 2 | 173983253 | 173984423 | 1170 | Gm10714 | CAST,MOLF,PJ | intron_variant,non_coding_transcript_variant |
| 3 | 4140392 | 4142765 | 2373 | - | CAST | intergenic_variant |
| 3 | 4217153 | 4220380 | 3227 | - | MOLF | intergenic_variant |
| 3 | 6274194 | 6276707 | 2513 | - | C58J,CE,DBA1J,DBA2J,FVB,MA/My,MRL,NOD,NOR,PL,SJL,SWR | intergenic_variant |
| 3 | 6424084 | 6431534 | 7450 | - | SPRET | intergenic_variant |
| 3 | 10654629 | 10656318 | 1689 | - | SPRET | intergenic_variant |
| 3 | 10854917 | 10861620 | 6703 | - | SPRET | intergenic_variant |
| 3 | 10988140 | 10991613 | 3473 | - | SPRET | intergenic_variant |
| 3 | 11505584 | 11506657 | 1073 | - | MOLF | intergenic_variant |
| 3 | 15269098 | 15270291 | 1193 | - | CAST,MOLF | intergenic_variant |
| 3 | 15297826 | 15300017 | 2191 | - | CE,DBA2J,LP | intergenic_variant |
| 3 | 18402343 | 18404439 | 2096 | Gm30667 | CAST,MOLF,SPRET | intron_variant,non_coding_transcript_variant |
| 3 | 20350851 | 20355041 | 4190 | - | SPRET | intergenic_variant |
| 3 | 22951995 | 22954663 | 2668 | - | MA/My,NOD,NOR,PL,SMJ,WSB | regulatory_region_variant,intergenic_variant |
| 3 | 25016629 | 25023643 | 7014 | - | SPRET | intergenic_variant |
| 3 | 26468513 | 26469790 | 1277 | - | SPRET | intergenic_variant |
| 3 | 26658970 | 26671078 | 12108 | - | MOLF | regulatory_region_variant,intergenic_variant |
| 3 | 26667003 | 26673058 | 6055 | - | MOLF | intergenic_variant |
| 3 | 27262040 | 27264304 | 2264 | Nceh1 | SPRET | intron_variant,regulatory_region_variant |
| 3 | 27525928 | 27529321 | 3393 | Fndc3b | AJ,AKR,C3H,CBA,CE,PL,RF | intron_variant,regulatory_region_variant |
| 3 | 27527909 | 27530369 | 2460 | Fndc3b | AJ,AKR,C3H,CAST,CBA,CE,MOLF,PL,RF,SPRET | intron_variant,regulatory_region_variant |
| 3 | 29508123 | 29510918 | 2795 | Egfem1 | SPRET | intron_variant |
| 3 | 29800550 | 29802143 | 1593 | - | MOLF | intergenic_variant |
| 3 | 33052867 | 33053874 | 1007 | Pex5l | SPRET | intron_variant |
| 3 | 33653812 | 33657484 | 3672 | - | WSB | intergenic_variant |
| 3 | 33750729 | 33757562 | 6833 | Gm43666 | SPRET | non_coding_transcript_exon_variant |
| 3 | 33778952 | 33780143 | 1191 | Gm43666 | CAST,MOLF | intron_variant,non_coding_transcript_variant,regulatory_region_variant |
| 3 | 35042058 | 35046185 | 4127 | - | SPRET | intergenic_variant |
| 3 | 35827056 | 35830836 | 3780 | Atp11b | DBA1J,RF | intron_variant,regulatory_region_variant |
| 3 | 40467275 | 40475809 | 8534 | - | CE | regulatory_region_variant,intergenic_variant |
| 3 | 42040451 | 42041553 | 1102 | - | CAST | intergenic_variant |
| 3 | 44757645 | 44761770 | 4125 | - | SPRET | regulatory_region_variant,intergenic_variant |
| 3 | 45730310 | 45732447 | 2137 | Gm38166,Gm32340 | SPRET | non_coding_transcript_exon_variant,non_coding_transcript_variant |
| 3 | 47530760 | 47532434 | 1674 | - | SPRET | intergenic_variant |
| 3 | 47577000 | 47581041 | 4041 | - | MOLF | intergenic_variant |
| 3 | 48654870 | 48657203 | 2333 | - | CE,SJL,SWR | intergenic_variant |
| 3 | 48655969 | 48657277 | 1308 | - | CE,SJL,SWR | intergenic_variant |
| 3 | 49273803 | 49282601 | 8798 | Gm20557 | SPRET | non_coding_transcript_exon_variant,intron_variant,regulatory_region_variant |
| 3 | 49274489 | 49282607 | 8118 | Gm20557 | SPRET | non_coding_transcript_exon_variant,intron_variant,regulatory_region_variant |
| 3 | 49308972 | 49315027 | 6055 | Gm20557 | AJ,AKR,BALB,BUB,C3H,CBA,CE,DBA1J,DBA2J,FVB,ILNJ2,MRL,NOD,NOR,NZB,PL,RF,RH2,SMJ,TH | non_coding_transcript_exon_variant,intron_variant |
| 3 | 54481975 | 54485272 | 3297 | - | SPRET | intergenic_variant |
| 3 | 54483215 | 54485441 | 2226 | - | SPRET | intergenic_variant |
| 3 | 57107557 | 57111400 | 3843 | - | CE,FVB,SJL,SMJ,SWR | regulatory_region_variant,intergenic_variant |
| 3 | 57107561 | 57109300 | 1739 | - | CE,FVB,SJL,SMJ,SWR | intergenic_variant |
| 3 | 57257044 | 57258647 | 1603 | Gm56807 | SPRET | intron_variant,non_coding_transcript_variant |
| 3 | 59794822 | 59797383 | 2561 | - | MOLF | intergenic_variant |
| 3 | 60775631 | 60778616 | 2985 | - | AKR | intergenic_variant |
| 3 | 62780416 | 62787413 | 6997 | - | SPRET | intergenic_variant |
| 3 | 67442649 | 67449164 | 6515 | - | SJL | intergenic_variant |
| 3 | 67629647 | 67632807 | 3160 | - | MOLF | intergenic_variant |
| 3 | 69477523 | 69479215 | 1692 | - | BUB,SEA | intergenic_variant |
| 3 | 69821873 | 69827579 | 5706 | - | SPRET | intergenic_variant |
| 3 | 71179575 | 71181625 | 2050 | - | SPRET | intergenic_variant |
| 3 | 71362763 | 71367606 | 4843 | - | SPRET | intergenic_variant |
| 3 | 71932412 | 71938143 | 5731 | - | MOLF | intergenic_variant |
| 3 | 72225318 | 72226596 | 1278 | - | MOLF,SPRET | intergenic_variant |
| 3 | 73877923 | 73879525 | 1602 | - | AKR,FVB,RF,SJL | intergenic_variant |
| 3 | 74467170 | 74469434 | 2264 | - | SPRET | intergenic_variant |
| 3 | 75835005 | 75840147 | 5142 | Golim4 | MOLF | intron_variant |
| 3 | 77182267 | 77184187 | 1920 | - | SPRET | intergenic_variant |

| | | | | | | |
|---|---|---|---|---|---|---|
| 3 | 77573997 | 77576342 | 2345 | - | 129S1,AKR,BALB,BTBR,BUB,C57L,CE,DBA1J,DBA2J,FVB,ILNJ2,KK,MA/My,MRL,NOD,NOR,NZB,NZO,NZW,PJ,PL,RF,RH2,SEA,SMJ,SWR,TH,WSB | intergenic_variant |
| 3 | 77574639 | 77582346 | 7707 | - | 129S1,AKR,BALB,BTBR,BUB,C57L,CE,DBA1J,DBA2J,FVB,ILNJ2,KK,MA/My,MRL,NOD,NOR,NZB,NZO,NZW,PJ,PL,RF,RH2,SEA,SMJ,SWR,TH,WSB | intergenic_variant |
| 3 | 78400887 | 78403687 | 2800 | - | B10D2,B10J,C58J,LP | regulatory_region_variant,intergenic_variant |
| 3 | 80197196 | 80203470 | 6274 | Gm37971 | SPRET | intron_variant,non_coding_transcript_variant |
| 3 | 83236047 | 83240286 | 4239 | Dchs2 | NZB,NZO,NZW | intron_variant |
| 3 | 91725467 | 91728167 | 2700 | - | MOLF | intergenic_variant |
| 3 | 92431401 | 92433954 | 2553 | - | NOD,NOR,WSB | regulatory_region_variant,intergenic_variant |
| 3 | 92431402 | 92435603 | 4201 | - | NOD,NOR,WSB | regulatory_region_variant,intergenic_variant |
| 3 | 93658104 | 93659520 | 1416 | - | CAST | intergenic_variant |
| 3 | 96145778 | 96154682 | 8904 | H2ac19,H2ac18,H3c15 | 129S1,AJ,BALB,C3H,C57L,CAST,CBA,CE,FVB,KK,MA/My,MOLF,NOR,NZB,NZO,NZW,PJ,RH2,SEA,SJL,SMJ,SWR,WSB | coding_transcript_variant,coding_sequence_variant,5_prime_UTR_variant,3_prime_UTR_variant,non_coding_transcript_variant,regulatory_region_variant |
| 3 | 96147881 | 96152570 | 4689 | H2ac19,H2ac18,Gm20( | 129S1,AJ,BALB,C3H,C57L,CAST,CBA,CE,FVB,KK,MA/My,MOLF,NOD,NOR,NZB,NZO,NZW,PJ,RH2,SEA,SJL,SMJ,SWR,WSB | upstream_gene_variant,non_coding_transcript_exon_variant,downstream_gene_variant,regulatory_region_variant |
| 3 | 96147882 | 96154684 | 6802 | H2ac19,H2ac18,H3c14 | AJ,BALB,C57L,CAST,CBA,CE,FVB,KK,NZO,NZW,SJL,SMJ,SWR | upstream_gene_variant,coding_transcript_variant,coding_sequence_variant,5_prime_UTR_variant,3_prime_UTR_variant,non_coding_transcript_variant,non_coding_transcript_exon_variant,downstream_gene_variant,regulatory_region_variant |
| 3 | 96361845 | 96363418 | 1573 | Gm9207 | 129S1,B10J,C57L,CBA,DBA2J,FVB,KK,NOR,NZO,NZW,SJL | non_coding_transcript_variant |
| 3 | 96394526 | 96395555 | 1029 | Gm25890,Gm22614 | 129S1,AJ,B10D2,BALB,C3H,C57L,CAST,CBA,CE,FVB,KK,MA/My,MOLF,NOD,NOR,NZB,NZO,NZW,PJ,RH2,SEA,SJL,SMJ,SWR,WSB | upstream_gene_variant,regulatory_region_variant |
| 3 | 99296164 | 99299520 | 3356 | - | SPRET | intergenic_variant |
| 3 | 100290630 | 100292278 | 1648 | - | SPRET | intergenic_variant |
| 3 | 106704233 | 106706350 | 2117 | - | SPRET | intergenic_variant |
| 3 | 106705344 | 106710887 | 5543 | - | SPRET | intergenic_variant |
| 3 | 108851201 | 108853666 | 2465 | Henmt1 | SPRET | intron_variant |
| 3 | 108851344 | 108854833 | 3489 | Henmt1 | SPRET | intron_variant |
| 3 | 109832361 | 109842112 | 9751 | Ntng1 | DBA2J | coding_sequence_variant,intron_variant,regulatory_region_variant |
| 3 | 110742646 | 110744301 | 1655 | - | SPRET | intergenic_variant |
| 3 | 112188470 | 112194973 | 6503 | - | SPRET | intergenic_variant |
| 3 | 112497920 | 112499427 | 1507 | Gm42526 | SPRET | non_coding_transcript_variant |
| 3 | 113142245 | 113143600 | 1355 | - | MOLF | intergenic_variant |
| 3 | 114147157 | 114149007 | 1850 | - | KK,MOLF | intergenic_variant |
| 3 | 114605192 | 114607475 | 2283 | - | SPRET | intergenic_variant |
| 3 | 118664586 | 118666109 | 1523 | Dpyd,Gm43410 | CAST | intron_variant,downstream_gene_variant |
| 3 | 122584760 | 122585875 | 1115 | Pde5a,4930447N08Rik | SPRET | intron_variant,intron_variant,non_coding_transcript_variant |
| 3 | 124917732 | 124919159 | 1427 | - | AJ,BALB,C3H,CAST,CBA,MOLF,NZB,RH2,SEA,SMJ,WSB | intergenic_variant |
| 3 | 126580362 | 126581723 | 1361 | Camk2d | 129S1,AJ,BALB,C3H,C57L,CBA,MOLF,NZB,RH2,SMJ | intron_variant |
| 3 | 129148426 | 129156367 | 7941 | Gm42629 | PJ,SEA,SPRET | non_coding_transcript_exon_variant,regulatory_region_variant |
| 3 | 133935516 | 133938223 | 2707 | Gm26691 | CAST | intron_variant,non_coding_transcript_variant,regulatory_region_variant |
| 3 | 133936163 | 133938270 | 2107 | Gm26691 | CAST | intron_variant,non_coding_transcript_variant,regulatory_region_variant |
| 3 | 134236670 | 134243441 | 6771 | Gm26820 | CAST,MOLF | intron_variant,non_coding_transcript_variant |
| 3 | 134597004 | 134599440 | 2436 | Tacr3 | SPRET | intron_variant |
| 3 | 134597015 | 134603008 | 5993 | Tacr3 | SPRET | intron_variant |
| 3 | 135283838 | 135286643 | 2805 | - | MOLF | regulatory_region_variant,intergenic_variant |
| 3 | 135661204 | 135664382 | 3178 | Gm31243 | CAST | intron_variant,non_coding_transcript_variant |
| 3 | 135811411 | 135812695 | 1284 | Bank1,Gm31243 | MOLF | intron_variant,intron_variant,non_coding_transcript_variant |
| 3 | 135811825 | 135813865 | 2040 | Bank1,Gm31243 | CAST,MOLF | intron_variant,intron_variant,non_coding_transcript_variant |
| 3 | 137234880 | 137238900 | 4020 | - | BUB,C58J,DBA2J,NZB,NZO,NZW,PL,RF,SMJ | regulatory_region_variant,intergenic_variant |
| 3 | 139092383 | 139093781 | 1398 | Stpg2 | CAST | intron_variant |
| 3 | 140021631 | 140022758 | 1127 | - | SPRET | intergenic_variant |
| 3 | 140390215 | 140392523 | 2308 | - | SPRET | intergenic_variant |
| 3 | 144651875 | 144654055 | 2180 | Gm16233 | SPRET | non_coding_transcript_exon_variant,intron_variant,regulatory_region_variant |
| 3 | 144743485 | 144750939 | 7454 | - | SPRET | intergenic_variant |
| 3 | 150900607 | 150902963 | 2356 | - | CAST | intergenic_variant |
| 3 | 151663788 | 151665646 | 1858 | Gm2939 | AJ,AKR,B10D2,B10J,BALB,BUB,C3H,C58J,DBA2J,FVB,ILNJ2,LP,MA/My,NOD,NOR,NZB,NZO,NZW,PJ,PL,RF,RH2,SEA,SJL,SWR,TH | non_coding_transcript_variant |
| 3 | 151717636 | 151724307 | 6671 | - | CAST | intergenic_variant |
| 3 | 151723184 | 151724316 | 1132 | - | CAST | intergenic_variant |
| 3 | 151754327 | 151758866 | 4539 | - | SPRET | intergenic_variant |
| 3 | 157117706 | 157120346 | 2640 | - | RF | intergenic_variant |
| 4 | 4310289 | 4319584 | 9295 | - | AJ,CAST,SPRET | regulatory_region_variant,intergenic_variant |
| 4 | 8121396 | 8122724 | 1328 | - | 129S1,C58J,LP,MA/My,MRL,RF,SJL | regulatory_region_variant,intergenic_variant |
| 4 | 8479535 | 8485899 | 6364 | - | AJ,KK,NOD,NOR | intergenic_variant |
| 4 | 12431211 | 12437975 | 6764 | - | BUB,FVB,MOLF,NZB,NZO,NZW,PJ,SEA,SJL,SWR,TH,WSB | intergenic_variant |
| 4 | 16798245 | 16800279 | 2034 | - | NOD,NOR | intergenic_variant |
| 4 | 21452313 | 21454677 | 2364 | - | SPRET | intergenic_variant |
| 4 | 22749880 | 22751463 | 1583 | - | TH | intergenic_variant |
| 4 | 24025452 | 24026477 | 1025 | - | MOLF | intergenic_variant |
| 4 | 28408135 | 28411948 | 3813 | - | SPRET | intergenic_variant |
| 4 | 30252025 | 30253667 | 1642 | - | SPRET | intergenic_variant |
| 4 | 30532738 | 30536423 | 3685 | - | SPRET | intergenic_variant |

| | | | | | | |
|---|---|---|---|---|---|---|
| 4 | 35616316 | 35619425 | 3109 | - | SPRET | intergenic_variant |
| 4 | 38582216 | 38584034 | 1818 | Gm12381 | CAST,SPRET | intron_variant,non_coding_transcript_variant,regulatory_region_variant |
| 4 | 53608064 | 53614549 | 6485 | Slc44a1 | SPRET | intron_variant |
| 4 | 59139859 | 59146269 | 6410 | Shoc1 | CAST | upstream_gene_variant |
| 4 | 59142636 | 59146270 | 3634 | - | CAST | intergenic_variant |
| 4 | 64530685 | 64533281 | 2596 | - | SPRET | intergenic_variant |
| 4 | 64532068 | 64533891 | 1823 | - | SPRET | intergenic_variant |
| 4 | 67029762 | 67037726 | 7964 | - | 129S1,C57L,PJ | intergenic_variant |
| 4 | 67052557 | 67054311 | 1754 | - | CAST | intergenic_variant |
| 4 | 67053400 | 67054614 | 1214 | - | 129S1,C57L,CAST,MOLF,PJ | intergenic_variant |
| 4 | 68052438 | 68053678 | 1240 | Gm11751 | SPRET | intron_variant,non_coding_transcript_variant |
| 4 | 68444331 | 68447468 | 3137 | - | MOLF | intergenic_variant |
| 4 | 69716847 | 69719018 | 2171 | - | SPRET | intergenic_variant |
| 4 | 69993376 | 69998561 | 5185 | - | 129S1,C57L,CAST,DBA1J,DBA2J,ILNJ2,KK,MOLF,NZB,NZO,NZW,PJ,SMJ,SPRET,WSB | intergenic_variant |
| 4 | 71833760 | 71838241 | 4481 | - | ILNJ2,PJ,WSB | intergenic_variant |
| 4 | 73161771 | 73164658 | 2887 | - | SPRET | intergenic_variant |
| 4 | 73972725 | 73978669 | 5944 | Frmd3 | SPRET | intron_variant |
| 4 | 77601804 | 77606915 | 5111 | - | SPRET | intergenic_variant |
| 4 | 78894866 | 78896396 | 1530 | - | SPRET | intergenic_variant |
| 4 | 79742400 | 79744124 | 1724 | - | SPRET | intergenic_variant |
| 4 | 81413530 | 81415976 | 2446 | - | 129S1,BTBR,C57L | intergenic_variant |
| 4 | 81413546 | 81418236 | 4690 | - | 129S1,BTBR,C57L | regulatory_region_variant,intergenic_variant |
| 4 | 81523402 | 81524998 | 1596 | - | SPRET | intergenic_variant |
| 4 | 81928730 | 81929857 | 1127 | - | MOLF | intergenic_variant |
| 4 | 81928733 | 81931417 | 2684 | - | AKR,B10D2,B10J,BUB,C3H,CBA,CE,DBA1J,DBA2J,FVB,KK,NOD,NOR,NZB,NZO,RF,SJL,TH | intergenic_variant |
| 4 | 81929192 | 81937576 | 8384 | - | 129S1,AJ,BALB,BTBR,C57L,RH2,SEA,SWR | regulatory_region_variant,intergenic_variant |
| 4 | 82844417 | 82846248 | 1831 | Frem1 | SPRET | intron_variant |
| 4 | 83086287 | 83088765 | 2478 | - | SPRET | regulatory_region_variant,intergenic_variant |
| 4 | 85390046 | 85393635 | 3589 | Sh3gl2 | B10D2,B10J,C58J,DBA1J,DBA2J,ILNJ2,PJ,RF | intron_variant |
| 4 | 90066706 | 90069596 | 2890 | - | SPRET | intergenic_variant |
| 4 | 91622375 | 91624556 | 2181 | - | MOLF | intergenic_variant |
| 4 | 91782677 | 91790003 | 7326 | - | CE,MA/My,NZB | intergenic_variant |
| 4 | 94224929 | 94228503 | 3574 | Gm12648 | SPRET | intron_variant,non_coding_transcript_variant |
| 4 | 94647929 | 94651467 | 3538 | Tek | CAST,MOLF,SPRET | intron_variant |
| 4 | 96190967 | 96197746 | 6779 | Cyp2j11 | CAST,MOLF | coding_sequence_variant,intron_variant |
| 4 | 97274443 | 97278991 | 4548 | - | SPRET | intergenic_variant |
| 4 | 97277171 | 97278988 | 1817 | - | SPRET | intergenic_variant |
| 4 | 98867062 | 98868066 | 1004 | Dock7 | 129S1,BTBR,C57L,MA/My,NOR,SJL,SWR,TH | intron_variant |
| 4 | 105491777 | 105494279 | 2502 | - | SPRET | intergenic_variant |
| 4 | 110984534 | 110986101 | 1567 | Agbl4 | AKR,KK,MRL,SWR | intron_variant |
| 4 | 112460259 | 112463366 | 3107 | Skint2 | AJ,BUB,C3H,CAST,CE,PL,SEA,SMJ,SPRET,SWR | intron_variant |
| 4 | 112974468 | 112978853 | 4385 | Skint6 | AJ | intron_variant |
| 4 | 112978183 | 112980675 | 2492 | Skint6 | AJ,DBA1J,FVB | intron_variant |
| 4 | 115124837 | 115131850 | 7013 | - | SPRET | intergenic_variant |
| 4 | 117792527 | 117793810 | 1283 | St3gal3 | AKR,ILNJ2,KK,MRL | intron_variant |
| 4 | 123788679 | 123791010 | 2331 | - | SPRET | intergenic_variant |
| 4 | 124281876 | 124285860 | 3984 | - | CAST | intergenic_variant |
| 4 | 129557150 | 129558407 | 1257 | Kpna6 | 129S1,AJ,AKR,BALB,BTBR,BUB,C3H,C57L,CBA,CE,DBA2J,FVB,ILNJ2,MA/My,MRL,NOD,NZO,NZW,RF,RH2,SJL,SWR,TH,WSB | upstream_gene_variant,intron_variant |
| 4 | 135787829 | 135792570 | 4741 | - | SPRET | intergenic_variant |
| 4 | 137066185 | 137067431 | 1246 | Cdc42 | C3H,CBA,NZW | intron_variant,regulatory_region_variant |
| 4 | 144478474 | 144487001 | 8527 | - | BTBR | regulatory_region_variant,intergenic_variant |
| 4 | 144498569 | 144499643 | 1074 | - | 129S1,BTBR,C57L | intergenic_variant |
| 4 | 152333926 | 152334987 | 1061 | Acot7 | SPRET | intron_variant |
| 5 | 3365861 | 3368771 | 2910 | - | BUB,MOLF,NOD,NOR | intergenic_variant |
| 5 | 5867926 | 5869997 | 2071 | - | CAST | intergenic_variant |
| 5 | 5869480 | 5870742 | 1262 | - | CAST | intergenic_variant |
| 5 | 6262520 | 6265739 | 3219 | - | C58J,MA/My,MRL,WSB | intergenic_variant |
| 5 | 8013404 | 8015361 | 1957 | Steap4 | 129S1,BALB,BTBR,C57L,CE,NZB,NZO,NZW,PJ,PL,RF,SEA,SJL,SWR,WSB | intron_variant,regulatory_region_variant |
| 5 | 8335754 | 8337065 | 1311 | Adam22 | SPRET | intron_variant |
| 5 | 9338913 | 9343521 | 4608 | Elapor2 | NOD,NOR | intron_variant |
| 5 | 12864046 | 12867600 | 3554 | - | CAST | intergenic_variant |
| 5 | 15014075 | 15020554 | 6479 | - | SPRET | regulatory_region_variant,intergenic_variant |
| 5 | 15663097 | 15664556 | 1459 | - | TH | regulatory_region_variant,intergenic_variant |
| 5 | 15726265 | 15727288 | 1023 | Gm21847 | NZO | intron_variant,non_coding_transcript_variant,upstream_gene_variant,regulatory_region_variant |
| 5 | 15829599 | 15832412 | 2813 | Speer4d,Gm21847 | FVB,MA/My,RF,SJL,SWR,WSB | downstream_gene_variant,intron_variant,non_coding_transcript_variant,regulatory_region_variant |
| 5 | 18249791 | 18251761 | 1970 | - | AKR,CAST,TH | intergenic_variant |
| 5 | 19029112 | 19035907 | 6795 | - | SPRET | intergenic_variant |
| 5 | 19185974 | 19188629 | 2655 | - | CAST,SPRET | regulatory_region_variant,intergenic_variant |
| 5 | 19314278 | 19316762 | 2484 | - | SPRET | intergenic_variant |
| 5 | 21913140 | 21914152 | 1012 | - | 129S1,BTBR,C57L,CAST,CE,DBA1J,DBA2J,MOLF,NOD,NOR,NZO,SPRET,WSB | intergenic_variant |
| 5 | 23223380 | 23224693 | 1313 | Lhfpl3 | 129S1,AKR,BALB,BTBR,BUB,C57L,C58J,CAST,CE,DBA1J,DBA2J,FVB,KK,MA/My,MRL,NOD,NOR,NZO,NZW,PJ,PL,RF,RH2,SEA,SJL,SMJ,SPRET,SWR,TH,WSB | intron_variant |
| 5 | 24155800 | 24156813 | 1013 | Hycc1 | CAST,SPRET | intron_variant |
| 5 | 28251160 | 28252722 | 1562 | Gm26608 | MOLF | intron_variant,non_coding_transcript_variant,regulatory_region_variant |
| 5 | 36585801 | 36587266 | 1465 | - | SPRET | intergenic_variant |
| 5 | 38286734 | 38292192 | 5458 | Stx18,Gm42858 | CAST,SPRET | intron_variant,non_coding_transcript_exon_variant |
| 5 | 43049306 | 43051506 | 2200 | - | MOLF | regulatory_region_variant,intergenic_variant |
| 5 | 44411353 | 44417732 | 6379 | - | SPRET | regulatory_region_variant,intergenic_variant |
| 5 | 47438629 | 47446518 | 7889 | - | FVB,SJL,SWR | intergenic_variant |

| | | | | | | |
|---|---|---|---|---|---|---|
| 5 | 49902527 | 49906049 | 3522 | - | NOR | intergenic_variant |
| 5 | 51112189 | 51115758 | 3569 | - | KK,MA/My,MOLF,NOD,NZB,NZO,NZW,PJ,SJL,SWR,TH,WSB | intergenic_variant |
| 5 | 51270921 | 51274527 | 3606 | - | SPRET | intergenic_variant |
| 5 | 54489325 | 54490379 | 1054 | - | CAST | intergenic_variant |
| 5 | 55538755 | 55541368 | 2613 | - | SPRET | intergenic_variant |
| 5 | 56439299 | 56445610 | 6311 | - | SPRET | intergenic_variant |
| 5 | 57030181 | 57031350 | 1169 | - | SPRET | intergenic_variant |
| 5 | 57907257 | 57909241 | 1984 | Pcdh7 | NZO | intron_variant |
| 5 | 58557564 | 58564500 | 6936 | - | MOLF | intergenic_variant |
| 5 | 59123699 | 59130291 | 6592 | - | CAST,MOLF | regulatory_region_variant,intergenic_variant |
| 5 | 60961199 | 60966384 | 5185 | Gm42860 | WSB | non_coding_transcript_exon_variant |
| 5 | 61140576 | 61144205 | 3629 | - | CAST | intergenic_variant |
| 5 | 61257422 | 61258924 | 1502 | - | SPRET | intergenic_variant |
| 5 | 62482446 | 62483562 | 1116 | - | TH | intergenic_variant |
| 5 | 66863982 | 66865055 | 1073 | - | BALB,RH2,SEA | intergenic_variant |
| 5 | 69691173 | 69693192 | 2019 | Yipf7 | BTBR,PL | intron_variant |
| 5 | 72391577 | 72393076 | 1499 | Atp10d | CAST,SPRET | intron_variant |
| 5 | 74660593 | 74664699 | 4106 | Scfd2 | SPRET | intron_variant |
| 5 | 74661433 | 74668001 | 6568 | Scfd2 | SPRET | intron_variant |
| 5 | 77915142 | 77916417 | 1275 | - | SWR | intergenic_variant |
| 5 | 79798289 | 79803794 | 5505 | - | 129S1,BALB,C57L,C58J,LP,NZO,NZW,RH2,SEA | intergenic_variant |
| 5 | 79957976 | 79963061 | 5085 | - | SPRET | intergenic_variant |
| 5 | 80876635 | 80879382 | 2747 | - | CAST | intergenic_variant |
| 5 | 80876638 | 80877808 | 1170 | - | CAST | intergenic_variant |
| 5 | 82704992 | 82706226 | 1234 | - | SPRET | intergenic_variant |
| 5 | 83309858 | 83311365 | 1507 | - | KK,NZB,NZO,NZW | intergenic_variant |
| 5 | 84889909 | 84891992 | 2083 | - | SPRET | intergenic_variant |
| 5 | 86846824 | 86849044 | 2220 | - | CAST | intergenic_variant |
| 5 | 91057574 | 91064025 | 6451 | - | CAST | intergenic_variant |
| 5 | 98466498 | 98470220 | 3722 | - | SPRET | intergenic_variant |
| 5 | 102497453 | 102498865 | 1412 | - | AKR,CAST,CE,ILNJ2,MOLF,MRL,NOD,NOR,NZO,NZW,PJ,PL,RF,SMJ,SPRET,TH,WSB | intergenic_variant |
| 5 | 103819475 | 103821328 | 1853 | - | MOLF | regulatory_region_variant,intergenic_variant |
| 5 | 104736052 | 104956008 | 219956 | Gm8258 | AJ,BALB,BUB,C3H,CAST,CBA,DBA1J,DBA2J,FVB,ILNJ2,MOLF,NZB,NZO,RH2,SEA,SJL,SWR,TH,WSB | non_coding_transcript_variant,regulatory_region_variant |
| 5 | 104752780 | 104993577 | 240797 | Gm8258,Zfp951 | SWR | non_coding_transcript_variant,coding_transcript_variant,coding_sequence_variant,3_prime_UTR_variant,intron_variant,regulatory_region_variant |
| 5 | 104922219 | 105005445 | 83226 | Gm8258,Zfp951 | BUB,NZO,RH2 | non_coding_transcript_variant,coding_transcript_variant,coding_sequence_variant,3_prime_UTR_variant,intron_variant,regulatory_region_variant |
| 5 | 105252845 | 105349836 | 96991 | Gbp8,Gbp9,Gbp4,Gm3 | SPRET | 5_prime_UTR_variant,intron_variant,coding_sequence_variant,5_prime_UTR_variant,intron_variant,coding_transcript_variant,non_coding_transcript_exon_variant,intron_variant,regulatory_region_variant |
| 5 | 107393133 | 107396597 | 3464 | Tgfbr3 | B10J | intron_variant,regulatory_region_variant |
| 5 | 107393835 | 107396633 | 2798 | Tgfbr3 | B10J | intron_variant,regulatory_region_variant |
| 5 | 116733726 | 116738148 | 4422 | - | SPRET | intergenic_variant |
| 5 | 121692042 | 121698525 | 6483 | - | SPRET | regulatory_region_variant,intergenic_variant |
| 5 | 121696946 | 121698524 | 1578 | - | SPRET | intergenic_variant |
| 5 | 130436391 | 130438163 | 1772 | Caln1 | MOLF | intron_variant |
| 5 | 135652140 | 135657838 | 5698 | - | AKR,BUB,ILNJ2,MRL | regulatory_region_variant,intergenic_variant |
| 5 | 138543038 | 138548236 | 5198 | - | SPRET | intergenic_variant |
| 5 | 148897740 | 148899710 | 1970 | Gm15409,1810059H22 | MOLF | non_coding_transcript_exon_variant,intron_variant,upstream_gene_variant,intron_variant,non_coding_transcript_variant |
| 6 | 3057218 | 3060516 | 3298 | - | SPRET | intergenic_variant |
| 6 | 6734331 | 6736612 | 2281 | Gm5873 | CAST | non_coding_transcript_variant |
| 6 | 6734498 | 6735519 | 1021 | Gm5873 | CAST | non_coding_transcript_exon_variant |
| 6 | 7390550 | 7392300 | 1750 | - | CAST | intergenic_variant |
| 6 | 7498542 | 7505412 | 6870 | - | SPRET | intergenic_variant |
| 6 | 11962991 | 11969930 | 6939 | Phf14 | CAST | intron_variant |
| 6 | 14440710 | 14446958 | 6248 | - | BALB,C3H,CBA,SWR | intergenic_variant |
| 6 | 19973397 | 19974812 | 1415 | - | SMJ | intergenic_variant |
| 6 | 21749708 | 21753787 | 4079 | - | CAST | intergenic_variant |
| 6 | 23668693 | 23675103 | 6410 | Cadps2 | SPRET | intron_variant |
| 6 | 24289418 | 24296196 | 6778 | - | SPRET | intergenic_variant |
| 6 | 26612486 | 26615408 | 2922 | - | CAST | intergenic_variant |
| 6 | 41474414 | 41475870 | 1456 | Gm37841 | SPRET | downstream_gene_variant |
| 6 | 44077405 | 44079385 | 1980 | - | SPRET | intergenic_variant |
| 6 | 45510748 | 45513411 | 2663 | Cntnap2 | SPRET | intron_variant |
| 6 | 46840800 | 46842422 | 1622 | Cntnap2 | CAST | intron_variant |
| 6 | 48123499 | 48129475 | 5976 | Gm16630 | SPRET | intron_variant,non_coding_transcript_variant |
| 6 | 56394390 | 56395571 | 1181 | Pde1c | SPRET | intron_variant |
| 6 | 57008619 | 57010073 | 1454 | Vmn1r8 | 129S1,CAST,KK,MOLF,NZO,NZW,PJ | 5_prime_UTR_variant,intron_variant |
| 6 | 59244126 | 59245913 | 1787 | - | SPRET | intergenic_variant |
| 6 | 59244774 | 59245871 | 1097 | - | SPRET | intergenic_variant |
| 6 | 60411944 | 60416634 | 4690 | - | CE,ILNJ2,PJ,SPRET,WSB | intergenic_variant |
| 6 | 60412577 | 60415981 | 3404 | - | CE,ILNJ2,PJ,SPRET,WSB | intergenic_variant |
| 6 | 61265685 | 61266785 | 1100 | Ccser1 | SPRET | intron_variant |
| 6 | 62843551 | 62844839 | 1288 | - | SPRET | intergenic_variant |
| 6 | 65192192 | 65199191 | 6999 | Gm44058 | SPRET | upstream_gene_variant |
| 6 | 65444756 | 65451849 | 7093 | - | CAST,KK | intergenic_variant |
| 6 | 65991376 | 65993770 | 2394 | - | SPRET | intergenic_variant |
| 6 | 66214446 | 66216044 | 1598 | - | SPRET | intergenic_variant |
| 6 | 66704751 | 66712554 | 7803 | Vmn1r37,Gm44261 | SPRET | coding_transcript_variant,non_coding_transcript_variant |
| 6 | 68911189 | 68913104 | 1915 | - | AKR,C58J,CAST,KK,MOLF,MRL,NOD,NOR,PL,RF | intergenic_variant |
| 6 | 69430420 | 69437100 | 6680 | - | WSB | intergenic_variant |
| 6 | 69531060 | 69534630 | 3570 | - | SPRET,WSB | intergenic_variant |

| | | | | | | |
|---|---|---|---|---|---|---|
| 6 | 69533170 | 69540124 | 6954 | - | SPRET,WSB | intergenic_variant |
| 6 | 73983969 | 73987185 | 3216 | - | C58J,KK,MOLF,NOD,NOR | intergenic_variant |
| 6 | 74215683 | 74217099 | 1416 | - | SPRET | intergenic_variant |
| 6 | 74313657 | 74318708 | 5051 | Gm26179 | AKR,BALB,MRL,PL,RF,RH2,SEA | non_coding_transcript_variant |
| 6 | 74316227 | 74318714 | 2487 | Gm26179 | AKR,BALB,MRL,PL,RF,RH2,SEA | non_coding_transcript_variant |
| 6 | 74353659 | 74355739 | 2080 | - | MOLF | intergenic_variant |
| 6 | 74895470 | 74896647 | 1177 | - | C58J,KK,MOLF,NOD,NOR | intergenic_variant |
| 6 | 76222119 | 76227552 | 5433 | - | SPRET | intergenic_variant |
| 6 | 76775272 | 76777835 | 2563 | - | SPRET | intergenic_variant |
| 6 | 78494361 | 78495806 | 1445 | - | SPRET | intergenic_variant |
| 6 | 79284022 | 79290271 | 6249 | - | KK,MOLF,NZB,NZO,NZW,TH | intergenic_variant |
| 6 | 80616300 | 80617733 | 1433 | Lrrtm4 | MOLF | intron_variant |
| 6 | 88438946 | 88440691 | 1745 | - | CAST,MOLF | intergenic_variant |
| 6 | 90953713 | 90954966 | 1253 | Iqsec1 | CE,MOLF,NZB,NZW,PJ,SEA | intron_variant |
| 6 | 93246633 | 93248754 | 2121 | 9530026P05Rik | SPRET | non_coding_transcript_exon_variant,intron_variant,regulatory_region_variant |
| 6 | 95388800 | 95402030 | 13230 | - | CE | intergenic_variant |
| 6 | 95389653 | 95402030 | 12377 | - | CE,MOLF | intergenic_variant |
| 6 | 95389745 | 95399166 | 9421 | - | CE,MOLF | intergenic_variant |
| 6 | 96738420 | 96745236 | 6816 | - | SPRET | intergenic_variant |
| 6 | 101567362 | 101577273 | 9911 | - | CAST | regulatory_region_variant,intergenic_variant |
| 6 | 101569547 | 101574612 | 5065 | - | CAST | regulatory_region_variant,intergenic_variant |
| 6 | 103454245 | 103459049 | 4804 | - | MOLF,NOD | intergenic_variant |
| 6 | 103466663 | 103469161 | 2498 | - | BUB,CAST,MOLF,NOD,NOR,SPRET | intergenic_variant |
| 6 | 104047473 | 104048919 | 1446 | - | SPRET | intergenic_variant |
| 6 | 105190962 | 105192877 | 1915 | - | SPRET | intergenic_variant |
| 6 | 105921660 | 105923210 | 1550 | Cntn4 | AJ,AKR,BALB,BUB,C3H,CAST,CBA,CE,DBA1J,ILNJ2,KK,MOLF,MRL,NZB,NZO,NZW,PJ,PL,RF,RH2,SEA,SPRET,WSB | intron_variant |
| 6 | 107116490 | 107123420 | 6930 | - | MOLF | regulatory_region_variant,intergenic_variant |
| 6 | 107149074 | 107150240 | 1166 | - | SPRET | intergenic_variant |
| 6 | 108816190 | 108817234 | 1044 | Edem1 | SPRET | intron_variant |
| 6 | 110386966 | 110400737 | 13771 | - | 129S1,AJ,BALB,BTBR,BUB,C57L,CE,DBA1J,DBA2J,FVB,NZB,NZO,NZW,PJ,RH2,SEA,SJL,SMJ,SWR,TH,WSB | intergenic_variant |
| 6 | 110667166 | 110668616 | 1450 | Grm7 | SPRET | intron_variant |
| 6 | 116008530 | 116011992 | 3462 | Tmcc1 | AKR,C3H,CBA,ILNJ2,MOLF,MRL,PL,RF,SPRET | intron_variant,regulatory_region_variant |
| 6 | 121563750 | 121571237 | 7487 | - | SPRET | intergenic_variant |
| 6 | 121674145 | 121681013 | 6868 | - | CAST,CE,MOLF,SPRET | regulatory_region_variant,intergenic_variant |
| 6 | 122520256 | 122521546 | 1290 | - | CAST,SPRET | intergenic_variant |
| 6 | 123788349 | 123790499 | 2150 | Vmn2r24 | SPRET | intron_variant |
| 6 | 123948582 | 123951944 | 3362 | A330044P14Rik | CE | intron_variant,non_coding_transcript_variant,regulatory_region_variant |
| 6 | 124539172 | 124540920 | 1748 | Gm18840 | SWR | non_coding_transcript_exon_variant,intron_variant |
| 6 | 126432422 | 126439710 | 7288 | - | CAST,MOLF | intergenic_variant |
| 6 | 130096234 | 130098813 | 2579 | Klra8 | WSB | intron_variant |
| 6 | 130726785 | 130733401 | 6616 | - | NZW | intergenic_variant |
| 6 | 131217016 | 131218210 | 1194 | Klra2 | AJ,AKR,BALB,C3H,C57L,CAST,CBA,CE,DBA1J,DBA2J,FVB,ILNJ2,KK,LP,MA/My,MOLF,MRL,NZB,NZO,NZW,PL,RF,RH2,SEA,SJL,SPRET,TH,WSB | intron_variant |
| 6 | 132245958 | 132247805 | 1847 | - | 129S1,AJ,AKR,BALB,BUB,C3H,C57L,CBA,DBA1J,DBA2J,ILNJ2,MOLF,MRL,NZB,PL,RF,RH2,SEA,SMJ | intergenic_variant |
| 6 | 132246016 | 132252885 | 6869 | - | 129S1,AJ,AKR,BALB,BUB,C3H,C57L,CBA,DBA1J,DBA2J,ILNJ2,MRL,NZB,PL,RF,RH2,SEA | intergenic_variant |
| 6 | 132246036 | 132250290 | 4254 | - | 129S1,AJ,AKR,BALB,BUB,C3H,C57L,CBA,DBA1J,DBA2J,ILNJ2,MOLF,MRL,NZB,PL,RF,RH2,SEA,SMJ | intergenic_variant |
| 6 | 133857156 | 133865968 | 8812 | - | SPRET | regulatory_region_variant,intergenic_variant |
| 6 | 149273884 | 149275377 | 1493 | - | SPRET | intergenic_variant |
| 7 | 4042770 | 4044738 | 1968 | Lair1 | SPRET | intron_variant |
| 7 | 6033982 | 6035282 | 1300 | - | CAST,MOLF,SPRET | intergenic_variant |
| 7 | 7219420 | 7226136 | 6716 | - | 129S1,AJ,BALB,BTBR,BUB,C57L,C58J,CBA,CE,ILNJ2,KK,LP,NOD,NOR,NZB,NZO,NZW,PL,RF,SJL,SWR,TH | regulatory_region_variant,intergenic_variant |
| 7 | 10439866 | 10441707 | 1841 | Nlrp4b | CAST,MOLF,SPRET | intron_variant |
| 7 | 11422409 | 11424014 | 1605 | Gm18850 | SPRET | upstream_gene_variant |
| 7 | 15010191 | 15017172 | 6981 | - | AJ,AKR,BTBR,ILNJ2,LP,MRL,RH2,SJL,WSB | intergenic_variant |
| 7 | 25281833 | 25283818 | 1985 | Gm38569 | SPRET | non_coding_transcript_exon_variant |
| 7 | 28333588 | 28334645 | 1057 | Gm44618 | SPRET | intron_variant,non_coding_transcript_variant,regulatory_region_variant |
| 7 | 31064904 | 31071721 | 6817 | - | 129S1,C58J,LP,MA/My,MRL,RF,SJL | intergenic_variant |
| 7 | 35194979 | 35196027 | 1048 | Tdrd12,Gm18558 | SPRET | intron_variant,downstream_gene_variant |
| 7 | 39264150 | 39266047 | 1897 | - | SPRET | intergenic_variant |
| 7 | 40655095 | 40658853 | 3758 | - | NZB,NZO,SMJ | intergenic_variant |
| 7 | 40657738 | 40660189 | 2451 | - | NZB,NZO,SMJ | intergenic_variant |
| 7 | 41396821 | 41400185 | 3364 | Vmn2r-ps56 | CAST | intron_variant,non_coding_transcript_variant |
| 7 | 42013287 | 42018512 | 5225 | - | WSB | regulatory_region_variant,intergenic_variant |
| 7 | 42491400 | 42499276 | 7876 | Gm2381 | CAST | intron_variant |
| 7 | 42493721 | 42496964 | 3243 | Gm2381 | CAST | intron_variant |
| 7 | 43549729 | 43562860 | 13131 | - | CAST | intergenic_variant |
| 7 | 43551188 | 43564130 | 12942 | - | AJ,BALB,C57L,CE,NZB,NZO,NZW,RH2,SEA,SJL,SMJ,SPRET,TH | intergenic_variant |
| 7 | 43553200 | 43562118 | 8918 | - | AJ,BALB,C57L,CAST,CE,DBA1J,DBA2J,ILNJ2,MOLF,NZB,NZO,NZW,PJ,RH2,SEA,SJL,SMJ,SPRET,TH | intergenic_variant |
| 7 | 46844210 | 46845732 | 1522 | Mrgpra6 | CAST,ILNJ2,MOLF,NZW,PJ,SJL | intron_variant |
| 7 | 47456469 | 47460815 | 4346 | Mrgprc7-ps | C57L | non_coding_transcript_variant |
| 7 | 47456589 | 47462701 | 6112 | Mrgprc7-ps | WSB | non_coding_transcript_variant |
| 7 | 48796518 | 48797708 | 1190 | Nav2,9130015G15Rik | CAST,ILNJ2,MOLF,NZW,PJ,SJL,SPRET | intron_variant,intron_variant,non_coding_transcript_variant,regulatory_region_variant |

| | | | | | | |
|---|---|---|---|---|---|---|
| 7 | 50292081 | 50293712 | 1631 | Nell1 | SPRET | intron_variant |
| 7 | 52418974 | 52425626 | 6652 | - | MA/My,SJL,TH | intergenic_variant |
| 7 | 52499454 | 52505885 | 6431 | - | MOLF | intergenic_variant |
| 7 | 52654009 | 52655405 | 1396 | - | SPRET | intergenic_variant |
| 7 | 52703647 | 52705428 | 1781 | - | SPRET | intergenic_variant |
| 7 | 54520511 | 54522579 | 2068 | Luzp2 | CAST,CE,ILNJ2,KK,MA/My,MOLF,NZB,NZO,NZW,PJ,SJL,TH,WSB | intron_variant |
| 7 | 54573208 | 54579924 | 6716 | Luzp2 | SPRET | intron_variant |
| 7 | 56209546 | 56212372 | 2826 | - | ILNJ2,KK,MOLF,NZW,PJ,SJL,SPRET | intergenic_variant |
| 7 | 56211710 | 56212821 | 1111 | - | ILNJ2,KK,MOLF,NZW,PJ,SJL,SPRET | intergenic_variant |
| 7 | 61038869 | 61041003 | 2134 | - | SPRET | intergenic_variant |
| 7 | 61091250 | 61092636 | 1386 | - | SPRET | intergenic_variant |
| 7 | 62170643 | 62181858 | 11215 | - | CAST | intergenic_variant |
| 7 | 62170644 | 62180145 | 9501 | - | CAST | intergenic_variant |
| 7 | 62172694 | 62179806 | 7112 | - | CAST | intergenic_variant |
| 7 | 62357533 | 62360013 | 2480 | - | SPRET | intergenic_variant |
| 7 | 62528034 | 62529598 | 1564 | - | CAST,KK | intergenic_variant |
| 7 | 62528034 | 62529598 | 1564 | - | CAST,KK | intergenic_variant |
| 7 | 70703806 | 70708637 | 4831 | Gm42397 | MOLF,NZB,NZO | intron_variant,non_coding_transcript_variant |
| 7 | 78889244 | 78890877 | 1633 | - | CAST,MOLF | intergenic_variant |
| 7 | 83398998 | 83400013 | 1015 | - | CAST | intergenic_variant |
| 7 | 84217241 | 84219512 | 2271 | Ctxnd1 | SPRET | intron_variant |
| 7 | 85009842 | 85018314 | 8472 | - | CAST,MOLF | regulatory_region_variant,intergenic_variant |
| 7 | 85015785 | 85018705 | 2920 | - | CAST,MOLF | intergenic_variant |
| 7 | 85453808 | 85455451 | 1643 | - | 129S1,BTBR,C3H,C57L,CAST,CBA,DBA1J,DBA2J,ILNJ2,KK,MOLF,NZB,NZO,NZW,PJ,SEA,SMJ | intergenic_variant |
| 7 | 85843847 | 85850470 | 6623 | Vmn2r76 | SPRET | downstream_gene_variant |
| 7 | 92837854 | 92844680 | 6826 | - | SPRET | intergenic_variant |
| 7 | 92946885 | 92949741 | 2856 | Gm44947 | SPRET | intron_variant,non_coding_transcript_variant |
| 7 | 93393172 | 93396471 | 3299 | - | SPRET | intergenic_variant |
| 7 | 102088700 | 102090015 | 1315 | Rrm1 | CAST | upstream_gene_variant,regulatory_region_variant |
| 7 | 102448218 | 102451884 | 3666 | Or51f23 | KK,NOD,SJL | upstream_gene_variant |
| 7 | 103172128 | 103173703 | 1575 | - | CAST | intergenic_variant |
| 7 | 104386571 | 104387828 | 1257 | - | SPRET | intergenic_variant |
| 7 | 105108322 | 105109497 | 1175 | Cckbr | BUB,FVB,NZO,SJL,TH | intron_variant |
| 7 | 108272067 | 108273346 | 1279 | - | MOLF | intergenic_variant |
| 7 | 108684248 | 108686144 | 1896 | - | NOD,SMJ,SWR | intergenic_variant |
| 7 | 111549136 | 111550348 | 1212 | - | CAST,NZO | intergenic_variant |
| 7 | 113637287 | 113638547 | 1260 | Spon1 | CE | intron_variant |
| 7 | 120317018 | 120319582 | 2564 | Mosmo | SPRET | intron_variant,regulatory_region_variant |
| 7 | 127827009 | 127828315 | 1306 | - | 129S1,BTBR,C57L,ILNJ2,KK,PJ,TH | intergenic_variant |
| 7 | 139444998 | 139446489 | 1491 | Adgra1 | SPRET | intron_variant |
| 7 | 140737759 | 140739107 | 1348 | Ptdss2 | MOLF,NZW | downstream_gene_variant |
| 7 | 142356731 | 142360182 | 3451 | - | SPRET | intergenic_variant |
| 8 | 4062270 | 4072815 | 10545 | Gm45048,Gm7397 | CAST | non_coding_transcript_variant |
| 8 | 5060036 | 5065949 | 5913 | - | SPRET | intergenic_variant |
| 8 | 5757490 | 5763570 | 6080 | - | CAST | intergenic_variant |
| 8 | 12316761 | 12319175 | 2414 | - | SPRET | regulatory_region_variant,intergenic_variant |
| 8 | 14840490 | 14842811 | 2321 | Dlgap2 | MOLF,NZB | intron_variant |
| 8 | 22457958 | 22459491 | 1533 | Ccdc70 | SPRET | upstream_gene_variant |
| 8 | 27852617 | 27854657 | 2040 | - | CAST,MOLF,PL,SPRET | regulatory_region_variant,intergenic_variant |
| 8 | 31717299 | 31722030 | 4731 | Fut10 | SPRET | intron_variant |
| 8 | 31717827 | 31720251 | 2424 | Fut10 | 129S1,AJ,AKR,BALB,BUB,C3H,C57L,CBA,DBA1J,DBA2J,FVB,MA/My,MRL,NOD,NOR,NZW,PL,RF,RH2,SJL,SMJ,SWR,TH,WSB | intron_variant |
| 8 | 31718762 | 31721375 | 2613 | Fut10 | SPRET | intron_variant |
| 8 | 31842247 | 31844294 | 2047 | - | 129S1,AJ,AKR,BALB,BUB,C3H,C57L,CAST,CBA,DBA1J,DBA2J,FVB,ILNJ2,MA/My,MRL,NOD,NOR,NZW,PL,RF,RH2,SJL,SMJ,SWR,TH,WSB | intergenic_variant |
| 8 | 34001855 | 34004140 | 2285 | - | CAST | regulatory_region_variant,intergenic_variant |
| 8 | 34229229 | 34230265 | 1036 | Gtf2e2 | SPRET | intron_variant |
| 8 | 36223018 | 36224163 | 1145 | Gm35021 | SPRET | non_coding_transcript_exon_variant |
| 8 | 42985202 | 42986222 | 1020 | - | SPRET | intergenic_variant |
| 8 | 45559151 | 45563265 | 4114 | - | SPRET | regulatory_region_variant,intergenic_variant |
| 8 | 45560739 | 45564059 | 3320 | - | SPRET | regulatory_region_variant,intergenic_variant |
| 8 | 45659053 | 45663100 | 4047 | - | SPRET | intergenic_variant |
| 8 | 47825560 | 47830273 | 4713 | Stox2 | SPRET | intron_variant |
| 8 | 50306061 | 50312788 | 6727 | - | SPRET | intergenic_variant |
| 8 | 56477864 | 56481907 | 4043 | Glra3 | CAST,MOLF | intron_variant |
| 8 | 60552222 | 60553420 | 1198 | - | AKR,BALB,BTBR,BUB,C57L,CBA,CE,DBA1J,DBA2J,ILNJ2,NZO,PJ,RF,SEA | intergenic_variant |
| 8 | 60777498 | 60778552 | 1054 | - | 129S1,BALB,C57L,CE,DBA2J,KK,MOLF,NZO,SMJ | intergenic_variant |
| 8 | 63750890 | 63754269 | 3379 | Spock3 | KK | intron_variant,regulatory_region_variant |
| 8 | 63824277 | 63828907 | 4630 | - | SPRET | intergenic_variant |
| 8 | 70305999 | 70307277 | 1278 | Pbx4 | WSB | intron_variant,regulatory_region_variant |
| 8 | 72857181 | 72862168 | 4987 | Or1ab2,Or2z9 | CAST | 5_prime_UTR_variant,intron_variant,downstream_gene_variant,regulatory_region_variant |
| 8 | 74611075 | 74612869 | 1794 | - | AKR,C57L,CE,MA/My,MRL,NOD,NOR,PL | intergenic_variant |
| 8 | 74765149 | 74766946 | 1797 | - | 129S1,BALB,BUB,C3H,CBA,DBA1J,DBA2J,FVB,ILNJ2,NZB,NZO,RF,SJL,SMJ,SPRET,SWR | intergenic_variant |
| 8 | 74914508 | 74922436 | 7928 | - | MOLF | intergenic_variant |
| 8 | 74918166 | 74924364 | 6198 | - | MOLF | intergenic_variant |
| 8 | 75751557 | 75753773 | 2216 | Hmgxb4,Gm45822 | CAST | intron_variant,intron_variant,non_coding_transcript_variant |
| 8 | 76782073 | 76786555 | 4482 | Gm45740 | SPRET | intron_variant,non_coding_transcript_variant |
| 8 | 79502973 | 79513473 | 10500 | Gm45446 | SPRET | non_coding_transcript_variant |
| 8 | 81866485 | 81868076 | 1591 | - | 129S1,AKR,BALB,BTBR,BUB,C57L,MRL,NOD,NOR,NZW,PJ,PL,RF,SEA,SJL | intergenic_variant |

| | | | | | | |
|---|---|---|---|---|---|---|
| 8 | 86323117 | 86327431 | 4314 | Gm45424 | 129S1,AJ,AKR,BALB,BTBR,C3H,C57L,CBA,CE,DBA1J,DBA2J,FVB,ILNJ2,KK,MA/My,MRL,PJ,PL,RF,RH2,SEA,SMJ | non_coding_transcript_exon_variant,regulatory_region_variant |
| 8 | 102757027 | 102758889 | 1862 | - | MOLF | intergenic_variant |
| 8 | 109992956 | 109997479 | 4523 | D030068K23Rik | MOLF | intron_variant,non_coding_transcript_variant |
| 8 | 109995028 | 109998474 | 3446 | D030068K23Rik | MOLF | intron_variant,non_coding_transcript_variant |
| 8 | 111873705 | 111878000 | 4295 | - | MOLF,SPRET | intergenic_variant |
| 8 | 113108924 | 113110120 | 1196 | - | BALB,CAST,MOLF,NZW,RH2,SEA,WSB | intergenic_variant |
| 8 | 129986989 | 129988195 | 1206 | - | KK,MOLF | intergenic_variant |
| 9 | 4182475 | 4188975 | 6500 | - | SPRET | regulatory_region_variant,intergenic_variant |
| 9 | 5170075 | 5172538 | 2463 | - | 129S1,AKR,CAST,CBA,MOLF,RF,SPRET,WSB | intergenic_variant |
| 9 | 5219529 | 5220757 | 1228 | - | CAST | intergenic_variant |
| 9 | 6742772 | 6749083 | 6311 | - | SPRET | intergenic_variant |
| 9 | 10343171 | 10344915 | 1744 | Cntn5 | CAST,WSB | intron_variant |
| 9 | 12373256 | 12379757 | 6501 | - | AKR,MRL,RF | intergenic_variant |
| 9 | 16744355 | 16751608 | 7253 | - | SPRET | intergenic_variant |
| 9 | 18981126 | 18984336 | 3210 | - | CAST | intergenic_variant |
| 9 | 22750874 | 22755891 | 5017 | Bbs9 | CAST | intron_variant |
| 9 | 24448106 | 24450535 | 2429 | - | WSB | intergenic_variant |
| 9 | 25571467 | 25573345 | 1878 | - | SPRET | intergenic_variant |
| 9 | 26880719 | 26883133 | 2414 | - | SPRET | intergenic_variant |
| 9 | 29477078 | 29480014 | 2936 | Ntm | CAST | intron_variant |
| 9 | 36087757 | 36089478 | 1721 | - | C57L,MOLF,NZO | intergenic_variant |
| 9 | 36284050 | 36288363 | 4313 | Gm48401 | C57L,MOLF,NZO | non_coding_transcript_exon_variant |
| 9 | 37743456 | 37746224 | 2768 | - | SPRET | intergenic_variant |
| 9 | 38175868 | 38176869 | 1001 | - | SPRET | intergenic_variant |
| 9 | 41394654 | 41395746 | 1092 | Mir100hg,Gm48742 | CE | intron_variant,non_coding_transcript_variant,regulatory_region_variant |
| 9 | 50838195 | 50841830 | 3635 | Sik2 | CAST | intron_variant,regulatory_region_variant |
| 9 | 57134250 | 57135483 | 1233 | - | MA/My,MOLF | regulatory_region_variant,intergenic_variant |
| 9 | 60459259 | 60460659 | 1400 | - | SPRET | intergenic_variant |
| 9 | 62544943 | 62546224 | 1281 | - | SPRET | regulatory_region_variant,intergenic_variant |
| 9 | 63343627 | 63347857 | 4230 | Iqch,Gm16759 | SPRET | intron_variant,intron_variant,non_coding_transcript_variant |
| 9 | 63345195 | 63348518 | 3323 | Iqch,Gm16759 | SPRET | intron_variant,intron_variant,non_coding_transcript_variant |
| 9 | 65753564 | 65755165 | 1601 | Trip4 | 129S1,AJ,AKR,B10D2,B10J,BALB,BTBR,BUB,C3H,C57L,CAST,CE,DBA1J,DBA2J,FVB,ILNJ2,KK,LP,MA/My,MOLF,MRL,NOD,NOR,NZB,NZO,PJ,PL,RF,RH2,SEA,SJL,SMJ,SPRET,SWR,TH | intron_variant,regulatory_region_variant |
| 9 | 66201246 | 66206979 | 5733 | - | SPRET | regulatory_region_variant,intergenic_variant |
| 9 | 68398173 | 68399900 | 1727 | 4930502A04Rik | AJ,BALB,BUB,C3H,CBA,DBA1J,DBA2J,FVB,ILNJ2,MA/My,NOD,NOR,NZO,PJ,RH2,SEA,SJL,SWR | intron_variant,non_coding_transcript_variant |
| 9 | 68398180 | 68401241 | 3061 | 4930502A04Rik | AJ,BALB,BUB,C3H,CBA,DBA1J,DBA2J,FVB,ILNJ2,MA/My,NOD,NOR,NZO,PJ,RH2,SEA,SJL,SWR | intron_variant,non_coding_transcript_variant |
| 9 | 72554701 | 72557436 | 2735 | - | SPRET | intergenic_variant |
| 9 | 78223497 | 78232046 | 8549 | Gm8074 | LP,NZW | non_coding_transcript_variant,regulatory_region_variant |
| 9 | 78223955 | 78231589 | 7634 | Gm8074 | SPRET | non_coding_transcript_variant,regulatory_region_variant |
| 9 | 78225554 | 78229768 | 4214 | Gm8074 | C58J,ILNJ2,LP,MOLF,NZW,SPRET | non_coding_transcript_variant,regulatory_region_variant |
| 9 | 79747426 | 79748845 | 1419 | Filip1 | 129S1,BTBR,C57L,C58J,CAST,ILNJ2,LP,MOLF,NZW,SPRET | intron_variant |
| 9 | 82704989 | 82708784 | 3795 | - | MOLF | intergenic_variant |
| 9 | 82704989 | 82708784 | 3795 | - | MOLF | intergenic_variant |
| 9 | 93176693 | 93178651 | 1958 | - | SPRET | regulatory_region_variant,intergenic_variant |
| 9 | 93487793 | 93489352 | 1559 | - | SPRET | intergenic_variant |
| 9 | 96234706 | 96236448 | 1742 | Atp1b3 | MRL,NZO,NZW | intron_variant,regulatory_region_variant |
| 9 | 96445462 | 96446537 | 1075 | Rasa2 | SPRET | intron_variant,regulatory_region_variant |
| 9 | 97291281 | 97293672 | 2391 | - | SPRET | intergenic_variant |
| 9 | 99937370 | 99938480 | 1110 | - | NOD,NOR,RF | intergenic_variant |
| 9 | 100694239 | 100695593 | 1354 | Stag1 | SPRET | intron_variant |
| 9 | 103558721 | 103561611 | 2890 | Tmem108 | SPRET | intron_variant,regulatory_region_variant |
| 9 | 105377767 | 105379197 | 1430 | Atp2c1 | 129S1,AJ,AKR,BALB,BTBR,BUB,C3H,C57L,CAST,CBA,CE,DBA1J,DBA2J,FVB,ILNJ2,KK,MOLF,MRL,NOD,NOR,NZB,NZO,NZW,PJ,PL,RF,RH2,SEA,SJL,SMJ,SPRET,SWR,TH,WSB | intron_variant |
| 9 | 119030479 | 119031637 | 1158 | Slc22a13 | SPRET | intron_variant |
| 9 | 122966093 | 122967773 | 1680 | Exosc7,Gm47159 | SPRET | downstream_gene_variant,non_coding_transcript_exon_variant,regulatory_region_variant |
| X | 5389200 | 5390927 | 1727 | - | MOLF | intergenic_variant |
| X | 5776993 | 5778346 | 1353 | - | PL | intergenic_variant |
| X | 15111212 | 15116617 | 5405 | - | SPRET | intergenic_variant |
| X | 15546613 | 15548560 | 1947 | - | SPRET | intergenic_variant |
| X | 15761377 | 15764108 | 2731 | Gm23516 | SPRET | non_coding_transcript_variant |
| X | 17907364 | 17908592 | 1228 | - | SPRET | intergenic_variant |
| X | 18048494 | 18053160 | 4666 | Kdm6a | SPRET | intron_variant,regulatory_region_variant |
| X | 21977119 | 21981560 | 4441 | - | SPRET | intergenic_variant |
| X | 23795973 | 23802149 | 6176 | - | SPRET | intergenic_variant |
| X | 32098874 | 32101935 | 3061 | - | C57L,KK,MOLF,NZB,NZO | intergenic_variant |
| X | 36667556 | 36672972 | 5416 | Rhox4c | SPRET | downstream_gene_variant,regulatory_region_variant |
| X | 37164419 | 37180018 | 15599 | Rhox11,Rhox11-ps3 | MOLF | coding_transcript_variant,non_coding_transcript_variant,regulatory_region_variant |
| X | 41850018 | 41855478 | 5460 | Tenm1 | SPRET | intron_variant |
| X | 41850267 | 41858185 | 7918 | Tenm1 | SPRET | intron_variant |
| X | 45394537 | 45396367 | 1830 | 4930515L19Rik | SPRET | intron_variant,non_coding_transcript_variant |
| X | 49300724 | 49307041 | 6317 | - | SPRET | intergenic_variant |
| X | 49326255 | 49328497 | 2242 | - | SPRET | intergenic_variant |
| X | 49326874 | 49329758 | 2884 | - | SPRET | intergenic_variant |
| X | 49477875 | 49481143 | 3268 | - | CAST | intergenic_variant |
| X | 49792793 | 49794872 | 2079 | - | SPRET | intergenic_variant |

| | | | | | | |
|---|---|---|---|---|---|---|
| X | 52660376 | 52666971 | 6595 | - | SPRET | intergenic_variant |
| X | 53273610 | 53275586 | 1976 | - | MOLF | intergenic_variant |
| X | 60383059 | 60389459 | 6400 | - | MOLF | intergenic_variant |
| X | 60510545 | 60515900 | 5355 | - | MOLF | intergenic_variant |
| X | 60511250 | 60515900 | 4650 | - | MOLF | intergenic_variant |
| X | 60991508 | 60999332 | 7824 | - | MOLF | intergenic_variant |
| X | 63566842 | 63568877 | 2035 | - | CAST | intergenic_variant |
| X | 63927322 | 63934356 | 7034 | - | SPRET | intergenic_variant |
| X | 66074977 | 66081279 | 6302 | - | MOLF | intergenic_variant |
| X | 66293937 | 66301467 | 7530 | - | MOLF | intergenic_variant |
| X | 66295963 | 66301497 | 5534 | - | MOLF | intergenic_variant |
| X | 69158921 | 69160986 | 2065 | 1700111N16Rik | SPRET | intron_variant,non_coding_transcript_variant |
| X | 69163571 | 69167247 | 3676 | 1700111N16Rik | SPRET | intron_variant,non_coding_transcript_variant |
| X | 70365641 | 70372606 | 6965 | - | MOLF | intergenic_variant |
| X | 71411368 | 71415524 | 4156 | - | SPRET | intergenic_variant |
| X | 71412542 | 71415570 | 3028 | - | SPRET | intergenic_variant |
| X | 74871492 | 74872836 | 1344 | Pls3 | SPRET | intron_variant,upstream_gene_variant |
| X | 77145043 | 77146912 | 1869 | - | CAST | intergenic_variant |
| X | 77467042 | 77468611 | 1569 | - | SPRET | regulatory_region_variant,intergenic_variant |
| X | 77711691 | 77713356 | 1665 | - | CAST | intergenic_variant |
| X | 78781471 | 78788948 | 7477 | - | SPRET | intergenic_variant |
| X | 79077783 | 79079972 | 2189 | - | MOLF | intergenic_variant |
| X | 79466032 | 79467495 | 1463 | - | AJ,AKR,BALB,BUB,C3H,CBA,DBA1J,DBA2J,KK,MRL,NZO,NZW,PJ,PL,RF,RH2,SEA,SJL,SMJ,WSB | intergenic_variant |
| X | 80192500 | 80194507 | 2007 | - | MOLF | intergenic_variant |
| X | 81153490 | 81154758 | 1268 | - | SPRET | intergenic_variant |
| X | 81632739 | 81634435 | 1696 | - | SPRET | intergenic_variant |
| X | 84856307 | 84857687 | 1380 | - | SPRET | intergenic_variant |
| X | 85810889 | 85812894 | 2005 | Il1rapl1 | CAST,MOLF,SPRET | intron_variant,coding_sequence_variant,3_prime_UTR_variant |
| X | 85935275 | 85942236 | 6961 | Il1rapl1 | CAST | intron_variant |
| X | 88970816 | 88971941 | 1125 | - | SPRET | intergenic_variant |
| X | 89257273 | 89258662 | 1389 | - | SPRET | intergenic_variant |
| X | 89257932 | 89259948 | 2016 | - | SPRET | intergenic_variant |
| X | 90068246 | 90070365 | 2119 | - | CAST | intergenic_variant |
| X | 91583584 | 91584927 | 1343 | Mageb18 | SPRET | intron_variant |
| X | 97026840 | 97028794 | 1954 | - | CAST | intergenic_variant |
| X | 100517022 | 100518453 | 1431 | - | MOLF | intergenic_variant |
| X | 101821339 | 101824078 | 2739 | Gm9119 | SPRET | non_coding_transcript_variant |
| X | 112254733 | 112255979 | 1246 | Dach2 | SPRET | intron_variant |
| X | 112687331 | 112688581 | 1250 | Dach2 | SPRET | intron_variant |
| X | 113900762 | 113907578 | 6816 | - | AJ,BALB,C3H,CBA,MRL | regulatory_region_variant,intergenic_variant |
| X | 116358549 | 116359647 | 1098 | - | AJ,BALB,C3H,CBA,MOLF,MRL,RH2 | intergenic_variant |
| X | 116416311 | 116421144 | 4833 | - | SPRET | intergenic_variant |
| X | 117072618 | 117075203 | 2585 | - | AJ,BALB,C3H,CBA,MRL | intergenic_variant |
| X | 117073645 | 117076032 | 2387 | - | RH2 | intergenic_variant |
| X | 118651675 | 118653655 | 1980 | - | SPRET | intergenic_variant |
| X | 120830933 | 120838952 | 8019 | - | SPRET | intergenic_variant |
| X | 133073592 | 133074731 | 1139 | - | MOLF,SPRET | intergenic_variant |
| X | 134878192 | 134879683 | 1491 | - | SPRET | intergenic_variant |
| X | 137171207 | 137172360 | 1153 | Il1rapl2 | SPRET | intron_variant |
| X | 138483419 | 138487522 | 4103 | Rnf128 | MOLF | intron_variant,regulatory_region_variant |
| X | 140375135 | 140376164 | 1029 | Col4a5 | SPRET | intron_variant |
| X | 141240408 | 141241738 | 1330 | - | MOLF | intergenic_variant |
| X | 145678269 | 145680422 | 2153 | - | CAST,MOLF | intergenic_variant |
| X | 145679192 | 145680445 | 1253 | - | CAST,MOLF,NZB | intergenic_variant |
| X | 146251290 | 146253017 | 1727 | Lrch2 | SPRET | downstream_gene_variant |
| X | 154011370 | 154012671 | 1301 | - | SPRET | intergenic_variant |
| X | 156845317 | 156846830 | 1513 | - | MOLF | intergenic_variant |
| X | 168856708 | 168859029 | 2321 | - | C58J,MA/My,PL | coding_transcript_variant |

**Table S10. Lists of high impact DUPs present in all 39 or in the 35 classical inbred strains.** The chromosome, starting and ending position, size, gene symbol, strains with the variant allele, and the predicted effect for each DUP are shown.

| CHR | START | END | SIZE | SYMBOL | STRAINS with VARIANT SV ALLELE | CONSEQUENCE |
|---|---|---|---|---|---|---|
| 1 | 16203253 | 16203495 | 242 | Rdh10 | LP | feature_elongation,3_prime_UTR_variant |
| 1 | 16203288 | 16203531 | 243 | Rdh10 | LP | feature_elongation,3_prime_UTR_variant |
| 1 | 38906629 | 38906704 | 75 | Chst10 | BALB,BTBR,C57L,DBA1J,DBA2J,FVB,ILNJ2,LP,MA/My,NOR,NZB,NZO,PL,SEA,SMJ,TH | feature_elongation,3_prime_UTR_variant |
| 1 | 57455110 | 57455178 | 68 | Maip1 | SMJ,SWR | feature_elongation,3_prime_UTR_variant |
| 1 | 59045982 | 59046081 | 99 | C2cd6 | 129S1,BALB,BTBR,BUB,C57L,MRL,NOD,NOR,PL,RH2,SEA,SJL,SWR | feature_elongation,coding_sequence_variant |
| 1 | 59046167 | 59046219 | 52 | C2cd6 | 129S1,AJ,AKR,B10D2,B10J,BALB,BTBR,BUB,C3H,C57L,C58J,CBA,CE,DBA1J,DBA2J,FVB,ILNJ2,KK,LP,MA/My,MRL,NOD,NOR,NZB,NZO,NZW,PJ,PL,RF,RH2,SEA,SJL,SMJ,SWR,TH | feature_elongation,coding_sequence_variant |
| 1 | 60109927 | 60109991 | 64 | Wdr12 | CE,LP | feature_elongation,3_prime_UTR_variant |
| 1 | 135363018 | 135363153 | 135 | Nav1 | 129S1,AJ,AKR,BTBR,C3H,CE,DBA1J,DBA2J,ILNJ2,KK,NZW,RF,RH2,SJL,SWR,TH | feature_elongation,3_prime_UTR_variant |
| 1 | 164021232 | 164021285 | 53 | F5 | DBA1J,DBA2J,ILNJ2 | feature_elongation,coding_sequence_variant |
| 1 | 165156624 | 165156685 | 61 | Dcaf6 | FVB | feature_elongation,3_prime_UTR_variant |
| 1 | 170708348 | 170708502 | 154 | Dusp12 | 129S1,BUB,C57L,FVB,MRL,NOD,NOR,NZB,NZO,RF,RH2,SWR,TH | feature_elongation,coding_sequence_variant,intron_variant |
| 1 | 173556632 | 173556875 | 243 | Ifi207 | AJ,BTBR,BUB,C58J,CE,DBA1J,DBA2J,FVB,KK,PL,RH2,SJL,SWR,TH | feature_elongation,coding_sequence_variant |
| 1 | 173556674 | 173557009 | 335 | Ifi207 | BTBR,KK | feature_elongation,coding_sequence_variant |
| 1 | 173587714 | 173587773 | 59 | Ifi204 | 129S1,AKR,BALB,BUB,C3H,C57L,C58J,CBA,CE,DBA1J,DBA2J,ILNJ2,MRL,NOD,NOR,NZB,NZO,PJ,RF,SEA,SJL,SMJ,SWR | feature_elongation,coding_sequence_variant |
| 1 | 173733716 | 173733775 | 59 | Ifi211 | 129S1,AKR,BALB,BUB,C3H,C57L,C58J,CBA,CE,DBA1J,DBA2J,ILNJ2,MRL,NOD,NOR,NZB,NZO,PJ,RF,SEA,SJL,SMJ,SWR | feature_elongation,coding_sequence_variant |
| 1 | 176850818 | 176851010 | 192 | Akt3 | SMJ | feature_elongation,3_prime_UTR_variant |
| 1 | 180056382 | 180056449 | 67 | Psen2 | CBA,CE,DBA1J,DBA2J,ILNJ2,PJ,SEA | feature_elongation,coding_sequence_variant,intron_variant |
| 1 | 180797691 | 180797800 | 109 | Tmem63a | AJ,BUB,C58J,CE,LP,NZB,RH2,SMJ | feature_elongation,coding_sequence_variant,intron_variant |
| 1 | 181632622 | 181632745 | 123 | Dnah14 | TH | feature_elongation,coding_sequence_variant,intron_variant |
| 10 | 7484379 | 7484450 | 71 | Lrp11 | MA/My | feature_elongation,coding_sequence_variant,intron_variant |
| 10 | 7690552 | 7690617 | 65 | Ppil4 | BUB,C3H,CBA,DBA1J,DBA2J,FVB,LP,MA/My,MRL,NOD,NOR,NZB,NZO,NZW,SWR | feature_elongation,coding_sequence_variant,intron_variant |
| 10 | 12424155 | 12424346 | 191 | Utrn | CE,DBA1J,DBA2J,LP,MA/My,TH | feature_elongation,5_prime_UTR_variant |
| 10 | 85887153 | 85887237 | 84 | Fbxo7 | C58J,LP,NZB | feature_elongation,3_prime_UTR_variant |
| 10 | 127578252 | 127578302 | 50 | Zbtb39 | AJ,BALB,FVB,NOD,NOR,PL | feature_elongation,coding_sequence_variant |
| 10 | 127878231 | 127878366 | 135 | Naca | SMJ | feature_elongation,coding_sequence_variant |
| 11 | 5673029 | 5673126 | 97 | Urgcp | B10D2,B10J,BUB,C58J,CE,FVB,KK,NOR,NZO,SJL,SWR | feature_elongation,coding_sequence_variant,intron_variant |
| 11 | 5854541 | 5854596 | 55 | Gck | AJ,AKR,B10D2,B10J,BALB,BTBR,BUB,C3H,C57L,C58J,CE,FVB,KK,MA/My,MRL,NOD,NOR,NZB,NZO,PJ,PL,RF,SEA,SJL,SWR,TH | feature_elongation,coding_sequence_variant,intron_variant |
| 11 | 29104848 | 29104938 | 90 | Pnpt1 | AJ,C3H,DBA1J,DBA2J,ILNJ2,MA/My,MRL,NZB,NZO,PL | feature_elongation,coding_sequence_variant,intron_variant |
| 11 | 40600846 | 40601032 | 186 | Hmmr | B10D2,B10J,BTBR,C58J,LP,MA/My | feature_elongation,coding_sequence_variant |
| 11 | 40600944 | 40601006 | 62 | Hmmr | BTBR | feature_elongation,coding_sequence_variant |
| 11 | 60267491 | 60267547 | 56 | Drc3 | AJ,SEA | feature_elongation,3_prime_UTR_variant |
| 11 | 64892697 | 64892770 | 73 | Elac2 | CE,ILNJ2 | feature_elongation,3_prime_UTR_variant |
| 11 | 77028564 | 77028622 | 58 | Efcab5 | TH | feature_elongation,coding_sequence_variant |
| 11 | 83193308 | 83193597 | 289 | Ap2b1 | KK | feature_elongation,5_prime_UTR_variant |
| 11 | 94430679 | 94430736 | 57 | Rsad1 | 129S1,BALB,BTBR,C57L,MA/My,NOD,NOR,PL,SJL,SMJ | feature_elongation,3_prime_UTR_variant |
| 11 | 95990842 | 95990945 | 103 | Calcoco2 | AJ,AKR,BUB,C3H,CBA,RH2 | feature_elongation,coding_sequence_variant |
| 11 | 99276823 | 99276881 | 58 | Krt10 | AKR,RF,RH2,SJL | feature_elongation,coding_sequence_variant |
| 11 | 113551867 | 113551918 | 51 | Cog1 | AJ,C3H,CBA,TH | feature_elongation,coding_sequence_variant,intron_variant |
| 12 | 71041499 | 71041992 | 493 | Psma3 | NOD,NOR | feature_elongation,3_prime_UTR_variant |
| 12 | 87994165 | 87994227 | 62 | Eif1ad11 | 129S1,AJ,AKR,BTBR,BUB,C3H,C57L,CBA,DBA1J,DBA2J,FVB,ILNJ2,MRL,NOR,PJ,RF,SEA,SJL,TH | feature_elongation,coding_sequence_variant,3_prime_UTR_variant |
| 12 | 112742722 | 112743238 | 516 | Ahnak2 | 129S1,B10D2,B10J,BALB,BTBR,BUB,C57L,C58J,DBA1J,DBA2J,ILNJ2,KK,LP,MA/My,MRL,NOD,RH2,SJL | feature_elongation,coding_sequence_variant |
| 12 | 112747707 | 112748270 | 563 | Ahnak2 | AJ | feature_elongation,coding_sequence_variant |
| 12 | 112819377 | 112819492 | 115 | Gpr132 | FVB,SWR | feature_elongation,coding_sequence_variant,5_prime_UTR_variant,intron_variant |
| 13 | 93233210 | 93233307 | 97 | Cmya5 | FVB,MRL,NZW,RF | feature_elongation,coding_sequence_variant |
| 14 | 45240799 | 45240866 | 67 | Ptger2 | C57L,NZO,PJ,PL | feature_elongation,3_prime_UTR_variant |
| 14 | 49457120 | 49457221 | 101 | Ccdc198 | AKR,BALB,CE,DBA1J,MRL,NOD,NZO,PJ,SMJ,SWR | feature_elongation,3_prime_UTR_variant |
| 14 | 49457149 | 49457217 | 68 | Ccdc198 | 129S1,AJ,AKR,BALB,C3H,CBA,CE,DBA1J,DBA2J,MRL,NZB,NZO,NZW,PJ,RF | feature_elongation,3_prime_UTR_variant |
| 14 | 55817708 | 55817799 | 91 | Psme1 | BUB,SJL | feature_elongation,coding_sequence_variant,intron_variant |
| 14 | 74992400 | 74992656 | 256 | Lrch1 | BUB,KK | feature_elongation,3_prime_UTR_variant |
| 15 | 38492530 | 38493454 | 924 | Azin1 | NZB,NZW | feature_elongation,coding_sequence_variant,intron_variant |
| 15 | 78660539 | 78660604 | 65 | Card10 | BALB,BUB,C3H,CBA,CE,DBA1J,DBA2J,PJ,PL | feature_elongation,3_prime_UTR_variant |
| 15 | 98748903 | 98748988 | 85 | Kmt2d | ILNJ2,NOD,NOR | feature_elongation,coding_sequence_variant |
| 15 | 99840402 | 99840708 | 306 | Fam186a | 129S1,C57L | feature_elongation,coding_sequence_variant |
| 15 | 101855780 | 101856043 | 263 | Krt78 | KK,RF,TH | feature_elongation,coding_sequence_variant |
| 16 | 32573538 | 32574279 | 741 | Muc4 | AJ,BALB,BTBR,BUB,C3H,C57L,CBA,CE,DBA1J,FVB,ILNJ2,KK,MRL,PL,RH2,SEA,SJL,SMJ,TH | feature_elongation,coding_sequence_variant |

| | | | | | | |
|---|---|---|---|---|---|---|
| 17 | 25314766 | 25314818 | 52 | Ift140 | 129S1,AJ,AKR,B10D2,BALB,BTBR,BUB,C3H,C57L,CBA,CE,DBA1J,DBA2J,FVB,ILNJ2,KK,MA/My,MRL,NOD,NOR,NZB,NZO,RF,RH2,SEA,SJL,SWR,TH | feature_elongation,3_prime_UTR_variant |
| 17 | 28115623 | 28115802 | 179 | Bltp3a | B10D2,MA/My,MRL,NZB,RF,SEA | feature_elongation,3_prime_UTR_variant |
| 17 | 31867576 | 31867645 | 69 | U2af1 | FVB,ILNJ2,PJ,SJL | feature_elongation,coding_sequence_variant,intron_variant |
| 17 | 34163412 | 34163476 | 64 | Wdr46 | PJ,SJL,TH | feature_elongation,coding_sequence_variant,intron_variant |
| 17 | 35931729 | 35931858 | 129 | Muc21 | PJ | feature_elongation,coding_sequence_variant |
| 17 | 45831665 | 45831734 | 69 | Aars2 | AKR,BALB,C3H,CBA,CE,PJ,RH2,SEA | feature_elongation,3_prime_UTR_variant |
| 17 | 46570366 | 46570481 | 115 | Tjap1 | BTBR,C57L,DBA1J,KK,LP,MA/My,NZO,RF | feature_elongation,coding_sequence_variant,intron_variant |
| 18 | 10595058 | 10595113 | 55 | Esco1 | 129S1,C57L | feature_elongation,coding_sequence_variant |
| 18 | 34402282 | 34402454 | 172 | Apc | AJ,AKR,BALB,C58J,CBA,DBA2J,ILNJ2,RH2,SEA | feature_elongation,coding_sequence_variant,intron_variant |
| 18 | 61328158 | 61328214 | 56 | Slc26a2 | KK,NZO,NZW | feature_elongation,3_prime_UTR_variant |
| 18 | 67926738 | 67926823 | 85 | Seh1l | AJ,BALB,ILNJ2,RH2,SMJ | feature_elongation,3_prime_UTR_variant |
| 18 | 75218780 | 75218954 | 174 | Dym | AKR,ILNJ2,RF | feature_elongation,coding_sequence_variant |
| 19 | 6171474 | 6171571 | 97 | Snx15 | TH | feature_elongation,coding_sequence_variant,intron_variant |
| 19 | 17729334 | 17729440 | 106 | Pcsk5 | 129S1,AKR,BALB,BTBR,C57L,C58J,FVB,ILNJ2,LP,NOD,NOR,NZB,NZO,NZW,PJ,PL,RF,SEA,SWR,TH | feature_elongation,coding_sequence_variant,intron_variant |
| 19 | 36096560 | 36096627 | 67 | Ankrd1 | AKR,RF,SJL,TH | feature_elongation,coding_sequence_variant,intron_variant |
| 2 | 3426224 | 3426427 | 203 | Dclre1c | MA/My,NZO,NZW | feature_elongation,coding_sequence_variant,intron_variant,feature_elongation,5_prime_UTR_variant,intron_variant |
| 2 | 26909695 | 26909825 | 130 | Cacfd1 | AJ,AKR,B10D2,B10J,BALB,C58J,CBA,DBA1J,DBA2J,ILNJ2,LP,NOR,NZB,NZW,PL,RH2,SEA,SWR | feature_elongation,3_prime_UTR_variant |
| 2 | 52305964 | 52306086 | 122 | Arl5a | 129S1,AJ,AKR,BUB,C3H,CBA,CE,DBA1J,DBA2J,ILNJ2,MRL,NOD,NOR,NZW,PJ,PL,RF,SEA,SMJ,SWR,TH | feature_elongation,coding_sequence_variant,intron_variant |
| 2 | 85671181 | 85671351 | 170 | Or5ar1 | KK | feature_elongation,coding_sequence_variant,3_prime_UTR_variant |
| 2 | 88483738 | 88484597 | 859 | Or4p4 | C3H,CBA,MA/My,NZW | feature_elongation,3_prime_UTR_variant |
| 2 | 88483764 | 88484706 | 942 | Or4p4 | C3H,CBA,CE,MA/My,NZW | feature_elongation,3_prime_UTR_variant |
| 2 | 119849813 | 119849887 | 74 | Mapkbp1 | AJ,AKR,BALB,CE,DBA1J,DBA2J,FVB,ILNJ2,PJ,PL,RF,RH2,SEA,SJL,SMJ,SWR,TH | feature_elongation,coding_sequence_variant,intron_variant |
| 2 | 152236668 | 152236787 | 119 | Sox12 | AJ,C58J,NOD,NZB,NZW,RF | feature_elongation,3_prime_UTR_variant |
| 2 | 154359700 | 154360115 | 415 | Cbfa2t2 | FVB,SWR | feature_elongation,coding_sequence_variant,intron_variant |
| 2 | 165133300 | 165133412 | 112 | Elmo2 | KK | feature_elongation,3_prime_UTR_variant |
| 2 | 165626713 | 165626793 | 80 | Zmynd8 | AKR,BTBR,BUB,C3H,C58J,DBA1J,DBA2J,FVB,ILNJ2,MA/My,NZB,NZO,PJ,RF,SJL,SMJ,SWR,TH | feature_elongation,3_prime_UTR_variant |
| 2 | 167820390 | 167820584 | 194 | Ptpn1 | 129S1,C57L,NZB | feature_elongation,3_prime_UTR_variant |
| 2 | 168590700 | 168590826 | 126 | Sall4 | B10D2 | feature_elongation,3_prime_UTR_variant |
| 2 | 172204807 | 172204907 | 100 | Aurka | FVB,SJL | feature_elongation,coding_sequence_variant,intron_variant |
| 3 | 36128569 | 36129037 | 468 | Acad9 | CE,DBA1J,NZB,NZO,NZW | feature_elongation,coding_sequence_variant,intron_variant |
| 3 | 68951947 | 68951999 | 52 | Trim59 | 129S1,C57L,FVB,SJL,SWR,TH | feature_elongation,5_prime_UTR_variant |
| 3 | 88020123 | 88020351 | 228 | Iqgap3 | NZB,NZO | feature_elongation,coding_sequence_variant,intron_variant |
| 3 | 92479161 | 92479239 | 78 | Ivl | NOD,NOR | feature_elongation,coding_sequence_variant |
| 3 | 93110593 | 93110665 | 72 | Flg2 | 129S1,AJ,BTBR,C3H,C57L,CBA,CE,ILNJ2,PJ,PL,SEA,TH | feature_elongation,coding_sequence_variant |
| 3 | 93113206 | 93113444 | 238 | Flg2 | NOD,NOR | feature_elongation,coding_sequence_variant |
| 3 | 93237575 | 93238112 | 537 | Hrnr | AJ,C3H,CBA,CE,MA/My | feature_elongation,coding_sequence_variant |
| 3 | 93304211 | 93304317 | 106 | Rptn | BALB,ILNJ2,KK,NOD,NOR,NZB,NZO,PJ,PL,RH2,SEA,TH | feature_elongation,coding_sequence_variant |
| 3 | 105893684 | 105893889 | 205 | Ovgp1 | AJ,C3H,CBA | feature_elongation,coding_sequence_variant |
| 3 | 107017328 | 107017422 | 94 | Kcna2 | AJ,AKR,BALB,C3H,CBA,DBA1J,DBA2J,ILNJ2,MA/My,MRL,NOD,NOR,NZB,NZO,NZW,PL,RF,RH2,TH | feature_elongation,3_prime_UTR_variant |
| 3 | 154283843 | 154284368 | 525 | Tyw3 | RF | feature_elongation,3_prime_UTR_variant |
| 4 | 70139206 | 70139390 | 184 | Cdk5rap2 | KK | feature_elongation,3_prime_UTR_variant |
| 4 | 88745704 | 88745762 | 58 | Ifna6 | CE | feature_elongation,coding_sequence_variant |
| 4 | 120804557 | 120804710 | 153 | Zfp69 | ILNJ2 | feature_elongation,coding_sequence_variant |
| 4 | 131950649 | 131950703 | 54 | Gmeb1 | DBA1J,KK,NZB,SMJ | feature_elongation,3_prime_UTR_variant |
| 4 | 133692302 | 133692479 | 177 | Hmgn2 | 129S1,BALB,BTBR,C57L,DBA1J,DBA2J,ILNJ2,NOD,PL,SMJ | feature_elongation,3_prime_UTR_variant |
| 4 | 136081060 | 136081117 | 57 | Hnrnpr | 129S1,B10D2,B10J,C57L,DBA1J,KK,NZO,PJ,SEA | feature_elongation,3_prime_UTR_variant |
| 4 | 145311899 | 145312126 | 227 | Zfp268 | 129S1,AKR,BTBR,BUB,C3H,C57L,C58J,CBA,CE,DBA1J,KK,LP,MA/My,MRL,NOD,NZB,NZO,PJ,PL,RF,SEA,SJL,SMJ,SWR | feature_elongation,5_prime_UTR_variant,intron_variant |
| 4 | 147958725 | 147958787 | 62 | Mfn2 | 129S1,AKR,BTBR,BUB,C3H,C57L,C58J,CBA,CE,DBA1J,DBA2J,KK,LP,MA/My,MRL,NOD,NZB,NZO,NZW,PJ,PL,RF,SEA,SJL,SMJ,SWR,TH | feature_elongation,3_prime_UTR_variant |
| 4 | 147958746 | 147958810 | 64 | Mfn2 | 129S1,AKR,BTBR,BUB,C3H,C57L,C58J,CBA,CE,DBA1J,DBA2J,KK,LP,MA/My,MRL,NOD,NZB,NZO,NZW,PJ,PL,RF,SEA,SJL,SMJ,SWR,TH | feature_elongation,3_prime_UTR_variant |
| 4 | 152101384 | 152101475 | 91 | Klhl21 | 129S1 | feature_elongation,3_prime_UTR_variant |
| 4 | 152101429 | 152101487 | 58 | Klhl21 | 129S1 | feature_elongation,3_prime_UTR_variant |
| 4 | 156302483 | 156302548 | 65 | Perm1 | AJ,BALB,C3H,C57L,C58J,DBA1J,DBA2J,MA/My,MRL,NOD,NOR,NZO,PL,RH2,SJL | feature_elongation,coding_sequence_variant |

| | | | | | | |
|---|---|---|---|---|---|---|
| 4 | 156302510 | 156302629 | 119 | Perm1 | AJ,BALB,C3H,C57L,C58J,DBA1J,DBA2J,ILNJ2,MA/My,MRL,NZO,PL,RH2,SJL,SMJ | feature_elongation,coding_sequence_variant |
| 5 | 8751924 | 8752102 | 178 | Abcb1a | BUB,NOD,NOR | feature_elongation,coding_sequence_variant,intron_variant |
| 5 | 76632060 | 76632135 | 75 | Exoc1l | AJ,AKR,BUB,FVB,PJ,PL,SJL,SMJ | feature_elongation,5_prime_UTR_variant |
| 5 | 115793879 | 115794000 | 121 | Bicdl1 | 129S1,AKR,CE,DBA1J,DBA2J,KK,MRL,NZB,NZO,RH2,SEA | feature_elongation,coding_sequence_variant,intron_variant |
| 5 | 118330475 | 118330533 | 58 | Rnft2 | 129S1,AJ,AKR,BALB,BUB,C3H,C57L,CBA,CE,DBA1J,DBA2J,ILNJ2,KK,NOR,NZB,NZO,PJ,PL,RH2,SEA,SJL,SMJ,TH | feature_elongation,3_prime_UTR_variant |
| 5 | 123710135 | 123710990 | 855 | Vps33a | 129S1,AJ,BALB,C57L,KK,MRL,NZW,PJ,RH2,SEA,SJL | feature_elongation,coding_sequence_variant,intron_variant |
| 5 | 125463413 | 125463640 | 227 | Ubc | SJL | feature_elongation,coding_sequence_variant |
| 5 | 129062020 | 129062140 | 120 | Stx2 | AJ,BALB,C58J,FVB,LP,MRL,NOD,NOR,NZB,NZO,NZW,RH2,SEA,SMJ,SWR,TH | feature_elongation,3_prime_UTR_variant |
| 5 | 137461977 | 137462074 | 97 | Zan | 129S1,BUB,LP,RF,SWR | feature_elongation,coding_sequence_variant |
| 6 | 5008455 | 5008608 | 153 | Ppp1r9a | PJ | feature_elongation,3_prime_UTR_variant |
| 6 | 40844545 | 40844663 | 118 | Mgam2-ps | BUB,FVB,LP,SJL,SWR,TH | feature_elongation,coding_sequence_variant |
| 6 | 84166818 | 84166905 | 87 | Dysf | NZB,NZO | feature_elongation,coding_sequence_variant,intron_variant |
| 6 | 90635724 | 90635820 | 96 | Iqsec1 | CE,PJ,SEA | feature_elongation,3_prime_UTR_variant |
| 6 | 92151525 | 92151724 | 199 | Mrps25 | CE,PJ,SEA | feature_elongation,3_prime_UTR_variant |
| 6 | 132549054 | 132549175 | 121 | Prh1 | AJ,AKR,BALB,BUB,C3H,C57L,CBA,DBA1J,DBA2J,ILNJ2,MRL,NZB,PL,RF,RH2,SEA | feature_elongation,coding_sequence_variant |
| 6 | 133083355 | 133083457 | 102 | Smim10l1 | 129S1,C58J,FVB,KK,LP,NOD,NOR,NZW,SJL,SMJ,TH | feature_elongation,3_prime_UTR_variant |
| 7 | 30451581 | 30451632 | 51 | Sbsn | DBA1J,DBA2J,NZB,NZO,SMJ | feature_elongation,coding_sequence_variant |
| 7 | 30452432 | 30452512 | 80 | Sbsn | NZW | feature_elongation,coding_sequence_variant |
| 7 | 85983550 | 85983603 | 53 | Or14a260 | 129S1,BTBR,C3H,C57L,CBA,DBA2J,ILNJ2,KK,NZB,NZO,NZW,PJ | feature_elongation,3_prime_UTR_variant |
| 7 | 98917972 | 98918107 | 135 | Map6 | C3H,CBA,DBA1J,DBA2J,ILNJ2,PJ,SEA | feature_elongation,coding_sequence_variant |
| 7 | 120545515 | 120546332 | 817 | Polr3e | C57L | feature_elongation,3_prime_UTR_variant |
| 7 | 140507818 | 140507872 | 54 | Nlrp6 | ILNJ2 | feature_elongation,3_prime_UTR_variant |
| 7 | 140681232 | 140681354 | 122 | Ano9 | NZW | feature_elongation,3_prime_UTR_variant |
| 7 | 140681253 | 140681314 | 61 | Ano9 | NZW | feature_elongation,3_prime_UTR_variant |
| 7 | 141224990 | 141225499 | 509 | Muc6 | NZW | feature_elongation,coding_sequence_variant |
| 7 | 141363058 | 141363152 | 94 | Muc5ac | 129S1,AKR,BALB,BTBR,BUB,C57L,C58J,CBA,FVB,KK,MA/My,MRL,NZB,PL,RF,RH2,SEA,SMJ,SWR | feature_elongation,coding_sequence_variant |
| 7 | 141412183 | 141413069 | 886 | Muc5b | SEA | feature_elongation,coding_sequence_variant |
| 7 | 141743689 | 141743890 | 201 | Gm10153 | BALB,MRL,RF,RH2,SEA | feature_elongation,coding_sequence_variant |
| 7 | 141755477 | 141755573 | 96 | Krtap5-3 | NZW | feature_elongation,coding_sequence_variant |
| 8 | 12917049 | 12917138 | 89 | Atp11a | C57L,ILNJ2,NZB,NZW,PJ,PL,SEA,SMJ,TH | feature_elongation,3_prime_UTR_variant |
| 8 | 13068280 | 13068376 | 96 | Mcf2l | 129S1,C57L,CE,ILNJ2,LP,NZB,NZO,NZW,PJ,PL,SEA,SMJ,TH | feature_elongation,3_prime_UTR_variant |
| 8 | 34110054 | 34110133 | 79 | Ubxn8 | 129S1,AJ,AKR,BALB,BUB,C3H,C57L,CBA,CE,DBA1J,DBA2J,FVB,MA/My,MRL,NOD,NOR,PL,RF,RH2,SJL,SMJ,SWR,TH | feature_elongation,3_prime_UTR_variant |
| 8 | 65194683 | 65194734 | 51 | Klhl2 | PJ,SEA | feature_elongation,3_prime_UTR_variant |
| 8 | 70360084 | 70360142 | 58 | Gatad2a | BALB,C57L,CBA,DBA2J,FVB,KK,RF,SJL,SMJ,TH | feature_elongation,3_prime_UTR_variant |
| 8 | 73005431 | 73005609 | 178 | Ap1m1 | 129S1,AKR,BALB,BUB,C3H,C57L,C58J,CBA,CE,DBA1J,DBA2J,FVB,ILNJ2,KK,MA/My,MRL,NZB,NZO,NZW,PJ,PL,RF,SEA,SJL,SMJ,SWR,TH | feature_elongation,coding_sequence_variant,intron_variant |
| 8 | 80366576 | 80366685 | 109 | Otud4 | CE | feature_elongation,coding_sequence_variant |
| 8 | 104991352 | 104991447 | 95 | Cklf | KK,PL | feature_elongation,3_prime_UTR_variant |
| 8 | 105076826 | 105076892 | 66 | Cmtm4 | KK,PL | feature_elongation,3_prime_UTR_variant |
| 8 | 125756560 | 125756698 | 138 | Tsnax | NZB | feature_elongation,3_prime_UTR_variant |
| 8 | 125756597 | 125756708 | 111 | Tsnax | NZB | feature_elongation,3_prime_UTR_variant |
| 9 | 96962198 | 96962273 | 75 | Slc25a36 | CE,NZB | feature_elongation,coding_sequence_variant,intron_variant |
| 9 | 99550276 | 99550348 | 72 | Dzip1l | NOD,NOR | feature_elongation,3_prime_UTR_variant |
| 9 | 110215211 | 110215277 | 66 | Ptpn23 | CBA | feature_elongation,coding_sequence_variant,intron_variant |
| 9 | 110815523 | 110815892 | 369 | Rtp3 | NZW | feature_elongation,coding_sequence_variant,intron_variant |
| 9 | 119607815 | 119607979 | 164 | Scn11a | BTBR,MRL | feature_elongation,coding_sequence_variant,intron_variant |
| X | 74423514 | 74423571 | 57 | F8 | 129S1,C57L,NZB | feature_elongation,5_prime_UTR_variant |
| X | 82530610 | 82530797 | 187 | Tsga8 | AJ,AKR,BALB,BUB,C3H,CBA,CE,DBA1J,DBA2J,ILNJ2,KK,MA/My,MRL,NZB,PJ,PL,RF,RH2,SEA,SMJ | feature_elongation,coding_sequence_variant |
| X | 82530695 | 82530874 | 179 | Tsga8 | MA/My,NZB | feature_elongation,coding_sequence_variant |
| X | 168772477 | 168772566 | 89 | Mid1 | B10D2,B10J,NOD,NOR,SJL,SMJ | feature_elongation,3_prime_UTR_variant |

| CHR | START | END | SIZE | SYMBOL | STRAINS with VARIANT SV ALLELE | CONSEQUENCE |
|---|---|---|---|---|---|---|
| 1 | 110816417 | 110831919 | 15502 | Cdh19 | NOD,NOR | feature_elongation,coding_sequence_variant,3_prime_UTR_variant,intron_variant |
| 1 | 117709473 | 118024554 | 315081 | Gm7145 | SWR | transcript_amplification |
| 1 | 151060176 | 151087731 | 27555 | 3110040M04Rik,Gm8947,Gm47996,Gm47995 | B10D2,B10J | transcript_amplification |
| 1 | 159901795 | 159962954 | 61159 | 4930523C07Rik | C3H | transcript_amplification |
| 1 | 173591669 | 173806055 | 214386 | Ifi211,Ifi203,Mndal | PJ,RF | transcript_amplification |
| 1 | 173737663 | 173806055 | 68392 | Ifi203 | AKR,BTBR,BUB,C58J,CBA,CE,DBA1J,DBA2J,ILNJ2,KK,NZB,NZO,PJ,PL,RF,SEA,SMJ,SWR,TH | transcript_amplification |
| 10 | 33340355 | 33344093 | 3738 | Trdn | KK | feature_elongation,coding_sequence_variant,intron_variant |
| 10 | 33340488 | 33342351 | 1863 | Trdn | KK | feature_elongation,coding_sequence_variant,intron_variant |
| 10 | 33342164 | 33344092 | 1928 | Trdn | KK | feature_elongation,coding_sequence_variant,intron_variant |
| 10 | 58056666 | 58061547 | 4881 | AW822073 | NZB | transcript_amplification |
| 10 | 58057567 | 58062450 | 4883 | AW822073 | 129S1,NOR | transcript_amplification |
| 10 | 93995634 | 94018245 | 22611 | Nr2c1 | ILNJ2 | feature_elongation,coding_sequence_variant,intron_variant |
| 11 | 99564024 | 99590225 | 26201 | Gm14190 | BALB,BUB,CBA,CE,MRL,NZB,PL,SEA | transcript_amplification |
| 11 | 107751339 | 107860972 | 109633 | Cacng5 | AKR | transcript_amplification |
| 12 | 31352598 | 31363544 | 10946 | Lamb1 | TH | feature_elongation,coding_sequence_variant,intron_variant |
| 12 | 37221052 | 37375832 | 154780 | Agmo | CBA | transcript_amplification |
| 12 | 108810263 | 108823579 | 13316 | Slc25a47 | AKR | transcript_amplification |
| 12 | 112742279 | 112744355 | 2076 | Ahnak2 | C58J,RH2 | feature_elongation,coding_sequence_variant |
| 12 | 112742280 | 112748427 | 6147 | Ahnak2 | C3H,PL,SEA | feature_elongation,coding_sequence_variant |
| 12 | 112743715 | 112748380 | 4665 | Ahnak2 | NZO | feature_elongation,coding_sequence_variant |
| 12 | 112744798 | 112748428 | 3630 | Ahnak2 | AJ | feature_elongation,coding_sequence_variant |
| 12 | 112745836 | 112748428 | 2592 | Ahnak2 | 129S1,B10J,BALB,BUB,DBA2J,NZO | feature_elongation,coding_sequence_variant |
| 13 | 66215801 | 66596564 | 380763 | Zfp998,Zfp999,Zfp997 | 129S1,AJ,AKR,BUB,CBA,FVB,KK,MRL,NOD,NZB,RF,SJL,TH | transcript_amplification |
| 13 | 113228363 | 113238135 | 9772 | Gzma | KK,NZB,NZW,RF | transcript_amplification |
| 13 | 120368879 | 120375129 | 6250 | Gm21761 | BTBR,C3H,C57L | transcript_amplification |
| 13 | 120368962 | 120375098 | 6136 | Gm21761 | 129S1,BTBR,C57L,KK,MRL | transcript_amplification |
| 13 | 120486361 | 120499016 | 12655 | Anxa2r2 | 129S1 | transcript_amplification |
| 14 | 15923561 | 16410495 | 486934 | Gm16440,Gm21103,Gm3476,Gm21560,Gm3594,Gm3411 | KK | transcript_amplification |
| 14 | 43765978 | 43855985 | 90007 | Gm10375,Gm5799 | C57L,NZO,PJ,PL | transcript_amplification |
| 14 | 52074876 | 52100002 | 25126 | Ear6 | 129S1,AJ,AKR,BALB,C3H,C57L,CBA,CE,KK,MRL,NZB,NZO,NZW,PJ,PL,RF,RH2,SEA,TH | transcript_amplification |
| 14 | 52076260 | 52101409 | 25149 | Ear6 | 129S1,AJ,AKR,BALB,C3H,C57L,CBA,CE,KK,MRL,NZB,NZO,NZW,PJ,PL,RF,RH2,SEA,TH | transcript_amplification |
| 15 | 3298353 | 3336085 | 37732 | Selenop,Ccdc152 | LP | transcript_amplification |
| 15 | 76922487 | 76946076 | 23589 | Apol6 | MA/My | transcript_amplification |
| 15 | 91730541 | 91734154 | 3613 | Smgc | CBA,PJ | feature_elongation,coding_sequence_variant,intron_variant |
| 16 | 88666970 | 88671824 | 4854 | Krtap19-2 | C58J | transcript_amplification |
| 17 | 7596967 | 7598125 | 1158 | Tcp10a | KK | feature_elongation,coding_sequence_variant,intron_variant |
| 17 | 30666348 | 31143185 | 476837 | Glo1,Dnah8,Btbd9 | 129S1,AJ,AKR,B10D2,C3H,C57L,CBA,CE,DBA1J,DBA2J,ILNJ2,MRL,PJ,RH2,SEA | transcript_amplification |
| 17 | 30725784 | 31111284 | 385500 | Glo1,Dnah8,Btbd9 | RF | transcript_amplification |
| 17 | 31217564 | 31250037 | 32473 | Umodl1 | PJ | transcript_amplification |
| 17 | 31441857 | 31499856 | 57999 | Rsph1 | C58J,ILNJ2,NZB,NZO,NZW | transcript_amplification |
| 17 | 31473829 | 31562278 | 88449 | Rsph1 | KK,NOR | transcript_amplification |
| 17 | 36387349 | 36449718 | 62369 | H2-T25,Gm19684,H2-Bl | BTBR | transcript_amplification |
| 18 | 61592541 | 61699963 | 107422 | Arhgef37 | NZW,SMJ | transcript_amplification |
| 2 | 32135487 | 32183245 | 47758 | Uck1,Swi5 | AKR | transcript_amplification |
| 2 | 118533766 | 118702353 | 168587 | Disp2,Ccdc9b,Plcb2,Knstrn,Phgr1,Inafm2 | BUB | transcript_amplification |
| 2 | 176402896 | 176769408 | 366512 | Gm14296,Gm14419,Gm14305,Gm14295,Gm14408 | PJ | transcript_amplification |
| 4 | 41883740 | 42247346 | 363606 | Il11ra3,Ccl27al,Ccl21f,Ccl21d,Spata31f1d,Gm20878 | NOD | transcript_amplification |
| 4 | 41886613 | 42247346 | 360733 | Il11ra3,Ccl27al,Ccl21f,Ccl21d,Spata31f1d,Gm20878 | AKR,NOR,NZB,PJ | transcript_amplification |
| 4 | 62007380 | 62077857 | 70477 | Mup21 | 129S1,AJ,AKR,BALB,BUB,C3H,C57L,CE,DBA1J,DBA2J,ILNJ2,MA/My,MRL,NOR,NZB,NZO,NZW,PJ,RF,RH2,SEA,SJL,SWR | transcript_amplification |
| 4 | 62413147 | 62442066 | 28919 | Alad,Hdhd3 | 129S1,AJ,AKR,B10D2,B10J,BALB,BUB,C3H,C57L,C58J,CBA,CE,DBA1J,DBA2J,FVB,ILNJ2,LP,MA/My,MRL,NOD,NOR,NZB,NZO,NZW,PJ,PL,RF,RH2,SEA,SJL,SWR,TH | transcript_amplification |
| 4 | 124746849 | 124899068 | 152219 | Cdca8,Epha10,Maneal,Airim | NOR | transcript_amplification |
| 4 | 143660682 | 143693357 | 32675 | Pramel16 | DBA2J | transcript_amplification |
| 4 | 143783909 | 143868185 | 84276 | Pramel4,Gm13102 | 129S1,BTBR,BUB,KK,MA/My,PJ,SJL,SMJ | transcript_amplification |
| 4 | 143820689 | 143852437 | 31748 | Gm13102 | C58J | transcript_amplification |
| 4 | 145158084 | 145310836 | 152752 | Zfp990 | DBA1J,DBA2J | transcript_amplification |
| 4 | 145159161 | 145311914 | 152753 | Zfp990 | DBA2J,LP | transcript_amplification |
| 5 | 11448768 | 11947739 | 498971 | Gm6460,Gm8906,Gm8897,Gm8922,Gm8926 | TH | transcript_amplification |

| | | | | | | |
|---|---|---|---|---|---|---|
| 5 | 87726730 | 87733196 | 6466 | Sult1e1 | BALB | feature_elongation,coding_sequence_variant,intron_variant |
| 5 | 138471917 | 138565066 | 93149 | 1700123K08Rik | AJ,RH2,SJL | transcript_amplification |
| 6 | 130020546 | 130239164 | 218618 | Klra8,Klra7,Klra9,Klra4 | KK,LP,NOD,NOR,NZB,NZW,SJL,SWR,TH | transcript_amplification |
| 7 | 9455229 | 9817901 | 362672 | Vmn2r50,Vmn2r49,Vmn2r48,Vmn2r47 | 129S1,BTBR,LP,NZB | transcript_amplification |
| 7 | 24419004 | 24436181 | 17177 | Lypd11 | BUB,C58J,MRL,PL | transcript_amplification |
| 7 | 30618519 | 30637774 | 19255 | Hamp2 | CE | transcript_amplification |
| 7 | 32516946 | 32869822 | 352876 | Scgb2b17,Scgb2b15,Scgb1b17,Scgb1b15 | NZB,NZW | transcript_amplification |
| 7 | 41685135 | 42087713 | 402578 | Vmn2r61,Vmn2r60 | ILNJ2,PJ | transcript_amplification |
| 7 | 101206784 | 101232156 | 25372 | Art2b | NZO | transcript_amplification |
| 7 | 104419177 | 104421446 | 2269 | Usp17le | KK | feature_elongation,coding_sequence_variant,intron_variant |
| 7 | 104419767 | 104424329 | 4562 | Usp17le | AJ,AKR,B10D2,C57L,CBA,DBA1J,LP,NOR,NZB,PJ,RF | feature_elongation,coding_sequence_variant,intron_variant |
| 7 | 108266621 | 108294904 | 28283 | Or5p81 | CE,KK,NOD,SMJ,SWR | transcript_amplification |
| 7 | 130194306 | 130269962 | 75656 | Tacc2 | B10D2 | feature_elongation,coding_sequence_variant,intron_variant |
| 7 | 139998710 | 140028824 | 30114 | Or13a21 | NZW,PL | transcript_amplification |
| 8 | 107142830 | 107147378 | 4548 | Zfp90 | MRL | feature_elongation,coding_sequence_variant,intron_variant |
| 9 | 18549616 | 18550919 | 1303 | Muc16 | NZB | feature_elongation,coding_sequence_variant |
| 9 | 56607833 | 56652136 | 44303 | Lingo1 | NOR | feature_elongation,5_prime_UTR_variant,intron_variant |
| 9 | 113809113 | 113849909 | 40796 | Fbxl2 | PL | feature_elongation,coding_sequence_variant,intron_variant |
| X | 7623820 | 7644895 | 21075 | Tfe3 | BALB | transcript_amplification |

| CHR | START | END | SIZE | SYMBOL | STRAINS with VARIANT SV ALLELE | CONSEQUENCE |
|---|---|---|---|---|---|---|
| 1 | 5163646 | 5163706 | 60 | Atp6v1h | CAST | feature_elongation,coding_sequence_variant,intron_variant |
| 1 | 16151971 | 16152208 | 237 | 4930444P10Rik | WSB | feature_elongation,5_prime_UTR_variant |
| 1 | 16203214 | 16203351 | 137 | Rdh10 | CAST,LP,MOLF,SPRET | feature_elongation,3_prime_UTR_variant |
| 1 | 16203288 | 16203531 | 243 | Rdh10 | LP | feature_elongation,3_prime_UTR_variant |
| 1 | 33822868 | 33822951 | 83 | Zfp451 | CAST,MOLF | feature_elongation,coding_sequence_variant,intron_variant |
| 1 | 52223506 | 52223587 | 81 | Gls | SPRET | feature_elongation,3_prime_UTR_variant |
| 1 | 54801370 | 54801453 | 83 | Ankrd44 | SPRET | feature_elongation,coding_sequence_variant,intron_variant |
| 1 | 57455110 | 57455178 | 68 | Maip1 | SMJ,SWR | feature_elongation,3_prime_UTR_variant |
| 1 | 59045982 | 59046081 | 99 | C2cd6 | 129S1,BALB,BTBR,BUB,C57L,MRL,NOD,NOR,PL,RH2,SEA,SJL,SWR,WSB | feature_elongation,coding_sequence_variant |
| 1 | 59046167 | 59046219 | 52 | C2cd6 | 129S1,AJ,AKR,B10D2,B10J,BALB,BTBR,BUB,C3H,C57L,C58J,CAST,CBA,CE,DBA1J,DBA2J,FVB,ILNJ2,KK,LP,MA/My,MOLF,MRL,NOD,NOR,NZB,NZO,NZW,PJ,PL,RF,RH2,SEA,SJL,SMJ,SWR,TH,WSB | feature_elongation,coding_sequence_variant |
| 1 | 60109927 | 60109991 | 64 | Wdr12 | CE,LP,MOLF,SPRET | feature_elongation,3_prime_UTR_variant |
| 1 | 78646556 | 78646653 | 97 | Utp14b | CAST | feature_elongation,3_prime_UTR_variant |
| 1 | 82430647 | 82430708 | 61 | Col4a4 | CAST,MOLF | feature_elongation,3_prime_UTR_variant |
| 1 | 135363018 | 135363153 | 135 | Nav1 | 129S1,AJ,AKR,BTBR,C3H,CE,DBA1J,DBA2J,ILNJ2,KK,NZW,RF,RH2,SJL,SPRET,SWR,TH,WSB | feature_elongation,3_prime_UTR_variant |
| 1 | 164006864 | 164006926 | 62 | F5 | CAST | feature_elongation,coding_sequence_variant,intron_variant |
| 1 | 164021231 | 164021284 | 53 | F5 | CAST,DBA1J,DBA2J,ILNJ2,MOLF,SPRET | feature_elongation,coding_sequence_variant |
| 1 | 165156624 | 165156685 | 61 | Dcaf6 | FVB | feature_elongation,3_prime_UTR_variant |
| 1 | 170708348 | 170708502 | 154 | Dusp12 | 129S1,BUB,C57L,FVB,MRL,NOD,NOR,NZB,NZO,RF,RH2,SWR,TH | feature_elongation,coding_sequence_variant,intron_variant |
| 1 | 173556688 | 173556770 | 82 | Ifi207 | AJ,BTBR,BUB,C58J,CE,DBA1J,DBA2J,FVB,KK,PL,RH2,SJL,SPRET,SWR,TH,WSB | feature_elongation,coding_sequence_variant |
| 1 | 173587714 | 173587773 | 59 | Ifi204 | 129S1,AKR,BALB,BUB,C3H,C57L,C58J,CBA,CE,DBA1J,DBA2J,ILNJ2,MRL,NOD,NOR,NZB,NZO,PJ,RF,SEA,SJL,SMJ,SWR,WSB | feature_elongation,coding_sequence_variant |
| 1 | 173733716 | 173733775 | 59 | Ifi211 | 129S1,AKR,BALB,BUB,C3H,C57L,C58J,CBA,CE,DBA1J,DBA2J,ILNJ2,MRL,NOD,NOR,NZB,NZO,PJ,RF,SEA,SJL,SMJ,SWR,WSB | feature_elongation,coding_sequence_variant |
| 1 | 175556504 | 175556616 | 112 | Wdr64 | SPRET | feature_elongation,coding_sequence_variant,intron_variant |
| 1 | 176850818 | 176851010 | 192 | Akt3 | SMJ | feature_elongation,3_prime_UTR_variant |
| 1 | 180056382 | 180056449 | 67 | Psen2 | CBA,CE,DBA1J,DBA2J,ILNJ2,PJ,SEA | feature_elongation,coding_sequence_variant,intron_variant |
| 1 | 180797691 | 180797800 | 109 | Tmem63a | AJ,BUB,C58J,CE,LP,NZB,RH2,SMJ | feature_elongation,coding_sequence_variant,intron_variant |
| 1 | 181632622 | 181632745 | 123 | Dnah14 | CAST,MOLF,TH | feature_elongation,coding_sequence_variant,intron_variant |
| 1 | 192527892 | 192527998 | 106 | Sertad4 | SPRET | feature_elongation,3_prime_UTR_variant |
| 10 | 7484379 | 7484450 | 71 | Lrp11 | MA/My | feature_elongation,coding_sequence_variant,intron_variant |
| 10 | 7690552 | 7690617 | 65 | Ppil4 | BUB,C3H,CAST,CBA,DBA1J,DBA2J,FVB,LP,MA/My,MRL,NOD,NOR,NZB,NZO,NZW,SWR | feature_elongation,coding_sequence_variant,intron_variant |
| 10 | 8600277 | 8600363 | 86 | Sash1 | CAST,MOLF | feature_elongation,3_prime_UTR_variant |
| 10 | 12424155 | 12424346 | 191 | Utrn | CE,DBA1J,DBA2J,LP,MA/My,TH | feature_elongation,5_prime_UTR_variant |
| 10 | 13083978 | 13084072 | 94 | Phactr2 | WSB | feature_elongation,3_prime_UTR_variant |
| 10 | 76320439 | 76320493 | 54 | Mcm3ap | CAST,MOLF | feature_elongation,coding_sequence_variant,intron_variant |
| 10 | 85887153 | 85887237 | 84 | Fbxo7 | C58J,LP,NZB | feature_elongation,3_prime_UTR_variant |
| 10 | 127578252 | 127578302 | 50 | Zbtb39 | AJ,BALB,FVB,NOD,NOR,PL | feature_elongation,coding_sequence_variant |
| 10 | 127878231 | 127878366 | 135 | Naca | CAST,MOLF,SMJ | feature_elongation,coding_sequence_variant |
| 11 | 5673029 | 5673126 | 97 | Urgcp | B10D2,B10J,BUB,C58J,CE,FVB,KK,MOLF,NOR,NZO,SJL,SWR | feature_elongation,coding_sequence_variant,intron_variant |
| 11 | 5854541 | 5854596 | 55 | Gck | AJ,AKR,B10D2,B10J,BALB,BTBR,BUB,C3H,C57L,C58J,CE,FVB,KK,MA/My,MOLF,MRL,NOD,NOR,NZB,NZO,PJ,PL,RF,SEA,SJL,SWR,TH,WSB | feature_elongation,coding_sequence_variant,intron_variant |
| 11 | 29104848 | 29104938 | 90 | Pnpt1 | AJ,C3H,CAST,DBA1J,DBA2J,ILNJ2,MA/My,MOLF,MRL,NZB,NZO,PL | feature_elongation,coding_sequence_variant,intron_variant |
| 11 | 40600783 | 40600906 | 123 | Hmmr | B10D2,B10J,BTBR,C58J,CAST,LP,MA/My,MOLF,SPRET | feature_elongation,coding_sequence_variant |
| 11 | 40600938 | 40601000 | 62 | Hmmr | BTBR,MOLF | feature_elongation,coding_sequence_variant |
| 11 | 60267491 | 60267546 | 55 | Drc3 | CAST | feature_elongation,3_prime_UTR_variant |
| 11 | 60692697 | 60692804 | 107 | Shmt1 | MOLF | feature_elongation,coding_sequence_variant,intron_variant |
| 11 | 64892697 | 64892770 | 73 | Elac2 | CE,ILNJ2 | feature_elongation,3_prime_UTR_variant |
| 11 | 77028569 | 77028686 | 117 | Efcab5 | CAST,MOLF,SPRET,TH | feature_elongation,coding_sequence_variant |
| 11 | 77028630 | 77028689 | 59 | Efcab5 | CAST,MOLF,TH | feature_elongation,coding_sequence_variant |
| 11 | 83193308 | 83193597 | 289 | Ap2b1 | KK | feature_elongation,5_prime_UTR_variant |
| 11 | 87413380 | 87413451 | 71 | Tex14 | SPRET | feature_elongation,coding_sequence_variant,intron_variant |
| 11 | 94430679 | 94430736 | 57 | Rsad1 | 129S1,BALB,BTBR,C57L,MA/My,NOD,NOR,PL,SJL,SMJ | feature_elongation,3_prime_UTR_variant |
| 11 | 95990842 | 95990945 | 103 | Calcoco2 | AJ,AKR,BUB,C3H,CAST,CBA,RH2,SPRET | feature_elongation,coding_sequence_variant |
| 11 | 97225110 | 97225297 | 187 | Gpr179 | SPRET | feature_elongation,coding_sequence_variant,3_prime_UTR_variant |
| 11 | 99276823 | 99276881 | 58 | Krt10 | AKR,RF,RH2,SJL | feature_elongation,coding_sequence_variant |
| 11 | 99549563 | 99549637 | 74 | Gm11563 | SPRET | feature_elongation,coding_sequence_variant |
| 11 | 99683648 | 99683721 | 73 | Gm11596 | CAST,MOLF | feature_elongation,coding_sequence_variant |
| 11 | 99705987 | 99706090 | 103 | Gm11564 | SPRET | feature_elongation,coding_sequence_variant |
| 11 | 99839625 | 99839824 | 199 | Krtap9-5 | SPRET | feature_elongation,coding_sequence_variant |
| 11 | 109464588 | 109464641 | 53 | Wipi1 | SPRET | feature_elongation,3_prime_UTR_variant |

| | | | | | | |
|---|---|---|---|---|---|---|
| 11 | 113551867 | 113551918 | 51 | Cog1 | AJ,C3H,CBA,TH | feature_elongation,coding_sequence_variant,intron_variant |
| 11 | 116348178 | 116348296 | 118 | Qrich2 | CAST,MOLF | feature_elongation,coding_sequence_variant |
| 12 | 3543825 | 3543916 | 91 | Asxl2 | MOLF | feature_elongation,coding_sequence_variant,intron_variant |
| 12 | 4159420 | 4159492 | 72 | Dnajc27 | CAST,MOLF | feature_elongation,coding_sequence_variant |
| 12 | 28917614 | 28917854 | 240 | Eipr1 | SPRET | feature_elongation,3_prime_UTR_variant |
| 12 | 71041499 | 71041992 | 493 | Psma3 | NOD,NOR | feature_elongation,3_prime_UTR_variant |
| 12 | 87994165 | 87994227 | 62 | Eif1ad11 | 129S1,AJ,AKR,BTBR,BUB,C3H,C57L,CBA,DBA1J,DBA2J,FVB,ILNJ2,MRL,NOR,PJ,RF,SEA,SJL,TH,WSB | feature_elongation,coding_sequence_variant,3_prime_UTR_variant |
| 12 | 112742722 | 112743238 | 516 | Ahnak2 | 129S1,B10D2,B10J,BALB,BTBR,BUB,C57L,C58J,DBA1J,DBA2J,ILNJ2,KK,LP,MA/My,MRL,NOD,RH2,SJL | feature_elongation,coding_sequence_variant |
| 12 | 112744476 | 112744992 | 516 | Ahnak2 | CAST | feature_elongation,coding_sequence_variant |
| 12 | 112747707 | 112748270 | 563 | Ahnak2 | AJ | feature_elongation,coding_sequence_variant |
| 12 | 112819377 | 112819492 | 115 | Gpr132 | FVB,SWR | feature_elongation,coding_sequence_variant,5_prime_UTR_variant,intron_variant |
| 13 | 24579156 | 24579238 | 82 | Cmah | SPRET | feature_elongation,5_prime_UTR_variant,intron_variant |
| 13 | 27387374 | 27387437 | 63 | Prl3c1 | SPRET | feature_elongation,coding_sequence_variant,intron_variant |
| 13 | 59953137 | 59953249 | 112 | Tut7 | SPRET | feature_elongation,coding_sequence_variant,intron_variant |
| 13 | 93233240 | 93233394 | 154 | Cmya5 | CAST,FVB,MOLF,MRL,NZW,RF | feature_elongation,coding_sequence_variant |
| 14 | 36641168 | 36641254 | 86 | Ccser2 | MOLF | feature_elongation,5_prime_UTR_variant,intron_variant |
| 14 | 38091647 | 38091741 | 94 | Nrg3 | MOLF | feature_elongation,3_prime_UTR_variant |
| 14 | 45240799 | 45240866 | 67 | Ptger2 | C57L,CAST,MOLF,NZO,PJ,PL | feature_elongation,3_prime_UTR_variant |
| 14 | 49457120 | 49457221 | 101 | Ccdc198 | AKR,BALB,CE,DBA1J,MRL,NOD,NZO,PJ,SMJ,SWR,WSB | feature_elongation,3_prime_UTR_variant |
| 14 | 49457149 | 49457217 | 68 | Ccdc198 | 129S1,AJ,AKR,BALB,C3H,CBA,CE,DBA1J,DBA2J,MRL,NZB,NZO,NZW,PJ,RF | feature_elongation,3_prime_UTR_variant |
| 14 | 51759277 | 51759705 | 428 | Gm7247 | SPRET | feature_elongation,coding_sequence_variant,intron_variant |
| 14 | 55817708 | 55817799 | 91 | Psme1 | BUB,SJL | feature_elongation,coding_sequence_variant,intron_variant |
| 14 | 56871597 | 56871707 | 110 | Parp4 | MOLF | feature_elongation,coding_sequence_variant,intron_variant |
| 14 | 62939929 | 62940055 | 126 | Ints6 | MOLF | feature_elongation,coding_sequence_variant,intron_variant |
| 14 | 72681392 | 72681469 | 77 | Gm9195 | CAST,SPRET | feature_elongation,coding_sequence_variant,intron_variant |
| 14 | 74992400 | 74992656 | 256 | Lrch1 | BUB,KK,MOLF | feature_elongation,3_prime_UTR_variant |
| 14 | 99297733 | 99297874 | 141 | Bora | CAST | feature_elongation,coding_sequence_variant,intron_variant |
| 15 | 5128216 | 5128274 | 58 | Card6 | MOLF | feature_elongation,coding_sequence_variant |
| 15 | 38492530 | 38493454 | 924 | Azin1 | NZB,NZW | feature_elongation,coding_sequence_variant,intron_variant |
| 15 | 58422163 | 58422228 | 65 | Fer1l6 | CAST | feature_elongation,coding_sequence_variant,intron_variant |
| 15 | 66792799 | 66792878 | 79 | Ccn4 | CAST | feature_elongation,3_prime_UTR_variant |
| 15 | 78660539 | 78660604 | 65 | Card10 | BALB,BUB,C3H,CBA,CE,DBA1J,DBA2J,PJ,PL | feature_elongation,3_prime_UTR_variant |
| 15 | 90453606 | 90453696 | 90 | Cpne8 | CAST,SPRET | feature_elongation,coding_sequence_variant,intron_variant |
| 15 | 98748903 | 98748988 | 85 | Kmt2d | ILNJ2,NOD,NOR | feature_elongation,coding_sequence_variant |
| 15 | 99840402 | 99840708 | 306 | Fam186a | 129S1,C57L | feature_elongation,coding_sequence_variant |
| 15 | 101325330 | 101325476 | 146 | Krt7 | MOLF | feature_elongation,coding_sequence_variant,intron_variant |
| 15 | 101855780 | 101856043 | 263 | Krt78 | KK,RF,TH | feature_elongation,coding_sequence_variant |
| 16 | 32573705 | 32574447 | 742 | Muc4 | AJ,BALB,BTBR,BUB,C3H,C57L,CBA,CE,DBA1J,FVB,ILNJ2,KK,MRL,PL,RH2,SEA,SJL,SMJ,TH,WSB | feature_elongation,coding_sequence_variant |
| 17 | 8642071 | 8642219 | 148 | T2 | SPRET | feature_elongation,3_prime_UTR_variant |
| 17 | 21807254 | 21807350 | 96 | Zfp948 | MOLF | feature_elongation,coding_sequence_variant |
| 17 | 25314802 | 25314853 | 51 | Ift140 | 129S1,AJ,AKR,B10D2,BALB,BTBR,BUB,C3H,C57L,CBA,CE,DBA1J,DBA2J,FVB,ILNJ2,KK,MA/My,MOLF,MRL,NOD,NOR,NZB,NZO,RF,RH2,SEA,SJL,SWR,TH,WSB | feature_elongation,3_prime_UTR_variant |
| 17 | 26663394 | 26663454 | 60 | Neurl1b | SPRET | feature_elongation,3_prime_UTR_variant |
| 17 | 28115621 | 28115800 | 179 | Bltp3a | B10D2,CAST,MA/My,MOLF,MRL,NZB,RF,SEA | feature_elongation,3_prime_UTR_variant |
| 17 | 31867576 | 31867645 | 69 | U2af1 | FVB,ILNJ2,PJ,SJL,SPRET | feature_elongation,coding_sequence_variant,intron_variant |
| 17 | 34163412 | 34163476 | 64 | Wdr46 | PJ,SJL,TH | feature_elongation,coding_sequence_variant,intron_variant |
| 17 | 35931530 | 35931843 | 313 | Muc21 | PJ,WSB | feature_elongation,coding_sequence_variant |
| 17 | 45831665 | 45831734 | 69 | Aars2 | AKR,BALB,C3H,CBA,CE,PJ,RH2,SEA | feature_elongation,3_prime_UTR_variant |
| 17 | 46570366 | 46570481 | 115 | Tjap1 | BTBR,C57L,DBA1J,KK,LP,MA/My,NZO,RF | feature_elongation,coding_sequence_variant,intron_variant |
| 17 | 56411925 | 56412121 | 196 | Plin4 | CAST,SPRET | feature_elongation,coding_sequence_variant |
| 18 | 3266980 | 3267033 | 53 | Crem | SPRET | feature_elongation,3_prime_UTR_variant |
| 18 | 3514499 | 3514551 | 52 | Bambi | SPRET | feature_elongation,3_prime_UTR_variant |
| 18 | 10595058 | 10595113 | 55 | Esco1 | 129S1,C57L,CAST,MOLF | feature_elongation,coding_sequence_variant |
| 18 | 20541369 | 20541667 | 298 | Dsg1b | SPRET | feature_elongation,coding_sequence_variant,intron_variant |
| 18 | 34402282 | 34402454 | 172 | Apc | AJ,AKR,BALB,C58J,CBA,DBA2J,ILNJ2,RH2,SEA | feature_elongation,coding_sequence_variant,intron_variant |
| 18 | 47038430 | 47038484 | 54 | Lvrn | CAST | feature_elongation,3_prime_UTR_variant |
| 18 | 61328158 | 61328214 | 56 | Slc26a2 | CAST,KK,MOLF,NZO,NZW | feature_elongation,3_prime_UTR_variant |
| 18 | 67926738 | 67926823 | 85 | Seh1l | AJ,BALB,ILNJ2,RH2,SMJ | feature_elongation,3_prime_UTR_variant |
| 18 | 75218780 | 75218954 | 174 | Dym | AKR,ILNJ2,MOLF,RF | feature_elongation,coding_sequence_variant |
| 19 | 6061448 | 6061509 | 61 | Capn1 | MOLF | feature_elongation,coding_sequence_variant,intron_variant |

| | | | | | | |
|---|---|---|---|---|---|---|
| 19 | 6171474 | 6171571 | 97 | Snx15 | TH | feature_elongation,coding_sequence_variant,intron_variant |
| 19 | 10581490 | 10581588 | 98 | Tkfc | WSB | feature_elongation,coding_sequence_variant,5_prime_UTR_variant,intron_variant |
| 19 | 11280599 | 11280673 | 74 | Ms4a14 | SPRET | feature_elongation,coding_sequence_variant |
| 19 | 17729334 | 17729440 | 106 | Pcsk5 | 129S1,AKR,BALB,BTBR,C57L,C58J,FVB,ILNJ2,LP,NOD,NOR,NZB,NZO,NZW,PJ,PL,RF,SEA,SWR,TH,WSB | feature_elongation,coding_sequence_variant,intron_variant |
| 19 | 18683752 | 18683807 | 55 | Carnmt1 | MOLF,NZB,NZO,NZW,PJ | feature_elongation,3_prime_UTR_variant |
| 19 | 36096560 | 36096627 | 67 | Ankrd1 | AKR,RF,SJL,TH | feature_elongation,coding_sequence_variant,intron_variant |
| 19 | 36713955 | 36714014 | 59 | Ppp1r3c | CAST | start_lost,feature_elongation,start_retained_variant,coding_sequence_variant,5_prime_UTR_variant |
| 19 | 41571696 | 41571819 | 123 | Lcor | SPRET | feature_elongation,coding_sequence_variant |
| 19 | 41872250 | 41872326 | 76 | Rrp12 | BALB,BUB,MOLF,NZO | feature_elongation,coding_sequence_variant,intron_variant |
| 19 | 42135929 | 42136001 | 72 | Marveld1 | MOLF | feature_elongation,5_prime_UTR_variant |
| 2 | 3426224 | 3426427 | 203 | Dclre1c | MA/My,NZO,NZW | feature_elongation,coding_sequence_variant,intron_variant,feature_elongation,5_prime_UTR_variant,intron_variant |
| 2 | 5803230 | 5803422 | 192 | Cdc123 | SPRET | feature_elongation,coding_sequence_variant,intron_variant |
| 2 | 26909695 | 26909825 | 130 | Cacfd1 | AJ,AKR,B10D2,B10J,BALB,C58J,CBA,DBA1J,DBA2J,ILNJ2,LP,NOR,NZB,NZW,PL,RH2,SEA,SWR,WSB | feature_elongation,3_prime_UTR_variant |
| 2 | 29051533 | 29051635 | 102 | Setx | MOLF | feature_elongation,coding_sequence_variant,intron_variant |
| 2 | 52305964 | 52306086 | 122 | Arl5a | 129S1,AJ,AKR,BUB,C3H,CBA,CE,DBA1J,DBA2J,ILNJ2,MRL,NOD,NOR,NZW,PJ,PL,RF,SEA,SMJ,SWR,TH,WSB | feature_elongation,coding_sequence_variant,intron_variant |
| 2 | 79163316 | 79163444 | 128 | Itga4 | MOLF | feature_elongation,3_prime_UTR_variant |
| 2 | 85671181 | 85671351 | 170 | Or5ar1 | KK | feature_elongation,coding_sequence_variant,3_prime_UTR_variant |
| 2 | 87901689 | 87901874 | 185 | Or5d36 | MOLF | feature_elongation,coding_sequence_variant,5_prime_UTR_variant,intron_variant |
| 2 | 87901728 | 87901864 | 136 | Or5d36 | MOLF | feature_elongation,5_prime_UTR_variant,intron_variant |
| 2 | 88483738 | 88484597 | 859 | Or4p4 | C3H,CBA,MA/My,NZW | feature_elongation,3_prime_UTR_variant |
| 2 | 88483764 | 88484706 | 942 | Or4p4 | C3H,CBA,CE,MA/My,NZW | feature_elongation,3_prime_UTR_variant |
| 2 | 119849813 | 119849887 | 74 | Mapkbp1 | AJ,AKR,BALB,CE,DBA1J,DBA2J,FVB,ILNJ2,PJ,PL,RF,RH2,SEA,SJL,SMJ,SWR,TH | feature_elongation,coding_sequence_variant,intron_variant |
| 2 | 120451349 | 120451427 | 78 | Haus2 | SPRET | feature_elongation,3_prime_UTR_variant |
| 2 | 125429459 | 125429539 | 80 | Cep152 | MOLF | feature_elongation,coding_sequence_variant,intron_variant |
| 2 | 152236668 | 152236787 | 119 | Sox12 | AJ,C58J,MOLF,NOD,NZB,NZW,RF | feature_elongation,3_prime_UTR_variant |
| 2 | 154359700 | 154360115 | 415 | Cbfa2t2 | FVB,SWR | feature_elongation,coding_sequence_variant,intron_variant |
| 2 | 155983989 | 155984073 | 84 | Nfs1 | WSB | feature_elongation,coding_sequence_variant,intron_variant,feature_elongation,coding_sequence_variant |
| 2 | 165133300 | 165133412 | 112 | Elmo2 | KK | feature_elongation,3_prime_UTR_variant |
| 2 | 165626713 | 165626793 | 80 | Zmynd8 | AKR,BTBR,BUB,C3H,C58J,DBA1J,DBA2J,FVB,ILNJ2,MA/My,NZB,NZO,PJ,RF,SJL,SMJ,SWR,TH,WSB | feature_elongation,3_prime_UTR_variant |
| 2 | 167820390 | 167820584 | 194 | Ptpn1 | 129S1,C57L,NZB | feature_elongation,3_prime_UTR_variant |
| 2 | 168590700 | 168590826 | 126 | Sall4 | B10D2 | feature_elongation,3_prime_UTR_variant |
| 2 | 172204807 | 172204907 | 100 | Aurka | FVB,SJL | feature_elongation,coding_sequence_variant,intron_variant |
| 3 | 9018638 | 9018748 | 110 | Tpd52 | MOLF,SPRET | feature_elongation,coding_sequence_variant,intron_variant |
| 3 | 35887025 | 35887150 | 125 | Atp11b | WSB | feature_elongation,coding_sequence_variant,intron_variant |
| 3 | 36128569 | 36129037 | 468 | Acad9 | CE,DBA1J,NZB,NZO,NZW | feature_elongation,coding_sequence_variant,intron_variant |
| 3 | 68951947 | 68951999 | 52 | Trim59 | 129S1,C57L,CAST,FVB,SJL,SWR,TH | feature_elongation,5_prime_UTR_variant |
| 3 | 88020123 | 88020351 | 228 | Iqgap3 | NZB,NZO | feature_elongation,coding_sequence_variant,intron_variant |
| 3 | 92479161 | 92479239 | 78 | Ivl | MOLF,NOD,NOR,SPRET | feature_elongation,coding_sequence_variant |
| 3 | 93110593 | 93110665 | 72 | Flg2 | 129S1,AJ,BTBR,C3H,C57L,CAST,CBA,CE,ILNJ2,PJ,PL,SEA,TH,WSB | feature_elongation,coding_sequence_variant |
| 3 | 93113206 | 93113444 | 238 | Flg2 | NOD,NOR | feature_elongation,coding_sequence_variant |
| 3 | 93231839 | 93232368 | 529 | Hrnr | CAST | feature_elongation,coding_sequence_variant |
| 3 | 93237575 | 93238112 | 537 | Hrnr | AJ,C3H,CBA,CE,MA/My | feature_elongation,coding_sequence_variant |
| 3 | 93304211 | 93304317 | 106 | Rptn | BALB,ILNJ2,KK,NOD,NOR,NZB,NZO,PJ,PL,RH2,SEA,SPRET,TH,WSB | feature_elongation,coding_sequence_variant |
| 3 | 105893684 | 105893742 | 58 | Ovgp1 | AJ,C3H,CBA,SPRET | feature_elongation,coding_sequence_variant |
| 3 | 107017328 | 107017422 | 94 | Kcna2 | AJ,AKR,BALB,C3H,CAST,CBA,DBA1J,DBA2J,ILNJ2,MA/My,MRL,NOD,NOR,NZB,NZO,NZW,PL,RF,RH2,TH,WSB | feature_elongation,3_prime_UTR_variant |
| 3 | 121156215 | 121156342 | 127 | Alg14 | SPRET | feature_elongation,3_prime_UTR_variant |
| 3 | 154283843 | 154284368 | 525 | Tyw3 | RF | feature_elongation,3_prime_UTR_variant |
| 3 | 154468185 | 154468247 | 62 | Erich3 | SPRET | feature_elongation,coding_sequence_variant |
| 4 | 10989522 | 10989604 | 82 | Plekhf2 | CAST | feature_elongation,3_prime_UTR_variant |
| 4 | 11171028 | 11171079 | 51 | Trp53inp1 | CAST | feature_elongation,3_prime_UTR_variant |
| 4 | 70139206 | 70139390 | 184 | Cdk5rap2 | KK | feature_elongation,3_prime_UTR_variant |
| 4 | 88745704 | 88745762 | 58 | Ifna6 | CE | feature_elongation,coding_sequence_variant |
| 4 | 101010497 | 101010563 | 66 | Jak1 | SPRET | feature_elongation,3_prime_UTR_variant |
| 4 | 120804557 | 120804710 | 153 | Zfp69 | ILNJ2 | feature_elongation,coding_sequence_variant |
| 4 | 131950649 | 131950703 | 54 | Gmeb1 | DBA1J,KK,MOLF,NZB,SMJ | feature_elongation,3_prime_UTR_variant |
| 4 | 133692302 | 133692479 | 177 | Hmgn2 | 129S1,BALB,BTBR,C57L,DBA1J,DBA2J,ILNJ2,NOD,PL,SMJ | feature_elongation,3_prime_UTR_variant |
| 4 | 136081060 | 136081117 | 57 | Hnrnpr | 129S1,B10D2,B10J,C57L,DBA1J,KK,MOLF,NZO,PJ,SEA | feature_elongation,3_prime_UTR_variant |
| 4 | 138016587 | 138016648 | 61 | Kif17 | CAST,SPRET | feature_elongation,coding_sequence_variant,intron_variant |

| | | | | | | |
|---|---|---|---|---|---|---|
| 4 | 145311899 | 145312126 | 227 | Zfp268 | 129S1,AKR,BTBR,BUB,C3H,C57L,C58J,CAST,CBA,CE,DBA1J,KK,LP,MA/My,MRL,NOD,NZB,NZO,PJ,PL,RF,SEA,SJL,SMJ,SWR,WSB | feature_elongation,5_prime_UTR_variant,intron_variant |
| 4 | 147958725 | 147958787 | 62 | Mfn2 | 129S1,AKR,BTBR,BUB,C3H,C57L,C58J,CBA,CE,DBA1J,DBA2J,KK,LP,MA/My,MRL,NOD,NZB,NZO,NZW,PJ,PL,RF,SEA,SJL,SMJ,SWR,TH,WSB | feature_elongation,3_prime_UTR_variant |
| 4 | 147958746 | 147958810 | 64 | Mfn2 | 129S1,AKR,BTBR,BUB,C3H,C57L,C58J,CBA,CE,DBA1J,DBA2J,KK,LP,MA/My,MRL,NOD,NZB,NZO,NZW,PJ,PL,RF,SEA,SJL,SMJ,SWR,TH,WSB | feature_elongation,3_prime_UTR_variant |
| 4 | 155526452 | 155526502 | 50 | Cfap74 | MOLF | feature_elongation,coding_sequence_variant,intron_variant |
| 4 | 156302483 | 156302548 | 65 | Perm1 | AJ,BALB,C3H,C57L,C58J,DBA1J,DBA2J,MA/My,MRL,NOD,NOR,NZO,PL,RH2,SJL | feature_elongation,coding_sequence_variant |
| 4 | 156302516 | 156302613 | 97 | Perm1 | AJ,BALB,C3H,C57L,C58J,DBA1J,DBA2J,ILNJ2,MA/My,MRL,NZO,PL,RH2,SJL,SMJ,WSB | feature_elongation,coding_sequence_variant |
| 5 | 8751924 | 8752102 | 178 | Abcb1a | BUB,NOD,NOR | feature_elongation,coding_sequence_variant,intron_variant |
| 5 | 33821832 | 33822116 | 284 | Tacc3 | CAST,SPRET | feature_elongation,coding_sequence_variant |
| 5 | 51614733 | 51614792 | 59 | Ppargc1a | MOLF | feature_elongation,3_prime_UTR_variant |
| 5 | 76632060 | 76632135 | 75 | Exoc1l | AJ,AKR,BUB,FVB,PJ,PL,SJL,SMJ | feature_elongation,5_prime_UTR_variant |
| 5 | 87823685 | 87823934 | 249 | Csn1s1 | CAST | feature_elongation,coding_sequence_variant,intron_variant |
| 5 | 88649969 | 88650065 | 96 | Enam | CAST | feature_elongation,coding_sequence_variant |
| 5 | 100216368 | 100216435 | 67 | Enoph1 | CAST,SPRET | feature_elongation,3_prime_UTR_variant |
| 5 | 107534399 | 107534477 | 78 | Brdt | CAST | feature_elongation,3_prime_UTR_variant |
| 5 | 108574751 | 108574818 | 67 | Pde6b | CAST | feature_elongation,coding_sequence_variant,intron_variant |
| 5 | 115793879 | 115794000 | 121 | Bicdl1 | 129S1,AKR,CE,DBA1J,DBA2J,KK,MRL,NZB,NZO,RH2,SEA | feature_elongation,coding_sequence_variant,intron_variant |
| 5 | 118330475 | 118330533 | 58 | Rnft2 | 129S1,AJ,AKR,BALB,BUB,C3H,C57L,CBA,CE,DBA1J,DBA2J,ILNJ2,KK,NOR,NZB,NZO,PJ,PL,RH2,SEA,SJL,SMJ,TH,WSB | feature_elongation,3_prime_UTR_variant |
| 5 | 123710135 | 123710990 | 855 | Vps33a | 129S1,AJ,BALB,C57L,KK,MRL,NZW,PJ,RH2,SEA,SJL,WSB | feature_elongation,coding_sequence_variant,intron_variant |
| 5 | 125463413 | 125463640 | 227 | Ubc | SJL | feature_elongation,coding_sequence_variant |
| 5 | 129062020 | 129062140 | 120 | Stx2 | AJ,BALB,C58J,FVB,LP,MOLF,MRL,NOD,NOR,NZB,NZO,NZW,RH2,SEA,SMJ,SWR,TH | feature_elongation,3_prime_UTR_variant |
| 5 | 137182292 | 137182380 | 88 | Muc17 | CAST | feature_elongation,coding_sequence_variant |
| 5 | 137461985 | 137462083 | 98 | Zan | LP | feature_elongation,coding_sequence_variant |
| 5 | 137462091 | 137462187 | 96 | Zan | 129S1,BUB,CAST,LP,MOLF,RF,SWR,WSB | feature_elongation,coding_sequence_variant |
| 5 | 147815835 | 147815939 | 104 | Slc46a3 | CAST,WSB | feature_elongation,3_prime_UTR_variant |
| 6 | 5008455 | 5008608 | 153 | Ppp1r9a | PJ | feature_elongation,3_prime_UTR_variant |
| 6 | 36502834 | 36502899 | 65 | Chrm2 | SPRET | feature_elongation,3_prime_UTR_variant |
| 6 | 40844009 | 40844127 | 118 | Mgam2-ps | MOLF | feature_elongation,coding_sequence_variant |
| 6 | 40844545 | 40844663 | 118 | Mgam2-ps | BUB,CAST,FVB,LP,SJL,SWR,TH,WSB | feature_elongation,coding_sequence_variant |
| 6 | 43093074 | 43093260 | 186 | Or2a54 | MOLF | feature_elongation,coding_sequence_variant |
| 6 | 83127300 | 83127361 | 61 | Wdr54 | CAST | feature_elongation,3_prime_UTR_variant |
| 6 | 84166818 | 84166905 | 87 | Dysf | MOLF,NZB,NZO | feature_elongation,coding_sequence_variant,intron_variant |
| 6 | 90635724 | 90635820 | 96 | Iqsec1 | CE,PJ,SEA | feature_elongation,3_prime_UTR_variant |
| 6 | 92151525 | 92151724 | 199 | Mrps25 | CE,PJ,SEA | feature_elongation,3_prime_UTR_variant |
| 6 | 132549054 | 132549175 | 121 | Prh1 | AJ,AKR,BALB,BUB,C3H,C57L,CBA,DBA1J,DBA2J,ILNJ2,MRL,NZB,PL,RF,RH2,SEA | feature_elongation,coding_sequence_variant |
| 6 | 133083404 | 133083456 | 52 | Smim10l1 | 129S1,C58J,FVB,KK,LP,NOD,NOR,NZW,SJL,SMJ,SPRET,TH,WSB | feature_elongation,3_prime_UTR_variant |
| 6 | 143113069 | 143113183 | 114 | Etnk1 | WSB | feature_elongation,5_prime_UTR_variant |
| 7 | 11889440 | 11889490 | 50 | Vmn1r78 | WSB | feature_elongation,3_prime_UTR_variant |
| 7 | 12508560 | 12508716 | 156 | Zscan18 | MOLF,SPRET | feature_elongation,coding_sequence_variant |
| 7 | 12508637 | 12508692 | 55 | Zscan18 | SPRET | feature_elongation,coding_sequence_variant |
| 7 | 30451470 | 30451546 | 76 | Sbsn | CAST,DBA1J,DBA2J,MOLF,NZB,NZO,SMJ | feature_elongation,coding_sequence_variant |
| 7 | 30451584 | 30451749 | 165 | Sbsn | NZW | feature_elongation,coding_sequence_variant |
| 7 | 30452385 | 30452437 | 52 | Sbsn | CAST,MOLF | feature_elongation,coding_sequence_variant |
| 7 | 30452432 | 30452512 | 80 | Sbsn | MOLF,NZW | feature_elongation,coding_sequence_variant |
| 7 | 30518388 | 30518454 | 66 | Ffar2 | SPRET | feature_elongation,3_prime_UTR_variant |
| 7 | 43326426 | 43326538 | 112 | Ctu1 | SPRET | feature_elongation,coding_sequence_variant |
| 7 | 43898363 | 43898474 | 111 | 2410002F23Rik | CAST | feature_elongation,5_prime_UTR_variant |
| 7 | 83547623 | 83547779 | 156 | Mesd | CAST | feature_elongation,3_prime_UTR_variant |
| 7 | 85983550 | 85983603 | 53 | Or14a260 | 129S1,BTBR,C3H,C57L,CBA,DBA2J,ILNJ2,KK,NZB,NZO,NZW,PJ | feature_elongation,3_prime_UTR_variant |
| 7 | 98917972 | 98918107 | 135 | Map6 | C3H,CBA,DBA1J,DBA2J,ILNJ2,MOLF,PJ,SEA | feature_elongation,coding_sequence_variant |
| 7 | 103882738 | 103882821 | 83 | Trim6 | CAST | feature_elongation,3_prime_UTR_variant |
| 7 | 119315944 | 119315996 | 52 | Thumpd1 | CAST | feature_elongation,coding_sequence_variant |
| 7 | 120545515 | 120546332 | 817 | Polr3e | C57L | feature_elongation,3_prime_UTR_variant |
| 7 | 128297801 | 128297852 | 51 | Inpp5f | SPRET | feature_elongation,3_prime_UTR_variant |
| 7 | 135300768 | 135301581 | 813 | Mki67 | SPRET | feature_elongation,coding_sequence_variant |
| 7 | 140443990 | 140444043 | 53 | Sirt3 | SPRET | feature_elongation,coding_sequence_variant,intron_variant |
| 7 | 140507818 | 140507872 | 54 | Nlrp6 | ILNJ2,WSB | feature_elongation,3_prime_UTR_variant |
| 7 | 140681232 | 140681354 | 122 | Ano9 | CAST,MOLF,NZW | feature_elongation,3_prime_UTR_variant |
| 7 | 140681257 | 140681318 | 61 | Ano9 | CAST,MOLF,NZW | feature_elongation,3_prime_UTR_variant |
| 7 | 141224990 | 141225499 | 509 | Muc6 | MOLF,NZW | feature_elongation,coding_sequence_variant |
| 7 | 141296808 | 141297097 | 289 | Muc2 | NZW,SPRET | feature_elongation,coding_sequence_variant |
| 7 | 141363058 | 141363152 | 94 | Muc5ac | 129S1,AKR,BALB,BTBR,BUB,C57L,C58J,CAST,CBA,FVB,KK,MA/My,MRL,NZB,PL,RF,RH2,SEA,SMJ,SPRET,SWR,WSB | feature_elongation,coding_sequence_variant |
| 7 | 141412183 | 141413069 | 886 | Muc5b | SEA | feature_elongation,coding_sequence_variant |
| 7 | 141743689 | 141743890 | 201 | Gm10153 | BALB,MRL,RF,RH2,SEA | feature_elongation,coding_sequence_variant |
| 7 | 141755477 | 141755573 | 96 | Krtap5-3 | NZW | feature_elongation,coding_sequence_variant |
| 7 | 141783413 | 141783468 | 55 | Krtap5-5 | CAST | feature_elongation,coding_sequence_variant |
| 7 | 141827607 | 141827718 | 111 | Gm4559 | MOLF | feature_elongation,coding_sequence_variant |

| | | | | | | |
|---|---|---|---|---|---|---|
| 7 | 141850232 | 141850345 | 113 | Krtap5-1 | MOLF,NZW,WSB | feature_elongation,coding_sequence_variant |
| 7 | 143190378 | 143190486 | 108 | Tnfrsf22 | CAST,MA/My,MOLF | feature_elongation,3_prime_UTR_variant |
| 8 | 3205261 | 3205330 | 69 | Insr | CAST | feature_elongation,3_prime_UTR_variant |
| 8 | 5134465 | 5134622 | 157 | Slc10a2 | SPRET | feature_elongation,3_prime_UTR_variant |
| 8 | 10476417 | 10476479 | 62 | Myo16 | CAST,MOLF,NZB | feature_elongation,coding_sequence_variant,intron_variant |
| 8 | 10620230 | 10620291 | 61 | Myo16 | CAST,MOLF | feature_elongation,coding_sequence_variant,intron_variant,feature_elongation,coding_sequence_variant |
| 8 | 12917049 | 12917138 | 89 | Atp11a | C57L,ILNJ2,NZB,NZW,PJ,PL,SEA,SMJ,TH | feature_elongation,3_prime_UTR_variant |
| 8 | 13068280 | 13068376 | 96 | Mcf2l | 129S1,C57L,CE,ILNJ2,LP,MOLF,NZB,NZO,NZW,PJ,PL,SEA,SMJ,TH | feature_elongation,3_prime_UTR_variant |
| 8 | 15076779 | 15076834 | 55 | Kbtbd11 | SPRET | feature_elongation,5_prime_UTR_variant |
| 8 | 34110054 | 34110133 | 79 | Ubxn8 | 129S1,AJ,AKR,BALB,BUB,C3H,C57L,CBA,CE,DBA1J,DBA2J,FVB,MA/My,MRL,NOD,NOR,PL,RF,RH2,SJL,SMJ,SWR,TH,WSB | feature_elongation,3_prime_UTR_variant |
| 8 | 46481335 | 46481387 | 52 | Lrp2bp | SPRET | feature_elongation,3_prime_UTR_variant |
| 8 | 55037559 | 55037704 | 145 | Asb5 | CAST | feature_elongation,coding_sequence_variant,intron_variant,feature_elongation,5_prime_UTR_variant,intron_variant |
| 8 | 65194683 | 65194734 | 51 | Klhl2 | PJ,SEA,WSB | feature_elongation,3_prime_UTR_variant |
| 8 | 70360084 | 70360142 | 58 | Gatad2a | BALB,C57L,CBA,DBA2J,FVB,KK,RF,SJL,SMJ,TH | feature_elongation,3_prime_UTR_variant |
| 8 | 72934873 | 72934988 | 115 | Rab8a | SPRET | feature_elongation,3_prime_UTR_variant |
| 8 | 72934906 | 72934967 | 61 | Rab8a | SPRET | feature_elongation,3_prime_UTR_variant |
| 8 | 73005431 | 73005609 | 178 | Ap1m1 | 129S1,AKR,BALB,BUB,C3H,C57L,C58J,CBA,CE,DBA1J,DBA2J,FVB,ILNJ2,KK,MA/My,MRL,NZB,NZO,NZW,PJ,PL,RF,SEA,SJL,SMJ,SWR,TH,WSB | feature_elongation,coding_sequence_variant,intron_variant |
| 8 | 79532286 | 79532338 | 52 | Lsm6 | SPRET | feature_elongation,3_prime_UTR_variant |
| 8 | 80366576 | 80366685 | 109 | Otud4 | CE | feature_elongation,coding_sequence_variant |
| 8 | 87568775 | 87568832 | 57 | N4bp1 | SPRET | feature_elongation,3_prime_UTR_variant |
| 8 | 104991331 | 104991428 | 97 | Cklf | KK,PL,SPRET,WSB | feature_elongation,3_prime_UTR_variant |
| 8 | 105076826 | 105076892 | 66 | Cmtm4 | KK,PL,WSB | feature_elongation,3_prime_UTR_variant |
| 8 | 125756560 | 125756698 | 138 | Tsnax | NZB | feature_elongation,3_prime_UTR_variant |
| 8 | 125756597 | 125756708 | 111 | Tsnax | NZB | feature_elongation,3_prime_UTR_variant |
| 9 | 14916731 | 14916846 | 115 | Panx1 | MOLF | feature_elongation,3_prime_UTR_variant |
| 9 | 19031907 | 19031958 | 51 | Or7g21 | MOLF | feature_elongation,5_prime_UTR_variant,intron_variant |
| 9 | 25188910 | 25189028 | 118 | Septin7 | MOLF,SPRET,WSB | feature_elongation,coding_sequence_variant,intron_variant |
| 9 | 45239778 | 45239842 | 64 | Tmprss13 | CAST | feature_elongation,coding_sequence_variant |
| 9 | 65395313 | 65395582 | 269 | Spg21 | CAST | feature_elongation,3_prime_UTR_variant |
| 9 | 72822674 | 72822768 | 94 | Prtg | SPRET | feature_elongation,3_prime_UTR_variant |
| 9 | 73010744 | 73010881 | 137 | Khdc3 | MOLF,SPRET | feature_elongation,coding_sequence_variant |
| 9 | 96962198 | 96962273 | 75 | Slc25a36 | CE,NZB | feature_elongation,coding_sequence_variant,intron_variant |
| 9 | 99550276 | 99550348 | 72 | Dzip1l | NOD,NOR | feature_elongation,3_prime_UTR_variant |
| 9 | 104167499 | 104167558 | 59 | Acp3 | SPRET | feature_elongation,3_prime_UTR_variant |
| 9 | 110215209 | 110215277 | 68 | Ptpn23 | CBA,MOLF | feature_elongation,coding_sequence_variant,intron_variant |
| 9 | 110815523 | 110815892 | 369 | Rtp3 | NZW | feature_elongation,coding_sequence_variant,intron_variant |
| 9 | 119607815 | 119607979 | 164 | Scn11a | BTBR,MRL | feature_elongation,coding_sequence_variant,intron_variant |
| X | 6227264 | 6227343 | 79 | Bmp15 | CAST | feature_elongation,3_prime_UTR_variant |
| X | 6690672 | 6690723 | 51 | Dgkk | CAST | feature_elongation,coding_sequence_variant |
| X | 74423514 | 74423571 | 57 | F8 | 129S1,C57L,NZB | feature_elongation,5_prime_UTR_variant |
| X | 82530610 | 82530797 | 187 | Tsga8 | AJ,AKR,BALB,BUB,C3H,CBA,CE,DBA1J,DBA2J,ILNJ2,KK,MA/My,MRL,NZB,PJ,PL,RF,RH2,SEA,SMJ,WSB | feature_elongation,coding_sequence_variant |
| X | 133852723 | 133852799 | 76 | Nxf2 | SPRET | feature_elongation,coding_sequence_variant |
| X | 134593573 | 134594091 | 518 | Tcp11x2 | SPRET | feature_elongation,coding_sequence_variant,5_prime_UTR_variant,intron_variant |
| X | 134698290 | 134698424 | 134 | Gprasp1 | CAST | feature_elongation,5_prime_UTR_variant,intron_variant |
| X | 134698468 | 134698683 | 215 | Gprasp1 | CAST,SPRET | feature_elongation,5_prime_UTR_variant,intron_variant |
| X | 134741200 | 134741423 | 223 | Gprasp2 | SPRET | feature_elongation,5_prime_UTR_variant,intron_variant |
| X | 143129710 | 143129851 | 141 | Alg13 | WSB | feature_elongation,coding_sequence_variant,intron_variant |
| X | 148587855 | 148587926 | 71 | Gm15097 | CAST | feature_elongation,coding_sequence_variant |
| X | 152177710 | 152177784 | 74 | Klf8 | MOLF | feature_elongation,3_prime_UTR_variant |
| X | 168772477 | 168772566 | 89 | Mid1 | B10D2,B10J,MOLF,NOD,NOR,SJL,SMJ | feature_elongation,3_prime_UTR_variant |

| CHR | START | END | SIZE | SYMBOL | STRAINS with VARIANT SV ALLELE | CONSEQUENCE |
|---|---|---|---|---|---|---|
| 1 | 58690090 | 58750341 | 60251 | Flacc1 | SPRET | transcript_amplification |
| 1 | 92417258 | 92435092 | 17834 | Or6b2b | BUB,C3H,CAST,CBA,DBA1J,DBA2J | transcript_amplification |
| 1 | 110816417 | 110831919 | 15502 | Cdh19 | NOD,NOR | feature_elongation,coding_sequence_variant,3_prime_UTR_variant,intron_variant |
| 1 | 117709473 | 118024554 | 315081 | Gm7145 | SWR | transcript_amplification |
| 1 | 139549434 | 139841799 | 292365 | Cfhr4,Cfhr2 | CAST,MOLF | transcript_amplification |
| 1 | 151060176 | 151087731 | 27555 | 3110040M04Rik,Gm8947,Gm47996,Gm47995 | B10D2,B10J | transcript_amplification |
| 1 | 159901795 | 159962954 | 61159 | 4930523C07Rik | C3H | transcript_amplification |
| 1 | 173591669 | 173806055 | 214386 | Ifi211,Ifi203,Mndal | PJ,RF,SPRET | transcript_amplification |
| 1 | 173737663 | 173806054 | 68391 | Ifi203 | AKR,BTBR,BUB,C58J,CBA,CE,DBA1J,DBA2J,ILNJ2,KK,NZB,NZO,PJ,PL,RF,SEA,SMJ,SPRET,SWR,TH | transcript_amplification |
| 1 | 194805605 | 194859147 | 53542 | Cr2 | SPRET | transcript_amplification |
| 10 | 7405164 | 7407158 | 1994 | Ulbp1 | MOLF | feature_elongation,coding_sequence_variant,intron_variant |
| 10 | 22059301 | 22248513 | 189212 | H60b | WSB | transcript_amplification |
| 10 | 33340355 | 33344093 | 3738 | Trdn | KK | feature_elongation,coding_sequence_variant,intron_variant |
| 10 | 33340488 | 33342351 | 1863 | Trdn | KK | feature_elongation,coding_sequence_variant,intron_variant |
| 10 | 33342164 | 33344092 | 1928 | Trdn | KK | feature_elongation,coding_sequence_variant,intron_variant |
| 10 | 58056666 | 58061547 | 4881 | AW822073 | NZB | transcript_amplification |
| 10 | 58057567 | 58062450 | 4883 | AW822073 | 129S1,NOR | transcript_amplification |
| 10 | 78348188 | 78421580 | 73392 | Ilvbl,Or10b1 | SPRET | transcript_amplification |
| 10 | 93995634 | 94018245 | 22611 | Nr2c1 | ILNJ2 | feature_elongation,coding_sequence_variant,intron_variant |
| 11 | 49414269 | 49520440 | 106171 | Or2y12,Or2y13,Or2y11,Or2y10 | SPRET | transcript_amplification |
| 11 | 59003366 | 59006771 | 3405 | Obscn | C3H,C57L,NOR,NZW,SEA,SJL | feature_elongation,coding_sequence_variant,intron_variant |
| 11 | 98496326 | 98531627 | 35301 | Gsdma3 | CAST | transcript_amplification |
| 11 | 99564024 | 99590225 | 26201 | Gm14190 | BALB,BUB,CBA,CE,MRL,NZB,PL,SEA | transcript_amplification |
| 11 | 99575616 | 99616203 | 40587 | Gm14190 | CAST | transcript_amplification |
| 11 | 107751339 | 107860972 | 109633 | Cacng5 | AKR | transcript_amplification |
| 11 | 121260518 | 121332370 | 71852 | Fn3krp | SPRET | transcript_amplification |
| 12 | 31352598 | 31363544 | 10946 | Lamb1 | TH | feature_elongation,coding_sequence_variant,intron_variant |
| 12 | 37221052 | 37375832 | 154780 | Agmo | CBA | transcript_amplification |
| 12 | 55125488 | 55352165 | 226677 | Srp54a,Ppp2r3c,Srp54b,Srp54c,Fam177a,Fam177a2 | SPRET | transcript_amplification |
| 12 | 108810263 | 108823579 | 13316 | Slc25a47 | AKR | transcript_amplification |
| 12 | 112742279 | 112744355 | 2076 | Ahnak2 | C58J,RH2 | feature_elongation,coding_sequence_variant |
| 12 | 112742280 | 112748427 | 6147 | Ahnak2 | C3H,MOLF,PL,SEA,SPRET,WSB | feature_elongation,coding_sequence_variant |
| 12 | 112743715 | 112748380 | 4665 | Ahnak2 | NZO | feature_elongation,coding_sequence_variant |
| 12 | 112744798 | 112748428 | 3630 | Ahnak2 | AJ | feature_elongation,coding_sequence_variant |
| 12 | 112745836 | 112748428 | 2592 | Ahnak2 | 129S1,B10J,BALB,BUB,DBA2J,NZO | feature_elongation,coding_sequence_variant |
| 13 | 35156836 | 35207729 | 50893 | Eci2 | SPRET | transcript_amplification |
| 13 | 66215801 | 66596564 | 380763 | Zfp998,Zfp999,Zfp997 | 129S1,AJ,AKR,BUB,CBA,FVB,KK,MRL,NOD,NZB,RF,SJL,SPRET,TH | transcript_amplification |
| 13 | 113228363 | 113238135 | 9772 | Gzma | KK,NZB,NZW,RF | transcript_amplification |
| 13 | 120072411 | 120119165 | 46754 | Ccl28 | SPRET | transcript_amplification |
| 13 | 120368879 | 120375129 | 6250 | Gm21761 | BTBR,C3H,C57L | transcript_amplification |
| 13 | 120368962 | 120375098 | 6136 | Gm21761 | 129S1,BTBR,C57L,KK,MRL | transcript_amplification |
| 13 | 120486361 | 120499016 | 12655 | Anxa2r2 | 129S1 | transcript_amplification |
| 14 | 15923561 | 16410495 | 486934 | Gm16440,Gm21103,Gm3476,Gm21560,Gm3594,Gm3411 | KK,MOLF | transcript_amplification |
| 14 | 20471026 | 20473838 | 2812 | Cfap70 | SPRET | feature_elongation,coding_sequence_variant,intron_variant |
| 14 | 43765978 | 43855985 | 90007 | Gm10375,Gm5799 | C57L,CAST,MOLF,NZO,PJ,PL | transcript_amplification |
| 14 | 52074876 | 52100002 | 25126 | Ear6 | 129S1,AJ,AKR,BALB,C3H,C57L,CBA,CE,KK,MRL,NZB,NZO,NZW,PJ,PL,RF,RH2,SEA,TH | transcript_amplification |
| 14 | 52076260 | 52101409 | 25149 | Ear6 | 129S1,AJ,AKR,BALB,C3H,C57L,CBA,CE,KK,MRL,NZB,NZO,NZW,PJ,PL,RF,RH2,SEA,TH | transcript_amplification |
| 14 | 56183147 | 56291138 | 107991 | Mcpt2,Mcpt1,Cma2,Mcpt9 | SPRET | transcript_amplification |
| 15 | 3298353 | 3336085 | 37732 | Selenop,Ccdc152 | LP,MOLF | transcript_amplification |
| 15 | 63692681 | 63730658 | 37977 | Gsdmc2 | MOLF | transcript_amplification |
| 15 | 76922487 | 76946076 | 23589 | Apol6 | MA/My | transcript_amplification |
| 15 | 91728970 | 91732538 | 3568 | Smgc | WSB | feature_elongation,coding_sequence_variant,intron_variant |
| 15 | 91730541 | 91734154 | 3613 | Smgc | CBA,PJ | feature_elongation,coding_sequence_variant,intron_variant |
| 16 | 32284135 | 32307408 | 23273 | Slc51a | MOLF | transcript_amplification |
| 16 | 32284178 | 32307325 | 23147 | Slc51a | MOLF | transcript_amplification |
| 16 | 88666970 | 88671824 | 4854 | Krtap19-2 | C58J | transcript_amplification |
| 17 | 7596967 | 7598125 | 1158 | Tcp10a | KK | feature_elongation,coding_sequence_variant,intron_variant |
| 17 | 9052729 | 9235028 | 182299 | Pde10a,1700010I14Rik | SPRET | transcript_amplification |
| 17 | 13447013 | 13456388 | 9375 | Smok2b | MOLF | transcript_amplification |
| 17 | 22314477 | 22407570 | 93093 | Zfp947 | SPRET | transcript_amplification |
| 17 | 30666348 | 31143185 | 476837 | Glo1,Dnah8,Btbd9 | 129S1,AJ,AKR,B10D2,C3H,C57L,CBA,CE,DBA1J,DBA2J,ILNJ2,MRL,PJ,RH2,SEA | transcript_amplification |
| 17 | 30725784 | 31111284 | 385500 | Glo1,Dnah8,Btbd9 | MOLF,RF | transcript_amplification |
| 17 | 31217564 | 31250037 | 32473 | Umodl1 | PJ | transcript_amplification |
| 17 | 31441857 | 31499856 | 57999 | Rsph1 | C58J,ILNJ2,NZB,NZO,NZW | transcript_amplification |

| | | | | | | |
|---|---|---|---|---|---|---|
| 17 | 31472596 | 31528746 | 56150 | Rsph1 | CAST | transcript_amplification |
| 17 | 31473829 | 31562278 | 88449 | Rsph1 | KK,MOLF,NOR | transcript_amplification |
| 17 | 36341360 | 36453492 | 112132 | H2-T25,H2-T22,H2-T27,Gm19684,H2-Bl | SPRET | transcript_amplification |
| 17 | 36387349 | 36449718 | 62369 | H2-T25,Gm19684,H2-Bl | BTBR | transcript_amplification |
| 17 | 38519624 | 38520991 | 1367 | Or2n1c | SPRET | feature_elongation,coding_sequence_variant,3_prime_UTR_variant |
| 18 | 61592541 | 61699963 | 107422 | Arhgef37 | NZW,SMJ | transcript_amplification |
| 19 | 39664393 | 39917625 | 253232 | Cyp2c68,Cyp2c40,Cyp2c69 | SPRET | transcript_amplification |
| 2 | 32135487 | 32183245 | 47758 | Uck1,Swi5 | AKR | transcript_amplification |
| 2 | 69502896 | 69568403 | 65507 | Fastkd1 | CAST | transcript_amplification |
| 2 | 77700000 | 77857702 | 157702 | Cwc22 | MOLF | transcript_amplification |
| 2 | 86614586 | 86616484 | 1898 | Or5t5 | MOLF | feature_elongation,coding_sequence_variant,5_prime_UTR_variant,intron_variant |
| 2 | 118533766 | 118702353 | 168587 | Disp2,Ccdc9b,Plcb2,Knstrn,Phgr1,Inafm2 | BUB | transcript_amplification |
| 2 | 151556595 | 151659420 | 102825 | Psmf1 | WSB | transcript_amplification |
| 2 | 176402896 | 176769408 | 366512 | Gm14296,Gm14419,Gm14305,Gm14295,Gm14408 | PJ | transcript_amplification |
| 2 | 176493556 | 176495075 | 1519 | Gm14295 | SPRET | feature_elongation,coding_sequence_variant,5_prime_UTR_variant,intron_variant |
| 3 | 40818698 | 40865657 | 46959 | Fundc2b,Abhd18 | SPRET | transcript_amplification,feature_elongation,coding_sequence_variant,5_prime_UTR_variant,intron_variant |
| 4 | 41883740 | 42247346 | 363606 | Il11ra3,Ccl27al,Ccl21f,Ccl21d,Spata31f1d,Gm20878 | NOD | transcript_amplification |
| 4 | 41886613 | 42247346 | 360733 | Il11ra3,Ccl27al,Ccl21f,Ccl21d,Spata31f1d,Gm20878 | AKR,NOR,NZB,PJ | transcript_amplification |
| 4 | 62007380 | 62077857 | 70477 | Mup21 | 129S1,AJ,AKR,BALB,BUB,C3H,C57L,CE,DBA1J,DBA2J,ILNJ2,MA/My,MOLF,MRL,NOR,NZB,NZO,NZW,PJ,RF,RH2,SEA,SJL,SWR,WSB | transcript_amplification |
| 4 | 62413147 | 62442066 | 28919 | Alad,Hdhd3 | 129S1,AJ,AKR,B10D2,B10J,BALB,BUB,C3H,C57L,C58J,CBA,CE,DBA1J,DBA2J,FVB,ILNJ2,LP,MA/My,MRL,NOD,NOR,NZB,NZO,NZW,PJ,PL,RF,RH2,SEA,SJL,SWR,TH | transcript_amplification |
| 4 | 124746849 | 124899068 | 152219 | Cdca8,Epha10,Maneal,Airim | NOR | transcript_amplification |
| 4 | 143660682 | 143693357 | 32675 | Pramel16 | DBA2J | transcript_amplification |
| 4 | 143783909 | 143868185 | 84276 | Pramel4,Gm13102 | 129S1,BTBR,BUB,KK,MA/My,PJ,SJL,SMJ,WSB | transcript_amplification |
| 4 | 143820689 | 143852437 | 31748 | Gm13102 | C58J | transcript_amplification |
| 4 | 145158084 | 145310836 | 152752 | Zfp990 | DBA1J,DBA2J | transcript_amplification |
| 4 | 145159161 | 145311914 | 152753 | Zfp990 | DBA2J,LP | transcript_amplification |
| 5 | 11448768 | 11947739 | 498971 | Gm6460,Gm8906,Gm8897,Gm8922,Gm8926 | TH | transcript_amplification |
| 5 | 29901773 | 29964489 | 62716 | Dnajb6 | CAST | transcript_amplification |
| 5 | 87726730 | 87733196 | 6466 | Sult1e1 | BALB | feature_elongation,coding_sequence_variant,intron_variant |
| 5 | 105154100 | 105215679 | 61579 | Gbp8 | SPRET | transcript_amplification |
| 5 | 138471917 | 138565066 | 93149 | 1700123K08Rik | AJ,RH2,SJL | transcript_amplification |
| 6 | 89902662 | 90135285 | 232623 | Vmn1r45,Vmn1r49,Vmn1r51,Vmn1r48,Vmn1r47,Vmn1r46,Vmn1r50 | CAST | transcript_amplification |
| 6 | 128864454 | 128947594 | 83140 | Clec2i | CAST,NZO | transcript_amplification |
| 6 | 130020546 | 130239164 | 218618 | Klra8,Klra7,Klra9,Klra4 | KK,LP,NOD,NOR,NZB,NZW,SJL,SWR,TH | transcript_amplification |
| 7 | 4575378 | 4576838 | 1460 | Ptprh | SPRET | feature_elongation,coding_sequence_variant,intron_variant |
| 7 | 4853380 | 4857442 | 4062 | Isoc2b | CAST | feature_elongation,coding_sequence_variant,5_prime_UTR_variant,intron_variant |
| 7 | 9455229 | 9817901 | 362672 | Vmn2r50,Vmn2r49,Vmn2r48,Vmn2r47 | 129S1,BTBR,LP,NZB | transcript_amplification |
| 7 | 24419004 | 24436181 | 17177 | Lypd11 | BUB,C58J,CAST,MOLF,MRL,PL | transcript_amplification |
| 7 | 30618519 | 30637774 | 19255 | Hamp2 | CE | transcript_amplification |
| 7 | 32516946 | 32869822 | 352876 | Scgb2b17,Scgb2b15,Scgb1b17,Scgb1b15 | NZB,NZW | transcript_amplification |
| 7 | 41685103 | 42087680 | 402577 | Vmn2r61,Vmn2r60 | CAST,ILNJ2,MOLF,PJ,WSB | transcript_amplification |
| 7 | 101206784 | 101232156 | 25372 | Art2b | NZO | transcript_amplification |
| 7 | 104419177 | 104421446 | 2269 | Usp17le | KK | feature_elongation,coding_sequence_variant,intron_variant |
| 7 | 104419767 | 104424329 | 4562 | Usp17le | AJ,AKR,B10D2,C57L,CBA,DBA1J,LP,NOR,NZB,PJ,RF | feature_elongation,coding_sequence_variant,intron_variant |
| 7 | 108266621 | 108294904 | 28283 | Or5p81 | CAST,CE,KK,MOLF,NOD,SMJ,SWR,WSB | transcript_amplification |
| 7 | 127317786 | 127320306 | 2520 | Ctf2 | CAST | feature_elongation,coding_sequence_variant,3_prime_UTR_variant,intron_variant |
| 7 | 127317821 | 127320294 | 2473 | Ctf2 | CAST | feature_elongation,coding_sequence_variant,3_prime_UTR_variant,intron_variant |
| 7 | 130194306 | 130269962 | 75656 | Tacc2 | B10D2 | feature_elongation,coding_sequence_variant,intron_variant |
| 7 | 130695013 | 130699432 | 4419 | Dmbt1 | CAST | feature_elongation,coding_sequence_variant,intron_variant |
| 7 | 130885662 | 130888990 | 3328 | Cdcp3 | CAST | feature_elongation,coding_sequence_variant,intron_variant |
| 7 | 139998710 | 140028824 | 30114 | Or13a21 | CAST,NZW,PL,SPRET | transcript_amplification |
| 8 | 72755702 | 72853912 | 98210 | Or2z8 | CAST | transcript_amplification |

| | | | | | | |
|---|---|---|---|---|---|---|
| 8 | 107142830 | 107147378 | 4548 | Zfp90 | MRL | feature_elongation,coding_sequence_variant,intron_variant |
| 8 | 111820177 | 111877683 | 57506 | Pdpr | SPRET | transcript_amplification |
| 9 | 18549616 | 18550919 | 1303 | Muc16 | NZB | feature_elongation,coding_sequence_variant |
| 9 | 18549684 | 18551576 | 1892 | Muc16 | MOLF | feature_elongation,coding_sequence_variant |
| 9 | 18549811 | 18551701 | 1890 | Muc16 | MOLF | feature_elongation,coding_sequence_variant |
| 9 | 39339307 | 39383012 | 43705 | Or8g35,Or8g34 | MOLF | transcript_amplification |
| 9 | 56607833 | 56652136 | 44303 | Lingo1 | NOR | feature_elongation,5_prime_UTR_variant,intron_variant |
| 9 | 113809113 | 113849909 | 40796 | Fbxl2 | PL | feature_elongation,coding_sequence_variant,intron_variant |
| X | 7623820 | 7644895 | 21075 | Tfe3 | BALB | transcript_amplification |

**Table S11. List of DELs present in all 39 or in the 35 classical inbred strains with CTCF binding sites.** The chromosome, starting and ending position of the CTCF recognition element and its sequence, size, gene symbol, strains with the variant allele for each DEL, the CTCF score (the strength of the motif match based on the position weight matrix), p-value and Q-value (p-value that is adjusted for the false discovery rate) are shown. These were determined using the CTCF (v0.99.11) R package in Bioconductor. All predicted sites have a P-value that is below the recommended cutoff (1 x $10^{-6}$). The number of peaks in the 42 ENCODE files with CTCF ChIP-Seq data obtained using mouse cells or tissues, which indicates that CTCF is bound to that site, and the data source for each peak is shown.

| DEL_NUM | DEL_CHR | DEL_START | DEL_END | DEL_LENGTH | STRAINS with VARIANT SV ALLELE | CTCF_CHR | CTCF_START | CTCF_END | CTCF_LENGTH | CTCF_STRAND | CTCF_SCORE | CTCF_PVALUE | CTCF_QVALUE | CTCF_SEQUENCE | # PEAKS | DATA_SOURCE |
|---|---|---|---|---|---|---|---|---|---|---|---|---|---|---|---|---|
| 1 | 1 | 4650900 | 4651097 | 197 | C3H.CBA.DBA1J.DBA2J.MRL.RF | 1 | 4650956 | 4650989 | 34 | + | 17.2439 | 5.79E-07 | 0.0995 | TAGCTGTCCTCAGATACTCCAGAAGAGGGCATCA | | |
| 2 | 1 | 6438536 | 6438750 | 214 | AJ,BALB,C3H,CBA,DBA1J,DBA2J,FVB,KK,MRL.NZO.RF.RH2.SEA.SMJ.TH | 1 | 6438609 | 6438642 | 34 | + | 16.4472 | 9.61E-07 | 0.125 | TAGCTGTCTTCAGACACACCAGAAGAGGGCGTCA | | |
| 3 | 1 | 13111148 | 13111363 | 215 | 129S1,AJ,AKR,BALB,BTBR,BUB,C3H,C57L,CBA,CE,DBA1J,DBA2J,FVB,ILNJ2,KK,LP,MA/My,MRL,NOD,NOR,NZB,NZO,NZW,PJ,PL,RF,RH2,SEA,SJL,SMJ,SWR,TH | 1 | 13111219 | 13111252 | 34 | + | 17.8293 | 3.95E-07 | 0.083 | TAGCTGTCTTCAGACCCTCCAGAAGAGGGCGTCA | | |
| 4 | 1 | 34041759 | 34041962 | 203 | CE.NOD.RF.SWR | 1 | 34041820 | 34041853 | 34 | + | 16.4634 | 9.51E-07 | 0.125 | TAGCTGTCTTCAGACGCACCAGAAGAGGGCATCA | | |
| 5 | 1 | 34129544 | 34129734 | 190 | DBA1J,DBA2J,ILNJ2,KK,MA/My,NOD,NOR,NZW,PJ,PL,SJL,SWR,TH | 1 | 34129587 | 34129620 | 34 | + | 18.5285 | 2.48E-07 | 0.0666 | CAGCTGTCTTCAGACACTCCAGAAGAGGGCGTCA | | |
| 6 | 1 | 39303819 | 39304362 | 543 | MA/Mv | 1 | 39304032 | 39304065 | 34 | + | 18.3089 | 2.87E-07 | 0.0691 | ATGCAGCTAGAGACACGAGCTCCAGGGGGTACTG | | |
| 7 | 1 | 47813238 | 47813450 | 212 | SWR | 1 | 47813306 | 47813339 | 34 | + | 18.3659 | 2.76E-07 | 0.069 | TAGCTGTCTTCAGACACTCCAGAAGAGGGCGCCA | | |
| 7 | 1 | 47813238 | 47813450 | 212 | SWR | 1 | 47813321 | 47813339 | 19 | + | 17.5574 | 6.32E-07 | 0.166 | ACTCCAGAAGAGGGCGCCA | | |
| 8 | 1 | 52343806 | 52343974 | 168 | BTBR.C57L.KK.NZO.NZW.SMJ.SWR | 1 | 52343849 | 52343867 | 19 | + | 17.8852 | 5.08E-07 | 0.151 | ACACCAGAAGAGGGCACCA | | |
| 9 | 1 | 52621206 | 52621399 | 193 | AKR,BALB,BUB,C58J,MRL,NOD,NOR,NZO,NZW.PL.RF.RH2.SEA.SMJ.SWR | 1 | 52621256 | 52621289 | 34 | + | 18.3659 | 2.76E-07 | 0.069 | TAGCTGTCTTCAGACACTCCAGAAGAGGGCGCCA | | |
| 9 | 1 | 52621206 | 52621399 | 193 | AKR,BALB,BUB,C58J,MRL,NOD,NOR,NZO,NZW,PL,RF,RH2,SEA,SMJ,SWR | 1 | 52621271 | 52621289 | 19 | + | 17.5574 | 6.32E-07 | 0.166 | ACTCCAGAAGAGGGCGCCA | | |
| 10 | 1 | 54942393 | 54942572 | 179 | ILNJ2.MA/Mv.NOD.NOR.PJ | 1 | 54942485 | 54942519 | 35 | + | 16.8807 | 5.80E-07 | 0.165 | CTGCTCTCAAGGCTTTGGAGCTGAAGGGGGCGTTG | | |
| 11 | 1 | 58189216 | 58189416 | 200 | BALB,BTBR,BUB,C57L,C58J,CBA,CE,DBA1J,DBA2J,MRL,PL,RH2,SEA,SJL | 1 | 58189272 | 58189305 | 34 | + | 16.4634 | 9.51E-07 | 0.125 | TAGCTGTCTTCAGACGCACCAGAAGAGGGCATCA | | |
| 12 | 1 | 59271285 | 59271491 | 206 | 129S1,BALB,BUB,CE,NOD,NOR,PL,RH2,SEA,SJL,SWR | 1 | 59271343 | 59271376 | 34 | + | 16.4472 | 9.61E-07 | 0.125 | TAGCTGTCTTCAGACACACCAGAAGAGGGCGTCA | | |
| 13 | 1 | 59823348 | 59823550 | 202 | AKR,BALB,BUB,ILNJ2,MA/My,NOD,NOR,NZB,NZO,NZW,PL,RH2,SEA,SJL,SMJ,SWR | 1 | 59823406 | 59823439 | 34 | + | 18.3659 | 2.76E-07 | 0.069 | TAGCTGTCTTCAGACACTCCAGAAGAGGGCGCCA | | |
| 13 | 1 | 59823348 | 59823550 | 202 | AKR,BALB,BUB,ILNJ2,MA/My,NOD,NOR,NZB,NZO,NZW,PL,RH2,SEA,SJL,SMJ,SWR | 1 | 59823421 | 59823439 | 19 | + | 17.5574 | 6.32E-07 | 0.166 | ACTCCAGAAGAGGGCGCCA | | |
| 14 | 1 | 69200181 | 69200898 | 717 | C58J | 1 | 69200651 | 69200685 | 35 | + | 15.9725 | 9.70E-07 | 0.212 | CTGCATGTACACCTACATGCCAGAATAGGGCATCA | | |
| 15 | 1 | 71683612 | 71683798 | 186 | AKR.BUB.PL.RF.SJL.SWR | 1 | 71683669 | 71683702 | 34 | + | 16.4472 | 9.61E-07 | 0.125 | TAGCTGTCTTCAGACACACCAGAAGAGGGCGTCA | | |
| 16 | 1 | 82772748 | 82772826 | 78 | RF | 1 | 82772782 | 82772815 | 34 | + | 16.7724 | 7.83E-07 | 0.115 | TAGCTGTCTTCAGATGCACCAGAAGAGGGCGTCA | | |
| 17 | 1 | 84949337 | 84949564 | 227 | AKR.BUB.CBA.MRL.NOD.PJ.PL.SMJ | 1 | 84949420 | 84949453 | 34 | + | 16.4634 | 9.51E-07 | 0.125 | TAGCTGTCTTCAGACGCACCAGAAGAGGGCATCA | | |
| 18 | 1 | 85194795 | 85194998 | 203 | AKR.BUB.CBA.MRL.NOD.SMJ | 1 | 85194854 | 85194887 | 34 | + | 16.4472 | 9.61E-07 | 0.125 | TAGCTGTCTTCAGACACACCAGAAGAGGGCGTCA | | |
| 19 | 1 | 99558814 | 99559114 | 300 | MRL.PL | 1 | 99558821 | 99558854 | 34 | + | 19.5041 | 1.26E-07 | 0.0441 | CTGCGATAACATTCGCCATCACAAGATGGCACCA | | |
| 20 | 1 | 100074793 | 100075329 | 536 | AKR.LP.MA/Mv.PJ.SEA | 1 | 100075293 | 100075326 | 34 | + | 16.8862 | 7.28E-07 | 0.112 | ATGCAGCTAGAGACATGAGCTCCGGGGGGCGGGG | | |
| 21 | 1 | 118652923 | 118653125 | 202 | AJ,BUB,C3H,CBA,CE,DBA1J,DBA2J,ILNJ2,KK,LP,MA/My,MRL,NOD,NOR,NZO,NZW,PJ,PL,RH2 | 1 | 118652982 | 118653016 | 35 | + | 18.1284 | 2.79E-07 | 0.109 | CTGTAGCTGTCTTCAGACACCAGAAGAGGGCATCA | | |
| 22 | 1 | 120021565 | 120021791 | 226 | 129S1,AJ,AKR,BALB,BTBR,BUB,C3H,C57L,CBA,CE,DBA1J,DBA2J,FVB,ILNJ2,KK,LP,MA/My,MRL,NOD,NOR,NZB,NZO,NZW,PJ,PL,RF,RH2,SEA,SJL,SMJ,SWR,TH | 1 | 120021645 | 120021678 | 34 | + | 18.374 | 2.75E-07 | 0.069 | TAGCTGTCTTCAGTCACTCCAGAAGAGGGCGCCA | 3 | ENCFF367UNR,ENCFF093ZRU |
| 22 | 1 | 120021565 | 120021791 | 226 | 129S1,AJ,AKR,BALB,BTBR,BUB,C3H,C57L,CBA,CE,DBA1J,DBA2J,FVB,ILNJ2,KK,LP,MA/My,MRL,NOD,NOR,NZB,NZO,NZW,PJ,PL,RF,RH2,SEA,SJL,SMJ,SWR,TH | 1 | 120021660 | 120021678 | 19 | + | 17.5574 | 6.32E-07 | 0.166 | ACTCCAGAAGAGGGCGCCA | 2 | ENCFF093ZRU,ENCFF367UNR |
| 23 | 1 | 127773686 | 127773912 | 226 | MRL.NZB.RF.SWR | 1 | 127773784 | 127773802 | 19 | + | 17.8852 | 5.08E-07 | 0.151 | ACACCAGAAGAGGGCACCA | | |
| 24 | 1 | 132752505 | 132752724 | 219 | AKR.ILNJ2.KK.NZB.NZW.SJL.SWR | 1 | 132752580 | 132752613 | 34 | + | 16.7724 | 7.83E-07 | 0.115 | TAGCTGTCTTCGGACACACCAGAAGAGGGCATCA | 1 | ENCFF093ZRU |
| 25 | 1 | 135334873 | 135335080 | 207 | 129S1,AJ,AKR,BTBR,BUB,C3H,CBA,CE,DBA1J,DBA2J,FVB,ILNJ2,KK,MA/My,MRL,NOD,NOR,NZB,NZO,NZW,PJ,PL,RF,RH2,SJL,SMJ,SWR,TH | 1 | 135334938 | 135334971 | 34 | + | 16.4634 | 9.51E-07 | 0.125 | TAGCTGTCTTCAGACGCACCAGAAGAGGGCATCA | | |
| 26 | 1 | 152763769 | 152763983 | 214 | AKR,BTBR,BUB,C57L,FVB,KK,NZB,NZO,NZW,RF,SJL,SMJ,SWR,TH | 1 | 152763842 | 152763875 | 34 | + | 16.4634 | 9.51E-07 | 0.125 | TAGCTGTCTTCAGACGCACCAGAAGAGGGCATCA | | |
| 27 | 1 | 153752071 | 153752274 | 203 | BUB,C57L,DBA1J,DBA2J,FVB,MA/My,RF,SJL,SWR,TH | 1 | 153752130 | 153752164 | 35 | + | 18.1284 | 2.79E-07 | 0.109 | CTGTAGCTGTCTTCAGACACCAGAAGAGGGCATCA | | |
| 28 | 1 | 153756202 | 153756697 | 495 | BUB,C57L,DBA1J,DBA2J,FVB,MA/My,RF,SJL,SWR,TH | 1 | 153756217 | 153756250 | 34 | + | 17.9106 | 3.74E-07 | 0.0805 | CCGCCCTCACATTCGCCGTCACAAGATGGCGCTG | | |
| 29 | 1 | 155320630 | 155320775 | 145 | SWR | 1 | 155320613 | 155320647 | 35 | + | 16.8624 | 5.86E-07 | 0.166 | GTGCTTTCTCTGCTTGTGGCCAGGCGGTGGTGGTG | | |
| 30 | 1 | 156350301 | 156350536 | 235 | 129S1,AJ,AKR,BALB,BTBR,BUB,C3H,C57L,C58J,CBA,CE,DBA1J,DBA2J,FVB,ILNJ2,KK,LP,MA/My,MRL,NOD,NOR,NZB,NZO,NZW,PJ,PL,RF,RH2,SEA,SWR,TH | 1 | 156350395 | 156350428 | 34 | + | 18.3659 | 2.76E-07 | 0.069 | TAGCTGTCTTCAGACACTCCAGAAGAGGGCGCCA | | |
| 30 | 1 | 156350301 | 156350536 | 235 | 129S1,AJ,AKR,BALB,BTBR,BUB,C3H,C57L,C58J,CBA,CE,DBA1J,DBA2J,FVB,ILNJ2,KK,LP,MA/My,MRL,NOD,NOR,NZB,NZO,NZW,PJ,PL,RF,RH2,SEA,SWR,TH | 1 | 156350410 | 156350428 | 19 | + | 17.5574 | 6.32E-07 | 0.166 | ACTCCAGAAGAGGGCGCCA | | |
| 31 | 1 | 160240285 | 160240519 | 234 | BALB.C58J.DBA1J.DBA2J.LP | 1 | 160240377 | 160240410 | 34 | + | 16.4634 | 9.51E-07 | 0.125 | TAGCTGTCTTCAGACGCACCAGAAGAGGGCATCA | | |
| 32 | 1 | 166094270 | 166094511 | 241 | BTBR,DBA1J,DBA2J,FVB,ILNJ2,KK,NOD,NOR,RF,SMJ,SWR | 1 | 166094354 | 166094387 | 34 | + | 17.7967 | 4.04E-07 | 0.0841 | CAGCTGTCCTCAGATACTCCAGAAGAGGGAGTCA | | |
| 33 | 1 | 167058302 | 167058527 | 225 | BTBR,FVB,ILNJ2,KK,NZB,NZO,NZW,RF,SMJ,SWR,TH | 1 | 167058377 | 167058410 | 34 | + | 17.2439 | 5.79E-07 | 0.0995 | TAGCTGTCCTCAGATACTCCAGAAGAGGGCATCA | | |
| 34 | 1 | 169345502 | 169345615 | 113 | 129S1,AJ,BTBR,BUB,C3H,C57L,CE,FVB,KK,MRL,NOD,NOR,NZB,NZO,NZW,PJ,PL,RF,RH2,SEA,SJL,SMJ,SWR,TH | 1 | 169345598 | 169345616 | 19 | + | 17.377 | 7.11E-07 | 0.174 | AGGCCAGCAGGGGGAGTCA | 5 | ENCFF821BIA,ENCFF024JOZ,ENCFF327LYS,ENCFF907XEG,ENCFF198SSR |
| 35 | 1 | 170757166 | 170757376 | 210 | 129S1,AJ,BTBR,BUB,C3H,C57L,CE,FVB,KK,MRL,NOD,NOR,NZB,NZO,NZW,PJ,PL,RF,RH2,SEA,SJL,SMJ,SWR,TH | 1 | 170757234 | 170757267 | 34 | + | 17.2439 | 5.79E-07 | 0.0995 | TAGCTGTCCTCAGATACTCCAGAAGAGGGCATCA | | |
| 36 | 1 | 171859494 | 171859700 | 206 | 129S1,AJ,AKR,BALB,BUB,C3H,C57L,CBA,CE,DBA1J,DBA2J,FVB,KK,MRL,NOD,NOR,NZB,NZO,NZW,PJ,PL,RF,RH2,SEA,SJL,SMJ,SWR,TH | 1 | 171859560 | 171859593 | 34 | + | 17.2358 | 5.82E-07 | 0.0997 | GAGCTGTCTTCAGATACACCAGAAGAGGGCGTCA | | |
| 37 | 1 | 171982743 | 171982946 | 203 | 129S1,AJ,AKR,BALB,BUB,C3H,C57L,CBA,CE,DBA1J,DBA2J,FVB,KK,MRL,NOD,NOR,NZB,NZO,NZW,PJ,PL,RF,RH2,SEA,SJL,SMJ,SWR,TH | 1 | 171982805 | 171982838 | 34 | + | 17.2439 | 5.79E-07 | 0.0995 | TAGCTGTCCTCAGATACTCCAGAAGAGGGCATCA | | |
| 38 | 1 | 173236114 | 173236314 | 200 | 129S1,AJ,AKR,BALB,BTBR,BUB,C3H,C57L,C58J,CBA,CE,DBA1J,DBA2J,FVB,ILNJ2,KK,MRL,NOD,NOR,NZB,NZO,PJ,PL,RF,RH2,SEA,SJL,SMJ,SWR,TH | 1 | 173236171 | 173236204 | 34 | + | 16.4634 | 9.51E-07 | 0.125 | TAGCTGTCTTCAGACGCACCAGAAGAGGGCATCA | 1 | ENCFF093ZRU |
| 39 | 1 | 177614791 | 177614872 | 81 | PL | 1 | 177614763 | 177614797 | 35 | + | 15.9266 | 9.95E-07 | 0.213 | ATGCACTGCCCCTGGGCTGCAAGAAGGGGGCCAAG | | |
| 40 | 1 | 179019252 | 179019473 | 221 | AKR | 1 | 179019347 | 179019365 | 19 | + | 17.9836 | 4.75E-07 | 0.151 | ACACCAGAAGAGGGCGGCA | | |
| 41 | 1 | 180627208 | 180627420 | 212 | AJ.BUB.CE.PJ.RH2.SEA.TH | 1 | 180627277 | 180627310 | 34 | + | 18.3659 | 2.76E-07 | 0.069 | TAGCTGTCTTCAGACACTCCAGAAGAGGGCGCCA | | |
| 41 | 1 | 180627208 | 180627420 | 212 | AJ.BUB.CE.PJ.RH2.SEA.TH | 1 | 180627292 | 180627310 | 19 | + | 17.5574 | 6.32E-07 | 0.166 | ACTCCAGAAGAGGGCGCCA | | |
| 42 | 1 | 181614406 | 181615076 | 670 | TH | 1 | 181614995 | 181615013 | 19 | + | 17.541 | 6.39E-07 | 0.167 | TTGCCTGCAGGTGGCAGAC | | |
| 43 | 1 | 182369734 | 182370226 | 492 | CE | 1 | 182370072 | 182370090 | 19 | + | 17.082 | 8.61E-07 | 0.188 | TGGCCACCAGGTGGAATAA | | |
| 44 | 1 | 183176095 | 183176331 | 236 | TH | 1 | 183176098 | 183176131 | 34 | + | 17.3496 | 5.41E-07 | 0.0962 | ATGTGGTGCTCACAGAGGCCAGGAGAGGGTGTCA | | |
| 45 | 1 | 185122912 | 185123183 | 271 | 129S1,AJ,AKR,BALB,BTBR,BUB,C3H,C57L,CE,FVB,ILNJ2,KK,MA/My,MRL,NOD,NOR,NZB,NZW,PJ,PL,RF,RH2,SEA,SJL,SWR,TH | 1 | 185123039 | 185123072 | 34 | + | 16.4634 | 9.51E-07 | 0.125 | TAGCTGTCTTCAGACGCACCAGAAGAGGGCATCA | | |
| 46 | 1 | 195027700 | 195027901 | 201 | NZO.NZW.RF | 1 | 195027759 | 195027792 | 34 | + | 16.4472 | 9.61E-07 | 0.125 | TAGCTGTCTTCAGACACACCAGAAGAGGGCGTCA | | |
| 47 | 1 | 16716263 | 16716466 | 203 | AJ,AKR,BALB,BUB,C3H,CBA,DBA1J,DBA2J,FVB,ILNJ2,KK,MRL,NOD,NOR,NZB,NZO,NZW,PJ,PL,RF,RH2,SEA,SJL,SMJ,SWR,TH | 1 | 16716389 | 16716407 | 19 | - | 17.8852 | 5.08E-07 | 0.151 | ACACCAGAAGAGGGCACCA | 1 | ENCFF093ZRU |
| 48 | 1 | 36005108 | 36005315 | 207 | RF | 1 | 36005232 | 36005265 | 34 | - | 17.8455 | 3.91E-07 | 0.0823 | TAGCTGTCCTCAGACACTCCAGAAGAGGGCGTCA | | |
| 49 | 1 | 36858706 | 36858904 | 198 | AJ,AKR,BUB,C3H,CBA,CE,FVB,KK,MRL,NOD,NZB,NZO,NZW,PJ,PL,RF,RH2,SJL,SMJ,SWR | 1 | 36858832 | 36858865 | 34 | - | 18.0732 | 3.36E-07 | 0.0757 | CAGCTGTCTTCAGACACTCCAGAAGAGGGCATCA | | |
| 50 | 1 | 38469541 | 38469667 | 126 | DBA1J.DBA2J.MA/Mv.NOR.RF | 1 | 38469529 | 38469563 | 35 | - | 15.9266 | 9.95E-07 | 0.213 | CTGCCCACCACTGGAGGGACCAGGAGGTGGTGCTC | | |
| 51 | 1 | 39391988 | 39392431 | 443 | MA/Mv | 1 | 39392018 | 39392036 | 19 | - | 20.3607 | 8.29E-08 | 0.0655 | AGGCCAGAAGAGGGCACCA | 5 | ENCFF099QNE.ENCFF821BIA.ENCFF502WDT.ENCFF367UNR.ENCFF093ZRU |
| 51 | 1 | 39391988 | 39392431 | 443 | MA/Mv | 1 | 39392018 | 39392052 | 35 | - | 27.2844 | 3.67E-10 | 0.00426 | CTGCATGTGTGCCTGCAGGCCAGAAGAGGGCACCA | 5 | ENCFF099QNE.ENCFF821BIA.ENCFF502WDT.ENCFF367UNR.ENCFF093ZRU |
| 52 | 1 | 45470542 | 45470747 | 205 | SWR.TH | 1 | 45470666 | 45470699 | 34 | - | 16.4472 | 9.61E-07 | 0.125 | TAGCTGTCTTCAGACACACCAGAAGAGGGCGTCA | | |
| 53 | 1 | 46921997 | 46922210 | 213 | 129S1,AJ,AKR,BTBR,BUB,C3H,C57L,CBA,CE,DBA1J,DBA2J,FVB,ILNJ2,KK,MA/My,MRL,NOD,NOR,NZO,NZW,PJ,PL,RF,RH2,SJL,SMJ.SWR.TH | 1 | 46922123 | 46922156 | 34 | - | 18.2439 | 3.00E-07 | 0.0708 | TAGCTGTCCTCAGATACTCCAGAAGGGGGCATCA | | |
| 54 | 1 | 52730551 | 52730765 | 214 | AKR,BALB,BUB,C58J,MRL,NOD,NOR,NZO,NZW,PL,RF,RH2,SEA,SMJ,SWR | 1 | 52730675 | 52730693 | 19 | - | 17.5574 | 6.32E-07 | 0.166 | ACTCCAGAAGAGGGCGCCA | | |
| 54 | 1 | 52730551 | 52730765 | 214 | AKR,BALB,BUB,C58J,MRL,NOD,NOR,NZO,NZW,PL,RF,RH2,SEA,SMJ,SWR | 1 | 52730675 | 52730708 | 34 | - | 18.3659 | 2.76E-07 | 0.069 | TAGCTGTCTTCAGACACTCCAGAAGAGGGCGCCA | | |
| 55 | 1 | 55201091 | 55201868 | 777 | SMJ | 1 | 55201269 | 55201287 | 19 | - | 19.0656 | 2.23E-07 | 0.105 | AGTCCAGAAGAGGGCGCCA | 27 | ENCFF805QBU,ENCFF093ZRU,ENCFF089GBC,ENCFF165LXO,ENCFF367UNR,ENCFF092NMJ,ENCFF198SSR,ENCFF951GOP,ENCFF201JUA,ENCFF065ODV,ENCFF136ZDK,ENCFF099QNE,ENCFF321AGQ,ENCFF097CAQ,ENCFF589ZBJ,ENCFF142CNG,ENCFF052KVO,ENCFF055HAP,ENCFF194HVL,ENCFF821BIA,ENCFF957SPH,ENCFF533APC,ENCFF024JOZ,ENCFF327LYS,ENCFF502WDT,ENCFF907XEG,ENCFF883UPM |
| 55 | 1 | 55201091 | 55201868 | 777 | SMJ | 1 | 55201269 | 55201303 | 35 | - | 19.5872 | 1.13E-07 | 0.0634 | GTACAGGCCCCCAAAGAGTCCAGAAGAGGGCGCCA | 27 | ENCFF805QBU,ENCFF093ZRU,ENCFF089GBC,ENCFF165LXO,ENCFF367UNR,ENCFF092NMJ,ENCFF198SSR,ENCFF951GOP,ENCFF201JUA,ENCFF065ODV,ENCFF136ZDK,ENCFF099QNE,ENCFF321AGQ,ENCFF097CAQ,ENCFF589ZBJ,ENCFF142CNG,ENCFF052KVO,ENCFF055HAP,ENCFF194HVL,ENCFF821BIA,ENCFF957SPH,ENCFF533APC,ENCFF024JOZ,ENCFF327LYS,ENCFF502WDT,ENCFF907XEG,ENCFF883UPM |
| 56 | 1 | 58599597 | 58599800 | 203 | CE | 1 | 58599721 | 58599739 | 19 | - | 17.5574 | 6.32E-07 | 0.166 | ACTCCAGAAGAGGGCGCCA | 4 | ENCFF821BIA.ENCFF201JUA.ENCFF093ZRU.ENCFF367UNR |
| 56 | 1 | 58599597 | 58599800 | 203 | CE | 1 | 58599721 | 58599754 | 34 | - | 20.252 | 7.40E-08 | 0.0316 | TAGCTGTCCTCAGACACTCCAGAAGAGGGCGCCA | 4 | ENCFF821BIA.ENCFF201JUA.ENCFF093ZRU.ENCFF367UNR |
| 57 | 1 | 58716964 | 58717166 | 202 | CE | 1 | 58716937 | 58716971 | 35 | - | 28.1835 | 1.64E-10 | 0.00426 | CTGCATGTACACCCACAGGCCAGAAGAGGGCACCA | 21 | ENCFF093ZRU,ENCFF367UNR,ENCFF136ZDK,ENCFF201JUA,ENCFF430PPJ,ENCFF099QNE,ENCFF327LYS,ENCFF821BIA,ENCFF907XEG,ENCFF024JOZ,ENCFF609MDN,ENCFF589ZBJ,ENCFF502WDT,ENCFF055HAP,ENCFF198SSR,ENCFF089GBC,ENCFF065ODV,ENCFF883UPM,ENCFF321AGQ,ENCFF092NMJ,ENCFF165LXO |
| 58 | 1 | 60206503 | 60206727 | 224 | CE.LP | 1 | 60206627 | 60206660 | 34 | - | 16.4634 | 9.51E-07 | 0.125 | TAGCTGTCTTCAGACGCACCAGAAGAGGGCATCA | | |
| 59 | 1 | 72485362 | 72485572 | 210 | AJ,AKR,BUB,C3H,C58J,MA/My,MRL,PJ,PL,RF,SJL,SWR,TH | 1 | 72485488 | 72485521 | 34 | - | 16.7724 | 7.83E-07 | 0.115 | TAGCTGTCTTCAGATGCACCAGAAGAGGGCGTCA | | |
| 60 | 1 | 74225488 | 74225705 | 217 | AJ.C3H.CBA.MRL.NOD.NZB.RH2.SMJ | 1 | 74225611 | 74225644 | 34 | - | 17.3902 | 5.27E-07 | 0.095 | TAGCTGTCCTCAGACACTCCAGAAGAGGGCATCA | 2 | ENCFF367UNR.ENCFF099QNE |
| 61 | 1 | 74722425 | 74722665 | 240 | 129S1,AJ,AKR,BTBR,BUB,C3H,C57L,CBA,DBA2J,ILNJ2,MA/My,MRL,NOD,NZB,NZO,PJ,PL,RF,RH2,SEA,SJL,SMJ,SWR | 1 | 74722550 | 74722583 | 34 | - | 18.561 | 2.42E-07 | 0.0656 | CAGCTGTCTTCAGACACACCAGAAGAGGGCATCA | | |
| 62 | 1 | 86210401 | 86210615 | 214 | 129S1,AJ,BTBR,BUB,C3H,C57L,C58J,CE,DBA1J,DBA2J,FVB,ILNJ2,KK,LP,NZB,NZO,NZW,RF,RH2,SJL,SMJ,SWR,TH | 1 | 86210526 | 86210560 | 35 | - | 18.1284 | 2.79E-07 | 0.109 | CTGTAGCTGTCTTCAGACACCAGAAGAGGGCATCA | | |
| 63 | 1 | 111824481 | 111824694 | 213 | 129S1,AJ,AKR,BALB,BTBR,BUB,C3H,C57L,CBA,CE,DBA1J,DBA2J,FVB,ILNJ2,KK,LP,MA/My,MRL,NOD,NOR,NZB,NZO,NZW,PJ,PL,RF,RH2,SEA,SJL,SMJ,SWR,TH | 1 | 111824594 | 111824627 | 34 | - | 16.4634 | 9.51E-07 | 0.125 | TAGCTGTCTTCAGACGCACCAGAAGAGGGCATCA | | |
| 64 | 1 | 134436504 | 134436705 | 201 | 129S1,AJ,AKR,BTBR,BUB,C3H,CBA,CE,DBA1J,DBA2J,FVB,ILNJ2,KK,MA/My,MRL,NOD,NOR,NZB,NZO,PJ,PL,RF,RH2,SJL,SMJ,SWR,TH | 1 | 134436621 | 134436655 | 35 | - | 15.9266 | 9.95E-07 | 0.213 | CTGTCTTCAGATGCACCAACCAGAAGAGGGCGTCA | | |
| 65 | 1 | 153741387 | 153741611 | 224 | BUB,C57L,DBA1J,DBA2J,FVB,MA/My,RF,SJL,SWR,TH | 1 | 153741527 | 153741560 | 34 | - | 16.4634 | 9.51E-07 | 0.125 | TAGCTGTCTTCAGACGCACCAGAAGAGGGCATCA | | |
| 66 | 1 | 156540039 | 156540240 | 201 | AJ,AKR,BALB,C3H,C58J,CBA,CE,DBA1J,DBA2J,FVB,ILNJ2,KK,LP,MA/My,PL,RF,RH2,TH | 1 | 156540165 | 156540183 | 19 | - | 17.5574 | 6.32E-07 | 0.166 | ACTCCAGAAGAGGGCGCCA | | |
| 66 | 1 | 156540039 | 156540240 | 201 | AJ,AKR,BALB,C3H,C58J,CBA,CE,DBA1J,DBA2J,FVB,ILNJ2,KK,LP,MA/My,PL,RF,RH2,TH | 1 | 156540165 | 156540198 | 34 | - | 18.3659 | 2.76E-07 | 0.069 | TAGCTGTCTTCAGACACTCCAGAAGAGGGCGCCA | | |
| 67 | 1 | 156706649 | 156706855 | 206 | AJ,AKR,BALB,C3H,C57L,C58J,CBA,CE,DBA1J,DBA2J,FVB,ILNJ2,LP,MA/My,MRL,NZB,NZO,PL,RF,RH2,TH | 1 | 156706776 | 156706794 | 19 | - | 17.5738 | 6.25E-07 | 0.166 | ACTCCAGAAGAGGGCGCTA | 1 | ENCFF367UNR |
| 67 | 1 | 156706649 | 156706855 | 206 | AJ,AKR,BALB,C3H,C57L,C58J,CBA,CE,DBA1J,DBA2J,FVB,ILNJ2,LP,MA/My,MRL,NZB,NZO,PL,RF,RH2,TH | 1 | 156706776 | 156706809 | 34 | - | 18.1789 | 3.13E-07 | 0.0725 | TAGCTGTCTTCAGACACTCCAGAAGAGGGCGCTA | 1 | ENCFF367UNR |
| 68 | 1 | 164024505 | 164024754 | 249 | DBA1J.DBA2J.ILNJ2 | 1 | 164024631 | 164024664 | 34 | - | 16.9187 | 7.13E-07 | 0.111 | TAGCTGTCTTCAGACGCACCAGAAGAGGGCGTCA | | |
| 69 | 1 | 169984133 | 169984345 | 212 | 129S1,AJ,BTBR,BUB,C3H,C57L,CE,FVB,KK,MRL,NOD,NOR,NZB,NZO,NZW,PJ,PL,RF,RH2,SEA,SJL,SMJ,SWR,TH | 1 | 169984258 | 169984291 | 34 | - | 16.4472 | 9.61E-07 | 0.125 | TAGCTGTCTTCAGACACACCAGAAGAGGGCGTCA | | |
| 70 | 1 | 170970781 | 170971000 | 219 | 129S1,AJ,BUB,C3H,C57L,CBA,CE,DBA1J,DBA2J,FVB,KK,MRL,NOD,NOR,NZB,NZO,NZW,PJ,PL,RH2,SEA,SJL,SMJ,SWR,TH | 1 | 170970906 | 170970939 | 34 | - | 17.8455 | 3.91E-07 | 0.0823 | TAGCTGTCCTCAGACACTCCAGAAGAGGGCGTCA | 1 | ENCFF367UNR |
| 71 | 1 | 171092141 | 171092503 | 362 | 129S1,AJ,BALB,BUB,C3H,C57L,CBA,CE,DBA1J,DBA2J,FVB,KK,MRL,NOD,NOR,NZB,NZO,NZW,PJ,PL,RF,RH2,SEA,SJL,SMJ,SWR,TH | 1 | 171092408 | 171092426 | 19 | - | 17.5574 | 6.32E-07 | 0.166 | ACTCCAGAAGAGGGCGCCA | 2 | ENCFF367UNR,ENCFF093ZRU |
| 71 | 1 | 171092141 | 171092503 | 362 | 129S1,AJ,BALB,BUB,C3H,C57L,CBA,CE,DBA1J,DBA2J,FVB,KK,MRL,NOD,NOR,NZB,NZO,NZW,PJ,PL,RF,RH2,SEA,SJL,SMJ,SWR,TH | 1 | 171092408 | 171092441 | 34 | - | 18.2358 | 3.02E-07 | 0.0709 | TAGCTGTCTTAAGACACTCCAGAAGAGGGCGCCA | 2 | ENCFF367UNR,ENCFF093ZRU |
| 72 | 1 | 171587537 | 171587801 | 264 | BUB.SWR | 1 | 171587529 | 171587563 | 35 | - | 15.9725 | 9.70E-07 | 0.212 | ATGCCATTCCTGGGCTTCTCCACAAGAAGGCCCAA | | |
| 73 | 1 | 171927823 | 171928048 | 225 | 129S1,AJ,AKR,BALB,BUB,C3H,C57L,CBA,CE,DBA1J,DBA2J,FVB,KK,MRL,NOD,NOR,NZB,NZO,NZW,PJ,PL,RF,RH2,SEA,SJL,SMJ,SWR,TH | 1 | 171927948 | 171927982 | 35 | - | 16.1193 | 8.93E-07 | 0.205 | TAGCTGTCTTCAGACAGCACCAGAAGAGGGCATCG | | |
| 74 | 1 | 179333294 | 179333530 | 236 | AKR | 1 | 179333418 | 179333451 | 34 | - | 17.2439 | 5.79E-07 | 0.0995 | TAGCTGTCCTCAGATACTCCAGAAGAGGGCATCA | | |
| 75 | 1 | 179528628 | 179528828 | 200 | AJ,AKR,BUB,C58J,CE,FVB,LP,NZW,PJ,RF,RH2,SEA,SJL,SWR,TH | 1 | 179528753 | 179528786 | 34 | - | 16.4472 | 9.61E-07 | 0.125 | TAGCTGTCTTCAGACACACCAGAAGAGGGCGTCA | | |
| 76 | 1 | 180378976 | 180379185 | 209 | TH | 1 | 180379110 | 180379143 | 34 | - | 17.8455 | 3.91E-07 | 0.0823 | TAGCTGTCCTCAGACACTCCAGAAGAGGGCGTCA | | |
| 77 | 1 | 180420708 | 180420917 | 209 | CBA.CE.DBA1J.DBA2J.PJ.SEA.TH | 1 | 180420832 | 180420865 | 34 | - | 17.1463 | 6.17E-07 | 0.103 | TAGCTGTCCTCAGACACTCCAGAAGAAGGCGCCA | | |
| 78 | 1 | 180893692 | 180893901 | 209 | TH | 1 | 180893827 | 180893860 | 34 | - | 16.6341 | 8.54E-07 | 0.12 | CTGTAGCTGTCAGACACTCCAGAAGAGGGAGTCA | | |
| 79 | 1 | 181834956 | 181835153 | 197 | TH | 1 | 181835082 | 181835115 | 34 | - | 16.4472 | 9.61E-07 | 0.125 | TAGCTGTCTTCAGACACACCAGAAGAGGGCGTCA | | |
| 80 | 1 | 184945846 | 184946111 | 265 | AKR.BTBR.C57L.ILNJ2.KK.MRL.NZW.PL | 1 | 184945970 | 184946003 | 34 | - | 17.6667 | 4.40E-07 | 0.0868 | TAGCTGTCTTCAGACACGCCAGAAGAGGGCGTCA | | |
| 81 | 1 | 190583486 | 190583994 | 508 | ILNJ2.SMJ.SWR | 1 | 190583569 | 190583603 | 35 | - | 20.5413 | 6.11E-08 | 0.046 | GTGCACTAGGGGGCTGTGTGCACTAGGGGGCTGTG | | |
| 81 | 1 | 190583486 | 190583994 | 508 | ILNJ2.SMJ.SWR | 1 | 190583744 | 190583778 | 35 | - | 17.3761 | 4.35E-07 | 0.137 | GTGCACGAGGGGGCTGTGTGCACTAGGGGGCTGTG | | |
| 81 | 1 | 190583486 | 190583994 | 508 | ILNJ2.SMJ.SWR | 1 | 190583776 | 190583810 | 35 | - | 17.1743 | 4.90E-07 | 0.148 | GTGCACGAGGGGGCTGTGTGCACTAGGGGGCCGTG | | |
| 81 | 1 | 190583486 | 190583994 | 508 | ILNJ2.SMJ.SWR | 1 | 190583823 | 190583857 | 35 | - | 18.2569 | 2.58E-07 | 0.106 | GTGCACTAGGGGGCTGTGTGCACGAGGGGGCTGTG | | |
| 81 | 1 | 190583486 | 190583994 | 508 | ILNJ2.SMJ.SWR | 1 | 190583887 | 190583921 | 35 | - | 18.2569 | 2.58E-07 | 0.106 | GTGCACTAGGGGGCTGTGTGCACGAGGGGGCTGTG | | |
| 81 | 1 | 190583486 | 190583994 | 508 | ILNJ2.SMJ.SWR | 1 | 190583936 | 190583970 | 35 | - | 18.2569 | 2.58E-07 | 0.106 | GTGCACTAGGGGGCTGTGTGCACGAGGGGGCTGTG | | |
| 81 | 1 | 190583486 | 190583994 | 508 | ILNJ2.SMJ.SWR | 1 | 190583983 | 190584017 | 35 | - | 20.5413 | 6.11E-08 | 0.046 | GTGCACTAGGGGGCTGTGTGCACTAGGGGGCTGTG | | |
| 82 | 2 | 4286762 | 4286962 | 200 | 129S1.C57L.FVB.MA/Mv.SMJ | 2 | 4286820 | 4286853 | 34 | + | 16.4472 | 9.61E-07 | 0.0987 | TAGCTGTCTTCAGACACACCAGAAGAGGGCGTCA | | |
| 83 | 2 | 20711335 | 20711531 | 196 | CE.FVB.SJL | 2 | 20711388 | 20711421 | 34 | + | 17.2439 | 5.79E-07 | 0.0793 | TAGCTGTCCTCAGATACTCCAGAAGAGGGCATCA | | |
| 84 | 2 | 22814131 | 22814358 | 227 | FVB.SJL | 2 | 22814214 | 22814247 | 34 | + | 16.4634 | 9.51E-07 | 0.0987 | TAGCTGTCTTCAGACGCACCAGAAGAGGGCATCA | | |
| 85 | 2 | 22816144 | 22816395 | 251 | FVB.SJL | 2 | 22816266 | 22816284 | 19 | + | 17.8852 | 5.08E-07 | 0.115 | ACACCAGAAGAGGGCACCA | | |
| 86 | 2 | 29572791 | 29573025 | 234 | 129S1.C57L.RF.SWR | 2 | 29572885 | 29572918 | 34 | + | 16.4634 | 9.51E-07 | 0.0987 | TAGCTGTCTTCAGACGCACCAGAAGAGGGCATCA | | |
| 87 | 2 | 29598902 | 29599113 | 211 | 129S1.C57L.RF.SWR | 2 | 29598987 | 29599005 | 19 | + | 17.8852 | 5.08E-07 | 0.115 | ACACCAGAAGAGGGCACCA | | |
| 88 | 2 | 29724135 | 29724359 | 224 | SWR | 2 | 29724217 | 29724250 | 34 | + | 16.4634 | 9.51E-07 | 0.0987 | TAGCTGTCTTCAGACGCACCAGAAGAGGGCATCA | | |
| 89 | 2 | 31623886 | 31624096 | 210 | NZO.SWR | 2 | 31623953 | 31623986 | 34 | + | 17.8455 | 3.91E-07 | 0.0641 | TAGCTGTCCTCAGACACTCCAGAAGAGGGCGTCA | 1 | ENCFF093ZRU |
| 90 | 2 | 32015896 | 32016109 | 213 | MA/Mv.SWR | 2 | 32015967 | 32016000 | 34 | + | 17.2439 | 5.79E-07 | 0.0793 | TAGCTGTCCTCAGATACTCCAGAAGAGGGCATCA | | |
| 91 | 2 | 32299901 | 32300071 | 170 | NZO.SWR | 2 | 32300009 | 32300042 | 34 | + | 16.4472 | 9.61E-07 | 0.0987 | TAGCTGTCTTCAGACACACCAGAAGAGGGCGTCA | | |
| 92 | 2 | 34721317 | 34721529 | 212 | NZO | 2 | 34721377 | 34721410 | 34 | + | 16.4634 | 9.51E-07 | 0.0987 | TAGCTGTCTTCAGACGCACCAGAAGAGGGCATCA | | |
| 93 | 2 | 39664924 | 39665237 | 313 | SMJ | 2 | 39664940 | 39664973 | 34 | + | 16.4878 | 9.37E-07 | 0.0987 | CTGCCCTCACAGTCGCCGTTACAAGATGGCGCTG | | |
| 94 | 2 | 43377366 | 43377593 | 227 | SMJ.TH | 2 | 43377450 | 43377484 | 35 | + | 19.6422 | 1.09E-07 | 0.0511 | CTGTAGCTATCCTCAGACACCAGAAGAGGGCATCA | | |

| | | | | | | | | | | | | | | | | |
|---|---|---|---|---|---|---|---|---|---|---|---|---|---|---|---|---|
| 95 | 2 | 44718289 | 44718512 | 223 | 129S1,AJ,AKR,BTBR,BUB,C3H,C58J,CBA,CE,DBA1J,DBA2J,FVB,ILNJ2,KK,MA/My,MRL,NOD,NOR,NZB,NZO,NZW,PJ,PL,RF,SEA,SJL,SMJ,SWR,TH | 2 | 44718368 | 44718401 | 34 | + | 17.6667 | 4.40E-07 | 0.0682 | TAGCTGTCTTCAGACACGCCAGAAGAGGGCGTCA | | |
| 96 | 2 | 45793996 | 45794198 | 202 | CE.KK.NZB.SMJ.TH | 2 | 45794056 | 45794089 | 34 | + | 16.4472 | 9.61E-07 | 0.0987 | TAGCTGTCTTCAGACACACCAGAAGAGGGCGTCA | | |
| 97 | 2 | 49907443 | 49907655 | 212 | 129S1.FVB.KK.SJL.TH | 2 | 49907512 | 49907545 | 34 | + | 18.3659 | 2.76E-07 | 0.0538 | TAGCTGTCTTCAGACACTCCAGAAGAGGGCGCCA | | |
| 97 | 2 | 49907443 | 49907655 | 212 | 129S1.FVB.KK.SJL.TH | 2 | 49907527 | 49907545 | 19 | + | 17.5574 | 6.32E-07 | 0.124 | ACTCCAGAAGAGGGCGCCA | | |
| 98 | 2 | 59532682 | 59532893 | 211 | 129S1.BTBR.C57L.DBA1J.TH | 2 | 59532750 | 59532783 | 34 | + | 18.3659 | 2.76E-07 | 0.0538 | TAGCTGTCTTCAGACACTCCAGAAGAGGGCGCCA | 2 | ENCFF367UNR.ENCFF093ZRU |
| 98 | 2 | 59532682 | 59532893 | 211 | 129S1.BTBR.C57L.DBA1J.TH | 2 | 59532765 | 59532783 | 19 | + | 17.5574 | 6.32E-07 | 0.124 | ACTCCAGAAGAGGGCGCCA | 2 | ENCFF367UNR.ENCFF093ZRU |
| 99 | 2 | 69737995 | 69738207 | 212 | AJ,AKR,BALB,BTBR,BUB,C3H,C57L,C58J,CBA,CE,DBA1J,DBA2J,FVB,ILNJ2,KK,MA/My,MRL,NOD,NOR,NZB,NZO,NZW,PJ,PL,RF,RH2,SEA,SJL,SMJ,SWR,TH | 2 | 69738060 | 69738093 | 34 | + | 18.3659 | 2.76E-07 | 0.0538 | TAGCTGTCTTCAGACACTCCAGAAGAGGGCGCCA | | |
| 99 | 2 | 69737995 | 69738207 | 212 | AJ,AKR,BALB,BTBR,BUB,C3H,C57L,C58J,CBA,CE,DBA1J,DBA2J,FVB,ILNJ2,KK,MA/My,MRL,NOD,NOR,NZB,NZO,NZW,PJ,PL,RF,RH2,SEA,SJL,SMJ,SWR,TH | 2 | 69738075 | 69738093 | 19 | + | 17.5574 | 6.32E-07 | 0.124 | ACTCCAGAAGAGGGCGCCA | | |
| 100 | 2 | 69751933 | 69752154 | 221 | AJ,AKR,BALB,BTBR,BUB,C3H,C57L,C58J,CBA,CE,DBA1J,DBA2J,FVB,ILNJ2,KK,MA/My,MRL,NOD,NOR,NZB,NZO,NZW,PJ,PL,RF,RH2,SEA,SJL,SMJ,SWR,TH | 2 | 69752026 | 69752044 | 19 | + | 17.8852 | 5.08E-07 | 0.115 | ACACCAGAAGAGGGCACCA | | |
| 101 | 2 | 71261769 | 71262032 | 263 | AJ,AKR,BUB,C3H,FVB,ILNJ2,MA/My,MRL,NOD,NOR,NZW,PL,RH2,SMJ | 2 | 71261890 | 71261923 | 34 | + | 18.3659 | 2.76E-07 | 0.0538 | TAGCTGTCTTCAGACACTCCAGAAGAGGGCGCCA | | |
| 101 | 2 | 71261769 | 71262032 | 263 | AJ,AKR,BUB,C3H,FVB,ILNJ2,MA/My,MRL,NOD.NOR.NZW.PL.RH2.SMJ | 2 | 71261905 | 71261923 | 19 | + | 17.5574 | 6.32E-07 | 0.124 | ACTCCAGAAGAGGGCGCCA | | |
| 102 | 2 | 71682700 | 71683032 | 332 | 129S1,AJ,AKR,BALB,BTBR,BUB,C57L,DBA1J,FVB,ILNJ2,MA/My,MRL,NOD,NOR,PL,RH2,SEA | 2 | 71682730 | 71682748 | 19 | + | 19.2951 | 1.88E-07 | 0.071 | CCGCCACTAGATGGTGCTA | 18 | ENCFF367UNR,ENCFF093ZRU,ENCFF201JUA,ENCFF099QNE,ENCFF502WDT,ENCFF821BIA,ENCFF907XEG,ENCFF609MDN,ENCFF136ZDK,ENCFF327LYS,ENCFF198SSR,ENCFF024JOZ,ENCFF089GBC,ENCFF589ZBJ,ENCFF430PPJ,ENCFF092NMJ,ENCFF165LXO |
| 103 | 2 | 71736187 | 71736588 | 401 | AJ,AKR,BALB,BTBR,BUB,C57L,DBA1J,FVB,ILNJ2,MA/My,MRL,NOD,NOR,PL,RH2,SEA | 2 | 71736531 | 71736565 | 35 | + | 24.9541 | 2.55E-09 | 0.00618 | CTGCATGTATGCCTGCACACCAGAAGAGGGCACCA | 2 | ENCFF093ZRU,ENCFF099QNE |
| 103 | 2 | 71736187 | 71736588 | 401 | AJ,AKR,BALB,BTBR,BUB,C57L,DBA1J,FVB,ILNJ2,MA/My,MRL,NOD,NOR,PL,RH2,SEA | 2 | 71736547 | 71736565 | 19 | + | 17.8852 | 5.08E-07 | 0.115 | ACACCAGAAGAGGGCACCA | 2 | ENCFF093ZRU,ENCFF099QNE |
| 104 | 2 | 84411853 | 84412062 | 209 | AJ,AKR,BALB,C3H,C57L,C58J,CBA,CE,DBA1J,DBA2J,FVB,ILNJ2,MA/My,MRL,NZW,PJ,RF,RH2,SEA,SJL,SMJ,SWR | 2 | 84411920 | 84411953 | 34 | + | 18.3659 | 2.76E-07 | 0.0538 | TAGCTGTCTTCAGACACTCCAGAAGAGGGCGCCA | | |
| 104 | 2 | 84411853 | 84412062 | 209 | AJ,AKR,BALB,C3H,C57L,C58J,CBA,CE,DBA1J,DBA2J,FVB,ILNJ2,MA/My,MRL,NZW,PJ,RF,RH2,SEA,SJL,SMJ,SWR | 2 | 84411935 | 84411953 | 19 | + | 17.5574 | 6.32E-07 | 0.124 | ACTCCAGAAGAGGGCGCCA | | |
| 105 | 2 | 91200289 | 91200503 | 214 | AJ,BALB,BUB,C57L,C58J,DBA1J,DBA2J,ILNJ2,KK,MA/My,NOD,NOR,NZB,NZO,NZW,PJ,PL,RH2,SEA,SJL,SMJ,SWR | 2 | 91200361 | 91200394 | 34 | + | 17.0081 | 6.74E-07 | 0.0859 | TAGCTGTCCCCAGACACTCCAGAAGAGGGAGTCA | | |
| 106 | 2 | 91578905 | 91579117 | 212 | AJ,BALB,C57L,C58J,DBA1J,DBA2J,FVB,ILNJ2,KK,MA/My,NOD,NOR,NZW,PJ,RH2,SEA,SJL,SMJ,SWR | 2 | 91578975 | 91579009 | 35 | + | 18.1284 | 2.79E-07 | 0.0894 | CTGTAGCTGTCTTCAGACACCAGAAGAGGGCATCA | | |
| 107 | 2 | 104858874 | 104859088 | 214 | 129S1,AJ,AKR,BALB,BTBR,BUB,C3H,C57L,CBA,CE,DBA1J,DBA2J,FVB,ILNJ2,KK,MRL,NOD,NOR,NZB,NZW,PJ,PL,RH2,SEA,SJL,SMJ,SWR | 2 | 104858947 | 104858980 | 34 | + | 17.8455 | 3.91E-07 | 0.0641 | TAGCTGTCCTCAGACACTCCAGAAGAGGGCGTCA | | |
| 108 | 2 | 107698529 | 107698739 | 210 | AJ,BALB,C3H,CBA,CE,DBA1J,DBA2J,FVB,ILNJ2,MA/My,RF,RH2 | 2 | 107698596 | 107698629 | 34 | + | 19.7236 | 1.08E-07 | 0.0317 | CTGTAGTCTTCAGATACACCAGAAGAGGGCATCA | | |
| 109 | 2 | 116731417 | 116731871 | 454 | 129S1,BTBR,BUB,C3H,C57L,C58J,CBA,NOD,NOR,PJ,SEA | 2 | 116731729 | 116731762 | 34 | + | 16.4472 | 9.61E-07 | 0.0987 | TAGCTGTCTTCAGACACACCAGAAGAGGGCGTCA | | |
| 110 | 2 | 120481519 | 120482365 | 846 | AJ,BALB,DBA1J,DBA2J,ILNJ2,PJ,PL,RF,RH2,SEA,SMJ,TH | 2 | 120481605 | 120481638 | 34 | + | 18.5854 | 2.38E-07 | 0.0516 | TTGCTGTCTTCAGACACACCAGAAGAGGGCATCG | | |
| 111 | 2 | 121569503 | 121569757 | 254 | 129S1,AJ,AKR,BALB,BTBR,BUB,C3H,C57L,C58J,CBA,CE,DBA1J,DBA2J,FVB,ILNJ2,KK,MA/My,NOD,NZB,NZO,NZW,PJ,PL,RF,RH2,SEA,SJL,SMJ,SWR,TH | 2 | 121569618 | 121569651 | 34 | + | 16.4634 | 9.51E-07 | 0.0987 | TAGCTGTCTTCAGACGCACCAGAAGAGGGCATCA | | |
| 112 | 2 | 126840760 | 126840964 | 204 | AJ,BALB,C3H,CBA,DBA2J,ILNJ2,KK,PJ,RH2,SEA,SMJ | 2 | 126840821 | 126840854 | 34 | + | 16.4472 | 9.61E-07 | 0.0987 | TAGCTGTCTTCAGACACACCAGAAGAGGGCGTCA | | |
| 113 | 2 | 126866298 | 126866519 | 221 | AJ,BALB,C3H,CBA,DBA2J,ILNJ2,KK,PJ,RH2,SEA,SMJ | 2 | 126866378 | 126866411 | 34 | + | 16.4634 | 9.51E-07 | 0.0987 | TAGCTGTCTTCAGACGCACCAGAAGAGGGCATCA | | |
| 114 | 2 | 127765871 | 127766059 | 188 | C3H.CBA.CE.DBA1J.DBA2J.ILNJ2.PJ.PL | 2 | 127765917 | 127765950 | 34 | + | 18.187 | 3.12E-07 | 0.0569 | TAGCTGTCTTCAGTCACTCCAGAAGAGGGCGCTA | | |
| 114 | 2 | 127765871 | 127766059 | 188 | C3H.CBA.CE.DBA1J.DBA2J.ILNJ2.PJ.PL | 2 | 127765932 | 127765950 | 19 | + | 17.5738 | 6.25E-07 | 0.124 | ACTCCAGAAGAGGGCGCTA | | |
| 115 | 2 | 140304111 | 140304532 | 421 | C58J.DBA1J.DBA2J.NZO.NZW | 2 | 140304127 | 140304160 | 34 | + | 19.748 | 1.06E-07 | 0.0313 | CCGCCCCCACATTCGCCGCCACAAGATGGCGCTG | | |
| 115 | 2 | 140304111 | 140304532 | 421 | C58J.DBA1J.DBA2J.NZO.NZW | 2 | 140304142 | 140304160 | 19 | + | 21.1148 | 4.45E-08 | 0.0407 | CCGCCACAAGATGGCGCTG | | |
| 116 | 2 | 141898627 | 141898826 | 199 | C58J.DBA1J.DBA2J.NZO.NZW | 2 | 141898684 | 141898717 | 34 | + | 16.4634 | 9.51E-07 | 0.0987 | TAGCTGTCTTCAGACGCACCAGAAGAGGGCATCA | | |
| 117 | 2 | 143708646 | 143709024 | 378 | 129S1,BTBR,C57L,C58J,CE,DBA1J,DBA2J,NZO,NZW | 2 | 143708885 | 143708918 | 34 | + | 16.4472 | 9.61E-07 | 0.0987 | TAGCTGTCTTCAGACACACCAGAAGAGGGCGTCA | | |
| 118 | 2 | 150730567 | 150730774 | 207 | 129S1.BUB.C57L.CE.NOD.NZB.NZW.RF | 2 | 150730646 | 150730664 | 19 | + | 17.8852 | 5.08E-07 | 0.115 | ACACCAGAAGAGGGCACCA | 1 | ENCFF093ZRU |
| 119 | 2 | 154495678 | 154495890 | 212 | AJ,BTBR,C58J,DBA1J,DBA2J,KK,NZB,NZO,NZW | 2 | 154495748 | 154495781 | 34 | + | 18.3659 | 2.76E-07 | 0.0538 | TAGCTGTCTTCAGACACTCCAGAAGAGGGCGCCA | 1 | ENCFF093ZRU |
| 119 | 2 | 154495678 | 154495890 | 212 | AJ,BTBR,C58J,DBA1J,DBA2J,KK,NZB,NZO,NZW | 2 | 154495763 | 154495781 | 19 | + | 17.5574 | 6.32E-07 | 0.124 | ACTCCAGAAGAGGGCGCCA | 1 | ENCFF093ZRU |
| 120 | 2 | 154896369 | 154896574 | 205 | 129S1,BALB,C3H,C57L,CBA,CE,FVB,MA/My,NOD,NZO,NZW,PL,SEA,SJL,SWR,TH | 2 | 154896428 | 154896462 | 35 | + | 18.1284 | 2.79E-07 | 0.0894 | CTGTAGCTGTCTTCAGACACCAGAAGAGGGCATCA | | |
| 121 | 2 | 155109406 | 155109656 | 250 | 129S1,BALB,C3H,C57L,CBA,CE,FVB,MA/My,NOD,NZO,NZW,PL,SEA,SJL,SMJ,SWR,TH | 2 | 155109532 | 155109565 | 34 | + | 16.4472 | 9.61E-07 | 0.0987 | TAGCTGTCTTCAGACACACCAGAAGAGGGCGTCA | | |
| 122 | 2 | 155972010 | 155972174 | 164 | 129S1.C57L | 2 | 155972055 | 155972088 | 34 | + | 19.0244 | 1.76E-07 | 0.0433 | TAGCTGTCCCCAGACACTCCAGAAGAGGGCATCA | 3 | ENCFF093ZRU.ENCFF099QNE.ENCFF367UNR |
| 123 | 2 | 164708088 | 164708289 | 201 | AJ,AKR,BALB,BTBR,BUB,C3H,C58J,CBA,DBA1J,DBA2J,FVB,ILNJ2,KK,MA/My,MRL,NOD,NOR,NZB,NZO,NZW,PJ,PL,RF,SEA,SJL,SMJ,SWR,TH | 2 | 164708146 | 164708179 | 34 | + | 17.2439 | 5.79E-07 | 0.0793 | TAGCTGTCCTCAGATACTCCAGAAGAGGGCATCA | | |
| 124 | 2 | 165758175 | 165758425 | 250 | KK | 2 | 165758242 | 165758275 | 34 | + | 17.1463 | 6.17E-07 | 0.0817 | TAGCTGTCCTCAGACACTCCAGAAGAAGGCGCCA | | |
| 125 | 2 | 165857809 | 165858028 | 219 | 129S1,AJ,AKR,BALB,BTBR,BUB,C3H,C57L,C58J,CBA,CE,DBA1J,DBA2J,FVB,ILNJ2,KK,MA/My,NOD,NOR,NZB,NZO,PJ,PL,RF,SEA,SJL,SMJ,SWR,TH | 2 | 165857886 | 165857919 | 34 | + | 16.4472 | 9.61E-07 | 0.0987 | TAGCTGTCTTCAGACACACCAGAAGAGGGCGTCA | | |
| 126 | 2 | 165919818 | 165920286 | 468 | KK | 2 | 165920020 | 165920053 | 34 | + | 19.2846 | 1.47E-07 | 0.0385 | TAGCTGTCTTCAGACTCGCCAGAAGAGGGCGTCA | 1 | ENCFF093ZRU |
| 126 | 2 | 165919818 | 165920286 | 468 | KK | 2 | 165920035 | 165920053 | 19 | + | 18 | 4.70E-07 | 0.113 | TCGCCAGAAGAGGGCGTCA | 1 | ENCFF093ZRU |
| 127 | 2 | 166595903 | 166596110 | 207 | CE.KK.TH | 2 | 166595967 | 166596000 | 34 | + | 16.4634 | 9.51E-07 | 0.0987 | TAGCTGTCTTCAGACGCACCAGAAGAGGGCATCA | | |
| 128 | 2 | 166822996 | 166823207 | 211 | 129S1.C57L.CE.KK | 2 | 166823048 | 166823081 | 34 | + | 16.4472 | 9.61E-07 | 0.0987 | TAGCTGTCTTCAGACACACCAGAAGAGGGCGTCA | 1 | ENCFF093ZRU |
| 129 | 2 | 168128156 | 168128226 | 70 | 129S1.C57L.NZB.TH | 2 | 168128154 | 168128188 | 35 | + | 17.633 | 3.74E-07 | 0.107 | GTACAGGTACCCATGGAGACCAGAAGAGGGCGTCA | 1 | ENCFF093ZRU |
| 130 | 2 | 168838895 | 168839092 | 197 | KK | 2 | 168838950 | 168838983 | 34 | + | 16.5447 | 9.04E-07 | 0.0987 | CAGCTGTCTTCAGACACACCAGAAGAGGGAGTCA | | |
| 131 | 2 | 17823736 | 17823949 | 213 | MA/Mv.RF | 2 | 17823859 | 17823892 | 34 | - | 18.7561 | 2.12E-07 | 0.0488 | TAGCTGCTCCCAGACACTCCAGAAGAGGGTGCCA | | |
| 132 | 2 | 18287500 | 18287703 | 203 | MA/Mv.RF | 2 | 18287628 | 18287661 | 34 | - | 17.2114 | 5.91E-07 | 0.0799 | TAGCTGTTTTCAGACACTCCAGAAGAGGGCATCA | | |
| 133 | 2 | 18417387 | 18417578 | 191 | MA/Mv.RF.SJL | 2 | 18417512 | 18417530 | 19 | - | 17.5574 | 6.32E-07 | 0.124 | ACTCCAGAAGAGGGCGCCA | | |
| 133 | 2 | 18417387 | 18417578 | 191 | MA/Mv.RF.SJL | 2 | 18417512 | 18417545 | 34 | - | 18.3659 | 2.76E-07 | 0.0538 | TAGCTGTCTTCAGACACTCCAGAAGAGGGCGCCA | | |
| 134 | 2 | 20462715 | 20462915 | 200 | FVB.SJL | 2 | 20462831 | 20462864 | 34 | - | 17.6667 | 4.40E-07 | 0.0682 | TAGCTGTCTTCAGACACGCCAGAAGAGGGCGTCA | | |
| 135 | 2 | 26123501 | 26123708 | 207 | MA/Mv.NZO | 2 | 26123626 | 26123659 | 34 | - | 16.7724 | 7.83E-07 | 0.0917 | TAGCTGTCTTCAGATGCACCAGAAGAGGGCGTCA | | |
| 136 | 2 | 29540278 | 29540389 | 111 | CE | 2 | 29540308 | 29540326 | 19 | - | 17.3934 | 7.03E-07 | 0.131 | TAGACTCCAGGGGGCAGCA | 20 | ENCFF367UNR,ENCFF092NMJ,ENCFF201JUA,ENCFF136ZDK,ENCFF093ZRU,ENCFF089GBC,ENCFF165LXO,ENCFF883UPM,ENCFF609MDN,ENCFF502WDT,ENCFF198SSR,ENCFF951GOP,ENCFF821BIA,ENCFF327LYS,ENCFF907XEG,ENCFF099QNE,ENCFF589ZBJ,ENCFF957SPH,ENCFF024JOZ,ENCFF055HAP |
| 137 | 2 | 32245375 | 32245572 | 197 | NZO.SWR | 2 | 32245486 | 32245504 | 19 | - | 17.5574 | 6.32E-07 | 0.124 | ACTCCAGAAGAGGGCGCCA | 2 | ENCFF093ZRU.ENCFF367UNR |
| 137 | 2 | 32245375 | 32245572 | 197 | NZO.SWR | 2 | 32245486 | 32245519 | 34 | - | 18.3659 | 2.76E-07 | 0.0538 | TAGCTGTCTTCAGACACTCCAGAAGAGGGCGCCA | 2 | ENCFF093ZRU.ENCFF367UNR |
| 138 | 2 | 37826868 | 37827078 | 210 | BUB.TH | 2 | 37826994 | 37827012 | 19 | - | 17.5574 | 6.32E-07 | 0.124 | ACTCCAGAAGAGGGCGCCA | 1 | ENCFF093ZRU |
| 138 | 2 | 37826868 | 37827078 | 210 | BUB.TH | 2 | 37826994 | 37827027 | 34 | - | 18.561 | 2.42E-07 | 0.0519 | AAGCTGTCTTCAGACACTCCAGAAGAGGGCGCCA | 1 | ENCFF093ZRU |
| 139 | 2 | 45026115 | 45026361 | 246 | 129S1,AJ,AKR,BTBR,BUB,C3H,C58J,CBA,CE,DBA1J,DBA2J,FVB,ILNJ2,KK,MA/My,MRL,NOD,NOR,NZB,NZO,NZW,PJ,PL,RF,SEA,SJL,SMJ,SWR,TH | 2 | 45026242 | 45026275 | 34 | - | 16.4472 | 9.61E-07 | 0.0987 | TAGCTGTCTTCAGACACACCAGAAGAGGGCGTCA | | |
| 140 | 2 | 50545989 | 50546196 | 207 | 129S1,AJ,AKR,BUB,C3H,C58J,CBA,CE,DBA1J,DBA2J,FVB,ILNJ2,KK,MRL,NOD,NOR,PJ,PL,RF,SEA,SJL,SMJ,SWR,TH | 2 | 50546111 | 50546144 | 34 | - | 16.4472 | 9.61E-07 | 0.0987 | TAGCTGTCTTCAGACACACCAGAAGAGGGCGTCA | | |
| 141 | 2 | 52688599 | 52688823 | 224 | NZB | 2 | 52688727 | 52688760 | 34 | - | 17.9431 | 3.66E-07 | 0.0623 | CAGCTGTCCTCAGACACTCCAGAAGAGGGAGTCA | | |
| 142 | 2 | 59682211 | 59682402 | 191 | 129S1.C57L.CE.KK.NOD.NOR.SJL | 2 | 59682310 | 59682328 | 19 | - | 17.5574 | 6.32E-07 | 0.124 | ACTCCAGAAGAGGGCGCCA | 4 | ENCFF093ZRU.ENCFF099QNE.ENCFF821BIA.ENCFF367UNR |
| 142 | 2 | 59682211 | 59682402 | 191 | 129S1.C57L.CE.KK.NOD.NOR.SJL | 2 | 59682310 | 59682343 | 34 | - | 18.3659 | 2.76E-07 | 0.0538 | TAGCTGTCTTCAGACACTCCAGAAGAGGGCGCCA | 4 | ENCFF093ZRU.ENCFF099QNE.ENCFF821BIA.ENCFF367UNR |
| 143 | 2 | 62877587 | 62877809 | 222 | 129S1,BTBR,DBA1J,DBA2J,FVB,KK,PJ,SEA.SJL.SWR.TH | 2 | 62877712 | 62877730 | 19 | - | 17.5574 | 6.32E-07 | 0.124 | ACTCCAGAAGAGGGCGCCA | | |
| 143 | 2 | 62877587 | 62877809 | 222 | 129S1,BTBR,DBA1J,DBA2J,FVB,KK,PJ,SEA.SJL.SWR.TH | 2 | 62877712 | 62877745 | 34 | - | 18.3659 | 2.76E-07 | 0.0538 | TAGCTGTCTTCAGACACTCCAGAAGAGGGCGCCA | | |
| 144 | 2 | 72472857 | 72473144 | 287 | MA/Mv.NZB | 2 | 72472999 | 72473017 | 19 | - | 18.7213 | 2.85E-07 | 0.0853 | GTGCCAGAAGAGGGCACTG | | |
| 144 | 2 | 72472857 | 72473144 | 287 | MA/Mv.NZB | 2 | 72472999 | 72473033 | 35 | - | 18.4404 | 2.31E-07 | 0.0809 | CTTCATGTATGCCTGTGTGCCAGAAGAGGGCACTG | | |
| 145 | 2 | 75691175 | 75691416 | 241 | NOD.NOR | 2 | 75691308 | 75691341 | 34 | - | 16.4634 | 9.51E-07 | 0.0987 | TAGCTGTCTTCAGACGCACCAGAAGAGGGCATCA | 1 | ENCFF093ZRU |
| 146 | 2 | 86294951 | 86295283 | 332 | AJ,AKR,BALB,BUB,C3H,C57L,CBA,CE,DBA1J,DBA2J,FVB,KK,MA/My,MRL,NOD,NOR,NZB,NZO,NZW,PJ,PL,RF,RH2,SEA,SMJ | 2 | 86295237 | 86295255 | 19 | - | 19.2951 | 1.88E-07 | 0.071 | CCGCCACTAGATGGTGCTA | 28 | ENCFF093ZRU,ENCFF367UNR,ENCFF805QBU,ENCFF099QNE,ENCFF883UPM,ENCFF821BIA,ENCFF601JVE,ENCFF316FYN,ENCFF327LYS,ENCFF907XEG,ENCFF024JOZ,ENCFF033UQX,ENCFF136ZDK,ENCFF589ZBJ,ENCFF097CAQ,ENCFF089GBC,ENCFF198SSR,ENCFF165LXO,ENCFF092NMJ,ENCFF052KVO,ENCFF321AGQ,ENCFF430PPJ,ENCFF281QMV,ENCFF957SPH,ENCFF194HVL,ENCFF951GOP,ENCFF533APC,ENCFF609MDN |
| 147 | 2 | 91968422 | 91968643 | 221 | AJ,AKR,BUB,C3H,C58J,CE,FVB,ILNJ2,MA/My,MRL,NZW,PL,RF,TH | 2 | 91968546 | 91968579 | 34 | - | 18.4797 | 2.56E-07 | 0.0524 | TTGCTGTCTTCAGACACACCAGAAGAGGGCATCA | | |
| 148 | 2 | 102031488 | 102031686 | 198 | AJ,BUB,FVB,ILNJ2,KK,NZB,NZO,NZW,RF,RH2,SJL,SMJ,SWR | 2 | 102031618 | 102031636 | 19 | - | 17.8852 | 5.08E-07 | 0.115 | ACACCAGAAGAGGGCACCA | 2 | ENCFF093ZRU |
| 149 | 2 | 103306113 | 103306966 | 853 | AJ,BALB,C57L,CBA,DBA1J,DBA2J,ILNJ2,RH2 | 2 | 103306701 | 103306734 | 34 | - | 16.6504 | 8.46E-07 | 0.0961 | TAGCTGTTTACAGACATACCAGAAGAGGGCATCA | 2 | ENCFF367UNR,ENCFF093ZRU |
| 150 | 2 | 104267716 | 104267905 | 189 | AJ,AKR,BALB,C3H,C57L,CBA,CE,DBA1J,DBA2J,FVB,ILNJ2,MRL,NZB,NZW,PL,RF,RH2,SMJ,SWR,TH | 2 | 104267840 | 104267873 | 34 | - | 16.4472 | 9.61E-07 | 0.0987 | TAGCTGTCTTCAGACACACCAGAAGAGGGCGTCA | | |
| 151 | 2 | 104438880 | 104439094 | 214 | 129S1,AJ,AKR,BALB,BTBR,BUB,C3H,C57L,CBA,CE,DBA1J,DBA2J,FVB,ILNJ2,KK,MRL,NOD,NOR,NZB,NZW,PJ,PL,RF,RH2,SEA,SJL,SWR,TH | 2 | 104439006 | 104439039 | 34 | - | 16.4472 | 9.61E-07 | 0.0987 | TAGCTGTCTTCAGACACACCAGAAGAGGGCGTCA | | |
| 152 | 2 | 108410493 | 108410722 | 229 | C3H.CE.RF | 2 | 108410621 | 108410654 | 34 | - | 16.6179 | 8.63E-07 | 0.0972 | TAGCTGTCTTCCGACACTCCAGAAGAGGGCGTCA | | |
| 153 | 2 | 119401873 | 119402089 | 216 | 129S1,AJ,AKR,BALB,BTBR,BUB,C3H,C57L,C58J,CBA,CE,DBA1J,DBA2J,FVB,ILNJ2,KK,MA/My,NOD,NZB,NZO,NZW,PJ,PL,RF,RH2,SEA,SJL,SMJ,SWR,TH | 2 | 119401998 | 119402016 | 19 | - | 17.5574 | 6.32E-07 | 0.124 | ACTCCAGAAGAGGGCGCCA | 2 | ENCFF093ZRU,ENCFF367UNR |
| 153 | 2 | 119401873 | 119402089 | 216 | 129S1,AJ,AKR,BALB,BTBR,BUB,C3H,C57L,C58J,CBA,CE,DBA1J,DBA2J,FVB,ILNJ2,KK,MA/My,NOD,NZB,NZO,NZW,PJ,PL,RF,RH2,SEA,SJL,SMJ,SWR,TH | 2 | 119401998 | 119402031 | 34 | - | 18.3659 | 2.76E-07 | 0.0538 | TAGCTGTCTTCAGACACTCCAGAAGAGGGCGCCA | 2 | ENCFF093ZRU,ENCFF367UNR |
| 154 | 2 | 121922117 | 121922193 | 76 | 129S1,AKR,BTBR,BUB,C57L,CE,FVB,MA/My,NZW,SJL,SWR | 2 | 121922155 | 121922188 | 34 | - | 19.9512 | 9.19E-08 | 0.0283 | GTGCTGTGCCGGCCTTGGCCTACAGAAGTCAGTG | | |
| 155 | 2 | 124933953 | 124934168 | 215 | 129S1,AKR,BTBR,BUB,C3H,C57L,C58J,CBA,CE,FVB,KK,NZB,NZO,NZW,RF,SJL,SWR,TH | 2 | 124934079 | 124934097 | 19 | - | 17.5574 | 6.32E-07 | 0.124 | ACTCCAGAAGAGGGCGCCA | 3 | ENCFF367UNR,ENCFF093ZRU,ENCFF099QNE |
| 155 | 2 | 124933953 | 124934168 | 215 | 129S1,AKR,BTBR,BUB,C3H,C57L,C58J,CBA,CE,FVB,KK,NZB,NZO,NZW,RF,SJL,SWR,TH | 2 | 124934079 | 124934112 | 34 | - | 18.3659 | 2.76E-07 | 0.0538 | TAGCTGTCTTCAGACACTCCAGAAGAGGGCGCCA | 3 | ENCFF367UNR,ENCFF093ZRU,ENCFF099QNE |
| 156 | 2 | 128482743 | 128482947 | 204 | C3H.CBA.DBA1J.DBA2J.ILNJ2.PJ.PL | 2 | 128482868 | 128482901 | 34 | - | 16.7724 | 7.83E-07 | 0.0917 | TAGCTGTCTTCAGATGCACCAGAAGAGGGCGTCA | | |
| 157 | 2 | 131910238 | 131910336 | 98 | ILNJ2.NOD.PJ | 2 | 131910326 | 131910360 | 35 | - | 20.1284 | 8.00E-08 | 0.0419 | CTGCATTTTAACCTGTGAGCCCCCAGGTGCCTGCA | | |
| 158 | 2 | 141730258 | 141730341 | 83 | C58J.DBA1J.DBA2J.NZO.NZW | 2 | 141730225 | 141730259 | 35 | - | 16.0642 | 9.21E-07 | 0.174 | CTTCATGTACACCTGCATGCCAGAAGAGGGCATGA | | |
| 159 | 2 | 144071185 | 144071436 | 251 | 129S1,BTBR,C57L,C58J,CE,DBA1J,DBA2J,NZO,NZW | 2 | 144071329 | 144071363 | 35 | - | 18.1835 | 2.70E-07 | 0.0882 | CTGTAGCTGTCTTCAGACACCAGAAGAGGGCATTG | | |
| 160 | 2 | 144402783 | 144403008 | 225 | 129S1,AJ,BTBR,C57L,C58J,CE,DBA1J,DBA2J.NZB.NZO.NZW.SMJ | 2 | 144402906 | 144402924 | 19 | - | 17.5574 | 6.32E-07 | 0.124 | ACTCCAGAAGAGGGCGCCA | 2 | ENCFF367UNR,ENCFF093ZRU |
| 160 | 2 | 144402783 | 144403008 | 225 | 129S1,AJ,BTBR,C57L,C58J,CE,DBA1J,DBA2J.NZB.NZO.NZW.SMJ | 2 | 144402906 | 144402939 | 34 | - | 18.3659 | 2.76E-07 | 0.0538 | TAGCTGTCTTCAGACACTCCAGAAGAGGGCGCCA | 2 | ENCFF367UNR,ENCFF093ZRU |
| 161 | 2 | 146118060 | 146118278 | 218 | AJ,BALB,BUB,C3H,CBA,FVB,ILNJ2,KK,MA/My,MRL,NOD,RF,SEA,SJL,SMJ,SWR,TH | 2 | 146118185 | 146118218 | 34 | - | 17.0325 | 6.63E-07 | 0.0853 | TAGCTGTCCTCAGACACTCCAGAAGAGGGAGGCA | | |
| 162 | 2 | 158140673 | 158141128 | 455 | KK.SJL | 2 | 158141089 | 158141122 | 34 | - | 17.0244 | 6.67E-07 | 0.0854 | GAGCAGGTGTGGCAGCAGTCCCAAGATGGCGCCC | | |
| 163 | 2 | 158211034 | 158211249 | 215 | KK.SJL | 2 | 158211160 | 158211178 | 19 | - | 17.5574 | 6.32E-07 | 0.124 | ACTCCAGAAGAGGGCGCCA | 1 | ENCFF093ZRU |
| 163 | 2 | 158211034 | 158211249 | 215 | KK.SJL | 2 | 158211160 | 158211193 | 34 | - | 18.3659 | 2.76E-07 | 0.0538 | TAGCTGTCTTCAGACACTCCAGAAGAGGGCGCCA | 1 | ENCFF093ZRU |
| 164 | 2 | 162885257 | 162885462 | 205 | KK | 2 | 162885382 | 162885415 | 34 | - | 16.4715 | 9.47E-07 | 0.0987 | TAGCTGTCTTCAGACACTCCAGAAGGGGGTGCCA | | |
| 165 | 2 | 164719690 | 164719896 | 206 | AJ,AKR,BALB,BTBR,BUB,C3H,C58J,CBA,DBA1J,DBA2J,FVB,ILNJ2,KK,MA/My,MRL,NOD,NOR,NZB,NZO,NZW,PJ,PL,RF,SEA,SJL,SMJ,SWR,TH | 2 | 164719814 | 164719832 | 19 | - | 17.5574 | 6.32E-07 | 0.124 | ACTCCAGAAGAGGGCGCCA | 5 | ENCFF093ZRU,ENCFF099QNE,ENCFF327LYS,ENCFF821BIA,ENCFF367UNR |
| 165 | 2 | 164719690 | 164719896 | 206 | AJ,AKR,BALB,BTBR,BUB,C3H,C58J,CBA,DBA1J,DBA2J,FVB,ILNJ2,KK,MA/My,MRL,NOD,NOR,NZB,NZO,NZW,PJ,PL,RF,SEA,SJL,SMJ,SWR,TH | 2 | 164719814 | 164719847 | 34 | - | 18.561 | 2.42E-07 | 0.0519 | AAGCTGTCTTCAGACACTCCAGAAGAGGGCGCCA | 5 | ENCFF093ZRU,ENCFF099QNE,ENCFF327LYS,ENCFF821BIA,ENCFF367UNR |
| 166 | 2 | 165620539 | 165620735 | 196 | KK | 2 | 165620666 | 165620699 | 34 | - | 16.4472 | 9.61E-07 | 0.0987 | TAGCTGTCTTCAGACACACCAGAAGAGGGCGTCA | | |
| 167 | 2 | 166745969 | 166746676 | 707 | 129S1.C57L.CE | 2 | 166746014 | 166746048 | 35 | - | 19.4954 | 1.20E-07 | 0.0535 | CCGCATGTATGCCTGCATGCCTGAAGAGGGCACCA | 4 | ENCFF089GBC.ENCFF055HAP.ENCFF502WDT.ENCFF093ZRU |
| 168 | 2 | 167289634 | 167289692 | 58 | CE | 2 | 167289629 | 167289662 | 34 | - | 20.0569 | 8.52E-08 | 0.027 | CTGCAGCTCACTCAGAGAACACCAGGTGCTACTA | | |
| 169 | 2 | 167981845 | 167982051 | 206 | 129S1.C57L.KK.NZB.TH | 2 | 167981967 | 167982000 | 34 | - | 17.8455 | 3.91E-07 | 0.0641 | TAGCTGTCCTCAGACACTCCAGAAGAGGGCGTCA | | |
| 170 | 2 | 168494045 | 168494249 | 204 | 129S1,AJ,AKR,BALB,BTBR,C3H,C57L,CBA,DBA1J,DBA2J,FVB,ILNJ2,MA/My,MRL,NOD,NOR,NZB,NZW,PJ,PL,RF,SEA,SJL,SMJ,SWR | 2 | 168494155 | 168494173 | 19 | - | 18.623 | 3.06E-07 | 0.0887 | ACACCAGAAGAGGGCGCCA | | |
| 170 | 2 | 168494045 | 168494249 | 204 | 129S1,AJ,AKR,BALB,BTBR,C3H,C57L,CBA,DBA1J,DBA2J,FVB,ILNJ2,MA/My,MRL,NOD,NOR,NZB,NZW,PJ,PL,RF,SEA,SJL,SMJ,SWR | 2 | 168494155 | 168494188 | 34 | - | 18.8537 | 1.98E-07 | 0.0465 | TAGCTGTCTTCAGACACACCAGAAGAGGGCGCCA | | |
| 171 | 2 | 180986188 | 180986461 | 273 | AJ,AKR,BUB,C3H,C58J,CE,DBA1J,DBA2J,KK,LP,MA/My,NZB,NZO,NZW,PJ,RF,RH2 | 2 | 180986342 | 180986375 | 34 | - | 17.4634 | 5.02E-07 | 0.0732 | GTGCACCACGACGCCCGGCCTGCAGGTGGATTTC | | |
| 172 | 2 | 181137837 | 181138042 | 205 | AJ,AKR,BALB,BUB,C3H,CE,DBA1J,DBA2J,KK,MA/My,NZB,PJ,RF,RH2,SEA | 2 | 181137965 | 181137998 | 34 | - | 17.6667 | 4.40E-07 | 0.0682 | TAGCTGTCTTCAGACACGCCAGAAGAGGGCGTCA | | |
| 173 | 3 | 15333748 | 15333856 | 108 | AJ,BALB,BUB,C3H,C58J,CBA,FVB,PL,SJL,SWR,TH | 3 | 15333749 | 15333782 | 34 | + | 16.561 | 8.95E-07 | 0.14 | CTGCCCTTGCAATCGCCATTACAAGATGGCGCTG | | |
| 174 | 3 | 27130519 | 27130731 | 212 | AJ.AKR.C3H.CBA.CE.PL.RF | 3 | 27130591 | 27130624 | 34 | + | 17.0488 | 6.56E-07 | 0.121 | TTGCTATCTTCAGATACACCAGAAGAGGGCATCA | | |
| 175 | 3 | 34817878 | 34818086 | 208 | AJ,BUB,C3H,CBA,CE,DBA1J,FVB,SJL,SWR | 3 | 34817944 | 34817977 | 34 | + | 17.9431 | 3.66E-07 | 0.0914 | CAGCTGTCCTCAGACACTCCAGAAGAGGGAGTCA | | |
| 176 | 3 | 51278907 | 51279060 | 153 | NZW | 3 | 51278910 | 51278943 | 34 | + | 18.3659 | 2.76E-07 | 0.0804 | TAGCTGTCTTCAGACACTCCAGAAGAGGGCGCCA | 3 | ENCFF367UNR.ENCFF093ZRU.ENCFF201JUA |
| 176 | 3 | 51278907 | 51279060 | 153 | NZW | 3 | 51278925 | 51278943 | 19 | + | 17.5574 | 6.32E-07 | 0.181 | ACTCCAGAAGAGGGCGCCA | 3 | ENCFF367UNR.ENCFF093ZRU.ENCFF201JUA |
| 177 | 3 | 54422208 | 54422497 | 289 | CE | 3 | 54422376 | 54422409 | 34 | + | 23.1545 | 7.96E-09 | 0.00773 | ATGTAGTGCCTGTGGATGCCAGCAGGAGGCATCA | | |
| 178 | 3 | 57646623 | 57646882 | 259 | AJ,CE,DBA1J,DBA2J,FVB,PJ,SEA,SJL,SMJ,SWR,TH | 3 | 57646740 | 57646773 | 34 | + | 16.4472 | 9.61E-07 | 0.141 | TAGCTGTCTTCAGACACACCAGAAGAGGGCGTCA | | |
| 179 | 3 | 65871941 | 65872016 | 75 | FVB.NOD.NOR.SWR | 3 | 65871965 | 65871998 | 34 | + | 16.6016 | 8.72E-07 | 0.138 | GTGCAGGTCCCCCTGTGGAAGCCAGAGGTCAGAT | | |
| 180 | 3 | 73305611 | 73305818 | 207 | AJ.CE.MA/Mv.NOD.NOR.RH2 | 3 | 73305676 | 73305709 | 34 | + | 16.4472 | 9.61E-07 | 0.141 | TAGCTGTCTTCAGACACACCAGAAGAGGGCGTCA | | |
| 181 | 3 | 87827833 | 87828033 | 200 | AJ,AKR,BALB,BTBR,BUB,C3H,CBA,CE,DBA1J,DBA2J,FVB,ILNJ2,KK,MA/My,MRL,NOD,NOR,NZB,NZO,NZW,PJ,PL,RH2,SEA,SJL,SMJ.SWR.TH | 3 | 87827905 | 87827923 | 19 | + | 17.8852 | 5.08E-07 | 0.167 | ACACCAGAAGAGGGCACCA | 6 | ENCFF367UNR,ENCFF907XEG,ENCFF327LYS,ENCFF883UPM,ENCFF198SSR,ENCFF093ZRU |
| 182 | 3 | 88193037 | 88193254 | 217 | MA/Mv.RF | 3 | 88193111 | 88193144 | 34 | + | 18.3659 | 2.76E-07 | 0.0804 | TAGCTGTCTTCAGACACTCCAGAAGAGGGCGCCA | 1 | ENCFF093ZRU |
| 182 | 3 | 88193037 | 88193254 | 217 | MA/Mv.RF | 3 | 88193126 | 88193144 | 19 | + | 17.5574 | 6.32E-07 | 0.181 | ACTCCAGAAGAGGGCGCCA | 1 | ENCFF093ZRU |
| 183 | 3 | 88591920 | 88592118 | 198 | AJ,AKR,BTBR,BUB,C3H,CBA,FVB,KK,MA/My,MRL,NZB,NZO,NZW,RF,SJL,SWR,TH | 3 | 88591974 | 88592007 | 34 | + | 18.3659 | 2.76E-07 | 0.0804 | TAGCTGTCTTCAGACACTCCAGAAGAGGGCGCCA | 10 | ENCFF821BIA,ENCFF907XEG,ENCFF099QNE,ENCFF502WDT,ENCFF367UNR,ENCFF024JOZ,ENCFF883UPM,ENCFF093ZRU,ENCFF055HAP |
| 183 | 3 | 88591920 | 88592118 | 198 | AJ,AKR,BTBR,BUB,C3H,CBA,FVB,KK,MA/My,MRL,NZB,NZO,NZW,RF,SJL,SWR,TH | 3 | 88591989 | 88592007 | 19 | + | 17.5574 | 6.32E-07 | 0.181 | ACTCCAGAAGAGGGCGCCA | 10 | ENCFF821BIA,ENCFF907XEG,ENCFF099QNE,ENCFF502WDT,ENCFF367UNR,ENCFF024JOZ,ENCFF883UPM,ENCFF093ZRU,ENCFF055HAP |
| 184 | 3 | 89069764 | 89070318 | 554 | KK | 3 | 89069769 | 89069802 | 34 | + | 16.4472 | 9.61E-07 | 0.141 | TAGCTGTCTTCAGACACACCAGAAGAGGGCGTCA | | |
| 185 | 3 | 93745376 | 93746044 | 668 | RF | 3 | 93745350 | 93745383 | 34 | + | 16.561 | 8.95E-07 | 0.14 | CTGCCCTTGCAATCGCCATTACAAGATGGCGCTG | | |
| 186 | 3 | 95120439 | 95120654 | 215 | KK.NOD.NOR.TH | 3 | 95120510 | 95120543 | 34 | + | 18.561 | 2.42E-07 | 0.079 | AAGCTGTCTTCAGACACTCCAGAAGAGGGCGCCA | 5 | ENCFF093ZRU.ENCFF367UNR.ENCFF821BIA.ENCFF201JUA |
| 186 | 3 | 95120439 | 95120654 | 215 | KK.NOD.NOR.TH | 3 | 95120525 | 95120543 | 19 | + | 17.5574 | 6.32E-07 | 0.181 | ACTCCAGAAGAGGGCGCCA | 5 | ENCFF093ZRU.ENCFF367UNR.ENCFF821BIA.ENCFF201JUA |

| | | | | | | | | | | | | | | | | |
|---|---|---|---|---|---|---|---|---|---|---|---|---|---|---|---|---|
| 187 | 3 | 95877937 | 95878146 | 209 | 129S1,AJ,BALB,C3H,C57L,CBA,CE,FVB,KK,MA/My,NOD,NOR,NZB,NZO,NZW,PJ,RH2,SEA.SJL.SMJ.SWR.TH | 3 | 95878005 | 95878038 | 34 | + | 17.8455 | 3.91E-07 | 0.0943 | TAGCTGTCCTCAGACACTCCAGAAGAGGGCGTCA | 1 | ENCFF055HAP |
| 188 | 3 | 96643107 | 96643318 | 211 | KK.MA/Mv.NZB.NZO.NZW.RF | 3 | 96643175 | 96643208 | 34 | + | 16.4634 | 9.51E-07 | 0.141 | TAGCTGTCTTCAGACGCACCAGAAGAGGGCATCA | 1 | ENCFF093ZRU |
| 189 | 3 | 107510464 | 107510674 | 210 | BALB.CE.MA/Mv.MRL.NZO.NZW.RH2.TH | 3 | 107510525 | 107510558 | 34 | + | 16.4634 | 9.51E-07 | 0.141 | TAGCTGTCTTCAGACGCACCAGAAGAGGGCATCA | | |
| 190 | 3 | 114945012 | 114945229 | 217 | AJ,AKR,BALB,C3H,CBA,DBA1J,DBA2J,FVB,ILNJ2,KK,MA/My,MRL,NZO,NZW,PJ,PL,RF,RH2.SEA.SJL.SMJ.SWR.TH | 3 | 114945088 | 114945121 | 34 | + | 18.3659 | 2.76E-07 | 0.0804 | TAGCTGTCTTCAGACACTCCAGAAGAGGGCGCCA | | |
| 190 | 3 | 114945012 | 114945229 | 217 | AJ,AKR,BALB,C3H,CBA,DBA1J,DBA2J,FVB,ILNJ2,KK,MA/My,MRL,NZO,NZW,PJ,PL,RF,RH2.SEA.SJL.SMJ.SWR.TH | 3 | 114945103 | 114945121 | 19 | + | 17.5574 | 6.32E-07 | 0.181 | ACTCCAGAAGAGGGCGCCA | | |
| 191 | 3 | 116546109 | 116546318 | 209 | 129S1,AJ,AKR,BALB,BUB,C3H,C57L,C58J,CBA,CE,DBA1J,DBA2J,FVB,ILNJ2,LP,MA/My,MRL,NOD,NOR,NZB,NZO,NZW,PL,RF,RH2,SJL.SMJ.SWR.TH | 3 | 116546176 | 116546209 | 34 | + | 17.8455 | 3.91E-07 | 0.0943 | TAGCTGTCCTCAGACACTCCAGAAGAGGGCGTCA | | |
| 192 | 3 | 122080634 | 122080834 | 200 | BTBR.DBA1J.DBA2J.SMJ.TH | 3 | 122080691 | 122080724 | 34 | + | 16.4472 | 9.61E-07 | 0.141 | TAGCTGTCTTCAGACACACCAGAAGAGGGCGTCA | | |
| 193 | 3 | 123347060 | 123347256 | 196 | AJ.BALB.C3H.CBA.NZB.RH2.SMJ | 3 | 123347113 | 123347146 | 34 | + | 16.4634 | 9.51E-07 | 0.141 | TAGCTGTCTTCAGACGCACCAGAAGAGGGCATCA | | |
| 194 | 3 | 125211965 | 125212171 | 206 | AJ.BALB.C3H.CBA.MA/Mv.NZB.RH2.SMJ | 3 | 125212045 | 125212063 | 19 | + | 17.8852 | 5.08E-07 | 0.167 | ACACCAGAAGAGGGCACCA | | |
| 195 | 3 | 127165356 | 127166351 | 995 | CE | 3 | 127165619 | 127165652 | 34 | + | 23.7073 | 5.04E-09 | 0.00582 | GTGCAGTGCTCACAGAGGCCAGACGAGGGAGCTA | 7 | ENCFF367UNR,ENCFF821BIA,ENCFF093ZRU,ENCFF089GBC,ENCFF201JUA,ENCFF502WDT,ENCFF099QNE |
| 195 | 3 | 127165356 | 127166351 | 995 | CE | 3 | 127165424 | 127165442 | 19 | - | 16.8525 | 9.96E-07 | 0.218 | TAACCAAAAGAGGGCACCA | 1 | ENCFF367UNR |
| 196 | 3 | 144189940 | 144190157 | 217 | 129S1,BTBR,BUB,C57L,C58J,CE,DBA2J,LP.NZB.NZO.RF | 3 | 144190015 | 144190048 | 34 | + | 17.4959 | 4.92E-07 | 0.106 | TAGCTGTCCCCAGACACACCAGAAGAGGGAGTCA | | |
| 197 | 3 | 153353444 | 153353503 | 59 | 129S1.BTBR.C57L | 3 | 153353434 | 153353468 | 35 | + | 16.7798 | 6.15E-07 | 0.191 | CAGCAGGTGGCTGCAGGTGGCTGCAGGTGGCAGCA | 5 | ENCFF907XEG.ENCFF136ZDK.ENCFF089GBC.ENCFF024JOZ.ENCFF327LYS |
| 197 | 3 | 153353444 | 153353503 | 59 | 129S1.BTBR.C57L | 3 | 153353444 | 153353478 | 35 | + | 20.4128 | 6.65E-08 | 0.0531 | CTGCAGGTGGCTGCAGGTGGCAGCAGGTGGCTGCA | 5 | ENCFF907XEG.ENCFF136ZDK.ENCFF089GBC.ENCFF024JOZ.ENCFF327LYS |
| 197 | 3 | 153353444 | 153353503 | 59 | 129S1.BTBR.C57L | 3 | 153353454 | 153353488 | 35 | + | 20.2569 | 7.36E-08 | 0.0562 | CTGCAGGTGGCAGCAGGTGGCTGCAGGTGGCAGCA | 5 | ENCFF907XEG.ENCFF136ZDK.ENCFF089GBC.ENCFF024JOZ.ENCFF327LYS |
| 197 | 3 | 153353444 | 153353503 | 59 | 129S1.BTBR.C57L | 3 | 153353464 | 153353498 | 35 | + | 17.3028 | 4.54E-07 | 0.168 | CAGCAGGTGGCTGCAGGTGGCAGCAGGTGGCTGCA | 5 | ENCFF907XEG.ENCFF136ZDK.ENCFF089GBC.ENCFF024JOZ.ENCFF327LYS |
| 197 | 3 | 153353444 | 153353503 | 59 | 129S1.BTBR.C57L | 3 | 153353474 | 153353508 | 35 | + | 20.2569 | 7.36E-08 | 0.0562 | CTGCAGGTGGCAGCAGGTGGCTGCAGGTGGCAGCA | 5 | ENCFF907XEG.ENCFF136ZDK.ENCFF089GBC.ENCFF024JOZ.ENCFF327LYS |
| 197 | 3 | 153353444 | 153353503 | 59 | 129S1.BTBR.C57L | 3 | 153353484 | 153353518 | 35 | + | 17.3028 | 4.54E-07 | 0.168 | CAGCAGGTGGCTGCAGGTGGCAGCAGGTGGCTGCA | 5 | ENCFF907XEG.ENCFF136ZDK.ENCFF089GBC.ENCFF024JOZ.ENCFF327LYS |
| 197 | 3 | 153353444 | 153353503 | 59 | 129S1.BTBR.C57L | 3 | 153353494 | 153353528 | 35 | + | 20.2569 | 7.36E-08 | 0.0562 | CTGCAGGTGGCAGCAGGTGGCTGCAGGTGGCAGCA | 5 | ENCFF907XEG.ENCFF136ZDK.ENCFF089GBC.ENCFF024JOZ.ENCFF327LYS |
| 198 | 3 | 10314329 | 10314661 | 332 | BUB,SMJ | 3 | 10314620 | 10314638 | 19 | - | 19.2951 | 1.88E-07 | 0.113 | CCGCCACTAGATGGTGCTA | 21 | ENCFF210BTO,ENCFF201JUA,ENCFF367UNR,ENCFF093ZRU,ENCFF099QNE,ENCFF024JOZ,ENCFF907XEG,ENCFF327LYS,ENCFF609MDN,ENCFF089GBC,ENCFF136ZDK,ENCFF198SSR,ENCFF883UPM,ENCFF821BIA,ENCFF321AGQ,ENCFF589ZBJ,ENCFF092NMJ,ENCFF165LXO,ENCFF502WDT,ENCFF052KVO,ENCFF430PPJ |
| 199 | 3 | 15610108 | 15610332 | 224 | AJ,BALB,BUB,C3H,C58J,CBA,CE,DBA2J,FVB,LP.PL.SJL.SWR.TH | 3 | 15610233 | 15610266 | 34 | - | 16.9187 | 7.13E-07 | 0.126 | TAGCTGTCTTCAGACGCACCAGAAGAGGGCGTCA | | |
| 200 | 3 | 23725635 | 23725840 | 205 | KK.MA/Mv.NOD.NOR.PJ.PL.SEA | 3 | 23725760 | 23725793 | 34 | - | 17.2439 | 5.79E-07 | 0.113 | TAGCTGTCCTCAGATACTCCAGAAGAGGGCATCA | | |
| 201 | 3 | 28792631 | 28793264 | 633 | DBA1J.DBA2J.ILNJ2 | 3 | 28793259 | 28793292 | 34 | - | 18.2195 | 3.05E-07 | 0.0836 | GTGCAATGTTCATGGAGGCCAGAAGAGAGCATCA | 1 | ENCFF201JUA |
| 202 | 3 | 38726235 | 38726671 | 436 | CE.SJL.SWR | 3 | 38726629 | 38726662 | 34 | - | 16.626 | 8.59E-07 | 0.137 | CTGCACTCACATTTGCCATTACAAGATGGCACTG | | |
| 203 | 3 | 51287969 | 51288181 | 212 | 129S1,BTBR,BUB,C3H,C57L,C58J,CBA,CE,FVB.LP.NZB.NZW.SJL.SMJ.SWR.TH | 3 | 51288095 | 51288113 | 19 | - | 17.5574 | 6.32E-07 | 0.181 | ACTCCAGAAGAGGGCGCCA | 3 | ENCFF367UNR,ENCFF093ZRU,ENCFF055HAP |
| 203 | 3 | 51287969 | 51288181 | 212 | 129S1,BTBR,BUB,C3H,C57L,C58J,CBA,CE,FVB.LP.NZB.NZW.SJL.SMJ.SWR.TH | 3 | 51288095 | 51288128 | 34 | - | 18.561 | 2.42E-07 | 0.079 | AAGCTGTCTTCAGACACTCCAGAAGAGGGCGCCA | 3 | ENCFF367UNR,ENCFF093ZRU,ENCFF055HAP |
| 204 | 3 | 54498103 | 54499012 | 909 | NOR | 3 | 54498766 | 54498784 | 19 | - | 17.2951 | 7.50E-07 | 0.193 | AGACCAGAAGAGGGCGACA | 18 | ENCFF198SSR,ENCFF165LXO,ENCFF883UPM,ENCFF502WDT,ENCFF099QNE,ENCFF589ZBJ,ENCFF136ZDK,ENCFF052KVO,ENCFF907XEG,ENCFF327LYS,ENCFF821BIA,ENCFF201JUA,ENCFF089GBC,ENCFF093ZRU,ENCFF024JOZ,ENCFF092NMJ,ENCFF367UNR,ENCFF055HAP |
| 204 | 3 | 54498103 | 54499012 | 909 | NOR | 3 | 54498766 | 54498799 | 34 | - | 28.8943 | 3.68E-11 | 0.00028 | GTGCAGTTCCCACAGAGACCAGAAGAGGGCGACA | 18 | ENCFF198SSR,ENCFF165LXO,ENCFF883UPM,ENCFF502WDT,ENCFF099QNE,ENCFF589ZBJ,ENCFF136ZDK,ENCFF052KVO,ENCFF907XEG,ENCFF327LYS,ENCFF821BIA,ENCFF201JUA,ENCFF089GBC,ENCFF093ZRU,ENCFF024JOZ,ENCFF092NMJ,ENCFF367UNR,ENCFF055HAP |
| 205 | 3 | 54903923 | 54904778 | 855 | AKR,BALB,BTBR,BUB,C3H,CBA,CE,ILNJ2,KK,MA/My,MRL,NOD,NOR,NZB,NZW,PL,RF,TH | 3 | 54904248 | 54904266 | 19 | - | 17.5574 | 6.32E-07 | 0.181 | TGGCCAGTTGGGGGCGCGC | | |
| 206 | 3 | 58454274 | 58454579 | 305 | AKR,BALB,BTBR,BUB,C3H,CBA,CE,DBA1J,DBA2J,ILNJ2,KK,MA/My,MRL,NOD,NOR,NZB.NZW.PJ.PL.RF.SEA.SJL.SMJ | 3 | 58454398 | 58454431 | 34 | - | 16.4472 | 9.61E-07 | 0.141 | TAGCTGTCTTCAGACACACCAGAAGAGGGCGTCA | | |
| 207 | 3 | 63040697 | 63040892 | 195 | DBA1J.DBA2J.FVB.SJL.SWR | 3 | 63040808 | 63040841 | 34 | - | 16.4472 | 9.61E-07 | 0.141 | TAGCTGTCTTCAGACACACCAGAAGAGGGCGTCA | | |
| 208 | 3 | 64138308 | 64138519 | 211 | SJL | 3 | 64138434 | 64138467 | 34 | - | 16.4472 | 9.61E-07 | 0.141 | TAGCTGTCTTCAGACACACCAGAAGAGGGCGTCA | | |
| 209 | 3 | 68358470 | 68358700 | 230 | CE | 3 | 68358582 | 68358615 | 34 | - | 18.935 | 1.88E-07 | 0.0693 | TTGCTGTCTTCAGACACACCAGAAGAGGGCGTCA | | |
| 210 | 3 | 75888649 | 75888947 | 298 | AJ,AKR,BTBR,DBA1J,DBA2J,NZB,NZO,NZW.PL.RF.SWR.TH | 3 | 75888829 | 75888862 | 34 | - | 18.3902 | 2.72E-07 | 0.0804 | GTGCAAGGCTAGATATGACCACTAGAGGCAGATG | 2 | ENCFF367UNR,ENCFF093ZRU |
| 211 | 3 | 84942247 | 84943062 | 815 | 129S1,AJ,C3H,C57L,CBA,CE,NZB,NZO,NZW.RF.SMJ | 3 | 84942601 | 84942619 | 19 | - | 17.2131 | 7.91E-07 | 0.196 | GGGCCAAAAGGGGGCAGTG | | |
| 211 | 3 | 84942247 | 84943062 | 815 | 129S1,AJ,C3H,C57L,CBA,CE,NZB,NZO,NZW.RF.SMJ | 3 | 84942601 | 84942635 | 35 | - | 16.4771 | 7.30E-07 | 0.211 | GTACAGGGGAACGCCAGGGCCAAAAGGGGGCAGTG | | |
| 212 | 3 | 86024561 | 86024772 | 211 | MA/Mv | 3 | 86024686 | 86024719 | 34 | - | 16.6341 | 8.54E-07 | 0.137 | TAGCTGTCTTCACACACTCCAGAAGAGGGCGTCA | 1 | ENCFF201JUA |
| 213 | 3 | 88186357 | 88186570 | 213 | MA/Mv | 3 | 88186484 | 88186517 | 34 | - | 16.4472 | 9.61E-07 | 0.141 | TAGCTGTCTTCAGACACACCAGAAGAGGGCGTCA | | |
| 214 | 3 | 88623391 | 88623589 | 198 | KK | 3 | 88623515 | 88623548 | 34 | - | 17.2439 | 5.79E-07 | 0.113 | TAGCTGTCCTCAGATACTCCAGAAGAGGGCATCA | | |
| 215 | 3 | 97788057 | 97788986 | 929 | BTBR,KK,MA/My,NOD,NOR,NZB,NZO,NZW | 3 | 97788730 | 97788763 | 34 | - | 22.0894 | 1.86E-08 | 0.0144 | GTGCAATGCCTGAAGATGTCAGAAGAGGGTGGTA | | |
| 216 | 3 | 104674062 | 104674260 | 198 | NZO.NZW | 3 | 104674189 | 104674207 | 19 | - | 17.5574 | 6.32E-07 | 0.181 | ACTCCAGAAGAGGGCGCCA | 2 | ENCFF093ZRU.ENCFF367UNR |
| 216 | 3 | 104674062 | 104674260 | 198 | NZO.NZW | 3 | 104674189 | 104674222 | 34 | - | 18.3659 | 2.76E-07 | 0.0804 | TAGCTGTCTTCAGACACTCCAGAAGAGGGCGCCA | 2 | ENCFF093ZRU.ENCFF367UNR |
| 217 | 3 | 115438271 | 115439173 | 902 | NZW | 3 | 115438570 | 115438603 | 34 | - | 21.3984 | 3.17E-08 | 0.0215 | CTGCTATCTCCAGACACACCAGAAGAGGGCATCA | 2 | ENCFF093ZRU.ENCFF367UNR |
| 218 | 3 | 121586611 | 121586983 | 372 | 129S1,AKR,BUB,C57L,C58J,CE,FVB,ILNJ2,LP,MRL,NOD,NOR,NZB,NZO,NZW,PJ,PL,RF,SEA.SJL.SWR.TH | 3 | 121586895 | 121586928 | 34 | - | 16.4472 | 9.61E-07 | 0.141 | TAGCTGTCTTCAGACACACCAGAAGAGGGCGTCA | | |
| 219 | 3 | 122209520 | 122209769 | 249 | AJ,BALB,BTBR,C3H,CBA,CE,DBA1J,DBA2J.KK.RH2.SMJ.TH | 3 | 122209702 | 122209735 | 34 | - | 17.0976 | 6.36E-07 | 0.119 | TAGCTGTCCTCAGACACCCCAGAAGAGGGCATCA | | |
| 220 | 3 | 127251475 | 127251695 | 220 | 129S1,AJ,BALB,C3H,C57L,CBA,NZB,PJ,RH2.SEA.SMJ | 3 | 127251608 | 127251641 | 34 | - | 18.8211 | 2.03E-07 | 0.0721 | TGGCTGTTTCCAGATACACCAGAAGGGGGCATCC | 8 | ENCFF367UNR,ENCFF201JUA,ENCFF099QNE,ENCFF327LYS,ENCFF024JOZ,ENCFF502WDT,ENCFF093ZRU,ENCFF055HAP |
| 221 | 3 | 146152281 | 146152506 | 225 | AJ,BALB,BUB,C3H,CBA,DBA1J,ILNJ2,MA/My,MRL,NOD,NOR,NZW,PJ,RF,RH2,SEA,SJL,SMJ | 3 | 146152404 | 146152437 | 34 | - | 19.7967 | 1.03E-07 | 0.0485 | TAGCTGTCCTCAGACACTCCAGAAGAGGGCACCA | 1 | ENCFF093ZRU |
| 222 | 3 | 151991341 | 151991559 | 218 | AJ.BALB.C3H.CE.ILNJ2.PL.RH2.SEA | 3 | 151991466 | 151991484 | 19 | - | 17.5574 | 6.32E-07 | 0.181 | ACTCCAGAAGAGGGCGCCA | 2 | ENCFF093ZRU.ENCFF367UNR |
| 222 | 3 | 151991341 | 151991559 | 218 | AJ.BALB.C3H.CE.ILNJ2.PL.RH2.SEA | 3 | 151991466 | 151991499 | 34 | - | 18.3659 | 2.76E-07 | 0.0804 | TAGCTGTCTTCAGACACTCCAGAAGAGGGCGCCA | 2 | ENCFF093ZRU.ENCFF367UNR |
| 223 | 4 | 42992514 | 42992726 | 212 | AKR,CE,KK,MRL,NOD,NOR,PJ,RF,SEA,SJL | 4 | 42992583 | 42992616 | 34 | + | 16.4228 | 9.76E-07 | 0.0938 | TAGCTGTCTTCAGATGCACCAGAAGAGGGCATCG | | |
| 224 | 4 | 52937050 | 52937266 | 216 | AJ.BALB.BUB.C3H.KK.MA/Mv.RH2 | 4 | 52937124 | 52937157 | 34 | + | 16.4634 | 9.51E-07 | 0.0931 | TAGCTGTCTTCAGACGCACCAGAAGAGGGCATCA | | |
| 225 | 4 | 53272922 | 53273122 | 200 | MA/Mv | 4 | 53272980 | 53273013 | 34 | + | 16.4472 | 9.61E-07 | 0.0931 | TAGCTGTCTTCAGACACACCAGAAGAGGGCGTCA | | |
| 226 | 4 | 54983434 | 54983674 | 240 | 129S1,AJ,AKR,BALB,BTBR,BUB,C3H,C57L,CBA,CE,DBA1J,DBA2J,FVB,ILNJ2,LP,MRL,NOD,NOR,NZB,NZO,NZW,PJ,PL,RF,RH2,SEA.SJL.SMJ.SWR.TH | 4 | 54983533 | 54983566 | 34 | + | 16.4634 | 9.51E-07 | 0.0931 | TAGCTGTCTTCAGACGCACCAGAAGAGGGCATCA | 1 | ENCFF093ZRU |
| 227 | 4 | 62180084 | 62180304 | 220 | BTBR.BUB | 4 | 62180162 | 62180195 | 34 | + | 17.8455 | 3.91E-07 | 0.0606 | TAGCTGTCCTCAGACACTCCAGAAGAGGGCGTCA | | |
| 228 | 4 | 99396359 | 99397068 | 709 | AJ,B10D2,B10J,BALB,BUB,C3H,C58J,CBA,KK.NZB.PJ.PL.RF.SEA | 4 | 99396698 | 99396731 | 34 | + | 20.3577 | 6.86E-08 | 0.0213 | CAGCCTTCTGCCTGCCGGCCAGAAGAGGGCACCA | | |
| 228 | 4 | 99396359 | 99397068 | 709 | AJ,B10D2,B10J,BALB,BUB,C3H,C58J,CBA,KK.NZB.PJ.PL.RF.SEA | 4 | 99396713 | 99396731 | 19 | + | 22.377 | 1.43E-08 | 0.0203 | CGGCCAGAAGAGGGCACCA | | |
| 229 | 4 | 101547401 | 101548101 | 700 | AKR.C58J.MA/Mv.MRL.PJ | 4 | 101547496 | 101547514 | 19 | + | 17.5246 | 6.45E-07 | 0.112 | AGGCCAGAAGAGGGCAACA | | |
| 230 | 4 | 106186410 | 106186673 | 263 | MA/My | 4 | 106186488 | 106186521 | 34 | + | 16.4634 | 9.51E-07 | 0.0931 | TAGCTGTCTTCAGACGCACCAGAAGAGGGCATCA | 10 | ENCFF093ZRU,ENCFF367UNR,ENCFF821BIA,ENCFF024JOZ,ENCFF136ZDK,ENCFF327LYS,ENCFF883UPM,ENCFF589ZBJ,ENCFF055HAP,ENCFF502WDT |
| 231 | 4 | 107626802 | 107627001 | 199 | AKR,BALB,BUB,C3H,CBA,CE,MRL,NOD,PL,RF.SEA.SJL.SMJ | 4 | 107626858 | 107626891 | 34 | + | 16.4472 | 9.61E-07 | 0.0931 | TAGCTGTCTTCAGACACACCAGAAGAGGGCGTCA | | |
| 232 | 4 | 108600337 | 108600534 | 197 | 129S1,AJ,AKR,BALB,BUB,C3H,C57L,C58J,CBA,CE,DBA1J,DBA2J,FVB,MRL,NOD,NZB,NZO,NZW,PJ,PL,RF,SEA,SJL,SWR,TH | 4 | 108600384 | 108600417 | 34 | + | 16.4472 | 9.61E-07 | 0.0931 | TAGCTGTCTTCAGACACACCAGAAGAGGGCGTCA | | |
| 233 | 4 | 115882689 | 115882951 | 262 | 129S1,AJ,AKR,B10D2,B10J,BTBR,C57L,C58J,CE,DBA1J,DBA2J,ILNJ2,KK,MA/My,MRL,NZB,NZO,NZW,PJ,PL,SEA,SMJ,SWR,TH | 4 | 115882754 | 115882787 | 34 | + | 16.4634 | 9.51E-07 | 0.0931 | TAGCTGTCTTCAGACGCACCAGAAGAGGGCATCA | 1 | ENCFF093ZRU |
| 234 | 4 | 116884796 | 116885031 | 235 | ILNJ2.TH | 4 | 116884873 | 116884907 | 35 | + | 16.0917 | 9.07E-07 | 0.15 | TAGCTGTCTTCAGACAGCACCAGAAGAGGGCATCA | 1 | ENCFF093ZRU |
| 235 | 4 | 117513241 | 117513470 | 229 | AJ,AKR,B10D2,B10J,BALB,BTBR,BUB,C3H,C58J,CBA,CE,DBA1J,DBA2J,FVB,ILNJ2,KK,MA/My,MRL,NOD,NOR,PL,RF,SJL,SWR,TH | 4 | 117513327 | 117513360 | 34 | + | 16.9593 | 6.95E-07 | 0.0815 | TAGCTGTCTTCAGACACTCCAGAAGGGGGCGTCA | | |
| 236 | 4 | 118453630 | 118453839 | 209 | 129S1,AJ,B10D2,B10J,BALB,BTBR,BUB,C3H,C57L,C58J,CBA,CE,DBA1J,DBA2J,FVB,ILNJ2,MA/My,NOD,NOR,NZB,PJ,PL,RF,SEA,SJL.SWR.TH | 4 | 118453697 | 118453730 | 34 | + | 16.4472 | 9.61E-07 | 0.0931 | TAGCTGTCTTCAGACACACCAGAAGAGGGCGTCA | | |
| 237 | 4 | 124708487 | 124708706 | 219 | 129S1,AJ,AKR,BALB,BTBR,BUB,C3H,C57L,C58J,CBA,CE,DBA1J,DBA2J,FVB,ILNJ2,KK,MA/My,MRL,NOD,NOR,NZO,NZW,PJ,PL,RF,SEA.SJL.SWR.TH | 4 | 124708565 | 124708598 | 34 | + | 18.3659 | 2.76E-07 | 0.0503 | TAGCTGTCTTCAGACACTCCAGAAGAGGGCGCCA | 2 | ENCFF367UNR,ENCFF093ZRU |
| 237 | 4 | 124708487 | 124708706 | 219 | 129S1,AJ,AKR,BALB,BTBR,BUB,C3H,C57L,C58J,CBA,CE,DBA1J,DBA2J,FVB,ILNJ2,KK,MA/My,MRL,NOD,NOR,NZO,NZW,PJ,PL,RF,SEA.SJL.SWR.TH | 4 | 124708580 | 124708598 | 19 | + | 17.5574 | 6.32E-07 | 0.111 | ACTCCAGAAGAGGGCGCCA | 2 | ENCFF367UNR,ENCFF093ZRU |
| 238 | 4 | 125057157 | 125057274 | 117 | AKR.MRL.TH | 4 | 125057198 | 125057231 | 34 | + | 17.4472 | 5.08E-07 | 0.0703 | CTGTAGTTCCTTCAGGTGTCACAGGGGAGCTCCC | | |
| 239 | 4 | 126339359 | 126339559 | 200 | B10D2.B10J.NZW | 4 | 126339418 | 126339451 | 34 | + | 17.2439 | 5.79E-07 | 0.0743 | TAGCTGTCCTCAGATACTCCAGAAGAGGGCATCA | | |
| 240 | 4 | 130410591 | 130410801 | 210 | FVB.KK.NOR.NZB.SJL.SWR | 4 | 130410659 | 130410692 | 34 | + | 18.3659 | 2.76E-07 | 0.0503 | TAGCTGTCTTCAGACACTCCAGAAGAGGGCGCCA | 1 | ENCFF093ZRU |
| 240 | 4 | 130410591 | 130410801 | 210 | FVB.KK.NOR.NZB.SJL.SWR | 4 | 130410674 | 130410692 | 19 | + | 17.5574 | 6.32E-07 | 0.111 | ACTCCAGAAGAGGGCGCCA | 1 | ENCFF093ZRU |
| 241 | 4 | 130969320 | 130969529 | 209 | 129S1,B10D2,B10J,BTBR,BUB,C57L,DBA1J,FVB,ILNJ2,KK,MA/My,NOR,NZB,PJ,PL,SEA.SJL.SMJ.SWR.TH | 4 | 130969386 | 130969419 | 34 | + | 16.4634 | 9.51E-07 | 0.0931 | TAGCTGTCTTCAGACGCACCAGAAGAGGGCATCA | | |
| 242 | 4 | 132507380 | 132507568 | 188 | DBA1J.KK.NZB.SMJ | 4 | 132507426 | 132507459 | 34 | + | 18.3333 | 2.82E-07 | 0.0507 | TAGCTGTCCTCAGACACACCAGAAGAGGGCGTCA | 1 | ENCFF093ZRU |
| 243 | 4 | 132630049 | 132630657 | 608 | DBA1J.KK.NZB.SMJ | 4 | 132630639 | 132630672 | 34 | + | 17.3902 | 5.27E-07 | 0.071 | TAGCTGTCCTCAGACACTCCAGAAGAGGGCATCA | | |
| 244 | 4 | 132893045 | 132893262 | 217 | 129S1,AJ,AKR,B10D2,B10J,BALB,BTBR,BUB,C3H,C57L,CBA,CE,DBA2J,FVB,ILNJ2,MA/My,MRL,NOD,NZW,PJ,PL,RF,RH2,SEA,SJL,SWR.TH | 4 | 132893125 | 132893158 | 34 | + | 16.4634 | 9.51E-07 | 0.0931 | TAGCTGTCTTCAGACGCACCAGAAGAGGGCATCA | | |
| 245 | 4 | 133763750 | 133764002 | 252 | 129S1,AJ,AKR,B10D2,B10J,BALB,BTBR,BUB,C3H,C57L,CBA,CE,DBA1J,DBA2J,FVB,ILNJ2,MA/My,MRL,NOD,NZW,PJ,PL,RF,RH2,SEA.SJL.SMJ.SWR.TH | 4 | 133763800 | 133763833 | 34 | + | 18.3659 | 2.76E-07 | 0.0503 | TAGCTGTCTTCAGACACTCCAGAAGAGGGCGCCA | | |
| 245 | 4 | 133763750 | 133764002 | 252 | 129S1,AJ,AKR,B10D2,B10J,BALB,BTBR,BUB,C3H,C57L,CBA,CE,DBA1J,DBA2J,FVB,ILNJ2,MA/My,MRL,NOD,NZW,PJ,PL,RF,RH2,SEA.SJL.SMJ.SWR.TH | 4 | 133763815 | 133763833 | 19 | + | 17.5574 | 6.32E-07 | 0.111 | ACTCCAGAAGAGGGCGCCA | | |
| 246 | 4 | 133966613 | 133966835 | 222 | 129S1,AJ,AKR,B10D2,B10J,BALB,BTBR,BUB,C3H,C57L,CBA,CE,DBA1J,DBA2J,FVB,ILNJ2,MA/My,MRL,NOD,NZW,PJ,PL,RF,RH2,SEA.SJL.SMJ.SWR.TH | 4 | 133966692 | 133966725 | 34 | + | 18.3659 | 2.76E-07 | 0.0503 | TAGCTGTCTTCAGACACTCCAGAAGAGGGCGCCA | 3 | ENCFF367UNR,ENCFF093ZRU,ENCFF099QNE |
| 246 | 4 | 133966613 | 133966835 | 222 | 129S1,AJ,AKR,B10D2,B10J,BALB,BTBR,BUB,C3H,C57L,CBA,CE,DBA1J,DBA2J,FVB,ILNJ2,MA/My,MRL,NOD,NZW,PJ,PL,RF,RH2,SEA.SJL.SMJ.SWR.TH | 4 | 133966707 | 133966725 | 19 | + | 17.5574 | 6.32E-07 | 0.111 | ACTCCAGAAGAGGGCGCCA | 3 | ENCFF367UNR,ENCFF093ZRU,ENCFF099QNE |
| 247 | 4 | 134137243 | 134137549 | 306 | 129S1,AJ,AKR,B10D2,B10J,BALB,BTBR,BUB,C3H,C57L,CBA,CE,DBA1J,DBA2J,FVB,ILNJ2,MA/My,MRL,NOD,NZW,PJ,PL,RF,RH2,SEA.SJL.SMJ.SWR.TH | 4 | 134137368 | 134137401 | 34 | + | 20.065 | 8.47E-08 | 0.0249 | ATGCAATGCCTGTGGAGGCCAGAACAGGGTACTG | | |
| 248 | 4 | 135562292 | 135562499 | 207 | B10D2.B10J.DBA1J.KK.NZO.PJ.SEA | 4 | 135562355 | 135562388 | 34 | + | 17.7805 | 4.08E-07 | 0.0623 | CTGTAGTCTGTAGACACTCCAGAAGAGGGCATCA | | |
| 249 | 4 | 137693725 | 137694100 | 375 | PJ | 4 | 137693980 | 137694014 | 35 | + | 17.7339 | 3.53E-07 | 0.0912 | CTGTACCCCCAGGCCCTGGCCTGCAGACGACGGTA | | |
| 250 | 4 | 137724808 | 137725016 | 208 | AJ,AKR,BALB,BUB,C3H,CBA,CE,DBA1J,DBA2J,FVB,ILNJ2,KK,MA/My,MRL,NZB,NZW,PL.RF.RH2.SJL.SMJ.SWR.TH | 4 | 137724864 | 137724897 | 34 | + | 17.8455 | 3.91E-07 | 0.0606 | TAGCTGTCCTCAGACACTCCAGAAGAGGGCGTCA | | |
| 251 | 4 | 143036628 | 143037583 | 955 | 129S1,BTBR,BUB,C57L,C58J,CE,DBA2J,FVB,KK,LP,MA/My,NOD,NZB,NZW,PJ,PL,RF,SEA.SJL.SMJ.SWR.TH | 4 | 143037575 | 143037608 | 34 | + | 16.5528 | 8.99E-07 | 0.0931 | TAGCTGTCTTCAGACACACCAGAAGAGGGCGTCG | | |
| 252 | 4 | 143089847 | 143090039 | 192 | 129S1,BTBR,BUB,C57L,C58J,CE,DBA2J,FVB,KK,LP,MA/My,MRL,NOD,NZB,NZW,PJ,PL,RF.SEA.SJL.SMJ.SWR.TH | 4 | 143089898 | 143089931 | 34 | + | 16.5203 | 9.18E-07 | 0.0931 | TAGCTGTCTTCCAACACTCCAGAAGAGGGCGTCA | | |
| 253 | 4 | 145208053 | 145208258 | 205 | 129S1,AKR,BTBR,BUB,C3H,C57L,CBA,CE,DBA1J,KK,MA/My,MRL,NOD,NZB,NZO,NZW,PL.RF.SJL.SMJ.SWR | 4 | 145208116 | 145208149 | 34 | + | 17.3902 | 5.27E-07 | 0.071 | TAGCTGTCCTCAGACACTCCAGAAGAGGGCATCA | | |
| 254 | 4 | 146617253 | 146617462 | 209 | 129S1,AKR,B10J,BALB,BTBR,BUB,C3H,C57L,C58J,CBA,CE,DBA1J,DBA2J,FVB,ILNJ2,KK,LP,MA/My,MRL,NOD,NZB,NZO,NZW,PJ,PL.RF.SEA.SJL.SMJ.SWR.TH | 4 | 146617318 | 146617351 | 34 | + | 16.4472 | 9.61E-07 | 0.0931 | TAGCTGTCTTCAGACACACCAGAAGAGGGCGTCA | | |
| 255 | 4 | 147009059 | 147009256 | 197 | 129S1,AKR,BTBR,BUB,C57L,DBA1J,KK,MA/My,MRL,NOD,NZB,NZW,PL,RF,SJL,SWR | 4 | 147009113 | 147009146 | 34 | + | 16.4472 | 9.61E-07 | 0.0931 | TAGCTGTCTTCAGACACACCAGAAGAGGGCGTCA | | |
| 256 | 4 | 148034970 | 148035154 | 184 | 129S1,AKR,BTBR,BUB,C3H,C57L,C58J,CBA,CE,DBA1J,DBA2J,KK,LP,MA/My,MRL,NOD,NZB,NZO,NZW,PJ,PL,RF,SEA,SJL,SMJ,SWR.TH | 4 | 148035063 | 148035096 | 34 | + | 16.4634 | 9.51E-07 | 0.0931 | TAGCTGTCTTCAGACGCACCAGAAGAGGGCATCA | | |
| 257 | 4 | 148626379 | 148626593 | 214 | 129S1,AJ,AKR,BALB,BTBR,BUB,C3H,C57L,C58J,CBA,CE,DBA1J,DBA2J,KK,LP,MA/My,MRL,NOD,NZB,NZO,NZW,PJ,PL,RF,SEA,SJL.SMJ.SWR.TH | 4 | 148626449 | 148626482 | 34 | + | 16.4634 | 9.51E-07 | 0.0931 | TAGCTGTCTTCAGACGCACCAGAAGAGGGCATCA | | |
| 258 | 4 | 149485040 | 149485236 | 196 | 129S1,AKR,BTBR,C3H,C57L,CBA,DBA1J,DBA2J,FVB,ILNJ2,MA/My,MRL,NOD,PL,SJL,SWR.TH | 4 | 149485088 | 149485121 | 34 | + | 16.4472 | 9.61E-07 | 0.0931 | TAGCTGTCTTCAGACACACCAGAAGAGGGCGTCA | | |
| 259 | 4 | 149507935 | 149508144 | 209 | 129S1,AKR,BTBR,C3H,C57L,C58J,CBA,CE,DBA1J,DBA2J,FVB,ILNJ2,LP,MA/My,MRL,NOD.NZO.PJ.PL.SEA.SJL.SWR.TH | 4 | 149508002 | 149508035 | 34 | + | 16.4472 | 9.61E-07 | 0.0931 | TAGCTGTCTTCAGACACACCAGAAGAGGGCGTCA | 1 | ENCFF093ZRU |
| 260 | 4 | 149685148 | 149685570 | 422 | FVB.ILNJ2 | 4 | 149685427 | 149685460 | 34 | + | 16.5366 | 9.09E-07 | 0.0931 | ATGTAGTGCCTATGTATGCCAGAAGGGGATGTTG | | |
| 261 | 4 | 150596892 | 150597663 | 771 | AJ.BALB.RH2 | 4 | 150597410 | 150597443 | 34 | + | 16.8537 | 7.43E-07 | 0.0844 | GAGCAGTGCCAGGAGGCAGCACTGGGTGCCACAT | | |
| 262 | 4 | 150764197 | 150764399 | 202 | AKR.C3H.CBA.MRL.NOD.NOR.NZO | 4 | 150764258 | 150764291 | 34 | + | 16.4472 | 9.61E-07 | 0.0931 | TAGCTGTCTTCAGACACACCAGAAGAGGGCGTCA | | |
| 263 | 4 | 154277883 | 154278259 | 376 | 129S1,AJ,AKR,C3H,CBA,CE,DBA2J,FVB,ILNJ2,NOD,NOR,RF,RH2,SJL,SMJ,SWR,TH | 4 | 154277897 | 154277930 | 34 | + | 16.4472 | 9.61E-07 | 0.0931 | TAGCTGTCTTCAGACACACCAGAAGAGGGCGTCA | | |
| 264 | 4 | 155719765 | 155720337 | 572 | CE.FVB.NOD.NOR.SJL.SWR.TH | 4 | 155719753 | 155719786 | 34 | + | 18.5854 | 2.38E-07 | 0.0492 | TTGCTGTCTTCAGACACACCAGAAGAGGGCATCG | 2 | ENCFF093ZRU.ENCFF367UNR |
| 265 | 4 | 155803163 | 155803380 | 217 | 129S1,AJ,AKR,BALB,BUB,C3H,CBA,CE,DBA1J,DBA2J,FVB,MRL,NOD,NOR,NZO,PL,RF,RH2.SJL.SWR | 4 | 155803237 | 155803270 | 34 | + | 16.4634 | 9.51E-07 | 0.0931 | TAGCTGTCTTCAGACGCACCAGAAGAGGGCATCA | | |
| 266 | 4 | 156163550 | 156163788 | 238 | 129S1,AJ,AKR,BALB,C3H,C57L,C58J,CBA,CE,DBA1J,DBA2J,FVB,ILNJ2,MA/My,MRL,NOD,NOR,NZO,PJ,PL,RF,RH2,SEA,SJL,SMJ,SWR.TH | 4 | 156163647 | 156163680 | 34 | + | 18.3333 | 2.82E-07 | 0.0507 | TAGCTGTCCTCAGACACACCAGAAGAGGGCGTCA | | |
| 267 | 4 | 4616551 | 4616742 | 191 | AJ | 4 | 4616675 | 4616708 | 34 | - | 16.4472 | 9.61E-07 | 0.0931 | TAGCTGTCTTCAGACACACCAGAAGAGGGCGTCA | | |
| 268 | 4 | 6226075 | 6226274 | 199 | 129S1,AKR,BALB,BTBR,BUB,C57L,FVB,ILNJ2,MA/My,NOD,NOR,NZW,PL,RF,RH2,SEA,SJL.SWR.TH | 4 | 6226201 | 6226234 | 34 | - | 17.7805 | 4.08E-07 | 0.0623 | CTGTAGTCTGTAGACACTCCAGAAGAGGGCATCA | | |
| 269 | 4 | 6542315 | 6543176 | 861 | BUB | 4 | 6542838 | 6542856 | 19 | - | 17.5902 | 6.18E-07 | 0.111 | ACACCAGAAGAGGGCACTG | | |
| 269 | 4 | 6542315 | 6543176 | 861 | BUB | 4 | 6542838 | 6542871 | 34 | - | 18.2195 | 3.05E-07 | 0.0526 | TAGCTGTCTTCAAACACACCAGAAGAGGGCACTG | | |
| 270 | 4 | 11121823 | 11122044 | 221 | AJ.NOD.NOR | 4 | 11121946 | 11121979 | 34 | - | 17.1382 | 6.20E-07 | 0.0772 | TAGCTGTCTCCAGACACTCCAGAAGAGGGCATCA | | |
| 271 | 4 | 44001313 | 44001508 | 195 | 129S1,BUB,C58J,CBA,DBA1J,DBA2J,FVB,ILNJ2.LP.PJ.SEA.SJL.SMJ.SWR.TH | 4 | 44001437 | 44001470 | 34 | - | 16.4472 | 9.61E-07 | 0.0931 | TAGCTGTCTTCAGACACACCAGAAGAGGGCGTCA | | |
| 272 | 4 | 45801866 | 45802068 | 202 | BUB,C58J,CBA,DBA1J,DBA2J,ILNJ2,LP,MA/My.NZW.PL.SMJ | 4 | 45801990 | 45802008 | 19 | - | 17.5574 | 6.32E-07 | 0.111 | ACTCCAGAAGAGGGCGCCA | 5 | ENCFF367UNR,ENCFF821BIA,ENCFF327LYS,ENCFF093ZRU,ENCFF099QNE |
| 272 | 4 | 45801866 | 45802068 | 202 | BUB,C58J,CBA,DBA1J,DBA2J,ILNJ2,LP,MA/My.NZW.PL.SMJ | 4 | 45801990 | 45802023 | 34 | - | 18.3659 | 2.76E-07 | 0.0503 | TAGCTGTCTTCAGACACTCCAGAAGAGGGCGCCA | 5 | ENCFF367UNR,ENCFF821BIA,ENCFF327LYS,ENCFF093ZRU,ENCFF099QNE |
| 273 | 4 | 48602148 | 48602480 | 332 | 129S1.BUB.C57L.KK.MA/Mv.NZW | 4 | 48602335 | 48602353 | 19 | - | 17.5574 | 6.32E-07 | 0.111 | ACTCCAGAAGAGGGCGCCA | 1 | ENCFF093ZRU |
| 273 | 4 | 48602148 | 48602480 | 332 | 129S1.BUB.C57L.KK.MA/Mv.NZW | 4 | 48602335 | 48602368 | 34 | - | 18.3659 | 2.76E-07 | 0.0503 | TAGCTGTCTTCAGACACTCCAGAAGAGGGCGCCA | 1 | ENCFF093ZRU |
| 274 | 4 | 49686463 | 49687059 | 596 | MA/Mv | 4 | 49686495 | 49686529 | 35 | - | 18.3119 | 2.49E-07 | 0.0757 | TAGCTGTGGGGGGGGGGTGCCAGCAGAGGGCAAGG | | |
| 275 | 4 | 51782138 | 51782351 | 213 | AJ,BALB,BUB,C3H,ILNJ2,KK,MA/My,NOD,NOR.PJ.RH2.SEA | 4 | 51782262 | 51782295 | 34 | - | 18.8618 | 1.97E-07 | 0.0437 | TAGCTGTCCTCAGATTCTCCAGAAGAGGGCATCA | | |
| 276 | 4 | 58164222 | 58164445 | 223 | 129S1,C57L,NOD,NOR,PJ,PL,SJL,SMJ,SWR.TH | 4 | 58164344 | 58164377 | 34 | - | 17.439 | 5.10E-07 | 0.0703 | TAGCTGTCTTCAGTCACTCCAGAAGATGGCGCTA | | |
| 277 | 4 | 59197770 | 59198025 | 255 | 129S1,AJ,AKR,BALB,BTBR,C3H,C57L,CBA,CE,DBA1J,DBA2J,FVB,ILNJ2,LP,MA/My,MRL,NOD,NOR,NZB,NZO,NZW,PJ,RF,RH2,SEA,SJL.SWR.TH | 4 | 59197895 | 59197928 | 34 | - | 17.9431 | 3.66E-07 | 0.0591 | CAGCTGTCCTCAGACACTCCAGAAGAGGGAGTCA | | |
| 278 | 4 | 65726184 | 65726807 | 623 | BTBR.CE.PJ | 4 | 65726293 | 65726326 | 34 | - | 17.6341 | 4.49E-07 | 0.0653 | CTGCACCTCAATCTGAGCCCTGTAGGAGGACCCG | | |

| | | | | | | | | | | | | | | | | |
|---|---|---|---|---|---|---|---|---|---|---|---|---|---|---|---|---|
| 279 | 4 | 83231895 | 83232131 | 236 | SMJ | 4 | 83232022 | 83232055 | 34 | - | 17.878 | 3.83E-07 | 0.06 | TAGCTGTCCTCAGACACACCAGAAGAGGGCATCA | | |
| 280 | 4 | 85681775 | 85681973 | 198 | 129S1,B10D2,B10J,BTBR,C57L,C58J,CE,DBA1J,DBA2J,ILNJ2,KK,NZO,PJ,PL,RF,RH2,SMJ | 4 | 85681901 | 85681935 | 35 | - | 18.1284 | 2.79E-07 | 0.0805 | CTGTAGCTGTCTTCAGACACCAGAAGAGGGCATCA | | |
| 281 | 4 | 87324876 | 87325083 | 207 | B10D2,B10J,C3H,C58J,CBA,DBA1J,DBA2J,ILNJ2,MA/My,NZO,PJ,PL,RF,SMJ | 4 | 87324998 | 87325031 | 34 | - | 18.8537 | 1.98E-07 | 0.0437 | CAGCTGTCTTCAGACACTCCAGTAGAGGGCATCA | 1 | ENCFF093ZRU |
| 282 | 4 | 92489080 | 92489300 | 220 | 129S1.C57L.RH2 | 4 | 92489206 | 92489239 | 34 | - | 17.6667 | 4.40E-07 | 0.0647 | TAGCTGTCTTCAGACACGCCAGAAGAGGGCGTCA | | |
| 283 | 4 | 99520761 | 99520975 | 214 | 129S1,AJ,BALB,BTBR,C3H,C57L,CBA,ILNJ2,KK,MA/My,NOD,NOR,NZO,RH2,SEA,SWR,TH | 4 | 99520887 | 99520905 | 19 | - | 17.5574 | 6.32E-07 | 0.111 | ACTCCAGAAGAGGGCGCCA | | |
| 283 | 4 | 99520761 | 99520975 | 214 | 129S1,AJ,BALB,BTBR,C3H,C57L,CBA,ILNJ2,KK,MA/My,NOD,NOR,NZO,RH2,SEA,SWR,TH | 4 | 99520887 | 99520920 | 34 | - | 18.2358 | 3.02E-07 | 0.0524 | TAGCTGTCTTAAGACACTCCAGAAGAGGGCGCCA | | |
| 284 | 4 | 100489654 | 100490283 | 629 | BUB,PL,RF | 4 | 100490179 | 100490212 | 34 | - | 19.4715 | 1.29E-07 | 0.0337 | CTGCAAAGATAAACGTGGCCAGAAGAGGTCCCTG | | |
| 285 | 4 | 103112995 | 103113237 | 242 | AJ,BALB,BUB,C3H,C58J,CBA,KK,NZW,PJ,PL,SEA,SMJ,TH | 4 | 103113120 | 103113153 | 34 | - | 16.4634 | 9.51E-07 | 0.0931 | TAGCTGTCTTCAGACGCACCAGAAGAGGGCATCA | | |
| 286 | 4 | 104220166 | 104220355 | 189 | 129S1,AJ,AKR,BALB,BTBR,BUB,C3H,C57L,C58J,CBA,CE,DBA1J,DBA2J,FVB,ILNJ2,KK,MA/My,MRL,NOD,NOR,NZB,NZO,NZW,PJ,PL,RF,SEA,SJL,SMJ,SWR,TH | 4 | 104220287 | 104220305 | 19 | - | 17.5574 | 6.32E-07 | 0.111 | ACTCCAGAAGAGGGCGCCA | 2 | ENCFF093ZRU,ENCFF099QNE |
| 286 | 4 | 104220166 | 104220355 | 189 | 129S1,AJ,AKR,BALB,BTBR,BUB,C3H,C57L,C58J,CBA,CE,DBA1J,DBA2J,FVB,ILNJ2,KK,MA/My,MRL,NOD,NOR,NZB,NZO,NZW,PJ,PL,RF,SEA,SJL,SMJ,SWR,TH | 4 | 104220287 | 104220320 | 34 | - | 18.561 | 2.42E-07 | 0.0493 | AAGCTGTCTTCAGACACTCCAGAAGAGGGCGCCA | 2 | ENCFF093ZRU,ENCFF099QNE |
| 287 | 4 | 104861973 | 104862054 | 81 | BALB,BTBR,C3H,C58J,CBA,CE,ILNJ2,KK,MA/My,NOD,NOR,PJ,RF,SEA,SJL,SMJ,SWR | 4 | 104861954 | 104861988 | 35 | - | 18.1927 | 2.68E-07 | 0.0793 | GGGCAGTGTGCAGCGCCTGCCTAGAGAGGGCAGTG | 18 | ENCFF055HAP,ENCFF065ODV,ENCFF136ZDK,ENCFF327LYS,ENCFF092NMJ,ENCFF089GBC,ENCFF430PPJ,ENCFF198SSR,ENCFF165LXO,ENCFF210BTO,ENCFF589ZBJ,ENCFF907XEG,ENCFF052KVO,ENCFF533APC,ENCFF609MDN,ENCFF957SPH,ENCFF194HVL,ENCFF821BIA |
| 287 | 4 | 104861973 | 104862054 | 81 | BALB,BTBR,C3H,C58J,CBA,CE,ILNJ2,KK,MA/My,NOD,NOR,PJ,RF,SEA,SJL,SMJ,SWR | 4 | 104861981 | 104862015 | 35 | - | 18.1927 | 2.68E-07 | 0.0793 | GGGCAGTGTGCAGCGCCTGCCTAGAGAGGGCAGTG | 20 | ENCFF055HAP,ENCFF065ODV,ENCFF136ZDK,ENCFF327LYS,ENCFF092NMJ,ENCFF089GBC,ENCFF430PPJ,ENCFF198SSR,ENCFF165LXO,ENCFF210BTO,ENCFF589ZBJ,ENCFF907XEG,ENCFF052KVO,ENCFF533APC,ENCFF609MDN,ENCFF957SPH,ENCFF194HVL,ENCFF099QNE,ENCFF821BIA,ENCFF093ZRU |
| 287 | 4 | 104861973 | 104862054 | 81 | BALB,BTBR,C3H,C58J,CBA,CE,ILNJ2,KK,MA/My,NOD,NOR,PJ,RF,SEA,SJL,SMJ,SWR | 4 | 104862008 | 104862042 | 35 | - | 18.1927 | 2.68E-07 | 0.0793 | GGGCAGTGTGCAGCGCCTGCCTAGAGAGGGCAGTG | 22 | ENCFF055HAP,ENCFF065ODV,ENCFF136ZDK,ENCFF327LYS,ENCFF092NMJ,ENCFF089GBC,ENCFF430PPJ,ENCFF198SSR,ENCFF165LXO,ENCFF210BTO,ENCFF589ZBJ,ENCFF907XEG,ENCFF052KVO,ENCFF533APC,ENCFF609MDN,ENCFF957SPH,ENCFF194HVL,ENCFF099QNE,ENCFF024JOZ,ENCFF367UNR,ENCFF821BIA,ENCFF093ZRU |
| 287 | 4 | 104861973 | 104862054 | 81 | BALB,BTBR,C3H,C58J,CBA,CE,ILNJ2,KK,MA/My,NOD,NOR,PJ,RF,SEA,SJL,SMJ,SWR | 4 | 104862035 | 104862069 | 35 | - | 17.2936 | 4.57E-07 | 0.106 | GGGCAGTGTGCAGCGTCTGCCTAGAGAGGGCAGTG | 24 | ENCFF055HAP,ENCFF065ODV,ENCFF136ZDK,ENCFF327LYS,ENCFF092NMJ,ENCFF089GBC,ENCFF430PPJ,ENCFF198SSR,ENCFF165LXO,ENCFF210BTO,ENCFF589ZBJ,ENCFF907XEG,ENCFF052KVO,ENCFF533APC,ENCFF609MDN,ENCFF957SPH,ENCFF194HVL,ENCFF099QNE,ENCFF024JOZ,ENCFF367UNR,ENCFF883UPM,ENCFF502WDT,ENCFF821BIA,ENCFF093ZRU |
| 288 | 4 | 107975161 | 107975376 | 215 | 129S1,AJ,AKR,BALB,BUB,C3H,C57L,CBA,CE,FVB,MRL,NOD,PL,RF,SEA,SJL,SMJ | 4 | 107975287 | 107975320 | 34 | - | 17.2439 | 5.79E-07 | 0.0743 | TAGCTGTCCTCAGATACTCCAGAAGAGGGCATCA | 4 | ENCFF093ZRU,ENCFF367UNR |
| 289 | 4 | 107976530 | 107976776 | 246 | 129S1,AJ,AKR,BALB,BUB,C3H,C57L,C58J,CBA,CE,DBA1J,DBA2J,FVB,MRL,NOD,NZB,NZO,NZW,PJ,PL,RF,SEA,SJL,SMJ,TH | 4 | 107976760 | 107976793 | 34 | - | 16.4472 | 9.61E-07 | 0.0931 | TAGCTGTCTTCAGACACACCAGAAGAGGGCGTCA | 1 | ENCFF093ZRU |
| 290 | 4 | 108502364 | 108502580 | 216 | 129S1,AJ,AKR,C57L,CE,FVB,MRL,SMJ,SWR | 4 | 108502491 | 108502509 | 19 | - | 17.5574 | 6.32E-07 | 0.111 | ACTCCAGAAGAGGGCGCCA | | |
| 290 | 4 | 108502364 | 108502580 | 216 | 129S1,AJ,AKR,C57L,CE,FVB,MRL,SMJ,SWR | 4 | 108502491 | 108502524 | 34 | - | 18.374 | 2.75E-07 | 0.0503 | TAGCTGTCTTCAGTCACTCCAGAAGAGGGCGCCA | | |
| 291 | 4 | 114029992 | 114030324 | 332 | 129S1,AJ,AKR,BALB,BTBR,BUB,C3H,C57L,CBA,CE,DBA1J,DBA2J,FVB,LP,MA/My,MRL,NOD,NOR,PL,RF,SEA,SJL,SMJ,SWR,TH | 4 | 114030284 | 114030302 | 19 | - | 19.2951 | 1.88E-07 | 0.0616 | CCGCCACTAGATGGTGCTA | 13 | ENCFF092NMJ,ENCFF201JUA,ENCFF367UNR,ENCFF024JOZ,ENCFF327LYS,ENCFF907XEG,ENCFF136ZDK,ENCFF093ZRU,ENCFF957SPH,ENCFF099QNE,ENCFF589ZBJ,ENCFF165LXO,ENCFF821BIA |
| 292 | 4 | 115899064 | 115899269 | 205 | MA/My | 4 | 115899175 | 115899208 | 34 | - | 19.2602 | 1.50E-07 | 0.0369 | CAGCTGTCTTCAGACAGACCAGAAGAGGGCATCA | | |
| 293 | 4 | 116520516 | 116520725 | 209 | ILNJ2.TH | 4 | 116520647 | 116520680 | 34 | - | 16.4472 | 9.61E-07 | 0.0931 | TAGCTGTCTTCAGACACACCAGAAGAGGGCGTCA | | |
| 294 | 4 | 116526566 | 116526934 | 368 | CE | 4 | 116526669 | 116526702 | 34 | - | 22.9675 | 9.26E-09 | 0.00502 | ATGCATTGCCCCTGAAGGCCAGAAGAGGGCATCT | 19 | ENCFF089GBC,ENCFF092NMJ,ENCFF367UNR,ENCFF093ZRU,ENCFF136ZDK,ENCFF883UPM,ENCFF165LXO,ENCFF327LYS,ENCFF907XEG,ENCFF821BIA,ENCFF099QNE,ENCFF024JOZ,ENCFF055HAP,ENCFF502WDT,ENCFF589ZBJ,ENCFF198SSR,ENCFF065ODV,ENCFF052KVO,ENCFF957SPH |
| 295 | 4 | 117931709 | 117931923 | 214 | 129S1,AJ,C57L,DBA1J,DBA2J,MA/My,NZB,NZO.NZW.PJ.SEA.SMJ.TH | 4 | 117931839 | 117931872 | 34 | - | 17.8943 | 3.78E-07 | 0.0599 | TAGCAGTCTTCAGACACTCCAGAAGAGGGCGTCA | | |
| 296 | 4 | 118528982 | 118529178 | 196 | ILNJ2.NZO.NZW | 4 | 118529106 | 118529140 | 35 | - | 18.1284 | 2.79E-07 | 0.0805 | CTGTAGCTGTCTTCAGACACCAGAAGAGGGCATCA | | |
| 297 | 4 | 119585380 | 119585646 | 266 | 129S1.C57L | 4 | 119585370 | 119585403 | 34 | - | 18.0894 | 3.33E-07 | 0.0556 | CAGCAAGTCCATCCCCTCCCAGGAGGAGGAAGAG | | |
| 298 | 4 | 123253072 | 123253293 | 221 | 129S1,BTBR,BUB,C57L,C58J,FVB,ILNJ2,NOD,NOR,NZB,NZW,SMJ,SWR,TH | 4 | 123253197 | 123253230 | 34 | - | 16.4634 | 9.51E-07 | 0.0931 | TAGCTGTCTTCAGACGCACCAGAAGAGGGCATCA | | |
| 299 | 4 | 125127243 | 125127602 | 359 | 129S1,BALB,BTBR,BUB,C3H,C57L,CBA,CE,DBA1J,DBA2J,FVB,ILNJ2,KK,MA/My,NOD,NOR,PJ,PL,RF,SEA,SJL,SWR | 4 | 125127529 | 125127562 | 34 | - | 17.2439 | 5.79E-07 | 0.0743 | TAGCTGTCCTCAGATACTCCAGAAGAGGGCATCA | | |
| 300 | 4 | 125499736 | 125499988 | 252 | 129S1,AJ,BALB,BTBR,BUB,C3H,C57L,C58J,CBA,CE,DBA1J,DBA2J,FVB,KK,MA/My,NOD,NOR,NZO,PJ,PL,RF,SEA,SJL,SWR,TH | 4 | 125499863 | 125499881 | 19 | - | 17.5574 | 6.32E-07 | 0.111 | ACTCCAGAAGAGGGCGCCA | 2 | ENCFF093ZRU |
| 300 | 4 | 125499736 | 125499988 | 252 | 129S1,AJ,BALB,BTBR,BUB,C3H,C57L,C58J,CBA,CE,DBA1J,DBA2J,FVB,KK,MA/My,NOD,NOR,NZO,PJ,PL,RF,SEA,SJL,SWR,TH | 4 | 125499863 | 125499896 | 34 | - | 18.3659 | 2.76E-07 | 0.0503 | TAGCTGTCTTCAGACACTCCAGAAGAGGGCGCCA | 2 | ENCFF093ZRU |
| 301 | 4 | 126822883 | 126823088 | 205 | B10D2.B10J | 4 | 126823009 | 126823042 | 34 | - | 17.8455 | 3.91E-07 | 0.0606 | TAGCTGTCCTCAGACACTCCAGAAGAGGGCGTCA | | |
| 302 | 4 | 126993272 | 126993488 | 216 | B10D2.B10J.ILNJ2.PJ.SEA | 4 | 126993399 | 126993417 | 19 | - | 17.5574 | 6.32E-07 | 0.111 | ACTCCAGAAGAGGGCGCCA | | |
| 302 | 4 | 126993272 | 126993488 | 216 | B10D2.B10J.ILNJ2.PJ.SEA | 4 | 126993399 | 126993432 | 34 | - | 18.3659 | 2.76E-07 | 0.0503 | TAGCTGTCTTCAGACACTCCAGAAGAGGGCGCCA | | |
| 303 | 4 | 127257041 | 127257592 | 551 | B10D2,B10J,KK,MA/My,NOR,PJ,SEA,SMJ | 4 | 127257312 | 127257330 | 19 | - | 19.8689 | 1.22E-07 | 0.0488 | GGACCTCTAGGGGGCGGTA | 16 | ENCFF092NMJ,ENCFF024JOZ,ENCFF093ZRU,ENCFF210BTO,ENCFF136ZDK,ENCFF089GBC,ENCFF198SSR,ENCFF367UNR,ENCFF589ZBJ,ENCFF099QNE,ENCFF052KVO,ENCFF327LYS,ENCFF907XEG,ENCFF097CAO,ENCFF821BIA,ENCFF533APC |
| 304 | 4 | 129366980 | 129367179 | 199 | 129S1,B10D2,B10J,BTBR,C57L,DBA1J,KK,NOR,NZB,PJ,PL,SEA,SMJ | 4 | 129367103 | 129367136 | 34 | - | 16.4472 | 9.61E-07 | 0.0931 | TAGCTGTCTTCAGACACACCAGAAGAGGGCGTCA | | |
| 305 | 4 | 129455317 | 129455519 | 202 | AJ,AKR,BALB,BTBR,C3H,C57L,CBA,CE,DBA2J,MRL,NZO,NZW,RF,RH2,TH | 4 | 129455439 | 129455472 | 34 | - | 16.4472 | 9.61E-07 | 0.0931 | TAGCTGTCTTCAGACACACCAGAAGAGGGCGTCA | | |
| 306 | 4 | 129513353 | 129513555 | 202 | 129S1,AJ,AKR,BALB,BTBR,BUB,C3H,C57L,CBA,CE,DBA2J,FVB,ILNJ2,MA/My,MRL,NOD,NZO,NZW,RF,RH2,SJL,SWR,TH | 4 | 129513479 | 129513512 | 34 | - | 16.4634 | 9.51E-07 | 0.0931 | TAGCTGTCTTCAGACGCACCAGAAGAGGGCATCA | 1 | ENCFF367UNR |
| 307 | 4 | 129531501 | 129531716 | 215 | 129S1,AJ,AKR,BALB,BTBR,BUB,C3H,C57L,CBA,CE,DBA2J,FVB,ILNJ2,MA/My,MRL,NOD,NZO,NZW,RF,RH2,SJL,SWR,TH | 4 | 129531628 | 129531661 | 34 | - | 16.4472 | 9.61E-07 | 0.0931 | TAGCTGTCTTCAGACACACCAGAAGAGGGCGTCA | 1 | ENCFF093ZRU |
| 308 | 4 | 132091453 | 132091651 | 198 | B10D2,B10J,DBA1J,KK,NZB,PJ,SEA,SJL,SMJ | 4 | 132091578 | 132091611 | 34 | - | 17.8455 | 3.91E-07 | 0.0606 | TAGCTGTCCTCAGACACTCCAGAAGAGGGCGTCA | 1 | ENCFF093ZRU |
| 309 | 4 | 132244335 | 132244626 | 291 | B10D2,B10J,DBA1J,KK,NZB,PJ,SEA,SMJ | 4 | 132244484 | 132244517 | 34 | - | 17.3902 | 5.27E-07 | 0.071 | TAGCTGTCCTCAGACACTCCAGAAGAGGGCATCA | | |
| 310 | 4 | 133508349 | 133508406 | 57 | PJ | 4 | 133508357 | 133508375 | 19 | - | 17.5246 | 6.45E-07 | 0.112 | TGGCCTCCAGGAGGCAGAG | | |
| 311 | 4 | 134418095 | 134418291 | 196 | ILNJ2.KK.NZB.NZO | 4 | 134418219 | 134418252 | 34 | - | 16.4472 | 9.61E-07 | 0.0931 | TAGCTGTCTTCAGACACACCAGAAGAGGGCGTCA | | |
| 312 | 4 | 134569787 | 134569996 | 209 | B10D2,B10J,BUB,KK,NZB,NZO,NZW,PJ,SEA | 4 | 134569914 | 134569947 | 34 | - | 16.4634 | 9.51E-07 | 0.0931 | TAGCTGTCTTCAGACGCACCAGAAGAGGGCATCA | | |
| 313 | 4 | 135619214 | 135619609 | 395 | SMJ | 4 | 135619608 | 135619641 | 34 | - | 20.4878 | 6.24E-08 | 0.0199 | GTGCAGTACTTGAGGAAGCCAGAAGGTGCAGCCT | | |
| 314 | 4 | 136007098 | 136007504 | 406 | 129S1,B10D2,B10J,C57L,DBA1J,KK,NZO,PJ,SEA | 4 | 136007426 | 136007459 | 34 | - | 17.2439 | 5.79E-07 | 0.0743 | TAGCTGTCCTCAGATACTCCAGAAGAGGGCATCA | | |
| 315 | 4 | 137637207 | 137637491 | 284 | AJ,AKR,BALB,BUB,C3H,CBA,CE,DBA1J,DBA2J,FVB,ILNJ2,KK,MA/My,MRL,NZB,NZW,PL,RF,RH2,SJL,SMJ,SWR,TH | 4 | 137637330 | 137637363 | 34 | - | 16.4634 | 9.51E-07 | 0.0931 | TAGCTGTCTTCAGACGCACCAGAAGAGGGCATCA | 1 | ENCFF093ZRU |
| 316 | 4 | 147973904 | 147974118 | 214 | 129S1,AKR,BTBR,BUB,C3H,C57L,C58J,CBA,CE,DBA1J,DBA2J,KK,LP,MA/My,MRL,NOD,NZB,NZO,NZW,PJ,PL,RF,SEA,SJL,SMJ,SWR,TH | 4 | 147974028 | 147974061 | 34 | - | 16.4472 | 9.61E-07 | 0.0931 | TAGCTGTCTTCAGACACACCAGAAGAGGGCGTCA | 1 | ENCFF367UNR |
| 317 | 4 | 148187928 | 148188132 | 204 | 129S1,AKR,BTBR,BUB,C3H,C57L,C58J,CBA,CE,DBA1J,DBA2J,KK,LP,MA/My,MRL,NOD,NZB,NZO,NZW,PJ,PL,RF,SEA,SJL,SMJ,SWR,TH | 4 | 148188052 | 148188085 | 34 | - | 17.2439 | 5.79E-07 | 0.0743 | TAGCTGTCCTCAGATACTCCAGAAGAGGGCATCA | | |
| 318 | 4 | 149077161 | 149077366 | 205 | 129S1,AJ,AKR,BALB,BTBR,BUB,C3H,C57L,C58J,CBA,CE,DBA1J,DBA2J,KK,LP,MA/My,MRL,NOD,NZB,NZO,NZW,PJ,PL,RF,SEA,SJL,SMJ,SWR,TH | 4 | 149077286 | 149077319 | 34 | - | 16.4472 | 9.61E-07 | 0.0931 | TAGCTGTCTTCAGACACACCAGAAGAGGGCGTCA | | |
| 319 | 4 | 149273378 | 149273602 | 224 | 129S1,BTBR,C57L,C58J,FVB,ILNJ2,LP,PJ,SEA,SWR | 4 | 149273454 | 149273487 | 34 | - | 16.4472 | 9.61E-07 | 0.0931 | TAGCTGTCTTCAGACACACCAGAAGAGGGCGTCA | 1 | ENCFF093ZRU |
| 320 | 4 | 155787642 | 155787782 | 140 | CE.ILNJ2.SMJ | 4 | 155787653 | 155787686 | 34 | - | 17.1545 | 6.13E-07 | 0.0768 | ATGCTCTCTTCAGACACACCAGAAGAGGGCATCA | 1 | ENCFF093ZRU |
| 321 | 5 | 3600883 | 3601075 | 192 | BUB.FVB.NOD.NOR.SJL.SMJ.SWR | 5 | 3600932 | 3600965 | 34 | + | 16.4472 | 9.61E-07 | 0.0776 | TAGCTGTCTTCAGACACACCAGAAGAGGGCGTCA | | |
| 322 | 5 | 8057604 | 8057830 | 226 | BUB.NOD.NOR | 5 | 8057686 | 8057719 | 34 | + | 17.6992 | 4.30E-07 | 0.0494 | TAGCTGTTTTCAGACACACCAGAAGAGGGCATCA | | |
| 323 | 5 | 30912183 | 30912401 | 218 | 129S1.AKR.BTBR.C57L.CE.SJL | 5 | 30912259 | 30912292 | 34 | + | 17.8455 | 3.91E-07 | 0.0466 | TAGCTGTCCTCAGACACTCCAGAAGAGGGCGTCA | | |
| 324 | 5 | 34546300 | 34546513 | 213 | BUB,C58J,FVB,MA/My,MRL,NOD,NOR,NZB,PL,SWR,TH | 5 | 34546370 | 34546403 | 34 | + | 18.561 | 2.42E-07 | 0.036 | AAGCTGTCTTCAGACACTCCAGAAGAGGGCGCCA | 2 | ENCFF093ZRU |
| 324 | 5 | 34546300 | 34546513 | 213 | BUB,C58J,FVB,MA/My,MRL,NOD,NOR,NZB,PL,SWR,TH | 5 | 34546385 | 34546403 | 19 | + | 17.5574 | 6.32E-07 | 0.0956 | ACTCCAGAAGAGGGCGCCA | 2 | ENCFF093ZRU |
| 325 | 5 | 60388722 | 60389537 | 815 | AJ,AKR,BALB,BTBR,BUB,C3H,C58J,CBA,CE,FVB,KK,MA/My,MRL,NOD,NZB,NZW,PL,RF.RH2.SEA.SJL.SMJ.SWR.TH | 5 | 60388744 | 60388778 | 35 | + | 20.7339 | 5.38E-08 | 0.0239 | GTGCAGGTCCCGGCCTGCGCCTGCGGAGGGCTCAG | | |
| 326 | 5 | 64505267 | 64505475 | 208 | AJ,AKR,BALB,BUB,C3H,C58J,CBA,CE,DBA1J,DBA2J,FVB,ILNJ2,KK,MA/My,MRL,NOD,NOR,NZB,NZO,NZW,PJ,PL,RF,RH2,SEA,SJL,SMJ,SWR,TH | 5 | 64505334 | 64505367 | 34 | + | 17.8211 | 3.97E-07 | 0.047 | TAGCTGTCTTCAATCACTCCAGAAGAGGGCACCA | 2 | ENCFF093ZRU,ENCFF367UNR |
| 327 | 5 | 73663695 | 73663812 | 117 | AJ,AKR,BALB,BUB,C3H,CBA,CE,DBA1J,DBA2J,FVB,ILNJ2,MA/My,MRL,NOD,NOR,NZO,NZW,PJ,PL,RF,RH2,SEA,SJL,SWR | 5 | 73663809 | 73663843 | 35 | + | 19.2477 | 1.40E-07 | 0.0457 | TTGCAGCTGTTGCACATTCCCACCAGGTGTTGGCT | | |
| 328 | 5 | 73664472 | 73664678 | 206 | C3H,CBA,DBA1J,DBA2J,ILNJ2,NOD,NOR | 5 | 73664534 | 73664568 | 35 | + | 18.0367 | 2.94E-07 | 0.0717 | CTGTAGCTGACTTCAGACACCAGAAGAGGGCATCA | 1 | ENCFF093ZRU |
| 329 | 5 | 76794536 | 76794785 | 249 | C3H,CBA,DBA1J,DBA2J,MA/My,NOD,NOR,PL,RF,SJL,SMJ,SWR | 5 | 76794642 | 76794675 | 34 | + | 16.4634 | 9.51E-07 | 0.0776 | TAGCTGTCTTCAGACGCACCAGAAGAGGGCATCA | | |
| 330 | 5 | 77241475 | 77241685 | 210 | AJ,BALB,MA/My,NOD,NOR,PL,RF,RH2,SEA,SJL,SMJ,SWR,TH | 5 | 77241541 | 77241574 | 34 | + | 16.4472 | 9.61E-07 | 0.0776 | TAGCTGTCTTCAGACACACCAGAAGAGGGCGTCA | 1 | ENCFF093ZRU |
| 331 | 5 | 80851008 | 80851217 | 209 | MA/My.RF.SMJ.SWR.TH | 5 | 80851075 | 80851108 | 34 | + | 16.4472 | 9.61E-07 | 0.0776 | TAGCTGTCTTCAGACACACCAGAAGAGGGCGTCA | | |
| 332 | 5 | 102729697 | 102729894 | 197 | AKR,CE,MRL,NOD,NOR,NZO,NZW,PJ,PL,RF,SMJ,TH | 5 | 102729752 | 102729785 | 34 | + | 18.1789 | 3.13E-07 | 0.0413 | TAGCTGTCTTCAGACACTCCAGAAGAGGGCGCTA | | |
| 332 | 5 | 102729697 | 102729894 | 197 | AKR,CE,MRL,NOD,NOR,NZO,NZW,PJ,PL,RF,SMJ,TH | 5 | 102729767 | 102729785 | 19 | + | 17.5738 | 6.25E-07 | 0.0956 | ACTCCAGAAGAGGGCGCTA | | |
| 333 | 5 | 103874398 | 103874620 | 222 | 129S1,AKR,BUB,C3H,C57L,CBA,CE,DBA1J,DBA2J,FVB,LP,MRL,NOD,NOR,NZB,NZO,NZW,PJ,PL,RF,SJL,SMJ,TH | 5 | 103874587 | 103874621 | 35 | + | 16.7156 | 6.38E-07 | 0.112 | GTGCAGCCCCACACAGGTGCCAGGTGGTGGCCTTG | | |
| 334 | 5 | 117360789 | 117360941 | 152 | CE.DBA1J.DBA2J.KK.NZB.NZO.NZW | 5 | 117360941 | 117360974 | 34 | + | 23.9268 | 4.19E-09 | 0.00154 | GCGCAGTGCCCTTAAAGGCCAGAAGAGGGCATCA | 2 | ENCFF502WDT.ENCFF136ZDK |
| 335 | 5 | 117450699 | 117450897 | 198 | 129S1,AJ,AKR,BALB,C57L,CE,DBA1J,DBA2J,KK,MRL,NZB,NZO,NZW,PJ,PL,RH2,SEA,SMJ,TH | 5 | 117450755 | 117450788 | 34 | + | 18.3659 | 2.76E-07 | 0.0384 | TAGCTGTCTTCAGACACTCCAGAAGAGGGCGCCA | 1 | ENCFF367UNR |
| 335 | 5 | 117450699 | 117450897 | 198 | 129S1,AJ,AKR,BALB,C57L,CE,DBA1J,DBA2J,KK,MRL,NZB,NZO,NZW,PJ,PL,RH2,SEA,SMJ,TH | 5 | 117450770 | 117450788 | 19 | + | 17.5574 | 6.32E-07 | 0.0956 | ACTCCAGAAGAGGGCGCCA | 1 | ENCFF367UNR |
| 336 | 5 | 118970204 | 118970414 | 210 | RF | 5 | 118970270 | 118970303 | 34 | + | 17.6667 | 4.40E-07 | 0.0501 | TAGCTGTCTTCAGACACGCCAGAAGAGGGCGTCA | | |
| 337 | 5 | 121209931 | 121210148 | 217 | AKR.KK.MA/My.PL.TH | 5 | 121210007 | 121210040 | 34 | + | 16.4634 | 9.51E-07 | 0.0776 | TAGCTGTCTTCAGACGCACCAGAAGAGGGCATCA | | |
| 338 | 5 | 121529444 | 121529645 | 201 | 129S1.AJ.BALB.C57L.ILNJ2.RH2.SEA | 5 | 121529616 | 121529649 | 34 | + | 16.4472 | 9.61E-07 | 0.0776 | TAGCTGTCTTCAGACACACCAGAAGAGGGCGTCA | | |
| 339 | 5 | 123460062 | 123460278 | 216 | ILNJ2 | 5 | 123460134 | 123460167 | 34 | + | 18.3659 | 2.76E-07 | 0.0384 | TAGCTGTCTTCAGACACTCCAGAAGAGGGCGCCA | | |
| 339 | 5 | 123460062 | 123460278 | 216 | ILNJ2 | 5 | 123460149 | 123460167 | 19 | + | 17.5574 | 6.32E-07 | 0.0956 | ACTCCAGAAGAGGGCGCCA | | |
| 340 | 5 | 124638339 | 124638566 | 227 | MA/My.NZW.PJ.PL | 5 | 124638425 | 124638458 | 34 | + | 18.3659 | 2.76E-07 | 0.0384 | TAGCTGTCTTCAGACACTCCAGAAGAGGGCGCCA | 1 | ENCFF093ZRU |
| 340 | 5 | 124638339 | 124638566 | 227 | MA/My.NZW.PJ.PL | 5 | 124638440 | 124638458 | 19 | + | 17.5574 | 6.32E-07 | 0.0956 | ACTCCAGAAGAGGGCGCCA | 1 | ENCFF093ZRU |
| 341 | 5 | 129463378 | 129463820 | 442 | 129S1,AJ,BALB,C57L,C58J,FVB,ILNJ2,KK,LP,MRL,NOD,NOR,NZB,NZO,NZW,RH2,SEA,SMJ,SWR,TH | 5 | 129463503 | 129463537 | 35 | + | 23.633 | 7.02E-09 | 0.00665 | CTACAGAGCTGCTGAGCTGCCAGCAGGGGGCACTG | | |
| 341 | 5 | 129463378 | 129463820 | 442 | 129S1,AJ,BALB,C57L,C58J,FVB,ILNJ2,KK,LP,MRL,NOD,NOR,NZB,NZO,NZW,RH2,SEA,SMJ,SWR,TH | 5 | 129463519 | 129463537 | 19 | + | 24.0164 | 2.47E-09 | 0.012 | CTGCCAGCAGGGGGCACTG | | |
| 342 | 5 | 129873224 | 129873594 | 370 | 129S1,AJ,BALB,C57L,FVB,ILNJ2,KK,MRL,NOD,NOR,NZW,RH2,SEA,SMJ,SWR | 5 | 129873573 | 129873606 | 34 | + | 17.2439 | 5.79E-07 | 0.0594 | TAGCTGTCCTCAGATACTCCAGAAGAGGGCATCA | | |
| 342 | 5 | 129873224 | 129873594 | 370 | 129S1,AJ,BALB,C57L,FVB,ILNJ2,KK,MRL,NOD,NOR,NZW,RH2,SEA,SMJ,SWR | 5 | 129873308 | 129873341 | 34 | - | 17.2439 | 5.79E-07 | 0.0594 | TAGCTGTCCTCAGATACTCCAGAAGAGGGCATCA | 1 | ENCFF093ZRU |
| 343 | 5 | 133655334 | 133655538 | 204 | KK.RF | 5 | 133655390 | 133655423 | 34 | + | 16.4472 | 9.61E-07 | 0.0776 | TAGCTGTCTTCAGACACACCAGAAGAGGGCGTCA | | |
| 344 | 5 | 134682194 | 134682425 | 231 | AJ,AKR,BUB,FVB,ILNJ2,MA/My,MRL,RF,SJL | 5 | 134682284 | 134682317 | 34 | + | 16.4634 | 9.51E-07 | 0.0776 | TAGCTGTCTTCAGACGCACCAGAAGAGGGCATCA | 1 | ENCFF093ZRU |
| 345 | 5 | 142831047 | 142831151 | 104 | 129S1,AKR,BTBR,BUB,C57L,CE,DBA1J,DBA2J,ILNJ2,KK,MA/My,PJ,SJL | 5 | 142831020 | 142831054 | 35 | + | 16.5963 | 6.83E-07 | 0.117 | GAGCAATGCCCCCAGCCCCTCACTAGGGGATTCTA | | |
| 346 | 5 | 144200658 | 144200761 | 103 | TH | 5 | 144200702 | 144200736 | 35 | + | 17.9174 | 3.16E-07 | 0.0742 | CTGCAGGTCGGGCAGGTGCTCTGTAGGTGGTGAGG | | |
| 347 | 5 | 144252698 | 144252889 | 191 | TH | 5 | 144252749 | 144252782 | 34 | + | 17.2439 | 5.79E-07 | 0.0594 | TAGCTGTCCTCAGATACTCCAGAAGAGGGCATCA | | |
| 348 | 5 | 146854382 | 146854590 | 208 | AKR,CBA,FVB,ILNJ2,LP,MA/My,MRL,NZB,NZO,NZW,RF,SJL,TH | 5 | 146854448 | 146854481 | 34 | + | 18.561 | 2.42E-07 | 0.036 | AAGCTGTCTTCAGACACTCCAGAAGAGGGCGCCA | 1 | ENCFF055HAP |
| 348 | 5 | 146854382 | 146854590 | 208 | AKR,CBA,FVB,ILNJ2,LP,MA/My,MRL,NZB,NZO,NZW,RF,SJL,TH | 5 | 146854463 | 146854481 | 19 | + | 17.5574 | 6.32E-07 | 0.0956 | ACTCCAGAAGAGGGCGCCA | 1 | ENCFF055HAP |
| 349 | 5 | 148672442 | 148673233 | 791 | CBA.CE.KK.RF.SEA | 5 | 148672973 | 148673006 | 34 | + | 24.2846 | 3.09E-09 | 0.00123 | GTGCAGTACCCAAGGAGGCCAGAAGAAGGCATCA | | |
| 350 | 5 | 5902507 | 5902884 | 377 | AJ,BUB,C3H,CBA,DBA1J,LP,NOD,NOR,PL,TH | 5 | 5902568 | 5902601 | 34 | - | 18.3089 | 2.87E-07 | 0.0389 | ATGCAGCTAGAGACACGAGCTCCAGGGGGTACTG | | |
| 351 | 5 | 14894861 | 14895427 | 566 | DBA1J,DBA2J,FVB,MA/My,RF,SJL,SWR | 5 | 14895071 | 14895105 | 35 | - | 22.8349 | 1.26E-08 | 0.00945 | GTGCAGTTGCCCATAGTGTCCAGAAGAGGGAGAAG | 3 | ENCFF093ZRU,ENCFF099QNE,ENCFF052KVO |
| 352 | 5 | 24021250 | 24021460 | 210 | 129S1,AKR,BALB,BTBR,BUB,C57L,C58J,CE,DBA1J,DBA2J,FVB,KK,MA/My,MRL,NOD,NOR,NZO,NZW,PJ,PL,RF,SEA,SJL,SMJ,SWR,TH | 5 | 24021377 | 24021410 | 34 | - | 16.4634 | 9.51E-07 | 0.0776 | TAGCTGTCTTCAGACGCACCAGAAGAGGGCATCA | | |
| 353 | 5 | 24602416 | 24602618 | 202 | DBA1J.DBA2J.KK.MRL.NZB.NZW.PJ | 5 | 24602536 | 24602554 | 19 | - | 17.5574 | 6.32E-07 | 0.0956 | ACTCCAGAAGAGGGCGCCA | | |
| 353 | 5 | 24602416 | 24602618 | 202 | DBA1J.DBA2J.KK.MRL.NZB.NZW.PJ | 5 | 24602536 | 24602569 | 34 | - | 18.3659 | 2.76E-07 | 0.0384 | TAGCTGTCTTCAGACACTCCAGAAGAGGGCGCCA | | |
| 354 | 5 | 62597231 | 62597444 | 213 | AJ,AKR,BUB,C58J,CBA,CE,FVB,KK,MA/My,MRL,NOD,NZB,NZW,RF,SJL,SMJ,SWR,TH | 5 | 62597357 | 62597375 | 19 | - | 17.5574 | 6.32E-07 | 0.0956 | ACTCCAGAAGAGGGCGCCA | | |
| 354 | 5 | 62597231 | 62597444 | 213 | AJ,AKR,BUB,C58J,CBA,CE,FVB,KK,MA/My,MRL,NOD,NZB,NZW,RF,SJL,SMJ,SWR,TH | 5 | 62597357 | 62597390 | 34 | - | 18.3659 | 2.76E-07 | 0.0384 | TAGCTGTCTTCAGACACTCCAGAAGAGGGCGCCA | | |
| 355 | 5 | 66420435 | 66420649 | 214 | 129S1,AJ,AKR,BALB,BTBR,BUB,C3H,C57L,CBA,CE,DBA1J,DBA2J,FVB,ILNJ2,KK,MA/My,MRL,NOD,NOR,NZB,NZO,PJ,PL,RF,RH2,SEA,SJL,SMJ,SWR | 5 | 66420568 | 66420601 | 34 | - | 17.2114 | 5.91E-07 | 0.06 | TAGCTGTCTTCGGACGCTCCAGAAGAGGGCGTCA | | |
| 356 | 5 | 86034776 | 86034989 | 213 | SMJ | 5 | 86034899 | 86034917 | 19 | - | 17.5574 | 6.32E-07 | 0.0956 | ACTCCAGAAGAGGGCGCCA | | |
| 356 | 5 | 86034776 | 86034989 | 213 | SMJ | 5 | 86034899 | 86034932 | 34 | - | 18.561 | 2.42E-07 | 0.036 | AAGCTGTCTTCAGACACTCCAGAAGAGGGCGCCA | | |
| 357 | 5 | 86826327 | 86826667 | 340 | BALB,C3H,CBA,CE,DBA1J,DBA2J,ILNJ2,NOD,NOR,PJ,RH2,SEA | 5 | 86826648 | 86826681 | 34 | - | 21.3984 | 3.17E-08 | 0.00778 | ATGCAGTGCCTGCAGAGGCCAGAAGATGATGTTG | | |
| 358 | 5 | 89274096 | 89274293 | 197 | SMJ | 5 | 89274220 | 89274253 | 34 | - | 16.4472 | 9.61E-07 | 0.0776 | TAGCTGTCTTCAGACACACCAGAAGAGGGCGTCA | 1 | ENCFF093ZRU |
| 359 | 5 | 93182561 | 93182783 | 222 | 129S1,AKR,BALB,BTBR,BUB,C3H,C57L,CBA,CE,DBA1J,DBA2J,FVB,MA/My,MRL,NOD,NOR,NZW,PJ,RF,RH2,SEA,SJL,SMJ,SWR,TH | 5 | 93182684 | 93182717 | 34 | - | 20.1301 | 8.09E-08 | 0.0159 | TTGCTGTTTTCAGACACGCCAGAAGAGGGCATAA | 6 | ENCFF093ZRU,ENCFF327LYS,ENCFF907XEG,ENCFF198SSR,ENCFF367UNR,ENCFF099QNE |
| 360 | 5 | 99470918 | 99471137 | 219 | 129S1,AJ,AKR,BALB,BUB,C3H,C57L,CBA,CE,DBA1J,DBA2J,FVB,ILNJ2,MA/My,MRL,NOD,NOR,NZB,NZO,NZW,PJ,PL,RF,RH2,SEA,SJL,SMJ,SWR,TH | 5 | 99471044 | 99471077 | 34 | - | 16.6341 | 8.54E-07 | 0.0745 | TAGCTGTCTTCACACACTCCAGAAGAGGGCGTCA | | |
| 361 | 5 | 104725116 | 104725328 | 212 | AJ,BALB,BUB,C3H,CBA,DBA1J,DBA2J,FVB,ILNJ2,KK,NZB,NZO,RH2,SEA,SJL,SWR,TH | 5 | 104725242 | 104725260 | 19 | - | 17.5574 | 6.32E-07 | 0.0956 | ACTCCAGAAGAGGGCGCCA | | |
| 361 | 5 | 104725116 | 104725328 | 212 | AJ,BALB,BUB,C3H,CBA,DBA1J,DBA2J,FVB,ILNJ2,KK,NZB,NZO,RH2,SEA,SJL,SWR,TH | 5 | 104725242 | 104725275 | 34 | - | 18.3659 | 2.76E-07 | 0.0384 | TAGCTGTCTTCAGACACTCCAGAAGAGGGCGCCA | | |
| 362 | 5 | 107043585 | 107043789 | 204 | BUB.C3H.CBA.FVB.PJ.PL.SJL.TH | 5 | 107043710 | 107043728 | 19 | - | 17.5574 | 6.32E-07 | 0.0956 | ACTCCAGAAGAGGGCGCCA | 2 | ENCFF093ZRU.ENCFF367UNR |
| 362 | 5 | 107043585 | 107043789 | 204 | BUB.C3H.CBA.FVB.PJ.PL.SJL.TH | 5 | 107043710 | 107043743 | 34 | - | 18.1789 | 3.13E-07 | 0.0413 | TAGCTGTCTTCAGCCACTCCAGAAGAGGGCGCCA | 2 | ENCFF093ZRU.ENCFF367UNR |
| 363 | 5 | 108634272 | 108634536 | 264 | BUB.C3H.CBA.FVB.PJ.PL.SJL.SWR.TH | 5 | 108634398 | 108634431 | 34 | - | 16.4472 | 9.61E-07 | 0.0776 | TAGCTGTCTTCAGACACACCAGAAGAGGGCGTCA | | |
| 364 | 5 | 115637943 | 115638135 | 192 | 129S1,AKR,BALB,C57L,CE,DBA1J,DBA2J,KK,MRL,NZB,NZO,PL,RH2,SEA | 5 | 115638067 | 115638100 | 34 | - | 16.4472 | 9.61E-07 | 0.0776 | TAGCTGTCTTCAGACACACCAGAAGAGGGCGTCA | | |
| 365 | 5 | 123815787 | 123816000 | 213 | 129S1,AJ,BALB,C57L,MRL,PL,RH2,SEA,SJL | 5 | 123815915 | 123815948 | 34 | - | 16.7724 | 7.83E-07 | 0.0704 | TAGCTGTCTTCAGATGCACCAGAAGAGGGCGTCA | | |

| | | | | | | | | | | | | | | | | |
|---|---|---|---|---|---|---|---|---|---|---|---|---|---|---|---|---|
| 366 | 5 | 125251357 | 125251564 | 207 | 129S1,AJ,BALB,BUB,C57L,KK,MRL,PL,RH2,SEA,SJL | 5 | 125251486 | 125251519 | 34 | - | 17.8455 | 3.91E-07 | 0.0466 | TAGCTGTCCTCAGACACTCCAGAAGAGGGCGTCA | | |
| 367 | 5 | 129055156 | 129055938 | 782 | C58J,FVB,LP,NOD,NOR,NZB,NZO,SMJ,TH | 5 | 129055325 | 129055343 | 19 | - | 18.7049 | 2.88E-07 | 0.0658 | GCACCAGAAGAGGGCGCCA | 2 | ENCFF093ZRU,ENCFF099QNE |
| 367 | 5 | 129055156 | 129055938 | 782 | C58J,FVB,LP,NOD,NOR,NZB,NZO,SMJ,TH | 5 | 129055325 | 129055358 | 34 | - | 18.7805 | 2.09E-07 | 0.0323 | CAGCTGACTTCAGATGCACCAGAAGAGGGCGCCA | 2 | ENCFF093ZRU,ENCFF099QNE |
| 367 | 5 | 129055156 | 129055938 | 782 | C58J,FVB,LP,NOD,NOR,NZB,NZO,SMJ,TH | 5 | 129055761 | 129055794 | 34 | - | 22.1789 | 1.74E-08 | 0.00476 | ATGCAGTGCCTGCATATACCAGGAGATGGTGTTG | | |
| 368 | 5 | 129919204 | 129919406 | 202 | AJ,BALB,MRL,RH2,SEA,SMJ | 5 | 129919334 | 129919352 | 19 | - | 17.5574 | 6.32E-07 | 0.0956 | ACTCCAGAAGAGGGCGCCA | | |
| 368 | 5 | 129919204 | 129919406 | 202 | AJ,BALB,MRL,RH2,SEA,SMJ | 5 | 129919334 | 129919367 | 34 | - | 18.561 | 2.42E-07 | 0.036 | AAGCTGTCTTCAGACACTCCAGAAGAGGGCGCCA | | |
| 369 | 5 | 130670175 | 130670377 | 202 | 129S1,C57L,FVB,ILNJ2,KK,LP,NZB,NZO,PL,TH | 5 | 130670299 | 130670332 | 34 | - | 17.2439 | 5.79E-07 | 0.0594 | TAGCTGTCCTCAGATACTCCAGAAGAGGGCATCA | | |
| 370 | 5 | 134767401 | 134768020 | 619 | KK | 5 | 134767394 | 134767427 | 34 | - | 16.4634 | 9.51E-07 | 0.0776 | TAGCTGTCTTCAGACGCACCAGAAGAGGGCATCA | | |
| 371 | 5 | 135842730 | 135842923 | 193 | AKR,BALB,BTBR,BUB,C57L,ILNJ2,MRL,PJ,PL,RH2,SEA,TH | 5 | 135842832 | 135842865 | 34 | - | 16.7724 | 7.83E-07 | 0.0704 | TAGCTGTCTTCAGATGCACCAGAAGAGGGCGTCA | | |
| 372 | 5 | 137708586 | 137708799 | 213 | 129S1,AKR,BUB,KK,LP,MA/My,MRL,NZB,NZO.NZW.RF.SWR | 5 | 137708713 | 137708746 | 34 | - | 16.4472 | 9.61E-07 | 0.0776 | TAGCTGTCTTCAGACACACCAGAAGAGGGCGTCA | | |
| 373 | 5 | 139789875 | 139789967 | 92 | ILNJ2 | 5 | 139789929 | 139789947 | 19 | - | 18.1967 | 4.11E-07 | 0.0795 | CAGCCAGCAGGGGGCTGCC | 7 | ENCFF367UNR,ENCFF136ZDK,ENCFF883UPM,ENCFF327LYS,ENCFF589ZBJ,ENCFF907XEG,ENCFF024JOZ |
| 374 | 5 | 140424475 | 140424541 | 66 | 129S1,C3H,CBA,CE,DBA1J,DBA2J,KK,MRL,NZB,NZO,NZW,PL,RF,SMJ,SWR,TH | 5 | 140424519 | 140424537 | 19 | - | 18 | 4.70E-07 | 0.0851 | TTCCCACCAGAGGGCAGGG | 3 | ENCFF198SSR,ENCFF327LYS,ENCFF907XEG |
| 375 | 5 | 140860908 | 140861484 | 576 | NZW | 5 | 140861031 | 140861064 | 34 | - | 24.2033 | 3.31E-09 | 0.00131 | GTGCAATGCCCACAGAGGTCAGAAGAGGGCATCA | | |
| 376 | 5 | 142965522 | 142965923 | 401 | 129S1,AKR,BTBR,BUB,C57L,CE,DBA1J,DBA2J,ILNJ2,KK,MA/My,NZB,NZW,PJ,SJL | 5 | 142965847 | 142965880 | 34 | - | 16.7724 | 7.83E-07 | 0.0704 | TAGCTGTCTTCAGATGCACCAGAAGAGGGCGTCA | | |
| 377 | 5 | 143399223 | 143399433 | 210 | CBA | 5 | 143399348 | 143399381 | 34 | - | 17.2439 | 5.79E-07 | 0.0594 | TAGCTGTCCTCAGATACTCCAGAAGAGGGCATCA | | |
| 378 | 5 | 144169069 | 144169275 | 206 | TH | 5 | 144169194 | 144169212 | 19 | - | 17.5574 | 6.32E-07 | 0.0956 | ACTCCAGAAGAGGGCGCCA | 3 | ENCFF367UNR.ENCFF099QNE.ENCFF093ZRU |
| 378 | 5 | 144169069 | 144169275 | 206 | TH | 5 | 144169194 | 144169227 | 34 | - | 18.3659 | 2.76E-07 | 0.0384 | TAGCTGTCTTCAGACACTCCAGAAGAGGGCGCCA | 3 | ENCFF367UNR.ENCFF099QNE.ENCFF093ZRU |
| 379 | 5 | 147874152 | 147874343 | 191 | CE | 5 | 147874275 | 147874308 | 34 | - | 16.4472 | 9.61E-07 | 0.0776 | TAGCTGTCTTCAGACACACCAGAAGAGGGCGTCA | | |
| 380 | 5 | 147946081 | 147946412 | 331 | 129S1,AJ,AKR,BALB,BTBR,BUB,C3H,C57L,ILNJ2,LP,MA/My,MRL,NOD,NOR,RH2,SEA,SMJ | 5 | 147946330 | 147946363 | 34 | - | 17.878 | 3.83E-07 | 0.0461 | TAGCTGTCCTCAGACACACCAGAAGAGGGCATCA | | |
| 381 | 6 | 12775576 | 12775770 | 194 | LP.PL | 6 | 12775681 | 12775715 | 35 | + | 18.6055 | 2.09E-07 | 0.0851 | CTGCGGACACCCCCAAGGTCCCGTAGAGGGCTCTC | | |
| 382 | 6 | 14419492 | 14419697 | 205 | AKR.FVB.MA/Mv.MRL.NZB.RH2 | 6 | 14419547 | 14419580 | 34 | + | 18.4797 | 2.56E-07 | 0.0559 | TTGCTGTCTTCAGACACACCAGAAGAGGGCATCA | | |
| 383 | 6 | 22434071 | 22434305 | 234 | C3H.CBA.CE.FVB.NOD.NOR.SJL.TH | 6 | 22434183 | 22434216 | 34 | + | 17.2439 | 5.79E-07 | 0.086 | TAGCTGTCCTCAGATACTCCAGAAGAGGGCATCA | | |
| 384 | 6 | 29766239 | 29766465 | 226 | 129S1,AJ,AKR,BALB,BTBR,BUB,C3H,C57L,CBA,DBA1J,DBA2J,FVB,MRL,NOD,NOR,NZO,PJ,PL,RF,RH2,SEA,SMJ | 6 | 29766323 | 29766356 | 34 | + | 18.3659 | 2.76E-07 | 0.0572 | TAGCTGTCTTCAGACACTCCAGAAGAGGGCGCCA | 7 | ENCFF099QNE,ENCFF821BIA,ENCFF327LYS,ENCFF089GBC,ENCFF907XEG,ENCFF093ZRU,ENCFF367UNR |
| 384 | 6 | 29766239 | 29766465 | 226 | 129S1,AJ,AKR,BALB,BTBR,BUB,C3H,C57L,CBA,DBA1J,DBA2J,FVB,MRL,NOD,NOR,NZO,PJ,PL,RF,RH2,SEA,SMJ | 6 | 29766338 | 29766356 | 19 | + | 17.5574 | 6.32E-07 | 0.138 | ACTCCAGAAGAGGGCGCCA | 7 | ENCFF099QNE,ENCFF821BIA,ENCFF327LYS,ENCFF089GBC,ENCFF907XEG,ENCFF093ZRU,ENCFF367UNR |
| 385 | 6 | 38251936 | 38252308 | 372 | KK | 6 | 38252153 | 38252186 | 34 | + | 17.9593 | 3.63E-07 | 0.0662 | TTGCTGTCTTTAGACACACCAGAAGAGGGCATCC | | |
| 386 | 6 | 38446690 | 38446986 | 296 | KK | 6 | 38446696 | 38446730 | 35 | + | 27.4679 | 3.12E-10 | 0.00222 | CTGCATGTATGCCTGCAGGCCAGAAGAGGGCACCA | 17 | ENCFF099QNE,ENCFF089GBC,ENCFF589ZBJ,ENCFF198SSR,ENCFF201JUA,ENCFF092NMJ,ENCFF883UPM,ENCFF055HAP,ENCFF821BIA,ENCFF502WDT,ENCFF327LYS,ENCFF024JOZ,ENCFF907XEG,ENCFF367UNR,ENCFF093ZRU,ENCFF052KVO,ENCFF136ZDK |
| 386 | 6 | 38446690 | 38446986 | 296 | KK | 6 | 38446712 | 38446730 | 19 | + | 20.3607 | 8.29E-08 | 0.0572 | AGGCCAGAAGAGGGCACCA | 18 | ENCFF099QNE,ENCFF089GBC,ENCFF589ZBJ,ENCFF198SSR,ENCFF201JUA,ENCFF092NMJ,ENCFF883UPM,ENCFF055HAP,ENCFF821BIA,ENCFF502WDT,ENCFF327LYS,ENCFF024JOZ,ENCFF907XEG,ENCFF065ODV,ENCFF367UNR,ENCFF093ZRU,ENCFF052KVO,ENCFF136ZDK |
| 387 | 6 | 38655403 | 38655621 | 218 | KK | 6 | 38655458 | 38655492 | 35 | + | 17.9266 | 3.14E-07 | 0.11 | CTGTAGTTGTCTTCAGACACCAGAAGAGGGAATCA | | |
| 388 | 6 | 39288054 | 39288260 | 206 | KK | 6 | 39288119 | 39288152 | 34 | + | 17.2439 | 5.79E-07 | 0.086 | TAGCTGTCCTCAGATACTCCAGAAGAGGGCATCA | | |
| 389 | 6 | 51578060 | 51578305 | 245 | AKR,CE,ILNJ2,KK,LP,MA/My,NOD,NOR,NZB,NZO,NZW,PJ,RF,TH | 6 | 51578161 | 51578194 | 34 | + | 16.4634 | 9.51E-07 | 0.112 | TAGCTGTCTTCAGACGCACCAGAAGAGGGCATCA | | |
| 390 | 6 | 60284496 | 60284710 | 214 | PJ | 6 | 60284567 | 60284600 | 34 | + | 16.4634 | 9.51E-07 | 0.112 | TAGCTGTCTTCAGACGCACCAGAAGAGGGCATCA | | |
| 391 | 6 | 67159845 | 67160042 | 197 | C58J,CE,ILNJ2,KK,NOD,NOR,NZB,NZO,PJ | 6 | 67159900 | 67159933 | 34 | + | 17.8455 | 3.91E-07 | 0.0687 | TAGCTGTCCTCAGACACTCCAGAAGAGGGCGTCA | | |
| 392 | 6 | 67229920 | 67230124 | 204 | C58J.KK.NOD.NOR.NZB.NZO | 6 | 67229983 | 67230016 | 34 | + | 17.2358 | 5.82E-07 | 0.0863 | TAGCTGTCCTCAGAAACTCCAGAAGAGGGCATCA | | |
| 393 | 6 | 67237874 | 67238086 | 212 | C58J.KK.MA/Mv.NOD.NOR.NZB.NZO | 6 | 67237944 | 67237977 | 34 | + | 18.3659 | 2.76E-07 | 0.0572 | TAGCTGTCTTCAGACACTCCAGAAGAGGGCGCCA | | |
| 393 | 6 | 67237874 | 67238086 | 212 | C58J.KK.MA/Mv.NOD.NOR.NZB.NZO | 6 | 67237959 | 67237977 | 19 | + | 17.5574 | 6.32E-07 | 0.138 | ACTCCAGAAGAGGGCGCCA | | |
| 394 | 6 | 67676116 | 67676448 | 332 | C58J,KK,MA/My,NOD,NOR,NZB,NZO | 6 | 67676146 | 67676164 | 19 | + | 19.2951 | 1.88E-07 | 0.0786 | CCGCCACTAGATGGTGCTA | 11 | ENCFF093ZRU,ENCFF367UNR,ENCFF430PPJ,ENCFF099QNE,ENCFF201JUA,ENCFF907XEG,ENCFF821BIA,ENCFF024JOZ.ENCFF089GBC.ENCFF327LYS.ENCFF092NMJ |
| 395 | 6 | 67900627 | 67900853 | 226 | CE.ILNJ2.PJ | 6 | 67900764 | 67900797 | 34 | + | 17.0325 | 6.63E-07 | 0.0937 | TAGCTGTCCTCAGACATCCCAGAAGAGGGCATCA | | |
| 396 | 6 | 72894129 | 72894304 | 175 | AKR.BALB.MRL.PL.RF.RH2.SEA | 6 | 72894210 | 72894243 | 34 | + | 17.748 | 4.17E-07 | 0.0716 | TTGCTGTCTTCAGACACACCAGGAGAGGGCATCA | | |
| 397 | 6 | 84053398 | 84053914 | 516 | NZB.NZO | 6 | 84053631 | 84053664 | 34 | + | 19.0163 | 1.77E-07 | 0.0463 | CAGCTGTCTTCAGACACACCAGAAGAGGGCGTCA | | |
| 398 | 6 | 85365886 | 85366071 | 185 | PJ.SEA | 6 | 85365930 | 85365963 | 34 | + | 17.2439 | 5.79E-07 | 0.086 | TAGCTGTCCTCAGATACTCCAGAAGAGGGCATCA | | |
| 399 | 6 | 86322228 | 86322279 | 51 | PJ.SEA | 6 | 86322247 | 86322280 | 34 | + | 17.5285 | 4.81E-07 | 0.0779 | CTGCTTTTGCACTCCTTTCCACTAGGTGCAGCAT | | |
| 400 | 6 | 86397596 | 86397652 | 56 | PJ,SEA | 6 | 86397622 | 86397656 | 35 | + | 21.1193 | 4.16E-08 | 0.0327 | CTGCTGTCACACACTTCAGCCTCCTGGAGGCGCTC | 20 | ENCFF201JUA,ENCFF092NMJ,ENCFF165LXO,ENCFF430PPJ,ENCFF502WDT,ENCFF055HAP,ENCFF589ZBJ,ENCFF093ZRU,ENCFF065ODV,ENCFF907XEG,ENCFF327LYS,ENCFF367UNR,ENCFF533APC,ENCFF198SSR,ENCFF883UPM,ENCFF052KVO,ENCFF099QNE,ENCFF142CNG,ENCFF089GBC,ENCFF024JOZ |
| 401 | 6 | 86693926 | 86694104 | 178 | 129S1,AJ,BTBR,C3H,C57L,C58J,CBA,DBA1J,DBA2J,FVB,NOD,NOR,PJ,RF,SEA,SJL,SMJ,SWR | 6 | 86693977 | 86694010 | 34 | + | 18.3659 | 2.76E-07 | 0.0572 | TAGCTGTCTTCAGACACTCCAGAAGAGGGCGCCA | | |
| 401 | 6 | 86693926 | 86694104 | 178 | 129S1,AJ,BTBR,C3H,C57L,C58J,CBA,DBA1J,DBA2J,FVB,NOD,NOR,PJ,RF,SEA,SJL,SMJ,SWR | 6 | 86693992 | 86694010 | 19 | + | 17.5574 | 6.32E-07 | 0.138 | ACTCCAGAAGAGGGCGCCA | | |
| 402 | 6 | 87010483 | 87010697 | 214 | PJ.SEA | 6 | 87010509 | 87010542 | 34 | + | 18.0813 | 3.34E-07 | 0.0635 | CAGCTGTCTTCAGAAACCCCAGAAGAGGGCGTCA | | |
| 403 | 6 | 87576521 | 87576723 | 202 | PJ.SEA | 6 | 87576580 | 87576613 | 34 | + | 16.4472 | 9.61E-07 | 0.112 | TAGCTGTCTTCAGACACACCAGAAGAGGGCGTCA | | |
| 404 | 6 | 97240759 | 97241016 | 257 | C58J,LP,MA/My | 6 | 97240918 | 97240952 | 35 | + | 17.0459 | 5.27E-07 | 0.143 | TTGCATGTACACTTGTTTGTCAGAAGAGGGCATCA | 9 | ENCFF099QNE,ENCFF201JUA,ENCFF821BIA,ENCFF367UNR,ENCFF089GBC,ENCFF907XEG,ENCFF055HAP,ENCFF093ZRU,ENCFF502WDT |
| 404 | 6 | 97240759 | 97241016 | 257 | C58J,LP,MA/My | 6 | 97240843 | 97240861 | 19 | - | 20.3607 | 8.29E-08 | 0.0572 | AGGCCAGAAGAGGGCACCA | 10 | ENCFF099QNE,ENCFF201JUA,ENCFF821BIA,ENCFF367UNR,ENCFF089GBC,ENCFF907XEG,ENCFF055HAP.ENCFF093ZRU.ENCFF024JOZ.ENCFF502WDT |
| 404 | 6 | 97240759 | 97241016 | 257 | C58J,LP,MA/My | 6 | 97240843 | 97240877 | 35 | - | 16.2844 | 8.14E-07 | 0.173 | TTTAAATGTACACCTCAGGCCAGAAGAGGGCACCA | 10 | ENCFF099QNE,ENCFF201JUA,ENCFF821BIA,ENCFF367UNR,ENCFF089GBC,ENCFF907XEG,ENCFF055HAP,ENCFF093ZRU,ENCFF024JOZ,ENCFF502WDT |
| 405 | 6 | 113552401 | 113552618 | 217 | CE.SMJ | 6 | 113552477 | 113552510 | 34 | + | 17.0813 | 6.43E-07 | 0.0919 | TAGCTGTTCTCAGACACTCCAGAAGAGGGAGTCA | | |
| 406 | 6 | 115754364 | 115754561 | 197 | AKR,C3H,CBA,ILNJ2,MRL,PL,RF | 6 | 115754420 | 115754453 | 34 | + | 18.878 | 1.95E-07 | 0.0488 | TAGCTGTCCCCAGATACTCCAGAAGAGGGCATCA | | |
| 407 | 6 | 115823861 | 115824076 | 215 | 129S1,AKR,BTBR,C3H,C57L,C58J,CBA,CE,DBA1J,DBA2J,ILNJ2,LP,MRL,NOD,NOR,NZB,NZO,NZW,PJ,PL,RF,RH2 | 6 | 115823932 | 115823965 | 34 | + | 18.1789 | 3.13E-07 | 0.0611 | TAGCTGTCTTCAGACACTCCAGAAGAGGGCGCTA | 1 | ENCFF805QBU |
| 407 | 6 | 115823861 | 115824076 | 215 | 129S1,AKR,BTBR,C3H,C57L,C58J,CBA,CE,DBA1J,DBA2J,ILNJ2,LP,MRL,NOD,NOR,NZB.NZO.NZW.PJ.PL.RF.RH2 | 6 | 115823947 | 115823965 | 19 | + | 17.5738 | 6.25E-07 | 0.138 | ACTCCAGAAGAGGGCGCTA | 2 | ENCFF805QBU,ENCFF210BTO |
| 408 | 6 | 117167033 | 117167258 | 225 | FVB.PJ.SMJ.TH | 6 | 117167115 | 117167148 | 34 | + | 16.4634 | 9.51E-07 | 0.112 | TAGCTGTCTTCAGACGCACCAGAAGAGGGCATCA | | |
| 409 | 6 | 120406728 | 120406932 | 204 | SJL.SWR | 6 | 120406788 | 120406821 | 34 | + | 16.4472 | 9.61E-07 | 0.112 | TAGCTGTCTTCAGACACACCAGAAGAGGGCGTCA | | |
| 410 | 6 | 123876009 | 123876171 | 162 | CE.FVB.NZO.PJ.SMJ.SWR | 6 | 123876073 | 123876091 | 19 | + | 17.5574 | 6.32E-07 | 0.138 | TCACCAGCAGGTGTCAGTC | | |
| 411 | 6 | 125042414 | 125042635 | 221 | SWR | 6 | 125042506 | 125042524 | 19 | + | 17.8852 | 5.08E-07 | 0.128 | ACACCAGAAGAGGGCACCA | | |
| 412 | 6 | 128822903 | 128823105 | 202 | 129S1,AJ,AKR,BALB,BUB,C3H,C57L,C58J,CBA,DBA1J,DBA2J,ILNJ2,MRL,NOD,NOR,NZO.PL.RF.SEA.SMJ.SWR | 6 | 128822963 | 128822996 | 34 | + | 18.561 | 2.42E-07 | 0.0549 | AAGCTGTCTTCAGACACTCCAGAAGAGGGCGCCA | | |
| 412 | 6 | 128822903 | 128823105 | 202 | 129S1,AJ,AKR,BALB,BUB,C3H,C57L,C58J,CBA,DBA1J,DBA2J,ILNJ2,MRL,NOD,NOR,NZO.PL.RF.SEA.SMJ.SWR | 6 | 128822978 | 128822996 | 19 | + | 17.5574 | 6.32E-07 | 0.138 | ACTCCAGAAGAGGGCGCCA | | |
| 413 | 6 | 129571429 | 129571631 | 202 | 129S1,AJ,AKR,BALB,BUB,C3H,C57L,C58J,CBA,CE,DBA1J,DBA2J,FVB,ILNJ2,LP,MA/My,MRL,NOD,NOR,NZO,PL,RF,RH2,SEA,SJL,SMJ,SWR,TH | 6 | 129571489 | 129571522 | 34 | + | 17.8455 | 3.91E-07 | 0.0687 | TAGCTGTCCTCAGACACTCCAGAAGAGGGCGTCA | | |
| 414 | 6 | 133195549 | 133195755 | 206 | NZO | 6 | 133195610 | 133195643 | 34 | + | 17.2439 | 5.79E-07 | 0.086 | TAGCTGTCCTCAGACCCTCCAGAAGAGGGAGTCA | | |
| 415 | 6 | 133683827 | 133684160 | 333 | AJ,AKR,BALB,BUB,C3H,C57L,CBA,DBA1J,DBA2J,ILNJ2,MRL,NZB,PL,RF,RH2,SEA | 6 | 133683857 | 133683875 | 19 | + | 19.2951 | 1.88E-07 | 0.0786 | CCGCCACTAGATGGTGCTA | 21 | ENCFF367UNR,ENCFF093ZRU,ENCFF099QNE,ENCFF821BIA,ENCFF609MDN,ENCFF907XEG,ENCFF327LYS,ENCFF089GBC,ENCFF033UQX,ENCFF052KVO,ENCFF024JOZ,ENCFF589ZBJ,ENCFF198SSR,ENCFF136ZDK,ENCFF165LXO,ENCFF430PPJ,ENCFF092NMJ,ENCFF951GOP,ENCFF883UPM,ENCFF957SPH,ENCFF281QMV |
| 416 | 6 | 135077717 | 135077887 | 170 | 129S1,AJ,AKR,BALB,BUB,C3H,C57L,CBA,CE,DBA1J,DBA2J,KK,MA/My,MRL,NOD,NOR,NZO.PL.RF.RH2.SEA.SJL.TH | 6 | 135077760 | 135077793 | 34 | + | 17.3902 | 5.27E-07 | 0.0822 | TAGCTGTCCTCAGACACTCCAGAAGAGGGCATCA | | |
| 417 | 6 | 138793203 | 138793359 | 156 | AJ,AKR,BALB,BUB,C3H,CBA,CE,DBA1J,DBA2J,ILNJ2,KK,MRL,NOD,NOR,PL,RF,RH2,SEA | 6 | 138793172 | 138793205 | 34 | + | 17.5203 | 4.84E-07 | 0.0781 | GTGAAGTGCCCTCATAGGCCAGCAGGTAGTATCA | | |
| 418 | 6 | 140585047 | 140585271 | 224 | BTBR.C57L.C58J.FVB.PJ.SMJ.SWR | 6 | 140585129 | 140585162 | 34 | + | 16.5691 | 8.90E-07 | 0.11 | TAGCTGTCTTCAGACGCACCAGAAGAGGGCATCG | | |
| 419 | 6 | 146718506 | 146718955 | 449 | 129S1,AJ,BALB,BTBR,C3H,C57L,CBA,CE,DBA1J,DBA2J,FVB,ILNJ2,MA/My,NOD,NOR,PJ,RF,SEA,SMJ,SWR,TH | 6 | 146718523 | 146718556 | 34 | + | 17.9106 | 3.74E-07 | 0.0672 | CCGCCCTCACATTCGCCGTCACAAGATGGCGCTG | | |
| 420 | 6 | 15883679 | 15883893 | 214 | 129S1,C57L,CE,FVB,ILNJ2,KK,SMJ,SWR,TH | 6 | 15883804 | 15883822 | 19 | - | 17.5574 | 6.32E-07 | 0.138 | ACTCCAGAAGAGGGCGCCA | | |
| 420 | 6 | 15883679 | 15883893 | 214 | 129S1,C57L,CE,FVB,ILNJ2,KK,SMJ,SWR,TH | 6 | 15883804 | 15883837 | 34 | - | 18.3659 | 2.76E-07 | 0.0572 | TAGCTGTCTTCAGACACTCCAGAAGAGGGCGCCA | | |
| 421 | 6 | 22275614 | 22275749 | 135 | BUB.SJL.SMJ.SWR | 6 | 22275726 | 22275759 | 34 | - | 16.4472 | 9.61E-07 | 0.112 | TAGCTGTCTTCAGACACACCAGAAGAGGGCGTCA | | |
| 422 | 6 | 29956156 | 29956369 | 213 | 129S1,AJ,AKR,BALB,BTBR,BUB,C3H,C57L,CBA,DBA1J,DBA2J,FVB,MRL,NOD,NOR,NZO,PJ,PL,RF,RH2,SEA,SMJ | 6 | 29956276 | 29956309 | 34 | - | 17.2439 | 5.79E-07 | 0.086 | TAGCTGTCCTCAGATACTCCAGAAGAGGGCATCA | | |
| 423 | 6 | 38034690 | 38034890 | 200 | BUB,C3H,CBA,CE,DBA1J,DBA2J,FVB,ILNJ2,MRL,NOD,NOR,PJ,PL,SEA,SJL,SMJ,SWR,TH | 6 | 38034804 | 38034822 | 19 | - | 17.5738 | 6.25E-07 | 0.138 | ACTCCAGAAGAGGGCGCTA | | |
| 423 | 6 | 38034690 | 38034890 | 200 | BUB,C3H,CBA,CE,DBA1J,DBA2J,FVB,ILNJ2,MRL,NOD,NOR,PJ,PL,SEA,SJL,SMJ,SWR,TH | 6 | 38034804 | 38034837 | 34 | - | 18.187 | 3.12E-07 | 0.0608 | TAGCTGTCTTCAGTCACTCCAGAAGAGGGCGCTA | | |
| 424 | 6 | 38193518 | 38193774 | 256 | BUB,C3H,CBA,CE,DBA1J,DBA2J,FVB,ILNJ2,KK,MRL,NOD,NOR,PJ,PL,SEA,SJL,SMJ,SWR.TH | 6 | 38193684 | 38193702 | 19 | - | 17.5574 | 6.32E-07 | 0.138 | ACTCCAGAAGAGGGCGCCA | | |
| 424 | 6 | 38193518 | 38193774 | 256 | BUB,C3H,CBA,CE,DBA1J,DBA2J,FVB,ILNJ2,KK,MRL,NOD,NOR,PJ,PL,SEA,SJL,SMJ,SWR.TH | 6 | 38193684 | 38193717 | 34 | - | 18.3659 | 2.76E-07 | 0.0572 | TAGCTGTCTTCAGACACTCCAGAAGAGGGCGCCA | | |
| 425 | 6 | 39368744 | 39368954 | 210 | KK | 6 | 39368873 | 39368891 | 19 | - | 17.5574 | 6.32E-07 | 0.138 | ACTCCAGAAGAGGGCGCCA | 2 | ENCFF367UNR.ENCFF093ZRU |
| 425 | 6 | 39368744 | 39368954 | 210 | KK | 6 | 39368873 | 39368906 | 34 | - | 18.3659 | 2.76E-07 | 0.0572 | TAGCTGTCTTCAGACACTCCAGAAGAGGGCGCCA | 2 | ENCFF367UNR.ENCFF093ZRU |
| 426 | 6 | 48727821 | 48728448 | 627 | CE.ILNJ2 | 6 | 48728348 | 48728381 | 34 | - | 16.6504 | 8.46E-07 | 0.107 | TAGCTGTCTTCAGATGCACCAGAAGAGGGCGACA | | |
| 427 | 6 | 51481038 | 51481234 | 196 | AKR,BALB,C58J,LP,MA/My,NOD,NOR,PJ,RF,RH2,SEA | 6 | 51481149 | 51481182 | 34 | - | 16.7724 | 7.83E-07 | 0.103 | TAGCTGTCTTCAGATGCACCAGAAGAGGGCGTCA | | |
| 428 | 6 | 56660466 | 56660661 | 195 | 129S1,C57L,KK,NZB,NZO,NZW | 6 | 56660585 | 56660618 | 34 | - | 16.4472 | 9.61E-07 | 0.112 | TAGCTGTCTTCAGACACACCAGAAGAGGGCGTCA | | |
| 429 | 6 | 67198988 | 67199206 | 218 | C58J,CE,ILNJ2,KK,NOD,NOR,NZB,NZO,PJ | 6 | 67199114 | 67199147 | 34 | - | 17.3902 | 5.27E-07 | 0.0822 | TAGCTGTCCTCAGACACTCCAGAAGAGGGCATCA | | |
| 430 | 6 | 72373495 | 72373690 | 195 | AKR,C3H,C58J,CBA,DBA1J,DBA2J,KK,MA/My,MRL,NOD,NOR,PL,RF,SMJ | 6 | 72373607 | 72373641 | 35 | - | 16.0917 | 9.07E-07 | 0.184 | TAGCTGTCTTCAGACAGCACCAGAAGAGGGCATCA | | |
| 431 | 6 | 72589221 | 72589438 | 217 | C3H,C58J,CBA,CE,DBA1J,DBA2J,ILNJ2,KK,MA/Mv.NOD.NOR.PJ.SMJ | 6 | 72589339 | 72589372 | 34 | - | 17.8455 | 3.91E-07 | 0.0687 | TAGCTGTCCTCAGACACTCCAGAAGAGGGCGTCA | | |
| 432 | 6 | 72768765 | 72769172 | 407 | AKR,BALB,C3H,C58J,CBA,DBA1J,DBA2J,KK,MA/My,MRL,NOD,NOR,PL,RF,RH2,SEA,SMJ | 6 | 72769092 | 72769125 | 34 | - | 16.4472 | 9.61E-07 | 0.112 | TAGCTGTCTTCAGACACACCAGAAGAGGGCGTCA | | |
| 433 | 6 | 76595416 | 76595610 | 194 | KK.PJ | 6 | 76595539 | 76595572 | 34 | - | 16.5447 | 9.04E-07 | 0.111 | CAGCTGTCTTCAGACACACCAGAAGAGGGAGTCA | | |
| 434 | 6 | 85503754 | 85503953 | 199 | PJ.SEA | 6 | 85503868 | 85503901 | 34 | - | 16.4472 | 9.61E-07 | 0.112 | TAGCTGTCTTCAGACACACCAGAAGAGGGCGTCA | | |
| 435 | 6 | 87043314 | 87043528 | 214 | 129S1,AJ,BTBR,C3H,C57L,C58J,CBA,DBA1J,DBA2J,FVB,NOD,NOR,PJ,RF,SEA,SJL,SMJ | 6 | 87043438 | 87043471 | 34 | - | 17.8455 | 3.91E-07 | 0.0687 | TAGCTGTCCTCAGACACTCCAGAAGAGGGCGTCA | | |
| 436 | 6 | 89350461 | 89350667 | 206 | 129S1,C3H,C57L,C58J,CBA,CE,DBA1J,DBA2J,FVB,KK,LP,MA/My,NOD,NOR,NZB,NZO,NZW,PJ,SEA,SMJ,SWR,TH | 6 | 89350587 | 89350620 | 34 | - | 16.4634 | 9.51E-07 | 0.112 | TAGCTGTCTTCAGACGCACCAGAAGAGGGCATCA | | |
| 437 | 6 | 95306402 | 95307243 | 841 | C58J.DBA1J.DBA2J.LP.MA/Mv | 6 | 95306799 | 95306832 | 34 | - | 18.3171 | 2.86E-07 | 0.0578 | TAGCTGTCTTCAGACCCCCCAGTAGAGGGCGTCA | | |
| 438 | 6 | 98733753 | 98733941 | 188 | SJL | 6 | 98733863 | 98733896 | 34 | - | 16.4472 | 9.61E-07 | 0.112 | TAGCTGTCTTCAGACACACCAGAAGAGGGCGTCA | | |
| 439 | 6 | 100326492 | 100326709 | 217 | BUB.NZB.NZO.NZW | 6 | 100326639 | 100326672 | 34 | - | 16.7724 | 7.83E-07 | 0.103 | TAGCTGTCTTCAGATGCACCAGAAGAGGGCGTCA | | |
| 440 | 6 | 113313661 | 113313897 | 236 | AKR,BTBR,C3H,C58J,CBA,CE,DBA1J,DBA2J,FVB,ILNJ2,LP,MRL,NOD,NOR,NZB,NZO,NZW,PJ,PL,RF,SJL,SMJ,SWR,TH | 6 | 113313789 | 113313822 | 34 | - | 16.4634 | 9.51E-07 | 0.112 | TAGCTGTCTTCAGACGCACCAGAAGAGGGCATCA | | |
| 441 | 6 | 114652335 | 114652396 | 61 | NZB.NZW | 6 | 114652367 | 114652401 | 35 | - | 16.7064 | 6.41E-07 | 0.153 | ATACAATTCCTTGTCCAGGCCAGAGGAAGGCACTG | | |
| 442 | 6 | 114652507 | 114652996 | 489 | 129S1,AKR,BTBR,C3H,C57L,C58J,CBA,CE,DBA1J,DBA2J,FVB,ILNJ2,LP,MRL,NOD,NOR,NZO,PJ,RF,SJL,SWR,TH | 6 | 114652489 | 114652523 | 35 | - | 16.7064 | 6.41E-07 | 0.153 | ATACAATTCCTTGTCCAGGCCAGAGGAAGGCACTG | | |
| 442 | 6 | 114652507 | 114652996 | 489 | 129S1,AKR,BTBR,C3H,C57L,C58J,CBA,CE,DBA1J,DBA2J,FVB,ILNJ2,LP,MRL,NOD,NOR,NZO,PJ,RF,SJL,SWR,TH | 6 | 114652550 | 114652584 | 35 | - | 18.0367 | 2.94E-07 | 0.105 | ATACAATTCCTCGTCCAGGCCAGAGGAAGGCACTG | | |
| 442 | 6 | 114652507 | 114652996 | 489 | 129S1,AKR,BTBR,C3H,C57L,C58J,CBA,CE,DBA1J,DBA2J,FVB,ILNJ2,LP,MRL,NOD,NOR,NZO,PJ,RF,SJL,SWR,TH | 6 | 114652611 | 114652645 | 35 | - | 16.7064 | 6.41E-07 | 0.153 | ATACAATTCCTTGTCCAGGCCAGAGGAAGGCACTG | | |
| 442 | 6 | 114652507 | 114652996 | 489 | 129S1,AKR,BTBR,C3H,C57L,C58J,CBA,CE,DBA1J,DBA2J,FVB,ILNJ2,LP,MRL,NOD,NOR,NZO,PJ,RF,SJL,SWR,TH | 6 | 114652672 | 114652706 | 35 | - | 16.7064 | 6.41E-07 | 0.153 | ATACAATTCCTTGTCCAGGCCAGAGGAAGGCACTG | | |
| 442 | 6 | 114652507 | 114652996 | 489 | 129S1,AKR,BTBR,C3H,C57L,C58J,CBA,CE,DBA1J,DBA2J,FVB,ILNJ2,LP,MRL,NOD,NOR,NZO,PJ,RF,SJL,SWR,TH | 6 | 114652733 | 114652767 | 35 | - | 17.2936 | 4.57E-07 | 0.132 | ATACAATTCCTCTTCCAGGCCAGAGGAAGGCACTG | | |
| 442 | 6 | 114652507 | 114652996 | 489 | 129S1,AKR,BTBR,C3H,C57L,C58J,CBA,CE,DBA1J,DBA2J,FVB,ILNJ2,LP,MRL,NOD,NOR,NZO,PJ,RF,SJL,SWR,TH | 6 | 114652794 | 114652828 | 35 | - | 16.7064 | 6.41E-07 | 0.153 | ATACAATTCCTTGTCCAGGCCAGAGGAAGGCACTG | | |
| 442 | 6 | 114652507 | 114652996 | 489 | 129S1,AKR,BTBR,C3H,C57L,C58J,CBA,CE,DBA1J,DBA2J,FVB,ILNJ2,LP,MRL,NOD,NOR,NZO,PJ,RF,SJL,SWR,TH | 6 | 114652855 | 114652889 | 35 | - | 16.7064 | 6.41E-07 | 0.153 | ATACAATTCCTTGTCCAGGCCAGAGGAAGGCACTG | | |
| 442 | 6 | 114652507 | 114652996 | 489 | 129S1,AKR,BTBR,C3H,C57L,C58J,CBA,CE,DBA1J,DBA2J,FVB,ILNJ2,LP,MRL,NOD,NOR,NZO,PJ,RF,SJL,SWR,TH | 6 | 114652916 | 114652950 | 35 | - | 16.7064 | 6.41E-07 | 0.153 | ATACAATTCCTTGTCCAGGCCAGAGGAAGGCACTG | | |
| 442 | 6 | 114652507 | 114652996 | 489 | 129S1,AKR,BTBR,C3H,C57L,C58J,CBA,CE,DBA1J,DBA2J,FVB,ILNJ2,LP,MRL,NOD,NOR.NZO.PJ.RF.SJL.SWR.TH | 6 | 114652977 | 114653011 | 35 | - | 17.2936 | 4.57E-07 | 0.132 | ATACAATTCCTCTTCCAGGCCAGAGGAAGGCACTG | | |
| 443 | 6 | 125275304 | 125275691 | 387 | AJ,AKR,BALB,C3H,C57L,C58J,CBA,DBA1J,DBA2J,FVB,MA/My,MRL,NOD,NOR,NZW,PJ,RF.SEA.SJL.TH | 6 | 125275615 | 125275648 | 34 | - | 17.9187 | 3.72E-07 | 0.0672 | TAGCTGTCCTCACATACTCCAGAAGAGGGCATCA | | |
| 444 | 6 | 133183035 | 133183380 | 345 | NOD,NOR,NZO,SWR | 6 | 133183338 | 133183356 | 19 | - | 19.2951 | 1.88E-07 | 0.0786 | CCGCCACTAGATGGTGCTA | 30 | ENCFF805QBU,ENCFF367UNR,ENCFF093ZRU,ENCFF099QNE,ENCFF327LYS,ENCFF316FYN,ENCFF089GBC,ENCFF821BIA,ENCFF907XEG,ENCFF201JUA,ENCFF024JOZ,ENCFF194HVL,ENCFF092NMJ,ENCFF589ZBJ,ENCFF430PPJ,ENCFF491RJK,ENCFF198SSR,ENCFF136ZDK,ENCFF957SPH,ENCFF321AGQ,ENCFF165LXO,ENCFF055HAP,ENCFF210BTO,ENCFF951GOP,ENCFF502WDT,ENCFF281QMV,ENCFF883UPM,ENCFF052KVO.ENCFF575MGI.ENCFF609MDN |
| 445 | 6 | 134257772 | 134257823 | 51 | CBA.NZB.NZO | 6 | 134257746 | 134257779 | 34 | - | 17.1463 | 6.17E-07 | 0.0894 | CAGCAGCAGCAGCAGCAGCAGCTTGGGGGAGCCA | | |
| 446 | 6 | 137640390 | 137640581 | 191 | 129S1,AJ,AKR,BTBR,BUB,C3H,C57L,CBA,CE,DBA1J,DBA2J,FVB,ILNJ2,MA/My,NOD,NOR,NZO,NZW,PL,RF,SJL,SWR,TH | 6 | 137640510 | 137640528 | 19 | - | 17.5574 | 6.32E-07 | 0.138 | ACTCCAGAAGAGGGCGCCA | 2 | ENCFF367UNR,ENCFF093ZRU |
| 446 | 6 | 137640390 | 137640581 | 191 | 129S1,AJ,AKR,BTBR,BUB,C3H,C57L,CBA,CE,DBA1J,DBA2J,FVB,ILNJ2,MA/My,NOD,NOR,NZO,NZW,PL,RF,SJL,SWR,TH | 6 | 137640510 | 137640543 | 34 | - | 18.561 | 2.42E-07 | 0.0549 | AAGCTGTCTTCAGACACTCCAGAAGAGGGCGCCA | 2 | ENCFF367UNR,ENCFF093ZRU |
| 447 | 6 | 141282831 | 141283746 | 915 | BTBR,C57L,C58J,FVB,NOD,NOR,PJ,SMJ,SWR | 6 | 141283635 | 141283669 | 35 | - | 21.2202 | 3.89E-08 | 0.0311 | ATGCAGAGGCCCACAGCGAACAGTAGGGGGAGCTC | 23 | ENCFF093ZRU,ENCFF092NMJ,ENCFF165LXO,ENCFF367UNR,ENCFF210BTO,ENCFF089GBC,ENCFF327LYS,ENCFF099QNE,ENCFF609MDN,ENCFF033UQX,ENCFF883UPM,ENCFF052KVO,ENCFF589ZBJ,ENCFF198SSR,ENCFF907XEG,ENCFF281QMV,ENCFF024JOZ,ENCFF194HVL,ENCFF136ZDK,ENCFF957SPH,ENCFF533APC.ENCFF491RJK.ENCFF821BIA |
| 448 | 6 | 142460856 | 142461291 | 435 | BTBR,C57L,C58J,FVB,KK,MA/My,NOD,NOR,NZB,PJ,SMJ,SWR | 6 | 142460869 | 142460887 | 19 | - | 17.5574 | 6.32E-07 | 0.138 | TGGCCAGTTGGGGGCGCGC | 20 | ENCFF093ZRU,ENCFF099QNE,ENCFF367UNR,ENCFF136ZDK,ENCFF430PPJ,ENCFF165LXO,ENCFF883UPM,ENCFF089GBC,ENCFF907XEG,ENCFF609MDN,ENCFF821BIA,ENCFF327LYS,ENCFF533APC,ENCFF198SSR,ENCFF024JOZ,ENCFF589ZBJ,ENCFF957SPH,ENCFF092NMJ,ENCFF281QMV,ENCFF194HVL |
| 449 | 6 | 145045771 | 145045958 | 187 | CBA.ILNJ2.MA/Mv.NOD.NOR.PJ.TH | 6 | 145045887 | 145045920 | 34 | - | 16.4634 | 9.51E-07 | 0.112 | TAGCTGTCTTCAGACGCACCAGAAGAGGGCATCA | | |
| 450 | 6 | 147084772 | 147085153 | 381 | 129S1,AJ,BALB,BTBR,C57L,CBA,FVB,ILNJ2,MA/My,NOD,NOR,PJ,RF,SEA,SMJ,SWR,TH | 6 | 147084969 | 147085003 | 35 | - | 21.1193 | 4.16E-08 | 0.0327 | CTGTATTGCCACCTCCTTCCCACAGGGTGGTATCC | | |
| 451 | 6 | 148748713 | 148748926 | 213 | KK.PL | 6 | 148748837 | 148748870 | 34 | - | 17.6667 | 4.40E-07 | 0.0735 | TAGCTGTTTTCAGACACTCCAGAAGAGGGCGTCA | | |
| 452 | 7 | 3455756 | 3455929 | 173 | AJ,AKR,KK,LP,MA/My,NOD,NOR,NZW,PL,RF,RH2,SJL,SWR,TH | 7 | 3455754 | 3455787 | 34 | + | 20.0976 | 8.28E-08 | 0.0223 | ATGCAGAGCTCATGATGACCAGAAGAGGGCCTCA | 3 | ENCFF099QNE,ENCFF092NMJ,ENCFF093ZRU |
| 453 | 7 | 6243937 | 6244022 | 85 | SMJ | 7 | 6243997 | 6244031 | 35 | + | 21.6239 | 2.95E-08 | 0.0191 | TGGCTATTCTGGAAAATGTCCAGCAGGTGGCAGCA | 30 | ENCFF092NMJ,ENCFF430PPJ,ENCFF201JUA,ENCFF093ZRU,ENCFF367UNR,ENCFF951GOP,ENCFF609MDN,ENCFF024JOZ,ENCFF327LYS,ENCFF316FYN,ENCFF907XEG,ENCFF099QNE,ENCFF821BIA,ENCFF533APC,ENCFF089GBC,ENCFF589ZBJ,ENCFF491RJK,ENCFF194HVL,ENCFF055HAP,ENCFF097CAQ,ENCFF142CNG,ENCFF136ZDK,ENCFF198SSR,ENCFF321AGQ,ENCFF281QMV,ENCFF502WDT,ENCFF957SPH,ENCFF883UPM,ENCFF210BTO,ENCFF165LXO |

| | | | | | | | | | | | | | | | | |
|---|---|---|---|---|---|---|---|---|---|---|---|---|---|---|---|---|
| 453 | 7 | 6243937 | 6244022 | 85 | SMJ | 7 | 6244013 | 6244031 | 19 | + | 22.5902 | 1.16E-08 | 0.0201 | TGTCCAGCAGGTGGCAGCA | 30 | ENCFF092NMJ,ENCFF430PPJ,ENCFF201JUA,ENCFF093ZRU,ENCFF367UNR,ENCFF951GOP,ENCFF609MDN,ENCFF024JOZ,ENCFF327LYS,ENCFF316FYN,ENCFF907XEG,ENCFF099QNE,ENCFF821BIA,ENCFF533APC,ENCFF089GBC,ENCFF589ZBJ,ENCFF491RJK,ENCFF194HVL,ENCFF055HAP,ENCFF097CAQ,ENCFF142CNG,ENCFF136ZDK,ENCFF198SSR,ENCFF321AGQ,ENCFF281QMV,ENCFF502WDT,ENCFF957SPH,ENCFF883UPM,ENCFF210BTO,ENCFF165LXO |
| 454 | 7 | 12177107 | 12177326 | 219 | AKR,MRL,RH2 | 7 | 12177184 | 12177217 | 34 | + | 17.2439 | 5.79E-07 | 0.0706 | TAGCTGTCCTCAGATACTCCAGAAGAGGGCATCA | | |
| 455 | 7 | 16025876 | 16026120 | 244 | 129S1,AJ,AKR,BALB,BTBR,C3H,C57L,C58J,CBA,DBA1J,DBA2J,ILNJ2,KK,LP,MA/My,MRL,NZB,NZO,NZW,PJ,RF,RH2,SEA,SJL,SMJ,TH | 7 | 16025987 | 16026020 | 34 | + | 18.1545 | 3.18E-07 | 0.0508 | TAGCTGTTTTCAGACACACCAGAAGAGGGCGTCA | | |
| 456 | 7 | 16381827 | 16382040 | 213 | 129S1,BTBR,BUB,C3H,C57L,CBA,CE,DBA1J,DBA2J,FVB,MA/My,NOD,NOR,NZB,PJ,RF,SEA,SMJ | 7 | 16381892 | 16381925 | 34 | + | 16.9919 | 6.81E-07 | 0.0775 | TCGCTGTCCTCAGACACACCAGAAGAGGGCATCA | | |
| 457 | 7 | 16687429 | 16687625 | 196 | 129S1,BTBR,C3H,C57L,CBA,FVB,NZB,NZW,PL,RF | 7 | 16687501 | 16687534 | 34 | + | 16.4634 | 9.51E-07 | 0.0901 | TAGCTGTCTTCAGACGCACCAGAAGAGGGCATCA | | |
| 458 | 7 | 18923391 | 18923619 | 228 | 129S1,AJ,BALB,BTBR,BUB,C3H,C57L,CBA,CE,DBA1J,DBA2J,FVB,ILNJ2,KK,LP,MA/My,MRL,NOD,NOR,NZB,NZW,PJ,PL,RF,SEA,SJL,SMJ | 7 | 18923467 | 18923500 | 34 | + | 18.3659 | 2.76E-07 | 0.0476 | TAGCTGTCTTCAGACACTCCAGAAGAGGGCGCCA | | |
| 458 | 7 | 18923391 | 18923619 | 228 | 129S1,AJ,BALB,BTBR,BUB,C3H,C57L,CBA,CE,DBA1J,DBA2J,FVB,ILNJ2,KK,LP,MA/My,MRL,NOD,NOR,NZB,NZW,PJ,PL,RF,SEA,SJL,SMJ | 7 | 18923482 | 18923500 | 19 | + | 17.5574 | 6.32E-07 | 0.106 | ACTCCAGAAGAGGGCGCCA | | |
| 459 | 7 | 19032923 | 19033153 | 230 | 129S1,BTBR,C3H,C57L,CBA,NOD,NOR,NZB,NZW,RF,SJL | 7 | 19032898 | 19032931 | 34 | + | 16.4797 | 9.42E-07 | 0.0901 | CTGCTATTGCCACTGCCCCCTCCATGAGGCACTT | | |
| 460 | 7 | 24479573 | 24479778 | 205 | BUB,NOD,NOR,SJL | 7 | 24479652 | 24479670 | 19 | + | 17.8852 | 5.08E-07 | 0.0973 | ACACCAGAAGAGGGCACCA | | |
| 461 | 7 | 25098674 | 25098895 | 221 | BUB,C58J,CE,ILNJ2,MRL,NOD,NOR,PL,SJL | 7 | 25098756 | 25098789 | 34 | + | 17.0488 | 6.56E-07 | 0.076 | TTGCTATCTTCAGATACACCAGAAGAGGGCATCA | 1 | ENCFF093ZRU |
| 462 | 7 | 27379616 | 27379838 | 222 | 129S1,ILNJ2,KK,LP,NZO,NZW,PJ,RF,SEA,SMJ | 7 | 27379696 | 27379729 | 34 | + | 18.1789 | 3.13E-07 | 0.0504 | TAGCTGTCTTCAGACACTCCAGAAGAGGGCGCTA | 1 | ENCFF093ZRU |
| 462 | 7 | 27379616 | 27379838 | 222 | 129S1,ILNJ2,KK,LP,NZO,NZW,PJ,RF,SEA,SMJ | 7 | 27379711 | 27379729 | 19 | + | 17.5738 | 6.25E-07 | 0.106 | ACTCCAGAAGAGGGCGCTA | 1 | ENCFF093ZRU |
| 463 | 7 | 27393103 | 27393303 | 200 | 129S1,LP,NZW | 7 | 27393161 | 27393194 | 34 | + | 16.4472 | 9.61E-07 | 0.0901 | TAGCTGTCTTCAGACACACCAGAAGAGGGCGTCA | | |
| 464 | 7 | 29687919 | 29688192 | 273 | AJ,AKR,BALB,BTBR,C57L,DBA1J,DBA2J,MA/My,MRL,NOD,NOR,NZB,NZO,PJ,PL,SEA,SMJ,TH | 7 | 29688049 | 29688082 | 34 | + | 18.1789 | 3.13E-07 | 0.0504 | TAGCTGTCTTCAGACACTCCAGAAGAGGGCGCTA | 1 | ENCFF093ZRU |
| 464 | 7 | 29687919 | 29688192 | 273 | AJ,AKR,BALB,BTBR,C57L,DBA1J,DBA2J,MA/My,MRL,NOD,NOR,NZB,NZO,PJ,PL,SEA,SMJ,TH | 7 | 29688064 | 29688082 | 19 | + | 17.5738 | 6.25E-07 | 0.106 | ACTCCAGAAGAGGGCGCTA | 1 | ENCFF093ZRU |
| 465 | 7 | 29746172 | 29746387 | 215 | 129S1,AJ,AKR,BALB,BTBR,C57L,DBA1J,DBA2J,LP,MA/My,MRL,NOD,NOR,NZB,NZO,NZW,PJ,PL,SEA,SJL,SMJ,TH | 7 | 29746242 | 29746275 | 34 | + | 16.5691 | 8.90E-07 | 0.0882 | TAGCTGTCTTCAGACGCACCAGAAGAGGGCATCG | | |
| 466 | 7 | 30505054 | 30505579 | 525 | DBA1J,DBA2J,NZB,NZO,NZW,SMJ | 7 | 30505511 | 30505544 | 34 | + | 18.3902 | 2.72E-07 | 0.0476 | GCGTCGAACACCCAATGCCCAGCAGGGGGCGCTG | 23 | ENCFF093ZRU,ENCFF805QBU,ENCFF165LXO,ENCFF099QNE,ENCFF367UNR,ENCFF327LYS,ENCFF033UQX,ENCFF321AGQ,ENCFF136ZDK,ENCFF907XEG,ENCFF821BIA,ENCFF201JUA,ENCFF589ZBJ,ENCFF089EWX,ENCFF052KVO,ENCFF024JOZ,ENCFF198SSR,ENCFF533APC,ENCFF883UPM,ENCFF957SPH,ENCFF210BTO,ENCFF097CAQ,ENCFF194HVL |
| 466 | 7 | 30505054 | 30505579 | 525 | DBA1J,DBA2J,NZB,NZO,NZW,SMJ | 7 | 30505526 | 30505544 | 19 | + | 23.3607 | 5.29E-09 | 0.0159 | TGCCCAGCAGGGGGCGCTG | 24 | ENCFF093ZRU,ENCFF805QBU,ENCFF165LXO,ENCFF099QNE,ENCFF367UNR,ENCFF327LYS,ENCFF033UQX,ENCFF321AGQ,ENCFF136ZDK,ENCFF907XEG,ENCFF821BIA,ENCFF201JUA,ENCFF589ZBJ,ENCFF089EWX,ENCFF052KVO,ENCFF024JOZ,ENCFF198SSR,ENCFF533APC,ENCFF883UPM,ENCFF957SPH,ENCFF210BTO,ENCFF097CAQ,ENCFF194HVL,ENCFF504JDS |
| 467 | 7 | 33882332 | 33882540 | 208 | BTBR,C57L,DBA1J,DBA2J,NZB,NZO,NZW,PJ,PL,SMJ,TH | 7 | 33882398 | 33882431 | 34 | + | 18.3659 | 2.76E-07 | 0.0476 | TAGCTGTCTTCAGACACTCCAGAAGAGGGCGCCA | | |
| 467 | 7 | 33882332 | 33882540 | 208 | BTBR,C57L,DBA1J,DBA2J,NZB,NZO,NZW,PJ,PL,SMJ,TH | 7 | 33882413 | 33882431 | 19 | + | 17.5574 | 6.32E-07 | 0.106 | ACTCCAGAAGAGGGCGCCA | | |
| 468 | 7 | 34705347 | 34705550 | 203 | C58J,MRL,NOD,NOR,NZB,NZO,NZW,RF,SJL | 7 | 34705408 | 34705441 | 34 | + | 17.2439 | 5.79E-07 | 0.0706 | TAGCTGTCCTCAGATACTCCAGAAGAGGGCATCA | | |
| 469 | 7 | 38318733 | 38319427 | 694 | 129S1,BTBR,BUB,C3H,C57L,CBA,KK,MRL,NZW | 7 | 38319111 | 38319129 | 19 | + | 17.459 | 6.74E-07 | 0.109 | TGTCCACTAGATGTCAGTG | | |
| 470 | 7 | 38634438 | 38635132 | 694 | 129S1,BTBR,BUB,C3H,C57L,CBA,KK,MRL,NZW,PL,SJL | 7 | 38634814 | 38634832 | 19 | + | 17.459 | 6.74E-07 | 0.109 | TGTCCACTAGATGTCAGTG | | |
| 471 | 7 | 44673252 | 44673464 | 212 | 129S1,AJ,AKR,BALB,C3H,CBA,DBA1J,DBA2J,ILNJ2,KK,MA/My,NZW,PJ,PL,RH2,SEA,SJL,SMJ | 7 | 44673321 | 44673354 | 34 | + | 19.6667 | 1.13E-07 | 0.0279 | TAGCAGTCTTCAGATGCACCAGCAGAGGGCATCA | 13 | ENCFF367UNR,ENCFF092NMJ,ENCFF089GBC,ENCFF136ZDK,ENCFF093ZRU,ENCFF198SSR,ENCFF589ZBJ,ENCFF099QNE,ENCFF907XEG,ENCFF821BIA,ENCFF024JOZ,ENCFF327LYS |
| 472 | 7 | 44974365 | 44975125 | 760 | ILNJ2,NZW,PJ,SJL | 7 | 44974927 | 44974961 | 35 | + | 24.3211 | 4.17E-09 | 0.00615 | CTGCTCAGAAGGCTGCAGGCCAGAAGGTGGCACCT | | |
| 472 | 7 | 44974365 | 44975125 | 760 | ILNJ2,NZW,PJ,SJL | 7 | 44974943 | 44974961 | 19 | + | 17.7869 | 5.43E-07 | 0.101 | AGGCCAGAAGGTGGCACCT | | |
| 473 | 7 | 45127703 | 45127909 | 206 | ILNJ2,NZW,PJ,SJL | 7 | 45127767 | 45127800 | 34 | + | 18.3659 | 2.76E-07 | 0.0476 | TAGCTGTCTTCAGACACTCCAGAAGAGGGCGCCA | 6 | ENCFF093ZRU,ENCFF367UNR,ENCFF821BIA,ENCFF024JOZ,ENCFF907XEG,ENCFF136ZDK |
| 473 | 7 | 45127703 | 45127909 | 206 | ILNJ2,NZW,PJ,SJL | 7 | 45127782 | 45127800 | 19 | + | 17.5574 | 6.32E-07 | 0.106 | ACTCCAGAAGAGGGCGCCA | 6 | ENCFF093ZRU,ENCFF367UNR,ENCFF821BIA,ENCFF024JOZ,ENCFF907XEG,ENCFF136ZDK |
| 474 | 7 | 46552375 | 46552576 | 201 | ILNJ2,NZW,PJ,SJL | 7 | 46552434 | 46552467 | 34 | + | 16.4472 | 9.61E-07 | 0.0901 | TAGCTGTCTTCAGACACACCAGAAGAGGGCGTCA | | |
| 475 | 7 | 63729548 | 63729713 | 165 | DBA1J,DBA2J,KK | 7 | 63729550 | 63729583 | 34 | + | 18.4309 | 2.64E-07 | 0.047 | CTGTGGTGTCACCGGGCTCCACTTGGTGTCACCG | | |
| 476 | 7 | 66192992 | 66193942 | 950 | DBA1J,DBA2J,ILNJ2,KK,PJ | 7 | 66193369 | 66193387 | 19 | + | 16.9672 | 9.26E-07 | 0.127 | ATGCCAGCAGAGGGCATTG | | |
| 477 | 7 | 73988652 | 73988862 | 210 | 129S1,C3H,C58J,CBA,CE,DBA1J,DBA2J,FVB,KK,MA/My,PJ,SMJ,SWR,TH | 7 | 73988720 | 73988753 | 34 | + | 17.3902 | 5.27E-07 | 0.0672 | TAGCTGTCCTCAGACACTCCAGAAGAGGGCATCA | | |
| 478 | 7 | 75171940 | 75172536 | 596 | C58J,CE | 7 | 75172148 | 75172181 | 34 | + | 21.0894 | 4.00E-08 | 0.0128 | ATGCAGTGGACACAGAGGCCAGAAGAGGGTGTCA | 10 | ENCFF367UNR,ENCFF327LYS,ENCFF055HAP,ENCFF198SSR,ENCFF883UPM,ENCFF502WDT,ENCFF093ZRU,ENCFF907XEG,ENCFF099QNE,ENCFF821BIA |
| 479 | 7 | 80483114 | 80483335 | 221 | AJ,AKR,BALB,BUB,FVB,MRL,NOD,NOR,NZB,NZO,NZW,PL,RF,RH2,SJL,SMJ,SWR,TH | 7 | 80483181 | 80483214 | 34 | + | 16.4634 | 9.51E-07 | 0.0901 | TAGCTGTCTTCAGACGCACCAGAAGAGGGCATCA | 2 | ENCFF367UNR,ENCFF055HAP |
| 480 | 7 | 80501124 | 80501327 | 203 | CE | 7 | 80501182 | 80501215 | 34 | + | 16.4472 | 9.61E-07 | 0.0901 | TAGCTGTCTTCAGACACACCAGAAGAGGGCGTCA | | |
| 481 | 7 | 82377239 | 82377988 | 749 | CE | 7 | 82377538 | 82377556 | 19 | + | 16.9836 | 9.17E-07 | 0.126 | GGAACAGTAGAGGGCACTA | 1 | ENCFF093ZRU |
| 482 | 7 | 93891792 | 93891994 | 202 | 129S1,AJ,AKR,BALB,BTBR,BUB,C3H,C57L,CBA,DBA1J,DBA2J,FVB,ILNJ2,KK,MA/My,MRL,NOD,NZB,NZO,NZW,PJ,PL,RF,RH2,SEA,SJL,SMJ,SWR,TH | 7 | 93891853 | 93891886 | 34 | + | 18.3659 | 2.76E-07 | 0.0476 | TAGCTGTCTTCAGACACTCCAGAAGAGGGCGCCA | | |
| 482 | 7 | 93891792 | 93891994 | 202 | 129S1,AJ,AKR,BALB,BTBR,BUB,C3H,C57L,CBA,DBA1J,DBA2J,FVB,ILNJ2,KK,MA/My,MRL,NOD,NZB,NZO,NZW,PJ,PL,RF,RH2,SEA,SJL,SMJ,SWR,TH | 7 | 93891868 | 93891886 | 19 | + | 17.5574 | 6.32E-07 | 0.106 | ACTCCAGAAGAGGGCGCCA | | |
| 483 | 7 | 97866704 | 97866790 | 86 | C3H,CBA,DBA1J,DBA2J,ILNJ2,PJ,SEA | 7 | 97866738 | 97866771 | 34 | + | 17.6504 | 4.44E-07 | 0.0613 | ATGAAGTGCCTGTGGAGGCCAGAAGAGGGTGTGT | | |
| 484 | 7 | 98426070 | 98426463 | 393 | C3H,CBA,DBA1J,DBA2J,ILNJ2,PJ,SEA | 7 | 98426393 | 98426427 | 35 | + | 17.578 | 3.87E-07 | 0.0917 | CTGCAGACCATGACCCCGCCCAGCAGAAGACATCT | | |
| 485 | 7 | 98667704 | 98667985 | 281 | NOD,NOR,SWR | 7 | 98667915 | 98667949 | 35 | + | 17.0092 | 5.39E-07 | 0.11 | CTGCAATCCCAGAACTCAGGAAGCAGAGGGAGGAA | | |
| 486 | 7 | 100187291 | 100187494 | 203 | 129S1,AJ,AKR,BALB,BTBR,BUB,C3H,C57L,CBA,DBA1J,DBA2J,FVB,ILNJ2,KK,MA/My,MRL,NZB,NZW,PJ,PL,RF,RH2,SEA,SJL,SMJ,TH | 7 | 100187352 | 100187385 | 34 | + | 16.5041 | 9.27E-07 | 0.0901 | TAGCTGTCTTCAGACGTACCAGAAGAGGGCATCG | 1 | ENCFF093ZRU |
| 487 | 7 | 105511289 | 105511505 | 216 | CE,NZO | 7 | 105511363 | 105511396 | 34 | + | 18.4472 | 2.62E-07 | 0.0467 | TTGCTGTCTTCAGACACTCCAGAAGAGGGCGTCA | | |
| 488 | 7 | 105722697 | 105722912 | 215 | CE,NZO | 7 | 105722768 | 105722801 | 34 | + | 18.4472 | 2.62E-07 | 0.0467 | TTGCTGTCTTCAGACACTCCAGAAGAGGGCGTCA | | |
| 489 | 7 | 115962272 | 115962483 | 211 | 129S1,BTBR,C57L,ILNJ2 | 7 | 115962341 | 115962374 | 34 | + | 16.4634 | 9.51E-07 | 0.0901 | TAGCTGTCTTCAGACGCACCAGAAGAGGGCATCA | | |
| 490 | 7 | 116119065 | 116119530 | 465 | 129S1,BTBR,C57L,ILNJ2 | 7 | 116119202 | 116119235 | 34 | + | 25.7317 | 8.56E-10 | 0.000754 | ATGCAGTGCCCACAGAGGCCAAAAGAGGGCATCA | 27 | ENCFF065ODV,ENCFF367UNR,ENCFF093ZRU,ENCFF136ZDK,ENCFF201JUA,ENCFF430PPJ,ENCFF589ZBJ,ENCFF099QNE,ENCFF609MDN,ENCFF327LYS,ENCFF198SSR,ENCFF951GOP,ENCFF907XEG,ENCFF821BIA,ENCFF024JOZ,ENCFF097CAQ,ENCFF089GBC,ENCFF194HVL,ENCFF055HAP,ENCFF491RJK,ENCFF502WDT,ENCFF092NMJ,ENCFF957SPH,ENCFF142CNG,ENCFF165LXO,ENCFF281QMV,ENCFF883UPM |
| 491 | 7 | 117906037 | 117907007 | 970 | BTBR,C3H,C57L,CBA,CE,DBA1J,NOR | 7 | 117906225 | 117906258 | 34 | + | 18.2114 | 3.07E-07 | 0.0499 | TAGCTGTTTTCAGACACTCCAGAAGGGGGCATCA | | |
| 492 | 7 | 118117181 | 118117252 | 71 | 129S1,C3H,C57L,CBA,DBA1J,DBA2J,NOR | 7 | 118117231 | 118117265 | 35 | + | 18.4404 | 2.31E-07 | 0.0698 | CAGCAGTGAGCAGCAGTGAGCAGCAGTGGGCAGTG | 21 | ENCFF821BIA,ENCFF367UNR,ENCFF327LYS,ENCFF430PPJ,ENCFF198SSR,ENCFF092NMJ,ENCFF907XEG,ENCFF089GBC,ENCFF609MDN,ENCFF589ZBJ,ENCFF136ZDK,ENCFF611HDQ,ENCFF210BTO,ENCFF099QNE,ENCFF024JOZ,ENCFF883UPM,ENCFF097CAQ,ENCFF957SPH,ENCFF165LXO,ENCFF194HVL |
| 493 | 7 | 120402070 | 120402368 | 298 | AJ,CE | 7 | 120402098 | 120402132 | 35 | + | 21.211 | 3.91E-08 | 0.0225 | TTGCATGTACATCTGCATGCCAGAAGAGGGCATCA | 4 | ENCFF093ZRU,ENCFF821BIA,ENCFF502WDT,ENCFF055HAP |
| 494 | 7 | 122663538 | 122663682 | 144 | AKR,BUB,C58J,MRL,NOD,NZW,PL,SWR,TH | 7 | 122663641 | 122663674 | 34 | + | 16.4472 | 9.61E-07 | 0.0901 | TAGCTGTCTTCAGACACACCAGAAGAGGGCGTCA | | |
| 495 | 7 | 125254058 | 125254109 | 51 | AKR,BALB,BUB,C58J,MRL,NZW,PL,RH2,SEA,SJL,SMJ,SWR,TH | 7 | 125254083 | 125254117 | 35 | + | 20.8073 | 5.13E-08 | 0.026 | CTCCAGCTCCCCCTGAGCACCTCTGGGTGGCGGCC | | |
| 496 | 7 | 125829363 | 125829582 | 219 | 129S1,AJ,C57L,CE,FVB,ILNJ2,KK,LP,MA/My,NOD,NZB,NZO,PJ,RF,SMJ,TH | 7 | 125829437 | 125829470 | 34 | + | 16.4472 | 9.61E-07 | 0.0901 | TAGCTGTCTTCAGACACACCAGAAGAGGGCGTCA | | |
| 497 | 7 | 125831479 | 125831685 | 206 | 129S1,AJ,C57L,CE,FVB,ILNJ2,KK,LP,MA/My,NZB,NZO,PJ,RF,SMJ,TH | 7 | 125831543 | 125831576 | 34 | + | 17.6992 | 4.30E-07 | 0.0601 | TAGCTGTTTTCAGACACACCAGAAGAGGGCATCA | | |
| 498 | 7 | 127058392 | 127058600 | 208 | TH | 7 | 127058458 | 127058491 | 34 | + | 16.4634 | 9.51E-07 | 0.0901 | TAGCTGTCTTCAGACGCACCAGAAGAGGGCATCA | | |
| 499 | 7 | 127291269 | 127291480 | 211 | TH | 7 | 127291338 | 127291371 | 34 | + | 16.4472 | 9.61E-07 | 0.0901 | TAGCTGTCTTCAGACACACCAGAAGAGGGCGTCA | | |
| 500 | 7 | 127562133 | 127562345 | 212 | 129S1,AJ,BTBR,C57L,CE,ILNJ2,KK,PJ,TH | 7 | 127562202 | 127562235 | 34 | + | 16.4472 | 9.61E-07 | 0.0901 | TAGCTGTCTTCAGACACACCAGAAGAGGGCGTCA | | |
| 501 | 7 | 128688347 | 128688560 | 213 | 129S1,BTBR,C57L,ILNJ2,LP,NZW,PJ | 7 | 128688420 | 128688453 | 34 | + | 16.4472 | 9.61E-07 | 0.0901 | TAGCTGTCTTCAGACACACCAGAAGAGGGCGTCA | | |
| 502 | 7 | 129087311 | 129087435 | 124 | LP | 7 | 129087413 | 129087431 | 19 | + | 18.7541 | 2.79E-07 | 0.0733 | AGACCAGAAGAGGGCAGCA | | |
| 503 | 7 | 139700857 | 139701023 | 166 | BUB,C58J,CBA,CE,FVB,ILNJ2,KK,LP,MA/My,NOD,NOR,PJ,SMJ,SWR,TH | 7 | 139700921 | 139700954 | 34 | + | 16.5691 | 8.90E-07 | 0.0882 | TAGCTGTCTTCAGACGCACCAGAAGAGGGCATCG | | |
| 504 | 7 | 140720666 | 140720899 | 233 | NZW | 7 | 140720659 | 140720692 | 34 | + | 17.2114 | 5.91E-07 | 0.0711 | TAGCTGTTTTCAGACACTCCAGAAGAGGGCATCA | 2 | ENCFF367UNR,ENCFF093ZRU |
| 505 | 7 | 141245456 | 141245685 | 229 | ILNJ2,NZW | 7 | 141245542 | 141245575 | 34 | + | 16.4634 | 9.51E-07 | 0.0901 | TAGCTGTCTTCAGACGCACCAGAAGAGGGCATCA | | |
| 506 | 7 | 142386113 | 142386335 | 222 | C58J,LP,MA/My,PJ,SMJ | 7 | 142386191 | 142386224 | 34 | + | 16.4634 | 9.51E-07 | 0.0901 | TAGCTGTCTTCAGACGCACCAGAAGAGGGCATCA | | |
| 507 | 7 | 143635221 | 143635295 | 74 | 129S1,C57L,KK | 7 | 143635264 | 143635282 | 19 | + | 20.1803 | 9.60E-08 | 0.0453 | TAGCCAGCAGAGGGTGCTG | 20 | ENCFF201JUA,ENCFF093ZRU,ENCFF430PPJ,ENCFF099QNE,ENCFF327LYS,ENCFF089GBC,ENCFF589ZBJ,ENCFF907XEG,ENCFF883UPM,ENCFF024JOZ,ENCFF198SSR,ENCFF092NMJ,ENCFF055HAP,ENCFF136ZDK,ENCFF502WDT,ENCFF957SPH,ENCFF609MDN,ENCFF575MGI,ENCFF210BTO,ENCFF165LXO |
| 508 | 7 | 6978904 | 6979134 | 230 | 129S1,AJ,BALB,BTBR,BUB,C57L,C58J,CBA,KK,LP,NOD,NOR,NZB,NZO,NZW,PL,RF,SJL,SWR,TH | 7 | 6979027 | 6979045 | 19 | - | 17.5574 | 6.32E-07 | 0.106 | ACTCCAGAAGAGGGCGCCA | 1 | ENCFF093ZRU |
| 508 | 7 | 6978904 | 6979134 | 230 | 129S1,AJ,BALB,BTBR,BUB,C57L,C58J,CBA,KK,LP,NOD,NOR,NZB,NZO,NZW,PL,RF,SJL,SWR,TH | 7 | 6979027 | 6979060 | 34 | - | 18.3659 | 2.76E-07 | 0.0476 | TAGCTGTCTTCAGACACTCCAGAAGAGGGCGCCA | 1 | ENCFF093ZRU |
| 509 | 7 | 9872445 | 9872657 | 212 | 129S1,AJ,BALB,BTBR,BUB,C57L,C58J,CBA,ILNJ2,KK,LP,NOD,NOR,NZB,NZO,NZW,PL,RF,SJL,SWR,TH | 7 | 9872569 | 9872587 | 19 | - | 17.8852 | 5.08E-07 | 0.0973 | ACACCAGAAGAGGGCACCA | | |
| 509 | 7 | 9872445 | 9872657 | 212 | 129S1,AJ,BALB,BTBR,BUB,C57L,C58J,CBA,ILNJ2,KK,LP,NOD,NOR,NZB,NZO,NZW,PL,RF,SJL,SWR,TH | 7 | 9872569 | 9872602 | 34 | - | 18.252 | 2.98E-07 | 0.049 | TAGCTGTCTTCAGATACACCAGAAGAGGGCACCA | | |
| 510 | 7 | 13066480 | 13067206 | 726 | C58J | 7 | 13066814 | 13066848 | 35 | - | 16.0367 | 9.36E-07 | 0.147 | ATGCACTGCCCCTGGGCCGCAAGAAGGGGGCCAAG | | |
| 511 | 7 | 15997031 | 15997131 | 100 | AKR,RH2,TH | 7 | 15997114 | 15997132 | 19 | - | 23.1311 | 6.72E-09 | 0.0163 | CCGCCACTAGAGGGCACCA | 33 | ENCFF201JUA,ENCFF093ZRU,ENCFF367UNR,ENCFF099QNE,ENCFF327LYS,ENCFF065ODV,ENCFF821BIA,ENCFF589ZBJ,ENCFF024JOZ,ENCFF089GBC,ENCFF321AGQ,ENCFF055HAP,ENCFF907XEG,ENCFF136ZDK,ENCFF198SSR,ENCFF092NMJ,ENCFF502WDT,ENCFF165LXO,ENCFF052KVO,ENCFF951GOP,ENCFF609MDN,ENCFF097CAQ,ENCFF430PPJ,ENCFF194HVL,ENCFF883UPM,ENCFF089EWX,ENCFF491RJK,ENCFF957SPH,ENCFF281QMV,ENCFF575MGI,ENCFF142CNG,ENCFF316FYN |
| 512 | 7 | 18737463 | 18737699 | 236 | 129S1,AJ,AKR,BALB,BTBR,BUB,C3H,C57L,C58J,CBA,DBA1J,DBA2J,FVB,ILNJ2,KK,LP,MA/My,MRL,NOD,NOR,NZB,NZO,NZW,PJ,PL,RF,RH2,SEA,SJL,SMJ,SWR,TH | 7 | 18737589 | 18737622 | 34 | - | 17.9431 | 3.66E-07 | 0.0552 | CAGCTGTCCTCAGACACTCCAGAAGAGGGAGTCA | | |
| 513 | 7 | 23886214 | 23886408 | 194 | 129S1,BTBR,BUB,C3H,C57L,CBA,NOD,NOR,NZB,NZW,SJL | 7 | 23886340 | 23886373 | 34 | - | 16.6341 | 8.54E-07 | 0.0866 | GTACACTCTCTTCAGACACCAGAAGAGGGCATCA | | |
| 514 | 7 | 27170467 | 27170563 | 96 | 129S1,BTBR,C57L,ILNJ2,LP,NZW | 7 | 27170501 | 27170534 | 34 | - | 17.9675 | 3.61E-07 | 0.0545 | CTGAATTACAGTCAATCCCCTGCAGAGGGCACGG | | |
| 514 | 7 | 27170467 | 27170563 | 96 | 129S1,BTBR,C57L,ILNJ2,LP,NZW | 7 | 27170549 | 27170582 | 34 | - | 17.9675 | 3.61E-07 | 0.0545 | CTGAATTACAGTCAATCCCCTGCAGAGGGCACGG | | |
| 515 | 7 | 29225674 | 29225896 | 222 | CE,FVB,ILNJ2 | 7 | 29225801 | 29225834 | 34 | - | 17.8455 | 3.91E-07 | 0.057 | TAGCTGTCCTCAGACACTCCAGAAGAGGGCGTCA | | |
| 516 | 7 | 29564831 | 29564955 | 124 | AJ,AKR,C57L,DBA1J,DBA2J,MA/My,SMJ | 7 | 29564918 | 29564951 | 34 | - | 16.8537 | 7.43E-07 | 0.0809 | GAGCAGGTGCAGCAGGGGACAGCAGGTGAAGCAG | | |
| 517 | 7 | 29853542 | 29854538 | 996 | AJ,AKR,BALB,BTBR,C57L,DBA1J,DBA2J,SMJ | 7 | 29853659 | 29853677 | 19 | - | 20.3607 | 8.29E-08 | 0.042 | AGGCCAGAAGAGGGCACCA | 3 | ENCFF367UNR,ENCFF093ZRU,ENCFF052KVO |
| 517 | 7 | 29853542 | 29854538 | 996 | AJ,AKR,BALB,BTBR,C57L,DBA1J,DBA2J,SMJ | 7 | 29853659 | 29853693 | 35 | - | 25.8716 | 1.22E-09 | 0.00341 | CTGCATGCATGCCTACAGGCCAGAAGAGGGCACCA | 3 | ENCFF367UNR,ENCFF093ZRU,ENCFF052KVO |
| 518 | 7 | 29907594 | 29907805 | 211 | ILNJ2 | 7 | 29907719 | 29907752 | 34 | - | 17.439 | 5.10E-07 | 0.0663 | TAGCAGTCTTCAGACACTCCAGAAGAGGGCATCA | | |
| 519 | 7 | 29914917 | 29915122 | 205 | AJ,AKR,BALB,BTBR,C57L,DBA1J,DBA2J,SMJ | 7 | 29915043 | 29915061 | 19 | - | 17.6393 | 5.98E-07 | 0.106 | GCTCCAGAAGAGGGCGCCA | 3 | ENCFF093ZRU,ENCFF821BIA,ENCFF367UNR |
| 519 | 7 | 29914917 | 29915122 | 205 | AJ,AKR,BALB,BTBR,C57L,DBA1J,DBA2J,SMJ | 7 | 29915043 | 29915076 | 34 | - | 18.8374 | 2.01E-07 | 0.04 | TAGCTGTCTTCAGACGCTCCAGAAGAGGGCGCCA | 3 | ENCFF093ZRU,ENCFF821BIA,ENCFF367UNR |
| 520 | 7 | 33812219 | 33812433 | 214 | DBA1J,DBA2J,NZB,NZO,NZW,SMJ | 7 | 33812345 | 33812378 | 34 | - | 17.6992 | 4.30E-07 | 0.0601 | TAGCTGTCCTCAGATACTCCAGAAGAGGGCGTCA | | |
| 521 | 7 | 35708388 | 35708500 | 112 | NZB,NZO | 7 | 35708384 | 35708402 | 19 | - | 16.9508 | 9.36E-07 | 0.127 | GGAGCAGCAGGGGGCAGCA | | |
| 521 | 7 | 35708388 | 35708500 | 112 | NZB,NZO | 7 | 35708418 | 35708436 | 19 | - | 16.9508 | 9.36E-07 | 0.127 | GGAGCAGCAGGGGGCAGCA | | |
| 522 | 7 | 43205618 | 43206076 | 458 | C57L,CE,DBA1J,DBA2J,ILNJ2,NZB,NZO,NZW,PJ,SJL,SMJ | 7 | 43205744 | 43205777 | 34 | - | 16.4472 | 9.61E-07 | 0.0901 | TAGCTGTCTTCAGACACACCAGAAGAGGGCGTCA | | |
| 523 | 7 | 45193765 | 45194552 | 787 | C57L,C58J | 7 | 45194132 | 45194150 | 19 | - | 22.0656 | 1.91E-08 | 0.0243 | AGGCCACAAGGGGGCACCA | 15 | ENCFF821BIA,ENCFF024JOZ,ENCFF136ZDK,ENCFF907XEG,ENCFF589ZBJ,ENCFF055HAP,ENCFF093ZRU,ENCFF198SSR,ENCFF367UNR,ENCFF883UPM,ENCFF052KVO,ENCFF065ODV,ENCFF502WDT,ENCFF142CNG,ENCFF099QNE |
| 523 | 7 | 45193765 | 45194552 | 787 | C57L,C58J | 7 | 45194132 | 45194166 | 35 | - | 26.7706 | 5.73E-10 | 0.00258 | TTGCAGGTGCATGTGGAGGCCACAAGGGGGCACCA | 15 | ENCFF821BIA,ENCFF024JOZ,ENCFF136ZDK,ENCFF907XEG,ENCFF589ZBJ,ENCFF055HAP,ENCFF093ZRU,ENCFF198SSR,ENCFF367UNR,ENCFF883UPM,ENCFF052KVO,ENCFF065ODV,ENCFF502WDT,ENCFF142CNG,ENCFF099QNE |
| 524 | 7 | 45227428 | 45227631 | 203 | 129S1,C3H,DBA1J,DBA2J,ILNJ2,NZW,PJ,SJL | 7 | 45227553 | 45227571 | 19 | - | 17.5574 | 6.32E-07 | 0.106 | ACTCCAGAAGAGGGCGCCA | 4 | ENCFF367UNR,ENCFF099QNE,ENCFF821BIA,ENCFF093ZRU |
| 524 | 7 | 45227428 | 45227631 | 203 | 129S1,C3H,DBA1J,DBA2J,ILNJ2,NZW,PJ,SJL | 7 | 45227553 | 45227586 | 34 | - | 18.3659 | 2.76E-07 | 0.0476 | TAGCTGTCTTCAGACACTCCAGAAGAGGGCGCCA | 4 | ENCFF367UNR,ENCFF099QNE,ENCFF821BIA,ENCFF093ZRU |
| 525 | 7 | 46474406 | 46474608 | 202 | C57L,CE,ILNJ2,NZW,PJ,SJL | 7 | 46474518 | 46474551 | 34 | - | 17.9106 | 3.74E-07 | 0.0556 | TAGCTGTCTTCAGACACTCCAGAAGAGGGCACCA | 1 | ENCFF093ZRU |
| 526 | 7 | 46499239 | 46499436 | 197 | ILNJ2,NZW,PJ,SJL | 7 | 46499363 | 46499396 | 34 | - | 17.374 | 5.32E-07 | 0.0676 | TAGCTGTCTTCAGACCCTCCAGAAGAGGGCATCA | | |
| 527 | 7 | 46512888 | 46513752 | 864 | CE | 7 | 46513033 | 46513066 | 34 | - | 19.626 | 1.16E-07 | 0.0284 | ACGCAGTGCCCACGGAGGTCAGAAGAGGGCATCT | | |
| 527 | 7 | 46512888 | 46513752 | 864 | CE | 7 | 46513429 | 46513462 | 34 | - | 17.6423 | 4.47E-07 | 0.0616 | TAGCTGTGTTCAGACACACCAGAAGAGGGCGTCA | | |
| 528 | 7 | 51911701 | 51912095 | 394 | ILNJ2,KK,NZW,PJ | 7 | 51911707 | 51911725 | 19 | - | 17.7705 | 5.49E-07 | 0.101 | TGGCCAGTTGAGGGCGCAA | | |
| 528 | 7 | 51911701 | 51912095 | 394 | ILNJ2,KK,NZW,PJ | 7 | 51911707 | 51911740 | 34 | - | 18.935 | 1.88E-07 | 0.0387 | CAGCATCGTTCTTCCTGGCCAGTTGAGGGCGCAA | | |
| 529 | 7 | 53455492 | 53455862 | 370 | 129S1,AJ,AKR,BALB,BTBR,BUB,C3H,C58J,CBA,DBA1J,DBA2J,FVB,ILNJ2,KK,MRL,NOD,NOR,NZB,NZO,NZW,PJ,PL,RF,SEA,SJL,SMJ,SWR | 7 | 53455809 | 53455842 | 34 | - | 16.5285 | 9.13E-07 | 0.0896 | GAGCGGGTGTGGCTGCCACCCCAAGATGGCGCCC | | |
| 530 | 7 | 58110186 | 58110385 | 199 | 129S1,AKR,BALB,BTBR,C3H,C57L,CBA,DBA1J,DBA2J,ILNJ2,KK,LP,NOD,NOR,NZW,PJ,PL,RF,SEA,SMJ,TH | 7 | 58110310 | 58110328 | 19 | - | 17.5574 | 6.32E-07 | 0.106 | ACTCCAGAAGAGGGCGCCA | | |
| 530 | 7 | 58110186 | 58110385 | 199 | 129S1,AKR,BALB,BTBR,C3H,C57L,CBA,DBA1J,DBA2J,ILNJ2,KK,LP,NOD,NOR,NZW,PJ,PL,RF,SEA,SMJ,TH | 7 | 58110310 | 58110343 | 34 | - | 18.3659 | 2.76E-07 | 0.0476 | TAGCTGTCTTCAGACACTCCAGAAGAGGGCGCCA | | |
| 531 | 7 | 64923743 | 64923874 | 131 | CE,DBA1J,DBA2J,ILNJ2,KK,NZW,PJ | 7 | 64923748 | 64923782 | 35 | - | 16.6055 | 6.79E-07 | 0.125 | CAGCAGAGGGAGGGCGGGGTCAGCAGAGGGAGGGC | | |
| 531 | 7 | 64923743 | 64923874 | 131 | CE,DBA1J,DBA2J,ILNJ2,KK,NZW,PJ | 7 | 64923804 | 64923838 | 35 | - | 16.1835 | 8.62E-07 | 0.142 | CAGCAGAGGGAGGGCGGGGTCAGCAGAGGGAGGGT | | |
| 532 | 7 | 71892084 | 71892273 | 189 | ILNJ2,NZB,NZO | 7 | 71892204 | 71892237 | 34 | - | 16.9593 | 6.95E-07 | 0.0781 | TTGCTCTCTTCAGACACACCAGAAGAGGGCATCA | | |
| 533 | 7 | 72493682 | 72494411 | 729 | 129S1,AJ,AKR,BALB,BUB,C3H,C57L,C58J,CBA,CE,DBA1J,DBA2J,FVB,ILNJ2,KK,MRL,NOD,NOR,NZB,NZO,NZW,PJ,PL,RF,RH2,SEA,SJL,SMJ,SWR,TH | 7 | 72493679 | 72493713 | 35 | - | 22.1284 | 2.08E-08 | 0.0153 | CTGCCGCCACACCCGCTCCCCACAAGAGGGTACTA | | |
| 534 | 7 | 78898442 | 78898672 | 230 | 129S1,C3H,CBA,CE,DBA1J,DBA2J,KK,NZB,NZO,NZW,PJ,SEA,SMJ,SWR,TH | 7 | 78898570 | 78898603 | 34 | - | 17.8455 | 3.91E-07 | 0.057 | TAGCTGTCCTCAGACACTCCAGAAGAGGGCGTCA | | |
| 535 | 7 | 78995801 | 78996628 | 827 | C58J,CE | 7 | 78995851 | 78995884 | 34 | - | 19.1789 | 1.58E-07 | 0.0349 | ATGCAGTAAGCACTCTGAACAGATGGGGGCGCTG | 26 | ENCFF093ZRU,ENCFF367UNR,ENCFF099QNE,ENCFF201JUA,ENCFF951GOP,ENCFF321AGQ,ENCFF065ODV,ENCFF601JVE,ENCFF821BIA,ENCFF907XEG,ENCFF327LYS,ENCFF055HAP,ENCFF957SPH,ENCFF609MDN,ENCFF089GBC,ENCFF024JOZ,ENCFF589ZBJ,ENCFF198SSR,ENCFF052KVO,ENCFF502WDT,ENCFF281QMV,ENCFF092NMJ,ENCFF136ZDK,ENCFF142CNG,ENCFF430PPJ,ENCFF165LXO |
| 536 | 7 | 80217359 | 80217562 | 203 | AJ,AKR,BALB,BUB,CE,FVB,MRL,NZB,NZO,NZW,PL,RF,RH2,SJL,SWR | 7 | 80217484 | 80217502 | 19 | - | 17.5574 | 6.32E-07 | 0.106 | ACTCCAGAAGAGGGCGCCA | 1 | ENCFF093ZRU |
| 536 | 7 | 80217359 | 80217562 | 203 | AJ,AKR,BALB,BUB,CE,FVB,MRL,NZB,NZO,NZW,PL,RF,RH2,SJL,SWR | 7 | 80217484 | 80217517 | 34 | - | 18.3659 | 2.76E-07 | 0.0476 | TAGCTGTCTTCAGACACTCCAGAAGAGGGCGCCA | 1 | ENCFF093ZRU |
| 537 | 7 | 92828155 | 92828874 | 719 | 129S1,SMJ | 7 | 92828469 | 92828503 | 35 | - | 16.0367 | 9.36E-07 | 0.147 | ATGCACTGCCCCTGGGCCGCAAGAAGGGGGCCAAG | | |
| 538 | 7 | 98800622 | 98800686 | 64 | C3H,CBA,DBA1J,DBA2J,ILNJ2,PJ,SEA,SMJ | 7 | 98800614 | 98800632 | 19 | - | 22.918 | 8.34E-09 | 0.0179 | CAGCCACAAGGGGGCGCCG | 38 | ENCFF093ZRU,ENCFF201JUA,ENCFF327LYS,ENCFF907XEG,ENCFF821BIA,ENCFF024JOZ,ENCFF092NMJ,ENCFF065ODV,ENCFF589ZBJ,ENCFF089GBC,ENCFF136ZDK,ENCFF055HAP,ENCFF198SSR,ENCFF165LXO,ENCFF502WDT,ENCFF430PPJ,ENCFF321AGQ,ENCFF883UPM,ENCFF951GOP,ENCFF210BTO,ENCFF957SPH,ENCFF575MGI,ENCFF609MDN,ENCFF281QMV,ENCFF316FYN,ENCFF194HVL,ENCFF142CNG,ENCFF089EWX,ENCFF533APC,ENCFF097CAQ,ENCFF052KVO,ENCFF491RJK,ENCFF601JVE,ENCFF349BUL,ENCFF033UQX,ENCFF805QBU,ENCFF784ASD |
| 538 | 7 | 98800622 | 98800686 | 64 | C3H,CBA,DBA1J,DBA2J,ILNJ2,PJ,SEA,SMJ | 7 | 98800614 | 98800648 | 35 | - | 18.055 | 2.91E-07 | 0.0783 | TTGTCTGAAGGCCTCACAGCCACAAGGGGGCGCCG | 39 | ENCFF093ZRU,ENCFF201JUA,ENCFF327LYS,ENCFF907XEG,ENCFF821BIA,ENCFF024JOZ,ENCFF092NMJ,ENCFF065ODV,ENCFF589ZBJ,ENCFF089GBC,ENCFF136ZDK,ENCFF055HAP,ENCFF198SSR,ENCFF165LXO,ENCFF502WDT,ENCFF430PPJ,ENCFF321AGQ,ENCFF883UPM,ENCFF951GOP,ENCFF210BTO,ENCFF957SPH,ENCFF575MGI,ENCFF609MDN,ENCFF281QMV,ENCFF316FYN,ENCFF194HVL,ENCFF142CNG,ENCFF089EWX,ENCFF533APC,ENCFF097CAQ,ENCFF052KVO,ENCFF491RJK,ENCFF601JVE,ENCFF349BUL,ENCFF033UQX,ENCFF805QBU,ENCFF784ASD,ENCFF086ZTV |

| | | | | | | | | | | | | | | | | |
|---|---|---|---|---|---|---|---|---|---|---|---|---|---|---|---|---|
| 538 | 7 | 98800622 | 98800686 | 64 | C3H,CBA,DBA1J,DBA2J,ILNJ2,PJ,SEA,SMJ | 7 | 98800678 | 98800696 | 19 | - | 21.8689 | 2.29E-08 | 0.0265 | CAGCCACAAGAGGGCGCCG | 40 | ENCFF093ZRU,ENCFF201JUA,ENCFF327LYS,ENCFF907XEG,ENCFF821BIA,ENCFF024JOZ,ENCFF092NMJ,ENCFF065ODV,ENCFF589ZBJ,ENCFF089GBC,ENCFF136ZDK,ENCFF055HAP,ENCFF198SSR,ENCFF165LXO,ENCFF502WDT,ENCFF430PPJ,ENCFF321AGQ,ENCFF883UPM,ENCFF951GOP,ENCFF210BTO,ENCFF957SPH,ENCFF575MGI,ENCFF609MDN,ENCFF281QMV,ENCFF316FYN,ENCFF194HVL,ENCFF142CNG,ENCFF089EWX,ENCFF533APC,ENCFF097CAQ,ENCFF052KVO,ENCFF491RJK,ENCFF601JVE,ENCFF349BUL,ENCFF033UQX,ENCFF805QBU,ENCFF784ASD,ENCFF086ZTV,ENCFF611HDQ |
| 538 | 7 | 98800622 | 98800686 | 64 | C3H,CBA,DBA1J,DBA2J,ILNJ2,PJ,SEA,SMJ | 7 | 98800678 | 98800711 | 34 | - | 17.6585 | 4.42E-07 | 0.0612 | AAGGCCTCACAGCCACAGCCACAAGAGGGCGCCG | 41 | ENCFF093ZRU,ENCFF201JUA,ENCFF327LYS,ENCFF907XEG,ENCFF821BIA,ENCFF024JOZ,ENCFF092NMJ,ENCFF065ODV,ENCFF589ZBJ,ENCFF089GBC,ENCFF136ZDK,ENCFF055HAP,ENCFF198SSR,ENCFF165LXO,ENCFF502WDT,ENCFF430PPJ,ENCFF321AGQ,ENCFF883UPM,ENCFF951GOP,ENCFF210BTO,ENCFF957SPH,ENCFF575MGI,ENCFF609MDN,ENCFF281QMV,ENCFF316FYN,ENCFF194HVL,ENCFF142CNG,ENCFF089EWX,ENCFF533APC,ENCFF097CAQ,ENCFF052KVO,ENCFF491RJK,ENCFF601JVE,ENCFF349BUL,ENCFF033UQX,ENCFF805QBU,ENCFF784ASD,ENCFF086ZTV,ENCFF611HDQ,ENCFF504JDS |
| 539 | 7 | 102723536 | 102723653 | 117 | TH | 7 | 102723633 | 102723666 | 34 | - | 16.6016 | 8.72E-07 | 0.0874 | CAGCCCTGGTGCCCGCCGCCGGGCGGGGGCACAA | | |
| 540 | 7 | 105238105 | 105238324 | 219 | BUB,CE,FVB,KK,NOD,NZW,SJL,SWR,TH | 7 | 105238214 | 105238247 | 34 | - | 17.9431 | 3.66E-07 | 0.0552 | CAGCTGTCCTCAGACACTCCAGAAGAGGGAGTCA | | |
| 541 | 7 | 108889853 | 108889956 | 103 | NOD,SWR | 7 | 108889883 | 108889917 | 35 | - | 16.0183 | 9.45E-07 | 0.148 | AGACAGTCACAGCCGGCTTCCAGCAGATGGCAGGC | 11 | ENCFF092NMJ,ENCFF883UPM,ENCFF367UNR,ENCFF609MDN,ENCFF093ZRU,ENCFF281QMV,ENCFF099QNE,ENCFF327LYS,ENCFF089GBC,ENCFF024JOZ,ENCFF907XEG |
| 542 | 7 | 109427832 | 109428044 | 212 | ILNJ2,NOD,SMJ,SWR | 7 | 109427959 | 109427992 | 34 | - | 16.9187 | 7.13E-07 | 0.0792 | TAGCTGTCTTCAGACGCACCAGAAGAGGGCGTCA | | |
| 543 | 7 | 109805736 | 109805939 | 203 | CE,ILNJ2,KK,NOD,SMJ,SWR | 7 | 109805861 | 109805879 | 19 | - | 17.3443 | 7.26E-07 | 0.112 | GCACCAGAAGAGGGCAGTA | | |
| 543 | 7 | 109805736 | 109805939 | 203 | CE,ILNJ2,KK,NOD,SMJ,SWR | 7 | 109805861 | 109805894 | 34 | - | 17.7886 | 4.06E-07 | 0.0584 | TAGCTGTCTTCAGATGCACCAGAAGAGGGCAGTA | | |
| 544 | 7 | 112442121 | 112442342 | 221 | 129S1,AJ,AKR,BALB,BUB,C3H,C57L,CBA,DBA1J,DBA2J,ILNJ2,MRL,NOD,NOR,NZB,NZO,NZW,PL,RH2,SJL,SMJ | 7 | 112442248 | 112442266 | 19 | - | 17.8852 | 5.08E-07 | 0.0973 | ACACCAGAAGAGGGCACCA | 2 | ENCFF093ZRU |
| 545 | 7 | 126682650 | 126682860 | 210 | AJ,AKR,BALB,BUB,C58J,MRL,NOD,NZB,NZO,NZW,PL,RH2,SEA,SJL,SWR,TH | 7 | 126682778 | 126682811 | 34 | - | 16.5203 | 9.18E-07 | 0.0899 | TAGCTGTCTTCCAACACTCCAGAAGAGGGCGTCA | | |
| 546 | 7 | 127057828 | 127058048 | 220 | TH | 7 | 127057954 | 127057987 | 34 | - | 17.6504 | 4.44E-07 | 0.0613 | CAGCTGTCCTCAGACACCCCAGAAGAGGGAGTCA | | |
| 547 | 7 | 127087491 | 127087700 | 209 | AJ,NOD,TH | 7 | 127087614 | 127087632 | 19 | - | 17.8852 | 5.08E-07 | 0.0973 | ACACCAGAAGAGGGCACCA | 3 | ENCFF367UNR,ENCFF093ZRU |
| 548 | 7 | 128191426 | 128191630 | 204 | 129S1,AJ,AKR,BALB,BTBR,BUB,C3H,C57L,C58J,CE,FVB,ILNJ2,KK,LP,MA/My,MRL,NOD,NZB,NZO,NZW,PJ,PL,RF,RH2,SEA,SJL,SWR,TH | 7 | 128191551 | 128191569 | 19 | - | 17.5574 | 6.32E-07 | 0.106 | ACTCCAGAAGAGGGCGCCA | 6 | ENCFF093ZRU,ENCFF367UNR,ENCFF055HAP,ENCFF821BIA,ENCFF502WDT |
| 548 | 7 | 128191426 | 128191630 | 204 | 129S1,AJ,AKR,BALB,BTBR,BUB,C3H,C57L,C58J,CE,FVB,ILNJ2,KK,LP,MA/My,MRL,NOD,NZB,NZO,NZW,PJ,PL,RF,RH2,SEA,SJL,SWR,TH | 7 | 128191551 | 128191584 | 34 | - | 18.561 | 2.42E-07 | 0.0451 | AAGCTGTCTTCAGACACTCCAGAAGAGGGCGCCA | 6 | ENCFF093ZRU,ENCFF367UNR,ENCFF055HAP,ENCFF821BIA,ENCFF502WDT |
| 549 | 7 | 128285722 | 128285919 | 197 | 129S1,BTBR,C57L,ILNJ2,PJ | 7 | 128285848 | 128285866 | 19 | - | 16.9016 | 9.66E-07 | 0.13 | ATTCCAGAAGAGGGCACCA | | |
| 549 | 7 | 128285722 | 128285919 | 197 | 129S1,BTBR,C57L,ILNJ2,PJ | 7 | 128285848 | 128285881 | 34 | - | 17.8455 | 3.91E-07 | 0.057 | TAGCTGTCTTCAGACATTCCAGAAGAGGGCACCA | | |
| 550 | 7 | 135404046 | 135404266 | 220 | ILNJ2 | 7 | 135404170 | 135404203 | 34 | - | 16.7724 | 7.83E-07 | 0.0827 | TAGCTGTCTTCAGATGCACCAGAAGAGGGCGTCA | | |
| 551 | 7 | 136409524 | 136410451 | 927 | ILNJ2 | 7 | 136409885 | 136409918 | 34 | - | 26.5203 | 4.09E-10 | 0.000451 | ATGCAGTACCAAAGGAGGCCAGAAGAGGGCATCA | 6 | ENCFF327LYS,ENCFF093ZRU,ENCFF052KVO,ENCFF367UNR,ENCFF821BIA,ENCFF099QNE |
| 552 | 7 | 138622828 | 138622924 | 96 | CBA,NZB,NZO | 7 | 138622913 | 138622946 | 34 | - | 18.1382 | 3.22E-07 | 0.0511 | CTTCAGTGAGTGCACAGGGCAGCAGGGGGCGTGG | | |
| 553 | 7 | 139078982 | 139079219 | 237 | AKR,BUB | 7 | 139079106 | 139079139 | 34 | - | 17.9187 | 3.72E-07 | 0.0556 | TAGCTGTCCTCACATACTCCAGAAGAGGGCATCA | | |
| 554 | 7 | 139237731 | 139237787 | 56 | 129S1,AJ,AKR,BALB,BTBR,BUB,C3H,C57L,C58J,CBA,CE,DBA1J,DBA2J,FVB,ILNJ2,LP,MA/My,NOD,NOR,PJ,PL,RF,SEA,SJL,SMJ,TH | 7 | 139237734 | 139237767 | 34 | - | 16.6016 | 8.72E-07 | 0.0874 | GTGCCCTGCTGATGGTGCCCTGCTGATGGTGCCC | | |
| 554 | 7 | 139237731 | 139237787 | 56 | 129S1,AJ,AKR,BALB,BTBR,BUB,C3H,C57L,C58J,CBA,CE,DBA1J,DBA2J,FVB,ILNJ2,LP,MA/My,NOD,NOR,PJ,PL,RF,SEA,SJL,SMJ,TH | 7 | 139237748 | 139237781 | 34 | - | 16.6016 | 8.72E-07 | 0.0874 | GTGCCCTGCTGATGGTGCCCTGCTGATGGTGCCC | | |
| 554 | 7 | 139237731 | 139237787 | 56 | 129S1,AJ,AKR,BALB,BTBR,BUB,C3H,C57L,C58J,CBA,CE,DBA1J,DBA2J,FVB,ILNJ2,LP,MA/My,NOD,NOR,PJ,PL,RF,SEA,SJL,SMJ,TH | 7 | 139237762 | 139237795 | 34 | - | 16.6016 | 8.72E-07 | 0.0874 | GTGCCCTGCTGATGGTGCCCTGCTGATGGTGCCC | | |
| 555 | 7 | 140859812 | 140860016 | 204 | CE,ILNJ2,NZW | 7 | 140859938 | 140859971 | 34 | - | 16.4472 | 9.61E-07 | 0.0901 | TAGCTGTCTTCAGACACACCAGAAGAGGGCGTCA | | |
| 556 | 7 | 143080490 | 143080704 | 214 | C58J,LP,MA/My,PJ,SMJ | 7 | 143080616 | 143080649 | 34 | - | 17.3902 | 5.27E-07 | 0.0672 | TAGCTGTCCTCAGACACTCCAGAAGAGGGCATCA | | |
| 557 | 8 | 4432324 | 4432536 | 212 | 129S1,NZB,NZO,NZW | 8 | 4432394 | 4432427 | 34 | + | 18.374 | 2.75E-07 | 0.0529 | TAGCTGTCTTCAGTCACTCCAGAAGAGGGCGCCA | | |
| 557 | 8 | 4432324 | 4432536 | 212 | 129S1,NZB,NZO,NZW | 8 | 4432409 | 4432427 | 19 | + | 17.5574 | 6.32E-07 | 0.125 | ACTCCAGAAGAGGGCGCCA | | |
| 558 | 8 | 4648391 | 4648596 | 205 | 129S1,NZB | 8 | 4648453 | 4648486 | 34 | + | 18.3659 | 2.76E-07 | 0.0529 | TAGCTGTCTTCAGACACTCCAGAAGAGGGCGCCA | | |
| 558 | 8 | 4648391 | 4648596 | 205 | 129S1,NZB | 8 | 4648468 | 4648486 | 19 | + | 17.5574 | 6.32E-07 | 0.125 | ACTCCAGAAGAGGGCGCCA | | |
| 559 | 8 | 10079713 | 10079922 | 209 | NZB | 8 | 10079780 | 10079813 | 34 | + | 16.4472 | 9.61E-07 | 0.0944 | TAGCTGTCTTCAGACACACCAGAAGAGGGCGTCA | | |
| 560 | 8 | 14099335 | 14099536 | 201 | C57L,ILNJ2,NZB,NZW,PJ,PL,SEA,SMJ,TH | 8 | 14099396 | 14099429 | 34 | + | 17.0569 | 6.53E-07 | 0.0808 | TAGCTGTCCTCAGATACTCCAGAAGAGGGCATTA | | |
| 561 | 8 | 26612106 | 26612313 | 207 | PL | 8 | 26612162 | 26612195 | 34 | + | 19.878 | 9.69E-08 | 0.0283 | TTGCTGTCCTCAGACACTCCAGAAGAGGGCATCA | 7 | ENCFF093ZRU,ENCFF327LYS,ENCFF089GBC,ENCFF502WDT,ENCFF024JOZ,ENCFF136ZDK,ENCFF883UPM |
| 562 | 8 | 26774528 | 26774641 | 113 | 129S1,AJ,AKR,BALB,BTBR,BUB,C3H,C57L,CBA,DBA1J,DBA2J,FVB,ILNJ2,KK,MA/My,MRL,NOD,NOR,NZB,NZO,NZW,RF,RH2,SJL,SMJ,SWR,TH | 8 | 26774624 | 26774657 | 34 | + | 23.0325 | 8.79E-09 | 0.005 | ATGCAATACTCACAGAAGCCAGAAGATGGCATCA | | |
| 563 | 8 | 34256951 | 34257173 | 222 | 129S1,AJ,AKR,BALB,BUB,C3H,C57L,CBA,CE,DBA1J,DBA2J,MA/My,MRL,NOD,NOR,PL,RF,RH2,SJL,SMJ,SWR,TH | 8 | 34257031 | 34257064 | 34 | + | 17.8455 | 3.91E-07 | 0.0622 | TAGCTGTCCTCAGACACTCCAGAAGAGGGCGTCA | 2 | ENCFF093ZRU,ENCFF367UNR |
| 564 | 8 | 34529462 | 34530078 | 616 | NOD,NOR,SJL,TH | 8 | 34529891 | 34529924 | 34 | + | 19.2846 | 1.47E-07 | 0.0367 | TAGCTGTCTCCAGACACACCAGAAGAGGGCAGCA | 2 | ENCFF099QNE,ENCFF093ZRU |
| 564 | 8 | 34529462 | 34530078 | 616 | NOD,NOR,SJL,TH | 8 | 34529906 | 34529924 | 19 | + | 17.2459 | 7.74E-07 | 0.137 | ACACCAGAAGAGGGCAGCA | 2 | ENCFF099QNE,ENCFF093ZRU |
| 565 | 8 | 43846469 | 43846695 | 226 | 129S1,AJ,AKR,BALB,BTBR,BUB,C3H,C57L,CBA,CE,DBA1J,DBA2J,FVB,ILNJ2,KK,MA/My,MRL,NOD,NOR,NZB,NZO,NZW,PJ,PL,RF,RH2,SEA,SJL,SMJ,SWR,TH | 8 | 43846552 | 43846585 | 34 | + | 18.3659 | 2.76E-07 | 0.0529 | TAGCTGTCTTCAGACACTCCAGAAGAGGGCGCCA | 1 | ENCFF093ZRU |
| 565 | 8 | 43846469 | 43846695 | 226 | 129S1,AJ,AKR,BALB,BTBR,BUB,C3H,C57L,CBA,CE,DBA1J,DBA2J,FVB,ILNJ2,KK,MA/My,MRL,NOD,NOR,NZB,NZO,NZW,PJ,PL,RF,RH2,SEA,SJL,SMJ,SWR,TH | 8 | 43846567 | 43846585 | 19 | + | 17.5574 | 6.32E-07 | 0.125 | ACTCCAGAAGAGGGCGCCA | 1 | ENCFF093ZRU |
| 566 | 8 | 48050905 | 48051104 | 199 | 129S1,C58J,ILNJ2,LP | 8 | 48050963 | 48050996 | 34 | + | 16.4146 | 9.81E-07 | 0.0956 | TAGCTGTCTTCAGACACACCAGAGGAGGGCGCCA | | |
| 567 | 8 | 59124965 | 59125219 | 254 | NZB,NZO | 8 | 59125079 | 59125112 | 34 | + | 16.4634 | 9.51E-07 | 0.0944 | TAGCTGTCTTCAGACGCACCAGAAGAGGGCATCA | | |
| 568 | 8 | 69444367 | 69444574 | 207 | 129S1,AJ,AKR,BALB,BTBR,BUB,C3H,C57L,C58J,CBA,CE,DBA1J,DBA2J,FVB,ILNJ2,KK,MA/My,MRL,NOD,NOR,NZB,NZO,NZW,PJ,PL,RF,RH2,SEA,SJL,SMJ,SWR,TH | 8 | 69444432 | 69444465 | 34 | + | 18.3659 | 2.76E-07 | 0.0529 | TAGCTGTCTTCAGACACTCCAGAAGAGGGCGCCA | 1 | ENCFF367UNR |
| 568 | 8 | 69444367 | 69444574 | 207 | 129S1,AJ,AKR,BALB,BTBR,BUB,C3H,C57L,C58J,CBA,CE,DBA1J,DBA2J,FVB,ILNJ2,KK,MA/My,MRL,NOD,NOR,NZB,NZO,NZW,PJ,PL,RF,RH2,SEA,SJL,SMJ,SWR,TH | 8 | 69444447 | 69444465 | 19 | + | 17.5574 | 6.32E-07 | 0.125 | ACTCCAGAAGAGGGCGCCA | 1 | ENCFF367UNR |
| 569 | 8 | 71698163 | 71698369 | 206 | 129S1,AKR,BALB,BUB,C3H,C57L,C58J,CBA,CE,DBA1J,DBA2J,FVB,ILNJ2,KK,MA/My,MRL,NZB,NZO,NZW,PJ,PL,RF,SEA,SJL,SMJ,SWR,TH | 8 | 71698225 | 71698258 | 34 | + | 16.4472 | 9.61E-07 | 0.0944 | TAGCTGTCTTCAGACACACCAGAAGAGGGCGTCA | | |
| 570 | 8 | 85700092 | 85700161 | 69 | 129S1,AJ,AKR,BALB,BTBR,C3H,C57L,C58J,CBA,CE,DBA1J,DBA2J,FVB,ILNJ2,KK,MA/My,MRL,NOD,NOR,NZW,PJ,PL,RF,RH2,SEA,SJL,SMJ,SWR | 8 | 85700118 | 85700152 | 35 | + | 16.5229 | 7.12E-07 | 0.145 | CTATAGTTCCAGCATCCGGGCAGCAGAGGAAGGAA | | |
| 571 | 8 | 88633549 | 88633760 | 211 | 129S1,AJ,AKR,BALB,BTBR,BUB,C3H,C57L,C58J,CBA,CE,DBA1J,DBA2J,FVB,ILNJ2,KK,MA/My,MRL,NZB,NZO,PJ,PL,RH2,SEA,SMJ,SWR | 8 | 88633618 | 88633651 | 34 | + | 18.561 | 2.42E-07 | 0.0509 | AAGCTGTCTTCAGACACTCCAGAAGAGGGCGCCA | 4 | ENCFF367UNR,ENCFF093ZRU |
| 571 | 8 | 88633549 | 88633760 | 211 | 129S1,AJ,AKR,BALB,BTBR,BUB,C3H,C57L,C58J,CBA,CE,DBA1J,DBA2J,FVB,ILNJ2,KK,MA/My,MRL,NZB,NZO,PJ,PL,RH2,SEA,SMJ,SWR | 8 | 88633633 | 88633651 | 19 | + | 17.5574 | 6.32E-07 | 0.125 | ACTCCAGAAGAGGGCGCCA | 4 | ENCFF367UNR,ENCFF093ZRU |
| 572 | 8 | 88761326 | 88761539 | 213 | AJ,C3H,CBA,CE,DBA1J,DBA2J,FVB,ILNJ2,MA/My,NZB,NZO,RH2,SJL | 8 | 88761398 | 88761431 | 34 | + | 16.7724 | 7.83E-07 | 0.088 | TAGCTGTCTTCAGATGCACCAGAAGAGGGCGTCA | | |
| 573 | 8 | 89829641 | 89829842 | 201 | 129S1,AKR,BALB,BTBR,BUB,C57L,C58J,CBA,CE,DBA1J,DBA2J,FVB,ILNJ2,KK,MA/My,MRL,NOD,NOR,NZB,NZO,NZW,PJ,PL,RF,SEA,SJL,SMJ,SWR,TH | 8 | 89829700 | 89829733 | 34 | + | 18.4472 | 2.62E-07 | 0.0526 | TTGCTGTCTTCAGACACTCCAGAAGAGGGCGTCA | | |
| 574 | 8 | 91290876 | 91291105 | 229 | 129S1,AKR,BALB,BUB,C57L,C58J,CBA,CE,DBA1J,DBA2J,FVB,ILNJ2,MA/My,MRL,NOD,NOR,NZB,NZO,NZW,PJ,PL,RF,SEA,SJL,SMJ,SWR,TH | 8 | 91290964 | 91290997 | 34 | + | 17.2439 | 5.79E-07 | 0.0757 | TAGCTGTCCTCAGATACTCCAGAAGAGGGCATCA | | |
| 575 | 8 | 95566171 | 95566415 | 244 | 129S1,AJ,AKR,B10D2,B10J,BALB,BTBR,BUB,C3H,C57L,CBA,CE,DBA1J,DBA2J,FVB,ILNJ2,KK,LP,MA/My,MRL,NOD,NOR,NZB,NZO,NZW,PJ,PL,RF,RH2,SEA,SJL,SMJ,SWR,TH | 8 | 95566272 | 95566305 | 34 | + | 16.4634 | 9.51E-07 | 0.0944 | TAGCTGTCTTCAGACGCACCAGAAGAGGGCATCA | 1 | ENCFF093ZRU |
| 576 | 8 | 96325489 | 96325700 | 211 | B10D2,B10J,BUB,C3H,CBA,DBA1J,DBA2J,LP,NOD,NOR,SWR | 8 | 96325558 | 96325591 | 34 | + | 17.8455 | 3.91E-07 | 0.0622 | TAGCTGTCCTCAGACACTCCAGAAGAGGGCGTCA | 1 | ENCFF093ZRU |
| 577 | 8 | 107379043 | 107379253 | 210 | 129S1,AJ,AKR,B10D2,B10J,BALB,BTBR,BUB,C3H,C57L,C58J,CBA,CE,DBA1J,DBA2J,FVB,ILNJ2,KK,LP,MA/My,MRL,NOD,NOR,NZB,NZO,NZW,PJ,PL,RF,RH2,SEA,SJL,SMJ,SWR,TH | 8 | 107379112 | 107379145 | 34 | + | 16.7724 | 7.83E-07 | 0.088 | TAGCTGTCTTCAGATGCACCAGAAGAGGGCGTCA | | |
| 578 | 8 | 107409254 | 107409464 | 210 | 129S1,AJ,AKR,B10D2,B10J,BALB,BTBR,C3H,C57L,C58J,CBA,CE,DBA1J,DBA2J,ILNJ2,KK,LP,MA/My,MRL,NZB,NZO,NZW,RH2,SEA,SMJ,TH | 8 | 107409320 | 107409353 | 34 | + | 18.1545 | 3.18E-07 | 0.0563 | TAGCTGTTTTCAGACACACCAGAAGAGGGCGTCA | 1 | ENCFF093ZRU |
| 579 | 8 | 107686861 | 107687072 | 211 | PL,RH2 | 8 | 107686926 | 107686959 | 34 | + | 17.2114 | 5.91E-07 | 0.0762 | TAGCTGTTTTCAGACACTCCAGAAGAGGGCATCA | | |
| 580 | 8 | 120855037 | 120855268 | 231 | NZB | 8 | 120855100 | 120855133 | 34 | + | 18.3496 | 2.79E-07 | 0.0534 | TAGCTGTCCTCAGACGCACCAGAAGAGGGCATCA | | |
| 581 | 8 | 4322215 | 4322435 | 220 | 129S1,NZB | 8 | 4322342 | 4322375 | 34 | - | 17.374 | 5.32E-07 | 0.0732 | TAGCTGTCTTCAGACCCTCCAGAAGAGGGCATCA | 1 | ENCFF093ZRU |
| 582 | 8 | 8797418 | 8797637 | 219 | NZB | 8 | 8797544 | 8797577 | 34 | - | 17.8455 | 3.91E-07 | 0.0622 | TAGCTGTCCTCAGACACTCCAGAAGAGGGCGTCA | | |
| 583 | 8 | 11594289 | 11594501 | 212 | 129S1,BTBR,C57L,ILNJ2,NZB,NZO,NZW,SMJ,TH | 8 | 11594414 | 11594447 | 34 | - | 17.2439 | 5.79E-07 | 0.0757 | TAGCTGTCCTCAGATACTCCAGAAGAGGGCATCA | 2 | ENCFF367UNR,ENCFF093ZRU |
| 584 | 8 | 13201984 | 13202201 | 217 | 129S1,C57L,CE,ILNJ2,LP,NZB,NZO,NZW,PJ,PL,SEA,SMJ,TH | 8 | 13202109 | 13202142 | 34 | - | 16.4634 | 9.51E-07 | 0.0944 | TAGCTGTCTTCAGACGCACCAGAAGAGGGCATCA | | |
| 585 | 8 | 23028135 | 23028189 | 54 | C57L,CE,ILNJ2,LP,NZB,PJ,SEA | 8 | 23028173 | 23028207 | 35 | - | 17.6697 | 3.66E-07 | 0.106 | GAGCACTGAGAGTGTCAGGCCGGCAGGGAGCACTG | | |
| 586 | 8 | 34254944 | 34255029 | 85 | 129S1,AJ,AKR,BALB,BUB,C3H,C57L,CBA,CE,DBA1J,DBA2J,MA/My,MRL,NOD,NOR,PL,RF,RH2,SJL,SMJ,SWR,TH | 8 | 34254918 | 34254951 | 34 | - | 18.8537 | 1.98E-07 | 0.0448 | TAGCGGCTACCGCCGCTGCCCCAAGATGGCACCC | | |
| 587 | 8 | 40042632 | 40043367 | 735 | 129S1,AJ,AKR,BALB,BTBR,BUB,C3H,C57L,CBA,CE,DBA1J,DBA2J,FVB,ILNJ2,KK,MA/My,MRL,NOD,NOR,NZB,NZO,NZW,PJ,PL,RF,RH2,SEA,SJL,SMJ,SWR,TH | 8 | 40042887 | 40042905 | 19 | - | 19.7377 | 1.35E-07 | 0.0604 | TGACCAGTAGGTGGCGAAA | 8 | ENCFF024JOZ,ENCFF093ZRU,ENCFF367UNR,ENCFF883UPM,ENCFF099QNE,ENCFF907XEG,ENCFF821BIA,ENCFF089GBC |
| 587 | 8 | 40042632 | 40043367 | 735 | 129S1,AJ,AKR,BALB,BTBR,BUB,C3H,C57L,CBA,CE,DBA1J,DBA2J,FVB,ILNJ2,KK,MA/My,MRL,NOD,NOR,NZB,NZO,NZW,PJ,PL,RF,RH2,SEA,SJL,SMJ,SWR,TH | 8 | 40042887 | 40042920 | 34 | - | 22.5285 | 1.32E-08 | 0.00669 | TTGCAGTTAAACTCTTGACCAGTAGGTGGCGAAA | 8 | ENCFF024JOZ,ENCFF093ZRU,ENCFF367UNR,ENCFF883UPM,ENCFF099QNE,ENCFF907XEG,ENCFF821BIA,ENCFF089GBC |
| 588 | 8 | 52818217 | 52818602 | 385 | B10D2,B10J,C58J,LP | 8 | 52818273 | 52818291 | 19 | - | 17.8852 | 5.08E-07 | 0.113 | CTGCCACCAGGGGGGGGCA | | |
| 589 | 8 | 58014846 | 58015045 | 199 | NZB,NZO | 8 | 58014971 | 58015004 | 34 | - | 16.4472 | 9.61E-07 | 0.0944 | TAGCTGTCTTCAGACACACCAGAAGAGGGCGTCA | | |
| 590 | 8 | 58092087 | 58092222 | 135 | NZB,NZO | 8 | 58092153 | 58092187 | 35 | - | 16.5046 | 7.19E-07 | 0.145 | CTGCAGAAGCAAGAGCCAGCCAGCAGGAGACTGGC | | |
| 591 | 8 | 63097579 | 63097788 | 209 | CE,KK | 8 | 63097707 | 63097725 | 19 | - | 17.5574 | 6.32E-07 | 0.125 | ACTCCAGAAGAGGGCGCCA | | |
| 591 | 8 | 63097579 | 63097788 | 209 | CE,KK | 8 | 63097707 | 63097740 | 34 | - | 18.3659 | 2.76E-07 | 0.0529 | TAGCTGTCTTCAGACACTCCAGAAGAGGGCGCCA | | |
| 592 | 8 | 72788991 | 72789812 | 821 | 129S1,AKR,BALB,BUB,C3H,C58J,CBA,DBA1J,DBA2J,FVB,ILNJ2,KK,NZB,NZO,NZW,RF,SJL,SMJ,SWR,TH | 8 | 72789174 | 72789192 | 19 | - | 18.5574 | 3.20E-07 | 0.0916 | CCTCCAGTAGAGGGCGCAC | 4 | ENCFF099QNE,ENCFF821BIA,ENCFF093ZRU,ENCFF367UNR |
| 592 | 8 | 72788991 | 72789812 | 821 | 129S1,AKR,BALB,BUB,C3H,C58J,CBA,DBA1J,DBA2J,FVB,ILNJ2,KK,NZB,NZO,NZW,RF,SJL,SMJ,SWR,TH | 8 | 72789174 | 72789208 | 35 | - | 18.2844 | 2.54E-07 | 0.0885 | ATGAGGTTTTTTCACCCCTCCAGTAGAGGGCGCAC | 4 | ENCFF099QNE,ENCFF821BIA,ENCFF093ZRU,ENCFF367UNR |
| 593 | 8 | 73031870 | 73032119 | 249 | 129S1,AKR,BALB,BUB,C3H,C57L,C58J,CBA,CE,DBA1J,DBA2J,FVB,ILNJ2,KK,MA/My,MRL,NZB,NZO,NZW,PJ,PL,RF,SEA,SJL,SMJ,SWR,TH | 8 | 73031982 | 73032015 | 34 | - | 16.4472 | 9.61E-07 | 0.0944 | TAGCTGTCTTCAGACACACCAGAAGAGGGCGTCA | | |
| 594 | 8 | 73052033 | 73052238 | 205 | 129S1,AKR,BALB,BUB,C3H,C57L,C58J,CBA,CE,DBA1J,DBA2J,FVB,ILNJ2,KK,MA/My,MRL,NZB,NZO,NZW,PJ,PL,RF,SEA,SJL,SMJ,SWR,TH | 8 | 73052156 | 73052189 | 34 | - | 17.2439 | 5.79E-07 | 0.0757 | TAGCTGTCCTCAGATACTCCAGAAGAGGGCATCA | | |
| 595 | 8 | 78623052 | 78623236 | 184 | C3H,C57L,KK,MA/My,NOD,NOR,NZB,NZO,PL,RF,SJL,TH | 8 | 78623166 | 78623199 | 34 | - | 16.4472 | 9.61E-07 | 0.0944 | TAGCTGTCTTCAGACACACCAGAAGAGGGCGTCA | | |
| 596 | 8 | 81536669 | 81536869 | 200 | C58J,SWR,TH | 8 | 81536794 | 81536827 | 34 | - | 17.2439 | 5.79E-07 | 0.0757 | TAGCTGTCCTCAGATACTCCAGAAGAGGGCATCA | | |
| 597 | 8 | 84734707 | 84734900 | 193 | 129S1,AJ,AKR,BALB,BTBR,C3H,C57L,C58J,CBA,CE,DBA1J,DBA2J,ILNJ2,KK,MA/My,MRL,NOD,NOR,NZW,PJ,PL,RF,RH2,SEA,SJL,SWR,TH | 8 | 84734829 | 84734862 | 34 | - | 17.2439 | 5.79E-07 | 0.0757 | TAGCTGTCCTCAGATACTCCAGAAGAGGGCATCA | | |
| 598 | 8 | 85839195 | 85839420 | 225 | 129S1,AJ,AKR,BALB,BTBR,C3H,C57L,C58J,CBA,CE,DBA1J,DBA2J,FVB,ILNJ2,KK,MA/My,MRL,NOD,NOR,NZW,PJ,PL,RF,RH2,SEA,SJL,SMJ,SWR | 8 | 85839322 | 85839355 | 34 | - | 16.4634 | 9.51E-07 | 0.0944 | TAGCTGTCTTCAGACGCACCAGAAGAGGGCATCA | | |
| 599 | 8 | 85875459 | 85875669 | 210 | 129S1,AJ,AKR,BALB,BTBR,C3H,C57L,C58J,CBA,CE,DBA1J,DBA2J,FVB,ILNJ2,KK,MA/My,MRL,NOD,NOR,NZW,PJ,PL,RF,RH2,SEA,SJL,SMJ,SWR | 8 | 85875584 | 85875617 | 34 | - | 16.4472 | 9.61E-07 | 0.0944 | TAGCTGTCTTCAGACACACCAGAAGAGGGCGTCA | | |
| 600 | 8 | 88704951 | 88705149 | 198 | AJ,BUB,C3H,C58J,RH2,SWR | 8 | 88705070 | 88705103 | 34 | - | 17.2439 | 5.79E-07 | 0.0757 | TAGCTGTCCTCAGATACTCCAGAAGAGGGCATCA | | |
| 601 | 8 | 88835742 | 88836237 | 495 | AJ,C3H,RH2 | 8 | 88836044 | 88836077 | 34 | - | 23.6341 | 5.36E-09 | 0.00345 | TTGCAGTGTTCCAGGAAGCCAGAAGAGGGCATCA | 3 | ENCFF367UNR,ENCFF052KVO,ENCFF093ZRU |
| 602 | 8 | 91354220 | 91354440 | 220 | BALB,CE,FVB,ILNJ2,MRL,NZB,NZO,NZW,PJ,SEA,TH | 8 | 91354335 | 91354353 | 19 | - | 17.5574 | 6.32E-07 | 0.125 | ACTCCAGAAGAGGGCGCCA | | |
| 602 | 8 | 91354220 | 91354440 | 220 | BALB,CE,FVB,ILNJ2,MRL,NZB,NZO,NZW,PJ,SEA,TH | 8 | 91354335 | 91354368 | 34 | - | 16.4228 | 9.76E-07 | 0.0953 | TAGCTGCCTTCAGACACTCCAGAAGAGGGCGCCA | | |
| 603 | 8 | 92428629 | 92428837 | 208 | 129S1,AKR,BALB,BUB,C57L,C58J,CBA,CE,DBA1J,DBA2J,FVB,ILNJ2,MA/My,MRL,NOD,NOR,NZB,NZO,NZW,PJ,PL,RF,SEA,SJL,SMJ,SWR,TH | 8 | 92428747 | 92428780 | 34 | - | 16.5447 | 9.04E-07 | 0.0944 | CAGCTGTCTTCAGACACACCAGAAGAGGGAGTCA | | |
| 604 | 8 | 105862123 | 105862340 | 217 | KK,PJ | 8 | 105862249 | 105862282 | 34 | - | 16.4472 | 9.61E-07 | 0.0944 | TAGCTGTCTTCAGACACACCAGAAGAGGGCGTCA | | |
| 605 | 8 | 107383846 | 107383989 | 143 | 129S1,AJ,AKR,B10D2,B10J,BALB,BTBR,BUB,C3H,C57L,C58J,CBA,CE,DBA1J,DBA2J,FVB,ILNJ2,KK,LP,MA/My,MRL,NOD,NOR,NZB,NZO,NZW,PJ,PL,RF,RH2,SEA,SJL,SMJ,SWR,TH | 8 | 107383820 | 107383854 | 35 | - | 17.211 | 4.79E-07 | 0.124 | TAGTAGATGCCCTTAAAGGCCAGAAGAGGGCATCA | 1 | ENCFF327LYS |
| 605 | 8 | 107383846 | 107383989 | 143 | 129S1,AJ,AKR,B10D2,B10J,BALB,BTBR,BUB,C3H,C57L,C58J,CBA,CE,DBA1J,DBA2J,FVB,ILNJ2,KK,LP,MA/My,MRL,NOD,NOR,NZB,NZO,NZW,PJ,PL,RF,RH2,SEA,SJL,SMJ,SWR,TH | 8 | 107383932 | 107383966 | 35 | - | 17.211 | 4.79E-07 | 0.124 | TAGTAGATGCCCTTAAAGGCCAGAAGAGGGCATCA | 1 | ENCFF327LYS |
| 606 | 8 | 112203061 | 112203481 | 420 | BALB,NZW,SEA | 8 | 112203185 | 112203219 | 35 | - | 18.1284 | 2.79E-07 | 0.0931 | CTGTAGCTGTCTTCAGACACCAGAAGAGGGCATCA | | |
| 607 | 8 | 114782557 | 114782781 | 224 | MA/My | 8 | 114782678 | 114782711 | 34 | - | 16.4472 | 9.61E-07 | 0.0944 | TAGCTGTCTTCAGACACACCAGAAGAGGGCGTCA | | |
| 608 | 8 | 122534540 | 122535391 | 851 | AJ,AKR,BALB,C3H,C58J,CE,ILNJ2,LP,MRL,NZB,PL,RF,RH2,SEA,SMJ | 8 | 122534969 | 122535002 | 34 | - | 22.1382 | 1.79E-08 | 0.0084 | ATGCGGTGTCTGTGGAAGCCAGAAGAGGGCGTCA | 5 | ENCFF093ZRU,ENCFF052KVO,ENCFF198SSR,ENCFF883UPM,ENCFF533APC |
| 609 | 8 | 124042117 | 124042289 | 172 | 129S1,AKR,BTBR,C3H,C57L,C58J,CBA,CE,DBA1J,DBA2J,LP,MA/My,MRL,PJ,RF,SMJ,TH | 8 | 124042193 | 124042226 | 34 | - | 17.2439 | 5.79E-07 | 0.0757 | TAGCTGTCCTCAGATACTCCAGAAGAGGGCATCA | 1 | ENCFF093ZRU |
| 610 | 8 | 125451361 | 125451567 | 206 | 129S1,AJ,BALB,BTBR,C57L,ILNJ2,NZB,RH2,SEA | 8 | 125451487 | 125451520 | 34 | - | 17.2439 | 5.79E-07 | 0.0757 | TAGCTGTCCTCAGATACTCCAGAAGAGGGCATCA | | |
| 611 | 9 | 21399770 | 21399984 | 214 | BTBR,C57L,C58J,FVB,MA/My,NOD,NOR,NZB,NZW,RF,SJL,SWR | 9 | 21399842 | 21399875 | 34 | + | 18.3659 | 2.76E-07 | 0.0438 | TAGCTGTCTTCAGACACTCCAGAAGAGGGCGCCA | 1 | ENCFF055HAP |
| 611 | 9 | 21399770 | 21399984 | 214 | BTBR,C57L,C58J,FVB,MA/My,NOD,NOR,NZB,NZW,RF,SJL,SWR | 9 | 21399857 | 21399875 | 19 | + | 17.5574 | 6.32E-07 | 0.112 | ACTCCAGAAGAGGGCGCCA | 1 | ENCFF055HAP |
| 612 | 9 | 22096827 | 22097557 | 730 | C57L,FVB,MA/My,NOD,NOR,NZB,NZW,RF,SJL,SWR | 9 | 22097012 | 22097030 | 19 | + | 18.5902 | 3.13E-07 | 0.0834 | CGGCCAGCAGATGTCAGCC | | |
| 613 | 9 | 28175690 | 28176147 | 457 | B10J | 9 | 28175963 | 28175996 | 34 | + | 17.3415 | 5.44E-07 | 0.0631 | CTGCTACTCCTGCAGCCCCCTGGAGGTAGTGGTC | | |
| 614 | 9 | 31329254 | 31329461 | 207 | CE,NZW | 9 | 31329319 | 31329352 | 34 | + | 16.4472 | 9.61E-07 | 0.0841 | TAGCTGTCTTCAGACACACCAGAAGAGGGCGTCA | | |
| 615 | 9 | 44068437 | 44068674 | 237 | 129S1,C57L,ILNJ2,SMJ | 9 | 44068532 | 44068565 | 34 | + | 17.8455 | 3.91E-07 | 0.0525 | TAGCTGTCCTCAGACACTCCAGAAGAGGGCGTCA | 1 | ENCFF093ZRU |
| 616 | 9 | 44269791 | 44270004 | 213 | 129S1,C57L | 9 | 44269862 | 44269895 | 34 | + | 16.4472 | 9.61E-07 | 0.0841 | TAGCTGTCTTCAGACACACCAGAAGAGGGCGTCA | | |
| 617 | 9 | 57241347 | 57242106 | 759 | CBA | 9 | 57241760 | 57241794 | 35 | + | 16.0367 | 9.36E-07 | 0.152 | ATGCACTGCCCCTGGGCCGCAAGAAGGGGGCCAAG | | |
| 618 | 9 | 57768529 | 57768763 | 234 | 129S1,C57L,MA/My,PL | 9 | 57768620 | 57768653 | 34 | + | 18.3659 | 2.76E-07 | 0.0438 | TAGCTGTCTTCAGACACTCCAGAAGAGGGCGCCA | | |
| 618 | 9 | 57768529 | 57768763 | 234 | 129S1,C57L,MA/My,PL | 9 | 57768635 | 57768653 | 19 | + | 17.5574 | 6.32E-07 | 0.112 | ACTCCAGAAGAGGGCGCCA | | |

| | | | | | | | | | | | | | | | | |
|---|---|---|---|---|---|---|---|---|---|---|---|---|---|---|---|---|
| 619 | 9 | 58170482 | 58170725 | 243 | 129S1,AJ,AKR,BTBR,BUB,C3H,C57L,CBA,CE,DBA1J,DBA2J,FVB,ILNJ2,KK,MA/My,MRL,NOD,NOR,NZB,NZO,NZW,PJ,PL,RF,RH2,SEA,SJL,SMJ,SWR,TH | 9 | 58170583 | 58170616 | 34 | + | 16.4472 | 9.61E-07 | 0.0841 | TAGCTGTCTTCAGACACACCAGAAGAGGGCGTCA | | |
| 620 | 9 | 62633509 | 62633735 | 226 | AJ,AKR,C3H,FVB,NOD,NOR,NZB,NZO,NZW,PL,RF,RH2,SJL,SWR,TH | 9 | 62633593 | 62633626 | 34 | + | 17.9431 | 3.66E-07 | 0.0511 | CAGCTGTCCTCAGACACTCCAGAAGAGGGAGTCA | | |
| 621 | 9 | 62835852 | 62836061 | 209 | AKR,CE,NZB,RF | 9 | 62835920 | 62835953 | 34 | + | 16.7724 | 7.83E-07 | 0.0768 | TAGCTGTCTTCAGATGCACCAGAAGAGGGCGTCA | | |
| 622 | 9 | 64847313 | 64847521 | 208 | CE,FVB,SJL | 9 | 64847380 | 64847413 | 34 | + | 18.6098 | 2.34E-07 | 0.0417 | TTGCTGTCTTCAGATGAACCAGAAGAGGGCATCA | 5 | ENCFF099QNE,ENCFF883UPM,ENCFF502WDT,ENCFF093ZRU,ENCFF055HAP |
| 623 | 9 | 65211259 | 65212206 | 947 | 129S1,BTBR,C57L,SMJ | 9 | 65212162 | 65212195 | 34 | + | 22.4878 | 1.36E-08 | 0.0055 | GTGCAGTCCCCACAAAACCCTGAAGAGGGCATTG | 2 | ENCFF367UNR,ENCFF093ZRU |
| 624 | 9 | 65275146 | 65275237 | 91 | FVB,SJL | 9 | 65275191 | 65275224 | 34 | + | 17.2439 | 5.79E-07 | 0.065 | TAGCTGTCCTCAGATACTCCAGAAGAGGGCATCA | 1 | ENCFF093ZRU |
| 625 | 9 | 65612376 | 65612632 | 256 | 129S1,AJ,AKR,B10D2,B10J,BALB,BTBR,BUB,C3H,C57L,CE,DBA1J,DBA2J,FVB,ILNJ2,KK,LP,MA/My,MRL,NOD,NOR,NZB,NZO,NZW,PJ,PL,RF,RH2,SEA,SJL,SMJ,SWR,TH | 9 | 65612490 | 65612523 | 34 | + | 16.4634 | 9.51E-07 | 0.0841 | TAGCTGTCTTCAGACGCACCAGAAGAGGGCATCA | | |
| 626 | 9 | 68755999 | 68756235 | 236 | 129S1,B10D2,B10J,BTBR,C57L,SMJ | 9 | 68756093 | 68756126 | 34 | + | 16.4634 | 9.51E-07 | 0.0841 | TAGCTGTCTTCAGACGCACCAGAAGAGGGCATCA | | |
| 627 | 9 | 72121537 | 72121742 | 205 | AJ,BALB,BUB,C3H,CBA,CE,DBA1J,DBA2J,FVB,ILNJ2,KK,MA/My,NOD,NOR,NZB,NZO,NZW,PJ,RH2,SEA,SJL,SMJ,SWR | 9 | 72121601 | 72121635 | 35 | + | 15.9633 | 9.75E-07 | 0.154 | CTGTAGCTGTCTTCAGACACCAGAAGAGGGTGACA | | |
| 628 | 9 | 72686700 | 72686924 | 224 | 129S1,AJ,AKR,BALB,BTBR,BUB,C3H,C57L,CBA,CE,DBA1J,DBA2J,FVB,ILNJ2,KK,MA/My,MRL,NOD,NOR,NZB,NZO,NZW,PJ,PL,RF,RH2,SEA,SJL,SMJ,SWR,TH | 9 | 72686785 | 72686818 | 34 | + | 17.9106 | 3.74E-07 | 0.0517 | TAGCTGTCTTCAGACACTCCAGAAGAGGGCACCA | | |
| 629 | 9 | 75354075 | 75354297 | 222 | AJ,AKR,BALB,BTBR,BUB,C3H,C58J,CBA,CE,DBA1J,DBA2J,FVB,ILNJ2,KK,LP,MA/My,MRL,NOD,NOR,NZB,NZO,NZW,PJ,PL,RF,RH2,SEA,SJL,SMJ,SWR,TH | 9 | 75354155 | 75354188 | 34 | + | 16.4797 | 9.42E-07 | 0.0841 | TAGCTGTCTTCAGACACTCCACAAGAGGGCGTCA | 1 | ENCFF093ZRU |
| 630 | 9 | 81459294 | 81459498 | 204 | AKR,BUB,C58J,CBA,ILNJ2,LP,MRL,NOD,NOR,NZW,PL,RF,SMJ | 9 | 81459356 | 81459389 | 34 | + | 16.4472 | 9.61E-07 | 0.0841 | TAGCTGTCTTCAGACACACCAGAAGAGGGCGTCA | | |
| 631 | 9 | 81743274 | 81743482 | 208 | AKR,BUB,C58J,CBA,ILNJ2,LP,MRL,NOD,NOR,NZW,PL,RF,SMJ | 9 | 81743340 | 81743373 | 34 | + | 17.8455 | 3.91E-07 | 0.0525 | TAGCTGTCCTCAGACACTCCAGAAGAGGGCGTCA | | |
| 632 | 9 | 84254494 | 84254590 | 96 | FVB,KK,NZO,NZW,SJL,SWR | 9 | 84254549 | 84254582 | 34 | + | 18.3659 | 2.76E-07 | 0.0438 | TAGCTGTCTTCAGACACTCCAGAAGAGGGCGCCA | 1 | ENCFF093ZRU |
| 632 | 9 | 84254494 | 84254590 | 96 | FVB,KK,NZO,NZW,SJL,SWR | 9 | 84254564 | 84254582 | 19 | + | 17.5574 | 6.32E-07 | 0.112 | ACTCCAGAAGAGGGCGCCA | 1 | ENCFF093ZRU |
| 633 | 9 | 89963839 | 89964035 | 196 | MRL,PJ,SEA,SMJ | 9 | 89963893 | 89963926 | 34 | + | 17.878 | 3.83E-07 | 0.0522 | TAGCTGTCCTCAGACACACCAGAAGAGGGCATCA | | |
| 634 | 9 | 90050101 | 90050295 | 194 | PJ,SEA | 9 | 90050153 | 90050186 | 34 | + | 18.3659 | 2.76E-07 | 0.0438 | TAGCTGTCTTCAGACACTCCAGAAGAGGGCGCCA | | |
| 634 | 9 | 90050101 | 90050295 | 194 | PJ,SEA | 9 | 90050168 | 90050186 | 19 | + | 17.5574 | 6.32E-07 | 0.112 | ACTCCAGAAGAGGGCGCCA | | |
| 635 | 9 | 96639537 | 96639627 | 90 | MRL,NZO,NZW | 9 | 96639549 | 96639582 | 34 | + | 22.9268 | 9.57E-09 | 0.0042 | GTGCAGGGCCCACAGAGGCCAGAAGAGGGTGTAG | | |
| 636 | 9 | 97047075 | 97047274 | 199 | MRL,NZO,NZW | 9 | 97047125 | 97047158 | 34 | + | 16.4472 | 9.61E-07 | 0.0841 | TAGCTGTCTTCAGACACACCAGAAGAGGGCGTCA | | |
| 637 | 9 | 103623033 | 103623964 | 931 | CBA | 9 | 103623482 | 103623516 | 35 | + | 16.0734 | 9.17E-07 | 0.15 | CAGCAATGTGGACATCAGCTCTCCAGAGGGGACTC | | |
| 638 | 9 | 104012716 | 104012930 | 214 | 129S1,AJ,AKR,BALB,BTBR,BUB,C3H,C57L,CBA,CE,DBA1J,DBA2J,FVB,ILNJ2,KK,MA/My,MRL,NOD,NOR,NZB,NZO,NZW,PJ,PL,RF,RH2,SEA,SJL,SMJ,SWR,TH | 9 | 104012787 | 104012820 | 34 | + | 18.3659 | 2.76E-07 | 0.0438 | TAGCTGTCTTCAGACACTCCAGAAGAGGGCGCCA | | |
| 638 | 9 | 104012716 | 104012930 | 214 | 129S1,AJ,AKR,BALB,BTBR,BUB,C3H,C57L,CBA,CE,DBA1J,DBA2J,FVB,ILNJ2,KK,MA/My,MRL,NOD,NOR,NZB,NZO,NZW,PJ,PL,RF,RH2,SEA,SJL,SMJ,SWR,TH | 9 | 104012802 | 104012820 | 19 | + | 17.5574 | 6.32E-07 | 0.112 | ACTCCAGAAGAGGGCGCCA | | |
| 639 | 9 | 106794248 | 106794448 | 200 | 129S1,AJ,AKR,BALB,BTBR,BUB,C3H,C57L,CE,FVB,ILNJ2,MA/My,MRL,NOD,NOR,NZB,NZO,NZW,PJ,PL,RF,RH2,SEA,SJL,SMJ,SWR,TH | 9 | 106794305 | 106794338 | 34 | + | 16.5691 | 8.90E-07 | 0.0831 | TAGCTGTCTTCAGACGCACCAGAAGAGGGCATCG | | |
| 640 | 9 | 108164987 | 108165192 | 205 | CBA | 9 | 108165050 | 108165083 | 34 | + | 17.9431 | 3.66E-07 | 0.0511 | CAGCTGTCCTCAGACACTCCAGAAGAGGGAGTCA | | |
| 641 | 9 | 109977312 | 109977518 | 206 | CBA | 9 | 109977376 | 109977409 | 34 | + | 16.4472 | 9.61E-07 | 0.0841 | TAGCTGTCTTCAGACACACCAGAAGAGGGCGTCA | | |
| 642 | 9 | 110130184 | 110130389 | 205 | CBA,NZW | 9 | 110130244 | 110130277 | 34 | + | 17.8455 | 3.91E-07 | 0.0525 | TAGCTGTCCTCAGACACTCCAGAAGAGGGCGTCA | | |
| 643 | 9 | 114575098 | 114575228 | 130 | AJ,BALB,C3H,CE,FVB,KK,MRL,PJ,PL,SEA | 9 | 114575094 | 114575128 | 35 | + | 17.4312 | 4.21E-07 | 0.0981 | CTGCTCTTCCAGAGGACCCACATCAGATGGCTCTG | | |
| 644 | 9 | 120405396 | 120405685 | 289 | BTBR,MRL | 9 | 120405517 | 120405550 | 34 | + | 18.0976 | 3.31E-07 | 0.0481 | GTGCACCACCATTGTCCGCCTCCAGGGGACTTCA | 11 | ENCFF099QNE,ENCFF052KVO,ENCFF165LXO,ENCFF092NMJ,ENCFF198SSR,ENCFF327LYS,ENCFF589ZBJ,ENCFF089GBC,ENCFF430PPJ,ENCFF093ZRU,ENCFF136ZDK |
| 645 | 9 | 121473629 | 121473707 | 78 | AKR,BUB,C58J,CBA,CE,DBA1J,DBA2J,MRL,NZW,RF,RH2,SMJ | 9 | 121473602 | 121473635 | 34 | + | 18.3333 | 2.82E-07 | 0.0442 | CTGCTCTCCTCTCTCTGGACTGGAGGGGGACCCC | | |
| 645 | 9 | 121473629 | 121473707 | 78 | AKR,BUB,C58J,CBA,CE,DBA1J,DBA2J,MRL,NZW,RF,RH2,SMJ | 9 | 121473641 | 121473674 | 34 | + | 18.1382 | 3.22E-07 | 0.0474 | CTGCTCTCCTCTCCCTGGACTGGAGGGGGACCCC | | |
| 646 | 9 | 121526776 | 121527001 | 225 | AKR,BUB,C58J,CBA,DBA1J,DBA2J,MRL,RF,RH2,SMJ | 9 | 121526751 | 121526784 | 34 | + | 16.6098 | 8.68E-07 | 0.0821 | GCCTTGCACTGCCAGTGGCCACCAGAGGGCACTG | 37 | ENCFF201JUA,ENCFF093ZRU,ENCFF367UNR,ENCFF327LYS,ENCFF136ZDK,ENCFF099QNE,ENCFF907XEG,ENCFF821BIA,ENCFF589ZBJ,ENCFF089GBC,ENCFF024JOZ,ENCFF321AGQ,ENCFF065ODV,ENCFF092NMJ,ENCFF502WDT,ENCFF198SSR,ENCFF430PPJ,ENCFF210BTO,ENCFF052KVO,ENCFF142CNG,ENCFF165LXO,ENCFF194HVL,ENCFF504JDS,ENCFF349BUL,ENCFF281QMV,ENCFF086ZTV,ENCFF575MGI,ENCFF601JVE,ENCFF805QBU,ENCFF491RJK,ENCFF611HDQ,ENCFF951GOP,ENCFF883UPM,ENCFF316FYN,ENCFF055HAP,ENCFF089EWX,ENCFF784ASD |
| 646 | 9 | 121526776 | 121527001 | 225 | AKR,BUB,C58J,CBA,DBA1J,DBA2J,MRL,RF,RH2,SMJ | 9 | 121526766 | 121526784 | 19 | + | 25.4262 | 3.18E-10 | 0.00481 | TGGCCACCAGAGGGCACTG | 39 | ENCFF201JUA,ENCFF093ZRU,ENCFF367UNR,ENCFF327LYS,ENCFF136ZDK,ENCFF099QNE,ENCFF907XEG,ENCFF821BIA,ENCFF589ZBJ,ENCFF089GBC,ENCFF024JOZ,ENCFF321AGQ,ENCFF065ODV,ENCFF092NMJ,ENCFF502WDT,ENCFF198SSR,ENCFF430PPJ,ENCFF210BTO,ENCFF052KVO,ENCFF142CNG,ENCFF165LXO,ENCFF194HVL,ENCFF504JDS,ENCFF349BUL,ENCFF281QMV,ENCFF086ZTV,ENCFF575MGI,ENCFF601JVE,ENCFF805QBU,ENCFF491RJK,ENCFF611HDQ,ENCFF951GOP,ENCFF883UPM,ENCFF316FYN,ENCFF055HAP,ENCFF089EWX,ENCFF784ASD,ENCFF097CAQ,ENCFF033UQX |
| 646 | 9 | 121526776 | 121527001 | 225 | AKR,BUB,C58J,CBA,DBA1J,DBA2J,MRL,RF,RH2,SMJ | 9 | 121526796 | 121526829 | 34 | + | 16.6098 | 8.68E-07 | 0.0821 | GCCTTGCACTGCCAGTGGCCACCAGAGGGCACTG | 39 | ENCFF201JUA,ENCFF093ZRU,ENCFF367UNR,ENCFF327LYS,ENCFF136ZDK,ENCFF099QNE,ENCFF907XEG,ENCFF821BIA,ENCFF589ZBJ,ENCFF089GBC,ENCFF024JOZ,ENCFF321AGQ,ENCFF065ODV,ENCFF092NMJ,ENCFF502WDT,ENCFF198SSR,ENCFF430PPJ,ENCFF210BTO,ENCFF052KVO,ENCFF142CNG,ENCFF165LXO,ENCFF194HVL,ENCFF504JDS,ENCFF349BUL,ENCFF281QMV,ENCFF086ZTV,ENCFF575MGI,ENCFF601JVE,ENCFF805QBU,ENCFF491RJK,ENCFF611HDQ,ENCFF951GOP,ENCFF883UPM,ENCFF316FYN,ENCFF055HAP,ENCFF089EWX,ENCFF784ASD,ENCFF097CAQ,ENCFF033UQX |
| 646 | 9 | 121526776 | 121527001 | 225 | AKR,BUB,C58J,CBA,DBA1J,DBA2J,MRL,RF,RH2,SMJ | 9 | 121526811 | 121526829 | 19 | + | 25.4262 | 3.18E-10 | 0.00481 | TGGCCACCAGAGGGCACTG | 39 | ENCFF201JUA,ENCFF093ZRU,ENCFF367UNR,ENCFF327LYS,ENCFF136ZDK,ENCFF099QNE,ENCFF907XEG,ENCFF821BIA,ENCFF589ZBJ,ENCFF089GBC,ENCFF024JOZ,ENCFF321AGQ,ENCFF065ODV,ENCFF092NMJ,ENCFF502WDT,ENCFF198SSR,ENCFF430PPJ,ENCFF210BTO,ENCFF052KVO,ENCFF142CNG,ENCFF165LXO,ENCFF194HVL,ENCFF504JDS,ENCFF349BUL,ENCFF281QMV,ENCFF086ZTV,ENCFF575MGI,ENCFF601JVE,ENCFF805QBU,ENCFF491RJK,ENCFF611HDQ,ENCFF951GOP,ENCFF883UPM,ENCFF316FYN,ENCFF055HAP,ENCFF089EWX,ENCFF784ASD,ENCFF097CAQ,ENCFF033UQX |
| 646 | 9 | 121526776 | 121527001 | 225 | AKR,BUB,C58J,CBA,DBA1J,DBA2J,MRL,RF,RH2,SMJ | 9 | 121526841 | 121526874 | 34 | + | 16.6098 | 8.68E-07 | 0.0821 | GCCTTGCACTGCCAGTGGCCACCAGAGGGCACTG | 34 | ENCFF201JUA,ENCFF093ZRU,ENCFF367UNR,ENCFF327LYS,ENCFF136ZDK,ENCFF099QNE,ENCFF907XEG,ENCFF821BIA,ENCFF589ZBJ,ENCFF089GBC,ENCFF024JOZ,ENCFF321AGQ,ENCFF065ODV,ENCFF092NMJ,ENCFF198SSR,ENCFF430PPJ,ENCFF210BTO,ENCFF052KVO,ENCFF165LXO,ENCFF504JDS,ENCFF349BUL,ENCFF601JVE,ENCFF951GOP,ENCFF883UPM,ENCFF316FYN,ENCFF055HAP,ENCFF089EWX,ENCFF784ASD,ENCFF097CAQ,ENCFF033UQX,ENCFF609MDN,ENCFF957SPH,ENCFF194HVL |
| 646 | 9 | 121526776 | 121527001 | 225 | AKR,BUB,C58J,CBA,DBA1J,DBA2J,MRL,RF,RH2,SMJ | 9 | 121526856 | 121526874 | 19 | + | 25.4262 | 3.18E-10 | 0.00481 | TGGCCACCAGAGGGCACTG | 32 | ENCFF201JUA,ENCFF093ZRU,ENCFF367UNR,ENCFF327LYS,ENCFF136ZDK,ENCFF099QNE,ENCFF907XEG,ENCFF821BIA,ENCFF589ZBJ,ENCFF089GBC,ENCFF024JOZ,ENCFF321AGQ,ENCFF092NMJ,ENCFF198SSR,ENCFF430PPJ,ENCFF210BTO,ENCFF165LXO,ENCFF504JDS,ENCFF601JVE,ENCFF951GOP,ENCFF883UPM,ENCFF316FYN,ENCFF055HAP,ENCFF089EWX,ENCFF784ASD,ENCFF097CAQ,ENCFF033UQX,ENCFF609MDN,ENCFF957SPH,ENCFF194HVL,ENCFF805QBU |
| 646 | 9 | 121526776 | 121527001 | 225 | AKR,BUB,C58J,CBA,DBA1J,DBA2J,MRL,RF,RH2,SMJ | 9 | 121526886 | 121526919 | 34 | + | 16.6098 | 8.68E-07 | 0.0821 | GCCTTGCACTGCCAGTGGCCACCAGAGGGCACTG | 32 | ENCFF201JUA,ENCFF093ZRU,ENCFF367UNR,ENCFF327LYS,ENCFF136ZDK,ENCFF099QNE,ENCFF907XEG,ENCFF821BIA,ENCFF589ZBJ,ENCFF089GBC,ENCFF024JOZ,ENCFF321AGQ,ENCFF092NMJ,ENCFF198SSR,ENCFF430PPJ,ENCFF165LXO,ENCFF601JVE,ENCFF951GOP,ENCFF883UPM,ENCFF316FYN,ENCFF055HAP,ENCFF089EWX,ENCFF784ASD,ENCFF097CAQ,ENCFF033UQX,ENCFF609MDN,ENCFF957SPH,ENCFF194HVL,ENCFF805QBU,ENCFF491RJK |
| 646 | 9 | 121526776 | 121527001 | 225 | AKR,BUB,C58J,CBA,DBA1J,DBA2J,MRL,RF,RH2,SMJ | 9 | 121526901 | 121526919 | 19 | + | 25.4262 | 3.18E-10 | 0.00481 | TGGCCACCAGAGGGCACTG | 32 | ENCFF201JUA,ENCFF093ZRU,ENCFF367UNR,ENCFF327LYS,ENCFF136ZDK,ENCFF099QNE,ENCFF907XEG,ENCFF821BIA,ENCFF589ZBJ,ENCFF089GBC,ENCFF024JOZ,ENCFF321AGQ,ENCFF092NMJ,ENCFF198SSR,ENCFF430PPJ,ENCFF165LXO,ENCFF601JVE,ENCFF951GOP,ENCFF883UPM,ENCFF316FYN,ENCFF055HAP,ENCFF089EWX,ENCFF784ASD,ENCFF097CAQ,ENCFF033UQX,ENCFF609MDN,ENCFF957SPH,ENCFF194HVL,ENCFF805QBU,ENCFF491RJK |
| 646 | 9 | 121526776 | 121527001 | 225 | AKR,BUB,C58J,CBA,DBA1J,DBA2J,MRL,RF,RH2,SMJ | 9 | 121526931 | 121526964 | 34 | + | 16.6098 | 8.68E-07 | 0.0821 | GCCTTGCACTGCCAGTGGCCACCAGAGGGCACTG | 32 | ENCFF093ZRU,ENCFF367UNR,ENCFF327LYS,ENCFF136ZDK,ENCFF099QNE,ENCFF907XEG,ENCFF821BIA,ENCFF589ZBJ,ENCFF089GBC,ENCFF024JOZ,ENCFF321AGQ,ENCFF092NMJ,ENCFF198SSR,ENCFF430PPJ,ENCFF165LXO,ENCFF601JVE,ENCFF951GOP,ENCFF883UPM,ENCFF316FYN,ENCFF055HAP,ENCFF784ASD,ENCFF097CAQ,ENCFF033UQX,ENCFF957SPH,ENCFF194HVL,ENCFF805QBU,ENCFF201JUA,ENCFF491RJK,ENCFF349BUL,ENCFF065ODV,ENCFF502WDT |
| 646 | 9 | 121526776 | 121527001 | 225 | AKR,BUB,C58J,CBA,DBA1J,DBA2J,MRL,RF,RH2,SMJ | 9 | 121526946 | 121526964 | 19 | + | 25.4262 | 3.18E-10 | 0.00481 | TGGCCACCAGAGGGCACTG | 30 | ENCFF093ZRU,ENCFF367UNR,ENCFF327LYS,ENCFF136ZDK,ENCFF099QNE,ENCFF907XEG,ENCFF821BIA,ENCFF589ZBJ,ENCFF089GBC,ENCFF024JOZ,ENCFF321AGQ,ENCFF092NMJ,ENCFF198SSR,ENCFF430PPJ,ENCFF165LXO,ENCFF601JVE,ENCFF951GOP,ENCFF883UPM,ENCFF316FYN,ENCFF055HAP,ENCFF033UQX,ENCFF957SPH,ENCFF194HVL,ENCFF805QBU,ENCFF201JUA,ENCFF491RJK,ENCFF349BUL,ENCFF065ODV,ENCFF502WDT |
| 646 | 9 | 121526776 | 121527001 | 225 | AKR,BUB,C58J,CBA,DBA1J,DBA2J,MRL,RF,RH2,SMJ | 9 | 121526976 | 121527009 | 34 | + | 17.6098 | 4.56E-07 | 0.057 | GCCTTGCACTGCCAGTGGCCACCAGGGGGCACTG | 28 | ENCFF093ZRU,ENCFF367UNR,ENCFF327LYS,ENCFF136ZDK,ENCFF099QNE,ENCFF907XEG,ENCFF821BIA,ENCFF589ZBJ,ENCFF089GBC,ENCFF024JOZ,ENCFF321AGQ,ENCFF092NMJ,ENCFF198SSR,ENCFF430PPJ,ENCFF165LXO,ENCFF601JVE,ENCFF883UPM,ENCFF055HAP,ENCFF957SPH,ENCFF194HVL,ENCFF805QBU,ENCFF201JUA,ENCFF491RJK,ENCFF349BUL,ENCFF065ODV,ENCFF502WDT,ENCFF504JDS |
| 646 | 9 | 121526776 | 121527001 | 225 | AKR,BUB,C58J,CBA,DBA1J,DBA2J,MRL,RF,RH2,SMJ | 9 | 121526991 | 121527009 | 19 | + | 26.4754 | 4.11E-11 | 0.00331 | TGGCCACCAGGGGGCACTG | 28 | ENCFF093ZRU,ENCFF367UNR,ENCFF327LYS,ENCFF136ZDK,ENCFF099QNE,ENCFF907XEG,ENCFF821BIA,ENCFF589ZBJ,ENCFF089GBC,ENCFF024JOZ,ENCFF321AGQ,ENCFF092NMJ,ENCFF198SSR,ENCFF430PPJ,ENCFF165LXO,ENCFF601JVE,ENCFF883UPM,ENCFF055HAP,ENCFF957SPH,ENCFF194HVL,ENCFF805QBU,ENCFF201JUA,ENCFF491RJK,ENCFF349BUL,ENCFF065ODV,ENCFF502WDT,ENCFF504JDS |
| 647 | 9 | 121672118 | 121672176 | 58 | AJ,AKR,BALB,BUB,C3H,C58J,CBA,CE,DBA1J,DBA2J,FVB,ILNJ2,LP,MA/My,MRL,NZW,PJ,RF,RH2,SEA,SMJ,TH | 9 | 121672120 | 121672138 | 19 | + | 18.2951 | 3.84E-07 | 0.0907 | ATGCCAGAAGAGGGCAGCA | 14 | ENCFF093ZRU,ENCFF821BIA,ENCFF092NMJ,ENCFF024JOZ,ENCFF883UPM,ENCFF099QNE,ENCFF367UNR,ENCFF430PPJ,ENCFF136ZDK,ENCFF907XEG,ENCFF589ZBJ,ENCFF198SSR,ENCFF327LYS,ENCFF089GBC |
| 648 | 9 | 7728093 | 7728287 | 194 | 129S1 | 9 | 7728213 | 7728246 | 34 | - | 19.0163 | 1.77E-07 | 0.0347 | CAGCTGTCTTCAGACACACCAGAAGAGGGCGTCA | | |
| 649 | 9 | 21654516 | 21654922 | 406 | BTBR,C57L,C58J,FVB,MA/My,NOD,NOR,NZB,NZW,RF,SJL,SWR | 9 | 21654843 | 21654876 | 34 | - | 17.878 | 3.83E-07 | 0.0522 | TAGCTGTCCTCAGACACACCAGAAGAGGGCATCA | | |
| 650 | 9 | 21979919 | 21980138 | 219 | BTBR,C57L,C58J,FVB,MA/My,NOD,NOR,NZB,NZW,RF,SJL,SWR | 9 | 21980044 | 21980077 | 34 | - | 16.4472 | 9.61E-07 | 0.0841 | TAGCTGTCTTCAGACACACCAGAAGAGGGCGTCA | | |
| 651 | 9 | 21987018 | 21987257 | 239 | BTBR,C57L,C58J,FVB,MA/My,NOD,NOR,NZB,NZW,RF,SJL,SWR | 9 | 21987137 | 21987170 | 34 | - | 16.4309 | 9.71E-07 | 0.0846 | TAGCTGTCTTCAGACGCTCCAGAAGAGGGCGTCA | | |
| 652 | 9 | 22053175 | 22053383 | 208 | BTBR,C57L,C58J,FVB,MA/My,NOD,NOR,NZB,NZW,RF,SJL,SWR | 9 | 22053300 | 22053333 | 34 | - | 17.3902 | 5.27E-07 | 0.0619 | TAGCTGTCCTCAGACACTCCAGAAGAGGGCATCA | | |
| 653 | 9 | 23346195 | 23346401 | 206 | 129S1,CBA,FVB,LP,NZO,RF,SJL,SWR,TH | 9 | 23346325 | 23346358 | 34 | - | 16.7724 | 7.83E-07 | 0.0768 | TAGCTGTCTTCAGATGCACCAGAAGAGGGCGTCA | | |
| 654 | 9 | 28171514 | 28171725 | 211 | BUB | 9 | 28171626 | 28171659 | 34 | - | 16.4472 | 9.61E-07 | 0.0841 | TAGCTGTCTTCAGACACACCAGAAGAGGGCGTCA | | |
| 655 | 9 | 31090670 | 31090883 | 213 | CBA,CE,DBA1J,LP,MA/My,MRL,NZO,NZW,TH | 9 | 31090795 | 31090813 | 19 | - | 17.5574 | 6.32E-07 | 0.112 | ACTCCAGAAGAGGGCGCCA | | |
| 655 | 9 | 31090670 | 31090883 | 213 | CBA,CE,DBA1J,LP,MA/My,MRL,NZO,NZW,TH | 9 | 31090795 | 31090828 | 34 | - | 18.2195 | 3.05E-07 | 0.0458 | TAGCTGTCTTCAGATACTCCAGAAGAGGGCGCCA | | |
| 656 | 9 | 32106328 | 32106527 | 199 | BTBR,C57L | 9 | 32106452 | 32106470 | 19 | - | 17.5574 | 6.32E-07 | 0.112 | ACTCCAGAAGAGGGCGCCA | 4 | ENCFF367UNR,ENCFF099QNE,ENCFF821BIA,ENCFF093ZRU |
| 657 | 9 | 32683283 | 32683607 | 324 | 129S1,AJ,AKR,BALB,BUB,C3H,DBA1J,FVB,MA/My,NOD,NOR,NZB,NZO,NZW,PL,RH2,SJL,SWR,TH | 9 | 32683565 | 32683598 | 34 | - | 16.9837 | 6.84E-07 | 0.0717 | CTGCCGTCACATTCGCTGTTACAAGATGGCGCTG | | |
| 658 | 9 | 35044852 | 35045057 | 205 | 129S1,AJ,AKR,BALB,BTBR,BUB,C3H,C57L,CE,DBA1J,DBA2J,FVB,KK,MA/My,MRL,NOD,NOR,NZB,NZO,NZW,PJ,PL,RF,RH2,SEA,SJL,SMJ,SWR,TH | 9 | 35044974 | 35045007 | 34 | - | 16.4472 | 9.61E-07 | 0.0841 | TAGCTGTCTTCAGACACACCAGAAGAGGGCGTCA | | |
| 659 | 9 | 37503035 | 37503239 | 204 | C57L,NZO | 9 | 37503158 | 37503191 | 34 | - | 16.7886 | 7.75E-07 | 0.0768 | CAGCTATCTTCAGACACTCCAGAAGAGGGCATCA | | |
| 660 | 9 | 40557987 | 40558226 | 239 | 129S1,AJ,AKR,BALB,BTBR,BUB,C3H,C57L,CBA,CE,DBA1J,DBA2J,FVB,ILNJ2,KK,MA/My,MRL,NOD,NOR,NZB,NZO,NZW,PJ,PL,RF,RH2,SEA,SJL,SMJ,SWR,TH | 9 | 40558119 | 40558152 | 34 | - | 19.252 | 1.51E-07 | 0.0314 | CTGCTCTCTTCAGAAACACCAGAAGAGGGCAACA | | |
| 661 | 9 | 45278410 | 45279232 | 822 | ILNJ2 | 9 | 45278468 | 45278502 | 35 | - | 17.0092 | 5.39E-07 | 0.113 | TTCCTATGCGCGCTGTCACCCAGGAGAGGGCAGCT | | |
| 662 | 9 | 45637505 | 45637723 | 218 | 129S1,C57L,KK | 9 | 45637667 | 45637700 | 34 | - | 17.3902 | 5.27E-07 | 0.0619 | TAGCTGTCCTCAGACACTCCAGAAGAGGGCATCA | | |
| 663 | 9 | 50540667 | 50540884 | 217 | AJ,BALB,BTBR,C3H,CBA,CE,DBA2J,ILNJ2,MRL,NOD,NOR,NZO,NZW,RH2,SJL | 9 | 50540792 | 50540825 | 34 | - | 16.4472 | 9.61E-07 | 0.0841 | TAGCTGTCTTCAGACACACCAGAAGAGGGCGTCA | | |
| 664 | 9 | 50901587 | 50902529 | 942 | 129S1,C57L | 9 | 50901837 | 50901855 | 19 | - | 17.5574 | 6.32E-07 | 0.112 | ACTCCAGAAGAGGGCGCCA | | |
| 664 | 9 | 50901587 | 50902529 | 942 | 129S1,C57L | 9 | 50901837 | 50901870 | 34 | - | 18.3659 | 2.76E-07 | 0.0438 | TAGCTGTCTTCAGACACTCCAGAAGAGGGCGCCA | | |
| 665 | 9 | 56915026 | 56915127 | 101 | MA/My | 9 | 56915036 | 56915054 | 19 | - | 18.0328 | 4.60E-07 | 0.1 | AGTCCAGAAGAGGGCACTG | | |
| 665 | 9 | 56915026 | 56915127 | 101 | MA/My | 9 | 56915036 | 56915070 | 35 | - | 24.8349 | 2.80E-09 | 0.00544 | GTGCAGGTACCTGTGGAGTCCAGAAGAGGGCACTG | | |
| 666 | 9 | 58835279 | 58835871 | 592 | 129S1,AJ,BUB,C3H,C57L,CE,DBA1J,DBA2J,MRL,PJ,RH2,SEA,SJL | 9 | 58835584 | 58835602 | 19 | - | 22.6885 | 1.05E-08 | 0.0194 | CAACCACCAGGTGGCACTG | 19 | ENCFF367UNR,ENCFF201JUA,ENCFF093ZRU,ENCFF099QNE,ENCFF821BIA,ENCFF327LYS,ENCFF089GBC,ENCFF609MDN,ENCFF024JOZ,ENCFF907XEG,ENCFF589ZBJ,ENCFF198SSR,ENCFF165LXO,ENCFF092NMJ,ENCFF136ZDK,ENCFF281QMV,ENCFF430PPJ,ENCFF883UPM |
| 666 | 9 | 58835279 | 58835871 | 592 | 129S1,AJ,BUB,C3H,C57L,CE,DBA1J,DBA2J,MRL,PJ,RH2,SEA,SJL | 9 | 58835584 | 58835617 | 34 | - | 25.9837 | 6.78E-10 | 0.000645 | CTGCATTTAAAAATTCAACCACCAGGTGGCACTG | 19 | ENCFF367UNR,ENCFF201JUA,ENCFF093ZRU,ENCFF099QNE,ENCFF821BIA,ENCFF327LYS,ENCFF089GBC,ENCFF609MDN,ENCFF024JOZ,ENCFF907XEG,ENCFF589ZBJ,ENCFF198SSR,ENCFF165LXO,ENCFF092NMJ,ENCFF136ZDK,ENCFF281QMV,ENCFF430PPJ,ENCFF883UPM |
| 667 | 9 | 59118352 | 59118557 | 205 | 129S1,AJ,AKR,BTBR,BUB,C3H,C57L,CBA,CE,DBA1J,DBA2J,FVB,ILNJ2,MA/My,MRL,NOD,NOR,NZB,PJ,PL,RF,RH2,SEA,SJL,SWR,TH | 9 | 59118477 | 59118495 | 19 | - | 17.5738 | 6.25E-07 | 0.112 | ACTCCAGAAGAGGGCGCTA | | |
| 667 | 9 | 59118352 | 59118557 | 205 | 129S1,AJ,AKR,BTBR,BUB,C3H,C57L,CBA,CE,DBA1J,DBA2J,FVB,ILNJ2,MA/My,MRL,NOD,NOR,NZB,PJ,PL,RF,RH2,SEA,SJL,SWR,TH | 9 | 59118477 | 59118510 | 34 | - | 18.1789 | 3.13E-07 | 0.0464 | TAGCTGTCTTCAGACACTCCAGAAGAGGGCGCTA | | |
| 668 | 9 | 60530828 | 60530923 | 95 | TH | 9 | 60530847 | 60530880 | 34 | - | 16.6423 | 8.50E-07 | 0.0811 | CTGCAGGTGGAGGCTGGCTCTGCAGGTGGAGGCA | | |
| 669 | 9 | 64182860 | 64183271 | 411 | AJ,BUB,C3H,CBA,DBA1J,DBA2J,FVB,ILNJ2,KK,MRL,NZB,NZO,NZW,PJ,RH2,SEA,SJL,SWR | 9 | 64183184 | 64183217 | 34 | - | 17.7236 | 4.24E-07 | 0.0552 | TAGCTGTCCTCAGACACTCCAGAAGAGGGCGACA | 1 | ENCFF093ZRU |
| 670 | 9 | 64223903 | 64224116 | 213 | AJ,BUB,C3H,CBA,CE,DBA1J,DBA2J,FVB,ILNJ2,KK,MRL,NZB,NZO,NZW,PJ,PL,RH2,SEA,SJL,SWR | 9 | 64224029 | 64224062 | 34 | - | 16.4146 | 9.81E-07 | 0.0852 | TAGCTGTCTTCTGACAGTCCAGAAGAGGGAGCCA | | |
| 671 | 9 | 65802980 | 65803183 | 203 | FVB,SJL | 9 | 65803106 | 65803139 | 34 | - | 17.8455 | 3.91E-07 | 0.0525 | TAGCTGTCCTCAGACACTCCAGAAGAGGGCGTCA | | |
| 672 | 9 | 69602146 | 69602753 | 607 | 129S1,BTBR,C57L,NOD,NOR,SMJ | 9 | 69602448 | 69602481 | 34 | - | 17.7967 | 4.04E-07 | 0.0536 | CTGCCCTTCCCTTTCTCTTCAGAAGAGGGTAACC | | |
| 673 | 9 | 73623558 | 73623834 | 276 | C58J,LP | 9 | 73623605 | 73623638 | 34 | - | 18.4797 | 2.56E-07 | 0.0428 | TTGCTGTCTTCAGACACACCAGAAGAGGGCATCA | | |
| 674 | 9 | 73778782 | 73778921 | 139 | BTBR,CE,MA/My,NZW | 9 | 73778772 | 73778805 | 34 | - | 25.1138 | 1.50E-09 | 0.00113 | CTGCAATGCTCACAAAGGCCAGAAGGAGGCATCA | 1 | ENCFF093ZRU |
| 675 | 9 | 78367169 | 78367378 | 209 | 129S1,BTBR,C57L,C58J,ILNJ2,LP,NZW,SMJ | 9 | 78367293 | 78367326 | 34 | - | 16.4472 | 9.61E-07 | 0.0841 | TAGCTGTCTTCAGACACACCAGAAGAGGGCGTCA | | |
| 676 | 9 | 78527028 | 78527271 | 243 | 129S1,AJ,AKR,BALB,BTBR,BUB,C3H,C57L,C58J,CBA,CE,DBA1J,DBA2J,FVB,ILNJ2,KK,LP,MRL,NOD,NOR,NZB,NZO,NZW,PJ,PL,RF,RH2,SEA,SJL,SMJ,SWR,TH | 9 | 78527153 | 78527186 | 34 | - | 17.9431 | 3.66E-07 | 0.0511 | CAGCTGTCCTCAGACACTCCAGAAGAGGGAGTCA | | |
| 677 | 9 | 82227019 | 82227278 | 259 | AKR,BUB,C58J,CBA,ILNJ2,LP,MRL,NOD,NOR,NZW,PL,RF | 9 | 82227165 | 82227198 | 34 | - | 18.7724 | 2.10E-07 | 0.0387 | GTGAACTGCCTGCGGAGGCCAGAAGAGGGTATTT | | |
| 678 | 9 | 100896766 | 100896990 | 224 | AJ,AKR,BALB,BUB,C3H,CBA,CE,DBA1J,DBA2J,FVB,ILNJ2,MRL,NOD,NOR,NZO,NZW,PJ,PL,RF,RH2,SEA,SJL,SMJ,SWR,TH | 9 | 100896894 | 100896927 | 34 | - | 16.4634 | 9.51E-07 | 0.0841 | TAGCTGTCTTCAGACGCACCAGAAGAGGGCATCA | | |
| 679 | 9 | 102840795 | 102840854 | 59 | AKR,MA/My,TH | 9 | 102840841 | 102840874 | 34 | - | 19.9106 | 9.47E-08 | 0.022 | CTGCACAGCCACACGTGCCCGTAAGGAGGCAGCA | | |
| 680 | 9 | 105469900 | 105470125 | 225 | 129S1,AJ,AKR,BALB,BTBR,BUB,C3H,C57L,CBA,CE,DBA1J,DBA2J,FVB,ILNJ2,KK,MA/My,MRL,NOD,NOR,NZB,NZO,NZW,PJ,PL,RF,RH2,SEA,SJL,SMJ,SWR,TH | 9 | 105470024 | 105470057 | 34 | - | 16.4634 | 9.51E-07 | 0.0841 | TAGCTGTCTTCAGACGCACCAGAAGAGGGCATCA | | |
| 681 | 9 | 108766825 | 108767030 | 205 | CBA,CE | 9 | 108766939 | 108766972 | 34 | - | 16.4634 | 9.51E-07 | 0.0841 | TAGCTGTCTTCAGACGCACCAGAAGAGGGCATCA | 1 | ENCFF093ZRU |
| 682 | 9 | 109905482 | 109905700 | 218 | CBA,CE,NZW | 9 | 109905611 | 109905629 | 19 | - | 17.5574 | 6.32E-07 | 0.112 | ACTCCAGAAGAGGGCGCCA | 3 | ENCFF093ZRU,ENCFF821BIA,ENCFF367UNR |
| 682 | 9 | 109905482 | 109905700 | 218 | CBA,CE,NZW | 9 | 109905611 | 109905644 | 34 | - | 18.561 | 2.42E-07 | 0.0423 | AAGCTGTCTTCAGACACTCCAGAAGAGGGCGCCA | 3 | ENCFF093ZRU,ENCFF821BIA,ENCFF367UNR |
| 683 | 9 | 110301435 | 110301690 | 255 | BTBR,CBA,FVB,NZB,NZO,NZW,SJL,SWR | 9 | 110301562 | 110301595 | 34 | - | 17.6667 | 4.40E-07 | 0.0557 | TAGCTGTTTTCAGACACTCCAGAAGAGGGCGTCA | | |
| 684 | 9 | 110639708 | 110639917 | 209 | CBA,FVB,NZB,NZO,PJ,SEA,SJL,SMJ,SWR | 9 | 110639823 | 110639856 | 34 | - | 17.3902 | 5.27E-07 | 0.0619 | TAGCTGTCCTCAGACACTCCAGAAGAGGGCATCA | | |
| 685 | 9 | 110766576 | 110766789 | 213 | 129S1,AJ,AKR,BALB,BTBR,BUB,C3H,C57L,CBA,CE,DBA1J,DBA2J,FVB,ILNJ2,KK,MA/My,MRL,NOD,NOR,NZB,NZO,NZW,PJ,PL,RF,RH2,SEA,SJL,SMJ,SWR,TH | 9 | 110766702 | 110766720 | 19 | - | 17.9672 | 4.81E-07 | 0.102 | GCACCAGAAGAGGGCACCA | 4 | ENCFF099QNE,ENCFF821BIA,ENCFF367UNR,ENCFF093ZRU |
| 685 | 9 | 110766576 | 110766789 | 213 | 129S1,AJ,AKR,BALB,BTBR,BUB,C3H,C57L,CBA,CE,DBA1J,DBA2J,FVB,ILNJ2,KK,MA/My,MRL,NOD,NOR,NZB,NZO,NZW,PJ,PL,RF,RH2,SEA,SJL,SMJ,SWR,TH | 9 | 110766702 | 110766735 | 34 | - | 17.252 | 5.76E-07 | 0.065 | TAGCTATCTTCAGCTGCACCAGAAGAGGGCACCA | 4 | ENCFF099QNE,ENCFF821BIA,ENCFF367UNR,ENCFF093ZRU |
| 686 | 9 | 110797604 | 110797814 | 210 | 129S1,AJ,AKR,BALB,BTBR,BUB,C3H,CBA,CE,DBA1J,DBA2J,FVB,ILNJ2,KK,MA/My,MRL,NOD,NOR,NZB,NZO,PJ,PL,RF,RH2,SEA,SJL,SMJ,SWR,TH | 9 | 110797729 | 110797762 | 34 | - | 22.9187 | 9.63E-09 | 0.00422 | TTGCTGTCCTCAGACGCCCCAGAAGAGGGCGCCA | 3 | ENCFF367UNR,ENCFF093ZRU,ENCFF821BIA |
| 687 | 9 | 114254713 | 114254853 | 140 | AJ,BALB,C3H,CE,FVB,MA/My | 9 | 114254833 | 114254866 | 34 | - | 23.5122 | 5.93E-09 | 0.00294 | GTGCAGTACCCAGGGAAGACAGAAGAGGGCATCA | 3 | ENCFF052KVO,ENCFF093ZRU,ENCFF327LYS |
| 688 | 9 | 114409647 | 114409861 | 214 | AJ,BALB,C3H,FVB,KK,PJ,SEA | 9 | 114409778 | 114409811 | 34 | - | 16.4146 | 9.81E-07 | 0.0852 | TAGCTGTCTTCTGACAGTCCAGAAGAGGGAGCCA | | |
| 689 | 9 | 118775277 | 118775711 | 434 | CE,ILNJ2 | 9 | 118775253 | 118775286 | 34 | - | 18.3415 | 2.81E-07 | 0.0442 | CAGCACACCCAGAGCCAGCCTGGAGGTGGAGACA | | |

| | | | | | | | | | | | | | | | | |
|---|---|---|---|---|---|---|---|---|---|---|---|---|---|---|---|---|
| 690 | 9 | 120673869 | 120674068 | 199 | BTBR | 9 | 120673997 | 120674030 | 34 | - | 16.4472 | 9.61E-07 | 0.0841 | TAGCTGTCTTCAGACACACCAGAAGAGGGCGTCA | | |
| 691 | 9 | 122027332 | 122028011 | 679 | ILNJ2.LP | 9 | 122027509 | 122027543 | 35 | - | 17.3211 | 4.50E-07 | 0.101 | TAGCACTCAGGACCAATAGCCCGGAGGTGGCACCA | | |
| 692 | 10 | 4301194 | 4301405 | 211 | NOD.NOR | 10 | 4301262 | 4301295 | 34 | + | 18.3659 | 2.76E-07 | 0.0597 | TAGCTGTCTTCAGACACTCCAGAAGAGGGCGCCA | | |
| 692 | 10 | 4301194 | 4301405 | 211 | NOD.NOR | 10 | 4301277 | 4301295 | 19 | + | 17.5574 | 6.32E-07 | 0.134 | ACTCCAGAAGAGGGCGCCA | | |
| 693 | 10 | 5732747 | 5732935 | 188 | BUB | 10 | 5732818 | 5732852 | 35 | + | 24.633 | 3.28E-09 | 0.0077 | CTGCATCTACACCTGAGTGCCAGAAGAGGGCATTA | | |
| 694 | 10 | 6125097 | 6125301 | 204 | BUB | 10 | 6125158 | 6125191 | 34 | + | 16.4634 | 9.51E-07 | 0.108 | TAGCTGTCTTCAGACGCACCAGAAGAGGGCATCA | | |
| 695 | 10 | 8245125 | 8245344 | 219 | BUB,C3H,CBA,CE,DBA1J,DBA2J,FVB,ILNJ2,KK,LP,MA/My,MRL,NOD,NOR,NZB,NZO,NZW.PJ.SEA.SWR.TH | 10 | 8245201 | 8245234 | 34 | + | 16.4634 | 9.51E-07 | 0.108 | TAGCTGTCTTCAGACGCACCAGAAGAGGGCATCA | 1 | ENCFF093ZRU |
| 696 | 10 | 8403667 | 8403875 | 208 | C3H,CBA,DBA1J,DBA2J,LP,MA/My,MRL,NZW | 10 | 8403734 | 8403767 | 34 | + | 16.4472 | 9.61E-07 | 0.108 | TAGCTGTCTTCAGACACACCAGAAGAGGGCGTCA | | |
| 697 | 10 | 12858582 | 12858748 | 166 | BUB.DBA1J.DBA2J | 10 | 12858646 | 12858679 | 34 | + | 17.878 | 3.83E-07 | 0.0708 | TAGCTGTCCTCAGACACACCAGAAGAGGGCATCA | | |
| 698 | 10 | 14468555 | 14468754 | 199 | DBA1J.DBA2J | 10 | 14468613 | 14468646 | 34 | + | 16.7724 | 7.83E-07 | 0.0988 | TAGCTGTCTTCAGATGCACCAGAAGAGGGCGTCA | | |
| 699 | 10 | 18635351 | 18635566 | 215 | SWR | 10 | 18635421 | 18635454 | 34 | + | 17.8455 | 3.91E-07 | 0.071 | TAGCTGTCCTCAGACACTCCAGAAGAGGGCGTCA | | |
| 700 | 10 | 28113448 | 28113651 | 203 | PJ.SEA | 10 | 28113509 | 28113542 | 34 | + | 17.8455 | 3.91E-07 | 0.071 | TAGCTGTCCTCAGACACTCCAGAAGAGGGCGTCA | | |
| 701 | 10 | 28244653 | 28244883 | 230 | PJ.SEA | 10 | 28244716 | 28244749 | 34 | + | 18.3659 | 2.76E-07 | 0.0597 | TAGCTGTCTTCAGACACTCCAGAAGAGGGCGCCA | | |
| 701 | 10 | 28244653 | 28244883 | 230 | PJ.SEA | 10 | 28244731 | 28244749 | 19 | + | 17.5574 | 6.32E-07 | 0.134 | ACTCCAGAAGAGGGCGCCA | | |
| 702 | 10 | 41588430 | 41588642 | 212 | 129S1.BTBR.C57L.KK | 10 | 41588498 | 41588531 | 34 | + | 16.4472 | 9.61E-07 | 0.108 | TAGCTGTCTTCAGACACACCAGAAGAGGGCGTCA | | |
| 703 | 10 | 62871079 | 62871333 | 254 | RF | 10 | 62871190 | 62871223 | 34 | + | 16.4634 | 9.51E-07 | 0.108 | TAGCTGTCTTCAGACGCACCAGAAGAGGGCATCA | | |
| 704 | 10 | 66158365 | 66158527 | 162 | NZW.PL | 10 | 66158374 | 66158407 | 34 | + | 16.4228 | 9.76E-07 | 0.109 | TAGCTGCCTTCAGACACTCCAGAAGAGGGCGCCA | | |
| 704 | 10 | 66158365 | 66158527 | 162 | NZW.PL | 10 | 66158389 | 66158407 | 19 | + | 17.5574 | 6.32E-07 | 0.134 | ACTCCAGAAGAGGGCGCCA | | |
| 705 | 10 | 67183906 | 67184247 | 341 | ILNJ2.NZW.SMJ | 10 | 67184049 | 67184083 | 35 | + | 27.4954 | 3.05E-10 | 0.00296 | CTGCATGTTTGCCTGCAGGCCAGAAGAGGGCACCA | 1 | ENCFF502WDT |
| 705 | 10 | 67183906 | 67184247 | 341 | ILNJ2.NZW.SMJ | 10 | 67184065 | 67184083 | 19 | + | 20.3607 | 8.29E-08 | 0.0512 | AGGCCAGAAGAGGGCACCA | 1 | ENCFF502WDT |
| 706 | 10 | 69238022 | 69238421 | 399 | 129S1.C3H.CBA.CE.NZB.NZO.NZW | 10 | 69238378 | 69238411 | 34 | + | 17.0244 | 6.67E-07 | 0.0928 | TAGCTGTCTTCAAACTCTCCAGAAGAGGGCATCA | | |
| 707 | 10 | 75792490 | 75792762 | 272 | BUB.RF | 10 | 75792619 | 75792653 | 35 | + | 16.0917 | 9.07E-07 | 0.18 | TAGCTGTCTTCAGACAGCACCAGAAGAGGGCATCA | | |
| 708 | 10 | 75908722 | 75908930 | 208 | BUB.RF | 10 | 75908774 | 75908807 | 34 | + | 16.4634 | 9.51E-07 | 0.108 | TAGCTGTCTTCAGACGCACCAGAAGAGGGCATCA | | |
| 709 | 10 | 81552995 | 81553194 | 199 | 129S1,AJ,AKR,BALB,C3H,C57L,CBA,CE,DBA1J,DBA2J,FVB,ILNJ2,KK,MA/My,MRL,NOD,NOR,NZB,NZO,NZW,PJ,PL,RF,RH2,SEA,SJL.SMJ.SWR.TH | 10 | 81553055 | 81553088 | 34 | + | 18.3659 | 2.76E-07 | 0.0597 | TAGCTGTCTTCAGACACTCCAGAAGAGGGCGCCA | | |
| 709 | 10 | 81552995 | 81553194 | 199 | 129S1,AJ,AKR,BALB,C3H,C57L,CBA,CE,DBA1J,DBA2J,FVB,ILNJ2,KK,MA/My,MRL,NOD,NOR,NZB,NZO,NZW,PJ,PL,RF,RH2,SEA,SJL.SMJ.SWR.TH | 10 | 81553070 | 81553088 | 19 | + | 17.5574 | 6.32E-07 | 0.134 | ACTCCAGAAGAGGGCGCCA | | |
| 710 | 10 | 82270216 | 82270432 | 216 | 129S1,AJ,AKR,BALB,BTBR,BUB,C3H,C57L,CBA,CE,DBA1J,DBA2J,FVB,ILNJ2,KK,MA/My,MRL,NOD,NOR,NZB,NZO,NZW,PJ,PL,RF,RH2.SEA.SJL.SMJ.SWR.TH | 10 | 82270305 | 82270323 | 19 | + | 17.8852 | 5.08E-07 | 0.121 | ACACCAGAAGAGGGCACCA | | |
| 711 | 10 | 87912384 | 87912587 | 203 | AJ,AKR,BALB,BTBR,BUB,C3H,C58J,CBA,CE,DBA1J,DBA2J,FVB,ILNJ2,KK,LP,MA/My,MRL,NOD,NOR,NZO,PJ,PL,RF,RH2,SEA,SWR,TH | 10 | 87912445 | 87912478 | 34 | + | 18.561 | 2.42E-07 | 0.0573 | AAGCTGTCTTCAGACACTCCAGAAGAGGGCGCCA | | |
| 711 | 10 | 87912384 | 87912587 | 203 | AJ,AKR,BALB,BTBR,BUB,C3H,C58J,CBA,CE,DBA1J,DBA2J,FVB,ILNJ2,KK,LP,MA/My,MRL,NOD,NOR,NZO,PJ,PL,RF,RH2,SEA,SWR,TH | 10 | 87912460 | 87912478 | 19 | + | 17.5574 | 6.32E-07 | 0.134 | ACTCCAGAAGAGGGCGCCA | | |
| 712 | 10 | 88454150 | 88454630 | 480 | BUB | 10 | 88454610 | 88454643 | 34 | + | 17.2602 | 5.73E-07 | 0.0854 | CCGCCACACCCGCTCCCCACACAAGGTGGTACTG | | |
| 713 | 10 | 102271123 | 102271455 | 332 | AKR,BUB,FVB,NOD,NOR,PL,RF | 10 | 102271153 | 102271171 | 19 | + | 19.2951 | 1.88E-07 | 0.0757 | CCGCCACTAGATGGTGCTA | 17 | ENCFF201JUA,ENCFF367UNR,ENCFF093ZRU,ENCFF099QNE,ENCFF327LYS,ENCFF821BIA,ENCFF907XEG,ENCFF024JOZ,ENCFF092NMJ,ENCFF089GBC,ENCFF609MDN,ENCFF198SSR,ENCFF589ZBJ,ENCFF165LXO.ENCFF430PPJ.ENCFF052KVO.ENCFF136ZDK |
| 714 | 10 | 109568707 | 109569050 | 343 | NZW.PL.SMJ | 10 | 109568771 | 109568804 | 34 | + | 16.4472 | 9.61E-07 | 0.108 | TAGCTGTCTTCAGACACACCAGAAGAGGGCGTCA | | |
| 715 | 10 | 121746484 | 121746687 | 203 | DBA1J.KK.SMJ | 10 | 121746550 | 121746583 | 34 | + | 17.2439 | 5.79E-07 | 0.0854 | TAGCTGTCCTCAGATACTCCAGAAGAGGGCATCA | | |
| 716 | 10 | 122789869 | 122790086 | 217 | AJ,BALB,BUB,CE,KK,MA/My,NOD,NOR,PL,RH2 | 10 | 122789944 | 122789977 | 34 | + | 19.0325 | 1.75E-07 | 0.0478 | CAGCTGTCTTCAGACGCACCAGAAGAGGGCATCA | 1 | ENCFF093ZRU |
| 717 | 10 | 124153915 | 124154353 | 438 | PL | 10 | 124154330 | 124154364 | 35 | + | 16.5505 | 7.01E-07 | 0.158 | ATGCATGTGTGTTCTGATGCCAGAAGAGGGCATCA | | |
| 718 | 10 | 127250434 | 127250660 | 226 | AJ,BALB,BUB,C3H,CBA,CE,FVB,NOD,NOR,PL,RH2 | 10 | 127250518 | 127250551 | 34 | + | 17.3902 | 5.27E-07 | 0.082 | TAGCTGTCCTCAGACACTCCAGAAGAGGGCATCA | | |
| 719 | 10 | 128168450 | 128168655 | 205 | BUB.C3H.CBA.FVB.KK.SMJ | 10 | 128168516 | 128168549 | 34 | + | 19.4797 | 1.28E-07 | 0.04 | TAGCTGTCCCCAGACACTCCAGAAGAGGGCGTCA | | |
| 720 | 10 | 128182651 | 128182859 | 208 | BUB.CE.FVB.KK.SMJ | 10 | 128182717 | 128182750 | 34 | + | 22.9187 | 9.63E-09 | 0.00615 | TTGCTGTCCTCAGACGCCCCAGAAGAGGGCGCCA | 5 | ENCFF093ZRU.ENCFF821BIA.ENCFF367UNR.ENCFF201JUA |
| 721 | 10 | 128734180 | 128734417 | 237 | BUB.CE.KK.RF.SMJ | 10 | 128734275 | 128734308 | 34 | + | 16.4634 | 9.51E-07 | 0.108 | TAGCTGTCTTCAGACGCACCAGAAGAGGGCATCA | | |
| 722 | 10 | 5766375 | 5766584 | 209 | 129S1,AJ,AKR,BALB,BTBR,BUB,C3H,C57L,CBA,CE,DBA1J,DBA2J,FVB,KK,LP,MA/My,MRL,NOD,NOR,NZB,NZO,NZW,PL,RF,RH2,SJL.SMJ.SWR.TH | 10 | 5766493 | 5766511 | 19 | - | 17.5574 | 6.32E-07 | 0.134 | ACTCCAGAAGAGGGCGCCA | | |
| 722 | 10 | 5766375 | 5766584 | 209 | 129S1,AJ,AKR,BALB,BTBR,BUB,C3H,C57L,CBA,CE,DBA1J,DBA2J,FVB,KK,LP,MA/My,MRL,NOD,NOR,NZB,NZO,NZW,PL,RF,RH2,SJL.SMJ.SWR.TH | 10 | 5766493 | 5766526 | 34 | - | 18.3659 | 2.76E-07 | 0.0597 | TAGCTGTCTTCAGACACTCCAGAAGAGGGCGCCA | | |
| 723 | 10 | 26874406 | 26874613 | 207 | 129S1,AJ,AKR,BALB,BTBR,BUB,C3H,C57L,CE,FVB,ILNJ2,KK,LP,MA/My,MRL,NOD,NZB,NZO,NZW,PJ,PL,RF,SEA,SJL,SWR,TH | 10 | 26874529 | 26874562 | 34 | - | 16.7724 | 7.83E-07 | 0.0988 | TAGCTGTCTTCAGATGCACCAGAAGAGGGCGTCA | | |
| 724 | 10 | 33254302 | 33254563 | 261 | KK | 10 | 33254431 | 33254449 | 19 | - | 17.8852 | 5.08E-07 | 0.121 | ACACCAGAAGAGGGCACCA | | |
| 725 | 10 | 42452279 | 42452492 | 213 | 129S1.BTBR.C57L.KK | 10 | 42452404 | 42452437 | 34 | - | 17.3008 | 5.58E-07 | 0.0844 | CAGCCGTCCTCAGACACTCCAGAAGAGGGAGTCA | | |
| 726 | 10 | 44348891 | 44349093 | 202 | ILNJ2 | 10 | 44349016 | 44349049 | 34 | - | 16.4472 | 9.61E-07 | 0.108 | TAGCTGTCTTCAGACACACCAGAAGAGGGCGTCA | | |
| 727 | 10 | 45822854 | 45823058 | 204 | TH | 10 | 45822978 | 45823011 | 34 | - | 17.9106 | 3.74E-07 | 0.07 | TAGCTGTCTTCAGACACTCCAGAAGAGGGCACCA | | |
| 728 | 10 | 62525683 | 62525903 | 220 | CE.RF | 10 | 62525808 | 62525841 | 34 | - | 17.2114 | 5.91E-07 | 0.0866 | TAGCTGTTTTCAGACACTCCAGAAGAGGGCATCA | | |
| 729 | 10 | 77768658 | 77768872 | 214 | BUB.TH | 10 | 77768784 | 77768817 | 34 | - | 17.8455 | 3.91E-07 | 0.071 | TAGCTGTCCTCAGACACTCCAGAAGAGGGCGTCA | | |
| 730 | 10 | 80047318 | 80047546 | 228 | BUB | 10 | 80047443 | 80047476 | 34 | - | 20.4146 | 6.58E-08 | 0.0258 | CAGCTGTCCTCAGACACTCCAGAAGAGGGCGTCA | | |
| 731 | 10 | 81840583 | 81840876 | 293 | BUB | 10 | 81840835 | 81840853 | 19 | - | 19.2951 | 1.88E-07 | 0.0757 | CCGCCACTAGATGGTGCTA | 13 | ENCFF821BIA,ENCFF136ZDK,ENCFF327LYS,ENCFF201JUA,ENCFF883UPM,ENCFF093ZRU,ENCFF502WDT,ENCFF367UNR,ENCFF099QNE,ENCFF065ODV,ENCFF055HAP,ENCFF052KVO,ENCFF589ZBJ |
| 732 | 10 | 95183504 | 95183706 | 202 | MRL | 10 | 95183630 | 95183664 | 35 | - | 18.1284 | 2.79E-07 | 0.0973 | CTGTAGCTGTCTTCAGACACCAGAAGAGGGCATCA | | |
| 733 | 10 | 108116899 | 108117117 | 218 | AKR.LP.NOD.NOR.PL.RF | 10 | 108117028 | 108117061 | 34 | - | 17.8455 | 3.91E-07 | 0.071 | TAGCTGTCCTCAGACACTCCAGAAGAGGGCGTCA | | |
| 734 | 10 | 111489624 | 111489834 | 210 | SMJ | 10 | 111489751 | 111489784 | 34 | - | 16.4634 | 9.51E-07 | 0.108 | TAGCTGTCTTCAGACGCACCAGAAGAGGGCATCA | | |
| 735 | 10 | 116817779 | 116817991 | 212 | C57L.C58J.NZW.SMJ | 10 | 116817904 | 116817922 | 19 | - | 17.5574 | 6.32E-07 | 0.134 | ACTCCAGAAGAGGGCGCCA | | |
| 735 | 10 | 116817779 | 116817991 | 212 | C57L.C58J.NZW.SMJ | 10 | 116817904 | 116817937 | 34 | - | 18.3659 | 2.76E-07 | 0.0597 | TAGCTGTCTTCAGACACTCCAGAAGAGGGCGCCA | | |
| 736 | 10 | 121237649 | 121237850 | 201 | AJ,BALB,BUB,CE,DBA1J,DBA2J,FVB,ILNJ2,KK.LP.MA/My.NOD.NOR.PL.RH2.SMJ | 10 | 121237774 | 121237807 | 34 | - | 16.4472 | 9.61E-07 | 0.108 | TAGCTGTCTTCAGACACACCAGAAGAGGGCGTCA | | |
| 737 | 10 | 121273312 | 121273520 | 208 | AJ,BALB,DBA1J,DBA2J,KK,MA/My,RH2,SMJ | 10 | 121273436 | 121273469 | 34 | - | 16.4472 | 9.61E-07 | 0.108 | TAGCTGTCTTCAGACACACCAGAAGAGGGCGTCA | | |
| 738 | 10 | 122376428 | 122376609 | 181 | DBA1J.KK | 10 | 122376545 | 122376563 | 19 | - | 17.5574 | 6.32E-07 | 0.134 | ACTCCAGAAGAGGGCGCCA | 1 | ENCFF093ZRU |
| 738 | 10 | 122376428 | 122376609 | 181 | DBA1J.KK | 10 | 122376545 | 122376578 | 34 | - | 18.3659 | 2.76E-07 | 0.0597 | TAGCTGTCTTCAGACACTCCAGAAGAGGGCGCCA | 1 | ENCFF093ZRU |
| 739 | 10 | 126515251 | 126515316 | 65 | DBA1J,DBA2J,ILNJ2,KK,NZW | 10 | 126515292 | 126515310 | 19 | - | 21.0984 | 4.51E-08 | 0.0444 | AGGCCAGAAGAGGGCGCCA | 19 | ENCFF367UNR,ENCFF093ZRU,ENCFF024JOZ,ENCFF327LYS,ENCFF430PPJ,ENCFF092NMJ,ENCFF821BIA,ENCFF136ZDK,ENCFF099QNE,ENCFF589ZBJ,ENCFF201JUA,ENCFF052KVO,ENCFF957SPH,ENCFF194HVL,ENCFF533APC.ENCFF033UOX.ENCFF907XEG.ENCFF281QMV.ENCFF089GBC |
| 739 | 10 | 126515251 | 126515316 | 65 | DBA1J,DBA2J,ILNJ2,KK,NZW | 10 | 126515292 | 126515326 | 35 | - | 28.055 | 1.84E-10 | 0.0029 | GTGCAGGTGCCGGTGGAGGCCAGAAGAGGGCGCCA | 19 | ENCFF367UNR,ENCFF093ZRU,ENCFF024JOZ,ENCFF327LYS,ENCFF430PPJ,ENCFF092NMJ,ENCFF821BIA,ENCFF136ZDK,ENCFF099QNE,ENCFF589ZBJ,ENCFF201JUA,ENCFF052KVO,ENCFF957SPH,ENCFF194HVL,ENCFF533APC.ENCFF033UOX.ENCFF907XEG.ENCFF281QMV.ENCFF089GBC |
| 740 | 10 | 126789754 | 126789818 | 64 | CE | 10 | 126789810 | 126789843 | 34 | - | 18.1545 | 3.18E-07 | 0.0637 | TAGCTGTTTTCAGACACACCAGAAGAGGGCGTCA | 2 | ENCFF367UNR.ENCFF093ZRU |
| 741 | 10 | 127200111 | 127200327 | 216 | AJ,BALB,BUB,C3H,CBA,CE,FVB,NOD,NOR,PL,RH2 | 10 | 127200236 | 127200269 | 34 | - | 16.4472 | 9.61E-07 | 0.108 | TAGCTGTCTTCAGACACACCAGAAGAGGGCGTCA | 1 | ENCFF093ZRU |
| 742 | 10 | 127303212 | 127303487 | 275 | AJ,BALB,BUB,C3H,CBA,CE,FVB,NOD,NOR,PL,RH2 | 10 | 127303337 | 127303370 | 34 | - | 17.8455 | 3.91E-07 | 0.071 | TAGCTGTCCTCAGACACTCCAGAAGAGGGCGTCA | | |
| 743 | 11 | 4838030 | 4838240 | 210 | 129S1,AJ,B10D2,B10J,BALB,BTBR,BUB,C57L,C58J,ILNJ2,KK,MA/My,NOD,NOR,NZW,PJ.PL.SEA.TH | 11 | 4838096 | 4838129 | 34 | + | 18.1789 | 3.13E-07 | 0.0375 | TAGCTGTCTTCAGACACTCCAGAAGAGGGCGCTA | 3 | ENCFF093ZRU,ENCFF502WDT,ENCFF055HAP |
| 743 | 11 | 4838030 | 4838240 | 210 | 129S1,AJ,B10D2,B10J,BALB,BTBR,BUB,C57L,C58J,ILNJ2,KK,MA/My,NOD,NOR,NZW,PJ.PL.SEA.TH | 11 | 4838111 | 4838129 | 19 | + | 17.5738 | 6.25E-07 | 0.0841 | ACTCCAGAAGAGGGCGCTA | 3 | ENCFF093ZRU,ENCFF502WDT,ENCFF055HAP |
| 744 | 11 | 17072653 | 17073442 | 789 | AJ,AKR,BALB,BTBR,BUB,C3H,C57L,CBA,DBA1J,DBA2J,ILNJ2,KK,MRL,NZB,NZO,NZW,PL.RF.SEA.SJL.SMJ | 11 | 17072770 | 17072788 | 19 | + | 18.7213 | 2.85E-07 | 0.0593 | CTACCACTAGGTGGTACTA | 2 | ENCFF099QNE,ENCFF093ZRU |
| 745 | 11 | 27738290 | 27738772 | 482 | AJ,C3H,DBA1J,DBA2J,ILNJ2,MRL,NZB,NZO,NZW.PL | 11 | 27738306 | 27738339 | 34 | + | 16.626 | 8.59E-07 | 0.065 | CTGCACTCACATTTGCCATTACAAGATGGCACTG | | |
| 746 | 11 | 32579330 | 32579542 | 212 | AJ,AKR,BALB,BTBR,BUB,C57L,CE,DBA1J,DBA2J,FVB,ILNJ2,KK,MA/My,NOD,NZB,NZO,NZW,PJ,PL,RF,RH2,SEA,SWR,TH | 11 | 32579401 | 32579434 | 34 | + | 18.3659 | 2.76E-07 | 0.0355 | TAGCTGTCTTCAGACACTCCAGAAGAGGGCGCCA | | |
| 746 | 11 | 32579330 | 32579542 | 212 | AJ,AKR,BALB,BTBR,BUB,C57L,CE,DBA1J,DBA2J,FVB,ILNJ2,KK,MA/My,NOD,NZB,NZO,NZW,PJ,PL,RF,RH2,SEA,SWR,TH | 11 | 32579416 | 32579434 | 19 | + | 17.5574 | 6.32E-07 | 0.0841 | ACTCCAGAAGAGGGCGCCA | | |
| 747 | 11 | 35733276 | 35733692 | 416 | MA/My.NOD.NZB.NZO.PJ.RF.SEA | 11 | 35733550 | 35733583 | 34 | + | 18.3659 | 2.76E-07 | 0.0355 | TAGCTGTCTTCAGACACTCCAGAAGAGGGCGCCA | 3 | ENCFF367UNR.ENCFF093ZRU.ENCFF055HAP |
| 747 | 11 | 35733276 | 35733692 | 416 | MA/My.NOD.NZB.NZO.PJ.RF.SEA | 11 | 35733565 | 35733583 | 19 | + | 17.5574 | 6.32E-07 | 0.0841 | ACTCCAGAAGAGGGCGCCA | 3 | ENCFF367UNR.ENCFF093ZRU.ENCFF055HAP |
| 748 | 11 | 37763550 | 37763747 | 197 | 129S1,AJ,B10D2,B10J,BALB,BTBR,BUB,C3H,C57L,CBA,DBA1J,DBA2J,LP,MA/My,MRL,NZB.NZO.PJ.PL.SEA | 11 | 37763618 | 37763636 | 19 | + | 17.8852 | 5.08E-07 | 0.0767 | ACACCAGAAGAGGGCACCA | | |
| 749 | 11 | 42137572 | 42137791 | 219 | B10D2,B10J,BTBR,C58J,DBA2J,LP,NZB,NZO.NZW | 11 | 42137649 | 42137682 | 34 | + | 17.8455 | 3.91E-07 | 0.0425 | TAGCTGTCCTCAGACACTCCAGAAGAGGGCGTCA | | |
| 750 | 11 | 44447425 | 44447633 | 208 | 129S1,AJ,AKR,B10D2,B10J,BALB,BTBR,BUB,C3H,C57L,C58J,CBA,CE,DBA1J,DBA2J,FVB,ILNJ2,KK,MA/My,MRL,NOD,NZB,NZO,NZW,PJ,PL,RF,RH2,SEA,SJL,SMJ,SWR,TH | 11 | 44447489 | 44447522 | 34 | + | 17.1382 | 6.20E-07 | 0.0547 | TAGCTGTCTTCACACGCACCAGAAGAGGGCATCA | 1 | ENCFF093ZRU |
| 751 | 11 | 46694584 | 46694796 | 212 | NZB | 11 | 46694655 | 46694688 | 34 | + | 17.3252 | 5.49E-07 | 0.052 | TAGCTGTCCTCAGACATTCCAGAAGAGGGCATCA | | |
| 752 | 11 | 48853378 | 48853455 | 77 | PJ.SEA | 11 | 48853367 | 48853400 | 34 | + | 17.252 | 5.76E-07 | 0.0529 | TAGCTGTCTTCCGACAGTCCAGAAGAGGGAGCCA | | |
| 753 | 11 | 50915627 | 50916611 | 984 | NZO | 11 | 50916291 | 50916325 | 35 | + | 20.4679 | 6.41E-08 | 0.0248 | AGGCAGGTGATTTTGGGGGCCAGTAGAGGGCACTA | | |
| 753 | 11 | 50915627 | 50916611 | 984 | NZO | 11 | 50916307 | 50916325 | 19 | + | 22.0656 | 1.91E-08 | 0.0183 | GGGCCAGTAGAGGGCACTA | | |
| 754 | 11 | 51680833 | 51681440 | 607 | NZO | 11 | 51681170 | 51681204 | 35 | + | 17.6881 | 3.62E-07 | 0.0746 | ATGTTGTCTGCCAAGGAGGCCAGAAGAGGGCATCT | | |
| 755 | 11 | 53269370 | 53269588 | 218 | B10D2,B10J,C58J,CE,KK,NOD,NOR,NZW | 11 | 53269444 | 53269477 | 34 | + | 16.4472 | 9.61E-07 | 0.067 | TAGCTGTCTTCAGACACACCAGAAGAGGGCGTCA | | |
| 756 | 11 | 57608749 | 57608951 | 202 | 129S1,AJ,AKR,BALB,BTBR,BUB,C3H,C57L,CBA,CE,DBA1J,DBA2J,FVB,ILNJ2,KK,LP,MA/My,MRL,NOD,NZW,PJ,PL,RF,RH2,SEA,SJL.SMJ.SWR.TH | 11 | 57608808 | 57608841 | 34 | + | 18.2195 | 3.05E-07 | 0.037 | TAGCTGTCTTCAGATACTCCAGAAGAGGGCGCCA | 3 | ENCFF093ZRU,ENCFF821BIA,ENCFF367UNR |
| 756 | 11 | 57608749 | 57608951 | 202 | 129S1,AJ,AKR,BALB,BTBR,BUB,C3H,C57L,CBA,CE,DBA1J,DBA2J,FVB,ILNJ2,KK,LP,MA/My,MRL,NOD,NZW,PJ,PL,RF,RH2,SEA,SJL.SMJ.SWR.TH | 11 | 57608823 | 57608841 | 19 | + | 17.5574 | 6.32E-07 | 0.0841 | ACTCCAGAAGAGGGCGCCA | 3 | ENCFF093ZRU,ENCFF821BIA,ENCFF367UNR |
| 757 | 11 | 59883055 | 59883279 | 224 | AJ,AKR,C3H,CBA,CE,LP,MA/My,MRL,NOD,PJ.PL.RF.SEA.SMJ | 11 | 59883135 | 59883168 | 34 | + | 16.9187 | 7.13E-07 | 0.0585 | TAGCTGTCTTCAGACGCACCAGAAGAGGGCGTCA | | |
| 758 | 11 | 62664365 | 62664559 | 194 | AJ,BUB,C3H,CBA,KK,MRL,NZB,NZO,PJ,SEA | 11 | 62664410 | 62664443 | 34 | + | 16.4309 | 9.71E-07 | 0.0673 | TAGCTGTCTTCAGACGCTCCAGAAGAGGGCGTCA | | |
| 759 | 11 | 68731430 | 68731633 | 203 | 129S1,AJ,AKR,BTBR,BUB,C57L,C58J,DBA1J,DBA2J,ILNJ2,KK,LP,NOR,NZB,NZO,NZW,PJ.PL.RF.SEA.SMJ | 11 | 68731508 | 68731526 | 19 | + | 17.8852 | 5.08E-07 | 0.0767 | ACACCAGAAGAGGGCACCA | 1 | ENCFF367UNR |
| 760 | 11 | 70372610 | 70372816 | 206 | DBA1J,DBA2J,FVB,NOR,NZW,PJ,SEA,SMJ | 11 | 70372674 | 70372707 | 34 | + | 17.2439 | 5.79E-07 | 0.0529 | TAGCTGTCCTCAGATACTCCAGAAGAGGGCATCA | | |
| 761 | 11 | 72469675 | 72469887 | 212 | CE | 11 | 72469744 | 72469777 | 34 | + | 16.4472 | 9.61E-07 | 0.067 | TAGCTGTCTTCAGACACACCAGAAGAGGGCGTCA | | |
| 762 | 11 | 76236826 | 76237046 | 220 | 129S1,AJ,AKR,BALB,BTBR,BUB,C3H,C57L,C58J,CBA,CE,DBA1J,DBA2J,FVB,ILNJ2,KK,MA/My,MRL,NOR,NZO,PJ,PL,RF,RH2,SEA,SJL.SMJ.SWR | 11 | 76236905 | 76236938 | 34 | + | 17.8455 | 3.91E-07 | 0.0425 | TAGCTGTCCTCAGACACTCCAGAAGAGGGCGTCA | | |
| 763 | 11 | 77680127 | 77680340 | 213 | CE | 11 | 77680199 | 77680232 | 34 | + | 16.5203 | 9.18E-07 | 0.067 | TAGCTGTCTTCCAACACTCCAGAAGAGGGCGTCA | | |
| 764 | 11 | 78797431 | 78797626 | 195 | CE.FVB.SMJ | 11 | 78797485 | 78797518 | 34 | + | 17.3902 | 5.27E-07 | 0.0504 | TAGCTGTCCTCAGACACTCCAGAAGAGGGCATCA | | |
| 765 | 11 | 79930254 | 79930432 | 178 | CE.TH | 11 | 79930290 | 79930323 | 34 | + | 17.8455 | 3.91E-07 | 0.0425 | TAGCTGTCCTCAGACACTCCAGAAGAGGGCGTCA | | |
| 766 | 11 | 80040036 | 80040224 | 188 | CE | 11 | 80040081 | 80040114 | 34 | + | 16.4472 | 9.61E-07 | 0.067 | TAGCTGTCTTCAGACACACCAGAAGAGGGCGTCA | | |
| 767 | 11 | 80514952 | 80515176 | 224 | AKR.BALB.C58J.NOR.RF.RH2.SMJ.TH | 11 | 80515034 | 80515067 | 34 | + | 18.3659 | 2.76E-07 | 0.0355 | TAGCTGTCTTCAGACACTCCAGAAGAGGGCGCCA | | |
| 767 | 11 | 80514952 | 80515176 | 224 | AKR.BALB.C58J.NOR.RF.RH2.SMJ.TH | 11 | 80515049 | 80515067 | 19 | + | 17.5574 | 6.32E-07 | 0.0841 | ACTCCAGAAGAGGGCGCCA | | |
| 768 | 11 | 80863039 | 80863304 | 265 | AKR.RF.TH | 11 | 80863163 | 80863196 | 34 | + | 16.7724 | 7.83E-07 | 0.0614 | TAGCTGTCTTCAGATGCACCAGAAGAGGGCGTCA | | |
| 769 | 11 | 85567317 | 85567517 | 200 | BALB | 11 | 85567463 | 85567481 | 19 | + | 17.9344 | 4.92E-07 | 0.0767 | AGGCCAGTAGAGGGAAGTG | 2 | ENCFF327LYS.ENCFF198SSR |
| 770 | 11 | 86524972 | 86525193 | 221 | 129S1,AJ,AKR,BTBR,C3H,C57L,C58J,CBA,DBA1J,DBA2J,KK,MRL,NZB,NZO,PL,RF,SJL,SWR | 11 | 86525049 | 86525082 | 34 | + | 16.4634 | 9.51E-07 | 0.067 | TAGCTGTCTTCAGACGCACCAGAAGAGGGCATCA | | |
| 771 | 11 | 86726154 | 86727029 | 875 | 129S1,AJ,AKR,BTBR,BUB,C3H,C57L,C58J,CBA,DBA1J,DBA2J,KK,MRL,NZW,PJ,PL,RF,SEA.SJL.SWR | 11 | 86726778 | 86726811 | 34 | + | 17.1951 | 5.97E-07 | 0.0536 | GTGCAGTGCACCTGAAGGCCAGAATGGGGAGTCA | | |
| 772 | 11 | 86734243 | 86734450 | 207 | 129S1,AJ,AKR,BTBR,BUB,C3H,C57L,C58J,CBA,DBA1J,DBA2J,KK,MRL,NZW,PJ,PL,RF,SEA.SJL.SWR | 11 | 86734308 | 86734341 | 34 | + | 16.4472 | 9.61E-07 | 0.067 | TAGCTGTCTTCAGACACACCAGAAGAGGGCGTCA | | |
| 773 | 11 | 87953735 | 87953981 | 246 | 129S1,AJ,AKR,BALB,BTBR,BUB,C3H,C57L,C58J,CBA,CE,DBA1J,DBA2J,FVB,ILNJ2,KK,MA/My,MRL,NOD,NOR,NZB,NZO,NZW,PJ,PL,RF,RH2,SEA,SJL,SMJ,SWR,TH | 11 | 87953838 | 87953871 | 34 | + | 16.4634 | 9.51E-07 | 0.067 | TAGCTGTCTTCAGACGCACCAGAAGAGGGCATCA | | |
| 774 | 11 | 88852579 | 88852780 | 201 | 129S1,AJ,AKR,BALB,BTBR,BUB,C3H,C57L,CBA,CE,DBA1J,DBA2J,FVB,ILNJ2,KK,MA/My,MRL,NOD,NOR,NZB,NZO,NZW,PJ,PL,RF,RH2.SEA.SJL.SMJ.SWR.TH | 11 | 88852638 | 88852671 | 34 | + | 17.6667 | 4.40E-07 | 0.0454 | TAGCTGTCTTCAGACACGCCAGAAGAGGGCGTCA | | |
| 775 | 11 | 89085554 | 89085769 | 215 | 129S1,AJ,AKR,BALB,BTBR,BUB,C3H,C57L,CBA,CE,DBA1J,DBA2J,FVB,ILNJ2,KK,MA/My,MRL,NOD,NOR,NZB,NZO,NZW,PJ,PL,RF,RH2.SEA.SJL.SMJ.SWR.TH | 11 | 89085627 | 89085660 | 34 | + | 17.3902 | 5.27E-07 | 0.0504 | TAGCTGTCCTCAGACACTCCAGAAGAGGGCATCA | | |
| 776 | 11 | 94975490 | 94975698 | 208 | AJ.AKR.BUB.C3H.CBA | 11 | 94975554 | 94975587 | 34 | + | 16.7724 | 7.83E-07 | 0.0614 | TAGCTGTCTTCAGATGCACCAGAAGAGGGCGTCA | | |
| 777 | 11 | 95624062 | 95624270 | 208 | AJ.AKR.BUB.C3H.CBA.RH2 | 11 | 95624128 | 95624161 | 34 | + | 17.878 | 3.83E-07 | 0.0424 | TAGCTGTCCTCAGACACACCAGAAGAGGGCATCA | | |
| 778 | 11 | 96654375 | 96654582 | 207 | AJ,AKR,BALB,BUB,CBA,MRL,NOD,NOR,NZW.PL.RH2.SJL.SMJ | 11 | 96654440 | 96654473 | 34 | + | 17.8455 | 3.91E-07 | 0.0425 | TAGCTGTCCTCAGACACTCCAGAAGAGGGCGTCA | 1 | ENCFF093ZRU |
| 779 | 11 | 97610099 | 97610455 | 356 | NZW.SWR | 11 | 97610285 | 97610319 | 35 | + | 16.4312 | 7.50E-07 | 0.111 | GTGCAGCGCCCAGCAGCCCCCGGGGGAAGGCAGCC | | |
| 780 | 11 | 98068810 | 98069020 | 210 | 129S1,BTBR,C3H,C57L,CE,DBA1J,DBA2J,FVB,ILNJ2,KK,MA/My,NOD,NOR,NZO,NZW,SWR | 11 | 98068879 | 98068912 | 34 | + | 17.2439 | 5.79E-07 | 0.0529 | TAGCTGTCCTCAGATACTCCAGAAGAGGGCATCA | | |
| 781 | 11 | 98911665 | 98911853 | 188 | CE.NZB | 11 | 98911752 | 98911786 | 35 | + | 17.3761 | 4.35E-07 | 0.0826 | ATCCATGTTTGCCTGCAGACCAGAAGAGGGCACCA | 4 | ENCFF093ZRU.ENCFF327LYS.ENCFF502WDT.ENCFF055HAP |
| 781 | 11 | 98911665 | 98911853 | 188 | CE.NZB | 11 | 98911768 | 98911786 | 19 | + | 19.3934 | 1.75E-07 | 0.0464 | AGACCAGAAGAGGGCACCA | 4 | ENCFF093ZRU.ENCFF327LYS.ENCFF502WDT.ENCFF055HAP |
| 782 | 11 | 101360474 | 101360697 | 223 | 129S1,AJ,BTBR,BUB,C3H,C57L,CE,DBA1J,DBA2J,ILNJ2,KK,MRL,NOD,NOR,NZB,NZO,SJL.TH | 11 | 101360551 | 101360584 | 34 | + | 17.2439 | 5.79E-07 | 0.0529 | TAGCTGTCCTCAGACCCTCCAGAAGAGGGAGTCA | | |
| 783 | 11 | 107016346 | 107016551 | 205 | BUB.ILNJ2.RF.SJL.TH | 11 | 107016409 | 107016442 | 34 | + | 17.8455 | 3.91E-07 | 0.0425 | TAGCTGTCCTCAGACACTCCAGAAGAGGGCGTCA | | |
| 784 | 11 | 107506814 | 107507028 | 214 | 129S1.BUB.CBA.NZW.SMJ.TH | 11 | 107506886 | 107506919 | 34 | + | 16.4634 | 9.51E-07 | 0.067 | TAGCTGTCTTCAGACGCACCAGAAGAGGGCATCA | | |
| 785 | 11 | 110083843 | 110084030 | 187 | 129S1,AJ,BTBR,C57L,CBA,CE,FVB,ILNJ2,MA/My,NOD,NOR,NZO,NZW,SJL,SMJ,SWR,TH | 11 | 110083888 | 110083921 | 34 | + | 17.2439 | 5.79E-07 | 0.0529 | TAGCTGTCCTCAGATACTCCAGAAGAGGGCATCA | | |
| 786 | 11 | 114497269 | 114497482 | 213 | 129S1,AJ,BALB,C3H,C57L,C58J,CBA,CE,ILNJ2,KK,MA/My,MRL,NOD,NOR,NZB,NZO,PL,RF.SEA.SMJ.TH | 11 | 114497341 | 114497374 | 34 | + | 18.3659 | 2.76E-07 | 0.0355 | TAGCTGTCTTCAGACACTCCAGAAGAGGGCGCCA | | |
| 786 | 11 | 114497269 | 114497482 | 213 | 129S1,AJ,BALB,C3H,C57L,C58J,CBA,CE,ILNJ2,KK,MA/My,MRL,NOD,NOR,NZB,NZO,PL,RF.SEA.SMJ.TH | 11 | 114497356 | 114497374 | 19 | + | 17.5574 | 6.32E-07 | 0.0841 | ACTCCAGAAGAGGGCGCCA | | |
| 787 | 11 | 114748153 | 114748599 | 446 | AJ.C3H.C57L.CBA.MA/My.PL | 11 | 114748484 | 114748502 | 19 | + | 17.4426 | 6.81E-07 | 0.0875 | AGGCCAGAAGGGGGCATTA | | |
| 788 | 11 | 115419818 | 115420037 | 219 | 129S1,BALB,C58J,DBA1J,DBA2J,KK,NOD,NOR,NZB,NZO,NZW,PL,RF,SEA,SMJ,TH | 11 | 115419894 | 115419927 | 34 | + | 17.2439 | 5.79E-07 | 0.0529 | TAGCTGTCCTCAGATACTCCAGAAGAGGGCATCA | | |
| 789 | 11 | 115966202 | 115966435 | 233 | AJ.C3H.C57L.CBA.CE.ILNJ2 | 11 | 115966291 | 115966324 | 34 | + | 16.4634 | 9.51E-07 | 0.067 | TAGCTGTCTTCAGACGCACCAGAAGAGGGCATCA | | |
| 790 | 11 | 115981177 | 115981386 | 209 | AJ.BTBR.C3H.C57L.CBA.CE.ILNJ2 | 11 | 115981244 | 115981277 | 34 | + | 16.8537 | 7.43E-07 | 0.0602 | TAGCTGTCCTTAGATACTCCAGAAGAGGGCATCA | | |
| 791 | 11 | 116177438 | 116177971 | 533 | AJ,BALB,BTBR,C3H,C57L,CBA,CE,DBA1J,DBA2J.ILNJ2.SEA | 11 | 116177579 | 116177613 | 35 | + | 15.9725 | 9.70E-07 | 0.125 | AGGCCTTGTGCCCTCCCTGCCTCCAGATGGTACAC | | |
| 792 | 11 | 116623774 | 116624003 | 229 | 129S1,AJ,AKR,BTBR,BUB,C3H,C57L,C58J,CBA,CE,FVB,ILNJ2,KK,MA/My,MRL,NOD,NOR,NZB,NZO,NZW,PL,RF,RH2,SJL,SMJ,SWR,TH | 11 | 116623839 | 116623873 | 35 | + | 17.2661 | 4.64E-07 | 0.0861 | CTGTAGCTATCCAGACACTCCAGAAGAGGGAGTCA | | |
| 793 | 11 | 119964213 | 119964281 | 68 | BALB,BTBR,C3H,C57L,C58J,CBA,DBA1J,DBA2J.ILNJ2.NZB.NZW.SEA.SJL | 11 | 119964269 | 119964303 | 35 | + | 16 | 9.55E-07 | 0.124 | CAGCTCTGTAGCACTTGGCCCAGCAGAGGGCCAGA | 11 | ENCFF093ZRU,ENCFF136ZDK,ENCFF198SSR,ENCFF883UPM,ENCFF589ZBJ,ENCFF165LXO,ENCFF327LYS,ENCFF089GBC.ENCFF024JOZ.ENCFF907XEG.ENCFF092NMJ |
| 794 | 11 | 120304680 | 120304890 | 210 | SJL | 11 | 120304748 | 120304781 | 34 | + | 18.3659 | 2.76E-07 | 0.0355 | TAGCTGTCTTCAGACACTCCAGAAGAGGGCGCCA | 4 | ENCFF055HAP.ENCFF367UNR.ENCFF093ZRU.ENCFF821BIA |
| 794 | 11 | 120304680 | 120304890 | 210 | SJL | 11 | 120304763 | 120304781 | 19 | + | 17.5574 | 6.32E-07 | 0.0841 | ACTCCAGAAGAGGGCGCCA | 4 | ENCFF055HAP.ENCFF367UNR.ENCFF093ZRU.ENCFF821BIA |

| | | | | | | | | | | | | | | | | |
|---|---|---|---|---|---|---|---|---|---|---|---|---|---|---|---|---|
| 795 | 11 | 3776948 | 3777149 | 201 | B10D2,B10J,C58J,KK | 11 | 3777060 | 3777093 | 34 | - | 17.1545 | 6.13E-07 | 0.0544 | TAGCTGTCTTCCAACAGTCCAGAAGAGGGAGCCA | 1 | ENCFF093ZRU |
| 796 | 11 | 4063836 | 4064041 | 205 | NZB | 11 | 4063961 | 4063994 | 34 | - | 18.0732 | 3.36E-07 | 0.0394 | CAGCTGTCTTCAGACACTCCAGAAGAGGGCATCA | | |
| 797 | 11 | 6374252 | 6374450 | 198 | AJ,AKR,B10D2,B10J,BALB,BTBR,C3H,C57L,C58J,MA/My,MRL,NOD,NOR,PL,RF,SEA,TH | 11 | 6374375 | 6374408 | 34 | - | 17.2439 | 5.79E-07 | 0.0529 | TAGCTGTCCTCAGATACTCCAGAAGAGGGCATCA | | |
| 798 | 11 | 11712655 | 11712862 | 207 | 129S1,BALB,BUB,C57L,C58J,CBA,CE,DBA1J,DBA2J,FVB,ILNJ2,MA/My,NZO,PJ,PL,SEA,SJL,SMJ,SWR,TH | 11 | 11712781 | 11712814 | 34 | - | 16.4634 | 9.51E-07 | 0.067 | TAGCTGTCTTCAGACGCACCAGAAGAGGGCATCA | | |
| 799 | 11 | 17899549 | 17899772 | 223 | 129S1,AJ,AKR,BALB,BTBR,BUB,C3H,C57L,C58J,CBA,CE,DBA1J,DBA2J,FVB,ILNJ2,KK,MA/My,MRL,NOD,NOR,NZB,NZO,NZW,PJ,PL,RF,SEA,SJL,SMJ,SWR,TH | 11 | 17899677 | 17899710 | 34 | - | 16.4634 | 9.51E-07 | 0.067 | TAGCTGTCTTCAGACGCACCAGAAGAGGGCATCA | | |
| 800 | 11 | 20020764 | 20021012 | 248 | AJ,AKR,BALB,C57L,CE,FVB,ILNJ2,KK,MA/My,NOD,NOR,NZB,NZO,NZW,PL,RF,RH2,SEA,SWR,TH | 11 | 20020891 | 20020924 | 34 | - | 16.4634 | 9.51E-07 | 0.067 | TAGCTGTCTTCAGACGCACCAGAAGAGGGCATCA | 1 | ENCFF093ZRU |
| 801 | 11 | 27013216 | 27013414 | 198 | AJ,BUB,C3H,C58J,CBA,DBA1J,DBA2J,ILNJ2,KK,MRL,NZB,NZO,NZW,PL,RF,SMJ,TH | 11 | 27013341 | 27013359 | 19 | - | 17.5574 | 6.32E-07 | 0.0841 | ACTCCAGAAGAGGGCGCCA | | |
| 801 | 11 | 27013216 | 27013414 | 198 | AJ,BUB,C3H,C58J,CBA,DBA1J,DBA2J,ILNJ2,KK,MRL,NZB,NZO,NZW,PL,RF,SMJ,TH | 11 | 27013341 | 27013374 | 34 | - | 18.3659 | 2.76E-07 | 0.0355 | TAGCTGTCTTCAGACACTCCAGAAGAGGGCGCCA | | |
| 802 | 11 | 30518373 | 30518594 | 221 | AJ,AKR,BALB,BTBR,BUB,C57L,CBA,CE,DBA1J,DBA2J,FVB,ILNJ2,NOD,NZB,NZO,PJ,PL,RF,RH2,SEA,SJL,SMJ,SWR,TH | 11 | 30518498 | 30518516 | 19 | - | 17.0328 | 8.88E-07 | 0.0977 | TCACCAGAAGAGGGCGTCA | | |
| 803 | 11 | 32547246 | 32547443 | 197 | AKR,BUB,CE,ILNJ2,KK,MA/My,NZW,PJ,PL,RF,SEA | 11 | 32547366 | 32547399 | 34 | - | 16.4472 | 9.61E-07 | 0.067 | TAGCTGTCTTCAGACACACCAGAAGAGGGCGTCA | | |
| 804 | 11 | 46273664 | 46273861 | 197 | 129S1,AJ,AKR,BALB,BTBR,BUB,C3H,C57L,CBA,CE,DBA1J,DBA2J,FVB,ILNJ2,KK,MA/My,MRL,NOD,NZB,NZW,PJ,PL,RF,RH2,SEA,SJL.SMJ.SWR.TH | 11 | 46273775 | 46273808 | 34 | - | 17.6667 | 4.40E-07 | 0.0454 | TAGCTGTTTTCAGACACTCCAGAAGAGGGCGTCA | | |
| 805 | 11 | 51581358 | 51581565 | 207 | BUB,CE,KK,MA/My,NOD,NZB,NZO,NZW,PL,SJL,SWR | 11 | 51581484 | 51581502 | 19 | - | 17.5574 | 6.32E-07 | 0.0841 | ACTCCAGAAGAGGGCGCCA | 3 | ENCFF093ZRU,ENCFF367UNR,ENCFF055HAP |
| 805 | 11 | 51581358 | 51581565 | 207 | BUB,CE,KK,MA/My,NOD,NZB,NZO,NZW,PL,SJL,SWR | 11 | 51581484 | 51581517 | 34 | - | 18.3659 | 2.76E-07 | 0.0355 | TAGCTGTCTTCAGACACTCCAGAAGAGGGCGCCA | 3 | ENCFF093ZRU,ENCFF367UNR,ENCFF055HAP |
| 806 | 11 | 55007301 | 55007509 | 208 | B10D2,B10J,C58J,KK,NOR,NZB,NZO,NZW | 11 | 55007425 | 55007443 | 19 | - | 17.5574 | 6.32E-07 | 0.0841 | ACTCCAGAAGAGGGCGCCA | | |
| 806 | 11 | 55007301 | 55007509 | 208 | B10D2,B10J,C58J,KK,NOR,NZB,NZO,NZW | 11 | 55007425 | 55007458 | 34 | - | 18.3659 | 2.76E-07 | 0.0355 | TAGCTGTCTTCAGACACTCCAGAAGAGGGCGCCA | | |
| 807 | 11 | 66412171 | 66412379 | 208 | 129S1,BALB,BTBR,C57L,CBA,CE,DBA1J,DBA2J,FVB,ILNJ2,LP,MA/My,NOR,NZB,NZW,PJ.RH2.SEA.SJL.SWR.TH | 11 | 66412299 | 66412332 | 34 | - | 18.5285 | 2.48E-07 | 0.0346 | CAGCTGTCTTCAGACACTCCAGAAGAGGGCGTCA | | |
| 808 | 11 | 68709676 | 68709771 | 95 | DBA1J.DBA2J.NOR.PJ.SEA.SMJ | 11 | 68709731 | 68709764 | 34 | - | 17.0081 | 6.74E-07 | 0.0572 | CTGCTGAGCCACCTGCCACCTCAGGGAGGACCAA | | |
| 809 | 11 | 70254661 | 70254981 | 320 | DBA1J.DBA2J.NOR.PJ.SEA.SMJ | 11 | 70254783 | 70254816 | 34 | - | 16.4472 | 9.61E-07 | 0.067 | TAGCTGTCTTCAGACACACCAGAAGAGGGCGTCA | | |
| 810 | 11 | 70686804 | 70687723 | 919 | 129S1,AKR,BALB,BTBR,C3H,C57L,C58J,CBA,CE,DBA1J,DBA2J,FVB,ILNJ2,KK,LP,MA/My,MRL,NOD,NOR,NZB,NZO,NZW,PJ,PL,RF,RH2,SEA,SJL,SMJ,SWR | 11 | 70687236 | 70687254 | 19 | - | 17.0164 | 8.98E-07 | 0.0984 | TTGCCAGGAGGTGGTGGCA | | |
| 811 | 11 | 72202854 | 72203141 | 287 | 129S1,BTBR,C57L,CE,DBA1J,DBA2J,NOR,PJ.SEA | 11 | 72202932 | 72202965 | 34 | - | 16.4146 | 9.81E-07 | 0.0678 | TAGCTGTCTTCTGACAGTCCAGAAGAGGGAGCCA | | |
| 812 | 11 | 76575390 | 76575614 | 224 | CE.NZO | 11 | 76575516 | 76575549 | 34 | - | 17.878 | 3.83E-07 | 0.0424 | TAGCTGTCCTCAGACACACCAGAAGAGGGCATCA | | |
| 813 | 11 | 77543219 | 77543448 | 229 | CE | 11 | 77543347 | 77543365 | 19 | - | 17.5738 | 6.25E-07 | 0.0841 | ACTCCAGAAGAGGGCGCTA | | |
| 813 | 11 | 77543219 | 77543448 | 229 | CE | 11 | 77543347 | 77543380 | 34 | - | 18.1789 | 3.13E-07 | 0.0375 | TAGCTGTCTTCAGACACTCCAGAAGAGGGCGCTA | | |
| 814 | 11 | 87360058 | 87360271 | 213 | BALB,CE,FVB,ILNJ2,KK,MA/My,NOD,NOR,NZB,NZO,NZW,RH2,SMJ,TH | 11 | 87360182 | 87360215 | 34 | - | 19.7154 | 1.09E-07 | 0.0208 | CAGCTGTCTTCAGACAGACCAGAAGAGGGCGTCA | 1 | ENCFF093ZRU |
| 815 | 11 | 87382208 | 87382417 | 209 | CE,NZW | 11 | 87382332 | 87382365 | 34 | - | 16.4472 | 9.61E-07 | 0.067 | TAGCTGTCTTCAGACACACCAGAAGAGGGCGTCA | | |
| 816 | 11 | 87929728 | 87929937 | 209 | 129S1,AJ,AKR,BALB,BTBR,BUB,C3H,C57L,C58J,CBA,CE,DBA1J,DBA2J,FVB,ILNJ2,KK,MA/My,MRL,NOD,NOR,NZB,NZO,NZW,PJ,PL,RF,RH2,SEA,SJL,SMJ,SWR,TH | 11 | 87929846 | 87929864 | 19 | - | 17.5574 | 6.32E-07 | 0.0841 | ACTCCAGAAGAGGGCGCCA | 1 | ENCFF093ZRU |
| 816 | 11 | 87929728 | 87929937 | 209 | 129S1,AJ,AKR,BALB,BTBR,BUB,C3H,C57L,C58J,CBA,CE,DBA1J,DBA2J,FVB,ILNJ2,KK,MA/My,MRL,NOD,NOR,NZB,NZO,NZW,PJ,PL,RF,RH2,SEA,SJL,SMJ,SWR,TH | 11 | 87929846 | 87929879 | 34 | - | 17.5041 | 4.89E-07 | 0.0482 | TAGCTGTCATCAGACACTCCAGAAGAGGGCGCCA | 1 | ENCFF093ZRU |
| 817 | 11 | 88121530 | 88121941 | 411 | BALB,CE,ILNJ2,KK,MA/My,NZB,NZO,RH2,SMJ | 11 | 88121643 | 88121676 | 34 | - | 17.4472 | 5.08E-07 | 0.0494 | ATTCAGTGTCTGTGGCAGCCAGAAGAGGACATCA | 1 | ENCFF502WDT |
| 818 | 11 | 92888287 | 92888467 | 180 | 129S1.BTBR.C57L.CE | 11 | 92888410 | 92888443 | 34 | - | 16.4472 | 9.61E-07 | 0.067 | TAGCTGTCTTCAGACACACCAGAAGAGGGCGTCA | | |
| 819 | 11 | 93582481 | 93582679 | 198 | AJ.AKR.C3H.CBA.CE.MRL.RF | 11 | 93582603 | 93582621 | 19 | - | 17.5574 | 6.32E-07 | 0.0841 | ACTCCAGAAGAGGGCGCCA | | |
| 819 | 11 | 93582481 | 93582679 | 198 | AJ.AKR.C3H.CBA.CE.MRL.RF | 11 | 93582603 | 93582636 | 34 | - | 18.3659 | 2.76E-07 | 0.0355 | TAGCTGTCTTCAGACACTCCAGAAGAGGGCGCCA | | |
| 820 | 11 | 95176149 | 95176341 | 192 | AJ.AKR.BUB.C3H.CBA.MA/My.RH2 | 11 | 95176259 | 95176277 | 19 | - | 19.4918 | 1.63E-07 | 0.0448 | ACACCAGTAGAGGGCACCA | 1 | ENCFF093ZRU |
| 821 | 11 | 95311633 | 95311855 | 222 | AJ.AKR.BUB.C3H.CBA.MA/My.RH2 | 11 | 95311754 | 95311787 | 34 | - | 17.8455 | 3.91E-07 | 0.0425 | TAGCTGTCCTCAGACACTCCAGAAGAGGGCGTCA | 1 | ENCFF367UNR |
| 822 | 11 | 96605925 | 96606153 | 228 | AJ,AKR,BALB,BUB,CBA,MRL,NOD,NOR,NZW,PL,RH2,SJL,SMJ | 11 | 96606049 | 96606082 | 34 | - | 17.9106 | 3.74E-07 | 0.0419 | TAGCTGTCTTCAGACACTCCAGAAGAGGGCACCA | | |
| 823 | 11 | 97212604 | 97213444 | 840 | NZW.SWR | 11 | 97213243 | 97213261 | 19 | - | 19.8525 | 1.24E-07 | 0.0393 | GAACCACTAGAGGGCACTG | 2 | ENCFF883UPM.ENCFF093ZRU |
| 824 | 11 | 101457599 | 101457805 | 206 | BUB.NZO.TH | 11 | 101457724 | 101457742 | 19 | - | 17.5574 | 6.32E-07 | 0.0841 | ACTCCAGAAGAGGGCGCCA | 3 | ENCFF093ZRU.ENCFF367UNR.ENCFF055HAP |
| 824 | 11 | 101457599 | 101457805 | 206 | BUB.NZO.TH | 11 | 101457724 | 101457757 | 34 | - | 18.3659 | 2.76E-07 | 0.0355 | TAGCTGTCTTCAGACACTCCAGAAGAGGGCGCCA | 3 | ENCFF093ZRU.ENCFF367UNR.ENCFF055HAP |
| 825 | 11 | 102173543 | 102174164 | 621 | 129S1,AKR,BALB,BTBR,BUB,C3H,C57L,CE,DBA1J,DBA2J,ILNJ2,MA/My,MRL,NOD,NOR,NZB,NZO,PL,RF,RH2,SEA,SJL,SWR | 11 | 102174014 | 102174047 | 34 | - | 17.6179 | 4.54E-07 | 0.046 | TAGCTGTCTTCAGACACCCCAGAAGAGGGCACCA | 1 | ENCFF093ZRU |
| 826 | 11 | 102983571 | 102984174 | 603 | NZW | 11 | 102983706 | 102983739 | 34 | - | 20.4553 | 6.39E-08 | 0.0144 | ATGCAGTACCCTCAGAGGCCGGCAGAGAGCATGG | | |
| 827 | 11 | 106333326 | 106333533 | 207 | CBA,NZB,NZO,NZW,RF,SJL,SMJ,SWR,TH | 11 | 106333451 | 106333469 | 19 | - | 17.8852 | 5.08E-07 | 0.0767 | ACACCAGAAGAGGGCACCA | 4 | ENCFF093ZRU,ENCFF367UNR,ENCFF883UPM,ENCFF502WDT |
| 828 | 11 | 106456654 | 106456889 | 235 | BUB.RF.SMJ | 11 | 106456779 | 106456812 | 34 | - | 16.9919 | 6.81E-07 | 0.0573 | TAGCTGTCCTCAGACTCTCCAGAAGAGGGAGTCA | 1 | ENCFF093ZRU |
| 829 | 11 | 113551313 | 113551509 | 196 | AJ.C3H.CBA.TH | 11 | 113551421 | 113551454 | 34 | - | 16.4472 | 9.61E-07 | 0.067 | TAGCTGTCTTCAGACACACCAGAAGAGGGCGTCA | 1 | ENCFF093ZRU |
| 830 | 11 | 114947659 | 114947853 | 194 | AJ,AKR,BUB,C3H,C57L,CBA,DBA1J,DBA2J,PL.RH2.SWR | 11 | 114947766 | 114947799 | 34 | - | 16.4472 | 9.61E-07 | 0.067 | TAGCTGTCTTCAGACACACCAGAAGAGGGCGTCA | | |
| 831 | 11 | 115662324 | 115662538 | 214 | 129S1,AJ,AKR,BALB,C3H,C57L,C58J,CBA,CE,DBA1J,DBA2J,ILNJ2,KK,MRL,NOD,NOR,NZB,NZO,NZW,PL,RF,RH2,SEA,TH | 11 | 115662449 | 115662482 | 34 | - | 16.7724 | 7.83E-07 | 0.0614 | TAGCTGTCTTCAGATGCACCAGAAGAGGGCGTCA | | |
| 832 | 11 | 117707068 | 117707173 | 105 | 129S1,AJ,AKR,BTBR,BUB,C3H,C57L,C58J,CBA,CE,DBA1J,DBA2J,ILNJ2,KK,MA/My,MRL,NZB,NZO,NZW,PL,RH2,SJL,SMJ | 11 | 117707090 | 117707108 | 19 | - | 17.623 | 6.05E-07 | 0.0841 | CTCCCACCAGGTGGAGGTG | | |
| 833 | 11 | 117724280 | 117724499 | 219 | TH | 11 | 117724406 | 117724439 | 34 | - | 16.4472 | 9.61E-07 | 0.067 | TAGCTGTCTTCAGACACACCAGAAGAGGGCGTCA | | |
| 834 | 11 | 119274143 | 119274333 | 190 | FVB.SWR | 11 | 119274258 | 119274291 | 34 | - | 18.0732 | 3.36E-07 | 0.0394 | CAGCTGTCTTCAGACACTCCAGAAGAGGGCATCA | | |
| 835 | 11 | 120787655 | 120787857 | 202 | SJL | 11 | 120787770 | 120787803 | 34 | - | 17.9106 | 3.74E-07 | 0.0419 | TAGCTGTCTTCAGACACTCCAGAAGAGGGCACCA | | |
| 836 | 12 | 17541134 | 17541346 | 212 | 129S1,AJ,AKR,BALB,BTBR,BUB,C3H,C57L,CBA,CE,DBA1J,DBA2J,FVB,ILNJ2,KK,MA/My,MRL,NOD,NOR,NZO,PJ,PL,RF,SEA,SJL,SMJ.SWR.TH | 12 | 17541207 | 17541240 | 34 | + | 18.3659 | 2.76E-07 | 0.0591 | TAGCTGTCTTCAGACACTCCAGAAGAGGGCGCCA | | |
| 836 | 12 | 17541134 | 17541346 | 212 | 129S1,AJ,AKR,BALB,BTBR,BUB,C3H,C57L,CBA,CE,DBA1J,DBA2J,FVB,ILNJ2,KK,MA/My,MRL,NOD,NOR,NZO,PJ,PL,RF,SEA,SJL,SMJ.SWR.TH | 12 | 17541222 | 17541240 | 19 | + | 17.5574 | 6.32E-07 | 0.128 | ACTCCAGAAGAGGGCGCCA | | |
| 837 | 12 | 18440677 | 18441250 | 573 | 129S1,BTBR,BUB,C3H,C57L,CBA,DBA2J,FVB,KK,MA/My,NOD,NOR,NZO,PL,SJL,SMJ,SWR.TH | 12 | 18440694 | 18440727 | 34 | + | 17.6748 | 4.37E-07 | 0.073 | CCGCCCCTACATTCGCCGTCACAAGATGGCGCTG | | |
| 838 | 12 | 18829354 | 18829852 | 498 | DBA2J | 12 | 18829780 | 18829813 | 34 | + | 18.374 | 2.75E-07 | 0.0591 | TAGCAACACTAAACGTGACCCCTAGATGGCAGCG | | |
| 839 | 12 | 21495295 | 21495494 | 199 | AJ,AKR,BALB,CE,DBA1J,ILNJ2,PJ,RF,SEA | 12 | 21495361 | 21495394 | 34 | + | 16.4634 | 9.51E-07 | 0.104 | TAGCTGTCTTCAGACGCACCAGAAGAGGGCATCA | | |
| 840 | 12 | 40629270 | 40629482 | 212 | CE.FVB.KK.NOD.NOR.SJL | 12 | 40629356 | 40629374 | 19 | + | 17.8852 | 5.08E-07 | 0.115 | ACACCAGAAGAGGGCACCA | 1 | ENCFF093ZRU |
| 841 | 12 | 56673684 | 56673915 | 231 | 129S1.BTBR.C57L.CE.FVB.NZW.PL | 12 | 56673772 | 56673805 | 34 | + | 16.5691 | 8.90E-07 | 0.102 | TAGCTGTCTTCAGACGCACCAGAAGAGGGCATCG | | |
| 842 | 12 | 59081758 | 59082613 | 855 | NZW | 12 | 59081797 | 59081830 | 34 | + | 18.065 | 3.38E-07 | 0.064 | TAGCTGTCTTCAGACTCACCAGAAGAGGGCGTCA | | |
| 842 | 12 | 59081758 | 59082613 | 855 | NZW | 12 | 59081812 | 59081830 | 19 | + | 17.0328 | 8.88E-07 | 0.147 | TCACCAGAAGAGGGCGTCA | | |
| 842 | 12 | 59081758 | 59082613 | 855 | NZW | 12 | 59081892 | 59081925 | 34 | + | 18.065 | 3.38E-07 | 0.064 | TAGCTGTCTTCAGACTCACCAGAAGAGGGCGTCA | | |
| 842 | 12 | 59081758 | 59082613 | 855 | NZW | 12 | 59081907 | 59081925 | 19 | + | 17.0328 | 8.88E-07 | 0.147 | TCACCAGAAGAGGGCGTCA | | |
| 842 | 12 | 59081758 | 59082613 | 855 | NZW | 12 | 59081987 | 59082020 | 34 | + | 18.065 | 3.38E-07 | 0.064 | TAGCTGTCTTCAGACTCACCAGAAGAGGGCGTCA | | |
| 842 | 12 | 59081758 | 59082613 | 855 | NZW | 12 | 59082002 | 59082020 | 19 | + | 17.0328 | 8.88E-07 | 0.147 | TCACCAGAAGAGGGCGTCA | | |
| 842 | 12 | 59081758 | 59082613 | 855 | NZW | 12 | 59082082 | 59082115 | 34 | + | 18.065 | 3.38E-07 | 0.064 | TAGCTGTCTTCAGACTCACCAGAAGAGGGCGTCA | | |
| 842 | 12 | 59081758 | 59082613 | 855 | NZW | 12 | 59082097 | 59082115 | 19 | + | 17.0328 | 8.88E-07 | 0.147 | TCACCAGAAGAGGGCGTCA | | |
| 842 | 12 | 59081758 | 59082613 | 855 | NZW | 12 | 59082177 | 59082210 | 34 | + | 18.065 | 3.38E-07 | 0.064 | TAGCTGTCTTCAGACTCACCAGAAGAGGGCGTCA | | |
| 842 | 12 | 59081758 | 59082613 | 855 | NZW | 12 | 59082192 | 59082210 | 19 | + | 17.0328 | 8.88E-07 | 0.147 | TCACCAGAAGAGGGCGTCA | | |
| 842 | 12 | 59081758 | 59082613 | 855 | NZW | 12 | 59082272 | 59082305 | 34 | + | 18.065 | 3.38E-07 | 0.064 | TAGCTGTCTTCAGACTCACCAGAAGAGGGCGTCA | | |
| 842 | 12 | 59081758 | 59082613 | 855 | NZW | 12 | 59082287 | 59082305 | 19 | + | 17.0328 | 8.88E-07 | 0.147 | TCACCAGAAGAGGGCGTCA | | |
| 842 | 12 | 59081758 | 59082613 | 855 | NZW | 12 | 59082367 | 59082400 | 34 | + | 18.065 | 3.38E-07 | 0.064 | TAGCTGTCTTCAGACTCACCAGAAGAGGGCGTCA | | |
| 842 | 12 | 59081758 | 59082613 | 855 | NZW | 12 | 59082382 | 59082400 | 19 | + | 17.0328 | 8.88E-07 | 0.147 | TCACCAGAAGAGGGCGTCA | | |
| 842 | 12 | 59081758 | 59082613 | 855 | NZW | 12 | 59082462 | 59082495 | 34 | + | 18.065 | 3.38E-07 | 0.064 | TAGCTGTCTTCAGACTCACCAGAAGAGGGCGTCA | | |
| 842 | 12 | 59081758 | 59082613 | 855 | NZW | 12 | 59082477 | 59082495 | 19 | + | 17.0328 | 8.88E-07 | 0.147 | TCACCAGAAGAGGGCGTCA | | |
| 842 | 12 | 59081758 | 59082613 | 855 | NZW | 12 | 59082557 | 59082590 | 34 | + | 18.065 | 3.38E-07 | 0.064 | TAGCTGTCTTCAGACTCACCAGAAGAGGGCGTCA | | |
| 842 | 12 | 59081758 | 59082613 | 855 | NZW | 12 | 59082572 | 59082590 | 19 | + | 17.0328 | 8.88E-07 | 0.147 | TCACCAGAAGAGGGCGTCA | | |
| 843 | 12 | 59081760 | 59081855 | 95 | B10J.MRL | 12 | 59081797 | 59081830 | 34 | + | 18.065 | 3.38E-07 | 0.064 | TAGCTGTCTTCAGACTCACCAGAAGAGGGCGTCA | | |
| 843 | 12 | 59081760 | 59081855 | 95 | B10J.MRL | 12 | 59081812 | 59081830 | 19 | + | 17.0328 | 8.88E-07 | 0.147 | TCACCAGAAGAGGGCGTCA | | |
| 844 | 12 | 59085423 | 59085660 | 237 | C57L.MRL | 12 | 59085407 | 59085440 | 34 | + | 18.065 | 3.38E-07 | 0.064 | TAGCTGTCTTCAGACTCACCAGAAGAGGGCGTCA | | |
| 844 | 12 | 59085423 | 59085660 | 237 | C57L.MRL | 12 | 59085422 | 59085440 | 19 | + | 17.0328 | 8.88E-07 | 0.147 | TCACCAGAAGAGGGCGTCA | | |
| 844 | 12 | 59085423 | 59085660 | 237 | C57L.MRL | 12 | 59085502 | 59085535 | 34 | + | 18.065 | 3.38E-07 | 0.064 | TAGCTGTCTTCAGACTCACCAGAAGAGGGCGTCA | | |
| 844 | 12 | 59085423 | 59085660 | 237 | C57L.MRL | 12 | 59085517 | 59085535 | 19 | + | 17.0328 | 8.88E-07 | 0.147 | TCACCAGAAGAGGGCGTCA | | |
| 845 | 12 | 71009574 | 71009797 | 223 | FVB.NZW.SJL.SWR | 12 | 71009648 | 71009681 | 34 | + | 18.1545 | 3.18E-07 | 0.0635 | TAGCTGTTTTCAGACACACCAGAAGAGGGCGTCA | | |
| 846 | 12 | 75326783 | 75327542 | 759 | FVB,SWR | 12 | 75327426 | 75327444 | 19 | + | 18.2623 | 3.93E-07 | 0.103 | TGGCCACTAGGTGGAGAAA | 20 | ENCFF367UNR,ENCFF093ZRU,ENCFF136ZDK,ENCFF430PPJ,ENCFF089GBC,ENCFF092NMJ,ENCFF821BIA,ENCFF099QNE,ENCFF907XEG,ENCFF024JOZ,ENCFF316FYN,ENCFF609MDN,ENCFF327LYS,ENCFF198SSR,ENCFF165LXO,ENCFF951GOP,ENCFF589ZBJ,ENCFF055HAP,ENCFF502WDT,ENCFF883UPM |
| 847 | 12 | 75675393 | 75676145 | 752 | CE.FVB.PJ.SEA.SMJ.SWR | 12 | 75676095 | 75676128 | 34 | + | 16.9593 | 6.95E-07 | 0.0904 | TTGCTCTCTTCAGACACACCAGAAGAGGGCATCA | | |
| 848 | 12 | 83681689 | 83681907 | 218 | KK.SWR | 12 | 83681765 | 83681798 | 34 | + | 16.4472 | 9.61E-07 | 0.104 | TAGCTGTCTTCAGACACACCAGAAGAGGGCGTCA | | |
| 849 | 12 | 84451168 | 84451400 | 232 | CE.NZO.PL.SJL.SWR | 12 | 84451260 | 84451293 | 34 | + | 18.374 | 2.75E-07 | 0.0591 | TAGCTGTCTTCAGTCACTCCAGAAGAGGGCGCCA | | |
| 849 | 12 | 84451168 | 84451400 | 232 | CE.NZO.PL.SJL.SWR | 12 | 84451275 | 84451293 | 19 | + | 17.5574 | 6.32E-07 | 0.128 | ACTCCAGAAGAGGGCGCCA | | |
| 850 | 12 | 85206961 | 85207164 | 203 | CE.SWR | 12 | 85207022 | 85207055 | 34 | + | 17.8455 | 3.91E-07 | 0.0684 | TAGCTGTCCTCAGACACTCCAGAAGAGGGCGTCA | | |
| 851 | 12 | 86478761 | 86478967 | 206 | CE.NOD.NOR.SWR | 12 | 86478827 | 86478860 | 34 | + | 18.2439 | 3.00E-07 | 0.0613 | TAGCTGTCCTCAGATACTCCAGAAGGGGGCATCA | 1 | ENCFF093ZRU |
| 852 | 12 | 86842730 | 86842806 | 76 | CE,NOD,NOR | 12 | 86842763 | 86842797 | 35 | + | 16.0826 | 9.12E-07 | 0.189 | TTGTTATAACATGTACTGTCCACTGGATGGTGCTG | 36 | ENCFF367UNR,ENCFF093ZRU,ENCFF099QNE,ENCFF024JOZ,ENCFF907XEG,ENCFF821BIA,ENCFF136ZDK,ENCFF589ZBJ,ENCFF321AGQ,ENCFF327LYS,ENCFF089GBC,ENCFF086ZTV,ENCFF198SSR,ENCFF092NMJ,ENCFF065ODV,ENCFF430PPJ,ENCFF165LXO,ENCFF883UPM,ENCFF052KVO,ENCFF951GOP,ENCFF601JVE,ENCFF504JDS,ENCFF281QMV,ENCFF957SPH,ENCFF210BTO,ENCFF575MGI,ENCFF609MDN,ENCFF316FYN,ENCFF142CNG,ENCFF194HVL,ENCFF533APC,ENCFF089EWX,ENCFF097CAQ,ENCFF491RJK,ENCFF611HDQ.ENCFF033UQX |
| 853 | 12 | 86887938 | 86888159 | 221 | BUB.FVB.NOD.NOR.NZB.SWR | 12 | 86888018 | 86888051 | 34 | + | 17.8455 | 3.91E-07 | 0.0684 | TAGCTGTCCTCAGACACTCCAGAAGAGGGCGTCA | | |
| 854 | 12 | 94063025 | 94063228 | 203 | BUB.NOD.PJ.SEA.SMJ | 12 | 94063085 | 94063118 | 34 | + | 18.3659 | 2.76E-07 | 0.0591 | TAGCTGTCTTCAGACACTCCAGAAGAGGGCGCCA | | |
| 854 | 12 | 94063025 | 94063228 | 203 | BUB.NOD.PJ.SEA.SMJ | 12 | 94063100 | 94063118 | 19 | + | 17.5574 | 6.32E-07 | 0.128 | ACTCCAGAAGAGGGCGCCA | | |
| 855 | 12 | 98146693 | 98147103 | 410 | CBA | 12 | 98146806 | 98146839 | 34 | + | 23.252 | 7.35E-09 | 0.00522 | ATGCGGTGCTCAAGGAGGCCAGAAGAGGGCGTCG | 3 | ENCFF367UNR.ENCFF052KVO.ENCFF093ZRU |
| 856 | 12 | 101855215 | 101855409 | 194 | 129S1,AKR,BALB,BTBR,BUB,C57L,CBA,CE,FVB,ILNJ2,KK,LP,MA/My,NOD,NOR,NZB,NZO,NZW,PJ,PL,RF,SEA,SJL,SWR,TH | 12 | 101855267 | 101855300 | 34 | + | 16.4472 | 9.61E-07 | 0.104 | TAGCTGTCTTCAGACACACCAGAAGAGGGCGTCA | | |
| 857 | 12 | 101959233 | 101959438 | 205 | AKR,BALB,BUB,CBA,CE,DBA1J,DBA2J,ILNJ2,KK,LP,MA/My,NOD,NOR,NZO,NZW,PL,RF.TH | 12 | 101959296 | 101959329 | 34 | + | 18.3659 | 2.76E-07 | 0.0591 | TAGCTGTCTTCAGACACTCCAGAAGAGGGCGCCA | 2 | ENCFF367UNR,ENCFF093ZRU |
| 857 | 12 | 101959233 | 101959438 | 205 | AKR,BALB,BUB,CBA,CE,DBA1J,DBA2J,ILNJ2,KK,LP,MA/My,NOD,NOR,NZO,NZW,PL,RF.TH | 12 | 101959311 | 101959329 | 19 | + | 17.5574 | 6.32E-07 | 0.128 | ACTCCAGAAGAGGGCGCCA | 2 | ENCFF367UNR,ENCFF093ZRU |
| 858 | 12 | 102719708 | 102719910 | 202 | CE.DBA1J.DBA2J.PJ.SEA | 12 | 102719768 | 102719801 | 34 | + | 17.7805 | 4.08E-07 | 0.0701 | CTGTAGTCTGTAGACACTCCAGAAGAGGGCATCA | | |
| 859 | 12 | 111029072 | 111029287 | 215 | KK.MA/My.RF | 12 | 111029145 | 111029178 | 34 | + | 16.4472 | 9.61E-07 | 0.104 | TAGCTGTCTTCAGACACACCAGAAGAGGGCGTCA | | |
| 860 | 12 | 111513573 | 111514384 | 811 | CE | 12 | 111514369 | 111514403 | 35 | + | 16.156 | 8.75E-07 | 0.185 | CTGTAACTGTCTTCAGACACCAGAAGAGGGCATCG | | |
| 861 | 12 | 118409499 | 118409727 | 228 | AJ.NOD.NOR.NZB.PL.RF | 12 | 118409584 | 118409617 | 34 | + | 17.6667 | 4.40E-07 | 0.0731 | TAGCTGTTTTCAGACACTCCAGAAGAGGGCGTCA | | |
| 862 | 12 | 12072994 | 12073735 | 741 | 129S1,AJ,AKR,BALB,BUB,C3H,C57L,CBA,CE,DBA1J,DBA2J,FVB,ILNJ2,KK,MA/My,MRL,NOD,NOR,PJ,PL,RF,SEA,SJL,SMJ,SWR,TH | 12 | 12073531 | 12073549 | 19 | - | 18.5902 | 3.13E-07 | 0.0897 | CGGCCAGCAGATGTCAGCC | | |
| 863 | 12 | 21334858 | 21335034 | 176 | 129S1,AJ,AKR,BALB,BTBR,BUB,C3H,C57L,CBA,CE,DBA1J,DBA2J,FVB,ILNJ2,KK,MA/My,MRL,NOD,NOR,NZO,PJ,PL,RF,SEA,SJL,SMJ.SWR.TH | 12 | 21334953 | 21334971 | 19 | - | 18.7049 | 2.88E-07 | 0.0872 | GCACCAGAAGAGGGCGCCA | 6 | ENCFF093ZRU,ENCFF367UNR,ENCFF136ZDK,ENCFF024JOZ,ENCFF089GBC,ENCFF883UPM |
| 863 | 12 | 21334858 | 21335034 | 176 | 129S1,AJ,AKR,BALB,BTBR,BUB,C3H,C57L,CBA,CE,DBA1J,DBA2J,FVB,ILNJ2,KK,MA/My,MRL,NOD,NOR,NZO,PJ,PL,RF,SEA,SJL,SMJ.SWR.TH | 12 | 21334953 | 21334986 | 34 | - | 19.1789 | 1.58E-07 | 0.0435 | TAGCTGTCTTCAGATGCACCAGAAGAGGGCGCCA | 6 | ENCFF093ZRU,ENCFF367UNR,ENCFF136ZDK,ENCFF024JOZ,ENCFF089GBC,ENCFF883UPM |
| 864 | 12 | 28874027 | 28874220 | 193 | 129S1,AJ,AKR,BALB,BTBR,BUB,C3H,C57L,C58J,CBA,CE,DBA1J,DBA2J,FVB,ILNJ2,KK,LP,MA/My,MRL,NOD,NOR,NZB,NZO,NZW,PJ,PL,RF,RH2,SEA,SJL,SMJ,SWR,TH | 12 | 28874152 | 28874185 | 34 | - | 17.6992 | 4.30E-07 | 0.0725 | TAGCTGTTTTCAGACACACCAGAAGAGGGCATCA | | |
| 865 | 12 | 29090511 | 29091363 | 852 | AKR RF | 12 | 29091009 | 29091042 | 34 | - | 18.6504 | 2.28E-07 | 0.055 | ATGCAGTGTCTGCAGAAGCCAGAAGAGAGTGTCA | | |
| 866 | 12 | 60631303 | 60632110 | 807 | MRL | 12 | 60631835 | 60631853 | 19 | - | 17.1475 | 8.25E-07 | 0.146 | CAACCACAAGGGGTCAGCA | 11 | ENCFF367UNR,ENCFF099QNE,ENCFF092NMJ,ENCFF089GBC,ENCFF198SSR,ENCFF093ZRU,ENCFF024JOZ,ENCFF907XEG,ENCFF502WDT,ENCFF327LYS,ENCFF055HAP |
| 866 | 12 | 60631303 | 60632110 | 807 | MRL | 12 | 60631835 | 60631869 | 35 | - | 18.1193 | 2.80E-07 | 0.097 | CAGCTTTCCCCTAATTCAACCACAAGGGGTCAGCA | 11 | ENCFF367UNR,ENCFF099QNE,ENCFF092NMJ,ENCFF089GBC,ENCFF198SSR,ENCFF093ZRU,ENCFF024JOZ,ENCFF907XEG,ENCFF502WDT,ENCFF327LYS,ENCFF055HAP |
| 867 | 12 | 69194549 | 69194777 | 228 | BTBR.C57L | 12 | 69194669 | 69194702 | 34 | - | 16.4634 | 9.51E-07 | 0.104 | TAGCTGTCTTCAGACGCACCAGAAGAGGGCATCA | | |
| 868 | 12 | 69749225 | 69749457 | 232 | SWR | 12 | 69749351 | 69749384 | 34 | - | 16.4634 | 9.51E-07 | 0.104 | TAGCTGTCTTCAGACGCACCAGAAGAGGGCATCA | 1 | ENCFF093ZRU |
| 869 | 12 | 70406728 | 70407498 | 770 | AJ,BALB,BUB,C3H,CBA,CE,DBA1J,DBA2J,FVB,ILNJ2,MRL,NZB,NZO,RH2,SEA,SJL,TH | 12 | 70407294 | 70407312 | 19 | - | 21.2131 | 4.09E-08 | 0.0421 | CAACCAGCAGGGGGCGGAG | 11 | ENCFF099QNE,ENCFF327LYS,ENCFF165LXO,ENCFF907XEG,ENCFF093ZRU,ENCFF194HVL,ENCFF136ZDK,ENCFF097CAQ,ENCFF589ZBJ,ENCFF883UPM,ENCFF198SSR |
| 870 | 12 | 76391797 | 76392008 | 211 | FVB.SWR | 12 | 76391917 | 76391950 | 34 | - | 17.8455 | 3.91E-07 | 0.0684 | TAGCTGTCCTCAGACACTCCAGAAGAGGGCGTCA | | |
| 871 | 12 | 76762132 | 76762552 | 420 | FVB.SWR | 12 | 76762254 | 76762287 | 34 | - | 18.4797 | 2.56E-07 | 0.0578 | TTGCTGTCTTCAGACACACCAGAAGAGGGCATCA | 1 | ENCFF093ZRU |
| 872 | 12 | 78902264 | 78903195 | 931 | AKR,BALB,C58J,KK,LP,MRL,NOD,NOR,NZO.PJ.PL.RF.SEA.SJL.SMJ.SWR.TH | 12 | 78902317 | 78902350 | 34 | - | 22.2602 | 1.63E-08 | 0.00935 | CAGCTGTTCTCAGACACACCAGAAGAGGGCATCG | | |
| 873 | 12 | 83449940 | 83450142 | 202 | CE.KK.SWR | 12 | 83450065 | 83450098 | 34 | - | 17.3902 | 5.27E-07 | 0.0795 | TAGCTGTCCTCAGACACTCCAGAAGAGGGCATCA | | |
| 874 | 12 | 91755280 | 91755484 | 204 | NOD | 12 | 91755406 | 91755439 | 34 | - | 16.7724 | 7.83E-07 | 0.0958 | TAGCTGTCTTCAGATGCACCAGAAGAGGGCGTCA | 5 | ENCFF093ZRU.ENCFF327LYS.ENCFF024JOZ.ENCFF136ZDK.ENCFF883UPM |
| 875 | 12 | 99805124 | 99805245 | 121 | SWR | 12 | 99805177 | 99805211 | 35 | - | 17.5413 | 3.95E-07 | 0.122 | GTGTAGTAGATCTCTTTGTCCAGTTGGTGGTGCTA | 14 | ENCFF093ZRU,ENCFF805QBU,ENCFF367UNR,ENCFF024JOZ,ENCFF327LYS,ENCFF092NMJ,ENCFF430PPJ,ENCFF055HAP,ENCFF883UPM,ENCFF099QNE,ENCFF821BIA,ENCFF089GBC,ENCFF907XEG,ENCFF502WDT |
| 876 | 12 | 100996573 | 100996756 | 183 | CE,PJ,SEA | 12 | 100996697 | 100996715 | 19 | - | 19.9836 | 1.12E-07 | 0.0594 | CTACCAGAAGAGGGCACCA | 7 | ENCFF093ZRU,ENCFF089GBC,ENCFF821BIA,ENCFF502WDT,ENCFF024JOZ,ENCFF099QNE,ENCFF367UNR |
| 877 | 12 | 108196976 | 108197191 | 215 | AJ,AKR,BUB,C3H,CBA,CE,DBA1J,DBA2J,FVB,ILNJ2,KK,MA/My,MRL,NOD,NOR,NZB,NZO,NZW,PJ,PL,RF,SEA,SJL,SMJ,SWR,TH | 12 | 108197101 | 108197134 | 34 | - | 17.8455 | 3.91E-07 | 0.0684 | TAGCTGTCCTCAGACACTCCAGAAGAGGGCGTCA | | |
| 878 | 12 | 108718974 | 108719201 | 227 | AJ,AKR,BUB,C3H,CBA,CE,DBA1J,DBA2J,FVB,ILNJ2,KK,MA/My,MRL,NOD,NOR,NZB,NZO,NZW,PJ,PL,RF,SEA,SJL,SMJ,SWR,TH | 12 | 108719101 | 108719134 | 34 | - | 16.4634 | 9.51E-07 | 0.104 | TAGCTGTCTTCAGACGCACCAGAAGAGGGCATCA | | |
| 879 | 12 | 110611339 | 110611551 | 212 | AKR,BUB,C3H,CBA,KK,MA/My,MRL,NOD,NOR,PJ,SEA,SJL | 12 | 110611465 | 110611498 | 34 | - | 16.4472 | 9.61E-07 | 0.104 | TAGCTGTCTTCAGACACACCAGAAGAGGGCGTCA | | |
| 880 | 12 | 110796942 | 110797151 | 209 | AKR,BUB,DBA1J,DBA2J,ILNJ2,KK,MA/My,MRL,NOD,NOR,NZB,NZW,PJ,RF,SEA,SJL | 12 | 110797068 | 110797086 | 19 | - | 17.8852 | 5.08E-07 | 0.115 | ACACCAGAAGAGGGCACCA | | |
| 881 | 12 | 110829111 | 110829325 | 214 | AKR,BUB,DBA1J,DBA2J,ILNJ2,KK,MA/My,MRL,NOD,NOR,NZB,NZW,PJ,RF,SEA,SJL | 12 | 110829229 | 110829247 | 19 | - | 17.5574 | 6.32E-07 | 0.128 | ACTCCAGAAGAGGGCGCCA | 2 | ENCFF093ZRU,ENCFF367UNR |
| 881 | 12 | 110829111 | 110829325 | 214 | AKR,BUB,DBA1J,DBA2J,ILNJ2,KK,MA/My,MRL,NOD,NOR,NZB,NZW,PJ,RF,SEA,SJL | 12 | 110829229 | 110829262 | 34 | - | 18.3659 | 2.76E-07 | 0.0591 | TAGCTGTCTTCAGACACTCCAGAAGAGGGCGCCA | 2 | ENCFF093ZRU,ENCFF367UNR |
| 882 | 12 | 111244368 | 111244786 | 418 | KK.MA/My.RF | 12 | 111244714 | 111244732 | 19 | - | 17.5738 | 6.25E-07 | 0.128 | ACTCCAGAAGAGGGCGCTA | 1 | ENCFF093ZRU |
| 882 | 12 | 111244368 | 111244786 | 418 | KK.MA/My.RF | 12 | 111244714 | 111244747 | 34 | - | 18.187 | 3.12E-07 | 0.0627 | TAGCTGTCTTCAGTCACTCCAGAAGAGGGCGCTA | 1 | ENCFF093ZRU |

| | | | | | | | | | | | | | | | | |
|---|---|---|---|---|---|---|---|---|---|---|---|---|---|---|---|---|
| 883 | 12 | 111689704 | 111689903 | 199 | AJ,AKR,C3H,CBA,CE,DBA1J,DBA2J,FVB,ILNJ2,MRL,NZB,NZO,NZW,PL,SMJ,SWR,TH | 12 | 111689829 | 111689862 | 34 | - | 17.2114 | 5.91E-07 | 0.0834 | TAGCTGTTTTCAGACACTCCAGAAGAGGGCATCA | | |
| 884 | 12 | 112713733 | 112713822 | 89 | AJ,AKR,C3H,CBA,CE,FVB,MRL,NZB,NZO,NZW,PJ,PL,SEA,SMJ,SWR,TH | 12 | 112713775 | 112713808 | 34 | - | 17.2114 | 5.91E-07 | 0.0834 | GAGCAACTGCCCTCTCATGCAGTAGAGGGCAGCT | 1 | ENCFF089GBC |
| 885 | 12 | 113171875 | 113172098 | 223 | 129S1,AJ,AKR,BALB,BUB,C58J,CE,FVB,KK,LP,MA/My,NZB,NZO,NZW,PJ,SEA,SWR | 12 | 113171998 | 113172032 | 35 | - | 19.5046 | 1.19E-07 | 0.0579 | GTGTACTGTCACTGACACACCAGAAGAGGGCATCA | | |
| 886 | 12 | 113759666 | 113760163 | 497 | DBA2J | 12 | 113759682 | 113759700 | 19 | - | 23.623 | 3.91E-09 | 0.0223 | CAACCAGCAGGGGGCGCTG | 10 | ENCFF821BIA,ENCFF024JOZ,ENCFF907XEG,ENCFF136ZDK,ENCFF589ZBJ,ENCFF883UPM,ENCFF367UNR,ENCFF533APC,ENCFF194HVL,ENCFF327LYS |
| 886 | 12 | 113759666 | 113760163 | 497 | DBA2J | 12 | 113759682 | 113759715 | 34 | - | 26.065 | 6.29E-10 | 0.000986 | CTGCAGGGATGATCACAACCAGCAGGGGGCGCTG | 10 | ENCFF821BIA,ENCFF024JOZ,ENCFF907XEG,ENCFF136ZDK,ENCFF589ZBJ,ENCFF883UPM,ENCFF367UNR,ENCFF533APC,ENCFF194HVL,ENCFF327LYS |
| 887 | 12 | 114074677 | 114074879 | 202 | FVB | 12 | 114074664 | 114074697 | 34 | - | 16.6667 | 8.37E-07 | 0.099 | CTACAGTGAATCTGTCATGCAGCAGGGGGTGCCA | | |
| 888 | 12 | 114283098 | 114283773 | 675 | 129S1,AJ,BALB,BUB,C58J,CE,DBA1J,DBA2J,FVB,ILNJ2,KK,LP,MA/My,PJ,RF,SEA,SMJ,SWR,TH | 12 | 114283574 | 114283592 | 19 | - | 20.2787 | 8.86E-08 | 0.0547 | GGGCCGGCAGGGGGCACTG | | |
| 888 | 12 | 114283098 | 114283773 | 675 | 129S1,AJ,BALB,BUB,C58J,CE,DBA1J,DBA2J,FVB,ILNJ2,KK,LP,MA/My,PJ,RF,SEA,SMJ,SWR,TH | 12 | 114283574 | 114283608 | 35 | - | 17.1743 | 4.90E-07 | 0.139 | CTCCACAAGGTACTTAGGGCCGGCAGGGGGCACTG | | |
| 889 | 12 | 114480188 | 114481029 | 841 | AJ,RF | 12 | 114480711 | 114480729 | 19 | - | 18.0656 | 4.49E-07 | 0.111 | TAACCAGCAGGGGGTACAA | | |
| 889 | 12 | 114480188 | 114481029 | 841 | AJ,RF | 12 | 114480711 | 114480744 | 34 | - | 20.1463 | 7.99E-08 | 0.0273 | CTGCAAGGGAGGTCATAACCAGCAGGGGGTACAA | | |
| 890 | 12 | 115250589 | 115251261 | 672 | FVB,SWR | 12 | 115250838 | 115250871 | 34 | - | 17.5691 | 4.69E-07 | 0.0751 | GAGCATGACCACAAGTCCACAGCAGGTGGCAACA | | |
| 891 | 12 | 115377650 | 115377916 | 266 | AKR,C3H,CBA,MRL,NZB,NZO,NZW,PL | 12 | 115377788 | 115377806 | 19 | - | 21.082 | 4.57E-08 | 0.0421 | GGACCAGCAGGGGGCACAA | | |
| 891 | 12 | 115377650 | 115377916 | 266 | AKR,C3H,CBA,MRL,NZB,NZO,NZW,PL | 12 | 115377788 | 115377821 | 34 | - | 26.813 | 3.09E-10 | 0.000662 | CTGCAGGTGCATTCAGGACCAGCAGGGGGCACAA | | |
| 892 | 12 | 115836299 | 115836552 | 253 | C3H,CBA,MRL,NZB,NZO,NZW,PL | 12 | 115836273 | 115836306 | 34 | - | 17.935 | 3.68E-07 | 0.0672 | ATGCATGGGAGCTCATGACCAGCAGGGGTCACCA | | |
| 893 | 12 | 116036212 | 116036412 | 200 | AJ,DBA1J,DBA2J,ILNJ2,RF | 12 | 116036329 | 116036362 | 34 | - | 16.7724 | 7.83E-07 | 0.0958 | TAGCTGTCTTCAGATGCACCAGAAGAGGGCGTCA | | |
| 894 | 12 | 117486933 | 117487149 | 216 | CE,MRL,PL | 12 | 117487056 | 117487074 | 19 | - | 17.5574 | 6.32E-07 | 0.128 | ACTCCAGAAGAGGGCGCCA | | |
| 894 | 12 | 117486933 | 117487149 | 216 | CE,MRL,PL | 12 | 117487056 | 117487089 | 34 | - | 18.3659 | 2.76E-07 | 0.0591 | TAGCTGTCTTCAGACACTCCAGAAGAGGGCGCCA | | |
| 895 | 12 | 117908038 | 117908250 | 212 | 129S1,BALB,BUB,C3H,C58J,CBA,DBA1J,DBA2J,LP,MA/My,MRL,NOD,NOR,PJ,PL,SEA,SJL | 12 | 117908167 | 117908200 | 34 | - | 18.1545 | 3.18E-07 | 0.0635 | TAGCTGTTTTCAGACACACCAGAAGAGGGCGTCA | | |
| 896 | 13 | 5429253 | 5429645 | 392 | C57L,KK,LP,NZB,NZO,NZW | 13 | 5429372 | 5429405 | 34 | + | 17.4146 | 5.18E-07 | 0.0915 | ATGCAGCTAGAGACACAAGCTCCAGGGGGTACTG | | |
| 897 | 13 | 10986009 | 10986431 | 422 | AKR,BTBR,C57L,ILNJ2,KK,LP,NOD,NOR,NZB,RF | 13 | 10986279 | 10986313 | 35 | + | 15.9266 | 9.95E-07 | 0.199 | GTACAGGGGAACACCAGGGCCAAAAGGGGGCAGTG | | |
| 897 | 13 | 10986009 | 10986431 | 422 | AKR,BTBR,C57L,ILNJ2,KK,LP,NOD,NOR,NZB,RF | 13 | 10986295 | 10986313 | 19 | + | 17.2131 | 7.91E-07 | 0.171 | GGGCCAAAAGGGGGCAGTG | | |
| 898 | 13 | 26676494 | 26677214 | 720 | KK,PJ,SEA | 13 | 26676886 | 26676920 | 35 | + | 16.0367 | 9.36E-07 | 0.193 | ATGCACTGCCCCTGGGCCGCAAGAAGGGGGCCAAG | | |
| 899 | 13 | 32167957 | 32168166 | 209 | LP | 13 | 32168025 | 32168058 | 34 | + | 16.4472 | 9.61E-07 | 0.118 | TAGCTGTCTTCAGACACACCAGAAGAGGGCGTCA | | |
| 900 | 13 | 33702837 | 33703032 | 195 | NZW | 13 | 33702888 | 33702921 | 34 | + | 17.8455 | 3.91E-07 | 0.0799 | TAGCTGTCCTCAGACACTCCAGAAGAGGGCGTCA | | |
| 901 | 13 | 34401688 | 34401948 | 260 | AJ,AKR,B10D2,B10J,BALB,BTBR,C3H,C57L,C58J,CBA,CE,DBA1J,DBA2J,ILNJ2,KK,LP,MA/My,MRL,NZB,NZO,NZW,PL,RF,RH2,SMJ,TH | 13 | 34401905 | 34401938 | 34 | + | 17.065 | 6.50E-07 | 0.103 | TAGCTGTTTTCAGATACTCCAGAAGAGGGCATCA | 1 | ENCFF093ZRU |
| 902 | 13 | 37565059 | 37565269 | 210 | 129S1,AJ,BALB,BTBR,BUB,C3H,C57L,CBA,FVB,NZW,RH2,SJL,SMJ,SWR,TH | 13 | 37565124 | 37565157 | 34 | + | 16.9187 | 7.13E-07 | 0.105 | TAGCTGTCTTCAGACGCACCAGAAGAGGGCGTCA | | |
| 903 | 13 | 37703561 | 37703788 | 227 | 129S1,AJ,AKR,BALB,BTBR,BUB,C3H,C57L,C58J,CBA,CE,FVB,ILNJ2,KK,LP,MRL,NZB,NZO,NZW,PJ,RF,RH2,SEA,SJL,SMJ,SWR,TH | 13 | 37703646 | 37703679 | 34 | + | 16.4634 | 9.51E-07 | 0.118 | TAGCTGTCTTCAGACGCACCAGAAGAGGGCATCA | | |
| 904 | 13 | 41041924 | 41042127 | 203 | NOD,NOR | 13 | 41041980 | 41042013 | 34 | + | 19.0163 | 1.77E-07 | 0.0535 | CAGCTGTCTTCAGACACACCAGAAGAGGGCGTCA | | |
| 905 | 13 | 44957435 | 44957644 | 209 | MA/My,MRL,NZO,PJ,SEA | 13 | 44957502 | 44957535 | 34 | + | 16.4634 | 9.51E-07 | 0.118 | TAGCTGTCTTCAGACGCACCAGAAGAGGGCATCA | 1 | ENCFF093ZRU |
| 906 | 13 | 53457814 | 53457988 | 174 | 129S1,AJ,AKR,BALB,BTBR,BUB,C3H,C57L,CBA,CE,DBA1J,DBA2J,FVB,ILNJ2,KK,MA/My,MRL,NOD,NOR,NZB,NZO,NZW,PJ,PL,RF,RH2,SEA,SJL,SMJ,SWR,TH | 13 | 53457846 | 53457879 | 34 | + | 17.9675 | 3.61E-07 | 0.0773 | TAGCTGTCTTCAGACCCACCAGAAGAGGGCATCG | | |
| 907 | 13 | 54235664 | 54235867 | 203 | AJ,BALB,C3H,CBA,CE | 13 | 54235724 | 54235757 | 34 | + | 16.4634 | 9.51E-07 | 0.118 | TAGCTGTCTTCAGACGCACCAGAAGAGGGCATCA | | |
| 908 | 13 | 54647937 | 54648136 | 199 | BTBR,C57L | 13 | 54647996 | 54648029 | 34 | + | 17.8293 | 3.95E-07 | 0.0803 | TAGCTGTCTTCAGACCCTCCAGAAGAGGGCGTCA | | |
| 909 | 13 | 55538000 | 55538394 | 394 | 129S1,AJ,BALB,BTBR,C3H,C57L,CBA,CE,DBA1J,DBA2J,ILNJ2,MRL,NZB,NZO,NZW,PJ,SEA,SWR | 13 | 55538071 | 55538104 | 34 | + | 18.561 | 2.42E-07 | 0.0646 | AAGCTGTCTTCAGACACTCCAGAAGAGGGCGCCA | | |
| 909 | 13 | 55538000 | 55538394 | 394 | 129S1,AJ,BALB,BTBR,C3H,C57L,CBA,CE,DBA1J,DBA2J,ILNJ2,MRL,NZB,NZO,NZW,PJ,SEA,SWR | 13 | 55538086 | 55538104 | 19 | + | 17.5574 | 6.32E-07 | 0.153 | ACTCCAGAAGAGGGCGCCA | | |
| 910 | 13 | 58493728 | 58493930 | 202 | BUB,FVB,MRL,SMJ,SWR | 13 | 58493787 | 58493820 | 34 | + | 18.3659 | 2.76E-07 | 0.0667 | TAGCTGTCTTCAGACACTCCAGAAGAGGGCGCCA | | |
| 910 | 13 | 58493728 | 58493930 | 202 | BUB,FVB,MRL,SMJ,SWR | 13 | 58493802 | 58493820 | 19 | + | 17.5574 | 6.32E-07 | 0.153 | ACTCCAGAAGAGGGCGCCA | | |
| 911 | 13 | 59642786 | 59642848 | 62 | AJ,BALB,PJ,SEA | 13 | 59642843 | 59642877 | 35 | + | 19.8349 | 9.67E-08 | 0.054 | ATGCAGTGCCATGCATCTGCCTGCTGAGGGAGGGC | | |
| 912 | 13 | 65159594 | 65159740 | 146 | 129S1,AJ,AKR,BALB,BTBR,BUB,C3H,C57L,C58J,CBA,CE,DBA1J,DBA2J,FVB,ILNJ2,KK,MRL,NOD,NOR,NZB,NZO,NZW,PJ,PL,RF,SEA,SJL,SMJ,SWR,TH | 13 | 65159715 | 65159749 | 35 | + | 19.789 | 9.95E-08 | 0.055 | TTGCAGGTGCCCCTAGAGACCAGCAGGAAGCAGTG | | |
| 913 | 13 | 70842683 | 70842893 | 210 | 129S1,BTBR,C57L,MRL,SWR,TH | 13 | 70842751 | 70842784 | 34 | + | 18.3659 | 2.76E-07 | 0.0667 | TAGCTGTCTTCAGACACTCCAGAAGAGGGCGCCA | | |
| 913 | 13 | 70842683 | 70842893 | 210 | 129S1,BTBR,C57L,MRL,SWR,TH | 13 | 70842766 | 70842784 | 19 | + | 17.5574 | 6.32E-07 | 0.153 | ACTCCAGAAGAGGGCGCCA | | |
| 914 | 13 | 74228665 | 74228774 | 109 | SWR | 13 | 74228665 | 74228698 | 34 | + | 16.4472 | 9.61E-07 | 0.118 | TAGCTGTCTTCAGACACACCAGAAGAGGGCGTCA | | |
| 915 | 13 | 75970433 | 75970637 | 204 | 129S1,AJ,AKR,BALB,BTBR,C3H,C57L,CBA,CE,DBA1J,DBA2J,FVB,ILNJ2,KK,MA/My,MRL,NZB,NZO,PJ,PL,RF,SEA,SJL,SMJ,SWR,TH | 13 | 75970493 | 75970526 | 34 | + | 18.3659 | 2.76E-07 | 0.0667 | TAGCTGTCTTCAGACACTCCAGAAGAGGGCGCCA | 1 | ENCFF093ZRU |
| 915 | 13 | 75970433 | 75970637 | 204 | 129S1,AJ,AKR,BALB,BTBR,C3H,C57L,CBA,CE,DBA1J,DBA2J,FVB,ILNJ2,KK,MA/My,MRL,NZB,NZO,PJ,PL,RF,SEA,SJL,SMJ,SWR,TH | 13 | 75970508 | 75970526 | 19 | + | 17.5574 | 6.32E-07 | 0.153 | ACTCCAGAAGAGGGCGCCA | 1 | ENCFF093ZRU |
| 916 | 13 | 82166887 | 82167137 | 250 | BALB,BUB,NZW,PJ,SEA | 13 | 82167094 | 82167127 | 34 | + | 16.6829 | 8.29E-07 | 0.112 | ATGCAGCTAGAGACACGAGCTCCAGGGGGTGGGG | | |
| 917 | 13 | 97129712 | 97130056 | 344 | 129S1,AKR,BTBR,C3H,C57L,C58J,CBA,CE,DBA1J,DBA2J,FVB,ILNJ2,KK,MA/My,MRL,NZB,NZO,PL,SJL,SMJ,SWR | 13 | 97129730 | 97129748 | 19 | + | 18.9672 | 2.39E-07 | 0.102 | TGGCCACCAGAGGACGGCT | | |
| 918 | 13 | 117351251 | 117351464 | 213 | AJ,AKR,BALB,CBA,DBA1J,FVB,ILNJ2,MA/My,NOD,NOR,RF,RH2,SJL,SMJ,SWR,TH | 13 | 117351321 | 117351354 | 34 | + | 16.4472 | 9.61E-07 | 0.118 | TAGCTGTCTTCAGACACACCAGAAGAGGGCGTCA | | |
| 919 | 13 | 117378214 | 117378420 | 206 | AJ,CBA,DBA1J,ILNJ2,MA/My,RH2,SMJ | 13 | 117378276 | 117378309 | 34 | + | 17.6992 | 4.30E-07 | 0.083 | TAGCTGTCCTCAGATACTCCAGAAGAGGGCGTCA | | |
| 920 | 13 | 117494412 | 117495255 | 843 | AJ,AKR,BALB,CBA,DBA1J,FVB,ILNJ2,MA/My,NOD,NOR,RF,RH2,SJL,SMJ,SWR,TH | 13 | 117495039 | 117495073 | 35 | + | 16.4587 | 7.38E-07 | 0.171 | ATGAGGTTTCTTCACCCCTCCAATAGAGGGCGCAC | | |
| 921 | 13 | 9103515 | 9103717 | 202 | AKR,BTBR,C57L,KK,LP,NOD,NOR,NZB,NZO,NZW,RF,TH | 13 | 9103645 | 9103678 | 34 | - | 16.4472 | 9.61E-07 | 0.118 | TAGCTGTCTTCAGACACACCAGAAGAGGGCGTCA | | |
| 922 | 13 | 9303882 | 9304098 | 216 | 129S1,AKR,BTBR,BUB,C57L,ILNJ2,KK,LP,MA/My,NOD,NOR,NZB,NZO,NZW,PJ,RF,SEA,SMJ,TH | 13 | 9304007 | 9304040 | 34 | - | 16.5691 | 8.90E-07 | 0.117 | TAGCTGTCTTCAGACGCACCAGAAGAGGGCATCG | | |
| 923 | 13 | 11942503 | 11942963 | 460 | AKR,BTBR,C57L,ILNJ2,KK,LP,NOD,NOR,NZB,RF | 13 | 11942792 | 11942810 | 19 | - | 17.7049 | 5.73E-07 | 0.149 | AGTCCAGAAGAGGGCAGTA | 11 | ENCFF327LYS,ENCFF089GBC,ENCFF367UNR,ENCFF883UPM,ENCFF502WDT,ENCFF589ZBJ,ENCFF093ZRU,ENCFF533APC,ENCFF099QNE,ENCFF052KVO,ENCFF136ZDK |
| 923 | 13 | 11942503 | 11942963 | 460 | AKR,BTBR,C57L,ILNJ2,KK,LP,NOD,NOR,NZB,RF | 13 | 11942792 | 11942826 | 35 | - | 27.055 | 4.49E-10 | 0.00272 | GTGCAGTGGCCTGTGGAGTCCAGAAGAGGGCAGTA | 11 | ENCFF327LYS,ENCFF089GBC,ENCFF367UNR,ENCFF883UPM,ENCFF502WDT,ENCFF589ZBJ,ENCFF093ZRU,ENCFF533APC,ENCFF099QNE,ENCFF052KVO,ENCFF136ZDK |
| 924 | 13 | 12389007 | 12389209 | 202 | AKR,ILNJ2,LP,RF | 13 | 12389132 | 12389165 | 34 | - | 16.4472 | 9.61E-07 | 0.118 | TAGCTGTCTTCAGACACACCAGAAGAGGGCGTCA | | |
| 925 | 13 | 17994320 | 17994532 | 212 | AJ,AKR,B10D2,B10J,BALB,BTBR,C3H,C57L,C58J,CBA,CE,DBA1J,DBA2J,ILNJ2,KK,LP,MA/My,MRL,NZB,PJ,PL,RF,RH2,SEA,SMJ | 13 | 17994444 | 17994477 | 34 | - | 18.0732 | 3.36E-07 | 0.0742 | TAGCTGTCCTCAGGTGTACCAGAAGAGGGCATCG | | |
| 926 | 13 | 20217537 | 20217736 | 199 | 129S1,AJ,AKR,B10D2,B10J,BALB,BTBR,BUB,C3H,C57L,C58J,CBA,DBA1J,DBA2J,FVB,ILNJ2,KK,LP,MA/My,MRL,NOD,NOR,NZB,PJ,PL,RF,RH2,SEA,SJL,SMJ,SWR | 13 | 20217657 | 20217691 | 35 | - | 15.9633 | 9.75E-07 | 0.197 | CTGTAGCTGTTCAGACACTCCAGAAGAGGGAGTCA | | |
| 927 | 13 | 22023451 | 22023990 | 539 | 129S1,ILNJ2 | 13 | 22023963 | 22023996 | 34 | - | 16.5447 | 9.04E-07 | 0.118 | CTGTACTTCCCATGGGCTCCTAGAGGTGGGACCC | | |
| 928 | 13 | 28987038 | 28988015 | 977 | KK,PJ,SEA | 13 | 28987559 | 28987592 | 34 | - | 17.2439 | 5.79E-07 | 0.0964 | TAGCTGTCCTCAGATACTCCAGAAGAGGGCATCA | | |
| 929 | 13 | 42626853 | 42626927 | 74 | CE | 13 | 42626824 | 42626858 | 35 | - | 26.1927 | 9.35E-10 | 0.00334 | CTGCACATACACCTGCAGGCCAGAAGAGGGCAGAA | | |
| 930 | 13 | 61950068 | 61950274 | 206 | BALB | 13 | 61950193 | 61950226 | 34 | - | 17.8455 | 3.91E-07 | 0.0799 | TAGCTGTCCTCAGACACTCCAGAAGAGGGCGTCA | | |
| 931 | 13 | 62146527 | 62146859 | 332 | 129S1,AJ,BUB,KK,MA/My,MRL,NOD,NOR,NZB,NZW,SMJ | 13 | 62146818 | 62146836 | 19 | - | 18.6557 | 2.99E-07 | 0.111 | CCGCCACTAGATGGTGGTA | 20 | ENCFF136ZDK,ENCFF089GBC,ENCFF367UNR,ENCFF201JUA,ENCFF093ZRU,ENCFF092NMJ,ENCFF165LXO,ENCFF198SSR,ENCFF055HAP,ENCFF883UPM,ENCFF502WDT,ENCFF024JOZ,ENCFF327LYS,ENCFF907XEG,ENCFF609MDN,ENCFF957SPH,ENCFF821BIA,ENCFF099QNE,ENCFF589ZBJ,ENCFF142CNG |
| 932 | 13 | 63191108 | 63191347 | 239 | 129S1,AJ,AKR,B10D2,B10J,BALB,BTBR,BUB,C3H,C57L,C58J,CBA,CE,DBA1J,DBA2J,FVB,ILNJ2,KK,LP,MA/My,MRL,NOD,NOR,NZB,NZO,NZW,PJ,PL,RF,RH2,SEA,SJL,SMJ,SWR,TH | 13 | 63191219 | 63191252 | 34 | - | 16.4472 | 9.61E-07 | 0.118 | TAGCTGTCTTCAGACACACCAGAAGAGGGCGTCA | | |
| 933 | 13 | 64352724 | 64352781 | 57 | 129S1,AJ,AKR,BALB,BTBR,BUB,C3H,C57L,C58J,CBA,CE,DBA1J,DBA2J,FVB,ILNJ2,KK,MRL,NOD,NOR,NZO,NZW,PJ,PL,RF,SEA,SJL,SMJ,SWR,TH | 13 | 64352747 | 64352765 | 19 | - | 19.6721 | 1.42E-07 | 0.0806 | TGTCCACCAGGGGGCTGTA | 16 | ENCFF821BIA,ENCFF136ZDK,ENCFF089GBC,ENCFF367UNR,ENCFF609MDN,ENCFF093ZRU,ENCFF907XEG,ENCFF024JOZ,ENCFF099QNE,ENCFF589ZBJ,ENCFF327LYS,ENCFF198SSR,ENCFF194HVL,ENCFF883UPM,ENCFF957SPH,ENCFF165LXO |
| 934 | 13 | 65562144 | 65562351 | 207 | C3H,C58J,CE,PJ,PL,SEA,SMJ | 13 | 65562265 | 65562298 | 34 | - | 17.2439 | 5.79E-07 | 0.0964 | TAGCTGTCCTCAGATACTCCAGAAGAGGGCATCA | | |
| 935 | 13 | 65704719 | 65704926 | 207 | C3H,C58J,CBA | 13 | 65704836 | 65704869 | 34 | - | 17.2439 | 5.79E-07 | 0.0964 | TAGCTGTCCTCAGATACTCCAGAAGAGGGCATCA | | |
| 936 | 13 | 67009400 | 67009602 | 202 | 129S1,AKR,MRL,NZW,RF | 13 | 67009523 | 67009556 | 34 | - | 17.2439 | 5.79E-07 | 0.0964 | TAGCTGTCCTCAGATACTCCAGAAGAGGGCATCA | | |
| 937 | 13 | 67499763 | 67499999 | 236 | 129S1,AJ,AKR,BALB,BTBR,BUB,C3H,C57L,C58J,CBA,CE,DBA1J,DBA2J,FVB,ILNJ2,KK,MRL,NOD,NOR,NZB,NZO,NZW,PJ,PL,RF,SEA,SJL,SMJ,SWR,TH | 13 | 67499887 | 67499920 | 34 | - | 16.9187 | 7.13E-07 | 0.105 | TAGCTGTCTTCAGACGCACCAGAAGAGGGCGTCA | | |
| 938 | 13 | 67550678 | 67550887 | 209 | 129S1,AJ,AKR,BALB,BTBR,BUB,C3H,C57L,C58J,CBA,CE,DBA1J,DBA2J,FVB,ILNJ2,KK,MRL,NOD,NOR,NZB,NZO,NZW,PJ,PL,RF,SEA,SJL,SMJ,SWR,TH | 13 | 67550801 | 67550834 | 34 | - | 17.6504 | 4.44E-07 | 0.0847 | CAGCTGTCCTCAGACACCCCAGAAGAGGGAGTCA | | |
| 939 | 13 | 69827560 | 69827760 | 200 | 129S1,AJ,AKR,BALB,BTBR,BUB,C3H,C57L,C58J,CBA,CE,DBA1J,DBA2J,FVB,ILNJ2,KK,MA/My,MRL,NOD,NOR,NZB,NZO,NZW,PJ,PL,RF,SEA,SJL,SMJ,SWR,TH | 13 | 69827672 | 69827705 | 34 | - | 16.4472 | 9.61E-07 | 0.118 | TAGCTGTCTTCAGACACACCAGAAGAGGGCGTCA | | |
| 940 | 13 | 70139139 | 70140037 | 898 | BUB,C58J,MA/My,MRL,NZW | 13 | 70139535 | 70139553 | 19 | - | 18.7869 | 2.72E-07 | 0.108 | CCAACACCAGGGGGCGCAA | 35 | ENCFF093ZRU,ENCFF367UNR,ENCFF805QBU,ENCFF201JUA,ENCFF099QNE,ENCFF210BTO,ENCFF092NMJ,ENCFF821BIA,ENCFF907XEG,ENCFF883UPM,ENCFF951GOP,ENCFF327LYS,ENCFF089GBC,ENCFF589ZBJ,ENCFF024JOZ,ENCFF055HAP,ENCFF430PPJ,ENCFF052KVO,ENCFF065ODV,ENCFF609MDN,ENCFF136ZDK,ENCFF502WDT,ENCFF198SSR,ENCFF165LXO,ENCFF033UQX,ENCFF097CAQ,ENCFF957SPH,ENCFF491RJK,ENCFF281QMV,ENCFF611HDQ,ENCFF533APC,ENCFF575MGI,ENCFF194HVL,ENCFF142CNG,ENCFF316FYN |
| 940 | 13 | 70139139 | 70140037 | 898 | BUB,C58J,MA/My,MRL,NZW | 13 | 70139535 | 70139568 | 34 | - | 24.6179 | 2.32E-09 | 0.0026 | GTGCTATCCCGACGTCCAACACCAGGGGGCGCAA | 35 | ENCFF093ZRU,ENCFF367UNR,ENCFF805QBU,ENCFF201JUA,ENCFF099QNE,ENCFF210BTO,ENCFF092NMJ,ENCFF821BIA,ENCFF907XEG,ENCFF883UPM,ENCFF951GOP,ENCFF327LYS,ENCFF089GBC,ENCFF589ZBJ,ENCFF024JOZ,ENCFF055HAP,ENCFF430PPJ,ENCFF052KVO,ENCFF065ODV,ENCFF609MDN,ENCFF136ZDK,ENCFF502WDT,ENCFF198SSR,ENCFF165LXO,ENCFF033UQX,ENCFF097CAQ,ENCFF957SPH,ENCFF491RJK,ENCFF281QMV,ENCFF611HDQ,ENCFF533APC,ENCFF575MGI,ENCFF194HVL,ENCFF142CNG,ENCFF316FYN |
| 941 | 13 | 70652814 | 70653022 | 208 | BUB,C58J,MA/My,MRL,NZW,SMJ,SWR | 13 | 70652940 | 70652973 | 34 | - | 16.4472 | 9.61E-07 | 0.118 | TAGCTGTCTTCAGACACACCAGAAGAGGGCGTCA | | |
| 942 | 13 | 92780607 | 92780826 | 219 | AJ,AKR,BUB,C3H,KK,MRL,NZB,NZO,NZW,PL,RF | 13 | 92780733 | 92780751 | 19 | - | 17.5574 | 6.32E-07 | 0.153 | ACTCCAGAAGAGGGCGCCA | 2 | ENCFF093ZRU,ENCFF367UNR |
| 942 | 13 | 92780607 | 92780826 | 219 | AJ,AKR,BUB,C3H,KK,MRL,NZB,NZO,NZW,PL,RF | 13 | 92780733 | 92780766 | 34 | - | 18.3659 | 2.76E-07 | 0.0667 | TAGCTGTCTTCAGACACTCCAGAAGAGGGCGCCA | 2 | ENCFF093ZRU,ENCFF367UNR |
| 943 | 13 | 102880857 | 102881824 | 967 | KK | 13 | 102881612 | 102881646 | 35 | - | 22.3578 | 1.77E-08 | 0.0175 | CTGCTGTAGCTAGGTGGATCCACTAGGTGGCAGGC | 34 | ENCFF367UNR,ENCFF093ZRU,ENCFF099QNE,ENCFF805QBU,ENCFF821BIA,ENCFF907XEG,ENCFF024JOZ,ENCFF065ODV,ENCFF055HAP,ENCFF327LYS,ENCFF502WDT,ENCFF052KVO,ENCFF092NMJ,ENCFF609MDN,ENCFF089GBC,ENCFF198SSR,ENCFF589ZBJ,ENCFF883UPM,ENCFF165LXO,ENCFF136ZDK,ENCFF321AGQ,ENCFF951GOP,ENCFF430PPJ,ENCFF097CAQ,ENCFF142CNG,ENCFF957SPH,ENCFF210BTO,ENCFF575MGI,ENCFF281QMV,ENCFF316FYN,ENCFF504JDS,ENCFF533APC,ENCFF491RJK,ENCFF194HVL |
| 944 | 13 | 103172698 | 103172889 | 191 | 129S1,C57L,NZW | 13 | 103172800 | 103172833 | 34 | - | 16.4472 | 9.61E-07 | 0.118 | TAGCTGTCTTCAGACACACCAGAAGAGGGCGTCA | | |
| 945 | 13 | 107067339 | 107067551 | 212 | CE,MA/My,NZO | 13 | 107067463 | 107067496 | 34 | - | 16.9431 | 7.02E-07 | 0.104 | TAGCTGTCTTCTCACACACCAGAAGAGGGCGTCA | | |
| 946 | 13 | 109343698 | 109343979 | 281 | CE,MA/My,NZO,TH | 13 | 109343896 | 109343914 | 19 | - | 17.5574 | 6.32E-07 | 0.153 | ACTCCAGAAGAGGGCGCCA | 1 | ENCFF093ZRU |
| 946 | 13 | 109343698 | 109343979 | 281 | CE,MA/My,NZO,TH | 13 | 109343896 | 109343929 | 34 | - | 18.3659 | 2.76E-07 | 0.0667 | TAGCTGTCTTCAGACACTCCAGAAGAGGGCGCCA | 1 | ENCFF093ZRU |
| 947 | 13 | 110666445 | 110666646 | 201 | CE,KK,MA/My,NOD,NOR,NZB,NZO,NZW,PL,SWR | 13 | 110666568 | 110666601 | 34 | - | 16.4472 | 9.61E-07 | 0.118 | TAGCTGTCTTCAGACACACCAGAAGAGGGCGTCA | | |
| 948 | 13 | 114135886 | 114136100 | 214 | AJ,BUB,CE,DBA1J,FVB,KK,MA/My,MRL,NZB,NZO,NZW,PL,RF,RH2,SJL,SWR | 13 | 114136012 | 114136045 | 34 | - | 16.6911 | 8.24E-07 | 0.112 | TAGCTGTCTTCAGACAGACCAGAAGAGGGCATCA | | |
| 949 | 13 | 117846610 | 117847056 | 446 | 129S1,BTBR,BUB,C3H,C57L,CBA,CE,DBA2J,MA/My,MRL,NZB,NZO,NZW,PL,SMJ,TH | 13 | 117846746 | 117846779 | 34 | - | 17.2439 | 5.79E-07 | 0.0964 | TAGCTGTCCTCAGATACTCCAGAAGAGGGCATCA | | |
| 950 | 13 | 120249513 | 120249721 | 208 | 129S1,BTBR,C3H,C57L,CBA,CE,DBA2J,KK,MA/My,NZO,SMJ | 13 | 120249637 | 120249655 | 19 | - | 17.5574 | 6.32E-07 | 0.153 | ACTCCAGAAGAGGGCGCCA | 2 | ENCFF093ZRU,ENCFF367UNR |
| 950 | 13 | 120249513 | 120249721 | 208 | 129S1,BTBR,C3H,C57L,CBA,CE,DBA2J,KK,MA/My,NZO,SMJ | 13 | 120249637 | 120249670 | 34 | - | 18.3659 | 2.76E-07 | 0.0667 | TAGCTGTCTTCAGACACTCCAGAAGAGGGCGCCA | 2 | ENCFF093ZRU,ENCFF367UNR |
| 951 | 14 | 9719254 | 9719452 | 198 | 129S1,C3H,C57L,CBA,DBA1J,DBA2J,KK,MRL,NZW,PJ,PL | 14 | 9719305 | 9719338 | 34 | + | 17.2439 | 5.79E-07 | 0.0851 | TAGCTGTCCTCAGATACTCCAGAAGAGGGCATCA | | |
| 952 | 14 | 10475334 | 10475536 | 202 | 129S1,C3H,C57L,CBA,DBA1J,DBA2J,KK,MRL,NZW,PJ,PL | 14 | 10475393 | 10475426 | 34 | + | 16.4472 | 9.61E-07 | 0.111 | TAGCTGTCTTCAGACACACCAGAAGAGGGCGTCA | | |
| 953 | 14 | 10505318 | 10505520 | 202 | 129S1,C3H,C57L,CBA,DBA1J,DBA2J,KK,MRL,NZB,NZO,NZW,PJ,PL | 14 | 10505378 | 10505411 | 34 | + | 16.4472 | 9.61E-07 | 0.111 | TAGCTGTCTTCAGACACACCAGAAGAGGGCGTCA | | |
| 954 | 14 | 13871991 | 13872939 | 948 | 129S1,C57L,KK,PL | 14 | 13872667 | 13872700 | 34 | + | 17.813 | 3.99E-07 | 0.0708 | TTGCTGTCTTCATGTACACCAGAAGAGGGCATCA | 1 | ENCFF093ZRU |
| 955 | 14 | 25792758 | 25792968 | 210 | NOD,NOR,PJ,PL,TH | 14 | 25792826 | 25792859 | 34 | + | 17.8455 | 3.91E-07 | 0.0699 | TAGCTGTCCTCAGACACTCCAGAAGAGGGCGTCA | | |
| 956 | 14 | 31832227 | 31832411 | 184 | PJ | 14 | 31832269 | 31832302 | 34 | + | 17.9106 | 3.74E-07 | 0.0681 | TAGCTGTCTTCAGACACTCCAGAAGAGGGCACCA | | |
| 957 | 14 | 34122424 | 34122620 | 196 | 129S1,C57L,PJ | 14 | 34122473 | 34122506 | 34 | + | 17.2439 | 5.79E-07 | 0.0851 | TAGCTGTCCTCAGATACTCCAGAAGAGGGCATCA | | |
| 958 | 14 | 34570036 | 34570608 | 572 | 129S1,C57L | 14 | 34570129 | 34570162 | 34 | + | 17.7073 | 4.28E-07 | 0.0732 | TAGCTGTCTTCAGATGTACCAGAAGGGGGCGTCA | 1 | ENCFF093ZRU |
| 959 | 14 | 45806904 | 45807114 | 210 | BUB,C57L,DBA1J,DBA2J,ILNJ2,KK,MA/My,MRL,NZB,NZO,PJ,PL,SJL,SMJ,SWR,TH | 14 | 45806974 | 45807007 | 34 | + | 17.3902 | 5.27E-07 | 0.0813 | TAGCTGTCCTCAGACACTCCAGAAGAGGGCATCA | | |
| 960 | 14 | 52406911 | 52407131 | 220 | 129S1,AJ,AKR,BALB,BTBR,BUB,C3H,C57L,CBA,CE,DBA1J,DBA2J,FVB,ILNJ2,KK,MA/My,MRL,NOD,NOR,NZB,NZO,NZW,PJ,PL,RF,RH2,SEA,SJL,SMJ,SWR,TH | 14 | 52406962 | 52406995 | 34 | + | 19.813 | 1.01E-07 | 0.0351 | TAGCTGTCTCCAGCCACTCCAGAAGAGGGCGCCA | 2 | ENCFF093ZRU,ENCFF367UNR |
| 960 | 14 | 52406911 | 52407131 | 220 | 129S1,AJ,AKR,BALB,BTBR,BUB,C3H,C57L,CBA,CE,DBA1J,DBA2J,FVB,ILNJ2,KK,MA/My,MRL,NOD,NOR,NZB,NZO,NZW,PJ,PL,RF,RH2,SEA,SJL,SMJ,SWR,TH | 14 | 52406977 | 52406995 | 19 | + | 17.5574 | 6.32E-07 | 0.158 | ACTCCAGAAGAGGGCGCCA | 2 | ENCFF093ZRU,ENCFF367UNR |
| 961 | 14 | 52423215 | 52423424 | 209 | 129S1,AJ,AKR,BALB,BTBR,BUB,C3H,C57L,CBA,CE,DBA1J,DBA2J,FVB,ILNJ2,KK,MA/My,MRL,NOD,NOR,NZB,NZO,NZW,PJ,PL,RF,RH2,SEA,SJL,SMJ,SWR,TH | 14 | 52423282 | 52423315 | 34 | + | 17.2439 | 5.79E-07 | 0.0851 | TAGCTGTCCTCAGATACTCCAGAAGAGGGCATCA | | |
| 962 | 14 | 57930794 | 57931001 | 207 | BTBR,BUB,DBA2J,KK | 14 | 57930840 | 57930873 | 34 | + | 16.4634 | 9.51E-07 | 0.111 | TAGCTGTCTTCAGACGCACCAGAAGAGGGCATCA | | |
| 963 | 14 | 58164418 | 58164630 | 212 | BTBR,BUB,DBA2J,NZB | 14 | 58164486 | 58164519 | 34 | + | 16.4472 | 9.61E-07 | 0.111 | TAGCTGTCTTCAGACACACCAGAAGAGGGCGTCA | | |
| 964 | 14 | 61774932 | 61775416 | 484 | 129S1,C3H,C57L,CE,DBA1J,DBA2J,LP | 14 | 61775405 | 61775438 | 34 | + | 16.4634 | 9.51E-07 | 0.111 | GAGTCTTTCACCTGGAGTCCAGAAGAGGGCACCA | 6 | ENCFF099QNE,ENCFF907XEG,ENCFF367UNR,ENCFF093ZRU,ENCFF821BIA,ENCFF327LYS |
| 965 | 14 | 61802426 | 61802481 | 55 | 129S1,C3H,C57L,CE,DBA1J,DBA2J,LP | 14 | 61802474 | 61802507 | 34 | + | 18.3659 | 2.76E-07 | 0.0595 | TAGCTGTCTTCAGACACTCCAGAAGAGGGCGCCA | 4 | ENCFF367UNR,ENCFF093ZRU,ENCFF502WDT,ENCFF055HAP |
| 966 | 14 | 68752725 | 68753464 | 739 | 129S1,C3H,C57L,CE,DBA1J,DBA2J,KK | 14 | 68753456 | 68753490 | 35 | + | 18.0183 | 2.98E-07 | 0.124 | ATGCAGGTACCCACATGGGCCTGAAGAGGGTATTG | | |
| 967 | 14 | 70059373 | 70059484 | 111 | 129S1,C57L,KK,RH2,SJL | 14 | 70059393 | 70059411 | 19 | + | 16.9836 | 9.17E-07 | 0.187 | GGGCCAGGAGGTGGCAGGA | | |
| 967 | 14 | 70059373 | 70059484 | 111 | 129S1,C57L,KK,RH2,SJL | 14 | 70059473 | 70059491 | 19 | + | 16.9836 | 9.17E-07 | 0.187 | GGGCCAGGAGGTGGCAGGA | | |
| 968 | 14 | 70914823 | 70915034 | 211 | 129S1,AJ,BALB,BUB,C3H,C57L,CE,DBA1J,DBA2J,LP,PL,SEA | 14 | 70914893 | 70914926 | 34 | + | 16.4634 | 9.51E-07 | 0.111 | TAGCTGTCTTCAGACGCACCAGAAGAGGGCATCA | | |
| 969 | 14 | 71062167 | 71062365 | 198 | 129S1,AJ,BALB,BUB,C57L,PL,SEA | 14 | 71062221 | 71062255 | 35 | + | 18.1284 | 2.79E-07 | 0.12 | CTGTAGCTGTCTTCAGACACCAGAAGAGGGCATCA | | |
| 970 | 14 | 72681872 | 72682080 | 208 | TH | 14 | 72681940 | 72681973 | 34 | + | 17.5528 | 4.74E-07 | 0.0767 | TAGCTGTCTTCGGATGCACCAGAAGAGGGCGTCA | | |
| 971 | 14 | 72764168 | 72764395 | 227 | 129S1,BUB,C57L,FVB,SJL,TH | 14 | 72764251 | 72764284 | 34 | + | 18.3659 | 2.76E-07 | 0.0595 | TAGCTGTCTTCAGACACTCCAGAAGAGGGCGCCA | | |
| 971 | 14 | 72764168 | 72764395 | 227 | 129S1,BUB,C57L,FVB,SJL,TH | 14 | 72764266 | 72764284 | 19 | + | 17.5574 | 6.32E-07 | 0.158 | ACTCCAGAAGAGGGCGCCA | | |
| 972 | 14 | 78468755 | 78468952 | 197 | NZB,NZO | 14 | 78468809 | 78468842 | 34 | + | 16.6341 | 8.54E-07 | 0.106 | TAGCTGTCTTCACACACTCCAGAAGAGGGCGTCA | | |
| 973 | 14 | 99308985 | 99309190 | 205 | AKR,CBA,CE,DBA1J,DBA2J,ILNJ2,MRL,PJ,PL,RF,RH2,SJL,SMJ,SWR,TH | 14 | 99309049 | 99309082 | 34 | + | 16.4634 | 9.51E-07 | 0.111 | TAGCTGTCTTCAGACGCACCAGAAGAGGGCATCA | | |
| 974 | 14 | 102967214 | 102967429 | 215 | C57L,C58J,ILNJ2,PJ | 14 | 102967287 | 102967320 | 34 | + | 16.4634 | 9.51E-07 | 0.111 | TAGCTGTCTTCAGACGCACCAGAAGAGGGCATCA | | |
| 975 | 14 | 110719314 | 110720152 | 838 | KK,PJ | 14 | 110719715 | 110719748 | 34 | + | 19.5366 | 1.23E-07 | 0.0379 | TTGCTGTCGCCAGACACACCAGAAGAGGGCATTG | | |
| 976 | 14 | 4272917 | 4273117 | 200 | 129S1,C3H,C57L,CBA,DBA1J,DBA2J,KK,MRL,NZW,PJ,PL | 14 | 4273043 | 4273061 | 19 | - | 17.5574 | 6.32E-07 | 0.158 | ACTCCAGAAGAGGGCGCCA | | |
| 976 | 14 | 4272917 | 4273117 | 200 | 129S1,C3H,C57L,CBA,DBA1J,DBA2J,KK,MRL,NZW,PJ,PL | 14 | 4273043 | 4273076 | 34 | - | 18.3659 | 2.76E-07 | 0.0595 | TAGCTGTCTTCAGACACTCCAGAAGAGGGCGCCA | | |
| 977 | 14 | 8255268 | 8255477 | 209 | 129S1,C3H,C57L,CBA,DBA1J,DBA2J,KK,MRL,NZW,PJ,PL | 14 | 8255394 | 8255427 | 34 | - | 17.2439 | 5.79E-07 | 0.0851 | TAGCTGTCCTCAGATACTCCAGAAGAGGGCATCA | | |
| 978 | 14 | 10309617 | 10309824 | 207 | 129S1,C3H,C57L,CBA,DBA1J,DBA2J,FVB,KK,MA/My,MRL,NOD,NOR,NZW,PJ,PL,SJL,SWR | 14 | 10309742 | 10309775 | 34 | - | 16.4472 | 9.61E-07 | 0.111 | TAGCTGTCTTCAGACACACCAGAAGAGGGCGTCA | | |

| | | | | | | | | | | | | | | | | |
|---|---|---|---|---|---|---|---|---|---|---|---|---|---|---|---|---|
| 979 | 14 | 10941884 | 10942081 | 197 | 129S1,C3H,C57L,CBA,DBA1J,DBA2J,KK,MRL,NZW,PJ,PL | 14 | 10941998 | 10942031 | 34 | - | 16.4472 | 9.61E-07 | 0.111 | TAGCTGTCTTCAGACACACCAGAAGAGGGCGTCA | | |
| 980 | 14 | 31809729 | 31809969 | 240 | PJ | 14 | 31809854 | 31809887 | 34 | - | 18.1057 | 3.29E-07 | 0.0647 | AAGCTGTCTTCAGACACTCCAGAAGAGGGCACCA | 2 | ENCFF093ZRU.ENCFF367UNR |
| 981 | 14 | 34374172 | 34374879 | 707 | 129S1.C57L | 14 | 34374799 | 34374832 | 34 | - | 18.1545 | 3.18E-07 | 0.0632 | TAGCTGTTTTCAGACACACCAGAAGAGGGCGTCA | | |
| 982 | 14 | 55073187 | 55073562 | 375 | 129S1,AJ,AKR,BALB,BTBR,BUB,C3H,C57L,CBA,CE,DBA1J,DBA2J,FVB,ILNJ2,KK,MA/My,MRL,NOD,NOR,NZB,NZO,NZW,PJ,PL,RF,RH2.SEA.SJL.SMJ.SWR.TH | 14 | 55073266 | 55073284 | 19 | - | 17.5574 | 6.32E-07 | 0.158 | ACTCCAGAAGAGGGCGCCA | 2 | ENCFF093ZRU,ENCFF367UNR |
| 982 | 14 | 55073187 | 55073562 | 375 | 129S1,AJ,AKR,BALB,BTBR,BUB,C3H,C57L,CBA,CE,DBA1J,DBA2J,FVB,ILNJ2,KK,MA/My,MRL,NOD,NOR,NZB,NZO,NZW,PJ,PL,RF,RH2.SEA.SJL.SMJ.SWR.TH | 14 | 55073266 | 55073299 | 34 | - | 18.3659 | 2.76E-07 | 0.0595 | TAGCTGTCTTCAGACACTCCAGAAGAGGGCGCCA | 2 | ENCFF093ZRU,ENCFF367UNR |
| 982 | 14 | 55073187 | 55073562 | 375 | 129S1,AJ,AKR,BALB,BTBR,BUB,C3H,C57L,CBA,CE,DBA1J,DBA2J,FVB,ILNJ2,KK,MA/My,MRL,NOD,NOR,NZB,NZO,NZW,PJ,PL,RF,RH2.SEA.SJL.SMJ.SWR.TH | 14 | 55073466 | 55073484 | 19 | - | 17.5574 | 6.32E-07 | 0.158 | ACTCCAGAAGAGGGCGCCA | 2 | ENCFF093ZRU,ENCFF367UNR |
| 982 | 14 | 55073187 | 55073562 | 375 | 129S1,AJ,AKR,BALB,BTBR,BUB,C3H,C57L,CBA,CE,DBA1J,DBA2J,FVB,ILNJ2,KK,MA/My,MRL,NOD,NOR,NZB,NZO,NZW,PJ,PL,RF,RH2.SEA.SJL.SMJ.SWR.TH | 14 | 55073466 | 55073499 | 34 | - | 18.3659 | 2.76E-07 | 0.0595 | TAGCTGTCTTCAGACACTCCAGAAGAGGGCGCCA | 2 | ENCFF093ZRU,ENCFF367UNR |
| 983 | 14 | 56371835 | 56372048 | 213 | BTBR.DBA2J.ILNJ2.SMJ | 14 | 56371960 | 56371993 | 34 | - | 17.8455 | 3.91E-07 | 0.0699 | TAGCTGTCCTCAGACACTCCAGAAGAGGGCGTCA | | |
| 984 | 14 | 58157612 | 58157839 | 227 | BTBR.BUB.DBA2J.NZB | 14 | 58157738 | 58157756 | 19 | - | 17.5738 | 6.25E-07 | 0.158 | ACTCCAGAAGAGGGCGCTA | 1 | ENCFF093ZRU |
| 984 | 14 | 58157612 | 58157839 | 227 | BTBR.BUB.DBA2J.NZB | 14 | 58157738 | 58157771 | 34 | - | 18.1789 | 3.13E-07 | 0.0629 | TAGCTGTCTTCAGACACTCCAGAAGAGGGCGCTA | 1 | ENCFF093ZRU |
| 985 | 14 | 71071418 | 71071519 | 101 | 129S1.AJ.BALB.BUB.C57L.PL.SEA | 14 | 71071489 | 71071507 | 19 | - | 18 | 4.70E-07 | 0.142 | TTGCCAGTTGGTGGCGCTG | 1 | ENCFF502WDT |
| 986 | 14 | 86821182 | 86821443 | 261 | AJ,AKR,BALB,BUB,C3H,CBA,DBA1J,DBA2J,FVB,NOD,NZO,PJ,RF,SEA,SJL,SMJ,SWR,TH | 14 | 86821309 | 86821342 | 34 | - | 17.0732 | 6.46E-07 | 0.0918 | TAGCTGTCTTCAGAGGCACCAGAAGAGGGCATCA | | |
| 987 | 14 | 95569359 | 95569809 | 450 | AJ,AKR,BALB,BTBR,BUB,C3H,CBA,CE,DBA1J,DBA2J,FVB,MRL,NOD,PJ,PL,RF,RH2,SEA.SJL.SMJ.SWR.TH | 14 | 95569766 | 95569799 | 34 | - | 17.9106 | 3.74E-07 | 0.0681 | CCGCCCTCACATTCGCCGTCACAAGATGGCGCTG | | |
| 988 | 14 | 105471353 | 105471577 | 224 | C57L.ILNJ2.KK | 14 | 105471494 | 105471527 | 34 | - | 16.4472 | 9.61E-07 | 0.111 | TAGCTGTCTTCAGACACACCAGAAGAGGGCGTCA | | |
| 989 | 14 | 120217434 | 120218364 | 930 | NZW | 14 | 120218362 | 120218380 | 19 | - | 17.7049 | 5.73E-07 | 0.153 | CTCCCAGCAGAGGGCGGGG | | |
| 990 | 15 | 8761077 | 8761323 | 246 | TH | 15 | 8761181 | 8761214 | 34 | + | 17.2439 | 5.79E-07 | 0.0855 | TAGCTGTCTTCGGACGCACCAGAAGAGGGCATCA | | |
| 991 | 15 | 20042070 | 20042280 | 210 | 129S1.BTBR.C57L | 15 | 20042135 | 20042168 | 34 | + | 18.3659 | 2.76E-07 | 0.0597 | TAGCTGTCTTCAGACACTCCAGAAGAGGGCGCCA | | |
| 991 | 15 | 20042070 | 20042280 | 210 | 129S1.BTBR.C57L | 15 | 20042150 | 20042168 | 19 | + | 17.5574 | 6.32E-07 | 0.126 | ACTCCAGAAGAGGGCGCCA | | |
| 992 | 15 | 34327615 | 34327806 | 191 | 129S1.BUB.C57L.NOD.NOR.PJ.SWR | 15 | 34327666 | 34327699 | 34 | + | 17.6992 | 4.30E-07 | 0.0743 | TAGCTGTTTTCAGACACACCAGAAGAGGGCATCA | 1 | ENCFF093ZRU |
| 993 | 15 | 34401319 | 34401527 | 208 | 129S1.BUB.C57L.NOD.NOR.PJ.SWR | 15 | 34401384 | 34401417 | 34 | + | 17.878 | 3.83E-07 | 0.0697 | TAGCTGTCCTCAGACACACCAGAAGAGGGCATCA | 2 | ENCFF093ZRU |
| 994 | 15 | 35071803 | 35071937 | 134 | 129S1,BUB,C57L,CE,KK,LP,NOD,NOR,PJ,SWR | 15 | 35071911 | 35071929 | 19 | + | 21.4262 | 3.39E-08 | 0.038 | CTGCCACTAGATGGCAGTG | 17 | ENCFF907XEG,ENCFF327LYS,ENCFF099QNE,ENCFF093ZRU,ENCFF821BIA,ENCFF092NMJ,ENCFF883UPM,ENCFF430PPJ,ENCFF165LXO,ENCFF609MDN,ENCFF198SSR,ENCFF589ZBJ,ENCFF367UNR,ENCFF136ZDK,ENCFF024JOZ,ENCFF055HAP,ENCFF502WDT |
| 995 | 15 | 38705673 | 38705898 | 225 | 129S1.C57L.CE | 15 | 38705756 | 38705789 | 34 | + | 18.9919 | 1.80E-07 | 0.0488 | TTGCTGTCTTCAGACACTCCAGAAGGGGGCATCA | | |
| 996 | 15 | 40306399 | 40306626 | 227 | 129S1.C57L | 15 | 40306483 | 40306516 | 34 | + | 19.8374 | 9.97E-08 | 0.0331 | TAGCTGTCCCCAGATGCACCAGAAGAGGGCATCA | | |
| 997 | 15 | 59179022 | 59179228 | 206 | ILNJ2 | 15 | 59179086 | 59179119 | 34 | + | 16.4472 | 9.61E-07 | 0.106 | TAGCTGTCTTCAGACACACCAGAAGAGGGCGTCA | 2 | ENCFF093ZRU |
| 998 | 15 | 63374806 | 63375024 | 218 | SMJ | 15 | 63374878 | 63374911 | 34 | + | 17.9106 | 3.74E-07 | 0.0689 | TAGCTGTCTTCAGACACTCCAGAAGAGGGCACCA | | |
| 999 | 15 | 71919977 | 71920649 | 672 | AJ,B10D2,B10J,BUB,C3H,CBA,DBA1J,DBA2J,FVB,ILNJ2,LP,MA/My,MRL,NOR,NZB,NZO,NZW,PL,SWR,TH | 15 | 71920159 | 71920192 | 34 | + | 17.9431 | 3.66E-07 | 0.0685 | GAACCATTACTCCTACTGGCACCAGGGGGCGCTC | | |
| 999 | 15 | 71919977 | 71920649 | 672 | AJ,B10D2,B10J,BUB,C3H,CBA,DBA1J,DBA2J,FVB,ILNJ2,LP,MA/My,MRL,NOR,NZB,NZO,NZW,PL,SWR,TH | 15 | 71920174 | 71920192 | 19 | + | 19.6066 | 1.49E-07 | 0.062 | CTGGCACCAGGGGGCGCTC | | |
| 1000 | 15 | 75807174 | 75807500 | 326 | ILNJ2.MA/My | 15 | 75807358 | 75807391 | 34 | + | 17.5041 | 4.89E-07 | 0.0783 | TAGCTGTACTCAGATACTCCAGAAGAGGGCATCA | | |
| 1001 | 15 | 76793414 | 76793610 | 196 | MA/My | 15 | 76793465 | 76793499 | 35 | + | 18.1284 | 2.79E-07 | 0.0951 | CTGTAGCTGTCTTCAGACACCAGAAGAGGGCATCA | | |
| 1002 | 15 | 79027128 | 79027563 | 435 | 129S1,AJ,AKR,BALB,C3H,C57L,C58J,CBA,CE,DBA1J,DBA2J,FVB,ILNJ2,KK,LP,MA/My,MRL,NOD,NOR,NZW,PJ,PL,SEA,SJL,SMJ,SWR.TH | 15 | 79027241 | 79027274 | 34 | + | 16.4472 | 9.61E-07 | 0.106 | TAGCTGTCTTCAGACACACCAGAAGAGGGCGTCA | | |
| 1002 | 15 | 79027128 | 79027563 | 435 | 129S1,AJ,AKR,BALB,C3H,C57L,C58J,CBA,CE,DBA1J,DBA2J,FVB,ILNJ2,KK,LP,MA/My,MRL,NOD,NOR,NZW,PJ,PL,SEA,SJL,SMJ,SWR.TH | 15 | 79027445 | 79027463 | 19 | + | 17.8852 | 5.08E-07 | 0.114 | ACACCAGAAGAGGGCACCA | | |
| 1003 | 15 | 79750969 | 79751172 | 203 | 129S1,AJ,AKR,BALB,C3H,C57L,C58J,CBA,CE,DBA1J,DBA2J,FVB,ILNJ2,KK,LP,MA/My,MRL,NOD,NOR,NZW,PJ,PL,SEA,SJL,SMJ,SWR.TH | 15 | 79751030 | 79751063 | 34 | + | 18.5285 | 2.48E-07 | 0.058 | CAGCTGTCTTCAGACACTCCAGAAGAGGGCGTCA | | |
| 1004 | 15 | 80909323 | 80909540 | 217 | DBA1J.DBA2J.ILNJ2 | 15 | 80909398 | 80909431 | 34 | + | 16.4472 | 9.61E-07 | 0.106 | TAGCTGTCTTCAGACACACCAGAAGAGGGCGTCA | | |
| 1005 | 15 | 81078503 | 81078725 | 222 | AJ,AKR,BALB,C3H,CBA,CE,DBA1J,DBA2J,FVB,ILNJ2,KK,MRL,NOD,NOR,NZW,PJ,PL,SEA,SJL,SMJ,SWR | 15 | 81078583 | 81078616 | 34 | + | 17.6179 | 4.54E-07 | 0.0756 | TAGCTGTCTTCAGACACCCCAGAAGAGGGCACCA | | |
| 1006 | 15 | 81626817 | 81627028 | 211 | AJ,AKR,BALB,BUB,C3H,CBA,CE,DBA1J,DBA2J,FVB,ILNJ2,KK,MA/My,MRL,NOD,NOR,NZB,NZO,NZW,PJ,PL,SEA | 15 | 81626887 | 81626920 | 34 | + | 17.6179 | 4.54E-07 | 0.0756 | TAGCTGTCTTCAGACACCCCAGAAGAGGGCACCA | 8 | ENCFF099QNE,ENCFF821BIA,ENCFF907XEG,ENCFF327LYS,ENCFF201JUA,ENCFF089GBC,ENCFF093ZRU,ENCFF367UNR |
| 1007 | 15 | 84938634 | 84938856 | 222 | AKR,BTBR,FVB,MRL,NOD,NOR,PL,SJL,SMJ.SWR.TH | 15 | 84938713 | 84938746 | 34 | + | 16.4472 | 9.61E-07 | 0.106 | TAGCTGTCTTCAGACACACCAGAAGAGGGCGTCA | | |
| 1008 | 15 | 88650856 | 88651067 | 211 | CE.FVB.ILNJ2 | 15 | 88650925 | 88650958 | 34 | + | 18.3659 | 2.76E-07 | 0.0597 | TAGCTGTCTTCAGACACTCCAGAAGAGGGCGCCA | | |
| 1008 | 15 | 88650856 | 88651067 | 211 | CE.FVB.ILNJ2 | 15 | 88650940 | 88650958 | 19 | + | 17.5574 | 6.32E-07 | 0.126 | ACTCCAGAAGAGGGCGCCA | | |
| 1009 | 15 | 88678875 | 88679090 | 215 | CE.FVB.ILNJ2.PL | 15 | 88678948 | 88678981 | 34 | + | 16.7724 | 7.83E-07 | 0.0991 | TAGCTGTCTTCAGATGCACCAGAAGAGGGCGTCA | | |
| 1010 | 15 | 96611858 | 96612059 | 201 | NZW.SMJ | 15 | 96611917 | 96611950 | 34 | + | 16.4146 | 9.81E-07 | 0.108 | TAGCTGTCTTCAGACACACCAGAGGAGGGCGCCA | | |
| 1011 | 15 | 97263461 | 97263669 | 208 | 129S1.C57L.C58J.KK | 15 | 97263515 | 97263548 | 34 | + | 16.4472 | 9.61E-07 | 0.106 | TAGCTGTCTTCAGACACACCAGAAGAGGGCGTCA | | |
| 1012 | 15 | 98041467 | 98041677 | 210 | AJ.BUB.C3H.CE.MA/My.RH2.SWR | 15 | 98041535 | 98041568 | 34 | + | 16.4472 | 9.61E-07 | 0.106 | TAGCTGTCTTCAGACACACCAGAAGAGGGCGTCA | 1 | ENCFF093ZRU |
| 1013 | 15 | 103239372 | 103239604 | 232 | AJ,C3H,CBA,CE,DBA1J,DBA2J,FVB,ILNJ2,LP,MRL,NOD,NOR,NZB,NZO,PJ,PL,RH2,SEA,SJL.SMJ | 15 | 103239461 | 103239494 | 34 | + | 16.4472 | 9.61E-07 | 0.106 | TAGCTGTCTTCAGACACACCAGAAGAGGGCGTCA | | |
| 1014 | 15 | 8214193 | 8214404 | 211 | AKR.LP.RF.TH | 15 | 8214317 | 8214350 | 34 | - | 17.2439 | 5.79E-07 | 0.0855 | TAGCTGTCCTCAGATACTCCAGAAGAGGGCATCA | | |
| 1015 | 15 | 10079399 | 10079603 | 204 | CE.ILNJ2.TH | 15 | 10079520 | 10079538 | 19 | - | 17.5574 | 6.32E-07 | 0.126 | ACTCCAGAAGAGGGCGCCA | | |
| 1015 | 15 | 10079399 | 10079603 | 204 | CE.ILNJ2.TH | 15 | 10079520 | 10079553 | 34 | - | 18.3659 | 2.76E-07 | 0.0597 | TAGCTGTCTTCAGACACTCCAGAAGAGGGCGCCA | | |
| 1016 | 15 | 19279137 | 19279340 | 203 | CE.ILNJ2.PJ.PL.SEA | 15 | 19279262 | 19279295 | 34 | - | 16.4634 | 9.51E-07 | 0.106 | TAGCTGTCTTCAGACGCACCAGAAGAGGGCATCA | | |
| 1017 | 15 | 25718101 | 25718325 | 224 | AJ,BUB,C3H,CBA,CE,DBA1J,DBA2J,FVB,MA/My,MRL,NZB,NZO,NZW,PL,RH2,SMJ,TH | 15 | 25718244 | 25718262 | 19 | - | 17.5574 | 6.32E-07 | 0.126 | ACTCCAGAAGAGGGCGCCA | 1 | ENCFF367UNR |
| 1017 | 15 | 25718101 | 25718325 | 224 | AJ,BUB,C3H,CBA,CE,DBA1J,DBA2J,FVB,MA/My,MRL,NZB,NZO,NZW,PL,RH2,SMJ,TH | 15 | 25718244 | 25718277 | 34 | - | 18.3659 | 2.76E-07 | 0.0597 | TAGCTGTCTTCAGACACTCCAGAAGAGGGCGCCA | 1 | ENCFF367UNR |
| 1018 | 15 | 33931150 | 33931426 | 276 | 129S1,AJ,BALB,BTBR,BUB,C57L,FVB,MA/My,MRL,NOD,NOR,PJ,RH2,SEA,SJL,SMJ,SWR.TH | 15 | 33931275 | 33931308 | 34 | - | 16.4472 | 9.61E-07 | 0.106 | TAGCTGTCTTCAGACACACCAGAAGAGGGCGTCA | | |
| 1019 | 15 | 36549090 | 36549308 | 218 | 129S1.C57L | 15 | 36549215 | 36549248 | 34 | - | 16.4472 | 9.61E-07 | 0.106 | TAGCTGTCTTCAGACACACCAGAAGAGGGCGTCA | | |
| 1020 | 15 | 36703165 | 36703529 | 364 | 129S1.C57L | 15 | 36703234 | 36703252 | 19 | - | 18.1475 | 4.25E-07 | 0.108 | TACCCAGAAGGGGGCACCC | | |
| 1021 | 15 | 37286120 | 37286363 | 243 | 129S1.C57L | 15 | 37286245 | 37286278 | 34 | - | 16.7724 | 7.83E-07 | 0.0991 | TAGCTGTCTTCGGACACACCAGAAGAGGGCATCA | | |
| 1022 | 15 | 39045839 | 39046043 | 204 | 129S1,C3H,C57L,CE,FVB,MA/My,MRL,NOD,NOR.PL.SWR | 15 | 39045965 | 39045983 | 19 | - | 17.5574 | 6.32E-07 | 0.126 | ACTCCAGAAGAGGGCGCCA | | |
| 1022 | 15 | 39045839 | 39046043 | 204 | 129S1,C3H,C57L,CE,FVB,MA/My,MRL,NOD,NOR.PL.SWR | 15 | 39045965 | 39045998 | 34 | - | 18.3659 | 2.76E-07 | 0.0597 | TAGCTGTCTTCAGACACTCCAGAAGAGGGCGCCA | | |
| 1023 | 15 | 41645394 | 41645602 | 208 | 129S1,AJ,AKR,BTBR,BUB,C3H,C57L,CBA,CE,DBA1J,DBA2J,FVB,ILNJ2,KK,LP,MA/My,MRL,NOD,NOR,NZB,NZO,NZW,PJ,PL,RH2,SMJ.SWR | 15 | 41645519 | 41645537 | 19 | - | 17.8852 | 5.08E-07 | 0.114 | ACACCAGAAGAGGGCACCA | 3 | ENCFF093ZRU,ENCFF099QNE,ENCFF367UNR |
| 1024 | 15 | 53264671 | 53264862 | 191 | 129S1,AJ,C57L,CE,DBA1J,DBA2J,FVB,ILNJ2,KK,MA/My,RH2,SJL,SMJ,TH | 15 | 53264796 | 53264829 | 34 | - | 16.4472 | 9.61E-07 | 0.106 | TAGCTGTCTTCAGACACACCAGAAGAGGGCGTCA | | |
| 1025 | 15 | 56577524 | 56577752 | 228 | AKR.BTBR.ILNJ2.NZB.NZO.NZW.SMJ | 15 | 56577650 | 56577683 | 34 | - | 16.4634 | 9.51E-07 | 0.106 | TAGCTGTCTTCAGACGCACCAGAAGAGGGCATCA | | |
| 1026 | 15 | 57954911 | 57955129 | 218 | BALB.ILNJ2.NOD.NOR.SEA.SMJ | 15 | 57955034 | 57955052 | 19 | - | 17.5574 | 6.32E-07 | 0.126 | ACTCCAGAAGAGGGCGCCA | | |
| 1026 | 15 | 57954911 | 57955129 | 218 | BALB.ILNJ2.NOD.NOR.SEA.SMJ | 15 | 57955034 | 57955067 | 34 | - | 18.3659 | 2.76E-07 | 0.0597 | TAGCTGTCTTCAGACACTCCAGAAGAGGGCGCCA | | |
| 1027 | 15 | 66393950 | 66394160 | 210 | BALB.C3H.CBA.CE.RF.SEA.SMJ.TH | 15 | 66394076 | 66394094 | 19 | - | 17.9672 | 4.81E-07 | 0.114 | GCACCAGAAGAGGGCACCA | | |
| 1027 | 15 | 66393950 | 66394160 | 210 | BALB.C3H.CBA.CE.RF.SEA.SMJ.TH | 15 | 66394076 | 66394109 | 34 | - | 18.8699 | 1.96E-07 | 0.051 | TAGCTGTCTTCAGACGCACCAGAAGAGGGCACCA | | |
| 1028 | 15 | 76826643 | 76826847 | 204 | MA/My | 15 | 76826770 | 76826788 | 19 | - | 17.8852 | 5.08E-07 | 0.114 | ACACCAGAAGAGGGCACCA | | |
| 1029 | 15 | 77219316 | 77219540 | 224 | MA/My.NOD.NOR.NZB.NZO | 15 | 77219435 | 77219453 | 19 | - | 16.9016 | 9.66E-07 | 0.156 | ATTCCAGAAGAGGGCACCA | 1 | ENCFF093ZRU |
| 1029 | 15 | 77219316 | 77219540 | 224 | MA/My.NOD.NOR.NZB.NZO | 15 | 77219435 | 77219468 | 34 | - | 17.8455 | 3.91E-07 | 0.0705 | TAGCTGTCTTCAGACATTCCAGAAGAGGGCACCA | 1 | ENCFF093ZRU |
| 1030 | 15 | 84601870 | 84602187 | 317 | 129S1,AJ,BALB,BUB,C3H,C57L,CBA,CE,DBA1J,DBA2J,ILNJ2,KK,MA/My,NZB,NZW,RF | 15 | 84601997 | 84602030 | 34 | - | 18.9512 | 1.85E-07 | 0.0495 | TTGCTGTCTTCAGACGCACCAGAAGAGGGCATCA | 5 | ENCFF093ZRU,ENCFF327LYS,ENCFF024JOZ,ENCFF367UNR,ENCFF907XEG |
| 1031 | 15 | 98992106 | 98992479 | 373 | SMJ | 15 | 98992083 | 98992116 | 34 | - | 17.6911 | 4.33E-07 | 0.0745 | GCGCAGTGTCACTCTAGTCCGATAGGAGGGCGCCA | 26 | ENCFF093ZRU,ENCFF086ZTV,ENCFF367UNR,ENCFF099QNE,ENCFF092NMJ,ENCFF136ZDK,ENCFF198SSR,ENCFF430PPJ,ENCFF165LXO,ENCFF316FYN,ENCFF821BIA,ENCFF065ODV,ENCFF055HAP,ENCFF327LYS,ENCFF024JOZ,ENCFF502WDT,ENCFF194HVL,ENCFF907XEG,ENCFF089GBC,ENCFF052KVO,ENCFF957SPH,ENCFF142CNG,ENCFF589ZBJ,ENCFF281QMV |
| 1032 | 15 | 99561481 | 99561681 | 200 | BTBR.SMJ | 15 | 99561607 | 99561640 | 34 | - | 18.1545 | 3.18E-07 | 0.0636 | TAGCTGTTTTCAGACACACCAGAAGAGGGCGTCA | 2 | ENCFF055HAP.ENCFF093ZRU |
| 1033 | 16 | 3387937 | 3388140 | 203 | SMJ | 16 | 3387995 | 3388028 | 34 | + | 18.561 | 2.42E-07 | 0.0684 | AAGCTGTCTTCAGACACTCCAGAAGAGGGCGCCA | | |
| 1033 | 16 | 3387937 | 3388140 | 203 | SMJ | 16 | 3388010 | 3388028 | 19 | + | 17.5574 | 6.32E-07 | 0.163 | ACTCCAGAAGAGGGCGCCA | | |
| 1034 | 16 | 3642482 | 3642701 | 219 | AJ.BUB.CE.MRL.PL.SMJ | 16 | 3642561 | 3642594 | 34 | + | 17.6179 | 4.54E-07 | 0.0884 | TAGCTGTCTTCAGACACCCCAGAAGAGGGCACCA | | |
| 1035 | 16 | 8566571 | 8566805 | 234 | AJ.AKR.PL | 16 | 8566664 | 8566697 | 34 | + | 17.0488 | 6.56E-07 | 0.107 | TTGCTATCTTCAGATACACCAGAAGAGGGCATCA | | |
| 1036 | 16 | 11241621 | 11241849 | 228 | 129S1,BTBR,C3H,C57L,CBA,DBA1J,DBA2J.ILNJ2.PJ | 16 | 11241625 | 11241659 | 35 | + | 23.1927 | 9.72E-09 | 0.0147 | TGGCACTGCAGCACCCTCACCACCAGGGGCCACTC | 3 | ENCFF093ZRU,ENCFF367UNR,ENCFF504JDS |
| 1036 | 16 | 11241621 | 11241849 | 228 | 129S1,BTBR,C3H,C57L,CBA,DBA1J,DBA2J.ILNJ2.PJ | 16 | 11241641 | 11241659 | 19 | + | 16.8525 | 9.96E-07 | 0.192 | TCACCACCAGGGGCCACTC | 3 | ENCFF093ZRU,ENCFF367UNR,ENCFF504JDS |
| 1036 | 16 | 11241621 | 11241849 | 228 | 129S1,BTBR,C3H,C57L,CBA,DBA1J,DBA2J.ILNJ2.PJ | 16 | 11241682 | 11241716 | 35 | + | 23.1927 | 9.72E-09 | 0.0147 | TGGCACTGCAGCACCCTCACCACCAGGGGCCACTC | 3 | ENCFF093ZRU,ENCFF367UNR,ENCFF504JDS |
| 1036 | 16 | 11241621 | 11241849 | 228 | 129S1,BTBR,C3H,C57L,CBA,DBA1J,DBA2J.ILNJ2.PJ | 16 | 11241698 | 11241716 | 19 | + | 16.8525 | 9.96E-07 | 0.192 | TCACCACCAGGGGCCACTC | 3 | ENCFF093ZRU,ENCFF367UNR,ENCFF504JDS |
| 1036 | 16 | 11241621 | 11241849 | 228 | 129S1,BTBR,C3H,C57L,CBA,DBA1J,DBA2J.ILNJ2.PJ | 16 | 11241739 | 11241773 | 35 | + | 23.1927 | 9.72E-09 | 0.0147 | TGGCACTGCAGCACCCTCACCACCAGGGGCCACTC | 3 | ENCFF093ZRU,ENCFF367UNR,ENCFF504JDS |
| 1036 | 16 | 11241621 | 11241849 | 228 | 129S1,BTBR,C3H,C57L,CBA,DBA1J,DBA2J.ILNJ2.PJ | 16 | 11241755 | 11241773 | 19 | + | 16.8525 | 9.96E-07 | 0.192 | TCACCACCAGGGGCCACTC | 3 | ENCFF093ZRU,ENCFF367UNR,ENCFF504JDS |
| 1036 | 16 | 11241621 | 11241849 | 228 | 129S1,BTBR,C3H,C57L,CBA,DBA1J,DBA2J.ILNJ2.PJ | 16 | 11241796 | 11241830 | 35 | + | 23.1927 | 9.72E-09 | 0.0147 | TGGCACTGCAGCACCCTCACCACCAGGGGCCACTC | 6 | ENCFF093ZRU,ENCFF367UNR,ENCFF201JUA,ENCFF099QNE |
| 1036 | 16 | 11241621 | 11241849 | 228 | 129S1,BTBR,C3H,C57L,CBA,DBA1J,DBA2J.ILNJ2.PJ | 16 | 11241812 | 11241830 | 19 | + | 16.8525 | 9.96E-07 | 0.192 | TCACCACCAGGGGCCACTC | 8 | ENCFF093ZRU,ENCFF367UNR,ENCFF201JUA,ENCFF099QNE,ENCFF821BIA,ENCFF024JOZ |
| 1037 | 16 | 28034310 | 28034481 | 171 | 129S1,BUB,C3H,C57L,CBA,DBA1J,DBA2J,ILNJ2.NZB.NZO.NZW | 16 | 28034299 | 28034332 | 34 | + | 20.561 | 5.91E-08 | 0.0278 | TTGGAGTCCCACCTGTGGCCAGCAGGATGCACTG | 1 | ENCFF093ZRU |
| 1038 | 16 | 32923392 | 32923622 | 230 | NOD.NOR.NZB.NZO.NZW | 16 | 32923480 | 32923513 | 34 | + | 18.3659 | 2.76E-07 | 0.07 | TAGCTGTCTTCAGACACTCCAGAAGAGGGCGCCA | | |
| 1038 | 16 | 32923392 | 32923622 | 230 | NOD.NOR.NZB.NZO.NZW | 16 | 32923495 | 32923513 | 19 | + | 17.5574 | 6.32E-07 | 0.163 | ACTCCAGAAGAGGGCGCCA | | |
| 1039 | 16 | 33721250 | 33721462 | 212 | SMJ | 16 | 33721318 | 33721351 | 34 | + | 16.4634 | 9.51E-07 | 0.126 | TAGCTGTCTTCAGACGCACCAGAAGAGGGCATCA | | |
| 1040 | 16 | 33827061 | 33827895 | 834 | 129S1.SMJ | 16 | 33827322 | 33827340 | 19 | + | 18.5902 | 3.13E-07 | 0.12 | CGGCCAGCAGATGTCAGCC | | |
| 1041 | 16 | 47037567 | 47037779 | 212 | 129S1.C3H.C58J.CBA.CE.SJL.TH | 16 | 47037635 | 47037668 | 34 | + | 16.4634 | 9.51E-07 | 0.126 | TAGCTGTCTTCAGACGCACCAGAAGAGGGCATCA | | |
| 1042 | 16 | 55771283 | 55772230 | 947 | CE | 16 | 55771497 | 55771530 | 34 | + | 24.8211 | 1.94E-09 | 0.00274 | CTGTAATATCACATTCTTCCAGAAGAGGGCGCCC | 29 | ENCFF201JUA,ENCFF099QNE,ENCFF821BIA,ENCFF907XEG,ENCFF024JOZ,ENCFF136ZDK,ENCFF065ODV,ENCFF327LYS,ENCFF589ZBJ,ENCFF316FYN,ENCFF502WDT,ENCFF198SSR,ENCFF055HAP,ENCFF092NMJ,ENCFF052KVO,ENCFF504JDS,ENCFF089GBC,ENCFF321AGQ,ENCFF430PPJ,ENCFF165LXO,ENCFF957SPH,ENCFF951GOP,ENCFF609MDN,ENCFF883UPM,ENCFF194HVL,ENCFF281QMV,ENCFF575MGI,ENCFF093ZRU.ENCFF367UNR |
| 1042 | 16 | 55771283 | 55772230 | 947 | CE | 16 | 55771512 | 55771530 | 19 | + | 18.7869 | 2.72E-07 | 0.113 | CTTCCAGAAGAGGGCGCCC | 29 | ENCFF201JUA,ENCFF099QNE,ENCFF821BIA,ENCFF907XEG,ENCFF024JOZ,ENCFF136ZDK,ENCFF065ODV,ENCFF327LYS,ENCFF589ZBJ,ENCFF316FYN,ENCFF502WDT,ENCFF198SSR,ENCFF055HAP,ENCFF092NMJ,ENCFF052KVO,ENCFF504JDS,ENCFF089GBC,ENCFF321AGQ,ENCFF430PPJ,ENCFF165LXO,ENCFF957SPH,ENCFF951GOP,ENCFF609MDN,ENCFF883UPM,ENCFF194HVL,ENCFF281QMV,ENCFF575MGI,ENCFF093ZRU.ENCFF367UNR |
| 1043 | 16 | 56446509 | 56446729 | 220 | 129S1,AJ,BALB,BTBR,BUB,C57L,C58J,CBA,CE,DBA1J,DBA2J,FVB,KK,LP,MRL,NZB,NZO,NZW,PL,RH2,SJL,SMJ,SWR | 16 | 56446587 | 56446620 | 34 | + | 16.4634 | 9.51E-07 | 0.126 | TAGCTGTCTTCAGACGCACCAGAAGAGGGCATCA | | |
| 1044 | 16 | 56936530 | 56936603 | 73 | CBA,NZW,SMJ | 16 | 56936602 | 56936636 | 35 | + | 16.9725 | 5.50E-07 | 0.169 | TTGCTACCTCTCAATAGCACCACAAGATGGCAGCC | 21 | ENCFF821BIA,ENCFF136ZDK,ENCFF430PPJ,ENCFF165LXO,ENCFF367UNR,ENCFF093ZRU,ENCFF907XEG,ENCFF327LYS,ENCFF609MDN,ENCFF055HAP,ENCFF957SPH,ENCFF589ZBJ,ENCFF024JOZ,ENCFF198SSR,ENCFF502WDT,ENCFF099QNE,ENCFF194HVL,ENCFF883UPM,ENCFF089GBC,ENCFF092NMJ,ENCFF951GOP |
| 1045 | 16 | 84565825 | 84566055 | 230 | BALB,BUB,C58J,CBA,CE,DBA1J,ILNJ2,KK,LP.MA/My.MRL.NZO.RH2.SMJ.TH | 16 | 84565914 | 84565947 | 34 | + | 16.4634 | 9.51E-07 | 0.126 | TAGCTGTCTTCAGACGCACCAGAAGAGGGCATCA | | |
| 1046 | 16 | 84679049 | 84679275 | 226 | BALB,BUB,C58J,CBA,CE,DBA1J,ILNJ2,LP,MRL.NZO.RH2 | 16 | 84679133 | 84679166 | 34 | + | 18.3659 | 2.76E-07 | 0.07 | TAGCTGTCTTCAGACACTCCAGAAGAGGGCGCCA | | |
| 1046 | 16 | 84679049 | 84679275 | 226 | BALB,BUB,C58J,CBA,CE,DBA1J,ILNJ2,LP,MRL.NZO.RH2 | 16 | 84679148 | 84679166 | 19 | + | 17.5574 | 6.32E-07 | 0.163 | ACTCCAGAAGAGGGCGCCA | | |
| 1047 | 16 | 90152683 | 90152802 | 119 | 129S1,BUB,C57L,C58J,DBA1J,DBA2J,FVB,MA/My,NOD,NOR,NZO,PJ,RF,SJL,SWR,TH | 16 | 90152774 | 90152808 | 35 | + | 17.0275 | 5.33E-07 | 0.166 | TAGCTGTCTTCAGACAGTACCACAAGAGGGCGTTA | | |
| 1048 | 16 | 8550249 | 8550461 | 212 | AJ.AKR.PL | 16 | 8550376 | 8550394 | 19 | - | 17.8852 | 5.08E-07 | 0.147 | ACACCAGAAGAGGGCACCA | 2 | ENCFF093ZRU |
| 1049 | 16 | 12874410 | 12874625 | 215 | KK | 16 | 12874535 | 12874553 | 19 | - | 17.5574 | 6.32E-07 | 0.163 | ACTCCAGAAGAGGGCGCCA | | |
| 1049 | 16 | 12874410 | 12874625 | 215 | KK | 16 | 12874535 | 12874568 | 34 | - | 18.3659 | 2.76E-07 | 0.07 | TAGCTGTCTTCAGACACTCCAGAAGAGGGCGCCA | | |
| 1050 | 16 | 13523815 | 13524189 | 374 | KK | 16 | 13524111 | 13524129 | 19 | - | 17.5574 | 6.32E-07 | 0.163 | ACTCCAGAAGAGGGCGCCA | 1 | ENCFF093ZRU |
| 1050 | 16 | 13523815 | 13524189 | 374 | KK | 16 | 13524111 | 13524144 | 34 | - | 18.3659 | 2.76E-07 | 0.07 | TAGCTGTCTTCAGACACTCCAGAAGAGGGCGCCA | 1 | ENCFF093ZRU |
| 1051 | 16 | 17016243 | 17016436 | 193 | ILNJ2 | 16 | 17016371 | 17016404 | 34 | - | 16.4472 | 9.61E-07 | 0.126 | TAGCTGTCTTCAGACACACCAGAAGAGGGCGTCA | 1 | ENCFF093ZRU |
| 1052 | 16 | 21532654 | 21532855 | 201 | PL.SMJ | 16 | 21532785 | 21532818 | 34 | - | 17.2114 | 5.91E-07 | 0.101 | TAGCTGTTTTCAGACACTCCAGAAGAGGGCATCA | 1 | ENCFF093ZRU |
| 1053 | 16 | 21841044 | 21841157 | 113 | TH | 16 | 21841064 | 21841097 | 34 | - | 26.5691 | 3.91E-10 | 0.00111 | ATGCAGTACCCATAGAGGCCAGAAGAGGGCATCA | 21 | ENCFF201JUA,ENCFF504JDS,ENCFF093ZRU,ENCFF367UNR,ENCFF092NMJ,ENCFF430PPJ,ENCFF099QNE,ENCFF821BIA,ENCFF327LYS,ENCFF883UPM,ENCFF065ODV,ENCFF589ZBJ,ENCFF907XEG,ENCFF089GBC,ENCFF198SSR,ENCFF136ZDK,ENCFF024JOZ,ENCFF321AGQ,ENCFF165LXO,ENCFF052KVO,ENCFF502WDT |
| 1054 | 16 | 22387451 | 22387651 | 200 | TH | 16 | 22387574 | 22387607 | 34 | - | 16.4472 | 9.61E-07 | 0.126 | TAGCTGTCTTCAGACACACCAGAAGAGGGCGTCA | 1 | ENCFF093ZRU |
| 1055 | 16 | 30110498 | 30110715 | 217 | 129S1,AJ,AKR,BALB,FVB,ILNJ2,KK,LP,MRL,NOD,NOR,NZB,NZO,NZW,PL,RF,RH2,SEA,TH | 16 | 30110624 | 30110642 | 19 | - | 17.5574 | 6.32E-07 | 0.163 | ACTCCAGAAGAGGGCGCCA | | |
| 1055 | 16 | 30110498 | 30110715 | 217 | 129S1,AJ,AKR,BALB,FVB,ILNJ2,KK,LP,MRL,NOD,NOR,NZB,NZO,NZW,PL,RF,RH2,SEA,TH | 16 | 30110624 | 30110657 | 34 | - | 18.3659 | 2.76E-07 | 0.07 | TAGCTGTCTTCAGACACTCCAGAAGAGGGCGCCA | | |
| 1056 | 16 | 38434781 | 38435003 | 222 | 129S1.C58J | 16 | 38434905 | 38434938 | 34 | - | 17.8455 | 3.91E-07 | 0.0823 | TAGCTGTCCTCAGACACTCCAGAAGAGGGCGTCA | 1 | ENCFF093ZRU |
| 1057 | 16 | 38753016 | 38753217 | 201 | AJ,AKR,BALB,BTBR,BUB,C3H,C57L,CBA,CE,DBA1J,DBA2J,FVB,ILNJ2,KK,MA/My,MRL,NOD,NOR,NZB,NZO,NZW,PL,RF,RH2,SEA,SJL,SMJ,SWR,TH | 16 | 38753140 | 38753173 | 34 | - | 16.4472 | 9.61E-07 | 0.126 | TAGCTGTCTTCAGACACACCAGAAGAGGGCGTCA | | |
| 1058 | 16 | 58015830 | 58016023 | 193 | AJ.BALB.DBA1J.DBA2J.MRL.RH2 | 16 | 58015956 | 58015989 | 34 | - | 16.4472 | 9.61E-07 | 0.126 | TAGCTGTCTTCAGACACACCAGAAGAGGGCGTCA | | |
| 1059 | 16 | 69825442 | 69825771 | 329 | CE | 16 | 69825730 | 69825763 | 34 | - | 20.065 | 8.47E-08 | 0.0354 | CCGCAATAGCATTCACCTTCACAAGGTGGCGCCG | 9 | ENCFF099QNE,ENCFF883UPM,ENCFF093ZRU,ENCFF907XEG,ENCFF142CNG,ENCFF327LYS,ENCFF502WDT,ENCFF055HAP,ENCFF024JOZ |
| 1060 | 16 | 74689541 | 74689943 | 402 | C58J.LP | 16 | 74689860 | 74689893 | 34 | - | 21.9431 | 2.09E-08 | 0.0144 | CTGCTGTATACCTGTGTGCCAGAAGAGGGCATCA | 1 | ENCFF093ZRU |
| 1061 | 16 | 78025149 | 78025489 | 340 | AJ,C58J,CBA,DBA1J,DBA2J,FVB,ILNJ2,KK,MA/My,NOD,NOR,NZB,NZO,NZW,PL,SJL,SWR | 16 | 78025173 | 78025191 | 19 | - | 17.2459 | 7.74E-07 | 0.177 | CAGCCAGGAGGGGGAAGTA | | |
| 1062 | 16 | 90781676 | 90781892 | 216 | 129S1,AKR,BUB,C57L,C58J,CBA,CE,DBA1J,DBA2J,FVB,ILNJ2,MA/My,NOD,NOR,NZB,NZO,NZW,PJ,PL,RF,SEA,SJL,SMJ,SWR,TH | 16 | 90781801 | 90781819 | 19 | - | 17.5574 | 6.32E-07 | 0.163 | ACTCCAGAAGAGGGCGCCA | | |
| 1062 | 16 | 90781676 | 90781892 | 216 | 129S1,AKR,BUB,C57L,C58J,CBA,CE,DBA1J,DBA2J,FVB,ILNJ2,MA/My,NOD,NOR,NZB,NZO,NZW,PJ,PL,RF,SEA,SJL,SMJ,SWR,TH | 16 | 90781801 | 90781834 | 34 | - | 18.3659 | 2.76E-07 | 0.07 | TAGCTGTCTTCAGACACTCCAGAAGAGGGCGCCA | | |
| 1063 | 16 | 91442215 | 91442426 | 211 | BUB,CBA,CE,DBA1J,FVB,MA/My,NZB,NZO,PL.SJL.SWR.TH | 16 | 91442338 | 91442371 | 34 | - | 16.4634 | 9.51E-07 | 0.126 | TAGCTGTCTTCAGACGCACCAGAAGAGGGCATCA | | |
| 1064 | 16 | 92796018 | 92796212 | 194 | CE.DBA1J.TH | 16 | 92796142 | 92796160 | 19 | - | 17.5574 | 6.32E-07 | 0.163 | ACTCCAGAAGAGGGCGCCA | | |
| 1064 | 16 | 92796018 | 92796212 | 194 | CE.DBA1J.TH | 16 | 92796142 | 92796175 | 34 | - | 18.3659 | 2.76E-07 | 0.07 | TAGCTGTCTTCAGACACTCCAGAAGAGGGCGCCA | | |
| 1065 | 16 | 93385602 | 93385797 | 195 | CE.DBA1J.NZW | 16 | 93385713 | 93385746 | 34 | - | 16.7724 | 7.83E-07 | 0.117 | TAGCTGTCTTCAGATGCACCAGAAGAGGGCGTCA | | |
| 1066 | 16 | 93655101 | 93655311 | 210 | NZW.TH | 16 | 93655226 | 93655244 | 19 | - | 17.5574 | 6.32E-07 | 0.163 | ACTCCAGAAGAGGGCGCCA | 4 | ENCFF093ZRU.ENCFF367UNR.ENCFF099QNE.ENCFF821BIA |
| 1066 | 16 | 93655101 | 93655311 | 210 | NZW.TH | 16 | 93655226 | 93655259 | 34 | - | 18.1789 | 3.13E-07 | 0.0746 | TAGCTGTCTTCAGCCACTCCAGAAGAGGGCGCCA | 4 | ENCFF093ZRU.ENCFF367UNR.ENCFF099QNE.ENCFF821BIA |
| 1067 | 17 | 3501501 | 3501571 | 70 | DBA2J,FVB,ILNJ2,MA/My,NOD,NOR,RF,SJL | 17 | 3501504 | 3501537 | 34 | + | 24.7317 | 2.10E-09 | 0.00133 | GTGCAGTGCCCAAGGAGGCCAGAAGAGGGAGTCT | 42 | ENCFF093ZRU,ENCFF201JUA,ENCFF367UNR,ENCFF099QNE,ENCFF065ODV,ENCFF504JDS,ENCFF821BIA,ENCFF327LYS,ENCFF321AGQ,ENCFF589ZBJ,ENCFF907XEG,ENCFF198SSR,ENCFF136ZDK,ENCFF052KVO,ENCFF024JOZ,ENCFF502WDT,ENCFF430PPJ,ENCFF055HAP,ENCFF883UPM,ENCFF210BTO,ENCFF957SPH,ENCFF165LXO,ENCFF092NMJ,ENCFF033UQX,ENCFF533APC,ENCFF089GBC,ENCFF194HVL,ENCFF784ASD,ENCFF575MGI,ENCFF951GOP,ENCFF089EWX,ENCFF316FYN,ENCFF491RJK,ENCFF349BUL,ENCFF611HDQ,ENCFF609MDN,ENCFF086ZTV,ENCFF281QMV,ENCFF097CAQ,ENCFF601JVE,ENCFF142CNG |
| 1068 | 17 | 13067260 | 13067484 | 224 | C3H,CBA,CE,ILNJ2,KK,MRL,NOD,NOR,PL,SJL.SMJ.SWR.TH | 17 | 13067340 | 13067373 | 34 | + | 16.4472 | 9.61E-07 | 0.0875 | TAGCTGTCTTCAGACACACCAGAAGAGGGCGTCA | | |
| 1069 | 17 | 17699068 | 17700031 | 963 | BUB.C3H.CBA.MRL.PJ.RF.SJL.SMJ.TH | 17 | 17699422 | 17699440 | 19 | + | 17.6721 | 5.85E-07 | 0.107 | ATACCAGAAGAGGGCACTG | | |

| | | | | | | | | | | | | | | | | |
|---|---|---|---|---|---|---|---|---|---|---|---|---|---|---|---|---|
| 1070 | 17 | 25186773 | 25187009 | 236 | 129S1,AJ,AKR,B10D2,BALB,BTBR,BUB,C3H,C57L,CBA,CE,DBA1J,DBA2J,FVB,ILNJ2,KK,MA/My,MRL,NOD,NOR,NZB,NZO,PJ,RF,RH2,SEA,SJL,SWR,TH | 17 | 25186866 | 25186899 | 34 | + | 18.3659 | 2.76E-07 | 0.0462 | TAGCTGTCTTCAGACACTCCAGAAGAGGGCGCCA | 3 | ENCFF367UNR,ENCFF821BIA,ENCFF093ZRU |
| 1070 | 17 | 25186773 | 25187009 | 236 | 129S1,AJ,AKR,B10D2,BALB,BTBR,BUB,C3H,C57L,CBA,CE,DBA1J,DBA2J,FVB,ILNJ2,KK,MA/My,MRL,NOD,NOR,NZB,NZO,PJ,RF,RH2,SEA,SJL,SWR,TH | 17 | 25186881 | 25186899 | 19 | + | 17.5574 | 6.32E-07 | 0.108 | ACTCCAGAAGAGGGCGCCA | 3 | ENCFF367UNR,ENCFF821BIA,ENCFF093ZRU |
| 1071 | 17 | 27296136 | 27296286 | 150 | KK,NZO | 17 | 27296254 | 27296288 | 35 | + | 25.7615 | 1.33E-09 | 0.0034 | GTGTAGTTTCTTCACCTCACCAGTAGAGGGCACAC | 14 | ENCFF099QNE,ENCFF502WDT,ENCFF198SSR,ENCFF055HAP,ENCFF092NMJ,ENCFF052KVO,ENCFF093ZRU,ENCFF327LYS,ENCFF136ZDK,ENCFF367UNR,ENCFF907XEG,ENCFF321AGQ,ENCFF065ODV,ENCFF589ZBJ |
| 1071 | 17 | 27296136 | 27296286 | 150 | KK,NZO | 17 | 27296270 | 27296288 | 19 | + | 19.2623 | 1.93E-07 | 0.0593 | TCACCAGTAGAGGGCACAC | 14 | ENCFF099QNE,ENCFF502WDT,ENCFF198SSR,ENCFF055HAP,ENCFF092NMJ,ENCFF052KVO,ENCFF093ZRU,ENCFF327LYS,ENCFF136ZDK,ENCFF367UNR,ENCFF907XEG,ENCFF321AGQ,ENCFF065ODV,ENCFF589ZBJ |
| 1072 | 17 | 28147362 | 28147577 | 215 | 129S1,AJ,AKR,B10D2,BALB,BTBR,C3H,C57L,CBA,CE,DBA1J,DBA2J,FVB,ILNJ2,KK,MA/My,MRL,NZB,NZW,RF,RH2,SEA,SJL,SMJ,TH | 17 | 28147435 | 28147468 | 34 | + | 16.4634 | 9.51E-07 | 0.0875 | TAGCTGTCTTCAGACGCACCAGAAGAGGGCATCA | | |
| 1073 | 17 | 28609790 | 28610196 | 406 | 129S1,AJ,AKR,B10D2,BALB,C3H,C57L,CBA,CE,DBA1J,DBA2J,FVB,ILNJ2,KK,MA/My,MRL,NOD,NOR,NZB,NZW,PJ,RF,RH2,SEA,SJL,SMJ,TH | 17 | 28609865 | 28609898 | 34 | + | 17.6992 | 4.30E-07 | 0.0594 | TAGCTGTTTTCAGACACACCAGAAGAGGGCATCA | | |
| 1074 | 17 | 28829573 | 28829634 | 61 | ILNJ2,KK,RF | 17 | 28829603 | 28829637 | 35 | + | 19.1835 | 1.46E-07 | 0.0519 | CTGCATGTACGCCTGCAGGTCAGAAGAAGGCAACC | | |
| 1075 | 17 | 29040625 | 29040679 | 54 | ILNJ2 | 17 | 29040656 | 29040690 | 35 | + | 19.2294 | 1.42E-07 | 0.0513 | GAGCACTCTCCCCTGCCAGCCATCAGGGAGCACTC | | |
| 1076 | 17 | 29758628 | 29758832 | 204 | KK,RF | 17 | 29758692 | 29758725 | 34 | + | 17.2439 | 5.79E-07 | 0.0713 | TAGCTGTCCTCAGATACTCCAGAAGAGGGCATCA | | |
| 1077 | 17 | 29759259 | 29759450 | 191 | KK,RF | 17 | 29759259 | 29759292 | 34 | + | 18.7398 | 2.14E-07 | 0.0412 | CAGAAATGCCTCCTCCGGCCAGGCGGTGGTGGCA | | |
| 1078 | 17 | 31526431 | 31526642 | 211 | AJ,AKR,BTBR,C3H,C58J,CBA,CE,DBA1J,FVB,ILNJ2,KK,MRL,NOD,NOR,NZB,NZO,NZW,PJ,PL,RF,SJL,SMJ,TH | 17 | 31526502 | 31526535 | 34 | + | 16.6911 | 8.24E-07 | 0.0833 | TAGCTGTCTTCAGACAGACCAGAAGAGGGCATCA | | |
| 1079 | 17 | 34200236 | 34200674 | 438 | AJ,AKR,B10D2,BALB,BUB,C3H,C58J,CBA,CE,DBA1J,DBA2J,FVB,ILNJ2,MA/My,MRL,NOD,NOR,NZB,NZO,NZW,PJ,PL,RF,RH2,SEA,SJL,SMJ,SWR,TH | 17 | 34200266 | 34200284 | 19 | + | 17.4918 | 6.59E-07 | 0.11 | CTGTCACAAGGTGGCGCTG | 22 | ENCFF093ZRU,ENCFF367UNR,ENCFF089GBC,ENCFF136ZDK,ENCFF099QNE,ENCFF430PPJ,ENCFF316FYN,ENCFF327LYS,ENCFF907XEG,ENCFF589ZBJ,ENCFF533APC,ENCFF097CAQ,ENCFF609MDN,ENCFF194HVL,ENCFF024JOZ,ENCFF092NMJ,ENCFF821BIA,ENCFF321AGQ,ENCFF281QMV,ENCFF957SPH,ENCFF198SSR,ENCFF165LXO |
| 1080 | 17 | 35254375 | 35254606 | 231 | AJ,AKR,B10D2,BALB,BUB,C3H,C58J,CBA,CE,DBA1J,DBA2J,FVB,ILNJ2,MA/My,MRL,NZB,NZO,NZW,PJ,PL,RF,RH2,SEA,SJL,SMJ,SWR,TH | 17 | 35254457 | 35254490 | 34 | + | 16.4472 | 9.61E-07 | 0.0875 | TAGCTGTCTTCAGACACACCAGAAGAGGGCGTCA | | |
| 1081 | 17 | 35680857 | 35681065 | 208 | FVB,NOD,NOR,NZB,PL,RH2,SWR | 17 | 35680833 | 35680867 | 35 | + | 16.1743 | 8.66E-07 | 0.138 | CTACAGAACAAGACCACAGCCAGCAGGGGCTGGTG | | |
| 1082 | 17 | 35772782 | 35772987 | 205 | AJ,B10D2,BALB,BUB,DBA2J,FVB,NOD,NOR,NZB,PL,RH2,SEA,SWR | 17 | 35772759 | 35772793 | 35 | + | 16.1743 | 8.66E-07 | 0.138 | CTACAGAACAAGACCACAGCCAGCAGGGGCTGGTG | | |
| 1083 | 17 | 36404377 | 36404961 | 584 | AJ,BUB,C58J,FVB,KK,NOD,NOR,NZB,PJ,PL,SJL,SWR | 17 | 36404654 | 36404688 | 35 | + | 18.0642 | 2.90E-07 | 0.0766 | CTGCTTTTCCCCAGGCAGGGCCCTAGGTGTCACAG | | |
| 1084 | 17 | 36563820 | 36564277 | 457 | ILNJ2 | 17 | 36563938 | 36563971 | 34 | + | 17.0894 | 6.39E-07 | 0.0742 | TAGTTGTCCTCAGACACACCAGAAGAGGGCACTG | | |
| 1084 | 17 | 36563820 | 36564277 | 457 | ILNJ2 | 17 | 36563953 | 36563971 | 19 | + | 17.5902 | 6.18E-07 | 0.108 | ACACCAGAAGAGGGCACTG | | |
| 1085 | 17 | 45988387 | 45988595 | 208 | NZW.SJL | 17 | 45988453 | 45988486 | 34 | + | 16.7398 | 7.99E-07 | 0.0823 | TAGCTGTCTTCGGACACTCCAGAAGAGGGCGTCA | | |
| 1086 | 17 | 47049087 | 47049288 | 201 | 129S1.NOD.NOR.NZW.SJL | 17 | 47049146 | 47049179 | 34 | + | 18.3659 | 2.76E-07 | 0.0462 | TAGCTGTCTTCAGACACTCCAGAAGAGGGCGCCA | 3 | ENCFF093ZRU.ENCFF367UNR |
| 1086 | 17 | 47049087 | 47049288 | 201 | 129S1.NOD.NOR.NZW.SJL | 17 | 47049161 | 47049179 | 19 | + | 17.5574 | 6.32E-07 | 0.108 | ACTCCAGAAGAGGGCGCCA | 3 | ENCFF093ZRU.ENCFF367UNR |
| 1087 | 17 | 47973872 | 47974100 | 228 | SJL.SWR | 17 | 47973957 | 47973990 | 34 | + | 16.4634 | 9.51E-07 | 0.0875 | TAGCTGTCTTCAGACGCACCAGAAGAGGGCATCA | 1 | ENCFF093ZRU |
| 1088 | 17 | 48460709 | 48461375 | 666 | TH | 17 | 48461322 | 48461355 | 34 | + | 22.0569 | 1.91E-08 | 0.00721 | GTGCAGTGGCCACAGAGGCCAGAAGAGGTTGCGG | | |
| 1089 | 17 | 57349188 | 57349469 | 281 | 129S1,C57L,C58J,FVB,LP,MA/My,NOD,NOR,SJL.SWR | 17 | 57349326 | 57349360 | 35 | + | 16.2477 | 8.31E-07 | 0.135 | TAGCTGTCTTCAGACACTCCCAGAAGAGGGCATCA | | |
| 1090 | 17 | 66111793 | 66111998 | 205 | AJ,BALB,BTBR,BUB,C58J,CE,DBA1J,DBA2J,FVB,KK,NOD,NOR,NZB,NZW,PJ,PL,RH2,SEA.SJL.SMJ.TH | 17 | 66111855 | 66111888 | 34 | + | 17.1382 | 6.20E-07 | 0.0734 | TAGCTGTCTCCAGACACTCCAGAAGAGGGCATCA | | |
| 1091 | 17 | 66809654 | 66809864 | 210 | TH | 17 | 66809719 | 66809752 | 34 | + | 16.5691 | 8.90E-07 | 0.0867 | TAGCTGTCTTCAGACGCACCAGAAGAGGGCATCG | | |
| 1092 | 17 | 74858714 | 74858927 | 213 | 129S1,AKR,BTBR,BUB,C57L,ILNJ2,KK,MA/My,NOD,NOR,NZB,NZO,NZW,SWR,TH | 17 | 74858784 | 74858817 | 34 | + | 16.4634 | 9.51E-07 | 0.0875 | TAGCTGTCTTCAGACGCACCAGAAGAGGGCATCA | | |
| 1093 | 17 | 82824212 | 82824420 | 208 | MA/Mv.NZW | 17 | 82824276 | 82824309 | 34 | + | 18.187 | 3.12E-07 | 0.049 | TAGCTGTCTTCAGTCACTCCAGAAGAGGGCGCTA | | |
| 1093 | 17 | 82824212 | 82824420 | 208 | MA/Mv.NZW | 17 | 82824291 | 82824309 | 19 | + | 17.5738 | 6.25E-07 | 0.108 | ACTCCAGAAGAGGGCGCTA | | |
| 1094 | 17 | 84750765 | 84750976 | 211 | DBA1J.DBA2J.ILNJ2 | 17 | 84750836 | 84750869 | 34 | + | 18.3984 | 2.70E-07 | 0.046 | TTGCTGTCTTCAGACACACCAGAAGAGGGCATTG | | |
| 1095 | 17 | 6383944 | 6384147 | 203 | AJ,AKR,BALB,C3H,C57L,CBA,DBA1J,DBA2J,ILNJ2,MRL,NZO,PJ,PL,RF,RH2,SEA,SMJ,TH | 17 | 6384069 | 6384102 | 34 | - | 16.4472 | 9.61E-07 | 0.0875 | TAGCTGTCTTCAGACACACCAGAAGAGGGCGTCA | | |
| 1096 | 17 | 12437641 | 12437820 | 179 | AJ,B10D2,C3H,CBA,DBA1J,DBA2J,MRL,PL,SMJ | 17 | 12437789 | 12437822 | 34 | - | 16.5691 | 8.90E-07 | 0.0867 | GCGCTCTGTCAACAGGAGCCACTAGGAGGCTGTT | 2 | ENCFF093ZRU,ENCFF092NMJ |
| 1097 | 17 | 14833823 | 14834644 | 821 | ILNJ2,NOD,NOR,NZW,SJL | 17 | 14833827 | 14833861 | 35 | - | 17.8899 | 3.21E-07 | 0.0817 | ATGAAGTTGTCACTGTTGCCCAGGAGATGCCACCA | | |
| 1098 | 17 | 27212481 | 27212706 | 225 | KK,NZO | 17 | 27212611 | 27212644 | 34 | - | 18.0244 | 3.47E-07 | 0.0522 | TAGCTGTTTTCAGATGCACCAGAAGAGGGCATCA | 1 | ENCFF093ZRU |
| 1099 | 17 | 29212554 | 29212761 | 207 | ILNJ2,NOD,NOR,PJ | 17 | 29212667 | 29212700 | 34 | - | 17.6667 | 4.40E-07 | 0.06 | TAGCTGTTTTCAGACACTCCAGAAGAGGGCGTCA | | |
| 1100 | 17 | 29242767 | 29243028 | 261 | ILNJ2,KK,NOD,NOR,PJ,RF | 17 | 29242892 | 29242925 | 34 | - | 17.8455 | 3.91E-07 | 0.0561 | TAGCTGTCCTCAGACACTCCAGAAGAGGGCGTCA | | |
| 1101 | 17 | 31682402 | 31682488 | 86 | KK,PJ | 17 | 31682405 | 31682438 | 34 | - | 16.9837 | 6.84E-07 | 0.0767 | CATCAGGCCCCTGGGCAGTCACTGGGTGGCACTG | | |
| 1102 | 17 | 32946225 | 32946432 | 207 | AJ,AKR,C3H,CBA,CE,ILNJ2,MA/My,MRL,NOD,NOR,PJ,PL,RF,SJL,TH | 17 | 32946350 | 32946383 | 34 | - | 16.6911 | 8.24E-07 | 0.0833 | TAGCTGTCTTCAGACAGACCAGAAGAGGGCATCA | | |
| 1103 | 17 | 35131173 | 35131273 | 100 | ILNJ2 | 17 | 35131242 | 35131260 | 19 | - | 20.8033 | 5.79E-08 | 0.0357 | CAGCCACTAGGGGTCGCTG | 41 | ENCFF093ZRU,ENCFF367UNR,ENCFF327LYS,ENCFF024JOZ,ENCFF907XEG,ENCFF589ZBJ,ENCFF805QBU,ENCFF086ZTV,ENCFF136ZDK,ENCFF198SSR,ENCFF165LXO,ENCFF089GBC,ENCFF099QNE,ENCFF821BIA,ENCFF321AGQ,ENCFF430PPJ,ENCFF092NMJ,ENCFF957SPH,ENCFF055HAP,ENCFF502WDT,ENCFF349BUL,ENCFF052KVO,ENCFF065ODV,ENCFF210BTO,ENCFF951GOP,ENCFF194HVL,ENCFF142CNG,ENCFF316FYN,ENCFF281QMV,ENCFF611HDQ,ENCFF883UPM,ENCFF784ASD,ENCFF601JVE,ENCFF491RJK,ENCFF533APC,ENCFF504JDS,ENCFF097CAQ,ENCFF575MGI,ENCFF089EWX,ENCFF609MDN,ENCFF033UQX |
| 1103 | 17 | 35131173 | 35131273 | 100 | ILNJ2 | 17 | 35131242 | 35131275 | 34 | - | 24.0325 | 3.83E-09 | 0.00206 | ATGCAGGACATGGTCCAGCCACTAGGGGTCGCTG | 41 | ENCFF093ZRU,ENCFF367UNR,ENCFF327LYS,ENCFF024JOZ,ENCFF907XEG,ENCFF589ZBJ,ENCFF805QBU,ENCFF086ZTV,ENCFF136ZDK,ENCFF198SSR,ENCFF165LXO,ENCFF089GBC,ENCFF099QNE,ENCFF821BIA,ENCFF321AGQ,ENCFF430PPJ,ENCFF092NMJ,ENCFF957SPH,ENCFF055HAP,ENCFF502WDT,ENCFF349BUL,ENCFF052KVO,ENCFF065ODV,ENCFF210BTO,ENCFF951GOP,ENCFF194HVL,ENCFF142CNG,ENCFF316FYN,ENCFF281QMV,ENCFF611HDQ,ENCFF883UPM,ENCFF784ASD,ENCFF601JVE,ENCFF491RJK,ENCFF533APC,ENCFF504JDS,ENCFF097CAQ,ENCFF575MGI,ENCFF089EWX,ENCFF609MDN,ENCFF033UQX |
| 1104 | 17 | 35443008 | 35443798 | 790 | AKR,BUB,C3H,C58J,CBA,CE,DBA1J,FVB,MA/My,MRL,RF,SWR | 17 | 35443030 | 35443063 | 34 | - | 17.374 | 5.32E-07 | 0.0676 | TAGCTGTTTTCAGACACCCCAGAAGAGGGCGTCA | 1 | ENCFF093ZRU |
| 1105 | 17 | 35493210 | 35493417 | 207 | AKR.C3H.C58J.CBA.CE.MA/Mv.MRL.RF | 17 | 35493339 | 35493372 | 34 | - | 17.9919 | 3.55E-07 | 0.0528 | TTGCTGTCTTCAGACACTCCAGAAGAGGGCATCA | 5 | ENCFF093ZRU.ENCFF821BIA.ENCFF367UNR.ENCFF502WDT.ENCFF055HAP |
| 1106 | 17 | 36108459 | 36108666 | 207 | 129S1,AJ,AKR,B10D2,BALB,BTBR,C3H,C57L,C58J,CBA,CE,DBA1J,DBA2J,LP,MA/My,MRL,NZB,NZO,NZW,PJ,PL,RF,RH2,SEA,SJL.SMJ.TH | 17 | 36108582 | 36108615 | 34 | - | 16.4472 | 9.61E-07 | 0.0875 | TAGCTGTCTTCAGACACACCAGAAGAGGGCGTCA | | |
| 1107 | 17 | 45700085 | 45700287 | 202 | 129S1,AJ,BTBR,BUB,C57L,C58J,DBA1J,DBA2J,FVB,ILNJ2,KK,LP,NOD,NOR,NZB,NZO,NZW,SJL,SMJ,SWR,TH | 17 | 45700211 | 45700244 | 34 | - | 16.4472 | 9.61E-07 | 0.0875 | TAGCTGTCTTCAGACACACCAGAAGAGGGCGTCA | | |
| 1108 | 17 | 46416574 | 46416783 | 209 | 129S1,BTBR,BUB,C57L,DBA1J,FVB,KK,LP,MA/My,NOD,NOR,NZB,NZO,NZW,RF,SJL,SWR | 17 | 46416698 | 46416731 | 34 | - | 16.6911 | 8.24E-07 | 0.0833 | TAGCTGTCTTCAGACAGACCAGAAGAGGGCATCA | | |
| 1109 | 17 | 48928111 | 48928579 | 468 | NZW.SJL | 17 | 48928108 | 48928126 | 19 | - | 21.0984 | 4.51E-08 | 0.0335 | AGGCCAGAAGAGGGCGCCA | 3 | ENCFF093ZRU.ENCFF052KVO.ENCFF033UQX |
| 1109 | 17 | 48928111 | 48928579 | 468 | NZW.SJL | 17 | 48928108 | 48928141 | 34 | - | 24.0732 | 3.70E-09 | 0.00201 | GTGCATATGCCCTGGAGGCCAGAAGAGGGCGCCA | 3 | ENCFF093ZRU.ENCFF052KVO.ENCFF033UQX |
| 1110 | 17 | 50249498 | 50250132 | 634 | NZW,SWR | 17 | 50249895 | 50249913 | 19 | - | 18.1475 | 4.25E-07 | 0.0913 | AGGCCTGCAGATGGCAGCA | 29 | ENCFF093ZRU,ENCFF024JOZ,ENCFF089GBC,ENCFF198SSR,ENCFF367UNR,ENCFF099QNE,ENCFF065ODV,ENCFF092NMJ,ENCFF821BIA,ENCFF136ZDK,ENCFF033UQX,ENCFF575MGI,ENCFF502WDT,ENCFF281QMV,ENCFF142CNG,ENCFF097CAQ,ENCFF327LYS,ENCFF883UPM,ENCFF907XEG,ENCFF055HAP,ENCFF957SPH,ENCFF165LXO,ENCFF201JUA,ENCFF504JDS,ENCFF589ZBJ,ENCFF533APC,ENCFF611HDQ,ENCFF491RJK.ENCFF194HVL |
| 1110 | 17 | 50249498 | 50250132 | 634 | NZW,SWR | 17 | 50249895 | 50249929 | 35 | - | 23.5138 | 7.67E-09 | 0.00814 | GTGCACAGGCATCCCCAGGCCTGCAGATGGCAGCA | 29 | ENCFF093ZRU,ENCFF024JOZ,ENCFF089GBC,ENCFF198SSR,ENCFF367UNR,ENCFF099QNE,ENCFF065ODV,ENCFF092NMJ,ENCFF821BIA,ENCFF136ZDK,ENCFF033UQX,ENCFF575MGI,ENCFF502WDT,ENCFF281QMV,ENCFF142CNG,ENCFF097CAQ,ENCFF327LYS,ENCFF883UPM,ENCFF907XEG,ENCFF055HAP,ENCFF957SPH,ENCFF165LXO,ENCFF201JUA,ENCFF504JDS,ENCFF589ZBJ,ENCFF533APC,ENCFF611HDQ,ENCFF491RJK.ENCFF194HVL |
| 1111 | 17 | 63745136 | 63745368 | 232 | BTBR.C58J.LP | 17 | 63745261 | 63745294 | 34 | - | 16.4634 | 9.51E-07 | 0.0875 | TAGCTGTCTTCAGACGCACCAGAAGAGGGCATCA | | |
| 1112 | 17 | 73117576 | 73117825 | 249 | 129S1,BALB,BTBR,C3H,C57L,CBA,DBA1J,DBA2J,FVB,MRL,NZW,PJ,RF,SEA,SJL,SMJ,TH | 17 | 73117701 | 73117734 | 34 | - | 16.4634 | 9.51E-07 | 0.0875 | TAGCTGTCTTCAGACGCACCAGAAGAGGGCATCA | | |
| 1113 | 17 | 73422765 | 73422988 | 223 | 129S1.AJ.AKR.BTBR.C57L.TH | 17 | 73422888 | 73422921 | 34 | - | 17.6992 | 4.30E-07 | 0.0594 | TAGCTGTCCTCAGATACTCCAGAAGAGGGCGTCA | | |
| 1114 | 17 | 74818039 | 74818243 | 204 | 129S1,BTBR,BUB,C57L,ILNJ2,KK,NOD,NOR.NZW.SWR | 17 | 74818167 | 74818200 | 34 | - | 16.4472 | 9.61E-07 | 0.0875 | TAGCTGTCTTCAGACACACCAGAAGAGGGCGTCA | | |
| 1115 | 17 | 75011349 | 75011557 | 208 | 129S1,AKR,BTBR,BUB,C57L,ILNJ2,KK,MA/My,NOD,NOR,NZB,NZO,NZW,SWR,TH | 17 | 75011477 | 75011510 | 34 | - | 19.9593 | 9.14E-08 | 0.0234 | CAGCTGTCCTCAGACACTCCAGAAGAGGGCATCA | | |
| 1116 | 17 | 79816051 | 79816123 | 72 | NZW | 17 | 79816035 | 79816069 | 35 | - | 16.945 | 5.59E-07 | 0.109 | GGGCAAGGAATGCCCAGGGCCACCAGGTGCCAGGA | | |
| 1117 | 17 | 80920252 | 80920447 | 195 | CE.TH | 17 | 80920381 | 80920414 | 34 | - | 16.4472 | 9.61E-07 | 0.0875 | TAGCTGTCTTCAGACACACCAGAAGAGGGCGTCA | | |
| 1118 | 17 | 85174859 | 85175031 | 172 | FVB.NOD.NOR.SJL.SWR | 17 | 85174954 | 85174972 | 19 | - | 17.5574 | 6.32E-07 | 0.108 | ACTCCAGAAGAGGGCGCCA | 4 | ENCFF093ZRU.ENCFF099QNE.ENCFF367UNR.ENCFF136ZDK |
| 1118 | 17 | 85174859 | 85175031 | 172 | FVB.NOD.NOR.SJL.SWR | 17 | 85174954 | 85174987 | 34 | - | 17.0813 | 6.43E-07 | 0.0744 | TAGCTATCTTCAGACACTCCAGAAGAGGGCGCCA | 4 | ENCFF093ZRU.ENCFF099QNE.ENCFF367UNR.ENCFF136ZDK |
| 1119 | 17 | 85746477 | 85746684 | 207 | 129S1.AKR.MRL.NZB.NZO.PJ.PL.RF | 17 | 85746601 | 85746634 | 34 | - | 16.4634 | 9.51E-07 | 0.0875 | TAGCTGTCTTCAGACGCACCAGAAGAGGGCATCA | | |
| 1120 | 17 | 86526229 | 86526750 | 521 | AJ,BALB,C3H,C57L,CBA,CE,LP,PJ,RH2,SEA | 17 | 86526260 | 86526293 | 34 | - | 16.5285 | 9.13E-07 | 0.0875 | ATGTAAAGCCTGCTGTGGCCTGCAGAGGTCTTTC | | |
| 1121 | 17 | 90033095 | 90033650 | 555 | 129S1,AJ,AKR,BALB,BTBR,BUB,C3H,C57L,CBA,CE,DBA1J,DBA2J,FVB,ILNJ2,KK,LP,MA/My,MRL,NOD,NOR,NZB,NZO,NZW,PJ,PL,RF,RH2,SEA,SJL,SMJ,SWR,TH | 17 | 90033166 | 90033199 | 34 | - | 27.4878 | 1.59E-10 | 0.000243 | ATGCAGTGCCCTGAGAGGCCAGAAGAGGGCATTA | 5 | ENCFF367UNR,ENCFF093ZRU,ENCFF099QNE,ENCFF024JOZ,ENCFF907XEG |
| 1122 | 18 | 33036769 | 33037332 | 563 | BALB,C3H,C58J,CE,MRL,PL,RF,RH2,SEA,TH | 18 | 33037126 | 33037160 | 35 | + | 24.2569 | 4.38E-09 | 0.0113 | CTGCAGTATTCTACCCTGACCACCAGAAGGCTCAG | 13 | ENCFF367UNR,ENCFF430PPJ,ENCFF093ZRU,ENCFF198SSR,ENCFF165LXO,ENCFF099QNE,ENCFF316FYN,ENCFF201JUA,ENCFF327LYS,ENCFF092NMJ,ENCFF907XEG,ENCFF089GBC,ENCFF024JOZ |
| 1123 | 18 | 35182798 | 35183042 | 244 | AJ,BALB,C58J,CBA,DBA2J,ILNJ2,RH2,SEA,TH | 18 | 35182900 | 35182933 | 34 | + | 16.4472 | 9.61E-07 | 0.128 | TAGCTGTCTTCAGACACACCAGAAGAGGGCGTCA | | |
| 1124 | 18 | 35897278 | 35897492 | 214 | AJ,BALB,C58J,CBA,DBA2J,ILNJ2,MRL,RH2,SEA,SMJ | 18 | 35897349 | 35897382 | 34 | + | 18.3659 | 2.76E-07 | 0.0797 | TAGCTGTCTTCAGACACTCCAGAAGAGGGCGCCA | | |
| 1124 | 18 | 35897278 | 35897492 | 214 | AJ,BALB,C58J,CBA,DBA2J,ILNJ2,MRL,RH2,SEA,SMJ | 18 | 35897364 | 35897382 | 19 | + | 17.5574 | 6.32E-07 | 0.164 | ACTCCAGAAGAGGGCGCCA | | |
| 1125 | 18 | 36850402 | 36850604 | 202 | 129S1,AJ,AKR,BALB,BUB,C3H,C57L,C58J,CBA,CE,DBA1J,DBA2J,FVB,ILNJ2,KK,MA/My,NOD,NOR,NZB,NZO,NZW,PL,RF,RH2,SEA,SJL,SMJ,SWR,TH | 18 | 36850463 | 36850496 | 34 | + | 17.9106 | 3.74E-07 | 0.0901 | TAGCTGTCTTCAGACACTCCAGAAGAGGGCACCA | | |
| 1126 | 18 | 53897352 | 53897558 | 206 | KK | 18 | 53897402 | 53897435 | 34 | + | 16.4634 | 9.51E-07 | 0.128 | TAGCTGTCTTCAGACGCACCAGAAGAGGGCATCA | | |
| 1127 | 18 | 60927156 | 60927365 | 209 | AJ,BALB,CBA,FVB,ILNJ2,NZB,RH2,SEA,SJL,SMJ | 18 | 60927225 | 60927258 | 34 | + | 17.4797 | 4.97E-07 | 0.101 | TAGCTGTCTCCAGATACACCAGAAGAGGGCATCA | | |
| 1128 | 18 | 62105918 | 62106299 | 381 | AJ,BALB,CBA,CE,FVB,ILNJ2,NZB,NZO,PL,RH2,SEA,SJL,SWR,TH | 18 | 62106025 | 62106058 | 34 | + | 18.3821 | 2.73E-07 | 0.0797 | TAGCAGTCTTCAGACACACCAGAAGAGGGCGTCA | | |
| 1129 | 18 | 65026986 | 65027207 | 221 | AJ,AKR,BALB,BTBR,BUB,C3H,C57L,CBA,CE,FVB,ILNJ2,KK,MRL,NOD,NOR,NZB,NZO,PL,RF,RH2,SJL,SMJ,SWR | 18 | 65027060 | 65027093 | 34 | + | 16.4472 | 9.61E-07 | 0.128 | TAGCTGTCTTCAGACACACCAGAAGAGGGCGTCA | | |
| 1130 | 18 | 65621248 | 65621452 | 204 | 129S1,AJ,AKR,BALB,BTBR,BUB,C3H,C57L,CBA,CE,FVB,ILNJ2,KK,MA/My,MRL,NOD,NOR,NZB,NZO,NZW,RF,RH2,SJL,SMJ,SWR,TH | 18 | 65621309 | 65621342 | 34 | + | 18.1789 | 3.13E-07 | 0.0837 | TAGCTGTCTTCAGACACTCCAGAAGAGGGCGCTA | | |
| 1130 | 18 | 65621248 | 65621452 | 204 | 129S1,AJ,AKR,BALB,BTBR,BUB,C3H,C57L,CBA,CE,FVB,ILNJ2,KK,MA/My,MRL,NOD,NOR,NZB,NZO,NZW,RF,RH2,SJL,SMJ,SWR,TH | 18 | 65621324 | 65621342 | 19 | + | 17.5738 | 6.25E-07 | 0.164 | ACTCCAGAAGAGGGCGCTA | | |
| 1131 | 18 | 65926153 | 65927144 | 991 | 129S1,AJ,BALB,BTBR,BUB,C3H,C57L,CBA,CE.DBA1J.DBA2J.ILNJ2.PJ.RH2.SEA | 18 | 65926121 | 65926155 | 35 | + | 16.1376 | 8.84E-07 | 0.212 | TTGCTCTCTTTCAGACACACCAGAAGAGGGCATCG | | |
| 1132 | 18 | 66564546 | 66564755 | 209 | 129S1,AKR,BTBR,BUB,C3H,C57L,CBA,CE,DBA1J,DBA2J,FVB,KK,MA/My,MRL,NOD,NOR.PJ.RF.SEA.SJL.SWR.TH | 18 | 66564612 | 66564645 | 34 | + | 17.2439 | 5.79E-07 | 0.107 | TAGCTGTCCTCAGATACTCCAGAAGAGGGCATCA | 1 | ENCFF883UPM |
| 1133 | 18 | 67935251 | 67935457 | 206 | 129S1,AJ,AKR,BALB,BTBR,BUB,C3H,C57L,CBA,CE,DBA1J,DBA2J,ILNJ2,KK,MA/My,MRL,PJ,PL,RF,RH2,SEA,SMJ | 18 | 67935316 | 67935349 | 34 | + | 16.4472 | 9.61E-07 | 0.128 | TAGCTGTCTTCAGACACACCAGAAGAGGGCGTCA | | |
| 1134 | 18 | 67958125 | 67958340 | 215 | 129S1,AJ,AKR,BALB,BTBR,BUB,C3H,C57L,CBA,CE,DBA1J,DBA2J,ILNJ2,KK,MA/My,MRL,PJ,PL,RF,RH2,SEA,SMJ | 18 | 67958199 | 67958232 | 34 | + | 17.2439 | 5.79E-07 | 0.107 | TAGCTGTCCTCAGATACTCCAGAAGAGGGCATCA | | |
| 1135 | 18 | 70708751 | 70708949 | 198 | RF | 18 | 70708807 | 70708840 | 34 | + | 16.4472 | 9.61E-07 | 0.128 | TAGCTGTCTTCAGACACACCAGAAGAGGGCGTCA | | |
| 1136 | 18 | 75591263 | 75591388 | 125 | AKR.ILNJ2.RF | 18 | 75591232 | 75591265 | 34 | + | 16.4146 | 9.81E-07 | 0.129 | CTGGTGTACACCCAGCCTTCAGGAGGTGGAGGCA | | |
| 1137 | 18 | 76008576 | 76008791 | 215 | 129S1,AJ,AKR,BALB,BTBR,BUB,C57L,FVB,ILNJ2.KK.MRL.NZW.RF.RH2 | 18 | 76008645 | 76008678 | 34 | + | 16.4472 | 9.61E-07 | 0.128 | TAGCTGTCTTCAGACACACCAGAAGAGGGCGTCA | | |
| 1138 | 18 | 76339281 | 76339499 | 218 | 129S1,AJ,AKR,BALB,BTBR,BUB,C3H,C57L,CBA,DBA1J,DBA2J,FVB,ILNJ2,KK,MRL,NZO.NZW.PJ.PL.RF.RH2.SEA.SMJ.TH | 18 | 76339356 | 76339389 | 34 | + | 18.1545 | 3.18E-07 | 0.0841 | TAGCTGTTTTCAGACACACCAGAAGAGGGCGTCA | | |
| 1139 | 18 | 76453914 | 76454118 | 204 | AJ,AKR,BALB,BUB,C3H,CBA,DBA1J,DBA2J,FVB,ILNJ2,KK,MRL,NZO,NZW,PJ,PL,RF,RH2.SEA.SMJ.TH | 18 | 76453970 | 76454003 | 34 | + | 16.4472 | 9.61E-07 | 0.128 | TAGCTGTCTTCAGACACACCAGAAGAGGGCGTCA | | |
| 1140 | 18 | 80026009 | 80026435 | 426 | NZO | 18 | 80026045 | 80026063 | 19 | + | 18.7705 | 2.75E-07 | 0.119 | AAGCCAGAAGAGGGCACCA | 6 | ENCFF093ZRU.ENCFF092NMJ.ENCFF883UPM.ENCFF089GBC.ENCFF055HAP.ENCFF502WDT |
| 1141 | 18 | 90596172 | 90596381 | 209 | KK.NZB.NZO.RF.TH | 18 | 90596240 | 90596273 | 34 | + | 17.8455 | 3.91E-07 | 0.0917 | TAGCTGTCCTCAGACACTCCAGAAGAGGGCGTCA | | |
| 1142 | 18 | 9473359 | 9473565 | 206 | 129S1,AJ,AKR,BALB,BTBR,C3H,C57L,CBA,ILNJ2,RF,RH2 | 18 | 9473484 | 9473502 | 19 | - | 17.5574 | 6.32E-07 | 0.164 | ACTCCAGAAGAGGGCGCCA | | |
| 1142 | 18 | 9473359 | 9473565 | 206 | 129S1,AJ,AKR,BALB,BTBR,C3H,C57L,CBA,ILNJ2,RF,RH2 | 18 | 9473484 | 9473517 | 34 | - | 18.374 | 2.75E-07 | 0.0797 | TAGCTGTCTTCAGTCACTCCAGAAGAGGGCGCCA | | |
| 1143 | 18 | 30500897 | 30501108 | 211 | 129S1.C57L.ILNJ2.SMJ | 18 | 30501023 | 30501056 | 34 | - | 17.6667 | 4.40E-07 | 0.0954 | TAGCTGTCTTCAGACACGCCAGAAGAGGGCGTCA | | |
| 1144 | 18 | 36845488 | 36845757 | 269 | 129S1,AJ,AKR,BALB,BUB,C3H,C57L,C58J,CBA,CE,DBA1J,DBA2J,FVB,ILNJ2,KK,MA/My,NOD,NOR,NZB,NZO,NZW,PL,RF,RH2,SEA.SJL.SMJ.SWR.TH | 18 | 36845612 | 36845645 | 34 | - | 16.4634 | 9.51E-07 | 0.128 | TAGCTGTCTTCAGACGCACCAGAAGAGGGCATCA | | |
| 1145 | 18 | 46650557 | 46650769 | 212 | KK | 18 | 46650684 | 46650717 | 34 | - | 16.4472 | 9.61E-07 | 0.128 | TAGCTGTCTTCAGACACACCAGAAGAGGGCGTCA | | |
| 1146 | 18 | 56813182 | 56813752 | 570 | CE | 18 | 56813466 | 56813499 | 34 | - | 17.065 | 6.50E-07 | 0.114 | TTGCTCTCTTCAGACACACCAGAAGAGGGCATCG | | |
| 1147 | 18 | 60795534 | 60795742 | 208 | AJ,BALB,CBA,FVB,ILNJ2,KK,NOD,NOR,NZB,RH2,SEA,SJL,SMJ | 18 | 60795660 | 60795693 | 34 | - | 16.4634 | 9.51E-07 | 0.128 | TAGCTGTCTTCAGACGCACCAGAAGAGGGCATCA | | |
| 1148 | 18 | 67280550 | 67280890 | 340 | AJ,AKR,BALB,FVB,ILNJ2,KK,NOD,NOR,NZB,NZO,NZW,PL,RH2,SJL,SMJ,SWR,TH | 18 | 67280717 | 67280751 | 35 | - | 22.789 | 1.30E-08 | 0.02 | CTGCATGTATGCCTGCATGCCAGGAGAGGGCATCA | | |
| 1149 | 18 | 67835027 | 67835242 | 215 | AJ,BALB,FVB,ILNJ2,KK,MRL,NOD,NOR,NZB,NZO,NZW,PL,RF,RH2,SJL,SMJ,SWR,TH | 18 | 67835148 | 67835181 | 34 | - | 17.8455 | 3.91E-07 | 0.0917 | TAGCTGTCCTCAGACACTCCAGAAGAGGGCGTCA | | |
| 1150 | 18 | 73682226 | 73682442 | 216 | 129S1,BTBR,C3H,C57L,CBA,DBA1J,DBA2J,PJ,PL,SEA | 18 | 73682350 | 73682383 | 34 | - | 17.3008 | 5.58E-07 | 0.106 | TAGCTGTCTTCAGATACACCAGAAGGGGGCGTCA | | |
| 1151 | 18 | 75110503 | 75110737 | 234 | AKR.ILNJ2.RF | 18 | 75110491 | 75110524 | 34 | - | 19.7805 | 1.04E-07 | 0.0445 | GAGCAGGTACACCTGTAGGCAGAAGAGGGCATCA | | |
| 1152 | 18 | 75732976 | 75733034 | 58 | AJ,AKR,BALB,ILNJ2,KK,MRL,NOD,NOR,NZB,NZW,RF,RH2 | 18 | 75732953 | 75732987 | 35 | - | 16.7982 | 6.08E-07 | 0.183 | GTGTTGATGCACTACTCTGCCACTAGAGGGTATCA | 38 | ENCFF367UNR,ENCFF093ZRU,ENCFF327LYS,ENCFF024JOZ,ENCFF907XEG,ENCFF092NMJ,ENCFF136ZDK,ENCFF089GBC,ENCFF198SSR,ENCFF430PPJ,ENCFF589ZBJ,ENCFF165LXO,ENCFF883UPM,ENCFF065ODV,ENCFF504JDS,ENCFF321AGQ,ENCFF055HAP,ENCFF502WDT,ENCFF951GOP,ENCFF210BTO,ENCFF575MGI,ENCFF609MDN,ENCFF957SPH,ENCFF052KVO,ENCFF316FYN,ENCFF601JVE,ENCFF281QMV,ENCFF491RJK,ENCFF533APC,ENCFF097CAQ,ENCFF033UQX,ENCFF611HDQ,ENCFF142CNG,ENCFF089EWX,ENCFF784ASD,ENCFF194HVL,ENCFF805QBU,ENCFF086ZTV |
| 1152 | 18 | 75732976 | 75733034 | 58 | AJ,AKR,BALB,ILNJ2,KK,MRL,NOD,NOR,NZB,NZW,RF,RH2 | 18 | 75733011 | 75733045 | 35 | - | 17.6972 | 3.60E-07 | 0.143 | GTGTTGATGCATCACTCTGCCACTAGAGGGTGTCA | 39 | ENCFF367UNR,ENCFF093ZRU,ENCFF327LYS,ENCFF024JOZ,ENCFF907XEG,ENCFF092NMJ,ENCFF136ZDK,ENCFF089GBC,ENCFF198SSR,ENCFF430PPJ,ENCFF589ZBJ,ENCFF165LXO,ENCFF883UPM,ENCFF065ODV,ENCFF504JDS,ENCFF321AGQ,ENCFF055HAP,ENCFF502WDT,ENCFF951GOP,ENCFF210BTO,ENCFF575MGI,ENCFF609MDN,ENCFF957SPH,ENCFF052KVO,ENCFF316FYN,ENCFF601JVE,ENCFF281QMV,ENCFF491RJK,ENCFF533APC,ENCFF097CAQ,ENCFF033UQX,ENCFF611HDQ,ENCFF142CNG,ENCFF089EWX,ENCFF784ASD,ENCFF194HVL,ENCFF805QBU,ENCFF086ZTV,ENCFF349BUL |
| 1153 | 18 | 77859046 | 77859198 | 152 | KK,TH | 18 | 77859043 | 77859077 | 35 | - | 16.6422 | 6.65E-07 | 0.185 | CTGCTACCCAAGACCTCTAACACCAGAGGCCACCC | | |
| 1153 | 18 | 77859046 | 77859198 | 152 | KK,TH | 18 | 77859095 | 77859129 | 35 | - | 16.6422 | 6.65E-07 | 0.185 | CTGCTACCCAAGACCTCTAACACCAGAGGCCACCC | | |
| 1153 | 18 | 77859046 | 77859198 | 152 | KK,TH | 18 | 77859146 | 77859180 | 35 | - | 16.6422 | 6.65E-07 | 0.185 | CTGCTACCCAAGACCTCTAACACCAGAGGCCACCC | | |
| 1153 | 18 | 77859046 | 77859198 | 152 | KK,TH | 18 | 77859197 | 77859231 | 35 | - | 16.6422 | 6.65E-07 | 0.185 | CTGCTACCCAAGACCTCTAACACCAGAGGCCACCC | | |
| 1154 | 18 | 82747749 | 82747802 | 53 | BUB,MA/My,NZB,SJL | 18 | 82747756 | 82747774 | 19 | - | 19.6393 | 1.45E-07 | 0.0832 | CAGCCACTAGAAGGCGGCA | | |
| 1154 | 18 | 82747749 | 82747802 | 53 | BUB,MA/My,NZB,SJL | 18 | 82747773 | 82747791 | 19 | - | 19.6393 | 1.45E-07 | 0.0832 | CAGCCACTAGAAGGCGGCA | | |
| 1154 | 18 | 82747749 | 82747802 | 53 | BUB,MA/My,NZB,SJL | 18 | 82747790 | 82747808 | 19 | - | 19.6393 | 1.45E-07 | 0.0832 | CAGCCACTAGAAGGCGGCA | | |
| 1155 | 18 | 84698206 | 84698411 | 205 | 129S1,BTBR,C57L,CE,NOD | 18 | 84698320 | 84698338 | 19 | - | 17.8852 | 5.08E-07 | 0.15 | ACACCAGAAGAGGGCACCA | | |
| 1155 | 18 | 84698206 | 84698411 | 205 | 129S1,BTBR,C57L,CE,NOD | 18 | 84698320 | 84698353 | 34 | - | 20.8862 | 4.65E-08 | 0.0264 | TTGCTGTCTTCAGACACACCAGAAGAGGGCACCA | | |
| 1156 | 18 | 84918691 | 84918951 | 260 | NOD | 18 | 84918814 | 84918847 | 34 | - | 16.4472 | 9.61E-07 | 0.128 | TAGCTGTCTTCAGACACACCAGAAGAGGGCGTCA | | |
| 1157 | 19 | 3881554 | 3881756 | 202 | FVB,SJL | 19 | 3881614 | 3881647 | 34 | + | 17.8455 | 3.91E-07 | 0.0605 | TAGCTGTCCTCAGACACTCCAGAAGAGGGCGTCA | | |
| 1158 | 19 | 4093330 | 4093537 | 207 | BUB,FVB,KK,NZW,SJL | 19 | 4093396 | 4093429 | 34 | + | 18.561 | 2.42E-07 | 0.0503 | AAGCTGTCTTCAGACACTCCAGAAGAGGGCGCCA | | |

| | | | | | | | | | | | | | | | | |
|---|---|---|---|---|---|---|---|---|---|---|---|---|---|---|---|---|
| 1158 | 19 | 4093330 | 4093537 | 207 | BUB,FVB,KK,NZW,SJL | 19 | 4093411 | 4093429 | 19 | + | 17.5574 | 6.32E-07 | 0.105 | ACTCCAGAAGAGGGCGCCA | | |
| 1159 | 19 | 5831218 | 5831420 | 202 | C58J,MRL,TH | 19 | 5831278 | 5831311 | 34 | + | 18.3659 | 2.76E-07 | 0.0518 | TAGCTGTCTTCAGACACTCCAGAAGAGGGCGCCA | 3 | ENCFF093ZRU,ENCFF201JUA,ENCFF367UNR |
| 1159 | 19 | 5831218 | 5831420 | 202 | C58J,MRL,TH | 19 | 5831293 | 5831311 | 19 | + | 17.5574 | 6.32E-07 | 0.105 | ACTCCAGAAGAGGGCGCCA | 3 | ENCFF093ZRU,ENCFF201JUA,ENCFF367UNR |
| 1160 | 19 | 5959320 | 5959517 | 197 | BUB,C58J,FVB,LP,MRL,SJL,TH | 19 | 5959375 | 5959408 | 34 | + | 16.4715 | 9.47E-07 | 0.0915 | TAGCTGTCCTCAGAGACACCAGAAGAGGGAGTCA | 2 | ENCFF093ZRU,ENCFF367UNR |
| 1161 | 19 | 6986583 | 6986793 | 210 | C58J,FVB,LP | 19 | 6986667 | 6986685 | 19 | + | 17.8852 | 5.08E-07 | 0.098 | ACACCAGAAGAGGGCACCA | | |
| 1162 | 19 | 7566552 | 7566755 | 203 | BUB,C58J,FVB,LP,MA/My | 19 | 7566615 | 7566648 | 34 | + | 18.3659 | 2.76E-07 | 0.0518 | TAGCTGTCTTCAGACACTCCAGAAGAGGGCGCCA | | |
| 1162 | 19 | 7566552 | 7566755 | 203 | BUB,C58J,FVB,LP,MA/My | 19 | 7566630 | 7566648 | 19 | + | 17.5574 | 6.32E-07 | 0.105 | ACTCCAGAAGAGGGCGCCA | | |
| 1163 | 19 | 8999241 | 8999451 | 210 | LP,TH | 19 | 8999307 | 8999340 | 34 | + | 18.1545 | 3.18E-07 | 0.0548 | TAGCTGTTTTCAGACACACCAGAAGAGGGCGTCA | | |
| 1164 | 19 | 15952888 | 15953170 | 282 | MA/My | 19 | 15952897 | 15952930 | 34 | + | 19.439 | 1.32E-07 | 0.0363 | ATGCAGGTGCCGTGGAGGGCAGAAGAAGGCACTG | | |
| 1165 | 19 | 20184297 | 20184497 | 200 | MRL,NZB,NZO,NZW,PJ | 19 | 20184355 | 20184388 | 34 | + | 17.2439 | 5.79E-07 | 0.0746 | TAGCTGTCCTCAGATACTCCAGAAGAGGGCATCA | | |
| 1166 | 19 | 22864591 | 22864699 | 108 | C3H,CBA,CE,DBA1J,DBA2J,ILNJ2,MRL,NZB,NZO,NZW,PJ | 19 | 22864644 | 22864677 | 34 | + | 20.6585 | 5.50E-08 | 0.0207 | CTGCAGTGCTCCCATCATCAAACAGGTGGTAGTC | | |
| 1166 | 19 | 22864591 | 22864699 | 108 | C3H,CBA,CE,DBA1J,DBA2J,ILNJ2,MRL,NZB,NZO,NZW,PJ | 19 | 22864680 | 22864713 | 34 | + | 20.6585 | 5.50E-08 | 0.0207 | CTGCAGTGCTCCCATCATCAAACAGGTGGTAGTC | | |
| 1167 | 19 | 23599180 | 23599687 | 507 | C3H,CBA,CE,DBA1J,DBA2J,MRL,NZO,NZW,PJ | 19 | 23599622 | 23599655 | 34 | + | 18.7642 | 2.11E-07 | 0.0466 | ATGCAGCTAGAGACACGAGCTCCGGGGGGCACTG | | |
| 1168 | 19 | 34952105 | 34952543 | 438 | NZW | 19 | 34952452 | 34952485 | 34 | + | 16.4228 | 9.76E-07 | 0.092 | TTGAAATTTTACCAGCTGGCACCAGAAGGAACTA | | |
| 1169 | 19 | 34959347 | 34959559 | 212 | AJ,BALB,C58J,LP,MRL,NZO,NZW,RH2,SWR,TH | 19 | 34959417 | 34959450 | 34 | + | 16.4472 | 9.61E-07 | 0.0915 | TAGCTGTCTTCAGACACACCAGAAGAGGGCGTCA | | |
| 1170 | 19 | 37035396 | 37035803 | 407 | 129S1,AJ,AKR,BALB,BTBR,BUB,C3H,C57L,CBA,CE,DBA1J,DBA2J,FVB,ILNJ2,KK,MA/My,MRL,NOD,NOR,NZB,NZO,NZW,PJ,PL,RF,RH2,SEA,SJL,SMJ,SWR,TH | 19 | 37035661 | 37035694 | 34 | + | 16.4472 | 9.61E-07 | 0.0915 | TAGCTGTCTTCAGACACACCAGAAGAGGGCGTCA | | |
| 1171 | 19 | 43701549 | 43701763 | 214 | 129S1,AJ,AKR,BALB,BUB,C3H,C57L,CBA,CE,DBA1J,DBA2J,FVB,ILNJ2,KK,MA/My,MRL,NOD,NOR,NZB,NZO,NZW,PJ,PL,RF,RH2,SEA,SJL,SMJ,SWR,TH | 19 | 43701619 | 43701653 | 35 | + | 18.3945 | 2.37E-07 | 0.074 | CTGTAGCTGTCTTGACGCACCAGAAGAGGGCATCA | | |
| 1172 | 19 | 44188278 | 44188506 | 228 | 129S1,BUB,C57L,FVB,KK,PJ,SEA,SMJ,SWR | 19 | 44188362 | 44188395 | 34 | + | 16.4634 | 9.51E-07 | 0.0915 | TAGCTGTCTTCAGACGCACCAGAAGAGGGCATCA | | |
| 1173 | 19 | 44997974 | 44998183 | 209 | C3H,CBA,CE,ILNJ2,MRL,PJ,SEA | 19 | 44998040 | 44998073 | 34 | + | 16.4472 | 9.61E-07 | 0.0915 | TAGCTGTCTTCAGACACACCAGAAGAGGGCGTCA | | |
| 1174 | 19 | 46943889 | 46944091 | 202 | 129S1,BTBR,C57L,CE | 19 | 46943947 | 46943980 | 34 | + | 18.3659 | 2.76E-07 | 0.0518 | TAGCTGTCTTCAGACACTCCAGAAGAGGGCGCCA | 3 | ENCFF093ZRU,ENCFF367UNR,ENCFF099QNE |
| 1174 | 19 | 46943889 | 46944091 | 202 | 129S1,BTBR,C57L,CE | 19 | 46943962 | 46943980 | 19 | + | 17.5574 | 6.32E-07 | 0.105 | ACTCCAGAAGAGGGCGCCA | 3 | ENCFF093ZRU,ENCFF367UNR,ENCFF099QNE |
| 1175 | 19 | 47146646 | 47147392 | 746 | AKR,BUB,C3H,CBA,DBA1J,FVB,ILNJ2,KK,MA/My,MRL,NOD,NOR,NZB,NZO,NZW,PJ,PL,RF,SEA,SJL,SMJ,SWR,TH | 19 | 47146634 | 47146652 | 19 | + | 17.1803 | 8.08E-07 | 0.121 | CTTCCAGCAGAGGGTACCG | | |
| 1176 | 19 | 51625316 | 51625537 | 221 | TH | 19 | 51625373 | 51625406 | 34 | + | 16.4472 | 9.61E-07 | 0.0915 | TAGCTGTCTTCAGACACACCAGAAGAGGGCGTCA | | |
| 1177 | 19 | 57855510 | 57855603 | 93 | BTBR,NZB,SMJ | 19 | 57855520 | 57855553 | 34 | + | 17.6504 | 4.44E-07 | 0.0644 | CTGCAGTGCTCCTGAGCTCCTGCAGTGGTCCCTG | | |
| 1178 | 19 | 61014015 | 61014217 | 202 | 129S1,AJ,AKR,BALB,BTBR,C3H,C57L,CBA,DBA1J,DBA2J,FVB,KK,MA/My,MRL,NOD,NOR,NZB,NZO,NZW,PJ,PL,RF,RH2,SEA,SMJ,SWR,TH | 19 | 61014064 | 61014097 | 34 | + | 18.9512 | 1.85E-07 | 0.044 | TTGCTCTCCTCAGACACACCAGAAGAGGGCATCG | | |
| 1179 | 19 | 3707476 | 3707679 | 203 | NZW | 19 | 3707599 | 3707632 | 34 | - | 16.4472 | 9.61E-07 | 0.0915 | TAGCTGTCTTCAGACACACCAGAAGAGGGCGTCA | | |
| 1180 | 19 | 4975003 | 4975230 | 227 | AJ,AKR,BALB,BUB,C3H,C58J,CBA,CE,DBA1J,DBA2J,FVB,ILNJ2,KK,LP,MA/My,MRL,NOD,NOR,NZW,PJ,PL,RF,RH2,SEA,SJL,SMJ,SWR | 19 | 4975131 | 4975149 | 19 | - | 17.5574 | 6.32E-07 | 0.105 | ACTCCAGAAGAGGGCGCCA | | |
| 1180 | 19 | 4975003 | 4975230 | 227 | AJ,AKR,BALB,BUB,C3H,C58J,CBA,CE,DBA1J,DBA2J,FVB,ILNJ2,KK,LP,MA/My,MRL,NOD,NOR,NZW,PJ,PL,RF,RH2,SEA,SJL,SMJ,SWR | 19 | 4975131 | 4975164 | 34 | - | 18.3659 | 2.76E-07 | 0.0518 | TAGCTGTCTTCAGACACTCCAGAAGAGGGCGCCA | | |
| 1181 | 19 | 5545017 | 5545243 | 226 | FVB,LP,SJL | 19 | 5545148 | 5545181 | 34 | - | 17.2439 | 5.79E-07 | 0.0746 | TAGCTGTCCTCAGACCCTCCAGAAGAGGGAGTCA | | |
| 1182 | 19 | 5827178 | 5827588 | 410 | C58J,MRL,TH | 19 | 5827510 | 5827543 | 34 | - | 16.6341 | 8.54E-07 | 0.0888 | TAGCTGTCTTCACACACTCCAGAAGAGGGCGTCA | 1 | ENCFF502WDT |
| 1183 | 19 | 8694037 | 8694264 | 227 | LP | 19 | 8694163 | 8694196 | 34 | - | 17.8455 | 3.91E-07 | 0.0605 | TAGCTGTCCTCAGACACTCCAGAAGAGGGCGTCA | | |
| 1184 | 19 | 10143554 | 10143752 | 198 | AJ,BALB,BUB,C3H,C58J,CBA,CE,DBA1J,DBA2J,FVB,ILNJ2,KK,LP,MA/My,MRL,PJ,RH2,SEA,SWR,TH | 19 | 10143678 | 10143711 | 34 | - | 17.8455 | 3.91E-07 | 0.0605 | TAGCTGTCCTCAGACACTCCAGAAGAGGGCGTCA | | |
| 1185 | 19 | 21730946 | 21731381 | 435 | MRL,NZB,NZO,NZW | 19 | 21731295 | 21731313 | 19 | - | 17.0164 | 8.98E-07 | 0.125 | AGGCCAAAAGAGGGCACCA | | |
| 1186 | 19 | 42098034 | 42098237 | 203 | 129S1,AJ,AKR,BALB,BUB,C3H,C57L,CBA,CE,DBA1J,DBA2J,FVB,ILNJ2,KK,MA/My,MRL,NOD,NOR,NZB,NZO,NZW,PJ,PL,RF,RH2,SEA,SJL,SMJ,SWR | 19 | 42098158 | 42098191 | 34 | - | 18.2358 | 3.02E-07 | 0.0537 | CAGCTGTCTTCAGACACCCCAGAAGAGGGCGTCA | | |
| 1187 | 19 | 45182180 | 45182440 | 260 | 129S1,AJ,AKR,BALB,BTBR,BUB,C57L,CE,DBA1J,DBA2J,FVB,KK,MA/My,NOD,NOR,NZB,NZO,NZW,PL,RF,RH2,SJL,SMJ,SWR,TH | 19 | 45182384 | 45182418 | 35 | - | 15.9725 | 9.70E-07 | 0.156 | CTGCAAGTGGAGGTTCACACCAGCAAGTGGAGGTT | | |
| 1188 | 19 | 51334196 | 51334697 | 501 | TH | 19 | 51334165 | 51334198 | 34 | - | 18.5122 | 2.50E-07 | 0.0509 | ATGCAGCTAGAGACATGAGCTCCAGGGGGTGCTG | | |
| 1189 | 19 | 55030903 | 55031030 | 127 | 129S1,AJ,AKR,BALB,BTBR,C3H,C57L,CBA,CE,DBA1J,DBA2J,FVB,ILNJ2,MA/My,MRL,NZB,NZW,PJ,PL,RH2,SEA,SWR,TH | 19 | 55030964 | 55030997 | 34 | - | 17.9512 | 3.64E-07 | 0.0589 | CAGCAGCACCATCACCCAGCAGCAGAGGCTGGCC | | |
| 1190 | X | 12066395 | 12066625 | 230 | SMJ | X | 12066483 | 12066516 | 34 | + | 17.8455 | 3.91E-07 | 0.127 | TAGCTGTCCTCAGACACTCCAGAAGAGGGCGTCA | | |
| 1191 | X | 45777945 | 45778667 | 722 | C58J,NZB,NZO | X | 45778033 | 45778066 | 34 | + | 17.0244 | 6.67E-07 | 0.164 | GAGCAGGTGTGGCAGCAGTCCCAAGATGGCGCCC | | |
| 1192 | X | 74782900 | 74783868 | 968 | B10D2,B10J,BALB,BUB,CBA,CE,DBA1J,DBA2J,KK,LP,MA/My,MRL,PL,RF,SMJ | X | 74783752 | 74783785 | 34 | + | 16.748 | 7.95E-07 | 0.174 | CTGTAGGGCAGCTGCCAGCCACTGGAGGGCCTGG | | |
| 1193 | X | 76438839 | 76439068 | 229 | CE | X | 76438927 | 76438960 | 34 | + | 17.8455 | 3.91E-07 | 0.127 | TAGCTGTCCTCAGACACTCCAGAAGAGGGCGTCA | | |
| 1194 | X | 96338262 | 96338951 | 689 | AJ,AKR,BALB,C3H,CBA,CE,FVB,KK,MA/My,MRL,NZW,RF,RH2,SEA,SMJ,SWR | X | 96338859 | 96338892 | 34 | + | 18.3333 | 2.82E-07 | 0.111 | CTGCATTGCCTTTGGAATTCAGTAGAGGGCTTCA | | |
| 1195 | X | 101776764 | 101777325 | 561 | CE | X | 101777059 | 101777077 | 19 | + | 16.8689 | 9.85E-07 | 0.291 | AGGCCACTAGAGAGCGCCA | | |
| 1196 | X | 105164505 | 105164710 | 205 | 129S1,AJ,BALB,C3H,C57L,CBA,CE,FVB,ILNJ2,KK,MRL,NZB,NZO,NZW,RH2,SJL,SWR | X | 105164568 | 105164601 | 34 | + | 18.3659 | 2.76E-07 | 0.111 | TAGCTGTCTTCAGACACTCCAGAAGAGGGCGCCA | 1 | ENCFF093ZRU |
| 1196 | X | 105164505 | 105164710 | 205 | 129S1,AJ,BALB,C3H,C57L,CBA,CE,FVB,ILNJ2,KK,MRL,NZB,NZO,NZW,RH2,SJL,SWR | X | 105164583 | 105164601 | 19 | + | 17.5574 | 6.32E-07 | 0.241 | ACTCCAGAAGAGGGCGCCA | 1 | ENCFF093ZRU |
| 1197 | X | 135108294 | 135108498 | 204 | 129S1,AKR,BTBR,BUB,C3H,C57L,C58J,CBA,CE,DBA1J,DBA2J,LP,MRL,NOD,NOR,NZB,NZO,NZW,PJ,PL,RF,SEA,SJL,SMJ,SWR | X | 135108356 | 135108389 | 34 | + | 16.4634 | 9.51E-07 | 0.182 | TAGCTGTCTTCAGACGCACCAGAAGAGGGCATCA | | |
| 1198 | X | 136038049 | 136038257 | 208 | BTBR,BUB,C3H,C58J,CBA,CE,DBA1J,DBA2J,LP,MRL,NZB,NZO,NZW,PJ,PL,SEA,SJL,SMJ,SWR | X | 136038103 | 136038136 | 34 | + | 17.8455 | 3.91E-07 | 0.127 | TAGCTGTCCTCAGACACTCCAGAAGAGGGCGTCA | | |
| 1199 | X | 143071882 | 143072078 | 196 | 129S1,AJ,BALB,C57L,FVB,KK,MA/My,MRL,NOD,NOR,NZB,NZW,RH2,SEA,SJL,TH | X | 143071934 | 143071967 | 34 | + | 16.6911 | 8.24E-07 | 0.177 | TAGCTGTCTTCAGACAGACCAGAAGAGGGCATCA | | |
| 1200 | X | 151885066 | 151885286 | 220 | 129S1,C57L,KK,SWR | X | 151885135 | 151885168 | 34 | + | 16.4634 | 9.51E-07 | 0.182 | TAGCTGTCTTCAGACGCACCAGAAGAGGGCATCA | | |
| 1201 | X | 161382114 | 161382431 | 317 | TH | X | 161382290 | 161382323 | 34 | + | 19.8455 | 9.92E-08 | 0.0627 | TAGCTGTTTTCAGACGCGCCAGAAGAGGGCGTCA | | |
| 1202 | X | 161635270 | 161635501 | 231 | AKR,BALB,RF,SEA,TH | X | 161635340 | 161635374 | 35 | + | 21.2752 | 3.74E-08 | 0.0444 | CTGTAGCTGTCTTCAGACACCAGAAGAGGGCGCCA | 1 | ENCFF093ZRU |
| 1202 | X | 161635270 | 161635501 | 231 | AKR,BALB,RF,SEA,TH | X | 161635356 | 161635374 | 19 | + | 18.623 | 3.06E-07 | 0.176 | ACACCAGAAGAGGGCGCCA | 1 | ENCFF093ZRU |
| 1203 | X | 51761546 | 51761743 | 197 | ILNJ2 | X | 51761668 | 51761701 | 34 | - | 17.5935 | 4.61E-07 | 0.138 | TAGCTGTCTCCAGACACTCCAGAAGAGGGCGTCA | | |
| 1204 | X | 71631910 | 71632159 | 249 | ILNJ2 | X | 71632036 | 71632069 | 34 | - | 16.4634 | 9.51E-07 | 0.182 | TAGCTGTCTTCAGACGCACCAGAAGAGGGCATCA | | |
| 1205 | X | 72981367 | 72981521 | 154 | CE | X | 72981474 | 72981507 | 34 | - | 16.4472 | 9.61E-07 | 0.182 | TAGCTGTCTTCAGACACACCAGAAGAGGGCGTCA | | |
| 1206 | X | 93173057 | 93173303 | 246 | CE,MA/My | X | 93173211 | 93173229 | 19 | - | 17.5574 | 0.000000632 | 0.241 | ACTCCAGAAGAGGGCGCCA | | |
| 1206 | X | 93173057 | 93173303 | 246 | CE,MA/My | X | 93173211 | 93173244 | 34 | - | 18.3659 | 0.000000276 | 0.111 | TAGCTGTCTTCAGACACTCCAGAAGAGGGCGCCA | | |
| 1207 | X | 94909554 | 94909779 | 225 | MA/My | X | 94909677 | 94909710 | 34 | - | 16.4634 | 0.000000951 | 0.182 | TAGCTGTCTTCAGACGCACCAGAAGAGGGCATCA | | |
| 1208 | X | 97939269 | 97939473 | 204 | AJ,AKR,BALB,C3H,CBA,FVB,KK,MA/My,MRL,NZW,RF,RH2,SEA,SMJ,SWR,TH | X | 97939392 | 97939425 | 34 | - | 16.4472 | 0.000000961 | 0.182 | TAGCTGTCTTCAGACACACCAGAAGAGGGCGTCA | | |
| 1209 | X | 142779858 | 142780354 | 496 | B10J | X | 142779945 | 142779963 | 19 | - | 19.6066 | 0.000000149 | 0.12 | GGGCCACAAGATGGCAGCA | 16 | ENCFF093ZRU,ENCFF092NMJ,ENCFF099QNE,ENCFF165LXO,ENCFF198SSR,ENCFF367UNR,ENCFF883UPM,ENCFF055HAP,ENCFF609MDN,ENCFF821BIA,ENCFF089GBC,ENCFF907XEG,ENCFF024JOZ,ENCFF430PPJ,ENCFF281QMV,ENCFF327LYS |
| 1210 | X | 145464952 | 145465181 | 229 | NZB | X | 145465081 | 145465114 | 34 | - | 17.8455 | 0.000000391 | 0.127 | TAGCTGTCCTCAGACACTCCAGAAGAGGGCGTCA | | |
| 1211 | X | 161500164 | 161500375 | 211 | TH | X | 161500288 | 161500321 | 34 | - | 16.4634 | 0.000000951 | 0.182 | TAGCTGTCTTCAGACGCACCAGAAGAGGGCATCA | | |
| 1212 | X | 162748235 | 162748354 | 119 | TH | X | 162748252 | 162748270 | 19 | - | 17.9672 | 0.000000481 | 0.219 | GCACCAGAAGAGGGCACCA | 1 | ENCFF093ZRU |
| 1212 | X | 162748235 | 162748354 | 119 | TH | X | 162748252 | 162748285 | 34 | - | 17.0894 | 0.000000639 | 0.161 | CGGTTGTCTTCAGACGCACCAGAAGAGGGCACCA | 1 | ENCFF093ZRU |
| 1213 | X | 168803516 | 168803837 | 321 | 129S1,C3H,C57L,DBA1J,MA/My,PL,SMJ | X | 168803837 | 168803855 | 19 | - | 21.4918 | 0.000000032 | 0.0593 | TGGCCACTGGGTGGCGCCA | 22 | ENCFF367UNR,ENCFF821BIA,ENCFF327LYS,ENCFF024JOZ,ENCFF907XEG,ENCFF589ZBJ,ENCFF430PPJ,ENCFF136ZDK,ENCFF089GBC,ENCFF092NMJ,ENCFF165LXO,ENCFF210BTO,ENCFF198SSR,ENCFF052KVO,ENCFF805QBU,ENCFF086ZTV,ENCFF281QMV,ENCFF609MDN,ENCFF089EWX,ENCFF033UQX,ENCFF316FYN,ENCFF097CAO |
| 1213 | X | 168803516 | 168803837 | 321 | 129S1,C3H,C57L,DBA1J,MA/My,PL,SMJ | X | 168803837 | 168803871 | 35 | - | 29.6697 | 3.96E-11 | 0.00118 | TTGCACTGACGTCCTCTGGCCACTGGGTGGCGCCA | 22 | ENCFF367UNR,ENCFF821BIA,ENCFF327LYS,ENCFF024JOZ,ENCFF907XEG,ENCFF589ZBJ,ENCFF430PPJ,ENCFF136ZDK,ENCFF089GBC,ENCFF092NMJ,ENCFF165LXO,ENCFF210BTO,ENCFF198SSR,ENCFF052KVO,ENCFF805QBU,ENCFF086ZTV,ENCFF281QMV,ENCFF609MDN,ENCFF089EWX,ENCFF033UQX,ENCFF316FYN,ENCFF097CAO |

| DEL_NUM | DEL_CHR | DEL_START | DEL_END | DEL_LENGTH | STRAINS with VARIANT SV ALLELE | CTCF CHR | CTCF START | CTCF END | CTCF LENGTH | CTCF STRAND | CTCF SCORE | CTCF PVALUE | CTCF QVALUE | CTCF SEQUENCE | # PEAKS | DATA SOURCE |
|---|---|---|---|---|---|---|---|---|---|---|---|---|---|---|---|---|
| 1 | 1 | 6574639 | 6578351 | 3712 | C3H,CBA,MRL,RF | 1 | 6577912 | 6577945 | 34 | + | 16.813 | 7.63E-07 | 0.115 | ATGCAGTTAGAGACACGAGCTCCGGAGGGTACTG | | |
| 2 | 1 | 7992966 | 8010088 | 17122 | DBA1J,DBA2J | 1 | 7999113 | 7999146 | 34 | + | 18.7154 | 2.18E-07 | 0.0621 | CTGCAACACCCCCAAGGTCCCCTAGAGGACTCTC | | |
| 3 | 1 | 7999065 | 8000377 | 1312 | AJ,BUB,C3H,CBA,CE,FVB,MRL,NZB,NZW,RF | 1 | 7999113 | 7999146 | 34 | + | 18.7154 | 2.18E-07 | 0.0621 | CTGCAACACCCCCAAGGTCCCCTAGAGGACTCTC | | |
| 4 | 1 | 7999067 | 8006330 | 7263 | 129S1,AJ,AKR,BALB,BTBR,BUB,C3H,C57L,CBA,FVB,ILNJ2,KK,LP,MA/My,NOD,NOR,NZO,PJ,PL,RF,RH2,SEA,SJL,SMJ,SWR,TH | 1 | 7999113 | 7999146 | 34 | + | 18.7154 | 2.18E-07 | 0.0621 | CTGCAACACCCCCAAGGTCCCCTAGAGGACTCTC | | |
| 5 | 1 | 9103386 | 9105463 | 2077 | C3H,CBA,MRL,RF | 1 | 9103973 | 9104006 | 34 | + | 16.5447 | 9.04E-07 | 0.124 | CAGCTGTCTTCGGACACACCAAAAGAGGGCATCA | | |
| 6 | 1 | 21434734 | 21449490 | 14756 | CE,MA/My,MRL,SMJ | 1 | 21439381 | 21439415 | 35 | + | 16.8165 | 6.02E-07 | 0.167 | CTGCATGTACAGCTGCCTGCGAGTAGAGGGCATAT | | |
| 6 | 1 | 21434734 | 21449490 | 14756 | CE,MA/My,MRL,SMJ | 1 | 21443544 | 21443578 | 35 | + | 21.0826 | 4.26E-08 | 0.0373 | CTGCTTGTATAGCTGCCTGCCAGAAGAGGGCATCT | | |
| 7 | 1 | 23821397 | 23825569 | 4172 | AJ,BTBR,C3H,C57L,CBA,CE,DBA1J,DBA2J,FVB,ILNJ2,MRL,NOD,NZB,NZO,NZW,PL,RF,RH2,SMJ,SWR | 1 | 23822174 | 23822207 | 34 | + | 17.6748 | 4.37E-07 | 0.0865 | GAGCATGACCACAAGTCCACAGCAGGTGGCAACG | | |
| 8 | 1 | 28067380 | 28069462 | 2082 | AKR,BALB,PJ,SEA | 1 | 28068862 | 28068895 | 34 | + | 18.1138 | 3.27E-07 | 0.0746 | GTCCAGTGCCTGTGGATACCAGAAAAGGGCACCA | | |
| 9 | 1 | 31769616 | 31773761 | 4145 | BUB,CE,FVB,NZB,NZO,PJ,RF,SJL,SWR | 1 | 31770394 | 31770427 | 34 | + | 17.6748 | 4.37E-07 | 0.0865 | GAGCATGACCACAAGTCCACAGCAGGTGGCAACG | | |
| 10 | 1 | 34815500 | 34818186 | 2686 | CE | 1 | 34815680 | 34815713 | 34 | + | 18.0081 | 3.51E-07 | 0.0774 | GCCTAGTACCCGAGGAAGCCAGAAGAGGGCGCTG | 1 | ENCFF093ZRU |
| 10 | 1 | 34815500 | 34818186 | 2686 | CE | 1 | 34815695 | 34815713 | 19 | + | 19.2131 | 2.00E-07 | 0.0983 | AAGCCAGAAGAGGGCGCTG | 1 | ENCFF093ZRU |
| 11 | 1 | 36886530 | 36887601 | 1071 | AJ,AKR,BUB,C3H,CBA,CE,FVB,KK,MRL,NOD,NZB,NZO,NZW,PJ,PL,RF,RH2,SJL,SMJ,SWR | 1 | 36887287 | 36887320 | 34 | + | 24.9106 | 1.79E-09 | 0.00232 | ATGCAGTGTCCACAGAGCCCAGAAGAGGGCGTTA | 8 | ENCFF099QNE,ENCFF821BIA,ENCFF052KVO,ENCFF136ZDK,ENCFF502WDT,ENCFF093ZRU,ENCFF367UNR,ENCFF055HAP |
| 12 | 1 | 36950335 | 36954018 | 3683 | PJ,PL,RF,SJL | 1 | 36951496 | 36951529 | 34 | + | 19.7805 | 1.04E-07 | 0.0393 | GTGTGGTGCCTGCAGAGGCCAGATGAGGGCATTA | 6 | ENCFF367UNR,ENCFF093ZRU,ENCFF821BIA,ENCFF589ZBJ,ENCFF502WDT,ENCFF055HAP |
| 13 | 1 | 48747500 | 48754530 | 7030 | AKR,BALB,BUB,C58J,MA/My,RF,RH2,SEA | 1 | 48747957 | 48747975 | 19 | + | 17.7213 | 5.67E-07 | 0.161 | CAAGCACCAGGGGGCACCC | 4 | ENCFF367UNR,ENCFF093ZRU,ENCFF099QNE,ENCFF821BIA |
| 14 | 1 | 48776956 | 48781128 | 4172 | AKR,BALB,BUB,C58J,MA/My,RF,RH2,SEA | 1 | 48777734 | 48777767 | 34 | + | 17.6748 | 4.37E-07 | 0.0865 | GAGCATGACCACAAGTCCACAGCAGGTGGCAACG | | |
| 15 | 1 | 60196947 | 60201458 | 4511 | CE,LP | 1 | 60197809 | 60197827 | 19 | + | 17.5082 | 6.52E-07 | 0.168 | CACCCAGCAGAGGGCAGAA | 25 | ENCFF201JUA,ENCFF367UNR,ENCFF093ZRU,ENCFF821BIA,ENCFF430PPJ,ENCFF099QNE,ENCFF907XEG,ENCFF327LYS,ENCFF024JOZ,ENCFF883UPM,ENCFF089GBC,ENCFF589ZBJ,ENCFF951GOP,ENCFF575MGI,ENCFF198SSR,ENCFF194HVL,ENCFF055HAP,ENCFF491RJK,ENCFF502WDT,ENCFF957SPH,ENCFF142CNG,ENCFF092NMJ,ENCFF609MDN,ENCFF281QMV,ENCFF165LXO |
| 16 | 1 | 68937631 | 68938988 | 1357 | CE,TH | 1 | 68937990 | 68938023 | 34 | + | 18.3089 | 2.87E-07 | 0.0691 | ATGCAGCTAGAGACACGAGCTCCAGGGGGTACTG | | |
| 17 | 1 | 70241038 | 70262271 | 21233 | PJ | 1 | 70259453 | 70259486 | 34 | + | 16.8943 | 7.24E-07 | 0.111 | TTGCTCTCTTCAGACATACCAGAAGAGGGCATCA | | |
| 18 | 1 | 79113299 | 79116721 | 3422 | LP | 1 | 79113329 | 79113363 | 35 | + | 20.6147 | 5.82E-08 | 0.0449 | GTGCTCTGCATGGTGGAGGCCAGAAGAGGGCCTTG | | |
| 19 | 1 | 91845388 | 91847169 | 1781 | PL | 1 | 91845666 | 91845684 | 19 | + | 18.5082 | 3.31E-07 | 0.127 | TGACCACCAGGTGGGGCCA | 1 | ENCFF367UNR |
| 20 | 1 | 101014532 | 101023802 | 9270 | AKR,LP,MA/My,PJ,SEA | 1 | 101017578 | 101017611 | 34 | + | 17.5691 | 4.69E-07 | 0.0895 | GAGCATGACCACAAGTCCACAGCAGGTGGCAACA | | |
| 21 | 1 | 111738581 | 111740196 | 1615 | AKR,DBA1J,DBA2J,FVB,KK,LP,MA/My,MRL,NZO,NZW,PJ,RF,SJL,SWR | 1 | 111739768 | 111739786 | 19 | + | 17.2295 | 7.83E-07 | 0.183 | CATGCACCAGGGGGCACTG | 1 | ENCFF093ZRU |
| 22 | 1 | 112367859 | 112372405 | 4546 | AKR,DBA1J,DBA2J,FVB,KK,LP,MA/My,MRL,NOD,NOR,NZO,NZW,PJ,RF,SJL,SMJ,SWR | 1 | 112368636 | 112368669 | 34 | + | 17.6748 | 4.37E-07 | 0.0865 | GAGCATGACCACAAGTCCACAGCAGGTGGCAACG | | |
| 23 | 1 | 114659383 | 114667534 | 8151 | FVB,SMJ,SWR | 1 | 114665422 | 114665456 | 35 | + | 19.4312 | 1.25E-07 | 0.0674 | GTGCAGGGGAATGCCAGGGCCAGGAGGGGGAGTG | | |
| 24 | 1 | 122917357 | 122922329 | 4972 | C57L | 1 | 122917534 | 122917552 | 19 | + | 17.8033 | 5.37E-07 | 0.156 | TGACCAGAAGAGGGCATCA | 1 | ENCFF093ZRU |
| 25 | 1 | 124366108 | 124390498 | 24390 | DBA1J,DBA2J,FVB,MA/My,NOD,NOR,NZB,PJ,SJL,TH | 1 | 124378041 | 124378074 | 34 | + | 21.3659 | 3.25E-08 | 0.0181 | ATGCAGTACCCAGAGAAACCAGATGAGGGCAACA | 12 | ENCFF093ZRU,ENCFF821BIA,ENCFF099QNE,ENCFF430PPJ,ENCFF092NMJ,ENCFF165LXO,ENCFF327LYS,ENCFF052KVO,ENCFF089GBC,ENCFF024JOZ,ENCFF367UNR,ENCFF907XEG |
| 26 | 1 | 125244494 | 125251709 | 7215 | AJ,AKR,C3H,CBA,CE,DBA1J,DBA2J,MRL,NZO,RH2,SWR | 1 | 125244480 | 125244514 | 35 | + | 18.578 | 2.12E-07 | 0.0938 | CTGCAGTGACATCTGTTTGCCTGTTGGGAGCCGCC | | |
| 27 | 1 | 127454751 | 127456902 | 2151 | 129S1,AJ,AKR,BALB,C3H,C57L,CE,DBA1J,DBA2J,FVB,ILNJ2,KK,MA/My,PJ,RH2,SEA,TH | 1 | 127455324 | 127455358 | 35 | + | 19.156 | 1.48E-07 | 0.0747 | GTGCAGATGACTGATGAGGCCAGAAGAGGGTGTCA | | |
| 28 | 1 | 131570298 | 131579356 | 9058 | 129S1,AKR,BALB,BTBR,BUB,CBA,DBA1J,DBA2J,FVB,ILNJ2,KK,MA/My,MRL,NZO,NZW,PJ,PL,RF,SEA,SJL,SMJ,SWR,TH | 1 | 131575768 | 131575786 | 19 | + | 17.1639 | 8.17E-07 | 0.187 | CCTCCACGAGGTGGCGGAA | | |
| 29 | 1 | 131572917 | 131579372 | 6455 | 129S1,AKR,BTBR,BUB,CBA,DBA1J,DBA2J,ILNJ2,MRL,PJ,PL,RF,SJL,SMJ,SWR,TH | 1 | 131575768 | 131575786 | 19 | + | 17.1639 | 8.17E-07 | 0.187 | CCTCCACGAGGTGGCGGAA | | |
| 30 | 1 | 142907046 | 142908327 | 1281 | MRL | 1 | 142907269 | 142907302 | 34 | + | 18.0569 | 3.40E-07 | 0.0757 | ATGCAGCTAGAGACATGAGCTCCAGGGGGTACTG | | |
| 31 | 1 | 146600623 | 146605232 | 4609 | 129S1,AKR,BTBR,BUB,C57L,DBA1J,DBA2J,KK,MA/My,MRL,NZB,NZO,NZW,RF,SJL,SMJ,TH | 1 | 146603056 | 146603090 | 35 | + | 17.1009 | 5.11E-07 | 0.152 | CTGCACTGTGGCCGGACCATCAGCAGGGGCAAGAA | | |
| 32 | 1 | 151252147 | 151254798 | 2651 | AKR,BTBR,C57L,SMJ | 1 | 151253269 | 151253287 | 19 | + | 20.377 | 8.18E-08 | 0.0655 | AGGCCAGAAGAGGGCACTA | 10 | ENCFF099QNE,ENCFF821BIA,ENCFF024JOZ,ENCFF327LYS,ENCFF907XEG,ENCFF502WDT,ENCFF883UPM,ENCFF093ZRU,ENCFF055HAP,ENCFF367UNR |
| 33 | 1 | 156292059 | 156298875 | 6816 | AKR | 1 | 156292961 | 156292995 | 35 | + | 16.5596 | 6.97E-07 | 0.181 | TAGCTGTCTTCAGACAGCACCAGAAGAGGGCGTCA | | |
| 33 | 1 | 156292059 | 156298875 | 6816 | AKR | 1 | 156293955 | 156293988 | 34 | + | 25.0569 | 1.57E-09 | 0.00213 | ATGCAGTGCTCTCAGAGGCCAGATGAGGGCAGCA | 4 | ENCFF093ZRU,ENCFF052KVO,ENCFF327LYS,ENCFF907XEG |
| 34 | 1 | 162825252 | 162827296 | 2044 | DBA1J,DBA2J | 1 | 162827211 | 162827244 | 34 | + | 17.4146 | 5.18E-07 | 0.0944 | ATGCAGCTAGAGACACAAGCTCCAGGGGGTACTG | | |
| 35 | 1 | 164051371 | 164057121 | 5750 | BTBR,BUB,DBA1J,DBA2J,FVB,ILNJ2,KK,NOD,NOR,NZB,NZO,PJ,RF,SEA,SJL,SMJ,SWR | 1 | 164055070 | 164055088 | 19 | + | 17.1639 | 8.17E-07 | 0.187 | CCTCCACGAGGTGGCGGAA | | |
| 36 | 1 | 171139023 | 171144086 | 5063 | AJ,NZB,NZO,RH2,SJL,SMJ | 1 | 171140221 | 171140255 | 35 | + | 16.5596 | 6.97E-07 | 0.181 | AAGAAGTTACACCCTGTTGCCAGTAGGTGGTTTTC | | |
| 37 | 1 | 171139990 | 171142349 | 2359 | 129S1,BALB,BUB,C3H,C57L,CBA,CE,DBA1J,DBA2J,FVB,KK,MRL,NOD,NOR,NZW,PJ,PL,RF,SEA,SWR,TH | 1 | 171140221 | 171140255 | 35 | + | 16.5596 | 6.97E-07 | 0.181 | AAGAAGTTACACCCTGTTGCCAGTAGGTGGTTTTC | | |
| 38 | 1 | 181223486 | 181226155 | 2669 | TH | 1 | 181225852 | 181225886 | 35 | + | 18.0459 | 2.93E-07 | 0.111 | GAGCTGGTGCCTGTGGGGGCCAGAAGAGGGCATTA | | |
| 39 | 1 | 182084680 | 182093665 | 8985 | AJ,AKR,BUB,C58J,CBA,CE,DBA1J,DBA2J,FVB,KK,LP,MRL,NOD,NOR,NZB,NZO,PL,RF,RH2,SMJ,SWR,TH | 1 | 182090097 | 182090115 | 19 | + | 17.1639 | 8.17E-07 | 0.187 | CCTCCACGAGGTGGCGGAA | | |
| 40 | 1 | 191665720 | 191674705 | 8985 | 129S1,AJ,AKR,BALB,BTBR,BUB,C3H,C57L,C58J,CBA,CE,DBA1J,DBA2J,FVB,ILNJ2,KK,MA/My,MRL,NOD,NOR,NZB,NZO,NZW,PJ,PL,RF,RH2,SEA,SJL,SMJ,SWR,TH | 1 | 191671137 | 191671155 | 19 | + | 17.1639 | 8.17E-07 | 0.187 | CCTCCACGAGGTGGCGGAA | | |
| 41 | 1 | 192890521 | 192892631 | 2110 | AKR,BALB,C58J,CE,RH2 | 1 | 192890661 | 192890694 | 34 | + | 17.5122 | 4.86E-07 | 0.0914 | CTGCCTACCTGTGTACAGCCACTGGGTGGTGCTG | 38 | ENCFF093ZRU,ENCFF201JUA,ENCFF367UNR,ENCFF099QNE,ENCFF805QBU,ENCFF052KVO,ENCFF321AGQ,ENCFF821BIA,ENCFF327LYS,ENCFF907XEG,ENCFF198SSR,ENCFF024JOZ,ENCFF136ZDK,ENCFF589ZBJ,ENCFF033UQX,ENCFF883UPM,ENCFF957SPH,ENCFF502WDT,ENCFF142CNG,ENCFF165LXO,ENCFF065ODV,ENCFF055HAP,ENCFF194HVL,ENCFF089GBC,ENCFF575MGI,ENCFF951GOP,ENCFF430PPJ,ENCFF281QMV,ENCFF533APC,ENCFF609MDN,ENCFF316FYN,ENCFF097CAQ,ENCFF210BTO,ENCFF092NMJ,ENCFF611HDQ,ENCFF349BUL,ENCFF491RJK,ENCFF089EWX |
| 42 | 1 | 195018272 | 195029899 | 11627 | NOD,NOR,SWR | 1 | 195027759 | 195027792 | 34 | + | 16.4472 | 9.61E-07 | 0.125 | TAGCTGTCTTCAGACACACCAGAAGAGGGCGTCA | | |
| 43 | 1 | 4053655 | 4058231 | 4576 | DBA1J,DBA2J | 1 | 4055563 | 4055597 | 35 | - | 16.8165 | 6.02E-07 | 0.167 | AAGAAGTTACACCCTGTTGCCAGCAGGTGGTTTTC | | |
| 44 | 1 | 18706495 | 18707789 | 1294 | AKR,DBA1J,DBA2J,NZB,NZO,TH | 1 | 18707121 | 18707139 | 19 | - | 20.1967 | 9.47E-08 | 0.0696 | TTGCCAGCAGGTGGTGGCA | | |
| 45 | 1 | 19353448 | 19356541 | 3093 | AKR,PL | 1 | 19353445 | 19353478 | 34 | - | 16.4553 | 9.56E-07 | 0.125 | CTGCCGCCACACCCGCTCCCCACAGATGGCATCA | | |
| 46 | 1 | 27198489 | 27205305 | 6816 | 129S1,AJ,AKR,BALB,BTBR,C3H,C57L,CBA,CE,DBA1J,DBA2J,FVB,ILNJ2,MRL,NOD,NZB,NZO,NZW,PJ,PL,RH2,SEA | 1 | 27198773 | 27198806 | 34 | - | 16.439 | 9.66E-07 | 0.125 | GGGCACCTGTCCTCCGGTCCAGAAGGTGGCAGGC | | |
| 47 | 1 | 46081034 | 46087369 | 6335 | TH | 1 | 46081099 | 46081133 | 35 | - | 19.4495 | 1.23E-07 | 0.067 | GTGCAGGCAACTATGACAATCAGTAGAGGGCACTA | | |
| 48 | 1 | 60047215 | 60049217 | 2002 | MA/My,SMJ,SWR | 1 | 60047907 | 60047925 | 19 | - | 17.9016 | 5.02E-07 | 0.151 | ACACCAGAAGAGGGCACTA | 2 | ENCFF099QNE,ENCFF093ZRU |
| 48 | 1 | 60047215 | 60049217 | 2002 | MA/My,SMJ,SWR | 1 | 60047907 | 60047940 | 34 | - | 21.3984 | 3.17E-08 | 0.0178 | TTGCTGTGGTCAGACACACCAGAAGAGGGCACTA | 2 | ENCFF099QNE,ENCFF093ZRU |
| 49 | 1 | 79079810 | 79086881 | 7071 | AJ,BALB,C3H,ILNJ2,RH2,SEA,SJL,SWR | 1 | 79086284 | 79086317 | 34 | - | 18.0569 | 3.40E-07 | 0.0757 | ATGCAGCTAGAGACATGAGCTCCAGGGGGTACTG | | |
| 50 | 1 | 80445362 | 80447379 | 2017 | LP | 1 | 80446058 | 80446092 | 35 | - | 16.3945 | 7.65E-07 | 0.188 | CAGCAGATTTGAATGATAGCCTCCAGGTAGCAGTA | | |
| 51 | 1 | 86406120 | 86410619 | 4499 | LP,RF | 1 | 86408613 | 86408646 | 34 | - | 18.2927 | 2.90E-07 | 0.0695 | CTGCTCTGCCCGGACTTATCTCAAGGGGCTGCTC | | |
| 52 | 1 | 99210287 | 99214834 | 4547 | 129S1,AKR,C57L,DBA1J,DBA2J,ILNJ2,LP,MA/My,NOD,NOR,PJ,SEA,SMJ,TH | 1 | 99214071 | 99214104 | 34 | - | 16.8618 | 7.40E-07 | 0.113 | CAGTTATGGCAACCCTATCCACAAGAAGGCAGCA | | |
| 53 | 1 | 100336705 | 100345239 | 8534 | KK | 1 | 100338136 | 100338170 | 35 | - | 19.9725 | 8.85E-08 | 0.0559 | GTGCTGCTACCTGGGCATGCCAGCAGAGGGTGTTG | | |
| 54 | 1 | 110212250 | 110216665 | 4415 | 129S1,AJ,AKR,BALB,BTBR,BUB,C3H,C57L,CBA,CE,DBA1J,DBA2J,FVB,ILNJ2,KK,LP,MA/My,MRL,PJ,PL,RF,RH2,SEA,SJL,SWR,TH | 1 | 110212716 | 110212750 | 35 | - | 19.8807 | 9.39E-08 | 0.0578 | CTGCAATTGGGGCCTCCATCCAATAGGTGACTGTG | | |
| 55 | 1 | 114139103 | 114143246 | 4143 | AKR,BUB,C3H,CBA,DBA1J,DBA2J,ILNJ2,KK,LP,MA/My,MRL,NOD,NOR,PJ,PL,RF,SJL | 1 | 114142443 | 114142476 | 34 | - | 17.6748 | 4.37E-07 | 0.0865 | GAGCATGACCACAAGTCCACAGCAGGTGGCAACG | | |
| 56 | 1 | 130202366 | 130204704 | 2338 | CBA,NOD,NOR,SMJ | 1 | 130202394 | 130202412 | 19 | - | 16.8852 | 9.76E-07 | 0.198 | TAGCCACTAGGTGGCATTT | 14 | ENCFF093ZRU,ENCFF099QNE,ENCFF092NMJ,ENCFF165LXO,ENCFF198SSR,ENCFF883UPM,ENCFF089GBC,ENCFF367UNR,ENCFF502WDT,ENCFF055HAP,ENCFF327LYS,ENCFF024JOZ,ENCFF907XEG,ENCFF430PPJ |
| 57 | 1 | 145991207 | 145992864 | 1657 | BTBR | 1 | 145992374 | 145992407 | 34 | - | 17.6829 | 4.35E-07 | 0.0865 | TTGCAGCTAGAGACATGAGCTCTGGGGGGCACTG | | |
| 58 | 1 | 151465125 | 151467115 | 1990 | 129S1,AJ,AKR,BTBR,C3H,C57L,CBA,CE,ILNJ2,MRL,PL,RH2,SMJ | 1 | 151466885 | 151466918 | 34 | - | 16.4878 | 9.37E-07 | 0.125 | CTGCAGACTCTACCTCCCCCACACGATGACACCA | | |
| 59 | 2 | 12001069 | 12003907 | 2838 | MA/My,NOD,NOR,SJL | 2 | 12001929 | 12001962 | 34 | + | 16.7886 | 7.75E-07 | 0.0917 | GAGCATGGACACAAGCCCACAGCAGATGGCAGCA | | |
| 60 | 2 | 26533496 | 26539651 | 6155 | MA/My | 2 | 26534083 | 26534116 | 34 | + | 24.2764 | 3.11E-09 | 0.00246 | CTGCACTTCCTGAAGTCACCAGGAGGGGGGTGTG | 25 | ENCFF093ZRU,ENCFF201JUA,ENCFF367UNR,ENCFF430PPJ,ENCFF099QNE,ENCFF065ODV,ENCFF327LYS,ENCFF321AGQ,ENCFF052KVO,ENCFF609MDN,ENCFF907XEG,ENCFF136ZDK,ENCFF589ZBJ,ENCFF504JDS,ENCFF055HAP,ENCFF024JOZ,ENCFF198SSR,ENCFF089GBC,ENCFF502WDT,ENCFF092NMJ,ENCFF165LXO,ENCFF821BIA,ENCFF883UPM,ENCFF957SPH,ENCFF281QMV |
| 60 | 2 | 26533496 | 26539651 | 6155 | MA/My | 2 | 26536977 | 26537011 | 35 | + | 21.367 | 3.52E-08 | 0.0261 | CTGCGGGGGCAAGATGTGGACAGTAGGTGGTGCTG | 41 | ENCFF327LYS,ENCFF367UNR,ENCFF093ZRU,ENCFF907XEG,ENCFF024JOZ,ENCFF089GBC,ENCFF201JUA,ENCFF099QNE,ENCFF589ZBJ,ENCFF092NMJ,ENCFF198SSR,ENCFF321AGQ,ENCFF430PPJ,ENCFF821BIA,ENCFF055HAP,ENCFF065ODV,ENCFF136ZDK,ENCFF165LXO,ENCFF502WDT,ENCFF883UPM,ENCFF504JDS,ENCFF957SPH,ENCFF951GOP,ENCFF609MDN,ENCFF142CNG,ENCFF281QMV,ENCFF210BTO,ENCFF316FYN,ENCFF349BUL,ENCFF194HVL,ENCFF601JVE,ENCFF575MGI,ENCFF611HDQ,ENCFF491RJK,ENCFF533APC,ENCFF052KVO,ENCFF089EWX,ENCFF033UQX,ENCFF097CAQ,ENCFF784ASD,ENCFF086ZTV |
| 60 | 2 | 26533496 | 26539651 | 6155 | MA/My | 2 | 26536993 | 26537011 | 19 | + | 17.1475 | 8.25E-07 | 0.142 | TGGACAGTAGGTGGTGCTG | 41 | ENCFF327LYS,ENCFF367UNR,ENCFF093ZRU,ENCFF907XEG,ENCFF024JOZ,ENCFF089GBC,ENCFF201JUA,ENCFF099QNE,ENCFF589ZBJ,ENCFF092NMJ,ENCFF198SSR,ENCFF321AGQ,ENCFF430PPJ,ENCFF821BIA,ENCFF055HAP,ENCFF065ODV,ENCFF136ZDK,ENCFF165LXO,ENCFF502WDT,ENCFF883UPM,ENCFF504JDS,ENCFF957SPH,ENCFF951GOP,ENCFF609MDN,ENCFF142CNG,ENCFF281QMV,ENCFF210BTO,ENCFF316FYN,ENCFF349BUL,ENCFF194HVL,ENCFF601JVE,ENCFF575MGI,ENCFF611HDQ,ENCFF491RJK,ENCFF533APC,ENCFF052KVO,ENCFF089EWX,ENCFF033UQX,ENCFF097CAQ,ENCFF784ASD,ENCFF086ZTV |
| 60 | 2 | 26533496 | 26539651 | 6155 | MA/My | 2 | 26537009 | 26537043 | 35 | + | 17.2661 | 4.64E-07 | 0.12 | CTGTAGGGACCATGTGTGGACAGTGGGTGGTGTG | 41 | ENCFF327LYS,ENCFF367UNR,ENCFF093ZRU,ENCFF907XEG,ENCFF024JOZ,ENCFF089GBC,ENCFF201JUA,ENCFF099QNE,ENCFF589ZBJ,ENCFF092NMJ,ENCFF198SSR,ENCFF321AGQ,ENCFF430PPJ,ENCFF821BIA,ENCFF055HAP,ENCFF065ODV,ENCFF136ZDK,ENCFF165LXO,ENCFF502WDT,ENCFF883UPM,ENCFF504JDS,ENCFF957SPH,ENCFF951GOP,ENCFF609MDN,ENCFF142CNG,ENCFF281QMV,ENCFF210BTO,ENCFF316FYN,ENCFF349BUL,ENCFF194HVL,ENCFF601JVE,ENCFF575MGI,ENCFF611HDQ,ENCFF491RJK,ENCFF533APC,ENCFF052KVO,ENCFF089EWX,ENCFF033UQX,ENCFF097CAQ,ENCFF784ASD,ENCFF086ZTV |
| 60 | 2 | 26533496 | 26539651 | 6155 | MA/My | 2 | 26537049 | 26537083 | 35 | + | 22.3853 | 1.74E-08 | 0.0175 | CTGTAGGGACCCTGCGTGGACAGTAGGTGGTGCTG | 42 | ENCFF327LYS,ENCFF367UNR,ENCFF093ZRU,ENCFF907XEG,ENCFF024JOZ,ENCFF089GBC,ENCFF201JUA,ENCFF099QNE,ENCFF589ZBJ,ENCFF092NMJ,ENCFF198SSR,ENCFF321AGQ,ENCFF430PPJ,ENCFF821BIA,ENCFF055HAP,ENCFF065ODV,ENCFF136ZDK,ENCFF165LXO,ENCFF502WDT,ENCFF883UPM,ENCFF504JDS,ENCFF957SPH,ENCFF951GOP,ENCFF609MDN,ENCFF142CNG,ENCFF281QMV,ENCFF210BTO,ENCFF316FYN,ENCFF349BUL,ENCFF194HVL,ENCFF601JVE,ENCFF575MGI,ENCFF611HDQ,ENCFF491RJK,ENCFF533APC,ENCFF052KVO,ENCFF089EWX,ENCFF033UQX,ENCFF097CAQ,ENCFF784ASD,ENCFF086ZTV,ENCFF805QBU |
| 60 | 2 | 26533496 | 26539651 | 6155 | MA/My | 2 | 26537065 | 26537083 | 19 | + | 17.1475 | 8.25E-07 | 0.142 | TGGACAGTAGGTGGTGCTG | 42 | ENCFF327LYS,ENCFF367UNR,ENCFF093ZRU,ENCFF907XEG,ENCFF024JOZ,ENCFF089GBC,ENCFF201JUA,ENCFF099QNE,ENCFF589ZBJ,ENCFF092NMJ,ENCFF198SSR,ENCFF321AGQ,ENCFF430PPJ,ENCFF821BIA,ENCFF055HAP,ENCFF065ODV,ENCFF136ZDK,ENCFF165LXO,ENCFF502WDT,ENCFF883UPM,ENCFF504JDS,ENCFF957SPH,ENCFF951GOP,ENCFF609MDN,ENCFF142CNG,ENCFF281QMV,ENCFF210BTO,ENCFF316FYN,ENCFF349BUL,ENCFF194HVL,ENCFF601JVE,ENCFF575MGI,ENCFF611HDQ,ENCFF491RJK,ENCFF533APC,ENCFF052KVO,ENCFF089EWX,ENCFF033UQX,ENCFF097CAQ,ENCFF784ASD,ENCFF086ZTV,ENCFF805QBU |
| 60 | 2 | 26533496 | 26539651 | 6155 | MA/My | 2 | 26537089 | 26537123 | 35 | + | 19.055 | 1.58E-07 | 0.0638 | CTGTAGGGACCCTGTGTGGACAGTGGGTGGTGCTG | 42 | ENCFF327LYS,ENCFF367UNR,ENCFF093ZRU,ENCFF907XEG,ENCFF024JOZ,ENCFF089GBC,ENCFF201JUA,ENCFF099QNE,ENCFF589ZBJ,ENCFF092NMJ,ENCFF198SSR,ENCFF321AGQ,ENCFF430PPJ,ENCFF821BIA,ENCFF055HAP,ENCFF065ODV,ENCFF136ZDK,ENCFF165LXO,ENCFF502WDT,ENCFF883UPM,ENCFF504JDS,ENCFF957SPH,ENCFF951GOP,ENCFF609MDN,ENCFF142CNG,ENCFF281QMV,ENCFF210BTO,ENCFF316FYN,ENCFF349BUL,ENCFF194HVL,ENCFF601JVE,ENCFF575MGI,ENCFF611HDQ,ENCFF491RJK,ENCFF533APC,ENCFF052KVO,ENCFF089EWX,ENCFF033UQX,ENCFF097CAQ,ENCFF784ASD,ENCFF086ZTV,ENCFF805QBU |
| 60 | 2 | 26533496 | 26539651 | 6155 | MA/My | 2 | 26537129 | 26537163 | 35 | + | 19.055 | 1.58E-07 | 0.0638 | CTGTAGGGACCCTGTGTGGACAGTGGGTGGTGCTG | 41 | ENCFF327LYS,ENCFF367UNR,ENCFF093ZRU,ENCFF907XEG,ENCFF024JOZ,ENCFF089GBC,ENCFF201JUA,ENCFF099QNE,ENCFF589ZBJ,ENCFF092NMJ,ENCFF198SSR,ENCFF321AGQ,ENCFF430PPJ,ENCFF821BIA,ENCFF055HAP,ENCFF065ODV,ENCFF136ZDK,ENCFF165LXO,ENCFF502WDT,ENCFF883UPM,ENCFF504JDS,ENCFF957SPH,ENCFF951GOP,ENCFF609MDN,ENCFF142CNG,ENCFF281QMV,ENCFF210BTO,ENCFF316FYN,ENCFF349BUL,ENCFF194HVL,ENCFF601JVE,ENCFF575MGI,ENCFF611HDQ,ENCFF491RJK,ENCFF533APC,ENCFF089EWX,ENCFF033UQX,ENCFF097CAQ,ENCFF784ASD,ENCFF086ZTV,ENCFF805QBU |
| 61 | 2 | 30056474 | 30059382 | 2908 | SWR | 2 | 30056884 | 30056918 | 35 | + | 27.789 | 2.34E-10 | 0.00232 | TTGCAGTAACAAGTGCCAACCAGAAGGTGGCGCTC | 18 | ENCFF024JOZ,ENCFF821BIA,ENCFF136ZDK,ENCFF092NMJ,ENCFF198SSR,ENCFF165LXO,ENCFF589ZBJ,ENCFF430PPJ,ENCFF093ZRU,ENCFF055HAP,ENCFF883UPM,ENCFF367UNR,ENCFF065ODV,ENCFF089GBC,ENCFF052KVO,ENCFF099QNE,ENCFF907XEG,ENCFF327LYS |
| 61 | 2 | 30056474 | 30059382 | 2908 | SWR | 2 | 30056900 | 30056918 | 19 | + | 19.9016 | 1.19E-07 | 0.0563 | CAACCAGAAGGTGGCGCTC | 19 | ENCFF024JOZ,ENCFF821BIA,ENCFF136ZDK,ENCFF092NMJ,ENCFF198SSR,ENCFF165LXO,ENCFF589ZBJ,ENCFF430PPJ,ENCFF093ZRU,ENCFF055HAP,ENCFF883UPM,ENCFF367UNR,ENCFF065ODV,ENCFF089GBC,ENCFF052KVO,ENCFF099QNE,ENCFF907XEG,ENCFF327LYS,ENCFF502WDT |
| 62 | 2 | 32214039 | 32217080 | 3041 | SWR | 2 | 32214508 | 32214541 | 34 | + | 23.5772 | 5.62E-09 | 0.00383 | ATACAGTGCCTCCAGAGGCCAGAAGAGGGCGTCC | 10 | ENCFF367UNR,ENCFF052KVO,ENCFF089GBC,ENCFF327LYS,ENCFF092NMJ,ENCFF093ZRU,ENCFF055HAP,ENCFF099QNE,ENCFF533APC,ENCFF502WDT |
| 63 | 2 | 40937266 | 40964021 | 26755 | TH | 2 | 40950950 | 40950968 | 19 | + | 16.918 | 9.56E-07 | 0.151 | CCACCTGCAGGTGGCACGG | 4 | ENCFF024JOZ,ENCFF093ZRU,ENCFF367UNR,ENCFF099QNE |
| 63 | 2 | 40937266 | 40964021 | 26755 | TH | 2 | 40950945 | 40950963 | 19 | - | 19.3934 | 1.75E-07 | 0.0686 | CCACCTGCAGGTGGCACTA | 4 | ENCFF024JOZ,ENCFF093ZRU,ENCFF367UNR,ENCFF099QNE |
| 64 | 2 | 47353781 | 47358263 | 4482 | FVB,NOD,NOR,TH | 2 | 47356240 | 47356273 | 34 | + | 26.0325 | 6.48E-10 | 0.000834 | CTGCTGTTTTCAGACTCACCAGAAGAGGGCAGCA | 11 | ENCFF052KVO,ENCFF093ZRU,ENCFF821BIA,ENCFF136ZDK,ENCFF907XEG,ENCFF327LYS,ENCFF367UNR,ENCFF099QNE,ENCFF055HAP,ENCFF502WDT,ENCFF065ODV |
| 64 | 2 | 47353781 | 47358263 | 4482 | FVB,NOD,NOR,TH | 2 | 47356255 | 47356273 | 19 | + | 19.6393 | 1.45E-07 | 0.0618 | TCACCAGAAGAGGGCAGCA | 10 | ENCFF052KVO,ENCFF093ZRU,ENCFF821BIA,ENCFF136ZDK,ENCFF907XEG,ENCFF327LYS,ENCFF367UNR,ENCFF099QNE,ENCFF055HAP,ENCFF065ODV |
| 65 | 2 | 47827408 | 47842021 | 14613 | KK,NZB | 2 | 47835268 | 47835286 | 19 | + | 17.4262 | 6.88E-07 | 0.129 | GAACCAGGAGGGGGCAGCA | 9 | ENCFF093ZRU,ENCFF367UNR,ENCFF821BIA,ENCFF024JOZ,ENCFF327LYS,ENCFF198SSR,ENCFF165LXO,ENCFF907XEG,ENCFF099QNE |
| 66 | 2 | 55954633 | 55959266 | 4633 | 129S1,AJ,AKR,BUB,C3H,C58J,CBA,CE,DBA1J,DBA2J,FVB,ILNJ2,KK,MRL,NOD,NOR,NZB,NZO,PJ,PL,RF,SEA,SJL,SMJ,SWR,TH | 2 | 55955488 | 55955521 | 34 | + | 17.6748 | 4.37E-07 | 0.0681 | GAGCATGACCACAAGTCCACAGCAGGTGGCAACG | | |
| 67 | 2 | 61693575 | 61699324 | 5749 | KK | 2 | 61698267 | 61698285 | 19 | + | 17.082 | 8.61E-07 | 0.144 | TGGCCACTGGGAGGCGCAA | 1 | ENCFF883UPM |
| 68 | 2 | 71234599 | 71236385 | 1786 | AJ,AKR,BUB,C3H,FVB,ILNJ2,MA/My,MRL,NOD,NOR,NZW,PL,RH2,SMJ | 2 | 71235587 | 71235621 | 35 | + | 17.8349 | 3.32E-07 | 0.0989 | CTGCATGTGCACTTGCAGGCCAGAAGAGGCCCTCA | | |
| 69 | 2 | 77827983 | 77834256 | 6273 | NZW | 2 | 77828429 | 77828447 | 19 | + | 17.0328 | 8.88E-07 | 0.145 | TTGCCAGCAGAGAGCGCTC | | |
| 70 | 2 | 86329424 | 86332537 | 3113 | AJ,AKR,BALB,BUB,C3H,C57L,CBA,CE,DBA1J,DBA2J,KK,MA/My,MRL,NOD,NOR,NZB,NZO,NZW,PJ,PL,RF,RH2,SEA,SMJ | 2 | 86332362 | 86332395 | 34 | + | 18.0976 | 3.31E-07 | 0.059 | GAGCAGATGCGGCCGCCACCCCAAGATGGTGCCC | | |
| 71 | 2 | 89997096 | 90079546 | 82450 | BUB,C58J,ILNJ2,KK,PL,SJL,SWR | 2 | 90008182 | 90008216 | 35 | + | 16.4587 | 7.38E-07 | 0.154 | CTGCAGTGGTGTAATGTGCTCAGCTGAGGGTATTG | | |
| 71 | 2 | 89997096 | 90079546 | 82450 | BUB,C58J,ILNJ2,KK,PL,SJL,SWR | 2 | 90025246 | 90025279 | 34 | + | 21.8537 | 2.23E-08 | 0.0103 | CCTCAGTTTTAGATGTGGCCACTAGATGGAGCCA | | |
| 71 | 2 | 89997096 | 90079546 | 82450 | BUB,C58J,ILNJ2,KK,PL,SJL,SWR | 2 | 90025261 | 90025279 | 19 | + | 21.8033 | 2.43E-08 | 0.0339 | TGGCCACTAGATGGAGCCA | | |
| 71 | 2 | 89997096 | 90079546 | 82450 | BUB,C58J,ILNJ2,KK,PL,SJL,SWR | 2 | 90041506 | 90041539 | 34 | + | 18.0407 | 3.43E-07 | 0.0602 | TAGCTGTCTTCAGAGTTACCAGAAGAGGGCTCCA | | |
| 71 | 2 | 89997096 | 90079546 | 82450 | BUB,C58J,ILNJ2,KK,PL,SJL,SWR | 2 | 90072049 | 90072082 | 34 | + | 20.2195 | 7.58E-08 | 0.025 | CTGCAAGCCTTGCAGTGGACACCAGAGGTTAGTG | | |
| 71 | 2 | 89997096 | 90079546 | 82450 | BUB,C58J,ILNJ2,KK,PL,SJL,SWR | 2 | 90072092 | 90072125 | 34 | + | 20.2195 | 7.58E-08 | 0.025 | CTGCAAGCCTTGCAGTGGACACCAGAGGTTAGTG | | |
| 71 | 2 | 89997096 | 90079546 | 82450 | BUB,C58J,ILNJ2,KK,PL,SJL,SWR | 2 | 90072135 | 90072168 | 34 | + | 20.2195 | 7.58E-08 | 0.025 | CTGCAAGCCTTGCAGTGGACACCAGAGGTTAGTG | | |
| 72 | 2 | 123355540 | 123367197 | 11657 | 129S1,AJ,AKR,BALB,BTBR,BUB,C3H,C57L,C58J,CBA,CE,DBA1J,DBA2J,FVB,ILNJ2,MA/My,NOD,NZB,NZO,NZW,PJ,PL,RF,RH2,SEA,SJL,SMJ,SWR,TH | 2 | 123355556 | 123355589 | 34 | + | 17.9106 | 3.74E-07 | 0.063 | CCGCCCTCACATTCGCCGTCACAAGATGGCGCTG | | |
| 72 | 2 | 123355540 | 123367197 | 11657 | 129S1,AJ,AKR,BALB,BTBR,BUB,C3H,C57L,C58J,CBA,CE,DBA1J,DBA2J,FVB,ILNJ2,MA/My,NOD,NZB,NZO,NZW,PJ,PL,RF,RH2,SEA,SJL,SMJ,SWR,TH | 2 | 123358366 | 123358399 | 34 | + | 17.9106 | 3.74E-07 | 0.063 | CCGCCCTCACATTCGCCGTCACAAGATGGCGCTG | | |
| 72 | 2 | 123355540 | 123367197 | 11657 | 129S1,AJ,AKR,BALB,BTBR,BUB,C3H,C57L,C58J,CBA,CE,DBA1J,DBA2J,FVB,ILNJ2,MA/My,NOD,NZB,NZO,NZW,PJ,PL,RF,RH2,SEA,SJL,SMJ,SWR,TH | 2 | 123361176 | 123361209 | 34 | + | 17.9106 | 3.74E-07 | 0.063 | CCGCCCTCACATTCGCCGTCACAAGATGGCGCTG | | |
| 72 | 2 | 123355540 | 123367197 | 11657 | 129S1,AJ,AKR,BALB,BTBR,BUB,C3H,C57L,C58J,CBA,CE,DBA1J,DBA2J,FVB,ILNJ2,MA/My,NOD,NZB,NZO,NZW,PJ,PL,RF,RH2,SEA,SJL,SMJ,SWR,TH | 2 | 123363954 | 123363987 | 34 | + | 17.9106 | 3.74E-07 | 0.063 | CCGCCCTCACATTCGCCGTCACAAGATGGCGCTG | | |
| 72 | 2 | 123355540 | 123367197 | 11657 | 129S1,AJ,AKR,BALB,BTBR,BUB,C3H,C57L,C58J,CBA,CE,DBA1J,DBA2J,FVB,ILNJ2,MA/My,NOD,NZB,NZO,NZW,PJ,PL,RF,RH2,SEA,SJL,SMJ,SWR,TH | 2 | 123366724 | 123366757 | 34 | + | 17.9106 | 3.74E-07 | 0.063 | CCGCCCTCACATTCGCCGTCACAAGATGGCGCTG | | |
| 73 | 2 | 150224221 | 150228022 | 3801 | NOD,NZB,NZW,RF | 2 | 150226615 | 150226648 | 34 | + | 17.6992 | 4.30E-07 | 0.0675 | TAGCTGTTTTCAGACACACCAGAAGAGGGCATCA | | |
| 74 | 2 | 157494162 | 157498335 | 4173 | 129S1,AJ,AKR,BTBR,BUB,C57L,C58J,DBA1J,DBA2J,ILNJ2,KK,MA/My,MRL,NZB,PJ,RF,SJL,TH | 2 | 157494939 | 157494972 | 34 | + | 17.6748 | 4.37E-07 | 0.0681 | GAGCATGACCACAAGTCCACAGCAGGTGGCAACG | | |
| 75 | 2 | 181561079 | 181569889 | 8810 | BUB,CE,PJ,RF | 2 | 181563701 | 181563719 | 19 | + | 17.8689 | 5.14E-07 | 0.116 | AAACCACTAGGTGGCAGAA | | |
| 76 | 2 | 9045436 | 9047482 | 2046 | ILNJ2 | 2 | 9046899 | 9046917 | 19 | - | 19.7377 | 1.35E-07 | 0.0603 | AGGCCAGAAGAGGGCAGTA | | |
| 76 | 2 | 9045436 | 9047482 | 2046 | ILNJ2 | 2 | 9046899 | 9046933 | 35 | - | 20.6422 | 5.72E-08 | 0.0344 | CTCCAGATAACTCTGGAGGCCAGAAGAGGGCAGTA | | |
| 77 | 2 | 15942107 | 15943149 | 1042 | SJL | 2 | 15942138 | 15942172 | 35 | - | 16.7706 | 6.18E-07 | 0.139 | CTGCAAAACCAACATTCAGGCAGTGGAGGGAGGGG | | |

| | | | | | | | | | | | | | | | | |
|---|---|---|---|---|---|---|---|---|---|---|---|---|---|---|---|---|
| 78 | 2 | 16025306 | 16034352 | 9046 | CE,PL,SJL | 2 | 16028873 | 16028891 | 19 | - | 17.1639 | 8.17E-07 | 0.141 | CCTCCACGAGGTGGCGGAA | | |
| 79 | 2 | 23561284 | 23565193 | 3909 | FVB,SJL | 2 | 23564268 | 23564301 | 34 | - | 18.1057 | 3.29E-07 | 0.0588 | AAGCTGTCTTCAGACACTCCAGAAGAGGGCACCA | | |
| 80 | 2 | 49101210 | 49106486 | 5276 | C58J,FVB,KK,PL,SJL,TH | 2 | 49102120 | 49102153 | 34 | - | 17.4715 | 5.00E-07 | 0.0729 | ACACAGTGCCCGAGGAGAACAGAAGAGGGCACTG | 4 | ENCFF093ZRU,ENCFF367UNR,ENCFF099QNE,ENCFF821BIA |
| 81 | 2 | 57104419 | 57113404 | 8985 | 129S1,AJ,AKR,BUB,C3H,C58J,CBA,CE,DBA1J,DBA2J,FVB,ILNJ2,KK,MRL,NOD,NOR,NZB,NZO,PJ,PL,RF,SEA,SJL,SMJ,SWR,TH | 2 | 57107974 | 57107992 | 19 | - | 17.1639 | 8.17E-07 | 0.141 | CCTCCACGAGGTGGCGGAA | | |
| 82 | 2 | 79608322 | 79609546 | 1224 | BTBR,KK,NZB,NZO | 2 | 79608330 | 79608348 | 19 | - | 17.5738 | 6.25E-07 | 0.124 | TGAACACTAGAGGGAGCCA | 18 | ENCFF327LYS,ENCFF821BIA,ENCFF092NMJ,ENCFF024JOZ,ENCFF165LXO,ENCFF052KVO,ENCFF589ZBJ,ENCFF065ODV,ENCFF430PPJ,ENCFF093ZRU,ENCFF883UPM,ENCFF198SSR,ENCFF367UNR,ENCFF907XEG,ENCFF502WDT,ENCFF055HAP,ENCFF099QNE,ENCFF089GBC |
| 83 | 2 | 99374632 | 99376981 | 2349 | NZO | 2 | 99376003 | 99376036 | 34 | - | 18.5447 | 2.45E-07 | 0.0522 | CCGCACCCACATTCGCCGTCACAAGATGGCGCTG | | |
| 84 | 2 | 99375563 | 99382994 | 7431 | 129S1,AJ,BTBR,BUB,FVB,ILNJ2,KK,NZB,NZW,SJL,SMJ,SWR | 2 | 99376003 | 99376036 | 34 | - | 18.5447 | 2.45E-07 | 0.0522 | CCGCACCCACATTCGCCGTCACAAGATGGCGCTG | | |
| 85 | 2 | 115161626 | 115165598 | 3972 | C3H,CBA,CE | 2 | 115161998 | 115162031 | 34 | - | 16.8211 | 7.59E-07 | 0.0913 | GTGCAGGTACCTTGTAGGTCAACAGAGGGCATTA | | |
| 86 | 2 | 132183235 | 132189870 | 6635 | ILNJ2,PJ | 2 | 132183886 | 132183919 | 34 | - | 16.3984 | 9.91E-07 | 0.0999 | TAGCTGTCTTCAGACGTACCAGAAGAGGGCATCA | | |
| 86 | 2 | 132183235 | 132189870 | 6635 | ILNJ2,PJ | 2 | 132184597 | 132184630 | 34 | - | 16.4065 | 9.86E-07 | 0.0997 | GGACAGTACCCATGGACACCAGAAGAGGGCGTCA | 1 | ENCFF093ZRU |
| 87 | 2 | 139875792 | 139880002 | 4210 | C58J,DBA1J,DBA2J,NZO,NZW | 2 | 139877940 | 139877973 | 34 | - | 17.7398 | 4.19E-07 | 0.0668 | GAGCAGTCACTGGGACTACAGGCAGGTGGCAGTG | 1 | ENCFF093ZRU |
| 88 | 2 | 150339866 | 150342833 | 2967 | BUB,MA/My,NOD,NZB,NZW,RF | 2 | 150340649 | 150340682 | 34 | - | 21.8293 | 2.28E-08 | 0.0105 | CTGCAATGCCCTTCACAGCAAGAAGGGGGTAGAA | | |
| 89 | 2 | 150936711 | 150944533 | 7822 | NOD,NZB,NZW,RF | 2 | 150943764 | 150943798 | 35 | - | 15.9817 | 9.65E-07 | 0.178 | CTGTAGTGAGCCTTCTGGACAAGCTGGGGGCAGAT | | |
| 90 | 2 | 154856866 | 154871571 | 14705 | 129S1,BALB,C3H,C57L,CBA,CE,FVB,MA/My,NOD,NZO,NZW,PL,SEA,SJL,SWR,TH | 2 | 154861841 | 154861859 | 19 | - | 18.1475 | 4.25E-07 | 0.107 | CAGCCAGCAGATGGCGTTC | 30 | ENCFF093ZRU,ENCFF367UNR,ENCFF327LYS,ENCFF099QNE,ENCFF907XEG,ENCFF024JOZ,ENCFF821BIA,ENCFF201JUA,ENCFF316FYN,ENCFF589ZBJ,ENCFF089GBC,ENCFF198SSR,ENCFF575MGI,ENCFF097CAQ,ENCFF055HAP,ENCFF609MDN,ENCFF136ZDK,ENCFF502WDT,ENCFF165LXO,ENCFF430PPJ,ENCFF092NMJ,ENCFF951GOP,ENCFF194HVL,ENCFF491RJK,ENCFF957SPH,ENCFF883UPM,ENCFF281QMV,ENCFF210BTO,ENCFF142CNG,ENCFF065ODV |
| 90 | 2 | 154856866 | 154871571 | 14705 | 129S1,BALB,C3H,C57L,CBA,CE,FVB,MA/My,NOD,NZO,NZW,PL,SEA,SJL,SWR,TH | 2 | 154861841 | 154861875 | 35 | - | 17.1101 | 5.08E-07 | 0.126 | AAGTTTTTCAAAGGTCCAGCCAGCAGATGGCGTTC | 30 | ENCFF093ZRU,ENCFF367UNR,ENCFF327LYS,ENCFF099QNE,ENCFF907XEG,ENCFF024JOZ,ENCFF821BIA,ENCFF201JUA,ENCFF316FYN,ENCFF589ZBJ,ENCFF089GBC,ENCFF198SSR,ENCFF575MGI,ENCFF097CAQ,ENCFF055HAP,ENCFF609MDN,ENCFF136ZDK,ENCFF502WDT,ENCFF165LXO,ENCFF430PPJ,ENCFF092NMJ,ENCFF951GOP,ENCFF194HVL,ENCFF491RJK,ENCFF957SPH,ENCFF883UPM,ENCFF281QMV,ENCFF210BTO,ENCFF142CNG,ENCFF065ODV |
| 91 | 2 | 154859783 | 154868599 | 8816 | BTBR | 2 | 154861841 | 154861859 | 19 | - | 18.1475 | 4.25E-07 | 0.107 | CAGCCAGCAGATGGCGTTC | 30 | ENCFF093ZRU,ENCFF367UNR,ENCFF327LYS,ENCFF099QNE,ENCFF907XEG,ENCFF024JOZ,ENCFF821BIA,ENCFF201JUA,ENCFF316FYN,ENCFF589ZBJ,ENCFF089GBC,ENCFF198SSR,ENCFF575MGI,ENCFF097CAQ,ENCFF055HAP,ENCFF609MDN,ENCFF136ZDK,ENCFF502WDT,ENCFF165LXO,ENCFF430PPJ,ENCFF092NMJ,ENCFF951GOP,ENCFF194HVL,ENCFF491RJK,ENCFF957SPH,ENCFF883UPM,ENCFF281QMV,ENCFF210BTO,ENCFF142CNG,ENCFF065ODV |
| 91 | 2 | 154859783 | 154868599 | 8816 | BTBR | 2 | 154861841 | 154861875 | 35 | - | 17.1101 | 5.08E-07 | 0.126 | AAGTTTTTCAAAGGTCCAGCCAGCAGATGGCGTTC | 30 | ENCFF093ZRU,ENCFF367UNR,ENCFF327LYS,ENCFF099QNE,ENCFF907XEG,ENCFF024JOZ,ENCFF821BIA,ENCFF201JUA,ENCFF316FYN,ENCFF589ZBJ,ENCFF089GBC,ENCFF198SSR,ENCFF575MGI,ENCFF097CAQ,ENCFF055HAP,ENCFF609MDN,ENCFF136ZDK,ENCFF502WDT,ENCFF165LXO,ENCFF430PPJ,ENCFF092NMJ,ENCFF951GOP,ENCFF194HVL,ENCFF491RJK,ENCFF957SPH,ENCFF883UPM,ENCFF281QMV,ENCFF210BTO,ENCFF142CNG,ENCFF065ODV |
| 92 | 2 | 158577454 | 158582054 | 4600 | NZB | 2 | 158578364 | 158578382 | 19 | - | 18.623 | 3.06E-07 | 0.0887 | ACACCAGAAGAGGGCGCCA | 4 | ENCFF821BIA,ENCFF367UNR,ENCFF093ZRU,ENCFF099QNE |
| 92 | 2 | 158577454 | 158582054 | 4600 | NZB | 2 | 158578364 | 158578397 | 34 | - | 20.4146 | 6.58E-08 | 0.0228 | TAGCTGTTTTCAGATACACCAGAAGAGGGCGCCA | 4 | ENCFF821BIA,ENCFF367UNR,ENCFF093ZRU,ENCFF099QNE |
| 93 | 2 | 163814229 | 163817940 | 3711 | KK | 2 | 163817939 | 163817973 | 35 | - | 23.4495 | 8.05E-09 | 0.0114 | CTGCATGTACACCTGCATACCAGAAGAGGGCATCA | | |
| 94 | 2 | 170514414 | 170515430 | 1016 | KK | 2 | 170514897 | 170514931 | 35 | - | 24.2844 | 4.29E-09 | 0.00792 | GTGCAGTTTCCAGGGGAGGCCAGATGAGGGCAGTA | 2 | ENCFF093ZRU,ENCFF052KVO |
| 95 | 2 | 172069292 | 172070535 | 1243 | CE | 2 | 172069683 | 172069716 | 34 | - | 16.7724 | 7.83E-07 | 0.0917 | TAGCTGTCTTCAGATGCACCAGAAGAGGGCGTCA | | |
| 95 | 2 | 172069292 | 172070535 | 1243 | CE | 2 | 172069923 | 172069941 | 19 | - | 17.0328 | 8.88E-07 | 0.145 | AGGCCACAAGAGGGCATCA | 20 | ENCFF024JOZ,ENCFF093ZRU,ENCFF092NMJ,ENCFF327LYS,ENCFF907XEG,ENCFF430PPJ,ENCFF165LXO,ENCFF883UPM,ENCFF033UQX,ENCFF136ZDK,ENCFF099QNE,ENCFF055HAP,ENCFF065ODV,ENCFF533APC,ENCFF281QMV,ENCFF052KVO,ENCFF142CNG,ENCFF589ZBJ,ENCFF089GBC,ENCFF502WDT |
| 95 | 2 | 172069292 | 172070535 | 1243 | CE | 2 | 172069923 | 172069956 | 34 | - | 28.2846 | 7.03E-11 | 0.000209 | GTGCAGTGCCATTGGAGGCCACAAGAGGGCATCA | 20 | ENCFF024JOZ,ENCFF093ZRU,ENCFF092NMJ,ENCFF327LYS,ENCFF907XEG,ENCFF430PPJ,ENCFF165LXO,ENCFF883UPM,ENCFF033UQX,ENCFF136ZDK,ENCFF099QNE,ENCFF055HAP,ENCFF065ODV,ENCFF533APC,ENCFF281QMV,ENCFF052KVO,ENCFF142CNG,ENCFF589ZBJ,ENCFF089GBC,ENCFF502WDT |
| 96 | 3 | 14733518 | 14739015 | 5497 | NZO,SMJ | 3 | 14737253 | 14737287 | 35 | + | 16.4404 | 7.46E-07 | 0.213 | AAGAAGCTGGACCCAGTGGCCAGCAGATGGCCCAC | | |
| 96 | 3 | 14733518 | 14739015 | 5497 | NZO,SMJ | 3 | 14737269 | 14737287 | 19 | + | 17.0328 | 8.88E-07 | 0.207 | TGGCCAGCAGATGGCCCAC | | |
| 97 | 3 | 24327722 | 24437946 | 110224 | NOD | 3 | 24375838 | 24375872 | 35 | + | 20.3303 | 7.02E-08 | 0.0549 | CAGCACAGCCTCATGGGTACCACAAGGGGGCATTC | 10 | ENCFF052KVO,ENCFF024JOZ,ENCFF327LYS,ENCFF092NMJ,ENCFF089GBC,ENCFF883UPM,ENCFF093ZRU,ENCFF502WDT,ENCFF142CNG,ENCFF907XEG |
| 97 | 3 | 24327722 | 24437946 | 110224 | NOD | 3 | 24382881 | 24382915 | 35 | + | 20.6881 | 5.55E-08 | 0.0469 | CTGCAGGTATCCAAGGAGGCCAGAAGAGGATGCTA | | |
| 97 | 3 | 24327722 | 24437946 | 110224 | NOD | 3 | 24410991 | 24411024 | 34 | + | 19.1463 | 1.62E-07 | 0.0645 | TTGCTGTCTTCGGACACACCAGAAGAGGGCTCCA | | |
| 98 | 3 | 37444733 | 37449592 | 4859 | 129S1,AJ,AKR,BTBR,C3H,C57L,CBA,CE,DBA1J,DBA2J,ILNJ2,MA/My,MRL,NOD,NOR,NZB,NZO,NZW,PL,RF,SJL,SMJ,SWR,TH | 3 | 37445319 | 37445352 | 34 | + | 16.4472 | 9.61E-07 | 0.141 | TAGCTGTCTTCAGACACACCAGAAGAGGGCGTCA | | |
| 99 | 3 | 42733346 | 42736195 | 2849 | NZW | 3 | 42733588 | 42733622 | 35 | + | 16.5229 | 7.12E-07 | 0.208 | CTGCAGAAAGCTCTGGCTACCACAAAGGGGCAGGG | | |
| 100 | 3 | 47208010 | 47211257 | 3247 | SJL,SWR | 3 | 47208972 | 47209005 | 34 | + | 21.2033 | 3.67E-08 | 0.0236 | ATGCAGCTAGAGACACGAGCTCCAGGGGGCACTG | | |
| 100 | 3 | 47208010 | 47211257 | 3247 | SJL,SWR | 3 | 47208987 | 47209005 | 19 | + | 16.8525 | 9.96E-07 | 0.218 | CGAGCTCCAGGGGGCACTG | | |
| 101 | 3 | 67003406 | 67011127 | 7721 | CE,SJL | 3 | 67007657 | 67007675 | 19 | + | 17.1639 | 8.17E-07 | 0.199 | CCTCCACGAGGTGGCGGAA | | |
| 102 | 3 | 69360956 | 69362295 | 1339 | AKR,BUB,CBA,CE,DBA1J,DBA2J,MRL,NZO,RF,SJL | 3 | 69361852 | 69361870 | 19 | + | 17.4918 | 6.59E-07 | 0.185 | AACCCACCAGGGGGCGACA | 4 | ENCFF093ZRU,ENCFF367UNR,ENCFF821BIA,ENCFF883UPM |
| 103 | 3 | 91035376 | 91134392 | 99016 | NOD,NOR,RF | 3 | 91130656 | 91130690 | 35 | + | 17.844 | 3.30E-07 | 0.139 | CGGCAGTATCCTGAGTCAAACAGCAGAGGGAGACA | 5 | ENCFF093ZRU,ENCFF052KVO,ENCFF089GBC,ENCFF092NMJ,ENCFF099QNE |
| 104 | 3 | 93807151 | 93812187 | 5036 | 129S1,AJ,AKR,C3H,CBA,DBA1J,DBA2J,FVB,KK,MRL,NOD,NOR,NZW,RF,SJL | 3 | 93807912 | 93807946 | 35 | + | 20.6972 | 5.51E-08 | 0.0467 | CTGTAGTAAAAAGTTACAACCAGAAGATGGTGGTC | | |
| 105 | 3 | 98303333 | 98318999 | 15666 | NOD,NOR,NZB,NZO,NZW | 3 | 98310981 | 98311015 | 35 | + | 24.4037 | 3.92E-09 | 0.01 | ATGCAGGTACCAACAGAGGCCAGGAGAGGGCACCA | | |
| 105 | 3 | 98303333 | 98318999 | 15666 | NOD,NOR,NZB,NZO,NZW | 3 | 98310997 | 98311015 | 19 | + | 19.4918 | 1.63E-07 | 0.107 | AGGCCAGGAGAGGGCACCA | | |
| 106 | 3 | 98309560 | 98318662 | 9102 | 129S1,BTBR,C57L,DBA1J,DBA2J,FVB,PJ,SEA,SJL,SMJ,SWR | 3 | 98310981 | 98311015 | 35 | + | 24.4037 | 3.92E-09 | 0.01 | ATGCAGGTACCAACAGAGGCCAGGAGAGGGCACCA | | |
| 106 | 3 | 98309560 | 98318662 | 9102 | 129S1,BTBR,C57L,DBA1J,DBA2J,FVB,PJ,SEA,SJL,SMJ,SWR | 3 | 98310997 | 98311015 | 19 | + | 19.4918 | 1.63E-07 | 0.107 | AGGCCAGGAGAGGGCACCA | | |
| 107 | 3 | 104658855 | 104660384 | 1529 | NZO,NZW | 3 | 104659367 | 104659400 | 34 | + | 16.561 | 8.95E-07 | 0.14 | ATGCTGGGGATCAGGGGGCCTGAAGAGGGAACTA | | |
| 107 | 3 | 104658855 | 104660384 | 1529 | NZO,NZW | 3 | 104660334 | 104660368 | 35 | + | 16.3211 | 7.98E-07 | 0.221 | GTGAGGCTTCTTCACACCGCCGGTAGAGGGCGCGC | 17 | ENCFF092NMJ,ENCFF089GBC,ENCFF093ZRU,ENCFF367UNR,ENCFF165LXO,ENCFF099QNE,ENCFF136ZDK,ENCFF821BIA,ENCFF033UQX,ENCFF907XEG,ENCFF327LYS,ENCFF609MDN,ENCFF589ZBJ,ENCFF957SPH,ENCFF194HVL,ENCFF198SSR,ENCFF024JOZ |
| 107 | 3 | 104658855 | 104660384 | 1529 | NZO,NZW | 3 | 104660350 | 104660368 | 19 | + | 16.9672 | 9.26E-07 | 0.212 | CCGCCGGTAGAGGGCGCGC | 17 | ENCFF092NMJ,ENCFF089GBC,ENCFF093ZRU,ENCFF367UNR,ENCFF165LXO,ENCFF099QNE,ENCFF136ZDK,ENCFF821BIA,ENCFF033UQX,ENCFF907XEG,ENCFF327LYS,ENCFF609MDN,ENCFF589ZBJ,ENCFF957SPH,ENCFF194HVL,ENCFF198SSR,ENCFF024JOZ |
| 108 | 3 | 116884656 | 116886383 | 1727 | NOD,NOR | 3 | 116885401 | 116885434 | 34 | + | 17.1301 | 6.23E-07 | 0.118 | CTGCAGTGCATGTGCAGGTCAGATGAGGGTGATG | | |
| 109 | 3 | 124663949 | 124670025 | 6076 | AJ,BALB,C3H,CBA,NZB,RH2,SMJ | 3 | 124670009 | 124670043 | 35 | + | 17.1651 | 4.92E-07 | 0.174 | CTGCAGTCTCAGGTTCCTGCCACCTGAGGTTTTGA | | |
| 110 | 3 | 127165543 | 127166576 | 1033 | 129S1,AJ,BALB,C3H,C57L,CBA,NZB,RH2,SMJ | 3 | 127165619 | 127165652 | 34 | + | 23.7073 | 5.04E-09 | 0.00582 | GTGCAGTGCTCACAGAGGCCAGACGAGGGAGCTA | 7 | ENCFF367UNR,ENCFF821BIA,ENCFF093ZRU,ENCFF089GBC,ENCFF201JUA,ENCFF502WDT,ENCFF099QNE |
| 111 | 3 | 133207572 | 133210278 | 2706 | NZB,NZO,NZW | 3 | 133208832 | 133208850 | 19 | + | 17.6066 | 6.11E-07 | 0.181 | AGGCCAGAAGAGGGCACTT | 5 | ENCFF821BIA,ENCFF907XEG,ENCFF327LYS,ENCFF367UNR,ENCFF093ZRU |
| 112 | 3 | 134793714 | 134795598 | 1884 | CE,DBA1J,DBA2J,FVB,KK,MA/My,NOD,NOR,PL,SJL,SMJ,SWR,TH | 3 | 134794727 | 134794761 | 35 | + | 16.4404 | 7.46E-07 | 0.213 | TTGCAATTGACGGCCCAGGACTGAAGGGGTCACGC | | |
| 113 | 3 | 139265775 | 139269940 | 4165 | 129S1,AJ,AKR,BALB,BTBR,BUB,C3H,C57L,CBA,DBA1J,DBA2J,FVB,ILNJ2,KK,MA/My,MRL,NOD,NOR,NZB,NZO,NZW,PJ,PL,RH2,SEA,SJL,SMJ,SWR,TH | 3 | 139266552 | 139266585 | 34 | + | 17.6748 | 4.37E-07 | 0.0998 | GAGCATGACCACAAGTCCACAGCAGGTGGCAACG | | |
| 114 | 3 | 145851498 | 145853085 | 1587 | 129S1,AJ,AKR,BALB,BTBR,BUB,C3H,C57L,CBA,DBA1J,ILNJ2,MA/My,NOD,NOR,NZB,NZO,PJ,RH2,SEA,SJL,SMJ | 3 | 145851473 | 145851506 | 34 | + | 19.6423 | 1.15E-07 | 0.0524 | CTGCAATTCCTAAAGGCTACAGTAGGTGGGGAGG | 4 | ENCFF502WDT,ENCFF327LYS,ENCFF883UPM,ENCFF055HAP |
| 115 | 3 | 152028772 | 152037819 | 9047 | 129S1,AJ,BALB,BTBR,C3H,C57L,CBA,CE,DBA1J,ILNJ2,KK,MRL,PL,RH2,SEA,SMJ | 3 | 152034233 | 152034251 | 19 | + | 17.1639 | 8.17E-07 | 0.199 | CCTCCACGAGGTGGCGGAA | | |
| 116 | 3 | 15946000 | 15952309 | 6309 | AJ,BALB,BUB,C3H,C58J,CBA,FVB,PL,SJL,SMJ,SWR,TH | 3 | 15951468 | 15951502 | 35 | - | 20.3945 | 6.73E-08 | 0.0532 | CTGCTGATCTCCTTGGCTATCAGCAGGTGGCAGAG | | |
| 117 | 3 | 17761787 | 17765886 | 4099 | SMJ | 3 | 17765885 | 17765919 | 35 | - | 17 | 5.42E-07 | 0.182 | GTGCACTTCTGCAACCCTAGCACCAGGGGAGGCAG | | |
| 118 | 3 | 19637793 | 19639492 | 1699 | SMJ | 3 | 19639464 | 19639497 | 34 | - | 17.4146 | 5.18E-07 | 0.109 | GAGCAGTGCGGGACTCTCCCTCCGGGTGGTACAC | | |
| 119 | 3 | 22318079 | 22321858 | 3779 | SMJ | 3 | 22318229 | 22318262 | 34 | - | 18.6992 | 2.20E-07 | 0.0753 | ATGCAGATCTCGCTGGGGCCTCCAGATGAAAGAA | | |
| 120 | 3 | 32424747 | 32430435 | 5688 | AJ,AKR,BUB,C3H,C58J,CBA,FVB,MA/My,NZB,PL,RF,SMJ,TH | 3 | 32426831 | 32426864 | 34 | - | 16.5122 | 9.23E-07 | 0.141 | CTGATCCTCCTGCCTCCACCTGCAGAGGGCTTGG | | |
| 121 | 3 | 32472327 | 32477425 | 5098 | AKR,BUB,C58J,DBA1J,DBA2J,FVB,ILNJ2,KK,MA/My,NZB,PL,RF,SMJ,SWR,TH | 3 | 32477069 | 32477102 | 34 | - | 16.7073 | 8.16E-07 | 0.134 | TTGCTCTCTTCAGGCACACCAGAAGAGGGCATTG | | |
| 122 | 3 | 32751846 | 32753860 | 2014 | SJL | 3 | 32752546 | 32752564 | 19 | - | 19.2787 | 1.91E-07 | 0.113 | TCGACACAAGGGGGCGCTG | 38 | ENCFF093ZRU,ENCFF201JUA,ENCFF367UNR,ENCFF327LYS,ENCFF907XEG,ENCFF805QBU,ENCFF821BIA,ENCFF089GBC,ENCFF198SSR,ENCFF024JOZ,ENCFF321AGQ,ENCFF589ZBJ,ENCFF055HAP,ENCFF502WDT,ENCFF065ODV,ENCFF092NMJ,ENCFF430PPJ,ENCFF136ZDK,ENCFF165LXO,ENCFF951GOP,ENCFF883UPM,ENCFF089EWX,ENCFF957SPH,ENCFF575MGI,ENCFF142CNG,ENCFF210BTO,ENCFF052KVO,ENCFF609MDN,ENCFF281QMV,ENCFF491RJK,ENCFF194HVL,ENCFF504JDS,ENCFF611HDQ,ENCFF533APC,ENCFF316FYN,ENCFF097CAQ,ENCFF033UQX,ENCFF601JVE |
| 122 | 3 | 32751846 | 32753860 | 2014 | SJL | 3 | 32752546 | 32752579 | 34 | - | 26.7886 | 3.16E-10 | 0.000932 | CTGCAGCCTCTGGCTTCGACACAAGGGGGCGCTG | 38 | ENCFF093ZRU,ENCFF201JUA,ENCFF367UNR,ENCFF327LYS,ENCFF907XEG,ENCFF805QBU,ENCFF821BIA,ENCFF089GBC,ENCFF198SSR,ENCFF024JOZ,ENCFF321AGQ,ENCFF589ZBJ,ENCFF055HAP,ENCFF502WDT,ENCFF065ODV,ENCFF092NMJ,ENCFF430PPJ,ENCFF136ZDK,ENCFF165LXO,ENCFF951GOP,ENCFF883UPM,ENCFF089EWX,ENCFF957SPH,ENCFF575MGI,ENCFF142CNG,ENCFF210BTO,ENCFF052KVO,ENCFF609MDN,ENCFF281QMV,ENCFF491RJK,ENCFF194HVL,ENCFF504JDS,ENCFF611HDQ,ENCFF533APC,ENCFF316FYN,ENCFF097CAQ,ENCFF033UQX,ENCFF601JVE |
| 123 | 3 | 36653835 | 36655412 | 1577 | AKR,MA/My,MRL,PL,RF,TH | 3 | 36654113 | 36654147 | 35 | - | 23.4954 | 7.78E-09 | 0.0141 | CTGCATGTATGCCTGCATGCCAGAAGAGGGCATCA | 7 | ENCFF093ZRU,ENCFF367UNR,ENCFF327LYS,ENCFF136ZDK,ENCFF589ZBJ,ENCFF883UPM,ENCFF099QNE |
| 124 | 3 | 39725069 | 39727372 | 2303 | AJ,ILNJ2,NZB | 3 | 39725308 | 39725342 | 35 | - | 20.0367 | 8.49E-08 | 0.0619 | GTGCAGGTGCCCAGGAAGGTCAGAAGAGGGCGTCA | | |
| 125 | 3 | 42137274 | 42141800 | 4526 | NZW | 3 | 42140980 | 42141013 | 34 | - | 17.6748 | 4.37E-07 | 0.0998 | GAGCATGACCACAAGTCCACAGCAGGTGGCAACG | | |
| 126 | 3 | 49246396 | 49250735 | 4339 | AJ,AKR,BALB,BUB,C3H,CBA,CE,DBA1J,DBA2J,FVB,ILNJ2,MRL,NOD,NOR,NZB,PL,RF,RH2,SJL,SMJ,SWR,TH | 3 | 49249763 | 49249797 | 35 | - | 16.4037 | 7.61E-07 | 0.216 | GAGCATTAAACACAAGTCCACAGCAGATGGCAGCA | | |
| 127 | 3 | 51008474 | 51012675 | 4201 | AJ,AKR,BALB,C3H,CBA,DBA1J,DBA2J,ILNJ2,KK,MA/My,MRL,NOD,NOR,NZO,NZW,PL,RF,RH2,SJL,SWR | 3 | 51011837 | 51011870 | 34 | - | 17.6748 | 4.37E-07 | 0.0998 | GAGCATGACCACAAGTCCACAGCAGGTGGCAACG | | |
| 128 | 3 | 65260037 | 65261181 | 1144 | PL | 3 | 65260089 | 65260107 | 19 | - | 20.2623 | 8.99E-08 | 0.0805 | TGGACACTAGAGGGCAGCA | 19 | ENCFF367UNR,ENCFF093ZRU,ENCFF201JUA,ENCFF099QNE,ENCFF821BIA,ENCFF092NMJ,ENCFF430PPJ,ENCFF589ZBJ,ENCFF165LXO,ENCFF198SSR,ENCFF055HAP,ENCFF502WDT,ENCFF907XEG,ENCFF089GBC,ENCFF194HVL,ENCFF024JOZ,ENCFF052KVO,ENCFF327LYS,ENCFF883UPM |
| 129 | 3 | 70479038 | 70483203 | 4165 | CE | 3 | 70482406 | 70482439 | 34 | - | 17.6748 | 4.37E-07 | 0.0998 | GAGCATGACCACAAGTCCACAGCAGGTGGCAACG | | |
| 130 | 3 | 87366327 | 87392732 | 26405 | FVB,NOD,NOR,SJL,SWR | 3 | 87388710 | 87388743 | 34 | - | 16.626 | 8.59E-07 | 0.137 | CTGCTGATCCTGACTTTCACACCAGGTGCTGACC | | |
| 131 | 3 | 124075843 | 124083592 | 7749 | AJ,BALB,C3H,CBA,NZB,RH2,SMJ | 3 | 124082749 | 124082767 | 19 | - | 17.4262 | 6.88E-07 | 0.186 | TTTCCTGTAGGTGGCAGCA | 4 | ENCFF099QNE,ENCFF502WDT,ENCFF055HAP,ENCFF093ZRU |
| 132 | 3 | 130481401 | 130482604 | 1203 | PJ,SEA | 3 | 130481479 | 130481512 | 34 | - | 17.7967 | 4.04E-07 | 0.0961 | TAGCTGTCCTCAGACACACCAGAAGAGGGCATTG | | |
| 133 | 3 | 152534995 | 152536279 | 1284 | TH | 3 | 152536152 | 152536186 | 35 | - | 16.945 | 5.59E-07 | 0.184 | CTGCAATGCCAGGCCCCAGCCCGTAGACAGCCCAG | | |
| 134 | 4 | 12113869 | 12124875 | 11006 | FVB,NZB,NZW | 4 | 12121471 | 12121504 | 34 | + | 16.9187 | 7.13E-07 | 0.0825 | TAGCTGTTTTCAGACACCCCAGAAGAGGGCATCA | | |
| 135 | 4 | 35070177 | 35077789 | 7612 | AJ,AKR,BTBR,BUB,C3H,CBA,DBA1J,DBA2J,FVB,ILNJ2,LP,MA/My,MRL,NOD,NOR,PJ,PL,RF,SEA,SJL,SMJ,SWR,TH | 4 | 35071286 | 35071320 | 35 | + | 16 | 9.55E-07 | 0.154 | TTGCAATTGAAGGCCCAGGACTGAAGGGGTCACGC | | |
| 136 | 4 | 35377121 | 35385412 | 8291 | MA/My,PJ,PL,RF,SEA,TH | 4 | 35385038 | 35385072 | 35 | + | 18.2752 | 2.55E-07 | 0.077 | CTGAAGGCCAGGTGTCCAGACACCAGAGGGCCCTT | | |
| 137 | 4 | 36465337 | 36591613 | 126276 | B10J | 4 | 36512178 | 36512211 | 34 | + | 17.878 | 3.83E-07 | 0.06 | ATGCAGCTAGAGACATGAGCTCTGGGGGGCACTG | | |
| 137 | 4 | 36465337 | 36591613 | 126276 | B10J | 4 | 36566674 | 36566708 | 35 | + | 18.844 | 1.80E-07 | 0.0629 | CTGCAGTGGTGAGGTGCAACCAACAGAGGTAGCAA | | |
| 138 | 4 | 53386784 | 53393102 | 6318 | CBA,DBA1J,DBA2J,MA/My,SMJ | 4 | 53393078 | 53393112 | 35 | + | 16 | 9.55E-07 | 0.154 | CAGCCGGGCGGGCACCTGTCCTCTAGGCGACGCTC | | |
| 139 | 4 | 88673191 | 88676167 | 2976 | 129S1,AKR,BALB,BTBR,CBA,DBA1J,DBA2J,ILNJ2,KK,MRL,NOR,PJ,RF,RH2,SEA,SJL,SWR,TH | 4 | 88674715 | 88674748 | 34 | + | 16.9675 | 6.91E-07 | 0.0815 | CACCCGTGTCCCTTCTGCCCAGCTGGTGGCAGTG | | |
| 139 | 4 | 88673191 | 88676167 | 2976 | 129S1,AKR,BALB,BTBR,CBA,DBA1J,DBA2J,ILNJ2,KK,MRL,NOR,PJ,RF,RH2,SEA,SJL,SWR,TH | 4 | 88675842 | 88675860 | 19 | - | 18.3115 | 3.79E-07 | 0.0887 | GGGCCACCGGAGGGCAGCA | 9 | ENCFF327LYS,ENCFF367UNR,ENCFF024JOZ,ENCFF093ZRU,ENCFF907XEG,ENCFF821BIA,ENCFF099QNE,ENCFF198SSR,ENCFF589ZBJ |
| 139 | 4 | 88673191 | 88676167 | 2976 | 129S1,AKR,BALB,BTBR,CBA,DBA1J,DBA2J,ILNJ2,KK,MRL,NOR,PJ,RF,RH2,SEA,SJL,SWR,TH | 4 | 88675842 | 88675876 | 35 | - | 22.7431 | 1.35E-08 | 0.0116 | CAGCAGCAGAGGGCGCGGGCCACCGGAGGGCAGCA | 10 | ENCFF327LYS,ENCFF367UNR,ENCFF024JOZ,ENCFF093ZRU,ENCFF907XEG,ENCFF821BIA,ENCFF099QNE,ENCFF198SSR,ENCFF589ZBJ,ENCFF089GBC |
| 139 | 4 | 88673191 | 88676167 | 2976 | 129S1,AKR,BALB,BTBR,CBA,DBA1J,DBA2J,ILNJ2,KK,MRL,NOR,PJ,RF,RH2,SEA,SJL,SWR,TH | 4 | 88676038 | 88676056 | 19 | - | 17.7049 | 5.73E-07 | 0.11 | TGGCCTGCAGGGGGAGCGC | 28 | ENCFF327LYS,ENCFF367UNR,ENCFF024JOZ,ENCFF093ZRU,ENCFF907XEG,ENCFF821BIA,ENCFF099QNE,ENCFF198SSR,ENCFF589ZBJ,ENCFF089GBC,ENCFF136ZDK,ENCFF165LXO,ENCFF092NMJ,ENCFF430PPJ,ENCFF033UQX,ENCFF210BTO,ENCFF491RJK,ENCFF883UPM,ENCFF957SPH,ENCFF951GOP,ENCFF281QMV,ENCFF194HVL,ENCFF575MGI,ENCFF611HDQ,ENCFF316FYN,ENCFF533APC,ENCFF097CAQ,ENCFF609MDN |
| 140 | 4 | 93839894 | 93845077 | 5183 | CE,MA/My,NZB,SWR | 4 | 93844615 | 93844648 | 34 | + | 19.3415 | 1.42E-07 | 0.0358 | CTGCCCCCACATTCGCCGTCACAAGATGGCGCTG | | |
| 141 | 4 | 100314955 | 100318686 | 3731 | SMJ | 4 | 100316606 | 100316624 | 19 | + | 17.2459 | 7.74E-07 | 0.123 | AGGCCACAAGGTGGAACCC | | |
| 142 | 4 | 102346746 | 102348654 | 1908 | AKR,CE,KK,MA/My,MRL,SMJ | 4 | 102348000 | 102348033 | 34 | + | 20.2683 | 7.32E-08 | 0.0224 | GTTCAGTGTCTGCAGAGACCAGAAGAGGGCATAG | | |
| 143 | 4 | 111923812 | 111925200 | 1388 | ILNJ2,LP,MA/My | 4 | 111924177 | 111924211 | 35 | + | 19.4587 | 1.23E-07 | 0.0492 | CTGCCATGAGGACTTCCATCCACCAGGCGTCACCT | | |
| 144 | 4 | 112502873 | 112504255 | 1382 | 129S1,AJ,AKR,BALB,BTBR,BUB,C3H,C57L,CBA,CE,DBA1J,DBA2J,FVB,MRL,NOD,NOR,PL,RF,SEA,SJL,SMJ,SWR,TH | 4 | 112503166 | 112503199 | 34 | + | 23.8537 | 4.46E-09 | 0.00288 | GTGTAGTGCCTGTGGAAGCCAGAAGATGGCGATA | | |
| 145 | 4 | 118615672 | 118683173 | 67501 | NZO,NZW | 4 | 118643041 | 118643074 | 34 | + | 17.1951 | 5.97E-07 | 0.0755 | CTGCTCTGGCACACAAGTTCTCAAGGTGGTGCTG | | |
| 146 | 4 | 118874076 | 118875461 | 1385 | ILNJ2 | 4 | 118874828 | 118874861 | 34 | + | 18.5285 | 2.48E-07 | 0.0494 | GTGCATTGCATACGGAGGCCAGAGGAGGGTGTCA | | |
| 147 | 4 | 119783624 | 119787795 | 4171 | BTBR | 4 | 119784401 | 119784434 | 34 | + | 17.6748 | 4.37E-07 | 0.0644 | GAGCATGACCACAAGTCCACAGCAGGTGGCAACG | | |
| 148 | 4 | 123962386 | 123964206 | 1820 | 129S1,AJ,AKR,BALB,BTBR,BUB,C3H,C57L,C58J,CBA,CE,DBA1J,DBA2J,FVB,ILNJ2,KK,MA/My,MRL,NOD,NOR,NZB,NZO,NZW,PJ,PL,RF,SEA,SJL,SMJ,SWR,TH | 4 | 123962467 | 123962500 | 34 | + | 18.4797 | 2.56E-07 | 0.0494 | TTGCTGTCTTCAGACACACCAGAAGAGGGCATCA | | |
| 149 | 4 | 124019753 | 124037387 | 17634 | RH2 | 4 | 124023435 | 124023469 | 35 | + | 18.1743 | 2.71E-07 | 0.0796 | CTGCATGTATACCTGCATATCAGAAGAGGGCATCA | | |
| 149 | 4 | 124019753 | 124037387 | 17634 | RH2 | 4 | 124036030 | 124036048 | 19 | + | 20.8361 | 5.63E-08 | 0.037 | TTGCCAGCAGGTGTCGCCA | 36 | ENCFF367UNR,ENCFF093ZRU,ENCFF099QNE,ENCFF201JUA,ENCFF821BIA,ENCFF601JVE,ENCFF327LYS,ENCFF089GBC,ENCFF907XEG,ENCFF092NMJ,ENCFF024JOZ,ENCFF430PPJ,ENCFF198SSR,ENCFF052KVO,ENCFF504JDS,ENCFF136ZDK,ENCFF165LXO,ENCFF589ZBJ,ENCFF065ODV,ENCFF055HAP,ENCFF033UQX,ENCFF951GOP,ENCFF502WDT,ENCFF321AGQ,ENCFF281QMV,ENCFF575MGI,ENCFF883UPM,ENCFF957SPH,ENCFF609MDN,ENCFF316FYN,ENCFF142CNG,ENCFF097CAQ,ENCFF194HVL,ENCFF491RJK,ENCFF210BTO,ENCFF533APC |
| 149 | 4 | 124019753 | 124037387 | 17634 | RH2 | 4 | 124024928 | 124024961 | 34 | - | 25.4878 | 1.07E-09 | 0.00107 | ATGCAGTACCAAAAAAAGCCAGAAGAGGGCGTCA | 18 | ENCFF136ZDK,ENCFF092NMJ,ENCFF504JDS,ENCFF821BIA,ENCFF024JOZ,ENCFF589ZBJ,ENCFF093ZRU,ENCFF327LYS,ENCFF907XEG,ENCFF089GBC,ENCFF165LXO,ENCFF883UPM,ENCFF367UNR,ENCFF099QNE,ENCFF033UQX,ENCFF533APC,ENCFF052KVO,ENCFF281QMV |
| 150 | 4 | 130105829 | 130107092 | 1263 | KK,NOR,NZB | 4 | 130106306 | 130106339 | 34 | + | 21.6748 | 2.56E-08 | 0.0107 | ATGCAGTGCTCACGGAGGCCAGAAGGCAGCGCTG | 2 | ENCFF055HAP,ENCFF502WDT |
| 151 | 4 | 133062379 | 133064528 | 2149 | 129S1,C57L,ILNJ2,SJL,SWR | 4 | 133062399 | 133062432 | 34 | + | 19.0163 | 1.77E-07 | 0.0412 | CAGCTGTCTTCAGACACACCAGAAGAGGGCGTCA | | |
| 152 | 4 | 137713848 | 137716193 | 2345 | AJ,AKR,PL,RF,RH2,TH | 4 | 137714913 | 137714947 | 35 | + | 17.2202 | 4.77E-07 | 0.108 | AGGCATGTGCCACCACTGCCCAGCAGAGGCCAGCC | | |
| 153 | 4 | 141492132 | 141493760 | 1628 | FVB,NOD | 4 | 141492936 | 141492969 | 34 | + | 16.4472 | 9.61E-07 | 0.0931 | TAGCTGTCTTCAGACACACCAGAAGAGGGCGTCA | | |
| 154 | 4 | 143852559 | 143861083 | 8524 | 129S1,BTBR,BUB,C57L,CE,DBA2J,KK,LP,MA/My,MRL,NOD,NZB,NZW,PJ,PL,RF,SEA,SJL,SMJ,TH | 4 | 143854585 | 143854618 | 34 | + | 17.0325 | 6.63E-07 | 0.0801 | TGGCAGGTCTCAGAGGAGCCGCTAGATGGCCCAC | | |
| 155 | 4 | 145442985 | 145451136 | 8151 | DBA2J,TH | 4 | 145450036 | 145450070 | 35 | + | 24.422 | 3.86E-09 | 0.00589 | CTGCAGTTACCCATGTACGCCATGAGAGGGCGTCA | | |
| 155 | 4 | 145442985 | 145451136 | 8151 | DBA2J,TH | 4 | 145447080 | 145447113 | 34 | - | 19.1301 | 1.64E-07 | 0.0392 | ATGCAGGGGAGCATCCTTCCAGGAGGTGGCACCT | | |
| 156 | 4 | 147490772 | 147492053 | 1281 | 129S1,AJ,AKR,B10D2,B10J,BALB,BTBR,BUB,C3H,C57L,C58J,CE,DBA1J,DBA2J,FVB,ILNJ2,KK,LP,MA/My,MRL,NOR,NZB,NZO,NZW,PJ,PL,RF,RH2,SEA,SJL,SMJ,SWR,TH | 4 | 147491241 | 147491259 | 19 | + | 18.8033 | 2.69E-07 | 0.0753 | TATCCACCAGAGGGTGCTG | 8 | ENCFF907XEG,ENCFF093ZRU,ENCFF092NMJ,ENCFF821BIA,ENCFF089GBC,ENCFF367UNR,ENCFF099QNE,ENCFF327LYS |
| 157 | 4 | 152523830 | 152525506 | 1676 | BUB,MA/My,MRL,NZB,NZO,NZW,SMJ | 4 | 152525044 | 152525077 | 34 | + | 17.4146 | 5.18E-07 | 0.0708 | CTGGGGGACTAACCTCACCCAGCAGGTGGCCCCC | | |
| 158 | 4 | 15236200 | 15242810 | 6610 | 129S1,AJ,AKR,BALB,BUB,C57L,ILNJ2,MA/My,NOD,NOR,NZW,PJ,PL,RF,RH2,SEA,SJL,SMJ | 4 | 15238091 | 15238109 | 19 | - | 17.1639 | 8.17E-07 | 0.127 | CCTCCACGAGGTGGCGGAA | | |
| 159 | 4 | 16319790 | 16327153 | 7363 | NZW | 4 | 16327046 | 16327080 | 35 | - | 20.9817 | 4.56E-08 | 0.0252 | ATGCAGGTGCCTATAGAGTCCAGAAGAGGGCTCTG | | |
| 160 | 4 | 23003638 | 23107598 | 103960 | TH | 4 | 23009510 | 23009528 | 19 | - | 17.459 | 6.74E-07 | 0.114 | CAGCCAGGAGGTGGAGCTG | | |
| 161 | 4 | 26970469 | 26972024 | 1555 | B10D2,B10J | 4 | 26971844 | 26971877 | 34 | - | 18.4797 | 2.56E-07 | 0.0494 | TTGCTGTCTTCAGACACACCAGAAGAGGGCATCA | | |
| 162 | 4 | 32790350 | 32797716 | 7366 | 129S1,AJ,AKR,BALB,BTBR,BUB,C3H,CBA,DBA1J,DBA2J,FVB,ILNJ2,MA/My,MRL,NOD,NOR,PJ,PL,RF,RH2,SEA,SJL,SMJ,SWR,TH | 4 | 32792239 | 32792257 | 19 | - | 17.1639 | 8.17E-07 | 0.127 | CCTCCACGAGGTGGCGGAA | | |
| 163 | 4 | 50177779 | 50182290 | 4511 | MA/My | 4 | 50180078 | 50180096 | 19 | - | 17.2951 | 7.50E-07 | 0.121 | CTGCCACTAGATGGTGCAG | 4 | ENCFF092NMJ,ENCFF907XEG,ENCFF093ZRU,ENCFF099QNE |
| 163 | 4 | 50177779 | 50182290 | 4511 | MA/My | 4 | 50180078 | 50180111 | 34 | - | 17.7805 | 4.08E-07 | 0.0623 | ATGTCCTTTACACAGCTGCCACTAGATGGTGCAG | 4 | ENCFF092NMJ,ENCFF907XEG,ENCFF093ZRU,ENCFF099QNE |
| 164 | 4 | 62323359 | 62326152 | 2793 | AJ,AKR,BALB,C3H,CBA,CE,DBA1J,DBA2J,ILNJ2,LP,MA/My,MRL,NZW,RF,RH2,SEA,SJL,SWR | 4 | 62325587 | 62325605 | 19 | - | 17.5574 | 6.32E-07 | 0.111 | ACTCCAGAAGAGGGCGCCA | 2 | ENCFF136ZDK,ENCFF821BIA |
| 164 | 4 | 62323359 | 62326152 | 2793 | AJ,AKR,BALB,C3H,CBA,CE,DBA1J,DBA2J,ILNJ2,LP,MA/My,MRL,NZW,RF,RH2,SEA,SJL,SWR | 4 | 62325587 | 62325620 | 34 | - | 18.3659 | 2.76E-07 | 0.0503 | TAGCTGTCTTCAGACACTCCAGAAGAGGGCGCCA | 2 | ENCFF136ZDK,ENCFF821BIA |
| 165 | 4 | 67280118 | 67284271 | 4153 | 129S1,C57L,CE,PJ | 4 | 67283468 | 67283501 | 34 | - | 17.6748 | 4.37E-07 | 0.0644 | GAGCATGACCACAAGTCCACAGCAGGTGGCAACG | | |
| 166 | 4 | 86871650 | 86873597 | 1947 | MA/My | 4 | 86872026 | 86872044 | 19 | - | 19.0328 | 2.28E-07 | 0.0693 | ATGCCAGCAGGGGGCAGAC | 3 | ENCFF099QNE,ENCFF024JOZ,ENCFF093ZRU |
| 167 | 4 | 89606946 | 89618459 | 11513 | AKR,BTBR,BUB,KK,MRL,SJL,SMJ,SWR,TH | 4 | 89614812 | 89614845 | 34 | - | 18.5203 | 2.49E-07 | 0.0494 | ATGCAGCTAGAGACACGAACTCCAGGGGGTACTG | | |
| 168 | 4 | 99498162 | 99503432 | 5270 | CE | 4 | 99499802 | 99499836 | 35 | - | 18.2294 | 2.62E-07 | 0.0784 | ATGCCGGTGCCTACAGAGGCCAGAAGAGGGCATTA | | |
| 169 | 4 | 100756127 | 100757362 | 1235 | BTBR,MA/My | 4 | 100756713 | 100756747 | 35 | - | 16.367 | 7.77E-07 | 0.139 | CTGCATTGCTAATTCAAGGCCAGAGGAAGACCCTG | | |
| 170 | 4 | 101720349 | 101729335 | 8986 | AJ,AKR,BALB,BUB,C3H,C58J,CBA,CE,KK,LP,MA/My,MRL,NZB,NZO,NZW,PJ,PL,RF,SEA,SJL,SMJ | 4 | 101723904 | 101723922 | 19 | - | 17.1639 | 8.17E-07 | 0.127 | CCTCCACGAGGTGGCGGAA | | |

| | | | | | | | | | | | | | | | | |
|---|---|---|---|---|---|---|---|---|---|---|---|---|---|---|---|---|
| 171 | 4 | 108003223 | 108012214 | 8991 | 129S1,AJ,AKR,BALB,BUB,C3H,C57L,C58J,CBA,CE,DBA1J,DBA2J,FVB,MRL,NOD,NZB,NZO,NZW,PJ,PL,RF,SEA,SJL,SMJ,SWR,TH | 4 | 108006782 | 108006800 | 19 | - | 17.1639 | 8.17E-07 | 0.127 | CCTCCACGAGGTGGCGGAA | | |
| 172 | 4 | 133560483 | 133561926 | 1443 | AJ,AKR,B10D2,B10J,BALB,BTBR,BUB,C3H,CBA,CE,DBA1J,DBA2J,NOD,NZW,PJ,RF,RH2,SEA,SMJ,TH | 4 | 133561354 | 133561388 | 35 | - | 24.0734 | 5.04E-09 | 0.0069 | CTGCATGTACACCTGTGAGCCAGAAGAGGGCATCA | | |
| 173 | 4 | 134121224 | 134122542 | 1318 | 129S1,AJ,AKR,B10D2,B10J,BALB,BTBR,BUB,C3H,C57L,CBA,CE,DBA1J,DBA2J,FVB,ILNJ2,MA/My,MRL,NOD,NZW,PJ,PL,RF,RH2,SEA,SJL,SMJ,SWR,TH | 4 | 134122079 | 134122112 | 34 | - | 16.5691 | 8.90E-07 | 0.0928 | TAGCTGTCTTCAGACGCACCAGAAGAGGGCATCG | 1 | ENCFF093ZRU |
| 174 | 4 | 135563475 | 135565159 | 1684 | 129S1,AKR,BUB,C3H,C57L,CBA,CE,ILNJ2,MRL,NZB,NZW,RF | 4 | 135564377 | 135564410 | 34 | - | 16.4634 | 9.51E-07 | 0.0931 | TAGCTGTCTTCAGACGCACCAGAAGAGGGCATCA | | |
| 175 | 4 | 139064418 | 139066349 | 1931 | FVB,ILNJ2,NZW | 4 | 139064586 | 139064604 | 19 | - | 21.9836 | 2.06E-08 | 0.0238 | CGACCTCCAGGTGGCGCTG | 38 | ENCFF093ZRU,ENCFF367UNR,ENCFF099QNE,ENCFF327LYS,ENCFF136ZDK,ENCFF086ZTV,ENCFF589ZBJ,ENCFF601JVE,ENCFF024JOZ,ENCFF821BIA,ENCFF907XEG,ENCFF321AGQ,ENCFF502WDT,ENCFF089GBC,ENCFF092NMJ,ENCFF165LXO,ENCFF198SSR,ENCFF210BTO,ENCFF957SPH,ENCFF951GOP,ENCFF430PPJ,ENCFF883UPM,ENCFF052KVO,ENCFF194HVL,ENCFF281QMV,ENCFF575MGI,ENCFF533APC,ENCFF316FYN,ENCFF349BUL,ENCFF611HDQ,ENCFF142CNG,ENCFF609MDN,ENCFF097CAQ,ENCFF033UQX,ENCFF504JDS,ENCFF491RJK,ENCFF089EWX,ENCFF055HAP |
| 176 | 4 | 142076271 | 142078885 | 2614 | AKR,B10D2,B10J,C3H,CBA,MA/My,MRL,NOR,SMJ | 4 | 142076420 | 142076438 | 19 | - | 19.1475 | 2.10E-07 | 0.0659 | CTACCACTAGGAGGCAGCA | 24 | ENCFF093ZRU,ENCFF367UNR,ENCFF430PPJ,ENCFF327LYS,ENCFF099QNE,ENCFF533APC,ENCFF821BIA,ENCFF033UQX,ENCFF907XEG,ENCFF024JOZ,ENCFF194HVL,ENCFF198SSR,ENCFF589ZBJ,ENCFF052KVO,ENCFF165LXO,ENCFF957SPH,ENCFF089GBC,ENCFF951GOP,ENCFF136ZDK,ENCFF092NMJ,ENCFF210BTO,ENCFF281QMV,ENCFF609MDN,ENCFF491RJK |
| 177 | 4 | 142282694 | 142285926 | 3232 | 129S1,AKR,B10D2,B10J,BTBR,BUB,C3H,C57L,CBA,CE,DBA1J,DBA2J,FVB,ILNJ2,KK,MA/My,MRL,NOD,NOR,NZO,NZW,PL,RF,SJL,SMJ,SWR,TH | 4 | 142282707 | 142282725 | 19 | - | 17.5574 | 6.32E-07 | 0.111 | TGGCCAGTTGGGGGCGCGC | | |
| 177 | 4 | 142282694 | 142285926 | 3232 | 129S1,AKR,B10D2,B10J,BTBR,BUB,C3H,C57L,CBA,CE,DBA1J,DBA2J,FVB,ILNJ2,KK,MA/My,MRL,NOD,NOR,NZO,NZW,PL,RF,SJL,SMJ,SWR,TH | 4 | 142282707 | 142282740 | 34 | - | 19.561 | 1.21E-07 | 0.0323 | CAGCATCGTTCTTCCTGGCCAGTTGGGGGCGCGC | | |
| 177 | 4 | 142282694 | 142285926 | 3232 | 129S1,AKR,B10D2,B10J,BTBR,BUB,C3H,C57L,CBA,CE,DBA1J,DBA2J,FVB,ILNJ2,KK,MA/My,MRL,NOD,NOR,NZO,NZW,PL,RF,SJL,SMJ,SWR,TH | 4 | 142285441 | 142285459 | 19 | - | 17.5574 | 6.32E-07 | 0.111 | TGGCCAGTTGGGGGCGCGC | | |
| 177 | 4 | 142282694 | 142285926 | 3232 | 129S1,AKR,B10D2,B10J,BTBR,BUB,C3H,C57L,CBA,CE,DBA1J,DBA2J,FVB,ILNJ2,KK,MA/My,MRL,NOD,NOR,NZO,NZW,PL,RF,SJL,SMJ,SWR,TH | 4 | 142285441 | 142285474 | 34 | - | 19.561 | 1.21E-07 | 0.0323 | CAGCATCGTTCTTCCTGGCCAGTTGGGGGCGCGC | | |
| 178 | 4 | 146687955 | 146689424 | 1469 | BTBR,C57L,C58J,LP,PJ | 4 | 146688398 | 146688416 | 19 | - | 16.9836 | 9.17E-07 | 0.133 | CGGCCGCCGGGGGGCGCGG | | |
| 179 | 4 | 149902732 | 149903779 | 1047 | SWR | 4 | 149903351 | 149903369 | 19 | - | 18.6066 | 3.09E-07 | 0.0804 | AGGCCAGAAGAGGGCACAA | | |
| 180 | 4 | 150825024 | 150827055 | 2031 | AKR,C3H,CBA,MRL,NOD,NOR,NZO | 4 | 150826242 | 150826260 | 19 | - | 18.082 | 4.44E-07 | 0.0983 | ATGCCAGGAGAGGGCACTA | | |
| 181 | 5 | 8646128 | 8648612 | 2484 | BUB,NOD,NOR,SMJ | 5 | 8647423 | 8647456 | 34 | + | 22.8211 | 1.04E-08 | 0.00317 | GTGTAGTGCCCACAGAAGCCAGAAGAGGGTGTCA | 1 | ENCFF093ZRU |
| 182 | 5 | 25435691 | 25444667 | 8976 | 129S1,AJ,AKR,BTBR,BUB,C3H,C57L,CBA,CE,DBA1J,DBA2J,ILNJ2,KK,MRL,NOD,NOR,NZB,NZO,NZW,PJ,RH2,SMJ | 5 | 25441152 | 25441170 | 19 | + | 17.1639 | 8.17E-07 | 0.11 | CCTCCACGAGGTGGCGGAA | | |
| 183 | 5 | 27294423 | 27298012 | 3589 | 129S1,BTBR,C57L,CE,NZB,NZW | 5 | 27296780 | 27296813 | 34 | + | 29.2683 | 2.44E-11 | 0.0000711 | GTGCAGTGCCCACAGAGGCCAGAAGAGGGCATCA | 1 | ENCFF093ZRU |
| 184 | 5 | 34894995 | 34896978 | 1983 | BUB,C58J,FVB,MA/My,MRL,NOD,NOR,NZB,PL,SWR,TH | 5 | 34895288 | 34895321 | 34 | + | 18.4797 | 2.56E-07 | 0.0369 | ATGCAGCTAGAGACATGAGCTCCAGGGGGAACTG | | |
| 185 | 5 | 43365280 | 43374325 | 9045 | BALB,BUB,DBA1J,DBA2J,FVB,ILNJ2,KK,LP,MA/My,NZB,NZO,PJ,PL,RH2,SEA,SWR,TH | 5 | 43370739 | 43370757 | 19 | + | 17.1639 | 8.17E-07 | 0.11 | CCTCCACGAGGTGGCGGAA | | |
| 186 | 5 | 44291697 | 44300683 | 8986 | 129S1,AJ,AKR,BALB,BTBR,BUB,C3H,C57L,CBA,CE,DBA1J,DBA2J,FVB,ILNJ2,KK,LP,MA/My,MRL,NOD,NOR,NZB,NZO,NZW,PJ,PL,RF,RH2,SEA,SJL,SWR,TH | 5 | 44297115 | 44297133 | 19 | + | 17.1639 | 8.17E-07 | 0.11 | CCTCCACGAGGTGGCGGAA | | |
| 187 | 5 | 73502368 | 73504088 | 1720 | C3H,CBA,DBA1J,DBA2J,ILNJ2,NOD,NOR,TH | 5 | 73503170 | 73503203 | 34 | + | 16.9024 | 7.21E-07 | 0.0673 | TTGCTGTCTTCAGACACACCAGAAGAGGGCATCT | 1 | ENCFF093ZRU |
| 188 | 5 | 75578187 | 75580079 | 1892 | 129S1,AKR,BALB,BUB,C57L,MA/My,NOD,NOR,PL,RF,RH2,SEA,SWR,TH | 5 | 75579194 | 75579228 | 35 | + | 15.9541 | 9.80E-07 | 0.141 | TTGCAATTGAAGGCCCAGGACTGAAGGGGTCACAC | | |
| 189 | 5 | 77078357 | 77087343 | 8986 | AJ,AKR,BUB,C3H,CBA,CE,DBA1J,DBA2J,FVB,KK,MA/My,MRL,NOD,NOR,PJ,PL,RF,SJL | 5 | 77083776 | 77083794 | 19 | + | 17.1639 | 8.17E-07 | 0.11 | CCTCCACGAGGTGGCGGAA | | |
| 190 | 5 | 88881297 | 88886977 | 5680 | NZO,RF | 5 | 88882842 | 88882860 | 19 | + | 21.2459 | 3.96E-08 | 0.0284 | TGGCCAGAAGAGGGCAGCC | 18 | ENCFF052KVO,ENCFF099QNE,ENCFF201JUA,ENCFF821BIA,ENCFF367UNR,ENCFF089GBC,ENCFF093ZRU,ENCFF502WDT,ENCFF883UPM,ENCFF198SSR,ENCFF957SPH,ENCFF589ZBJ,ENCFF097CAQ,ENCFF142CNG,ENCFF055HAP,ENCFF327LYS,ENCFF136ZDK,ENCFF907XEG |
| 191 | 5 | 112080455 | 112082414 | 1959 | 129S1,AKR,C57L,NZW,RF | 5 | 112080860 | 112080878 | 19 | + | 17.1148 | 8.43E-07 | 0.11 | CTTCCAGCAGAGGGCGTCC | 2 | ENCFF093ZRU,ENCFF099QNE |
| 192 | 5 | 122330904 | 122338270 | 7366 | 129S1,AJ,BALB,BUB,C57L,ILNJ2,KK,MRL,NOD,NOR,NZW,PL,RF,RH2,SEA | 5 | 122336367 | 122336385 | 19 | + | 17.1639 | 8.17E-07 | 0.11 | CCTCCACGAGGTGGCGGAA | | |
| 193 | 5 | 123404716 | 123406370 | 1654 | 129S1,AJ,BALB,C57L,MRL,RH2,SEA,SJL | 5 | 123405151 | 123405169 | 19 | + | 18.2787 | 3.88E-07 | 0.077 | TGGCCTCTGGGGGGCACTG | 12 | ENCFF821BIA,ENCFF201JUA,ENCFF024JOZ,ENCFF327LYS,ENCFF430PPJ,ENCFF198SSR,ENCFF907XEG,ENCFF055HAP,ENCFF883UPM,ENCFF093ZRU,ENCFF367UNR,ENCFF099QNE |
| 194 | 5 | 126364341 | 126369157 | 4816 | AKR,C3H,CBA,FVB,ILNJ2,NOD,NOR,SMJ,SWR,TH | 5 | 126366477 | 126366510 | 34 | + | 18.878 | 1.95E-07 | 0.0309 | GTGCAGTGCCCATGGAGTCCAGAAGAGAGTGTCA | | |
| 195 | 5 | 133565691 | 133567154 | 1463 | NZW,PL,RF | 5 | 133566387 | 133566421 | 35 | + | 17.945 | 3.11E-07 | 0.0739 | GTACAGTTCCCTGTGGTGGCCAGAAGAGGATGCAG | | |
| 196 | 5 | 135632340 | 135634416 | 2076 | BALB,BTBR,C57L,MA/My,NZW,PL,RF,RH2,SEA,TH | 5 | 135633903 | 135633936 | 34 | + | 16.4472 | 9.61E-07 | 0.0776 | TAGCTGTCTTCAGACACACCAGAAGAGGGCGTCA | | |
| 197 | 5 | 144606644 | 144615629 | 8985 | 129S1,AJ,AKR,BALB,BTBR,BUB,C3H,C57L,CBA,CE,DBA1J,DBA2J,FVB,ILNJ2,KK,LP,MA/My,MRL,NOD,NOR,NZB,NZO,NZW,PJ,PL,RF,RH2,SEA,SJL,SMJ,SWR,TH | 5 | 144612061 | 144612079 | 19 | + | 17.1639 | 8.17E-07 | 0.11 | CCTCCACGAGGTGGCGGAA | | |
| 198 | 5 | 149840264 | 149841314 | 1050 | AKR,BUB,CE,MRL,PJ,SEA,SJL | 5 | 149841005 | 149841038 | 34 | + | 27.9268 | 1.02E-10 | 0.000125 | CTGCAGTATCCAAGGTGACCAATAGAGGGCACCA | 17 | ENCFF093ZRU,ENCFF367UNR,ENCFF099QNE,ENCFF089GBC,ENCFF430PPJ,ENCFF092NMJ,ENCFF198SSR,ENCFF589ZBJ,ENCFF165LXO,ENCFF821BIA,ENCFF055HAP,ENCFF609MDN,ENCFF201JUA,ENCFF024JOZ,ENCFF281QMV,ENCFF907XEG,ENCFF327LYS |
| 198 | 5 | 149840264 | 149841314 | 1050 | AKR,BUB,CE,MRL,PJ,SEA,SJL | 5 | 149841020 | 149841038 | 19 | + | 20.0492 | 1.06E-07 | 0.0377 | TGACCAATAGAGGGCACCA | 17 | ENCFF093ZRU,ENCFF367UNR,ENCFF099QNE,ENCFF089GBC,ENCFF430PPJ,ENCFF092NMJ,ENCFF198SSR,ENCFF589ZBJ,ENCFF165LXO,ENCFF821BIA,ENCFF055HAP,ENCFF609MDN,ENCFF201JUA,ENCFF024JOZ,ENCFF281QMV,ENCFF907XEG,ENCFF327LYS |
| 199 | 5 | 11830147 | 11834655 | 4508 | 129S1,BTBR,C57L,CE,NZB,NZO,NZW | 5 | 11830206 | 11830224 | 19 | - | 17.2131 | 7.91E-07 | 0.109 | CAGCCTCCAGATGGCGGTT | | |
| 199 | 5 | 11830147 | 11834655 | 4508 | 129S1,BTBR,C57L,CE,NZB,NZO,NZW | 5 | 11830206 | 11830240 | 35 | - | 17.4862 | 4.08E-07 | 0.0856 | TGGTGCTGCGGCCGTACAGCCTCCAGATGGCGGTT | | |
| 200 | 5 | 16872418 | 16879719 | 7301 | DBA1J,DBA2J | 5 | 16872746 | 16872779 | 34 | - | 19.3415 | 1.42E-07 | 0.0244 | CTGCCCCCACATTCGCCGTCACAAGATGGCGCTG | | |
| 201 | 5 | 21623456 | 21628025 | 4569 | BUB,TH | 5 | 21624940 | 21624974 | 35 | - | 18.5505 | 2.16E-07 | 0.0596 | CTGCATATACGCTGGCATGCCAGAGGAGGGCATCA | 1 | ENCFF502WDT |
| 201 | 5 | 21623456 | 21628025 | 4569 | BUB,TH | 5 | 21625798 | 21625816 | 19 | - | 18.5574 | 3.20E-07 | 0.0687 | TGGACAGCAGAGGGCAGAA | 17 | ENCFF093ZRU,ENCFF367UNR,ENCFF430PPJ,ENCFF099QNE,ENCFF165LXO,ENCFF024JOZ,ENCFF327LYS,ENCFF821BIA,ENCFF055HAP,ENCFF907XEG,ENCFF502WDT,ENCFF089GBC,ENCFF609MDN,ENCFF883UPM,ENCFF092NMJ,ENCFF589ZBJ,ENCFF198SSR |
| 202 | 5 | 22563552 | 22569432 | 5880 | NZO | 5 | 22564216 | 22564250 | 35 | - | 17.8257 | 3.34E-07 | 0.0763 | CTGCTGGGTCATGAGTGCATCAGTAGGTGGCAGAG | | |
| 203 | 5 | 41552619 | 41559806 | 7187 | 129S1,BALB,BTBR,BUB,C57L,DBA1J,DBA2J,FVB,ILNJ2,KK,LP,MA/My,NZB,NZO,NZW,PJ,PL,RH2,SEA,SWR,TH | 5 | 41552948 | 41552981 | 34 | - | 19.3415 | 1.42E-07 | 0.0244 | CTGCCCCCACATTCGCCGTCACAAGATGGCGCTG | | |
| 204 | 5 | 46412889 | 46547285 | 134396 | NOD,NOR | 5 | 46504932 | 46504950 | 19 | - | 17.8197 | 5.31E-07 | 0.0885 | TGGCCACACGGGGGCAGCA | 32 | ENCFF367UNR,ENCFF201JUA,ENCFF093ZRU,ENCFF099QNE,ENCFF907XEG,ENCFF089GBC,ENCFF024JOZ,ENCFF327LYS,ENCFF430PPJ,ENCFF086ZTV,ENCFF821BIA,ENCFF092NMJ,ENCFF055HAP,ENCFF198SSR,ENCFF589ZBJ,ENCFF502WDT,ENCFF210BTO,ENCFF533APC,ENCFF165LXO,ENCFF142CNG,ENCFF883UPM,ENCFF089EWX,ENCFF491RJK,ENCFF951GOP,ENCFF281QMV,ENCFF957SPH,ENCFF575MGI,ENCFF065ODV,ENCFF194HVL,ENCFF609MDN,ENCFF136ZDK,ENCFF316FYN |
| 205 | 5 | 92081179 | 92082898 | 1719 | 129S1,C57L | 5 | 92082707 | 92082740 | 34 | - | 18.4797 | 2.56E-07 | 0.0369 | TTGCTGTCTTCAGACACACCAGAAGAGGGCATCA | | |
| 206 | 5 | 106362998 | 106364840 | 1842 | DBA1J,DBA2J | 5 | 106364631 | 106364664 | 34 | - | 19.9024 | 9.52E-08 | 0.0182 | CTGCAGCTCCTTCTCGATTCCCCAGAGGGCGTCT | | |
| 207 | 5 | 110048410 | 110055880 | 7470 | 129S1,AJ,AKR,BALB,BTBR,BUB,C3H,C57L,CBA,CE,DBA1J,DBA2J,FVB,ILNJ2,KK,LP,MA/My,MRL,NOD,NOR,NZB,NZO,NZW,PJ,PL,RF,RH2,SEA,SJL,SMJ,SWR,TH | 5 | 110050403 | 110050421 | 19 | - | 17.1639 | 8.17E-07 | 0.11 | CCTCCACGAGGTGGCGGAA | | |
| 208 | 6 | 20047095 | 20048709 | 1614 | 129S1,C57L,NZB | 6 | 20047452 | 20047485 | 34 | + | 16.439 | 9.66E-07 | 0.112 | ATGCAGCTAGAGACAAGAGCTCCAGGGGGTACTG | | |
| 209 | 6 | 72061617 | 72064986 | 3369 | AKR,C58J,KK,MRL,NOD,NOR,PL,RF | 6 | 72064190 | 72064223 | 34 | + | 16.5935 | 8.77E-07 | 0.109 | CTGGACTTGAACCTAGGTCCTCTTGAGGGCAGCC | | |
| 210 | 6 | 75293336 | 75298897 | 5561 | 129S1,AJ,BTBR,BUB,C3H,C57L,C58J,CBA,CE,DBA1J,DBA2J,FVB,ILNJ2,MA/My,PJ,SMJ,TH | 6 | 75293489 | 75293522 | 34 | + | 18.3089 | 2.87E-07 | 0.0578 | ATGCAGCTAGAGACACGAGCTCCAGGGGGTACTG | | |
| 211 | 6 | 87187825 | 87190105 | 2280 | AKR,BALB,MRL,PL,RH2 | 6 | 87188708 | 87188741 | 34 | + | 25.8943 | 7.37E-10 | 0.000921 | GTGCAGTACCCACAGAGGCCAGAAGAGGGTGTCA | 4 | ENCFF052KVO,ENCFF907XEG,ENCFF198SSR,ENCFF093ZRU |
| 212 | 6 | 87998841 | 88002494 | 3653 | PJ,SEA | 6 | 88002159 | 88002177 | 19 | + | 19.6557 | 1.43E-07 | 0.0702 | TGGACAGCAGGTGGCAGTC | 1 | ENCFF093ZRU |
| 213 | 6 | 110311190 | 110315356 | 4166 | 129S1,AJ,BALB,BTBR,BUB,C57L,CE,DBA1J,DBA2J,FVB,KK,NZB,NZO,NZW,PJ,RH2,SEA,SJL,SMJ,SWR,TH | 6 | 110311968 | 110312001 | 34 | + | 17.6748 | 4.37E-07 | 0.0735 | GAGCATGACCACAAGTCCACAGCAGGTGGCAACG | | |
| 214 | 6 | 116023487 | 116026395 | 2908 | AKR,C3H,CBA,ILNJ2,MRL,PL,RF | 6 | 116026077 | 116026111 | 35 | + | 19.2661 | 1.39E-07 | 0.068 | GTGCAGCTGCCTGTGGCATCCAGAGGAGGGCATTC | | |
| 215 | 6 | 116225897 | 116227824 | 1927 | 129S1,BTBR,C57L,C58J,DBA1J,DBA2J,FVB,LP,NOD,NOR,NZO,RH2,SJL,SWR | 6 | 116227262 | 116227295 | 34 | + | 19.252 | 1.51E-07 | 0.0415 | GCCCAGTGCCCACAGAAGCCAGAAGAGGGCATTA | | |
| 216 | 6 | 117926810 | 117928342 | 1532 | CE | 6 | 117928130 | 117928163 | 34 | + | 24.0894 | 3.65E-09 | 0.00278 | CTGCAGTGACCACAGGGGCCGGAAGAGGGTGTTA | | |
| 217 | 6 | 119880567 | 119882567 | 2000 | 129S1,AKR,C57L,DBA1J,DBA2J,MRL,RF | 6 | 119882280 | 119882314 | 35 | + | 18.844 | 1.80E-07 | 0.0783 | AAGGAGTTCCTGCCACCTGCCACCAGAAGGCTGTG | | |
| 218 | 6 | 131117693 | 131119801 | 2108 | CE,FVB,KK,LP,MA/My,NOD,NOR,NZB,NZW,SJL,SWR,TH | 6 | 131119486 | 131119520 | 35 | + | 17.0183 | 5.36E-07 | 0.144 | GTGCAGACACACCCAAGGTCCACTAGAGGAATCCC | | |
| 219 | 6 | 133701362 | 133702521 | 1159 | NZO | 6 | 133701756 | 133701789 | 34 | + | 20.3171 | 7.06E-08 | 0.0241 | ATGCAGGGGAGCATCCTTCCAGAAGGTGGCGCCT | 4 | ENCFF093ZRU,ENCFF883UPM,ENCFF052KVO,ENCFF533APC |
| 219 | 6 | 133701362 | 133702521 | 1159 | NZO | 6 | 133701771 | 133701789 | 19 | + | 17.082 | 8.61E-07 | 0.161 | CTTCCAGAAGGTGGCGCCT | 4 | ENCFF093ZRU,ENCFF883UPM,ENCFF052KVO,ENCFF533APC |
| 220 | 6 | 134265471 | 134266990 | 1519 | AJ,AKR,BALB,BUB,C3H,C58J,CBA,CE,DBA1J,DBA2J,FVB,KK,MA/My,MRL,NOD,NOR,NZB,NZO,NZW,PL,RF,RH2,SEA,SJL,SWR,TH | 6 | 134266950 | 134266983 | 34 | + | 16.7398 | 7.99E-07 | 0.104 | GTTCAGTGCCTGAGGAGGTCAGAAGAGGGTATTA | | |
| 221 | 6 | 137536163 | 137543921 | 7758 | LP | 6 | 137541151 | 137541184 | 34 | + | 18.5041 | 2.52E-07 | 0.0559 | GTGCAGTGCCCATGGAGGGCAGCGGAAGGGGGCA | | |
| 222 | 6 | 142651900 | 142653817 | 1917 | BTBR,C57L,C58J,FVB,KK,MA/My,NOD,NOR,NZB,PJ,SMJ,SWR | 6 | 142652601 | 142652634 | 34 | + | 26.0081 | 6.63E-10 | 0.000854 | CTGCAGTGCCCTTGCAGACCAGAAGAGGGCTCCT | 19 | ENCFF065ODV,ENCFF093ZRU,ENCFF024JOZ,ENCFF367UNR,ENCFF165LXO,ENCFF907XEG,ENCFF198SSR,ENCFF589ZBJ,ENCFF821BIA,ENCFF052KVO,ENCFF533APC,ENCFF194HVL,ENCFF099QNE,ENCFF142CNG,ENCFF055HAP,ENCFF502WDT,ENCFF136ZDK,ENCFF883UPM,ENCFF957SPH |
| 222 | 6 | 142651900 | 142653817 | 1917 | BTBR,C57L,C58J,FVB,KK,MA/My,NOD,NOR,NZB,PJ,SMJ,SWR | 6 | 142653605 | 142653638 | 34 | + | 17.0894 | 6.39E-07 | 0.0917 | GTGCTCTTCCCTAAGTGAGCCCCAGAGGCCTCTG | | |
| 223 | 6 | 33759109 | 33760608 | 1499 | 129S1,C57L,FVB,SWR,TH | 6 | 33759120 | 33759153 | 34 | - | 17.6016 | 4.59E-07 | 0.0756 | ATGCAGCTAGAGAAACAAGCTCCAGGGGGTACTG | | |
| 224 | 6 | 49168598 | 49169823 | 1225 | CE,ILNJ2 | 6 | 49169064 | 49169097 | 34 | - | 20.5447 | 5.98E-08 | 0.0216 | ATGCAGTGGCTGAGGGAGCCAGAAGTGGGCATCA | | |
| 225 | 6 | 51315681 | 51318580 | 2899 | FVB,SJL,SWR | 6 | 51318241 | 51318274 | 34 | - | 19.9593 | 9.14E-08 | 0.0291 | GTGTAGTGCCTTCAGAGGCCAGAAGAAGGTGTTG | | |
| 226 | 6 | 68549897 | 68597377 | 47480 | MA/My | 6 | 68583210 | 68583228 | 19 | - | 18.9016 | 2.51E-07 | 0.0903 | TGACCAGCAGAGGGAGACA | 7 | ENCFF089GBC,ENCFF092NMJ,ENCFF093ZRU,ENCFF099QNE,ENCFF502WDT,ENCFF055HAP,ENCFF883UPM |
| 227 | 6 | 73263771 | 73271134 | 7363 | 129S1,AJ,AKR,BALB,BTBR,BUB,C3H,C57L,C58J,CBA,CE,DBA1J,FVB,ILNJ2,KK,MA/My,MRL,NOD,NOR,NZB,NZO,NZW,PJ,PL,RF,RH2,SEA,SJL,SWR,TH | 6 | 73265660 | 73265678 | 19 | - | 17.1639 | 8.17E-07 | 0.157 | CCTCCACGAGGTGGCGGAA | | |
| 228 | 6 | 97084487 | 97085635 | 1148 | SEA | 6 | 97085384 | 97085402 | 19 | - | 18.9836 | 2.36E-07 | 0.0872 | CAGCCAGGAGAGGGCAGTG | 3 | ENCFF024JOZ,ENCFF093ZRU,ENCFF099QNE |
| 229 | 6 | 108540388 | 108542659 | 2271 | AJ,AKR,BALB,C3H,C58J,CBA,CE,DBA1J,DBA2J,ILNJ2,LP,MRL,NZB,NZO,NZW,PL,RF,RH2,SEA,SWR,TH | 6 | 108541110 | 108541144 | 35 | - | 17.2018 | 4.82E-07 | 0.136 | CTGCTGTGTACTGATGTGGCAGCCAGAGGGCTGGA | | |
| 230 | 6 | 117562052 | 117569379 | 7327 | FVB,NZB,SJL,SWR | 6 | 117562576 | 117562609 | 34 | - | 19.3415 | 1.42E-07 | 0.0398 | CTGCCCCCACATTCGCCGTCACAAGATGGCGCTG | | |
| 231 | 6 | 126454104 | 126455789 | 1685 | NOD,NOR | 6 | 126455341 | 126455359 | 19 | - | 17 | 9.07E-07 | 0.164 | ATACCACCAGAGGGTGGTG | | |
| 232 | 6 | 134603615 | 134606964 | 3349 | 129S1,C57L,SMJ | 6 | 134606483 | 134606516 | 34 | - | 21.2114 | 3.65E-08 | 0.0146 | ATGCATTTCCTGAGAAGGCCAGAAGAGGCCATCA | | |
| 233 | 6 | 136073294 | 136075054 | 1760 | 129S1,AJ,AKR,BALB,BTBR,BUB,C3H,C57L,CBA,CE,DBA1J,DBA2J,FVB,KK,MA/My,MRL,NOD,NOR,NZB,NZW,PJ,PL,RF,RH2,SEA,SJL,SWR,TH | 6 | 136073456 | 136073489 | 34 | - | 24.4472 | 2.69E-09 | 0.00227 | ATGCAGTGCTCCTGAAGGCCAGCAGAGGGTGACA | 20 | ENCFF430PPJ,ENCFF024JOZ,ENCFF093ZRU,ENCFF136ZDK,ENCFF092NMJ,ENCFF589ZBJ,ENCFF883UPM,ENCFF099QNE,ENCFF533APC,ENCFF327LYS,ENCFF907XEG,ENCFF281QMV,ENCFF575MGI,ENCFF194HVL,ENCFF198SSR,ENCFF165LXO,ENCFF089GBC,ENCFF957SPH,ENCFF367UNR,ENCFF821BIA |
| 234 | 7 | 9652668 | 9662040 | 9372 | 129S1,AJ,AKR,BALB,BTBR,BUB,C57L,C58J,CBA,CE,FVB,ILNJ2,KK,LP,MRL,NOD,NOR,NZB,NZO,NZW,PL,RF,RH2,SJL,SWR,TH | 7 | 9656605 | 9656638 | 34 | + | 19.626 | 1.16E-07 | 2.84E-02 | GAGCAGGAGCCCACCAGGCCTCCTGGGGGCGGTG | | |
| 235 | 7 | 23006921 | 23015447 | 8526 | C3H,LP,MA/My,PJ,SEA,SMJ | 7 | 23010822 | 23010840 | 19 | + | 17.1475 | 8.25E-07 | 0.121 | CTGCCAGTAGATGGCCCCA | | |
| 235 | 7 | 23006921 | 23015447 | 8526 | C3H,LP,MA/My,PJ,SEA,SMJ | 7 | 23014370 | 23014388 | 19 | - | 20.5902 | 6.88E-08 | 0.0415 | TGGCCACCAGAGGGAGCCT | | |
| 235 | 7 | 23006921 | 23015447 | 8526 | C3H,LP,MA/My,PJ,SEA,SMJ | 7 | 23014370 | 23014404 | 35 | - | 16.422 | 7.54E-07 | 0.133 | CACCTTTGGAGTCAGTTGGCCACCAGAGGGAGCCT | | |
| 236 | 7 | 29314626 | 29323329 | 8703 | 129S1,AJ,AKR,BALB,BTBR,BUB,C3H,C57L,CE,DBA1J,DBA2J,ILNJ2,LP,MRL,NOD,NOR,NZB,NZO,NZW,PJ,PL,RF,SEA,SJL,SMJ,TH | 7 | 29320044 | 29320062 | 19 | + | 17.1639 | 8.17E-07 | 0.12 | CCTCCACGAGGTGGCGGAA | | |
| 237 | 7 | 30388060 | 30397045 | 8985 | 129S1,AJ,AKR,BALB,BTBR,C57L,C58J,CE,DBA1J,DBA2J,ILNJ2,LP,MA/My,MRL,NOD,NOR,NZB,NZO,NZW,PJ,PL,RF,SEA,SJL,SMJ,TH | 7 | 30393477 | 30393495 | 19 | + | 17.1639 | 8.17E-07 | 0.12 | CCTCCACGAGGTGGCGGAA | | |
| 238 | 7 | 30936693 | 30958743 | 22050 | DBA1J,DBA2J,NZB,NZO,NZW,SMJ | 7 | 30946472 | 30946505 | 34 | + | 18.3089 | 2.87E-07 | 0.0481 | ATGCAGCTAGAGACACGAGCTCCAGGGGGTACTG | | |
| 238 | 7 | 30936693 | 30958743 | 22050 | DBA1J,DBA2J,NZB,NZO,NZW,SMJ | 7 | 30957664 | 30957698 | 35 | + | 17.1101 | 5.08E-07 | 0.107 | ATGCTATTGTGAGGACTGCTCACCAGAGGGTACCA | | |
| 239 | 7 | 37879981 | 37882428 | 2447 | AJ,BALB,FVB,MA/My,NOD,NOR,RH2,SEA | 7 | 37882122 | 37882155 | 34 | + | 24.1057 | 3.60E-09 | 0.00208 | GTCCGGTTCTCGACCTTGCCACCAGAGGGCGCTG | 39 | ENCFF367UNR,ENCFF093ZRU,ENCFF201JUA,ENCFF099QNE,ENCFF805QBU,ENCFF821BIA,ENCFF327LYS,ENCFF907XEG,ENCFF589ZBJ,ENCFF136ZDK,ENCFF198SSR,ENCFF055HAP,ENCFF024JOZ,ENCFF089GBC,ENCFF092NMJ,ENCFF065ODV,ENCFF165LXO,ENCFF052KVO,ENCFF502WDT,ENCFF951GOP,ENCFF210BTO,ENCFF883UPM,ENCFF957SPH,ENCFF430PPJ,ENCFF142CNG,ENCFF281QMV,ENCFF194HVL,ENCFF491RJK,ENCFF349BUL,ENCFF504JDS,ENCFF611HDQ,ENCFF533APC,ENCFF089EWX,ENCFF316FYN,ENCFF609MDN,ENCFF575MGI,ENCFF033UQX,ENCFF086ZTV,ENCFF097CAQ |
| 239 | 7 | 37879981 | 37882428 | 2447 | AJ,BALB,FVB,MA/My,NOD,NOR,RH2,SEA | 7 | 37882137 | 37882155 | 19 | + | 24.7377 | 9.44E-10 | 0.00739 | TTGCCACCAGAGGGCGCTG | 39 | ENCFF367UNR,ENCFF093ZRU,ENCFF201JUA,ENCFF099QNE,ENCFF805QBU,ENCFF821BIA,ENCFF327LYS,ENCFF907XEG,ENCFF589ZBJ,ENCFF136ZDK,ENCFF198SSR,ENCFF055HAP,ENCFF024JOZ,ENCFF089GBC,ENCFF092NMJ,ENCFF065ODV,ENCFF165LXO,ENCFF052KVO,ENCFF502WDT,ENCFF951GOP,ENCFF210BTO,ENCFF883UPM,ENCFF957SPH,ENCFF430PPJ,ENCFF142CNG,ENCFF281QMV,ENCFF194HVL,ENCFF491RJK,ENCFF349BUL,ENCFF504JDS,ENCFF611HDQ,ENCFF533APC,ENCFF089EWX,ENCFF316FYN,ENCFF609MDN,ENCFF575MGI,ENCFF033UQX,ENCFF086ZTV,ENCFF097CAQ |
| 240 | 7 | 44903706 | 44904721 | 1015 | ILNJ2,NZW,PJ,SJL | 7 | 44903753 | 44903786 | 34 | + | 18.0894 | 3.33E-07 | 0.0521 | CAGCTGTCTTTAGACACACCAGAAGAGGGCATTG | | |
| 241 | 7 | 46803949 | 46809133 | 5184 | C57L | 7 | 46805315 | 46805348 | 34 | + | 25.3496 | 1.21E-09 | 0.000926 | GTGCAGTACCCAAAGCTGCCAGATGAGGGCATCA | 11 | ENCFF821BIA,ENCFF092NMJ,ENCFF089GBC,ENCFF327LYS,ENCFF136ZDK,ENCFF093ZRU,ENCFF099QNE,ENCFF502WDT,ENCFF367UNR,ENCFF052KVO,ENCFF065ODV |
| 242 | 7 | 47117678 | 47131601 | 13923 | ILNJ2,SJL | 7 | 47123827 | 47123845 | 19 | + | 16.9016 | 9.66E-07 | 0.13 | TTACCACTAGGAGGCAGAC | | |
| 243 | 7 | 47968414 | 47978239 | 9825 | C57L,FVB,NOD,NOR,SWR | 7 | 47976345 | 47976379 | 35 | + | 22.7982 | 1.29E-08 | 0.0119 | GTGCAGCTGCATGTGGAAGCCAGAAGAGGGCATTG | | |
| 244 | 7 | 49385468 | 49387086 | 1618 | ILNJ2,NZW,PJ,SJL | 7 | 49386064 | 49386098 | 35 | + | 23.3761 | 8.49E-09 | 0.00913 | CTGCATATATGCCTGCATGCCAGAAGAGGGTGCCA | | |
| 245 | 7 | 56091157 | 56113386 | 22229 | AKR | 7 | 56098850 | 56098883 | 34 | + | 18.0569 | 3.40E-07 | 0.0528 | ATGCAGCTAGAGACATGAGCTCCAGGGGGTACTG | | |
| 246 | 7 | 60345708 | 60373558 | 27850 | CE | 7 | 60360207 | 60360240 | 34 | + | 18.2439 | 3.00E-07 | 0.0492 | TTGCAGGTCCTGAAAAGGCCAGAAGAGTGCTGCA | | |
| 247 | 7 | 63776305 | 63785350 | 9045 | 129S1,AKR,BALB,BTBR,C3H,CBA,CE,DBA1J,DBA2J,ILNJ2,KK,LP,NOD,NOR,NZB,NZO,NZW,PJ,PL,RF,SEA,SMJ | 7 | 63781765 | 63781783 | 19 | + | 17.1639 | 8.17E-07 | 0.12 | CCTCCACGAGGTGGCGGAA | | |
| 248 | 7 | 69619037 | 69620466 | 1429 | SMJ | 7 | 69619521 | 69619555 | 35 | + | 16.0459 | 9.31E-07 | 0.147 | TTGCACATACCTTTCGAAGCGAGCAGGGGGCACCT | | |
| 249 | 7 | 71950589 | 71951904 | 1315 | BUB,C58J,CE,FVB,MA/My,NOD,NOR,NZW,SWR | 7 | 71951588 | 71951622 | 35 | + | 16.8257 | 5.99E-07 | 0.116 | CTGTTTGAGCACCTGGTCTCCAGTTGGTGGCGCTG | 2 | ENCFF093ZRU,ENCFF052KVO |
| 250 | 7 | 75473062 | 75476355 | 3293 | C58J,CE,FVB,SMJ,SWR,TH | 7 | 75475456 | 75475489 | 34 | + | 17.1463 | 6.17E-07 | 0.0728 | TAGCTGTCTTTAGATGCGCCAGAAGAGGGCATCA | | |
| 251 | 7 | 76619084 | 76635354 | 16270 | ILNJ2,MA/My | 7 | 76621155 | 76621188 | 34 | + | 16.813 | 7.63E-07 | 0.0822 | ATTCGATTTCTCCAACTGCCACTAGTTGGCGCTG | 17 | ENCFF093ZRU,ENCFF367UNR,ENCFF504JDS,ENCFF165LXO,ENCFF430PPJ,ENCFF198SSR,ENCFF099QNE,ENCFF589ZBJ,ENCFF821BIA,ENCFF883UPM,ENCFF609MDN,ENCFF024JOZ,ENCFF907XEG,ENCFF092NMJ,ENCFF327LYS,ENCFF089GBC,ENCFF281QMV |
| 251 | 7 | 76619084 | 76635354 | 16270 | ILNJ2,MA/My | 7 | 76621170 | 76621188 | 19 | + | 17.6066 | 6.11E-07 | 0.106 | CTGCCACTAGTTGGCGCTG | 17 | ENCFF093ZRU,ENCFF367UNR,ENCFF504JDS,ENCFF165LXO,ENCFF430PPJ,ENCFF198SSR,ENCFF099QNE,ENCFF589ZBJ,ENCFF821BIA,ENCFF883UPM,ENCFF609MDN,ENCFF024JOZ,ENCFF907XEG,ENCFF092NMJ,ENCFF327LYS,ENCFF089GBC,ENCFF281QMV |
| 252 | 7 | 97858780 | 97860271 | 1491 | C3H,CBA,DBA1J,DBA2J,ILNJ2,PJ,SEA | 7 | 97859659 | 97859693 | 35 | + | 16.1651 | 8.71E-07 | 0.143 | TTGCAATTGAAGGCCCAGGACGGAAGGGGTCACGC | | |
| 253 | 7 | 102938893 | 102943444 | 4551 | NZO,TH | 7 | 102939668 | 102939701 | 34 | + | 17.6748 | 4.37E-07 | 0.0607 | GAGCATGACCACAAGTCCACAGCAGGTGGCAACG | | |
| 254 | 7 | 105968059 | 105976874 | 8815 | CE,NZO | 7 | 105970676 | 105970694 | 19 | + | 17.8689 | 5.14E-07 | 0.0979 | AAACCACTAGGTGGCAGAA | | |
| 255 | 7 | 106664831 | 106666847 | 2016 | NOD,SMJ,SWR | 7 | 106666508 | 106666542 | 35 | + | 16.4128 | 7.57E-07 | 0.133 | CAGCTCAGCCACGTTCCCATCACCAGGGGCCAGCT | | |
| 256 | 7 | 112387750 | 112406429 | 18679 | SJL | 7 | 112405866 | 112405899 | 34 | + | 18.0244 | 3.47E-07 | 0.0534 | TTGCAGCTTCTTCTGGAGACTACAGGAGGCACCA | | |
| 257 | 7 | 116031339 | 116037017 | 5678 | 129S1,AKR,BALB,BTBR,C3H,C57L,CBA,CE,DBA1J,DBA2J,ILNJ2,KK,MRL,NOR,NZW,PL,RH2,SJL | 7 | 116035155 | 116035173 | 19 | + | 17.1639 | 8.17E-07 | 0.12 | CCTCCACGAGGTGGCGGAA | | |
| 258 | 7 | 127233514 | 127235214 | 1700 | AJ,NOD | 7 | 127234624 | 127234642 | 19 | + | 18.8033 | 2.69E-07 | 0.0728 | GTGCCAGAAGAGGGCGGCG | | |
| 259 | 7 | 3158490 | 3161965 | 3475 | MRL | 7 | 3159528 | 3159562 | 35 | - | 16 | 9.55E-07 | 0.148 | CTGCTGTGTCCTGACTTTGCCCGCAGGGGTATTTG | | |
| 260 | 7 | 3719960 | 3739682 | 19722 | MRL | 7 | 3728545 | 3728563 | 19 | - | 18.6885 | 2.92E-07 | 0.0743 | TAACCAGGAGAGGGCAGCA | | |
| 261 | 7 | 14815373 | 14818589 | 3216 | NZO | 7 | 14816472 | 14816490 | 19 | - | 17.918 | 4.97E-07 | 0.0973 | AAGCCAGGAGAGGGCACTA | | |
| 261 | 7 | 14815373 | 14818589 | 3216 | NZO | 7 | 14816472 | 14816506 | 35 | - | 26.5321 | 7.03E-10 | 0.00258 | TTGCATTTATGCCTGCAAGCCAGGAGAGGGCACTA | | |
| 262 | 7 | 20910453 | 20916877 | 6424 | CE | 7 | 20913466 | 20913500 | 35 | - | 19.9266 | 9.11E-08 | 0.0385 | ATGCAATTCATAGTGGAGACCAGAAGATGGCCTTA | | |
| 263 | 7 | 27561165 | 27586055 | 24890 | KK,NZO | 7 | 27563112 | 27563145 | 34 | - | 17.4553 | 5.05E-07 | 0.0659 | CTACAGTGGCCATGGAGACCAGAAGAAGGCATCA | | |
| 264 | 7 | 27633164 | 27641490 | 8326 | 129S1,BUB,CE,ILNJ2,KK,LP,NZB,NZO,NZW,PJ,SEA,SJL,SMJ | 7 | 27633455 | 27633473 | 19 | - | 19.2951 | 1.88E-07 | 0.0611 | CCGCCACTAGATGGTGCTA | | |
| 264 | 7 | 27633164 | 27641490 | 8326 | 129S1,BUB,CE,ILNJ2,KK,LP,NZB,NZO,NZW,PJ,SEA,SJL,SMJ | 7 | 27641449 | 27641467 | 19 | - | 19.2951 | 1.88E-07 | 0.0611 | CCGCCACTAGATGGTGCTA | 15 | ENCFF092NMJ,ENCFF201JUA,ENCFF093ZRU,ENCFF430PPJ,ENCFF367UNR,ENCFF099QNE,ENCFF327LYS,ENCFF065ODV,ENCFF055HAP,ENCFF907XEG,ENCFF136ZDK,ENCFF589ZBJ,ENCFF089GBC,ENCFF821BIA,ENCFF198SSR |

| | | | | | | | | | | | | | | | | |
|---|---|---|---|---|---|---|---|---|---|---|---|---|---|---|---|---|
| 265 | 7 | 27718631 | 27727443 | 8812 | 129S1,AJ,BALB,BUB,CE,FVB,ILNJ2,KK,LP,MRL,NZB,NZO,NZW,SJL,SMJ | 7 | 27725284 | 27725317 | 34 | - | 17.0325 | 6.63E-07 | 0.0765 | TGGCAGGTCTCAGAGGAGCCGCTAGATGGCCCAC | | |
| 266 | 7 | 39154714 | 39165031 | 10317 | 129S1,BTBR,BUB,C3H,C57L,CBA,CE,KK,MRL,NZW,PL,SJL,SMJ | 7 | 39155367 | 39155400 | 34 | - | 17.1301 | 6.23E-07 | 0.0735 | CAGCTATCTTCAGATACACCAGAAGAGGGCATCA | | |
| 267 | 7 | 46512747 | 46514141 | 1394 | ILNJ2.NZW.PJ.SJL | 7 | 46513033 | 46513066 | 34 | - | 19.626 | 1.16E-07 | 0.0284 | ACGCAGTGCCCACGGAGGTCAGAAGAGGGCATCT | | |
| 267 | 7 | 46512747 | 46514141 | 1394 | ILNJ2.NZW.PJ.SJL | 7 | 46513429 | 46513462 | 34 | - | 17.6423 | 4.47E-07 | 0.0616 | TAGCTGTGTTCAGACACACCAGAAGAGGGCGTCA | | |
| 268 | 7 | 51625875 | 51629913 | 4038 | ILNJ2,KK,NZW,PJ | 7 | 51629203 | 51629237 | 35 | - | 22.4495 | 1.66E-08 | 0.0134 | CTGCAGTTACTGTGTCTGAGCAGTAGGCGGCAGAA | 24 | ENCFF093ZRU,ENCFF367UNR,ENCFF099QNE,ENCFF430PPJ,ENCFF201JUA,ENCFF327LYS,ENCFF609MDN,ENCFF089GBC,ENCFF055HAP,ENCFF321AGQ,ENCFF907XEG,ENCFF024JOZ,ENCFF821BIA,ENCFF589ZBJ,ENCFF502WDT,ENCFF575MGI,ENCFF957SPH,ENCFF316FYN,ENCFF092NMJ,ENCFF065ODV,ENCFF136ZDK,ENCFF198SSR,ENCFF165LXO,ENCFF281QMV |
| 269 | 7 | 63252709 | 63254294 | 1585 | CE | 7 | 63252783 | 63252817 | 35 | - | 16.9725 | 5.50E-07 | 0.111 | CAGCAGTTCCCACACCTGACAGAAAGATGGCTGTG | | |
| 270 | 7 | 72466892 | 72468373 | 1481 | 129S1,AJ,AKR,BALB,BUB,C3H,C58J,CBA,CE,FVB,KK,MRL,NOD,NOR,NZB,NZO,RF,RH2,SJL,SMJ,SWR,TH | 7 | 72467534 | 72467568 | 35 | - | 16 | 9.55E-07 | 0.148 | TTGCAATTGAAGGCCCAGGACTGAAGGGGTCACGC | | |
| 271 | 7 | 84709089 | 84710391 | 1302 | 129S1,AJ,AKR,BALB,BTBR,BUB,C3H,C57L,CBA,DBA1J,DBA2J,FVB,ILNJ2,KK,MA/My,MRL,NOD,NOR,NZB,NZO,NZW,PJ,PL,RF,RH2,SEA,SJL,SMJ,SWR,TH | 7 | 84709367 | 84709400 | 34 | - | 17.2846 | 5.64E-07 | 0.0699 | CTGCAAGAGGAGAGCTTGCCTGCAGAGAGCACTC | | |
| 272 | 7 | 88463086 | 88464661 | 1575 | SMJ | 7 | 88464587 | 88464620 | 34 | - | 18.0569 | 3.40E-07 | 0.0528 | ATGCAGCTAGAGACATGAGCTCCAGGGGGTACTG | | |
| 273 | 7 | 94920123 | 94976361 | 56238 | SMJ | 7 | 94969281 | 94969314 | 34 | - | 20.187 | 7.76E-08 | 0.0212 | TTGCTGTTTTCAGACACACCAGAAGAGGGCATCA | | |
| 274 | 7 | 102173101 | 102174345 | 1244 | KK.NOD.SJL.SWR | 7 | 102173660 | 102173694 | 35 | - | 17.6606 | 3.68E-07 | 0.0897 | GGGAAATGCCATCTCAGACACACCAGAGGGCACTG | | |
| 275 | 7 | 102499908 | 102503117 | 3209 | KK.NOD.SJL.SWR | 7 | 102502504 | 102502522 | 19 | - | 17.0656 | 8.70E-07 | 0.124 | AGGCCACAAGAGGGAACTC | | |
| 276 | 7 | 109710735 | 109712782 | 2047 | NOD.SMJ.SWR | 7 | 109712572 | 109712605 | 34 | - | 18.813 | 2.04E-07 | 0.0402 | TAGCTGTCCCCAGATATTCCAGAAGAGGGCATCA | | |
| 277 | 7 | 112866154 | 112870131 | 3977 | C3H,CBA,DBA1J,DBA2J,ILNJ2,NOR,NZO,SMJ,TH | 7 | 112867716 | 112867734 | 19 | - | 17.9836 | 4.75E-07 | 0.0964 | GTTCCAGCAGGTGGCAGCC | 30 | ENCFF093ZRU,ENCFF367UNR,ENCFF099QNE,ENCFF201JUA,ENCFF821BIA,ENCFF210BTO,ENCFF327LYS,ENCFF907XEG,ENCFF024JOZ,ENCFF089GBC,ENCFF198SSR,ENCFF092NMJ,ENCFF589ZBJ,ENCFF430PPJ,ENCFF951GOP,ENCFF533APC,ENCFF033UQX,ENCFF165LXO,ENCFF957SPH,ENCFF136ZDK,ENCFF097CAQ,ENCFF321AGQ,ENCFF052KVO,ENCFF609MDN,ENCFF575MGI,ENCFF491RJK,ENCFF281QMV,ENCFF316FYN,ENCFF883UPM,ENCFF194HVL |
| 277 | 7 | 112866154 | 112870131 | 3977 | C3H,CBA,DBA1J,DBA2J,ILNJ2,NOR,NZO,SMJ,TH | 7 | 112867716 | 112867750 | 35 | - | 23.2202 | 9.52E-09 | 0.00969 | GTGCCATTACAACAACGTTCCAGCAGGTGGCAGCC | 30 | ENCFF093ZRU,ENCFF367UNR,ENCFF099QNE,ENCFF201JUA,ENCFF821BIA,ENCFF210BTO,ENCFF327LYS,ENCFF907XEG,ENCFF024JOZ,ENCFF089GBC,ENCFF198SSR,ENCFF092NMJ,ENCFF589ZBJ,ENCFF430PPJ,ENCFF951GOP,ENCFF533APC,ENCFF033UQX,ENCFF165LXO,ENCFF957SPH,ENCFF136ZDK,ENCFF097CAQ,ENCFF321AGQ,ENCFF052KVO,ENCFF609MDN,ENCFF575MGI,ENCFF491RJK,ENCFF281QMV,ENCFF316FYN,ENCFF883UPM,ENCFF194HVL |
| 277 | 7 | 112866154 | 112870131 | 3977 | C3H,CBA,DBA1J,DBA2J,ILNJ2,NOR,NZO,SMJ,TH | 7 | 112869315 | 112869348 | 34 | - | 16.7236 | 8.07E-07 | 0.084 | CTGCACTGCCAGCTGCAAACACAGCAGGGCCTCA | | |
| 278 | 7 | 122558697 | 122564120 | 5423 | MRL | 7 | 122564036 | 122564069 | 34 | - | 17.9756 | 3.59E-07 | 0.0544 | TTGCACAGTCTCAGCCCTGCAACAGAGGGCACCG | | |
| 279 | 7 | 122669754 | 122676858 | 7104 | FVB.SJL | 7 | 122674680 | 122674713 | 34 | - | 19.6504 | 1.14E-07 | 0.0282 | ATGCAGTTCCTATGAAGGCCAGAAGAGGATATCG | 2 | ENCFF099QNE.ENCFF093ZRU |
| 280 | 7 | 133840225 | 133853081 | 12856 | DBA1J.DBA2J | 7 | 133840774 | 133840792 | 19 | - | 17.3607 | 7.18E-07 | 0.112 | AGGCCAGAAGAGGGTGCCA | 1 | ENCFF052KVO |
| 280 | 7 | 133840225 | 133853081 | 12856 | DBA1J.DBA2J | 7 | 133840774 | 133840808 | 35 | - | 24.1376 | 4.80E-09 | 0.0065 | GTGCAGGTACCCTAAGAGGCCAGAAGAGGGTGCCA | 1 | ENCFF052KVO |
| 281 | 7 | 138400694 | 138402588 | 1894 | NZB.NZO | 7 | 138401683 | 138401701 | 19 | - | 17.9836 | 4.75E-07 | 0.0964 | GCACCAGAAGAGGGCACTA | | |
| 282 | 8 | 5659784 | 5666072 | 6288 | 129S1.NZB | 8 | 5660465 | 5660498 | 34 | + | 17.813 | 3.99E-07 | 0.063 | CAGCACTCCCAACTTCGCCCAGTCCAGGGCACCC | | |
| 283 | 8 | 7799142 | 7807592 | 8450 | KK.SMJ | 8 | 7805087 | 7805121 | 35 | + | 17.4037 | 4.28E-07 | 0.117 | ATGCAGTGCCCCCACAGGGGCCATAGGGGACATCA | | |
| 284 | 8 | 19444791 | 19447672 | 2881 | 129S1,AJ,BALB,BTBR,BUB,C3H,CBA,DBA1J,DBA2J,FVB,KK,MRL,NOD,NOR,NZB,NZW,PL.RH2.SJL.SMJ.SWR.TH | 8 | 19447648 | 19447682 | 35 | + | 16.1101 | 8.98E-07 | 0.163 | CTGCTGCCACACCCGCTCCCCACAGGGTGTCATCT | | |
| 285 | 8 | 19833367 | 19834900 | 1533 | MA/My,RF | 8 | 19834560 | 19834593 | 34 | + | 21.6992 | 2.52E-08 | 0.0107 | GTGAGGTTCATATAGGGGCCACCAGAGGGCGCGA | 12 | ENCFF201JUA,ENCFF093ZRU,ENCFF367UNR,ENCFF099QNE,ENCFF430PPJ,ENCFF821BIA,ENCFF092NMJ,ENCFF198SSR,ENCFF589ZBJ,ENCFF327LYS,ENCFF907XEG,ENCFF089GBC |
| 285 | 8 | 19833367 | 19834900 | 1533 | MA/My,RF | 8 | 19834575 | 19834593 | 19 | + | 22 | 2.03E-08 | 0.0275 | GGGCCACCAGAGGGCGCGA | 13 | ENCFF201JUA,ENCFF093ZRU,ENCFF367UNR,ENCFF099QNE,ENCFF430PPJ,ENCFF821BIA,ENCFF092NMJ,ENCFF198SSR,ENCFF589ZBJ,ENCFF327LYS,ENCFF907XEG,ENCFF089GBC,ENCFF136ZDK |
| 286 | 8 | 32165219 | 32171803 | 6584 | 129S1,AJ,AKR,BALB,BUB,C3H,C57L,CBA,DBA1J,DBA2J,FVB,MA/My,MRL,NOD,NOR,NZW,PL,RF,RH2,SJL,SMJ,SWR,TH | 8 | 32169807 | 32169841 | 35 | + | 18.1284 | 2.79E-07 | 0.0931 | CTGTAGCTGTCTTCAGACACCAGAAGAGGGCATCA | | |
| 287 | 8 | 34570509 | 34571667 | 1158 | 129S1,AJ,AKR,BALB,BUB,C3H,C57L,CBA,CE,DBA1J,DBA2J,MA/My,MRL,NOD,NOR,PL,RF,RH2.SJL.SMJ.SWR.TH | 8 | 34570716 | 34570749 | 34 | + | 21.9756 | 2.03E-08 | 0.00915 | CTGCTGTCTTCAGTCACACCAGAAGGGGGCATTG | 1 | ENCFF093ZRU |
| 288 | 8 | 41627138 | 41628522 | 1384 | 129S1,AJ,AKR,BALB,BTBR,BUB,C3H,C57L,CBA,CE,DBA1J,DBA2J,FVB,ILNJ2,KK,MA/My,MRL,NOD,NOR,PJ,PL,RF,RH2,SEA,SJL,SMJ,SWR,TH | 8 | 41627232 | 41627250 | 19 | + | 17.2787 | 7.58E-07 | 0.136 | CTTCCAGTAGAGGGCGTCA | 1 | ENCFF821BIA |
| 289 | 8 | 52184613 | 52188769 | 4156 | 129S1,AJ,B10D2,B10J,BUB,C58J,CBA,FVB,ILNJ2,KK,LP,MA/My,MRL,NZB,NZO,NZW,PL,RH2,SJL,SMJ,SWR | 8 | 52185390 | 52185423 | 34 | + | 17.6748 | 4.37E-07 | 0.066 | GAGCATGACCACAAGTCCACAGCAGGTGGCAACG | | |
| 290 | 8 | 64385349 | 64388640 | 3291 | AKR,BALB,C3H,C57L,CBA,CE,DBA1J,DBA2J,MRL,NZB,NZO,PJ,PL,RF,SEA,SMJ | 8 | 64388574 | 64388608 | 35 | + | 23.9266 | 5.63E-09 | 0.00912 | CTGCATGTATACCTTCATGCCAGAAGAGGGCCCTA | | |
| 291 | 8 | 65362517 | 65378480 | 15963 | C58J.LP.PJ.SEA | 8 | 65366314 | 65366347 | 34 | + | 16.8211 | 7.59E-07 | 0.0873 | CAGCTGTCTTCAGACACACCTGAAGAGGGCATCG | | |
| 292 | 8 | 67816635 | 67837561 | 20926 | C58J.PJ.SEA.TH | 8 | 67829169 | 67829202 | 34 | + | 22.252 | 1.64E-08 | 0.00794 | GAGCAGGTGCAGCCACCGCCCCAAGATGGCGCCC | | |
| 293 | 8 | 69811986 | 69814316 | 2330 | NZB.NZO | 8 | 69813175 | 69813208 | 34 | + | 16.9268 | 7.10E-07 | 0.0845 | ATGCAGTTAGAGACATGAGCTCTGGGGGGTACTG | | |
| 294 | 8 | 69912663 | 69914766 | 2103 | AKR.MA/My.NZB.RF | 8 | 69912826 | 69912859 | 34 | + | 17.3902 | 5.27E-07 | 0.0727 | TAGCTGTCCTCAGACACTCCAGAAGAGGGCATCA | | |
| 295 | 8 | 70094207 | 70102187 | 7980 | 129S1,AKR,BALB,BTBR,BUB,C3H,C57L,C58J,CBA,CE,DBA1J,DBA2J,FVB,ILNJ2,KK,MA/My,MRL,NZB,NZO,NZW,PJ,PL,RF,SEA,SJL,SMJ,SWR,TH | 8 | 70096875 | 70096908 | 34 | + | 17.6748 | 4.37E-07 | 0.066 | GAGCATGACCACAAGTCCACAGCAGGTGGCAACG | | |
| 296 | 8 | 70151239 | 70152818 | 1579 | 129S1,BALB,BTBR,FVB,ILNJ2,KK,NZW,RF,SJL.SWR.TH | 8 | 70151240 | 70151273 | 34 | + | 18.5691 | 2.41E-07 | 0.0509 | TAGCTGTCTTCATACACACCACAAGAGGGCACCA | | |
| 296 | 8 | 70151239 | 70152818 | 1579 | 129S1,BALB,BTBR,FVB,ILNJ2,KK,NZW,RF,SJL.SWR.TH | 8 | 70151255 | 70151273 | 19 | + | 18.541 | 3.24E-07 | 0.092 | ACACCACAAGAGGGCACCA | | |
| 297 | 8 | 83788780 | 83790841 | 2061 | 129S1,AJ,AKR,BALB,BTBR,C3H,C57L,CBA,CE,DBA1J,DBA2J,ILNJ2,KK,MA/My,MRL,NOD,NOR,NZW,PJ,PL,RF,RH2,SEA,SJL | 8 | 83790314 | 83790347 | 34 | + | 19.0569 | 1.72E-07 | 0.0405 | CAGCTGTCTTCTCACACACCAGAAGAGGGCATCA | 1 | ENCFF821BIA |
| 298 | 8 | 86957834 | 86960871 | 3037 | 129S1,AKR,BALB,BTBR,C57L,MRL,PJ,PL,RF,SEA | 8 | 86960511 | 86960545 | 35 | + | 21.9817 | 2.31E-08 | 0.0209 | CTGCTGTGTCACATGTACATCAGTAGAGGGCAGAG | 4 | ENCFF093ZRU,ENCFF367UNR,ENCFF327LYS,ENCFF883UPM |
| 299 | 8 | 89130672 | 89132551 | 1879 | 129S1,AKR,BALB,BTBR,BUB,C57L,C58J,CBA,CE,DBA1J,DBA2J,FVB,ILNJ2,KK,MA/My,MRL,NOD,NOR,NZB,NZO,NZW,PJ,PL,RF,SEA,SJL,SMJ,SWR,TH | 8 | 89131683 | 89131717 | 35 | + | 16 | 9.55E-07 | 0.168 | TTGCAATTGAAGGCCCAGGACTGAAGGGGTCACGC | | |
| 300 | 8 | 93307104 | 93308797 | 1693 | C58J.CBA.DBA1J.DBA2J.MA/My.SMJ | 8 | 93307102 | 93307136 | 35 | + | 16.1376 | 8.84E-07 | 0.162 | ATGCAAGTGCCCTCAAGGTCCAGAAGAAGGCATCA | | |
| 301 | 8 | 122559579 | 122568003 | 8424 | 129S1,BTBR,C57L,FVB,ILNJ2,NOD,NOR,NZB,SJL,SWR | 8 | 122563720 | 122563738 | 19 | + | 18.0984 | 4.40E-07 | 0.107 | TGGCCAGTAGATGGCCCTC | | |
| 302 | 8 | 122862058 | 122864866 | 2808 | 129S1,BTBR,C57L,FVB,NOD,NOR,SJL,SWR | 8 | 122862392 | 122862426 | 35 | + | 25.367 | 1.84E-09 | 0.00549 | CTGCACTGGCAGGGAGGAGCCAGAGGGGGGCACTC | 10 | ENCFF093ZRU,ENCFF092NMJ,ENCFF089GBC,ENCFF907XEG,ENCFF327LYS,ENCFF367UNR,ENCFF883UPM.ENCFF099QNE.ENCFF533APC.ENCFF821BIA |
| 303 | 8 | 123891450 | 123898816 | 7366 | 129S1,AJ,AKR,BALB,BTBR,C3H,C57L,C58J,CE,ILNJ2,LP,MA/My,MRL,NZB,PJ,PL,RF,RH2,SEA,SMJ,TH | 8 | 123896912 | 123896930 | 19 | + | 17.1639 | 8.17E-07 | 0.141 | CCTCCACGAGGTGGCGGAA | | |
| 304 | 8 | 126825819 | 126841407 | 15588 | NZB | 8 | 126826791 | 126826824 | 34 | + | 16.5772 | 8.86E-07 | 0.0942 | TTGCAACATGCTTTCTAACCACCAGGGGTCTCCA | 8 | ENCFF821BIA,ENCFF093ZRU,ENCFF024JOZ,ENCFF201JUA,ENCFF327LYS,ENCFF883UPM,ENCFF367UNR,ENCFF099QNE |
| 305 | 8 | 5337936 | 5340543 | 2607 | 129S1.NZB | 8 | 5339918 | 5339951 | 34 | - | 16.5366 | 9.09E-07 | 0.0944 | TAGCTGTCTTCAGAAAGACCAGAAGAGGGCATCA | | |
| 306 | 8 | 13592370 | 13594401 | 2031 | C57L,ILNJ2,NZB,NZW,PJ,PL,SEA,SMJ,TH | 8 | 13592688 | 13592722 | 35 | - | 16.5229 | 7.12E-07 | 0.145 | CTGCAGGTCCCCGGAAAAGACACAAGGTGCTCCTT | | |
| 307 | 8 | 13760404 | 13765757 | 5353 | AKR,BALB,C57L,CE,ILNJ2,LP,MRL,NZB,NZW,PJ,PL,RF,RH2,SEA,SMJ,TH | 8 | 13760378 | 13760411 | 34 | - | 16.4959 | 9.32E-07 | 0.0944 | ATCTGGTGCCTGAGGAGGCCAGAAGAGGGCGTCA | | |
| 308 | 8 | 30113636 | 30149023 | 35387 | 129S1,AJ,AKR,BALB,BTBR,BUB,C3H,C57L,CBA,CE,DBA1J,DBA2J,FVB,KK,MA/My,MRL,NOD,NOR,NZB,NZO,NZW,RF,RH2,SJL,SMJ,SWR,TH | 8 | 30128974 | 30129007 | 34 | - | 19.6748 | 1.12E-07 | 0.0314 | CTGCAAGAGGAGAGCTTGCCTGCAGAGGGTACTG | | |
| 308 | 8 | 30113636 | 30149023 | 35387 | 129S1,AJ,AKR,BALB,BTBR,BUB,C3H,C57L,CBA,CE,DBA1J,DBA2J,FVB,KK,MA/My,MRL,NOD,NOR,NZB,NZO,NZW,RF,RH2,SJL,SMJ,SWR,TH | 8 | 30129120 | 30129138 | 19 | - | 17.5902 | 6.18E-07 | 0.125 | CAGCCAGAAGGTGGAACTG | | |
| 309 | 8 | 30937812 | 30940317 | 2505 | 129S1,AJ,AKR,BALB,BTBR,BUB,C3H,C57L,CBA,DBA1J,DBA2J,FVB,KK,MA/My,MRL,NOD,NOR,NZW,RF,RH2,SJL,SMJ,SWR,TH | 8 | 30938561 | 30938594 | 34 | - | 17.2439 | 5.79E-07 | 0.0757 | CTGCAGTCCCCACTCTCACCTCCTGGAAGCTGTC | | |
| 310 | 8 | 33679371 | 33682361 | 2990 | 129S1,AJ,AKR,BALB,BUB,C57L,CBA,FVB,MA/My,MRL,NOD,NOR,PL,RF,RH2,SJL,SMJ,SWR,TH | 8 | 33679685 | 33679718 | 34 | - | 17.3171 | 5.52E-07 | 0.0747 | ATGAAGTGTCCATGACAGCCAGAAGAGGGGGTCA | | |
| 311 | 8 | 33721334 | 33722796 | 1462 | AJ,AKR,BALB,BUB,C3H,CBA,DBA1J,DBA2J,FVB,MA/My,MRL,NOD,NOR,PL,RF,RH2,SJL,SMJ,SWR,TH | 8 | 33721613 | 33721646 | 34 | - | 17.1057 | 6.33E-07 | 0.0794 | TAGCTGTCTCCAGACACTCCAGAAGAGGGTGCCA | | |
| 312 | 8 | 41899353 | 41902713 | 3360 | 129S1,AJ,AKR,BALB,BTBR,BUB,C57L,CBA,CE,FVB,ILNJ2,KK,MA/My,MRL,NOD,NOR,NZB,NZO,NZW,PJ,PL,RF,RH2,SEA,SJL,SMJ,SWR,TH | 8 | 41901617 | 41901651 | 35 | - | 17.6055 | 3.80E-07 | 0.109 | CTGCACTCCCACCTTCTTACCTGTTGGGGCATCTT | | |
| 313 | 8 | 61451998 | 61459097 | 7099 | KK | 8 | 61459055 | 61459088 | 34 | - | 19.3415 | 1.42E-07 | 0.036 | CTGCCCCCACATTCGCCGTCACAAGATGGCGCTG | | |
| 314 | 8 | 64402735 | 64404140 | 1405 | AKR,BALB,C3H,C57L,CBA,DBA1J,DBA2J,MRL.PL.RF.SMJ | 8 | 64403904 | 64403937 | 34 | - | 18.3333 | 2.82E-07 | 0.0534 | CAGCCTTTCCCTAATCAAACAACAGGGGGCAGCT | | |
| 315 | 8 | 72778628 | 72786928 | 8300 | 129S1,BALB,BUB,C58J,CBA,DBA1J,FVB,NZO,PJ,RF,SEA,SMJ,SWR | 8 | 72778911 | 72778929 | 19 | - | 19.2951 | 1.88E-07 | 0.0709 | CCGCCACTAGATGGTGCTA | 4 | ENCFF367UNR,ENCFF099QNE,ENCFF093ZRU,ENCFF821BIA |
| 315 | 8 | 72778628 | 72786928 | 8300 | 129S1,BALB,BUB,C58J,CBA,DBA1J,FVB,NZO,PJ,RF,SEA,SMJ,SWR | 8 | 72786885 | 72786903 | 19 | - | 19.2951 | 1.88E-07 | 0.0709 | CCGCCACTAGATGGTGCTA | 13 | ENCFF367UNR,ENCFF093ZRU,ENCFF430PPJ,ENCFF198SSR,ENCFF099QNE,ENCFF907XEG,ENCFF589ZBJ,ENCFF136ZDK,ENCFF821BIA,ENCFF327LYS,ENCFF089GBC,ENCFF165LXO,ENCFF092NMJ |
| 316 | 8 | 85863834 | 85868848 | 5014 | 129S1,AJ,AKR,BALB,BTBR,C3H,C57L,C58J,CBA,CE,DBA1J,DBA2J,FVB,ILNJ2,KK,MA/My,MRL,NOD,NOR,NZW,PJ,PL,RF,RH2,SEA,SJL,SMJ,SWR | 8 | 85867676 | 85867710 | 35 | - | 17.3761 | 4.35E-07 | 0.118 | GTGCGGGTCAAGCCCACTACCAGTCGGTGGAGGTC | | |
| 317 | 8 | 85865510 | 85868834 | 3324 | 129S1,AKR,BALB,BTBR,C57L,MRL,PJ,PL,SEA | 8 | 85867676 | 85867710 | 35 | - | 17.3761 | 4.35E-07 | 0.118 | GTGCGGGTCAAGCCCACTACCAGTCGGTGGAGGTC | | |
| 318 | 8 | 86901570 | 86906928 | 5358 | AJ,C3H,C58J,CBA,CE,DBA1J,DBA2J,FVB,ILNJ2,KK,MA/My,RH2,SMJ,TH | 8 | 86906797 | 86906830 | 34 | - | 16.9919 | 6.81E-07 | 0.083 | TAGCTGTCTTCAGACACACCAGAAGGGGGCATCA | | |
| 319 | 8 | 116320524 | 116321770 | 1246 | NZB | 8 | 116321464 | 116321498 | 35 | - | 16.1284 | 8.89E-07 | 0.162 | ATGACCTCCCATCTGCAGGCCAGCAGAGAGCGGTC | | |
| 320 | 8 | 120616300 | 120623780 | 7480 | NZB | 8 | 120619854 | 120619872 | 19 | - | 17.1639 | 8.17E-07 | 0.141 | CCTCCACGAGGTGGCGGAA | | |
| 321 | 8 | 124035438 | 124037028 | 1590 | 129S1,AKR,BTBR,C3H,C57L,C58J,CBA,CE,DBA1J,DBA2J,LP,MA/My,MRL,PJ,RF,SMJ,TH | 8 | 124035699 | 124035733 | 35 | - | 24.0275 | 5.22E-09 | 0.00885 | CTGCATGTACACCTGCACACCAGAAGAGGGCGTCA | 4 | ENCFF093ZRU,ENCFF089GBC,ENCFF136ZDK,ENCFF883UPM |
| 322 | 8 | 125533216 | 125537813 | 4597 | AJ.BALB.ILNJ2.RH2.SEA | 8 | 125533688 | 125533721 | 34 | - | 16.7724 | 7.83E-07 | 0.088 | TAGCTGTCTTCAGATGCACCAGAAGAGGGCGTCA | | |
| 322 | 8 | 125533216 | 125537813 | 4597 | AJ,BALB,ILNJ2,RH2,SEA | 8 | 125537137 | 125537155 | 19 | - | 19.5738 | 1.53E-07 | 0.0643 | CCACCACCAGAGGGTGCTG | 19 | ENCFF201JUA,ENCFF092NMJ,ENCFF136ZDK,ENCFF367UNR,ENCFF093ZRU,ENCFF089GBC,ENCFF165LXO,ENCFF589ZBJ,ENCFF198SSR,ENCFF883UPM,ENCFF502WDT,ENCFF327LYS,ENCFF194HVL,ENCFF099QNE,ENCFF907XEG,ENCFF821BIA,ENCFF024JOZ,ENCFF055HAP,ENCFF957SPH |
| 323 | 9 | 5098696 | 5175487 | 76791 | MA/My | 9 | 5146773 | 5146806 | 34 | + | 17.7154 | 4.26E-07 | 0.0552 | ATGGGCCTACATCTGCAGCCAGCAGGTGGAGCTG | 2 | ENCFF883UPM.ENCFF327LYS |
| 323 | 9 | 5098696 | 5175487 | 76791 | MA/My | 9 | 5146788 | 5146806 | 19 | + | 20.6393 | 6.61E-08 | 0.0435 | CAGCCAGCAGGTGGAGCTG | 2 | ENCFF883UPM.ENCFF327LYS |
| 324 | 9 | 28751227 | 28754396 | 3169 | NZO | 9 | 28753692 | 28753726 | 35 | + | 16.4771 | 7.30E-07 | 0.133 | GTGCAGGTGCCCCATTAAGCCATCAGTTGGCTCTT | | |
| 325 | 9 | 46279685 | 46280711 | 1026 | KK | 9 | 46280487 | 46280521 | 35 | + | 20.2661 | 7.32E-08 | 0.0342 | GTGCAGGTGCTTTTGGAAGCCAGAAGAGGGTGGTG | | |
| 326 | 9 | 84186186 | 84189692 | 3506 | 129S1,AJ,BALB,BTBR,C3H,C57L,CE,MRL,NOD,NOR,NZB,PJ,RH2,SEA,TH | 9 | 84186589 | 84186622 | 34 | + | 20.878 | 4.68E-08 | 0.0133 | CGCCAGTACCTGTGGAGGCCAGAAGGGGGCATCA | 12 | ENCFF367UNR,ENCFF821BIA,ENCFF198SSR,ENCFF089GBC,ENCFF907XEG,ENCFF055HAP,ENCFF589ZBJ,ENCFF327LYS,ENCFF883UPM,ENCFF093ZRU,ENCFF099QNE,ENCFF502WDT |
| 326 | 9 | 84186186 | 84189692 | 3506 | 129S1,AJ,BALB,BTBR,C3H,C57L,CE,MRL,NOD,NOR,NZB,PJ,RH2,SEA,TH | 9 | 84186604 | 84186622 | 19 | + | 17.4262 | 6.88E-07 | 0.118 | AGGCCAGAAGGGGGCATCA | 12 | ENCFF367UNR,ENCFF821BIA,ENCFF198SSR,ENCFF089GBC,ENCFF907XEG,ENCFF055HAP,ENCFF589ZBJ,ENCFF327LYS,ENCFF883UPM,ENCFF093ZRU,ENCFF099QNE,ENCFF502WDT |
| 327 | 9 | 88474227 | 88512163 | 37936 | CBA,DBA1J,DBA2J,FVB,MA/My,MRL,NZO,NZW.PL.RF.SJL.SWR | 9 | 88475350 | 88475384 | 35 | + | 21.1743 | 4.01E-08 | 0.0238 | TTGCATGCACACCTGCATGCCAGAAGAGGGCATCA | | |
| 327 | 9 | 88474227 | 88512163 | 37936 | CBA,DBA1J,DBA2J,FVB,MA/My,MRL,NZO,NZW,PL,RF,SJL,SWR | 9 | 88483475 | 88483509 | 35 | + | 16 | 9.55E-07 | 0.152 | TAGCTGTCTAAACAACACCCCAGAAGAGGGCATCA | | |
| 327 | 9 | 88474227 | 88512163 | 37936 | CBA,DBA1J,DBA2J,FVB,MA/My,MRL,NZO,NZW,PL,RF,SJL,SWR | 9 | 88493910 | 88493943 | 34 | + | 18.4797 | 2.56E-07 | 0.0428 | TTGCTGTCTTCAGACACACCAGAAGAGGGCATCA | | |
| 328 | 9 | 98775914 | 98791030 | 15116 | AJ,BALB,C3H,CBA,MRL,NOD,NOR,NZO,NZW,PJ,RH2,SJL,SWR | 9 | 98778903 | 98778937 | 35 | + | 23.9083 | 5.71E-09 | 0.00787 | CTGCAGGAACAAGAACCAACCAGAAGGTGGCACGC | | |
| 328 | 9 | 98775914 | 98791030 | 15116 | AJ,BALB,C3H,CBA,MRL,NOD,NOR,NZO,NZW,PJ,RH2,SJL,SWR | 9 | 98778919 | 98778937 | 19 | + | 17 | 9.07E-07 | 0.131 | CAACCAGAAGGTGGCACGC | | |
| 329 | 9 | 102792273 | 102795819 | 3546 | SMJ | 9 | 102794916 | 102794934 | 19 | + | 18.7049 | 2.88E-07 | 0.0804 | TGACCTGCAGAGGGAGCCA | 9 | ENCFF367UNR,ENCFF089GBC,ENCFF907XEG,ENCFF502WDT,ENCFF055HAP,ENCFF093ZRU,ENCFF198SSR,ENCFF024JOZ,ENCFF327LYS |
| 330 | 9 | 103463291 | 103464850 | 1559 | 129S1,AJ,BALB,BTBR,BUB,C3H,C57L,CBA,FVB,ILNJ2,NOD,NOR,NZB,NZW,PL,RH2,SMJ | 9 | 103463698 | 103463731 | 34 | + | 16.6585 | 8.41E-07 | 0.0807 | GAGCAAATCCTGCCCCTGGCGGCAGGGGTTGGCC | | |
| 331 | 9 | 105091140 | 105093238 | 2098 | 129S1,AJ,AKR,BALB,BTBR,BUB,C3H,C57L,CBA,CE,DBA1J,DBA2J,FVB,ILNJ2,KK,MRL,NOD,NOR,NZB,NZO,NZW,PJ,PL,RF,RH2,SEA.SJL.SMJ.SWR.TH | 9 | 105091878 | 105091911 | 34 | + | 17.7317 | 4.21E-07 | 0.055 | CTGGAGTTTCTGCAAAAGCCAGAAGAGGACAAGC | 11 | ENCFF093ZRU,ENCFF198SSR,ENCFF092NMJ,ENCFF430PPJ,ENCFF099QNE,ENCFF883UPM,ENCFF575MGI,ENCFF327LYS,ENCFF491RJK,ENCFF907XEG,ENCFF089GBC |
| 332 | 9 | 107684620 | 107687707 | 3087 | AKR.DBA1J.DBA2J.KK.NZB.NZO.RF | 9 | 107686862 | 107686895 | 34 | + | 18.8537 | 1.98E-07 | 0.0374 | GTGCAGTATCCTTGGAGGTCAGAAGAGGGTGTCA | | |
| 333 | 9 | 115698240 | 115706809 | 8569 | CE | 9 | 115701742 | 115701775 | 34 | + | 24.3984 | 2.80E-09 | 0.00168 | ATGCAACATCCAAGGAGGCCAGAAGAGGGCGCCA | 13 | ENCFF367UNR,ENCFF821BIA,ENCFF024JOZ,ENCFF327LYS,ENCFF907XEG,ENCFF883UPM,ENCFF589ZBJ,ENCFF093ZRU,ENCFF957SPH,ENCFF052KVO,ENCFF033UQX,ENCFF099QNE,ENCFF136ZDK |
| 333 | 9 | 115698240 | 115706809 | 8569 | CE | 9 | 115701757 | 115701775 | 19 | + | 21.0984 | 4.51E-08 | 0.0361 | AGGCCAGAAGAGGGCGCCA | 14 | ENCFF367UNR,ENCFF821BIA,ENCFF024JOZ,ENCFF327LYS,ENCFF907XEG,ENCFF883UPM,ENCFF589ZBJ,ENCFF093ZRU,ENCFF957SPH,ENCFF052KVO,ENCFF033UQX,ENCFF099QNE,ENCFF136ZDK,ENCFF533APC |
| 333 | 9 | 115698240 | 115706809 | 8569 | CE | 9 | 115699436 | 115699454 | 19 | - | 18.9344 | 2.45E-07 | 0.075 | TTGTCAGTAGGGGGCACTG | 14 | ENCFF821BIA,ENCFF093ZRU,ENCFF092NMJ,ENCFF024JOZ,ENCFF089GBC,ENCFF430PPJ,ENCFF165LXO,ENCFF883UPM,ENCFF198SSR,ENCFF099QNE,ENCFF327LYS,ENCFF575MGI,ENCFF907XEG,ENCFF367UNR |
| 334 | 9 | 123725884 | 123727491 | 1607 | 129S1,C57L,DBA1J,DBA2J | 9 | 123726341 | 123726374 | 34 | + | 24.3252 | 2.98E-09 | 0.00176 | CTGCAATTTCCGTTTTTCCCACCAGAGGGAGACT | 19 | ENCFF367UNR,ENCFF093ZRU,ENCFF099QNE,ENCFF957SPH,ENCFF821BIA,ENCFF089GBC,ENCFF024JOZ,ENCFF907XEG,ENCFF092NMJ,ENCFF327LYS,ENCFF198SSR,ENCFF136ZDK,ENCFF609MDN,ENCFF430PPJ,ENCFF589ZBJ,ENCFF281QMV,ENCFF883UPM,ENCFF165LXO,ENCFF951GOP |
| 335 | 9 | 8287923 | 8299496 | 11573 | AKR.MRL.RF | 9 | 8292742 | 8292775 | 34 | - | 18.0569 | 3.40E-07 | 0.0487 | ATGCAGCTAGAGACATGAGCTCCAGGGGGTACTG | | |
| 336 | 9 | 12129514 | 12139476 | 9962 | AKR.MRL.RF | 9 | 12138704 | 12138722 | 19 | - | 20.5902 | 6.88E-08 | 0.0447 | TGGCCACCAGAGGGAGCCT | | |
| 337 | 9 | 16689217 | 16712607 | 23390 | RF | 9 | 16710670 | 16710704 | 35 | - | 18.5229 | 2.19E-07 | 0.0679 | CTGCAGGTATCCACAGAAGCCAGAAGAGGACATTT | | |
| 338 | 9 | 38324381 | 38329200 | 4819 | 129S1,AJ,AKR,BALB,BTBR,BUB,C3H,CBA,DBA2J,NOD,NOR,PL,RF,RH2,SJL,SMJ,SWR,TH | 9 | 38328402 | 38328435 | 34 | - | 16.5691 | 8.90E-07 | 0.0831 | GAGCATGACCACAAGTCCACAGCAGATGGCAACA | | |
| 339 | 9 | 40543217 | 40544565 | 1348 | 129S1,AJ,BALB,BTBR,BUB,C3H,C57L,CBA,CE,DBA1J,FVB,ILNJ2,KK,MA/My,NZB,NZO,NZW,PJ,PL,RH2,SEA,SJL,SMJ,SWR,TH | 9 | 40544368 | 40544401 | 34 | - | 16.4472 | 9.61E-07 | 0.0841 | TAGCTGTCTTCAGACACACCAGAAGAGGGCGTCA | | |
| 340 | 9 | 42079372 | 42080794 | 1422 | BTBR.DBA1J.KK.MA/My.NZO | 9 | 42079964 | 42079997 | 34 | - | 16.4959 | 9.32E-07 | 0.0841 | CTGCCATTTTAAATAGGTCCAACAGAGGACACGG | | |
| 341 | 9 | 50537573 | 50543309 | 5736 | 129S1.C57L | 9 | 50540792 | 50540825 | 34 | - | 16.4472 | 9.61E-07 | 0.0841 | TAGCTGTCTTCAGACACACCAGAAGAGGGCGTCA | | |
| 342 | 9 | 66047483 | 66048500 | 1017 | FVB.SJL | 9 | 66048462 | 66048495 | 34 | - | 16.7724 | 7.83E-07 | 0.0768 | TAGCTGTCTTCAGATGCACCAGAAGAGGGCGTCA | | |
| 343 | 9 | 90809291 | 90824213 | 14922 | AJ,AKR,BALB,C3H,CE,FVB,KK,NZW,RH2,SJL.SWR.TH | 9 | 90820203 | 90820236 | 34 | - | 16.561 | 8.95E-07 | 0.0832 | TGGCAGGTCTCAGAGAAGCCGCTAGATGGCCCAC | | |
| 344 | 9 | 99437904 | 99442426 | 4522 | NOD.NOR | 9 | 99439557 | 99439590 | 34 | - | 21.0488 | 4.12E-08 | 0.0123 | CTGCTGTCTTCAGACACACCAGAAGAGGGCATCA | 1 | ENCFF093ZRU |
| 345 | 9 | 105359744 | 105363341 | 3597 | FVB.SJL.SWR | 9 | 105360222 | 105360240 | 19 | - | 17.3607 | 7.18E-07 | 0.12 | AGGCCAGAAGAGGGTGCCA | | |
| 346 | 9 | 109029516 | 109031200 | 1684 | CBA.KK.NZB.NZO.NZW | 9 | 109030439 | 109030472 | 34 | - | 19.6504 | 1.14E-07 | 0.0255 | TAGCTGTCCTCAGATACTCCAGAAGAGGGCACCA | | |
| 347 | 9 | 117816612 | 117825965 | 9353 | 129S1,AKR,BUB,C57L,KK,LP,MA/My,MRL,NZW,RF,RH2 | 9 | 117824608 | 117824642 | 35 | - | 17.2385 | 4.72E-07 | 0.104 | GTGCACCTTCCCTGCCAGGCCTGCAGAGAGTGCTC | | |
| 348 | 9 | 117904826 | 117906636 | 1810 | 129S1,AJ,BALB,C3H,C57L,FVB,ILNJ2,KK,NZW,PJ,SEA,SJL,SWR,TH | 9 | 117906615 | 117906633 | 19 | - | 19.7213 | 1.37E-07 | 0.056 | CTGCCAGCAGGGGGAAGTC | | |
| 349 | 9 | 122016531 | 122018498 | 1967 | CE | 9 | 122017972 | 122018005 | 34 | - | 21.7317 | 2.46E-08 | 0.00852 | CAGCAGTGCCTCTCTAGGCCACCAGGGGAAGTGC | 19 | ENCFF024JOZ,ENCFF136ZDK,ENCFF821BIA,ENCFF089GBC,ENCFF092NMJ,ENCFF367UNR,ENCFF093ZRU,ENCFF165LXO,ENCFF201JUA,ENCFF430PPJ,ENCFF055HAP,ENCFF589ZBJ,ENCFF502WDT,ENCFF052KVO,ENCFF907XEG,ENCFF099QNE,ENCFF327LYS,ENCFF065ODV,ENCFF198SSR |
| 350 | 9 | 123275284 | 123276751 | 1467 | 129S1,C57L | 9 | 123276362 | 123276380 | 19 | - | 21.1148 | 4.45E-08 | 0.0361 | AGGCCAGAAGGGGGCACTG | 10 | ENCFF327LYS,ENCFF092NMJ,ENCFF907XEG,ENCFF089GBC,ENCFF821BIA,ENCFF198SSR,ENCFF093ZRU,ENCFF367UNR,ENCFF055HAP,ENCFF502WDT |
| 350 | 9 | 123275284 | 123276751 | 1467 | 129S1,C57L | 9 | 123276362 | 123276395 | 34 | - | 27.8618 | 1.09E-10 | 0.000241 | GTGCAGTGTGCACAGAGGCCAGAAGGGGGCACTG | 10 | ENCFF327LYS,ENCFF092NMJ,ENCFF907XEG,ENCFF089GBC,ENCFF821BIA,ENCFF198SSR,ENCFF093ZRU,ENCFF367UNR.ENCFF055HAP.ENCFF502WDT |
| 351 | 10 | 5667712 | 5676755 | 9043 | 129S1,AJ,AKR,BALB,BTBR,BUB,C3H,C57L,CBA,CE,DBA1J,DBA2J,FVB,ILNJ2,KK,LP,MA/My,MRL,NOD,NOR,NZB,NZO,NZW,PJ,PL,RF,RH2,SEA,SJL,SMJ,SWR,TH | 10 | 5673168 | 5673186 | 19 | + | 17.1639 | 8.17E-07 | 0.153 | CCTCCACGAGGTGGCGGAA | | |
| 352 | 10 | 6119591 | 6125951 | 6360 | C3H,CBA,CE,DBA1J,DBA2J,FVB,LP,MA/My,MRL,NZB,NZO,NZW,SWR | 10 | 6125158 | 6125191 | 34 | + | 16.4634 | 9.51E-07 | 0.108 | TAGCTGTCTTCAGACGCACCAGAAGAGGGCATCA | | |
| 353 | 10 | 7447953 | 7449580 | 1627 | BUB | 10 | 7449049 | 7449082 | 34 | + | 17.3008 | 5.58E-07 | 0.0844 | CTGTAGTCCCAGCACTGGGCACATGGAGGTAGGA | | |
| 354 | 10 | 20247050 | 20249241 | 2191 | FVB,NOD,NOR,TH | 10 | 20247939 | 20247973 | 35 | + | 20.8899 | 4.85E-08 | 0.0343 | GTGAAGTTACTCACAGAGACCACAAGAGGGTGCTG | 9 | ENCFF367UNR,ENCFF093ZRU,ENCFF907XEG,ENCFF055HAP,ENCFF821BIA,ENCFF099QNE,ENCFF533APC,ENCFF052KVO,ENCFF502WDT |
| 355 | 10 | 22007795 | 22015276 | 7481 | LP | 10 | 22010389 | 22010423 | 35 | + | 16.1743 | 8.66E-07 | 0.176 | TGGCAGTTCAGTTCATAGGTCAGCAGAGGCAGCTC | | |
| 356 | 10 | 22381714 | 22393564 | 11850 | AJ,BTBR,C57L,FVB,NOD,NZO | 10 | 22387247 | 22387280 | 34 | + | 17.748 | 4.17E-07 | 0.0738 | GTGCTGTTTCTGATTGGTCCACAGGGTGAAGCTC | | |
| 357 | 10 | 22394322 | 22397009 | 2687 | 129S1,AJ,BALB,BTBR,BUB,C57L,FVB,KK,MA/My,NOD,NOR,NZO,PL,RH2,SJL | 10 | 22395937 | 22395970 | 34 | + | 18.6748 | 2.24E-07 | 0.0551 | ATGCTGTCTTCAGACACACCAGAAGAGGGCATCA | | |

| | | | | | | | | | | | | | | | | |
|---|---|---|---|---|---|---|---|---|---|---|---|---|---|---|---|---|
| 358 | 10 | 22610173 | 22619222 | 9049 | 129S1,AJ,AKR,BALB,BTBR,BUB,C57L,FVB,ILNJ2,KK,LP,MA/My,NOD,NOR,NZB,NZO,NZW.PL.RF.RH2.SJL.SWR.TH | 10 | 22615634 | 22615652 | 19 | + | 17.1639 | 8.17E-07 | 0.153 | CCTCCACGAGGTGGCGGAA | | |
| 359 | 10 | 23153987 | 23155187 | 1200 | 129S1,AKR,BTBR,C3H,C57L,FVB,ILNJ2,KK,LP,MRL,NOD,NZB,NZO,NZW,RF,SMJ,SWR,TH | 10 | 23154880 | 23154914 | 35 | + | 15.9908 | 9.60E-07 | 0.184 | CTGCACTGACCCCCTATGCCCACCTGGGTGGTCTT | | |
| 360 | 10 | 24724192 | 24730560 | 6368 | 129S1.BTBR.C57L.NZO.NZW | 10 | 24726443 | 24726477 | 35 | + | 17.4679 | 4.13E-07 | 0.121 | TGGCAGTGTGGCCAGCTACCCAGCAGATGCCTACC | | |
| 361 | 10 | 41312639 | 41321624 | 8985 | 129S1.BTBR.C57L.KK.MA/Mv | 10 | 41318056 | 41318074 | 19 | + | 17.1639 | 8.17E-07 | 0.153 | CCTCCACGAGGTGGCGGAA | | |
| 362 | 10 | 41928524 | 41931471 | 2947 | 129S1.BTBR.C57L.KK | 10 | 41929030 | 41929063 | 34 | + | 17.561 | 4.71E-07 | 0.0776 | CTGCCAAACCAGCACGCGCCAGGGGTGGGCACCA | | |
| 363 | 10 | 76065594 | 76121703 | 56109 | DBA1J | 10 | 76088758 | 76088792 | 35 | + | 16.3486 | 7.85E-07 | 0.168 | TTGCATGTACACCTCTACACTAGAAGGGGGCACCA | | |
| 363 | 10 | 76065594 | 76121703 | 56109 | DBA1J | 10 | 76107256 | 76107274 | 19 | + | 19.4918 | 1.63E-07 | 0.0697 | CGGCCAGCAGAGGGAGACA | 31 | ENCFF093ZRU,ENCFF367UNR,ENCFF430PPJ,ENCFF099QNE,ENCFF327LYS,ENCFF907XEG,ENCFF951GOP,ENCFF609MDN,ENCFF575MGI,ENCFF198SSR,ENCFF089GBC,ENCFF136ZDK,ENCFF092NMJ,ENCFF821BIA,ENCFF024JOZ,ENCFF165LXO,ENCFF097CAQ,ENCFF502WDT,ENCFF055HAP,ENCFF194HVL,ENCFF491RJK,ENCFF611HDQ,ENCFF957SPH,ENCFF533APC,ENCFF589ZBJ,ENCFF281QMV,ENCFF883UPM,ENCFF210BTO.ENCFF349BUL.ENCFF052KVO.ENCFF142CNG |
| 363 | 10 | 76065594 | 76121703 | 56109 | DBA1J | 10 | 76065646 | 76065664 | 19 | - | 20.3607 | 8.29E-08 | 0.0512 | AGGCCAGAAGAGGGCACCA | 15 | ENCFF093ZRU,ENCFF092NMJ,ENCFF367UNR,ENCFF089GBC,ENCFF589ZBJ,ENCFF099QNE,ENCFF198SSR,ENCFF327LYS,ENCFF065ODV,ENCFF024JOZ,ENCFF907XEG,ENCFF055HAP,ENCFF821BIA,ENCFF502WDT.ENCFF883UPM |
| 363 | 10 | 76065594 | 76121703 | 56109 | DBA1J | 10 | 76065646 | 76065680 | 35 | - | 22.7706 | 1.32E-08 | 0.0157 | GTGCTTGTACACCAGTAGGCCAGAAGAGGGCACCA | 16 | ENCFF093ZRU,ENCFF092NMJ,ENCFF367UNR,ENCFF089GBC,ENCFF589ZBJ,ENCFF099QNE,ENCFF198SSR,ENCFF327LYS,ENCFF065ODV,ENCFF024JOZ,ENCFF907XEG,ENCFF055HAP,ENCFF821BIA,ENCFF502WDT.ENCFF883UPM.ENCFF052KVO |
| 363 | 10 | 76065594 | 76121703 | 56109 | DBA1J | 10 | 76075414 | 76075447 | 34 | - | 18.2602 | 2.97E-07 | 0.0608 | TTCCAGTGCCCATGGAGGCCAGATGAGGGCATCA | 2 | ENCFF367UNR.ENCFF198SSR |
| 363 | 10 | 76065594 | 76121703 | 56109 | DBA1J | 10 | 76096696 | 76096729 | 34 | - | 17.7886 | 4.06E-07 | 0.0726 | CAGCCACTCACATTATGGACAGCAGAGGGAGCAA | 14 | ENCFF093ZRU,ENCFF821BIA,ENCFF367UNR,ENCFF883UPM,ENCFF099QNE,ENCFF089GBC,ENCFF609MDN,ENCFF327LYS,ENCFF281QMV,ENCFF907XEG,ENCFF024JOZ,ENCFF430PPJ,ENCFF092NMJ,ENCFF198SSR |
| 363 | 10 | 76065594 | 76121703 | 56109 | DBA1J | 10 | 76106573 | 76106606 | 34 | - | 17.1707 | 6.07E-07 | 0.0883 | CAGCACTGTTGTGTCAGGGCAGCAGAGGCCAGCA | | |
| 364 | 10 | 93446669 | 93452954 | 6285 | 129S1,AJ,AKR,BALB,BTBR,BUB,C3H,C57L,C58J,CBA,CE,DBA1J,DBA2J,FVB,ILNJ2,KK,LP,MA/My,NOD,NOR,NZO,NZW,PJ,PL,RF,RH2,SEA,SJL,SMJ,SWR,TH | 10 | 93447419 | 93447453 | 35 | + | 19.211 | 1.43E-07 | 0.0651 | GTTCACTGACAGCTAATCAACACTAGATGGCGCTC | 29 | ENCFF367UNR,ENCFF093ZRU,ENCFF099QNE,ENCFF201JUA,ENCFF821BIA,ENCFF210BTO,ENCFF089GBC,ENCFF024JOZ,ENCFF907XEG,ENCFF327LYS,ENCFF198SSR,ENCFF589ZBJ,ENCFF092NMJ,ENCFF136ZDK,ENCFF430PPJ,ENCFF165LXO,ENCFF504JDS,ENCFF089EWX,ENCFF194HVL,ENCFF491RJK,ENCFF097CAQ,ENCFF951GOP,ENCFF611HDQ,ENCFF281QMV,ENCFF575MGI,ENCFF957SPH,ENCFF883UPM,ENCFF609MDN.ENCFF316FYN |
| 364 | 10 | 93446669 | 93452954 | 6285 | 129S1,AJ,AKR,BALB,BTBR,BUB,C3H,C57L,C58J,CBA,CE,DBA1J,DBA2J,FVB,ILNJ2,KK,LP,MA/My,NOD,NOR,NZO,NZW,PJ,PL,RF,RH2,SEA,SJL,SMJ,SWR,TH | 10 | 93447435 | 93447453 | 19 | + | 17.459 | 6.74E-07 | 0.14 | TCAACACTAGATGGCGCTC | 29 | ENCFF367UNR,ENCFF093ZRU,ENCFF099QNE,ENCFF201JUA,ENCFF821BIA,ENCFF210BTO,ENCFF089GBC,ENCFF024JOZ,ENCFF907XEG,ENCFF327LYS,ENCFF198SSR,ENCFF589ZBJ,ENCFF092NMJ,ENCFF136ZDK,ENCFF430PPJ,ENCFF165LXO,ENCFF504JDS,ENCFF089EWX,ENCFF194HVL,ENCFF491RJK,ENCFF097CAQ,ENCFF951GOP,ENCFF611HDQ,ENCFF281QMV,ENCFF575MGI,ENCFF957SPH,ENCFF883UPM,ENCFF609MDN.ENCFF316FYN |
| 365 | 10 | 99975145 | 99980410 | 5265 | AKR,BUB,C3H,CBA,FVB,MA/My,NOD,NOR,NZB.PL.RF.SMJ.SWR.TH | 10 | 99977809 | 99977842 | 34 | + | 18.935 | 1.88E-07 | 0.0499 | CAGCTGTCTTCAAACGCACCAGAAGAGGGCATCA | | |
| 366 | 10 | 110661782 | 110692028 | 30246 | NZB | 10 | 110667962 | 110667996 | 35 | + | 16.3211 | 7.98E-07 | 0.169 | CTGCAGCATTAGGACGGTTACTCTAGAGGGAGGTC | | |
| 367 | 10 | 116598096 | 116599983 | 1887 | SMJ | 10 | 116599104 | 116599138 | 35 | + | 17.4862 | 4.08E-07 | 0.12 | TTGCAATTGAAGGTCCAGGCCTGAAGGGGTCATGC | | |
| 368 | 10 | 3516435 | 3519398 | 2963 | ILNJ2 | 10 | 3518009 | 3518027 | 19 | - | 18.0492 | 4.54E-07 | 0.12 | GGGCCAGAAGAGGGCGATG | 19 | ENCFF024JOZ,ENCFF821BIA,ENCFF198SSR,ENCFF092NMJ,ENCFF165LXO,ENCFF589ZBJ,ENCFF907XEG,ENCFF883UPM,ENCFF093ZRU,ENCFF502WDT,ENCFF367UNR,ENCFF194HVL,ENCFF033UQX,ENCFF957SPH.ENCFF052KVO.ENCFF327LYS.ENCFF533APC.ENCFF136ZDK.ENCFF099QNE |
| 368 | 10 | 3516435 | 3519398 | 2963 | ILNJ2 | 10 | 3518009 | 3518042 | 34 | - | 26.3984 | 4.60E-10 | 0.000687 | GTGCAGTACTCATAGGGGCCAGAAGAGGGCGATG | 19 | ENCFF024JOZ,ENCFF821BIA,ENCFF198SSR,ENCFF092NMJ,ENCFF165LXO,ENCFF589ZBJ,ENCFF907XEG,ENCFF883UPM,ENCFF093ZRU,ENCFF502WDT,ENCFF367UNR,ENCFF194HVL,ENCFF033UQX,ENCFF957SPH.ENCFF052KVO.ENCFF327LYS.ENCFF533APC.ENCFF136ZDK.ENCFF099QNE |
| 369 | 10 | 8419894 | 8427032 | 7138 | 129S1,AJ,AKR,BALB,BTBR,BUB,C3H,C57L,CBA,CE,DBA1J,DBA2J,FVB,ILNJ2,KK,LP,MA/My,MRL,NOD,NOR,NZB,NZO,NZW,PJ,PL,RF,RH2,SEA,SJL,SMJ,SWR,TH | 10 | 8421638 | 8421656 | 19 | - | 17.1639 | 8.17E-07 | 0.153 | CCTCCACGAGGTGGCGGAA | | |
| 370 | 10 | 11362943 | 11368756 | 5813 | AJ,AKR,BALB,BTBR,C3H,C57L,CBA,CE,KK,MRL.PJ.PL.RF.RH2.SEA.SJL.SMJ | 10 | 11363201 | 11363234 | 34 | - | 16.8862 | 7.28E-07 | 0.096 | CCGCGATAACATTTGCCATCACAAGATGGCGCCG | 2 | ENCFF821BIA,ENCFF367UNR |
| 371 | 10 | 14706214 | 14710819 | 4605 | DBA1J.DBA2J | 10 | 14709800 | 14709833 | 34 | - | 16.439 | 9.66E-07 | 0.108 | ATGCAGCTAGAGACAAGAGCTCCAGGGGGTACTG | | |
| 372 | 10 | 85658030 | 85673227 | 15197 | BUB | 10 | 85665250 | 85665283 | 34 | - | 17.1138 | 6.30E-07 | 0.0904 | CTCCAGAACCAGGACCGACCTCCAGCGGGCAGGA | 1 | ENCFF093ZRU |
| 372 | 10 | 85658030 | 85673227 | 15197 | BUB | 10 | 85672418 | 85672436 | 19 | - | 19.8361 | 1.25E-07 | 0.0608 | CAACCAGAAGAGGGCACTA | 1 | ENCFF093ZRU |
| 372 | 10 | 85658030 | 85673227 | 15197 | BUB | 10 | 85672418 | 85672452 | 35 | - | 23.2385 | 9.40E-09 | 0.013 | GTGCAGGCACCTATGACAACCAGAAGAGGGCACTA | 1 | ENCFF093ZRU |
| 373 | 10 | 87908050 | 87909941 | 1891 | BUB.FVB | 10 | 87908907 | 87908941 | 35 | - | 15.9541 | 9.80E-07 | 0.186 | TTGCAATTGAAGGCCCAGGACTGAAGGGGTCACAC | | |
| 374 | 10 | 92586396 | 92590013 | 3617 | 129S1,AJ,AKR,BALB,BTBR,BUB,C3H,C57L,C58J,CBA,CE,DBA1J,DBA2J,FVB,ILNJ2,KK,LP,MA/My,MRL,NOD,NOR,NZO,NZW,PJ,PL,RF,RH2,SEA,SJL,SMJ,SWR,TH | 10 | 92586696 | 92586729 | 34 | - | 17.9106 | 3.74E-07 | 0.07 | CCGCCCTCACATTCGCCGTCACAAGATGGCGCTG | | |
| 374 | 10 | 92586396 | 92590013 | 3617 | 129S1,AJ,AKR,BALB,BTBR,BUB,C3H,C57L,C58J,CBA,CE,DBA1J,DBA2J,FVB,ILNJ2,KK,LP,MA/My,MRL,NOD,NOR,NZO,NZW,PJ,PL,RF,RH2,SEA,SJL,SMJ,SWR,TH | 10 | 92589971 | 92590004 | 34 | - | 17.9106 | 3.74E-07 | 0.07 | CCGCCCTCACATTCGCCGTCACAAGATGGCGCTG | | |
| 375 | 10 | 94389306 | 94393572 | 4266 | PL | 10 | 94391227 | 94391260 | 34 | - | 17.0569 | 6.53E-07 | 0.0917 | ATGCAGGTCCCCCGAGGTCTAGGAGGTGCCACTG | | |
| 376 | 10 | 116964985 | 116977667 | 12682 | SMJ | 10 | 116968787 | 116968820 | 34 | - | 18.122 | 3.25E-07 | 0.0647 | TAGCTGTTTTCAGAGACACCAGAAGAGGGCATTA | 1 | ENCFF093ZRU |
| 377 | 11 | 7791883 | 7795001 | 3118 | KK | 11 | 7792329 | 7792363 | 35 | + | 16.2752 | 8.19E-07 | 0.115 | ATGCAATCAGACGTGCCTTCCACAGGATGCCAGCA | | |
| 378 | 11 | 8870293 | 8879277 | 8984 | 129S1,AJ,AKR,BALB,BTBR,BUB,C3H,C57L,C58J,CBA,CE,DBA1J,DBA2J,FVB,ILNJ2,KK,MA/My,MRL,NOD,NZB,NZO,PJ,PL,RF,SEA,SJL.SMJ.SWR.TH | 11 | 8875709 | 8875727 | 19 | + | 17.1639 | 8.17E-07 | 0.0957 | CCTCCACGAGGTGGCGGAA | | |
| 379 | 11 | 15748458 | 15750981 | 2523 | AKR.C3H.MA/Mv.MRL.NZB.NZW.RF | 11 | 15750578 | 15750596 | 19 | + | 17.459 | 6.74E-07 | 0.087 | CAGCCAGGAGGTGGAGCTG | | |
| 380 | 11 | 28375579 | 28434583 | 59004 | PL | 11 | 28382537 | 28382571 | 35 | + | 16.5596 | 6.97E-07 | 0.107 | CTGTAGGGTCTCCATACAACCAGCAGAAGTAGGTT | | |
| 380 | 11 | 28375579 | 28434583 | 59004 | PL | 11 | 28427283 | 28427317 | 35 | - | 17.7615 | 3.47E-07 | 0.073 | CTGCACTGCCTTCTCAAACCCTGCAGGGAGCCTCA | | |
| 381 | 11 | 43340607 | 43345457 | 4850 | FVB,SJL,SWR | 11 | 43343279 | 43343297 | 19 | + | 17.4098 | 6.96E-07 | 0.0881 | TAACCAGTAGGTGTCACTC | 14 | ENCFF367UNR,ENCFF093ZRU,ENCFF092NMJ,ENCFF165LXO,ENCFF430PPJ,ENCFF099QNE,ENCFF327LYS,ENCFF907XEG,ENCFF821BIA,ENCFF055HAP,ENCFF089GBC,ENCFF024JOZ,ENCFF502WDT,ENCFF198SSR |
| 382 | 11 | 48716159 | 48717570 | 1411 | CE.NZO.NZW.PJ.SEA | 11 | 48717462 | 48717496 | 35 | + | 22.7248 | 1.36E-08 | 0.00983 | GTGCTGGGCCGGGGCCGGACCGGGAGGGGGCGGCG | | |
| 383 | 11 | 50336005 | 50339169 | 3164 | B10D2,B10J,BUB,C58J,CE,DBA2J,KK,MA/My,NOD,NOR,NZB,NZO,NZW,PJ,PL,SEA,SJL,SWR.TH | 11 | 50339131 | 50339164 | 34 | + | 19.561 | 1.21E-07 | 0.0223 | CAGCATCGTTCTTCCTGGCCAGTTGGGGGCGCGC | 5 | ENCFF092NMJ,ENCFF093ZRU,ENCFF907XEG,ENCFF099QNE,ENCFF821BIA |
| 383 | 11 | 50336005 | 50339169 | 3164 | B10D2,B10J,BUB,C58J,CE,DBA2J,KK,MA/My,NOD,NOR,NZB,NZO,NZW,PJ,PL,SEA,SJL,SWR.TH | 11 | 50339146 | 50339164 | 19 | + | 17.5574 | 6.32E-07 | 0.0841 | TGGCCAGTTGGGGGCGCGC | 5 | ENCFF092NMJ,ENCFF093ZRU,ENCFF907XEG,ENCFF099QNE,ENCFF821BIA |
| 384 | 11 | 51555030 | 51561912 | 6882 | B10D2,B10J,BUB,C58J,CE,DBA2J,KK,MA/My.NOD.NOR.NZB.PJ.SEA.SWR.TH | 11 | 51555560 | 51555594 | 35 | + | 17.2294 | 4.74E-07 | 0.0871 | CTGCACGTATACCTGCATGTCAGAAGAGGGTATCA | | |
| 385 | 11 | 58580417 | 58587147 | 6730 | AJ,BTBR,BUB,C3H,C57L,CBA,DBA1J,DBA2J,FVB,ILNJ2,KK,MA/My,MRL,NOR,NZB,NZO,NZW.PJ.SEA.SJL.SWR.TH | 11 | 58580433 | 58580466 | 34 | + | 17.5041 | 4.89E-07 | 0.0482 | GAGCAGATGTGGCCGCGGCCCAAAGATGGCACCC | | |
| 386 | 11 | 60532890 | 60548505 | 15615 | C57L | 11 | 60536080 | 60536113 | 34 | + | 18.4797 | 2.56E-07 | 0.0347 | TTGCTGTCTTCAGACACACCAGAAGAGGGCATCA | | |
| 387 | 11 | 61764061 | 61765118 | 1057 | DBA1J.DBA2J.NOR | 11 | 61764065 | 61764098 | 34 | + | 16.4472 | 9.61E-07 | 0.067 | TAGCTGTCTTCAGACACACCAGAAGAGGGCGTCA | | |
| 388 | 11 | 76439631 | 76448680 | 9049 | 129S1,AJ,AKR,BALB,BTBR,BUB,C3H,C57L,C58J,CBA,CE,DBA1J,DBA2J,FVB,ILNJ2,KK,MA/My,MRL,NOD,NOR,NZB,NZO,NZW,PJ,PL,RF,RH2,SEA,SJL,SMJ,SWR,TH | 11 | 76445091 | 76445109 | 19 | + | 17.1639 | 8.17E-07 | 0.0957 | CCTCCACGAGGTGGCGGAA | | |
| 389 | 11 | 86773137 | 86782182 | 9045 | 129S1,AJ,AKR,BTBR,BUB,C3H,C57L,C58J,CBA,CE,DBA1J,DBA2J,MRL,NZB,NZO,NZW.PJ.PL.RF.SEA.SJL.SWR | 11 | 86778597 | 86778615 | 19 | + | 17.1639 | 8.17E-07 | 0.0957 | CCTCCACGAGGTGGCGGAA | | |
| 390 | 11 | 95839915 | 95842682 | 2767 | KK.MA/Mv.MRL.RF.SWR.TH | 11 | 95842532 | 95842565 | 34 | + | 17.8455 | 3.91E-07 | 0.0425 | TAGCTGTCCTCAGACACTCCAGAAGAGGGCGTCA | | |
| 391 | 11 | 102974483 | 102983530 | 9047 | 129S1,AJ,AKR,BALB,BTBR,BUB,C3H,C57L,CBA,CE,DBA1J,DBA2J,FVB,ILNJ2,KK,MRL,NOD,NOR,NZO,NZW,RF,RH2,SEA,SJL,SMJ | 11 | 102979945 | 102979963 | 19 | + | 17.1639 | 8.17E-07 | 0.0957 | CCTCCACGAGGTGGCGGAA | | |
| 392 | 11 | 103399750 | 103420209 | 20459 | BALB,CE,FVB,ILNJ2,KK,MRL,NOD,NOR,NZB.SEA.SJL.SMJ | 11 | 103411552 | 103411585 | 34 | + | 18.6423 | 2.29E-07 | 0.0335 | ATGCAGTGCCCATAGAGGCTAGAAGAAGGCATCA | | |
| 393 | 11 | 104292216 | 104293342 | 1126 | AKR.NZO.NZW.RF.RH2 | 11 | 104292883 | 104292901 | 19 | + | 16.9508 | 9.36E-07 | 0.1 | ACACCAGAAGAGGGCAGTG | 2 | ENCFF093ZRU.ENCFF099QNE |
| 394 | 11 | 111094965 | 111117212 | 22247 | AKR.RH2 | 11 | 111096172 | 111096205 | 34 | + | 18.3089 | 2.87E-07 | 0.0357 | ATGCAGCTAGAGACACGAGCTCCAGGGGGTACTG | | |
| 395 | 11 | 117299837 | 117302699 | 2862 | SMJ | 11 | 117301581 | 117301615 | 35 | + | 17.6239 | 3.76E-07 | 0.0758 | CTGTAGCCGTCTTCACACACCAGAAGAGGGCATCA | 1 | ENCFF093ZRU |
| 396 | 11 | 6694896 | 6709348 | 14452 | KK | 11 | 6702961 | 6702979 | 19 | - | 17.1639 | 8.17E-07 | 0.0957 | CCTCCACGAGGTGGCGGAA | | |
| 397 | 11 | 6699417 | 6708391 | 8974 | 129S1,AJ,AKR,B10D2,B10J,BALB,BTBR,C3H,C57L,C58J,FVB,MA/My,MRL,NOD,NOR,NZB.NZW.PL.RF.SEA.TH | 11 | 6702961 | 6702979 | 19 | - | 17.1639 | 8.17E-07 | 0.0957 | CCTCCACGAGGTGGCGGAA | | |
| 398 | 11 | 8220598 | 8224260 | 3662 | AJ,AKR,B10D2,B10J,BALB,BTBR,BUB,C3H,C57L,C58J,CE,KK,MA/My,MRL,NOD,NOR,NZB.PJ.PL.RF.SEA.SJL.SWR | 11 | 8223457 | 8223490 | 34 | - | 17.6748 | 4.37E-07 | 0.0453 | GAGCATGACCACAAGTCCACAGCAGGTGGCAACG | | |
| 399 | 11 | 10148082 | 10151004 | 2922 | MA/Mv.NZB.NZO | 11 | 10148078 | 10148112 | 35 | - | 19.4679 | 1.22E-07 | 0.0371 | CTGCCGCCACACCCGCTCCCCACAAGAGGGAAGGC | | |
| 400 | 11 | 31640225 | 31642354 | 2129 | AKR,BUB,C3H,CBA,CE,FVB,ILNJ2,KK,MRL,NOD.PJ.PL.SEA.SJL.SMJ.SWR.TH | 11 | 31642250 | 31642284 | 35 | - | 17.211 | 4.79E-07 | 0.0877 | CTGCAGTTCTATGGGGGGTCCTGTGGAGGTGGCTG | | |
| 401 | 11 | 40013869 | 40014938 | 1069 | 129S1,AJ,AKR,BALB,BUB,C3H,C57L,CBA,DBA1J.MRL.PL | 11 | 40013895 | 40013928 | 34 | - | 16.9268 | 7.10E-07 | 0.0585 | ATGCAGTTAGAGACATGAGCTCTGGGGGGTACTG | | |
| 402 | 11 | 45615252 | 45623164 | 7912 | 129S1,AJ,AKR,B10D2,B10J,BALB,BTBR,BUB,C3H,C57L,C58J,CBA,CE,DBA1J,DBA2J,FVB,ILNJ2,KK,MA/My,MRL,NOD,NZB,NZO,NZW,PJ,PL,RF,RH2,SEA,SJL,SMJ,SWR,TH | 11 | 45615601 | 45615635 | 35 | - | 16.2569 | 8.27E-07 | 0.116 | CGGCGGGTCCCGGACCGGTCCGGAAGGTGGCCGGC | | |
| 403 | 11 | 58990668 | 58994253 | 3585 | CE,NZB,NZO,SMJ | 11 | 58993343 | 58993376 | 34 | - | 28.0569 | 8.91E-11 | 0.000177 | ATGCAGTGCCCACAGAGGCCATAAGAGGGCGCCA | 18 | ENCFF099QNE,ENCFF201JUA,ENCFF136ZDK,ENCFF589ZBJ,ENCFF327LYS,ENCFF198SSR,ENCFF907XEG,ENCFF367UNR,ENCFF093ZRU,ENCFF055HAP,ENCFF883UPM,ENCFF533APC,ENCFF033UQX,ENCFF821BIA.ENCFF065ODV.ENCFF142CNG.ENCFF052KVO.ENCFF502WDT |
| 404 | 11 | 59032525 | 59035385 | 2860 | CE.SMJ | 11 | 59034656 | 59034689 | 34 | - | 20.1951 | 7.72E-08 | 0.0164 | CAGCTGTCTCCAGACACACCAGAAGAGGGCATCA | | |
| 405 | 11 | 60469470 | 60476834 | 7364 | 129S1,AKR,BTBR,BUB,C57L,CE,FVB,ILNJ2,LP,MA/My,NOD,NZW,PL,RF,SEA,SMJ,SWR | 11 | 60471361 | 60471379 | 19 | - | 17.1639 | 8.17E-07 | 0.0957 | CCTCCACGAGGTGGCGGAA | | |
| 406 | 11 | 69178082 | 69180400 | 2318 | AKR.PL.RF | 11 | 69179734 | 69179768 | 35 | - | 23.2294 | 9.46E-09 | 0.00768 | CTGCATGTAAGCCTACATGCCAGAAGAGGGCATTG | | |
| 407 | 11 | 95831825 | 95833551 | 1726 | DBA1J.DBA2J.ILNJ2.NZB.NZO.PJ.SEA | 11 | 95832638 | 95832671 | 34 | - | 17.7317 | 4.21E-07 | 0.0444 | TAGCTGTCCTCAGATACACCAGAAGAGGGCATCA | | |
| 408 | 11 | 102507104 | 102510538 | 3434 | AJ,CBA | 11 | 102510471 | 102510489 | 19 | - | 17.8033 | 5.37E-07 | 0.0792 | AGGCCAGTAGAGGGCGAAG | 14 | ENCFF093ZRU,ENCFF367UNR,ENCFF907XEG,ENCFF136ZDK,ENCFF821BIA,ENCFF327LYS,ENCFF589ZBJ,ENCFF052KVO,ENCFF198SSR,ENCFF502WDT,ENCFF883UPM,ENCFF055HAP,ENCFF065ODV,ENCFF142CNG |
| 408 | 11 | 102507104 | 102510538 | 3434 | AJ,CBA | 11 | 102510471 | 102510504 | 34 | - | 23.2602 | 7.30E-09 | 0.00289 | AGGCAGTACCTGAGGAGGCCAGTAGAGGGCGAAG | 14 | ENCFF093ZRU,ENCFF367UNR,ENCFF907XEG,ENCFF136ZDK,ENCFF821BIA,ENCFF327LYS,ENCFF589ZBJ,ENCFF052KVO,ENCFF198SSR,ENCFF502WDT,ENCFF883UPM,ENCFF055HAP,ENCFF065ODV,ENCFF142CNG |
| 409 | 11 | 112636523 | 112637629 | 1106 | NOD.NOR | 11 | 112636789 | 112636822 | 34 | - | 17.6667 | 4.40E-07 | 0.0454 | TAGCTGTCTTCAGACACGCCAGAAGAGGGCGTCA | | |
| 410 | 11 | 115662353 | 115664762 | 2409 | 129S1,AJ,AKR,BALB,C3H,C57L,C58J,CBA,CE,DBA1J,DBA2J,ILNJ2,KK,MRL,NOD,NOR,NZB,NZO,NZW,PL,RF,RH2,SEA,SMJ,TH | 11 | 115662449 | 115662482 | 34 | - | 16.7724 | 7.83E-07 | 0.0614 | TAGCTGTCTTCAGATGCACCAGAAGAGGGCGTCA | | |
| 411 | 11 | 116447580 | 116448778 | 1198 | 129S1,BALB,C58J,DBA1J,DBA2J,FVB,KK,MA/My,NOD,NOR,NZB,NZO,NZW,PL,RF,SEA,SJL.SMJ.SWR.TH | 11 | 116448596 | 116448630 | 35 | - | 17.4404 | 4.19E-07 | 0.081 | CTGCTGACCCCAGCCTGTGACAGTAGAAGGCGGGC | | |
| 412 | 11 | 121017059 | 121018285 | 1226 | SJL | 11 | 121017423 | 121017456 | 34 | - | 21.9593 | 2.06E-08 | 0.00614 | GTGTAGTTCCCAAAGAGGCCACTAGAGAGCATTA | 18 | ENCFF201JUA,ENCFF092NMJ,ENCFF504JDS,ENCFF821BIA,ENCFF165LXO,ENCFF093ZRU,ENCFF065ODV,ENCFF367UNR,ENCFF327LYS,ENCFF198SSR,ENCFF502WDT,ENCFF055HAP,ENCFF907XEG,ENCFF099QNE.ENCFF589ZBJ.ENCFF024JOZ.ENCFF052KVO.ENCFF883UPM |
| 413 | 12 | 4194672 | 4222791 | 28119 | B10D2 | 12 | 4208707 | 4208740 | 34 | + | 17.3902 | 5.27E-07 | 0.0795 | TAGCTGTCCTCAGACACTCCAGAAGAGGGCATCA | | |
| 413 | 12 | 4194672 | 4222791 | 28119 | B10D2 | 12 | 4214420 | 4214453 | 34 | + | 17.5528 | 4.74E-07 | 0.0755 | TAGCTGTTTTCAGATACACCAGAAGAGGGCATCA | 1 | ENCFF093ZRU |
| 413 | 12 | 4194672 | 4222791 | 28119 | B10D2 | 12 | 4215333 | 4215351 | 19 | + | 17.4754 | 6.66E-07 | 0.132 | CCACCTCCAGGGGGCAGCT | | |
| 413 | 12 | 4194672 | 4222791 | 28119 | B10D2 | 12 | 4203958 | 4203991 | 34 | - | 16.7073 | 8.16E-07 | 0.0978 | GAGATCTGCCTGCCTCTGCCTCCAGAGGGCTATG | | |
| 414 | 12 | 5663770 | 5667937 | 4167 | AJ.BALB.ILNJ2.MRL.NZB.SEA.SMJ | 12 | 5664546 | 5664579 | 34 | + | 17.6748 | 4.37E-07 | 0.073 | GAGCATGACCACAAGTCCACAGCAGGTGGCAACG | | |
| 415 | 12 | 18784494 | 18789572 | 5078 | 129S1,BTBR,BUB,C3H,CBA,DBA2J,FVB,MA/My.NOD.NOR.PL.SJL.SMJ.SWR.TH | 12 | 18786288 | 18786321 | 34 | + | 17.2439 | 5.79E-07 | 0.0827 | TAGCTGTCCTCAGATACTCCAGAAGAGGGCATCA | | |
| 416 | 12 | 19039121 | 19045787 | 6666 | AJ,AKR,BALB,C57L,CE,DBA1J,ILNJ2,PJ,RF.SEA | 12 | 19045481 | 19045499 | 19 | + | 18.8525 | 2.59E-07 | 0.0823 | TCTACACTAGGGGGCGCTG | | |
| 417 | 12 | 19293147 | 19299715 | 6568 | 129S1,BTBR,BUB,C3H,C57L,CBA,DBA2J,FVB,KK,MA/My,MRL,NOD,NOR,NZO,PL,SJL,SMJ.SWR.TH | 12 | 19297709 | 19297727 | 19 | + | 17.1639 | 8.17E-07 | 0.145 | CCTCCACGAGGTGGCGGAA | | |
| 418 | 12 | 20055455 | 20058152 | 2697 | AJ.CE.DBA1J.PJ | 12 | 20056798 | 20056816 | 19 | + | 18.1148 | 4.35E-07 | 0.109 | TCTACACTAGGGGGCACTG | | |
| 419 | 12 | 20624757 | 20630333 | 5576 | DBA2J | 12 | 20627189 | 20627222 | 34 | + | 18.3659 | 2.76E-07 | 0.0591 | CTGAAGGCCCAGCCTTAGCCTTCAGGTGGCTGCC | | |
| 420 | 12 | 20744352 | 20757157 | 12805 | MRL | 12 | 20747473 | 20747506 | 34 | + | 21.9593 | 2.06E-08 | 0.011 | ATGCTATGTTCAAAGTAGCCACAAGAGGGCTCTA | 1 | ENCFF367UNR |
| 420 | 12 | 20744352 | 20757157 | 12805 | MRL | 12 | 20747488 | 20747506 | 19 | + | 17.3934 | 7.03E-07 | 0.134 | TAGCCACAAGAGGGCTCTA | 1 | ENCFF367UNR |
| 420 | 12 | 20744352 | 20757157 | 12805 | MRL | 12 | 20753456 | 20753474 | 19 | + | 17.1639 | 8.17E-07 | 0.145 | CCTCCACGAGGTGGCGGAA | | |
| 421 | 12 | 20748888 | 20754653 | 5765 | 129S1,AJ,AKR,BALB,BTBR,BUB,C3H,C57L,CBA,CE,DBA1J,FVB,ILNJ2,KK,MA/My,NOD,NOR,NZO,PJ,PL,RF,SEA,SJL,SMJ,SWR,TH | 12 | 20753456 | 20753474 | 19 | + | 17.1639 | 8.17E-07 | 0.145 | CCTCCACGAGGTGGCGGAA | | |
| 422 | 12 | 20991097 | 20993450 | 2353 | MRL | 12 | 20993034 | 20993067 | 34 | + | 21.6667 | 2.58E-08 | 0.0127 | ATGCTATGTTCAAAGCAGCCACGAGAGGGCTCCA | 2 | ENCFF821BIA.ENCFF367UNR |
| 423 | 12 | 23656010 | 23659599 | 3589 | MRL | 12 | 23656749 | 23656783 | 35 | + | 23.9633 | 5.48E-09 | 0.0077 | GTGCAGGTGCCCATGGAGGCCTGAAGAGGGCACCT | | |
| 424 | 12 | 29163923 | 29167447 | 3524 | 129S1,AJ,AKR,BALB,BUB,C3H,CBA,CE,DBA1J,DBA2J,KK,LP,MRL,NOD,NOR,NZO,NZW.PJ.PL.RF.RH2.SEA.SMJ | 12 | 29165332 | 29165365 | 34 | + | 18.9431 | 1.87E-07 | 0.0482 | CTGGCATCCCAGAGGCTTCCAACAGAGGGCAACA | | |
| 425 | 12 | 44893398 | 44897080 | 3682 | NZO,NZW | 12 | 44895654 | 44895687 | 34 | + | 17.7073 | 4.28E-07 | 0.0725 | GCGCACTCGTGTGAATAACCACAAGGGGGTGCCC | 22 | ENCFF093ZRU,ENCFF367UNR,ENCFF201JUA,ENCFF099QNE,ENCFF821BIA,ENCFF136ZDK,ENCFF327LYS,ENCFF907XEG,ENCFF024JOZ,ENCFF089GBC,ENCFF575MGI,ENCFF165LXO,ENCFF589ZBJ,ENCFF198SSR,ENCFF092NMJ,ENCFF052KVO,ENCFF430PPJ,ENCFF502WDT,ENCFF055HAP,ENCFF951GOP,ENCFF281QMV.ENCFF609MDN |
| 425 | 12 | 44893398 | 44897080 | 3682 | NZO,NZW | 12 | 44895669 | 44895687 | 19 | + | 18.0328 | 4.60E-07 | 0.113 | TAACCACAAGGGGGTGCCC | 22 | ENCFF093ZRU,ENCFF367UNR,ENCFF201JUA,ENCFF099QNE,ENCFF821BIA,ENCFF136ZDK,ENCFF327LYS,ENCFF907XEG,ENCFF024JOZ,ENCFF089GBC,ENCFF575MGI,ENCFF165LXO,ENCFF589ZBJ,ENCFF198SSR,ENCFF092NMJ,ENCFF052KVO,ENCFF430PPJ,ENCFF502WDT,ENCFF055HAP,ENCFF951GOP,ENCFF281QMV.ENCFF609MDN |
| 426 | 12 | 51347640 | 51356509 | 8869 | 129S1.C57L.NZB | 12 | 51355062 | 51355095 | 34 | + | 16.4309 | 9.71E-07 | 0.104 | TAGCTGTCTTCAGGGACACCAGAAGAGGGCATCA | | |
| 427 | 12 | 54821004 | 54829990 | 8986 | AJ,AKR,BALB,C3H,CBA,CE,DBA1J,DBA2J,FVB,ILNJ2,KK,NOD,NOR,NZW,PJ,RF,RH2,SEA.SJL.SMJ.SWR.TH | 12 | 54826422 | 54826440 | 19 | + | 17.1639 | 8.17E-07 | 0.145 | CCTCCACGAGGTGGCGGAA | | |
| 428 | 12 | 55451509 | 55452656 | 1147 | PJ | 12 | 55451601 | 55451635 | 35 | + | 16.6147 | 6.75E-07 | 0.161 | GTGTAGGTGCCCACAGTGTACAGAAGAGGGCAATA | | |
| 429 | 12 | 56672966 | 56676585 | 3619 | AJ,AKR,BALB,C3H,CBA,DBA1J,DBA2J,ILNJ2,KK,MRL,NOD,NOR,RF,SEA,SJL,SMJ,SWR | 12 | 56673772 | 56673805 | 34 | + | 16.5691 | 8.90E-07 | 0.102 | TAGCTGTCTTCAGACGCACCAGAAGAGGGCATCG | | |
| 430 | 12 | 59081758 | 59084703 | 2945 | AJ,BALB,C3H,CBA,DBA1J,DBA2J,ILNJ2,KK,MA/Mv.RH2.SEA | 12 | 59081797 | 59081830 | 34 | + | 18.065 | 3.38E-07 | 0.064 | TAGCTGTCTTCAGACTCACCAGAAGAGGGCGTCA | | |
| 430 | 12 | 59081758 | 59083753 | 1995 | NZO | 12 | 59081797 | 59081830 | 34 | + | 18.065 | 3.38E-07 | 0.064 | TAGCTGTCTTCAGACTCACCAGAAGAGGGCGTCA | | |
| 430 | 12 | 59081758 | 59084703 | 2945 | AJ,BALB,C3H,CBA,DBA1J,DBA2J,ILNJ2,KK,MA/Mv.RH2.SEA | 12 | 59081812 | 59081830 | 19 | + | 17.0328 | 8.88E-07 | 0.147 | TCACCAGAAGAGGGCGTCA | | |
| 430 | 12 | 59081758 | 59083753 | 1995 | NZO | 12 | 59081812 | 59081830 | 19 | + | 17.0328 | 8.88E-07 | 0.147 | TCACCAGAAGAGGGCGTCA | | |
| 430 | 12 | 59081758 | 59084703 | 2945 | AJ,BALB,C3H,CBA,DBA1J,DBA2J,ILNJ2,KK,MA/Mv.RH2.SEA | 12 | 59081892 | 59081925 | 34 | + | 18.065 | 3.38E-07 | 0.064 | TAGCTGTCTTCAGACTCACCAGAAGAGGGCGTCA | | |
| 430 | 12 | 59081758 | 59083753 | 1995 | NZO | 12 | 59081892 | 59081925 | 34 | + | 18.065 | 3.38E-07 | 0.064 | TAGCTGTCTTCAGACTCACCAGAAGAGGGCGTCA | | |
| 430 | 12 | 59081758 | 59084703 | 2945 | AJ,BALB,C3H,CBA,DBA1J,DBA2J,ILNJ2,KK,MA/Mv.RH2.SEA | 12 | 59081907 | 59081925 | 19 | + | 17.0328 | 8.88E-07 | 0.147 | TCACCAGAAGAGGGCGTCA | | |
| 430 | 12 | 59081758 | 59083753 | 1995 | NZO | 12 | 59081907 | 59081925 | 19 | + | 17.0328 | 8.88E-07 | 0.147 | TCACCAGAAGAGGGCGTCA | | |
| 430 | 12 | 59081758 | 59084703 | 2945 | AJ,BALB,C3H,CBA,DBA1J,DBA2J,ILNJ2,KK,MA/Mv.RH2.SEA | 12 | 59081987 | 59082020 | 34 | + | 18.065 | 3.38E-07 | 0.064 | TAGCTGTCTTCAGACTCACCAGAAGAGGGCGTCA | | |
| 430 | 12 | 59081758 | 59083753 | 1995 | NZO | 12 | 59081987 | 59082020 | 34 | + | 18.065 | 3.38E-07 | 0.064 | TAGCTGTCTTCAGACTCACCAGAAGAGGGCGTCA | | |
| 430 | 12 | 59081758 | 59084703 | 2945 | AJ,BALB,C3H,CBA,DBA1J,DBA2J,ILNJ2,KK,MA/Mv.RH2.SEA | 12 | 59082002 | 59082020 | 19 | + | 17.0328 | 8.88E-07 | 0.147 | TCACCAGAAGAGGGCGTCA | | |
| 430 | 12 | 59081758 | 59083753 | 1995 | NZO | 12 | 59082002 | 59082020 | 19 | + | 17.0328 | 8.88E-07 | 0.147 | TCACCAGAAGAGGGCGTCA | | |
| 430 | 12 | 59081758 | 59084703 | 2945 | AJ,BALB,C3H,CBA,DBA1J,DBA2J,ILNJ2,KK,MA/Mv.RH2.SEA | 12 | 59082082 | 59082115 | 34 | + | 18.065 | 3.38E-07 | 0.064 | TAGCTGTCTTCAGACTCACCAGAAGAGGGCGTCA | | |
| 430 | 12 | 59081758 | 59083753 | 1995 | NZO | 12 | 59082082 | 59082115 | 34 | + | 18.065 | 3.38E-07 | 0.064 | TAGCTGTCTTCAGACTCACCAGAAGAGGGCGTCA | | |
| 430 | 12 | 59081758 | 59084703 | 2945 | AJ,BALB,C3H,CBA,DBA1J,DBA2J,ILNJ2,KK,MA/Mv.RH2.SEA | 12 | 59082097 | 59082115 | 19 | + | 17.0328 | 8.88E-07 | 0.147 | TCACCAGAAGAGGGCGTCA | | |
| 430 | 12 | 59081758 | 59083753 | 1995 | NZO | 12 | 59082097 | 59082115 | 19 | + | 17.0328 | 8.88E-07 | 0.147 | TCACCAGAAGAGGGCGTCA | | |
| 430 | 12 | 59081758 | 59084703 | 2945 | AJ,BALB,C3H,CBA,DBA1J,DBA2J,ILNJ2,KK,MA/Mv.RH2.SEA | 12 | 59082177 | 59082210 | 34 | + | 18.065 | 3.38E-07 | 0.064 | TAGCTGTCTTCAGACTCACCAGAAGAGGGCGTCA | | |
| 430 | 12 | 59081758 | 59083753 | 1995 | NZO | 12 | 59082177 | 59082210 | 34 | + | 18.065 | 3.38E-07 | 0.064 | TAGCTGTCTTCAGACTCACCAGAAGAGGGCGTCA | | |
| 430 | 12 | 59081758 | 59084703 | 2945 | AJ,BALB,C3H,CBA,DBA1J,DBA2J,ILNJ2,KK,MA/Mv.RH2.SEA | 12 | 59082192 | 59082210 | 19 | + | 17.0328 | 8.88E-07 | 0.147 | TCACCAGAAGAGGGCGTCA | | |
| 430 | 12 | 59081758 | 59083753 | 1995 | NZO | 12 | 59082192 | 59082210 | 19 | + | 17.0328 | 8.88E-07 | 0.147 | TCACCAGAAGAGGGCGTCA | | |
| 430 | 12 | 59081758 | 59084703 | 2945 | AJ,BALB,C3H,CBA,DBA1J,DBA2J,ILNJ2,KK,MA/Mv.RH2.SEA | 12 | 59082272 | 59082305 | 34 | + | 18.065 | 3.38E-07 | 0.064 | TAGCTGTCTTCAGACTCACCAGAAGAGGGCGTCA | | |
| 430 | 12 | 59081758 | 59083753 | 1995 | NZO | 12 | 59082272 | 59082305 | 34 | + | 18.065 | 3.38E-07 | 0.064 | TAGCTGTCTTCAGACTCACCAGAAGAGGGCGTCA | | |
| 430 | 12 | 59081758 | 59084703 | 2945 | AJ,BALB,C3H,CBA,DBA1J,DBA2J,ILNJ2,KK,MA/Mv.RH2.SEA | 12 | 59082287 | 59082305 | 19 | + | 17.0328 | 8.88E-07 | 0.147 | TCACCAGAAGAGGGCGTCA | | |
| 430 | 12 | 59081758 | 59083753 | 1995 | NZO | 12 | 59082287 | 59082305 | 19 | + | 17.0328 | 8.88E-07 | 0.147 | TCACCAGAAGAGGGCGTCA | | |

| | | | | | | | | | | | | | | | | |
|---|---|---|---|---|---|---|---|---|---|---|---|---|---|---|---|---|
| 430 | 12 | 59081758 | 59084703 | 2945 | AJ,BALB,C3H,CBA,DBA1J,DBA2J,ILNJ2,KK,MA/Mv.RH2.SEA | 12 | 59082367 | 59082400 | 34 | + | 18.065 | 3.38E-07 | 0.064 | TAGCTGTCTTCAGACTCACCAGAAGAGGGCGTCA | | |
| 430 | 12 | 59081758 | 59083753 | 1995 | NZO | 12 | 59082367 | 59082400 | 34 | + | 18.065 | 3.38E-07 | 0.064 | TAGCTGTCTTCAGACTCACCAGAAGAGGGCGTCA | | |
| 430 | 12 | 59081758 | 59084703 | 2945 | AJ,BALB,C3H,CBA,DBA1J,DBA2J,ILNJ2,KK,MA/Mv.RH2.SEA | 12 | 59082382 | 59082400 | 19 | + | 17.0328 | 8.88E-07 | 0.147 | TCACCAGAAGAGGGCGTCA | | |
| 430 | 12 | 59081758 | 59083753 | 1995 | NZO | 12 | 59082382 | 59082400 | 19 | + | 17.0328 | 8.88E-07 | 0.147 | TCACCAGAAGAGGGCGTCA | | |
| 430 | 12 | 59081758 | 59084703 | 2945 | AJ,BALB,C3H,CBA,DBA1J,DBA2J,ILNJ2,KK,MA/Mv.RH2.SEA | 12 | 59082462 | 59082495 | 34 | + | 18.065 | 3.38E-07 | 0.064 | TAGCTGTCTTCAGACTCACCAGAAGAGGGCGTCA | | |
| 430 | 12 | 59081758 | 59083753 | 1995 | NZO | 12 | 59082462 | 59082495 | 34 | + | 18.065 | 3.38E-07 | 0.064 | TAGCTGTCTTCAGACTCACCAGAAGAGGGCGTCA | | |
| 430 | 12 | 59081758 | 59084703 | 2945 | AJ,BALB,C3H,CBA,DBA1J,DBA2J,ILNJ2,KK,MA/Mv.RH2.SEA | 12 | 59082477 | 59082495 | 19 | + | 17.0328 | 8.88E-07 | 0.147 | TCACCAGAAGAGGGCGTCA | | |
| 430 | 12 | 59081758 | 59083753 | 1995 | NZO | 12 | 59082477 | 59082495 | 19 | + | 17.0328 | 8.88E-07 | 0.147 | TCACCAGAAGAGGGCGTCA | | |
| 430 | 12 | 59081758 | 59084703 | 2945 | AJ,BALB,C3H,CBA,DBA1J,DBA2J,ILNJ2,KK,MA/Mv.RH2.SEA | 12 | 59082557 | 59082590 | 34 | + | 18.065 | 3.38E-07 | 0.064 | TAGCTGTCTTCAGACTCACCAGAAGAGGGCGTCA | | |
| 430 | 12 | 59081758 | 59083753 | 1995 | NZO | 12 | 59082557 | 59082590 | 34 | + | 18.065 | 3.38E-07 | 0.064 | TAGCTGTCTTCAGACTCACCAGAAGAGGGCGTCA | | |
| 430 | 12 | 59081758 | 59084703 | 2945 | AJ,BALB,C3H,CBA,DBA1J,DBA2J,ILNJ2,KK,MA/Mv.RH2.SEA | 12 | 59082572 | 59082590 | 19 | + | 17.0328 | 8.88E-07 | 0.147 | TCACCAGAAGAGGGCGTCA | | |
| 430 | 12 | 59081758 | 59083753 | 1995 | NZO | 12 | 59082572 | 59082590 | 19 | + | 17.0328 | 8.88E-07 | 0.147 | TCACCAGAAGAGGGCGTCA | | |
| 430 | 12 | 59081758 | 59084703 | 2945 | AJ,BALB,C3H,CBA,DBA1J,DBA2J,ILNJ2,KK,MA/Mv.RH2.SEA | 12 | 59082652 | 59082685 | 34 | + | 18.065 | 3.38E-07 | 0.064 | TAGCTGTCTTCAGACTCACCAGAAGAGGGCGTCA | | |
| 430 | 12 | 59081758 | 59083753 | 1995 | NZO | 12 | 59082652 | 59082685 | 34 | + | 18.065 | 3.38E-07 | 0.064 | TAGCTGTCTTCAGACTCACCAGAAGAGGGCGTCA | | |
| 430 | 12 | 59081758 | 59084703 | 2945 | AJ,BALB,C3H,CBA,DBA1J,DBA2J,ILNJ2,KK,MA/Mv.RH2.SEA | 12 | 59082667 | 59082685 | 19 | + | 17.0328 | 8.88E-07 | 0.147 | TCACCAGAAGAGGGCGTCA | | |
| 430 | 12 | 59081758 | 59083753 | 1995 | NZO | 12 | 59082667 | 59082685 | 19 | + | 17.0328 | 8.88E-07 | 0.147 | TCACCAGAAGAGGGCGTCA | | |
| 430 | 12 | 59081758 | 59084703 | 2945 | AJ,BALB,C3H,CBA,DBA1J,DBA2J,ILNJ2,KK,MA/Mv.RH2.SEA | 12 | 59082747 | 59082780 | 34 | + | 18.065 | 3.38E-07 | 0.064 | TAGCTGTCTTCAGACTCACCAGAAGAGGGCGTCA | | |
| 430 | 12 | 59081758 | 59083753 | 1995 | NZO | 12 | 59082747 | 59082780 | 34 | + | 18.065 | 3.38E-07 | 0.064 | TAGCTGTCTTCAGACTCACCAGAAGAGGGCGTCA | | |
| 430 | 12 | 59081758 | 59084703 | 2945 | AJ,BALB,C3H,CBA,DBA1J,DBA2J,ILNJ2,KK,MA/Mv.RH2.SEA | 12 | 59082762 | 59082780 | 19 | + | 17.0328 | 8.88E-07 | 0.147 | TCACCAGAAGAGGGCGTCA | | |
| 430 | 12 | 59081758 | 59083753 | 1995 | NZO | 12 | 59082762 | 59082780 | 19 | + | 17.0328 | 8.88E-07 | 0.147 | TCACCAGAAGAGGGCGTCA | | |
| 430 | 12 | 59081758 | 59084703 | 2945 | AJ,BALB,C3H,CBA,DBA1J,DBA2J,ILNJ2,KK,MA/Mv.RH2.SEA | 12 | 59082842 | 59082875 | 34 | + | 18.065 | 3.38E-07 | 0.064 | TAGCTGTCTTCAGACTCACCAGAAGAGGGCGTCA | | |
| 430 | 12 | 59081758 | 59083753 | 1995 | NZO | 12 | 59082842 | 59082875 | 34 | + | 18.065 | 3.38E-07 | 0.064 | TAGCTGTCTTCAGACTCACCAGAAGAGGGCGTCA | | |
| 430 | 12 | 59081758 | 59084703 | 2945 | AJ,BALB,C3H,CBA,DBA1J,DBA2J,ILNJ2,KK,MA/Mv.RH2.SEA | 12 | 59082857 | 59082875 | 19 | + | 17.0328 | 8.88E-07 | 0.147 | TCACCAGAAGAGGGCGTCA | | |
| 430 | 12 | 59081758 | 59083753 | 1995 | NZO | 12 | 59082857 | 59082875 | 19 | + | 17.0328 | 8.88E-07 | 0.147 | TCACCAGAAGAGGGCGTCA | | |
| 430 | 12 | 59081758 | 59084703 | 2945 | AJ,BALB,C3H,CBA,DBA1J,DBA2J,ILNJ2,KK,MA/Mv.RH2.SEA | 12 | 59082937 | 59082970 | 34 | + | 18.065 | 3.38E-07 | 0.064 | TAGCTGTCTTCAGACTCACCAGAAGAGGGCGTCA | | |
| 430 | 12 | 59081758 | 59083753 | 1995 | NZO | 12 | 59082937 | 59082970 | 34 | + | 18.065 | 3.38E-07 | 0.064 | TAGCTGTCTTCAGACTCACCAGAAGAGGGCGTCA | | |
| 430 | 12 | 59081758 | 59084703 | 2945 | AJ,BALB,C3H,CBA,DBA1J,DBA2J,ILNJ2,KK,MA/Mv.RH2.SEA | 12 | 59082952 | 59082970 | 19 | + | 17.0328 | 8.88E-07 | 0.147 | TCACCAGAAGAGGGCGTCA | | |
| 430 | 12 | 59081758 | 59083753 | 1995 | NZO | 12 | 59082952 | 59082970 | 19 | + | 17.0328 | 8.88E-07 | 0.147 | TCACCAGAAGAGGGCGTCA | | |
| 430 | 12 | 59081758 | 59084703 | 2945 | AJ,BALB,C3H,CBA,DBA1J,DBA2J,ILNJ2,KK,MA/Mv.RH2.SEA | 12 | 59083032 | 59083065 | 34 | + | 18.065 | 3.38E-07 | 0.064 | TAGCTGTCTTCAGACTCACCAGAAGAGGGCGTCA | | |
| 430 | 12 | 59081758 | 59083753 | 1995 | NZO | 12 | 59083032 | 59083065 | 34 | + | 18.065 | 3.38E-07 | 0.064 | TAGCTGTCTTCAGACTCACCAGAAGAGGGCGTCA | | |
| 430 | 12 | 59081758 | 59084703 | 2945 | AJ,BALB,C3H,CBA,DBA1J,DBA2J,ILNJ2,KK,MA/Mv.RH2.SEA | 12 | 59083047 | 59083065 | 19 | + | 17.0328 | 8.88E-07 | 0.147 | TCACCAGAAGAGGGCGTCA | | |
| 430 | 12 | 59081758 | 59083753 | 1995 | NZO | 12 | 59083047 | 59083065 | 19 | + | 17.0328 | 8.88E-07 | 0.147 | TCACCAGAAGAGGGCGTCA | | |
| 430 | 12 | 59081758 | 59084703 | 2945 | AJ,BALB,C3H,CBA,DBA1J,DBA2J,ILNJ2,KK,MA/Mv.RH2.SEA | 12 | 59083127 | 59083160 | 34 | + | 18.065 | 3.38E-07 | 0.064 | TAGCTGTCTTCAGACTCACCAGAAGAGGGCGTCA | | |
| 430 | 12 | 59081758 | 59083753 | 1995 | NZO | 12 | 59083127 | 59083160 | 34 | + | 18.065 | 3.38E-07 | 0.064 | TAGCTGTCTTCAGACTCACCAGAAGAGGGCGTCA | | |
| 430 | 12 | 59081758 | 59084703 | 2945 | AJ,BALB,C3H,CBA,DBA1J,DBA2J,ILNJ2,KK,MA/Mv.RH2.SEA | 12 | 59083142 | 59083160 | 19 | + | 17.0328 | 8.88E-07 | 0.147 | TCACCAGAAGAGGGCGTCA | | |
| 430 | 12 | 59081758 | 59083753 | 1995 | NZO | 12 | 59083142 | 59083160 | 19 | + | 17.0328 | 8.88E-07 | 0.147 | TCACCAGAAGAGGGCGTCA | | |
| 430 | 12 | 59081758 | 59084703 | 2945 | AJ,BALB,C3H,CBA,DBA1J,DBA2J,ILNJ2,KK,MA/Mv.RH2.SEA | 12 | 59083222 | 59083255 | 34 | + | 18.065 | 3.38E-07 | 0.064 | TAGCTGTCTTCAGACTCACCAGAAGAGGGCGTCA | | |
| 430 | 12 | 59081758 | 59083753 | 1995 | NZO | 12 | 59083222 | 59083255 | 34 | + | 18.065 | 3.38E-07 | 0.064 | TAGCTGTCTTCAGACTCACCAGAAGAGGGCGTCA | | |
| 430 | 12 | 59081758 | 59084703 | 2945 | AJ,BALB,C3H,CBA,DBA1J,DBA2J,ILNJ2,KK,MA/Mv.RH2.SEA | 12 | 59083237 | 59083255 | 19 | + | 17.0328 | 8.88E-07 | 0.147 | TCACCAGAAGAGGGCGTCA | | |
| 430 | 12 | 59081758 | 59083753 | 1995 | NZO | 12 | 59083237 | 59083255 | 19 | + | 17.0328 | 8.88E-07 | 0.147 | TCACCAGAAGAGGGCGTCA | | |
| 430 | 12 | 59081758 | 59084703 | 2945 | AJ,BALB,C3H,CBA,DBA1J,DBA2J,ILNJ2,KK,MA/Mv.RH2.SEA | 12 | 59083317 | 59083350 | 34 | + | 18.065 | 3.38E-07 | 0.064 | TAGCTGTCTTCAGACTCACCAGAAGAGGGCGTCA | | |
| 430 | 12 | 59081758 | 59083753 | 1995 | NZO | 12 | 59083317 | 59083350 | 34 | + | 18.065 | 3.38E-07 | 0.064 | TAGCTGTCTTCAGACTCACCAGAAGAGGGCGTCA | | |
| 430 | 12 | 59081758 | 59084703 | 2945 | AJ,BALB,C3H,CBA,DBA1J,DBA2J,ILNJ2,KK,MA/Mv.RH2.SEA | 12 | 59083332 | 59083350 | 19 | + | 17.0328 | 8.88E-07 | 0.147 | TCACCAGAAGAGGGCGTCA | | |
| 430 | 12 | 59081758 | 59083753 | 1995 | NZO | 12 | 59083332 | 59083350 | 19 | + | 17.0328 | 8.88E-07 | 0.147 | TCACCAGAAGAGGGCGTCA | | |
| 430 | 12 | 59081758 | 59084703 | 2945 | AJ,BALB,C3H,CBA,DBA1J,DBA2J,ILNJ2,KK,MA/Mv.RH2.SEA | 12 | 59083412 | 59083445 | 34 | + | 18.065 | 3.38E-07 | 0.064 | TAGCTGTCTTCAGACTCACCAGAAGAGGGCGTCA | | |
| 430 | 12 | 59081758 | 59083753 | 1995 | NZO | 12 | 59083412 | 59083445 | 34 | + | 18.065 | 3.38E-07 | 0.064 | TAGCTGTCTTCAGACTCACCAGAAGAGGGCGTCA | | |
| 430 | 12 | 59081758 | 59084703 | 2945 | AJ,BALB,C3H,CBA,DBA1J,DBA2J,ILNJ2,KK,MA/Mv.RH2.SEA | 12 | 59083427 | 59083445 | 19 | + | 17.0328 | 8.88E-07 | 0.147 | TCACCAGAAGAGGGCGTCA | | |
| 430 | 12 | 59081758 | 59083753 | 1995 | NZO | 12 | 59083427 | 59083445 | 19 | + | 17.0328 | 8.88E-07 | 0.147 | TCACCAGAAGAGGGCGTCA | | |
| 430 | 12 | 59081758 | 59084703 | 2945 | AJ,BALB,C3H,CBA,DBA1J,DBA2J,ILNJ2,KK,MA/Mv.RH2.SEA | 12 | 59083507 | 59083540 | 34 | + | 18.065 | 3.38E-07 | 0.064 | TAGCTGTCTTCAGACTCACCAGAAGAGGGCGTCA | | |
| 430 | 12 | 59081758 | 59083753 | 1995 | NZO | 12 | 59083507 | 59083540 | 34 | + | 18.065 | 3.38E-07 | 0.064 | TAGCTGTCTTCAGACTCACCAGAAGAGGGCGTCA | | |
| 430 | 12 | 59081758 | 59084703 | 2945 | AJ,BALB,C3H,CBA,DBA1J,DBA2J,ILNJ2,KK,MA/Mv.RH2.SEA | 12 | 59083522 | 59083540 | 19 | + | 17.0328 | 8.88E-07 | 0.147 | TCACCAGAAGAGGGCGTCA | | |
| 430 | 12 | 59081758 | 59083753 | 1995 | NZO | 12 | 59083522 | 59083540 | 19 | + | 17.0328 | 8.88E-07 | 0.147 | TCACCAGAAGAGGGCGTCA | | |
| 430 | 12 | 59081758 | 59084703 | 2945 | AJ,BALB,C3H,CBA,DBA1J,DBA2J,ILNJ2,KK,MA/Mv.RH2.SEA | 12 | 59083602 | 59083635 | 34 | + | 18.065 | 3.38E-07 | 0.064 | TAGCTGTCTTCAGACTCACCAGAAGAGGGCGTCA | | |
| 430 | 12 | 59081758 | 59083753 | 1995 | NZO | 12 | 59083602 | 59083635 | 34 | + | 18.065 | 3.38E-07 | 0.064 | TAGCTGTCTTCAGACTCACCAGAAGAGGGCGTCA | | |
| 430 | 12 | 59081758 | 59084703 | 2945 | AJ,BALB,C3H,CBA,DBA1J,DBA2J,ILNJ2,KK,MA/Mv.RH2.SEA | 12 | 59083617 | 59083635 | 19 | + | 17.0328 | 8.88E-07 | 0.147 | TCACCAGAAGAGGGCGTCA | | |
| 430 | 12 | 59081758 | 59083753 | 1995 | NZO | 12 | 59083617 | 59083635 | 19 | + | 17.0328 | 8.88E-07 | 0.147 | TCACCAGAAGAGGGCGTCA | | |
| 430 | 12 | 59081758 | 59084703 | 2945 | AJ,BALB,C3H,CBA,DBA1J,DBA2J,ILNJ2,KK,MA/Mv.RH2.SEA | 12 | 59083697 | 59083730 | 34 | + | 18.065 | 3.38E-07 | 0.064 | TAGCTGTCTTCAGACTCACCAGAAGAGGGCGTCA | | |
| 430 | 12 | 59081758 | 59083753 | 1995 | NZO | 12 | 59083697 | 59083730 | 34 | + | 18.065 | 3.38E-07 | 0.064 | TAGCTGTCTTCAGACTCACCAGAAGAGGGCGTCA | | |
| 430 | 12 | 59081758 | 59084703 | 2945 | AJ,BALB,C3H,CBA,DBA1J,DBA2J,ILNJ2,KK,MA/Mv.RH2.SEA | 12 | 59083712 | 59083730 | 19 | + | 17.0328 | 8.88E-07 | 0.147 | TCACCAGAAGAGGGCGTCA | | |
| 430 | 12 | 59081758 | 59083753 | 1995 | NZO | 12 | 59083712 | 59083730 | 19 | + | 17.0328 | 8.88E-07 | 0.147 | TCACCAGAAGAGGGCGTCA | | |
| 430 | 12 | 59081758 | 59084703 | 2945 | AJ,BALB,C3H,CBA,DBA1J,DBA2J,ILNJ2,KK,MA/Mv.RH2.SEA | 12 | 59083792 | 59083825 | 34 | + | 18.065 | 3.38E-07 | 0.064 | TAGCTGTCTTCAGACTCACCAGAAGAGGGCGTCA | | |
| 430 | 12 | 59081758 | 59084703 | 2945 | AJ,BALB,C3H,CBA,DBA1J,DBA2J,ILNJ2,KK,MA/Mv.RH2.SEA | 12 | 59083807 | 59083825 | 19 | + | 17.0328 | 8.88E-07 | 0.147 | TCACCAGAAGAGGGCGTCA | | |
| 430 | 12 | 59081758 | 59084703 | 2945 | AJ,BALB,C3H,CBA,DBA1J,DBA2J,ILNJ2,KK,MA/Mv.RH2.SEA | 12 | 59083887 | 59083920 | 34 | + | 18.065 | 3.38E-07 | 0.064 | TAGCTGTCTTCAGACTCACCAGAAGAGGGCGTCA | | |
| 430 | 12 | 59081758 | 59084703 | 2945 | AJ,BALB,C3H,CBA,DBA1J,DBA2J,ILNJ2,KK,MA/Mv.RH2.SEA | 12 | 59083902 | 59083920 | 19 | + | 17.0328 | 8.88E-07 | 0.147 | TCACCAGAAGAGGGCGTCA | | |
| 430 | 12 | 59081758 | 59084703 | 2945 | AJ,BALB,C3H,CBA,DBA1J,DBA2J,ILNJ2,KK,MA/Mv.RH2.SEA | 12 | 59083982 | 59084015 | 34 | + | 18.065 | 3.38E-07 | 0.064 | TAGCTGTCTTCAGACTCACCAGAAGAGGGCGTCA | | |
| 430 | 12 | 59081758 | 59084703 | 2945 | AJ,BALB,C3H,CBA,DBA1J,DBA2J,ILNJ2,KK,MA/Mv.RH2.SEA | 12 | 59083997 | 59084015 | 19 | + | 17.0328 | 8.88E-07 | 0.147 | TCACCAGAAGAGGGCGTCA | | |
| 430 | 12 | 59081758 | 59084703 | 2945 | AJ,BALB,C3H,CBA,DBA1J,DBA2J,ILNJ2,KK,MA/Mv.RH2.SEA | 12 | 59084077 | 59084110 | 34 | + | 18.065 | 3.38E-07 | 0.064 | TAGCTGTCTTCAGACTCACCAGAAGAGGGCGTCA | | |
| 430 | 12 | 59081758 | 59084703 | 2945 | AJ,BALB,C3H,CBA,DBA1J,DBA2J,ILNJ2,KK,MA/Mv.RH2.SEA | 12 | 59084092 | 59084110 | 19 | + | 17.0328 | 8.88E-07 | 0.147 | TCACCAGAAGAGGGCGTCA | | |
| 430 | 12 | 59081758 | 59084703 | 2945 | AJ,BALB,C3H,CBA,DBA1J,DBA2J,ILNJ2,KK,MA/Mv.RH2.SEA | 12 | 59084172 | 59084205 | 34 | + | 18.065 | 3.38E-07 | 0.064 | TAGCTGTCTTCAGACTCACCAGAAGAGGGCGTCA | | |
| 430 | 12 | 59081758 | 59084703 | 2945 | AJ,BALB,C3H,CBA,DBA1J,DBA2J,ILNJ2,KK,MA/Mv.RH2.SEA | 12 | 59084187 | 59084205 | 19 | + | 17.0328 | 8.88E-07 | 0.147 | TCACCAGAAGAGGGCGTCA | | |
| 430 | 12 | 59081758 | 59084703 | 2945 | AJ,BALB,C3H,CBA,DBA1J,DBA2J,ILNJ2,KK,MA/Mv.RH2.SEA | 12 | 59084267 | 59084300 | 34 | + | 18.065 | 3.38E-07 | 0.064 | TAGCTGTCTTCAGACTCACCAGAAGAGGGCGTCA | | |
| 430 | 12 | 59081758 | 59084703 | 2945 | AJ,BALB,C3H,CBA,DBA1J,DBA2J,ILNJ2,KK,MA/Mv.RH2.SEA | 12 | 59084282 | 59084300 | 19 | + | 17.0328 | 8.88E-07 | 0.147 | TCACCAGAAGAGGGCGTCA | | |
| 430 | 12 | 59081758 | 59084703 | 2945 | AJ,BALB,C3H,CBA,DBA1J,DBA2J,ILNJ2,KK,MA/Mv.RH2.SEA | 12 | 59084362 | 59084395 | 34 | + | 18.065 | 3.38E-07 | 0.064 | TAGCTGTCTTCAGACTCACCAGAAGAGGGCGTCA | | |
| 430 | 12 | 59081758 | 59084703 | 2945 | AJ,BALB,C3H,CBA,DBA1J,DBA2J,ILNJ2,KK,MA/Mv.RH2.SEA | 12 | 59084377 | 59084395 | 19 | + | 17.0328 | 8.88E-07 | 0.147 | TCACCAGAAGAGGGCGTCA | | |
| 430 | 12 | 59081758 | 59084703 | 2945 | AJ,BALB,C3H,CBA,DBA1J,DBA2J,ILNJ2,KK,MA/Mv.RH2.SEA | 12 | 59084457 | 59084490 | 34 | + | 18.065 | 3.38E-07 | 0.064 | TAGCTGTCTTCAGACTCACCAGAAGAGGGCGTCA | | |
| 430 | 12 | 59081758 | 59084703 | 2945 | AJ,BALB,C3H,CBA,DBA1J,DBA2J,ILNJ2,KK,MA/Mv.RH2.SEA | 12 | 59084472 | 59084490 | 19 | + | 17.0328 | 8.88E-07 | 0.147 | TCACCAGAAGAGGGCGTCA | | |
| 430 | 12 | 59081758 | 59084703 | 2945 | AJ,BALB,C3H,CBA,DBA1J,DBA2J,ILNJ2,KK,MA/Mv.RH2.SEA | 12 | 59084552 | 59084585 | 34 | + | 18.065 | 3.38E-07 | 0.064 | TAGCTGTCTTCAGACTCACCAGAAGAGGGCGTCA | | |
| 430 | 12 | 59081758 | 59084703 | 2945 | AJ,BALB,C3H,CBA,DBA1J,DBA2J,ILNJ2,KK,MA/Mv.RH2.SEA | 12 | 59084567 | 59084585 | 19 | + | 17.0328 | 8.88E-07 | 0.147 | TCACCAGAAGAGGGCGTCA | | |
| 430 | 12 | 59081758 | 59084703 | 2945 | AJ,BALB,C3H,CBA,DBA1J,DBA2J,ILNJ2,KK,MA/Mv.RH2.SEA | 12 | 59084647 | 59084680 | 34 | + | 18.065 | 3.38E-07 | 0.064 | TAGCTGTCTTCAGACTCACCAGAAGAGGGCGTCA | | |
| 430 | 12 | 59081758 | 59084703 | 2945 | AJ,BALB,C3H,CBA,DBA1J,DBA2J,ILNJ2,KK,MA/Mv.RH2.SEA | 12 | 59084662 | 59084680 | 19 | + | 17.0328 | 8.88E-07 | 0.147 | TCACCAGAAGAGGGCGTCA | | |
| 431 | 12 | 59081831 | 59085915 | 4084 | BUB,FVB | 12 | 59081892 | 59081925 | 34 | + | 18.065 | 3.38E-07 | 0.064 | TAGCTGTCTTCAGACTCACCAGAAGAGGGCGTCA | | |
| 431 | 12 | 59081831 | 59085915 | 4084 | BUB,FVB | 12 | 59081907 | 59081925 | 19 | + | 17.0328 | 8.88E-07 | 0.147 | TCACCAGAAGAGGGCGTCA | | |
| 431 | 12 | 59081831 | 59085915 | 4084 | BUB,FVB | 12 | 59081987 | 59082020 | 34 | + | 18.065 | 3.38E-07 | 0.064 | TAGCTGTCTTCAGACTCACCAGAAGAGGGCGTCA | | |
| 431 | 12 | 59081831 | 59085915 | 4084 | BUB,FVB | 12 | 59082002 | 59082020 | 19 | + | 17.0328 | 8.88E-07 | 0.147 | TCACCAGAAGAGGGCGTCA | | |
| 431 | 12 | 59081831 | 59085915 | 4084 | BUB,FVB | 12 | 59082082 | 59082115 | 34 | + | 18.065 | 3.38E-07 | 0.064 | TAGCTGTCTTCAGACTCACCAGAAGAGGGCGTCA | | |
| 431 | 12 | 59081831 | 59085915 | 4084 | BUB,FVB | 12 | 59082097 | 59082115 | 19 | + | 17.0328 | 8.88E-07 | 0.147 | TCACCAGAAGAGGGCGTCA | | |
| 431 | 12 | 59081831 | 59085915 | 4084 | BUB,FVB | 12 | 59082177 | 59082210 | 34 | + | 18.065 | 3.38E-07 | 0.064 | TAGCTGTCTTCAGACTCACCAGAAGAGGGCGTCA | | |
| 431 | 12 | 59081831 | 59085915 | 4084 | BUB,FVB | 12 | 59082192 | 59082210 | 19 | + | 17.0328 | 8.88E-07 | 0.147 | TCACCAGAAGAGGGCGTCA | | |
| 431 | 12 | 59081831 | 59085915 | 4084 | BUB,FVB | 12 | 59082272 | 59082305 | 34 | + | 18.065 | 3.38E-07 | 0.064 | TAGCTGTCTTCAGACTCACCAGAAGAGGGCGTCA | | |
| 431 | 12 | 59081831 | 59085915 | 4084 | BUB,FVB | 12 | 59082287 | 59082305 | 19 | + | 17.0328 | 8.88E-07 | 0.147 | TCACCAGAAGAGGGCGTCA | | |
| 431 | 12 | 59081831 | 59085915 | 4084 | BUB,FVB | 12 | 59082367 | 59082400 | 34 | + | 18.065 | 3.38E-07 | 0.064 | TAGCTGTCTTCAGACTCACCAGAAGAGGGCGTCA | | |
| 431 | 12 | 59081831 | 59085915 | 4084 | BUB,FVB | 12 | 59082382 | 59082400 | 19 | + | 17.0328 | 8.88E-07 | 0.147 | TCACCAGAAGAGGGCGTCA | | |
| 431 | 12 | 59081831 | 59085915 | 4084 | BUB,FVB | 12 | 59082462 | 59082495 | 34 | + | 18.065 | 3.38E-07 | 0.064 | TAGCTGTCTTCAGACTCACCAGAAGAGGGCGTCA | | |
| 431 | 12 | 59081831 | 59085915 | 4084 | BUB,FVB | 12 | 59082477 | 59082495 | 19 | + | 17.0328 | 8.88E-07 | 0.147 | TCACCAGAAGAGGGCGTCA | | |
| 431 | 12 | 59081831 | 59085915 | 4084 | BUB,FVB | 12 | 59082557 | 59082590 | 34 | + | 18.065 | 3.38E-07 | 0.064 | TAGCTGTCTTCAGACTCACCAGAAGAGGGCGTCA | | |
| 431 | 12 | 59081831 | 59085915 | 4084 | BUB,FVB | 12 | 59082572 | 59082590 | 19 | + | 17.0328 | 8.88E-07 | 0.147 | TCACCAGAAGAGGGCGTCA | | |
| 431 | 12 | 59081831 | 59085915 | 4084 | BUB,FVB | 12 | 59082652 | 59082685 | 34 | + | 18.065 | 3.38E-07 | 0.064 | TAGCTGTCTTCAGACTCACCAGAAGAGGGCGTCA | | |
| 431 | 12 | 59081831 | 59085915 | 4084 | BUB,FVB | 12 | 59082667 | 59082685 | 19 | + | 17.0328 | 8.88E-07 | 0.147 | TCACCAGAAGAGGGCGTCA | | |
| 431 | 12 | 59081831 | 59085915 | 4084 | BUB,FVB | 12 | 59082747 | 59082780 | 34 | + | 18.065 | 3.38E-07 | 0.064 | TAGCTGTCTTCAGACTCACCAGAAGAGGGCGTCA | | |
| 431 | 12 | 59081831 | 59085915 | 4084 | BUB,FVB | 12 | 59082762 | 59082780 | 19 | + | 17.0328 | 8.88E-07 | 0.147 | TCACCAGAAGAGGGCGTCA | | |
| 431 | 12 | 59081831 | 59085915 | 4084 | BUB,FVB | 12 | 59082842 | 59082875 | 34 | + | 18.065 | 3.38E-07 | 0.064 | TAGCTGTCTTCAGACTCACCAGAAGAGGGCGTCA | | |
| 431 | 12 | 59081831 | 59085915 | 4084 | BUB,FVB | 12 | 59082857 | 59082875 | 19 | + | 17.0328 | 8.88E-07 | 0.147 | TCACCAGAAGAGGGCGTCA | | |
| 431 | 12 | 59081831 | 59085915 | 4084 | BUB,FVB | 12 | 59082937 | 59082970 | 34 | + | 18.065 | 3.38E-07 | 0.064 | TAGCTGTCTTCAGACTCACCAGAAGAGGGCGTCA | | |
| 431 | 12 | 59081831 | 59085915 | 4084 | BUB,FVB | 12 | 59082952 | 59082970 | 19 | + | 17.0328 | 8.88E-07 | 0.147 | TCACCAGAAGAGGGCGTCA | | |
| 431 | 12 | 59081831 | 59085915 | 4084 | BUB,FVB | 12 | 59083032 | 59083065 | 34 | + | 18.065 | 3.38E-07 | 0.064 | TAGCTGTCTTCAGACTCACCAGAAGAGGGCGTCA | | |
| 431 | 12 | 59081831 | 59085915 | 4084 | BUB,FVB | 12 | 59083047 | 59083065 | 19 | + | 17.0328 | 8.88E-07 | 0.147 | TCACCAGAAGAGGGCGTCA | | |
| 431 | 12 | 59081831 | 59085915 | 4084 | BUB,FVB | 12 | 59083127 | 59083160 | 34 | + | 18.065 | 3.38E-07 | 0.064 | TAGCTGTCTTCAGACTCACCAGAAGAGGGCGTCA | | |
| 431 | 12 | 59081831 | 59085915 | 4084 | BUB,FVB | 12 | 59083142 | 59083160 | 19 | + | 17.0328 | 8.88E-07 | 0.147 | TCACCAGAAGAGGGCGTCA | | |
| 431 | 12 | 59081831 | 59085915 | 4084 | BUB,FVB | 12 | 59083222 | 59083255 | 34 | + | 18.065 | 3.38E-07 | 0.064 | TAGCTGTCTTCAGACTCACCAGAAGAGGGCGTCA | | |
| 431 | 12 | 59081831 | 59085915 | 4084 | BUB,FVB | 12 | 59083237 | 59083255 | 19 | + | 17.0328 | 8.88E-07 | 0.147 | TCACCAGAAGAGGGCGTCA | | |
| 431 | 12 | 59081831 | 59085915 | 4084 | BUB,FVB | 12 | 59083317 | 59083350 | 34 | + | 18.065 | 3.38E-07 | 0.064 | TAGCTGTCTTCAGACTCACCAGAAGAGGGCGTCA | | |
| 431 | 12 | 59081831 | 59085915 | 4084 | BUB,FVB | 12 | 59083332 | 59083350 | 19 | + | 17.0328 | 8.88E-07 | 0.147 | TCACCAGAAGAGGGCGTCA | | |
| 431 | 12 | 59081831 | 59085915 | 4084 | BUB,FVB | 12 | 59083412 | 59083445 | 34 | + | 18.065 | 3.38E-07 | 0.064 | TAGCTGTCTTCAGACTCACCAGAAGAGGGCGTCA | | |
| 431 | 12 | 59081831 | 59085915 | 4084 | BUB,FVB | 12 | 59083427 | 59083445 | 19 | + | 17.0328 | 8.88E-07 | 0.147 | TCACCAGAAGAGGGCGTCA | | |
| 431 | 12 | 59081831 | 59085915 | 4084 | BUB,FVB | 12 | 59083507 | 59083540 | 34 | + | 18.065 | 3.38E-07 | 0.064 | TAGCTGTCTTCAGACTCACCAGAAGAGGGCGTCA | | |
| 431 | 12 | 59081831 | 59085915 | 4084 | BUB,FVB | 12 | 59083522 | 59083540 | 19 | + | 17.0328 | 8.88E-07 | 0.147 | TCACCAGAAGAGGGCGTCA | | |
| 431 | 12 | 59081831 | 59085915 | 4084 | BUB,FVB | 12 | 59083602 | 59083635 | 34 | + | 18.065 | 3.38E-07 | 0.064 | TAGCTGTCTTCAGACTCACCAGAAGAGGGCGTCA | | |
| 431 | 12 | 59081831 | 59085915 | 4084 | BUB,FVB | 12 | 59083617 | 59083635 | 19 | + | 17.0328 | 8.88E-07 | 0.147 | TCACCAGAAGAGGGCGTCA | | |
| 431 | 12 | 59081831 | 59085915 | 4084 | BUB,FVB | 12 | 59083697 | 59083730 | 34 | + | 18.065 | 3.38E-07 | 0.064 | TAGCTGTCTTCAGACTCACCAGAAGAGGGCGTCA | | |
| 431 | 12 | 59081831 | 59085915 | 4084 | BUB,FVB | 12 | 59083712 | 59083730 | 19 | + | 17.0328 | 8.88E-07 | 0.147 | TCACCAGAAGAGGGCGTCA | | |
| 431 | 12 | 59081831 | 59085915 | 4084 | BUB,FVB | 12 | 59083792 | 59083825 | 34 | + | 18.065 | 3.38E-07 | 0.064 | TAGCTGTCTTCAGACTCACCAGAAGAGGGCGTCA | | |
| 431 | 12 | 59081831 | 59085915 | 4084 | BUB,FVB | 12 | 59083807 | 59083825 | 19 | + | 17.0328 | 8.88E-07 | 0.147 | TCACCAGAAGAGGGCGTCA | | |
| 431 | 12 | 59081831 | 59085915 | 4084 | BUB,FVB | 12 | 59083887 | 59083920 | 34 | + | 18.065 | 3.38E-07 | 0.064 | TAGCTGTCTTCAGACTCACCAGAAGAGGGCGTCA | | |
| 431 | 12 | 59081831 | 59085915 | 4084 | BUB,FVB | 12 | 59083902 | 59083920 | 19 | + | 17.0328 | 8.88E-07 | 0.147 | TCACCAGAAGAGGGCGTCA | | |
| 431 | 12 | 59081831 | 59085915 | 4084 | BUB,FVB | 12 | 59083982 | 59084015 | 34 | + | 18.065 | 3.38E-07 | 0.064 | TAGCTGTCTTCAGACTCACCAGAAGAGGGCGTCA | | |
| 431 | 12 | 59081831 | 59085915 | 4084 | BUB,FVB | 12 | 59083997 | 59084015 | 19 | + | 17.0328 | 8.88E-07 | 0.147 | TCACCAGAAGAGGGCGTCA | | |
| 431 | 12 | 59081831 | 59085915 | 4084 | BUB,FVB | 12 | 59084077 | 59084110 | 34 | + | 18.065 | 3.38E-07 | 0.064 | TAGCTGTCTTCAGACTCACCAGAAGAGGGCGTCA | | |
| 431 | 12 | 59081831 | 59085915 | 4084 | BUB,FVB | 12 | 59084092 | 59084110 | 19 | + | 17.0328 | 8.88E-07 | 0.147 | TCACCAGAAGAGGGCGTCA | | |
| 431 | 12 | 59081831 | 59085915 | 4084 | BUB,FVB | 12 | 59084172 | 59084205 | 34 | + | 18.065 | 3.38E-07 | 0.064 | TAGCTGTCTTCAGACTCACCAGAAGAGGGCGTCA | | |
| 431 | 12 | 59081831 | 59085915 | 4084 | BUB,FVB | 12 | 59084187 | 59084205 | 19 | + | 17.0328 | 8.88E-07 | 0.147 | TCACCAGAAGAGGGCGTCA | | |
| 431 | 12 | 59081831 | 59085915 | 4084 | BUB,FVB | 12 | 59084267 | 59084300 | 34 | + | 18.065 | 3.38E-07 | 0.064 | TAGCTGTCTTCAGACTCACCAGAAGAGGGCGTCA | | |
| 431 | 12 | 59081831 | 59085915 | 4084 | BUB,FVB | 12 | 59084282 | 59084300 | 19 | + | 17.0328 | 8.88E-07 | 0.147 | TCACCAGAAGAGGGCGTCA | | |
| 431 | 12 | 59081831 | 59085915 | 4084 | BUB,FVB | 12 | 59084362 | 59084395 | 34 | + | 18.065 | 3.38E-07 | 0.064 | TAGCTGTCTTCAGACTCACCAGAAGAGGGCGTCA | | |
| 431 | 12 | 59081831 | 59085915 | 4084 | BUB,FVB | 12 | 59084377 | 59084395 | 19 | + | 17.0328 | 8.88E-07 | 0.147 | TCACCAGAAGAGGGCGTCA | | |
| 431 | 12 | 59081831 | 59085915 | 4084 | BUB,FVB | 12 | 59084457 | 59084490 | 34 | + | 18.065 | 3.38E-07 | 0.064 | TAGCTGTCTTCAGACTCACCAGAAGAGGGCGTCA | | |
| 431 | 12 | 59081831 | 59085915 | 4084 | BUB,FVB | 12 | 59084472 | 59084490 | 19 | + | 17.0328 | 8.88E-07 | 0.147 | TCACCAGAAGAGGGCGTCA | | |
| 431 | 12 | 59081831 | 59085915 | 4084 | BUB,FVB | 12 | 59084552 | 59084585 | 34 | + | 18.065 | 3.38E-07 | 0.064 | TAGCTGTCTTCAGACTCACCAGAAGAGGGCGTCA | | |
| 431 | 12 | 59081831 | 59085915 | 4084 | BUB,FVB | 12 | 59084567 | 59084585 | 19 | + | 17.0328 | 8.88E-07 | 0.147 | TCACCAGAAGAGGGCGTCA | | |
| 431 | 12 | 59081831 | 59085915 | 4084 | BUB,FVB | 12 | 59084647 | 59084680 | 34 | + | 18.065 | 3.38E-07 | 0.064 | TAGCTGTCTTCAGACTCACCAGAAGAGGGCGTCA | | |
| 431 | 12 | 59081831 | 59085915 | 4084 | BUB,FVB | 12 | 59084662 | 59084680 | 19 | + | 17.0328 | 8.88E-07 | 0.147 | TCACCAGAAGAGGGCGTCA | | |
| 431 | 12 | 59081831 | 59085915 | 4084 | BUB,FVB | 12 | 59084742 | 59084775 | 34 | + | 18.065 | 3.38E-07 | 0.064 | TAGCTGTCTTCAGACTCACCAGAAGAGGGCGTCA | | |
| 431 | 12 | 59081831 | 59085915 | 4084 | BUB,FVB | 12 | 59084757 | 59084775 | 19 | + | 17.0328 | 8.88E-07 | 0.147 | TCACCAGAAGAGGGCGTCA | | |
| 431 | 12 | 59081831 | 59085915 | 4084 | BUB,FVB | 12 | 59084837 | 59084870 | 34 | + | 18.065 | 3.38E-07 | 0.064 | TAGCTGTCTTCAGACTCACCAGAAGAGGGCGTCA | | |
| 431 | 12 | 59081831 | 59085915 | 4084 | BUB,FVB | 12 | 59084852 | 59084870 | 19 | + | 17.0328 | 8.88E-07 | 0.147 | TCACCAGAAGAGGGCGTCA | | |
| 431 | 12 | 59081831 | 59085915 | 4084 | BUB,FVB | 12 | 59084932 | 59084965 | 34 | + | 18.065 | 3.38E-07 | 0.064 | TAGCTGTCTTCAGACTCACCAGAAGAGGGCGTCA | | |
| 431 | 12 | 59081831 | 59085915 | 4084 | BUB,FVB | 12 | 59084947 | 59084965 | 19 | + | 17.0328 | 8.88E-07 | 0.147 | TCACCAGAAGAGGGCGTCA | | |
| 431 | 12 | 59081831 | 59085915 | 4084 | BUB,FVB | 12 | 59085027 | 59085060 | 34 | + | 18.065 | 3.38E-07 | 0.064 | TAGCTGTCTTCAGACTCACCAGAAGAGGGCGTCA | | |
| 431 | 12 | 59081831 | 59085915 | 4084 | BUB,FVB | 12 | 59085042 | 59085060 | 19 | + | 17.0328 | 8.88E-07 | 0.147 | TCACCAGAAGAGGGCGTCA | | |
| 431 | 12 | 59081831 | 59085915 | 4084 | BUB,FVB | 12 | 59085122 | 59085155 | 34 | + | 18.065 | 3.38E-07 | 0.064 | TAGCTGTCTTCAGACTCACCAGAAGAGGGCGTCA | | |
| 431 | 12 | 59081831 | 59085915 | 4084 | BUB,FVB | 12 | 59085137 | 59085155 | 19 | + | 17.0328 | 8.88E-07 | 0.147 | TCACCAGAAGAGGGCGTCA | | |
| 431 | 12 | 59081831 | 59085915 | 4084 | BUB,FVB | 12 | 59085217 | 59085250 | 34 | + | 18.065 | 3.38E-07 | 0.064 | TAGCTGTCTTCAGACTCACCAGAAGAGGGCGTCA | | |
| 431 | 12 | 59081831 | 59085915 | 4084 | BUB,FVB | 12 | 59085232 | 59085250 | 19 | + | 17.0328 | 8.88E-07 | 0.147 | TCACCAGAAGAGGGCGTCA | | |
| 431 | 12 | 59081831 | 59085915 | 4084 | BUB,FVB | 12 | 59085312 | 59085345 | 34 | + | 18.065 | 3.38E-07 | 0.064 | TAGCTGTCTTCAGACTCACCAGAAGAGGGCGTCA | | |
| 431 | 12 | 59081831 | 59085915 | 4084 | BUB,FVB | 12 | 59085327 | 59085345 | 19 | + | 17.0328 | 8.88E-07 | 0.147 | TCACCAGAAGAGGGCGTCA | | |
| 431 | 12 | 59081831 | 59085915 | 4084 | BUB,FVB | 12 | 59085407 | 59085440 | 34 | + | 18.065 | 3.38E-07 | 0.064 | TAGCTGTCTTCAGACTCACCAGAAGAGGGCGTCA | | |
| 431 | 12 | 59081831 | 59085915 | 4084 | BUB,FVB | 12 | 59085422 | 59085440 | 19 | + | 17.0328 | 8.88E-07 | 0.147 | TCACCAGAAGAGGGCGTCA | | |
| 431 | 12 | 59081831 | 59085915 | 4084 | BUB,FVB | 12 | 59085502 | 59085535 | 34 | + | 18.065 | 3.38E-07 | 0.064 | TAGCTGTCTTCAGACTCACCAGAAGAGGGCGTCA | | |
| 431 | 12 | 59081831 | 59085915 | 4084 | BUB,FVB | 12 | 59085517 | 59085535 | 19 | + | 17.0328 | 8.88E-07 | 0.147 | TCACCAGAAGAGGGCGTCA | | |
| 431 | 12 | 59081831 | 59085915 | 4084 | BUB,FVB | 12 | 59085692 | 59085725 | 34 | + | 18.065 | 3.38E-07 | 0.064 | TAGCTGTCTTCAGACTCACCAGAAGAGGGCGTCA | | |
| 431 | 12 | 59081831 | 59085915 | 4084 | BUB,FVB | 12 | 59085707 | 59085725 | 19 | + | 17.0328 | 8.88E-07 | 0.147 | TCACCAGAAGAGGGCGTCA | | |
| 431 | 12 | 59081831 | 59085915 | 4084 | BUB,FVB | 12 | 59085787 | 59085820 | 34 | + | 16.4472 | 9.61E-07 | 0.104 | TAGCTGTCTTCAGACACACCAGAAGAGGGCGTCA | 2 | ENCFF024JOZ,ENCFF883UPM |
| 432 | 12 | 59082294 | 59085144 | 2850 | AJ,BTBR,CBA,KK,SJL,SWR | 12 | 59082272 | 59082305 | 34 | + | 18.065 | 3.38E-07 | 0.064 | TAGCTGTCTTCAGACTCACCAGAAGAGGGCGTCA | | |
| 432 | 12 | 59082294 | 59085144 | 2850 | AJ,BTBR,CBA,KK,SJL,SWR | 12 | 59082287 | 59082305 | 19 | + | 17.0328 | 8.88E-07 | 0.147 | TCACCAGAAGAGGGCGTCA | | |
| 432 | 12 | 59082294 | 59085144 | 2850 | AJ,BTBR,CBA,KK,SJL,SWR | 12 | 59082367 | 59082400 | 34 | + | 18.065 | 3.38E-07 | 0.064 | TAGCTGTCTTCAGACTCACCAGAAGAGGGCGTCA | | |
| 432 | 12 | 59082294 | 59085144 | 2850 | AJ,BTBR,CBA,KK,SJL,SWR | 12 | 59082382 | 59082400 | 19 | + | 17.0328 | 8.88E-07 | 0.147 | TCACCAGAAGAGGGCGTCA | | |
| 432 | 12 | 59082294 | 59085144 | 2850 | AJ,BTBR,CBA,KK,SJL,SWR | 12 | 59082462 | 59082495 | 34 | + | 18.065 | 3.38E-07 | 0.064 | TAGCTGTCTTCAGACTCACCAGAAGAGGGCGTCA | | |
| 432 | 12 | 59082294 | 59085144 | 2850 | AJ,BTBR,CBA,KK,SJL,SWR | 12 | 59082477 | 59082495 | 19 | + | 17.0328 | 8.88E-07 | 0.147 | TCACCAGAAGAGGGCGTCA | | |
| 432 | 12 | 59082294 | 59085144 | 2850 | AJ,BTBR,CBA,KK,SJL,SWR | 12 | 59082557 | 59082590 | 34 | + | 18.065 | 3.38E-07 | 0.064 | TAGCTGTCTTCAGACTCACCAGAAGAGGGCGTCA | | |
| 432 | 12 | 59082294 | 59085144 | 2850 | AJ,BTBR,CBA,KK,SJL,SWR | 12 | 59082572 | 59082590 | 19 | + | 17.0328 | 8.88E-07 | 0.147 | TCACCAGAAGAGGGCGTCA | | |
| 432 | 12 | 59082294 | 59085144 | 2850 | AJ,BTBR,CBA,KK,SJL,SWR | 12 | 59082652 | 59082685 | 34 | + | 18.065 | 3.38E-07 | 0.064 | TAGCTGTCTTCAGACTCACCAGAAGAGGGCGTCA | | |
| 432 | 12 | 59082294 | 59085144 | 2850 | AJ,BTBR,CBA,KK,SJL,SWR | 12 | 59082667 | 59082685 | 19 | + | 17.0328 | 8.88E-07 | 0.147 | TCACCAGAAGAGGGCGTCA | | |
| 432 | 12 | 59082294 | 59085144 | 2850 | AJ,BTBR,CBA,KK,SJL,SWR | 12 | 59082747 | 59082780 | 34 | + | 18.065 | 3.38E-07 | 0.064 | TAGCTGTCTTCAGACTCACCAGAAGAGGGCGTCA | | |
| 432 | 12 | 59082294 | 59085144 | 2850 | AJ,BTBR,CBA,KK,SJL,SWR | 12 | 59082762 | 59082780 | 19 | + | 17.0328 | 8.88E-07 | 0.147 | TCACCAGAAGAGGGCGTCA | | |
| 432 | 12 | 59082294 | 59085144 | 2850 | AJ,BTBR,CBA,KK,SJL,SWR | 12 | 59082842 | 59082875 | 34 | + | 18.065 | 3.38E-07 | 0.064 | TAGCTGTCTTCAGACTCACCAGAAGAGGGCGTCA | | |
| 432 | 12 | 59082294 | 59085144 | 2850 | AJ,BTBR,CBA,KK,SJL,SWR | 12 | 59082857 | 59082875 | 19 | + | 17.0328 | 8.88E-07 | 0.147 | TCACCAGAAGAGGGCGTCA | | |
| 432 | 12 | 59082294 | 59085144 | 2850 | AJ,BTBR,CBA,KK,SJL,SWR | 12 | 59082937 | 59082970 | 34 | + | 18.065 | 3.38E-07 | 0.064 | TAGCTGTCTTCAGACTCACCAGAAGAGGGCGTCA | | |
| 432 | 12 | 59082294 | 59085144 | 2850 | AJ,BTBR,CBA,KK,SJL,SWR | 12 | 59082952 | 59082970 | 19 | + | 17.0328 | 8.88E-07 | 0.147 | TCACCAGAAGAGGGCGTCA | | |
| 432 | 12 | 59082294 | 59085144 | 2850 | AJ,BTBR,CBA,KK,SJL,SWR | 12 | 59083032 | 59083065 | 34 | + | 18.065 | 3.38E-07 | 0.064 | TAGCTGTCTTCAGACTCACCAGAAGAGGGCGTCA | | |
| 432 | 12 | 59082294 | 59085144 | 2850 | AJ,BTBR,CBA,KK,SJL,SWR | 12 | 59083047 | 59083065 | 19 | + | 17.0328 | 8.88E-07 | 0.147 | TCACCAGAAGAGGGCGTCA | | |
| 432 | 12 | 59082294 | 59085144 | 2850 | AJ,BTBR,CBA,KK,SJL,SWR | 12 | 59083127 | 59083160 | 34 | + | 18.065 | 3.38E-07 | 0.064 | TAGCTGTCTTCAGACTCACCAGAAGAGGGCGTCA | | |
| 432 | 12 | 59082294 | 59085144 | 2850 | AJ,BTBR,CBA,KK,SJL,SWR | 12 | 59083142 | 59083160 | 19 | + | 17.0328 | 8.88E-07 | 0.147 | TCACCAGAAGAGGGCGTCA | | |

| | | | | | | | | | | | | | | | | |
|---|---|---|---|---|---|---|---|---|---|---|---|---|---|---|---|---|
| 432 | 12 | 59082294 | 59085144 | 2850 | AJ,BTBR,CBA,KK,SJL,SWR | 12 | 59083222 | 59083255 | 34 | + | 18.065 | 3.38E-07 | 0.064 | TAGCTGTCTTCAGACTCACCAGAAGAGGGCGTCA | | |
| 432 | 12 | 59082294 | 59085144 | 2850 | AJ,BTBR,CBA,KK,SJL,SWR | 12 | 59083237 | 59083255 | 19 | + | 17.0328 | 8.88E-07 | 0.147 | TCACCAGAAGAGGGCGTCA | | |
| 432 | 12 | 59082294 | 59085144 | 2850 | AJ,BTBR,CBA,KK,SJL,SWR | 12 | 59083317 | 59083350 | 34 | + | 18.065 | 3.38E-07 | 0.064 | TAGCTGTCTTCAGACTCACCAGAAGAGGGCGTCA | | |
| 432 | 12 | 59082294 | 59085144 | 2850 | AJ,BTBR,CBA,KK,SJL,SWR | 12 | 59083332 | 59083350 | 19 | + | 17.0328 | 8.88E-07 | 0.147 | TCACCAGAAGAGGGCGTCA | | |
| 432 | 12 | 59082294 | 59085144 | 2850 | AJ,BTBR,CBA,KK,SJL,SWR | 12 | 59083412 | 59083445 | 34 | + | 18.065 | 3.38E-07 | 0.064 | TAGCTGTCTTCAGACTCACCAGAAGAGGGCGTCA | | |
| 432 | 12 | 59082294 | 59085144 | 2850 | AJ,BTBR,CBA,KK,SJL,SWR | 12 | 59083427 | 59083445 | 19 | + | 17.0328 | 8.88E-07 | 0.147 | TCACCAGAAGAGGGCGTCA | | |
| 432 | 12 | 59082294 | 59085144 | 2850 | AJ,BTBR,CBA,KK,SJL,SWR | 12 | 59083507 | 59083540 | 34 | + | 18.065 | 3.38E-07 | 0.064 | TAGCTGTCTTCAGACTCACCAGAAGAGGGCGTCA | | |
| 432 | 12 | 59082294 | 59085144 | 2850 | AJ,BTBR,CBA,KK,SJL,SWR | 12 | 59083522 | 59083540 | 19 | + | 17.0328 | 8.88E-07 | 0.147 | TCACCAGAAGAGGGCGTCA | | |
| 432 | 12 | 59082294 | 59085144 | 2850 | AJ,BTBR,CBA,KK,SJL,SWR | 12 | 59083602 | 59083635 | 34 | + | 18.065 | 3.38E-07 | 0.064 | TAGCTGTCTTCAGACTCACCAGAAGAGGGCGTCA | | |
| 432 | 12 | 59082294 | 59085144 | 2850 | AJ,BTBR,CBA,KK,SJL,SWR | 12 | 59083617 | 59083635 | 19 | + | 17.0328 | 8.88E-07 | 0.147 | TCACCAGAAGAGGGCGTCA | | |
| 432 | 12 | 59082294 | 59085144 | 2850 | AJ,BTBR,CBA,KK,SJL,SWR | 12 | 59083697 | 59083730 | 34 | + | 18.065 | 3.38E-07 | 0.064 | TAGCTGTCTTCAGACTCACCAGAAGAGGGCGTCA | | |
| 432 | 12 | 59082294 | 59085144 | 2850 | AJ,BTBR,CBA,KK,SJL,SWR | 12 | 59083712 | 59083730 | 19 | + | 17.0328 | 8.88E-07 | 0.147 | TCACCAGAAGAGGGCGTCA | | |
| 432 | 12 | 59082294 | 59085144 | 2850 | AJ,BTBR,CBA,KK,SJL,SWR | 12 | 59083792 | 59083825 | 34 | + | 18.065 | 3.38E-07 | 0.064 | TAGCTGTCTTCAGACTCACCAGAAGAGGGCGTCA | | |
| 432 | 12 | 59082294 | 59085144 | 2850 | AJ,BTBR,CBA,KK,SJL,SWR | 12 | 59083807 | 59083825 | 19 | + | 17.0328 | 8.88E-07 | 0.147 | TCACCAGAAGAGGGCGTCA | | |
| 432 | 12 | 59082294 | 59085144 | 2850 | AJ,BTBR,CBA,KK,SJL,SWR | 12 | 59083887 | 59083920 | 34 | + | 18.065 | 3.38E-07 | 0.064 | TAGCTGTCTTCAGACTCACCAGAAGAGGGCGTCA | | |
| 432 | 12 | 59082294 | 59085144 | 2850 | AJ,BTBR,CBA,KK,SJL,SWR | 12 | 59083902 | 59083920 | 19 | + | 17.0328 | 8.88E-07 | 0.147 | TCACCAGAAGAGGGCGTCA | | |
| 432 | 12 | 59082294 | 59085144 | 2850 | AJ,BTBR,CBA,KK,SJL,SWR | 12 | 59083982 | 59084015 | 34 | + | 18.065 | 3.38E-07 | 0.064 | TAGCTGTCTTCAGACTCACCAGAAGAGGGCGTCA | | |
| 432 | 12 | 59082294 | 59085144 | 2850 | AJ,BTBR,CBA,KK,SJL,SWR | 12 | 59083997 | 59084015 | 19 | + | 17.0328 | 8.88E-07 | 0.147 | TCACCAGAAGAGGGCGTCA | | |
| 432 | 12 | 59082294 | 59085144 | 2850 | AJ,BTBR,CBA,KK,SJL,SWR | 12 | 59084077 | 59084110 | 34 | + | 18.065 | 3.38E-07 | 0.064 | TAGCTGTCTTCAGACTCACCAGAAGAGGGCGTCA | | |
| 432 | 12 | 59082294 | 59085144 | 2850 | AJ,BTBR,CBA,KK,SJL,SWR | 12 | 59084092 | 59084110 | 19 | + | 17.0328 | 8.88E-07 | 0.147 | TCACCAGAAGAGGGCGTCA | | |
| 432 | 12 | 59082294 | 59085144 | 2850 | AJ,BTBR,CBA,KK,SJL,SWR | 12 | 59084172 | 59084205 | 34 | + | 18.065 | 3.38E-07 | 0.064 | TAGCTGTCTTCAGACTCACCAGAAGAGGGCGTCA | | |
| 432 | 12 | 59082294 | 59085144 | 2850 | AJ,BTBR,CBA,KK,SJL,SWR | 12 | 59084187 | 59084205 | 19 | + | 17.0328 | 8.88E-07 | 0.147 | TCACCAGAAGAGGGCGTCA | | |
| 432 | 12 | 59082294 | 59085144 | 2850 | AJ,BTBR,CBA,KK,SJL,SWR | 12 | 59084267 | 59084300 | 34 | + | 18.065 | 3.38E-07 | 0.064 | TAGCTGTCTTCAGACTCACCAGAAGAGGGCGTCA | | |
| 432 | 12 | 59082294 | 59085144 | 2850 | AJ,BTBR,CBA,KK,SJL,SWR | 12 | 59084282 | 59084300 | 19 | + | 17.0328 | 8.88E-07 | 0.147 | TCACCAGAAGAGGGCGTCA | | |
| 432 | 12 | 59082294 | 59085144 | 2850 | AJ,BTBR,CBA,KK,SJL,SWR | 12 | 59084362 | 59084395 | 34 | + | 18.065 | 3.38E-07 | 0.064 | TAGCTGTCTTCAGACTCACCAGAAGAGGGCGTCA | | |
| 432 | 12 | 59082294 | 59085144 | 2850 | AJ,BTBR,CBA,KK,SJL,SWR | 12 | 59084377 | 59084395 | 19 | + | 17.0328 | 8.88E-07 | 0.147 | TCACCAGAAGAGGGCGTCA | | |
| 432 | 12 | 59082294 | 59085144 | 2850 | AJ,BTBR,CBA,KK,SJL,SWR | 12 | 59084457 | 59084490 | 34 | + | 18.065 | 3.38E-07 | 0.064 | TAGCTGTCTTCAGACTCACCAGAAGAGGGCGTCA | | |
| 432 | 12 | 59082294 | 59085144 | 2850 | AJ,BTBR,CBA,KK,SJL,SWR | 12 | 59084472 | 59084490 | 19 | + | 17.0328 | 8.88E-07 | 0.147 | TCACCAGAAGAGGGCGTCA | | |
| 432 | 12 | 59082294 | 59085144 | 2850 | AJ,BTBR,CBA,KK,SJL,SWR | 12 | 59084552 | 59084585 | 34 | + | 18.065 | 3.38E-07 | 0.064 | TAGCTGTCTTCAGACTCACCAGAAGAGGGCGTCA | | |
| 432 | 12 | 59082294 | 59085144 | 2850 | AJ,BTBR,CBA,KK,SJL,SWR | 12 | 59084567 | 59084585 | 19 | + | 17.0328 | 8.88E-07 | 0.147 | TCACCAGAAGAGGGCGTCA | | |
| 432 | 12 | 59082294 | 59085144 | 2850 | AJ,BTBR,CBA,KK,SJL,SWR | 12 | 59084647 | 59084680 | 34 | + | 18.065 | 3.38E-07 | 0.064 | TAGCTGTCTTCAGACTCACCAGAAGAGGGCGTCA | | |
| 432 | 12 | 59082294 | 59085144 | 2850 | AJ,BTBR,CBA,KK,SJL,SWR | 12 | 59084662 | 59084680 | 19 | + | 17.0328 | 8.88E-07 | 0.147 | TCACCAGAAGAGGGCGTCA | | |
| 432 | 12 | 59082294 | 59085144 | 2850 | AJ,BTBR,CBA,KK,SJL,SWR | 12 | 59084742 | 59084775 | 34 | + | 18.065 | 3.38E-07 | 0.064 | TAGCTGTCTTCAGACTCACCAGAAGAGGGCGTCA | | |
| 432 | 12 | 59082294 | 59085144 | 2850 | AJ,BTBR,CBA,KK,SJL,SWR | 12 | 59084757 | 59084775 | 19 | + | 17.0328 | 8.88E-07 | 0.147 | TCACCAGAAGAGGGCGTCA | | |
| 432 | 12 | 59082294 | 59085144 | 2850 | AJ,BTBR,CBA,KK,SJL,SWR | 12 | 59084837 | 59084870 | 34 | + | 18.065 | 3.38E-07 | 0.064 | TAGCTGTCTTCAGACTCACCAGAAGAGGGCGTCA | | |
| 432 | 12 | 59082294 | 59085144 | 2850 | AJ,BTBR,CBA,KK,SJL,SWR | 12 | 59084852 | 59084870 | 19 | + | 17.0328 | 8.88E-07 | 0.147 | TCACCAGAAGAGGGCGTCA | | |
| 432 | 12 | 59082294 | 59085144 | 2850 | AJ,BTBR,CBA,KK,SJL,SWR | 12 | 59084932 | 59084965 | 34 | + | 18.065 | 3.38E-07 | 0.064 | TAGCTGTCTTCAGACTCACCAGAAGAGGGCGTCA | | |
| 432 | 12 | 59082294 | 59085144 | 2850 | AJ,BTBR,CBA,KK,SJL,SWR | 12 | 59084947 | 59084965 | 19 | + | 17.0328 | 8.88E-07 | 0.147 | TCACCAGAAGAGGGCGTCA | | |
| 432 | 12 | 59082294 | 59085144 | 2850 | AJ,BTBR,CBA,KK,SJL,SWR | 12 | 59085027 | 59085060 | 34 | + | 18.065 | 3.38E-07 | 0.064 | TAGCTGTCTTCAGACTCACCAGAAGAGGGCGTCA | | |
| 432 | 12 | 59082294 | 59085144 | 2850 | AJ,BTBR,CBA,KK,SJL,SWR | 12 | 59085042 | 59085060 | 19 | + | 17.0328 | 8.88E-07 | 0.147 | TCACCAGAAGAGGGCGTCA | | |
| 432 | 12 | 59082294 | 59085144 | 2850 | AJ,BTBR,CBA,KK,SJL,SWR | 12 | 59085122 | 59085155 | 34 | + | 18.065 | 3.38E-07 | 0.064 | TAGCTGTCTTCAGACTCACCAGAAGAGGGCGTCA | | |
| 432 | 12 | 59082294 | 59085144 | 2850 | AJ,BTBR,CBA,KK,SJL,SWR | 12 | 59085137 | 59085155 | 19 | + | 17.0328 | 8.88E-07 | 0.147 | TCACCAGAAGAGGGCGTCA | | |
| 433 | 12 | 59082858 | 59085898 | 3040 | SMJ | 12 | 59082842 | 59082875 | 34 | + | 18.065 | 3.38E-07 | 0.064 | TAGCTGTCTTCAGACTCACCAGAAGAGGGCGTCA | | |
| 433 | 12 | 59082858 | 59085898 | 3040 | SMJ | 12 | 59082857 | 59082875 | 19 | + | 17.0328 | 8.88E-07 | 0.147 | TCACCAGAAGAGGGCGTCA | | |
| 433 | 12 | 59082858 | 59085898 | 3040 | SMJ | 12 | 59082937 | 59082970 | 34 | + | 18.065 | 3.38E-07 | 0.064 | TAGCTGTCTTCAGACTCACCAGAAGAGGGCGTCA | | |
| 433 | 12 | 59082858 | 59085898 | 3040 | SMJ | 12 | 59082952 | 59082970 | 19 | + | 17.0328 | 8.88E-07 | 0.147 | TCACCAGAAGAGGGCGTCA | | |
| 433 | 12 | 59082858 | 59085898 | 3040 | SMJ | 12 | 59083032 | 59083065 | 34 | + | 18.065 | 3.38E-07 | 0.064 | TAGCTGTCTTCAGACTCACCAGAAGAGGGCGTCA | | |
| 433 | 12 | 59082858 | 59085898 | 3040 | SMJ | 12 | 59083047 | 59083065 | 19 | + | 17.0328 | 8.88E-07 | 0.147 | TCACCAGAAGAGGGCGTCA | | |
| 433 | 12 | 59082858 | 59085898 | 3040 | SMJ | 12 | 59083127 | 59083160 | 34 | + | 18.065 | 3.38E-07 | 0.064 | TAGCTGTCTTCAGACTCACCAGAAGAGGGCGTCA | | |
| 433 | 12 | 59082858 | 59085898 | 3040 | SMJ | 12 | 59083142 | 59083160 | 19 | + | 17.0328 | 8.88E-07 | 0.147 | TCACCAGAAGAGGGCGTCA | | |
| 433 | 12 | 59082858 | 59085898 | 3040 | SMJ | 12 | 59083222 | 59083255 | 34 | + | 18.065 | 3.38E-07 | 0.064 | TAGCTGTCTTCAGACTCACCAGAAGAGGGCGTCA | | |
| 433 | 12 | 59082858 | 59085898 | 3040 | SMJ | 12 | 59083237 | 59083255 | 19 | + | 17.0328 | 8.88E-07 | 0.147 | TCACCAGAAGAGGGCGTCA | | |
| 433 | 12 | 59082858 | 59085898 | 3040 | SMJ | 12 | 59083317 | 59083350 | 34 | + | 18.065 | 3.38E-07 | 0.064 | TAGCTGTCTTCAGACTCACCAGAAGAGGGCGTCA | | |
| 433 | 12 | 59082858 | 59085898 | 3040 | SMJ | 12 | 59083332 | 59083350 | 19 | + | 17.0328 | 8.88E-07 | 0.147 | TCACCAGAAGAGGGCGTCA | | |
| 433 | 12 | 59082858 | 59085898 | 3040 | SMJ | 12 | 59083412 | 59083445 | 34 | + | 18.065 | 3.38E-07 | 0.064 | TAGCTGTCTTCAGACTCACCAGAAGAGGGCGTCA | | |
| 433 | 12 | 59082858 | 59085898 | 3040 | SMJ | 12 | 59083427 | 59083445 | 19 | + | 17.0328 | 8.88E-07 | 0.147 | TCACCAGAAGAGGGCGTCA | | |
| 433 | 12 | 59082858 | 59085898 | 3040 | SMJ | 12 | 59083507 | 59083540 | 34 | + | 18.065 | 3.38E-07 | 0.064 | TAGCTGTCTTCAGACTCACCAGAAGAGGGCGTCA | | |
| 433 | 12 | 59082858 | 59085898 | 3040 | SMJ | 12 | 59083522 | 59083540 | 19 | + | 17.0328 | 8.88E-07 | 0.147 | TCACCAGAAGAGGGCGTCA | | |
| 433 | 12 | 59082858 | 59085898 | 3040 | SMJ | 12 | 59083602 | 59083635 | 34 | + | 18.065 | 3.38E-07 | 0.064 | TAGCTGTCTTCAGACTCACCAGAAGAGGGCGTCA | | |
| 433 | 12 | 59082858 | 59085898 | 3040 | SMJ | 12 | 59083617 | 59083635 | 19 | + | 17.0328 | 8.88E-07 | 0.147 | TCACCAGAAGAGGGCGTCA | | |
| 433 | 12 | 59082858 | 59085898 | 3040 | SMJ | 12 | 59083697 | 59083730 | 34 | + | 18.065 | 3.38E-07 | 0.064 | TAGCTGTCTTCAGACTCACCAGAAGAGGGCGTCA | | |
| 433 | 12 | 59082858 | 59085898 | 3040 | SMJ | 12 | 59083712 | 59083730 | 19 | + | 17.0328 | 8.88E-07 | 0.147 | TCACCAGAAGAGGGCGTCA | | |
| 433 | 12 | 59082858 | 59085898 | 3040 | SMJ | 12 | 59083792 | 59083825 | 34 | + | 18.065 | 3.38E-07 | 0.064 | TAGCTGTCTTCAGACTCACCAGAAGAGGGCGTCA | | |
| 433 | 12 | 59082858 | 59085898 | 3040 | SMJ | 12 | 59083807 | 59083825 | 19 | + | 17.0328 | 8.88E-07 | 0.147 | TCACCAGAAGAGGGCGTCA | | |
| 433 | 12 | 59082858 | 59085898 | 3040 | SMJ | 12 | 59083887 | 59083920 | 34 | + | 18.065 | 3.38E-07 | 0.064 | TAGCTGTCTTCAGACTCACCAGAAGAGGGCGTCA | | |
| 433 | 12 | 59082858 | 59085898 | 3040 | SMJ | 12 | 59083902 | 59083920 | 19 | + | 17.0328 | 8.88E-07 | 0.147 | TCACCAGAAGAGGGCGTCA | | |
| 433 | 12 | 59082858 | 59085898 | 3040 | SMJ | 12 | 59083982 | 59084015 | 34 | + | 18.065 | 3.38E-07 | 0.064 | TAGCTGTCTTCAGACTCACCAGAAGAGGGCGTCA | | |
| 433 | 12 | 59082858 | 59085898 | 3040 | SMJ | 12 | 59083997 | 59084015 | 19 | + | 17.0328 | 8.88E-07 | 0.147 | TCACCAGAAGAGGGCGTCA | | |
| 433 | 12 | 59082858 | 59085898 | 3040 | SMJ | 12 | 59084077 | 59084110 | 34 | + | 18.065 | 3.38E-07 | 0.064 | TAGCTGTCTTCAGACTCACCAGAAGAGGGCGTCA | | |
| 433 | 12 | 59082858 | 59085898 | 3040 | SMJ | 12 | 59084092 | 59084110 | 19 | + | 17.0328 | 8.88E-07 | 0.147 | TCACCAGAAGAGGGCGTCA | | |
| 433 | 12 | 59082858 | 59085898 | 3040 | SMJ | 12 | 59084172 | 59084205 | 34 | + | 18.065 | 3.38E-07 | 0.064 | TAGCTGTCTTCAGACTCACCAGAAGAGGGCGTCA | | |
| 433 | 12 | 59082858 | 59085898 | 3040 | SMJ | 12 | 59084187 | 59084205 | 19 | + | 17.0328 | 8.88E-07 | 0.147 | TCACCAGAAGAGGGCGTCA | | |
| 433 | 12 | 59082858 | 59085898 | 3040 | SMJ | 12 | 59084267 | 59084300 | 34 | + | 18.065 | 3.38E-07 | 0.064 | TAGCTGTCTTCAGACTCACCAGAAGAGGGCGTCA | | |
| 433 | 12 | 59082858 | 59085898 | 3040 | SMJ | 12 | 59084282 | 59084300 | 19 | + | 17.0328 | 8.88E-07 | 0.147 | TCACCAGAAGAGGGCGTCA | | |
| 433 | 12 | 59082858 | 59085898 | 3040 | SMJ | 12 | 59084362 | 59084395 | 34 | + | 18.065 | 3.38E-07 | 0.064 | TAGCTGTCTTCAGACTCACCAGAAGAGGGCGTCA | | |
| 433 | 12 | 59082858 | 59085898 | 3040 | SMJ | 12 | 59084377 | 59084395 | 19 | + | 17.0328 | 8.88E-07 | 0.147 | TCACCAGAAGAGGGCGTCA | | |
| 433 | 12 | 59082858 | 59085898 | 3040 | SMJ | 12 | 59084457 | 59084490 | 34 | + | 18.065 | 3.38E-07 | 0.064 | TAGCTGTCTTCAGACTCACCAGAAGAGGGCGTCA | | |
| 433 | 12 | 59082858 | 59085898 | 3040 | SMJ | 12 | 59084472 | 59084490 | 19 | + | 17.0328 | 8.88E-07 | 0.147 | TCACCAGAAGAGGGCGTCA | | |
| 433 | 12 | 59082858 | 59085898 | 3040 | SMJ | 12 | 59084552 | 59084585 | 34 | + | 18.065 | 3.38E-07 | 0.064 | TAGCTGTCTTCAGACTCACCAGAAGAGGGCGTCA | | |
| 433 | 12 | 59082858 | 59085898 | 3040 | SMJ | 12 | 59084567 | 59084585 | 19 | + | 17.0328 | 8.88E-07 | 0.147 | TCACCAGAAGAGGGCGTCA | | |
| 433 | 12 | 59082858 | 59085898 | 3040 | SMJ | 12 | 59084647 | 59084680 | 34 | + | 18.065 | 3.38E-07 | 0.064 | TAGCTGTCTTCAGACTCACCAGAAGAGGGCGTCA | | |
| 433 | 12 | 59082858 | 59085898 | 3040 | SMJ | 12 | 59084662 | 59084680 | 19 | + | 17.0328 | 8.88E-07 | 0.147 | TCACCAGAAGAGGGCGTCA | | |
| 433 | 12 | 59082858 | 59085898 | 3040 | SMJ | 12 | 59084742 | 59084775 | 34 | + | 18.065 | 3.38E-07 | 0.064 | TAGCTGTCTTCAGACTCACCAGAAGAGGGCGTCA | | |
| 433 | 12 | 59082858 | 59085898 | 3040 | SMJ | 12 | 59084757 | 59084775 | 19 | + | 17.0328 | 8.88E-07 | 0.147 | TCACCAGAAGAGGGCGTCA | | |
| 433 | 12 | 59082858 | 59085898 | 3040 | SMJ | 12 | 59084837 | 59084870 | 34 | + | 18.065 | 3.38E-07 | 0.064 | TAGCTGTCTTCAGACTCACCAGAAGAGGGCGTCA | | |
| 433 | 12 | 59082858 | 59085898 | 3040 | SMJ | 12 | 59084852 | 59084870 | 19 | + | 17.0328 | 8.88E-07 | 0.147 | TCACCAGAAGAGGGCGTCA | | |
| 433 | 12 | 59082858 | 59085898 | 3040 | SMJ | 12 | 59084932 | 59084965 | 34 | + | 18.065 | 3.38E-07 | 0.064 | TAGCTGTCTTCAGACTCACCAGAAGAGGGCGTCA | | |
| 433 | 12 | 59082858 | 59085898 | 3040 | SMJ | 12 | 59084947 | 59084965 | 19 | + | 17.0328 | 8.88E-07 | 0.147 | TCACCAGAAGAGGGCGTCA | | |
| 433 | 12 | 59082858 | 59085898 | 3040 | SMJ | 12 | 59085027 | 59085060 | 34 | + | 18.065 | 3.38E-07 | 0.064 | TAGCTGTCTTCAGACTCACCAGAAGAGGGCGTCA | | |
| 433 | 12 | 59082858 | 59085898 | 3040 | SMJ | 12 | 59085042 | 59085060 | 19 | + | 17.0328 | 8.88E-07 | 0.147 | TCACCAGAAGAGGGCGTCA | | |
| 433 | 12 | 59082858 | 59085898 | 3040 | SMJ | 12 | 59085122 | 59085155 | 34 | + | 18.065 | 3.38E-07 | 0.064 | TAGCTGTCTTCAGACTCACCAGAAGAGGGCGTCA | | |
| 433 | 12 | 59082858 | 59085898 | 3040 | SMJ | 12 | 59085137 | 59085155 | 19 | + | 17.0328 | 8.88E-07 | 0.147 | TCACCAGAAGAGGGCGTCA | | |
| 433 | 12 | 59082858 | 59085898 | 3040 | SMJ | 12 | 59085217 | 59085250 | 34 | + | 18.065 | 3.38E-07 | 0.064 | TAGCTGTCTTCAGACTCACCAGAAGAGGGCGTCA | | |
| 433 | 12 | 59082858 | 59085898 | 3040 | SMJ | 12 | 59085232 | 59085250 | 19 | + | 17.0328 | 8.88E-07 | 0.147 | TCACCAGAAGAGGGCGTCA | | |
| 433 | 12 | 59082858 | 59085898 | 3040 | SMJ | 12 | 59085312 | 59085345 | 34 | + | 18.065 | 3.38E-07 | 0.064 | TAGCTGTCTTCAGACTCACCAGAAGAGGGCGTCA | | |
| 433 | 12 | 59082858 | 59085898 | 3040 | SMJ | 12 | 59085327 | 59085345 | 19 | + | 17.0328 | 8.88E-07 | 0.147 | TCACCAGAAGAGGGCGTCA | | |
| 433 | 12 | 59082858 | 59085898 | 3040 | SMJ | 12 | 59085407 | 59085440 | 34 | + | 18.065 | 3.38E-07 | 0.064 | TAGCTGTCTTCAGACTCACCAGAAGAGGGCGTCA | | |
| 433 | 12 | 59082858 | 59085898 | 3040 | SMJ | 12 | 59085422 | 59085440 | 19 | + | 17.0328 | 8.88E-07 | 0.147 | TCACCAGAAGAGGGCGTCA | | |
| 433 | 12 | 59082858 | 59085898 | 3040 | SMJ | 12 | 59085502 | 59085535 | 34 | + | 18.065 | 3.38E-07 | 0.064 | TAGCTGTCTTCAGACTCACCAGAAGAGGGCGTCA | | |
| 433 | 12 | 59082858 | 59085898 | 3040 | SMJ | 12 | 59085517 | 59085535 | 19 | + | 17.0328 | 8.88E-07 | 0.147 | TCACCAGAAGAGGGCGTCA | | |
| 433 | 12 | 59082858 | 59085898 | 3040 | SMJ | 12 | 59085692 | 59085725 | 34 | + | 18.065 | 3.38E-07 | 0.064 | TAGCTGTCTTCAGACTCACCAGAAGAGGGCGTCA | | |
| 433 | 12 | 59082858 | 59085898 | 3040 | SMJ | 12 | 59085707 | 59085725 | 19 | + | 17.0328 | 8.88E-07 | 0.147 | TCACCAGAAGAGGGCGTCA | | |
| 433 | 12 | 59082858 | 59085898 | 3040 | SMJ | 12 | 59085787 | 59085820 | 34 | + | 16.4472 | 9.61E-07 | 0.104 | TAGCTGTCTTCAGACACACCAGAAGAGGGCGTCA | 2 | ENCFF024JOZ,ENCFF883UPM |
| 434 | 12 | 59083722 | 59085622 | 1900 | AKR,C57L,CE,NOD,NOR,RF | 12 | 59083697 | 59083730 | 34 | + | 18.065 | 3.38E-07 | 0.064 | TAGCTGTCTTCAGACTCACCAGAAGAGGGCGTCA | | |
| 434 | 12 | 59083722 | 59085622 | 1900 | AKR,C57L,CE,NOD,NOR,RF | 12 | 59083712 | 59083730 | 19 | + | 17.0328 | 8.88E-07 | 0.147 | TCACCAGAAGAGGGCGTCA | | |
| 434 | 12 | 59083722 | 59085622 | 1900 | AKR,C57L,CE,NOD,NOR,RF | 12 | 59083792 | 59083825 | 34 | + | 18.065 | 3.38E-07 | 0.064 | TAGCTGTCTTCAGACTCACCAGAAGAGGGCGTCA | | |
| 434 | 12 | 59083722 | 59085622 | 1900 | AKR,C57L,CE,NOD,NOR,RF | 12 | 59083807 | 59083825 | 19 | + | 17.0328 | 8.88E-07 | 0.147 | TCACCAGAAGAGGGCGTCA | | |
| 434 | 12 | 59083722 | 59085622 | 1900 | AKR,C57L,CE,NOD,NOR,RF | 12 | 59083887 | 59083920 | 34 | + | 18.065 | 3.38E-07 | 0.064 | TAGCTGTCTTCAGACTCACCAGAAGAGGGCGTCA | | |
| 434 | 12 | 59083722 | 59085622 | 1900 | AKR,C57L,CE,NOD,NOR,RF | 12 | 59083902 | 59083920 | 19 | + | 17.0328 | 8.88E-07 | 0.147 | TCACCAGAAGAGGGCGTCA | | |
| 434 | 12 | 59083722 | 59085622 | 1900 | AKR,C57L,CE,NOD,NOR,RF | 12 | 59083982 | 59084015 | 34 | + | 18.065 | 3.38E-07 | 0.064 | TAGCTGTCTTCAGACTCACCAGAAGAGGGCGTCA | | |
| 434 | 12 | 59083722 | 59085622 | 1900 | AKR,C57L,CE,NOD,NOR,RF | 12 | 59083997 | 59084015 | 19 | + | 17.0328 | 8.88E-07 | 0.147 | TCACCAGAAGAGGGCGTCA | | |
| 434 | 12 | 59083722 | 59085622 | 1900 | AKR,C57L,CE,NOD,NOR,RF | 12 | 59084077 | 59084110 | 34 | + | 18.065 | 3.38E-07 | 0.064 | TAGCTGTCTTCAGACTCACCAGAAGAGGGCGTCA | | |
| 434 | 12 | 59083722 | 59085622 | 1900 | AKR,C57L,CE,NOD,NOR,RF | 12 | 59084092 | 59084110 | 19 | + | 17.0328 | 8.88E-07 | 0.147 | TCACCAGAAGAGGGCGTCA | | |
| 434 | 12 | 59083722 | 59085622 | 1900 | AKR,C57L,CE,NOD,NOR,RF | 12 | 59084172 | 59084205 | 34 | + | 18.065 | 3.38E-07 | 0.064 | TAGCTGTCTTCAGACTCACCAGAAGAGGGCGTCA | | |
| 434 | 12 | 59083722 | 59085622 | 1900 | AKR,C57L,CE,NOD,NOR,RF | 12 | 59084187 | 59084205 | 19 | + | 17.0328 | 8.88E-07 | 0.147 | TCACCAGAAGAGGGCGTCA | | |
| 434 | 12 | 59083722 | 59085622 | 1900 | AKR,C57L,CE,NOD,NOR,RF | 12 | 59084267 | 59084300 | 34 | + | 18.065 | 3.38E-07 | 0.064 | TAGCTGTCTTCAGACTCACCAGAAGAGGGCGTCA | | |
| 434 | 12 | 59083722 | 59085622 | 1900 | AKR,C57L,CE,NOD,NOR,RF | 12 | 59084282 | 59084300 | 19 | + | 17.0328 | 8.88E-07 | 0.147 | TCACCAGAAGAGGGCGTCA | | |
| 434 | 12 | 59083722 | 59085622 | 1900 | AKR,C57L,CE,NOD,NOR,RF | 12 | 59084362 | 59084395 | 34 | + | 18.065 | 3.38E-07 | 0.064 | TAGCTGTCTTCAGACTCACCAGAAGAGGGCGTCA | | |
| 434 | 12 | 59083722 | 59085622 | 1900 | AKR,C57L,CE,NOD,NOR,RF | 12 | 59084377 | 59084395 | 19 | + | 17.0328 | 8.88E-07 | 0.147 | TCACCAGAAGAGGGCGTCA | | |
| 434 | 12 | 59083722 | 59085622 | 1900 | AKR,C57L,CE,NOD,NOR,RF | 12 | 59084457 | 59084490 | 34 | + | 18.065 | 3.38E-07 | 0.064 | TAGCTGTCTTCAGACTCACCAGAAGAGGGCGTCA | | |
| 434 | 12 | 59083722 | 59085622 | 1900 | AKR,C57L,CE,NOD,NOR,RF | 12 | 59084472 | 59084490 | 19 | + | 17.0328 | 8.88E-07 | 0.147 | TCACCAGAAGAGGGCGTCA | | |
| 434 | 12 | 59083722 | 59085622 | 1900 | AKR,C57L,CE,NOD,NOR,RF | 12 | 59084552 | 59084585 | 34 | + | 18.065 | 3.38E-07 | 0.064 | TAGCTGTCTTCAGACTCACCAGAAGAGGGCGTCA | | |
| 434 | 12 | 59083722 | 59085622 | 1900 | AKR,C57L,CE,NOD,NOR,RF | 12 | 59084567 | 59084585 | 19 | + | 17.0328 | 8.88E-07 | 0.147 | TCACCAGAAGAGGGCGTCA | | |
| 434 | 12 | 59083722 | 59085622 | 1900 | AKR,C57L,CE,NOD,NOR,RF | 12 | 59084647 | 59084680 | 34 | + | 18.065 | 3.38E-07 | 0.064 | TAGCTGTCTTCAGACTCACCAGAAGAGGGCGTCA | | |
| 434 | 12 | 59083722 | 59085622 | 1900 | AKR,C57L,CE,NOD,NOR,RF | 12 | 59084662 | 59084680 | 19 | + | 17.0328 | 8.88E-07 | 0.147 | TCACCAGAAGAGGGCGTCA | | |
| 434 | 12 | 59083722 | 59085622 | 1900 | AKR,C57L,CE,NOD,NOR,RF | 12 | 59084742 | 59084775 | 34 | + | 18.065 | 3.38E-07 | 0.064 | TAGCTGTCTTCAGACTCACCAGAAGAGGGCGTCA | | |
| 434 | 12 | 59083722 | 59085622 | 1900 | AKR,C57L,CE,NOD,NOR,RF | 12 | 59084757 | 59084775 | 19 | + | 17.0328 | 8.88E-07 | 0.147 | TCACCAGAAGAGGGCGTCA | | |
| 434 | 12 | 59083722 | 59085622 | 1900 | AKR,C57L,CE,NOD,NOR,RF | 12 | 59084837 | 59084870 | 34 | + | 18.065 | 3.38E-07 | 0.064 | TAGCTGTCTTCAGACTCACCAGAAGAGGGCGTCA | | |
| 434 | 12 | 59083722 | 59085622 | 1900 | AKR,C57L,CE,NOD,NOR,RF | 12 | 59084852 | 59084870 | 19 | + | 17.0328 | 8.88E-07 | 0.147 | TCACCAGAAGAGGGCGTCA | | |
| 434 | 12 | 59083722 | 59085622 | 1900 | AKR,C57L,CE,NOD,NOR,RF | 12 | 59084932 | 59084965 | 34 | + | 18.065 | 3.38E-07 | 0.064 | TAGCTGTCTTCAGACTCACCAGAAGAGGGCGTCA | | |
| 434 | 12 | 59083722 | 59085622 | 1900 | AKR,C57L,CE,NOD,NOR,RF | 12 | 59084947 | 59084965 | 19 | + | 17.0328 | 8.88E-07 | 0.147 | TCACCAGAAGAGGGCGTCA | | |
| 434 | 12 | 59083722 | 59085622 | 1900 | AKR,C57L,CE,NOD,NOR,RF | 12 | 59085027 | 59085060 | 34 | + | 18.065 | 3.38E-07 | 0.064 | TAGCTGTCTTCAGACTCACCAGAAGAGGGCGTCA | | |
| 434 | 12 | 59083722 | 59085622 | 1900 | AKR,C57L,CE,NOD,NOR,RF | 12 | 59085042 | 59085060 | 19 | + | 17.0328 | 8.88E-07 | 0.147 | TCACCAGAAGAGGGCGTCA | | |
| 434 | 12 | 59083722 | 59085622 | 1900 | AKR,C57L,CE,NOD,NOR,RF | 12 | 59085122 | 59085155 | 34 | + | 18.065 | 3.38E-07 | 0.064 | TAGCTGTCTTCAGACTCACCAGAAGAGGGCGTCA | | |
| 434 | 12 | 59083722 | 59085622 | 1900 | AKR,C57L,CE,NOD,NOR,RF | 12 | 59085137 | 59085155 | 19 | + | 17.0328 | 8.88E-07 | 0.147 | TCACCAGAAGAGGGCGTCA | | |
| 434 | 12 | 59083722 | 59085622 | 1900 | AKR,C57L,CE,NOD,NOR,RF | 12 | 59085217 | 59085250 | 34 | + | 18.065 | 3.38E-07 | 0.064 | TAGCTGTCTTCAGACTCACCAGAAGAGGGCGTCA | | |
| 434 | 12 | 59083722 | 59085622 | 1900 | AKR,C57L,CE,NOD,NOR,RF | 12 | 59085232 | 59085250 | 19 | + | 17.0328 | 8.88E-07 | 0.147 | TCACCAGAAGAGGGCGTCA | | |
| 434 | 12 | 59083722 | 59085622 | 1900 | AKR,C57L,CE,NOD,NOR,RF | 12 | 59085312 | 59085345 | 34 | + | 18.065 | 3.38E-07 | 0.064 | TAGCTGTCTTCAGACTCACCAGAAGAGGGCGTCA | | |
| 434 | 12 | 59083722 | 59085622 | 1900 | AKR,C57L,CE,NOD,NOR,RF | 12 | 59085327 | 59085345 | 19 | + | 17.0328 | 8.88E-07 | 0.147 | TCACCAGAAGAGGGCGTCA | | |
| 434 | 12 | 59083722 | 59085622 | 1900 | AKR,C57L,CE,NOD,NOR,RF | 12 | 59085407 | 59085440 | 34 | + | 18.065 | 3.38E-07 | 0.064 | TAGCTGTCTTCAGACTCACCAGAAGAGGGCGTCA | | |
| 434 | 12 | 59083722 | 59085622 | 1900 | AKR,C57L,CE,NOD,NOR,RF | 12 | 59085422 | 59085440 | 19 | + | 17.0328 | 8.88E-07 | 0.147 | TCACCAGAAGAGGGCGTCA | | |
| 434 | 12 | 59083722 | 59085622 | 1900 | AKR,C57L,CE,NOD,NOR,RF | 12 | 59085502 | 59085535 | 34 | + | 18.065 | 3.38E-07 | 0.064 | TAGCTGTCTTCAGACTCACCAGAAGAGGGCGTCA | | |
| 434 | 12 | 59083722 | 59085622 | 1900 | AKR,C57L,CE,NOD,NOR,RF | 12 | 59085517 | 59085535 | 19 | + | 17.0328 | 8.88E-07 | 0.147 | TCACCAGAAGAGGGCGTCA | | |
| 435 | 12 | 59084291 | 59085621 | 1330 | 129S1 | 12 | 59084267 | 59084300 | 34 | + | 18.065 | 3.38E-07 | 0.064 | TAGCTGTCTTCAGACTCACCAGAAGAGGGCGTCA | | |
| 435 | 12 | 59084291 | 59085621 | 1330 | 129S1 | 12 | 59084282 | 59084300 | 19 | + | 17.0328 | 8.88E-07 | 0.147 | TCACCAGAAGAGGGCGTCA | | |
| 435 | 12 | 59084291 | 59085621 | 1330 | 129S1 | 12 | 59084362 | 59084395 | 34 | + | 18.065 | 3.38E-07 | 0.064 | TAGCTGTCTTCAGACTCACCAGAAGAGGGCGTCA | | |
| 435 | 12 | 59084291 | 59085621 | 1330 | 129S1 | 12 | 59084377 | 59084395 | 19 | + | 17.0328 | 8.88E-07 | 0.147 | TCACCAGAAGAGGGCGTCA | | |
| 435 | 12 | 59084291 | 59085621 | 1330 | 129S1 | 12 | 59084457 | 59084490 | 34 | + | 18.065 | 3.38E-07 | 0.064 | TAGCTGTCTTCAGACTCACCAGAAGAGGGCGTCA | | |
| 435 | 12 | 59084291 | 59085621 | 1330 | 129S1 | 12 | 59084472 | 59084490 | 19 | + | 17.0328 | 8.88E-07 | 0.147 | TCACCAGAAGAGGGCGTCA | | |
| 435 | 12 | 59084291 | 59085621 | 1330 | 129S1 | 12 | 59084552 | 59084585 | 34 | + | 18.065 | 3.38E-07 | 0.064 | TAGCTGTCTTCAGACTCACCAGAAGAGGGCGTCA | | |
| 435 | 12 | 59084291 | 59085621 | 1330 | 129S1 | 12 | 59084567 | 59084585 | 19 | + | 17.0328 | 8.88E-07 | 0.147 | TCACCAGAAGAGGGCGTCA | | |
| 435 | 12 | 59084291 | 59085621 | 1330 | 129S1 | 12 | 59084647 | 59084680 | 34 | + | 18.065 | 3.38E-07 | 0.064 | TAGCTGTCTTCAGACTCACCAGAAGAGGGCGTCA | | |
| 435 | 12 | 59084291 | 59085621 | 1330 | 129S1 | 12 | 59084662 | 59084680 | 19 | + | 17.0328 | 8.88E-07 | 0.147 | TCACCAGAAGAGGGCGTCA | | |
| 435 | 12 | 59084291 | 59085621 | 1330 | 129S1 | 12 | 59084742 | 59084775 | 34 | + | 18.065 | 3.38E-07 | 0.064 | TAGCTGTCTTCAGACTCACCAGAAGAGGGCGTCA | | |
| 435 | 12 | 59084291 | 59085621 | 1330 | 129S1 | 12 | 59084757 | 59084775 | 19 | + | 17.0328 | 8.88E-07 | 0.147 | TCACCAGAAGAGGGCGTCA | | |
| 435 | 12 | 59084291 | 59085621 | 1330 | 129S1 | 12 | 59084837 | 59084870 | 34 | + | 18.065 | 3.38E-07 | 0.064 | TAGCTGTCTTCAGACTCACCAGAAGAGGGCGTCA | | |
| 435 | 12 | 59084291 | 59085621 | 1330 | 129S1 | 12 | 59084852 | 59084870 | 19 | + | 17.0328 | 8.88E-07 | 0.147 | TCACCAGAAGAGGGCGTCA | | |
| 435 | 12 | 59084291 | 59085621 | 1330 | 129S1 | 12 | 59084932 | 59084965 | 34 | + | 18.065 | 3.38E-07 | 0.064 | TAGCTGTCTTCAGACTCACCAGAAGAGGGCGTCA | | |
| 435 | 12 | 59084291 | 59085621 | 1330 | 129S1 | 12 | 59084947 | 59084965 | 19 | + | 17.0328 | 8.88E-07 | 0.147 | TCACCAGAAGAGGGCGTCA | | |
| 435 | 12 | 59084291 | 59085621 | 1330 | 129S1 | 12 | 59085027 | 59085060 | 34 | + | 18.065 | 3.38E-07 | 0.064 | TAGCTGTCTTCAGACTCACCAGAAGAGGGCGTCA | | |
| 435 | 12 | 59084291 | 59085621 | 1330 | 129S1 | 12 | 59085042 | 59085060 | 19 | + | 17.0328 | 8.88E-07 | 0.147 | TCACCAGAAGAGGGCGTCA | | |
| 435 | 12 | 59084291 | 59085621 | 1330 | 129S1 | 12 | 59085122 | 59085155 | 34 | + | 18.065 | 3.38E-07 | 0.064 | TAGCTGTCTTCAGACTCACCAGAAGAGGGCGTCA | | |
| 435 | 12 | 59084291 | 59085621 | 1330 | 129S1 | 12 | 59085137 | 59085155 | 19 | + | 17.0328 | 8.88E-07 | 0.147 | TCACCAGAAGAGGGCGTCA | | |
| 435 | 12 | 59084291 | 59085621 | 1330 | 129S1 | 12 | 59085217 | 59085250 | 34 | + | 18.065 | 3.38E-07 | 0.064 | TAGCTGTCTTCAGACTCACCAGAAGAGGGCGTCA | | |
| 435 | 12 | 59084291 | 59085621 | 1330 | 129S1 | 12 | 59085232 | 59085250 | 19 | + | 17.0328 | 8.88E-07 | 0.147 | TCACCAGAAGAGGGCGTCA | | |
| 435 | 12 | 59084291 | 59085621 | 1330 | 129S1 | 12 | 59085312 | 59085345 | 34 | + | 18.065 | 3.38E-07 | 0.064 | TAGCTGTCTTCAGACTCACCAGAAGAGGGCGTCA | | |
| 435 | 12 | 59084291 | 59085621 | 1330 | 129S1 | 12 | 59085327 | 59085345 | 19 | + | 17.0328 | 8.88E-07 | 0.147 | TCACCAGAAGAGGGCGTCA | | |
| 435 | 12 | 59084291 | 59085621 | 1330 | 129S1 | 12 | 59085407 | 59085440 | 34 | + | 18.065 | 3.38E-07 | 0.064 | TAGCTGTCTTCAGACTCACCAGAAGAGGGCGTCA | | |
| 435 | 12 | 59084291 | 59085621 | 1330 | 129S1 | 12 | 59085422 | 59085440 | 19 | + | 17.0328 | 8.88E-07 | 0.147 | TCACCAGAAGAGGGCGTCA | | |
| 435 | 12 | 59084291 | 59085621 | 1330 | 129S1 | 12 | 59085502 | 59085535 | 34 | + | 18.065 | 3.38E-07 | 0.064 | TAGCTGTCTTCAGACTCACCAGAAGAGGGCGTCA | | |
| 435 | 12 | 59084291 | 59085621 | 1330 | 129S1 | 12 | 59085517 | 59085535 | 19 | + | 17.0328 | 8.88E-07 | 0.147 | TCACCAGAAGAGGGCGTCA | | |
| 436 | 12 | 60730399 | 60733428 | 3029 | PJ | 12 | 60730981 | 60731014 | 34 | + | 20.8211 | 4.88E-08 | 0.0195 | ATGTACTACTTACTGAGGCCAGAAGAGGGCAACA | 3 | ENCFF055HAP,ENCFF093ZRU,ENCFF502WDT |
| 436 | 12 | 60730399 | 60733428 | 3029 | PJ | 12 | 60730996 | 60731014 | 19 | + | 17.5246 | 6.45E-07 | 0.13 | AGGCCAGAAGAGGGCAACA | 3 | ENCFF055HAP,ENCFF093ZRU,ENCFF502WDT |
| 437 | 12 | 69240468 | 69241582 | 1114 | 129S1,BTBR,C57L,CE,NZB,NZO,PJ | 12 | 69241419 | 69241452 | 34 | + | 16.7724 | 7.83E-07 | 0.0958 | TAGCTGTCTTCGGACACACCAGAAGAGGGCATCA | 1 | ENCFF093ZRU |
| 438 | 12 | 69593575 | 69602620 | 9045 | AJ,AKR,BALB,BTBR,BUB,C3H,C57L,CBA,CE,FVB,KK,MA/My,MRL,NOD,NOR,NZW,PL,RF.RH2.SEA.SJL.SMJ.SWR.TH | 12 | 69599035 | 69599053 | 19 | + | 17.1639 | 8.17E-07 | 0.145 | CCTCCACGAGGTGGCGGAA | | |
| 439 | 12 | 82579407 | 82580533 | 1126 | CE.SWR | 12 | 82580278 | 82580311 | 34 | + | 20.6992 | 5.34E-08 | 0.0206 | ATGCAAAGCTAACTGCGGCCAGCAGAGGCCCCGG | 1 | ENCFF093ZRU |
| 440 | 12 | 87593448 | 87988990 | 395542 | CE.NOD.SWR | 12 | 87672777 | 87672810 | 34 | + | 17.9675 | 3.61E-07 | 0.0662 | ATGCAGTAATAAGAACAACCATAAGGTGGTGCTC | | |
| 440 | 12 | 87593448 | 87988990 | 395542 | CE.NOD.SWR | 12 | 87697366 | 87697384 | 19 | + | 17.6557 | 5.92E-07 | 0.126 | TGCCCAACAGAGGGCACCC | | |
| 440 | 12 | 87593448 | 87988990 | 395542 | CE.NOD.SWR | 12 | 87790496 | 87790529 | 34 | + | 17.9675 | 3.61E-07 | 0.0662 | ATGCAGTAATAAGAACAACCATAAGGTGGTGCTC | | |
| 440 | 12 | 87593448 | 87988990 | 395542 | CE.NOD.SWR | 12 | 87815101 | 87815119 | 19 | + | 17.6557 | 5.92E-07 | 0.126 | TGCCCAACAGAGGGCACCC | | |
| 440 | 12 | 87593448 | 87988990 | 395542 | CE,NOD,SWR | 12 | 87950885 | 87950919 | 35 | + | 20.5321 | 6.15E-08 | 0.0389 | CTGAAGTAACAAGAACAAACCAGAAGGTGGTGCTC | 7 | ENCFF092NMJ,ENCFF024JOZ,ENCFF089GBC,ENCFF099QNE,ENCFF502WDT,ENCFF093ZRU,ENCFF907XEG |
| 440 | 12 | 87593448 | 87988990 | 395542 | CE,NOD,SWR | 12 | 87950972 | 87951005 | 34 | + | 17.9675 | 3.61E-07 | 0.0662 | ATGCAGTAATAAGAACAACCATAAGGTGGTGCTC | 7 | ENCFF092NMJ,ENCFF024JOZ,ENCFF089GBC,ENCFF099QNE,ENCFF502WDT,ENCFF093ZRU,ENCFF907XEG |
| 440 | 12 | 87593448 | 87988990 | 395542 | CE.NOD.SWR | 12 | 87968631 | 87968649 | 19 | + | 17.6557 | 5.92E-07 | 0.126 | TGCCCAACAGAGGGCACCC | | |
| 440 | 12 | 87593448 | 87988990 | 395542 | CE.NOD.SWR | 12 | 87623305 | 87623339 | 35 | - | 21.3028 | 3.68E-08 | 0.0274 | TTGCAGGTAAAGGTATGTGCCAGAAGATGGAGCTC | | |
| 440 | 12 | 87593448 | 87988990 | 395542 | CE.NOD.SWR | 12 | 87770625 | 87770658 | 34 | - | 18.2358 | 3.02E-07 | 0.0615 | ATGCAGCTAGAGACACGAGCTCCTGGGGGCGCTA | | |
| 440 | 12 | 87593448 | 87988990 | 395542 | CE.NOD.SWR | 12 | 87867060 | 87867094 | 35 | - | 19.7523 | 1.02E-07 | 0.053 | ATGCAGTAAATAAGAACAACCAGAAGGTGGTGCTC | | |
| 440 | 12 | 87593448 | 87988990 | 395542 | CE.NOD.SWR | 12 | 87867146 | 87867180 | 35 | - | 19.6697 | 1.07E-07 | 0.0544 | CTGAACTAACACGAATCAACCAGAAGGTGGTGCTC | | |
| 440 | 12 | 87593448 | 87988990 | 395542 | CE.NOD.SWR | 12 | 87867262 | 87867296 | 35 | - | 20.5321 | 6.15E-08 | 0.0389 | CTGAAGTAACAAGAACAAACCAGAAGGTGGTGCTC | | |
| 440 | 12 | 87593448 | 87988990 | 395542 | CE.NOD.SWR | 12 | 87875841 | 87875874 | 34 | - | 16.9431 | 7.02E-07 | 0.0905 | ATGCAGCTAGAGACAGGAACTCTGGGGGGCACTG | | |
| 441 | 12 | 88114738 | 88138497 | 23759 | CE | 12 | 88124882 | 88124915 | 34 | + | 16.9431 | 7.02E-07 | 0.0905 | ATGCAGCTAGAGACAGGAACTCTGGGGGGCACTG | | |
| 441 | 12 | 88114738 | 88138497 | 23759 | CE | 12 | 88133462 | 88133496 | 35 | + | 20.5321 | 6.15E-08 | 0.0389 | CTGAAGTAACAAGAACAAACCAGAAGGTGGTGCTC | | |
| 441 | 12 | 88114738 | 88138497 | 23759 | CE | 12 | 88133578 | 88133612 | 35 | + | 19.6697 | 1.07E-07 | 0.0544 | CTGAACTAACACGAATCAACCAGAAGGTGGTGCTC | | |
| 441 | 12 | 88114738 | 88138497 | 23759 | CE | 12 | 88133664 | 88133698 | 35 | + | 19.7523 | 1.02E-07 | 0.053 | ATGCAGTAAATAAGAACAACCAGAAGGTGGTGCTC | | |
| 442 | 12 | 93505347 | 93509518 | 4171 | AJ,BUB,C3H,C58J,DBA1J,DBA2J,FVB,ILNJ2,KK,NOD,NZO,NZW,PJ,PL,RF,SEA,SJL,SMJ.SWR.TH | 12 | 93506123 | 93506156 | 34 | + | 17.6748 | 4.37E-07 | 0.073 | GAGCATGACCACAAGTCCACAGCAGGTGGCAACG | | |
| 443 | 12 | 99099864 | 99106207 | 6343 | 129S1,BTBR,C57L,FVB,KK,NZB,NZW,SWR | 12 | 99105995 | 99106028 | 34 | + | 17.2114 | 5.91E-07 | 0.0834 | TTGCTGGGCAAGCAGTTCTCAGCAGGGGACACCC | | |
| 444 | 12 | 109087950 | 109109567 | 21617 | AJ.AKR.C3H.CBA.TH | 12 | 109108884 | 109108918 | 35 | + | 16.7156 | 6.38E-07 | 0.158 | CTCCTGTCATCTCCGCATGGCACCAGGTGGCCCTC | | |
| 445 | 12 | 10202122 | 10210229 | 8107 | 129S1,AJ,AKR,BALB,BTBR,BUB,C3H,C57L,C58J,CBA,CE,DBA1J,DBA2J,FVB,ILNJ2,KK,MA/My,MRL,NOD,NOR,NZB,NZO,NZW,PJ,PL,RF.SEA.SJL.SMJ.SWR.TH | 12 | 10203885 | 10203918 | 34 | - | 19.8455 | 9.92E-08 | 0.0321 | CTGCAGCTCTAGCAGAGGCAAGGAGGGGTCTCTC | | |
| 446 | 12 | 16163283 | 16164419 | 1136 | NZB | 12 | 16163649 | 16163667 | 19 | - | 18.7049 | 2.88E-07 | 0.0872 | AGGCCAGCAGAGGGCATTA | 3 | ENCFF367UNR.ENCFF821BIA.ENCFF093ZRU |
| 446 | 12 | 16163283 | 16164419 | 1136 | NZB | 12 | 16163649 | 16163683 | 35 | - | 25.367 | 1.84E-09 | 0.00483 | GTGCAGGTGCCTGTAGAGGCCAGCAGAGGGCATTA | 3 | ENCFF367UNR.ENCFF821BIA.ENCFF093ZRU |
| 447 | 12 | 17814509 | 17819957 | 5448 | AKR | 12 | 17817059 | 17817093 | 35 | - | 24.5138 | 3.60E-09 | 0.00636 | GTGCAGGTGCCCGTGGAGGCCTGAAGAGGGCACCT | | |
| 448 | 12 | 22831229 | 22836104 | 4875 | 129S1,AJ,AKR,BALB,BTBR,BUB,C3H,C57L,CBA,CE,DBA1J,DBA2J,FVB,ILNJ2,KK,MA/My,NOD,NOR,NZO,PJ,PL,RF,SEA,SJL,SMJ,SWR.TH | 12 | 22833318 | 22833351 | 34 | - | 19.2602 | 1.50E-07 | 0.0419 | CTGAAGGCCCAGCCTTGGCCTTCAGGTGGCTGCC | | |
| 449 | 12 | 23647683 | 23656009 | 8326 | 129S1,BTBR,BUB,C3H,CBA,DBA2J,FVB,MA/My,NOD,NOR,PL,SJL,SWR,TH | 12 | 23647975 | 23647993 | 19 | - | 19.2951 | 1.88E-07 | 0.0758 | CCGCCACTAGATGGTGCTA | | |
| 449 | 12 | 23647683 | 23656009 | 8326 | 129S1,BTBR,BUB,C3H,CBA,DBA2J,FVB,MA/My,NOD,NOR,PL,SJL,SWR,TH | 12 | 23655969 | 23655987 | 19 | - | 19.2951 | 1.88E-07 | 0.0758 | CCGCCACTAGATGGTGCTA | | |
| 450 | 12 | 28813380 | 28814490 | 1110 | BTBR.C57L.FVB.MA/Mv.SJL.SWR | 12 | 28813686 | 28813704 | 19 | - | 17.2295 | 7.83E-07 | 0.142 | AGGCCAGAAGAGGGCAATG | 5 | ENCFF367UNR.ENCFF099QNE.ENCFF821BIA.ENCFF502WDT.ENCFF093ZRU |
| 450 | 12 | 28813380 | 28814490 | 1110 | BTBR.C57L.FVB.MA/Mv.SJL.SWR | 12 | 28813686 | 28813719 | 34 | - | 26.7398 | 3.32E-10 | 0.000691 | ATGCAGTGCCTATGGAGGCCAGAAGAGGGCAATG | 5 | ENCFF367UNR.ENCFF099QNE.ENCFF821BIA.ENCFF502WDT.ENCFF093ZRU |
| 451 | 12 | 45240779 | 45244396 | 3617 | KK.NZO.NZW | 12 | 45242311 | 45242344 | 34 | - | 16.439 | 9.66E-07 | 0.104 | CCCCAGTGTTCCCCTCCTCCACTGGGGGCCGCTG | | |
| 452 | 12 | 55287456 | 55300490 | 13034 | CE | 12 | 55295347 | 55295380 | 34 | - | 16.8943 | 7.24E-07 | 0.0917 | TTGCTCTCTTCAGACATACCAGAAGAGGGCATCA | | |
| 453 | 12 | 71851709 | 71858416 | 6707 | FVB.SWR | 12 | 71856527 | 71856560 | 34 | - | 24.2439 | 3.20E-09 | 0.0028 | GCGCAGTGCCTGTGGAGACCAGAAGAGGGCGTTG | 4 | ENCFF093ZRU.ENCFF367UNR.ENCFF099QNE.ENCFF052KVO |
| 454 | 12 | 83334886 | 83336451 | 1565 | C58J.CE.KK.PL.SJL.SWR | 12 | 83335584 | 83335618 | 35 | - | 16.578 | 6.90E-07 | 0.162 | CTGCAATTAATGTCCCAGGACAGAAGGGGTCATGT | | |
| 455 | 12 | 90227474 | 90230259 | 2785 | SWR | 12 | 90229161 | 90229179 | 19 | - | 18.8197 | 2.66E-07 | 0.0837 | TGACCAGGAGGGGGCAACG | 1 | ENCFF093ZRU |
| 456 | 12 | 90902496 | 90903583 | 1087 | BUB.SWR | 12 | 90902657 | 90902675 | 19 | - | 17.2131 | 7.91E-07 | 0.143 | GTACCAGTAGGAGGCACTA | | |

| | | | | | | | | | | | | | | | | |
|---|---|---|---|---|---|---|---|---|---|---|---|---|---|---|---|---|
| 457 | 12 | 105070280 | 105073289 | 3009 | CE | 12 | 105070495 | 105070513 | 19 | - | 20.0164 | 1.09E-07 | 0.0593 | GAACCACCAGGGGGCGCAC | 36 | ENCFF093ZRU,ENCFF201JUA,ENCFF367UNR,ENCFF099QNE,ENCFF024JOZ,ENCFF065ODV,ENCFF821BIA,ENCFF805QBU,ENCFF327LYS,ENCFF089GBC,ENCFF907XEG,ENCFF198SSR,ENCFF092NMJ,ENCFF136ZDK,ENCFF589ZBJ,ENCFF609MDN,ENCFF430PPJ,ENCFF033UQX,ENCFF165LXO,ENCFF321AGQ,ENCFF883UPM,ENCFF142CNG,ENCFF491RJK,ENCFF502WDT,ENCFF097CAQ,ENCFF281QMV,ENCFF055HAP,ENCFF210BTO,ENCFF052KVO,ENCFF957SPH,ENCFF533APC,ENCFF194HVL,ENCFF089EWX,ENCFF575MGI,ENCFF601JVE.ENCFF951GOP |
| 458 | 12 | 110504408 | 110508319 | 3911 | DBA1J.DBA2J.ILNJ2 | 12 | 110504898 | 110504932 | 35 | - | 16.4495 | 7.42E-07 | 0.17 | GTGCCCTTGCCTGTGGAGGCCAGAGGAGGACACTG | | |
| 459 | 12 | 113799961 | 113806063 | 6102 | 129S1,AJ,BALB,BUB,C58J,CE,DBA1J,DBA2J,ILNJ2,KK,LP,MA/My,PJ,RF,SEA,SMJ,TH | 12 | 113802072 | 113802090 | 19 | - | 17.8033 | 5.37E-07 | 0.119 | GAACCAGCAGAGGGCGATG | 2 | ENCFF367UNR,ENCFF883UPM |
| 460 | 12 | 113870577 | 113884203 | 13626 | AJ,CE,DBA1J,DBA2J,ILNJ2,RF | 12 | 113875577 | 113875595 | 19 | - | 17.6066 | 6.11E-07 | 0.128 | GGACCAACAGGGGGCGCAC | 10 | ENCFF367UNR,ENCFF589ZBJ,ENCFF883UPM,ENCFF201JUA,ENCFF033UQX,ENCFF194HVL,ENCFF957SPH,ENCFF052KVO.ENCFF327LYS.ENCFF533APC |
| 460 | 12 | 113870577 | 113884203 | 13626 | AJ,CE,DBA1J,DBA2J,ILNJ2,RF | 12 | 113875577 | 113875610 | 34 | - | 22.2764 | 1.61E-08 | 0.00928 | TTGCAGGGCTGCTCAGGACCAACAGGGGGCGCAC | 10 | ENCFF367UNR,ENCFF589ZBJ,ENCFF883UPM,ENCFF201JUA,ENCFF033UQX,ENCFF194HVL,ENCFF957SPH,ENCFF052KVO.ENCFF327LYS.ENCFF533APC |
| 461 | 12 | 114032177 | 114042890 | 10713 | 129S1,AJ,BALB,BUB,C58J,KK,LP,MA/My,PJ,SEA.SMJ.TH | 12 | 114032357 | 114032375 | 19 | - | 18.459 | 3.43E-07 | 0.0947 | GGACCAGCAGGGGACACTG | | |
| 462 | 12 | 114437524 | 114451059 | 13535 | 129S1,BALB,FVB,KK,LP,MA/My,PJ,SEA,SWR | 12 | 114442550 | 114442568 | 19 | - | 19.3934 | 1.75E-07 | 0.073 | TGACCAGCAGGGGGTGCTT | | |
| 462 | 12 | 114437524 | 114451059 | 13535 | 129S1,BALB,FVB,KK,LP,MA/My,PJ,SEA,SWR | 12 | 114442550 | 114442583 | 34 | - | 17.0894 | 6.39E-07 | 0.0874 | CAGTTCAGGCACCCATGACCAGCAGGGGGTGCTT | | |
| 462 | 12 | 114437524 | 114451059 | 13535 | 129S1,BALB,FVB,KK,LP,MA/My,PJ,SEA,SWR | 12 | 114449667 | 114449700 | 34 | - | 17.2439 | 5.79E-07 | 0.0827 | CTGGTATGACAAGTTTGTTCAGAAGGGGGCACAG | | |
| 463 | 12 | 114457611 | 114468277 | 10666 | BALB.FVB.KK.LP.PJ.SEA.SWR.TH | 12 | 114459548 | 114459566 | 19 | - | 21.5246 | 3.11E-08 | 0.0421 | TGACCAGCAGGAGGCGCTC | 3 | ENCFF052KVO.ENCFF502WDT.ENCFF055HAP |
| 464 | 12 | 114500133 | 114501768 | 1635 | AJ.RF | 12 | 114501022 | 114501055 | 34 | - | 17.2439 | 5.79E-07 | 0.0827 | CTGGTATGACAAGTTTGTTCAGAAGGGGGCACAG | | |
| 465 | 12 | 115063524 | 115070903 | 7379 | AJ.DBA2J.FVB.RF.SWR | 12 | 115063684 | 115063717 | 34 | - | 17.5772 | 4.66E-07 | 0.0751 | CTGCAGTCTCAGGTCTGCGCAATTGGGGCCGGCG | | |
| 466 | 12 | 115232326 | 115243936 | 11610 | 129S1,BALB,BUB,C58J,KK,LP,MA/My,PJ,SMJ.TH | 12 | 115240554 | 115240572 | 19 | - | 21.082 | 4.57E-08 | 0.0421 | GGACCAGCAGGGGGCACAA | | |
| 466 | 12 | 115232326 | 115243936 | 11610 | 129S1,BALB,BUB,C58J,KK,LP,MA/My,PJ,SMJ.TH | 12 | 115240554 | 115240587 | 34 | - | 26.2439 | 5.32E-10 | 0.000914 | CTGCAGGTGCGTTCAGGACCAGCAGGGGGCACAA | | |
| 467 | 12 | 115246831 | 115251582 | 4751 | 129S1,AKR,BALB,BUB,C3H,C58J,CBA,FVB,KK,LP,MA/My,MRL,NZB,NZO,NZW,PJ,PL,SEA.SMJ.TH | 12 | 115250838 | 115250871 | 34 | - | 17.5691 | 4.69E-07 | 0.0751 | GAGCATGACCACAAGTCCACAGCAGGTGGCAACA | | |
| 468 | 12 | 115842562 | 115862128 | 19566 | C3H.MRL.NZB.NZW.PL | 12 | 115854486 | 115854504 | 19 | - | 17.4262 | 6.88E-07 | 0.132 | TGACCAGCAGGGGTCATCA | | |
| 469 | 12 | 115849768 | 115855995 | 6227 | 129S1,AJ,BALB,BUB,C58J,FVB,KK,LP,MA/My,PJ,SEA,SMJ,SWR,TH | 12 | 115854486 | 115854504 | 19 | - | 17.4262 | 6.88E-07 | 0.132 | TGACCAGCAGGGGTCATCA | | |
| 470 | 12 | 116675766 | 116677331 | 1565 | C3H,CBA,DBA1J,DBA2J,ILNJ2,MRL,PL,RF | 12 | 116677008 | 116677041 | 34 | - | 17.5854 | 4.64E-07 | 0.0751 | GTGCTGAACATAGAGGGGCCTCCAGAGAGCACTG | | |
| 471 | 13 | 21987842 | 21990110 | 2268 | 129S1.C57L.ILNJ2 | 13 | 21988286 | 21988319 | 34 | + | 16.5447 | 9.04E-07 | 0.118 | CTGTACTTCCCATGGGCTCCTAGAGGTGGGACCC | | |
| 472 | 13 | 21996577 | 22004435 | 7858 | 129S1,BTBR,C57L,CE,ILNJ2,KK,LP,PJ,SEA | 13 | 22002035 | 22002053 | 19 | + | 17.1639 | 8.17E-07 | 0.174 | CCTCCACGAGGTGGCGGAA | | |
| 473 | 13 | 22516446 | 22519482 | 3036 | BTBR,C57L,KK,LP,NZB,NZO,NZW | 13 | 22518884 | 22518918 | 35 | + | 16.6147 | 6.75E-07 | 0.163 | CTGCATTTGCCTCCATTCACCTCTGGAGGAAACTT | | |
| 474 | 13 | 24125793 | 24127042 | 1249 | BTBR,C57L,KK,LP,NOD,NOR,NZB,PJ,SEA,SMJ | 13 | 24127007 | 24127040 | 34 | + | 17.4553 | 5.05E-07 | 0.0902 | TTAACGTACCTGCAGAGGCCAGAAGAGGGCACTG | 11 | ENCFF093ZRU,ENCFF092NMJ,ENCFF089GBC,ENCFF589ZBJ,ENCFF198SSR,ENCFF883UPM,ENCFF327LYS,ENCFF957SPH.ENCFF533APC.ENCFF024JOZ.ENCFF907XEG |
| 474 | 13 | 24125793 | 24127042 | 1249 | BTBR,C57L,KK,LP,NOD,NOR,NZB,PJ,SEA,SMJ | 13 | 24127022 | 24127040 | 19 | + | 20.0656 | 1.05E-07 | 0.0686 | AGGCCAGAAGAGGGCACTG | 11 | ENCFF093ZRU,ENCFF092NMJ,ENCFF089GBC,ENCFF589ZBJ,ENCFF198SSR,ENCFF883UPM,ENCFF327LYS,ENCFF957SPH.ENCFF533APC.ENCFF024JOZ.ENCFF907XEG |
| 475 | 13 | 25191372 | 25199698 | 8326 | ILNJ2,KK,LP,NOD,NOR,NZB,PJ,SEA,SMJ,TH | 13 | 25191406 | 25191424 | 19 | + | 19.2951 | 1.88E-07 | 0.091 | CCGCCACTAGATGGTGCTA | 16 | ENCFF093ZRU,ENCFF367UNR,ENCFF099QNE,ENCFF502WDT,ENCFF201JUA,ENCFF065ODV,ENCFF089GBC,ENCFF327LYS,ENCFF821BIA,ENCFF907XEG,ENCFF589ZBJ,ENCFF092NMJ,ENCFF136ZDK,ENCFF198SSR.ENCFF430PPJ.ENCFF055HAP |
| 475 | 13 | 25191372 | 25199698 | 8326 | ILNJ2,KK,LP,NOD,NOR,NZB,PJ,SEA,SMJ,TH | 13 | 25199400 | 25199418 | 19 | + | 19.2951 | 1.88E-07 | 0.091 | CCGCCACTAGATGGTGCTA | | |
| 476 | 13 | 40070107 | 40071627 | 1520 | 129S1,AJ,AKR,B10D2,B10J,BALB,BTBR,BUB,C3H,C57L,C58J,CBA,CE,DBA1J,DBA2J,FVB,ILNJ2,KK,LP,MA/My,MRL,NOD,NOR,NZB,NZO,NZW,PJ,PL,RF,RH2,SEA,SJL,SMJ,SWR.TH | 13 | 40070812 | 40070846 | 35 | + | 16.367 | 7.77E-07 | 0.175 | GTGCATGTGTCTATGGGATCCAGAAGAGGGCATCA | | |
| 477 | 13 | 66045246 | 66046451 | 1205 | 129S1,AKR,BUB,C3H,CBA,MRL,NZO,NZW,RF | 13 | 66045624 | 66045657 | 34 | + | 17.2439 | 5.79E-07 | 0.0964 | TAGCTGTCCTCAGATACTCCAGAAGAGGGCATCA | | |
| 478 | 13 | 68237879 | 68251890 | 14011 | 129S1,AJ,AKR,BALB,BTBR,BUB,C3H,C57L,C58J,CBA,CE,DBA1J,DBA2J,FVB,ILNJ2,KK,MA/My,MRL,NOD,NOR,NZB,NZO,NZW,PJ,PL,RF,SEA,SJL,SMJ,SWR,TH | 13 | 68244407 | 68244425 | 19 | + | 19.9672 | 1.13E-07 | 0.0716 | TGGCCACCAGAGGGTGCTT | | |
| 479 | 13 | 70411741 | 70417889 | 6148 | MRL | 13 | 70411945 | 70411978 | 34 | + | 17.5041 | 4.89E-07 | 0.0884 | CTCTAGGGCTTTGCCAGACCACCAGGGGGTGCCA | 6 | ENCFF327LYS.ENCFF907XEG.ENCFF367UNR.ENCFF093ZRU.ENCFF099QNE.ENCFF024JOZ |
| 479 | 13 | 70411741 | 70417889 | 6148 | MRL | 13 | 70411960 | 70411978 | 19 | + | 20.4098 | 7.98E-08 | 0.0634 | AGACCACCAGGGGGTGCCA | 6 | ENCFF327LYS.ENCFF907XEG.ENCFF367UNR.ENCFF093ZRU.ENCFF099QNE.ENCFF024JOZ |
| 480 | 13 | 75970323 | 75971418 | 1095 | BUB.NOD.NOR.NZW | 13 | 75970493 | 75970526 | 34 | + | 18.3659 | 2.76E-07 | 0.0667 | TAGCTGTCTTCAGACACTCCAGAAGAGGGCGCCA | 1 | ENCFF093ZRU |
| 480 | 13 | 75970323 | 75971418 | 1095 | BUB.NOD.NOR.NZW | 13 | 75970508 | 75970526 | 19 | + | 17.5574 | 6.32E-07 | 0.153 | ACTCCAGAAGAGGGCGCCA | 1 | ENCFF093ZRU |
| 481 | 13 | 76598303 | 76603571 | 5268 | ILNJ2 | 13 | 76599744 | 76599778 | 35 | + | 16.7615 | 6.21E-07 | 0.156 | TAGCAGTTGCCACCTGTGTGCTCTTGGGGGCGGGG | | |
| 482 | 13 | 82163306 | 82172180 | 8874 | SWR.TH | 13 | 82167094 | 82167127 | 34 | + | 16.6829 | 8.29E-07 | 0.112 | ATGCAGCTAGAGACACGAGCTCCAGGGGGTGGGG | | |
| 483 | 13 | 105929624 | 105936088 | 6464 | AJ,AKR,BUB,CE,DBA1J,MA/My,NOD,NOR,NZB.NZO | 13 | 105930278 | 105930296 | 19 | + | 17.0328 | 8.88E-07 | 0.181 | TTGCCAGCAGAGAGCGCTC | | |
| 484 | 13 | 106563529 | 106565171 | 1642 | CE.MA/My.NZO | 13 | 106563753 | 106563786 | 34 | + | 19.0976 | 1.68E-07 | 0.0517 | GAGCAGGTGCAGCTGCCACCCCAAGATGGTGCCC | | |
| 485 | 13 | 116904425 | 116910505 | 6080 | 129S1,BTBR,BUB,C3H,C57L,CE,DBA2J,KK,MRL.NZB.NZO.NZW.PL | 13 | 116909082 | 116909100 | 19 | + | 17.2295 | 7.83E-07 | 0.17 | CTGCCAGAAGAGGGAAGTA | | |
| 486 | 13 | 116907587 | 116909876 | 2289 | AKR.BALB.CBA.RF.SMJ.TH | 13 | 116909082 | 116909100 | 19 | + | 17.2295 | 7.83E-07 | 0.17 | CTGCCAGAAGAGGGAAGTA | | |
| 487 | 13 | 120664931 | 120675600 | 10669 | BTBR.C57L | 13 | 120667830 | 120667863 | 34 | + | 19.2276 | 1.53E-07 | 0.0486 | TTGCTTTGCCTATGGAGTCCTGCAGGGGGAGGAC | | |
| 488 | 13 | 18042027 | 18051875 | 9848 | AKR.ILNJ2.LP.PJ.RF.SEA | 13 | 18044584 | 18044618 | 35 | - | 19.4587 | 1.23E-07 | 0.0632 | CAGCAGGGGCCAGCAGGGACCAGTAGGGGCCAGAA | | |
| 488 | 13 | 18042027 | 18051875 | 9848 | AKR.ILNJ2.LP.PJ.RF.SEA | 13 | 18044604 | 18044638 | 35 | - | 17.5872 | 3.85E-07 | 0.12 | TAGCAGGGACTATCAGGGACCAGCAGGGGCCAGCA | | |
| 489 | 13 | 23064254 | 23140550 | 76296 | NOD.NOR | 13 | 23105720 | 23105754 | 35 | - | 19 | 1.64E-07 | 0.0747 | CAGCAGTTGCCACCTGTGTGCTCTTGGGGGCGGGG | | |
| 490 | 13 | 31515462 | 31534163 | 18701 | BUB | 13 | 31526531 | 31526549 | 19 | - | 19.0984 | 2.17E-07 | 0.098 | AGACCAGAAGAGGGCACTG | 2 | ENCFF327LYS.ENCFF907XEG |
| 490 | 13 | 31515462 | 31534163 | 18701 | BUB | 13 | 31526531 | 31526565 | 35 | - | 17.7064 | 3.58E-07 | 0.115 | GTACAGGTATCTGTGTAGACCAGAAGAGGGCACTG | 2 | ENCFF327LYS.ENCFF907XEG |
| 491 | 13 | 40505647 | 40506926 | 1279 | 129S1,AJ,AKR,B10D2,B10J,BALB,BTBR,BUB,C3H,C57L,C58J,CBA,CE,DBA1J,DBA2J,FVB,ILNJ2,KK,LP,MA/My,MRL,NOD,NOR,NZB,NZO,NZW,PJ,PL,RF,RH2,SEA,SJL,SMJ,SWR.TH | 13 | 40506924 | 40506958 | 35 | - | 15.9541 | 9.80E-07 | 0.198 | CTGCTTTACTACTTCAAAGGCACTTGAGGGCAGGT | | |
| 492 | 13 | 60971665 | 60973516 | 1851 | PJ.SEA | 13 | 60972610 | 60972644 | 35 | - | 17.1468 | 4.98E-07 | 0.137 | TTGCAATTGAAGGCCCAGGCCTGAAGGGGTCATGC | | |
| 493 | 13 | 68590110 | 68598851 | 8741 | AJ,AKR,BALB,BUB,C3H,CBA,DBA1J,DBA2J,FVB,ILNJ2,KK,MRL,NOD,NOR,NZB,NZO,NZW,PJ,RF,SEA,SJL,TH | 13 | 68596292 | 68596310 | 19 | - | 17.8689 | 5.14E-07 | 0.139 | AAACCACTAGGTGGCAGAA | | |
| 494 | 13 | 76641139 | 76654006 | 12867 | 129S1,AJ,AKR,BALB,BTBR,C3H,C57L,CBA,CE,DBA1J,DBA2J,FVB,KK,MA/My,MRL,NZB,NZO,NZW,PJ,RF,SEA,SJL,SMJ,TH | 13 | 76652934 | 76652967 | 34 | - | 19.3171 | 1.44E-07 | 0.0472 | CTGCAAGAGGACAGCTTGCCTGCAGAGGGTACTC | | |
| 495 | 13 | 76642026 | 76653329 | 11303 | BUB.ILNJ2.NOD.NOR.SWR | 13 | 76652934 | 76652967 | 34 | - | 19.3171 | 1.44E-07 | 0.0472 | CTGCAAGAGGACAGCTTGCCTGCAGAGGGTACTC | | |
| 496 | 13 | 93335115 | 93339256 | 4141 | 129S1,AJ,AKR,BALB,BTBR,C3H,C57L,C58J,CBA,CE,DBA1J,DBA2J,FVB,ILNJ2,KK,MA/My,MRL,NZB,NZO,NZW,PJ,PL,RF,SEA,SJL,SMJ.SWR | 13 | 93338453 | 93338486 | 34 | - | 17.6748 | 4.37E-07 | 0.0839 | GAGCATGACCACAAGTCCACAGCAGGTGGCAACG | | |
| 497 | 13 | 99122778 | 99131763 | 8985 | 129S1,AJ,AKR,BALB,BTBR,BUB,C3H,C57L,C58J,CBA,CE,DBA1J,DBA2J,FVB,ILNJ2,KK,MA/My,MRL,NOD,NOR,NZB,NZO,PL,SJL,SMJ.SWR | 13 | 99126333 | 99126351 | 19 | - | 17.1639 | 8.17E-07 | 0.174 | CCTCCACGAGGTGGCGGAA | | |
| 498 | 13 | 112283888 | 112286234 | 2346 | AJ,DBA1J,KK,MA/My,NZO,PL,RF,RH2,SWR,TH | 13 | 112284156 | 112284189 | 34 | - | 17.3252 | 5.49E-07 | 0.0944 | CTGCATTTTATGAGGCTGCCAGAAGGAGATGGTC | | |
| 499 | 13 | 116010250 | 116017170 | 6920 | AJ.DBA1J.MA/My.RH2 | 13 | 116015957 | 116015991 | 35 | - | 21.8899 | 2.46E-08 | 0.0216 | CTGCATTTATAACAATTTCCCAGTTGATGGCAGTC | 1 | ENCFF093ZRU |
| 500 | 14 | 4871219 | 4875491 | 4272 | 129S1,C3H,C57L,CBA,DBA1J,DBA2J,KK,MRL.NZW.PJ.PL | 14 | 4871235 | 4871268 | 34 | + | 19.3415 | 1.42E-07 | 0.0418 | CTGCCCCCACATTCGCCGTCACAAGATGGCGCTG | | |
| 501 | 14 | 5013070 | 5033506 | 20436 | 129S1,C3H,C57L,CBA,DBA1J,DBA2J,KK,MRL.NZW.PJ.PL | 14 | 5017881 | 5017914 | 34 | + | 17.9756 | 3.59E-07 | 0.0664 | ATGCAGCTAGAGACATGAACTCCGGGTGGCACTG | | |
| 502 | 14 | 6916484 | 6922010 | 5526 | 129S1,C3H,C57L,CBA,DBA1J,DBA2J,KK,MRL.NZW.PJ.PL | 14 | 6920257 | 6920291 | 35 | + | 16.4404 | 7.46E-07 | 0.2 | AAGAAGCTGGACCCAGTGGCCAGCAGATGGCCCAC | | |
| 502 | 14 | 6916484 | 6922010 | 5526 | 129S1,C3H,C57L,CBA,DBA1J,DBA2J,KK,MRL.NZW.PJ.PL | 14 | 6920273 | 6920291 | 19 | + | 17.0328 | 8.88E-07 | 0.184 | TGGCCAGCAGATGGCCCAC | | |
| 503 | 14 | 15552829 | 15559931 | 7102 | C57L.PL | 14 | 15558759 | 15558792 | 34 | + | 20.626 | 5.64E-08 | 0.0235 | CTGGAGGGGCAGCCTTGGCCTTCAGGTGGCAGCC | | |
| 504 | 14 | 16578683 | 16580837 | 2154 | 129S1.C57L.KK.PL | 14 | 16580306 | 16580339 | 34 | + | 18.0488 | 3.42E-07 | 0.0647 | TTGCTGTATGCGTAGAGGCCAGAAGAGGGTGTCA | | |
| 505 | 14 | 22494653 | 22568513 | 73860 | PJ | 14 | 22514564 | 22514597 | 34 | + | 26.6016 | 3.79E-10 | 0.000974 | ATGCAGTGCCCATGGAGGCCAGGAGAGGGCATCA | 5 | ENCFF093ZRU.ENCFF052KVO.ENCFF055HAP.ENCFF367UNR.ENCFF502WDT |
| 506 | 14 | 28970837 | 28977767 | 6930 | 129S1,AJ,AKR,BALB,BTBR,BUB,C3H,C57L,CBA,CE,FVB,KK,MA/My,MRL,NOD,NOR,NZO,NZW,PJ,PL,RF,RH2,SEA,SJL,SMJ,SWR,TH | 14 | 28973330 | 28973348 | 19 | + | 18.5738 | 3.17E-07 | 0.117 | AAACCACCAGGTGGCAGAA | | |
| 507 | 14 | 32612656 | 32614149 | 1493 | DBA1J.DBA2J.ILNJ2.NZB.PJ | 14 | 32613896 | 32613929 | 34 | + | 19.6667 | 1.13E-07 | 0.0368 | GTGCAGTGCCCTCGAAGGCCAGTGGAGGGTTTGG | | |
| 508 | 14 | 35526970 | 35528584 | 1614 | BUB.MA/My.NZB.PJ.SWR | 14 | 35527353 | 35527386 | 34 | + | 18.3902 | 2.72E-07 | 0.0594 | CAGCTGGGCCTCTTTGAGTCACTGGGGGGCAGAG | | |
| 509 | 14 | 37830579 | 37839837 | 9258 | BTBR.C57L.NZO.PJ.PL | 14 | 37832715 | 37832748 | 34 | + | 17.0325 | 6.63E-07 | 0.0931 | TGGCAGGTCTCAGAGGAGCCGCTAGATGGCCCAC | | |
| 510 | 14 | 38781300 | 38936510 | 155210 | LP | 14 | 38847603 | 38847621 | 19 | + | 17.1639 | 8.17E-07 | 0.178 | TCCCCAGCAGGTGGAACTG | | |
| 511 | 14 | 44125853 | 44255959 | 130106 | CBA,DBA1J,DBA2J,ILNJ2 | 14 | 44162639 | 44162657 | 19 | + | 19.2951 | 1.88E-07 | 0.0904 | CCGCCACTAGATGGTGCTA | 30 | ENCFF367UNR,ENCFF093ZRU,ENCFF099QNE,ENCFF201JUA,ENCFF327LYS,ENCFF821BIA,ENCFF907XEG,ENCFF589ZBJ,ENCFF089GBC,ENCFF198SSR,ENCFF430PPJ,ENCFF024JOZ,ENCFF092NMJ,ENCFF321AGQ,ENCFF136ZDK,ENCFF165LXO,ENCFF316FYN,ENCFF033UQX,ENCFF097CAQ,ENCFF052KVO,ENCFF951GOP,ENCFF957SPH,ENCFF281QMV,ENCFF883UPM,ENCFF210BTO,ENCFF194HVL,ENCFF609MDN,ENCFF611HDQ.ENCFF491RJK.ENCFF575MGI |
| 511 | 14 | 44125853 | 44255959 | 130106 | CBA,DBA1J,DBA2J,ILNJ2 | 14 | 44170621 | 44170639 | 19 | + | 18.6557 | 2.99E-07 | 0.113 | CCGCCACTAGATGGTGGTA | 23 | ENCFF093ZRU,ENCFF367UNR,ENCFF052KVO,ENCFF099QNE,ENCFF201JUA,ENCFF821BIA,ENCFF024JOZ,ENCFF327LYS,ENCFF907XEG,ENCFF089GBC,ENCFF092NMJ,ENCFF609MDN,ENCFF136ZDK,ENCFF589ZBJ,ENCFF430PPJ,ENCFF198SSR,ENCFF055HAP,ENCFF165LXO,ENCFF502WDT,ENCFF883UPM,ENCFF951GOP.ENCFF957SPH.ENCFF281QMV |
| 511 | 14 | 44125853 | 44255959 | 130106 | CBA.DBA1J.DBA2J.ILNJ2 | 14 | 44254995 | 44255013 | 19 | + | 16.9672 | 9.26E-07 | 0.188 | AGGCCAGAAGAGGGAACTG | | |
| 511 | 14 | 44125853 | 44255959 | 130106 | CBA.DBA1J.DBA2J.ILNJ2 | 14 | 44183356 | 44183374 | 19 | - | 16.9672 | 9.26E-07 | 0.188 | AGGCCAGAAGAGGGAACTG | | |
| 511 | 14 | 44125853 | 44255959 | 130106 | CBA,DBA1J,DBA2J,ILNJ2 | 14 | 44221550 | 44221568 | 19 | - | 24.2295 | 1.90E-09 | 0.0188 | TGGCCAGCAGGGGGCGCAC | 29 | ENCFF093ZRU,ENCFF367UNR,ENCFF201JUA,ENCFF099QNE,ENCFF327LYS,ENCFF907XEG,ENCFF589ZBJ,ENCFF024JOZ,ENCFF198SSR,ENCFF951GOP,ENCFF821BIA,ENCFF136ZDK,ENCFF321AGQ,ENCFF097CAQ,ENCFF092NMJ,ENCFF089GBC,ENCFF033UQX,ENCFF165LXO,ENCFF491RJK,ENCFF430PPJ,ENCFF957SPH,ENCFF281QMV,ENCFF883UPM,ENCFF210BTO,ENCFF194HVL,ENCFF609MDN,ENCFF533APC,ENCFF611HDQ.ENCFF575MGI |
| 511 | 14 | 44125853 | 44255959 | 130106 | CBA,DBA1J,DBA2J,ILNJ2 | 14 | 44221550 | 44221583 | 34 | - | 23.561 | 5.69E-09 | 0.00511 | AGGGAGTTGCTGCTTTGGCCAGCAGGGGGCGCAC | 29 | ENCFF093ZRU,ENCFF367UNR,ENCFF201JUA,ENCFF099QNE,ENCFF327LYS,ENCFF907XEG,ENCFF589ZBJ,ENCFF024JOZ,ENCFF198SSR,ENCFF951GOP,ENCFF821BIA,ENCFF136ZDK,ENCFF321AGQ,ENCFF097CAQ,ENCFF092NMJ,ENCFF089GBC,ENCFF033UQX,ENCFF165LXO,ENCFF491RJK,ENCFF430PPJ,ENCFF957SPH,ENCFF281QMV,ENCFF883UPM,ENCFF210BTO,ENCFF194HVL,ENCFF609MDN,ENCFF533APC,ENCFF611HDQ.ENCFF575MGI |
| 511 | 14 | 44125853 | 44255959 | 130106 | CBA,DBA1J,DBA2J,ILNJ2 | 14 | 44225063 | 44225081 | 19 | - | 18.2787 | 3.88E-07 | 0.128 | AGGCCAGAAGAGGGCGATA | 28 | ENCFF093ZRU,ENCFF367UNR,ENCFF052KVO,ENCFF099QNE,ENCFF201JUA,ENCFF136ZDK,ENCFF327LYS,ENCFF033UQX,ENCFF821BIA,ENCFF907XEG,ENCFF024JOZ,ENCFF198SSR,ENCFF092NMJ,ENCFF589ZBJ,ENCFF097CAQ,ENCFF089GBC,ENCFF321AGQ,ENCFF165LXO,ENCFF491RJK,ENCFF957SPH,ENCFF883UPM,ENCFF281QMV,ENCFF533APC,ENCFF430PPJ,ENCFF609MDN,ENCFF951GOP,ENCFF194HVL,ENCFF575MGI |
| 511 | 14 | 44125853 | 44255959 | 130106 | CBA,DBA1J,DBA2J,ILNJ2 | 14 | 44225063 | 44225096 | 34 | - | 25.0163 | 1.63E-09 | 0.00214 | GCGCAGTGCCCATAGAGGCCAGAAGAGGGCGATA | 28 | ENCFF093ZRU,ENCFF367UNR,ENCFF052KVO,ENCFF099QNE,ENCFF201JUA,ENCFF136ZDK,ENCFF327LYS,ENCFF033UQX,ENCFF821BIA,ENCFF907XEG,ENCFF024JOZ,ENCFF198SSR,ENCFF092NMJ,ENCFF589ZBJ,ENCFF097CAQ,ENCFF089GBC,ENCFF321AGQ,ENCFF165LXO,ENCFF491RJK,ENCFF957SPH,ENCFF883UPM,ENCFF281QMV,ENCFF533APC,ENCFF430PPJ,ENCFF609MDN,ENCFF951GOP,ENCFF194HVL,ENCFF575MGI |
| 512 | 14 | 44879155 | 44887080 | 7925 | C57L.NZO.PJ.PL | 14 | 44880161 | 44880194 | 34 | + | 24.7236 | 2.11E-09 | 0.00247 | ATGCAATTTCTCCAGAGGCCAGAAGAGGGCATGG | | |
| 513 | 14 | 49652511 | 49653639 | 1128 | 129S1,AJ,AKR,BALB,BUB,C3H,CBA,CE,DBA1J,DBA2J,FVB,ILNJ2,MA/My,MRL,NOD,NOR,NZB,NZO,NZW,PJ,PL,RF,RH2,SEA,SJL,SMJ.SWR.TH | 14 | 49653113 | 49653146 | 34 | + | 16.4472 | 9.61E-07 | 0.111 | TAGCTGTCTTCAGACACACCAGAAGAGGGCGTCA | | |
| 514 | 14 | 51599291 | 51600361 | 1070 | 129S1,AJ,AKR,BALB,BUB,C3H,C57L,CBA,CE,DBA1J,DBA2J,FVB,ILNJ2,KK,MA/My,MRL,NOD,NOR,NZB,NZO,NZW,PJ,PL,RF,RH2,SEA,SJL,SMJ,SWR,TH | 14 | 51599790 | 51599823 | 34 | + | 17.9756 | 3.59E-07 | 0.0664 | CTGAAGGGCCAACCTTGGCCTTCAGGGGCAGCCA | | |
| 515 | 14 | 51619535 | 51623466 | 3931 | 129S1,AJ,AKR,BALB,BUB,C3H,C57L,CBA,CE,DBA1J,DBA2J,KK,MRL,NZB,NZO,NZW,PJ,PL,RF,RH2,SEA,TH | 14 | 51620067 | 51620100 | 34 | + | 18.3171 | 2.86E-07 | 0.0602 | GGGCAGAGTCTCCAGATGACGGCAGGGGGCGGGG | | |
| 516 | 14 | 53157579 | 53166165 | 8586 | DBA1J.DBA2J.NZW | 14 | 53157817 | 53157850 | 34 | + | 20.8049 | 4.94E-08 | 0.0235 | ATTTGGTTCCTCCTATGGCCTCCAGAGGGCGGTG | | |
| 516 | 14 | 53157579 | 53166165 | 8586 | DBA1J.DBA2J.NZW | 14 | 53157832 | 53157850 | 19 | + | 22.4918 | 1.28E-08 | 0.0351 | TGGCCTCCAGAGGGCGGTG | | |
| 517 | 14 | 53216805 | 53220988 | 4183 | ILNJ2.NOD | 14 | 53220594 | 53220628 | 35 | + | 18.1009 | 2.83E-07 | 0.121 | CATTTGTTTCCTCCTATGGCCTCCAGAGGGCGCTG | | |
| 517 | 14 | 53216805 | 53220988 | 4183 | ILNJ2.NOD | 14 | 53220595 | 53220628 | 34 | + | 18.8699 | 1.96E-07 | 0.0508 | ATTTGTTTCCTCCTATGGCCTCCAGAGGGCGCTG | | |
| 517 | 14 | 53216805 | 53220988 | 4183 | ILNJ2.NOD | 14 | 53220610 | 53220628 | 19 | + | 23.1311 | 6.72E-09 | 0.0294 | TGGCCTCCAGAGGGCGCTG | | |
| 518 | 14 | 53909278 | 53913728 | 4450 | FVB,ILNJ2,MA/My,NOD,NOR,NZB,PL,SJL,SMJ.SWR | 14 | 53913334 | 53913368 | 35 | + | 18.1009 | 2.83E-07 | 0.121 | CATTTGTTTCCTCCTATGGCCTCCAGAGGGCGCTG | 1 | ENCFF093ZRU |
| 518 | 14 | 53909278 | 53913728 | 4450 | FVB,ILNJ2,MA/My,NOD,NOR,NZB,PL,SJL,SMJ.SWR | 14 | 53913335 | 53913368 | 34 | + | 18.8699 | 1.96E-07 | 0.0508 | ATTTGTTTCCTCCTATGGCCTCCAGAGGGCGCTG | 1 | ENCFF093ZRU |
| 518 | 14 | 53909278 | 53913728 | 4450 | FVB,ILNJ2,MA/My,NOD,NOR,NZB,PL,SJL,SMJ.SWR | 14 | 53913350 | 53913368 | 19 | + | 23.1311 | 6.72E-09 | 0.0294 | TGGCCTCCAGAGGGCGCTG | 1 | ENCFF093ZRU |
| 519 | 14 | 60011763 | 60015164 | 3401 | AJ,AKR,BALB,C3H,CBA,CE,DBA1J,DBA2J,MRL,NZB,PJ,PL,RH2,SEA,SMJ | 14 | 60012248 | 60012281 | 34 | + | 19.561 | 1.21E-07 | 0.0376 | CAGCATCGTTCTTCCTGGCCAGTTGGGGGCGCGC | | |
| 519 | 14 | 60011763 | 60015164 | 3401 | AJ,AKR,BALB,C3H,CBA,CE,DBA1J,DBA2J,MRL,NZB,PJ,PL,RH2,SEA,SMJ | 14 | 60012263 | 60012281 | 19 | + | 17.5574 | 6.32E-07 | 0.158 | TGGCCAGTTGGGGGCGCGC | | |
| 519 | 14 | 60011763 | 60015164 | 3401 | AJ,AKR,BALB,C3H,CBA,CE,DBA1J,DBA2J,MRL,NZB,PJ,PL,RH2,SEA,SMJ | 14 | 60014777 | 60014810 | 34 | + | 19.561 | 1.21E-07 | 0.0376 | CAGCATCGTTCTTCCTGGCCAGTTGGGGGCGCGC | | |
| 519 | 14 | 60011763 | 60015164 | 3401 | AJ,AKR,BALB,C3H,CBA,CE,DBA1J,DBA2J,MRL,NZB,PJ,PL,RH2,SEA,SMJ | 14 | 60014792 | 60014810 | 19 | + | 17.5574 | 6.32E-07 | 0.158 | TGGCCAGTTGGGGGCGCGC | | |
| 520 | 14 | 69047722 | 69050461 | 2739 | 129S1.BTBR.C57L.KK.LP.TH | 14 | 69048113 | 69048131 | 19 | + | 17.5574 | 6.32E-07 | 0.158 | TGGCCAGTTGGGGGCGCGC | | |
| 521 | 14 | 90975758 | 90977224 | 1466 | NZO | 14 | 90976900 | 90976933 | 34 | + | 21.2683 | 3.49E-08 | 0.0184 | ATGCAGTGCCTCAAACATCCAAAAGAAGGCACTT | | |
| 522 | 14 | 19427725 | 19429889 | 2164 | 129S1.C57L.KK.SMJ | 14 | 19428222 | 19428255 | 34 | - | 18.5935 | 2.37E-07 | 0.0571 | GTGCTGTATGTGTAGAGGCCAGAAGAGGGTGTCA | | |
| 523 | 14 | 25912256 | 25913989 | 1733 | AJ.BALB.NZW.RH2.SEA | 14 | 25913556 | 25913589 | 34 | - | 16.4472 | 9.61E-07 | 0.111 | TAGCTGTCTTCAGACACACCAGAAGAGGGCGTCA | | |
| 524 | 14 | 33594943 | 33597797 | 2854 | BTBR | 14 | 33596497 | 33596530 | 34 | - | 16.8374 | 7.51E-07 | 0.0977 | CAGCAGGTCTTGACACTTCCTGGAGGAGTCACCC | | |
| 525 | 14 | 43050984 | 43055970 | 4986 | CBA.DBA1J.DBA2J | 14 | 43051941 | 43051974 | 34 | - | 25.3984 | 1.16E-09 | 0.00172 | ATGCAATTTCTCCAGAGGCCAGTAGAGGGCATGA | 4 | ENCFF367UNR.ENCFF201JUA.ENCFF099QNE.ENCFF093ZRU |
| 526 | 14 | 44041824 | 44050145 | 8321 | BTBR,C57L,FVB,KK,NOD,NOR,NZB,NZO,NZW,PJ,PL | 14 | 44042116 | 44042134 | 19 | - | 19.2951 | 1.88E-07 | 0.0904 | CCGCCACTAGATGGTGCTA | 13 | ENCFF092NMJ,ENCFF136ZDK,ENCFF165LXO,ENCFF198SSR,ENCFF093ZRU,ENCFF367UNR,ENCFF099QNE,ENCFF821BIA,ENCFF327LYS,ENCFF907XEG,ENCFF024JOZ,ENCFF089GBC,ENCFF589ZBJ |
| 526 | 14 | 44041824 | 44050145 | 8321 | BTBR,C57L,FVB,KK,NOD,NOR,NZB,NZO,NZW,PJ,PL | 14 | 44050104 | 44050122 | 19 | - | 19.2951 | 1.88E-07 | 0.0904 | CCGCCACTAGATGGTGCTA | 28 | ENCFF201JUA,ENCFF093ZRU,ENCFF367UNR,ENCFF883UPM,ENCFF609MDN,ENCFF099QNE,ENCFF097CAQ,ENCFF194HVL,ENCFF907XEG,ENCFF136ZDK,ENCFF327LYS,ENCFF821BIA,ENCFF089GBC,ENCFF491RJK,ENCFF589ZBJ,ENCFF321AGQ,ENCFF052KVO,ENCFF055HAP,ENCFF065ODV,ENCFF957SPH,ENCFF281QMV,ENCFF142CNG,ENCFF092NMJ,ENCFF198SSR,ENCFF165LXO,ENCFF430PPJ,ENCFF502WDT,ENCFF024JOZ |
| 527 | 14 | 52067424 | 52086384 | 18960 | BUB,C57L,DBA1J,DBA2J,FVB,ILNJ2,MA/My,NOD,NOR,RF,SJL,SMJ,SWR | 14 | 52076111 | 52076129 | 19 | - | 17.5246 | 6.45E-07 | 0.16 | TGGTCAGCAGAGGGCAGGA | 9 | ENCFF367UNR,ENCFF092NMJ,ENCFF089GBC,ENCFF907XEG,ENCFF883UPM,ENCFF093ZRU,ENCFF099QNE,ENCFF327LYS.ENCFF024JOZ |
| 527 | 14 | 52067424 | 52086384 | 18960 | BUB,C57L,DBA1J,DBA2J,FVB,ILNJ2,MA/My,NOD,NOR,RF,SJL,SMJ,SWR | 14 | 52076111 | 52076145 | 35 | - | 18.3303 | 2.47E-07 | 0.112 | CTGTAAGAGCTTCTTGTGGTCAGCAGAGGGCAGGA | 9 | ENCFF367UNR,ENCFF092NMJ,ENCFF089GBC,ENCFF907XEG,ENCFF883UPM,ENCFF093ZRU,ENCFF099QNE,ENCFF327LYS.ENCFF024JOZ |
| 528 | 14 | 52113662 | 52120861 | 7199 | 129S1,AJ,AKR,BALB,C3H,C57L,CBA,CE,DBA1J,DBA2J,KK,MRL,NZB,NZO,NZW,PJ,PL,RF,RH2,SEA,TH | 14 | 52118709 | 52118743 | 35 | - | 16.1009 | 9.03E-07 | 0.221 | CTACTGTTCGTCACCAGGATCTCCAGAGGGCAGCT | | |
| 529 | 14 | 53012170 | 53013603 | 1433 | BUB | 14 | 53013487 | 53013520 | 34 | - | 17.6992 | 4.30E-07 | 0.0732 | CTGCAATACCAGACTGAAGCTGAAGGTGTCTCCG | 4 | ENCFF089GBC.ENCFF883UPM.ENCFF093ZRU.ENCFF099QNE |
| 530 | 14 | 71406924 | 71409672 | 2748 | BTBR | 14 | 71409615 | 71409648 | 34 | - | 18.0569 | 3.40E-07 | 0.0647 | ATGCAGCTAGAGTCACGAGCTCCAGGGGGTACTG | | |
| 531 | 14 | 76170441 | 76173026 | 2585 | SWR | 14 | 76171985 | 76172018 | 34 | - | 27.8943 | 1.05E-10 | 0.000521 | GTGCAGTGCCCTTGGAGGCCAGAAGAGGGCATCA | 20 | ENCFF024JOZ,ENCFF089GBC,ENCFF093ZRU,ENCFF367UNR,ENCFF198SSR,ENCFF201JUA,ENCFF883UPM,ENCFF065ODV,ENCFF136ZDK,ENCFF194HVL,ENCFF327LYS,ENCFF589ZBJ,ENCFF052KVO,ENCFF957SPH,ENCFF321AGQ,ENCFF821BIA,ENCFF502WDT,ENCFF099QNE,ENCFF907XEG,ENCFF055HAP |
| 532 | 14 | 76525327 | 76526557 | 1230 | NZB.NZO | 14 | 76526258 | 76526291 | 34 | - | 17.1382 | 6.20E-07 | 0.089 | TAGCTGTCTTCAAGCACACCAGCAGAGGGCATCA | | |
| 533 | 14 | 77777140 | 77778495 | 1355 | AJ.BALB.NZB.NZO.SEA.SJL | 14 | 77778142 | 77778176 | 35 | - | 22.2569 | 1.90E-08 | 0.0271 | CTGCTGTGTCACATGTGCATCAGTAGGTGGCAGAG | 2 | ENCFF821BIA.ENCFF367UNR |
| 534 | 14 | 94168361 | 94174974 | 6613 | AJ,AKR,BALB,BTBR,BUB,C3H,CBA,CE,DBA1J,DBA2J,FVB,MRL,NOD,PJ,PL,RF,RH2,SEA,SJL,SMJ,SWR,TH | 14 | 94174171 | 94174204 | 34 | - | 16.9106 | 7.17E-07 | 0.0962 | GAGCATGACCACAAATCCACAGCAGGTGGCAACG | | |
| 535 | 14 | 99330219 | 99331758 | 1539 | AKR,CBA,CE,DBA1J,DBA2J,ILNJ2,MRL,PJ,PL,RF,RH2,SJL,SMJ,SWR,TH | 14 | 99330761 | 99330795 | 35 | - | 16.6789 | 6.51E-07 | 0.189 | GTGCAGTGTGCGCCATTCAGCTAAAGAGGGCTCCC | | |
| 536 | 14 | 105129975 | 105196398 | 66423 | NZB | 14 | 105143349 | 105143367 | 19 | - | 21.082 | 4.57E-08 | 0.0571 | AAGCCAGCAGAGGGCACCA | 2 | ENCFF089GBC.ENCFF052KVO |
| 536 | 14 | 105129975 | 105196398 | 66423 | NZB | 14 | 105143349 | 105143383 | 35 | - | 25.211 | 2.08E-09 | 0.00868 | TTGTAGTTATCTATGGAAGCCAGCAGAGGGCACCA | 2 | ENCFF089GBC.ENCFF052KVO |
| 537 | 14 | 120489443 | 120490691 | 1248 | BTBR.C57L | 14 | 120490579 | 120490613 | 35 | - | 18.7064 | 1.96E-07 | 0.0995 | CAGCACTCCTAGCTCAGGCCCTGCAGAGGTCATTG | | |
| 538 | 15 | 8224437 | 8226850 | 2413 | LP.TH | 15 | 8225786 | 8225819 | 34 | + | 16.4634 | 9.51E-07 | 0.106 | TAGCTGTCTTCAGACGCACCAGAAGAGGGCATCA | | |
| 539 | 15 | 15784854 | 15789357 | 4503 | PL | 15 | 15785631 | 15785664 | 34 | + | 17.6748 | 4.37E-07 | 0.0745 | GAGCATGACCACAAGTCCACAGCAGGTGGCAACG | | |
| 540 | 15 | 29125046 | 29131360 | 6314 | C3H.CBA.DBA1J.DBA2J.MA/My.MRL | 15 | 29128130 | 29128163 | 34 | + | 16.5041 | 9.27E-07 | 0.106 | ATGCAGCTAGAGACACGAGCTCTGGGGGGCGGGG | | |
| 541 | 15 | 38951626 | 38952650 | 1024 | C3H.MRL.NOD.NOR.PL.SWR | 15 | 38952166 | 38952200 | 35 | + | 16.4037 | 7.61E-07 | 0.163 | CTGCATGTACACCTGCATGCCAGAAGAGAGTATTA | | |
| 542 | 15 | 41750645 | 41752728 | 2083 | 129S1.AKR.BUB.C3H.C57L.KK.PL.SWR | 15 | 41752251 | 41752269 | 19 | + | 17.459 | 6.74E-07 | 0.13 | CAGCCAGGAGGTGGAGCTG | | |
| 543 | 15 | 59033542 | 59034868 | 1326 | 129S1.C57L.MRL | 15 | 59033558 | 59033576 | 19 | + | 17.5738 | 6.25E-07 | 0.126 | CCACCAGCAGGCGGCACTG | 2 | ENCFF093ZRU.ENCFF367UNR |
| 544 | 15 | 59033550 | 59039293 | 5743 | 129S1,BALB,C57L,ILNJ2,MRL,NOD,NOR,SEA | 15 | 59033558 | 59033576 | 19 | + | 17.5738 | 6.25E-07 | 0.126 | CCACCAGCAGGCGGCACTG | 2 | ENCFF093ZRU,ENCFF367UNR |
| 545 | 15 | 68993767 | 69001031 | 7264 | BUB,DBA1J,DBA2J,ILNJ2,KK,LP,NZB,NZO,PL,TH | 15 | 68994062 | 68994096 | 35 | + | 15.9817 | 9.65E-07 | 0.183 | CTGCTCTGACCACTGAAACTCAGAGGAGGGAGCTA | | |

| | | | | | | | | | | | | | | | | |
|---|---|---|---|---|---|---|---|---|---|---|---|---|---|---|---|---|
| 546 | 15 | 71919987 | 71921209 | 1222 | AJ,B10D2,B10J,BALB,BTBR,BUB,C3H,C57L,CBA,CE,DBA1J,DBA2J,FVB,ILNJ2,KK,LP,MA/My,MRL,NOD,NOR,NZB,NZO,NZW,PJ,PL,RF,SEA,SJL,SMJ,SWR,TH | 15 | 71920159 | 71920192 | 34 | + | 17.9431 | 3.66E-07 | 0.0685 | GAACCATTACTCCTACTGGCACCAGGGGGCGCTC | | |
| 546 | 15 | 71919987 | 71921209 | 1222 | AJ,B10D2,B10J,BALB,BTBR,BUB,C3H,C57L,CBA,CE,DBA1J,DBA2J,FVB,ILNJ2,KK,LP,MA/My,MRL,NOD,NOR,NZB,NZO,NZW,PJ,PL,RF,SEA,SJL,SMJ,SWR,TH | 15 | 71920174 | 71920192 | 19 | + | 19.6066 | 1.49E-07 | 0.062 | CTGGCACCAGGGGGCGCTC | | |
| 547 | 15 | 75036685 | 75047588 | 10903 | AJ,BALB,C3H,CBA,CE,NOD,NOR,NZB,NZO,SEA,TH | 15 | 75047549 | 75047567 | 19 | + | 19.7049 | 1.38E-07 | 0.0593 | TGGCCAGCAGGTGGCAGGT | 35 | ENCFF201JUA,ENCFF093ZRU,ENCFF367UNR,ENCFF099QNE,ENCFF327LYS,ENCFF502WDT,ENCFF024JOZ,ENCFF907XEG,ENCFF951GOP,ENCFF316FYN,ENCFF033UQX,ENCFF821BIA,ENCFF198SSR,ENCFF165LXO,ENCFF589ZBJ,ENCFF092NMJ,ENCFF575MGI,ENCFF052KVO,ENCFF957SPH,ENCFF097CAQ,ENCFF136ZDK,ENCFF883UPM,ENCFF089EWX,ENCFF210BTO,ENCFF611HDQ,ENCFF065ODV,ENCFF089GBC,ENCFF055HAP,ENCFF281QMV,ENCFF504JDS,ENCFF533APC,ENCFF142CNG,ENCFF491RJK,ENCFF194HVL,ENCFF430PPJ |
| 548 | 15 | 77464039 | 77486565 | 22526 | MA/Mv | 15 | 77485627 | 77485645 | 19 | + | 17.1967 | 7.99E-07 | 0.144 | GGGCCAGAAGAGGGCGTCA | | |
| 549 | 15 | 14476678 | 14485306 | 8628 | LP.SMJ | 15 | 14484237 | 14484270 | 34 | - | 17.6748 | 4.37E-07 | 0.0745 | GAGCATGACCACAAGTCCACAGCAGGTGGCAACG | | |
| 550 | 15 | 14480550 | 14484986 | 4436 | BALB,C3H,CBA,DBA2J,MRL,NOD,NOR,NZB,NZO,PL,SJL,SWR | 15 | 14484237 | 14484270 | 34 | - | 17.6748 | 4.37E-07 | 0.0745 | GAGCATGACCACAAGTCCACAGCAGGTGGCAACG | | |
| 551 | 15 | 14845014 | 14853191 | 8177 | BUB.CE.ILNJ2.MRL.PL | 15 | 14845280 | 14845313 | 34 | - | 17.0407 | 6.60E-07 | 0.0918 | CTGTGATAACATTTGCCATCACAAGATGGCGCCA | 2 | ENCFF055HAP.ENCFF502WDT |
| 552 | 15 | 19495443 | 19499482 | 4039 | 129S1,BTBR,C3H,C57L,CBA,CE,ILNJ2,KK,LP.MA/Mv.NZW.PL.SWR | 15 | 19498735 | 19498768 | 34 | - | 17.6748 | 4.37E-07 | 0.0745 | GAGCATGACCACAAGTCCACAGCAGGTGGCAACG | | |
| 553 | 15 | 20647297 | 20650504 | 3207 | MRL.PL.SWR | 15 | 20650022 | 20650055 | 34 | - | 16.8943 | 7.24E-07 | 0.0956 | ATGCAGTCCCCATGGAGGCCAAAACATGGCATCG | | |
| 554 | 15 | 36481361 | 36483947 | 2586 | 129S1.C57L | 15 | 36481793 | 36481826 | 34 | - | 22.8293 | 1.04E-08 | 0.00719 | GTACAGTTCCAACAGAGGCCAGAAGAGGGCATAA | 2 | ENCFF093ZRU.ENCFF327LYS |
| 555 | 15 | 57899641 | 57903272 | 3631 | ILNJ2 | 15 | 57902792 | 57902825 | 34 | - | 21.374 | 3.23E-08 | 0.0158 | GTGCAGGTACAGAAGAGGCCAGAAAAGGGCACCA | | |
| 556 | 15 | 65495252 | 65496497 | 1245 | SWR | 15 | 65495222 | 65495256 | 35 | - | 16.5046 | 7.19E-07 | 0.158 | TTGTAGGTAGAGGTATAGGCCTGAAGATGGAGCTT | | |
| 557 | 15 | 76444012 | 76451202 | 7190 | AJ,BALB,C3H,C57L,CBA,CE,FVB,MA/My,MRL,NOD,NOR,NZB,NZO,PL,RF,SEA | 15 | 76447567 | 76447585 | 19 | - | 17.1639 | 8.17E-07 | 0.145 | CCTCCACGAGGTGGCGGAA | | |
| 558 | 15 | 77568069 | 77576340 | 8271 | AJ,BALB,C3H,C57L,CBA,CE,FVB,MA/My,MRL,PL,RF,SEA,TH | 15 | 77568356 | 77568374 | 19 | - | 19.2951 | 1.88E-07 | 0.0694 | CCGCCACTAGATGGTGCTA | 26 | ENCFF367UNR,ENCFF093ZRU,ENCFF201JUA,ENCFF099QNE,ENCFF821BIA,ENCFF327LYS,ENCFF024JOZ,ENCFF907XEG,ENCFF136ZDK,ENCFF097CAQ,ENCFF589ZBJ,ENCFF609MDN,ENCFF165LXO,ENCFF092NMJ,ENCFF198SSR,ENCFF055HAP,ENCFF491RJK,ENCFF089GBC,ENCFF430PPJ,ENCFF951GOP,ENCFF281QMV,ENCFF502WDT,ENCFF142CNG,ENCFF883UPM,ENCFF957SPH,ENCFF194HVL |
| 559 | 15 | 88473285 | 88477436 | 4151 | C58J.KK.LP.NOD.NOR.PL.SWR.TH | 15 | 88476636 | 88476669 | 34 | - | 17.6748 | 4.37E-07 | 0.0745 | GAGCATGACCACAAGTCCACAGCAGGTGGCAACG | | |
| 560 | 15 | 88920563 | 88924461 | 3898 | KK.NOD.NOR.SWR | 15 | 88921019 | 88921052 | 34 | - | 18.5447 | 2.45E-07 | 0.0579 | CCGCACCCACATTCGCCGTCACAAGATGGCGCTG | | |
| 561 | 16 | 3146833 | 3155603 | 8770 | AJ,AKR,B10J,C3H,ILNJ2,KK,NOD,NOR,NZB,PJ,RF,SJL,SMJ | 16 | 3150976 | 3151010 | 35 | + | 16.156 | 8.75E-07 | 0.21 | AAGAAACTGGACCCTGTGGCCAGTGGATGGCGCTC | | |
| 561 | 16 | 3146833 | 3155603 | 8770 | AJ,AKR,B10J,C3H,ILNJ2,KK,NOD,NOR,NZB,PJ,RF,SJL,SMJ | 16 | 3150992 | 3151010 | 19 | + | 18.9344 | 2.45E-07 | 0.108 | TGGCCAGTGGATGGCGCTC | | |
| 562 | 16 | 25181459 | 25184935 | 3476 | BUB | 16 | 25183390 | 25183423 | 34 | + | 18.1951 | 3.10E-07 | 0.0743 | TTGCTGTCTTCAACCACACCAGAAGAGGGCATCA | 1 | ENCFF093ZRU |
| 563 | 16 | 27019888 | 27021582 | 1694 | ILNJ2.NZW | 16 | 27020856 | 27020874 | 19 | + | 17.0984 | 8.52E-07 | 0.181 | AGACCAGCAGAGGGAAGCC | | |
| 564 | 16 | 55758881 | 55763593 | 4712 | CBA.NZW.SMJ | 16 | 55761350 | 55761383 | 34 | + | 27.9919 | 9.53E-11 | 0.000516 | ATGCAGTGCCTACAGAGGCCAGAAGAGGGAACTG | 2 | ENCFF883UPM.ENCFF502WDT |
| 564 | 16 | 55758881 | 55763593 | 4712 | CBA.NZW.SMJ | 16 | 55761365 | 55761383 | 19 | + | 16.9672 | 9.26E-07 | 0.189 | AGGCCAGAAGAGGGAACTG | 2 | ENCFF883UPM.ENCFF502WDT |
| 565 | 16 | 57557229 | 57565509 | 8280 | CBA.NZW.SMJ | 16 | 57560706 | 57560740 | 35 | + | 17.1376 | 5.00E-07 | 0.161 | CTGAAGTGCCCTTAGCCAGCCAGCAAGTGCCAGAG | | |
| 566 | 16 | 68419265 | 68423830 | 4565 | BUB.CE.FVB.NOD.NOR.SWR | 16 | 68420025 | 68420058 | 34 | + | 17.6748 | 4.37E-07 | 0.0869 | GAGCATGACCACAAGTCCACAGCAGGTGGCAACG | | |
| 567 | 16 | 80812498 | 80816661 | 4163 | CE.DBA1J.MA/Mv.NZB.NZO.NZW | 16 | 80813271 | 80813304 | 34 | + | 17.6748 | 4.37E-07 | 0.0869 | GAGCATGACCACAAGTCCACAGCAGGTGGCAACG | | |
| 568 | 16 | 13064660 | 13065719 | 1059 | KK | 16 | 13065419 | 13065453 | 35 | - | 19.945 | 9.01E-08 | 0.0577 | CTGTAATCCCAGCACTGAAGCAGCAGGTGGGGGCA | | |
| 569 | 16 | 45973323 | 45975447 | 2124 | 129S1,C3H,C58J,CBA,CE,FVB,SJL,SWR,TH | 16 | 45974269 | 45974302 | 34 | - | 17.2114 | 5.91E-07 | 0.101 | GAGCAACTCCTTCACAAACCAGGTGATGGCACAG | | |
| 570 | 16 | 46147201 | 46149617 | 2416 | C3H.C58J.CBA | 16 | 46148113 | 46148146 | 34 | - | 17.3089 | 5.55E-07 | 0.0993 | ATGCAGCTAGAGACACAAGCTCCAGGGGGTACTA | | |
| 571 | 16 | 53496551 | 53498570 | 2019 | C3H.FVB.SWR.TH | 16 | 53496778 | 53496811 | 34 | - | 17.9106 | 3.74E-07 | 0.0805 | CCGCCCTCACATTCGCCGTCACAAGATGGCGCTG | | |
| 572 | 16 | 55143094 | 55146239 | 3145 | 129S1.C58J.CBA.LP.NZW.SJL | 16 | 55144653 | 55144686 | 34 | - | 16.5122 | 9.23E-07 | 0.126 | ATGCATCCTCAGCAGCCACCTGTAGGTGGACCTT | | |
| 573 | 16 | 55728126 | 55730825 | 2699 | 129S1,BTBR,C57L,C58J,CE,FVB,KK,LP,MRL.SJL.SWR | 16 | 55730163 | 55730196 | 34 | - | 18.8049 | 2.05E-07 | 0.062 | TTGCTGTCTTCAGATGCACCAGAAGAGGGCATCA | 2 | ENCFF093ZRU,ENCFF099QNE |
| 574 | 16 | 56142527 | 56155583 | 13056 | C58J.LP | 16 | 56142863 | 56142896 | 34 | - | 16.5691 | 8.90E-07 | 0.126 | GAGCAGGTGTGGCAGCAGTCCCAAGATGGCACCC | | |
| 575 | 16 | 75931926 | 75940912 | 8986 | AJ,C58J,CE,DBA2J,LP,NZB,NZO,NZW,SMJ | 16 | 75935481 | 75935499 | 19 | - | 17.1639 | 8.17E-07 | 0.181 | CCTCCACGAGGTGGCGGAA | | |
| 576 | 16 | 82311038 | 82413162 | 102124 | MA/My | 16 | 82318653 | 82318671 | 19 | - | 16.8525 | 9.96E-07 | 0.192 | CATCCACTAGGTGGTAGTA | 7 | ENCFF093ZRU,ENCFF099QNE,ENCFF092NMJ,ENCFF430PPJ,ENCFF089GBC,ENCFF907XEG,ENCFF024JOZ |
| 577 | 16 | 90138425 | 90146760 | 8335 | 129S1,BUB,C57L,C58J,DBA1J,DBA2J,FVB,MA/My,NOD,NOR,NZO,PJ,RF,SJL,SWR,TH | 16 | 90138716 | 90138734 | 19 | - | 19.2951 | 1.88E-07 | 0.0958 | CCGCCACTAGATGGTGCTA | | |
| 577 | 16 | 90138425 | 90146760 | 8335 | 129S1,BUB,C57L,C58J,DBA1J,DBA2J,FVB,MA/My,NOD,NOR,NZO,PJ,RF,SJL,SWR,TH | 16 | 90146719 | 90146737 | 19 | - | 19.2951 | 1.88E-07 | 0.0958 | CCGCCACTAGATGGTGCTA | 14 | ENCFF367UNR,ENCFF093ZRU,ENCFF099QNE,ENCFF024JOZ,ENCFF883UPM,ENCFF089GBC,ENCFF821BIA,ENCFF907XEG,ENCFF589ZBJ,ENCFF136ZDK,ENCFF165LXO,ENCFF198SSR,ENCFF430PPJ,ENCFF092NMJ |
| 578 | 16 | 93420107 | 93438493 | 18386 | TH | 16 | 93423823 | 93423841 | 19 | - | 17.3443 | 7.26E-07 | 0.173 | TGTCCAGGAGGTGGCAGAG | 26 | ENCFF201JUA,ENCFF093ZRU,ENCFF367UNR,ENCFF136ZDK,ENCFF430PPJ,ENCFF099QNE,ENCFF165LXO,ENCFF951GOP,ENCFF883UPM,ENCFF502WDT,ENCFF907XEG,ENCFF024JOZ,ENCFF327LYS,ENCFF821BIA,ENCFF575MGI,ENCFF609MDN,ENCFF589ZBJ,ENCFF281QMV,ENCFF957SPH,ENCFF198SSR,ENCFF194HVL,ENCFF491RJK,ENCFF055HAP,ENCFF089GBC,ENCFF210BTO,ENCFF092NMJ |
| 578 | 16 | 93420107 | 93438493 | 18386 | TH | 16 | 93423823 | 93423856 | 34 | - | 17 | 6.77E-07 | 0.109 | TTCCTATCCCACTGCTGTCCAGGAGGTGGCAGAG | 26 | ENCFF201JUA,ENCFF093ZRU,ENCFF367UNR,ENCFF136ZDK,ENCFF430PPJ,ENCFF099QNE,ENCFF165LXO,ENCFF951GOP,ENCFF883UPM,ENCFF502WDT,ENCFF907XEG,ENCFF024JOZ,ENCFF327LYS,ENCFF821BIA,ENCFF575MGI,ENCFF609MDN,ENCFF589ZBJ,ENCFF281QMV,ENCFF957SPH,ENCFF198SSR,ENCFF194HVL,ENCFF491RJK,ENCFF055HAP,ENCFF089GBC,ENCFF210BTO,ENCFF092NMJ |
| 578 | 16 | 93420107 | 93438493 | 18386 | TH | 16 | 93425950 | 93425968 | 19 | - | 17.7869 | 5.43E-07 | 0.153 | AGGCCTCCAGGTGGCAGAG | | |
| 579 | 16 | 93494339 | 93503324 | 8985 | CE.DBA1J.NZB.NZW.TH | 16 | 93497895 | 93497913 | 19 | - | 17.1639 | 8.17E-07 | 0.181 | CCTCCACGAGGTGGCGGAA | | |
| 580 | 16 | 93494344 | 93502628 | 8284 | 129S1,AJ,AKR,BALB,BUB,C3H,C57L,C58J,CBA,DBA2J,FVB,ILNJ2,KK,LP,MA/My,MRL,NOD,NOR,NZO,PJ,PL,RF,RH2,SEA,SJL,SMJ,SWR | 16 | 93497895 | 93497913 | 19 | - | 17.1639 | 8.17E-07 | 0.181 | CCTCCACGAGGTGGCGGAA | | |
| 581 | 17 | 5766826 | 5768611 | 1785 | CE | 17 | 5768142 | 5768175 | 34 | + | 16.935 | 7.06E-07 | 0.0775 | CTGTGGTGAGAGCCATGGCCTGGGGGGGGCGGGG | | |
| 582 | 17 | 7779608 | 7782187 | 2579 | AJ,BALB,BUB,C3H,CBA,DBA1J,DBA2J,FVB,KK,MA/My,MRL,NOD,NOR,RH2,SEA,SMJ | 17 | 7781046 | 7781079 | 34 | + | 20.3984 | 6.66E-08 | 0.0187 | TAGCTGTCTTCAGATGCGCCAGAAGAGGGCGCCA | 5 | ENCFF093ZRU,ENCFF367UNR,ENCFF099QNE,ENCFF821BIA,ENCFF907XEG |
| 582 | 17 | 7779608 | 7782187 | 2579 | AJ,BALB,BUB,C3H,CBA,DBA1J,DBA2J,FVB,KK,MA/My,MRL,NOD,NOR,RH2,SEA,SMJ | 17 | 7781061 | 7781079 | 19 | + | 19.6721 | 1.42E-07 | 0.05 | GCGCCAGAAGAGGGCGCCA | 5 | ENCFF093ZRU,ENCFF367UNR,ENCFF099QNE,ENCFF821BIA,ENCFF907XEG |
| 583 | 17 | 22472369 | 22475611 | 3242 | 129S1,AJ,BALB,C57L,CE,DBA1J,DBA2J,NZB,NZO,RH2,SEA,SJL,SMJ,TH | 17 | 22472795 | 22472813 | 19 | + | 19.623 | 1.47E-07 | 0.0507 | GGACCACCAGAGGGCAATA | | |
| 584 | 17 | 24129763 | 24132897 | 3134 | 129S1,AJ,AKR,B10D2,BTBR,BUB,CE,DBA1J,DBA2J,FVB,ILNJ2,KK,MA/My,NOD,NOR,NZW.PJ.RF.RH2.SEA.SJL | 17 | 24130248 | 24130281 | 34 | + | 17.0244 | 6.67E-07 | 0.0758 | GAGCAGGTGTGGCAGCAGTCCCAAGATGGCGCCC | | |
| 584 | 17 | 24129763 | 24132897 | 3134 | 129S1,AJ,AKR,B10D2,BTBR,BUB,CE,DBA1J,DBA2J,FVB,ILNJ2,KK,MA/My,NOD,NOR,NZW.PJ.RF.RH2.SEA.SJL | 17 | 24131471 | 24131504 | 34 | + | 16.561 | 8.95E-07 | 0.087 | GAGCTGCTTTCAGACACACCAGAAGAGGGCATCC | | |
| 585 | 17 | 27881997 | 27884128 | 2131 | 129S1.BALB.C57L.ILNJ2.RH2.SEA | 17 | 27882741 | 27882775 | 35 | + | 19.5596 | 1.15E-07 | 0.0458 | CTGCATGTGCATCTGCACCCCAGAAGAGGGCATAG | 1 | ENCFF093ZRU |
| 586 | 17 | 29684972 | 29686324 | 1352 | 129S1,B10D2,BALB,C57L,DBA1J,DBA2J,RH2,SEA | 17 | 29685426 | 29685459 | 34 | + | 16.9919 | 6.81E-07 | 0.0767 | TAGCTGTCTTCAGACACACCAGAAGGGGGCATCA | | |
| 587 | 17 | 36043641 | 36047726 | 4085 | 129S1,AJ,AKR,B10D2,BALB,BTBR,C3H,C57L,C58J,CBA,CE,DBA2J,LP,MA/My,MRL,NZB,NZO,NZW,PL,RF,RH2,SEA,SJL,SMJ,TH | 17 | 36047365 | 36047399 | 35 | + | 19.4771 | 1.21E-07 | 0.047 | CTACAGCTCCCCCTCCTGGCCAGCTGAGGTCCCAG | 31 | ENCFF093ZRU,ENCFF367UNR,ENCFF805QBU,ENCFF099QNE,ENCFF327LYS,ENCFF201JUA,ENCFF907XEG,ENCFF821BIA,ENCFF024JOZ,ENCFF136ZDK,ENCFF589ZBJ,ENCFF165LXO,ENCFF198SSR,ENCFF089GBC,ENCFF033UQX,ENCFF092NMJ,ENCFF210BTO,ENCFF957SPH,ENCFF951GOP,ENCFF281QMV,ENCFF089EWX,ENCFF097CAQ,ENCFF533APC,ENCFF611HDQ,ENCFF430PPJ,ENCFF194HVL,ENCFF575MGI,ENCFF349BUL.ENCFF609MDN.ENCFF316FYN.ENCFF491RJK |
| 587 | 17 | 36043641 | 36047726 | 4085 | 129S1,AJ,AKR,B10D2,BALB,BTBR,C3H,C57L,C58J,CBA,CE,DBA2J,LP,MA/My,MRL,NZB,NZO,NZW,PL,RF,RH2,SEA,SJL,SMJ,TH | 17 | 36044171 | 36044204 | 34 | - | 18.7398 | 2.14E-07 | 0.0412 | ATGCAGTGGTCCCAGAGGCCAGAAGAGGCCTTTG | | |
| 587 | 17 | 36043641 | 36047726 | 4085 | 129S1,AJ,AKR,B10D2,BALB,BTBR,C3H,C57L,C58J,CBA,CE,DBA2J,LP,MA/My,MRL,NZB,NZO,NZW,PL,RF,RH2,SEA,SJL,SMJ,TH | 17 | 36047368 | 36047386 | 19 | - | 20.2787 | 8.86E-08 | 0.0417 | TGGCCAGGAGGGGGAGCTG | 31 | ENCFF093ZRU,ENCFF367UNR,ENCFF805QBU,ENCFF099QNE,ENCFF327LYS,ENCFF201JUA,ENCFF907XEG,ENCFF821BIA,ENCFF024JOZ,ENCFF136ZDK,ENCFF589ZBJ,ENCFF165LXO,ENCFF198SSR,ENCFF089GBC,ENCFF033UQX,ENCFF092NMJ,ENCFF210BTO,ENCFF957SPH,ENCFF951GOP,ENCFF281QMV,ENCFF089EWX,ENCFF097CAQ,ENCFF533APC,ENCFF611HDQ,ENCFF430PPJ,ENCFF194HVL,ENCFF575MGI,ENCFF349BUL.ENCFF609MDN.ENCFF316FYN.ENCFF491RJK |
| 587 | 17 | 36043641 | 36047726 | 4085 | 129S1,AJ,AKR,B10D2,BALB,BTBR,C3H,C57L,C58J,CBA,CE,DBA2J,LP,MA/My,MRL,NZB,NZO,NZW,PL,RF,RH2,SEA,SJL,SMJ,TH | 17 | 36047368 | 36047402 | 35 | - | 19.3853 | 1.29E-07 | 0.0483 | ACGCTGGGACCTCAGCTGGCCAGGAGGGGGAGCTG | 31 | ENCFF093ZRU,ENCFF367UNR,ENCFF805QBU,ENCFF099QNE,ENCFF327LYS,ENCFF201JUA,ENCFF907XEG,ENCFF821BIA,ENCFF024JOZ,ENCFF136ZDK,ENCFF589ZBJ,ENCFF165LXO,ENCFF198SSR,ENCFF089GBC,ENCFF033UQX,ENCFF092NMJ,ENCFF210BTO,ENCFF957SPH,ENCFF951GOP,ENCFF281QMV,ENCFF089EWX,ENCFF097CAQ,ENCFF533APC,ENCFF611HDQ,ENCFF430PPJ,ENCFF194HVL,ENCFF575MGI,ENCFF349BUL.ENCFF609MDN.ENCFF316FYN.ENCFF491RJK |
| 588 | 17 | 36483020 | 36484765 | 1745 | ILNJ2 | 17 | 36483009 | 36483043 | 35 | + | 17.4679 | 4.13E-07 | 0.0933 | TTGTAGCTGTCCAGACACACCAGAAGAGGGCATCC | | |
| 589 | 17 | 37201352 | 37205593 | 4241 | PJ | 17 | 37203392 | 37203426 | 35 | + | 20.2202 | 7.54E-08 | 0.0346 | TAGCATTGTGCCTTCACGGACAGTAGGTGGTGCTG | 29 | ENCFF093ZRU,ENCFF201JUA,ENCFF367UNR,ENCFF099QNE,ENCFF327LYS,ENCFF821BIA,ENCFF907XEG,ENCFF065ODV,ENCFF089GBC,ENCFF589ZBJ,ENCFF055HAP,ENCFF316FYN,ENCFF430PPJ,ENCFF321AGQ,ENCFF502WDT,ENCFF092NMJ,ENCFF198SSR,ENCFF024JOZ,ENCFF165LXO,ENCFF951GOP,ENCFF136ZDK,ENCFF957SPH,ENCFF883UPM,ENCFF281QMV,ENCFF142CNG,ENCFF575MGI,ENCFF609MDN,ENCFF194HVL.ENCFF052KVO |
| 590 | 17 | 38614972 | 38688148 | 73176 | FVB.TH | 17 | 38625166 | 38625184 | 19 | + | 18.7049 | 2.88E-07 | 0.0728 | CAGCCAGCAGGTGGCAAAA | | |
| 591 | 17 | 45773031 | 45774937 | 1906 | NZW.SJL | 17 | 45773600 | 45773633 | 34 | + | 26.0569 | 6.34E-10 | 0.000552 | ATGCAGTGCCCATGGAGGCCAGAAGAGGGCATAA | 2 | ENCFF093ZRU.ENCFF502WDT |
| 592 | 17 | 50555814 | 50560277 | 4463 | NZW.SWR | 17 | 50556593 | 50556626 | 34 | + | 17.6748 | 4.37E-07 | 0.0599 | GAGCATGACCACAAGTCCACAGCAGGTGGCAACG | | |
| 593 | 17 | 53912047 | 53917681 | 5634 | 129S1,BALB,C3H,C57L,CBA,DBA1J,DBA2J,FVB,LP,MA/Mv,MRL,PJ,PL,SMJ | 17 | 53912503 | 53912536 | 34 | + | 16.4634 | 9.51E-07 | 0.0875 | TAGCTGTCTTCAGACGCACCAGAAGAGGGCATCA | | |
| 593 | 17 | 53912047 | 53917681 | 5634 | 129S1,BALB,C3H,C57L,CBA,DBA1J,DBA2J,FVB,LP,MA/Mv,MRL,PJ,PL,SMJ | 17 | 53915352 | 53915385 | 34 | + | 16.4634 | 9.51E-07 | 0.0875 | TAGCTGTCTTCAGACGCACCAGAAGAGGGCATCA | | |
| 594 | 17 | 74858727 | 74860295 | 1568 | 129S1,BTBR,BUB,C57L,ILNJ2,KK,NOD,NOR.NZW.SWR | 17 | 74858784 | 74858817 | 34 | + | 16.4634 | 9.51E-07 | 0.0875 | TAGCTGTCTTCAGACGCACCAGAAGAGGGCATCA | | |
| 595 | 17 | 76073723 | 76077535 | 3812 | CE | 17 | 76075835 | 76075869 | 35 | + | 18.9908 | 1.65E-07 | 0.0555 | TTGCAGTTCCTCCTGCCTACCACTAGGGTCTGACA | 9 | ENCFF821BIA,ENCFF589ZBJ,ENCFF089GBC,ENCFF165LXO,ENCFF093ZRU,ENCFF367UNR,ENCFF052KVO,ENCFF099QNE.ENCFF136ZDK |
| 596 | 17 | 78798950 | 78800983 | 2033 | C3H.CBA.DBA1J.DBA2J | 17 | 78799505 | 78799523 | 19 | + | 17.1639 | 8.17E-07 | 0.123 | GTACCACCAGAGGGTGCTC | | |
| 597 | 17 | 79524290 | 79526198 | 1908 | 129S1,AJ,AKR,BALB,BTBR,BUB,C3H,C57L,C58J,CBA,CE,DBA1J,DBA2J,FVB,ILNJ2,KK,LP,MA/My,MRL,NOD,NOR,NZB,NZO,NZW,PJ,PL,RF,RH2,SEA,SJL,SMJ,SWR | 17 | 79525261 | 79525295 | 35 | + | 17.0826 | 5.16E-07 | 0.104 | GTGCAGATCCCCTTGGATATCAGAAGAGGGTGTTG | | |
| 598 | 17 | 80863880 | 80865744 | 1864 | CE | 17 | 80865058 | 80865092 | 35 | + | 17.3486 | 4.42E-07 | 0.0965 | ATGTGAGTGCCCGCAGTGTCCAGAAGAGGGCACCA | 11 | ENCFF367UNR,ENCFF099QNE,ENCFF065ODV,ENCFF327LYS,ENCFF136ZDK,ENCFF194HVL,ENCFF093ZRU.ENCFF142CNG.ENCFF052KVO.ENCFF502WDT.ENCFF055HAP |
| 598 | 17 | 80863880 | 80865744 | 1864 | CE | 17 | 80865059 | 80865092 | 34 | + | 18.0894 | 3.33E-07 | 0.0512 | TGTGAGTGCCCGCAGTGTCCAGAAGAGGGCACCA | 11 | ENCFF367UNR,ENCFF099QNE,ENCFF065ODV,ENCFF327LYS,ENCFF136ZDK,ENCFF194HVL,ENCFF093ZRU.ENCFF142CNG.ENCFF052KVO.ENCFF502WDT.ENCFF055HAP |
| 598 | 17 | 80863880 | 80865744 | 1864 | CE | 17 | 80865074 | 80865092 | 19 | + | 20.7213 | 6.18E-08 | 0.0363 | TGTCCAGAAGAGGGCACCA | 11 | ENCFF367UNR,ENCFF099QNE,ENCFF065ODV,ENCFF327LYS,ENCFF136ZDK,ENCFF194HVL,ENCFF093ZRU.ENCFF142CNG.ENCFF052KVO.ENCFF502WDT.ENCFF055HAP |
| 599 | 17 | 92376319 | 92377383 | 1064 | 129S1,AJ,AKR,BALB,BTBR,BUB,C3H,C57L,CBA,CE,DBA1J,DBA2J,FVB,ILNJ2,KK,LP,MA/My,MRL,NOD,NOR,NZB,NZO,NZW,PL,RF,RH2,SEA,SJL,SMJ,SWR,TH | 17 | 92377083 | 92377101 | 19 | + | 17.5082 | 6.52E-07 | 0.109 | TAGCCACCAGGGGTTGGTG | | |
| 600 | 17 | 92377355 | 92378425 | 1070 | AJ,BALB,CBA,FVB,NOD,NOR,PJ,SJL | 17 | 92378121 | 92378139 | 19 | + | 17.5082 | 6.52E-07 | 0.109 | TAGCCACCAGGGGTTGGTG | | |
| 601 | 17 | 15037907 | 15041019 | 3112 | PI | 17 | 15038681 | 15038715 | 35 | - | 16.9174 | 5.68E-07 | 0.109 | ATGCATTTATCTCCTGTAGCCACCAGTGGCCACAG | | |
| 602 | 17 | 15823104 | 15830259 | 7155 | PI | 17 | 15823420 | 15823453 | 34 | - | 17.9106 | 3.74E-07 | 0.0548 | CCGCCCTCACATTCGCCGTCACAAGATGGCGCTG | | |
| 602 | 17 | 15823104 | 15830259 | 7155 | PI | 17 | 15830217 | 15830250 | 34 | - | 17.9106 | 3.74E-07 | 0.0548 | CCGCCCTCACATTCGCCGTCACAAGATGGCGCTG | | |
| 603 | 17 | 22433896 | 22441403 | 7507 | 129S1,AJ,BALB,C57L,CE,DBA1J,DBA2J,NZB,NZO,RH2,SEA,SJL,SMJ,TH | 17 | 22436073 | 22436091 | 19 | - | 16.9672 | 9.26E-07 | 0.13 | CTGCCAGAAGAGGGCATCA | | |
| 604 | 17 | 26510423 | 26517276 | 6853 | AJ,B10D2,BTBR,CE,DBA1J,DBA2J,FVB,ILNJ2,MA/My,NOD,NOR,NZO,NZW,PJ,RF,SJL | 17 | 26510637 | 26510670 | 34 | - | 16.5691 | 8.90E-07 | 0.0867 | GAGCAGGTGTGGCAGCAGTCCCAAGATGGCACCC | | |
| 605 | 17 | 29570509 | 29573938 | 3429 | KK.RF | 17 | 29572271 | 29572304 | 34 | - | 16.9593 | 6.95E-07 | 0.0768 | TTGCTCTCTTCAGACACACCAGAAGAGGGCATCA | | |
| 606 | 17 | 32545606 | 32549712 | 4106 | MA/Mv.SJL | 17 | 32548692 | 32548725 | 34 | - | 17.0488 | 6.56E-07 | 0.0752 | TTGCTATCTTCAGATACACCAGAAGAGGGCATCA | 4 | ENCFF367UNR.ENCFF821BIA.ENCFF093ZRU.ENCFF099QNE |
| 606 | 17 | 32545606 | 32549712 | 4106 | MA/Mv.SJL | 17 | 32549507 | 32549541 | 35 | - | 16.4771 | 7.30E-07 | 0.126 | ATGCAGGTGTTCCTGGAAGCCAAAAGAGGGCTACA | | |
| 607 | 17 | 35497854 | 35499443 | 1589 | SJL.TH | 17 | 35498595 | 35498628 | 34 | - | 20.4146 | 6.58E-08 | 0.0185 | TTGCAGTCTTCAGACACACCAGAAGAGGGCATCA | 2 | ENCFF367UNR.ENCFF093ZRU |
| 608 | 17 | 35526624 | 35530945 | 4321 | PJ | 17 | 35529549 | 35529583 | 35 | - | 18.0642 | 2.90E-07 | 0.0766 | CTGCTTTTCCCCAGGCAGGGCCCTAGGTGTCACAG | | |
| 609 | 17 | 36518879 | 36521909 | 3030 | 129S1.BTBR.C57L.ILNJ2.LP | 17 | 36521891 | 36521925 | 35 | - | 17.5688 | 3.89E-07 | 0.0913 | CTGCAGGGGATGGGACCTTCCAGAAGTGGGCAGCT | | |
| 610 | 17 | 43515298 | 43517402 | 2104 | 129S1.BTBR.C57L.LP | 17 | 43515647 | 43515681 | 35 | - | 17.0367 | 5.30E-07 | 0.105 | TTGCTGTTGTTTTTTCTGGACAGTGGATGGAACTG | | |
| 611 | 17 | 45434942 | 45436779 | 1837 | NZW.SJL | 17 | 45435063 | 45435081 | 19 | - | 20.0656 | 1.05E-07 | 0.043 | AGGCCAGAAGAGGGCACTG | | |
| 611 | 17 | 45434942 | 45436779 | 1837 | NZW.SJL | 17 | 45436718 | 45436752 | 35 | - | 19.8807 | 9.39E-08 | 0.0405 | CTGAATGTATGCCTGCAGGCCAGAAGAGGGCATCA | | |
| 612 | 17 | 47552077 | 47558544 | 6467 | DBA2J | 17 | 47557648 | 47557681 | 34 | - | 16.9512 | 6.99E-07 | 0.077 | CTGCTGATGACAGAATGGCGAGAAGAGGGCAGTG | | |
| 613 | 17 | 49748216 | 49755734 | 7518 | 129S1,AJ,AKR,BALB,BTBR,BUB,C3H,C57L,CBA,CE,DBA1J,DBA2J,FVB,ILNJ2,KK,LP,MA/My,MRL,NOD,NOR,NZB,NZO,NZW,PJ,PL,RF,RH2,SEA,SJL,SMJ,SWR,TH | 17 | 49748197 | 49748230 | 34 | - | 22.0163 | 1.97E-08 | 0.00736 | CTGTGGCACCCATGAAGGCCAGAAGAGGGCACTG | 2 | ENCFF093ZRU,ENCFF089GBC |
| 614 | 17 | 77687073 | 77708739 | 21666 | C58J | 17 | 77705360 | 77705393 | 34 | - | 16.5935 | 8.77E-07 | 0.0863 | GCCCAGTGCCCAAGGAGGCCAGAAGGGGACGTCA | | |
| 615 | 17 | 82023635 | 82030270 | 6635 | CE.MA/Mv.NZW | 17 | 82023759 | 82023777 | 19 | - | 17.8852 | 5.08E-07 | 0.0979 | ACACCAGAAGAGGGCACCA | | |
| 616 | 17 | 88924865 | 88928139 | 3274 | 129S1,AJ,AKR,BALB,C3H,CBA,CE,MRL,NOD,NOR,PJ,RF,RH2,SEA | 17 | 88927654 | 88927672 | 19 | - | 17.5574 | 6.32E-07 | 0.108 | TGGCCAGTTGGGGGCGCGC | | |
| 616 | 17 | 88924865 | 88928139 | 3274 | 129S1,AJ,AKR,BALB,C3H,CBA,CE,MRL,NOD,NOR,PJ,RF,RH2,SEA | 17 | 88927654 | 88927687 | 34 | - | 19.561 | 1.21E-07 | 0.029 | CAGCATCGTTCTTCCTGGCCAGTTGGGGGCGCGC | | |
| 617 | 18 | 25452308 | 25454707 | 2399 | CE | 18 | 25454253 | 25454286 | 34 | + | 24.1138 | 3.58E-09 | 0.00429 | CTGCAGTTCCATATCCTGCCAATAGGAGGGAGCA | 15 | ENCFF024JOZ,ENCFF093ZRU,ENCFF821BIA,ENCFF198SSR,ENCFF367UNR,ENCFF136ZDK,ENCFF099QNE,ENCFF052KVO,ENCFF055HAP,ENCFF883UPM,ENCFF065ODV,ENCFF589ZBJ,ENCFF502WDT,ENCFF327LYS.ENCFF907XEG |
| 618 | 18 | 30144539 | 30149093 | 4554 | BALB,C3H,C58J,CE,ILNJ2,KK,MRL,NZB,NZO,PL,RF,RH2,SEA | 18 | 30145316 | 30145349 | 34 | + | 16.7398 | 7.99E-07 | 0.121 | TAGCATGACCACAAGTCCACAGCAGGTGGCAACG | | |
| 619 | 18 | 30399560 | 30405253 | 5693 | 129S1,AKR,BALB,BUB,C3H,C57L,C58J,CE,FVB,ILNJ2,KK,MA/My,MRL,NZB,NZO,NZW,PL,RF,RH2,SEA,SJL,SMJ,SWR,TH | 18 | 30403404 | 30403422 | 19 | + | 17.1639 | 8.17E-07 | 0.188 | CCTCCACGAGGTGGCGGAA | | |
| 620 | 18 | 48333828 | 48353963 | 20135 | KK | 18 | 48339455 | 48339488 | 34 | + | 17.3089 | 5.55E-07 | 0.106 | ATGCAGCTAGAGACACGAGCTCCAGAGGGTACTG | | |
| 621 | 18 | 51195398 | 51197141 | 1743 | KK | 18 | 51195965 | 51195998 | 34 | + | 17.9756 | 3.59E-07 | 0.0887 | CTGCTTTCTTCAGACACACCAAAAGAGGGCACCA | | |
| 622 | 18 | 61446801 | 61448445 | 1644 | CE | 18 | 61447202 | 61447220 | 19 | + | 17.2131 | 7.91E-07 | 0.185 | TGACCAGCAGGGGATGCTC | 40 | ENCFF093ZRU,ENCFF099QNE,ENCFF321AGQ,ENCFF821BIA,ENCFF327LYS,ENCFF065ODV,ENCFF092NMJ,ENCFF907XEG,ENCFF024JOZ,ENCFF504JDS,ENCFF136ZDK,ENCFF089GBC,ENCFF165LXO,ENCFF430PPJ,ENCFF589ZBJ,ENCFF198SSR,ENCFF055HAP,ENCFF052KVO,ENCFF601JVE,ENCFF502WDT,ENCFF951GOP,ENCFF575MGI,ENCFF609MDN,ENCFF957SPH,ENCFF883UPM,ENCFF210BTO,ENCFF142CNG,ENCFF281QMV,ENCFF316FYN,ENCFF533APC,ENCFF805QBU,ENCFF194HVL,ENCFF491RJK,ENCFF089EWX,ENCFF086ZTV,ENCFF611HDQ,ENCFF349BUL,ENCFF784ASD,ENCFF097CAQ,ENCFF033UQX |
| 623 | 18 | 65695235 | 65696608 | 1373 | NZB.NZO.NZW | 18 | 65695868 | 65695901 | 34 | + | 16.8537 | 7.43E-07 | 0.117 | ATGCAGGACCCACTGCAGTCAGAAAAGGGCATCA | | |
| 624 | 18 | 75968843 | 75971972 | 3129 | PJ | 18 | 75970863 | 75970897 | 35 | + | 16.7523 | 6.24E-07 | 0.183 | GTGCTTCCTCTTTCTCTTGCCTGCAGGGGGAGCCA | 29 | ENCFF201JUA,ENCFF367UNR,ENCFF093ZRU,ENCFF821BIA,ENCFF099QNE,ENCFF430PPJ,ENCFF327LYS,ENCFF907XEG,ENCFF024JOZ,ENCFF198SSR,ENCFF316FYN,ENCFF589ZBJ,ENCFF883UPM,ENCFF089GBC,ENCFF575MGI,ENCFF194HVL,ENCFF097CAQ,ENCFF052KVO,ENCFF957SPH,ENCFF504JDS,ENCFF165LXO,ENCFF092NMJ,ENCFF136ZDK,ENCFF210BTO,ENCFF951GOP,ENCFF281QMV,ENCFF533APC,ENCFF491RJK.ENCFF609MDN |
| 624 | 18 | 75968843 | 75971972 | 3129 | PJ | 18 | 75970879 | 75970897 | 19 | + | 19.2951 | 1.88E-07 | 0.0952 | TTGCCTGCAGGGGGAGCCA | 29 | ENCFF201JUA,ENCFF367UNR,ENCFF093ZRU,ENCFF821BIA,ENCFF099QNE,ENCFF430PPJ,ENCFF327LYS,ENCFF907XEG,ENCFF024JOZ,ENCFF198SSR,ENCFF316FYN,ENCFF589ZBJ,ENCFF883UPM,ENCFF089GBC,ENCFF575MGI,ENCFF194HVL,ENCFF097CAQ,ENCFF052KVO,ENCFF957SPH,ENCFF504JDS,ENCFF165LXO,ENCFF092NMJ,ENCFF136ZDK,ENCFF210BTO,ENCFF951GOP,ENCFF281QMV,ENCFF533APC,ENCFF491RJK.ENCFF609MDN |
| 625 | 18 | 78947770 | 78955313 | 7543 | KK | 18 | 78953414 | 78953447 | 34 | + | 16.9024 | 7.21E-07 | 0.117 | TTGCTGTCTTCAGACACACCAGAAGAGGGCATCT | 1 | ENCFF093ZRU |
| 626 | 18 | 82012680 | 82017213 | 4533 | BTBR | 18 | 82016447 | 82016465 | 19 | + | 21.623 | 2.86E-08 | 0.0404 | CAGCCAGCAGATGGCAGTA | 21 | ENCFF093ZRU,ENCFF099QNE,ENCFF201JUA,ENCFF367UNR,ENCFF589ZBJ,ENCFF316FYN,ENCFF089GBC,ENCFF821BIA,ENCFF907XEG,ENCFF055HAP,ENCFF327LYS,ENCFF430PPJ,ENCFF092NMJ,ENCFF024JOZ,ENCFF281QMV,ENCFF165LXO,ENCFF502WDT,ENCFF575MGI,ENCFF142CNG,ENCFF883UPM,ENCFF198SSR |
| 627 | 18 | 82711670 | 82720554 | 8884 | 129S1,BTBR,BUB,C57L,MA/Mv,NZB,SJL | 18 | 82716987 | 82717005 | 19 | + | 17.1639 | 8.17E-07 | 0.188 | CCTCCACGAGGTGGCGGAA | | |
| 628 | 18 | 16983427 | 16987983 | 4556 | 129S1,BTBR,C57L,C58J,CE,FVB,KK,MRL,PJ,RF,SEA,SJL,SMJ,SWR | 18 | 16987181 | 16987214 | 34 | - | 16.9106 | 7.17E-07 | 0.116 | GAGCATGACCACAAATCCACAGCAGGTGGCAACG | | |
| 629 | 19 | 7625617 | 7630337 | 4720 | C58J.LP | 19 | 7630253 | 7630287 | 35 | + | 16.4679 | 7.34E-07 | 0.137 | TTGCTATGCCACCTTTACCCCAGCTGGAGGCAAGA | | |
| 630 | 19 | 8496891 | 8501647 | 4756 | BUB.C58J.FVB.LP.MA/Mv.TH | 19 | 8501370 | 8501403 | 34 | + | 17.6829 | 4.35E-07 | 0.0635 | CAGCATTATGCCTGCAACCCAGAAGATGGCACCA | 1 | ENCFF093ZRU |
| 631 | 19 | 12339345 | 12344228 | 4883 | AJ,BALB,CE,MA/My,NOD,NOR,PL,RH2,SEA | 19 | 12339361 | 12339394 | 34 | + | 19.3415 | 1.42E-07 | 0.0379 | CTGCCCCCACATTCGCCGTCACAAGATGGCGCTG | | |
| 632 | 19 | 20060209 | 20065174 | 4965 | MRL.NZB.NZO.NZW.PJ | 19 | 20064281 | 20064314 | 34 | + | 17.4715 | 5.00E-07 | 0.0688 | GTGCAGCTTGAGACACGAGCTCCGGGGGTACTG | | |
| 633 | 19 | 38363465 | 38370531 | 7066 | 129S1,AJ,AKR,BALB,BUB,C3H,C57L,CBA,CE,DBA1J,DBA2J,FVB,ILNJ2,KK,MRL,NOD,NOR,NZB,NZO,NZW,PJ,PL,RF,RH2,SEA,SJL,SMJ,SWR,TH | 19 | 38368882 | 38368900 | 19 | + | 17.1639 | 8.17E-07 | 0.122 | CCTCCACGAGGTGGCGGAA | | |
| 634 | 19 | 9129625 | 9139091 | 9466 | C58J,KK,LP | 19 | 9129735 | 9129753 | 19 | - | 18.3115 | 3.79E-07 | 0.0839 | CAGCCAGGAGGGGGAGCTG | | |
| 635 | 19 | 9775232 | 9831269 | 56037 | KK | 19 | 9806496 | 9806514 | 19 | - | 19.2951 | 1.88E-07 | 0.0596 | CCGCCACTAGATGGTGCTA | | |
| 635 | 19 | 9775232 | 9831269 | 56037 | KK | 19 | 9814487 | 9814505 | 19 | - | 19.2951 | 1.88E-07 | 0.0596 | CCGCCACTAGATGGTGCTA | 19 | ENCFF201JUA,ENCFF367UNR,ENCFF093ZRU,ENCFF099QNE,ENCFF327LYS,ENCFF024JOZ,ENCFF136ZDK,ENCFF907XEG,ENCFF089GBC,ENCFF589ZBJ,ENCFF821BIA,ENCFF502WDT,ENCFF165LXO,ENCFF609MDN.ENCFF198SSR.ENCFF430PPJ.ENCFF092NMJ.ENCFF065ODV.ENCFF055HAP |
| 636 | 19 | 9806205 | 9814528 | 8323 | BUB,ILNJ2,LP | 19 | 9806496 | 9806514 | 19 | - | 19.2951 | 1.88E-07 | 0.0596 | CCGCCACTAGATGGTGCTA | | |
| 636 | 19 | 9806205 | 9814528 | 8323 | BUB,ILNJ2,LP | 19 | 9814487 | 9814505 | 19 | - | 19.2951 | 1.88E-07 | 0.0596 | CCGCCACTAGATGGTGCTA | 19 | ENCFF201JUA,ENCFF367UNR,ENCFF093ZRU,ENCFF099QNE,ENCFF327LYS,ENCFF024JOZ,ENCFF136ZDK,ENCFF907XEG,ENCFF089GBC,ENCFF589ZBJ,ENCFF821BIA,ENCFF502WDT,ENCFF165LXO,ENCFF609MDN.ENCFF198SSR.ENCFF430PPJ.ENCFF092NMJ.ENCFF065ODV.ENCFF055HAP |

| | | | | | | | | | | | | | | | |
|---|---|---|---|---|---|---|---|---|---|---|---|---|---|---|---|
| 637 | 19 | 18897511 | 18901666 | 4155 | 129S1,AKR,BALB,BTBR,C57L,C58J,FVB,LP,MA/My,NZB,NZO,NZW,PJ,RF,SEA,SMJ,SWR.TH | 19 | 18900863 | 18900896 | 34 | - | 17.6748 | 4.37E-07 | 0.0638 | GAGCATGACCACAAGTCCACAGCAGGTGGCAACG | | |
| 638 | 19 | 23760109 | 23762086 | 1977 | ILNJ2.MA/Mv.NOD.NOR | 19 | 23762035 | 23762068 | 34 | - | 17.4228 | 5.16E-07 | 0.0699 | ATACAGTCCCTGCAGAGGCCAGAAAAGGGCGTCA | | |
| 639 | 19 | 25889730 | 25942646 | 52916 | FVB | 19 | 25932182 | 25932216 | 35 | - | 16.5688 | 6.93E-07 | 0.133 | CTGCAGTGATGAGGCTGGGTCAGAGGGTGGGTCTG | | |
| 640 | 19 | 35316835 | 35324489 | 7654 | C58J.LP | 19 | 35321316 | 35321334 | 19 | - | 17.459 | 6.74E-07 | 0.109 | CAGCCAGGAGGTGGAGCTG | | |
| 640 | 19 | 35316835 | 35324489 | 7654 | C58J.LP | 19 | 35321316 | 35321349 | 34 | - | 19.0407 | 1.74E-07 | 0.0424 | TCGCCCTTTCATCCTCAGCCAGGAGGTGGAGCTG | | |
| 641 | 19 | 36750143 | 36753392 | 3249 | NZW | 19 | 36750817 | 36750850 | 34 | - | 16.5122 | 9.23E-07 | 0.0915 | CTGCCTTGTCTGCACCTAGCAGCAGAGGAAGCAG | | |
| 642 | 19 | 42411534 | 42412960 | 1426 | NZW | 19 | 42411819 | 42411837 | 19 | - | 17.3607 | 7.18E-07 | 0.111 | GGACCAGCAGAGGGAGCAG | | |
| 643 | 19 | 51728011 | 51731054 | 3043 | TH | 19 | 51730125 | 51730158 | 34 | - | 18.3984 | 2.70E-07 | 0.0518 | TTGCTGTCTTCAGACACACCAGAAGAGGGCATTG | | |
| 644 | X | 9505391 | 9507522 | 2131 | DBA1J | X | 9506503 | 9506536 | 34 | + | 17.0569 | 6.53E-07 | 0.163 | CAGCTTTGTCTGTAGGTCCCTGTAGATGGTGCTG | 5 | ENCFF367UNR.ENCFF093ZRU.ENCFF055HAP.ENCFF502WDT.ENCFF099QNE |
| 645 | X | 15340282 | 15349267 | 8985 | AKR.SMJ | X | 15345699 | 15345717 | 19 | + | 17.1639 | 8.17E-07 | 0.276 | CCTCCACGAGGTGGCGGAA | | |
| 646 | X | 45749358 | 45755295 | 5937 | C58J | X | 45753611 | 45753644 | 34 | + | 20.2846 | 7.23E-08 | 0.0524 | ATGCATTGGTTTCAGAAGCCAGAAGAGGGCATTG | | |
| 647 | X | 53911929 | 53913128 | 1199 | 129S1,AJ,AKR,B10D2,B10J,BALB,BTBR,BUB,C3H,C57L,C58J,CBA,CE,DBA1J,DBA2J,FVB,ILNJ2,KK,LP,MA/My,MRL,NOD,NOR,NZB,NZO,NZW,PJ,PL,RF,RH2,SEA,SJL,SMJ,SWR.TH | X | 53912721 | 53912739 | 19 | + | 17.7541 | 5.54E-07 | 0.229 | GCTCCAGCAGGTGGCGGAA | | |
| 648 | X | 89388721 | 89434202 | 45481 | BUB.DBA1J.DBA2J.PJ | X | 89419892 | 89419925 | 34 | + | 19.1382 | 1.63E-07 | 0.0885 | GAGTAGTGACAGTGGTTGCCACTGGGGGGTGGAG | | |
| 649 | X | 100543794 | 100550473 | 6679 | NZB.NZO | X | 100547253 | 100547271 | 19 | + | 18.6393 | 3.02E-07 | 0.176 | ATGCCAGAAGAGGGCACTG | | |
| 649 | X | 100543794 | 100550473 | 6679 | NZB,NZO | X | 100548407 | 100548441 | 35 | + | 27.5505 | 2.90E-10 | 0.00367 | CTGCATGTACATCCGCAGGCCAGAAGAGGGCGCCA | 23 | ENCFF201JUA,ENCFF367UNR,ENCFF093ZRU,ENCFF099QNE,ENCFF092NMJ,ENCFF821BIA,ENCFF883UPM,ENCFF165LXO,ENCFF055HAP,ENCFF907XEG,ENCFF281QMV,ENCFF194HVL,ENCFF024JOZ,ENCFF589ZBJ,ENCFF136ZDK,ENCFF052KVO,ENCFF198SSR,ENCFF327LYS,ENCFF065ODV,ENCFF957SPH,ENCFF502WDT,ENCFF089GBC,ENCFF142CNG |
| 649 | X | 100543794 | 100550473 | 6679 | NZB,NZO | X | 100548423 | 100548441 | 19 | + | 21.0984 | 4.51E-08 | 0.0714 | AGGCCAGAAGAGGGCGCCA | 23 | ENCFF201JUA,ENCFF367UNR,ENCFF093ZRU,ENCFF099QNE,ENCFF092NMJ,ENCFF821BIA,ENCFF883UPM,ENCFF165LXO,ENCFF055HAP,ENCFF907XEG,ENCFF281QMV,ENCFF194HVL,ENCFF024JOZ,ENCFF589ZBJ,ENCFF136ZDK,ENCFF052KVO,ENCFF198SSR,ENCFF327LYS,ENCFF065ODV,ENCFF957SPH,ENCFF502WDT,ENCFF089GBC,ENCFF142CNG |
| 649 | X | 100543794 | 100550473 | 6679 | NZB.NZO | X | 100545105 | 100545138 | 34 | - | 17.065 | 6.50E-07 | 0.163 | TTGCTCTCTTCAGACACACCAGAAGAGGGCATCG | | |
| 649 | X | 100543794 | 100550473 | 6679 | NZB.NZO | X | 100548256 | 100548289 | 34 | - | 19.813 | 1.01E-07 | 0.064 | TAGCAGTCCTCAGACACACCAGAAGAGGGCATCA | 4 | ENCFF201JUA.ENCFF367UNR.ENCFF093ZRU.ENCFF099QNE |
| 649 | X | 100543794 | 100550473 | 6679 | NZB.NZO | X | 100549498 | 100549532 | 35 | - | 16.2936 | 8.10E-07 | 0.272 | CTGCATCCATTTCCAACAGCCACTGGATGGATCCT | | |
| 650 | X | 120085327 | 120087588 | 2261 | AJ.BALB.C3H.CBA.MRL.RH2 | X | 120086901 | 120086934 | 34 | + | 16.4634 | 9.51E-07 | 0.182 | TTGCAGTTCCACACCCCCTTAGCAGGTGGTGATG | | |
| 650 | X | 120085327 | 120087588 | 2261 | AJ.BALB.C3H.CBA.MRL.RH2 | X | 120085983 | 120086016 | 34 | - | 18.1789 | 3.13E-07 | 0.117 | CTGCAGTGACTAGAATTTCCCCCTGGGGACACTT | | |
| 651 | X | 140682743 | 140685747 | 3004 | CE | X | 140683658 | 140683692 | 35 | + | 17.6606 | 3.68E-07 | 0.181 | ATGCAATCAGACGTGCCTTCCACAAGATGCCAACA | | |
| 652 | X | 145282896 | 145284496 | 1600 | NZB | X | 145283283 | 145283316 | 34 | + | 17.626 | 4.52E-07 | 0.136 | ATGCAGCTAGAGACACAAACTCCAGGGGGTACTG | | |
| 653 | X | 149212433 | 149214652 | 2219 | NZB | X | 149212579 | 149212612 | 34 | + | 21.8537 | 2.23E-08 | 0.0245 | TTGTAGTTCCAATAGAGGCCAGAAGAGAGCACTG | | |
| 654 | X | 25868498 | 25886343 | 17845 | C58J | X | 25873720 | 25873738 | 19 | - | 22.3115 | 1.52E-08 | 0.0382 | CTACCACCAGAGGGCAGCA | | |
| 654 | X | 25868498 | 25886343 | 17845 | C58J | X | 25873720 | 25873754 | 35 | - | 19.5138 | 1.19E-07 | 0.0914 | ATGTCTTCTAACCCGCCTACCACCAGAGGGCAGCA | | |
| 655 | X | 57817028 | 57819402 | 2374 | C58J.CE.ILNJ2.LP.PJ | X | 57819318 | 57819336 | 19 | - | 17.5574 | 6.32E-07 | 0.241 | TGGCCAGTTGGGGGCGCGC | | |
| 656 | X | 77467075 | 77468905 | 1830 | CE.ILNJ2 | X | 77468675 | 77468693 | 19 | - | 17.3934 | 7.03E-07 | 0.255 | CTACCAGCAGATGTCACCG | | |
| 657 | X | 77700667 | 77712405 | 11738 | NZB | X | 77706194 | 77706227 | 34 | - | 19.252 | 1.51E-07 | 0.0835 | CTGCAGTTCTAGCACTCAGGAGATGGGGGCAGCA | 1 | ENCFF093ZRU |
| 658 | X | 81929953 | 81932678 | 2725 | NZB | X | 81931627 | 81931660 | 34 | - | 16.7317 | 8.03E-07 | 0.174 | CTGAAGTGCCTTATTGGGCCTCAGGGGGAAAGTA | | |
| 659 | X | 95905680 | 95915455 | 9775 | FVB.MRL.NZW.SWR | X | 95906342 | 95906375 | 34 | - | 16.439 | 9.66E-07 | 0.182 | CTCAAGTGCTCGAGTGGACCAGAAGGGGGTACCT | | |
| 660 | X | 112874751 | 112882934 | 8183 | AJ.BALB.C3H.CBA.MRL.RH2 | X | 112876040 | 112876058 | 19 | - | 22.1967 | 1.68E-08 | 0.0409 | TGGCCAGTAGGGGGAGCTC | 1 | ENCFF367UNR |
| 660 | X | 112874751 | 112882934 | 8183 | AJ.BALB.C3H.CBA.MRL.RH2 | X | 112876040 | 112876074 | 35 | - | 21.6147 | 2.97E-08 | 0.0407 | AAGCTGTAGGTCATTCTGGCCAGTAGGGGGAGCTC | 1 | ENCFF367UNR |
| 661 | X | 113011737 | 113018247 | 6510 | AJ.BALB.C3H.CBA.MRL.RH2 | X | 113017999 | 113018032 | 34 | - | 18.0569 | 3.40E-07 | 0.119 | ATGCAGCTAGAGACATGAGCTCCAGGGGGTACTG | | |
| 662 | X | 116226546 | 116231732 | 5186 | AJ.BALB.C3H.CBA.MRL.RH2 | X | 116228572 | 116228606 | 35 | - | 16.5596 | 6.97E-07 | 0.257 | AAGAAGTTACACCCTGTTGCCAGTAGGTGGTTTTC | | |
| 663 | X | 117773297 | 117836311 | 63014 | AJ.BALB.C3H.CBA.MRL.RH2 | X | 117798779 | 117798812 | 34 | - | 17.9837 | 3.57E-07 | 0.123 | GAGCGGGTGCTGCCGCTGCCCCAAGATGGCGCCT | | |
| 664 | X | 145950498 | 145951747 | 1249 | NZB | X | 145950955 | 145950973 | 19 | - | 17.3443 | 7.26E-07 | 0.258 | TTGCCAGAAGAGGGCATCA | 2 | ENCFF055HAP.ENCFF502WDT |
| 664 | X | 145950498 | 145951747 | 1249 | NZB | X | 145950955 | 145950989 | 35 | - | 24.7248 | 3.05E-09 | 0.0103 | CTGCATGTACACCTAATTGCCAGAAGAGGGCATCA | 2 | ENCFF055HAP.ENCFF502WDT |

| DEL_NUM | DEL_CHR | DEL_START | DEL_END | DEL_LENGTH | STRAINS with VARIANT SV ALLELE | CTCF CHR | CTCF START | CTCF END | CTCF LENGTH | CTCF STRAND | CTCF SCORE | CTCF PVALUE | CTCF QVALUE | CTCF SEQUENCE | # PEAKS | DATA SOURCE |
|---|---|---|---|---|---|---|---|---|---|---|---|---|---|---|---|---|
| 1 | 1 | 4650900 | 4651097 | 197 | C3H,CAST,CBA,DBA1J,DBA2J,MRL,RF,SPRET.WSB | 1 | 4650956 | 4650989 | 34 | + | 17.2439 | 5.79E-07 | 0.0995 | TAGCTGTCCTCAGATACTCCAGAAGAGGGCATCA | | |
| 2 | 1 | 6417255 | 6417531 | 276 | SPRET | 1 | 6417381 | 6417414 | 34 | + | 17.2439 | 5.79E-07 | 0.0995 | TAGCTGTCCTCAGATACTCCAGAAGAGGGCATCA | | |
| 3 | 1 | 6438536 | 6438750 | 214 | AJ,BALB,C3H,CAST,CBA,DBA1J,DBA2J,FVB,KK,MOLF,MRL,NZO,RF,RH2,SEA,SMJ,SPRET.TH.WSB | 1 | 6438609 | 6438642 | 34 | + | 16.4472 | 9.61E-07 | 0.125 | TAGCTGTCTTCAGACACACCAGAAGAGGGCGTCA | | |
| 4 | 1 | 9611657 | 9611713 | 56 | CAST,SPRET | 1 | 9611635 | 9611669 | 35 | + | 16.4954 | 7.23E-07 | 0.182 | ATACAGTGACACCTACTTGCCACTAGGAGAGGCTG | 32 | ENCFF093ZRU,ENCFF201JUA,ENCFF099QNE,ENCFF821BIA,ENCFF601JVE,ENCFF327LYS,ENCFF089GBC,ENCFF024JOZ,ENCFF907XEG,ENCFF589ZBJ,ENCFF136ZDK,ENCFF055HAP,ENCFF430PPJ,ENCFF502WDT,ENCFF092NMJ,ENCFF883UPM,ENCFF321AGQ,ENCFF198SSR,ENCFF367UNR,ENCFF210BTO,ENCFF052KVO,ENCFF575MGI,ENCFF165LXO,ENCFF957SPH,ENCFF533APC,ENCFF065ODV,ENCFF281QMV,ENCFF504JDS,ENCFF194HVL,ENCFF142CNG,ENCFF609MDN,ENCFF951GOP |
| 4 | 1 | 9611657 | 9611713 | 56 | CAST,SPRET | 1 | 9611691 | 9611725 | 35 | + | 18.789 | 1.86E-07 | 0.0879 | ATACAGTGACACCTACTTGCCAGTAGGAGGTTCTT | 28 | ENCFF093ZRU,ENCFF201JUA,ENCFF099QNE,ENCFF821BIA,ENCFF601JVE,ENCFF327LYS,ENCFF089GBC,ENCFF024JOZ,ENCFF907XEG,ENCFF589ZBJ,ENCFF136ZDK,ENCFF055HAP,ENCFF430PPJ,ENCFF502WDT,ENCFF092NMJ,ENCFF883UPM,ENCFF321AGQ,ENCFF198SSR,ENCFF367UNR,ENCFF052KVO,ENCFF575MGI,ENCFF165LXO,ENCFF957SPH,ENCFF065ODV,ENCFF281QMV,ENCFF504JDS,ENCFF609MDN,ENCFF951GOP |
| 4 | 1 | 9611657 | 9611713 | 56 | CAST,SPRET | 1 | 9611638 | 9611672 | 35 | - | 17.6881 | 3.62E-07 | 0.124 | CCGCAGCCTCTCCTAGTGGCAAGTAGGTGTCACTG | 32 | ENCFF093ZRU,ENCFF201JUA,ENCFF099QNE,ENCFF821BIA,ENCFF601JVE,ENCFF327LYS,ENCFF089GBC,ENCFF024JOZ,ENCFF907XEG,ENCFF589ZBJ,ENCFF136ZDK,ENCFF055HAP,ENCFF430PPJ,ENCFF502WDT,ENCFF092NMJ,ENCFF883UPM,ENCFF321AGQ,ENCFF198SSR,ENCFF367UNR,ENCFF210BTO,ENCFF052KVO,ENCFF575MGI,ENCFF165LXO,ENCFF957SPH,ENCFF533APC,ENCFF065ODV,ENCFF281QMV,ENCFF504JDS,ENCFF194HVL,ENCFF142CNG,ENCFF609MDN,ENCFF951GOP |
| 5 | 1 | 13070920 | 13071126 | 206 | SPRET | 1 | 13070983 | 13071016 | 34 | + | 16.4472 | 9.61E-07 | 0.125 | TAGCTGTCTTCAGACACACCAGAAGAGGGCGTCA | | |
| 6 | 1 | 13111148 | 13111363 | 215 | 129S1,AJ,AKR,BALB,BTBR,BUB,C3H,C57L,CAST,CBA,CE,DBA1J,DBA2J,FVB,ILNJ2,KK,LP,MA/My,MOLF,MRL,NOD,NOR,NZB,NZO,NZW,PJ,PL,RF,RH2,SEA,SJL,SMJ,SPRET,SWR.TH.WSB | 1 | 13111219 | 13111252 | 34 | + | 17.8293 | 3.95E-07 | 0.083 | TAGCTGTCTTCAGACCCTCCAGAAGAGGGCGTCA | | |
| 7 | 1 | 13260703 | 13261118 | 415 | SPRET | 1 | 13260972 | 13261005 | 34 | + | 17.1138 | 6.30E-07 | 0.104 | TAGCTATCTTCAGACACACCAGAAGAGGGCACCA | 1 | ENCFF093ZRU |
| 7 | 1 | 13260703 | 13261118 | 415 | SPRET | 1 | 13260987 | 13261005 | 19 | + | 17.8852 | 5.08E-07 | 0.151 | ACACCAGAAGAGGGCACCA | 1 | ENCFF093ZRU |
| 8 | 1 | 20622804 | 20623208 | 404 | SPRET | 1 | 20622817 | 20622850 | 34 | + | 17.5447 | 4.76E-07 | 0.0903 | GAGCGGGTGTGGCAGCAGCCCCAAGATGGCGCCC | | |
| 9 | 1 | 22579690 | 22579881 | 191 | SPRET | 1 | 22579739 | 22579772 | 34 | + | 16.4634 | 9.51E-07 | 0.125 | TAGCTGTCTTCAGACGCACCAGAAGAGGGCATCA | | |
| 10 | 1 | 34041759 | 34041962 | 203 | CAST,CE,MOLF,NOD,RF,SPRET,SWR,WSB | 1 | 34041820 | 34041853 | 34 | + | 16.4634 | 9.51E-07 | 0.125 | TAGCTGTCTTCAGACGCACCAGAAGAGGGCATCA | | |
| 11 | 1 | 34129544 | 34129734 | 190 | DBA1J,DBA2J,ILNJ2,KK,MA/My,NOD,NOR,NZW,PJ,PL,SJL,SPRET,SWR,TH | 1 | 34129587 | 34129620 | 34 | + | 18.5285 | 2.48E-07 | 0.0666 | CAGCTGTCTTCAGACACTCCAGAAGAGGGCGTCA | | |
| 12 | 1 | 35746625 | 35747540 | 915 | MOLF | 1 | 35747436 | 35747470 | 35 | + | 19.7523 | 1.02E-07 | 0.0599 | ATGCAGATGCACTCAATGGCCAGTAGGGAGCATTG | 5 | ENCFF367UNR.ENCFF093ZRU.ENCFF907XEG.ENCFF883UPM.ENCFF099QNE |
| 13 | 1 | 37458652 | 37458746 | 94 | SPRET | 1 | 37458686 | 37458719 | 34 | + | 19.0407 | 1.74E-07 | 0.0542 | GTGCAGTACCTCCTACTCCCAGTGGGGGCAGTAC | 39 | ENCFF367UNR,ENCFF093ZRU,ENCFF201JUA,ENCFF327LYS,ENCFF136ZDK,ENCFF089GBC,ENCFF099QNE,ENCFF198SSR,ENCFF092NMJ,ENCFF321AGQ,ENCFF065ODV,ENCFF907XEG,ENCFF589ZBJ,ENCFF024JOZ,ENCFF883UPM,ENCFF430PPJ,ENCFF805QBU,ENCFF502WDT,ENCFF055HAP,ENCFF165LXO,ENCFF601JVE,ENCFF052KVO,ENCFF611HDQ,ENCFF957SPH,ENCFF504JDS,ENCFF349BUL,ENCFF609MDN,ENCFF210BTO,ENCFF951GOP,ENCFF491RJK,ENCFF281QMV,ENCFF194HVL,ENCFF533APC,ENCFF033UQX,ENCFF316FYN,ENCFF575MGI,ENCFF097CAQ,ENCFF089EWX,ENCFF142CNG |
| 13 | 1 | 37458652 | 37458746 | 94 | SPRET | 1 | 37458733 | 37458766 | 34 | + | 19.0407 | 1.74E-07 | 0.0542 | GTGCAGTACCTCCTACTCCCAGTGGGGGCAGTAC | 37 | ENCFF367UNR,ENCFF093ZRU,ENCFF201JUA,ENCFF327LYS,ENCFF136ZDK,ENCFF089GBC,ENCFF099QNE,ENCFF198SSR,ENCFF092NMJ,ENCFF321AGQ,ENCFF065ODV,ENCFF907XEG,ENCFF589ZBJ,ENCFF024JOZ,ENCFF883UPM,ENCFF430PPJ,ENCFF805QBU,ENCFF502WDT,ENCFF055HAP,ENCFF165LXO,ENCFF601JVE,ENCFF052KVO,ENCFF957SPH,ENCFF504JDS,ENCFF349BUL,ENCFF609MDN,ENCFF210BTO,ENCFF951GOP,ENCFF491RJK,ENCFF281QMV,ENCFF194HVL,ENCFF533APC,ENCFF033UQX,ENCFF316FYN,ENCFF575MGI.ENCFF097CAQ.ENCFF089EWX |
| 14 | 1 | 39303819 | 39304362 | 543 | MA/Mv.MOLF | 1 | 39304032 | 39304065 | 34 | + | 18.3089 | 2.87E-07 | 0.0691 | ATGCAGCTAGAGACACGAGCTCCAGGGGGTACTG | | |
| 15 | 1 | 39520112 | 39520306 | 194 | SPRET | 1 | 39520163 | 39520196 | 34 | + | 16.4472 | 9.61E-07 | 0.125 | TAGCTGTCTTCAGACACACCAGAAGAGGGCGTCA | | |
| 16 | 1 | 45708127 | 45708325 | 198 | SPRET | 1 | 45708182 | 45708215 | 34 | + | 17.8455 | 3.91E-07 | 0.0823 | TAGCTGTCCTCAGACACTCCAGAAGAGGGCGTCA | | |
| 17 | 1 | 47813238 | 47813450 | 212 | CAST.MOLF.SPRET.SWR | 1 | 47813306 | 47813339 | 34 | + | 18.3659 | 2.76E-07 | 0.069 | TAGCTGTCTTCAGACACTCCAGAAGAGGGCGCCA | | |
| 17 | 1 | 47813238 | 47813450 | 212 | CAST.MOLF.SPRET.SWR | 1 | 47813321 | 47813339 | 19 | + | 17.5574 | 6.32E-07 | 0.166 | ACTCCAGAAGAGGGCGCCA | | |
| 18 | 1 | 52343806 | 52343974 | 168 | BTBR,C57L,CAST,KK,MOLF,NZO,NZW,SMJ,SPRET.SWR.WSB | 1 | 52343849 | 52343867 | 19 | + | 17.8852 | 5.08E-07 | 0.151 | ACACCAGAAGAGGGCACCA | | |
| 19 | 1 | 52621206 | 52621399 | 193 | AKR,BALB,BUB,C58J,CAST,MOLF,MRL,NOD,NOR,NZO,NZW,PL,RF,RH2,SEA,SMJ,SPRET.SWR.WSB | 1 | 52621256 | 52621289 | 34 | + | 18.3659 | 2.76E-07 | 0.069 | TAGCTGTCTTCAGACACTCCAGAAGAGGGCGCCA | | |
| 19 | 1 | 52621206 | 52621399 | 193 | AKR,BALB,BUB,C58J,CAST,MOLF,MRL,NOD,NOR,NZO,NZW,PL,RF,RH2,SEA,SMJ,SPRET.SWR.WSB | 1 | 52621271 | 52621289 | 19 | + | 17.5574 | 6.32E-07 | 0.166 | ACTCCAGAAGAGGGCGCCA | | |
| 20 | 1 | 54942393 | 54942572 | 179 | ILNJ2.MA/Mv.NOD.NOR.PJ | 1 | 54942485 | 54942519 | 35 | + | 16.8807 | 5.80E-07 | 0.165 | CTGCTCTCAAGGCTTTGGAGCTGAAGGGGGCGTTG | | |
| 21 | 1 | 55145657 | 55145856 | 199 | CAST.MOLF | 1 | 55145713 | 55145746 | 34 | + | 16.4472 | 9.61E-07 | 0.125 | TAGCTGTCTTCAGACACACCAGAAGAGGGCGTCA | | |
| 22 | 1 | 58189216 | 58189416 | 200 | BALB,BTBR,BUB,C57L,C58J,CAST,CBA,CE,DBA1J,DBA2J,MOLF,MRL,PL,RH2,SEA,SJL,SPRET.WSB | 1 | 58189272 | 58189305 | 34 | + | 16.4634 | 9.51E-07 | 0.125 | TAGCTGTCTTCAGACGCACCAGAAGAGGGCATCA | | |
| 23 | 1 | 59271285 | 59271491 | 206 | 129S1,BALB,BUB,CAST,CE,MOLF,NOD,NOR,PL,RH2,SEA,SJL,SPRET,SWR,WSB | 1 | 59271343 | 59271376 | 34 | + | 16.4472 | 9.61E-07 | 0.125 | TAGCTGTCTTCAGACACACCAGAAGAGGGCGTCA | | |
| 24 | 1 | 59823348 | 59823550 | 202 | AKR,BALB,BUB,CAST,ILNJ2,MA/My,MOLF,NOD,NOR,NZB,NZO,NZW,PL,RH2,SEA,SJL,SMJ.SPRET.SWR.WSB | 1 | 59823406 | 59823439 | 34 | + | 18.3659 | 2.76E-07 | 0.069 | TAGCTGTCTTCAGACACTCCAGAAGAGGGCGCCA | | |
| 24 | 1 | 59823348 | 59823550 | 202 | AKR,BALB,BUB,CAST,ILNJ2,MA/My,MOLF,NOD,NOR,NZB,NZO,NZW,PL,RH2,SEA,SJL,SMJ.SPRET.SWR.WSB | 1 | 59823421 | 59823439 | 19 | + | 17.5574 | 6.32E-07 | 0.166 | ACTCCAGAAGAGGGCGCCA | | |
| 25 | 1 | 65143093 | 65143326 | 233 | CAST.MOLF.SPRET | 1 | 65143184 | 65143217 | 34 | + | 17.2439 | 5.79E-07 | 0.0995 | TAGCTGTCCTCAGATACTCCAGAAGAGGGCATCA | | |
| 26 | 1 | 69200181 | 69200898 | 717 | C58J | 1 | 69200651 | 69200685 | 35 | + | 15.9725 | 9.70E-07 | 0.212 | CTGCATGTACACCTACATGCCAGAATAGGGCATCA | | |
| 27 | 1 | 71683612 | 71683798 | 186 | AKR,BUB,CAST,PL,RF,SJL,SPRET,SWR,WSB | 1 | 71683669 | 71683702 | 34 | + | 16.4472 | 9.61E-07 | 0.125 | TAGCTGTCTTCAGACACACCAGAAGAGGGCGTCA | | |
| 28 | 1 | 74750870 | 74751615 | 745 | SPRET | 1 | 74751299 | 74751332 | 34 | + | 19.0325 | 1.75E-07 | 0.0544 | CAGCTGTCTTCAGACGCACCAGAAGAGGGCATCA | 8 | ENCFF099QNE,ENCFF821BIA,ENCFF327LYS,ENCFF093ZRU,ENCFF367UNR,ENCFF055HAP,ENCFF502WDT,ENCFF883UPM |
| 29 | 1 | 74796099 | 74796302 | 203 | SPRET | 1 | 74796161 | 74796194 | 34 | + | 18.187 | 3.12E-07 | 0.0723 | TAGCTGTCTTCAGTCACTCCAGAAGAGGGCGCTA | 1 | ENCFF093ZRU |
| 29 | 1 | 74796099 | 74796302 | 203 | SPRET | 1 | 74796176 | 74796194 | 19 | + | 17.5738 | 6.25E-07 | 0.166 | ACTCCAGAAGAGGGCGCTA | 1 | ENCFF093ZRU |
| 30 | 1 | 74912586 | 74912803 | 217 | SPRET | 1 | 74912662 | 74912695 | 34 | + | 17.9756 | 3.59E-07 | 0.0784 | TAGCTGTCTTTAGACACTCCAGAAGAGGGCGCCA | 1 | ENCFF093ZRU |
| 30 | 1 | 74912586 | 74912803 | 217 | SPRET | 1 | 74912677 | 74912695 | 19 | + | 17.5574 | 6.32E-07 | 0.166 | ACTCCAGAAGAGGGCGCCA | 1 | ENCFF093ZRU |
| 31 | 1 | 75793093 | 75793291 | 198 | SPRET | 1 | 75793152 | 75793185 | 34 | + | 18.187 | 3.12E-07 | 0.0723 | TAGCTGTCTTCTGACACTCCAGAAGAGGGCGCCA | 1 | ENCFF093ZRU |
| 31 | 1 | 75793093 | 75793291 | 198 | SPRET | 1 | 75793167 | 75793185 | 19 | + | 17.5574 | 6.32E-07 | 0.166 | ACTCCAGAAGAGGGCGCCA | 1 | ENCFF093ZRU |
| 32 | 1 | 77776634 | 77776955 | 321 | SPRET | 1 | 77776670 | 77776704 | 35 | + | 24.1743 | 4.67E-09 | 0.0113 | GTGCAGGGGCCCAAGGAGGCCAGAAGAGGGCATTG | | |
| 33 | 1 | 82772748 | 82772826 | 78 | MOLF.RF | 1 | 82772782 | 82772815 | 34 | + | 16.7724 | 7.83E-07 | 0.115 | TAGCTGTCTTCAGATGCACCAGAAGAGGGCGTCA | | |
| 34 | 1 | 84949337 | 84949564 | 227 | AKR,BUB,CAST,CBA,MRL,NOD,PJ,PL,SMJ,SPRET | 1 | 84949420 | 84949453 | 34 | + | 16.4634 | 9.51E-07 | 0.125 | TAGCTGTCTTCAGACGCACCAGAAGAGGGCATCA | | |
| 35 | 1 | 85194795 | 85194998 | 203 | AKR.BUB.CBA.MRL.NOD.SMJ | 1 | 85194854 | 85194887 | 34 | + | 16.4472 | 9.61E-07 | 0.125 | TAGCTGTCTTCAGACACACCAGAAGAGGGCGTCA | | |
| 36 | 1 | 89496454 | 89496770 | 316 | SPRET | 1 | 89496447 | 89496481 | 35 | + | 19.2018 | 1.44E-07 | 0.0731 | CTGCCTTGCTGCCTTCCAGGCTGGAGGTGGAGCCT | | |
| 37 | 1 | 90505197 | 90505521 | 324 | SPRET | 1 | 90505311 | 90505344 | 34 | + | 16.4959 | 9.32E-07 | 0.125 | AAGAAGTTCCAGGCATTACCAGAAGGCGGGGCCA | 5 | ENCFF052KVO.ENCFF367UNR.ENCFF093ZRU.ENCFF821BIA.ENCFF099QNE |
| 38 | 1 | 92884514 | 92884845 | 331 | SPRET | 1 | 92884542 | 92884560 | 19 | + | 19.2951 | 1.88E-07 | 0.096 | CCGCCACTAGATGGTGCTA | 21 | ENCFF367UNR,ENCFF093ZRU,ENCFF052KVO,ENCFF201JUA,ENCFF099QNE,ENCFF821BIA,ENCFF502WDT,ENCFF883UPM,ENCFF327LYS,ENCFF907XEG,ENCFF136ZDK,ENCFF609MDN,ENCFF089GBC,ENCFF024JOZ,ENCFF198SSR,ENCFF589ZBJ,ENCFF165LXO,ENCFF430PPJ,ENCFF194HVL,ENCFF092NMJ,ENCFF055HAP |
| 39 | 1 | 99558814 | 99559114 | 300 | CAST.MOLF.MRL.PL.SPRET | 1 | 99558821 | 99558854 | 34 | + | 19.5041 | 1.26E-07 | 0.0441 | CTGCGATAACATTCGCCATCACAAGATGGCACCA | | |
| 40 | 1 | 100074793 | 100075329 | 536 | AKR.LP.MA/Mv.PJ.SEA | 1 | 100075293 | 100075326 | 34 | + | 16.8862 | 7.28E-07 | 0.112 | ATGCAGCTAGAGACATGAGCTCCGGGGGGCGGGG | | |
| 41 | 1 | 113174834 | 113175692 | 858 | MOLF | 1 | 113175254 | 113175272 | 19 | + | 21.459 | 3.29E-08 | 0.052 | GGGCCACCAGGGGTCGCCA | 6 | ENCFF092NMJ.ENCFF907XEG.ENCFF093ZRU.ENCFF502WDT.ENCFF099QNE.ENCFF327LYS |
| 42 | 1 | 117424489 | 117424689 | 200 | CAST.MOLF.SPRET | 1 | 117424545 | 117424578 | 34 | + | 16.4634 | 9.51E-07 | 0.125 | TAGCTGTCTTCAGACGCACCAGAAGAGGGCATCA | | |
| 43 | 1 | 118652923 | 118653125 | 202 | AJ,BUB,C3H,CAST,CBA,CE,DBA1J,DBA2J,ILNJ2,KK,LP,MA/My,MOLF,MRL,NOD,NOR,NZO,NZW,PJ,PL,RH2,SPRET,WSB | 1 | 118652982 | 118653016 | 35 | + | 18.1284 | 2.79E-07 | 0.109 | CTGTAGCTGTCTTCAGACACCAGAAGAGGGCATCA | | |
| 44 | 1 | 119545512 | 119546395 | 883 | MOLF | 1 | 119546115 | 119546149 | 35 | + | 23.1743 | 9.85E-09 | 0.0169 | CTGCATGTATACCTACATGCCAGAAGAGGGCATCA | 9 | ENCFF099QNE,ENCFF024JOZ,ENCFF907XEG,ENCFF055HAP,ENCFF502WDT,ENCFF367UNR,ENCFF821BIA.ENCFF093ZRU.ENCFF327LYS |
| 45 | 1 | 119987857 | 119988271 | 414 | SPRET | 1 | 119987912 | 119987946 | 35 | + | 18.1284 | 2.79E-07 | 0.109 | CTGTAGCTGTCTTCAGACACCAGAAGAGGGCATCA | | |
| 46 | 1 | 120021565 | 120021787 | 222 | 129S1,AJ,AKR,BALB,BTBR,BUB,C3H,C57L,CAST,CBA,CE,DBA1J,DBA2J,FVB,ILNJ2,KK,LP,MA/My,MOLF,MRL,NOD,NOR,NZB,NZO,NZW,PJ,PL,RF,RH2,SEA,SJL,SMJ,SPRET,SWR.TH.WSB | 1 | 120021645 | 120021678 | 34 | + | 18.374 | 2.75E-07 | 0.069 | TAGCTGTCTTCAGTCACTCCAGAAGAGGGCGCCA | 3 | ENCFF367UNR,ENCFF093ZRU |
| 46 | 1 | 120021565 | 120021787 | 222 | 129S1,AJ,AKR,BALB,BTBR,BUB,C3H,C57L,CAST,CBA,CE,DBA1J,DBA2J,FVB,ILNJ2,KK,LP,MA/My,MOLF,MRL,NOD,NOR,NZB,NZO,NZW,PJ,PL,RF,RH2,SEA,SJL,SMJ,SPRET,SWR.TH.WSB | 1 | 120021660 | 120021678 | 19 | + | 17.5574 | 6.32E-07 | 0.166 | ACTCCAGAAGAGGGCGCCA | 2 | ENCFF093ZRU,ENCFF367UNR |
| 47 | 1 | 122076804 | 122077207 | 403 | SPRET | 1 | 122076859 | 122076892 | 34 | + | 18.9431 | 1.87E-07 | 0.0559 | TTGCAACTGTCTTTGACACCAGAAGAGGGCGTCA | | |
| 48 | 1 | 125760530 | 125760603 | 73 | SPRET | 1 | 125760596 | 125760614 | 19 | + | 18.8852 | 2.53E-07 | 0.112 | TTGCCAGCAGGTGACACCC | | |
| 49 | 1 | 127773686 | 127773912 | 226 | CAST,MOLF,MRL,NZB,RF,SPRET,SWR,WSB | 1 | 127773784 | 127773802 | 19 | + | 17.8852 | 5.08E-07 | 0.151 | ACACCAGAAGAGGGCACCA | | |
| 50 | 1 | 131085566 | 131085764 | 198 | MOLF.SPRET | 1 | 131085623 | 131085656 | 34 | + | 19.065 | 1.71E-07 | 0.0538 | CTGTAGCTGTAGCTGACACCAGAAGAGGGAGTCA | | |
| 51 | 1 | 131493793 | 131494117 | 324 | SPRET | 1 | 131493762 | 131493795 | 34 | + | 18.3089 | 2.87E-07 | 0.0691 | ATGCAGCTAGAGACACGAGCTCCAGGGGGTACTG | | |
| 52 | 1 | 131830171 | 131830366 | 195 | CAST.MOLF | 1 | 131830224 | 131830257 | 34 | + | 19.0163 | 1.77E-07 | 0.0544 | CAGCTGTCTTCAGACACACCAGAAGAGGGCGTCA | | |
| 53 | 1 | 132152903 | 132153099 | 196 | CAST | 1 | 132153010 | 132153043 | 34 | + | 16.6098 | 8.68E-07 | 0.122 | TAGCTGTCTTCAAATGTACCAGAAGAGGGCGTCA | 2 | ENCFF093ZRU.ENCFF502WDT |
| 54 | 1 | 132752505 | 132752724 | 219 | AKR,ILNJ2,KK,MOLF,NZB,NZW,SJL,SPRET,SWR | 1 | 132752580 | 132752613 | 34 | + | 16.7724 | 7.83E-07 | 0.115 | TAGCTGTCTTCGGACACACCAGAAGAGGGCATCA | 1 | ENCFF093ZRU |
| 55 | 1 | 132953534 | 132953907 | 373 | CAST | 1 | 132953900 | 132953934 | 35 | + | 16.0367 | 9.36E-07 | 0.209 | ATGCACTGCCCCTGGGCCGCAAGAAGGGGGCCAAG | | |
| 56 | 1 | 135334873 | 135335080 | 207 | 129S1,AJ,AKR,BTBR,BUB,C3H,CAST,CBA,CE,DBA1J,DBA2J,FVB,ILNJ2,KK,MA/My,MOLF,MRL,NOD,NOR,NZB,NZO,NZW,PJ,PL,RF,RH2,SJL,SMJ,SPRET,SWR,TH,WSB | 1 | 135334938 | 135334971 | 34 | + | 16.4634 | 9.51E-07 | 0.125 | TAGCTGTCTTCAGACGCACCAGAAGAGGGCATCA | | |
| 57 | 1 | 137843970 | 137844469 | 499 | SPRET | 1 | 137844055 | 137844088 | 34 | + | 18.3089 | 2.87E-07 | 0.0691 | ATGCAGCTAGAGACACGAGCTCCAGGGGGTACTG | | |
| 58 | 1 | 152763769 | 152763983 | 214 | AKR,BTBR,BUB,C57L,CAST,FVB,KK,MOLF,NZB,NZO,NZW,RF,SJL,SMJ,SPRET,SWR,TH.WSB | 1 | 152763842 | 152763875 | 34 | + | 16.4634 | 9.51E-07 | 0.125 | TAGCTGTCTTCAGACGCACCAGAAGAGGGCATCA | | |
| 59 | 1 | 153752071 | 153752274 | 203 | BUB,C57L,CAST,DBA1J,DBA2J,FVB,MA/My,RF.SJL.SPRET.SWR.TH.WSB | 1 | 153752130 | 153752164 | 35 | + | 18.1284 | 2.79E-07 | 0.109 | CTGTAGCTGTCTTCAGACACCAGAAGAGGGCATCA | | |
| 60 | 1 | 153756202 | 153756697 | 495 | BUB,C57L,CAST,DBA1J,DBA2J,FVB,MA/My,RF.SJL.SPRET.SWR.TH.WSB | 1 | 153756217 | 153756250 | 34 | + | 17.9106 | 3.74E-07 | 0.0805 | CCGCCCTCACATTCGCCGTCACAAGATGGCGCTG | | |
| 61 | 1 | 155320630 | 155320775 | 145 | SPRET.SWR.WSB | 1 | 155320613 | 155320647 | 35 | + | 16.8624 | 5.86E-07 | 0.166 | GTGCTTTCTCTGCTTGTGGCCAGGCGGTGGTGGTG | | |
| 62 | 1 | 156038223 | 156038426 | 203 | SPRET | 1 | 156038284 | 156038317 | 34 | + | 16.4472 | 9.61E-07 | 0.125 | TAGCTGTCTTCAGACACACCAGAAGAGGGCGTCA | | |
| 63 | 1 | 156350301 | 156350536 | 235 | 129S1,AJ,AKR,BALB,BTBR,BUB,C3H,C57L,C58J,CAST,CBA,CE,DBA1J,DBA2J,FVB,ILNJ2,KK,LP,MA/My,MOLF,MRL,NOD,NOR,NZB,NZO,NZW,PJ,PL,RF,RH2,SEA,SWR,TH,WSB | 1 | 156350395 | 156350428 | 34 | + | 18.3659 | 2.76E-07 | 0.069 | TAGCTGTCTTCAGACACTCCAGAAGAGGGCGCCA | | |
| 63 | 1 | 156350301 | 156350536 | 235 | 129S1,AJ,AKR,BALB,BTBR,BUB,C3H,C57L,C58J,CAST,CBA,CE,DBA1J,DBA2J,FVB,ILNJ2,KK,LP,MA/My,MOLF,MRL,NOD,NOR,NZB,NZO,NZW,PJ,PL,RF,RH2,SEA,SWR,TH,WSB | 1 | 156350410 | 156350428 | 19 | + | 17.5574 | 6.32E-07 | 0.166 | ACTCCAGAAGAGGGCGCCA | | |
| 64 | 1 | 160240285 | 160240519 | 234 | BALB.C58J.DBA1J.DBA2J.LP.MOLF | 1 | 160240377 | 160240410 | 34 | + | 16.4634 | 9.51E-07 | 0.125 | TAGCTGTCTTCAGACGCACCAGAAGAGGGCATCA | | |
| 65 | 1 | 165558346 | 165558413 | 67 | SPRET | 1 | 165558395 | 165558429 | 35 | + | 23.1284 | 1.02E-08 | 0.0171 | CTGCCATGAGACTCTGGCACCAGCAGATGGCACTT | 5 | ENCFF093ZRU.ENCFF024JOZ.ENCFF198SSR.ENCFF907XEG.ENCFF367UNR |
| 66 | 1 | 166094270 | 166094511 | 241 | BTBR,CAST,DBA1J,DBA2J,FVB,ILNJ2,KK,MOLF,NOD,NOR,RF,SMJ,SPRET,SWR,WSB | 1 | 166094354 | 166094387 | 34 | + | 17.7967 | 4.04E-07 | 0.0841 | CAGCTGTCCTCAGATACTCCAGAAGAGGGAGTCA | | |
| 67 | 1 | 167058302 | 167058527 | 225 | BTBR,FVB,ILNJ2,KK,NZB,NZO,NZW,RF,SMJ.SPRET.SWR.TH | 1 | 167058377 | 167058410 | 34 | + | 17.2439 | 5.79E-07 | 0.0995 | TAGCTGTCCTCAGATACTCCAGAAGAGGGCATCA | | |
| 68 | 1 | 169345502 | 169345615 | 113 | 129S1,AJ,BTBR,BUB,C3H,C57L,CE,FVB,KK,MOLF,MRL,NOD,NOR,NZB,NZO,NZW,PJ,PL,RF,RH2,SEA,SJL,SMJ,SPRET,SWR,TH,WSB | 1 | 169345598 | 169345616 | 19 | + | 17.377 | 7.11E-07 | 0.174 | AGGCCAGCAGGGGGAGTCA | 5 | ENCFF821BIA,ENCFF024JOZ,ENCFF327LYS,ENCFF907XEG,ENCFF198SSR |
| 69 | 1 | 170757166 | 170757376 | 210 | 129S1,AJ,BTBR,BUB,C3H,C57L,CAST,CE,FVB,KK,MRL,NOD,NOR,NZB,NZO,NZW,PJ,PL,RF,RH2,SEA,SJL,SMJ,SPRET,SWR,TH,WSB | 1 | 170757234 | 170757267 | 34 | + | 17.2439 | 5.79E-07 | 0.0995 | TAGCTGTCCTCAGATACTCCAGAAGAGGGCATCA | | |
| 70 | 1 | 171859494 | 171859700 | 206 | 129S1,AJ,AKR,BALB,BUB,C3H,C57L,CBA,CE,DBA1J,DBA2J,FVB,KK,MRL,NOD,NOR,NZB,NZO,NZW,PJ,PL,RF,RH2,SEA,SJL,SMJ,SWR.TH.WSB | 1 | 171859560 | 171859593 | 34 | + | 17.2358 | 5.82E-07 | 0.0997 | GAGCTGTCTTCAGATACACCAGAAGAGGGCGTCA | | |
| 71 | 1 | 171982743 | 171982946 | 203 | 129S1,AJ,AKR,BALB,BUB,C3H,C57L,CAST,CBA,CE,DBA1J,DBA2J,FVB,KK,MRL,NOD,NOR,NZB,NZO,NZW,PJ,PL,RF,RH2,SEA,SJL,SMJ.SPRET.SWR.TH.WSB | 1 | 171982805 | 171982838 | 34 | + | 17.2439 | 5.79E-07 | 0.0995 | TAGCTGTCCTCAGATACTCCAGAAGAGGGCATCA | | |
| 72 | 1 | 172882436 | 172882681 | 245 | SPRET | 1 | 172882539 | 172882572 | 34 | + | 16.4634 | 9.51E-07 | 0.125 | TAGCTGTCTTCAGACGCACCAGAAGAGGGCATCA | | |
| 73 | 1 | 173236114 | 173236314 | 200 | 129S1,AJ,AKR,BALB,BTBR,BUB,C3H,C57L,C58J,CBA,CE,DBA1J,DBA2J,FVB,ILNJ2,KK,MRL,NOD,NOR,NZB,NZO,PJ,PL,RF,RH2,SEA,SJL,SMJ,SPRET,SWR,TH,WSB | 1 | 173236171 | 173236204 | 34 | + | 16.4634 | 9.51E-07 | 0.125 | TAGCTGTCTTCAGACGCACCAGAAGAGGGCATCA | 1 | ENCFF093ZRU |
| 74 | 1 | 177614791 | 177614872 | 81 | PL | 1 | 177614763 | 177614797 | 35 | + | 15.9266 | 9.95E-07 | 0.213 | ATGCACTGCCCCTGGGCTGCAAGAAGGGGGCCAAG | | |
| 75 | 1 | 179019252 | 179019473 | 221 | AKR.CAST.MOLF.SPRET | 1 | 179019347 | 179019365 | 19 | + | 17.9836 | 4.75E-07 | 0.151 | ACACCAGAAGAGGGCGGCA | | |
| 76 | 1 | 179530319 | 179531227 | 908 | SPRET | 1 | 179530567 | 179530600 | 34 | + | 18.4146 | 2.67E-07 | 0.0688 | TTGCTGTCTTCAGACATACCAGAAGAGGGCATCA | 4 | ENCFF093ZRU.ENCFF883UPM.ENCFF055HAP.ENCFF502WDT |
| 77 | 1 | 180627208 | 180627420 | 212 | AJ,BUB,CAST,CE,MOLF,PJ,RH2,SEA,SPRET.TH.WSB | 1 | 180627277 | 180627310 | 34 | + | 18.3659 | 2.76E-07 | 0.069 | TAGCTGTCTTCAGACACTCCAGAAGAGGGCGCCA | | |
| 77 | 1 | 180627208 | 180627420 | 212 | AJ,BUB,CAST,CE,MOLF,PJ,RH2,SEA,SPRET.TH.WSB | 1 | 180627292 | 180627310 | 19 | + | 17.5574 | 6.32E-07 | 0.166 | ACTCCAGAAGAGGGCGCCA | | |
| 78 | 1 | 181614406 | 181615076 | 670 | MOLF.TH | 1 | 181614995 | 181615013 | 19 | + | 17.541 | 6.39E-07 | 0.167 | TTGCCTGCAGGTGGCAGAC | | |
| 79 | 1 | 182369734 | 182370226 | 492 | CE | 1 | 182370072 | 182370090 | 19 | + | 17.082 | 8.61E-07 | 0.188 | TGGCCACCAGGTGGAATAA | | |
| 80 | 1 | 183176095 | 183176331 | 236 | MOLF.SPRET.TH | 1 | 183176098 | 183176131 | 34 | + | 17.3496 | 5.41E-07 | 0.0962 | ATGTGGTGCTCACAGAGGCCAGGAGAGGGTGTCA | | |
| 81 | 1 | 183892033 | 183892204 | 171 | SPRET | 1 | 183892195 | 183892228 | 34 | + | 25.5041 | 1.05E-09 | 0.00163 | GTGCAATGCCCAAAGAGTCCAGAAGAGGGCATCA | 2 | ENCFF093ZRU.ENCFF052KVO |
| 82 | 1 | 185122912 | 185123183 | 271 | 129S1,AJ,AKR,BALB,BTBR,BUB,C3H,C57L,CAST,CE,FVB,ILNJ2,KK,MA/My,MOLF,MRL,NOD,NOR,NZB,NZW,PJ,PL,RF,RH2,SEA,SJL.SPRET.SWR.TH.WSB | 1 | 185123039 | 185123072 | 34 | + | 16.4634 | 9.51E-07 | 0.125 | TAGCTGTCTTCAGACGCACCAGAAGAGGGCATCA | | |
| 83 | 1 | 186534856 | 186535080 | 224 | SPRET | 1 | 186534937 | 186534970 | 34 | + | 16.4472 | 9.61E-07 | 0.125 | TAGCTGTCTTCAGACACACCAGAAGAGGGCGTCA | | |
| 84 | 1 | 195027700 | 195027901 | 201 | NZO.NZW.RF | 1 | 195027759 | 195027792 | 34 | + | 16.4472 | 9.61E-07 | 0.125 | TAGCTGTCTTCAGACACACCAGAAGAGGGCGTCA | | |
| 85 | 1 | 10314970 | 10315066 | 96 | SPRET | 1 | 10314943 | 10314976 | 34 | - | 17.0569 | 6.53E-07 | 0.106 | CAGAGGTGCCCTCAGAGGCCAGAAGAGGTCATCA | | |
| 86 | 1 | 10677545 | 10677854 | 309 | SPRET | 1 | 10677597 | 10677631 | 35 | - | 16.0642 | 9.21E-07 | 0.207 | TAGCTGTACTTCAGACACTCCAGAAGAGGGCGTCT | | |
| 87 | 1 | 16716263 | 16716466 | 203 | AJ,AKR,BALB,BUB,C3H,CAST,CBA,DBA1J,DBA2J,FVB,ILNJ2,KK,MOLF,MRL,NOD,NOR,NZB,NZO,NZW,PJ,PL,RF,RH2,SEA,SJL,SMJ.SPRET.SWR.TH | 1 | 16716389 | 16716407 | 19 | - | 17.8852 | 5.08E-07 | 0.151 | ACACCAGAAGAGGGCACCA | 1 | ENCFF093ZRU |
| 88 | 1 | 17262599 | 17263327 | 728 | SPRET | 1 | 17263263 | 17263296 | 34 | - | 16.5447 | 9.04E-07 | 0.124 | CAGCTGTCTTCAGACACACCAGAAGAGGGAGTCA | | |
| 89 | 1 | 17263146 | 17263350 | 204 | CAST.MOLF | 1 | 17263263 | 17263296 | 34 | - | 16.5447 | 9.04E-07 | 0.124 | CAGCTGTCTTCAGACACACCAGAAGAGGGAGTCA | | |
| 90 | 1 | 19246828 | 19247046 | 218 | CAST.MOLF | 1 | 19246952 | 19246985 | 34 | - | 18.0732 | 3.36E-07 | 0.0757 | CAGCTGTCTTCAGACACTCCAGAAGAGGGCATCA | | |
| 91 | 1 | 36005108 | 36005315 | 207 | CAST.MOLF.RF.SPRET | 1 | 36005232 | 36005265 | 34 | - | 17.8455 | 3.91E-07 | 0.0823 | TAGCTGTCCTCAGACACTCCAGAAGAGGGCGTCA | | |
| 92 | 1 | 36858706 | 36858904 | 198 | AJ,AKR,BUB,C3H,CBA,CE,FVB,KK,MRL,NOD,NZB,NZO,NZW,PJ,PL,RF,RH2,SJL,SMJ,SPRET.SWR.WSB | 1 | 36858832 | 36858865 | 34 | - | 18.0732 | 3.36E-07 | 0.0757 | CAGCTGTCTTCAGACACTCCAGAAGAGGGCATCA | | |
| 93 | 1 | 37275426 | 37275633 | 207 | CAST | 1 | 37275549 | 37275582 | 34 | - | 17.3496 | 5.41E-07 | 0.0962 | TAGCTGTCCTCAGATACTCCAGAAGAGGGCATCG | | |
| 94 | 1 | 37693221 | 37693771 | 550 | SPRET | 1 | 37693279 | 37693297 | 19 | - | 21.0984 | 4.51E-08 | 0.0555 | AGGCCAGAAGAGGGCGCCA | 14 | ENCFF907XEG,ENCFF327LYS,ENCFF093ZRU,ENCFF198SSR,ENCFF165LXO,ENCFF589ZBJ,ENCFF367UNR,ENCFF033UQX,ENCFF065ODV,ENCFF052KVO,ENCFF099QNE,ENCFF136ZDK,ENCFF142CNG,ENCFF821BIA |
| 94 | 1 | 37693221 | 37693771 | 550 | SPRET | 1 | 37693279 | 37693312 | 34 | - | 29.0813 | 3.00E-11 | 0.000226 | ATGCAATGCCCATAGAGGCCAGAAGAGGGCGCCA | 14 | ENCFF907XEG,ENCFF327LYS,ENCFF093ZRU,ENCFF198SSR,ENCFF165LXO,ENCFF589ZBJ,ENCFF367UNR,ENCFF033UQX,ENCFF065ODV,ENCFF052KVO,ENCFF099QNE,ENCFF136ZDK,ENCFF142CNG,ENCFF821BIA |
| 95 | 1 | 38469541 | 38469667 | 126 | DBA1J.DBA2J.MA/Mv.MOLF.NOR.RF | 1 | 38469529 | 38469563 | 35 | - | 15.9266 | 9.95E-07 | 0.213 | CTGCCCACCACTGGAGGGACCAGGAGGTGGTGCTC | | |
| 96 | 1 | 38644657 | 38644781 | 124 | CAST | 1 | 38644713 | 38644731 | 19 | - | 17.9016 | 5.02E-07 | 0.151 | TGGCCAGCGGAGGGCAGAG | 4 | ENCFF821BIA.ENCFF502WDT.ENCFF883UPM.ENCFF055HAP |
| 97 | 1 | 39391988 | 39392431 | 443 | MA/Mv.MOLF.SPRET | 1 | 39392018 | 39392036 | 19 | - | 20.3607 | 8.29E-08 | 0.0655 | AGGCCAGAAGAGGGCACCA | 5 | ENCFF099QNE.ENCFF821BIA.ENCFF502WDT.ENCFF367UNR.ENCFF093ZRU |
| 97 | 1 | 39391988 | 39392431 | 443 | MA/Mv.MOLF.SPRET | 1 | 39392018 | 39392052 | 35 | - | 27.2844 | 3.67E-10 | 0.00426 | CTGCATGTGTGCCTGCAGGCCAGAAGAGGGCACCA | 5 | ENCFF099QNE.ENCFF821BIA.ENCFF502WDT.ENCFF367UNR.ENCFF093ZRU |
| 98 | 1 | 43932151 | 43932356 | 205 | CAST.MOLF.SPRET | 1 | 43932275 | 43932308 | 34 | - | 16.7724 | 7.83E-07 | 0.115 | TAGCTGTCTTCAGATGCACCAGAAGAGGGCGTCA | | |
| 99 | 1 | 45470542 | 45470747 | 205 | CAST.MOLF.SPRET.SWR.TH | 1 | 45470666 | 45470699 | 34 | - | 16.4472 | 9.61E-07 | 0.125 | TAGCTGTCTTCAGACACACCAGAAGAGGGCGTCA | | |

| | | | | | | | | | | | | | | | | |
|---|---|---|---|---|---|---|---|---|---|---|---|---|---|---|---|---|
| 100 | 1 | 46921997 | 46922210 | 213 | 129S1,AJ,AKR,BTBR,BUB,C3H,C57L,CAST,CBA,CE,DBA1J,DBA2J,FVB,ILNJ2,KK,MA/My,MOLF,MRL,NOD,NOR,NZO,NZW,PJ,PL,RF,RH2,SJL,SMJ,SPRET,SWR,TH,WSB | 1 | 46922123 | 46922156 | 34 | - | 18.2439 | 3.00E-07 | 0.0708 | TAGCTGTCCTCAGATACTCCAGAAGGGGGCATCA | | |
| 101 | 1 | 51181542 | 51181722 | 180 | CAST.SPRET | 1 | 51181648 | 51181666 | 19 | - | 17.0328 | 8.88E-07 | 0.19 | AGACCAGAAGAGGGAGCCA | 1 | ENCFF093ZRU |
| 102 | 1 | 52730551 | 52730765 | 214 | AKR,BALB,BUB,C58J,CAST,MOLF,MRL,NOD,NOR,NZO,NZW,PL,RF,RH2,SEA,SMJ,SPRFT.SWR.WSB | 1 | 52730675 | 52730693 | 19 | - | 17.5574 | 6.32E-07 | 0.166 | ACTCCAGAAGAGGGCGCCA | | |
| 102 | 1 | 52730551 | 52730765 | 214 | AKR,BALB,BUB,C58J,CAST,MOLF,MRL,NOD,NOR,NZO,NZW,PL,RF,RH2,SEA,SMJ,SPRFT.SWR.WSB | 1 | 52730675 | 52730708 | 34 | - | 18.3659 | 2.76E-07 | 0.069 | TAGCTGTCTTCAGACACTCCAGAAGAGGGCGCCA | | |
| 103 | 1 | 55201091 | 55201868 | 777 | SMJ | 1 | 55201269 | 55201287 | 19 | - | 19.0656 | 2.23E-07 | 0.105 | AGTCCAGAAGAGGGCGCCA | 27 | ENCFF805QBU,ENCFF093ZRU,ENCFF089GBC,ENCFF165LXO,ENCFF367UNR,ENCFF092NMJ,ENCFF198SSR,ENCFF951GOP,ENCFF201JUA,ENCFF065ODV,ENCFF136ZDK,ENCFF099QNE,ENCFF321AGQ,ENCFF097CAQ,ENCFF589ZBJ,ENCFF142CNG,ENCFF052KVO,ENCFF055HAP,ENCFF194HVL,ENCFF821BIA,ENCFF957SPH,ENCFF533APC,ENCFF024JOZ,ENCFF327LYS,ENCFF502WDT,ENCFF907XEG,ENCFF883UPM |
| 103 | 1 | 55201091 | 55201868 | 777 | SMJ | 1 | 55201269 | 55201303 | 35 | - | 19.5872 | 1.13E-07 | 0.0634 | GTACAGGCCCCCAAAGAGTCCAGAAGAGGGCGCCA | 27 | ENCFF805QBU,ENCFF093ZRU,ENCFF089GBC,ENCFF165LXO,ENCFF367UNR,ENCFF092NMJ,ENCFF198SSR,ENCFF951GOP,ENCFF201JUA,ENCFF065ODV,ENCFF136ZDK,ENCFF099QNE,ENCFF321AGQ,ENCFF097CAQ,ENCFF589ZBJ,ENCFF142CNG,ENCFF052KVO,ENCFF055HAP,ENCFF194HVL,ENCFF821BIA,ENCFF957SPH,ENCFF533APC,ENCFF024JOZ,ENCFF327LYS,ENCFF502WDT,ENCFF907XEG,ENCFF883UPM |
| 104 | 1 | 58716964 | 58717166 | 202 | CE | 1 | 58716937 | 58716971 | 35 | - | 28.1835 | 1.64E-10 | 0.00426 | CTGCATGTACACCCACAGGCCAGAAGAGGGCACCA | 21 | ENCFF093ZRU,ENCFF367UNR,ENCFF136ZDK,ENCFF201JUA,ENCFF430PPJ,ENCFF099QNE,ENCFF327LYS,ENCFF821BIA,ENCFF907XEG,ENCFF024JOZ,ENCFF609MDN,ENCFF589ZBJ,ENCFF502WDT,ENCFF055HAP,ENCFF198SSR,ENCFF089GBC,ENCFF065ODV,ENCFF883UPM,ENCFF321AGQ,ENCFF092NMJ,ENCFF165LXO |
| 105 | 1 | 60206503 | 60206727 | 224 | CE.LP.MOLF | 1 | 60206627 | 60206660 | 34 | - | 16.4634 | 9.51E-07 | 0.125 | TAGCTGTCTTCAGACGCACCAGAAGAGGGCATCA | | |
| 106 | 1 | 64675932 | 64676136 | 204 | CAST.MOLF | 1 | 64675936 | 64675954 | 19 | - | 18.3279 | 3.75E-07 | 0.135 | AGTCCAGAAGAGGGCACCA | | |
| 106 | 1 | 64675932 | 64676136 | 204 | CAST.MOLF | 1 | 64675936 | 64675969 | 34 | - | 18.6098 | 2.34E-07 | 0.0647 | TAGCTGTCTTCAGACAGTCCAGAAGAGGGCACCA | | |
| 107 | 1 | 65274734 | 65275432 | 698 | CAST.MOLF | 1 | 65275271 | 65275289 | 19 | - | 16.9016 | 9.66E-07 | 0.197 | CAACCAGCAGATGACACTG | | |
| 107 | 1 | 65274734 | 65275432 | 698 | CAST.MOLF | 1 | 65275271 | 65275305 | 35 | - | 16.8165 | 6.02E-07 | 0.167 | ATGTACTTGAATATAACAACCAGCAGATGACACTG | | |
| 108 | 1 | 71509875 | 71510288 | 413 | SPRET | 1 | 71510244 | 71510277 | 34 | - | 25.8943 | 7.37E-10 | 0.00133 | GTGCAGTACCCACAGAGGCCAGAGGAGGGCATCA | 2 | ENCFF367UNR.ENCFF052KVO |
| 109 | 1 | 72126974 | 72127712 | 738 | CAST.MOLF.SPRET | 1 | 72127489 | 72127507 | 19 | - | 18.5902 | 3.13E-07 | 0.123 | CGGCCAGCAGATGTCAGCC | 1 | ENCFF093ZRU |
| 110 | 1 | 72485362 | 72485572 | 210 | AJ,AKR,BUB,C3H,C58J,CAST,MA/My,MRL,PJ,PL,RF,SJL,SPRET,SWR,TH | 1 | 72485488 | 72485521 | 34 | - | 16.7724 | 7.83E-07 | 0.115 | TAGCTGTCTTCAGATGCACCAGAAGAGGGCGTCA | | |
| 111 | 1 | 74225488 | 74225705 | 217 | AJ,C3H,CAST,CBA,MOLF,MRL,NOD,NZB,RH2,SMJ,SPRET | 1 | 74225611 | 74225644 | 34 | - | 17.3902 | 5.27E-07 | 0.095 | TAGCTGTCCTCAGACACTCCAGAAGAGGGCATCA | 2 | ENCFF367UNR,ENCFF099QNE |
| 112 | 1 | 74366091 | 74366332 | 241 | CAST | 1 | 74366281 | 74366299 | 19 | - | 17.5574 | 6.32E-07 | 0.166 | ACTCCAGAAGAGGGCGCCA | 13 | ENCFF201JUA,ENCFF327LYS,ENCFF821BIA,ENCFF055HAP,ENCFF136ZDK,ENCFF502WDT,ENCFF883UPM,ENCFF024JOZ,ENCFF907XEG,ENCFF589ZBJ,ENCFF367UNR,ENCFF093ZRU,ENCFF099QNE |
| 112 | 1 | 74366091 | 74366332 | 241 | CAST | 1 | 74366281 | 74366314 | 34 | - | 20.935 | 4.49E-08 | 0.0226 | CAGCTGTCTTCAGACACTCCAGAAGAGGGCGCCA | 13 | ENCFF201JUA,ENCFF327LYS,ENCFF821BIA,ENCFF055HAP,ENCFF136ZDK,ENCFF502WDT,ENCFF883UPM,ENCFF024JOZ,ENCFF907XEG,ENCFF589ZBJ,ENCFF367UNR,ENCFF093ZRU,ENCFF099QNE |
| 113 | 1 | 74722425 | 74722665 | 240 | 129S1,AJ,AKR,BTBR,BUB,C3H,C57L,CAST,CBA,DBA2J,ILNJ2,MA/My,MOLF,MRL,NOD,NZB,NZO,PJ,PL,RF,RH2,SEA,SJL,SMJ,SPRFT.SWR.WSB | 1 | 74722550 | 74722583 | 34 | - | 18.561 | 2.42E-07 | 0.0656 | CAGCTGTCTTCAGACACACCAGAAGAGGGCATCA | | |
| 114 | 1 | 74766765 | 74767597 | 832 | CAST | 1 | 74766940 | 74766973 | 34 | - | 17.2276 | 5.85E-07 | 0.0999 | TAGCTGTCTTCAGACACACCAGTAGAGGGCGTCA | | |
| 115 | 1 | 75089412 | 75089620 | 208 | SPRET | 1 | 75089533 | 75089566 | 34 | - | 17.8455 | 3.91E-07 | 0.0823 | TAGCTGTCCTCAGACACTCCAGAAGAGGGCGTCA | | |
| 116 | 1 | 84742731 | 84742926 | 195 | SPRET | 1 | 84742855 | 84742888 | 34 | - | 16.4634 | 9.51E-07 | 0.125 | TAGCTGTCTTCAGACGCACCAGAAGAGGGCATCA | | |
| 117 | 1 | 85275566 | 85275758 | 192 | SPRET | 1 | 85275729 | 85275762 | 34 | - | 19.0163 | 1.77E-07 | 0.0544 | GTGCAGTAACCCAGGAGACCAGGAGAGGACATTG | | |
| 118 | 1 | 86059707 | 86059902 | 195 | SPRET | 1 | 86059831 | 86059864 | 34 | - | 16.4472 | 9.61E-07 | 0.125 | TAGCTGTCTTCAGACACACCAGAAGAGGGCGTCA | 1 | ENCFF093ZRU |
| 119 | 1 | 86210401 | 86210615 | 214 | 129S1,AJ,BTBR,BUB,C3H,C57L,C58J,CAST,CE,DBA1J,DBA2J,FVB,ILNJ2,KK,LP,NZB,NZO,NZW,RF,RH2,SJL,SMJ,SPRET,SWR,TH,WSB | 1 | 86210526 | 86210560 | 35 | - | 18.1284 | 2.79E-07 | 0.109 | CTGTAGCTGTCTTCAGACACCAGAAGAGGGCATCA | | |
| 120 | 1 | 87008617 | 87009203 | 586 | SPRET | 1 | 87008641 | 87008674 | 34 | - | 17.4472 | 5.08E-07 | 0.0936 | CAGTTGTGACCTCTGCTGTCAGTAGATGGAGGCA | | |
| 121 | 1 | 90517599 | 90518062 | 463 | SPRET | 1 | 90517949 | 90517982 | 34 | - | 17.7886 | 4.06E-07 | 0.0843 | ATGCAGTGGCCATAGAGGACAGAAGAGGACATCA | | |
| 122 | 1 | 106538207 | 106538385 | 178 | SPRET | 1 | 106538232 | 106538266 | 35 | - | 19.4404 | 1.24E-07 | 0.0671 | TGGCAATCCGCATCTCTGCCCTCCAGGTGGCAGGC | | |
| 123 | 1 | 111824481 | 111824694 | 213 | 129S1,AJ,AKR,BALB,BTBR,BUB,C3H,C57L,CAST,CBA,CE,DBA1J,DBA2J,FVB,ILNJ2,KK,LP,MA/My,MOLF,MRL,NOD,NOR,NZB,NZO,NZW,PJ,PL,RF,RH2,SEA,SJL,SMJ,SPRET,SWR.TH | 1 | 111824594 | 111824627 | 34 | - | 16.4634 | 9.51E-07 | 0.125 | TAGCTGTCTTCAGACGCACCAGAAGAGGGCATCA | | |
| 124 | 1 | 118560315 | 118560510 | 195 | CAST.MOLF.SPRET | 1 | 118560441 | 118560475 | 35 | - | 16.844 | 5.92E-07 | 0.166 | TTGCAGCTGTCTTTAGACACCAGAAGAGGGTGTCA | | |
| 125 | 1 | 120627713 | 120627922 | 209 | SPRET | 1 | 120627837 | 120627870 | 34 | - | 16.6016 | 8.72E-07 | 0.122 | TAGCTGTCTTCAGAGACACCAGAAGAGGGCATCA | | |
| 126 | 1 | 121286254 | 121286973 | 719 | CAST.MOLF | 1 | 121286641 | 121286674 | 34 | - | 16.9106 | 7.17E-07 | 0.111 | CAGCAGGGGCGGGAGGGGCAGCAGGGGGGCAGCA | | |
| 127 | 1 | 131927366 | 131927574 | 208 | CAST.MOLF.SPRET | 1 | 131927479 | 131927512 | 34 | - | 16.4472 | 9.61E-07 | 0.125 | TAGCTGTCTTCAGACACACCAGAAGAGGGCGTCA | | |
| 128 | 1 | 132028835 | 132029546 | 711 | SPRET | 1 | 132028871 | 132028904 | 34 | - | 24.122 | 3.55E-09 | 0.00389 | ATGCAGTGTCTGCAGAGACCAGCAGAAGGCATCA | | |
| 129 | 1 | 132420846 | 132421043 | 197 | SPRET | 1 | 132420973 | 132421006 | 34 | - | 16.7724 | 7.83E-07 | 0.115 | TAGCTGTCTTCAGATGCACCAGAAGAGGGCGTCA | 2 | ENCFF093ZRU.ENCFF367UNR |
| 130 | 1 | 134436504 | 134436705 | 201 | 129S1,AJ,AKR,BTBR,BUB,C3H,CAST,CBA,CE,DBA1J,DBA2J,FVB,ILNJ2,KK,MA/My,MRL,NOD,NOR,NZB,NZO,PJ,PL,RF,RH2,SJL,SMJ.SPRET.SWR.TH | 1 | 134436621 | 134436655 | 35 | - | 15.9266 | 9.95E-07 | 0.213 | CTGTCTTCAGATGCACCAACCAGAAGAGGGCGTCA | | |
| 131 | 1 | 134531319 | 134531527 | 208 | SPRET | 1 | 134531443 | 134531476 | 34 | - | 16.4472 | 9.61E-07 | 0.125 | TAGCTGTCTTCAGACACACCAGAAGAGGGCGTCA | 3 | ENCFF367UNR.ENCFF093ZRU.ENCFF821BIA |
| 132 | 1 | 136987580 | 136988183 | 603 | CAST | 1 | 136988073 | 136988106 | 34 | - | 19.3984 | 1.36E-07 | 0.0461 | ATGCAACACCTGCAGAGGTCAGAAGAGGGCATCT | | |
| 133 | 1 | 152637974 | 152638158 | 184 | SPRET | 1 | 152638085 | 152638103 | 19 | - | 18.623 | 3.06E-07 | 0.122 | ACACCAGAAGAGGGCGCCA | 1 | ENCFF093ZRU |
| 133 | 1 | 152637974 | 152638158 | 184 | SPRET | 1 | 152638085 | 152638118 | 34 | - | 17.6179 | 4.54E-07 | 0.0884 | TGGCTGTCTTCAGACACACCAGAAGAGGGCGCCA | 1 | ENCFF093ZRU |
| 134 | 1 | 153277599 | 153277820 | 221 | BUB,C57L,CAST,DBA1J,DBA2J,FVB,NOD,NOR,RF,SJL,SPRET,SWR,TH,WSB | 1 | 153277728 | 153277762 | 35 | - | 19.5413 | 1.17E-07 | 0.0645 | GTGTAGCTCCTGCATCTGACCACACGGCGGCACTG | 39 | ENCFF367UNR,ENCFF093ZRU,ENCFF805QBU,ENCFF349BUL,ENCFF099QNE,ENCFF201JUA,ENCFF327LYS,ENCFF086ZTV,ENCFF024JOZ,ENCFF907XEG,ENCFF821BIA,ENCFF198SSR,ENCFF092NMJ,ENCFF089GBC,ENCFF589ZBJ,ENCFF165LXO,ENCFF321AGQ,ENCFF052KVO,ENCFF430PPJ,ENCFF136ZDK,ENCFF033UQX,ENCFF601JVE,ENCFF951GOP,ENCFF097CAQ,ENCFF957SPH,ENCFF491RJK,ENCFF281QMV,ENCFF065ODV,ENCFF194HVL,ENCFF609MDN,ENCFF575MGI,ENCFF504JDS,ENCFF210BTO,ENCFF883UPM,ENCFF142CNG,ENCFF055HAP,ENCFF533APC,ENCFF316FYN,ENCFF502WDT |
| 135 | 1 | 153326066 | 153326263 | 197 | SPRET | 1 | 153326188 | 153326221 | 34 | - | 17.2439 | 5.79E-07 | 0.0995 | TAGCTGTCCTCAGATACTCCAGAAGAGGGCATCA | | |
| 136 | 1 | 153741387 | 153741611 | 224 | BUB,C57L,CAST,DBA1J,DBA2J,FVB,MA/My,RF.SJL.SPRET.SWR.TH.WSB | 1 | 153741527 | 153741560 | 34 | - | 16.4634 | 9.51E-07 | 0.125 | TAGCTGTCTTCAGACGCACCAGAAGAGGGCATCA | | |
| 137 | 1 | 155957998 | 155958194 | 196 | SPRET | 1 | 155958120 | 155958153 | 34 | - | 17.3252 | 5.49E-07 | 0.0971 | TAGCTGTCCTCAGACATTCCAGAAGAGGGCATCA | | |
| 138 | 1 | 156540039 | 156540240 | 201 | AJ,AKR,BALB,C3H,C58J,CAST,CBA,CE,DBA1J,DBA2J,FVB,ILNJ2,KK,LP,MA/My,MOLF,PL.RF.RH2.SPRET.TH | 1 | 156540165 | 156540183 | 19 | - | 17.5574 | 6.32E-07 | 0.166 | ACTCCAGAAGAGGGCGCCA | | |
| 138 | 1 | 156540039 | 156540240 | 201 | AJ,AKR,BALB,C3H,C58J,CAST,CBA,CE,DBA1J,DBA2J,FVB,ILNJ2,KK,LP,MA/My,MOLF,PL.RF.RH2.SPRET.TH | 1 | 156540165 | 156540198 | 34 | - | 18.3659 | 2.76E-07 | 0.069 | TAGCTGTCTTCAGACACTCCAGAAGAGGGCGCCA | | |
| 139 | 1 | 156706649 | 156706855 | 206 | AJ,AKR,BALB,C3H,C57L,C58J,CAST,CBA,CE,DBA1J,DBA2J,FVB,ILNJ2,LP,MA/My,MOLF,MRL,NZB,NZO,PL,RF,RH2,SPRET,TH,WSB | 1 | 156706776 | 156706794 | 19 | - | 17.5738 | 6.25E-07 | 0.166 | ACTCCAGAAGAGGGCGCTA | 1 | ENCFF367UNR |
| 139 | 1 | 156706649 | 156706855 | 206 | AJ,AKR,BALB,C3H,C57L,C58J,CAST,CBA,CE,DBA1J,DBA2J,FVB,ILNJ2,LP,MA/My,MOLF,MRL,NZB,NZO,PL,RF,RH2,SPRET,TH,WSB | 1 | 156706776 | 156706809 | 34 | - | 18.1789 | 3.13E-07 | 0.0725 | TAGCTGTCTTCAGACACTCCAGAAGAGGGCGCTA | 1 | ENCFF367UNR |
| 140 | 1 | 160609796 | 160609993 | 197 | MOLF.SPRET | 1 | 160609922 | 160609955 | 34 | - | 17.1626 | 6.10E-07 | 0.102 | TAGCTGTCTTTGGATGCACCAGAAGAGGGCGTCA | | |
| 141 | 1 | 160749482 | 160749903 | 421 | SPRET | 1 | 160749702 | 160749720 | 19 | - | 17.0656 | 8.70E-07 | 0.189 | AGGCCAGAAGAGGGTGCTG | 18 | ENCFF099QNE,ENCFF198SSR,ENCFF201JUA,ENCFF907XEG,ENCFF367UNR,ENCFF883UPM,ENCFF055HAP,ENCFF052KVO,ENCFF093ZRU,ENCFF136ZDK,ENCFF957SPH,ENCFF097CAQ,ENCFF589ZBJ,ENCFF821BIA.ENCFF327LYS.ENCFF142CNG.ENCFF502WDT |
| 141 | 1 | 160749482 | 160749903 | 421 | SPRET | 1 | 160749702 | 160749736 | 35 | - | 20.2018 | 7.63E-08 | 0.0519 | CTGCTAGTGCATACTGAGGCCAGAAGAGGGTGCTG | 18 | ENCFF099QNE,ENCFF198SSR,ENCFF201JUA,ENCFF907XEG,ENCFF367UNR,ENCFF883UPM,ENCFF055HAP,ENCFF052KVO,ENCFF093ZRU,ENCFF136ZDK,ENCFF957SPH,ENCFF097CAQ,ENCFF589ZBJ,ENCFF821BIA.ENCFF327LYS.ENCFF142CNG.ENCFF502WDT |
| 142 | 1 | 161004065 | 161004141 | 76 | SPRET | 1 | 161004125 | 161004158 | 34 | - | 19.5447 | 1.23E-07 | 0.0433 | TTGCTGTCTTTAGACACACCAGAAGGGGGCGTCA | | |
| 143 | 1 | 163846831 | 163847035 | 204 | SPRET | 1 | 163846957 | 163846975 | 19 | - | 17.5574 | 6.32E-07 | 0.166 | ACTCCAGAAGAGGGCGCCA | 3 | ENCFF093ZRU.ENCFF367UNR.ENCFF099QNE |
| 143 | 1 | 163846831 | 163847035 | 204 | SPRET | 1 | 163846957 | 163846990 | 34 | - | 18.3659 | 2.76E-07 | 0.069 | TAGCTGTCTTCAGACACTCCAGAAGAGGGCGCCA | 3 | ENCFF093ZRU.ENCFF367UNR.ENCFF099QNE |
| 144 | 1 | 164024505 | 164024754 | 249 | CAST,DBA1J,DBA2J,ILNJ2,MOLF,SPRET | 1 | 164024631 | 164024664 | 34 | - | 16.9187 | 7.13E-07 | 0.111 | TAGCTGTCTTCAGACGCACCAGAAGAGGGCGTCA | | |
| 145 | 1 | 169984133 | 169984345 | 212 | 129S1,AJ,BTBR,BUB,C3H,C57L,CAST,CE,FVB,KK,MOLF,MRL,NOD,NOR,NZB,NZO,NZW,PJ,PL,RF,RH2,SEA,SJL,SMJ,SPRET,SWR.TH.WSB | 1 | 169984258 | 169984291 | 34 | - | 16.4472 | 9.61E-07 | 0.125 | TAGCTGTCTTCAGACACACCAGAAGAGGGCGTCA | | |
| 146 | 1 | 170970781 | 170971000 | 219 | 129S1,AJ,BUB,C3H,C57L,CBA,CE,DBA1J,DBA2J,FVB,KK,MOLF,MRL,NOD,NOR,NZB,NZO,NZW,PJ,PL,RH2,SEA,SJL,SMJ,SPRET,SWR.TH.WSB | 1 | 170970906 | 170970939 | 34 | - | 17.8455 | 3.91E-07 | 0.0823 | TAGCTGTCCTCAGACACTCCAGAAGAGGGCGTCA | 1 | ENCFF367UNR |
| 147 | 1 | 171092141 | 171092503 | 362 | 129S1,AJ,BALB,BUB,C3H,C57L,CBA,CE,DBA1J,DBA2J,FVB,KK,MRL,NOD,NOR,NZB,NZO,NZW,PJ,PL,RF,RH2,SEA,SJL,SMJ,SPRET.SWR.TH.WSB | 1 | 171092408 | 171092426 | 19 | - | 17.5574 | 6.32E-07 | 0.166 | ACTCCAGAAGAGGGCGCCA | 2 | ENCFF367UNR,ENCFF093ZRU |
| 147 | 1 | 171092141 | 171092503 | 362 | 129S1,AJ,BALB,BUB,C3H,C57L,CBA,CE,DBA1J,DBA2J,FVB,KK,MRL,NOD,NOR,NZB,NZO,NZW,PJ,PL,RF,RH2,SEA,SJL,SMJ,SPRET.SWR.TH.WSB | 1 | 171092408 | 171092441 | 34 | - | 18.2358 | 3.02E-07 | 0.0709 | TAGCTGTCTTAAGACACTCCAGAAGAGGGCGCCA | 2 | ENCFF367UNR,ENCFF093ZRU |
| 148 | 1 | 171191203 | 171191407 | 204 | SPRET | 1 | 171191221 | 171191255 | 35 | - | 18.3211 | 2.48E-07 | 0.104 | CTGTAGCTGTCTTCAGACACCAGAAGGTGGCATCA | | |
| 149 | 1 | 171261337 | 171261533 | 196 | SPRET | 1 | 171261441 | 171261474 | 34 | - | 16.6585 | 8.41E-07 | 0.119 | TAGCTGTCTTCAGACAGTCCAGAAGAGGGCGTCA | | |
| 150 | 1 | 171587537 | 171587801 | 264 | BUB.SWR | 1 | 171587529 | 171587563 | 35 | - | 15.9725 | 9.70E-07 | 0.212 | ATGCCATTCCTGGGCTTCTCCACAAGAAGGCCCAA | | |
| 151 | 1 | 171927823 | 171928048 | 225 | 129S1,AJ,AKR,BALB,BUB,C3H,C57L,CAST,CBA,CE,DBA1J,DBA2J,FVB,KK,MRL,NOD,NOR,NZB,NZO,NZW,PJ,PL,RF,RH2,SEA,SJL,SMJ.SPRET.SWR.TH.WSB | 1 | 171927948 | 171927982 | 35 | - | 16.1193 | 8.93E-07 | 0.205 | TAGCTGTCTTCAGACAGCACCAGAAGAGGGCATCG | | |
| 152 | 1 | 172436265 | 172436460 | 195 | MOLF.SPRET | 1 | 172436385 | 172436403 | 19 | - | 17.2459 | 7.74E-07 | 0.182 | ACACCAGAAGAGGGCAGCA | | |
| 153 | 1 | 179333294 | 179333530 | 236 | AKR.CAST.MOLF.SPRET | 1 | 179333418 | 179333451 | 34 | - | 17.2439 | 5.79E-07 | 0.0995 | TAGCTGTCCTCAGATACTCCAGAAGAGGGCATCA | | |
| 154 | 1 | 179528628 | 179528828 | 200 | AJ,AKR,BUB,C58J,CAST,CE,FVB,LP,MOLF,NZW,PJ,RF,RH2,SEA,SJL,SPRET,SWR,TH,WSB | 1 | 179528753 | 179528786 | 34 | - | 16.4472 | 9.61E-07 | 0.125 | TAGCTGTCTTCAGACACACCAGAAGAGGGCGTCA | | |
| 155 | 1 | 180378976 | 180379185 | 209 | CAST.MOLF.SPRET.TH | 1 | 180379110 | 180379143 | 34 | - | 17.8455 | 3.91E-07 | 0.0823 | TAGCTGTCCTCAGACACTCCAGAAGAGGGCGTCA | | |
| 156 | 1 | 180420708 | 180420917 | 209 | CAST,CBA,CE,DBA1J,DBA2J,MOLF,PJ,SEA.SPRET.TH | 1 | 180420832 | 180420865 | 34 | - | 17.1463 | 6.17E-07 | 0.103 | TAGCTGTCCTCAGACACTCCAGAAGAAGGCGCCA | | |
| 157 | 1 | 180893692 | 180893901 | 209 | CAST.MOLF.SPRET.TH | 1 | 180893827 | 180893860 | 34 | - | 16.6341 | 8.54E-07 | 0.12 | CTGTAGCTGTCAGACACTCCAGAAGAGGGAGTCA | | |
| 158 | 1 | 181834956 | 181835153 | 197 | MOLF.SPRET.TH | 1 | 181835082 | 181835115 | 34 | - | 16.4472 | 9.61E-07 | 0.125 | TAGCTGTCTTCAGACACACCAGAAGAGGGCGTCA | | |
| 159 | 1 | 184945846 | 184946111 | 265 | AKR,BTBR,C57L,CAST,ILNJ2,KK,MOLF,MRL.NZW.PL.SPRET | 1 | 184945970 | 184946003 | 34 | - | 17.6667 | 4.40E-07 | 0.0868 | TAGCTGTCTTCAGACACGCCAGAAGAGGGCGTCA | | |
| 160 | 1 | 189672692 | 189672905 | 213 | CAST.MOLF.SPRET | 1 | 189672799 | 189672832 | 34 | - | 16.4472 | 9.61E-07 | 0.125 | TAGCTGTCTTCAGACACACCAGAAGAGGGCGTCA | | |
| 161 | 1 | 190583278 | 190584105 | 827 | SPRET | 1 | 190583283 | 190583317 | 35 | - | 20.3394 | 6.98E-08 | 0.0494 | GTGCACTAGGGGGCTGTGTGCACTAGGGGGCCGTG | | |
| 161 | 1 | 190583278 | 190584105 | 827 | SPRET | 1 | 190583569 | 190583603 | 35 | - | 20.5413 | 6.11E-08 | 0.046 | GTGCACTAGGGGGCTGTGTGCACTAGGGGGCTGTG | | |
| 161 | 1 | 190583278 | 190584105 | 827 | SPRET | 1 | 190583744 | 190583778 | 35 | - | 17.3761 | 4.35E-07 | 0.137 | GTGCACGAGGGGGCTGTGTGCACTAGGGGGCTGTG | | |
| 161 | 1 | 190583278 | 190584105 | 827 | SPRET | 1 | 190583776 | 190583810 | 35 | - | 17.1743 | 4.90E-07 | 0.148 | GTGCACGAGGGGGCTGTGTGCACTAGGGGGCCGTG | | |
| 161 | 1 | 190583278 | 190584105 | 827 | SPRET | 1 | 190583823 | 190583857 | 35 | - | 18.2569 | 2.58E-07 | 0.106 | GTGCACTAGGGGGCTGTGTGCACGAGGGGGCTGTG | | |
| 161 | 1 | 190583278 | 190584105 | 827 | SPRET | 1 | 190583887 | 190583921 | 35 | - | 18.2569 | 2.58E-07 | 0.106 | GTGCACTAGGGGGCTGTGTGCACGAGGGGGCTGTG | | |
| 161 | 1 | 190583278 | 190584105 | 827 | SPRET | 1 | 190583936 | 190583970 | 35 | - | 18.2569 | 2.58E-07 | 0.106 | GTGCACTAGGGGGCTGTGTGCACGAGGGGGCTGTG | | |
| 161 | 1 | 190583278 | 190584105 | 827 | SPRET | 1 | 190583983 | 190584017 | 35 | - | 20.5413 | 6.11E-08 | 0.046 | GTGCACTAGGGGGCTGTGTGCACTAGGGGGCTGTG | | |
| 161 | 1 | 190583278 | 190584105 | 827 | SPRET | 1 | 190584015 | 190584049 | 35 | - | 18.5138 | 2.21E-07 | 0.0962 | GTGCACTAGGGGGCCGTGTGCACGAGGGGGCTGTG | | |
| 161 | 1 | 190583278 | 190584105 | 827 | SPRET | 1 | 190584079 | 190584112 | 34 | - | 19.8049 | 1.02E-07 | 0.0389 | GTGCACTGGGGGCTGTGTGCACAAGGGGGCTGTG | | |
| 162 | 1 | 190583486 | 190583994 | 508 | CAST,ILNJ2,MOLF,SMJ,SPRET,SWR,WSB | 1 | 190583569 | 190583603 | 35 | - | 20.5413 | 6.11E-08 | 0.046 | GTGCACTAGGGGGCTGTGTGCACTAGGGGGCTGTG | | |
| 162 | 1 | 190583486 | 190583994 | 508 | CAST,ILNJ2,MOLF,SMJ,SPRET,SWR,WSB | 1 | 190583744 | 190583778 | 35 | - | 17.3761 | 4.35E-07 | 0.137 | GTGCACGAGGGGGCTGTGTGCACTAGGGGGCTGTG | | |
| 162 | 1 | 190583486 | 190583994 | 508 | CAST,ILNJ2,MOLF,SMJ,SPRET,SWR,WSB | 1 | 190583776 | 190583810 | 35 | - | 17.1743 | 4.90E-07 | 0.148 | GTGCACGAGGGGGCTGTGTGCACTAGGGGGCCGTG | | |
| 162 | 1 | 190583486 | 190583994 | 508 | CAST,ILNJ2,MOLF,SMJ,SPRET,SWR,WSB | 1 | 190583823 | 190583857 | 35 | - | 18.2569 | 2.58E-07 | 0.106 | GTGCACTAGGGGGCTGTGTGCACGAGGGGGCTGTG | | |
| 162 | 1 | 190583486 | 190583994 | 508 | CAST,ILNJ2,MOLF,SMJ,SPRET,SWR,WSB | 1 | 190583887 | 190583921 | 35 | - | 18.2569 | 2.58E-07 | 0.106 | GTGCACTAGGGGGCTGTGTGCACGAGGGGGCTGTG | | |
| 162 | 1 | 190583486 | 190583994 | 508 | CAST,ILNJ2,MOLF,SMJ,SPRET,SWR,WSB | 1 | 190583936 | 190583970 | 35 | - | 18.2569 | 2.58E-07 | 0.106 | GTGCACTAGGGGGCTGTGTGCACGAGGGGGCTGTG | | |
| 162 | 1 | 190583486 | 190583994 | 508 | CAST,ILNJ2,MOLF,SMJ,SPRET,SWR,WSB | 1 | 190583983 | 190584017 | 35 | - | 20.5413 | 6.11E-08 | 0.046 | GTGCACTAGGGGGCTGTGTGCACTAGGGGGCTGTG | | |
| 163 | 1 | 191257515 | 191257676 | 161 | SPRET | 1 | 191257603 | 191257636 | 34 | - | 16.626 | 8.59E-07 | 0.121 | TAGCTATCTTCAGACACTCCAGAAGAGGGCACCA | | |
| 164 | 1 | 192936425 | 192936652 | 227 | SPRET | 1 | 192936533 | 192936567 | 35 | - | 16.0917 | 9.07E-07 | 0.206 | TAGCTGTCTTCAGACAGCACCAGAAGAGGGCATCA | 1 | ENCFF093ZRU |
| 165 | 2 | 4286762 | 4286962 | 200 | 129S1,C57L,CAST,FVB,MA/My,SMJ,SPRET | 2 | 4286820 | 4286853 | 34 | + | 16.4472 | 9.61E-07 | 0.0987 | TAGCTGTCTTCAGACACACCAGAAGAGGGCGTCA | | |
| 166 | 2 | 4742950 | 4743166 | 216 | CAST.SPRET | 2 | 4743024 | 4743057 | 34 | + | 17.8455 | 3.91E-07 | 0.0641 | TAGCTGTCCTCAGACACTCCAGAAGAGGGCGTCA | | |
| 167 | 2 | 4787811 | 4787919 | 108 | CAST | 2 | 4787803 | 4787837 | 35 | + | 16.0275 | 9.40E-07 | 0.177 | GTGCACGGCGCAGTGATGTCCTGTAGAAGTCAGTG | | |
| 168 | 2 | 6605267 | 6605491 | 224 | SPRET | 2 | 6605340 | 6605373 | 34 | + | 16.4634 | 9.51E-07 | 0.0987 | TAGCTGTCTTCAGACGCACCAGAAGAGGGCATCA | | |
| 169 | 2 | 20711335 | 20711531 | 196 | CAST.CE.FVB.SJL.SPRET.WSB | 2 | 20711388 | 20711421 | 34 | + | 17.2439 | 5.79E-07 | 0.0793 | TAGCTGTCCTCAGATACTCCAGAAGAGGGCATCA | | |
| 170 | 2 | 22814131 | 22814358 | 227 | CAST.FVB.SJL.SPRET.WSB | 2 | 22814214 | 22814247 | 34 | + | 16.4634 | 9.51E-07 | 0.0987 | TAGCTGTCTTCAGACGCACCAGAAGAGGGCATCA | | |
| 171 | 2 | 22816144 | 22816395 | 251 | CAST.FVB.SJL.SPRET.WSB | 2 | 22816266 | 22816284 | 19 | + | 17.8852 | 5.08E-07 | 0.115 | ACACCAGAAGAGGGCACCA | | |
| 172 | 2 | 22865667 | 22865875 | 208 | CAST.SPRET | 2 | 22865733 | 22865766 | 34 | + | 17.878 | 3.83E-07 | 0.0635 | TAGCTGTCCTCAGACACACCAGAAGAGGGCATCA | 1 | ENCFF093ZRU |
| 173 | 2 | 29266603 | 29266825 | 222 | CAST.MOLF.SPRET | 2 | 29266682 | 29266715 | 34 | + | 16.4634 | 9.51E-07 | 0.0987 | TAGCTGTCTTCAGACGCACCAGAAGAGGGCATCA | | |
| 174 | 2 | 29487710 | 29487913 | 203 | CAST.MOLF.SPRET | 2 | 29487770 | 29487803 | 34 | + | 18.3659 | 2.76E-07 | 0.0538 | TAGCTGTCTTCAGACACTCCAGAAGAGGGCGCCA | | |
| 174 | 2 | 29487710 | 29487913 | 203 | CAST.MOLF.SPRET | 2 | 29487785 | 29487803 | 19 | + | 17.5574 | 6.32E-07 | 0.124 | ACTCCAGAAGAGGGCGCCA | | |
| 175 | 2 | 29572791 | 29573025 | 234 | 129S1,C57L,CAST,MOLF,RF,SPRET,SWR,WSB | 2 | 29572885 | 29572918 | 34 | + | 16.4634 | 9.51E-07 | 0.0987 | TAGCTGTCTTCAGACGCACCAGAAGAGGGCATCA | | |
| 176 | 2 | 29598902 | 29599113 | 211 | 129S1,C57L,CAST,MOLF,RF,SPRET,SWR,WSB | 2 | 29598987 | 29599005 | 19 | + | 17.8852 | 5.08E-07 | 0.115 | ACACCAGAAGAGGGCACCA | | |
| 177 | 2 | 29724135 | 29724359 | 224 | MOLF.SPRET.SWR.WSB | 2 | 29724217 | 29724250 | 34 | + | 16.4634 | 9.51E-07 | 0.0987 | TAGCTGTCTTCAGACGCACCAGAAGAGGGCATCA | | |
| 178 | 2 | 30121845 | 30122051 | 206 | SPRET | 2 | 30121907 | 30121940 | 34 | + | 16.7724 | 7.83E-07 | 0.0917 | TAGCTGTCTTCAGATGCACCAGAAGAGGGCGTCA | | |
| 179 | 2 | 30834389 | 30834605 | 216 | CAST | 2 | 30834464 | 30834497 | 34 | + | 17.2439 | 5.79E-07 | 0.0793 | TAGCTGTCCTCAGATACTCCAGAAGAGGGCATCA | | |
| 180 | 2 | 31623886 | 31624096 | 210 | MOLF.NZO.SPRET.SWR | 2 | 31623953 | 31623986 | 34 | + | 17.8455 | 3.91E-07 | 0.0641 | TAGCTGTCCTCAGACACTCCAGAAGAGGGCGTCA | 1 | ENCFF093ZRU |
| 181 | 2 | 32015895 | 32016107 | 212 | CAST.MA/My.MOLF.SPRET.SWR | 2 | 32015967 | 32016000 | 34 | + | 17.2439 | 5.79E-07 | 0.0793 | TAGCTGTCCTCAGATACTCCAGAAGAGGGCATCA | | |
| 182 | 2 | 32300060 | 32300229 | 169 | MOLF.NZO.SPRET.SWR | 2 | 32300179 | 32300212 | 34 | + | 16.4472 | 9.61E-07 | 0.0987 | TAGCTGTCTTCAGACACACCAGAAGAGGGCGTCA | 1 | ENCFF367UNR |
| 183 | 2 | 34721317 | 34721529 | 212 | CAST.NZO.SPRET | 2 | 34721377 | 34721410 | 34 | + | 16.4634 | 9.51E-07 | 0.0987 | TAGCTGTCTTCAGACGCACCAGAAGAGGGCATCA | | |
| 184 | 2 | 38255178 | 38255389 | 211 | CAST.SPRET | 2 | 38255245 | 38255278 | 34 | + | 16.4472 | 9.61E-07 | 0.0987 | TAGCTGTCTTCAGACACACCAGAAGAGGGCGTCA | | |
| 185 | 2 | 38343290 | 38343494 | 204 | SPRET | 2 | 38343350 | 38343383 | 34 | + | 16.9187 | 7.13E-07 | 0.0879 | TAGCTGTCTTCAGACGCACCAGAAGAGGGCGTCA | 2 | ENCFF093ZRU.ENCFF367UNR |
| 186 | 2 | 38777992 | 38778193 | 201 | CAST.MOLF.SPRET | 2 | 38778051 | 38778084 | 34 | + | 18.3659 | 2.76E-07 | 0.0538 | TAGCTGTCTTCAGACACTCCAGAAGAGGGCGCCA | 1 | ENCFF093ZRU |
| 186 | 2 | 38777992 | 38778193 | 201 | CAST.MOLF.SPRET | 2 | 38778066 | 38778084 | 19 | + | 17.5574 | 6.32E-07 | 0.124 | ACTCCAGAAGAGGGCGCCA | 1 | ENCFF093ZRU |
| 187 | 2 | 39664924 | 39665237 | 313 | SMJ.SPRET | 2 | 39664940 | 39664973 | 34 | + | 16.4878 | 9.37E-07 | 0.0987 | CTGCCCTCACAGTCGCCGTTACAAGATGGCGCTG | | |
| 188 | 2 | 43377366 | 43377593 | 227 | SMJ.SPRET.TH | 2 | 43377450 | 43377484 | 35 | + | 19.6422 | 1.09E-07 | 0.0511 | CTGTAGCTATCCTCAGACACCAGAAGAGGGCATCA | | |
| 189 | 2 | 44718289 | 44718512 | 223 | 129S1,AJ,AKR,BTBR,BUB,C3H,C58J,CAST,CBA,CE,DBA1J,DBA2J,FVB,ILNJ2,KK,MA/My,MOLF,MRL,NOD,NOR,NZB,NZO,NZW,PJ,PL,RF,SEA,SJL,SMJ,SPRET,SWR,TH,WSB | 2 | 44718368 | 44718401 | 34 | + | 17.6667 | 4.40E-07 | 0.0682 | TAGCTGTCTTCAGACACGCCAGAAGAGGGCGTCA | | |
| 190 | 2 | 45793996 | 45794198 | 202 | CE.KK.NZB.SMJ.SPRET.TH | 2 | 45794056 | 45794089 | 34 | + | 16.4472 | 9.61E-07 | 0.0987 | TAGCTGTCTTCAGACACACCAGAAGAGGGCGTCA | | |
| 191 | 2 | 49907443 | 49907655 | 212 | 129S1,CAST,FVB,KK,MOLF,SJL,SPRET,TH,WSB | 2 | 49907512 | 49907545 | 34 | + | 18.3659 | 2.76E-07 | 0.0538 | TAGCTGTCTTCAGACACTCCAGAAGAGGGCGCCA | | |
| 191 | 2 | 49907443 | 49907655 | 212 | 129S1,CAST,FVB,KK,MOLF,SJL,SPRET,TH,WSB | 2 | 49907527 | 49907545 | 19 | + | 17.5574 | 6.32E-07 | 0.124 | ACTCCAGAAGAGGGCGCCA | | |
| 192 | 2 | 55195512 | 55195714 | 202 | CAST.MOLF.SPRET | 2 | 55195573 | 55195606 | 34 | + | 17.8455 | 3.91E-07 | 0.0641 | TAGCTGTCCTCAGACACTCCAGAAGAGGGCGTCA | | |
| 193 | 2 | 59532682 | 59532893 | 211 | 129S1,BTBR,C57L,CAST,DBA1J,MOLF,SPRET.TH | 2 | 59532750 | 59532783 | 34 | + | 18.3659 | 2.76E-07 | 0.0538 | TAGCTGTCTTCAGACACTCCAGAAGAGGGCGCCA | 2 | ENCFF367UNR,ENCFF093ZRU |
| 193 | 2 | 59532682 | 59532893 | 211 | 129S1,BTBR,C57L,CAST,DBA1J,MOLF,SPRET.TH | 2 | 59532765 | 59532783 | 19 | + | 17.5574 | 6.32E-07 | 0.124 | ACTCCAGAAGAGGGCGCCA | 2 | ENCFF367UNR,ENCFF093ZRU |
| 194 | 2 | 59847380 | 59847580 | 200 | SPRET | 2 | 59847438 | 59847471 | 34 | + | 16.4472 | 9.61E-07 | 0.0987 | TAGCTGTCTTCAGACACACCAGAAGAGGGCGTCA | | |
| 195 | 2 | 69737990 | 69738203 | 213 | AJ,AKR,BALB,BTBR,BUB,C3H,C57L,C58J,CAST,CBA,CE,DBA1J,DBA2J,FVB,ILNJ2,KK,MA/My,MOLF,MRL,NOD,NOR,NZB,NZO,NZW,PJ,PL,RF,RH2,SEA,SJL,SMJ,SPRET,SWR.TH.WSB | 2 | 69738060 | 69738093 | 34 | + | 18.3659 | 2.76E-07 | 0.0538 | TAGCTGTCTTCAGACACTCCAGAAGAGGGCGCCA | | |
| 195 | 2 | 69737990 | 69738203 | 213 | AJ,AKR,BALB,BTBR,BUB,C3H,C57L,C58J,CAST,CBA,CE,DBA1J,DBA2J,FVB,ILNJ2,KK,MA/My,MOLF,MRL,NOD,NOR,NZB,NZO,NZW,PJ,PL,RF,RH2,SEA,SJL,SMJ,SPRET,SWR.TH.WSB | 2 | 69738075 | 69738093 | 19 | + | 17.5574 | 6.32E-07 | 0.124 | ACTCCAGAAGAGGGCGCCA | | |

| | | | | | | | | | | | | | | | | |
|---|---|---|---|---|---|---|---|---|---|---|---|---|---|---|---|---|
| 196 | 2 | 69751933 | 69752154 | 221 | AJ,AKR,BALB,BTBR,BUB,C3H,C57L,C58J,CAST,CBA,CE,DBA1J,DBA2J,FVB,ILNJ2,KK,MA/My,MOLF,MRL,NOD,NOR,NZB,NZO,NZW,PJ,PL,RF,RH2,SEA,SJL,SMJ,SPRET,SWR.TH.WSB | 2 | 69752026 | 69752044 | 19 | + | 17.8852 | 5.08E-07 | 0.115 | ACACCAGAAGAGGGCACCA | | |
| 197 | 2 | 71261769 | 71262032 | 263 | AJ,AKR,BUB,C3H,CAST,FVB,ILNJ2,MA/My,MOLF,MRL,NOD,NOR,NZW,PL,RH2,SMJ,SPRET | 2 | 71261890 | 71261923 | 34 | + | 18.3659 | 2.76E-07 | 0.0538 | TAGCTGTCTTCAGACACTCCAGAAGAGGGCGCCA | | |
| 197 | 2 | 71261769 | 71262032 | 263 | AJ,AKR,BUB,C3H,CAST,FVB,ILNJ2,MA/My,MOLF,MRL,NOD,NOR,NZW,PL,RH2,SMJ,SPRET | 2 | 71261905 | 71261923 | 19 | + | 17.5574 | 6.32E-07 | 0.124 | ACTCCAGAAGAGGGCGCCA | | |
| 198 | 2 | 71682700 | 71683032 | 332 | 129S1,AJ,AKR,BALB,BTBR,BUB,C57L,CAST,DBA1J,FVB,ILNJ2,MA/My,MOLF,MRL,NOD,NOR,PL,RH2,SEA,SPRET | 2 | 71682730 | 71682748 | 19 | + | 19.2951 | 1.88E-07 | 0.071 | CCGCCACTAGATGGTGCTA | 18 | ENCFF367UNR,ENCFF093ZRU,ENCFF201JUA,ENCFF099QNE,ENCFF502WDT,ENCFF821BIA,ENCFF907XEG,ENCFF609MDN,ENCFF136ZDK,ENCFF327LYS,ENCFF198SSR,ENCFF024JOZ,ENCFF089GBC,ENCFF589ZBJ,ENCFF430PPJ,ENCFF092NMJ,ENCFF165LXO |
| 199 | 2 | 71736187 | 71736588 | 401 | AJ,AKR,BALB,BTBR,BUB,C57L,CAST,DBA1J,FVB,ILNJ2,MA/My,MOLF,MRL,NOD,NOR,PL,RH2,SEA | 2 | 71736531 | 71736565 | 35 | + | 24.9541 | 2.55E-09 | 0.00618 | CTGCATGTATGCCTGCACACCAGAAGAGGGCACCA | 2 | ENCFF093ZRU,ENCFF099QNE |
| 199 | 2 | 71736187 | 71736588 | 401 | AJ,AKR,BALB,BTBR,BUB,C57L,CAST,DBA1J,FVB,ILNJ2,MA/My,MOLF,MRL,NOD,NOR,PL,RH2,SEA | 2 | 71736547 | 71736565 | 19 | + | 17.8852 | 5.08E-07 | 0.115 | ACACCAGAAGAGGGCACCA | 2 | ENCFF093ZRU,ENCFF099QNE |
| 200 | 2 | 74069403 | 74069619 | 216 | SPRET | 2 | 74069476 | 74069509 | 34 | + | 16.4472 | 9.61E-07 | 0.0987 | TAGCTGTCTTCAGACACACCAGAAGAGGGCGTCA | | |
| 201 | 2 | 75873206 | 75874050 | 844 | SPRET | 2 | 75873897 | 75873930 | 34 | + | 17.8618 | 3.87E-07 | 0.0639 | TAGCTGTCTTCAGACCCACCAGAAGAGGGCATCA | 1 | ENCFF093ZRU |
| 202 | 2 | 84411853 | 84412062 | 209 | AJ,AKR,BALB,C3H,C57L,C58J,CAST,CBA,CE,DBA1J,DBA2J,FVB,ILNJ2,MA/My,MOLF,MRL,NZW,PJ,RF,RH2,SEA,SJL,SMJ,SPRET,SWR,WSB | 2 | 84411920 | 84411953 | 34 | + | 18.3659 | 2.76E-07 | 0.0538 | TAGCTGTCTTCAGACACTCCAGAAGAGGGCGCCA | | |
| 202 | 2 | 84411853 | 84412062 | 209 | AJ,AKR,BALB,C3H,C57L,C58J,CAST,CBA,CE,DBA1J,DBA2J,FVB,ILNJ2,MA/My,MOLF,MRL,NZW,PJ,RF,RH2,SEA,SJL,SMJ,SPRET,SWR,WSB | 2 | 84411935 | 84411953 | 19 | + | 17.5574 | 6.32E-07 | 0.124 | ACTCCAGAAGAGGGCGCCA | | |
| 203 | 2 | 84514518 | 84514694 | 176 | SPRET | 2 | 84514575 | 84514608 | 34 | + | 16.4472 | 9.61E-07 | 0.0987 | TAGCTGTCTTCAGACACACCAGAAGAGGGCGTCA | | |
| 204 | 2 | 85111833 | 85112027 | 194 | MOLF.SPRET | 2 | 85111872 | 85111905 | 34 | + | 17.9106 | 3.74E-07 | 0.063 | TAGCTGTCTTCAGACACTCCAGAAGAGGGCACCA | 1 | ENCFF367UNR |
| 205 | 2 | 90973599 | 90973806 | 207 | SPRET | 2 | 90973665 | 90973698 | 34 | + | 16.7724 | 7.83E-07 | 0.0917 | TAGCTGTCTTCAGATGCACCAGAAGAGGGCGTCA | 2 | ENCFF093ZRU.ENCFF367UNR |
| 206 | 2 | 91200289 | 91200503 | 214 | AJ,BALB,BUB,C57L,C58J,CAST,DBA1J,DBA2J,ILNJ2,KK,MA/My,MOLF,NOD,NOR,NZB,NZO,NZW,PJ,PL,RH2,SEA,SJL,SMJ,SPRET,SWR.WSB | 2 | 91200361 | 91200394 | 34 | + | 17.0081 | 6.74E-07 | 0.0859 | TAGCTGTCCCCAGACACTCCAGAAGAGGGAGTCA | | |
| 207 | 2 | 91578905 | 91579117 | 212 | AJ,BALB,C57L,C58J,CAST,DBA1J,DBA2J,FVB,ILNJ2,KK,MA/My,MOLF,NOD,NOR,NZW,PJ,RH2,SEA,SJL,SMJ,SPRET,SWR,WSB | 2 | 91578975 | 91579009 | 35 | + | 18.1284 | 2.79E-07 | 0.0894 | CTGTAGCTGTCTTCAGACACCAGAAGAGGGCATCA | | |
| 208 | 2 | 93065566 | 93066037 | 471 | SPRET | 2 | 93065765 | 93065798 | 34 | + | 17.0813 | 6.43E-07 | 0.0837 | CTGCAGTCACCAGTGCAGTCTCTAGGTGTGAGCC | | |
| 209 | 2 | 97866502 | 97866614 | 112 | CAST.SPRET | 2 | 97866500 | 97866533 | 34 | + | 17.0244 | 6.67E-07 | 0.0854 | GAGCAGGTGTGGCAGCAGTCCCAAGATGGCGCCC | | |
| 210 | 2 | 104858874 | 104859088 | 214 | 129S1,AJ,AKR,BALB,BTBR,BUB,C3H,C57L,CAST,CBA,CE,DBA1J,DBA2J,FVB,ILNJ2,KK,MOLF,MRL,NOD,NOR,NZB,NZW,PJ,PL,RH2,SEA,SJL,SMJ,SPRET,SWR,WSB | 2 | 104858947 | 104858980 | 34 | + | 17.8455 | 3.91E-07 | 0.0641 | TAGCTGTCCTCAGACACTCCAGAAGAGGGCGTCA | | |
| 211 | 2 | 107698529 | 107698739 | 210 | AJ,BALB,C3H,CAST,CBA,CE,DBA1J,DBA2J,FVB,ILNJ2,MA/My,MOLF,RF,RH2,SPRET | 2 | 107698596 | 107698629 | 34 | + | 19.7236 | 1.08E-07 | 0.0317 | CTGTAGTCTTCAGATACACCAGAAGAGGGCATCA | | |
| 212 | 2 | 112278346 | 112278534 | 188 | SPRET | 2 | 112278392 | 112278425 | 34 | + | 17.2114 | 5.91E-07 | 0.0799 | TAGCTGTTTTCAGACACTCCAGAAGAGGGCATCA | | |
| 213 | 2 | 116007258 | 116007448 | 190 | SPRET | 2 | 116007299 | 116007332 | 34 | + | 17.2114 | 5.91E-07 | 0.0799 | TAGCTGTTTTCAGACACTCCAGAAGAGGGCATCA | | |
| 214 | 2 | 116731417 | 116731871 | 454 | 129S1,BTBR,BUB,C3H,C57L,C58J,CAST,CBA,MOLF,NOD,NOR,PJ,SEA,SPRET | 2 | 116731729 | 116731762 | 34 | + | 16.4472 | 9.61E-07 | 0.0987 | TAGCTGTCTTCAGACACACCAGAAGAGGGCGTCA | | |
| 215 | 2 | 120481519 | 120482365 | 846 | AJ,BALB,DBA1J,DBA2J,ILNJ2,PJ,PL,RF,RH2,SEA,SMJ,TH | 2 | 120481605 | 120481638 | 34 | + | 18.5854 | 2.38E-07 | 0.0516 | TTGCTGTCTTCAGACACACCAGAAGAGGGCATCG | | |
| 216 | 2 | 121569503 | 121569757 | 254 | 129S1,AJ,AKR,BALB,BTBR,BUB,C3H,C57L,C58J,CAST,CBA,CE,DBA1J,DBA2J,FVB,ILNJ2,KK,MA/My,NOD,NZB,NZO,NZW,PJ,PL,RF,RH2,SEA,SJL,SMJ,SPRET,SWR,TH,WSB | 2 | 121569618 | 121569651 | 34 | + | 16.4634 | 9.51E-07 | 0.0987 | TAGCTGTCTTCAGACGCACCAGAAGAGGGCATCA | | |
| 217 | 2 | 125679525 | 125679736 | 211 | MOLF.SPRET | 2 | 125679577 | 125679610 | 34 | + | 16.6179 | 8.63E-07 | 0.0972 | CTGTAGCTGTCTTCAGACACACCAGAGGGCATCA | | |
| 218 | 2 | 126536974 | 126537174 | 200 | SPRET | 2 | 126537020 | 126537053 | 34 | + | 18.1545 | 3.18E-07 | 0.0576 | TAGCTGTTTTCAGACACACCAGAAGAGGGCGTCA | | |
| 219 | 2 | 126840760 | 126840964 | 204 | AJ,BALB,C3H,CAST,CBA,DBA2J,ILNJ2,KK,MOLF,PJ,RH2,SEA,SMJ,SPRET,WSB | 2 | 126840821 | 126840854 | 34 | + | 16.4472 | 9.61E-07 | 0.0987 | TAGCTGTCTTCAGACACACCAGAAGAGGGCGTCA | | |
| 220 | 2 | 126866298 | 126866519 | 221 | AJ,BALB,C3H,CAST,CBA,DBA2J,ILNJ2,KK,MOLF,PJ,RH2,SEA,SMJ,SPRET,WSB | 2 | 126866378 | 126866411 | 34 | + | 16.4634 | 9.51E-07 | 0.0987 | TAGCTGTCTTCAGACGCACCAGAAGAGGGCATCA | | |
| 221 | 2 | 127765871 | 127766059 | 188 | C3H,CBA,CE,DBA1J,DBA2J,ILNJ2,MOLF,PJ,PL.SPRET | 2 | 127765917 | 127765950 | 34 | + | 18.187 | 3.12E-07 | 0.0569 | TAGCTGTCTTCAGTCACTCCAGAAGAGGGCGCTA | | |
| 221 | 2 | 127765871 | 127766059 | 188 | C3H,CBA,CE,DBA1J,DBA2J,ILNJ2,MOLF,PJ,PL.SPRET | 2 | 127765932 | 127765950 | 19 | + | 17.5738 | 6.25E-07 | 0.124 | ACTCCAGAAGAGGGCGCTA | | |
| 222 | 2 | 128665230 | 128665435 | 205 | SPRET | 2 | 128665284 | 128665317 | 34 | + | 16.4472 | 9.61E-07 | 0.0987 | TAGCTGTCTTCAGACACACCAGAAGAGGGCGTCA | 1 | ENCFF093ZRU |
| 223 | 2 | 131971640 | 131971846 | 206 | SPRET | 2 | 131971705 | 131971738 | 34 | + | 17.6992 | 4.30E-07 | 0.0675 | TAGCTGTTTTCAGACACACCAGAAGAGGGCATCA | | |
| 224 | 2 | 140304111 | 140304532 | 421 | C58J,CAST,DBA1J,DBA2J,MOLF,NZO,NZW,SPRET,WSB | 2 | 140304127 | 140304160 | 34 | + | 19.748 | 1.06E-07 | 0.0313 | CCGCCCCCACATTCGCCGCCACAAGATGGCGCTG | | |
| 224 | 2 | 140304111 | 140304532 | 421 | C58J,CAST,DBA1J,DBA2J,MOLF,NZO,NZW,SPRET,WSB | 2 | 140304142 | 140304160 | 19 | + | 21.1148 | 4.45E-08 | 0.0407 | CCGCCACAAGATGGCGCTG | | |
| 225 | 2 | 141898627 | 141898826 | 199 | C58J,CAST,DBA1J,DBA2J,MOLF,NZO,NZW,SPRET | 2 | 141898684 | 141898717 | 34 | + | 16.4634 | 9.51E-07 | 0.0987 | TAGCTGTCTTCAGACGCACCAGAAGAGGGCATCA | | |
| 226 | 2 | 143708646 | 143709024 | 378 | 129S1,BTBR,C57L,C58J,CAST,CE,DBA1J,DBA2J,MOLF,NZO,NZW,SPRET | 2 | 143708885 | 143708918 | 34 | + | 16.4472 | 9.61E-07 | 0.0987 | TAGCTGTCTTCAGACACACCAGAAGAGGGCGTCA | | |
| 227 | 2 | 150730567 | 150730774 | 207 | 129S1,BUB,C57L,CAST,CE,MOLF,NOD,NZB,NZW,RF,SPRET,WSB | 2 | 150730646 | 150730664 | 19 | + | 17.8852 | 5.08E-07 | 0.115 | ACACCAGAAGAGGGCACCA | 1 | ENCFF093ZRU |
| 228 | 2 | 153201930 | 153202668 | 738 | MOLF | 2 | 153202266 | 153202299 | 34 | + | 16.4472 | 9.61E-07 | 0.0987 | TAGCTGTCTTCAGACACACCAGAAGAGGGCGTCA | | |
| 229 | 2 | 154495678 | 154495890 | 212 | AJ,BTBR,C58J,CAST,DBA1J,DBA2J,KK,MOLF.NZB.NZO.NZW.SPRET.WSB | 2 | 154495748 | 154495781 | 34 | + | 18.3659 | 2.76E-07 | 0.0538 | TAGCTGTCTTCAGACACTCCAGAAGAGGGCGCCA | 1 | ENCFF093ZRU |
| 229 | 2 | 154495678 | 154495890 | 212 | AJ,BTBR,C58J,CAST,DBA1J,DBA2J,KK,MOLF.NZB.NZO.NZW.SPRET.WSB | 2 | 154495763 | 154495781 | 19 | + | 17.5574 | 6.32E-07 | 0.124 | ACTCCAGAAGAGGGCGCCA | 1 | ENCFF093ZRU |
| 230 | 2 | 154608963 | 154609893 | 930 | CAST | 2 | 154609752 | 154609785 | 34 | + | 16.7724 | 7.83E-07 | 0.0917 | TAGCTGTCTTCAGATGCACCAGAAGAGGGCGTCA | | |
| 231 | 2 | 154770780 | 154770984 | 204 | CAST.SPRET | 2 | 154770842 | 154770875 | 34 | + | 16.4472 | 9.61E-07 | 0.0987 | TAGCTGTCTTCAGACACACCAGAAGAGGGCGTCA | | |
| 232 | 2 | 154896369 | 154896574 | 205 | 129S1,BALB,C3H,C57L,CAST,CBA,CE,FVB,MA/My,NOD,NZO,NZW,PL,SEA,SJL,SPRET,SWR.TH.WSB | 2 | 154896428 | 154896462 | 35 | + | 18.1284 | 2.79E-07 | 0.0894 | CTGTAGCTGTCTTCAGACACCAGAAGAGGGCATCA | | |
| 233 | 2 | 155109406 | 155109656 | 250 | 129S1,BALB,C3H,C57L,CAST,CBA,CE,FVB,MA/My,NOD,NZO,NZW,PL,SEA,SJL,SMJ,SPRET.SWR.TH.WSB | 2 | 155109532 | 155109565 | 34 | + | 16.4472 | 9.61E-07 | 0.0987 | TAGCTGTCTTCAGACACACCAGAAGAGGGCGTCA | | |
| 234 | 2 | 155972010 | 155972174 | 164 | 129S1.C57L | 2 | 155972055 | 155972088 | 34 | + | 19.0244 | 1.76E-07 | 0.0433 | TAGCTGTCCCCAGACACTCCAGAAGAGGGCATCA | 3 | ENCFF093ZRU.ENCFF099QNE.ENCFF367UNR |
| 235 | 2 | 156047519 | 156047726 | 207 | SPRET | 2 | 156047581 | 156047614 | 34 | + | 16.4472 | 9.61E-07 | 0.0987 | TAGCTGTCTTCAGACACACCAGAAGAGGGCGTCA | | |
| 236 | 2 | 156478930 | 156479671 | 741 | SPRET | 2 | 156479102 | 156479135 | 34 | + | 16.8049 | 7.67E-07 | 0.0917 | TTGCTCTCTTCAGAAACACCAGAAGAGGGCATCA | | |
| 237 | 2 | 156666964 | 156667168 | 204 | SPRET | 2 | 156667026 | 156667059 | 34 | + | 16.4472 | 9.61E-07 | 0.0987 | TAGCTGTCTTCAGACACACCAGAAGAGGGCGTCA | | |
| 238 | 2 | 156835199 | 156835399 | 200 | SPRET | 2 | 156835255 | 156835288 | 34 | + | 16.4472 | 9.61E-07 | 0.0987 | TAGCTGTCTTCAGACACACCAGAAGAGGGCGTCA | | |
| 239 | 2 | 163158058 | 163158220 | 162 | WSB | 2 | 163158216 | 163158249 | 34 | + | 17.2358 | 5.82E-07 | 0.0796 | GTGTGGCCTCTCTGAGGGCCTCCAGGTGGCAGGG | | |
| 240 | 2 | 164708088 | 164708289 | 201 | AJ,AKR,BALB,BTBR,BUB,C3H,C58J,CAST,CBA,DBA1J,DBA2J,FVB,ILNJ2,KK,MA/My,MRL,NOD,NOR,NZB,NZO,NZW,PJ,PL,RF,SEA,SJL,SMJ,SPRET,SWR,TH,WSB | 2 | 164708146 | 164708179 | 34 | + | 17.2439 | 5.79E-07 | 0.0793 | TAGCTGTCCTCAGATACTCCAGAAGAGGGCATCA | | |
| 241 | 2 | 165758175 | 165758425 | 250 | CAST.KK.SPRET | 2 | 165758242 | 165758275 | 34 | + | 17.1463 | 6.17E-07 | 0.0817 | TAGCTGTCCTCAGACACTCCAGAAGAAGGCGCCA | | |
| 242 | 2 | 165857809 | 165858028 | 219 | 129S1,AJ,AKR,BALB,BTBR,BUB,C3H,C57L,C58J,CAST,CBA,CE,DBA1J,DBA2J,FVB,ILNJ2,KK,MA/My,NOD,NOR,NZB,NZO,PJ,PL,RF,SEA,SJL,SMJ,SPRET,SWR,TH,WSB | 2 | 165857886 | 165857919 | 34 | + | 16.4472 | 9.61E-07 | 0.0987 | TAGCTGTCTTCAGACACACCAGAAGAGGGCGTCA | | |
| 243 | 2 | 165919818 | 165920286 | 468 | KK | 2 | 165920020 | 165920053 | 34 | + | 19.2846 | 1.47E-07 | 0.0385 | TAGCTGTCTTCAGACTCGCCAGAAGAGGGCGTCA | 1 | ENCFF093ZRU |
| 243 | 2 | 165919818 | 165920286 | 468 | KK | 2 | 165920035 | 165920053 | 19 | + | 18 | 4.70E-07 | 0.113 | TCGCCAGAAGAGGGCGTCA | 1 | ENCFF093ZRU |
| 244 | 2 | 166595903 | 166596110 | 207 | CAST.CE.KK.SPRET.TH | 2 | 166595967 | 166596000 | 34 | + | 16.4634 | 9.51E-07 | 0.0987 | TAGCTGTCTTCAGACGCACCAGAAGAGGGCATCA | | |
| 245 | 2 | 166822996 | 166823207 | 211 | 129S1.C57L.CAST.CE.KK.SPRET.WSB | 2 | 166823048 | 166823081 | 34 | + | 16.4472 | 9.61E-07 | 0.0987 | TAGCTGTCTTCAGACACACCAGAAGAGGGCGTCA | 1 | ENCFF093ZRU |
| 246 | 2 | 167743704 | 167743921 | 217 | SPRET | 2 | 167743779 | 167743812 | 34 | + | 16.4472 | 9.61E-07 | 0.0987 | TAGCTGTCTTCAGACACACCAGAAGAGGGCGTCA | | |
| 247 | 2 | 168128156 | 168128226 | 70 | 129S1.C57L.NZB.TH | 2 | 168128154 | 168128188 | 35 | + | 17.633 | 3.74E-07 | 0.107 | GTACAGGTACCCATGGAGACCAGAAGAGGGCGTCA | 1 | ENCFF093ZRU |
| 248 | 2 | 168518524 | 168518737 | 213 | SPRET | 2 | 168518594 | 168518627 | 34 | + | 18.5447 | 2.45E-07 | 0.0522 | CAGCTGTCTTCAGACGCTCCAGAAGAGGGCATCA | | |
| 249 | 2 | 168838895 | 168839092 | 197 | KK.SPRET | 2 | 168838950 | 168838983 | 34 | + | 16.5447 | 9.04E-07 | 0.0987 | CAGCTGTCTTCAGACACACCAGAAGAGGGAGTCA | | |
| 250 | 2 | 174978710 | 174978926 | 216 | BUB.KK.NOD.NOR.SPRET | 2 | 174978783 | 174978816 | 34 | + | 18.3659 | 2.76E-07 | 0.0538 | TAGCTGTCTTCAGACACTCCAGAAGAGGGCGCCA | | |
| 250 | 2 | 174978710 | 174978926 | 216 | BUB.KK.NOD.NOR.SPRET | 2 | 174978798 | 174978816 | 19 | + | 17.5574 | 6.32E-07 | 0.124 | ACTCCAGAAGAGGGCGCCA | | |
| 251 | 2 | 181267330 | 181267529 | 199 | SPRET | 2 | 181267384 | 181267418 | 35 | + | 18.1284 | 2.79E-07 | 0.0894 | CTGTAGCTGTCTTCAGACACCAGAAGAGGGCATCA | 1 | ENCFF093ZRU |
| 252 | 2 | 10973281 | 10973404 | 123 | SPRET | 2 | 10973254 | 10973288 | 35 | - | 15.9266 | 9.95E-07 | 0.181 | GTGTTGTGTACCTGTAACCCCAGCAGATGGAGACA | | |
| 253 | 2 | 13414079 | 13414210 | 131 | SPRET | 2 | 13414142 | 13414175 | 34 | - | 16.4472 | 9.61E-07 | 0.0987 | TAGCTGTCTTCAGACACACCAGAAGAGGGCGTCA | | |
| 254 | 2 | 15943762 | 15944296 | 534 | SPRET | 2 | 15944155 | 15944189 | 35 | - | 18.6055 | 2.09E-07 | 0.0757 | TTGTATGTATCCCTGCAGGCCAGAAGACGGCACTG | | |
| 255 | 2 | 17823736 | 17823949 | 213 | CAST.MA/My.MOLF.RF.SPRET | 2 | 17823859 | 17823892 | 34 | - | 18.7561 | 2.12E-07 | 0.0488 | TAGCTGCTCCCAGACACTCCAGAAGAGGGTGCCA | | |
| 256 | 2 | 18287500 | 18287703 | 203 | CAST.MA/My.MOLF.RF.SPRET | 2 | 18287628 | 18287661 | 34 | - | 17.2114 | 5.91E-07 | 0.0799 | TAGCTGTTTTCAGACACTCCAGAAGAGGGCATCA | | |
| 257 | 2 | 18417387 | 18417578 | 191 | CAST.MA/My.MOLF.RF.SJL.SPRET | 2 | 18417512 | 18417530 | 19 | - | 17.5574 | 6.32E-07 | 0.124 | ACTCCAGAAGAGGGCGCCA | | |
| 257 | 2 | 18417387 | 18417578 | 191 | CAST.MA/My.MOLF.RF.SJL.SPRET | 2 | 18417512 | 18417545 | 34 | - | 18.3659 | 2.76E-07 | 0.0538 | TAGCTGTCTTCAGACACTCCAGAAGAGGGCGCCA | | |
| 258 | 2 | 20462715 | 20462915 | 200 | CAST.FVB.SJL.SPRET | 2 | 20462831 | 20462864 | 34 | - | 17.6667 | 4.40E-07 | 0.0682 | TAGCTGTCTTCAGACACGCCAGAAGAGGGCGTCA | | |
| 259 | 2 | 20691711 | 20691908 | 197 | SPRET | 2 | 20691825 | 20691858 | 34 | - | 18.9919 | 1.80E-07 | 0.0438 | TTGCTGTCTTCAGACACTCCAGAAGGGGGCATCA | | |
| 260 | 2 | 22810762 | 22811696 | 934 | SPRET | 2 | 22810893 | 22810926 | 34 | - | 16.4634 | 9.51E-07 | 0.0987 | TAGCTGTCTTCAGACGCACCAGAAGAGGGCATCA | | |
| 261 | 2 | 25079339 | 25079547 | 208 | CAST | 2 | 25079461 | 25079494 | 34 | - | 17.2195 | 5.88E-07 | 0.0799 | TAGCTGTCTTCCAACAGTCCAGAAGAGGGCGTCA | | |
| 262 | 2 | 26123501 | 26123708 | 207 | MA/My.MOLF.NZO.SPRET | 2 | 26123626 | 26123659 | 34 | - | 16.7724 | 7.83E-07 | 0.0917 | TAGCTGTCTTCAGATGCACCAGAAGAGGGCGTCA | | |
| 263 | 2 | 27796309 | 27796530 | 221 | SPRET | 2 | 27796433 | 27796466 | 34 | - | 17.6016 | 4.59E-07 | 0.0697 | TAGCTGTCTTCAGACATGCCAGAAGAGGGCGTCA | | |
| 264 | 2 | 28495248 | 28495492 | 244 | CAST.MOLF.SPRET | 2 | 28495372 | 28495390 | 19 | - | 17.5574 | 6.32E-07 | 0.124 | ACTCCAGAAGAGGGCGCCA | | |
| 264 | 2 | 28495248 | 28495492 | 244 | CAST.MOLF.SPRET | 2 | 28495372 | 28495405 | 34 | - | 18.3659 | 2.76E-07 | 0.0538 | TAGCTGTCTTCAGACACTCCAGAAGAGGGCGCCA | | |
| 265 | 2 | 28523391 | 28523590 | 199 | MOLF.SPRET | 2 | 28523519 | 28523552 | 34 | - | 16.4472 | 9.61E-07 | 0.0987 | TAGCTGTCTTCAGACACACCAGAAGAGGGCGTCA | 1 | ENCFF093ZRU |
| 266 | 2 | 28934207 | 28934428 | 221 | CAST.SPRET | 2 | 28934332 | 28934365 | 34 | - | 17.878 | 3.83E-07 | 0.0635 | TAGCTGTCCTCAGACACACCAGAAGAGGGCATCA | | |
| 267 | 2 | 28949247 | 28949449 | 202 | CAST.MOLF.SPRET | 2 | 28949370 | 28949403 | 34 | - | 17.5935 | 4.61E-07 | 0.0698 | TAGCTGTCTCCAGACACTCCAGAAGAGGGCGTCA | | |
| 268 | 2 | 29801061 | 29801270 | 209 | SPRET | 2 | 29801181 | 29801199 | 19 | - | 17.5574 | 6.32E-07 | 0.124 | ACTCCAGAAGAGGGCGCCA | 3 | ENCFF093ZRU.ENCFF821BIA.ENCFF367UNR |
| 268 | 2 | 29801061 | 29801270 | 209 | SPRET | 2 | 29801181 | 29801214 | 34 | - | 18.3659 | 2.76E-07 | 0.0538 | TAGCTGTCTTCAGACACTCCAGAAGAGGGCGCCA | 3 | ENCFF093ZRU.ENCFF821BIA.ENCFF367UNR |
| 269 | 2 | 30763433 | 30763876 | 443 | SPRET | 2 | 30763472 | 30763505 | 34 | - | 16.4472 | 9.61E-07 | 0.0987 | TAGCTGTCTTCAGACACACCAGAAGAGGGCGTCA | | |
| 270 | 2 | 31558403 | 31558602 | 199 | MOLF.SPRET | 2 | 31558527 | 31558560 | 34 | - | 16.4472 | 9.61E-07 | 0.0987 | TAGCTGTCTTCAGACACACCAGAAGAGGGCGTCA | | |
| 271 | 2 | 31571130 | 31571337 | 207 | MOLF,SPRET | 2 | 31571253 | 31571271 | 19 | - | 17.5574 | 6.32E-07 | 0.124 | ACTCCAGAAGAGGGCGCCA | 9 | ENCFF327LYS,ENCFF092NMJ,ENCFF024JOZ,ENCFF089GBC,ENCFF821BIA,ENCFF093ZRU,ENCFF367UNR,ENCFF099QNE,ENCFF055HAP |
| 271 | 2 | 31571130 | 31571337 | 207 | MOLF,SPRET | 2 | 31571253 | 31571286 | 34 | - | 18.561 | 2.42E-07 | 0.0519 | AAGCTGTCTTCAGACACTCCAGAAGAGGGCGCCA | 9 | ENCFF327LYS,ENCFF092NMJ,ENCFF024JOZ,ENCFF089GBC,ENCFF821BIA,ENCFF093ZRU,ENCFF367UNR,ENCFF099QNE,ENCFF055HAP |
| 272 | 2 | 31576666 | 31577045 | 379 | SPRET | 2 | 31576781 | 31576814 | 34 | - | 16.7398 | 7.99E-07 | 0.0927 | TAGCTGTCTTCAGACACACCAGTAGAGGGTGCCA | 2 | ENCFF093ZRU.ENCFF099QNE |
| 273 | 2 | 32245375 | 32245575 | 200 | CAST.MOLF.NZO.SPRET.SWR | 2 | 32245486 | 32245504 | 19 | - | 17.5574 | 6.32E-07 | 0.124 | ACTCCAGAAGAGGGCGCCA | 2 | ENCFF093ZRU.ENCFF367UNR |
| 273 | 2 | 32245375 | 32245575 | 200 | CAST.MOLF.NZO.SPRET.SWR | 2 | 32245486 | 32245519 | 34 | - | 18.3659 | 2.76E-07 | 0.0538 | TAGCTGTCTTCAGACACTCCAGAAGAGGGCGCCA | 2 | ENCFF093ZRU.ENCFF367UNR |
| 274 | 2 | 32697191 | 32697387 | 196 | SPRET | 2 | 32697313 | 32697346 | 34 | - | 17.2439 | 5.79E-07 | 0.0793 | TAGCTGTCCTCAGATACTCCAGAAGAGGGCATCA | 1 | ENCFF093ZRU |
| 275 | 2 | 32926717 | 32926920 | 203 | SPRET | 2 | 32926842 | 32926875 | 34 | - | 16.4472 | 9.61E-07 | 0.0987 | TAGCTGTCTTCAGACACACCAGAAGAGGGCGTCA | | |
| 276 | 2 | 37826869 | 37827070 | 201 | BUB.CAST.MOLF.SPRET.TH.WSB | 2 | 37826994 | 37827012 | 19 | - | 17.5574 | 6.32E-07 | 0.124 | ACTCCAGAAGAGGGCGCCA | 1 | ENCFF093ZRU |
| 276 | 2 | 37826869 | 37827070 | 201 | BUB.CAST.MOLF.SPRET.TH.WSB | 2 | 37826994 | 37827027 | 34 | - | 18.561 | 2.42E-07 | 0.0519 | AAGCTGTCTTCAGACACTCCAGAAGAGGGCGCCA | 1 | ENCFF093ZRU |
| 277 | 2 | 38740385 | 38740585 | 200 | SPRET | 2 | 38740510 | 38740543 | 34 | - | 17.0813 | 6.43E-07 | 0.0837 | TAGCTGTTCTCAGACACTCCAGAAGAGGGAGTCA | | |
| 278 | 2 | 45026115 | 45026361 | 246 | 129S1,AJ,AKR,BTBR,BUB,C3H,C58J,CAST,CBA,CE,DBA1J,DBA2J,FVB,ILNJ2,KK,MA/My,MOLF,MRL,NOD,NOR,NZB,NZO,NZW,PJ,PL,RF,SEA,SJL,SMJ,SPRET,SWR,TH | 2 | 45026242 | 45026275 | 34 | - | 16.4472 | 9.61E-07 | 0.0987 | TAGCTGTCTTCAGACACACCAGAAGAGGGCGTCA | | |
| 279 | 2 | 50180965 | 50181057 | 92 | SPRET | 2 | 50180950 | 50180984 | 35 | - | 16.2385 | 8.36E-07 | 0.165 | CTGTAATTCTACTACTTGACCACTAGGAGAAAGAT | 18 | ENCFF093ZRU,ENCFF367UNR,ENCFF065ODV,ENCFF024JOZ,ENCFF165LXO,ENCFF099QNE,ENCFF089GBC,ENCFF430PPJ,ENCFF092NMJ,ENCFF055HAP,ENCFF883UPM,ENCFF502WDT,ENCFF907XEG,ENCFF821BIA,ENCFF327LYS,ENCFF052KVO,ENCFF589ZBJ,ENCFF198SSR |
| 280 | 2 | 50545989 | 50546196 | 207 | 129S1,AJ,AKR,BUB,C3H,C58J,CAST,CBA,CE,DBA1J,DBA2J,FVB,ILNJ2,KK,MOLF,MRL,NOD,NOR,PJ,PL,RF,SEA,SJL,SMJ,SPRET,SWR.TH.WSB | 2 | 50546111 | 50546144 | 34 | - | 16.4472 | 9.61E-07 | 0.0987 | TAGCTGTCTTCAGACACACCAGAAGAGGGCGTCA | | |
| 281 | 2 | 52688599 | 52688823 | 224 | CAST.MOLF.NZB.SPRET.WSB | 2 | 52688727 | 52688760 | 34 | - | 17.9431 | 3.66E-07 | 0.0623 | CAGCTGTCCTCAGACACTCCAGAAGAGGGAGTCA | | |
| 282 | 2 | 53060871 | 53061006 | 135 | SPRET | 2 | 53060892 | 53060925 | 34 | - | 17.8943 | 3.78E-07 | 0.0633 | GTGCTCTCTTCAGACACACCAGAAGAGGGCATCA | | |
| 283 | 2 | 59382258 | 59382710 | 452 | SPRET | 2 | 59382348 | 59382381 | 34 | - | 17.5203 | 4.84E-07 | 0.0717 | TAGCTGTTTTCTGACACACCAGAAGAGGGCATCA | 1 | ENCFF093ZRU |
| 284 | 2 | 59414947 | 59415152 | 205 | CAST.MOLF.SPRET.WSB | 2 | 59415071 | 59415089 | 19 | - | 17.5574 | 6.32E-07 | 0.124 | ACTCCAGAAGAGGGCGCCA | | |
| 284 | 2 | 59414947 | 59415152 | 205 | CAST.MOLF.SPRET.WSB | 2 | 59415071 | 59415104 | 34 | - | 18.3659 | 2.76E-07 | 0.0538 | TAGCTGTCTTCAGACACTCCAGAAGAGGGCGCCA | | |
| 285 | 2 | 59682211 | 59682402 | 191 | 129S1,C57L,CAST,CE,KK,MOLF,NOD,NOR,SJL.SPRET.WSB | 2 | 59682310 | 59682328 | 19 | - | 17.5574 | 6.32E-07 | 0.124 | ACTCCAGAAGAGGGCGCCA | 4 | ENCFF093ZRU,ENCFF099QNE,ENCFF821BIA,ENCFF367UNR |
| 285 | 2 | 59682211 | 59682402 | 191 | 129S1,C57L,CAST,CE,KK,MOLF,NOD,NOR,SJL.SPRET.WSB | 2 | 59682310 | 59682343 | 34 | - | 18.3659 | 2.76E-07 | 0.0538 | TAGCTGTCTTCAGACACTCCAGAAGAGGGCGCCA | 4 | ENCFF093ZRU,ENCFF099QNE,ENCFF821BIA,ENCFF367UNR |
| 286 | 2 | 60665314 | 60665818 | 504 | MOLF.SPRET | 2 | 60665724 | 60665757 | 34 | - | 16.4472 | 9.61E-07 | 0.0987 | TAGCTGTCTTCAGACACACCAGAAGAGGGCGTCA | | |
| 287 | 2 | 62877587 | 62877809 | 222 | 129S1,BTBR,CAST,DBA1J,DBA2J,FVB,KK,MOLF,PJ,SEA,SJL,SPRET,SWR,TH,WSB | 2 | 62877712 | 62877730 | 19 | - | 17.5574 | 6.32E-07 | 0.124 | ACTCCAGAAGAGGGCGCCA | | |
| 287 | 2 | 62877587 | 62877809 | 222 | 129S1,BTBR,CAST,DBA1J,DBA2J,FVB,KK,MOLF,PJ,SEA,SJL,SPRET,SWR,TH,WSB | 2 | 62877712 | 62877745 | 34 | - | 18.3659 | 2.76E-07 | 0.0538 | TAGCTGTCTTCAGACACTCCAGAAGAGGGCGCCA | | |
| 288 | 2 | 67717821 | 67718597 | 776 | SPRET | 2 | 67718200 | 67718233 | 34 | - | 18.5528 | 2.44E-07 | 0.0521 | TTACAGTACCTGCAGAGGCCCAAAGAGGGCACCA | 2 | ENCFF052KVO.ENCFF093ZRU |
| 289 | 2 | 70326734 | 70326945 | 211 | SPRET | 2 | 70326859 | 70326892 | 34 | - | 17.6667 | 4.40E-07 | 0.0682 | TAGCTGTCTTCAGACACGCCAGAAGAGGGCGTCA | | |
| 290 | 2 | 71203237 | 71203416 | 179 | SPRET | 2 | 71203361 | 71203394 | 34 | - | 16.8293 | 7.55E-07 | 0.091 | TTGCTCTCTTAAGACACACCAGAAGAGGGCATCA | | |
| 291 | 2 | 72472857 | 72473144 | 287 | MA/My.MOLF.NZB | 2 | 72472999 | 72473017 | 19 | - | 18.7213 | 2.85E-07 | 0.0853 | GTGCCAGAAGAGGGCACTG | | |
| 291 | 2 | 72472857 | 72473144 | 287 | MA/My.MOLF.NZB | 2 | 72472999 | 72473033 | 35 | - | 18.4404 | 2.31E-07 | 0.0809 | CTTCATGTATGCCTGTGTGCCAGAAGAGGGCACTG | | |
| 292 | 2 | 73817571 | 73817779 | 208 | CAST.SPRET | 2 | 73817694 | 73817727 | 34 | - | 19.5122 | 1.26E-07 | 0.035 | CTGTAGTACTCAGACACTCCAGAAGAGGGAGTCA | | |
| 293 | 2 | 75691175 | 75691416 | 241 | CAST.MOLF.NOD.NOR.SPRET | 2 | 75691308 | 75691341 | 34 | - | 16.4634 | 9.51E-07 | 0.0987 | TAGCTGTCTTCAGACGCACCAGAAGAGGGCATCA | 1 | ENCFF093ZRU |
| 294 | 2 | 82786935 | 82787884 | 949 | SPRET | 2 | 82787858 | 82787892 | 35 | - | 16.7248 | 6.34E-07 | 0.141 | ATGCAATTTGGGCATATATCCAGAAGATGCCCCAA | | |
| 295 | 2 | 84627497 | 84627700 | 203 | SPRET | 2 | 84627620 | 84627638 | 19 | - | 17.8852 | 5.08E-07 | 0.115 | ACACCAGAAGAGGGCACCA | | |
| 295 | 2 | 84627497 | 84627700 | 203 | SPRET | 2 | 84627620 | 84627653 | 34 | - | 17.1138 | 6.30E-07 | 0.0826 | TAGCTATCTTCAGACACACCAGAAGAGGGCACCA | | |
| 296 | 2 | 86294951 | 86295283 | 332 | AJ,AKR,BALB,BUB,C3H,C57L,CBA,CE,DBA1J,DBA2J,FVB,KK,MA/My,MRL,NOD,NOR,NZB,NZO,NZW,PJ,PL,RF,RH2,SEA,SMJ,SPRET,WSB | 2 | 86295237 | 86295255 | 19 | - | 19.2951 | 1.88E-07 | 0.071 | CCGCCACTAGATGGTGCTA | 28 | ENCFF093ZRU,ENCFF367UNR,ENCFF805QBU,ENCFF099QNE,ENCFF883UPM,ENCFF821BIA,ENCFF601JVE,ENCFF316FYN,ENCFF327LYS,ENCFF907XEG,ENCFF024JOZ,ENCFF033UQX,ENCFF136ZDK,ENCFF589ZBJ,ENCFF097CAQ,ENCFF089GBC,ENCFF198SSR,ENCFF165LXO,ENCFF092NMJ,ENCFF052KVO,ENCFF321AGQ,ENCFF430PPJ,ENCFF281QMV,ENCFF957SPH,ENCFF194HVL,ENCFF951GOP,ENCFF533APC,ENCFF609MDN |
| 297 | 2 | 91968422 | 91968643 | 221 | AJ,AKR,BUB,C3H,C58J,CAST,CE,FVB,ILNJ2,MA/My,MOLF,MRL,NZW,PL,RF,SPRET,TH,WSB | 2 | 91968546 | 91968579 | 34 | - | 18.4797 | 2.56E-07 | 0.0524 | TTGCTGTCTTCAGACACACCAGAAGAGGGCATCA | | |
| 298 | 2 | 93620769 | 93621111 | 342 | SPRET | 2 | 93620935 | 93620969 | 35 | - | 16.2569 | 8.27E-07 | 0.164 | CTGCTGTGATTTCAGGGTACCAGGTGATGGCAGAT | | |
| 299 | 2 | 94184621 | 94184823 | 202 | SPRET | 2 | 94184747 | 94184765 | 19 | - | 17.5574 | 6.32E-07 | 0.124 | ACTCCAGAAGAGGGCGCCA | | |
| 299 | 2 | 94184621 | 94184823 | 202 | SPRET | 2 | 94184747 | 94184780 | 34 | - | 18.3659 | 2.76E-07 | 0.0538 | TAGCTGTCTTCAGACACTCCAGAAGAGGGCGCCA | | |
| 300 | 2 | 102031488 | 102031686 | 198 | AJ,BUB,CAST,FVB,ILNJ2,KK,MOLF,NZB,NZO,NZW,RF,RH2,SJL,SMJ,SPRET,SWR,WSB | 2 | 102031618 | 102031636 | 19 | - | 17.8852 | 5.08E-07 | 0.115 | ACACCAGAAGAGGGCACCA | 2 | ENCFF093ZRU |
| 301 | 2 | 103306113 | 103306966 | 853 | AJ,BALB,C57L,CBA,DBA1J,DBA2J,ILNJ2,RH2 | 2 | 103306701 | 103306734 | 34 | - | 16.6504 | 8.46E-07 | 0.0961 | TAGCTGTTTACAGACATACCAGAAGAGGGCATCA | 2 | ENCFF367UNR,ENCFF093ZRU |
| 302 | 2 | 104267716 | 104267907 | 191 | AJ,AKR,BALB,C3H,C57L,CAST,CBA,CE,DBA1J,DBA2J,FVB,ILNJ2,MOLF,MRL,NZB,NZW,PL,RF,RH2,SMJ,SPRET,SWR,TH,WSB | 2 | 104267840 | 104267873 | 34 | - | 16.4472 | 9.61E-07 | 0.0987 | TAGCTGTCTTCAGACACACCAGAAGAGGGCGTCA | | |
| 303 | 2 | 104438880 | 104439094 | 214 | 129S1,AJ,AKR,BALB,BTBR,BUB,C3H,C57L,CAST,CBA,CE,DBA1J,DBA2J,FVB,ILNJ2,KK,MOLF,MRL,NOD,NOR,NZB,NZW,PJ,PL,RF,RH2,SEA,SJL,SPRET,SWR,TH,WSB | 2 | 104439006 | 104439039 | 34 | - | 16.4472 | 9.61E-07 | 0.0987 | TAGCTGTCTTCAGACACACCAGAAGAGGGCGTCA | | |
| 304 | 2 | 108410493 | 108410722 | 229 | C3H.CAST.CE.MOLF.RF.SPRET | 2 | 108410621 | 108410654 | 34 | - | 16.6179 | 8.63E-07 | 0.0972 | TAGCTGTCTTCCGACACTCCAGAAGAGGGCGTCA | | |
| 305 | 2 | 112293855 | 112294077 | 222 | SPRET | 2 | 112293977 | 112294010 | 34 | - | 16.4634 | 9.51E-07 | 0.0987 | TAGCTGTCTTCAGACGCACCAGAAGAGGGCATCA | | |
| 306 | 2 | 119111535 | 119111774 | 239 | MOLF.SPRET | 2 | 119111659 | 119111693 | 35 | - | 16.2477 | 8.31E-07 | 0.164 | TAGCTGTCTTCAGACACTCCCAGAAGAGGGCATCA | | |
| 307 | 2 | 119401873 | 119402089 | 216 | 129S1,AJ,AKR,BALB,BTBR,BUB,C3H,C57L,C58J,CAST,CBA,CE,DBA1J,DBA2J,FVB,ILNJ2,KK,MA/My,NOD,NZB,NZO,NZW,PJ,PL,RF,RH2,SEA,SJL,SMJ,SPRET,SWR,TH,WSB | 2 | 119401998 | 119402016 | 19 | - | 17.5574 | 6.32E-07 | 0.124 | ACTCCAGAAGAGGGCGCCA | 2 | ENCFF093ZRU,ENCFF367UNR |

| | | | | | | | | | | | | | | | | |
|---|---|---|---|---|---|---|---|---|---|---|---|---|---|---|---|---|
| 307 | 2 | 119401873 | 119402089 | 216 | 129S1,AJ,AKR,BALB,BTBR,BUB,C3H,C57L,C58J,CAST,CBA,CE,DBA1J,DBA2J,FVB,ILNJ2,KK,MA/My,NOD,NZB,NZO,NZW,PJ,PL,RF,RH2,SEA,SJL,SMJ,SPRET,SWR,TH,WSB | 2 | 119401998 | 119402031 | 34 | - | 18.3659 | 2.76E-07 | 0.0538 | TAGCTGTCTTCAGACACTCCAGAAGAGGGCGCCA | 2 | ENCFF093ZRU,ENCFF367UNR |
| 308 | 2 | 121239602 | 121239796 | 194 | SPRET | 2 | 121239720 | 121239753 | 34 | - | 16.4634 | 9.51E-07 | 0.0987 | TAGCTGTCTTCAGACGCACCAGAAGAGGGCATCA | 1 | ENCFF093ZRU |
| 309 | 2 | 121560387 | 121560604 | 217 | SPRET | 2 | 121560509 | 121560542 | 34 | - | 17.7805 | 4.08E-07 | 0.0656 | TAGCTGTCCTCAAACACACCAGAAGAGGGCATCA | | |
| 310 | 2 | 121922117 | 121922193 | 76 | 129S1,AKR,BTBR,BUB,C57L,CE,FVB,MA/Mv.NZW.SJL.SWR.WSB | 2 | 121922155 | 121922188 | 34 | - | 19.9512 | 9.19E-08 | 0.0283 | GTGCTGTGCCGGCCTTGGCCTACAGAAGTCAGTG | | |
| 311 | 2 | 124933953 | 124934168 | 215 | 129S1,AKR,BTBR,BUB,C3H,C57L,C58J,CAST,CBA,CE,FVB,KK,MOLF,NZB,NZO,NZW,RF,SJL,SPRET,SWR,TH | 2 | 124934079 | 124934097 | 19 | - | 17.5574 | 6.32E-07 | 0.124 | ACTCCAGAAGAGGGCGCCA | 3 | ENCFF367UNR,ENCFF093ZRU,ENCFF099QNE |
| 311 | 2 | 124933953 | 124934168 | 215 | 129S1,AKR,BTBR,BUB,C3H,C57L,C58J,CAST,CBA,CE,FVB,KK,MOLF,NZB,NZO,NZW,RF,SJL,SPRET,SWR,TH | 2 | 124934079 | 124934112 | 34 | - | 18.3659 | 2.76E-07 | 0.0538 | TAGCTGTCTTCAGACACTCCAGAAGAGGGCGCCA | 3 | ENCFF367UNR,ENCFF093ZRU,ENCFF099QNE |
| 312 | 2 | 126873331 | 126873530 | 199 | SPRET | 2 | 126873441 | 126873474 | 34 | - | 16.4472 | 9.61E-07 | 0.0987 | TAGCTGTCTTCAGACACACCAGAAGAGGGCGTCA | 1 | ENCFF093ZRU |
| 313 | 2 | 128482743 | 128482947 | 204 | C3H,CBA,DBA1J,DBA2J,ILNJ2,PJ,PL,SPRET | 2 | 128482868 | 128482901 | 34 | - | 16.7724 | 7.83E-07 | 0.0917 | TAGCTGTCTTCAGATGCACCAGAAGAGGGCGTCA | | |
| 314 | 2 | 131910313 | 131910437 | 124 | CAST.ILNJ2.NOD.PJ.SPRET.WSB | 2 | 131910326 | 131910360 | 35 | - | 20.1284 | 8.00E-08 | 0.0419 | CTGCATTTTAACCTGTGAGCCCCCAGGTGCCTGCA | | |
| 314 | 2 | 131910313 | 131910437 | 124 | CAST.ILNJ2.NOD.PJ.SPRET.WSB | 2 | 131910356 | 131910390 | 35 | - | 20.1284 | 8.00E-08 | 0.0419 | CTGCATTTTAACCTGTGAGCCCCCAGGTGCCTGCA | | |
| 315 | 2 | 132183231 | 132184230 | 999 | CAST | 2 | 132183886 | 132183919 | 34 | - | 16.3984 | 9.91E-07 | 0.0999 | TAGCTGTCTTCAGACGTACCAGAAGAGGGCATCA | | |
| 316 | 2 | 141730258 | 141730341 | 83 | C58J.CAST.DBA1J.DBA2J.NZO.NZW | 2 | 141730225 | 141730259 | 35 | - | 16.0642 | 9.21E-07 | 0.174 | CTTCATGTACACCTGCATGCCAGAAGAGGGCATGA | | |
| 317 | 2 | 144071185 | 144071436 | 251 | 129S1,BTBR,C57L,C58J,CE,DBA1J,DBA2J,MOLF.NZO.NZW | 2 | 144071329 | 144071363 | 35 | - | 18.1835 | 2.70E-07 | 0.0882 | CTGTAGCTGTCTTCAGACACCAGAAGAGGGCATTG | | |
| 318 | 2 | 144286867 | 144286935 | 68 | SPRET | 2 | 144286842 | 144286876 | 35 | - | 16.7156 | 6.38E-07 | 0.141 | GTGGTGTCGTCCACACAGGCCAGAAGGGGGCATCA | | |
| 319 | 2 | 144402783 | 144403008 | 225 | 129S1,AJ,BTBR,C57L,C58J,CAST,CE,DBA1J,DBA2J,MOLF,NZB,NZO,NZW,SMJ,SPRET | 2 | 144402906 | 144402924 | 19 | - | 17.5574 | 6.32E-07 | 0.124 | ACTCCAGAAGAGGGCGCCA | 2 | ENCFF367UNR,ENCFF093ZRU |
| 319 | 2 | 144402783 | 144403008 | 225 | 129S1,AJ,BTBR,C57L,C58J,CAST,CE,DBA1J,DBA2J,MOLF,NZB,NZO,NZW,SMJ,SPRET | 2 | 144402906 | 144402939 | 34 | - | 18.3659 | 2.76E-07 | 0.0538 | TAGCTGTCTTCAGACACTCCAGAAGAGGGCGCCA | 2 | ENCFF367UNR,ENCFF093ZRU |
| 320 | 2 | 146118060 | 146118278 | 218 | AJ,BALB,BUB,C3H,CAST,CBA,FVB,ILNJ2,KK,MA/My,MRL,NOD,RF,SEA,SJL,SMJ,SPRET.SWR.TH.WSB | 2 | 146118185 | 146118218 | 34 | - | 17.0325 | 6.63E-07 | 0.0853 | TAGCTGTCCTCAGACACTCCAGAAGAGGGAGGCA | | |
| 321 | 2 | 150203548 | 150203777 | 229 | SPRET | 2 | 150203672 | 150203705 | 34 | - | 17.3902 | 5.27E-07 | 0.075 | TAGCTGTCCTCAGACACTCCAGAAGAGGGCATCA | | |
| 322 | 2 | 153692983 | 153693447 | 464 | SPRET | 2 | 153693121 | 153693139 | 19 | - | 25.2459 | 4.18E-10 | 0.0181 | CGGCCACCAGGTGGCACTG | 26 | ENCFF093ZRU,ENCFF367UNR,ENCFF099QNE,ENCFF821BIA,ENCFF907XEG,ENCFF024JOZ,ENCFF327LYS,ENCFF951GOP,ENCFF316FYN,ENCFF502WDT,ENCFF055HAP,ENCFF609MDN,ENCFF575MGI,ENCFF165LXO,ENCFF194HVL,ENCFF097CAQ,ENCFF883UPM,ENCFF092NMJ,ENCFF089GBC,ENCFF589ZBJ,ENCFF136ZDK,ENCFF957SPH,ENCFF491RJK,ENCFF198SSR,ENCFF142CNG,ENCFF281QMV |
| 323 | 2 | 154136578 | 154136911 | 333 | SPRET | 2 | 154136870 | 154136888 | 19 | - | 19.2951 | 1.88E-07 | 0.071 | CCGCCACTAGATGGTGCTA | 22 | ENCFF367UNR,ENCFF201JUA,ENCFF093ZRU,ENCFF099QNE,ENCFF316FYN,ENCFF024JOZ,ENCFF821BIA,ENCFF907XEG,ENCFF327LYS,ENCFF136ZDK,ENCFF589ZBJ,ENCFF198SSR,ENCFF089GBC,ENCFF165LXO,ENCFF092NMJ,ENCFF430PPJ,ENCFF052KVO,ENCFF883UPM,ENCFF957SPH,ENCFF281QMV,ENCFF194HVL,ENCFF609MDN |
| 324 | 2 | 156253224 | 156253438 | 214 | SPRET | 2 | 156253350 | 156253368 | 19 | - | 17.5574 | 6.32E-07 | 0.124 | ACTCCAGAAGAGGGCGCCA | 3 | ENCFF367UNR,ENCFF099QNE,ENCFF821BIA |
| 324 | 2 | 156253224 | 156253438 | 214 | SPRET | 2 | 156253350 | 156253383 | 34 | - | 18.3659 | 2.76E-07 | 0.0538 | TAGCTGTCTTCAGACACTCCAGAAGAGGGCGCCA | 3 | ENCFF367UNR,ENCFF099QNE,ENCFF821BIA |
| 325 | 2 | 158140673 | 158141128 | 455 | KK,MOLF,SJL,SPRET | 2 | 158141089 | 158141122 | 34 | - | 17.0244 | 6.67E-07 | 0.0854 | GAGCAGGTGTGGCAGCAGTCCCAAGATGGCGCCC | | |
| 326 | 2 | 158211034 | 158211249 | 215 | CAST,KK,MOLF,SJL,SPRET | 2 | 158211160 | 158211178 | 19 | - | 17.5574 | 6.32E-07 | 0.124 | ACTCCAGAAGAGGGCGCCA | 1 | ENCFF093ZRU |
| 326 | 2 | 158211034 | 158211249 | 215 | CAST,KK,MOLF,SJL,SPRET | 2 | 158211160 | 158211193 | 34 | - | 18.3659 | 2.76E-07 | 0.0538 | TAGCTGTCTTCAGACACTCCAGAAGAGGGCGCCA | 1 | ENCFF093ZRU |
| 327 | 2 | 160509038 | 160509251 | 213 | CAST,MOLF,SPRET | 2 | 160509164 | 160509197 | 34 | - | 17.2439 | 5.79E-07 | 0.0793 | TAGCTGTCCTCAGATACTCCAGAAGAGGGCATCA | | |
| 328 | 2 | 162885257 | 162885462 | 205 | KK,MOLF,SPRET | 2 | 162885382 | 162885415 | 34 | - | 16.4715 | 9.47E-07 | 0.0987 | TAGCTGTCTTCAGACACTCCAGAAGGGGGTGCCA | | |
| 329 | 2 | 164719690 | 164719896 | 206 | AJ,AKR,BALB,BTBR,BUB,C3H,C58J,CAST,CBA,DBA1J,DBA2J,FVB,ILNJ2,KK,MA/My,MRL,NOD,NOR,NZB,NZO,NZW,PJ,PL,RF,SEA,SJL,SMJ,SPRET,SWR,TH,WSB | 2 | 164719814 | 164719832 | 19 | - | 17.5574 | 6.32E-07 | 0.124 | ACTCCAGAAGAGGGCGCCA | 5 | ENCFF093ZRU,ENCFF099QNE,ENCFF327LYS,ENCFF821BIA,ENCFF367UNR |
| 329 | 2 | 164719690 | 164719896 | 206 | AJ,AKR,BALB,BTBR,BUB,C3H,C58J,CAST,CBA,DBA1J,DBA2J,FVB,ILNJ2,KK,MA/My,MRL,NOD,NOR,NZB,NZO,NZW,PJ,PL,RF,SEA,SJL,SMJ,SPRET,SWR,TH,WSB | 2 | 164719814 | 164719847 | 34 | - | 18.561 | 2.42E-07 | 0.0519 | AAGCTGTCTTCAGACACTCCAGAAGAGGGCGCCA | 5 | ENCFF093ZRU,ENCFF099QNE,ENCFF327LYS,ENCFF821BIA,ENCFF367UNR |
| 330 | 2 | 165620539 | 165620735 | 196 | KK.SPRET | 2 | 165620666 | 165620699 | 34 | - | 16.4472 | 9.61E-07 | 0.0987 | TAGCTGTCTTCAGACACACCAGAAGAGGGCGTCA | | |
| 331 | 2 | 165926667 | 165926889 | 222 | CAST.SPRET | 2 | 165926876 | 165926894 | 19 | - | 17.3279 | 7.34E-07 | 0.133 | GGACCACAAGGGGTCGCTC | 3 | ENCFF093ZRU.ENCFF367UNR.ENCFF099QNE |
| 332 | 2 | 166262534 | 166262645 | 111 | CAST | 2 | 166262508 | 166262541 | 34 | - | 16.3902 | 9.96E-07 | 0.1 | CTTCAGTGCTGGGAGCTGCCTCCGGGAGCCAGGC | | |
| 333 | 2 | 166745969 | 166746676 | 707 | 129S1.C57L.CE.WSB | 2 | 166746014 | 166746048 | 35 | - | 19.4954 | 1.20E-07 | 0.0535 | CCGCATGTATGCCTGCATGCCTGAAGAGGGCACCA | 4 | ENCFF089GBC.ENCFF055HAP.ENCFF502WDT.ENCFF093ZRU |
| 334 | 2 | 167289634 | 167289692 | 58 | CAST.CE | 2 | 167289629 | 167289662 | 34 | - | 20.0569 | 8.52E-08 | 0.027 | CTGCAGCTCACTCAGAGAACACCAGGTGCTACTA | | |
| 335 | 2 | 167913303 | 167913516 | 213 | CAST.SPRET | 2 | 167913434 | 167913467 | 34 | - | 16.4634 | 9.51E-07 | 0.0987 | TAGCTGTCTTCAGACGCACCAGAAGAGGGCATCA | | |
| 336 | 2 | 167981845 | 167982051 | 206 | 129S1.C57L.KK.NZB.SPRET.TH.WSB | 2 | 167981967 | 167982000 | 34 | - | 17.8455 | 3.91E-07 | 0.0641 | TAGCTGTCCTCAGACACTCCAGAAGAGGGCGTCA | | |
| 337 | 2 | 168494045 | 168494249 | 204 | 129S1,AJ,AKR,BALB,BTBR,C3H,C57L,CAST,CBA,DBA1J,DBA2J,FVB,ILNJ2,MA/My,MRL,NOD,NOR,NZB,NZW,PJ,PL,RF,SEA,SJL,SMJ.SPRET.SWR.WSB | 2 | 168494155 | 168494173 | 19 | - | 18.623 | 3.06E-07 | 0.0887 | ACACCAGAAGAGGGCGCCA | | |
| 337 | 2 | 168494045 | 168494249 | 204 | 129S1,AJ,AKR,BALB,BTBR,C3H,C57L,CAST,CBA,DBA1J,DBA2J,FVB,ILNJ2,MA/My,MRL,NOD,NOR,NZB,NZW,PJ,PL,RF,SEA,SJL,SMJ.SPRET.SWR.WSB | 2 | 168494155 | 168494188 | 34 | - | 18.8537 | 1.98E-07 | 0.0465 | TAGCTGTCTTCAGACACACCAGAAGAGGGCGCCA | | |
| 338 | 2 | 168527516 | 168527902 | 386 | SPRET | 2 | 168527628 | 168527661 | 34 | - | 18.1545 | 3.18E-07 | 0.0576 | TAGCTGTTTTCAGACACACCAGAAGAGGGCGTCA | 1 | ENCFF093ZRU |
| 339 | 2 | 172657515 | 172657821 | 306 | CAST | 2 | 172657628 | 172657662 | 35 | - | 20.3853 | 6.77E-08 | 0.0381 | GTGCAGGTGCTCATGGTGACCAGAAGAAGGTGCTA | | |
| 340 | 2 | 176939843 | 176940048 | 205 | SPRET | 2 | 176939968 | 176940001 | 34 | - | 16.4472 | 9.61E-07 | 0.0987 | TAGCTGTCTTCAGACACACCAGAAGAGGGCGTCA | | |
| 341 | 2 | 180986188 | 180986461 | 273 | AJ,AKR,BUB,C3H,C58J,CAST,CE,DBA1J,DBA2J,KK,LP,MA/My,MOLF,NZB,NZO,NZW,PJ,RF,RH2,SPRET | 2 | 180986342 | 180986375 | 34 | - | 17.4634 | 5.02E-07 | 0.0732 | GTGCACCACGACGCCCGGCCTGCAGGTGGATTTC | | |
| 342 | 2 | 181137837 | 181138042 | 205 | AJ,AKR,BALB,BUB,C3H,CAST,CE,DBA1J,DBA2J,KK,MA/My,MOLF,NZB,PJ,RF,RH2,SEA,SPRET | 2 | 181137965 | 181137998 | 34 | - | 17.6667 | 4.40E-07 | 0.0682 | TAGCTGTCTTCAGACACGCCAGAAGAGGGCGTCA | | |
| 343 | 3 | 3101915 | 3102116 | 201 | CAST.MOLF | 3 | 3101973 | 3102006 | 34 | + | 18.561 | 2.42E-07 | 0.079 | AAGCTGTCTTCAGACACTCCAGAAGAGGGCGCCA | | |
| 343 | 3 | 3101915 | 3102116 | 201 | CAST.MOLF | 3 | 3101988 | 3102006 | 19 | + | 17.5574 | 6.32E-07 | 0.181 | ACTCCAGAAGAGGGCGCCA | | |
| 344 | 3 | 7125983 | 7126484 | 501 | SPRET | 3 | 7125999 | 7126032 | 34 | + | 17.0813 | 6.43E-07 | 0.12 | CTGCACTCACATTTGCCATTACAAGATGGCGCTG | | |
| 345 | 3 | 8587560 | 8587773 | 213 | CAST.MOLF.SPRET | 3 | 8587631 | 8587664 | 34 | + | 17.8455 | 3.91E-07 | 0.0943 | TAGCTGTCCTCAGACACTCCAGAAGAGGGCGTCA | | |
| 346 | 3 | 10596600 | 10596745 | 145 | SPRET | 3 | 10596603 | 10596636 | 34 | + | 18.8049 | 2.05E-07 | 0.0723 | ATGCAGTTCTGAATCCTCCCTGTAGGAGGGGGTC | | |
| 347 | 3 | 15333732 | 15334083 | 351 | AJ,BALB,BUB,C3H,C58J,CAST,CBA,FVB,MOLF,PL,SJL,SPRET,SWR,TH | 3 | 15333749 | 15333782 | 34 | + | 16.561 | 8.95E-07 | 0.14 | CTGCCCTTGCAATCGCCATTACAAGATGGCGCTG | | |
| 348 | 3 | 16079337 | 16079669 | 332 | MOLF,SPRET | 3 | 16079368 | 16079386 | 19 | + | 19.2951 | 1.88E-07 | 0.113 | CCGCCACTAGATGGTGCTA | 20 | ENCFF093ZRU,ENCFF201JUA,ENCFF367UNR,ENCFF052KVO,ENCFF430PPJ,ENCFF198SSR,ENCFF136ZDK,ENCFF092NMJ,ENCFF099QNE,ENCFF821BIA,ENCFF883UPM,ENCFF024JOZ,ENCFF327LYS,ENCFF907XEG,ENCFF089GBC,ENCFF589ZBJ,ENCFF165LXO,ENCFF055HAP,ENCFF502WDT |
| 349 | 3 | 24410923 | 24411313 | 390 | SPRET | 3 | 24410991 | 24411024 | 34 | + | 19.1463 | 1.62E-07 | 0.0645 | TTGCTGTCTTCGGACACACCAGAAGAGGGCTCCA | | |
| 350 | 3 | 26287563 | 26287781 | 218 | SPRET | 3 | 26287640 | 26287673 | 34 | + | 17.626 | 4.52E-07 | 0.101 | TAGCTGTCTCCAGACACACCAGAAGAGGGCATCA | | |
| 351 | 3 | 27130519 | 27130731 | 212 | AJ,AKR,C3H,CAST,CBA,CE,MOLF,PL,RF,SPRET | 3 | 27130591 | 27130624 | 34 | + | 17.0488 | 6.56E-07 | 0.121 | TTGCTATCTTCAGATACACCAGAAGAGGGCATCA | | |
| 352 | 3 | 27534894 | 27534952 | 58 | MOLF | 3 | 27534865 | 27534898 | 34 | + | 17.2195 | 5.88E-07 | 0.114 | CTGCTGTATCAGCTCCTGCCTCCAGGTTTCTGTC | | |
| 353 | 3 | 28216303 | 28216468 | 165 | SPRET | 3 | 28216316 | 28216334 | 19 | + | 17.2459 | 7.74E-07 | 0.195 | GAGTCACCAGGGGGCACCC | 1 | ENCFF093ZRU |
| 353 | 3 | 28216303 | 28216468 | 165 | SPRET | 3 | 28216349 | 28216367 | 19 | + | 17.2459 | 7.74E-07 | 0.195 | GAGTCACCAGGGGGCACCC | 1 | ENCFF093ZRU |
| 353 | 3 | 28216303 | 28216468 | 165 | SPRET | 3 | 28216415 | 28216433 | 19 | + | 17.2459 | 7.74E-07 | 0.195 | GAGTCACCAGGGGGCACCC | 1 | ENCFF093ZRU |
| 354 | 3 | 34817878 | 34818086 | 208 | AJ,BUB,C3H,CAST,CBA,CE,DBA1J,FVB,MOLF.SJL.SPRET.SWR.WSB | 3 | 34817944 | 34817977 | 34 | + | 17.9431 | 3.66E-07 | 0.0914 | CAGCTGTCCTCAGACACTCCAGAAGAGGGAGTCA | | |
| 355 | 3 | 34826304 | 34826602 | 298 | SPRET | 3 | 34826360 | 34826393 | 34 | + | 24.0894 | 3.65E-09 | 0.00457 | CTGCAGTTCCTGTGGAGGCCAGAAGAGGACCTCA | 2 | ENCFF052KVO.ENCFF533APC |
| 356 | 3 | 35816000 | 35816203 | 203 | SPRET | 3 | 35816062 | 35816095 | 34 | + | 18.3659 | 2.76E-07 | 0.0804 | TAGCTGTCTTCAGACACTCCAGAAGAGGGCGCCA | 3 | ENCFF367UNR.ENCFF093ZRU.ENCFF201JUA |
| 356 | 3 | 35816000 | 35816203 | 203 | SPRET | 3 | 35816077 | 35816095 | 19 | + | 17.5574 | 6.32E-07 | 0.181 | ACTCCAGAAGAGGGCGCCA | 3 | ENCFF367UNR.ENCFF093ZRU.ENCFF201JUA |
| 357 | 3 | 37445241 | 37445504 | 263 | CAST.ILNJ2.MOLF.TH | 3 | 37445319 | 37445352 | 34 | + | 16.4472 | 9.61E-07 | 0.141 | TAGCTGTCTTCAGACACACCAGAAGAGGGCGTCA | | |
| 358 | 3 | 45027727 | 45027918 | 191 | SPRET | 3 | 45027781 | 45027814 | 34 | + | 16.4472 | 9.61E-07 | 0.141 | TAGCTGTCTTCAGACACACCAGAAGAGGGCGTCA | | |
| 359 | 3 | 51278907 | 51279060 | 153 | CAST.MOLF.NZW | 3 | 51278910 | 51278943 | 34 | + | 18.3659 | 2.76E-07 | 0.0804 | TAGCTGTCTTCAGACACTCCAGAAGAGGGCGCCA | 3 | ENCFF367UNR.ENCFF093ZRU.ENCFF201JUA |
| 359 | 3 | 51278907 | 51279060 | 153 | CAST.MOLF.NZW | 3 | 51278925 | 51278943 | 19 | + | 17.5574 | 6.32E-07 | 0.181 | ACTCCAGAAGAGGGCGCCA | 3 | ENCFF367UNR.ENCFF093ZRU.ENCFF201JUA |
| 360 | 3 | 51333797 | 51334035 | 238 | SPRET | 3 | 51333892 | 51333925 | 34 | + | 16.4634 | 9.51E-07 | 0.141 | TAGCTGTCTTCAGACGCACCAGAAGAGGGCATCA | 1 | ENCFF093ZRU |
| 361 | 3 | 54422208 | 54422497 | 289 | CE.WSB | 3 | 54422376 | 54422409 | 34 | + | 23.1545 | 7.96E-09 | 0.00773 | ATGTAGTGCCTGTGGATGCCAGCAGGAGGCATCA | | |
| 362 | 3 | 56490068 | 56490647 | 579 | MOLF | 3 | 56490587 | 56490620 | 34 | + | 16.8862 | 7.28E-07 | 0.127 | TTGCTCTCTTCAGTCACACCAGAAGAGGGCATTG | | |
| 362 | 3 | 56490068 | 56490647 | 579 | MOLF | 3 | 56490362 | 56490395 | 34 | - | 18.3659 | 2.76E-07 | 0.0804 | ATGCAGTGCCTATGGATGCCAGAAGTAGGCATTG | | |
| 363 | 3 | 56950315 | 56950368 | 53 | CAST.MOLF | 3 | 56950322 | 56950355 | 34 | + | 16.9919 | 6.81E-07 | 0.123 | TAGCTGTCTTAAGACTCTCCAGAAGAGGGCATCA | | |
| 364 | 3 | 57646623 | 57646882 | 259 | AJ,CAST,CE,DBA1J,DBA2J,FVB,MOLF,PJ,SEA,SJL,SMJ,SPRET,SWR,TH,WSB | 3 | 57646740 | 57646773 | 34 | + | 16.4472 | 9.61E-07 | 0.141 | TAGCTGTCTTCAGACACACCAGAAGAGGGCGTCA | | |
| 365 | 3 | 61102502 | 61102695 | 193 | CAST | 3 | 61102565 | 61102599 | 35 | + | 17.0183 | 5.36E-07 | 0.18 | TTGCACATAGAGGTATGGACCAGAAGATGGAGCAC | | |
| 366 | 3 | 62053247 | 62053846 | 599 | MOLF | 3 | 62053278 | 62053311 | 34 | + | 17.0325 | 6.63E-07 | 0.122 | ATGCAGATAGAGACATGAGCTCCAGGGGGTACTG | | |
| 367 | 3 | 65871941 | 65872016 | 75 | FVB.NOD.NOR.SWR | 3 | 65871965 | 65871998 | 34 | + | 16.6016 | 8.72E-07 | 0.138 | GTGCAGGTCCCCCTGTGGAAGCCAGAGGTCAGAT | | |
| 368 | 3 | 68754138 | 68754373 | 235 | CAST.MOLF.SPRET | 3 | 68754196 | 68754229 | 34 | + | 18.3659 | 2.76E-07 | 0.0804 | TAGCTGTCTTCAGACACTCCAGAAGAGGGCGCCA | | |
| 368 | 3 | 68754138 | 68754373 | 235 | CAST.MOLF.SPRET | 3 | 68754211 | 68754229 | 19 | + | 17.5574 | 6.32E-07 | 0.181 | ACTCCAGAAGAGGGCGCCA | | |
| 369 | 3 | 73305611 | 73305818 | 207 | AJ,CAST,CE,MA/My,MOLF,NOD,NOR,RH2,SPRET.WSB | 3 | 73305676 | 73305709 | 34 | + | 16.4472 | 9.61E-07 | 0.141 | TAGCTGTCTTCAGACACACCAGAAGAGGGCGTCA | | |
| 370 | 3 | 87827833 | 87828033 | 200 | AJ,AKR,BALB,BTBR,BUB,C3H,CAST,CBA,CE,DBA1J,DBA2J,FVB,ILNJ2,KK,MA/My,MOLF,MRL,NOD,NOR,NZB,NZO,NZW,PJ,PL,RH2,SEA,SJL,SMJ,SPRET,SWR,TH | 3 | 87827905 | 87827923 | 19 | + | 17.8852 | 5.08E-07 | 0.167 | ACACCAGAAGAGGGCACCA | 6 | ENCFF367UNR,ENCFF907XEG,ENCFF327LYS,ENCFF883UPM,ENCFF198SSR,ENCFF093ZRU |
| 371 | 3 | 88061331 | 88061594 | 263 | CAST.SPRET | 3 | 88061427 | 88061460 | 34 | + | 17.2439 | 5.79E-07 | 0.113 | TAGCTGTCCTCAGATACTCCAGAAGAGGGCATCA | | |
| 372 | 3 | 88193037 | 88193254 | 217 | CAST,MA/My,MOLF,RF,SPRET,WSB | 3 | 88193111 | 88193144 | 34 | + | 18.3659 | 2.76E-07 | 0.0804 | TAGCTGTCTTCAGACACTCCAGAAGAGGGCGCCA | 1 | ENCFF093ZRU |
| 372 | 3 | 88193037 | 88193254 | 217 | CAST,MA/My,MOLF,RF,SPRET,WSB | 3 | 88193126 | 88193144 | 19 | + | 17.5574 | 6.32E-07 | 0.181 | ACTCCAGAAGAGGGCGCCA | 1 | ENCFF093ZRU |
| 373 | 3 | 88591919 | 88592117 | 198 | AJ,AKR,BTBR,BUB,C3H,CAST,CBA,FVB,KK,MA/My,MOLF,MRL,NZB,NZO,NZW,RF,SJL,SPRET,SWR,TH | 3 | 88591974 | 88592007 | 34 | + | 18.3659 | 2.76E-07 | 0.0804 | TAGCTGTCTTCAGACACTCCAGAAGAGGGCGCCA | 10 | ENCFF821BIA,ENCFF907XEG,ENCFF099QNE,ENCFF502WDT,ENCFF367UNR,ENCFF024JOZ,ENCFF883UPM,ENCFF093ZRU,ENCFF055HAP |
| 373 | 3 | 88591919 | 88592117 | 198 | AJ,AKR,BTBR,BUB,C3H,CAST,CBA,FVB,KK,MA/My,MOLF,MRL,NZB,NZO,NZW,RF,SJL,SPRET.SWR.TH | 3 | 88591989 | 88592007 | 19 | + | 17.5574 | 6.32E-07 | 0.181 | ACTCCAGAAGAGGGCGCCA | 10 | ENCFF821BIA,ENCFF907XEG,ENCFF099QNE,ENCFF502WDT,ENCFF367UNR,ENCFF024JOZ,ENCFF883UPM,ENCFF093ZRU,ENCFF055HAP |
| 374 | 3 | 89069764 | 89070318 | 554 | KK.MOLF | 3 | 89069769 | 89069802 | 34 | + | 16.4472 | 9.61E-07 | 0.141 | TAGCTGTCTTCAGACACACCAGAAGAGGGCGTCA | | |
| 375 | 3 | 89111000 | 89111194 | 194 | SPRET | 3 | 89111047 | 89111080 | 34 | + | 16.4472 | 9.61E-07 | 0.141 | TAGCTGTCTTCAGACACACCAGAAGAGGGCGTCA | 1 | ENCFF093ZRU |
| 376 | 3 | 89714039 | 89714877 | 838 | SPRET | 3 | 89714258 | 89714291 | 34 | + | 23.1463 | 8.01E-09 | 0.00776 | ATGCACTACCCAAGAAGACCAGAAGAGGGCATCA | 15 | ENCFF099QNE,ENCFF327LYS,ENCFF201JUA,ENCFF136ZDK,ENCFF198SSR,ENCFF052KVO,ENCFF093ZRU,ENCFF367UNR,ENCFF065ODV,ENCFF502WDT,ENCFF055HAP,ENCFF194HVL,ENCFF589ZBJ,ENCFF821BIA.ENCFF907XEG |
| 377 | 3 | 90062441 | 90062651 | 210 | CAST.MOLF.SPRET | 3 | 90062509 | 90062542 | 34 | + | 18.7642 | 2.11E-07 | 0.0734 | TAGCTGTCCTCAGATACGCCAGAAGAGGGCATTA | | |
| 378 | 3 | 90188327 | 90188852 | 525 | MOLF | 3 | 90188571 | 90188604 | 34 | + | 22.1301 | 1.80E-08 | 0.014 | AAGCAGTACCCACAGAGGCCAGAAGAGGGTATTG | 1 | ENCFF093ZRU |
| 379 | 3 | 90212358 | 90212561 | 203 | CAST.SPRET | 3 | 90212405 | 90212438 | 34 | + | 16.4634 | 9.51E-07 | 0.141 | TAGCTGTCTTCAGACGCACCAGAAGAGGGCATCA | | |
| 380 | 3 | 93745376 | 93746044 | 668 | CAST.RF | 3 | 93745350 | 93745383 | 34 | + | 16.561 | 8.95E-07 | 0.14 | CTGCCCTTGCAATCGCCATTACAAGATGGCGCTG | | |
| 381 | 3 | 95057403 | 95057644 | 241 | SPRET | 3 | 95057457 | 95057490 | 34 | + | 16.4472 | 9.61E-07 | 0.141 | TAGCTGTCTTCAGACACACCAGAAGAGGGCGTCA | | |
| 382 | 3 | 95120439 | 95120654 | 215 | CAST,KK,MOLF,NOD,NOR,SPRET,TH,WSB | 3 | 95120510 | 95120543 | 34 | + | 18.561 | 2.42E-07 | 0.079 | AAGCTGTCTTCAGACACTCCAGAAGAGGGCGCCA | 5 | ENCFF093ZRU,ENCFF367UNR,ENCFF821BIA,ENCFF201JUA |
| 382 | 3 | 95120439 | 95120654 | 215 | CAST,KK,MOLF,NOD,NOR,SPRET,TH,WSB | 3 | 95120525 | 95120543 | 19 | + | 17.5574 | 6.32E-07 | 0.181 | ACTCCAGAAGAGGGCGCCA | 5 | ENCFF093ZRU,ENCFF367UNR,ENCFF821BIA,ENCFF201JUA |
| 383 | 3 | 95255739 | 95255942 | 203 | MOLF,SPRET | 3 | 95255800 | 95255833 | 34 | + | 18.3659 | 2.76E-07 | 0.0804 | TAGCTGTCTTCAGACACTCCAGAAGAGGGCGCCA | 8 | ENCFF201JUA,ENCFF502WDT,ENCFF821BIA,ENCFF093ZRU,ENCFF367UNR,ENCFF055HAP,ENCFF099QNE |
| 383 | 3 | 95255739 | 95255942 | 203 | MOLF,SPRET | 3 | 95255815 | 95255833 | 19 | + | 17.5574 | 6.32E-07 | 0.181 | ACTCCAGAAGAGGGCGCCA | 8 | ENCFF201JUA,ENCFF502WDT,ENCFF821BIA,ENCFF093ZRU,ENCFF367UNR,ENCFF055HAP,ENCFF099QNE |
| 384 | 3 | 95291152 | 95291422 | 270 | CAST.SPRET | 3 | 95291280 | 95291313 | 34 | + | 17.2439 | 5.79E-07 | 0.113 | TAGCTGTCCTCAGATACTCCAGAAGAGGGCATCA | | |
| 385 | 3 | 95474270 | 95474445 | 175 | CAST.SPRET | 3 | 95474325 | 95474358 | 34 | + | 18.3984 | 2.70E-07 | 0.0804 | TAGCTGTCTTCAGACACACCAGAAGAGGGCACCA | 1 | ENCFF093ZRU |
| 385 | 3 | 95474270 | 95474445 | 175 | CAST.SPRET | 3 | 95474340 | 95474358 | 19 | + | 17.8852 | 5.08E-07 | 0.167 | ACACCAGAAGAGGGCACCA | 1 | ENCFF093ZRU |
| 386 | 3 | 95877937 | 95878146 | 209 | 129S1,AJ,BALB,C3H,C57L,CAST,CBA,CE,FVB,KK,MA/My,MOLF,NOD,NOR,NZB,NZO,NZW,PJ,RH2,SEA,SJL,SMJ,SPRET,SWR,TH,WSB | 3 | 95878005 | 95878038 | 34 | + | 17.8455 | 3.91E-07 | 0.0943 | TAGCTGTCCTCAGACACTCCAGAAGAGGGCGTCA | 1 | ENCFF055HAP |
| 387 | 3 | 96643107 | 96643318 | 211 | KK,MA/My,MOLF,NZB,NZO,NZW,RF,SPRET | 3 | 96643175 | 96643208 | 34 | + | 16.4634 | 9.51E-07 | 0.141 | TAGCTGTCTTCAGACGCACCAGAAGAGGGCATCA | 1 | ENCFF093ZRU |
| 388 | 3 | 104659686 | 104660623 | 937 | SPRET | 3 | 104660334 | 104660368 | 35 | + | 16.3211 | 7.98E-07 | 0.221 | GTGAGGCTTCTTCACACCGCCGGTAGAGGGCGCGC | 17 | ENCFF092NMJ,ENCFF089GBC,ENCFF093ZRU,ENCFF367UNR,ENCFF165LXO,ENCFF099QNE,ENCFF136ZDK,ENCFF821BIA,ENCFF033UQX,ENCFF907XEG,ENCFF327LYS,ENCFF609MDN,ENCFF589ZBJ,ENCFF957SPH,ENCFF194HVL,ENCFF198SSR,ENCFF024JOZ |
| 388 | 3 | 104659686 | 104660623 | 937 | SPRET | 3 | 104660350 | 104660368 | 19 | + | 16.9672 | 9.26E-07 | 0.212 | CCGCCGGTAGAGGGCGCGC | 17 | ENCFF092NMJ,ENCFF089GBC,ENCFF093ZRU,ENCFF367UNR,ENCFF165LXO,ENCFF099QNE,ENCFF136ZDK,ENCFF821BIA,ENCFF033UQX,ENCFF907XEG,ENCFF327LYS,ENCFF609MDN,ENCFF589ZBJ,ENCFF957SPH,ENCFF194HVL,ENCFF198SSR,ENCFF024JOZ |
| 389 | 3 | 107510464 | 107510674 | 210 | BALB,CAST,CE,MA/My,MOLF,MRL,NZO,NZW.RH2.SPRET.TH.WSB | 3 | 107510525 | 107510558 | 34 | + | 16.4634 | 9.51E-07 | 0.141 | TAGCTGTCTTCAGACGCACCAGAAGAGGGCATCA | | |
| 390 | 3 | 109821097 | 109821491 | 394 | SPRET | 3 | 109821389 | 109821407 | 19 | + | 18.6393 | 3.02E-07 | 0.132 | TGGCCTGCAGGAGGCAGTG | | |
| 391 | 3 | 114136741 | 114136935 | 194 | CAST | 3 | 114136798 | 114136832 | 35 | + | 19.3486 | 1.32E-07 | 0.0815 | CTCCAGTGGGAGTCCTCCTACAGTAGAGGGCACTT | | |
| 392 | 3 | 114945012 | 114945229 | 217 | AJ,AKR,BALB,C3H,CAST,CBA,DBA1J,DBA2J,FVB,ILNJ2,KK,MA/My,MOLF,MRL,NZO,NZW,PJ,PL,RF,RH2,SEA,SJL,SMJ,SWR,TH,WSB | 3 | 114945088 | 114945121 | 34 | + | 18.3659 | 2.76E-07 | 0.0804 | TAGCTGTCTTCAGACACTCCAGAAGAGGGCGCCA | | |
| 392 | 3 | 114945012 | 114945229 | 217 | AJ,AKR,BALB,C3H,CAST,CBA,DBA1J,DBA2J,FVB,ILNJ2,KK,MA/My,MOLF,MRL,NZO,NZW,PJ,PL,RF,RH2,SEA,SJL,SMJ,SWR,TH,WSB | 3 | 114945103 | 114945121 | 19 | + | 17.5574 | 6.32E-07 | 0.181 | ACTCCAGAAGAGGGCGCCA | | |
| 393 | 3 | 116481376 | 116481552 | 176 | CAST.MOLF.SPRET | 3 | 116481417 | 116481450 | 34 | + | 17.2439 | 5.79E-07 | 0.113 | TAGCTGTCCTCAGATACTCCAGAAGAGGGCATCA | | |
| 394 | 3 | 116546109 | 116546318 | 209 | 129S1,AJ,AKR,BALB,BUB,C3H,C57L,C58J,CAST,CBA,CE,DBA1J,DBA2J,FVB,ILNJ2,LP,MA/My,MOLF,MRL,NOD,NOR,NZB,NZO,NZW,PL,RF,RH2,SJL,SMJ,SPRET,SWR,TH,WSB | 3 | 116546176 | 116546209 | 34 | + | 17.8455 | 3.91E-07 | 0.0943 | TAGCTGTCCTCAGACACTCCAGAAGAGGGCGTCA | | |
| 395 | 3 | 116560724 | 116561021 | 297 | MOLF,SPRET | 3 | 116560768 | 116560802 | 35 | + | 24.6239 | 3.30E-09 | 0.00966 | TTGCAGTGACAACCTGTCGCCACCTGGTGGCGTGC | 24 | ENCFF201JUA,ENCFF093ZRU,ENCFF092NMJ,ENCFF024JOZ,ENCFF367UNR,ENCFF089GBC,ENCFF165LXO,ENCFF907XEG,ENCFF883UPM,ENCFF099QNE,ENCFF821BIA,ENCFF136ZDK,ENCFF052KVO,ENCFF316FYN,ENCFF142CNG,ENCFF589ZBJ,ENCFF502WDT,ENCFF097CAQ,ENCFF491RJK,ENCFF055HAP,ENCFF194HVL,ENCFF957SPH,ENCFF198SSR,ENCFF327LYS |
| 395 | 3 | 116560724 | 116561021 | 297 | MOLF,SPRET | 3 | 116560782 | 116560800 | 19 | - | 19.918 | 1.18E-07 | 0.091 | ACGCCACCAGGTGGCGACA | 24 | ENCFF201JUA,ENCFF093ZRU,ENCFF092NMJ,ENCFF024JOZ,ENCFF367UNR,ENCFF089GBC,ENCFF165LXO,ENCFF907XEG,ENCFF883UPM,ENCFF099QNE,ENCFF821BIA,ENCFF136ZDK,ENCFF052KVO,ENCFF316FYN,ENCFF142CNG,ENCFF589ZBJ,ENCFF502WDT,ENCFF097CAQ,ENCFF491RJK,ENCFF055HAP,ENCFF194HVL,ENCFF957SPH,ENCFF198SSR,ENCFF327LYS |
| 396 | 3 | 117699915 | 117700245 | 330 | MOLF | 3 | 117700002 | 117700036 | 35 | + | 17.0826 | 5.16E-07 | 0.177 | CTGCACCTTCCACAGGAGAGCACTGGGTGGTCCCA | | |
| 397 | 3 | 122080634 | 122080834 | 200 | BTBR,CAST,DBA1J,DBA2J,MOLF,SMJ,SPRET.TH | 3 | 122080691 | 122080724 | 34 | + | 16.4472 | 9.61E-07 | 0.141 | TAGCTGTCTTCAGACACACCAGAAGAGGGCGTCA | | |
| 398 | 3 | 123347060 | 123347256 | 196 | AJ,BALB,C3H,CAST,CBA,MOLF,NZB,RH2,SMJ.SPRET | 3 | 123347113 | 123347146 | 34 | + | 16.4634 | 9.51E-07 | 0.141 | TAGCTGTCTTCAGACGCACCAGAAGAGGGCATCA | | |
| 399 | 3 | 125211965 | 125212171 | 206 | AJ,BALB,C3H,CAST,CBA,MA/My,MOLF,NZB,RH2,SMJ,SPRET | 3 | 125212045 | 125212063 | 19 | + | 17.8852 | 5.08E-07 | 0.167 | ACACCAGAAGAGGGCACCA | | |
| 400 | 3 | 127165356 | 127166351 | 995 | CE | 3 | 127165619 | 127165652 | 34 | + | 23.7073 | 5.04E-09 | 0.00582 | GTGCAGTGCTCACAGAGGCCAGACGAGGGAGCTA | 7 | ENCFF367UNR,ENCFF821BIA,ENCFF093ZRU,ENCFF089GBC,ENCFF201JUA,ENCFF502WDT,ENCFF099QNE |
| 400 | 3 | 127165356 | 127166351 | 995 | CE | 3 | 127165424 | 127165442 | 19 | - | 16.8525 | 9.96E-07 | 0.218 | TAACCAAAAGAGGGCACCA | 1 | ENCFF367UNR |
| 401 | 3 | 134417213 | 134417541 | 328 | CAST.MOLF.SPRET | 3 | 134417437 | 134417470 | 34 | + | 18.3089 | 2.87E-07 | 0.0804 | ATGCAGCTAGAGACACGAGCTCCAGGGGGTACTG | | |
| 402 | 3 | 135236387 | 135236997 | 610 | SPRET | 3 | 135236896 | 135236929 | 34 | + | 18.0244 | 3.47E-07 | 0.0884 | TAGCTGTTTTCAGATGCACCAGAAGAGGGCATCA | 1 | ENCFF502WDT |
| 403 | 3 | 135293085 | 135293597 | 512 | CAST | 3 | 135293516 | 135293534 | 19 | + | 19.6885 | 1.40E-07 | 0.101 | AGGCCTCCAGGGGGCGGAA | | |
| 404 | 3 | 138168818 | 138169123 | 305 | CAST | 3 | 138169032 | 138169050 | 19 | + | 19.8689 | 1.22E-07 | 0.0926 | TGGCCACCAGAGGGCAAGC | 26 | ENCFF201JUA,ENCFF093ZRU,ENCFF367UNR,ENCFF099QNE,ENCFF883UPM,ENCFF052KVO,ENCFF089GBC,ENCFF327LYS,ENCFF907XEG,ENCFF092NMJ,ENCFF024JOZ,ENCFF609MDN,ENCFF821BIA,ENCFF951GOP,ENCFF055HAP,ENCFF589ZBJ,ENCFF430PPJ,ENCFF502WDT,ENCFF198SSR,ENCFF097CAQ,ENCFF210BTO,ENCFF165LXO,ENCFF957SPH,ENCFF136ZDK,ENCFF281QMV,ENCFF194HVL |
| 405 | 3 | 142876606 | 142877437 | 831 | SPRET | 3 | 142876643 | 142876677 | 35 | + | 16.4587 | 7.38E-07 | 0.212 | CTGCAATCTCCCACCTCCAACAGTAGGTGCCTGGT | | |
| 406 | 3 | 144189940 | 144190157 | 217 | 129S1,BTBR,BUB,C57L,C58J,CAST,CE,DBA2J,LP,MOLF,NZB,NZO,RF,SPRET | 3 | 144190015 | 144190048 | 34 | + | 17.4959 | 4.92E-07 | 0.106 | TAGCTGTCCCCAGACACACCAGAAGAGGGAGTCA | | |
| 407 | 3 | 146097612 | 146098290 | 678 | CAST.MOLF | 3 | 146098280 | 146098313 | 34 | + | 16.4146 | 9.81E-07 | 0.142 | TAGCTGTCTTCAGACACACCAGAGGAGGGCGCCA | 1 | ENCFF093ZRU |
| 408 | 3 | 152269767 | 152269866 | 99 | SPRET | 3 | 152269740 | 152269773 | 34 | + | 18.1382 | 3.22E-07 | 0.0855 | CTCCAGTTGTTAGAGCACCCTCCAGGTGGTGGAG | | |
| 409 | 3 | 153353444 | 153353503 | 59 | 129S1.BTBR.C57L.CAST | 3 | 153353434 | 153353468 | 35 | + | 16.7798 | 6.15E-07 | 0.191 | CAGCAGGTGGCTGCAGGTGGCTGCAGGTGGCAGCA | 5 | ENCFF907XEG.ENCFF136ZDK.ENCFF089GBC.ENCFF024JOZ.ENCFF327LYS |
| 409 | 3 | 153353444 | 153353503 | 59 | 129S1.BTBR.C57L.CAST | 3 | 153353444 | 153353478 | 35 | + | 20.4128 | 6.65E-08 | 0.0531 | CTGCAGGTGGCTGCAGGTGGCAGCAGGTGGCTGCA | 5 | ENCFF907XEG.ENCFF136ZDK.ENCFF089GBC.ENCFF024JOZ.ENCFF327LYS |
| 409 | 3 | 153353444 | 153353503 | 59 | 129S1.BTBR.C57L.CAST | 3 | 153353454 | 153353488 | 35 | + | 20.2569 | 7.36E-08 | 0.0562 | CTGCAGGTGGCAGCAGGTGGCTGCAGGTGGCAGCA | 5 | ENCFF907XEG.ENCFF136ZDK.ENCFF089GBC.ENCFF024JOZ.ENCFF327LYS |
| 409 | 3 | 153353444 | 153353503 | 59 | 129S1.BTBR.C57L.CAST | 3 | 153353464 | 153353498 | 35 | + | 17.3028 | 4.54E-07 | 0.168 | CAGCAGGTGGCTGCAGGTGGCAGCAGGTGGCTGCA | 5 | ENCFF907XEG.ENCFF136ZDK.ENCFF089GBC.ENCFF024JOZ.ENCFF327LYS |
| 409 | 3 | 153353444 | 153353503 | 59 | 129S1.BTBR.C57L.CAST | 3 | 153353474 | 153353508 | 35 | + | 20.2569 | 7.36E-08 | 0.0562 | CTGCAGGTGGCAGCAGGTGGCTGCAGGTGGCAGCA | 5 | ENCFF907XEG.ENCFF136ZDK.ENCFF089GBC.ENCFF024JOZ.ENCFF327LYS |
| 409 | 3 | 153353444 | 153353503 | 59 | 129S1.BTBR.C57L.CAST | 3 | 153353484 | 153353518 | 35 | + | 17.3028 | 4.54E-07 | 0.168 | CAGCAGGTGGCTGCAGGTGGCAGCAGGTGGCTGCA | 5 | ENCFF907XEG.ENCFF136ZDK.ENCFF089GBC.ENCFF024JOZ.ENCFF327LYS |
| 409 | 3 | 153353444 | 153353503 | 59 | 129S1.BTBR.C57L.CAST | 3 | 153353494 | 153353528 | 35 | + | 20.2569 | 7.36E-08 | 0.0562 | CTGCAGGTGGCAGCAGGTGGCTGCAGGTGGCAGCA | 5 | ENCFF907XEG.ENCFF136ZDK.ENCFF089GBC.ENCFF024JOZ.ENCFF327LYS |
| 410 | 3 | 156546724 | 156546945 | 221 | CAST.MOLF.SPRET | 3 | 156546807 | 156546840 | 34 | + | 16.9919 | 6.81E-07 | 0.123 | CAGCTGTCTTCAGACATTCCAGAAGGGGGAGTCA | | |
| 411 | 3 | 156694498 | 156694706 | 208 | CAST.MOLF.SPRET.WSB | 3 | 156694556 | 156694589 | 34 | + | 16.6341 | 8.54E-07 | 0.137 | CTGTAGCTGTCAGACACTCCAGAAGAGGGAGTCA | | |

| | | | | | | | | | | | | | | | | |
|---|---|---|---|---|---|---|---|---|---|---|---|---|---|---|---|---|
| 412 | 3 | 10314329 | 10314661 | 332 | BUB,CAST,MOLF,SMJ,SPRET,WSB | 3 | 10314620 | 10314638 | 19 | - | 19.2951 | 1.88E-07 | 0.113 | CCGCCACTAGATGGTGCTA | 21 | ENCFF210BTO,ENCFF201JUA,ENCFF367UNR,ENCFF093ZRU,ENCFF099QNE,ENCFF024JOZ,ENCFF907XEG,ENCFF327LYS,ENCFF609MDN,ENCFF089GBC,ENCFF136ZDK,ENCFF198SSR,ENCFF883UPM,ENCFF821BIA,ENCFF321AGQ,ENCFF589ZBJ,ENCFF092NMJ,ENCFF165LXO,ENCFF502WDT,ENCFF052KVO,ENCFF430PPJ |
| 413 | 3 | 15610108 | 15610332 | 224 | AJ,BALB,BUB,C3H,C58J,CAST,CBA,CE,DBA2J.FVB.LP.PL.SJL.SWR.TH | 3 | 15610233 | 15610266 | 34 | - | 16.9187 | 7.13E-07 | 0.126 | TAGCTGTCTTCAGACGCACCAGAAGAGGGCGTCA | | |
| 414 | 3 | 16928820 | 16929026 | 206 | SPRET | 3 | 16928945 | 16928978 | 34 | - | 16.7724 | 7.83E-07 | 0.131 | TAGCTGTCTTCAGATGCACCAGAAGAGGGCGTCA | | |
| 415 | 3 | 20129686 | 20129891 | 205 | CAST.SPRET.WSB | 3 | 20129811 | 20129844 | 34 | - | 16.4472 | 9.61E-07 | 0.141 | TAGCTGTCTTCAGACACACCAGAAGAGGGCGTCA | | |
| 416 | 3 | 23725635 | 23725840 | 205 | KK,MA/My,MOLF,NOD,NOR,PJ,PL,SEA,SPRET | 3 | 23725760 | 23725793 | 34 | - | 17.2439 | 5.79E-07 | 0.113 | TAGCTGTCCTCAGATACTCCAGAAGAGGGCATCA | | |
| 417 | 3 | 24620518 | 24620723 | 205 | MOLF.SPRET | 3 | 24620644 | 24620677 | 34 | - | 18.1545 | 3.18E-07 | 0.0853 | TAGCTGTTTTCAGACACACCAGAAGAGGGCGTCA | | |
| 418 | 3 | 28105540 | 28105797 | 257 | MOLF | 3 | 28105681 | 28105714 | 34 | - | 16.7724 | 7.83E-07 | 0.131 | CTGCAGGTTGTCCTCTGACCTCCACATGGAGCAG | | |
| 419 | 3 | 28792631 | 28793264 | 633 | CAST.DBA1J.DBA2J.ILNJ2 | 3 | 28793259 | 28793292 | 34 | - | 18.2195 | 3.05E-07 | 0.0836 | GTGCAATGTTCATGGAGGCCAGAAGAGAGCATCA | 1 | ENCFF201JUA |
| 420 | 3 | 32399123 | 32399358 | 235 | SPRET | 3 | 32399172 | 32399205 | 34 | - | 18.8049 | 2.05E-07 | 0.0723 | GGGCTTTTTTCCCAGCAGCCACAAGGTGGCTCAC | 4 | ENCFF092NMJ.ENCFF430PPJ.ENCFF093ZRU.ENCFF089GBC |
| 421 | 3 | 34140180 | 34140398 | 218 | CAST.MOLF | 3 | 34140246 | 34140279 | 34 | - | 20.8049 | 4.94E-08 | 0.0293 | GTGCAGTACCCATTGTGCCCAGGAAAGGGCATTA | | |
| 422 | 3 | 38726235 | 38726671 | 436 | CAST.CE.SJL.SPRET.SWR.WSB | 3 | 38726629 | 38726662 | 34 | - | 16.626 | 8.59E-07 | 0.137 | CTGCACTCACATTTGCCATTACAAGATGGCACTG | | |
| 423 | 3 | 43876242 | 43876556 | 314 | SPRET | 3 | 43876330 | 43876348 | 19 | - | 17.9508 | 4.86E-07 | 0.167 | TTATCAGTAGGGGGCACCG | 1 | ENCFF093ZRU |
| 423 | 3 | 43876242 | 43876556 | 314 | SPRET | 3 | 43876330 | 43876363 | 34 | - | 18.8211 | 2.03E-07 | 0.0721 | GTACAATACTCATGGTTATCAGTAGGGGGCACCG | 1 | ENCFF093ZRU |
| 424 | 3 | 51287969 | 51288181 | 212 | 129S1,BTBR,BUB,C3H,C57L,C58J,CAST,CBA,CE,FVB,LP,MOLF,NZB,NZW,SJL,SMJ,SPRET.SWR.TH.WSB | 3 | 51288095 | 51288113 | 19 | - | 17.5574 | 6.32E-07 | 0.181 | ACTCCAGAAGAGGGCGCCA | 3 | ENCFF367UNR,ENCFF093ZRU,ENCFF055HAP |
| 424 | 3 | 51287969 | 51288181 | 212 | 129S1,BTBR,BUB,C3H,C57L,C58J,CAST,CBA,CE,FVB,LP,MOLF,NZB,NZW,SJL,SMJ,SPRET.SWR.TH.WSB | 3 | 51288095 | 51288128 | 34 | - | 18.561 | 2.42E-07 | 0.079 | AAGCTGTCTTCAGACACTCCAGAAGAGGGCGCCA | 3 | ENCFF367UNR,ENCFF093ZRU,ENCFF055HAP |
| 425 | 3 | 52974288 | 52975185 | 897 | MOLF | 3 | 52974860 | 52974893 | 34 | - | 22.813 | 1.05E-08 | 0.0094 | ATGCAGTGCTCACAAAGGCCAGATGAGGGCATCA | | |
| 426 | 3 | 53599759 | 53600337 | 578 | CAST.MOLF | 3 | 53599986 | 53600020 | 35 | - | 17.2844 | 4.59E-07 | 0.169 | CTGCACATCAAACTCCGGGCCACAGGAGGAAACAG | | |
| 427 | 3 | 54498103 | 54499012 | 909 | NOR | 3 | 54498766 | 54498784 | 19 | - | 17.2951 | 7.50E-07 | 0.193 | AGACCAGAAGAGGGCGACA | 18 | ENCFF198SSR,ENCFF165LXO,ENCFF883UPM,ENCFF502WDT,ENCFF099QNE,ENCFF589ZBJ,ENCFF136ZDK,ENCFF052KVO,ENCFF907XEG,ENCFF327LYS,ENCFF821BIA,ENCFF201JUA,ENCFF089GBC,ENCFF093ZRU.ENCFF024JOZ.ENCFF092NMJ.ENCFF367UNR.ENCFF055HAP |
| 427 | 3 | 54498103 | 54499012 | 909 | NOR | 3 | 54498766 | 54498799 | 34 | - | 28.8943 | 3.68E-11 | 0.00028 | GTGCAGTTCCCACAGAGACCAGAAGAGGGCGACA | 18 | ENCFF198SSR,ENCFF165LXO,ENCFF883UPM,ENCFF502WDT,ENCFF099QNE,ENCFF589ZBJ,ENCFF136ZDK,ENCFF052KVO,ENCFF907XEG,ENCFF327LYS,ENCFF821BIA,ENCFF201JUA,ENCFF089GBC,ENCFF093ZRU.ENCFF024JOZ.ENCFF092NMJ.ENCFF367UNR.ENCFF055HAP |
| 428 | 3 | 54522944 | 54523815 | 871 | SPRET | 3 | 54522934 | 54522967 | 34 | - | 17.0244 | 6.67E-07 | 0.122 | TAGCTGTCTTCAAACTCTCCAGAAGAGGGCATCA | | |
| 429 | 3 | 54903875 | 54904775 | 900 | AKR,BALB,BTBR,BUB,C3H,CAST,CBA,CE,ILNJ2,KK,MA/My,MOLF,MRL,NOD,NOR,NZB,NZW,PL,RF,SPRET.TH.WSB | 3 | 54904248 | 54904266 | 19 | - | 17.5574 | 6.32E-07 | 0.181 | TGGCCAGTTGGGGGCGCGC | | |
| 430 | 3 | 58454274 | 58454579 | 305 | AKR,BALB,BTBR,BUB,C3H,CAST,CBA,CE,DBA1J,DBA2J,ILNJ2,KK,MA/My,MOLF,MRL,NOD,NOR,NZB,NZW,PJ,PL,RF,SEA,SJL,SMJ.SPRET | 3 | 58454398 | 58454431 | 34 | - | 16.4472 | 9.61E-07 | 0.141 | TAGCTGTCTTCAGACACACCAGAAGAGGGCGTCA | | |
| 431 | 3 | 63040697 | 63040892 | 195 | CAST,DBA1J,DBA2J,FVB,MOLF,SJL,SPRET.SWR | 3 | 63040808 | 63040841 | 34 | - | 16.4472 | 9.61E-07 | 0.141 | TAGCTGTCTTCAGACACACCAGAAGAGGGCGTCA | | |
| 432 | 3 | 64138308 | 64138519 | 211 | CAST.SJL.SPRET | 3 | 64138434 | 64138467 | 34 | - | 16.4472 | 9.61E-07 | 0.141 | TAGCTGTCTTCAGACACACCAGAAGAGGGCGTCA | | |
| 433 | 3 | 64476805 | 64477772 | 967 | CAST.SPRET | 3 | 64477495 | 64477528 | 34 | - | 17.4146 | 5.18E-07 | 0.109 | ATGCAGCTAGAGACACAAGCTCCAGGGGGTACTG | | |
| 434 | 3 | 65369687 | 65370635 | 948 | CAST | 3 | 65370145 | 65370178 | 34 | - | 18.2927 | 2.90E-07 | 0.0809 | TTGCTGTCTTCAGACACACCAGAAGAGGGCATTA | | |
| 435 | 3 | 68358485 | 68358694 | 209 | CAST.CE.MOLF.SPRET | 3 | 68358582 | 68358615 | 34 | - | 18.935 | 1.88E-07 | 0.0693 | TTGCTGTCTTCAGACACACCAGAAGAGGGCGTCA | | |
| 436 | 3 | 70553823 | 70554446 | 623 | SPRET | 3 | 70553926 | 70553960 | 35 | - | 18.2661 | 2.56E-07 | 0.121 | GTGCAGGGGAACGCCAGGGCCAAAAGGGGGGAGTG | | |
| 437 | 3 | 74068515 | 74068714 | 199 | CAST.MOLF | 3 | 74068633 | 74068666 | 34 | - | 16.4472 | 9.61E-07 | 0.141 | TAGCTGTCTTCAGACACACCAGAAGAGGGCGTCA | | |
| 438 | 3 | 75888399 | 75888699 | 300 | AJ,AKR,BTBR,CAST,DBA1J,DBA2J,MOLF,NZB,NZO,NZW,PL,RF,SWR,TH | 3 | 75888531 | 75888564 | 34 | - | 18.3902 | 2.72E-07 | 0.0804 | GTGCAAGGCTAGATATGACCACTAGAGGCAGATG | 14 | ENCFF093ZRU,ENCFF367UNR,ENCFF907XEG,ENCFF024JOZ,ENCFF589ZBJ,ENCFF136ZDK,ENCFF198SSR,ENCFF092NMJ,ENCFF099QNE,ENCFF821BIA,ENCFF327LYS,ENCFF502WDT,ENCFF055HAP,ENCFF089GBC |
| 439 | 3 | 80004576 | 80004996 | 420 | CAST | 3 | 80004859 | 80004892 | 34 | - | 18.1057 | 3.29E-07 | 0.0863 | ATGCACTGCCCAAGAAGACCATAAGAGGGTGATG | | |
| 440 | 3 | 84942247 | 84943062 | 815 | 129S1,AJ,C3H,C57L,CBA,CE,NZB,NZO,NZW,RF,SMJ | 3 | 84942601 | 84942619 | 19 | - | 17.2131 | 7.91E-07 | 0.196 | GGGCCAAAAGGGGGCAGTG | | |
| 440 | 3 | 84942247 | 84943062 | 815 | 129S1,AJ,C3H,C57L,CBA,CE,NZB,NZO,NZW,RF,SMJ | 3 | 84942601 | 84942635 | 35 | - | 16.4771 | 7.30E-07 | 0.211 | GTACAGGGGAACGCCAGGGCCAAAAGGGGGCAGTG | | |
| 441 | 3 | 86024561 | 86024772 | 211 | CAST.MA/My.MOLF.SPRET.WSB | 3 | 86024686 | 86024719 | 34 | - | 16.6341 | 8.54E-07 | 0.137 | TAGCTGTCTTCACACACTCCAGAAGAGGGCGTCA | 1 | ENCFF201JUA |
| 442 | 3 | 87489504 | 87490256 | 752 | SPRET | 3 | 87490036 | 87490070 | 35 | - | 23.4771 | 7.88E-09 | 0.0141 | CTGCTGTGTCACATGTGTGTCAGTAGGTGGCAGAG | 4 | ENCFF821BIA.ENCFF093ZRU.ENCFF367UNR.ENCFF099QNE |
| 443 | 3 | 88037897 | 88038107 | 210 | SPRET | 3 | 88038030 | 88038063 | 34 | - | 16.4634 | 9.51E-07 | 0.141 | TAGCTGTCTTCAGACGCACCAGAAGAGGGCATCA | | |
| 444 | 3 | 88186357 | 88186570 | 213 | CAST.MA/My.MOLF.SPRET | 3 | 88186484 | 88186517 | 34 | - | 16.4472 | 9.61E-07 | 0.141 | TAGCTGTCTTCAGACACACCAGAAGAGGGCGTCA | | |
| 445 | 3 | 88623391 | 88623589 | 198 | CAST.KK.MOLF.SPRET | 3 | 88623515 | 88623548 | 34 | - | 17.2439 | 5.79E-07 | 0.113 | TAGCTGTCCTCAGATACTCCAGAAGAGGGCATCA | | |
| 446 | 3 | 88940904 | 88941159 | 255 | SPRET | 3 | 88941033 | 88941066 | 34 | - | 16.4634 | 9.51E-07 | 0.141 | TAGCTGTCTTCAGACGCACCAGAAGAGGGCATCA | 3 | ENCFF099QNE.ENCFF093ZRU.ENCFF367UNR |
| 447 | 3 | 90101670 | 90101870 | 200 | SPRET | 3 | 90101795 | 90101828 | 34 | - | 17.2439 | 5.79E-07 | 0.113 | TAGCTGTCCTCAGATACTCCAGAAGAGGGCATCA | | |
| 448 | 3 | 91870035 | 91870277 | 242 | CAST.MOLF.SPRET | 3 | 91870157 | 91870190 | 34 | - | 17.9431 | 3.66E-07 | 0.0914 | CAGCTGTCCTCAGACACTCCAGAAGAGGGAGTCA | | |
| 449 | 3 | 93394059 | 93394382 | 323 | SPRET | 3 | 93394189 | 93394222 | 34 | - | 20.1057 | 8.23E-08 | 0.0417 | CAGCTGTCTTCAGATGCGCCAGAAGAGGGCATCA | | |
| 450 | 3 | 94727620 | 94727812 | 192 | SPRET | 3 | 94727741 | 94727774 | 34 | - | 16.4472 | 9.61E-07 | 0.141 | TAGCTGTCTTCAGACACACCAGAAGAGGGCGTCA | 1 | ENCFF093ZRU |
| 451 | 3 | 95715036 | 95715238 | 202 | CAST.SPRET | 3 | 95715160 | 95715178 | 19 | - | 17.5574 | 6.32E-07 | 0.181 | ACTCCAGAAGAGGGCGCCA | 5 | ENCFF093ZRU.ENCFF367UNR.ENCFF201JUA.ENCFF055HAP.ENCFF502WDT |
| 451 | 3 | 95715036 | 95715238 | 202 | CAST.SPRET | 3 | 95715160 | 95715193 | 34 | - | 18.3659 | 2.76E-07 | 0.0804 | TAGCTGTCTTCAGACACTCCAGAAGAGGGCGCCA | 5 | ENCFF093ZRU.ENCFF367UNR.ENCFF201JUA.ENCFF055HAP.ENCFF502WDT |
| 452 | 3 | 97049074 | 97049145 | 71 | SPRET | 3 | 97049073 | 97049107 | 35 | - | 19.6972 | 1.06E-07 | 0.0707 | CTGCAGAGGCTGGTCTGGCCCTGGAGGGGGCTCAG | | |
| 453 | 3 | 97238881 | 97239804 | 923 | SPRET | 3 | 97239686 | 97239704 | 19 | - | 18.7049 | 2.88E-07 | 0.131 | GCACCAGAAGAGGGCGCCA | 17 | ENCFF201JUA,ENCFF430PPJ,ENCFF092NMJ,ENCFF093ZRU,ENCFF367UNR,ENCFF099QNE,ENCFF589ZBJ,ENCFF198SSR,ENCFF883UPM,ENCFF907XEG,ENCFF327LYS,ENCFF821BIA,ENCFF089GBC,ENCFF055HAP.ENCFF502WDT.ENCFF142CNG.ENCFF024JOZ |
| 453 | 3 | 97238881 | 97239804 | 923 | SPRET | 3 | 97239686 | 97239719 | 34 | - | 19.3252 | 1.43E-07 | 0.0598 | TAGCTGTCTTCAGACGCACCAGAAGAGGGCGCCA | 17 | ENCFF201JUA,ENCFF430PPJ,ENCFF092NMJ,ENCFF093ZRU,ENCFF367UNR,ENCFF099QNE,ENCFF589ZBJ,ENCFF198SSR,ENCFF883UPM,ENCFF907XEG,ENCFF327LYS,ENCFF821BIA,ENCFF089GBC,ENCFF055HAP.ENCFF502WDT.ENCFF142CNG.ENCFF024JOZ |
| 454 | 3 | 97788057 | 97788986 | 929 | BTBR,KK,MA/My,MOLF,NOD,NOR,NZB,NZO,NZW | 3 | 97788730 | 97788763 | 34 | - | 22.0894 | 1.86E-08 | 0.0144 | GTGCAATGCCTGAAGATGTCAGAAGAGGGTGGTA | | |
| 455 | 3 | 100839236 | 100839972 | 736 | SPRET | 3 | 100839564 | 100839582 | 19 | - | 17.377 | 7.11E-07 | 0.188 | TAGCCGCTAGGGGGTGCCC | 11 | ENCFF136ZDK,ENCFF093ZRU,ENCFF198SSR,ENCFF367UNR,ENCFF883UPM,ENCFF099QNE,ENCFF533APC.ENCFF907XEG.ENCFF052KVO.ENCFF821BIA.ENCFF327LYS |
| 456 | 3 | 103097442 | 103097646 | 204 | SPRET | 3 | 103097562 | 103097595 | 34 | - | 17.2439 | 5.79E-07 | 0.113 | TAGCTGTCCTCAGATACTCCAGAAGAGGGCATCA | | |
| 457 | 3 | 104674062 | 104674260 | 198 | CAST.MOLF.NZO.NZW.SPRET | 3 | 104674189 | 104674207 | 19 | - | 17.5574 | 6.32E-07 | 0.181 | ACTCCAGAAGAGGGCGCCA | 2 | ENCFF093ZRU.ENCFF367UNR |
| 457 | 3 | 104674062 | 104674260 | 198 | CAST.MOLF.NZO.NZW.SPRET | 3 | 104674189 | 104674222 | 34 | - | 18.3659 | 2.76E-07 | 0.0804 | TAGCTGTCTTCAGACACTCCAGAAGAGGGCGCCA | 2 | ENCFF093ZRU.ENCFF367UNR |
| 458 | 3 | 105800480 | 105800551 | 71 | SPRET | 3 | 105800463 | 105800497 | 35 | - | 20.578 | 5.97E-08 | 0.0492 | GTGCAGAGCCAGCTCTTCGTCTGCAGGGGGAAATC | 1 | ENCFF367UNR |
| 459 | 3 | 115438271 | 115439173 | 902 | NZW | 3 | 115438570 | 115438603 | 34 | - | 21.3984 | 3.17E-08 | 0.0215 | CTGCTATCTCCAGACACACCAGAAGAGGGCATCA | 2 | ENCFF093ZRU.ENCFF367UNR |
| 460 | 3 | 115721324 | 115722067 | 743 | MOLF | 3 | 115721817 | 115721850 | 34 | - | 16.5854 | 8.81E-07 | 0.139 | TTACAGGTCCCCATGAAGGCAGAAGAGGGAGCTG | | |
| 461 | 3 | 116351834 | 116352481 | 647 | MOLF | 3 | 116352416 | 116352449 | 34 | - | 19.1626 | 1.60E-07 | 0.0641 | ATGCAGCTAGAGACACGACCTCCGGGGGGTACTG | | |
| 462 | 3 | 121586611 | 121586983 | 372 | 129S1,AKR,BUB,C57L,C58J,CAST,CE,FVB,ILNJ2,LP,MRL,NOD,NOR,NZB,NZO,NZW,PJ,PL,RF,SEA,SJL,SPRET,SWR,TH,WSB | 3 | 121586895 | 121586928 | 34 | - | 16.4472 | 9.61E-07 | 0.141 | TAGCTGTCTTCAGACACACCAGAAGAGGGCGTCA | | |
| 463 | 3 | 121613021 | 121613181 | 160 | SPRET | 3 | 121613153 | 121613171 | 19 | - | 17 | 9.07E-07 | 0.209 | CAGCCAGGAGATGGCACAG | | |
| 463 | 3 | 121613021 | 121613181 | 160 | SPRET | 3 | 121613153 | 121613187 | 35 | - | 17.1835 | 4.87E-07 | 0.173 | TTACTGTCCTCTCAGACAGCCAGGAGATGGCACAG | | |
| 464 | 3 | 122209520 | 122209769 | 249 | AJ,BALB,BTBR,C3H,CAST,CBA,CE,DBA1J,DBA2J,KK,MOLF,RH2,SMJ,SPRET,TH,WSB | 3 | 122209702 | 122209735 | 34 | - | 17.0976 | 6.36E-07 | 0.119 | TAGCTGTCCTCAGACACCCCAGAAGAGGGCATCA | | |
| 465 | 3 | 123300348 | 123300856 | 508 | SPRET | 3 | 123300548 | 123300582 | 35 | - | 17.4495 | 4.17E-07 | 0.161 | CTGAACTTACTTGAAGGTACCAGTAGATGGCGTAC | 21 | ENCFF201JUA,ENCFF367UNR,ENCFF093ZRU,ENCFF430PPJ,ENCFF099QNE,ENCFF165LXO,ENCFF052KVO,ENCFF327LYS,ENCFF024JOZ,ENCFF883UPM,ENCFF609MDN,ENCFF589ZBJ,ENCFF502WDT,ENCFF907XEG,ENCFF821BIA,ENCFF198SSR,ENCFF055HAP,ENCFF089GBC,ENCFF092NMJ,ENCFF065ODV,ENCFF136ZDK |
| 466 | 3 | 127251475 | 127251695 | 220 | 129S1,AJ,BALB,C3H,C57L,CAST,CBA,NZB,PJ.RH2.SEA.SMJ.SPRET | 3 | 127251608 | 127251641 | 34 | - | 18.8211 | 2.03E-07 | 0.0721 | TGGCTGTTTCCAGATACACCAGAAGGGGGCATCC | 8 | ENCFF367UNR,ENCFF201JUA,ENCFF099QNE,ENCFF327LYS,ENCFF024JOZ,ENCFF502WDT,ENCFF093ZRU.ENCFF055HAP |
| 467 | 3 | 130829909 | 130830095 | 186 | SPRET | 3 | 130830023 | 130830041 | 19 | - | 17.5574 | 6.32E-07 | 0.181 | ACTCCAGAAGAGGGCGCCA | | |
| 467 | 3 | 130829909 | 130830095 | 186 | SPRET | 3 | 130830023 | 130830056 | 34 | - | 17.9756 | 3.59E-07 | 0.0905 | TAGCTGTCTTTAGACACTCCAGAAGAGGGCGCCA | | |
| 468 | 3 | 130870519 | 130870762 | 243 | MOLF.SPRET | 3 | 130870644 | 130870677 | 34 | - | 17.2439 | 5.79E-07 | 0.113 | TAGCTGTCCTCAGATACTCCAGAAGAGGGCATCA | | |
| 469 | 3 | 132071533 | 132071730 | 197 | CAST.MOLF.SPRET | 3 | 132071656 | 132071689 | 34 | - | 17.4472 | 5.08E-07 | 0.108 | TAGCTGTCTTCAGACACACCAGAAGGGGGCGTCA | | |
| 470 | 3 | 136247333 | 136247773 | 440 | CAST | 3 | 136247381 | 136247415 | 35 | - | 20.055 | 8.39E-08 | 0.0613 | ATGCAGGTTCCTGCAAAATCCAGAAGAGGGAGCCG | 1 | ENCFF052KVO |
| 471 | 3 | 142477649 | 142478306 | 657 | CAST | 3 | 142477810 | 142477828 | 19 | - | 18.623 | 3.06E-07 | 0.132 | ACACCAGAAGAGGGCGCCA | 2 | ENCFF093ZRU.ENCFF367UNR |
| 471 | 3 | 142477649 | 142478306 | 657 | CAST | 3 | 142477810 | 142477843 | 34 | - | 18.4634 | 2.59E-07 | 0.0804 | TAGCTGTCTTTAGACACACCAGAAGAGGGCGCCA | 2 | ENCFF093ZRU.ENCFF367UNR |
| 472 | 3 | 142554232 | 142554435 | 203 | SPRET | 3 | 142554355 | 142554388 | 34 | - | 16.4146 | 9.81E-07 | 0.142 | TAGCTGTCTTCAGACACACCAGAGGAGGGCGCCA | | |
| 473 | 3 | 145272641 | 145272989 | 348 | CAST.MOLF | 3 | 145272947 | 145272980 | 34 | - | 16.561 | 8.95E-07 | 0.14 | CTGCCCTTGCAATCGCCATTACAAGATGGCGCTG | | |
| 474 | 3 | 146152281 | 146152506 | 225 | AJ,BALB,BUB,C3H,CAST,CBA,DBA1J,ILNJ2,MA/My,MOLF,MRL,NOD,NOR,NZW,PJ,RF,RH2.SEA.SJL.SMJ.SPRET.WSB | 3 | 146152404 | 146152437 | 34 | - | 19.7967 | 1.03E-07 | 0.0485 | TAGCTGTCCTCAGACACTCCAGAAGAGGGCACCA | 1 | ENCFF093ZRU |
| 475 | 3 | 151674607 | 151674724 | 117 | CAST | 3 | 151674682 | 151674716 | 35 | - | 15.9541 | 9.80E-07 | 0.242 | GTGCAGTGTGGGCAATGGATCAGAAGAAGGAAGAG | | |
| 476 | 3 | 151991341 | 151991559 | 218 | AJ,BALB,C3H,CAST,CE,ILNJ2,MOLF,PL,RH2.SEA.SPRET.WSB | 3 | 151991466 | 151991484 | 19 | - | 17.5574 | 6.32E-07 | 0.181 | ACTCCAGAAGAGGGCGCCA | 2 | ENCFF093ZRU,ENCFF367UNR |
| 476 | 3 | 151991341 | 151991559 | 218 | AJ,BALB,C3H,CAST,CE,ILNJ2,MOLF,PL,RH2.SEA.SPRET.WSB | 3 | 151991466 | 151991499 | 34 | - | 18.3659 | 2.76E-07 | 0.0804 | TAGCTGTCTTCAGACACTCCAGAAGAGGGCGCCA | 2 | ENCFF093ZRU,ENCFF367UNR |
| 477 | 3 | 152044101 | 152044308 | 207 | CAST.SPRET | 3 | 152044208 | 152044241 | 34 | - | 16.4472 | 9.61E-07 | 0.141 | TAGCTGTCTTCAGACACACCAGAAGAGGGCGTCA | | |
| 478 | 3 | 153525927 | 153526757 | 830 | SPRET | 3 | 153526700 | 153526733 | 34 | - | 16.7805 | 7.79E-07 | 0.131 | CAGCTCTACCGTGAGAAACCACCCGGGGTCACAC | | |
| 479 | 4 | 7576419 | 7577178 | 759 | CAST | 4 | 7577160 | 7577194 | 35 | + | 18.5872 | 2.11E-07 | 0.0689 | GTGCAAGTGCTAACAGAGACCAGAAGAGGGCATTG | | |
| 480 | 4 | 9455167 | 9455243 | 76 | SPRET | 4 | 9455141 | 9455175 | 35 | + | 17.9817 | 3.04E-07 | 0.0845 | ATGCAGTAGAGGGAAGGGGCCAGTAGGTGGTTAAT | 1 | ENCFF093ZRU |
| 481 | 4 | 10412243 | 10412552 | 309 | SPRET | 4 | 10412259 | 10412292 | 34 | + | 17.065 | 6.50E-07 | 0.0792 | CTGCGATAACATTCGCCATCACAAGATGGTGCCA | | |
| 482 | 4 | 10483863 | 10484069 | 206 | SPRET | 4 | 10483925 | 10483958 | 34 | + | 17.8455 | 3.91E-07 | 0.0606 | TAGCTGTCCTCAGACACTCCAGAAGAGGGCGTCA | | |
| 483 | 4 | 11396446 | 11396641 | 195 | SPRET | 4 | 11396497 | 11396530 | 34 | + | 17.3902 | 5.27E-07 | 0.071 | TAGCTGTCCTCAGACACTCCAGAAGAGGGCATCA | | |
| 484 | 4 | 14112938 | 14113573 | 635 | SPRET | 4 | 14113208 | 14113226 | 19 | + | 17.5902 | 6.18E-07 | 0.111 | CTACCTGAAGGTGGCGCTG | 3 | ENCFF093ZRU.ENCFF089GBC.ENCFF502WDT |
| 485 | 4 | 16316679 | 16316779 | 100 | SPRET | 4 | 16316694 | 16316727 | 34 | + | 17.5203 | 4.84E-07 | 0.0682 | ATGTACTGCAAGGGAAGACCAGAAGAGGGCATCT | | |
| 486 | 4 | 21724969 | 21725182 | 213 | CAST.SPRET.WSB | 4 | 21725040 | 21725073 | 34 | + | 16.4472 | 9.61E-07 | 0.0931 | TAGCTGTCTTCAGACACACCAGAAGAGGGCGTCA | | |
| 487 | 4 | 31826773 | 31827772 | 999 | SPRET | 4 | 31827446 | 31827464 | 19 | + | 19.5902 | 1.51E-07 | 0.0547 | TCACCGGTAGGTGGCGCCA | 19 | ENCFF367UNR,ENCFF093ZRU,ENCFF136ZDK,ENCFF327LYS,ENCFF165LXO,ENCFF589ZBJ,ENCFF099QNE,ENCFF821BIA,ENCFF609MDN,ENCFF052KVO,ENCFF504JDS,ENCFF024JOZ,ENCFF907XEG,ENCFF883UPM,ENCFF142CNG,ENCFF198SSR,ENCFF055HAP,ENCFF957SPH,ENCFF502WDT |
| 488 | 4 | 33011422 | 33011626 | 204 | MOLF.SPRET | 4 | 33011484 | 33011517 | 34 | + | 17.7317 | 4.21E-07 | 0.0636 | TAGCTGTCCTCAGATACACCAGAAGAGGGCATCA | 3 | ENCFF093ZRU.ENCFF367UNR.ENCFF821BIA |
| 489 | 4 | 34780735 | 34780951 | 216 | NZB.NZO.NZW.SPRET.TH.WSB | 4 | 34780809 | 34780842 | 34 | + | 16.4472 | 9.61E-07 | 0.0931 | TAGCTGTCTTCAGACACACCAGAAGAGGGCGTCA | | |
| 490 | 4 | 38928845 | 38929269 | 424 | CAST.SPRET | 4 | 38929232 | 38929265 | 34 | + | 16.5772 | 8.86E-07 | 0.0924 | TTGCAGCGTCGTTCTTGGCCAGTTGGGCGCGCGC | | |
| 491 | 4 | 41146195 | 41146386 | 191 | SPRET | 4 | 41146180 | 41146213 | 34 | + | 16.7642 | 7.87E-07 | 0.0862 | TAGCTGTCTTCAGAAGCACCAGAAGAGGGCGTCA | | |
| 492 | 4 | 42992514 | 42992726 | 212 | AKR,CAST,CE,KK,MRL,NOD,NOR,PJ,RF,SEA.SJL.SPRET | 4 | 42992583 | 42992616 | 34 | + | 16.4228 | 9.76E-07 | 0.0938 | TAGCTGTCTTCAGATGCACCAGAAGAGGGCATCG | | |
| 493 | 4 | 43703231 | 43703652 | 421 | CAST | 4 | 43703578 | 43703611 | 34 | + | 25.2195 | 1.36E-09 | 0.00124 | GTGTAGTGTTCATGGAGGCCAGAAGGGGGCGCTG | 21 | ENCFF367UNR,ENCFF821BIA,ENCFF099QNE,ENCFF092NMJ,ENCFF024JOZ,ENCFF165LXO,ENCFF327LYS,ENCFF430PPJ,ENCFF089GBC,ENCFF052KVO,ENCFF589ZBJ,ENCFF136ZDK,ENCFF907XEG,ENCFF093ZRU,ENCFF502WDT,ENCFF065ODV,ENCFF533APC,ENCFF198SSR,ENCFF055HAP,ENCFF321AGQ,ENCFF033UOX |
| 493 | 4 | 43703231 | 43703652 | 421 | CAST | 4 | 43703593 | 43703611 | 19 | + | 21.8525 | 2.33E-08 | 0.0254 | AGGCCAGAAGGGGGCGCTG | 21 | ENCFF367UNR,ENCFF821BIA,ENCFF099QNE,ENCFF092NMJ,ENCFF024JOZ,ENCFF165LXO,ENCFF327LYS,ENCFF430PPJ,ENCFF089GBC,ENCFF052KVO,ENCFF589ZBJ,ENCFF136ZDK,ENCFF907XEG,ENCFF093ZRU,ENCFF502WDT,ENCFF065ODV,ENCFF533APC,ENCFF198SSR,ENCFF055HAP,ENCFF321AGQ,ENCFF033UOX |
| 494 | 4 | 43830146 | 43830206 | 60 | CAST | 4 | 43830198 | 43830232 | 35 | + | 16.3394 | 7.89E-07 | 0.14 | CATCAGACCCCCTTGGAGGCCAGAAGAGGGCATCA | | |
| 495 | 4 | 45773441 | 45774101 | 660 | SPRET | 4 | 45773658 | 45773691 | 34 | + | 18.1707 | 3.15E-07 | 0.0539 | CAGCTGTCTTTAGACACACCAGAAGAGGGCATCA | | |
| 496 | 4 | 46773089 | 46773632 | 543 | CAST,MOLF | 4 | 46773407 | 46773441 | 35 | + | 18.3394 | 2.45E-07 | 0.0747 | GTGTGGATACTCATGGTGACCAGCAGAGGGCATCA | 13 | ENCFF093ZRU,ENCFF430PPJ,ENCFF024JOZ,ENCFF092NMJ,ENCFF907XEG,ENCFF089GBC,ENCFF165LXO,ENCFF055HAP,ENCFF502WDT,ENCFF136ZDK,ENCFF142CNG,ENCFF327LYS,ENCFF589ZBJ |
| 496 | 4 | 46773089 | 46773632 | 543 | CAST,MOLF | 4 | 46773423 | 46773441 | 19 | + | 20.1148 | 1.01E-07 | 0.0443 | TGACCAGCAGAGGGCATCA | 13 | ENCFF093ZRU,ENCFF430PPJ,ENCFF024JOZ,ENCFF092NMJ,ENCFF907XEG,ENCFF089GBC,ENCFF165LXO,ENCFF055HAP,ENCFF502WDT,ENCFF136ZDK,ENCFF142CNG,ENCFF327LYS,ENCFF589ZBJ |
| 497 | 4 | 49230812 | 49231016 | 204 | CAST.SPRET | 4 | 49230871 | 49230904 | 34 | + | 18.561 | 2.42E-07 | 0.0493 | CAGCTGTCTTCAGACACACCAGAAGAGGGCATCA | | |
| 498 | 4 | 52937050 | 52937266 | 216 | AJ,BALB,BUB,C3H,CAST,KK,MA/My,MOLF,RH2.SPRET | 4 | 52937124 | 52937157 | 34 | + | 16.4634 | 9.51E-07 | 0.0931 | TAGCTGTCTTCAGACGCACCAGAAGAGGGCATCA | | |
| 499 | 4 | 53272922 | 53273122 | 200 | CAST.MA/My.MOLF.SPRET | 4 | 53272980 | 53273013 | 34 | + | 16.4472 | 9.61E-07 | 0.0931 | TAGCTGTCTTCAGACACACCAGAAGAGGGCGTCA | | |
| 500 | 4 | 54983434 | 54983674 | 240 | 129S1,AJ,AKR,BALB,BTBR,BUB,C3H,C57L,CAST,CBA,CE,DBA1J,DBA2J,FVB,ILNJ2,LP,MOLF,MRL,NOD,NOR,NZB,NZO,NZW,PJ,PL,RF,RH2,SEA,SJL,SMJ,SPRET,SWR,TH,WSB | 4 | 54983533 | 54983566 | 34 | + | 16.4634 | 9.51E-07 | 0.0931 | TAGCTGTCTTCAGACGCACCAGAAGAGGGCATCA | 1 | ENCFF093ZRU |
| 501 | 4 | 57317673 | 57318639 | 966 | SPRET | 4 | 57317929 | 57317963 | 35 | + | 16.5413 | 7.04E-07 | 0.133 | ATACAGTCCAGGATTCCAGCCAGGGGATGGTGCCA | | |
| 502 | 4 | 59001033 | 59001243 | 210 | SPRET | 4 | 59001081 | 59001114 | 34 | + | 16.4634 | 9.51E-07 | 0.0931 | TAGCTGTCTTCAGACGCACCAGAAGAGGGCATCA | | |
| 503 | 4 | 59278987 | 59279197 | 210 | MOLF.SPRET | 4 | 59279054 | 59279087 | 34 | + | 18.8293 | 2.02E-07 | 0.0441 | CAGCTGTCTTCAGACACACCAGAAGAGGGCGTTA | | |
| 504 | 4 | 62180084 | 62180304 | 220 | BTBR.BUB.CAST.MOLF.SPRET.WSB | 4 | 62180162 | 62180195 | 34 | + | 17.8455 | 3.91E-07 | 0.0606 | TAGCTGTCCTCAGACACTCCAGAAGAGGGCGTCA | | |
| 505 | 4 | 74224847 | 74225060 | 213 | SPRET | 4 | 74224913 | 74224946 | 34 | + | 19.1382 | 1.63E-07 | 0.0392 | CTGTAGCTGTCAGACACACCAGAAGAGGGCATCA | | |
| 506 | 4 | 81289409 | 81289545 | 136 | CAST | 4 | 81289505 | 81289538 | 34 | + | 17.1545 | 6.13E-07 | 0.0768 | GTGTTATGCCTGTGGAGCCCAGAAGAGGGTATTG | | |
| 507 | 4 | 83438170 | 83438817 | 647 | CAST | 4 | 83438740 | 83438758 | 19 | + | 21.0984 | 4.51E-08 | 0.0333 | AGGCCAGAAGAGGGCGCCA | 11 | ENCFF093ZRU,ENCFF367UNR,ENCFF024JOZ,ENCFF136ZDK,ENCFF327LYS,ENCFF589ZBJ,ENCFF883UPM,ENCFF142CNG.ENCFF052KVO.ENCFF502WDT.ENCFF055HAP |
| 508 | 4 | 94012525 | 94012848 | 323 | CAST | 4 | 94012543 | 94012576 | 34 | + | 20.3171 | 7.06E-08 | 0.0218 | CCGCAATAGCATTCGCCATCACAAGATGGCGCCG | 14 | ENCFF052KVO,ENCFF089GBC,ENCFF367UNR,ENCFF165LXO,ENCFF198SSR,ENCFF502WDT,ENCFF957SPH,ENCFF821BIA,ENCFF194HVL,ENCFF907XEG,ENCFF024JOZ,ENCFF093ZRU,ENCFF142CNG,ENCFF327LYS |
| 509 | 4 | 95812629 | 95813000 | 371 | SPRET | 4 | 95812673 | 95812691 | 19 | + | 17.2131 | 7.91E-07 | 0.124 | TGTCCACTAGAGGGTGCCT | 20 | ENCFF210BTO,ENCFF093ZRU,ENCFF430PPJ,ENCFF099QNE,ENCFF367UNR,ENCFF907XEG,ENCFF327LYS,ENCFF024JOZ,ENCFF609MDN,ENCFF575MGI,ENCFF821BIA,ENCFF089GBC,ENCFF198SSR,ENCFF589ZBJ,ENCFF883UPM,ENCFF092NMJ,ENCFF957SPH,ENCFF281QMV,ENCFF165LXO,ENCFF951GOP |
| 510 | 4 | 96009186 | 96009335 | 149 | MOLF.SPRET | 4 | 96009212 | 96009245 | 34 | + | 16.4634 | 9.51E-07 | 0.0931 | TAGCTGTCTTCAGACGCACCAGAAGAGGGCATCA | 1 | ENCFF093ZRU |
| 511 | 4 | 99396359 | 99397068 | 709 | AJ,B10D2,B10J,BALB,BUB,C3H,C58J,CBA,KK.NZB.PJ.PL.RF.SEA.WSB | 4 | 99396698 | 99396731 | 34 | + | 20.3577 | 6.86E-08 | 0.0213 | CAGCCTTCTGCCTGCCGGCCAGAAGAGGGCACCA | | |
| 511 | 4 | 99396359 | 99397068 | 709 | AJ,B10D2,B10J,BALB,BUB,C3H,C58J,CBA,KK.NZB.PJ.PL.RF.SEA.WSB | 4 | 99396713 | 99396731 | 19 | + | 22.377 | 1.43E-08 | 0.0203 | CGGCCAGAAGAGGGCACCA | | |
| 512 | 4 | 100528589 | 100529197 | 608 | SPRET | 4 | 100528711 | 100528744 | 34 | + | 16.6179 | 8.63E-07 | 0.0909 | GAGCAGTGGCTGGGGGAGTCAGGAGATGGCAAGA | | |
| 513 | 4 | 100630386 | 100630599 | 213 | CAST.SPRET | 4 | 100630456 | 100630489 | 34 | + | 17.9431 | 3.66E-07 | 0.0591 | CAGCTGTCCTCAGACACTCCAGAAGAGGGAGTCA | | |
| 514 | 4 | 101547401 | 101548101 | 700 | AKR.C58J.MA/My.MRL.PJ | 4 | 101547496 | 101547514 | 19 | + | 17.5246 | 6.45E-07 | 0.112 | AGGCCAGAAGAGGGCAACA | | |
| 515 | 4 | 106186410 | 106186673 | 263 | CAST,MA/My,MOLF,SPRET | 4 | 106186488 | 106186521 | 34 | + | 16.4634 | 9.51E-07 | 0.0931 | TAGCTGTCTTCAGACGCACCAGAAGAGGGCATCA | 10 | ENCFF093ZRU,ENCFF367UNR,ENCFF821BIA,ENCFF024JOZ,ENCFF136ZDK,ENCFF327LYS,ENCFF883UPM,ENCFF589ZBJ.ENCFF055HAP.ENCFF502WDT |
| 516 | 4 | 107626802 | 107627001 | 199 | AKR,BALB,BUB,C3H,CAST,CBA,CE,MOLF,MRL,NOD,PL,RF,SEA,SJL,SMJ,SPRET,WSB | 4 | 107626858 | 107626891 | 34 | + | 16.4472 | 9.61E-07 | 0.0931 | TAGCTGTCTTCAGACACACCAGAAGAGGGCGTCA | | |
| 517 | 4 | 108019659 | 108019873 | 214 | SPRET | 4 | 108019731 | 108019764 | 34 | + | 16.4472 | 9.61E-07 | 0.0931 | TAGCTGTCTTCAGACACACCAGAAGAGGGCGTCA | | |
| 518 | 4 | 108163750 | 108163951 | 201 | SPRET | 4 | 108163804 | 108163837 | 34 | + | 16.5691 | 8.90E-07 | 0.0928 | TAGCTGTCTTCAGACGCACCAGAAGAGGGCATCG | | |
| 519 | 4 | 108532027 | 108532227 | 200 | CAST.SPRET | 4 | 108532076 | 108532109 | 34 | + | 17.2439 | 5.79E-07 | 0.0743 | TAGCTGTCCTCAGATACTCCAGAAGAGGGCATCA | | |
| 520 | 4 | 108600337 | 108600534 | 197 | 129S1,AJ,AKR,BALB,BUB,C3H,C57L,C58J,CAST,CBA,CE,DBA1J,DBA2J,FVB,MRL,NOD,NZB,NZO,NZW,PJ,PL,RF,SEA,SJL,SPRET.SWR.TH.WSB | 4 | 108600384 | 108600417 | 34 | + | 16.4472 | 9.61E-07 | 0.0931 | TAGCTGTCTTCAGACACACCAGAAGAGGGCGTCA | | |
| 521 | 4 | 109313352 | 109313556 | 204 | CAST.MOLF.SPRET | 4 | 109313413 | 109313446 | 34 | + | 16.4634 | 9.51E-07 | 0.0931 | TAGCTGTCTTCAGACGCACCAGAAGAGGGCATCA | | |
| 522 | 4 | 111412043 | 111412139 | 96 | SPRET | 4 | 111412013 | 111412046 | 34 | + | 17.3415 | 5.44E-07 | 0.0723 | CTGCTGTCCTCCAACCATACTCCAGAGGTTGGTA | | |
| 523 | 4 | 114872793 | 114872983 | 190 | SPRET | 4 | 114872840 | 114872873 | 34 | + | 16.4472 | 9.61E-07 | 0.0931 | TAGCTGTCTTCAGACACACCAGAAGAGGGCGTCA | 1 | ENCFF367UNR |
| 524 | 4 | 115882689 | 115882951 | 262 | 129S1,AJ,AKR,B10D2,B10J,BTBR,C57L,C58J,CE,DBA1J,DBA2J,ILNJ2,KK,MA/My,MOLF,MRL,NZB,NZO,NZW,PJ,PL,SEA,SMJ,SPRET.SWR.TH.WSB | 4 | 115882754 | 115882787 | 34 | + | 16.4634 | 9.51E-07 | 0.0931 | TAGCTGTCTTCAGACGCACCAGAAGAGGGCATCA | 1 | ENCFF093ZRU |
| 525 | 4 | 115898291 | 115898865 | 574 | SPRET | 4 | 115898725 | 115898758 | 34 | + | 20.7236 | 5.25E-08 | 0.0177 | CAGCTGTTTTCAGACACACCAGAAGAGGGCGTCA | 1 | ENCFF093ZRU |
| 526 | 4 | 116057211 | 116057471 | 260 | MOLF | 4 | 116057207 | 116057241 | 35 | + | 17.3394 | 4.45E-07 | 0.104 | GTGCAGCTGCCTGCAGCGTCCAGCAGAAGACATCA | | |
| 527 | 4 | 116178545 | 116178736 | 191 | SPRET | 4 | 116178596 | 116178629 | 34 | + | 16.4472 | 9.61E-07 | 0.0931 | TAGCTGTCTTCAGACACACCAGAAGAGGGCGTCA | 1 | ENCFF093ZRU |
| 528 | 4 | 116884788 | 116885017 | 229 | CAST.ILNJ2.MOLF.SPRET.TH | 4 | 116884873 | 116884907 | 35 | + | 16.0917 | 9.07E-07 | 0.15 | TAGCTGTCTTCAGACAGCACCAGAAGAGGGCATCA | 1 | ENCFF093ZRU |
| 529 | 4 | 117027832 | 117028163 | 331 | SPRET | 4 | 117027868 | 117027901 | 34 | + | 17.1951 | 5.97E-07 | 0.0755 | TTGCTATCTTCAGACACACCAGAAGAGGGCATCA | 1 | ENCFF093ZRU |
| 530 | 4 | 117513241 | 117513470 | 229 | AJ,AKR,B10D2,B10J,BALB,BTBR,BUB,C3H,C58J,CAST,CBA,CE,DBA1J,DBA2J,FVB,ILNJ2,KK,MA/My,MOLF,MRL,NOD,NOR,PL,RF,SJL.SPRET.SWR.TH.WSB | 4 | 117513327 | 117513360 | 34 | + | 16.9593 | 6.95E-07 | 0.0815 | TAGCTGTCTTCAGACACTCCAGAAGGGGGCGTCA | | |
| 531 | 4 | 118403501 | 118403713 | 212 | CAST.SPRET | 4 | 118403570 | 118403603 | 34 | + | 16.4472 | 9.61E-07 | 0.0931 | TAGCTGTCTTCAGACACACCAGAAGAGGGCGTCA | | |
| 532 | 4 | 118453630 | 118453839 | 209 | 129S1,AJ,B10D2,B10J,BALB,BTBR,BUB,C3H,C57L,C58J,CAST,CBA,CE,DBA1J,DBA2J,FVB,ILNJ2,MA/My,MOLF,NOD,NOR,NZB,PJ,PL,RF,SEA,SJL,SPRET,SWR,TH,WSB | 4 | 118453697 | 118453730 | 34 | + | 16.4472 | 9.61E-07 | 0.0931 | TAGCTGTCTTCAGACACACCAGAAGAGGGCGTCA | | |
| 533 | 4 | 119365912 | 119366739 | 827 | SPRET | 4 | 119366240 | 119366274 | 35 | + | 16.4587 | 7.38E-07 | 0.136 | CTGCATGTATGCCTGCAGACCAGAAGAAGCCATTG | | |
| 534 | 4 | 119815578 | 119815778 | 200 | SPRET | 4 | 119815635 | 119815668 | 34 | + | 18.1545 | 3.18E-07 | 0.0543 | TAGCTGTTTTCAGACACACCAGAAGAGGGCGTCA | 1 | ENCFF093ZRU |
| 535 | 4 | 120755109 | 120755302 | 193 | SPRET | 4 | 120755159 | 120755192 | 34 | + | 17.2439 | 5.79E-07 | 0.0743 | TAGCTGTCCTCAGATACTCCAGAAGAGGGCATCA | | |
| 536 | 4 | 124656269 | 124656474 | 205 | SPRET | 4 | 124656329 | 124656363 | 35 | + | 18.1284 | 2.79E-07 | 0.0805 | CTGTAGCTGTCTTCAGACACCAGAAGAGGGCATCA | | |
| 537 | 4 | 124708487 | 124708706 | 219 | 129S1,AJ,AKR,BALB,BTBR,BUB,C3H,C57L,C58J,CAST,CBA,CE,DBA1J,DBA2J,FVB,ILNJ2,KK,MA/My,MOLF,MRL,NOD,NOR,NZO,NZW,PJ,PL,RF,SEA,SJL,SPRET,SWR,TH,WSB | 4 | 124708565 | 124708598 | 34 | + | 18.3659 | 2.76E-07 | 0.0503 | TAGCTGTCTTCAGACACTCCAGAAGAGGGCGCCA | 2 | ENCFF367UNR,ENCFF093ZRU |

| | | | | | | | | | | | | | | | | |
|---|---|---|---|---|---|---|---|---|---|---|---|---|---|---|---|---|
| 537 | 4 | 124708487 | 124708706 | 219 | 129S1,AJ,AKR,BALB,BTBR,BUB,C3H,C57L,C58J,CAST,CBA,CE,DBA1J,DBA2J,FVB,ILNJ2,KK,MA/My,MOLF,MRL,NOD,NOR,NZO,NZW,PJ,PL,RF,SEA,SJL,SPRET,SWR,TH,WSB | 4 | 124708580 | 124708598 | 19 | + | 17.5574 | 6.32E-07 | 0.111 | ACTCCAGAAGAGGGCGCCA | 2 | ENCFF367UNR,ENCFF093ZRU |
| 538 | 4 | 124956873 | 124957064 | 191 | SPRET | 4 | 124956920 | 124956953 | 34 | + | 17.6016 | 4.59E-07 | 0.066 | TAGCTGTCTTCAGACATGCCAGAAGAGGGCGTCA | | |
| 539 | 4 | 125057157 | 125057274 | 117 | AKR.MOLF.MRL.TH | 4 | 125057198 | 125057231 | 34 | + | 17.4472 | 5.08E-07 | 0.0703 | CTGTAGTTCCTTCAGGTGTCACAGGGGAGCTCCC | | |
| 540 | 4 | 125165621 | 125165829 | 208 | SPRET | 4 | 125165687 | 125165720 | 34 | + | 18.0244 | 3.47E-07 | 0.0571 | TAGCTGTTTTCAGATGCACCAGAAGAGGGCATCA | 2 | ENCFF093ZRU |
| 541 | 4 | 126106251 | 126106470 | 219 | CAST.SPRET | 4 | 126106326 | 126106359 | 34 | + | 16.4472 | 9.61E-07 | 0.0931 | TAGCTGTCTTCAGACACACCAGAAGAGGGCGTCA | | |
| 542 | 4 | 126339359 | 126339559 | 200 | B10D2.B10J.CAST.MOLF.NZW.SPRET | 4 | 126339418 | 126339451 | 34 | + | 17.2439 | 5.79E-07 | 0.0743 | TAGCTGTCCTCAGATACTCCAGAAGAGGGCATCA | | |
| 543 | 4 | 130191175 | 130191311 | 136 | CAST | 4 | 130191161 | 130191195 | 35 | + | 23.3119 | 8.90E-09 | 0.00927 | GTGCAATGCTTCCTCTTTAACAGAAGGGGGCGCGT | 36 | ENCFF093ZRU,ENCFF367UNR,ENCFF136ZDK,ENCFF099QNE,ENCFF065ODV,ENCFF821BIA,ENCFF327LYS,ENCFF055HAP,ENCFF589ZBJ,ENCFF907XEG,ENCFF024JOZ,ENCFF502WDT,ENCFF092NMJ,ENCFF052KVO,ENCFF089GBC,ENCFF198SSR,ENCFF165LXO,ENCFF033UQX,ENCFF321AGQ,ENCFF430PPJ,ENCFF957SPH,ENCFF142CNG,ENCFF281QMV,ENCFF210BTO,ENCFF601JVE,ENCFF883UPM,ENCFF316FYN,ENCFF951GOP,ENCFF504JDS,ENCFF533APC,ENCFF609MDN,ENCFF491RJK,ENCFF611HDQ,ENCFF575MGI,ENCFF097CAQ.ENCFF194HVL |
| 544 | 4 | 130303894 | 130304089 | 195 | SPRET | 4 | 130303956 | 130303974 | 19 | + | 17.5574 | 6.32E-07 | 0.111 | ACTCCAGAAGAGGGCGCCA | 1 | ENCFF093ZRU |
| 545 | 4 | 130312458 | 130312684 | 226 | SPRET | 4 | 130312542 | 130312575 | 34 | + | 16.4472 | 9.61E-07 | 0.0931 | TAGCTGTCTTCAGACACACCAGAAGAGGGCGTCA | 1 | ENCFF093ZRU |
| 546 | 4 | 130410591 | 130410801 | 210 | CAST,FVB,KK,MOLF,NOR,NZB,SJL,SPRET,SWR | 4 | 130410659 | 130410692 | 34 | + | 18.3659 | 2.76E-07 | 0.0503 | TAGCTGTCTTCAGACACTCCAGAAGAGGGCGCCA | 1 | ENCFF093ZRU |
| 546 | 4 | 130410591 | 130410801 | 210 | CAST,FVB,KK,MOLF,NOR,NZB,SJL,SPRET,SWR | 4 | 130410674 | 130410692 | 19 | + | 17.5574 | 6.32E-07 | 0.111 | ACTCCAGAAGAGGGCGCCA | 1 | ENCFF093ZRU |
| 547 | 4 | 130969320 | 130969529 | 209 | 129S1,B10D2,B10J,BTBR,BUB,C57L,CAST,DBA1J,FVB,ILNJ2,KK,MA/My,MOLF,NOR,NZB,PJ,PL,SEA,SJL,SMJ,SPRET,SWR,TH,WSB | 4 | 130969386 | 130969419 | 34 | + | 16.4634 | 9.51E-07 | 0.0931 | TAGCTGTCTTCAGACGCACCAGAAGAGGGCATCA | | |
| 548 | 4 | 132497309 | 132497438 | 129 | SPRET | 4 | 132497338 | 132497371 | 34 | + | 18.0732 | 3.36E-07 | 0.056 | CAGCTGTCTTCAGACACTCCAGAAGAGGGCATCA | | |
| 549 | 4 | 132500651 | 132500848 | 197 | CAST.SPRET | 4 | 132500704 | 132500737 | 34 | + | 16.7724 | 7.83E-07 | 0.086 | TAGCTGTCTTCAGATGCACCAGAAGAGGGCGTCA | | |
| 550 | 4 | 132507380 | 132507568 | 188 | CAST,DBA1J,KK,MOLF,NZB,SMJ,SPRET | 4 | 132507426 | 132507459 | 34 | + | 18.3333 | 2.82E-07 | 0.0507 | TAGCTGTCCTCAGACACACCAGAAGAGGGCGTCA | 1 | ENCFF093ZRU |
| 551 | 4 | 132513693 | 132513895 | 202 | SPRET | 4 | 132513754 | 132513787 | 34 | + | 18.935 | 1.88E-07 | 0.0423 | TTGCTGTCTTCAGACACACCAGAAGAGGGCGTCA | | |
| 552 | 4 | 132526245 | 132526455 | 210 | SPRET | 4 | 132526291 | 132526324 | 34 | + | 17.2439 | 5.79E-07 | 0.0743 | TAGCTGTCCTCAGATACTCCAGAAGAGGGCATCA | 1 | ENCFF093ZRU |
| 553 | 4 | 132630049 | 132630657 | 608 | CAST.DBA1J.KK.MOLF.NZB.SMJ | 4 | 132630639 | 132630672 | 34 | + | 17.3902 | 5.27E-07 | 0.071 | TAGCTGTCCTCAGACACTCCAGAAGAGGGCATCA | | |
| 554 | 4 | 132893045 | 132893262 | 217 | 129S1,AJ,AKR,B10D2,B10J,BALB,BTBR,BUB,C3H,C57L,CAST,CBA,CE,DBA2J,FVB,ILNJ2,MA/My,MRL,NOD,NZW,PJ,PL,RF,RH2,SEA,SJL,SPRET,SWR,TH,WSB | 4 | 132893125 | 132893158 | 34 | + | 16.4634 | 9.51E-07 | 0.0931 | TAGCTGTCTTCAGACGCACCAGAAGAGGGCATCA | | |
| 555 | 4 | 133339317 | 133339885 | 568 | CAST,MOLF,SPRET | 4 | 133339302 | 133339335 | 34 | + | 29.0407 | 3.14E-11 | 0.000147 | GTGCAGTACCAGAGGAGACCAGAAGAGGGCACTA | 14 | ENCFF821BIA,ENCFF092NMJ,ENCFF136ZDK,ENCFF367UNR,ENCFF165LXO,ENCFF093ZRU,ENCFF502WDT,ENCFF052KVO,ENCFF089GBC,ENCFF024JOZ,ENCFF907XEG,ENCFF198SSR,ENCFF327LYS,ENCFF589ZBJ |
| 555 | 4 | 133339317 | 133339885 | 568 | CAST,MOLF,SPRET | 4 | 133339317 | 133339335 | 19 | + | 19.4098 | 1.73E-07 | 0.0591 | AGACCAGAAGAGGGCACTA | 14 | ENCFF821BIA,ENCFF092NMJ,ENCFF136ZDK,ENCFF367UNR,ENCFF165LXO,ENCFF093ZRU,ENCFF502WDT,ENCFF052KVO,ENCFF089GBC,ENCFF024JOZ,ENCFF907XEG,ENCFF198SSR,ENCFF327LYS,ENCFF589ZBJ |
| 555 | 4 | 133339317 | 133339885 | 568 | CAST,MOLF,SPRET | 4 | 133339871 | 133339904 | 34 | + | 27.5122 | 1.55E-10 | 0.00034 | GTGCAGTACTAGAGGAGACCAGAAGAGGGCACTG | 9 | ENCFF052KVO,ENCFF089GBC,ENCFF907XEG,ENCFF198SSR,ENCFF092NMJ,ENCFF136ZDK,ENCFF099QNE.ENCFF367UNR.ENCFF093ZRU |
| 556 | 4 | 133763750 | 133764002 | 252 | 129S1,AJ,AKR,B10D2,B10J,BALB,BTBR,BUB,C3H,C57L,CAST,CBA,CE,DBA1J,DBA2J,FVB,ILNJ2,MA/My,MOLF,MRL,NOD,NZW,PJ,PL,RF,RH2,SEA,SJL,SMJ,SPRET,SWR,TH,WSB | 4 | 133763800 | 133763833 | 34 | + | 18.3659 | 2.76E-07 | 0.0503 | TAGCTGTCTTCAGACACTCCAGAAGAGGGCGCCA | | |
| 556 | 4 | 133763750 | 133764002 | 252 | 129S1,AJ,AKR,B10D2,B10J,BALB,BTBR,BUB,C3H,C57L,CAST,CBA,CE,DBA1J,DBA2J,FVB,ILNJ2,MA/My,MOLF,MRL,NOD,NZW,PJ,PL,RF,RH2,SEA,SJL,SMJ,SPRET,SWR,TH,WSB | 4 | 133763815 | 133763833 | 19 | + | 17.5574 | 6.32E-07 | 0.111 | ACTCCAGAAGAGGGCGCCA | | |
| 557 | 4 | 133966613 | 133966835 | 222 | 129S1,AJ,AKR,B10D2,B10J,BALB,BTBR,BUB,C3H,C57L,CAST,CBA,CE,DBA1J,DBA2J,FVB,ILNJ2,MA/My,MOLF,MRL,NOD,NZW,PJ,PL,RF,RH2,SEA,SJL,SMJ,SPRET,SWR,TH,WSB | 4 | 133966692 | 133966725 | 34 | + | 18.3659 | 2.76E-07 | 0.0503 | TAGCTGTCTTCAGACACTCCAGAAGAGGGCGCCA | 3 | ENCFF367UNR,ENCFF093ZRU,ENCFF099QNE |
| 557 | 4 | 133966613 | 133966835 | 222 | 129S1,AJ,AKR,B10D2,B10J,BALB,BTBR,BUB,C3H,C57L,CAST,CBA,CE,DBA1J,DBA2J,FVB,ILNJ2,MA/My,MOLF,MRL,NOD,NZW,PJ,PL,RF,RH2,SEA,SJL,SMJ,SPRET,SWR,TH,WSB | 4 | 133966707 | 133966725 | 19 | + | 17.5574 | 6.32E-07 | 0.111 | ACTCCAGAAGAGGGCGCCA | 3 | ENCFF367UNR,ENCFF093ZRU,ENCFF099QNE |
| 558 | 4 | 134137243 | 134137549 | 306 | 129S1,AJ,AKR,B10D2,B10J,BALB,BTBR,BUB,C3H,C57L,CBA,CE,DBA1J,DBA2J,FVB,ILNJ2,MA/My,MRL,NOD,NZW,PJ,PL,RF,RH2,SEA,SJL,SMJ,SWR,TH,WSB | 4 | 134137368 | 134137401 | 34 | + | 20.065 | 8.47E-08 | 0.0249 | ATGCAATGCCTGTGGAGGCCAGAACAGGGTACTG | | |
| 559 | 4 | 134317754 | 134318534 | 780 | SPRET | 4 | 134318014 | 134318048 | 35 | + | 19.422 | 1.26E-07 | 0.0499 | TTGCAGTTACAGGTTATGGCCAGCAGGGCGGCCCC | 25 | ENCFF367UNR,ENCFF093ZRU,ENCFF821BIA,ENCFF099QNE,ENCFF210BTO,ENCFF052KVO,ENCFF430PPJ,ENCFF092NMJ,ENCFF165LXO,ENCFF024JOZ,ENCFF065ODV,ENCFF198SSR,ENCFF883UPM,ENCFF136ZDK,ENCFF033UQX,ENCFF316FYN,ENCFF589ZBJ,ENCFF327LYS,ENCFF907XEG,ENCFF194HVL,ENCFF957SPH.ENCFF055HAP.ENCFF142CNG.ENCFF089GBC.ENCFF502WDT |
| 560 | 4 | 134441901 | 134442124 | 223 | SPRET | 4 | 134441957 | 134441990 | 34 | + | 16.4472 | 9.61E-07 | 0.0931 | TAGCTGTCTTCAGACACACCAGAAGAGGGCGTCA | | |
| 561 | 4 | 134553136 | 134553331 | 195 | SPRET | 4 | 134553189 | 134553222 | 34 | + | 18.3659 | 2.76E-07 | 0.0503 | TAGCTGTCTTCAGACACTCCAGAAGAGGGCGCCA | 3 | ENCFF093ZRU.ENCFF367UNR.ENCFF821BIA |
| 561 | 4 | 134553136 | 134553331 | 195 | SPRET | 4 | 134553204 | 134553222 | 19 | + | 17.5574 | 6.32E-07 | 0.111 | ACTCCAGAAGAGGGCGCCA | 3 | ENCFF093ZRU.ENCFF367UNR.ENCFF821BIA |
| 562 | 4 | 134952939 | 134953150 | 211 | SPRET | 4 | 134953003 | 134953036 | 34 | + | 16.4472 | 9.61E-07 | 0.0931 | TAGCTGTCTTCAGACACACCAGAAGAGGGCGTCA | | |
| 563 | 4 | 135246832 | 135247012 | 180 | SPRET | 4 | 135246870 | 135246903 | 34 | + | 19.1057 | 1.67E-07 | 0.0396 | CTGTAGCTGTCAGACACTCCAGAAGAGGGCGTCA | 1 | ENCFF093ZRU |
| 564 | 4 | 135562292 | 135562499 | 207 | B10D2,B10J,CAST,DBA1J,KK,MOLF,NZO,PJ,SEA,SPRET | 4 | 135562355 | 135562388 | 34 | + | 17.7805 | 4.08E-07 | 0.0623 | CTGTAGTCTGTAGACACTCCAGAAGAGGGCATCA | | |
| 565 | 4 | 135705382 | 135705771 | 389 | SPRET | 4 | 135705630 | 135705663 | 34 | + | 16.4472 | 9.61E-07 | 0.0931 | TAGCTGTCTTCAGACACACCAGAAGAGGGCGTCA | | |
| 566 | 4 | 137398651 | 137398736 | 85 | CAST | 4 | 137398621 | 137398654 | 34 | + | 17.065 | 6.50E-07 | 0.0792 | TTGCTCTCTTCAGACACACCAGAAGAGGGCATCG | | |
| 567 | 4 | 137693725 | 137694100 | 375 | PJ | 4 | 137693980 | 137694014 | 35 | + | 17.7339 | 3.53E-07 | 0.0912 | CTGTACCCCCAGGCCCTGGCCTGCAGACGACGGTA | | |
| 568 | 4 | 137724808 | 137725016 | 208 | AJ,AKR,BALB,BUB,C3H,CAST,CBA,CE,DBA1J,DBA2J,FVB,ILNJ2,KK,MA/My,MRL,NZB,NZW,PL,RF,RH2,SJL,SMJ,SPRET,SWR,TH,WSB | 4 | 137724864 | 137724897 | 34 | + | 17.8455 | 3.91E-07 | 0.0606 | TAGCTGTCCTCAGACACTCCAGAAGAGGGCGTCA | | |
| 569 | 4 | 139430891 | 139431297 | 406 | SPRET | 4 | 139430940 | 139430973 | 34 | + | 27.3821 | 1.77E-10 | 0.000352 | ATGCAATTCCCTTGGAGGCCAGAAGGGGGCATCA | 13 | ENCFF024JOZ,ENCFF092NMJ,ENCFF198SSR,ENCFF327LYS,ENCFF883UPM,ENCFF367UNR,ENCFF093ZRU,ENCFF052KVO,ENCFF099QNE,ENCFF821BIA,ENCFF907XEG,ENCFF136ZDK,ENCFF589ZBJ |
| 569 | 4 | 139430891 | 139431297 | 406 | SPRET | 4 | 139430955 | 139430973 | 19 | + | 17.4262 | 6.88E-07 | 0.115 | AGGCCAGAAGGGGGCATCA | 13 | ENCFF024JOZ,ENCFF092NMJ,ENCFF198SSR,ENCFF327LYS,ENCFF883UPM,ENCFF367UNR,ENCFF093ZRU,ENCFF052KVO,ENCFF099QNE,ENCFF821BIA,ENCFF907XEG,ENCFF136ZDK,ENCFF589ZBJ |
| 570 | 4 | 141480755 | 141481065 | 310 | CAST | 4 | 141480783 | 141480816 | 34 | + | 16.4472 | 9.61E-07 | 0.0931 | TAGCTGTCTTCAGACACACCAGAAGAGGGCGTCA | | |
| 571 | 4 | 141660929 | 141661131 | 202 | CAST.MOLF.SPRET | 4 | 141660989 | 141661022 | 34 | + | 17.2439 | 5.79E-07 | 0.0743 | TAGCTGTCCTCAGATACTCCAGAAGAGGGCATCA | | |
| 572 | 4 | 142962694 | 142963274 | 580 | SPRET | 4 | 142962953 | 142962986 | 34 | + | 18.7724 | 2.10E-07 | 0.0451 | TAGCAATCTTCAGACACACCAGGAGAGGGCGCCA | 3 | ENCFF367UNR.ENCFF093ZRU.ENCFF821BIA |
| 572 | 4 | 142962694 | 142963274 | 580 | SPRET | 4 | 142962968 | 142962986 | 19 | + | 17.7541 | 5.54E-07 | 0.109 | ACACCAGGAGAGGGCGCCA | 3 | ENCFF367UNR.ENCFF093ZRU.ENCFF821BIA |
| 573 | 4 | 143036628 | 143037583 | 955 | 129S1,BTBR,BUB,C57L,C58J,CE,DBA2J,FVB,KK,LP,MA/My,NOD,NZB,NZW,PJ,PL,RF,SEA.SJL.SMJ.SWR.TH.WSB | 4 | 143037575 | 143037608 | 34 | + | 16.5528 | 8.99E-07 | 0.0931 | TAGCTGTCTTCAGACACACCAGAAGAGGGCGTCG | | |
| 574 | 4 | 143089847 | 143090039 | 192 | 129S1,BTBR,BUB,C57L,C58J,CE,DBA2J,FVB,KK,LP,MA/My,MOLF,MRL,NOD,NZB,NZW,PJ,PL,RF,SEA,SJL,SMJ,SPRET,SWR,TH,WSB | 4 | 143089898 | 143089931 | 34 | + | 16.5203 | 9.18E-07 | 0.0931 | TAGCTGTCTTCCAACACTCCAGAAGAGGGCGTCA | | |
| 575 | 4 | 144917499 | 144917715 | 216 | SPRET | 4 | 144917711 | 144917745 | 35 | + | 16.3303 | 7.94E-07 | 0.141 | ATGCAGGAGCCTGCGGGGGCCAGAGGAGGGCATAA | | |
| 576 | 4 | 145208053 | 145208258 | 205 | 129S1,AKR,BTBR,BUB,C3H,C57L,CAST,CBA,CE,DBA1J,KK,MA/My,MRL,NOD,NZB,NZO,NZW,PL,RF,SJL,SMJ,SPRET,SWR,WSB | 4 | 145208116 | 145208149 | 34 | + | 17.3902 | 5.27E-07 | 0.071 | TAGCTGTCCTCAGACACTCCAGAAGAGGGCATCA | | |
| 577 | 4 | 146617253 | 146617462 | 209 | 129S1,AKR,B10J,BALB,BTBR,BUB,C3H,C57L,C58J,CAST,CBA,CE,DBA1J,DBA2J,FVB,ILNJ2,KK,LP,MA/My,MOLF,MRL,NOD,NZB,NZO,NZW,PJ,PL,RF,SEA,SJL,SMJ,SPRET,SWR.TH.WSB | 4 | 146617318 | 146617351 | 34 | + | 16.4472 | 9.61E-07 | 0.0931 | TAGCTGTCTTCAGACACACCAGAAGAGGGCGTCA | | |
| 578 | 4 | 147009059 | 147009256 | 197 | 129S1,AKR,BTBR,BUB,C57L,DBA1J,KK,MA/My,MRL,NOD,NZB,NZW,PL,RF,SJL,SWR | 4 | 147009113 | 147009146 | 34 | + | 16.4472 | 9.61E-07 | 0.0931 | TAGCTGTCTTCAGACACACCAGAAGAGGGCGTCA | | |
| 579 | 4 | 148034970 | 148035154 | 184 | 129S1,AKR,BTBR,BUB,C3H,C57L,C58J,CBA,CE,DBA1J,DBA2J,KK,LP,MA/My,MRL,NOD,NZB,NZO,NZW,PJ,PL,RF,SEA,SJL,SMJ,SPRET.SWR.TH.WSB | 4 | 148035063 | 148035096 | 34 | + | 16.4634 | 9.51E-07 | 0.0931 | TAGCTGTCTTCAGACGCACCAGAAGAGGGCATCA | | |
| 580 | 4 | 148200733 | 148200839 | 106 | SPRET | 4 | 148200803 | 148200836 | 34 | + | 16.4472 | 9.61E-07 | 0.0931 | TAGCTGTCTTCAGACACACCAGAAGAGGGCGTCA | | |
| 581 | 4 | 148625767 | 148626704 | 937 | CAST | 4 | 148626449 | 148626482 | 34 | + | 16.4634 | 9.51E-07 | 0.0931 | TAGCTGTCTTCAGACGCACCAGAAGAGGGCATCA | | |
| 582 | 4 | 148626379 | 148626593 | 214 | 129S1,AJ,AKR,BALB,BTBR,BUB,C3H,C57L,C58J,CBA,CE,DBA1J,DBA2J,KK,LP,MA/My,MOLF,MRL,NOD,NZB,NZO,NZW,PJ,PL,RF,SEA,SJL,SMJ,SPRET,SWR,TH,WSB | 4 | 148626449 | 148626482 | 34 | + | 16.4634 | 9.51E-07 | 0.0931 | TAGCTGTCTTCAGACGCACCAGAAGAGGGCATCA | | |
| 583 | 4 | 148953146 | 148953391 | 245 | SPRET | 4 | 148953247 | 148953280 | 34 | + | 16.4472 | 9.61E-07 | 0.0931 | TAGCTGTCTTCAGACACACCAGAAGAGGGCGTCA | 1 | ENCFF367UNR |
| 584 | 4 | 149485040 | 149485236 | 196 | 129S1,AKR,BTBR,C3H,C57L,CAST,CBA,DBA1J,DBA2J,FVB,ILNJ2,MA/My,MOLF,MRL,NOD,PL,SJL,SPRET,SWR,TH | 4 | 149485088 | 149485121 | 34 | + | 16.4472 | 9.61E-07 | 0.0931 | TAGCTGTCTTCAGACACACCAGAAGAGGGCGTCA | | |
| 585 | 4 | 149507935 | 149508144 | 209 | 129S1,AKR,BTBR,C3H,C57L,C58J,CAST,CBA,CE,DBA1J,DBA2J,FVB,ILNJ2,LP,MA/My,MOLF,MRL,NOD,NZO,PJ,PL,SEA,SJL,SPRET,SWR,TH,WSB | 4 | 149508002 | 149508035 | 34 | + | 16.4472 | 9.61E-07 | 0.0931 | TAGCTGTCTTCAGACACACCAGAAGAGGGCGTCA | 1 | ENCFF093ZRU |
| 586 | 4 | 149685148 | 149685570 | 422 | CAST,FVB,ILNJ2,MOLF,SPRET | 4 | 149685427 | 149685460 | 34 | + | 16.5366 | 9.09E-07 | 0.0931 | ATGTAGTGCCTATGTATGCCAGAAGGGGATGTTG | | |
| 587 | 4 | 150595608 | 150595932 | 324 | SPRET | 4 | 150595841 | 150595874 | 34 | + | 16.4472 | 9.61E-07 | 0.0931 | TAGCTGTCTTCAGACACACCAGAAGAGGGCGTCA | 2 | ENCFF367UNR,ENCFF093ZRU |
| 588 | 4 | 150596892 | 150597663 | 771 | AJ,BALB,RH2 | 4 | 150597410 | 150597443 | 34 | + | 16.8537 | 7.43E-07 | 0.0844 | GAGCAGTGCCAGGAGGCAGCACTGGGTGCCACAT | | |
| 589 | 4 | 150622997 | 150623196 | 199 | CAST,MOLF,SPRET | 4 | 150623056 | 150623089 | 34 | + | 17.3659 | 5.35E-07 | 0.0717 | TAGCTATTCTCAGAAACCCCAGAAGAGGGCATCA | | |
| 590 | 4 | 150764197 | 150764399 | 202 | AKR,C3H,CAST,CBA,MRL,NOD,NOR,NZO,SPRET | 4 | 150764258 | 150764291 | 34 | + | 16.4472 | 9.61E-07 | 0.0931 | TAGCTGTCTTCAGACACACCAGAAGAGGGCGTCA | | |
| 591 | 4 | 152063061 | 152063255 | 194 | MOLF,SPRET | 4 | 152063102 | 152063135 | 34 | + | 17.8455 | 3.91E-07 | 0.0606 | TAGCTGTCCTCAGACACTCCAGAAGAGGGCGTCA | | |
| 592 | 4 | 152120800 | 152120996 | 196 | CAST.MOLF.SPRET.WSB | 4 | 152120854 | 152120887 | 34 | + | 18.8699 | 1.96E-07 | 0.0436 | TAGCTGTCTTCAGACGCACCAGAAGAGGGCACCA | | |
| 592 | 4 | 152120800 | 152120996 | 196 | CAST.MOLF.SPRET.WSB | 4 | 152120869 | 152120887 | 19 | + | 17.9672 | 4.81E-07 | 0.103 | GCACCAGAAGAGGGCACCA | | |
| 593 | 4 | 152165445 | 152165663 | 218 | CAST.SPRET.WSB | 4 | 152165522 | 152165555 | 34 | + | 16.4634 | 9.51E-07 | 0.0931 | TAGCTGTCTTCAGACGCACCAGAAGAGGGCATCA | | |
| 594 | 4 | 154277898 | 154278273 | 375 | 129S1,AJ,AKR,C3H,CAST,CBA,CE,DBA2J,FVB,ILNJ2,MOLF,NOD,NOR,RF,RH2,SJL,SMJ,SPRET,SWR,TH | 4 | 154277897 | 154277930 | 34 | + | 16.4472 | 9.61E-07 | 0.0931 | TAGCTGTCTTCAGACACACCAGAAGAGGGCGTCA | | |
| 594 | 4 | 154277898 | 154278273 | 375 | 129S1,AJ,AKR,C3H,CAST,CBA,CE,DBA2J,FVB,ILNJ2,MOLF,NOD,NOR,RF,RH2,SJL,SMJ,SPRET,SWR,TH | 4 | 154278273 | 154278306 | 34 | + | 18.3984 | 2.70E-07 | 0.0503 | TAGCTGTCTTCAGACACACCAGAAGAGGGCACCA | | |
| 595 | 4 | 155581452 | 155581584 | 132 | CAST,MOLF,SPRET | 4 | 155581442 | 155581475 | 34 | + | 17.2764 | 5.67E-07 | 0.0739 | TAGCTGTCTTTAGACACGCCAGAAGAGGGCGTCA | | |
| 596 | 4 | 155601743 | 155601957 | 214 | SPRET | 4 | 155601895 | 155601928 | 34 | + | 17.6911 | 4.33E-07 | 0.0642 | CTGCATGCATCCTGCAGGGCAGAAGAGGGCACTA | | |
| 597 | 4 | 155719765 | 155720337 | 572 | CE,FVB,NOD,NOR,SJL,SWR,TH,WSB | 4 | 155719753 | 155719786 | 34 | + | 18.5854 | 2.38E-07 | 0.0492 | TTGCTGTCTTCAGACACACCAGAAGAGGGCATCG | 2 | ENCFF093ZRU,ENCFF367UNR |
| 598 | 4 | 155803163 | 155803380 | 217 | 129S1,AJ,AKR,BALB,BUB,C3H,CAST,CBA,CE,DBA1J,DBA2J,FVB,MOLF,MRL,NOD,NOR,NZO,PL,RF,RH2,SJL,SPRET,SWR,WSB | 4 | 155803237 | 155803270 | 34 | + | 16.4634 | 9.51E-07 | 0.0931 | TAGCTGTCTTCAGACGCACCAGAAGAGGGCATCA | | |
| 599 | 4 | 156163550 | 156163788 | 238 | 129S1,AJ,AKR,BALB,C3H,C57L,C58J,CAST,CBA,CE,DBA1J,DBA2J,FVB,ILNJ2,MA/My,MOLF,MRL,NOD,NOR,NZO,PJ,PL,RF,RH2,SEA,SJL,SMJ,SPRET,SWR,TH,WSB | 4 | 156163647 | 156163680 | 34 | + | 18.3333 | 2.82E-07 | 0.0507 | TAGCTGTCCTCAGACACACCAGAAGAGGGCGTCA | | |
| 600 | 4 | 4616551 | 4616742 | 191 | AJ.CAST.SPRET | 4 | 4616675 | 4616708 | 34 | - | 16.4472 | 9.61E-07 | 0.0931 | TAGCTGTCTTCAGACACACCAGAAGAGGGCGTCA | | |
| 601 | 4 | 6226075 | 6226274 | 199 | 129S1,AKR,BALB,BTBR,BUB,C57L,CAST,FVB,ILNJ2,MA/My,MOLF,NOD,NOR,NZW,PL,RF,RH2,SEA,SJL,SPRET,SWR,TH,WSB | 4 | 6226201 | 6226234 | 34 | - | 17.7805 | 4.08E-07 | 0.0623 | CTGTAGTCTGTAGACACTCCAGAAGAGGGCATCA | | |
| 602 | 4 | 6542315 | 6543176 | 861 | BUB | 4 | 6542838 | 6542856 | 19 | - | 17.5902 | 6.18E-07 | 0.111 | ACACCAGAAGAGGGCACTG | | |
| 602 | 4 | 6542315 | 6543176 | 861 | BUB | 4 | 6542838 | 6542871 | 34 | - | 18.2195 | 3.05E-07 | 0.0526 | TAGCTGTCTTCAAACACACCAGAAGAGGGCACTG | | |
| 603 | 4 | 11121823 | 11122044 | 221 | AJ.CAST.NOD.NOR.SPRET | 4 | 11121946 | 11121979 | 34 | - | 17.1382 | 6.20E-07 | 0.0772 | TAGCTGTCTCCAGACACTCCAGAAGAGGGCATCA | | |
| 604 | 4 | 11210188 | 11210392 | 204 | CAST.SPRET | 4 | 11210311 | 11210329 | 19 | - | 17.5574 | 6.32E-07 | 0.111 | ACTCCAGAAGAGGGCGCCA | 2 | ENCFF093ZRU.ENCFF367UNR |
| 604 | 4 | 11210188 | 11210392 | 204 | CAST.SPRET | 4 | 11210311 | 11210344 | 34 | - | 18.3659 | 2.76E-07 | 0.0503 | TAGCTGTCTTCAGACACTCCAGAAGAGGGCGCCA | 2 | ENCFF093ZRU.ENCFF367UNR |
| 605 | 4 | 18619866 | 18620073 | 207 | CAST.SPRET | 4 | 18619990 | 18620008 | 19 | - | 17.5574 | 6.32E-07 | 0.111 | ACTCCAGAAGAGGGCGCCA | 1 | ENCFF093ZRU |
| 605 | 4 | 18619866 | 18620073 | 207 | CAST.SPRET | 4 | 18619990 | 18620023 | 34 | - | 18.3659 | 2.76E-07 | 0.0503 | TAGCTGTCTTCAGACACTCCAGAAGAGGGCGCCA | 1 | ENCFF093ZRU |
| 606 | 4 | 19289455 | 19289654 | 199 | SPRET | 4 | 19289574 | 19289592 | 19 | - | 17.5574 | 6.32E-07 | 0.111 | ACTCCAGAAGAGGGCGCCA | 1 | ENCFF093ZRU |
| 606 | 4 | 19289455 | 19289654 | 199 | SPRET | 4 | 19289574 | 19289607 | 34 | - | 18.3659 | 2.76E-07 | 0.0503 | TAGCTGTCTTCAGACACTCCAGAAGAGGGCGCCA | 1 | ENCFF093ZRU |
| 607 | 4 | 23276622 | 23276849 | 227 | SPRET | 4 | 23276750 | 23276783 | 34 | - | 18.4634 | 2.59E-07 | 0.0496 | TTGCTGTCTTCAGATGTACCAGGAGAGGGCGTCA | | |
| 608 | 4 | 24934035 | 24934602 | 567 | SPRET | 4 | 24934216 | 24934249 | 34 | - | 18.0325 | 3.45E-07 | 0.0569 | GTACAGTGTTCCAGGAGGCCAGAAGAGGGCAATT | | |
| 609 | 4 | 44001313 | 44001508 | 195 | 129S1,BUB,C58J,CBA,DBA1J,DBA2J,FVB,ILNJ2,LP,MOLF,PJ,SEA,SJL,SMJ,SPRET,SWR,TH,WSB | 4 | 44001437 | 44001470 | 34 | - | 16.4472 | 9.61E-07 | 0.0931 | TAGCTGTCTTCAGACACACCAGAAGAGGGCGTCA | | |
| 610 | 4 | 44861872 | 44862068 | 196 | SPRET | 4 | 44861996 | 44862029 | 34 | - | 18.9187 | 1.90E-07 | 0.0426 | CAGCTGTCTTCAAACACACCAGAAGAGGGCGTCA | 1 | ENCFF093ZRU |
| 611 | 4 | 45801866 | 45802068 | 202 | BUB,C58J,CAST,CBA,DBA1J,DBA2J,ILNJ2,LP,MA/My,MOLF,NZW,PL,SMJ,SPRET | 4 | 45801990 | 45802008 | 19 | - | 17.5574 | 6.32E-07 | 0.111 | ACTCCAGAAGAGGGCGCCA | 5 | ENCFF367UNR,ENCFF821BIA,ENCFF327LYS,ENCFF093ZRU,ENCFF099QNE |
| 611 | 4 | 45801866 | 45802068 | 202 | BUB,C58J,CAST,CBA,DBA1J,DBA2J,ILNJ2,LP,MA/My,MOLF,NZW,PL,SMJ,SPRET | 4 | 45801990 | 45802023 | 34 | - | 18.3659 | 2.76E-07 | 0.0503 | TAGCTGTCTTCAGACACTCCAGAAGAGGGCGCCA | 5 | ENCFF367UNR,ENCFF821BIA,ENCFF327LYS,ENCFF093ZRU,ENCFF099QNE |
| 612 | 4 | 48602148 | 48602480 | 332 | 129S1,BUB,C57L,CAST,KK,MA/My,MOLF,NZW.SPRET | 4 | 48602335 | 48602353 | 19 | - | 17.5574 | 6.32E-07 | 0.111 | ACTCCAGAAGAGGGCGCCA | 1 | ENCFF093ZRU |
| 612 | 4 | 48602148 | 48602480 | 332 | 129S1,BUB,C57L,CAST,KK,MA/My,MOLF,NZW.SPRET | 4 | 48602335 | 48602368 | 34 | - | 18.3659 | 2.76E-07 | 0.0503 | TAGCTGTCTTCAGACACTCCAGAAGAGGGCGCCA | 1 | ENCFF093ZRU |
| 613 | 4 | 48697044 | 48697248 | 204 | SPRET.WSB | 4 | 48697167 | 48697185 | 19 | - | 17.8852 | 5.08E-07 | 0.103 | ACACCAGAAGAGGGCACCA | | |
| 614 | 4 | 49686463 | 49687059 | 596 | MA/My.MOLF | 4 | 49686495 | 49686529 | 35 | - | 18.3119 | 2.49E-07 | 0.0757 | TAGCTGTGGGGGGGGGGTGCCAGCAGAGGGCAAGG | | |
| 615 | 4 | 51782138 | 51782351 | 213 | AJ,BALB,BUB,C3H,CAST,ILNJ2,KK,MA/My,MOLF,NOD,NOR,PJ,RH2,SEA,SPRET | 4 | 51782262 | 51782295 | 34 | - | 18.8618 | 1.97E-07 | 0.0437 | TAGCTGTCCTCAGATTCTCCAGAAGAGGGCATCA | | |
| 616 | 4 | 57294341 | 57294545 | 204 | CAST.MOLF.SPRET | 4 | 57294465 | 57294498 | 34 | - | 16.4472 | 9.61E-07 | 0.0931 | TAGCTGTCTTCAGACACACCAGAAGAGGGCGTCA | | |
| 617 | 4 | 58164222 | 58164445 | 223 | 129S1,C57L,CAST,MOLF,NOD,NOR,PJ,PL,SJL,SMJ,SPRET,SWR,TH | 4 | 58164344 | 58164377 | 34 | - | 17.439 | 5.10E-07 | 0.0703 | TAGCTGTCTTCAGTCACTCCAGAAGATGGCGCTA | | |
| 618 | 4 | 59197770 | 59198025 | 255 | 129S1,AJ,AKR,BALB,BTBR,C3H,C57L,CBA,CE,DBA1J,DBA2J,FVB,ILNJ2,LP,MA/My,MOLF,MRL,NOD,NOR,NZB,NZO,NZW,PJ,RF,RH2,SEA,SJL,SPRET,SWR,TH,WSB | 4 | 59197895 | 59197928 | 34 | - | 17.9431 | 3.66E-07 | 0.0591 | CAGCTGTCCTCAGACACTCCAGAAGAGGGAGTCA | | |
| 619 | 4 | 61956073 | 61956182 | 109 | MOLF | 4 | 61956050 | 61956083 | 34 | - | 25.2439 | 1.33E-09 | 0.00122 | GTGCAGCACTCAGCCTGGCCAGTAGAGGGCAATC | | |
| 620 | 4 | 65726149 | 65726366 | 217 | BTBR.CAST.CE.MOLF.PJ.SPRET.WSB | 4 | 65726293 | 65726326 | 34 | - | 17.6341 | 4.49E-07 | 0.0653 | CTGCACCTCAATCTGAGCCCTGTAGGAGGACCCG | | |
| 621 | 4 | 78846445 | 78846643 | 198 | SPRET | 4 | 78846571 | 78846604 | 34 | - | 16.4634 | 9.51E-07 | 0.0931 | TTGCTGTCTTCAGACACACCAGAAGAGGGAGTCA | | |
| 622 | 4 | 83231895 | 83232131 | 236 | CAST.MOLF.SMJ.SPRET | 4 | 83232022 | 83232055 | 34 | - | 17.878 | 3.83E-07 | 0.06 | TAGCTGTCCTCAGACACACCAGAAGAGGGCATCA | | |
| 623 | 4 | 85191804 | 85191910 | 106 | SPRET | 4 | 85191905 | 85191939 | 35 | - | 16.3945 | 7.65E-07 | 0.138 | CTGCCACCTCCTCTGTCCACAAGCAGAGGGCAGTG | 8 | ENCFF092NMJ,ENCFF089GBC,ENCFF024JOZ,ENCFF093ZRU,ENCFF907XEG,ENCFF316FYN,ENCFF099QNE.ENCFF327LYS |
| 624 | 4 | 85681775 | 85681973 | 198 | 129S1,B10D2,B10J,BTBR,C57L,C58J,CAST,CE,DBA1J,DBA2J,ILNJ2,KK,MOLF,NZO,PJ,PL,RF,RH2,SMJ,SPRET,WSB | 4 | 85681901 | 85681935 | 35 | - | 18.1284 | 2.79E-07 | 0.0805 | CTGTAGCTGTCTTCAGACACCAGAAGAGGGCATCA | | |
| 625 | 4 | 87324876 | 87325083 | 207 | B10D2,B10J,C3H,C58J,CAST,CBA,DBA1J,DBA2J,ILNJ2,MA/My,MOLF,NZO,PJ,PL,RF,SMJ,SPRET,WSB | 4 | 87324998 | 87325031 | 34 | - | 18.8537 | 1.98E-07 | 0.0437 | CAGCTGTCTTCAGACACTCCAGTAGAGGGCATCA | 1 | ENCFF093ZRU |
| 626 | 4 | 87513583 | 87513715 | 132 | SPRET | 4 | 87513686 | 87513719 | 34 | - | 16.9268 | 7.10E-07 | 0.0825 | TAGCTGTTTTCAGTCACCCCAGAAGAGGGCATCA | | |
| 627 | 4 | 89614571 | 89615541 | 970 | SPRET | 4 | 89614812 | 89614845 | 34 | - | 18.5203 | 2.49E-07 | 0.0494 | ATGCAGCTAGAGACACGAACTCCAGGGGGTACTG | | |
| 628 | 4 | 92489080 | 92489300 | 220 | 129S1.C57L.CAST.MOLF.RH2.SPRET | 4 | 92489206 | 92489239 | 34 | - | 17.6667 | 4.40E-07 | 0.0647 | TAGCTGTCTTCAGACACGCCAGAAGAGGGCGTCA | | |
| 629 | 4 | 94487598 | 94487811 | 213 | SPRET | 4 | 94487707 | 94487725 | 19 | - | 18.5574 | 3.20E-07 | 0.082 | CCTCCAGTAGAGGGCGCAC | 3 | ENCFF367UNR.ENCFF093ZRU.ENCFF136ZDK |
| 629 | 4 | 94487598 | 94487811 | 213 | SPRET | 4 | 94487707 | 94487741 | 35 | - | 19.6606 | 1.08E-07 | 0.0448 | ATGAGGTTTCTTCACCCCTCCAGTAGAGGGCGCAC | 3 | ENCFF367UNR.ENCFF093ZRU.ENCFF136ZDK |
| 630 | 4 | 99520761 | 99520975 | 214 | 129S1,AJ,BALB,BTBR,C3H,C57L,CAST,CBA,ILNJ2,KK,MA/My,MOLF,NOD,NOR,NZO,RH2,SEA,SPRET,SWR,TH,WSB | 4 | 99520887 | 99520905 | 19 | - | 17.5574 | 6.32E-07 | 0.111 | ACTCCAGAAGAGGGCGCCA | | |
| 630 | 4 | 99520761 | 99520975 | 214 | 129S1,AJ,BALB,BTBR,C3H,C57L,CAST,CBA,ILNJ2,KK,MA/My,MOLF,NOD,NOR,NZO,RH2,SEA,SPRET,SWR,TH,WSB | 4 | 99520887 | 99520920 | 34 | - | 18.2358 | 3.02E-07 | 0.0524 | TAGCTGTCTTAAGACACTCCAGAAGAGGGCGCCA | | |
| 631 | 4 | 100489654 | 100490283 | 629 | BUB.PL.RF | 4 | 100490179 | 100490212 | 34 | - | 19.4715 | 1.29E-07 | 0.0337 | CTGCAAAGATAAACGTGGCCAGAAGAGGTCCCTG | | |
| 632 | 4 | 102041013 | 102041852 | 839 | CAST.MOLF | 4 | 102041135 | 102041168 | 34 | - | 17.7398 | 4.19E-07 | 0.0633 | CTGCAAGAGGAGAGCTTGCCTGCAGAGAGCGCTC | | |
| 633 | 4 | 103049830 | 103050012 | 182 | SPRET | 4 | 103049813 | 103049831 | 19 | - | 20.3115 | 8.63E-08 | 0.0415 | CAGCCAGTAGAGGGCGATA | 13 | ENCFF821BIA,ENCFF099QNE,ENCFF327LYS,ENCFF092NMJ,ENCFF589ZBJ,ENCFF198SSR,ENCFF907XEG,ENCFF055HAP,ENCFF052KVO,ENCFF367UNR,ENCFF093ZRU,ENCFF136ZDK,ENCFF502WDT |

| | | | | | | | | | | | | | | | | |
|---|---|---|---|---|---|---|---|---|---|---|---|---|---|---|---|---|
| 633 | 4 | 103049830 | 103050012 | 182 | SPRET | 4 | 103049813 | 103049847 | 35 | - | 20.156 | 7.86E-08 | 0.0364 | TAGCGGTACTGAACATCAGCCAGTAGAGGGCGATA | 13 | ENCFF821BIA,ENCFF099QNE,ENCFF327LYS,ENCFF092NMJ,ENCFF589ZBJ,ENCFF198SSR,ENCFF907XEG,ENCFF055HAP,ENCFF052KVO,ENCFF367UNR,ENCFF093ZRU,ENCFF136ZDK,ENCFF502WDT |
| 634 | 4 | 103112995 | 103113237 | 242 | AJ,BALB,BUB,C3H,C58J,CAST,CBA,KK,MOLF,NZW,PJ,PL,SEA,SMJ,SPRET,TH,WSB | 4 | 103113120 | 103113153 | 34 | - | 16.4634 | 9.51E-07 | 0.0931 | TAGCTGTCTTCAGACGCACCAGAAGAGGGCATCA | | |
| 635 | 4 | 104220166 | 104220355 | 189 | 129S1,AJ,AKR,BALB,BTBR,BUB,C3H,C57L,C58J,CAST,CBA,CE,DBA1J,DBA2J,FVB,ILNJ2,KK,MA/My,MOLF,MRL,NOD,NOR,NZB,NZO,NZW,PJ,PL,RF,SEA,SJL,SMJ,SPRET,SWR.TH.WSB | 4 | 104220287 | 104220305 | 19 | - | 17.5574 | 6.32E-07 | 0.111 | ACTCCAGAAGAGGGCGCCA | 2 | ENCFF093ZRU,ENCFF099QNE |
| 635 | 4 | 104220166 | 104220355 | 189 | 129S1,AJ,AKR,BALB,BTBR,BUB,C3H,C57L,C58J,CAST,CBA,CE,DBA1J,DBA2J,FVB,ILNJ2,KK,MA/My,MOLF,MRL,NOD,NOR,NZB,NZO,NZW,PJ,PL,RF,SEA,SJL,SMJ,SPRET,SWR.TH.WSB | 4 | 104220287 | 104220320 | 34 | - | 18.561 | 2.42E-07 | 0.0493 | AAGCTGTCTTCAGACACTCCAGAAGAGGGCGCCA | 2 | ENCFF093ZRU,ENCFF099QNE |
| 636 | 4 | 104664106 | 104664351 | 245 | CAST.MOLF.SPRET | 4 | 104664260 | 104664278 | 19 | - | 17.5574 | 6.32E-07 | 0.111 | ACTCCAGAAGAGGGCGCCA | | |
| 636 | 4 | 104664106 | 104664351 | 245 | CAST.MOLF.SPRET | 4 | 104664260 | 104664293 | 34 | - | 18.3659 | 2.76E-07 | 0.0503 | TAGCTGTCTTCAGACACTCCAGAAGAGGGCGCCA | | |
| 637 | 4 | 104861912 | 104862047 | 135 | BALB,BTBR,C3H,C58J,CAST,CBA,CE,ILNJ2,KK,MA/My,MOLF,NOD,NOR,PJ,RF,SEA,SJL.SMJ.SPRET.SWR.WSB | 4 | 104861927 | 104861961 | 35 | - | 18.1927 | 2.68E-07 | 0.0793 | GGGCAGTGTGCAGCGCCTGCCTAGAGAGGGCAGTG | 18 | ENCFF055HAP,ENCFF065ODV,ENCFF136ZDK,ENCFF327LYS,ENCFF092NMJ,ENCFF089GBC,ENCFF430PPJ,ENCFF198SSR,ENCFF165LXO,ENCFF210BTO,ENCFF589ZBJ,ENCFF907XEG,ENCFF052KVO,ENCFF533APC.ENCFF609MDN.ENCFF957SPH.ENCFF194HVL.ENCFF821BIA |
| 637 | 4 | 104861912 | 104862047 | 135 | BALB,BTBR,C3H,C58J,CAST,CBA,CE,ILNJ2,KK,MA/My,MOLF,NOD,NOR,PJ,RF,SEA,SJL.SMJ.SPRET.SWR.WSB | 4 | 104861954 | 104861988 | 35 | - | 18.1927 | 2.68E-07 | 0.0793 | GGGCAGTGTGCAGCGCCTGCCTAGAGAGGGCAGTG | 18 | ENCFF055HAP,ENCFF065ODV,ENCFF136ZDK,ENCFF327LYS,ENCFF092NMJ,ENCFF089GBC,ENCFF430PPJ,ENCFF198SSR,ENCFF165LXO,ENCFF210BTO,ENCFF589ZBJ,ENCFF907XEG,ENCFF052KVO,ENCFF533APC.ENCFF609MDN.ENCFF957SPH.ENCFF194HVL.ENCFF821BIA |
| 637 | 4 | 104861912 | 104862047 | 135 | BALB,BTBR,C3H,C58J,CAST,CBA,CE,ILNJ2,KK,MA/My,MOLF,NOD,NOR,PJ,RF,SEA,SJL,SMJ,SPRET,SWR,WSB | 4 | 104861981 | 104862015 | 35 | - | 18.1927 | 2.68E-07 | 0.0793 | GGGCAGTGTGCAGCGCCTGCCTAGAGAGGGCAGTG | 20 | ENCFF055HAP,ENCFF065ODV,ENCFF136ZDK,ENCFF327LYS,ENCFF092NMJ,ENCFF089GBC,ENCFF430PPJ,ENCFF198SSR,ENCFF165LXO,ENCFF210BTO,ENCFF589ZBJ,ENCFF907XEG,ENCFF052KVO,ENCFF533APC,ENCFF609MDN,ENCFF957SPH,ENCFF194HVL,ENCFF099QNE,ENCFF821BIA,ENCFF093ZRU |
| 637 | 4 | 104861912 | 104862047 | 135 | BALB,BTBR,C3H,C58J,CAST,CBA,CE,ILNJ2,KK,MA/My,MOLF,NOD,NOR,PJ,RF,SEA,SJL,SMJ,SPRET,SWR,WSB | 4 | 104862008 | 104862042 | 35 | - | 18.1927 | 2.68E-07 | 0.0793 | GGGCAGTGTGCAGCGCCTGCCTAGAGAGGGCAGTG | 22 | ENCFF055HAP,ENCFF065ODV,ENCFF136ZDK,ENCFF327LYS,ENCFF092NMJ,ENCFF089GBC,ENCFF430PPJ,ENCFF198SSR,ENCFF165LXO,ENCFF210BTO,ENCFF589ZBJ,ENCFF907XEG,ENCFF052KVO,ENCFF533APC,ENCFF609MDN,ENCFF957SPH,ENCFF194HVL,ENCFF099QNE,ENCFF024JOZ,ENCFF367UNR,ENCFF821BIA.ENCFF093ZRU |
| 637 | 4 | 104861912 | 104862047 | 135 | BALB,BTBR,C3H,C58J,CAST,CBA,CE,ILNJ2,KK,MA/My,MOLF,NOD,NOR,PJ,RF,SEA,SJL,SMJ,SPRET,SWR,WSB | 4 | 104862035 | 104862069 | 35 | - | 17.2936 | 4.57E-07 | 0.106 | GGGCAGTGTGCAGCGTCTGCCTAGAGAGGGCAGTG | 24 | ENCFF055HAP,ENCFF065ODV,ENCFF136ZDK,ENCFF327LYS,ENCFF092NMJ,ENCFF089GBC,ENCFF430PPJ,ENCFF198SSR,ENCFF165LXO,ENCFF210BTO,ENCFF589ZBJ,ENCFF907XEG,ENCFF052KVO,ENCFF533APC,ENCFF609MDN,ENCFF957SPH,ENCFF194HVL,ENCFF099QNE,ENCFF024JOZ,ENCFF367UNR,ENCFF883UPM.ENCFF502WDT.ENCFF821BIA.ENCFF093ZRU |
| 638 | 4 | 106392773 | 106392952 | 179 | CAST.MOLF | 4 | 106392898 | 106392931 | 34 | - | 18.5447 | 2.45E-07 | 0.0494 | TTGCTGTCTTTAGACACACCAGAAGAGGGCGTCA | | |
| 639 | 4 | 107975161 | 107975376 | 215 | 129S1,AJ,AKR,BALB,BUB,C3H,C57L,CAST,CBA,CE,FVB,MOLF,MRL,NOD,PL,RF,SEA,SJL.SMJ.SPRET.WSB | 4 | 107975287 | 107975320 | 34 | - | 17.2439 | 5.79E-07 | 0.0743 | TAGCTGTCCTCAGATACTCCAGAAGAGGGCATCA | 4 | ENCFF093ZRU,ENCFF367UNR |
| 640 | 4 | 107976530 | 107976776 | 246 | 129S1,AJ,AKR,BALB,BUB,C3H,C57L,C58J,CAST,CBA,CE,DBA1J,DBA2J,FVB,MOLF,MRL,NOD,NZB,NZO,NZW,PJ,PL,RF,SEA,SJL,SMJ.SPRET.TH.WSB | 4 | 107976760 | 107976793 | 34 | - | 16.4472 | 9.61E-07 | 0.0931 | TAGCTGTCTTCAGACACACCAGAAGAGGGCGTCA | 1 | ENCFF093ZRU |
| 641 | 4 | 108095428 | 108095602 | 174 | SPRET | 4 | 108095444 | 108095478 | 35 | - | 16.7339 | 6.31E-07 | 0.125 | CTGTATATACACCTGAATGCCACTGGAGGGTATCA | | |
| 642 | 4 | 108178631 | 108178832 | 201 | SPRET | 4 | 108178755 | 108178788 | 34 | - | 17.2439 | 5.79E-07 | 0.0743 | TAGCTGTCCTCAGATACTCCAGAAGAGGGCATCA | | |
| 643 | 4 | 108181653 | 108181848 | 195 | MOLF.SPRET | 4 | 108181776 | 108181809 | 34 | - | 18.935 | 1.88E-07 | 0.0423 | TTGCTGTCTTCAGACACACCAGAAGAGGGCGTCA | 1 | ENCFF093ZRU |
| 644 | 4 | 108502364 | 108502580 | 216 | 129S1,AJ,AKR,C57L,CAST,CE,FVB,MOLF,MRL.SMJ.SPRET.SWR.WSB | 4 | 108502491 | 108502509 | 19 | - | 17.5574 | 6.32E-07 | 0.111 | ACTCCAGAAGAGGGCGCCA | | |
| 644 | 4 | 108502364 | 108502580 | 216 | 129S1,AJ,AKR,C57L,CAST,CE,FVB,MOLF,MRL.SMJ.SPRET.SWR.WSB | 4 | 108502491 | 108502524 | 34 | - | 18.374 | 2.75E-07 | 0.0503 | TAGCTGTCTTCAGTCACTCCAGAAGAGGGCGCCA | | |
| 645 | 4 | 108760793 | 108760990 | 197 | CAST | 4 | 108760919 | 108760953 | 35 | - | 16.844 | 5.92E-07 | 0.121 | TTGCAGCTGTCTTTAGACACCAGAAGAGGGTGTCA | | |
| 646 | 4 | 114029993 | 114030325 | 332 | 129S1,AJ,AKR,BALB,BTBR,BUB,C3H,C57L,CAST,CBA,CE,DBA1J,DBA2J,FVB,LP,MA/My,MOLF,MRL,NOD,NOR,PL,RF,SEA,SJL,SMJ.SWR.TH.WSB | 4 | 114030284 | 114030302 | 19 | - | 19.2951 | 1.88E-07 | 0.0616 | CCGCCACTAGATGGTGCTA | 13 | ENCFF092NMJ,ENCFF201JUA,ENCFF367UNR,ENCFF024JOZ,ENCFF327LYS,ENCFF907XEG,ENCFF136ZDK,ENCFF093ZRU,ENCFF957SPH,ENCFF099QNE,ENCFF589ZBJ,ENCFF165LXO,ENCFF821BIA |
| 647 | 4 | 115899064 | 115899269 | 205 | MA/My.WSB | 4 | 115899175 | 115899208 | 34 | - | 19.2602 | 1.50E-07 | 0.0369 | CAGCTGTCTTCAGACAGACCAGAAGAGGGCATCA | | |
| 648 | 4 | 116520516 | 116520725 | 209 | CAST.ILNJ2.MOLF.SPRET.TH | 4 | 116520647 | 116520680 | 34 | - | 16.4472 | 9.61E-07 | 0.0931 | TAGCTGTCTTCAGACACACCAGAAGAGGGCGTCA | | |
| 649 | 4 | 116526566 | 116526934 | 368 | CE | 4 | 116526669 | 116526702 | 34 | - | 22.9675 | 9.26E-09 | 0.00502 | ATGCATTGCCCCTGAAGGCCAGAAGAGGGCATCT | 19 | ENCFF089GBC,ENCFF092NMJ,ENCFF367UNR,ENCFF093ZRU,ENCFF136ZDK,ENCFF883UPM,ENCFF165LXO,ENCFF327LYS,ENCFF907XEG,ENCFF821BIA,ENCFF099QNE,ENCFF024JOZ,ENCFF055HAP,ENCFF502WDT.ENCFF589ZBJ.ENCFF198SSR.ENCFF065ODV.ENCFF052KVO.ENCFF957SPH |
| 650 | 4 | 117931709 | 117931923 | 214 | 129S1,AJ,C57L,DBA1J,DBA2J,MA/My,NZB,NZO.NZW.PJ.SEA.SMJ.SPRET.TH | 4 | 117931839 | 117931872 | 34 | - | 17.8943 | 3.78E-07 | 0.0599 | TAGCAGTCTTCAGACACTCCAGAAGAGGGCGTCA | | |
| 651 | 4 | 118528982 | 118529178 | 196 | CAST.ILNJ2.MOLF.NZO.NZW.SPRET | 4 | 118529106 | 118529140 | 35 | - | 18.1284 | 2.79E-07 | 0.0805 | CTGTAGCTGTCTTCAGACACCAGAAGAGGGCATCA | | |
| 652 | 4 | 118922466 | 118922863 | 397 | SPRET | 4 | 118922613 | 118922646 | 34 | - | 17.439 | 5.10E-07 | 0.0703 | TAGCTGTCTCCAGACACACCAGAAGAGGGCATTA | | |
| 653 | 4 | 119124593 | 119124796 | 203 | SPRET | 4 | 119124713 | 119124746 | 34 | - | 17.2439 | 5.79E-07 | 0.0743 | TAGCTGTCCTCAGATACTCCAGAAGAGGGCATCA | 1 | ENCFF093ZRU |
| 654 | 4 | 119539489 | 119540145 | 656 | SPRET | 4 | 119540083 | 119540117 | 35 | - | 16.4312 | 7.50E-07 | 0.136 | TTGCCAGCCCCACCCCCTGCCCCCAGGAGGCAGAG | | |
| 655 | 4 | 119585380 | 119585646 | 266 | 129S1.C57L | 4 | 119585370 | 119585403 | 34 | - | 18.0894 | 3.33E-07 | 0.0556 | CAGCAAGTCCATCCCCTCCCAGGAGGAGGAAGAG | | |
| 656 | 4 | 119906572 | 119906817 | 245 | CAST.MOLF.SPRET | 4 | 119906696 | 119906729 | 34 | - | 17.2439 | 5.79E-07 | 0.0743 | TAGCTGTCCTCAGATACTCCAGAAGAGGGCATCA | | |
| 657 | 4 | 121043579 | 121043772 | 193 | SPRET | 4 | 121043703 | 121043721 | 19 | - | 17.5574 | 6.32E-07 | 0.111 | ACTCCAGAAGAGGGCGCCA | 3 | ENCFF093ZRU.ENCFF367UNR.ENCFF821BIA |
| 657 | 4 | 121043579 | 121043772 | 193 | SPRET | 4 | 121043703 | 121043736 | 34 | - | 18.3659 | 2.76E-07 | 0.0503 | TAGCTGTCTTCAGACACTCCAGAAGAGGGCGCCA | 3 | ENCFF093ZRU.ENCFF367UNR.ENCFF821BIA |
| 658 | 4 | 123187876 | 123188069 | 193 | SPRET | 4 | 123188001 | 123188019 | 19 | - | 17.4754 | 6.66E-07 | 0.114 | TGTCCAGAAGAGGGCGTCA | 3 | ENCFF093ZRU.ENCFF367UNR.ENCFF099QNE |
| 659 | 4 | 123253072 | 123253293 | 221 | 129S1,BTBR,BUB,C57L,C58J,CAST,FVB,ILNJ2,MOLF,NOD,NOR,NZB,NZW,SMJ,SPRET.SWR.TH.WSB | 4 | 123253197 | 123253230 | 34 | - | 16.4634 | 9.51E-07 | 0.0931 | TAGCTGTCTTCAGACGCACCAGAAGAGGGCATCA | | |
| 660 | 4 | 123651314 | 123651512 | 198 | CAST.MOLF.SPRET | 4 | 123651434 | 123651467 | 34 | - | 16.4472 | 9.61E-07 | 0.0931 | TAGCTGTCTTCAGACACACCAGAAGAGGGCGTCA | | |
| 661 | 4 | 125127243 | 125127602 | 359 | 129S1,BALB,BTBR,BUB,C3H,C57L,CAST,CBA,CE,DBA1J,DBA2J,FVB,ILNJ2,KK,MA/My,NOD,NOR,PJ,PL,RF,SEA,SJL,SPRET,SWR | 4 | 125127529 | 125127562 | 34 | - | 17.2439 | 5.79E-07 | 0.0743 | TAGCTGTCCTCAGATACTCCAGAAGAGGGCATCA | | |
| 662 | 4 | 125499736 | 125499988 | 252 | 129S1,AJ,BALB,BTBR,BUB,C3H,C57L,C58J,CAST,CBA,CE,DBA1J,DBA2J,FVB,KK,MA/My,MOLF,NOD,NOR,NZO,PJ,PL,RF,SEA,SJL.SPRET.SWR.TH | 4 | 125499863 | 125499881 | 19 | - | 17.5574 | 6.32E-07 | 0.111 | ACTCCAGAAGAGGGCGCCA | 2 | ENCFF093ZRU |
| 662 | 4 | 125499736 | 125499988 | 252 | 129S1,AJ,BALB,BTBR,BUB,C3H,C57L,C58J,CAST,CBA,CE,DBA1J,DBA2J,FVB,KK,MA/My,MOLF,NOD,NOR,NZO,PJ,PL,RF,SEA,SJL.SPRET.SWR.TH | 4 | 125499863 | 125499896 | 34 | - | 18.3659 | 2.76E-07 | 0.0503 | TAGCTGTCTTCAGACACTCCAGAAGAGGGCGCCA | 2 | ENCFF093ZRU |
| 663 | 4 | 126822883 | 126823088 | 205 | B10D2.B10J.MOLF.SPRET | 4 | 126823009 | 126823042 | 34 | - | 17.8455 | 3.91E-07 | 0.0606 | TAGCTGTCCTCAGACACTCCAGAAGAGGGCGTCA | | |
| 664 | 4 | 126993272 | 126993492 | 220 | B10D2,B10J,CAST,ILNJ2,MOLF,PJ,SEA,SPRET.WSB | 4 | 126993399 | 126993417 | 19 | - | 17.5574 | 6.32E-07 | 0.111 | ACTCCAGAAGAGGGCGCCA | | |
| 664 | 4 | 126993272 | 126993492 | 220 | B10D2,B10J,CAST,ILNJ2,MOLF,PJ,SEA,SPRET.WSB | 4 | 126993399 | 126993432 | 34 | - | 18.3659 | 2.76E-07 | 0.0503 | TAGCTGTCTTCAGACACTCCAGAAGAGGGCGCCA | | |
| 665 | 4 | 127257041 | 127257592 | 551 | B10D2,B10J,KK,MA/My,NOR,PJ,SEA,SMJ | 4 | 127257312 | 127257330 | 19 | - | 19.8689 | 1.22E-07 | 0.0488 | GGACCTCTAGGGGGCGGTA | 16 | ENCFF092NMJ,ENCFF024JOZ,ENCFF093ZRU,ENCFF210BTO,ENCFF136ZDK,ENCFF089GBC,ENCFF198SSR,ENCFF367UNR,ENCFF589ZBJ,ENCFF099QNE,ENCFF052KVO,ENCFF327LYS,ENCFF907XEG,ENCFF097CAO.ENCFF821BIA.ENCFF533APC |
| 666 | 4 | 128463505 | 128464091 | 586 | SPRET | 4 | 128463571 | 128463604 | 34 | - | 19.0488 | 1.73E-07 | 0.0407 | ATGCAGTATCCTCAGAATCCAGAAGAGGACGTTG | | |
| 667 | 4 | 129366980 | 129367179 | 199 | 129S1,B10D2,B10J,BTBR,C57L,DBA1J,KK,MOLF,NOR,NZB,PJ,PL,SEA,SMJ,SPRET,WSB | 4 | 129367103 | 129367136 | 34 | - | 16.4472 | 9.61E-07 | 0.0931 | TAGCTGTCTTCAGACACACCAGAAGAGGGCGTCA | | |
| 668 | 4 | 129455317 | 129455519 | 202 | AJ,AKR,BALB,BTBR,C3H,C57L,CAST,CBA,CE,DBA2J,MRL,NZO,NZW,RF,RH2,SPRET,TH.WSB | 4 | 129455439 | 129455472 | 34 | - | 16.4472 | 9.61E-07 | 0.0931 | TAGCTGTCTTCAGACACACCAGAAGAGGGCGTCA | | |
| 669 | 4 | 129513353 | 129513555 | 202 | 129S1,AJ,AKR,BALB,BTBR,BUB,C3H,C57L,CAST,CBA,CE,DBA2J,FVB,ILNJ2,MA/My,MRL,NOD,NZO,NZW,RF,RH2,SJL,SPRET,SWR.TH.WSB | 4 | 129513479 | 129513512 | 34 | - | 16.4634 | 9.51E-07 | 0.0931 | TAGCTGTCTTCAGACGCACCAGAAGAGGGCATCA | 1 | ENCFF367UNR |
| 670 | 4 | 129531501 | 129531716 | 215 | 129S1,AJ,AKR,BALB,BTBR,BUB,C3H,C57L,CAST,CBA,CE,DBA2J,FVB,ILNJ2,MA/My,MRL,NOD,NZO,NZW,RF,RH2,SJL,SPRET,SWR.TH.WSB | 4 | 129531628 | 129531661 | 34 | - | 16.4472 | 9.61E-07 | 0.0931 | TAGCTGTCTTCAGACACACCAGAAGAGGGCGTCA | 1 | ENCFF093ZRU |
| 671 | 4 | 131891160 | 131891500 | 340 | MOLF | 4 | 131891244 | 131891262 | 19 | - | 17.8852 | 5.08E-07 | 0.103 | ACACCAGAAGAGGGCACCA | 1 | ENCFF093ZRU |
| 671 | 4 | 131891160 | 131891500 | 340 | MOLF | 4 | 131891244 | 131891277 | 34 | - | 18.3984 | 2.70E-07 | 0.0503 | TAGCTGTCTTCAGACACACCAGAAGAGGGCACCA | 1 | ENCFF093ZRU |
| 672 | 4 | 132091453 | 132091651 | 198 | B10D2,B10J,CAST,DBA1J,KK,MOLF,NZB,PJ.SEA.SJL.SMJ.SPRET | 4 | 132091578 | 132091611 | 34 | - | 17.8455 | 3.91E-07 | 0.0606 | TAGCTGTCCTCAGACACTCCAGAAGAGGGCGTCA | 1 | ENCFF093ZRU |
| 673 | 4 | 132244335 | 132244626 | 291 | B10D2,B10J,CAST,DBA1J,KK,MOLF,NZB,PJ.SEA.SMJ.SPRET.WSB | 4 | 132244484 | 132244517 | 34 | - | 17.3902 | 5.27E-07 | 0.071 | TAGCTGTCCTCAGACACTCCAGAAGAGGGCATCA | | |
| 674 | 4 | 132550376 | 132551318 | 942 | SPRET | 4 | 132551063 | 132551081 | 19 | - | 17.541 | 6.39E-07 | 0.111 | AGGCCAGAAGAGGGCAATA | 23 | ENCFF367UNR,ENCFF093ZRU,ENCFF099QNE,ENCFF055HAP,ENCFF502WDT,ENCFF907XEG,ENCFF589ZBJ,ENCFF024JOZ,ENCFF327LYS,ENCFF821BIA,ENCFF136ZDK,ENCFF321AGQ,ENCFF883UPM,ENCFF609MDN,ENCFF089GBC,ENCFF052KVO,ENCFF198SSR,ENCFF165LXO,ENCFF092NMJ,ENCFF142CNG,ENCFF951GOP.ENCFF575MGI.ENCFF430PPJ |
| 674 | 4 | 132550376 | 132551318 | 942 | SPRET | 4 | 132551063 | 132551096 | 34 | - | 29.2276 | 2.55E-11 | 0.00013 | GTGCAGTGCCTCCAGAGGCCAGAAGAGGGCAATA | 23 | ENCFF367UNR,ENCFF093ZRU,ENCFF099QNE,ENCFF055HAP,ENCFF502WDT,ENCFF907XEG,ENCFF589ZBJ,ENCFF024JOZ,ENCFF327LYS,ENCFF821BIA,ENCFF136ZDK,ENCFF321AGQ,ENCFF883UPM,ENCFF609MDN,ENCFF089GBC,ENCFF052KVO,ENCFF198SSR,ENCFF165LXO,ENCFF092NMJ,ENCFF142CNG,ENCFF951GOP.ENCFF575MGI.ENCFF430PPJ |
| 675 | 4 | 133110219 | 133110540 | 321 | SPRET | 4 | 133110311 | 133110344 | 34 | - | 16.7724 | 7.83E-07 | 0.086 | TAGCTGTCTTCAGATGCACCAGAAGAGGGCGTCA | | |
| 676 | 4 | 133508349 | 133508406 | 57 | PL | 4 | 133508357 | 133508375 | 19 | - | 17.5246 | 6.45E-07 | 0.112 | TGGCCTCCAGGAGGCAGAG | | |
| 677 | 4 | 134018509 | 134018707 | 198 | SPRET | 4 | 134018633 | 134018666 | 34 | - | 16.4472 | 9.61E-07 | 0.0931 | TAGCTGTCTTCAGACACACCAGAAGAGGGCGTCA | | |
| 678 | 4 | 134418095 | 134418291 | 196 | ILNJ2.KK.MOLF.NZB.NZO.SPRET | 4 | 134418219 | 134418252 | 34 | - | 16.4472 | 9.61E-07 | 0.0931 | TAGCTGTCTTCAGACACACCAGAAGAGGGCGTCA | | |
| 679 | 4 | 134569787 | 134569996 | 209 | B10D2,B10J,BUB,CAST,KK,MOLF,NZB,NZO,NZW.PJ.SEA.SPRET | 4 | 134569914 | 134569947 | 34 | - | 16.4634 | 9.51E-07 | 0.0931 | TAGCTGTCTTCAGACGCACCAGAAGAGGGCATCA | | |
| 680 | 4 | 135174764 | 135174956 | 192 | CAST.MOLF.SPRET.WSB | 4 | 135174879 | 135174912 | 34 | - | 16.7724 | 7.83E-07 | 0.086 | TAGCTGTCTTCAGATGCACCAGAAGAGGGCGTCA | | |
| 681 | 4 | 135358665 | 135358935 | 270 | SPRET | 4 | 135358675 | 135358709 | 35 | - | 19.1101 | 1.53E-07 | 0.0564 | GTGCAGTGCCCAGTGAGAGTAAGAAGGTGGCACAA | | |
| 682 | 4 | 135619214 | 135619609 | 395 | SMJ.SPRET | 4 | 135619608 | 135619641 | 34 | - | 20.4878 | 6.24E-08 | 0.0199 | GTGCAGTACTTGAGGAAGCCAGAAGGTGCAGCCT | | |
| 683 | 4 | 136007098 | 136007504 | 406 | 129S1,B10D2,B10J,C57L,DBA1J,KK,MOLF,NZO.PJ.SEA.SPRET | 4 | 136007426 | 136007459 | 34 | - | 17.2439 | 5.79E-07 | 0.0743 | TAGCTGTCCTCAGATACTCCAGAAGAGGGCATCA | | |
| 684 | 4 | 136117747 | 136117987 | 240 | SPRET | 4 | 136117857 | 136117890 | 34 | - | 16.8537 | 7.43E-07 | 0.0844 | TAGCTGTCCTTAGATACTCCAGAAGAGGGCATCA | | |
| 685 | 4 | 137118357 | 137118874 | 517 | CAST | 4 | 137118809 | 137118827 | 19 | - | 18.9344 | 2.45E-07 | 0.072 | ATGCCAGAAGAGGGCACCA | 16 | ENCFF136ZDK,ENCFF092NMJ,ENCFF089GBC,ENCFF198SSR,ENCFF367UNR,ENCFF093ZRU,ENCFF883UPM,ENCFF327LYS,ENCFF099QNE,ENCFF821BIA,ENCFF907XEG,ENCFF589ZBJ,ENCFF024JOZ,ENCFF055HAP.ENCFF052KVO.ENCFF502WDT |
| 685 | 4 | 137118357 | 137118874 | 517 | CAST | 4 | 137118809 | 137118843 | 35 | - | 24.8165 | 2.84E-09 | 0.00507 | ATGCATGTACACCTGCATGCCAGAAGAGGGCACCA | 16 | ENCFF136ZDK,ENCFF092NMJ,ENCFF089GBC,ENCFF198SSR,ENCFF367UNR,ENCFF093ZRU,ENCFF883UPM,ENCFF327LYS,ENCFF099QNE,ENCFF821BIA,ENCFF907XEG,ENCFF589ZBJ,ENCFF024JOZ,ENCFF055HAP.ENCFF052KVO.ENCFF502WDT |
| 686 | 4 | 137154002 | 137154191 | 189 | CAST.SPRET | 4 | 137154120 | 137154153 | 34 | - | 16.5122 | 9.23E-07 | 0.0931 | TAGCTGTCTTCAGACACACCACAAGAGGGCATCA | 1 | ENCFF093ZRU |
| 687 | 4 | 137362877 | 137363070 | 193 | SPRET | 4 | 137363004 | 137363037 | 34 | - | 17.6667 | 4.40E-07 | 0.0647 | TAGCTGTCTTCAGACACGCCAGAAGAGGGCGTCA | 1 | ENCFF093ZRU |
| 688 | 4 | 137637207 | 137637491 | 284 | AJ,AKR,BALB,BUB,C3H,CAST,CBA,CE,DBA1J,DBA2J,FVB,ILNJ2,KK,MA/My,MOLF,MRL,NZB,NZW,PL,RF,RH2,SJL,SMJ,SPRET,SWR.TH.WSB | 4 | 137637330 | 137637363 | 34 | - | 16.4634 | 9.51E-07 | 0.0931 | TAGCTGTCTTCAGACGCACCAGAAGAGGGCATCA | 1 | ENCFF093ZRU |
| 689 | 4 | 138274513 | 138274598 | 85 | SPRET | 4 | 138274547 | 138274565 | 19 | - | 21.8525 | 2.33E-08 | 0.0254 | AGGCCAGAAGGGGGCGCTG | 16 | ENCFF024JOZ,ENCFF093ZRU,ENCFF589ZBJ,ENCFF092NMJ,ENCFF165LXO,ENCFF367UNR,ENCFF198SSR,ENCFF099QNE,ENCFF883UPM,ENCFF957SPH,ENCFF052KVO,ENCFF089GBC,ENCFF327LYS,ENCFF907XEG.ENCFF821BIA.ENCFF430PPJ |
| 689 | 4 | 138274513 | 138274598 | 85 | SPRET | 4 | 138274547 | 138274580 | 34 | - | 17.3902 | 5.27E-07 | 0.071 | AGCTAGTACCTATGGAGGCCAGAAGGGGGCGCTG | 16 | ENCFF024JOZ,ENCFF093ZRU,ENCFF589ZBJ,ENCFF092NMJ,ENCFF165LXO,ENCFF367UNR,ENCFF198SSR,ENCFF099QNE,ENCFF883UPM,ENCFF957SPH,ENCFF052KVO,ENCFF089GBC,ENCFF327LYS,ENCFF907XEG.ENCFF821BIA.ENCFF430PPJ |
| 689 | 4 | 138274513 | 138274598 | 85 | SPRET | 4 | 138274547 | 138274581 | 35 | - | 19.0183 | 1.62E-07 | 0.0589 | TAGCTAGTACCTATGGAGGCCAGAAGGGGGCGCTG | 16 | ENCFF024JOZ,ENCFF093ZRU,ENCFF589ZBJ,ENCFF092NMJ,ENCFF165LXO,ENCFF367UNR,ENCFF198SSR,ENCFF099QNE,ENCFF883UPM,ENCFF957SPH,ENCFF052KVO,ENCFF089GBC,ENCFF327LYS,ENCFF907XEG.ENCFF821BIA.ENCFF430PPJ |
| 690 | 4 | 140620430 | 140620721 | 291 | SPRET | 4 | 140620553 | 140620586 | 34 | - | 17.1382 | 6.20E-07 | 0.0772 | TAGCTGTCTCCAGACACTCCAGAAGAGGGCATCA | | |
| 691 | 4 | 140923534 | 140924433 | 899 | SPRET | 4 | 140923709 | 140923742 | 34 | - | 22.2033 | 1.70E-08 | 0.00777 | CAGCTGTTTTCAGATGTGCCAGAAGAGGGCATCA | 2 | ENCFF367UNR.ENCFF093ZRU |
| 692 | 4 | 141292246 | 141292446 | 200 | SPRET | 4 | 141292370 | 141292403 | 34 | - | 16.4472 | 9.61E-07 | 0.0931 | TAGCTGTCTTCAGACACACCAGAAGAGGGCGTCA | 1 | ENCFF367UNR |
| 693 | 4 | 142943122 | 142943313 | 191 | CAST.SPRET | 4 | 142943237 | 142943270 | 34 | - | 16.4472 | 9.61E-07 | 0.0931 | TAGCTGTCTTCAGACACACCAGAAGAGGGCGTCA | | |
| 694 | 4 | 147973903 | 147974117 | 214 | 129S1,AKR,BTBR,BUB,C3H,C57L,C58J,CBA,CE,DBA1J,DBA2J,KK,LP,MA/My,MRL,NOD,NZB,NZO,NZW,PJ,PL,RF,SEA,SJL,SMJ,SPRET.SWR.TH.WSB | 4 | 147974028 | 147974061 | 34 | - | 16.4472 | 9.61E-07 | 0.0931 | TAGCTGTCTTCAGACACACCAGAAGAGGGCGTCA | 1 | ENCFF367UNR |
| 695 | 4 | 148187918 | 148188120 | 202 | 129S1,AKR,BTBR,BUB,C3H,C57L,C58J,CBA,CE,DBA1J,DBA2J,KK,LP,MA/My,MRL,NOD,NZB,NZO,NZW,PJ,PL,RF,SEA,SJL,SMJ,SPRET.SWR.TH.WSB | 4 | 148188052 | 148188085 | 34 | - | 17.2439 | 5.79E-07 | 0.0743 | TAGCTGTCCTCAGATACTCCAGAAGAGGGCATCA | | |
| 696 | 4 | 149077161 | 149077366 | 205 | 129S1,AJ,AKR,BALB,BTBR,BUB,C3H,C57L,C58J,CAST,CBA,CE,DBA1J,DBA2J,KK,LP,MA/My,MRL,NOD,NZB,NZO,NZW,PJ,PL,RF,SEA,SJL,SMJ,SPRET,SWR,TH,WSB | 4 | 149077286 | 149077319 | 34 | - | 16.4472 | 9.61E-07 | 0.0931 | TAGCTGTCTTCAGACACACCAGAAGAGGGCGTCA | | |
| 697 | 4 | 149273328 | 149273540 | 212 | 129S1,BTBR,C57L,C58J,CAST,FVB,ILNJ2,LP.MOLF.PJ.SEA.SPRET.SWR.WSB | 4 | 149273454 | 149273487 | 34 | - | 16.4472 | 9.61E-07 | 0.0931 | TAGCTGTCTTCAGACACACCAGAAGAGGGCGTCA | 1 | ENCFF093ZRU |
| 698 | 4 | 149295948 | 149296143 | 195 | SPRET | 4 | 149296075 | 149296093 | 19 | - | 17.5574 | 6.32E-07 | 0.111 | ACTCCAGAAGAGGGCGCCA | 4 | ENCFF367UNR.ENCFF093ZRU.ENCFF099QNE.ENCFF821BIA |
| 698 | 4 | 149295948 | 149296143 | 195 | SPRET | 4 | 149296075 | 149296108 | 34 | - | 20.252 | 7.40E-08 | 0.0225 | TAGCTGTCCTCAGACACTCCAGAAGAGGGCGCCA | 4 | ENCFF367UNR.ENCFF093ZRU.ENCFF099QNE.ENCFF821BIA |
| 699 | 4 | 149755489 | 149755672 | 183 | SPRET | 4 | 149755610 | 149755643 | 34 | - | 16.7724 | 7.83E-07 | 0.086 | TAGCTGTCTTCAGATGCACCAGAAGAGGGCGTCA | | |
| 700 | 4 | 155149781 | 155149954 | 173 | MOLF | 4 | 155149765 | 155149798 | 34 | - | 18.8455 | 1.99E-07 | 0.0439 | TTGCTCTCCTCAGACACACCAGAAGAGGGCATCA | 1 | ENCFF093ZRU |
| 701 | 4 | 155746913 | 155747126 | 213 | SPRET | 4 | 155747037 | 155747070 | 34 | - | 17.878 | 3.83E-07 | 0.06 | TAGCTGTCCTCAGACACACCAGAAGAGGGCATCA | | |
| 702 | 4 | 155787642 | 155787782 | 140 | CE.ILNJ2.SMJ | 4 | 155787653 | 155787686 | 34 | - | 17.1545 | 6.13E-07 | 0.0768 | ATGCTCTCTTCAGACACACCAGAAGAGGGCATCA | 1 | ENCFF093ZRU |
| 703 | 4 | 156130367 | 156130582 | 215 | SPRET | 4 | 156130493 | 156130527 | 35 | - | 19.6422 | 1.09E-07 | 0.0452 | TTGCAGCTGTCTTTAGACACCAGAAGAGGGCGTCA | | |
| 704 | 4 | 156193374 | 156193574 | 200 | CAST.MOLF.SPRET | 4 | 156193499 | 156193532 | 34 | - | 17.5528 | 4.74E-07 | 0.0674 | TAGCTGTCCTCAGACACCCCAGAAGAGGGCGTCA | 6 | ENCFF024JOZ.ENCFF327LYS.ENCFF089GBC.ENCFF055HAP.ENCFF093ZRU.ENCFF367UNR |
| 705 | 5 | 3600883 | 3601075 | 192 | BUB,CAST,FVB,MOLF,NOD,NOR,SJL,SMJ,SPRET.SWR | 5 | 3600932 | 3600965 | 34 | + | 16.4472 | 9.61E-07 | 0.0776 | TAGCTGTCTTCAGACACACCAGAAGAGGGCGTCA | | |
| 706 | 5 | 4101317 | 4101760 | 443 | SPRET | 5 | 4101336 | 4101369 | 34 | + | 17.0813 | 6.43E-07 | 0.0633 | CTGCACTCACATTTGCCATTACAAGATGGCGCTG | | |
| 707 | 5 | 5402789 | 5402993 | 204 | SPRET | 5 | 5402844 | 5402877 | 34 | + | 16.4472 | 9.61E-07 | 0.0776 | TAGCTGTCTTCAGACACACCAGAAGAGGGCGTCA | | |
| 708 | 5 | 8057604 | 8057830 | 226 | BUB.MOLF.NOD.NOR.SPRET | 5 | 8057686 | 8057719 | 34 | + | 17.6992 | 4.30E-07 | 0.0494 | TAGCTGTTTTCAGACACACCAGAAGAGGGCATCA | | |
| 709 | 5 | 8511120 | 8511329 | 209 | CAST.MOLF.SPRET | 5 | 8511187 | 8511220 | 34 | + | 16.4472 | 9.61E-07 | 0.0776 | TAGCTGTCTTCAGACACACCAGAAGAGGGCGTCA | | |
| 710 | 5 | 8593054 | 8593917 | 863 | SPRET | 5 | 8593611 | 8593629 | 19 | + | 17.0328 | 8.88E-07 | 0.113 | TGTCCAGTAGATGGCTCCA | 3 | ENCFF093ZRU.ENCFF821BIA.ENCFF367UNR |
| 711 | 5 | 8805115 | 8805324 | 209 | SPRET | 5 | 8805183 | 8805216 | 34 | + | 18.4959 | 2.53E-07 | 0.0369 | CAGCTGTCTTCAGACATACCAGAAGAGGGCATCA | | |
| 712 | 5 | 14995762 | 14996686 | 924 | FVB.RF.SPRET.WSB | 5 | 14996651 | 14996684 | 34 | + | 20.3902 | 6.70E-08 | 0.0138 | CTGCCTCACAAGCTGACTCCACCAGAGGGCCCTG | | |
| 713 | 5 | 21730049 | 21730103 | 54 | MOLF | 5 | 21730074 | 21730107 | 34 | + | 17.8049 | 4.02E-07 | 0.0474 | TAGCTGTTTTCAGACACACCAGAAGAGGGCATCG | | |
| 714 | 5 | 24601363 | 24601567 | 204 | SPRET | 5 | 24601432 | 24601465 | 34 | + | 20.252 | 7.40E-08 | 0.015 | TAGCTGTCCTCAGACACTCCAGAAGAGGGCGCCA | 7 | ENCFF093ZRU,ENCFF367UNR,ENCFF821BIA,ENCFF136ZDK,ENCFF099QNE,ENCFF327LYS,ENCFF055HAP |
| 714 | 5 | 24601363 | 24601567 | 204 | SPRET | 5 | 24601447 | 24601465 | 19 | + | 17.5574 | 6.32E-07 | 0.0956 | ACTCCAGAAGAGGGCGCCA | 7 | ENCFF093ZRU,ENCFF367UNR,ENCFF821BIA,ENCFF136ZDK,ENCFF099QNE,ENCFF327LYS,ENCFF055HAP |
| 715 | 5 | 26635602 | 26635653 | 51 | CAST.SPRET | 5 | 26635592 | 26635626 | 35 | + | 21.578 | 3.05E-08 | 0.0168 | CTGCAGAGTTCCAAGATGGCCACCAGGAAGCAGTG | | |
| 716 | 5 | 29753527 | 29753742 | 215 | CAST | 5 | 29753599 | 29753632 | 34 | + | 16.4634 | 9.51E-07 | 0.0776 | TAGCTGTCTTCAGACGCACCAGAAGAGGGCATCA | | |
| 717 | 5 | 30122555 | 30122781 | 226 | MOLF.SPRET | 5 | 30122639 | 30122672 | 34 | + | 16.4472 | 9.61E-07 | 0.0776 | TAGCTGTCTTCAGACACACCAGAAGAGGGCGTCA | | |
| 718 | 5 | 30912183 | 30912401 | 218 | 129S1,AKR,BTBR,C57L,CAST,CE,MOLF,SJL.SPRET.WSB | 5 | 30912259 | 30912292 | 34 | + | 17.8455 | 3.91E-07 | 0.0466 | TAGCTGTCCTCAGACACTCCAGAAGAGGGCGTCA | | |
| 719 | 5 | 31179350 | 31179560 | 210 | CAST.MOLF.SPRET | 5 | 31179436 | 31179454 | 19 | + | 17.8852 | 5.08E-07 | 0.0862 | ACACCAGAAGAGGGCACCA | 1 | ENCFF093ZRU |
| 720 | 5 | 31299636 | 31299852 | 216 | CAST.MOLF.SPRET | 5 | 31299712 | 31299745 | 34 | + | 16.4634 | 9.51E-07 | 0.0776 | TAGCTGTCTTCAGACGCACCAGAAGAGGGCATCA | | |
| 721 | 5 | 31376527 | 31376593 | 66 | SPRET | 5 | 31376565 | 31376598 | 34 | + | 17.8862 | 3.81E-07 | 0.0461 | CTGTCATTCCAGAGCTAACCACAGGGAGGAGCGA | 16 | ENCFF093ZRU,ENCFF367UNR,ENCFF052KVO,ENCFF024JOZ,ENCFF092NMJ,ENCFF430PPJ,ENCFF165LXO,ENCFF198SSR,ENCFF099QNE,ENCFF821BIA,ENCFF502WDT,ENCFF327LYS,ENCFF089GBC,ENCFF055HAP.ENCFF589ZBJ.ENCFF907XEG |
| 722 | 5 | 31713680 | 31713881 | 201 | SPRET | 5 | 31713723 | 31713756 | 34 | + | 18.2927 | 2.90E-07 | 0.0392 | TTGCTGTCTTCAGACACACCAGAAGAGGGCATTA | | |
| 723 | 5 | 32721649 | 32721882 | 233 | SPRET | 5 | 32721741 | 32721774 | 34 | + | 16.4472 | 9.61E-07 | 0.0776 | TAGCTGTCTTCAGACACACCAGAAGAGGGCGTCA | | |
| 724 | 5 | 33896265 | 33896475 | 210 | SPRET | 5 | 33896332 | 33896365 | 34 | + | 16.4634 | 9.51E-07 | 0.0776 | TAGCTGTCTTCAGACGCACCAGAAGAGGGCATCA | | |
| 725 | 5 | 34546300 | 34546513 | 213 | BUB,C58J,CAST,FVB,MA/My,MOLF,MRL,NOD,NOR,NZB,PL,SPRET,SWR,TH,WSB | 5 | 34546370 | 34546403 | 34 | + | 18.561 | 2.42E-07 | 0.036 | AAGCTGTCTTCAGACACTCCAGAAGAGGGCGCCA | 2 | ENCFF093ZRU |
| 725 | 5 | 34546300 | 34546513 | 213 | BUB,C58J,CAST,FVB,MA/My,MOLF,MRL,NOD,NOR,NZB,PL,SPRET,SWR,TH,WSB | 5 | 34546385 | 34546403 | 19 | + | 17.5574 | 6.32E-07 | 0.0956 | ACTCCAGAAGAGGGCGCCA | 2 | ENCFF093ZRU |
| 726 | 5 | 38577543 | 38577737 | 194 | SPRET | 5 | 38577601 | 38577634 | 34 | + | 16.6911 | 8.24E-07 | 0.0728 | CTGTCTCTTCAGACAGCACCAGAAGAGGGCATCA | 1 | ENCFF093ZRU |
| 727 | 5 | 38858229 | 38858394 | 165 | SPRET | 5 | 38858319 | 38858352 | 34 | + | 17.2439 | 5.79E-07 | 0.0594 | TAGCTGTCCTCAGATACTCCAGAAGAGGGCATCA | | |
| 728 | 5 | 46896670 | 46897133 | 463 | SPRET | 5 | 46897094 | 46897127 | 34 | + | 16.9675 | 6.91E-07 | 0.066 | CTGCATCGTTCTTGCTGGCAAGTTGGGGGCGCGC | | |
| 729 | 5 | 48570000 | 48570211 | 211 | SPRET | 5 | 48570069 | 48570102 | 34 | + | 20.3659 | 6.82E-08 | 0.0139 | TTGCTGTCCTCAGACACACCAGAAGAGGGCATCA | 1 | ENCFF093ZRU |
| 730 | 5 | 53758093 | 53758304 | 211 | SPRET | 5 | 53758162 | 53758195 | 34 | + | 17.8455 | 3.91E-07 | 0.0466 | TAGCTGTCCTCAGACACTCCAGAAGAGGGCGTCA | | |
| 731 | 5 | 60388722 | 60389537 | 815 | AJ,AKR,BALB,BTBR,BUB,C3H,C58J,CAST,CBA,CE,FVB,KK,MA/My,MOLF,MRL,NOD,NZB,NZW,PL,RF,RH2,SEA,SJL,SMJ,SPRET,SWR.TH.WSB | 5 | 60388744 | 60388778 | 35 | + | 20.7339 | 5.38E-08 | 0.0239 | GTGCAGGTCCCGGCCTGCGCCTGCGGAGGGCTCAG | | |
| 732 | 5 | 63494006 | 63494708 | 702 | CAST | 5 | 63494490 | 63494523 | 34 | + | 18.0569 | 3.40E-07 | 0.043 | ATGCAGCTAGAGACATGAGCTCCAGGGGGTACTG | | |
| 733 | 5 | 64505267 | 64505475 | 208 | AJ,AKR,BALB,BUB,C3H,C58J,CAST,CBA,CE,DBA1J,DBA2J,FVB,ILNJ2,KK,MA/My,MOLF,MRL,NOD,NOR,NZB,NZO,NZW,PJ,PL,RF,RH2,SEA,SJL,SMJ,SPRET,SWR,TH,WSB | 5 | 64505334 | 64505367 | 34 | + | 17.8211 | 3.97E-07 | 0.047 | TAGCTGTCTTCAATCACTCCAGAAGAGGGCACCA | 2 | ENCFF093ZRU,ENCFF367UNR |
| 734 | 5 | 65900226 | 65900435 | 209 | MOLF.SPRET | 5 | 65900295 | 65900328 | 34 | + | 17.9187 | 3.72E-07 | 0.0456 | TAGCTGTCCTCACATACTCCAGAAGAGGGCATCA | | |

| | | | | | | | | | | | | | | | | |
|---|---|---|---|---|---|---|---|---|---|---|---|---|---|---|---|---|
| 735 | 5 | 67441629 | 67441830 | 201 | SPRET | 5 | 67441689 | 67441722 | 34 | + | 18.3659 | 2.76E-07 | 0.0384 | TAGCTGTCTTCAGACACTCCAGAAGAGGGCGCCA | 2 | ENCFF367UNR,ENCFF093ZRU |
| 735 | 5 | 67441629 | 67441830 | 201 | SPRET | 5 | 67441704 | 67441722 | 19 | + | 17.5574 | 6.32E-07 | 0.0956 | ACTCCAGAAGAGGGCGCCA | 2 | ENCFF367UNR,ENCFF093ZRU |
| 736 | 5 | 69999662 | 69999715 | 53 | SPRET | 5 | 69999694 | 69999712 | 19 | + | 21.0328 | 4.77E-08 | 0.0293 | TGACCAGGAGGGGGCAGTG | | |
| 737 | 5 | 73060964 | 73061167 | 203 | SPRET | 5 | 73061026 | 73061059 | 34 | + | 16.626 | 8.59E-07 | 0.0748 | TAGCTGTCTTCGGATACACCAGAAGAGGGCATCA | 2 | ENCFF093ZRU,ENCFF367UNR |
| 738 | 5 | 73663695 | 73663812 | 117 | AJ,AKR,BALB,BUB,C3H,CAST,CBA,CE,DBA1J,DBA2J,FVB,ILNJ2,MA/My,MOLF,MRL,NOD,NOR,NZO,NZW,PJ,PL,RF,RH2,SEA,SJL,SPRET,SWR,WSB | 5 | 73663809 | 73663843 | 35 | + | 19.2477 | 1.40E-07 | 0.0457 | TTGCAGCTGTTGCACATTCCCACCAGGTGTTGGCT | | |
| 739 | 5 | 73664472 | 73664678 | 206 | C3H,CBA,DBA1J,DBA2J,ILNJ2,NOD,NOR,SPRET | 5 | 73664534 | 73664568 | 35 | + | 18.0367 | 2.94E-07 | 0.0717 | CTGTAGCTGACTTCAGACACCAGAAGAGGGCATCA | 1 | ENCFF093ZRU |
| 740 | 5 | 76551033 | 76551261 | 228 | SPRET | 5 | 76551119 | 76551152 | 34 | + | 16.8211 | 7.59E-07 | 0.0699 | TAGCTGTCCTCAGACCCTCCAGAAGAGGGTGTCA | | |
| 741 | 5 | 76794536 | 76794785 | 249 | C3H,CAST,CBA,DBA1J,DBA2J,MA/My,NOD,NOR,PL,RF,SJL,SMJ,SPRET,SWR,WSB | 5 | 76794642 | 76794675 | 34 | + | 16.4634 | 9.51E-07 | 0.0776 | TAGCTGTCTTCAGACGCACCAGAAGAGGGCATCA | | |
| 742 | 5 | 77241475 | 77241685 | 210 | AJ,BALB,CAST,MA/My,NOD,NOR,PL,RF,RH2,SEA,SJL,SMJ,SPRET,SWR,TH,WSB | 5 | 77241541 | 77241574 | 34 | + | 16.4472 | 9.61E-07 | 0.0776 | TAGCTGTCTTCAGACACACCAGAAGAGGGCGTCA | 1 | ENCFF093ZRU |
| 743 | 5 | 77289716 | 77289919 | 203 | CAST.SPRET | 5 | 77289770 | 77289803 | 34 | + | 17.6667 | 4.40E-07 | 0.0501 | TAGCTGTCTTCAGACACGCCAGAAGAGGGCGTCA | | |
| 744 | 5 | 77569567 | 77569948 | 381 | SPRET | 5 | 77569661 | 77569694 | 34 | + | 17.6098 | 4.56E-07 | 0.0512 | TAGCTGTCTTCAGACTCACCAGAAGAGGGCATCA | | |
| 745 | 5 | 80851008 | 80851217 | 209 | CAST,MA/My,MOLF,RF,SMJ,SPRET,SWR,TH,WSB | 5 | 80851075 | 80851108 | 34 | + | 16.4472 | 9.61E-07 | 0.0776 | TAGCTGTCTTCAGACACACCAGAAGAGGGCGTCA | | |
| 746 | 5 | 85185127 | 85185388 | 261 | SPRET | 5 | 85185292 | 85185325 | 34 | + | 16.4634 | 9.51E-07 | 0.0776 | TAGCTGTCTTCAGACGCACCAGAAGAGGGCATCA | | |
| 747 | 5 | 92789172 | 92789369 | 197 | CAST.SPRET | 5 | 92789227 | 92789260 | 34 | + | 18.1545 | 3.18E-07 | 0.0415 | TAGCTGTTTTCAGACACACCAGAAGAGGGCGTCA | | |
| 748 | 5 | 102729697 | 102729894 | 197 | AKR,CAST,CE,MOLF,MRL,NOD,NOR,NZO,NZW,PJ,PL,RF,SMJ,SPRET,TH,WSB | 5 | 102729752 | 102729785 | 34 | + | 18.1789 | 3.13E-07 | 0.0413 | TAGCTGTCTTCAGACACTCCAGAAGAGGGCGCTA | | |
| 748 | 5 | 102729697 | 102729894 | 197 | AKR,CAST,CE,MOLF,MRL,NOD,NOR,NZO,NZW,PJ,PL,RF,SMJ,SPRET,TH,WSB | 5 | 102729767 | 102729785 | 19 | + | 17.5738 | 6.25E-07 | 0.0956 | ACTCCAGAAGAGGGCGCTA | | |
| 749 | 5 | 103223488 | 103223888 | 400 | SPRET | 5 | 103223487 | 103223520 | 34 | + | 16.5285 | 9.13E-07 | 0.0776 | GTGCTGCCCCCTACTGGAGCAGAGGATGGCGCAC | 7 | ENCFF367UNR,ENCFF052KVO,ENCFF093ZRU,ENCFF883UPM,ENCFF099QNE,ENCFF821BIA,ENCFF533APC |
| 749 | 5 | 103223488 | 103223888 | 400 | SPRET | 5 | 103223488 | 103223506 | 19 | - | 18.918 | 2.48E-07 | 0.0608 | GCTCCAGTAGGGGGCAGCA | 7 | ENCFF367UNR,ENCFF052KVO,ENCFF093ZRU,ENCFF883UPM,ENCFF099QNE,ENCFF821BIA,ENCFF533APC |
| 749 | 5 | 103223488 | 103223888 | 400 | SPRET | 5 | 103223488 | 103223522 | 35 | - | 17.3028 | 4.54E-07 | 0.0915 | GTGTGCGCCATCCTCTGCTCCAGTAGGGGGCAGCA | 7 | ENCFF367UNR,ENCFF052KVO,ENCFF093ZRU,ENCFF883UPM,ENCFF099QNE,ENCFF821BIA,ENCFF533APC |
| 750 | 5 | 103874398 | 103874620 | 222 | 129S1,AKR,BUB,C3H,C57L,CAST,CBA,CE,DBA1J,DBA2J,FVB,LP,MRL,NOD,NOR,NZB,NZO,NZW,PJ,PL,RF,SJL,SMJ,SPRET,TH,WSB | 5 | 103874587 | 103874621 | 35 | + | 16.7156 | 6.38E-07 | 0.112 | GTGCAGCCCCACACAGGTGCCAGGTGGTGGCCTTG | | |
| 751 | 5 | 104062779 | 104062940 | 161 | SPRET | 5 | 104062868 | 104062902 | 35 | + | 23.789 | 6.25E-09 | 0.00624 | CTGCATGTATAACTGCATACCAGAAGAGGGCACCA | 20 | ENCFF367UNR,ENCFF093ZRU,ENCFF201JUA,ENCFF052KVO,ENCFF092NMJ,ENCFF089GBC,ENCFF430PPJ,ENCFF198SSR,ENCFF883UPM,ENCFF165LXO,ENCFF099QNE,ENCFF136ZDK,ENCFF821BIA,ENCFF907XEG,ENCFF055HAP,ENCFF327LYS,ENCFF065ODV,ENCFF589ZBJ,ENCFF024JOZ,ENCFF502WDT |
| 751 | 5 | 104062779 | 104062940 | 161 | SPRET | 5 | 104062884 | 104062902 | 19 | + | 17.9672 | 4.81E-07 | 0.0853 | ATACCAGAAGAGGGCACCA | 20 | ENCFF367UNR,ENCFF093ZRU,ENCFF201JUA,ENCFF052KVO,ENCFF092NMJ,ENCFF089GBC,ENCFF430PPJ,ENCFF198SSR,ENCFF883UPM,ENCFF165LXO,ENCFF099QNE,ENCFF136ZDK,ENCFF821BIA,ENCFF907XEG,ENCFF055HAP,ENCFF327LYS,ENCFF065ODV,ENCFF589ZBJ,ENCFF024JOZ,ENCFF502WDT |
| 752 | 5 | 108238851 | 108239044 | 193 | SPRET | 5 | 108238821 | 108238854 | 34 | + | 17.5691 | 4.69E-07 | 0.052 | TTGCTCTCTTCAGAGACACCAGAAGAGGGCATCA | | |
| 753 | 5 | 108548366 | 108548577 | 211 | SPRET | 5 | 108548434 | 108548467 | 34 | + | 16.4472 | 9.61E-07 | 0.0776 | TAGCTGTCTTCAGACACACCAGAAGAGGGCGTCA | | |
| 754 | 5 | 111586951 | 111587340 | 389 | SPRET | 5 | 111587132 | 111587150 | 19 | + | 18.2951 | 3.84E-07 | 0.0766 | TGGCCTGCAGAGGTCACCA | | |
| 755 | 5 | 114297412 | 114297617 | 205 | CAST.MOLF.SPRET | 5 | 114297476 | 114297509 | 34 | + | 18.3659 | 2.76E-07 | 0.0384 | TAGCTGTCTTCAGACACTCCAGAAGAGGGCGCCA | 3 | ENCFF093ZRU.ENCFF821BIA.ENCFF367UNR |
| 755 | 5 | 114297412 | 114297617 | 205 | CAST.MOLF.SPRET | 5 | 114297491 | 114297509 | 19 | + | 17.5574 | 6.32E-07 | 0.0956 | ACTCCAGAAGAGGGCGCCA | 3 | ENCFF093ZRU.ENCFF821BIA.ENCFF367UNR |
| 756 | 5 | 114656971 | 114657173 | 202 | CAST.SPRET | 5 | 114657030 | 114657063 | 34 | + | 17.2439 | 5.79E-07 | 0.0594 | TAGCTGTCCTCAGATACTCCAGAAGAGGGCATCA | | |
| 757 | 5 | 115287160 | 115287378 | 218 | CAST.MOLF.SPRET | 5 | 115287235 | 115287268 | 34 | + | 18.3659 | 2.76E-07 | 0.0384 | TAGCTGTCTTCAGACACTCCAGAAGAGGGCGCCA | | |
| 757 | 5 | 115287160 | 115287378 | 218 | CAST.MOLF.SPRET | 5 | 115287250 | 115287268 | 19 | + | 17.5574 | 6.32E-07 | 0.0956 | ACTCCAGAAGAGGGCGCCA | | |
| 758 | 5 | 115872767 | 115872834 | 67 | CAST | 5 | 115872804 | 115872838 | 35 | + | 26.2385 | 8.99E-10 | 0.00242 | ATGCAGTTGCCCAAGGAAGCCAGAAGAGGGCACGG | 8 | ENCFF093ZRU,ENCFF052KVO,ENCFF092NMJ,ENCFF430PPJ,ENCFF055HAP,ENCFF502WDT,ENCFF201JUA,ENCFF089GBC |
| 759 | 5 | 116444949 | 116445086 | 137 | MOLF | 5 | 116444980 | 116445013 | 34 | + | 20.9919 | 4.30E-08 | 0.0099 | GTGCAGGAGCCAGCAAGGCCAGCAGAGGGAGTCA | | |
| 760 | 5 | 117360789 | 117360941 | 152 | CE,DBA1J,DBA2J,KK,NZB,NZO,NZW,WSB | 5 | 117360941 | 117360974 | 34 | + | 23.9268 | 4.19E-09 | 0.00154 | GCGCAGTGCCCTTAAAGGCCAGAAGAGGGCATCA | 2 | ENCFF502WDT,ENCFF136ZDK |
| 761 | 5 | 117450699 | 117450897 | 198 | 129S1,AJ,AKR,BALB,C57L,CAST,CE,DBA1J,DBA2J,KK,MOLF,MRL,NZB,NZO,NZW,PJ,PL,RH2,SEA,SMJ,SPRET,TH,WSB | 5 | 117450755 | 117450788 | 34 | + | 18.3659 | 2.76E-07 | 0.0384 | TAGCTGTCTTCAGACACTCCAGAAGAGGGCGCCA | 1 | ENCFF367UNR |
| 761 | 5 | 117450699 | 117450897 | 198 | 129S1,AJ,AKR,BALB,C57L,CAST,CE,DBA1J,DBA2J,KK,MOLF,MRL,NZB,NZO,NZW,PJ,PL,RH2,SEA,SMJ,SPRET,TH,WSB | 5 | 117450770 | 117450788 | 19 | + | 17.5574 | 6.32E-07 | 0.0956 | ACTCCAGAAGAGGGCGCCA | 1 | ENCFF367UNR |
| 762 | 5 | 118970204 | 118970414 | 210 | MOLF.RF.SPRET | 5 | 118970270 | 118970303 | 34 | + | 17.6667 | 4.40E-07 | 0.0501 | TAGCTGTCTTCAGACACGCCAGAAGAGGGCGTCA | | |
| 763 | 5 | 120413462 | 120413790 | 328 | SPRET | 5 | 120413700 | 120413733 | 34 | + | 23.6667 | 5.21E-09 | 0.00184 | GTGCAGTGTCTGTGGAAGCCAGAAGAGGGAGTCA | 1 | ENCFF093ZRU |
| 764 | 5 | 121209931 | 121210148 | 217 | AKR,CAST,KK,MA/My,MOLF,PL,SPRET,TH | 5 | 121210007 | 121210040 | 34 | + | 16.4634 | 9.51E-07 | 0.0776 | TAGCTGTCTTCAGACGCACCAGAAGAGGGCATCA | | |
| 765 | 5 | 121309342 | 121309545 | 203 | CAST.MOLF.SPRET | 5 | 121309404 | 121309437 | 34 | + | 16.9919 | 6.81E-07 | 0.0655 | TAGCTGTCTTCAGACACACCAGAAGGGGGCATCA | 1 | ENCFF093ZRU |
| 766 | 5 | 121529444 | 121529645 | 201 | 129S1,AJ,BALB,C57L,ILNJ2,RH2,SEA,SPRET,WSB | 5 | 121529616 | 121529649 | 34 | + | 16.4472 | 9.61E-07 | 0.0776 | TAGCTGTCTTCAGACACACCAGAAGAGGGCGTCA | | |
| 767 | 5 | 123052827 | 123053266 | 439 | SPRET | 5 | 123052928 | 123052962 | 35 | + | 17.7156 | 3.56E-07 | 0.0793 | AAGCATTTCAAGCAATCGAGCAACAGGGGGAGCCA | | |
| 768 | 5 | 123337058 | 123337715 | 657 | CAST | 5 | 123337132 | 123337166 | 35 | + | 27.4679 | 3.12E-10 | 0.00145 | CTGCATGTATGCCTGCAGGCCAGAAGAGGGCACCA | 2 | ENCFF136ZDK.ENCFF589ZBJ |
| 768 | 5 | 123337058 | 123337715 | 657 | CAST | 5 | 123337148 | 123337166 | 19 | + | 20.3607 | 8.29E-08 | 0.0341 | AGGCCAGAAGAGGGCACCA | 2 | ENCFF136ZDK.ENCFF589ZBJ |
| 769 | 5 | 123460062 | 123460278 | 216 | CAST.ILNJ2.MOLF.SPRET | 5 | 123460134 | 123460167 | 34 | + | 18.3659 | 2.76E-07 | 0.0384 | TAGCTGTCTTCAGACACTCCAGAAGAGGGCGCCA | | |
| 769 | 5 | 123460062 | 123460278 | 216 | CAST.ILNJ2.MOLF.SPRET | 5 | 123460149 | 123460167 | 19 | + | 17.5574 | 6.32E-07 | 0.0956 | ACTCCAGAAGAGGGCGCCA | | |
| 770 | 5 | 124638339 | 124638566 | 227 | CAST,MA/My,MOLF,NZW,PJ,PL,SPRET,WSB | 5 | 124638425 | 124638458 | 34 | + | 18.3659 | 2.76E-07 | 0.0384 | TAGCTGTCTTCAGACACTCCAGAAGAGGGCGCCA | 1 | ENCFF093ZRU |
| 770 | 5 | 124638339 | 124638566 | 227 | CAST,MA/My,MOLF,NZW,PJ,PL,SPRET,WSB | 5 | 124638440 | 124638458 | 19 | + | 17.5574 | 6.32E-07 | 0.0956 | ACTCCAGAAGAGGGCGCCA | 1 | ENCFF093ZRU |
| 771 | 5 | 125291561 | 125291764 | 203 | SPRET | 5 | 125291621 | 125291655 | 35 | + | 15.9633 | 9.75E-07 | 0.14 | CTGTAGCTGTCTTCAGACACCAGAAGAGGGTGACA | | |
| 772 | 5 | 127599877 | 127600792 | 915 | SPRET | 5 | 127599933 | 127599967 | 35 | + | 24.6881 | 3.14E-09 | 0.00406 | GGGCCGTTCCCCACCTCAGCCACTAGGGGGTGGTG | 24 | ENCFF093ZRU,ENCFF367UNR,ENCFF099QNE,ENCFF502WDT,ENCFF821BIA,ENCFF609MDN,ENCFF805QBU,ENCFF052KVO,ENCFF907XEG,ENCFF024JOZ,ENCFF327LYS,ENCFF575MGI,ENCFF092NMJ,ENCFF194HVL,ENCFF198SSR,ENCFF589ZBJ,ENCFF491RJK,ENCFF957SPH,ENCFF165LXO,ENCFF089GBC,ENCFF281QMV,ENCFF136ZDK,ENCFF316FYN,ENCFF430PPJ |
| 772 | 5 | 127599877 | 127600792 | 915 | SPRET | 5 | 127599949 | 127599967 | 19 | + | 20.1639 | 9.73E-08 | 0.037 | CAGCCACTAGGGGGTGGTG | 24 | ENCFF093ZRU,ENCFF367UNR,ENCFF099QNE,ENCFF502WDT,ENCFF821BIA,ENCFF609MDN,ENCFF805QBU,ENCFF052KVO,ENCFF907XEG,ENCFF024JOZ,ENCFF327LYS,ENCFF575MGI,ENCFF092NMJ,ENCFF194HVL,ENCFF198SSR,ENCFF589ZBJ,ENCFF491RJK,ENCFF957SPH,ENCFF165LXO,ENCFF089GBC,ENCFF281QMV,ENCFF136ZDK,ENCFF316FYN,ENCFF430PPJ |
| 773 | 5 | 129463378 | 129463820 | 442 | 129S1,AJ,BALB,C57L,C58J,CAST,FVB,ILNJ2,KK,LP,MOLF,MRL,NOD,NOR,NZB,NZO,NZW,RH2,SEA,SMJ,SPRET,SWR,TH,WSB | 5 | 129463503 | 129463537 | 35 | + | 23.633 | 7.02E-09 | 0.00665 | CTACAGAGCTGCTGAGCTGCCAGCAGGGGGCACTG | | |
| 773 | 5 | 129463378 | 129463820 | 442 | 129S1,AJ,BALB,C57L,C58J,CAST,FVB,ILNJ2,KK,LP,MOLF,MRL,NOD,NOR,NZB,NZO,NZW,RH2,SEA,SMJ,SPRET,SWR,TH,WSB | 5 | 129463519 | 129463537 | 19 | + | 24.0164 | 2.47E-09 | 0.012 | CTGCCAGCAGGGGGCACTG | | |
| 774 | 5 | 129873224 | 129873594 | 370 | 129S1,AJ,BALB,C57L,CAST,FVB,ILNJ2,KK,MOLF,MRL,NOD,NOR,NZW,RH2,SEA,SMJ,SPRET,SWR | 5 | 129873573 | 129873606 | 34 | + | 17.2439 | 5.79E-07 | 0.0594 | TAGCTGTCCTCAGATACTCCAGAAGAGGGCATCA | | |
| 774 | 5 | 129873224 | 129873594 | 370 | 129S1,AJ,BALB,C57L,CAST,FVB,ILNJ2,KK,MOLF,MRL,NOD,NOR,NZW,RH2,SEA,SMJ,SPRET,SWR | 5 | 129873308 | 129873341 | 34 | - | 17.2439 | 5.79E-07 | 0.0594 | TAGCTGTCCTCAGATACTCCAGAAGAGGGCATCA | 1 | ENCFF093ZRU |
| 775 | 5 | 129880706 | 129880960 | 254 | CAST | 5 | 129880818 | 129880851 | 34 | + | 17.0976 | 6.36E-07 | 0.063 | TAGCTGTCCTCAGACACTCCAGAAGATGGCGTCA | | |
| 776 | 5 | 129921418 | 129921629 | 211 | SPRET | 5 | 129921487 | 129921520 | 34 | + | 18.3333 | 2.82E-07 | 0.0388 | TAGCTGTCCTCAGACACACCAGAAGAGGGCGTCA | | |
| 777 | 5 | 129981805 | 129982011 | 206 | CAST.MOLF.SPRET | 5 | 129981883 | 129981901 | 19 | + | 17.8852 | 5.08E-07 | 0.0862 | ACACCAGAAGAGGGCACCA | | |
| 778 | 5 | 130864676 | 130864981 | 305 | SPRET | 5 | 130864684 | 130864702 | 19 | + | 16.9836 | 9.17E-07 | 0.115 | CCACCTGCAGGGGGCGTCA | | |
| 779 | 5 | 133655334 | 133655538 | 204 | CAST.KK.MOLF.RF.SPRET.WSB | 5 | 133655390 | 133655423 | 34 | + | 16.4472 | 9.61E-07 | 0.0776 | TAGCTGTCTTCAGACACACCAGAAGAGGGCGTCA | | |
| 780 | 5 | 134682194 | 134682425 | 231 | AJ,AKR,BUB,CAST,FVB,ILNJ2,MA/My,MOLF,MRL,RF,SJL,SPRET,WSB | 5 | 134682284 | 134682317 | 34 | + | 16.4634 | 9.51E-07 | 0.0776 | TAGCTGTCTTCAGACGCACCAGAAGAGGGCATCA | 1 | ENCFF093ZRU |
| 781 | 5 | 135401425 | 135401635 | 210 | CAST.MOLF.SPRET | 5 | 135401493 | 135401526 | 34 | + | 16.4472 | 9.61E-07 | 0.0776 | TAGCTGTCTTCAGACACACCAGAAGAGGGCGTCA | | |
| 782 | 5 | 137772178 | 137772537 | 359 | SPRET | 5 | 137772243 | 137772276 | 34 | + | 16.4472 | 9.61E-07 | 0.0776 | TAGCTGTCTTCAGACACACCAGAAGAGGGCGTCA | | |
| 783 | 5 | 139416152 | 139416538 | 386 | SPRET | 5 | 139416295 | 139416328 | 34 | + | 20.2114 | 7.63E-08 | 0.0153 | CTGCTGTCCTGAACACGTACACCAGGGAGCAGCC | 1 | ENCFF093ZRU |
| 784 | 5 | 142631247 | 142631464 | 217 | CAST.MOLF.SPRET | 5 | 142631321 | 142631354 | 34 | + | 16.4634 | 9.51E-07 | 0.0776 | TAGCTGTCTTCAGACGCACCAGAAGAGGGCATCA | | |
| 785 | 5 | 142831047 | 142831151 | 104 | 129S1,AKR,BTBR,BUB,C57L,CAST,CE,DBA1J,DBA2J,ILNJ2,KK,MA/My,MOLF,PJ,SJL,SPRET | 5 | 142831020 | 142831054 | 35 | + | 16.5963 | 6.83E-07 | 0.117 | GAGCAATGCCCCCAGCCCCTCACTAGGGGATTCTA | | |
| 786 | 5 | 143322460 | 143322586 | 126 | CAST | 5 | 143322453 | 143322487 | 35 | + | 22.5688 | 1.52E-08 | 0.0109 | ATGCAGATGCTCACAGAGGCCAGAAGGGGGCGTTC | 16 | ENCFF201JUA,ENCFF367UNR,ENCFF099QNE,ENCFF052KVO,ENCFF589ZBJ,ENCFF093ZRU,ENCFF198SSR,ENCFF055HAP,ENCFF883UPM,ENCFF502WDT,ENCFF089GBC,ENCFF327LYS,ENCFF907XEG,ENCFF092NMJ,ENCFF430PPJ |
| 786 | 5 | 143322460 | 143322586 | 126 | CAST | 5 | 143322469 | 143322487 | 19 | + | 17.3115 | 7.42E-07 | 0.105 | AGGCCAGAAGGGGGCGTTC | 17 | ENCFF201JUA,ENCFF367UNR,ENCFF099QNE,ENCFF052KVO,ENCFF589ZBJ,ENCFF093ZRU,ENCFF198SSR,ENCFF055HAP,ENCFF883UPM,ENCFF502WDT,ENCFF089GBC,ENCFF327LYS,ENCFF907XEG,ENCFF092NMJ,ENCFF430PPJ,ENCFF024JOZ |
| 787 | 5 | 143877765 | 143878620 | 855 | MOLF | 5 | 143878149 | 143878183 | 35 | + | 20.5505 | 6.07E-08 | 0.0261 | GTGCAGATGCCCATGGAGACCAGAAGGGGGTATTG | 1 | ENCFF093ZRU |
| 788 | 5 | 144200658 | 144200761 | 103 | MOLF.TH | 5 | 144200702 | 144200736 | 35 | + | 17.9174 | 3.16E-07 | 0.0742 | CTGCAGGTCGGGCAGGTGCTCTGTAGGTGGTGAGG | | |
| 789 | 5 | 144252698 | 144252889 | 191 | MOLF.TH | 5 | 144252749 | 144252782 | 34 | + | 17.2439 | 5.79E-07 | 0.0594 | TAGCTGTCCTCAGATACTCCAGAAGAGGGCATCA | | |
| 790 | 5 | 144953798 | 144954122 | 324 | SPRET | 5 | 144953765 | 144953799 | 35 | + | 18.9541 | 1.68E-07 | 0.052 | ATGGCCTTTCCCCACTGGGCCACTAGATGGCTCTG | 4 | ENCFF093ZRU.ENCFF099QNE.ENCFF367UNR.ENCFF055HAP |
| 790 | 5 | 144953798 | 144954122 | 324 | SPRET | 5 | 144953766 | 144953799 | 34 | + | 19.6341 | 1.15E-07 | 0.0211 | TGGCCTTTCCCCACTGGGCCACTAGATGGCTCTG | 4 | ENCFF093ZRU.ENCFF099QNE.ENCFF367UNR.ENCFF055HAP |
| 790 | 5 | 144953798 | 144954122 | 324 | SPRET | 5 | 144953781 | 144953799 | 19 | + | 17.1148 | 8.43E-07 | 0.11 | GGGCCACTAGATGGCTCTG | 4 | ENCFF093ZRU.ENCFF099QNE.ENCFF367UNR.ENCFF055HAP |
| 791 | 5 | 146301974 | 146302116 | 142 | SPRET | 5 | 146302072 | 146302105 | 34 | + | 16.4797 | 9.42E-07 | 0.0776 | TAGCTGTTCTCAGACACACCAGAAGAAGGCATCA | | |
| 792 | 5 | 146604325 | 146604854 | 529 | CAST | 5 | 146604709 | 146604742 | 34 | + | 28 | 9.45E-11 | 0.000121 | ATGCAGTACCCATAGAAGCCAGAAGAGGGCACTG | 9 | ENCFF821BIA,ENCFF052KVO,ENCFF136ZDK,ENCFF367UNR,ENCFF093ZRU,ENCFF589ZBJ,ENCFF502WDT,ENCFF201JUA,ENCFF099QNE |
| 792 | 5 | 146604325 | 146604854 | 529 | CAST | 5 | 146604724 | 146604742 | 19 | + | 18.4754 | 3.39E-07 | 0.0713 | AAGCCAGAAGAGGGCACTG | 9 | ENCFF821BIA,ENCFF052KVO,ENCFF136ZDK,ENCFF367UNR,ENCFF093ZRU,ENCFF589ZBJ,ENCFF502WDT,ENCFF201JUA,ENCFF099QNE |
| 793 | 5 | 146854382 | 146854590 | 208 | AKR,CAST,CBA,FVB,ILNJ2,LP,MA/My,MOLF,MRL,NZB,NZO,NZW,RF,SJL,TH,WSB | 5 | 146854448 | 146854481 | 34 | + | 18.561 | 2.42E-07 | 0.036 | AAGCTGTCTTCAGACACTCCAGAAGAGGGCGCCA | 1 | ENCFF055HAP |
| 793 | 5 | 146854382 | 146854590 | 208 | AKR,CAST,CBA,FVB,ILNJ2,LP,MA/My,MOLF,MRL,NZB,NZO,NZW,RF,SJL,TH,WSB | 5 | 146854463 | 146854481 | 19 | + | 17.5574 | 6.32E-07 | 0.0956 | ACTCCAGAAGAGGGCGCCA | 1 | ENCFF055HAP |
| 794 | 5 | 146882139 | 146882838 | 699 | SPRET | 5 | 146882285 | 146882318 | 34 | + | 16.4634 | 9.51E-07 | 0.0776 | TAGCTGTCTTCAGACGCACCAGAAGAGGGCATCA | | |
| 795 | 5 | 147270735 | 147270928 | 193 | SPRET | 5 | 147270777 | 147270810 | 34 | + | 17.2439 | 5.79E-07 | 0.0594 | TAGCTGTCCTCAGATACTCCAGAAGAGGGCATCA | | |
| 796 | 5 | 147344596 | 147344786 | 190 | SPRET | 5 | 147344645 | 147344678 | 34 | + | 18.1463 | 3.20E-07 | 0.0416 | TTGCTGTCTTCAGATACACCAGAAGAGGGCATTA | | |
| 797 | 5 | 148672442 | 148673233 | 791 | CAST.CBA.CE.KK.MOLF.RF.SEA | 5 | 148672973 | 148673006 | 34 | + | 24.2846 | 3.09E-09 | 0.00123 | GTGCAGTACCCAAGGAGGCCAGAAGAAGGCATCA | | |
| 798 | 5 | 5902507 | 5902884 | 377 | AJ,BUB,C3H,CBA,DBA1J,LP,NOD,NOR,PL,TH,WSB | 5 | 5902568 | 5902601 | 34 | - | 18.3089 | 2.87E-07 | 0.0389 | ATGCAGCTAGAGACACGAGCTCCAGGGGGTACTG | | |
| 799 | 5 | 11074526 | 11074746 | 220 | CAST | 5 | 11074649 | 11074682 | 34 | - | 17.3902 | 5.27E-07 | 0.0561 | TAGCTGTCCTCAGACACTCCAGAAGAGGGCATCA | | |
| 800 | 5 | 14894861 | 14895427 | 566 | DBA1J,DBA2J,FVB,MA/My,RF,SJL,SWR,WSB | 5 | 14895071 | 14895105 | 35 | - | 22.8349 | 1.26E-08 | 0.00945 | GTGCAGTTGCCCATAGTGTCCAGAAGAGGGAGAAG | 3 | ENCFF093ZRU,ENCFF099QNE,ENCFF052KVO |
| 801 | 5 | 22171441 | 22171663 | 222 | CAST.MOLF.SPRET | 5 | 22171558 | 22171591 | 34 | - | 17.6098 | 4.56E-07 | 0.0512 | TAGCTGTCTTCAGACTCACCAGAAGAGGGCATCA | | |
| 802 | 5 | 22235293 | 22235495 | 202 | CAST.SPRET | 5 | 22235419 | 22235452 | 34 | - | 16.7724 | 7.83E-07 | 0.0704 | TAGCTGTCTTCAGATGCACCAGAAGAGGGCGTCA | | |
| 803 | 5 | 22723288 | 22724100 | 812 | SPRET | 5 | 22724074 | 22724107 | 34 | - | 16.561 | 8.95E-07 | 0.0769 | TAGCTGTCTTAGGACACACCAGAAGAGGGCATTG | | |
| 804 | 5 | 24021250 | 24021460 | 210 | 129S1,AKR,BALB,BTBR,BUB,C57L,C58J,CAST,CE,DBA1J,DBA2J,FVB,KK,MA/My,MOLF,MRL,NOD,NOR,NZO,NZW,PJ,PL,RF,SEA,SJL,SMJ,SPRET,SWR,TH,WSB | 5 | 24021377 | 24021410 | 34 | - | 16.4634 | 9.51E-07 | 0.0776 | TAGCTGTCTTCAGACGCACCAGAAGAGGGCATCA | | |
| 805 | 5 | 24602416 | 24602618 | 202 | CAST,DBA1J,DBA2J,KK,MOLF,MRL,NZB,NZW,PJ,SPRET | 5 | 24602536 | 24602554 | 19 | - | 17.5574 | 6.32E-07 | 0.0956 | ACTCCAGAAGAGGGCGCCA | | |
| 805 | 5 | 24602416 | 24602618 | 202 | CAST,DBA1J,DBA2J,KK,MOLF,MRL,NZB,NZW,PJ,SPRET | 5 | 24602536 | 24602569 | 34 | - | 18.3659 | 2.76E-07 | 0.0384 | TAGCTGTCTTCAGACACTCCAGAAGAGGGCGCCA | | |
| 806 | 5 | 29542289 | 29542492 | 203 | MOLF.SPRET | 5 | 29542415 | 29542448 | 34 | - | 19.0163 | 1.77E-07 | 0.0289 | CAGCTGTCTTCAGACACACCAGAAGAGGGCGTCA | | |
| 807 | 5 | 29823265 | 29823519 | 254 | SPRET | 5 | 29823394 | 29823412 | 19 | - | 17.8852 | 5.08E-07 | 0.0862 | ACACCAGAAGAGGGCACCA | 3 | ENCFF367UNR.ENCFF099QNE.ENCFF093ZRU |
| 807 | 5 | 29823265 | 29823519 | 254 | SPRET | 5 | 29823394 | 29823427 | 34 | - | 22.5935 | 1.25E-08 | 0.00364 | TTGCTGTTTTCAGACACACCAGAAGAGGGCACCA | 3 | ENCFF367UNR.ENCFF099QNE.ENCFF093ZRU |
| 808 | 5 | 30228056 | 30228305 | 249 | CAST.MOLF.SPRET.WSB | 5 | 30228183 | 30228216 | 34 | - | 16.4634 | 9.51E-07 | 0.0776 | TAGCTGTCTTCAGACGCACCAGAAGAGGGCATCA | | |
| 809 | 5 | 30516238 | 30516369 | 131 | SPRET | 5 | 30516268 | 30516301 | 34 | - | 18.6179 | 2.33E-07 | 0.0351 | CTGCCGTCTCAGGAACACCCACAAGATGTTGCAG | | |
| 810 | 5 | 30575026 | 30575228 | 202 | SPRET | 5 | 30575201 | 30575234 | 34 | - | 17.1626 | 6.10E-07 | 0.0613 | CAGCCATGAAAGCAGCTTCCCCCAGAGGGAAGCG | | |
| 811 | 5 | 30986976 | 30987176 | 200 | SPRET | 5 | 30987089 | 30987122 | 34 | - | 16.4472 | 9.61E-07 | 0.0776 | TAGCTGTCTTCAGACACACCAGAAGAGGGCGTCA | | |
| 812 | 5 | 32952025 | 32952269 | 244 | SPRET | 5 | 32952150 | 32952183 | 34 | - | 16.9187 | 7.13E-07 | 0.067 | TAGCTGTCTTCAGACGCACCAGAAGAGGGCGTCA | 2 | ENCFF093ZRU.ENCFF099QNE |
| 813 | 5 | 37511400 | 37512048 | 648 | MOLF | 5 | 37511925 | 37511943 | 19 | - | 17.0984 | 8.52E-07 | 0.111 | AGTCCAGCAGGGGGCAAAA | 24 | ENCFF201JUA,ENCFF093ZRU,ENCFF367UNR,ENCFF099QNE,ENCFF065ODV,ENCFF092NMJ,ENCFF821BIA,ENCFF430PPJ,ENCFF907XEG,ENCFF327LYS,ENCFF198SSR,ENCFF609MDN,ENCFF024JOZ,ENCFF883UPM,ENCFF589ZBJ,ENCFF052KVO,ENCFF504JDS,ENCFF957SPH,ENCFF136ZDK,ENCFF165LXO,ENCFF951GOP.ENCFF281QMV.ENCFF089GBC.ENCFF575MGI |
| 813 | 5 | 37511400 | 37512048 | 648 | MOLF | 5 | 37511925 | 37511958 | 34 | - | 24.065 | 3.73E-09 | 0.00141 | CTGTAATGCCATGAAAGTCCAGCAGGGGGCAAAA | 24 | ENCFF201JUA,ENCFF093ZRU,ENCFF367UNR,ENCFF099QNE,ENCFF065ODV,ENCFF092NMJ,ENCFF821BIA,ENCFF430PPJ,ENCFF907XEG,ENCFF327LYS,ENCFF198SSR,ENCFF609MDN,ENCFF024JOZ,ENCFF883UPM,ENCFF589ZBJ,ENCFF052KVO,ENCFF504JDS,ENCFF957SPH,ENCFF136ZDK,ENCFF165LXO,ENCFF951GOP,ENCFF281QMV,ENCFF089GBC,ENCFF575MGI |
| 814 | 5 | 53818180 | 53818342 | 162 | SPRET | 5 | 53818273 | 53818291 | 19 | - | 17.5902 | 6.18E-07 | 0.0956 | ACACCAGAAGAGGGCACTG | | |
| 815 | 5 | 62597231 | 62597444 | 213 | AJ,AKR,BUB,C58J,CAST,CBA,CE,FVB,KK,MA/My,MOLF,MRL,NOD,NZB,NZW,RF,SJL,SMJ,SPRET,SWR,TH,WSB | 5 | 62597357 | 62597375 | 19 | - | 17.5574 | 6.32E-07 | 0.0956 | ACTCCAGAAGAGGGCGCCA | | |
| 815 | 5 | 62597231 | 62597444 | 213 | AJ,AKR,BUB,C58J,CAST,CBA,CE,FVB,KK,MA/My,MOLF,MRL,NOD,NZB,NZW,RF,SJL,SMJ,SPRET,SWR,TH,WSB | 5 | 62597357 | 62597390 | 34 | - | 18.3659 | 2.76E-07 | 0.0384 | TAGCTGTCTTCAGACACTCCAGAAGAGGGCGCCA | | |
| 816 | 5 | 66420435 | 66420649 | 214 | 129S1,AJ,AKR,BALB,BTBR,BUB,C3H,C57L,CAST,CBA,CE,DBA1J,DBA2J,FVB,ILNJ2,KK,MA/My,MOLF,MRL,NOD,NOR,NZB,NZO,PJ,PL,RF,RH2,SEA,SJL,SMJ,SPRET,SWR,WSB | 5 | 66420568 | 66420601 | 34 | - | 17.2114 | 5.91E-07 | 0.06 | TAGCTGTCTTCGGACGCTCCAGAAGAGGGCGTCA | | |
| 817 | 5 | 76713403 | 76713491 | 88 | SPRET | 5 | 76713383 | 76713416 | 34 | - | 23.2764 | 7.20E-09 | 0.00238 | GTGTAGTGCCTCTGTAGGCCAGAAGAGGGCAGAT | 39 | ENCFF201JUA,ENCFF093ZRU,ENCFF099QNE,ENCFF065ODV,ENCFF821BIA,ENCFF136ZDK,ENCFF327LYS,ENCFF086ZTV,ENCFF805QBU,ENCFF321AGQ,ENCFF089GBC,ENCFF024JOZ,ENCFF883UPM,ENCFF907XEG,ENCFF198SSR,ENCFF589ZBJ,ENCFF430PPJ,ENCFF052KVO,ENCFF092NMJ,ENCFF165LXO,ENCFF504JDS,ENCFF957SPH,ENCFF951GOP,ENCFF142CNG,ENCFF210BTO,ENCFF533APC,ENCFF194HVL,ENCFF097CAQ,ENCFF491RJK,ENCFF575MGI,ENCFF281QMV,ENCFF601JVE,ENCFF349BUL,ENCFF611HDQ,ENCFF609MDN,ENCFF089EWX,ENCFF033UQX,ENCFF316FYN,ENCFF784ASD |
| 818 | 5 | 86034776 | 86034989 | 213 | CAST.SMJ.SPRET | 5 | 86034899 | 86034917 | 19 | - | 17.5574 | 6.32E-07 | 0.0956 | ACTCCAGAAGAGGGCGCCA | | |
| 818 | 5 | 86034776 | 86034989 | 213 | CAST.SMJ.SPRET | 5 | 86034899 | 86034932 | 34 | - | 18.561 | 2.42E-07 | 0.036 | AAGCTGTCTTCAGACACTCCAGAAGAGGGCGCCA | | |
| 819 | 5 | 86826327 | 86826667 | 340 | BALB,C3H,CBA,CE,DBA1J,DBA2J,ILNJ2,NOD.NOR.PJ.RH2.SEA | 5 | 86826648 | 86826681 | 34 | - | 21.3984 | 3.17E-08 | 0.00778 | ATGCAGTGCCTGCAGAGGCCAGAAGATGATGTTG | | |
| 820 | 5 | 89274096 | 89274294 | 198 | CAST.SMJ.SPRET.WSB | 5 | 89274220 | 89274253 | 34 | - | 16.4472 | 9.61E-07 | 0.0776 | TAGCTGTCTTCAGACACACCAGAAGAGGGCGTCA | 1 | ENCFF093ZRU |
| 821 | 5 | 93182561 | 93182783 | 222 | 129S1,AKR,BALB,BTBR,BUB,C3H,C57L,CBA,CE,DBA1J,DBA2J,FVB,MA/My,MRL,NOD,NOR,NZW,PJ,RF,RH2,SEA,SJL,SMJ,SPRET,SWR,TH,WSB | 5 | 93182684 | 93182717 | 34 | - | 20.1301 | 8.09E-08 | 0.0159 | TTGCTGTTTTCAGACACGCCAGAAGAGGGCATAA | 6 | ENCFF093ZRU,ENCFF327LYS,ENCFF907XEG,ENCFF198SSR,ENCFF367UNR,ENCFF099QNE |
| 822 | 5 | 97201355 | 97201576 | 221 | SPRET | 5 | 97201479 | 97201512 | 34 | - | 16.748 | 7.95E-07 | 0.0712 | TAGCCGTCCTCAGACACTCCAGAAGAGGGCATCA | | |
| 823 | 5 | 99280514 | 99281301 | 787 | SPRET | 5 | 99280705 | 99280739 | 35 | - | 20.2385 | 7.45E-08 | 0.0297 | GTGCATTTCTCCATGGCATCCAGAAGAGGTCATTA | 1 | ENCFF327LYS |
| 824 | 5 | 99470918 | 99471137 | 219 | 129S1,AJ,AKR,BALB,BUB,C3H,C57L,CAST,CBA,CE,DBA1J,DBA2J,FVB,ILNJ2,MA/My,MOLF,MRL,NOD,NOR,NZB,NZO,NZW,PJ,PL,RF,RH2,SEA,SJL,SMJ,SPRET,SWR,TH,WSB | 5 | 99471044 | 99471077 | 34 | - | 16.6341 | 8.54E-07 | 0.0745 | TAGCTGTCTTCACACACTCCAGAAGAGGGCGTCA | | |
| 825 | 5 | 100166253 | 100167203 | 950 | SPRET | 5 | 100166541 | 100166574 | 34 | - | 17.4715 | 5.00E-07 | 0.0542 | TAGCTGTCTTCAGATGTGCCAGAAGAGGGCATCA | | |
| 826 | 5 | 103482552 | 103482791 | 239 | CAST.SPRET | 5 | 103482674 | 103482707 | 34 | - | 16.4472 | 9.61E-07 | 0.0776 | TAGCTGTCTTCAGACACACCAGAAGAGGGCGTCA | | |
| 827 | 5 | 104084907 | 104085281 | 374 | CAST | 5 | 104085257 | 104085291 | 35 | - | 22.7982 | 1.29E-08 | 0.00955 | GTGTGGTGCCAGGGGCCAGGCACTAGGGGGCGATG | 34 | ENCFF504JDS,ENCFF201JUA,ENCFF805QBU,ENCFF093ZRU,ENCFF430PPJ,ENCFF367UNR,ENCFF099QNE,ENCFF821BIA,ENCFF065ODV,ENCFF327LYS,ENCFF052KVO,ENCFF907XEG,ENCFF316FYN,ENCFF502WDT,ENCFF198SSR,ENCFF033UQX,ENCFF575MGI,ENCFF609MDN,ENCFF055HAP,ENCFF589ZBJ,ENCFF097CAQ,ENCFF142CNG,ENCFF024JOZ,ENCFF883UPM,ENCFF957SPH,ENCFF165LXO,ENCFF951GOP,ENCFF194HVL,ENCFF210BTO,ENCFF281QMV,ENCFF136ZDK,ENCFF491RJK,ENCFF089GBC,ENCFF092NMJ |
| 828 | 5 | 104725116 | 104725328 | 212 | AJ,BALB,BUB,C3H,CBA,DBA1J,DBA2J,FVB,ILNJ2,KK,MOLF,NZB,NZO,RH2,SEA,SJL,SWR,TH,WSB | 5 | 104725242 | 104725260 | 19 | - | 17.5574 | 6.32E-07 | 0.0956 | ACTCCAGAAGAGGGCGCCA | | |
| 828 | 5 | 104725116 | 104725328 | 212 | AJ,BALB,BUB,C3H,CBA,DBA1J,DBA2J,FVB,ILNJ2,KK,MOLF,NZB,NZO,RH2,SEA,SJL,SWR,TH,WSB | 5 | 104725242 | 104725275 | 34 | - | 18.3659 | 2.76E-07 | 0.0384 | TAGCTGTCTTCAGACACTCCAGAAGAGGGCGCCA | | |
| 829 | 5 | 107043585 | 107043789 | 204 | BUB,C3H,CBA,FVB,PJ,PL,SJL,SPRET,TH,WSB | 5 | 107043710 | 107043728 | 19 | - | 17.5574 | 6.32E-07 | 0.0956 | ACTCCAGAAGAGGGCGCCA | 2 | ENCFF093ZRU,ENCFF367UNR |
| 829 | 5 | 107043585 | 107043789 | 204 | BUB,C3H,CBA,FVB,PJ,PL,SJL,SPRET,TH,WSB | 5 | 107043710 | 107043743 | 34 | - | 18.1789 | 3.13E-07 | 0.0413 | TAGCTGTCTTCAGCCACTCCAGAAGAGGGCGCCA | 2 | ENCFF093ZRU,ENCFF367UNR |

| | | | | | | | | | | | | | | | | |
|---|---|---|---|---|---|---|---|---|---|---|---|---|---|---|---|---|
| 830 | 5 | 107132879 | 107133079 | 200 | SPRET | 5 | 107133002 | 107133035 | 34 | - | 17.6016 | 4.59E-07 | 0.0514 | TTGCTGTCTTCGGACACACCAGTAGAGGGTGTCA | 2 | ENCFF367UNR,ENCFF093ZRU |
| 831 | 5 | 108330271 | 108330490 | 219 | SPRET | 5 | 108330400 | 108330418 | 19 | - | 17.5574 | 6.32E-07 | 0.0956 | ACTCCAGAAGAGGGCGCCA | 3 | ENCFF821BIA,ENCFF093ZRU,ENCFF367UNR |
| 831 | 5 | 108330271 | 108330490 | 219 | SPRET | 5 | 108330400 | 108330433 | 34 | - | 16.4228 | 9.76E-07 | 0.0782 | TAGCTGCCTTCAGACACTCCAGAAGAGGGCGCCA | 3 | ENCFF821BIA,ENCFF093ZRU,ENCFF367UNR |
| 832 | 5 | 108634272 | 108634536 | 264 | BUB,C3H,CAST,CBA,FVB,MOLF,PJ,PL,SJL,SPRET,SWR,TH,WSB | 5 | 108634398 | 108634431 | 34 | - | 16.4472 | 9.61E-07 | 0.0776 | TAGCTGTCTTCAGACACACCAGAAGAGGGCGTCA | | |
| 833 | 5 | 110094119 | 110094643 | 524 | SPRET | 5 | 110094179 | 110094197 | 19 | - | 18.9344 | 2.45E-07 | 0.0604 | ATGCCAGAAGAGGGCACCA | | |
| 834 | 5 | 110398742 | 110398939 | 197 | CAST.SPRET | 5 | 110398864 | 110398897 | 34 | - | 16.4472 | 9.61E-07 | 0.0776 | TAGCTGTCTTCAGACACACCAGAAGAGGGCGTCA | | |
| 835 | 5 | 110468592 | 110468811 | 219 | CAST.SPRET | 5 | 110468732 | 110468765 | 34 | - | 17.9431 | 3.66E-07 | 0.0451 | CAGCTGTCCTCAGACACTCCAGAAGAGGGAGTCA | | |
| 836 | 5 | 111097374 | 111097587 | 213 | CAST.SPRET | 5 | 111097501 | 111097519 | 19 | - | 17.5574 | 6.32E-07 | 0.0956 | ACTCCAGAAGAGGGCGCCA | 2 | ENCFF367UNR.ENCFF093ZRU |
| 836 | 5 | 111097374 | 111097587 | 213 | CAST.SPRET | 5 | 111097501 | 111097534 | 34 | - | 18.3659 | 2.76E-07 | 0.0384 | TAGCTGTCTTCAGACACTCCAGAAGAGGGCGCCA | 2 | ENCFF367UNR.ENCFF093ZRU |
| 837 | 5 | 113949312 | 113949519 | 207 | CAST.MOLF.SPRET | 5 | 113949422 | 113949440 | 19 | - | 17.5574 | 6.32E-07 | 0.0956 | ACTCCAGAAGAGGGCGCCA | | |
| 837 | 5 | 113949312 | 113949519 | 207 | CAST.MOLF.SPRET | 5 | 113949422 | 113949455 | 34 | - | 18.561 | 2.42E-07 | 0.036 | AAGCTGTCTTCAGACACTCCAGAAGAGGGCGCCA | | |
| 838 | 5 | 114085501 | 114086035 | 534 | SPRET | 5 | 114085546 | 114085564 | 19 | - | 17.9836 | 4.75E-07 | 0.0853 | ACACCAGAAGAGGGCGGCA | 9 | ENCFF367UNR,ENCFF024JOZ,ENCFF099QNE,ENCFF821BIA,ENCFF907XEG,ENCFF052KVO,ENCFF093ZRU,ENCFF055HAP,ENCFF502WDT |
| 838 | 5 | 114085501 | 114086035 | 534 | SPRET | 5 | 114085546 | 114085579 | 34 | - | 17.9593 | 3.63E-07 | 0.0449 | TAGCTGTCTTCAGATACACCAGAAGAGGGCGGCA | 9 | ENCFF367UNR,ENCFF024JOZ,ENCFF099QNE,ENCFF821BIA,ENCFF907XEG,ENCFF052KVO,ENCFF093ZRU.ENCFF055HAP.ENCFF502WDT |
| 839 | 5 | 115637943 | 115638135 | 192 | 129S1,AKR,BALB,C57L,CAST,CE,DBA1J,DBA2J,KK,MOLF,MRL,NZB,NZO,PL,RH2,SEA | 5 | 115638067 | 115638100 | 34 | - | 16.4472 | 9.61E-07 | 0.0776 | TAGCTGTCTTCAGACACACCAGAAGAGGGCGTCA | | |
| 840 | 5 | 115972312 | 115972517 | 205 | CAST.MOLF.SPRET | 5 | 115972437 | 115972455 | 19 | - | 17.5574 | 6.32E-07 | 0.0956 | ACTCCAGAAGAGGGCGCCA | | |
| 840 | 5 | 115972312 | 115972517 | 205 | CAST.MOLF.SPRET | 5 | 115972437 | 115972470 | 34 | - | 18.3659 | 2.76E-07 | 0.0384 | TAGCTGTCTTCAGACACTCCAGAAGAGGGCGCCA | | |
| 841 | 5 | 118059672 | 118059871 | 199 | SPRET | 5 | 118059797 | 118059830 | 34 | - | 17.2114 | 5.91E-07 | 0.06 | TAGCTGTTTTCAGACACTCCAGAAGAGGGCATCA | | |
| 842 | 5 | 118190666 | 118190861 | 195 | CAST.SPRET | 5 | 118190790 | 118190823 | 34 | - | 16.4472 | 9.61E-07 | 0.0776 | TAGCTGTCTTCAGACACACCAGAAGAGGGCGTCA | | |
| 843 | 5 | 118392646 | 118392878 | 232 | SPRET | 5 | 118392759 | 118392792 | 34 | - | 16.4634 | 9.51E-07 | 0.0776 | TAGCTGTCTTCAGACGCACCAGAAGAGGGCATCA | | |
| 844 | 5 | 123261829 | 123262440 | 611 | SPRET | 5 | 123261958 | 123261991 | 34 | - | 17.3008 | 5.58E-07 | 0.0584 | ATCCAGCCCCAGAAGACTCCAGAAGAGGGCATCA | | |
| 845 | 5 | 123815787 | 123816000 | 213 | 129S1,AJ,BALB,C57L,CAST,MOLF,MRL,PL,RH2,SEA,SJL,SPRET | 5 | 123815915 | 123815948 | 34 | - | 16.7724 | 7.83E-07 | 0.0704 | TAGCTGTCTTCAGATGCACCAGAAGAGGGCGTCA | | |
| 846 | 5 | 124418837 | 124419040 | 203 | SPRET | 5 | 124418962 | 124418995 | 34 | - | 16.4472 | 9.61E-07 | 0.0776 | TAGCTGTCTTCAGACACACCAGAAGAGGGCGTCA | | |
| 847 | 5 | 124437464 | 124437831 | 367 | SPRET | 5 | 124437557 | 124437575 | 19 | - | 18.7705 | 2.75E-07 | 0.0643 | AAGCCAGAAGAGGGCACCA | 32 | ENCFF367UNR,ENCFF093ZRU,ENCFF201JUA,ENCFF099QNE,ENCFF210BTO,ENCFF883UPM,ENCFF821BIA,ENCFF907XEG,ENCFF327LYS,ENCFF316FYN,ENCFF609MDN,ENCFF575MGI,ENCFF089GBC,ENCFF024JOZ,ENCFF198SSR,ENCFF136ZDK,ENCFF055HAP,ENCFF589ZBJ,ENCFF092NMJ,ENCFF194HVL,ENCFF097CAQ,ENCFF165LXO,ENCFF491RJK,ENCFF957SPH,ENCFF502WDT,ENCFF430PPJ,ENCFF321AGQ,ENCFF065ODV,ENCFF951GOP,ENCFF281QMV,ENCFF052KVO,ENCFF533APC |
| 847 | 5 | 124437464 | 124437831 | 367 | SPRET | 5 | 124437557 | 124437591 | 35 | - | 27.0459 | 4.52E-10 | 0.00161 | CTGCAGGTATGCTTGCAAGCCAGAAGAGGGCACCA | 32 | ENCFF367UNR,ENCFF093ZRU,ENCFF201JUA,ENCFF099QNE,ENCFF210BTO,ENCFF883UPM,ENCFF821BIA,ENCFF907XEG,ENCFF327LYS,ENCFF316FYN,ENCFF609MDN,ENCFF575MGI,ENCFF089GBC,ENCFF024JOZ,ENCFF198SSR,ENCFF136ZDK,ENCFF055HAP,ENCFF589ZBJ,ENCFF092NMJ,ENCFF194HVL,ENCFF097CAQ,ENCFF165LXO,ENCFF491RJK,ENCFF957SPH,ENCFF502WDT,ENCFF430PPJ,ENCFF321AGQ,ENCFF065ODV,ENCFF951GOP,ENCFF281QMV,ENCFF052KVO,ENCFF533APC |
| 848 | 5 | 124946078 | 124946479 | 401 | SPRET | 5 | 124946259 | 124946277 | 19 | - | 17.0492 | 8.79E-07 | 0.113 | GCACCTCCAGAGGGCACTC | 29 | ENCFF201JUA,ENCFF093ZRU,ENCFF367UNR,ENCFF099QNE,ENCFF210BTO,ENCFF907XEG,ENCFF327LYS,ENCFF024JOZ,ENCFF821BIA,ENCFF089GBC,ENCFF092NMJ,ENCFF430PPJ,ENCFF609MDN,ENCFF198SSR,ENCFF055HAP,ENCFF097CAQ,ENCFF589ZBJ,ENCFF957SPH,ENCFF575MGI,ENCFF883UPM,ENCFF502WDT,ENCFF316FYN,ENCFF142CNG,ENCFF165LXO,ENCFF951GOP,ENCFF281QMV,ENCFF491RJK,ENCFF089FWX,ENCFF136ZDK |
| 849 | 5 | 125121883 | 125122136 | 253 | CAST | 5 | 125122069 | 125122102 | 34 | - | 16.9268 | 7.10E-07 | 0.0669 | GTGCCGTGTCCACAGAGGCCTGAGGAAGGAGGCC | | |
| 850 | 5 | 125251357 | 125251564 | 207 | 129S1,AJ,BALB,BUB,C57L,CAST,KK,MOLF,MRL,PL,RH2,SEA,SJL,SPRET,WSB | 5 | 125251486 | 125251519 | 34 | - | 17.8455 | 3.91E-07 | 0.0466 | TAGCTGTCCTCAGACACTCCAGAAGAGGGCGTCA | | |
| 851 | 5 | 125444533 | 125444753 | 220 | SPRET | 5 | 125444642 | 125444675 | 34 | - | 17.6992 | 4.30E-07 | 0.0494 | TAGCTGTCCTCAGATACTCCAGAAGAGGGCGTCA | | |
| 852 | 5 | 125582706 | 125582824 | 118 | CAST.MOLF.SPRET | 5 | 125582753 | 125582786 | 34 | - | 17.8455 | 3.91E-07 | 0.0466 | TAGCTGTCCTCAGACACTCCAGAAGAGGGCGTCA | 1 | ENCFF093ZRU |
| 853 | 5 | 129055156 | 129055938 | 782 | C58J,FVB,LP,MOLF,NOD,NOR,NZB,NZO,SMJ,TH,WSB | 5 | 129055325 | 129055343 | 19 | - | 18.7049 | 2.88E-07 | 0.0658 | GCACCAGAAGAGGGCGCCA | 2 | ENCFF093ZRU,ENCFF099QNE |
| 853 | 5 | 129055156 | 129055938 | 782 | C58J,FVB,LP,MOLF,NOD,NOR,NZB,NZO,SMJ.TH.WSB | 5 | 129055325 | 129055358 | 34 | - | 18.7805 | 2.09E-07 | 0.0323 | CAGCTGACTTCAGATGCACCAGAAGAGGGCGCCA | 2 | ENCFF093ZRU,ENCFF099QNE |
| 853 | 5 | 129055156 | 129055938 | 782 | C58J,FVB,LP,MOLF,NOD,NOR,NZB,NZO,SMJ,TH,WSB | 5 | 129055761 | 129055794 | 34 | - | 22.1789 | 1.74E-08 | 0.00476 | ATGCAGTGCCTGCATATACCAGGAGATGGTGTTG | | |
| 854 | 5 | 129711672 | 129711889 | 217 | CAST.MOLF.SPRET | 5 | 129711792 | 129711825 | 34 | - | 21.3577 | 3.27E-08 | 0.00799 | TAGCTGTTCTCAGATACTCCAGAAGAGGGCACCA | 5 | ENCFF099QNE,ENCFF055HAP,ENCFF201JUA,ENCFF093ZRU,ENCFF367UNR |
| 855 | 5 | 129919204 | 129919406 | 202 | AJ,BALB,CAST,MOLF,MRL,RH2,SEA,SMJ,SPRET | 5 | 129919334 | 129919352 | 19 | - | 17.5574 | 6.32E-07 | 0.0956 | ACTCCAGAAGAGGGCGCCA | | |
| 855 | 5 | 129919204 | 129919406 | 202 | AJ,BALB,CAST,MOLF,MRL,RH2,SEA,SMJ,SPRET | 5 | 129919334 | 129919367 | 34 | - | 18.561 | 2.42E-07 | 0.036 | AAGCTGTCTTCAGACACTCCAGAAGAGGGCGCCA | | |
| 856 | 5 | 129951473 | 129951667 | 194 | MOLF.SPRET | 5 | 129951597 | 129951630 | 34 | - | 17.2439 | 5.79E-07 | 0.0594 | TAGCTGTCCTCAGATACTCCAGAAGAGGGCATCA | | |
| 857 | 5 | 130095172 | 130095371 | 199 | SPRET | 5 | 130095300 | 130095318 | 19 | - | 18.2295 | 4.01E-07 | 0.0781 | CCACCAGCAGAGGGCATCA | 5 | ENCFF821BIA.ENCFF055HAP.ENCFF367UNR.ENCFF093ZRU.ENCFF099QNE |
| 858 | 5 | 130316752 | 130317086 | 334 | CAST | 5 | 130316759 | 130316793 | 35 | - | 21.1468 | 4.08E-08 | 0.0205 | GTGCAGATGACCCCAGGGGCCGGAAGAGGGCATCA | | |
| 859 | 5 | 130670175 | 130670377 | 202 | 129S1,C57L,CAST,FVB,ILNJ2,KK,LP,NZB,NZO,PL,SPRET,TH | 5 | 130670299 | 130670332 | 34 | - | 17.2439 | 5.79E-07 | 0.0594 | TAGCTGTCCTCAGATACTCCAGAAGAGGGCATCA | | |
| 860 | 5 | 134767268 | 134767474 | 206 | SPRET | 5 | 134767394 | 134767427 | 34 | - | 16.4634 | 9.51E-07 | 0.0776 | TAGCTGTCTTCAGACGCACCAGAAGAGGGCATCA | | |
| 861 | 5 | 134767401 | 134768020 | 619 | KK | 5 | 134767394 | 134767427 | 34 | - | 16.4634 | 9.51E-07 | 0.0776 | TAGCTGTCTTCAGACGCACCAGAAGAGGGCATCA | | |
| 862 | 5 | 135125669 | 135125911 | 242 | SPRET | 5 | 135125785 | 135125818 | 34 | - | 17.2439 | 5.79E-07 | 0.0594 | TAGCTGTCCTCAGATACTCCAGAAGAGGGCATCA | | |
| 863 | 5 | 135286603 | 135286664 | 61 | CAST,MOLF | 5 | 135286591 | 135286609 | 19 | - | 16.9836 | 9.17E-07 | 0.115 | TGAACTGCAGGGGGCGCAA | 10 | ENCFF367UNR,ENCFF821BIA,ENCFF327LYS,ENCFF089GBC,ENCFF052KVO,ENCFF136ZDK,ENCFF502WDT,ENCFF093ZRU,ENCFF533APC,ENCFF033UOX |
| 864 | 5 | 135842730 | 135842923 | 193 | AKR,BALB,BTBR,BUB,C57L,CAST,ILNJ2,MOLF,MRL,PJ,PL,RH2,SEA,SPRET,TH,WSB | 5 | 135842832 | 135842865 | 34 | - | 16.7724 | 7.83E-07 | 0.0704 | TAGCTGTCTTCAGATGCACCAGAAGAGGGCGTCA | | |
| 865 | 5 | 137028851 | 137029054 | 203 | CAST.SPRET | 5 | 137028975 | 137029008 | 34 | - | 17.5935 | 4.61E-07 | 0.0515 | TAGCTGTCTCCAGACACTCCAGAAGAGGGCGTCA | | |
| 866 | 5 | 137689715 | 137690086 | 371 | CAST | 5 | 137689727 | 137689760 | 34 | - | 16.7724 | 7.83E-07 | 0.0704 | TAGCTGTCTTCAGATGCACCAGAAGAGGGCGTCA | | |
| 867 | 5 | 137708586 | 137708799 | 213 | 129S1,AKR,BUB,CAST,KK,LP,MA/My,MOLF,MRL,NZB,NZO,NZW,RF,SPRET,SWR,WSB | 5 | 137708713 | 137708746 | 34 | - | 16.4472 | 9.61E-07 | 0.0776 | TAGCTGTCTTCAGACACACCAGAAGAGGGCGTCA | | |
| 868 | 5 | 139789875 | 139789967 | 92 | ILNJ2 | 5 | 139789929 | 139789947 | 19 | - | 18.1967 | 4.11E-07 | 0.0795 | CAGCCAGCAGGGGGCTGCC | 7 | ENCFF367UNR,ENCFF136ZDK,ENCFF883UPM,ENCFF327LYS,ENCFF589ZBJ,ENCFF907XEG,ENCFF024JOZ |
| 869 | 5 | 140424433 | 140424961 | 528 | CAST.MOLF.NZW.SPRET.WSB | 5 | 140424453 | 140424471 | 19 | - | 18 | 4.70E-07 | 0.0851 | TTCCCACCAGAGGGCAGGG | 3 | ENCFF198SSR.ENCFF327LYS.ENCFF907XEG |
| 869 | 5 | 140424433 | 140424961 | 528 | CAST.MOLF.NZW.SPRET.WSB | 5 | 140424519 | 140424537 | 19 | - | 18 | 4.70E-07 | 0.0851 | TTCCCACCAGAGGGCAGGG | 3 | ENCFF198SSR.ENCFF327LYS.ENCFF907XEG |
| 869 | 5 | 140424433 | 140424961 | 528 | CAST.MOLF.NZW.SPRET.WSB | 5 | 140424585 | 140424603 | 19 | - | 18 | 4.70E-07 | 0.0851 | TTCCCACCAGAGGGCAGGG | 3 | ENCFF198SSR.ENCFF327LYS.ENCFF907XEG |
| 869 | 5 | 140424433 | 140424961 | 528 | CAST.MOLF.NZW.SPRET.WSB | 5 | 140424651 | 140424669 | 19 | - | 18 | 4.70E-07 | 0.0851 | TTCCCACCAGAGGGCAGGG | 2 | ENCFF327LYS.ENCFF907XEG |
| 869 | 5 | 140424433 | 140424961 | 528 | CAST.MOLF.NZW.SPRET.WSB | 5 | 140424717 | 140424735 | 19 | - | 18 | 4.70E-07 | 0.0851 | TTCCCACCAGAGGGCAGGG | 2 | ENCFF327LYS.ENCFF907XEG |
| 869 | 5 | 140424433 | 140424961 | 528 | CAST.MOLF.NZW.SPRET.WSB | 5 | 140424783 | 140424801 | 19 | - | 18 | 4.70E-07 | 0.0851 | TTCCCACCAGAGGGCAGGG | | |
| 869 | 5 | 140424433 | 140424961 | 528 | CAST.MOLF.NZW.SPRET.WSB | 5 | 140424849 | 140424867 | 19 | - | 18 | 4.70E-07 | 0.0851 | TTCCCACCAGAGGGCAGGG | | |
| 869 | 5 | 140424433 | 140424961 | 528 | CAST.MOLF.NZW.SPRET.WSB | 5 | 140424915 | 140424933 | 19 | - | 18 | 4.70E-07 | 0.0851 | TTCCCACCAGAGGGCAGGG | 5 | ENCFF093ZRU.ENCFF089GBC.ENCFF092NMJ.ENCFF430PPJ.ENCFF327LYS |
| 870 | 5 | 140424475 | 140424541 | 66 | 129S1,C3H,CBA,CE,DBA1J,DBA2J,KK,MRL,NZB,NZO,PL,RF,SMJ,SWR,TH | 5 | 140424519 | 140424537 | 19 | - | 18 | 4.70E-07 | 0.0851 | TTCCCACCAGAGGGCAGGG | 3 | ENCFF198SSR,ENCFF327LYS,ENCFF907XEG |
| 871 | 5 | 140860908 | 140861484 | 576 | MOLF.NZW | 5 | 140861031 | 140861064 | 34 | - | 24.2033 | 3.31E-09 | 0.00131 | GTGCAATGCCCACAGAGGTCAGAAGAGGGCATCA | | |
| 872 | 5 | 142282796 | 142283229 | 433 | CAST.MOLF.SPRET | 5 | 142282910 | 142282943 | 34 | - | 18.2927 | 2.90E-07 | 0.0392 | TAGCTGTTTTCAGATGCACCAGAAGAGGGCGTTA | | |
| 873 | 5 | 142937522 | 142937723 | 201 | SPRET | 5 | 142937647 | 142937680 | 34 | - | 16.4472 | 9.61E-07 | 0.0776 | TAGCTGTCTTCAGACACACCAGAAGAGGGCGTCA | | |
| 874 | 5 | 142965522 | 142965923 | 401 | 129S1,AKR,BTBR,BUB,C57L,CAST,CE,DBA1J,DBA2J,ILNJ2,KK,MA/My,MOLF,NZB,NZW,PJ,SJL,WSB | 5 | 142965847 | 142965880 | 34 | - | 16.7724 | 7.83E-07 | 0.0704 | TAGCTGTCTTCAGATGCACCAGAAGAGGGCGTCA | | |
| 875 | 5 | 143010278 | 143010498 | 220 | CAST.MOLF.SPRET | 5 | 143010406 | 143010439 | 34 | - | 16.4634 | 9.51E-07 | 0.0776 | TAGCTGTCTTCAGACGCACCAGAAGAGGGCATCA | | |
| 876 | 5 | 143092335 | 143092694 | 359 | SPRET | 5 | 143092368 | 143092401 | 34 | - | 22.3577 | 1.51E-08 | 0.00422 | GTACAGTGCCTACAGAAGCCAGAAGAGGGCATTG | 1 | ENCFF093ZRU |
| 877 | 5 | 143263305 | 143264285 | 980 | SPRET | 5 | 143264118 | 143264151 | 34 | - | 20.3659 | 6.82E-08 | 0.0139 | TTGCTGTCCTCAGACACACCAGAAGAGGGCATCA | | |
| 878 | 5 | 143399223 | 143399433 | 210 | CAST.CBA.SPRET.WSB | 5 | 143399348 | 143399381 | 34 | - | 17.2439 | 5.79E-07 | 0.0594 | TAGCTGTCCTCAGATACTCCAGAAGAGGGCATCA | | |
| 879 | 5 | 143425598 | 143425799 | 201 | CAST.MOLF.SPRET | 5 | 143425710 | 143425743 | 34 | - | 16.4472 | 9.61E-07 | 0.0776 | TAGCTGTCTTCAGACACACCAGAAGAGGGCGTCA | | |
| 880 | 5 | 144118854 | 144119196 | 342 | SPRET | 5 | 144119079 | 144119097 | 19 | - | 17.8852 | 5.08E-07 | 0.0862 | ACACCAGAAGAGGGCACCA | | |
| 880 | 5 | 144118854 | 144119196 | 342 | SPRET | 5 | 144119079 | 144119112 | 34 | - | 18.3008 | 2.89E-07 | 0.039 | TAGCTGTCTTCAAACACACCAGAAGAGGGCACCA | | |
| 881 | 5 | 144169070 | 144169276 | 206 | MOLF.SPRET.TH | 5 | 144169194 | 144169212 | 19 | - | 17.5574 | 6.32E-07 | 0.0956 | ACTCCAGAAGAGGGCGCCA | 3 | ENCFF367UNR.ENCFF099QNE.ENCFF093ZRU |
| 881 | 5 | 144169070 | 144169276 | 206 | MOLF.SPRET.TH | 5 | 144169194 | 144169227 | 34 | - | 18.3659 | 2.76E-07 | 0.0384 | TAGCTGTCTTCAGACACTCCAGAAGAGGGCGCCA | 3 | ENCFF367UNR.ENCFF099QNE.ENCFF093ZRU |
| 882 | 5 | 145152966 | 145153146 | 180 | SPRET | 5 | 145152962 | 145152996 | 35 | - | 17.1835 | 4.87E-07 | 0.0955 | CTTCATATACAGCTGCATGCCAGAAGAGGGCATCA | | |
| 883 | 5 | 147178873 | 147179246 | 373 | NZO.SPRET | 5 | 147178904 | 147178937 | 34 | - | 16.7724 | 7.83E-07 | 0.0704 | TAGCTGTCTTCAGATGCACCAGAAGAGGGCGTCA | | |
| 884 | 5 | 147806520 | 147806596 | 76 | MOLF.SPRET | 5 | 147806521 | 147806539 | 19 | - | 17.5246 | 6.45E-07 | 0.0966 | CTGCCTGCAGAGGGCAGAG | 2 | ENCFF907XEG.ENCFF327LYS |
| 885 | 5 | 147874165 | 147874392 | 227 | CE.SPRET | 5 | 147874275 | 147874308 | 34 | - | 16.4472 | 9.61E-07 | 0.0776 | TAGCTGTCTTCAGACACACCAGAAGAGGGCGTCA | | |
| 886 | 5 | 147946081 | 147946412 | 331 | 129S1,AJ,AKR,BALB,BTBR,BUB,C3H,C57L,CAST,ILNJ2,LP,MA/My,MOLF,MRL,NOD,NOR,RH2,SEA,SMJ,SPRET,WSB | 5 | 147946330 | 147946363 | 34 | - | 17.878 | 3.83E-07 | 0.0461 | TAGCTGTCCTCAGACACACCAGAAGAGGGCATCA | | |
| 887 | 6 | 5208486 | 5208988 | 502 | SPRET | 6 | 5208552 | 5208585 | 34 | + | 16.7317 | 8.03E-07 | 0.104 | CTGCAGGGGCTAAGGGACACACCAAGGGGCCCTG | | |
| 888 | 6 | 12775593 | 12775787 | 194 | LP.PL.SPRET.WSB | 6 | 12775681 | 12775715 | 35 | + | 18.6055 | 2.09E-07 | 0.0851 | CTGCGGACACCCCCAAGGTCCCGTAGAGGGCTCTC | | |
| 889 | 6 | 14419492 | 14419697 | 205 | AKR.FVB.MA/My.MRL.NZB.RH2 | 6 | 14419547 | 14419580 | 34 | + | 18.4797 | 2.56E-07 | 0.0559 | TTGCTGTCTTCAGACACACCAGAAGAGGGCATCA | | |
| 890 | 6 | 17315338 | 17315533 | 195 | CAST.MOLF.SPRET | 6 | 17315391 | 17315424 | 34 | + | 18.8537 | 1.98E-07 | 0.0489 | TAGCTGTCTTCAGACACACCAGAAGAGGGCGCCA | 4 | ENCFF093ZRU.ENCFF821BIA.ENCFF367UNR.ENCFF201JUA |
| 890 | 6 | 17315338 | 17315533 | 195 | CAST.MOLF.SPRET | 6 | 17315406 | 17315424 | 19 | + | 18.623 | 3.06E-07 | 0.0981 | ACACCAGAAGAGGGCGCCA | 4 | ENCFF093ZRU.ENCFF821BIA.ENCFF367UNR.ENCFF201JUA |
| 891 | 6 | 22434071 | 22434305 | 234 | C3H,CBA,CE,FVB,NOD,NOR,SJL,SPRET,TH,WSB | 6 | 22434183 | 22434216 | 34 | + | 17.2439 | 5.79E-07 | 0.086 | TAGCTGTCCTCAGATACTCCAGAAGAGGGCATCA | | |
| 892 | 6 | 23584187 | 23585137 | 950 | MOLF | 6 | 23584655 | 23584689 | 35 | + | 16.3211 | 7.98E-07 | 0.171 | GTGCATTCCTTCTAAAGGCCCTGTGGGTGGAGCTT | | |
| 893 | 6 | 25263993 | 25264344 | 351 | SPRET | 6 | 25264006 | 25264039 | 34 | + | 21.7073 | 2.50E-08 | 0.0111 | GAGCAGGTGTGGCTGCGGCCCCAAGATGGCGCCC | | |
| 894 | 6 | 28323111 | 28323931 | 820 | MOLF | 6 | 28323895 | 28323913 | 19 | + | 17.6557 | 5.92E-07 | 0.138 | TGGGCACCAGGGGTCGCCC | | |
| 895 | 6 | 29511439 | 29511651 | 212 | CAST.SPRET | 6 | 29511525 | 29511543 | 19 | + | 17.8852 | 5.08E-07 | 0.128 | ACACCAGAAGAGGGCACCA | | |
| 896 | 6 | 29766239 | 29766465 | 226 | 129S1,AJ,AKR,BALB,BTBR,BUB,C3H,C57L,CAST,CBA,DBA1J,DBA2J,FVB,MOLF,MRL,NOD,NOR,NZO,PJ,PL,RF,RH2,SEA,SMJ,SPRET | 6 | 29766323 | 29766356 | 34 | + | 18.3659 | 2.76E-07 | 0.0572 | TAGCTGTCTTCAGACACTCCAGAAGAGGGCGCCA | 7 | ENCFF099QNE,ENCFF821BIA,ENCFF327LYS,ENCFF089GBC,ENCFF907XEG,ENCFF093ZRU,ENCFF367UNR |
| 896 | 6 | 29766239 | 29766465 | 226 | 129S1,AJ,AKR,BALB,BTBR,BUB,C3H,C57L,CAST,CBA,DBA1J,DBA2J,FVB,MOLF,MRL,NOD,NOR,NZO,PJ,PL,RF,RH2,SEA,SMJ,SPRET | 6 | 29766338 | 29766356 | 19 | + | 17.5574 | 6.32E-07 | 0.138 | ACTCCAGAAGAGGGCGCCA | 7 | ENCFF099QNE,ENCFF821BIA,ENCFF327LYS,ENCFF089GBC,ENCFF907XEG,ENCFF093ZRU,ENCFF367UNR |
| 897 | 6 | 31678699 | 31678960 | 261 | SPRET | 6 | 31678694 | 31678727 | 34 | + | 19.5285 | 1.24E-07 | 0.0363 | CTGCAGTCCTACCTCTCTGCTGCAGAGGGCTGGT | | |
| 898 | 6 | 36988364 | 36989215 | 851 | CAST | 6 | 36988703 | 36988721 | 19 | + | 18.9344 | 2.45E-07 | 0.0886 | ATGCCAGAAGAGGGCACCA | 6 | ENCFF089GBC.ENCFF907XEG.ENCFF198SSR.ENCFF883UPM.ENCFF093ZRU.ENCFF024JOZ |
| 899 | 6 | 36995299 | 36995501 | 202 | CAST.SPRET | 6 | 36995355 | 36995388 | 34 | + | 16.4472 | 9.61E-07 | 0.112 | TAGCTGTCTTCAGACACACCAGAAGAGGGCGTCA | | |
| 900 | 6 | 37973532 | 37973759 | 227 | CAST.MOLF.SPRET | 6 | 37973608 | 37973641 | 34 | + | 16.5854 | 8.81E-07 | 0.11 | TAGCTGTCCCCAGACACTCCAGAAGAGGGTGTCA | | |
| 901 | 6 | 38251936 | 38252308 | 372 | CAST.KK.MOLF.SPRET | 6 | 38252153 | 38252186 | 34 | + | 17.9593 | 3.63E-07 | 0.0662 | TTGCTGTCTTTAGACACACCAGAAGAGGGCATCC | | |
| 902 | 6 | 38342448 | 38342657 | 209 | SPRET | 6 | 38342496 | 38342529 | 34 | + | 16.4146 | 9.81E-07 | 0.113 | TAGCTGTCTTCAGACACACCAGAGGAGGGCGCCA | | |
| 903 | 6 | 38355973 | 38356212 | 239 | CAST.SPRET | 6 | 38356071 | 38356104 | 34 | + | 17.3496 | 5.41E-07 | 0.0831 | TAGCTGTCCTCAGATACTCCAGAAGAGGGCATCG | | |
| 904 | 6 | 38446690 | 38446986 | 296 | KK,MOLF | 6 | 38446696 | 38446730 | 35 | + | 27.4679 | 3.12E-10 | 0.00222 | CTGCATGTATGCCTGCAGGCCAGAAGAGGGCACCA | 17 | ENCFF099QNE,ENCFF089GBC,ENCFF589ZBJ,ENCFF198SSR,ENCFF201JUA,ENCFF092NMJ,ENCFF883UPM,ENCFF055HAP,ENCFF821BIA,ENCFF502WDT,ENCFF327LYS,ENCFF024JOZ,ENCFF907XEG,ENCFF367UNR.ENCFF093ZRU.ENCFF052KVO.ENCFF136ZDK |
| 904 | 6 | 38446690 | 38446986 | 296 | KK,MOLF | 6 | 38446712 | 38446730 | 19 | + | 20.3607 | 8.29E-08 | 0.0572 | AGGCCAGAAGAGGGCACCA | 18 | ENCFF099QNE,ENCFF089GBC,ENCFF589ZBJ,ENCFF198SSR,ENCFF201JUA,ENCFF092NMJ,ENCFF883UPM,ENCFF055HAP,ENCFF821BIA,ENCFF502WDT,ENCFF327LYS,ENCFF024JOZ,ENCFF907XEG,ENCFF065ODV,ENCFF367UNR,ENCFF093ZRU,ENCFF052KVO,ENCFF136ZDK |
| 905 | 6 | 38655396 | 38655614 | 218 | CAST.KK.MOLF.SPRET | 6 | 38655458 | 38655492 | 35 | + | 17.9266 | 3.14E-07 | 0.11 | CTGTAGTTGTCTTCAGACACCAGAAGAGGGAATCA | | |
| 906 | 6 | 38662525 | 38662750 | 225 | CAST.SPRET | 6 | 38662596 | 38662629 | 34 | + | 16.4472 | 9.61E-07 | 0.112 | TAGCTGTCTTCAGACACACCAGAAGAGGGCGTCA | | |
| 907 | 6 | 39215370 | 39215577 | 207 | SPRET | 6 | 39215436 | 39215469 | 34 | + | 16.4472 | 9.61E-07 | 0.112 | TAGCTGTCTTCAGACACACCAGAAGAGGGCGTCA | | |
| 908 | 6 | 39288054 | 39288260 | 206 | KK.MOLF.SPRET | 6 | 39288119 | 39288152 | 34 | + | 17.2439 | 5.79E-07 | 0.086 | TAGCTGTCCTCAGATACTCCAGAAGAGGGCATCA | | |
| 909 | 6 | 41159830 | 41160207 | 377 | SPRET | 6 | 41160207 | 41160240 | 34 | + | 18.6179 | 2.33E-07 | 0.0538 | ATGCAGTGCTTATGGAGGCCATAAGGTGGTGTAG | | |
| 910 | 6 | 47834635 | 47834843 | 208 | CAST.MOLF.SPRET | 6 | 47834701 | 47834734 | 34 | + | 17.2439 | 5.79E-07 | 0.086 | TAGCTGTCCTCAGATACTCCAGAAGAGGGCATCA | | |
| 911 | 6 | 51578060 | 51578305 | 245 | AKR,CAST,CE,ILNJ2,KK,LP,MA/My,MOLF,NOD,NOR,NZB,NZO,NZW,PJ,RF,SPRET,TH | 6 | 51578161 | 51578194 | 34 | + | 16.4634 | 9.51E-07 | 0.112 | TAGCTGTCTTCAGACGCACCAGAAGAGGGCATCA | | |
| 912 | 6 | 52099696 | 52099911 | 215 | CAST.MOLF.SPRET | 6 | 52099768 | 52099801 | 34 | + | 18.3659 | 2.76E-07 | 0.0572 | TAGCTGTCTTCAGACACTCCAGAAGAGGGCGCCA | 2 | ENCFF367UNR.ENCFF821BIA |
| 912 | 6 | 52099696 | 52099911 | 215 | CAST.MOLF.SPRET | 6 | 52099783 | 52099801 | 19 | + | 17.5574 | 6.32E-07 | 0.138 | ACTCCAGAAGAGGGCGCCA | 2 | ENCFF367UNR.ENCFF821BIA |
| 913 | 6 | 52271388 | 52271601 | 213 | SPRET | 6 | 52271458 | 52271491 | 34 | + | 17.9187 | 3.72E-07 | 0.0672 | TAGCTGTCCTCACATACTCCAGAAGAGGGCATCA | | |
| 914 | 6 | 54964042 | 54964283 | 241 | CAST.MOLF.SPRET | 6 | 54964140 | 54964173 | 34 | + | 16.9187 | 7.13E-07 | 0.0978 | TAGCTGTCTTCAGACGCACCAGAAGAGGGCGTCA | 1 | ENCFF093ZRU |
| 915 | 6 | 60284495 | 60284709 | 214 | CAST.PJ.SPRET.WSB | 6 | 60284567 | 60284600 | 34 | + | 16.4634 | 9.51E-07 | 0.112 | TAGCTGTCTTCAGACGCACCAGAAGAGGGCATCA | | |
| 916 | 6 | 60689283 | 60689933 | 650 | CAST.MOLF.SPRET | 6 | 60689473 | 60689506 | 34 | + | 17.6992 | 4.30E-07 | 0.0729 | TAGCTGTTTTCAGACACACCAGAAGAGGGCATCA | | |
| 917 | 6 | 67159845 | 67160042 | 197 | C58J,CAST,CE,ILNJ2,KK,MOLF,NOD,NOR,NZB.NZO.PJ.SPRET.WSB | 6 | 67159900 | 67159933 | 34 | + | 17.8455 | 3.91E-07 | 0.0687 | TAGCTGTCCTCAGACACTCCAGAAGAGGGCGTCA | | |
| 918 | 6 | 67229920 | 67230124 | 204 | C58J,KK,MOLF,NOD,NOR,NZB,NZO,SPRET | 6 | 67229983 | 67230016 | 34 | + | 17.2358 | 5.82E-07 | 0.0863 | TAGCTGTCCTCAGAAACTCCAGAAGAGGGCATCA | | |
| 919 | 6 | 67237874 | 67238086 | 212 | C58J,CAST,KK,MA/My,MOLF,NOD,NOR,NZB,NZO,SPRET | 6 | 67237944 | 67237977 | 34 | + | 18.3659 | 2.76E-07 | 0.0572 | TAGCTGTCTTCAGACACTCCAGAAGAGGGCGCCA | | |
| 919 | 6 | 67237874 | 67238086 | 212 | C58J,CAST,KK,MA/My,MOLF,NOD,NOR,NZB,NZO,SPRET | 6 | 67237959 | 67237977 | 19 | + | 17.5574 | 6.32E-07 | 0.138 | ACTCCAGAAGAGGGCGCCA | | |
| 920 | 6 | 67676116 | 67676448 | 332 | C58J,CAST,KK,MA/My,MOLF,NOD,NOR,NZB,NZO,SPRET,WSB | 6 | 67676146 | 67676164 | 19 | + | 19.2951 | 1.88E-07 | 0.0786 | CCGCCACTAGATGGTGCTA | 11 | ENCFF093ZRU,ENCFF367UNR,ENCFF430PPJ,ENCFF099QNE,ENCFF201JUA,ENCFF907XEG,ENCFF821BIA,ENCFF024JOZ,ENCFF089GBC,ENCFF327LYS,ENCFF092NMJ |
| 921 | 6 | 67900627 | 67900853 | 226 | CE,ILNJ2,PJ | 6 | 67900764 | 67900797 | 34 | + | 17.0325 | 6.63E-07 | 0.0937 | TAGCTGTCCTCAGACATCCCAGAAGAGGGCATCA | | |
| 922 | 6 | 72894129 | 72894304 | 175 | AKR,BALB,MRL,PL,RF,RH2,SEA | 6 | 72894210 | 72894243 | 34 | + | 17.748 | 4.17E-07 | 0.0716 | TTGCTGTCTTCAGACACACCAGGAGAGGGCATCA | | |
| 923 | 6 | 82124805 | 82125595 | 790 | SPRET | 6 | 82124884 | 82124917 | 34 | + | 19.065 | 1.71E-07 | 0.0454 | TTGCAGTGCCCATGGAGGCCAACAGAGGATGTCA | | |
| 924 | 6 | 84053398 | 84053914 | 516 | NZB,NZO | 6 | 84053631 | 84053664 | 34 | + | 19.0163 | 1.77E-07 | 0.0463 | CAGCTGTCTTCAGACACACCAGAAGAGGGCGTCA | | |
| 925 | 6 | 85365886 | 85366071 | 185 | PJ,SEA,SPRET | 6 | 85365930 | 85365963 | 34 | + | 17.2439 | 5.79E-07 | 0.086 | TAGCTGTCCTCAGATACTCCAGAAGAGGGCATCA | | |
| 926 | 6 | 86322228 | 86322279 | 51 | MOLF,PJ,SEA | 6 | 86322247 | 86322280 | 34 | + | 17.5285 | 4.81E-07 | 0.0779 | CTGCTTTTGCACTCCTTTCCACTAGGTGCAGCAT | | |
| 927 | 6 | 86386014 | 86386513 | 499 | CAST | 6 | 86386378 | 86386412 | 35 | + | 19.3853 | 1.29E-07 | 0.0651 | CTGTAGCTGTCTTTACACACCAGAAGAGGGCATTG | | |
| 928 | 6 | 86397596 | 86397652 | 56 | CAST,MOLF,PJ,SEA,SPRET | 6 | 86397622 | 86397656 | 35 | + | 21.1193 | 4.16E-08 | 0.0327 | CTGCTGTCACACACTTCAGCCTCCTGGAGGCGCTC | 20 | ENCFF201JUA,ENCFF092NMJ,ENCFF165LXO,ENCFF430PPJ,ENCFF502WDT,ENCFF055HAP,ENCFF589ZBJ,ENCFF093ZRU,ENCFF065ODV,ENCFF907XEG,ENCFF327LYS,ENCFF367UNR,ENCFF533APC,ENCFF198SSR,ENCFF883UPM,ENCFF052KVO,ENCFF099QNE,ENCFF142CNG,ENCFF089GBC,ENCFF024JOZ |
| 929 | 6 | 86506559 | 86506718 | 159 | SPRET | 6 | 86506594 | 86506627 | 34 | + | 16.4634 | 9.51E-07 | 0.112 | TAGCTGTCTTCAGACGCACCAGAAGAGGGCATCA | 1 | ENCFF367UNR |
| 930 | 6 | 86693926 | 86694104 | 178 | 129S1,AJ,BTBR,C3H,C57L,C58J,CAST,CBA,DBA1J,DBA2J,FVB,MOLF,NOD,NOR,PJ,RF,SEA,SJL,SMJ,SPRET,SWR | 6 | 86693977 | 86694010 | 34 | + | 18.3659 | 2.76E-07 | 0.0572 | TAGCTGTCTTCAGACACTCCAGAAGAGGGCGCCA | | |
| 930 | 6 | 86693926 | 86694104 | 178 | 129S1,AJ,BTBR,C3H,C57L,C58J,CAST,CBA,DBA1J,DBA2J,FVB,MOLF,NOD,NOR,PJ,RF,SEA,SJL,SMJ,SPRET,SWR | 6 | 86693992 | 86694010 | 19 | + | 17.5574 | 6.32E-07 | 0.138 | ACTCCAGAAGAGGGCGCCA | | |
| 931 | 6 | 87010483 | 87010697 | 214 | CAST.MOLF.PJ.SEA.SPRET | 6 | 87010509 | 87010542 | 34 | + | 18.0813 | 3.34E-07 | 0.0635 | CAGCTGTCTTCAGAAACCCCAGAAGAGGGCGTCA | | |
| 932 | 6 | 87576521 | 87576723 | 202 | CAST.MOLF.PJ.SEA.SPRET | 6 | 87576580 | 87576613 | 34 | + | 16.4472 | 9.61E-07 | 0.112 | TAGCTGTCTTCAGACACACCAGAAGAGGGCGTCA | | |
| 933 | 6 | 94269206 | 94269871 | 665 | MOLF | 6 | 94269825 | 94269859 | 35 | + | 21.367 | 3.52E-08 | 0.0299 | CTGTATATACACCTGCATGCCAGAAGAGGGCATCA | | |
| 934 | 6 | 97240759 | 97241016 | 257 | C58J,CAST,LP,MA/My | 6 | 97240918 | 97240952 | 35 | + | 17.0459 | 5.27E-07 | 0.143 | TTGCATGTACACTTGTTTGTCAGAAGAGGGCATCA | 9 | ENCFF099QNE,ENCFF201JUA,ENCFF821BIA,ENCFF367UNR,ENCFF089GBC,ENCFF907XEG,ENCFF055HAP,ENCFF093ZRU,ENCFF502WDT |
| 934 | 6 | 97240759 | 97241016 | 257 | C58J,CAST,LP,MA/My | 6 | 97240843 | 97240861 | 19 | - | 20.3607 | 8.29E-08 | 0.0572 | AGGCCAGAAGAGGGCACCA | 10 | ENCFF099QNE,ENCFF201JUA,ENCFF821BIA,ENCFF367UNR,ENCFF089GBC,ENCFF907XEG,ENCFF055HAP,ENCFF093ZRU,ENCFF024JOZ,ENCFF502WDT |
| 934 | 6 | 97240759 | 97241016 | 257 | C58J,CAST,LP,MA/My | 6 | 97240843 | 97240877 | 35 | - | 16.2844 | 8.14E-07 | 0.173 | TTTAAATGTACACCTCAGGCCAGAAGAGGGCACCA | 10 | ENCFF099QNE,ENCFF201JUA,ENCFF821BIA,ENCFF367UNR,ENCFF089GBC,ENCFF907XEG,ENCFF055HAP,ENCFF093ZRU,ENCFF024JOZ,ENCFF502WDT |
| 935 | 6 | 98520394 | 98520579 | 185 | CAST.MOLF.SPRET.WSB | 6 | 98520436 | 98520469 | 34 | + | 16.4634 | 9.51E-07 | 0.112 | TAGCTGTCTTCAGACGCACCAGAAGAGGGCATCA | | |
| 936 | 6 | 103995417 | 103996076 | 659 | SPRET | 6 | 103995544 | 103995577 | 34 | + | 17.2683 | 5.70E-07 | 0.0858 | TTGCTCTCTTCAGATACACCAGAAGAGGGCGTCA | 1 | ENCFF093ZRU |
| 937 | 6 | 112134934 | 112135197 | 263 | MOLF | 6 | 112135057 | 112135090 | 34 | + | 17.8537 | 3.89E-07 | 0.0686 | CTGCCGTGTTCAAAGGAGCTAGAAGAGGGCATCA | | |
| 938 | 6 | 112263112 | 112263274 | 162 | CAST | 6 | 112263162 | 112263196 | 35 | + | 19.4862 | 1.21E-07 | 0.0624 | ATGCAGATGCTTGCAAAGACCAGAAGAGGGCATCA | | |
| 939 | 6 | 112349953 | 112350015 | 62 | CAST | 6 | 112349957 | 112349991 | 35 | + | 16.211 | 8.49E-07 | 0.177 | CTGCAGCCCAGCCTCTTGGCCTCCAGGTAGGTGGT | | |
| 940 | 6 | 113309790 | 113309951 | 161 | SPRET | 6 | 113309828 | 113309861 | 34 | + | 16.4634 | 9.51E-07 | 0.112 | TAGCTGTCTTCAGACGCACCAGAAGAGGGCATCA | 6 | ENCFF099QNE.ENCFF024JOZ.ENCFF907XEG.ENCFF327LYS.ENCFF093ZRU.ENCFF367UNR |
| 941 | 6 | 113552401 | 113552618 | 217 | CAST.CE.MOLF.SMJ.SPRET | 6 | 113552477 | 113552510 | 34 | + | 17.0813 | 6.43E-07 | 0.0919 | TAGCTGTTCTCAGACACTCCAGAAGAGGGAGTCA | | |
| 942 | 6 | 115661031 | 115661236 | 205 | SPRET | 6 | 115661096 | 115661129 | 34 | + | 18.4797 | 2.56E-07 | 0.0559 | TTGCTGTCTTCAGACACACCAGAAGAGGGCATCA | | |
| 943 | 6 | 115703817 | 115704015 | 198 | SPRET | 6 | 115703874 | 115703907 | 34 | + | 16.4472 | 9.61E-07 | 0.112 | TAGCTGTCTTCAGACACACCAGAAGAGGGCGTCA | | |
| 944 | 6 | 115754364 | 115754561 | 197 | AKR,C3H,CBA,ILNJ2,MOLF,MRL,PL,RF,SPRET | 6 | 115754420 | 115754453 | 34 | + | 18.878 | 1.95E-07 | 0.0488 | TAGCTGTCCCCAGATACTCCAGAAGAGGGCATCA | | |
| 945 | 6 | 115823861 | 115824076 | 215 | 129S1,AKR,BTBR,C3H,C57L,C58J,CAST,CBA,CE,DBA1J,DBA2J,ILNJ2,LP,MOLF,MRL,NOD,NOR,NZB,NZO,NZW,PJ,PL,RF,RH2,SPRET,WSB | 6 | 115823932 | 115823965 | 34 | + | 18.1789 | 3.13E-07 | 0.0611 | TAGCTGTCTTCAGACACTCCAGAAGAGGGCGCTA | 1 | ENCFF805QBU |
| 945 | 6 | 115823861 | 115824076 | 215 | 129S1,AKR,BTBR,C3H,C57L,C58J,CAST,CBA,CE,DBA1J,DBA2J,ILNJ2,LP,MOLF,MRL,NOD,NOR,NZB,NZO,NZW,PJ,PL,RF,RH2,SPRET,WSB | 6 | 115823947 | 115823965 | 19 | + | 17.5738 | 6.25E-07 | 0.138 | ACTCCAGAAGAGGGCGCTA | 2 | ENCFF805QBU,ENCFF210BTO |
| 946 | 6 | 117167033 | 117167258 | 225 | CAST,FVB.MOLF.PJ.SMJ.TH | 6 | 117167115 | 117167148 | 34 | + | 16.4634 | 9.51E-07 | 0.112 | TAGCTGTCTTCAGACGCACCAGAAGAGGGCATCA | | |
| 947 | 6 | 117900078 | 117900285 | 207 | SPRET | 6 | 117900137 | 117900170 | 34 | + | 18.3659 | 2.76E-07 | 0.0572 | TAGCTGTCTTCAGACACTCCAGAAGAGGGCGCCA | 2 | ENCFF099QNE.ENCFF367UNR |
| 947 | 6 | 117900078 | 117900285 | 207 | SPRET | 6 | 117900152 | 117900170 | 19 | + | 17.5574 | 6.32E-07 | 0.138 | ACTCCAGAAGAGGGCGCCA | 2 | ENCFF099QNE.ENCFF367UNR |
| 948 | 6 | 117963407 | 117963818 | 411 | SPRET | 6 | 117963676 | 117963709 | 34 | + | 16.4472 | 9.61E-07 | 0.112 | TAGCTGTCTTCAGACACACCAGAAGAGGGCGTCA | | |
| 949 | 6 | 120406728 | 120406932 | 204 | CAST.MOLF.SJL.SPRET.SWR | 6 | 120406788 | 120406821 | 34 | + | 16.4472 | 9.61E-07 | 0.112 | TAGCTGTCTTCAGACACACCAGAAGAGGGCGTCA | | |
| 950 | 6 | 120534180 | 120534394 | 214 | CAST.MOLF.SPRET | 6 | 120534266 | 120534284 | 19 | + | 17.8852 | 5.08E-07 | 0.128 | ACACCAGAAGAGGGCACCA | | |
| 951 | 6 | 123876009 | 123876171 | 162 | CAST,CE,FVB,NZO,PJ,SMJ,SPRET,SWR,WSB | 6 | 123876073 | 123876091 | 19 | + | 17.5574 | 6.32E-07 | 0.138 | TCACCAGCAGGTGTCAGTC | | |
| 952 | 6 | 123908089 | 123908421 | 332 | SPRET | 6 | 123908118 | 123908136 | 19 | + | 19.2951 | 1.88E-07 | 0.0786 | CCGCCACTAGATGGTGCTA | 19 | ENCFF367UNR,ENCFF093ZRU,ENCFF099QNE,ENCFF092NMJ,ENCFF430PPJ,ENCFF883UPM,ENCFF201JUA,ENCFF821BIA,ENCFF609MDN,ENCFF907XEG,ENCFF024JOZ,ENCFF089GBC,ENCFF502WDT,ENCFF327LYS.ENCFF589ZBJ.ENCFF055HAP.ENCFF165LXO.ENCFF136ZDK.ENCFF198SSR |
| 953 | 6 | 125022354 | 125022570 | 216 | SPRET | 6 | 125022427 | 125022460 | 34 | + | 16.4472 | 9.61E-07 | 0.112 | TAGCTGTCTTCAGACACACCAGAAGAGGGCGTCA | | |
| 954 | 6 | 125042414 | 125042635 | 221 | CAST.SPRET.SWR | 6 | 125042506 | 125042524 | 19 | + | 17.8852 | 5.08E-07 | 0.128 | ACACCAGAAGAGGGCACCA | | |
| 955 | 6 | 125135259 | 125135466 | 207 | CAST.MOLF.SPRET | 6 | 125135323 | 125135356 | 34 | + | 16.4472 | 9.61E-07 | 0.112 | TAGCTGTCTTCAGACACACCAGAAGAGGGCGTCA | | |
| 956 | 6 | 127413442 | 127414164 | 722 | SPRET | 6 | 127413569 | 127413602 | 34 | + | 16.6748 | 8.33E-07 | 0.107 | TTGCTGTCTTCAGACACACCGGAAGAGGGCATCG | | |
| 957 | 6 | 128443942 | 128444151 | 209 | SPRET | 6 | 128444010 | 128444043 | 34 | + | 16.4634 | 9.51E-07 | 0.112 | TAGCTGTCTTCAGACGCACCAGAAGAGGGCATCA | | |

| | | | | | | | | | | | | | | | | |
|---|---|---|---|---|---|---|---|---|---|---|---|---|---|---|---|---|
| 958 | 6 | 128822903 | 128823105 | 202 | 129S1,AJ,AKR,BALB,BUB,C3H,C57L,C58J,CAST,CBA,DBA1J,DBA2J,ILNJ2,MOLF,MRL,NOD,NOR,NZO,PL,RF,SEA,SMJ,SPRET,SWR,WSB | 6 | 128822963 | 128822996 | 34 | + | 18.561 | 2.42E-07 | 0.0549 | AAGCTGTCTTCAGACACTCCAGAAGAGGGCGCCA | | |
| 958 | 6 | 128822903 | 128823105 | 202 | 129S1,AJ,AKR,BALB,BUB,C3H,C57L,C58J,CAST,CBA,DBA1J,DBA2J,ILNJ2,MOLF,MRL,NOD,NOR,NZO,PL,RF,SEA,SMJ,SPRET,SWR,WSB | 6 | 128822978 | 128822996 | 19 | + | 17.5574 | 6.32E-07 | 0.138 | ACTCCAGAAGAGGGCGCCA | | |
| 959 | 6 | 129571429 | 129571631 | 202 | 129S1,AJ,AKR,BALB,BUB,C3H,C57L,C58J,CAST,CBA,CE,DBA1J,DBA2J,FVB,ILNJ2,LP,MA/My,MOLF,MRL,NOD,NOR,NZO,PL,RF,RH2,SEA,SJL,SMJ,SPRET,SWR,TH,WSB | 6 | 129571489 | 129571522 | 34 | + | 17.8455 | 3.91E-07 | 0.0687 | TAGCTGTCCTCAGACACTCCAGAAGAGGGCGTCA | | |
| 960 | 6 | 133195547 | 133195754 | 207 | NZO,SPRET | 6 | 133195610 | 133195643 | 34 | + | 17.2439 | 5.79E-07 | 0.086 | TAGCTGTCCTCAGACCCTCCAGAAGAGGGAGTCA | | |
| 961 | 6 | 133683827 | 133684160 | 333 | AJ,AKR,BALB,BUB,C3H,C57L,CAST,CBA,DBA1J,DBA2J,ILNJ2,MOLF,MRL,NZB,PL,RF,RH2,SEA,SPRET | 6 | 133683857 | 133683875 | 19 | + | 19.2951 | 1.88E-07 | 0.0786 | CCGCCACTAGATGGTGCTA | 21 | ENCFF367UNR,ENCFF093ZRU,ENCFF099QNE,ENCFF821BIA,ENCFF609MDN,ENCFF907XEG,ENCFF327LYS,ENCFF089GBC,ENCFF033UQX,ENCFF052KVO,ENCFF024JOZ,ENCFF589ZBJ,ENCFF198SSR,ENCFF136ZDK,ENCFF165LXO,ENCFF430PPJ,ENCFF092NMJ,ENCFF951GOP,ENCFF883UPM,ENCFF957SPH,ENCFF281QMV |
| 962 | 6 | 134883099 | 134883315 | 216 | SPRET | 6 | 134883170 | 134883203 | 34 | + | 17.2439 | 5.79E-07 | 0.086 | TAGCTGTCCTCAGATACTCCAGAAGAGGGCATCA | 1 | ENCFF367UNR |
| 963 | 6 | 135077717 | 135077887 | 170 | 129S1,AJ,AKR,BALB,BUB,C3H,C57L,CAST,CBA,CE,DBA1J,DBA2J,KK,MA/My,MRL,NOD,NOR,NZO,PL,RF,RH2,SEA,SJL,SPRET,TH,WSB | 6 | 135077760 | 135077793 | 34 | + | 17.3902 | 5.27E-07 | 0.0822 | TAGCTGTCCTCAGACACTCCAGAAGAGGGCATCA | | |
| 964 | 6 | 135485236 | 135485609 | 373 | SPRET | 6 | 135485567 | 135485600 | 34 | + | 19.8618 | 9.80E-08 | 0.0306 | GTGCGGTGCCTACAGAGGCTAGAAGAGGGCATCA | | |
| 965 | 6 | 138793203 | 138793359 | 156 | AJ,AKR,BALB,BUB,C3H,CBA,CE,DBA1J,DBA2J,ILNJ2,KK,MOLF,MRL,NOD,NOR,PL,RF,RH2,SEA | 6 | 138793172 | 138793205 | 34 | + | 17.5203 | 4.84E-07 | 0.0781 | GTGAAGTGCCCTCATAGGCCAGCAGGTAGTATCA | | |
| 966 | 6 | 140585047 | 140585271 | 224 | BTBR,C57L,C58J,CAST,FVB,MOLF,PJ,SMJ,SPRET,SWR,WSB | 6 | 140585129 | 140585162 | 34 | + | 16.5691 | 8.90E-07 | 0.11 | TAGCTGTCTTCAGACGCACCAGAAGAGGGCATCG | | |
| 967 | 6 | 141777544 | 141778432 | 888 | CAST | 6 | 141778399 | 141778432 | 34 | + | 18.1138 | 3.27E-07 | 0.0627 | GTGCAGATGCCCAGAGAGCCGGAAAAGGGCACCC | | |
| 968 | 6 | 142653565 | 142653637 | 72 | SPRET | 6 | 142653605 | 142653638 | 34 | + | 17.0894 | 6.39E-07 | 0.0917 | GTGCTCTTCCCTAAGTGAGCCCCAGAGGCCTCTG | | |
| 969 | 6 | 144494924 | 144494999 | 75 | SPRET | 6 | 144494902 | 144494936 | 35 | + | 19.2569 | 1.39E-07 | 0.0681 | CTGAAATGAACTCCTCCCACCAGATGGGGGCAGTC | 1 | ENCFF093ZRU |
| 970 | 6 | 145317858 | 145318013 | 155 | CAST,MOLF,SPRET | 6 | 145317936 | 145317969 | 34 | + | 16.7154 | 8.12E-07 | 0.105 | GAGAACTGACTACAGCTACCAGCAGAGGCCAGCA | 8 | ENCFF024JOZ,ENCFF907XEG,ENCFF198SSR,ENCFF883UPM,ENCFF609MDN,ENCFF367UNR,ENCFF093ZRU,ENCFF327LYS |
| 971 | 6 | 146718506 | 146718955 | 449 | 129S1,AJ,BALB,BTBR,C3H,C57L,CAST,CBA,CE,DBA1J,DBA2J,FVB,ILNJ2,MA/My,MOLF,NOD,NOR,PJ,RF,SEA,SMJ,SPRET,SWR,TH,WSB | 6 | 146718523 | 146718556 | 34 | + | 17.9106 | 3.74E-07 | 0.0672 | CCGCCCTCACATTCGCCGTCACAAGATGGCGCTG | | |
| 972 | 6 | 148813106 | 148813304 | 198 | CAST,MOLF | 6 | 148813162 | 148813195 | 34 | + | 16.4472 | 9.61E-07 | 0.112 | TAGCTGTCTTCAGACACACCAGAAGAGGGCGTCA | | |
| 973 | 6 | 149026002 | 149026650 | 648 | SPRET | 6 | 149026648 | 149026682 | 35 | + | 17.2294 | 4.74E-07 | 0.135 | CTGCATGTACACCTGCATGCCAGAAGAAGACATCA | | |
| 974 | 6 | 3116181 | 3116513 | 332 | SPRET | 6 | 3116473 | 3116491 | 19 | - | 18.6557 | 2.99E-07 | 0.0973 | CCGCCACTAGATGGTGGTA | | |
| 975 | 6 | 10100356 | 10100639 | 283 | SPRET | 6 | 10100561 | 10100594 | 34 | - | 17.878 | 3.83E-07 | 0.0681 | CAGCTGTCCTCAGACATTCCAGAAGAGGGAGTCA | | |
| 976 | 6 | 14641164 | 14642109 | 945 | SPRET | 6 | 14641716 | 14641734 | 19 | - | 17.5902 | 6.18E-07 | 0.138 | ACACCAGAAGAGGGCACTG | | |
| 976 | 6 | 14641164 | 14642109 | 945 | SPRET | 6 | 14641716 | 14641750 | 35 | - | 16.3211 | 7.98E-07 | 0.171 | CTACATGTACTTTTGCACACCAGAAGAGGGCACTG | | |
| 977 | 6 | 15883679 | 15883893 | 214 | 129S1,C57L,CAST,CE,FVB,ILNJ2,KK,MOLF,SMJ,SPRET,SWR,TH,WSB | 6 | 15883804 | 15883822 | 19 | - | 17.5574 | 6.32E-07 | 0.138 | ACTCCAGAAGAGGGCGCCA | | |
| 977 | 6 | 15883679 | 15883893 | 214 | 129S1,C57L,CAST,CE,FVB,ILNJ2,KK,MOLF,SMJ,SPRET,SWR,TH,WSB | 6 | 15883804 | 15883837 | 34 | - | 18.3659 | 2.76E-07 | 0.0572 | TAGCTGTCTTCAGACACTCCAGAAGAGGGCGCCA | | |
| 978 | 6 | 22275614 | 22275749 | 135 | BUB,MOLF,SJL,SMJ,SPRET,SWR | 6 | 22275726 | 22275759 | 34 | - | 16.4472 | 9.61E-07 | 0.112 | TAGCTGTCTTCAGACACACCAGAAGAGGGCGTCA | | |
| 979 | 6 | 24911448 | 24911639 | 191 | CAST,MOLF,SPRET | 6 | 24911564 | 24911598 | 35 | - | 17.1284 | 5.03E-07 | 0.139 | TAGCTGTCTTCAGCCCGCACCAGAAGAGGGCATCA | | |
| 980 | 6 | 28412800 | 28413000 | 200 | CAST,SPRET | 6 | 28412916 | 28412934 | 19 | - | 16.9016 | 9.66E-07 | 0.17 | ATTCCAGAAGAGGGCACCA | 1 | ENCFF093ZRU |
| 980 | 6 | 28412800 | 28413000 | 200 | CAST,SPRET | 6 | 28412916 | 28412949 | 34 | - | 17.748 | 4.17E-07 | 0.0716 | TAGCTGTCTTCAAACATTCCAGAAGAGGGCACCA | 1 | ENCFF093ZRU |
| 981 | 6 | 29840873 | 29841076 | 203 | SPRET | 6 | 29841001 | 29841035 | 35 | - | 16.8624 | 5.86E-07 | 0.15 | GTGTACTCCAGTAGACACTCCAGAAGAGGGCATCA | | |
| 982 | 6 | 29956156 | 29956369 | 213 | 129S1,AJ,AKR,BALB,BTBR,BUB,C3H,C57L,CAST,CBA,DBA1J,DBA2J,FVB,MOLF,MRL,NOD,NOR,NZO,PJ,PL,RF,RH2,SEA,SMJ,SPRET,WSB | 6 | 29956276 | 29956309 | 34 | - | 17.2439 | 5.79E-07 | 0.086 | TAGCTGTCCTCAGATACTCCAGAAGAGGGCATCA | | |
| 983 | 6 | 35124393 | 35124604 | 211 | CAST,MOLF,SPRET | 6 | 35124522 | 35124555 | 34 | - | 16.4634 | 9.51E-07 | 0.112 | TAGCTGTCTTCAGACGCACCAGAAGAGGGCATCA | | |
| 984 | 6 | 37826077 | 37826280 | 203 | CAST,MOLF,SPRET | 6 | 37826200 | 37826218 | 19 | - | 17.5574 | 6.32E-07 | 0.138 | ACTCCAGAAGAGGGCGCCA | | |
| 984 | 6 | 37826077 | 37826280 | 203 | CAST,MOLF,SPRET | 6 | 37826200 | 37826233 | 34 | - | 18.3659 | 2.76E-07 | 0.0572 | TAGCTGTCTTCAGACACTCCAGAAGAGGGCGCCA | | |
| 985 | 6 | 38034690 | 38034890 | 200 | BUB,C3H,CAST,CBA,CE,DBA1J,DBA2J,FVB,ILNJ2,MOLF,MRL,NOD,NOR,PJ,PL,SEA,SJL,SMJ,SPRET,SWR,TH | 6 | 38034804 | 38034822 | 19 | - | 17.5738 | 6.25E-07 | 0.138 | ACTCCAGAAGAGGGCGCTA | | |
| 985 | 6 | 38034690 | 38034890 | 200 | BUB,C3H,CAST,CBA,CE,DBA1J,DBA2J,FVB,ILNJ2,MOLF,MRL,NOD,NOR,PJ,PL,SEA,SJL,SMJ,SPRET,SWR,TH | 6 | 38034804 | 38034837 | 34 | - | 18.187 | 3.12E-07 | 0.0608 | TAGCTGTCTTCAGTCACTCCAGAAGAGGGCGCTA | | |
| 986 | 6 | 38193518 | 38193774 | 256 | BUB,C3H,CAST,CBA,CE,DBA1J,DBA2J,FVB,ILNJ2,KK,MOLF,MRL,NOD,NOR,PJ,PL,SEA,SJL,SMJ,SPRET,SWR,TH,WSB | 6 | 38193684 | 38193702 | 19 | - | 17.5574 | 6.32E-07 | 0.138 | ACTCCAGAAGAGGGCGCCA | | |
| 986 | 6 | 38193518 | 38193774 | 256 | BUB,C3H,CAST,CBA,CE,DBA1J,DBA2J,FVB,ILNJ2,KK,MOLF,MRL,NOD,NOR,PJ,PL,SEA,SJL,SMJ,SPRET,SWR,TH,WSB | 6 | 38193684 | 38193717 | 34 | - | 18.3659 | 2.76E-07 | 0.0572 | TAGCTGTCTTCAGACACTCCAGAAGAGGGCGCCA | | |
| 987 | 6 | 39368744 | 39368954 | 210 | KK,MOLF,SPRET | 6 | 39368873 | 39368891 | 19 | - | 17.5574 | 6.32E-07 | 0.138 | ACTCCAGAAGAGGGCGCCA | 2 | ENCFF367UNR,ENCFF093ZRU |
| 987 | 6 | 39368744 | 39368954 | 210 | KK,MOLF,SPRET | 6 | 39368873 | 39368906 | 34 | - | 18.3659 | 2.76E-07 | 0.0572 | TAGCTGTCTTCAGACACTCCAGAAGAGGGCGCCA | 2 | ENCFF367UNR,ENCFF093ZRU |
| 988 | 6 | 46329520 | 46329705 | 185 | CAST,MOLF,SPRET | 6 | 46329633 | 46329666 | 34 | - | 17.9919 | 3.55E-07 | 0.0653 | TTGCTGTCTTCAGACACTCCAGAAGAGGGCATCA | | |
| 989 | 6 | 47938094 | 47938950 | 856 | CAST | 6 | 47938371 | 47938389 | 19 | - | 18.4426 | 3.47E-07 | 0.105 | CCGCCTCCAGAGGGAGCCA | 4 | ENCFF093ZRU,ENCFF327LYS,ENCFF055HAP,ENCFF502WDT |
| 990 | 6 | 48727821 | 48728448 | 627 | CAST,CE,ILNJ2,SPRET | 6 | 48728348 | 48728381 | 34 | - | 16.6504 | 8.46E-07 | 0.107 | TAGCTGTCTTCAGATGCACCAGAAGAGGGCGACA | | |
| 991 | 6 | 50503816 | 50503938 | 122 | MOLF | 6 | 50503861 | 50503879 | 19 | - | 17.541 | 6.39E-07 | 0.139 | CTACCACAAGGTGTCGCTG | 34 | ENCFF093ZRU,ENCFF367UNR,ENCFF099QNE,ENCFF201JUA,ENCFF327LYS,ENCFF907XEG,ENCFF821BIA,ENCFF024JOZ,ENCFF089GBC,ENCFF504JDS,ENCFF065ODV,ENCFF055HAP,ENCFF589ZBJ,ENCFF430PPJ,ENCFF092NMJ,ENCFF165LXO,ENCFF321AGQ,ENCFF052KVO,ENCFF198SSR,ENCFF502WDT,ENCFF136ZDK,ENCFF883UPM,ENCFF033UQX,ENCFF609MDN,ENCFF097CAQ,ENCFF951GOP,ENCFF142CNG,ENCFF601JVE,ENCFF281QMV,ENCFF491RJK,ENCFF575MGI,ENCFF957SPH,ENCFF194HVL,ENCFF316FYN |
| 991 | 6 | 50503816 | 50503938 | 122 | MOLF | 6 | 50503861 | 50503894 | 34 | - | 22.0407 | 1.93E-08 | 0.00917 | GTGCCATTCACCTCACTACCACAAGGTGTCGCTG | 34 | ENCFF093ZRU,ENCFF367UNR,ENCFF099QNE,ENCFF201JUA,ENCFF327LYS,ENCFF907XEG,ENCFF821BIA,ENCFF024JOZ,ENCFF089GBC,ENCFF504JDS,ENCFF065ODV,ENCFF055HAP,ENCFF589ZBJ,ENCFF430PPJ,ENCFF092NMJ,ENCFF165LXO,ENCFF321AGQ,ENCFF052KVO,ENCFF198SSR,ENCFF502WDT,ENCFF136ZDK,ENCFF883UPM,ENCFF033UQX,ENCFF609MDN,ENCFF097CAQ,ENCFF951GOP,ENCFF142CNG,ENCFF601JVE,ENCFF281QMV,ENCFF491RJK,ENCFF575MGI,ENCFF957SPH,ENCFF194HVL,ENCFF316FYN |
| 992 | 6 | 51481038 | 51481234 | 196 | AKR,BALB,C58J,CAST,LP,MA/My,MOLF,NOD,NOR,PJ,RF,RH2,SEA,SPRET | 6 | 51481149 | 51481182 | 34 | - | 16.7724 | 7.83E-07 | 0.103 | TAGCTGTCTTCAGATGCACCAGAAGAGGGCGTCA | | |
| 993 | 6 | 54534618 | 54535334 | 716 | SPRET | 6 | 54534908 | 54534941 | 34 | - | 21.5203 | 2.89E-08 | 0.0123 | GTGCAGTGCCATCAGAAACCAGAGAAGGGCACTG | | |
| 994 | 6 | 56660466 | 56660661 | 195 | 129S1,C57L,CAST,KK,MOLF,NZB,NZO,NZW | 6 | 56660585 | 56660618 | 34 | - | 16.4472 | 9.61E-07 | 0.112 | TAGCTGTCTTCAGACACACCAGAAGAGGGCGTCA | | |
| 995 | 6 | 64977874 | 64978036 | 162 | CAST,SPRET | 6 | 64977954 | 64977987 | 34 | - | 16.4146 | 9.81E-07 | 0.113 | TAGCTGTCTTCAGACACACCAGAGGAGGGCGCCA | | |
| 996 | 6 | 65002542 | 65002756 | 214 | SPRET | 6 | 65002666 | 65002699 | 34 | - | 16.4634 | 9.51E-07 | 0.112 | TAGCTGTCTTCAGACGCACCAGAAGAGGGCATCA | | |
| 997 | 6 | 66940699 | 66940952 | 253 | SPRET | 6 | 66940821 | 66940839 | 19 | - | 17.5574 | 6.32E-07 | 0.138 | ACTCCAGAAGAGGGCGCCA | 8 | ENCFF024JOZ,ENCFF099QNE,ENCFF136ZDK,ENCFF821BIA,ENCFF201JUA,ENCFF055HAP,ENCFF367UNR,ENCFF093ZRU |
| 997 | 6 | 66940699 | 66940952 | 253 | SPRET | 6 | 66940821 | 66940854 | 34 | - | 18.3659 | 2.76E-07 | 0.0572 | TAGCTGTCTTCAGACACTCCAGAAGAGGGCGCCA | 8 | ENCFF024JOZ,ENCFF099QNE,ENCFF136ZDK,ENCFF821BIA,ENCFF201JUA,ENCFF055HAP,ENCFF367UNR,ENCFF093ZRU |
| 998 | 6 | 67198988 | 67199206 | 218 | C58J,CE,ILNJ2,KK,MOLF,NOD,NOR,NZB,NZO,PJ,SPRET | 6 | 67199114 | 67199147 | 34 | - | 17.3902 | 5.27E-07 | 0.0822 | TAGCTGTCCTCAGACACTCCAGAAGAGGGCATCA | | |
| 999 | 6 | 72373495 | 72373690 | 195 | AKR,C3H,C58J,CAST,CBA,DBA1J,DBA2J,KK,MA/My,MOLF,MRL,NOD,NOR,PL,RF,SMJ,SPRET,WSB | 6 | 72373607 | 72373641 | 35 | - | 16.0917 | 9.07E-07 | 0.184 | TAGCTGTCTTCAGACAGCACCAGAAGAGGGCATCA | | |
| 1000 | 6 | 72589221 | 72589438 | 217 | C3H,C58J,CAST,CBA,CE,DBA1J,DBA2J,ILNJ2,KK,MA/My,MOLF,NOD,NOR,PJ,SMJ,SPRET,WSB | 6 | 72589339 | 72589372 | 34 | - | 17.8455 | 3.91E-07 | 0.0687 | TAGCTGTCCTCAGACACTCCAGAAGAGGGCGTCA | | |
| 1001 | 6 | 72768765 | 72769172 | 407 | AKR,BALB,C3H,C58J,CAST,CBA,DBA1J,DBA2J,KK,MA/My,MOLF,MRL,NOD,NOR,PL,RF,RH2,SEA,SMJ,SPRET | 6 | 72769092 | 72769125 | 34 | - | 16.4472 | 9.61E-07 | 0.112 | TAGCTGTCTTCAGACACACCAGAAGAGGGCGTCA | | |
| 1002 | 6 | 76595416 | 76595610 | 194 | CAST,KK,MOLF,PJ,SPRET | 6 | 76595539 | 76595572 | 34 | - | 16.5447 | 9.04E-07 | 0.111 | CAGCTGTCTTCAGACACACCAGAAGAGGGAGTCA | | |
| 1003 | 6 | 81414749 | 81414955 | 206 | SPRET | 6 | 81414861 | 81414895 | 35 | - | 16.0917 | 9.07E-07 | 0.184 | TAGCTGTCTTCAGACAGCACCAGAAGAGGGCATCA | | |
| 1004 | 6 | 85503754 | 85503953 | 199 | PJ,SEA,SPRET | 6 | 85503868 | 85503901 | 34 | - | 16.4472 | 9.61E-07 | 0.112 | TAGCTGTCTTCAGACACACCAGAAGAGGGCGTCA | | |
| 1005 | 6 | 86868310 | 86869155 | 845 | SPRET | 6 | 86868698 | 86868716 | 19 | - | 17.8852 | 5.08E-07 | 0.128 | ACACCAGAAGAGGGCACCA | | |
| 1006 | 6 | 87043314 | 87043528 | 214 | 129S1,AJ,BTBR,C3H,C57L,C58J,CAST,CBA,DBA1J,DBA2J,FVB,MOLF,NOD,NOR,PJ,RF,SEA,SJL,SMJ,SPRET | 6 | 87043438 | 87043471 | 34 | - | 17.8455 | 3.91E-07 | 0.0687 | TAGCTGTCCTCAGACACTCCAGAAGAGGGCGTCA | | |
| 1007 | 6 | 89350461 | 89350667 | 206 | 129S1,C3H,C57L,C58J,CAST,CBA,CE,DBA1J,DBA2J,FVB,KK,LP,MA/My,MOLF,NOD,NOR,NZB,NZO,NZW,PJ,SEA,SMJ,SPRET,SWR,TH,WSB | 6 | 89350587 | 89350620 | 34 | - | 16.4634 | 9.51E-07 | 0.112 | TAGCTGTCTTCAGACGCACCAGAAGAGGGCATCA | | |
| 1008 | 6 | 89635214 | 89635548 | 334 | CAST,SPRET | 6 | 89635303 | 89635336 | 34 | - | 16.7886 | 7.75E-07 | 0.103 | CTGGAGCACTGTAGCCATGCACTAGGTGGCTCTG | | |
| 1008 | 6 | 89635214 | 89635548 | 334 | CAST,SPRET | 6 | 89635334 | 89635367 | 34 | - | 16.7886 | 7.75E-07 | 0.103 | CTGGAGCACTGTAGCCATGCACTAGGTGGCTCTG | 1 | ENCFF093ZRU |
| 1008 | 6 | 89635214 | 89635548 | 334 | CAST,SPRET | 6 | 89635457 | 89635490 | 34 | - | 16.7886 | 7.75E-07 | 0.103 | CTGGAGCACTGTAGCCATGCACTAGGTGGCTCTG | 6 | ENCFF093ZRU,ENCFF089GBC,ENCFF092NMJ,ENCFF367UNR,ENCFF883UPM,ENCFF907XEG |
| 1008 | 6 | 89635214 | 89635548 | 334 | CAST,SPRET | 6 | 89635488 | 89635521 | 34 | - | 16.5691 | 8.90E-07 | 0.11 | CTGGAGCACTGTGACCATTCACTAGGTGGCTCTG | 6 | ENCFF093ZRU,ENCFF089GBC,ENCFF092NMJ,ENCFF367UNR,ENCFF883UPM,ENCFF907XEG |
| 1009 | 6 | 95306402 | 95307243 | 841 | C58J,DBA1J,DBA2J,LP,MA/My | 6 | 95306799 | 95306832 | 34 | - | 18.3171 | 2.86E-07 | 0.0578 | TAGCTGTCTTCAGACCCCCCAGTAGAGGGCGTCA | | |
| 1010 | 6 | 97690337 | 97690536 | 199 | SPRET | 6 | 97690459 | 97690492 | 34 | - | 18.0976 | 3.31E-07 | 0.0631 | TAGCTGTCTCCAGACGCACCAGAAGAGGGCATCA | | |
| 1011 | 6 | 98733753 | 98733941 | 188 | CAST,MOLF,SJL,SPRET | 6 | 98733863 | 98733896 | 34 | - | 16.4472 | 9.61E-07 | 0.112 | TAGCTGTCTTCAGACACACCAGAAGAGGGCGTCA | | |
| 1012 | 6 | 99966379 | 99966581 | 202 | SPRET | 6 | 99966490 | 99966523 | 34 | - | 17.252 | 5.76E-07 | 0.086 | TAGCTGTCTTCCGACAGTCCAGAAGAGGGAGCCA | 1 | ENCFF093ZRU |
| 1013 | 6 | 100212016 | 100212269 | 253 | SPRET | 6 | 100211997 | 100212031 | 35 | - | 19.0642 | 1.57E-07 | 0.0734 | CTGAAGTTCCAGCATGAGGCCACTTGGGTGCCCTT | 1 | ENCFF093ZRU |
| 1014 | 6 | 100326492 | 100326709 | 217 | BUB,CAST,MOLF,NZB,NZO,NZW,SPRET,WSB | 6 | 100326639 | 100326672 | 34 | - | 16.7724 | 7.83E-07 | 0.103 | TAGCTGTCTTCAGATGCACCAGAAGAGGGCGTCA | | |
| 1015 | 6 | 102123118 | 102123562 | 444 | MOLF | 6 | 102123456 | 102123490 | 35 | - | 18.9817 | 1.66E-07 | 0.075 | CTGCATCTGACTGTTGAGTCCTCCAGAAGGCAGTC | | |
| 1016 | 6 | 113313661 | 113313897 | 236 | AKR,BTBR,C3H,C58J,CAST,CBA,CE,DBA1J,DBA2J,FVB,ILNJ2,LP,MOLF,MRL,NOD,NOR,NZB,NZO,NZW,PJ,PL,RF,SJL,SMJ,SPRET,SWR,TH,WSB | 6 | 113313789 | 113313822 | 34 | - | 16.4634 | 9.51E-07 | 0.112 | TAGCTGTCTTCAGACGCACCAGAAGAGGGCATCA | | |
| 1017 | 6 | 113588137 | 113588575 | 438 | CAST,MOLF | 6 | 113588153 | 113588186 | 34 | - | 17.3659 | 5.35E-07 | 0.0827 | TTGCTGTCCTCAGACACACCAGAAGAAGGCATCG | | |
| 1018 | 6 | 114652335 | 114652396 | 61 | NZB,NZW | 6 | 114652367 | 114652401 | 35 | - | 16.7064 | 6.41E-07 | 0.153 | ATACAATTCCTTGTCCAGGCCAGAGGAAGGCACTG | | |
| 1019 | 6 | 114652512 | 114653000 | 488 | 129S1,AKR,BTBR,C3H,C57L,C58J,CAST,CBA,CE,DBA1J,DBA2J,FVB,ILNJ2,LP,MOLF,MRL,NOD,NOR,NZO,PJ,RF,SJL,SPRET,SWR,TH,WSB | 6 | 114652489 | 114652523 | 35 | - | 16.7064 | 6.41E-07 | 0.153 | ATACAATTCCTTGTCCAGGCCAGAGGAAGGCACTG | | |
| 1019 | 6 | 114652512 | 114653000 | 488 | 129S1,AKR,BTBR,C3H,C57L,C58J,CAST,CBA,CE,DBA1J,DBA2J,FVB,ILNJ2,LP,MOLF,MRL,NOD,NOR,NZO,PJ,RF,SJL,SPRET,SWR,TH,WSB | 6 | 114652550 | 114652584 | 35 | - | 18.0367 | 2.94E-07 | 0.105 | ATACAATTCCTCGTCCAGGCCAGAGGAAGGCACTG | | |
| 1019 | 6 | 114652512 | 114653000 | 488 | 129S1,AKR,BTBR,C3H,C57L,C58J,CAST,CBA,CE,DBA1J,DBA2J,FVB,ILNJ2,LP,MOLF,MRL,NOD,NOR,NZO,PJ,RF,SJL,SPRET,SWR,TH,WSB | 6 | 114652611 | 114652645 | 35 | - | 16.7064 | 6.41E-07 | 0.153 | ATACAATTCCTTGTCCAGGCCAGAGGAAGGCACTG | | |
| 1019 | 6 | 114652512 | 114653000 | 488 | 129S1,AKR,BTBR,C3H,C57L,C58J,CAST,CBA,CE,DBA1J,DBA2J,FVB,ILNJ2,LP,MOLF,MRL,NOD,NOR,NZO,PJ,RF,SJL,SPRET,SWR,TH,WSB | 6 | 114652672 | 114652706 | 35 | - | 16.7064 | 6.41E-07 | 0.153 | ATACAATTCCTTGTCCAGGCCAGAGGAAGGCACTG | | |
| 1019 | 6 | 114652512 | 114653000 | 488 | 129S1,AKR,BTBR,C3H,C57L,C58J,CAST,CBA,CE,DBA1J,DBA2J,FVB,ILNJ2,LP,MOLF,MRL,NOD,NOR,NZO,PJ,RF,SJL,SPRET,SWR,TH,WSB | 6 | 114652733 | 114652767 | 35 | - | 17.2936 | 4.57E-07 | 0.132 | ATACAATTCCTCTTCCAGGCCAGAGGAAGGCACTG | | |
| 1019 | 6 | 114652512 | 114653000 | 488 | 129S1,AKR,BTBR,C3H,C57L,C58J,CAST,CBA,CE,DBA1J,DBA2J,FVB,ILNJ2,LP,MOLF,MRL,NOD,NOR,NZO,PJ,RF,SJL,SPRET,SWR.TH.WSB | 6 | 114652794 | 114652828 | 35 | - | 16.7064 | 6.41E-07 | 0.153 | ATACAATTCCTTGTCCAGGCCAGAGGAAGGCACTG | | |
| 1019 | 6 | 114652512 | 114653000 | 488 | 129S1,AKR,BTBR,C3H,C57L,C58J,CAST,CBA,CE,DBA1J,DBA2J,FVB,ILNJ2,LP,MOLF,MRL,NOD,NOR,NZO,PJ,RF,SJL,SPRET,SWR,TH,WSB | 6 | 114652855 | 114652889 | 35 | - | 16.7064 | 6.41E-07 | 0.153 | ATACAATTCCTTGTCCAGGCCAGAGGAAGGCACTG | | |
| 1019 | 6 | 114652512 | 114653000 | 488 | 129S1,AKR,BTBR,C3H,C57L,C58J,CAST,CBA,CE,DBA1J,DBA2J,FVB,ILNJ2,LP,MOLF,MRL,NOD,NOR,NZO,PJ,RF,SJL,SPRET,SWR,TH,WSB | 6 | 114652916 | 114652950 | 35 | - | 16.7064 | 6.41E-07 | 0.153 | ATACAATTCCTTGTCCAGGCCAGAGGAAGGCACTG | | |
| 1019 | 6 | 114652512 | 114653000 | 488 | 129S1,AKR,BTBR,C3H,C57L,C58J,CAST,CBA,CE,DBA1J,DBA2J,FVB,ILNJ2,LP,MOLF,MRL,NOD,NOR,NZO,PJ,RF,SJL,SPRET,SWR,TH,WSB | 6 | 114652977 | 114653011 | 35 | - | 17.2936 | 4.57E-07 | 0.132 | ATACAATTCCTCTTCCAGGCCAGAGGAAGGCACTG | | |
| 1020 | 6 | 116145166 | 116145370 | 204 | SPRET | 6 | 116145293 | 116145326 | 34 | - | 18.1707 | 3.15E-07 | 0.0613 | TAGCTGTTTTCAGACGCACCAGAAGAGGGCATCA | | |
| 1021 | 6 | 116483185 | 116484001 | 816 | SPRET | 6 | 116483292 | 116483325 | 34 | - | 16.6179 | 8.63E-07 | 0.108 | CAGCCATGGCCTGGGCTTCCAGTGGAGAGCGCTG | | |
| 1022 | 6 | 119858448 | 119858642 | 194 | SPRET | 6 | 119858572 | 119858605 | 34 | - | 16.4472 | 9.61E-07 | 0.112 | TAGCTGTCTTCAGACACACCAGAAGAGGGCGTCA | 2 | ENCFF093ZRU,ENCFF367UNR |
| 1023 | 6 | 120586955 | 120587268 | 313 | CAST,MOLF,SPRET | 6 | 120586968 | 120587001 | 34 | - | 19.6911 | 1.11E-07 | 0.0333 | TTGCTGTTGTCAGACACACCAGAAGAGGGCATCA | 1 | ENCFF093ZRU |
| 1024 | 6 | 124500566 | 124500815 | 249 | MOLF,SPRET | 6 | 124500681 | 124500714 | 34 | - | 17.2439 | 5.79E-07 | 0.086 | TAGCTGTCCTCAGATACTCCAGAAGAGGGCATCA | | |
| 1025 | 6 | 124762422 | 124762624 | 202 | MOLF,SPRET | 6 | 124762548 | 124762581 | 34 | - | 16.4472 | 9.61E-07 | 0.112 | TAGCTGTCTTCAGACACACCAGAAGAGGGCGTCA | | |
| 1026 | 6 | 124765132 | 124766018 | 886 | MOLF | 6 | 124765959 | 124765992 | 34 | - | 16.7886 | 7.75E-07 | 0.103 | CTGCAGCTGTCTTCACACACTGCAGAAGGCACAA | | |
| 1027 | 6 | 125275304 | 125275691 | 387 | AJ,AKR,BALB,C3H,C57L,C58J,CAST,CBA,DBA1J,DBA2J,FVB,MA/My,MOLF,MRL,NOD,NOR,NZW,PJ,RF,SEA,SJL,SPRET,TH | 6 | 125275615 | 125275648 | 34 | - | 17.9187 | 3.72E-07 | 0.0672 | TAGCTGTCCTCACATACTCCAGAAGAGGGCATCA | | |
| 1028 | 6 | 125433523 | 125433722 | 199 | MOLF,SPRET | 6 | 125433648 | 125433681 | 34 | - | 16.4472 | 9.61E-07 | 0.112 | TAGCTGTCTTCAGACACACCAGAAGAGGGCGTCA | 1 | ENCFF093ZRU |
| 1029 | 6 | 125587890 | 125588113 | 223 | MOLF | 6 | 125587887 | 125587905 | 19 | - | 17.082 | 8.61E-07 | 0.161 | ACACCACCAGGGGGAGCGG | 3 | ENCFF367UNR,ENCFF099QNE,ENCFF093ZRU |
| 1030 | 6 | 126875047 | 126875258 | 211 | SPRET | 6 | 126875159 | 126875192 | 34 | - | 18.5203 | 2.49E-07 | 0.0559 | CAGCTCTCTTCAGATGTGCCAGAAGAGGGCATCA | | |
| 1031 | 6 | 128063144 | 128063284 | 140 | CAST,MOLF,SPRET | 6 | 128063180 | 128063214 | 35 | - | 16.5963 | 6.83E-07 | 0.158 | CTGCAGGGGCCCAGCCCTGGCTGCAGGGGTCCATC | | |
| 1031 | 6 | 128063144 | 128063284 | 140 | CAST,MOLF,SPRET | 6 | 128063280 | 128063314 | 35 | - | 16.5963 | 6.83E-07 | 0.158 | CTGCAGGGGCCCAGCCCTGGCTGCAGGGGTCCATC | | |
| 1032 | 6 | 133183035 | 133183380 | 345 | NOD,NOR,NZO,SWR | 6 | 133183338 | 133183356 | 19 | - | 19.2951 | 1.88E-07 | 0.0786 | CCGCCACTAGATGGTGCTA | 30 | ENCFF805QBU,ENCFF367UNR,ENCFF093ZRU,ENCFF099QNE,ENCFF327LYS,ENCFF316FYN,ENCFF089GBC,ENCFF821BIA,ENCFF907XEG,ENCFF201JUA,ENCFF024JOZ,ENCFF194HVL,ENCFF092NMJ,ENCFF589ZBJ,ENCFF430PPJ,ENCFF491RJK,ENCFF198SSR,ENCFF136ZDK,ENCFF957SPH,ENCFF321AGQ,ENCFF165LXO,ENCFF055HAP,ENCFF210BTO,ENCFF951GOP,ENCFF502WDT,ENCFF281QMV,ENCFF883UPM,ENCFF052KVO,ENCFF575MGI,ENCFF609MDN |
| 1033 | 6 | 134257767 | 134257821 | 54 | CAST,CBA,NZB,NZO,SPRET,WSB | 6 | 134257746 | 134257779 | 34 | - | 17.1463 | 6.17E-07 | 0.0894 | CAGCAGCAGCAGCAGCAGCAGCTTGGGGGAGCCA | | |
| 1034 | 6 | 137640390 | 137640581 | 191 | 129S1,AJ,AKR,BTBR,BUB,C3H,C57L,CAST,CBA,CE,DBA1J,DBA2J,FVB,ILNJ2,MA/My,NOD,NOR,NZO,NZW,PL,RF,SJL,SPRET,SWR,TH,WSB | 6 | 137640510 | 137640528 | 19 | - | 17.5574 | 6.32E-07 | 0.138 | ACTCCAGAAGAGGGCGCCA | 2 | ENCFF367UNR,ENCFF093ZRU |
| 1034 | 6 | 137640390 | 137640581 | 191 | 129S1,AJ,AKR,BTBR,BUB,C3H,C57L,CAST,CBA,CE,DBA1J,DBA2J,FVB,ILNJ2,MA/My,NOD,NOR,NZO,NZW,PL,RF,SJL,SPRET,SWR,TH,WSB | 6 | 137640510 | 137640543 | 34 | - | 18.561 | 2.42E-07 | 0.0549 | AAGCTGTCTTCAGACACTCCAGAAGAGGGCGCCA | 2 | ENCFF367UNR,ENCFF093ZRU |
| 1035 | 6 | 141282831 | 141283746 | 915 | BTBR,C57L,C58J,FVB,NOD,NOR,PJ,SMJ,SWR,WSB | 6 | 141283635 | 141283669 | 35 | - | 21.2202 | 3.89E-08 | 0.0311 | ATGCAGAGGCCCACAGCGAACAGTAGGGGGAGCTC | 23 | ENCFF093ZRU,ENCFF092NMJ,ENCFF165LXO,ENCFF367UNR,ENCFF210BTO,ENCFF089GBC,ENCFF327LYS,ENCFF099QNE,ENCFF609MDN,ENCFF033UQX,ENCFF883UPM,ENCFF052KVO,ENCFF589ZBJ,ENCFF198SSR,ENCFF907XEG,ENCFF281QMV,ENCFF024JOZ,ENCFF194HVL,ENCFF136ZDK,ENCFF957SPH,ENCFF533APC,ENCFF491RJK,ENCFF821BIA |
| 1036 | 6 | 142460856 | 142461291 | 435 | BTBR,C57L,C58J,FVB,KK,MA/My,MOLF,NOD,NOR,NZB,PJ,SMJ,SPRET,SWR,WSB | 6 | 142460869 | 142460887 | 19 | - | 17.5574 | 6.32E-07 | 0.138 | TGGCCAGTTGGGGGCGCGC | 20 | ENCFF093ZRU,ENCFF099QNE,ENCFF367UNR,ENCFF136ZDK,ENCFF430PPJ,ENCFF165LXO,ENCFF883UPM,ENCFF089GBC,ENCFF907XEG,ENCFF609MDN,ENCFF821BIA,ENCFF327LYS,ENCFF533APC,ENCFF198SSR,ENCFF024JOZ,ENCFF589ZBJ,ENCFF957SPH,ENCFF092NMJ,ENCFF281QMV,ENCFF194HVL |
| 1037 | 6 | 145045771 | 145045958 | 187 | CAST,CBA,ILNJ2,MA/My,NOD,NOR,PJ,SPRET,TH | 6 | 145045887 | 145045920 | 34 | - | 16.4634 | 9.51E-07 | 0.112 | TAGCTGTCTTCAGACGCACCAGAAGAGGGCATCA | | |
| 1038 | 6 | 146158464 | 146158936 | 472 | CAST,SPRET | 6 | 146158652 | 146158685 | 34 | - | 21.122 | 3.90E-08 | 0.0153 | TTGCAATACCCACATTGCCCTGGAGGGGGGGGCC | 15 | ENCFF089GBC,ENCFF024JOZ,ENCFF198SSR,ENCFF165LXO,ENCFF883UPM,ENCFF327LYS,ENCFF367UNR,ENCFF093ZRU,ENCFF907XEG,ENCFF136ZDK,ENCFF589ZBJ,ENCFF099QNE,ENCFF502WDT,ENCFF052KVO,ENCFF055HAP |
| 1039 | 6 | 146440672 | 146440871 | 199 | SPRET | 6 | 146440790 | 146440823 | 34 | - | 16.4472 | 9.61E-07 | 0.112 | TAGCTGTCTTCAGACACACCAGAAGAGGGCGTCA | 1 | ENCFF093ZRU |
| 1040 | 6 | 147084772 | 147085153 | 381 | 129S1,AJ,BALB,BTBR,C57L,CBA,FVB,ILNJ2,MA/My,NOD,NOR,PJ,RF,SEA,SMJ,SPRET,SWR,TH,WSB | 6 | 147084969 | 147085003 | 35 | - | 21.1193 | 4.16E-08 | 0.0327 | CTGTATTGCCACCTCCTTCCCACAGGGTGGTATCC | | |
| 1041 | 6 | 148748713 | 148748926 | 213 | KK,PL,SPRET | 6 | 148748837 | 148748870 | 34 | - | 17.6667 | 4.40E-07 | 0.0735 | TAGCTGTTTTCAGACACTCCAGAAGAGGGCGTCA | | |
| 1042 | 6 | 148831212 | 148831399 | 187 | SPRET | 6 | 148831323 | 148831341 | 19 | - | 17.0328 | 8.88E-07 | 0.162 | AGACCAGAAGAGGGAGCCA | | |
| 1043 | 7 | 3223061 | 3223863 | 802 | CAST,MOLF | 7 | 3223422 | 3223455 | 34 | + | 18.3659 | 2.76E-07 | 0.0476 | TAGCTGTCTTCAGACACTCCAGAAGAGGGCGCCA | | |
| 1043 | 7 | 3223061 | 3223863 | 802 | CAST,MOLF | 7 | 3223437 | 3223455 | 19 | + | 17.5574 | 6.32E-07 | 0.106 | ACTCCAGAAGAGGGCGCCA | | |
| 1044 | 7 | 3314032 | 3314511 | 479 | CAST,MOLF | 7 | 3314231 | 3314265 | 35 | + | 17.5505 | 3.93E-07 | 0.0924 | GTGCTGGGTAGTGCAGCGGCCAGCAGATGGCCAGA | 6 | ENCFF821BIA,ENCFF327LYS,ENCFF907XEG,ENCFF367UNR,ENCFF194HVL,ENCFF198SSR |
| 1045 | 7 | 3455756 | 3455929 | 173 | AJ,AKR,KK,LP,MA/My,NOD,NOR,NZW,PL,RF,RH2,SJL,SWR,TH | 7 | 3455754 | 3455787 | 34 | + | 20.0976 | 8.28E-08 | 0.0223 | ATGCAGAGCTCATGATGACCAGAAGAGGGCCTCA | 3 | ENCFF099QNE,ENCFF092NMJ,ENCFF093ZRU |
| 1046 | 7 | 3595651 | 3595853 | 202 | MOLF | 7 | 3595712 | 3595745 | 34 | + | 16.4228 | 9.76E-07 | 0.0903 | TAGCTGTCTTCAGATGCACCAGAAGAGGGCATCG | | |

| | | | | | | | | | | | | | | | | |
|---|---|---|---|---|---|---|---|---|---|---|---|---|---|---|---|---|
| 1047 | 7 | 6243937 | 6244022 | 85 | SMJ | 7 | 6243997 | 6244031 | 35 | + | 21.6239 | 2.95E-08 | 0.0191 | TGGCTATTCTGGAAAATGTCCAGCAGGTGGCAGCA | 30 | ENCFF092NMJ,ENCFF430PPJ,ENCFF201JUA,ENCFF093ZRU,ENCFF367UNR,ENCFF951GOP,ENCFF609MDN,ENCFF024JOZ,ENCFF327LYS,ENCFF316FYN,ENCFF907XEG,ENCFF099QNE,ENCFF821BIA,ENCFF533APC,ENCFF089GBC,ENCFF589ZBJ,ENCFF491RJK,ENCFF194HVL,ENCFF055HAP,ENCFF097CAQ,ENCFF142CNG,ENCFF136ZDK,ENCFF198SSR,ENCFF321AGQ,ENCFF281QMV,ENCFF502WDT,ENCFF957SPH,ENCFF883UPM,ENCFF210BTO,ENCFF165LXO |
| 1047 | 7 | 6243937 | 6244022 | 85 | SMJ | 7 | 6244013 | 6244031 | 19 | + | 22.5902 | 1.16E-08 | 0.0201 | TGTCCAGCAGGTGGCAGCA | 30 | ENCFF092NMJ,ENCFF430PPJ,ENCFF201JUA,ENCFF093ZRU,ENCFF367UNR,ENCFF951GOP,ENCFF609MDN,ENCFF024JOZ,ENCFF327LYS,ENCFF316FYN,ENCFF907XEG,ENCFF099QNE,ENCFF821BIA,ENCFF533APC,ENCFF089GBC,ENCFF589ZBJ,ENCFF491RJK,ENCFF194HVL,ENCFF055HAP,ENCFF097CAQ,ENCFF142CNG,ENCFF136ZDK,ENCFF198SSR,ENCFF321AGQ,ENCFF281QMV,ENCFF502WDT,ENCFF957SPH,ENCFF883UPM,ENCFF210BTO,ENCFF165LXO |
| 1048 | 7 | 6387999 | 6388657 | 658 | SPRET | 7 | 6388595 | 6388629 | 35 | + | 16.4128 | 7.57E-07 | 0.133 | CTGCTCTTTTCTCAGAAGTCCACAAGGTGTCCTTG | | |
| 1049 | 7 | 12177107 | 12177326 | 219 | AKR,CAST,MOLF,MRL,RH2,SPRET,WSB | 7 | 12177184 | 12177217 | 34 | + | 17.2439 | 5.79E-07 | 0.0706 | TAGCTGTCCTCAGATACTCCAGAAGAGGGCATCA | | |
| 1050 | 7 | 16025876 | 16026120 | 244 | 129S1,AJ,AKR,BALB,BTBR,C3H,C57L,C58J,CBA,DBA1J,DBA2J,ILNJ2,KK,LP,MA/My,MOLF,MRL,NZB,NZO,NZW,PJ,RF,RH2,SEA,SJL,SMJ,SPRET,TH | 7 | 16025987 | 16026020 | 34 | + | 18.1545 | 3.18E-07 | 0.0508 | TAGCTGTTTTCAGACACACCAGAAGAGGGCGTCA | | |
| 1051 | 7 | 16216935 | 16217570 | 635 | SPRET | 7 | 16217565 | 16217598 | 34 | + | 17.8374 | 3.93E-07 | 0.0571 | TTGCTGTCTTCAGACACACCAGAAGATGGCATCG | | |
| 1052 | 7 | 16381827 | 16382040 | 213 | 129S1,BTBR,BUB,C3H,C57L,CAST,CBA,CE,DBA1J,DBA2J,FVB,MA/My,MOLF,NOD,NOR,NZB,PJ,RF,SEA,SMJ,SPRET,WSB | 7 | 16381892 | 16381925 | 34 | + | 16.9919 | 6.81E-07 | 0.0775 | TCGCTGTCCTCAGACACACCAGAAGAGGGCATCA | | |
| 1053 | 7 | 16687429 | 16687625 | 196 | 129S1,BTBR,C3H,C57L,CAST,CBA,FVB,MOLF,NZB,NZW,PL,RF,SPRET | 7 | 16687501 | 16687534 | 34 | + | 16.4634 | 9.51E-07 | 0.0901 | TAGCTGTCTTCAGACGCACCAGAAGAGGGCATCA | | |
| 1054 | 7 | 18923396 | 18923606 | 210 | 129S1,AJ,BALB,BTBR,BUB,C3H,C57L,CAST,CBA,CE,DBA1J,DBA2J,FVB,ILNJ2,KK,LP,MA/My,MOLF,MRL,NOD,NOR,NZB,NZW,PJ,PL,RF,SEA,SJL,SMJ,SPRET | 7 | 18923467 | 18923500 | 34 | + | 18.3659 | 2.76E-07 | 0.0476 | TAGCTGTCTTCAGACACTCCAGAAGAGGGCGCCA | | |
| 1054 | 7 | 18923396 | 18923606 | 210 | 129S1,AJ,BALB,BTBR,BUB,C3H,C57L,CAST,CBA,CE,DBA1J,DBA2J,FVB,ILNJ2,KK,LP,MA/My,MOLF,MRL,NOD,NOR,NZB,NZW,PJ,PL,RF,SEA,SJL,SMJ,SPRET | 7 | 18923482 | 18923500 | 19 | + | 17.5574 | 6.32E-07 | 0.106 | ACTCCAGAAGAGGGCGCCA | | |
| 1055 | 7 | 19032913 | 19033643 | 730 | 129S1,BTBR,C3H,C57L,CAST,CBA,MOLF,NOD,NOR,NZB,NZW,RF,SJL | 7 | 19032898 | 19032931 | 34 | + | 16.4797 | 9.42E-07 | 0.0901 | CTGCTATTGCCACTGCCCCCTCCATGAGGCACTT | | |
| 1056 | 7 | 24110910 | 24111126 | 216 | SPRET | 7 | 24110971 | 24111004 | 34 | + | 17.2439 | 5.79E-07 | 0.0706 | TAGCTGTCCTCAGATACTCCAGAAGAGGGCATCA | | |
| 1057 | 7 | 24215615 | 24216200 | 585 | SPRET | 7 | 24215910 | 24215943 | 34 | + | 20.2358 | 7.49E-08 | 0.0208 | GTGTAGTGACCAAGGAGGCCACGAGAGGGAGTCA | 18 | ENCFF024JOZ,ENCFF327LYS,ENCFF093ZRU,ENCFF821BIA,ENCFF589ZBJ,ENCFF883UPM,ENCFF099QNE,ENCFF502WDT,ENCFF957SPH,ENCFF089GBC,ENCFF052KVO,ENCFF533APC,ENCFF055HAP,ENCFF367UNR,ENCFF092NMJ,ENCFF142CNG,ENCFF430PPJ,ENCFF907XEG |
| 1058 | 7 | 24479573 | 24479778 | 205 | BUB,CAST,MOLF,NOD,NOR,SJL,SPRET,WSB | 7 | 24479652 | 24479670 | 19 | + | 17.8852 | 5.08E-07 | 0.0973 | ACACCAGAAGAGGGCACCA | | |
| 1059 | 7 | 25018887 | 25019093 | 206 | MOLF,SPRET | 7 | 25018951 | 25018984 | 34 | + | 18.3008 | 2.89E-07 | 0.0482 | TAGCAGTCTTCAGACACACCAGAAGAGGGCGTTG | 1 | ENCFF093ZRU |
| 1060 | 7 | 25098674 | 25098895 | 221 | BUB,C58J,CAST,CE,ILNJ2,MOLF,MRL,NOD,NOR,PL,SJL,SPRET,WSB | 7 | 25098756 | 25098789 | 34 | + | 17.0488 | 6.56E-07 | 0.076 | TTGCTATCTTCAGATACACCAGAAGAGGGCATCA | 1 | ENCFF093ZRU |
| 1061 | 7 | 26475451 | 26475513 | 62 | CAST | 7 | 26475471 | 26475505 | 35 | + | 19.1009 | 1.54E-07 | 0.0537 | CTCAAGTAGTCGTTTCTGACCAGTAGATGGCAGTA | 18 | ENCFF367UNR,ENCFF052KVO,ENCFF092NMJ,ENCFF136ZDK,ENCFF430PPJ,ENCFF907XEG,ENCFF327LYS,ENCFF093ZRU,ENCFF099QNE,ENCFF024JOZ,ENCFF194HVL,ENCFF089GBC,ENCFF589ZBJ,ENCFF198SSR,ENCFF883UPM,ENCFF957SPH,ENCFF165LXO,ENCFF533APC |
| 1061 | 7 | 26475451 | 26475513 | 62 | CAST | 7 | 26475487 | 26475505 | 19 | + | 21.918 | 2.19E-08 | 0.0261 | TGACCAGTAGATGGCAGTA | 18 | ENCFF367UNR,ENCFF052KVO,ENCFF092NMJ,ENCFF136ZDK,ENCFF430PPJ,ENCFF907XEG,ENCFF327LYS,ENCFF093ZRU,ENCFF099QNE,ENCFF024JOZ,ENCFF194HVL,ENCFF089GBC,ENCFF589ZBJ,ENCFF198SSR,ENCFF883UPM,ENCFF957SPH,ENCFF165LXO,ENCFF533APC |
| 1062 | 7 | 27344310 | 27344514 | 204 | SPRET | 7 | 27344372 | 27344405 | 34 | + | 17.8455 | 3.91E-07 | 0.057 | TAGCTGTCTTCAGACATTCCAGAAGAGGGCACCA | | |
| 1062 | 7 | 27344310 | 27344514 | 204 | SPRET | 7 | 27344387 | 27344405 | 19 | + | 16.9016 | 9.66E-07 | 0.13 | ATTCCAGAAGAGGGCACCA | | |
| 1063 | 7 | 27379616 | 27379838 | 222 | 129S1,CAST,ILNJ2,KK,LP,MOLF,NZO,NZW,PJ,RF,SEA,SMJ,SPRET | 7 | 27379696 | 27379729 | 34 | + | 18.1789 | 3.13E-07 | 0.0504 | TAGCTGTCTTCAGACACTCCAGAAGAGGGCGCTA | 1 | ENCFF093ZRU |
| 1063 | 7 | 27379616 | 27379838 | 222 | 129S1,CAST,ILNJ2,KK,LP,MOLF,NZO,NZW,PJ,RF,SEA,SMJ,SPRET | 7 | 27379711 | 27379729 | 19 | + | 17.5738 | 6.25E-07 | 0.106 | ACTCCAGAAGAGGGCGCTA | 1 | ENCFF093ZRU |
| 1064 | 7 | 27393103 | 27393303 | 200 | 129S1,CAST,LP,MOLF,NZW,SPRET | 7 | 27393161 | 27393194 | 34 | + | 16.4472 | 9.61E-07 | 0.0901 | TAGCTGTCTTCAGACACACCAGAAGAGGGCGTCA | | |
| 1065 | 7 | 27974998 | 27975072 | 74 | MOLF | 7 | 27975025 | 27975058 | 34 | + | 18.7886 | 2.07E-07 | 0.0405 | CAGCAGCAGCAGCAGGGAGCAGCAGGGAGCAGCA | | |
| 1065 | 7 | 27974998 | 27975072 | 74 | MOLF | 7 | 27975070 | 27975103 | 34 | + | 17.1545 | 6.13E-07 | 0.0727 | GAGCAGCAGCAGCAGGGAGCAGCAGGGAGCAGCA | | |
| 1066 | 7 | 28267749 | 28267859 | 110 | MOLF | 7 | 28267817 | 28267850 | 34 | + | 22.7317 | 1.12E-08 | 0.00473 | CAGCAGATATCTCATATACCAGCAGGGGGCAGCA | 34 | ENCFF093ZRU,ENCFF367UNR,ENCFF099QNE,ENCFF821BIA,ENCFF805QBU,ENCFF321AGQ,ENCFF065ODV,ENCFF327LYS,ENCFF907XEG,ENCFF136ZDK,ENCFF024JOZ,ENCFF589ZBJ,ENCFF165LXO,ENCFF089GBC,ENCFF198SSR,ENCFF210BTO,ENCFF055HAP,ENCFF092NMJ,ENCFF609MDN,ENCFF502WDT,ENCFF086ZTV,ENCFF883UPM,ENCFF430PPJ,ENCFF142CNG,ENCFF281QMV,ENCFF957SPH,ENCFF194HVL,ENCFF491RJK,ENCFF052KVO,ENCFF097CAQ,ENCFF951GOP,ENCFF316FYN,ENCFF575MGI,ENCFF533APC |
| 1066 | 7 | 28267749 | 28267859 | 110 | MOLF | 7 | 28267832 | 28267850 | 19 | + | 20.6885 | 6.34E-08 | 0.0403 | ATACCAGCAGGGGGCAGCA | 34 | ENCFF093ZRU,ENCFF367UNR,ENCFF099QNE,ENCFF821BIA,ENCFF805QBU,ENCFF321AGQ,ENCFF065ODV,ENCFF327LYS,ENCFF907XEG,ENCFF136ZDK,ENCFF024JOZ,ENCFF589ZBJ,ENCFF165LXO,ENCFF089GBC,ENCFF198SSR,ENCFF210BTO,ENCFF055HAP,ENCFF092NMJ,ENCFF609MDN,ENCFF502WDT,ENCFF086ZTV,ENCFF883UPM,ENCFF430PPJ,ENCFF142CNG,ENCFF281QMV,ENCFF957SPH,ENCFF194HVL,ENCFF491RJK,ENCFF052KVO,ENCFF097CAQ,ENCFF951GOP,ENCFF316FYN,ENCFF575MGI,ENCFF533APC |
| 1067 | 7 | 28714514 | 28714616 | 102 | MOLF | 7 | 28714574 | 28714607 | 34 | + | 28.3659 | 6.46E-11 | 0.00019 | GTGCAGTACTTTCGGAGGCCAGAAGAGGGCACCA | 15 | ENCFF367UNR,ENCFF099QNE,ENCFF327LYS,ENCFF907XEG,ENCFF055HAP,ENCFF821BIA,ENCFF883UPM,ENCFF093ZRU,ENCFF052KVO,ENCFF589ZBJ,ENCFF033UQX,ENCFF065ODV,ENCFF502WDT,ENCFF136ZDK,ENCFF533APC |
| 1067 | 7 | 28714514 | 28714616 | 102 | MOLF | 7 | 28714589 | 28714607 | 19 | + | 20.3607 | 8.29E-08 | 0.042 | AGGCCAGAAGAGGGCACCA | 15 | ENCFF367UNR,ENCFF099QNE,ENCFF327LYS,ENCFF907XEG,ENCFF055HAP,ENCFF821BIA,ENCFF883UPM,ENCFF093ZRU,ENCFF052KVO,ENCFF589ZBJ,ENCFF033UQX,ENCFF065ODV,ENCFF502WDT,ENCFF136ZDK,ENCFF533APC |
| 1068 | 7 | 29687919 | 29688192 | 273 | AJ,AKR,BALB,BTBR,C57L,CAST,DBA1J,DBA2J,MA/My,MOLF,MRL,NOD,NOR,NZB,NZO,PJ,PL,SEA,SMJ,SPRET,TH | 7 | 29688049 | 29688082 | 34 | + | 18.1789 | 3.13E-07 | 0.0504 | TAGCTGTCTTCAGACACTCCAGAAGAGGGCGCTA | 1 | ENCFF093ZRU |
| 1068 | 7 | 29687919 | 29688192 | 273 | AJ,AKR,BALB,BTBR,C57L,CAST,DBA1J,DBA2J,MA/My,MOLF,MRL,NOD,NOR,NZB,NZO,PJ,PL,SEA,SMJ,SPRET,TH | 7 | 29688064 | 29688082 | 19 | + | 17.5738 | 6.25E-07 | 0.106 | ACTCCAGAAGAGGGCGCTA | 1 | ENCFF093ZRU |
| 1069 | 7 | 29746172 | 29746387 | 215 | 129S1,AJ,AKR,BALB,BTBR,C57L,CAST,DBA1J,DBA2J,LP,MA/My,MOLF,MRL,NOD,NOR,NZB,NZO,NZW,PJ,PL,SEA,SJL,SMJ,SPRET,TH | 7 | 29746242 | 29746275 | 34 | + | 16.5691 | 8.90E-07 | 0.0882 | TAGCTGTCTTCAGACGCACCAGAAGAGGGCATCG | | |
| 1070 | 7 | 30505054 | 30505579 | 525 | DBA1J,DBA2J,MOLF,NZB,NZO,NZW,SMJ | 7 | 30505511 | 30505544 | 34 | + | 18.3902 | 2.72E-07 | 0.0476 | GCGTCGAACACCCAATGCCCAGCAGGGGGCGCTG | 23 | ENCFF093ZRU,ENCFF805QBU,ENCFF165LXO,ENCFF099QNE,ENCFF367UNR,ENCFF327LYS,ENCFF033UQX,ENCFF321AGQ,ENCFF136ZDK,ENCFF907XEG,ENCFF821BIA,ENCFF201JUA,ENCFF589ZBJ,ENCFF089EWX,ENCFF052KVO,ENCFF024JOZ,ENCFF198SSR,ENCFF533APC,ENCFF883UPM,ENCFF957SPH,ENCFF210BTO,ENCFF097CAQ,ENCFF194HVL |
| 1070 | 7 | 30505054 | 30505579 | 525 | DBA1J,DBA2J,MOLF,NZB,NZO,NZW,SMJ | 7 | 30505526 | 30505544 | 19 | + | 23.3607 | 5.29E-09 | 0.0159 | TGCCCAGCAGGGGGCGCTG | 24 | ENCFF093ZRU,ENCFF805QBU,ENCFF165LXO,ENCFF099QNE,ENCFF367UNR,ENCFF327LYS,ENCFF033UQX,ENCFF321AGQ,ENCFF136ZDK,ENCFF907XEG,ENCFF821BIA,ENCFF201JUA,ENCFF589ZBJ,ENCFF089EWX,ENCFF052KVO,ENCFF024JOZ,ENCFF198SSR,ENCFF533APC,ENCFF883UPM,ENCFF957SPH,ENCFF210BTO,ENCFF097CAQ,ENCFF194HVL,ENCFF504JDS |
| 1071 | 7 | 33681867 | 33682199 | 332 | SPRET | 7 | 33681896 | 33681914 | 19 | + | 19.2951 | 1.88E-07 | 0.0611 | CCGCCACTAGATGGTGCTA | 17 | ENCFF201JUA,ENCFF092NMJ,ENCFF367UNR,ENCFF093ZRU,ENCFF883UPM,ENCFF099QNE,ENCFF821BIA,ENCFF327LYS,ENCFF024JOZ,ENCFF907XEG,ENCFF198SSR,ENCFF136ZDK,ENCFF165LXO,ENCFF589ZBJ,ENCFF957SPH,ENCFF194HVL,ENCFF491RJK |
| 1072 | 7 | 33882332 | 33882540 | 208 | BTBR,C57L,CAST,DBA1J,DBA2J,MOLF,NZB,NZO,NZW,PJ,PL,SMJ,SPRET,TH | 7 | 33882398 | 33882431 | 34 | + | 18.3659 | 2.76E-07 | 0.0476 | TAGCTGTCTTCAGACACTCCAGAAGAGGGCGCCA | | |
| 1072 | 7 | 33882332 | 33882540 | 208 | BTBR,C57L,CAST,DBA1J,DBA2J,MOLF,NZB,NZO,NZW,PJ,PL,SMJ,SPRET,TH | 7 | 33882413 | 33882431 | 19 | + | 17.5574 | 6.32E-07 | 0.106 | ACTCCAGAAGAGGGCGCCA | | |
| 1073 | 7 | 34705347 | 34705550 | 203 | C58J,MRL,NOD,NOR,NZB,NZO,NZW,RF,SJL,SPRET | 7 | 34705408 | 34705441 | 34 | + | 17.2439 | 5.79E-07 | 0.0706 | TAGCTGTCCTCAGATACTCCAGAAGAGGGCATCA | | |
| 1074 | 7 | 38318735 | 38319429 | 694 | 129S1,BTBR,BUB,C3H,C57L,CBA,KK,MRL,NZW,WSB | 7 | 38319111 | 38319129 | 19 | + | 17.459 | 6.74E-07 | 0.109 | TGTCCACTAGATGTCAGTG | | |
| 1075 | 7 | 38634438 | 38635132 | 694 | 129S1,BTBR,BUB,C3H,C57L,CBA,KK,MRL,NZW,PL,SJL,WSB | 7 | 38634814 | 38634832 | 19 | + | 17.459 | 6.74E-07 | 0.109 | TGTCCACTAGATGTCAGTG | | |
| 1076 | 7 | 42392351 | 42392514 | 163 | CAST,SPRET | 7 | 42392400 | 42392433 | 34 | + | 16.4472 | 9.61E-07 | 0.0901 | TAGCTGTCTTCAGACACACCAGAAGAGGGCGTCA | | |
| 1077 | 7 | 44673252 | 44673464 | 212 | 129S1,AJ,AKR,BALB,C3H,CAST,CBA,DBA1J,DBA2J,ILNJ2,KK,MA/My,MOLF,NZW,PJ,PL,RH2,SEA,SJL,SMJ,SPRET,WSB | 7 | 44673321 | 44673354 | 34 | + | 19.6667 | 1.13E-07 | 0.0279 | TAGCAGTCTTCAGATGCACCAGCAGAGGGCATCA | 13 | ENCFF367UNR,ENCFF092NMJ,ENCFF089GBC,ENCFF136ZDK,ENCFF093ZRU,ENCFF198SSR,ENCFF589ZBJ,ENCFF099QNE,ENCFF907XEG,ENCFF821BIA,ENCFF024JOZ,ENCFF327LYS |
| 1078 | 7 | 44974365 | 44975125 | 760 | CAST.ILNJ2.MOLF.NZW.PJ.SJL.SPRET | 7 | 44974927 | 44974961 | 35 | + | 24.3211 | 4.17E-09 | 0.00615 | CTGCTCAGAAGGCTGCAGGCCAGAAGGTGGCACCT | | |
| 1078 | 7 | 44974365 | 44975125 | 760 | CAST.ILNJ2.MOLF.NZW.PJ.SJL.SPRET | 7 | 44974943 | 44974961 | 19 | + | 17.7869 | 5.43E-07 | 0.101 | AGGCCAGAAGGTGGCACCT | | |
| 1079 | 7 | 45127703 | 45127909 | 206 | CAST.ILNJ2.MOLF.NZW.PJ.SJL.SPRET | 7 | 45127767 | 45127800 | 34 | + | 18.3659 | 2.76E-07 | 0.0476 | TAGCTGTCTTCAGACACTCCAGAAGAGGGCGCCA | 6 | ENCFF093ZRU.ENCFF367UNR.ENCFF821BIA.ENCFF024JOZ.ENCFF907XEG.ENCFF136ZDK |
| 1079 | 7 | 45127703 | 45127909 | 206 | CAST.ILNJ2.MOLF.NZW.PJ.SJL.SPRET | 7 | 45127782 | 45127800 | 19 | + | 17.5574 | 6.32E-07 | 0.106 | ACTCCAGAAGAGGGCGCCA | 6 | ENCFF093ZRU.ENCFF367UNR.ENCFF821BIA.ENCFF024JOZ.ENCFF907XEG.ENCFF136ZDK |
| 1080 | 7 | 46552375 | 46552576 | 201 | CAST.ILNJ2.MOLF.NZW.PJ.SJL.SPRET | 7 | 46552434 | 46552467 | 34 | + | 16.4472 | 9.61E-07 | 0.0901 | TAGCTGTCTTCAGACACACCAGAAGAGGGCGTCA | | |
| 1081 | 7 | 63729548 | 63729713 | 165 | CAST.DBA1J.DBA2J.KK.MOLF.SPRET | 7 | 63729550 | 63729583 | 34 | + | 18.4309 | 2.64E-07 | 0.047 | CTGTGGTGTCACCGGGCTCCACTTGGTGTCACCG | | |
| 1082 | 7 | 63831977 | 63832592 | 615 | SPRET | 7 | 63832561 | 63832579 | 19 | + | 18.0492 | 4.54E-07 | 0.0941 | TGTCCTCAAGAGGGCACTG | 5 | ENCFF093ZRU.ENCFF099QNE.ENCFF907XEG.ENCFF089GBC.ENCFF821BIA |
| 1082 | 7 | 63831977 | 63832592 | 615 | SPRET | 7 | 63831972 | 63832005 | 34 | - | 19.4472 | 1.31E-07 | 0.0308 | CTGCTACTCCTGAGACTGGCAGCAGGGGTAGTTC | | |
| 1083 | 7 | 64615605 | 64615839 | 234 | CAST.SPRET | 7 | 64615697 | 64615730 | 34 | + | 18.3659 | 2.76E-07 | 0.0476 | TAGCTGTCTTCAGACACTCCAGAAGAGGGCGCCA | | |
| 1083 | 7 | 64615605 | 64615839 | 234 | CAST.SPRET | 7 | 64615712 | 64615730 | 19 | + | 17.5574 | 6.32E-07 | 0.106 | ACTCCAGAAGAGGGCGCCA | | |
| 1084 | 7 | 65346095 | 65346613 | 518 | SPRET | 7 | 65346215 | 65346248 | 34 | + | 18.9837 | 1.81E-07 | 0.0381 | ATGCAGTCTAACTGAGAGCCACCTGATGGCAGGA | 1 | ENCFF093ZRU |
| 1084 | 7 | 65346095 | 65346613 | 518 | SPRET | 7 | 65346387 | 65346420 | 34 | + | 18.2195 | 3.05E-07 | 0.0497 | ATGTGGTACCTCTGGTAACCAGAAGAGGGCATCT | 19 | ENCFF093ZRU,ENCFF430PPJ,ENCFF367UNR,ENCFF092NMJ,ENCFF052KVO,ENCFF165LXO,ENCFF099QNE,ENCFF907XEG,ENCFF327LYS,ENCFF821BIA,ENCFF589ZBJ,ENCFF198SSR,ENCFF024JOZ,ENCFF089GBC,ENCFF055HAP,ENCFF136ZDK,ENCFF502WDT,ENCFF321AGQ,ENCFF065ODV |
| 1084 | 7 | 65346095 | 65346613 | 518 | SPRET | 7 | 65346228 | 65346261 | 34 | - | 17.6911 | 4.33E-07 | 0.0602 | GTACAGCTCTGAGTCCTGCCATCAGGTGGCTCTC | 4 | ENCFF093ZRU.ENCFF430PPJ.ENCFF367UNR.ENCFF092NMJ |
| 1085 | 7 | 66192992 | 66193942 | 950 | DBA1J.DBA2J.ILNJ2.KK.MOLF.PJ | 7 | 66193369 | 66193387 | 19 | + | 16.9672 | 9.26E-07 | 0.127 | ATGCCAGCAGAGGGCATTG | | |
| 1086 | 7 | 73988652 | 73988862 | 210 | 129S1,C3H,C58J,CAST,CBA,CE,DBA1J,DBA2J,FVB,KK,MA/My,MOLF,PJ,SMJ,SPRET,SWR,TH,WSB | 7 | 73988720 | 73988753 | 34 | + | 17.3902 | 5.27E-07 | 0.0672 | TAGCTGTCCTCAGACACTCCAGAAGAGGGCATCA | | |
| 1087 | 7 | 74013900 | 74014026 | 126 | SPRET | 7 | 74013987 | 74014021 | 35 | + | 17.9725 | 3.06E-07 | 0.08 | ATGCAAGTGCCTGAGAAGGCCGGAAGAGGGCATCA | | |
| 1088 | 7 | 75171940 | 75172536 | 596 | C58J,CE,WSB | 7 | 75172148 | 75172181 | 34 | + | 21.0894 | 4.00E-08 | 0.0128 | ATGCAGTGGACACAGAGGCCAGAAGAGGGTGTCA | 10 | ENCFF367UNR,ENCFF327LYS,ENCFF055HAP,ENCFF198SSR,ENCFF883UPM,ENCFF502WDT,ENCFF093ZRU,ENCFF907XEG,ENCFF099QNE,ENCFF821BIA |
| 1089 | 7 | 75475084 | 75475855 | 771 | SPRET | 7 | 75475456 | 75475489 | 34 | + | 17.1463 | 6.17E-07 | 0.0728 | TAGCTGTCTTTAGATGCGCCAGAAGAGGGCATCA | | |
| 1090 | 7 | 79217948 | 79218138 | 190 | SPRET | 7 | 79218080 | 79218114 | 35 | + | 18.1284 | 2.79E-07 | 0.0768 | TTGCAAGTCTCAGATCTGGCCAGCAGGGGAAGAAA | 2 | ENCFF093ZRU.ENCFF092NMJ |
| 1091 | 7 | 80483114 | 80483335 | 221 | AJ,AKR,BALB,BUB,CAST,FVB,MRL,NOD,NOR,NZB,NZO,NZW,PL,RF,RH2,SJL,SMJ,SPRET,SWR,TH,WSB | 7 | 80483181 | 80483214 | 34 | + | 16.4634 | 9.51E-07 | 0.0901 | TAGCTGTCTTCAGACGCACCAGAAGAGGGCATCA | 2 | ENCFF367UNR,ENCFF055HAP |
| 1092 | 7 | 80501124 | 80501327 | 203 | CAST.CE.MOLF | 7 | 80501182 | 80501215 | 34 | + | 16.4472 | 9.61E-07 | 0.0901 | TAGCTGTCTTCAGACACACCAGAAGAGGGCGTCA | | |
| 1093 | 7 | 81230225 | 81230874 | 649 | SPRET | 7 | 81230753 | 81230786 | 34 | + | 16.4472 | 9.61E-07 | 0.0901 | TAGCTGTCTTCAGACACACCAGAAGAGGGCGTCA | | |
| 1094 | 7 | 82377239 | 82377988 | 749 | CE | 7 | 82377538 | 82377556 | 19 | + | 16.9836 | 9.17E-07 | 0.126 | GGAACAGTAGAGGGCACTA | 1 | ENCFF093ZRU |
| 1095 | 7 | 86930390 | 86930587 | 197 | CAST.MOLF.SPRET | 7 | 86930444 | 86930477 | 34 | + | 16.4472 | 9.61E-07 | 0.0901 | TAGCTGTCTTCAGACACACCAGAAGAGGGCGTCA | | |
| 1096 | 7 | 93891792 | 93891994 | 202 | 129S1,AJ,AKR,BALB,BTBR,BUB,C3H,C57L,CAST,CBA,DBA1J,DBA2J,FVB,ILNJ2,KK,MA/My,MOLF,MRL,NOD,NZB,NZO,NZW,PJ,PL,RF,RH2,SEA,SJL,SMJ,SWR,TH,WSB | 7 | 93891853 | 93891886 | 34 | + | 18.3659 | 2.76E-07 | 0.0476 | TAGCTGTCTTCAGACACTCCAGAAGAGGGCGCCA | | |
| 1096 | 7 | 93891792 | 93891994 | 202 | 129S1,AJ,AKR,BALB,BTBR,BUB,C3H,C57L,CAST,CBA,DBA1J,DBA2J,FVB,ILNJ2,KK,MA/My,MOLF,MRL,NOD,NZB,NZO,NZW,PJ,PL,RF,RH2,SEA,SJL,SMJ,SWR,TH,WSB | 7 | 93891868 | 93891886 | 19 | + | 17.5574 | 6.32E-07 | 0.106 | ACTCCAGAAGAGGGCGCCA | | |
| 1097 | 7 | 97866770 | 97866826 | 56 | C3H,CBA,DBA1J,DBA2J,ILNJ2,MOLF,PJ,SEA,SPRET | 7 | 97866738 | 97866771 | 34 | + | 17.6504 | 4.44E-07 | 0.0613 | ATGAAGTGCCTGTGGAGGCCAGAAGAGGGTGTGT | | |
| 1098 | 7 | 98426070 | 98426463 | 393 | C3H,CBA,DBA1J,DBA2J,ILNJ2,MOLF,PJ,SEA | 7 | 98426393 | 98426427 | 35 | + | 17.578 | 3.87E-07 | 0.0917 | CTGCAGACCATGACCCCGCCCAGCAGAAGACATCT | | |
| 1099 | 7 | 98667704 | 98667985 | 281 | NOD,NOR,SWR | 7 | 98667915 | 98667949 | 35 | + | 17.0092 | 5.39E-07 | 0.11 | CTGCAATCCCAGAACTCAGGAAGCAGAGGAGGAA | | |
| 1100 | 7 | 99212591 | 99212957 | 366 | SPRET | 7 | 99212628 | 99212661 | 34 | + | 17.6341 | 4.49E-07 | 0.0617 | CTCCAGAGACAGCTCCGGCCTGGTGATGGCGCCC | 8 | ENCFF099QNE,ENCFF092NMJ,ENCFF198SSR,ENCFF089GBC,ENCFF907XEG,ENCFF316FYN,ENCFF093ZRU,ENCFF821BIA |
| 1101 | 7 | 100187291 | 100187494 | 203 | 129S1,AJ,AKR,BALB,BTBR,BUB,C3H,C57L,CAST,CBA,DBA1J,DBA2J,FVB,ILNJ2,KK,MA/My,MOLF,MRL,NZB,NZW,PJ,PL,RF,RH2,SEA,SJL,SMJ,SPRET,TH | 7 | 100187352 | 100187385 | 34 | + | 16.5041 | 9.27E-07 | 0.0901 | TAGCTGTCTTCAGACGTACCAGAAGAGGGCATCG | 1 | ENCFF093ZRU |
| 1102 | 7 | 101791852 | 101792052 | 200 | CAST.MOLF.SMJ.SPRET | 7 | 101791908 | 101791941 | 34 | + | 16.4634 | 9.51E-07 | 0.0901 | TAGCTGTCTTCAGACGCACCAGAAGAGGGCATCA | | |
| 1103 | 7 | 105102771 | 105102969 | 198 | SPRET | 7 | 105102827 | 105102861 | 35 | + | 16.1468 | 8.80E-07 | 0.143 | TAGCTGTACTTCAGACACTCCAGAAGAGGGCATCA | | |
| 1104 | 7 | 105511289 | 105511505 | 216 | CAST.CE.MOLF.NZO.SPRET.WSB | 7 | 105511363 | 105511396 | 34 | + | 18.4472 | 2.62E-07 | 0.0467 | TTGCTGTCTTCAGACACTCCAGAAGAGGGCGTCA | | |
| 1105 | 7 | 105722697 | 105722912 | 215 | CE.NZO.SPRET | 7 | 105722768 | 105722801 | 34 | + | 18.4472 | 2.62E-07 | 0.0467 | TTGCTGTCTTCAGACACTCCAGAAGAGGGCGTCA | | |
| 1106 | 7 | 108547685 | 108547888 | 203 | CAST.SPRET | 7 | 108547858 | 108547891 | 34 | + | 17.9837 | 3.57E-07 | 0.0542 | CTGGAGTTACAAGTGAGGCCACTAGGTGAGAGTG | | |
| 1107 | 7 | 115962272 | 115962483 | 211 | 129S1,BTBR,C57L,CAST,ILNJ2,MOLF,SPRET | 7 | 115962341 | 115962374 | 34 | + | 16.4634 | 9.51E-07 | 0.0901 | TAGCTGTCTTCAGACGCACCAGAAGAGGGCATCA | | |
| 1108 | 7 | 116119065 | 116119530 | 465 | 129S1,BTBR,C57L,CAST,ILNJ2,MOLF | 7 | 116119202 | 116119235 | 34 | + | 25.7317 | 8.56E-10 | 0.000754 | ATGCAGTGCCCACAGAGGCCAAAAGAGGGCATCA | 27 | ENCFF065ODV,ENCFF367UNR,ENCFF093ZRU,ENCFF136ZDK,ENCFF201JUA,ENCFF430PPJ,ENCFF589ZBJ,ENCFF099QNE,ENCFF609MDN,ENCFF327LYS,ENCFF198SSR,ENCFF951GOP,ENCFF907XEG,ENCFF821BIA,ENCFF024JOZ,ENCFF097CAQ,ENCFF089GBC,ENCFF194HVL,ENCFF055HAP,ENCFF491RJK,ENCFF502WDT,ENCFF092NMJ,ENCFF957SPH,ENCFF142CNG,ENCFF165LXO,ENCFF281QMV,ENCFF883UPM |
| 1109 | 7 | 117906037 | 117907007 | 970 | BTBR,C3H,C57L,CAST,CBA,CE,DBA1J,MOLF,NOR,WSB | 7 | 117906225 | 117906258 | 34 | + | 18.2114 | 3.07E-07 | 0.0499 | TAGCTGTTTTCAGACACTCCAGAAGGGGGCATCA | | |
| 1110 | 7 | 118117181 | 118117252 | 71 | 129S1,C3H,C57L,CAST,CBA,DBA1J,DBA2J,MOLF,NOR,SPRET | 7 | 118117231 | 118117265 | 35 | + | 18.4404 | 2.31E-07 | 0.0698 | CAGCAGTGAGCAGCAGTGAGCAGCAGTGGGCAGTG | 21 | ENCFF821BIA,ENCFF367UNR,ENCFF327LYS,ENCFF430PPJ,ENCFF198SSR,ENCFF092NMJ,ENCFF907XEG,ENCFF089GBC,ENCFF609MDN,ENCFF589ZBJ,ENCFF136ZDK,ENCFF611HDQ,ENCFF210BTO,ENCFF099QNE,ENCFF024JOZ,ENCFF883UPM,ENCFF097CAQ,ENCFF957SPH,ENCFF165LXO,ENCFF194HVL |
| 1111 | 7 | 120402070 | 120402368 | 298 | AJ.CE.MOLF | 7 | 120402098 | 120402132 | 35 | + | 21.211 | 3.91E-08 | 0.0225 | TTGCATGTACATCTGCATGCCAGAAGAGGGCATCA | 4 | ENCFF093ZRU.ENCFF821BIA.ENCFF502WDT.ENCFF055HAP |
| 1112 | 7 | 120729952 | 120730011 | 59 | SPRET | 7 | 120729927 | 120729960 | 34 | + | 27.0407 | 2.48E-10 | 0.000329 | ATGCAGTACCTGAGGAGGCCAGAAGAGGGCATCA | | |
| 1113 | 7 | 122663538 | 122663682 | 144 | AKR,BUB,C58J,MRL,NOD,NZW,PL,SWR,TH,WSB | 7 | 122663641 | 122663674 | 34 | + | 16.4472 | 9.61E-07 | 0.0901 | TAGCTGTCTTCAGACACACCAGAAGAGGGCGTCA | | |
| 1114 | 7 | 125254058 | 125254109 | 51 | AKR,BALB,BUB,C58J,CAST,MOLF,MRL,NZW,PL,RH2,SEA,SJL,SMJ,SWR,TH | 7 | 125254083 | 125254117 | 35 | + | 20.8073 | 5.13E-08 | 0.026 | CTCCAGCTCCCCCTGAGCACCTCTGGGTGGCGGCC | | |
| 1115 | 7 | 125829363 | 125829582 | 219 | 129S1,AJ,C57L,CAST,CE,FVB,ILNJ2,KK,LP,MA/My,MOLF,NOD,NZB,NZO,PJ,RF,SMJ,SPRET,TH,WSB | 7 | 125829437 | 125829470 | 34 | + | 16.4472 | 9.61E-07 | 0.0901 | TAGCTGTCTTCAGACACACCAGAAGAGGGCGTCA | | |
| 1116 | 7 | 125831479 | 125831685 | 206 | 129S1,AJ,C57L,CAST,CE,FVB,ILNJ2,KK,LP,MA/My,MOLF,NZB,NZO,PJ,RF,SMJ,SPRET,TH,WSB | 7 | 125831543 | 125831576 | 34 | + | 17.6992 | 4.30E-07 | 0.0601 | TAGCTGTTTTCAGACACACCAGAAGAGGGCATCA | | |
| 1117 | 7 | 127058392 | 127058600 | 208 | SPRET.TH | 7 | 127058458 | 127058491 | 34 | + | 16.4634 | 9.51E-07 | 0.0901 | TAGCTGTCTTCAGACGCACCAGAAGAGGGCATCA | | |
| 1118 | 7 | 127230152 | 127230355 | 203 | MOLF.SPRET | 7 | 127230212 | 127230246 | 35 | + | 18.1284 | 2.79E-07 | 0.0768 | CTGTAGCTGTCTTCAGACACCAGAAGAGGGCATCA | | |
| 1119 | 7 | 127274124 | 127274352 | 228 | MOLF | 7 | 127274314 | 127274332 | 19 | + | 23.6066 | 4.01E-09 | 0.0145 | TCGCCAGTAGAGGGCGCTA | 39 | ENCFF367UNR,ENCFF093ZRU,ENCFF201JUA,ENCFF099QNE,ENCFF821BIA,ENCFF327LYS,ENCFF805QBU,ENCFF907XEG,ENCFF089GBC,ENCFF589ZBJ,ENCFF065ODV,ENCFF024JOZ,ENCFF092NMJ,ENCFF136ZDK,ENCFF430PPJ,ENCFF198SSR,ENCFF321AGQ,ENCFF055HAP,ENCFF165LXO,ENCFF502WDT,ENCFF883UPM,ENCFF504JDS,ENCFF052KVO,ENCFF951GOP,ENCFF281QMV,ENCFF957SPH,ENCFF575MGI,ENCFF609MDN,ENCFF142CNG,ENCFF210BTO,ENCFF601JVE,ENCFF316FYN,ENCFF611HDQ,ENCFF533APC,ENCFF097CAQ,ENCFF089EWX,ENCFF491RJK,ENCFF194HVL,ENCFF033UQX |
| 1120 | 7 | 127291279 | 127291486 | 207 | CAST.MOLF.SPRET.TH | 7 | 127291338 | 127291371 | 34 | + | 16.4472 | 9.61E-07 | 0.0901 | TAGCTGTCTTCAGACACACCAGAAGAGGGCGTCA | | |
| 1121 | 7 | 127513018 | 127513429 | 411 | SPRET | 7 | 127513082 | 127513115 | 34 | + | 16.9919 | 6.81E-07 | 0.0775 | TAGCTGTCCTCAGACTCTCCAGAAGAGGGAGTCA | | |
| 1122 | 7 | 127562133 | 127562345 | 212 | 129S1,AJ,BTBR,C57L,CAST,CE,ILNJ2,KK,MOLF,PJ,SPRET,TH | 7 | 127562202 | 127562235 | 34 | + | 16.4472 | 9.61E-07 | 0.0901 | TAGCTGTCTTCAGACACACCAGAAGAGGGCGTCA | | |
| 1123 | 7 | 127642335 | 127642585 | 250 | CAST.SPRET | 7 | 127642375 | 127642408 | 34 | + | 18.3984 | 2.70E-07 | 0.0475 | TAGCTGTCTTCAGACACACCAGAAGAGGGCACCA | | |
| 1123 | 7 | 127642335 | 127642585 | 250 | CAST.SPRET | 7 | 127642390 | 127642408 | 19 | + | 17.8852 | 5.08E-07 | 0.0973 | ACACCAGAAGAGGGCACCA | | |
| 1124 | 7 | 128688347 | 128688560 | 213 | 129S1,BTBR,C57L,ILNJ2,LP,MOLF,NZW,PJ,SPRET,WSB | 7 | 128688420 | 128688453 | 34 | + | 16.4472 | 9.61E-07 | 0.0901 | TAGCTGTCTTCAGACACACCAGAAGAGGGCGTCA | | |
| 1125 | 7 | 129087311 | 129087435 | 124 | LP | 7 | 129087413 | 129087431 | 19 | + | 18.7541 | 2.79E-07 | 0.0733 | AGACCAGAAGAGGGCAGCA | | |
| 1126 | 7 | 129833895 | 129834080 | 185 | CAST.MOLF.SPRET | 7 | 129833929 | 129833962 | 34 | + | 16.4472 | 9.61E-07 | 0.0901 | TAGCTGTCTTCAGACACACCAGAAGAGGGCGTCA | | |
| 1127 | 7 | 130656464 | 130656822 | 358 | CAST,MOLF | 7 | 130656695 | 130656729 | 35 | + | 16.8624 | 5.86E-07 | 0.115 | ATGATCATTCAGAGAGTGGCCAGCAGAGGGCAGTA | 19 | ENCFF093ZRU,ENCFF201JUA,ENCFF821BIA,ENCFF052KVO,ENCFF430PPJ,ENCFF367UNR,ENCFF099QNE,ENCFF589ZBJ,ENCFF883UPM,ENCFF609MDN,ENCFF327LYS,ENCFF089GBC,ENCFF024JOZ,ENCFF907XEG,ENCFF198SSR,ENCFF165LXO,ENCFF092NMJ,ENCFF957SPH,ENCFF281QMV |
| 1127 | 7 | 130656464 | 130656822 | 358 | CAST,MOLF | 7 | 130656711 | 130656729 | 19 | + | 24.4426 | 1.40E-09 | 0.00895 | TGGCCAGCAGAGGGCAGTA | 19 | ENCFF093ZRU,ENCFF201JUA,ENCFF821BIA,ENCFF052KVO,ENCFF430PPJ,ENCFF367UNR,ENCFF099QNE,ENCFF589ZBJ,ENCFF883UPM,ENCFF609MDN,ENCFF327LYS,ENCFF089GBC,ENCFF024JOZ,ENCFF907XEG,ENCFF198SSR,ENCFF165LXO,ENCFF092NMJ,ENCFF957SPH,ENCFF281QMV |
| 1128 | 7 | 139700857 | 139701023 | 166 | BUB,C58J,CAST,CBA,CE,FVB,ILNJ2,KK,LP,MA/My,MOLF,NOD,NOR,PJ,SMJ,SPRET,SWR,TH | 7 | 139700921 | 139700954 | 34 | + | 16.5691 | 8.90E-07 | 0.0882 | TAGCTGTCTTCAGACGCACCAGAAGAGGGCATCG | | |
| 1129 | 7 | 140720666 | 140720899 | 233 | CAST.MOLF.NZW | 7 | 140720659 | 140720692 | 34 | + | 17.2114 | 5.91E-07 | 0.0711 | TAGCTGTTTTCAGACACTCCAGAAGAGGGCATCA | 2 | ENCFF367UNR.ENCFF093ZRU |
| 1130 | 7 | 141245456 | 141245685 | 229 | CAST.ILNJ2.MOLF.NZW.SPRET.WSB | 7 | 141245542 | 141245575 | 34 | + | 16.4634 | 9.51E-07 | 0.0901 | TAGCTGTCTTCAGACGCACCAGAAGAGGGCATCA | | |
| 1131 | 7 | 142386113 | 142386335 | 222 | C58J,CAST,LP,MA/My,MOLF,PJ,SMJ,SPRET | 7 | 142386191 | 142386224 | 34 | + | 16.4634 | 9.51E-07 | 0.0901 | TAGCTGTCTTCAGACGCACCAGAAGAGGGCATCA | | |
| 1132 | 7 | 143635288 | 143635360 | 72 | 129S1,C57L,KK,SPRET | 7 | 143635301 | 143635319 | 19 | + | 20.1803 | 9.60E-08 | 0.0453 | TAGCCAGCAGAGGGTGCTG | 23 | ENCFF201JUA,ENCFF093ZRU,ENCFF430PPJ,ENCFF099QNE,ENCFF327LYS,ENCFF089GBC,ENCFF589ZBJ,ENCFF907XEG,ENCFF883UPM,ENCFF024JOZ,ENCFF198SSR,ENCFF092NMJ,ENCFF055HAP,ENCFF136ZDK,ENCFF502WDT,ENCFF957SPH,ENCFF609MDN,ENCFF575MGI,ENCFF210BTO,ENCFF165LXO,ENCFF194HVL,ENCFF367UNR |

| | | | | | | | | | | | | | | | | |
|---|---|---|---|---|---|---|---|---|---|---|---|---|---|---|---|---|
| 1132 | 7 | 143635288 | 143635360 | 72 | 129S1,C57L,KK,SPRET | 7 | 143635357 | 143635391 | 35 | + | 16 | 9.55E-07 | 0.148 | GAGTCTGTCTCAGACCTAGCCAGCAGGGGGTGCTG | 20 | ENCFF093ZRU,ENCFF430PPJ,ENCFF099QNE,ENCFF327LYS,ENCFF089GBC,ENCFF589ZBJ,ENCFF907XEG,ENCFF883UPM,ENCFF024JOZ,ENCFF198SSR,ENCFF092NMJ,ENCFF055HAP,ENCFF136ZDK,ENCFF502WDT,ENCFF609MDN,ENCFF575MGI,ENCFF165LXO,ENCFF194HVL,ENCFF367UNR,ENCFF201JUA |
| 1132 | 7 | 143635288 | 143635360 | 72 | 129S1,C57L,KK,SPRET | 7 | 143635358 | 143635391 | 34 | + | 16.7805 | 7.79E-07 | 0.0827 | AGTCTGTCTCAGACCTAGCCAGCAGGGGGTGCTG | 20 | ENCFF093ZRU,ENCFF430PPJ,ENCFF099QNE,ENCFF327LYS,ENCFF089GBC,ENCFF589ZBJ,ENCFF907XEG,ENCFF883UPM,ENCFF024JOZ,ENCFF198SSR,ENCFF092NMJ,ENCFF055HAP,ENCFF136ZDK,ENCFF502WDT,ENCFF609MDN,ENCFF575MGI,ENCFF165LXO,ENCFF194HVL,ENCFF367UNR,ENCFF201JUA |
| 1133 | 7 | 3610833 | 3611161 | 328 | SPRET | 7 | 3610975 | 3611008 | 34 | - | 19.2602 | 1.50E-07 | 0.0336 | TTGCTGTCTTCAGATGCACCAGAAGAGGGCGTCA | | |
| 1134 | 7 | 4682563 | 4682773 | 210 | CAST,MOLF,SPRET | 7 | 4682680 | 4682713 | 34 | - | 17.6992 | 4.30E-07 | 0.0601 | TAGCTGTCCTCAGATACTCCAGAAGAGGGCGTCA | | |
| 1135 | 7 | 4964562 | 4965067 | 505 | SPRET | 7 | 4964791 | 4964809 | 19 | - | 18.2623 | 3.93E-07 | 0.0868 | AGGCCAGAAGAGGGCGACA | | |
| 1135 | 7 | 4964562 | 4965067 | 505 | SPRET | 7 | 4964791 | 4964824 | 34 | - | 21.8293 | 2.28E-08 | 0.00817 | GTTCAGTGTCCAAAGAGGCCAGAAGAGGGCGACA | | |
| 1136 | 7 | 5214401 | 5214802 | 401 | SPRET | 7 | 5214744 | 5214778 | 35 | - | 19.9541 | 8.95E-08 | 0.0382 | CTGCTCTGGTGGCCGGAGCCCTCCAGGAGGCGACC | | |
| 1137 | 7 | 6978904 | 6979134 | 230 | 129S1,AJ,BALB,BTBR,BUB,C57L,C58J,CBA,KK,LP,NOD,NOR,NZB,NZO,NZW,PL,RF,SJL,SPRET,SWR,TH | 7 | 6979027 | 6979045 | 19 | - | 17.5574 | 6.32E-07 | 0.106 | ACTCCAGAAGAGGGCGCCA | 1 | ENCFF093ZRU |
| 1137 | 7 | 6978904 | 6979134 | 230 | 129S1,AJ,BALB,BTBR,BUB,C57L,C58J,CBA,KK,LP,NOD,NOR,NZB,NZO,NZW,PL,RF,SJL,SPRET,SWR,TH | 7 | 6979027 | 6979060 | 34 | - | 18.3659 | 2.76E-07 | 0.0476 | TAGCTGTCTTCAGACACTCCAGAAGAGGGCGCCA | 1 | ENCFF093ZRU |
| 1138 | 7 | 9872445 | 9872657 | 212 | 129S1,AJ,BALB,BTBR,BUB,C57L,C58J,CAST,CBA,ILNJ2,KK,LP,MOLF,NOD,NOR,NZB,NZO,NZW,PL,RF,SJL,SPRET,SWR,TH | 7 | 9872569 | 9872587 | 19 | - | 17.8852 | 5.08E-07 | 0.0973 | ACACCAGAAGAGGGCACCA | | |
| 1138 | 7 | 9872445 | 9872657 | 212 | 129S1,AJ,BALB,BTBR,BUB,C57L,C58J,CAST,CBA,ILNJ2,KK,LP,MOLF,NOD,NOR,NZB,NZO,NZW,PL,RF,SJL,SPRET,SWR,TH | 7 | 9872569 | 9872602 | 34 | - | 18.252 | 2.98E-07 | 0.049 | TAGCTGTCTTCAGATACACCAGAAGAGGGCACCA | | |
| 1139 | 7 | 12549565 | 12549773 | 208 | CAST,SPRET | 7 | 12549692 | 12549725 | 34 | - | 19.9837 | 8.98E-08 | 0.0236 | TAGCTGTCCCCAGACGCACCAGAAGAGGGCATCA | | |
| 1140 | 7 | 12687802 | 12687991 | 189 | SPRET | 7 | 12687920 | 12687953 | 34 | - | 21.8943 | 2.17E-08 | 0.00792 | CAGCCGTCCCCAGACACACCAGAAGAGGGCGTCA | | |
| 1141 | 7 | 13066480 | 13067206 | 726 | C58J,CAST,MOLF,SPRET | 7 | 13066814 | 13066848 | 35 | - | 16.0367 | 9.36E-07 | 0.147 | ATGCACTGCCCCTGGGCCGCAAGAAGGGGGCCAAG | | |
| 1142 | 7 | 15987164 | 15987395 | 231 | SPRET | 7 | 15987288 | 15987321 | 34 | - | 16.4472 | 9.61E-07 | 0.0901 | TAGCTGTCTTCAGACACACCAGAAGAGGGCGTCA | | |
| 1143 | 7 | 15997031 | 15997131 | 100 | AKR,RH2,TH | 7 | 15997114 | 15997132 | 19 | - | 23.1311 | 6.72E-09 | 0.0163 | CCGCCACTAGAGGGCACCA | 33 | ENCFF201JUA,ENCFF093ZRU,ENCFF367UNR,ENCFF099QNE,ENCFF327LYS,ENCFF065ODV,ENCFF821BIA,ENCFF589ZBJ,ENCFF024JOZ,ENCFF089GBC,ENCFF321AGQ,ENCFF055HAP,ENCFF907XEG,ENCFF136ZDK,ENCFF198SSR,ENCFF092NMJ,ENCFF502WDT,ENCFF165LXO,ENCFF052KVO,ENCFF951GOP,ENCFF609MDN,ENCFF097CAQ,ENCFF430PPJ,ENCFF194HVL,ENCFF883UPM,ENCFF089EWX,ENCFF491RJK,ENCFF957SPH,ENCFF281QMV,ENCFF575MGI,ENCFF142CNG,ENCFF316FYN |
| 1144 | 7 | 18737463 | 18737699 | 236 | 129S1,AJ,AKR,BALB,BTBR,BUB,C3H,C57L,C58J,CAST,CBA,DBA1J,DBA2J,FVB,ILNJ2,KK,LP,MA/My,MOLF,MRL,NOD,NOR,NZB,NZO,NZW,PJ,PL,RF,RH2,SEA,SJL,SMJ,SPRET,SWR,TH,WSB | 7 | 18737589 | 18737622 | 34 | - | 17.9431 | 3.66E-07 | 0.0552 | CAGCTGTCCTCAGACACTCCAGAAGAGGGAGTCA | | |
| 1145 | 7 | 21184101 | 21184259 | 158 | MOLF | 7 | 21184103 | 21184121 | 19 | - | 16.8689 | 9.85E-07 | 0.131 | ATACCAGCAGAGGGCAGCT | | |
| 1145 | 7 | 21184101 | 21184259 | 158 | MOLF | 7 | 21184103 | 21184136 | 34 | - | 27.3902 | 1.75E-10 | 0.000279 | CTGCAGCTCCGCAGCATACCAGCAGAGGGCAGCT | | |
| 1146 | 7 | 23886214 | 23886408 | 194 | 129S1,BTBR,BUB,C3H,C57L,CAST,CBA,MOLF,NOD,NOR,NZB,NZW,SJL,SPRET | 7 | 23886340 | 23886373 | 34 | - | 16.6341 | 8.54E-07 | 0.0866 | GTACACTCTCTTCAGACACCAGAAGAGGGCATCA | | |
| 1147 | 7 | 24358648 | 24358702 | 54 | SPRET | 7 | 24358662 | 24358680 | 19 | - | 17.9672 | 4.81E-07 | 0.0964 | AGACCTGTAGAGGGCACCA | 2 | ENCFF367UNR,ENCFF502WDT |
| 1147 | 7 | 24358648 | 24358702 | 54 | SPRET | 7 | 24358662 | 24358696 | 35 | - | 21.6606 | 2.88E-08 | 0.0187 | GTGCAGGTGTCATCAGAGACCTGTAGAGGGCACCA | 2 | ENCFF367UNR,ENCFF502WDT |
| 1148 | 7 | 26947852 | 26948054 | 202 | CAST,SPRET | 7 | 26947981 | 26948014 | 34 | - | 16.4472 | 9.61E-07 | 0.0901 | TAGCTGTCTTCAGACACACCAGAAGAGGGCGTCA | | |
| 1149 | 7 | 27170467 | 27170563 | 96 | 129S1,BTBR,C57L,CAST,ILNJ2,LP,MOLF,NZW,SPRET | 7 | 27170501 | 27170534 | 34 | - | 17.9675 | 3.61E-07 | 0.0545 | CTGAATTACAGTCAATCCCCTGCAGAGGGCACGG | | |
| 1149 | 7 | 27170467 | 27170563 | 96 | 129S1,BTBR,C57L,CAST,ILNJ2,LP,MOLF,NZW,SPRET | 7 | 27170549 | 27170582 | 34 | - | 17.9675 | 3.61E-07 | 0.0545 | CTGAATTACAGTCAATCCCCTGCAGAGGGCACGG | | |
| 1150 | 7 | 27948593 | 27948958 | 365 | SPRET | 7 | 27948575 | 27948593 | 19 | - | 19.7213 | 1.37E-07 | 0.053 | AGGCCAGAAGAGGGCAGCA | 1 | ENCFF093ZRU |
| 1150 | 7 | 27948593 | 27948958 | 365 | SPRET | 7 | 27948575 | 27948608 | 34 | - | 21.878 | 2.19E-08 | 0.00797 | GTGAAGGACCCAGGGAGGCCAGAAGAGGGCAGCA | 1 | ENCFF093ZRU |
| 1151 | 7 | 28128163 | 28128381 | 218 | SPRET | 7 | 28128278 | 28128311 | 34 | - | 16.4634 | 9.51E-07 | 0.0901 | TAGCTGTCTTCAGACGCACCAGAAGAGGGCATCA | 1 | ENCFF093ZRU |
| 1152 | 7 | 28876988 | 28877189 | 201 | SPRET | 7 | 28877113 | 28877146 | 34 | - | 16.4472 | 9.61E-07 | 0.0901 | TAGCTGTCTTCAGACACACCAGAAGAGGGCGTCA | | |
| 1153 | 7 | 28967158 | 28968149 | 991 | SPRET | 7 | 28967222 | 28967255 | 34 | - | 17.7398 | 4.19E-07 | 0.0596 | TTGCTCTCTTCAGACACACCAGTAGAGGGCATCA | 2 | ENCFF093ZRU,ENCFF327LYS |
| 1154 | 7 | 29206849 | 29207053 | 204 | SPRET | 7 | 29206967 | 29207001 | 35 | - | 18.5963 | 2.10E-07 | 0.0658 | CTGTAGCTGTCTTCAGACACCAGAAGAGGGCGTCA | | |
| 1155 | 7 | 29225674 | 29225896 | 222 | CAST,CE,FVB,ILNJ2,MOLF,SPRET,WSB | 7 | 29225801 | 29225834 | 34 | - | 17.8455 | 3.91E-07 | 0.057 | TAGCTGTCCTCAGACACTCCAGAAGAGGGCGTCA | | |
| 1156 | 7 | 29296576 | 29296907 | 331 | SPRET | 7 | 29296866 | 29296884 | 19 | - | 19.2951 | 1.88E-07 | 0.0611 | CCGCCACTAGATGGTGCTA | 26 | ENCFF093ZRU,ENCFF367UNR,ENCFF201JUA,ENCFF099QNE,ENCFF821BIA,ENCFF327LYS,ENCFF024JOZ,ENCFF907XEG,ENCFF609MDN,ENCFF136ZDK,ENCFF165LXO,ENCFF321AGQ,ENCFF089GBC,ENCFF194HVL,ENCFF589ZBJ,ENCFF491RJK,ENCFF055HAP,ENCFF198SSR,ENCFF065ODV,ENCFF092NMJ,ENCFF957SPH,ENCFF052KVO,ENCFF502WDT,ENCFF142CNG,ENCFF430PPJ,ENCFF883UPM |
| 1157 | 7 | 29564830 | 29564933 | 103 | AJ,AKR,C57L,DBA1J,DBA2J,MA/My,MOLF,SMJ | 7 | 29564918 | 29564951 | 34 | - | 16.8537 | 7.43E-07 | 0.0809 | GAGCAGGTGCAGCAGGGGACAGCAGGTGAAGCAG | | |
| 1158 | 7 | 29853542 | 29854538 | 996 | AJ,AKR,BALB,BTBR,C57L,DBA1J,DBA2J,SMJ | 7 | 29853659 | 29853677 | 19 | - | 20.3607 | 8.29E-08 | 0.042 | AGGCCAGAAGAGGGCACCA | 3 | ENCFF367UNR,ENCFF093ZRU,ENCFF052KVO |
| 1158 | 7 | 29853542 | 29854538 | 996 | AJ,AKR,BALB,BTBR,C57L,DBA1J,DBA2J,SMJ | 7 | 29853659 | 29853693 | 35 | - | 25.8716 | 1.22E-09 | 0.00341 | CTGCATGCATGCCTACAGGCCAGAAGAGGGCACCA | 3 | ENCFF367UNR,ENCFF093ZRU,ENCFF052KVO |
| 1159 | 7 | 29907594 | 29907805 | 211 | CAST,ILNJ2,SPRET,WSB | 7 | 29907719 | 29907752 | 34 | - | 17.439 | 5.10E-07 | 0.0663 | TAGCAGTCTTCAGACACTCCAGAAGAGGGCATCA | | |
| 1160 | 7 | 29914917 | 29915122 | 205 | AJ,AKR,BALB,BTBR,C57L,CAST,DBA1J,DBA2J,MOLF,SMJ,SPRET | 7 | 29915043 | 29915061 | 19 | - | 17.6393 | 5.98E-07 | 0.106 | GCTCCAGAAGAGGGCGCCA | 3 | ENCFF093ZRU,ENCFF821BIA,ENCFF367UNR |
| 1160 | 7 | 29914917 | 29915122 | 205 | AJ,AKR,BALB,BTBR,C57L,CAST,DBA1J,DBA2J,MOLF,SMJ,SPRET | 7 | 29915043 | 29915076 | 34 | - | 18.8374 | 2.01E-07 | 0.04 | TAGCTGTCTTCAGACGCTCCAGAAGAGGGCGCCA | 3 | ENCFF093ZRU,ENCFF821BIA,ENCFF367UNR |
| 1161 | 7 | 30443885 | 30444359 | 474 | SPRET | 7 | 30443973 | 30444006 | 34 | - | 16.7967 | 7.71E-07 | 0.0827 | CTGCAGTACATAAAAAGGTGAGAAGAGGGCAGTA | 2 | ENCFF093ZRU,ENCFF198SSR |
| 1162 | 7 | 33812219 | 33812433 | 214 | CAST,DBA1J,DBA2J,MOLF,NZB,NZO,NZW,SMJ,SPRET | 7 | 33812345 | 33812378 | 34 | - | 17.6992 | 4.30E-07 | 0.0601 | TAGCTGTCCTCAGATACTCCAGAAGAGGGCGTCA | | |
| 1163 | 7 | 35448473 | 35448575 | 102 | SPRET | 7 | 35448463 | 35448497 | 35 | - | 16.9725 | 5.50E-07 | 0.111 | GTGCAGTGCAGTGGTACCACCACATGATGGAGTAG | 1 | ENCFF093ZRU |
| 1163 | 7 | 35448473 | 35448575 | 102 | SPRET | 7 | 35448557 | 35448591 | 35 | - | 16.9725 | 5.50E-07 | 0.111 | GTGCAGTGCAGTGGTACCACCACATGATGGAGTAG | 1 | ENCFF093ZRU |
| 1164 | 7 | 35708388 | 35708500 | 112 | NZB,NZO | 7 | 35708384 | 35708402 | 19 | - | 16.9508 | 9.36E-07 | 0.127 | GGAGCAGCAGGGGGCAGCA | | |
| 1164 | 7 | 35708388 | 35708500 | 112 | NZB,NZO | 7 | 35708418 | 35708436 | 19 | - | 16.9508 | 9.36E-07 | 0.127 | GGAGCAGCAGGGGGCAGCA | | |
| 1165 | 7 | 42127413 | 42127866 | 453 | SPRET | 7 | 42127515 | 42127548 | 34 | - | 16.4634 | 9.51E-07 | 0.0901 | TAGCTGTCTTCAGACGCACCAGAAGAGGGCATCA | 1 | ENCFF093ZRU |
| 1166 | 7 | 42937025 | 42937928 | 903 | CAST,SPRET | 7 | 42937136 | 42937169 | 34 | - | 18.3821 | 2.73E-07 | 0.0476 | GTACAGTTACTGAGGAGGCCAGAAGAGGGAATTG | | |
| 1167 | 7 | 43205618 | 43206076 | 458 | C57L,CAST,CE,DBA1J,DBA2J,ILNJ2,MOLF,NZB,NZO,NZW,PJ,SJL,SMJ,SPRET,WSB | 7 | 43205744 | 43205777 | 34 | - | 16.4472 | 9.61E-07 | 0.0901 | TAGCTGTCTTCAGACACACCAGAAGAGGGCGTCA | | |
| 1168 | 7 | 44944325 | 44944548 | 223 | CAST,MOLF,SPRET | 7 | 44944450 | 44944483 | 34 | - | 16.7073 | 8.16E-07 | 0.0846 | TAGCTGTCTTCAGATGCACCAGAAGAGGGAGCCA | 5 | ENCFF093ZRU,ENCFF883UPM,ENCFF502WDT,ENCFF907XEG,ENCFF055HAP |
| 1169 | 7 | 45193765 | 45194552 | 787 | C57L,C58J | 7 | 45194132 | 45194150 | 19 | - | 22.0656 | 1.91E-08 | 0.0243 | AGGCCACAAGGGGGCACCA | 15 | ENCFF821BIA,ENCFF024JOZ,ENCFF136ZDK,ENCFF907XEG,ENCFF589ZBJ,ENCFF055HAP,ENCFF093ZRU,ENCFF198SSR,ENCFF367UNR,ENCFF883UPM,ENCFF052KVO,ENCFF065ODV,ENCFF502WDT,ENCFF142CNG,ENCFF099QNE |
| 1169 | 7 | 45193765 | 45194552 | 787 | C57L,C58J | 7 | 45194132 | 45194166 | 35 | - | 26.7706 | 5.73E-10 | 0.00258 | TTGCAGGTGCATGTGGAGGCCACAAGGGGGCACCA | 15 | ENCFF821BIA,ENCFF024JOZ,ENCFF136ZDK,ENCFF907XEG,ENCFF589ZBJ,ENCFF055HAP,ENCFF093ZRU,ENCFF198SSR,ENCFF367UNR,ENCFF883UPM,ENCFF052KVO,ENCFF065ODV,ENCFF502WDT,ENCFF142CNG,ENCFF099QNE |
| 1170 | 7 | 45227428 | 45227631 | 203 | 129S1,C3H,CAST,DBA1J,DBA2J,ILNJ2,MOLF,NZW,PJ,SJL,SPRET | 7 | 45227553 | 45227571 | 19 | - | 17.5574 | 6.32E-07 | 0.106 | ACTCCAGAAGAGGGCGCCA | 4 | ENCFF367UNR,ENCFF099QNE,ENCFF821BIA,ENCFF093ZRU |
| 1170 | 7 | 45227428 | 45227631 | 203 | 129S1,C3H,CAST,DBA1J,DBA2J,ILNJ2,MOLF,NZW,PJ,SJL,SPRET | 7 | 45227553 | 45227586 | 34 | - | 18.3659 | 2.76E-07 | 0.0476 | TAGCTGTCTTCAGACACTCCAGAAGAGGGCGCCA | 4 | ENCFF367UNR,ENCFF099QNE,ENCFF821BIA,ENCFF093ZRU |
| 1171 | 7 | 46474406 | 46474608 | 202 | C57L,CAST,CE,ILNJ2,MOLF,NZW,PJ,SJL,WSB | 7 | 46474518 | 46474551 | 34 | - | 17.9106 | 3.74E-07 | 0.0556 | TAGCTGTCTTCAGACACTCCAGAAGAGGGCACCA | 1 | ENCFF093ZRU |
| 1172 | 7 | 46499239 | 46499436 | 197 | CAST,ILNJ2,MOLF,NZW,PJ,SJL,SPRET | 7 | 46499363 | 46499396 | 34 | - | 17.374 | 5.32E-07 | 0.0676 | TAGCTGTCTTCAGACCCTCCAGAAGAGGGCATCA | | |
| 1173 | 7 | 51911701 | 51912095 | 394 | CAST,ILNJ2,KK,MOLF,NZW,PJ,SPRET | 7 | 51911707 | 51911725 | 19 | - | 17.7705 | 5.49E-07 | 0.101 | TGGCCAGTTGAGGGCGCAA | | |
| 1173 | 7 | 51911701 | 51912095 | 394 | CAST,ILNJ2,KK,MOLF,NZW,PJ,SPRET | 7 | 51911707 | 51911740 | 34 | - | 18.935 | 1.88E-07 | 0.0387 | CAGCATCGTTCTTCCTGGCCAGTTGAGGGCGCAA | | |
| 1174 | 7 | 52217735 | 52218006 | 271 | SPRET | 7 | 52217858 | 52217892 | 35 | - | 20.0917 | 8.19E-08 | 0.0359 | CTGCATTTCTGAGTTCACTCCACTAGTAGGCAGCC | | |
| 1175 | 7 | 53455492 | 53455862 | 370 | 129S1,AJ,AKR,BALB,BTBR,BUB,C3H,C58J,CBA,DBA1J,DBA2J,FVB,ILNJ2,KK,MOLF,MRL,NOD,NOR,NZB,NZO,NZW,PJ,PL,RF,SEA,SJL,SMJ,SWR,WSB | 7 | 53455809 | 53455842 | 34 | - | 16.5285 | 9.13E-07 | 0.0896 | GAGCGGGTGTGGCTGCCACCCCAAGATGGCGCCC | | |
| 1176 | 7 | 58110186 | 58110385 | 199 | 129S1,AKR,BALB,BTBR,C3H,C57L,CAST,CBA,DBA1J,DBA2J,ILNJ2,KK,LP,MOLF,NOD,NOR,NZW,PJ,PL,RF,SEA,SMJ,TH | 7 | 58110310 | 58110328 | 19 | - | 17.5574 | 6.32E-07 | 0.106 | ACTCCAGAAGAGGGCGCCA | | |
| 1176 | 7 | 58110186 | 58110385 | 199 | 129S1,AKR,BALB,BTBR,C3H,C57L,CAST,CBA,DBA1J,DBA2J,ILNJ2,KK,LP,MOLF,NOD,NOR,NZW,PJ,PL,RF,SEA,SMJ,TH | 7 | 58110310 | 58110343 | 34 | - | 18.3659 | 2.76E-07 | 0.0476 | TAGCTGTCTTCAGACACTCCAGAAGAGGGCGCCA | | |
| 1177 | 7 | 59309561 | 59310296 | 735 | SPRET | 7 | 59310202 | 59310235 | 34 | - | 18.7398 | 2.14E-07 | 0.0413 | TTGCTGTCTTCAGATGTACCAGAAGAGGGCATCA | | |
| 1178 | 7 | 64923743 | 64923874 | 131 | CE,DBA1J,DBA2J,ILNJ2,KK,MOLF,NZW,PJ,SPRET,WSB | 7 | 64923748 | 64923782 | 35 | - | 16.6055 | 6.79E-07 | 0.125 | CAGCAGAGGGAGGGCGGGGTCAGCAGAGGGAGGGC | | |
| 1178 | 7 | 64923743 | 64923874 | 131 | CE,DBA1J,DBA2J,ILNJ2,KK,MOLF,NZW,PJ,SPRET,WSB | 7 | 64923804 | 64923838 | 35 | - | 16.1835 | 8.62E-07 | 0.142 | CAGCAGAGGGAGGGCGGGGTCAGCAGAGGGAGGGT | | |
| 1179 | 7 | 71892084 | 71892273 | 189 | CAST,ILNJ2,MOLF,NZB,NZO | 7 | 71892204 | 71892237 | 34 | - | 16.9593 | 6.95E-07 | 0.0781 | TTGCTCTCTTCAGACACACCAGAAGAGGGCATCA | | |
| 1180 | 7 | 72493682 | 72494411 | 729 | 129S1,AJ,AKR,BALB,BUB,C3H,C57L,C58J,CAST,CBA,CE,DBA1J,DBA2J,FVB,ILNJ2,KK,MOLF,MRL,NOD,NOR,NZB,NZO,NZW,PJ,PL,RF,RH2,SEA,SJL,SMJ,SPRET,SWR,TH,WSB | 7 | 72493679 | 72493713 | 35 | - | 22.1284 | 2.08E-08 | 0.0153 | CTGCCGCCACACCCGCTCCCCACAAGAGGGTACTA | | |
| 1181 | 7 | 78898442 | 78898672 | 230 | 129S1,C3H,CAST,CBA,CE,DBA1J,DBA2J,KK,MOLF,NZB,NZO,NZW,PJ,SEA,SMJ,SPRET,SWR,TH,WSB | 7 | 78898570 | 78898603 | 34 | - | 17.8455 | 3.91E-07 | 0.057 | TAGCTGTCCTCAGACACTCCAGAAGAGGGCGTCA | | |
| 1182 | 7 | 78995801 | 78996628 | 827 | C58J,CE,WSB | 7 | 78995851 | 78995884 | 34 | - | 19.1789 | 1.58E-07 | 0.0349 | ATGCAGTAAGCACTCTGAACAGATGGGGGCGCTG | 26 | ENCFF093ZRU,ENCFF367UNR,ENCFF099QNE,ENCFF201JUA,ENCFF951GOP,ENCFF321AGQ,ENCFF065ODV,ENCFF601JVE,ENCFF821BIA,ENCFF907XEG,ENCFF327LYS,ENCFF055HAP,ENCFF957SPH,ENCFF609MDN,ENCFF089GBC,ENCFF024JOZ,ENCFF589ZBJ,ENCFF198SSR,ENCFF052KVO,ENCFF502WDT,ENCFF281QMV,ENCFF092NMJ,ENCFF136ZDK,ENCFF142CNG,ENCFF430PPJ,ENCFF165LXO |
| 1183 | 7 | 80217359 | 80217562 | 203 | AJ,AKR,BALB,BUB,CE,FVB,MRL,NZB,NZO,NZW,PL,RF,RH2,SJL,SWR,WSB | 7 | 80217484 | 80217502 | 19 | - | 17.5574 | 6.32E-07 | 0.106 | ACTCCAGAAGAGGGCGCCA | 1 | ENCFF093ZRU |
| 1183 | 7 | 80217359 | 80217562 | 203 | AJ,AKR,BALB,BUB,CE,FVB,MRL,NZB,NZO,NZW,PL,RF,RH2,SJL,SWR,WSB | 7 | 80217484 | 80217517 | 34 | - | 18.3659 | 2.76E-07 | 0.0476 | TAGCTGTCTTCAGACACTCCAGAAGAGGGCGCCA | 1 | ENCFF093ZRU |
| 1184 | 7 | 81437214 | 81437429 | 215 | SPRET | 7 | 81437327 | 81437360 | 34 | - | 19.0325 | 1.75E-07 | 0.0373 | CAGCTGTCTTCAGACGCACCAGAAGAGGGCATCA | 1 | ENCFF093ZRU |
| 1185 | 7 | 88877905 | 88878126 | 221 | CAST,MOLF,SPRET | 7 | 88878030 | 88878063 | 34 | - | 17.9106 | 3.74E-07 | 0.0556 | TAGCTGTCTTCAGACACTCCAGAAGAGGGCACCA | | |
| 1186 | 7 | 92828155 | 92828874 | 719 | 129S1,CAST,MOLF,SMJ,SPRET | 7 | 92828469 | 92828503 | 35 | - | 16.0367 | 9.36E-07 | 0.147 | ATGCACTGCCCCTGGGCCGCAAGAAGGGGGCCAAG | | |
| 1187 | 7 | 98800622 | 98800686 | 64 | C3H,CAST,CBA,DBA1J,DBA2J,ILNJ2,MOLF,PJ,SEA,SMJ | 7 | 98800614 | 98800632 | 19 | - | 22.918 | 8.34E-09 | 0.0179 | CAGCCACAAGGGGGCGCCG | 38 | ENCFF093ZRU,ENCFF201JUA,ENCFF327LYS,ENCFF907XEG,ENCFF821BIA,ENCFF024JOZ,ENCFF092NMJ,ENCFF065ODV,ENCFF589ZBJ,ENCFF089GBC,ENCFF136ZDK,ENCFF055HAP,ENCFF198SSR,ENCFF165LXO,ENCFF502WDT,ENCFF430PPJ,ENCFF321AGQ,ENCFF883UPM,ENCFF951GOP,ENCFF210BTO,ENCFF957SPH,ENCFF575MGI,ENCFF609MDN,ENCFF281QMV,ENCFF316FYN,ENCFF194HVL,ENCFF142CNG,ENCFF089EWX,ENCFF533APC,ENCFF097CAQ,ENCFF052KVO,ENCFF491RJK,ENCFF601JVE,ENCFF349BUL,ENCFF033UQX,ENCFF805QBU,ENCFF784ASD |
| 1187 | 7 | 98800622 | 98800686 | 64 | C3H,CAST,CBA,DBA1J,DBA2J,ILNJ2,MOLF,PJ,SEA,SMJ | 7 | 98800614 | 98800648 | 35 | - | 18.055 | 2.91E-07 | 0.0783 | TTGTCTGAAGGCCTCACAGCCACAAGGGGGCGCCG | 39 | ENCFF093ZRU,ENCFF201JUA,ENCFF327LYS,ENCFF907XEG,ENCFF821BIA,ENCFF024JOZ,ENCFF092NMJ,ENCFF065ODV,ENCFF589ZBJ,ENCFF089GBC,ENCFF136ZDK,ENCFF055HAP,ENCFF198SSR,ENCFF165LXO,ENCFF502WDT,ENCFF430PPJ,ENCFF321AGQ,ENCFF883UPM,ENCFF951GOP,ENCFF210BTO,ENCFF957SPH,ENCFF575MGI,ENCFF609MDN,ENCFF281QMV,ENCFF316FYN,ENCFF194HVL,ENCFF142CNG,ENCFF089EWX,ENCFF533APC,ENCFF097CAQ,ENCFF052KVO,ENCFF491RJK,ENCFF601JVE,ENCFF349BUL,ENCFF033UQX,ENCFF805QBU,ENCFF784ASD,ENCFF086ZTV |
| 1187 | 7 | 98800622 | 98800686 | 64 | C3H,CAST,CBA,DBA1J,DBA2J,ILNJ2,MOLF,PJ,SEA,SMJ | 7 | 98800678 | 98800696 | 19 | - | 21.8689 | 2.29E-08 | 0.0265 | CAGCCACAAGAGGGCGCCG | 40 | ENCFF093ZRU,ENCFF201JUA,ENCFF327LYS,ENCFF907XEG,ENCFF821BIA,ENCFF024JOZ,ENCFF092NMJ,ENCFF065ODV,ENCFF589ZBJ,ENCFF089GBC,ENCFF136ZDK,ENCFF055HAP,ENCFF198SSR,ENCFF165LXO,ENCFF502WDT,ENCFF430PPJ,ENCFF321AGQ,ENCFF883UPM,ENCFF951GOP,ENCFF210BTO,ENCFF957SPH,ENCFF575MGI,ENCFF609MDN,ENCFF281QMV,ENCFF316FYN,ENCFF194HVL,ENCFF142CNG,ENCFF089EWX,ENCFF533APC,ENCFF097CAQ,ENCFF052KVO,ENCFF491RJK,ENCFF601JVE,ENCFF349BUL,ENCFF033UQX,ENCFF805QBU,ENCFF784ASD,ENCFF086ZTV,ENCFF611HDQ |
| 1187 | 7 | 98800622 | 98800686 | 64 | C3H,CAST,CBA,DBA1J,DBA2J,ILNJ2,MOLF,PJ,SEA,SMJ | 7 | 98800678 | 98800711 | 34 | - | 17.6585 | 4.42E-07 | 0.0612 | AAGGCCTCACAGCCACAGCCACAAGAGGGCGCCG | 41 | ENCFF093ZRU,ENCFF201JUA,ENCFF327LYS,ENCFF907XEG,ENCFF821BIA,ENCFF024JOZ,ENCFF092NMJ,ENCFF065ODV,ENCFF589ZBJ,ENCFF089GBC,ENCFF136ZDK,ENCFF055HAP,ENCFF198SSR,ENCFF165LXO,ENCFF502WDT,ENCFF430PPJ,ENCFF321AGQ,ENCFF883UPM,ENCFF951GOP,ENCFF210BTO,ENCFF957SPH,ENCFF575MGI,ENCFF609MDN,ENCFF281QMV,ENCFF316FYN,ENCFF194HVL,ENCFF142CNG,ENCFF089EWX,ENCFF533APC,ENCFF097CAQ,ENCFF052KVO,ENCFF491RJK,ENCFF601JVE,ENCFF349BUL,ENCFF033UQX,ENCFF805QBU,ENCFF784ASD,ENCFF086ZTV,ENCFF611HDQ,ENCFF504JDS |
| 1188 | 7 | 102723536 | 102723653 | 117 | CAST,TH,WSB | 7 | 102723633 | 102723666 | 34 | - | 16.6016 | 8.72E-07 | 0.0874 | CAGCCCTGGTGCCCGCCGCCGGGCGGGGGCACAA | | |
| 1189 | 7 | 105238105 | 105238324 | 219 | BUB,CAST,CE,FVB,KK,MOLF,NOD,NZW,SJL,SPRET,SWR,TH | 7 | 105238214 | 105238247 | 34 | - | 17.9431 | 3.66E-07 | 0.0552 | CAGCTGTCCTCAGACACTCCAGAAGAGGGAGTCA | | |
| 1190 | 7 | 108668185 | 108668847 | 662 | SPRET | 7 | 108668544 | 108668577 | 34 | - | 18.0976 | 3.31E-07 | 0.052 | TTGCAATGCTCGTGGAAGCCAGAAAGGGGTGCTT | | |
| 1191 | 7 | 108889853 | 108889956 | 103 | NOD,SWR | 7 | 108889883 | 108889917 | 35 | - | 16.0183 | 9.45E-07 | 0.148 | AGACAGTCACAGCCGGCTTCCAGCAGATGGCAGGC | 11 | ENCFF092NMJ,ENCFF883UPM,ENCFF367UNR,ENCFF609MDN,ENCFF093ZRU,ENCFF281QMV,ENCFF099QNE,ENCFF327LYS,ENCFF089GBC,ENCFF024JOZ,ENCFF907XEG |
| 1192 | 7 | 109282919 | 109283723 | 804 | CAST | 7 | 109283308 | 109283341 | 34 | - | 17.6341 | 4.49E-07 | 0.0617 | ATACAGTACCTGTAGAGGTCAGAAGAGGGCATCA | 1 | ENCFF093ZRU |
| 1193 | 7 | 109427832 | 109428044 | 212 | CAST,ILNJ2,MOLF,NOD,SMJ,SPRET,SWR | 7 | 109427959 | 109427992 | 34 | - | 16.9187 | 7.13E-07 | 0.0792 | TAGCTGTCTTCAGACGCACCAGAAGAGGGCGTCA | | |
| 1194 | 7 | 109805736 | 109805939 | 203 | CAST,CE,ILNJ2,KK,MOLF,NOD,SMJ,SPRET,SWR,WSB | 7 | 109805861 | 109805879 | 19 | - | 17.3443 | 7.26E-07 | 0.112 | GCACCAGAAGAGGGCAGTA | | |
| 1194 | 7 | 109805736 | 109805939 | 203 | CAST,CE,ILNJ2,KK,MOLF,NOD,SMJ,SPRET,SWR,WSB | 7 | 109805861 | 109805894 | 34 | - | 17.7886 | 4.06E-07 | 0.0584 | TAGCTGTCTTCAGATGCACCAGAAGAGGGCAGTA | | |
| 1195 | 7 | 112442123 | 112442345 | 222 | 129S1,AJ,AKR,BALB,BUB,C3H,C57L,CAST,CBA,DBA1J,DBA2J,ILNJ2,MOLF,MRL,NOD,NOR,NZB,NZO,NZW,PL,RH2,SJL,SMJ,SPRET | 7 | 112442248 | 112442266 | 19 | - | 17.8852 | 5.08E-07 | 0.0973 | ACACCAGAAGAGGGCACCA | 2 | ENCFF093ZRU |
| 1196 | 7 | 124203005 | 124203198 | 193 | SPRET | 7 | 124203129 | 124203147 | 19 | - | 17.5574 | 6.32E-07 | 0.106 | ACTCCAGAAGAGGGCGCCA | 3 | ENCFF367UNR,ENCFF099QNE,ENCFF093ZRU |
| 1196 | 7 | 124203005 | 124203198 | 193 | SPRET | 7 | 124203129 | 124203162 | 34 | - | 18.3659 | 2.76E-07 | 0.0476 | TAGCTGTCTTCAGACACTCCAGAAGAGGGCGCCA | 3 | ENCFF367UNR,ENCFF099QNE,ENCFF093ZRU |
| 1197 | 7 | 124985399 | 124985806 | 407 | MOLF | 7 | 124985767 | 124985800 | 34 | - | 17.0244 | 6.67E-07 | 0.0768 | GAGCAGGTGTGGCAGCAGTCCCAAGATGGCGCCC | | |
| 1198 | 7 | 126682650 | 126682860 | 210 | AJ,AKR,BALB,BUB,C58J,CAST,MOLF,MRL,NOD,NZB,NZO,NZW,PL,RH2,SEA,SJL,SPRET,SWR,TH,WSB | 7 | 126682778 | 126682811 | 34 | - | 16.5203 | 9.18E-07 | 0.0899 | TAGCTGTCTTCCAACACTCCAGAAGAGGGCGTCA | | |
| 1199 | 7 | 127057828 | 127058048 | 220 | MOLF,SPRET,TH | 7 | 127057954 | 127057987 | 34 | - | 17.6504 | 4.44E-07 | 0.0613 | CAGCTGTCCTCAGACACCCCAGAAGAGGGAGTCA | | |
| 1200 | 7 | 127087491 | 127087700 | 209 | AJ,MOLF,NOD,SPRET,TH,WSB | 7 | 127087614 | 127087632 | 19 | - | 17.8852 | 5.08E-07 | 0.0973 | ACACCAGAAGAGGGCACCA | 3 | ENCFF367UNR,ENCFF093ZRU |
| 1201 | 7 | 127250696 | 127250913 | 217 | MOLF,SPRET | 7 | 127250815 | 127250848 | 34 | - | 17.8455 | 3.91E-07 | 0.057 | TAGCTGTCCTCAGACACTCCAGAAGAGGGCGTCA | | |
| 1202 | 7 | 128191426 | 128191630 | 204 | 129S1,AJ,AKR,BALB,BTBR,BUB,C3H,C57L,C58J,CAST,CE,FVB,ILNJ2,KK,LP,MA/My,MOLF,MRL,NOD,NZB,NZO,NZW,PJ,PL,RF,RH2,SEA,SJL,SPRET,SWR,TH,WSB | 7 | 128191551 | 128191569 | 19 | - | 17.5574 | 6.32E-07 | 0.106 | ACTCCAGAAGAGGGCGCCA | 6 | ENCFF093ZRU,ENCFF367UNR,ENCFF055HAP,ENCFF821BIA,ENCFF502WDT |
| 1202 | 7 | 128191426 | 128191630 | 204 | 129S1,AJ,AKR,BALB,BTBR,BUB,C3H,C57L,C58J,CAST,CE,FVB,ILNJ2,KK,LP,MA/My,MOLF,MRL,NOD,NZB,NZO,NZW,PJ,PL,RF,RH2,SEA,SJL,SPRET,SWR,TH,WSB | 7 | 128191551 | 128191584 | 34 | - | 18.561 | 2.42E-07 | 0.0451 | AAGCTGTCTTCAGACACTCCAGAAGAGGGCGCCA | 6 | ENCFF093ZRU,ENCFF367UNR,ENCFF055HAP,ENCFF821BIA,ENCFF502WDT |
| 1203 | 7 | 128285722 | 128285919 | 197 | 129S1,BTBR,C57L,CAST,ILNJ2,MOLF,PJ,SPRET | 7 | 128285848 | 128285866 | 19 | - | 16.9016 | 9.66E-07 | 0.13 | ATTCCAGAAGAGGGCACCA | | |
| 1203 | 7 | 128285722 | 128285919 | 197 | 129S1,BTBR,C57L,CAST,ILNJ2,MOLF,PJ,SPRET | 7 | 128285848 | 128285881 | 34 | - | 17.8455 | 3.91E-07 | 0.057 | TAGCTGTCTTCAGACATTCCAGAAGAGGGCACCA | | |
| 1204 | 7 | 130080794 | 130080989 | 195 | MOLF,SPRET,WSB | 7 | 130080918 | 130080936 | 19 | - | 17.5574 | 6.32E-07 | 0.106 | ACTCCAGAAGAGGGCGCCA | 4 | ENCFF821BIA,ENCFF367UNR,ENCFF093ZRU,ENCFF099QNE |
| 1204 | 7 | 130080794 | 130080989 | 195 | MOLF,SPRET,WSB | 7 | 130080918 | 130080951 | 34 | - | 18.3659 | 2.76E-07 | 0.0476 | TAGCTGTCTTCAGACACTCCAGAAGAGGGCGCCA | 4 | ENCFF821BIA,ENCFF367UNR,ENCFF093ZRU,ENCFF099QNE |
| 1205 | 7 | 131429510 | 131429878 | 368 | SPRET | 7 | 131429807 | 131429840 | 34 | - | 24.2033 | 3.31E-09 | 0.00197 | CTGCAGCGCCCTCCACAGGCAGCAGAAGGCAGAA | | |
| 1206 | 7 | 133277922 | 133277976 | 54 | SPRET | 7 | 133277933 | 133277951 | 19 | - | 17.2787 | 7.58E-07 | 0.115 | GCTCCAGCAGGGGGTGCTA | 17 | ENCFF099QNE,ENCFF052KVO,ENCFF092NMJ,ENCFF089GBC,ENCFF198SSR,ENCFF136ZDK,ENCFF024JOZ,ENCFF883UPM,ENCFF093ZRU,ENCFF957SPH,ENCFF327LYS,ENCFF055HAP,ENCFF367UNR,ENCFF502WDT,ENCFF589ZBJ,ENCFF907XEG,ENCFF533APC |
| 1207 | 7 | 135404046 | 135404266 | 220 | CAST,ILNJ2,MOLF,SPRET | 7 | 135404170 | 135404203 | 34 | - | 16.7724 | 7.83E-07 | 0.0827 | TAGCTGTCTTCAGATGCACCAGAAGAGGGCGTCA | | |
| 1208 | 7 | 135424592 | 135424696 | 104 | CAST | 7 | 135424629 | 135424662 | 34 | - | 27.2602 | 1.99E-10 | 0.000297 | GTGCAGTTCCCACAGCAGCCAGAAGAGGTTGCTG | | |
| 1209 | 7 | 136409524 | 136410451 | 927 | ILNJ2 | 7 | 136409885 | 136409918 | 34 | - | 26.5203 | 4.09E-10 | 0.000451 | ATGCAGTACCAAAGGAGGCCAGAAGAGGGCATCA | 6 | ENCFF327LYS,ENCFF093ZRU,ENCFF052KVO,ENCFF367UNR,ENCFF821BIA,ENCFF099QNE |
| 1210 | 7 | 138622828 | 138622924 | 96 | CBA,MOLF,NZB,NZO | 7 | 138622913 | 138622946 | 34 | - | 18.1382 | 3.22E-07 | 0.0511 | CTTCAGTGAGTGCACAGGGCAGCAGGGGGCGTGG | | |
| 1211 | 7 | 139078982 | 139079219 | 237 | AKR,BUB,CAST,MOLF,SPRET | 7 | 139079106 | 139079139 | 34 | - | 17.9187 | 3.72E-07 | 0.0556 | TAGCTGTCCTCACATACTCCAGAAGAGGGCATCA | | |
| 1212 | 7 | 139237731 | 139237787 | 56 | 129S1,AJ,AKR,BALB,BTBR,BUB,C3H,C57L,C58J,CBA,CE,DBA1J,DBA2J,FVB,ILNJ2,LP,MA/My,MOLF,NOD,NOR,PJ,PL,RF,SEA,SJL,SMJ,TH,WSB | 7 | 139237734 | 139237767 | 34 | - | 16.6016 | 8.72E-07 | 0.0874 | GTGCCCTGCTGATGGTGCCCTGCTGATGGTGCCC | | |
| 1212 | 7 | 139237731 | 139237787 | 56 | 129S1,AJ,AKR,BALB,BTBR,BUB,C3H,C57L,C58J,CBA,CE,DBA1J,DBA2J,FVB,ILNJ2,LP,MA/My,MOLF,NOD,NOR,PJ,PL,RF,SEA,SJL,SMJ,TH,WSB | 7 | 139237748 | 139237781 | 34 | - | 16.6016 | 8.72E-07 | 0.0874 | GTGCCCTGCTGATGGTGCCCTGCTGATGGTGCCC | | |
| 1212 | 7 | 139237731 | 139237787 | 56 | 129S1,AJ,AKR,BALB,BTBR,BUB,C3H,C57L,C58J,CBA,CE,DBA1J,DBA2J,FVB,ILNJ2,LP,MA/My,MOLF,NOD,NOR,PJ,PL,RF,SEA,SJL,SMJ,TH,WSB | 7 | 139237762 | 139237795 | 34 | - | 16.6016 | 8.72E-07 | 0.0874 | GTGCCCTGCTGATGGTGCCCTGCTGATGGTGCCC | | |
| 1213 | 7 | 140859812 | 140860016 | 204 | CAST,CE,ILNJ2,MOLF,NZW,SPRET | 7 | 140859938 | 140859971 | 34 | - | 16.4472 | 9.61E-07 | 0.0901 | TAGCTGTCTTCAGACACACCAGAAGAGGGCGTCA | | |

| | | | | | | | | | | | | | | | | |
|---|---|---|---|---|---|---|---|---|---|---|---|---|---|---|---|---|
| 1214 | 7 | 143080490 | 143080704 | 214 | C58J,CAST,LP,MA/My,MOLF,PJ,SMJ,SPRET | 7 | 143080616 | 143080649 | 34 | - | 17.3902 | 5.27E-07 | 0.0672 | TAGCTGTCCTCAGACACTCCAGAAGAGGGCATCA | | |
| 1215 | 7 | 144018218 | 144018434 | 216 | SPRET | 7 | 144018378 | 144018411 | 34 | - | 17.8537 | 3.89E-07 | 0.057 | TTGCAGTGCCCACGAGGGCCCAGAGAAGGCATTG | | |
| 1216 | 7 | 144137572 | 144137644 | 72 | SPRET | 7 | 144137557 | 144137591 | 35 | - | 28.055 | 1.84E-10 | 0.00248 | CTGTAGTGCCCAGCAGAGACCAGCAGAGGGCCCCT | 29 | ENCFF093ZRU,ENCFF367UNR,ENCFF099QNE,ENCFF052KVO,ENCFF327LYS,ENCFF089GBC,ENCFF907XEG,ENCFF821BIA,ENCFF065ODV,ENCFF316FYN,ENCFF024JOZ,ENCFF589ZBJ,ENCFF321AGQ,ENCFF609MDN,ENCFF092NMJ,ENCFF136ZDK,ENCFF055HAP,ENCFF430PPJ,ENCFF165LXO,ENCFF097CAQ,ENCFF198SSR,ENCFF502WDT,ENCFF575MGI,ENCFF194HVL,ENCFF951GOP,ENCFF142CNG,ENCFF281QMV,ENCFF957SPH.ENCFF883UPM |
| 1217 | 8 | 4432324 | 4432536 | 212 | 129S1,CAST,NZB,NZO,NZW,SPRET,WSB | 8 | 4432394 | 4432427 | 34 | + | 18.374 | 2.75E-07 | 0.0529 | TAGCTGTCTTCAGTCACTCCAGAAGAGGGCGCCA | | |
| 1217 | 8 | 4432324 | 4432536 | 212 | 129S1,CAST,NZB,NZO,NZW,SPRET,WSB | 8 | 4432409 | 4432427 | 19 | + | 17.5574 | 6.32E-07 | 0.125 | ACTCCAGAAGAGGGCGCCA | | |
| 1218 | 8 | 4648391 | 4648596 | 205 | 129S1.CAST.NZB.SPRET | 8 | 4648453 | 4648486 | 34 | + | 18.3659 | 2.76E-07 | 0.0529 | TAGCTGTCTTCAGACACTCCAGAAGAGGGCGCCA | | |
| 1218 | 8 | 4648391 | 4648596 | 205 | 129S1.CAST.NZB.SPRET | 8 | 4648468 | 4648486 | 19 | + | 17.5574 | 6.32E-07 | 0.125 | ACTCCAGAAGAGGGCGCCA | | |
| 1219 | 8 | 10079713 | 10079922 | 209 | CAST.MOLF.NZB.SPRET | 8 | 10079780 | 10079813 | 34 | + | 16.4472 | 9.61E-07 | 0.0944 | TAGCTGTCTTCAGACACACCAGAAGAGGGCGTCA | | |
| 1220 | 8 | 14099335 | 14099536 | 201 | C57L,CAST,ILNJ2,MOLF,NZB,NZW,PJ,PL,SEA,SMJ.SPRET.TH | 8 | 14099396 | 14099429 | 34 | + | 17.0569 | 6.53E-07 | 0.0808 | TAGCTGTCCTCAGATACTCCAGAAGAGGGCATTA | | |
| 1221 | 8 | 24727002 | 24727206 | 204 | SPRET | 8 | 24727062 | 24727095 | 34 | + | 17.3902 | 5.27E-07 | 0.0727 | TAGCTGTCCTCAGACACTCCAGAAGAGGGCATCA | | |
| 1222 | 8 | 25921209 | 25921404 | 195 | SPRET | 8 | 25921263 | 25921296 | 34 | + | 17.5122 | 4.86E-07 | 0.07 | TAGCTGTTTTCAGACACACCAGAAGAGGGCATTA | | |
| 1223 | 8 | 26612106 | 26612313 | 207 | CAST,MOLF,PL | 8 | 26612162 | 26612195 | 34 | + | 19.878 | 9.69E-08 | 0.0283 | TTGCTGTCCTCAGACACTCCAGAAGAGGGCATCA | 7 | ENCFF093ZRU,ENCFF327LYS,ENCFF089GBC,ENCFF502WDT,ENCFF024JOZ,ENCFF136ZDK,ENCFF883UPM |
| 1224 | 8 | 26774528 | 26774641 | 113 | 129S1,AJ,AKR,BALB,BTBR,BUB,C3H,C57L,CAST,CBA,DBA1J,DBA2J,FVB,ILNJ2,KK,MA/My,MRL,NOD,NOR,NZB,NZO,NZW,RF,RH2,SJL,SMJ,SWR,TH | 8 | 26774624 | 26774657 | 34 | + | 23.0325 | 8.79E-09 | 0.005 | ATGCAATACTCACAGAAGCCAGAAGATGGCATCA | | |
| 1225 | 8 | 27636887 | 27637122 | 235 | SPRET | 8 | 27637116 | 27637150 | 35 | + | 17.1376 | 5.00E-07 | 0.127 | GTGTATATGACCATGGAGGCCAGAAGAGGGCATTA | | |
| 1226 | 8 | 34256951 | 34257173 | 222 | 129S1,AJ,AKR,BALB,BUB,C3H,C57L,CAST,CBA,CE,DBA1J,DBA2J,MA/My,MOLF,MRL,NOD,NOR,PL,RF,RH2,SJL,SMJ,SPRET,SWR,TH,WSB | 8 | 34257031 | 34257064 | 34 | + | 17.8455 | 3.91E-07 | 0.0622 | TAGCTGTCCTCAGACACTCCAGAAGAGGGCGTCA | 2 | ENCFF093ZRU,ENCFF367UNR |
| 1227 | 8 | 34529462 | 34530078 | 616 | NOD,NOR,SJL,TH | 8 | 34529891 | 34529924 | 34 | + | 19.2846 | 1.47E-07 | 0.0367 | TAGCTGTCTCCAGACACACCAGAAGAGGGCAGCA | 2 | ENCFF099QNE,ENCFF093ZRU |
| 1227 | 8 | 34529462 | 34530078 | 616 | NOD,NOR,SJL,TH | 8 | 34529906 | 34529924 | 19 | + | 17.2459 | 7.74E-07 | 0.137 | ACACCAGAAGAGGGCAGCA | 2 | ENCFF099QNE,ENCFF093ZRU |
| 1228 | 8 | 42394512 | 42394716 | 204 | SPRET | 8 | 42394573 | 42394606 | 34 | + | 16.4472 | 9.61E-07 | 0.0944 | TAGCTGTCTTCAGACACACCAGAAGAGGGCGTCA | | |
| 1229 | 8 | 43631416 | 43632182 | 766 | SPRET | 8 | 43631475 | 43631509 | 35 | + | 18.1468 | 2.76E-07 | 0.0931 | AAGCAAGCGACTTTGGAGGCCACAAGGGGGCACCG | 6 | ENCFF367UNR,ENCFF024JOZ,ENCFF093ZRU,ENCFF099QNE,ENCFF327LYS,ENCFF533APC |
| 1229 | 8 | 43631416 | 43632182 | 766 | SPRET | 8 | 43631491 | 43631509 | 19 | + | 21.7541 | 2.54E-08 | 0.0301 | AGGCCACAAGGGGGCACCG | 7 | ENCFF367UNR,ENCFF024JOZ,ENCFF093ZRU,ENCFF099QNE,ENCFF327LYS,ENCFF533APC,ENCFF907XEG |
| 1230 | 8 | 43846469 | 43846695 | 226 | 129S1,AJ,AKR,BALB,BTBR,BUB,C3H,C57L,CBA,CE,DBA1J,DBA2J,FVB,ILNJ2,KK,MA/My,MRL,NOD,NOR,NZB,NZO,NZW,PJ,PL,RF,RH2,SEA,SJL,SMJ,SPRET,SWR,TH,WSB | 8 | 43846552 | 43846585 | 34 | + | 18.3659 | 2.76E-07 | 0.0529 | TAGCTGTCTTCAGACACTCCAGAAGAGGGCGCCA | 1 | ENCFF093ZRU |
| 1230 | 8 | 43846469 | 43846695 | 226 | 129S1,AJ,AKR,BALB,BTBR,BUB,C3H,C57L,CBA,CE,DBA1J,DBA2J,FVB,ILNJ2,KK,MA/My,MRL,NOD,NOR,NZB,NZO,NZW,PJ,PL,RF,RH2,SEA,SJL,SMJ,SPRET,SWR,TH,WSB | 8 | 43846567 | 43846585 | 19 | + | 17.5574 | 6.32E-07 | 0.125 | ACTCCAGAAGAGGGCGCCA | 1 | ENCFF093ZRU |
| 1231 | 8 | 47151867 | 47152062 | 195 | SPRET | 8 | 47151922 | 47151955 | 34 | + | 16.7724 | 7.83E-07 | 0.088 | TAGCTGTCTTCAGATGCACCAGAAGAGGGCGTCA | | |
| 1232 | 8 | 47157658 | 47157864 | 206 | SPRET | 8 | 47157710 | 47157743 | 34 | + | 17.6992 | 4.30E-07 | 0.0654 | TAGCTGTCCTCAGATACTCCAGAAGAGGGCGTCA | | |
| 1233 | 8 | 47826070 | 47826243 | 173 | SPRET | 8 | 47826108 | 47826141 | 34 | + | 16.9106 | 7.17E-07 | 0.0846 | TAGCTATCTTCAGGCACTCCAGAAGAGGGCGCCA | 4 | ENCFF099QNE.ENCFF821BIA.ENCFF367UNR.ENCFF093ZRU |
| 1233 | 8 | 47826070 | 47826243 | 173 | SPRET | 8 | 47826123 | 47826141 | 19 | + | 17.5574 | 6.32E-07 | 0.125 | ACTCCAGAAGAGGGCGCCA | 4 | ENCFF099QNE.ENCFF821BIA.ENCFF367UNR.ENCFF093ZRU |
| 1234 | 8 | 48050905 | 48051104 | 199 | 129S1,C58J,CAST,ILNJ2,LP,MOLF,SPRET | 8 | 48050963 | 48050996 | 34 | + | 16.4146 | 9.81E-07 | 0.0956 | TAGCTGTCTTCAGACACACCAGAGGAGGGCGCCA | | |
| 1235 | 8 | 48085259 | 48085853 | 594 | SPRET | 8 | 48085661 | 48085694 | 34 | + | 18.1382 | 3.22E-07 | 0.0568 | CTGCAGCGCGCACGATCATCTCCAGGGGACGACC | | |
| 1236 | 8 | 59124965 | 59125219 | 254 | CAST.MOLF.NZB.NZO.SPRET | 8 | 59125079 | 59125112 | 34 | + | 16.4634 | 9.51E-07 | 0.0944 | TAGCTGTCTTCAGACGCACCAGAAGAGGGCATCA | | |
| 1237 | 8 | 65338666 | 65338993 | 327 | SPRET | 8 | 65338702 | 65338735 | 34 | + | 27.3902 | 1.75E-10 | 0.000352 | ATGCAGTACCAGAAGAGGCCAGAAGAGGGCATTG | 16 | ENCFF099QNE,ENCFF367UNR,ENCFF430PPJ,ENCFF089GBC,ENCFF024JOZ,ENCFF092NMJ,ENCFF136ZDK,ENCFF201JUA,ENCFF165LXO,ENCFF589ZBJ,ENCFF883UPM,ENCFF821BIA,ENCFF052KVO,ENCFF327LYS,ENCFF093ZRU,ENCFF907XEG |
| 1238 | 8 | 68259840 | 68260038 | 198 | CAST.MOLF.SPRET | 8 | 68259897 | 68259930 | 34 | + | 17.8455 | 3.91E-07 | 0.0622 | TAGCTGTCCTCAGACACTCCAGAAGAGGGCGTCA | | |
| 1239 | 8 | 69444367 | 69444574 | 207 | 129S1,AJ,AKR,BALB,BTBR,BUB,C3H,C57L,C58J,CAST,CBA,CE,DBA1J,DBA2J,FVB,ILNJ2,KK,MA/My,MRL,NOD,NOR,NZB,NZO,NZW,PJ,PL,RF,RH2,SEA,SJL,SMJ,SPRET,SWR,TH.WSB | 8 | 69444432 | 69444465 | 34 | + | 18.3659 | 2.76E-07 | 0.0529 | TAGCTGTCTTCAGACACTCCAGAAGAGGGCGCCA | 1 | ENCFF367UNR |
| 1239 | 8 | 69444367 | 69444574 | 207 | 129S1,AJ,AKR,BALB,BTBR,BUB,C3H,C57L,C58J,CAST,CBA,CE,DBA1J,DBA2J,FVB,ILNJ2,KK,MA/My,MRL,NOD,NOR,NZB,NZO,NZW,PJ,PL,RF,RH2,SEA,SJL,SMJ,SPRET,SWR,TH.WSB | 8 | 69444447 | 69444465 | 19 | + | 17.5574 | 6.32E-07 | 0.125 | ACTCCAGAAGAGGGCGCCA | 1 | ENCFF367UNR |
| 1240 | 8 | 70463074 | 70463905 | 831 | SPRET | 8 | 70463542 | 70463560 | 19 | + | 17.4098 | 6.96E-07 | 0.13 | TGGCCACCAGGCGGAACCC | | |
| 1240 | 8 | 70463074 | 70463905 | 831 | SPRET | 8 | 70463758 | 70463791 | 34 | + | 17.3902 | 5.27E-07 | 0.0727 | TAGCTGTCCTCAGACACTCCAGAAGAGGGCATCA | | |
| 1241 | 8 | 71698163 | 71698369 | 206 | 129S1,AKR,BALB,BUB,C3H,C57L,C58J,CAST,CBA,CE,DBA1J,DBA2J,FVB,ILNJ2,KK,MA/My,MRL,NZB,NZO,NZW,PJ,PL,RF,SEA,SJL,SMJ,SWR,TH,WSB | 8 | 71698225 | 71698258 | 34 | + | 16.4472 | 9.61E-07 | 0.0944 | TAGCTGTCTTCAGACACACCAGAAGAGGGCGTCA | | |
| 1242 | 8 | 84437555 | 84437754 | 199 | SPRET | 8 | 84437612 | 84437645 | 34 | + | 16.4472 | 9.61E-07 | 0.0944 | TAGCTGTCTTCAGACACACCAGAAGAGGGCGTCA | | |
| 1243 | 8 | 85632785 | 85633006 | 221 | SPRET | 8 | 85632863 | 85632896 | 34 | + | 18.4065 | 2.69E-07 | 0.0529 | CTGTTGTCTTCAGACGCACCAGAAGAGGGCATCA | 1 | ENCFF093ZRU |
| 1244 | 8 | 85700092 | 85700161 | 69 | 129S1,AJ,AKR,BALB,BTBR,C3H,C57L,C58J,CBA,CE,DBA1J,DBA2J,FVB,ILNJ2,KK,MA/My,MRL,NOD,NOR,NZW,PJ,PL,RF,RH2,SEA,SJL.SMJ.SWR.WSB | 8 | 85700118 | 85700152 | 35 | + | 16.5229 | 7.12E-07 | 0.145 | CTATAGTTCCAGCATCCGGGCAGCAGAGGAAGGAA | | |
| 1245 | 8 | 88633549 | 88633760 | 211 | 129S1,AJ,AKR,BALB,BTBR,BUB,C3H,C57L,C58J,CAST,CBA,CE,DBA1J,DBA2J,FVB,ILNJ2,KK,MA/My,MOLF,MRL,NZB,NZO,PJ,PL,RH2,SEA,SMJ,SPRET,SWR,WSB | 8 | 88633618 | 88633651 | 34 | + | 18.561 | 2.42E-07 | 0.0509 | AAGCTGTCTTCAGACACTCCAGAAGAGGGCGCCA | 4 | ENCFF367UNR,ENCFF093ZRU |
| 1245 | 8 | 88633549 | 88633760 | 211 | 129S1,AJ,AKR,BALB,BTBR,BUB,C3H,C57L,C58J,CAST,CBA,CE,DBA1J,DBA2J,FVB,ILNJ2,KK,MA/My,MOLF,MRL,NZB,NZO,PJ,PL,RH2,SEA,SMJ,SPRET,SWR,WSB | 8 | 88633633 | 88633651 | 19 | + | 17.5574 | 6.32E-07 | 0.125 | ACTCCAGAAGAGGGCGCCA | 4 | ENCFF367UNR,ENCFF093ZRU |
| 1246 | 8 | 88761326 | 88761539 | 213 | AJ,C3H,CAST,CBA,CE,DBA1J,DBA2J,FVB,ILNJ2,MA/My,MOLF,NZB,NZO,RH2,SJL,SPRET.WSB | 8 | 88761398 | 88761431 | 34 | + | 16.7724 | 7.83E-07 | 0.088 | TAGCTGTCTTCAGATGCACCAGAAGAGGGCGTCA | | |
| 1247 | 8 | 89829641 | 89829842 | 201 | 129S1,AKR,BALB,BTBR,BUB,C57L,C58J,CAST,CBA,CE,DBA1J,DBA2J,FVB,ILNJ2,KK,MA/My,MRL,NOD,NOR,NZB,NZO,NZW,PJ,PL,RF,SEA,SJL,SMJ,SWR,TH,WSB | 8 | 89829700 | 89829733 | 34 | + | 18.4472 | 2.62E-07 | 0.0526 | TTGCTGTCTTCAGACACTCCAGAAGAGGGCGTCA | | |
| 1248 | 8 | 91290876 | 91291105 | 229 | 129S1,AKR,BALB,BUB,C57L,C58J,CAST,CBA,CE,DBA1J,DBA2J,FVB,ILNJ2,MA/My,MRL,NOD,NOR,NZB,NZO,NZW,PJ,PL,RF,SEA,SJL,SMJ,SPRET,SWR,TH,WSB | 8 | 91290964 | 91290997 | 34 | + | 17.2439 | 5.79E-07 | 0.0757 | TAGCTGTCCTCAGATACTCCAGAAGAGGGCATCA | | |
| 1249 | 8 | 91519150 | 91519593 | 443 | CAST | 8 | 91519373 | 91519406 | 34 | + | 16.4553 | 9.56E-07 | 0.0944 | ATGCAGCTAGAGACACGGGCTCTGGGGGGTACTG | | |
| 1250 | 8 | 91673715 | 91673906 | 191 | CAST,SPRET | 8 | 91673764 | 91673797 | 34 | + | 16.4472 | 9.61E-07 | 0.0944 | TAGCTGTCTTCAGACACACCAGAAGAGGGCGTCA | | |
| 1251 | 8 | 94925310 | 94925854 | 544 | CAST,SPRET | 8 | 94925341 | 94925374 | 34 | + | 16.4472 | 9.61E-07 | 0.0944 | TAGCTGTCTTCAGACACACCAGAAGAGGGCGTCA | 1 | ENCFF093ZRU |
| 1252 | 8 | 95566171 | 95566415 | 244 | 129S1,AJ,AKR,B10D2,B10J,BALB,BTBR,BUB,C3H,C57L,CAST,CBA,CE,DBA1J,DBA2J,FVB,ILNJ2,KK,LP,MA/My,MOLF,MRL,NOD,NOR,NZB,NZO,NZW,PJ,PL,RF,RH2,SEA,SJL,SMJ.SPRET.SWR.TH.WSB | 8 | 95566272 | 95566305 | 34 | + | 16.4634 | 9.51E-07 | 0.0944 | TAGCTGTCTTCAGACGCACCAGAAGAGGGCATCA | 1 | ENCFF093ZRU |
| 1253 | 8 | 96077347 | 96077551 | 204 | SPRET | 8 | 96077409 | 96077442 | 34 | + | 16.6341 | 8.54E-07 | 0.0925 | CTGTAGCTGTCAGACACTCCAGAAGAGGGAGTCA | 1 | ENCFF093ZRU |
| 1254 | 8 | 96325489 | 96325700 | 211 | B10D2,B10J,BUB,C3H,CBA,DBA1J,DBA2J,LP.MOLF.NOD.NOR.SPRET.SWR | 8 | 96325558 | 96325591 | 34 | + | 17.8455 | 3.91E-07 | 0.0622 | TAGCTGTCCTCAGACACTCCAGAAGAGGGCGTCA | 1 | ENCFF093ZRU |
| 1255 | 8 | 96560875 | 96561288 | 413 | SPRET | 8 | 96561140 | 96561174 | 35 | + | 20.6789 | 5.58E-08 | 0.0352 | CTGCATGTGTGCCTATAGGCCAGAAGAGGGTATCA | | |
| 1256 | 8 | 96643548 | 96643759 | 211 | CAST.MOLF.SPRET.WSB | 8 | 96643616 | 96643649 | 34 | + | 18.3659 | 2.76E-07 | 0.0529 | TAGCTGTCTTCAGACACTCCAGAAGAGGGCGCCA | | |
| 1256 | 8 | 96643548 | 96643759 | 211 | CAST.MOLF.SPRET.WSB | 8 | 96643631 | 96643649 | 19 | + | 17.5574 | 6.32E-07 | 0.125 | ACTCCAGAAGAGGGCGCCA | | |
| 1257 | 8 | 103643468 | 103644271 | 803 | MOLF | 8 | 103643681 | 103643699 | 19 | + | 17.6721 | 5.85E-07 | 0.122 | ATACCAGAAGAGGGCACTG | | |
| 1258 | 8 | 106028229 | 106028746 | 517 | SPRET | 8 | 106028397 | 106028430 | 34 | + | 16.4472 | 9.61E-07 | 0.0944 | TAGCTGTCTTCAGACACACCAGAAGAGGGCGTCA | | |
| 1259 | 8 | 106637325 | 106637529 | 204 | MOLF.SPRET | 8 | 106637386 | 106637419 | 34 | + | 16.4634 | 9.51E-07 | 0.0944 | TAGCTGTCTTCAGACGCACCAGAAGAGGGCATCA | 1 | ENCFF093ZRU |
| 1260 | 8 | 106655678 | 106655741 | 63 | MOLF | 8 | 106655735 | 106655769 | 35 | + | 17.4037 | 4.28E-07 | 0.117 | CTCCTGTTCCATCTCTCCTCCATTGGATGGCACCT | | |
| 1261 | 8 | 106917095 | 106917322 | 227 | SPRET | 8 | 106917157 | 106917190 | 34 | + | 16.4634 | 9.51E-07 | 0.0944 | TAGCTGTCTTCAGACGCACCAGAAGAGGGCATCA | | |
| 1262 | 8 | 107379043 | 107379253 | 210 | 129S1,AJ,AKR,B10D2,B10J,BALB,BTBR,BUB,C3H,C57L,C58J,CAST,CBA,CE,DBA1J,DBA2J,FVB,ILNJ2,KK,LP,MA/My,MOLF,MRL,NOD,NOR,NZB,NZO,NZW,PJ,PL,RF,RH2,SEA.SJL.SMJ.SPRET.SWR.TH | 8 | 107379112 | 107379145 | 34 | + | 16.7724 | 7.83E-07 | 0.088 | TAGCTGTCTTCAGATGCACCAGAAGAGGGCGTCA | | |
| 1263 | 8 | 107409254 | 107409464 | 210 | 129S1,AJ,AKR,B10D2,B10J,BALB,BTBR,C3H,C57L,C58J,CAST,CBA,CE,DBA1J,DBA2J,ILNJ2,KK,LP,MA/My,MOLF,MRL,NZB,NZO,NZW,RH2,SEA,SMJ,SPRET,TH | 8 | 107409320 | 107409353 | 34 | + | 18.1545 | 3.18E-07 | 0.0563 | TAGCTGTTTTCAGACACACCAGAAGAGGGCGTCA | 1 | ENCFF093ZRU |
| 1264 | 8 | 107686861 | 107687072 | 211 | CAST,MOLF,PL,RH2,SPRET | 8 | 107686926 | 107686959 | 34 | + | 17.2114 | 5.91E-07 | 0.0762 | TAGCTGTTTTCAGACACTCCAGAAGAGGGCATCA | | |
| 1265 | 8 | 107707031 | 107707233 | 202 | CAST,MOLF,SPRET,WSB | 8 | 107707091 | 107707124 | 34 | + | 18.3659 | 2.76E-07 | 0.0529 | TAGCTGTCTTCAGACACTCCAGAAGAGGGCGCCA | 2 | ENCFF093ZRU,ENCFF367UNR |
| 1265 | 8 | 107707031 | 107707233 | 202 | CAST,MOLF,SPRET,WSB | 8 | 107707106 | 107707124 | 19 | + | 17.5574 | 6.32E-07 | 0.125 | ACTCCAGAAGAGGGCGCCA | 2 | ENCFF093ZRU,ENCFF367UNR |
| 1266 | 8 | 107880973 | 107881178 | 205 | SPRET | 8 | 107881036 | 107881069 | 34 | + | 16.4472 | 9.61E-07 | 0.0944 | TAGCTGTCTTCAGACACACCAGAAGAGGGCGTCA | | |
| 1267 | 8 | 110298575 | 110298779 | 204 | SPRET | 8 | 110298635 | 110298668 | 34 | + | 16.4634 | 9.51E-07 | 0.0944 | TAGCTGTCTTCAGACGCACCAGAAGAGGGCATCA | 2 | ENCFF099QNE,ENCFF093ZRU |
| 1268 | 8 | 110604102 | 110604311 | 209 | CAST,MOLF,SPRET | 8 | 110604169 | 110604202 | 34 | + | 16.4472 | 9.61E-07 | 0.0944 | TAGCTGTCTTCAGACACACCAGAAGAGGGCGTCA | | |
| 1269 | 8 | 111211229 | 111211433 | 204 | MOLF,SPRET | 8 | 111211292 | 111211325 | 34 | + | 17.2439 | 5.79E-07 | 0.0757 | TAGCTGTCCTCAGATACTCCAGAAGAGGGCATCA | | |
| 1270 | 8 | 114195187 | 114195386 | 199 | CAST | 8 | 114195243 | 114195276 | 34 | + | 19.0163 | 1.77E-07 | 0.0412 | CAGCTGTCTTCAGACACACCAGAAGAGGGCAGCA | | |
| 1271 | 8 | 119626093 | 119626728 | 635 | CAST | 8 | 119626131 | 119626164 | 34 | + | 23.5366 | 5.81E-09 | 0.0037 | CTGTAATACCCCTTGTGGCCAGAAGAAGGCATTA | 1 | ENCFF052KVO |
| 1271 | 8 | 119626093 | 119626728 | 635 | CAST | 8 | 119626133 | 119626167 | 35 | - | 15.9908 | 9.60E-07 | 0.168 | ATGTAATGCCTTCTTCTGGCCACAAGGGGTATTAC | 1 | ENCFF052KVO |
| 1271 | 8 | 119626093 | 119626728 | 635 | CAST | 8 | 119626384 | 119626417 | 34 | - | 18.3008 | 2.89E-07 | 0.0536 | GTGCTCCTCCCAGGGAGCCCACCAGAGAGCAACA | | |
| 1272 | 8 | 120855037 | 120855268 | 231 | CAST,MOLF,NZB,SPRET,WSB | 8 | 120855100 | 120855133 | 34 | + | 18.3496 | 2.79E-07 | 0.0534 | TAGCTGTCCTCAGACGCACCAGAAGAGGGCATCA | | |
| 1273 | 8 | 120971802 | 120972022 | 220 | CAST,SPRET | 8 | 120971880 | 120971913 | 34 | + | 16.4228 | 9.76E-07 | 0.0953 | TAGCTGTCTTCAGATGCACCAGAAGAGGGCATCG | | |
| 1274 | 8 | 121156273 | 121156483 | 210 | SPRET | 8 | 121156341 | 121156374 | 34 | + | 17.8455 | 3.91E-07 | 0.0622 | TAGCTGTCCTCAGACACTCCAGAAGAGGGCGTCA | 1 | ENCFF093ZRU |
| 1275 | 8 | 125104076 | 125104294 | 218 | SPRET | 8 | 125104154 | 125104187 | 34 | + | 16.7724 | 7.83E-07 | 0.088 | TAGCTGTCTTCAGATGCACCAGAAGAGGGCGTCA | 1 | ENCFF093ZRU |
| 1276 | 8 | 127186534 | 127187331 | 797 | SPRET | 8 | 127187199 | 127187217 | 19 | + | 23.623 | 3.91E-09 | 0.0175 | CAGCCACCAGAGGGCGCCC | 34 | ENCFF093ZRU,ENCFF367UNR,ENCFF805QBU,ENCFF092NMJ,ENCFF099QNE,ENCFF821BIA,ENCFF430PPJ,ENCFF316FYN,ENCFF321AGQ,ENCFF327LYS,ENCFF033UQX,ENCFF589ZBJ,ENCFF907XEG,ENCFF065ODV,ENCFF609MDN,ENCFF136ZDK,ENCFF198SSR,ENCFF052KVO,ENCFF142CNG,ENCFF281QMV,ENCFF024JOZ,ENCFF089GBC,ENCFF097CAQ,ENCFF349BUL,ENCFF165LXO,ENCFF194HVL,ENCFF502WDT,ENCFF957SPH,ENCFF491RJK,ENCFF533APC,ENCFF210BTO,ENCFF055HAP,ENCFF883UPM,ENCFF089EWX |
| 1277 | 8 | 129667882 | 129668214 | 332 | SPRET | 8 | 129667915 | 129667933 | 19 | + | 19.2951 | 1.88E-07 | 0.0709 | CCGCCACTAGATGGTGCTA | 20 | ENCFF093ZRU,ENCFF367UNR,ENCFF099QNE,ENCFF201JUA,ENCFF502WDT,ENCFF883UPM,ENCFF821BIA,ENCFF327LYS,ENCFF907XEG,ENCFF165LXO,ENCFF024JOZ,ENCFF055HAP,ENCFF089GBC,ENCFF589ZBJ,ENCFF136ZDK,ENCFF092NMJ,ENCFF430PPJ,ENCFF198SSR |
| 1278 | 8 | 4322215 | 4322435 | 220 | 129S1.NZB.SPRET | 8 | 4322342 | 4322375 | 34 | - | 17.374 | 5.32E-07 | 0.0732 | TAGCTGTCTTCAGACCCTCCAGAAGAGGGCATCA | 1 | ENCFF093ZRU |
| 1279 | 8 | 8797418 | 8797637 | 219 | CAST.MOLF.NZB.SPRET.WSB | 8 | 8797544 | 8797577 | 34 | - | 17.8455 | 3.91E-07 | 0.0622 | TAGCTGTCCTCAGACACTCCAGAAGAGGGCGTCA | | |
| 1280 | 8 | 11538442 | 11538645 | 203 | SPRET | 8 | 11538562 | 11538595 | 34 | - | 17.9919 | 3.55E-07 | 0.0594 | TTGCTGTCTTCAGACACTCCAGAAGAGGGCATCA | 1 | ENCFF093ZRU |
| 1281 | 8 | 11594289 | 11594501 | 212 | 129S1,BTBR,C57L,CAST,ILNJ2,MOLF,NZB,NZO,NZW,SMJ,SPRET,TH | 8 | 11594414 | 11594447 | 34 | - | 17.2439 | 5.79E-07 | 0.0757 | TAGCTGTCCTCAGATACTCCAGAAGAGGGCATCA | 2 | ENCFF367UNR,ENCFF093ZRU |
| 1282 | 8 | 12695543 | 12696089 | 546 | CAST | 8 | 12695770 | 12695804 | 35 | - | 17.1927 | 4.84E-07 | 0.125 | CTGCTCTGCCTCCTTCCCTACTGTAGATGCAGCCT | | |
| 1283 | 8 | 13201984 | 13202201 | 217 | 129S1,C57L,CAST,CE,ILNJ2,LP,MOLF,NZB,NZO,NZW,PJ,PL,SEA,SMJ,SPRET,TH,WSB | 8 | 13202109 | 13202142 | 34 | - | 16.4634 | 9.51E-07 | 0.0944 | TAGCTGTCTTCAGACGCACCAGAAGAGGGCATCA | | |
| 1284 | 8 | 14020319 | 14020526 | 207 | SPRET | 8 | 14020445 | 14020478 | 34 | - | 17.3902 | 5.27E-07 | 0.0727 | TAGCTGTCCTCAGACACTCCAGAAGAGGGCATCA | | |
| 1285 | 8 | 14941568 | 14941657 | 89 | SPRET | 8 | 14941604 | 14941637 | 34 | - | 20.2114 | 7.63E-08 | 0.0242 | CTGCAGTCCCAGATGTGCCCAGGAGGCAGAGGTG | 3 | ENCFF093ZRU.ENCFF589ZBJ.ENCFF136ZDK |
| 1286 | 8 | 22913631 | 22913840 | 209 | SPRET | 8 | 22913760 | 22913793 | 34 | - | 16.9187 | 7.13E-07 | 0.0845 | TAGCTGTCTTCAGACGCACCAGAAGAGGGCGTCA | 1 | ENCFF093ZRU |
| 1287 | 8 | 23028135 | 23028189 | 54 | C57L,CAST,CE,ILNJ2,LP,MOLF,NZB,PJ,SEA | 8 | 23028173 | 23028207 | 35 | - | 17.6697 | 3.66E-07 | 0.106 | GAGCACTGAGAGTGTCAGGCCGGCAGGGAGCACTG | | |
| 1288 | 8 | 27695237 | 27695427 | 190 | SPRET | 8 | 27695361 | 27695394 | 34 | - | 16.7724 | 7.83E-07 | 0.088 | TAGCTGTCTTCAGATGCACCAGAAGAGGGCGTCA | 3 | ENCFF502WDT.ENCFF093ZRU.ENCFF055HAP |
| 1289 | 8 | 33835947 | 33836506 | 559 | CAST.SPRET | 8 | 33836123 | 33836156 | 34 | - | 17.9919 | 3.55E-07 | 0.0594 | CTGTCATTGTTCCCATCTCCTGCAGGAGGCAGTG | 1 | ENCFF052KVO |
| 1290 | 8 | 34254944 | 34255029 | 85 | 129S1,AJ,AKR,BALB,BUB,C3H,C57L,CBA,CE,DBA1J,DBA2J,MA/My,MRL,NOD,NOR,PL,RF,RH2,SJL,SMJ,SWR,TH,WSB | 8 | 34254918 | 34254951 | 34 | - | 18.8537 | 1.98E-07 | 0.0448 | TAGCGGCTACCGCCGCTGCCCCAAGATGGCACCC | | |
| 1291 | 8 | 34437729 | 34438580 | 851 | CAST | 8 | 34438324 | 34438357 | 34 | - | 26.3659 | 4.74E-10 | 0.000598 | CTGCAGTGCCTGAGGAGCCCAGAGGAGGGCGTCA | 1 | ENCFF052KVO |
| 1292 | 8 | 43099450 | 43099639 | 189 | SPRET | 8 | 43099544 | 43099577 | 34 | - | 19.4878 | 1.28E-07 | 0.034 | CCGCTGTTTTCAGACACACCAGAAGAGGGCATCG | | |
| 1293 | 8 | 44711261 | 44711367 | 106 | SPRET | 8 | 44711330 | 44711363 | 34 | - | 17.0976 | 6.36E-07 | 0.0797 | ACACAATTCCCGCAGAGGCCAGAAGAGGGTGTTG | | |
| 1294 | 8 | 48429411 | 48429649 | 238 | SPRET | 8 | 48429549 | 48429582 | 34 | - | 16.4634 | 9.51E-07 | 0.0944 | TAGCTGTCTTCAGACGCACCAGAAGAGGGCATCA | 1 | ENCFF093ZRU |
| 1295 | 8 | 52818217 | 52818602 | 385 | B10D2.B10J.C58J.LP.WSB | 8 | 52818273 | 52818291 | 19 | - | 17.8852 | 5.08E-07 | 0.113 | CTGCCACCAGGGGGGGGCA | | |
| 1296 | 8 | 57048188 | 57048398 | 210 | SPRET | 8 | 57048308 | 57048341 | 34 | - | 16.4634 | 9.51E-07 | 0.0944 | TAGCTGTCTTCAGACGCACCAGAAGAGGGCATCA | | |
| 1297 | 8 | 58014846 | 58015045 | 199 | CAST.MOLF.NZB.NZO.SPRET | 8 | 58014971 | 58015004 | 34 | - | 16.4472 | 9.61E-07 | 0.0944 | TAGCTGTCTTCAGACACACCAGAAGAGGGCGTCA | | |
| 1298 | 8 | 58092087 | 58092222 | 135 | MOLF.NZB.NZO | 8 | 58092153 | 58092187 | 35 | - | 16.5046 | 7.19E-07 | 0.145 | CTGCAGAAGCAAGAGCCAGCCAGCAGGAGACTGGC | | |
| 1299 | 8 | 63097579 | 63097788 | 209 | CAST.CE.KK.MOLF.SPRET | 8 | 63097707 | 63097725 | 19 | - | 17.5574 | 6.32E-07 | 0.125 | ACTCCAGAAGAGGGCGCCA | | |
| 1299 | 8 | 63097579 | 63097788 | 209 | CAST.CE.KK.MOLF.SPRET | 8 | 63097707 | 63097740 | 34 | - | 18.3659 | 2.76E-07 | 0.0529 | TAGCTGTCTTCAGACACTCCAGAAGAGGGCGCCA | | |
| 1300 | 8 | 64987739 | 64988116 | 377 | SPRET | 8 | 64988040 | 64988073 | 34 | - | 16.7967 | 7.71E-07 | 0.088 | CTGCGCCATCTGTTGCTGCCAGAGGATGTCAGCC | | |
| 1301 | 8 | 68744645 | 68745214 | 569 | SPRET | 8 | 68744904 | 68744937 | 34 | - | 17.8211 | 3.97E-07 | 0.0628 | ATGCAGCTAGAGACACGTGCTCCAGGGGGTACTG | | |
| 1302 | 8 | 70807731 | 70807934 | 203 | SPRET | 8 | 70807857 | 70807875 | 19 | - | 17.5574 | 6.32E-07 | 0.125 | ACTCCAGAAGAGGGCGCCA | 2 | ENCFF093ZRU.ENCFF367UNR |
| 1302 | 8 | 70807731 | 70807934 | 203 | SPRET | 8 | 70807857 | 70807890 | 34 | - | 18.3659 | 2.76E-07 | 0.0529 | TAGCTGTCTTCAGACACTCCAGAAGAGGGCGCCA | 2 | ENCFF093ZRU.ENCFF367UNR |
| 1303 | 8 | 71431062 | 71431207 | 145 | 129S1,AKR,BALB,BUB,C3H,C57L,C58J,CBA,CE,DBA1J,DBA2J,FVB,ILNJ2,KK,MA/My,MRL,NZB,NZW,PJ,RF,SEA,SJL,SMJ,SPRET,SWR,TH,WSB | 8 | 71431173 | 71431206 | 34 | - | 16.3984 | 9.91E-07 | 0.0962 | CTGGAATTACAAATGTGTCCACTAGGGCCCACCA | | |
| 1304 | 8 | 71920408 | 71920608 | 200 | SPRET | 8 | 71920535 | 71920568 | 34 | - | 16.4472 | 9.61E-07 | 0.0944 | TAGCTGTCTTCAGACACACCAGAAGAGGGCGTCA | 1 | ENCFF093ZRU |
| 1305 | 8 | 72269064 | 72269273 | 209 | CAST | 8 | 72269187 | 72269220 | 34 | - | 17.3902 | 5.27E-07 | 0.0727 | TAGCTGTCCTCAGACACTCCAGAAGAGGGCATCA | | |
| 1306 | 8 | 72570278 | 72570483 | 205 | CAST,MOLF | 8 | 72570401 | 72570434 | 34 | - | 17.3902 | 5.27E-07 | 0.0727 | TAGCTGTCCTCAGACACTCCAGAAGAGGGCATCA | | |
| 1307 | 8 | 72788991 | 72789812 | 821 | 129S1,AKR,BALB,BUB,C3H,C58J,CBA,DBA1J,DBA2J,FVB,ILNJ2,KK,NZB,NZO,NZW,RF,SJL,SMJ,SPRET,SWR,TH,WSB | 8 | 72789174 | 72789192 | 19 | - | 18.5574 | 3.20E-07 | 0.0916 | CCTCCAGTAGAGGGCGCAC | 4 | ENCFF099QNE,ENCFF821BIA,ENCFF093ZRU,ENCFF367UNR |
| 1307 | 8 | 72788991 | 72789812 | 821 | 129S1,AKR,BALB,BUB,C3H,C58J,CBA,DBA1J,DBA2J,FVB,ILNJ2,KK,NZB,NZO,NZW,RF,SJL.SMJ.SPRET.SWR.TH.WSB | 8 | 72789174 | 72789208 | 35 | - | 18.2844 | 2.54E-07 | 0.0885 | ATGAGGTTTTTTCACCCCTCCAGTAGAGGGCGCAC | 4 | ENCFF099QNE,ENCFF821BIA,ENCFF093ZRU,ENCFF367UNR |
| 1308 | 8 | 73031870 | 73032119 | 249 | 129S1,AKR,BALB,BUB,C3H,C57L,C58J,CBA,CE,DBA1J,DBA2J,FVB,ILNJ2,KK,MA/My,MRL,NZB,NZO,NZW,PJ,PL,RF,SEA,SJL,SMJ,SPRET,SWR,TH,WSB | 8 | 73031982 | 73032015 | 34 | - | 16.4472 | 9.61E-07 | 0.0944 | TAGCTGTCTTCAGACACACCAGAAGAGGGCGTCA | | |
| 1309 | 8 | 73052033 | 73052238 | 205 | 129S1,AKR,BALB,BUB,C3H,C57L,C58J,CBA,CE,DBA1J,DBA2J,FVB,ILNJ2,KK,MA/My,MRL,NZB,NZO,NZW,PJ,PL,RF,SEA,SJL,SMJ,SPRET,SWR,TH,WSB | 8 | 73052156 | 73052189 | 34 | - | 17.2439 | 5.79E-07 | 0.0757 | TAGCTGTCCTCAGATACTCCAGAAGAGGGCATCA | | |
| 1310 | 8 | 73410658 | 73410850 | 192 | CAST,MOLF | 8 | 73410654 | 73410687 | 34 | - | 16.5447 | 9.04E-07 | 0.0944 | CAGCTGTCTTCAGACACACCAGAAGAGGGAGTCA | | |
| 1311 | 8 | 75672199 | 75672401 | 202 | SPRET | 8 | 75672323 | 75672356 | 34 | - | 17.2439 | 5.79E-07 | 0.0757 | TAGCTGTCCTCAGATACTCCAGAAGAGGGCATCA | | |
| 1312 | 8 | 75899913 | 75900111 | 198 | MOLF,SPRET,WSB | 8 | 75900038 | 75900071 | 34 | - | 16.4634 | 9.51E-07 | 0.0944 | TAGCTGTCTTCAGACGCACCAGAAGAGGGCATCA | | |
| 1313 | 8 | 78269094 | 78269292 | 198 | SPRET | 8 | 78269204 | 78269237 | 34 | - | 17.3902 | 5.27E-07 | 0.0727 | TAGCTGTCCTCAGACACTCCAGAAGAGGGCATCA | | |
| 1314 | 8 | 78623052 | 78623236 | 184 | C3H,C57L,KK,MA/My,MOLF,NOD,NOR,NZB,NZO,PL,RF,SJL,SPRET,TH,WSB | 8 | 78623166 | 78623199 | 34 | - | 16.4472 | 9.61E-07 | 0.0944 | TAGCTGTCTTCAGACACACCAGAAGAGGGCGTCA | | |
| 1315 | 8 | 79141486 | 79141681 | 195 | SPRET | 8 | 79141607 | 79141640 | 34 | - | 16.4472 | 9.61E-07 | 0.0944 | TAGCTGTCTTCAGACACACCAGAAGAGGGCGTCA | | |
| 1316 | 8 | 79200799 | 79201373 | 574 | SPRET | 8 | 79200888 | 79200922 | 35 | - | 16.0092 | 9.50E-07 | 0.167 | CTGCAATGGCTACGACATTCCAGAAGATTGCATTT | | |
| 1317 | 8 | 81536669 | 81536869 | 200 | C58J.CAST.MOLF.SPRET.SWR.TH | 8 | 81536794 | 81536827 | 34 | - | 17.2439 | 5.79E-07 | 0.0757 | TAGCTGTCCTCAGATACTCCAGAAGAGGGCATCA | | |
| 1318 | 8 | 84734707 | 84734900 | 193 | 129S1,AJ,AKR,BALB,BTBR,C3H,C57L,C58J,CBA,CE,DBA1J,DBA2J,ILNJ2,KK,MA/My,MRL,NOD,NOR,NZW,PJ,PL,RF,RH2,SEA,SJL,SPRET.SWR.TH.WSB | 8 | 84734829 | 84734862 | 34 | - | 17.2439 | 5.79E-07 | 0.0757 | TAGCTGTCCTCAGATACTCCAGAAGAGGGCATCA | | |
| 1319 | 8 | 85601217 | 85601390 | 173 | CAST | 8 | 85601322 | 85601355 | 34 | - | 17.4228 | 5.16E-07 | 0.0726 | TAGCTGTTTTCAGACACACCAGGAGAGGGCGTCA | | |
| 1320 | 8 | 85839195 | 85839420 | 225 | 129S1,AJ,AKR,BALB,BTBR,C3H,C57L,C58J,CAST,CBA,CE,DBA1J,DBA2J,FVB,ILNJ2,KK,MA/My,MOLF,MRL,NOD,NOR,NZW,PJ,PL,RF,RH2,SEA,SJL,SMJ,SPRET,SWR,WSB | 8 | 85839322 | 85839355 | 34 | - | 16.4634 | 9.51E-07 | 0.0944 | TAGCTGTCTTCAGACGCACCAGAAGAGGGCATCA | | |
| 1321 | 8 | 85875459 | 85875669 | 210 | 129S1,AJ,AKR,BALB,BTBR,C3H,C57L,C58J,CAST,CBA,CE,DBA1J,DBA2J,FVB,ILNJ2,KK,MA/My,MOLF,MRL,NOD,NOR,NZW,PJ,PL,RF,RH2,SEA,SJL,SMJ,SPRET,SWR,WSB | 8 | 85875584 | 85875617 | 34 | - | 16.4472 | 9.61E-07 | 0.0944 | TAGCTGTCTTCAGACACACCAGAAGAGGGCGTCA | | |

| | | | | | | | | | | | | | | | | |
|---|---|---|---|---|---|---|---|---|---|---|---|---|---|---|---|---|
| 1322 | 8 | 88704945 | 88705144 | 199 | AJ,BUB,C3H,C58J,CAST,MOLF,RH2,SPRET,SWR | 8 | 88705070 | 88705103 | 34 | - | 17.2439 | 5.79E-07 | 0.0757 | TAGCTGTCCTCAGATACTCCAGAAGAGGGCATCA | | |
| 1323 | 8 | 88835742 | 88836237 | 495 | AJ.C3H.MOLF.RH2 | 8 | 88836044 | 88836077 | 34 | - | 23.6341 | 5.36E-09 | 0.00345 | TTGCAGTGTTCCAGGAAGCCAGAAGAGGGCATCA | 3 | ENCFF367UNR.ENCFF052KVO.ENCFF093ZRU |
| 1324 | 8 | 91354220 | 91354440 | 220 | BALB,CE,FVB,ILNJ2,MRL,NZB,NZO,NZW,PJ.SEA.SPRET.TH | 8 | 91354335 | 91354353 | 19 | - | 17.5574 | 6.32E-07 | 0.125 | ACTCCAGAAGAGGGCGCCA | | |
| 1324 | 8 | 91354220 | 91354440 | 220 | BALB,CE,FVB,ILNJ2,MRL,NZB,NZO,NZW,PJ.SEA.SPRET.TH | 8 | 91354335 | 91354368 | 34 | - | 16.4228 | 9.76E-07 | 0.0953 | TAGCTGCCTTCAGACACTCCAGAAGAGGGCGCCA | | |
| 1325 | 8 | 92428629 | 92428837 | 208 | 129S1,AKR,BALB,BUB,C57L,C58J,CAST,CBA,CE,DBA1J,DBA2J,FVB,ILNJ2,MA/My,MOLF,MRL,NOD,NOR,NZB,NZO,NZW,PJ,PL,RF,SEA,SJL,SMJ,SPRET,SWR,TH,WSB | 8 | 92428747 | 92428780 | 34 | - | 16.5447 | 9.04E-07 | 0.0944 | CAGCTGTCTTCAGACACACCAGAAGAGGGAGTCA | | |
| 1326 | 8 | 96686996 | 96687193 | 197 | CAST.MOLF.SPRET | 8 | 96687120 | 96687153 | 34 | - | 16.4472 | 9.61E-07 | 0.0944 | TAGCTGTCTTCAGACACACCAGAAGAGGGCGTCA | | |
| 1327 | 8 | 105862123 | 105862340 | 217 | CAST.KK.MOLF.PJ.SPRET | 8 | 105862249 | 105862282 | 34 | - | 16.4472 | 9.61E-07 | 0.0944 | TAGCTGTCTTCAGACACACCAGAAGAGGGCGTCA | | |
| 1328 | 8 | 107383846 | 107383989 | 143 | 129S1,AJ,AKR,B10D2,B10J,BALB,BTBR,BUB,C3H,C57L,C58J,CAST,CBA,CE,DBA1J,DBA2J,FVB,ILNJ2,KK,LP,MA/My,MOLF,MRL,NOD,NOR,NZB,NZO,NZW,PJ,PL,RF,RH2,SEA.SJL.SMJ.SPRET.SWR.TH | 8 | 107383820 | 107383854 | 35 | - | 17.211 | 4.79E-07 | 0.124 | TAGTAGATGCCCTTAAAGGCCAGAAGAGGGCATCA | 1 | ENCFF327LYS |
| 1328 | 8 | 107383846 | 107383989 | 143 | 129S1,AJ,AKR,B10D2,B10J,BALB,BTBR,BUB,C3H,C57L,C58J,CAST,CBA,CE,DBA1J,DBA2J,FVB,ILNJ2,KK,LP,MA/My,MOLF,MRL,NOD,NOR,NZB,NZO,NZW,PJ,PL,RF,RH2,SEA.SJL.SMJ.SPRET.SWR.TH | 8 | 107383932 | 107383966 | 35 | - | 17.211 | 4.79E-07 | 0.124 | TAGTAGATGCCCTTAAAGGCCAGAAGAGGGCATCA | 1 | ENCFF327LYS |
| 1329 | 8 | 107954088 | 107954669 | 581 | CAST | 8 | 107954089 | 107954123 | 35 | - | 19.3486 | 1.32E-07 | 0.06 | CAGCAGTGTCCCCTCTTTCACAGCAGGTGAAAGCA | | |
| 1330 | 8 | 111585341 | 111585432 | 91 | SPRET | 8 | 111585312 | 111585345 | 34 | - | 17.6667 | 4.40E-07 | 0.0661 | CGGCTGCTCCCCGGCTCACCTGCTGGTGGTAGTC | | |
| 1331 | 8 | 111917233 | 111917450 | 217 | MOLF.SPRET | 8 | 111917356 | 111917389 | 34 | - | 17.8455 | 3.91E-07 | 0.0622 | TAGCTGTCCTCAGACACTCCAGAAGAGGGCGTCA | | |
| 1332 | 8 | 111993025 | 111993229 | 204 | SPRET | 8 | 111993148 | 111993181 | 34 | - | 18.0976 | 3.31E-07 | 0.0574 | TAGCTGTCTCCAGACGCACCAGAAGAGGGCATCA | | |
| 1333 | 8 | 112203061 | 112203481 | 420 | BALB,CAST,MOLF,NZW,SEA,SPRET,WSB | 8 | 112203185 | 112203219 | 35 | - | 18.1284 | 2.79E-07 | 0.0931 | CTGTAGCTGTCTTCAGACACCAGAAGAGGGCATCA | | |
| 1334 | 8 | 114782557 | 114782781 | 224 | CAST.MA/Mv.MOLF.SPRET | 8 | 114782678 | 114782711 | 34 | - | 16.4472 | 9.61E-07 | 0.0944 | TAGCTGTCTTCAGACACACCAGAAGAGGGCGTCA | | |
| 1335 | 8 | 119906923 | 119907131 | 208 | CAST.MOLF.SPRET | 8 | 119907047 | 119907080 | 34 | - | 17.2439 | 5.79E-07 | 0.0757 | TAGCTGTCCTCAGATACTCCAGAAGAGGGCATCA | | |
| 1336 | 8 | 120118811 | 120119029 | 218 | CAST.MOLF.SPRET.WSB | 8 | 120118934 | 120118967 | 34 | - | 17.8455 | 3.91E-07 | 0.0622 | TAGCTGTCCTCAGACACTCCAGAAGAGGGCGTCA | | |
| 1337 | 8 | 120464048 | 120464260 | 212 | CAST.MOLF.SPRET.WSB | 8 | 120464173 | 120464206 | 34 | - | 17.1382 | 6.20E-07 | 0.0787 | TAGCTGTCTCCAGACACTCCAGAAGAGGGCATCA | | |
| 1338 | 8 | 120496166 | 120496370 | 204 | CAST.MOLF.SPRET | 8 | 120496289 | 120496322 | 34 | - | 16.4146 | 9.81E-07 | 0.0956 | TAGCTGTCTTCAGACACACCAGAGGAGGGCGCCA | | |
| 1339 | 8 | 122534540 | 122535391 | 851 | AJ,AKR,BALB,C3H,C58J,CAST,CE,ILNJ2,LP,MOLF,MRL,NZB,PL,RF,RH2,SEA,SMJ,WSB | 8 | 122534969 | 122535002 | 34 | - | 22.1382 | 1.79E-08 | 0.0084 | ATGCGGTGTCTGTGGAAGCCAGAAGAGGGCGTCA | 5 | ENCFF093ZRU,ENCFF052KVO,ENCFF198SSR,ENCFF883UPM,ENCFF533APC |
| 1340 | 8 | 124042117 | 124042289 | 172 | 129S1,AKR,BTBR,C3H,C57L,C58J,CBA,CE,DBA1J,DBA2J,LP,MA/My,MRL,PJ,RF,SMJ,SPRET.TH | 8 | 124042193 | 124042226 | 34 | - | 17.2439 | 5.79E-07 | 0.0757 | TAGCTGTCCTCAGATACTCCAGAAGAGGGCATCA | 1 | ENCFF093ZRU |
| 1341 | 8 | 124527557 | 124528387 | 830 | SPRET | 8 | 124527813 | 124527831 | 19 | - | 19.7213 | 1.37E-07 | 0.0604 | AGGCCAGAAGAGGGCAGCA | 5 | ENCFF093ZRU.ENCFF136ZDK.ENCFF055HAP.ENCFF533APC.ENCFF502WDT |
| 1341 | 8 | 124527557 | 124528387 | 830 | SPRET | 8 | 124527813 | 124527847 | 35 | - | 24.5872 | 3.40E-09 | 0.00735 | CTGCAAGTGACCTTGAAGGCCAGAAGAGGGCAGCA | 5 | ENCFF093ZRU.ENCFF136ZDK.ENCFF055HAP.ENCFF533APC.ENCFF502WDT |
| 1342 | 8 | 125140725 | 125140815 | 90 | SPRET | 8 | 125140747 | 125140781 | 35 | - | 18.9908 | 1.65E-07 | 0.0678 | CTGCTGTGCCGGATGCTTGCCTGGAGGGAGCATGA | | |
| 1343 | 8 | 125451361 | 125451567 | 206 | 129S1,AJ,BALB,BTBR,C57L,CAST,ILNJ2,MOLF.NZB.RH2.SEA.SPRET.WSB | 8 | 125451487 | 125451520 | 34 | - | 17.2439 | 5.79E-07 | 0.0757 | TAGCTGTCCTCAGATACTCCAGAAGAGGGCATCA | | |
| 1344 | 8 | 125533563 | 125533790 | 227 | SPRET | 8 | 125533688 | 125533721 | 34 | - | 16.7724 | 7.83E-07 | 0.088 | TAGCTGTCTTCAGATGCACCAGAAGAGGGCGTCA | | |
| 1345 | 9 | 3122502 | 3123374 | 872 | CAST.MOLF.SPRET | 9 | 3122970 | 3123003 | 34 | + | 16.439 | 9.66E-07 | 0.0843 | ATGCAGCTAGAGACAAGAGCTCCAGGGGGTACTG | | |
| 1346 | 9 | 3342119 | 3342317 | 198 | SPRET | 9 | 3342173 | 3342206 | 34 | + | 16.4634 | 9.51E-07 | 0.0841 | TAGCTGTCTTCAGACGCACCAGAAGAGGGCATCA | | |
| 1347 | 9 | 5614293 | 5614507 | 214 | SPRET | 9 | 5614353 | 5614386 | 34 | + | 17.8618 | 3.87E-07 | 0.0524 | TAGCTGTCTTCAGACCCACCAGAAGAGGGCATCA | | |
| 1348 | 9 | 6237797 | 6238030 | 233 | CAST.MOLF.SPRET | 9 | 6237865 | 6237898 | 34 | + | 17.2439 | 5.79E-07 | 0.065 | TAGCTGTCCTCAGATACTCCAGAAGAGGGCATCA | | |
| 1349 | 9 | 13657804 | 13658018 | 214 | SPRET | 9 | 13657780 | 13657813 | 34 | + | 16.626 | 8.59E-07 | 0.0816 | CTGTAGCTGTCTTCAGACACACCAGAGGGCGCAT | | |
| 1350 | 9 | 13702385 | 13702600 | 215 | CAST.MOLF.SPRET.WSB | 9 | 13702472 | 13702490 | 19 | + | 17.8852 | 5.08E-07 | 0.102 | ACACCAGAAGAGGGCACCA | 1 | ENCFF093ZRU |
| 1351 | 9 | 13866508 | 13866726 | 218 | SPRET | 9 | 13866585 | 13866618 | 34 | + | 17.2439 | 5.79E-07 | 0.065 | TAGCTGTCCTCAGATACTCCAGAAGAGGGCATCA | | |
| 1352 | 9 | 15213414 | 15213609 | 195 | CAST,MOLF,SPRET | 9 | 15213467 | 15213500 | 34 | + | 18.3659 | 2.76E-07 | 0.0438 | TAGCTGTCTTCAGACACTCCAGAAGAGGGCGCCA | 12 | ENCFF024JOZ,ENCFF065ODV,ENCFF089GBC,ENCFF327LYS,ENCFF198SSR,ENCFF201JUA,ENCFF367UNR,ENCFF907XEG,ENCFF883UPM,ENCFF093ZRU,ENCFF099QNE,ENCFF821BIA |
| 1352 | 9 | 15213414 | 15213609 | 195 | CAST,MOLF,SPRET | 9 | 15213482 | 15213500 | 19 | + | 17.5574 | 6.32E-07 | 0.112 | ACTCCAGAAGAGGGCGCCA | 12 | ENCFF024JOZ,ENCFF065ODV,ENCFF089GBC,ENCFF327LYS,ENCFF198SSR,ENCFF201JUA,ENCFF367UNR,ENCFF907XEG,ENCFF883UPM,ENCFF093ZRU,ENCFF099QNE,ENCFF821BIA |
| 1353 | 9 | 20312250 | 20312468 | 218 | CAST.MOLF | 9 | 20312326 | 20312359 | 34 | + | 16.5528 | 8.99E-07 | 0.0834 | TAGCTGTCTTCAGACACACCAGAAGAGGGCGTCG | | |
| 1354 | 9 | 20922445 | 20922662 | 217 | CAST.MOLF.SPRET | 9 | 20922505 | 20922539 | 35 | + | 15.9817 | 9.65E-07 | 0.153 | TAGCTGTCTTCAGACAGTACCAGAAGAGGGCATCA | 3 | ENCFF093ZRU.ENCFF367UNR.ENCFF055HAP |
| 1355 | 9 | 20945306 | 20945523 | 217 | CAST.MOLF.SPRET.WSB | 9 | 20945381 | 20945414 | 34 | + | 16.4472 | 9.61E-07 | 0.0841 | TAGCTGTCTTCAGACACACCAGAAGAGGGCGTCA | 2 | ENCFF093ZRU |
| 1356 | 9 | 21194359 | 21194598 | 239 | SPRET | 9 | 21194451 | 21194484 | 34 | + | 16.4472 | 9.61E-07 | 0.0841 | TAGCTGTCTTCAGACACACCAGAAGAGGGCGTCA | 1 | ENCFF093ZRU |
| 1357 | 9 | 21399770 | 21399984 | 214 | BTBR,C57L,C58J,CAST,FVB,MA/My,MOLF,NOD,NOR,NZB,NZW,RF,SJL,SPRET,SWR,WSB | 9 | 21399842 | 21399875 | 34 | + | 18.3659 | 2.76E-07 | 0.0438 | TAGCTGTCTTCAGACACTCCAGAAGAGGGCGCCA | 1 | ENCFF055HAP |
| 1357 | 9 | 21399770 | 21399984 | 214 | BTBR,C57L,C58J,CAST,FVB,MA/My,MOLF,NOD,NOR,NZB,NZW,RF,SJL,SPRET,SWR,WSB | 9 | 21399857 | 21399875 | 19 | + | 17.5574 | 6.32E-07 | 0.112 | ACTCCAGAAGAGGGCGCCA | 1 | ENCFF055HAP |
| 1358 | 9 | 22096827 | 22097557 | 730 | C57L,CAST,FVB,MA/My,MOLF,NOD,NOR,NZB,NZW,RF,SJL,SPRET,SWR | 9 | 22097012 | 22097030 | 19 | + | 18.5902 | 3.13E-07 | 0.0834 | CGGCCAGCAGATGTCAGCC | | |
| 1359 | 9 | 22140117 | 22140311 | 194 | SPRET | 9 | 22140161 | 22140194 | 34 | + | 16.813 | 7.63E-07 | 0.076 | CTGTAGCTGTCTTTAGACACACCAGAGGGCATCA | | |
| 1360 | 9 | 22319096 | 22319309 | 213 | CAST.MOLF.SPRET | 9 | 22319167 | 22319200 | 34 | + | 18.3659 | 2.76E-07 | 0.0438 | TAGCTGTCTTCAGACACTCCAGAAGAGGGCGCCA | 2 | ENCFF093ZRU.ENCFF055HAP |
| 1360 | 9 | 22319096 | 22319309 | 213 | CAST.MOLF.SPRET | 9 | 22319182 | 22319200 | 19 | + | 17.5574 | 6.32E-07 | 0.112 | ACTCCAGAAGAGGGCGCCA | 2 | ENCFF093ZRU.ENCFF055HAP |
| 1361 | 9 | 27607727 | 27607929 | 202 | SPRET | 9 | 27607787 | 27607820 | 34 | + | 16.4472 | 9.61E-07 | 0.0841 | TAGCTGTCTTCAGACACACCAGAAGAGGGCGTCA | | |
| 1362 | 9 | 28175690 | 28176147 | 457 | B10J | 9 | 28175963 | 28175996 | 34 | + | 17.3415 | 5.44E-07 | 0.0631 | CTGCTACTCCTGCAGCCCCCTGGAGGTAGTGGTC | | |
| 1363 | 9 | 28268347 | 28268589 | 242 | SPRET | 9 | 28268394 | 28268427 | 34 | + | 17.878 | 3.83E-07 | 0.0522 | TAGCTGTCCTCAGACACACCAGAAGAGGGCATCA | 1 | ENCFF093ZRU |
| 1364 | 9 | 31329254 | 31329461 | 207 | CAST.CE.MOLF.NZW.SPRET | 9 | 31329319 | 31329352 | 34 | + | 16.4472 | 9.61E-07 | 0.0841 | TAGCTGTCTTCAGACACACCAGAAGAGGGCGTCA | | |
| 1365 | 9 | 40735885 | 40736105 | 220 | SPRET | 9 | 40735963 | 40735996 | 34 | + | 16.9919 | 6.81E-07 | 0.0715 | TAGCTGTCTTCAGACACACCAGAAGGGGCATCA | 1 | ENCFF093ZRU |
| 1366 | 9 | 43361318 | 43361710 | 392 | MOLF.SPRET | 9 | 43361619 | 43361652 | 34 | + | 19.9106 | 9.47E-08 | 0.022 | GTTCAGTTCCCTGCACCTACACCAGGTGGCTCAC | 2 | ENCFF327LYS.ENCFF907XEG |
| 1367 | 9 | 44068437 | 44068674 | 237 | 129S1,C57L,CAST,ILNJ2,MOLF,SMJ,SPRET | 9 | 44068532 | 44068565 | 34 | + | 17.8455 | 3.91E-07 | 0.0525 | TAGCTGTCCTCAGACACTCCAGAAGAGGGCGTCA | 1 | ENCFF093ZRU |
| 1368 | 9 | 44269791 | 44270004 | 213 | 129S1.C57L.CAST.MOLF.SPRET | 9 | 44269862 | 44269895 | 34 | + | 16.4472 | 9.61E-07 | 0.0841 | TAGCTGTCTTCAGACACACCAGAAGAGGGCGTCA | | |
| 1369 | 9 | 44682430 | 44682632 | 202 | SPRET | 9 | 44682490 | 44682523 | 34 | + | 16.4472 | 9.61E-07 | 0.0841 | TAGCTGTCTTCAGACACACCAGAAGAGGGCGTCA | | |
| 1370 | 9 | 49272690 | 49273259 | 569 | SPRET | 9 | 49273026 | 49273059 | 34 | + | 25.2602 | 1.31E-09 | 0.00104 | GTGCAGTGTCTGTGGGGAACAGAAGAGGGCACCG | | |
| 1371 | 9 | 53507635 | 53507928 | 293 | CAST | 9 | 53507778 | 53507811 | 34 | + | 20.4146 | 6.58E-08 | 0.0169 | TTGCAGTCTTCAGACACACCAGAAGAGGGCATCA | | |
| 1372 | 9 | 53574114 | 53574328 | 214 | CAST.MOLF.SPRET | 9 | 53574171 | 53574204 | 34 | + | 16.4634 | 9.51E-07 | 0.0841 | TAGCTGTCTTCAGACGCACCAGAAGAGGGCATCA | 1 | ENCFF367UNR |
| 1373 | 9 | 57005820 | 57006017 | 197 | SPRET | 9 | 57005874 | 57005907 | 34 | + | 18.6992 | 2.20E-07 | 0.0402 | TTGCTGTCTTCAGACACTCCAGAAGGTGGCGTCA | 1 | ENCFF093ZRU |
| 1374 | 9 | 57241347 | 57242106 | 759 | CAST.CBA | 9 | 57241760 | 57241794 | 35 | + | 16.0367 | 9.36E-07 | 0.152 | ATGCACTGCCCCTGGGCCGCAAGAAGGGGGCCAAG | | |
| 1375 | 9 | 57768529 | 57768763 | 234 | 129S1,C57L,CAST,MA/My,MOLF,PL,SPRET | 9 | 57768620 | 57768653 | 34 | + | 18.3659 | 2.76E-07 | 0.0438 | TAGCTGTCTTCAGACACTCCAGAAGAGGGCGCCA | | |
| 1375 | 9 | 57768529 | 57768763 | 234 | 129S1,C57L,CAST,MA/My,MOLF,PL,SPRET | 9 | 57768635 | 57768653 | 19 | + | 17.5574 | 6.32E-07 | 0.112 | ACTCCAGAAGAGGGCGCCA | | |
| 1376 | 9 | 57891741 | 57891943 | 202 | CAST.SPRET | 9 | 57891798 | 57891831 | 34 | + | 17.2114 | 5.91E-07 | 0.066 | TAGCTGTTTTCAGACACTCCAGAAGAGGGCATCA | | |
| 1377 | 9 | 58170482 | 58170725 | 243 | 129S1,AJ,AKR,BTBR,BUB,C3H,C57L,CAST,CBA,CE,DBA1J,DBA2J,FVB,ILNJ2,KK,MA/My,MOLF,MRL,NOD,NOR,NZB,NZO,NZW,PJ,PL,RF,RH2,SEA,SJL,SMJ,SPRET,SWR,TH,WSB | 9 | 58170583 | 58170616 | 34 | + | 16.4472 | 9.61E-07 | 0.0841 | TAGCTGTCTTCAGACACACCAGAAGAGGGCGTCA | | |
| 1378 | 9 | 62633509 | 62633735 | 226 | AJ,AKR,C3H,CAST,FVB,MOLF,NOD,NOR,NZB,NZO,NZW,PL,RF,RH2,SJL,SPRET,SWR,TH | 9 | 62633593 | 62633626 | 34 | + | 17.9431 | 3.66E-07 | 0.0511 | CAGCTGTCCTCAGACACTCCAGAAGAGGGAGTCA | | |
| 1379 | 9 | 62827108 | 62827311 | 203 | SPRET | 9 | 62827166 | 62827199 | 34 | + | 16.4472 | 9.61E-07 | 0.0841 | TAGCTGTCTTCAGACACACCAGAAGAGGGCGTCA | 1 | ENCFF093ZRU |
| 1380 | 9 | 62835852 | 62836061 | 209 | AKR.CAST.CE.MOLF.NZB.RF.SPRET | 9 | 62835920 | 62835953 | 34 | + | 16.7724 | 7.83E-07 | 0.0768 | TAGCTGTCTTCAGATGCACCAGAAGAGGGCGTCA | | |
| 1381 | 9 | 63511800 | 63511861 | 61 | SPRET | 9 | 63511798 | 63511831 | 34 | + | 18.3659 | 2.76E-07 | 0.0438 | CTCCCCTTCCCCCATCCCCCACCAGGGAGTGCTC | | |
| 1381 | 9 | 63511800 | 63511861 | 61 | SPRET | 9 | 63511829 | 63511862 | 34 | + | 18.5122 | 2.50E-07 | 0.0428 | CTCCCCTTCCCCCACCCCCCACCAGGGAGTGCTC | | |
| 1381 | 9 | 63511800 | 63511861 | 61 | SPRET | 9 | 63511860 | 63511893 | 34 | + | 18.3659 | 2.76E-07 | 0.0438 | CTCCCCTTCCCCCATCCCCCACCAGGGAGTGCTC | | |
| 1382 | 9 | 64046323 | 64046651 | 328 | SPRET | 9 | 64046354 | 64046372 | 19 | + | 19.2951 | 1.88E-07 | 0.0659 | CCGCCACTAGATGGTGCTA | 32 | ENCFF093ZRU,ENCFF099QNE,ENCFF201JUA,ENCFF821BIA,ENCFF024JOZ,ENCFF907XEG,ENCFF327LYS,ENCFF052KVO,ENCFF609MDN,ENCFF089GBC,ENCFF589ZBJ,ENCFF092NMJ,ENCFF136ZDK,ENCFF065ODV,ENCFF316FYN,ENCFF097CAQ,ENCFF055HAP,ENCFF198SSR,ENCFF430PPJ,ENCFF165LXO,ENCFF210BTO,ENCFF502WDT,ENCFF321AGQ,ENCFF281QMV,ENCFF957SPH,ENCFF951GOP,ENCFF533APC,ENCFF883UPM.ENCFF194HVL.ENCFF491RJK.ENCFF575MGI |
| 1383 | 9 | 64847313 | 64847521 | 208 | CAST.CE.FVB.MOLF.SJL | 9 | 64847380 | 64847413 | 34 | + | 18.6098 | 2.34E-07 | 0.0417 | TTGCTGTCTTCAGATGAACCAGAAGAGGGCATCA | 5 | ENCFF099QNE.ENCFF883UPM.ENCFF502WDT.ENCFF093ZRU.ENCFF055HAP |
| 1384 | 9 | 65211259 | 65212206 | 947 | 129S1.BTBR.C57L.SMJ | 9 | 65212162 | 65212195 | 34 | + | 22.4878 | 1.36E-08 | 0.0055 | GTGCAGTCCCCACAAAACCCTGAAGAGGGCATTG | 2 | ENCFF367UNR.ENCFF093ZRU |
| 1385 | 9 | 65275146 | 65275237 | 91 | CAST.FVB.MOLF.SJL.SPRET | 9 | 65275191 | 65275224 | 34 | + | 17.2439 | 5.79E-07 | 0.065 | TAGCTGTCCTCAGATACTCCAGAAGAGGGCATCA | 1 | ENCFF093ZRU |
| 1386 | 9 | 65612376 | 65612632 | 256 | 129S1,AJ,AKR,B10D2,B10J,BALB,BTBR,BUB,C3H,C57L,CAST,CE,DBA1J,DBA2J,FVB,ILNJ2,KK,LP,MA/My,MOLF,MRL,NOD,NOR,NZB,NZO,NZW,PJ,PL,RF,RH2,SEA,SJL,SMJ,SPRET.SWR.TH | 9 | 65612490 | 65612523 | 34 | + | 16.4634 | 9.51E-07 | 0.0841 | TAGCTGTCTTCAGACGCACCAGAAGAGGGCATCA | | |
| 1387 | 9 | 65763212 | 65763407 | 195 | SPRET | 9 | 65763265 | 65763298 | 34 | + | 18.3659 | 2.76E-07 | 0.0438 | TAGCTGTCTTCAGACACTCCAGAAGAGGGCGCCA | 4 | ENCFF821BIA.ENCFF093ZRU.ENCFF367UNR.ENCFF099QNE |
| 1387 | 9 | 65763212 | 65763407 | 195 | SPRET | 9 | 65763280 | 65763298 | 19 | + | 17.5574 | 6.32E-07 | 0.112 | ACTCCAGAAGAGGGCGCCA | 4 | ENCFF821BIA.ENCFF093ZRU.ENCFF367UNR.ENCFF099QNE |
| 1388 | 9 | 68755999 | 68756235 | 236 | 129S1,B10D2,B10J,BTBR,C57L,CAST,MOLF.SMJ.SPRET.WSB | 9 | 68756093 | 68756126 | 34 | + | 16.4634 | 9.51E-07 | 0.0841 | TAGCTGTCTTCAGACGCACCAGAAGAGGGCATCA | | |
| 1389 | 9 | 69706187 | 69706949 | 762 | CAST | 9 | 69706297 | 69706331 | 35 | + | 20.633 | 5.75E-08 | 0.0295 | TTGCACTTGCCTGTGGAGGCCAGAAGAGGCCATCA | | |
| 1390 | 9 | 71550787 | 71550985 | 198 | CAST | 9 | 71550844 | 71550877 | 34 | + | 16.7724 | 7.83E-07 | 0.0768 | TAGCTGTCTTCAGATGCACCAGAAGAGGGCGTCA | | |
| 1391 | 9 | 72121537 | 72121742 | 205 | AJ,BALB,BUB,C3H,CAST,CBA,CE,DBA1J,DBA2J,FVB,ILNJ2,KK,MA/My,MOLF,NOD,NOR,NZB,NZO,NZW,PJ,RH2,SEA,SJL,SMJ,SPRET.SWR | 9 | 72121601 | 72121635 | 35 | + | 15.9633 | 9.75E-07 | 0.154 | CTGTAGCTGTCTTCAGACACCAGAAGAGGGTGACA | | |
| 1392 | 9 | 72573872 | 72574093 | 221 | SPRET | 9 | 72573941 | 72573974 | 34 | + | 16.4634 | 9.51E-07 | 0.0841 | TAGCTGTCTTCAGACGCACCAGAAGAGGGCATCA | 2 | ENCFF367UNR.ENCFF093ZRU |
| 1393 | 9 | 72591103 | 72591320 | 217 | SPRET | 9 | 72591161 | 72591194 | 34 | + | 16.4472 | 9.61E-07 | 0.0841 | TAGCTGTCTTCAGACACACCAGAAGAGGGCGTCA | 1 | ENCFF093ZRU |
| 1394 | 9 | 72686700 | 72686924 | 224 | 129S1,AJ,AKR,BALB,BTBR,BUB,C3H,C57L,CAST,CBA,CE,DBA1J,DBA2J,FVB,ILNJ2,KK,MA/My,MOLF,MRL,NOD,NOR,NZB,NZO,NZW,PJ,PL,RF,RH2,SEA,SJL,SMJ,SPRET,SWR.TH | 9 | 72686785 | 72686818 | 34 | + | 17.9106 | 3.74E-07 | 0.0517 | TAGCTGTCTTCAGACACTCCAGAAGAGGGCACCA | | |
| 1395 | 9 | 72878423 | 72878638 | 215 | CAST.MOLF.SPRET | 9 | 72878493 | 72878526 | 34 | + | 17.8455 | 3.91E-07 | 0.0525 | TAGCTGTCCTCAGACACTCCAGAAGAGGGCGTCA | | |
| 1396 | 9 | 75354075 | 75354297 | 222 | AJ,AKR,BALB,BTBR,BUB,C3H,C58J,CAST,CBA,CE,DBA1J,DBA2J,FVB,ILNJ2,KK,LP,MA/My,MOLF,MRL,NOD,NOR,NZB,NZO,NZW,PJ,PL,RF,RH2,SEA,SJL,SMJ,SPRET,SWR,TH | 9 | 75354155 | 75354188 | 34 | + | 16.4797 | 9.42E-07 | 0.0841 | TAGCTGTCTTCAGACACTCCACAAGAGGGCGTCA | 1 | ENCFF093ZRU |
| 1397 | 9 | 81459294 | 81459498 | 204 | AKR,BUB,C58J,CAST,CBA,ILNJ2,LP,MOLF,MRL,NOD,NOR,NZW,PL,RF,SMJ,SPRET,WSB | 9 | 81459356 | 81459389 | 34 | + | 16.4472 | 9.61E-07 | 0.0841 | TAGCTGTCTTCAGACACACCAGAAGAGGGCGTCA | | |
| 1398 | 9 | 81743274 | 81743482 | 208 | AKR,BUB,C58J,CAST,CBA,ILNJ2,LP,MOLF,MRL,NOD,NOR,NZW,PL,RF,SMJ,SPRET,WSB | 9 | 81743340 | 81743373 | 34 | + | 17.8455 | 3.91E-07 | 0.0525 | TAGCTGTCCTCAGACACTCCAGAAGAGGGCGTCA | | |
| 1399 | 9 | 84254494 | 84254590 | 96 | CAST,FVB,KK,NZO,NZW,SJL,SPRET,SWR | 9 | 84254549 | 84254582 | 34 | + | 18.3659 | 2.76E-07 | 0.0438 | TAGCTGTCTTCAGACACTCCAGAAGAGGGCGCCA | 1 | ENCFF093ZRU |
| 1399 | 9 | 84254494 | 84254590 | 96 | CAST,FVB,KK,NZO,NZW,SJL,SPRET,SWR | 9 | 84254564 | 84254582 | 19 | + | 17.5574 | 6.32E-07 | 0.112 | ACTCCAGAAGAGGGCGCCA | 1 | ENCFF093ZRU |
| 1400 | 9 | 88620840 | 88621058 | 218 | SPRET | 9 | 88620894 | 88620927 | 34 | + | 17.6829 | 4.35E-07 | 0.0554 | TAGCTGTCTTCAGACGCGCCAGAAGAGGGCATCA | | |
| 1401 | 9 | 89963839 | 89964035 | 196 | CAST.MRL.PJ.SEA.SMJ.WSB | 9 | 89963893 | 89963926 | 34 | + | 17.878 | 3.83E-07 | 0.0522 | TAGCTGTCCTCAGACACACCAGAAGAGGGCATCA | | |
| 1402 | 9 | 90050101 | 90050295 | 194 | CAST.PJ.SEA.SPRET | 9 | 90050153 | 90050186 | 34 | + | 18.3659 | 2.76E-07 | 0.0438 | TAGCTGTCTTCAGACACTCCAGAAGAGGGCGCCA | | |
| 1402 | 9 | 90050101 | 90050295 | 194 | CAST.PJ.SEA.SPRET | 9 | 90050168 | 90050186 | 19 | + | 17.5574 | 6.32E-07 | 0.112 | ACTCCAGAAGAGGGCGCCA | | |
| 1403 | 9 | 96639537 | 96639627 | 90 | MRL.NZO.NZW | 9 | 96639549 | 96639582 | 34 | + | 22.9268 | 9.57E-09 | 0.0042 | GTGCAGGGCCCACAGAGGCCAGAAGAGGGTGTAG | | |
| 1404 | 9 | 97047074 | 97047273 | 199 | MRL.NZO.NZW.SPRET | 9 | 97047125 | 97047158 | 34 | + | 16.4472 | 9.61E-07 | 0.0841 | TAGCTGTCTTCAGACACACCAGAAGAGGGCGTCA | | |
| 1405 | 9 | 99326959 | 99327561 | 602 | SPRET | 9 | 99327062 | 99327095 | 34 | + | 16.7398 | 7.99E-07 | 0.0779 | AAGGAGTGTGAATAGAGTCCAGAAGGTGGCGCTC | 25 | ENCFF093ZRU,ENCFF367UNR,ENCFF099QNE,ENCFF821BIA,ENCFF316FYN,ENCFF065ODV,ENCFF055HAP,ENCFF327LYS,ENCFF609MDN,ENCFF907XEG,ENCFF502WDT,ENCFF089GBC,ENCFF092NMJ,ENCFF198SSR,ENCFF052KVO,ENCFF957SPH,ENCFF136ZDK,ENCFF024JOZ,ENCFF589ZBJ,ENCFF533APC,ENCFF883UPM.ENCFF142CNG.ENCFF281QMV.ENCFF430PPJ.ENCFF165LXO |
| 1405 | 9 | 99326959 | 99327561 | 602 | SPRET | 9 | 99327077 | 99327095 | 19 | + | 18.4098 | 3.55E-07 | 0.0872 | AGTCCAGAAGGTGGCGCTC | 25 | ENCFF093ZRU,ENCFF367UNR,ENCFF099QNE,ENCFF821BIA,ENCFF316FYN,ENCFF065ODV,ENCFF055HAP,ENCFF327LYS,ENCFF609MDN,ENCFF907XEG,ENCFF502WDT,ENCFF089GBC,ENCFF092NMJ,ENCFF198SSR,ENCFF052KVO,ENCFF957SPH,ENCFF136ZDK,ENCFF024JOZ,ENCFF589ZBJ,ENCFF533APC,ENCFF883UPM.ENCFF142CNG.ENCFF281QMV.ENCFF430PPJ.ENCFF165LXO |
| 1406 | 9 | 103623033 | 103623964 | 931 | CBA | 9 | 103623482 | 103623516 | 35 | + | 16.0734 | 9.17E-07 | 0.15 | CAGCAATGTGGACATCAGCTCTCCAGAGGGGACTC | | |
| 1407 | 9 | 104012716 | 104012930 | 214 | 129S1,AJ,AKR,BALB,BTBR,BUB,C3H,C57L,CAST,CBA,CE,DBA1J,DBA2J,FVB,ILNJ2,KK,MA/My,MOLF,MRL,NOD,NOR,NZB,NZO,NZW,PJ,PL,RF,RH2,SEA,SJL,SMJ,SPRET,SWR.TH.WSB | 9 | 104012787 | 104012820 | 34 | + | 18.3659 | 2.76E-07 | 0.0438 | TAGCTGTCTTCAGACACTCCAGAAGAGGGCGCCA | | |
| 1407 | 9 | 104012716 | 104012930 | 214 | 129S1,AJ,AKR,BALB,BTBR,BUB,C3H,C57L,CAST,CBA,CE,DBA1J,DBA2J,FVB,ILNJ2,KK,MA/My,MOLF,MRL,NOD,NOR,NZB,NZO,NZW,PJ,PL,RF,RH2,SEA,SJL,SMJ,SPRET,SWR.TH.WSB | 9 | 104012802 | 104012820 | 19 | + | 17.5574 | 6.32E-07 | 0.112 | ACTCCAGAAGAGGGCGCCA | | |
| 1408 | 9 | 106794248 | 106794448 | 200 | 129S1,AJ,AKR,BALB,BTBR,BUB,C3H,C57L,CAST,CE,FVB,ILNJ2,MA/My,MRL,NOD,NOR,NZB,NZO,NZW,PJ,PL,RF,RH2,SEA,SJL,SMJ,SPRET.SWR.TH.WSB | 9 | 106794305 | 106794338 | 34 | + | 16.5691 | 8.90E-07 | 0.0831 | TAGCTGTCTTCAGACGCACCAGAAGAGGGCATCG | | |
| 1409 | 9 | 107658793 | 107659178 | 385 | SPRET | 9 | 107659056 | 107659074 | 19 | + | 17.2295 | 7.83E-07 | 0.125 | AGGCCAGAAGAGGGCAATG | | |
| 1410 | 9 | 108075251 | 108075448 | 197 | SPRET | 9 | 108075301 | 108075334 | 34 | + | 16.4472 | 9.61E-07 | 0.0841 | TAGCTGTCTTCAGACACACCAGAAGAGGGCGTCA | 1 | ENCFF093ZRU |
| 1411 | 9 | 108164987 | 108165192 | 205 | CAST.CBA.MOLF.SPRET.WSB | 9 | 108165050 | 108165083 | 34 | + | 17.9431 | 3.66E-07 | 0.0511 | CAGCTGTCCTCAGACACTCCAGAAGAGGGAGTCA | | |
| 1412 | 9 | 109737668 | 109737901 | 233 | SPRET | 9 | 109737760 | 109737793 | 34 | + | 17.2439 | 5.79E-07 | 0.065 | TAGCTGTCCTCAGATACTCCAGAAGAGGGCATCA | | |
| 1413 | 9 | 109977312 | 109977518 | 206 | CBA,MOLF.SPRET | 9 | 109977376 | 109977409 | 34 | + | 16.4472 | 9.61E-07 | 0.0841 | TAGCTGTCTTCAGACACACCAGAAGAGGGCGTCA | | |
| 1414 | 9 | 110130180 | 110130386 | 206 | CAST.CBA.MOLF.NZW.SPRET | 9 | 110130244 | 110130277 | 34 | + | 17.8455 | 3.91E-07 | 0.0525 | TAGCTGTCCTCAGACACTCCAGAAGAGGGCGTCA | | |
| 1415 | 9 | 113829697 | 113829896 | 199 | CAST | 9 | 113829753 | 113829786 | 34 | + | 17.2439 | 5.79E-07 | 0.065 | TAGCTGTCCTCAGATACTCCAGAAGAGGGCATCA | | |
| 1416 | 9 | 114794703 | 114794892 | 189 | CAST | 9 | 114794811 | 114794844 | 34 | + | 18.5854 | 2.38E-07 | 0.0421 | TTGCTGTCTTCAGACACACCAGAAGAGGGCATCG | 1 | ENCFF093ZRU |
| 1417 | 9 | 114917407 | 114917647 | 240 | SPRET | 9 | 114917480 | 114917514 | 35 | + | 17.6881 | 3.62E-07 | 0.0909 | CAACATCTGTAGCTAGTGCCCACTAGATGGCGGTA | 31 | ENCFF093ZRU,ENCFF367UNR,ENCFF099QNE,ENCFF201JUA,ENCFF052KVO,ENCFF024JOZ,ENCFF821BIA,ENCFF907XEG,ENCFF327LYS,ENCFF165LXO,ENCFF198SSR,ENCFF089GBC,ENCFF589ZBJ,ENCFF092NMJ,ENCFF033UQX,ENCFF957SPH,ENCFF086ZTV,ENCFF575MGI,ENCFF430PPJ,ENCFF136ZDK,ENCFF951GOP,ENCFF281QMV,ENCFF097CAQ,ENCFF533APC,ENCFF210BTO,ENCFF194HVL,ENCFF611HDQ,ENCFF609MDN.ENCFF316FYN.ENCFF491RJK.ENCFF883UPM |
| 1417 | 9 | 114917407 | 114917647 | 240 | SPRET | 9 | 114917496 | 114917514 | 19 | + | 21.082 | 4.57E-08 | 0.0361 | TGCCCACTAGATGGCGGTA | 31 | ENCFF093ZRU,ENCFF367UNR,ENCFF099QNE,ENCFF201JUA,ENCFF052KVO,ENCFF024JOZ,ENCFF821BIA,ENCFF907XEG,ENCFF327LYS,ENCFF165LXO,ENCFF198SSR,ENCFF089GBC,ENCFF589ZBJ,ENCFF092NMJ,ENCFF033UQX,ENCFF957SPH,ENCFF086ZTV,ENCFF575MGI,ENCFF430PPJ,ENCFF136ZDK,ENCFF951GOP,ENCFF281QMV,ENCFF097CAQ,ENCFF533APC,ENCFF210BTO,ENCFF194HVL,ENCFF611HDQ,ENCFF609MDN.ENCFF316FYN.ENCFF491RJK.ENCFF883UPM |
| 1418 | 9 | 115659525 | 115659968 | 443 | CAST.SPRET | 9 | 115659826 | 115659859 | 34 | + | 18.4959 | 2.53E-07 | 0.0428 | CAGCTGTCTTCAGACATACCAGAAGAGGGCATCA | | |
| 1419 | 9 | 118482316 | 118482425 | 109 | CAST | 9 | 118482307 | 118482340 | 34 | + | 26.6179 | 3.73E-10 | 0.000467 | GTGCAGTGTCCACAAAGGCCAGAAGAGGGCATCA | | |
| 1420 | 9 | 120405396 | 120405685 | 289 | BTBR,MOLF,MRL,SPRET,WSB | 9 | 120405517 | 120405550 | 34 | + | 18.0976 | 3.31E-07 | 0.0481 | GTGCACCACCATTGTCCGCCTCCAGGGGACTTCA | 11 | ENCFF099QNE,ENCFF052KVO,ENCFF165LXO,ENCFF092NMJ,ENCFF198SSR,ENCFF327LYS,ENCFF589ZBJ,ENCFF089GBC.ENCFF430PPJ.ENCFF093ZRU.ENCFF136ZDK |
| 1421 | 9 | 120516370 | 120516559 | 189 | BTBR.CAST.MOLF.SPRET | 9 | 120516422 | 120516455 | 34 | + | 16.4634 | 9.51E-07 | 0.0841 | TAGCTGTCTTCAGACGCACCAGAAGAGGGCATCA | 3 | ENCFF093ZRU.ENCFF099QNE.ENCFF367UNR |
| 1422 | 9 | 120621906 | 120622353 | 447 | SPRET | 9 | 120622150 | 120622183 | 34 | + | 28.1707 | 7.92E-11 | 0.000215 | ATGCAGTGCCTGAAGCAACCAGAAGAGGGCGTTA | 4 | ENCFF821BIA.ENCFF052KVO.ENCFF093ZRU.ENCFF367UNR |
| 1423 | 9 | 121473629 | 121473707 | 78 | AKR,BUB,C58J,CAST,CBA,CE,DBA1J,DBA2J.MRL.NZW.RF.RH2.SMJ.SPRET | 9 | 121473602 | 121473635 | 34 | + | 18.3333 | 2.82E-07 | 0.0442 | CTGCTCTCCTCTCTCTGGACTGGAGGGGGACCCC | | |
| 1423 | 9 | 121473629 | 121473707 | 78 | AKR,BUB,C58J,CAST,CBA,CE,DBA1J,DBA2J.MRL.NZW.RF.RH2.SMJ.SPRET | 9 | 121473641 | 121473674 | 34 | + | 18.1382 | 3.22E-07 | 0.0474 | CTGCTCTCCTCTCCCTGGACTGGAGGGGGACCCC | | |
| 1424 | 9 | 121526776 | 121527001 | 225 | AKR,BUB,C58J,CAST,CBA,DBA1J,DBA2J,MOLF,MRL,RF,RH2,SMJ,SPRET | 9 | 121526751 | 121526784 | 34 | + | 16.6098 | 8.68E-07 | 0.0821 | GCCTTGCACTGCCAGTGGCCACCAGAGGGCACTG | 37 | ENCFF201JUA,ENCFF093ZRU,ENCFF367UNR,ENCFF327LYS,ENCFF136ZDK,ENCFF099QNE,ENCFF907XEG,ENCFF821BIA,ENCFF589ZBJ,ENCFF089GBC,ENCFF024JOZ,ENCFF321AGQ,ENCFF065ODV,ENCFF092NMJ,ENCFF502WDT,ENCFF198SSR,ENCFF430PPJ,ENCFF210BTO,ENCFF052KVO,ENCFF142CNG,ENCFF165LXO,ENCFF194HVL,ENCFF504JDS,ENCFF349BUL,ENCFF281QMV,ENCFF086ZTV,ENCFF575MGI,ENCFF601JVE,ENCFF805QBU,ENCFF491RJK,ENCFF611HDQ,ENCFF951GOP,ENCFF883UPM,ENCFF316FYN,ENCFF055HAP,ENCFF089EWX,ENCFF784ASD |
| 1424 | 9 | 121526776 | 121527001 | 225 | AKR,BUB,C58J,CAST,CBA,DBA1J,DBA2J,MOLF,MRL,RF,RH2,SMJ,SPRET | 9 | 121526766 | 121526784 | 19 | + | 25.4262 | 3.18E-10 | 0.00481 | TGGCCACCAGAGGGCACTG | 39 | ENCFF201JUA,ENCFF093ZRU,ENCFF367UNR,ENCFF327LYS,ENCFF136ZDK,ENCFF099QNE,ENCFF907XEG,ENCFF821BIA,ENCFF589ZBJ,ENCFF089GBC,ENCFF024JOZ,ENCFF321AGQ,ENCFF065ODV,ENCFF092NMJ,ENCFF502WDT,ENCFF198SSR,ENCFF430PPJ,ENCFF210BTO,ENCFF052KVO,ENCFF142CNG,ENCFF165LXO,ENCFF194HVL,ENCFF504JDS,ENCFF349BUL,ENCFF281QMV,ENCFF086ZTV,ENCFF575MGI,ENCFF601JVE,ENCFF805QBU,ENCFF491RJK,ENCFF611HDQ,ENCFF951GOP,ENCFF883UPM,ENCFF316FYN,ENCFF055HAP,ENCFF089EWX,ENCFF784ASD,ENCFF097CAQ,ENCFF033UQX |
| 1424 | 9 | 121526776 | 121527001 | 225 | AKR,BUB,C58J,CAST,CBA,DBA1J,DBA2J,MOLF,MRL,RF,RH2,SMJ,SPRET | 9 | 121526796 | 121526829 | 34 | + | 16.6098 | 8.68E-07 | 0.0821 | GCCTTGCACTGCCAGTGGCCACCAGAGGGCACTG | 39 | ENCFF201JUA,ENCFF093ZRU,ENCFF367UNR,ENCFF327LYS,ENCFF136ZDK,ENCFF099QNE,ENCFF907XEG,ENCFF821BIA,ENCFF589ZBJ,ENCFF089GBC,ENCFF024JOZ,ENCFF321AGQ,ENCFF065ODV,ENCFF092NMJ,ENCFF502WDT,ENCFF198SSR,ENCFF430PPJ,ENCFF210BTO,ENCFF052KVO,ENCFF142CNG,ENCFF165LXO,ENCFF194HVL,ENCFF504JDS,ENCFF349BUL,ENCFF281QMV,ENCFF086ZTV,ENCFF575MGI,ENCFF601JVE,ENCFF805QBU,ENCFF491RJK,ENCFF611HDQ,ENCFF951GOP,ENCFF883UPM,ENCFF316FYN,ENCFF055HAP,ENCFF089EWX,ENCFF784ASD,ENCFF097CAQ,ENCFF033UQX |

| | | | | | | | | | | | | | | | | |
|---|---|---|---|---|---|---|---|---|---|---|---|---|---|---|---|---|
| 1424 | 9 | 121526776 | 121527001 | 225 | AKR,BUB,C58J,CAST,CBA,DBA1J,DBA2J,MOLF,MRL,RF,RH2,SMJ,SPRET | 9 | 121526811 | 121526829 | 19 | + | 25.4262 | 3.18E-10 | 0.00481 | TGGCCACCAGAGGGCACTG | 39 | ENCFF201JUA,ENCFF093ZRU,ENCFF367UNR,ENCFF327LYS,ENCFF136ZDK,ENCFF099QNE,ENCFF907XEG,ENCFF821BIA,ENCFF589ZBJ,ENCFF089GBC,ENCFF024JOZ,ENCFF321AGQ,ENCFF065ODV,ENCFF092NMJ,ENCFF502WDT,ENCFF198SSR,ENCFF430PPJ,ENCFF210BTO,ENCFF052KVO,ENCFF142CNG,ENCFF165LXO,ENCFF194HVL,ENCFF504JDS,ENCFF349BUL,ENCFF281QMV,ENCFF086ZTV,ENCFF575MGI,ENCFF601JVE,ENCFF805QBU,ENCFF491RJK,ENCFF611HDQ,ENCFF951GOP,ENCFF883UPM,ENCFF316FYN,ENCFF055HAP,ENCFF089EWX,ENCFF784ASD,ENCFF097CAQ,ENCFF033UQX |
| 1424 | 9 | 121526776 | 121527001 | 225 | AKR,BUB,C58J,CAST,CBA,DBA1J,DBA2J,MOLF,MRL,RF,RH2,SMJ,SPRET | 9 | 121526841 | 121526874 | 34 | + | 16.6098 | 8.68E-07 | 0.0821 | GCCTTGCACTGCCAGTGGCCACCAGAGGGCACTG | 34 | ENCFF201JUA,ENCFF093ZRU,ENCFF367UNR,ENCFF327LYS,ENCFF136ZDK,ENCFF099QNE,ENCFF907XEG,ENCFF821BIA,ENCFF589ZBJ,ENCFF089GBC,ENCFF024JOZ,ENCFF321AGQ,ENCFF065ODV,ENCFF092NMJ,ENCFF198SSR,ENCFF430PPJ,ENCFF210BTO,ENCFF052KVO,ENCFF165LXO,ENCFF504JDS,ENCFF349BUL,ENCFF601JVE,ENCFF951GOP,ENCFF883UPM,ENCFF316FYN,ENCFF055HAP,ENCFF089EWX,ENCFF784ASD,ENCFF097CAQ,ENCFF033UQX,ENCFF609MDN,ENCFF957SPH,ENCFF194HVL |
| 1424 | 9 | 121526776 | 121527001 | 225 | AKR,BUB,C58J,CAST,CBA,DBA1J,DBA2J,MOLF,MRL,RF,RH2,SMJ,SPRET | 9 | 121526856 | 121526874 | 19 | + | 25.4262 | 3.18E-10 | 0.00481 | TGGCCACCAGAGGGCACTG | 32 | ENCFF201JUA,ENCFF093ZRU,ENCFF367UNR,ENCFF327LYS,ENCFF136ZDK,ENCFF099QNE,ENCFF907XEG,ENCFF821BIA,ENCFF589ZBJ,ENCFF089GBC,ENCFF024JOZ,ENCFF321AGQ,ENCFF092NMJ,ENCFF198SSR,ENCFF430PPJ,ENCFF210BTO,ENCFF165LXO,ENCFF504JDS,ENCFF601JVE,ENCFF951GOP,ENCFF883UPM,ENCFF316FYN,ENCFF055HAP,ENCFF089EWX,ENCFF784ASD,ENCFF097CAQ,ENCFF033UQX,ENCFF609MDN.ENCFF957SPH.ENCFF194HVL.ENCFF805QBU |
| 1424 | 9 | 121526776 | 121527001 | 225 | AKR,BUB,C58J,CAST,CBA,DBA1J,DBA2J,MOLF,MRL,RF,RH2,SMJ,SPRET | 9 | 121526886 | 121526919 | 34 | + | 16.6098 | 8.68E-07 | 0.0821 | GCCTTGCACTGCCAGTGGCCACCAGAGGGCACTG | 32 | ENCFF201JUA,ENCFF093ZRU,ENCFF367UNR,ENCFF327LYS,ENCFF136ZDK,ENCFF099QNE,ENCFF907XEG,ENCFF821BIA,ENCFF589ZBJ,ENCFF089GBC,ENCFF024JOZ,ENCFF321AGQ,ENCFF092NMJ,ENCFF198SSR,ENCFF430PPJ,ENCFF165LXO,ENCFF601JVE,ENCFF951GOP,ENCFF883UPM,ENCFF316FYN,ENCFF055HAP,ENCFF089EWX,ENCFF784ASD,ENCFF097CAQ,ENCFF033UQX,ENCFF609MDN,ENCFF957SPH,ENCFF194HVL.ENCFF805QBU.ENCFF491RJK |
| 1424 | 9 | 121526776 | 121527001 | 225 | AKR,BUB,C58J,CAST,CBA,DBA1J,DBA2J,MOLF,MRL,RF,RH2,SMJ,SPRET | 9 | 121526901 | 121526919 | 19 | + | 25.4262 | 3.18E-10 | 0.00481 | TGGCCACCAGAGGGCACTG | 32 | ENCFF201JUA,ENCFF093ZRU,ENCFF367UNR,ENCFF327LYS,ENCFF136ZDK,ENCFF099QNE,ENCFF907XEG,ENCFF821BIA,ENCFF589ZBJ,ENCFF089GBC,ENCFF024JOZ,ENCFF321AGQ,ENCFF092NMJ,ENCFF198SSR,ENCFF430PPJ,ENCFF165LXO,ENCFF601JVE,ENCFF951GOP,ENCFF883UPM,ENCFF316FYN,ENCFF055HAP,ENCFF089EWX,ENCFF784ASD,ENCFF097CAQ,ENCFF033UQX,ENCFF609MDN,ENCFF957SPH,ENCFF194HVL.ENCFF805QBU.ENCFF491RJK |
| 1424 | 9 | 121526776 | 121527001 | 225 | AKR,BUB,C58J,CAST,CBA,DBA1J,DBA2J,MOLF,MRL,RF,RH2,SMJ,SPRET | 9 | 121526931 | 121526964 | 34 | + | 16.6098 | 8.68E-07 | 0.0821 | GCCTTGCACTGCCAGTGGCCACCAGAGGGCACTG | 32 | ENCFF093ZRU,ENCFF367UNR,ENCFF327LYS,ENCFF136ZDK,ENCFF099QNE,ENCFF907XEG,ENCFF821BIA,ENCFF589ZBJ,ENCFF089GBC,ENCFF024JOZ,ENCFF321AGQ,ENCFF092NMJ,ENCFF198SSR,ENCFF430PPJ,ENCFF165LXO,ENCFF601JVE,ENCFF951GOP,ENCFF883UPM,ENCFF316FYN,ENCFF055HAP,ENCFF784ASD,ENCFF097CAQ,ENCFF033UQX,ENCFF957SPH,ENCFF194HVL,ENCFF805QBU,ENCFF201JUA,ENCFF491RJK.ENCFF349BUL.ENCFF065ODV.ENCFF502WDT |
| 1424 | 9 | 121526776 | 121527001 | 225 | AKR,BUB,C58J,CAST,CBA,DBA1J,DBA2J,MOLF,MRL,RF,RH2,SMJ,SPRET | 9 | 121526946 | 121526964 | 19 | + | 25.4262 | 3.18E-10 | 0.00481 | TGGCCACCAGAGGGCACTG | 30 | ENCFF093ZRU,ENCFF367UNR,ENCFF327LYS,ENCFF136ZDK,ENCFF099QNE,ENCFF907XEG,ENCFF821BIA,ENCFF589ZBJ,ENCFF089GBC,ENCFF024JOZ,ENCFF321AGQ,ENCFF092NMJ,ENCFF198SSR,ENCFF430PPJ,ENCFF165LXO,ENCFF601JVE,ENCFF951GOP,ENCFF883UPM,ENCFF316FYN,ENCFF055HAP,ENCFF033UQX,ENCFF957SPH,ENCFF194HVL,ENCFF805QBU,ENCFF201JUA,ENCFF491RJK,ENCFF349BUL,ENCFF065ODV.ENCFF502WDT |
| 1424 | 9 | 121526776 | 121527001 | 225 | AKR,BUB,C58J,CAST,CBA,DBA1J,DBA2J,MOLF,MRL,RF,RH2,SMJ,SPRET | 9 | 121526976 | 121527009 | 34 | + | 17.6098 | 4.56E-07 | 0.057 | GCCTTGCACTGCCAGTGGCCACCAGGGGGCACTG | 28 | ENCFF093ZRU,ENCFF367UNR,ENCFF327LYS,ENCFF136ZDK,ENCFF099QNE,ENCFF907XEG,ENCFF821BIA,ENCFF589ZBJ,ENCFF089GBC,ENCFF024JOZ,ENCFF321AGQ,ENCFF092NMJ,ENCFF198SSR,ENCFF430PPJ,ENCFF165LXO,ENCFF601JVE,ENCFF883UPM,ENCFF055HAP,ENCFF957SPH,ENCFF194HVL,ENCFF805QBU,ENCFF201JUA,ENCFF491RJK,ENCFF349BUL,ENCFF065ODV,ENCFF502WDT,ENCFF504JDS |
| 1424 | 9 | 121526776 | 121527001 | 225 | AKR,BUB,C58J,CAST,CBA,DBA1J,DBA2J,MOLF,MRL,RF,RH2,SMJ,SPRET | 9 | 121526991 | 121527009 | 19 | + | 26.4754 | 4.11E-11 | 0.00331 | TGGCCACCAGGGGGCACTG | 28 | ENCFF093ZRU,ENCFF367UNR,ENCFF327LYS,ENCFF136ZDK,ENCFF099QNE,ENCFF907XEG,ENCFF821BIA,ENCFF589ZBJ,ENCFF089GBC,ENCFF024JOZ,ENCFF321AGQ,ENCFF092NMJ,ENCFF198SSR,ENCFF430PPJ,ENCFF165LXO,ENCFF601JVE,ENCFF883UPM,ENCFF055HAP,ENCFF957SPH,ENCFF194HVL,ENCFF805QBU,ENCFF201JUA,ENCFF491RJK,ENCFF349BUL,ENCFF065ODV,ENCFF502WDT,ENCFF504JDS |
| 1425 | 9 | 121672118 | 121672176 | 58 | AJ,AKR,BALB,BUB,C3H,C58J,CAST,CBA,CE,DBA1J,DBA2J,FVB,ILNJ2,LP,MA/My,MOLF,MRL,NZW,PJ,RF,RH2,SEA,SMJ,SPRET,TH.WSB | 9 | 121672120 | 121672138 | 19 | + | 18.2951 | 3.84E-07 | 0.0907 | ATGCCAGAAGAGGGCAGCA | 14 | ENCFF093ZRU,ENCFF821BIA,ENCFF092NMJ,ENCFF024JOZ,ENCFF883UPM,ENCFF099QNE,ENCFF367UNR,ENCFF430PPJ,ENCFF136ZDK,ENCFF907XEG,ENCFF589ZBJ,ENCFF198SSR,ENCFF327LYS,ENCFF089GBC |
| 1426 | 9 | 121889384 | 121889857 | 473 | MOLF | 9 | 121889485 | 121889503 | 19 | + | 17.2131 | 7.91E-07 | 0.126 | AGGCCAGTGGAGGGCACTG | | |
| 1427 | 9 | 123307627 | 123307703 | 76 | SPRET,WSB | 9 | 123307593 | 123307627 | 35 | + | 16.9083 | 5.71E-07 | 0.117 | TCGGCGTGCGCATCACTGGCCTGCAGGGGGCGCCC | 23 | ENCFF367UNR,ENCFF093ZRU,ENCFF092NMJ,ENCFF210BTO,ENCFF099QNE,ENCFF089GBC,ENCFF821BIA,ENCFF883UPM,ENCFF327LYS,ENCFF321AGQ,ENCFF033UQX,ENCFF589ZBJ,ENCFF957SPH,ENCFF136ZDK,ENCFF907XEG,ENCFF165LXO,ENCFF198SSR,ENCFF502WDT,ENCFF024JOZ,ENCFF430PPJ,ENCFF055HAP.ENCFF142CNG.ENCFF281QMV |
| 1427 | 9 | 123307627 | 123307703 | 76 | SPRET,WSB | 9 | 123307594 | 123307627 | 34 | + | 18.9268 | 1.89E-07 | 0.0363 | CGGCGTGCGCATCACTGGCCTGCAGGGGGCGCCC | 23 | ENCFF367UNR,ENCFF093ZRU,ENCFF092NMJ,ENCFF210BTO,ENCFF099QNE,ENCFF089GBC,ENCFF821BIA,ENCFF883UPM,ENCFF327LYS,ENCFF321AGQ,ENCFF033UQX,ENCFF589ZBJ,ENCFF957SPH,ENCFF136ZDK,ENCFF907XEG,ENCFF165LXO,ENCFF198SSR,ENCFF502WDT,ENCFF024JOZ,ENCFF430PPJ,ENCFF055HAP.ENCFF142CNG.ENCFF281QMV |
| 1427 | 9 | 123307627 | 123307703 | 76 | SPRET,WSB | 9 | 123307609 | 123307627 | 19 | + | 22.9508 | 8.07E-09 | 0.0177 | TGGCCTGCAGGGGGCGCCC | 23 | ENCFF367UNR,ENCFF093ZRU,ENCFF092NMJ,ENCFF210BTO,ENCFF099QNE,ENCFF089GBC,ENCFF821BIA,ENCFF883UPM,ENCFF327LYS,ENCFF321AGQ,ENCFF033UQX,ENCFF589ZBJ,ENCFF957SPH,ENCFF136ZDK,ENCFF907XEG,ENCFF165LXO,ENCFF198SSR,ENCFF502WDT,ENCFF024JOZ,ENCFF430PPJ,ENCFF055HAP.ENCFF142CNG.ENCFF281QMV |
| 1428 | 9 | 6057428 | 6057634 | 206 | CAST.MOLF.SPRET | 9 | 6057553 | 6057586 | 34 | - | 17.2439 | 5.79E-07 | 0.065 | TAGCTGTCCTCAGATACTCCAGAAGAGGGCATCA | | |
| 1429 | 9 | 7728093 | 7728287 | 194 | 129S1.CAST.MOLF.SPRET | 9 | 7728213 | 7728246 | 34 | - | 19.0163 | 1.77E-07 | 0.0347 | CAGCTGTCTTCAGACACACCAGAAGAGGGCGTCA | | |
| 1430 | 9 | 20322717 | 20323045 | 328 | MOLF | 9 | 20322985 | 20323003 | 19 | - | 17.3443 | 7.26E-07 | 0.121 | TTGCCAGAAGAGGGCATCA | 4 | ENCFF367UNR.ENCFF502WDT.ENCFF142CNG.ENCFF055HAP |
| 1430 | 9 | 20322717 | 20323045 | 328 | MOLF | 9 | 20322985 | 20323019 | 35 | - | 24.422 | 3.86E-09 | 0.00637 | CTGCACGTGCACTTGCTTGCCAGAAGAGGGCATCA | 4 | ENCFF367UNR.ENCFF502WDT.ENCFF142CNG.ENCFF055HAP |
| 1431 | 9 | 20920655 | 20920831 | 176 | CAST | 9 | 20920693 | 20920711 | 19 | - | 17.5902 | 6.18E-07 | 0.112 | ACACCAGAAGAGGGCACTG | | |
| 1432 | 9 | 21267624 | 21267976 | 352 | MOLF | 9 | 21267757 | 21267790 | 34 | - | 18 | 3.53E-07 | 0.0499 | CAGCTGTGTTCAGATGTACCAGAAGAGGGAGTCA | 1 | ENCFF502WDT |
| 1433 | 9 | 21321397 | 21321497 | 100 | MOLF | 9 | 21321370 | 21321403 | 34 | - | 19.9919 | 8.93E-08 | 0.0211 | CCGTATCTGCCACCTCTGCCGGCAGAGGGCACCA | 30 | ENCFF093ZRU,ENCFF367UNR,ENCFF805QBU,ENCFF099QNE,ENCFF327LYS,ENCFF065ODV,ENCFF907XEG,ENCFF024JOZ,ENCFF089GBC,ENCFF821BIA,ENCFF092NMJ,ENCFF198SSR,ENCFF589ZBJ,ENCFF136ZDK,ENCFF430PPJ,ENCFF165LXO,ENCFF883UPM,ENCFF097CAQ,ENCFF089EWX,ENCFF194HVL,ENCFF957SPH,ENCFF210BTO,ENCFF951GOP,ENCFF575MGI,ENCFF281QMV,ENCFF052KVO,ENCFF533APC,ENCFF321AGQ.ENCFF491RJK.ENCFF316FYN |
| 1434 | 9 | 21654516 | 21654922 | 406 | BTBR,C57L,C58J,CAST,FVB,MA/My,MOLF,NOD,NOR,NZB,NZW,RF,SJL,SPRET,SWR,WSB | 9 | 21654843 | 21654876 | 34 | - | 17.878 | 3.83E-07 | 0.0522 | TAGCTGTCCTCAGACACACCAGAAGAGGGCATCA | | |
| 1435 | 9 | 21979919 | 21980138 | 219 | BTBR,C57L,C58J,CAST,FVB,MA/My,MOLF,NOD,NOR,NZB,NZW,RF,SJL,SPRET,SWR,WSB | 9 | 21980044 | 21980077 | 34 | - | 16.4472 | 9.61E-07 | 0.0841 | TAGCTGTCTTCAGACACACCAGAAGAGGGCGTCA | | |
| 1436 | 9 | 21987018 | 21987257 | 239 | BTBR,C57L,C58J,CAST,FVB,MA/My,MOLF,NOD,NOR,NZB,NZW,RF,SJL,SPRET,SWR,WSB | 9 | 21987137 | 21987170 | 34 | - | 16.4309 | 9.71E-07 | 0.0846 | TAGCTGTCTTCAGACGCTCCAGAAGAGGGCGTCA | | |
| 1437 | 9 | 22053175 | 22053383 | 208 | BTBR,C57L,C58J,CAST,FVB,MA/My,MOLF,NOD,NOR,NZB,NZW,RF,SJL,SPRET,SWR,WSB | 9 | 22053300 | 22053333 | 34 | - | 17.3902 | 5.27E-07 | 0.0619 | TAGCTGTCCTCAGACACTCCAGAAGAGGGCATCA | | |
| 1438 | 9 | 23346195 | 23346401 | 206 | 129S1,CAST,CBA,FVB,LP,MOLF,NZO,RF,SJL,SPRET,SWR,TH | 9 | 23346325 | 23346358 | 34 | - | 16.7724 | 7.83E-07 | 0.0768 | TAGCTGTCTTCAGATGCACCAGAAGAGGGCGTCA | | |
| 1439 | 9 | 28171514 | 28171725 | 211 | BUB.CAST.MOLF.SPRET | 9 | 28171626 | 28171659 | 34 | - | 16.4472 | 9.61E-07 | 0.0841 | TAGCTGTCTTCAGACACACCAGAAGAGGGCGTCA | | |
| 1440 | 9 | 31076810 | 31077009 | 199 | SPRET | 9 | 31076933 | 31076951 | 19 | - | 17.5574 | 6.32E-07 | 0.112 | ACTCCAGAAGAGGGCGCCA | 5 | ENCFF093ZRU.ENCFF367UNR.ENCFF055HAP.ENCFF821BIA.ENCFF099QNE |
| 1440 | 9 | 31076810 | 31077009 | 199 | SPRET | 9 | 31076933 | 31076966 | 34 | - | 18.3659 | 2.76E-07 | 0.0438 | TAGCTGTCTTCAGACACTCCAGAAGAGGGCGCCA | 5 | ENCFF093ZRU.ENCFF367UNR.ENCFF055HAP.ENCFF821BIA.ENCFF099QNE |
| 1441 | 9 | 31090670 | 31090883 | 213 | CAST,CBA,CE,DBA1J,LP,MA/My,MOLF,MRL,NZO,NZW,SPRET,TH,WSB | 9 | 31090795 | 31090813 | 19 | - | 17.5574 | 6.32E-07 | 0.112 | ACTCCAGAAGAGGGCGCCA | | |
| 1441 | 9 | 31090670 | 31090883 | 213 | CAST,CBA,CE,DBA1J,LP,MA/My,MOLF,MRL,NZO,NZW,SPRET,TH,WSB | 9 | 31090795 | 31090828 | 34 | - | 18.2195 | 3.05E-07 | 0.0458 | TAGCTGTCTTCAGATACTCCAGAAGAGGGCGCCA | | |
| 1442 | 9 | 31144561 | 31145419 | 858 | SPRET | 9 | 31145256 | 31145289 | 34 | - | 16.4472 | 9.61E-07 | 0.0841 | TAGCTGTCTTCAGACACACCAGAAGAGGGCGTCA | | |
| 1443 | 9 | 31280108 | 31280515 | 407 | SPRET | 9 | 31280440 | 31280473 | 34 | - | 16.4634 | 9.51E-07 | 0.0841 | TAGCTGTCTTCAGACGCACCAGAAGAGGGCATCA | | |
| 1444 | 9 | 31281233 | 31281422 | 189 | SPRET | 9 | 31281351 | 31281384 | 34 | - | 17.1463 | 6.17E-07 | 0.0675 | TAGCTGTCCTCAAATACTCCAGAAGAGGGCATCA | | |
| 1445 | 9 | 32106328 | 32106527 | 199 | BTBR.C57L.MOLF.SPRET | 9 | 32106452 | 32106470 | 19 | - | 17.5574 | 6.32E-07 | 0.112 | ACTCCAGAAGAGGGCGCCA | 4 | ENCFF367UNR.ENCFF099QNE.ENCFF821BIA.ENCFF093ZRU |
| 1446 | 9 | 32683283 | 32683607 | 324 | 129S1,AJ,AKR,BALB,BUB,C3H,CAST,DBA1J,FVB,MA/My,MOLF,NOD,NOR,NZB,NZO,NZW,PL,RH2,SJL,SPRET,SWR,TH,WSB | 9 | 32683565 | 32683598 | 34 | - | 16.9837 | 6.84E-07 | 0.0717 | CTGCCGTCACATTCGCTGTTACAAGATGGCGCTG | | |
| 1447 | 9 | 35044852 | 35045057 | 205 | 129S1,AJ,AKR,BALB,BTBR,BUB,C3H,C57L,CAST,CE,DBA1J,DBA2J,FVB,KK,MA/My,MOLF,MRL,NOD,NOR,NZB,NZO,NZW,PJ,PL,RF,RH2,SEA,SJL,SMJ,SPRET,SWR,TH,WSB | 9 | 35044974 | 35045007 | 34 | - | 16.4472 | 9.61E-07 | 0.0841 | TAGCTGTCTTCAGACACACCAGAAGAGGGCGTCA | | |
| 1448 | 9 | 35107719 | 35108185 | 466 | SPRET | 9 | 35107850 | 35107883 | 34 | - | 17.5041 | 4.89E-07 | 0.0594 | TTGCTGTCTTCAGACACTCCAGAAGAGGGTACCA | | |
| 1449 | 9 | 37503035 | 37503239 | 204 | C57L.CAST.MOLF.NZO.SPRET | 9 | 37503158 | 37503191 | 34 | - | 16.7886 | 7.75E-07 | 0.0768 | CAGCTATCTTCAGACACTCCAGAAGAGGGCATCA | | |
| 1450 | 9 | 37862211 | 37862960 | 749 | MOLF | 9 | 37862930 | 37862964 | 35 | - | 18.7339 | 1.93E-07 | 0.064 | TTGCAGTTACCCCTTTTTGCCTGGAGATGAAGATA | | |
| 1451 | 9 | 40544335 | 40544815 | 480 | WSB | 9 | 40544368 | 40544401 | 34 | - | 16.4472 | 9.61E-07 | 0.0841 | TAGCTGTCTTCAGACACACCAGAAGAGGGCGTCA | | |
| 1452 | 9 | 40557987 | 40558226 | 239 | 129S1,AJ,AKR,BALB,BTBR,BUB,C3H,C57L,CAST,CBA,CE,DBA1J,DBA2J,FVB,ILNJ2,KK,MA/My,MOLF,MRL,NOD,NOR,NZB,NZO,NZW,PJ,PL,RF,RH2,SEA,SJL,SMJ,SWR,TH,WSB | 9 | 40558119 | 40558152 | 34 | - | 19.252 | 1.51E-07 | 0.0314 | CTGCTCTCTTCAGAAACACCAGAAGAGGGCAACA | | |
| 1453 | 9 | 43689540 | 43690531 | 991 | SPRET | 9 | 43690171 | 43690205 | 35 | - | 16.6422 | 6.65E-07 | 0.126 | CTGTAATCACAGAACACCTCCAGCAGGGGATGGGG | | |
| 1454 | 9 | 43965030 | 43965208 | 178 | SPRET | 9 | 43965125 | 43965158 | 34 | - | 17.9106 | 3.74E-07 | 0.0517 | TAGCTGTCTTCAGACACTCCAGAAGAGGGCACCA | | |
| 1455 | 9 | 44273509 | 44273708 | 199 | SPRET | 9 | 44273629 | 44273662 | 34 | - | 16.4472 | 9.61E-07 | 0.0841 | TAGCTGTCTTCAGACACACCAGAAGAGGGCGTCA | 3 | ENCFF093ZRU.ENCFF821BIA |
| 1456 | 9 | 44472559 | 44472774 | 215 | SPRET | 9 | 44472668 | 44472701 | 34 | - | 17.2439 | 5.79E-07 | 0.065 | TAGCTGTCCTCAGATACTCCAGAAGAGGGCATCA | 1 | ENCFF093ZRU |
| 1457 | 9 | 44649718 | 44649923 | 205 | SPRET | 9 | 44649841 | 44649874 | 34 | - | 20.7967 | 4.97E-08 | 0.0139 | CTGTAGTCTTCAGACGCACCAGAAGAGGGCGTCA | 2 | ENCFF093ZRU.ENCFF099QNE |
| 1458 | 9 | 45278410 | 45279232 | 822 | ILNJ2.WSB | 9 | 45278468 | 45278502 | 35 | - | 17.0092 | 5.39E-07 | 0.113 | TTCCTATGCGCGCTGTCACCCAGGAGAGGGCAGCT | | |
| 1459 | 9 | 45637550 | 45637735 | 185 | 129S1.C57L.CAST.KK.MOLF | 9 | 45637667 | 45637700 | 34 | - | 17.3902 | 5.27E-07 | 0.0619 | TAGCTGTCCTCAGACACTCCAGAAGAGGGCATCA | | |
| 1460 | 9 | 50505078 | 50505305 | 227 | SPRET | 9 | 50505045 | 50505078 | 34 | - | 28.5203 | 5.49E-11 | 0.000184 | ATGCAGTGCCCATGGAGACCAGAAGAGGGCACCA | 29 | ENCFF883UPM,ENCFF089GBC,ENCFF201JUA,ENCFF210BTO,ENCFF093ZRU,ENCFF367UNR,ENCFF092NMJ,ENCFF805QBU,ENCFF327LYS,ENCFF165LXO,ENCFF430PPJ,ENCFF099QNE,ENCFF065ODV,ENCFF907XEG,ENCFF321AGQ,ENCFF589ZBJ,ENCFF533APC,ENCFF097CAQ,ENCFF055HAP,ENCFF609MDN,ENCFF502WDT,ENCFF821BIA,ENCFF136ZDK,ENCFF194HVL,ENCFF024JOZ,ENCFF198SSR,ENCFF142CNG,ENCFF052KVO.ENCFF957SPH |
| 1461 | 9 | 50540667 | 50540884 | 217 | AJ,BALB,BTBR,C3H,CBA,CE,DBA2J,ILNJ2,MRL,NOD,NOR,NZO,NZW,RH2,SJL,WSB | 9 | 50540792 | 50540825 | 34 | - | 16.4472 | 9.61E-07 | 0.0841 | TAGCTGTCTTCAGACACACCAGAAGAGGGCGTCA | | |
| 1462 | 9 | 50901715 | 50901968 | 253 | CAST.SPRET | 9 | 50901837 | 50901855 | 19 | - | 17.5574 | 6.32E-07 | 0.112 | ACTCCAGAAGAGGGCGCCA | | |
| 1462 | 9 | 50901715 | 50901968 | 253 | CAST.SPRET | 9 | 50901837 | 50901870 | 34 | - | 18.3659 | 2.76E-07 | 0.0438 | TAGCTGTCTTCAGACACTCCAGAAGAGGGCGCCA | | |
| 1463 | 9 | 50901587 | 50902529 | 942 | 129S1.C57L.MOLF | 9 | 50901837 | 50901855 | 19 | - | 17.5574 | 6.32E-07 | 0.112 | ACTCCAGAAGAGGGCGCCA | | |
| 1463 | 9 | 50901587 | 50902529 | 942 | 129S1.C57L.MOLF | 9 | 50901837 | 50901870 | 34 | - | 18.3659 | 2.76E-07 | 0.0438 | TAGCTGTCTTCAGACACTCCAGAAGAGGGCGCCA | | |
| 1464 | 9 | 56828226 | 56828412 | 186 | SPRET | 9 | 56828337 | 56828370 | 34 | - | 16.4472 | 9.61E-07 | 0.0841 | TAGCTGTCTTCAGACACACCAGAAGAGGGCGTCA | | |
| 1465 | 9 | 56915026 | 56915127 | 101 | CAST.MA/Mv.MOLF | 9 | 56915036 | 56915054 | 19 | - | 18.0328 | 4.60E-07 | 0.1 | AGTCCAGAAGAGGGCACTG | | |
| 1465 | 9 | 56915026 | 56915127 | 101 | CAST.MA/Mv.MOLF | 9 | 56915036 | 56915070 | 35 | - | 24.8349 | 2.80E-09 | 0.00544 | GTGCAGGTACCTGTGGAGTCCAGAAGAGGGCACTG | | |
| 1466 | 9 | 57569799 | 57570043 | 244 | SPRET | 9 | 57569910 | 57569943 | 34 | - | 17.8455 | 3.91E-07 | 0.0525 | TAGCTGTCCTCAGACACTCCAGAAGAGGGCGTCA | | |
| 1467 | 9 | 57754866 | 57755396 | 530 | SPRET | 9 | 57755312 | 57755345 | 34 | - | 17.5528 | 4.74E-07 | 0.0583 | TAGCTGTCTTCGGATGCACCAGAAGAGGGCGTCA | | |
| 1468 | 9 | 58835279 | 58835871 | 592 | 129S1,AJ,BUB,C3H,C57L,CE,DBA1J,DBA2J,MRL,PJ,RH2,SEA,SJL,WSB | 9 | 58835584 | 58835602 | 19 | - | 22.6885 | 1.05E-08 | 0.0194 | CAACCACCAGGTGGCACTG | 19 | ENCFF367UNR,ENCFF201JUA,ENCFF093ZRU,ENCFF099QNE,ENCFF821BIA,ENCFF327LYS,ENCFF089GBC,ENCFF609MDN,ENCFF024JOZ,ENCFF907XEG,ENCFF589ZBJ,ENCFF198SSR,ENCFF165LXO,ENCFF092NMJ,ENCFF136ZDK.ENCFF281QMV.ENCFF430PPJ.ENCFF883UPM |
| 1468 | 9 | 58835279 | 58835871 | 592 | 129S1,AJ,BUB,C3H,C57L,CE,DBA1J,DBA2J,MRL,PJ,RH2,SEA,SJL,WSB | 9 | 58835584 | 58835617 | 34 | - | 25.9837 | 6.78E-10 | 0.000645 | CTGCATTTAAAAATTCAACCACCAGGTGGCACTG | 19 | ENCFF367UNR,ENCFF201JUA,ENCFF093ZRU,ENCFF099QNE,ENCFF821BIA,ENCFF327LYS,ENCFF089GBC,ENCFF609MDN,ENCFF024JOZ,ENCFF907XEG,ENCFF589ZBJ,ENCFF198SSR,ENCFF165LXO,ENCFF092NMJ,ENCFF136ZDK.ENCFF281QMV.ENCFF430PPJ.ENCFF883UPM |
| 1469 | 9 | 59118352 | 59118557 | 205 | 129S1,AJ,AKR,BTBR,BUB,C3H,C57L,CAST,CBA,CE,DBA1J,DBA2J,FVB,ILNJ2,MA/My,MOLF,MRL,NOD,NOR,NZB,PJ,PL,RF,RH2,SEA.SJL.SPRET.SWR.TH | 9 | 59118477 | 59118495 | 19 | - | 17.5738 | 6.25E-07 | 0.112 | ACTCCAGAAGAGGGCGCTA | | |
| 1469 | 9 | 59118352 | 59118557 | 205 | 129S1,AJ,AKR,BTBR,BUB,C3H,C57L,CAST,CBA,CE,DBA1J,DBA2J,FVB,ILNJ2,MA/My,MOLF,MRL,NOD,NOR,NZB,PJ,PL,RF,RH2,SEA.SJL.SPRET.SWR.TH | 9 | 59118477 | 59118510 | 34 | - | 18.1789 | 3.13E-07 | 0.0464 | TAGCTGTCTTCAGACACTCCAGAAGAGGGCGCTA | | |
| 1470 | 9 | 59332259 | 59332453 | 194 | SPRET | 9 | 59332375 | 59332408 | 34 | - | 16.7724 | 7.83E-07 | 0.0768 | TAGCTGTCTTCAGATGCACCAGAAGAGGGCGTCA | 2 | ENCFF367UNR.ENCFF093ZRU |
| 1471 | 9 | 59439615 | 59439833 | 218 | SPRET | 9 | 59439749 | 59439782 | 34 | - | 19.5285 | 1.24E-07 | 0.0274 | CTGCTCTCTTCAGACACACCAGAAGAGGGCATCA | 3 | ENCFF093ZRU.ENCFF883UPM.ENCFF502WDT |
| 1472 | 9 | 60530829 | 60530905 | 76 | MOLF.SPRET.TH | 9 | 60530847 | 60530880 | 34 | - | 16.6423 | 8.50E-07 | 0.0811 | CTGCAGGTGGAGGCTGGCTCTGCAGGTGGAGGCA | | |
| 1473 | 9 | 64182857 | 64183264 | 407 | AJ,BUB,C3H,CAST,CBA,DBA1J,DBA2J,FVB,ILNJ2,KK,MOLF,MRL,NZB,NZO,NZW,PJ,RH2.SEA.SJL.SPRET.SWR | 9 | 64183184 | 64183217 | 34 | - | 17.7236 | 4.24E-07 | 0.0552 | TAGCTGTCCTCAGACACTCCAGAAGAGGGCGACA | 1 | ENCFF093ZRU |
| 1474 | 9 | 64223903 | 64224116 | 213 | AJ,BUB,C3H,CAST,CBA,CE,DBA1J,DBA2J,FVB,ILNJ2,KK,MOLF,MRL,NZB,NZO,NZW,PJ,PL,RH2,SEA,SJL,SPRET,SWR,WSB | 9 | 64224029 | 64224062 | 34 | - | 16.4146 | 9.81E-07 | 0.0852 | TAGCTGTCTTCTGACAGTCCAGAAGAGGGAGCCA | | |
| 1475 | 9 | 65802980 | 65803183 | 203 | CAST.FVB.MOLF.SJL.SPRET | 9 | 65803106 | 65803139 | 34 | - | 17.8455 | 3.91E-07 | 0.0525 | TAGCTGTCCTCAGACACTCCAGAAGAGGGCGTCA | | |
| 1476 | 9 | 66696628 | 66696818 | 190 | MOLF | 9 | 66696733 | 66696751 | 19 | - | 18.5574 | 3.20E-07 | 0.0843 | ACGCCAGAAGAGGGCACTG | | |
| 1477 | 9 | 69602146 | 69602753 | 607 | 129S1,BTBR,C57L,CAST,NOD,NOR,SMJ,WSB | 9 | 69602448 | 69602481 | 34 | - | 17.7967 | 4.04E-07 | 0.0536 | CTGCCCTTCCCTTTCTCTTCAGAAGAGGGTAACC | | |
| 1478 | 9 | 71702251 | 71703122 | 871 | MOLF.SPRET | 9 | 71702745 | 71702779 | 35 | - | 16.3761 | 7.73E-07 | 0.137 | GTACAGTCCCGGATCCCAGCCAGGGGATGGTTCCA | | |
| 1479 | 9 | 73065976 | 73066086 | 110 | CAST | 9 | 73065996 | 73066029 | 34 | - | 24.1707 | 3.41E-09 | 0.00194 | CTGCAGTGTCTGCAGAGGCCAGAAGAGAGCATTA | | |
| 1480 | 9 | 73623558 | 73623834 | 276 | C58J.CAST.LP.MOLF.SPRET | 9 | 73623605 | 73623638 | 34 | - | 18.4797 | 2.56E-07 | 0.0428 | TTGCTGTCTTCAGACACACCAGAAGAGGGCATCA | | |
| 1481 | 9 | 73778782 | 73778921 | 139 | BTBR,CAST,CE,MA/My,MOLF,NZW,SPRET | 9 | 73778772 | 73778805 | 34 | - | 25.1138 | 1.50E-09 | 0.00113 | CTGCAATGCTCACAAAGGCCAGAAGGAGGCATCA | 1 | ENCFF093ZRU |
| 1482 | 9 | 76233877 | 76234347 | 470 | SPRET | 9 | 76234305 | 76234338 | 34 | - | 17.0813 | 6.43E-07 | 0.0691 | CTGCACTCACATTTGCCATTACAAGATGGCGCTG | | |
| 1483 | 9 | 76793687 | 76794218 | 531 | SPRET | 9 | 76793671 | 76793689 | 19 | - | 18.1803 | 4.16E-07 | 0.0944 | CAGCCAGGAGGGGGCAGGA | 1 | ENCFF093ZRU |
| 1484 | 9 | 77287101 | 77287250 | 149 | SPRET | 9 | 77287084 | 77287102 | 19 | - | 24.623 | 1.10E-09 | 0.00915 | TGGCCAGCAGGGGGCAGTC | 30 | ENCFF093ZRU,ENCFF367UNR,ENCFF099QNE,ENCFF089GBC,ENCFF907XEG,ENCFF024JOZ,ENCFF327LYS,ENCFF821BIA,ENCFF316FYN,ENCFF055HAP,ENCFF092NMJ,ENCFF198SSR,ENCFF430PPJ,ENCFF033UQX,ENCFF502WDT,ENCFF883UPM,ENCFF589ZBJ,ENCFF142CNG,ENCFF097CAQ,ENCFF165LXO,ENCFF136ZDK,ENCFF575MGI,ENCFF491RJK,ENCFF281QMV,ENCFF210BTO,ENCFF957SPH,ENCFF533APC,ENCFF951GOP.ENCFF609MDN.ENCFF194HVL |
| 1484 | 9 | 77287101 | 77287250 | 149 | SPRET | 9 | 77287084 | 77287117 | 34 | - | 26.6016 | 3.79E-10 | 0.000467 | CTGCATGGCAGAAGGTGGCCAGCAGGGGGCAGTC | 30 | ENCFF093ZRU,ENCFF367UNR,ENCFF099QNE,ENCFF089GBC,ENCFF907XEG,ENCFF024JOZ,ENCFF327LYS,ENCFF821BIA,ENCFF316FYN,ENCFF055HAP,ENCFF092NMJ,ENCFF198SSR,ENCFF430PPJ,ENCFF033UQX,ENCFF502WDT,ENCFF883UPM,ENCFF589ZBJ,ENCFF142CNG,ENCFF097CAQ,ENCFF165LXO,ENCFF136ZDK,ENCFF575MGI,ENCFF491RJK,ENCFF281QMV,ENCFF210BTO,ENCFF957SPH,ENCFF533APC,ENCFF951GOP.ENCFF609MDN.ENCFF194HVL |
| 1485 | 9 | 78367169 | 78367378 | 209 | 129S1,BTBR,C57L,C58J,CAST,ILNJ2,LP,MOLF.NZW.SMJ.SPRET | 9 | 78367293 | 78367326 | 34 | - | 16.4472 | 9.61E-07 | 0.0841 | TAGCTGTCTTCAGACACACCAGAAGAGGGCGTCA | | |
| 1486 | 9 | 78378356 | 78378556 | 200 | SPRET | 9 | 78378470 | 78378488 | 19 | - | 17.5738 | 6.25E-07 | 0.112 | ACTCCAGAAGAGGGCGCTA | 5 | ENCFF367UNR.ENCFF099QNE.ENCFF821BIA.ENCFF201JUA.ENCFF055HAP |
| 1486 | 9 | 78378356 | 78378556 | 200 | SPRET | 9 | 78378470 | 78378503 | 34 | - | 18.1789 | 3.13E-07 | 0.0464 | TAGCTGTCTTCAGACACTCCAGAAGAGGGCGCTA | 5 | ENCFF367UNR.ENCFF099QNE.ENCFF821BIA.ENCFF201JUA.ENCFF055HAP |
| 1487 | 9 | 78400878 | 78401735 | 857 | SPRET | 9 | 78401406 | 78401439 | 34 | - | 17.2439 | 5.79E-07 | 0.065 | TAGCTGTCCTCAGATACTCCAGAAGAGGGCATCA | | |
| 1488 | 9 | 78527028 | 78527271 | 243 | 129S1,AJ,AKR,BALB,BTBR,BUB,C3H,C57L,C58J,CAST,CBA,CE,DBA1J,DBA2J,FVB,ILNJ2,KK,LP,MOLF,MRL,NOD,NOR,NZB,NZO,NZW,PJ,PL,RF,RH2,SEA,SJL,SMJ,SPRET,SWR.TH | 9 | 78527153 | 78527186 | 34 | - | 17.9431 | 3.66E-07 | 0.0511 | CAGCTGTCCTCAGACACTCCAGAAGAGGGAGTCA | | |
| 1489 | 9 | 82227019 | 82227278 | 259 | AKR,BUB,C58J,CBA,ILNJ2,LP,MOLF,MRL,NOD,NOR,NZW,PL,RF | 9 | 82227165 | 82227198 | 34 | - | 18.7724 | 2.10E-07 | 0.0387 | GTGAACTGCCTGCGGAGGCCAGAAGAGGGTATTT | | |
| 1490 | 9 | 99052643 | 99052843 | 200 | SPRET | 9 | 99052770 | 99052803 | 34 | - | 21.3008 | 3.41E-08 | 0.0108 | GTGCTGTCCTCAGACACACCAGAAGAGGGCATCA | | |
| 1491 | 9 | 100896766 | 100896990 | 224 | AJ,AKR,BALB,BUB,C3H,CAST,CBA,CE,DBA1J,DBA2J,FVB,ILNJ2,MOLF,MRL,NOD,NOR,NZO,NZW,PJ,PL,RF,RH2,SEA,SJL,SMJ,SPRET,SWR,TH,WSB | 9 | 100896894 | 100896927 | 34 | - | 16.4634 | 9.51E-07 | 0.0841 | TAGCTGTCTTCAGACGCACCAGAAGAGGGCATCA | | |
| 1492 | 9 | 102466706 | 102466909 | 203 | SPRET | 9 | 102466909 | 102466943 | 35 | - | 16.7156 | 6.38E-07 | 0.124 | TAGCTGTTTTCAGACAATACCAGAAGAGGGCATCT | | |
| 1493 | 9 | 102840795 | 102840854 | 59 | AKR.MA/Mv.MOLF.TH | 9 | 102840841 | 102840874 | 34 | - | 19.9106 | 9.47E-08 | 0.022 | CTGCACAGCCACACGTGCCCGTAAGGAGGCAGCA | | |
| 1494 | 9 | 105469900 | 105470125 | 225 | 129S1,AJ,AKR,BALB,BTBR,BUB,C3H,C57L,CAST,CBA,CE,DBA1J,DBA2J,FVB,ILNJ2,KK,MA/My,MOLF,MRL,NOD,NOR,NZB,NZO,NZW,PJ,PL,RF,RH2,SEA,SJL,SMJ,SPRET,SWR.TH.WSB | 9 | 105470024 | 105470057 | 34 | - | 16.4634 | 9.51E-07 | 0.0841 | TAGCTGTCTTCAGACGCACCAGAAGAGGGCATCA | | |
| 1495 | 9 | 106416321 | 106416712 | 391 | CAST.MOLF | 9 | 106416543 | 106416561 | 19 | - | 16.8525 | 9.96E-07 | 0.137 | AGGCCAGAAGAAGGCACCA | | |
| 1496 | 9 | 108010081 | 108010291 | 210 | MOLF.SPRET | 9 | 108010189 | 108010222 | 34 | - | 16.5691 | 8.90E-07 | 0.0831 | TAGCTGTCTTCAGACGCACCAGAAGAGGGCATCG | | |
| 1497 | 9 | 108073674 | 108073871 | 197 | CAST.MOLF.SPRET | 9 | 108073798 | 108073831 | 34 | - | 16.4634 | 9.51E-07 | 0.0841 | TAGCTGTCTTCAGACGCACCAGAAGAGGGCATCA | 4 | ENCFF201JUA.ENCFF093ZRU.ENCFF367UNR.ENCFF099QNE |
| 1498 | 9 | 108766825 | 108767030 | 205 | CAST.CBA.CE.MOLF.SPRET | 9 | 108766939 | 108766972 | 34 | - | 16.4634 | 9.51E-07 | 0.0841 | TAGCTGTCTTCAGACGCACCAGAAGAGGGCATCA | 1 | ENCFF093ZRU |
| 1499 | 9 | 109473241 | 109473437 | 196 | SPRET | 9 | 109473356 | 109473389 | 34 | - | 16.4472 | 9.61E-07 | 0.0841 | TAGCTGTCTTCAGACACACCAGAAGAGGGCGTCA | | |
| 1500 | 9 | 109905482 | 109905700 | 218 | CAST.CBA.CE.MOLF.NZW.SPRET | 9 | 109905611 | 109905629 | 19 | - | 17.5574 | 6.32E-07 | 0.112 | ACTCCAGAAGAGGGCGCCA | 3 | ENCFF093ZRU.ENCFF821BIA.ENCFF367UNR |
| 1500 | 9 | 109905482 | 109905700 | 218 | CAST.CBA.CE.MOLF.NZW.SPRET | 9 | 109905611 | 109905644 | 34 | - | 18.561 | 2.42E-07 | 0.0423 | AAGCTGTCTTCAGACACTCCAGAAGAGGGCGCCA | 3 | ENCFF093ZRU.ENCFF821BIA.ENCFF367UNR |
| 1501 | 9 | 110301435 | 110301690 | 255 | BTBR,CAST,CBA,FVB,MOLF,NZB,NZO,NZW,SJL,SPRET,SWR,WSB | 9 | 110301562 | 110301595 | 34 | - | 17.6667 | 4.40E-07 | 0.0557 | TAGCTGTTTTCAGACACTCCAGAAGAGGGCGTCA | | |
| 1502 | 9 | 110639708 | 110639917 | 209 | CAST,CBA,FVB,MOLF,NZB,NZO,PJ,SEA,SJL,SMJ,SPRET,SWR | 9 | 110639823 | 110639856 | 34 | - | 17.3902 | 5.27E-07 | 0.0619 | TAGCTGTCCTCAGACACTCCAGAAGAGGGCATCA | | |
| 1503 | 9 | 110766576 | 110766786 | 210 | 129S1,AJ,AKR,BALB,BTBR,BUB,C3H,C57L,CAST,CBA,CE,DBA1J,DBA2J,FVB,ILNJ2,KK,MA/My,MOLF,MRL,NOD,NOR,NZB,NZO,NZW,PJ,PL,RF,RH2,SEA,SJL,SMJ,SPRET,SWR.TH.WSB | 9 | 110766702 | 110766720 | 19 | - | 17.9672 | 4.81E-07 | 0.102 | GCACCAGAAGAGGGCACCA | 4 | ENCFF099QNE,ENCFF821BIA,ENCFF367UNR,ENCFF093ZRU |
| 1503 | 9 | 110766576 | 110766786 | 210 | 129S1,AJ,AKR,BALB,BTBR,BUB,C3H,C57L,CAST,CBA,CE,DBA1J,DBA2J,FVB,ILNJ2,KK,MA/My,MOLF,MRL,NOD,NOR,NZB,NZO,NZW,PJ,PL,RF,RH2,SEA,SJL,SMJ,SPRET,SWR.TH.WSB | 9 | 110766702 | 110766735 | 34 | - | 17.252 | 5.76E-07 | 0.065 | TAGCTATCTTCAGCTGCACCAGAAGAGGGCACCA | 4 | ENCFF099QNE,ENCFF821BIA,ENCFF367UNR,ENCFF093ZRU |

| | | | | | | | | | | | | | | | | |
|---|---|---|---|---|---|---|---|---|---|---|---|---|---|---|---|---|
| 1504 | 9 | 110797604 | 110797815 | 211 | 129S1,AJ,AKR,BALB,BTBR,BUB,C3H,C57L,CAST,CBA,CE,DBA1J,DBA2J,FVB,ILNJ2,KK,MA/My,MOLF,MRL,NOD,NOR,NZB,NZO,PJ,PL,RF,RH2,SEA,SJL,SMJ,SPRET,SWR,TH,WSB | 9 | 110797729 | 110797762 | 34 | - | 22.9187 | 9.63E-09 | 0.00422 | TTGCTGTCCTCAGACGCCCCAGAAGAGGGCGCCA | 3 | ENCFF367UNR,ENCFF093ZRU,ENCFF821BIA |
| 1505 | 9 | 113310305 | 113310442 | 137 | CAST | 9 | 113310301 | 113310334 | 34 | - | 21.3659 | 3.25E-08 | 0.0104 | CAGCAGTGAACCCGGCAACCATTAGGTGGCGTTG | | |
| 1506 | 9 | 113512163 | 113512718 | 555 | SPRET | 9 | 113512347 | 113512380 | 34 | - | 18.6829 | 2.23E-07 | 0.0405 | TTGCTCTCTTCAGACCTACCAGAAGAGGGCATTG | | |
| 1507 | 9 | 113584035 | 113584285 | 250 | CAST,MOLF,SPRET,WSB | 9 | 113584146 | 113584179 | 34 | - | 16.4309 | 9.71E-07 | 0.0846 | TAGCTGTCTTCAGACGCTCCAGAAGAGGGCGTCA | | |
| 1508 | 9 | 113733649 | 113733872 | 223 | CAST,SPRET | 9 | 113733775 | 113733808 | 34 | - | 17.5285 | 4.81E-07 | 0.0587 | TAGCTGTCCTCAGGTACTCCAGAAGAGGGCGTCA | 1 | ENCFF093ZRU |
| 1509 | 9 | 114210909 | 114211124 | 215 | SPRET | 9 | 114211029 | 114211047 | 19 | - | 17.5738 | 6.25E-07 | 0.112 | ACTCCAGAAGAGGGCGCTA | 5 | ENCFF821BIA,ENCFF093ZRU,ENCFF367UNR,ENCFF099QNE,ENCFF055HAP |
| 1509 | 9 | 114210909 | 114211124 | 215 | SPRET | 9 | 114211029 | 114211062 | 34 | - | 18.187 | 3.12E-07 | 0.0463 | TAGCTGTCTTCAGTCACTCCAGAAGAGGGCGCTA | 5 | ENCFF821BIA,ENCFF093ZRU,ENCFF367UNR,ENCFF099QNE,ENCFF055HAP |
| 1510 | 9 | 114254713 | 114254853 | 140 | AJ,BALB,C3H,CE,FVB,MA/My,WSB | 9 | 114254833 | 114254866 | 34 | - | 23.5122 | 5.93E-09 | 0.00294 | GTGCAGTACCCAGGGAAGACAGAAGAGGGCATCA | 3 | ENCFF052KVO,ENCFF093ZRU,ENCFF327LYS |
| 1511 | 9 | 114323075 | 114323428 | 353 | SPRET | 9 | 114323273 | 114323306 | 34 | - | 16.6341 | 8.54E-07 | 0.0813 | TAGCTGTCTTCACACACTCCAGAAGAGGGCGTCA | | |
| 1512 | 9 | 114409647 | 114409861 | 214 | AJ,BALB,C3H,CAST,FVB,KK,MOLF,PJ,SEA,SPRET | 9 | 114409778 | 114409811 | 34 | - | 16.4146 | 9.81E-07 | 0.0852 | TAGCTGTCTTCTGACAGTCCAGAAGAGGGAGCCA | | |
| 1513 | 9 | 116209156 | 116209647 | 491 | MOLF | 9 | 116209222 | 116209256 | 35 | - | 20.5963 | 5.89E-08 | 0.0297 | GTGCAGGTGTCTGAGGAGACCAGAAGAGGGCATTG | 3 | ENCFF093ZRU,ENCFF367UNR,ENCFF883UPM |
| 1514 | 9 | 118775277 | 118775711 | 434 | CE,ILNJ2,WSB | 9 | 118775253 | 118775286 | 34 | - | 18.3415 | 2.81E-07 | 0.0442 | CAGCACACCCAGAGCCAGCCTGGAGGTGGAGACA | | |
| 1515 | 9 | 120074080 | 120074866 | 786 | SPRET | 9 | 120074701 | 120074719 | 19 | - | 17.0492 | 8.79E-07 | 0.129 | AGGCCACAAGAGGGCATTA | 14 | ENCFF136ZDK,ENCFF327LYS,ENCFF092NMJ,ENCFF165LXO,ENCFF089GBC,ENCFF907XEG,ENCFF198SSR,ENCFF883UPM,ENCFF502WDT,ENCFF367UNR,ENCFF093ZRU,ENCFF052KVO,ENCFF055HAP,ENCFF099QNE |
| 1515 | 9 | 120074080 | 120074866 | 786 | SPRET | 9 | 120074701 | 120074735 | 35 | - | 24.6422 | 3.26E-09 | 0.0058 | GTGCAATTACTGAAGTAGGCCACAAGAGGGCATTA | 14 | ENCFF136ZDK,ENCFF327LYS,ENCFF092NMJ,ENCFF165LXO,ENCFF089GBC,ENCFF907XEG,ENCFF198SSR,ENCFF883UPM,ENCFF502WDT,ENCFF367UNR,ENCFF093ZRU,ENCFF052KVO,ENCFF055HAP,ENCFF099QNE |
| 1516 | 9 | 120673869 | 120674068 | 199 | BTBR,CAST,MOLF,SPRET | 9 | 120673997 | 120674030 | 34 | - | 16.4472 | 9.61E-07 | 0.0841 | TAGCTGTCTTCAGACACACCAGAAGAGGGCGTCA | | |
| 1517 | 9 | 121198146 | 121198362 | 216 | SPRET | 9 | 121198273 | 121198291 | 19 | - | 17.5574 | 6.32E-07 | 0.112 | ACTCCAGAAGAGGGCGCCA | 4 | ENCFF093ZRU,ENCFF367UNR |
| 1517 | 9 | 121198146 | 121198362 | 216 | SPRET | 9 | 121198273 | 121198306 | 34 | - | 18.1789 | 3.13E-07 | 0.0464 | TAGCTGTCTTCAGCCACTCCAGAAGAGGGCGCCA | 4 | ENCFF093ZRU,ENCFF367UNR |
| 1518 | 9 | 122027332 | 122028011 | 679 | ILNJ2,LP | 9 | 122027509 | 122027543 | 35 | - | 17.3211 | 4.50E-07 | 0.101 | TAGCACTCAGGACCAATAGCCCGGAGGTGGCACCA | | |
| 1519 | 9 | 123276283 | 123276525 | 242 | SPRET | 9 | 123276362 | 123276380 | 19 | - | 21.1148 | 4.45E-08 | 0.0361 | AGGCCAGAAGGGGGCACTG | 10 | ENCFF327LYS,ENCFF092NMJ,ENCFF907XEG,ENCFF089GBC,ENCFF821BIA,ENCFF198SSR,ENCFF093ZRU,ENCFF367UNR,ENCFF055HAP,ENCFF502WDT |
| 1519 | 9 | 123276283 | 123276525 | 242 | SPRET | 9 | 123276362 | 123276395 | 34 | - | 27.8618 | 1.09E-10 | 0.000241 | GTGCAGTGTGCACAGAGGCCAGAAGGGGGCACTG | 10 | ENCFF327LYS,ENCFF092NMJ,ENCFF907XEG,ENCFF089GBC,ENCFF821BIA,ENCFF198SSR,ENCFF093ZRU,ENCFF367UNR,ENCFF055HAP,ENCFF502WDT |
| 1520 | 10 | 4301194 | 4301405 | 211 | CAST,MOLF,NOD,NOR,SPRET | 10 | 4301262 | 4301295 | 34 | + | 18.3659 | 2.76E-07 | 0.0597 | TAGCTGTCTTCAGACACTCCAGAAGAGGGCGCCA | | |
| 1520 | 10 | 4301194 | 4301405 | 211 | CAST,MOLF,NOD,NOR,SPRET | 10 | 4301277 | 4301295 | 19 | + | 17.5574 | 6.32E-07 | 0.134 | ACTCCAGAAGAGGGCGCCA | | |
| 1521 | 10 | 5732747 | 5732935 | 188 | BUB | 10 | 5732818 | 5732852 | 35 | + | 24.633 | 3.28E-09 | 0.0077 | CTGCATCTACACCTGAGTGCCAGAAGAGGGCATTA | | |
| 1522 | 10 | 6125097 | 6125301 | 204 | BUB | 10 | 6125158 | 6125191 | 34 | + | 16.4634 | 9.51E-07 | 0.108 | TAGCTGTCTTCAGACGCACCAGAAGAGGGCATCA | | |
| 1523 | 10 | 8245125 | 8245344 | 219 | BUB,C3H,CAST,CBA,CE,DBA1J,DBA2J,FVB,ILNJ2,KK,LP,MA/My,MOLF,MRL,NOD,NOR,NZB,NZO,NZW,PJ,SEA,SPRET,SWR,TH,WSB | 10 | 8245201 | 8245234 | 34 | + | 16.4634 | 9.51E-07 | 0.108 | TAGCTGTCTTCAGACGCACCAGAAGAGGGCATCA | 1 | ENCFF093ZRU |
| 1524 | 10 | 8403667 | 8403875 | 208 | C3H,CAST,CBA,DBA1J,DBA2J,LP,MA/My,MOLF,MRL,NZW,SPRET | 10 | 8403734 | 8403767 | 34 | + | 16.4472 | 9.61E-07 | 0.108 | TAGCTGTCTTCAGACACACCAGAAGAGGGCGTCA | | |
| 1525 | 10 | 11110646 | 11110849 | 203 | CAST,MOLF,SPRET | 10 | 11110706 | 11110739 | 34 | + | 17.6667 | 4.40E-07 | 0.0748 | TAGCTGTCTTCAGACACGCCAGAAGAGGGCGTCA | | |
| 1526 | 10 | 12858582 | 12858748 | 166 | BUB,DBA1J,DBA2J,MOLF,SPRET,WSB | 10 | 12858646 | 12858679 | 34 | + | 17.878 | 3.83E-07 | 0.0708 | TAGCTGTCCTCAGACACACCAGAAGAGGGCATCA | | |
| 1527 | 10 | 14468555 | 14468754 | 199 | CAST,DBA1J,DBA2J,MOLF,SPRET | 10 | 14468613 | 14468646 | 34 | + | 16.7724 | 7.83E-07 | 0.0988 | TAGCTGTCTTCAGATGCACCAGAAGAGGGCGTCA | | |
| 1528 | 10 | 17658382 | 17658591 | 209 | CAST,SPRET | 10 | 17658448 | 17658481 | 34 | + | 16.6911 | 8.24E-07 | 0.102 | TAGCTGTCTTCAGACAGACCAGAAGAGGGCATCA | | |
| 1529 | 10 | 18635351 | 18635566 | 215 | CAST,MOLF,SPRET,SWR | 10 | 18635421 | 18635454 | 34 | + | 17.8455 | 3.91E-07 | 0.071 | TAGCTGTCCTCAGACACTCCAGAAGAGGGCGTCA | | |
| 1530 | 10 | 23687692 | 23688227 | 535 | SPRET | 10 | 23688225 | 23688258 | 34 | + | 18.2764 | 2.93E-07 | 0.0604 | AGGCAATACCACAGTAAACCACAAGGTGGTGCTT | 29 | ENCFF093ZRU,ENCFF367UNR,ENCFF201JUA,ENCFF099QNE,ENCFF821BIA,ENCFF907XEG,ENCFF065ODV,ENCFF052KVO,ENCFF089GBC,ENCFF024JOZ,ENCFF589ZBJ,ENCFF327LYS,ENCFF198SSR,ENCFF092NMJ,ENCFF430PPJ,ENCFF321AGQ,ENCFF055HAP,ENCFF165LXO,ENCFF316FYN,ENCFF601JVE,ENCFF504JDS,ENCFF502WDT,ENCFF136ZDK,ENCFF951GOP,ENCFF281QMV,ENCFF575MGI,ENCFF957SPH,ENCFF883UPM,ENCFF609MDN |
| 1531 | 10 | 23812420 | 23812642 | 222 | CAST,MOLF,SPRET,WSB | 10 | 23812498 | 23812532 | 35 | + | 16.0917 | 9.07E-07 | 0.18 | TAGCTGTCTTCAGACAGCACCAGAAGAGGGCATCA | | |
| 1532 | 10 | 23850967 | 23851422 | 455 | SPRET | 10 | 23851374 | 23851392 | 19 | + | 20.1148 | 1.01E-07 | 0.055 | TTGCCAGCAGGTGTCACTA | 2 | ENCFF093ZRU,ENCFF883UPM |
| 1533 | 10 | 24021428 | 24021627 | 199 | CAST,MOLF,SPRET | 10 | 24021481 | 24021514 | 34 | + | 17.1057 | 6.33E-07 | 0.0907 | TAGCTGTGTTCAGACACACCAGAAGAGGGCATTG | | |
| 1534 | 10 | 25849498 | 25850248 | 750 | CAST,MOLF,SPRET | 10 | 25850120 | 25850154 | 35 | + | 16.7615 | 6.21E-07 | 0.148 | TAGCAGTTGCCACCTGTGTGCTCTTGGGGGCGGGG | | |
| 1535 | 10 | 28113448 | 28113651 | 203 | CAST,MOLF,PJ,SEA,SPRET | 10 | 28113509 | 28113542 | 34 | + | 17.8455 | 3.91E-07 | 0.071 | TAGCTGTCCTCAGACACTCCAGAAGAGGGCGTCA | | |
| 1536 | 10 | 28244653 | 28244883 | 230 | CAST,MOLF,PJ,SEA,SPRET | 10 | 28244716 | 28244749 | 34 | + | 18.3659 | 2.76E-07 | 0.0597 | TAGCTGTCTTCAGACACTCCAGAAGAGGGCGCCA | | |
| 1536 | 10 | 28244653 | 28244883 | 230 | CAST,MOLF,PJ,SEA,SPRET | 10 | 28244731 | 28244749 | 19 | + | 17.5574 | 6.32E-07 | 0.134 | ACTCCAGAAGAGGGCGCCA | | |
| 1537 | 10 | 30764189 | 30764923 | 734 | CAST | 10 | 30764667 | 30764700 | 34 | + | 22.2602 | 1.63E-08 | 0.00949 | GTGCAGTGCCCATGAAGGCCAGAAGAGAGCATCA | | |
| 1538 | 10 | 41588430 | 41588642 | 212 | 129S1,BTBR,C57L,CAST,KK,MOLF,SPRET,WSB | 10 | 41588498 | 41588531 | 34 | + | 16.4472 | 9.61E-07 | 0.108 | TAGCTGTCTTCAGACACACCAGAAGAGGGCGTCA | | |
| 1539 | 10 | 41706774 | 41707043 | 269 | WSB | 10 | 41707041 | 41707075 | 35 | + | 25.1743 | 2.14E-09 | 0.00677 | CAGCAGTTAGCTGCTGTGAGCACTAGGTGGCACTC | 32 | ENCFF093ZRU,ENCFF201JUA,ENCFF367UNR,ENCFF099QNE,ENCFF210BTO,ENCFF052KVO,ENCFF821BIA,ENCFF089GBC,ENCFF327LYS,ENCFF024JOZ,ENCFF907XEG,ENCFF065ODV,ENCFF092NMJ,ENCFF194HVL,ENCFF136ZDK,ENCFF589ZBJ,ENCFF502WDT,ENCFF055HAP,ENCFF198SSR,ENCFF321AGQ,ENCFF430PPJ,ENCFF951GOP,ENCFF165LXO,ENCFF491RJK,ENCFF883UPM,ENCFF957SPH,ENCFF281QMV,ENCFF575MGI,ENCFF142CNG,ENCFF316FYN,ENCFF609MDN,ENCFF089EWX |
| 1540 | 10 | 57342487 | 57342706 | 219 | CAST,MOLF,SPRET,WSB | 10 | 57342563 | 57342596 | 34 | + | 16.4634 | 9.51E-07 | 0.108 | TAGCTGTCTTCAGACGCACCAGAAGAGGGCATCA | | |
| 1541 | 10 | 59829858 | 59830079 | 221 | CAST,MOLF,SPRET,WSB | 10 | 59829936 | 59829969 | 34 | + | 18.3659 | 2.76E-07 | 0.0597 | TAGCTGTCTTCAGACACTCCAGAAGAGGGCGCCA | 2 | ENCFF367UNR |
| 1541 | 10 | 59829858 | 59830079 | 221 | CAST,MOLF,SPRET,WSB | 10 | 59829951 | 59829969 | 19 | + | 17.5574 | 6.32E-07 | 0.134 | ACTCCAGAAGAGGGCGCCA | 2 | ENCFF367UNR |
| 1542 | 10 | 61273514 | 61273830 | 316 | SPRET | 10 | 61273517 | 61273550 | 34 | + | 17.2439 | 5.79E-07 | 0.0854 | TAGCTGTCCTCAGATACTCCAGAAGAGGGCATCA | | |
| 1543 | 10 | 61486183 | 61486386 | 203 | SPRET,WSB | 10 | 61486244 | 61486277 | 34 | + | 16.4472 | 9.61E-07 | 0.108 | TAGCTGTCTTCAGACACACCAGAAGAGGGCGTCA | 1 | ENCFF093ZRU |
| 1544 | 10 | 62871079 | 62871333 | 254 | MOLF,RF,SPRET | 10 | 62871190 | 62871223 | 34 | + | 16.4634 | 9.51E-07 | 0.108 | TAGCTGTCTTCAGACGCACCAGAAGAGGGCATCA | | |
| 1545 | 10 | 63440844 | 63441763 | 919 | CAST | 10 | 63441241 | 63441275 | 35 | + | 16.4954 | 7.23E-07 | 0.161 | ATGCAGGGACATTCAGAGGCCAGAAGAGGGACATC | | |
| 1546 | 10 | 66158365 | 66158527 | 162 | CAST,NZW,PL,SPRET | 10 | 66158374 | 66158407 | 34 | + | 16.4228 | 9.76E-07 | 0.109 | TAGCTGCCTTCAGACACTCCAGAAGAGGGCGCCA | | |
| 1546 | 10 | 66158365 | 66158527 | 162 | CAST,NZW,PL,SPRET | 10 | 66158389 | 66158407 | 19 | + | 17.5574 | 6.32E-07 | 0.134 | ACTCCAGAAGAGGGCGCCA | | |
| 1547 | 10 | 67183906 | 67184247 | 341 | ILNJ2,MOLF,NZW,SMJ | 10 | 67184049 | 67184083 | 35 | + | 27.4954 | 3.05E-10 | 0.00296 | CTGCATGTTTGCCTGCAGGCCAGAAGAGGGCACCA | 1 | ENCFF502WDT |
| 1547 | 10 | 67183906 | 67184247 | 341 | ILNJ2,MOLF,NZW,SMJ | 10 | 67184065 | 67184083 | 19 | + | 20.3607 | 8.29E-08 | 0.0512 | AGGCCAGAAGAGGGCACCA | 1 | ENCFF502WDT |
| 1548 | 10 | 67834123 | 67834201 | 78 | SPRET | 10 | 67834123 | 67834141 | 19 | + | 20.7213 | 6.18E-08 | 0.0483 | TTGCCACTAGATGGCGCTT | 37 | ENCFF093ZRU,ENCFF367UNR,ENCFF201JUA,ENCFF099QNE,ENCFF210BTO,ENCFF907XEG,ENCFF089GBC,ENCFF821BIA,ENCFF092NMJ,ENCFF327LYS,ENCFF951GOP,ENCFF430PPJ,ENCFF052KVO,ENCFF198SSR,ENCFF883UPM,ENCFF502WDT,ENCFF165LXO,ENCFF024JOZ,ENCFF055HAP,ENCFF136ZDK,ENCFF316FYN,ENCFF589ZBJ,ENCFF065ODV,ENCFF349BUL,ENCFF089EWX,ENCFF533APC,ENCFF097CAQ,ENCFF281QMV,ENCFF957SPH,ENCFF611HDQ,ENCFF609MDN,ENCFF033UQX,ENCFF491RJK,ENCFF504JDS,ENCFF142CNG,ENCFF575MGI,ENCFF194HVL |
| 1548 | 10 | 67834123 | 67834201 | 78 | SPRET | 10 | 67834185 | 67834219 | 35 | + | 24.6789 | 3.17E-09 | 0.00758 | CTTCCGTACCAACCCCCTGCCACTAGATGGCGCTT | 35 | ENCFF093ZRU,ENCFF367UNR,ENCFF201JUA,ENCFF099QNE,ENCFF210BTO,ENCFF907XEG,ENCFF089GBC,ENCFF821BIA,ENCFF092NMJ,ENCFF327LYS,ENCFF951GOP,ENCFF430PPJ,ENCFF052KVO,ENCFF198SSR,ENCFF883UPM,ENCFF502WDT,ENCFF165LXO,ENCFF024JOZ,ENCFF055HAP,ENCFF136ZDK,ENCFF316FYN,ENCFF589ZBJ,ENCFF065ODV,ENCFF349BUL,ENCFF089EWX,ENCFF533APC,ENCFF097CAQ,ENCFF281QMV,ENCFF611HDQ,ENCFF609MDN,ENCFF033UQX,ENCFF491RJK,ENCFF504JDS,ENCFF142CNG,ENCFF805QBU |
| 1548 | 10 | 67834123 | 67834201 | 78 | SPRET | 10 | 67834201 | 67834219 | 19 | + | 20.3443 | 8.41E-08 | 0.0513 | CTGCCACTAGATGGCGCTT | 35 | ENCFF093ZRU,ENCFF367UNR,ENCFF201JUA,ENCFF099QNE,ENCFF210BTO,ENCFF907XEG,ENCFF089GBC,ENCFF821BIA,ENCFF092NMJ,ENCFF327LYS,ENCFF951GOP,ENCFF430PPJ,ENCFF052KVO,ENCFF198SSR,ENCFF883UPM,ENCFF502WDT,ENCFF165LXO,ENCFF024JOZ,ENCFF055HAP,ENCFF136ZDK,ENCFF316FYN,ENCFF589ZBJ,ENCFF065ODV,ENCFF349BUL,ENCFF089EWX,ENCFF533APC,ENCFF097CAQ,ENCFF281QMV,ENCFF611HDQ,ENCFF609MDN,ENCFF033UQX,ENCFF491RJK,ENCFF504JDS,ENCFF142CNG,ENCFF805QBU |
| 1549 | 10 | 68050990 | 68051825 | 835 | SPRET | 10 | 68051524 | 68051558 | 35 | + | 15.9541 | 9.80E-07 | 0.186 | ATGCATGTGCCCATGGAGTTCAGAAGAGGGAGCCA | | |
| 1550 | 10 | 69238022 | 69238421 | 399 | 129S1,C3H,CBA,CE,NZB,NZO,NZW,SPRET | 10 | 69238378 | 69238411 | 34 | + | 17.0244 | 6.67E-07 | 0.0928 | TAGCTGTCTTCAAACTCTCCAGAAGAGGGCATCA | | |
| 1551 | 10 | 71152738 | 71152935 | 197 | CAST,MOLF,SPRET | 10 | 71152793 | 71152826 | 34 | + | 16.4634 | 9.51E-07 | 0.108 | TAGCTGTCTTCAGACGCACCAGAAGAGGGCATCA | 1 | ENCFF093ZRU |
| 1552 | 10 | 75792490 | 75792762 | 272 | BUB,CAST,RF,SPRET,WSB | 10 | 75792619 | 75792653 | 35 | + | 16.0917 | 9.07E-07 | 0.18 | TAGCTGTCTTCAGACAGCACCAGAAGAGGGCATCA | | |
| 1553 | 10 | 75883964 | 75884882 | 918 | CAST,MOLF | 10 | 75884397 | 75884431 | 35 | + | 22.7156 | 1.37E-08 | 0.0158 | TTGCATGTATGCCTGCACACCAGAAGAGGGCACCA | 3 | ENCFF089GBC,ENCFF883UPM,ENCFF093ZRU |
| 1553 | 10 | 75883964 | 75884882 | 918 | CAST,MOLF | 10 | 75884413 | 75884431 | 19 | + | 17.8852 | 5.08E-07 | 0.121 | ACACCAGAAGAGGGCACCA | 3 | ENCFF089GBC,ENCFF883UPM,ENCFF093ZRU |
| 1554 | 10 | 75908722 | 75908930 | 208 | BUB,CAST,MOLF,RF,SPRET,WSB | 10 | 75908774 | 75908807 | 34 | + | 16.4634 | 9.51E-07 | 0.108 | TAGCTGTCTTCAGACGCACCAGAAGAGGGCATCA | | |
| 1555 | 10 | 76289608 | 76289808 | 200 | SPRET | 10 | 76289667 | 76289700 | 34 | + | 16.4472 | 9.61E-07 | 0.108 | TAGCTGTCTTCAGACACACCAGAAGAGGGCGTCA | | |
| 1556 | 10 | 80108682 | 80108847 | 165 | CAST,MOLF,SPRET | 10 | 80108702 | 80108736 | 35 | + | 18.0183 | 2.98E-07 | 0.101 | CTGTAGCTGTCTTCAGATACCAGAAGAGGGCATCA | | |
| 1557 | 10 | 81455637 | 81455826 | 189 | SPRET | 10 | 81455707 | 81455725 | 19 | + | 17.7213 | 5.67E-07 | 0.129 | AGACCAGCAGAGGGCATCA | 1 | ENCFF093ZRU |
| 1558 | 10 | 81552995 | 81553194 | 199 | 129S1,AJ,AKR,BALB,C3H,C57L,CBA,CE,DBA1J,DBA2J,FVB,ILNJ2,KK,MA/My,MRL,NOD,NOR,NZB,NZO,NZW,PJ,PL,RF,RH2,SEA,SJL,SMJ,SWR,TH | 10 | 81553055 | 81553088 | 34 | + | 18.3659 | 2.76E-07 | 0.0597 | TAGCTGTCTTCAGACACTCCAGAAGAGGGCGCCA | | |
| 1558 | 10 | 81552995 | 81553194 | 199 | 129S1,AJ,AKR,BALB,C3H,C57L,CBA,CE,DBA1J,DBA2J,FVB,ILNJ2,KK,MA/My,MRL,NOD,NOR,NZB,NZO,NZW,PJ,PL,RF,RH2,SEA,SJL,SMJ,SWR,TH | 10 | 81553070 | 81553088 | 19 | + | 17.5574 | 6.32E-07 | 0.134 | ACTCCAGAAGAGGGCGCCA | | |
| 1559 | 10 | 81688575 | 81689127 | 552 | SPRET | 10 | 81688958 | 81688991 | 34 | + | 17.8537 | 3.89E-07 | 0.071 | CTGCAGGGTCTTGGGCCGCCGGAAGGTGGGTGAC | | |
| 1560 | 10 | 82270254 | 82270432 | 178 | 129S1,AJ,AKR,BALB,BTBR,BUB,C3H,C57L,CAST,CBA,CE,DBA1J,DBA2J,FVB,ILNJ2,KK,MA/My,MOLF,MRL,NOD,NOR,NZB,NZO,NZW,PJ,PL,RF,RH2,SEA,SJL,SMJ,SPRET,SWR,TH,WSB | 10 | 82270305 | 82270323 | 19 | + | 17.8852 | 5.08E-07 | 0.121 | ACACCAGAAGAGGGCACCA | | |
| 1561 | 10 | 84633653 | 84633854 | 201 | SPRET | 10 | 84633711 | 84633744 | 34 | + | 17.4715 | 5.00E-07 | 0.0795 | TAGCTGTCTTCAGATGTGCCAGAAGAGGGCATCA | | |
| 1562 | 10 | 87912384 | 87912587 | 203 | AJ,AKR,BALB,BTBR,BUB,C3H,C58J,CAST,CBA,CE,DBA1J,DBA2J,FVB,ILNJ2,KK,LP,MA/My,MOLF,MRL,NOD,NOR,NZO,PJ,PL,RF,RH2,SEA,SPRET,SWR,TH | 10 | 87912445 | 87912478 | 34 | + | 18.561 | 2.42E-07 | 0.0573 | AAGCTGTCTTCAGACACTCCAGAAGAGGGCGCCA | | |
| 1562 | 10 | 87912384 | 87912587 | 203 | AJ,AKR,BALB,BTBR,BUB,C3H,C58J,CAST,CBA,CE,DBA1J,DBA2J,FVB,ILNJ2,KK,LP,MA/My,MOLF,MRL,NOD,NOR,NZO,PJ,PL,RF,RH2,SEA,SPRET,SWR,TH | 10 | 87912460 | 87912478 | 19 | + | 17.5574 | 6.32E-07 | 0.134 | ACTCCAGAAGAGGGCGCCA | | |
| 1563 | 10 | 88454150 | 88454630 | 480 | BUB,CAST,MOLF,SPRET,WSB | 10 | 88454610 | 88454643 | 34 | + | 17.2602 | 5.73E-07 | 0.0854 | CCGCCACACCCGCTCCCCACACAAGGTGGTACTG | | |
| 1564 | 10 | 90159085 | 90159140 | 55 | SPRET | 10 | 90159121 | 90159155 | 35 | + | 16.5229 | 7.12E-07 | 0.16 | CTCCTTGTACACCTGCATGCCAGAAGGGGGCATCA | | |
| 1565 | 10 | 95371766 | 95371974 | 208 | SPRET | 10 | 95371830 | 95371863 | 34 | + | 18.3659 | 2.76E-07 | 0.0597 | TAGCTGTCTTCAGACACTCCAGAAGAGGGCGCCA | 4 | ENCFF055HAP,ENCFF367UNR,ENCFF201JUA,ENCFF099QNE |
| 1565 | 10 | 95371766 | 95371974 | 208 | SPRET | 10 | 95371845 | 95371863 | 19 | + | 17.5574 | 6.32E-07 | 0.134 | ACTCCAGAAGAGGGCGCCA | 4 | ENCFF055HAP,ENCFF367UNR,ENCFF201JUA,ENCFF099QNE |
| 1566 | 10 | 102271123 | 102271455 | 332 | AKR,BUB,CAST,FVB,NOD,NOR,PL,RF,SPRET,WSB | 10 | 102271153 | 102271171 | 19 | + | 19.2951 | 1.88E-07 | 0.0757 | CCGCCACTAGATGGTGCTA | 17 | ENCFF201JUA,ENCFF367UNR,ENCFF093ZRU,ENCFF099QNE,ENCFF327LYS,ENCFF821BIA,ENCFF907XEG,ENCFF024JOZ,ENCFF092NMJ,ENCFF089GBC,ENCFF609MDN,ENCFF198SSR,ENCFF589ZBJ,ENCFF165LXO,ENCFF430PPJ,ENCFF052KVO,ENCFF136ZDK |
| 1567 | 10 | 108130023 | 108130218 | 195 | SPRET | 10 | 108130076 | 108130109 | 34 | + | 16.4472 | 9.61E-07 | 0.108 | TAGCTGTCTTCAGACACACCAGAAGAGGGCGTCA | | |
| 1568 | 10 | 108402112 | 108402303 | 191 | CAST,SPRET | 10 | 108402162 | 108402195 | 34 | + | 17.3902 | 5.27E-07 | 0.082 | TAGCTGTCCTCAGACACTCCAGAAGAGGGCATCA | | |
| 1569 | 10 | 109568707 | 109569050 | 343 | CAST,MOLF,NZW,PL,SMJ,SPRET,WSB | 10 | 109568771 | 109568804 | 34 | + | 16.4472 | 9.61E-07 | 0.108 | TAGCTGTCTTCAGACACACCAGAAGAGGGCGTCA | | |
| 1570 | 10 | 118209884 | 118210079 | 195 | SPRET | 10 | 118209939 | 118209972 | 34 | + | 17.2439 | 5.79E-07 | 0.0854 | TAGCTGTCCTCAGATACTCCAGAAGAGGGCATCA | | |
| 1571 | 10 | 119141918 | 119142372 | 454 | MOLF | 10 | 119142065 | 119142098 | 34 | + | 26.1951 | 5.57E-10 | 0.000748 | CTGCTGTTCCCAGACACACCAGAAGAGGGCATTG | | |
| 1572 | 10 | 120102114 | 120102315 | 201 | CAST,MOLF,SPRET | 10 | 120102171 | 120102205 | 35 | + | 18.1284 | 2.79E-07 | 0.0973 | CTGTAGCTGTCTTCAGACACCAGAAGAGGGCATCA | | |
| 1573 | 10 | 120390334 | 120390534 | 200 | SPRET | 10 | 120390393 | 120390426 | 34 | + | 16.4472 | 9.61E-07 | 0.108 | TAGCTGTCTTCAGACACACCAGAAGAGGGCGTCA | | |
| 1574 | 10 | 121746484 | 121746687 | 203 | CAST,DBA1J,KK,MOLF,SMJ | 10 | 121746550 | 121746583 | 34 | + | 17.2439 | 5.79E-07 | 0.0854 | TAGCTGTCCTCAGATACTCCAGAAGAGGGCATCA | | |
| 1575 | 10 | 122789869 | 122790086 | 217 | AJ,BALB,BUB,CAST,CE,KK,MA/My,MOLF,NOD,NOR,PL,RH2,SPRET | 10 | 122789944 | 122789977 | 34 | + | 19.0325 | 1.75E-07 | 0.0478 | CAGCTGTCTTCAGACGCACCAGAAGAGGGCATCA | 1 | ENCFF093ZRU |
| 1576 | 10 | 124153915 | 124154353 | 438 | PL | 10 | 124154330 | 124154364 | 35 | + | 16.5505 | 7.01E-07 | 0.158 | ATGCATGTGTGTTCTGATGCCAGAAGAGGGCATCA | | |
| 1577 | 10 | 127250434 | 127250660 | 226 | AJ,BALB,BUB,C3H,CAST,CBA,CE,FVB,MOLF,NOD,NOR,PL,RH2,SPRET,WSB | 10 | 127250518 | 127250551 | 34 | + | 17.3902 | 5.27E-07 | 0.082 | TAGCTGTCCTCAGACACTCCAGAAGAGGGCATCA | | |
| 1578 | 10 | 127510713 | 127510838 | 125 | SPRET | 10 | 127510717 | 127510751 | 35 | + | 20.2018 | 7.63E-08 | 0.0446 | ATGTAGTACCCTGTGGAGACCAGAAGAGGGTGACA | 31 | ENCFF093ZRU,ENCFF367UNR,ENCFF099QNE,ENCFF907XEG,ENCFF327LYS,ENCFF024JOZ,ENCFF089GBC,ENCFF065ODV,ENCFF589ZBJ,ENCFF430PPJ,ENCFF055HAP,ENCFF136ZDK,ENCFF198SSR,ENCFF321AGQ,ENCFF502WDT,ENCFF092NMJ,ENCFF165LXO,ENCFF052KVO,ENCFF504JDS,ENCFF883UPM,ENCFF601JVE,ENCFF951GOP,ENCFF575MGI,ENCFF957SPH,ENCFF609MDN,ENCFF210BTO,ENCFF097CAQ,ENCFF805QBU,ENCFF033UQX,ENCFF316FYN,ENCFF784ASD |
| 1578 | 10 | 127510713 | 127510838 | 125 | SPRET | 10 | 127510778 | 127510811 | 34 | + | 19.3252 | 1.43E-07 | 0.0426 | ATGTAGTACCCTTGGAGACCAGAAGAGGGTGACA | 35 | ENCFF093ZRU,ENCFF367UNR,ENCFF099QNE,ENCFF907XEG,ENCFF327LYS,ENCFF024JOZ,ENCFF089GBC,ENCFF065ODV,ENCFF589ZBJ,ENCFF430PPJ,ENCFF055HAP,ENCFF136ZDK,ENCFF198SSR,ENCFF321AGQ,ENCFF502WDT,ENCFF092NMJ,ENCFF165LXO,ENCFF052KVO,ENCFF504JDS,ENCFF883UPM,ENCFF601JVE,ENCFF951GOP,ENCFF575MGI,ENCFF957SPH,ENCFF609MDN,ENCFF210BTO,ENCFF097CAQ,ENCFF805QBU,ENCFF033UQX,ENCFF316FYN,ENCFF784ASD,ENCFF281QMV,ENCFF533APC,ENCFF142CNG,ENCFF194HVL |
| 1579 | 10 | 128108930 | 128109134 | 204 | SPRET | 10 | 128108992 | 128109025 | 34 | + | 18.3659 | 2.76E-07 | 0.0597 | TAGCTGTCTTCAGACACTCCAGAAGAGGGCGCCA | 7 | ENCFF093ZRU,ENCFF099QNE,ENCFF821BIA,ENCFF367UNR |
| 1579 | 10 | 128108930 | 128109134 | 204 | SPRET | 10 | 128109007 | 128109025 | 19 | + | 17.5574 | 6.32E-07 | 0.134 | ACTCCAGAAGAGGGCGCCA | 7 | ENCFF093ZRU,ENCFF099QNE,ENCFF821BIA,ENCFF367UNR |
| 1580 | 10 | 128168450 | 128168655 | 205 | BUB,C3H,CAST,CBA,FVB,KK,MOLF,SMJ,SPRET | 10 | 128168516 | 128168549 | 34 | + | 19.4797 | 1.28E-07 | 0.04 | TAGCTGTCCCCAGACACTCCAGAAGAGGGCGTCA | | |
| 1581 | 10 | 128182651 | 128182859 | 208 | BUB,CAST,CE,FVB,KK,MOLF,SMJ,SPRET,WSB | 10 | 128182717 | 128182750 | 34 | + | 22.9187 | 9.63E-09 | 0.00615 | TTGCTGTCCTCAGACGCCCCAGAAGAGGGCGCCA | 5 | ENCFF093ZRU,ENCFF821BIA,ENCFF367UNR,ENCFF201JUA |
| 1582 | 10 | 128734180 | 128734417 | 237 | BUB,CAST,CE,KK,MOLF,RF,SMJ,SPRET,WSB | 10 | 128734275 | 128734308 | 34 | + | 16.4634 | 9.51E-07 | 0.108 | TAGCTGTCTTCAGACGCACCAGAAGAGGGCATCA | | |
| 1583 | 10 | 3517533 | 3518153 | 620 | SPRET | 10 | 3518009 | 3518027 | 19 | - | 18.0492 | 4.54E-07 | 0.12 | GGGCCAGAAGAGGGCGATG | 19 | ENCFF024JOZ,ENCFF821BIA,ENCFF198SSR,ENCFF092NMJ,ENCFF165LXO,ENCFF589ZBJ,ENCFF907XEG,ENCFF883UPM,ENCFF093ZRU,ENCFF502WDT,ENCFF367UNR,ENCFF194HVL,ENCFF033UQX,ENCFF957SPH,ENCFF052KVO,ENCFF327LYS,ENCFF533APC,ENCFF136ZDK,ENCFF099QNE |
| 1583 | 10 | 3517533 | 3518153 | 620 | SPRET | 10 | 3518009 | 3518042 | 34 | - | 26.3984 | 4.60E-10 | 0.000687 | GTGCAGTACTCATAGGGGCCAGAAGAGGGCGATG | 19 | ENCFF024JOZ,ENCFF821BIA,ENCFF198SSR,ENCFF092NMJ,ENCFF165LXO,ENCFF589ZBJ,ENCFF907XEG,ENCFF883UPM,ENCFF093ZRU,ENCFF502WDT,ENCFF367UNR,ENCFF194HVL,ENCFF033UQX,ENCFF957SPH,ENCFF052KVO,ENCFF327LYS,ENCFF533APC,ENCFF136ZDK,ENCFF099QNE |
| 1584 | 10 | 5766375 | 5766584 | 209 | 129S1,AJ,AKR,BALB,BTBR,BUB,C3H,C57L,CAST,CBA,CE,DBA1J,DBA2J,FVB,KK,LP,MA/My,MOLF,MRL,NOD,NOR,NZB,NZO,NZW,PL,RF,RH2,SJL,SMJ,SPRET,SWR,TH | 10 | 5766493 | 5766511 | 19 | - | 17.5574 | 6.32E-07 | 0.134 | ACTCCAGAAGAGGGCGCCA | | |
| 1584 | 10 | 5766375 | 5766584 | 209 | 129S1,AJ,AKR,BALB,BTBR,BUB,C3H,C57L,CAST,CBA,CE,DBA1J,DBA2J,FVB,KK,LP,MA/My,MOLF,MRL,NOD,NOR,NZB,NZO,NZW,PL,RF,RH2,SJL,SMJ,SPRET,SWR,TH | 10 | 5766493 | 5766526 | 34 | - | 18.3659 | 2.76E-07 | 0.0597 | TAGCTGTCTTCAGACACTCCAGAAGAGGGCGCCA | | |
| 1585 | 10 | 12913930 | 12914055 | 125 | SPRET | 10 | 12913898 | 12913931 | 34 | - | 20.6829 | 5.41E-08 | 0.0225 | ATGTAGTGCCCAAGGAAGCCAGAAGAGGGTATCA | | |
| 1586 | 10 | 19622215 | 19622847 | 632 | SPRET | 10 | 19622588 | 19622622 | 35 | - | 16 | 9.55E-07 | 0.184 | TTGCAATTGAAGGCCCAGGACTGAAGGGGTCACGC | | |
| 1587 | 10 | 20842102 | 20842166 | 64 | SPRET,WSB | 10 | 20842128 | 20842146 | 19 | - | 21.9672 | 2.09E-08 | 0.0336 | TGACCACCAGAGGGCAGGA | 39 | ENCFF093ZRU,ENCFF367UNR,ENCFF201JUA,ENCFF099QNE,ENCFF821BIA,ENCFF907XEG,ENCFF055HAP,ENCFF327LYS,ENCFF024JOZ,ENCFF089GBC,ENCFF198SSR,ENCFF589ZBJ,ENCFF136ZDK,ENCFF065ODV,ENCFF092NMJ,ENCFF883UPM,ENCFF349BUL,ENCFF052KVO,ENCFF165LXO,ENCFF430PPJ,ENCFF951GOP,ENCFF957SPH,ENCFF086ZTV,ENCFF033UQX,ENCFF575MGI,ENCFF142CNG,ENCFF491RJK,ENCFF210BTO,ENCFF784ASD,ENCFF281QMV,ENCFF097CAQ,ENCFF609MDN,ENCFF194HVL,ENCFF805QBU,ENCFF611HDQ,ENCFF533APC,ENCFF504JDS,ENCFF316FYN,ENCFF089EWX |
| 1588 | 10 | 21581423 | 21582187 | 764 | CAST,MOLF | 10 | 21582105 | 21582139 | 35 | - | 17.789 | 3.41E-07 | 0.108 | TGGCAGTCTCCACAGCTTTCCTCTAGGGGCCAGGG | | |
| 1589 | 10 | 22039501 | 22039833 | 332 | MOLF,TH | 10 | 22039791 | 22039809 | 19 | - | 19.2951 | 1.88E-07 | 0.0757 | CCGCCACTAGATGGTGCTA | 22 | ENCFF201JUA,ENCFF367UNR,ENCFF093ZRU,ENCFF099QNE,ENCFF821BIA,ENCFF327LYS,ENCFF024JOZ,ENCFF089GBC,ENCFF907XEG,ENCFF883UPM,ENCFF198SSR,ENCFF136ZDK,ENCFF430PPJ,ENCFF502WDT,ENCFF589ZBJ,ENCFF609MDN,ENCFF165LXO,ENCFF052KVO,ENCFF092NMJ,ENCFF055HAP,ENCFF194HVL,ENCFF957SPH |
| 1590 | 10 | 25006888 | 25007767 | 879 | CAST | 10 | 25007608 | 25007642 | 35 | - | 18.7706 | 1.89E-07 | 0.0777 | GTGCATTCAGGTGTGGAGGCCAGAAGAGGGCTGAT | | |
| 1591 | 10 | 25063248 | 25063540 | 292 | CAST,MOLF,SPRET | 10 | 25063437 | 25063471 | 35 | - | 16.9908 | 5.44E-07 | 0.138 | CAGCAAGGAGCAGTGAGGGGCAGTGGGGGGCAGCA | | |
| 1591 | 10 | 25063248 | 25063540 | 292 | CAST,MOLF,SPRET | 10 | 25063538 | 25063572 | 35 | - | 16.9908 | 5.44E-07 | 0.138 | CAGCAAGGAGCAGTGAGGGGCAGTGGGGGGCAGCA | | |
| 1592 | 10 | 25160736 | 25160937 | 201 | SPRET | 10 | 25160854 | 25160887 | 34 | - | 18.5528 | 2.44E-07 | 0.0575 | TAGCTGTTTTCAGATACACCAGAAGGGGGCATCA | 8 | ENCFF821BIA,ENCFF367UNR,ENCFF093ZRU,ENCFF198SSR,ENCFF055HAP,ENCFF883UPM,ENCFF502WDT,ENCFF099QNE |
| 1593 | 10 | 26874406 | 26874613 | 207 | 129S1,AJ,AKR,BALB,BTBR,BUB,C3H,C57L,CAST,CE,FVB,ILNJ2,KK,LP,MA/My,MOLF,MRL,NOD,NZB,NZO,NZW,PJ,PL,RF,SEA,SJL,SPRET,SWR,TH,WSB | 10 | 26874529 | 26874562 | 34 | - | 16.7724 | 7.83E-07 | 0.0988 | TAGCTGTCTTCAGATGCACCAGAAGAGGGCGTCA | | |
| 1594 | 10 | 30062007 | 30062202 | 195 | SPRET | 10 | 30062130 | 30062163 | 34 | - | 16.9187 | 7.13E-07 | 0.0951 | TAGCTGTCTTCAGACGCACCAGAAGAGGGCGTCA | | |
| 1595 | 10 | 33254302 | 33254563 | 261 | CAST,KK,MOLF,SPRET,WSB | 10 | 33254431 | 33254449 | 19 | - | 17.8852 | 5.08E-07 | 0.121 | ACACCAGAAGAGGGCACCA | | |
| 1596 | 10 | 36110868 | 36111075 | 207 | CAST,MOLF,SPRET,WSB | 10 | 36110993 | 36111026 | 34 | - | 16.4472 | 9.61E-07 | 0.108 | TAGCTGTCTTCAGACACACCAGAAGAGGGCGTCA | | |
| 1597 | 10 | 41197149 | 41197395 | 246 | MOLF,SPRET | 10 | 41197274 | 41197307 | 34 | - | 16.4634 | 9.51E-07 | 0.108 | TAGCTGTCTTCAGACGCACCAGAAGAGGGCATCA | | |
| 1598 | 10 | 41612706 | 41612951 | 245 | SPRET | 10 | 41612766 | 41612800 | 35 | - | 21.0826 | 4.26E-08 | 0.0317 | GTGTAGGTTCCCAAATAGGCCAGAAGAGGGCATTG | | |
| 1599 | 10 | 42452279 | 42452492 | 213 | 129S1,BTBR,C57L,CAST,KK,MOLF,SPRET,WSB | 10 | 42452404 | 42452437 | 34 | - | 17.3008 | 5.58E-07 | 0.0844 | CAGCCGTCCTCAGACACTCCAGAAGAGGGAGTCA | | |
| 1600 | 10 | 42800626 | 42801154 | 528 | SPRET | 10 | 42800803 | 42800837 | 35 | - | 22.1927 | 1.99E-08 | 0.0194 | CTGCATGTCCACCTGCATGCCAGAAGAGGACATCA | | |
| 1601 | 10 | 44348891 | 44349093 | 202 | ILNJ2,SPRET | 10 | 44349016 | 44349049 | 34 | - | 16.4472 | 9.61E-07 | 0.108 | TAGCTGTCTTCAGACACACCAGAAGAGGGCGTCA | | |
| 1602 | 10 | 45545329 | 45546048 | 719 | SPRET | 10 | 45545799 | 45545832 | 34 | - | 20.0813 | 8.38E-08 | 0.0304 | GTCCAGTGCCCAATGAAGCCAGCAGAGGTCATCA | | |
| 1603 | 10 | 45822854 | 45823058 | 204 | SPRET,TH | 10 | 45822978 | 45823011 | 34 | - | 17.9106 | 3.74E-07 | 0.07 | TAGCTGTCTTCAGACACTCCAGAAGAGGGCACCA | | |
| 1604 | 10 | 49461584 | 49462136 | 552 | CAST,MOLF,SPRET,TH | 10 | 49462113 | 49462131 | 19 | - | 16.8525 | 9.96E-07 | 0.168 | CGAGCTCCAGGGGGCACTG | | |
| 1604 | 10 | 49461584 | 49462136 | 552 | CAST,MOLF,SPRET,TH | 10 | 49462113 | 49462146 | 34 | - | 21.2033 | 3.67E-08 | 0.0174 | ATGCAGCTAGAGACACGAGCTCCAGGGGGCACTG | | |
| 1605 | 10 | 56230496 | 56231212 | 716 | CAST,SPRET | 10 | 56230787 | 56230821 | 35 | - | 16.0367 | 9.36E-07 | 0.182 | ATGCACTGCCCCTGGGCCGCAAGAAGGGGGCCAAG | | |
| 1606 | 10 | 57085135 | 57085566 | 431 | CAST,MOLF | 10 | 57085315 | 57085348 | 34 | - | 17.3496 | 5.41E-07 | 0.0834 | ATGCAGCTAGAGACACGAGCTTCAGGGGGCACTG | | |
| 1607 | 10 | 58288195 | 58288837 | 642 | SPRET | 10 | 58288769 | 58288787 | 19 | - | 19.5902 | 1.51E-07 | 0.0671 | AGACCAGAAGGTGGCACCA | 7 | ENCFF367UNR,ENCFF093ZRU,ENCFF099QNE,ENCFF136ZDK,ENCFF821BIA,ENCFF502WDT,ENCFF883UPM |
| 1608 | 10 | 59651046 | 59651253 | 207 | CAST,MOLF,SPRET | 10 | 59651171 | 59651204 | 34 | - | 16.4634 | 9.51E-07 | 0.108 | TAGCTGTCTTCAGACGCACCAGAAGAGGGCATCA | | |
| 1609 | 10 | 59858974 | 59859176 | 202 | CAST,MOLF,SPRET | 10 | 59859099 | 59859117 | 19 | - | 17.5738 | 6.25E-07 | 0.134 | ACTCCAGAAGAGGGCGCTA | 3 | ENCFF093ZRU,ENCFF367UNR,ENCFF821BIA |
| 1609 | 10 | 59858974 | 59859176 | 202 | CAST,MOLF,SPRET | 10 | 59859099 | 59859132 | 34 | - | 20.065 | 8.47E-08 | 0.0305 | TAGCTGTCCTCAGACACTCCAGAAGAGGGCGCTA | 3 | ENCFF093ZRU,ENCFF367UNR,ENCFF821BIA |
| 1610 | 10 | 61253241 | 61253441 | 200 | SPRET,WSB | 10 | 61253366 | 61253384 | 19 | - | 17.5574 | 6.32E-07 | 0.134 | ACTCCAGAAGAGGGCGCCA | 3 | ENCFF821BIA,ENCFF093ZRU,ENCFF367UNR |
| 1610 | 10 | 61253241 | 61253441 | 200 | SPRET,WSB | 10 | 61253366 | 61253399 | 34 | - | 18.3659 | 2.76E-07 | 0.0597 | TAGCTGTCTTCAGACACTCCAGAAGAGGGCGCCA | 3 | ENCFF821BIA,ENCFF093ZRU,ENCFF367UNR |
| 1611 | 10 | 61358511 | 61359229 | 718 | SPRET | 10 | 61358633 | 61358651 | 19 | - | 17.5574 | 6.32E-07 | 0.134 | ACTCCAGAAGAGGGCGCCA | 1 | ENCFF093ZRU |
| 1611 | 10 | 61358511 | 61359229 | 718 | SPRET | 10 | 61358633 | 61358666 | 34 | - | 18.3659 | 2.76E-07 | 0.0597 | TAGCTGTCTTCAGACACTCCAGAAGAGGGCGCCA | 1 | ENCFF093ZRU |

| | | | | | | | | | | | | | | | | |
|---|---|---|---|---|---|---|---|---|---|---|---|---|---|---|---|---|
| 1612 | 10 | 62328119 | 62328306 | 187 | CAST.SPRET | 10 | 62328237 | 62328270 | 34 | - | 16.5691 | 8.90E-07 | 0.107 | TAGCTGTCTTCAGACGCACCAGAAGAGGGCATCG | | |
| 1613 | 10 | 62525683 | 62525903 | 220 | CAST.CE.MOLF.RF.SPRET | 10 | 62525808 | 62525841 | 34 | - | 17.2114 | 5.91E-07 | 0.0866 | TAGCTGTTTTCAGACACTCCAGAAGAGGGCATCA | | |
| 1614 | 10 | 62528927 | 62529143 | 216 | SPRET | 10 | 62529064 | 62529082 | 19 | - | 18.623 | 3.06E-07 | 0.0973 | ACACCAGAAGAGGGCGCCA | | |
| 1614 | 10 | 62528927 | 62529143 | 216 | SPRET | 10 | 62529064 | 62529097 | 34 | - | 18.8537 | 1.98E-07 | 0.0514 | TAGCTGTCTTCAGACACACCAGAAGAGGGCGCCA | | |
| 1615 | 10 | 62733553 | 62733760 | 207 | CAST.SPRET | 10 | 62733675 | 62733708 | 34 | - | 16.4634 | 9.51E-07 | 0.108 | TAGCTGTCTTCAGACGCACCAGAAGAGGGCATCA | | |
| 1616 | 10 | 66356318 | 66356545 | 227 | SPRET | 10 | 66356441 | 66356474 | 34 | - | 16.4634 | 9.51E-07 | 0.108 | TAGCTGTCTTCAGACGCACCAGAAGAGGGCATCA | | |
| 1617 | 10 | 68848690 | 68849495 | 805 | CAST.MOLF | 10 | 68849310 | 68849343 | 34 | - | 18.7073 | 2.19E-07 | 0.0543 | GTGCAGTATCATTAGGAACCAGAGGATGGCATCT | | |
| 1618 | 10 | 77589331 | 77589540 | 209 | CAST | 10 | 77589513 | 77589546 | 34 | - | 20.4715 | 6.31E-08 | 0.0252 | TTGCTGTACTCAGAAACACCAGAAGAGGGCATCA | 7 | ENCFF093ZRU,ENCFF821BIA,ENCFF883UPM,ENCFF052KVO,ENCFF367UNR,ENCFF055HAP,ENCFF502WDT |
| 1619 | 10 | 77591480 | 77591680 | 200 | SPRET | 10 | 77591543 | 77591577 | 35 | - | 21.1101 | 4.19E-08 | 0.0313 | CTGCATGTACACCTGCATGCCAGAAGAAGGCATCT | | |
| 1620 | 10 | 77768658 | 77768872 | 214 | BUB.CAST.MOLF.SPRET.TH | 10 | 77768784 | 77768817 | 34 | - | 17.8455 | 3.91E-07 | 0.071 | TAGCTGTCCTCAGACACTCCAGAAGAGGGCGTCA | | |
| 1621 | 10 | 77799197 | 77799391 | 194 | SPRET | 10 | 77799281 | 77799314 | 34 | - | 17.065 | 6.50E-07 | 0.0917 | TTGCTGTCTTCAGACACACCAGAAGAGGGCATGG | | |
| 1622 | 10 | 79265042 | 79265251 | 209 | CAST.MOLF.SPRET | 10 | 79265168 | 79265201 | 34 | - | 18.935 | 1.88E-07 | 0.0499 | TTGCTGTCTTCAGACACACCAGAAGAGGGCGTCA | | |
| 1623 | 10 | 80047318 | 80047546 | 228 | BUB.CAST.MOLF.SPRET.WSB | 10 | 80047443 | 80047476 | 34 | - | 20.4146 | 6.58E-08 | 0.0258 | CAGCTGTCCTCAGACACTCCAGAAGAGGGCGTCA | | |
| 1624 | 10 | 80080888 | 80081001 | 113 | CAST,MOLF,SPRET | 10 | 80080906 | 80080939 | 34 | - | 17.6667 | 4.40E-07 | 0.0748 | TAGCTGTCTTCAGACACGCCAGAAGAGGGCGTCA | 17 | ENCFF136ZDK,ENCFF092NMJ,ENCFF093ZRU,ENCFF430PPJ,ENCFF883UPM,ENCFF502WDT,ENCFF367UNR,ENCFF957SPH,ENCFF099QNE,ENCFF327LYS,ENCFF907XEG,ENCFF089GBC,ENCFF589ZBJ,ENCFF055HAP,ENCFF024JOZ,ENCFF821BIA,ENCFF198SSR |
| 1625 | 10 | 81117424 | 81118000 | 576 | SPRET | 10 | 81117837 | 81117870 | 34 | - | 17.9106 | 3.74E-07 | 0.07 | TAGCTGTCTTCAGACACTCCAGAAGAGGGCACCA | 1 | ENCFF093ZRU |
| 1626 | 10 | 81210813 | 81211019 | 206 | SPRET | 10 | 81210954 | 81210987 | 34 | - | 17.1463 | 6.17E-07 | 0.089 | TAGCTGTCCTCAGACACTCCAGAAGAAGGCGCCA | | |
| 1627 | 10 | 95183504 | 95183706 | 202 | CAST.MOLF.MRL.SPRET | 10 | 95183630 | 95183664 | 35 | - | 18.1284 | 2.79E-07 | 0.0973 | CTGTAGCTGTCTTCAGACACCAGAAGAGGGCATCA | | |
| 1628 | 10 | 97099901 | 97099955 | 54 | SPRET | 10 | 97099870 | 97099904 | 35 | - | 19.2661 | 1.39E-07 | 0.0639 | GTGCACATCCTCTCAGAGGCCAGAAGAAGGCAATA | | |
| 1629 | 10 | 98286647 | 98287188 | 541 | SPRET | 10 | 98287100 | 98287133 | 34 | - | 17.8455 | 3.91E-07 | 0.071 | CTGCTGTCTTCATACACACCAGAAGAGGGCATAT | 1 | ENCFF093ZRU |
| 1630 | 10 | 108116899 | 108117117 | 218 | AKR,CAST,LP,NOD,NOR,PL,RF,SPRET,WSB | 10 | 108117028 | 108117061 | 34 | - | 17.8455 | 3.91E-07 | 0.071 | TAGCTGTCCTCAGACACTCCAGAAGAGGGCGTCA | | |
| 1631 | 10 | 111489624 | 111489834 | 210 | CAST.MOLF.SMJ.SPRET | 10 | 111489751 | 111489784 | 34 | - | 16.4634 | 9.51E-07 | 0.108 | TAGCTGTCTTCAGACGCACCAGAAGAGGGCATCA | | |
| 1632 | 10 | 111941599 | 111941930 | 331 | CAST,SPRET | 10 | 111941889 | 111941907 | 19 | - | 19.2951 | 1.88E-07 | 0.0757 | CCGCCACTAGATGGTGCTA | 33 | ENCFF201JUA,ENCFF367UNR,ENCFF093ZRU,ENCFF099QNE,ENCFF821BIA,ENCFF316FYN,ENCFF033UQX,ENCFF327LYS,ENCFF907XEG,ENCFF097CAQ,ENCFF136ZDK,ENCFF024JOZ,ENCFF589ZBJ,ENCFF089GBC,ENCFF052KVO,ENCFF165LXO,ENCFF055HAP,ENCFF430PPJ,ENCFF065ODV,ENCFF092NMJ,ENCFF502WDT,ENCFF198SSR,ENCFF321AGQ,ENCFF951GOP,ENCFF957SPH,ENCFF883UPM,ENCFF575MGI,ENCFF194HVL,ENCFF281QMV,ENCFF533APC,ENCFF142CNG,ENCFF491RJK,ENCFF609MDN |
| 1633 | 10 | 116817779 | 116817991 | 212 | C57L,C58J,CAST,MOLF,NZW,SMJ,SPRET | 10 | 116817904 | 116817922 | 19 | - | 17.5574 | 6.32E-07 | 0.134 | ACTCCAGAAGAGGGCGCCA | | |
| 1633 | 10 | 116817779 | 116817991 | 212 | C57L,C58J,CAST,MOLF,NZW,SMJ,SPRET | 10 | 116817904 | 116817937 | 34 | - | 18.3659 | 2.76E-07 | 0.0597 | TAGCTGTCTTCAGACACTCCAGAAGAGGGCGCCA | | |
| 1634 | 10 | 119947716 | 119947897 | 181 | SPRET | 10 | 119947726 | 119947760 | 35 | - | 22.7064 | 1.38E-08 | 0.0158 | CTGCATCTATACCTGCATACCAGAAGAGGGCATCA | 5 | ENCFF367UNR.ENCFF099QNE.ENCFF907XEG.ENCFF055HAP.ENCFF093ZRU |
| 1635 | 10 | 120330583 | 120331258 | 675 | WSB | 10 | 120330572 | 120330605 | 34 | - | 16.5854 | 8.81E-07 | 0.107 | TAGCTGTCCCCAGACACTCCAGAAGAGGGTGTCA | | |
| 1636 | 10 | 121237649 | 121237850 | 201 | AJ,BALB,BUB,CAST,CE,DBA1J,DBA2J,FVB,ILNJ2,KK,LP,MA/My,MOLF,NOD,NOR,PL,RH2,SMJ,SPRET | 10 | 121237774 | 121237807 | 34 | - | 16.4472 | 9.61E-07 | 0.108 | TAGCTGTCTTCAGACACACCAGAAGAGGGCGTCA | | |
| 1637 | 10 | 121273312 | 121273520 | 208 | AJ,BALB,CAST,DBA1J,DBA2J,KK,MA/My,MOLF,RH2,SMJ,SPRET | 10 | 121273436 | 121273469 | 34 | - | 16.4472 | 9.61E-07 | 0.108 | TAGCTGTCTTCAGACACACCAGAAGAGGGCGTCA | | |
| 1638 | 10 | 122376428 | 122376609 | 181 | CAST.DBA1J.KK.MOLF | 10 | 122376545 | 122376563 | 19 | - | 17.5574 | 6.32E-07 | 0.134 | ACTCCAGAAGAGGGCGCCA | 1 | ENCFF093ZRU |
| 1638 | 10 | 122376428 | 122376609 | 181 | CAST.DBA1J.KK.MOLF | 10 | 122376545 | 122376578 | 34 | - | 18.3659 | 2.76E-07 | 0.0597 | TAGCTGTCTTCAGACACTCCAGAAGAGGGCGCCA | 1 | ENCFF093ZRU |
| 1639 | 10 | 126515251 | 126515316 | 65 | DBA1J,DBA2J,ILNJ2,KK,NZW,WSB | 10 | 126515292 | 126515310 | 19 | - | 21.0984 | 4.51E-08 | 0.0444 | AGGCCAGAAGAGGGCGCCA | 19 | ENCFF367UNR,ENCFF093ZRU,ENCFF024JOZ,ENCFF327LYS,ENCFF430PPJ,ENCFF092NMJ,ENCFF821BIA,ENCFF136ZDK,ENCFF099QNE,ENCFF589ZBJ,ENCFF201JUA,ENCFF052KVO,ENCFF957SPH,ENCFF194HVL,ENCFF533APC,ENCFF033UQX,ENCFF907XEG,ENCFF281QMV,ENCFF089GBC |
| 1639 | 10 | 126515251 | 126515316 | 65 | DBA1J,DBA2J,ILNJ2,KK,NZW,WSB | 10 | 126515292 | 126515326 | 35 | - | 28.055 | 1.84E-10 | 0.0029 | GTGCAGGTGCCGGTGGAGGCCAGAAGAGGGCGCCA | 19 | ENCFF367UNR,ENCFF093ZRU,ENCFF024JOZ,ENCFF327LYS,ENCFF430PPJ,ENCFF092NMJ,ENCFF821BIA,ENCFF136ZDK,ENCFF099QNE,ENCFF589ZBJ,ENCFF201JUA,ENCFF052KVO,ENCFF957SPH,ENCFF194HVL,ENCFF533APC,ENCFF033UQX,ENCFF907XEG,ENCFF281QMV,ENCFF089GBC |
| 1640 | 10 | 126731956 | 126732305 | 349 | CAST.MOLF | 10 | 126731953 | 126731986 | 34 | - | 19.3171 | 1.44E-07 | 0.0428 | GTGTGTTGCCCACAGAAGCCAGAAGAGGGCATCA | 4 | ENCFF093ZRU.ENCFF055HAP.ENCFF502WDT.ENCFF533APC |
| 1641 | 10 | 126789754 | 126789818 | 64 | CE | 10 | 126789810 | 126789843 | 34 | - | 18.1545 | 3.18E-07 | 0.0637 | TAGCTGTTTTCAGACACACCAGAAGAGGGCGTCA | 2 | ENCFF367UNR.ENCFF093ZRU |
| 1642 | 10 | 126921917 | 126922150 | 233 | CAST.MOLF.SPRET | 10 | 126922036 | 126922069 | 34 | - | 17.6667 | 4.40E-07 | 0.0748 | TAGCTGTTTTCAGACACTCCAGAAGAGGGCGTCA | | |
| 1643 | 10 | 127200111 | 127200327 | 216 | AJ,BALB,BUB,C3H,CAST,CBA,CE,FVB,MOLF,NOD,NOR,PL,RH2,SPRET,WSB | 10 | 127200236 | 127200269 | 34 | - | 16.4472 | 9.61E-07 | 0.108 | TAGCTGTCTTCAGACACACCAGAAGAGGGCGTCA | 1 | ENCFF093ZRU |
| 1644 | 10 | 127303212 | 127303487 | 275 | AJ,BALB,BUB,C3H,CAST,CBA,CE,FVB,MOLF.NOD.NOR.PL.RH2.SPRET.WSB | 10 | 127303337 | 127303370 | 34 | - | 17.8455 | 3.91E-07 | 0.071 | TAGCTGTCCTCAGACACTCCAGAAGAGGGCGTCA | | |
| 1645 | 11 | 3893063 | 3893255 | 192 | B10D2.B10J.C58J.KK.SPRET | 11 | 3893113 | 3893146 | 34 | + | 16.4472 | 9.61E-07 | 0.067 | TAGCTGTCTTCAGACACACCAGAAGAGGGCGTCA | | |
| 1646 | 11 | 4660359 | 4660556 | 197 | SPRET | 11 | 4660410 | 4660443 | 34 | + | 17.2114 | 5.91E-07 | 0.0533 | TAGCTGTTTTCAGACACTCCAGAAGAGGGCATCA | 2 | ENCFF099QNE.ENCFF093ZRU |
| 1647 | 11 | 4838030 | 4838240 | 210 | 129S1,AJ,B10D2,B10J,BALB,BTBR,BUB,C57L,C58J,CAST,ILNJ2,KK,MA/My,NOD,NOR,NZW,PJ,PL,SEA,SPRET,TH,WSB | 11 | 4838096 | 4838129 | 34 | + | 18.1789 | 3.13E-07 | 0.0375 | TAGCTGTCTTCAGACACTCCAGAAGAGGGCGCTA | 3 | ENCFF093ZRU,ENCFF502WDT,ENCFF055HAP |
| 1647 | 11 | 4838030 | 4838240 | 210 | 129S1,AJ,B10D2,B10J,BALB,BTBR,BUB,C57L,C58J,CAST,ILNJ2,KK,MA/My,NOD,NOR,NZW,PJ,PL,SEA,SPRET,TH,WSB | 11 | 4838111 | 4838129 | 19 | + | 17.5738 | 6.25E-07 | 0.0841 | ACTCCAGAAGAGGGCGCTA | 3 | ENCFF093ZRU,ENCFF502WDT,ENCFF055HAP |
| 1648 | 11 | 5901163 | 5901358 | 195 | SPRET | 11 | 5901216 | 5901249 | 34 | + | 18.3659 | 2.76E-07 | 0.0355 | TAGCTGTCTTCAGACACTCCAGAAGAGGGCGCCA | 4 | ENCFF099QNE.ENCFF821BIA.ENCFF093ZRU.ENCFF367UNR |
| 1648 | 11 | 5901163 | 5901358 | 195 | SPRET | 11 | 5901231 | 5901249 | 19 | + | 17.5574 | 6.32E-07 | 0.0841 | ACTCCAGAAGAGGGCGCCA | 4 | ENCFF099QNE.ENCFF821BIA.ENCFF093ZRU.ENCFF367UNR |
| 1649 | 11 | 6284707 | 6284935 | 228 | SPRET | 11 | 6284793 | 6284826 | 34 | + | 18.8537 | 1.98E-07 | 0.0308 | TAGCTGTCTTCAGACACACCAGAAGAGGGCGCCA | 4 | ENCFF099QNE.ENCFF367UNR.ENCFF821BIA.ENCFF093ZRU |
| 1649 | 11 | 6284707 | 6284935 | 228 | SPRET | 11 | 6284808 | 6284826 | 19 | + | 18.623 | 3.06E-07 | 0.0609 | ACACCAGAAGAGGGCGCCA | 4 | ENCFF099QNE.ENCFF367UNR.ENCFF821BIA.ENCFF093ZRU |
| 1650 | 11 | 17072653 | 17073442 | 789 | AJ,AKR,BALB,BTBR,BUB,C3H,C57L,CBA,DBA1J,DBA2J,ILNJ2,KK,MRL,NZB,NZO,NZW,PL,RF,SEA,SJL,SMJ | 11 | 17072770 | 17072788 | 19 | + | 18.7213 | 2.85E-07 | 0.0593 | CTACCACTAGGTGGTACTA | 2 | ENCFF099QNE,ENCFF093ZRU |
| 1651 | 11 | 17228526 | 17228728 | 202 | CAST.SPRET | 11 | 17228567 | 17228600 | 34 | + | 17.2439 | 5.79E-07 | 0.0529 | TAGCTGTCCTCAGATACTCCAGAAGAGGGCATCA | | |
| 1652 | 11 | 18718566 | 18719504 | 938 | SPRET | 11 | 18718891 | 18718924 | 34 | + | 18.3089 | 2.87E-07 | 0.0357 | ATGCACTGTCTATTAAGGACAGAAGGGGGTGACA | | |
| 1653 | 11 | 23255382 | 23255598 | 216 | SPRET | 11 | 23255456 | 23255489 | 34 | + | 16.4634 | 9.51E-07 | 0.067 | TAGCTGTCTTCAGACGCACCAGAAGAGGGCATCA | 5 | ENCFF093ZRU.ENCFF367UNR.ENCFF099QNE.ENCFF821BIA.ENCFF055HAP |
| 1654 | 11 | 27738290 | 27738772 | 482 | AJ,C3H,DBA1J,DBA2J,ILNJ2,MOLF,MRL,NZB.NZO.NZW.PL.SPRET | 11 | 27738306 | 27738339 | 34 | + | 16.626 | 8.59E-07 | 0.065 | CTGCACTCACATTTGCCATTACAAGATGGCACTG | | |
| 1655 | 11 | 32579330 | 32579542 | 212 | AJ,AKR,BALB,BTBR,BUB,C57L,CAST,CE,DBA1J,DBA2J,FVB,ILNJ2,KK,MA/My,MOLF,NOD,NZB,NZO,NZW,PJ,PL,RF,RH2,SEA,SPRET,SWR,TH,WSB | 11 | 32579401 | 32579434 | 34 | + | 18.3659 | 2.76E-07 | 0.0355 | TAGCTGTCTTCAGACACTCCAGAAGAGGGCGCCA | | |
| 1655 | 11 | 32579330 | 32579542 | 212 | AJ,AKR,BALB,BTBR,BUB,C57L,CAST,CE,DBA1J,DBA2J,FVB,ILNJ2,KK,MA/My,MOLF,NOD,NZB,NZO,NZW,PJ,PL,RF,RH2,SEA,SPRET,SWR,TH,WSB | 11 | 32579416 | 32579434 | 19 | + | 17.5574 | 6.32E-07 | 0.0841 | ACTCCAGAAGAGGGCGCCA | | |
| 1656 | 11 | 32757124 | 32757873 | 749 | SPRET | 11 | 32757766 | 32757800 | 35 | + | 22.211 | 1.96E-08 | 0.0122 | ATGCAGGTTCCCGTGGAGGCCAGAAGAGGGTGTCA | 1 | ENCFF093ZRU |
| 1657 | 11 | 35733276 | 35733692 | 416 | CAST,MA/My,MOLF,NOD,NZB,NZO,PJ,RF,SEA,SPRET | 11 | 35733550 | 35733583 | 34 | + | 18.3659 | 2.76E-07 | 0.0355 | TAGCTGTCTTCAGACACTCCAGAAGAGGGCGCCA | 3 | ENCFF367UNR,ENCFF093ZRU,ENCFF055HAP |
| 1657 | 11 | 35733276 | 35733692 | 416 | CAST,MA/My,MOLF,NOD,NZB,NZO,PJ,RF,SEA,SPRET | 11 | 35733565 | 35733583 | 19 | + | 17.5574 | 6.32E-07 | 0.0841 | ACTCCAGAAGAGGGCGCCA | 3 | ENCFF367UNR,ENCFF093ZRU,ENCFF055HAP |
| 1658 | 11 | 37763540 | 37763744 | 204 | 129S1,AJ,B10D2,B10J,BALB,BTBR,BUB,C3H,C57L,CAST,CBA,DBA1J,DBA2J,LP,MA/My,MOLF,MRL,NZB,NZO,PJ,PL,SEA,SPRET,WSB | 11 | 37763618 | 37763636 | 19 | + | 17.8852 | 5.08E-07 | 0.0767 | ACACCAGAAGAGGGCACCA | | |
| 1659 | 11 | 42137572 | 42137791 | 219 | B10D2,B10J,BTBR,C58J,CAST,DBA2J,LP,MOLF,NZB,NZO,NZW,SPRET,WSB | 11 | 42137649 | 42137682 | 34 | + | 17.8455 | 3.91E-07 | 0.0425 | TAGCTGTCCTCAGACACTCCAGAAGAGGGCGTCA | | |
| 1660 | 11 | 44447425 | 44447633 | 208 | 129S1,AJ,AKR,B10D2,B10J,BALB,BTBR,BUB,C3H,C57L,C58J,CAST,CBA,CE,DBA1J,DBA2J,FVB,ILNJ2,KK,MA/My,MRL,NOD,NZB,NZO,NZW,PJ,PL,RF,RH2,SEA,SJL,SMJ,SWR.TH | 11 | 44447489 | 44447522 | 34 | + | 17.1382 | 6.20E-07 | 0.0547 | TAGCTGTCTTCACACGCACCAGAAGAGGGCATCA | 1 | ENCFF093ZRU |
| 1661 | 11 | 46694584 | 46694796 | 212 | CAST.MOLF.NZB.SPRET.WSB | 11 | 46694655 | 46694688 | 34 | + | 17.3252 | 5.49E-07 | 0.052 | TAGCTGTCCTCAGACATTCCAGAAGAGGGCATCA | | |
| 1662 | 11 | 48853378 | 48853455 | 77 | PJ.SEA | 11 | 48853367 | 48853400 | 34 | + | 17.252 | 5.76E-07 | 0.0529 | TAGCTGTCTTCCGACAGTCCAGAAGAGGGAGCCA | | |
| 1663 | 11 | 49701490 | 49701705 | 215 | SPRET | 11 | 49701549 | 49701582 | 34 | + | 17.2439 | 5.79E-07 | 0.0529 | TAGCTGTCCTCAGATACTCCAGAAGAGGGCATCA | | |
| 1664 | 11 | 50915627 | 50916611 | 984 | CAST.MOLF.NZO.SPRET | 11 | 50916291 | 50916325 | 35 | + | 20.4679 | 6.41E-08 | 0.0248 | AGGCAGGTGATTTTGGGGGCCAGTAGAGGGCACTA | | |
| 1664 | 11 | 50915627 | 50916611 | 984 | CAST.MOLF.NZO.SPRET | 11 | 50916307 | 50916325 | 19 | + | 22.0656 | 1.91E-08 | 0.0183 | GGGCCAGTAGAGGGCACTA | | |
| 1665 | 11 | 51555353 | 51555607 | 254 | SPRET | 11 | 51555560 | 51555594 | 35 | + | 17.2294 | 4.74E-07 | 0.0871 | CTGCACGTATACCTGCATGTCAGAAGAGGGTATCA | | |
| 1666 | 11 | 51609643 | 51610141 | 498 | SPRET | 11 | 51609745 | 51609778 | 34 | + | 16.9187 | 7.13E-07 | 0.0585 | CAGCAGTGCCCCTATGGTCCTGCAGAAGCAGGGG | | |
| 1667 | 11 | 51680833 | 51681440 | 607 | MOLF.NZO | 11 | 51681170 | 51681204 | 35 | + | 17.6881 | 3.62E-07 | 0.0746 | ATGTTGTCTGCCAAGGAGGCCAGAAGAGGGCATCT | | |
| 1668 | 11 | 53228923 | 53229131 | 208 | SPRET | 11 | 53228984 | 53229017 | 34 | + | 16.4472 | 9.61E-07 | 0.067 | TAGCTGTCTTCAGACACACCAGAAGAGGGCGTCA | | |
| 1669 | 11 | 53269370 | 53269588 | 218 | B10D2,B10J,C58J,CAST,CE,KK,MOLF,NOD,NOR,NZW,SPRET | 11 | 53269444 | 53269477 | 34 | + | 16.4472 | 9.61E-07 | 0.067 | TAGCTGTCTTCAGACACACCAGAAGAGGGCGTCA | | |
| 1670 | 11 | 53329737 | 53329929 | 192 | SPRET | 11 | 53329788 | 53329821 | 34 | + | 18.0732 | 3.36E-07 | 0.0394 | TAGCTGTCCTCAGGTGTACCAGAAGAGGGCATCG | | |
| 1671 | 11 | 53737507 | 53737704 | 197 | SPRET | 11 | 53737558 | 53737592 | 35 | + | 18.3945 | 2.37E-07 | 0.057 | CTGTAGCTGTCTTGACGCACCAGAAGAGGGCATCA | 2 | ENCFF367UNR.ENCFF883UPM |
| 1672 | 11 | 57608749 | 57608951 | 202 | 129S1,AJ,AKR,BALB,BTBR,BUB,C3H,C57L,CAST,CBA,CE,DBA1J,DBA2J,FVB,ILNJ2,KK,LP,MA/My,MRL,NOD,NZW,PJ,PL,RF,RH2,SEA,SJL,SMJ,SPRET,SWR,TH,WSB | 11 | 57608808 | 57608841 | 34 | + | 18.2195 | 3.05E-07 | 0.037 | TAGCTGTCTTCAGATACTCCAGAAGAGGGCGCCA | 3 | ENCFF093ZRU,ENCFF821BIA,ENCFF367UNR |
| 1672 | 11 | 57608749 | 57608951 | 202 | 129S1,AJ,AKR,BALB,BTBR,BUB,C3H,C57L,CAST,CBA,CE,DBA1J,DBA2J,FVB,ILNJ2,KK,LP,MA/My,MRL,NOD,NZW,PJ,PL,RF,RH2,SEA,SJL,SMJ,SPRET,SWR,TH,WSB | 11 | 57608823 | 57608841 | 19 | + | 17.5574 | 6.32E-07 | 0.0841 | ACTCCAGAAGAGGGCGCCA | 3 | ENCFF093ZRU,ENCFF821BIA,ENCFF367UNR |
| 1673 | 11 | 59883055 | 59883279 | 224 | AJ,AKR,C3H,CAST,CBA,CE,LP,MA/My,MOLF,MRL,NOD,PJ,PL,RF,SEA,SMJ,SPRET | 11 | 59883135 | 59883168 | 34 | + | 16.9187 | 7.13E-07 | 0.0585 | TAGCTGTCTTCAGACGCACCAGAAGAGGGCGTCA | | |
| 1674 | 11 | 62664365 | 62664558 | 193 | AJ,BUB,C3H,CAST,CBA,KK,MOLF,MRL,NZB,NZO,PJ,SEA | 11 | 62664410 | 62664443 | 34 | + | 16.4309 | 9.71E-07 | 0.0673 | TAGCTGTCTTCAGACGCTCCAGAAGAGGGCGTCA | | |
| 1675 | 11 | 68072580 | 68072781 | 201 | CAST.MOLF.SPRET | 11 | 68072639 | 68072672 | 34 | + | 16.4472 | 9.61E-07 | 0.067 | TAGCTGTCTTCAGACACACCAGAAGAGGGCGTCA | | |
| 1676 | 11 | 68731434 | 68731638 | 204 | 129S1,AJ,AKR,BTBR,BUB,C57L,C58J,CAST,DBA1J,DBA2J,ILNJ2,KK,LP,MOLF,NOR,NZB,NZO,NZW,PJ,PL,RF,SEA,SMJ,SPRET | 11 | 68731508 | 68731526 | 19 | + | 17.8852 | 5.08E-07 | 0.0767 | ACACCAGAAGAGGGCACCA | 1 | ENCFF367UNR |
| 1677 | 11 | 69083735 | 69084355 | 620 | SPRET | 11 | 69083816 | 69083850 | 35 | + | 24.1468 | 4.77E-09 | 0.00544 | TTGCAATTAAAGTGTTCTGCCACCAGGTGGTGGCC | 28 | ENCFF367UNR,ENCFF093ZRU,ENCFF201JUA,ENCFF099QNE,ENCFF821BIA,ENCFF024JOZ,ENCFF089GBC,ENCFF907XEG,ENCFF052KVO,ENCFF327LYS,ENCFF092NMJ,ENCFF055HAP,ENCFF198SSR,ENCFF502WDT,ENCFF430PPJ,ENCFF589ZBJ,ENCFF165LXO,ENCFF883UPM,ENCFF951GOP,ENCFF504JDS,ENCFF065ODV,ENCFF609MDN,ENCFF575MGI,ENCFF957SPH,ENCFF281QMV,ENCFF316FYN,ENCFF136ZDK,ENCFF194HVL |
| 1677 | 11 | 69083735 | 69084355 | 620 | SPRET | 11 | 69083832 | 69083850 | 19 | + | 19.6066 | 1.49E-07 | 0.0436 | CTGCCACCAGGTGGTGGCC | 29 | ENCFF367UNR,ENCFF093ZRU,ENCFF201JUA,ENCFF099QNE,ENCFF821BIA,ENCFF024JOZ,ENCFF089GBC,ENCFF907XEG,ENCFF052KVO,ENCFF327LYS,ENCFF092NMJ,ENCFF055HAP,ENCFF198SSR,ENCFF502WDT,ENCFF430PPJ,ENCFF589ZBJ,ENCFF165LXO,ENCFF883UPM,ENCFF951GOP,ENCFF504JDS,ENCFF065ODV,ENCFF609MDN,ENCFF575MGI,ENCFF957SPH,ENCFF281QMV,ENCFF316FYN,ENCFF136ZDK,ENCFF194HVL.ENCFF142CNG |
| 1678 | 11 | 69141628 | 69142047 | 419 | SPRET | 11 | 69141977 | 69142011 | 35 | + | 18.4862 | 2.24E-07 | 0.0548 | GGGCAGGTACCTCTGGAAGCCAGAAGAGGACACTG | | |
| 1679 | 11 | 70372610 | 70372816 | 206 | CAST,DBA1J,DBA2J,FVB,MOLF,NOR,NZW,PJ,SEA,SMJ,SPRET | 11 | 70372674 | 70372707 | 34 | + | 17.2439 | 5.79E-07 | 0.0529 | TAGCTGTCCTCAGATACTCCAGAAGAGGGCATCA | | |
| 1680 | 11 | 72469676 | 72469887 | 211 | CE.SPRET | 11 | 72469744 | 72469777 | 34 | + | 16.4472 | 9.61E-07 | 0.067 | TAGCTGTCTTCAGACACACCAGAAGAGGGCGTCA | | |
| 1681 | 11 | 76236826 | 76237046 | 220 | 129S1,AJ,AKR,BALB,BTBR,BUB,C3H,C57L,C58J,CAST,CBA,CE,DBA1J,DBA2J,FVB,ILNJ2,KK,MA/My,MOLF,MRL,NOR,NZO,PJ,PL,RF,RH2,SEA,SJL,SMJ,SPRET,SWR,WSB | 11 | 76236905 | 76236938 | 34 | + | 17.8455 | 3.91E-07 | 0.0425 | TAGCTGTCCTCAGACACTCCAGAAGAGGGCGTCA | | |
| 1682 | 11 | 76848711 | 76848876 | 165 | SPRET | 11 | 76848873 | 76848906 | 34 | + | 22.4715 | 1.38E-08 | 0.00459 | GTGCAGTGTCTGAGGACACCAGAAGAGGGTGTCA | 1 | ENCFF093ZRU |
| 1683 | 11 | 77136810 | 77137012 | 202 | CAST.MOLF.SPRET | 11 | 77136870 | 77136903 | 34 | + | 17.8455 | 3.91E-07 | 0.0425 | TAGCTGTCCTCAGACACTCCAGAAGAGGGCGTCA | | |
| 1684 | 11 | 77665843 | 77666027 | 184 | CAST.SPRET | 11 | 77665877 | 77665910 | 34 | + | 17.2439 | 5.79E-07 | 0.0529 | TAGCTGTCCTCAGATACTCCAGAAGAGGGCATCA | 1 | ENCFF093ZRU |
| 1685 | 11 | 77680127 | 77680340 | 213 | CAST.CE.MOLF.SPRET | 11 | 77680199 | 77680232 | 34 | + | 16.5203 | 9.18E-07 | 0.067 | TAGCTGTCTTCCAACACTCCAGAAGAGGGCGTCA | | |
| 1686 | 11 | 78193945 | 78194775 | 830 | SPRET | 11 | 78194387 | 78194420 | 34 | + | 18.813 | 2.04E-07 | 0.0312 | TAGCTGTTTTCAGTCACACCAGAAGGGGGCATCG | 1 | ENCFF093ZRU |
| 1687 | 11 | 78350527 | 78351105 | 578 | CAST | 11 | 78350698 | 78350731 | 34 | + | 16.7724 | 7.83E-07 | 0.0614 | TAGCTGTCTTCAGATGCACCAGAAGAGGGCGTCA | | |
| 1688 | 11 | 78456063 | 78456273 | 210 | CAST.MOLF.SPRET | 11 | 78456130 | 78456163 | 34 | + | 16.4634 | 9.51E-07 | 0.067 | TAGCTGTCTTCAGACGCACCAGAAGAGGGCATCA | | |
| 1689 | 11 | 78797431 | 78797626 | 195 | CAST.CE.FVB.MOLF.SMJ.SPRET.WSB | 11 | 78797485 | 78797518 | 34 | + | 17.3902 | 5.27E-07 | 0.0504 | TAGCTGTCCTCAGACACTCCAGAAGAGGGCATCA | | |
| 1690 | 11 | 79817296 | 79817500 | 204 | SPRET | 11 | 79817358 | 79817391 | 34 | + | 16.4472 | 9.61E-07 | 0.067 | TAGCTGTCTTCAGACACACCAGAAGAGGGCGTCA | | |
| 1691 | 11 | 79930254 | 79930432 | 178 | CAST.CE.MOLF.SPRET.TH | 11 | 79930290 | 79930323 | 34 | + | 17.8455 | 3.91E-07 | 0.0425 | TAGCTGTCCTCAGACACTCCAGAAGAGGGCGTCA | | |
| 1692 | 11 | 80015549 | 80015755 | 206 | SPRET | 11 | 80015613 | 80015646 | 34 | + | 16.4634 | 9.51E-07 | 0.067 | TAGCTGTCTTCAGACGCACCAGAAGAGGGCATCA | | |
| 1693 | 11 | 80019020 | 80019252 | 232 | CAST.MOLF.SPRET | 11 | 80019107 | 80019140 | 34 | + | 17.2439 | 5.79E-07 | 0.0529 | TAGCTGTCCTCAGATACTCCAGAAGAGGGCATCA | | |
| 1694 | 11 | 80040042 | 80040224 | 182 | CAST.CE.MOLF.SPRET | 11 | 80040081 | 80040114 | 34 | + | 16.4472 | 9.61E-07 | 0.067 | TAGCTGTCTTCAGACACACCAGAAGAGGGCGTCA | | |
| 1695 | 11 | 80514952 | 80515176 | 224 | AKR,BALB,C58J,CAST,MOLF,NOR,RF,RH2,SMJ.SPRET.TH.WSB | 11 | 80515034 | 80515067 | 34 | + | 18.3659 | 2.76E-07 | 0.0355 | TAGCTGTCTTCAGACACTCCAGAAGAGGGCGCCA | | |
| 1695 | 11 | 80514952 | 80515176 | 224 | AKR,BALB,C58J,CAST,MOLF,NOR,RF,RH2,SMJ.SPRET.TH.WSB | 11 | 80515049 | 80515067 | 19 | + | 17.5574 | 6.32E-07 | 0.0841 | ACTCCAGAAGAGGGCGCCA | | |
| 1696 | 11 | 80863039 | 80863304 | 265 | AKR.CAST.MOLF.RF.SPRET.TH.WSB | 11 | 80863163 | 80863196 | 34 | + | 16.7724 | 7.83E-07 | 0.0614 | TAGCTGTCTTCAGATGCACCAGAAGAGGGCGTCA | | |
| 1697 | 11 | 82443715 | 82443889 | 174 | AKR,C58J,CAST,MA/My,MOLF,RF,SPRET | 11 | 82443747 | 82443780 | 34 | + | 16.4472 | 9.61E-07 | 0.067 | TAGCTGTCTTCAGACACACCAGAAGAGGGCGTCA | | |
| 1698 | 11 | 84668923 | 84669656 | 733 | CAST | 11 | 84669501 | 84669535 | 35 | + | 17.6239 | 3.76E-07 | 0.0758 | CTGTATATATGCCTGCATGCCAGAAGAGAGCGCCA | | |
| 1699 | 11 | 85499990 | 85500098 | 108 | SPRET | 11 | 85500031 | 85500064 | 34 | + | 20.813 | 4.91E-08 | 0.0119 | GTGCAGTACCATCAGAGCCCAGAAAAGGGCATCA | 2 | ENCFF367UNR.ENCFF093ZRU |
| 1700 | 11 | 86145711 | 86145916 | 205 | CAST.MOLF.SPRET | 11 | 86145774 | 86145807 | 34 | + | 17.2439 | 5.79E-07 | 0.0529 | TAGCTGTCCTCAGATACTCCAGAAGAGGGCATCA | | |
| 1701 | 11 | 86524972 | 86525193 | 221 | 129S1,AJ,AKR,BTBR,C3H,C57L,C58J,CAST,CBA,DBA1J,DBA2J,KK,MOLF,MRL,NZB,NZO.PL.RF.SJL.SPRET.SWR | 11 | 86525049 | 86525082 | 34 | + | 16.4634 | 9.51E-07 | 0.067 | TAGCTGTCTTCAGACGCACCAGAAGAGGGCATCA | | |
| 1702 | 11 | 86726154 | 86727029 | 875 | 129S1,AJ,AKR,BTBR,BUB,C3H,C57L,C58J,CBA,DBA1J,DBA2J,KK,MOLF,MRL,NZW,PJ,PL,RF,SEA,SJL,SWR | 11 | 86726778 | 86726811 | 34 | + | 17.1951 | 5.97E-07 | 0.0536 | GTGCAGTGCACCTGAAGGCCAGAATGGGGAGTCA | | |
| 1703 | 11 | 86734243 | 86734450 | 207 | 129S1,AJ,AKR,BTBR,BUB,C3H,C57L,C58J,CAST,CBA,DBA1J,DBA2J,KK,MRL,NZW,PJ,PL,RF,SEA,SJL,SPRET,SWR | 11 | 86734308 | 86734341 | 34 | + | 16.4472 | 9.61E-07 | 0.067 | TAGCTGTCTTCAGACACACCAGAAGAGGGCGTCA | | |
| 1704 | 11 | 87436657 | 87436886 | 229 | MOLF.SPRET | 11 | 87436695 | 87436728 | 34 | + | 17.8455 | 3.91E-07 | 0.0425 | TAGCTGTCCTCAGACACTCCAGAAGAGGGCGTCA | | |
| 1705 | 11 | 87953735 | 87953981 | 246 | 129S1,AJ,AKR,BALB,BTBR,BUB,C3H,C57L,C58J,CBA,CE,DBA1J,DBA2J,FVB,ILNJ2,KK,MA/My,MRL,NOD,NOR,NZB,NZO,NZW,PJ,PL,RF,RH2,SEA,SJL,SMJ,SPRET,SWR,TH,WSB | 11 | 87953838 | 87953871 | 34 | + | 16.4634 | 9.51E-07 | 0.067 | TAGCTGTCTTCAGACGCACCAGAAGAGGGCATCA | | |
| 1706 | 11 | 88300082 | 88300278 | 196 | SPRET | 11 | 88300136 | 88300169 | 34 | + | 16.7724 | 7.83E-07 | 0.0614 | TAGCTGTCTTCAGATGCACCAGAAGAGGGCGTCA | | |
| 1707 | 11 | 88390176 | 88390370 | 194 | CAST.SPRET | 11 | 88390228 | 88390261 | 34 | + | 17.8293 | 3.95E-07 | 0.0428 | TAGCTGTCTTCAGACCCTCCAGAAGAGGGCGTCA | 3 | ENCFF367UNR.ENCFF093ZRU.ENCFF024JOZ |
| 1708 | 11 | 88852579 | 88852780 | 201 | 129S1,AJ,AKR,BALB,BTBR,BUB,C3H,C57L,CBA,CE,DBA1J,DBA2J,FVB,ILNJ2,KK,MA/My,MOLF,MRL,NOD,NOR,NZB,NZO,NZW,PJ,PL,RF,RH2,SEA,SJL,SMJ,SPRET,SWR,TH,WSB | 11 | 88852638 | 88852671 | 34 | + | 17.6667 | 4.40E-07 | 0.0454 | TAGCTGTCTTCAGACACGCCAGAAGAGGGCGTCA | | |
| 1709 | 11 | 89085554 | 89085769 | 215 | 129S1,AJ,AKR,BALB,BTBR,BUB,C3H,C57L,CBA,CE,DBA1J,DBA2J,FVB,ILNJ2,KK,MA/My,MRL,NOD,NOR,NZB,NZO,NZW,PJ,PL,RF,RH2,SEA,SJL,SMJ,SPRET,SWR,TH,WSB | 11 | 89085627 | 89085660 | 34 | + | 17.3902 | 5.27E-07 | 0.0504 | TAGCTGTCCTCAGACACTCCAGAAGAGGGCATCA | | |
| 1710 | 11 | 94975490 | 94975698 | 208 | AJ,AKR,BUB,C3H,CAST,CBA,MOLF,SPRET | 11 | 94975554 | 94975587 | 34 | + | 16.7724 | 7.83E-07 | 0.0614 | TAGCTGTCTTCAGATGCACCAGAAGAGGGCGTCA | | |
| 1711 | 11 | 95218800 | 95218970 | 170 | SPRET | 11 | 95218835 | 95218868 | 34 | + | 17.6992 | 4.30E-07 | 0.0449 | TAGCTGTCCTCAGATACTCCAGAAGAGGGCGTCA | 1 | ENCFF367UNR |
| 1712 | 11 | 95624062 | 95624270 | 208 | AJ,AKR,BUB,C3H,CBA,MOLF,RH2,SPRET | 11 | 95624128 | 95624161 | 34 | + | 17.878 | 3.83E-07 | 0.0424 | TAGCTGTCCTCAGACACACCAGAAGAGGGCATCA | | |
| 1713 | 11 | 95922564 | 95922784 | 220 | MOLF | 11 | 95922663 | 95922696 | 34 | + | 18.7398 | 2.14E-07 | 0.0321 | CAGCTCTACTCCCAGCAACCACATGGTGGCTCAC | 14 | ENCFF367UNR,ENCFF093ZRU,ENCFF089GBC,ENCFF092NMJ,ENCFF136ZDK,ENCFF589ZBJ,ENCFF502WDT,ENCFF198SSR,ENCFF883UPM,ENCFF055HAP,ENCFF907XEG,ENCFF024JOZ,ENCFF533APC,ENCFF327LYS |
| 1714 | 11 | 96654375 | 96654582 | 207 | AJ,AKR,BALB,BUB,CAST,CBA,MOLF,MRL,NOD,NOR,NZW,PL,RH2,SJL,SMJ,SPRET,WSB | 11 | 96654440 | 96654473 | 34 | + | 17.8455 | 3.91E-07 | 0.0425 | TAGCTGTCCTCAGACACTCCAGAAGAGGGCGTCA | 1 | ENCFF093ZRU |
| 1715 | 11 | 97321431 | 97322161 | 730 | SPRET | 11 | 97321620 | 97321653 | 34 | + | 18.187 | 3.12E-07 | 0.0374 | TTGCTGTCTCCAGACACACCTGAAGAGGGCATTG | | |
| 1716 | 11 | 97605281 | 97605514 | 233 | SPRET | 11 | 97605394 | 97605427 | 34 | + | 18.8537 | 1.98E-07 | 0.0308 | ACGCAAGTATGCCTGTGGCCAGAAGAGGGCATCA | 16 | ENCFF821BIA,ENCFF089GBC,ENCFF092NMJ,ENCFF907XEG,ENCFF024JOZ,ENCFF430PPJ,ENCFF093ZRU,ENCFF589ZBJ,ENCFF136ZDK,ENCFF198SSR,ENCFF367UNR,ENCFF883UPM,ENCFF099QNE,ENCFF055HAP,ENCFF502WDT,ENCFF327LYS |
| 1716 | 11 | 97605281 | 97605514 | 233 | SPRET | 11 | 97605409 | 97605427 | 19 | + | 18.7705 | 2.75E-07 | 0.0587 | TGGCCAGAAGAGGGCATCA | 16 | ENCFF821BIA,ENCFF089GBC,ENCFF092NMJ,ENCFF907XEG,ENCFF024JOZ,ENCFF430PPJ,ENCFF093ZRU,ENCFF589ZBJ,ENCFF136ZDK,ENCFF198SSR,ENCFF367UNR,ENCFF883UPM,ENCFF099QNE,ENCFF055HAP,ENCFF502WDT,ENCFF327LYS |
| 1717 | 11 | 97610099 | 97610455 | 356 | NZW.SWR | 11 | 97610285 | 97610319 | 35 | + | 16.4312 | 7.50E-07 | 0.111 | GTGCAGCGCCCAGCAGCCCCCGGGGGAAGGCAGCC | | |
| 1718 | 11 | 98068810 | 98069020 | 210 | 129S1,BTBR,C3H,C57L,CAST,CE,DBA1J,DBA2J,FVB,ILNJ2,KK,MA/My,MOLF,NOD,NOR,NZO,NZW,SPRET,SWR,WSB | 11 | 98068879 | 98068912 | 34 | + | 17.2439 | 5.79E-07 | 0.0529 | TAGCTGTCCTCAGATACTCCAGAAGAGGGCATCA | | |
| 1719 | 11 | 98100719 | 98100916 | 197 | SPRET | 11 | 98100764 | 98100797 | 34 | + | 18.3659 | 2.76E-07 | 0.0355 | TAGCTGTCTTCAGACACTCCAGAAGAGGGCGCCA | 2 | ENCFF093ZRU |
| 1719 | 11 | 98100719 | 98100916 | 197 | SPRET | 11 | 98100779 | 98100797 | 19 | + | 17.5574 | 6.32E-07 | 0.0841 | ACTCCAGAAGAGGGCGCCA | 2 | ENCFF093ZRU |
| 1720 | 11 | 98133839 | 98134051 | 212 | SPRET | 11 | 98133908 | 98133941 | 34 | + | 18.3659 | 2.76E-07 | 0.0355 | TAGCTGTCTTCAGACACTCCAGAAGAGGGCGCCA | 5 | ENCFF821BIA.ENCFF093ZRU.ENCFF367UNR.ENCFF201JUA.ENCFF099QNE |
| 1720 | 11 | 98133839 | 98134051 | 212 | SPRET | 11 | 98133923 | 98133941 | 19 | + | 17.5574 | 6.32E-07 | 0.0841 | ACTCCAGAAGAGGGCGCCA | 5 | ENCFF821BIA.ENCFF093ZRU.ENCFF367UNR.ENCFF201JUA.ENCFF099QNE |
| 1721 | 11 | 98575478 | 98575693 | 215 | SPRET | 11 | 98575498 | 98575531 | 34 | + | 20.561 | 5.91E-08 | 0.0136 | GTGCAGTGCCCACCAAGGCCAAAAGAGGTCATCT | 3 | ENCFF093ZRU.ENCFF883UPM.ENCFF502WDT |
| 1722 | 11 | 99626493 | 99626846 | 353 | MOLF | 11 | 99626703 | 99626736 | 34 | + | 17.8293 | 3.95E-07 | 0.0428 | ATGCAGCTACAGACATGAGCTCTGGGGGGTACTG | | |

| | | | | | | | | | | | | | | | | |
|---|---|---|---|---|---|---|---|---|---|---|---|---|---|---|---|---|
| 1723 | 11 | 101360474 | 101360697 | 223 | 129S1,AJ,BTBR,BUB,C3H,C57L,CAST,CE,DBA1J,DBA2J,ILNJ2,KK,MOLF,MRL,NOD,NOR.NZB.NZO.SJL.SPRET.TH.WSB | 11 | 101360551 | 101360584 | 34 | + | 17.2439 | 5.79E-07 | 0.0529 | TAGCTGTCCTCAGACCCTCCAGAAGAGGGAGTCA | | |
| 1724 | 11 | 103855628 | 103855850 | 222 | SPRET | 11 | 103855708 | 103855741 | 34 | + | 16.4634 | 9.51E-07 | 0.067 | TAGCTGTCTTCAGACGCACCAGAAGAGGGCATCA | | |
| 1725 | 11 | 104151342 | 104151545 | 203 | CAST.SPRET | 11 | 104151397 | 104151430 | 34 | + | 16.626 | 8.59E-07 | 0.065 | TAGCTGTCTTCAGACACCCCAGCAGAGGGCATCA | | |
| 1726 | 11 | 106504836 | 106504888 | 52 | SPRET | 11 | 106504857 | 106504875 | 19 | + | 19.4426 | 1.69E-07 | 0.0459 | ATTCCAGTAGGGGGCGCTC | 18 | ENCFF093ZRU,ENCFF367UNR,ENCFF136ZDK,ENCFF327LYS,ENCFF805QBU,ENCFF099QNE,ENCFF907XEG,ENCFF089GBC,ENCFF198SSR,ENCFF210BTO,ENCFF883UPM,ENCFF589ZBJ,ENCFF065ODV,ENCFF055HAP.ENCFF821BIA.ENCFF502WDT.ENCFF052KVO.ENCFF142CNG |
| 1727 | 11 | 106656583 | 106657373 | 790 | SPRET | 11 | 106656676 | 106656709 | 34 | + | 17.2276 | 5.85E-07 | 0.0531 | TAGCTGTCTTCAGACACACCAGAAGAGGGCACAG | | |
| 1728 | 11 | 106856057 | 106856408 | 351 | CAST | 11 | 106856285 | 106856318 | 34 | + | 16.7724 | 7.83E-07 | 0.0614 | TAGCTGTATTCAGACACACCACAAGAGGGCATCA | | |
| 1729 | 11 | 107016346 | 107016551 | 205 | BUB.CAST.ILNJ2.RF.SJL.SPRET.TH | 11 | 107016409 | 107016442 | 34 | + | 17.8455 | 3.91E-07 | 0.0425 | TAGCTGTCCTCAGACACTCCAGAAGAGGGCGTCA | | |
| 1730 | 11 | 107500050 | 107500257 | 207 | SPRET | 11 | 107500107 | 107500140 | 34 | + | 16.5691 | 8.90E-07 | 0.0664 | TAGCTGTCTTCAGACGCACCAGAAGAGGGCATCG | | |
| 1731 | 11 | 107506814 | 107507028 | 214 | 129S1,BUB,CAST,CBA,NZW,SMJ,SPRET,TH.WSB | 11 | 107506886 | 107506919 | 34 | + | 16.4634 | 9.51E-07 | 0.067 | TAGCTGTCTTCAGACGCACCAGAAGAGGGCATCA | | |
| 1732 | 11 | 110083843 | 110084030 | 187 | 129S1,AJ,BTBR,C57L,CBA,CE,FVB,ILNJ2,MA/My,MOLF,NOD,NOR,NZO,NZW,SJL,SMJ,SPRET.SWR.TH | 11 | 110083888 | 110083921 | 34 | + | 17.2439 | 5.79E-07 | 0.0529 | TAGCTGTCCTCAGATACTCCAGAAGAGGGCATCA | | |
| 1733 | 11 | 111095926 | 111096506 | 580 | SPRET | 11 | 111096172 | 111096205 | 34 | + | 18.3089 | 2.87E-07 | 0.0357 | ATGCAGCTAGAGACACGAGCTCCAGGGGGTACTG | | |
| 1734 | 11 | 114497269 | 114497482 | 213 | 129S1,AJ,BALB,C3H,C57L,C58J,CAST,CBA,CE,ILNJ2,KK,MA/My,MOLF,MRL,NOD,NOR,NZB,NZO,PL,RF,SEA,SMJ,SPRET,TH,WSB | 11 | 114497341 | 114497374 | 34 | + | 18.3659 | 2.76E-07 | 0.0355 | TAGCTGTCTTCAGACACTCCAGAAGAGGGCGCCA | | |
| 1734 | 11 | 114497269 | 114497482 | 213 | 129S1,AJ,BALB,C3H,C57L,C58J,CAST,CBA,CE,ILNJ2,KK,MA/My,MOLF,MRL,NOD,NOR,NZB,NZO,PL,RF,SEA,SMJ,SPRET,TH,WSB | 11 | 114497356 | 114497374 | 19 | + | 17.5574 | 6.32E-07 | 0.0841 | ACTCCAGAAGAGGGCGCCA | | |
| 1735 | 11 | 114748153 | 114748599 | 446 | AJ.C3H.C57L.CBA.MA/My.PL | 11 | 114748484 | 114748502 | 19 | + | 17.4426 | 6.81E-07 | 0.0875 | AGGCCAGAAGGGGGCATTA | | |
| 1736 | 11 | 115419818 | 115420037 | 219 | 129S1,BALB,C58J,DBA1J,DBA2J,KK,MOLF,NOD,NOR,NZB,NZO,NZW,PL,RF,SEA,SMJ,SPRET.TH | 11 | 115419894 | 115419927 | 34 | + | 17.2439 | 5.79E-07 | 0.0529 | TAGCTGTCCTCAGATACTCCAGAAGAGGGCATCA | | |
| 1737 | 11 | 115966202 | 115966435 | 233 | AJ,C3H,C57L,CAST,CBA,CE,ILNJ2,MOLF,SPRET.WSB | 11 | 115966291 | 115966324 | 34 | + | 16.4634 | 9.51E-07 | 0.067 | TAGCTGTCTTCAGACGCACCAGAAGAGGGCATCA | | |
| 1738 | 11 | 115981177 | 115981386 | 209 | AJ,BTBR,C3H,C57L,CAST,CBA,CE,ILNJ2,MOLF.SPRET.WSB | 11 | 115981244 | 115981277 | 34 | + | 16.8537 | 7.43E-07 | 0.0602 | TAGCTGTCCTTAGATACTCCAGAAGAGGGCATCA | | |
| 1739 | 11 | 116177438 | 116177971 | 533 | AJ,BALB,BTBR,C3H,C57L,CAST,CBA,CE,DBA1J,DBA2J,ILNJ2,MOLF,SEA,SPRET | 11 | 116177579 | 116177613 | 35 | + | 15.9725 | 9.70E-07 | 0.125 | AGGCCTTGTGCCCTCCCTGCCTCCAGATGGTACAC | | |
| 1740 | 11 | 116623779 | 116624008 | 229 | 129S1,AJ,AKR,BTBR,BUB,C3H,C57L,C58J,CBA,CE,FVB,ILNJ2,KK,MA/My,MRL,NOD,NOR,NZB,NZO,NZW,PL,RF,RH2,SJL,SMJ,SPRET.SWR.TH.WSB | 11 | 116623839 | 116623873 | 35 | + | 17.2661 | 4.64E-07 | 0.0861 | CTGTAGCTATCCAGACACTCCAGAAGAGGGAGTCA | | |
| 1741 | 11 | 116690786 | 116690997 | 211 | CAST.MOLF.SPRET | 11 | 116690855 | 116690888 | 34 | + | 16.4472 | 9.61E-07 | 0.067 | TAGCTGTCTTCAGACACACCAGAAGAGGGCGTCA | | |
| 1742 | 11 | 116929635 | 116929850 | 215 | CAST | 11 | 116929708 | 116929741 | 34 | + | 18.3659 | 2.76E-07 | 0.0355 | TAGCTGTCTTCAGACACTCCAGAAGAGGGCGCCA | 9 | ENCFF367UNR,ENCFF099QNE,ENCFF024JOZ,ENCFF055HAP,ENCFF327LYS,ENCFF093ZRU,ENCFF821BIA,ENCFF907XEG |
| 1742 | 11 | 116929635 | 116929850 | 215 | CAST | 11 | 116929723 | 116929741 | 19 | + | 17.5574 | 6.32E-07 | 0.0841 | ACTCCAGAAGAGGGCGCCA | 9 | ENCFF367UNR,ENCFF099QNE,ENCFF024JOZ,ENCFF055HAP,ENCFF327LYS,ENCFF093ZRU,ENCFF821BIA,ENCFF907XEG |
| 1743 | 11 | 116944895 | 116945095 | 200 | SPRET | 11 | 116944954 | 116944987 | 34 | + | 17.2439 | 5.79E-07 | 0.0529 | TAGCTGTCCTCAGATACTCCAGAAGAGGGCATCA | | |
| 1744 | 11 | 116964584 | 116964797 | 213 | SPRET | 11 | 116964655 | 116964688 | 34 | + | 17.2439 | 5.79E-07 | 0.0529 | TAGCTGTCCTCAGATACTCCAGAAGAGGGCATCA | | |
| 1745 | 11 | 117307938 | 117308137 | 199 | CAST,MOLF | 11 | 117307962 | 117307995 | 34 | + | 25.3008 | 1.27E-09 | 0.000812 | ATGCAGTTTCTAAGGAAGCCAGAAGAGGGCGTTG | 10 | ENCFF093ZRU,ENCFF089GBC,ENCFF092NMJ,ENCFF907XEG,ENCFF430PPJ,ENCFF327LYS,ENCFF502WDT.ENCFF055HAP.ENCFF065ODV.ENCFF052KVO |
| 1746 | 11 | 119964213 | 119964281 | 68 | BALB,BTBR,C3H,C57L,C58J,CAST,CBA,DBA1J,DBA2J,ILNJ2,MOLF,NZB,NZW,SEA,SJL.SPRET.WSB | 11 | 119964269 | 119964303 | 35 | + | 16 | 9.55E-07 | 0.124 | CAGCTCTGTAGCACTTGGCCCAGCAGAGGGCCAGA | 11 | ENCFF093ZRU,ENCFF136ZDK,ENCFF198SSR,ENCFF883UPM,ENCFF589ZBJ,ENCFF165LXO,ENCFF327LYS,ENCFF089GBC,ENCFF024JOZ,ENCFF907XEG,ENCFF092NMJ |
| 1747 | 11 | 120304680 | 120304890 | 210 | CAST.MOLF.SJL.SPRET | 11 | 120304748 | 120304781 | 34 | + | 18.3659 | 2.76E-07 | 0.0355 | TAGCTGTCTTCAGACACTCCAGAAGAGGGCGCCA | 4 | ENCFF055HAP.ENCFF367UNR.ENCFF093ZRU.ENCFF821BIA |
| 1747 | 11 | 120304680 | 120304890 | 210 | CAST.MOLF.SJL.SPRET | 11 | 120304763 | 120304781 | 19 | + | 17.5574 | 6.32E-07 | 0.0841 | ACTCCAGAAGAGGGCGCCA | 4 | ENCFF055HAP.ENCFF367UNR.ENCFF093ZRU.ENCFF821BIA |
| 1748 | 11 | 3776948 | 3777149 | 201 | B10D2.B10J.C58J.CAST.KK.SPRET | 11 | 3777060 | 3777093 | 34 | - | 17.1545 | 6.13E-07 | 0.0544 | TAGCTGTCTTCCAACAGTCCAGAAGAGGGAGCCA | 1 | ENCFF093ZRU |
| 1749 | 11 | 4063836 | 4064041 | 205 | NZB.SPRET | 11 | 4063961 | 4063994 | 34 | - | 18.0732 | 3.36E-07 | 0.0394 | CAGCTGTCTTCAGACACTCCAGAAGAGGGCATCA | | |
| 1750 | 11 | 6157813 | 6158017 | 204 | SPRET | 11 | 6157938 | 6157971 | 34 | - | 16.4472 | 9.61E-07 | 0.067 | TAGCTGTCTTCAGACACACCAGAAGAGGGCGTCA | | |
| 1751 | 11 | 6374252 | 6374450 | 198 | AJ,AKR,B10D2,B10J,BALB,BTBR,C3H,C57L,C58J,CAST,MA/My,MRL,NOD,NOR,PL,RF,SEA.SPRET.TH.WSB | 11 | 6374375 | 6374408 | 34 | - | 17.2439 | 5.79E-07 | 0.0529 | TAGCTGTCCTCAGATACTCCAGAAGAGGGCATCA | | |
| 1752 | 11 | 7394518 | 7394820 | 302 | SPRET | 11 | 7394686 | 7394719 | 34 | - | 27.5447 | 1.50E-10 | 0.000227 | CTGCAGTGCCCATAGAAGCCAGAAGGGGGTGTCA | 2 | ENCFF093ZRU.ENCFF052KVO |
| 1753 | 11 | 11712655 | 11712862 | 207 | 129S1,BALB,BUB,C57L,C58J,CAST,CBA,CE,DBA1J,DBA2J,FVB,ILNJ2,MA/My,NZO,PJ,PL,SEA,SJL,SMJ,SPRET,SWR,TH,WSB | 11 | 11712781 | 11712814 | 34 | - | 16.4634 | 9.51E-07 | 0.067 | TAGCTGTCTTCAGACGCACCAGAAGAGGGCATCA | | |
| 1754 | 11 | 12373868 | 12374081 | 213 | SPRET | 11 | 12373995 | 12374013 | 19 | - | 17.5738 | 6.25E-07 | 0.0841 | ACTCCAGAAGAGGGCGCTA | 1 | ENCFF093ZRU |
| 1754 | 11 | 12373868 | 12374081 | 213 | SPRET | 11 | 12373995 | 12374028 | 34 | - | 18.187 | 3.12E-07 | 0.0374 | TAGCTGTCTTCAGTCACTCCAGAAGAGGGCGCTA | 1 | ENCFF093ZRU |
| 1755 | 11 | 16013595 | 16013790 | 195 | SPRET | 11 | 16013715 | 16013748 | 34 | - | 17.8455 | 3.91E-07 | 0.0425 | TAGCTGTCCTCAGACACTCCAGAAGAGGGCGTCA | | |
| 1756 | 11 | 16436602 | 16437347 | 745 | MOLF | 11 | 16437146 | 16437179 | 34 | - | 20.252 | 7.40E-08 | 0.0159 | ATGCAGTTAGAGACACGAGCTCCAGGGGGTACTG | | |
| 1757 | 11 | 17899549 | 17899772 | 223 | 129S1,AJ,AKR,BALB,BTBR,BUB,C3H,C57L,C58J,CAST,CBA,CE,DBA1J,DBA2J,FVB,ILNJ2,KK,MA/My,MOLF,MRL,NOD,NOR,NZB,NZO,NZW,PJ,PL,RF,SEA,SJL,SMJ,SPRET,SWR.TH.WSB | 11 | 17899677 | 17899710 | 34 | - | 16.4634 | 9.51E-07 | 0.067 | TAGCTGTCTTCAGACGCACCAGAAGAGGGCATCA | | |
| 1758 | 11 | 19755515 | 19756011 | 496 | SPRET | 11 | 19755813 | 19755847 | 35 | - | 22.0826 | 2.15E-08 | 0.0126 | GTGCAGGTGCCGATGGAGGCCAGAAGAGGGTGTTG | | |
| 1759 | 11 | 20020764 | 20021012 | 248 | AJ,AKR,BALB,C57L,CAST,CE,FVB,ILNJ2,KK,MA/My,MOLF,NOD,NOR,NZB,NZO,NZW,PL,RF,RH2,SEA,SPRET,SWR,TH,WSB | 11 | 20020891 | 20020924 | 34 | - | 16.4634 | 9.51E-07 | 0.067 | TAGCTGTCTTCAGACGCACCAGAAGAGGGCATCA | 1 | ENCFF093ZRU |
| 1760 | 11 | 23773895 | 23774105 | 210 | CAST.MOLF.SPRET.WSB | 11 | 23774019 | 23774037 | 19 | - | 17.5574 | 6.32E-07 | 0.0841 | ACTCCAGAAGAGGGCGCCA | | |
| 1760 | 11 | 23773895 | 23774105 | 210 | CAST.MOLF.SPRET.WSB | 11 | 23774019 | 23774052 | 34 | - | 18.3659 | 2.76E-07 | 0.0355 | TAGCTGTCTTCAGACACTCCAGAAGAGGGCGCCA | | |
| 1761 | 11 | 27013216 | 27013414 | 198 | AJ,BUB,C3H,C58J,CAST,CBA,DBA1J,DBA2J,ILNJ2,KK,MOLF,MRL,NZB,NZO,NZW,PL,RF.SMJ.SPRET.TH | 11 | 27013341 | 27013359 | 19 | - | 17.5574 | 6.32E-07 | 0.0841 | ACTCCAGAAGAGGGCGCCA | | |
| 1761 | 11 | 27013216 | 27013414 | 198 | AJ,BUB,C3H,C58J,CAST,CBA,DBA1J,DBA2J,ILNJ2,KK,MOLF,MRL,NZB,NZO,NZW,PL,RF.SMJ.SPRET.TH | 11 | 27013341 | 27013374 | 34 | - | 18.3659 | 2.76E-07 | 0.0355 | TAGCTGTCTTCAGACACTCCAGAAGAGGGCGCCA | | |
| 1762 | 11 | 30518373 | 30518594 | 221 | AJ,AKR,BALB,BTBR,BUB,C57L,CAST,CBA,CE,DBA1J,DBA2J,FVB,ILNJ2,NOD,NZB,NZO,PJ,PL,RF,RH2,SEA,SJL,SMJ,SPRET,SWR.TH.WSB | 11 | 30518498 | 30518516 | 19 | - | 17.0328 | 8.88E-07 | 0.0977 | TCACCAGAAGAGGGCGTCA | | |
| 1763 | 11 | 32122666 | 32122876 | 210 | SPRET | 11 | 32122790 | 32122823 | 34 | - | 16.4472 | 9.61E-07 | 0.067 | TAGCTGTCTTCAGACACACCAGAAGAGGGCGTCA | | |
| 1764 | 11 | 32547246 | 32547443 | 197 | AKR,BUB,CE,ILNJ2,KK,MA/My,NZW,PJ,PL,RF.SEA.SPRET | 11 | 32547366 | 32547399 | 34 | - | 16.4472 | 9.61E-07 | 0.067 | TAGCTGTCTTCAGACACACCAGAAGAGGGCGTCA | | |
| 1765 | 11 | 32552533 | 32552825 | 292 | SPRET | 11 | 32552727 | 32552760 | 34 | - | 17.2439 | 5.79E-07 | 0.0529 | TAGCTGTCCTCAGATACTCCAGAAGAGGGCATCA | 2 | ENCFF367UNR.ENCFF093ZRU |
| 1766 | 11 | 43536096 | 43536992 | 896 | WSB | 11 | 43536310 | 43536343 | 34 | - | 21 | 4.27E-08 | 0.0108 | ATGCAATGCCCTTGGAGACCAGTAGGGGGGATAG | 16 | ENCFF504JDS,ENCFF327LYS,ENCFF821BIA,ENCFF367UNR,ENCFF092NMJ,ENCFF907XEG,ENCFF136ZDK,ENCFF589ZBJ,ENCFF198SSR,ENCFF093ZRU,ENCFF883UPM,ENCFF957SPH,ENCFF052KVO,ENCFF099QNE.ENCFF065ODV.ENCFF502WDT |
| 1767 | 11 | 43560752 | 43560818 | 66 | CAST | 11 | 43560778 | 43560811 | 34 | - | 16.4309 | 9.71E-07 | 0.0673 | TAGCTGTCTTCAGACGCTCCAGAAGAGGGCGTCA | | |
| 1768 | 11 | 43710883 | 43711100 | 217 | SPRET | 11 | 43711012 | 43711045 | 34 | - | 16.4634 | 9.51E-07 | 0.067 | TAGCTGTCTTCAGACGCACCAGAAGAGGGCATCA | | |
| 1769 | 11 | 46273664 | 46273861 | 197 | 129S1,AJ,AKR,BALB,BTBR,BUB,C3H,C57L,CAST,CBA,CE,DBA1J,DBA2J,FVB,ILNJ2,KK,MA/My,MOLF,MRL,NOD,NZB,NZW,PJ,PL,RF,RH2,SEA,SJL,SMJ,SPRET,SWR,TH | 11 | 46273775 | 46273808 | 34 | - | 17.6667 | 4.40E-07 | 0.0454 | TAGCTGTTTTCAGACACTCCAGAAGAGGGCGTCA | | |
| 1770 | 11 | 46448100 | 46448296 | 196 | MOLF | 11 | 46448220 | 46448253 | 34 | - | 16.5447 | 9.04E-07 | 0.067 | TAGCTGTCCTCAGATACTCCAGAAGAAGGCACCA | | |
| 1771 | 11 | 51581358 | 51581565 | 207 | BUB,CAST,CE,KK,MA/My,MOLF,NOD,NZB,NZO.NZW.PL.SJL.SPRET.SWR.WSB | 11 | 51581484 | 51581502 | 19 | - | 17.5574 | 6.32E-07 | 0.0841 | ACTCCAGAAGAGGGCGCCA | 3 | ENCFF093ZRU,ENCFF367UNR,ENCFF055HAP |
| 1771 | 11 | 51581358 | 51581565 | 207 | BUB,CAST,CE,KK,MA/My,MOLF,NOD,NZB,NZO.NZW.PL.SJL.SPRET.SWR.WSB | 11 | 51581484 | 51581517 | 34 | - | 18.3659 | 2.76E-07 | 0.0355 | TAGCTGTCTTCAGACACTCCAGAAGAGGGCGCCA | 3 | ENCFF093ZRU,ENCFF367UNR,ENCFF055HAP |
| 1772 | 11 | 51588304 | 51588645 | 341 | SPRET | 11 | 51588638 | 51588672 | 35 | - | 16.8257 | 5.99E-07 | 0.0984 | CTGCATGTATATCTGTGTGCCAGAAGAGGACATCA | | |
| 1773 | 11 | 51907525 | 51908454 | 929 | SPRET | 11 | 51908075 | 51908108 | 34 | - | 17.2358 | 5.82E-07 | 0.053 | TAGCTGTCTCTAGACACACCAGAAGAGGGCATCA | | |
| 1774 | 11 | 54765083 | 54765216 | 133 | SPRET | 11 | 54765122 | 54765140 | 19 | - | 20.4426 | 7.76E-08 | 0.0322 | AGACCAGAAGGGGGCACCA | 5 | ENCFF093ZRU.ENCFF367UNR.ENCFF907XEG.ENCFF099QNE.ENCFF502WDT |
| 1774 | 11 | 54765083 | 54765216 | 133 | SPRET | 11 | 54765122 | 54765156 | 35 | - | 18.3394 | 2.45E-07 | 0.0583 | ATGCCTGGTACCATGGAGACCAGAAGGGGGCACCA | 5 | ENCFF093ZRU.ENCFF367UNR.ENCFF907XEG.ENCFF099QNE.ENCFF502WDT |
| 1775 | 11 | 54767439 | 54767584 | 145 | SPRET | 11 | 54767499 | 54767532 | 34 | - | 18.065 | 3.38E-07 | 0.0394 | TAGCTGTCCTCACACACTCCAGAAGAGGGCATCA | | |
| 1776 | 11 | 54856888 | 54857068 | 180 | SPRET | 11 | 54857000 | 54857033 | 34 | - | 16.4309 | 9.71E-07 | 0.0673 | TAGCTGTCTTCAGACGCTCCAGAAGAGGGCGTCA | | |
| 1777 | 11 | 55007301 | 55007509 | 208 | B10D2,B10J,C58J,CAST,KK,MOLF,NOR,NZB.NZO.NZW.SPRET | 11 | 55007425 | 55007443 | 19 | - | 17.5574 | 6.32E-07 | 0.0841 | ACTCCAGAAGAGGGCGCCA | | |
| 1777 | 11 | 55007301 | 55007509 | 208 | B10D2,B10J,C58J,CAST,KK,MOLF,NOR,NZB.NZO.NZW.SPRET | 11 | 55007425 | 55007458 | 34 | - | 18.3659 | 2.76E-07 | 0.0355 | TAGCTGTCTTCAGACACTCCAGAAGAGGGCGCCA | | |
| 1778 | 11 | 57811251 | 57811499 | 248 | SPRET | 11 | 57811293 | 57811326 | 34 | - | 26.7642 | 3.24E-10 | 0.000338 | ATGCAGTGCCTGGAGAGGCCAGAAGAGGGGGCTG | 16 | ENCFF367UNR,ENCFF099QNE,ENCFF821BIA,ENCFF136ZDK,ENCFF198SSR,ENCFF502WDT,ENCFF055HAP,ENCFF957SPH,ENCFF327LYS,ENCFF907XEG,ENCFF589ZBJ,ENCFF533APC,ENCFF052KVO,ENCFF065ODV.ENCFF883UPM.ENCFF093ZRU |
| 1779 | 11 | 58093132 | 58093250 | 118 | CAST | 11 | 58093211 | 58093229 | 19 | - | 17.0164 | 8.98E-07 | 0.0984 | TTGCCAGGAGGTGGTGGCA | | |
| 1780 | 11 | 60172414 | 60172784 | 370 | CAST | 11 | 60172743 | 60172776 | 34 | - | 17.9756 | 3.59E-07 | 0.0407 | CAGCTGTCTTCAAACACTCCAGAAGAGGGCATCA | | |
| 1781 | 11 | 60715262 | 60715480 | 218 | SPRET | 11 | 60715381 | 60715414 | 34 | - | 17.5935 | 4.61E-07 | 0.0464 | TAGCTGTCTCCAGACACTCCAGAAGAGGGCGTCA | | |
| 1782 | 11 | 66412171 | 66412379 | 208 | 129S1,BALB,BTBR,C57L,CAST,CBA,CE,DBA1J,DBA2J,FVB,ILNJ2,LP,MA/My,MOLF,NOR,NZB,NZW,PJ,RH2,SEA,SJL,SPRET,SWR,TH | 11 | 66412299 | 66412332 | 34 | - | 18.5285 | 2.48E-07 | 0.0346 | CAGCTGTCTTCAGACACTCCAGAAGAGGGCGTCA | | |
| 1783 | 11 | 68709676 | 68709771 | 95 | DBA1J.DBA2J.MOLF.NOR.PJ.SEA.SMJ | 11 | 68709731 | 68709764 | 34 | - | 17.0081 | 6.74E-07 | 0.0572 | CTGCTGAGCCACCTGCCACCTCAGGGAGGACCAA | | |
| 1784 | 11 | 70254661 | 70254981 | 320 | CAST,DBA1J,DBA2J,MOLF,NOR,PJ,SEA,SMJ.SPRET | 11 | 70254783 | 70254816 | 34 | - | 16.4472 | 9.61E-07 | 0.067 | TAGCTGTCTTCAGACACACCAGAAGAGGGCGTCA | | |
| 1785 | 11 | 70349136 | 70349352 | 216 | SPRET | 11 | 70349263 | 70349296 | 34 | - | 19.7967 | 1.03E-07 | 0.0199 | TAGCTGTCCTCAGACACTCCAGAAGAGGGCACCA | 2 | ENCFF093ZRU.ENCFF367UNR |
| 1786 | 11 | 70615343 | 70615407 | 64 | CAST | 11 | 70615325 | 70615358 | 34 | - | 17.7642 | 4.12E-07 | 0.0438 | CTGCTTATCCAGGTGTGGACCCCAGGGAGCACCT | | |
| 1786 | 11 | 70615343 | 70615407 | 64 | CAST | 11 | 70615357 | 70615390 | 34 | - | 18.1057 | 3.29E-07 | 0.0388 | CTGCTGATCCAGGTGTGTACCCTAGGGAGCACCT | | |
| 1787 | 11 | 70686804 | 70687723 | 919 | 129S1,AKR,BALB,BTBR,C3H,C57L,C58J,CAST,CBA,CE,DBA1J,DBA2J,FVB,ILNJ2,KK,LP,MA/My,MOLF,MRL,NOD,NOR,NZB,NZO,NZW,PJ,PL,RF,RH2,SEA,SJL,SMJ,SPRET,SWR.WSB | 11 | 70687236 | 70687254 | 19 | - | 17.0164 | 8.98E-07 | 0.0984 | TTGCCAGGAGGTGGTGGCA | | |
| 1788 | 11 | 72202854 | 72203141 | 287 | 129S1,BTBR,C57L,CAST,CE,DBA1J,DBA2J.MOLF.NOR.PJ.SEA.SPRET | 11 | 72202932 | 72202965 | 34 | - | 16.4146 | 9.81E-07 | 0.0678 | TAGCTGTCTTCTGACAGTCCAGAAGAGGGAGCCA | | |
| 1789 | 11 | 76282472 | 76282685 | 213 | CAST | 11 | 76282440 | 76282473 | 34 | - | 21.2276 | 3.60E-08 | 0.00954 | CTGCAGATGTCAGACACACCAGAAGAGGGCATCA | 4 | ENCFF367UNR.ENCFF821BIA.ENCFF093ZRU.ENCFF201JUA |
| 1790 | 11 | 76575390 | 76575614 | 224 | CAST.CE.MOLF.NZO.SPRET | 11 | 76575516 | 76575549 | 34 | - | 17.878 | 3.83E-07 | 0.0424 | TAGCTGTCCTCAGACACACCAGAAGAGGGCATCA | | |
| 1791 | 11 | 77530891 | 77531111 | 220 | MOLF.SPRET | 11 | 77531015 | 77531048 | 34 | - | 18.3252 | 2.84E-07 | 0.0357 | CTTCAGACGCACCAGGCACCAGAAGAGGGCATCA | | |
| 1792 | 11 | 77543219 | 77543448 | 229 | CAST.CE.MOLF.SPRET.WSB | 11 | 77543347 | 77543365 | 19 | - | 17.5738 | 6.25E-07 | 0.0841 | ACTCCAGAAGAGGGCGCTA | | |
| 1792 | 11 | 77543219 | 77543448 | 229 | CAST.CE.MOLF.SPRET.WSB | 11 | 77543347 | 77543380 | 34 | - | 18.1789 | 3.13E-07 | 0.0375 | TAGCTGTCTTCAGACACTCCAGAAGAGGGCGCTA | | |
| 1793 | 11 | 77554155 | 77554378 | 223 | SPRET | 11 | 77554296 | 77554329 | 34 | - | 16.4472 | 9.61E-07 | 0.067 | TAGCTGTCTTCAGACACACCAGAAGAGGGCGTCA | | |
| 1794 | 11 | 78143667 | 78143888 | 221 | CAST.MOLF.SPRET | 11 | 78143794 | 78143827 | 34 | - | 16.7724 | 7.83E-07 | 0.0614 | TAGCTGTCTTCAGATGCACCAGAAGAGGGCGTCA | | |
| 1795 | 11 | 78218674 | 78218888 | 214 | CAST,SPRET | 11 | 78218799 | 78218832 | 34 | - | 16.3984 | 9.91E-07 | 0.0682 | TAGCTGTCTTCAGACGTACCAGAAGAGGGCATCA | 9 | ENCFF099QNE,ENCFF092NMJ,ENCFF821BIA,ENCFF093ZRU,ENCFF089GBC,ENCFF327LYS,ENCFF907XEG.ENCFF055HAP.ENCFF883UPM |
| 1796 | 11 | 78319987 | 78320200 | 213 | SPRET | 11 | 78320111 | 78320145 | 35 | - | 18.1101 | 2.82E-07 | 0.0643 | CTACATATACACCTGCATGCCAGAAGAGGGCATCA | | |
| 1797 | 11 | 79957101 | 79957471 | 370 | SPRET | 11 | 79957398 | 79957416 | 19 | - | 17.5574 | 6.32E-07 | 0.0841 | ACTCCAGAAGAGGGCGCCA | 7 | ENCFF821BIA,ENCFF136ZDK,ENCFF367UNR,ENCFF093ZRU,ENCFF502WDT,ENCFF099QNE,ENCFF055HAP |
| 1797 | 11 | 79957101 | 79957471 | 370 | SPRET | 11 | 79957398 | 79957431 | 34 | - | 20.252 | 7.40E-08 | 0.0159 | TAGCTGTCCTCAGACACTCCAGAAGAGGGCGCCA | 7 | ENCFF821BIA,ENCFF136ZDK,ENCFF367UNR,ENCFF093ZRU,ENCFF502WDT,ENCFF099QNE,ENCFF055HAP |
| 1798 | 11 | 80181984 | 80182935 | 951 | SPRET | 11 | 80182032 | 80182050 | 19 | - | 19.1311 | 2.12E-07 | 0.0522 | AAGCCACAAGAGGGCACTG | 11 | ENCFF089GBC,ENCFF327LYS,ENCFF092NMJ,ENCFF907XEG,ENCFF055HAP,ENCFF093ZRU,ENCFF883UPM.ENCFF052KVO.ENCFF502WDT.ENCFF099QNE.ENCFF367UNR |
| 1798 | 11 | 80181984 | 80182935 | 951 | SPRET | 11 | 80182032 | 80182066 | 35 | - | 20.5963 | 5.89E-08 | 0.0236 | GTGTTGGTACCCACAGAAGCCACAAGAGGGCACTG | 11 | ENCFF089GBC,ENCFF327LYS,ENCFF092NMJ,ENCFF907XEG,ENCFF055HAP,ENCFF093ZRU,ENCFF883UPM.ENCFF052KVO.ENCFF502WDT.ENCFF099QNE.ENCFF367UNR |
| 1798 | 11 | 80181984 | 80182935 | 951 | SPRET | 11 | 80182395 | 80182428 | 34 | - | 21.0976 | 3.97E-08 | 0.0102 | TTGCAGTGCCTTTGGAGATCAAAAGAGGGCACTG | | |
| 1799 | 11 | 87322977 | 87323227 | 250 | SPRET | 11 | 87323098 | 87323131 | 34 | - | 17.2439 | 5.79E-07 | 0.0529 | TAGCTGTCCTCAGATACTCCAGAAGAGGGCATCA | | |
| 1800 | 11 | 87360058 | 87360271 | 213 | BALB,CAST,CE,FVB,ILNJ2,KK,MA/My,MOLF,NOD,NOR,NZB,NZO,NZW,RH2,SMJ,SPRET.TH.WSB | 11 | 87360182 | 87360215 | 34 | - | 19.7154 | 1.09E-07 | 0.0208 | CAGCTGTCTTCAGACAGACCAGAAGAGGGCGTCA | 1 | ENCFF093ZRU |
| 1801 | 11 | 87382209 | 87382418 | 209 | CAST.CE.MOLF.NZW.SPRET | 11 | 87382332 | 87382365 | 34 | - | 16.4472 | 9.61E-07 | 0.067 | TAGCTGTCTTCAGACACACCAGAAGAGGGCGTCA | | |
| 1802 | 11 | 87929728 | 87929937 | 209 | 129S1,AJ,AKR,BALB,BTBR,BUB,C3H,C57L,C58J,CAST,CBA,CE,DBA1J,DBA2J,FVB,ILNJ2,KK,MA/My,MOLF,MRL,NOD,NOR,NZB,NZO,NZW,PJ,PL,RF,RH2,SEA,SJL,SMJ,SPRET.SWR.TH.WSB | 11 | 87929846 | 87929864 | 19 | - | 17.5574 | 6.32E-07 | 0.0841 | ACTCCAGAAGAGGGCGCCA | 1 | ENCFF093ZRU |
| 1802 | 11 | 87929728 | 87929937 | 209 | 129S1,AJ,AKR,BALB,BTBR,BUB,C3H,C57L,C58J,CAST,CBA,CE,DBA1J,DBA2J,FVB,ILNJ2,KK,MA/My,MOLF,MRL,NOD,NOR,NZB,NZO,NZW,PJ,PL,RF,RH2,SEA,SJL,SMJ,SPRET.SWR.TH.WSB | 11 | 87929846 | 87929879 | 34 | - | 17.5041 | 4.89E-07 | 0.0482 | TAGCTGTCATCAGACACTCCAGAAGAGGGCGCCA | 1 | ENCFF093ZRU |
| 1803 | 11 | 88121530 | 88121941 | 411 | BALB,CE,ILNJ2,KK,MA/My,NZB,NZO,RH2,SMJ.WSB | 11 | 88121643 | 88121676 | 34 | - | 17.4472 | 5.08E-07 | 0.0494 | ATTCAGTGTCTGTGGCAGCCAGAAGAGGACATCA | 1 | ENCFF502WDT |
| 1804 | 11 | 88149213 | 88149680 | 467 | SPRET | 11 | 88149237 | 88149255 | 19 | - | 19.3279 | 1.84E-07 | 0.0476 | AGGCCAGAAGGGGGCGATA | 17 | ENCFF093ZRU,ENCFF099QNE,ENCFF327LYS,ENCFF821BIA,ENCFF024JOZ,ENCFF589ZBJ,ENCFF089GBC,ENCFF883UPM,ENCFF165LXO,ENCFF033UQX,ENCFF367UNR,ENCFF065ODV,ENCFF502WDT,ENCFF136ZDK.ENCFF055HAP.ENCFF052KVO.ENCFF142CNG |
| 1804 | 11 | 88149213 | 88149680 | 467 | SPRET | 11 | 88149237 | 88149271 | 35 | - | 28.156 | 1.68E-10 | 0.00145 | GTGCAGTTGCCCTCAGAGGCCAGAAGGGGGCGATA | 17 | ENCFF093ZRU,ENCFF099QNE,ENCFF327LYS,ENCFF821BIA,ENCFF024JOZ,ENCFF589ZBJ,ENCFF089GBC,ENCFF883UPM,ENCFF165LXO,ENCFF033UQX,ENCFF367UNR,ENCFF065ODV,ENCFF502WDT,ENCFF136ZDK.ENCFF055HAP.ENCFF052KVO.ENCFF142CNG |
| 1805 | 11 | 92888286 | 92888465 | 179 | 129S1.BTBR.C57L.CAST.CE.SPRET | 11 | 92888410 | 92888443 | 34 | - | 16.4472 | 9.61E-07 | 0.067 | TAGCTGTCTTCAGACACACCAGAAGAGGGCGTCA | | |
| 1806 | 11 | 93582481 | 93582679 | 198 | AJ,AKR,C3H,CAST,CBA,CE,MOLF,MRL,RF,SPRET | 11 | 93582603 | 93582621 | 19 | - | 17.5574 | 6.32E-07 | 0.0841 | ACTCCAGAAGAGGGCGCCA | | |
| 1806 | 11 | 93582481 | 93582679 | 198 | AJ,AKR,C3H,CAST,CBA,CE,MOLF,MRL,RF,SPRET | 11 | 93582603 | 93582636 | 34 | - | 18.3659 | 2.76E-07 | 0.0355 | TAGCTGTCTTCAGACACTCCAGAAGAGGGCGCCA | | |
| 1807 | 11 | 94584735 | 94584946 | 211 | SPRET | 11 | 94584859 | 94584892 | 34 | - | 16.7724 | 7.83E-07 | 0.0614 | TAGCTGTCTTCAGATGCACCAGAAGAGGGCGTCA | | |
| 1808 | 11 | 94761214 | 94762208 | 994 | SPRET | 11 | 94761356 | 94761389 | 34 | - | 20.374 | 6.78E-08 | 0.015 | GTGCAGAACACGTTCTGGACAGCAGAGGGGGCTG | 33 | ENCFF367UNR,ENCFF093ZRU,ENCFF327LYS,ENCFF099QNE,ENCFF907XEG,ENCFF430PPJ,ENCFF821BIA,ENCFF024JOZ,ENCFF065ODV,ENCFF165LXO,ENCFF589ZBJ,ENCFF198SSR,ENCFF316FYN,ENCFF033UQX,ENCFF097CAQ,ENCFF194HVL,ENCFF136ZDK,ENCFF052KVO,ENCFF491RJK,ENCFF055HAP,ENCFF089GBC,ENCFF502WDT,ENCFF142CNG,ENCFF883UPM,ENCFF957SPH,ENCFF504JDS,ENCFF092NMJ,ENCFF210BTO,ENCFF533APC,ENCFF281QMV,ENCFF951GOP,ENCFF609MDN,ENCFF321AGQ |
| 1809 | 11 | 95176149 | 95176341 | 192 | AJ,AKR,BUB,C3H,CAST,CBA,MA/My,MOLF,RH2.SPRET.WSB | 11 | 95176259 | 95176277 | 19 | - | 19.4918 | 1.63E-07 | 0.0448 | ACACCAGTAGAGGGCACCA | 1 | ENCFF093ZRU |
| 1810 | 11 | 95311633 | 95311855 | 222 | AJ,AKR,BUB,C3H,CAST,CBA,MA/My,MOLF,RH2.SPRET.WSB | 11 | 95311754 | 95311787 | 34 | - | 17.8455 | 3.91E-07 | 0.0425 | TAGCTGTCCTCAGACACTCCAGAAGAGGGCGTCA | 1 | ENCFF367UNR |
| 1811 | 11 | 95648341 | 95648550 | 209 | SPRET | 11 | 95648460 | 95648493 | 34 | - | 17.2439 | 5.79E-07 | 0.0529 | TAGCTGTCCTCAGATACTCCAGAAGAGGGCATCA | | |
| 1812 | 11 | 96013210 | 96013395 | 185 | SPRET | 11 | 96013329 | 96013347 | 19 | - | 17.5738 | 6.25E-07 | 0.0841 | ACTCCAGAAGAGGGCGCTA | 1 | ENCFF055HAP |
| 1812 | 11 | 96013210 | 96013395 | 185 | SPRET | 11 | 96013329 | 96013362 | 34 | - | 18.187 | 3.12E-07 | 0.0374 | TAGCTGTCTTCAGTCACTCCAGAAGAGGGCGCTA | 1 | ENCFF055HAP |
| 1813 | 11 | 96605925 | 96606153 | 228 | AJ,AKR,BALB,BUB,CAST,CBA,MOLF,MRL,NOD,NOR,NZW,PL,RH2,SJL,SMJ,SPRET | 11 | 96606049 | 96606082 | 34 | - | 17.9106 | 3.74E-07 | 0.0419 | TAGCTGTCTTCAGACACTCCAGAAGAGGGCACCA | | |
| 1814 | 11 | 96756069 | 96756414 | 345 | SPRET | 11 | 96756229 | 96756263 | 35 | - | 16.3578 | 7.81E-07 | 0.112 | GTGCATATTTGTATGCACCCCAGAAGAGGGCATTG | | |
| 1815 | 11 | 97212583 | 97213436 | 853 | MOLF.NZW.SPRET.SWR | 11 | 97213243 | 97213261 | 19 | - | 19.8525 | 1.24E-07 | 0.0393 | GAACCACTAGAGGGCACTG | 2 | ENCFF883UPM.ENCFF093ZRU |
| 1816 | 11 | 97537576 | 97537798 | 222 | SPRET | 11 | 97537699 | 97537732 | 34 | - | 16.9919 | 6.81E-07 | 0.0573 | TAGCTGTCTTCAGACACACCAGAAGGGGGCATCA | | |
| 1817 | 11 | 98898898 | 98899091 | 193 | SPRET | 11 | 98899020 | 98899053 | 34 | - | 17.2439 | 5.79E-07 | 0.0529 | TAGCTGTCCTCAGATACTCCAGAAGAGGGCATCA | 1 | ENCFF093ZRU |
| 1818 | 11 | 99760927 | 99761181 | 254 | SPRET | 11 | 99761054 | 99761087 | 34 | - | 16.7724 | 7.83E-07 | 0.0614 | TAGCTGTCTTCAGATGCACCAGAAGAGGGCGTCA | | |
| 1819 | 11 | 101457599 | 101457805 | 206 | BUB.CAST.MOLF.NZO.SPRET.TH | 11 | 101457724 | 101457742 | 19 | - | 17.5574 | 6.32E-07 | 0.0841 | ACTCCAGAAGAGGGCGCCA | 3 | ENCFF093ZRU.ENCFF367UNR.ENCFF055HAP |
| 1819 | 11 | 101457599 | 101457805 | 206 | BUB.CAST.MOLF.NZO.SPRET.TH | 11 | 101457724 | 101457757 | 34 | - | 18.3659 | 2.76E-07 | 0.0355 | TAGCTGTCTTCAGACACTCCAGAAGAGGGCGCCA | 3 | ENCFF093ZRU.ENCFF367UNR.ENCFF055HAP |
| 1820 | 11 | 101601427 | 101601626 | 199 | CAST | 11 | 101601552 | 101601585 | 34 | - | 16.5122 | 9.23E-07 | 0.067 | TAGCTGTCTTCGGTTACACCAGAAGAGGGCAACA | | |
| 1821 | 11 | 102173543 | 102174164 | 621 | 129S1,AKR,BALB,BTBR,BUB,C3H,C57L,CE,DBA1J,DBA2J,ILNJ2,MA/My,MRL,NOD,NOR,NZB,NZO,PL,RF,RH2,SEA,SJL,SWR | 11 | 102174014 | 102174047 | 34 | - | 17.6179 | 4.54E-07 | 0.046 | TAGCTGTCTTCAGACACCCCAGAAGAGGGCACCA | 1 | ENCFF093ZRU |
| 1822 | 11 | 102983571 | 102984174 | 603 | CAST.MOLF.NZW.SPRET | 11 | 102983706 | 102983739 | 34 | - | 20.4553 | 6.39E-08 | 0.0144 | ATGCAGTACCCTCAGAGGCCGGCAGAGAGCATGG | | |
| 1823 | 11 | 104994510 | 104994728 | 218 | CAST.SPRET | 11 | 104994636 | 104994669 | 34 | - | 16.9187 | 7.13E-07 | 0.0585 | TAGCTGTCTTCAGACGCACCAGAAGAGGGCGTCA | | |
| 1824 | 11 | 106062534 | 106062732 | 198 | CAST.SPRET | 11 | 106062662 | 106062695 | 34 | - | 16.4472 | 9.61E-07 | 0.067 | TAGCTGTCTTCAGACACACCAGAAGAGGGCGTCA | | |
| 1825 | 11 | 106333326 | 106333533 | 207 | CAST,CBA,NZB,NZO,NZW,RF,SJL,SMJ,SPRET.SWR.TH.WSB | 11 | 106333451 | 106333469 | 19 | - | 17.8852 | 5.08E-07 | 0.0767 | ACACCAGAAGAGGGCACCA | 4 | ENCFF093ZRU,ENCFF367UNR,ENCFF883UPM,ENCFF502WDT |
| 1826 | 11 | 106344769 | 106344974 | 205 | CAST.SPRET | 11 | 106344895 | 106344928 | 34 | - | 16.4472 | 9.61E-07 | 0.067 | TAGCTGTCTTCAGACACACCAGAAGAGGGCGTCA | 1 | ENCFF093ZRU |
| 1827 | 11 | 106456654 | 106456889 | 235 | BUB.CAST.RF.SMJ.SPRET.WSB | 11 | 106456779 | 106456812 | 34 | - | 16.9919 | 6.81E-07 | 0.0573 | TAGCTGTCCTCAGACTCTCCAGAAGAGGGAGTCA | 1 | ENCFF093ZRU |
| 1828 | 11 | 106876475 | 106876685 | 210 | SPRET | 11 | 106876588 | 106876621 | 34 | - | 17.8618 | 3.87E-07 | 0.0425 | TTGCTGTCCTCAGACACTCCAGAAGAGGGAGTCA | 2 | ENCFF093ZRU.ENCFF367UNR |
| 1829 | 11 | 107314710 | 107314788 | 78 | SPRET | 11 | 107314684 | 107314717 | 34 | - | 16.4472 | 9.61E-07 | 0.067 | TAGCTGTCTTCAGACACGCCAGGAGAGGGTGCCA | | |
| 1830 | 11 | 107397875 | 107398075 | 200 | SPRET | 11 | 107397999 | 107398032 | 34 | - | 16.4472 | 9.61E-07 | 0.067 | TAGCTGTCTTCAGACACACCAGAAGAGGGCGTCA | | |
| 1831 | 11 | 113551313 | 113551509 | 196 | AJ.C3H.CAST.CBA.MOLF.SPRET.TH | 11 | 113551421 | 113551454 | 34 | - | 16.4472 | 9.61E-07 | 0.067 | TAGCTGTCTTCAGACACACCAGAAGAGGGCGTCA | 1 | ENCFF093ZRU |
| 1832 | 11 | 113561646 | 113561853 | 207 | SPRET | 11 | 113561765 | 113561798 | 34 | - | 16.3984 | 9.91E-07 | 0.0682 | TTGCTATCTTCAGATGCACCAGAAGAGGGCGTCT | 6 | ENCFF821BIA.ENCFF093ZRU.ENCFF502WDT.ENCFF099QNE.ENCFF367UNR.ENCFF055HAP |
| 1833 | 11 | 114038934 | 114039620 | 686 | SPRET | 11 | 114039386 | 114039419 | 34 | - | 17.1545 | 6.13E-07 | 0.0544 | GTGTAATGCCCATGAAGTCCAGAAGGGAGCATCA | | |

| | | | | | | | | | | | | | | | | |
|---|---|---|---|---|---|---|---|---|---|---|---|---|---|---|---|---|
| 1834 | 11 | 114456485 | 114457223 | 738 | SPRET | 11 | 114456812 | 114456845 | 34 | - | 16.4472 | 9.61E-07 | 0.067 | TAGCTGTCTTCAGACACACCAGAAGAGGGCGTCA | | |
| 1835 | 11 | 114465462 | 114466109 | 647 | MOLF | 11 | 114465797 | 114465831 | 35 | - | 17.4404 | 4.19E-07 | 0.081 | ATGTATACCTACCTGCATGCCAGAAGATGGCATCA | | |
| 1836 | 11 | 114658567 | 114658704 | 137 | SPRET | 11 | 114658696 | 114658729 | 34 | - | 16.4472 | 9.61E-07 | 0.067 | TAGCTGTCTTCAGACACACCAGAAGAGGGCGTCA | | |
| 1837 | 11 | 114947643 | 114947834 | 191 | AJ,AKR,BUB,C3H,C57L,CAST,CBA,DBA1J,DBA2J,MOLF,PL,RH2,SPRET,SWR,WSB | 11 | 114947766 | 114947799 | 34 | - | 16.4472 | 9.61E-07 | 0.067 | TAGCTGTCTTCAGACACACCAGAAGAGGGCGTCA | | |
| 1838 | 11 | 115662324 | 115662538 | 214 | 129S1,AJ,AKR,BALB,C3H,C57L,C58J,CAST,CBA,CE,DBA1J,DBA2J,ILNJ2,KK,MOLF,MRL,NOD,NOR,NZB,NZO,NZW,PL,RF,RH2,SEA,SPRET,TH,WSB | 11 | 115662449 | 115662482 | 34 | - | 16.7724 | 7.83E-07 | 0.0614 | TAGCTGTCTTCAGATGCACCAGAAGAGGGCGTCA | | |
| 1839 | 11 | 115754789 | 115754969 | 180 | CAST,MOLF,SPRET | 11 | 115754896 | 115754914 | 19 | - | 17.5574 | 6.32E-07 | 0.0841 | ACTCCAGAAGAGGGCGCCA | 11 | ENCFF136ZDK,ENCFF821BIA,ENCFF093ZRU,ENCFF883UPM,ENCFF367UNR,ENCFF327LYS,ENCFF099QNE,ENCFF024JOZ,ENCFF055HAP,ENCFF907XEG,ENCFF502WDT |
| 1839 | 11 | 115754789 | 115754969 | 180 | CAST,MOLF,SPRET | 11 | 115754896 | 115754929 | 34 | - | 18.374 | 2.75E-07 | 0.0355 | TAGCTGTCTTCAGTCACTCCAGAAGAGGGCGCCA | 11 | ENCFF136ZDK,ENCFF821BIA,ENCFF093ZRU,ENCFF883UPM,ENCFF367UNR,ENCFF327LYS,ENCFF099QNE,ENCFF024JOZ,ENCFF055HAP,ENCFF907XEG,ENCFF502WDT |
| 1840 | 11 | 116365549 | 116365747 | 198 | SPRET | 11 | 116365662 | 116365695 | 34 | - | 16.4472 | 9.61E-07 | 0.067 | TAGCTGTCTTCAGACACACCAGAAGAGGGCGTCA | | |
| 1841 | 11 | 116582637 | 116582882 | 245 | CAST,MOLF,SPRET | 11 | 116582757 | 116582775 | 19 | - | 17.5574 | 6.32E-07 | 0.0841 | ACTCCAGAAGAGGGCGCCA | 4 | ENCFF093ZRU,ENCFF099QNE,ENCFF055HAP,ENCFF367UNR |
| 1841 | 11 | 116582637 | 116582882 | 245 | CAST,MOLF,SPRET | 11 | 116582757 | 116582790 | 34 | - | 18.3659 | 2.76E-07 | 0.0355 | TAGCTGTCTTCAGACACTCCAGAAGAGGGCGCCA | 4 | ENCFF093ZRU,ENCFF099QNE,ENCFF055HAP,ENCFF367UNR |
| 1842 | 11 | 117484928 | 117485134 | 206 | CAST,MOLF,SPRET | 11 | 117485029 | 117485062 | 34 | - | 17.2439 | 5.79E-07 | 0.0529 | TAGCTGTCCTCAGATACTCCAGAAGAGGGCATCA | | |
| 1843 | 11 | 117707066 | 117707173 | 107 | 129S1,AJ,AKR,BTBR,BUB,C3H,C57L,C58J,CAST,CBA,CE,DBA1J,DBA2J,ILNJ2,KK,MA/My,MOLF,MRL,NZB,NZO,NZW,PL,RH2,SJL,SMJ,SPRET | 11 | 117707090 | 117707108 | 19 | - | 17.623 | 6.05E-07 | 0.0841 | CTCCCACCAGGTGGAGGTG | | |
| 1844 | 11 | 117724280 | 117724499 | 219 | CAST.MOLF.SPRET.TH | 11 | 117724406 | 117724439 | 34 | - | 16.4472 | 9.61E-07 | 0.067 | TAGCTGTCTTCAGACACACCAGAAGAGGGCGTCA | | |
| 1845 | 11 | 117993865 | 117993920 | 55 | SPRET | 11 | 117993880 | 117993913 | 34 | - | 17.7154 | 4.26E-07 | 0.0448 | ATGCCTTACCACACCATGCCACATGGGGGAGCAC | 41 | ENCFF093ZRU,ENCFF367UNR,ENCFF099QNE,ENCFF327LYS,ENCFF907XEG,ENCFF136ZDK,ENCFF065ODV,ENCFF821BIA,ENCFF024JOZ,ENCFF321AGQ,ENCFF201JUA,ENCFF092NMJ,ENCFF198SSR,ENCFF589ZBJ,ENCFF165LXO,ENCFF055HAP,ENCFF089GBC,ENCFF430PPJ,ENCFF883UPM,ENCFF957SPH,ENCFF504JDS,ENCFF951GOP,ENCFF210BTO,ENCFF601JVE,ENCFF281QMV,ENCFF194HVL,ENCFF316FYN,ENCFF575MGI,ENCFF533APC,ENCFF142CNG,ENCFF349BUL,ENCFF611HDQ,ENCFF609MDN,ENCFF491RJK,ENCFF052KVO,ENCFF097CAQ,ENCFF086ZTV,ENCFF784ASD,ENCFF089EWX,ENCFF033UQX,ENCFF805QBU |
| 1846 | 11 | 119274143 | 119274333 | 190 | FVB.SPRET.SWR | 11 | 119274258 | 119274291 | 34 | - | 18.0732 | 3.36E-07 | 0.0394 | CAGCTGTCTTCAGACACTCCAGAAGAGGGCATCA | | |
| 1847 | 11 | 120397448 | 120397656 | 208 | CAST | 11 | 120397524 | 120397558 | 35 | - | 16.0092 | 9.50E-07 | 0.124 | CTGTAGGGGTCACCCCTATGCTGTAGGGGTCACCT | | |
| 1848 | 11 | 120787655 | 120787857 | 202 | MOLF.SJL.SPRET | 11 | 120787770 | 120787803 | 34 | - | 17.9106 | 3.74E-07 | 0.0419 | TAGCTGTCTTCAGACACTCCAGAAGAGGGCACCA | | |
| 1849 | 11 | 121444862 | 121445052 | 190 | SPRET | 11 | 121444959 | 121444977 | 19 | - | 17.5738 | 6.25E-07 | 0.0841 | ACTCCAGAAGAGGGCGCTA | 3 | ENCFF367UNR.ENCFF099QNE.ENCFF093ZRU |
| 1849 | 11 | 121444862 | 121445052 | 190 | SPRET | 11 | 121444959 | 121444992 | 34 | - | 18.187 | 3.12E-07 | 0.0374 | TAGCTGTCTTCAGTCACTCCAGAAGAGGGCGCTA | 3 | ENCFF367UNR.ENCFF099QNE.ENCFF093ZRU |
| 1850 | 12 | 3484868 | 3485073 | 205 | SPRET | 12 | 3484910 | 3484943 | 34 | + | 16.4634 | 9.51E-07 | 0.104 | TAGCTGTCTTCAGACGCACCAGAAGAGGGCATCA | | |
| 1851 | 12 | 4224481 | 4224675 | 194 | CAST.MOLF.SPRET | 12 | 4224523 | 4224556 | 34 | + | 17.3902 | 5.27E-07 | 0.0795 | TAGCTGTCCTCAGACACTCCAGAAGAGGGCATCA | | |
| 1852 | 12 | 8718419 | 8718637 | 218 | SPRET | 12 | 8718493 | 8718526 | 34 | + | 16.5691 | 8.90E-07 | 0.102 | TAGCTGTCTTCAGACGCACCAGAAGAGGGCATCG | | |
| 1853 | 12 | 16769105 | 16769176 | 71 | CAST | 12 | 16769094 | 16769112 | 19 | + | 22.2131 | 1.66E-08 | 0.0362 | CCTCCACCAGGGGGCAGCA | 38 | ENCFF367UNR,ENCFF093ZRU,ENCFF201JUA,ENCFF099QNE,ENCFF504JDS,ENCFF805QBU,ENCFF327LYS,ENCFF907XEG,ENCFF821BIA,ENCFF089GBC,ENCFF165LXO,ENCFF092NMJ,ENCFF210BTO,ENCFF024JOZ,ENCFF198SSR,ENCFF321AGQ,ENCFF589ZBJ,ENCFF065ODV,ENCFF033UQX,ENCFF430PPJ,ENCFF136ZDK,ENCFF055HAP,ENCFF052KVO,ENCFF502WDT,ENCFF089EWX,ENCFF957SPH,ENCFF883UPM,ENCFF142CNG,ENCFF609MDN,ENCFF951GOP,ENCFF281QMV,ENCFF194HVL,ENCFF533APC,ENCFF575MGI,ENCFF491RJK,ENCFF316FYN,ENCFF097CAQ,ENCFF611HDQ |
| 1854 | 12 | 17541134 | 17541346 | 212 | 129S1,AJ,AKR,BALB,BTBR,BUB,C3H,C57L,CBA,CE,DBA1J,DBA2J,FVB,ILNJ2,KK,MA/My,MRL,NOD,NOR,NZO,PJ,PL,RF,SEA,SJL,SMJ.SPRET.SWR.TH.WSB | 12 | 17541207 | 17541240 | 34 | + | 18.3659 | 2.76E-07 | 0.0591 | TAGCTGTCTTCAGACACTCCAGAAGAGGGCGCCA | | |
| 1854 | 12 | 17541134 | 17541346 | 212 | 129S1,AJ,AKR,BALB,BTBR,BUB,C3H,C57L,CBA,CE,DBA1J,DBA2J,FVB,ILNJ2,KK,MA/My,MRL,NOD,NOR,NZO,PJ,PL,RF,SEA,SJL,SMJ.SPRET.SWR.TH.WSB | 12 | 17541222 | 17541240 | 19 | + | 17.5574 | 6.32E-07 | 0.128 | ACTCCAGAAGAGGGCGCCA | | |
| 1855 | 12 | 18440677 | 18441250 | 573 | 129S1,BTBR,BUB,C3H,C57L,CBA,DBA2J,FVB,KK,MA/My,NOD,NOR,NZO,PL,SJL,SMJ,SPRET,SWR,TH | 12 | 18440694 | 18440727 | 34 | + | 17.6748 | 4.37E-07 | 0.073 | CCGCCCCTACATTCGCCGTCACAAGATGGCGCTG | | |
| 1856 | 12 | 18829354 | 18829852 | 498 | DBA2J | 12 | 18829780 | 18829813 | 34 | + | 18.374 | 2.75E-07 | 0.0591 | TAGCAACACTAAACGTGACCCCTAGATGGCAGCG | | |
| 1857 | 12 | 21294259 | 21294484 | 225 | SPRET | 12 | 21294284 | 21294318 | 35 | + | 15.9541 | 9.80E-07 | 0.196 | CTGAAGAGGGCACTCGGGACCTGCAGGAGGAACTC | | |
| 1858 | 12 | 21495295 | 21495494 | 199 | AJ,AKR,BALB,CAST,CE,DBA1J,ILNJ2,PJ,RF,SEA,SPRET | 12 | 21495361 | 21495394 | 34 | + | 16.4634 | 9.51E-07 | 0.104 | TAGCTGTCTTCAGACGCACCAGAAGAGGGCATCA | | |
| 1859 | 12 | 29229009 | 29229258 | 249 | SPRET | 12 | 29229059 | 29229092 | 34 | + | 16.4472 | 9.61E-07 | 0.104 | TAGCTGTCTTCAGACACACCAGAAGAGGGCGTCA | | |
| 1860 | 12 | 36119619 | 36119801 | 182 | CAST.MOLF.SPRET | 12 | 36119667 | 36119700 | 34 | + | 16.4472 | 9.61E-07 | 0.104 | TAGCTGTCTTCAGACACACCAGAAGAGGGCGTCA | | |
| 1861 | 12 | 36279119 | 36279357 | 238 | SPRET | 12 | 36279233 | 36279251 | 19 | + | 17.2459 | 7.74E-07 | 0.142 | ACACCAGAAGAGGGCAGCA | | |
| 1862 | 12 | 40035122 | 40036089 | 967 | CAST.MOLF.SPRET | 12 | 40035763 | 40035797 | 35 | + | 18.3211 | 2.48E-07 | 0.0911 | CAGCTCTCCTGCATCCCTGCCACCAGGGTGAGCTC | 4 | ENCFF327LYS.ENCFF136ZDK.ENCFF589ZBJ.ENCFF883UPM |
| 1863 | 12 | 40184886 | 40185087 | 201 | CAST.MOLF.SPRET | 12 | 40184944 | 40184977 | 34 | + | 18.3659 | 2.76E-07 | 0.0591 | TAGCTGTCTTCAGACACTCCAGAAGAGGGCGCCA | 2 | ENCFF093ZRU.ENCFF367UNR |
| 1863 | 12 | 40184886 | 40185087 | 201 | CAST.MOLF.SPRET | 12 | 40184959 | 40184977 | 19 | + | 17.5574 | 6.32E-07 | 0.128 | ACTCCAGAAGAGGGCGCCA | 2 | ENCFF093ZRU.ENCFF367UNR |
| 1864 | 12 | 40629270 | 40629482 | 212 | CAST,CE,FVB,KK,MOLF,NOD,NOR,SJL,SPRET | 12 | 40629356 | 40629374 | 19 | + | 17.8852 | 5.08E-07 | 0.115 | ACACCAGAAGAGGGCACCA | 1 | ENCFF093ZRU |
| 1865 | 12 | 40815323 | 40815412 | 89 | SPRET | 12 | 40815373 | 40815407 | 35 | + | 18.0642 | 2.90E-07 | 0.0986 | GTGCACTTACCCACAGAGGCCAGAAGAGGATATCA | | |
| 1866 | 12 | 45779439 | 45779995 | 556 | SPRET | 12 | 45779892 | 45779926 | 35 | + | 17.1651 | 4.92E-07 | 0.139 | CTGCACTTCCGGTTATGATACAGCAGGGGCAAGAT | | |
| 1867 | 12 | 55243398 | 55243656 | 258 | SPRET | 12 | 55243478 | 55243511 | 34 | + | 17.5041 | 4.89E-07 | 0.0768 | CTGTAGCTCCAGTTCCCAGCACCACATGGCAGCT | | |
| 1867 | 12 | 55243398 | 55243656 | 258 | SPRET | 12 | 55243564 | 55243597 | 34 | + | 17.5041 | 4.89E-07 | 0.0768 | CTGTAGCTCCAGTTCCCAGCACCACATGGCAGCT | | |
| 1867 | 12 | 55243398 | 55243656 | 258 | SPRET | 12 | 55243607 | 55243640 | 34 | + | 17.5041 | 4.89E-07 | 0.0768 | CTGTAGCTCCAGTTCCCAGCACCACATGGCAGCT | | |
| 1868 | 12 | 55317601 | 55317687 | 86 | SPRET | 12 | 55317638 | 55317671 | 34 | + | 17.5041 | 4.89E-07 | 0.0768 | CTGTAGCTCCAGTTCCCAGCACCACATGGCAGCT | | |
| 1869 | 12 | 56673684 | 56673915 | 231 | 129S1,BTBR,C57L,CAST,CE,FVB,MOLF,NZW,PL,SPRET,WSB | 12 | 56673772 | 56673805 | 34 | + | 16.5691 | 8.90E-07 | 0.102 | TAGCTGTCTTCAGACGCACCAGAAGAGGGCATCG | | |
| 1870 | 12 | 57257001 | 57257194 | 193 | SPRET | 12 | 57257059 | 57257092 | 34 | + | 17.3902 | 5.27E-07 | 0.0795 | TAGCTGTCCTCAGACACTCCAGAAGAGGGCATCA | | |
| 1871 | 12 | 59081760 | 59081855 | 95 | B10J.MRL | 12 | 59081797 | 59081830 | 34 | + | 18.065 | 3.38E-07 | 0.064 | TAGCTGTCTTCAGACTCACCAGAAGAGGGCGTCA | | |
| 1871 | 12 | 59081760 | 59081855 | 95 | B10J.MRL | 12 | 59081812 | 59081830 | 19 | + | 17.0328 | 8.88E-07 | 0.147 | TCACCAGAAGAGGGCGTCA | | |
| 1872 | 12 | 59081758 | 59082613 | 855 | NZW | 12 | 59081797 | 59081830 | 34 | + | 18.065 | 3.38E-07 | 0.064 | TAGCTGTCTTCAGACTCACCAGAAGAGGGCGTCA | | |
| 1872 | 12 | 59081758 | 59082613 | 855 | NZW | 12 | 59081812 | 59081830 | 19 | + | 17.0328 | 8.88E-07 | 0.147 | TCACCAGAAGAGGGCGTCA | | |
| 1872 | 12 | 59081758 | 59082613 | 855 | NZW | 12 | 59081892 | 59081925 | 34 | + | 18.065 | 3.38E-07 | 0.064 | TAGCTGTCTTCAGACTCACCAGAAGAGGGCGTCA | | |
| 1872 | 12 | 59081758 | 59082613 | 855 | NZW | 12 | 59081907 | 59081925 | 19 | + | 17.0328 | 8.88E-07 | 0.147 | TCACCAGAAGAGGGCGTCA | | |
| 1872 | 12 | 59081758 | 59082613 | 855 | NZW | 12 | 59081987 | 59082020 | 34 | + | 18.065 | 3.38E-07 | 0.064 | TAGCTGTCTTCAGACTCACCAGAAGAGGGCGTCA | | |
| 1872 | 12 | 59081758 | 59082613 | 855 | NZW | 12 | 59082002 | 59082020 | 19 | + | 17.0328 | 8.88E-07 | 0.147 | TCACCAGAAGAGGGCGTCA | | |
| 1872 | 12 | 59081758 | 59082613 | 855 | NZW | 12 | 59082082 | 59082115 | 34 | + | 18.065 | 3.38E-07 | 0.064 | TAGCTGTCTTCAGACTCACCAGAAGAGGGCGTCA | | |
| 1872 | 12 | 59081758 | 59082613 | 855 | NZW | 12 | 59082097 | 59082115 | 19 | + | 17.0328 | 8.88E-07 | 0.147 | TCACCAGAAGAGGGCGTCA | | |
| 1872 | 12 | 59081758 | 59082613 | 855 | NZW | 12 | 59082177 | 59082210 | 34 | + | 18.065 | 3.38E-07 | 0.064 | TAGCTGTCTTCAGACTCACCAGAAGAGGGCGTCA | | |
| 1872 | 12 | 59081758 | 59082613 | 855 | NZW | 12 | 59082192 | 59082210 | 19 | + | 17.0328 | 8.88E-07 | 0.147 | TCACCAGAAGAGGGCGTCA | | |
| 1872 | 12 | 59081758 | 59082613 | 855 | NZW | 12 | 59082272 | 59082305 | 34 | + | 18.065 | 3.38E-07 | 0.064 | TAGCTGTCTTCAGACTCACCAGAAGAGGGCGTCA | | |
| 1872 | 12 | 59081758 | 59082613 | 855 | NZW | 12 | 59082287 | 59082305 | 19 | + | 17.0328 | 8.88E-07 | 0.147 | TCACCAGAAGAGGGCGTCA | | |
| 1872 | 12 | 59081758 | 59082613 | 855 | NZW | 12 | 59082367 | 59082400 | 34 | + | 18.065 | 3.38E-07 | 0.064 | TAGCTGTCTTCAGACTCACCAGAAGAGGGCGTCA | | |
| 1872 | 12 | 59081758 | 59082613 | 855 | NZW | 12 | 59082382 | 59082400 | 19 | + | 17.0328 | 8.88E-07 | 0.147 | TCACCAGAAGAGGGCGTCA | | |
| 1872 | 12 | 59081758 | 59082613 | 855 | NZW | 12 | 59082462 | 59082495 | 34 | + | 18.065 | 3.38E-07 | 0.064 | TAGCTGTCTTCAGACTCACCAGAAGAGGGCGTCA | | |
| 1872 | 12 | 59081758 | 59082613 | 855 | NZW | 12 | 59082477 | 59082495 | 19 | + | 17.0328 | 8.88E-07 | 0.147 | TCACCAGAAGAGGGCGTCA | | |
| 1872 | 12 | 59081758 | 59082613 | 855 | NZW | 12 | 59082557 | 59082590 | 34 | + | 18.065 | 3.38E-07 | 0.064 | TAGCTGTCTTCAGACTCACCAGAAGAGGGCGTCA | | |
| 1872 | 12 | 59081758 | 59082613 | 855 | NZW | 12 | 59082572 | 59082590 | 19 | + | 17.0328 | 8.88E-07 | 0.147 | TCACCAGAAGAGGGCGTCA | | |
| 1873 | 12 | 59085423 | 59085660 | 237 | C57L.MRL.WSB | 12 | 59085407 | 59085440 | 34 | + | 18.065 | 3.38E-07 | 0.064 | TAGCTGTCTTCAGACTCACCAGAAGAGGGCGTCA | | |
| 1873 | 12 | 59085423 | 59085660 | 237 | C57L.MRL.WSB | 12 | 59085422 | 59085440 | 19 | + | 17.0328 | 8.88E-07 | 0.147 | TCACCAGAAGAGGGCGTCA | | |
| 1873 | 12 | 59085423 | 59085660 | 237 | C57L.MRL.WSB | 12 | 59085502 | 59085535 | 34 | + | 18.065 | 3.38E-07 | 0.064 | TAGCTGTCTTCAGACTCACCAGAAGAGGGCGTCA | | |
| 1873 | 12 | 59085423 | 59085660 | 237 | C57L.MRL.WSB | 12 | 59085517 | 59085535 | 19 | + | 17.0328 | 8.88E-07 | 0.147 | TCACCAGAAGAGGGCGTCA | | |
| 1874 | 12 | 59147401 | 59147574 | 173 | MOLF.SPRET | 12 | 59147432 | 59147465 | 34 | + | 17.2439 | 5.79E-07 | 0.0827 | TAGCTGTCCTCAGATACTCCAGAAGAGGGCATCA | | |
| 1875 | 12 | 61256843 | 61256936 | 93 | SPRET | 12 | 61256925 | 61256959 | 35 | + | 16.945 | 5.59E-07 | 0.148 | GTGTAATTTTCTGAAGAGGCCAGCAGAGGGTCATA | | |
| 1876 | 12 | 65103719 | 65103896 | 177 | SPRET | 12 | 65103753 | 65103786 | 34 | + | 16.4472 | 9.61E-07 | 0.104 | TAGCTGTCTTCAGACACACCAGAAGAGGGCGTCA | | |
| 1877 | 12 | 68200712 | 68200913 | 201 | MOLF.SPRET | 12 | 68200772 | 68200805 | 34 | + | 17.2439 | 5.79E-07 | 0.0827 | TAGCTGTCCTCAGATACTCCAGAAGAGGGCATCA | | |
| 1878 | 12 | 71009574 | 71009797 | 223 | CAST,FVB,MOLF,NZW,SJL,SPRET,SWR,WSB | 12 | 71009648 | 71009681 | 34 | + | 18.1545 | 3.18E-07 | 0.0635 | TAGCTGTTTTCAGACACACCAGAAGAGGGCGTCA | | |
| 1879 | 12 | 71023306 | 71024141 | 835 | MOLF | 12 | 71023700 | 71023733 | 34 | + | 17.6585 | 4.42E-07 | 0.0733 | TTGCTGTCTTCAGACACACCAGAAGAGGGCGTAA | 3 | ENCFF367UNR.ENCFF099QNE.ENCFF093ZRU |
| 1880 | 12 | 73037474 | 73037676 | 202 | SPRET | 12 | 73037535 | 73037568 | 34 | + | 16.4472 | 9.61E-07 | 0.104 | TAGCTGTCTTCAGACACACCAGAAGAGGGCGTCA | | |
| 1881 | 12 | 73182083 | 73182499 | 416 | SPRET | 12 | 73182357 | 73182390 | 34 | + | 16.4634 | 9.51E-07 | 0.104 | TAGCTGTCTTCAGACGCACCAGAAGAGGGCATCA | | |
| 1882 | 12 | 75326783 | 75327542 | 759 | FVB,SWR | 12 | 75327426 | 75327444 | 19 | + | 18.2623 | 3.93E-07 | 0.103 | TGGCCACTAGGTGGAGAAA | 20 | ENCFF367UNR,ENCFF093ZRU,ENCFF136ZDK,ENCFF430PPJ,ENCFF089GBC,ENCFF092NMJ,ENCFF821BIA,ENCFF099QNE,ENCFF907XEG,ENCFF024JOZ,ENCFF316FYN,ENCFF609MDN,ENCFF327LYS,ENCFF198SSR,ENCFF165LXO,ENCFF951GOP,ENCFF589ZBJ,ENCFF055HAP,ENCFF502WDT,ENCFF883UPM |
| 1883 | 12 | 75484696 | 75485324 | 628 | SPRET | 12 | 75484725 | 75484758 | 34 | + | 20.0813 | 8.38E-08 | 0.0282 | CTGCACTGTTCACAGCAGCCAGGAAATGGAACCA | | |
| 1884 | 12 | 75675393 | 75676145 | 752 | CAST,CE,FVB,PJ,SEA,SMJ,SWR | 12 | 75676095 | 75676128 | 34 | + | 16.9593 | 6.95E-07 | 0.0904 | TTGCTCTCTTCAGACACACCAGAAGAGGGCATCA | | |
| 1885 | 12 | 76826249 | 76826455 | 206 | SPRET | 12 | 76826315 | 76826348 | 34 | + | 16.4959 | 9.32E-07 | 0.104 | GTACAGTGTAGCTGTCATCCAGAAGAGGGCATCA | | |
| 1886 | 12 | 80425230 | 80426032 | 802 | CAST,MOLF | 12 | 80425516 | 80425534 | 19 | + | 22.459 | 1.32E-08 | 0.0324 | TGGCCAGAAGAGGGCACTG | 2 | ENCFF093ZRU.ENCFF089GBC |
| 1887 | 12 | 80530177 | 80530469 | 292 | SPRET | 12 | 80530354 | 80530387 | 34 | + | 19.6911 | 1.11E-07 | 0.0343 | GTGCAGTGCCTGAGGAGGCCAGAGGGGGCATCAG | 4 | ENCFF367UNR.ENCFF093ZRU.ENCFF821BIA.ENCFF052KVO |
| 1888 | 12 | 80964193 | 80964393 | 200 | CAST,MOLF,SPRET | 12 | 80964252 | 80964285 | 34 | + | 16.7724 | 7.83E-07 | 0.0958 | TAGCTGTCTTCAGATGCACCAGAAGAGGGCGTCA | | |
| 1889 | 12 | 83681689 | 83681907 | 218 | CAST,KK,MOLF,SPRET,SWR,WSB | 12 | 83681765 | 83681798 | 34 | + | 16.4472 | 9.61E-07 | 0.104 | TAGCTGTCTTCAGACACACCAGAAGAGGGCGTCA | | |
| 1890 | 12 | 83798154 | 83798384 | 230 | SPRET | 12 | 83798243 | 83798276 | 34 | + | 16.4634 | 9.51E-07 | 0.104 | TAGCTGTCTTCAGACGCACCAGAAGAGGGCATCA | 2 | ENCFF093ZRU.ENCFF055HAP |
| 1891 | 12 | 84451168 | 84451400 | 232 | CAST,CE,MOLF,NZO,PL,SJL,SPRET,SWR,WSB | 12 | 84451260 | 84451293 | 34 | + | 18.374 | 2.75E-07 | 0.0591 | TAGCTGTCTTCAGTCACTCCAGAAGAGGGCGCCA | | |
| 1891 | 12 | 84451168 | 84451400 | 232 | CAST,CE,MOLF,NZO,PL,SJL,SPRET,SWR,WSB | 12 | 84451275 | 84451293 | 19 | + | 17.5574 | 6.32E-07 | 0.128 | ACTCCAGAAGAGGGCGCCA | | |
| 1892 | 12 | 84665006 | 84665202 | 196 | CAST.SPRET | 12 | 84665055 | 84665088 | 34 | + | 16.4472 | 9.61E-07 | 0.104 | TAGCTGTCTTCAGACACACCAGAAGAGGGCGTCA | | |
| 1893 | 12 | 85206961 | 85207164 | 203 | CAST.CE.MOLF.SPRET.SWR.WSB | 12 | 85207022 | 85207055 | 34 | + | 17.8455 | 3.91E-07 | 0.0684 | TAGCTGTCCTCAGACACTCCAGAAGAGGGCGTCA | | |
| 1894 | 12 | 86478761 | 86478967 | 206 | CAST,CE,MOLF,NOD,NOR,SPRET,SWR,WSB | 12 | 86478827 | 86478860 | 34 | + | 18.2439 | 3.00E-07 | 0.0613 | TAGCTGTCCTCAGATACTCCAGAAGGGGGCATCA | 1 | ENCFF093ZRU |
| 1895 | 12 | 86842730 | 86842806 | 76 | CE,NOD,NOR | 12 | 86842763 | 86842797 | 35 | + | 16.0826 | 9.12E-07 | 0.189 | TTGTTATAACATGTACTGTCCACTGGATGGTGCTG | 36 | ENCFF367UNR,ENCFF093ZRU,ENCFF099QNE,ENCFF024JOZ,ENCFF907XEG,ENCFF821BIA,ENCFF136ZDK,ENCFF589ZBJ,ENCFF321AGQ,ENCFF327LYS,ENCFF089GBC,ENCFF086ZTV,ENCFF198SSR,ENCFF092NMJ,ENCFF065ODV,ENCFF430PPJ,ENCFF165LXO,ENCFF883UPM,ENCFF052KVO,ENCFF951GOP,ENCFF601JVE,ENCFF504JDS,ENCFF281QMV,ENCFF957SPH,ENCFF210BTO,ENCFF575MGI,ENCFF609MDN,ENCFF316FYN,ENCFF142CNG,ENCFF194HVL,ENCFF533APC,ENCFF089EWX,ENCFF097CAQ,ENCFF491RJK,ENCFF611HDQ,ENCFF033UQX |
| 1896 | 12 | 86887938 | 86888159 | 221 | BUB,CAST,FVB,MOLF,NOD,NOR,NZB,SPRET,SWR | 12 | 86888018 | 86888051 | 34 | + | 17.8455 | 3.91E-07 | 0.0684 | TAGCTGTCCTCAGACACTCCAGAAGAGGGCGTCA | | |
| 1897 | 12 | 87569342 | 87569661 | 319 | CAST.MOLF | 12 | 87569492 | 87569525 | 34 | + | 16.7317 | 8.03E-07 | 0.0971 | ATGCAGCTAGAGACAGGAGCTCTGGGGGGCACTG | | |
| 1898 | 12 | 89751774 | 89752699 | 925 | SPRET | 12 | 89752080 | 89752114 | 35 | + | 16.5321 | 7.08E-07 | 0.165 | TTGTCGTTATTCCTAAATTCCAGCAGGTGGTGATG | | |
| 1899 | 12 | 91979732 | 91979797 | 65 | SPRET | 12 | 91979724 | 91979758 | 35 | + | 17.633 | 3.74E-07 | 0.118 | CTGCTGACAGCTTCCGGTGCCAGCAGAGGCCAGCC | | |
| 1900 | 12 | 94063025 | 94063228 | 203 | BUB,CAST,MOLF,NOD,PJ,SEA,SMJ,WSB | 12 | 94063085 | 94063118 | 34 | + | 18.3659 | 2.76E-07 | 0.0591 | TAGCTGTCTTCAGACACTCCAGAAGAGGGCGCCA | | |
| 1900 | 12 | 94063025 | 94063228 | 203 | BUB,CAST,MOLF,NOD,PJ,SEA,SMJ,WSB | 12 | 94063100 | 94063118 | 19 | + | 17.5574 | 6.32E-07 | 0.128 | ACTCCAGAAGAGGGCGCCA | | |
| 1901 | 12 | 95948604 | 95948800 | 196 | CAST.MOLF.SPRET | 12 | 95948659 | 95948692 | 34 | + | 18.8455 | 1.99E-07 | 0.05 | TAGCTGTCTTCAGTCGCTCCAGAAGAGGGCGCCA | 1 | ENCFF052KVO |
| 1901 | 12 | 95948604 | 95948800 | 196 | CAST.MOLF.SPRET | 12 | 95948674 | 95948692 | 19 | + | 17.6393 | 5.98E-07 | 0.127 | GCTCCAGAAGAGGGCGCCA | 1 | ENCFF052KVO |
| 1902 | 12 | 98146693 | 98147103 | 410 | CBA | 12 | 98146806 | 98146839 | 34 | + | 23.252 | 7.35E-09 | 0.00522 | ATGCGGTGCTCAAGGAGGCCAGAAGAGGGCGTCG | 3 | ENCFF367UNR.ENCFF052KVO.ENCFF093ZRU |
| 1903 | 12 | 99479004 | 99479110 | 106 | CAST | 12 | 99479035 | 99479068 | 34 | + | 16.8943 | 7.24E-07 | 0.0917 | CAGCACCTGTTCTTCCTGCCAGCAGGTGTCCTTC | | |
| 1904 | 12 | 101855215 | 101855409 | 194 | 129S1,AKR,BALB,BTBR,BUB,C57L,CAST,CBA,CE,FVB,ILNJ2,KK,LP,MA/My,MOLF,NOD,NOR,NZB,NZO,NZW,PJ,PL,RF,SEA,SJL,SPRET,SWR,TH | 12 | 101855267 | 101855300 | 34 | + | 16.4472 | 9.61E-07 | 0.104 | TAGCTGTCTTCAGACACACCAGAAGAGGGCGTCA | | |
| 1905 | 12 | 101906295 | 101906507 | 212 | CAST.SPRET | 12 | 101906364 | 101906397 | 34 | + | 18.3659 | 2.76E-07 | 0.0591 | TAGCTGTCTTCAGACACTCCAGAAGAGGGCGCCA | 1 | ENCFF367UNR |
| 1905 | 12 | 101906295 | 101906507 | 212 | CAST.SPRET | 12 | 101906379 | 101906397 | 19 | + | 17.5574 | 6.32E-07 | 0.128 | ACTCCAGAAGAGGGCGCCA | 1 | ENCFF367UNR |
| 1906 | 12 | 101959233 | 101959438 | 205 | AKR,BALB,BUB,CAST,CBA,CE,DBA1J,DBA2J,ILNJ2,KK,LP,MA/My,MOLF,NOD,NOR,NZO,NZW,PL,RF,SPRET,TH,WSB | 12 | 101959296 | 101959329 | 34 | + | 18.3659 | 2.76E-07 | 0.0591 | TAGCTGTCTTCAGACACTCCAGAAGAGGGCGCCA | 2 | ENCFF367UNR,ENCFF093ZRU |
| 1906 | 12 | 101959233 | 101959438 | 205 | AKR,BALB,BUB,CAST,CBA,CE,DBA1J,DBA2J,ILNJ2,KK,LP,MA/My,MOLF,NOD,NOR,NZO,NZW,PL,RF,SPRET,TH,WSB | 12 | 101959311 | 101959329 | 19 | + | 17.5574 | 6.32E-07 | 0.128 | ACTCCAGAAGAGGGCGCCA | 2 | ENCFF367UNR,ENCFF093ZRU |
| 1907 | 12 | 102719708 | 102719910 | 202 | CAST,CE,DBA1J,DBA2J,MOLF,PJ,SEA,SPRET,WSB | 12 | 102719768 | 102719801 | 34 | + | 17.7805 | 4.08E-07 | 0.0701 | CTGTAGTCTGTAGACACTCCAGAAGAGGGCATCA | | |
| 1908 | 12 | 103054156 | 103054263 | 107 | MOLF.SPRET | 12 | 103054216 | 103054249 | 34 | + | 17.6748 | 4.37E-07 | 0.073 | CCGCTGTCTTCAGACACACCAGAAGAGGGCATCA | 1 | ENCFF093ZRU |
| 1909 | 12 | 103486456 | 103486663 | 207 | CAST.MOLF.SPRET | 12 | 103486521 | 103486554 | 34 | + | 17.2439 | 5.79E-07 | 0.0827 | TAGCTGTCCTCAGATACTCCAGAAGAGGGCATCA | | |
| 1910 | 12 | 109561377 | 109561443 | 66 | SPRET | 12 | 109561400 | 109561434 | 35 | + | 24.8532 | 2.76E-09 | 0.00581 | TTGCATTTCCTGGTGGGGGCCACTGGATGGCTCCT | 2 | ENCFF099QNE.ENCFF327LYS |
| 1911 | 12 | 111029072 | 111029287 | 215 | CAST.KK.MA/My.MOLF.RF.SPRET | 12 | 111029145 | 111029178 | 34 | + | 16.4472 | 9.61E-07 | 0.104 | TAGCTGTCTTCAGACACACCAGAAGAGGGCGTCA | | |
| 1912 | 12 | 111513573 | 111514384 | 811 | CE | 12 | 111514369 | 111514403 | 35 | + | 16.156 | 8.75E-07 | 0.185 | CTGTAACTGTCTTCAGACACCAGAAGAGGGCATCG | | |
| 1913 | 12 | 118409499 | 118409727 | 228 | AJ,CAST,MOLF,NOD,NOR,NZB,PL,RF,SPRET | 12 | 118409584 | 118409617 | 34 | + | 17.6667 | 4.40E-07 | 0.0731 | TAGCTGTTTTCAGACACTCCAGAAGAGGGCGTCA | | |
| 1914 | 12 | 11387071 | 11387143 | 72 | CAST,MOLF,SPRET | 12 | 11387059 | 11387077 | 19 | - | 21.3607 | 3.59E-08 | 0.0421 | CTGCCAGCAGGTGGCGACA | 23 | ENCFF201JUA,ENCFF367UNR,ENCFF093ZRU,ENCFF321AGQ,ENCFF099QNE,ENCFF136ZDK,ENCFF883UPM,ENCFF821BIA,ENCFF327LYS,ENCFF052KVO,ENCFF907XEG,ENCFF024JOZ,ENCFF065ODV,ENCFF502WDT,ENCFF055HAP,ENCFF609MDN,ENCFF589ZBJ,ENCFF198SSR,ENCFF089GBC,ENCFF165LXO,ENCFF092NMJ,ENCFF430PPJ,ENCFF951GOP |
| 1914 | 12 | 11387071 | 11387143 | 72 | CAST,MOLF,SPRET | 12 | 11387059 | 11387093 | 35 | - | 22.7431 | 1.35E-08 | 0.0142 | CTGAAGCACATACATACTGCCAGCAGGTGGCGACA | 23 | ENCFF201JUA,ENCFF367UNR,ENCFF093ZRU,ENCFF321AGQ,ENCFF099QNE,ENCFF136ZDK,ENCFF883UPM,ENCFF821BIA,ENCFF327LYS,ENCFF052KVO,ENCFF907XEG,ENCFF024JOZ,ENCFF065ODV,ENCFF502WDT,ENCFF055HAP,ENCFF609MDN,ENCFF589ZBJ,ENCFF198SSR,ENCFF089GBC,ENCFF165LXO,ENCFF092NMJ.ENCFF430PPJ.ENCFF951GOP |
| 1915 | 12 | 12072994 | 12073735 | 741 | 129S1,AJ,AKR,BALB,BUB,C3H,C57L,CAST,CBA,CE,DBA1J,DBA2J,FVB,ILNJ2,KK,MA/My,MOLF,MRL,NOD,NOR,PJ,PL,RF,SEA,SJL,SMJ,SPRET,SWR,TH | 12 | 12073531 | 12073549 | 19 | - | 18.5902 | 3.13E-07 | 0.0897 | CGGCCAGCAGATGTCAGCC | | |
| 1916 | 12 | 21334858 | 21335034 | 176 | 129S1,AJ,AKR,BALB,BTBR,BUB,C3H,C57L,CAST,CBA,CE,DBA1J,DBA2J,FVB,ILNJ2,KK,MA/My,MRL,NOD,NOR,NZO,PJ,PL,RF,SEA,SJL,SMJ,SPRET,SWR,TH,WSB | 12 | 21334953 | 21334971 | 19 | - | 18.7049 | 2.88E-07 | 0.0872 | GCACCAGAAGAGGGCGCCA | 6 | ENCFF093ZRU,ENCFF367UNR,ENCFF136ZDK,ENCFF024JOZ,ENCFF089GBC,ENCFF883UPM |
| 1916 | 12 | 21334858 | 21335034 | 176 | 129S1,AJ,AKR,BALB,BTBR,BUB,C3H,C57L,CAST,CBA,CE,DBA1J,DBA2J,FVB,ILNJ2,KK,MA/My,MRL,NOD,NOR,NZO,PJ,PL,RF,SEA,SJL,SMJ,SPRET,SWR,TH,WSB | 12 | 21334953 | 21334986 | 34 | - | 19.1789 | 1.58E-07 | 0.0435 | TAGCTGTCTTCAGATGCACCAGAAGAGGGCGCCA | 6 | ENCFF093ZRU,ENCFF367UNR,ENCFF136ZDK,ENCFF024JOZ,ENCFF089GBC,ENCFF883UPM |
| 1917 | 12 | 28874027 | 28874220 | 193 | 129S1,AJ,AKR,BALB,BTBR,BUB,C3H,C57L,C58J,CAST,CBA,CE,DBA1J,DBA2J,FVB,ILNJ2,KK,LP,MA/My,MRL,NOD,NOR,NZB,NZO,NZW,PJ,PL,RF,RH2,SEA,SJL,SMJ,SPRET,SWR.TH.WSB | 12 | 28874152 | 28874185 | 34 | - | 17.6992 | 4.30E-07 | 0.0725 | TAGCTGTTTTCAGACACACCAGAAGAGGGCATCA | | |
| 1918 | 12 | 29090511 | 29091363 | 852 | AKR.RF.WSB | 12 | 29091009 | 29091042 | 34 | - | 18.6504 | 2.28E-07 | 0.055 | ATGCAGTGTCTGCAGAAGCCAGAAGAGAGTGTCA | | |
| 1919 | 12 | 31745912 | 31746128 | 216 | SPRET | 12 | 31746037 | 31746070 | 34 | - | 19.0325 | 1.75E-07 | 0.0463 | CAGCTGTCTTCAGACGCACCAGAAGAGGGCATCA | 1 | ENCFF093ZRU |
| 1920 | 12 | 51831421 | 51832193 | 772 | CAST,MOLF | 12 | 51831817 | 51831850 | 34 | - | 27.1626 | 2.20E-10 | 0.000529 | ATGCAGTGCCCATGCAGGCCACAAGAGGGCATTG | 34 | ENCFF201JUA,ENCFF367UNR,ENCFF093ZRU,ENCFF805QBU,ENCFF430PPJ,ENCFF951GOP,ENCFF065ODV,ENCFF589ZBJ,ENCFF136ZDK,ENCFF883UPM,ENCFF321AGQ,ENCFF609MDN,ENCFF907XEG,ENCFF033UQX,ENCFF821BIA,ENCFF099QNE,ENCFF327LYS,ENCFF052KVO,ENCFF097CAQ,ENCFF502WDT,ENCFF089GBC,ENCFF142CNG,ENCFF601JVE,ENCFF957SPH,ENCFF491RJK,ENCFF055HAP,ENCFF198SSR,ENCFF533APC,ENCFF194HVL,ENCFF092NMJ,ENCFF281QMV,ENCFF165LXO,ENCFF210BTO,ENCFF024JOZ |
| 1921 | 12 | 51953482 | 51953676 | 194 | CAST.MOLF.SPRET | 12 | 51953606 | 51953639 | 34 | - | 17.8455 | 3.91E-07 | 0.0684 | TAGCTGTCCTCAGACACTCCAGAAGAGGGCGTCA | | |
| 1922 | 12 | 52719548 | 52719663 | 115 | CAST.MOLF | 12 | 52719598 | 52719632 | 35 | - | 16.0275 | 9.40E-07 | 0.192 | ATGCCACTCACCAGTGCAGTCACCAGGCGGCACCG | | |
| 1923 | 12 | 57306385 | 57306580 | 195 | SPRET | 12 | 57306499 | 57306532 | 34 | - | 16.4472 | 9.61E-07 | 0.104 | TAGCTGTCTTCAGACACACCAGAAGAGGGCGTCA | | |
| 1924 | 12 | 60631303 | 60632110 | 807 | MRL | 12 | 60631835 | 60631853 | 19 | - | 17.1475 | 8.25E-07 | 0.146 | CAACCACAAGGGGTCAGCA | 11 | ENCFF367UNR,ENCFF099QNE,ENCFF092NMJ,ENCFF089GBC,ENCFF198SSR,ENCFF093ZRU,ENCFF024JOZ,ENCFF907XEG,ENCFF502WDT,ENCFF327LYS,ENCFF055HAP |
| 1924 | 12 | 60631303 | 60632110 | 807 | MRL | 12 | 60631835 | 60631869 | 35 | - | 18.1193 | 2.80E-07 | 0.097 | CAGCTTTCCCCTAATTCAACCACAAGGGGTCAGCA | 11 | ENCFF367UNR,ENCFF099QNE,ENCFF092NMJ,ENCFF089GBC,ENCFF198SSR,ENCFF093ZRU,ENCFF024JOZ.ENCFF907XEG.ENCFF502WDT.ENCFF327LYS.ENCFF055HAP |
| 1925 | 12 | 61769968 | 61770229 | 261 | SPRET | 12 | 61770144 | 61770162 | 19 | - | 17.459 | 6.74E-07 | 0.132 | CAGCCAGGAGGTGGAGCTG | 11 | ENCFF052KVO,ENCFF092NMJ,ENCFF093ZRU,ENCFF907XEG,ENCFF089GBC,ENCFF533APC,ENCFF099QNE,ENCFF142CNG,ENCFF502WDT,ENCFF055HAP,ENCFF024JOZ |
| 1926 | 12 | 63976839 | 63977714 | 875 | MOLF | 12 | 63977325 | 63977359 | 35 | - | 16.5596 | 6.97E-07 | 0.163 | CTGTAGTACAGCTTTAGGTCCACCAGAGGTTCTTT | | |
| 1927 | 12 | 66818546 | 66818762 | 216 | CAST.MOLF.SPRET | 12 | 66818671 | 66818704 | 34 | - | 16.4634 | 9.51E-07 | 0.104 | TAGCTGTCTTCAGACGCACCAGAAGAGGGCATCA | | |
| 1928 | 12 | 69194549 | 69194777 | 228 | BTBR.C57L.CAST.MOLF.SPRET | 12 | 69194669 | 69194702 | 34 | - | 16.4634 | 9.51E-07 | 0.104 | TAGCTGTCTTCAGACGCACCAGAAGAGGGCATCA | | |
| 1929 | 12 | 69699298 | 69699480 | 182 | SPRET | 12 | 69699422 | 69699456 | 35 | - | 17.1927 | 4.84E-07 | 0.138 | CTGTAGCTGTCCTCAGACACCAGAAGAGGGAGTCA | | |
| 1930 | 12 | 69749225 | 69749457 | 232 | SPRET.SWR | 12 | 69749351 | 69749384 | 34 | - | 16.4634 | 9.51E-07 | 0.104 | TAGCTGTCTTCAGACGCACCAGAAGAGGGCATCA | 1 | ENCFF093ZRU |
| 1931 | 12 | 70406728 | 70407498 | 770 | AJ,BALB,BUB,C3H,CBA,CE,DBA1J,DBA2J,FVB,ILNJ2,MRL,NZB,NZO,RH2,SEA,SJL,TH | 12 | 70407294 | 70407312 | 19 | - | 21.2131 | 4.09E-08 | 0.0421 | CAACCAGCAGGGGGCGGAG | 11 | ENCFF099QNE,ENCFF327LYS,ENCFF165LXO,ENCFF907XEG,ENCFF093ZRU,ENCFF194HVL,ENCFF136ZDK,ENCFF097CAQ,ENCFF589ZBJ,ENCFF883UPM,ENCFF198SSR |
| 1932 | 12 | 71050452 | 71050673 | 221 | CAST.MOLF.SPRET | 12 | 71050578 | 71050611 | 34 | - | 16.4634 | 9.51E-07 | 0.104 | TAGCTGTCTTCAGACGCACCAGAAGAGGGCATCA | 1 | ENCFF093ZRU |
| 1933 | 12 | 71269306 | 71269523 | 217 | CAST.MOLF.SPRET | 12 | 71269431 | 71269449 | 19 | - | 17.5574 | 6.32E-07 | 0.128 | ACTCCAGAAGAGGGCGCCA | 1 | ENCFF093ZRU |
| 1933 | 12 | 71269306 | 71269523 | 217 | CAST.MOLF.SPRET | 12 | 71269431 | 71269464 | 34 | - | 18.3659 | 2.76E-07 | 0.0591 | TAGCTGTCTTCAGACACTCCAGAAGAGGGCGCCA | | |
| 1934 | 12 | 73072826 | 73073082 | 256 | SPRET | 12 | 73072951 | 73072984 | 34 | - | 17.9106 | 3.74E-07 | 0.0677 | TAGCTGTCTTCAGACACTCCAGAAGAGGGCACCA | | |
| 1935 | 12 | 76391797 | 76392008 | 211 | CAST.FVB.MOLF.SPRET.SWR | 12 | 76391917 | 76391950 | 34 | - | 17.8455 | 3.91E-07 | 0.0684 | TAGCTGTCCTCAGACACTCCAGAAGAGGGCGTCA | | |
| 1936 | 12 | 76762132 | 76762552 | 420 | FVB.SWR | 12 | 76762254 | 76762287 | 34 | - | 18.4797 | 2.56E-07 | 0.0578 | TTGCTGTCTTCAGACACACCAGAAGAGGGCATCA | 1 | ENCFF093ZRU |
| 1937 | 12 | 76824131 | 76824562 | 431 | MOLF.SPRET | 12 | 76824383 | 76824416 | 34 | - | 17.2439 | 5.79E-07 | 0.0827 | TAGCTGTCCTCAGATACTCCAGAAGAGGGCATCA | | |
| 1938 | 12 | 76915920 | 76916137 | 217 | CAST.MOLF.SPRET | 12 | 76916062 | 76916095 | 34 | - | 17 | 6.77E-07 | 0.0898 | CTGCCTCTGCCGAGATTATCAGCAGGTGCCACCA | | |
| 1939 | 12 | 77280378 | 77280579 | 201 | SPRET | 12 | 77280502 | 77280535 | 34 | - | 17.6423 | 4.47E-07 | 0.0738 | TAGCTGTGTTCAGACACACCAGAAGAGGGCGTCA | | |
| 1940 | 12 | 78845058 | 78845902 | 844 | SPRET | 12 | 78845886 | 78845919 | 34 | - | 18.5854 | 2.38E-07 | 0.0566 | TTGCTGTCTTCAGACACACCAGAAGAGGGCATCG | | |

| | | | | | | | | | | | | | | | | |
|---|---|---|---|---|---|---|---|---|---|---|---|---|---|---|---|---|
| 1941 | 12 | 78845810 | 78845902 | 92 | SPRET | 12 | 78845886 | 78845919 | 34 | - | 18.5854 | 2.38E-07 | 0.0566 | TTGCTGTCTTCAGACACACCAGAAGAGGGCATCG | | |
| 1942 | 12 | 78902264 | 78903195 | 931 | AKR,BALB,C58J,KK,LP,MRL,NOD,NOR,NZO,PJ,PL,RF,SEA,SJL,SMJ,SWR,TH,WSB | 12 | 78902317 | 78902350 | 34 | - | 22.2602 | 1.63E-08 | 0.00935 | CAGCTGTTCTCAGACACACCAGAAGAGGGCATCG | | |
| 1943 | 12 | 79828421 | 79828812 | 391 | SPRET | 12 | 79828751 | 79828784 | 34 | - | 25.8293 | 7.82E-10 | 0.00113 | ATGCAGTACCTGTGGAGGCCAGAAGAGGGCGTGG | 15 | ENCFF099QNE,ENCFF092NMJ,ENCFF024JOZ,ENCFF089GBC,ENCFF136ZDK,ENCFF198SSR,ENCFF907XEG,ENCFF055HAP,ENCFF165LXO,ENCFF327LYS,ENCFF065ODV,ENCFF093ZRU,ENCFF502WDT,ENCFF589ZBJ,ENCFF367UNR |
| 1944 | 12 | 80207471 | 80207675 | 204 | SPRET | 12 | 80207596 | 80207629 | 34 | - | 17.8455 | 3.91E-07 | 0.0684 | TAGCTGTCCTCAGACACTCCAGAAGAGGGCGTCA | | |
| 1945 | 12 | 80219328 | 80219418 | 90 | SPRET | 12 | 80219335 | 80219369 | 35 | - | 20.4587 | 6.45E-08 | 0.0398 | CTGCAGTGTTACCATGCAGCCACTTGAGACCACCG | | |
| 1946 | 12 | 81577580 | 81577951 | 371 | SPRET | 12 | 81577707 | 81577725 | 19 | - | 17.5574 | 6.32E-07 | 0.128 | ACTCCAGAAGAGGGCGCCA | 1 | ENCFF093ZRU |
| 1946 | 12 | 81577580 | 81577951 | 371 | SPRET | 12 | 81577707 | 81577740 | 34 | - | 18.561 | 2.42E-07 | 0.0569 | AAGCTGTCTTCAGACACTCCAGAAGAGGGCGCCA | 1 | ENCFF093ZRU |
| 1947 | 12 | 81833575 | 81834277 | 702 | CAST,MOLF,SPRET | 12 | 81834234 | 81834267 | 34 | - | 27.7967 | 1.16E-10 | 0.000409 | GTGCAGTGCCTGCCATGTCCAGAAGAGGGCGTAA | 23 | ENCFF099QNE,ENCFF210BTO,ENCFF092NMJ,ENCFF821BIA,ENCFF024JOZ,ENCFF327LYS,ENCFF136ZDK,ENCFF430PPJ,ENCFF201JUA,ENCFF502WDT,ENCFF198SSR,ENCFF055HAP,ENCFF883UPM,ENCFF093ZRU,ENCFF033UQX,ENCFF065ODV,ENCFF089GBC,ENCFF589ZBJ,ENCFF142CNG,ENCFF052KVO,ENCFF907XEG,ENCFF533APC,ENCFF367UNR |
| 1948 | 12 | 83449940 | 83450142 | 202 | CAST,CE,KK,MOLF,SPRET,SWR,WSB | 12 | 83450065 | 83450098 | 34 | - | 17.3902 | 5.27E-07 | 0.0795 | TAGCTGTCCTCAGACACTCCAGAAGAGGGCATCA | | |
| 1949 | 12 | 84323235 | 84323944 | 709 | MOLF | 12 | 84323754 | 84323788 | 35 | - | 26.0734 | 1.03E-09 | 0.00465 | CTGCATGTACACCTGCAGGCCAGAAGAGGGCATCA | 12 | ENCFF099QNE,ENCFF821BIA,ENCFF136ZDK,ENCFF327LYS,ENCFF052KVO,ENCFF907XEG,ENCFF093ZRU,ENCFF589ZBJ,ENCFF883UPM,ENCFF502WDT,ENCFF055HAP,ENCFF367UNR |
| 1950 | 12 | 84706777 | 84706999 | 222 | MOLF | 12 | 84706897 | 84706930 | 34 | - | 16.4472 | 9.61E-07 | 0.104 | TAGCTGTCTTCAGACACACCAGAAGAGGGCGTCA | | |
| 1951 | 12 | 85033800 | 85034003 | 203 | CAST,MOLF,SPRET | 12 | 85033924 | 85033942 | 19 | - | 17.8852 | 5.08E-07 | 0.115 | ACACCAGAAGAGGGCACCA | 1 | ENCFF502WDT |
| 1952 | 12 | 85115415 | 85115655 | 240 | SPRET | 12 | 85115536 | 85115569 | 34 | - | 17.6829 | 4.35E-07 | 0.073 | TAGCTGTCTTCAGACGCGCCAGAAGAGGGCATCA | 1 | ENCFF093ZRU |
| 1953 | 12 | 89178524 | 89178676 | 152 | MOLF | 12 | 89178571 | 89178589 | 19 | - | 19.1475 | 2.10E-07 | 0.0777 | TGACCACTGGGTGGCAGCA | 11 | ENCFF024JOZ,ENCFF165LXO,ENCFF327LYS,ENCFF092NMJ,ENCFF089GBC,ENCFF198SSR,ENCFF907XEG,ENCFF093ZRU,ENCFF883UPM,ENCFF099QNE,ENCFF367UNR |
| 1953 | 12 | 89178524 | 89178676 | 152 | MOLF | 12 | 89178571 | 89178605 | 35 | - | 16.055 | 9.26E-07 | 0.191 | GGGCGCTTAAAATGCATGACCACTGGGTGGCAGCA | 11 | ENCFF024JOZ,ENCFF165LXO,ENCFF327LYS,ENCFF092NMJ,ENCFF089GBC,ENCFF198SSR,ENCFF907XEG,ENCFF093ZRU,ENCFF883UPM,ENCFF099QNE,ENCFF367UNR |
| 1954 | 12 | 91755280 | 91755484 | 204 | CAST,MOLF,NOD,SPRET | 12 | 91755406 | 91755439 | 34 | - | 16.7724 | 7.83E-07 | 0.0958 | TAGCTGTCTTCAGATGCACCAGAAGAGGGCGTCA | 5 | ENCFF093ZRU,ENCFF327LYS,ENCFF024JOZ,ENCFF136ZDK,ENCFF883UPM |
| 1955 | 12 | 99805124 | 99805245 | 121 | SWR | 12 | 99805177 | 99805211 | 35 | - | 17.5413 | 3.95E-07 | 0.122 | GTGTAGTAGATCTCTTTGTCCAGTTGGTGGTGCTA | 14 | ENCFF093ZRU,ENCFF805QBU,ENCFF367UNR,ENCFF024JOZ,ENCFF327LYS,ENCFF092NMJ,ENCFF430PPJ,ENCFF055HAP,ENCFF883UPM,ENCFF099QNE,ENCFF821BIA,ENCFF089GBC,ENCFF907XEG,ENCFF502WDT |
| 1956 | 12 | 100568873 | 100569066 | 193 | CAST,MOLF,SPRET | 12 | 100568989 | 100569022 | 34 | - | 18.1545 | 3.18E-07 | 0.0635 | TAGCTGTTTTCAGACACACCAGAAGAGGGCGTCA | | |
| 1957 | 12 | 100996573 | 100996756 | 183 | CAST,CE,MOLF,PJ,SEA,SPRET,WSB | 12 | 100996697 | 100996715 | 19 | - | 19.9836 | 1.12E-07 | 0.0594 | CTACCAGAAGAGGGCACCA | 7 | ENCFF093ZRU,ENCFF089GBC,ENCFF821BIA,ENCFF502WDT,ENCFF024JOZ,ENCFF099QNE,ENCFF367UNR |
| 1958 | 12 | 102003316 | 102003700 | 384 | CAST | 12 | 102003508 | 102003541 | 34 | - | 17.878 | 3.83E-07 | 0.0683 | TAGCTGTCCTCAGACACACCAGAAGAGGGCATCA | 1 | ENCFF093ZRU |
| 1959 | 12 | 107320122 | 107320729 | 607 | SPRET | 12 | 107320343 | 107320376 | 34 | - | 18.3089 | 2.87E-07 | 0.0597 | ATGCAGCTAGAGACACGAGCTCCAGGGGGTACTG | | |
| 1960 | 12 | 108196976 | 108197191 | 215 | AJ,AKR,BUB,C3H,CAST,CBA,CE,DBA1J,DBA2J,FVB,ILNJ2,KK,MA/My,MOLF,MRL,NOD,NOR,NZB,NZO,NZW,PJ,PL,RF,SEA,SJL,SMJ,SPRET,SWR,TH | 12 | 108197101 | 108197134 | 34 | - | 17.8455 | 3.91E-07 | 0.0684 | TAGCTGTCCTCAGACACTCCAGAAGAGGGCGTCA | | |
| 1961 | 12 | 108304978 | 108305039 | 61 | SPRET | 12 | 108305018 | 108305036 | 19 | - | 20.8197 | 5.71E-08 | 0.0463 | TCTCCAGCAGGGGGCACAA | 29 | ENCFF093ZRU,ENCFF367UNR,ENCFF327LYS,ENCFF805QBU,ENCFF099QNE,ENCFF907XEG,ENCFF086ZTV,ENCFF024JOZ,ENCFF089GBC,ENCFF092NMJ,ENCFF198SSR,ENCFF609MDN,ENCFF821BIA,ENCFF589ZBJ,ENCFF165LXO,ENCFF033UQX,ENCFF097CAQ,ENCFF430PPJ,ENCFF136ZDK,ENCFF281QMV,ENCFF957SPH,ENCFF210BTO,ENCFF491RJK,ENCFF951GOP,ENCFF052KVO,ENCFF575MGI,ENCFF533APC,ENCFF316FYN,ENCFF194HVL |
| 1961 | 12 | 108304978 | 108305039 | 61 | SPRET | 12 | 108305018 | 108305051 | 34 | - | 24.9431 | 1.74E-09 | 0.00191 | GGGAAGTTCCCCCTGTCTCCAGCAGGGGGCACAA | 29 | ENCFF093ZRU,ENCFF367UNR,ENCFF327LYS,ENCFF805QBU,ENCFF099QNE,ENCFF907XEG,ENCFF086ZTV,ENCFF024JOZ,ENCFF089GBC,ENCFF092NMJ,ENCFF198SSR,ENCFF609MDN,ENCFF821BIA,ENCFF589ZBJ,ENCFF165LXO,ENCFF033UQX,ENCFF097CAQ,ENCFF430PPJ,ENCFF136ZDK,ENCFF281QMV,ENCFF957SPH,ENCFF210BTO,ENCFF491RJK,ENCFF951GOP,ENCFF052KVO,ENCFF575MGI,ENCFF533APC,ENCFF316FYN,ENCFF194HVL |
| 1962 | 12 | 108718974 | 108719201 | 227 | AJ,AKR,BUB,C3H,CAST,CBA,CE,DBA1J,DBA2J,FVB,ILNJ2,KK,MA/My,MOLF,MRL,NOD,NOR,NZB,NZO,NZW,PJ,PL,RF,SEA,SJL,SMJ,SPRET,SWR,TH,WSB | 12 | 108719101 | 108719134 | 34 | - | 16.4634 | 9.51E-07 | 0.104 | TAGCTGTCTTCAGACGCACCAGAAGAGGGCATCA | | |
| 1963 | 12 | 110611339 | 110611551 | 212 | AKR,BUB,C3H,CAST,CBA,KK,MA/My,MOLF,MRL,NOD,NOR,PJ,SEA,SJL,SPRET | 12 | 110611465 | 110611498 | 34 | - | 16.4472 | 9.61E-07 | 0.104 | TAGCTGTCTTCAGACACACCAGAAGAGGGCGTCA | | |
| 1964 | 12 | 110646704 | 110647014 | 310 | SPRET | 12 | 110646896 | 110646929 | 34 | - | 16.7724 | 7.83E-07 | 0.0958 | TAGCTGTCTTCAGATGCACCAGAAGAGGGCGTCA | | |
| 1965 | 12 | 110733027 | 110733230 | 203 | SPRET | 12 | 110733148 | 110733181 | 34 | - | 16.4634 | 9.51E-07 | 0.104 | TAGCTGTCTTCAGACGCACCAGAAGAGGGCATCA | 1 | ENCFF093ZRU |
| 1966 | 12 | 110796942 | 110797151 | 209 | AKR,BUB,CAST,DBA1J,DBA2J,ILNJ2,KK,MA/My,MOLF,MRL,NOD,NOR,NZB,NZW,PJ,RF,SEA,SJL,SPRET | 12 | 110797068 | 110797086 | 19 | - | 17.8852 | 5.08E-07 | 0.115 | ACACCAGAAGAGGGCACCA | | |
| 1967 | 12 | 110829111 | 110829325 | 214 | AKR,BUB,CAST,DBA1J,DBA2J,ILNJ2,KK,MA/My,MOLF,MRL,NOD,NOR,NZB,NZW,PJ,RF,SEA,SJL,SPRET | 12 | 110829229 | 110829247 | 19 | - | 17.5574 | 6.32E-07 | 0.128 | ACTCCAGAAGAGGGCGCCA | 2 | ENCFF093ZRU,ENCFF367UNR |
| 1967 | 12 | 110829111 | 110829325 | 214 | AKR,BUB,CAST,DBA1J,DBA2J,ILNJ2,KK,MA/My,MOLF,MRL,NOD,NOR,NZB,NZW,PJ,RF,SEA,SJL,SPRET | 12 | 110829229 | 110829262 | 34 | - | 18.3659 | 2.76E-07 | 0.0591 | TAGCTGTCTTCAGACACTCCAGAAGAGGGCGCCA | 2 | ENCFF093ZRU,ENCFF367UNR |
| 1968 | 12 | 111244368 | 111244786 | 418 | CAST,KK,MA/My,MOLF,RF,SPRET | 12 | 111244714 | 111244732 | 19 | - | 17.5738 | 6.25E-07 | 0.128 | ACTCCAGAAGAGGGCGCTA | 1 | ENCFF093ZRU |
| 1968 | 12 | 111244368 | 111244786 | 418 | CAST,KK,MA/My,MOLF,RF,SPRET | 12 | 111244714 | 111244747 | 34 | - | 18.187 | 3.12E-07 | 0.0627 | TAGCTGTCTTCAGTCACTCCAGAAGAGGGCGCTA | 1 | ENCFF093ZRU |
| 1969 | 12 | 111689704 | 111689903 | 199 | AJ,AKR,C3H,CAST,CBA,CE,DBA1J,DBA2J,FVB,ILNJ2,MOLF,MRL,NZB,NZO,NZW,PL,SMJ,SPRET,SWR,TH,WSB | 12 | 111689829 | 111689862 | 34 | - | 17.2114 | 5.91E-07 | 0.0834 | TAGCTGTTTTCAGACACTCCAGAAGAGGGCATCA | | |
| 1970 | 12 | 112713733 | 112713822 | 89 | AJ,AKR,C3H,CAST,CBA,CE,FVB,MRL,NZB,NZO,NZW,PJ,PL,SEA,SMJ,SPRET,SWR,TH,WSB | 12 | 112713775 | 112713808 | 34 | - | 17.2114 | 5.91E-07 | 0.0834 | GAGCAACTGCCCTCTCATGCAGTAGAGGGCAGCT | 1 | ENCFF089GBC |
| 1971 | 12 | 113171875 | 113172098 | 223 | 129S1,AJ,AKR,BALB,BUB,C58J,CE,FVB,KK,LP,MA/My,NZB,NZO,NZW,PJ,SEA,SWR,WSB | 12 | 113171998 | 113172032 | 35 | - | 19.5046 | 1.19E-07 | 0.0579 | GTGTACTGTCACTGACACACCAGAAGAGGGCATCA | | |
| 1972 | 12 | 113759666 | 113760163 | 497 | DBA2J | 12 | 113759682 | 113759700 | 19 | - | 23.623 | 3.91E-09 | 0.0223 | CAACCAGCAGGGGGCGCTG | 10 | ENCFF821BIA,ENCFF024JOZ,ENCFF907XEG,ENCFF136ZDK,ENCFF589ZBJ,ENCFF883UPM,ENCFF367UNR,ENCFF533APC,ENCFF194HVL,ENCFF327LYS |
| 1972 | 12 | 113759666 | 113760163 | 497 | DBA2J | 12 | 113759682 | 113759715 | 34 | - | 26.065 | 6.29E-10 | 0.000986 | CTGCAGGGATGATCACAACCAGCAGGGGGCGCTG | 10 | ENCFF821BIA,ENCFF024JOZ,ENCFF907XEG,ENCFF136ZDK,ENCFF589ZBJ,ENCFF883UPM,ENCFF367UNR,ENCFF533APC,ENCFF194HVL,ENCFF327LYS |
| 1973 | 12 | 114074677 | 114074879 | 202 | FVB | 12 | 114074664 | 114074697 | 34 | - | 16.6667 | 8.37E-07 | 0.099 | CTACAGTGAATCTGTCATGCAGCAGGGGGTGCCA | | |
| 1974 | 12 | 114283098 | 114283773 | 675 | 129S1,AJ,BALB,BUB,C58J,CE,DBA1J,DBA2J,FVB,ILNJ2,KK,LP,MA/My,PJ,RF,SEA,SMJ,SPRET,SWR,TH,WSB | 12 | 114283574 | 114283592 | 19 | - | 20.2787 | 8.86E-08 | 0.0547 | GGGCCGGCAGGGGGCACTG | | |
| 1974 | 12 | 114283098 | 114283773 | 675 | 129S1,AJ,BALB,BUB,C58J,CE,DBA1J,DBA2J,FVB,ILNJ2,KK,LP,MA/My,PJ,RF,SEA,SMJ,SPRET,SWR,TH,WSB | 12 | 114283574 | 114283608 | 35 | - | 17.1743 | 4.90E-07 | 0.139 | CTCCACAAGGTACTTAGGGCCGGCAGGGGGCACTG | | |
| 1975 | 12 | 114480188 | 114481029 | 841 | AJ,RF,SPRET | 12 | 114480711 | 114480729 | 19 | - | 18.0656 | 4.49E-07 | 0.111 | TAACCAGCAGGGGGTACAA | | |
| 1975 | 12 | 114480188 | 114481029 | 841 | AJ,RF,SPRET | 12 | 114480711 | 114480744 | 34 | - | 20.1463 | 7.99E-08 | 0.0273 | CTGCAAGGGAGGTCATAACCAGCAGGGGGTACAA | | |
| 1976 | 12 | 115250589 | 115251261 | 672 | FVB,SWR | 12 | 115250838 | 115250871 | 34 | - | 17.5691 | 4.69E-07 | 0.0751 | GAGCATGACCACAAGTCCACAGCAGGTGGCAACA | | |
| 1977 | 12 | 115377650 | 115377916 | 266 | AKR,C3H,CBA,MRL,NZB,NZO,NZW,PL | 12 | 115377788 | 115377806 | 19 | - | 21.082 | 4.57E-08 | 0.0421 | GGACCAGCAGGGGGCACAA | | |
| 1977 | 12 | 115377650 | 115377916 | 266 | AKR,C3H,CBA,MRL,NZB,NZO,NZW,PL | 12 | 115377788 | 115377821 | 34 | - | 26.813 | 3.09E-10 | 0.000662 | CTGCAGGTGCATTCAGGACCAGCAGGGGGCACAA | | |
| 1978 | 12 | 115836299 | 115836552 | 253 | C3H,CBA,MRL,NZB,NZO,NZW,PL | 12 | 115836273 | 115836306 | 34 | - | 17.935 | 3.68E-07 | 0.0672 | ATGCATGGGAGCTCATGACCAGCAGGGGTCACCA | | |
| 1979 | 12 | 116036212 | 116036412 | 200 | AJ,DBA1J,DBA2J,ILNJ2,RF,SPRET | 12 | 116036329 | 116036362 | 34 | - | 16.7724 | 7.83E-07 | 0.0958 | TAGCTGTCTTCAGATGCACCAGAAGAGGGCGTCA | | |
| 1980 | 12 | 117486933 | 117487149 | 216 | CAST,CE,MOLF,MRL,PL,SPRET,WSB | 12 | 117487056 | 117487074 | 19 | - | 17.5574 | 6.32E-07 | 0.128 | ACTCCAGAAGAGGGCGCCA | | |
| 1980 | 12 | 117486933 | 117487149 | 216 | CAST,CE,MOLF,MRL,PL,SPRET,WSB | 12 | 117487056 | 117487089 | 34 | - | 18.3659 | 2.76E-07 | 0.0591 | TAGCTGTCTTCAGACACTCCAGAAGAGGGCGCCA | | |
| 1981 | 12 | 117908038 | 117908250 | 212 | 129S1,BALB,BUB,C3H,C58J,CAST,CBA,DBA1J,DBA2J,LP,MA/My,MOLF,MRL,NOD,NOR,PJ,PL,SEA,SJL,SPRET | 12 | 117908167 | 117908200 | 34 | - | 18.1545 | 3.18E-07 | 0.0635 | TAGCTGTTTTCAGACACACCAGAAGAGGGCGTCA | | |
| 1982 | 12 | 118518608 | 118518801 | 193 | CAST,SPRET | 12 | 118518724 | 118518757 | 34 | - | 17.8455 | 3.91E-07 | 0.0684 | TAGCTGTCCTCAGACACTCCAGAAGAGGGCGTCA | | |
| 1983 | 12 | 118839896 | 118840088 | 192 | MOLF,SPRET | 12 | 118840020 | 118840053 | 34 | - | 16.5528 | 8.99E-07 | 0.102 | TAGCTGTCTTCAGACACACCAGAAGAGGGCGTCG | | |
| 1984 | 13 | 5429253 | 5429645 | 392 | C57L,KK,LP,NZB,NZO,NZW,SPRET | 13 | 5429372 | 5429405 | 34 | + | 17.4146 | 5.18E-07 | 0.0915 | ATGCAGCTAGAGACACAAGCTCCAGGGGGTACTG | | |
| 1985 | 13 | 10986009 | 10986431 | 422 | AKR,BTBR,C57L,ILNJ2,KK,LP,MOLF,NOD,NOR,NZB,RF,SPRET | 13 | 10986279 | 10986313 | 35 | + | 15.9266 | 9.95E-07 | 0.199 | GTACAGGGGAACACCAGGGCCAAAAGGGGGCAGTG | | |
| 1985 | 13 | 10986009 | 10986431 | 422 | AKR,BTBR,C57L,ILNJ2,KK,LP,MOLF,NOD,NOR,NZB,RF,SPRET | 13 | 10986295 | 10986313 | 19 | + | 17.2131 | 7.91E-07 | 0.171 | GGGCCAAAAGGGGGCAGTG | | |
| 1986 | 13 | 20114399 | 20115239 | 840 | CAST | 13 | 20115121 | 20115154 | 34 | + | 18.2276 | 3.03E-07 | 0.0698 | CTGGAGGGCTTTACCTGGGCACCTGGGGGAGCCA | | |
| 1987 | 13 | 20481672 | 20482244 | 572 | SPRET | 13 | 20482170 | 20482188 | 19 | + | 22.3443 | 1.47E-08 | 0.038 | TAACCACCAGGTGGCGCAA | 32 | ENCFF093ZRU,ENCFF367UNR,ENCFF099QNE,ENCFF052KVO,ENCFF821BIA,ENCFF327LYS,ENCFF024JOZ,ENCFF907XEG,ENCFF089GBC,ENCFF430PPJ,ENCFF502WDT,ENCFF055HAP,ENCFF589ZBJ,ENCFF198SSR,ENCFF065ODV,ENCFF165LXO,ENCFF092NMJ,ENCFF136ZDK,ENCFF883UPM,ENCFF281QMV,ENCFF097CAQ,ENCFF957SPH,ENCFF210BTO,ENCFF142CNG,ENCFF575MGI,ENCFF951GOP,ENCFF316FYN,ENCFF194HVL,ENCFF504JDS,ENCFF533APC,ENCFF609MDN,ENCFF491RJK |
| 1988 | 13 | 26676494 | 26677214 | 720 | CAST,KK,MOLF,PJ,SEA,SPRET | 13 | 26676886 | 26676920 | 35 | + | 16.0367 | 9.36E-07 | 0.193 | ATGCACTGCCCCTGGGCCGCAAGAAGGGGGCCAAG | | |
| 1989 | 13 | 31566773 | 31567439 | 666 | SPRET | 13 | 31566844 | 31566877 | 34 | + | 19.439 | 1.32E-07 | 0.0449 | ATGCAGTGCTTGTGGAAGCTAGAAGAGGGCATCA | | |
| 1990 | 13 | 32167957 | 32168166 | 209 | CAST,LP,MOLF,SPRET,WSB | 13 | 32168025 | 32168058 | 34 | + | 16.4472 | 9.61E-07 | 0.118 | TAGCTGTCTTCAGACACACCAGAAGAGGGCGTCA | | |
| 1991 | 13 | 32897160 | 32897397 | 237 | SPRET | 13 | 32897253 | 32897286 | 34 | + | 16.4634 | 9.51E-07 | 0.118 | TAGCTGTCTTCAGACGCACCAGAAGAGGGCATCA | | |
| 1992 | 13 | 33702837 | 33703032 | 195 | CAST,MOLF,NZW,SPRET | 13 | 33702888 | 33702921 | 34 | + | 17.8455 | 3.91E-07 | 0.0799 | TAGCTGTCCTCAGACACTCCAGAAGAGGGCGTCA | | |
| 1993 | 13 | 34401688 | 34401948 | 260 | AJ,AKR,B10D2,B10J,BALB,BTBR,C3H,C57L,C58J,CBA,CE,DBA1J,DBA2J,ILNJ2,KK,LP,MA/My,MOLF,MRL,NZB,NZO,NZW,PL,RF,RH2,SMJ,TH,WSB | 13 | 34401905 | 34401938 | 34 | + | 17.065 | 6.50E-07 | 0.103 | TAGCTGTTTTCAGATACTCCAGAAGAGGGCATCA | 1 | ENCFF093ZRU |
| 1994 | 13 | 36091042 | 36091516 | 474 | SPRET | 13 | 36091054 | 36091087 | 34 | + | 17.1138 | 6.30E-07 | 0.101 | TTGCTATCTTCAGACACACCAGAAGAGGGCATTG | 2 | ENCFF093ZRU,ENCFF367UNR |
| 1995 | 13 | 37565055 | 37565266 | 211 | 129S1,AJ,BALB,BTBR,BUB,C3H,C57L,CAST,CBA,FVB,MOLF,NZW,RH2,SJL,SMJ,SPRET,SWR,TH,WSB | 13 | 37565124 | 37565157 | 34 | + | 16.9187 | 7.13E-07 | 0.105 | TAGCTGTCTTCAGACGCACCAGAAGAGGGCGTCA | | |
| 1996 | 13 | 37703561 | 37703788 | 227 | 129S1,AJ,AKR,BALB,BTBR,BUB,C3H,C57L,C58J,CAST,CBA,CE,FVB,ILNJ2,KK,LP,MOLF,MRL,NZB,NZO,NZW,PJ,RF,RH2,SEA,SJL,SMJ,SPRET,SWR,TH,WSB | 13 | 37703646 | 37703679 | 34 | + | 16.4634 | 9.51E-07 | 0.118 | TAGCTGTCTTCAGACGCACCAGAAGAGGGCATCA | | |
| 1997 | 13 | 40959214 | 40959416 | 202 | CAST,MOLF,SPRET | 13 | 40959273 | 40959306 | 34 | + | 17.2927 | 5.61E-07 | 0.0957 | TAGCTGTCCTCAAACACTCCAGAAGAGGGCATCA | | |
| 1998 | 13 | 41041924 | 41042127 | 203 | CAST,MOLF,NOD,NOR,SPRET,WSB | 13 | 41041980 | 41042013 | 34 | + | 19.0163 | 1.77E-07 | 0.0535 | CAGCTGTCTTCAGACACACCAGAAGAGGGCGTCA | | |
| 1999 | 13 | 41419920 | 41420119 | 199 | CAST,MOLF,SPRET | 13 | 41419977 | 41420010 | 34 | + | 17.6667 | 4.40E-07 | 0.0842 | TAGCTGTTTTCAGACACTCCAGAAGAGGGCGTCA | | |
| 2000 | 13 | 43332668 | 43332746 | 78 | SPRET | 13 | 43332676 | 43332709 | 34 | + | 20.0081 | 8.83E-08 | 0.0337 | CTGCAGTGCCCAGTGCCTTCAGAGGATGACAGCC | | |
| 2001 | 13 | 43739848 | 43740205 | 357 | MOLF,SPRET | 13 | 43740066 | 43740100 | 35 | + | 17.2844 | 4.59E-07 | 0.133 | TAGCTGTACTTCCGATGCACCAGAAGAGGGCATTC | | |
| 2002 | 13 | 44957435 | 44957644 | 209 | CAST,MA/My,MOLF,MRL,NZO,PJ,SEA,SPRET,WSB | 13 | 44957502 | 44957535 | 34 | + | 16.4634 | 9.51E-07 | 0.118 | TAGCTGTCTTCAGACGCACCAGAAGAGGGCATCA | 1 | ENCFF093ZRU |
| 2003 | 13 | 45028140 | 45028334 | 194 | SPRET | 13 | 45028192 | 45028225 | 34 | + | 16.4634 | 9.51E-07 | 0.118 | TAGCTGTCTTCAGACGCACCAGAAGAGGGCATCA | | |
| 2004 | 13 | 47148578 | 47148756 | 178 | MOLF | 13 | 47148623 | 47148656 | 34 | + | 16.4472 | 9.61E-07 | 0.118 | TAGCTGTCTTCAGACACACCAGAAGAGGGCGTCA | | |
| 2005 | 13 | 48801803 | 48801886 | 83 | SPRET | 13 | 48801812 | 48801846 | 35 | + | 19.6422 | 1.09E-07 | 0.058 | CTGCATCCCTAGCCCTTGCCCAGGTGAGGGCTCTA | | |
| 2006 | 13 | 49175967 | 49176165 | 198 | CAST,SPRET | 13 | 49176023 | 49176056 | 34 | + | 16.4309 | 9.71E-07 | 0.118 | TAGCTGTCTTCAGACGCTCCAGAAGAGGGCGTCA | | |
| 2007 | 13 | 51181904 | 51182456 | 552 | CAST | 13 | 51181931 | 51181965 | 35 | + | 17.0092 | 5.39E-07 | 0.145 | CTGTAGCTGTTTTCAGACACCAGAAGAGGGCATCA | | |
| 2008 | 13 | 51880607 | 51880867 | 260 | CAST,MOLF,SPRET | 13 | 51880723 | 51880757 | 35 | + | 18.1284 | 2.79E-07 | 0.1 | CTGTAGCTGTCTTCAGACACCAGAAGAGGGCATCA | | |
| 2009 | 13 | 53457818 | 53457988 | 170 | 129S1,AJ,AKR,BALB,BTBR,BUB,C3H,C57L,CAST,CBA,CE,DBA1J,DBA2J,FVB,ILNJ2,KK,MA/My,MRL,NOD,NOR,NZB,NZO,NZW,PJ,PL,RF,RH2,SEA,SJL,SMJ,SPRET,SWR,TH,WSB | 13 | 53457846 | 53457879 | 34 | + | 17.9675 | 3.61E-07 | 0.0773 | TAGCTGTCTTCAGACCCACCAGAAGAGGGCATCG | | |
| 2010 | 13 | 53681629 | 53681883 | 254 | CAST,MOLF,SPRET | 13 | 53681740 | 53681773 | 34 | + | 16.4472 | 9.61E-07 | 0.118 | TAGCTGTCTTCAGACACACCAGAAGAGGGCGTCA | | |
| 2011 | 13 | 54235664 | 54235867 | 203 | AJ,BALB,C3H,CBA,CE,SPRET | 13 | 54235724 | 54235757 | 34 | + | 16.4634 | 9.51E-07 | 0.118 | TAGCTGTCTTCAGACGCACCAGAAGAGGGCATCA | | |
| 2012 | 13 | 54647937 | 54648136 | 199 | BTBR,C57L,CAST,MOLF,SPRET | 13 | 54647996 | 54648029 | 34 | + | 17.8293 | 3.95E-07 | 0.0803 | TAGCTGTCTTCAGACCCTCCAGAAGAGGGCGTCA | | |
| 2013 | 13 | 54888829 | 54889030 | 201 | CAST,MOLF,SPRET | 13 | 54888887 | 54888920 | 34 | + | 16.4472 | 9.61E-07 | 0.118 | TAGCTGTCTTCAGACACACCAGAAGAGGGCGTCA | | |
| 2014 | 13 | 55538000 | 55538394 | 394 | 129S1,AJ,BALB,BTBR,C3H,C57L,CAST,CBA,CE,DBA1J,DBA2J,ILNJ2,MOLF,MRL,NZB,NZO,NZW,PJ,SEA,SPRET,SWR,WSB | 13 | 55538071 | 55538104 | 34 | + | 18.561 | 2.42E-07 | 0.0646 | AAGCTGTCTTCAGACACTCCAGAAGAGGGCGCCA | | |
| 2014 | 13 | 55538000 | 55538394 | 394 | 129S1,AJ,BALB,BTBR,C3H,C57L,CAST,CBA,CE,DBA1J,DBA2J,ILNJ2,MOLF,MRL,NZB,NZO,NZW,PJ,SEA,SPRET,SWR,WSB | 13 | 55538086 | 55538104 | 19 | + | 17.5574 | 6.32E-07 | 0.153 | ACTCCAGAAGAGGGCGCCA | | |
| 2015 | 13 | 56647440 | 56648286 | 846 | SPRET | 13 | 56648102 | 56648135 | 34 | + | 19.2927 | 1.46E-07 | 0.0473 | TGGCTGATCCAGCACTTCTCACTAGGTGGCACCA | 18 | ENCFF367UNR,ENCFF093ZRU,ENCFF821BIA,ENCFF092NMJ,ENCFF201JUA,ENCFF198SSR,ENCFF609MDN,ENCFF024JOZ,ENCFF089GBC,ENCFF327LYS,ENCFF099QNE,ENCFF907XEG,ENCFF575MGI,ENCFF957SPH,ENCFF430PPJ,ENCFF951GOP,ENCFF281QMV,ENCFF165LXO |
| 2016 | 13 | 56725420 | 56726317 | 897 | SPRET | 13 | 56726044 | 56726077 | 34 | + | 17.2764 | 5.67E-07 | 0.0964 | TTGCCCTTCTGTCTTCAGCCAAGTGAGGGCAGCA | 8 | ENCFF093ZRU,ENCFF024JOZ,ENCFF165LXO,ENCFF198SSR,ENCFF609MDN,ENCFF367UNR,ENCFF907XEG,ENCFF327LYS |
| 2017 | 13 | 57024926 | 57025078 | 152 | SPRET | 13 | 57024898 | 57024932 | 35 | + | 16.8165 | 6.02E-07 | 0.153 | CTGCAGTGGGGAGGAGAGTGCAGATGGAGGCACTG | | |
| 2018 | 13 | 58446347 | 58446563 | 216 | MOLF,SPRET | 13 | 58446422 | 58446455 | 34 | + | 17.9431 | 3.66E-07 | 0.0779 | CAGCTGTCCTCAGACACTCCAGAAGAGGGAGTCA | | |
| 2019 | 13 | 58493728 | 58493930 | 202 | BUB,CAST,FVB,MOLF,MRL,SMJ,SPRET,SWR,WSB | 13 | 58493787 | 58493820 | 34 | + | 18.3659 | 2.76E-07 | 0.0667 | TAGCTGTCTTCAGACACTCCAGAAGAGGGCGCCA | | |
| 2019 | 13 | 58493728 | 58493930 | 202 | BUB,CAST,FVB,MOLF,MRL,SMJ,SPRET,SWR,WSB | 13 | 58493802 | 58493820 | 19 | + | 17.5574 | 6.32E-07 | 0.153 | ACTCCAGAAGAGGGCGCCA | | |
| 2020 | 13 | 58681183 | 58681410 | 227 | CAST,MOLF,SPRET | 13 | 58681268 | 58681302 | 35 | + | 17.422 | 4.24E-07 | 0.128 | TAGCTGTCTTCAGACAGCGCCAGAAGAGGGCATCA | | |
| 2021 | 13 | 59642786 | 59642848 | 62 | AJ,BALB,MOLF,PJ,SEA,WSB | 13 | 59642843 | 59642877 | 35 | + | 19.8349 | 9.67E-08 | 0.054 | ATGCAGTGCCATGCATCTGCCTGCTGAGGGAGGGC | | |
| 2022 | 13 | 59825079 | 59825595 | 516 | MOLF | 13 | 59825432 | 59825466 | 35 | + | 16.0917 | 9.07E-07 | 0.19 | TAGCTGTCTTCAGACAGCACCAGAAGAGGGCATCA | 1 | ENCFF367UNR |
| 2023 | 13 | 62138322 | 62138522 | 200 | NOD,SPRET | 13 | 62138381 | 62138414 | 34 | + | 17.2439 | 5.79E-07 | 0.0964 | TAGCTGTCCTCAGATACTCCAGAAGAGGGCATCA | | |
| 2024 | 13 | 62923626 | 62923831 | 205 | SPRET | 13 | 62923688 | 62923721 | 34 | + | 16.4634 | 9.51E-07 | 0.118 | TAGCTGTCTTCAGACGCACCAGAAGAGGGCATCA | | |
| 2025 | 13 | 65159594 | 65159740 | 146 | 129S1,AJ,AKR,BALB,BTBR,BUB,C3H,C57L,C58J,CBA,CE,DBA1J,DBA2J,FVB,ILNJ2,KK,MRL,NOD,NOR,NZB,NZO,NZW,PJ,PL,RF,SEA,SJL,SMJ,SWR,TH,WSB | 13 | 65159715 | 65159749 | 35 | + | 19.789 | 9.95E-08 | 0.055 | TTGCAGGTGCCCCTAGAGACCAGCAGGAAGCAGTG | | |
| 2026 | 13 | 70842683 | 70842893 | 210 | 129S1,BTBR,C57L,CAST,MOLF,MRL,SPRET,SWR,TH,WSB | 13 | 70842751 | 70842784 | 34 | + | 18.3659 | 2.76E-07 | 0.0667 | TAGCTGTCTTCAGACACTCCAGAAGAGGGCGCCA | | |
| 2026 | 13 | 70842683 | 70842893 | 210 | 129S1,BTBR,C57L,CAST,MOLF,MRL,SPRET,SWR,TH,WSB | 13 | 70842766 | 70842784 | 19 | + | 17.5574 | 6.32E-07 | 0.153 | ACTCCAGAAGAGGGCGCCA | | |
| 2027 | 13 | 73800485 | 73800691 | 206 | MOLF,SPRET | 13 | 73800548 | 73800581 | 34 | + | 18.3659 | 2.76E-07 | 0.0667 | TAGCTGTCTTCAGACACTCCAGAAGAGGGCGCCA | | |
| 2027 | 13 | 73800485 | 73800691 | 206 | MOLF,SPRET | 13 | 73800563 | 73800581 | 19 | + | 17.5574 | 6.32E-07 | 0.153 | ACTCCAGAAGAGGGCGCCA | | |
| 2028 | 13 | 73874665 | 73874896 | 231 | SPRET | 13 | 73874750 | 73874783 | 34 | + | 16.5528 | 8.99E-07 | 0.117 | CAGCTGTCTTCAGACACACCAGAAGAGGGAGGGC | | |
| 2029 | 13 | 74228665 | 74228774 | 109 | SPRET,SWR | 13 | 74228665 | 74228698 | 34 | + | 16.4472 | 9.61E-07 | 0.118 | TAGCTGTCTTCAGACACACCAGAAGAGGGCGTCA | | |
| 2030 | 13 | 75970433 | 75970637 | 204 | 129S1,AJ,AKR,BALB,BTBR,C3H,C57L,CBA,CE,DBA1J,DBA2J,FVB,ILNJ2,KK,MA/My,MRL,NZB,NZO,PJ,PL,RF,SEA,SJL,SMJ,SWR,TH,WSB | 13 | 75970493 | 75970526 | 34 | + | 18.3659 | 2.76E-07 | 0.0667 | TAGCTGTCTTCAGACACTCCAGAAGAGGGCGCCA | 1 | ENCFF093ZRU |
| 2030 | 13 | 75970433 | 75970637 | 204 | 129S1,AJ,AKR,BALB,BTBR,C3H,C57L,CBA,CE,DBA1J,DBA2J,FVB,ILNJ2,KK,MA/My,MRL,NZB,NZO,PJ,PL,RF,SEA,SJL,SMJ,SWR,TH,WSB | 13 | 75970508 | 75970526 | 19 | + | 17.5574 | 6.32E-07 | 0.153 | ACTCCAGAAGAGGGCGCCA | 1 | ENCFF093ZRU |
| 2031 | 13 | 82166887 | 82167137 | 250 | BALB,BUB,CAST,MOLF,NZW,PJ,SEA | 13 | 82167094 | 82167127 | 34 | + | 16.6829 | 8.29E-07 | 0.112 | ATGCAGCTAGAGACACGAGCTCCAGGGGGTGGGG | | |
| 2032 | 13 | 86304036 | 86304228 | 192 | CAST,SPRET | 13 | 86304091 | 86304124 | 34 | + | 16.9593 | 6.95E-07 | 0.104 | TTGCTCTCTTCAGACACACCAGAAGAGGGCATCA | | |
| 2033 | 13 | 92320810 | 92321648 | 838 | WSB | 13 | 92320809 | 92320843 | 35 | + | 18.5321 | 2.18E-07 | 0.0887 | CTGCATATATGCATGCATGCCAGAAGAGAGCACCA | | |
| 2034 | 13 | 97129712 | 97130056 | 344 | 129S1,AKR,BTBR,C3H,C57L,C58J,CBA,CE,DBA1J,DBA2J,FVB,ILNJ2,KK,MA/My,MOLF,MRL,NZB,NZO,PL,SJL,SMJ,SWR | 13 | 97129730 | 97129748 | 19 | + | 18.9672 | 2.39E-07 | 0.102 | TGGCCACCAGAGGACGGCT | | |
| 2035 | 13 | 97202695 | 97202889 | 194 | SPRET | 13 | 97202823 | 97202857 | 35 | + | 17.5505 | 3.93E-07 | 0.122 | CTGTATGTATGCCTGCAGGCCAGAAAATGGCACCA | | |
| 2036 | 13 | 100717571 | 100717780 | 209 | CAST,MOLF,SPRET,WSB | 13 | 100717637 | 100717670 | 34 | + | 17.8455 | 3.91E-07 | 0.0799 | TAGCTGTCCTCAGACACTCCAGAAGAGGGCGTCA | | |
| 2037 | 13 | 100801788 | 100802009 | 221 | SPRET | 13 | 100801858 | 100801891 | 34 | + | 16.4472 | 9.61E-07 | 0.118 | TAGCTGTCTTCAGACACACCAGAAGAGGGCGTCA | | |
| 2038 | 13 | 100957308 | 100957503 | 195 | SPRET | 13 | 100957362 | 100957395 | 34 | + | 18.0813 | 3.34E-07 | 0.074 | TAGCTGTCTCCAGACACACCAGAAGAGGGCGTCA | 1 | ENCFF093ZRU |
| 2039 | 13 | 103223065 | 103223277 | 212 | SPRET | 13 | 103223134 | 103223167 | 34 | + | 21.1463 | 3.83E-08 | 0.0191 | TTGCTGTCCTCGGACACACCAGAAGAGGGCATCA | 1 | ENCFF093ZRU |
| 2040 | 13 | 104410721 | 104410907 | 186 | SPRET | 13 | 104410765 | 104410798 | 34 | + | 16.4472 | 9.61E-07 | 0.118 | TAGCTGTCTTCAGACACACCAGAAGAGGGCGTCA | | |
| 2041 | 13 | 107104470 | 107104666 | 196 | SPRET | 13 | 107104524 | 107104557 | 34 | + | 16.6341 | 8.54E-07 | 0.114 | CTGTAGCTGTCAGACACTCCAGAAGAGGGAGTCA | | |
| 2042 | 13 | 111494293 | 111494492 | 199 | SPRET | 13 | 111494346 | 111494379 | 34 | + | 18.935 | 1.88E-07 | 0.0557 | TTGCTGTCTTCAGACACACCAGAAGAGGGCGTCA | | |
| 2043 | 13 | 111772273 | 111772488 | 215 | SPRET | 13 | 111772326 | 111772359 | 34 | + | 17.8455 | 3.91E-07 | 0.0799 | TAGCTGTCCTCAGACACTCCAGAAGAGGGCGTCA | | |
| 2044 | 13 | 111806221 | 111806681 | 460 | SPRET | 13 | 111806676 | 111806709 | 34 | + | 16.9593 | 6.95E-07 | 0.104 | TTGCTCTCTTCAGACACACCAGAAGAGGGCATCA | | |
| 2045 | 13 | 114264850 | 114265796 | 946 | SPRET | 13 | 114265061 | 114265094 | 34 | + | 23.4878 | 6.05E-09 | 0.00529 | GTGCAGTACCCATAGACACCAGGAGAGGGCATAG | 12 | ENCFF821BIA,ENCFF052KVO,ENCFF065ODV,ENCFF367UNR,ENCFF327LYS,ENCFF024JOZ,ENCFF907XEG,ENCFF198SSR,ENCFF093ZRU,ENCFF055HAP,ENCFF502WDT,ENCFF099QNE |
| 2046 | 13 | 117351251 | 117351464 | 213 | AJ,AKR,BALB,CAST,CBA,DBA1J,FVB,ILNJ2,MA/My,MOLF,NOD,NOR,RF,RH2,SJL,SMJ,SPRET,SWR,TH,WSB | 13 | 117351321 | 117351354 | 34 | + | 16.4472 | 9.61E-07 | 0.118 | TAGCTGTCTTCAGACACACCAGAAGAGGGCGTCA | | |
| 2047 | 13 | 117378214 | 117378420 | 206 | AJ,CAST,CBA,DBA1J,ILNJ2,MA/My,MOLF,RH2,SMJ,SPRET | 13 | 117378276 | 117378309 | 34 | + | 17.6992 | 4.30E-07 | 0.083 | TAGCTGTCCTCAGATACTCCAGAAGAGGGCGTCA | | |
| 2048 | 13 | 117384582 | 117384780 | 198 | SPRET | 13 | 117384638 | 117384671 | 34 | + | 17.2439 | 5.79E-07 | 0.0964 | TAGCTGTCCTCAGATACTCCAGAAGAGGGCATCA | | |
| 2049 | 13 | 117494412 | 117495255 | 843 | AJ,AKR,BALB,CAST,CBA,DBA1J,FVB,ILNJ2,MA/My,MOLF,NOD,NOR,RF,RH2,SJL,SMJ,SPRET,SWR,TH,WSB | 13 | 117495039 | 117495073 | 35 | + | 16.4587 | 7.38E-07 | 0.171 | ATGAGGTTTCTTCACCCCTCCAATAGAGGGCGCAC | | |
| 2050 | 13 | 3646978 | 3647185 | 207 | SPRET | 13 | 3647105 | 3647138 | 34 | - | 16.4472 | 9.61E-07 | 0.118 | TAGCTGTCTTCAGACACACCAGAAGAGGGCGTCA | 1 | ENCFF093ZRU |
| 2051 | 13 | 9103515 | 9103717 | 202 | AKR,BTBR,C57L,CAST,KK,LP,MOLF,NOD,NOR,NZB,NZO,NZW,RF,SPRET,TH | 13 | 9103645 | 9103678 | 34 | - | 16.4472 | 9.61E-07 | 0.118 | TAGCTGTCTTCAGACACACCAGAAGAGGGCGTCA | | |
| 2052 | 13 | 9303882 | 9304098 | 216 | 129S1,AKR,BTBR,BUB,C57L,CAST,ILNJ2,KK,LP,MA/My,MOLF,NOD,NOR,NZB,NZO,NZW,PJ,RF,SEA,SMJ,SPRET,TH,WSB | 13 | 9304007 | 9304040 | 34 | - | 16.5691 | 8.90E-07 | 0.117 | TAGCTGTCTTCAGACGCACCAGAAGAGGGCATCG | | |
| 2053 | 13 | 10767788 | 10768155 | 367 | SPRET | 13 | 10768104 | 10768137 | 34 | - | 25.6829 | 8.95E-10 | 0.00151 | GTGCAGTGCCAACAGAGCTCAGAAGAGGGCACTG | | |
| 2054 | 13 | 11942503 | 11942963 | 460 | AKR,BTBR,C57L,ILNJ2,KK,LP,MOLF,NOD,NOR,NZB,RF | 13 | 11942792 | 11942810 | 19 | - | 17.7049 | 5.73E-07 | 0.149 | AGTCCAGAAGAGGGCAGTA | 11 | ENCFF327LYS,ENCFF089GBC,ENCFF367UNR,ENCFF883UPM,ENCFF502WDT,ENCFF589ZBJ,ENCFF093ZRU,ENCFF533APC,ENCFF099QNE,ENCFF052KVO,ENCFF136ZDK |

| | | | | | | | | | | | | | | | | |
|---|---|---|---|---|---|---|---|---|---|---|---|---|---|---|---|---|
| 2054 | 13 | 11942503 | 11942963 | 460 | AKR,BTBR,C57L,ILNJ2,KK,LP,MOLF,NOD,N OR.NZB.RF | 13 | 11942792 | 11942826 | 35 | - | 27.055 | 4.49E-10 | 0.00272 | GTGCAGTGGCCTGTGGAGTCCAGAAGAGGGCAGTA | 11 | ENCFF327LYS,ENCFF089GBC,ENCFF367UNR,ENCFF883UPM,ENCFF502WDT,ENCFF589ZBJ,ENCFF093ZR U,ENCFF533APC,ENCFF099QNE,ENCFF052KVO,ENCFF136ZDK |
| 2055 | 13 | 12221837 | 12222023 | 186 | CAST.SPRET | 13 | 12221954 | 12221987 | 34 | - | 17.2439 | 5.79E-07 | 0.0964 | TAGCTGTCCTCAGATACTCCAGAAGAGGGCATCA | | |
| 2056 | 13 | 12389007 | 12389209 | 202 | AKR.CAST.ILNJ2.LP.MOLF.RF.SPRET | 13 | 12389132 | 12389165 | 34 | - | 16.4472 | 9.61E-07 | 0.118 | TAGCTGTCTTCAGACACACCAGAAGAGGGCGTCA | | |
| 2057 | 13 | 17994320 | 17994532 | 212 | AJ,AKR,B10D2,B10J,BALB,BTBR,C3H,C57L ,C58J,CAST,CBA,CE,DBA1J,DBA2J,ILNJ2, KK,LP,MA/My,MOLF,MRL,NZB,PJ,PL,RF,RH 2.SEA.SMJ.SPRET | 13 | 17994444 | 17994477 | 34 | - | 18.0732 | 3.36E-07 | 0.0742 | TAGCTGTCCTCAGGTGTACCAGAAGAGGGCATCG | | |
| 2058 | 13 | 20217537 | 20217736 | 199 | 129S1,AJ,AKR,B10D2,B10J,BALB,BTBR,BU B,C3H,C57L,C58J,CAST,CBA,DBA1J,DBA2 J,FVB,ILNJ2,KK,LP,MA/My,MOLF,MRL,NOD, NOR,NZB,PJ,PL,RF,RH2,SEA,SJL,SMJ,SPR ET.SWR.WSB | 13 | 20217657 | 20217691 | 35 | - | 15.9633 | 9.75E-07 | 0.197 | CTGTAGCTGTTCAGACACTCCAGAAGAGGGAGTCA | | |
| 2059 | 13 | 21108593 | 21108650 | 57 | SPRET | 13 | 21108644 | 21108678 | 35 | - | 16.2477 | 8.31E-07 | 0.182 | TAGCTGTCTTCAGACACTCCCAGAAGAGGGCATCA | | |
| 2060 | 13 | 22023451 | 22023990 | 539 | 129S1.ILNJ2 | 13 | 22023963 | 22023996 | 34 | - | 16.5447 | 9.04E-07 | 0.118 | CTGTACTTCCCATGGGCTCCTAGAGGTGGGACCC | | |
| 2061 | 13 | 28987038 | 28988015 | 977 | CAST.KK.MOLF.PJ.SEA.SPRET | 13 | 28987559 | 28987592 | 34 | - | 17.2439 | 5.79E-07 | 0.0964 | TAGCTGTCCTCAGATACTCCAGAAGAGGGCATCA | | |
| 2062 | 13 | 34975323 | 34975532 | 209 | CAST.MOLF.SPRET.WSB | 13 | 34975448 | 34975481 | 34 | - | 16.4634 | 9.51E-07 | 0.118 | TAGCTGTCTTCAGACGCACCAGAAGAGGGCATCA | | |
| 2063 | 13 | 39820408 | 39820612 | 204 | SPRET | 13 | 39820534 | 39820567 | 34 | - | 17.6016 | 4.59E-07 | 0.0853 | TAGCTGTCTTCAGACATGCCAGAAGAGGGCGTCA | | |
| 2064 | 13 | 40906148 | 40906604 | 456 | MOLF.SPRET | 13 | 40906239 | 40906272 | 34 | - | 17.6992 | 4.30E-07 | 0.083 | TAGCTGTTTTCAGACACACCAGAAGAGGGCATCA | | |
| 2065 | 13 | 41003431 | 41003702 | 271 | CAST.MOLF.SPRET | 13 | 41003555 | 41003573 | 19 | - | 17.5574 | 6.32E-07 | 0.153 | ACTCCAGAAGAGGGCGCCA | 2 | ENCFF093ZRU.ENCFF367UNR |
| 2065 | 13 | 41003431 | 41003702 | 271 | CAST.MOLF.SPRET | 13 | 41003555 | 41003588 | 34 | - | 18.3659 | 2.76E-07 | 0.0667 | TAGCTGTCTTCAGACACTCCAGAAGAGGGCGCCA | 2 | ENCFF093ZRU.ENCFF367UNR |
| 2066 | 13 | 41823196 | 41823286 | 90 | CAST | 13 | 41823204 | 41823222 | 19 | - | 17.377 | 7.11E-07 | 0.162 | AGGCCAGAAGAGGGTGCTA | | |
| 2067 | 13 | 42626853 | 42626927 | 74 | CAST.CE.MOLF | 13 | 42626824 | 42626858 | 35 | - | 26.1927 | 9.35E-10 | 0.00334 | CTGCACATACACCTGCAGGCCAGAAGAGGGCAGAA | | |
| 2068 | 13 | 45114566 | 45114660 | 94 | SPRET | 13 | 45114617 | 45114651 | 35 | - | 16.0092 | 9.50E-07 | 0.194 | GTGCTGTTCCTGCTCTCAGCAGCTAGATGGAAAAC | 3 | ENCFF092NMJ.ENCFF093ZRU.ENCFF099QNE |
| 2069 | 13 | 45160175 | 45160242 | 67 | CAST,MOLF,SPRET | 13 | 45160154 | 45160187 | 34 | - | 22.1057 | 1.84E-08 | 0.0111 | ATGTATTCCTAGAAGAGGCCAGAAGAGGGCACTG | 27 | ENCFF589ZBJ,ENCFF024JOZ,ENCFF136ZDK,ENCFF327LYS,ENCFF281QMV,ENCFF089GBC,ENCFF052KV O,ENCFF097CAQ,ENCFF321AGQ,ENCFF907XEG,ENCFF883UPM,ENCFF142CNG,ENCFF198SSR,ENCFF957 SPH,ENCFF165LXO,ENCFF533APC,ENCFF504JDS,ENCFF201JUA,ENCFF821BIA,ENCFF502WDT,ENCFF43 0PPJ,ENCFF065ODV,ENCFF805QBU,ENCFF092NMJ,ENCFF093ZRU,ENCFF367UNR,ENCFF099QNE |
| 2069 | 13 | 45160175 | 45160242 | 67 | CAST,MOLF,SPRET | 13 | 45160215 | 45160233 | 19 | - | 20.8033 | 5.79E-08 | 0.0616 | AGGCCAGAAGAGGGCGCTG | 27 | ENCFF589ZBJ,ENCFF024JOZ,ENCFF136ZDK,ENCFF327LYS,ENCFF281QMV,ENCFF089GBC,ENCFF052KV O,ENCFF097CAQ,ENCFF321AGQ,ENCFF907XEG,ENCFF883UPM,ENCFF142CNG,ENCFF198SSR,ENCFF957 SPH,ENCFF165LXO,ENCFF533APC,ENCFF504JDS,ENCFF201JUA,ENCFF821BIA,ENCFF502WDT,ENCFF43 0PPJ,ENCFF065ODV,ENCFF805QBU,ENCFF092NMJ,ENCFF093ZRU,ENCFF367UNR,ENCFF099QNE |
| 2069 | 13 | 45160175 | 45160242 | 67 | CAST,MOLF,SPRET | 13 | 45160215 | 45160248 | 34 | - | 19.6098 | 1.17E-07 | 0.0413 | GCCTAGTGCCTACAGAGGCCAGAAGAGGGCGCTG | 27 | ENCFF589ZBJ,ENCFF024JOZ,ENCFF136ZDK,ENCFF327LYS,ENCFF281QMV,ENCFF089GBC,ENCFF052KV O,ENCFF097CAQ,ENCFF321AGQ,ENCFF907XEG,ENCFF883UPM,ENCFF142CNG,ENCFF198SSR,ENCFF957 SPH,ENCFF165LXO,ENCFF533APC,ENCFF504JDS,ENCFF201JUA,ENCFF821BIA,ENCFF502WDT,ENCFF43 0PPJ,ENCFF065ODV,ENCFF805QBU,ENCFF092NMJ,ENCFF093ZRU,ENCFF367UNR,ENCFF099QNE |
| 2070 | 13 | 46856601 | 46856902 | 301 | SPRET | 13 | 46856822 | 46856855 | 34 | - | 16.813 | 7.63E-07 | 0.109 | TTGCAATGCCCATGGAGGTTGGAAGAGGGCACCA | 16 | ENCFF210BTO,ENCFF024JOZ,ENCFF136ZDK,ENCFF092NMJ,ENCFF821BIA,ENCFF089GBC,ENCFF589ZBJ, ENCFF055HAP,ENCFF327LYS,ENCFF502WDT,ENCFF367UNR,ENCFF198SSR,ENCFF194HVL,ENCFF093ZR U,ENCFF957SPH,ENCFF907XEG |
| 2071 | 13 | 59988726 | 59988799 | 73 | SPRET | 13 | 59988741 | 59988775 | 35 | - | 18.8532 | 1.79E-07 | 0.0792 | ATGCAATGCATACAGCTTAACAGCAGAGGGGGCAA | 41 | ENCFF093ZRU,ENCFF367UNR,ENCFF201JUA,ENCFF099QNE,ENCFF589ZBJ,ENCFF327LYS,ENCFF821BIA, ENCFF907XEG,ENCFF065ODV,ENCFF089GBC,ENCFF136ZDK,ENCFF024JOZ,ENCFF198SSR,ENCFF092NM J,ENCFF805QBU,ENCFF321AGQ,ENCFF430PPJ,ENCFF502WDT,ENCFF504JDS,ENCFF349BUL,ENCFF883U PM,ENCFF165LXO,ENCFF951GOP,ENCFF957SPH,ENCFF052KVO,ENCFF601JVE,ENCFF609MDN,ENCFF57 5MGI,ENCFF281QMV,ENCFF210BTO,ENCFF316FYN,ENCFF142CNG,ENCFF097CAQ,ENCFF491RJK,ENCFF6 11HDQ,ENCFF194HVL,ENCFF533APC,ENCFF089EWX,ENCFF784ASD,ENCFF033UQX,ENCFF086ZTV |
| 2072 | 13 | 61950068 | 61950274 | 206 | BALB.CAST.MOLF.WSB | 13 | 61950193 | 61950226 | 34 | - | 17.8455 | 3.91E-07 | 0.0799 | TAGCTGTCCTCAGACACTCCAGAAGAGGGCGTCA | | |
| 2073 | 13 | 62146528 | 62146860 | 332 | 129S1,AJ,BUB,KK,MA/My,MRL,NOD,NOR,N ZB,NZW,SMJ,SPRET | 13 | 62146818 | 62146836 | 19 | - | 18.6557 | 2.99E-07 | 0.111 | CCGCCACTAGATGGTGGTA | 20 | ENCFF136ZDK,ENCFF089GBC,ENCFF367UNR,ENCFF201JUA,ENCFF093ZRU,ENCFF092NMJ,ENCFF165LX O,ENCFF198SSR,ENCFF055HAP,ENCFF883UPM,ENCFF502WDT,ENCFF024JOZ,ENCFF327LYS,ENCFF907X EG,ENCFF609MDN,ENCFF957SPH,ENCFF821BIA,ENCFF099QNE,ENCFF589ZBJ,ENCFF142CNG |
| 2074 | 13 | 62554119 | 62554334 | 215 | CAST.MOLF.SPRET | 13 | 62554244 | 62554262 | 19 | - | 17.5574 | 6.32E-07 | 0.153 | ACTCCAGAAGAGGGCGCCA | | |
| 2074 | 13 | 62554119 | 62554334 | 215 | CAST.MOLF.SPRET | 13 | 62554244 | 62554277 | 34 | - | 18.3659 | 2.76E-07 | 0.0667 | TAGCTGTCTTCAGACACTCCAGAAGAGGGCGCCA | | |
| 2075 | 13 | 62628918 | 62629123 | 205 | SPRET | 13 | 62629044 | 62629077 | 34 | - | 16.4634 | 9.51E-07 | 0.118 | TAGCTGTCTTCAGACGCACCAGAAGAGGGCATCA | | |
| 2076 | 13 | 64352724 | 64352781 | 57 | 129S1,AJ,AKR,BALB,BTBR,BUB,C3H,C57L, C58J,CBA,CE,DBA1J,DBA2J,FVB,ILNJ2,KK, MRL,NOD,NOR,NZO,NZW,PJ,PL,RF,SEA,SJ L.SMJ.SWR.TH.WSB | 13 | 64352747 | 64352765 | 19 | - | 19.6721 | 1.42E-07 | 0.0806 | TGTCCACCAGGGGGCTGTA | 16 | ENCFF821BIA,ENCFF136ZDK,ENCFF089GBC,ENCFF367UNR,ENCFF609MDN,ENCFF093ZRU,ENCFF907XE G,ENCFF024JOZ,ENCFF099QNE,ENCFF589ZBJ,ENCFF327LYS,ENCFF198SSR,ENCFF194HVL,ENCFF883UP M,ENCFF957SPH,ENCFF165LXO |
| 2077 | 13 | 65562144 | 65562351 | 207 | C3H,C58J,CE,PJ,PL,SEA,SMJ,SPRET,WSB | 13 | 65562265 | 65562298 | 34 | - | 17.2439 | 5.79E-07 | 0.0964 | TAGCTGTCCTCAGATACTCCAGAAGAGGGCATCA | | |
| 2078 | 13 | 65704714 | 65704921 | 207 | C3H.C58J.CBA.SPRET.WSB | 13 | 65704836 | 65704869 | 34 | - | 17.2439 | 5.79E-07 | 0.0964 | TAGCTGTCCTCAGATACTCCAGAAGAGGGCATCA | | |
| 2079 | 13 | 67009400 | 67009602 | 202 | 129S1.AKR.MRL.NZW.RF | 13 | 67009523 | 67009556 | 34 | - | 17.2439 | 5.79E-07 | 0.0964 | TAGCTGTCCTCAGATACTCCAGAAGAGGGCATCA | | |
| 2080 | 13 | 67499763 | 67499999 | 236 | 129S1,AJ,AKR,BALB,BTBR,BUB,C3H,C57L, C58J,CAST,CBA,CE,DBA1J,DBA2J,FVB,ILN J2,KK,MOLF,MRL,NOD,NOR,NZB,NZO,NZW ,PJ,PL,RF,SEA,SJL,SMJ,SPRET,SWR,TH,W SB | 13 | 67499887 | 67499920 | 34 | - | 16.9187 | 7.13E-07 | 0.105 | TAGCTGTCTTCAGACGCACCAGAAGAGGGCGTCA | | |
| 2081 | 13 | 67550678 | 67550887 | 209 | 129S1,AJ,AKR,BALB,BTBR,BUB,C3H,C57L, C58J,CAST,CBA,CE,DBA1J,DBA2J,FVB,ILN J2,KK,MOLF,MRL,NOD,NOR,NZB,NZO,NZW ,PJ,PL,RF,SEA,SJL,SMJ,SPRET,SWR,TH,W SB | 13 | 67550801 | 67550834 | 34 | - | 17.6504 | 4.44E-07 | 0.0847 | CAGCTGTCCTCAGACACCCCAGAAGAGGGAGTCA | | |
| 2082 | 13 | 69827549 | 69827749 | 200 | 129S1,AJ,AKR,BALB,BTBR,BUB,C3H,C57L, C58J,CBA,CE,DBA1J,DBA2J,FVB,ILNJ2,KK, MA/My,MOLF,MRL,NOD,NOR,NZB,NZO,NZ W,PJ,PL,RF,SEA,SJL,SMJ,SPRET,SWR,TH, WSB | 13 | 69827672 | 69827705 | 34 | - | 16.4472 | 9.61E-07 | 0.118 | TAGCTGTCTTCAGACACACCAGAAGAGGGCGTCA | | |
| 2083 | 13 | 70139139 | 70140037 | 898 | BUB,C58J,MA/My,MRL,NZW | 13 | 70139535 | 70139553 | 19 | - | 18.7869 | 2.72E-07 | 0.108 | CCAACACCAGGGGGCGCAA | 35 | ENCFF093ZRU,ENCFF367UNR,ENCFF805QBU,ENCFF201JUA,ENCFF099QNE,ENCFF210BTO,ENCFF092NM J,ENCFF821BIA,ENCFF907XEG,ENCFF883UPM,ENCFF951GOP,ENCFF327LYS,ENCFF089GBC,ENCFF589Z BJ,ENCFF024JOZ,ENCFF055HAP,ENCFF430PPJ,ENCFF052KVO,ENCFF065ODV,ENCFF609MDN,ENCFF136 ZDK,ENCFF502WDT,ENCFF198SSR,ENCFF165LXO,ENCFF033UQX,ENCFF097CAQ,ENCFF957SPH,ENCFF4 91RJK,ENCFF281QMV,ENCFF611HDQ,ENCFF533APC,ENCFF575MGI,ENCFF194HVL,ENCFF142CNG,ENCFF 316FYN |
| 2083 | 13 | 70139139 | 70140037 | 898 | BUB,C58J,MA/My,MRL,NZW | 13 | 70139535 | 70139568 | 34 | - | 24.6179 | 2.32E-09 | 0.0026 | GTGCTATCCCGACGTCCAACACCAGGGGGCGCAA | 35 | ENCFF093ZRU,ENCFF367UNR,ENCFF805QBU,ENCFF201JUA,ENCFF099QNE,ENCFF210BTO,ENCFF092NM J,ENCFF821BIA,ENCFF907XEG,ENCFF883UPM,ENCFF951GOP,ENCFF327LYS,ENCFF089GBC,ENCFF589Z BJ,ENCFF024JOZ,ENCFF055HAP,ENCFF430PPJ,ENCFF052KVO,ENCFF065ODV,ENCFF609MDN,ENCFF136 ZDK,ENCFF502WDT,ENCFF198SSR,ENCFF165LXO,ENCFF033UQX,ENCFF097CAQ,ENCFF957SPH,ENCFF4 91RJK,ENCFF281QMV,ENCFF611HDQ,ENCFF533APC,ENCFF575MGI,ENCFF194HVL,ENCFF142CNG,ENCFF 316FYN |
| 2084 | 13 | 70652814 | 70653022 | 208 | BUB,C58J,CAST,MA/My,MOLF,MRL,NZW,S MJ.SPRET.SWR.WSB | 13 | 70652940 | 70652973 | 34 | - | 16.4472 | 9.61E-07 | 0.118 | TAGCTGTCTTCAGACACACCAGAAGAGGGCGTCA | | |
| 2085 | 13 | 73714578 | 73715286 | 708 | SPRET | 13 | 73714883 | 73714916 | 34 | - | 16.4797 | 9.42E-07 | 0.118 | CAGCCTCTTTAAGGCTGGACACCAGGTGGCTGCA | | |
| 2086 | 13 | 74709751 | 74709957 | 206 | SPRET | 13 | 74709876 | 74709909 | 34 | - | 17.0081 | 6.74E-07 | 0.104 | TAGCTGTCCCCAGACACTCCAGAAGAGGGAGTCA | | |
| 2087 | 13 | 84882579 | 84883039 | 460 | SPRET | 13 | 84882997 | 84883030 | 34 | - | 17.0813 | 6.43E-07 | 0.102 | CTGCACTCACATTTGCCATTACAAGATGGCGCTG | | |
| 2088 | 13 | 91484131 | 91484340 | 209 | CAST.MOLF.SPRET | 13 | 91484271 | 91484304 | 34 | - | 16.9919 | 6.81E-07 | 0.104 | TAGCTGTCTTCAGACACACCAGAAGGGGGCATCA | | |
| 2089 | 13 | 92780607 | 92780826 | 219 | AJ,AKR,BUB,C3H,CAST,KK,MOLF,MRL,NZB .NZO.NZW.PL.RF.SPRET.WSB | 13 | 92780733 | 92780751 | 19 | - | 17.5574 | 6.32E-07 | 0.153 | ACTCCAGAAGAGGGCGCCA | 2 | ENCFF093ZRU,ENCFF367UNR |
| 2089 | 13 | 92780607 | 92780826 | 219 | AJ,AKR,BUB,C3H,CAST,KK,MOLF,MRL,NZB .NZO.NZW.PL.RF.SPRET.WSB | 13 | 92780733 | 92780766 | 34 | - | 18.3659 | 2.76E-07 | 0.0667 | TAGCTGTCTTCAGACACTCCAGAAGAGGGCGCCA | 2 | ENCFF093ZRU,ENCFF367UNR |
| 2090 | 13 | 93706997 | 93707193 | 196 | SPRET | 13 | 93707122 | 93707155 | 34 | - | 16.4472 | 9.61E-07 | 0.118 | TAGCTGTCTTCAGACACACCAGAAGAGGGCGTCA | | |
| 2091 | 13 | 94622033 | 94622228 | 195 | CAST.SPRET | 13 | 94622154 | 94622187 | 34 | - | 16.7724 | 7.83E-07 | 0.109 | TAGCTGTCTTCAGATGCACCAGAAGAGGGCGTCA | | |
| 2092 | 13 | 98594993 | 98595478 | 485 | WSB | 13 | 98595163 | 98595181 | 19 | - | 17.0328 | 8.88E-07 | 0.181 | AGGCCACAAGAGGGCATCA | | |
| 2093 | 13 | 98962876 | 98963115 | 239 | CAST.SPRET | 13 | 98963077 | 98963111 | 35 | - | 19.9908 | 8.75E-08 | 0.0502 | CTGCATGTATGCCTGAAGGCCAGAAAAGGGCACCA | | |
| 2094 | 13 | 98964621 | 98964892 | 271 | SPRET | 13 | 98964853 | 98964886 | 34 | - | 18.2114 | 3.07E-07 | 0.0699 | TTGCTGTATACAGACACACCAGAAGAGGGCGTCA | 2 | ENCFF093ZRU.ENCFF367UNR |
| 2095 | 13 | 102880857 | 102881824 | 967 | KK | 13 | 102881612 | 102881646 | 35 | - | 22.3578 | 1.77E-08 | 0.0175 | CTGCTGTAGCTAGGTGGATCCACTAGGTGGCAGGC | 34 | ENCFF367UNR,ENCFF093ZRU,ENCFF099QNE,ENCFF805QBU,ENCFF821BIA,ENCFF907XEG,ENCFF024JOZ ,ENCFF065ODV,ENCFF055HAP,ENCFF327LYS,ENCFF502WDT,ENCFF052KVO,ENCFF092NMJ,ENCFF609M DN,ENCFF089GBC,ENCFF198SSR,ENCFF589ZBJ,ENCFF883UPM,ENCFF165LXO,ENCFF136ZDK,ENCFF321 AGQ,ENCFF951GOP,ENCFF430PPJ,ENCFF097CAQ,ENCFF142CNG,ENCFF957SPH,ENCFF210BTO,ENCFF5 75MGI,ENCFF281QMV,ENCFF316FYN,ENCFF504JDS,ENCFF533APC,ENCFF491RJK,ENCFF194HVL |
| 2096 | 13 | 103172698 | 103172889 | 191 | 129S1.C57L.MOLF.NZW.SPRET | 13 | 103172800 | 103172833 | 34 | - | 16.4472 | 9.61E-07 | 0.118 | TAGCTGTCTTCAGACACACCAGAAGAGGGCGTCA | | |
| 2097 | 13 | 107067339 | 107067551 | 212 | CAST.CE.MA/My.MOLF.NZO.SPRET | 13 | 107067463 | 107067496 | 34 | - | 16.9431 | 7.02E-07 | 0.104 | TAGCTGTCTTCTCACACACCAGAAGAGGGCGTCA | | |
| 2098 | 13 | 107258204 | 107258581 | 377 | SPRET | 13 | 107258253 | 107258286 | 34 | - | 17.6016 | 4.59E-07 | 0.0853 | CAGGACCGCTACGGACCTCCACCAGGTGGCCGAG | | |
| 2099 | 13 | 107408135 | 107408362 | 227 | SPRET | 13 | 107408256 | 107408289 | 34 | - | 16.5691 | 8.90E-07 | 0.117 | TAGCTGTCTTCAGACGCACCAGAAGAGGGCATCG | | |
| 2100 | 13 | 109343698 | 109343979 | 281 | CAST.CE.MA/My.MOLF.NZO.SPRET.TH | 13 | 109343896 | 109343914 | 19 | - | 17.5574 | 6.32E-07 | 0.153 | ACTCCAGAAGAGGGCGCCA | 1 | ENCFF093ZRU |
| 2100 | 13 | 109343698 | 109343979 | 281 | CAST.CE.MA/My.MOLF.NZO.SPRET.TH | 13 | 109343896 | 109343929 | 34 | - | 18.3659 | 2.76E-07 | 0.0667 | TAGCTGTCTTCAGACACTCCAGAAGAGGGCGCCA | 1 | ENCFF093ZRU |
| 2101 | 13 | 110666445 | 110666646 | 201 | CAST,CE,KK,MA/My,MOLF,NOD,NOR,NZB,N ZO.NZW.PL.SPRET.SWR | 13 | 110666568 | 110666601 | 34 | - | 16.4472 | 9.61E-07 | 0.118 | TAGCTGTCTTCAGACACACCAGAAGAGGGCGTCA | | |
| 2102 | 13 | 111712033 | 111712256 | 223 | SPRET | 13 | 111712156 | 111712189 | 34 | - | 16.4472 | 9.61E-07 | 0.118 | TAGCTGTCTTCAGACACACCAGAAGAGGGCGTCA | | |
| 2103 | 13 | 113658880 | 113659082 | 202 | SPRET | 13 | 113659006 | 113659039 | 34 | - | 16.4472 | 9.61E-07 | 0.118 | TAGCTGTCTTCAGACACACCAGAAGAGGGCGTCA | | |
| 2104 | 13 | 114135886 | 114136100 | 214 | AJ,BUB,CAST,CE,DBA1J,FVB,KK,MA/My,M OLF,MRL,NZB,NZO,NZW,PL,RF,RH2,SJL,S PRET.SWR.WSB | 13 | 114136012 | 114136045 | 34 | - | 16.6911 | 8.24E-07 | 0.112 | TAGCTGTCTTCAGACAGACCAGAAGAGGGCATCA | | |
| 2105 | 13 | 117846610 | 117847056 | 446 | 129S1,BTBR,BUB,C3H,C57L,CAST,CBA,CE ,DBA2J,MA/My,MOLF,MRL,NZB,NZO,NZW,P L.SMJ.TH | 13 | 117846746 | 117846779 | 34 | - | 17.2439 | 5.79E-07 | 0.0964 | TAGCTGTCCTCAGATACTCCAGAAGAGGGCATCA | | |
| 2106 | 13 | 119402051 | 119402255 | 204 | SPRET | 13 | 119402176 | 119402209 | 34 | - | 16.4472 | 9.61E-07 | 0.118 | TAGCTGTCTTCAGACACACCAGAAGAGGGCGTCA | | |
| 2107 | 13 | 120233833 | 120234008 | 175 | SPRET | 13 | 120233837 | 120233870 | 34 | - | 18.6992 | 2.20E-07 | 0.0613 | CAGCTGTCTTCAGACACACCAGAAGAGGGCGTTC | 1 | ENCFF093ZRU |
| 2108 | 13 | 120249513 | 120249721 | 208 | 129S1,BTBR,C3H,C57L,CAST,CBA,CE,DBA 2J.KK.MA/My.NZO.SMJ.SPRET | 13 | 120249637 | 120249655 | 19 | - | 17.5574 | 6.32E-07 | 0.153 | ACTCCAGAAGAGGGCGCCA | 2 | ENCFF093ZRU,ENCFF367UNR |
| 2108 | 13 | 120249513 | 120249721 | 208 | 129S1,BTBR,C3H,C57L,CAST,CBA,CE,DBA 2J.KK.MA/My.NZO.SMJ.SPRET | 13 | 120249637 | 120249670 | 34 | - | 18.3659 | 2.76E-07 | 0.0667 | TAGCTGTCTTCAGACACTCCAGAAGAGGGCGCCA | 2 | ENCFF093ZRU,ENCFF367UNR |
| 2109 | 13 | 120268882 | 120269112 | 230 | SPRET | 13 | 120269012 | 120269030 | 19 | - | 16.9344 | 9.46E-07 | 0.187 | AGACCAGAAGAGGGCACGG | | |
| 2110 | 14 | 9719254 | 9719452 | 198 | 129S1,C3H,C57L,CBA,DBA1J,DBA2J,KK,M OLF.MRL.NZW.PJ.PL.SPRET.WSB | 14 | 9719305 | 9719338 | 34 | + | 17.2439 | 5.79E-07 | 0.0851 | TAGCTGTCCTCAGATACTCCAGAAGAGGGCATCA | | |
| 2111 | 14 | 10475334 | 10475536 | 202 | 129S1,C3H,C57L,CBA,DBA1J,DBA2J,KK,M OLF.MRL.NZW.PJ.PL.SPRET | 14 | 10475393 | 10475426 | 34 | + | 16.4472 | 9.61E-07 | 0.111 | TAGCTGTCTTCAGACACACCAGAAGAGGGCGTCA | | |
| 2112 | 14 | 10505318 | 10505520 | 202 | 129S1,C3H,C57L,CAST,CBA,DBA1J,DBA2J ,KK,MOLF,MRL,NZB,NZO,NZW,PJ,PL,SPRE T.WSB | 14 | 10505378 | 10505411 | 34 | + | 16.4472 | 9.61E-07 | 0.111 | TAGCTGTCTTCAGACACACCAGAAGAGGGCGTCA | | |
| 2113 | 14 | 11805963 | 11806015 | 52 | SPRET | 14 | 11806011 | 11806045 | 35 | + | 16.0642 | 9.21E-07 | 0.223 | ATGCAGAGGGCTGAGGAGGGCTGCAGGTGGCTGTG | | |
| 2114 | 14 | 13829574 | 13829750 | 176 | SPRET | 14 | 13829661 | 13829694 | 34 | + | 17.6098 | 4.56E-07 | 0.0753 | CAGCAGTTGTAGGTGCTGACACCTGGTGCTGCTC | | |
| 2115 | 14 | 13871991 | 13872939 | 948 | 129S1.C57L.KK.MOLF.PL | 14 | 13872667 | 13872700 | 34 | + | 17.813 | 3.99E-07 | 0.0708 | TTGCTGTCTTCATGTACACCAGAAGAGGGCATCA | 1 | ENCFF093ZRU |
| 2116 | 14 | 14424924 | 14425216 | 292 | SPRET | 14 | 14425206 | 14425240 | 35 | + | 22.9908 | 1.13E-08 | 0.0197 | CTGTATGTACACTTACATGCCAGAAGAGGGCGCCA | 21 | ENCFF093ZRU,ENCFF367UNR,ENCFF201JUA,ENCFF099QNE,ENCFF136ZDK,ENCFF052KVO,ENCFF165LX O,ENCFF430PPJ,ENCFF821BIA,ENCFF024JOZ,ENCFF327LYS,ENCFF609MDN,ENCFF907XEG,ENCFF065O DV,ENCFF589ZBJ,ENCFF089GBC,ENCFF198SSR,ENCFF883UPM,ENCFF055HAP,ENCFF092NMJ,ENCFF502 WDT |
| 2117 | 14 | 14553417 | 14553626 | 209 | CAST.SPRET | 14 | 14553477 | 14553511 | 35 | + | 17.0092 | 5.39E-07 | 0.169 | CTGTAGCTGTTTTCAGACACCAGAAGAGGGCATCA | | |
| 2118 | 14 | 16340200 | 16340736 | 536 | CAST | 14 | 16340440 | 16340473 | 34 | + | 20.626 | 5.64E-08 | 0.0235 | CTGGAGGGGGCAGCCTTGGCCTTCAGGTGGCAGCC | | |
| 2119 | 14 | 20352203 | 20352408 | 205 | SPRET | 14 | 20352250 | 20352283 | 34 | + | 17.6667 | 4.40E-07 | 0.0741 | TAGCTGTCTTCAGACACGCCAGAAGAGGGCGTCA | | |
| 2120 | 14 | 21838676 | 21838877 | 201 | SPRET | 14 | 21838734 | 21838767 | 34 | + | 16.4634 | 9.51E-07 | 0.111 | TAGCTGTCTTCAGACGCACCAGAAGAGGGCATCA | 1 | ENCFF093ZRU |
| 2121 | 14 | 24360221 | 24360670 | 449 | SPRET | 14 | 24360238 | 24360271 | 34 | + | 17.0813 | 6.43E-07 | 0.0915 | CTGCACTCACATTTGCCATTACAAGATGGCGCTG | | |
| 2122 | 14 | 24361337 | 24361437 | 100 | SPRET | 14 | 24361354 | 24361387 | 34 | + | 26.6179 | 3.73E-10 | 0.000974 | ATGCAACTCCCTTAGTGGCCAGAAGAGGGAGCCA | 33 | ENCFF201JUA,ENCFF093ZRU,ENCFF367UNR,ENCFF099QNE,ENCFF210BTO,ENCFF821BIA,ENCFF327LYS, ENCFF089GBC,ENCFF907XEG,ENCFF321AGQ,ENCFF024JOZ,ENCFF504JDS,ENCFF065ODV,ENCFF136ZD K,ENCFF052KVO,ENCFF092NMJ,ENCFF165LXO,ENCFF316FYN,ENCFF589ZBJ,ENCFF198SSR,ENCFF055H AP,ENCFF194HVL,ENCFF097CAQ,ENCFF502WDT,ENCFF883UPM,ENCFF957SPH,ENCFF430PPJ,ENCFF575 MGI,ENCFF601JVE,ENCFF142CNG,ENCFF281QMV,ENCFF951GOP,ENCFF609MDN |
| 2122 | 14 | 24361337 | 24361437 | 100 | SPRET | 14 | 24361369 | 24361387 | 19 | + | 20.3934 | 8.07E-08 | 0.0613 | TGGCCAGAAGAGGGAGCCA | 33 | ENCFF201JUA,ENCFF093ZRU,ENCFF367UNR,ENCFF099QNE,ENCFF210BTO,ENCFF821BIA,ENCFF327LYS, ENCFF089GBC,ENCFF907XEG,ENCFF321AGQ,ENCFF024JOZ,ENCFF504JDS,ENCFF065ODV,ENCFF136ZD K,ENCFF052KVO,ENCFF092NMJ,ENCFF165LXO,ENCFF316FYN,ENCFF589ZBJ,ENCFF198SSR,ENCFF055H AP,ENCFF194HVL,ENCFF097CAQ,ENCFF502WDT,ENCFF883UPM,ENCFF957SPH,ENCFF430PPJ,ENCFF575 MGI,ENCFF601JVE,ENCFF142CNG,ENCFF281QMV,ENCFF951GOP,ENCFF609MDN |
| 2123 | 14 | 25792758 | 25792968 | 210 | CAST,MOLF,NOD,NOR,PJ,PL,SPRET,TH | 14 | 25792826 | 25792859 | 34 | + | 17.8455 | 3.91E-07 | 0.0699 | TAGCTGTCCTCAGACACTCCAGAAGAGGGCGTCA | | |
| 2124 | 14 | 26388681 | 26388749 | 68 | MOLF | 14 | 26388707 | 26388740 | 34 | + | 18.4797 | 2.56E-07 | 0.0581 | TTGCTGTCTTCAGACACACCAGAAGAGGGCATCA | 3 | ENCFF099QNE.ENCFF089GBC.ENCFF093ZRU |
| 2125 | 14 | 26439674 | 26439886 | 212 | CAST.MOLF.SPRET.WSB | 14 | 26439743 | 26439776 | 34 | + | 18.3659 | 2.76E-07 | 0.0595 | TAGCTGTCTTCAGACACTCCAGAAGAGGGCGCCA | | |
| 2125 | 14 | 26439674 | 26439886 | 212 | CAST.MOLF.SPRET.WSB | 14 | 26439758 | 26439776 | 19 | + | 17.5574 | 6.32E-07 | 0.158 | ACTCCAGAAGAGGGCGCCA | | |
| 2126 | 14 | 27526036 | 27526177 | 141 | CAST,FVB,MOLF,SPRET | 14 | 27526050 | 27526084 | 35 | + | 17.5963 | 3.82E-07 | 0.141 | AGTCAGTTCCACCTATCCCCCACTAGGGGGTGGAG | 27 | ENCFF093ZRU,ENCFF367UNR,ENCFF201JUA,ENCFF099QNE,ENCFF210BTO,ENCFF907XEG,ENCFF327LY S,ENCFF089GBC,ENCFF821BIA,ENCFF024JOZ,ENCFF136ZDK,ENCFF883UPM,ENCFF092NMJ,ENCFF165LX O,ENCFF589ZBJ,ENCFF430PPJ,ENCFF198SSR,ENCFF502WDT,ENCFF504JDS,ENCFF055HAP,ENCFF321A GQ,ENCFF052KVO,ENCFF951GOP,ENCFF316FYN,ENCFF609MDN,ENCFF065ODV,ENCFF957SPH |
| 2127 | 14 | 30533675 | 30533981 | 306 | CAST.MOLF.SPRET | 14 | 30533839 | 30533872 | 34 | + | 16.4472 | 9.61E-07 | 0.111 | TAGCTGTCTTCAGACACACCAGAAGAGGGCGTCA | | |
| 2128 | 14 | 31832227 | 31832411 | 184 | MOLF.PJ | 14 | 31832269 | 31832302 | 34 | + | 17.9106 | 3.74E-07 | 0.0681 | TAGCTGTCTTCAGACACTCCAGAAGAGGGCACCA | | |
| 2129 | 14 | 34122424 | 34122630 | 206 | 129S1,C57L,CAST,MOLF,PJ,SPRET,WSB | 14 | 34122473 | 34122506 | 34 | + | 17.2439 | 5.79E-07 | 0.0851 | TAGCTGTCCTCAGATACTCCAGAAGAGGGCATCA | | |
| 2130 | 14 | 34570036 | 34570608 | 572 | 129S1.C57L | 14 | 34570129 | 34570162 | 34 | + | 17.7073 | 4.28E-07 | 0.0732 | TAGCTGTCTTCAGATGTACCAGAAGGGGGCGTCA | 1 | ENCFF093ZRU |
| 2131 | 14 | 45806904 | 45807114 | 210 | BUB,C57L,CAST,DBA1J,DBA2J,ILNJ2,KK,M A/My,MOLF,MRL,NZB,NZO,PJ,PL,SJL,SMJ,S PRET.SWR.TH.WSB | 14 | 45806974 | 45807007 | 34 | + | 17.3902 | 5.27E-07 | 0.0813 | TAGCTGTCCTCAGACACTCCAGAAGAGGGCATCA | | |
| 2132 | 14 | 48200723 | 48200933 | 210 | SPRET | 14 | 48200805 | 48200838 | 34 | + | 17.4228 | 5.16E-07 | 0.0804 | TAGCTGTTTTCAGACACACCAGGAGAGGGCGTCA | | |
| 2133 | 14 | 51142148 | 51142356 | 208 | SPRET | 14 | 51142216 | 51142249 | 34 | + | 18.3659 | 2.76E-07 | 0.0595 | TAGCTGTCTTCAGACACTCCAGAAGAGGGCGCCA | 3 | ENCFF821BIA.ENCFF367UNR.ENCFF093ZRU |
| 2133 | 14 | 51142148 | 51142356 | 208 | SPRET | 14 | 51142231 | 51142249 | 19 | + | 17.5574 | 6.32E-07 | 0.158 | ACTCCAGAAGAGGGCGCCA | 3 | ENCFF821BIA.ENCFF367UNR.ENCFF093ZRU |
| 2134 | 14 | 52103183 | 52103376 | 193 | SPRET | 14 | 52103235 | 52103268 | 34 | + | 16.4472 | 9.61E-07 | 0.111 | TAGCTGTCTTCAGACACACCAGAAGAGGGCGTCA | | |
| 2135 | 14 | 52406911 | 52407134 | 223 | 129S1,AJ,AKR,BALB,BTBR,BUB,C3H,C57L, CAST,CBA,CE,DBA1J,DBA2J,FVB,ILNJ2,K K,MA/My,MOLF,MRL,NOD,NOR,NZB,NZO,NZ W,PJ,PL,RF,RH2,SEA,SJL,SMJ,SPRET,SW R.TH.WSB | 14 | 52406962 | 52406995 | 34 | + | 19.813 | 1.01E-07 | 0.0351 | TAGCTGTCTCCAGCCACTCCAGAAGAGGGCGCCA | 2 | ENCFF093ZRU,ENCFF367UNR |
| 2135 | 14 | 52406911 | 52407134 | 223 | 129S1,AJ,AKR,BALB,BTBR,BUB,C3H,C57L, CAST,CBA,CE,DBA1J,DBA2J,FVB,ILNJ2,K K,MA/My,MOLF,MRL,NOD,NOR,NZB,NZO,NZ W,PJ,PL,RF,RH2,SEA,SJL,SMJ,SPRET,SW R.TH.WSB | 14 | 52406977 | 52406995 | 19 | + | 17.5574 | 6.32E-07 | 0.158 | ACTCCAGAAGAGGGCGCCA | 2 | ENCFF093ZRU,ENCFF367UNR |
| 2136 | 14 | 52423215 | 52423424 | 209 | 129S1,AJ,AKR,BALB,BTBR,BUB,C3H,C57L, CAST,CBA,CE,DBA1J,DBA2J,FVB,ILNJ2,K K,MA/My,MRL,NOD,NOR,NZB,NZO,NZW,PJ, PL,RF,RH2,SEA,SJL,SMJ,SPRET,SWR,TH, WSB | 14 | 52423282 | 52423315 | 34 | + | 17.2439 | 5.79E-07 | 0.0851 | TAGCTGTCCTCAGATACTCCAGAAGAGGGCATCA | | |
| 2137 | 14 | 52592246 | 52592464 | 218 | SPRET | 14 | 52592323 | 52592356 | 34 | + | 17.4228 | 5.16E-07 | 0.0804 | TAGCTGTTTTCAGACGCACCAGAAGATGGCATCA | | |
| 2138 | 14 | 57929877 | 57930072 | 195 | CAST.MOLF.SPRET | 14 | 57929930 | 57929963 | 34 | + | 16.4472 | 9.61E-07 | 0.111 | TAGCTGTCTTCAGACACACCAGAAGAGGGCGTCA | | |
| 2139 | 14 | 57930795 | 57931001 | 206 | BTBR,BUB,CAST,DBA2J,KK,MOLF,SPRET, WSB | 14 | 57930840 | 57930873 | 34 | + | 16.4634 | 9.51E-07 | 0.111 | TAGCTGTCTTCAGACGCACCAGAAGAGGGCATCA | | |
| 2140 | 14 | 58164418 | 58164630 | 212 | BTBR,BUB,CAST,DBA2J,MOLF,NZB,SPRET .WSB | 14 | 58164486 | 58164519 | 34 | + | 16.4472 | 9.61E-07 | 0.111 | TAGCTGTCTTCAGACACACCAGAAGAGGGCGTCA | | |
| 2141 | 14 | 61352951 | 61353301 | 350 | SPRET | 14 | 61353185 | 61353218 | 34 | + | 18.3089 | 2.87E-07 | 0.0602 | ATGCAGCTAGAGACACGAGCTCCAGGGGGTACTG | | |
| 2142 | 14 | 61774932 | 61775416 | 484 | 129S1,C3H,C57L,CAST,CE,DBA1J,DBA2J,L P.MOLF.SPRET | 14 | 61775405 | 61775438 | 34 | + | 16.4634 | 9.51E-07 | 0.111 | GAGTCTTTCACCTGGAGTCCAGAAGAGGGCACCA | 6 | ENCFF099QNE,ENCFF907XEG,ENCFF367UNR,ENCFF093ZRU,ENCFF821BIA,ENCFF327LYS |
| 2143 | 14 | 61802426 | 61802481 | 55 | 129S1,C3H,C57L,CAST,CE,DBA1J,DBA2J,L P.MOLF.SPRET | 14 | 61802474 | 61802507 | 34 | + | 18.3659 | 2.76E-07 | 0.0595 | TAGCTGTCTTCAGACACTCCAGAAGAGGGCGCCA | 4 | ENCFF367UNR,ENCFF093ZRU,ENCFF502WDT,ENCFF055HAP |
| 2144 | 14 | 63321619 | 63321820 | 201 | SPRET | 14 | 63321676 | 63321709 | 34 | + | 16.4634 | 9.51E-07 | 0.111 | TAGCTGTCTTCAGACGCACCAGAAGAGGGCATCA | | |
| 2145 | 14 | 67282156 | 67282253 | 97 | SPRET | 14 | 67282173 | 67282207 | 35 | + | 16.0826 | 9.12E-07 | 0.222 | ATGCCCTGCAAGTCACTAACCACTAGGGGATGACA | 13 | ENCFF367UNR,ENCFF024JOZ,ENCFF093ZRU,ENCFF165LXO,ENCFF609MDN,ENCFF099QNE,ENCFF198SS R,ENCFF089GBC,ENCFF821BIA,ENCFF327LYS,ENCFF430PPJ,ENCFF092NMJ,ENCFF907XEG |
| 2146 | 14 | 68752725 | 68753464 | 739 | 129S1,C3H,C57L,CE,DBA1J,DBA2J,KK,WS B | 14 | 68753456 | 68753490 | 35 | + | 18.0183 | 2.98E-07 | 0.124 | ATGCAGGTACCCACATGGGCCTGAAGAGGGTATTG | | |
| 2147 | 14 | 70059373 | 70059484 | 111 | 129S1.C57L.KK.MOLF.RH2.SJL.SPRET | 14 | 70059393 | 70059411 | 19 | + | 16.9836 | 9.17E-07 | 0.187 | GGGCCAGGAGGTGGCAGGA | | |
| 2147 | 14 | 70059373 | 70059484 | 111 | 129S1.C57L.KK.MOLF.RH2.SJL.SPRET | 14 | 70059473 | 70059491 | 19 | + | 16.9836 | 9.17E-07 | 0.187 | GGGCCAGGAGGTGGCAGGA | | |
| 2148 | 14 | 70914823 | 70915034 | 211 | 129S1,AJ,BALB,BUB,C3H,C57L,CAST,CE,D BA1J,DBA2J,LP,MOLF,PL,SEA,SPRET | 14 | 70914893 | 70914926 | 34 | + | 16.4634 | 9.51E-07 | 0.111 | TAGCTGTCTTCAGACGCACCAGAAGAGGGCATCA | | |
| 2149 | 14 | 71062167 | 71062365 | 198 | 129S1,AJ,BALB,BUB,C57L,CAST,MOLF,PL, SEA.SPRET | 14 | 71062221 | 71062255 | 35 | + | 18.1284 | 2.79E-07 | 0.12 | CTGTAGCTGTCTTCAGACACCAGAAGAGGGCATCA | | |
| 2150 | 14 | 72681872 | 72682080 | 208 | TH | 14 | 72681940 | 72681973 | 34 | + | 17.5528 | 4.74E-07 | 0.0767 | TAGCTGTCTTCGGATGCACCAGAAGAGGGCGTCA | | |
| 2151 | 14 | 72764168 | 72764395 | 227 | 129S1,BUB,C57L,CAST,FVB,MOLF,SJL,SP RET.TH | 14 | 72764251 | 72764284 | 34 | + | 18.3659 | 2.76E-07 | 0.0595 | TAGCTGTCTTCAGACACTCCAGAAGAGGGCGCCA | | |
| 2151 | 14 | 72764168 | 72764395 | 227 | 129S1,BUB,C57L,CAST,FVB,MOLF,SJL,SP RET.TH | 14 | 72764266 | 72764284 | 19 | + | 17.5574 | 6.32E-07 | 0.158 | ACTCCAGAAGAGGGCGCCA | | |

| | | | | | | | | | | | | | | | | |
|---|---|---|---|---|---|---|---|---|---|---|---|---|---|---|---|---|
| 2152 | 14 | 73598641 | 73599113 | 472 | SPRET | 14 | 73599058 | 73599092 | 35 | + | 21.422 | 3.39E-08 | 0.0404 | GTGCAGAGGCCAGTGGAGGCCAGAAGAGGGTGTTA | 2 | ENCFF093ZRU,ENCFF367UNR |
| 2153 | 14 | 73921967 | 73922164 | 197 | SPRET | 14 | 73922004 | 73922037 | 34 | + | 16.8943 | 7.24E-07 | 0.0965 | TTGCTCTCTTCAGACATACCAGAAGAGGGCATCA | | |
| 2154 | 14 | 78468755 | 78468952 | 197 | CAST.MOLF.NZB.NZO | 14 | 78468809 | 78468842 | 34 | + | 16.6341 | 8.54E-07 | 0.106 | TAGCTGTCTTCACACACTCCAGAAGAGGGCGTCA | | |
| 2155 | 14 | 81412083 | 81412778 | 695 | CAST.SPRET | 14 | 81412097 | 81412130 | 34 | + | 16.4472 | 9.61E-07 | 0.111 | TAGCTGTCTTCAGACACACCAGAAGAGGGCGTCA | | |
| 2156 | 14 | 82408849 | 82409263 | 414 | SPRET | 14 | 82408855 | 82408888 | 34 | + | 16.5691 | 8.90E-07 | 0.108 | GAGCAGGTGTGGCAGCAGTCCCAAGATGGCACCC | | |
| 2157 | 14 | 99308985 | 99309190 | 205 | AKR,CAST,CBA,CE,DBA1J,DBA2J,ILNJ2,MRL,PJ,PL,RF,RH2,SJL,SMJ,SPRET,SWR,TH,WSB | 14 | 99309049 | 99309082 | 34 | + | 16.4634 | 9.51E-07 | 0.111 | TAGCTGTCTTCAGACGCACCAGAAGAGGGCATCA | | |
| 2158 | 14 | 101941300 | 101941498 | 198 | CAST.SPRET | 14 | 101941353 | 101941386 | 34 | + | 17.2114 | 5.91E-07 | 0.0861 | TAGCTGTTTTCAGACACTCCAGAAGAGGGCATCA | | |
| 2159 | 14 | 102967214 | 102967429 | 215 | C57L,C58J,CAST,ILNJ2,MOLF,PJ,SPRET | 14 | 102967287 | 102967320 | 34 | + | 16.4634 | 9.51E-07 | 0.111 | TAGCTGTCTTCAGACGCACCAGAAGAGGGCATCA | | |
| 2160 | 14 | 109079019 | 109079477 | 458 | CAST.SPRET | 14 | 109079308 | 109079342 | 35 | + | 16.789 | 6.11E-07 | 0.182 | GTGCAGGGGAACGCCAGGGCCAAAAGGGGGAGTGG | | |
| 2161 | 14 | 110719314 | 110720152 | 838 | KK.PJ | 14 | 110719715 | 110719748 | 34 | + | 19.5366 | 1.23E-07 | 0.0379 | TTGCTGTCGCCAGACACACCAGAAGAGGGCATTG | | |
| 2162 | 14 | 114101579 | 114102255 | 676 | SPRET | 14 | 114101791 | 114101825 | 35 | + | 20.3119 | 7.10E-08 | 0.0558 | CTGCAGTGCGCTCTCCATGCCACTGGGTGAGCGTC | | |
| 2163 | 14 | 118614928 | 118615417 | 489 | CAST | 14 | 118615370 | 118615403 | 34 | + | 16.4634 | 9.51E-07 | 0.111 | TTGCTGTCTTCAGACACACCAGAAGAGGGAGTCA | | |
| 2164 | 14 | 121054548 | 121054753 | 205 | SPRET | 14 | 121054610 | 121054643 | 34 | + | 17.6667 | 4.40E-07 | 0.0741 | TAGCTGTTTTCAGACACTCCAGAAGAGGGCGTCA | | |
| 2165 | 14 | 121216335 | 121217224 | 889 | SPRET | 14 | 121217172 | 121217190 | 19 | + | 20.1803 | 9.60E-08 | 0.0652 | TGACCACCAGGGGGAGGAC | 25 | ENCFF093ZRU,ENCFF367UNR,ENCFF136ZDK,ENCFF099QNE,ENCFF201JUA,ENCFF502WDT,ENCFF024JOZ,ENCFF907XEG,ENCFF089GBC,ENCFF883UPM,ENCFF821BIA,ENCFF092NMJ,ENCFF327LYS,ENCFF609MDN,ENCFF430PPJ,ENCFF198SSR,ENCFF194HVL,ENCFF165LXO,ENCFF589ZBJ,ENCFF957SPH,ENCFF281QMV,ENCFF951GOP,ENCFF575MGI,ENCFF052KVO,ENCFF533APC |
| 2166 | 14 | 4272917 | 4273117 | 200 | 129S1,C3H,C57L,CBA,DBA1J,DBA2J,KK,MOLF,MRL,NZW,PJ,PL,SPRET | 14 | 4273043 | 4273061 | 19 | - | 17.5574 | 6.32E-07 | 0.158 | ACTCCAGAAGAGGGCGCCA | | |
| 2166 | 14 | 4272917 | 4273117 | 200 | 129S1,C3H,C57L,CBA,DBA1J,DBA2J,KK,MOLF,MRL,NZW,PJ,PL,SPRET | 14 | 4273043 | 4273076 | 34 | - | 18.3659 | 2.76E-07 | 0.0595 | TAGCTGTCTTCAGACACTCCAGAAGAGGGCGCCA | | |
| 2167 | 14 | 8255268 | 8255477 | 209 | 129S1,C3H,C57L,CAST,CBA,DBA1J,DBA2J,KK,MOLF,MRL,NZW,PJ,PL,SPRET | 14 | 8255394 | 8255427 | 34 | - | 17.2439 | 5.79E-07 | 0.0851 | TAGCTGTCCTCAGATACTCCAGAAGAGGGCATCA | | |
| 2168 | 14 | 10309618 | 10309825 | 207 | 129S1,C3H,C57L,CAST,CBA,DBA1J,DBA2J,FVB,KK,MA/My,MOLF,MRL,NOD,NOR,NZW,PJ,PL,SJL,SPRET,SWR | 14 | 10309742 | 10309775 | 34 | - | 16.4472 | 9.61E-07 | 0.111 | TAGCTGTCTTCAGACACACCAGAAGAGGGCGTCA | | |
| 2169 | 14 | 10941884 | 10942081 | 197 | 129S1,C3H,C57L,CBA,DBA1J,DBA2J,KK,MOLF.MRL.NZW.PJ.PL.SPRET | 14 | 10941998 | 10942031 | 34 | - | 16.4472 | 9.61E-07 | 0.111 | TAGCTGTCTTCAGACACACCAGAAGAGGGCGTCA | | |
| 2170 | 14 | 25849065 | 25849284 | 219 | CAST.MOLF.SPRET.WSB | 14 | 25849193 | 25849211 | 19 | - | 17.5574 | 6.32E-07 | 0.158 | ACTCCAGAAGAGGGCGCCA | 2 | ENCFF367UNR.ENCFF093ZRU |
| 2170 | 14 | 25849065 | 25849284 | 219 | CAST.MOLF.SPRET.WSB | 14 | 25849193 | 25849226 | 34 | - | 18.3659 | 2.76E-07 | 0.0595 | TAGCTGTCTTCAGACACTCCAGAAGAGGGCGCCA | 2 | ENCFF367UNR.ENCFF093ZRU |
| 2171 | 14 | 25915180 | 25915377 | 197 | CAST.MOLF.SPRET | 14 | 25915290 | 25915323 | 34 | - | 16.4472 | 9.61E-07 | 0.111 | TAGCTGTCTTCAGACACACCAGAAGAGGGCGTCA | | |
| 2172 | 14 | 31171605 | 31171784 | 179 | SPRET | 14 | 31171645 | 31171678 | 34 | - | 18.4797 | 2.56E-07 | 0.0581 | TTGCTGTCTTCAGACACACCAGAAGAGGGCATCA | 3 | ENCFF093ZRU.ENCFF821BIA.ENCFF367UNR |
| 2173 | 14 | 31809729 | 31809969 | 240 | CAST.MOLF.PJ.SPRET | 14 | 31809854 | 31809887 | 34 | - | 18.1057 | 3.29E-07 | 0.0647 | AAGCTGTCTTCAGACACTCCAGAAGAGGGCACCA | 2 | ENCFF093ZRU.ENCFF367UNR |
| 2174 | 14 | 34374172 | 34374879 | 707 | 129S1.C57L.SPRET | 14 | 34374799 | 34374832 | 34 | - | 18.1545 | 3.18E-07 | 0.0632 | TAGCTGTTTTCAGACACACCAGAAGAGGGCGTCA | | |
| 2175 | 14 | 34475157 | 34475232 | 75 | SPRET | 14 | 34475126 | 34475159 | 34 | - | 19.2195 | 1.54E-07 | 0.0444 | TAGCTGTTTTTTCACACACCAGCAGAGGGCATCA | 4 | ENCFF093ZRU.ENCFF883UPM.ENCFF055HAP.ENCFF502WDT |
| 2176 | 14 | 51905472 | 51905638 | 166 | SPRET | 14 | 51905550 | 51905583 | 34 | - | 27 | 2.58E-10 | 0.00085 | ATGCAATTCCCCCAGAGGCCAGAAGAGGGCATGG | 2 | ENCFF093ZRU.ENCFF052KVO |
| 2177 | 14 | 52347222 | 52347464 | 242 | SPRET | 14 | 52347346 | 52347379 | 34 | - | 16.4472 | 9.61E-07 | 0.111 | TAGCTGTCTTCAGACACACCAGAAGAGGGCGTCA | | |
| 2178 | 14 | 52372531 | 52372763 | 232 | SPRET | 14 | 52372657 | 52372691 | 35 | - | 16.0917 | 9.07E-07 | 0.222 | TAGCTGTCTTCAGACAGCACCAGAAGAGGGCATCA | | |
| 2179 | 14 | 54556375 | 54556572 | 197 | SPRET | 14 | 54556497 | 54556530 | 34 | - | 17.2114 | 5.91E-07 | 0.0861 | TAGCTGTTTTCAGACACTCCAGAAGAGGGCATCA | | |
| 2180 | 14 | 55073334 | 55073533 | 199 | 129S1,AJ,AKR,BALB,BTBR,BUB,C3H,C57L,CAST,CBA,CE,DBA1J,DBA2J,FVB,ILNJ2,KK,MA/My,MOLF,MRL,NOD,NOR,NZB,NZO,NZW,PJ,PL,RF,RH2,SEA,SJL,SMJ,SPRET,SWR.TH.WSB | 14 | 55073466 | 55073484 | 19 | - | 17.5574 | 6.32E-07 | 0.158 | ACTCCAGAAGAGGGCGCCA | 2 | ENCFF093ZRU,ENCFF367UNR |
| 2180 | 14 | 55073334 | 55073533 | 199 | 129S1,AJ,AKR,BALB,BTBR,BUB,C3H,C57L,CAST,CBA,CE,DBA1J,DBA2J,FVB,ILNJ2,KK,MA/My,MOLF,MRL,NOD,NOR,NZB,NZO,NZW,PJ,PL,RF,RH2,SEA,SJL,SMJ,SPRET,SWR.TH.WSB | 14 | 55073466 | 55073499 | 34 | - | 18.3659 | 2.76E-07 | 0.0595 | TAGCTGTCTTCAGACACTCCAGAAGAGGGCGCCA | 2 | ENCFF093ZRU,ENCFF367UNR |
| 2181 | 14 | 55997507 | 55997839 | 332 | SPRET | 14 | 55997798 | 55997816 | 19 | - | 19.2951 | 1.88E-07 | 0.0904 | CCGCCACTAGATGGTGCTA | 33 | ENCFF093ZRU,ENCFF367UNR,ENCFF099QNE,ENCFF327LYS,ENCFF907XEG,ENCFF821BIA,ENCFF136ZDK,ENCFF316FYN,ENCFF024JOZ,ENCFF589ZBJ,ENCFF033UQX,ENCFF089GBC,ENCFF198SSR,ENCFF092NMJ,ENCFF097CAQ,ENCFF321AGQ,ENCFF491RJK,ENCFF065ODV,ENCFF055HAP,ENCFF430PPJ,ENCFF165LXO,ENCFF502WDT,ENCFF957SPH,ENCFF883UPM,ENCFF210BTO,ENCFF281QMV,ENCFF575MGI,ENCFF142CNG,ENCFF951GOP,ENCFF052KVO,ENCFF194HVL,ENCFF533APC,ENCFF609MDN |
| 2182 | 14 | 56115808 | 56116029 | 221 | CAST,SPRET | 14 | 56115933 | 56115966 | 34 | - | 17.2439 | 5.79E-07 | 0.0851 | TAGCTGTCCTCAGACCCTCCAGAAGAGGGAGTCA | 8 | ENCFF093ZRU,ENCFF821BIA,ENCFF883UPM,ENCFF089GBC,ENCFF024JOZ,ENCFF367UNR,ENCFF198SSR,ENCFF907XEG |
| 2183 | 14 | 56371835 | 56372048 | 213 | BTBR,CAST,DBA2J,ILNJ2,MOLF,SMJ,SPRET.WSB | 14 | 56371960 | 56371993 | 34 | - | 17.8455 | 3.91E-07 | 0.0699 | TAGCTGTCCTCAGACACTCCAGAAGAGGGCGTCA | | |
| 2184 | 14 | 57684375 | 57684576 | 201 | CAST.MOLF | 14 | 57684500 | 57684533 | 34 | - | 16.7724 | 7.83E-07 | 0.0992 | TAGCTGTCTTCAGATGCACCAGAAGAGGGCGTCA | | |
| 2185 | 14 | 58157612 | 58157839 | 227 | BTBR,BUB,CAST,DBA2J,MOLF,NZB,WSB | 14 | 58157738 | 58157756 | 19 | - | 17.5738 | 6.25E-07 | 0.158 | ACTCCAGAAGAGGGCGCTA | 1 | ENCFF093ZRU |
| 2185 | 14 | 58157612 | 58157839 | 227 | BTBR,BUB,CAST,DBA2J,MOLF,NZB,WSB | 14 | 58157738 | 58157771 | 34 | - | 18.1789 | 3.13E-07 | 0.0629 | TAGCTGTCTTCAGACACTCCAGAAGAGGGCGCTA | 1 | ENCFF093ZRU |
| 2186 | 14 | 65304707 | 65304916 | 209 | SPRET | 14 | 65304891 | 65304925 | 35 | - | 18.633 | 2.05E-07 | 0.102 | CTGCAGGTGTGATGTCCACCCTGCAGGTGGTGATG | | |
| 2187 | 14 | 65947884 | 65948076 | 192 | CAST.SPRET | 14 | 65948008 | 65948041 | 34 | - | 17.2439 | 5.79E-07 | 0.0851 | TAGCTGTCCTCAGATACTCCAGAAGAGGGCATCA | | |
| 2188 | 14 | 67795357 | 67795568 | 211 | CAST.MOLF.SPRET | 14 | 67795483 | 67795501 | 19 | - | 17.5574 | 6.32E-07 | 0.158 | ACTCCAGAAGAGGGCGCCA | | |
| 2188 | 14 | 67795357 | 67795568 | 211 | CAST.MOLF.SPRET | 14 | 67795483 | 67795516 | 34 | - | 18.1789 | 3.13E-07 | 0.0629 | TAGCTGTCTTCAGCCACTCCAGAAGAGGGCGCCA | | |
| 2189 | 14 | 71071418 | 71071519 | 101 | 129S1,AJ,BALB,BUB,C57L,MOLF,PL,SEA,SPRET | 14 | 71071489 | 71071507 | 19 | - | 18 | 4.70E-07 | 0.142 | TTGCCAGTTGGTGGCGCTG | 1 | ENCFF502WDT |
| 2190 | 14 | 72725200 | 72725597 | 397 | CAST | 14 | 72725372 | 72725390 | 19 | - | 17.1148 | 8.43E-07 | 0.18 | TTTCCAGGAGGGGGCGGGG | | |
| 2191 | 14 | 73503823 | 73503945 | 122 | SPRET | 14 | 73503803 | 73503836 | 34 | - | 27.9024 | 1.04E-10 | 0.000521 | ATGCAGTACCTGCAGAGGCCAGAAGAGGGCATTG | 18 | ENCFF367UNR,ENCFF024JOZ,ENCFF327LYS,ENCFF092NMJ,ENCFF093ZRU,ENCFF907XEG,ENCFF198SSR,ENCFF055HAP,ENCFF883UPM,ENCFF957SPH,ENCFF136ZDK,ENCFF099QNE,ENCFF589ZBJ,ENCFF052KVO,ENCFF142CNG,ENCFF502WDT,ENCFF821BIA,ENCFF065ODV |
| 2192 | 14 | 76062190 | 76062663 | 473 | SPRET | 14 | 76062657 | 76062675 | 19 | - | 17.0656 | 8.70E-07 | 0.184 | AGGCCAGAAGAGGGTGCTG | 2 | ENCFF093ZRU.ENCFF367UNR |
| 2192 | 14 | 76062190 | 76062663 | 473 | SPRET | 14 | 76062657 | 76062690 | 34 | - | 20.3008 | 7.15E-08 | 0.0276 | ATGTGGTACCTGTGGAGGCCAGAAGAGGGTGCTG | 2 | ENCFF093ZRU.ENCFF367UNR |
| 2193 | 14 | 76097973 | 76098180 | 207 | SPRET | 14 | 76098103 | 76098136 | 34 | - | 16.7398 | 7.99E-07 | 0.101 | TAGCTGTCTTCGGACACTCCAGAAGAGGGCGTCA | | |
| 2194 | 14 | 86799121 | 86799329 | 208 | SPRET | 14 | 86799245 | 86799278 | 34 | - | 17.2439 | 5.79E-07 | 0.0851 | TAGCTGTCCTCAGATACTCCAGAAGAGGGCATCA | | |
| 2195 | 14 | 86821182 | 86821443 | 261 | AJ,AKR,BALB,BUB,C3H,CAST,CBA,DBA1J,DBA2J,FVB,NOD,NZO,PJ,RF,SEA,SJL,SMJ,SPRET.SWR.TH.WSB | 14 | 86821309 | 86821342 | 34 | - | 17.0732 | 6.46E-07 | 0.0918 | TAGCTGTCTTCAGAGGCACCAGAAGAGGGCATCA | | |
| 2196 | 14 | 87369255 | 87369307 | 52 | SPRET | 14 | 87369245 | 87369278 | 34 | - | 17.6179 | 4.54E-07 | 0.0752 | TAGCTGTTTTCAGACACACCAGAAGAGGGCATTG | | |
| 2197 | 14 | 95569359 | 95569809 | 450 | AJ,AKR,BALB,BTBR,BUB,C3H,CAST,CBA,CE,DBA1J,DBA2J,FVB,MRL,NOD,PJ,PL,RF,RH2.SEA.SJL.SMJ.SWR.TH.WSB | 14 | 95569766 | 95569799 | 34 | - | 17.9106 | 3.74E-07 | 0.0681 | CCGCCCTCACATTCGCCGTCACAAGATGGCGCTG | | |
| 2198 | 14 | 99487121 | 99487338 | 217 | CAST.SPRET | 14 | 99487246 | 99487279 | 34 | - | 16.4634 | 9.51E-07 | 0.111 | TAGCTGTCTTCAGACGCACCAGAAGAGGGCATCA | | |
| 2199 | 14 | 101995282 | 101995478 | 196 | CAST.SPRET | 14 | 101995403 | 101995437 | 35 | - | 17.6422 | 3.72E-07 | 0.139 | TTGCAACTGTCTTTAGACACCAGAAGAGGGCGTCA | | |
| 2200 | 14 | 105471353 | 105471577 | 224 | C57L.ILNJ2.KK.SPRET | 14 | 105471494 | 105471527 | 34 | - | 16.4472 | 9.61E-07 | 0.111 | TAGCTGTCTTCAGACACACCAGAAGAGGGCGTCA | | |
| 2201 | 14 | 111392223 | 111392356 | 133 | SPRET | 14 | 111392282 | 111392315 | 34 | - | 18 | 3.53E-07 | 0.0659 | TTGTTATGCTTCATCTTCCCAGCAGATGGTGGCA | | |
| 2202 | 14 | 113498864 | 113498935 | 71 | MOLF.SPRET | 14 | 113498915 | 113498949 | 35 | - | 17.9541 | 3.09E-07 | 0.126 | CTGCTTTGATTCTCCACATCCACTGGGTGGCTCAC | 3 | ENCFF093ZRU.ENCFF099QNE.ENCFF502WDT |
| 2203 | 14 | 120217434 | 120218364 | 930 | NZW | 14 | 120218362 | 120218380 | 19 | - | 17.7049 | 5.73E-07 | 0.153 | CTCCCAGCAGAGGGCGGGG | | |
| 2204 | 14 | 122491872 | 122492068 | 196 | SPRET | 14 | 122491996 | 122492030 | 35 | - | 18.1284 | 2.79E-07 | 0.12 | CTGTAGCTGTCTTCAGACACCAGAAGAGGGCATCA | | |
| 2205 | 15 | 6197637 | 6197834 | 197 | SPRET | 15 | 6197690 | 6197723 | 34 | + | 16.4472 | 9.61E-07 | 0.106 | TAGCTGTCTTCAGACACACCAGAAGAGGGCGTCA | | |
| 2206 | 15 | 8225726 | 8225928 | 202 | SPRET | 15 | 8225786 | 8225819 | 34 | + | 16.4634 | 9.51E-07 | 0.106 | TAGCTGTCTTCAGACGCACCAGAAGAGGGCATCA | | |
| 2207 | 15 | 8760894 | 8761469 | 575 | CAST.MOLF.SPRET.TH | 15 | 8761181 | 8761214 | 34 | + | 17.2439 | 5.79E-07 | 0.0855 | TAGCTGTCTTCGGACGCACCAGAAGAGGGCATCA | | |
| 2208 | 15 | 12063493 | 12063918 | 425 | CAST | 15 | 12063523 | 12063557 | 35 | + | 21.578 | 3.05E-08 | 0.0255 | CTGCATATATGCCTACATGTCAGAAGAGGGCACCA | | |
| 2208 | 15 | 12063493 | 12063918 | 425 | CAST | 15 | 12063817 | 12063835 | 19 | + | 17.4262 | 6.88E-07 | 0.131 | TCTACACCAGAGGGCAGCA | | |
| 2209 | 15 | 20042070 | 20042280 | 210 | 129S1.BTBR.C57L.MOLF.SPRET | 15 | 20042135 | 20042168 | 34 | + | 18.3659 | 2.76E-07 | 0.0597 | TAGCTGTCTTCAGACACTCCAGAAGAGGGCGCCA | | |
| 2209 | 15 | 20042070 | 20042280 | 210 | 129S1.BTBR.C57L.MOLF.SPRET | 15 | 20042150 | 20042168 | 19 | + | 17.5574 | 6.32E-07 | 0.126 | ACTCCAGAAGAGGGCGCCA | | |
| 2210 | 15 | 29575520 | 29575627 | 107 | SPRET | 15 | 29575500 | 29575533 | 34 | + | 17.8862 | 3.81E-07 | 0.0697 | ATGCAGCTAGAGACACGAACTCTAGGGGGTACTG | | |
| 2211 | 15 | 34327615 | 34327806 | 191 | 129S1,BUB,C57L,NOD,NOR,PJ,SPRET,SWR | 15 | 34327666 | 34327699 | 34 | + | 17.6992 | 4.30E-07 | 0.0743 | TAGCTGTTTTCAGACACACCAGAAGAGGGCATCA | 1 | ENCFF093ZRU |
| 2212 | 15 | 34401319 | 34401527 | 208 | 129S1,BUB,C57L,NOD,NOR,PJ,SPRET,SWR | 15 | 34401384 | 34401417 | 34 | + | 17.878 | 3.83E-07 | 0.0697 | TAGCTGTCCTCAGACACACCAGAAGAGGGCATCA | 2 | ENCFF093ZRU |
| 2213 | 15 | 35071805 | 35071937 | 132 | 129S1,BUB,C57L,CE,KK,LP,MOLF,NOD,NOR,PJ,SPRET,SWR | 15 | 35071911 | 35071929 | 19 | + | 21.4262 | 3.39E-08 | 0.038 | CTGCCACTAGATGGCAGTG | 17 | ENCFF907XEG,ENCFF327LYS,ENCFF099QNE,ENCFF093ZRU,ENCFF821BIA,ENCFF092NMJ,ENCFF883UPM,ENCFF430PPJ,ENCFF165LXO,ENCFF609MDN,ENCFF198SSR,ENCFF589ZBJ,ENCFF367UNR,ENCFF136ZDK,ENCFF024JOZ,ENCFF055HAP,ENCFF502WDT |
| 2214 | 15 | 37943481 | 37944040 | 559 | MOLF.SPRET | 15 | 37943878 | 37943911 | 34 | + | 16.6341 | 8.54E-07 | 0.104 | CTGCAGGGTCTTGGGCCACCGGAAGGTGGGTGAC | | |
| 2215 | 15 | 37954715 | 37954917 | 202 | SPRET | 15 | 37954774 | 37954807 | 34 | + | 16.9919 | 6.81E-07 | 0.093 | TAGCTGTCCTCAGACTCTCCAGAAGAGGGAGTCA | 1 | ENCFF093ZRU |
| 2216 | 15 | 38705673 | 38705898 | 225 | 129S1.C57L.CAST.CE.MOLF.SPRET | 15 | 38705756 | 38705789 | 34 | + | 18.9919 | 1.80E-07 | 0.0488 | TTGCTGTCTTCAGACACTCCAGAAGAGGGGCATCA | | |
| 2217 | 15 | 40306399 | 40306626 | 227 | 129S1.C57L.CAST.MOLF | 15 | 40306483 | 40306516 | 34 | + | 19.8374 | 9.97E-08 | 0.0331 | TAGCTGTCCCCAGATGCACCAGAAGAGGGCATCA | | |
| 2218 | 15 | 41887302 | 41887448 | 146 | SPRET | 15 | 41887337 | 41887370 | 34 | + | 16.6748 | 8.33E-07 | 0.103 | CTGTAACTCCAGCAGTGGACAGGAGAGATCAGTG | | |
| 2219 | 15 | 49332280 | 49332485 | 205 | CAST.MOLF | 15 | 49332305 | 49332338 | 34 | + | 17.3333 | 5.46E-07 | 0.0834 | CGGGAGTGCTTTGCCTGGGCACCTGGGGGAGCCA | | |
| 2220 | 15 | 52340941 | 52341767 | 826 | CAST.SPRET | 15 | 52341119 | 52341137 | 19 | + | 18.5902 | 3.13E-07 | 0.0907 | CGGCCAGCAGATGTCAGCC | | |
| 2221 | 15 | 54567167 | 54567377 | 210 | CAST.MOLF.SPRET | 15 | 54567235 | 54567268 | 34 | + | 16.4634 | 9.51E-07 | 0.106 | TAGCTGTCTTCAGACGCACCAGAAGAGGGCATCA | | |
| 2222 | 15 | 55371088 | 55371292 | 204 | CAST.MOLF.SPRET | 15 | 55371142 | 55371175 | 34 | + | 17.8455 | 3.91E-07 | 0.0705 | TAGCTGTCCTCAGACACTCCAGAAGAGGGCGTCA | | |
| 2223 | 15 | 58002353 | 58002614 | 261 | SPRET | 15 | 58002474 | 58002507 | 34 | + | 16.4634 | 9.51E-07 | 0.106 | TAGCTGTCTTCAGACGCACCAGAAGAGGGCATCA | 4 | ENCFF367UNR.ENCFF099QNE.ENCFF055HAP.ENCFF093ZRU |
| 2224 | 15 | 59179022 | 59179228 | 206 | CAST.ILNJ2.MOLF.SPRET | 15 | 59179086 | 59179119 | 34 | + | 16.4472 | 9.61E-07 | 0.106 | TAGCTGTCTTCAGACACACCAGAAGAGGGCGTCA | 2 | ENCFF093ZRU |
| 2225 | 15 | 63374806 | 63375024 | 218 | CAST.MOLF.SMJ.SPRET | 15 | 63374878 | 63374911 | 34 | + | 17.9106 | 3.74E-07 | 0.0689 | TAGCTGTCTTCAGACACTCCAGAAGAGGGCACCA | | |
| 2226 | 15 | 63700937 | 63701014 | 77 | SPRET | 15 | 63700992 | 63701025 | 34 | + | 17.8211 | 3.97E-07 | 0.0712 | CTGCAAGTCACATTCTGAACGCTAGGGGACGCGC | 22 | ENCFF201JUA,ENCFF367UNR,ENCFF093ZRU,ENCFF099QNE,ENCFF136ZDK,ENCFF089GBC,ENCFF092NMJ,ENCFF589ZBJ,ENCFF165LXO,ENCFF198SSR,ENCFF504JDS,ENCFF821BIA,ENCFF609MDN,ENCFF502WDT,ENCFF055HAP,ENCFF907XEG,ENCFF883UPM,ENCFF327LYS,ENCFF024JOZ,ENCFF142CNG,ENCFF065ODV,ENCFF052KVO |
| 2227 | 15 | 71920001 | 71920752 | 751 | AJ,B10D2,B10J,BUB,C3H,CAST,CBA,DBA1J,DBA2J,FVB,ILNJ2,LP,MA/My,MOLF,MRL,NOR,NZB,NZO,NZW,PL,SPRET,SWR,TH,WSB | 15 | 71920159 | 71920192 | 34 | + | 17.9431 | 3.66E-07 | 0.0685 | GAACCATTACTCCTACTGGCACCAGGGGGCGCTC | | |
| 2227 | 15 | 71920001 | 71920752 | 751 | AJ,B10D2,B10J,BUB,C3H,CAST,CBA,DBA1J,DBA2J,FVB,ILNJ2,LP,MA/My,MOLF,MRL,NOR,NZB,NZO,NZW,PL,SPRET,SWR,TH,WSB | 15 | 71920174 | 71920192 | 19 | + | 19.6066 | 1.49E-07 | 0.062 | CTGGCACCAGGGGGCGCTC | | |
| 2228 | 15 | 74102515 | 74102610 | 95 | SPRET | 15 | 74102503 | 74102536 | 34 | + | 17.1057 | 6.33E-07 | 0.0898 | CTGCAGGTCCCAGCTCAGCCTGTAGGTCCCAGCT | | |
| 2229 | 15 | 75807174 | 75807500 | 326 | ILNJ2.MA/My.MOLF.SPRET | 15 | 75807358 | 75807391 | 34 | + | 17.5041 | 4.89E-07 | 0.0783 | TAGCTGTACTCAGATACTCCAGAAGAGGGCATCA | | |
| 2230 | 15 | 76779751 | 76780169 | 418 | SPRET | 15 | 76779770 | 76779803 | 34 | + | 16.7724 | 7.83E-07 | 0.0991 | TAGCTGTTTTCAGATACCCCAGAAGAGGGCATCA | 1 | ENCFF093ZRU |
| 2231 | 15 | 76793414 | 76793610 | 196 | CAST.MA/My.MOLF | 15 | 76793465 | 76793499 | 35 | + | 18.1284 | 2.79E-07 | 0.0951 | CTGTAGCTGTCTTCAGACACCAGAAGAGGGCATCA | | |
| 2232 | 15 | 77244690 | 77244906 | 216 | MOLF.SPRET | 15 | 77244764 | 77244797 | 34 | + | 16.9919 | 6.81E-07 | 0.093 | TAGCTGTCTTCAGACACACCAGAAGGGGGCATCA | 1 | ENCFF093ZRU |
| 2233 | 15 | 77256768 | 77257170 | 402 | MOLF.SPRET | 15 | 77256981 | 77257014 | 34 | + | 17.6829 | 4.35E-07 | 0.0745 | TAGCTGTCTTCAGACGCGCCAGAAGAGGGCATCA | | |
| 2234 | 15 | 79027128 | 79027563 | 435 | 129S1,AJ,AKR,BALB,C3H,C57L,C58J,CAST,CBA,CE,DBA1J,DBA2J,FVB,ILNJ2,KK,LP,MA/My,MOLF,MRL,NOD,NOR,NZW,PJ,PL,SEA,SJL,SMJ.SPRET.SWR.TH | 15 | 79027241 | 79027274 | 34 | + | 16.4472 | 9.61E-07 | 0.106 | TAGCTGTCTTCAGACACACCAGAAGAGGGCGTCA | | |
| 2234 | 15 | 79027128 | 79027563 | 435 | 129S1,AJ,AKR,BALB,C3H,C57L,C58J,CAST,CBA,CE,DBA1J,DBA2J,FVB,ILNJ2,KK,LP,MA/My,MOLF,MRL,NOD,NOR,NZW,PJ,PL,SEA,SJL,SMJ.SPRET.SWR.TH | 15 | 79027445 | 79027463 | 19 | + | 17.8852 | 5.08E-07 | 0.114 | ACACCAGAAGAGGGCACCA | | |
| 2235 | 15 | 79750969 | 79751172 | 203 | 129S1,AJ,AKR,BALB,C3H,C57L,C58J,CAST,CBA,CE,DBA1J,DBA2J,FVB,ILNJ2,KK,LP,MA/My,MRL,NOD,NOR,NZW,PJ,PL,SEA,SJL,SMJ.SPRET.SWR.TH | 15 | 79751030 | 79751063 | 34 | + | 18.5285 | 2.48E-07 | 0.058 | CAGCTGTCTTCAGACACTCCAGAAGAGGGCGTCA | | |
| 2236 | 15 | 80739319 | 80739983 | 664 | SPRET | 15 | 80739899 | 80739933 | 35 | + | 17.0092 | 5.39E-07 | 0.138 | ATGCAGTGCTTTCCCTTTACCACTACAGGGCATGT | | |
| 2237 | 15 | 80836083 | 80836318 | 235 | CAST.MOLF.SPRET | 15 | 80836180 | 80836213 | 34 | + | 16.4472 | 9.61E-07 | 0.106 | TAGCTGTCTTCAGACACACCAGAAGAGGGCGTCA | | |
| 2238 | 15 | 80885208 | 80885415 | 207 | CAST.MOLF.SPRET | 15 | 80885272 | 80885305 | 34 | + | 16.4634 | 9.51E-07 | 0.106 | TAGCTGTCTTCAGACGCACCAGAAGAGGGCATCA | | |
| 2239 | 15 | 80909323 | 80909540 | 217 | CAST,DBA1J,DBA2J,ILNJ2,MOLF,SPRET,WSB | 15 | 80909398 | 80909431 | 34 | + | 16.4472 | 9.61E-07 | 0.106 | TAGCTGTCTTCAGACACACCAGAAGAGGGCGTCA | | |
| 2240 | 15 | 81078503 | 81078725 | 222 | AJ,AKR,BALB,C3H,CAST,CBA,CE,DBA1J,DBA2J,FVB,ILNJ2,KK,MOLF,MRL,NOD,NOR,NZW,PJ,PL,SEA,SJL,SMJ,SPRET,SWR,WSB | 15 | 81078583 | 81078616 | 34 | + | 17.6179 | 4.54E-07 | 0.0756 | TAGCTGTCTTCAGACACCCCAGAAGAGGGCACCA | | |
| 2241 | 15 | 81554862 | 81555081 | 219 | SPRET | 15 | 81554936 | 81554969 | 34 | + | 18.3333 | 2.82E-07 | 0.0603 | TAGCTGTCCTCAGACACACCAGAAGAGGGCGTCA | | |
| 2242 | 15 | 81585676 | 81585886 | 210 | MOLF | 15 | 81585737 | 81585771 | 35 | + | 24.8073 | 2.86E-09 | 0.00686 | CTGCATGTATGCCTACGCACCAGAAGAGGGCACCA | 8 | ENCFF821BIA,ENCFF055HAP,ENCFF089GBC,ENCFF093ZRU,ENCFF502WDT,ENCFF099QNE,ENCFF367UNR,ENCFF052KVO |
| 2242 | 15 | 81585676 | 81585886 | 210 | MOLF | 15 | 81585753 | 81585771 | 19 | + | 17.9672 | 4.81E-07 | 0.114 | GCACCAGAAGAGGGCACCA | 8 | ENCFF821BIA,ENCFF055HAP,ENCFF089GBC,ENCFF093ZRU,ENCFF502WDT,ENCFF099QNE,ENCFF367UNR,ENCFF052KVO |
| 2243 | 15 | 81626817 | 81627028 | 211 | AJ,AKR,BALB,BUB,C3H,CAST,CBA,CE,DBA1J,DBA2J,FVB,ILNJ2,KK,MA/My,MOLF,MRL,NOD,NOR,NZB,NZO,NZW,PJ,PL,SEA,SPRET.WSB | 15 | 81626887 | 81626920 | 34 | + | 17.6179 | 4.54E-07 | 0.0756 | TAGCTGTCTTCAGACACCCCAGAAGAGGGCACCA | 8 | ENCFF099QNE,ENCFF821BIA,ENCFF907XEG,ENCFF327LYS,ENCFF201JUA,ENCFF089GBC,ENCFF093ZRU,ENCFF367UNR |
| 2244 | 15 | 84258986 | 84259434 | 448 | SPRET | 15 | 84259013 | 84259046 | 34 | + | 25.5285 | 1.03E-09 | 0.00136 | GTGCAGTACCTATGGAGACCAGAAGAGGGCATCA | 3 | ENCFF367UNR.ENCFF052KVO.ENCFF093ZRU |
| 2245 | 15 | 84545380 | 84545572 | 192 | CAST.MOLF.SPRET | 15 | 84545418 | 84545451 | 34 | + | 16.4472 | 9.61E-07 | 0.106 | TAGCTGTCTTCAGACACACCAGAAGAGGGCGTCA | | |
| 2246 | 15 | 84938634 | 84938856 | 222 | AKR,BTBR,CAST,FVB,MOLF,MRL,NOD,NOR,PL,SJL,SMJ.SPRET.SWR.TH | 15 | 84938713 | 84938746 | 34 | + | 16.4472 | 9.61E-07 | 0.106 | TAGCTGTCTTCAGACACACCAGAAGAGGGCGTCA | | |
| 2247 | 15 | 85474972 | 85475248 | 276 | MOLF | 15 | 85475222 | 85475256 | 35 | + | 16.2294 | 8.40E-07 | 0.171 | TTGCACTGTGAGGATATGGCCACTAGGGTGTGTGT | 2 | ENCFF093ZRU.ENCFF052KVO |
| 2248 | 15 | 88650856 | 88651067 | 211 | CE.FVB.ILNJ2.MOLF.SPRET | 15 | 88650925 | 88650958 | 34 | + | 18.3659 | 2.76E-07 | 0.0597 | TAGCTGTCTTCAGACACTCCAGAAGAGGGCGCCA | | |
| 2248 | 15 | 88650856 | 88651067 | 211 | CE.FVB.ILNJ2.MOLF.SPRET | 15 | 88650940 | 88650958 | 19 | + | 17.5574 | 6.32E-07 | 0.126 | ACTCCAGAAGAGGGCGCCA | | |
| 2249 | 15 | 88678875 | 88679090 | 215 | CAST.CE.FVB.ILNJ2.MOLF.PL.SPRET | 15 | 88678948 | 88678981 | 34 | + | 16.7724 | 7.83E-07 | 0.0991 | TAGCTGTCTTCAGATGCACCAGAAGAGGGCGTCA | | |
| 2250 | 15 | 93272957 | 93273147 | 190 | SPRET | 15 | 93273035 | 93273068 | 34 | + | 17.065 | 6.50E-07 | 0.0909 | TTGCTCTCTTCAGACACACCAGAAGAGGGCATCG | | |
| 2251 | 15 | 93401958 | 93402165 | 207 | SPRET | 15 | 93402016 | 93402049 | 34 | + | 16.4472 | 9.61E-07 | 0.106 | TAGCTGTCTTCAGACACACCAGAAGAGGGCGTCA | | |
| 2252 | 15 | 94884419 | 94884740 | 321 | SPRET | 15 | 94884395 | 94884429 | 35 | + | 18.0642 | 2.90E-07 | 0.0972 | CTGCCCTGTGCCCAGCCAGGCCCCTGGTGGCACCA | | |
| 2253 | 15 | 95353815 | 95353899 | 84 | MOLF | 15 | 95353817 | 95353850 | 34 | + | 17.9106 | 3.74E-07 | 0.0689 | TAGCTGTCTTCAGACACTCCAGAAGAGGGCACCA | 1 | ENCFF093ZRU |
| 2254 | 15 | 96611858 | 96612059 | 201 | CAST.MOLF.NZW.SMJ.SPRET.WSB | 15 | 96611917 | 96611950 | 34 | + | 16.4146 | 9.81E-07 | 0.108 | TAGCTGTCTTCAGACACACCAGAGGAGGGCGCCA | | |
| 2255 | 15 | 97263461 | 97263669 | 208 | 129S1.C57L.C58J.CAST.KK.MOLF | 15 | 97263515 | 97263548 | 34 | + | 16.4472 | 9.61E-07 | 0.106 | TAGCTGTCTTCAGACACACCAGAAGAGGGCGTCA | | |
| 2256 | 15 | 97666927 | 97667538 | 611 | MOLF | 15 | 97667291 | 97667324 | 34 | + | 16.561 | 8.95E-07 | 0.106 | CTGCCTTTAGCCCATCAGCCACCAGAGGTAGTGG | | |
| 2257 | 15 | 97763967 | 97764221 | 254 | CAST,MOLF | 15 | 97764079 | 97764112 | 34 | + | 26.2114 | 5.48E-10 | 0.000884 | ATGCAGTACCTGTGGAGGCCACATGAGGGCACTA | 16 | ENCFF093ZRU,ENCFF821BIA,ENCFF136ZDK,ENCFF907XEG,ENCFF024JOZ,ENCFF165LXO,ENCFF198SSR,ENCFF883UPM,ENCFF367UNR,ENCFF052KVO,ENCFF065ODV,ENCFF055HAP,ENCFF142CNG,ENCFF327LYS,ENCFF502WDT,ENCFF589ZBJ |
| 2258 | 15 | 98041467 | 98041677 | 210 | AJ,BUB,C3H,CAST,CE,MA/My,MOLF,RH2,SPRET.SWR.WSB | 15 | 98041535 | 98041568 | 34 | + | 16.4472 | 9.61E-07 | 0.106 | TAGCTGTCTTCAGACACACCAGAAGAGGGCGTCA | 1 | ENCFF093ZRU |
| 2259 | 15 | 98393264 | 98393466 | 202 | MOLF.SPRET | 15 | 98393326 | 98393359 | 34 | + | 17.6667 | 4.40E-07 | 0.0746 | TAGCTGTTTTCAGACACTCCAGAAGAGGGCGTCA | | |
| 2260 | 15 | 98458489 | 98458682 | 193 | SPRET | 15 | 98458541 | 98458574 | 34 | + | 17.7154 | 4.26E-07 | 0.0739 | TAGCTGTCCTCAGATGCTCCAGAAGAGGGCATCA | | |
| 2261 | 15 | 98788185 | 98788403 | 218 | CAST.MOLF.SPRET | 15 | 98788261 | 98788294 | 34 | + | 17.2439 | 5.79E-07 | 0.0855 | TAGCTGTCCTCAGATACTCCAGAAGAGGGCATCA | | |
| 2262 | 15 | 98918523 | 98918918 | 395 | CAST.MOLF | 15 | 98918581 | 98918614 | 34 | + | 16.4634 | 9.51E-07 | 0.106 | TAGCTGTCTTCAGACGCACCAGAAGAGGGCATCA | | |
| 2263 | 15 | 99001377 | 99001573 | 196 | CAST.MOLF.SPRET | 15 | 99001426 | 99001459 | 34 | + | 18.4797 | 2.56E-07 | 0.058 | TTGCTGTCTTCAGACACACCAGAAGAGGGCATCA | | |
| 2264 | 15 | 100205755 | 100206076 | 321 | SPRET | 15 | 100205750 | 100205783 | 34 | + | 21.3902 | 3.19E-08 | 0.0157 | TAGCCGTCTTCGGACTTACCACTAGAGGGCACCA | 21 | ENCFF092NMJ,ENCFF430PPJ,ENCFF093ZRU,ENCFF367UNR,ENCFF883UPM,ENCFF201JUA,ENCFF099QNE,ENCFF609MDN,ENCFF907XEG,ENCFF821BIA,ENCFF589ZBJ,ENCFF089GBC,ENCFF327LYS,ENCFF024JOZ,ENCFF065ODV,ENCFF136ZDK,ENCFF055HAP,ENCFF198SSR,ENCFF165LXO,ENCFF951GOP,ENCFF502WDT |
| 2264 | 15 | 100205755 | 100206076 | 321 | SPRET | 15 | 100205765 | 100205783 | 19 | + | 22.623 | 1.13E-08 | 0.0255 | TTACCACTAGAGGGCACCA | 22 | ENCFF092NMJ,ENCFF430PPJ,ENCFF093ZRU,ENCFF367UNR,ENCFF883UPM,ENCFF201JUA,ENCFF099QNE,ENCFF609MDN,ENCFF907XEG,ENCFF821BIA,ENCFF589ZBJ,ENCFF089GBC,ENCFF327LYS,ENCFF024JOZ,ENCFF065ODV,ENCFF136ZDK,ENCFF055HAP,ENCFF198SSR,ENCFF165LXO,ENCFF951GOP,ENCFF502WDT,ENCFF957SPH |
| 2265 | 15 | 100522067 | 100522279 | 212 | CAST.MOLF.SPRET | 15 | 100522137 | 100522170 | 34 | + | 20.3659 | 6.82E-08 | 0.0257 | TTGCTGTCCTCAGACACACCAGAAGAGGGCATCA | | |
| 2266 | 15 | 102296009 | 102296220 | 211 | SPRET | 15 | 102296071 | 102296104 | 34 | + | 18.1707 | 3.15E-07 | 0.0634 | TAGCTGTTTTCAGACGCACCAGAAGAGGGCATCA | 1 | ENCFF093ZRU |
| 2267 | 15 | 103239372 | 103239604 | 232 | AJ,C3H,CAST,CBA,CE,DBA1J,DBA2J,FVB,ILNJ2,LP,MOLF,MRL,NOD,NOR,NZB,NZO,PJ,PL,RH2,SEA,SJL,SMJ,SPRET,WSB | 15 | 103239461 | 103239494 | 34 | + | 16.4472 | 9.61E-07 | 0.106 | TAGCTGTCTTCAGACACACCAGAAGAGGGCGTCA | | |
| 2268 | 15 | 103313976 | 103314182 | 206 | CAST.MOLF.SPRET | 15 | 103314054 | 103314072 | 19 | + | 17.8852 | 5.08E-07 | 0.114 | ACACCAGAAGAGGGCACCA | | |
| 2269 | 15 | 103797268 | 103797600 | 332 | CAST,MOLF,SPRET | 15 | 103797297 | 103797315 | 19 | + | 19.2951 | 1.88E-07 | 0.0694 | CCGCCACTAGATGGTGCTA | 31 | ENCFF093ZRU,ENCFF201JUA,ENCFF367UNR,ENCFF052KVO,ENCFF099QNE,ENCFF327LYS,ENCFF907XEG,ENCFF316FYN,ENCFF951GOP,ENCFF821BIA,ENCFF589ZBJ,ENCFF024JOZ,ENCFF321AGQ,ENCFF136ZDK,ENCFF092NMJ,ENCFF089GBC,ENCFF609MDN,ENCFF165LXO,ENCFF430PPJ,ENCFF198SSR,ENCFF097CAQ,ENCFF504JDS,ENCFF883UPM,ENCFF194HVL,ENCFF957SPH,ENCFF281QMV,ENCFF491RJK,ENCFF055HAP.ENCFF533APC.ENCFF575MGI.ENCFF502WDT |
| 2270 | 15 | 4519520 | 4519732 | 212 | SPRET | 15 | 4519639 | 4519672 | 34 | - | 16.4472 | 9.61E-07 | 0.106 | TAGCTGTCTTCAGACACACCAGAAGAGGGCGTCA | | |
| 2271 | 15 | 8098069 | 8098236 | 167 | CAST | 15 | 8098120 | 8098153 | 34 | - | 16.4634 | 9.51E-07 | 0.106 | TAGCTGTCTTCAGACGCACCAGAAGAGGGCATCA | | |
| 2272 | 15 | 8214193 | 8214404 | 211 | AKR.CAST.LP.MOLF.RF.SPRET.TH | 15 | 8214317 | 8214350 | 34 | - | 17.2439 | 5.79E-07 | 0.0855 | TAGCTGTCCTCAGATACTCCAGAAGAGGGCATCA | | |
| 2273 | 15 | 8571113 | 8571950 | 837 | SPRET | 15 | 8571597 | 8571630 | 34 | - | 16.7724 | 7.83E-07 | 0.0991 | TAGCTGTCTTCAGACACACCAGTAGAGGGCATCA | | |
| 2274 | 15 | 10079399 | 10079603 | 204 | CAST.CE.ILNJ2.MOLF.SPRET.TH | 15 | 10079520 | 10079538 | 19 | - | 17.5574 | 6.32E-07 | 0.126 | ACTCCAGAAGAGGGCGCCA | | |
| 2274 | 15 | 10079399 | 10079603 | 204 | CAST.CE.ILNJ2.MOLF.SPRET.TH | 15 | 10079520 | 10079553 | 34 | - | 18.3659 | 2.76E-07 | 0.0597 | TAGCTGTCTTCAGACACTCCAGAAGAGGGCGCCA | | |
| 2275 | 15 | 10991667 | 10991854 | 187 | SPRET | 15 | 10991848 | 10991866 | 19 | - | 17.2459 | 7.74E-07 | 0.142 | ACACCAGAAGAGGGCAGCA | 8 | ENCFF907XEG,ENCFF024JOZ,ENCFF327LYS,ENCFF093ZRU,ENCFF099QNE,ENCFF367UNR,ENCFF821BIA,ENCFF883UPM |
| 2275 | 15 | 10991667 | 10991854 | 187 | SPRET | 15 | 10991848 | 10991881 | 34 | - | 17.6504 | 4.44E-07 | 0.0749 | TAGCTGTCTTCAGACACACCAGAAGAGGGCAGCA | 8 | ENCFF907XEG,ENCFF024JOZ,ENCFF327LYS,ENCFF093ZRU,ENCFF099QNE,ENCFF367UNR,ENCFF821BIA,ENCFF883UPM |
| 2276 | 15 | 19279137 | 19279340 | 203 | CE.ILNJ2.PJ.PL.SEA.SPRET | 15 | 19279262 | 19279295 | 34 | - | 16.4634 | 9.51E-07 | 0.106 | TAGCTGTCTTCAGACGCACCAGAAGAGGGCATCA | | |
| 2277 | 15 | 25718101 | 25718323 | 222 | AJ,BUB,C3H,CAST,CBA,CE,DBA1J,DBA2J,FVB,MA/My,MOLF,MRL,NZB,NZO,NZW,PL,RH2.SMJ.SPRET.TH.WSB | 15 | 25718244 | 25718262 | 19 | - | 17.5574 | 6.32E-07 | 0.126 | ACTCCAGAAGAGGGCGCCA | 1 | ENCFF367UNR |
| 2277 | 15 | 25718101 | 25718323 | 222 | AJ,BUB,C3H,CAST,CBA,CE,DBA1J,DBA2J,FVB,MA/My,MOLF,MRL,NZB,NZO,NZW,PL,RH2.SMJ.SPRET.TH.WSB | 15 | 25718244 | 25718277 | 34 | - | 18.3659 | 2.76E-07 | 0.0597 | TAGCTGTCTTCAGACACTCCAGAAGAGGGCGCCA | 1 | ENCFF367UNR |

| | | | | | | | | | | | | | | | | |
|---|---|---|---|---|---|---|---|---|---|---|---|---|---|---|---|---|
| 2278 | 15 | 33931150 | 33931426 | 276 | 129S1,AJ,BALB,BTBR,BUB,C57L,CAST,FVB,MA/My,MOLF,MRL,NOD,NOR,PJ,RH2,SEA.SJL.SMJ.SPRET.SWR.TH.WSB | 15 | 33931275 | 33931308 | 34 | - | 16.4472 | 9.61E-07 | 0.106 | TAGCTGTCTTCAGACACACCAGAAGAGGGCGTCA | | |
| 2279 | 15 | 34155820 | 34155897 | 77 | SPRET | 15 | 34155836 | 34155854 | 19 | - | 23.6066 | 4.01E-09 | 0.0207 | CGGCCACCAGGGGGCAGGA | 24 | ENCFF367UNR,ENCFF504JDS,ENCFF136ZDK,ENCFF089GBC,ENCFF093ZRU,ENCFF092NMJ,ENCFF099QNE,ENCFF165LXO,ENCFF821BIA,ENCFF065ODV,ENCFF327LYS,ENCFF609MDN,ENCFF201JUA,ENCFF052KVO,ENCFF951GOP,ENCFF055HAP,ENCFF907XEG,ENCFF024JOZ,ENCFF502WDT,ENCFF957SPH,ENCFF883UPM,ENCFF589ZBJ,ENCFF198SSR,ENCFF142CNG |
| 2280 | 15 | 34469980 | 34470069 | 89 | SPRET | 15 | 34470023 | 34470041 | 19 | - | 18.0656 | 4.49E-07 | 0.112 | GAGCCTGTAGGTGGCGCTG | 18 | ENCFF136ZDK,ENCFF092NMJ,ENCFF093ZRU,ENCFF367UNR,ENCFF907XEG,ENCFF099QNE,ENCFF055HAP,ENCFF327LYS,ENCFF821BIA,ENCFF502WDT,ENCFF142CNG,ENCFF024JOZ,ENCFF589ZBJ,ENCFF198SSR,ENCFF281QMV,ENCFF089GBC,ENCFF883UPM,ENCFF957SPH |
| 2280 | 15 | 34469980 | 34470069 | 89 | SPRET | 15 | 34470023 | 34470057 | 35 | - | 19.9174 | 9.17E-08 | 0.0508 | CTGGAGTTAGATTGAGGAGCCTGTAGGTGGCGCTG | 18 | ENCFF136ZDK,ENCFF092NMJ,ENCFF093ZRU,ENCFF367UNR,ENCFF907XEG,ENCFF099QNE,ENCFF055HAP,ENCFF327LYS,ENCFF821BIA,ENCFF502WDT,ENCFF142CNG,ENCFF024JOZ,ENCFF589ZBJ,ENCFF198SSR,ENCFF281QMV,ENCFF089GBC,ENCFF883UPM,ENCFF957SPH |
| 2281 | 15 | 35543103 | 35543302 | 199 | SPRET | 15 | 35543213 | 35543246 | 34 | - | 16.6341 | 8.54E-07 | 0.104 | TAGCTGTCTTCACACACTCCAGAAGAGGGCGTCA | | |
| 2282 | 15 | 36549090 | 36549308 | 218 | 129S1.C57L.CAST.SPRET | 15 | 36549215 | 36549248 | 34 | - | 16.4472 | 9.61E-07 | 0.106 | TAGCTGTCTTCAGACACACCAGAAGAGGGCGTCA | | |
| 2283 | 15 | 36703165 | 36703529 | 364 | 129S1.C57L.CAST | 15 | 36703234 | 36703252 | 19 | - | 18.1475 | 4.25E-07 | 0.108 | TACCCAGAAGGGGGCACCC | | |
| 2284 | 15 | 37125934 | 37126410 | 476 | CAST | 15 | 37126286 | 37126319 | 34 | - | 19.9512 | 9.19E-08 | 0.0314 | GTGTAGTCCCTATCATGGCCAGAAGAGGGGATCA | 1 | ENCFF052KVO |
| 2285 | 15 | 37286120 | 37286363 | 243 | 129S1.C57L.CAST.MOLF.SPRET | 15 | 37286245 | 37286278 | 34 | - | 16.7724 | 7.83E-07 | 0.0991 | TAGCTGTCTTCGGACACACCAGAAGAGGGCATCA | | |
| 2286 | 15 | 39045839 | 39046043 | 204 | 129S1,C3H,C57L,CAST,CE,FVB,MA/My,MOLF,MRL,NOD,NOR,PL,SPRET,SWR,WSB | 15 | 39045965 | 39045983 | 19 | - | 17.5574 | 6.32E-07 | 0.126 | ACTCCAGAAGAGGGCGCCA | | |
| 2286 | 15 | 39045839 | 39046043 | 204 | 129S1,C3H,C57L,CAST,CE,FVB,MA/My,MOLF,MRL,NOD,NOR,PL,SPRET,SWR,WSB | 15 | 39045965 | 39045998 | 34 | - | 18.3659 | 2.76E-07 | 0.0597 | TAGCTGTCTTCAGACACTCCAGAAGAGGGCGCCA | | |
| 2287 | 15 | 41645394 | 41645602 | 208 | 129S1,AJ,AKR,BTBR,BUB,C3H,C57L,CBA,CE,DBA1J,DBA2J,FVB,ILNJ2,KK,LP,MA/My,MOLF,MRL,NOD,NOR,NZB,NZO,NZW,PJ,PL,RH2,SMJ,SPRET,SWR,WSB | 15 | 41645519 | 41645537 | 19 | - | 17.8852 | 5.08E-07 | 0.114 | ACACCAGAAGAGGGCACCA | 3 | ENCFF093ZRU,ENCFF099QNE,ENCFF367UNR |
| 2288 | 15 | 53264671 | 53264862 | 191 | 129S1,AJ,C57L,CE,DBA1J,DBA2J,FVB,ILNJ2,KK,MA/My,RH2,SJL,SMJ,SPRET,TH,WSB | 15 | 53264796 | 53264829 | 34 | - | 16.4472 | 9.61E-07 | 0.106 | TAGCTGTCTTCAGACACACCAGAAGAGGGCGTCA | | |
| 2289 | 15 | 55013550 | 55013785 | 235 | SPRET | 15 | 55013682 | 55013700 | 19 | - | 17.8852 | 5.08E-07 | 0.114 | ACACCAGAAGAGGGCACCA | 5 | ENCFF093ZRU.ENCFF821BIA.ENCFF099QNE.ENCFF883UPM.ENCFF367UNR |
| 2289 | 15 | 55013550 | 55013785 | 235 | SPRET | 15 | 55013682 | 55013715 | 34 | - | 22.5935 | 1.25E-08 | 0.00818 | TTGCTGTTTTCAGACACACCAGAAGAGGGCACCA | 5 | ENCFF093ZRU.ENCFF821BIA.ENCFF099QNE.ENCFF883UPM.ENCFF367UNR |
| 2290 | 15 | 55167036 | 55167414 | 378 | MOLF.SPRET | 15 | 55167334 | 55167367 | 34 | - | 16.4634 | 9.51E-07 | 0.106 | TAGCTGTCTTCAGACGCACCAGAAGAGGGCATCA | | |
| 2291 | 15 | 56577524 | 56577752 | 228 | AKR,BTBR,CAST,ILNJ2,MOLF,NZB,NZO,NZW.SMJ.SPRET.WSB | 15 | 56577650 | 56577683 | 34 | - | 16.4634 | 9.51E-07 | 0.106 | TAGCTGTCTTCAGACGCACCAGAAGAGGGCATCA | | |
| 2292 | 15 | 57954911 | 57955129 | 218 | BALB,CAST,ILNJ2,MOLF,NOD,NOR,SEA,SMJ.SPRET | 15 | 57955034 | 57955052 | 19 | - | 17.5574 | 6.32E-07 | 0.126 | ACTCCAGAAGAGGGCGCCA | | |
| 2292 | 15 | 57954911 | 57955129 | 218 | BALB,CAST,ILNJ2,MOLF,NOD,NOR,SEA,SMJ.SPRET | 15 | 57955034 | 57955067 | 34 | - | 18.3659 | 2.76E-07 | 0.0597 | TAGCTGTCTTCAGACACTCCAGAAGAGGGCGCCA | | |
| 2293 | 15 | 58638304 | 58638500 | 196 | CAST.SPRET | 15 | 58638427 | 58638460 | 34 | - | 16.9919 | 6.81E-07 | 0.093 | TAGCTGTCTTCAGACACACCAGAAGGGGGCATCA | | |
| 2294 | 15 | 63968803 | 63968998 | 195 | SPRET | 15 | 63968927 | 63968960 | 34 | - | 16.4472 | 9.61E-07 | 0.106 | TAGCTGTCTTCAGACACACCAGAAGAGGGCGTCA | | |
| 2295 | 15 | 65494995 | 65495237 | 242 | CAST | 15 | 65495222 | 65495256 | 35 | - | 16.5046 | 7.19E-07 | 0.158 | TTGTAGGTAGAGGTATAGGCCTGAAGATGGAGCTT | | |
| 2296 | 15 | 66393950 | 66394160 | 210 | BALB,C3H,CAST,CBA,CE,MOLF,RF,SEA,SMJ.SPRET.TH | 15 | 66394076 | 66394094 | 19 | - | 17.9672 | 4.81E-07 | 0.114 | GCACCAGAAGAGGGCACCA | | |
| 2296 | 15 | 66393950 | 66394160 | 210 | BALB,C3H,CAST,CBA,CE,MOLF,RF,SEA,SMJ.SPRET.TH | 15 | 66394076 | 66394109 | 34 | - | 18.8699 | 1.96E-07 | 0.051 | TAGCTGTCTTCAGACGCACCAGAAGAGGGCACCA | | |
| 2297 | 15 | 70656986 | 70657189 | 203 | CAST.MOLF.SPRET | 15 | 70657113 | 70657146 | 34 | - | 17.3902 | 5.27E-07 | 0.0815 | TAGCTGTCCTCAGACACTCCAGAAGAGGGCATCA | | |
| 2298 | 15 | 73906228 | 73906285 | 57 | SPRET | 15 | 73906220 | 73906238 | 19 | - | 17.7049 | 5.73E-07 | 0.122 | TGGGCACAAGGGGGCAGCC | 1 | ENCFF093ZRU |
| 2299 | 15 | 76826643 | 76826847 | 204 | CAST.MA/My.MOLF.SPRET | 15 | 76826770 | 76826788 | 19 | - | 17.8852 | 5.08E-07 | 0.114 | ACACCAGAAGAGGGCACCA | | |
| 2300 | 15 | 77219316 | 77219540 | 224 | MA/My.NOD.NOR.NZB.NZO.SPRET | 15 | 77219435 | 77219453 | 19 | - | 16.9016 | 9.66E-07 | 0.156 | ATTCCAGAAGAGGGCACCA | 1 | ENCFF093ZRU |
| 2300 | 15 | 77219316 | 77219540 | 224 | MA/My.NOD.NOR.NZB.NZO.SPRET | 15 | 77219435 | 77219468 | 34 | - | 17.8455 | 3.91E-07 | 0.0705 | TAGCTGTCTTCAGACATTCCAGAAGAGGGCACCA | 1 | ENCFF093ZRU |
| 2301 | 15 | 80753013 | 80753217 | 204 | SPRET | 15 | 80753139 | 80753172 | 34 | - | 17.878 | 3.83E-07 | 0.0697 | TAGCTGTCCTCAGACACACCAGAAGAGGGCATCA | | |
| 2302 | 15 | 83916269 | 83916737 | 468 | SPRET | 15 | 83916590 | 83916623 | 34 | - | 17.5203 | 4.84E-07 | 0.0777 | GGGCAGTGCCTGCAGAAGCCAGAAGAGAGAATCA | | |
| 2303 | 15 | 84157902 | 84158130 | 228 | CAST.MOLF.SPRET | 15 | 84158027 | 84158060 | 34 | - | 17.252 | 5.76E-07 | 0.0855 | TAGCTGTCTTCCGACAGTCCAGAAGAGGGAGCCA | | |
| 2304 | 15 | 84601870 | 84602187 | 317 | 129S1,AJ,BALB,BUB,C3H,C57L,CAST,CBA,CE,DBA1J,DBA2J,ILNJ2,KK,MA/My,MOLF,NZB.NZW.RF.SPRET.WSB | 15 | 84601997 | 84602030 | 34 | - | 18.9512 | 1.85E-07 | 0.0495 | TTGCTGTCTTCAGACGCACCAGAAGAGGGCATCA | 5 | ENCFF093ZRU,ENCFF327LYS,ENCFF024JOZ,ENCFF367UNR,ENCFF907XEG |
| 2305 | 15 | 85771201 | 85772049 | 848 | MOLF | 15 | 85771302 | 85771335 | 34 | - | 17.5691 | 4.69E-07 | 0.0764 | TTGCTCTCTTCAGAGACACCAGAAGAGGGCATCA | | |
| 2306 | 15 | 86534647 | 86534785 | 138 | SPRET | 15 | 86534633 | 86534651 | 19 | - | 18.918 | 2.48E-07 | 0.0799 | TGTACAGCAGAGGGCACCA | | |
| 2307 | 15 | 92836977 | 92837184 | 207 | CAST.MOLF.SPRET | 15 | 92837103 | 92837136 | 34 | - | 16.4472 | 9.61E-07 | 0.106 | TAGCTGTCTTCAGACACACCAGAAGAGGGCGTCA | | |
| 2308 | 15 | 94371902 | 94372758 | 856 | CAST | 15 | 94372244 | 94372277 | 34 | - | 24.0081 | 3.91E-09 | 0.00358 | GTGCAGTAACCAGGGAGGCCAGGAGAGGGCATCA | 5 | ENCFF093ZRU.ENCFF052KVO.ENCFF502WDT.ENCFF055HAP.ENCFF142CNG |
| 2309 | 15 | 98992106 | 98992479 | 373 | CAST,MOLF,SMJ,SPRET | 15 | 98992083 | 98992116 | 34 | - | 17.6911 | 4.33E-07 | 0.0745 | GCGCAGTGTCACTCTAGTCCGATAGGAGGCGCCA | 26 | ENCFF093ZRU,ENCFF086ZTV,ENCFF367UNR,ENCFF099QNE,ENCFF092NMJ,ENCFF136ZDK,ENCFF198SSR,ENCFF430PPJ,ENCFF165LXO,ENCFF316FYN,ENCFF821BIA,ENCFF065ODV,ENCFF055HAP,ENCFF327LYS,ENCFF024JOZ,ENCFF502WDT,ENCFF194HVL,ENCFF907XEG,ENCFF089GBC,ENCFF052KVO,ENCFF957SPH,ENCFF142CNG,ENCFF589ZBJ,ENCFF281QMV |
| 2310 | 15 | 99561481 | 99561681 | 200 | BTBR.CAST.MOLF.SMJ.SPRET.WSB | 15 | 99561607 | 99561640 | 34 | - | 18.1545 | 3.18E-07 | 0.0636 | TAGCTGTTTTCAGACACACCAGAAGAGGGCGTCA | 2 | ENCFF055HAP.ENCFF093ZRU |
| 2311 | 15 | 99947955 | 99948191 | 236 | CAST.MOLF.SPRET | 15 | 99948078 | 99948111 | 34 | - | 17.3902 | 5.27E-07 | 0.0815 | TAGCTGTCCTCAGACACTCCAGAAGAGGGCATCA | | |
| 2312 | 15 | 101897297 | 101897454 | 157 | CAST | 15 | 101897352 | 101897370 | 19 | - | 17.1311 | 8.34E-07 | 0.146 | AGGCCAGAAGAGGGCGTTA | 9 | ENCFF907XEG,ENCFF089GBC,ENCFF198SSR,ENCFF533APC,ENCFF055HAP,ENCFF327LYS,ENCFF065ODV.ENCFF142CNG.ENCFF502WDT |
| 2312 | 15 | 101897297 | 101897454 | 157 | CAST | 15 | 101897352 | 101897385 | 34 | - | 22.813 | 1.05E-08 | 0.00719 | ATGTGGTGCCTGCAGAGGCCAGAAGAGGGCGTTA | 9 | ENCFF907XEG,ENCFF089GBC,ENCFF198SSR,ENCFF533APC,ENCFF055HAP,ENCFF327LYS,ENCFF065ODV.ENCFF142CNG.ENCFF502WDT |
| 2313 | 15 | 102592701 | 102593176 | 475 | SPRET | 15 | 102593022 | 102593055 | 34 | - | 17.8455 | 3.91E-07 | 0.0705 | TAGCTGTCCTCAGACACTCCAGAAGAGGGCGTCA | 1 | ENCFF093ZRU |
| 2314 | 15 | 102981751 | 102981948 | 197 | SPRET | 15 | 102981869 | 102981902 | 34 | - | 17.3902 | 5.27E-07 | 0.0815 | TAGCTGTCCTCAGACACTCCAGAAGAGGGCATCA | | |
| 2315 | 16 | 3387937 | 3388140 | 203 | CAST.MOLF.SMJ.SPRET | 16 | 3387995 | 3388028 | 34 | + | 18.561 | 2.42E-07 | 0.0684 | AAGCTGTCTTCAGACACTCCAGAAGAGGGCGCCA | | |
| 2315 | 16 | 3387937 | 3388140 | 203 | CAST.MOLF.SMJ.SPRET | 16 | 3388010 | 3388028 | 19 | + | 17.5574 | 6.32E-07 | 0.163 | ACTCCAGAAGAGGGCGCCA | | |
| 2316 | 16 | 3484299 | 3485024 | 725 | CAST.MOLF | 16 | 3484445 | 3484478 | 34 | + | 20.9512 | 4.43E-08 | 0.0236 | ATGCAGCTAGAGACATGAGCTCCAGGGGGCACTG | | |
| 2316 | 16 | 3484299 | 3485024 | 725 | CAST.MOLF | 16 | 3484460 | 3484478 | 19 | + | 17.2295 | 7.83E-07 | 0.178 | TGAGCTCCAGGGGGCACTG | | |
| 2317 | 16 | 3642482 | 3642701 | 219 | AJ,BUB,CAST,CE,MOLF,MRL,PL,SMJ,SPRET.WSB | 16 | 3642561 | 3642594 | 34 | + | 17.6179 | 4.54E-07 | 0.0884 | TAGCTGTCTTCAGACACCCCAGAAGAGGGCACCA | | |
| 2318 | 16 | 8566571 | 8566805 | 234 | AJ.AKR.CAST.MOLF.PL.SPRET.WSB | 16 | 8566664 | 8566697 | 34 | + | 17.0488 | 6.56E-07 | 0.107 | TTGCTATCTTCAGATACACCAGAAGAGGGCATCA | | |
| 2319 | 16 | 11241621 | 11241849 | 228 | 129S1,BTBR,C3H,C57L,CBA,DBA1J,DBA2J.ILNJ2.PJ.SPRET.WSB | 16 | 11241625 | 11241659 | 35 | + | 23.1927 | 9.72E-09 | 0.0147 | TGGCACTGCAGCACCCTCACCACCAGGGGCCACTC | 3 | ENCFF093ZRU,ENCFF367UNR,ENCFF504JDS |
| 2319 | 16 | 11241621 | 11241849 | 228 | 129S1,BTBR,C3H,C57L,CBA,DBA1J,DBA2J.ILNJ2.PJ.SPRET.WSB | 16 | 11241641 | 11241659 | 19 | + | 16.8525 | 9.96E-07 | 0.192 | TCACCACCAGGGGCCACTC | 3 | ENCFF093ZRU,ENCFF367UNR,ENCFF504JDS |
| 2319 | 16 | 11241621 | 11241849 | 228 | 129S1,BTBR,C3H,C57L,CBA,DBA1J,DBA2J.ILNJ2.PJ.SPRET.WSB | 16 | 11241682 | 11241716 | 35 | + | 23.1927 | 9.72E-09 | 0.0147 | TGGCACTGCAGCACCCTCACCACCAGGGGCCACTC | 3 | ENCFF093ZRU,ENCFF367UNR,ENCFF504JDS |
| 2319 | 16 | 11241621 | 11241849 | 228 | 129S1,BTBR,C3H,C57L,CBA,DBA1J,DBA2J.ILNJ2.PJ.SPRET.WSB | 16 | 11241698 | 11241716 | 19 | + | 16.8525 | 9.96E-07 | 0.192 | TCACCACCAGGGGCCACTC | 3 | ENCFF093ZRU,ENCFF367UNR,ENCFF504JDS |
| 2319 | 16 | 11241621 | 11241849 | 228 | 129S1,BTBR,C3H,C57L,CBA,DBA1J,DBA2J.ILNJ2.PJ.SPRET.WSB | 16 | 11241739 | 11241773 | 35 | + | 23.1927 | 9.72E-09 | 0.0147 | TGGCACTGCAGCACCCTCACCACCAGGGGCCACTC | 3 | ENCFF093ZRU,ENCFF367UNR,ENCFF504JDS |
| 2319 | 16 | 11241621 | 11241849 | 228 | 129S1,BTBR,C3H,C57L,CBA,DBA1J,DBA2J.ILNJ2.PJ.SPRET.WSB | 16 | 11241755 | 11241773 | 19 | + | 16.8525 | 9.96E-07 | 0.192 | TCACCACCAGGGGCCACTC | 3 | ENCFF093ZRU,ENCFF367UNR,ENCFF504JDS |
| 2319 | 16 | 11241621 | 11241849 | 228 | 129S1,BTBR,C3H,C57L,CBA,DBA1J,DBA2J.ILNJ2.PJ.SPRET.WSB | 16 | 11241796 | 11241830 | 35 | + | 23.1927 | 9.72E-09 | 0.0147 | TGGCACTGCAGCACCCTCACCACCAGGGGCCACTC | 6 | ENCFF093ZRU,ENCFF367UNR,ENCFF201JUA,ENCFF099QNE |
| 2319 | 16 | 11241621 | 11241849 | 228 | 129S1,BTBR,C3H,C57L,CBA,DBA1J,DBA2J.ILNJ2.PJ.SPRET.WSB | 16 | 11241812 | 11241830 | 19 | + | 16.8525 | 9.96E-07 | 0.192 | TCACCACCAGGGGCCACTC | 8 | ENCFF093ZRU,ENCFF367UNR,ENCFF201JUA,ENCFF099QNE,ENCFF821BIA,ENCFF024JOZ |
| 2320 | 16 | 11488213 | 11488297 | 84 | SPRET | 16 | 11488187 | 11488220 | 34 | + | 17.3659 | 5.35E-07 | 0.0971 | CTGTCTGTCTCTCAGCAGCCTGCAGGGGACAGTG | 22 | ENCFF201JUA,ENCFF093ZRU,ENCFF321AGQ,ENCFF883UPM,ENCFF136ZDK,ENCFF092NMJ,ENCFF052KVO,ENCFF089GBC,ENCFF327LYS,ENCFF907XEG,ENCFF430PPJ,ENCFF033UQX,ENCFF589ZBJ,ENCFF198SSR,ENCFF194HVL,ENCFF504JDS,ENCFF502WDT,ENCFF099QNE,ENCFF805QBU,ENCFF821BIA,ENCFF210BTO,ENCFF086ZTV |
| 2320 | 16 | 11488213 | 11488297 | 84 | SPRET | 16 | 11488229 | 11488262 | 34 | + | 21.4472 | 3.05E-08 | 0.0181 | CTGTCTGTCTCTCGGCAGCCTGCAGGGGGCAGTG | 25 | ENCFF201JUA,ENCFF093ZRU,ENCFF321AGQ,ENCFF883UPM,ENCFF136ZDK,ENCFF092NMJ,ENCFF052KVO,ENCFF089GBC,ENCFF327LYS,ENCFF907XEG,ENCFF430PPJ,ENCFF033UQX,ENCFF589ZBJ,ENCFF198SSR,ENCFF194HVL,ENCFF504JDS,ENCFF502WDT,ENCFF099QNE,ENCFF805QBU,ENCFF821BIA,ENCFF210BTO,ENCFF086ZTV,ENCFF024JOZ,ENCFF165LXO,ENCFF316FYN |
| 2320 | 16 | 11488213 | 11488297 | 84 | SPRET | 16 | 11488244 | 11488262 | 19 | + | 20.1803 | 9.60E-08 | 0.0682 | CAGCCTGCAGGGGGCAGTG | 26 | ENCFF201JUA,ENCFF093ZRU,ENCFF321AGQ,ENCFF883UPM,ENCFF136ZDK,ENCFF092NMJ,ENCFF052KVO,ENCFF089GBC,ENCFF327LYS,ENCFF907XEG,ENCFF430PPJ,ENCFF033UQX,ENCFF589ZBJ,ENCFF198SSR,ENCFF194HVL,ENCFF504JDS,ENCFF502WDT,ENCFF099QNE,ENCFF805QBU,ENCFF821BIA,ENCFF210BTO,ENCFF086ZTV,ENCFF024JOZ,ENCFF165LXO,ENCFF316FYN,ENCFF609MDN |
| 2320 | 16 | 11488213 | 11488297 | 84 | SPRET | 16 | 11488271 | 11488304 | 34 | + | 18.3415 | 2.81E-07 | 0.07 | CTGTCTGTCTCTCGGCAGCCTGCAGGAGGCAGTG | 28 | ENCFF093ZRU,ENCFF321AGQ,ENCFF883UPM,ENCFF136ZDK,ENCFF092NMJ,ENCFF052KVO,ENCFF089GBC,ENCFF327LYS,ENCFF907XEG,ENCFF430PPJ,ENCFF033UQX,ENCFF589ZBJ,ENCFF198SSR,ENCFF194HVL,ENCFF502WDT,ENCFF099QNE,ENCFF805QBU,ENCFF821BIA,ENCFF210BTO,ENCFF086ZTV,ENCFF024JOZ,ENCFF165LXO,ENCFF316FYN,ENCFF609MDN,ENCFF957SPH,ENCFF533APC,ENCFF504JDS,ENCFF055HAP |
| 2321 | 16 | 15462386 | 15462584 | 198 | SPRET | 16 | 15462443 | 15462476 | 34 | + | 18.0813 | 3.34E-07 | 0.0764 | CAGCTGTCTTCAGAAACCCCAGAAGAGGGCGTCA | | |
| 2322 | 16 | 15476985 | 15477173 | 188 | CAST.MOLF.SPRET | 16 | 15477030 | 15477063 | 34 | + | 17.0813 | 6.43E-07 | 0.106 | TAGCTGTTCTCAGACACTCCAGAAGAGGGAGTCA | | |
| 2323 | 16 | 16805673 | 16805887 | 214 | CAST.MOLF.SPRET | 16 | 16805743 | 16805776 | 34 | + | 18.1789 | 3.13E-07 | 0.0746 | TAGCTGTCTTCAGACACTCCAGAAGAGGGCGCTA | 1 | ENCFF093ZRU |
| 2323 | 16 | 16805673 | 16805887 | 214 | CAST.MOLF.SPRET | 16 | 16805758 | 16805776 | 19 | + | 17.5738 | 6.25E-07 | 0.163 | ACTCCAGAAGAGGGCGCTA | 1 | ENCFF093ZRU |
| 2324 | 16 | 17002039 | 17002229 | 190 | CAST.SPRET | 16 | 17002087 | 17002120 | 34 | + | 16.4472 | 9.61E-07 | 0.126 | TAGCTGTCTTCAGACACACCAGAAGAGGGCGTCA | | |
| 2325 | 16 | 17049623 | 17049836 | 213 | CAST.MOLF.SPRET | 16 | 17049692 | 17049725 | 34 | + | 17.2439 | 5.79E-07 | 0.1 | TAGCTGTCCTCAGATACTCCAGAAGAGGGCATCA | | |
| 2326 | 16 | 17885484 | 17885710 | 226 | CAST.MOLF.SPRET | 16 | 17885568 | 17885601 | 34 | + | 16.4472 | 9.61E-07 | 0.126 | TAGCTGTCTTCAGACACACCAGAAGAGGGCGTCA | 1 | ENCFF093ZRU |
| 2327 | 16 | 18339664 | 18339858 | 194 | CAST.MOLF.SPRET.WSB | 16 | 18339712 | 18339745 | 34 | + | 17.8455 | 3.91E-07 | 0.0823 | TAGCTGTCCTCAGACACTCCAGAAGAGGGCGTCA | | |
| 2328 | 16 | 18606256 | 18606509 | 253 | CAST.MOLF.SPRET.WSB | 16 | 18606361 | 18606394 | 34 | + | 16.4472 | 9.61E-07 | 0.126 | TAGCTGTCTTCAGACACACCAGAAGAGGGCGTCA | | |
| 2329 | 16 | 20301600 | 20301807 | 207 | CAST | 16 | 20301673 | 20301706 | 34 | + | 16.4472 | 9.61E-07 | 0.126 | TAGCTGTCTTCAGACACACCAGAAGAGGGCGTCA | | |
| 2330 | 16 | 20308829 | 20309028 | 199 | CAST.MOLF.SPRET | 16 | 20308885 | 20308918 | 34 | + | 18.1545 | 3.18E-07 | 0.0752 | TAGCTGTTTTCAGACACACCAGAAGAGGGCGTCA | | |
| 2331 | 16 | 20594757 | 20594971 | 214 | MOLF.SPRET | 16 | 20594829 | 20594862 | 34 | + | 17.6667 | 4.40E-07 | 0.0869 | TAGCTGTCTTCAGACACGCCAGAAGAGGGCGTCA | | |
| 2332 | 16 | 28034310 | 28034481 | 171 | 129S1,BUB,C3H,C57L,CBA,DBA1J,DBA2J,ILNJ2.NZB.NZO.NZW | 16 | 28034299 | 28034332 | 34 | + | 20.561 | 5.91E-08 | 0.0278 | TTGGAGTCCCACCTGTGGCCAGCAGGATGCACTG | 1 | ENCFF093ZRU |
| 2333 | 16 | 30999358 | 30999556 | 198 | SPRET | 16 | 30999421 | 30999454 | 34 | + | 16.4472 | 9.61E-07 | 0.126 | TAGCTGTCTTCAGACACACCAGAAGAGGGCGTCA | 1 | ENCFF093ZRU |
| 2334 | 16 | 31849861 | 31850053 | 192 | MOLF | 16 | 31849904 | 31849937 | 34 | + | 20.4228 | 6.54E-08 | 0.0298 | CTGCTGTCTTCAGACACACCAGGAGAGGGCATCG | 1 | ENCFF093ZRU |
| 2335 | 16 | 32432111 | 32432308 | 197 | SPRET | 16 | 32432169 | 32432202 | 34 | + | 16.4472 | 9.61E-07 | 0.126 | TAGCTGTCTTCAGACACACCAGAAGAGGGCGTCA | 6 | ENCFF367UNR.ENCFF099QNE.ENCFF821BIA.ENCFF136ZDK.ENCFF883UPM.ENCFF093ZRU |
| 2336 | 16 | 32468923 | 32469125 | 202 | SPRET | 16 | 32468983 | 32469016 | 34 | + | 16.4472 | 9.61E-07 | 0.126 | TAGCTGTCTTCAGACACACCAGAAGAGGGCGTCA | | |
| 2337 | 16 | 32788669 | 32789155 | 486 | CAST.MOLF.SPRET | 16 | 32788770 | 32788803 | 34 | + | 17.2439 | 5.79E-07 | 0.1 | TAGCTGTCCTCAGATACTCCAGAAGAGGGCATCA | | |
| 2338 | 16 | 32923392 | 32923622 | 230 | CAST,MOLF,NOD,NOR,NZB,NZO,NZW,SPRET | 16 | 32923480 | 32923513 | 34 | + | 18.3659 | 2.76E-07 | 0.07 | TAGCTGTCTTCAGACACTCCAGAAGAGGGCGCCA | | |
| 2338 | 16 | 32923392 | 32923622 | 230 | CAST,MOLF,NOD,NOR,NZB,NZO,NZW,SPRET | 16 | 32923495 | 32923513 | 19 | + | 17.5574 | 6.32E-07 | 0.163 | ACTCCAGAAGAGGGCGCCA | | |
| 2339 | 16 | 33721250 | 33721462 | 212 | CAST.MOLF.SMJ.SPRET | 16 | 33721318 | 33721351 | 34 | + | 16.4634 | 9.51E-07 | 0.126 | TAGCTGTCTTCAGACGCACCAGAAGAGGGCATCA | | |
| 2340 | 16 | 33826997 | 33827987 | 990 | 129S1.CAST.MOLF.SMJ.SPRET.WSB | 16 | 33827322 | 33827340 | 19 | + | 18.5902 | 3.13E-07 | 0.12 | CGGCCAGCAGATGTCAGCC | | |
| 2341 | 16 | 34546840 | 34547026 | 186 | CAST.MOLF.SPRET | 16 | 34546905 | 34546938 | 34 | + | 16.4309 | 9.71E-07 | 0.127 | TAGCTGTCTTCAGACGCTCCAGAAGAGGGCGTCA | | |
| 2342 | 16 | 35509406 | 35509615 | 209 | CAST.MOLF | 16 | 35509470 | 35509503 | 34 | + | 17.2114 | 5.91E-07 | 0.101 | TAGCTGTTTTCAGACACTCCAGAAGAGGGCATCA | | |
| 2343 | 16 | 37043652 | 37043987 | 335 | MOLF.SPRET | 16 | 37043721 | 37043754 | 34 | + | 16.5691 | 8.90E-07 | 0.126 | TAGCTGTCTTCAGACGCACCAGAAGAGGGCATCG | 1 | ENCFF093ZRU |
| 2343 | 16 | 37043652 | 37043987 | 335 | MOLF.SPRET | 16 | 37043845 | 37043878 | 34 | + | 16.5691 | 8.90E-07 | 0.126 | TAGCTGTCTTCAGACGCACCAGAAGAGGGCATCG | 1 | ENCFF093ZRU |
| 2344 | 16 | 37939971 | 37940171 | 200 | SPRET | 16 | 37940032 | 37940065 | 34 | + | 16.4634 | 9.51E-07 | 0.126 | TAGCTGTCTTCAGACGCACCAGAAGAGGGCATCA | 2 | ENCFF093ZRU.ENCFF367UNR |
| 2345 | 16 | 46549662 | 46549859 | 197 | SPRET | 16 | 46549717 | 46549750 | 34 | + | 16.7724 | 7.83E-07 | 0.117 | TAGCTGTCTTCAGATGCACCAGAAGAGGGCGTCA | | |
| 2346 | 16 | 47037602 | 47037653 | 51 | 129S1,C3H,C58J,CAST,CBA,CE,MOLF,SJL,SPRET.TH | 16 | 47037635 | 47037668 | 34 | + | 16.4634 | 9.51E-07 | 0.126 | TAGCTGTCTTCAGACGCACCAGAAGAGGGCATCA | | |
| 2347 | 16 | 47494710 | 47494914 | 204 | CAST.SPRET | 16 | 47494774 | 47494807 | 34 | + | 16.4472 | 9.61E-07 | 0.126 | TAGCTGTCTTCAGACACACCAGAAGAGGGCGTCA | | |
| 2348 | 16 | 55771283 | 55772230 | 947 | CE | 16 | 55771497 | 55771530 | 34 | + | 24.8211 | 1.94E-09 | 0.00274 | CTGTAATATCACATTCTTCCAGAAGAGGGCGCCC | 29 | ENCFF201JUA,ENCFF099QNE,ENCFF821BIA,ENCFF907XEG,ENCFF024JOZ,ENCFF136ZDK,ENCFF065ODV,ENCFF327LYS,ENCFF589ZBJ,ENCFF316FYN,ENCFF502WDT,ENCFF198SSR,ENCFF055HAP,ENCFF092NMJ,ENCFF052KVO,ENCFF504JDS,ENCFF089GBC,ENCFF321AGQ,ENCFF430PPJ,ENCFF165LXO,ENCFF957SPH,ENCFF951GOP,ENCFF609MDN,ENCFF883UPM,ENCFF194HVL,ENCFF281QMV,ENCFF575MGI,ENCFF093ZRU.ENCFF367UNR |
| 2348 | 16 | 55771283 | 55772230 | 947 | CE | 16 | 55771512 | 55771530 | 19 | + | 18.7869 | 2.72E-07 | 0.113 | CTTCCAGAAGAGGGCGCCC | 29 | ENCFF201JUA,ENCFF099QNE,ENCFF821BIA,ENCFF907XEG,ENCFF024JOZ,ENCFF136ZDK,ENCFF065ODV,ENCFF327LYS,ENCFF589ZBJ,ENCFF316FYN,ENCFF502WDT,ENCFF198SSR,ENCFF055HAP,ENCFF092NMJ,ENCFF052KVO,ENCFF504JDS,ENCFF089GBC,ENCFF321AGQ,ENCFF430PPJ,ENCFF165LXO,ENCFF957SPH,ENCFF951GOP,ENCFF609MDN,ENCFF883UPM,ENCFF194HVL,ENCFF281QMV,ENCFF575MGI,ENCFF093ZRU.ENCFF367UNR |
| 2349 | 16 | 55875445 | 55875717 | 272 | SPRET | 16 | 55875519 | 55875552 | 34 | + | 16.4472 | 9.61E-07 | 0.126 | TAGCTGTCTTCAGACACACCAGAAGAGGGCGTCA | | |
| 2350 | 16 | 56446509 | 56446729 | 220 | 129S1,AJ,BALB,BTBR,BUB,C57L,C58J,CAST,CBA,CE,DBA1J,DBA2J,FVB,KK,LP,MOLF,MRL,NZB,NZO,NZW,PL,RH2,SJL,SMJ,SPRET.SWR.WSB | 16 | 56446587 | 56446620 | 34 | + | 16.4634 | 9.51E-07 | 0.126 | TAGCTGTCTTCAGACGCACCAGAAGAGGGCATCA | | |
| 2351 | 16 | 56936530 | 56936603 | 73 | CAST,CBA,MOLF,NZW,SMJ | 16 | 56936602 | 56936636 | 35 | + | 16.9725 | 5.50E-07 | 0.169 | TTGCTACCTCTCAATAGCACCACAAGATGGCAGCC | 21 | ENCFF821BIA,ENCFF136ZDK,ENCFF430PPJ,ENCFF165LXO,ENCFF367UNR,ENCFF093ZRU,ENCFF907XEG,ENCFF327LYS,ENCFF609MDN,ENCFF055HAP,ENCFF957SPH,ENCFF589ZBJ,ENCFF024JOZ,ENCFF198SSR,ENCFF502WDT,ENCFF099QNE,ENCFF194HVL,ENCFF883UPM,ENCFF089GBC,ENCFF092NMJ,ENCFF951GOP |
| 2352 | 16 | 59764426 | 59764920 | 494 | SPRET | 16 | 59764742 | 59764776 | 35 | + | 17.1284 | 5.03E-07 | 0.161 | CAGCTGTCCCGCATCTGTACTACCAGGGGGAACTA | | |
| 2353 | 16 | 73607014 | 73608002 | 988 | CAST.MOLF.SPRET | 16 | 73607950 | 73607983 | 34 | + | 16.4715 | 9.47E-07 | 0.126 | CTGCCTTACAAGCTGACTCCACCAGAGTGCCCTG | | |
| 2354 | 16 | 80033614 | 80033926 | 312 | SPRET | 16 | 80033636 | 80033669 | 34 | + | 18.8537 | 1.98E-07 | 0.0609 | CTGCAGTTAGAGATACAAGCTCTGGGGGGTACTG | | |
| 2355 | 16 | 84679049 | 84679275 | 226 | BALB,BUB,C58J,CAST,CBA,CE,DBA1J,ILNJ2,LP,MOLF,MRL,NZO,RH2,SPRET | 16 | 84679133 | 84679166 | 34 | + | 18.3659 | 2.76E-07 | 0.07 | TAGCTGTCTTCAGACACTCCAGAAGAGGGCGCCA | | |
| 2355 | 16 | 84679049 | 84679275 | 226 | BALB,BUB,C58J,CAST,CBA,CE,DBA1J,ILNJ2,LP,MOLF,MRL,NZO,RH2,SPRET | 16 | 84679148 | 84679166 | 19 | + | 17.5574 | 6.32E-07 | 0.163 | ACTCCAGAAGAGGGCGCCA | | |
| 2356 | 16 | 86718871 | 86719069 | 198 | CAST.MOLF | 16 | 86718927 | 86718960 | 34 | + | 16.7561 | 7.91E-07 | 0.117 | TCGCTGTGTTCAGACACACCAGAAGAGGGCGTCA | | |
| 2357 | 16 | 87624266 | 87624474 | 208 | CAST.MOLF.SPRET | 16 | 87624339 | 87624372 | 34 | + | 16.4634 | 9.51E-07 | 0.126 | TAGCTGTCTTCAGACGCACCAGAAGAGGGCATCA | | |
| 2358 | 16 | 88192439 | 88192648 | 209 | CAST.MOLF.SPRET | 16 | 88192505 | 88192538 | 34 | + | 16.7724 | 7.83E-07 | 0.117 | TAGCTGTCTTCAGATGCACCAGAAGAGGGCGTCA | | |
| 2359 | 16 | 89178547 | 89178701 | 154 | MOLF | 16 | 89178572 | 89178605 | 34 | + | 18.3089 | 2.87E-07 | 0.07 | ATGCAGCTAGAGACACGAGCTCCAGGGGGTACTG | | |
| 2360 | 16 | 89912432 | 89912638 | 206 | SPRET | 16 | 89912485 | 89912518 | 34 | + | 17.9919 | 3.55E-07 | 0.0786 | TTGCTGTCTTCAGACACTCCAGAAGAGGGCATCA | 1 | ENCFF093ZRU |
| 2361 | 16 | 90152689 | 90152802 | 113 | 129S1,BUB,C57L,C58J,CAST,DBA1J,DBA2J,FVB,MA/My,NOD,NOR,NZO,PJ,RF,SJL,SWR.TH | 16 | 90152774 | 90152808 | 35 | + | 17.0275 | 5.33E-07 | 0.166 | TAGCTGTCTTCAGACAGTACCACAAGAGGGCGTTA | | |
| 2362 | 16 | 91031893 | 91031968 | 75 | SPRET | 16 | 91031895 | 91031929 | 35 | + | 18.8716 | 1.77E-07 | 0.0891 | ATGCAGGTGACTCTGCTAGCCACTGGGTGACACAC | | |
| 2363 | 16 | 95618097 | 95618300 | 203 | CAST.SPRET | 16 | 95618086 | 95618120 | 35 | + | 17.1835 | 4.87E-07 | 0.158 | CTGCAAGGGATGCCCACAGCCACCAGAAGCTGCAA | | |
| 2363 | 16 | 95618097 | 95618300 | 203 | CAST.SPRET | 16 | 95618115 | 95618149 | 35 | + | 16.0092 | 9.50E-07 | 0.217 | CTGCAAGGGATTCCCACAGCCACCAGAAGCTGCAA | | |
| 2363 | 16 | 95618097 | 95618300 | 203 | CAST.SPRET | 16 | 95618144 | 95618178 | 35 | + | 16.0092 | 9.50E-07 | 0.217 | CTGCAAGGGATTCCCACAGCCACCAGAAGCTGCAA | | |
| 2363 | 16 | 95618097 | 95618300 | 203 | CAST.SPRET | 16 | 95618202 | 95618236 | 35 | + | 16.0092 | 9.50E-07 | 0.217 | CTGCAAGGGATTCCCACAGCCACCAGAAGCTGCAA | | |
| 2363 | 16 | 95618097 | 95618300 | 203 | CAST.SPRET | 16 | 95618231 | 95618265 | 35 | + | 16.0092 | 9.50E-07 | 0.217 | CTGCAAGGGATTCCCACAGCCACCAGAAGCTGCAA | | |
| 2363 | 16 | 95618097 | 95618300 | 203 | CAST.SPRET | 16 | 95618289 | 95618323 | 35 | + | 17.7982 | 3.39E-07 | 0.129 | CTGCAAGGGATTCCCACAGCCACCAGAAGCTGCCA | | |
| 2364 | 16 | 96522938 | 96523775 | 837 | MOLF | 16 | 96523521 | 96523539 | 19 | + | 21.1148 | 4.45E-08 | 0.0538 | AGGCCAGAAGGGGGCACTG | 8 | ENCFF093ZRU,ENCFF024JOZ,ENCFF089GBC,ENCFF589ZBJ,ENCFF198SSR,ENCFF099QNE,ENCFF883UPM.ENCFF327LYS |
| 2365 | 16 | 8550246 | 8550459 | 213 | AJ.AKR.CAST.MOLF.PL.SPRET.WSB | 16 | 8550376 | 8550394 | 19 | - | 17.8852 | 5.08E-07 | 0.147 | ACACCAGAAGAGGGCACCA | 2 | ENCFF093ZRU |
| 2366 | 16 | 11099892 | 11100094 | 202 | SPRET | 16 | 11099996 | 11100014 | 19 | - | 17.5574 | 6.32E-07 | 0.163 | ACTCCAGAAGAGGGCGCCA | 1 | ENCFF093ZRU |
| 2366 | 16 | 11099892 | 11100094 | 202 | SPRET | 16 | 11099996 | 11100029 | 34 | - | 20.252 | 7.40E-08 | 0.0323 | TAGCTGTCCTCAGACACTCCAGAAGAGGGCGCCA | 1 | ENCFF093ZRU |
| 2367 | 16 | 12874410 | 12874625 | 215 | CAST.KK.MOLF.SPRET | 16 | 12874535 | 12874553 | 19 | - | 17.5574 | 6.32E-07 | 0.163 | ACTCCAGAAGAGGGCGCCA | | |
| 2367 | 16 | 12874410 | 12874625 | 215 | CAST.KK.MOLF.SPRET | 16 | 12874535 | 12874568 | 34 | - | 18.3659 | 2.76E-07 | 0.07 | TAGCTGTCTTCAGACACTCCAGAAGAGGGCGCCA | | |
| 2368 | 16 | 13523815 | 13524189 | 374 | CAST.KK.MOLF.SPRET | 16 | 13524111 | 13524129 | 19 | - | 17.5574 | 6.32E-07 | 0.163 | ACTCCAGAAGAGGGCGCCA | 1 | ENCFF093ZRU |
| 2368 | 16 | 13523815 | 13524189 | 374 | CAST.KK.MOLF.SPRET | 16 | 13524111 | 13524144 | 34 | - | 18.3659 | 2.76E-07 | 0.07 | TAGCTGTCTTCAGACACTCCAGAAGAGGGCGCCA | 1 | ENCFF093ZRU |
| 2369 | 16 | 14109328 | 14109432 | 104 | SPRET | 16 | 14109334 | 14109352 | 19 | - | 21.0328 | 4.77E-08 | 0.0545 | TCACCTGCAGGGGGCGCCG | 30 | ENCFF367UNR,ENCFF093ZRU,ENCFF327LYS,ENCFF430PPJ,ENCFF099QNE,ENCFF907XEG,ENCFF024JOZ,ENCFF821BIA,ENCFF198SSR,ENCFF136ZDK,ENCFF033UQX,ENCFF165LXO,ENCFF609MDN,ENCFF052KVO,ENCFF589ZBJ,ENCFF142CNG,ENCFF097CAQ,ENCFF089GBC,ENCFF092NMJ,ENCFF533APC,ENCFF194HVL,ENCFF957SPH,ENCFF055HAP,ENCFF281QMV,ENCFF883UPM,ENCFF502WDT,ENCFF210BTO,ENCFF321AGQ.ENCFF201JUA.ENCFF065ODV |
| 2369 | 16 | 14109328 | 14109432 | 104 | SPRET | 16 | 14109334 | 14109367 | 34 | - | 23.4797 | 6.09E-09 | 0.00589 | GTGCGGCTCTGTTCCTCACCTGCAGGGGGCGCCG | 30 | ENCFF367UNR,ENCFF093ZRU,ENCFF327LYS,ENCFF430PPJ,ENCFF099QNE,ENCFF907XEG,ENCFF024JOZ,ENCFF821BIA,ENCFF198SSR,ENCFF136ZDK,ENCFF033UQX,ENCFF165LXO,ENCFF609MDN,ENCFF052KVO,ENCFF589ZBJ,ENCFF142CNG,ENCFF097CAQ,ENCFF089GBC,ENCFF092NMJ,ENCFF533APC,ENCFF194HVL,ENCFF957SPH,ENCFF055HAP,ENCFF281QMV,ENCFF883UPM,ENCFF502WDT,ENCFF210BTO,ENCFF321AGQ.ENCFF201JUA.ENCFF065ODV |
| 2370 | 16 | 14366846 | 14367181 | 335 | SPRET | 16 | 14366885 | 14366919 | 35 | - | 18.1284 | 2.79E-07 | 0.115 | CTGTAGCTGTCTTCAGACACCAGAAGAGGGCATCA | | |
| 2371 | 16 | 16921498 | 16922063 | 565 | SPRET | 16 | 16921643 | 16921676 | 34 | - | 16.8699 | 7.36E-07 | 0.114 | GTGCAGTGCCTACGGGAGCTGGGAGAGGACACAG | | |
| 2372 | 16 | 16924480 | 16924677 | 197 | SPRET | 16 | 16924599 | 16924632 | 34 | - | 16.4634 | 9.51E-07 | 0.126 | TAGCTGTCTTCAGACGCACCAGAAGAGGGCATCA | 1 | ENCFF093ZRU |
| 2373 | 16 | 17016243 | 17016436 | 193 | ILNJ2.SPRET | 16 | 17016371 | 17016404 | 34 | - | 16.4472 | 9.61E-07 | 0.126 | TAGCTGTCTTCAGACACACCAGAAGAGGGCGTCA | 1 | ENCFF093ZRU |
| 2374 | 16 | 17250040 | 17250912 | 872 | SPRET | 16 | 17250609 | 17250627 | 19 | - | 17.9672 | 4.81E-07 | 0.147 | ATACCAGAAGAGGGCACCA | 22 | ENCFF201JUA,ENCFF092NMJ,ENCFF093ZRU,ENCFF052KVO,ENCFF430PPJ,ENCFF099QNE,ENCFF165LXO,ENCFF065ODV,ENCFF907XEG,ENCFF327LYS,ENCFF609MDN,ENCFF821BIA,ENCFF589ZBJ,ENCFF055HAP,ENCFF502WDT,ENCFF089GBC,ENCFF198SSR,ENCFF024JOZ,ENCFF883UPM,ENCFF951GOP,ENCFF136ZDK,ENCFF367UNR |
| 2374 | 16 | 17250040 | 17250912 | 872 | SPRET | 16 | 17250609 | 17250643 | 35 | - | 22.7431 | 1.35E-08 | 0.018 | CTGCATGTATTTCTGCATACCAGAAGAGGGCACCA | 22 | ENCFF201JUA,ENCFF092NMJ,ENCFF093ZRU,ENCFF052KVO,ENCFF430PPJ,ENCFF099QNE,ENCFF165LXO,ENCFF065ODV,ENCFF907XEG,ENCFF327LYS,ENCFF609MDN,ENCFF821BIA,ENCFF589ZBJ,ENCFF055HAP,ENCFF502WDT,ENCFF089GBC,ENCFF198SSR,ENCFF024JOZ,ENCFF883UPM,ENCFF951GOP,ENCFF136ZDK,ENCFF367UNR |
| 2375 | 16 | 17402056 | 17402253 | 197 | SPRET | 16 | 17402145 | 17402178 | 34 | - | 16.4634 | 9.51E-07 | 0.126 | TAGCTGTCTTCAGACGCACCAGAAGAGGGCATCA | | |
| 2376 | 16 | 17419257 | 17419802 | 545 | SPRET | 16 | 17419253 | 17419271 | 19 | - | 17.2951 | 7.50E-07 | 0.176 | AGGACAGAAGGGGGCACCA | 15 | ENCFF367UNR,ENCFF821BIA,ENCFF136ZDK,ENCFF198SSR,ENCFF165LXO,ENCFF093ZRU,ENCFF055HAP,ENCFF502WDT,ENCFF883UPM,ENCFF327LYS,ENCFF957SPH,ENCFF024JOZ,ENCFF099QNE,ENCFF142CNG,ENCFF907XEG |
| 2376 | 16 | 17419257 | 17419802 | 545 | SPRET | 16 | 17419253 | 17419287 | 35 | - | 24.2202 | 4.51E-09 | 0.011 | CTGCATGTATGCCTGCAGGACAGAAGGGGGCACCA | 15 | ENCFF367UNR,ENCFF821BIA,ENCFF136ZDK,ENCFF198SSR,ENCFF165LXO,ENCFF093ZRU,ENCFF055HAP,ENCFF502WDT,ENCFF883UPM,ENCFF327LYS,ENCFF957SPH,ENCFF024JOZ,ENCFF099QNE,ENCFF142CNG,ENCFF907XEG |
| 2377 | 16 | 18183257 | 18183569 | 312 | CAST.MOLF.SPRET | 16 | 18183386 | 18183419 | 34 | - | 16.4472 | 9.61E-07 | 0.126 | TAGCTGTCTTCAGACACACCAGAAGAGGGCGTCA | | |
| 2378 | 16 | 18342025 | 18342290 | 265 | MOLF.SPRET | 16 | 18342206 | 18342239 | 34 | - | 16.5122 | 9.23E-07 | 0.126 | TAGCTGTCTTCAGACACACCACAAGAGGGCATCA | 2 | ENCFF367UNR.ENCFF093ZRU |
| 2379 | 16 | 20170928 | 20171137 | 209 | CAST.MOLF.SPRET | 16 | 20171049 | 20171082 | 34 | - | 17.2439 | 5.79E-07 | 0.1 | TAGCTGTCCTCAGATACTCCAGAAGAGGGCATCA | | |
| 2380 | 16 | 21532654 | 21532855 | 201 | CAST.MOLF.PL.SMJ.SPRET.WSB | 16 | 21532785 | 21532818 | 34 | - | 17.2114 | 5.91E-07 | 0.101 | TAGCTGTTTTCAGACACTCCAGAAGAGGGCATCA | 1 | ENCFF093ZRU |

| | | | | | | | | | | | | | | | | |
|---|---|---|---|---|---|---|---|---|---|---|---|---|---|---|---|---|
| 2381 | 16 | 21841044 | 21841157 | 113 | MOLF,TH | 16 | 21841064 | 21841097 | 34 | - | 26.5691 | 3.91E-10 | 0.00111 | ATGCAGTACCCATAGAGGCCAGAAGAGGGCATCA | 21 | ENCFF201JUA,ENCFF504JDS,ENCFF093ZRU,ENCFF367UNR,ENCFF092NMJ,ENCFF430PPJ,ENCFF099QNE,ENCFF821BIA,ENCFF327LYS,ENCFF883UPM,ENCFF065ODV,ENCFF589ZBJ,ENCFF907XEG,ENCFF089GBC,ENCFF198SSR,ENCFF136ZDK,ENCFF024JOZ,ENCFF321AGQ,ENCFF165LXO,ENCFF052KVO,ENCFF502WDT |
| 2382 | 16 | 22029507 | 22029708 | 201 | CAST.SPRET | 16 | 22029635 | 22029668 | 34 | - | 18.1545 | 3.18E-07 | 0.0752 | TAGCTGTTTTCAGACACACCAGAAGAGGGCGTCA | | |
| 2383 | 16 | 22387451 | 22387651 | 200 | CAST.MOLF.SPRET.TH | 16 | 22387574 | 22387607 | 34 | - | 16.4472 | 9.61E-07 | 0.126 | TAGCTGTCTTCAGACACACCAGAAGAGGGCGTCA | 1 | ENCFF093ZRU |
| 2384 | 16 | 24650603 | 24650817 | 214 | SPRET | 16 | 24650732 | 24650765 | 34 | - | 16.4634 | 9.51E-07 | 0.126 | TAGCTGTCTTCAGACGCACCAGAAGAGGGCATCA | 1 | ENCFF093ZRU |
| 2385 | 16 | 28109360 | 28109556 | 196 | CAST.MOLF.SPRET | 16 | 28109483 | 28109516 | 34 | - | 16.4472 | 9.61E-07 | 0.126 | TAGCTGTCTTCAGACACACCAGAAGAGGGCGTCA | | |
| 2386 | 16 | 30110498 | 30110715 | 217 | 129S1,AJ,AKR,BALB,CAST,FVB,ILNJ2,KK,LP,MOLF,MRL,NOD,NOR,NZB,NZO,NZW,PL,RF,RH2,SEA,SPRET,TH | 16 | 30110624 | 30110642 | 19 | - | 17.5574 | 6.32E-07 | 0.163 | ACTCCAGAAGAGGGCGCCA | | |
| 2386 | 16 | 30110498 | 30110715 | 217 | 129S1,AJ,AKR,BALB,CAST,FVB,ILNJ2,KK,LP,MOLF,MRL,NOD,NOR,NZB,NZO,NZW,PL,RF,RH2,SEA,SPRET,TH | 16 | 30110624 | 30110657 | 34 | - | 18.3659 | 2.76E-07 | 0.07 | TAGCTGTCTTCAGACACTCCAGAAGAGGGCGCCA | | |
| 2387 | 16 | 30161224 | 30162157 | 933 | SPRET | 16 | 30161281 | 30161315 | 35 | - | 18.2752 | 2.55E-07 | 0.108 | CTGTAGTGCAGGATCCCAGCCAGAGAATGGTGCCA | | |
| 2388 | 16 | 31803766 | 31803822 | 56 | MOLF | 16 | 31803806 | 31803839 | 34 | - | 16.4634 | 9.51E-07 | 0.126 | TAGCTGTCTTCAGACGCACCAGAAGAGGGCATCA | | |
| 2389 | 16 | 32885120 | 32885191 | 71 | CAST.MOLF.SPRET | 16 | 32885190 | 32885223 | 34 | - | 16.8943 | 7.24E-07 | 0.112 | CTGCAGTCTTCCTCCTCACAGGCAGGTGACAGCG | 5 | ENCFF327LYS.ENCFF136ZDK.ENCFF589ZBJ.ENCFF883UPM.ENCFF194HVL |
| 2390 | 16 | 33041378 | 33042230 | 852 | WSB | 16 | 33042073 | 33042091 | 19 | - | 17.8033 | 5.37E-07 | 0.152 | AAACCAGAAGAGGGCACCA | 1 | ENCFF367UNR |
| 2391 | 16 | 35348324 | 35348521 | 197 | CAST.MOLF.SPRET | 16 | 35348449 | 35348482 | 34 | - | 16.9187 | 7.13E-07 | 0.112 | TAGCTGTCTTCAGACGCACCAGAAGAGGGCGTCA | | |
| 2392 | 16 | 35676561 | 35676787 | 226 | CAST.MOLF.SPRET | 16 | 35676688 | 35676706 | 19 | - | 17.5738 | 6.25E-07 | 0.163 | ACTCCAGAAGAGGGCGCTA | | |
| 2392 | 16 | 35676561 | 35676787 | 226 | CAST.MOLF.SPRET | 16 | 35676688 | 35676721 | 34 | - | 18.1789 | 3.13E-07 | 0.0746 | TAGCTGTCTTCAGACACTCCAGAAGAGGGCGCTA | | |
| 2393 | 16 | 37602026 | 37602469 | 443 | MOLF.SPRET | 16 | 37602389 | 37602422 | 34 | - | 16.7724 | 7.83E-07 | 0.117 | TAGCTGTCTTCAGATGCACCAGAAGAGGGCGTCA | | |
| 2394 | 16 | 38434781 | 38435003 | 222 | 129S1.C58J.MOLF.SPRET | 16 | 38434905 | 38434938 | 34 | - | 17.8455 | 3.91E-07 | 0.0823 | TAGCTGTCCTCAGACACTCCAGAAGAGGGCGTCA | 1 | ENCFF093ZRU |
| 2395 | 16 | 38753016 | 38753217 | 201 | AJ,AKR,BALB,BTBR,BUB,C3H,C57L,CAST,CBA,CE,DBA1J,DBA2J,FVB,ILNJ2,KK,MA/My,MRL,NOD,NOR,NZB,NZO,NZW,PL,RF,RH2.SEA.SJL.SMJ.SWR.TH.WSB | 16 | 38753140 | 38753173 | 34 | - | 16.4472 | 9.61E-07 | 0.126 | TAGCTGTCTTCAGACACACCAGAAGAGGGCGTCA | | |
| 2396 | 16 | 46746523 | 46746784 | 261 | SPRET | 16 | 46746579 | 46746612 | 34 | - | 18.0976 | 3.31E-07 | 0.076 | ATGCAAAGCCAACTTCTGCGGCCAGAGGGTGCTG | 9 | ENCFF093ZRU,ENCFF024JOZ,ENCFF907XEG,ENCFF821BIA,ENCFF198SSR,ENCFF367UNR,ENCFF194HVL,ENCFF327LYS.ENCFF957SPH |
| 2397 | 16 | 48797525 | 48798152 | 627 | CAST | 16 | 48798028 | 48798061 | 34 | - | 18.4797 | 2.56E-07 | 0.0693 | TTGCTGTCTTCAGACACACCAGAAGAGGGCATCA | | |
| 2398 | 16 | 50729412 | 50730075 | 663 | SPRET | 16 | 50729862 | 50729895 | 34 | - | 18.3089 | 2.87E-07 | 0.07 | ATGCAGCTAGAGACACGAGCTCCAGGGGGTACTG | | |
| 2399 | 16 | 54970056 | 54970261 | 205 | CAST.SPRET | 16 | 54970180 | 54970213 | 34 | - | 17.2439 | 5.79E-07 | 0.1 | TAGCTGTCCTCAGATACTCCAGAAGAGGGCATCA | | |
| 2400 | 16 | 56358343 | 56358674 | 331 | CAST.SPRET | 16 | 56358635 | 56358668 | 34 | - | 21.6829 | 2.55E-08 | 0.0162 | GAGCAGGTGCGGCCACCGCCCCAAGATGGCGCCC | | |
| 2401 | 16 | 58015830 | 58016023 | 193 | AJ,BALB,DBA1J,DBA2J,MOLF,MRL,RH2,SPRET | 16 | 58015956 | 58015989 | 34 | - | 16.4472 | 9.61E-07 | 0.126 | TAGCTGTCTTCAGACACACCAGAAGAGGGCGTCA | | |
| 2402 | 16 | 69825442 | 69825771 | 329 | CE | 16 | 69825730 | 69825763 | 34 | - | 20.065 | 8.47E-08 | 0.0354 | CCGCAATAGCATTCACCTTCACAAGGTGGCGCCG | 9 | ENCFF099QNE,ENCFF883UPM,ENCFF093ZRU,ENCFF907XEG,ENCFF142CNG,ENCFF327LYS,ENCFF502WDT.ENCFF055HAP.ENCFF024JOZ |
| 2403 | 16 | 74689541 | 74689943 | 402 | C58J.LP | 16 | 74689860 | 74689893 | 34 | - | 21.9431 | 2.09E-08 | 0.0144 | CTGCTGTATACCTGTGTGCCAGAAGAGGGCATCA | 1 | ENCFF093ZRU |
| 2404 | 16 | 78025149 | 78025489 | 340 | AJ,C58J,CAST,CBA,DBA1J,DBA2J,FVB,ILNJ2,KK,MA/My,MOLF,NOD,NOR,NZB,NZO,NZW,PL,SJL,SPRET,SWR,WSB | 16 | 78025173 | 78025191 | 19 | - | 17.2459 | 7.74E-07 | 0.177 | CAGCCAGGAGGGGGAAGTA | | |
| 2405 | 16 | 85582372 | 85582580 | 208 | CAST.MOLF | 16 | 85582499 | 85582533 | 35 | - | 16.0917 | 9.07E-07 | 0.214 | TAGCTGTCTTCAGACAGCACCAGAAGAGGGCATCA | | |
| 2406 | 16 | 90781676 | 90781892 | 216 | 129S1,AKR,BUB,C57L,C58J,CAST,CBA,CE,DBA1J,DBA2J,FVB,ILNJ2,MA/My,MOLF,NOD,NOR,NZB,NZO,NZW,PJ,PL,RF,SEA,SJL,SMJ.SPRET.SWR.TH.WSB | 16 | 90781801 | 90781819 | 19 | - | 17.5574 | 6.32E-07 | 0.163 | ACTCCAGAAGAGGGCGCCA | | |
| 2406 | 16 | 90781676 | 90781892 | 216 | 129S1,AKR,BUB,C57L,C58J,CAST,CBA,CE,DBA1J,DBA2J,FVB,ILNJ2,MA/My,MOLF,NOD,NOR,NZB,NZO,NZW,PJ,PL,RF,SEA,SJL,SMJ.SPRET.SWR.TH.WSB | 16 | 90781801 | 90781834 | 34 | - | 18.3659 | 2.76E-07 | 0.07 | TAGCTGTCTTCAGACACTCCAGAAGAGGGCGCCA | | |
| 2407 | 16 | 91279855 | 91280058 | 203 | SPRET | 16 | 91279977 | 91280010 | 34 | - | 16.4472 | 9.61E-07 | 0.126 | TAGCTGTCTTCAGACACACCAGAAGAGGGCGTCA | 1 | ENCFF093ZRU |
| 2408 | 16 | 91442215 | 91442426 | 211 | BUB,CAST,CBA,CE,DBA1J,FVB,MA/My,MOLF,NZB,NZO,PL,SJL,SPRET,SWR,TH | 16 | 91442338 | 91442371 | 34 | - | 16.4634 | 9.51E-07 | 0.126 | TAGCTGTCTTCAGACGCACCAGAAGAGGGCATCA | | |
| 2409 | 16 | 92796018 | 92796212 | 194 | CAST.CE.DBA1J.MOLF.SPRET.TH | 16 | 92796142 | 92796160 | 19 | - | 17.5574 | 6.32E-07 | 0.163 | ACTCCAGAAGAGGGCGCCA | | |
| 2409 | 16 | 92796018 | 92796212 | 194 | CAST.CE.DBA1J.MOLF.SPRET.TH | 16 | 92796142 | 92796175 | 34 | - | 18.3659 | 2.76E-07 | 0.07 | TAGCTGTCTTCAGACACTCCAGAAGAGGGCGCCA | | |
| 2410 | 16 | 93385602 | 93385797 | 195 | CE.DBA1J.MOLF.NZW.SPRET | 16 | 93385713 | 93385746 | 34 | - | 16.7724 | 7.83E-07 | 0.117 | TAGCTGTCTTCAGATGCACCAGAAGAGGGCGTCA | | |
| 2411 | 16 | 93655110 | 93655316 | 206 | MOLF.NZW.SPRET.TH | 16 | 93655226 | 93655244 | 19 | - | 17.5574 | 6.32E-07 | 0.163 | ACTCCAGAAGAGGGCGCCA | 4 | ENCFF093ZRU.ENCFF367UNR.ENCFF099QNE.ENCFF821BIA |
| 2411 | 16 | 93655110 | 93655316 | 206 | MOLF.NZW.SPRET.TH | 16 | 93655226 | 93655259 | 34 | - | 18.1789 | 3.13E-07 | 0.0746 | TAGCTGTCTTCAGCCACTCCAGAAGAGGGCGCCA | 4 | ENCFF093ZRU.ENCFF367UNR.ENCFF099QNE.ENCFF821BIA |
| 2412 | 16 | 93816193 | 93816402 | 209 | SPRET | 16 | 93816317 | 93816350 | 34 | - | 16.4634 | 9.51E-07 | 0.126 | TAGCTGTCTTCAGACGCACCAGAAGAGGGCATCA | 1 | ENCFF093ZRU |
| 2413 | 16 | 95744113 | 95744303 | 190 | CAST.MOLF.SPRET.WSB | 16 | 95744227 | 95744260 | 34 | - | 16.4634 | 9.51E-07 | 0.126 | TAGCTGTCTTCAGACGCACCAGAAGAGGGCATCA | | |
| 2414 | 16 | 96886224 | 96886874 | 650 | SPRET | 16 | 96886485 | 96886518 | 34 | - | 17.8049 | 4.02E-07 | 0.084 | GTGAAGTGCCCAAGGAGGCCAAAAGATGGTGTCA | | |
| 2415 | 17 | 3501501 | 3501571 | 70 | CAST,DBA2J,FVB,ILNJ2,MA/My,MOLF,NOD,NOR,RF,SJL,SPRET | 17 | 3501504 | 3501537 | 34 | + | 24.7317 | 2.10E-09 | 0.00133 | GTGCAGTGCCCAAGGAGGCCAGAAGAGGGAGTCT | 42 | ENCFF093ZRU,ENCFF201JUA,ENCFF367UNR,ENCFF099QNE,ENCFF065ODV,ENCFF504JDS,ENCFF821BIA,ENCFF327LYS,ENCFF321AGQ,ENCFF589ZBJ,ENCFF907XEG,ENCFF198SSR,ENCFF136ZDK,ENCFF052KVO,ENCFF024JOZ,ENCFF502WDT,ENCFF430PPJ,ENCFF055HAP,ENCFF883UPM,ENCFF210BTO,ENCFF957SPH,ENCFF165LXO,ENCFF092NMJ,ENCFF033UQX,ENCFF533APC,ENCFF089GBC,ENCFF194HVL,ENCFF784ASD,ENCFF575MGI,ENCFF951GOP,ENCFF089EWX,ENCFF316FYN,ENCFF491RJK,ENCFF349BUL,ENCFF611HDQ,ENCFF609MDN,ENCFF086ZTV,ENCFF281QMV,ENCFF097CAQ,ENCFF601JVE,ENCFF142CNG |
| 2416 | 17 | 6150177 | 6150930 | 753 | SPRET | 17 | 6150485 | 6150519 | 35 | + | 18.1284 | 2.79E-07 | 0.0764 | CTGTAGCTGTCTTCAGACACCAGAAGAGGGCATCA | | |
| 2417 | 17 | 13067260 | 13067484 | 224 | C3H,CAST,CBA,CE,ILNJ2,KK,MOLF,MRL,NOD,NOR,PL,SJL,SMJ,SPRET,SWR,TH | 17 | 13067340 | 13067373 | 34 | + | 16.4472 | 9.61E-07 | 0.0875 | TAGCTGTCTTCAGACACACCAGAAGAGGGCGTCA | | |
| 2418 | 17 | 13916787 | 13917347 | 560 | SPRET | 17 | 13917034 | 13917067 | 34 | + | 16.5528 | 8.99E-07 | 0.0873 | TAGCTGTCTTCCAACACACCAGAAGAGGGCATCA | 2 | ENCFF093ZRU.ENCFF367UNR |
| 2419 | 17 | 15194780 | 15194983 | 203 | SPRET | 17 | 15194845 | 15194878 | 34 | + | 16.9593 | 6.95E-07 | 0.0768 | TTGCTCTCTTCAGACACACCAGAAGAGGGCATCA | | |
| 2420 | 17 | 15257501 | 15257704 | 203 | SPRET | 17 | 15257566 | 15257599 | 34 | + | 16.9593 | 6.95E-07 | 0.0768 | TTGCTCTCTTCAGACACACCAGAAGAGGGCATCA | | |
| 2421 | 17 | 17699068 | 17700031 | 963 | BUB.C3H.CBA.MRL.PJ.RF.SJL.SMJ.TH | 17 | 17699422 | 17699440 | 19 | + | 17.6721 | 5.85E-07 | 0.107 | ATACCAGAAGAGGGCACTG | | |
| 2422 | 17 | 25186801 | 25187009 | 208 | 129S1,AJ,AKR,B10D2,BALB,BTBR,BUB,C3H,C57L,CAST,CBA,CE,DBA1J,DBA2J,FVB,ILNJ2,KK,MA/My,MRL,NOD,NOR,NZB,NZO,PJ,RF,RH2,SEA,SJL,SPRET,SWR,TH,WSB | 17 | 25186866 | 25186899 | 34 | + | 18.3659 | 2.76E-07 | 0.0462 | TAGCTGTCTTCAGACACTCCAGAAGAGGGCGCCA | 3 | ENCFF367UNR,ENCFF821BIA,ENCFF093ZRU |
| 2422 | 17 | 25186801 | 25187009 | 208 | 129S1,AJ,AKR,B10D2,BALB,BTBR,BUB,C3H,C57L,CAST,CBA,CE,DBA1J,DBA2J,FVB,ILNJ2,KK,MA/My,MRL,NOD,NOR,NZB,NZO,PJ,RF,RH2,SEA,SJL,SPRET,SWR,TH,WSB | 17 | 25186881 | 25186899 | 19 | + | 17.5574 | 6.32E-07 | 0.108 | ACTCCAGAAGAGGGCGCCA | 3 | ENCFF367UNR,ENCFF821BIA,ENCFF093ZRU |
| 2423 | 17 | 27296136 | 27296286 | 150 | CAST,KK,NZO | 17 | 27296254 | 27296288 | 35 | + | 25.7615 | 1.33E-09 | 0.0034 | GTGTAGTTTCTTCACCTCACCAGTAGAGGGCACAC | 14 | ENCFF099QNE,ENCFF502WDT,ENCFF198SSR,ENCFF055HAP,ENCFF092NMJ,ENCFF052KVO,ENCFF093ZRU,ENCFF327LYS,ENCFF136ZDK,ENCFF367UNR,ENCFF907XEG,ENCFF321AGQ,ENCFF065ODV,ENCFF589ZBJ |
| 2423 | 17 | 27296136 | 27296286 | 150 | CAST,KK,NZO | 17 | 27296270 | 27296288 | 19 | + | 19.2623 | 1.93E-07 | 0.0593 | TCACCAGTAGAGGGCACAC | 14 | ENCFF099QNE,ENCFF502WDT,ENCFF198SSR,ENCFF055HAP,ENCFF092NMJ,ENCFF052KVO,ENCFF093ZRU,ENCFF327LYS,ENCFF136ZDK,ENCFF367UNR,ENCFF907XEG,ENCFF321AGQ,ENCFF065ODV,ENCFF589ZBJ |
| 2424 | 17 | 28147362 | 28147577 | 215 | 129S1,AJ,AKR,B10D2,BALB,BTBR,C3H,C57L,CAST,CBA,CE,DBA1J,DBA2J,FVB,ILNJ2,KK,MA/My,MOLF,MRL,NZB,NZW,RF,RH2,SEA,SJL,SMJ,SPRET,TH,WSB | 17 | 28147435 | 28147468 | 34 | + | 16.4634 | 9.51E-07 | 0.0875 | TAGCTGTCTTCAGACGCACCAGAAGAGGGCATCA | | |
| 2425 | 17 | 28609798 | 28610016 | 218 | 129S1,AJ,AKR,B10D2,BALB,C3H,C57L,CAST,CBA,CE,DBA1J,DBA2J,FVB,ILNJ2,KK,MA/My,MOLF,MRL,NOD,NOR,NZB,NZW,PJ,RF,RH2,SEA,SJL,SMJ,SPRET,TH,WSB | 17 | 28609865 | 28609898 | 34 | + | 17.6992 | 4.30E-07 | 0.0594 | TAGCTGTTTTCAGACACACCAGAAGAGGGCATCA | | |
| 2426 | 17 | 28750481 | 28750857 | 376 | SPRET | 17 | 28750480 | 28750513 | 34 | + | 18.3821 | 2.73E-07 | 0.0462 | GTGCAGTGCCCACAGAGGCTAGAAGAGAGCATTG | | |
| 2427 | 17 | 28829543 | 28829644 | 101 | CAST.ILNJ2.KK.MOLF.RF.SPRET | 17 | 28829603 | 28829637 | 35 | + | 19.1835 | 1.46E-07 | 0.0519 | CTGCATGTACGCCTGCAGGTCAGAAGAAGGCAACC | | |
| 2428 | 17 | 29040624 | 29040732 | 108 | ILNJ2.SPRET | 17 | 29040656 | 29040690 | 35 | + | 19.2294 | 1.42E-07 | 0.0513 | GAGCACTCTCCCCTGCCAGCCATCAGGGAGCACTC | | |
| 2428 | 17 | 29040624 | 29040732 | 108 | ILNJ2.SPRET | 17 | 29040683 | 29040717 | 35 | + | 19.2294 | 1.42E-07 | 0.0513 | GAGCACTCTCCCCTGCCAGCCATCAGGGAGCACTC | | |
| 2428 | 17 | 29040624 | 29040732 | 108 | ILNJ2.SPRET | 17 | 29040710 | 29040744 | 35 | + | 19.2294 | 1.42E-07 | 0.0513 | GAGCACTCTCCCCTGCCAGCCATCAGGGAGCACTC | | |
| 2429 | 17 | 29519527 | 29519732 | 205 | SPRET | 17 | 29519581 | 29519614 | 34 | + | 18.3659 | 2.76E-07 | 0.0462 | TAGCTGTCTTCAGACACTCCAGAAGAGGGCGCCA | | |
| 2429 | 17 | 29519527 | 29519732 | 205 | SPRET | 17 | 29519596 | 29519614 | 19 | + | 17.5574 | 6.32E-07 | 0.108 | ACTCCAGAAGAGGGCGCCA | | |
| 2430 | 17 | 29531130 | 29531281 | 151 | SPRET | 17 | 29531159 | 29531192 | 34 | + | 18.3984 | 2.70E-07 | 0.046 | ATGCAATGCCCATAGAGGCCAAAAGAGGCCAGAT | | |
| 2431 | 17 | 29734996 | 29735130 | 134 | CAST.SPRET | 17 | 29735081 | 29735114 | 34 | + | 17.1789 | 6.04E-07 | 0.0725 | TAGCTGTCCTCAGATATTCCAGAAGAGGGCATCA | | |
| 2432 | 17 | 29758628 | 29758832 | 204 | CAST.KK.MOLF.RF | 17 | 29758692 | 29758725 | 34 | + | 17.2439 | 5.79E-07 | 0.0713 | TAGCTGTCCTCAGATACTCCAGAAGAGGGCATCA | | |
| 2433 | 17 | 29759259 | 29759450 | 191 | CAST.KK.MOLF.RF | 17 | 29759259 | 29759292 | 34 | + | 18.7398 | 2.14E-07 | 0.0412 | CAGAAATGCCTCCTCCGGCCAGGCGGTGGTGGCA | | |
| 2434 | 17 | 31526431 | 31526642 | 211 | AJ,AKR,BTBR,C3H,C58J,CAST,CBA,CE,DBA1J,FVB,ILNJ2,KK,MOLF,MRL,NOD,NOR,NZB,NZO,NZW,PJ,PL,RF,SJL,SMJ,SPRET,TH,WSB | 17 | 31526502 | 31526535 | 34 | + | 16.6911 | 8.24E-07 | 0.0833 | TAGCTGTCTTCAGACAGACCAGAAGAGGGCATCA | | |
| 2435 | 17 | 34200236 | 34200674 | 438 | AJ,AKR,B10D2,BALB,BUB,C3H,C58J,CAST,CBA,CE,DBA1J,DBA2J,FVB,ILNJ2,MA/My,MOLF,MRL,NOD,NOR,NZB,NZO,NZW,PJ,PL,RF,RH2,SEA,SJL,SMJ,SPRET,SWR,TH,WSB | 17 | 34200266 | 34200284 | 19 | + | 17.4918 | 6.59E-07 | 0.11 | CTGTCACAAGGTGGCGCTG | 22 | ENCFF093ZRU,ENCFF367UNR,ENCFF089GBC,ENCFF136ZDK,ENCFF099QNE,ENCFF430PPJ,ENCFF316FYN,ENCFF327LYS,ENCFF907XEG,ENCFF589ZBJ,ENCFF533APC,ENCFF097CAQ,ENCFF609MDN,ENCFF194HVL,ENCFF024JOZ,ENCFF092NMJ,ENCFF821BIA,ENCFF321AGQ,ENCFF281QMV,ENCFF957SPH,ENCFF198SSR,ENCFF165LXO |
| 2436 | 17 | 35254375 | 35254606 | 231 | AJ,AKR,B10D2,BALB,BUB,C3H,C58J,CAST,CBA,CE,DBA1J,DBA2J,FVB,ILNJ2,MA/My,MOLF,MRL,NZB,NZO,NZW,PJ,PL,RF,RH2,SEA,SJL,SMJ,SPRET,SWR,TH,WSB | 17 | 35254457 | 35254490 | 34 | + | 16.4472 | 9.61E-07 | 0.0875 | TAGCTGTCTTCAGACACACCAGAAGAGGGCGTCA | | |
| 2437 | 17 | 35635649 | 35635857 | 208 | SPRET | 17 | 35635627 | 35635661 | 35 | + | 16.1743 | 8.66E-07 | 0.138 | CTACAGAACAAGACCACAGCCAGCAGGGGCTGGTG | | |
| 2438 | 17 | 35680857 | 35681065 | 208 | FVB.NOD.NOR.NZB.PL.RH2.SWR | 17 | 35680833 | 35680867 | 35 | + | 16.1743 | 8.66E-07 | 0.138 | CTACAGAACAAGACCACAGCCAGCAGGGGCTGGTG | | |
| 2439 | 17 | 35772782 | 35772987 | 205 | AJ,B10D2,BALB,BUB,CAST,DBA2J,FVB,MOLF,NOD,NOR,NZB,PL,RH2,SEA,SPRET,SWR | 17 | 35772759 | 35772793 | 35 | + | 16.1743 | 8.66E-07 | 0.138 | CTACAGAACAAGACCACAGCCAGCAGGGGCTGGTG | | |
| 2440 | 17 | 36030832 | 36031282 | 450 | MOLF | 17 | 36031040 | 36031074 | 35 | + | 26.6514 | 6.35E-10 | 0.00239 | CTGCATGTCCACCTGCATGCCAGAAGAGGGCATTA | 14 | ENCFF201JUA,ENCFF821BIA,ENCFF089GBC,ENCFF165LXO,ENCFF093ZRU,ENCFF092NMJ,ENCFF367UNR,ENCFF609MDN,ENCFF099QNE,ENCFF327LYS,ENCFF907XEG,ENCFF024JOZ,ENCFF055HAP,ENCFF198SSR |
| 2441 | 17 | 36404377 | 36404855 | 478 | AJ,BUB,C58J,FVB,KK,NOD,NOR,NZB,PJ,PL,SJL,SWR,WSB | 17 | 36404654 | 36404688 | 35 | + | 18.0642 | 2.90E-07 | 0.0766 | CTGCTTTTCCCCAGGCAGGGCCCTAGGTGTCACAG | | |
| 2442 | 17 | 43317099 | 43317266 | 167 | SPRET | 17 | 43317152 | 43317185 | 34 | + | 17.9512 | 3.64E-07 | 0.0538 | ATGTGGTGCCCATGGGAGCCACAAGAAGGCATCA | | |
| 2443 | 17 | 45777983 | 45778168 | 185 | SPRET | 17 | 45778020 | 45778053 | 34 | + | 16.7724 | 7.83E-07 | 0.0815 | TAGCTGTCTTCAGATGCACCAGAAGAGGGCGTCA | 2 | ENCFF093ZRU.ENCFF367UNR |
| 2444 | 17 | 45988387 | 45988595 | 208 | MOLF.NZW.SJL.SPRET | 17 | 45988453 | 45988486 | 34 | + | 16.7398 | 7.99E-07 | 0.0823 | TAGCTGTCTTCGGACACTCCAGAAGAGGGCGTCA | | |
| 2445 | 17 | 47049087 | 47049288 | 201 | 129S1,CAST,MOLF,NOD,NOR,NZW,SJL,SPRET,WSB | 17 | 47049146 | 47049179 | 34 | + | 18.3659 | 2.76E-07 | 0.0462 | TAGCTGTCTTCAGACACTCCAGAAGAGGGCGCCA | 3 | ENCFF093ZRU,ENCFF367UNR |
| 2445 | 17 | 47049087 | 47049288 | 201 | 129S1,CAST,MOLF,NOD,NOR,NZW,SJL,SPRET.WSB | 17 | 47049161 | 47049179 | 19 | + | 17.5574 | 6.32E-07 | 0.108 | ACTCCAGAAGAGGGCGCCA | 3 | ENCFF093ZRU,ENCFF367UNR |
| 2446 | 17 | 47973872 | 47974101 | 229 | CAST.MOLF.SJL.SWR | 17 | 47973957 | 47973990 | 34 | + | 16.4634 | 9.51E-07 | 0.0875 | TAGCTGTCTTCAGACGCACCAGAAGAGGGCATCA | 1 | ENCFF093ZRU |
| 2447 | 17 | 48253606 | 48253772 | 166 | SPRET | 17 | 48253624 | 48253658 | 35 | + | 16.9174 | 5.68E-07 | 0.109 | CTGTCCTGTCCTAGAAGGGCCACCAGAGGATGCCC | | |
| 2448 | 17 | 48460709 | 48461375 | 666 | TH | 17 | 48461322 | 48461355 | 34 | + | 22.0569 | 1.91E-08 | 0.00721 | GTGCAGTGGCCACAGAGGCCAGAAGAGGTTGCGG | | |
| 2449 | 17 | 52217253 | 52217457 | 204 | SPRET | 17 | 52217318 | 52217351 | 34 | + | 17.2439 | 5.79E-07 | 0.0713 | TAGCTGTCCTCAGATACTCCAGAAGAGGGCATCA | | |
| 2450 | 17 | 53464670 | 53464866 | 196 | SPRET | 17 | 53464720 | 53464753 | 34 | + | 17.1463 | 6.17E-07 | 0.0733 | TAGCTGTCCTCAAATACTCCAGAAGAGGGCATCA | | |
| 2451 | 17 | 53909591 | 53909799 | 208 | CAST.MOLF.SPRET | 17 | 53909657 | 53909690 | 34 | + | 16.4634 | 9.51E-07 | 0.0875 | TAGCTGTCTTCAGACGCACCAGAAGAGGGCATCA | | |
| 2452 | 17 | 53915286 | 53915494 | 208 | CAST.MOLF | 17 | 53915352 | 53915385 | 34 | + | 16.4634 | 9.51E-07 | 0.0875 | TAGCTGTCTTCAGACGCACCAGAAGAGGGCATCA | | |
| 2453 | 17 | 56372834 | 56373035 | 201 | MOLF,SPRET | 17 | 56372886 | 56372919 | 34 | + | 18.7886 | 2.07E-07 | 0.0401 | TAGCTGTCTTCAGACATACCAGAAGAGGGCGCCA | 8 | ENCFF055HAP,ENCFF093ZRU,ENCFF367UNR,ENCFF099QNE,ENCFF024JOZ,ENCFF907XEG,ENCFF089GBC.ENCFF052KVO |
| 2453 | 17 | 56372834 | 56373035 | 201 | MOLF,SPRET | 17 | 56372901 | 56372919 | 19 | + | 18.7049 | 2.88E-07 | 0.0728 | ATACCAGAAGAGGGCGCCA | 8 | ENCFF055HAP,ENCFF093ZRU,ENCFF367UNR,ENCFF099QNE,ENCFF024JOZ,ENCFF907XEG,ENCFF089GBC.ENCFF052KVO |
| 2454 | 17 | 57349188 | 57349469 | 281 | 129S1,C57L,C58J,CAST,FVB,LP,MA/My,MOLF.NOD.NOR.SJL.SPRET.SWR | 17 | 57349326 | 57349360 | 35 | + | 16.2477 | 8.31E-07 | 0.135 | TAGCTGTCTTCAGACACTCCCAGAAGAGGGCATCA | | |
| 2455 | 17 | 57607186 | 57607424 | 238 | CAST.MOLF.SPRET | 17 | 57607257 | 57607290 | 34 | + | 18.1789 | 3.13E-07 | 0.0491 | TAGCTGTCTTCAGACACTCCAGAAGAGGGCGCTA | 1 | ENCFF055HAP |
| 2455 | 17 | 57607186 | 57607424 | 238 | CAST.MOLF.SPRET | 17 | 57607272 | 57607290 | 19 | + | 17.5738 | 6.25E-07 | 0.108 | ACTCCAGAAGAGGGCGCTA | 1 | ENCFF055HAP |
| 2456 | 17 | 58982164 | 58983136 | 972 | CAST.MOLF | 17 | 58982651 | 58982685 | 35 | + | 17.7156 | 3.56E-07 | 0.0868 | GTGTACTTGTCTCCAGAGGCCAGAAGATAGCACCA | | |
| 2457 | 17 | 59345864 | 59346117 | 253 | CAST.MOLF.SPRET | 17 | 59345975 | 59346008 | 34 | + | 17.2439 | 5.79E-07 | 0.0713 | TAGCTGTCCTCAGATACTCCAGAAGAGGGCATCA | | |
| 2458 | 17 | 60083136 | 60083560 | 424 | CAST | 17 | 60083292 | 60083325 | 34 | + | 18.0569 | 3.40E-07 | 0.0516 | ATGCAGCTAGAGACATGAGCTCCAGGGGGTACTG | | |
| 2459 | 17 | 61268553 | 61269039 | 486 | CAST.SPRET | 17 | 61268546 | 61268579 | 34 | + | 16.6585 | 8.41E-07 | 0.0841 | GTGCAGATGTACCGAAAGACTCCAGGAGGAGCCC | | |
| 2460 | 17 | 62807399 | 62807594 | 195 | CAST.MOLF.SPRET | 17 | 62807452 | 62807485 | 34 | + | 17.122 | 6.26E-07 | 0.0736 | TAGCTGTCCTCAGATACTCCAGAAGAGGGCAACA | | |
| 2461 | 17 | 66032534 | 66033439 | 905 | CAST,MOLF | 17 | 66033386 | 66033419 | 34 | + | 19.5691 | 1.21E-07 | 0.0289 | CTGTAGTTCTGCCGATGACCACAAGGGGAAGAGC | 26 | ENCFF367UNR,ENCFF093ZRU,ENCFF092NMJ,ENCFF089GBC,ENCFF024JOZ,ENCFF165LXO,ENCFF099QNE,ENCFF210BTO,ENCFF883UPM,ENCFF052KVO,ENCFF821BIA,ENCFF327LYS,ENCFF065ODV,ENCFF033UQX,ENCFF136ZDK,ENCFF055HAP,ENCFF533APC,ENCFF194HVL,ENCFF198SSR,ENCFF907XEG,ENCFF491RJK,ENCFF502WDT,ENCFF281QMV,ENCFF142CNG,ENCFF957SPH,ENCFF589ZBJ |
| 2462 | 17 | 66111793 | 66111998 | 205 | AJ,BALB,BTBR,BUB,C58J,CAST,CE,DBA1J,DBA2J,FVB,KK,MOLF,NOD,NOR,NZB,NZW,PJ,PL,RH2,SEA,SJL,SMJ,SPRET,TH | 17 | 66111855 | 66111888 | 34 | + | 17.1382 | 6.20E-07 | 0.0734 | TAGCTGTCTCCAGACACTCCAGAAGAGGGCATCA | | |
| 2463 | 17 | 66262264 | 66262470 | 206 | CAST.MOLF.SPRET.WSB | 17 | 66262329 | 66262362 | 34 | + | 16.4472 | 9.61E-07 | 0.0875 | TAGCTGTCTTCAGACACACCAGAAGAGGGCGTCA | | |
| 2464 | 17 | 66461994 | 66462051 | 57 | SPRET | 17 | 66462024 | 66462058 | 35 | + | 21.2752 | 3.74E-08 | 0.022 | GTGCAGGCAATTATGATGACCAGTAGAGGGCTCTG | 1 | ENCFF502WDT |
| 2464 | 17 | 66461994 | 66462051 | 57 | SPRET | 17 | 66462040 | 66462058 | 19 | + | 18.6557 | 2.99E-07 | 0.0748 | TGACCAGTAGAGGGCTCTG | 1 | ENCFF502WDT |
| 2465 | 17 | 66809654 | 66809864 | 210 | CAST.MOLF.SPRET.TH.WSB | 17 | 66809719 | 66809752 | 34 | + | 16.5691 | 8.90E-07 | 0.0867 | TAGCTGTCTTCAGACGCACCAGAAGAGGGCATCG | | |
| 2466 | 17 | 71823249 | 71823835 | 586 | CAST.MOLF | 17 | 71823674 | 71823707 | 34 | + | 17.3902 | 5.27E-07 | 0.0672 | CAGCTGTCTTTAGACACCCCAGAAGAGGGCATCA | | |
| 2467 | 17 | 74745188 | 74745662 | 474 | SPRET | 17 | 74745505 | 74745538 | 34 | + | 16.7642 | 7.87E-07 | 0.0816 | CCGCTGCTCCTCTGGCGGCCGCTTGAGGATGCCC | | |
| 2468 | 17 | 74746280 | 74746486 | 206 | SPRET | 17 | 74746344 | 74746377 | 34 | + | 16.4472 | 9.61E-07 | 0.0875 | TAGCTGTCTTCAGACACACCAGAAGAGGGCGTCA | | |
| 2469 | 17 | 74858714 | 74858927 | 213 | 129S1,AKR,BTBR,BUB,C57L,CAST,ILNJ2,KK,MA/My,MOLF,NOD,NOR,NZB,NZO,NZW,SPRET.SWR.TH | 17 | 74858784 | 74858817 | 34 | + | 16.4634 | 9.51E-07 | 0.0875 | TAGCTGTCTTCAGACGCACCAGAAGAGGGCATCA | | |
| 2470 | 17 | 82824212 | 82824420 | 208 | CAST.MA/My.MOLF.NZW.SPRET.WSB | 17 | 82824276 | 82824309 | 34 | + | 18.187 | 3.12E-07 | 0.049 | TAGCTGTCTTCAGTCACTCCAGAAGAGGGCGCTA | | |
| 2470 | 17 | 82824212 | 82824420 | 208 | CAST.MA/My.MOLF.NZW.SPRET.WSB | 17 | 82824291 | 82824309 | 19 | + | 17.5738 | 6.25E-07 | 0.108 | ACTCCAGAAGAGGGCGCTA | | |
| 2471 | 17 | 82843248 | 82843451 | 203 | SPRET | 17 | 82843306 | 82843339 | 34 | + | 18.0732 | 3.36E-07 | 0.0515 | CAGCTGTCTTCAGACACTCCAGAAGAGGGCATCA | | |
| 2472 | 17 | 84750765 | 84750976 | 211 | CAST,DBA1J,DBA2J,ILNJ2,MOLF,SPRET | 17 | 84750836 | 84750869 | 34 | + | 18.3984 | 2.70E-07 | 0.046 | TTGCTGTCTTCAGACACACCAGAAGAGGGCATTG | | |
| 2473 | 17 | 85465272 | 85465503 | 231 | CAST.MOLF | 17 | 85465360 | 85465394 | 35 | + | 21.4404 | 3.35E-08 | 0.0209 | CTGCTGTCTTCAGACAGCACCAGAAGAGGGCATCA | | |
| 2474 | 17 | 88473901 | 88474397 | 496 | SPRET | 17 | 88474107 | 88474141 | 35 | + | 26.4954 | 7.25E-10 | 0.00239 | GTGCAGGTGCCCAAGGTGACCAGAAGAGGGCACTG | | |
| 2474 | 17 | 88473901 | 88474397 | 496 | SPRET | 17 | 88474123 | 88474141 | 19 | + | 21.4918 | 3.20E-08 | 0.0305 | TGACCAGAAGAGGGCACTG | | |
| 2475 | 17 | 88496023 | 88496361 | 338 | SPRET | 17 | 88496282 | 88496316 | 35 | + | 18.8991 | 1.74E-07 | 0.0578 | ATGCAGATGCTGGAGGAGCACAGAAGAGGGCACCA | | |
| 2476 | 17 | 88575204 | 88575907 | 703 | SPRET | 17 | 88575804 | 88575838 | 35 | + | 23.1743 | 9.85E-09 | 0.00958 | GTGCAGGTGCCCTCGGAGACCAGAAGAGGGCGTTG | 3 | ENCFF052KVO.ENCFF093ZRU.ENCFF533APC |
| 2477 | 17 | 88719209 | 88719304 | 95 | CAST | 17 | 88719195 | 88719228 | 34 | + | 20.0407 | 8.62E-08 | 0.0225 | CAGCTGTCTTCAGATGTGCCAGAAGAGGGCATCA | | |
| 2478 | 17 | 3102080 | 3102417 | 337 | SPRET | 17 | 3102091 | 3102124 | 34 | - | 17.561 | 4.71E-07 | 0.0623 | ATGCAGTTAGAGACATGAGCTCCGGGGGGTACTG | | |
| 2479 | 17 | 6383944 | 6384147 | 203 | AJ,AKR,BALB,C3H,C57L,CAST,CBA,DBA1J,DBA2J,ILNJ2,MRL,NZO,PJ,PL,RF,RH2,SEA,SMJ,TH | 17 | 6384069 | 6384102 | 34 | - | 16.4472 | 9.61E-07 | 0.0875 | TAGCTGTCTTCAGACACACCAGAAGAGGGCGTCA | | |
| 2480 | 17 | 7768146 | 7768342 | 196 | SPRET | 17 | 7768270 | 7768303 | 34 | - | 17.6667 | 4.40E-07 | 0.06 | TAGCTGTCTTCAGACACGCCAGAAGAGGGCGTCA | | |
| 2481 | 17 | 12137339 | 12137498 | 159 | MOLF | 17 | 12137415 | 12137449 | 35 | - | 19.7064 | 1.05E-07 | 0.0437 | CCGCCTTCCTGGCTTCTGTCCACTAGATGCCAGTA | 23 | ENCFF093ZRU,ENCFF099QNE,ENCFF052KVO,ENCFF198SSR,ENCFF821BIA,ENCFF201JUA,ENCFF089GBC,ENCFF883UPM,ENCFF092NMJ,ENCFF609MDN,ENCFF575MGI,ENCFF024JOZ,ENCFF907XEG,ENCFF533APC,ENCFF430PPJ,ENCFF367UNR,ENCFF281QMV,ENCFF327LYS,ENCFF055HAP,ENCFF165LXO,ENCFF316FYN.ENCFF502WDT.ENCFF142CNG |
| 2482 | 17 | 12437641 | 12437820 | 179 | AJ,B10D2,C3H,CBA,DBA1J,DBA2J,MOLF,MRL.PL.SMJ | 17 | 12437789 | 12437822 | 34 | - | 16.5691 | 8.90E-07 | 0.0867 | GCGCTCTGTCAACAGGAGCCACTAGGAGGCTGTT | 2 | ENCFF093ZRU,ENCFF092NMJ |
| 2483 | 17 | 14833823 | 14834644 | 821 | ILNJ2.NOD.NOR.NZW.SJL.WSB | 17 | 14833827 | 14833861 | 35 | - | 17.8899 | 3.21E-07 | 0.0817 | ATGAAGTTGTCACTGTTGCCCAGGAGATGCCACCA | | |
| 2484 | 17 | 15798599 | 15798801 | 202 | SPRET | 17 | 15798723 | 15798741 | 19 | - | 17.8852 | 5.08E-07 | 0.0979 | ACACCAGAAGAGGGCACCA | | |
| 2484 | 17 | 15798599 | 15798801 | 202 | SPRET | 17 | 15798723 | 15798756 | 34 | - | 19.0813 | 1.70E-07 | 0.0357 | TAGCTGTCTTCGAACACACCAGAAGAGGGCACCA | | |
| 2485 | 17 | 23418804 | 23419153 | 349 | CAST,SPRET | 17 | 23419007 | 23419041 | 35 | - | 24.367 | 4.03E-09 | 0.0054 | CTGCAATTTCCCGAGCTGAACAGCAGAGGGAGACA | 22 | ENCFF093ZRU,ENCFF367UNR,ENCFF136ZDK,ENCFF099QNE,ENCFF883UPM,ENCFF907XEG,ENCFF821BIA,ENCFF609MDN,ENCFF951GOP,ENCFF327LYS,ENCFF024JOZ,ENCFF089GBC,ENCFF198SSR,ENCFF092NMJ,ENCFF052KVO,ENCFF589ZBJ,ENCFF055HAP,ENCFF165LXO,ENCFF430PPJ,ENCFF957SPH,ENCFF502WDT.ENCFF281QMV |
| 2486 | 17 | 23790827 | 23791034 | 207 | SPRET | 17 | 23790983 | 23791016 | 34 | - | 16.9919 | 6.81E-07 | 0.0767 | TAGCTGTCTTCACATGCACCAGAAGAGGGCATCA | | |
| 2487 | 17 | 24483536 | 24483744 | 208 | CAST | 17 | 24483645 | 24483678 | 34 | - | 16.4472 | 9.61E-07 | 0.0875 | TAGCTGTCTTCAGACACACCAGAAGAGGGCGTCA | | |
| 2488 | 17 | 24524725 | 24524928 | 203 | CAST | 17 | 24524770 | 24524803 | 34 | - | 18.4797 | 2.56E-07 | 0.0444 | TTGCTGTCTTCAGACACACCAGAAGAGGGCATCA | 1 | ENCFF093ZRU |
| 2489 | 17 | 24968595 | 24968809 | 214 | SPRET | 17 | 24968720 | 24968753 | 34 | - | 16.4634 | 9.51E-07 | 0.0875 | TAGCTGTCTTCAGACGCACCAGAAGAGGGCATCA | | |
| 2490 | 17 | 25895146 | 25895343 | 197 | CAST.MOLF.SPRET | 17 | 25895264 | 25895282 | 19 | - | 17.5574 | 6.32E-07 | 0.108 | ACTCCAGAAGAGGGCGCCA | | |
| 2490 | 17 | 25895146 | 25895343 | 197 | CAST.MOLF.SPRET | 17 | 25895264 | 25895297 | 34 | - | 18.3659 | 2.76E-07 | 0.0462 | TAGCTGTCTTCAGACACTCCAGAAGAGGGCGCCA | | |
| 2491 | 17 | 27212481 | 27212706 | 225 | CAST.KK.NZO | 17 | 27212611 | 27212644 | 34 | - | 18.0244 | 3.47E-07 | 0.0522 | TAGCTGTTTTCAGATGCACCAGAAGAGGGCATCA | 1 | ENCFF093ZRU |
| 2492 | 17 | 28438357 | 28438584 | 227 | SPRET | 17 | 28438482 | 28438515 | 34 | - | 17.4797 | 4.97E-07 | 0.0646 | TAGCTGTTTTCAGACACTCCAGAAGAGGGCGTTA | | |
| 2493 | 17 | 28745733 | 28746321 | 588 | SPRET | 17 | 28745804 | 28745822 | 19 | - | 17.0328 | 8.88E-07 | 0.128 | GAACCAGAAGATGGCACCA | 1 | ENCFF093ZRU |
| 2494 | 17 | 28822620 | 28822861 | 241 | MOLF.SPRET | 17 | 28822747 | 28822765 | 19 | - | 17.9672 | 4.81E-07 | 0.0964 | GCACCAGAAGAGGGCACCA | 1 | ENCFF093ZRU |
| 2494 | 17 | 28822620 | 28822861 | 241 | MOLF.SPRET | 17 | 28822747 | 28822780 | 34 | - | 18.8699 | 1.96E-07 | 0.0389 | TAGCTGTCTTCAGACGCACCAGAAGAGGGCACCA | 1 | ENCFF093ZRU |
| 2495 | 17 | 29212554 | 29212761 | 207 | CAST.ILNJ2.NOD.NOR.PJ.SPRET | 17 | 29212667 | 29212700 | 34 | - | 17.6667 | 4.40E-07 | 0.06 | TAGCTGTTTTCAGACACTCCAGAAGAGGGCGTCA | | |
| 2496 | 17 | 29242767 | 29243028 | 261 | CAST,ILNJ2,KK,MOLF,NOD,NOR,PJ,RF,SPRET | 17 | 29242892 | 29242925 | 34 | - | 17.8455 | 3.91E-07 | 0.0561 | TAGCTGTCCTCAGACACTCCAGAAGAGGGCGTCA | | |
| 2497 | 17 | 31682403 | 31682489 | 86 | CAST.KK.MOLF.PL | 17 | 31682405 | 31682438 | 34 | - | 16.9837 | 6.84E-07 | 0.0767 | CATCAGGCCCCTGGGCAGTCACTGGGTGGCACTG | | |
| 2498 | 17 | 32946225 | 32946432 | 207 | AJ,AKR,C3H,CAST,CBA,CE,ILNJ2,MA/My,MOLF,MRL,NOD,NOR,PJ,PL,RF,SJL,SPRET,TH.WSB | 17 | 32946350 | 32946383 | 34 | - | 16.6911 | 8.24E-07 | 0.0833 | TAGCTGTCTTCAGACAGACCAGAAGAGGGCATCA | | |
| 2499 | 17 | 34885403 | 34885606 | 203 | SPRET | 17 | 34885523 | 34885556 | 34 | - | 16.4472 | 9.61E-07 | 0.0875 | TAGCTGTCTTCAGACACACCAGAAGAGGGCGTCA | | |

| | | | | | | | | | | | | | | | | |
|---|---|---|---|---|---|---|---|---|---|---|---|---|---|---|---|---|
| 2500 | 17 | 35131173 | 35131273 | 100 | ILNJ2 | 17 | 35131242 | 35131260 | 19 | - | 20.8033 | 5.79E-08 | 0.0357 | CAGCCACTAGGGGTCGCTG | 41 | ENCFF093ZRU,ENCFF367UNR,ENCFF327LYS,ENCFF024JOZ,ENCFF907XEG,ENCFF589ZBJ,ENCFF805QBU,ENCFF086ZTV,ENCFF136ZDK,ENCFF198SSR,ENCFF165LXO,ENCFF089GBC,ENCFF099QNE,ENCFF821BIA,ENCFF321AGQ,ENCFF430PPJ,ENCFF092NMJ,ENCFF957SPH,ENCFF055HAP,ENCFF502WDT,ENCFF349BUL,ENCFF052KVO,ENCFF065ODV,ENCFF210BTO,ENCFF951GOP,ENCFF194HVL,ENCFF142CNG,ENCFF316FYN,ENCFF281QMV,ENCFF611HDQ,ENCFF883UPM,ENCFF784ASD,ENCFF601JVE,ENCFF491RJK,ENCFF533APC,ENCFF504JDS,ENCFF097CAQ,ENCFF575MGI,ENCFF089EWX,ENCFF609MDN,ENCFF033UQX |
| 2500 | 17 | 35131173 | 35131273 | 100 | ILNJ2 | 17 | 35131242 | 35131275 | 34 | - | 24.0325 | 3.83E-09 | 0.00206 | ATGCAGGACATGGTCCAGCCACTAGGGGTCGCTG | 41 | ENCFF093ZRU,ENCFF367UNR,ENCFF327LYS,ENCFF024JOZ,ENCFF907XEG,ENCFF589ZBJ,ENCFF805QBU,ENCFF086ZTV,ENCFF136ZDK,ENCFF198SSR,ENCFF165LXO,ENCFF089GBC,ENCFF099QNE,ENCFF821BIA,ENCFF321AGQ,ENCFF430PPJ,ENCFF092NMJ,ENCFF957SPH,ENCFF055HAP,ENCFF502WDT,ENCFF349BUL,ENCFF052KVO,ENCFF065ODV,ENCFF210BTO,ENCFF951GOP,ENCFF194HVL,ENCFF142CNG,ENCFF316FYN,ENCFF281QMV,ENCFF611HDQ,ENCFF883UPM,ENCFF784ASD,ENCFF601JVE,ENCFF491RJK,ENCFF533APC,ENCFF504JDS,ENCFF097CAQ,ENCFF575MGI,ENCFF089EWX,ENCFF609MDN,ENCFF033UQX |
| 2501 | 17 | 35306519 | 35306724 | 205 | SPRET | 17 | 35306644 | 35306677 | 34 | - | 16.4146 | 9.81E-07 | 0.0882 | TAGCTGTCTTCAGACACACCAGAGGAGGGCGCCA | 1 | ENCFF093ZRU |
| 2502 | 17 | 35443008 | 35443798 | 790 | AKR,BUB,C3H,C58J,CBA,CE,DBA1J,FVB,MA/Mv.MRL.RF.SWR | 17 | 35443030 | 35443063 | 34 | - | 17.374 | 5.32E-07 | 0.0676 | TAGCTGTTTTCAGACACCCCAGAAGAGGGCGTCA | 1 | ENCFF093ZRU |
| 2503 | 17 | 35493210 | 35493417 | 207 | AKR,C3H,C58J,CBA,CE,MA/My,MRL,RF,WSB | 17 | 35493339 | 35493372 | 34 | - | 17.9919 | 3.55E-07 | 0.0528 | TTGCTGTCTTCAGACACTCCAGAAGAGGGCATCA | 5 | ENCFF093ZRU,ENCFF821BIA,ENCFF367UNR,ENCFF502WDT,ENCFF055HAP |
| 2504 | 17 | 35797370 | 35797566 | 196 | SPRET | 17 | 35797485 | 35797518 | 34 | - | 16.4472 | 9.61E-07 | 0.0875 | TAGCTGTCTTCAGACACACCAGAAGAGGGCGTCA | | |
| 2505 | 17 | 36097603 | 36097969 | 366 | SPRET | 17 | 36097811 | 36097844 | 34 | - | 16.4472 | 9.61E-07 | 0.0875 | TAGCTGTCTTCAGACACACCAGAAGAGGGCGTCA | 1 | ENCFF367UNR |
| 2506 | 17 | 36108459 | 36108666 | 207 | 129S1,AJ,AKR,B10D2,BALB,BTBR,C3H,C57L,C58J,CAST,CBA,CE,DBA1J,DBA2J,LP,MA/My,MOLF,MRL,NZB,NZO,NZW,PJ,PL,RF,RH2,SEA,SJL,SMJ,SPRET,TH,WSB | 17 | 36108582 | 36108615 | 34 | - | 16.4472 | 9.61E-07 | 0.0875 | TAGCTGTCTTCAGACACACCAGAAGAGGGCGTCA | | |
| 2507 | 17 | 36186515 | 36186768 | 253 | MOLF.SPRET | 17 | 36186640 | 36186673 | 34 | - | 17.2276 | 5.85E-07 | 0.0716 | TAGCTGTCTTCAGACACACCAGTAGAGGGCGTCA | 3 | ENCFF367UNR.ENCFF093ZRU |
| 2508 | 17 | 36778072 | 36778274 | 202 | SPRET | 17 | 36778196 | 36778229 | 34 | - | 16.4472 | 9.61E-07 | 0.0875 | TAGCTGTCTTCAGACACACCAGAAGAGGGCGTCA | | |
| 2509 | 17 | 45700085 | 45700287 | 202 | 129S1,AJ,BTBR,BUB,C57L,C58J,CAST,DBA1J,DBA2J,FVB,ILNJ2,KK,LP,MOLF,NOD,NOR,NZB,NZO,NZW,SJL,SMJ,SPRET,SWR,TH.WSB | 17 | 45700211 | 45700244 | 34 | - | 16.4472 | 9.61E-07 | 0.0875 | TAGCTGTCTTCAGACACACCAGAAGAGGGCGTCA | | |
| 2510 | 17 | 46416574 | 46416783 | 209 | 129S1,BTBR,BUB,C57L,CAST,DBA1J,FVB,KK,LP,MA/My,MOLF,NOD,NOR,NZB,NZO,NZW.RF.SJL.SPRET.SWR.WSB | 17 | 46416698 | 46416731 | 34 | - | 16.6911 | 8.24E-07 | 0.0833 | TAGCTGTCTTCAGACAGACCAGAAGAGGGCATCA | | |
| 2511 | 17 | 47186114 | 47187070 | 956 | MOLF | 17 | 47186756 | 47186774 | 19 | - | 21.0984 | 4.51E-08 | 0.0335 | AGGCCAGAAGAGGGCGCCA | 19 | ENCFF210BTO,ENCFF367UNR,ENCFF093ZRU,ENCFF024JOZ,ENCFF198SSR,ENCFF097CAQ,ENCFF327LYS,ENCFF502WDT,ENCFF136ZDK,ENCFF055HAP,ENCFF821BIA,ENCFF589ZBJ,ENCFF142CNG,ENCFF099QNE,ENCFF883UPM,ENCFF907XEG,ENCFF533APC,ENCFF957SPH,ENCFF052KVO |
| 2511 | 17 | 47186114 | 47187070 | 956 | MOLF | 17 | 47186756 | 47186790 | 35 | - | 20.7064 | 5.48E-08 | 0.028 | GTACACCTGTACCTGCAGGCCAGAAGAGGGCGCCA | 18 | ENCFF210BTO,ENCFF367UNR,ENCFF093ZRU,ENCFF024JOZ,ENCFF198SSR,ENCFF097CAQ,ENCFF327LYS,ENCFF502WDT,ENCFF136ZDK,ENCFF055HAP,ENCFF821BIA,ENCFF589ZBJ,ENCFF142CNG,ENCFF099QNE.ENCFF883UPM.ENCFF907XEG.ENCFF533APC.ENCFF052KVO |
| 2512 | 17 | 48407648 | 48407837 | 189 | SPRET | 17 | 48407701 | 48407719 | 19 | - | 16.9344 | 9.46E-07 | 0.132 | GGGCCACCAGGGGGAGGAT | 16 | ENCFF430PPJ,ENCFF367UNR,ENCFF093ZRU,ENCFF907XEG,ENCFF327LYS,ENCFF198SSR,ENCFF609MDN,ENCFF099QNE,ENCFF024JOZ,ENCFF097CAQ,ENCFF194HVL,ENCFF821BIA,ENCFF089GBC,ENCFF165LXO.ENCFF589ZBJ.ENCFF281QMV |
| 2513 | 17 | 48928111 | 48928579 | 468 | MOLF.NZW.SJL | 17 | 48928108 | 48928126 | 19 | - | 21.0984 | 4.51E-08 | 0.0335 | AGGCCAGAAGAGGGCGCCA | 3 | ENCFF093ZRU.ENCFF052KVO.ENCFF033UQX |
| 2513 | 17 | 48928111 | 48928579 | 468 | MOLF.NZW.SJL | 17 | 48928108 | 48928141 | 34 | - | 24.0732 | 3.70E-09 | 0.00201 | GTGCATATGCCCTGGAGGCCAGAAGAGGGCGCCA | 3 | ENCFF093ZRU.ENCFF052KVO.ENCFF033UQX |
| 2514 | 17 | 50249498 | 50250132 | 634 | MOLF,NZW,SWR | 17 | 50249895 | 50249913 | 19 | - | 18.1475 | 4.25E-07 | 0.0913 | AGGCCTGCAGATGGCAGCA | 29 | ENCFF093ZRU,ENCFF024JOZ,ENCFF089GBC,ENCFF198SSR,ENCFF367UNR,ENCFF099QNE,ENCFF065ODV,ENCFF092NMJ,ENCFF821BIA,ENCFF136ZDK,ENCFF033UQX,ENCFF575MGI,ENCFF502WDT,ENCFF281QMV,ENCFF142CNG,ENCFF097CAQ,ENCFF327LYS,ENCFF883UPM,ENCFF907XEG,ENCFF055HAP,ENCFF957SPH,ENCFF165LXO,ENCFF201JUA,ENCFF504JDS,ENCFF589ZBJ,ENCFF533APC,ENCFF611HDQ,ENCFF491RJK.ENCFF194HVL |
| 2514 | 17 | 50249498 | 50250132 | 634 | MOLF,NZW,SWR | 17 | 50249895 | 50249929 | 35 | - | 23.5138 | 7.67E-09 | 0.00814 | GTGCACAGGCATCCCCAGGCCTGCAGATGGCAGCA | 29 | ENCFF093ZRU,ENCFF024JOZ,ENCFF089GBC,ENCFF198SSR,ENCFF367UNR,ENCFF099QNE,ENCFF065ODV,ENCFF092NMJ,ENCFF821BIA,ENCFF136ZDK,ENCFF033UQX,ENCFF575MGI,ENCFF502WDT,ENCFF281QMV,ENCFF142CNG,ENCFF097CAQ,ENCFF327LYS,ENCFF883UPM,ENCFF907XEG,ENCFF055HAP,ENCFF957SPH,ENCFF165LXO,ENCFF201JUA,ENCFF504JDS,ENCFF589ZBJ,ENCFF533APC,ENCFF611HDQ,ENCFF491RJK.ENCFF194HVL |
| 2515 | 17 | 63745136 | 63745368 | 232 | BTBR.C58J.CAST.LP.MOLF.SPRET | 17 | 63745261 | 63745294 | 34 | - | 16.4634 | 9.51E-07 | 0.0875 | TAGCTGTCTTCAGACGCACCAGAAGAGGGCATCA | | |
| 2516 | 17 | 66069058 | 66069412 | 354 | CAST.MOLF | 17 | 66069230 | 66069264 | 35 | - | 16.9908 | 5.44E-07 | 0.107 | CTGCTGAGCCATCTCGCCAGCCCCAGAAGGCAGCT | | |
| 2517 | 17 | 66085933 | 66086145 | 212 | SPRET | 17 | 66086059 | 66086092 | 34 | - | 18.5285 | 2.48E-07 | 0.0444 | CAGCTGTCTTCAGACACTCCAGAAGAGGGCGTCA | | |
| 2518 | 17 | 73117576 | 73117825 | 249 | 129S1,BALB,BTBR,C3H,C57L,CAST,CBA,DBA1J,DBA2J,FVB,MOLF,MRL,NZW,PJ,RF,SEA.SJL.SMJ.SPRET.TH | 17 | 73117701 | 73117734 | 34 | - | 16.4634 | 9.51E-07 | 0.0875 | TAGCTGTCTTCAGACGCACCAGAAGAGGGCATCA | | |
| 2519 | 17 | 73422599 | 73422988 | 389 | 129S1,AJ,AKR,BTBR,C57L,CAST,MOLF,SPRET.TH.WSB | 17 | 73422888 | 73422921 | 34 | - | 17.6992 | 4.30E-07 | 0.0594 | TAGCTGTCCTCAGATACTCCAGAAGAGGGCGTCA | | |
| 2520 | 17 | 74447812 | 74448031 | 219 | SPRET | 17 | 74447926 | 74447944 | 19 | - | 17.5574 | 6.32E-07 | 0.108 | ACTCCAGAAGAGGGCGCCA | 1 | ENCFF367UNR |
| 2520 | 17 | 74447812 | 74448031 | 219 | SPRET | 17 | 74447926 | 74447959 | 34 | - | 18.3659 | 2.76E-07 | 0.0462 | TAGCTGTCTTCAGACACTCCAGAAGAGGGCGCCA | 1 | ENCFF367UNR |
| 2521 | 17 | 74770909 | 74771135 | 226 | SPRET | 17 | 74771018 | 74771051 | 34 | - | 16.7724 | 7.83E-07 | 0.0815 | TAGCTGTCTTCAGATGCACCAGAAGAGGGCGTCA | 1 | ENCFF093ZRU |
| 2522 | 17 | 74818039 | 74818243 | 204 | 129S1,BTBR,BUB,C57L,CAST,ILNJ2,KK,MOLF.NOD.NOR.NZW.SWR | 17 | 74818167 | 74818200 | 34 | - | 16.4472 | 9.61E-07 | 0.0875 | TAGCTGTCTTCAGACACACCAGAAGAGGGCGTCA | | |
| 2523 | 17 | 75011349 | 75011557 | 208 | 129S1,AKR,BTBR,BUB,C57L,CAST,ILNJ2,KK,MA/My,MOLF,NOD,NOR,NZB,NZO,NZW,SPRET.SWR.TH | 17 | 75011477 | 75011510 | 34 | - | 19.9593 | 9.14E-08 | 0.0234 | CAGCTGTCCTCAGACACTCCAGAAGAGGGCATCA | | |
| 2524 | 17 | 80498389 | 80498591 | 202 | SPRET | 17 | 80498503 | 80498536 | 34 | - | 16.8699 | 7.36E-07 | 0.0792 | TAGCTGTCTTCAGACACCCCAGAAGAGGGCAGCA | 3 | ENCFF367UNR.ENCFF099QNE.ENCFF093ZRU |
| 2525 | 17 | 80920252 | 80920447 | 195 | CAST.CE.SPRET.TH | 17 | 80920381 | 80920414 | 34 | - | 16.4472 | 9.61E-07 | 0.0875 | TAGCTGTCTTCAGACACACCAGAAGAGGGCGTCA | | |
| 2526 | 17 | 85174859 | 85175031 | 172 | FVB.NOD.NOR.SJL.SPRET.SWR | 17 | 85174954 | 85174972 | 19 | - | 17.5574 | 6.32E-07 | 0.108 | ACTCCAGAAGAGGGCGCCA | 4 | ENCFF093ZRU.ENCFF099QNE.ENCFF367UNR.ENCFF136ZDK |
| 2526 | 17 | 85174859 | 85175031 | 172 | FVB.NOD.NOR.SJL.SPRET.SWR | 17 | 85174954 | 85174987 | 34 | - | 17.0813 | 6.43E-07 | 0.0744 | TAGCTATCTTCAGACACTCCAGAAGAGGGCGCCA | 4 | ENCFF093ZRU.ENCFF099QNE.ENCFF367UNR.ENCFF136ZDK |
| 2527 | 17 | 85628359 | 85628439 | 80 | CAST.MOLF | 17 | 85628393 | 85628427 | 35 | - | 16.7523 | 6.24E-07 | 0.115 | GTGCAGTTGCCCAATGAGGACAGAAGAGGACCTCA | | |
| 2528 | 17 | 85746477 | 85746684 | 207 | 129S1,AKR,CAST,MOLF,MRL,NZB,NZO,PJ,PL.RF.SPRET | 17 | 85746601 | 85746634 | 34 | - | 16.4634 | 9.51E-07 | 0.0875 | TAGCTGTCTTCAGACGCACCAGAAGAGGGCATCA | | |
| 2529 | 17 | 86526229 | 86526750 | 521 | AJ,BALB,C3H,C57L,CBA,CE,LP,PJ,RH2,SEA | 17 | 86526260 | 86526293 | 34 | - | 16.5285 | 9.13E-07 | 0.0875 | ATGTAAAGCCTGCTGTGGCCTGCAGAGGTCTTTC | | |
| 2530 | 17 | 87322908 | 87323605 | 697 | CAST | 17 | 87323465 | 87323483 | 19 | - | 17.3443 | 7.26E-07 | 0.116 | TTGCCAGAAGAGGGCATCA | | |
| 2530 | 17 | 87322908 | 87323605 | 697 | CAST | 17 | 87323465 | 87323499 | 35 | - | 19.6972 | 1.06E-07 | 0.0438 | CTACACATACACCTGCTTGCCAGAAGAGGGCATCA | | |
| 2531 | 17 | 87731573 | 87731766 | 193 | SPRET | 17 | 87731696 | 87731714 | 19 | - | 17.8852 | 5.08E-07 | 0.0979 | ACACCAGAAGAGGGCACCA | 2 | ENCFF093ZRU.ENCFF099QNE |
| 2531 | 17 | 87731573 | 87731766 | 193 | SPRET | 17 | 87731696 | 87731729 | 34 | - | 18.3984 | 2.70E-07 | 0.046 | TAGCTGTCTTCAGACACACCAGAAGAGGGCACCA | 2 | ENCFF093ZRU.ENCFF099QNE |
| 2532 | 17 | 90033095 | 90033650 | 555 | 129S1,AJ,AKR,BALB,BTBR,BUB,C3H,C57L,CAST,CBA,CE,DBA1J,DBA2J,FVB,ILNJ2,KK,LP,MA/My,MRL,NOD,NOR,NZB,NZO,NZW,PJ,PL,RF,RH2,SEA,SJL,SMJ,SWR,TH,WSB | 17 | 90033166 | 90033199 | 34 | - | 27.4878 | 1.59E-10 | 0.000243 | ATGCAGTGCCCTGAGAGGCCAGAAGAGGGCATTA | 5 | ENCFF367UNR,ENCFF093ZRU,ENCFF099QNE,ENCFF024JOZ,ENCFF907XEG |
| 2533 | 18 | 6296293 | 6296496 | 203 | MOLF.SPRET | 18 | 6296354 | 6296387 | 34 | + | 18.3902 | 2.72E-07 | 0.0797 | CAGCTGTCGTCAGATGCACCAGAAGAGGGCATCA | | |
| 2534 | 18 | 21837300 | 21837507 | 207 | CAST.MOLF.SPRET | 18 | 21837349 | 21837382 | 34 | + | 17.878 | 3.83E-07 | 0.091 | TAGCTGTCCTCAGACACACCAGAAGAGGGCATCA | | |
| 2535 | 18 | 33036769 | 33037332 | 563 | BALB,C3H,C58J,CE,MRL,PL,RF,RH2,SEA,TH | 18 | 33037126 | 33037160 | 35 | + | 24.2569 | 4.38E-09 | 0.0113 | CTGCAGTATTCTACCCTGACCACCAGAAGGCTCAG | 13 | ENCFF367UNR,ENCFF430PPJ,ENCFF093ZRU,ENCFF198SSR,ENCFF165LXO,ENCFF099QNE,ENCFF316FYN,ENCFF201JUA,ENCFF327LYS,ENCFF092NMJ,ENCFF907XEG,ENCFF089GBC,ENCFF024JOZ |
| 2536 | 18 | 34807037 | 34807242 | 205 | CAST.MOLF.SPRET | 18 | 34807100 | 34807133 | 34 | + | 17.7805 | 4.08E-07 | 0.0934 | CTGTAGTCTGTAGACACTCCAGAAGAGGGCATCA | | |
| 2537 | 18 | 34834214 | 34834950 | 736 | CAST | 18 | 34834428 | 34834461 | 34 | + | 19.3902 | 1.37E-07 | 0.0538 | TTGCACTTTCCAGGGTTTGCAGAGGGTGGCACAA | | |
| 2538 | 18 | 35092637 | 35093403 | 766 | MOLF | 18 | 35093223 | 35093257 | 35 | + | 17.9358 | 3.13E-07 | 0.133 | CTGCCTGTTCAGATCCAATCCAGGAGGTGGCCCCA | 1 | ENCFF093ZRU |
| 2539 | 18 | 35182798 | 35183042 | 244 | AJ,BALB,C58J,CAST,CBA,DBA2J,ILNJ2,MOLF.RH2.SEA.SPRET.TH.WSB | 18 | 35182900 | 35182933 | 34 | + | 16.4472 | 9.61E-07 | 0.128 | TAGCTGTCTTCAGACACACCAGAAGAGGGCGTCA | | |
| 2540 | 18 | 35300767 | 35300978 | 211 | SPRET | 18 | 35300837 | 35300870 | 34 | + | 16.4472 | 9.61E-07 | 0.128 | TAGCTGTCTTCAGACACACCAGAAGAGGGCGTCA | | |
| 2541 | 18 | 35728619 | 35728812 | 193 | SPRET | 18 | 35728681 | 35728714 | 34 | + | 16.5691 | 8.90E-07 | 0.127 | TAGCTGTCTTCAGACGCACCAGAAGAGGGCATCG | 1 | ENCFF093ZRU |
| 2542 | 18 | 35897294 | 35897504 | 210 | AJ,BALB,C58J,CAST,CBA,DBA2J,ILNJ2,MOLF.MRL.RH2.SEA.SMJ.SPRET.WSB | 18 | 35897349 | 35897382 | 34 | + | 18.3659 | 2.76E-07 | 0.0797 | TAGCTGTCTTCAGACACTCCAGAAGAGGGCGCCA | | |
| 2542 | 18 | 35897294 | 35897504 | 210 | AJ,BALB,C58J,CAST,CBA,DBA2J,ILNJ2,MOLF.MRL.RH2.SEA.SMJ.SPRET.WSB | 18 | 35897364 | 35897382 | 19 | + | 17.5574 | 6.32E-07 | 0.164 | ACTCCAGAAGAGGGCGCCA | | |
| 2543 | 18 | 36223453 | 36223667 | 214 | CAST.MOLF.SPRET | 18 | 36223525 | 36223558 | 34 | + | 16.4634 | 9.51E-07 | 0.128 | TAGCTGTCTTCAGACGCACCAGAAGAGGGCATCA | | |
| 2544 | 18 | 36850402 | 36850604 | 202 | 129S1,AJ,AKR,BALB,BUB,C3H,C57L,C58J,CAST,CBA,CE,DBA1J,DBA2J,FVB,ILNJ2,KK,MA/My,MOLF,NOD,NOR,NZB,NZO,NZW,PL,RF,RH2,SEA,SJL,SMJ,SPRET,SWR,TH,WSB | 18 | 36850463 | 36850496 | 34 | + | 17.9106 | 3.74E-07 | 0.0901 | TAGCTGTCTTCAGACACTCCAGAAGAGGGCACCA | | |
| 2545 | 18 | 36918166 | 36918368 | 202 | CAST.MOLF.SPRET | 18 | 36918226 | 36918259 | 34 | + | 17.2439 | 5.79E-07 | 0.107 | TAGCTGTCCTCAGATACTCCAGAAGAGGGCATCA | | |
| 2546 | 18 | 42651937 | 42652131 | 194 | CAST.MOLF.SPRET | 18 | 42651984 | 42652017 | 34 | + | 16.9675 | 6.91E-07 | 0.115 | TAGCTGTTTTCAGACACACCAGGAGAGGGCATCA | 1 | ENCFF093ZRU |
| 2547 | 18 | 47552389 | 47552588 | 199 | SPRET | 18 | 47552446 | 47552479 | 34 | + | 16.4472 | 9.61E-07 | 0.128 | TAGCTGTCTTCAGACACACCAGAAGAGGGCGTCA | | |
| 2548 | 18 | 53897352 | 53897558 | 206 | CAST.KK.MOLF.SPRET | 18 | 53897402 | 53897435 | 34 | + | 16.4634 | 9.51E-07 | 0.128 | TAGCTGTCTTCAGACGCACCAGAAGAGGGCATCA | | |
| 2549 | 18 | 56715366 | 56715574 | 208 | CAST.MOLF.SPRET | 18 | 56715432 | 56715465 | 34 | + | 17.2439 | 5.79E-07 | 0.107 | TAGCTGTCCTCAGATACTCCAGAAGAGGGCATCA | 1 | ENCFF093ZRU |
| 2550 | 18 | 56876405 | 56876614 | 209 | SPRET | 18 | 56876473 | 56876506 | 34 | + | 16.8537 | 7.43E-07 | 0.117 | TAGCTGTTTTCAGACACTCCAGAAGAGGGAGGCA | | |
| 2551 | 18 | 60069296 | 60069491 | 195 | SPRET | 18 | 60069349 | 60069382 | 34 | + | 18.2358 | 3.02E-07 | 0.0822 | TAGCTGTCTTAAGACACTCCAGAAGAGGGCGCCA | | |
| 2551 | 18 | 60069296 | 60069491 | 195 | SPRET | 18 | 60069364 | 60069382 | 19 | + | 17.5574 | 6.32E-07 | 0.164 | ACTCCAGAAGAGGGCGCCA | | |
| 2552 | 18 | 60927156 | 60927365 | 209 | AJ,BALB,CAST,CBA,FVB,ILNJ2,NZB,RH2,SEA.SJL.SMJ | 18 | 60927225 | 60927258 | 34 | + | 17.4797 | 4.97E-07 | 0.101 | TAGCTGTCTCCAGATACACCAGAAGAGGGCATCA | | |
| 2553 | 18 | 62105918 | 62106299 | 381 | AJ,BALB,CBA,CE,FVB,ILNJ2,NZB,NZO,PL,RH2.SEA.SJL.SWR.TH | 18 | 62106025 | 62106058 | 34 | + | 18.3821 | 2.73E-07 | 0.0797 | TAGCAGTCTTCAGACACACCAGAAGAGGGCGTCA | | |
| 2554 | 18 | 65026986 | 65027207 | 221 | AJ,AKR,BALB,BTBR,BUB,C3H,C57L,CAST,CBA,CE,FVB,ILNJ2,KK,MOLF,MRL,NOD,NOR,NZB,NZO,PL,RF,RH2,SJL,SMJ,SPRET,SWR.WSB | 18 | 65027060 | 65027093 | 34 | + | 16.4472 | 9.61E-07 | 0.128 | TAGCTGTCTTCAGACACACCAGAAGAGGGCGTCA | | |
| 2555 | 18 | 65621248 | 65621451 | 203 | 129S1,AJ,AKR,BALB,BTBR,BUB,C3H,C57L,CAST,CBA,CE,FVB,ILNJ2,KK,MA/My,MOLF,MRL,NOD,NOR,NZB,NZO,NZW,RF,RH2,SJL,SMJ.SPRET.SWR.TH.WSB | 18 | 65621309 | 65621342 | 34 | + | 18.1789 | 3.13E-07 | 0.0837 | TAGCTGTCTTCAGACACTCCAGAAGAGGGCGCTA | | |
| 2555 | 18 | 65621248 | 65621451 | 203 | 129S1,AJ,AKR,BALB,BTBR,BUB,C3H,C57L,CAST,CBA,CE,FVB,ILNJ2,KK,MA/My,MOLF,MRL,NOD,NOR,NZB,NZO,NZW,RF,RH2,SJL,SMJ.SPRET.SWR.TH.WSB | 18 | 65621324 | 65621342 | 19 | + | 17.5738 | 6.25E-07 | 0.164 | ACTCCAGAAGAGGGCGCTA | | |
| 2556 | 18 | 65926153 | 65927144 | 991 | 129S1,AJ,BALB,BTBR,BUB,C3H,C57L,CBA,CE.DBA1J.DBA2J.ILNJ2.PJ.RH2.SEA | 18 | 65926121 | 65926155 | 35 | + | 16.1376 | 8.84E-07 | 0.212 | TTGCTCTCTTTCAGACACACCAGAAGAGGGCATCG | | |
| 2557 | 18 | 66564546 | 66564755 | 209 | 129S1,AKR,BTBR,BUB,C3H,C57L,CAST,CBA,CE,DBA1J,DBA2J,FVB,KK,MA/My,MOLF,MRL,NOD,NOR,PJ,RF,SEA,SJL,SPRET,SWR.TH | 18 | 66564612 | 66564645 | 34 | + | 17.2439 | 5.79E-07 | 0.107 | TAGCTGTCCTCAGATACTCCAGAAGAGGGCATCA | 1 | ENCFF883UPM |
| 2558 | 18 | 67486370 | 67486479 | 109 | CAST.MOLF.WSB | 18 | 67486446 | 67486479 | 34 | + | 21.1789 | 3.74E-08 | 0.0224 | ATGTAGTACCTAAAGATGTCAGAAGAGGGCACCA | 1 | ENCFF052KVO |
| 2559 | 18 | 67935251 | 67935457 | 206 | 129S1,AJ,AKR,BALB,BTBR,BUB,C3H,C57L,CAST,CBA,CE,DBA1J,DBA2J,ILNJ2,KK,MA/My,MOLF,MRL,PJ,PL,RF,RH2,SEA,SMJ,SPRET.WSB | 18 | 67935316 | 67935349 | 34 | + | 16.4472 | 9.61E-07 | 0.128 | TAGCTGTCTTCAGACACACCAGAAGAGGGCGTCA | | |
| 2560 | 18 | 67958125 | 67958340 | 215 | 129S1,AJ,AKR,BALB,BTBR,BUB,C3H,C57L,CAST,CBA,CE,DBA1J,DBA2J,ILNJ2,KK,MA/My,MOLF,MRL,PJ,PL,RF,RH2,SEA,SMJ,SPRET.WSB | 18 | 67958199 | 67958232 | 34 | + | 17.2439 | 5.79E-07 | 0.107 | TAGCTGTCCTCAGATACTCCAGAAGAGGGCATCA | | |
| 2561 | 18 | 70708751 | 70708949 | 198 | CAST.MOLF.RF.SPRET | 18 | 70708807 | 70708840 | 34 | + | 16.4472 | 9.61E-07 | 0.128 | TAGCTGTCTTCAGACACACCAGAAGAGGGCGTCA | | |
| 2562 | 18 | 74835279 | 74835474 | 195 | SPRET | 18 | 74835331 | 74835364 | 34 | + | 16.7724 | 7.83E-07 | 0.12 | TAGCTGTCTTCAGATGCACCAGAAGAGGGCGTCA | | |
| 2563 | 18 | 75591263 | 75591388 | 125 | AKR.CAST.ILNJ2.MOLF.RF | 18 | 75591232 | 75591265 | 34 | + | 16.4146 | 9.81E-07 | 0.129 | CTGGTGTACACCCAGCCTTCAGGAGGTGGAGGCA | | |
| 2564 | 18 | 76008576 | 76008791 | 215 | 129S1,AJ,AKR,BALB,BTBR,BUB,C57L,CAST,FVB,ILNJ2,KK,MOLF,MRL,NZW,RF,RH2,SPRET | 18 | 76008645 | 76008678 | 34 | + | 16.4472 | 9.61E-07 | 0.128 | TAGCTGTCTTCAGACACACCAGAAGAGGGCGTCA | | |
| 2565 | 18 | 76339281 | 76339499 | 218 | 129S1,AJ,AKR,BALB,BTBR,BUB,C3H,C57L,CAST,CBA,DBA1J,DBA2J,FVB,ILNJ2,KK,MOLF,MRL,NZO,NZW,PJ,PL,RF,RH2,SEA,SMJ.SPRET.TH.WSB | 18 | 76339356 | 76339389 | 34 | + | 18.1545 | 3.18E-07 | 0.0841 | TAGCTGTTTTCAGACACACCAGAAGAGGGCGTCA | | |
| 2566 | 18 | 76453914 | 76454118 | 204 | AJ,AKR,BALB,BUB,C3H,CAST,CBA,DBA1J,DBA2J,FVB,ILNJ2,KK,MOLF,MRL,NZO,NZW,PJ,PL,RF,RH2,SEA,SMJ,SPRET,TH,WSB | 18 | 76453970 | 76454003 | 34 | + | 16.4472 | 9.61E-07 | 0.128 | TAGCTGTCTTCAGACACACCAGAAGAGGGCGTCA | | |
| 2567 | 18 | 76989936 | 76990150 | 214 | SPRET | 18 | 76989996 | 76990029 | 34 | + | 16.4634 | 9.51E-07 | 0.128 | TAGCTGTCTTCAGACGCACCAGAAGAGGGCATCA | 4 | ENCFF821BIA.ENCFF089GBC.ENCFF092NMJ.ENCFF136ZDK |
| 2568 | 18 | 77765409 | 77765629 | 220 | SPRET | 18 | 77765491 | 77765524 | 34 | + | 17.3902 | 5.27E-07 | 0.104 | TAGCTGTCCTCAGACACTCCAGAAGAGGGCATCA | | |
| 2569 | 18 | 77892888 | 77893086 | 198 | SPRET | 18 | 77893078 | 77893111 | 34 | + | 19.813 | 1.01E-07 | 0.0439 | CAGCTGTCCTCAGATACTCCAGAAGAGGGCATCA | | |
| 2570 | 18 | 80026009 | 80026435 | 426 | NZO | 18 | 80026045 | 80026063 | 19 | + | 18.7705 | 2.75E-07 | 0.119 | AAGCCAGAAGAGGGCACCA | 6 | ENCFF093ZRU.ENCFF092NMJ.ENCFF883UPM.ENCFF089GBC.ENCFF055HAP.ENCFF502WDT |
| 2571 | 18 | 90596172 | 90596381 | 209 | CAST.KK.MOLF.NZB.NZO.RF.TH.WSB | 18 | 90596240 | 90596273 | 34 | + | 17.8455 | 3.91E-07 | 0.0917 | TAGCTGTCCTCAGACACTCCAGAAGAGGGCGTCA | | |
| 2572 | 18 | 4255040 | 4255249 | 209 | CAST.SPRET | 18 | 4255064 | 4255098 | 35 | - | 16.6789 | 6.51E-07 | 0.185 | CTGCACAGTGGGGGGCTGCACAGTGGGGGGCTGCA | | |
| 2572 | 18 | 4255040 | 4255249 | 209 | CAST.SPRET | 18 | 4255108 | 4255142 | 35 | - | 16.6789 | 6.51E-07 | 0.185 | CTGCACAGTGGGGGGCTGCACAGTGGGGGGCTGCA | | |
| 2572 | 18 | 4255040 | 4255249 | 209 | CAST.SPRET | 18 | 4255123 | 4255157 | 35 | - | 16.6789 | 6.51E-07 | 0.185 | CTGCACAGTGGGGGGCTGCACAGTGGGGGGCTGCA | | |
| 2572 | 18 | 4255040 | 4255249 | 209 | CAST.SPRET | 18 | 4255138 | 4255172 | 35 | - | 16.6789 | 6.51E-07 | 0.185 | CTGCACAGTGGGGGGCTGCACAGTGGGGGGCTGCA | | |
| 2573 | 18 | 9473359 | 9473565 | 206 | 129S1,AJ,AKR,BALB,BTBR,C3H,C57L,CAST.CBA.ILNJ2.MOLF.RF.RH2.SPRET | 18 | 9473484 | 9473502 | 19 | - | 17.5574 | 6.32E-07 | 0.164 | ACTCCAGAAGAGGGCGCCA | | |
| 2573 | 18 | 9473359 | 9473565 | 206 | 129S1,AJ,AKR,BALB,BTBR,C3H,C57L,CAST.CBA.ILNJ2.MOLF.RF.RH2.SPRET | 18 | 9473484 | 9473517 | 34 | - | 18.374 | 2.75E-07 | 0.0797 | TAGCTGTCTTCAGTCACTCCAGAAGAGGGCGCCA | | |
| 2574 | 18 | 10204304 | 10204514 | 210 | SPRET | 18 | 10204414 | 10204432 | 19 | - | 17.7213 | 5.67E-07 | 0.159 | AGACCAGCAGAGGGCATCA | | |
| 2575 | 18 | 11594437 | 11594818 | 381 | SPRET | 18 | 11594723 | 11594756 | 34 | - | 22.0813 | 1.87E-08 | 0.014 | ATGTAGTCCCCACAGAGGCCAGAAGGAGGTGCCA | 2 | ENCFF093ZRU.ENCFF367UNR |
| 2576 | 18 | 12249944 | 12250154 | 210 | CAST | 18 | 12250081 | 12250114 | 34 | - | 17.3902 | 5.27E-07 | 0.104 | TAGCTGTCCTCAGACACTCCAGAAGAGGGCATCA | | |
| 2577 | 18 | 17708095 | 17708437 | 342 | CAST.MOLF.SPRET | 18 | 17708395 | 17708428 | 34 | - | 16.6098 | 8.68E-07 | 0.125 | CCGCTCCCACATTCGCCGTCACAAGATGGCGCTG | | |
| 2578 | 18 | 21173119 | 21173316 | 197 | SPRET | 18 | 21173242 | 21173275 | 34 | - | 16.4472 | 9.61E-07 | 0.128 | TAGCTGTCTTCAGACACACCAGAAGAGGGCGTCA | | |
| 2579 | 18 | 24327513 | 24327696 | 183 | SPRET | 18 | 24327634 | 24327652 | 19 | - | 17.0328 | 8.88E-07 | 0.193 | TCACCAGAAGAGGGCGTCA | 2 | ENCFF093ZRU.ENCFF367UNR |
| 2579 | 18 | 24327513 | 24327696 | 183 | SPRET | 18 | 24327634 | 24327667 | 34 | - | 16.8049 | 7.67E-07 | 0.12 | TAGCTGATTTCAGACTCACCAGAAGAGGGCGTCA | 2 | ENCFF093ZRU.ENCFF367UNR |
| 2580 | 18 | 24578839 | 24579056 | 217 | CAST.MOLF.SPRET.WSB | 18 | 24578965 | 24578998 | 34 | - | 17.2439 | 5.79E-07 | 0.107 | TAGCTGTCCTCAGATACTCCAGAAGAGGGCATCA | | |
| 2581 | 18 | 29420135 | 29420334 | 199 | CAST.MOLF.SPRET | 18 | 29420257 | 29420290 | 34 | - | 18.3333 | 2.82E-07 | 0.0799 | TAGCTGTCCTCAGACACACCAGAAGAGGGCGTCA | | |
| 2582 | 18 | 29585440 | 29586106 | 666 | SPRET | 18 | 29585880 | 29585913 | 34 | - | 16.439 | 9.66E-07 | 0.128 | ATGCAGCTAGAGACAAGAGCTCCAGGGGGTACTG | | |
| 2583 | 18 | 30500897 | 30501108 | 211 | 129S1,C57L,CAST,ILNJ2,MOLF,SMJ,SPRET | 18 | 30501023 | 30501056 | 34 | - | 17.6667 | 4.40E-07 | 0.0954 | TAGCTGTCTTCAGACACGCCAGAAGAGGGCGTCA | | |
| 2584 | 18 | 34030046 | 34030268 | 222 | SPRET | 18 | 34030170 | 34030203 | 34 | - | 17.813 | 3.99E-07 | 0.0927 | GAGCTGTCGTCAGACCCTCCAGAAGAGGGCATCA | | |
| 2585 | 18 | 34795414 | 34795661 | 247 | SPRET | 18 | 34795576 | 34795609 | 34 | - | 16.4472 | 9.61E-07 | 0.128 | TAGCTGTCTTCAGACACACCAGAAGAGGGCGTCA | | |
| 2586 | 18 | 36845488 | 36845757 | 269 | 129S1,AJ,AKR,BALB,BUB,C3H,C57L,C58J,CAST,CBA,CE,DBA1J,DBA2J,FVB,ILNJ2,KK,MA/My,MOLF,NOD,NOR,NZB,NZO,NZW,PL,RF,RH2,SEA,SJL,SMJ,SPRET,SWR,TH,WSB | 18 | 36845612 | 36845645 | 34 | - | 16.4634 | 9.51E-07 | 0.128 | TAGCTGTCTTCAGACGCACCAGAAGAGGGCATCA | | |
| 2587 | 18 | 38444299 | 38444500 | 201 | SPRET | 18 | 38444424 | 38444442 | 19 | - | 17.5574 | 6.32E-07 | 0.164 | ACTCCAGAAGAGGGCGCCA | 6 | ENCFF201JUA.ENCFF367UNR.ENCFF093ZRU.ENCFF099QNE.ENCFF092NMJ.ENCFF198SSR |
| 2587 | 18 | 38444299 | 38444500 | 201 | SPRET | 18 | 38444424 | 38444457 | 34 | - | 18.3659 | 2.76E-07 | 0.0797 | TAGCTGTCTTCAGACACTCCAGAAGAGGGCGCCA | 7 | ENCFF201JUA,ENCFF367UNR,ENCFF093ZRU,ENCFF099QNE,ENCFF092NMJ,ENCFF198SSR,ENCFF883UPM |
| 2588 | 18 | 42384759 | 42384992 | 233 | SPRET | 18 | 42384871 | 42384904 | 34 | - | 16.9919 | 6.81E-07 | 0.115 | TAGCTGTCTCCAGATACTCCAGAAGAGGGCATCA | | |
| 2589 | 18 | 46650557 | 46650769 | 212 | CAST.KK.MOLF.SPRET | 18 | 46650684 | 46650717 | 34 | - | 16.4472 | 9.61E-07 | 0.128 | TAGCTGTCTTCAGACACACCAGAAGAGGGCGTCA | | |
| 2590 | 18 | 47562456 | 47562618 | 162 | CAST | 18 | 47562569 | 47562602 | 34 | - | 17.2439 | 5.79E-07 | 0.107 | TAGCTGTCCTCAGATACTCCAGAAGAGGGCATCA | | |
| 2591 | 18 | 56813084 | 56813672 | 588 | CE.MOLF | 18 | 56813466 | 56813499 | 34 | - | 17.065 | 6.50E-07 | 0.114 | TTGCTCTCTTCAGACACACCAGAAGAGGGCATCG | | |
| 2592 | 18 | 58466585 | 58466997 | 412 | SPRET | 18 | 58466610 | 58466643 | 34 | - | 16.439 | 9.66E-07 | 0.128 | ATGCAGCTAGAGACAAGAGCTCCAGGGGGTACTG | | |
| 2593 | 18 | 60795534 | 60795742 | 208 | AJ,BALB,CAST,CBA,FVB,ILNJ2,KK,MOLF,NOD,NOR,NZB,RH2,SEA,SJL,SMJ,SPRET,WSB | 18 | 60795660 | 60795693 | 34 | - | 16.4634 | 9.51E-07 | 0.128 | TAGCTGTCTTCAGACGCACCAGAAGAGGGCATCA | | |
| 2594 | 18 | 64461395 | 64461601 | 206 | MOLF.SPRET | 18 | 64461523 | 64461556 | 34 | - | 16.6911 | 8.24E-07 | 0.122 | TAGCTGTCTTCAGACAGACCAGAAGAGGGCATCA | | |
| 2595 | 18 | 65616746 | 65616951 | 205 | MOLF | 18 | 65616870 | 65616903 | 34 | - | 16.4634 | 9.51E-07 | 0.128 | TAGCTGTCTTCAGACGCACCAGAAGAGGGCATCA | | |
| 2596 | 18 | 66055045 | 66055139 | 94 | SPRET | 18 | 66055085 | 66055103 | 19 | - | 17.9508 | 4.86E-07 | 0.15 | TGACCACCAGAGGGAAACC | 37 | ENCFF201JUA,ENCFF093ZRU,ENCFF327LYS,ENCFF504JDS,ENCFF089GBC,ENCFF367UNR,ENCFF024JOZ,ENCFF099QNE,ENCFF907XEG,ENCFF092NMJ,ENCFF165LXO,ENCFF430PPJ,ENCFF198SSR,ENCFF589ZBJ,ENCFF609MDN,ENCFF055HAP,ENCFF821BIA,ENCFF316FYN,ENCFF136ZDK,ENCFF502WDT,ENCFF951GOP,ENCFF533APC,ENCFF883UPM,ENCFF097CAQ,ENCFF281QMV,ENCFF033UQX,ENCFF089EWX,ENCFF575MGI,ENCFF142CNG,ENCFF957SPH,ENCFF210BTO,ENCFF052KVO,ENCFF491RJK,ENCFF321AGQ,ENCFF065ODV.ENCFF086ZTV.ENCFF194HVL |
| 2596 | 18 | 66055045 | 66055139 | 94 | SPRET | 18 | 66055085 | 66055118 | 34 | - | 23.1707 | 7.85E-09 | 0.00775 | GTGAAGTTCCAAAGCTGACCACCAGAGGGAAACC | 37 | ENCFF201JUA,ENCFF093ZRU,ENCFF327LYS,ENCFF504JDS,ENCFF089GBC,ENCFF367UNR,ENCFF024JOZ,ENCFF099QNE,ENCFF907XEG,ENCFF092NMJ,ENCFF165LXO,ENCFF430PPJ,ENCFF198SSR,ENCFF589ZBJ,ENCFF609MDN,ENCFF055HAP,ENCFF821BIA,ENCFF316FYN,ENCFF136ZDK,ENCFF502WDT,ENCFF951GOP,ENCFF533APC,ENCFF883UPM,ENCFF097CAQ,ENCFF281QMV,ENCFF033UQX,ENCFF089EWX,ENCFF575MGI,ENCFF142CNG,ENCFF957SPH,ENCFF210BTO,ENCFF052KVO,ENCFF491RJK,ENCFF321AGQ,ENCFF065ODV.ENCFF086ZTV.ENCFF194HVL |
| 2597 | 18 | 67280550 | 67280887 | 337 | AJ,AKR,BALB,FVB,ILNJ2,KK,NOD,NOR,NZB,NZO,NZW,PL,RH2,SJL,SMJ,SWR,TH,WSB | 18 | 67280717 | 67280751 | 35 | - | 22.789 | 1.30E-08 | 0.02 | CTGCATGTATGCCTGCATGCCAGGAGAGGGCATCA | | |
| 2598 | 18 | 67689292 | 67689492 | 200 | CAST.SPRET | 18 | 67689412 | 67689445 | 34 | - | 18.0732 | 3.36E-07 | 0.0867 | CAGCTGTCTTCAGACACTCCAGAAGAGGGCATCA | | |
| 2599 | 18 | 67835027 | 67835242 | 215 | AJ,BALB,CAST,FVB,ILNJ2,KK,MOLF,MRL,NOD,NOR,NZB,NZO,NZW,PL,RF,RH2,SJL,SMJ.SPRET.SWR.TH | 18 | 67835148 | 67835181 | 34 | - | 17.8455 | 3.91E-07 | 0.0917 | TAGCTGTCCTCAGACACTCCAGAAGAGGGCGTCA | | |
| 2600 | 18 | 68083120 | 68083206 | 86 | MOLF.SPRET | 18 | 68083157 | 68083191 | 35 | - | 17.1927 | 4.84E-07 | 0.169 | CTGGAGAGGCAGCTCTGGTCCTGGAGAGGGCAGCT | | |
| 2601 | 18 | 73682226 | 73682442 | 216 | 129S1,BTBR,C3H,C57L,CAST,CBA,DBA1J,DBA2J.MOLF.PJ.PL.SEA.SPRET | 18 | 73682350 | 73682383 | 34 | - | 17.3008 | 5.58E-07 | 0.106 | TAGCTGTCTTCAGATACACCAGAAGGGGGCGTCA | | |

| | | | | | | | | | | | | | | | | |
|---|---|---|---|---|---|---|---|---|---|---|---|---|---|---|---|---|
| 2602 | 18 | 75110503 | 75110737 | 234 | AKR,ILNJ2,MOLF,RF | 18 | 75110491 | 75110524 | 34 | - | 19.7805 | 1.04E-07 | 0.0445 | GAGCAGGTACACCTGTAGGCAGAAGAGGGCATCA | | |
| 2603 | 18 | 75732976 | 75733034 | 58 | AJ,AKR,BALB,ILNJ2,KK,MOLF,MRL,NOD,NOR,NZB,NZW,RF,RH2 | 18 | 75732953 | 75732987 | 35 | - | 16.7982 | 6.08E-07 | 0.183 | GTGTTGATGCACTACTCTGCCACTAGAGGGTATCA | 38 | ENCFF367UNR,ENCFF093ZRU,ENCFF327LYS,ENCFF024JOZ,ENCFF907XEG,ENCFF092NMJ,ENCFF136ZDK,ENCFF089GBC,ENCFF198SSR,ENCFF430PPJ,ENCFF589ZBJ,ENCFF165LXO,ENCFF883UPM,ENCFF065ODV,ENCFF504JDS,ENCFF321AGQ,ENCFF055HAP,ENCFF502WDT,ENCFF951GOP,ENCFF210BTO,ENCFF575MGI,ENCFF609MDN,ENCFF957SPH,ENCFF052KVO,ENCFF316FYN,ENCFF601JVE,ENCFF281QMV,ENCFF491RJK,ENCFF533APC,ENCFF097CAQ,ENCFF033UQX,ENCFF611HDQ,ENCFF142CNG,ENCFF089EWX,ENCFF784ASD,ENCFF194HVL,ENCFF805QBU,ENCFF086ZTV |
| 2603 | 18 | 75732976 | 75733034 | 58 | AJ,AKR,BALB,ILNJ2,KK,MOLF,MRL,NOD,NOR,NZB,NZW,RF,RH2 | 18 | 75733011 | 75733045 | 35 | - | 17.6972 | 3.60E-07 | 0.143 | GTGTTGATGCATCACTCTGCCACTAGAGGGTGTCA | 39 | ENCFF367UNR,ENCFF093ZRU,ENCFF327LYS,ENCFF024JOZ,ENCFF907XEG,ENCFF092NMJ,ENCFF136ZDK,ENCFF089GBC,ENCFF198SSR,ENCFF430PPJ,ENCFF589ZBJ,ENCFF165LXO,ENCFF883UPM,ENCFF065ODV,ENCFF504JDS,ENCFF321AGQ,ENCFF055HAP,ENCFF502WDT,ENCFF951GOP,ENCFF210BTO,ENCFF575MGI,ENCFF609MDN,ENCFF957SPH,ENCFF052KVO,ENCFF316FYN,ENCFF601JVE,ENCFF281QMV,ENCFF491RJK,ENCFF533APC,ENCFF097CAQ,ENCFF033UQX,ENCFF611HDQ,ENCFF142CNG,ENCFF089EWX,ENCFF784ASD,ENCFF194HVL,ENCFF805QBU,ENCFF086ZTV,ENCFF349BUL |
| 2604 | 18 | 77105832 | 77106036 | 204 | SPRET | 18 | 77105959 | 77105992 | 34 | - | 17.2114 | 5.91E-07 | 0.109 | TAGCTGTTTTCAGACACTCCAGAAGAGGGCATCA | | |
| 2605 | 18 | 77850742 | 77850964 | 222 | CAST | 18 | 77850897 | 77850915 | 19 | - | 17.4918 | 6.59E-07 | 0.168 | TAGCCAGGAGGTGGAGGCA | | |
| 2606 | 18 | 77858969 | 77859328 | 359 | KK.MOLF.SPRET.TH | 18 | 77858991 | 77859025 | 35 | - | 16.6422 | 6.65E-07 | 0.185 | CTGCTACCCAAGACCTCTAACACCAGAGGCCACCC | | |
| 2606 | 18 | 77858969 | 77859328 | 359 | KK.MOLF.SPRET.TH | 18 | 77859043 | 77859077 | 35 | - | 16.6422 | 6.65E-07 | 0.185 | CTGCTACCCAAGACCTCTAACACCAGAGGCCACCC | | |
| 2606 | 18 | 77858969 | 77859328 | 359 | KK.MOLF.SPRET.TH | 18 | 77859095 | 77859129 | 35 | - | 16.6422 | 6.65E-07 | 0.185 | CTGCTACCCAAGACCTCTAACACCAGAGGCCACCC | | |
| 2606 | 18 | 77858969 | 77859328 | 359 | KK.MOLF.SPRET.TH | 18 | 77859146 | 77859180 | 35 | - | 16.6422 | 6.65E-07 | 0.185 | CTGCTACCCAAGACCTCTAACACCAGAGGCCACCC | | |
| 2606 | 18 | 77858969 | 77859328 | 359 | KK.MOLF.SPRET.TH | 18 | 77859197 | 77859231 | 35 | - | 16.6422 | 6.65E-07 | 0.185 | CTGCTACCCAAGACCTCTAACACCAGAGGCCACCC | | |
| 2606 | 18 | 77858969 | 77859328 | 359 | KK.MOLF.SPRET.TH | 18 | 77859248 | 77859282 | 35 | - | 16.6422 | 6.65E-07 | 0.185 | CTGCTACCCAAGACCTCTAACACCAGAGGCCACCC | | |
| 2606 | 18 | 77858969 | 77859328 | 359 | KK.MOLF.SPRET.TH | 18 | 77859299 | 77859333 | 35 | - | 16.6422 | 6.65E-07 | 0.185 | CTGCTACCCAAGACCTCTAACACCAGAGGCCACCC | | |
| 2607 | 18 | 81915964 | 81916316 | 352 | CAST.SPRET.WSB | 18 | 81915980 | 81916013 | 34 | - | 20.0813 | 8.38E-08 | 0.0386 | GTGCAGTGCTCACCACAGCCTCCAGAGTTCAGTG | | |
| 2608 | 18 | 82587126 | 82587188 | 62 | SPRET | 18 | 82587163 | 82587181 | 19 | - | 23.5246 | 4.39E-09 | 0.022 | TCACCAGCAGGGGGCGCTC | 36 | ENCFF093ZRU,ENCFF201JUA,ENCFF099QNE,ENCFF136ZDK,ENCFF327LYS,ENCFF907XEG,ENCFF024JOZ,ENCFF589ZBJ,ENCFF089GBC,ENCFF065ODV,ENCFF210BTO,ENCFF198SSR,ENCFF165LXO,ENCFF883UPM,ENCFF092NMJ,ENCFF321AGQ,ENCFF430PPJ,ENCFF142CNG,ENCFF957SPH,ENCFF805QBU,ENCFF033UQX,ENCFF951GOP,ENCFF281QMV,ENCFF504JDS,ENCFF609MDN,ENCFF086ZTV,ENCFF089EWX,ENCFF533APC,ENCFF316FYN,ENCFF349BUL,ENCFF784ASD,ENCFF194HVL,ENCFF052KVO,ENCFF575MGI,ENCFF097CAQ.ENCFF491RJK |
| 2608 | 18 | 82587126 | 82587188 | 62 | SPRET | 18 | 82587163 | 82587196 | 34 | - | 18.1057 | 3.29E-07 | 0.0854 | GGGATTTTTCCAACCTCACCAGCAGGGGGCGCTC | 36 | ENCFF093ZRU,ENCFF201JUA,ENCFF099QNE,ENCFF136ZDK,ENCFF327LYS,ENCFF907XEG,ENCFF024JOZ,ENCFF589ZBJ,ENCFF089GBC,ENCFF065ODV,ENCFF210BTO,ENCFF198SSR,ENCFF165LXO,ENCFF883UPM,ENCFF092NMJ,ENCFF321AGQ,ENCFF430PPJ,ENCFF142CNG,ENCFF957SPH,ENCFF805QBU,ENCFF033UQX,ENCFF951GOP,ENCFF281QMV,ENCFF504JDS,ENCFF609MDN,ENCFF086ZTV,ENCFF089EWX,ENCFF533APC,ENCFF316FYN,ENCFF349BUL,ENCFF784ASD,ENCFF194HVL,ENCFF052KVO,ENCFF575MGI,ENCFF097CAQ.ENCFF491RJK |
| 2608 | 18 | 82587126 | 82587188 | 62 | SPRET | 18 | 82587163 | 82587197 | 35 | - | 18.1009 | 2.83E-07 | 0.126 | AGGGATTTTTCCAACCTCACCAGCAGGGGGCGCTC | 36 | ENCFF093ZRU,ENCFF201JUA,ENCFF099QNE,ENCFF136ZDK,ENCFF327LYS,ENCFF907XEG,ENCFF024JOZ,ENCFF589ZBJ,ENCFF089GBC,ENCFF065ODV,ENCFF210BTO,ENCFF198SSR,ENCFF165LXO,ENCFF883UPM,ENCFF092NMJ,ENCFF321AGQ,ENCFF430PPJ,ENCFF142CNG,ENCFF957SPH,ENCFF805QBU,ENCFF033UQX,ENCFF951GOP,ENCFF281QMV,ENCFF504JDS,ENCFF609MDN,ENCFF086ZTV,ENCFF089EWX,ENCFF533APC,ENCFF316FYN,ENCFF349BUL,ENCFF784ASD,ENCFF194HVL,ENCFF052KVO,ENCFF575MGI,ENCFF097CAQ.ENCFF491RJK |
| 2609 | 18 | 82747749 | 82747802 | 53 | BUB,CAST,MA/My,NZB,SJL,SPRET,WSB | 18 | 82747756 | 82747774 | 19 | - | 19.6393 | 1.45E-07 | 0.0832 | CAGCCACTAGAAGGCGGCA | | |
| 2609 | 18 | 82747749 | 82747802 | 53 | BUB,CAST,MA/My,NZB,SJL,SPRET,WSB | 18 | 82747773 | 82747791 | 19 | - | 19.6393 | 1.45E-07 | 0.0832 | CAGCCACTAGAAGGCGGCA | | |
| 2609 | 18 | 82747749 | 82747802 | 53 | BUB,CAST,MA/My,NZB,SJL,SPRET,WSB | 18 | 82747790 | 82747808 | 19 | - | 19.6393 | 1.45E-07 | 0.0832 | CAGCCACTAGAAGGCGGCA | | |
| 2610 | 18 | 84698206 | 84698411 | 205 | 129S1,BTBR,C57L,CAST,CE,MOLF,NOD,WSB | 18 | 84698320 | 84698338 | 19 | - | 17.8852 | 5.08E-07 | 0.15 | ACACCAGAAGAGGGCACCA | | |
| 2610 | 18 | 84698206 | 84698411 | 205 | 129S1,BTBR,C57L,CAST,CE,MOLF,NOD,WSB | 18 | 84698320 | 84698353 | 34 | - | 20.8862 | 4.65E-08 | 0.0264 | TTGCTGTCTTCAGACACACCAGAAGAGGGCACCA | | |
| 2611 | 18 | 84918690 | 84918950 | 260 | NOD.SPRET | 18 | 84918814 | 84918847 | 34 | - | 16.4472 | 9.61E-07 | 0.128 | TAGCTGTCTTCAGACACACCAGAAGAGGGCGTCA | | |
| 2612 | 19 | 3497470 | 3497565 | 95 | SPRET | 19 | 3497497 | 3497531 | 35 | + | 24.5229 | 3.57E-09 | 0.0059 | ATGCAGGTATCTGAGGAGGCCAGAAGAGGGCGCTG | 5 | ENCFF367UNR.ENCFF052KVO.ENCFF136ZDK.ENCFF327LYS.ENCFF502WDT |
| 2612 | 19 | 3497470 | 3497565 | 95 | SPRET | 19 | 3497513 | 3497531 | 19 | + | 20.8033 | 5.79E-08 | 0.0346 | AGGCCAGAAGAGGGCGCTG | 5 | ENCFF367UNR.ENCFF052KVO.ENCFF136ZDK.ENCFF327LYS.ENCFF502WDT |
| 2613 | 19 | 3881554 | 3881756 | 202 | FVB.MOLF.SJL.SPRET | 19 | 3881614 | 3881647 | 34 | + | 17.8455 | 3.91E-07 | 0.0605 | TAGCTGTCCTCAGACACTCCAGAAGAGGGCGTCA | | |
| 2614 | 19 | 4093364 | 4093543 | 179 | BUB,CAST,FVB,KK,MOLF,NZW,SJL,SPRET,WSB | 19 | 4093396 | 4093429 | 34 | + | 18.561 | 2.42E-07 | 0.0503 | AAGCTGTCTTCAGACACTCCAGAAGAGGGCGCCA | | |
| 2614 | 19 | 4093364 | 4093543 | 179 | BUB,CAST,FVB,KK,MOLF,NZW,SJL,SPRET,WSB | 19 | 4093411 | 4093429 | 19 | + | 17.5574 | 6.32E-07 | 0.105 | ACTCCAGAAGAGGGCGCCA | | |
| 2615 | 19 | 4734461 | 4734661 | 200 | SPRET | 19 | 4734520 | 4734553 | 34 | + | 16.4472 | 9.61E-07 | 0.0915 | TAGCTGTCTTCAGACACACCAGAAGAGGGCGTCA | | |
| 2616 | 19 | 5672567 | 5672790 | 223 | CAST.MOLF.SPRET | 19 | 5672648 | 5672681 | 34 | + | 18.1789 | 3.13E-07 | 0.0547 | TAGCTGTCTTCAGACACTCCAGAAGAGGGCGCTA | | |
| 2616 | 19 | 5672567 | 5672790 | 223 | CAST.MOLF.SPRET | 19 | 5672663 | 5672681 | 19 | + | 17.5738 | 6.25E-07 | 0.105 | ACTCCAGAAGAGGGCGCTA | | |
| 2617 | 19 | 5831218 | 5831420 | 202 | C58J.CAST.MOLF.MRL.SPRET.TH | 19 | 5831278 | 5831311 | 34 | + | 18.3659 | 2.76E-07 | 0.0518 | TAGCTGTCTTCAGACACTCCAGAAGAGGGCGCCA | 3 | ENCFF093ZRU.ENCFF201JUA.ENCFF367UNR |
| 2617 | 19 | 5831218 | 5831420 | 202 | C58J.CAST.MOLF.MRL.SPRET.TH | 19 | 5831293 | 5831311 | 19 | + | 17.5574 | 6.32E-07 | 0.105 | ACTCCAGAAGAGGGCGCCA | 3 | ENCFF093ZRU.ENCFF201JUA.ENCFF367UNR |
| 2618 | 19 | 5959320 | 5959517 | 197 | BUB,C58J,CAST,FVB,LP,MOLF,MRL,SJL,SPRET,TH | 19 | 5959375 | 5959408 | 34 | + | 16.4715 | 9.47E-07 | 0.0915 | TAGCTGTCCTCAGAGACACCAGAAGAGGGAGTCA | 2 | ENCFF093ZRU,ENCFF367UNR |
| 2619 | 19 | 6087171 | 6087357 | 186 | BUB.SPRET | 19 | 6087226 | 6087259 | 34 | + | 17.6992 | 4.30E-07 | 0.0634 | TAGCTGTTTTCAGACACACCAGAAGAGGGCATCA | | |
| 2620 | 19 | 6445379 | 6445595 | 216 | BUB,C58J,CAST,CBA,DBA1J,DBA2J,LP,MA/My.MOLF.NZW.SPRET.TH.WSB | 19 | 6445453 | 6445486 | 34 | + | 16.4634 | 9.51E-07 | 0.0915 | TAGCTGTCTTCAGACGCACCAGAAGAGGGCATCA | | |
| 2621 | 19 | 6484541 | 6484749 | 208 | CAST.MOLF.SPRET | 19 | 6484595 | 6484628 | 34 | + | 18.3659 | 2.76E-07 | 0.0518 | TAGCTGTCTTCAGACACTCCAGAAGAGGGCGCCA | 1 | ENCFF093ZRU |
| 2621 | 19 | 6484541 | 6484749 | 208 | CAST.MOLF.SPRET | 19 | 6484610 | 6484628 | 19 | + | 17.5574 | 6.32E-07 | 0.105 | ACTCCAGAAGAGGGCGCCA | 1 | ENCFF093ZRU |
| 2622 | 19 | 6986583 | 6986793 | 210 | C58J.CAST.FVB.LP.MOLF.SPRET | 19 | 6986667 | 6986685 | 19 | + | 17.8852 | 5.08E-07 | 0.098 | ACACCAGAAGAGGGCACCA | | |
| 2623 | 19 | 7566552 | 7566755 | 203 | BUB,C58J,CAST,FVB,LP,MA/My,MOLF,SPRET,WSB | 19 | 7566615 | 7566648 | 34 | + | 18.3659 | 2.76E-07 | 0.0518 | TAGCTGTCTTCAGACACTCCAGAAGAGGGCGCCA | | |
| 2623 | 19 | 7566552 | 7566755 | 203 | BUB,C58J,CAST,FVB,LP,MA/My,MOLF,SPRET,WSB | 19 | 7566630 | 7566648 | 19 | + | 17.5574 | 6.32E-07 | 0.105 | ACTCCAGAAGAGGGCGCCA | | |
| 2624 | 19 | 7645609 | 7646514 | 905 | CAST.MOLF | 19 | 7646060 | 7646093 | 34 | + | 17.8455 | 3.91E-07 | 0.0605 | CTGCACTGCTCCATGCTGGCAGAAGACGCCATCA | | |
| 2625 | 19 | 8999241 | 8999451 | 210 | CAST.LP.MOLF.SPRET.TH.WSB | 19 | 8999307 | 8999340 | 34 | + | 18.1545 | 3.18E-07 | 0.0548 | TAGCTGTTTTCAGACACACCAGAAGAGGGCGTCA | | |
| 2626 | 19 | 10119963 | 10120195 | 232 | MOLF.SPRET | 19 | 10120052 | 10120085 | 34 | + | 16.4959 | 9.32E-07 | 0.0915 | TAGCTGTCTTCAGACACGCCAGAAGAGGGCGTAG | 1 | ENCFF093ZRU |
| 2627 | 19 | 10535412 | 10535585 | 173 | SPRET | 19 | 10535535 | 10535553 | 19 | + | 19.0164 | 2.30E-07 | 0.0672 | GTGCCAGAAGAGGGCACCA | 6 | ENCFF367UNR.ENCFF201JUA.ENCFF093ZRU.ENCFF502WDT.ENCFF055HAP.ENCFF327LYS |
| 2628 | 19 | 15952888 | 15953170 | 282 | MA/My.WSB | 19 | 15952897 | 15952930 | 34 | + | 19.439 | 1.32E-07 | 0.0363 | ATGCAGGTGCCGTGGAGGGCAGAAGAAGGCACTG | | |
| 2629 | 19 | 20184297 | 20184497 | 200 | MOLF.MRL.NZB.NZO.NZW.PJ.SPRET | 19 | 20184355 | 20184388 | 34 | + | 17.2439 | 5.79E-07 | 0.0746 | TAGCTGTCCTCAGATACTCCAGAAGAGGGCATCA | | |
| 2630 | 19 | 22864530 | 22864702 | 172 | C3H,CBA,CE,DBA1J,DBA2J,ILNJ2,MOLF,MRL.NZB.NZO.NZW.PJ.SPRET | 19 | 22864644 | 22864677 | 34 | + | 20.6585 | 5.50E-08 | 0.0207 | CTGCAGTGCTCCCATCATCAAACAGGTGGTAGTC | | |
| 2630 | 19 | 22864530 | 22864702 | 172 | C3H,CBA,CE,DBA1J,DBA2J,ILNJ2,MOLF,MRL.NZB.NZO.NZW.PJ.SPRET | 19 | 22864680 | 22864713 | 34 | + | 20.6585 | 5.50E-08 | 0.0207 | CTGCAGTGCTCCCATCATCAAACAGGTGGTAGTC | | |
| 2631 | 19 | 23599180 | 23599687 | 507 | C3H,CBA,CE,DBA1J,DBA2J,MOLF,MRL,NZO.NZW.PJ.SPRET | 19 | 23599622 | 23599655 | 34 | + | 18.7642 | 2.11E-07 | 0.0466 | ATGCAGCTAGAGACACGAGCTCCGGGGGGCACTG | | |
| 2632 | 19 | 29304190 | 29304551 | 361 | SPRET | 19 | 29304399 | 29304432 | 34 | + | 18.0813 | 3.34E-07 | 0.0566 | GTGTAGTGCCTATTGAAGCCATAAGAGGGTGTCA | 1 | ENCFF055HAP |
| 2633 | 19 | 29502804 | 29503213 | 409 | CAST.MOLF.SPRET.WSB | 19 | 29503070 | 29503103 | 34 | + | 18.2927 | 2.90E-07 | 0.0526 | TAGCTGTCTTCGGTTACACCAGAAGAGGGCAGCA | 4 | ENCFF821BIA.ENCFF055HAP.ENCFF367UNR.ENCFF093ZRU |
| 2633 | 19 | 29502804 | 29503213 | 409 | CAST.MOLF.SPRET.WSB | 19 | 29503085 | 29503103 | 19 | + | 17.2459 | 7.74E-07 | 0.116 | ACACCAGAAGAGGGCAGCA | 4 | ENCFF821BIA.ENCFF055HAP.ENCFF367UNR.ENCFF093ZRU |
| 2634 | 19 | 29717130 | 29717197 | 67 | MOLF | 19 | 29717163 | 29717196 | 34 | + | 19.9024 | 9.52E-08 | 0.0296 | TGGGAGTGCCTATGGAGGCCAGAAGGTGGCACAA | 12 | ENCFF093ZRU,ENCFF367UNR,ENCFF821BIA,ENCFF201JUA,ENCFF907XEG,ENCFF198SSR,ENCFF883UPM,ENCFF957SPH,ENCFF502WDT,ENCFF327LYS,ENCFF055HAP,ENCFF142CNG |
| 2634 | 19 | 29717130 | 29717197 | 67 | MOLF | 19 | 29717178 | 29717196 | 19 | + | 18.8033 | 2.69E-07 | 0.0716 | AGGCCAGAAGGTGGCACAA | 12 | ENCFF093ZRU,ENCFF367UNR,ENCFF821BIA,ENCFF201JUA,ENCFF907XEG,ENCFF198SSR,ENCFF883UPM,ENCFF957SPH,ENCFF502WDT,ENCFF327LYS,ENCFF055HAP,ENCFF142CNG |
| 2635 | 19 | 34952105 | 34952543 | 438 | NZW | 19 | 34952452 | 34952485 | 34 | + | 16.4228 | 9.76E-07 | 0.092 | TTGAAATTTTACCAGCTGGCACCAGAAGGAACTA | | |
| 2636 | 19 | 34959347 | 34959559 | 212 | AJ,BALB,C58J,CAST,LP,MOLF,MRL,NZO,NZW,RH2,SPRET,SWR,TH | 19 | 34959417 | 34959450 | 34 | + | 16.4472 | 9.61E-07 | 0.0915 | TAGCTGTCTTCAGACACACCAGAAGAGGGCGTCA | | |
| 2637 | 19 | 36919686 | 36919891 | 205 | SPRET | 19 | 36919749 | 36919782 | 34 | + | 18.3659 | 2.76E-07 | 0.0518 | TAGCTGTCTTCAGACACTCCAGAAGAGGGCGCCA | 2 | ENCFF093ZRU.ENCFF367UNR |
| 2637 | 19 | 36919686 | 36919891 | 205 | SPRET | 19 | 36919764 | 36919782 | 19 | + | 17.5574 | 6.32E-07 | 0.105 | ACTCCAGAAGAGGGCGCCA | 2 | ENCFF093ZRU.ENCFF367UNR |
| 2638 | 19 | 37035396 | 37035803 | 407 | 129S1,AJ,AKR,BALB,BTBR,BUB,C3H,C57L,CAST,CBA,CE,DBA1J,DBA2J,FVB,ILNJ2,KK,MA/My,MOLF,MRL,NOD,NOR,NZB,NZO,NZW,PJ,PL,RF,RH2,SEA,SJL,SMJ,SPRET,SWR.TH.WSB | 19 | 37035661 | 37035694 | 34 | + | 16.4472 | 9.61E-07 | 0.0915 | TAGCTGTCTTCAGACACACCAGAAGAGGGCGTCA | | |
| 2639 | 19 | 37160509 | 37160718 | 209 | SPRET | 19 | 37160536 | 37160569 | 34 | + | 16.4472 | 9.61E-07 | 0.0915 | TAGCTGTCTTCAGACACACCAGAAGAGGGCGTCA | | |
| 2640 | 19 | 40963770 | 40964707 | 937 | CAST | 19 | 40964421 | 40964454 | 34 | + | 16.6341 | 8.54E-07 | 0.0888 | TAGCTGTCTTCACACACTCCAGAAGAGGGCGTCA | | |
| 2641 | 19 | 41411560 | 41411761 | 201 | CAST.MOLF.SPRET | 19 | 41411618 | 41411651 | 34 | + | 18.1789 | 3.13E-07 | 0.0547 | TAGCTGTCTTCAGCCACTCCAGAAGAGGGCGCCA | | |
| 2641 | 19 | 41411560 | 41411761 | 201 | CAST.MOLF.SPRET | 19 | 41411633 | 41411651 | 19 | + | 17.5574 | 6.32E-07 | 0.105 | ACTCCAGAAGAGGGCGCCA | | |
| 2642 | 19 | 43701549 | 43701763 | 214 | 129S1,AJ,AKR,BALB,BUB,C3H,C57L,CAST,CBA,CE,DBA1J,DBA2J,FVB,ILNJ2,KK,MA/My,MOLF,MRL,NOD,NOR,NZB,NZO,NZW,PJ,PL,RF,RH2,SEA,SJL,SMJ,SPRET,SWR,TH,WSB | 19 | 43701619 | 43701653 | 35 | + | 18.3945 | 2.37E-07 | 0.074 | CTGTAGCTGTCTTGACGCACCAGAAGAGGGCATCA | | |
| 2643 | 19 | 44188278 | 44188506 | 228 | 129S1,BUB,C57L,CAST,FVB,KK,MOLF,PJ,SEA,SMJ,SPRET,SWR | 19 | 44188362 | 44188395 | 34 | + | 16.4634 | 9.51E-07 | 0.0915 | TAGCTGTCTTCAGACGCACCAGAAGAGGGCATCA | | |
| 2644 | 19 | 44997974 | 44998183 | 209 | C3H,CAST,CBA,CE,ILNJ2,MOLF,MRL,PJ,SEA,SPRET,WSB | 19 | 44998040 | 44998073 | 34 | + | 16.4472 | 9.61E-07 | 0.0915 | TAGCTGTCTTCAGACACACCAGAAGAGGGCGTCA | | |
| 2645 | 19 | 45403521 | 45404140 | 619 | SPRET | 19 | 45403752 | 45403770 | 19 | + | 17.3934 | 7.03E-07 | 0.111 | CAGGCAGCAGGGGGCAGCC | 2 | ENCFF093ZRU.ENCFF367UNR |
| 2646 | 19 | 45658525 | 45658798 | 273 | CAST | 19 | 45658660 | 45658693 | 34 | + | 18.3577 | 2.78E-07 | 0.0519 | TTGCTGTCTTCAGACACACCAGAAGAGGGCAACA | | |
| 2647 | 19 | 45883589 | 45883791 | 202 | CAST.MOLF.SPRET.WSB | 19 | 45883649 | 45883682 | 34 | + | 18.3659 | 2.76E-07 | 0.0518 | TAGCTGTCTTCAGACACTCCAGAAGAGGGCGCCA | 2 | ENCFF093ZRU.ENCFF055HAP |
| 2647 | 19 | 45883589 | 45883791 | 202 | CAST.MOLF.SPRET.WSB | 19 | 45883664 | 45883682 | 19 | + | 17.5574 | 6.32E-07 | 0.105 | ACTCCAGAAGAGGGCGCCA | 2 | ENCFF093ZRU.ENCFF055HAP |
| 2648 | 19 | 46684990 | 46685560 | 570 | SPRET | 19 | 46684987 | 46685021 | 35 | + | 17.1009 | 5.11E-07 | 0.115 | CTGCAGAAGCCGCCAGAAGCCGCCAGAAGCTGCCC | | |
| 2649 | 19 | 46943889 | 46944091 | 202 | 129S1,BTBR,C57L,CAST,CE,MOLF,SPRET,WSB | 19 | 46943947 | 46943980 | 34 | + | 18.3659 | 2.76E-07 | 0.0518 | TAGCTGTCTTCAGACACTCCAGAAGAGGGCGCCA | 3 | ENCFF093ZRU,ENCFF367UNR,ENCFF099QNE |
| 2649 | 19 | 46943889 | 46944091 | 202 | 129S1,BTBR,C57L,CAST,CE,MOLF,SPRET,WSB | 19 | 46943962 | 46943980 | 19 | + | 17.5574 | 6.32E-07 | 0.105 | ACTCCAGAAGAGGGCGCCA | 3 | ENCFF093ZRU,ENCFF367UNR,ENCFF099QNE |
| 2650 | 19 | 47146640 | 47147355 | 715 | AKR,BUB,C3H,CBA,DBA1J,FVB,ILNJ2,KK,MA/My,MRL,NOD,NOR,NZB,NZO,NZW,PJ,PL,RF,SEA,SJL,SMJ,SWR,TH,WSB | 19 | 47146634 | 47146652 | 19 | + | 17.1803 | 8.08E-07 | 0.121 | CTTCCAGCAGAGGGTACCG | | |
| 2651 | 19 | 50468853 | 50469049 | 196 | SPRET | 19 | 50468907 | 50468940 | 34 | + | 16.4634 | 9.51E-07 | 0.0915 | TAGCTGTCTTCAGACGCACCAGAAGAGGGCATCA | | |
| 2652 | 19 | 51610478 | 51610625 | 147 | SPRET | 19 | 51610549 | 51610582 | 34 | + | 16.748 | 7.95E-07 | 0.0859 | CTGCAGTTACTCCAAATTCCTCTATGTGGCTCAC | | |
| 2653 | 19 | 51625316 | 51625537 | 221 | CAST.MOLF.SPRET.TH | 19 | 51625373 | 51625406 | 34 | + | 16.4472 | 9.61E-07 | 0.0915 | TAGCTGTCTTCAGACACACCAGAAGAGGGCGTCA | | |
| 2654 | 19 | 53256741 | 53256947 | 206 | SPRET | 19 | 53256811 | 53256844 | 34 | + | 16.4472 | 9.61E-07 | 0.0915 | TAGCTGTCTTCAGACACACCAGAAGAGGGCGTCA | 3 | ENCFF093ZRU.ENCFF821BIA.ENCFF367UNR |
| 2655 | 19 | 53597968 | 53598170 | 202 | MOLF.SPRET.WSB | 19 | 53598026 | 53598059 | 34 | + | 16.4472 | 9.61E-07 | 0.0915 | TAGCTGTCTTCAGACACACCAGAAGAGGGCGTCA | | |
| 2656 | 19 | 53632532 | 53632765 | 233 | MOLF,SPRET | 19 | 53632610 | 53632643 | 34 | + | 18.1545 | 3.18E-07 | 0.0548 | TAGCTGTCTTCCAACAGTCCAGAAGGGGGAGCCA | 7 | ENCFF821BIA,ENCFF093ZRU,ENCFF198SSR,ENCFF367UNR,ENCFF055HAP,ENCFF201JUA,ENCFF099QNE |
| 2656 | 19 | 53632532 | 53632765 | 233 | MOLF,SPRET | 19 | 53632625 | 53632643 | 19 | + | 17.0164 | 8.98E-07 | 0.125 | AGTCCAGAAGGGGGAGCCA | 7 | ENCFF821BIA,ENCFF093ZRU,ENCFF198SSR,ENCFF367UNR,ENCFF055HAP,ENCFF201JUA,ENCFF099QNE |
| 2657 | 19 | 56968626 | 56968773 | 147 | SPRET | 19 | 56968688 | 56968721 | 34 | + | 16.7642 | 7.87E-07 | 0.0853 | CTGTGTGGTCACCCAGGTCCACTAGGGGTCACCC | | |
| 2657 | 19 | 56968626 | 56968773 | 147 | SPRET | 19 | 56968703 | 56968721 | 19 | + | 17.1148 | 8.43E-07 | 0.122 | GGTCCACTAGGGGTCACCC | | |
| 2658 | 19 | 61014015 | 61014217 | 202 | 129S1,AJ,AKR,BALB,BTBR,C3H,C57L,CAST,CBA,DBA1J,DBA2J,FVB,KK,MA/My,MRL,NOD,NOR,NZB,NZO,NZW,PJ,PL,RF,RH2,SEA,SMJ,SPRET,SWR,TH | 19 | 61014064 | 61014097 | 34 | + | 18.9512 | 1.85E-07 | 0.044 | TTGCTCTCCTCAGACACACCAGAAGAGGGCATCG | | |
| 2659 | 19 | 3707476 | 3707679 | 203 | CAST.MOLF.NZW.SPRET.WSB | 19 | 3707599 | 3707632 | 34 | - | 16.4472 | 9.61E-07 | 0.0915 | TAGCTGTCTTCAGACACACCAGAAGAGGGCGTCA | | |
| 2660 | 19 | 4099091 | 4099479 | 388 | CAST,MOLF | 19 | 4099262 | 4099296 | 35 | - | 24.8073 | 2.86E-09 | 0.00546 | CTGCATGTACACCTGCATGCCAGAAGAGGGCATCG | 11 | ENCFF099QNE,ENCFF024JOZ,ENCFF201JUA,ENCFF327LYS,ENCFF957SPH,ENCFF093ZRU,ENCFF367UNR,ENCFF589ZBJ,ENCFF055HAP,ENCFF907XEG,ENCFF502WDT |
| 2661 | 19 | 4481674 | 4481822 | 148 | CAST.MOLF.SPRET | 19 | 4481731 | 4481764 | 34 | - | 17.1626 | 6.10E-07 | 0.0767 | CTGTAACTTCAGCGGGGGGCAGTGGGGGGGACAG | | |
| 2662 | 19 | 4975003 | 4975230 | 227 | AJ,AKR,BALB,BUB,C3H,C58J,CAST,CBA,CE,DBA1J,DBA2J,FVB,ILNJ2,KK,LP,MA/My,MOLF,MRL,NOD,NOR,NZW,PJ,PL,RF,RH2,SEA,SJL,SMJ,SPRET,SWR,WSB | 19 | 4975131 | 4975149 | 19 | - | 17.5574 | 6.32E-07 | 0.105 | ACTCCAGAAGAGGGCGCCA | | |
| 2662 | 19 | 4975003 | 4975230 | 227 | AJ,AKR,BALB,BUB,C3H,C58J,CAST,CBA,CE,DBA1J,DBA2J,FVB,ILNJ2,KK,LP,MA/My,MOLF,MRL,NOD,NOR,NZW,PJ,PL,RF,RH2,SEA,SJL,SMJ,SPRET,SWR,WSB | 19 | 4975131 | 4975164 | 34 | - | 18.3659 | 2.76E-07 | 0.0518 | TAGCTGTCTTCAGACACTCCAGAAGAGGGCGCCA | | |
| 2663 | 19 | 5545017 | 5545243 | 226 | CAST.FVB.LP.MOLF.SJL.SPRET | 19 | 5545148 | 5545181 | 34 | - | 17.2439 | 5.79E-07 | 0.0746 | TAGCTGTCCTCAGACCCTCCAGAAGAGGGAGTCA | | |
| 2664 | 19 | 5827178 | 5827588 | 410 | C58J.CAST.MOLF.MRL.SPRET.TH | 19 | 5827510 | 5827543 | 34 | - | 16.6341 | 8.54E-07 | 0.0888 | TAGCTGTCTTCACACACTCCAGAAGAGGGCGTCA | 1 | ENCFF502WDT |
| 2665 | 19 | 6142731 | 6143089 | 358 | SPRET | 19 | 6143004 | 6143037 | 34 | - | 18.4797 | 2.56E-07 | 0.0512 | TTGCTGTCTTCAGACACACCAGAAGAGGGCATCA | | |
| 2666 | 19 | 7476508 | 7476708 | 200 | SPRET | 19 | 7476633 | 7476666 | 34 | - | 16.4472 | 9.61E-07 | 0.0915 | TAGCTGTCTTCAGACACACCAGAAGAGGGCGTCA | | |
| 2667 | 19 | 8694037 | 8694264 | 227 | CAST.LP.MOLF.SPRET | 19 | 8694163 | 8694196 | 34 | - | 17.8455 | 3.91E-07 | 0.0605 | TAGCTGTCCTCAGACACTCCAGAAGAGGGCGTCA | | |
| 2668 | 19 | 9843243 | 9843459 | 216 | SPRET | 19 | 9843368 | 9843401 | 34 | - | 17.8293 | 3.95E-07 | 0.0608 | TAGCTGTCTTCAGACCCTCCAGAAGAGGGCGTCA | | |
| 2669 | 19 | 9989498 | 9989709 | 211 | MOLF.SPRET | 19 | 9989624 | 9989657 | 34 | - | 17.6341 | 4.49E-07 | 0.0646 | TAGCTGTCTCCAGACACACCAGAAGAGGGCAGGC | | |
| 2670 | 19 | 10143554 | 10143752 | 198 | AJ,BALB,BUB,C3H,C58J,CAST,CBA,CE,DBA1J,DBA2J,FVB,ILNJ2,KK,LP,MA/My,MOLF,MRL,PJ,RH2,SEA,SPRET,SWR,TH,WSB | 19 | 10143678 | 10143711 | 34 | - | 17.8455 | 3.91E-07 | 0.0605 | TAGCTGTCCTCAGACACTCCAGAAGAGGGCGTCA | | |
| 2671 | 19 | 13130437 | 13130634 | 197 | CAST.MOLF | 19 | 13130562 | 13130580 | 19 | - | 17.5574 | 6.32E-07 | 0.105 | ACTCCAGAAGAGGGCGCCA | | |
| 2671 | 19 | 13130437 | 13130634 | 197 | CAST.MOLF | 19 | 13130562 | 13130595 | 34 | - | 18.3659 | 2.76E-07 | 0.0518 | TAGCTGTCTTCAGACACTCCAGAAGAGGGCGCCA | | |
| 2672 | 19 | 16012609 | 16012811 | 202 | SPRET | 19 | 16012740 | 16012773 | 34 | - | 16.4634 | 9.51E-07 | 0.0915 | TAGCTGTCTTCAGACGCACCAGAAGAGGGCATCA | | |
| 2673 | 19 | 21417457 | 21417686 | 229 | CAST.SPRET | 19 | 21417589 | 21417622 | 34 | - | 17.2439 | 5.79E-07 | 0.0746 | TAGCTGTCCTCAGATACTCCAGAAGAGGGCATCA | | |
| 2674 | 19 | 21730946 | 21731381 | 435 | CAST.MOLF.MRL.NZB.NZO.NZW | 19 | 21731295 | 21731313 | 19 | - | 17.0164 | 8.98E-07 | 0.125 | AGGCCAAAAGAGGGCACCA | | |
| 2675 | 19 | 24945586 | 24945777 | 191 | SPRET | 19 | 24945702 | 24945720 | 19 | - | 17.9836 | 4.75E-07 | 0.098 | ACACCAGAAGAGGGCGGCA | | |
| 2676 | 19 | 25195880 | 25196052 | 172 | SPRET | 19 | 25195994 | 25196027 | 34 | - | 17.122 | 6.26E-07 | 0.0776 | TAGCTGTCTTCAGACTCTCCAGAAGAGGGCATCA | | |
| 2677 | 19 | 29043043 | 29043752 | 709 | CAST | 19 | 29043306 | 29043340 | 35 | - | 20.4312 | 6.57E-08 | 0.0338 | CTGCATACACACCTGCATGCCGGAAGAGGGCATCA | | |
| 2678 | 19 | 36952805 | 36953009 | 204 | CAST.MOLF.SPRET | 19 | 36952932 | 36952965 | 34 | - | 16.4634 | 9.51E-07 | 0.0915 | TAGCTGTCTTCAGACGCACCAGAAGAGGGCATCA | | |
| 2679 | 19 | 42098034 | 42098237 | 203 | 129S1,AJ,AKR,BALB,BUB,C3H,C57L,CAST,CBA,CE,DBA1J,DBA2J,FVB,ILNJ2,KK,MA/My,MOLF,MRL,NOD,NOR,NZB,NZO,NZW,PJ,PL,RF,RH2,SEA,SJL,SMJ,SPRET,SWR,WSB | 19 | 42098158 | 42098191 | 34 | - | 18.2358 | 3.02E-07 | 0.0537 | CAGCTGTCTTCAGACACCCCAGAAGAGGGCGTCA | | |
| 2680 | 19 | 45797346 | 45797539 | 193 | MOLF.SPRET | 19 | 45797440 | 45797473 | 34 | - | 16.9675 | 6.91E-07 | 0.0817 | TAGCTGTCCTCAGACATTCCAGAAGAGGGAGGCA | 1 | ENCFF093ZRU |
| 2681 | 19 | 46000357 | 46000561 | 204 | CAST.SPRET | 19 | 46000480 | 46000513 | 34 | - | 16.7073 | 8.16E-07 | 0.0868 | TAGCTGTCTTCAGATGTACCAGAAGAGGGCGTCA | | |
| 2682 | 19 | 46721132 | 46721336 | 204 | MOLF,SPRET | 19 | 46721257 | 46721275 | 19 | - | 17.5574 | 6.32E-07 | 0.105 | ACTCCAGAAGAGGGCGCCA | 9 | ENCFF136ZDK,ENCFF907XEG,ENCFF589ZBJ,ENCFF055HAP,ENCFF367UNR,ENCFF502WDT,ENCFF093ZRU,ENCFF821BIA,ENCFF099QNE |
| 2683 | 19 | 51334196 | 51334697 | 501 | CAST.MOLF.SPRET.TH | 19 | 51334165 | 51334198 | 34 | - | 18.5122 | 2.50E-07 | 0.0509 | ATGCAGCTAGAGACATGAGCTCCAGGGGGTGCTG | | |
| 2684 | 19 | 55030903 | 55031030 | 127 | 129S1,AJ,AKR,BALB,BTBR,C3H,C57L,CAST,CBA,CE,DBA1J,DBA2J,FVB,ILNJ2,MA/My,MOLF,MRL,NZB,NZW,PJ,PL,RH2,SEA,SWR,TH.WSB | 19 | 55030964 | 55030997 | 34 | - | 17.9512 | 3.64E-07 | 0.0589 | CAGCAGCACCATCACCCAGCAGCAGAGGCTGGCC | | |
| 2685 | 19 | 55663814 | 55664022 | 208 | CAST.MOLF.SPRET | 19 | 55663926 | 55663959 | 34 | - | 17.2439 | 5.79E-07 | 0.0746 | TAGCTGTCCTCAGATACTCCAGAAGAGGGCATCA | | |
| 2686 | 19 | 56190225 | 56190466 | 241 | MOLF | 19 | 56190394 | 56190412 | 19 | - | 18.1803 | 4.16E-07 | 0.0893 | GTGCCACAAGATGGCAGCA | 31 | ENCFF093ZRU,ENCFF201JUA,ENCFF367UNR,ENCFF052KVO,ENCFF099QNE,ENCFF024JOZ,ENCFF327LYS,ENCFF198SSR,ENCFF907XEG,ENCFF821BIA,ENCFF033UQX,ENCFF055HAP,ENCFF589ZBJ,ENCFF609MDN,ENCFF502WDT,ENCFF089GBC,ENCFF065ODV,ENCFF136ZDK,ENCFF092NMJ,ENCFF575MGI,ENCFF142CNG,ENCFF097CAQ,ENCFF883UPM,ENCFF281QMV,ENCFF430PPJ,ENCFF957SPH,ENCFF533APC,ENCFF165LXO.ENCFF194HVL.ENCFF611HDQ.ENCFF491RJK |
| 2686 | 19 | 56190225 | 56190466 | 241 | MOLF | 19 | 56190394 | 56190428 | 35 | - | 21.5321 | 3.14E-08 | 0.0216 | TTGCTCTGGCCTGGAGGTGCCACAAGATGGCAGCA | 31 | ENCFF093ZRU,ENCFF201JUA,ENCFF367UNR,ENCFF052KVO,ENCFF099QNE,ENCFF024JOZ,ENCFF327LYS,ENCFF198SSR,ENCFF907XEG,ENCFF821BIA,ENCFF033UQX,ENCFF055HAP,ENCFF589ZBJ,ENCFF609MDN,ENCFF502WDT,ENCFF089GBC,ENCFF065ODV,ENCFF136ZDK,ENCFF092NMJ,ENCFF575MGI,ENCFF142CNG,ENCFF097CAQ,ENCFF883UPM,ENCFF281QMV,ENCFF430PPJ,ENCFF957SPH,ENCFF533APC,ENCFF165LXO.ENCFF194HVL.ENCFF611HDQ.ENCFF491RJK |
| 2687 | 19 | 56203214 | 56203760 | 546 | SPRET | 19 | 56203526 | 56203559 | 34 | - | 16.439 | 9.66E-07 | 0.0916 | ATGCAGCTAGAGACAAGAGCTCCAGGGGGTACTG | | |
| 2688 | 19 | 58725839 | 58726513 | 674 | CAST | 19 | 58725828 | 58725861 | 34 | - | 17.2439 | 5.79E-07 | 0.0746 | CTGCTATTCCTCCAGTCTGCACTGGGGGTGACTT | | |
| 2689 | 19 | 60325317 | 60325518 | 201 | SPRET | 19 | 60325432 | 60325450 | 19 | - | 17.5574 | 6.32E-07 | 0.105 | ACTCCAGAAGAGGGCGCCA | 2 | ENCFF093ZRU.ENCFF099QNE |
| 2689 | 19 | 60325317 | 60325518 | 201 | SPRET | 19 | 60325432 | 60325465 | 34 | - | 18.3659 | 2.76E-07 | 0.0518 | TAGCTGTCTTCAGACACTCCAGAAGAGGGCGCCA | 2 | ENCFF093ZRU.ENCFF099QNE |
| 2690 | X | 6091627 | 6092616 | 989 | SPRET | X | 6092184 | 6092217 | 34 | + | 16.4878 | 9.37E-07 | 0.182 | CAGCCTTTCCCTAATTCAACCCCAGGGGTCAGCA | | |
| 2691 | X | 7821122 | 7821324 | 202 | SPRET | X | 7821182 | 7821215 | 34 | + | 20.0407 | 8.62E-08 | 0.0579 | CAGCTGTCTTCAGATGTGCCAGAAGAGGGCATCA | 1 | ENCFF502WDT |
| 2692 | X | 12066395 | 12066625 | 230 | CAST.MOLF.SMJ.SPRET | X | 12066483 | 12066516 | 34 | + | 17.8455 | 3.91E-07 | 0.127 | TAGCTGTCCTCAGACACTCCAGAAGAGGGCGTCA | | |
| 2693 | X | 12749452 | 12749673 | 221 | CAST.MOLF.SPRET | X | 12749528 | 12749561 | 34 | + | 18.3659 | 2.76E-07 | 0.111 | TAGCTGTCTTCAGACACTCCAGAAGAGGGCGCCA | | |
| 2693 | X | 12749452 | 12749673 | 221 | CAST.MOLF.SPRET | X | 12749543 | 12749561 | 19 | + | 17.5574 | 6.32E-07 | 0.241 | ACTCCAGAAGAGGGCGCCA | | |
| 2694 | X | 13094236 | 13094439 | 203 | MOLF.SPRET | X | 13094298 | 13094331 | 34 | + | 17.2439 | 5.79E-07 | 0.156 | TAGCTGTCCTCAGATACTCCAGAAGAGGGCATCA | | |
| 2695 | X | 15746242 | 15746315 | 73 | SPRET | X | 15746282 | 15746316 | 35 | + | 16.422 | 7.54E-07 | 0.266 | ATGCAGTGAAAGTGTTAACACTGTAGATGGCACAA | | |
| 2696 | X | 20849064 | 20849417 | 353 | CAST.MOLF.SPRET | X | 20849199 | 20849232 | 34 | + | 19.5528 | 1.22E-07 | 0.0713 | TAGCAGTTTTCAGACACACCAGAAGAGGGCATTG | | |
| 2697 | X | 38780774 | 38781019 | 245 | CAST.MOLF.SPRET | X | 38780860 | 38780893 | 34 | + | 18.3496 | 2.79E-07 | 0.111 | TAGCTGTCCTCAGACGCACCAGAAGAGGGCATCA | | |
| 2698 | X | 41681307 | 41681519 | 212 | MOLF.SPRET | X | 41681369 | 41681402 | 34 | + | 18.1545 | 3.18E-07 | 0.117 | TAGCTGTTTTCAGACACACCAGAAGAGGGCGTCA | | |
| 2699 | X | 45777945 | 45778667 | 722 | C58J.CAST.NZB.NZO.WSB | X | 45778033 | 45778066 | 34 | + | 17.0244 | 6.67E-07 | 0.164 | GAGCAGGTGTGGCAGCAGTCCCAAGATGGCGCCC | | |
| 2700 | X | 47068253 | 47068455 | 202 | SPRET | X | 47068311 | 47068344 | 34 | + | 16.4472 | 9.61E-07 | 0.182 | TAGCTGTCTTCAGACACACCAGAAGAGGGCGTCA | 1 | ENCFF093ZRU |
| 2701 | X | 47596325 | 47596535 | 210 | SPRET | X | 47596392 | 47596425 | 34 | + | 18.3659 | 2.76E-07 | 0.111 | TAGCTGTCTTCAGACACTCCAGAAGAGGGCGCCA | 1 | ENCFF093ZRU |
| 2701 | X | 47596325 | 47596535 | 210 | SPRET | X | 47596407 | 47596425 | 19 | + | 17.5574 | 6.32E-07 | 0.241 | ACTCCAGAAGAGGGCGCCA | 1 | ENCFF093ZRU |
| 2702 | X | 47758223 | 47758430 | 207 | CAST.MOLF.SPRET | X | 47758287 | 47758320 | 34 | + | 18.3659 | 2.76E-07 | 0.111 | TAGCTGTCTTCAGACACTCCAGAAGAGGGCGCCA | | |
| 2702 | X | 47758223 | 47758430 | 207 | CAST.MOLF.SPRET | X | 47758302 | 47758320 | 19 | + | 17.5574 | 6.32E-07 | 0.241 | ACTCCAGAAGAGGGCGCCA | | |
| 2703 | X | 51779307 | 51779506 | 199 | SPRET | X | 51779364 | 51779397 | 34 | + | 16.4472 | 9.61E-07 | 0.182 | TAGCTGTCTTCAGACACACCAGAAGAGGGCGTCA | | |
| 2704 | X | 54765139 | 54765250 | 111 | CAST.MOLF.SPRET | X | 54765241 | 54765259 | 19 | + | 16.8689 | 9.85E-07 | 0.291 | TAGCCACCAGATGTAGCCG | | |
| 2705 | X | 55426554 | 55426743 | 189 | SPRET | X | 55426597 | 55426630 | 34 | + | 16.4472 | 9.61E-07 | 0.182 | TAGCTGTCTTCAGACACACCAGAAGAGGGCGTCA | | |
| 2706 | X | 56681924 | 56682363 | 439 | SPRET | X | 56682073 | 56682106 | 34 | + | 17.6992 | 4.30E-07 | 0.134 | TAGCTGTTTTCAGACACACCAGAAGAGGGCATCA | | |
| 2707 | X | 62977341 | 62977544 | 203 | SPRET | X | 62977403 | 62977436 | 34 | + | 17.9756 | 3.59E-07 | 0.123 | TAGCTGTCTTTAGACACTCCAGAAGAGGGCGCCA | 1 | ENCFF367UNR |
| 2707 | X | 62977341 | 62977544 | 203 | SPRET | X | 62977418 | 62977436 | 19 | + | 17.5574 | 6.32E-07 | 0.241 | ACTCCAGAAGAGGGCGCCA | 1 | ENCFF367UNR |

| | | | | | | | | | | | | | | | | |
|---|---|---|---|---|---|---|---|---|---|---|---|---|---|---|---|---|
| 2708 | X | 74782900 | 74783868 | 968 | B10D2,B10J,BALB,BUB,CAST,CBA,CE,DBA1J,DBA2J,KK,LP,MA/My,MRL,PL,RF,SMJ | X | 74783752 | 74783785 | 34 | + | 16.748 | 7.95E-07 | 0.174 | CTGTAGGGCAGCTGCCAGCCACTGGAGGGCCTGG | | |
| 2709 | X | 74802066 | 74803034 | 968 | MOLF | X | 74802180 | 74802213 | 34 | + | 16.748 | 7.95E-07 | 0.174 | CTGTAGGGCAGCTGCCAGCCACTGGAGGGCCTGG | | |
| 2710 | X | 76438855 | 76439319 | 464 | CAST.CE.MOLF.SPRET | X | 76438927 | 76438960 | 34 | + | 17.8455 | 3.91E-07 | 0.127 | TAGCTGTCCTCAGACACTCCAGAAGAGGGCGTCA | | |
| 2711 | X | 87803042 | 87803926 | 884 | CAST.SPRET | X | 87803747 | 87803780 | 34 | + | 16.439 | 9.66E-07 | 0.182 | GAGCAGCAGCTTTGGCATCCAGAAGGTGGTCCTG | | |
| 2712 | X | 91050828 | 91051026 | 198 | SPRET | X | 91050881 | 91050915 | 35 | + | 18.1284 | 2.79E-07 | 0.156 | CTGTAGCTGTCTTCAGACACCAGAAGAGGGCATCA | | |
| 2713 | X | 93289435 | 93289806 | 371 | SPRET | X | 93289495 | 93289528 | 34 | + | 16.4472 | 9.61E-07 | 0.182 | TAGCTGTCTTCAGACACACCAGAAGAGGGCGTCA | | |
| 2714 | X | 96338290 | 96338932 | 642 | AJ,AKR,BALB,C3H,CAST,CBA,CE,FVB,KK,MA/My,MOLF,MRL,NZW,RF,RH2,SEA,SMJ,SPRET.SWR.WSB | X | 96338859 | 96338892 | 34 | + | 18.3333 | 2.82E-07 | 0.111 | CTGCATTGCCTTTGGAATTCAGTAGAGGGCTTCA | | |
| 2715 | X | 101453984 | 101454188 | 204 | SPRET | X | 101454044 | 101454077 | 34 | + | 16.4472 | 9.61E-07 | 0.182 | TAGCTGTCTTCAGACACACCAGAAGAGGGCGTCA | | |
| 2716 | X | 101776708 | 101777266 | 558 | CE.SPRET | X | 101777059 | 101777077 | 19 | + | 16.8689 | 9.85E-07 | 0.291 | AGGCCACTAGAGAGCGCCA | | |
| 2717 | X | 104328080 | 104328278 | 198 | CAST.MOLF.SPRET | X | 104328125 | 104328158 | 34 | + | 18.0732 | 3.36E-07 | 0.119 | TAGCTGTCTTCAGACACCCCAGAAGAGGGCGCCA | | |
| 2718 | X | 105164505 | 105164710 | 205 | 129S1,AJ,BALB,C3H,C57L,CAST,CBA,CE,FVB,ILNJ2,KK,MOLF,MRL,NZB,NZO,NZW,RH2,SJL,SPRET,SWR | X | 105164568 | 105164601 | 34 | + | 18.3659 | 2.76E-07 | 0.111 | TAGCTGTCTTCAGACACTCCAGAAGAGGGCGCCA | 1 | ENCFF093ZRU |
| 2718 | X | 105164505 | 105164710 | 205 | 129S1,AJ,BALB,C3H,C57L,CAST,CBA,CE,FVB,ILNJ2,KK,MOLF,MRL,NZB,NZO,NZW,RH2,SJL,SPRET,SWR | X | 105164583 | 105164601 | 19 | + | 17.5574 | 6.32E-07 | 0.241 | ACTCCAGAAGAGGGCGCCA | 1 | ENCFF093ZRU |
| 2719 | X | 106503562 | 106503659 | 97 | SPRET | X | 106503586 | 106503619 | 34 | + | 18.4959 | 2.53E-07 | 0.11 | ATGCAGCTAGAGAAACGAGCTCCAGGGGGTACTG | | |
| 2720 | X | 109172221 | 109172775 | 554 | CE.MOLF.SPRET | X | 109172655 | 109172688 | 34 | + | 16.5691 | 8.90E-07 | 0.182 | TAGCTGTCTTCAGACGCACCAGAAGAGGGCATCG | | |
| 2721 | X | 111661155 | 111661334 | 179 | SPRET | X | 111661192 | 111661225 | 34 | + | 16.9512 | 6.99E-07 | 0.164 | TAGCTGTCCTCAGATACCCCAGAAGAGGGCATCA | | |
| 2722 | X | 128660334 | 128660523 | 189 | MOLF.SPRET.WSB | X | 128660380 | 128660413 | 34 | + | 17.2439 | 5.79E-07 | 0.156 | TAGCTGTCCTCAGATACTCCAGAAGAGGGCATCA | | |
| 2723 | X | 128876138 | 128876397 | 259 | MOLF.SPRET | X | 128876255 | 128876288 | 34 | + | 17.9431 | 3.66E-07 | 0.124 | CAGCTGTCCTCAGACACTCCAGAAGAGGGAGTCA | | |
| 2724 | X | 133415951 | 133416155 | 204 | MOLF.SPRET | X | 133416013 | 133416046 | 34 | + | 17.878 | 3.83E-07 | 0.126 | TAGCTGTCCTCAGACACACCAGAAGAGGGCATCA | | |
| 2725 | X | 135108294 | 135108498 | 204 | 129S1,AKR,BTBR,BUB,C3H,C57L,C58J,CAST,CBA,CE,DBA1J,DBA2J,LP,MOLF,MRL,NOD,NOR,NZB,NZO,NZW,PJ,PL,RF,SEA,SJL,SMJ,SPRET,SWR | X | 135108356 | 135108389 | 34 | + | 16.4634 | 9.51E-07 | 0.182 | TAGCTGTCTTCAGACGCACCAGAAGAGGGCATCA | | |
| 2726 | X | 136038049 | 136038257 | 208 | BTBR,BUB,C3H,C58J,CAST,CBA,CE,DBA1J,DBA2J,LP,MOLF,MRL,NZB,NZO,NZW,PJ,PL,SEA,SJL,SMJ,SPRET,SWR | X | 136038103 | 136038136 | 34 | + | 17.8455 | 3.91E-07 | 0.127 | TAGCTGTCCTCAGACACTCCAGAAGAGGGCGTCA | | |
| 2727 | X | 143071882 | 143072078 | 196 | 129S1,AJ,BALB,C57L,CAST,FVB,KK,MA/My,MOLF,MRL,NOD,NOR,NZB,NZW,RH2,SEA,SJL,SPRET,TH,WSB | X | 143071934 | 143071967 | 34 | + | 16.6911 | 8.24E-07 | 0.177 | TAGCTGTCTTCAGACAGACCAGAAGAGGGCATCA | | |
| 2728 | X | 150358184 | 150358784 | 600 | SPRET | X | 150358335 | 150358368 | 34 | + | 24.1301 | 3.53E-09 | 0.00689 | ATGCAGTGCCTAAAGAGACCAGAAGAGGGAGTCA | 7 | ENCFF821BIA,ENCFF093ZRU,ENCFF367UNR,ENCFF065ODV,ENCFF055HAP,ENCFF099QNE,ENCFF502WDT |
| 2729 | X | 151885066 | 151885286 | 220 | 129S1.C57L.KK.MOLF.SPRET.SWR | X | 151885135 | 151885168 | 34 | + | 16.4634 | 9.51E-07 | 0.182 | TAGCTGTCTTCAGACGCACCAGAAGAGGGCATCA | | |
| 2730 | X | 155252797 | 155253784 | 987 | SPRET | X | 155253512 | 155253545 | 34 | + | 16.4146 | 9.81E-07 | 0.183 | TAGCTGTCTTCTGACAGTCCAGAAGAGGGAGCCA | | |
| 2731 | X | 155253451 | 155253654 | 203 | CAST.MOLF | X | 155253512 | 155253545 | 34 | + | 16.4146 | 9.81E-07 | 0.183 | TAGCTGTCTTCTGACAGTCCAGAAGAGGGAGCCA | | |
| 2732 | X | 158141028 | 158141230 | 202 | MOLF.SPRET | X | 158141090 | 158141123 | 34 | + | 17.2439 | 5.79E-07 | 0.156 | TAGCTGTCCTCAGATACTCCAGAAGAGGGCATCA | | |
| 2733 | X | 161382128 | 161382445 | 317 | CAST.MOLF.SPRET.TH | X | 161382290 | 161382323 | 34 | + | 19.8455 | 9.92E-08 | 0.0627 | TAGCTGTTTTCAGACGCGCCAGAAGAGGGCGTCA | | |
| 2734 | X | 161635270 | 161635501 | 231 | AKR,BALB,CAST,MOLF,RF,SEA,SPRET,TH,WSB | X | 161635340 | 161635374 | 35 | + | 21.2752 | 3.74E-08 | 0.0444 | CTGTAGCTGTCTTCAGACACCAGAAGAGGGCGCCA | 1 | ENCFF093ZRU |
| 2734 | X | 161635270 | 161635501 | 231 | AKR,BALB,CAST,MOLF,RF,SEA,SPRET,TH,WSB | X | 161635356 | 161635374 | 19 | + | 18.623 | 3.06E-07 | 0.176 | ACACCAGAAGAGGGCGCCA | 1 | ENCFF093ZRU |
| 2735 | X | 7366293 | 7366733 | 440 | MOLF | X | 7366733 | 7366766 | 34 | - | 19.1382 | 1.63E-07 | 0.0885 | CTGTAGCTGTCAGACACCAGAAGAGGGCATCA | | |
| 2736 | X | 7891140 | 7891289 | 149 | SPRET | X | 7891222 | 7891240 | 19 | - | 16.9672 | 9.26E-07 | 0.287 | CTGCCAGAAGAGGGCATCA | 5 | ENCFF367UNR.ENCFF093ZRU.ENCFF065ODV.ENCFF055HAP.ENCFF502WDT |
| 2736 | X | 7891140 | 7891289 | 149 | SPRET | X | 7891222 | 7891256 | 35 | - | 23.8257 | 6.08E-09 | 0.0156 | CTGCATATACATCTACCTGCCAGAAGAGGGCATCA | 5 | ENCFF367UNR.ENCFF093ZRU.ENCFF065ODV.ENCFF055HAP.ENCFF502WDT |
| 2737 | X | 12187259 | 12187464 | 205 | SPRET | X | 12187384 | 12187417 | 34 | - | 16.4472 | 9.61E-07 | 0.182 | TAGCTGTCTTCAGACACACCAGAAGAGGGCGTCA | | |
| 2738 | X | 12729679 | 12729892 | 213 | CAST.MOLF.SPRET | X | 12729805 | 12729823 | 19 | - | 17.5574 | 6.32E-07 | 0.241 | ACTCCAGAAGAGGGCGCCA | 3 | ENCFF821BIA.ENCFF093ZRU.ENCFF367UNR |
| 2738 | X | 12729679 | 12729892 | 213 | CAST.MOLF.SPRET | X | 12729805 | 12729838 | 34 | - | 18.3659 | 2.76E-07 | 0.111 | TAGCTGTCTTCAGACACTCCAGAAGAGGGCGCCA | 3 | ENCFF821BIA.ENCFF093ZRU.ENCFF367UNR |
| 2739 | X | 15978321 | 15979209 | 888 | SPRET | X | 15978339 | 15978372 | 34 | - | 16.9593 | 6.95E-07 | 0.164 | TTGCTCTCTTCAGACACACCAGAAGAGGGCATCA | | |
| 2740 | X | 19525742 | 19526546 | 804 | SPRET | X | 19526143 | 19526176 | 34 | - | 18.935 | 1.88E-07 | 0.0965 | AAGCACTTCCCTGGGGCCACTCTAGATGGCAGCC | | |
| 2741 | X | 20268569 | 20268774 | 205 | CAST.MOLF.SPRET | X | 20268696 | 20268729 | 34 | - | 16.4472 | 9.61E-07 | 0.182 | TAGCTGTCTTCAGACACACCAGAAGAGGGCGTCA | | |
| 2742 | X | 41947360 | 41947718 | 358 | SPRET | X | 41947461 | 41947494 | 34 | - | 16.6341 | 8.54E-07 | 0.18 | ATGCAGCTAGAAACACAAGCTCCAGGGGGTACTG | | |
| 2743 | X | 47567900 | 47568109 | 209 | MOLF.SPRET | X | 47568031 | 47568049 | 19 | - | 17.8852 | 5.08E-07 | 0.222 | ACACCAGAAGAGGGCACCA | 2 | ENCFF093ZRU.ENCFF367UNR |
| 2743 | X | 47567900 | 47568109 | 209 | MOLF.SPRET | X | 47568031 | 47568064 | 34 | - | 22.5935 | 1.25E-08 | 0.0172 | TTGCTGTTTTCAGACACACCAGAAGAGGGCACCA | 2 | ENCFF093ZRU.ENCFF367UNR |
| 2744 | X | 51761546 | 51761743 | 197 | CAST.ILNJ2.MOLF.SPRET.WSB | X | 51761668 | 51761701 | 34 | - | 17.5935 | 4.61E-07 | 0.138 | TAGCTGTCTCCAGACACTCCAGAAGAGGGCGTCA | | |
| 2745 | X | 53293797 | 53294129 | 332 | CAST.MOLF.SPRET | X | 53294088 | 53294106 | 19 | - | 19.2951 | 1.88E-07 | 0.139 | CCGCCACTAGATGGTGCTA | | |
| 2746 | X | 53992597 | 53992928 | 331 | SPRET | X | 53992886 | 53992904 | 19 | - | 19.2951 | 1.88E-07 | 0.139 | CCGCCACTAGATGGTGCTA | | |
| 2747 | X | 65768065 | 65768258 | 193 | CAST.MOLF.SPRET | X | 65768180 | 65768213 | 34 | - | 16.4472 | 0.000000961 | 0.182 | TAGCTGTCTTCAGACACACCAGAAGAGGGCGTCA | | |
| 2748 | X | 71631910 | 71632159 | 249 | CAST.ILNJ2.MOLF.WSB | X | 71632036 | 71632069 | 34 | - | 16.4634 | 0.000000951 | 0.182 | TAGCTGTCTTCAGACGCACCAGAAGAGGGCATCA | | |
| 2749 | X | 72746568 | 72746689 | 121 | SPRET | X | 72746545 | 72746578 | 34 | - | 18.3984 | 0.00000027 | 0.111 | TAGCTGTCTTCAGACACACCAGAAGAGGGCACCA | | |
| 2750 | X | 72981361 | 72981550 | 189 | CAST.CE.MOLF.SPRET.WSB | X | 72981474 | 72981507 | 34 | - | 16.4472 | 0.000000961 | 0.182 | TAGCTGTCTTCAGACACACCAGAAGAGGGCGTCA | | |
| 2751 | X | 74317317 | 74317519 | 202 | SPRET | X | 74317432 | 74317450 | 19 | - | 17.5574 | 0.000000632 | 0.241 | ACTCCAGAAGAGGGCGCCA | | |
| 2751 | X | 74317317 | 74317519 | 202 | SPRET | X | 74317432 | 74317465 | 34 | - | 18.3659 | 0.000000276 | 0.111 | TAGCTGTCTTCAGACACTCCAGAAGAGGGCGCCA | | |
| 2752 | X | 87877879 | 87878092 | 213 | CAST.MOLF.SPRET | X | 87878004 | 87878022 | 19 | - | 17.6557 | 0.000000592 | 0.238 | ATTCCAGAAGAGGGCGCTA | | |
| 2752 | X | 87877879 | 87878092 | 213 | CAST.MOLF.SPRET | X | 87878004 | 87878037 | 34 | - | 18.122 | 0.000000325 | 0.119 | TAGCTGTCTTCAGTCATTCCAGAAGAGGGCGCTA | | |
| 2753 | X | 88321528 | 88321834 | 306 | MOLF | X | 88321532 | 88321566 | 35 | - | 16.2661 | 0.000000823 | 0.273 | CTGCATCTACTCCTCTCCTCCAGGAGGAGCAACCC | | |
| 2754 | X | 89642925 | 89643924 | 999 | SPRET | X | 89642938 | 89642971 | 34 | - | 16.9512 | 0.000000699 | 0.164 | TTGCTGTTCTCAGGTACACCGGAAGAGGGTATCA | | |
| 2755 | X | 90385404 | 90385921 | 517 | CAST.MOLF.SPRET | X | 90385518 | 90385552 | 35 | - | 18.1284 | 0.000000279 | 0.156 | CTGTAGCTGTCTTCAGACACCAGAAGAGGGCATCA | | |
| 2756 | X | 93173027 | 93173303 | 276 | CAST.CE.MA/Mv.MOLF.SPRET.WSB | X | 93173211 | 93173229 | 19 | - | 17.5574 | 0.000000632 | 0.241 | ACTCCAGAAGAGGGCGCCA | | |
| 2756 | X | 93173027 | 93173303 | 276 | CAST.CE.MA/Mv.MOLF.SPRET.WSB | X | 93173211 | 93173244 | 34 | - | 18.3659 | 0.000000276 | 0.111 | TAGCTGTCTTCAGACACTCCAGAAGAGGGCGCCA | | |
| 2757 | X | 94909554 | 94909779 | 225 | CAST.MA/Mv.MOLF.SPRET | X | 94909677 | 94909710 | 34 | - | 16.4634 | 0.000000951 | 0.182 | TAGCTGTCTTCAGACGCACCAGAAGAGGGCATCA | | |
| 2758 | X | 97749616 | 97750081 | 465 | SPRET | X | 97749993 | 97750026 | 34 | - | 17.878 | 0.000000383 | 0.126 | TAGCTGTCCTCAGACACACCAGAAGAGGGCATCA | | |
| 2759 | X | 97939269 | 97939473 | 204 | AJ,AKR,BALB,C3H,CAST,CBA,FVB,KK,MA/My,MOLF,MRL,NZW,RF,RH2,SEA,SMJ,SPRET.SWR.TH | X | 97939392 | 97939425 | 34 | - | 16.4472 | 0.000000961 | 0.182 | TAGCTGTCTTCAGACACACCAGAAGAGGGCGTCA | | |
| 2760 | X | 100868015 | 100868908 | 893 | SPRET | X | 100868206 | 100868224 | 19 | - | 18.5574 | 0.00000032 | 0.18 | CCTCCAGTAGAGGGCGCAC | | |
| 2760 | X | 100868015 | 100868908 | 893 | SPRET | X | 100868206 | 100868240 | 35 | - | 19.6606 | 0.000000108 | 0.0892 | ATGAGGTTTCTTCACCCCTCCAGTAGAGGGCGCAC | | |
| 2761 | X | 101269240 | 101269422 | 182 | SPRET | X | 101269396 | 101269429 | 34 | - | 16.4634 | 0.000000951 | 0.182 | TAGCTGTCTTCAGACGCACCAGAAGAGGGCATCA | | |
| 2762 | X | 105945120 | 105945714 | 594 | MOLF | X | 105945237 | 105945255 | 19 | - | 17.5738 | 0.000000625 | 0.241 | TGGCCTGTAGGTGGTGGCG | | |
| 2763 | X | 110212741 | 110212951 | 210 | CAST.SPRET | X | 110212865 | 110212898 | 34 | - | 17.8618 | 0.000000387 | 0.126 | TAGCTGTCTTCAGACCCACCAGAAGAGGGCATCA | | |
| 2764 | X | 116151445 | 116152048 | 603 | SPRET | X | 116151803 | 116151837 | 35 | - | 17.9541 | 0.000000309 | 0.162 | CTGCATCTCTTTGCATCAGCCACCAGGTGAATCCT | | |
| 2765 | X | 134735477 | 134735674 | 197 | SPRET | X | 134735602 | 134735635 | 34 | - | 17.6667 | 0.00000044 | 0.135 | TAGCTGTCTTCAGACACGCCAGAAGAGGGCGTCA | | |
| 2766 | X | 142779858 | 142780354 | 496 | B10J | X | 142779945 | 142779963 | 19 | - | 19.6066 | 0.000000149 | 0.12 | GGGCCACAAGATGGCAGCA | 16 | ENCFF093ZRU,ENCFF092NMJ,ENCFF099QNE,ENCFF165LXO,ENCFF198SSR,ENCFF367UNR,ENCFF883UPM,ENCFF055HAP,ENCFF609MDN,ENCFF821BIA,ENCFF089GBC,ENCFF907XEG,ENCFF024JOZ,ENCFF430PPJ,ENCFF281QMV,ENCFF327LYS |
| 2767 | X | 143026541 | 143026747 | 206 | SPRET | X | 143026667 | 143026700 | 34 | - | 17.0813 | 0.000000643 | 0.162 | TAGCTGTTCTCAGACACTCCAGAAGAGGGAGTCA | | |
| 2768 | X | 143271716 | 143271927 | 211 | SPRET | X | 143271831 | 143271864 | 34 | - | 17.9106 | 0.000000374 | 0.125 | TAGCTGTCTTCAGACACTCCAGAAGAGGGCACCA | | |
| 2769 | X | 145464952 | 145465181 | 229 | CAST.MOLF.NZB | X | 145465081 | 145465114 | 34 | - | 17.8455 | 0.000000391 | 0.127 | TAGCTGTCCTCAGACACTCCAGAAGAGGGCGTCA | | |
| 2770 | X | 150096825 | 150097049 | 224 | MOLF | X | 150096933 | 150096951 | 19 | - | 20.2459 | 9.11E-08 | 0.0913 | AGGCCAGGAGAGGGCGCTA | 9 | ENCFF093ZRU,ENCFF136ZDK,ENCFF052KVO,ENCFF589ZBJ,ENCFF883UPM,ENCFF065ODV,ENCFF502WDT,ENCFF142CNG,ENCFF055HAP |
| 2770 | X | 150096825 | 150097049 | 224 | MOLF | X | 150096933 | 150096966 | 34 | - | 19.0244 | 0.000000176 | 0.0925 | ATGTATTATTCATGGAGGCCAGGAGAGGGCGCTA | 9 | ENCFF093ZRU,ENCFF136ZDK,ENCFF052KVO,ENCFF589ZBJ,ENCFF883UPM,ENCFF065ODV,ENCFF502WDT,ENCFF142CNG,ENCFF055HAP |
| 2771 | X | 150741935 | 150742143 | 208 | SPRET | X | 150742055 | 150742088 | 34 | - | 17.2114 | 0.000000591 | 0.157 | TAGCTGTCTTCAGACACGCCAGAAGAGGGCATCA | | |
| 2772 | X | 161500163 | 161500375 | 212 | MOLF.SPRET.TH | X | 161500288 | 161500321 | 34 | - | 16.4634 | 0.000000951 | 0.182 | TAGCTGTCTTCAGACGCACCAGAAGAGGGCATCA | | |
| 2773 | X | 161503738 | 161503925 | 187 | SPRET | X | 161503852 | 161503885 | 34 | - | 16.4472 | 0.000000961 | 0.182 | TAGCTGTCTTCAGACACACCAGAAGAGGGCGTCA | | |
| 2774 | X | 161586049 | 161587035 | 986 | SPRET | X | 161586425 | 161586459 | 35 | - | 21.2844 | 3.72E-08 | 0.0443 | CTGCATATACACCTACATGCCAGAAGAGGGTATCA | | |
| 2775 | X | 162748235 | 162748354 | 119 | MOLF.TH | X | 162748252 | 162748270 | 19 | - | 17.9672 | 0.000000481 | 0.219 | GCACCAGAAGAGGGCACCA | 1 | ENCFF093ZRU |
| 2775 | X | 162748235 | 162748354 | 119 | MOLF.TH | X | 162748252 | 162748285 | 34 | - | 17.0894 | 0.000000639 | 0.161 | CGGTTGTCTTCAGACGCACCAGAAGAGGGCACCA | 1 | ENCFF093ZRU |
| 2776 | X | 168762246 | 168762498 | 252 | SPRET | X | 168762286 | 168762304 | 19 | - | 16.8852 | 0.000000976 | 0.291 | TCACCACCGGGTGGCGACA | 2 | ENCFF055HAP.ENCFF502WDT |
| 2776 | X | 168762246 | 168762498 | 252 | SPRET | X | 168762286 | 168762319 | 34 | - | 21.374 | 3.23E-08 | 0.0307 | CCGCATTCTCCTCTGTCACCACCGGGTGGCGACA | 2 | ENCFF055HAP.ENCFF502WDT |
| 2777 | X | 168803157 | 168803854 | 697 | 129S1,C3H,C57L,DBA1J,MA/My,MOLF,PL,SMJ | X | 168803139 | 168803157 | 19 | - | 21.4918 | 0.000000032 | 0.0593 | TGGCCACTGGGTGGCGCCA | 22 | ENCFF367UNR,ENCFF099QNE,ENCFF805QBU,ENCFF327LYS,ENCFF907XEG,ENCFF609MDN,ENCFF024JOZ,ENCFF316FYN,ENCFF089GBC,ENCFF589ZBJ,ENCFF430PPJ,ENCFF165LXO,ENCFF136ZDK,ENCFF198SSR,ENCFF092NMJ,ENCFF502WDT,ENCFF065ODV,ENCFF321AGQ,ENCFF052KVO,ENCFF055HAP,ENCFF210BTO.ENCFF281QMV |
| 2777 | X | 168803157 | 168803854 | 697 | 129S1,C3H,C57L,DBA1J,MA/My,MOLF,PL,SMJ | X | 168803139 | 168803173 | 35 | - | 30.8073 | 1.23E-11 | 0.000675 | TTGCAGTGACGTCCTGTGGCCACTGGGTGGCGCCA | 22 | ENCFF367UNR,ENCFF099QNE,ENCFF805QBU,ENCFF327LYS,ENCFF907XEG,ENCFF609MDN,ENCFF024JOZ,ENCFF316FYN,ENCFF089GBC,ENCFF589ZBJ,ENCFF430PPJ,ENCFF165LXO,ENCFF136ZDK,ENCFF198SSR,ENCFF092NMJ,ENCFF502WDT,ENCFF065ODV,ENCFF321AGQ,ENCFF052KVO,ENCFF055HAP,ENCFF210BTO,ENCFF281QMV |
| 2777 | X | 168803157 | 168803854 | 697 | 129S1,C3H,C57L,DBA1J,MA/My,MOLF,PL,SMJ | X | 168803837 | 168803855 | 19 | - | 21.4918 | 0.000000032 | 0.0593 | TGGCCACTGGGTGGCGCCA | 22 | ENCFF367UNR,ENCFF821BIA,ENCFF327LYS,ENCFF024JOZ,ENCFF907XEG,ENCFF589ZBJ,ENCFF430PPJ,ENCFF136ZDK,ENCFF089GBC,ENCFF092NMJ,ENCFF165LXO,ENCFF210BTO,ENCFF198SSR,ENCFF052KVO,ENCFF805QBU,ENCFF086ZTV,ENCFF281QMV,ENCFF609MDN,ENCFF089EWX,ENCFF033UQX,ENCFF316FYN,ENCFF097CAO |
| 2777 | X | 168803157 | 168803854 | 697 | 129S1,C3H,C57L,DBA1J,MA/My,MOLF,PL,SMJ | X | 168803837 | 168803871 | 35 | - | 29.6697 | 3.96E-11 | 0.00118 | TTGCACTGACGTCCTCTGGCCACTGGGTGGCGCCA | 22 | ENCFF367UNR,ENCFF821BIA,ENCFF327LYS,ENCFF024JOZ,ENCFF907XEG,ENCFF589ZBJ,ENCFF430PPJ,ENCFF136ZDK,ENCFF089GBC,ENCFF092NMJ,ENCFF165LXO,ENCFF210BTO,ENCFF198SSR,ENCFF052KVO,ENCFF805QBU,ENCFF086ZTV,ENCFF281QMV,ENCFF609MDN,ENCFF089EWX,ENCFF033UQX,ENCFF316FYN,ENCFF097CAO |

| DEL_NUM | DEL_CHR | DEL_START | DEL_END | DEL_LENGTH | STRAINS with VARIANT SV ALLELE | CTCF CHR | CTCF START | CTCF END | CTCF LENGTH | CTCF STRAND | CTCF SCORE | CTCF PVALUE | CTCF QVALUE | CTCF SEQUENCE | # PEAKS | DATA SOURCE |
|---|---|---|---|---|---|---|---|---|---|---|---|---|---|---|---|---|
| 1 | 1 | 6574639 | 6578351 | 3712 | C3H,CBA,MRL,RF | 1 | 6577912 | 6577945 | 34 | + | 16.813 | 7.63E-07 | 0.115 | ATGCAGTTAGAGACACGAGCTCCGGAGGGTACTG | | |
| 2 | 1 | 7247898 | 7250015 | 2117 | CAST,SPRET | 1 | 7249036 | 7249069 | 34 | + | 17.2114 | 5.91E-07 | 0.101 | GAGCAACTCCTTCACAAACCAGGTGATGGCACAG | | |
| 3 | 1 | 7992966 | 8010088 | 17122 | DBA1J,DBA2J | 1 | 7999113 | 7999146 | 34 | + | 18.7154 | 2.18E-07 | 0.0621 | CTGCAACACCCCCAAGGTCCCCTAGAGGACTCTC | | |
| 4 | 1 | 7999065 | 8000377 | 1312 | AJ,BUB,C3H,CBA,CE,FVB,MRL,NZB,NZW,RF | 1 | 7999113 | 7999146 | 34 | + | 18.7154 | 2.18E-07 | 0.0621 | CTGCAACACCCCCAAGGTCCCCTAGAGGACTCTC | | |
| 5 | 1 | 7999067 | 8006330 | 7263 | 129S1,AJ,AKR,BALB,BTBR,BUB,C3H,C57L,CBA,FVB,ILNJ2,KK,LP,MA/My,MOLF,NOD,NOR,NZO,PJ,PL,RF,RH2,SEA,SJL,SMJ,SWR,TH | 1 | 7999113 | 7999146 | 34 | + | 18.7154 | 2.18E-07 | 0.0621 | CTGCAACACCCCCAAGGTCCCCTAGAGGACTCTC | | |
| 6 | 1 | 8604799 | 8606083 | 1284 | CAST | 1 | 8605461 | 8605494 | 34 | + | 18.5041 | 2.52E-07 | 0.0667 | CTGCTGCTGTGAATGAGACCACCTGGGGGTGGAG | | |
| 6 | 1 | 8604799 | 8606083 | 1284 | CAST | 1 | 8605464 | 8605497 | 34 | + | 17.7317 | 4.21E-07 | 0.0858 | CTGCTGTGAATGAGACCACCTGGGGGTGGAGCCC | | |
| 7 | 1 | 9103386 | 9105463 | 2077 | C3H,CBA,MRL,RF | 1 | 9103973 | 9104006 | 34 | + | 16.5447 | 9.04E-07 | 0.124 | CAGCTGTCTTCGGACACACCAAAAGAGGGCATCA | | |
| 8 | 1 | 15827886 | 15830971 | 3085 | SPRET | 1 | 15828084 | 15828118 | 35 | + | 17.3853 | 4.33E-07 | 0.137 | CTACTGTGTCACATGTGCGTCAGTAGGTGGCAGAG | | |
| 9 | 1 | 17535831 | 17550810 | 14979 | SPRET | 1 | 17535860 | 17535893 | 34 | + | 17.439 | 5.10E-07 | 0.0938 | CTGCACTCCAACCCATAGGCTGTGGAAGGCAGTA | | |
| 10 | 1 | 18023067 | 18036879 | 13812 | CAST,WSB | 1 | 18033809 | 18033842 | 34 | + | 19.3415 | 1.42E-07 | 0.0472 | CTGCCCCCACATTCGCCGTCACAAGATGGCGCTG | | |
| 11 | 1 | 18885174 | 18891345 | 6171 | SPRET | 1 | 18885249 | 18885282 | 34 | + | 16.7724 | 7.83E-07 | 0.115 | TTGAGGTTTCTGCCTCAGGCTCCAGAGGGAGCCA | | |
| 12 | 1 | 20642679 | 20644237 | 1558 | CAST | 1 | 20642873 | 20642906 | 34 | + | 21.3415 | 3.31E-08 | 0.0184 | GTGCCGTGCTTACAGAGGCCAGGAGATGACACCA | | |
| 13 | 1 | 21434734 | 21449490 | 14756 | CE,MA/My,MRL,SMJ | 1 | 21439381 | 21439415 | 35 | + | 16.8165 | 6.02E-07 | 0.167 | CTGCATGTACAGCTGCCTGCGAGTAGAGGGCATAT | | |
| 13 | 1 | 21434734 | 21449490 | 14756 | CE,MA/My,MRL,SMJ | 1 | 21443544 | 21443578 | 35 | + | 21.0826 | 4.26E-08 | 0.0373 | CTGCTTGTATAGCTGCCTGCCAGAAGAGGGCATCT | | |
| 14 | 1 | 22218201 | 22219712 | 1511 | CAST | 1 | 22218426 | 22218459 | 34 | + | 16.9756 | 6.88E-07 | 0.109 | ATGCAGCTAGAGACACGAGCTCCAGGGGGTACGG | | |
| 15 | 1 | 23821397 | 23825569 | 4172 | AJ,BTBR,C3H,C57L,CAST,CBA,CE,DBA1J,DBA2J,FVB,ILNJ2,MOLF,MRL,NOD,NZB,NZO,NZW,PL,RF,RH2,SMJ,SPRET,SWR,WSB | 1 | 23822174 | 23822207 | 34 | + | 17.6748 | 4.37E-07 | 0.0865 | GAGCATGACCACAAGTCCACAGCAGGTGGCAACG | | |
| 16 | 1 | 24840799 | 24851399 | 10600 | CAST,SPRET | 1 | 24843802 | 24843835 | 34 | + | 17.5691 | 4.69E-07 | 0.0895 | GAGCATGACCACAAGTCCACAGCAGGTGGCAACA | | |
| 17 | 1 | 24842762 | 24846950 | 4188 | MOLF | 1 | 24843802 | 24843835 | 34 | + | 17.5691 | 4.69E-07 | 0.0895 | GAGCATGACCACAAGTCCACAGCAGGTGGCAACA | | |
| 18 | 1 | 27767167 | 27782007 | 14840 | SPRET | 1 | 27771019 | 27771037 | 19 | + | 18.5738 | 3.17E-07 | 0.124 | AAACCACCAGGTGGCAGAA | | |
| 19 | 1 | 28066386 | 28105144 | 38758 | CAST | 1 | 28068862 | 28068895 | 34 | + | 18.1138 | 3.27E-07 | 0.0746 | GTCCAGTGCCTGTGGATACCAGAAAAGGGCACCA | | |
| 20 | 1 | 28067380 | 28069462 | 2082 | AKR,BALB,MOLF,PJ,SEA,SPRET,WSB | 1 | 28068862 | 28068895 | 34 | + | 18.1138 | 3.27E-07 | 0.0746 | GTCCAGTGCCTGTGGATACCAGAAAAGGGCACCA | | |
| 21 | 1 | 30998964 | 31007205 | 8241 | SPRET | 1 | 31003107 | 31003140 | 34 | + | 17.4472 | 5.08E-07 | 0.0936 | TTGCAATGCCACTGAAAGACTCCGGGAGGAGCCC | | |
| 22 | 1 | 31769616 | 31773761 | 4145 | BUB,CAST,CE,FVB,MOLF,NZB,NZO,PJ,RF,SJL,SPRET,SWR | 1 | 31770394 | 31770427 | 34 | + | 17.6748 | 4.37E-07 | 0.0865 | GAGCATGACCACAAGTCCACAGCAGGTGGCAACG | | |
| 23 | 1 | 34815500 | 34818186 | 2686 | CE,MOLF | 1 | 34815680 | 34815713 | 34 | + | 18.0081 | 3.51E-07 | 0.0774 | GCCTAGTACCCGAGGAAGCCAGAAGAGGGCGCTG | 1 | ENCFF093ZRU |
| 23 | 1 | 34815500 | 34818186 | 2686 | CE,MOLF | 1 | 34815695 | 34815713 | 19 | + | 19.2131 | 2.00E-07 | 0.0983 | AAGCCAGAAGAGGGCGCTG | 1 | ENCFF093ZRU |
| 24 | 1 | 36886530 | 36887601 | 1071 | AJ,AKR,BUB,C3H,CBA,CE,FVB,KK,MRL,NOD,NZB,NZO,NZW,PJ,PL,RF,RH2,SJL,SMJ,SWR,WSB | 1 | 36887287 | 36887320 | 34 | + | 24.9106 | 1.79E-09 | 0.00232 | ATGCAGTGTCCACAGAGCCCAGAAGAGGGCGTTA | 8 | ENCFF099QNE,ENCFF821BIA,ENCFF052KVO,ENCFF136ZDK,ENCFF502WDT,ENCFF093ZRU,ENCFF367UNR,ENCFF055HAP |
| 25 | 1 | 36950335 | 36954018 | 3683 | PJ,PL,RF,SJL | 1 | 36951496 | 36951529 | 34 | + | 19.7805 | 1.04E-07 | 0.0393 | GTGTGGTGCCTGCAGAGGCCAGATGAGGGCATTA | 6 | ENCFF367UNR,ENCFF093ZRU,ENCFF821BIA,ENCFF589ZBJ,ENCFF502WDT,ENCFF055HAP |
| 26 | 1 | 37086715 | 37087861 | 1146 | CAST | 1 | 37087517 | 37087550 | 34 | + | 18.3984 | 2.70E-07 | 0.069 | TAGCTGTCTTCAGACACACCAGAAGAGGGCACCA | | |
| 26 | 1 | 37086715 | 37087861 | 1146 | CAST | 1 | 37087532 | 37087550 | 19 | + | 17.8852 | 5.08E-07 | 0.151 | ACACCAGAAGAGGGCACCA | | |
| 27 | 1 | 48747500 | 48754530 | 7030 | AKR,BALB,BUB,C58J,CAST,MA/My,MOLF,RF,RH2,SEA,SPRET | 1 | 48747957 | 48747975 | 19 | + | 17.7213 | 5.67E-07 | 0.161 | CAAGCACCAGGGGGCACCC | 4 | ENCFF367UNR,ENCFF093ZRU,ENCFF099QNE,ENCFF821BIA |
| 28 | 1 | 48776956 | 48781128 | 4172 | AKR,BALB,BUB,C58J,CAST,MA/My,MOLF,RF,RH2,SEA,SPRET | 1 | 48777734 | 48777767 | 34 | + | 17.6748 | 4.37E-07 | 0.0865 | GAGCATGACCACAAGTCCACAGCAGGTGGCAACG | | |
| 29 | 1 | 49360189 | 49368561 | 8372 | CAST,MOLF,SPRET | 1 | 49362722 | 49362740 | 19 | + | 18.5738 | 3.17E-07 | 0.124 | AAACCACCAGGTGGCAGAA | | |
| 30 | 1 | 50332804 | 50336708 | 3904 | SPRET | 1 | 50336405 | 50336439 | 35 | + | 17.7064 | 3.58E-07 | 0.123 | GTGCATGGCCTGCGGGGGGACAGGAGAGGACGCTG | | |
| 31 | 1 | 53991907 | 54002673 | 10766 | SPRET | 1 | 53992850 | 53992883 | 34 | + | 16.4309 | 9.71E-07 | 0.125 | CTGCTATCATAAAAACATACAGCAGATGGTGGTG | | |
| 32 | 1 | 55314183 | 55321845 | 7662 | SPRET | 1 | 55320270 | 55320304 | 35 | + | 16.3578 | 7.81E-07 | 0.19 | ATGCTGGGTACCGTGTTGGCCATAAGAGGCCACCA | | |
| 33 | 1 | 60196947 | 60201458 | 4511 | CE,LP,MOLF | 1 | 60197809 | 60197827 | 19 | + | 17.5082 | 6.52E-07 | 0.168 | CACCCAGCAGAGGGCAGAA | 25 | ENCFF201JUA,ENCFF367UNR,ENCFF093ZRU,ENCFF821BIA,ENCFF430PPJ,ENCFF099QNE,ENCFF907XEG,ENCFF327LYS,ENCFF024JOZ,ENCFF883UPM,ENCFF089GBC,ENCFF589ZBJ,ENCFF951GOP,ENCFF575MGI,ENCFF198SSR,ENCFF194HVL,ENCFF055HAP,ENCFF491RJK,ENCFF502WDT,ENCFF957SPH,ENCFF142CNG,ENCFF092NMJ,ENCFF609MDN,ENCFF281QMV,ENCFF165LXO |
| 34 | 1 | 68937631 | 68938988 | 1357 | CE,SPRET,TH | 1 | 68937990 | 68938023 | 34 | + | 18.3089 | 2.87E-07 | 0.0691 | ATGCAGCTAGAGACACGAGCTCCAGGGGGTACTG | | |
| 35 | 1 | 70241038 | 70262271 | 21233 | PJ | 1 | 70259453 | 70259486 | 34 | + | 16.8943 | 7.24E-07 | 0.111 | TTGCTCTCTTCAGACATACCAGAAGAGGGCATCA | | |
| 36 | 1 | 74251880 | 74253617 | 1737 | CAST | 1 | 74252567 | 74252600 | 34 | + | 17.9268 | 3.70E-07 | 0.0802 | GTGCAATGACCACAGAGACCAAAAGAAGGCATTA | | |
| 37 | 1 | 79113477 | 79116897 | 3420 | LP,MOLF | 1 | 79116750 | 79116784 | 35 | + | 20.6147 | 5.82E-08 | 0.0449 | GTGCTCTGCATGGTGGAGGCCAGAAGAGGGCCTTG | | |
| 38 | 1 | 91845388 | 91847169 | 1781 | PL | 1 | 91845666 | 91845684 | 19 | + | 18.5082 | 3.31E-07 | 0.127 | TGACCACCAGGTGGGGCCA | 1 | ENCFF367UNR |
| 39 | 1 | 101014532 | 101023802 | 9270 | AKR,LP,MA/My,PJ,SEA | 1 | 101017578 | 101017611 | 34 | + | 17.5691 | 4.69E-07 | 0.0895 | GAGCATGACCACAAGTCCACAGCAGGTGGCAACA | | |
| 40 | 1 | 101017060 | 101021216 | 4156 | CAST,MOLF,SPRET | 1 | 101017578 | 101017611 | 34 | + | 17.5691 | 4.69E-07 | 0.0895 | GAGCATGACCACAAGTCCACAGCAGGTGGCAACA | | |
| 41 | 1 | 103322446 | 103337860 | 15414 | SPRET | 1 | 103326228 | 103326262 | 35 | + | 17.0459 | 5.27E-07 | 0.156 | CAGCAGTTTGTTCAAGTGACCATCAGCTGGCACTT | | |
| 42 | 1 | 111165717 | 111167805 | 2088 | SPRET | 1 | 111167523 | 111167556 | 34 | + | 16.4715 | 9.47E-07 | 0.125 | CAGCCTTTCCCTAATTCAACAACAGGGGTCAGCA | | |
| 43 | 1 | 111738581 | 111740196 | 1615 | AKR,DBA1J,DBA2J,FVB,KK,LP,MA/My,MRL,NZO,NZW,PJ,RF,SJL,SWR | 1 | 111739768 | 111739786 | 19 | + | 17.2295 | 7.83E-07 | 0.183 | CATGCACCAGGGGGCACTG | 1 | ENCFF093ZRU |
| 44 | 1 | 112367859 | 112372405 | 4546 | AKR,DBA1J,DBA2J,FVB,KK,LP,MA/My,MOLF,MRL,NOD,NOR,NZO,NZW,PJ,RF,SJL,SMJ,SWR | 1 | 112368636 | 112368669 | 34 | + | 17.6748 | 4.37E-07 | 0.0865 | GAGCATGACCACAAGTCCACAGCAGGTGGCAACG | | |
| 45 | 1 | 114659383 | 114667534 | 8151 | CAST,FVB,MOLF,SMJ,SPRET,SWR,WSB | 1 | 114665422 | 114665456 | 35 | + | 19.4312 | 1.25E-07 | 0.0674 | GTGCAGGGGAATGCCAGGGCCAGGAGGGGGGAGTG | | |
| 46 | 1 | 122917357 | 122922329 | 4972 | C57L | 1 | 122917534 | 122917552 | 19 | + | 17.8033 | 5.37E-07 | 0.156 | TGACCAGAAGAGGGCATCA | 1 | ENCFF093ZRU |
| 47 | 1 | 124366108 | 124390498 | 24390 | DBA1J,DBA2J,FVB,MA/My,NOD,NOR,NZB,PJ,SJL,TH | 1 | 124378041 | 124378074 | 34 | + | 21.3659 | 3.25E-08 | 0.0181 | ATGCAGTACCCAGAGAAACCAGATGAGGGCAACA | 12 | ENCFF093ZRU,ENCFF821BIA,ENCFF099QNE,ENCFF430PPJ,ENCFF092NMJ,ENCFF165LXO,ENCFF327LYS,ENCFF052KVO,ENCFF089GBC,ENCFF024JOZ,ENCFF367UNR,ENCFF907XEG |
| 48 | 1 | 125244494 | 125251709 | 7215 | AJ,AKR,C3H,CAST,CBA,CE,DBA1J,DBA2J,MOLF,MRL,NZO,RH2,SPRET,SWR | 1 | 125244480 | 125244514 | 35 | + | 18.578 | 2.12E-07 | 0.0938 | CTGCAGTGACATCTGTTTGCCTGTTGGGAGCCGCC | | |
| 49 | 1 | 127454751 | 127456902 | 2151 | 129S1,AJ,AKR,BALB,C3H,C57L,CE,DBA1J,DBA2J,FVB,ILNJ2,KK,MA/My,PJ,RH2,SEA,TH | 1 | 127455324 | 127455358 | 35 | + | 19.156 | 1.48E-07 | 0.0747 | GTGCAGATGACTGATGAGGCCAGAAGAGGGTGTCA | | |
| 50 | 1 | 127526402 | 127529760 | 3358 | CAST,MOLF | 1 | 127529256 | 127529290 | 35 | + | 18.1835 | 2.70E-07 | 0.108 | GTGCAGGTGTTCACAGAGGCCAAAAGAGGGTACTG | | |
| 51 | 1 | 128062643 | 128064163 | 1520 | SPRET | 1 | 128063354 | 128063387 | 34 | + | 20.0244 | 8.72E-08 | 0.035 | TTGCTGTCCTCAGAAACACCAGAAGAGGGCATTA | | |
| 52 | 1 | 131570298 | 131579356 | 9058 | 129S1,AKR,BALB,BTBR,BUB,CAST,CBA,DBA1J,DBA2J,FVB,ILNJ2,KK,MA/My,MOLF,MRL,NZO,NZW,PJ,PL,RF,SEA,SJL,SMJ,SPRET,SWR,TH,WSB | 1 | 131575768 | 131575786 | 19 | + | 17.1639 | 8.17E-07 | 0.187 | CCTCCACGAGGTGGCGGAA | | |
| 53 | 1 | 131572917 | 131579372 | 6455 | 129S1,AKR,BTBR,BUB,CBA,DBA1J,DBA2J,ILNJ2,MRL,PJ,PL,RF,SJL,SMJ,SWR,TH,WSB | 1 | 131575768 | 131575786 | 19 | + | 17.1639 | 8.17E-07 | 0.187 | CCTCCACGAGGTGGCGGAA | | |
| 54 | 1 | 133882433 | 133886545 | 4112 | SPRET | 1 | 133885556 | 133885589 | 34 | + | 17.1951 | 5.97E-07 | 0.101 | TTGCTATCTTCAGACACACCAGAAGAGGGCATCA | 1 | ENCFF093ZRU |
| 55 | 1 | 142907046 | 142908327 | 1281 | CAST,MOLF,MRL,SPRET | 1 | 142907269 | 142907302 | 34 | + | 18.0569 | 3.40E-07 | 0.0757 | ATGCAGCTAGAGACATGAGCTCCAGGGGGTACTG | | |
| 56 | 1 | 146600623 | 146605232 | 4609 | 129S1,AKR,BTBR,BUB,C57L,CAST,DBA1J,DBA2J,KK,MA/My,MOLF,MRL,NZB,NZO,NZW,RF,SJL,SMJ,SPRET,TH | 1 | 146603056 | 146603090 | 35 | + | 17.1009 | 5.11E-07 | 0.152 | CTGCACTGTGGCCGGACCATCAGCAGGGGCAAGAA | | |
| 57 | 1 | 151252149 | 151254800 | 2651 | AKR,BTBR,C57L,MOLF,SMJ,SPRET | 1 | 151253269 | 151253287 | 19 | + | 20.377 | 8.18E-08 | 0.0655 | AGGCCAGAAGAGGGCACTA | 10 | ENCFF099QNE,ENCFF821BIA,ENCFF024JOZ,ENCFF327LYS,ENCFF907XEG,ENCFF502WDT,ENCFF883UPM,ENCFF093ZRU,ENCFF055HAP,ENCFF367UNR |
| 58 | 1 | 151334144 | 151340269 | 6125 | WSB | 1 | 151339959 | 151339993 | 35 | + | 21.4954 | 3.22E-08 | 0.0314 | TTGCATATATGACTAAAAGCCAGAAGAGGGCACCA | 5 | ENCFF367UNR,ENCFF055HAP,ENCFF502WDT,ENCFF093ZRU,ENCFF099QNE |
| 58 | 1 | 151334144 | 151340269 | 6125 | WSB | 1 | 151339975 | 151339993 | 19 | + | 18.7705 | 2.75E-07 | 0.117 | AAGCCAGAAGAGGGCACCA | 5 | ENCFF367UNR,ENCFF055HAP,ENCFF502WDT,ENCFF093ZRU,ENCFF099QNE |
| 59 | 1 | 151830581 | 151833073 | 2492 | SPRET | 1 | 151831507 | 151831525 | 19 | + | 17.5738 | 6.25E-07 | 0.166 | TGGGCAGCAGGGGGAGGTG | | |
| 60 | 1 | 156292059 | 156298875 | 6816 | AKR | 1 | 156292961 | 156292995 | 35 | + | 16.5596 | 6.97E-07 | 0.181 | TAGCTGTCTTCAGACAGCACCAGAAGAGGGCGTCA | | |
| 60 | 1 | 156292059 | 156298875 | 6816 | AKR | 1 | 156293955 | 156293988 | 34 | + | 25.0569 | 1.57E-09 | 0.00213 | ATGCAGTGCTCTCAGAGGCCAGATGAGGGCAGCA | 4 | ENCFF093ZRU,ENCFF052KVO,ENCFF327LYS,ENCFF907XEG |
| 61 | 1 | 160239336 | 160240649 | 1313 | SPRET | 1 | 160240377 | 160240410 | 34 | + | 16.4634 | 9.51E-07 | 0.125 | TAGCTGTCTTCAGACGCACCAGAAGAGGGCATCA | | |
| 62 | 1 | 162825252 | 162827296 | 2044 | DBA1J,DBA2J | 1 | 162827211 | 162827244 | 34 | + | 17.4146 | 5.18E-07 | 0.0944 | ATGCAGCTAGAGACACAAGCTCCAGGGGGTACTG | | |
| 63 | 1 | 164051371 | 164057121 | 5750 | BTBR,BUB,CAST,DBA1J,DBA2J,FVB,ILNJ2,KK,MOLF,NOD,NOR,NZB,NZO,PJ,RF,SEA,SJL,SMJ,SPRET,SWR,WSB | 1 | 164055070 | 164055088 | 19 | + | 17.1639 | 8.17E-07 | 0.187 | CCTCCACGAGGTGGCGGAA | | |
| 64 | 1 | 165207611 | 165211704 | 4093 | CAST | 1 | 165209461 | 165209495 | 35 | + | 18.8073 | 1.84E-07 | 0.0873 | CAGCTGTCTTCAGATACACCCAGAAGAGGGCGTCA | | |
| 65 | 1 | 167166711 | 167170912 | 4201 | WSB | 1 | 167170599 | 167170617 | 19 | + | 19.2131 | 2.00E-07 | 0.0983 | AAGCCAGAAGAGGGCGCTG | 1 | ENCFF052KVO |
| 66 | 1 | 171139023 | 171144086 | 5063 | AJ,NZB,NZO,RH2,SJL,SMJ | 1 | 171140221 | 171140255 | 35 | + | 16.5596 | 6.97E-07 | 0.181 | AAGAAGTTACACCCTGTTGCCAGTAGGTGGTTTTC | | |
| 67 | 1 | 171139990 | 171142349 | 2359 | 129S1,BALB,BUB,C3H,C57L,CAST,CBA,CE,DBA1J,DBA2J,FVB,KK,MRL,NOD,NOR,NZW,PJ,PL,RF,SEA,SPRET,SWR,TH,WSB | 1 | 171140221 | 171140255 | 35 | + | 16.5596 | 6.97E-07 | 0.181 | AAGAAGTTACACCCTGTTGCCAGTAGGTGGTTTTC | | |
| 68 | 1 | 177573476 | 177576283 | 2807 | SPRET | 1 | 177574953 | 177574971 | 19 | + | 19.541 | 1.57E-07 | 0.0873 | CCCCCAGCAGGGGGCAGTC | | |
| 69 | 1 | 181223486 | 181226155 | 2669 | CAST,MOLF,TH | 1 | 181225852 | 181225886 | 35 | + | 18.0459 | 2.93E-07 | 0.111 | GAGCTGGTGCCTGTGGGGGCCAGAAGAGGGCATTA | | |
| 70 | 1 | 181975324 | 181984270 | 8946 | CAST | 1 | 181977698 | 181977731 | 34 | + | 26.6748 | 3.53E-10 | 0.000876 | ATGCAATACTCATGGCAGCCAGAAGAGGGCACTA | 26 | ENCFF367UNR,ENCFF201JUA,ENCFF093ZRU,ENCFF430PPJ,ENCFF589ZBJ,ENCFF052KVO,ENCFF055HAP,ENCFF024JOZ,ENCFF065ODV,ENCFF327LYS,ENCFF907XEG,ENCFF609MDN,ENCFF198SSR,ENCFF502WDT,ENCFF821BIA,ENCFF099QNE,ENCFF136ZDK,ENCFF165LXO,ENCFF321AGQ,ENCFF089GBC,ENCFF194HVL,ENCFF142CNG,ENCFF883UPM,ENCFF092NMJ,ENCFF957SPH,ENCFF951GOP |
| 70 | 1 | 181975324 | 181984270 | 8946 | CAST | 1 | 181977713 | 181977731 | 19 | + | 20.8033 | 5.79E-08 | 0.062 | CAGCCAGAAGAGGGCACTA | 26 | ENCFF367UNR,ENCFF201JUA,ENCFF093ZRU,ENCFF430PPJ,ENCFF589ZBJ,ENCFF052KVO,ENCFF055HAP,ENCFF024JOZ,ENCFF065ODV,ENCFF327LYS,ENCFF907XEG,ENCFF609MDN,ENCFF198SSR,ENCFF502WDT,ENCFF821BIA,ENCFF099QNE,ENCFF136ZDK,ENCFF165LXO,ENCFF321AGQ,ENCFF089GBC,ENCFF194HVL,ENCFF142CNG,ENCFF883UPM,ENCFF092NMJ,ENCFF957SPH,ENCFF951GOP |
| 71 | 1 | 182084680 | 182093665 | 8985 | AJ,AKR,BUB,C58J,CAST,CBA,CE,DBA1J,DBA2J,FVB,KK,LP,MOLF,MRL,NOD,NOR,NZB,NZO,PL,RF,RH2,SMJ,SPRET,SWR,TH,WSB | 1 | 182090097 | 182090115 | 19 | + | 17.1639 | 8.17E-07 | 0.187 | CCTCCACGAGGTGGCGGAA | | |
| 72 | 1 | 189695768 | 189698406 | 2638 | SPRET | 1 | 189696126 | 189696160 | 35 | + | 18.3578 | 2.43E-07 | 0.103 | CTGCAGCTCCAAGCAGTGGCATCCAGGTGGCCAGA | | |
| 73 | 1 | 191665720 | 191674705 | 8985 | 129S1,AJ,AKR,BALB,BTBR,BUB,C3H,C57L,C58J,CAST,CBA,CE,DBA1J,DBA2J,FVB,ILNJ2,KK,MA/My,MOLF,MRL,NOD,NOR,NZB,NZO,NZW,PJ,PL,RF,RH2,SEA,SJL,SMJ,SPRET,SWR,TH,WSB | 1 | 191671137 | 191671155 | 19 | + | 17.1639 | 8.17E-07 | 0.187 | CCTCCACGAGGTGGCGGAA | | |
| 74 | 1 | 192890521 | 192892631 | 2110 | AKR,BALB,C58J,CE,RH2 | 1 | 192890661 | 192890694 | 34 | + | 17.5122 | 4.86E-07 | 0.0914 | CTGCCTACCTGTGTACAGCCACTGGGTGGTGCTG | 38 | ENCFF093ZRU,ENCFF201JUA,ENCFF367UNR,ENCFF099QNE,ENCFF805QBU,ENCFF052KVO,ENCFF321AGQ,ENCFF821BIA,ENCFF327LYS,ENCFF907XEG,ENCFF198SSR,ENCFF024JOZ,ENCFF136ZDK,ENCFF589ZBJ,ENCFF033UQX,ENCFF883UPM,ENCFF957SPH,ENCFF502WDT,ENCFF142CNG,ENCFF165LXO,ENCFF065ODV,ENCFF055HAP,ENCFF194HVL,ENCFF089GBC,ENCFF575MGI,ENCFF951GOP,ENCFF430PPJ,ENCFF281QMV,ENCFF533APC,ENCFF609MDN,ENCFF316FYN,ENCFF097CAQ,ENCFF210BTO,ENCFF092NMJ,ENCFF611HDQ,ENCFF349BUL,ENCFF491RJK,ENCFF089EWX |
| 75 | 1 | 194916386 | 194922552 | 6166 | MOLF | 1 | 194916698 | 194916731 | 34 | + | 19.3333 | 1.42E-07 | 0.0473 | GTGCAGGTCCCAAGCAAGGCAGCCGGTGGCAGAG | | |
| 76 | 1 | 195018272 | 195029899 | 11627 | NOD,NOR,SWR | 1 | 195027759 | 195027792 | 34 | + | 16.4472 | 9.61E-07 | 0.125 | TAGCTGTCTTCAGACACACCAGAAGAGGGCGTCA | | |
| 77 | 1 | 4053655 | 4058231 | 4576 | DBA1J,DBA2J,SPRET | 1 | 4055563 | 4055597 | 35 | - | 16.8165 | 6.02E-07 | 0.167 | AAGAAGTTACACCCTGTTGCCAGCAGGTGGTTTTC | | |
| 78 | 1 | 18706495 | 18707789 | 1294 | AKR,DBA1J,DBA2J,NZB,NZO,TH | 1 | 18707121 | 18707139 | 19 | - | 20.1967 | 9.47E-08 | 0.0696 | TTGCCAGCAGGTGGTGGCA | | |
| 79 | 1 | 19259169 | 19260731 | 1562 | MOLF,SPRET | 1 | 19259423 | 19259441 | 19 | - | 16.8525 | 9.96E-07 | 0.2 | CCACCAGTAGGTGGAACAA | | |
| 80 | 1 | 19353448 | 19356541 | 3093 | AKR,CAST,MOLF,PL,SPRET,WSB | 1 | 19353445 | 19353478 | 34 | - | 16.4553 | 9.56E-07 | 0.125 | CTGCCGCCACACCCGCTCCCCACAGATGGCATCA | | |
| 81 | 1 | 24114265 | 24120725 | 6460 | CAST,MOLF,SPRET | 1 | 24115480 | 24115498 | 19 | - | 16.9016 | 9.66E-07 | 0.197 | TGAGCAGCAGGGGTCGCCA | | |
| 82 | 1 | 27198489 | 27205305 | 6816 | 129S1,AJ,AKR,BALB,BTBR,C3H,C57L,CAST,CBA,CE,DBA1J,DBA2J,FVB,ILNJ2,MOLF,MRL,NOD,NZB,NZO,NZW,PJ,PL,RH2,SEA,SPRET,WSB | 1 | 27198773 | 27198806 | 34 | - | 16.439 | 9.66E-07 | 0.125 | GGGCACCTGTCCTCCGGTCCAGAAGGTGGCAGGC | | |
| 83 | 1 | 33888096 | 33889614 | 1518 | CE,MOLF | 1 | 33888656 | 33888689 | 34 | - | 19.561 | 1.21E-07 | 0.043 | CTGCTATCTTCAGACACACCAGAAGAGGGCAATG | 1 | ENCFF093ZRU |
| 84 | 1 | 37267077 | 37268295 | 1218 | SPRET | 1 | 37267608 | 37267641 | 34 | - | 17.878 | 3.83E-07 | 0.0813 | TAGCTGTCCTCAGACACACCAGAAGAGGGCATCA | | |
| 85 | 1 | 42957636 | 42961368 | 3732 | CAST | 1 | 42958312 | 42958345 | 34 | - | 18.1057 | 3.29E-07 | 0.0749 | CTGCATTTGTTAGCACAAACAGCAAGGGGCAGCC | | |
| 86 | 1 | 43032873 | 43036534 | 3661 | SPRET | 1 | 43033142 | 43033160 | 19 | - | 20.3607 | 8.29E-08 | 0.0655 | AGGCCAGAAGAGGGCACCA | 1 | ENCFF093ZRU |
| 86 | 1 | 43032873 | 43036534 | 3661 | SPRET | 1 | 43033142 | 43033176 | 35 | - | 27.4679 | 3.12E-10 | 0.00426 | CTGCATGTATGCCTGCAGGCCAGAAGAGGGCACCA | 1 | ENCFF093ZRU |
| 87 | 1 | 45994846 | 46004108 | 9262 | CAST | 1 | 45995813 | 45995846 | 34 | - | 21.0325 | 4.17E-08 | 0.0215 | ATGCAGCTAAAGGCACAACCTCCAGGGGGTACTG | | |
| 88 | 1 | 46081034 | 46087369 | 6335 | TH | 1 | 46081099 | 46081133 | 35 | - | 19.4495 | 1.23E-07 | 0.067 | GTGCAGGCAACTATGACAATCAGTAGAGGGCACTA | | |
| 89 | 1 | 60047215 | 60049217 | 2002 | MA/My,SMJ,SWR | 1 | 60047907 | 60047925 | 19 | - | 17.9016 | 5.02E-07 | 0.151 | ACACCAGAAGAGGGCACTA | 2 | ENCFF099QNE,ENCFF093ZRU |
| 89 | 1 | 60047215 | 60049217 | 2002 | MA/My,SMJ,SWR | 1 | 60047907 | 60047940 | 34 | - | 21.3984 | 3.17E-08 | 0.0178 | TTGCTGTGGTCAGACACACCAGAAGAGGGCACTA | 2 | ENCFF099QNE,ENCFF093ZRU |
| 90 | 1 | 65809323 | 65812982 | 3659 | WSB | 1 | 65809560 | 65809594 | 35 | - | 17.1284 | 5.03E-07 | 0.151 | ATGCAGAGACCTCAGCCAAACATTAGGTGGAGCTC | | |
| 91 | 1 | 70988395 | 70993315 | 4920 | SPRET | 1 | 70990845 | 70990878 | 34 | - | 18.0732 | 3.36E-07 | 0.0757 | CAGCTGTCCTTAGACACTCCACAAGAGGGAGTCA | | |
| 92 | 1 | 71518396 | 71521712 | 3316 | SPRET | 1 | 71519555 | 71519588 | 34 | - | 16.6504 | 8.46E-07 | 0.12 | TAGCTGTCTTCCGACACACCAGAAGAGGGCATCA | | |
| 93 | 1 | 71972539 | 71973798 | 1259 | CAST | 1 | 71973078 | 71973111 | 34 | - | 22.8943 | 9.83E-09 | 0.00768 | ATGCAGTGCCTGCAGAAGCCAGGAGAAGGGGCCG | | |
| 94 | 1 | 79079810 | 79086881 | 7071 | AJ,BALB,C3H,ILNJ2,RH2,SEA,SJL,SWR | 1 | 79086284 | 79086317 | 34 | - | 18.0569 | 3.40E-07 | 0.0757 | ATGCAGCTAGAGACATGAGCTCCAGGGGGTACTG | | |
| 95 | 1 | 80445362 | 80447379 | 2017 | LP,MOLF | 1 | 80446058 | 80446092 | 35 | - | 16.3945 | 7.65E-07 | 0.188 | CAGCAGATTTGAATGATAGCCTCCAGGTAGCAGTA | | |
| 96 | 1 | 81719284 | 81720742 | 1458 | SPRET | 1 | 81720075 | 81720108 | 34 | - | 18.6423 | 2.29E-07 | 0.0639 | CTGCTGCTTCCCCTGCTGCCTCCAGGAGATAGAA | | |
| 97 | 1 | 82553591 | 82555463 | 1872 | SPRET | 1 | 82554081 | 82554114 | 34 | - | 18.1301 | 3.24E-07 | 0.0741 | ATGCAGTGCCTGAAGTGGCCAGAAGGGCACATCAG | 5 | ENCFF198SSR,ENCFF502WDT,ENCFF055HAP,ENCFF093ZRU,ENCFF367UNR |
| 98 | 1 | 86406120 | 86410619 | 4499 | LP,RF | 1 | 86408613 | 86408646 | 34 | - | 18.2927 | 2.90E-07 | 0.0695 | CTGCTCTGCCCGGACTTATCTCAAGGGGCTGCTC | | |
| 99 | 1 | 90118246 | 90124454 | 6208 | MOLF | 1 | 90121169 | 90121187 | 19 | - | 21.0984 | 4.51E-08 | 0.0555 | AGGCCAGAAGAGGGCGCCA | | |
| 99 | 1 | 90118246 | 90124454 | 6208 | MOLF | 1 | 90121169 | 90121202 | 34 | - | 22.6179 | 1.23E-08 | 0.00902 | GAGAAGCACCTGCAGAGGCCAGAAGAGGGCGCCA | | |
| 100 | 1 | 99210287 | 99214834 | 4547 | 129S1,AKR,C57L,DBA1J,DBA2J,ILNJ2,LP,MA/My,NOD,NOR,PJ,SEA,SMJ,TH | 1 | 99214071 | 99214104 | 34 | - | 16.8618 | 7.40E-07 | 0.113 | CAGTTATGGCAACCCTATCCACAAGAAGGCAGCA | | |
| 101 | 1 | 100336705 | 100345239 | 8534 | KK | 1 | 100338136 | 100338170 | 35 | - | 19.9725 | 8.85E-08 | 0.0559 | GTGCTGCTACCTGGGCATGCCAGCAGAGGGTGTTG | | |
| 102 | 1 | 101345342 | 101348772 | 3430 | CAST,MOLF | 1 | 101346363 | 101346397 | 35 | - | 16.5321 | 7.08E-07 | 0.182 | CTGCAGTGGGGAAATGAAACCTGAAGAAGCCACTT | | |
| 103 | 1 | 110212250 | 110216665 | 4415 | 129S1,AJ,AKR,BALB,BTBR,BUB,C3H,C57L,CBA,CE,DBA1J,DBA2J,FVB,ILNJ2,KK,LP,MA/My,MRL,PJ,PL,RF,RH2,SEA,SJL,SWR,TH | 1 | 110212716 | 110212750 | 35 | - | 19.8807 | 9.39E-08 | 0.0578 | CTGCAATTGGGGCCTCCATCCAATAGGTGACTGTG | | |
| 104 | 1 | 110886664 | 110888748 | 2084 | SPRET | 1 | 110887284 | 110887302 | 19 | - | 17.9672 | 4.81E-07 | 0.151 | AAGCCACCAGGTGGAACCC | | |
| 105 | 1 | 111681398 | 111683191 | 1793 | SPRET | 1 | 111682370 | 111682404 | 35 | - | 20.6606 | 5.65E-08 | 0.044 | CAGCAGTGACAGCACACAGCCTCAAGATGCTGGTG | | |
| 106 | 1 | 114139103 | 114143246 | 4143 | AKR,BUB,C3H,CAST,CBA,DBA1J,DBA2J,ILNJ2,KK,LP,MA/My,MOLF,MRL,NOD,NOR,PJ,PL,RF,SJL,SPRET,WSB | 1 | 114142443 | 114142476 | 34 | - | 17.6748 | 4.37E-07 | 0.0865 | GAGCATGACCACAAGTCCACAGCAGGTGGCAACG | | |
| 107 | 1 | 114871479 | 114873316 | 1837 | CAST | 1 | 114872420 | 114872453 | 34 | - | 17.748 | 4.17E-07 | 0.0854 | CTGCAATGTTTCTTCTCACCAGAACGTGGCCCAA | | |
| 108 | 1 | 115034345 | 115039574 | 5229 | SPRET | 1 | 115039434 | 115039467 | 34 | - | 16.8943 | 7.24E-07 | 0.111 | TTGCTCTCTTCAGACATACCAGAAGAGGGCATCA | | |
| 109 | 1 | 119784920 | 119786439 | 1519 | SPRET | 1 | 119785314 | 119785332 | 19 | - | 16.9672 | 9.26E-07 | 0.193 | AGGCCAGAAGAGGGAACTG | | |
| 110 | 1 | 127753561 | 127754828 | 1267 | SPRET | 1 | 127754149 | 127754182 | 34 | - | 18.4553 | 2.60E-07 | 0.0675 | ATACAGTGCCCATGGAAGCCAGAAGAGGGTGTCA | 3 | ENCFF093ZRU,ENCFF502WDT,ENCFF055HAP |
| 111 | 1 | 128156095 | 128158034 | 1939 | MOLF | 1 | 128157005 | 128157023 | 19 | - | 19.9836 | 1.12E-07 | 0.0736 | TCACCAGAAGAGGGCACTG | 1 | ENCFF093ZRU |
| 112 | 1 | 130202366 | 130204704 | 2338 | CBA,NOD,NOR,SMJ | 1 | 130202394 | 130202412 | 19 | - | 16.8852 | 9.76E-07 | 0.198 | TAGCCACTAGGTGGCATTT | 14 | ENCFF093ZRU,ENCFF099QNE,ENCFF092NMJ,ENCFF165LXO,ENCFF198SSR,ENCFF883UPM,ENCFF089GBC,ENCFF367UNR,ENCFF502WDT,ENCFF055HAP,ENCFF327LYS,ENCFF024JOZ,ENCFF907XEG,ENCFF430PPJ |
| 113 | 1 | 130255549 | 130257399 | 1850 | SPRET | 1 | 130256923 | 130256956 | 34 | - | 19.4065 | 1.35E-07 | 0.046 | CTGCAGCCTCTGCTCGAGGCAGCAGGAGGGACCA | | |
| 114 | 1 | 130841681 | 130843578 | 1897 | SPRET | 1 | 130842541 | 130842575 | 35 | - | 16 | 9.55E-07 | 0.211 | TTGCAATTGAAGGCCCAGGACTGAAGGGGTCACGC | | |
| 115 | 1 | 132035312 | 132036635 | 1323 | SPRET | 1 | 132036592 | 132036625 | 34 | - | 17.6992 | 4.30E-07 | 0.0861 | CTGCGTTCTCTCTGGTGGTCTCCAGATGGCTCTC | | |
| 116 | 1 | 140424592 | 140431252 | 6660 | WSB | 1 | 140425323 | 140425341 | 19 | - | 17.1475 | 8.25E-07 | 0.187 | TATCCACTAGATGGAACTG | | |
| 117 | 1 | 142539774 | 142543491 | 3717 | CAST,MOLF,SPRET | 1 | 142543025 | 142543058 | 34 | - | 17.6748 | 4.37E-07 | 0.0865 | GAGCATGACCACAAGTCCACAGCAGGTGGCAACG | | |
| 118 | 1 | 144822122 | 144824532 | 2410 | SPRET | 1 | 144822270 | 144822304 | 35 | - | 17.6789 | 3.64E-07 | 0.124 | CTGTAAGGCTGTCTATGTTCCAGTAGATGGCCTTA | | |
| 119 | 1 | 145991206 | 145992863 | 1657 | BTBR,CAST,MOLF | 1 | 145992374 | 145992407 | 34 | - | 17.6829 | 4.35E-07 | 0.0865 | TTGCAGCTAGAGACATGAGCTCTGGGGGGCACTG | | |
| 120 | 1 | 151465125 | 151467115 | 1990 | 129S1,AJ,AKR,BTBR,C3H,C57L,CAST,CBA,CE,ILNJ2,MOLF,MRL,PL,RH2,SMJ,SPRET | 1 | 151466885 | 151466918 | 34 | - | 16.4878 | 9.37E-07 | 0.125 | CTGCAGACTCTACCTCCCCCACACGATGACACCA | | |
| 121 | 1 | 151508379 | 151509405 | 1026 | SPRET | 1 | 151508533 | 151508567 | 35 | - | 24.8073 | 2.86E-09 | 0.00883 | GTGCAGGTACCCACTGAGACCAGTAGAGGGCATCG | 7 | ENCFF093ZRU,ENCFF821BIA,ENCFF907XEG,ENCFF367UNR,ENCFF502WDT,ENCFF065ODV,ENCFF052KVO |
| 122 | 1 | 160190004 | 160196431 | 6427 | MOLF | 1 | 160192546 | 160192579 | 34 | - | 18.5041 | 2.52E-07 | 0.0667 | TAGCTGTTCACAAACACACCAGAAGAGGGCATCA | 2 | ENCFF093ZRU,ENCFF099QNE |
| 123 | 1 | 161185106 | 161187909 | 2803 | SPRET | 1 | 161185910 | 161185928 | 19 | - | 17.5082 | 6.52E-07 | 0.168 | TGGCCTCTAGGAGGCGCTT | 2 | ENCFF093ZRU,ENCFF502WDT |
| 124 | 1 | 166287668 | 166292439 | 4771 | SPRET | 1 | 166287966 | 166287999 | 34 | - | 17.2276 | 5.85E-07 | 0.0999 | GTGCAGGGACCATGCAAGTCAGAAGAGGGCATTG | | |
| 125 | 1 | 166549752 | 166551428 | 1676 | WSB | 1 | 166550522 | 166550556 | 35 | - | 20.5413 | 6.11E-08 | 0.046 | GTGTAGGTACCCGTGGAGTCCAGAAGAGGGCATTG | 1 | ENCFF093ZRU |
| 125 | 1 | 166549752 | 166551428 | 1676 | WSB | 1 | 166551329 | 166551363 | 35 | - | 16.5138 | 7.15E-07 | 0.182 | TAGCTTTTCCTCTTTTTCTCCAGTAGATGGAGATT | 2 | ENCFF099QNE,ENCFF093ZRU |
| 126 | 1 | 171004246 | 171007625 | 3379 | SPRET | 1 | 171006627 | 171006660 | 34 | - | 16.7724 | 7.83E-07 | 0.115 | TAGCTGTCTTCAGATGCACCAGAAGAGGGCGTCA | | |
| 127 | 1 | 188305180 | 188307300 | 2120 | MOLF | 1 | 188305513 | 188305547 | 35 | - | 16.3945 | 7.65E-07 | 0.188 | GTGCAGAAGCCAGTGGATACCAGAAGAAGGCATTG | | |
| 128 | 1 | 188788846 | 188794982 | 6136 | CAST,MOLF | 1 | 188789817 | 188789835 | 19 | - | 17.1967 | 7.99E-07 | 0.185 | ACGCCAGAAGGGGGCAGAG | 15 | ENCFF821BIA,ENCFF327LYS,ENCFF589ZBJ,ENCFF198SSR,ENCFF883UPM,ENCFF065ODV,ENCFF367UNR,ENCFF033UQX,ENCFF052KVO,ENCFF502WDT,ENCFF136ZDK,ENCFF099QNE,ENCFF142CNG,ENCFF093ZRU,ENCFF055HAP |
| 128 | 1 | 188788846 | 188794982 | 6136 | CAST,MOLF | 1 | 188789817 | 188789850 | 34 | - | 28.0976 | 8.54E-11 | 0.000437 | GTGCAGTACCCATAGACGCCAGAAGGGGGCAGAG | 15 | ENCFF821BIA,ENCFF327LYS,ENCFF589ZBJ,ENCFF198SSR,ENCFF883UPM,ENCFF065ODV,ENCFF367UNR,ENCFF033UQX,ENCFF052KVO,ENCFF502WDT,ENCFF136ZDK,ENCFF099QNE,ENCFF142CNG,ENCFF093ZRU,ENCFF055HAP |
| 129 | 2 | 3551071 | 3559484 | 8413 | SPRET | 2 | 3556057 | 3556075 | 19 | + | 17.4262 | 6.88E-07 | 0.129 | AGGCCAGAAGGGGGCATCA | | |
| 130 | 2 | 7954282 | 7971589 | 17307 | CAST | 2 | 7965314 | 7965348 | 35 | + | 19.9266 | 9.11E-08 | 0.0456 | CTGCACCTGCTCCCATCAGCCACCAGAGGAAGTCT | | |
| 131 | 2 | 8380605 | 8383407 | 2802 | SPRET | 2 | 8380622 | 8380655 | 34 | + | 20.4065 | 6.62E-08 | 0.0229 | CTGCACTCACATTTGCCACTACAAGATGGCACTG | | |
| 132 | 2 | 9239803 | 9242428 | 2625 | SPRET | 2 | 9240665 | 9240698 | 34 | + | 16.7886 | 7.75E-07 | 0.0917 | GAGCATGGACACAAGCCCACAGCAGATGGCAGCA | | |
| 133 | 2 | 12001069 | 12003907 | 2838 | CAST,MA/My,MOLF,NOD,NOR,SJL,SPRET | 2 | 12001929 | 12001962 | 34 | + | 16.7886 | 7.75E-07 | 0.0917 | GAGCATGGACACAAGCCCACAGCAGATGGCAGCA | | |
| 134 | 2 | 18426899 | 18429134 | 2235 | SPRET | 2 | 18427103 | 18427136 | 34 | + | 16.5447 | 9.04E-07 | 0.0987 | CAGCTATCTCCACACGAGTCAGCAGGAGGAACCA | | |
| 135 | 2 | 21837483 | 21841966 | 4483 | CAST,SPRET | 2 | 21838260 | 21838293 | 34 | + | 17.6748 | 4.37E-07 | 0.0681 | GAGCATGACCACAAGTCCACAGCAGGTGGCAACG | | |
| 136 | 2 | 23622950 | 23626712 | 3762 | SPRET | 2 | 23624482 | 23624515 | 34 | + | 18.1545 | 3.18E-07 | 0.0576 | CTGCACTTCTTAGTGTTGCAGCCAGGGAGTGGCA | | |
| 137 | 2 | 25625033 | 25626147 | 1114 | CAST | 2 | 25625556 | 25625590 | 35 | + | 17.3211 | 4.50E-07 | 0.118 | AAGCAGTGGCCTTCCTCTCCCACCAGGTGGTTGAT | | |
| 138 | 2 | 25940011 | 25941509 | 1498 | CAST | 2 | 25941388 | 25941421 | 34 | + | 25.0732 | 1.55E-09 | 0.00154 | CTGCAGTGCCTGCAGAAGCCAGAAAAGGGTGCTG | | |
| 139 | 2 | 26532638 | 26538794 | 6156 | CAST,MA/My,MOLF | 2 | 26534083 | 26534116 | 34 | + | 24.2764 | 3.11E-09 | 0.00246 | CTGCACTTCCTGAAGTCACCAGGAGGGGGGTGTG | 25 | ENCFF093ZRU,ENCFF201JUA,ENCFF367UNR,ENCFF430PPJ,ENCFF099QNE,ENCFF065ODV,ENCFF327LYS,ENCFF321AGQ,ENCFF052KVO,ENCFF609MDN,ENCFF907XEG,ENCFF136ZDK,ENCFF589ZBJ,ENCFF504JDS,ENCFF055HAP,ENCFF024JOZ,ENCFF198SSR,ENCFF089GBC,ENCFF502WDT,ENCFF092NMJ,ENCFF165LXO,ENCFF821BIA,ENCFF883UPM,ENCFF957SPH,ENCFF281QMV |

| | | | | | | | | | | | | | | | | |
|---|---|---|---|---|---|---|---|---|---|---|---|---|---|---|---|---|
| 139 | 2 | 26532638 | 26538794 | 6156 | CAST,MA/My,MOLF | 2 | 26536977 | 26537011 | 35 | + | 21.367 | 3.52E-08 | 0.0261 | CTGCGGGGGCAAGATGTGGACAGTAGGTGGTGCTG | 41 | ENCFF327LYS,ENCFF367UNR,ENCFF093ZRU,ENCFF907XEG,ENCFF024JOZ,ENCFF089GBC,ENCFF201JUA,ENCFF099QNE,ENCFF589ZBJ,ENCFF092NMJ,ENCFF198SSR,ENCFF321AGQ,ENCFF430PPJ,ENCFF821BIA,ENCFF055HAP,ENCFF065ODV,ENCFF136ZDK,ENCFF165LXO,ENCFF502WDT,ENCFF883UPM,ENCFF504JDS,ENCFF957SPH,ENCFF951GOP,ENCFF609MDN,ENCFF142CNG,ENCFF281QMV,ENCFF210BTO,ENCFF316FYN,ENCFF349BUL,ENCFF194HVL,ENCFF601JVE,ENCFF575MGI,ENCFF611HDQ,ENCFF491RJK,ENCFF533APC,ENCFF052KVO,ENCFF089EWX,ENCFF033UQX,ENCFF097CAQ,ENCFF784ASD,ENCFF086ZTV |
| 139 | 2 | 26532638 | 26538794 | 6156 | CAST,MA/My,MOLF | 2 | 26536993 | 26537011 | 19 | + | 17.1475 | 8.25E-07 | 0.142 | TGGACAGTAGGTGGTGCTG | 41 | ENCFF327LYS,ENCFF367UNR,ENCFF093ZRU,ENCFF907XEG,ENCFF024JOZ,ENCFF089GBC,ENCFF201JUA,ENCFF099QNE,ENCFF589ZBJ,ENCFF092NMJ,ENCFF198SSR,ENCFF321AGQ,ENCFF430PPJ,ENCFF821BIA,ENCFF055HAP,ENCFF065ODV,ENCFF136ZDK,ENCFF165LXO,ENCFF502WDT,ENCFF883UPM,ENCFF504JDS,ENCFF957SPH,ENCFF951GOP,ENCFF609MDN,ENCFF142CNG,ENCFF281QMV,ENCFF210BTO,ENCFF316FYN,ENCFF349BUL,ENCFF194HVL,ENCFF601JVE,ENCFF575MGI,ENCFF611HDQ,ENCFF491RJK,ENCFF533APC,ENCFF052KVO,ENCFF089EWX,ENCFF033UQX,ENCFF097CAQ,ENCFF784ASD,ENCFF086ZTV |
| 139 | 2 | 26532638 | 26538794 | 6156 | CAST,MA/My,MOLF | 2 | 26537009 | 26537043 | 35 | + | 17.2661 | 4.64E-07 | 0.12 | CTGTAGGGACCATGTGTGGACAGTGGGTGGTGGTG | 41 | ENCFF327LYS,ENCFF367UNR,ENCFF093ZRU,ENCFF907XEG,ENCFF024JOZ,ENCFF089GBC,ENCFF201JUA,ENCFF099QNE,ENCFF589ZBJ,ENCFF092NMJ,ENCFF198SSR,ENCFF321AGQ,ENCFF430PPJ,ENCFF821BIA,ENCFF055HAP,ENCFF065ODV,ENCFF136ZDK,ENCFF165LXO,ENCFF502WDT,ENCFF883UPM,ENCFF504JDS,ENCFF957SPH,ENCFF951GOP,ENCFF609MDN,ENCFF142CNG,ENCFF281QMV,ENCFF210BTO,ENCFF316FYN,ENCFF349BUL,ENCFF194HVL,ENCFF601JVE,ENCFF575MGI,ENCFF611HDQ,ENCFF491RJK,ENCFF533APC,ENCFF052KVO,ENCFF089EWX,ENCFF033UQX,ENCFF097CAQ,ENCFF784ASD,ENCFF086ZTV |
| 139 | 2 | 26532638 | 26538794 | 6156 | CAST,MA/My,MOLF | 2 | 26537049 | 26537083 | 35 | + | 22.3853 | 1.74E-08 | 0.0175 | CTGTAGGGACCCTGCGTGGACAGTAGGTGGTGCTG | 42 | ENCFF327LYS,ENCFF367UNR,ENCFF093ZRU,ENCFF907XEG,ENCFF024JOZ,ENCFF089GBC,ENCFF201JUA,ENCFF099QNE,ENCFF589ZBJ,ENCFF092NMJ,ENCFF198SSR,ENCFF321AGQ,ENCFF430PPJ,ENCFF821BIA,ENCFF055HAP,ENCFF065ODV,ENCFF136ZDK,ENCFF165LXO,ENCFF502WDT,ENCFF883UPM,ENCFF504JDS,ENCFF957SPH,ENCFF951GOP,ENCFF609MDN,ENCFF142CNG,ENCFF281QMV,ENCFF210BTO,ENCFF316FYN,ENCFF349BUL,ENCFF194HVL,ENCFF601JVE,ENCFF575MGI,ENCFF611HDQ,ENCFF491RJK,ENCFF533APC,ENCFF052KVO,ENCFF089EWX,ENCFF033UQX,ENCFF097CAQ,ENCFF784ASD,ENCFF086ZTV,ENCFF805QBU |
| 139 | 2 | 26532638 | 26538794 | 6156 | CAST,MA/My,MOLF | 2 | 26537065 | 26537083 | 19 | + | 17.1475 | 8.25E-07 | 0.142 | TGGACAGTAGGTGGTGCTG | 42 | ENCFF327LYS,ENCFF367UNR,ENCFF093ZRU,ENCFF907XEG,ENCFF024JOZ,ENCFF089GBC,ENCFF201JUA,ENCFF099QNE,ENCFF589ZBJ,ENCFF092NMJ,ENCFF198SSR,ENCFF321AGQ,ENCFF430PPJ,ENCFF821BIA,ENCFF055HAP,ENCFF065ODV,ENCFF136ZDK,ENCFF165LXO,ENCFF502WDT,ENCFF883UPM,ENCFF504JDS,ENCFF957SPH,ENCFF951GOP,ENCFF609MDN,ENCFF142CNG,ENCFF281QMV,ENCFF210BTO,ENCFF316FYN,ENCFF349BUL,ENCFF194HVL,ENCFF601JVE,ENCFF575MGI,ENCFF611HDQ,ENCFF491RJK,ENCFF533APC,ENCFF052KVO,ENCFF089EWX,ENCFF033UQX,ENCFF097CAQ,ENCFF784ASD,ENCFF086ZTV,ENCFF805QBU |
| 139 | 2 | 26532638 | 26538794 | 6156 | CAST,MA/My,MOLF | 2 | 26537089 | 26537123 | 35 | + | 19.055 | 1.58E-07 | 0.0638 | CTGTAGGGACCCTGTGTGGACAGTGGGTGGTGCTG | 42 | ENCFF327LYS,ENCFF367UNR,ENCFF093ZRU,ENCFF907XEG,ENCFF024JOZ,ENCFF089GBC,ENCFF201JUA,ENCFF099QNE,ENCFF589ZBJ,ENCFF092NMJ,ENCFF198SSR,ENCFF321AGQ,ENCFF430PPJ,ENCFF821BIA,ENCFF055HAP,ENCFF065ODV,ENCFF136ZDK,ENCFF165LXO,ENCFF502WDT,ENCFF883UPM,ENCFF504JDS,ENCFF957SPH,ENCFF951GOP,ENCFF609MDN,ENCFF142CNG,ENCFF281QMV,ENCFF210BTO,ENCFF316FYN,ENCFF349BUL,ENCFF194HVL,ENCFF601JVE,ENCFF575MGI,ENCFF611HDQ,ENCFF491RJK,ENCFF533APC,ENCFF052KVO,ENCFF089EWX,ENCFF033UQX,ENCFF097CAQ,ENCFF784ASD,ENCFF086ZTV,ENCFF805QBU |
| 139 | 2 | 26532638 | 26538794 | 6156 | CAST,MA/My,MOLF | 2 | 26537129 | 26537163 | 35 | + | 19.055 | 1.58E-07 | 0.0638 | CTGTAGGGACCCTGTGTGGACAGTGGGTGGTGCTG | 41 | ENCFF327LYS,ENCFF367UNR,ENCFF093ZRU,ENCFF907XEG,ENCFF024JOZ,ENCFF089GBC,ENCFF201JUA,ENCFF099QNE,ENCFF589ZBJ,ENCFF092NMJ,ENCFF198SSR,ENCFF321AGQ,ENCFF430PPJ,ENCFF821BIA,ENCFF055HAP,ENCFF065ODV,ENCFF136ZDK,ENCFF165LXO,ENCFF502WDT,ENCFF883UPM,ENCFF504JDS,ENCFF957SPH,ENCFF951GOP,ENCFF609MDN,ENCFF142CNG,ENCFF281QMV,ENCFF210BTO,ENCFF316FYN,ENCFF349BUL,ENCFF194HVL,ENCFF601JVE,ENCFF575MGI,ENCFF611HDQ,ENCFF491RJK,ENCFF533APC,ENCFF089EWX,ENCFF033UQX,ENCFF097CAQ,ENCFF784ASD,ENCFF086ZTV,ENCFF805QBU |
| 140 | 2 | 30056474 | 30059382 | 2908 | SWR | 2 | 30056884 | 30056918 | 35 | + | 27.789 | 2.34E-10 | 0.00232 | TTGCAGTAACAAGTGCCAACCAGAAGGTGGCGCTC | 18 | ENCFF024JOZ,ENCFF821BIA,ENCFF136ZDK,ENCFF092NMJ,ENCFF198SSR,ENCFF165LXO,ENCFF589ZBJ,ENCFF430PPJ,ENCFF093ZRU,ENCFF055HAP,ENCFF883UPM,ENCFF367UNR,ENCFF065ODV,ENCFF089GBC.ENCFF052KVO.ENCFF099QNE.ENCFF907XEG.ENCFF327LYS |
| 140 | 2 | 30056474 | 30059382 | 2908 | SWR | 2 | 30056900 | 30056918 | 19 | + | 19.9016 | 1.19E-07 | 0.0563 | CAACCAGAAGGTGGCGCTC | 19 | ENCFF024JOZ,ENCFF821BIA,ENCFF136ZDK,ENCFF092NMJ,ENCFF198SSR,ENCFF165LXO,ENCFF589ZBJ,ENCFF430PPJ,ENCFF093ZRU,ENCFF055HAP,ENCFF883UPM,ENCFF367UNR,ENCFF065ODV,ENCFF089GBC,ENCFF052KVO,ENCFF099QNE,ENCFF907XEG,ENCFF327LYS,ENCFF502WDT |
| 141 | 2 | 32060055 | 32062073 | 2018 | MOLF | 2 | 32061837 | 32061871 | 35 | + | 17.3211 | 4.50E-07 | 0.118 | GTGCAGGTGTCTACAGAGGCCAGAAGAGGGCTTAG | 2 | ENCFF093ZRU.ENCFF821BIA |
| 142 | 2 | 32214039 | 32217080 | 3041 | SWR | 2 | 32214508 | 32214541 | 34 | + | 23.5772 | 5.62E-09 | 0.00383 | ATACAGTGCCTCCAGAGGCCAGAAGAGGGCGTCC | 10 | ENCFF367UNR,ENCFF052KVO,ENCFF089GBC,ENCFF327LYS,ENCFF092NMJ,ENCFF093ZRU,ENCFF055HAP.ENCFF099QNE.ENCFF533APC.ENCFF502WDT |
| 143 | 2 | 40937266 | 40964021 | 26755 | TH | 2 | 40950950 | 40950968 | 19 | + | 16.918 | 9.56E-07 | 0.151 | CCACCTGCAGGTGGCACGG | 4 | ENCFF024JOZ.ENCFF093ZRU.ENCFF367UNR.ENCFF099QNE |
| 143 | 2 | 40937266 | 40964021 | 26755 | TH | 2 | 40950945 | 40950963 | 19 | - | 19.3934 | 1.75E-07 | 0.0686 | CCACCTGCAGGTGGCACTA | 4 | ENCFF024JOZ.ENCFF093ZRU.ENCFF367UNR.ENCFF099QNE |
| 144 | 2 | 47353781 | 47358263 | 4482 | FVB,NOD,NOR,TH | 2 | 47356240 | 47356273 | 34 | + | 26.0325 | 6.48E-10 | 0.000834 | CTGCTGTTTTCAGACTCACCAGAAGAGGGCAGCA | 11 | ENCFF052KVO,ENCFF093ZRU,ENCFF821BIA,ENCFF136ZDK,ENCFF907XEG,ENCFF327LYS,ENCFF367UNR.ENCFF099QNE.ENCFF055HAP.ENCFF502WDT.ENCFF065ODV |
| 144 | 2 | 47353781 | 47358263 | 4482 | FVB,NOD,NOR,TH | 2 | 47356255 | 47356273 | 19 | + | 19.6393 | 1.45E-07 | 0.0618 | TCACCAGAAGAGGGCAGCA | 10 | ENCFF052KVO,ENCFF093ZRU,ENCFF821BIA,ENCFF136ZDK,ENCFF907XEG,ENCFF327LYS,ENCFF367UNR.ENCFF099QNE.ENCFF055HAP.ENCFF065ODV |
| 145 | 2 | 47827408 | 47842021 | 14613 | KK,NZB | 2 | 47835268 | 47835286 | 19 | + | 17.4262 | 6.88E-07 | 0.129 | GAACCAGGAGGGGGCAGCA | 9 | ENCFF093ZRU,ENCFF367UNR,ENCFF821BIA,ENCFF024JOZ,ENCFF327LYS,ENCFF198SSR,ENCFF165LXO,ENCFF907XEG.ENCFF099QNE |
| 146 | 2 | 55954633 | 55959266 | 4633 | 129S1,AJ,AKR,BUB,C3H,C58J,CAST,CBA,CE,DBA1J,DBA2J,FVB,ILNJ2,KK,MOLF,MRL,NOD,NOR,NZB,NZO,PJ,PL,RF,SEA,SJL,SMJ.SPRET.SWR.TH.WSB | 2 | 55955488 | 55955521 | 34 | + | 17.6748 | 4.37E-07 | 0.0681 | GAGCATGACCACAAGTCCACAGCAGGTGGCAACG | | |
| 147 | 2 | 61693575 | 61699324 | 5749 | KK | 2 | 61698267 | 61698285 | 19 | + | 17.082 | 8.61E-07 | 0.144 | TGGCCACTGGGAGGCGCAA | 1 | ENCFF883UPM |
| 148 | 2 | 66144396 | 66146671 | 2275 | SPRET | 2 | 66144507 | 66144540 | 34 | + | 16.4472 | 9.61E-07 | 0.0987 | TAGCTGTCTTCAGACACACCAGAAGAGGGCGTCA | | |
| 149 | 2 | 71234599 | 71236385 | 1786 | AJ,AKR,BUB,C3H,CAST,FVB,ILNJ2,MA/My,MOLF,MRL,NOD,NOR,NZW,PL,RH2,SMJ | 2 | 71235587 | 71235621 | 35 | + | 17.8349 | 3.32E-07 | 0.0989 | CTGCATGTGCACTTGCAGGCCAGAAGAGGCCCTCA | | |
| 150 | 2 | 77827983 | 77834256 | 6273 | CAST.MOLF.NZW.SPRET | 2 | 77828429 | 77828447 | 19 | + | 17.0328 | 8.88E-07 | 0.145 | TTGCCAGCAGAGAGCGCTC | | |
| 151 | 2 | 86329424 | 86332537 | 3113 | AJ,AKR,BALB,BUB,C3H,C57L,CBA,CE,DBA1J,DBA2J,KK,MA/My,MRL,NOD,NOR,NZB,NZO.NZW.PJ.PL.RF.RH2.SEA.SMJ | 2 | 86332362 | 86332395 | 34 | + | 18.0976 | 3.31E-07 | 0.059 | GAGCAGATGCGGCCGCCACCCCAAGATGGTGCCC | | |
| 152 | 2 | 89255055 | 89257513 | 2458 | MOLF.SPRET | 2 | 89255867 | 89255901 | 35 | + | 16.1284 | 8.89E-07 | 0.172 | GTGTAGATGTCCATCGAGTCCAGAAGAGGGAGTTG | | |
| 153 | 2 | 89997115 | 90079547 | 82432 | BUB,C58J,ILNJ2,KK,MOLF,PL,SJL,SWR,WSB | 2 | 90008182 | 90008216 | 35 | + | 16.4587 | 7.38E-07 | 0.154 | CTGCAGTGGTGTAATGTGCTCAGCTGAGGGTATTG | | |
| 153 | 2 | 89997115 | 90079547 | 82432 | BUB,C58J,ILNJ2,KK,MOLF,PL,SJL,SWR,WSB | 2 | 90025246 | 90025279 | 34 | + | 21.8537 | 2.23E-08 | 0.0103 | CCTCAGTTTTAGATGTGGCCACTAGATGGAGCCA | | |
| 153 | 2 | 89997115 | 90079547 | 82432 | BUB,C58J,ILNJ2,KK,MOLF,PL,SJL,SWR,WSB | 2 | 90025261 | 90025279 | 19 | + | 21.8033 | 2.43E-08 | 0.0339 | TGGCCACTAGATGGAGCCA | | |
| 153 | 2 | 89997115 | 90079547 | 82432 | BUB,C58J,ILNJ2,KK,MOLF,PL,SJL,SWR,WSB | 2 | 90041506 | 90041539 | 34 | + | 18.0407 | 3.43E-07 | 0.0602 | TAGCTGTCTTCAGAGTTACCAGAAGAGGGCTCCA | | |
| 153 | 2 | 89997115 | 90079547 | 82432 | BUB,C58J,ILNJ2,KK,MOLF,PL,SJL,SWR,WSB | 2 | 90072049 | 90072082 | 34 | + | 20.2195 | 7.58E-08 | 0.025 | CTGCAAGCCTTGCAGTGGACACCAGAGGTTAGTG | | |
| 153 | 2 | 89997115 | 90079547 | 82432 | BUB,C58J,ILNJ2,KK,MOLF,PL,SJL,SWR,WSB | 2 | 90072092 | 90072125 | 34 | + | 20.2195 | 7.58E-08 | 0.025 | CTGCAAGCCTTGCAGTGGACACCAGAGGTTAGTG | | |
| 153 | 2 | 89997115 | 90079547 | 82432 | BUB,C58J,ILNJ2,KK,MOLF,PL,SJL,SWR,WSB | 2 | 90072135 | 90072168 | 34 | + | 20.2195 | 7.58E-08 | 0.025 | CTGCAAGCCTTGCAGTGGACACCAGAGGTTAGTG | | |
| 154 | 2 | 91116652 | 91118678 | 2026 | MOLF | 2 | 91118643 | 91118676 | 34 | + | 18.2033 | 3.08E-07 | 0.0565 | TAGCTGTCCTCAGATGCACCAGAAGAGGGCATCA | 3 | ENCFF093ZRU.ENCFF367UNR.ENCFF099QNE |
| 155 | 2 | 103793716 | 103795418 | 1702 | SPRET | 2 | 103794927 | 103794960 | 34 | + | 16.4715 | 9.47E-07 | 0.0987 | ATGCAGTCCTTGGGAAGACCACTGGGTGTTGTTG | | |
| 156 | 2 | 114098969 | 114235273 | 136304 | MOLF | 2 | 114223143 | 114223176 | 34 | + | 16.6504 | 8.46E-07 | 0.0961 | CTGCAGCAGCAGCAGCAGAAGGAAGGGGGAGGAG | | |
| 156 | 2 | 114098969 | 114235273 | 136304 | MOLF | 2 | 114114405 | 114114439 | 35 | - | 17.0367 | 5.30E-07 | 0.129 | CTGTAGCTAAACAGGACTTCCAGTGGATGGAGGTA | | |
| 157 | 2 | 118211969 | 118215514 | 3545 | SPRET | 2 | 118214479 | 118214512 | 34 | + | 18.3984 | 2.70E-07 | 0.0538 | TAGCTGTCTTCAGACACACCAGAAGAGGGCACCA | | |
| 157 | 2 | 118211969 | 118215514 | 3545 | SPRET | 2 | 118214494 | 118214512 | 19 | + | 17.8852 | 5.08E-07 | 0.115 | ACACCAGAAGAGGGCACCA | | |
| 158 | 2 | 119495569 | 119496633 | 1064 | SPRET | 2 | 119496398 | 119496432 | 35 | + | 19.6606 | 1.08E-07 | 0.0506 | ATGAGGTTTCTTCACCCCTCCAGTAGAGGGCGCAC | 6 | ENCFF093ZRU.ENCFF821BIA.ENCFF327LYS.ENCFF589ZBJ.ENCFF367UNR.ENCFF136ZDK |
| 158 | 2 | 119495569 | 119496633 | 1064 | SPRET | 2 | 119496414 | 119496432 | 19 | + | 18.5574 | 3.20E-07 | 0.0909 | CCTCCAGTAGAGGGCGCAC | 6 | ENCFF093ZRU.ENCFF821BIA.ENCFF327LYS.ENCFF589ZBJ.ENCFF367UNR.ENCFF136ZDK |
| 159 | 2 | 121820019 | 121823343 | 3324 | SPRET | 2 | 121821733 | 121821766 | 34 | + | 18.3089 | 2.87E-07 | 0.0543 | ATGCAGCTAGAGACACGAGCTCCAGGGGGTACTG | | |
| 160 | 2 | 122046959 | 122048424 | 1465 | SPRET | 2 | 122046928 | 122046961 | 34 | + | 18.1707 | 3.15E-07 | 0.0573 | CAGCTGTCTTTAGACACACCAGAAGAGGGCATCA | | |
| 161 | 2 | 123355540 | 123367197 | 11657 | 129S1,AJ,AKR,BALB,BTBR,BUB,C3H,C57L,C58J,CAST,CBA,CE,DBA1J,DBA2J,FVB,ILNJ2,MA/My,MOLF,NOD,NZB,NZO,NZW,PJ,PL,RF,RH2,SEA,SJL,SMJ,SPRET,SWR,TH,WSB | 2 | 123355556 | 123355589 | 34 | + | 17.9106 | 3.74E-07 | 0.063 | CCGCCCTCACATTCGCCGTCACAAGATGGCGCTG | | |
| 161 | 2 | 123355540 | 123367197 | 11657 | 129S1,AJ,AKR,BALB,BTBR,BUB,C3H,C57L,C58J,CAST,CBA,CE,DBA1J,DBA2J,FVB,ILNJ2,MA/My,MOLF,NOD,NZB,NZO,NZW,PJ,PL,RF,RH2,SEA,SJL,SMJ,SPRET,SWR,TH,WSB | 2 | 123358366 | 123358399 | 34 | + | 17.9106 | 3.74E-07 | 0.063 | CCGCCCTCACATTCGCCGTCACAAGATGGCGCTG | | |
| 161 | 2 | 123355540 | 123367197 | 11657 | 129S1,AJ,AKR,BALB,BTBR,BUB,C3H,C57L,C58J,CAST,CBA,CE,DBA1J,DBA2J,FVB,ILNJ2,MA/My,MOLF,NOD,NZB,NZO,NZW,PJ,PL,RF,RH2,SEA,SJL,SMJ,SPRET,SWR,TH,WSB | 2 | 123361176 | 123361209 | 34 | + | 17.9106 | 3.74E-07 | 0.063 | CCGCCCTCACATTCGCCGTCACAAGATGGCGCTG | | |
| 161 | 2 | 123355540 | 123367197 | 11657 | 129S1,AJ,AKR,BALB,BTBR,BUB,C3H,C57L,C58J,CAST,CBA,CE,DBA1J,DBA2J,FVB,ILNJ2,MA/My,MOLF,NOD,NZB,NZO,NZW,PJ,PL,RF,RH2,SEA,SJL,SMJ,SPRET,SWR,TH,WSB | 2 | 123363954 | 123363987 | 34 | + | 17.9106 | 3.74E-07 | 0.063 | CCGCCCTCACATTCGCCGTCACAAGATGGCGCTG | | |
| 161 | 2 | 123355540 | 123367197 | 11657 | 129S1,AJ,AKR,BALB,BTBR,BUB,C3H,C57L,C58J,CAST,CBA,CE,DBA1J,DBA2J,FVB,ILNJ2,MA/My,MOLF,NOD,NZB,NZO,NZW,PJ,PL,RF,RH2,SEA,SJL,SMJ,SPRET,SWR,TH,WSB | 2 | 123366724 | 123366757 | 34 | + | 17.9106 | 3.74E-07 | 0.063 | CCGCCCTCACATTCGCCGTCACAAGATGGCGCTG | | |
| 162 | 2 | 125799092 | 125803640 | 4548 | C3H.CBA.DBA2J.ILNJ2.PJ.SPRET | 2 | 125799863 | 125799896 | 34 | + | 17.9675 | 3.61E-07 | 0.0616 | GAGCATGACCACAAGTCTACAGCAGGTGGCAACG | | |
| 163 | 2 | 128533119 | 128534508 | 1389 | SPRET | 2 | 128533121 | 128533139 | 19 | + | 17.5902 | 6.18E-07 | 0.124 | ACACCAGAAGAGGGCACTG | | |
| 164 | 2 | 130230554 | 130231611 | 1057 | SPRET | 2 | 130230890 | 130230923 | 34 | + | 19.1301 | 1.64E-07 | 0.0413 | TTGTAGTGCTCTTGGAGGACAGAAGAGGGCATCA | 18 | ENCFF052KVO,ENCFF099QNE,ENCFF430PPJ,ENCFF821BIA,ENCFF092NMJ,ENCFF089GBC,ENCFF093ZRU,ENCFF367UNR,ENCFF907XEG,ENCFF502WDT,ENCFF327LYS,ENCFF055HAP,ENCFF024JOZ,ENCFF065ODV.ENCFF589ZBJ.ENCFF198SSR.ENCFF165LXO.ENCFF883UPM |
| 165 | 2 | 137903626 | 137909412 | 5786 | SPRET | 2 | 137903640 | 137903673 | 34 | + | 16.6423 | 8.50E-07 | 0.0962 | ATGCAGCTATAGATACGAGCTCTGGGGGGTACTG | | |
| 166 | 2 | 149025912 | 149068124 | 42212 | SPRET | 2 | 149037816 | 149037850 | 35 | + | 18.7798 | 1.87E-07 | 0.0703 | ATGTACTGCCCCAATTTTGACTCTAGGGGTCACTA | 2 | ENCFF052KVO.ENCFF502WDT |
| 167 | 2 | 150224221 | 150228022 | 3801 | CAST.MOLF.NOD.NZB.NZW.RF.SPRET | 2 | 150226615 | 150226648 | 34 | + | 17.6992 | 4.30E-07 | 0.0675 | TAGCTGTTTTCAGACACACCAGAAGAGGGCATCA | | |
| 168 | 2 | 150926159 | 150938775 | 12616 | CAST | 2 | 150930637 | 150930671 | 35 | + | 15.9817 | 9.65E-07 | 0.178 | CTGTAGTGAGCCTTCTGGACAAGCTGGGGGCAGAT | | |
| 169 | 2 | 152693660 | 152694740 | 1080 | SPRET | 2 | 152694058 | 152694091 | 34 | + | 17.3821 | 5.29E-07 | 0.0753 | TTGCTATCTTCAGACACACCAGGAGGGGGCATTG | 1 | ENCFF093ZRU |
| 170 | 2 | 155971128 | 155973271 | 2143 | SPRET | 2 | 155972055 | 155972088 | 34 | + | 19.0244 | 1.76E-07 | 0.0433 | TAGCTGTCCCCAGACACTCCAGAAGAGGGCATCA | 3 | ENCFF093ZRU.ENCFF099QNE.ENCFF367UNR |
| 170 | 2 | 155971128 | 155973271 | 2143 | SPRET | 2 | 155972231 | 155972264 | 34 | + | 16.4472 | 9.61E-07 | 0.0987 | TAGCTGTCTTCAGACACACCAGAAGAGGGCGTCA | 2 | ENCFF093ZRU.ENCFF099QNE |
| 170 | 2 | 155971128 | 155973271 | 2143 | SPRET | 2 | 155971377 | 155971410 | 34 | - | 18.3008 | 2.89E-07 | 0.0546 | GTGCAATGTTCGTGAAGACCAGAAGATGACACAG | | |
| 171 | 2 | 157494162 | 157498335 | 4173 | 129S1,AJ,AKR,BTBR,BUB,C57L,C58J,DBA1J,DBA2J,ILNJ2,KK,MA/My,MOLF,MRL,NZB,PJ.RF.SJL.SPRET.TH | 2 | 157494939 | 157494972 | 34 | + | 17.6748 | 4.37E-07 | 0.0681 | GAGCATGACCACAAGTCCACAGCAGGTGGCAACG | | |
| 172 | 2 | 164592725 | 164595427 | 2702 | SPRET | 2 | 164592915 | 164592948 | 34 | + | 16.6911 | 8.24E-07 | 0.0945 | TAGCTGTCTTCAGACACACCAGTAGAGGGCATTG | | |
| 173 | 2 | 167249818 | 167255940 | 6122 | SPRET | 2 | 167255733 | 167255766 | 34 | + | 21.9187 | 2.13E-08 | 0.00999 | ATGCAGTTTACTCAGAGGCCAGAAGAGGGTGTCA | | |
| 174 | 2 | 170194921 | 170205924 | 11003 | WSB | 2 | 170196945 | 170196978 | 34 | + | 18.5285 | 2.48E-07 | 0.0524 | GCGTCATTCTAGGAAGAGACACCAGGGGGCACTC | 34 | ENCFF093ZRU,ENCFF367UNR,ENCFF201JUA,ENCFF099QNE,ENCFF821BIA,ENCFF907XEG,ENCFF327LYS,ENCFF089GBC,ENCFF024JOZ,ENCFF052KVO,ENCFF165LXO,ENCFF589ZBJ,ENCFF198SSR,ENCFF805QBU,ENCFF504JDS,ENCFF430PPJ,ENCFF092NMJ,ENCFF502WDT,ENCFF055HAP,ENCFF951GOP,ENCFF883UPM,ENCFF136ZDK,ENCFF033UQX,ENCFF065ODV,ENCFF194HVL,ENCFF609MDN,ENCFF097CAQ,ENCFF281QMV,ENCFF957SPH,ENCFF575MGI,ENCFF142CNG,ENCFF316FYN,ENCFF210BTO,ENCFF533APC |
| 174 | 2 | 170194921 | 170205924 | 11003 | WSB | 2 | 170196960 | 170196978 | 19 | + | 17.9016 | 5.02E-07 | 0.115 | GAGACACCAGGGGGCACTC | 34 | ENCFF093ZRU,ENCFF367UNR,ENCFF201JUA,ENCFF099QNE,ENCFF821BIA,ENCFF907XEG,ENCFF327LYS,ENCFF089GBC,ENCFF024JOZ,ENCFF052KVO,ENCFF165LXO,ENCFF589ZBJ,ENCFF198SSR,ENCFF805QBU,ENCFF504JDS,ENCFF430PPJ,ENCFF092NMJ,ENCFF502WDT,ENCFF055HAP,ENCFF951GOP,ENCFF883UPM,ENCFF136ZDK,ENCFF033UQX,ENCFF065ODV,ENCFF194HVL,ENCFF609MDN,ENCFF097CAQ,ENCFF281QMV,ENCFF957SPH,ENCFF575MGI,ENCFF142CNG,ENCFF316FYN,ENCFF210BTO,ENCFF533APC |
| 174 | 2 | 170194921 | 170205924 | 11003 | WSB | 2 | 170197546 | 170197580 | 35 | - | 16.055 | 9.26E-07 | 0.175 | CTGCAATCCCAGCACTTAGGAGGCAGAGGGAGGAA | | |
| 174 | 2 | 170194921 | 170205924 | 11003 | WSB | 2 | 170198763 | 170198796 | 34 | - | 18.5285 | 2.48E-07 | 0.0524 | CTGTTGAGGAAGCCACTGGCTCTAGGGGGCGCTA | 16 | ENCFF327LYS,ENCFF024JOZ,ENCFF367UNR,ENCFF092NMJ,ENCFF093ZRU,ENCFF089GBC,ENCFF883UPM,ENCFF589ZBJ,ENCFF821BIA,ENCFF099QNE,ENCFF097CAQ,ENCFF491RJK,ENCFF052KVO,ENCFF907XEG.ENCFF957SPH.ENCFF198SSR |
| 175 | 2 | 178636041 | 178639185 | 3144 | CAST | 2 | 178636593 | 178636627 | 35 | + | 18.2018 | 2.67E-07 | 0.088 | ATGCAGATGCCTTCTGGGGCCAGCAGATGGTTTCA | | |
| 176 | 2 | 181561079 | 181569889 | 8810 | BUB.CAST.CE.MOLF.PJ.RF | 2 | 181563701 | 181563719 | 19 | + | 17.8689 | 5.14E-07 | 0.116 | AAACCACTAGGTGGCAGAA | | |
| 177 | 2 | 9045436 | 9047482 | 2046 | CAST.ILNJ2.MOLF.SPRET | 2 | 9046899 | 9046917 | 19 | - | 19.7377 | 1.35E-07 | 0.0603 | AGGCCAGAAGAGGGCAGTA | | |
| 177 | 2 | 9045436 | 9047482 | 2046 | CAST.ILNJ2.MOLF.SPRET | 2 | 9046899 | 9046933 | 35 | - | 20.6422 | 5.72E-08 | 0.0344 | CTCCAGATAACTCTGGAGGCCAGAAGAGGGCAGTA | | |
| 178 | 2 | 15942107 | 15943149 | 1042 | SJL | 2 | 15942138 | 15942172 | 35 | - | 16.7706 | 6.18E-07 | 0.139 | CTGCAAAACCAACATTCAGGCAGTGGAGGGAGGGG | | |
| 179 | 2 | 16025306 | 16034352 | 9046 | CAST.CE.MOLF.PL.SJL.SPRET | 2 | 16028873 | 16028891 | 19 | - | 17.1639 | 8.17E-07 | 0.141 | CCTCCACGAGGTGGCGGAA | | |
| 180 | 2 | 17213074 | 17214916 | 1842 | CAST | 2 | 17213587 | 17213620 | 34 | - | 20.9675 | 4.38E-08 | 0.0172 | CTGCTGTCTTCAGACACACCAGAAGAGGGCATTG | | |
| 181 | 2 | 22196359 | 22201705 | 5346 | SPRET | 2 | 22200154 | 22200188 | 35 | - | 16.7615 | 6.21E-07 | 0.139 | TAGCAGTTGCCACCTGTGTGCTCTTGGGGGCGGGG | | |
| 182 | 2 | 23561284 | 23565193 | 3909 | FVB.SJL | 2 | 23564268 | 23564301 | 34 | - | 18.1057 | 3.29E-07 | 0.0588 | AAGCTGTCTTCAGACACTCCAGAAGAGGGCACCA | | |
| 183 | 2 | 28993499 | 29006418 | 12919 | SPRET | 2 | 28996883 | 28996916 | 34 | - | 19.748 | 1.06E-07 | 0.0313 | GTGCAGTGCCCAAGACATCCAAGAAAGGGCAGCA | | |
| 184 | 2 | 30820322 | 30821486 | 1164 | SPRET | 2 | 30821077 | 30821110 | 34 | - | 16.7967 | 7.71E-07 | 0.0917 | ATGCAGGGCACTCACTGGCCAAGAGTGGGCGGAG | | |
| 185 | 2 | 49101210 | 49106486 | 5276 | C58J.FVB.KK.PL.SJL.TH | 2 | 49102120 | 49102153 | 34 | - | 17.4715 | 5.00E-07 | 0.0729 | ACACAGTGCCCGAGGAGAACAGAAGAGGGCACTG | 4 | ENCFF093ZRU.ENCFF367UNR.ENCFF099QNE.ENCFF821BIA |
| 186 | 2 | 57104419 | 57113404 | 8985 | 129S1,AJ,AKR,BUB,C3H,C58J,CAST,CBA,CE,DBA1J,DBA2J,FVB,ILNJ2,KK,MOLF,MRL,NOD,NOR,NZB,NZO,PJ,PL,RF,SEA,SJL,SMJ.SPRET.SWR.TH.WSB | 2 | 57107974 | 57107992 | 19 | - | 17.1639 | 8.17E-07 | 0.141 | CCTCCACGAGGTGGCGGAA | | |
| 187 | 2 | 71290350 | 71291766 | 1416 | SPRET | 2 | 71290669 | 71290702 | 34 | - | 17.1382 | 6.20E-07 | 0.0819 | CTGTCCATCCCTTCTTTGCCCCCAGGTGGCTCCT | | |
| 188 | 2 | 71386387 | 71387509 | 1122 | SPRET | 2 | 71386607 | 71386625 | 19 | - | 21.2131 | 4.09E-08 | 0.0407 | AGGCCAGAAGGGGGCGGTG | 5 | ENCFF093ZRU.ENCFF821BIA.ENCFF136ZDK.ENCFF502WDT.ENCFF052KVO |
| 188 | 2 | 71386387 | 71387509 | 1122 | SPRET | 2 | 71386607 | 71386641 | 35 | - | 21.0917 | 4.24E-08 | 0.0295 | ATGTAGGAGCTCATGAAGGCCAGAAGGGGGCGGTG | 5 | ENCFF093ZRU.ENCFF821BIA.ENCFF136ZDK.ENCFF502WDT.ENCFF052KVO |
| 189 | 2 | 72784752 | 72787050 | 2298 | WSB | 2 | 72786942 | 72786975 | 34 | - | 17.8618 | 3.87E-07 | 0.0639 | TAGCTGTCTTCAGACCCACCAGAAGAGGGCATCA | 5 | ENCFF367UNR.ENCFF821BIA.ENCFF055HAP.ENCFF502WDT.ENCFF093ZRU |
| 190 | 2 | 79608322 | 79609546 | 1224 | BTBR,KK,NZB,NZO | 2 | 79608330 | 79608348 | 19 | - | 17.5738 | 6.25E-07 | 0.124 | TGAACACTAGAGGGAGCCA | 18 | ENCFF327LYS,ENCFF821BIA,ENCFF092NMJ,ENCFF024JOZ,ENCFF165LXO,ENCFF052KVO,ENCFF589ZBJ,ENCFF065ODV,ENCFF430PPJ,ENCFF093ZRU,ENCFF883UPM,ENCFF198SSR,ENCFF367UNR,ENCFF907XEG.ENCFF502WDT.ENCFF055HAP.ENCFF099QNE.ENCFF089GBC |
| 191 | 2 | 83063735 | 83066498 | 2763 | MOLF | 2 | 83065149 | 83065183 | 35 | - | 16.1101 | 8.98E-07 | 0.172 | CTGCATCTCTTTCCATTGGCTACTGGGTGGAGCCT | | |
| 192 | 2 | 86377777 | 86378838 | 1061 | SPRET | 2 | 86378815 | 86378833 | 19 | - | 18.3607 | 3.67E-07 | 0.0983 | TCTCCAGAAGATGGCACCA | | |
| 193 | 2 | 86475324 | 86481482 | 6158 | SPRET | 2 | 86475423 | 86475457 | 35 | - | 18.2661 | 2.56E-07 | 0.0863 | GTGCAGGGGAACGCCAGGGCCAAAAGGGGGGAGTG | | |
| 194 | 2 | 86689698 | 86692361 | 2663 | SPRET | 2 | 86691778 | 86691811 | 34 | - | 17.5691 | 4.69E-07 | 0.0703 | ATGCAGCTAGAGACATGATCTCCAGGGGGTACTG | | |
| 195 | 2 | 90206605 | 90211464 | 4859 | CAST | 2 | 90211164 | 90211197 | 34 | - | 20.2195 | 7.58E-08 | 0.025 | CTGCAAGCCTTGCAGTGGACACCAGAGGTTAGTG | | |
| 195 | 2 | 90206605 | 90211464 | 4859 | CAST | 2 | 90211207 | 90211240 | 34 | - | 20.2195 | 7.58E-08 | 0.025 | CTGCAAGCCTTGCAGTGGACACCAGAGGTTAGTG | | |
| 195 | 2 | 90206605 | 90211464 | 4859 | CAST | 2 | 90211250 | 90211283 | 34 | - | 20.2195 | 7.58E-08 | 0.025 | CTGCAAGCCTTGCAGTGGACACCAGAGGTTAGTG | | |
| 196 | 2 | 92275010 | 92278638 | 3628 | SPRET | 2 | 92277888 | 92277906 | 19 | - | 16.918 | 9.56E-07 | 0.151 | AGGCCAGCAGAGGGTGCTT | 7 | ENCFF089GBC,ENCFF092NMJ,ENCFF093ZRU,ENCFF055HAP,ENCFF099QNE,ENCFF327LYS,ENCFF367UNR |
| 196 | 2 | 92275010 | 92278638 | 3628 | SPRET | 2 | 92277888 | 92277921 | 34 | - | 18.626 | 2.32E-07 | 0.0513 | GCCCAGTGCCAATGGAGGCCAGCAGAGGGTGCTT | 7 | ENCFF089GBC,ENCFF092NMJ,ENCFF093ZRU,ENCFF055HAP,ENCFF099QNE,ENCFF327LYS,ENCFF367UNR |
| 197 | 2 | 97288730 | 97291486 | 2756 | CAST | 2 | 97289980 | 97290013 | 34 | - | 18.3089 | 2.87E-07 | 0.0543 | ATGCAGCTAGAGACACGAGCTCCAGGGGGTACTG | | |
| 198 | 2 | 98126174 | 98130194 | 4020 | MOLF | 2 | 98130093 | 98130126 | 34 | - | 20.2114 | 7.63E-08 | 0.0251 | ATGCAGCTAGAGACATGAGCTCTAGGGGGCACTA | | |
| 199 | 2 | 99374632 | 99376981 | 2349 | NZO | 2 | 99376003 | 99376036 | 34 | - | 18.5447 | 2.45E-07 | 0.0522 | CCGCACCCACATTCGCCGTCACAAGATGGCGCTG | | |
| 200 | 2 | 99375563 | 99382994 | 7431 | 129S1,AJ,BTBR,BUB,CAST,FVB,ILNJ2,KK,MOLF,NZB,NZW,SJL,SMJ,SPRET,SWR | 2 | 99376003 | 99376036 | 34 | - | 18.5447 | 2.45E-07 | 0.0522 | CCGCACCCACATTCGCCGTCACAAGATGGCGCTG | | |
| 201 | 2 | 102791267 | 102792953 | 1686 | SPRET | 2 | 102792726 | 102792759 | 34 | - | 18.3089 | 2.87E-07 | 0.0543 | ATGCAGCTAGAGACACGAGCTCCAGGGGGTACTG | | |
| 202 | 2 | 104878875 | 104882288 | 3413 | SPRET | 2 | 104881673 | 104881707 | 35 | - | 18.2936 | 2.52E-07 | 0.0853 | GTGCATGTGCACCACCTGACCAGCAGAAGATGCCA | | |
| 203 | 2 | 115161626 | 115165598 | 3972 | C3H.CBA.CE | 2 | 115161998 | 115162031 | 34 | - | 16.8211 | 7.59E-07 | 0.0913 | GTGCAGGTACCTTGTAGGTCAACAGAGGGCATTA | | |
| 204 | 2 | 118483741 | 118484852 | 1111 | SPRET | 2 | 118484290 | 118484323 | 34 | - | 16.6667 | 8.37E-07 | 0.0956 | CTGCAACTCTCTTCAGACTCACCAGAAGGAGGCA | | |
| 205 | 2 | 119498599 | 119503375 | 4776 | SPRET | 2 | 119499725 | 119499759 | 35 | - | 18.3394 | 2.45E-07 | 0.0834 | ATGAGGTTTCTTCACTCCTCCAGTAGAGGGCGCAT | 22 | ENCFF210BTO,ENCFF093ZRU,ENCFF367UNR,ENCFF430PPJ,ENCFF092NMJ,ENCFF089GBC,ENCFF327LYS,ENCFF136ZDK,ENCFF052KVO,ENCFF099QNE,ENCFF883UPM,ENCFF907XEG,ENCFF097CAQ,ENCFF609MDN,ENCFF589ZBJ,ENCFF024JOZ,ENCFF821BIA,ENCFF198SSR,ENCFF575MGI,ENCFF281QMV,ENCFF957SPH.ENCFF165LXO |
| 206 | 2 | 121651474 | 121653056 | 1582 | CAST | 2 | 121651658 | 121651691 | 34 | - | 17.122 | 6.26E-07 | 0.0825 | TAGCTGTCTTCAGACACACCAGAAGAGGGCACAA | | |
| 207 | 2 | 125437118 | 125438228 | 1110 | SPRET | 2 | 125437319 | 125437337 | 19 | - | 18.7705 | 2.75E-07 | 0.0841 | TGGCCAGAAGAGGGCATCA | 1 | ENCFF367UNR |
| 208 | 2 | 127291931 | 127295543 | 3612 | CAST | 2 | 127295031 | 127295049 | 19 | - | 18.7049 | 2.88E-07 | 0.0857 | CAGACACCAGAGGGCAGTG | | |
| 209 | 2 | 131564341 | 131567208 | 2867 | SPRET | 2 | 131565257 | 131565290 | 34 | - | 19 | 1.79E-07 | 0.0436 | ATGCAGTGTCACAAACCATCACCAGGGAGCACGT | | |
| 209 | 2 | 131564341 | 131567208 | 2867 | SPRET | 2 | 131565297 | 131565330 | 34 | - | 19 | 1.79E-07 | 0.0436 | ATGCAGTGTCACAAACCATCACCAGGGAGCACGT | | |
| 210 | 2 | 132183235 | 132189870 | 6635 | ILNJ2.MOLF.PJ | 2 | 132183886 | 132183919 | 34 | - | 16.3984 | 9.91E-07 | 0.0999 | TAGCTGTCTTCAGACGTACCAGAAGAGGGCATCA | | |
| 210 | 2 | 132183235 | 132189870 | 6635 | ILNJ2.MOLF.PJ | 2 | 132184597 | 132184630 | 34 | - | 16.4065 | 9.86E-07 | 0.0997 | GGACAGTACCCATGGACACCAGAAGAGGGCGTCA | 1 | ENCFF093ZRU |
| 211 | 2 | 132723584 | 132725394 | 1810 | CAST | 2 | 132724408 | 132724441 | 34 | - | 17.3984 | 5.24E-07 | 0.0749 | TAGCTGTTTTCAGGCACACCAGAAGAGGGCATCC | | |
| 212 | 2 | 139875792 | 139880002 | 4210 | C58J.DBA1J.DBA2J.MOLF.NZO.NZW | 2 | 139877940 | 139877973 | 34 | - | 17.7398 | 4.19E-07 | 0.0668 | GAGCAGTCACTGGGACTACAGGCAGGTGGCAGTG | 1 | ENCFF093ZRU |
| 213 | 2 | 150339866 | 150342833 | 2967 | BUB,CAST,MA/My,MOLF,NOD,NZB,NZW,RF.SPRET | 2 | 150340649 | 150340682 | 34 | - | 21.8293 | 2.28E-08 | 0.0105 | CTGCAATGCCCTTCACAGCAAGAAGGGGTAGAA | | |
| 214 | 2 | 150936711 | 150944533 | 7822 | NOD.NZB.NZW.RF | 2 | 150943764 | 150943798 | 35 | - | 15.9817 | 9.65E-07 | 0.178 | CTGTAGTGAGCCTTCTGGACAAGCTGGGGGCAGAT | | |
| 215 | 2 | 154514164 | 154515454 | 1290 | WSB | 2 | 154515222 | 154515255 | 34 | - | 16.8943 | 7.24E-07 | 0.0886 | TTGCTCTCTTCAGACATACCAGAAGAGGGCATCA | | |
| 216 | 2 | 154856866 | 154871571 | 14705 | 129S1,BALB,C3H,C57L,CAST,CBA,CE,FVB,MA/My,NOD,NZO,NZW,PL,SEA,SJL,SPRET,SWR,TH,WSB | 2 | 154861841 | 154861859 | 19 | - | 18.1475 | 4.25E-07 | 0.107 | CAGCCAGCAGATGGCGTTC | 30 | ENCFF093ZRU,ENCFF367UNR,ENCFF327LYS,ENCFF099QNE,ENCFF907XEG,ENCFF024JOZ,ENCFF821BIA,ENCFF201JUA,ENCFF316FYN,ENCFF589ZBJ,ENCFF089GBC,ENCFF198SSR,ENCFF575MGI,ENCFF097CAQ,ENCFF055HAP,ENCFF609MDN,ENCFF136ZDK,ENCFF502WDT,ENCFF165LXO,ENCFF430PPJ,ENCFF092NMJ,ENCFF951GOP,ENCFF194HVL,ENCFF491RJK,ENCFF957SPH,ENCFF883UPM,ENCFF281QMV,ENCFF210BTO.ENCFF142CNG.ENCFF065ODV |

| | | | | | | | | | | | | | | | | |
|---|---|---|---|---|---|---|---|---|---|---|---|---|---|---|---|---|
| 216 | 2 | 154856866 | 154871571 | 14705 | 129S1,BALB,C3H,C57L,CAST,CBA,CE,FVB,MA/My,NOD,NZO,NZW,PL,SEA,SJL,SPRET,SWR,TH,WSB | 2 | 154861841 | 154861875 | 35 | - | 17.1101 | 5.08E-07 | 0.126 | AAGTTTTTCAAAGGTCCAGCCAGCAGATGGCGTTC | 30 | ENCFF093ZRU,ENCFF367UNR,ENCFF327LYS,ENCFF099QNE,ENCFF907XEG,ENCFF024JOZ,ENCFF821BIA,ENCFF201JUA,ENCFF316FYN,ENCFF589ZBJ,ENCFF089GBC,ENCFF198SSR,ENCFF575MGI,ENCFF097CAQ,ENCFF055HAP,ENCFF609MDN,ENCFF136ZDK,ENCFF502WDT,ENCFF165LXO,ENCFF430PPJ,ENCFF092NMJ,ENCFF951GOP,ENCFF194HVL,ENCFF491RJK,ENCFF957SPH,ENCFF883UPM,ENCFF281QMV,ENCFF210BTO.ENCFF142CNG.ENCFF065ODV |
| 217 | 2 | 154859783 | 154868599 | 8816 | BTBR | 2 | 154861841 | 154861859 | 19 | - | 18.1475 | 4.25E-07 | 0.107 | CAGCCAGCAGATGGCGTTC | 30 | ENCFF093ZRU,ENCFF367UNR,ENCFF327LYS,ENCFF099QNE,ENCFF907XEG,ENCFF024JOZ,ENCFF821BIA,ENCFF201JUA,ENCFF316FYN,ENCFF589ZBJ,ENCFF089GBC,ENCFF198SSR,ENCFF575MGI,ENCFF097CAQ,ENCFF055HAP,ENCFF609MDN,ENCFF136ZDK,ENCFF502WDT,ENCFF165LXO,ENCFF430PPJ,ENCFF092NMJ,ENCFF951GOP,ENCFF194HVL,ENCFF491RJK,ENCFF957SPH,ENCFF883UPM,ENCFF281QMV,ENCFF210BTO.ENCFF142CNG.ENCFF065ODV |
| 217 | 2 | 154859783 | 154868599 | 8816 | BTBR | 2 | 154861841 | 154861875 | 35 | - | 17.1101 | 5.08E-07 | 0.126 | AAGTTTTTCAAAGGTCCAGCCAGCAGATGGCGTTC | 30 | ENCFF093ZRU,ENCFF367UNR,ENCFF327LYS,ENCFF099QNE,ENCFF907XEG,ENCFF024JOZ,ENCFF821BIA,ENCFF201JUA,ENCFF316FYN,ENCFF589ZBJ,ENCFF089GBC,ENCFF198SSR,ENCFF575MGI,ENCFF097CAQ,ENCFF055HAP,ENCFF609MDN,ENCFF136ZDK,ENCFF502WDT,ENCFF165LXO,ENCFF430PPJ,ENCFF092NMJ,ENCFF951GOP,ENCFF194HVL,ENCFF491RJK,ENCFF957SPH,ENCFF883UPM,ENCFF281QMV,ENCFF210BTO.ENCFF142CNG.ENCFF065ODV |
| 218 | 2 | 155421926 | 155423080 | 1154 | CAST | 2 | 155423015 | 155423049 | 35 | - | 17.3761 | 4.35E-07 | 0.117 | CTGGAGTTGTATCACTCACCCTCCAGAGGGGACTG | | |
| 219 | 2 | 155608519 | 155610591 | 2072 | MOLF | 2 | 155609299 | 155609332 | 34 | - | 21.2114 | 3.65E-08 | 0.0149 | GTGCAGTGCCTGAGGGGGCCAGAGGAGGGTGTGT | | |
| 220 | 2 | 156641184 | 156651433 | 10249 | SPRET | 2 | 156649405 | 156649438 | 34 | - | 16.5366 | 9.09E-07 | 0.0987 | TAGCTGTCTTCAGAGATACCAGAAGAGGGCATCA | | |
| 221 | 2 | 158577454 | 158582054 | 4600 | NZB | 2 | 158578364 | 158578382 | 19 | - | 18.623 | 3.06E-07 | 0.0887 | ACACCAGAAGAGGGCGCCA | 4 | ENCFF821BIA.ENCFF367UNR.ENCFF093ZRU.ENCFF099QNE |
| 221 | 2 | 158577454 | 158582054 | 4600 | NZB | 2 | 158578364 | 158578397 | 34 | - | 20.4146 | 6.58E-08 | 0.0228 | TAGCTGTTTTCAGATACACCAGAAGAGGGCGCCA | 4 | ENCFF821BIA.ENCFF367UNR.ENCFF093ZRU.ENCFF099QNE |
| 222 | 2 | 163814229 | 163817940 | 3711 | KK | 2 | 163817939 | 163817973 | 35 | - | 23.4495 | 8.05E-09 | 0.0114 | CTGCATGTACACCTGCATACCAGAAGAGGGCATCA | | |
| 223 | 2 | 170514414 | 170515430 | 1016 | KK | 2 | 170514897 | 170514931 | 35 | - | 24.2844 | 4.29E-09 | 0.00792 | GTGCAGTTTCCAGGGGAGGCCAGATGAGGGCAGTA | 2 | ENCFF093ZRU.ENCFF052KVO |
| 224 | 2 | 172069292 | 172070535 | 1243 | CE | 2 | 172069683 | 172069716 | 34 | - | 16.7724 | 7.83E-07 | 0.0917 | TAGCTGTCTTCAGATGCACCAGAAGAGGGCGTCA | | |
| 224 | 2 | 172069292 | 172070535 | 1243 | CE | 2 | 172069923 | 172069941 | 19 | - | 17.0328 | 8.88E-07 | 0.145 | AGGCCACAAGAGGGCATCA | 20 | ENCFF024JOZ,ENCFF093ZRU,ENCFF092NMJ,ENCFF327LYS,ENCFF907XEG,ENCFF430PPJ,ENCFF165LXO,ENCFF883UPM,ENCFF033UQX,ENCFF136ZDK,ENCFF099QNE,ENCFF055HAP,ENCFF065ODV,ENCFF533APC,ENCFF281QMV,ENCFF052KVO,ENCFF142CNG,ENCFF589ZBJ,ENCFF089GBC,ENCFF502WDT |
| 224 | 2 | 172069292 | 172070535 | 1243 | CE | 2 | 172069923 | 172069956 | 34 | - | 28.2846 | 7.03E-11 | 0.000209 | GTGCAGTGCCATTGGAGGCCACAAGAGGGCATCA | 20 | ENCFF024JOZ,ENCFF093ZRU,ENCFF092NMJ,ENCFF327LYS,ENCFF907XEG,ENCFF430PPJ,ENCFF165LXO,ENCFF883UPM,ENCFF033UQX,ENCFF136ZDK,ENCFF099QNE,ENCFF055HAP,ENCFF065ODV,ENCFF533APC,ENCFF281QMV,ENCFF052KVO,ENCFF142CNG,ENCFF589ZBJ,ENCFF089GBC,ENCFF502WDT |
| 225 | 3 | 3181454 | 3193756 | 12302 | CAST.MOLF | 3 | 3181871 | 3181904 | 34 | + | 18.3659 | 2.76E-07 | 0.0804 | TAGCTGTCTTCAGACACTCCAGAAGAGGGCGCCA | | |
| 225 | 3 | 3181454 | 3193756 | 12302 | CAST.MOLF | 3 | 3181886 | 3181904 | 19 | + | 17.5574 | 6.32E-07 | 0.181 | ACTCCAGAAGAGGGCGCCA | | |
| 226 | 3 | 10177986 | 10179884 | 1898 | CAST | 3 | 10178744 | 10178777 | 34 | + | 18.1626 | 3.17E-07 | 0.0852 | CTTCAGTTCAAATTCCTGGCTCAAGAGGGAACTA | | |
| 227 | 3 | 10286015 | 10288693 | 2678 | SPRET | 3 | 10286862 | 10286895 | 34 | + | 16.7886 | 7.75E-07 | 0.131 | GAGCATGGACACAAGCCCACAGCAGATGGCAGCA | | |
| 228 | 3 | 10594242 | 10596653 | 2411 | MOLF.SPRET | 3 | 10596603 | 10596636 | 34 | + | 18.8049 | 2.05E-07 | 0.0723 | ATGCAGTTCTGAATCCTCCCTGTAGGAGGGGGTC | | |
| 229 | 3 | 10595192 | 10597066 | 1874 | MOLF | 3 | 10596603 | 10596636 | 34 | + | 18.8049 | 2.05E-07 | 0.0723 | ATGCAGTTCTGAATCCTCCCTGTAGGAGGGGGTC | | |
| 230 | 3 | 12111701 | 12113864 | 2163 | SPRET | 3 | 12113552 | 12113585 | 34 | + | 17.3496 | 5.41E-07 | 0.111 | CTGCCCTCTCAGTCGCCGTTACAAGATGGCGCTG | | |
| 231 | 3 | 12501565 | 12502999 | 1434 | SPRET | 3 | 12502927 | 12502961 | 35 | + | 16.3119 | 8.02E-07 | 0.221 | GTGAAGCTGGCAGTCCCAGGCACCAGGTGGAGACA | | |
| 232 | 3 | 14733518 | 14739016 | 5498 | CAST.MOLF.NZO.SMJ.SPRET | 3 | 14737253 | 14737287 | 35 | + | 16.4404 | 7.46E-07 | 0.213 | AAGAAGCTGGACCCAGTGGCCAGCAGATGGCCCAC | | |
| 232 | 3 | 14733518 | 14739016 | 5498 | CAST.MOLF.NZO.SMJ.SPRET | 3 | 14737269 | 14737287 | 19 | + | 17.0328 | 8.88E-07 | 0.207 | TGGCCAGCAGATGGCCCAC | | |
| 233 | 3 | 24327722 | 24437946 | 110224 | NOD | 3 | 24375838 | 24375872 | 35 | + | 20.3303 | 7.02E-08 | 0.0549 | CAGCACAGCCTCATGGGTACCACAAGGGGGCATTC | 10 | ENCFF052KVO,ENCFF024JOZ,ENCFF327LYS,ENCFF092NMJ,ENCFF089GBC,ENCFF883UPM,ENCFF093ZRU.ENCFF502WDT.ENCFF142CNG.ENCFF907XEG |
| 233 | 3 | 24327722 | 24437946 | 110224 | NOD | 3 | 24382881 | 24382915 | 35 | + | 20.6881 | 5.55E-08 | 0.0469 | CTGCAGGTATCCAAGGAGGCCAGAAGAGGATGCTA | | |
| 233 | 3 | 24327722 | 24437946 | 110224 | NOD | 3 | 24410991 | 24411024 | 34 | + | 19.1463 | 1.62E-07 | 0.0645 | TTGCTGTCTTCGGACACACCAGAAGAGGGCTCCA | | |
| 234 | 3 | 26145866 | 26148355 | 2489 | MOLF | 3 | 26147272 | 26147306 | 35 | + | 15.9358 | 9.90E-07 | 0.244 | ATGCAGTGGCACACACTGCCCACTTGTTGGCCCAT | | |
| 235 | 3 | 27337895 | 27339047 | 1152 | SPRET | 3 | 27338547 | 27338581 | 35 | + | 27.6881 | 2.57E-10 | 0.00321 | GTGCAGATCCACCGTAAGGCCAGAAGAGGGCGCAA | 16 | ENCFF024JOZ,ENCFF821BIA,ENCFF907XEG,ENCFF430PPJ,ENCFF201JUA,ENCFF367UNR,ENCFF092NMJ,ENCFF589ZBJ,ENCFF198SSR,ENCFF093ZRU,ENCFF502WDT,ENCFF052KVO,ENCFF033UQX,ENCFF099QNE.ENCFF327LYS.ENCFF089GBC |
| 235 | 3 | 27337895 | 27339047 | 1152 | SPRET | 3 | 27338563 | 27338581 | 19 | + | 19.3443 | 1.82E-07 | 0.111 | AGGCCAGAAGAGGGCGCAA | 16 | ENCFF024JOZ,ENCFF821BIA,ENCFF907XEG,ENCFF430PPJ,ENCFF201JUA,ENCFF367UNR,ENCFF092NMJ,ENCFF589ZBJ,ENCFF198SSR,ENCFF093ZRU,ENCFF502WDT,ENCFF052KVO,ENCFF033UQX,ENCFF099QNE.ENCFF327LYS.ENCFF089GBC |
| 236 | 3 | 28906501 | 28912013 | 5512 | MOLF | 3 | 28911123 | 28911156 | 34 | + | 19.5285 | 1.24E-07 | 0.0556 | CCCCACTTCACAGGGCGTCCTCCAGGTGGCAGCC | | |
| 236 | 3 | 28906501 | 28912013 | 5512 | MOLF | 3 | 28911138 | 28911156 | 19 | + | 18.9672 | 2.39E-07 | 0.122 | CGTCCTCCAGGTGGCAGCC | | |
| 237 | 3 | 37444733 | 37449592 | 4859 | 129S1,AJ,AKR,BTBR,C3H,C57L,CAST,CBA,CE,DBA1J,DBA2J,ILNJ2,MA/My,MRL,NOD,NOR,NZB,NZO,NZW,PL,RF,SJL,SMJ,SPRET,SWR,TH,WSB | 3 | 37445319 | 37445352 | 34 | + | 16.4472 | 9.61E-07 | 0.141 | TAGCTGTCTTCAGACACACCAGAAGAGGGCGTCA | | |
| 238 | 3 | 42733346 | 42736195 | 2849 | MOLF.NZW | 3 | 42733588 | 42733622 | 35 | + | 16.5229 | 7.12E-07 | 0.208 | CTGCAGAAAGCTCTGGCTACCACAAAGGGGCAGGG | | |
| 239 | 3 | 44549501 | 44555372 | 5871 | SPRET | 3 | 44553995 | 44554013 | 19 | + | 19.6885 | 1.40E-07 | 0.101 | TGACCTGGAGGGGGCGCTA | | |
| 240 | 3 | 45451014 | 45455023 | 4009 | CAST | 3 | 45452446 | 45452480 | 35 | + | 18.1193 | 2.80E-07 | 0.126 | CTGCAGACACCCCCAAGGTCCCCTAGAGGACTCTC | | |
| 241 | 3 | 47208010 | 47211257 | 3247 | SJL.SWR | 3 | 47208972 | 47209005 | 34 | + | 21.2033 | 3.67E-08 | 0.0236 | ATGCAGCTAGAGACACGAGCTCCAGGGGGCACTG | | |
| 241 | 3 | 47208010 | 47211257 | 3247 | SJL.SWR | 3 | 47208987 | 47209005 | 19 | + | 16.8525 | 9.96E-07 | 0.218 | CGAGCTCCAGGGGGCACTG | | |
| 242 | 3 | 59670841 | 59680801 | 9960 | WSB | 3 | 59679833 | 59679867 | 35 | + | 16.9083 | 5.71E-07 | 0.185 | CTGTAGTTGACGGCCCAGGACTGAAGGGGTCAGGC | | |
| 243 | 3 | 62915214 | 62925746 | 10532 | SPRET | 3 | 62923972 | 62924005 | 34 | + | 17.374 | 5.32E-07 | 0.109 | GTGCAGTGTCCACAGATGCATGAAGTGGGCACTG | | |
| 244 | 3 | 67003406 | 67011127 | 7721 | CAST.CE.MOLF.SJL.SPRET.WSB | 3 | 67007657 | 67007675 | 19 | + | 17.1639 | 8.17E-07 | 0.199 | CCTCCACGAGGTGGCGGAA | | |
| 245 | 3 | 67541336 | 67542448 | 1112 | MOLF | 3 | 67541733 | 67541766 | 34 | + | 17.9593 | 3.63E-07 | 0.091 | CTGGAGGGCTTTGCCAGAGCACCTGGGGGCGCCA | | |
| 245 | 3 | 67541336 | 67542448 | 1112 | MOLF | 3 | 67541721 | 67541754 | 34 | - | 16.6748 | 8.33E-07 | 0.135 | GTGCTCTGGCAAAGCCCTCCAGGACGGGGCACAC | | |
| 246 | 3 | 69360956 | 69362295 | 1339 | AKR,BUB,CBA,CE,DBA1J,DBA2J,MRL,NZO,RF,SJL | 3 | 69361852 | 69361870 | 19 | + | 17.4918 | 6.59E-07 | 0.185 | AACCCACCAGGGGGCGACA | 4 | ENCFF093ZRU,ENCFF367UNR,ENCFF821BIA,ENCFF883UPM |
| 247 | 3 | 74039020 | 74042728 | 3708 | SPRET | 3 | 74039756 | 74039789 | 34 | + | 17.122 | 6.26E-07 | 0.118 | ATGCAGTGCTCAAGGAGGTCAGAAGAGAGCATCA | | |
| 248 | 3 | 87915117 | 87921671 | 6554 | MOLF | 3 | 87918095 | 87918128 | 34 | + | 16.5203 | 9.18E-07 | 0.141 | CGTTAATTCTCAGAGGATCCAGAAGAGGGCGCTG | 31 | ENCFF201JUA,ENCFF367UNR,ENCFF093ZRU,ENCFF099QNE,ENCFF821BIA,ENCFF805QBU,ENCFF065ODV,ENCFF092NMJ,ENCFF052KVO,ENCFF055HAP,ENCFF609MDN,ENCFF327LYS,ENCFF430PPJ,ENCFF502WDT,ENCFF316FYN,ENCFF907XEG,ENCFF589ZBJ,ENCFF198SSR,ENCFF033UQX,ENCFF024JOZ,ENCFF089GBC,ENCFF194HVL,ENCFF136ZDK,ENCFF142CNG,ENCFF491RJK,ENCFF951GOP,ENCFF883UPM,ENCFF210BTO.ENCFF957SPH.ENCFF281QMV.ENCFF533APC |
| 248 | 3 | 87915117 | 87921671 | 6554 | MOLF | 3 | 87918110 | 87918128 | 19 | + | 17.2623 | 7.66E-07 | 0.195 | GATCCAGAAGAGGGCGCTG | 31 | ENCFF201JUA,ENCFF367UNR,ENCFF093ZRU,ENCFF099QNE,ENCFF821BIA,ENCFF805QBU,ENCFF065ODV,ENCFF092NMJ,ENCFF052KVO,ENCFF055HAP,ENCFF609MDN,ENCFF327LYS,ENCFF430PPJ,ENCFF502WDT,ENCFF316FYN,ENCFF907XEG,ENCFF589ZBJ,ENCFF198SSR,ENCFF033UQX,ENCFF024JOZ,ENCFF089GBC,ENCFF194HVL,ENCFF136ZDK,ENCFF142CNG,ENCFF491RJK,ENCFF951GOP,ENCFF883UPM,ENCFF210BTO.ENCFF957SPH.ENCFF281QMV.ENCFF533APC |
| 248 | 3 | 87915117 | 87921671 | 6554 | MOLF | 3 | 87918388 | 87918421 | 34 | - | 16.8537 | 7.43E-07 | 0.129 | GTGCATCACCATGCCCAGCCTCTAGGGGGACTTT | 5 | ENCFF201JUA.ENCFF367UNR.ENCFF093ZRU.ENCFF099QNE.ENCFF024JOZ |
| 249 | 3 | 88179646 | 88182234 | 2588 | SPRET | 3 | 88181484 | 88181517 | 34 | + | 23.6179 | 5.43E-09 | 0.00602 | GTGCAGTAACCCTGCAGGCCAGAAGAGGGCGTAA | 19 | ENCFF907XEG,ENCFF327LYS,ENCFF136ZDK,ENCFF198SSR,ENCFF089GBC,ENCFF589ZBJ,ENCFF092NMJ,ENCFF201JUA,ENCFF065ODV,ENCFF883UPM,ENCFF093ZRU,ENCFF502WDT,ENCFF099QNE,ENCFF367UNR,ENCFF142CNG,ENCFF533APC,ENCFF052KVO,ENCFF821BIA,ENCFF055HAP |
| 250 | 3 | 90033749 | 90034914 | 1165 | MOLF | 3 | 90034783 | 90034816 | 34 | + | 17.3496 | 5.41E-07 | 0.111 | TTGCTGTACCAGAAGAGGGCAACAGGTGTCATTA | | |
| 251 | 3 | 90048748 | 90050325 | 1577 | SPRET | 3 | 90049424 | 90049457 | 34 | + | 16.4553 | 9.56E-07 | 0.141 | CAGCACTCACAGAGGCAGGCAGAGGAAGGCAGCA | | |
| 252 | 3 | 91035376 | 91134392 | 99016 | NOD.NOR.RF | 3 | 91130656 | 91130690 | 35 | + | 17.844 | 3.30E-07 | 0.139 | CGGCAGTATCCTGAGTCAAACAGCAGAGGGAGACA | 5 | ENCFF093ZRU.ENCFF052KVO.ENCFF089GBC.ENCFF092NMJ.ENCFF099QNE |
| 253 | 3 | 92469625 | 92470962 | 1337 | CAST.MOLF.SPRET | 3 | 92469958 | 92469991 | 34 | + | 23.8943 | 4.31E-09 | 0.00518 | GTGCAGTGCCCACAAGTGCCAAAAGAGGGCATGA | | |
| 254 | 3 | 93744648 | 93757424 | 12776 | MOLF | 3 | 93745350 | 93745383 | 34 | + | 16.561 | 8.95E-07 | 0.14 | CTGCCCTTGCAATCGCCATTACAAGATGGCGCTG | | |
| 255 | 3 | 93800269 | 93807745 | 7476 | C3H,CBA,DBA1J,DBA2J,FVB,KK,MOLF,MRL | 3 | 93800436 | 93800470 | 35 | + | 21.2752 | 3.74E-08 | 0.0363 | CTGTAGTAACAAGAAACAACCAGAAGATGGTGGTC | | |
| 256 | 3 | 93807151 | 93812187 | 5036 | 129S1,AJ,AKR,C3H,CBA,DBA1J,DBA2J,FVB,KK,MRL,NOD,NOR,NZW,RF,SJL,SPRET,WSB | 3 | 93807912 | 93807946 | 35 | + | 20.6972 | 5.51E-08 | 0.0467 | CTGTAGTAAAAAGTTACAACCAGAAGATGGTGGTC | | |
| 257 | 3 | 96994099 | 96996810 | 2711 | MOLF,SPRET | 3 | 96994834 | 96994852 | 19 | + | 25.6066 | 2.33E-10 | 0.00962 | TGGCCACCAGAGGGCGCTC | 20 | ENCFF201JUA,ENCFF024JOZ,ENCFF367UNR,ENCFF092NMJ,ENCFF093ZRU,ENCFF165LXO,ENCFF065ODV,ENCFF099QNE,ENCFF198SSR,ENCFF089GBC,ENCFF502WDT,ENCFF055HAP,ENCFF327LYS,ENCFF907XEG,ENCFF142CNG,ENCFF883UPM,ENCFF589ZBJ,ENCFF136ZDK,ENCFF957SPH,ENCFF821BIA |
| 257 | 3 | 96994099 | 96996810 | 2711 | MOLF.SPRET | 3 | 96995790 | 96995824 | 35 | + | 16.8991 | 5.74E-07 | 0.185 | TTGACCCTATGTCAGAGGGCCAGCAGGTGGCGGTG | 1 | ENCFF883UPM |
| 257 | 3 | 96994099 | 96996810 | 2711 | MOLF.SPRET | 3 | 96995806 | 96995824 | 19 | + | 22.7541 | 9.85E-09 | 0.0386 | GGGCCAGCAGGTGGCGGTG | 1 | ENCFF883UPM |
| 258 | 3 | 97262713 | 97264443 | 1730 | SPRET | 3 | 97263524 | 97263557 | 34 | + | 18.1545 | 3.18E-07 | 0.0853 | ATGCTGTCTTCATGCACACCAGAAGAGGGCATCA | | |
| 259 | 3 | 98303333 | 98318999 | 15666 | CAST,MOLF,NOD,NOR,NZB,NZO,NZW,SPRET | 3 | 98310981 | 98311015 | 35 | + | 24.4037 | 3.92E-09 | 0.01 | ATGCAGGTACCAACAGAGGCCAGGAGAGGGCACCA | | |
| 259 | 3 | 98303333 | 98318999 | 15666 | CAST,MOLF,NOD,NOR,NZB,NZO,NZW,SPRET | 3 | 98310997 | 98311015 | 19 | + | 19.4918 | 1.63E-07 | 0.107 | AGGCCAGGAGAGGGCACCA | | |
| 260 | 3 | 98309576 | 98318661 | 9085 | 129S1,BTBR,C57L,DBA1J,DBA2J,FVB,PJ,SEA,SJL,SMJ,SWR,WSB | 3 | 98310981 | 98311015 | 35 | + | 24.4037 | 3.92E-09 | 0.01 | ATGCAGGTACCAACAGAGGCCAGGAGAGGGCACCA | | |
| 260 | 3 | 98309576 | 98318661 | 9085 | 129S1,BTBR,C57L,DBA1J,DBA2J,FVB,PJ,SEA,SJL,SMJ,SWR,WSB | 3 | 98310997 | 98311015 | 19 | + | 19.4918 | 1.63E-07 | 0.107 | AGGCCAGGAGAGGGCACCA | | |
| 261 | 3 | 104658855 | 104660384 | 1529 | NZO.NZW | 3 | 104659367 | 104659400 | 34 | + | 16.561 | 8.95E-07 | 0.14 | ATGCTGGGGATCAGGGGGCCTGAAGAGGGAACTA | | |
| 261 | 3 | 104658855 | 104660384 | 1529 | NZO,NZW | 3 | 104660334 | 104660368 | 35 | + | 16.3211 | 7.98E-07 | 0.221 | GTGAGGCTTCTTCACACCGCCGGTAGAGGGCGCGC | 17 | ENCFF092NMJ,ENCFF089GBC,ENCFF093ZRU,ENCFF367UNR,ENCFF165LXO,ENCFF099QNE,ENCFF136ZDK,ENCFF821BIA,ENCFF033UQX,ENCFF907XEG,ENCFF327LYS,ENCFF609MDN,ENCFF589ZBJ,ENCFF957SPH.ENCFF194HVL.ENCFF198SSR.ENCFF024JOZ |
| 261 | 3 | 104658855 | 104660384 | 1529 | NZO,NZW | 3 | 104660350 | 104660368 | 19 | + | 16.9672 | 9.26E-07 | 0.212 | CCGCCGGTAGAGGGCGCGC | 17 | ENCFF092NMJ,ENCFF089GBC,ENCFF093ZRU,ENCFF367UNR,ENCFF165LXO,ENCFF099QNE,ENCFF136ZDK,ENCFF821BIA,ENCFF033UQX,ENCFF907XEG,ENCFF327LYS,ENCFF609MDN,ENCFF589ZBJ,ENCFF957SPH.ENCFF194HVL.ENCFF198SSR.ENCFF024JOZ |
| 262 | 3 | 106151170 | 106201848 | 50678 | SPRET | 3 | 106164953 | 106164987 | 35 | + | 16.1101 | 8.98E-07 | 0.233 | GTGAAGCTGGAAATCTCAGGCACCAGGTGGAGGCA | | |
| 263 | 3 | 116884656 | 116886383 | 1727 | NOD.NOR | 3 | 116885401 | 116885434 | 34 | + | 17.1301 | 6.23E-07 | 0.118 | CTGCAGTGCATGTGCAGGTCAGATGAGGGTGATG | | |
| 264 | 3 | 124595937 | 124598433 | 2496 | SPRET | 3 | 124596146 | 124596179 | 34 | + | 17.3089 | 5.55E-07 | 0.112 | ATGCAGCTAGAGACACGAGCTCCAGAGGGTACTG | | |
| 265 | 3 | 124663949 | 124670025 | 6076 | AJ,BALB,C3H,CBA,MOLF,NZB,RH2,SMJ,SPRET | 3 | 124670009 | 124670043 | 35 | + | 17.1651 | 4.92E-07 | 0.174 | CTGCAGTCTCAGGTTCCTGCCACCTGAGGTTTTGA | | |
| 266 | 3 | 126014210 | 126019598 | 5388 | SPRET | 3 | 126014552 | 126014585 | 34 | + | 16.4309 | 9.71E-07 | 0.142 | CTGCAAGAGGGGAGCTGGCCAGCAGAGAGTACTC | | |
| 267 | 3 | 127165543 | 127166576 | 1033 | 129S1,AJ,BALB,C3H,C57L,CBA,NZB,RH2,SMJ | 3 | 127165619 | 127165652 | 34 | + | 23.7073 | 5.04E-09 | 0.00582 | GTGCAGTGCTCACAGAGGCCAGACGAGGGAGCTA | 7 | ENCFF367UNR,ENCFF821BIA,ENCFF093ZRU,ENCFF089GBC,ENCFF201JUA,ENCFF502WDT,ENCFF099QNE |
| 268 | 3 | 131249997 | 131253715 | 3718 | CAST.MOLF.SPRET | 3 | 131250013 | 131250046 | 34 | + | 17.9106 | 3.74E-07 | 0.0926 | CCGCCCTCACATTCGCCGTCACAAGATGGCGCTG | | |
| 268 | 3 | 131249997 | 131253715 | 3718 | CAST.MOLF.SPRET | 3 | 131253318 | 131253351 | 34 | + | 17.9106 | 3.74E-07 | 0.0926 | CCGCCCTCACATTCGCCGTCACAAGATGGCGCTG | | |
| 269 | 3 | 132685181 | 132686256 | 1075 | SPRET | 3 | 132685994 | 132686028 | 35 | + | 23.9266 | 5.63E-09 | 0.0124 | CTGCAGGTGCTCTCAGGGGGCAGAAGAGGGCGCTC | 4 | ENCFF093ZRU.ENCFF052KVO.ENCFF883UPM.ENCFF533APC |
| 270 | 3 | 133207572 | 133210278 | 2706 | NZB.NZO.NZW | 3 | 133208832 | 133208850 | 19 | + | 17.6066 | 6.11E-07 | 0.181 | AGGCCAGAAGAGGGCACTT | 5 | ENCFF821BIA.ENCFF907XEG.ENCFF327LYS.ENCFF367UNR.ENCFF093ZRU |
| 271 | 3 | 134793714 | 134795598 | 1884 | CAST,CE,DBA1J,DBA2J,FVB,KK,MA/My,MOLF,NOD,NOR,PL,SJL,SMJ,SPRET,SWR,TH | 3 | 134794727 | 134794761 | 35 | + | 16.4404 | 7.46E-07 | 0.213 | TTGCAATTGACGGCCCAGGACTGAAGGGGTCACGC | | |
| 272 | 3 | 139265775 | 139269940 | 4165 | 129S1,AJ,AKR,BALB,BTBR,BUB,C3H,C57L,CAST,CBA,DBA1J,DBA2J,FVB,ILNJ2,KK,MA/My,MOLF,MRL,NOD,NOR,NZB,NZO,NZW,PJ,PL,RH2,SEA,SJL,SMJ,SPRET,SWR,TH,WSB | 3 | 139266552 | 139266585 | 34 | + | 17.6748 | 4.37E-07 | 0.0998 | GAGCATGACCACAAGTCCACAGCAGGTGGCAACG | | |
| 273 | 3 | 145851498 | 145853085 | 1587 | 129S1,AJ,AKR,BALB,BTBR,BUB,C3H,C57L,CBA,DBA1J,ILNJ2,MA/My,NOD,NOR,NZB,NZO,PJ,RH2,SEA,SJL,SMJ | 3 | 145851473 | 145851506 | 34 | + | 19.6423 | 1.15E-07 | 0.0524 | CTGCAATTCCTAAAGGCTACAGTAGGTGGGGAGG | 4 | ENCFF502WDT,ENCFF327LYS,ENCFF883UPM,ENCFF055HAP |
| 274 | 3 | 152028772 | 152037819 | 9047 | 129S1,AJ,AKR,BALB,BTBR,C3H,C57L,CAST,CBA,CE,DBA1J,ILNJ2,KK,MOLF,MRL,PL,RH2,SEA,SMJ,SPRET,WSB | 3 | 152034233 | 152034251 | 19 | + | 17.1639 | 8.17E-07 | 0.199 | CCTCCACGAGGTGGCGGAA | | |
| 275 | 3 | 154052135 | 154053350 | 1215 | MOLF | 3 | 154052448 | 154052482 | 35 | + | 16.6422 | 6.65E-07 | 0.199 | GTGCAATTGCCCACAGAGGCCAGAAAGGGGTGTCC | | |
| 276 | 3 | 156194087 | 156198183 | 4096 | SPRET | 3 | 156195779 | 156195797 | 19 | + | 18.5246 | 3.27E-07 | 0.135 | TAGCCACTAGAGGGCAAAG | | |
| 277 | 3 | 159530474 | 159532919 | 2445 | SPRET | 3 | 159532852 | 159532885 | 34 | + | 21.3252 | 3.35E-08 | 0.0223 | GTGCAGTGCCCATGGAATCCAGAAGAGGACATCG | | |
| 278 | 3 | 3054105 | 3055688 | 1583 | CAST.MOLF | 3 | 3055321 | 3055354 | 34 | - | 16.5041 | 9.27E-07 | 0.141 | CTGTGGTGTCACAGTGCTCCACTGGGTGTCACAG | | |
| 279 | 3 | 4127873 | 4132298 | 4425 | CAST | 3 | 4131556 | 4131589 | 34 | - | 19.2033 | 1.56E-07 | 0.0632 | CAGCATGACCACAAGTCCACAGCAGGTGGCAACA | | |
| 280 | 3 | 14294257 | 14297501 | 3244 | SPRET | 3 | 14296164 | 14296197 | 34 | - | 18.3333 | 2.82E-07 | 0.0804 | GAGCAGGTGCGGCCGCCACCCCAAGATGGTGCCC | | |
| 281 | 3 | 14369646 | 14375807 | 6161 | CAST | 3 | 14375420 | 14375453 | 34 | - | 16.7724 | 7.83E-07 | 0.131 | CTGCAAGACGAGAGCTTGCCTGCAGAGAGTACTC | | |
| 282 | 3 | 15946000 | 15952309 | 6309 | AJ,BALB,BUB,C3H,C58J,CBA,FVB,MOLF,PL,SJL,SMJ,SWR,TH | 3 | 15951468 | 15951502 | 35 | - | 20.3945 | 6.73E-08 | 0.0532 | CTGCTGATCTCCTTGGCTATCAGCAGGTGGCAGAG | | |
| 283 | 3 | 17761787 | 17765886 | 4099 | SMJ.SPRET | 3 | 17765885 | 17765919 | 35 | - | 17 | 5.42E-07 | 0.182 | GTGCACTTCTGCAACCCTAGCACCAGGGGAGGCAG | | |
| 284 | 3 | 19597233 | 19599518 | 2285 | SPRET | 3 | 19598669 | 19598703 | 35 | - | 16.1101 | 8.98E-07 | 0.233 | CTATAATAGCCGGTTTCCCCAGTGGATGGCACTG | | |
| 285 | 3 | 19637793 | 19639492 | 1699 | SMJ.WSB | 3 | 19639464 | 19639497 | 34 | - | 17.4146 | 5.18E-07 | 0.109 | GAGCAGTGCGGGACTCTCCCTCCGGGTGGTACAC | | |
| 286 | 3 | 22318079 | 22321858 | 3779 | CAST.SMJ | 3 | 22318229 | 22318262 | 34 | - | 18.6992 | 2.20E-07 | 0.0753 | ATGCAGATCTCGCTGGGGCCTCCAGATGAAAGAA | | |
| 287 | 3 | 26947353 | 26952027 | 4674 | MOLF | 3 | 26950093 | 26950111 | 19 | - | 20.0656 | 1.05E-07 | 0.085 | AGGCCAGAAGAGGGCACTG | | |
| 288 | 3 | 29837225 | 29840301 | 3076 | SPRET | 3 | 29837231 | 29837264 | 34 | - | 18.3089 | 2.87E-07 | 0.0804 | ATGCAGCTAGAGACACGAGCTCCAGGGGGTACTG | | |
| 289 | 3 | 30890235 | 30894213 | 3978 | CAST | 3 | 30893788 | 30893822 | 35 | - | 21.5505 | 3.11E-08 | 0.0328 | CTGCTGAGCTGCCCACACTCCAGTGGAGGGCTCCA | | |
| 290 | 3 | 30893604 | 30895040 | 1436 | SPRET | 3 | 30893788 | 30893822 | 35 | - | 21.5505 | 3.11E-08 | 0.0328 | CTGCTGAGCTGCCCACACTCCAGTGGAGGGCTCCA | | |
| 291 | 3 | 32424747 | 32430435 | 5688 | AJ,AKR,BUB,C3H,C58J,CBA,FVB,MA/My,NZB,PL,RF,SMJ,TH | 3 | 32426831 | 32426864 | 34 | - | 16.5122 | 9.23E-07 | 0.141 | CTGATCCTCCTGCCTCCACCTGCAGAGGGCTTGG | | |
| 292 | 3 | 32472327 | 32477425 | 5098 | AKR,BUB,C58J,DBA1J,DBA2J,FVB,ILNJ2,KK,MA/My,NZB,PL,RF,SMJ,SWR,TH | 3 | 32477069 | 32477102 | 34 | - | 16.7073 | 8.16E-07 | 0.134 | TTGCTCTCTTCAGGCACACCAGAAGAGGGCATTG | | |
| 293 | 3 | 32751846 | 32753860 | 2014 | SJL | 3 | 32752546 | 32752564 | 19 | - | 19.2787 | 1.91E-07 | 0.113 | TCGACACAAGGGGGCGCTG | 38 | ENCFF093ZRU,ENCFF201JUA,ENCFF367UNR,ENCFF327LYS,ENCFF907XEG,ENCFF805QBU,ENCFF821BIA,ENCFF089GBC,ENCFF198SSR,ENCFF024JOZ,ENCFF321AGQ,ENCFF589ZBJ,ENCFF055HAP,ENCFF502WDT,ENCFF065ODV,ENCFF092NMJ,ENCFF430PPJ,ENCFF136ZDK,ENCFF165LXO,ENCFF951GOP,ENCFF883UPM,ENCFF089EWX,ENCFF957SPH,ENCFF575MGI,ENCFF142CNG,ENCFF210BTO,ENCFF052KVO,ENCFF609MDN,ENCFF281QMV,ENCFF491RJK,ENCFF194HVL,ENCFF504JDS,ENCFF611HDQ,ENCFF533APC,ENCFF316FYN,ENCFF097CAQ,ENCFF033UQX,ENCFF601JVE |
| 293 | 3 | 32751846 | 32753860 | 2014 | SJL | 3 | 32752546 | 32752579 | 34 | - | 26.7886 | 3.16E-10 | 0.000932 | CTGCAGCCTCTGGCTTCGACACAAGGGGGCGCTG | 38 | ENCFF093ZRU,ENCFF201JUA,ENCFF367UNR,ENCFF327LYS,ENCFF907XEG,ENCFF805QBU,ENCFF821BIA,ENCFF089GBC,ENCFF198SSR,ENCFF024JOZ,ENCFF321AGQ,ENCFF589ZBJ,ENCFF055HAP,ENCFF502WDT,ENCFF065ODV,ENCFF092NMJ,ENCFF430PPJ,ENCFF136ZDK,ENCFF165LXO,ENCFF951GOP,ENCFF883UPM,ENCFF089EWX,ENCFF957SPH,ENCFF575MGI,ENCFF142CNG,ENCFF210BTO,ENCFF052KVO,ENCFF609MDN,ENCFF281QMV,ENCFF491RJK,ENCFF194HVL,ENCFF504JDS,ENCFF611HDQ,ENCFF533APC,ENCFF316FYN,ENCFF097CAQ,ENCFF033UQX,ENCFF601JVE |
| 294 | 3 | 32816449 | 32819992 | 3543 | SPRET | 3 | 32819739 | 32819773 | 35 | - | 16.844 | 5.92E-07 | 0.188 | CTGCCATGCCTGGCTCGTACCAGGCGATGGTGGCA | | |
| 295 | 3 | 36653835 | 36655412 | 1577 | AKR,MA/My,MRL,PL,RF,TH | 3 | 36654113 | 36654147 | 35 | - | 23.4954 | 7.78E-09 | 0.0141 | CTGCATGTATGCCTGCATGCCAGAAGAGGGCATCA | 7 | ENCFF093ZRU,ENCFF367UNR,ENCFF327LYS,ENCFF136ZDK,ENCFF589ZBJ,ENCFF883UPM,ENCFF099QNE |
| 296 | 3 | 39725069 | 39727372 | 2303 | AJ.ILNJ2.NZB | 3 | 39725308 | 39725342 | 35 | - | 20.0367 | 8.49E-08 | 0.0619 | GTGCAGGTGCCCAGGAAGGTCAGAAGAGGGCGTCA | | |
| 297 | 3 | 42137274 | 42141800 | 4526 | CAST.MOLF.NZW.SPRET | 3 | 42140980 | 42141013 | 34 | - | 17.6748 | 4.37E-07 | 0.0998 | GAGCATGACCACAAGTCCACAGCAGGTGGCAACG | | |
| 298 | 3 | 43819107 | 43822420 | 3313 | SPRET | 3 | 43822306 | 43822324 | 19 | - | 17.082 | 8.61E-07 | 0.205 | TTTCCTCCAGGTGGAGCTA | | |
| 299 | 3 | 44496909 | 44499790 | 2881 | SPRET | 3 | 44498270 | 44498303 | 34 | - | 17.4146 | 5.18E-07 | 0.109 | ATGCAGCTAGAGACACAAGCTCCAGGGGGTACTG | | |
| 300 | 3 | 49246396 | 49250735 | 4339 | AJ,AKR,BALB,BUB,C3H,CAST,CBA,CE,DBA1J,DBA2J,FVB,ILNJ2,MOLF,MRL,NOD,NOR,NZB,PL,RF,RH2,SJL,SMJ,SPRET,SWR,TH,WSB | 3 | 49249763 | 49249797 | 35 | - | 16.4037 | 7.61E-07 | 0.216 | GAGCATTAAACACAAGTCCACAGCAGATGGCAGCA | | |
| 301 | 3 | 51008474 | 51012675 | 4201 | AJ,AKR,BALB,C3H,CAST,CBA,DBA1J,DBA2J,ILNJ2,KK,MA/My,MOLF,MRL,NOD,NOR,NZO,NZW,PL,RF,RH2,SJL,SPRET,SWR,WSB | 3 | 51011837 | 51011870 | 34 | - | 17.6748 | 4.37E-07 | 0.0998 | GAGCATGACCACAAGTCCACAGCAGGTGGCAACG | | |
| 302 | 3 | 54137504 | 54139577 | 2073 | SPRET | 3 | 54139549 | 54139582 | 34 | - | 22.9106 | 9.70E-09 | 0.00895 | ATGCAGTGATGCTGGAGGCCAGAAGAGGGCATCA | 3 | ENCFF099QNE.ENCFF052KVO.ENCFF093ZRU |
| 303 | 3 | 54404719 | 54407415 | 2696 | MOLF | 3 | 54406745 | 54406778 | 34 | - | 16.4472 | 9.61E-07 | 0.141 | TAGCTGTCTTCAGACACACCAGAAGAGGGCGTCA | | |
| 304 | 3 | 65260037 | 65261181 | 1144 | PL | 3 | 65260089 | 65260107 | 19 | - | 20.2623 | 8.99E-08 | 0.0805 | TGGACACTAGAGGGCAGCA | 19 | ENCFF367UNR,ENCFF093ZRU,ENCFF201JUA,ENCFF099QNE,ENCFF821BIA,ENCFF092NMJ,ENCFF430PPJ,ENCFF589ZBJ,ENCFF165LXO,ENCFF198SSR,ENCFF055HAP,ENCFF502WDT,ENCFF907XEG,ENCFF089GBC.ENCFF194HVL.ENCFF024JOZ.ENCFF052KVO.ENCFF327LYS.ENCFF883UPM |
| 305 | 3 | 69851640 | 69858125 | 6485 | SPRET | 3 | 69851753 | 69851771 | 19 | - | 17.2131 | 7.91E-07 | 0.196 | GGGCCAAAAGGGGGCAGTG | | |
| 305 | 3 | 69851640 | 69858125 | 6485 | SPRET | 3 | 69851753 | 69851787 | 35 | - | 15.9266 | 9.95E-07 | 0.244 | GTACAGGGGAACACCAGGGCCAAAAGGGGGCAGTG | | |
| 306 | 3 | 70479038 | 70483203 | 4165 | CAST.CE.MOLF.SPRET | 3 | 70482406 | 70482439 | 34 | - | 17.6748 | 4.37E-07 | 0.0998 | GAGCATGACCACAAGTCCACAGCAGGTGGCAACG | | |
| 307 | 3 | 73781753 | 73784074 | 2321 | SPRET | 3 | 73783985 | 73784018 | 34 | - | 17.9106 | 3.74E-07 | 0.0926 | TAGCTGTCTTCAGACACTCCAGAAGAGGGCACCA | | |
| 308 | 3 | 84942496 | 84944573 | 2077 | CAST.MOLF.SPRET | 3 | 84942601 | 84942619 | 19 | - | 17.2131 | 7.91E-07 | 0.196 | GGGCCAAAAGGGGGCAGTG | | |
| 308 | 3 | 84942496 | 84944573 | 2077 | CAST.MOLF.SPRET | 3 | 84942601 | 84942635 | 35 | - | 16.4771 | 7.30E-07 | 0.211 | GTACAGGGGAACGCCAGGGCCAAAAGGGGGCAGTG | | |
| 309 | 3 | 87192551 | 87194830 | 2279 | WSB | 3 | 87194777 | 87194810 | 34 | - | 23.1626 | 7.90E-09 | 0.00773 | GTGCATTGCCTGGGGAGGCCAGATGATGGCGCTT | | |
| 310 | 3 | 87366327 | 87392732 | 26405 | FVB.NOD.NOR.SJL.SWR | 3 | 87388710 | 87388743 | 34 | - | 16.626 | 8.59E-07 | 0.137 | CTGCTGATCCTGACTTTCACACCAGGTGCTGACC | | |
| 311 | 3 | 96361784 | 96362963 | 1179 | MOLF | 3 | 96362445 | 96362478 | 34 | - | 16.8293 | 7.55E-07 | 0.13 | CCGGACACCCCCCTACCACCACCAGATGCCACCC | | |
| 312 | 3 | 104592036 | 104596565 | 4529 | SPRET | 3 | 104593234 | 104593268 | 35 | - | 21.578 | 3.05E-08 | 0.0324 | TTGCATGTACAACTGCATGCCAGAAGAGGGCATCA | | |
| 313 | 3 | 108513368 | 108515845 | 2477 | CAST | 3 | 108514237 | 108514270 | 34 | - | 18.8618 | 1.97E-07 | 0.0709 | ATGCAGTACCTGAGGACTCCAGAAGAGGACATGA | 1 | ENCFF093ZRU |
| 314 | 3 | 116330647 | 116332426 | 1779 | SPRET | 3 | 116331378 | 116331412 | 35 | - | 16.3303 | 7.94E-07 | 0.221 | GTGCAGAGACCACAGCGACACACTAGGTGGAAAGA | | |
| 315 | 3 | 116349710 | 116353282 | 3572 | SPRET | 3 | 116352416 | 116352449 | 34 | - | 19.1626 | 1.60E-07 | 0.0641 | ATGCAGCTAGAGACACGACCTCCGGGGGGTACTG | | |
| 315 | 3 | 116349710 | 116353282 | 3572 | SPRET | 3 | 116353145 | 116353178 | 34 | - | 23.2683 | 7.25E-09 | 0.00731 | CTGCGGTGCCCAGAGAGGTCAGAAGATGGCACCT | | |
| 316 | 3 | 116542023 | 116543148 | 1125 | MOLF | 3 | 116542665 | 116542698 | 34 | - | 23.4959 | 6.01E-09 | 0.00639 | ATACAGTTCCCACAGAGGCCAGAAGAGGGCATCA | 33 | ENCFF201JUA,ENCFF805QBU,ENCFF367UNR,ENCFF093ZRU,ENCFF210BTO,ENCFF099QNE,ENCFF327LYS,ENCFF821BIA,ENCFF024JOZ,ENCFF502WDT,ENCFF907XEG,ENCFF589ZBJ,ENCFF055HAP,ENCFF089GBC,ENCFF136ZDK,ENCFF198SSR,ENCFF491RJK,ENCFF609MDN,ENCFF321AGQ,ENCFF065ODV,ENCFF430PPJ,ENCFF165LXO,ENCFF097CAQ,ENCFF194HVL,ENCFF092NMJ,ENCFF957SPH,ENCFF052KVO,ENCFF533APC,ENCFF142CNG,ENCFF883UPM,ENCFF575MGI,ENCFF281QMV,ENCFF951GOP |
| 317 | 3 | 124075843 | 124083592 | 7749 | AJ,BALB,C3H,CAST,CBA,MOLF,NZB,RH2,SMJ | 3 | 124082749 | 124082767 | 19 | - | 17.4262 | 6.88E-07 | 0.186 | TTTCCTGTAGGTGGCAGCA | 4 | ENCFF099QNE,ENCFF502WDT,ENCFF055HAP,ENCFF093ZRU |
| 318 | 3 | 130481401 | 130482604 | 1203 | CAST.PJ.SEA | 3 | 130481479 | 130481512 | 34 | - | 17.7967 | 4.04E-07 | 0.0961 | TAGCTGTCCTCAGACACACCAGAAGAGGGCATTG | | |
| 319 | 3 | 134949660 | 134951704 | 2044 | CAST | 3 | 134951685 | 134951718 | 34 | - | 17.9431 | 3.66E-07 | 0.0914 | TTGCACTGTCCTTCCTTTCCACATGGTGGGGCTA | 20 | ENCFF201JUA,ENCFF089GBC,ENCFF821BIA,ENCFF052KVO,ENCFF883UPM,ENCFF430PPJ,ENCFF165LXO,ENCFF327LYS,ENCFF093ZRU,ENCFF609MDN,ENCFF055HAP,ENCFF907XEG,ENCFF024JOZ,ENCFF099QNE,ENCFF502WDT,ENCFF198SSR,ENCFF367UNR,ENCFF589ZBJ,ENCFF065ODV,ENCFF092NMJ |
| 320 | 3 | 135242158 | 135245049 | 2891 | SPRET | 3 | 135243508 | 135243541 | 34 | - | 17.6179 | 4.54E-07 | 0.101 | TAGCTGTCTTCAGACACCCCAGAAGAGGGCACCA | 4 | ENCFF093ZRU.ENCFF367UNR.ENCFF821BIA.ENCFF883UPM |
| 321 | 3 | 151448106 | 151449980 | 1874 | SPRET | 3 | 151449489 | 151449522 | 34 | - | 18.4797 | 2.56E-07 | 0.0799 | TTGCTGTCTTCAGACACACCAGAAGAGGGCATCA | | |
| 322 | 3 | 152534995 | 152536279 | 1284 | TH | 3 | 152536152 | 152536186 | 35 | - | 16.945 | 5.59E-07 | 0.184 | CTGCAATGCCAGGCCCCAGCCCGTAGACAGCCCAG | | |
| 323 | 4 | 5460810 | 5465373 | 4563 | CAST.SPRET | 4 | 5461594 | 5461627 | 34 | + | 16.7398 | 7.99E-07 | 0.0873 | TAGCATGACCACAAGTCCACAGCAGGTGGCAACG | | |
| 324 | 4 | 7924800 | 7926775 | 1975 | CAST | 4 | 7926052 | 7926085 | 34 | + | 17.6911 | 4.33E-07 | 0.0642 | CAGCCTTTCCCTAATTCAACAGCAGGGGTCAGCT | | |
| 325 | 4 | 12113869 | 12124875 | 11006 | FVB.NZB.NZW | 4 | 12121471 | 12121504 | 34 | + | 16.9187 | 7.13E-07 | 0.0825 | TAGCTGTTTTCAGACACCCCAGAAGAGGGCATCA | | |

| | | | | | | | | | | | | | | | | |
|---|---|---|---|---|---|---|---|---|---|---|---|---|---|---|---|---|
| 326 | 4 | 19561347 | 19563261 | 1914 | SPRET | 4 | 19562100 | 19562133 | 34 | + | 26.2033 | 5.53E-10 | 0.000726 | ATGCACTGCCTGTGGAGGCCAGAAGAGGGCATCA | 23 | ENCFF367UNR,ENCFF089GBC,ENCFF024JOZ,ENCFF093ZRU,ENCFF198SSR,ENCFF165LXO,ENCFF210BTO,ENCFF883UPM,ENCFF099QNE,ENCFF052KVO,ENCFF821BIA,ENCFF533APC,ENCFF065ODV,ENCFF907XEG,ENCFF055HAP,ENCFF194HVL,ENCFF327LYS,ENCFF502WDT,ENCFF142CNG,ENCFF321AGQ,ENCFF589ZBJ,ENCFF957SPH,ENCFF136ZDK |
| 327 | 4 | 21788120 | 21789426 | 1306 | SPRET | 4 | 21788405 | 21788438 | 34 | + | 16.4472 | 9.61E-07 | 0.0931 | TAGCTGTCTTCAGACACACCAGAAGAGGGCGTCA | | |
| 328 | 4 | 32970036 | 32971281 | 1245 | MOLF | 4 | 32970347 | 32970381 | 35 | + | 19.844 | 9.61E-08 | 0.0418 | CTGCATGTACACCTGTGTGCCTGAAGAGGGCCCAG | | |
| 329 | 4 | 35070177 | 35077789 | 7612 | AJ,AKR,BTBR,BUB,C3H,CAST,CBA,DBA1J,DBA2J,FVB,ILNJ2,LP,MA/My,MOLF,MRL,NOD,NOR,PJ,PL,RF,SEA,SJL,SMJ,SPRET,SWR,TH,WSB | 4 | 35071286 | 35071320 | 35 | + | 16 | 9.55E-07 | 0.154 | TTGCAATTGAAGGCCCAGGACTGAAGGGGTCACGC | | |
| 330 | 4 | 35268642 | 35269655 | 1013 | SPRET | 4 | 35269465 | 35269499 | 35 | + | 17.633 | 3.74E-07 | 0.0942 | GTGCTGTGCACGGGTCCTGCCAGAAGAGAGCTGGT | | |
| 331 | 4 | 35377121 | 35385412 | 8291 | MA/My,PJ,PL,RF,SEA,TH | 4 | 35385038 | 35385072 | 35 | + | 18.2752 | 2.55E-07 | 0.077 | CTGAAGGCCAGGTGTCCAGACACCAGAGGGCCCTT | | |
| 332 | 4 | 36465337 | 36591613 | 126276 | B10J | 4 | 36512178 | 36512211 | 34 | + | 17.878 | 3.83E-07 | 0.06 | ATGCAGCTAGAGACATGAGCTCTGGGGGGCACTG | | |
| 332 | 4 | 36465337 | 36591613 | 126276 | B10J | 4 | 36566674 | 36566708 | 35 | + | 18.844 | 1.80E-07 | 0.0629 | CTGCAGTGGTGAGGTGCAACCAACAGAGGTAGCAA | | |
| 333 | 4 | 36635705 | 36641111 | 5406 | CAST | 4 | 36637692 | 36637725 | 34 | + | 17.3089 | 5.55E-07 | 0.0732 | GTGCAGTCTTGTGCCAGGCCAGATGAAGGCAGCC | | |
| 334 | 4 | 37443950 | 37449388 | 5438 | SPRET | 4 | 37445438 | 37445471 | 34 | + | 18.0894 | 3.33E-07 | 0.0556 | CTAAAATGTGAAGCTTGACCAGCAGGGGGCACTA | 6 | ENCFF093ZRU,ENCFF907XEG,ENCFF367UNR,ENCFF502WDT,ENCFF099QNE,ENCFF055HAP |
| 334 | 4 | 37443950 | 37449388 | 5438 | SPRET | 4 | 37445453 | 37445471 | 19 | + | 25.1639 | 4.83E-10 | 0.00638 | TGACCAGCAGGGGGCACTA | 6 | ENCFF093ZRU,ENCFF907XEG,ENCFF367UNR,ENCFF502WDT,ENCFF099QNE,ENCFF055HAP |
| 335 | 4 | 39598186 | 39599313 | 1127 | CAST | 4 | 39598520 | 39598554 | 35 | + | 20.8807 | 4.88E-08 | 0.0263 | CTGCATGTATGCCTGCAGGCCAGAAAAGGGCACCA | | |
| 336 | 4 | 40294801 | 40298430 | 3629 | SPRET | 4 | 40297041 | 40297074 | 34 | + | 21.2927 | 3.43E-08 | 0.0131 | GTGTAGTGCCCTTAGATGCCAGAAGAGGGTATCA | | |
| 336 | 4 | 40294801 | 40298430 | 3629 | SPRET | 4 | 40297591 | 40297624 | 34 | + | 17.5528 | 4.74E-07 | 0.0674 | TAGCTGTCTTCAGATGCACCAGTAGAGGGCGTCA | 1 | ENCFF093ZRU |
| 337 | 4 | 40296780 | 40298156 | 1376 | CAST | 4 | 40297041 | 40297074 | 34 | + | 21.2927 | 3.43E-08 | 0.0131 | GTGTAGTGCCCTTAGATGCCAGAAGAGGGTATCA | | |
| 337 | 4 | 40296780 | 40298156 | 1376 | CAST | 4 | 40297591 | 40297624 | 34 | + | 17.5528 | 4.74E-07 | 0.0674 | TAGCTGTCTTCAGATGCACCAGTAGAGGGCGTCA | 1 | ENCFF093ZRU |
| 338 | 4 | 43527278 | 43529136 | 1858 | MOLF | 4 | 43528669 | 43528702 | 34 | + | 16.4228 | 9.76E-07 | 0.0938 | TAGCTGCCTTCAGACACTCCAGAAGAGGGCGCCA | 13 | ENCFF099QNE,ENCFF024JOZ,ENCFF052KVO,ENCFF089GBC,ENCFF367UNR,ENCFF589ZBJ,ENCFF502WDT,ENCFF198SSR,ENCFF907XEG,ENCFF136ZDK,ENCFF327LYS,ENCFF093ZRU,ENCFF821BIA |
| 338 | 4 | 43527278 | 43529136 | 1858 | MOLF | 4 | 43528684 | 43528702 | 19 | + | 17.5574 | 6.32E-07 | 0.111 | ACTCCAGAAGAGGGCGCCA | 13 | ENCFF099QNE,ENCFF024JOZ,ENCFF052KVO,ENCFF089GBC,ENCFF367UNR,ENCFF589ZBJ,ENCFF502WDT,ENCFF198SSR,ENCFF907XEG,ENCFF136ZDK,ENCFF327LYS,ENCFF093ZRU,ENCFF821BIA |
| 338 | 4 | 43527278 | 43529136 | 1858 | MOLF | 4 | 43528063 | 43528096 | 34 | - | 24.5285 | 2.50E-09 | 0.00189 | GTGAAGTACCTGCAGAGGCCAGAGGAGGGCGCTG | 25 | ENCFF092NMJ,ENCFF089GBC,ENCFF321AGQ,ENCFF430PPJ,ENCFF883UPM,ENCFF367UNR,ENCFF165LXO,ENCFF093ZRU,ENCFF327LYS,ENCFF502WDT,ENCFF033UQX,ENCFF055HAP,ENCFF821BIA,ENCFF957SPH,ENCFF194HVL,ENCFF907XEG,ENCFF099QNE,ENCFF142CNG,ENCFF052KVO,ENCFF024JOZ,ENCFF533APC,ENCFF065ODV,ENCFF198SSR,ENCFF136ZDK,ENCFF589ZBJ |
| 339 | 4 | 45115109 | 45116975 | 1866 | SPRET | 4 | 45116670 | 45116703 | 34 | + | 17.5691 | 4.69E-07 | 0.0668 | TAGCTGTCTTCAGACACACCAGAAGAGGGCAGTG | 1 | ENCFF093ZRU |
| 339 | 4 | 45115109 | 45116975 | 1866 | SPRET | 4 | 45116685 | 45116703 | 19 | + | 16.9508 | 9.36E-07 | 0.135 | ACACCAGAAGAGGGCAGTG | 1 | ENCFF093ZRU |
| 340 | 4 | 52299418 | 52300892 | 1474 | SPRET | 4 | 52299401 | 52299419 | 19 | + | 19.0328 | 2.28E-07 | 0.0693 | TGCCCTGCAGGGGGCACTC | 6 | ENCFF502WDT,ENCFF093ZRU,ENCFF957SPH,ENCFF194HVL,ENCFF099QNE,ENCFF367UNR |
| 341 | 4 | 53386784 | 53393102 | 6318 | CAST,CBA,DBA1J,DBA2J,MA/My,MOLF,SMJ,SPRET | 4 | 53393078 | 53393112 | 35 | + | 16 | 9.55E-07 | 0.154 | CAGCCGGGCGGGCACCTGTCCTCTAGGCGACGCTC | | |
| 342 | 4 | 61134288 | 61141873 | 7585 | CAST | 4 | 61139173 | 61139206 | 34 | + | 20.439 | 6.46E-08 | 0.0204 | GCGCAACACTCAGCTTGGCCAGTAGAGGGCAATC | 5 | ENCFF367UNR,ENCFF092NMJ,ENCFF089GBC,ENCFF907XEG,ENCFF327LYS |
| 342 | 4 | 61134288 | 61141873 | 7585 | CAST | 4 | 61139188 | 61139206 | 19 | + | 20.6721 | 6.43E-08 | 0.038 | TGGCCAGTAGAGGGCAATC | 5 | ENCFF367UNR,ENCFF092NMJ,ENCFF089GBC,ENCFF907XEG,ENCFF327LYS |
| 343 | 4 | 61229445 | 61233649 | 4204 | MOLF | 4 | 61229793 | 61229826 | 34 | + | 19.9106 | 9.47E-08 | 0.0268 | GAGCAAAACTAAGCTTGGCCAGTAGGGGGCAACT | | |
| 343 | 4 | 61229445 | 61233649 | 4204 | MOLF | 4 | 61229808 | 61229826 | 19 | + | 19.8033 | 1.28E-07 | 0.0499 | TGGCCAGTAGGGGGCAACT | | |
| 344 | 4 | 71621047 | 71628286 | 7239 | SPRET | 4 | 71622314 | 71622347 | 34 | + | 17.5935 | 4.61E-07 | 0.0662 | CTGTAGTAGGAGAGCTTGCCAGCAGAGAGTGCTC | | |
| 345 | 4 | 73175663 | 73190235 | 14572 | SPRET | 4 | 73180261 | 73180279 | 19 | + | 17.1148 | 8.43E-07 | 0.128 | CTGCCAGTAGGGGGAAGTT | | |
| 346 | 4 | 73494769 | 73509067 | 14298 | WSB | 4 | 73508448 | 73508481 | 34 | + | 17.6992 | 4.30E-07 | 0.064 | ATGCAGTGCAAGCTGAGACCAGAGAAGGGCATCA | | |
| 347 | 4 | 74798472 | 74801964 | 3492 | SPRET | 4 | 74800158 | 74800191 | 34 | + | 16.7886 | 7.75E-07 | 0.086 | ATGCACCTAGAGACACGAGCTCCAGGGGGTACTG | | |
| 348 | 4 | 79333924 | 79336572 | 2648 | SPRET | 4 | 79334781 | 79334814 | 34 | + | 16.9106 | 7.17E-07 | 0.0827 | GAGCATGGACAGAAGCCCACAGCAGATGGCAGCA | | |
| 349 | 4 | 85333129 | 85334329 | 1200 | SPRET | 4 | 85334231 | 85334249 | 19 | + | 17.2131 | 7.91E-07 | 0.124 | TGCCCAGAAGAGGGAGCTA | 1 | ENCFF093ZRU |
| 350 | 4 | 88673191 | 88676167 | 2976 | 129S1,AKR,BALB,BTBR,CBA,DBA1J,DBA2J,ILNJ2,KK,MRL,NOR,PJ,RF,RH2,SEA,SJL,SWR,TH | 4 | 88674715 | 88674748 | 34 | + | 16.9675 | 6.91E-07 | 0.0815 | CACCCGTGTCCCTTCTGCCCAGCTGGTGGCAGTG | | |
| 350 | 4 | 88673191 | 88676167 | 2976 | 129S1,AKR,BALB,BTBR,CBA,DBA1J,DBA2J,ILNJ2,KK,MRL,NOR,PJ,RF,RH2,SEA,SJL,SWR,TH | 4 | 88675842 | 88675860 | 19 | - | 18.3115 | 3.79E-07 | 0.0887 | GGGCCACCGGAGGGCAGCA | 9 | ENCFF327LYS,ENCFF367UNR,ENCFF024JOZ,ENCFF093ZRU,ENCFF907XEG,ENCFF821BIA,ENCFF099QNE,ENCFF198SSR,ENCFF589ZBJ |
| 350 | 4 | 88673191 | 88676167 | 2976 | 129S1,AKR,BALB,BTBR,CBA,DBA1J,DBA2J,ILNJ2,KK,MRL,NOR,PJ,RF,RH2,SEA,SJL,SWR,TH | 4 | 88675842 | 88675876 | 35 | - | 22.7431 | 1.35E-08 | 0.0116 | CAGCAGCAGAGGGCGCGGGCCACCGGAGGGCAGCA | 10 | ENCFF327LYS,ENCFF367UNR,ENCFF024JOZ,ENCFF093ZRU,ENCFF907XEG,ENCFF821BIA,ENCFF099QNE,ENCFF198SSR,ENCFF589ZBJ,ENCFF089GBC |
| 350 | 4 | 88673191 | 88676167 | 2976 | 129S1,AKR,BALB,BTBR,CBA,DBA1J,DBA2J,ILNJ2,KK,MRL,NOR,PJ,RF,RH2,SEA,SJL,SWR,TH | 4 | 88676038 | 88676056 | 19 | - | 17.7049 | 5.73E-07 | 0.11 | TGGCCTGCAGGGGGAGCGC | 28 | ENCFF327LYS,ENCFF367UNR,ENCFF024JOZ,ENCFF093ZRU,ENCFF907XEG,ENCFF821BIA,ENCFF099QNE,ENCFF198SSR,ENCFF589ZBJ,ENCFF089GBC,ENCFF136ZDK,ENCFF165LXO,ENCFF092NMJ,ENCFF430PPJ,ENCFF033UQX,ENCFF210BTO,ENCFF491RJK,ENCFF883UPM,ENCFF957SPH,ENCFF951GOP,ENCFF281QMV,ENCFF194HVL,ENCFF575MGI,ENCFF611HDQ,ENCFF316FYN,ENCFF533APC,ENCFF097CAQ,ENCFF609MDN |
| 351 | 4 | 93839894 | 93845077 | 5183 | CAST,CE,MA/My,MOLF,NZB,SPRET,SWR | 4 | 93844615 | 93844648 | 34 | + | 19.3415 | 1.42E-07 | 0.0358 | CTGCCCCCACATTCGCCGTCACAAGATGGCGCTG | | |
| 352 | 4 | 96400426 | 96402849 | 2423 | CAST,MOLF,WSB | 4 | 96402718 | 96402751 | 34 | + | 17.6992 | 4.30E-07 | 0.064 | TAGCTGTCCTCTGACACACCAGAAGAGGGCATCA | | |
| 353 | 4 | 98635155 | 98640475 | 5320 | SPRET | 4 | 98635245 | 98635279 | 35 | + | 16.0642 | 9.21E-07 | 0.151 | GTGCAGGTACCTGAAATAACCAGAAGAGAGCGTTT | | |
| 354 | 4 | 100314955 | 100318686 | 3731 | SMJ,WSB | 4 | 100316606 | 100316624 | 19 | + | 17.2459 | 7.74E-07 | 0.123 | AGGCCACAAGGTGGAACCC | | |
| 355 | 4 | 102346746 | 102348654 | 1908 | AKR,CE,KK,MA/My,MRL,SMJ | 4 | 102348000 | 102348033 | 34 | + | 20.2683 | 7.32E-08 | 0.0224 | GTTCAGTGTCTGCAGAGACCAGAAGAGGGCATAG | | |
| 356 | 4 | 108176653 | 108178228 | 1575 | SPRET | 4 | 108177164 | 108177197 | 34 | + | 26.6341 | 3.67E-10 | 0.000551 | TTGCAGTGCCCCAAAACGCCAGAAGGGGGCGTCT | 21 | ENCFF024JOZ,ENCFF092NMJ,ENCFF165LXO,ENCFF089GBC,ENCFF883UPM,ENCFF198SSR,ENCFF093ZRU,ENCFF367UNR,ENCFF052KVO,ENCFF033UQX,ENCFF065ODV,ENCFF821BIA,ENCFF589ZBJ,ENCFF136ZDK,ENCFF504JDS,ENCFF327LYS,ENCFF907XEG,ENCFF099QNE,ENCFF321AGQ,ENCFF502WDT,ENCFF055HAP |
| 357 | 4 | 111923812 | 111925200 | 1388 | ILNJ2,LP,MA/My,MOLF | 4 | 111924177 | 111924211 | 35 | + | 19.4587 | 1.23E-07 | 0.0492 | CTGCCATGAGGACTTCCATCCACCAGGCGTCACCT | | |
| 358 | 4 | 112502873 | 112504255 | 1382 | 129S1,AJ,AKR,BALB,BTBR,BUB,C3H,C57L,CAST,CBA,CE,DBA1J,DBA2J,FVB,MOLF,MRL,NOD,NOR,PL,RF,SEA,SJL,SMJ,SWR,TH,WSB | 4 | 112503166 | 112503199 | 34 | + | 23.8537 | 4.46E-09 | 0.00288 | GTGTAGTGCCTGTGGAAGCCAGAAGATGGCGATA | | |
| 359 | 4 | 118615672 | 118683173 | 67501 | NZO,NZW | 4 | 118643041 | 118643074 | 34 | + | 17.1951 | 5.97E-07 | 0.0755 | CTGCTCTGGCACACAAGTTCTCAAGGTGGTGCTG | | |
| 360 | 4 | 118874076 | 118875461 | 1385 | ILNJ2,MOLF | 4 | 118874828 | 118874861 | 34 | + | 18.5285 | 2.48E-07 | 0.0494 | GTGCATTGCATACGGAGGCCAGAGGAGGGTGTCA | | |
| 361 | 4 | 119783624 | 119787795 | 4171 | BTBR,CAST,MOLF,SPRET,WSB | 4 | 119784401 | 119784434 | 34 | + | 17.6748 | 4.37E-07 | 0.0644 | GAGCATGACCACAAGTCCACAGCAGGTGGCAACG | | |
| 362 | 4 | 123962386 | 123964206 | 1820 | 129S1,AJ,AKR,BALB,BTBR,BUB,C3H,C57L,C58J,CAST,CBA,CE,DBA1J,DBA2J,FVB,ILNJ2,KK,MA/My,MRL,NOD,NOR,NZB,NZO,NZW,PJ,PL,RF,SEA,SJL,SMJ,SPRET,SWR,TH,WSB | 4 | 123962467 | 123962500 | 34 | + | 18.4797 | 2.56E-07 | 0.0494 | TTGCTGTCTTCAGACACACCAGAAGAGGGCATCA | | |
| 363 | 4 | 124019753 | 124037387 | 17634 | RH2 | 4 | 124023435 | 124023469 | 35 | + | 18.1743 | 2.71E-07 | 0.0796 | CTGCATGTATACCTGCATATCAGAAGAGGGCATCA | | |
| 363 | 4 | 124019753 | 124037387 | 17634 | RH2 | 4 | 124036030 | 124036048 | 19 | + | 20.8361 | 5.63E-08 | 0.037 | TTGCCAGCAGGTGTCGCCA | 36 | ENCFF367UNR,ENCFF093ZRU,ENCFF099QNE,ENCFF201JUA,ENCFF821BIA,ENCFF601JVE,ENCFF327LYS,ENCFF089GBC,ENCFF907XEG,ENCFF092NMJ,ENCFF024JOZ,ENCFF430PPJ,ENCFF198SSR,ENCFF052KVO,ENCFF504JDS,ENCFF136ZDK,ENCFF165LXO,ENCFF589ZBJ,ENCFF065ODV,ENCFF055HAP,ENCFF033UQX,ENCFF951GOP,ENCFF502WDT,ENCFF321AGQ,ENCFF281QMV,ENCFF575MGI,ENCFF883UPM,ENCFF957SPH,ENCFF609MDN,ENCFF316FYN,ENCFF142CNG,ENCFF097CAQ,ENCFF194HVL,ENCFF491RJK,ENCFF210BTO,ENCFF533APC |
| 363 | 4 | 124019753 | 124037387 | 17634 | RH2 | 4 | 124024928 | 124024961 | 34 | - | 25.4878 | 1.07E-09 | 0.00107 | ATGCAGTACCAAAAAAAGCCAGAAGAGGGCGTCA | 18 | ENCFF136ZDK,ENCFF092NMJ,ENCFF504JDS,ENCFF821BIA,ENCFF024JOZ,ENCFF589ZBJ,ENCFF093ZRU,ENCFF327LYS,ENCFF907XEG,ENCFF089GBC,ENCFF165LXO,ENCFF883UPM,ENCFF367UNR,ENCFF099QNE,ENCFF033UQX,ENCFF533APC,ENCFF052KVO,ENCFF281QMV |
| 364 | 4 | 126931746 | 126935251 | 3505 | SPRET | 4 | 126934679 | 126934712 | 34 | + | 17.9431 | 3.66E-07 | 0.0591 | TAGCTGTCCTCAGATAGTCCAGAAGAGGGCATCA | 2 | ENCFF099QNE,ENCFF093ZRU |
| 365 | 4 | 130105829 | 130107092 | 1263 | KK,MOLF,NOR,NZB | 4 | 130106306 | 130106339 | 34 | + | 21.6748 | 2.56E-08 | 0.0107 | ATGCAGTGCTCACGGAGGCCAGAAGGCAGCGCTG | 2 | ENCFF055HAP,ENCFF502WDT |
| 366 | 4 | 130345556 | 130346914 | 1358 | SPRET | 4 | 130346405 | 130346438 | 34 | + | 18.4797 | 2.56E-07 | 0.0494 | TTGCTGTCTTCAGACACACCAGAAGAGGGCATCA | | |
| 367 | 4 | 132355360 | 132356644 | 1284 | SPRET | 4 | 132356229 | 132356262 | 34 | + | 19.9512 | 9.19E-08 | 0.0265 | CAGCTGTCAGTCATCCCCACAGCAGGGGGCGCCA | 29 | ENCFF367UNR,ENCFF201JUA,ENCFF093ZRU,ENCFF092NMJ,ENCFF024JOZ,ENCFF089GBC,ENCFF883UPM,ENCFF136ZDK,ENCFF430PPJ,ENCFF951GOP,ENCFF589ZBJ,ENCFF327LYS,ENCFF907XEG,ENCFF609MDN,ENCFF198SSR,ENCFF055HAP,ENCFF065ODV,ENCFF533APC,ENCFF097CAQ,ENCFF142CNG,ENCFF194HVL,ENCFF821BIA,ENCFF491RJK,ENCFF957SPH,ENCFF502WDT,ENCFF099QNE,ENCFF165LXO,ENCFF210BTO,ENCFF052KVO |
| 367 | 4 | 132355360 | 132356644 | 1284 | SPRET | 4 | 132356244 | 132356262 | 19 | + | 17.6557 | 5.92E-07 | 0.111 | CCCACAGCAGGGGGCGCCA | 29 | ENCFF367UNR,ENCFF201JUA,ENCFF093ZRU,ENCFF092NMJ,ENCFF024JOZ,ENCFF089GBC,ENCFF883UPM,ENCFF136ZDK,ENCFF430PPJ,ENCFF951GOP,ENCFF589ZBJ,ENCFF327LYS,ENCFF907XEG,ENCFF609MDN,ENCFF198SSR,ENCFF055HAP,ENCFF065ODV,ENCFF533APC,ENCFF097CAQ,ENCFF142CNG,ENCFF194HVL,ENCFF821BIA,ENCFF491RJK,ENCFF957SPH,ENCFF502WDT,ENCFF099QNE,ENCFF165LXO,ENCFF210BTO,ENCFF052KVO |
| 368 | 4 | 133034365 | 133035999 | 1634 | SPRET | 4 | 133034823 | 133034856 | 34 | + | 23.4472 | 6.26E-09 | 0.0037 | CTGCACTACACAGATCCTCCAGAAGGGGGCATGG | 37 | ENCFF093ZRU,ENCFF367UNR,ENCFF099QNE,ENCFF086ZTV,ENCFF821BIA,ENCFF327LYS,ENCFF065ODV,ENCFF024JOZ,ENCFF907XEG,ENCFF321AGQ,ENCFF136ZDK,ENCFF589ZBJ,ENCFF609MDN,ENCFF033UQX,ENCFF055HAP,ENCFF089GBC,ENCFF502WDT,ENCFF198SSR,ENCFF957SPH,ENCFF092NMJ,ENCFF575MGI,ENCFF097CAQ,ENCFF165LXO,ENCFF194HVL,ENCFF142CNG,ENCFF611HDQ,ENCFF281QMV,ENCFF533APC,ENCFF504JDS,ENCFF052KVO,ENCFF951GOP,ENCFF430PPJ,ENCFF316FYN,ENCFF601JVE,ENCFF210BTO,ENCFF491RJK,ENCFF883UPM |
| 369 | 4 | 133062379 | 133064528 | 2149 | 129S1,C57L,ILNJ2,SJL,SWR | 4 | 133062399 | 133062432 | 34 | + | 19.0163 | 1.77E-07 | 0.0412 | CAGCTGTCTTCAGACACACCAGAAGAGGGCGTCA | | |
| 370 | 4 | 137713848 | 137716193 | 2345 | AJ,AKR,PL,RF,RH2,TH | 4 | 137714913 | 137714947 | 35 | + | 17.2202 | 4.77E-07 | 0.108 | AGGCATGTGCCACCACTGCCCAGCAGAGGCCAGCC | | |
| 371 | 4 | 141492132 | 141493760 | 1628 | FVB,NOD | 4 | 141492936 | 141492969 | 34 | + | 16.4472 | 9.61E-07 | 0.0931 | TAGCTGTCTTCAGACACACCAGAAGAGGGCGTCA | | |
| 372 | 4 | 143401446 | 143423863 | 22417 | SPRET | 4 | 143408198 | 143408216 | 19 | + | 18.5738 | 3.17E-07 | 0.0816 | AAACCACCAGGTGGCAGAA | | |
| 373 | 4 | 143852435 | 143861238 | 8803 | 129S1,BTBR,BUB,C57L,CAST,CE,DBA2J,KK,LP,MA/My,MOLF,MRL,NOD,NZB,NZW,PJ,PL,RF,SEA,SJL,SMJ,TH,WSB | 4 | 143854585 | 143854618 | 34 | + | 17.0325 | 6.63E-07 | 0.0801 | TGGCAGGTCTCAGAGGAGCCGCTAGATGGCCCAC | | |
| 374 | 4 | 145442985 | 145451136 | 8151 | DBA2J,TH | 4 | 145450036 | 145450070 | 35 | + | 24.422 | 3.86E-09 | 0.00589 | CTGCAGTTACCCATGTACGCCATGAGAGGGCGTCA | | |
| 374 | 4 | 145442985 | 145451136 | 8151 | DBA2J,TH | 4 | 145447080 | 145447113 | 34 | - | 19.1301 | 1.64E-07 | 0.0392 | ATGCAGGGGAGCATCCTTCCAGGAGGTGGCACCT | | |
| 375 | 4 | 147490771 | 147492052 | 1281 | 129S1,AJ,AKR,B10D2,B10J,BALB,BTBR,BUB,C3H,C57L,C58J,CAST,CE,DBA1J,DBA2J,FVB,ILNJ2,KK,LP,MA/My,MRL,NOR,NZB,NZO,NZW,PJ,PL,RF,RH2,SEA,SJL,SMJ,SPRET,SWR,TH,WSB | 4 | 147491241 | 147491259 | 19 | + | 18.8033 | 2.69E-07 | 0.0753 | TATCCACCAGAGGGTGCTG | 8 | ENCFF907XEG,ENCFF093ZRU,ENCFF092NMJ,ENCFF821BIA,ENCFF089GBC,ENCFF367UNR,ENCFF099QNE,ENCFF327LYS |
| 376 | 4 | 150672892 | 150679192 | 6300 | MOLF | 4 | 150675207 | 150675240 | 34 | + | 19.0325 | 1.75E-07 | 0.0409 | GTGCACTGCAAGTGGAGACCAGAAGAGGACATTG | | |
| 377 | 4 | 152165196 | 152166495 | 1299 | MOLF | 4 | 152165522 | 152165555 | 34 | + | 16.4634 | 9.51E-07 | 0.0931 | TAGCTGTCTTCAGACGCACCAGAAGAGGGCATCA | | |
| 378 | 4 | 152523830 | 152525506 | 1676 | BUB,MA/My,MRL,NZB,NZO,NZW,SMJ | 4 | 152525044 | 152525077 | 34 | + | 17.4146 | 5.18E-07 | 0.0708 | CTGGGGGACTAACCTCACCCAGCAGGTGGCCCCC | | |
| 379 | 4 | 152605826 | 152607898 | 2072 | CAST | 4 | 152607797 | 152607815 | 19 | + | 20.8033 | 5.79E-08 | 0.037 | AGGCCAGAAGAGGGCGCTG | 1 | ENCFF093ZRU |
| 380 | 4 | 156593953 | 156599643 | 5690 | ILNJ2,WSB | 4 | 156594623 | 156594656 | 34 | + | 17.6016 | 4.59E-07 | 0.066 | TAGCTGTCTTCAGACATGCCAGAAGAGGGCGTCA | | |
| 381 | 4 | 3889494 | 3891346 | 1852 | SPRET | 4 | 3890435 | 3890468 | 34 | - | 24.065 | 3.73E-09 | 0.00253 | CTGCAGTGCCCACCACGACCACAAGAAGCCAACA | | |
| 382 | 4 | 8664934 | 8666549 | 1615 | SPRET | 4 | 8665594 | 8665627 | 34 | - | 19.7967 | 1.03E-07 | 0.0285 | TAGCTGTCCTCAGACACTCCAGAAGAGGGCACCA | 4 | ENCFF099QNE,ENCFF821BIA,ENCFF093ZRU,ENCFF367UNR |
| 383 | 4 | 15236200 | 15242810 | 6610 | 129S1,AJ,AKR,BALB,BUB,C57L,CAST,ILNJ2,MA/My,NOD,NOR,NZW,PJ,PL,RF,RH2,SEA,SJL,SMJ,SPRET | 4 | 15238091 | 15238109 | 19 | - | 17.1639 | 8.17E-07 | 0.127 | CCTCCACGAGGTGGCGGAA | | |
| 384 | 4 | 16319790 | 16327153 | 7363 | CAST,NZW,SPRET | 4 | 16327046 | 16327080 | 35 | - | 20.9817 | 4.56E-08 | 0.0252 | ATGCAGGTGCCTATAGAGTCCAGAAGAGGGCTCTG | | |
| 385 | 4 | 23003638 | 23107598 | 103960 | TH | 4 | 23009510 | 23009528 | 19 | - | 17.459 | 6.74E-07 | 0.114 | CAGCCAGGAGGTGGAGCTG | | |
| 386 | 4 | 26970469 | 26972024 | 1555 | B10D2,B10J | 4 | 26971844 | 26971877 | 34 | - | 18.4797 | 2.56E-07 | 0.0494 | TTGCTGTCTTCAGACACACCAGAAGAGGGCATCA | | |
| 387 | 4 | 30169975 | 30172782 | 2807 | SPRET | 4 | 30172552 | 30172585 | 34 | - | 17.4146 | 5.18E-07 | 0.0708 | ATGCAGCTAGAGACACAAGCTCCAGGGGGTACTG | | |
| 388 | 4 | 32790350 | 32797716 | 7366 | 129S1,AJ,AKR,BALB,BTBR,BUB,C3H,CAST,CBA,DBA1J,DBA2J,FVB,ILNJ2,MA/My,MOLF,MRL,NOD,NOR,PJ,PL,RF,RH2,SEA,SJL,SMJ,SPRET,SWR,TH,WSB | 4 | 32792239 | 32792257 | 19 | - | 17.1639 | 8.17E-07 | 0.127 | CCTCCACGAGGTGGCGGAA | | |
| 389 | 4 | 40435312 | 40443984 | 8672 | CAST | 4 | 40443899 | 40443932 | 34 | - | 17.3252 | 5.49E-07 | 0.0728 | GAGCGGGTGTGGCTGCGGCCCCTAGATGGCACCT | | |
| 390 | 4 | 45835768 | 45846657 | 10889 | CAST,MOLF | 4 | 45844489 | 45844522 | 34 | - | 18.1138 | 3.27E-07 | 0.0551 | CAGTCATTCTGGCTTTGTCCACATGGTGGCGCTT | 29 | ENCFF093ZRU,ENCFF367UNR,ENCFF099QNE,ENCFF052KVO,ENCFF907XEG,ENCFF327LYS,ENCFF201JUA,ENCFF089GBC,ENCFF024JOZ,ENCFF065ODV,ENCFF430PPJ,ENCFF821BIA,ENCFF055HAP,ENCFF502WDT,ENCFF092NMJ,ENCFF589ZBJ,ENCFF609MDN,ENCFF316FYN,ENCFF198SSR,ENCFF194HVL,ENCFF321AGQ,ENCFF142CNG,ENCFF136ZDK,ENCFF165LXO,ENCFF957SPH,ENCFF281QMV,ENCFF951GOP,ENCFF575MGI,ENCFF883UPM |
| 391 | 4 | 46831653 | 46834946 | 3293 | MOLF | 4 | 46833335 | 46833369 | 35 | - | 16.6881 | 6.48E-07 | 0.126 | CCGCCCTTGCTTCCTGCTGCCACATGGTGGCGCAA | 28 | ENCFF805QBU,ENCFF024JOZ,ENCFF349BUL,ENCFF092NMJ,ENCFF165LXO,ENCFF430PPJ,ENCFF093ZRU,ENCFF052KVO,ENCFF198SSR,ENCFF065ODV,ENCFF099QNE,ENCFF316FYN,ENCFF821BIA,ENCFF055HAP,ENCFF502WDT,ENCFF367UNR,ENCFF033UQX,ENCFF589ZBJ,ENCFF327LYS,ENCFF142CNG,ENCFF136ZDK,ENCFF194HVL,ENCFF281QMV,ENCFF097CAQ,ENCFF883UPM,ENCFF089GBC,ENCFF533APC,ENCFF907XEG |
| 392 | 4 | 47430622 | 47432871 | 2249 | SPRET | 4 | 47430761 | 47430795 | 35 | - | 22.055 | 2.19E-08 | 0.0164 | CTGCACTTCCTGTCTTTATCCAGTTGGGGCCAGCG | | |
| 393 | 4 | 48513775 | 48518954 | 5179 | SPRET | 4 | 48516588 | 48516621 | 34 | - | 17.8455 | 3.91E-07 | 0.0606 | TAGCTGTCCTCAGACACTCCAGAAGAGGGCGTCA | | |
| 393 | 4 | 48513775 | 48518954 | 5179 | SPRET | 4 | 48517590 | 48517624 | 35 | - | 16.7615 | 6.21E-07 | 0.125 | TAGCAGTTGCCACCTGTGTGCTCTTGGGGGCGGGG | | |
| 394 | 4 | 50177779 | 50182290 | 4511 | MA/My,MOLF | 4 | 50180078 | 50180096 | 19 | - | 17.2951 | 7.50E-07 | 0.121 | CTGCCACTAGATGGTGCAG | 4 | ENCFF092NMJ,ENCFF907XEG,ENCFF093ZRU,ENCFF099QNE |
| 394 | 4 | 50177779 | 50182290 | 4511 | MA/My,MOLF | 4 | 50180078 | 50180111 | 34 | - | 17.7805 | 4.08E-07 | 0.0623 | ATGTCCTTTACACAGCTGCCACTAGATGGTGCAG | 4 | ENCFF092NMJ,ENCFF907XEG,ENCFF093ZRU,ENCFF099QNE |
| 395 | 4 | 50341491 | 50344272 | 2781 | SPRET | 4 | 50342769 | 50342787 | 19 | - | 17.9016 | 5.02E-07 | 0.103 | TGGCCAGCAGATGGCCCAA | | |
| 395 | 4 | 50341491 | 50344272 | 2781 | SPRET | 4 | 50342769 | 50342803 | 35 | - | 16.5688 | 6.93E-07 | 0.132 | AAGAAGCTGGACCCAGTGGCCAGCAGATGGCCCAA | | |
| 396 | 4 | 59601932 | 59603739 | 1807 | SPRET | 4 | 59602912 | 59602946 | 35 | - | 22.156 | 2.04E-08 | 0.0156 | CTGCACATATGCCTACATACCAGAAGAGGGCAACA | 15 | ENCFF136ZDK,ENCFF092NMJ,ENCFF821BIA,ENCFF589ZBJ,ENCFF327LYS,ENCFF367UNR,ENCFF089GBC,ENCFF093ZRU,ENCFF198SSR,ENCFF883UPM,ENCFF502WDT,ENCFF024JOZ,ENCFF055HAP,ENCFF099QNE,ENCFF907XEG |
| 397 | 4 | 62323359 | 62326152 | 2793 | AJ,AKR,BALB,C3H,CBA,CE,DBA1J,DBA2J,ILNJ2,LP,MA/My,MRL,NZW,RF,RH2,SEA,SJL,SWR | 4 | 62325587 | 62325605 | 19 | - | 17.5574 | 6.32E-07 | 0.111 | ACTCCAGAAGAGGGCGCCA | 2 | ENCFF136ZDK,ENCFF821BIA |
| 397 | 4 | 62323359 | 62326152 | 2793 | AJ,AKR,BALB,C3H,CBA,CE,DBA1J,DBA2J,ILNJ2,LP,MA/My,MRL,NZW,RF,RH2,SEA,SJL,SWR | 4 | 62325587 | 62325620 | 34 | - | 18.3659 | 2.76E-07 | 0.0503 | TAGCTGTCTTCAGACACTCCAGAAGAGGGCGCCA | 2 | ENCFF136ZDK,ENCFF821BIA |
| 398 | 4 | 67280118 | 67284271 | 4153 | 129S1,C57L,CAST,CE,MOLF,PJ,SPRET | 4 | 67283468 | 67283501 | 34 | - | 17.6748 | 4.37E-07 | 0.0644 | GAGCATGACCACAAGTCCACAGCAGGTGGCAACG | | |
| 399 | 4 | 72823333 | 72824601 | 1268 | SPRET | 4 | 72824514 | 72824548 | 35 | - | 22.0734 | 2.16E-08 | 0.0163 | CTGCTGTGTCACATATGCAACAGTAGGTGGCAGAA | 3 | ENCFF024JOZ,ENCFF093ZRU,ENCFF099QNE |
| 400 | 4 | 86871650 | 86873597 | 1947 | MA/My | 4 | 86872026 | 86872044 | 19 | - | 19.0328 | 2.28E-07 | 0.0693 | ATGCCAGCAGGGGGCAGAC | 3 | ENCFF099QNE,ENCFF024JOZ,ENCFF093ZRU |
| 401 | 4 | 88107260 | 88111202 | 3942 | SPRET | 4 | 88107922 | 88107955 | 34 | - | 18.7154 | 2.18E-07 | 0.0464 | GTGTAGTGCAGGTCACTGTCAACAGAGGGCACAA | 7 | ENCFF198SSR,ENCFF883UPM,ENCFF055HAP,ENCFF502WDT,ENCFF093ZRU,ENCFF367UNR,ENCFF099QNE |
| 402 | 4 | 89606946 | 89618459 | 11513 | AKR,BTBR,BUB,KK,MOLF,MRL,SJL,SMJ,SWR,TH | 4 | 89614812 | 89614845 | 34 | - | 18.5203 | 2.49E-07 | 0.0494 | ATGCAGCTAGAGACACGAACTCCAGGGGGTACTG | | |
| 403 | 4 | 94870239 | 94871463 | 1224 | SPRET | 4 | 94871124 | 94871142 | 19 | - | 20.3607 | 8.29E-08 | 0.0406 | AGGCCAGAAGAGGGCACCA | | |
| 403 | 4 | 94870239 | 94871463 | 1224 | SPRET | 4 | 94871124 | 94871158 | 35 | - | 20.4954 | 6.30E-08 | 0.0311 | CTTCATGTATGCCTGCAGGCCAGAAGAGGGCACCA | | |
| 404 | 4 | 98386008 | 98387148 | 1140 | SPRET | 4 | 98386430 | 98386448 | 19 | - | 18.1475 | 4.25E-07 | 0.0953 | TGGCCAGGAGGTGGTGGTG | | |
| 405 | 4 | 99404200 | 99405932 | 1732 | CAST,MOLF | 4 | 99405305 | 99405323 | 19 | - | 18.7213 | 2.85E-07 | 0.0776 | AGGCCAGTAGAGGGCGTCA | 8 | ENCFF093ZRU,ENCFF099QNE,ENCFF327LYS,ENCFF089GBC,ENCFF907XEG,ENCFF033UQX,ENCFF533APC,ENCFF052KVO |
| 405 | 4 | 99404200 | 99405932 | 1732 | CAST,MOLF | 4 | 99405305 | 99405338 | 34 | - | 29.2114 | 2.60E-11 | 0.00013 | ATGCAGTGCCTGAGGAGGCCAGTAGAGGGCGTCA | 8 | ENCFF093ZRU,ENCFF099QNE,ENCFF327LYS,ENCFF089GBC,ENCFF907XEG,ENCFF033UQX,ENCFF533APC,ENCFF052KVO |
| 406 | 4 | 99498162 | 99503432 | 5270 | CE | 4 | 99499802 | 99499836 | 35 | - | 18.2294 | 2.62E-07 | 0.0784 | ATGCCGGTGCCTACAGAGGCCAGAAGAGGGCATTA | | |
| 407 | 4 | 100756127 | 100757362 | 1235 | BTBR,MA/My,WSB | 4 | 100756713 | 100756747 | 35 | - | 16.367 | 7.77E-07 | 0.139 | CTGCATTGCTAATTCAAGGCCAGAGGAAGACCCTG | | |
| 408 | 4 | 101720349 | 101729335 | 8986 | AJ,AKR,BALB,BUB,C3H,C58J,CAST,CBA,CE,KK,LP,MA/My,MOLF,MRL,NZB,NZO,NZW,PJ,PL,RF,SEA,SJL,SMJ,SPRET,WSB | 4 | 101723904 | 101723922 | 19 | - | 17.1639 | 8.17E-07 | 0.127 | CCTCCACGAGGTGGCGGAA | | |
| 409 | 4 | 108003223 | 108012214 | 8991 | 129S1,AJ,AKR,BALB,BUB,C3H,C57L,C58J,CAST,CBA,CE,DBA1J,DBA2J,FVB,MOLF,MRL,NOD,NZB,NZO,NZW,PJ,PL,RF,SEA,SJL,SMJ,SPRET,SWR,TH,WSB | 4 | 108006782 | 108006800 | 19 | - | 17.1639 | 8.17E-07 | 0.127 | CCTCCACGAGGTGGCGGAA | | |
| 410 | 4 | 123746581 | 123748958 | 2377 | SPRET | 4 | 123748619 | 123748652 | 34 | - | 16.9024 | 7.21E-07 | 0.0829 | CTGCACTTTCACCCATAGCCTCTACTGGGCTCTC | | |
| 411 | 4 | 124769705 | 124770952 | 1247 | SPRET | 4 | 124770549 | 124770582 | 34 | - | 17.3902 | 5.27E-07 | 0.071 | TAGCTGTCCTCAGACACTCCAGAAGAGGGCATCA | 1 | ENCFF093ZRU |
| 412 | 4 | 131090948 | 131092678 | 1730 | SPRET | 4 | 131092105 | 131092123 | 19 | - | 17.1311 | 8.34E-07 | 0.128 | TGACCACAAGTGGGCGCTC | 21 | ENCFF093ZRU,ENCFF367UNR,ENCFF089GBC,ENCFF092NMJ,ENCFF136ZDK,ENCFF165LXO,ENCFF099QNE,ENCFF327LYS,ENCFF210BTO,ENCFF821BIA,ENCFF609MDN,ENCFF907XEG,ENCFF052KVO,ENCFF316FYN,ENCFF024JOZ,ENCFF198SSR,ENCFF430PPJ,ENCFF589ZBJ,ENCFF883UPM,ENCFF281QMV,ENCFF957SPH |
| 412 | 4 | 131090948 | 131092678 | 1730 | SPRET | 4 | 131092105 | 131092138 | 34 | - | 19.6179 | 1.17E-07 | 0.0314 | CTGACCTTTCAGATGTGACCACAAGTGGGCGCTC | 21 | ENCFF093ZRU,ENCFF367UNR,ENCFF089GBC,ENCFF092NMJ,ENCFF136ZDK,ENCFF165LXO,ENCFF099QNE,ENCFF327LYS,ENCFF210BTO,ENCFF821BIA,ENCFF609MDN,ENCFF907XEG,ENCFF052KVO,ENCFF316FYN,ENCFF024JOZ,ENCFF198SSR,ENCFF430PPJ,ENCFF589ZBJ,ENCFF883UPM,ENCFF281QMV,ENCFF957SPH |
| 413 | 4 | 133560483 | 133561926 | 1443 | AJ,AKR,B10D2,B10J,BALB,BTBR,BUB,C3H,CBA,CE,DBA1J,DBA2J,NOD,NZW,PJ,RF,RH2,SEA,SMJ,TH | 4 | 133561354 | 133561388 | 35 | - | 24.0734 | 5.04E-09 | 0.0069 | CTGCATGTACACCTGTGAGCCAGAAGAGGGCATCA | | |
| 414 | 4 | 134121224 | 134122542 | 1318 | 129S1,AJ,AKR,B10D2,B10J,BALB,BTBR,BUB,C3H,C57L,CBA,CE,DBA1J,DBA2J,FVB,ILNJ2,MA/My,MRL,NOD,NZW,PJ,PL,RF,RH2,SEA,SJL,SMJ,SWR,TH,WSB | 4 | 134122079 | 134122112 | 34 | - | 16.5691 | 8.90E-07 | 0.0928 | TAGCTGTCTTCAGACGCACCAGAAGAGGGCATCG | 1 | ENCFF093ZRU |
| 415 | 4 | 135563475 | 135565159 | 1684 | 129S1,AKR,BUB,C3H,C57L,CBA,CE,ILNJ2,MRL,NZB,NZW,RF,SPRET | 4 | 135564377 | 135564410 | 34 | - | 16.4634 | 9.51E-07 | 0.0931 | TAGCTGTCTTCAGACGCACCAGAAGAGGGCATCA | | |
| 416 | 4 | 139064418 | 139066349 | 1931 | FVB,ILNJ2,NZW | 4 | 139064586 | 139064604 | 19 | - | 21.9836 | 2.06E-08 | 0.0238 | CGACCTCCAGGTGGCGCTG | 38 | ENCFF093ZRU,ENCFF367UNR,ENCFF099QNE,ENCFF327LYS,ENCFF136ZDK,ENCFF086ZTV,ENCFF589ZBJ,ENCFF601JVE,ENCFF024JOZ,ENCFF821BIA,ENCFF907XEG,ENCFF321AGQ,ENCFF502WDT,ENCFF089GBC,ENCFF092NMJ,ENCFF165LXO,ENCFF198SSR,ENCFF210BTO,ENCFF957SPH,ENCFF951GOP,ENCFF430PPJ,ENCFF883UPM,ENCFF052KVO,ENCFF194HVL,ENCFF281QMV,ENCFF575MGI,ENCFF533APC,ENCFF316FYN,ENCFF349BUL,ENCFF611HDQ,ENCFF142CNG,ENCFF609MDN,ENCFF097CAQ,ENCFF033UQX,ENCFF504JDS,ENCFF491RJK,ENCFF089EWX,ENCFF055HAP |
| 417 | 4 | 142076271 | 142078885 | 2614 | AKR,B10D2,B10J,C3H,CBA,MA/My,MOLF,MRL,NOR,SMJ | 4 | 142076420 | 142076438 | 19 | - | 19.1475 | 2.10E-07 | 0.0659 | CTACCACTAGGAGGCAGCA | 24 | ENCFF093ZRU,ENCFF367UNR,ENCFF430PPJ,ENCFF327LYS,ENCFF099QNE,ENCFF533APC,ENCFF821BIA,ENCFF033UQX,ENCFF907XEG,ENCFF024JOZ,ENCFF194HVL,ENCFF198SSR,ENCFF589ZBJ,ENCFF052KVO,ENCFF165LXO,ENCFF957SPH,ENCFF089GBC,ENCFF951GOP,ENCFF136ZDK,ENCFF092NMJ,ENCFF210BTO,ENCFF281QMV,ENCFF609MDN,ENCFF491RJK |
| 418 | 4 | 142282694 | 142285926 | 3232 | 129S1,AKR,B10D2,B10J,BTBR,BUB,C3H,C57L,CAST,CBA,CE,DBA1J,DBA2J,FVB,ILNJ2,KK,MA/My,MOLF,MRL,NOD,NOR,NZO,NZW,PL,RF,SJL,SMJ,SPRET,SWR,TH,WSB | 4 | 142282707 | 142282725 | 19 | - | 17.5574 | 6.32E-07 | 0.111 | TGGCCAGTTGGGGGCGCGC | | |

| | | | | | | | | | | | | | | | | |
|---|---|---|---|---|---|---|---|---|---|---|---|---|---|---|---|---|
| 418 | 4 | 142282694 | 142285926 | 3232 | 129S1,AKR,B10D2,B10J,BTBR,BUB,C3H,C57L,CAST,CBA,CE,DBA1J,DBA2J,FVB,ILNJ2,KK,MA/My,MOLF,MRL,NOD,NOR,NZO,NZW,PL,RF,SJL,SMJ,SPRET,SWR,TH,WSB | 4 | 142282707 | 142282740 | 34 | - | 19.561 | 1.21E-07 | 0.0323 | CAGCATCGTTCTTCCTGGCCAGTTGGGGGCGCGC | | |
| 418 | 4 | 142282694 | 142285926 | 3232 | 129S1,AKR,B10D2,B10J,BTBR,BUB,C3H,C57L,CAST,CBA,CE,DBA1J,DBA2J,FVB,ILNJ2,KK,MA/My,MOLF,MRL,NOD,NOR,NZO,NZW,PL,RF,SJL,SMJ,SPRET,SWR,TH,WSB | 4 | 142285441 | 142285459 | 19 | - | 17.5574 | 6.32E-07 | 0.111 | TGGCCAGTTGGGGGCGCGC | | |
| 418 | 4 | 142282694 | 142285926 | 3232 | 129S1,AKR,B10D2,B10J,BTBR,BUB,C3H,C57L,CAST,CBA,CE,DBA1J,DBA2J,FVB,ILNJ2,KK,MA/My,MOLF,MRL,NOD,NOR,NZO,NZW,PL,RF,SJL,SMJ,SPRET,SWR,TH,WSB | 4 | 142285441 | 142285474 | 34 | - | 19.561 | 1.21E-07 | 0.0323 | CAGCATCGTTCTTCCTGGCCAGTTGGGGGCGCGC | | |
| 419 | 4 | 142922370 | 142923593 | 1223 | MOLF | 4 | 142922340 | 142922374 | 35 | - | 19.3761 | 1.29E-07 | 0.0508 | CTGCATGTGCACCTGTGTGCCAGAAGAGGACATTA | | |
| 420 | 4 | 146687955 | 146689424 | 1469 | BTBR.C57L.C58J.LP.PJ | 4 | 146688398 | 146688416 | 19 | - | 16.9836 | 9.17E-07 | 0.133 | CGGCCGCCGGGGGGCGCGG | | |
| 421 | 4 | 149902732 | 149903779 | 1047 | SWR | 4 | 149903351 | 149903369 | 19 | - | 18.6066 | 3.09E-07 | 0.0804 | AGGCCAGAAGAGGGCACAA | | |
| 422 | 4 | 150825024 | 150827055 | 2031 | AKR.C3H.CBA.MRL.NOD.NOR.NZO | 4 | 150826242 | 150826260 | 19 | - | 18.082 | 4.44E-07 | 0.0983 | ATGCCAGGAGAGGGCACTA | | |
| 423 | 5 | 5770855 | 5774029 | 3174 | SPRET | 5 | 5771855 | 5771888 | 34 | + | 18.3984 | 2.70E-07 | 0.0382 | TTGCTGTCTTCAGACACACCAGAAGAGGGCATTG | 1 | ENCFF093ZRU |
| 424 | 5 | 8646128 | 8648612 | 2484 | BUB.NOD.NOR.SMJ | 5 | 8647423 | 8647456 | 34 | + | 22.8211 | 1.04E-08 | 0.00317 | GTGTAGTGCCCACAGAAGCCAGAAGAGGGTGTCA | | |
| 425 | 5 | 9898650 | 9907462 | 8812 | SPRET | 5 | 9901257 | 9901275 | 19 | + | 18.5738 | 3.17E-07 | 0.0683 | AAACCACCAGGTGGCAGAA | | |
| 426 | 5 | 17145433 | 17163002 | 17569 | SPRET | 5 | 17152593 | 17152611 | 19 | + | 19.2295 | 1.98E-07 | 0.0533 | GGGCCACTAGGGGGCACAT | 19 | ENCFF367UNR,ENCFF093ZRU,ENCFF201JUA,ENCFF099QNE,ENCFF821BIA,ENCFF907XEG,ENCFF327LYS,ENCFF024JOZ,ENCFF609MDN,ENCFF502WDT,ENCFF055HAP,ENCFF198SSR,ENCFF589ZBJ,ENCFF089GBC,ENCFF165LXO,ENCFF092NMJ,ENCFF281QMV,ENCFF883UPM,ENCFF430PPJ |
| 427 | 5 | 17182017 | 17187716 | 5699 | SPRET | 5 | 17182501 | 17182535 | 35 | + | 17.0092 | 5.39E-07 | 0.101 | TTGCAATTGATGGCCCAGGGCTGAAGGGGCCACCA | | |
| 428 | 5 | 23446637 | 23448692 | 2055 | SPRET | 5 | 23447674 | 23447708 | 35 | + | 19.7706 | 1.01E-07 | 0.0366 | CAGCAGTTCAGTTCACAGGTCAGCAGAGGCAGCTC | | |
| 429 | 5 | 25435691 | 25444667 | 8976 | 129S1,AJ,AKR,BTBR,BUB,C3H,C57L,CAST,CBA,CE,DBA1J,DBA2J,ILNJ2,KK,MRL,NOD,NOR,NZB,NZO,NZW,PJ,RH2,SMJ,SPRET,WSB | 5 | 25441152 | 25441170 | 19 | + | 17.1639 | 8.17E-07 | 0.11 | CCTCCACGAGGTGGCGGAA | | |
| 430 | 5 | 26166046 | 26182147 | 16101 | 129S1.BTBR.C57L.CE.MOLF.NZW.WSB | 5 | 26169218 | 26169251 | 34 | + | 19.0976 | 1.68E-07 | 0.0278 | GAGCATTGATCACTGTGACCGCTAGGAGGCGCAG | | |
| 430 | 5 | 26166046 | 26182147 | 16101 | 129S1.BTBR.C57L.CE.MOLF.NZW.WSB | 5 | 26169233 | 26169251 | 19 | + | 17.0328 | 8.88E-07 | 0.113 | TGACCGCTAGGAGGCGCAG | | |
| 431 | 5 | 27294423 | 27298012 | 3589 | 129S1.BTBR.C57L.CE.NZB.NZW | 5 | 27296780 | 27296813 | 34 | + | 29.2683 | 2.44E-11 | 0.0000711 | GTGCAGTGCCCACAGAGGCCAGAAGAGGGCATCA | 1 | ENCFF093ZRU |
| 432 | 5 | 31613658 | 31615551 | 1893 | CAST | 5 | 31613638 | 31613671 | 34 | + | 17.2195 | 5.88E-07 | 0.0599 | TAGCTGTTTCCAGATGCACCAGAAGAGGGTGTCA | | |
| 433 | 5 | 34894995 | 34896978 | 1983 | BUB,C58J,FVB,MA/My,MRL,NOD,NOR,NZB,PL,SWR,TH | 5 | 34895288 | 34895321 | 34 | + | 18.4797 | 2.56E-07 | 0.0369 | ATGCAGCTAGAGACATGAGCTCCAGGGGGAACTG | | |
| 434 | 5 | 43365280 | 43374325 | 9045 | BALB,BUB,CAST,DBA1J,DBA2J,FVB,ILNJ2,KK,LP,MA/My,MOLF,NZB,NZO,PJ,PL,RH2,SEA,SPRET,SWR,TH,WSB | 5 | 43370739 | 43370757 | 19 | + | 17.1639 | 8.17E-07 | 0.11 | CCTCCACGAGGTGGCGGAA | | |
| 435 | 5 | 44291697 | 44300683 | 8986 | 129S1,AJ,AKR,BALB,BTBR,BUB,C3H,C57L,CAST,CBA,CE,DBA1J,DBA2J,FVB,ILNJ2,KK,LP,MA/My,MOLF,MRL,NOD,NOR,NZB,NZO,NZW,PJ,PL,RF,RH2,SEA,SJL,SPRET,SWR,TH.WSB | 5 | 44297115 | 44297133 | 19 | + | 17.1639 | 8.17E-07 | 0.11 | CCTCCACGAGGTGGCGGAA | | |
| 436 | 5 | 60831803 | 60833018 | 1215 | SPRET | 5 | 60832640 | 60832674 | 35 | + | 19.3853 | 1.29E-07 | 0.0432 | CTGCGAAGCTCGTCACTTACCAGGAGAGGGTGCTG | | |
| 437 | 5 | 64292059 | 64294498 | 2439 | WSB | 5 | 64293845 | 64293878 | 34 | + | 28.1057 | 8.47E-11 | 0.000118 | GTGCATTGCCTGTGGAGGCCAGAAGAGGGCACTG | 29 | ENCFF201JUA,ENCFF367UNR,ENCFF093ZRU,ENCFF052KVO,ENCFF099QNE,ENCFF821BIA,ENCFF092NMJ,ENCFF210BTO,ENCFF430PPJ,ENCFF065ODV,ENCFF951GOP,ENCFF609MDN,ENCFF327LYS,ENCFF907XEG,ENCFF024JOZ,ENCFF502WDT,ENCFF055HAP,ENCFF589ZBJ,ENCFF097CAQ,ENCFF136ZDK,ENCFF198SSR,ENCFF194HVL,ENCFF142CNG,ENCFF491RJK,ENCFF957SPH,ENCFF883UPM,ENCFF533APC,ENCFF165LXO.ENCFF089GBC |
| 437 | 5 | 64292059 | 64294498 | 2439 | WSB | 5 | 64293860 | 64293878 | 19 | + | 20.0656 | 1.05E-07 | 0.0375 | AGGCCAGAAGAGGGCACTG | 29 | ENCFF201JUA,ENCFF367UNR,ENCFF093ZRU,ENCFF052KVO,ENCFF099QNE,ENCFF821BIA,ENCFF092NMJ,ENCFF210BTO,ENCFF430PPJ,ENCFF065ODV,ENCFF951GOP,ENCFF609MDN,ENCFF327LYS,ENCFF907XEG,ENCFF024JOZ,ENCFF502WDT,ENCFF055HAP,ENCFF589ZBJ,ENCFF097CAQ,ENCFF136ZDK,ENCFF198SSR,ENCFF194HVL,ENCFF142CNG,ENCFF491RJK,ENCFF957SPH,ENCFF883UPM,ENCFF533APC,ENCFF165LXO,ENCFF089GBC |
| 438 | 5 | 65703225 | 65704949 | 1724 | MOLF | 5 | 65704248 | 65704281 | 34 | + | 17.3902 | 5.27E-07 | 0.0561 | TAGCTGTCCTCAGACACTCCAGAAGAGGGCATCA | 2 | ENCFF367UNR,ENCFF093ZRU |
| 439 | 5 | 73502368 | 73504088 | 1720 | C3H,CBA,DBA1J,DBA2J,ILNJ2,MOLF,NOD,NOR,TH | 5 | 73503170 | 73503203 | 34 | + | 16.9024 | 7.21E-07 | 0.0673 | TTGCTGTCTTCAGACACACCAGAAGAGGGCATCT | 1 | ENCFF093ZRU |
| 440 | 5 | 73810235 | 73811629 | 1394 | SPRET | 5 | 73810415 | 73810448 | 34 | + | 21.1789 | 3.74E-08 | 0.00889 | GTGCAGTGCTTGCAGAGATCAGAAGAGGACACCA | | |
| 441 | 5 | 75578187 | 75580079 | 1892 | 129S1,AKR,BALB,BUB,C57L,CAST,MA/My,NOD,NOR,PL,RF,RH2,SEA,SPRET,SWR,TH,WSB | 5 | 75579194 | 75579228 | 35 | + | 15.9541 | 9.80E-07 | 0.141 | TTGCAATTGAAGGCCCAGGACTGAAGGGGTCACAC | | |
| 442 | 5 | 77078156 | 77089005 | 10849 | SPRET | 5 | 77083776 | 77083794 | 19 | + | 17.1639 | 8.17E-07 | 0.11 | CCTCCACGAGGTGGCGGAA | | |
| 443 | 5 | 77078357 | 77087343 | 8986 | AJ,AKR,BUB,C3H,CAST,CBA,CE,DBA1J,DBA2J,FVB,KK,MA/My,MOLF,MRL,NOD,NOR,PJ,PL,RF,SJL,WSB | 5 | 77083776 | 77083794 | 19 | + | 17.1639 | 8.17E-07 | 0.11 | CCTCCACGAGGTGGCGGAA | | |
| 444 | 5 | 77437892 | 77439740 | 1848 | SPRET | 5 | 77438590 | 77438624 | 35 | + | 27.4679 | 3.12E-10 | 0.00145 | CTGCATGTATGCCTGCAGGCCAGAAGAGGGCACCA | 5 | ENCFF367UNR.ENCFF093ZRU.ENCFF099QNE.ENCFF502WDT.ENCFF055HAP |
| 444 | 5 | 77437892 | 77439740 | 1848 | SPRET | 5 | 77438606 | 77438624 | 19 | + | 20.3607 | 8.29E-08 | 0.0341 | AGGCCAGAAGAGGGCACCA | 5 | ENCFF367UNR.ENCFF093ZRU.ENCFF099QNE.ENCFF502WDT.ENCFF055HAP |
| 445 | 5 | 78621707 | 78627135 | 5428 | SPRET.WSB | 5 | 78622332 | 78622366 | 35 | + | 16.367 | 7.77E-07 | 0.126 | CTCCAGGTTTGACCAGATGGCAGCAGGGGGCCCCC | | |
| 446 | 5 | 88881297 | 88886977 | 5680 | NZO,RF | 5 | 88882842 | 88882860 | 19 | + | 21.2459 | 3.96E-08 | 0.0284 | TGGCCAGAAGAGGGCAGCC | 18 | ENCFF052KVO,ENCFF099QNE,ENCFF201JUA,ENCFF821BIA,ENCFF367UNR,ENCFF089GBC,ENCFF093ZRU,ENCFF502WDT,ENCFF883UPM,ENCFF198SSR,ENCFF957SPH,ENCFF589ZBJ,ENCFF097CAQ,ENCFF142CNG,ENCFF055HAP,ENCFF327LYS,ENCFF136ZDK,ENCFF907XEG |
| 447 | 5 | 103808565 | 103809960 | 1395 | SPRET | 5 | 103808788 | 103808806 | 19 | + | 17.3279 | 7.34E-07 | 0.104 | CTTCCACCAGGTGGAGGAA | | |
| 448 | 5 | 106939755 | 106940958 | 1203 | CAST | 5 | 106940488 | 106940521 | 34 | + | 17.2033 | 5.94E-07 | 0.0603 | TAGCTGTTGTCAGACACACCAGAAGAGGGCATCA | | |
| 449 | 5 | 111513297 | 111514694 | 1397 | SPRET | 5 | 111513576 | 111513609 | 34 | + | 19.065 | 1.71E-07 | 0.0282 | CTGCTGTCTTCAGACACACCAGAGGAGGGCGTCA | | |
| 450 | 5 | 112080455 | 112082414 | 1959 | 129S1.AKR.C57L.NZW.RF | 5 | 112080860 | 112080878 | 19 | + | 17.1148 | 8.43E-07 | 0.11 | CTTCCAGCAGAGGGCGTCC | 2 | ENCFF093ZRU.ENCFF099QNE |
| 451 | 5 | 114953245 | 114963642 | 10397 | MOLF | 5 | 114962329 | 114962362 | 34 | + | 20.9919 | 4.30E-08 | 0.0099 | TGGCAGTGCTCATAGAGGCCAGAAGAGGGCGTGG | 18 | ENCFF201JUA,ENCFF093ZRU,ENCFF367UNR,ENCFF099QNE,ENCFF907XEG,ENCFF327LYS,ENCFF055HAP,ENCFF024JOZ,ENCFF502WDT,ENCFF589ZBJ,ENCFF198SSR,ENCFF136ZDK,ENCFF086ZTV,ENCFF821BIA,ENCFF533APC,ENCFF210BTO,ENCFF805QBU,ENCFF349BUI |
| 451 | 5 | 114953245 | 114963642 | 10397 | MOLF | 5 | 114962549 | 114962582 | 34 | + | 25.8699 | 7.54E-10 | 0.000444 | GTGCAGTGCCTGTGGAGGTCACTAGAGGGTGCTG | 34 | ENCFF201JUA,ENCFF093ZRU,ENCFF367UNR,ENCFF099QNE,ENCFF907XEG,ENCFF327LYS,ENCFF055HAP,ENCFF024JOZ,ENCFF502WDT,ENCFF589ZBJ,ENCFF198SSR,ENCFF136ZDK,ENCFF086ZTV,ENCFF821BIA,ENCFF210BTO,ENCFF033UQX,ENCFF065ODV,ENCFF609MDN,ENCFF321AGQ,ENCFF092NMJ,ENCFF052KVO,ENCFF165LXO,ENCFF089GBC,ENCFF430PPJ,ENCFF957SPH,ENCFF575MGI,ENCFF491RJK,ENCFF883UPM,ENCFF504JDS,ENCFF281QMV,ENCFF142CNG,ENCFF951GOP,ENCFF097CAQ,ENCFF316FYN |
| 452 | 5 | 120661798 | 120665365 | 3567 | CAST | 5 | 120663968 | 120664002 | 35 | + | 17.4587 | 4.15E-07 | 0.0865 | CAGTGGCTCAGTCCTGCTGCCTGCAGGTGGAGCTA | 24 | ENCFF430PPJ,ENCFF165LXO,ENCFF316FYN,ENCFF327LYS,ENCFF609MDN,ENCFF093ZRU,ENCFF367UNR,ENCFF907XEG,ENCFF589ZBJ,ENCFF024JOZ,ENCFF194HVL,ENCFF136ZDK,ENCFF821BIA,ENCFF097CAQ,ENCFF099QNE,ENCFF055HAP,ENCFF198SSR,ENCFF089GBC,ENCFF957SPH,ENCFF502WDT,ENCFF533APC,ENCFF142CNG,ENCFF092NMJ,ENCFF883UPM |
| 452 | 5 | 120661798 | 120665365 | 3567 | CAST | 5 | 120663984 | 120664002 | 19 | + | 18.082 | 4.44E-07 | 0.0836 | CTGCCTGCAGGTGGAGCTA | 24 | ENCFF430PPJ,ENCFF165LXO,ENCFF316FYN,ENCFF327LYS,ENCFF609MDN,ENCFF093ZRU,ENCFF367UNR,ENCFF907XEG,ENCFF589ZBJ,ENCFF024JOZ,ENCFF194HVL,ENCFF136ZDK,ENCFF821BIA,ENCFF097CAQ,ENCFF099QNE,ENCFF055HAP,ENCFF198SSR,ENCFF089GBC,ENCFF957SPH,ENCFF502WDT,ENCFF533APC,ENCFF142CNG,ENCFF092NMJ,ENCFF883UPM |
| 452 | 5 | 120661798 | 120665365 | 3567 | CAST | 5 | 120662671 | 120662704 | 34 | - | 17.6992 | 4.30E-07 | 0.0494 | TAGCTGTTTTCAGACACACCAGAAGAGGGCATCA | 1 | ENCFF093ZRU |
| 453 | 5 | 122330904 | 122338270 | 7366 | 129S1,AJ,BALB,BUB,C57L,CAST,ILNJ2,KK,MOLF,MRL,NOD,NOR,NZW,PL,RF,RH2,SEA,SPRET,WSB | 5 | 122336367 | 122336385 | 19 | + | 17.1639 | 8.17E-07 | 0.11 | CCTCCACGAGGTGGCGGAA | | |
| 454 | 5 | 123335970 | 123342488 | 6518 | MOLF | 5 | 123337132 | 123337166 | 35 | + | 27.4679 | 3.12E-10 | 0.00145 | CTGCATGTATGCCTGCAGGCCAGAAGAGGGCACCA | 2 | ENCFF136ZDK.ENCFF589ZBJ |
| 454 | 5 | 123335970 | 123342488 | 6518 | MOLF | 5 | 123337148 | 123337166 | 19 | + | 20.3607 | 8.29E-08 | 0.0341 | AGGCCAGAAGAGGGCACCA | 2 | ENCFF136ZDK.ENCFF589ZBJ |
| 454 | 5 | 123335970 | 123342488 | 6518 | MOLF | 5 | 123341001 | 123341019 | 19 | + | 17.5738 | 6.25E-07 | 0.0956 | GTGCCACTGGAGGGCGCTA | 7 | ENCFF093ZRU,ENCFF065ODV,ENCFF136ZDK,ENCFF052KVO,ENCFF502WDT,ENCFF055HAP,ENCFF589ZBJ |
| 455 | 5 | 123404716 | 123406370 | 1654 | 129S1,AJ,BALB,C57L,CAST,MRL,RH2,SEA,SJL | 5 | 123405151 | 123405169 | 19 | + | 18.2787 | 3.88E-07 | 0.077 | TGGCCTCTGGGGGGCACTG | 12 | ENCFF821BIA,ENCFF201JUA,ENCFF024JOZ,ENCFF327LYS,ENCFF430PPJ,ENCFF198SSR,ENCFF907XEG,ENCFF055HAP.ENCFF883UPM.ENCFF093ZRU.ENCFF367UNR.ENCFF099QNE |
| 456 | 5 | 126364341 | 126369157 | 4816 | AKR,C3H,CBA,FVB,ILNJ2,NOD,NOR,SMJ,SWR,TH | 5 | 126366477 | 126366510 | 34 | + | 18.878 | 1.95E-07 | 0.0309 | GTGCAGTGCCCATGGAGTCCAGAAGAGAGTGTCA | | |
| 457 | 5 | 129036857 | 129037949 | 1092 | WSB | 5 | 129037741 | 129037759 | 19 | + | 16.9508 | 9.36E-07 | 0.116 | ACTCCAGCAGATGGCGCAG | | |
| 457 | 5 | 129036857 | 129037949 | 1092 | WSB | 5 | 129037229 | 129037247 | 19 | - | 20.8689 | 5.48E-08 | 0.0313 | TCTCCAGCAGGTGGCACTC | | |
| 458 | 5 | 129495665 | 129496973 | 1308 | SPRET | 5 | 129496099 | 129496117 | 19 | + | 19.5902 | 1.51E-07 | 0.0464 | TTCCCAGCAGAGGGCACTC | | |
| 459 | 5 | 133565691 | 133567154 | 1463 | CAST.NZW.PL.RF.WSB | 5 | 133566387 | 133566421 | 35 | + | 17.945 | 3.11E-07 | 0.0739 | GTACAGTTCCCTGTGGTGGCCAGAAGAGGATGCAG | 1 | ENCFF367UNR |
| 460 | 5 | 135632340 | 135634416 | 2076 | BALB,BTBR,C57L,CAST,MA/My,NZW,PL,RF,RH2,SEA,SPRET,TH,WSB | 5 | 135633903 | 135633936 | 34 | + | 16.4472 | 9.61E-07 | 0.0776 | TAGCTGTCTTCAGACACACCAGAAGAGGGCGTCA | | |
| 461 | 5 | 135633935 | 135636005 | 2070 | MOLF.NOD.SJL | 5 | 135633903 | 135633936 | 34 | + | 16.4472 | 9.61E-07 | 0.0776 | TAGCTGTCTTCAGACACACCAGAAGAGGGCGTCA | | |
| 462 | 5 | 138573750 | 138575157 | 1407 | SPRET | 5 | 138573899 | 138573932 | 34 | + | 22.6179 | 1.23E-08 | 0.00358 | AGGCAGTGTCCAAAGGGGCCAGAAGAGGGCATCA | | |
| 463 | 5 | 141585006 | 141591199 | 6193 | SPRET | 5 | 141585226 | 141585259 | 34 | + | 18.3089 | 2.87E-07 | 0.0389 | ATGCAGCTAGAGACACGAGCTCCAGGGGGTACTG | | |
| 464 | 5 | 141723554 | 141726030 | 2476 | SPRET | 5 | 141724641 | 141724674 | 34 | + | 23.8293 | 4.55E-09 | 0.00166 | ATGCAGTACCAGCAGAGGGCAGAAGAGGGCGTCT | 11 | ENCFF099QNE,ENCFF052KVO,ENCFF065ODV,ENCFF821BIA,ENCFF093ZRU,ENCFF089GBC,ENCFF589ZBJ,ENCFF136ZDK,ENCFF907XEG,ENCFF367UNR,ENCFF327LYS |
| 464 | 5 | 141723554 | 141726030 | 2476 | SPRET | 5 | 141724646 | 141724664 | 19 | + | 17.9672 | 4.81E-07 | 0.0853 | GTACCAGCAGAGGGCAGAA | 11 | ENCFF099QNE,ENCFF052KVO,ENCFF065ODV,ENCFF821BIA,ENCFF093ZRU,ENCFF089GBC,ENCFF589ZBJ.ENCFF136ZDK.ENCFF907XEG.ENCFF367UNR.ENCFF327LYS |
| 465 | 5 | 144606644 | 144615629 | 8985 | 129S1,AJ,AKR,BALB,BTBR,BUB,C3H,C57L,CAST,CBA,CE,DBA1J,DBA2J,FVB,ILNJ2,KK,LP,MA/My,MOLF,MRL,NOD,NOR,NZB,NZO,NZW,PJ,PL,RF,RH2,SEA,SJL,SMJ,SPRET,SWR.TH.WSB | 5 | 144612061 | 144612079 | 19 | + | 17.1639 | 8.17E-07 | 0.11 | CCTCCACGAGGTGGCGGAA | | |
| 466 | 5 | 149434120 | 149437328 | 3208 | MOLF | 5 | 149434810 | 149434843 | 34 | + | 23.9756 | 4.02E-09 | 0.0015 | CTGGTGTGCCCACAGAACCCAGAAGAGGGCACCA | 1 | ENCFF093ZRU |
| 467 | 5 | 149840264 | 149841314 | 1050 | AKR,BUB,CAST,CE,MRL,PJ,SEA,SJL | 5 | 149841005 | 149841038 | 34 | + | 27.9268 | 1.02E-10 | 0.000125 | CTGCAGTATCCAAGGTGACCAATAGAGGGCACCA | 17 | ENCFF093ZRU,ENCFF367UNR,ENCFF099QNE,ENCFF089GBC,ENCFF430PPJ,ENCFF092NMJ,ENCFF198SSR,ENCFF589ZBJ,ENCFF165LXO,ENCFF821BIA,ENCFF055HAP,ENCFF609MDN,ENCFF201JUA,ENCFF024JOZ,ENCFF281QMV,ENCFF907XEG,ENCFF327LYS |
| 467 | 5 | 149840264 | 149841314 | 1050 | AKR,BUB,CAST,CE,MRL,PJ,SEA,SJL | 5 | 149841020 | 149841038 | 19 | + | 20.0492 | 1.06E-07 | 0.0377 | TGACCAATAGAGGGCACCA | 17 | ENCFF093ZRU,ENCFF367UNR,ENCFF099QNE,ENCFF089GBC,ENCFF430PPJ,ENCFF092NMJ,ENCFF198SSR,ENCFF589ZBJ,ENCFF165LXO,ENCFF821BIA,ENCFF055HAP,ENCFF609MDN,ENCFF201JUA,ENCFF024JOZ,ENCFF281QMV,ENCFF907XEG,ENCFF327LYS |
| 468 | 5 | 150917371 | 150919643 | 2272 | SPRET | 5 | 150918730 | 150918763 | 34 | + | 27.7642 | 1.20E-10 | 0.00014 | ATGCAGTGCCAATGGAGGCCAGAAGAGGGCGTCG | 5 | ENCFF821BIA.ENCFF052KVO.ENCFF093ZRU.ENCFF099QNE.ENCFF367UNR |
| 469 | 5 | 151525371 | 151526405 | 1034 | CAST | 5 | 151525881 | 151525914 | 34 | + | 18.8699 | 1.96E-07 | 0.031 | GTGCTGTGTCCACCATAGCCAGCAGAGGTGAACA | | |
| 470 | 5 | 11830149 | 11834657 | 4508 | 129S1,BTBR,C57L,CE,NZB,NZO,NZW,WSB | 5 | 11830206 | 11830224 | 19 | - | 17.2131 | 7.91E-07 | 0.109 | CAGCCTCCAGATGGCGGTT | | |
| 470 | 5 | 11830149 | 11834657 | 4508 | 129S1,BTBR,C57L,CE,NZB,NZO,NZW,WSB | 5 | 11830206 | 11830240 | 35 | - | 17.4862 | 4.08E-07 | 0.0856 | TGGTGCTGCGGCCGTACAGCCTCCAGATGGCGGTT | | |
| 471 | 5 | 15704189 | 15708248 | 4059 | CAST,WSB | 5 | 15704267 | 15704285 | 19 | - | 19.2951 | 1.88E-07 | 0.0516 | CCGCCACTAGATGGTGCTA | 13 | ENCFF136ZDK,ENCFF883UPM,ENCFF093ZRU,ENCFF367UNR,ENCFF099QNE,ENCFF327LYS,ENCFF194HVL,ENCFF821BIA,ENCFF024JOZ,ENCFF907XEG,ENCFF589ZBJ,ENCFF957SPH |
| 472 | 5 | 16872418 | 16879719 | 7301 | CAST.DBA1J.DBA2J.MOLF.SPRET | 5 | 16872746 | 16872779 | 34 | - | 19.3415 | 1.42E-07 | 0.0244 | CTGCCCCCACATTCGCCGTCACAAGATGGCGCTG | | |
| 473 | 5 | 17517842 | 17524153 | 6311 | MOLF.SPRET | 5 | 17521790 | 17521808 | 19 | - | 18.5082 | 3.31E-07 | 0.0703 | TTGCCAGCAGATGGCATTG | | |
| 474 | 5 | 21623456 | 21628025 | 4569 | BUB.TH | 5 | 21624940 | 21624974 | 35 | - | 18.5505 | 2.16E-07 | 0.0596 | CTGCATATACGCTGGCATGCCAGAGGAGGGCATCA | 1 | ENCFF502WDT |
| 474 | 5 | 21623456 | 21628025 | 4569 | BUB,TH | 5 | 21625798 | 21625816 | 19 | - | 18.5574 | 3.20E-07 | 0.0687 | TGGACAGCAGAGGGCAGAA | 17 | ENCFF093ZRU,ENCFF367UNR,ENCFF430PPJ,ENCFF099QNE,ENCFF165LXO,ENCFF024JOZ,ENCFF327LYS,ENCFF821BIA,ENCFF055HAP,ENCFF907XEG,ENCFF502WDT,ENCFF089GBC,ENCFF609MDN,ENCFF883UPM,ENCFF092NMJ,ENCFF589ZBJ,ENCFF198SSR |
| 475 | 5 | 22563552 | 22569432 | 5880 | NZO.WSB | 5 | 22564216 | 22564250 | 35 | - | 17.8257 | 3.34E-07 | 0.0763 | CTGCTGGGTCATGAGTGCATCAGTAGGTGGCAGAG | | |
| 476 | 5 | 22722566 | 22726795 | 4229 | MOLF | 5 | 22724074 | 22724107 | 34 | - | 16.561 | 8.95E-07 | 0.0769 | TAGCTGTCTTAGGACACACCAGAAGAGGGCATTG | | |
| 477 | 5 | 27848664 | 27849933 | 1269 | MOLF | 5 | 27849093 | 27849126 | 34 | - | 23.7967 | 4.68E-09 | 0.00169 | ATGCAGTGCCCATGAAGGCCAGAAGAGAGCACCC | | |
| 478 | 5 | 31486772 | 31491400 | 4628 | CAST.SPRET | 5 | 31488036 | 31488054 | 19 | - | 17.0328 | 8.88E-07 | 0.113 | TGGCCAGCAGATGGCCCAC | | |
| 478 | 5 | 31486772 | 31491400 | 4628 | CAST.SPRET | 5 | 31488036 | 31488070 | 35 | - | 16.6239 | 6.72E-07 | 0.116 | AAGAAGCTAGACCCAGTGGCCAGCAGATGGCCCAC | | |
| 479 | 5 | 32700757 | 32702898 | 2141 | CAST.MOLF | 5 | 32701730 | 32701763 | 34 | - | 19.9024 | 9.52E-08 | 0.0182 | GTGTGGTGCCTGCAGAGGCCAGGAGAAGGCGTCA | | |
| 480 | 5 | 32729153 | 32730886 | 1733 | SPRET | 5 | 32729408 | 32729426 | 19 | - | 17.0328 | 8.88E-07 | 0.113 | AAGCCAGCAGAGGTCACCG | 3 | ENCFF093ZRU.ENCFF367UNR.ENCFF327LYS |
| 481 | 5 | 37511788 | 37513130 | 1342 | SPRET | 5 | 37511925 | 37511943 | 19 | - | 17.0984 | 8.52E-07 | 0.111 | AGTCCAGCAGGGGGCAAAA | 24 | ENCFF201JUA,ENCFF093ZRU,ENCFF367UNR,ENCFF099QNE,ENCFF065ODV,ENCFF092NMJ,ENCFF821BIA,ENCFF430PPJ,ENCFF907XEG,ENCFF327LYS,ENCFF198SSR,ENCFF609MDN,ENCFF024JOZ,ENCFF883UPM,ENCFF589ZBJ,ENCFF052KVO,ENCFF504JDS,ENCFF957SPH,ENCFF136ZDK,ENCFF165LXO,ENCFF951GOP,ENCFF281QMV,ENCFF089GBC,ENCFF575MGI |
| 481 | 5 | 37511788 | 37513130 | 1342 | SPRET | 5 | 37511925 | 37511958 | 34 | - | 24.065 | 3.73E-09 | 0.00141 | CTGTAATGCCATGAAAGTCCAGCAGGGGGCAAAA | 24 | ENCFF201JUA,ENCFF093ZRU,ENCFF367UNR,ENCFF099QNE,ENCFF065ODV,ENCFF092NMJ,ENCFF821BIA,ENCFF430PPJ,ENCFF907XEG,ENCFF327LYS,ENCFF198SSR,ENCFF609MDN,ENCFF024JOZ,ENCFF883UPM,ENCFF589ZBJ,ENCFF052KVO,ENCFF504JDS,ENCFF957SPH,ENCFF136ZDK,ENCFF165LXO,ENCFF951GOP,ENCFF281QMV,ENCFF089GBC,ENCFF575MGI |
| 482 | 5 | 38601308 | 38602665 | 1357 | SPRET | 5 | 38602449 | 38602482 | 34 | - | 17.4146 | 5.18E-07 | 0.0557 | ATGCAGCTAGAGACACAAGCTCCAGGGGGTACTG | | |
| 483 | 5 | 41552619 | 41559806 | 7187 | 129S1,BALB,BTBR,BUB,C57L,CAST,DBA1J,DBA2J,FVB,ILNJ2,KK,LP,MA/My,MOLF,NZB,NZO,NZW,PJ,PL,RH2,SEA,SPRET,SWR,TH,WSB | 5 | 41552948 | 41552981 | 34 | - | 19.3415 | 1.42E-07 | 0.0244 | CTGCCCCCACATTCGCCGTCACAAGATGGCGCTG | | |
| 484 | 5 | 46412889 | 46547285 | 134396 | NOD,NOR | 5 | 46504932 | 46504950 | 19 | - | 17.8197 | 5.31E-07 | 0.0885 | TGGCCACACGGGGGCAGCA | 32 | ENCFF367UNR,ENCFF201JUA,ENCFF093ZRU,ENCFF099QNE,ENCFF907XEG,ENCFF089GBC,ENCFF024JOZ,ENCFF327LYS,ENCFF430PPJ,ENCFF086ZTV,ENCFF821BIA,ENCFF092NMJ,ENCFF055HAP,ENCFF198SSR,ENCFF589ZBJ,ENCFF502WDT,ENCFF210BTO,ENCFF533APC,ENCFF165LXO,ENCFF142CNG,ENCFF883UPM,ENCFF089EWX,ENCFF491RJK,ENCFF951GOP,ENCFF281QMV,ENCFF957SPH,ENCFF575MGI,ENCFF065ODV,ENCFF194HVL,ENCFF609MDN,ENCFF136ZDK,ENCFF316FYN |
| 485 | 5 | 63134451 | 63138621 | 4170 | MOLF.SPRET | 5 | 63137818 | 63137851 | 34 | - | 17.6748 | 4.37E-07 | 0.05 | GAGCATGACCACAAGTCCACAGCAGGTGGCAACG | | |
| 486 | 5 | 66392338 | 66397948 | 5610 | MOLF | 5 | 66393382 | 66393400 | 19 | - | 20.8033 | 5.79E-08 | 0.0313 | AGGCCAGAAGAGGGCGCTG | 14 | ENCFF367UNR,ENCFF136ZDK,ENCFF198SSR,ENCFF589ZBJ,ENCFF907XEG,ENCFF883UPM,ENCFF033UQX,ENCFF097CAQ,ENCFF327LYS,ENCFF052KVO,ENCFF194HVL,ENCFF533APC,ENCFF093ZRU,ENCFF957SPH |
| 486 | 5 | 66392338 | 66397948 | 5610 | MOLF | 5 | 66393382 | 66393416 | 35 | - | 26.5505 | 6.92E-10 | 0.00207 | GTGCAGGTACCTGCAGAGGCCAGAAGAGGGCGCTG | 14 | ENCFF367UNR,ENCFF136ZDK,ENCFF198SSR,ENCFF589ZBJ,ENCFF907XEG,ENCFF883UPM,ENCFF033UQX,ENCFF097CAQ,ENCFF327LYS,ENCFF052KVO,ENCFF194HVL,ENCFF533APC,ENCFF093ZRU,ENCFF957SPH |
| 487 | 5 | 78108224 | 78114539 | 6315 | SPRET | 5 | 78108326 | 78108344 | 19 | - | 17.2131 | 7.91E-07 | 0.109 | GGGCCAAAAGGGGGCAGTG | | |
| 487 | 5 | 78108224 | 78114539 | 6315 | SPRET | 5 | 78108326 | 78108360 | 35 | - | 16.4771 | 7.30E-07 | 0.121 | GTACAGGGGAACGCCAGGGCCAAAAGGGGGCAGTG | | |
| 488 | 5 | 78173028 | 78178480 | 5452 | SPRET | 5 | 78173867 | 78173900 | 34 | - | 21.0894 | 4.00E-08 | 0.00936 | CTGCTGTTACAGAGAGTGACAGCAGGTGGTGATC | | |
| 489 | 5 | 90539217 | 90541419 | 2202 | SPRET | 5 | 90540268 | 90540302 | 35 | - | 17.0917 | 5.14E-07 | 0.0988 | CTGCCTTTCAAGTGCACCACCACTGGGTGGAATTT | | |
| 490 | 5 | 92081179 | 92082898 | 1719 | 129S1.C57L.MOLF | 5 | 92082707 | 92082740 | 34 | - | 18.4797 | 2.56E-07 | 0.0369 | TTGCTGTCTTCAGACACACCAGAAGAGGGCATCA | | |
| 491 | 5 | 93856172 | 93885101 | 28929 | CAST | 5 | 93872557 | 93872590 | 34 | - | 20.252 | 7.40E-08 | 0.015 | ATGCAGTTAGAGACACGAGCTCCAGGGGGTACTG | | |
| 492 | 5 | 100054895 | 100062001 | 7106 | CAST.SPRET | 5 | 100054907 | 100054940 | 34 | - | 16.4309 | 9.71E-07 | 0.078 | CAGCGTTGTTCTTCCTGGCCAGTCGAGGGCGCGC | | |
| 493 | 5 | 102120526 | 102122566 | 2040 | SPRET | 5 | 102120618 | 102120636 | 19 | - | 17.5574 | 6.32E-07 | 0.0956 | ACTCCAGAAGAGGGCGCCA | 12 | ENCFF024JOZ,ENCFF327LYS,ENCFF589ZBJ,ENCFF093ZRU,ENCFF201JUA,ENCFF821BIA,ENCFF198SSR,ENCFF367UNR,ENCFF502WDT,ENCFF055HAP,ENCFF099QNE,ENCFF907XEG |
| 493 | 5 | 102120526 | 102122566 | 2040 | SPRET | 5 | 102120618 | 102120651 | 34 | - | 18.3659 | 2.76E-07 | 0.0384 | TAGCTGTCTTCAGACACTCCAGAAGAGGGCGCCA | 12 | ENCFF024JOZ,ENCFF327LYS,ENCFF589ZBJ,ENCFF093ZRU,ENCFF201JUA,ENCFF821BIA,ENCFF198SSR,ENCFF367UNR,ENCFF502WDT,ENCFF055HAP,ENCFF099QNE,ENCFF907XEG |
| 494 | 5 | 106362998 | 106364840 | 1842 | DBA1J.DBA2J | 5 | 106364631 | 106364664 | 34 | - | 19.9024 | 9.52E-08 | 0.0182 | CTGCAGCTCCTTCTCGATTCCCCAGAGGGCGTCT | | |
| 495 | 5 | 107970138 | 107973969 | 3831 | CAST | 5 | 107971099 | 107971117 | 19 | - | 17.9836 | 4.75E-07 | 0.0853 | ACACCAGAAGAGGGCGGCA | | |
| 495 | 5 | 107970138 | 107973969 | 3831 | CAST | 5 | 107971099 | 107971132 | 34 | - | 18.1057 | 3.29E-07 | 0.0424 | TAGCTGTCTTCAGACACACCAGAAGAGGGCGGCA | | |
| 496 | 5 | 110048410 | 110055880 | 7470 | 129S1,AJ,AKR,BALB,BTBR,BUB,C3H,C57L,CAST,CBA,CE,DBA1J,DBA2J,FVB,ILNJ2,KK,LP,MA/My,MOLF,MRL,NOD,NOR,NZB,NZO,NZW,PJ,PL,RF,RH2,SEA,SJL,SMJ,SPRET,SWR.TH.WSB | 5 | 110050403 | 110050421 | 19 | - | 17.1639 | 8.17E-07 | 0.11 | CCTCCACGAGGTGGCGGAA | | |
| 497 | 5 | 113409179 | 113411009 | 1830 | SPRET | 5 | 113410675 | 113410708 | 34 | - | 16.7724 | 7.83E-07 | 0.0704 | TAGCTGTCTTCAGATGCACCAGAAGAGGGCGTCA | | |
| 498 | 5 | 113805321 | 113808610 | 3289 | MOLF | 5 | 113805785 | 113805818 | 34 | - | 17.5691 | 4.69E-07 | 0.052 | TTGCTGTCTTCAGACACCCCAGAAGAGGGCATCC | | |
| 499 | 5 | 115076163 | 115079549 | 3386 | SPRET | 5 | 115079401 | 115079419 | 19 | - | 18.623 | 3.06E-07 | 0.0668 | ACACCAGAAGAGGGCGCCA | 12 | ENCFF093ZRU,ENCFF092NMJ,ENCFF821BIA,ENCFF024JOZ,ENCFF089GBC,ENCFF907XEG,ENCFF883UPM,ENCFF099QNE,ENCFF055HAP,ENCFF367UNR,ENCFF327LYS,ENCFF502WDT |
| 499 | 5 | 115076163 | 115079549 | 3386 | SPRET | 5 | 115079401 | 115079434 | 34 | - | 18.8537 | 1.98E-07 | 0.0312 | TAGCTGTCTTCAGACACACCAGAAGAGGGCGCCA | 12 | ENCFF093ZRU,ENCFF092NMJ,ENCFF821BIA,ENCFF024JOZ,ENCFF089GBC,ENCFF907XEG,ENCFF883UPM,ENCFF099QNE,ENCFF055HAP,ENCFF367UNR,ENCFF327LYS,ENCFF502WDT |
| 500 | 5 | 116987395 | 116988727 | 1332 | SPRET | 5 | 116988723 | 116988756 | 34 | - | 18.4146 | 2.67E-07 | 0.0381 | TTGCTGTCTTCAGACATACCAGAAGAGGGCATCA | 1 | ENCFF093ZRU |
| 501 | 5 | 122623031 | 122625059 | 2028 | SPRET | 5 | 122624497 | 122624530 | 34 | - | 21.9756 | 2.03E-08 | 0.00544 | CTGCTGTCTTCAGACGCACCAGAAGAGGGCGTCA | 4 | ENCFF367UNR.ENCFF093ZRU.ENCFF502WDT.ENCFF099QNE |
| 502 | 5 | 122993487 | 122994694 | 1207 | MOLF | 5 | 122994075 | 122994109 | 35 | - | 21.5046 | 3.20E-08 | 0.0173 | CTGCATATACACCTATACTCCAGAAGAGGGCATTA | 10 | ENCFF201JUA,ENCFF589ZBJ,ENCFF907XEG,ENCFF821BIA,ENCFF502WDT,ENCFF883UPM,ENCFF367UNR,ENCFF093ZRU,ENCFF055HAP,ENCFF099QNE |
| 503 | 5 | 125429722 | 125431532 | 1810 | CAST | 5 | 125430148 | 125430166 | 19 | - | 18.8689 | 2.56E-07 | 0.0627 | AGGCCAGAAGAGGGCAGTC | 9 | ENCFF367UNR,ENCFF093ZRU,ENCFF065ODV,ENCFF907XEG,ENCFF589ZBJ,ENCFF052KVO,ENCFF502WDT,ENCFF033UQX,ENCFF136ZDK |
| 503 | 5 | 125429722 | 125431532 | 1810 | CAST | 5 | 125430148 | 125430181 | 34 | - | 29.9756 | 1.10E-11 | 0.0000481 | GTGCAGTGCCTGTAGAGGCCAGAAGAGGGCAGTC | 9 | ENCFF367UNR,ENCFF093ZRU,ENCFF065ODV,ENCFF907XEG,ENCFF589ZBJ,ENCFF052KVO,ENCFF502WDT,ENCFF033UQX,ENCFF136ZDK |
| 504 | 5 | 126112500 | 126117528 | 5028 | SPRET | 5 | 126117297 | 126117330 | 34 | - | 26.374 | 4.70E-10 | 0.000319 | GTGCAGTTTCAGTACAGGCCAGATGAGGGCGCTG | 3 | ENCFF093ZRU.ENCFF089GBC.ENCFF052KVO |
| 505 | 5 | 130541910 | 130543955 | 2045 | MOLF | 5 | 130542323 | 130542356 | 34 | - | 17.9675 | 3.61E-07 | 0.0447 | CTGCCACTCCACTGCCAGGCACCACAGGGCCCTC | | |
| 506 | 5 | 134662665 | 134664547 | 1882 | CAST | 5 | 134662647 | 134662680 | 34 | - | 18.1707 | 3.15E-07 | 0.0413 | GTGCGCCACCACCACCCGGCAGCAGGTGGATCTC | | |
| 507 | 5 | 135615481 | 135618180 | 2699 | SPRET | 5 | 135617958 | 135617991 | 34 | - | 28.7967 | 4.08E-11 | 0.0000838 | GTGCAGTGCCCACAGAGGCCAGGAGAGGGAGCCC | 3 | ENCFF367UNR.ENCFF821BIA.ENCFF093ZRU |
| 508 | 5 | 137030063 | 137032342 | 2279 | CAST | 5 | 137030550 | 137030583 | 34 | - | 16.9593 | 6.95E-07 | 0.066 | TTGCTCTCTTCAGACACACCAGAAGAGGGCATCA | | |
| 508 | 5 | 137030063 | 137032342 | 2279 | CAST | 5 | 137032253 | 137032287 | 35 | - | 16.5229 | 7.12E-07 | 0.119 | CTGAACTGCATCCCTCTAGACACTAGGGGTTTGTG | | |
| 509 | 5 | 138480203 | 138494647 | 14444 | SPRET | 5 | 138483525 | 138483558 | 34 | - | 16.4472 | 9.61E-07 | 0.0776 | ATGCAATTGCTGATTCAGCCACCAGAGGCATATT | 1 | ENCFF093ZRU |
| 510 | 5 | 141194963 | 141198012 | 3049 | WSB | 5 | 141195297 | 141195330 | 34 | - | 24.6667 | 2.22E-09 | 0.000949 | GTGCAGTACCCTCAGAGACCAGAAGAGGGCCTCA | 2 | ENCFF093ZRU.ENCFF052KVO |
| 511 | 5 | 146414581 | 146417115 | 2534 | CAST.MOLF.SPRET.WSB | 5 | 146414877 | 146414910 | 34 | - | 18 | 3.53E-07 | 0.0441 | GTGCTGTGTGCACGGAAGCCAGAAGAGTGCACTG | | |
| 511 | 5 | 146414581 | 146417115 | 2534 | CAST,MOLF,SPRET,WSB | 5 | 146416540 | 146416558 | 19 | - | 23.2295 | 6.09E-09 | 0.015 | TAGCCACTAGAGGGCGGTG | 23 | ENCFF093ZRU,ENCFF316FYN,ENCFF367UNR,ENCFF201JUA,ENCFF052KVO,ENCFF099QNE,ENCFF065ODV,ENCFF327LYS,ENCFF136ZDK,ENCFF089GBC,ENCFF907XEG,ENCFF821BIA,ENCFF883UPM,ENCFF024JOZ,ENCFF055HAP,ENCFF092NMJ,ENCFF430PPJ,ENCFF609MDN,ENCFF502WDT,ENCFF589ZBJ,ENCFF957SPH.ENCFF198SSR.ENCFF165LXO |
| 511 | 5 | 146414581 | 146417115 | 2534 | CAST,MOLF,SPRET,WSB | 5 | 146416540 | 146416574 | 35 | - | 20.156 | 7.86E-08 | 0.0307 | ATGATGCTGCATCGTGTAGCCACTAGAGGGCGGTG | 23 | ENCFF093ZRU,ENCFF316FYN,ENCFF367UNR,ENCFF201JUA,ENCFF052KVO,ENCFF099QNE,ENCFF065ODV,ENCFF327LYS,ENCFF136ZDK,ENCFF089GBC,ENCFF907XEG,ENCFF821BIA,ENCFF883UPM,ENCFF024JOZ,ENCFF055HAP,ENCFF092NMJ,ENCFF430PPJ,ENCFF609MDN,ENCFF502WDT,ENCFF589ZBJ,ENCFF957SPH,ENCFF198SSR,ENCFF165LXO |
| 512 | 5 | 147331236 | 147333515 | 2279 | SPRET | 5 | 147332061 | 147332079 | 19 | - | 21.0984 | 4.51E-08 | 0.0288 | AGGCCAGAAGAGGGCGCCA | 3 | ENCFF052KVO.ENCFF502WDT.ENCFF093ZRU |
| 512 | 5 | 147331236 | 147333515 | 2279 | SPRET | 5 | 147332061 | 147332094 | 34 | - | 27.626 | 1.38E-10 | 0.000153 | GTGCAGTGTACACAGAGGCCAGAAGAGGGCGCCA | 3 | ENCFF052KVO.ENCFF502WDT.ENCFF093ZRU |
| 513 | 5 | 148393190 | 148395742 | 2552 | SPRET | 5 | 148394291 | 148394309 | 19 | - | 17.2131 | 7.91E-07 | 0.109 | AGGCCAGAAGAGGGCAACG | | |

| | | | | | | | | | | | | | | | | |
|---|---|---|---|---|---|---|---|---|---|---|---|---|---|---|---|---|
| 514 | 5 | 149663913 | 149665357 | 1444 | SPRET | 5 | 149664369 | 149664387 | 19 | - | 20.3607 | 8.29E-08 | 0.0341 | AGGCCAGAAGAGGGCACCA | 13 | ENCFF136ZDK,ENCFF092NMJ,ENCFF065ODV,ENCFF821BIA,ENCFF907XEG,ENCFF089GBC,ENCFF093ZRU,ENCFF099QNE,ENCFF052KVO,ENCFF502WDT,ENCFF142CNG,ENCFF367UNR,ENCFF055HAP |
| 514 | 5 | 149663913 | 149665357 | 1444 | SPRET | 5 | 149664369 | 149664402 | 34 | - | 30.8699 | 3.80E-12 | 0.000034 | GTGCAGTGCCTGTGGAGGCCAGAAGAGGGCACCA | 13 | ENCFF136ZDK,ENCFF092NMJ,ENCFF065ODV,ENCFF821BIA,ENCFF907XEG,ENCFF089GBC,ENCFF093ZRU,ENCFF099QNE,ENCFF052KVO,ENCFF502WDT,ENCFF142CNG,ENCFF367UNR,ENCFF055HAP |
| 515 | 6 | 11822853 | 11864716 | 41863 | CAST | 6 | 11846142 | 11846160 | 19 | + | 17.8852 | 5.08E-07 | 0.128 | ACACCAGAAGAGGGCACCA | | |
| 516 | 6 | 20047089 | 20048712 | 1623 | 129S1.C57L.MOLF.NZB | 6 | 20047452 | 20047485 | 34 | + | 16.439 | 9.66E-07 | 0.112 | ATGCAGCTAGAGACAAGAGCTCCAGGGGGTACTG | | |
| 517 | 6 | 25540348 | 25543465 | 3117 | CAST.MOLF.SPRET | 6 | 25540802 | 25540820 | 19 | + | 17.5574 | 6.32E-07 | 0.138 | TGGCCAGTTGGGGGCGCGC | | |
| 517 | 6 | 25540348 | 25543465 | 3117 | CAST.MOLF.SPRET | 6 | 25543441 | 25543459 | 19 | + | 17.5574 | 6.32E-07 | 0.138 | TGGCCAGTTGGGGGCGCGC | | |
| 518 | 6 | 26012217 | 26016058 | 3841 | SPRET | 6 | 26015920 | 26015953 | 34 | + | 16.4715 | 9.47E-07 | 0.112 | CTGCAAGACGACAGCTTGCCTGTAGAGAGTGCTC | | |
| 519 | 6 | 26040672 | 26042857 | 2185 | MOLF.SPRET | 6 | 26040856 | 26040889 | 34 | + | 17.9187 | 3.72E-07 | 0.0672 | GTGCAGTTAGAGACACGAGCTCTGGGGGGTACTG | | |
| 520 | 6 | 37420827 | 37424397 | 3570 | MOLF | 6 | 37421097 | 37421115 | 19 | + | 17.541 | 6.39E-07 | 0.139 | TGACCTCCAGGTGTCAGCA | | |
| 521 | 6 | 42909500 | 42965037 | 55537 | CAST | 6 | 42948304 | 42948337 | 34 | + | 16.7642 | 7.87E-07 | 0.103 | ATGCAGCTAGAAACACTAGCTCCAGGGGGTACTG | | |
| 522 | 6 | 42943254 | 42974604 | 31350 | SPRET | 6 | 42948304 | 42948337 | 34 | + | 16.7642 | 7.87E-07 | 0.103 | ATGCAGCTAGAAACACTAGCTCCAGGGGGTACTG | | |
| 523 | 6 | 50483868 | 50485204 | 1336 | SPRET | 6 | 50484652 | 50484686 | 35 | + | 18.6789 | 1.99E-07 | 0.0828 | CTGTAGGTATTCCCAGACGCCAGAAGAGGGTGTTA | | |
| 524 | 6 | 60666737 | 60673182 | 6445 | CAST.MOLF.SPRET | 6 | 60668914 | 60668947 | 34 | + | 17.0325 | 6.63E-07 | 0.0937 | TGGCAGGTCTCAGAGGAGCCGCTAGATGGCCCAC | | |
| 525 | 6 | 67194374 | 67198297 | 3923 | SPRET | 6 | 67196987 | 67197020 | 34 | + | 19.5772 | 1.20E-07 | 0.0354 | GAGCTGGTGTGGCCGCGGCCCCAAGATGGCGCCC | | |
| 526 | 6 | 72061617 | 72064986 | 3369 | AKR.C58J.KK.MRL.NOD.NOR.PL.RF | 6 | 72064190 | 72064223 | 34 | + | 16.5935 | 8.77E-07 | 0.109 | CTGGACTTGAACCTAGGTCCTCTTGAGGGCAGCC | | |
| 527 | 6 | 72828220 | 72830391 | 2171 | SPRET | 6 | 72828837 | 72828870 | 34 | + | 18.7236 | 2.17E-07 | 0.0516 | AGGTAATCTTGGCAGCTGCCAGGAGGGGGCAGTG | 30 | ENCFF093ZRU,ENCFF367UNR,ENCFF210BTO,ENCFF099QNE,ENCFF201JUA,ENCFF609MDN,ENCFF907XEG,ENCFF198SSR,ENCFF327LYS,ENCFF024JOZ,ENCFF097CAQ,ENCFF089GBC,ENCFF589ZBJ,ENCFF194HVL,ENCFF502WDT,ENCFF491RJK,ENCFF821BIA,ENCFF883UPM,ENCFF165LXO,ENCFF136ZDK,ENCFF092NMJ,ENCFF142CNG,ENCFF052KVO,ENCFF281QMV,ENCFF055HAP,ENCFF430PPJ,ENCFF533APC,ENCFF321AGQ.ENCFF957SPH.ENCFF065ODV |
| 527 | 6 | 72828220 | 72830391 | 2171 | SPRET | 6 | 72828852 | 72828870 | 19 | + | 20.1967 | 9.47E-08 | 0.0605 | CTGCCAGGAGGGGGCAGTG | 30 | ENCFF093ZRU,ENCFF367UNR,ENCFF210BTO,ENCFF099QNE,ENCFF201JUA,ENCFF609MDN,ENCFF907XEG,ENCFF198SSR,ENCFF327LYS,ENCFF024JOZ,ENCFF097CAQ,ENCFF089GBC,ENCFF589ZBJ,ENCFF194HVL,ENCFF502WDT,ENCFF491RJK,ENCFF821BIA,ENCFF883UPM,ENCFF165LXO,ENCFF136ZDK,ENCFF092NMJ,ENCFF142CNG,ENCFF052KVO,ENCFF281QMV,ENCFF055HAP,ENCFF430PPJ,ENCFF533APC,ENCFF321AGQ.ENCFF957SPH.ENCFF065ODV |
| 528 | 6 | 75293336 | 75298897 | 5561 | 129S1,AJ,BTBR,BUB,C3H,C57L,C58J,CBA,CE,DBA1J,DBA2J,FVB,ILNJ2,MA/My,PJ,SMJ.TH | 6 | 75293489 | 75293522 | 34 | + | 18.3089 | 2.87E-07 | 0.0578 | ATGCAGCTAGAGACACGAGCTCCAGGGGGTACTG | | |
| 529 | 6 | 82370967 | 82374514 | 3547 | SPRET | 6 | 82372683 | 82372716 | 34 | + | 22.9756 | 9.20E-09 | 0.00543 | TTGCTGTGCCCATATGAACCAGAAGAGGGCATCA | | |
| 530 | 6 | 82372433 | 82375323 | 2890 | CAST.MOLF | 6 | 82372683 | 82372716 | 34 | + | 22.9756 | 9.20E-09 | 0.00543 | TTGCTGTGCCCATATGAACCAGAAGAGGGCATCA | | |
| 531 | 6 | 85544902 | 85545940 | 1038 | SPRET | 6 | 85545864 | 85545897 | 34 | + | 16.6829 | 8.29E-07 | 0.106 | TTGCTCTCTCCAGACACACCGGAAGAGGGCATCA | | |
| 532 | 6 | 86253653 | 86255396 | 1743 | SPRET | 6 | 86254787 | 86254820 | 34 | + | 21.0407 | 4.15E-08 | 0.016 | CTGCTTCTCCTCACCTTGCCACTTGGGGGCCTTG | 5 | ENCFF093ZRU.ENCFF327LYS.ENCFF502WDT.ENCFF883UPM.ENCFF533APC |
| 533 | 6 | 87187825 | 87190105 | 2280 | AKR.BALB.MRL.PL.RH2.WSB | 6 | 87188708 | 87188741 | 34 | + | 25.8943 | 7.37E-10 | 0.000921 | GTGCAGTACCCACAGAGGCCAGAAGAGGGTGTCA | 4 | ENCFF052KVO.ENCFF907XEG.ENCFF198SSR.ENCFF093ZRU |
| 534 | 6 | 87998841 | 88002494 | 3653 | MOLF.PJ.SEA | 6 | 88002159 | 88002177 | 19 | + | 19.6557 | 1.43E-07 | 0.0702 | TGGACAGCAGGTGGCAGTC | 1 | ENCFF093ZRU |
| 535 | 6 | 88118423 | 88120184 | 1761 | MOLF | 6 | 88118897 | 88118931 | 35 | + | 19.2477 | 1.40E-07 | 0.0684 | ATGCAGGAACTTGAAGAGGCCAGAAGAGGGCATCT | | |
| 536 | 6 | 93219683 | 93220789 | 1106 | SPRET | 6 | 93220355 | 93220389 | 35 | + | 17.9083 | 3.18E-07 | 0.11 | CTGCAGTGCATAGCTCCTTCCAGGAGAGATCTCCA | | |
| 537 | 6 | 101432652 | 101434340 | 1688 | MOLF | 6 | 101433872 | 101433906 | 35 | + | 16.8165 | 6.02E-07 | 0.15 | ATGCAGGCCCAGCTCATGGGCCCTTGGAGGCACTT | | |
| 538 | 6 | 107694333 | 107696718 | 2385 | CAST | 6 | 107695388 | 107695422 | 35 | + | 24.211 | 4.54E-09 | 0.00886 | GTGCAGGTACCCAAGGAGGCCAGAAGAGGGCATCA | | |
| 539 | 6 | 110311190 | 110315356 | 4166 | 129S1,AJ,BALB,BTBR,BUB,C57L,CAST,CE,DBA1J,DBA2J,FVB,KK,MOLF,NZB,NZO,NZW,PJ,RH2,SEA,SJL,SMJ,SPRET,SWR,TH,WSB | 6 | 110311968 | 110312001 | 34 | + | 17.6748 | 4.37E-07 | 0.0735 | GAGCATGACCACAAGTCCACAGCAGGTGGCAACG | | |
| 540 | 6 | 116023487 | 116026395 | 2908 | AKR.C3H.CBA.ILNJ2.MOLF.MRL.PL.RF | 6 | 116026077 | 116026111 | 35 | + | 19.2661 | 1.39E-07 | 0.068 | GTGCAGCTGCCTGTGGCATCCAGAGGAGGGCATTC | | |
| 541 | 6 | 116225897 | 116227824 | 1927 | 129S1,BTBR,C57L,C58J,CAST,DBA1J,DBA2J,FVB,LP,MOLF,NOD,NOR,NZO,RH2,SJL,SWR | 6 | 116227262 | 116227295 | 34 | + | 19.252 | 1.51E-07 | 0.0415 | GCCCAGTGCCCACAGAAGCCAGAAGAGGGCATTA | | |
| 542 | 6 | 117720445 | 117729977 | 9532 | CAST | 6 | 117729467 | 117729485 | 19 | + | 18.4262 | 3.51E-07 | 0.105 | CAGCCAGGAGATGGCAGCA | 6 | ENCFF024JOZ.ENCFF907XEG.ENCFF089GBC.ENCFF093ZRU.ENCFF099QNE.ENCFF367UNR |
| 543 | 6 | 117926810 | 117928342 | 1532 | CE | 6 | 117928130 | 117928163 | 34 | + | 24.0894 | 3.65E-09 | 0.00278 | CTGCAGTGACCACAGGGGCCGGAAGAGGGTGTTA | | |
| 544 | 6 | 119880567 | 119882567 | 2000 | 129S1.AKR.C57L.DBA1J.DBA2J.MRL.RF | 6 | 119882280 | 119882314 | 35 | + | 18.844 | 1.80E-07 | 0.0783 | AAGGAGTTCCTGCCACCTGCCACCAGAAGGCTGTG | | |
| 545 | 6 | 124542731 | 124555928 | 13197 | SPRET | 6 | 124547160 | 124547194 | 35 | + | 27.1009 | 4.31E-10 | 0.00234 | CAGCCATTCCCCACATTGACCACCAGATGGCGCCA | 39 | ENCFF201JUA,ENCFF093ZRU,ENCFF367UNR,ENCFF099QNE,ENCFF805QBU,ENCFF821BIA,ENCFF065ODV,ENCFF907XEG,ENCFF024JOZ,ENCFF092NMJ,ENCFF327LYS,ENCFF089GBC,ENCFF055HAP,ENCFF052KVO,ENCFF136ZDK,ENCFF033UQX,ENCFF589ZBJ,ENCFF430PPJ,ENCFF165LXO,ENCFF502WDT,ENCFF321AGQ,ENCFF198SSR,ENCFF210BTO,ENCFF194HVL,ENCFF089EWX,ENCFF142CNG,ENCFF097CAQ,ENCFF504JDS,ENCFF883UPM,ENCFF957SPH,ENCFF951GOP,ENCFF281QMV,ENCFF601JVE,ENCFF611HDQ,ENCFF533APC,ENCFF316FYN,ENCFF575MGI,ENCFF491RJK,ENCFF609MDN |
| 545 | 6 | 124542731 | 124555928 | 13197 | SPRET | 6 | 124547161 | 124547194 | 34 | + | 19.0569 | 1.72E-07 | 0.0455 | AGCCATTCCCCACATTGACCACCAGATGGCGCCA | 39 | ENCFF201JUA,ENCFF093ZRU,ENCFF367UNR,ENCFF099QNE,ENCFF805QBU,ENCFF821BIA,ENCFF065ODV,ENCFF907XEG,ENCFF024JOZ,ENCFF092NMJ,ENCFF327LYS,ENCFF089GBC,ENCFF055HAP,ENCFF052KVO,ENCFF136ZDK,ENCFF033UQX,ENCFF589ZBJ,ENCFF430PPJ,ENCFF165LXO,ENCFF502WDT,ENCFF321AGQ,ENCFF198SSR,ENCFF210BTO,ENCFF194HVL,ENCFF089EWX,ENCFF142CNG,ENCFF097CAQ,ENCFF504JDS,ENCFF883UPM,ENCFF957SPH,ENCFF951GOP,ENCFF281QMV,ENCFF601JVE,ENCFF611HDQ,ENCFF533APC,ENCFF316FYN,ENCFF575MGI,ENCFF491RJK,ENCFF609MDN |
| 545 | 6 | 124542731 | 124555928 | 13197 | SPRET | 6 | 124547176 | 124547194 | 19 | + | 24.6393 | 1.08E-09 | 0.0175 | TGACCACCAGATGGCGCCA | 39 | ENCFF201JUA,ENCFF093ZRU,ENCFF367UNR,ENCFF099QNE,ENCFF805QBU,ENCFF821BIA,ENCFF065ODV,ENCFF907XEG,ENCFF024JOZ,ENCFF092NMJ,ENCFF327LYS,ENCFF089GBC,ENCFF055HAP,ENCFF052KVO,ENCFF136ZDK,ENCFF033UQX,ENCFF589ZBJ,ENCFF430PPJ,ENCFF165LXO,ENCFF502WDT,ENCFF321AGQ,ENCFF198SSR,ENCFF210BTO,ENCFF194HVL,ENCFF089EWX,ENCFF142CNG,ENCFF097CAQ,ENCFF504JDS,ENCFF883UPM,ENCFF957SPH,ENCFF951GOP,ENCFF281QMV,ENCFF601JVE,ENCFF611HDQ,ENCFF533APC,ENCFF316FYN,ENCFF575MGI,ENCFF491RJK,ENCFF609MDN |
| 546 | 6 | 126550531 | 126554034 | 3503 | CAST | 6 | 126553150 | 126553183 | 34 | + | 17.5528 | 4.74E-07 | 0.0774 | CTGCTCTTCCTGCTCAGCCCACTCGAGGTAAGAG | | |
| 547 | 6 | 131117693 | 131119801 | 2108 | CE,FVB,KK,LP,MA/My,NOD,NOR,NZB,NZW,SJL,SWR,TH,WSB | 6 | 131119486 | 131119520 | 35 | + | 17.0183 | 5.36E-07 | 0.144 | GTGCAGACACACCCAAGGTCCACTAGAGGAATCCC | | |
| 548 | 6 | 133055073 | 133057618 | 2545 | SPRET | 6 | 133056686 | 133056719 | 34 | + | 19.2358 | 1.52E-07 | 0.0418 | CTGCAGCTCCCTGAACATTCAGCAGATGTCTCAA | | |
| 549 | 6 | 133701362 | 133702521 | 1159 | NZO | 6 | 133701756 | 133701789 | 34 | + | 20.3171 | 7.06E-08 | 0.0241 | ATGCAGGGGAGCATCCTTCCAGAAGGTGGCGCCT | 4 | ENCFF093ZRU.ENCFF883UPM.ENCFF052KVO.ENCFF533APC |
| 549 | 6 | 133701362 | 133702521 | 1159 | NZO | 6 | 133701771 | 133701789 | 19 | + | 17.082 | 8.61E-07 | 0.161 | CTTCCAGAAGGTGGCGCCT | 4 | ENCFF093ZRU.ENCFF883UPM.ENCFF052KVO.ENCFF533APC |
| 550 | 6 | 133892240 | 133893865 | 1625 | SPRET | 6 | 133893430 | 133893464 | 35 | + | 17.1193 | 5.06E-07 | 0.14 | TGGCTGGTGCCTGTGAAGGCCAGAAGAGGGAACTG | | |
| 550 | 6 | 133892240 | 133893865 | 1625 | SPRET | 6 | 133893446 | 133893464 | 19 | + | 16.9672 | 9.26E-07 | 0.165 | AGGCCAGAAGAGGGAACTG | | |
| 551 | 6 | 134265471 | 134266990 | 1519 | AJ,AKR,BALB,BUB,C3H,C58J,CAST,CBA,CE,DBA1J,DBA2J,FVB,KK,MA/My,MRL,NOD,NOR,NZB,NZO,NZW,PL,RF,RH2,SEA,SJL,SWR,TH,WSB | 6 | 134266950 | 134266983 | 34 | + | 16.7398 | 7.99E-07 | 0.104 | GTTCAGTGCCTGAGGAGGTCAGAAGAGGGTATTA | | |
| 552 | 6 | 136011961 | 136016184 | 4223 | CAST | 6 | 136015346 | 136015380 | 35 | + | 16.1101 | 8.98E-07 | 0.183 | CTGTAATCCCAGCACTTGGGAGGCAGAGGGAGGCG | | |
| 553 | 6 | 137536163 | 137543921 | 7758 | LP | 6 | 137541151 | 137541184 | 34 | + | 18.5041 | 2.52E-07 | 5.59E-02 | GTGCAGTGCCCATGGAGGGCAGCGGAAGGGGGCA | | |
| 554 | 6 | 142651900 | 142653817 | 1917 | BTBR,C57L,C58J,FVB,KK,MA/My,NOD,NOR,NZB,PJ,SMJ,SWR,WSB | 6 | 142652601 | 142652634 | 34 | + | 26.0081 | 6.63E-10 | 0.000854 | CTGCAGTGCCCTTGCAGACCAGAAGAGGGCTCCT | 19 | ENCFF065ODV,ENCFF093ZRU,ENCFF024JOZ,ENCFF367UNR,ENCFF165LXO,ENCFF907XEG,ENCFF198SSR,ENCFF589ZBJ,ENCFF821BIA,ENCFF052KVO,ENCFF533APC,ENCFF194HVL,ENCFF099QNE,ENCFF142CNG,ENCFF055HAP,ENCFF502WDT,ENCFF136ZDK,ENCFF883UPM,ENCFF957SPH |
| 554 | 6 | 142651900 | 142653817 | 1917 | BTBR,C57L,C58J,FVB,KK,MA/My,NOD,NOR,NZB,PJ,SMJ,SWR,WSB | 6 | 142653605 | 142653638 | 34 | + | 17.0894 | 6.39E-07 | 0.0917 | GTGCTCTTCCCTAAGTGAGCCCCAGAGGCCTCTG | | |
| 555 | 6 | 149031730 | 149035638 | 3908 | CAST | 6 | 149033960 | 149033978 | 19 | + | 18.3607 | 3.67E-07 | 0.108 | TTACCACTGGGTGGCACCA | 29 | ENCFF367UNR,ENCFF093ZRU,ENCFF201JUA,ENCFF052KVO,ENCFF099QNE,ENCFF327LYS,ENCFF821BIA,ENCFF907XEG,ENCFF024JOZ,ENCFF502WDT,ENCFF065ODV,ENCFF589ZBJ,ENCFF055HAP,ENCFF089GBC,ENCFF165LXO,ENCFF316FYN,ENCFF198SSR,ENCFF883UPM,ENCFF194HVL,ENCFF575MGI,ENCFF092NMJ,ENCFF951GOP,ENCFF491RJK,ENCFF136ZDK,ENCFF142CNG,ENCFF957SPH,ENCFF281QMV,ENCFF609MDN,ENCFF430PPJ |
| 555 | 6 | 149031730 | 149035638 | 3908 | CAST | 6 | 149033296 | 149033330 | 35 | - | 17.0826 | 5.16E-07 | 0.141 | TAGCTGTCCTCAGACACTTTCAGAAGAGGGCACCA | | |
| 556 | 6 | 3115443 | 3116581 | 1138 | CAST | 6 | 3116473 | 3116491 | 19 | - | 18.6557 | 2.99E-07 | 0.0973 | CCGCCACTAGATGGTGGTA | | |
| 557 | 6 | 11974138 | 11976269 | 2131 | SPRET | 6 | 11975711 | 11975744 | 34 | - | 19.439 | 1.32E-07 | 0.0379 | CTGCAAGAGGAGAGCTTGCCTGCAGAGGGTACTC | | |
| 558 | 6 | 12780271 | 12781779 | 1508 | SPRET | 6 | 12781676 | 12781709 | 34 | - | 16.3902 | 9.96E-07 | 0.115 | AAGCAGCTAGAGACACGAGCTCCAGGGGGCATCA | | |
| 559 | 6 | 13117201 | 13119777 | 2576 | SPRET | 6 | 13118899 | 13118932 | 34 | - | 16.7886 | 7.75E-07 | 0.103 | GAGCATGGACACAAGCCCACAGCAGATGGCAGCA | | |
| 560 | 6 | 29668231 | 29669316 | 1085 | CAST.MOLF | 6 | 29668533 | 29668566 | 34 | - | 17.3171 | 5.52E-07 | 0.0844 | TAGCTGTCTCCAGACGCCCCAGAAGAGGGCATCA | | |
| 561 | 6 | 33759109 | 33760608 | 1499 | 129S1.C57L.FVB.SWR.TH.WSB | 6 | 33759120 | 33759153 | 34 | - | 17.6016 | 4.59E-07 | 0.0756 | ATGCAGCTAGAGAAACAAGCTCCAGGGGGTACTG | | |
| 562 | 6 | 35102364 | 35103434 | 1070 | SPRET | 6 | 35102343 | 35102376 | 34 | - | 18.935 | 1.88E-07 | 0.0476 | TTGCTGTCTTCAGACACACCAGAAGAGGGCGTCA | 2 | ENCFF099QNE.ENCFF093ZRU |
| 563 | 6 | 38186305 | 38188486 | 2181 | CAST | 6 | 38186709 | 38186742 | 34 | - | 27.3008 | 1.92E-10 | 0.000448 | CTGCAGTGCCTGTAGAGGTCAGAAGGGGGCATTA | | |
| 564 | 6 | 38551661 | 38552931 | 1270 | CAST | 6 | 38551871 | 38551904 | 34 | - | 17.2114 | 5.91E-07 | 0.087 | TAGCTGTTTTCAGACACTCCAGAAGAGGGCATCA | | |
| 565 | 6 | 42977119 | 42987294 | 10175 | WSB | 6 | 42983298 | 42983331 | 34 | - | 16.4146 | 9.81E-07 | 0.113 | CTGGAATGGCTCCACGCCCCAATGGATGGCGCAC | | |
| 566 | 6 | 49168598 | 49169823 | 1225 | CAST.CE.ILNJ2 | 6 | 49169064 | 49169097 | 34 | - | 20.5447 | 5.98E-08 | 0.0216 | ATGCAGTGGCTGAGGGAGCCAGAAGTGGGCATCA | | |
| 567 | 6 | 51315681 | 51318580 | 2899 | FVB.SJL.SWR | 6 | 51318241 | 51318274 | 34 | - | 19.9593 | 9.14E-08 | 0.0291 | GTGTAGTGCCTTCAGAGGCCAGAAGAAGGTGTTG | | |
| 568 | 6 | 58517853 | 58521527 | 3674 | SPRET | 6 | 58520031 | 58520049 | 19 | - | 19.541 | 1.57E-07 | 0.0734 | CTTCCAGAAGGTGGCGCCG | 20 | ENCFF065ODV,ENCFF093ZRU,ENCFF136ZDK,ENCFF092NMJ,ENCFF052KVO,ENCFF430PPJ,ENCFF367UNR,ENCFF165LXO,ENCFF883UPM,ENCFF327LYS,ENCFF099QNE,ENCFF957SPH,ENCFF907XEG,ENCFF055HAP,ENCFF821BIA,ENCFF024JOZ,ENCFF089GBC,ENCFF502WDT,ENCFF589ZBJ,ENCFF198SSR |
| 568 | 6 | 58517853 | 58521527 | 3674 | SPRET | 6 | 58520031 | 58520064 | 34 | - | 22 | 2.00E-08 | 0.00938 | ATGCAGGGGAGCATCCTTCCAGAAGGTGGCGCCG | 20 | ENCFF065ODV,ENCFF093ZRU,ENCFF136ZDK,ENCFF092NMJ,ENCFF052KVO,ENCFF430PPJ,ENCFF367UNR,ENCFF165LXO,ENCFF883UPM,ENCFF327LYS,ENCFF099QNE,ENCFF957SPH,ENCFF907XEG,ENCFF055HAP,ENCFF821BIA,ENCFF024JOZ,ENCFF089GBC,ENCFF502WDT,ENCFF589ZBJ,ENCFF198SSR |
| 569 | 6 | 65036402 | 65039156 | 2754 | SPRET | 6 | 65036817 | 65036835 | 19 | - | 17.459 | 6.74E-07 | 0.143 | TAACCAGTGGGGGGCAGTG | | |
| 570 | 6 | 68549897 | 68597377 | 47480 | MA/My,SPRET,WSB | 6 | 68583210 | 68583228 | 19 | - | 18.9016 | 2.51E-07 | 0.0903 | TGACCAGCAGAGGGAGACA | 7 | ENCFF089GBC,ENCFF092NMJ,ENCFF093ZRU,ENCFF099QNE,ENCFF502WDT,ENCFF055HAP,ENCFF883UPM |
| 571 | 6 | 69729660 | 69733813 | 4153 | WSB | 6 | 69731099 | 69731132 | 34 | - | 17.3659 | 5.35E-07 | 0.0827 | ATGCAGCTAGAGACAGGAGCTCCAGGGGGTGCTG | | |
| 572 | 6 | 71134567 | 71139713 | 5146 | SPRET | 6 | 71139411 | 71139429 | 19 | - | 17.9508 | 4.86E-07 | 0.128 | AGGCCACCTGGGGGCACTC | 14 | ENCFF136ZDK,ENCFF099QNE,ENCFF821BIA,ENCFF210BTO,ENCFF327LYS,ENCFF198SSR,ENCFF367UNR,ENCFF907XEG,ENCFF093ZRU,ENCFF957SPH,ENCFF883UPM,ENCFF589ZBJ,ENCFF194HVL,ENCFF024JOZ |
| 572 | 6 | 71134567 | 71139713 | 5146 | SPRET | 6 | 71139411 | 71139445 | 35 | - | 23.3853 | 8.44E-09 | 0.0122 | GGGCAGTCCTTTACCCAGGCCACCTGGGGGCACTC | 14 | ENCFF136ZDK,ENCFF099QNE,ENCFF821BIA,ENCFF210BTO,ENCFF327LYS,ENCFF198SSR,ENCFF367UNR,ENCFF907XEG,ENCFF093ZRU,ENCFF957SPH,ENCFF883UPM,ENCFF589ZBJ,ENCFF194HVL,ENCFF024JOZ |
| 573 | 6 | 73263771 | 73271134 | 7363 | 129S1,AJ,AKR,BALB,BTBR,BUB,C3H,C57L,C58J,CAST,CBA,CE,DBA1J,FVB,ILNJ2,KK,MA/My,MOLF,MRL,NOD,NOR,NZB,NZO,NZW,PJ,PL,RF,RH2,SEA,SJL,SPRET,SWR,TH,WSB | 6 | 73265660 | 73265678 | 19 | - | 17.1639 | 8.17E-07 | 0.157 | CCTCCACGAGGTGGCGGAA | | |
| 574 | 6 | 77910840 | 77912003 | 1163 | SPRET | 6 | 77911347 | 77911380 | 34 | - | 21.5935 | 2.73E-08 | 0.0118 | TTGCAGTGGCCACAGAGGCCAGCTGAGGGCCTTA | | |
| 575 | 6 | 85815027 | 85819857 | 4830 | MOLF | 6 | 85816470 | 85816503 | 34 | - | 16.5691 | 8.90E-07 | 0.11 | TAGCTGTCTTCAGATGCACCAGAAGGTGGCATCA | 2 | ENCFF055HAP.ENCFF093ZRU |
| 576 | 6 | 86284544 | 86287260 | 2716 | SPRET | 6 | 86287120 | 86287154 | 35 | - | 17.9633 | 3.08E-07 | 0.108 | CAGCAGTGTTAAGTCAGGCACAGAGGAGGGCAGTA | | |
| 577 | 6 | 97084487 | 97085635 | 1148 | SEA | 6 | 97085384 | 97085402 | 19 | - | 18.9836 | 2.36E-07 | 0.0872 | CAGCCAGGAGAGGGCAGTG | 3 | ENCFF024JOZ.ENCFF093ZRU.ENCFF099QNE |
| 578 | 6 | 100746877 | 100749459 | 2582 | SPRET | 6 | 100747649 | 100747682 | 34 | - | 17.7805 | 4.08E-07 | 0.0707 | CACCTGTGCCCATAGAAACCAGAAGAGGGCATCA | | |
| 579 | 6 | 108540388 | 108542659 | 2271 | AJ,AKR,BALB,C3H,C58J,CAST,CBA,CE,DBA1J,DBA2J,ILNJ2,LP,MOLF,MRL,NZB,NZO,NZW,PL,RF,RH2,SEA,SPRET,SWR,TH | 6 | 108541110 | 108541144 | 35 | - | 17.2018 | 4.82E-07 | 0.136 | CTGCTGTGTACTGATGTGGCAGCCAGAGGGCTGGA | | |
| 580 | 6 | 114848172 | 114849607 | 1435 | SPRET | 6 | 114848431 | 114848449 | 19 | - | 17.1148 | 8.43E-07 | 0.159 | AGGCCAGAAGAGGGCGTCA | 4 | ENCFF093ZRU.ENCFF327LYS.ENCFF502WDT.ENCFF052KVO |
| 580 | 6 | 114848172 | 114849607 | 1435 | SPRET | 6 | 114848431 | 114848464 | 34 | - | 21.065 | 4.07E-08 | 0.0158 | GTATAGTGCCCACAGAGGCCAGAAGAGGGCGTCA | 4 | ENCFF093ZRU.ENCFF327LYS.ENCFF502WDT.ENCFF052KVO |
| 581 | 6 | 117562052 | 117569379 | 7327 | CAST.FVB.MOLF.NZB.SJL.SPRET.SWR | 6 | 117562576 | 117562609 | 34 | - | 19.3415 | 1.42E-07 | 0.0398 | CTGCCCCCACATTCGCCGTCACAAGATGGCGCTG | | |
| 582 | 6 | 126454104 | 126455789 | 1685 | NOD.NOR | 6 | 126455341 | 126455359 | 19 | - | 17 | 9.07E-07 | 0.164 | ATACCACCAGAGGGTGGTG | | |
| 583 | 6 | 130506277 | 130507358 | 1081 | SPRET | 6 | 130506814 | 130506847 | 34 | - | 17.6667 | 4.40E-07 | 0.0735 | TAGCTGTCTTCAGACACGCCAGAAGAGGGCGTCA | | |
| 584 | 6 | 134603615 | 134606964 | 3349 | 129S1.C57L.SMJ | 6 | 134606483 | 134606516 | 34 | - | 21.2114 | 3.65E-08 | 0.0146 | ATGCATTTCCTGAGAAGGCCAGAAGAGGCCATCA | | |
| 585 | 6 | 136073294 | 136075054 | 1760 | 129S1,AJ,AKR,BALB,BTBR,BUB,C3H,C57L,CBA,CE,DBA1J,DBA2J,FVB,KK,MA/My,MRL,NOD,NOR,NZB,NZW,PJ,PL,RF,RH2,SEA,SJL,SWR,TH,WSB | 6 | 136073456 | 136073489 | 34 | - | 24.4472 | 2.69E-09 | 0.00227 | ATGCAGTGCTCCTGAAGGCCAGCAGAGGGTGACA | 20 | ENCFF430PPJ,ENCFF024JOZ,ENCFF093ZRU,ENCFF136ZDK,ENCFF092NMJ,ENCFF589ZBJ,ENCFF883UPM,ENCFF099QNE,ENCFF533APC,ENCFF327LYS,ENCFF907XEG,ENCFF281QMV,ENCFF575MGI,ENCFF194HVL,ENCFF198SSR,ENCFF165LXO,ENCFF089GBC,ENCFF957SPH,ENCFF367UNR,ENCFF821BIA |
| 586 | 6 | 142676290 | 142677951 | 1661 | SPRET | 6 | 142677441 | 142677459 | 19 | - | 20.0656 | 1.05E-07 | 0.0608 | AGGCCAGAAGAGGGCACTG | 12 | ENCFF024JOZ,ENCFF907XEG,ENCFF327LYS,ENCFF821BIA,ENCFF198SSR,ENCFF883UPM,ENCFF957SPH,ENCFF367UNR,ENCFF055HAP,ENCFF093ZRU,ENCFF502WDT,ENCFF099QNE |
| 586 | 6 | 142676290 | 142677951 | 1661 | SPRET | 6 | 142677441 | 142677474 | 34 | - | 17.0813 | 6.43E-07 | 0.0919 | GTCTGGAGCCTGCAGAGGCCAGAAGAGGGCACTG | 12 | ENCFF024JOZ,ENCFF907XEG,ENCFF327LYS,ENCFF821BIA,ENCFF198SSR,ENCFF883UPM,ENCFF957SPH,ENCFF367UNR,ENCFF055HAP,ENCFF093ZRU,ENCFF502WDT,ENCFF099QNE |
| 587 | 6 | 148027474 | 148032663 | 5189 | SPRET | 6 | 148028724 | 148028742 | 19 | - | 17.0328 | 8.88E-07 | 0.162 | TGGCCAGCAGATGGCCCAC | | |
| 587 | 6 | 148027474 | 148032663 | 5189 | SPRET | 6 | 148028724 | 148028758 | 35 | - | 16.4404 | 7.46E-07 | 0.165 | AAGAAGCTGGACCCAGTGGCCAGCAGATGGCCCAC | | |
| 588 | 7 | 3240450 | 3242592 | 2142 | SPRET | 7 | 3240529 | 3240562 | 34 | + | 18.3496 | 2.79E-07 | 0.0479 | TTGCTGTCTTCAGACACACCAGAAGAGGGCATCC | | |
| 589 | 7 | 4420966 | 4425909 | 4943 | SPRET | 7 | 4424503 | 4424537 | 35 | + | 17.1101 | 5.08E-07 | 0.107 | ATGCACTACCTCCATGTTGTCTCAAGGAGGTGCCA | 4 | ENCFF093ZRU.ENCFF367UNR.ENCFF907XEG.ENCFF502WDT |
| 590 | 7 | 9652668 | 9662040 | 9372 | 129S1,AJ,AKR,BALB,BTBR,BUB,C57L,C58J,CBA,CE,FVB,ILNJ2,KK,LP,MOLF,MRL,NOD,NOR,NZB,NZO,NZW,PL,RF,RH2,SJL,SWR,TH | 7 | 9656605 | 9656638 | 34 | + | 19.626 | 1.16E-07 | 0.0284 | GAGCAGGAGCCCACCAGGCCTCCTGGGGGCGGTG | | |
| 591 | 7 | 16538675 | 16539999 | 1324 | SPRET | 7 | 16538776 | 16538809 | 34 | + | 18.8537 | 1.98E-07 | 0.0397 | GAACTGTTCCTCATATGTCCACAAGAGGGCATCC | 26 | ENCFF093ZRU,ENCFF367UNR,ENCFF099QNE,ENCFF821BIA,ENCFF201JUA,ENCFF210BTO,ENCFF136ZDK,ENCFF316FYN,ENCFF065ODV,ENCFF321AGQ,ENCFF907XEG,ENCFF055HAP,ENCFF024JOZ,ENCFF502WDT,ENCFF327LYS,ENCFF198SSR,ENCFF089GBC,ENCFF589ZBJ,ENCFF165LXO,ENCFF609MDN,ENCFF092NMJ,ENCFF951GOP,ENCFF430PPJ,ENCFF281QMV,ENCFF052KVO,ENCFF883UPM |
| 592 | 7 | 18273438 | 18276254 | 2816 | SPRET | 7 | 18274299 | 18274332 | 34 | + | 16.5935 | 8.77E-07 | 0.0876 | GAGCATGGACACAAGCACACAGCAGATGGCAGCA | | |
| 593 | 7 | 18803347 | 18804550 | 1203 | SPRET | 7 | 18803588 | 18803622 | 35 | + | 24.1376 | 4.80E-09 | 0.0065 | ATGCAGGCACCTTTGACAGCCAGAAGAGGGCGCTG | 31 | ENCFF201JUA,ENCFF093ZRU,ENCFF367UNR,ENCFF430PPJ,ENCFF165LXO,ENCFF210BTO,ENCFF805QBU,ENCFF099QNE,ENCFF951GOP,ENCFF065ODV,ENCFF821BIA,ENCFF907XEG,ENCFF055HAP,ENCFF609MDN,ENCFF033UQX,ENCFF502WDT,ENCFF327LYS,ENCFF024JOZ,ENCFF589ZBJ,ENCFF136ZDK,ENCFF142CNG,ENCFF097CAQ,ENCFF194HVL,ENCFF281QMV,ENCFF089GBC,ENCFF198SSR,ENCFF533APC,ENCFF957SPH.ENCFF092NMJ.ENCFF052KVO.ENCFF883UPM |
| 593 | 7 | 18803347 | 18804550 | 1203 | SPRET | 7 | 18803604 | 18803622 | 19 | + | 21.2295 | 4.03E-08 | 0.0332 | CAGCCAGAAGAGGGCGCTG | 31 | ENCFF201JUA,ENCFF093ZRU,ENCFF367UNR,ENCFF430PPJ,ENCFF165LXO,ENCFF210BTO,ENCFF805QBU,ENCFF099QNE,ENCFF951GOP,ENCFF065ODV,ENCFF821BIA,ENCFF907XEG,ENCFF055HAP,ENCFF609MDN,ENCFF033UQX,ENCFF502WDT,ENCFF327LYS,ENCFF024JOZ,ENCFF589ZBJ,ENCFF136ZDK,ENCFF142CNG,ENCFF097CAQ,ENCFF194HVL,ENCFF281QMV,ENCFF089GBC,ENCFF198SSR,ENCFF533APC,ENCFF957SPH.ENCFF092NMJ.ENCFF052KVO.ENCFF883UPM |
| 594 | 7 | 19107618 | 19108934 | 1316 | SPRET | 7 | 19108405 | 19108438 | 34 | + | 23.7886 | 4.71E-09 | 0.00249 | GTGCAGCGCCCTCTGAGGCCAGAAGGGGACTCCA | | |
| 595 | 7 | 23006921 | 23015447 | 8526 | C3H.LP.MA/My.PJ.SEA.SMJ.SPRET | 7 | 23010822 | 23010840 | 19 | + | 17.1475 | 8.25E-07 | 0.121 | CTGCCAGTAGATGGCCCCA | | |
| 595 | 7 | 23006921 | 23015447 | 8526 | C3H.LP.MA/My.PJ.SEA.SMJ.SPRET | 7 | 23014370 | 23014388 | 19 | - | 20.5902 | 6.88E-08 | 0.0415 | TGGCCACCAGAGGGAGCCT | | |
| 595 | 7 | 23006921 | 23015447 | 8526 | C3H.LP.MA/My.PJ.SEA.SMJ.SPRET | 7 | 23014370 | 23014404 | 35 | - | 16.422 | 7.54E-07 | 0.133 | CACCTTTGGAGTCAGTTGGCCACCAGAGGGAGCCT | | |
| 596 | 7 | 23537036 | 23549090 | 12054 | SPRET | 7 | 23545861 | 23545895 | 35 | + | 16.0183 | 9.45E-07 | 0.148 | CTGCATCTACCACAGTCAGCCTCCAGGTAGACCCT | | |
| 597 | 7 | 25104927 | 25107036 | 2109 | SPRET | 7 | 25106819 | 25106852 | 34 | + | 19.1951 | 1.57E-07 | 0.0346 | ATGCAGTGTCCACAAAGGCCAGAAGGAGTCAGAT | | |
| 598 | 7 | 26710242 | 26714093 | 3851 | CAST | 7 | 26713905 | 26713938 | 34 | + | 17.7967 | 4.04E-07 | 0.0582 | TAGCAGTCTTCAGACACACCAGAAGAGGGCATCC | | |
| 599 | 7 | 27654192 | 27658208 | 4016 | MOLF | 7 | 27654932 | 27654965 | 34 | + | 17.2114 | 5.91E-07 | 0.0711 | TAGCTGTTTTCAGACACTCCAGAAGAGGGCATCA | | |
| 600 | 7 | 28735781 | 28737466 | 1685 | SPRET | 7 | 28736842 | 28736875 | 34 | + | 29.7805 | 1.37E-11 | 0.000102 | GTGCAGTGCTCTCAGAGGCCAGAAGAGGGCACTG | 13 | ENCFF093ZRU,ENCFF099QNE,ENCFF821BIA,ENCFF327LYS,ENCFF052KVO,ENCFF136ZDK,ENCFF089GBC,ENCFF589ZBJ,ENCFF055HAP,ENCFF883UPM,ENCFF065ODV,ENCFF502WDT,ENCFF367UNR |
| 600 | 7 | 28735781 | 28737466 | 1685 | SPRET | 7 | 28736857 | 28736875 | 19 | + | 20.0656 | 1.05E-07 | 0.0453 | AGGCCAGAAGAGGGCACTG | 13 | ENCFF093ZRU,ENCFF099QNE,ENCFF821BIA,ENCFF327LYS,ENCFF052KVO,ENCFF136ZDK,ENCFF089GBC,ENCFF589ZBJ,ENCFF055HAP,ENCFF883UPM,ENCFF065ODV,ENCFF502WDT,ENCFF367UNR |
| 601 | 7 | 29314626 | 29323329 | 8703 | 129S1,AJ,AKR,BALB,BTBR,BUB,C3H,C57L,CAST,CE,DBA1J,DBA2J,ILNJ2,LP,MOLF,MRL,NOD,NOR,NZB,NZO,NZW,PJ,PL,RF,SEA,SJL,SMJ,SPRET,TH | 7 | 29320044 | 29320062 | 19 | + | 17.1639 | 8.17E-07 | 0.12 | CCTCCACGAGGTGGCGGAA | | |
| 602 | 7 | 29519333 | 29521472 | 2139 | SPRET | 7 | 29521388 | 29521421 | 34 | + | 17.5528 | 4.74E-07 | 0.0635 | CTGACACACCCTCTGCTGCCCCCAGGGAGCACCA | | |
| 603 | 7 | 29657120 | 29658278 | 1158 | CAST.MOLF | 7 | 29657119 | 29657153 | 35 | + | 20.4862 | 6.34E-08 | 0.0299 | GTGCTGCTGCCCACACAGGCCAGAAGAGGGAGTCA | | |
| 604 | 7 | 30388060 | 30397045 | 8985 | 129S1,AJ,AKR,BALB,BTBR,C57L,C58J,CAST,CE,DBA1J,DBA2J,ILNJ2,LP,MA/My,MOLF,MRL,NOD,NOR,NZB,NZO,NZW,PJ,PL,RF,SEA.SJL.SMJ.SPRET.TH.WSB | 7 | 30393477 | 30393495 | 19 | + | 17.1639 | 8.17E-07 | 0.12 | CCTCCACGAGGTGGCGGAA | | |
| 605 | 7 | 30888997 | 30967905 | 78908 | SPRET | 7 | 30946472 | 30946505 | 34 | + | 18.3089 | 2.87E-07 | 0.0481 | ATGCAGCTAGAGACACGAGCTCCAGGGGGTACTG | | |
| 605 | 7 | 30888997 | 30967905 | 78908 | SPRET | 7 | 30957664 | 30957698 | 35 | + | 17.1101 | 5.08E-07 | 0.107 | ATGCTATTGTGAGGACTGCTCACCAGAGGGTACCA | | |
| 606 | 7 | 30936691 | 30958742 | 22051 | DBA1J,DBA2J,MOLF,NZB,NZO,NZW,SMJ | 7 | 30946472 | 30946505 | 34 | + | 18.3089 | 2.87E-07 | 0.0481 | ATGCAGCTAGAGACACGAGCTCCAGGGGGTACTG | | |
| 606 | 7 | 30936691 | 30958742 | 22051 | DBA1J,DBA2J,MOLF,NZB,NZO,NZW,SMJ | 7 | 30957664 | 30957698 | 35 | + | 17.1101 | 5.08E-07 | 0.107 | ATGCTATTGTGAGGACTGCTCACCAGAGGGTACCA | | |
| 607 | 7 | 35365806 | 35366908 | 1102 | SPRET | 7 | 35365815 | 35365849 | 35 | + | 26.2661 | 8.79E-10 | 0.00297 | GTGCAGTGCCAGGGCCTGTCCTGAAGATGGCATTG | 3 | ENCFF093ZRU.ENCFF092NMJ.ENCFF089GBC |
| 608 | 7 | 37879981 | 37882428 | 2447 | AJ,BALB,FVB,MA/My,NOD,NOR,RH2,SEA | 7 | 37882122 | 37882155 | 34 | + | 24.1057 | 3.60E-09 | 0.00208 | GTCCGGTTCTCGACCTTGCCACCAGAGGGCGCTG | 39 | ENCFF367UNR,ENCFF093ZRU,ENCFF201JUA,ENCFF099QNE,ENCFF805QBU,ENCFF821BIA,ENCFF327LYS,ENCFF907XEG,ENCFF589ZBJ,ENCFF136ZDK,ENCFF198SSR,ENCFF055HAP,ENCFF024JOZ,ENCFF089GBC,ENCFF092NMJ,ENCFF065ODV,ENCFF165LXO,ENCFF052KVO,ENCFF502WDT,ENCFF951GOP,ENCFF210BTO,ENCFF883UPM,ENCFF957SPH,ENCFF430PPJ,ENCFF142CNG,ENCFF281QMV,ENCFF194HVL,ENCFF491RJK,ENCFF349BUL,ENCFF504JDS,ENCFF611HDQ,ENCFF533APC,ENCFF089EWX,ENCFF316FYN,ENCFF609MDN,ENCFF575MGI,ENCFF033UQX,ENCFF086ZTV,ENCFF097CAQ |
| 608 | 7 | 37879981 | 37882428 | 2447 | AJ,BALB,FVB,MA/My,NOD,NOR,RH2,SEA | 7 | 37882137 | 37882155 | 19 | + | 24.7377 | 9.44E-10 | 0.00739 | TTGCCACCAGAGGGCGCTG | 39 | ENCFF367UNR,ENCFF093ZRU,ENCFF201JUA,ENCFF099QNE,ENCFF805QBU,ENCFF821BIA,ENCFF327LYS,ENCFF907XEG,ENCFF589ZBJ,ENCFF136ZDK,ENCFF198SSR,ENCFF055HAP,ENCFF024JOZ,ENCFF089GBC,ENCFF092NMJ,ENCFF065ODV,ENCFF165LXO,ENCFF052KVO,ENCFF502WDT,ENCFF951GOP,ENCFF210BTO,ENCFF883UPM,ENCFF957SPH,ENCFF430PPJ,ENCFF142CNG,ENCFF281QMV,ENCFF194HVL,ENCFF491RJK,ENCFF349BUL,ENCFF504JDS,ENCFF611HDQ,ENCFF533APC,ENCFF089EWX,ENCFF316FYN,ENCFF609MDN,ENCFF575MGI,ENCFF033UQX,ENCFF086ZTV,ENCFF097CAQ |
| 609 | 7 | 41652882 | 41653890 | 1008 | SPRET | 7 | 41653024 | 41653058 | 35 | + | 23.4495 | 8.05E-09 | 0.00885 | CTGCTCCTGCCACCCCAAGCCAACAGGTGGCAGTA | 1 | ENCFF052KVO |
| 609 | 7 | 41652882 | 41653890 | 1008 | SPRET | 7 | 41653040 | 41653058 | 19 | + | 17.3115 | 7.42E-07 | 0.114 | AAGCCAACAGGTGGCAGTA | 1 | ENCFF052KVO |
| 610 | 7 | 42765489 | 42803176 | 37687 | CAST | 7 | 42787510 | 42787528 | 19 | + | 17.5902 | 6.18E-07 | 0.106 | GGACCACTAGATGGTGCTG | | |
| 611 | 7 | 42818002 | 42819480 | 1478 | SPRET | 7 | 42819121 | 42819154 | 34 | + | 16.8455 | 7.47E-07 | 0.0812 | CTGTAGCTTTCATACACACCAGAAGAGGGTGTCA | | |
| 612 | 7 | 44903706 | 44904721 | 1015 | ILNJ2,NZW,PJ,SJL | 7 | 44903753 | 44903786 | 34 | + | 18.0894 | 3.33E-07 | 0.0521 | CAGCTGTCTTTAGACACACCAGAAGAGGGCATTG | | |

| | | | | | | | | | | | | | | | | |
|---|---|---|---|---|---|---|---|---|---|---|---|---|---|---|---|---|
| 613 | 7 | 46803949 | 46809133 | 5184 | C57L,SPRET | 7 | 46805315 | 46805348 | 34 | + | 25.3496 | 1.21E-09 | 0.000926 | GTGCAGTACCCAAAGCTGCCAGATGAGGGCATCA | 11 | ENCFF821BIA,ENCFF092NMJ,ENCFF089GBC,ENCFF327LYS,ENCFF136ZDK,ENCFF093ZRU,ENCFF099QNE,ENCFF502WDT,ENCFF367UNR,ENCFF052KVO,ENCFF065ODV |
| 614 | 7 | 47117678 | 47131601 | 13923 | ILNJ2.MOLF.SJL | 7 | 47123827 | 47123845 | 19 | + | 16.9016 | 9.66E-07 | 0.13 | TTACCACTAGGAGGCAGAC | | |
| 615 | 7 | 47968418 | 47978336 | 9918 | C57L.FVB.NOD.NOR.SPRET.SWR | 7 | 47976345 | 47976379 | 35 | + | 22.7982 | 1.29E-08 | 0.0119 | GTGCAGCTGCATGTGGAAGCCAGAAGAGGGCATTG | | |
| 616 | 7 | 48052651 | 48059866 | 7215 | WSB | 7 | 48059499 | 48059533 | 35 | + | 22.7982 | 1.29E-08 | 0.0119 | GTGCAGCTGCATGTGGAAGCCAGAAGAGGGCATTG | | |
| 617 | 7 | 49385468 | 49387086 | 1618 | ILNJ2.MOLF.NZW.PJ.SJL | 7 | 49386064 | 49386098 | 35 | + | 23.3761 | 8.49E-09 | 0.00913 | CTGCATATATGCCTGCATGCCAGAAGAGGGTGCCA | | |
| 618 | 7 | 56091157 | 56113386 | 22229 | AKR | 7 | 56098850 | 56098883 | 34 | + | 18.0569 | 3.40E-07 | 0.0528 | ATGCAGCTAGAGACATGAGCTCCAGGGGGTACTG | | |
| 619 | 7 | 60345708 | 60373558 | 27850 | CE | 7 | 60360207 | 60360240 | 34 | + | 18.2439 | 3.00E-07 | 0.0492 | TTGCAGGTCCTGAAAAGGCCAGAAGAGTGCTGCA | | |
| 620 | 7 | 63776305 | 63785350 | 9045 | 129S1,AKR,BALB,BTBR,C3H,CAST,CBA,CE,DBA1J,DBA2J,ILNJ2,KK,LP,MOLF,NOD,NOR,NZB,NZO,NZW,PJ,PL,RF,SEA,SMJ,SPRET.WSB | 7 | 63781765 | 63781783 | 19 | + | 17.1639 | 8.17E-07 | 0.12 | CCTCCACGAGGTGGCGGAA | | |
| 621 | 7 | 65170284 | 65172410 | 2126 | CAST | 7 | 65172364 | 65172397 | 34 | + | 24.8374 | 1.91E-09 | 0.00129 | CAGCAGCTCTACCATCCTCCACCAGGTGGAGGCT | 8 | ENCFF089GBC,ENCFF327LYS,ENCFF092NMJ,ENCFF099QNE,ENCFF093ZRU,ENCFF907XEG,ENCFF024JOZ,ENCFF198SSR |
| 622 | 7 | 65492887 | 65494996 | 2109 | SPRET | 7 | 65493129 | 65493162 | 34 | + | 30.3496 | 7.10E-12 | 0.0000748 | GTGCAGTACCATCAGAGGCCAGAAGAGGGCACTG | 14 | ENCFF099QNE,ENCFF821BIA,ENCFF024JOZ,ENCFF089GBC,ENCFF092NMJ,ENCFF327LYS,ENCFF907XEG,ENCFF502WDT,ENCFF589ZBJ,ENCFF367UNR,ENCFF052KVO,ENCFF033UQX,ENCFF093ZRU,ENCFF136ZDK |
| 622 | 7 | 65492887 | 65494996 | 2109 | SPRET | 7 | 65493144 | 65493162 | 19 | + | 20.0656 | 1.05E-07 | 0.0453 | AGGCCAGAAGAGGGCACTG | 14 | ENCFF099QNE,ENCFF821BIA,ENCFF024JOZ,ENCFF089GBC,ENCFF092NMJ,ENCFF327LYS,ENCFF907XEG,ENCFF502WDT,ENCFF589ZBJ,ENCFF367UNR,ENCFF052KVO,ENCFF033UQX,ENCFF093ZRU,ENCFF136ZDK |
| 623 | 7 | 69619037 | 69620466 | 1429 | SMJ | 7 | 69619521 | 69619555 | 35 | + | 16.0459 | 9.31E-07 | 0.147 | TTGCACATACCTTTCGAAGCGAGCAGGGGGCACCT | | |
| 624 | 7 | 71950589 | 71951904 | 1315 | BUB,C58J,CE,FVB,MA/My,NOD,NOR,NZW,SWR | 7 | 71951588 | 71951622 | 35 | + | 16.8257 | 5.99E-07 | 0.116 | CTGTTTGAGCACCTGGTCTCCAGTTGGTGGCGCTG | 2 | ENCFF093ZRU,ENCFF052KVO |
| 625 | 7 | 75473062 | 75476355 | 3293 | C58J.CE.FVB.SMJ.SWR.TH.WSB | 7 | 75475456 | 75475489 | 34 | + | 17.1463 | 6.17E-07 | 0.0728 | TAGCTGTCTTTAGATGCGCCAGAAGAGGGCATCA | | |
| 626 | 7 | 76619084 | 76635354 | 16270 | ILNJ2,MA/My | 7 | 76621155 | 76621188 | 34 | + | 16.813 | 7.63E-07 | 0.0822 | ATTCGATTTCTCCAACTGCCACTAGTTGGCGCTG | 17 | ENCFF093ZRU,ENCFF367UNR,ENCFF504JDS,ENCFF165LXO,ENCFF430PPJ,ENCFF198SSR,ENCFF099QNE,ENCFF589ZBJ,ENCFF821BIA,ENCFF883UPM,ENCFF609MDN,ENCFF024JOZ,ENCFF907XEG,ENCFF092NMJ,ENCFF327LYS,ENCFF089GBC,ENCFF281QMV |
| 626 | 7 | 76619084 | 76635354 | 16270 | ILNJ2,MA/My | 7 | 76621170 | 76621188 | 19 | + | 17.6066 | 6.11E-07 | 0.106 | CTGCCACTAGTTGGCGCTG | 17 | ENCFF093ZRU,ENCFF367UNR,ENCFF504JDS,ENCFF165LXO,ENCFF430PPJ,ENCFF198SSR,ENCFF099QNE,ENCFF589ZBJ,ENCFF821BIA,ENCFF883UPM,ENCFF609MDN,ENCFF024JOZ,ENCFF907XEG,ENCFF092NMJ,ENCFF327LYS,ENCFF089GBC,ENCFF281QMV |
| 627 | 7 | 82645934 | 82652446 | 6512 | CAST | 7 | 82649488 | 82649521 | 34 | + | 17.3577 | 5.38E-07 | 0.068 | ATGAAGTACTTGCAGAGGCCAATAGAGGGTGTCA | | |
| 628 | 7 | 83553272 | 83559286 | 6014 | CAST | 7 | 83556665 | 83556698 | 34 | + | 19.5528 | 1.22E-07 | 0.0292 | ATGCACATCCCCCAGAGTCCAGAAGGAGACACCA | | |
| 629 | 7 | 84138996 | 84140946 | 1950 | CAST.MOLF | 7 | 84140248 | 84140281 | 34 | + | 17.4146 | 5.18E-07 | 0.0667 | ATGCAGCTAGAGACACAAGCTCCAGGGGGTACTG | | |
| 630 | 7 | 89137743 | 89141287 | 3544 | CAST.MOLF | 7 | 89139232 | 89139265 | 34 | + | 22.5772 | 1.27E-08 | 0.00516 | ATGCAATGCCCTTGGAGGCCAGAAGAAGGCATTA | | |
| 631 | 7 | 93890842 | 93897958 | 7116 | SPRET | 7 | 93891853 | 93891886 | 34 | + | 18.3659 | 2.76E-07 | 0.0476 | TAGCTGTCTTCAGACACTCCAGAAGAGGGCGCCA | | |
| 631 | 7 | 93890842 | 93897958 | 7116 | SPRET | 7 | 93891868 | 93891886 | 19 | + | 17.5574 | 6.32E-07 | 0.106 | ACTCCAGAAGAGGGCGCCA | | |
| 632 | 7 | 97858780 | 97860271 | 1491 | C3H,CBA,DBA1J,DBA2J,ILNJ2,MOLF,PJ,SEA | 7 | 97859659 | 97859693 | 35 | + | 16.1651 | 8.71E-07 | 0.143 | TTGCAATTGAAGGCCCAGGACGGAAGGGGTCACGC | | |
| 633 | 7 | 102938893 | 102943444 | 4551 | MOLF.NZO.TH.WSB | 7 | 102939668 | 102939701 | 34 | + | 17.6748 | 4.37E-07 | 0.0607 | GAGCATGACCACAAGTCCACAGCAGGTGGCAACG | | |
| 634 | 7 | 104358271 | 104371144 | 12873 | SPRET | 7 | 104362500 | 104362534 | 35 | + | 16.7615 | 6.21E-07 | 0.119 | TAGCAGTTGCCACCTGTGTGCTCTTGGGGGCGGGG | | |
| 635 | 7 | 105968059 | 105976874 | 8815 | CE.NZO.SPRET | 7 | 105970676 | 105970694 | 19 | + | 17.8689 | 5.14E-07 | 0.0979 | AAACCACTAGGTGGCAGAA | | |
| 636 | 7 | 106664831 | 106666847 | 2016 | NOD.SMJ.SWR | 7 | 106666508 | 106666542 | 35 | + | 16.4128 | 7.57E-07 | 0.133 | CAGCTCAGCCACGTTCCCATCACCAGGGGCCAGCT | | |
| 637 | 7 | 108417747 | 108424464 | 6717 | MOLF | 7 | 108419958 | 108419991 | 34 | + | 20.9431 | 4.46E-08 | 0.014 | GTGTAGTGGCCACAGAGGCCAAAAGAGGGTGCCA | | |
| 638 | 7 | 110428403 | 110430366 | 1963 | CAST | 7 | 110429273 | 110429306 | 34 | + | 18.3171 | 2.86E-07 | 0.0481 | TAGCTGTCTTCAGACACACCAGAAGAGGGCACTG | | |
| 638 | 7 | 110428403 | 110430366 | 1963 | CAST | 7 | 110429288 | 110429306 | 19 | + | 17.5902 | 6.18E-07 | 0.106 | ACACCAGAAGAGGGCACTG | | |
| 639 | 7 | 112387750 | 112406429 | 18679 | SJL | 7 | 112405866 | 112405899 | 34 | + | 18.0244 | 3.47E-07 | 0.0534 | TTGCAGCTTCTTCTGGAGACTACAGGAGGCACCA | | |
| 640 | 7 | 114722497 | 114724268 | 1771 | SPRET | 7 | 114722713 | 114722731 | 19 | + | 20.2951 | 8.74E-08 | 0.0431 | TCCCCAGTAGGTGGCGCTG | 17 | ENCFF367UNR,ENCFF093ZRU,ENCFF821BIA,ENCFF052KVO,ENCFF092NMJ,ENCFF589ZBJ,ENCFF136ZDK,ENCFF024JOZ,ENCFF089GBC,ENCFF165LXO,ENCFF198SSR,ENCFF883UPM,ENCFF502WDT,ENCFF033UQX,ENCFF327LYS,ENCFF533APC,ENCFF907XEG |
| 641 | 7 | 116031339 | 116037017 | 5678 | 129S1,AKR,BALB,BTBR,C3H,C57L,CAST,CBA,CE,DBA1J,DBA2J,ILNJ2,KK,MOLF,MRL,NOR,NZW,PL,RH2,SJL,SPRET,WSB | 7 | 116035155 | 116035173 | 19 | + | 17.1639 | 8.17E-07 | 0.12 | CCTCCACGAGGTGGCGGAA | | |
| 642 | 7 | 119330659 | 119334382 | 3723 | SPRET | 7 | 119330660 | 119330678 | 19 | + | 25.2131 | 4.47E-10 | 0.007 | TGGCCACTAGGGGGCACTC | 17 | ENCFF024JOZ,ENCFF136ZDK,ENCFF165LXO,ENCFF052KVO,ENCFF821BIA,ENCFF089GBC,ENCFF093ZRU,ENCFF055HAP,ENCFF883UPM,ENCFF198SSR,ENCFF957SPH,ENCFF099QNE,ENCFF327LYS,ENCFF907XEG,ENCFF502WDT,ENCFF367UNR,ENCFF589ZBJ |
| 643 | 7 | 121788205 | 121789300 | 1095 | CAST | 7 | 121788884 | 121788917 | 34 | + | 27.5122 | 1.55E-10 | 2.69E-04 | ATGCAGTACCAGCAGGGGCCAGAAGAGGGTGCAG | 35 | ENCFF093ZRU,ENCFF201JUA,ENCFF805QBU,ENCFF367UNR,ENCFF065ODV,ENCFF099QNE,ENCFF821BIA,ENCFF907XEG,ENCFF327LYS,ENCFF089GBC,ENCFF092NMJ,ENCFF024JOZ,ENCFF136ZDK,ENCFF321AGQ,ENCFF316FYN,ENCFF589ZBJ,ENCFF052KVO,ENCFF198SSR,ENCFF055HAP,ENCFF165LXO,ENCFF430PPJ,ENCFF033UQX,ENCFF502WDT,ENCFF575MGI,ENCFF210BTO,ENCFF504JDS,ENCFF097CAQ,ENCFF142CNG,ENCFF281QMV,ENCFF957SPH,ENCFF883UPM,ENCFF491RJK,ENCFF194HVL,ENCFF601JVE,ENCFF533APC |
| 644 | 7 | 123264632 | 123266393 | 1761 | CAST | 7 | 123264847 | 123264881 | 35 | + | 19.3119 | 1.35E-07 | 0.0493 | CTGTAATTGCGCTGAGTGCCCACTAGGAGGTGATA | 8 | ENCFF093ZRU,ENCFF089GBC,ENCFF327LYS,ENCFF430PPJ,ENCFF907XEG,ENCFF883UPM,ENCFF502WDT,ENCFF052KVO |
| 645 | 7 | 124568255 | 124569772 | 1517 | CAST | 7 | 124569476 | 124569509 | 34 | + | 18.7236 | 2.17E-07 | 0.0418 | ATGCAGAGCCCAGGGAGGCCAGAAGAGTGCATCC | | |
| 646 | 7 | 126281621 | 126284304 | 2683 | CAST | 7 | 126283381 | 126283414 | 34 | + | 19.1138 | 1.66E-07 | 0.036 | CTGCAACTCTGTTTTCTGCCTCCCGGAGGCAGAC | | |
| 647 | 7 | 127233514 | 127235214 | 1700 | AJ.NOD.WSB | 7 | 127234624 | 127234642 | 19 | + | 18.8033 | 2.69E-07 | 7.28E-02 | GTGCCAGAAGAGGGCGGCG | | |
| 648 | 7 | 135084952 | 135086448 | 1496 | SPRET | 7 | 135085264 | 135085298 | 35 | + | 16.9083 | 5.71E-07 | 0.114 | ATGTATGCATGCCTAGACACCAGAAGAGGGCACCA | 1 | ENCFF093ZRU |
| 648 | 7 | 135084952 | 135086448 | 1496 | SPRET | 7 | 135085280 | 135085298 | 19 | + | 17.8852 | 5.08E-07 | 0.0973 | ACACCAGAAGAGGGCACCA | 1 | ENCFF093ZRU |
| 649 | 7 | 137393454 | 137395482 | 2028 | SPRET | 7 | 137395453 | 137395487 | 35 | + | 18.4495 | 2.29E-07 | 0.0697 | GGGTTGTGGGAGGGGCTGACCAGGAGGGGGCAGTG | | |
| 649 | 7 | 137393454 | 137395482 | 2028 | SPRET | 7 | 137395469 | 137395487 | 19 | + | 21.0328 | 4.77E-08 | 0.0346 | TGACCAGGAGGGGGCAGTG | | |
| 650 | 7 | 137394197 | 137395521 | 1324 | MOLF.WSB | 7 | 137395453 | 137395487 | 35 | + | 18.4495 | 2.29E-07 | 0.0697 | GGGTTGTGGGAGGGGCTGACCAGGAGGGGGCAGTG | | |
| 650 | 7 | 137394197 | 137395521 | 1324 | MOLF.WSB | 7 | 137395469 | 137395487 | 19 | + | 21.0328 | 4.77E-08 | 0.0346 | TGACCAGGAGGGGGCAGTG | | |
| 651 | 7 | 140630036 | 140631567 | 1531 | WSB | 7 | 140630105 | 140630138 | 34 | + | 17.0894 | 6.39E-07 | 0.0746 | GTACACTCTCTTCAGACACCAGAAGAGGGCGTCA | 1 | ENCFF502WDT |
| 652 | 7 | 142322548 | 142339219 | 16671 | SPRET | 7 | 142325524 | 142325557 | 34 | + | 16.6911 | 8.24E-07 | 0.0848 | CTGCAAGAGGAGAGCTTGCCAGCAGAGAGTGCTC | | |
| 653 | 7 | 3158490 | 3161965 | 3475 | CAST.MOLF.MRL.SPRET | 7 | 3159528 | 3159562 | 35 | - | 16 | 9.55E-07 | 0.148 | CTGCTGTGTCCTGACTTTGCCCGCAGGGGTATTTG | | |
| 654 | 7 | 3719960 | 3739682 | 19722 | MRL | 7 | 3728545 | 3728563 | 19 | - | 18.6885 | 2.92E-07 | 0.0743 | TAACCAGGAGAGGGCAGCA | | |
| 655 | 7 | 11887754 | 11892685 | 4931 | MOLF | 7 | 11887990 | 11888023 | 34 | - | 16.9837 | 6.84E-07 | 0.0778 | GTTCTGTTCCCAGCAAGTACACCAGGTGGCTCAC | | |
| 656 | 7 | 12735989 | 12737781 | 1792 | CAST | 7 | 12737527 | 12737560 | 34 | - | 16.7724 | 7.83E-07 | 0.0827 | TAGCTGTCTTCAGATGCACCAGAAGAGGGCGTCA | 1 | ENCFF093ZRU |
| 657 | 7 | 14815373 | 14818589 | 3216 | NZO | 7 | 14816472 | 14816490 | 19 | - | 17.918 | 4.97E-07 | 0.0973 | AAGCCAGGAGAGGGCACTA | | |
| 657 | 7 | 14815373 | 14818589 | 3216 | NZO | 7 | 14816472 | 14816506 | 35 | - | 26.5321 | 7.03E-10 | 0.00258 | TTGCATTTATGCCTGCAAGCCAGGAGAGGGCACTA | | |
| 658 | 7 | 14858495 | 14867114 | 8619 | SPRET | 7 | 14867032 | 14867065 | 34 | - | 16.4309 | 9.71E-07 | 0.0901 | TAGCAACTTTACAAGAGTCCCCAGGTGGTGCTG | | |
| 659 | 7 | 17749509 | 17751002 | 1493 | SPRET | 7 | 17750302 | 17750336 | 35 | - | 15.9541 | 9.80E-07 | 0.15 | TTGCAATTGAAGGCCCAGGACTGAAGGGGTCACAC | | |
| 660 | 7 | 18022346 | 18026099 | 3753 | SPRET | 7 | 18022831 | 18022864 | 34 | - | 18.8049 | 2.05E-07 | 0.0402 | ATGCAGTTCTGAATCCTCCCTGTAGGAGGGGGTC | | |
| 661 | 7 | 18061973 | 18069787 | 7814 | SPRET | 7 | 18065384 | 18065417 | 34 | - | 17.0081 | 6.74E-07 | 0.0773 | ATGCAGTTAGAGACATGAGCTCTGGGGGGTACCA | | |
| 662 | 7 | 19662458 | 19682558 | 20100 | SPRET | 7 | 19673059 | 19673093 | 35 | - | 18.2844 | 2.54E-07 | 0.0739 | TTGCGTTCCTCCCACTGGTCCACCAGAGGACATCC | | |
| 663 | 7 | 20910453 | 20916877 | 6424 | CE | 7 | 20913466 | 20913500 | 35 | - | 19.9266 | 9.11E-08 | 0.0385 | ATGCAATTCATAGTGGAGACCAGAAGATGGCCTTA | | |
| 664 | 7 | 23491820 | 23500880 | 9060 | SPRET | 7 | 23494641 | 23494674 | 34 | - | 16.9106 | 7.17E-07 | 0.0793 | ATGCAGCTAGAGACAGGAGCTCCAGGGGGTACTG | | |
| 665 | 7 | 27561165 | 27586055 | 24890 | KK.NZO | 7 | 27563112 | 27563145 | 34 | - | 17.4553 | 5.05E-07 | 0.0659 | CTACAGTGGCCATGGAGACCAGAAGAAGGCATCA | | |
| 666 | 7 | 27633164 | 27641490 | 8326 | 129S1,BUB,CAST,CE,ILNJ2,KK,LP,MOLF,NZB,NZO,NZW,PJ,SEA,SJL,SMJ,SPRET,WSB | 7 | 27633455 | 27633473 | 19 | - | 19.2951 | 1.88E-07 | 0.0611 | CCGCCACTAGATGGTGCTA | | |
| 666 | 7 | 27633164 | 27641490 | 8326 | 129S1,BUB,CAST,CE,ILNJ2,KK,LP,MOLF,NZB,NZO,NZW,PJ,SEA,SJL,SMJ,SPRET,WSB | 7 | 27641449 | 27641467 | 19 | - | 19.2951 | 1.88E-07 | 0.0611 | CCGCCACTAGATGGTGCTA | 15 | ENCFF092NMJ,ENCFF201JUA,ENCFF093ZRU,ENCFF430PPJ,ENCFF367UNR,ENCFF099QNE,ENCFF327LYS,ENCFF065ODV,ENCFF055HAP,ENCFF907XEG,ENCFF136ZDK,ENCFF589ZBJ,ENCFF089GBC,ENCFF821BIA,ENCFF198SSR |
| 667 | 7 | 27718631 | 27727443 | 8812 | 129S1,AJ,BALB,BUB,CAST,CE,FVB,ILNJ2,KK,LP,MOLF,MRL,NZB,NZO,NZW,SJL,SMJ,SPRET.WSB | 7 | 27725284 | 27725317 | 34 | - | 17.0325 | 6.63E-07 | 0.0765 | TGGCAGGTCTCAGAGGAGCCGCTAGATGGCCCAC | | |
| 668 | 7 | 27743670 | 27746166 | 2496 | MOLF | 7 | 27745355 | 27745388 | 34 | - | 17.1626 | 6.10E-07 | 0.0725 | GTGCAGACTCATGGGTAGGCTGCAGATGCCGCTA | | |
| 669 | 7 | 34928856 | 34930885 | 2029 | CAST | 7 | 34929933 | 34929966 | 34 | - | 25 | 1.66E-09 | 0.00115 | ATGCAGTACCCATGGAGGCCAGAAGAGGGGGCCA | 21 | ENCFF327LYS,ENCFF136ZDK,ENCFF093ZRU,ENCFF821BIA,ENCFF907XEG,ENCFF165LXO,ENCFF198SSR,ENCFF589ZBJ,ENCFF883UPM,ENCFF957SPH,ENCFF089GBC,ENCFF367UNR,ENCFF092NMJ,ENCFF065ODV,ENCFF052KVO,ENCFF024JOZ,ENCFF099QNE,ENCFF430PPJ,ENCFF055HAP,ENCFF502WDT,ENCFF281QMV |
| 670 | 7 | 39154714 | 39165031 | 10317 | 129S1,BTBR,BUB,C3H,C57L,CBA,CE,KK,MRL.NZW.PL.SJL.SMJ.WSB | 7 | 39155367 | 39155400 | 34 | - | 17.1301 | 6.23E-07 | 0.0735 | CAGCTATCTTCAGATACACCAGAAGAGGGCATCA | | |
| 671 | 7 | 46512747 | 46514141 | 1394 | ILNJ2.MOLF.NZW.PJ.SJL | 7 | 46513033 | 46513066 | 34 | - | 19.626 | 1.16E-07 | 0.0284 | ACGCAGTGCCCACGGAGGTCAGAAGAGGGCATCT | | |
| 671 | 7 | 46512747 | 46514141 | 1394 | ILNJ2.MOLF.NZW.PJ.SJL | 7 | 46513429 | 46513462 | 34 | - | 17.6423 | 4.47E-07 | 0.0616 | TAGCTGTGTTCAGACACACCAGAAGAGGGCGTCA | | |
| 672 | 7 | 51625875 | 51629913 | 4038 | ILNJ2,KK,NZW,PJ | 7 | 51629203 | 51629237 | 35 | - | 22.4495 | 1.66E-08 | 0.0134 | CTGCAGTTACTGTGTCTGAGCAGTAGGCGGCAGAA | 24 | ENCFF093ZRU,ENCFF367UNR,ENCFF099QNE,ENCFF430PPJ,ENCFF201JUA,ENCFF327LYS,ENCFF609MDN,ENCFF089GBC,ENCFF055HAP,ENCFF321AGQ,ENCFF907XEG,ENCFF024JOZ,ENCFF821BIA,ENCFF589ZBJ,ENCFF502WDT,ENCFF575MGI,ENCFF957SPH,ENCFF316FYN,ENCFF092NMJ,ENCFF065ODV,ENCFF136ZDK,ENCFF198SSR,ENCFF165LXO,ENCFF281QMV |
| 673 | 7 | 55358136 | 55371212 | 13076 | CAST | 7 | 55360820 | 55360838 | 19 | - | 21.3934 | 3.48E-08 | 0.0315 | TTGCCACTAGGTGGCAGAA | 35 | ENCFF201JUA,ENCFF093ZRU,ENCFF504JDS,ENCFF367UNR,ENCFF099QNE,ENCFF210BTO,ENCFF327LYS,ENCFF092NMJ,ENCFF089GBC,ENCFF907XEG,ENCFF024JOZ,ENCFF055HAP,ENCFF589ZBJ,ENCFF321AGQ,ENCFF821BIA,ENCFF430PPJ,ENCFF165LXO,ENCFF502WDT,ENCFF198SSR,ENCFF065ODV,ENCFF883UPM,ENCFF136ZDK,ENCFF033UQX,ENCFF951GOP,ENCFF575MGI,ENCFF281QMV,ENCFF609MDN,ENCFF097CAQ,ENCFF491RJK,ENCFF194HVL,ENCFF957SPH,ENCFF611HDQ,ENCFF316FYN,ENCFF142CNG,ENCFF052KVO |
| 673 | 7 | 55358136 | 55371212 | 13076 | CAST | 7 | 55360820 | 55360853 | 34 | - | 22.2846 | 1.60E-08 | 0.00624 | ATGCAGCGTGTATCTTTGCCACTAGGTGGCAGAA | 35 | ENCFF201JUA,ENCFF093ZRU,ENCFF504JDS,ENCFF367UNR,ENCFF099QNE,ENCFF210BTO,ENCFF327LYS,ENCFF092NMJ,ENCFF089GBC,ENCFF907XEG,ENCFF024JOZ,ENCFF055HAP,ENCFF589ZBJ,ENCFF321AGQ,ENCFF821BIA,ENCFF430PPJ,ENCFF165LXO,ENCFF502WDT,ENCFF198SSR,ENCFF065ODV,ENCFF883UPM,ENCFF136ZDK,ENCFF033UQX,ENCFF951GOP,ENCFF575MGI,ENCFF281QMV,ENCFF609MDN,ENCFF097CAQ,ENCFF491RJK,ENCFF194HVL,ENCFF957SPH,ENCFF611HDQ,ENCFF316FYN,ENCFF142CNG,ENCFF052KVO |
| 673 | 7 | 55358136 | 55371212 | 13076 | CAST | 7 | 55370618 | 55370636 | 19 | - | 18.2295 | 4.01E-07 | 0.088 | AGACCAGGAGAGGGCACTG | | |
| 673 | 7 | 55358136 | 55371212 | 13076 | CAST | 7 | 55370618 | 55370652 | 35 | - | 22.1835 | 2.00E-08 | 0.0149 | CTGTGGTTTTCCACAAAGACCAGGAGAGGGCACTG | | |
| 674 | 7 | 55613157 | 55615897 | 2740 | SPRET | 7 | 55615703 | 55615737 | 35 | - | 19.4954 | 1.20E-07 | 4.60E-02 | CTGCATGTACACCCATGTGTCAGAAGAGGGCATCA | | |
| 675 | 7 | 58109118 | 58110738 | 1620 | SPRET | 7 | 58110310 | 58110328 | 19 | - | 17.5574 | 6.32E-07 | 0.106 | ACTCCAGAAGAGGGCGCCA | | |
| 675 | 7 | 58109118 | 58110738 | 1620 | SPRET | 7 | 58110310 | 58110343 | 34 | - | 18.3659 | 2.76E-07 | 0.0476 | TAGCTGTCTTCAGACACTCCAGAAGAGGGCGCCA | | |
| 676 | 7 | 63252709 | 63254294 | 1585 | CE | 7 | 63252783 | 63252817 | 35 | - | 16.9725 | 5.50E-07 | 0.111 | CAGCAGTTCCCACACCTGACAGAAAGATGGCTGTG | | |
| 677 | 7 | 64515546 | 64517287 | 1741 | SPRET | 7 | 64516443 | 64516477 | 35 | - | 17.9266 | 3.14E-07 | 0.0817 | GTGAAGTGCTTCCTCCAACACTCTAGGTGGCAGAG | | |
| 678 | 7 | 64922535 | 64924168 | 1633 | CAST | 7 | 64923748 | 64923782 | 35 | - | 16.6055 | 6.79E-07 | 0.125 | CAGCAGAGGGAGGGCGGGGTCAGCAGAGGGAGGGC | | |
| 678 | 7 | 64922535 | 64924168 | 1633 | CAST | 7 | 64923804 | 64923838 | 35 | - | 16.1835 | 8.62E-07 | 0.142 | CAGCAGAGGGAGGGCGGGGTCAGCAGAGGGAGGGT | | |
| 679 | 7 | 72466892 | 72468373 | 1481 | 129S1,AJ,AKR,BALB,BUB,C3H,C58J,CBA,CE,FVB,KK,MOLF,MRL,NOD,NOR,NZB,NZO,RF,RH2,SJL,SMJ,SPRET,SWR,TH | 7 | 72467534 | 72467568 | 35 | - | 16 | 9.55E-07 | 0.148 | TTGCAATTGAAGGCCCAGGACTGAAGGGGTCACGC | | |
| 680 | 7 | 74731313 | 74732438 | 1125 | SPRET | 7 | 74731596 | 74731614 | 19 | - | 17.5902 | 6.18E-07 | 0.106 | CTGCCAGTAGGTGGCAGAT | 6 | ENCFF367UNR.ENCFF093ZRU.ENCFF092NMJ.ENCFF907XEG.ENCFF089GBC.ENCFF099QNE |
| 681 | 7 | 75890589 | 75899884 | 9295 | MOLF.SPRET | 7 | 75890860 | 75890878 | 19 | - | 17.2131 | 7.91E-07 | 0.118 | GGGCCAAAAGGGGGCAGTG | | |
| 681 | 7 | 75890589 | 75899884 | 9295 | MOLF.SPRET | 7 | 75890860 | 75890894 | 35 | - | 16.4771 | 7.30E-07 | 0.13 | GTACAGGGGAACGCCAGGGCCAAAAGGGGGCAGTG | | |
| 682 | 7 | 80737931 | 80740104 | 2173 | SPRET | 7 | 80738879 | 80738897 | 19 | - | 20.082 | 1.04E-07 | 0.0453 | TGTCCAGAAGAGGGCAGCA | 16 | ENCFF136ZDK,ENCFF024JOZ,ENCFF589ZBJ,ENCFF907XEG,ENCFF052KVO,ENCFF367UNR,ENCFF883UPM,ENCFF198SSR,ENCFF821BIA,ENCFF093ZRU,ENCFF957SPH,ENCFF327LYS,ENCFF065ODV,ENCFF502WDT,ENCFF055HAP,ENCFF099QNE |
| 682 | 7 | 80737931 | 80740104 | 2173 | SPRET | 7 | 80738879 | 80738912 | 34 | - | 22.8374 | 1.03E-08 | 0.00445 | GTGGAGGGCCCACAATGTCCAGAAGAGGGCAGCA | 16 | ENCFF136ZDK,ENCFF024JOZ,ENCFF589ZBJ,ENCFF907XEG,ENCFF052KVO,ENCFF367UNR,ENCFF883UPM,ENCFF198SSR,ENCFF821BIA,ENCFF093ZRU,ENCFF957SPH,ENCFF327LYS,ENCFF065ODV,ENCFF502WDT,ENCFF055HAP,ENCFF099QNE |
| 683 | 7 | 82343027 | 82344114 | 1087 | SPRET | 7 | 82343389 | 82343422 | 34 | - | 16.878 | 7.32E-07 | 0.0801 | TTGCTCTCTTCAGACACACCAGAAGAGGGCATTG | | |
| 684 | 7 | 84709089 | 84710391 | 1302 | 129S1,AJ,AKR,BALB,BTBR,BUB,C3H,C57L,CBA,DBA1J,DBA2J,FVB,ILNJ2,KK,MA/My,MRL,NOD,NOR,NZB,NZO,NZW,PJ,PL,RF,RH2,SEA,SJL,SMJ,SWR,TH | 7 | 84709367 | 84709400 | 34 | - | 17.2846 | 5.64E-07 | 0.0699 | CTGCAAGAGGAGAGCTTGCCTGCAGAGAGCACTC | | |
| 685 | 7 | 88463086 | 88464661 | 1575 | SMJ | 7 | 88464587 | 88464620 | 34 | - | 18.0569 | 3.40E-07 | 0.0528 | ATGCAGCTAGAGACATGAGCTCCAGGGGGTACTG | | |
| 686 | 7 | 94920123 | 94976361 | 56238 | SMJ | 7 | 94969281 | 94969314 | 34 | - | 20.187 | 7.76E-08 | 0.0212 | TTGCTGTTTTCAGACACACCAGAAGAGGGCATCA | | |
| 687 | 7 | 99006822 | 99008895 | 2073 | SPRET | 7 | 99007255 | 99007288 | 34 | - | 17.4228 | 5.16E-07 | 0.0666 | TAGCTGTTTTCAGATACACCAGAAGAGGGCATCC | 3 | ENCFF093ZRU.ENCFF367UNR.ENCFF099QNE |
| 688 | 7 | 99970795 | 99975214 | 4419 | SPRET | 7 | 99974853 | 99974871 | 19 | - | 21.0164 | 4.84E-08 | 0.0349 | TTGCCTGCAGGGGGCAGCA | 34 | ENCFF093ZRU,ENCFF367UNR,ENCFF099QNE,ENCFF327LYS,ENCFF210BTO,ENCFF089GBC,ENCFF201JUA,ENCFF821BIA,ENCFF907XEG,ENCFF951GOP,ENCFF024JOZ,ENCFF316FYN,ENCFF589ZBJ,ENCFF198SSR,ENCFF065ODV,ENCFF165LXO,ENCFF092NMJ,ENCFF430PPJ,ENCFF055HAP,ENCFF136ZDK,ENCFF502WDT,ENCFF097CAQ,ENCFF194HVL,ENCFF142CNG,ENCFF957SPH,ENCFF281QMV,ENCFF052KVO,ENCFF533APC,ENCFF575MGI,ENCFF883UPM,ENCFF321AGQ,ENCFF491RJK,ENCFF609MDN,ENCFF611HDQ |
| 688 | 7 | 99970795 | 99975214 | 4419 | SPRET | 7 | 99974853 | 99974886 | 34 | - | 21.9593 | 2.06E-08 | 0.0076 | CCGCTGCATACCCTGTTGCCTGCAGGGGGCAGCA | 34 | ENCFF093ZRU,ENCFF367UNR,ENCFF099QNE,ENCFF327LYS,ENCFF210BTO,ENCFF089GBC,ENCFF201JUA,ENCFF821BIA,ENCFF907XEG,ENCFF951GOP,ENCFF024JOZ,ENCFF316FYN,ENCFF589ZBJ,ENCFF198SSR,ENCFF065ODV,ENCFF165LXO,ENCFF092NMJ,ENCFF430PPJ,ENCFF055HAP,ENCFF136ZDK,ENCFF502WDT,ENCFF097CAQ,ENCFF194HVL,ENCFF142CNG,ENCFF957SPH,ENCFF281QMV,ENCFF052KVO,ENCFF533APC,ENCFF575MGI,ENCFF883UPM,ENCFF321AGQ,ENCFF491RJK,ENCFF609MDN,ENCFF611HDQ |
| 689 | 7 | 102173101 | 102174345 | 1244 | KK.NOD.SJL.SPRET.SWR | 7 | 102173660 | 102173694 | 35 | - | 17.6606 | 3.68E-07 | 0.0897 | GGGAAATGCCATCTCAGACACACCAGAGGGCACTG | | |
| 690 | 7 | 102499908 | 102503117 | 3209 | KK.NOD.SJL.SPRET.SWR | 7 | 102502504 | 102502522 | 19 | - | 17.0656 | 8.70E-07 | 0.124 | AGGCCACAAGAGGGAACTC | | |
| 691 | 7 | 109710735 | 109712782 | 2047 | CAST.MOLF.NOD.SMJ.SPRET.SWR | 7 | 109712572 | 109712605 | 34 | - | 18.813 | 2.04E-07 | 0.0402 | TAGCTGTCCCCAGATATTCCAGAAGAGGGCATCA | | |
| 692 | 7 | 112866154 | 112870131 | 3977 | C3H,CBA,DBA1J,DBA2J,ILNJ2,MOLF,NOR,NZO,SMJ,TH | 7 | 112867716 | 112867734 | 19 | - | 17.9836 | 4.75E-07 | 0.0964 | GTTCCAGCAGGTGGCAGCC | 30 | ENCFF093ZRU,ENCFF367UNR,ENCFF099QNE,ENCFF201JUA,ENCFF821BIA,ENCFF210BTO,ENCFF327LYS,ENCFF907XEG,ENCFF024JOZ,ENCFF089GBC,ENCFF198SSR,ENCFF092NMJ,ENCFF589ZBJ,ENCFF430PPJ,ENCFF951GOP,ENCFF533APC,ENCFF033UQX,ENCFF165LXO,ENCFF957SPH,ENCFF136ZDK,ENCFF097CAQ,ENCFF321AGQ,ENCFF052KVO,ENCFF609MDN,ENCFF575MGI,ENCFF491RJK,ENCFF281QMV,ENCFF316FYN.ENCFF883UPM.ENCFF194HVL |
| 692 | 7 | 112866154 | 112870131 | 3977 | C3H,CBA,DBA1J,DBA2J,ILNJ2,MOLF,NOR,NZO,SMJ,TH | 7 | 112867716 | 112867750 | 35 | - | 23.2202 | 9.52E-09 | 0.00969 | GTGCCATTACAACAACGTTCCAGCAGGTGGCAGCC | 30 | ENCFF093ZRU,ENCFF367UNR,ENCFF099QNE,ENCFF201JUA,ENCFF821BIA,ENCFF210BTO,ENCFF327LYS,ENCFF907XEG,ENCFF024JOZ,ENCFF089GBC,ENCFF198SSR,ENCFF092NMJ,ENCFF589ZBJ,ENCFF430PPJ,ENCFF951GOP,ENCFF533APC,ENCFF033UQX,ENCFF165LXO,ENCFF957SPH,ENCFF136ZDK,ENCFF097CAQ,ENCFF321AGQ,ENCFF052KVO,ENCFF609MDN,ENCFF575MGI,ENCFF491RJK,ENCFF281QMV,ENCFF316FYN.ENCFF883UPM.ENCFF194HVL |
| 692 | 7 | 112866154 | 112870131 | 3977 | C3H,CBA,DBA1J,DBA2J,ILNJ2,MOLF,NOR,NZO,SMJ,TH | 7 | 112869315 | 112869348 | 34 | - | 16.7236 | 8.07E-07 | 0.084 | CTGCACTGCCAGCTGCAAACACAGCAGGGCCTCA | | |
| 693 | 7 | 115926063 | 115929685 | 3622 | 129S1.BTBR.C57L.ILNJ2.MOLF | 7 | 115926548 | 115926581 | 34 | - | 18.3984 | 2.70E-07 | 0.0475 | TTGCTGTCTTCAGACACACCAGAAGAGGGCATTG | 1 | ENCFF093ZRU |
| 694 | 7 | 122558697 | 122564120 | 5423 | MRL | 7 | 122564036 | 122564069 | 34 | - | 17.9756 | 3.59E-07 | 0.0544 | TTGCACAGTCTCAGCCCTGCAACAGAGGGCACCG | | |
| 695 | 7 | 122669754 | 122676858 | 7104 | FVB.SJL | 7 | 122674680 | 122674713 | 34 | - | 19.6504 | 1.14E-07 | 0.0282 | ATGCAGTTCCTATGAAGGCCAGAAGAGGATATCG | 2 | ENCFF099QNE.ENCFF093ZRU |
| 696 | 7 | 125445214 | 125448255 | 3041 | WSB | 7 | 125446706 | 125446739 | 34 | - | 16.6016 | 8.72E-07 | 0.0874 | CTGCTATTCTCCTTCACCCCACCAGATGTCTTCT | 1 | ENCFF099QNE |
| 697 | 7 | 133840225 | 133853081 | 12856 | DBA1J.DBA2J | 7 | 133840774 | 133840792 | 19 | - | 17.3607 | 7.18E-07 | 0.112 | AGGCCAGAAGAGGGTGCCA | 1 | ENCFF052KVO |
| 697 | 7 | 133840225 | 133853081 | 12856 | DBA1J.DBA2J | 7 | 133840774 | 133840808 | 35 | - | 24.1376 | 4.80E-09 | 0.0065 | GTGCAGGTACCCTAAGAGGCCAGAAGAGGGTGCCA | 1 | ENCFF052KVO |
| 698 | 7 | 138400694 | 138402588 | 1894 | MOLF.NZB.NZO | 7 | 138401683 | 138401701 | 19 | - | 17.9836 | 4.75E-07 | 0.0964 | GCACCAGAAGAGGGCACTA | | |
| 699 | 8 | 3132050 | 3165147 | 33097 | SPRET | 8 | 3142227 | 3142245 | 19 | + | 17.377 | 7.11E-07 | 0.132 | TGTCCAGCAGGGGACACAC | | |
| 699 | 8 | 3132050 | 3165147 | 33097 | SPRET | 8 | 3146189 | 3146222 | 34 | + | 20.3008 | 7.15E-08 | 0.0231 | GGGCAGTTCCCTCAGAGGCCAGAAGAAGGAATCA | | |
| 700 | 8 | 4595818 | 4602409 | 6591 | SPRET | 8 | 4596392 | 4596425 | 34 | + | 16.7724 | 7.83E-07 | 0.088 | CTGCAAGACGAGAGCTTGCCTGCAGAGAGTACTC | | |
| 701 | 8 | 5659784 | 5666072 | 6288 | 129S1.CAST.NZB.SPRET | 8 | 5660465 | 5660498 | 34 | + | 17.813 | 3.99E-07 | 0.063 | CAGCACTCCCAACTTCGCCCAGTCCAGGGCACCC | | |
| 702 | 8 | 7799142 | 7807592 | 8450 | KK.SMJ | 8 | 7805087 | 7805121 | 35 | + | 17.4037 | 4.28E-07 | 0.117 | ATGCAGTGCCCCCACAGGGGCCATAGGGACATCA | | |
| 703 | 8 | 19444790 | 19447671 | 2881 | 129S1,AJ,BALB,BTBR,BUB,C3H,CAST,CBA,DBA1J,DBA2J,FVB,KK,MOLF,MRL,NOD,NOR,NZB,NZW,PL,RH2,SJL,SMJ,SPRET,SWR,TH | 8 | 19447648 | 19447682 | 35 | + | 16.1101 | 8.98E-07 | 0.163 | CTGCTGCCACACCCGCTCCCCACAGGGTGTCATCT | | |
| 704 | 8 | 19549274 | 19562479 | 13205 | CAST | 8 | 19556267 | 19556285 | 19 | + | 17.7213 | 5.67E-07 | 0.12 | AGGACAGCAGATGGCACTA | 2 | ENCFF502WDT,ENCFF055HAP |
| 705 | 8 | 19833367 | 19834900 | 1533 | MA/My,RF | 8 | 19834560 | 19834593 | 34 | + | 21.6992 | 2.52E-08 | 0.0107 | GTGAGGTTCATATAGGGGCCACCAGAGGGCGCGA | 12 | ENCFF201JUA,ENCFF093ZRU,ENCFF367UNR,ENCFF099QNE,ENCFF430PPJ,ENCFF821BIA,ENCFF092NMJ,ENCFF198SSR,ENCFF589ZBJ,ENCFF327LYS,ENCFF907XEG,ENCFF089GBC |
| 705 | 8 | 19833367 | 19834900 | 1533 | MA/My,RF | 8 | 19834575 | 19834593 | 19 | + | 22 | 2.03E-08 | 0.0275 | GGGCCACCAGAGGGCGCGA | 13 | ENCFF201JUA,ENCFF093ZRU,ENCFF367UNR,ENCFF099QNE,ENCFF430PPJ,ENCFF821BIA,ENCFF092NMJ,ENCFF198SSR,ENCFF589ZBJ,ENCFF327LYS,ENCFF907XEG,ENCFF089GBC,ENCFF136ZDK |
| 706 | 8 | 21748114 | 22122932 | 374818 | BTBR.CAST.MOLF.NOR.NZB.NZW | 8 | 21889112 | 21889145 | 34 | + | 16.7805 | 7.79E-07 | 0.088 | ATGCAGCTAGAGACACAAGCTCTAGGGGGTACTG | | |
| 706 | 8 | 21748114 | 22122932 | 374818 | BTBR.CAST.MOLF.NOR.NZB.NZW | 8 | 22026318 | 22026351 | 34 | + | 16.7805 | 7.79E-07 | 0.088 | ATGCAGCTAGAGACACAAGCTCTAGGGGGTACTG | | |
| 706 | 8 | 21748114 | 22122932 | 374818 | BTBR.CAST.MOLF.NOR.NZB.NZW | 8 | 21780469 | 21780487 | 19 | - | 17.7213 | 5.67E-07 | 0.12 | AGGACAGCAGATGGCACTA | | |
| 706 | 8 | 21748114 | 22122932 | 374818 | BTBR.CAST.MOLF.NOR.NZB.NZW | 8 | 21992420 | 21992438 | 19 | - | 17.4918 | 6.59E-07 | 0.128 | AGAACAGCAGATGGCGCTA | | |
| 707 | 8 | 25018753 | 25021632 | 2879 | SPRET | 8 | 25020353 | 25020387 | 35 | + | 18.3303 | 2.47E-07 | 0.0869 | GTCCTGGGCAGCCTCACAACCACAAGGGGGCGATG | 7 | ENCFF093ZRU,ENCFF367UNR,ENCFF327LYS,ENCFF821BIA,ENCFF024JOZ,ENCFF907XEG,ENCFF198SSR |
| 707 | 8 | 25018753 | 25021632 | 2879 | SPRET | 8 | 25020369 | 25020387 | 19 | + | 19.1311 | 2.12E-07 | 0.0758 | CAACCACAAGGGGGCGATG | 7 | ENCFF093ZRU,ENCFF367UNR,ENCFF327LYS,ENCFF821BIA,ENCFF024JOZ,ENCFF907XEG,ENCFF198SSR |
| 707 | 8 | 25018753 | 25021632 | 2879 | SPRET | 8 | 25019051 | 25019084 | 34 | - | 18.5041 | 2.52E-07 | 0.052 | CTGCAGGACCTGCATCGACATCCAGATGACACGG | | |
| 707 | 8 | 25018753 | 25021632 | 2879 | SPRET | 8 | 25019906 | 25019939 | 34 | - | 16.5041 | 9.27E-07 | 0.0944 | CTGCCCTAATCCCAGCACTCAGGAGGTGGAGGCA | | |
| 708 | 8 | 27544054 | 27545085 | 1031 | CAST | 8 | 27544953 | 27544986 | 34 | + | 19.9512 | 9.19E-08 | 0.0275 | ATGAAATTCTTTTGATTGCCTCTAGGTGGCGACA | 30 | ENCFF201JUA,ENCFF093ZRU,ENCFF367UNR,ENCFF099QNE,ENCFF821BIA,ENCFF907XEG,ENCFF089GBC,ENCFF024JOZ,ENCFF327LYS,ENCFF321AGQ,ENCFF198SSR,ENCFF065ODV,ENCFF165LXO,ENCFF502WDT,ENCFF055HAP,ENCFF430PPJ,ENCFF589ZBJ,ENCFF092NMJ,ENCFF136ZDK,ENCFF052KVO,ENCFF883UPM,ENCFF504JDS,ENCFF951GOP,ENCFF194HVL,ENCFF609MDN,ENCFF281QMV,ENCFF957SPH,ENCFF316FYN.ENCFF575MGI.ENCFF142CNG |
| 708 | 8 | 27544054 | 27545085 | 1031 | CAST | 8 | 27544968 | 27544986 | 19 | + | 18.6557 | 2.99E-07 | 0.0888 | TTGCCTCTAGGTGGCGACA | 30 | ENCFF201JUA,ENCFF093ZRU,ENCFF367UNR,ENCFF099QNE,ENCFF821BIA,ENCFF907XEG,ENCFF089GBC,ENCFF024JOZ,ENCFF327LYS,ENCFF321AGQ,ENCFF198SSR,ENCFF065ODV,ENCFF165LXO,ENCFF502WDT,ENCFF055HAP,ENCFF430PPJ,ENCFF589ZBJ,ENCFF092NMJ,ENCFF136ZDK,ENCFF052KVO,ENCFF883UPM,ENCFF504JDS,ENCFF951GOP,ENCFF194HVL,ENCFF609MDN,ENCFF281QMV,ENCFF957SPH,ENCFF316FYN.ENCFF575MGI.ENCFF142CNG |
| 709 | 8 | 27966240 | 27969045 | 2805 | SPRET | 8 | 27966996 | 27967030 | 35 | + | 16.6972 | 6.44E-07 | 0.138 | ATGAAAGTACCAGTGGAGGCCAGAAGAGGGCATTG | | |
| 710 | 8 | 32165219 | 32171803 | 6584 | 129S1,AJ,AKR,BALB,BUB,C3H,C57L,CAST,CBA,DBA1J,DBA2J,FVB,MA/My,MRL,NOD,NOR,NZW,PL,RF,RH2,SJL,SMJ,SPRET,SWR,TH.WSB | 8 | 32169807 | 32169841 | 35 | + | 18.1284 | 2.79E-07 | 0.0931 | CTGTAGCTGTCTTCAGACACCAGAAGAGGGCATCA | | |
| 711 | 8 | 34570509 | 34571667 | 1158 | 129S1,AJ,AKR,BALB,BUB,C3H,C57L,CBA,CE,DBA1J,DBA2J,MA/My,MRL,NOD,NOR,PL,RF,RH2,SJL,SMJ,SWR,TH,WSB | 8 | 34570716 | 34570749 | 34 | + | 21.9756 | 2.03E-08 | 0.00915 | CTGCTGTCTTCAGTCACACCAGAAGGGGGCATTG | 1 | ENCFF093ZRU |

| | | | | | | | | | | | | | | | | |
|---|---|---|---|---|---|---|---|---|---|---|---|---|---|---|---|---|
| 712 | 8 | 41627138 | 41628522 | 1384 | 129S1,AJ,AKR,BALB,BTBR,BUB,C3H,C57L,CBA,CE,DBA1J,DBA2J,FVB,ILNJ2,KK,MA/My,MRL,NOD,NOR,PJ,PL,RF,RH2,SEA,SJL,SMJ,SWR,TH | 8 | 41627232 | 41627250 | 19 | + | 17.2787 | 7.58E-07 | 0.136 | CTTCCAGTAGAGGGCGTCA | 1 | ENCFF821BIA |
| 713 | 8 | 45557214 | 45558325 | 1111 | SPRET | 8 | 45557323 | 45557356 | 34 | + | 20.6748 | 5.44E-08 | 0.0191 | CTGCAAAGCTAATTGTGTCCAGCAGATGCCCCTG | 7 | ENCFF024JOZ,ENCFF327LYS,ENCFF092NMJ,ENCFF907XEG,ENCFF089GBC,ENCFF093ZRU,ENCFF099QNE |
| 714 | 8 | 48086132 | 48087139 | 1007 | SPRET | 8 | 48086996 | 48087029 | 34 | + | 19.5285 | 1.24E-07 | 0.0333 | ATGCAGTGCCTGCAGAGGCCAGAAGAAAGTAGCA | | |
| 715 | 8 | 48258313 | 48259398 | 1085 | SPRET | 8 | 48259013 | 48259047 | 35 | + | 17.0826 | 5.16E-07 | 0.128 | CTGCATATATGGTTACAAGCCAGAAAAGGGCACCA | | |
| 716 | 8 | 52184613 | 52188769 | 4156 | 129S1,AJ,B10D2,B10J,BUB,C58J,CAST,CBA,FVB,ILNJ2,KK,LP,MA/My,MOLF,MRL,NZB,NZO,NZW,PL,RH2,SJL,SMJ,SPRET,SWR,WSB | 8 | 52185390 | 52185423 | 34 | + | 17.6748 | 4.37E-07 | 0.066 | GAGCATGACCACAAGTCCACAGCAGGTGGCAACG | | |
| 717 | 8 | 64385349 | 64388640 | 3291 | AKR,BALB,C3H,C57L,CAST,CBA,CE,DBA1J,DBA2J,MRL,NZB,NZO,PJ,PL,RF,SEA,SMJ | 8 | 64388574 | 64388608 | 35 | + | 23.9266 | 5.63E-09 | 0.00912 | CTGCATGTATACCTTCATGCCAGAAGAGGGCCCTA | | |
| 718 | 8 | 64698279 | 64703946 | 5667 | CAST | 8 | 64700258 | 64700291 | 34 | + | 18.0569 | 3.40E-07 | 0.0578 | ATGCAGCTAGAGACATGAGCTCCAGGGGGTACTG | | |
| 719 | 8 | 65362517 | 65378480 | 15963 | C58J.LP.PJ.SEA | 8 | 65366314 | 65366347 | 34 | + | 16.8211 | 7.59E-07 | 0.0873 | CAGCTGTCTTCAGACACACCTGAAGAGGGCATCG | | |
| 720 | 8 | 67816635 | 67837561 | 20926 | C58J.PJ.SEA.TH.WSB | 8 | 67829169 | 67829202 | 34 | + | 22.252 | 1.64E-08 | 0.00794 | GAGCAGGTGCAGCCACCGCCCCAAGATGGCGCCC | | |
| 721 | 8 | 69811986 | 69814316 | 2330 | NZB.NZO | 8 | 69813175 | 69813208 | 34 | + | 16.9268 | 7.10E-07 | 0.0845 | ATGCAGTTAGAGACATGAGCTCTGGGGGGTACTG | | |
| 722 | 8 | 69907761 | 69915596 | 7835 | SPRET | 8 | 69912826 | 69912859 | 34 | + | 17.3902 | 5.27E-07 | 0.0727 | TAGCTGTCCTCAGACACTCCAGAAGAGGGCATCA | | |
| 723 | 8 | 69912662 | 69914765 | 2103 | AKR.CAST.MA/Mv.MOLF.NZB.RF.WSB | 8 | 69912826 | 69912859 | 34 | + | 17.3902 | 5.27E-07 | 0.0727 | TAGCTGTCCTCAGACACTCCAGAAGAGGGCATCA | | |
| 724 | 8 | 69982260 | 69984363 | 2103 | CAST.MOLF | 8 | 69982424 | 69982457 | 34 | + | 17.3902 | 5.27E-07 | 0.0727 | TAGCTGTCCTCAGACACTCCAGAAGAGGGCATCA | | |
| 725 | 8 | 70094207 | 70102187 | 7980 | 129S1,AKR,BALB,BTBR,BUB,C3H,C57L,C58J,CAST,CBA,CE,DBA1J,DBA2J,FVB,ILNJ2,KK,MA/My,MOLF,MRL,NZB,NZO,NZW,PJ,PL,RF,SEA,SJL,SMJ,SWR,TH,WSB | 8 | 70096875 | 70096908 | 34 | + | 17.6748 | 4.37E-07 | 0.066 | GAGCATGACCACAAGTCCACAGCAGGTGGCAACG | | |
| 726 | 8 | 70096098 | 70098978 | 2880 | SPRET | 8 | 70096875 | 70096908 | 34 | + | 17.6748 | 4.37E-07 | 0.066 | GAGCATGACCACAAGTCCACAGCAGGTGGCAACG | | |
| 727 | 8 | 70144215 | 70153249 | 9034 | CAST.MOLF | 8 | 70151240 | 70151273 | 34 | + | 18.5691 | 2.41E-07 | 0.0509 | TAGCTGTCTTCATACACACCACAAGAGGGCACCA | | |
| 727 | 8 | 70144215 | 70153249 | 9034 | CAST.MOLF | 8 | 70151255 | 70151273 | 19 | + | 18.541 | 3.24E-07 | 0.092 | ACACCACAAGAGGGCACCA | | |
| 728 | 8 | 70151241 | 70152819 | 1578 | 129S1,BALB,BTBR,FVB,ILNJ2,KK,NZW,RF,SJL,SWR,TH,WSB | 8 | 70151240 | 70151273 | 34 | + | 18.5691 | 2.41E-07 | 0.0509 | TAGCTGTCTTCATACACACCACAAGAGGGCACCA | | |
| 728 | 8 | 70151241 | 70152819 | 1578 | 129S1,BALB,BTBR,FVB,ILNJ2,KK,NZW,RF,SJL,SWR,TH,WSB | 8 | 70151255 | 70151273 | 19 | + | 18.541 | 3.24E-07 | 0.092 | ACACCACAAGAGGGCACCA | | |
| 729 | 8 | 71696006 | 71698945 | 2939 | SPRET | 8 | 71698225 | 71698258 | 34 | + | 16.4472 | 9.61E-07 | 0.0944 | TAGCTGTCTTCAGACACACCAGAAGAGGGCGTCA | | |
| 730 | 8 | 72239133 | 72244758 | 5625 | SPRET | 8 | 72241993 | 72242027 | 35 | + | 16.8807 | 5.80E-07 | 0.136 | CCGACGAGCCCAGAGTCCCCCAGCAGATGGCGCTC | 7 | ENCFF821BIA.ENCFF093ZRU.ENCFF367UNR.ENCFF099QNE |
| 730 | 8 | 72239133 | 72244758 | 5625 | SPRET | 8 | 72242009 | 72242027 | 19 | + | 19.0164 | 2.30E-07 | 0.08 | CCCCCAGCAGATGGCGCTC | 7 | ENCFF821BIA.ENCFF093ZRU.ENCFF367UNR.ENCFF099QNE |
| 731 | 8 | 73410655 | 73413640 | 2985 | SPRET | 8 | 73412349 | 73412382 | 34 | + | 24.626 | 2.30E-09 | 0.00188 | GTGCAGTGTCACCAGAGGCCAAAAGAGGGTACCA | | |
| 731 | 8 | 73410655 | 73413640 | 2985 | SPRET | 8 | 73410654 | 73410687 | 34 | - | 16.5447 | 9.04E-07 | 0.0944 | CAGCTGTCTTCAGACACACCAGAAGAGGGAGTCA | | |
| 732 | 8 | 74090969 | 74101816 | 10847 | SPRET | 8 | 74099498 | 74099532 | 35 | + | 17.9817 | 3.04E-07 | 0.0977 | AAGAAGTTAGACCCTGTTGCCAGCAGGTGGCCTTC | | |
| 733 | 8 | 83788780 | 83790841 | 2061 | 129S1,AJ,AKR,BALB,BTBR,C3H,C57L,CBA,CE,DBA1J,DBA2J,ILNJ2,KK,MA/My,MRL,NOD,NOR,NZW,PJ,PL,RF,RH2,SEA,SJL,WSB | 8 | 83790314 | 83790347 | 34 | + | 19.0569 | 1.72E-07 | 0.0405 | CAGCTGTCTTCTCACACACCAGAAGAGGGCATCA | 1 | ENCFF821BIA |
| 734 | 8 | 86957834 | 86960871 | 3037 | 129S1,AKR,BALB,BTBR,C57L,MRL,PJ,PL,RF,SEA | 8 | 86960511 | 86960545 | 35 | + | 21.9817 | 2.31E-08 | 0.0209 | CTGCTGTGTCACATGTACATCAGTAGAGGGCAGAG | 4 | ENCFF093ZRU,ENCFF367UNR,ENCFF327LYS,ENCFF883UPM |
| 735 | 8 | 89130672 | 89132551 | 1879 | 129S1,AKR,BALB,BTBR,BUB,C57L,C58J,CBA,CE,DBA1J,DBA2J,FVB,ILNJ2,KK,MA/My,MRL,NOD,NOR,NZB,NZO,NZW,PJ,PL,RF,SEA.SJL.SMJ.SWR.TH.WSB | 8 | 89131683 | 89131717 | 35 | + | 16 | 9.55E-07 | 0.168 | TTGCAATTGAAGGCCCAGGACTGAAGGGGTCACGC | | |
| 736 | 8 | 89130713 | 89131844 | 1131 | SPRET | 8 | 89131683 | 89131717 | 35 | + | 16 | 9.55E-07 | 0.168 | TTGCAATTGAAGGCCCAGGACTGAAGGGGTCACGC | | |
| 737 | 8 | 93307104 | 93308797 | 1693 | C58J.CBA.DBA1J.DBA2J.MA/Mv.SMJ | 8 | 93307102 | 93307136 | 35 | + | 16.1376 | 8.84E-07 | 0.162 | ATGCAAGTGCCCTCAAGGTCCAGAAGAAGGCATCA | | |
| 738 | 8 | 96508464 | 96509750 | 1286 | SPRET | 8 | 96509395 | 96509429 | 35 | + | 16.7523 | 6.24E-07 | 0.137 | TTGGATTGTAGTATACCAACCACTGGGTGGCACTG | 4 | ENCFF367UNR.ENCFF198SSR.ENCFF821BIA.ENCFF093ZRU |
| 738 | 8 | 96508464 | 96509750 | 1286 | SPRET | 8 | 96509411 | 96509429 | 19 | + | 17.5246 | 6.45E-07 | 0.126 | CAACCACTGGGTGGCACTG | 4 | ENCFF367UNR.ENCFF198SSR.ENCFF821BIA.ENCFF093ZRU |
| 739 | 8 | 105584309 | 105591165 | 6856 | CAST.MOLF.SPRET | 8 | 105586679 | 105586712 | 34 | + | 18.3659 | 2.76E-07 | 0.0529 | TAGCTGTCTTCAGACACTCCAGAAGAGGGCGCCA | | |
| 739 | 8 | 105584309 | 105591165 | 6856 | CAST.MOLF.SPRET | 8 | 105586694 | 105586712 | 19 | + | 17.5574 | 6.32E-07 | 0.125 | ACTCCAGAAGAGGGCGCCA | | |
| 740 | 8 | 110066405 | 110067541 | 1136 | CAST.MOLF.SPRET | 8 | 110066510 | 110066543 | 34 | + | 23.8699 | 4.40E-09 | 0.00301 | GTGCAGTGTCCAAGGAGGCCAGAAGAGGGTGTCC | | |
| 741 | 8 | 112362773 | 112365114 | 2341 | MOLF | 8 | 112363347 | 112363381 | 35 | + | 21.6055 | 2.99E-08 | 0.0243 | CTGCATATATGACTGCAGGCCAGAAGAGGACACCA | | |
| 742 | 8 | 120311712 | 120313301 | 1589 | SPRET | 8 | 120312678 | 120312711 | 34 | + | 17.5285 | 4.81E-07 | 0.0695 | AGGCAGTTCTCACAGCTGCCAACTGGGGGGGGGG | | |
| 743 | 8 | 120399230 | 120401088 | 1858 | CAST | 8 | 120399682 | 120399700 | 19 | + | 19.1967 | 2.02E-07 | 0.0739 | CAGCCACCAGGTGGAGGAA | | |
| 743 | 8 | 120399230 | 120401088 | 1858 | CAST | 8 | 120400875 | 120400908 | 34 | + | 27 | 2.58E-10 | 0.000434 | ATGCAGTGCCTATAGAAGCCAGTAGAGGGCATCA | 20 | ENCFF099QNE,ENCFF821BIA,ENCFF165LXO,ENCFF198SSR,ENCFF092NMJ,ENCFF589ZBJ,ENCFF055HAP,ENCFF052KVO,ENCFF093ZRU,ENCFF533APC,ENCFF033UQX,ENCFF327LYS,ENCFF065ODV,ENCFF907XEG,ENCFF024JOZ,ENCFF502WDT,ENCFF089GBC,ENCFF367UNR,ENCFF136ZDK,ENCFF142CNG |
| 744 | 8 | 122559579 | 122568003 | 8424 | 129S1,BTBR,C57L,CAST,FVB,ILNJ2,MOLF,NOD,NOR,NZB,SJL,SPRET,SWR,WSB | 8 | 122563720 | 122563738 | 19 | + | 18.0984 | 4.40E-07 | 0.107 | TGGCCAGTAGATGGCCCTC | | |
| 745 | 8 | 122862058 | 122864866 | 2808 | 129S1,BTBR,C57L,FVB,NOD,NOR,SJL,SWR | 8 | 122862392 | 122862426 | 35 | + | 25.367 | 1.84E-09 | 0.00549 | CTGCACTGGCAGGGAGGAGCCAGAGGGGGGCACTC | 10 | ENCFF093ZRU,ENCFF092NMJ,ENCFF089GBC,ENCFF907XEG,ENCFF327LYS,ENCFF367UNR,ENCFF883UPM,ENCFF099QNE,ENCFF533APC,ENCFF821BIA |
| 746 | 8 | 123891450 | 123898816 | 7366 | 129S1,AJ,AKR,BALB,BTBR,C3H,C57L,C58J,CAST,CE,ILNJ2,LP,MA/My,MOLF,MRL,NZB,PJ,PL,RF,RH2,SEA,SMJ,SPRET,TH,WSB | 8 | 123896912 | 123896930 | 19 | + | 17.1639 | 8.17E-07 | 0.141 | CCTCCACGAGGTGGCGGAA | | |
| 747 | 8 | 126825819 | 126841407 | 15588 | NZB | 8 | 126826791 | 126826824 | 34 | + | 16.5772 | 8.86E-07 | 0.0942 | TTGCAACATGCTTTCTAACCACCAGGGGTCTCCA | 8 | ENCFF821BIA,ENCFF093ZRU,ENCFF024JOZ,ENCFF201JUA,ENCFF327LYS,ENCFF883UPM,ENCFF367UNR,ENCFF099QNE |
| 748 | 8 | 3949469 | 3951518 | 2049 | SPRET | 8 | 3951232 | 3951265 | 34 | - | 16.8943 | 7.24E-07 | 0.085 | ATGCAGCTAGAGACAAGAGCTCCGGGGGGCACTG | | |
| 749 | 8 | 5337936 | 5340543 | 2607 | 129S1.NZB | 8 | 5339918 | 5339951 | 34 | - | 16.5366 | 9.09E-07 | 0.0944 | TAGCTGTCTTCAGAAAGACCAGAAGAGGGCATCA | | |
| 750 | 8 | 7244130 | 7246974 | 2844 | SPRET | 8 | 7246786 | 7246819 | 34 | - | 17.5041 | 4.89E-07 | 0.0702 | TTGCTGTCTTCAAACACACCAGAAGATGGCATCC | | |
| 751 | 8 | 9799577 | 9801196 | 1619 | SPRET | 8 | 9800611 | 9800644 | 34 | - | 18.4715 | 2.57E-07 | 0.0522 | GTGCAGTGCCCAAGAAGGCCAAAGGAGGGTGTGG | 1 | ENCFF093ZRU |
| 752 | 8 | 11639416 | 11645962 | 6546 | SPRET | 8 | 11641717 | 11641735 | 19 | - | 17.8852 | 5.08E-07 | 0.113 | ACACCAGAAGAGGGCACCA | 2 | ENCFF502WDT.ENCFF883UPM |
| 752 | 8 | 11639416 | 11645962 | 6546 | SPRET | 8 | 11641717 | 11641750 | 34 | - | 18.3984 | 2.70E-07 | 0.0529 | TAGCTGTCTTCAGACACACCAGAAGAGGGCACCA | 2 | ENCFF502WDT.ENCFF883UPM |
| 753 | 8 | 13592370 | 13594401 | 2031 | C57L,ILNJ2,MOLF,NZB,NZW,PJ,PL,SEA,SMJ,TH | 8 | 13592688 | 13592722 | 35 | - | 16.5229 | 7.12E-07 | 0.145 | CTGCAGGTCCCCGGAAAAGACACAAGGTGCTCCTT | | |
| 754 | 8 | 13760404 | 13765757 | 5353 | AKR,BALB,C57L,CAST,CE,ILNJ2,LP,MOLF,MRL,NZB,NZW,PJ,PL,RF,RH2,SEA,SMJ,SPRET,TH,WSB | 8 | 13760378 | 13760411 | 34 | - | 16.4959 | 9.32E-07 | 0.0944 | ATCTGGTGCCTGAGGAGGCCAGAAGAGGGCGTCA | | |
| 755 | 8 | 18071688 | 18073078 | 1390 | WSB | 8 | 18072336 | 18072354 | 19 | - | 17.9508 | 4.86E-07 | 0.113 | TGGCCTCTAGAGGGTACTG | | |
| 756 | 8 | 26185085 | 26187174 | 2089 | SPRET | 8 | 26186233 | 26186266 | 34 | - | 17.4878 | 4.94E-07 | 0.0707 | TAGCTGTCCTTAGACACACCAGAAGAGGGCATCA | | |
| 757 | 8 | 26310845 | 26312704 | 1859 | CAST | 8 | 26311536 | 26311570 | 35 | - | 22.1743 | 2.02E-08 | 0.0192 | CTGCATGTACATCTGCATACCAGAAGAGGGCATTG | 3 | ENCFF367UNR.ENCFF055HAP.ENCFF093ZRU |
| 758 | 8 | 30113636 | 30149023 | 35387 | 129S1,AJ,AKR,BALB,BTBR,BUB,C3H,C57L,CAST,CBA,CE,DBA1J,DBA2J,FVB,KK,MA/My,MOLF,MRL,NOD,NOR,NZB,NZO,NZW,RF,RH2,SJL,SMJ,SPRET,SWR,TH,WSB | 8 | 30128974 | 30129007 | 34 | - | 19.6748 | 1.12E-07 | 0.0314 | CTGCAAGAGGAGAGCTTGCCTGCAGAGGGTACTG | | |
| 758 | 8 | 30113636 | 30149023 | 35387 | 129S1,AJ,AKR,BALB,BTBR,BUB,C3H,C57L,CAST,CBA,CE,DBA1J,DBA2J,FVB,KK,MA/My,MOLF,MRL,NOD,NOR,NZB,NZO,NZW,RF,RH2,SJL,SMJ,SPRET,SWR,TH,WSB | 8 | 30129120 | 30129138 | 19 | - | 17.5902 | 6.18E-07 | 0.125 | CAGCCAGAAGGTGGAACTG | | |
| 759 | 8 | 30937812 | 30940317 | 2505 | 129S1,AJ,AKR,BALB,BTBR,BUB,C3H,C57L,CBA,DBA1J,DBA2J,FVB,KK,MA/My,MRL,NOD,NOR,NZW,RF,RH2,SJL,SMJ,SWR,TH,WSB | 8 | 30938561 | 30938594 | 34 | - | 17.2439 | 5.79E-07 | 0.0757 | CTGCAGTCCCCACTCTCACCTCCTGGAAGCTGTC | | |
| 760 | 8 | 31432520 | 31435161 | 2641 | SPRET | 8 | 31432574 | 31432607 | 34 | - | 23.0407 | 8.73E-09 | 0.00499 | ATGCAGTGTCTATGAAAGCCAGGAGAGGGCGTTG | 1 | ENCFF093ZRU |
| 761 | 8 | 33679371 | 33682361 | 2990 | 129S1,AJ,AKR,BALB,BUB,C57L,CAST,CBA,FVB,MA/My,MOLF,MRL,NOD,NOR,PL,RF,RH2,SJL,SMJ,SPRET,SWR,TH,WSB | 8 | 33679685 | 33679718 | 34 | - | 17.3171 | 5.52E-07 | 0.0747 | ATGAAGTGTCCATGACAGCCAGAAGAGGGGGTCA | | |
| 762 | 8 | 33721334 | 33722796 | 1462 | AJ,AKR,BALB,BUB,C3H,CBA,DBA1J,DBA2J,FVB,MA/My,MRL,NOD,NOR,PL,RF,RH2,SJL,SMJ.SWR.TH | 8 | 33721613 | 33721646 | 34 | - | 17.1057 | 6.33E-07 | 0.0794 | TAGCTGTCTCCAGACACTCCAGAAGAGGGTGCCA | | |
| 763 | 8 | 34057198 | 34059014 | 1816 | WSB | 8 | 34057771 | 34057805 | 35 | - | 16.3211 | 7.98E-07 | 0.153 | CTGTATGTACATTTACACATCAGAAGGTGGCACTG | | |
| 764 | 8 | 41899353 | 41902713 | 3360 | 129S1,AJ,AKR,BALB,BTBR,BUB,C57L,CAST,CBA,CE,FVB,ILNJ2,KK,MA/My,MOLF,MRL,NOD,NOR,NZB,NZO,NZW,PJ,PL,RF,RH2,SEA,SJL,SMJ,SPRET,SWR,TH,WSB | 8 | 41901617 | 41901651 | 35 | - | 17.6055 | 3.80E-07 | 0.109 | CTGCACTCCCACCTTCTTACCTGTTGGGGCATCTT | | |
| 765 | 8 | 44073189 | 44098987 | 25798 | MOLF | 8 | 44076507 | 44076540 | 34 | - | 17.1301 | 6.23E-07 | 0.0789 | CAGCTGTCCTCAGACATCCCAGAAGAGGGAATCA | | |
| 766 | 8 | 55387673 | 55393568 | 5895 | SPRET | 8 | 55393240 | 55393273 | 34 | - | 17.4472 | 5.08E-07 | 0.0719 | CAGCACTCCCACTTTCGCCCAGTCCAGGGCACCC | | |
| 767 | 8 | 58884948 | 58895372 | 10424 | SPRET | 8 | 58890047 | 58890080 | 34 | - | 16.8699 | 7.36E-07 | 0.0858 | ATGCAACTAGAGGCATGAGCTCCAGGGGGTACTG | | |
| 768 | 8 | 61451998 | 61459097 | 7099 | CAST.KK.MOLF.SPRET | 8 | 61459055 | 61459088 | 34 | - | 19.3415 | 1.42E-07 | 0.036 | CTGCCCCCACATTCGCCGTCACAAGATGGCGCTG | | |
| 769 | 8 | 64402735 | 64404140 | 1405 | AKR,BALB,C3H,C57L,CBA,DBA1J,DBA2J,MRL,PL,RF,SMJ,SPRET | 8 | 64403904 | 64403937 | 34 | - | 18.3333 | 2.82E-07 | 0.0534 | CAGCCTTTCCCTAATCAAACAACAGGGGGCAGCT | | |
| 770 | 8 | 65403291 | 65404549 | 1258 | SPRET | 8 | 65403641 | 65403659 | 19 | - | 19.7213 | 1.37E-07 | 0.0604 | AGGCCAGAAGAGGGCAGCA | 1 | ENCFF502WDT |
| 770 | 8 | 65403291 | 65404549 | 1258 | SPRET | 8 | 65403641 | 65403675 | 35 | - | 17.7982 | 3.39E-07 | 0.103 | ATGGAGGTGCTCAATGAGGCCAGAAGAGGGCAGCA | 1 | ENCFF502WDT |
| 771 | 8 | 66710028 | 66713777 | 3749 | SPRET | 8 | 66713691 | 66713724 | 34 | - | 18.6098 | 2.34E-07 | 0.0504 | CTGCTGTCTTCAGACACACCAGAAGAGGGTGTCA | | |
| 772 | 8 | 72778628 | 72786928 | 8300 | 129S1,BALB,BUB,C58J,CAST,CBA,DBA1J,FVB.NZO.PJ.RF.SEA.SMJ.SWR.WSB | 8 | 72778911 | 72778929 | 19 | - | 19.2951 | 1.88E-07 | 0.0709 | CCGCCACTAGATGGTGCTA | 4 | ENCFF367UNR,ENCFF099QNE,ENCFF093ZRU,ENCFF821BIA |
| 772 | 8 | 72778628 | 72786928 | 8300 | 129S1,BALB,BUB,C58J,CAST,CBA,DBA1J,FVB,NZO,PJ,RF,SEA,SMJ,SWR,WSB | 8 | 72786885 | 72786903 | 19 | - | 19.2951 | 1.88E-07 | 0.0709 | CCGCCACTAGATGGTGCTA | 13 | ENCFF367UNR,ENCFF093ZRU,ENCFF430PPJ,ENCFF198SSR,ENCFF099QNE,ENCFF907XEG,ENCFF589ZBJ,ENCFF136ZDK,ENCFF821BIA,ENCFF327LYS,ENCFF089GBC,ENCFF165LXO,ENCFF092NMJ |
| 773 | 8 | 79669197 | 79671255 | 2058 | SPRET | 8 | 79670432 | 79670466 | 35 | - | 19.3303 | 1.33E-07 | 0.0605 | CAGCAGTGAGTAGGAGCTGCCAGCAGGTGGATTCC | | |
| 774 | 8 | 85375149 | 85376950 | 1801 | SPRET | 8 | 85376274 | 85376307 | 34 | - | 16.7073 | 8.16E-07 | 0.09 | TAGCTGTATTCAGACACACCAGAAGAGGGCGTCA | | |
| 775 | 8 | 85863834 | 85868848 | 5014 | 129S1,AJ,AKR,BALB,BTBR,C3H,C57L,C58J,CAST,CBA,CE,DBA1J,DBA2J,FVB,ILNJ2,KK,MA/My,MOLF,MRL,NOD,NOR,NZW,PJ,PL,RF,RH2,SEA,SJL,SMJ,SPRET,SWR,WSB | 8 | 85867676 | 85867710 | 35 | - | 17.3761 | 4.35E-07 | 0.118 | GTGCGGGTCAAGCCCACTACCAGTCGGTGGAGGTC | | |
| 776 | 8 | 85865458 | 85868451 | 2993 | 129S1,AKR,BALB,BTBR,C57L,MRL,PJ,PL,SEA,WSB | 8 | 85867676 | 85867710 | 35 | - | 17.3761 | 4.35E-07 | 0.118 | GTGCGGGTCAAGCCCACTACCAGTCGGTGGAGGTC | | |
| 777 | 8 | 86901570 | 86906928 | 5358 | AJ,C3H,C58J,CAST,CBA,CE,DBA1J,DBA2J,FVB,ILNJ2,KK,MA/My,MOLF,RH2,SMJ,SPRET,TH | 8 | 86906797 | 86906830 | 34 | - | 16.9919 | 6.81E-07 | 0.083 | TAGCTGTCTTCAGACACACCAGAAGGGGGCATCA | | |
| 778 | 8 | 92889861 | 92893242 | 3381 | SPRET | 8 | 92892150 | 92892183 | 34 | - | 17.1463 | 6.17E-07 | 0.0784 | GTGCAGTGTGCTCAGAGCACACTCGAGGGCGGGG | | |
| 779 | 8 | 96521062 | 96523990 | 2928 | SPRET | 8 | 96522471 | 96522505 | 35 | - | 21.7523 | 2.70E-08 | 0.0227 | CTGCATGTACAACTGTGGGCCAGAAGGAGGCATCA | 1 | ENCFF093ZRU |
| 780 | 8 | 116320524 | 116321770 | 1246 | NZB | 8 | 116321464 | 116321498 | 35 | - | 16.1284 | 8.89E-07 | 0.162 | ATGACCTCCCATCTGCAGGCCAGCAGAGAGCGGTC | | |
| 781 | 8 | 120616300 | 120623780 | 7480 | CAST.MOLF.NZB.SPRET.WSB | 8 | 120619854 | 120619872 | 19 | - | 17.1639 | 8.17E-07 | 0.141 | CCTCCACGAGGTGGCGGAA | | |
| 782 | 8 | 122544625 | 122545633 | 1008 | WSB | 8 | 122545257 | 122545290 | 34 | - | 16.7724 | 7.83E-07 | 0.088 | CAGCTGGGTTCTCACCGGCCTCTGGGTGTCAGTG | 1 | ENCFF093ZRU |
| 783 | 8 | 124035438 | 124037028 | 1590 | 129S1,AKR,BTBR,C3H,C57L,C58J,CBA,CE,DBA1J,DBA2J,LP,MA/My,MRL,PJ,RF,SMJ,SPRET,TH,WSB | 8 | 124035699 | 124035733 | 35 | - | 24.0275 | 5.22E-09 | 0.00885 | CTGCATGTACACCTGCACACCAGAAGAGGGCGTCA | 4 | ENCFF093ZRU,ENCFF089GBC,ENCFF136ZDK,ENCFF883UPM |
| 784 | 8 | 125193638 | 125196446 | 2808 | CAST | 8 | 125193771 | 125193804 | 34 | - | 17.6829 | 4.35E-07 | 0.0658 | ATGCAGTCATTCATGCAACCACAAGAGAGCATCA | 1 | ENCFF093ZRU |
| 785 | 8 | 125495743 | 125506407 | 10664 | SPRET | 8 | 125502682 | 125502700 | 19 | - | 19.6885 | 1.40E-07 | 0.0613 | AGCCCAGTAGGGGGCGCCC | 35 | ENCFF093ZRU,ENCFF367UNR,ENCFF099QNE,ENCFF033UQX,ENCFF327LYS,ENCFF805QBU,ENCFF024JOZ,ENCFF907XEG,ENCFF201JUA,ENCFF136ZDK,ENCFF821BIA,ENCFF089GBC,ENCFF065ODV,ENCFF951GOP,ENCFF165LXO,ENCFF609MDN,ENCFF198SSR,ENCFF589ZBJ,ENCFF097CAQ,ENCFF430PPJ,ENCFF055HAP,ENCFF092NMJ,ENCFF142CNG,ENCFF349BUL,ENCFF281QMV,ENCFF957SPH,ENCFF502WDT,ENCFF210BTO,ENCFF883UPM,ENCFF491RJK,ENCFF194HVL,ENCFF533APC,ENCFF316FYN,ENCFF575MGI,ENCFF052KVO |
| 785 | 8 | 125495743 | 125506407 | 10664 | SPRET | 8 | 125502682 | 125502715 | 34 | - | 23.9756 | 4.02E-09 | 0.00283 | CAGCATCTTCCTAAGAGCCCAGTAGGGGGCGCCC | 35 | ENCFF093ZRU,ENCFF367UNR,ENCFF099QNE,ENCFF033UQX,ENCFF327LYS,ENCFF805QBU,ENCFF024JOZ,ENCFF907XEG,ENCFF201JUA,ENCFF136ZDK,ENCFF821BIA,ENCFF089GBC,ENCFF065ODV,ENCFF951GOP,ENCFF165LXO,ENCFF609MDN,ENCFF198SSR,ENCFF589ZBJ,ENCFF097CAQ,ENCFF430PPJ,ENCFF055HAP,ENCFF092NMJ,ENCFF142CNG,ENCFF349BUL,ENCFF281QMV,ENCFF957SPH,ENCFF502WDT,ENCFF210BTO,ENCFF883UPM,ENCFF491RJK,ENCFF194HVL,ENCFF533APC,ENCFF316FYN,ENCFF575MGI,ENCFF052KVO |
| 786 | 8 | 125533216 | 125537813 | 4597 | AJ.BALB.ILNJ2.MOLF.RH2.SEA | 8 | 125533688 | 125533721 | 34 | - | 16.7724 | 7.83E-07 | 0.088 | TAGCTGTCTTCAGATGCACCAGAAGAGGGCGTCA | | |
| 786 | 8 | 125533216 | 125537813 | 4597 | AJ,BALB,ILNJ2,MOLF,RH2,SEA | 8 | 125537137 | 125537155 | 19 | - | 19.5738 | 1.53E-07 | 0.0643 | CCACCACCAGAGGGTGCTG | 19 | ENCFF201JUA,ENCFF092NMJ,ENCFF136ZDK,ENCFF367UNR,ENCFF093ZRU,ENCFF089GBC,ENCFF165LXO,ENCFF589ZBJ,ENCFF198SSR,ENCFF883UPM,ENCFF502WDT,ENCFF327LYS,ENCFF194HVL,ENCFF099QNE.ENCFF907XEG.ENCFF821BIA.ENCFF024JOZ.ENCFF055HAP.ENCFF957SPH |
| 787 | 9 | 5098696 | 5175487 | 76791 | MA/Mv | 9 | 5146773 | 5146806 | 34 | + | 17.7154 | 4.26E-07 | 0.0552 | ATGGGCCTACATCTGCAGCCAGCAGGTGGAGCTG | 2 | ENCFF883UPM.ENCFF327LYS |
| 787 | 9 | 5098696 | 5175487 | 76791 | MA/Mv | 9 | 5146788 | 5146806 | 19 | + | 20.6393 | 6.61E-08 | 0.0435 | CAGCCAGCAGGTGGAGCTG | 2 | ENCFF883UPM.ENCFF327LYS |
| 788 | 9 | 6657611 | 6664119 | 6508 | CAST.MOLF.SPRET.WSB | 9 | 6663711 | 6663729 | 19 | + | 18.4262 | 3.51E-07 | 0.0872 | GAGCCAGAAGATGGCGCCG | | |
| 789 | 9 | 8063513 | 8068095 | 4582 | CAST | 9 | 8067811 | 8067845 | 35 | + | 21.1468 | 4.08E-08 | 0.024 | CTGCTGTGCTGCTAACCTGCCACCAGAGGCCTAAT | | |
| 790 | 9 | 8659859 | 8667584 | 7725 | SPRET | 9 | 8660728 | 8660746 | 19 | + | 17.459 | 6.74E-07 | 0.116 | CAGCCAGGAGGTGGAGCTG | | |
| 791 | 9 | 13665525 | 13669697 | 4172 | CAST.MOLF.SPRET.WSB | 9 | 13666302 | 13666335 | 34 | + | 17.6748 | 4.37E-07 | 0.0556 | GAGCATGACCACAAGTCCACAGCAGGTGGCAACG | | |
| 792 | 9 | 17012843 | 17014481 | 1638 | SPRET | 9 | 17014181 | 17014214 | 34 | + | 21.6016 | 2.71E-08 | 0.00913 | CTTCAGTCTCCTGGGAGGCCAGAAGAGGGCATTG | | |
| 793 | 9 | 18995799 | 18997763 | 1964 | SPRET | 9 | 18996466 | 18996484 | 19 | + | 18.1803 | 4.16E-07 | 0.0944 | ACACCACCAGGGGTCACTA | 3 | ENCFF502WDT.ENCFF533APC.ENCFF055HAP |
| 794 | 9 | 19776248 | 19780347 | 4099 | SPRET | 9 | 19776952 | 19776985 | 34 | + | 17.5691 | 4.69E-07 | 0.058 | GAGCATGACCACAAGTCCACAGCAGGTGGCAACA | | |
| 795 | 9 | 22231248 | 22232771 | 1523 | CAST | 9 | 22231293 | 22231326 | 34 | + | 17.2114 | 5.91E-07 | 0.066 | TAGCTGTTTTCAGACACTCCAGAAGAGGGCATCA | 1 | ENCFF093ZRU |
| 796 | 9 | 25720147 | 25721637 | 1490 | CAST | 9 | 25721031 | 25721064 | 34 | + | 24.6585 | 2.24E-09 | 0.00145 | ATGCAGTGCCTGTAGAGACCAGAAGAGGGAGTCA | 2 | ENCFF093ZRU.ENCFF052KVO |
| 797 | 9 | 28751227 | 28754396 | 3169 | NZO | 9 | 28753692 | 28753726 | 35 | + | 16.4771 | 7.30E-07 | 0.133 | GTGCAGGTGCCCCATTAAGCCATCAGTTGGCTCTT | | |
| 798 | 9 | 37673166 | 37678573 | 5407 | SPRET | 9 | 37673830 | 37673848 | 19 | + | 17.459 | 6.74E-07 | 0.116 | CAGCCAGGAGGTGGAGCTG | | |
| 798 | 9 | 37673166 | 37678573 | 5407 | SPRET | 9 | 37674023 | 37674041 | 19 | + | 17.6721 | 5.85E-07 | 0.11 | CAGCCAGGAGGGGGTGCTG | 6 | ENCFF024JOZ.ENCFF327LYS.ENCFF055HAP.ENCFF502WDT.ENCFF093ZRU.ENCFF883UPM |
| 799 | 9 | 39965630 | 39966732 | 1102 | MOLF | 9 | 39966029 | 39966063 | 35 | + | 18.156 | 2.74E-07 | 0.0779 | TTGCCTGTGTCCATGGAAGCCAGAAGAGGGCACTG | | |
| 799 | 9 | 39965630 | 39966732 | 1102 | MOLF | 9 | 39966045 | 39966063 | 19 | + | 18.4754 | 3.39E-07 | 0.0863 | AAGCCAGAAGAGGGCACTG | | |
| 800 | 9 | 40138136 | 40142860 | 4724 | CAST | 9 | 40139563 | 40139581 | 19 | + | 17.3934 | 7.03E-07 | 0.119 | TGGACACAAGGTGGCGACA | 24 | ENCFF093ZRU,ENCFF201JUA,ENCFF367UNR,ENCFF099QNE,ENCFF821BIA,ENCFF327LYS,ENCFF907XEG,ENCFF430PPJ,ENCFF092NMJ,ENCFF089GBC,ENCFF024JOZ,ENCFF198SSR,ENCFF589ZBJ,ENCFF165LXO,ENCFF883UPM,ENCFF951GOP,ENCFF055HAP,ENCFF609MDN,ENCFF502WDT,ENCFF575MGI,ENCFF136ZDK,ENCFF281QMV,ENCFF052KVO,ENCFF504JDS |
| 801 | 9 | 46279685 | 46280711 | 1026 | KK | 9 | 46280487 | 46280521 | 35 | + | 20.2661 | 7.32E-08 | 0.0342 | GTGCAGGTGCTTTTGGAAGCCAGAAGAGGGTGGTG | | |
| 802 | 9 | 51876986 | 51880228 | 3242 | SPRET | 9 | 51877848 | 51877866 | 19 | + | 19.6393 | 1.45E-07 | 0.0584 | TCACCAGAAGAGGGCAGCA | 14 | ENCFF201JUA,ENCFF367UNR,ENCFF093ZRU,ENCFF089GBC,ENCFF883UPM,ENCFF099QNE,ENCFF136ZDK,ENCFF198SSR,ENCFF055HAP,ENCFF502WDT,ENCFF821BIA,ENCFF907XEG,ENCFF024JOZ,ENCFF327LYS |
| 803 | 9 | 53507569 | 53509256 | 1687 | MOLF | 9 | 53507778 | 53507811 | 34 | + | 20.4146 | 6.58E-08 | 0.0169 | TTGCAGTCTTCAGACACACCAGAAGAGGGCATCA | | |
| 803 | 9 | 53507569 | 53509256 | 1687 | MOLF | 9 | 53508079 | 53508112 | 34 | + | 20.4146 | 6.58E-08 | 0.0169 | TTGCAGTCTTCAGACACACCAGAAGAGGGCATCA | | |
| 804 | 9 | 54737926 | 54739518 | 1592 | MOLF | 9 | 54738092 | 54738110 | 19 | + | 17.082 | 8.61E-07 | 0.129 | TGGCCTCTGGGTGGCAGCA | 34 | ENCFF093ZRU,ENCFF201JUA,ENCFF367UNR,ENCFF052KVO,ENCFF099QNE,ENCFF024JOZ,ENCFF327LYS,ENCFF907XEG,ENCFF089GBC,ENCFF821BIA,ENCFF092NMJ,ENCFF589ZBJ,ENCFF065ODV,ENCFF136ZDK,ENCFF033UQX,ENCFF055HAP,ENCFF198SSR,ENCFF609MDN,ENCFF430PPJ,ENCFF502WDT,ENCFF957SPH,ENCFF142CNG,ENCFF097CAQ,ENCFF210BTO,ENCFF883UPM,ENCFF611HDQ,ENCFF165LXO,ENCFF491RJK,ENCFF533APC,ENCFF951GOP,ENCFF281QMV,ENCFF575MGI,ENCFF194HVL,ENCFF316FYN |
| 805 | 9 | 55700686 | 55705671 | 4985 | MOLF | 9 | 55704373 | 55704407 | 35 | + | 16.0092 | 9.50E-07 | 0.152 | GTGCAATGCTGCCTACTGTGCTGAAGAGGGACTCA | | |
| 806 | 9 | 57594100 | 57602718 | 8618 | SPRET | 9 | 57595526 | 57595559 | 34 | + | 27.6911 | 1.30E-10 | 0.000272 | GTGCAGCACCCACGGGGGCCAGAAGGGGGCATCA | 23 | ENCFF099QNE,ENCFF052KVO,ENCFF024JOZ,ENCFF165LXO,ENCFF367UNR,ENCFF093ZRU,ENCFF883UPM,ENCFF533APC,ENCFF821BIA,ENCFF491RJK,ENCFF194HVL,ENCFF055HAP,ENCFF502WDT,ENCFF907XEG,ENCFF136ZDK,ENCFF097CAQ,ENCFF065ODV,ENCFF327LYS,ENCFF198SSR,ENCFF589ZBJ,ENCFF210BTO,ENCFF957SPH,ENCFF142CNG |
| 806 | 9 | 57594100 | 57602718 | 8618 | SPRET | 9 | 57595541 | 57595559 | 19 | + | 17.5082 | 6.52E-07 | 0.114 | GGGCCAGAAGGGGGCATCA | 23 | ENCFF099QNE,ENCFF052KVO,ENCFF024JOZ,ENCFF165LXO,ENCFF367UNR,ENCFF093ZRU,ENCFF883UPM,ENCFF533APC,ENCFF821BIA,ENCFF491RJK,ENCFF194HVL,ENCFF055HAP,ENCFF502WDT,ENCFF907XEG,ENCFF136ZDK,ENCFF097CAQ,ENCFF065ODV,ENCFF327LYS,ENCFF198SSR,ENCFF589ZBJ,ENCFF210BTO,ENCFF957SPH,ENCFF142CNG |
| 807 | 9 | 58473517 | 58475642 | 2125 | SPRET | 9 | 58475133 | 58475166 | 34 | + | 20.7642 | 5.09E-08 | 0.014 | ATGCAATACCTGAAGAGGCCAGAAGAGAGTGCTG | 19 | ENCFF367UNR,ENCFF093ZRU,ENCFF089GBC,ENCFF092NMJ,ENCFF430PPJ,ENCFF165LXO,ENCFF198SSR,ENCFF099QNE,ENCFF052KVO,ENCFF907XEG,ENCFF055HAP,ENCFF821BIA,ENCFF589ZBJ,ENCFF502WDT,ENCFF065ODV,ENCFF327LYS,ENCFF883UPM,ENCFF024JOZ,ENCFF136ZDK |
| 808 | 9 | 67331274 | 67332593 | 1319 | CAST | 9 | 67332366 | 67332399 | 34 | + | 18.3984 | 2.70E-07 | 0.0438 | TTGCTGTCTTCAGACACACCAGAAGAGGGCATTG | | |
| 809 | 9 | 67605804 | 67610641 | 4837 | CAST | 9 | 67610020 | 67610054 | 35 | + | 16.0092 | 9.50E-07 | 0.152 | CTGCATGTACACCTGCACACCAGAAGAAGACATCA | | |
| 810 | 9 | 73256196 | 73259933 | 3737 | SPRET | 9 | 73259675 | 73259708 | 34 | + | 16.935 | 7.06E-07 | 0.0726 | CTTCGGTACCTTCCGATGCCAGAAGAAGGTGCCA | | |
| 811 | 9 | 84186186 | 84189692 | 3506 | 129S1,AJ,BALB,BTBR,C3H,C57L,CE,MRL,NOD,NOR,NZB,PJ,RH2,SEA,TH | 9 | 84186589 | 84186622 | 34 | + | 20.878 | 4.68E-08 | 0.0133 | CGCCAGTACCTGTGGAGGCCAGAAGGGGGCATCA | 12 | ENCFF367UNR,ENCFF821BIA,ENCFF198SSR,ENCFF089GBC,ENCFF907XEG,ENCFF055HAP,ENCFF589ZBJ,ENCFF327LYS,ENCFF883UPM,ENCFF093ZRU,ENCFF099QNE,ENCFF502WDT |
| 811 | 9 | 84186186 | 84189692 | 3506 | 129S1,AJ,BALB,BTBR,C3H,C57L,CE,MRL,NOD,NOR,NZB,PJ,RH2,SEA,TH | 9 | 84186604 | 84186622 | 19 | + | 17.4262 | 6.88E-07 | 0.118 | AGGCCAGAAGGGGGCATCA | 12 | ENCFF367UNR,ENCFF821BIA,ENCFF198SSR,ENCFF089GBC,ENCFF907XEG,ENCFF055HAP,ENCFF589ZBJ,ENCFF327LYS,ENCFF883UPM,ENCFF093ZRU,ENCFF099QNE,ENCFF502WDT |
| 812 | 9 | 85671193 | 85674693 | 3500 | SPRET | 9 | 85672374 | 85672407 | 34 | + | 18.3171 | 2.86E-07 | 0.0443 | CTGCAGTTCCCCCAGCAGCAGGTGGTGGGCAGGT | | |
| 813 | 9 | 86352592 | 86357632 | 5040 | SPRET | 9 | 86356432 | 86356466 | 35 | + | 17.7798 | 3.43E-07 | 0.0879 | CTGTTGCTTTCTTCAGACACCAGAAGAGGGCACCT | | |
| 814 | 9 | 87342032 | 87345185 | 3153 | CAST | 9 | 87343292 | 87343326 | 35 | + | 19.4037 | 1.27E-07 | 0.0494 | GTGTAGATACATGAAGAGGCCAGAAGAGGGCATTG | | |
| 815 | 9 | 88474227 | 88512163 | 37936 | CBA,DBA1J,DBA2J,FVB,MA/My,MOLF,MRL,NZO,NZW,PL,RF,SJL,SWR,WSB | 9 | 88475350 | 88475384 | 35 | + | 21.1743 | 4.01E-08 | 0.0238 | TTGCATGCACACCTGCATGCCAGAAGAGGGCATCA | | |
| 815 | 9 | 88474227 | 88512163 | 37936 | CBA,DBA1J,DBA2J,FVB,MA/My,MOLF,MRL,NZO,NZW,PL,RF,SJL,SWR,WSB | 9 | 88483475 | 88483509 | 35 | + | 16 | 9.55E-07 | 0.152 | TAGCTGTCTAAACAACACCCCAGAAGAGGGCATCA | | |
| 815 | 9 | 88474227 | 88512163 | 37936 | CBA,DBA1J,DBA2J,FVB,MA/My,MOLF,MRL,NZO,NZW,PL,RF,SJL,SWR,WSB | 9 | 88493910 | 88493943 | 34 | + | 18.4797 | 2.56E-07 | 0.0428 | TTGCTGTCTTCAGACACACCAGAAGAGGGCATCA | | |
| 816 | 9 | 98775914 | 98791030 | 15116 | AJ,BALB,C3H,CBA,MOLF,MRL,NOD,NOR,NZO.NZW.PJ.RH2.SJL.SPRET.SWR | 9 | 98778903 | 98778937 | 35 | + | 23.9083 | 5.71E-09 | 0.00787 | CTGCAGGAACAAGAACCAACCAGAAGGTGGCACGC | | |
| 816 | 9 | 98775914 | 98791030 | 15116 | AJ,BALB,C3H,CBA,MOLF,MRL,NOD,NOR,NZO,NZW,PJ,RH2,SJL,SPRET,SWR | 9 | 98778919 | 98778937 | 19 | + | 17 | 9.07E-07 | 0.131 | CAACCAGAAGGTGGCACGC | | |

| | | | | | | | | | | | | | | | | |
|---|---|---|---|---|---|---|---|---|---|---|---|---|---|---|---|---|
| 817 | 9 | 99128443 | 99131204 | 2761 | SPRET | 9 | 99129526 | 99129559 | 34 | + | 19.0407 | 1.74E-07 | 0.0344 | GAGCGGCTTCCTGTGTGGCCAGAAGAGGGGACTG | | |
| 817 | 9 | 99128443 | 99131204 | 2761 | SPRET | 9 | 99130599 | 99130633 | 35 | + | 16.3761 | 7.73E-07 | 0.137 | CTGCTGTGGTCTAGAAAGGCCTCCAGATGCCTGGG | | |
| 818 | 9 | 99427414 | 99429695 | 2281 | SPRET | 9 | 99428467 | 99428501 | 35 | + | 18.9633 | 1.67E-07 | 0.0583 | TTGCATGTACACCTACATGTCACAAGAGGGCATCA | 6 | ENCFF821BIA,ENCFF883UPM,ENCFF367UNR,ENCFF055HAP,ENCFF502WDT,ENCFF093ZRU |
| 819 | 9 | 102792273 | 102795819 | 3546 | SMJ | 9 | 102794916 | 102794934 | 19 | + | 18.7049 | 2.88E-07 | 0.0804 | TGACCTGCAGAGGGAGCCA | 9 | ENCFF367UNR,ENCFF089GBC,ENCFF907XEG,ENCFF502WDT,ENCFF055HAP,ENCFF093ZRU,ENCFF198SSR,ENCFF024JOZ,ENCFF327LYS |
| 820 | 9 | 103463291 | 103464850 | 1559 | 129S1,AJ,BALB,BTBR,BUB,C3H,C57L,CAST,CBA,FVB,ILNJ2,NOD,NOR,NZB,NZW,PL,RH2,SMJ | 9 | 103463698 | 103463731 | 34 | + | 16.6585 | 8.41E-07 | 0.0807 | GAGCAAATCCTGCCCCTGGCGGCAGGGGTTGGCC | | |
| 821 | 9 | 105091140 | 105093238 | 2098 | 129S1,AJ,AKR,BALB,BTBR,BUB,C3H,C57L,CBA,CE,DBA1J,DBA2J,FVB,ILNJ2,KK,MRL,NOD,NOR,NZB,NZO,NZW,PJ,PL,RF,RH2,SEA,SJL,SMJ,SWR,TH,WSB | 9 | 105091878 | 105091911 | 34 | + | 17.7317 | 4.21E-07 | 0.055 | CTGGAGTTTCTGCAAAAGCCAGAAGAGGACAAGC | 11 | ENCFF093ZRU,ENCFF198SSR,ENCFF092NMJ,ENCFF430PPJ,ENCFF099QNE,ENCFF883UPM,ENCFF575MGI,ENCFF327LYS,ENCFF491RJK,ENCFF907XEG,ENCFF089GBC |
| 822 | 9 | 105735295 | 105737220 | 1925 | SPRET | 9 | 105736082 | 105736115 | 34 | + | 16.5935 | 8.77E-07 | 0.0825 | CTGATCATCCTGCCTTGACCTCCAGAGGGCTTAA | | |
| 823 | 9 | 107684620 | 107687707 | 3087 | AKR,DBA1J,DBA2J,KK,NZB,NZO,RF | 9 | 107686862 | 107686895 | 34 | + | 18.8537 | 1.98E-07 | 0.0374 | GTGCAGTATCCTTGGAGGTCAGAAGAGGGTGTCA | | |
| 824 | 9 | 108579417 | 108580656 | 1239 | SPRET | 9 | 108579603 | 108579636 | 34 | + | 17.6992 | 4.30E-07 | 0.0552 | TAGCTGTTTTCAGACACACCAGAAGAGGGCATCA | | |
| 825 | 9 | 109233553 | 109237606 | 4053 | SPRET | 9 | 109235578 | 109235611 | 34 | + | 16.5854 | 8.81E-07 | 0.0825 | TTGGTCTCTTCAGACTCACCAGAAGAGGGCACCG | 2 | ENCFF093ZRU,ENCFF099QNE |
| 825 | 9 | 109233553 | 109237606 | 4053 | SPRET | 9 | 109235593 | 109235611 | 19 | + | 19.9672 | 1.13E-07 | 0.0516 | TCACCAGAAGAGGGCACCG | 2 | ENCFF093ZRU,ENCFF099QNE |
| 826 | 9 | 114254265 | 114256660 | 2395 | SPRET | 9 | 114254638 | 114254672 | 35 | + | 20.1927 | 7.68E-08 | 0.0356 | GTACAGTTGCAGGTGAAGGCCAGAAGAGGGGGCTG | 1 | ENCFF093ZRU |
| 826 | 9 | 114254265 | 114256660 | 2395 | SPRET | 9 | 114254833 | 114254866 | 34 | - | 23.5122 | 5.93E-09 | 0.00294 | GTGCAGTACCCAGGGAAGACAGAAGAGGGCATCA | 3 | ENCFF052KVO,ENCFF093ZRU,ENCFF327LYS |
| 827 | 9 | 115698240 | 115706809 | 8569 | CE | 9 | 115701742 | 115701775 | 34 | + | 24.3984 | 2.80E-09 | 0.00168 | ATGCAACATCCAAGGAGGCCAGAAGAGGGCGCCA | 13 | ENCFF367UNR,ENCFF821BIA,ENCFF024JOZ,ENCFF327LYS,ENCFF907XEG,ENCFF883UPM,ENCFF589ZBJ,ENCFF093ZRU,ENCFF957SPH,ENCFF052KVO,ENCFF033UQX,ENCFF099QNE,ENCFF136ZDK |
| 827 | 9 | 115698240 | 115706809 | 8569 | CE | 9 | 115701757 | 115701775 | 19 | + | 21.0984 | 4.51E-08 | 0.0361 | AGGCCAGAAGAGGGCGCCA | 14 | ENCFF367UNR,ENCFF821BIA,ENCFF024JOZ,ENCFF327LYS,ENCFF907XEG,ENCFF883UPM,ENCFF589ZBJ,ENCFF093ZRU,ENCFF957SPH,ENCFF052KVO,ENCFF033UQX,ENCFF099QNE,ENCFF136ZDK,ENCFF533APC |
| 827 | 9 | 115698240 | 115706809 | 8569 | CE | 9 | 115699436 | 115699454 | 19 | - | 18.9344 | 2.45E-07 | 0.075 | TTGTCAGTAGGGGGCACTG | 14 | ENCFF821BIA,ENCFF093ZRU,ENCFF092NMJ,ENCFF024JOZ,ENCFF089GBC,ENCFF430PPJ,ENCFF165LXO,ENCFF883UPM,ENCFF198SSR,ENCFF099QNE,ENCFF327LYS,ENCFF575MGI,ENCFF907XEG,ENCFF367UNR |
| 828 | 9 | 123725884 | 123727491 | 1607 | 129S1,C57L,DBA1J,DBA2J,MOLF | 9 | 123726341 | 123726374 | 34 | + | 24.3252 | 2.98E-09 | 0.00176 | CTGCAATTTCCGTTTTTCCCACCAGAGGGAGACT | 19 | ENCFF367UNR,ENCFF093ZRU,ENCFF099QNE,ENCFF957SPH,ENCFF821BIA,ENCFF089GBC,ENCFF024JOZ,ENCFF907XEG,ENCFF092NMJ,ENCFF327LYS,ENCFF198SSR,ENCFF136ZDK,ENCFF609MDN,ENCFF430PPJ,ENCFF589ZBJ,ENCFF281QMV,ENCFF883UPM,ENCFF165LXO,ENCFF951GOP |
| 829 | 9 | 124215413 | 124231407 | 15994 | SPRET | 9 | 124222290 | 124222323 | 34 | + | 20.0976 | 8.28E-08 | 0.02 | GTGTCATGCCCTGTGTGGCCAGTAGATGTCAGAA | 27 | ENCFF367UNR,ENCFF504JDS,ENCFF093ZRU,ENCFF201JUA,ENCFF052KVO,ENCFF099QNE,ENCFF430PPJ,ENCFF907XEG,ENCFF327LYS,ENCFF198SSR,ENCFF024JOZ,ENCFF589ZBJ,ENCFF165LXO,ENCFF821BIA,ENCFF055HAP,ENCFF092NMJ,ENCFF951GOP,ENCFF194HVL,ENCFF136ZDK,ENCFF957SPH,ENCFF065ODV,ENCFF089GBC,ENCFF321AGQ,ENCFF609MDN,ENCFF502WDT,ENCFF883UPM,ENCFF281QMV |
| 829 | 9 | 124215413 | 124231407 | 15994 | SPRET | 9 | 124222305 | 124222323 | 19 | + | 17.377 | 7.11E-07 | 0.12 | TGGCCAGTAGATGTCAGAA | 27 | ENCFF367UNR,ENCFF504JDS,ENCFF093ZRU,ENCFF201JUA,ENCFF052KVO,ENCFF099QNE,ENCFF430PPJ,ENCFF907XEG,ENCFF327LYS,ENCFF198SSR,ENCFF024JOZ,ENCFF589ZBJ,ENCFF165LXO,ENCFF821BIA,ENCFF055HAP,ENCFF092NMJ,ENCFF951GOP,ENCFF194HVL,ENCFF136ZDK,ENCFF957SPH,ENCFF065ODV,ENCFF089GBC,ENCFF321AGQ,ENCFF609MDN,ENCFF502WDT,ENCFF883UPM,ENCFF281QMV |
| 829 | 9 | 124215413 | 124231407 | 15994 | SPRET | 9 | 124223478 | 124223512 | 35 | - | 19.4771 | 1.21E-07 | 0.048 | CTGTGGTTCCCTGCTGTGGCCAGAAGAAGGTATCA | | |
| 830 | 9 | 8287923 | 8299496 | 11573 | AKR,MRL,RF | 9 | 8292742 | 8292775 | 34 | - | 18.0569 | 3.40E-07 | 0.0487 | ATGCAGCTAGAGACATGAGCTCCAGGGGGTACTG | | |
| 831 | 9 | 11811907 | 11826559 | 14652 | SPRET | 9 | 11813239 | 11813257 | 19 | - | 17.082 | 8.61E-07 | 0.129 | CGGCCACTCGGTGGCACAA | | |
| 832 | 9 | 12127434 | 12144036 | 16602 | SPRET | 9 | 12138704 | 12138722 | 19 | - | 20.5902 | 6.88E-08 | 4.47E-02 | TGGCCACCAGAGGGAGCCT | | |
| 833 | 9 | 12129514 | 12139476 | 9962 | AKR,MRL,RF,WSB | 9 | 12138704 | 12138722 | 19 | - | 20.5902 | 6.88E-08 | 0.0447 | TGGCCACCAGAGGGAGCCT | | |
| 834 | 9 | 16689217 | 16712607 | 23390 | RF | 9 | 16710670 | 16710704 | 35 | - | 18.5229 | 2.19E-07 | 0.0679 | CTGCAGGTATCCACAGAAGCCAGAAGAGGACATTT | | |
| 835 | 9 | 30470019 | 30472022 | 2003 | SPRET | 9 | 30470000 | 30470033 | 34 | - | 18.4065 | 2.69E-07 | 0.0438 | TAGCTGTCTCCAGATGCACCAGAAGAGGGCGTCA | | |
| 836 | 9 | 38324381 | 38329200 | 4819 | 129S1,AJ,AKR,BALB,BTBR,BUB,C3H,CAST,CBA,DBA2J,NOD,NOR,PL,RF,RH2,SJL,SMJ,SWR,TH,WSB | 9 | 38328402 | 38328435 | 34 | - | 16.5691 | 8.90E-07 | 0.0831 | GAGCATGACCACAAGTCCACAGCAGATGGCAACA | | |
| 837 | 9 | 39098325 | 39101524 | 3199 | SPRET | 9 | 39098481 | 39098515 | 35 | - | 16.4128 | 7.57E-07 | 0.136 | CTGCTGAACTAGAGTGGATCCAGCGGGGGTCACTG | | |
| 838 | 9 | 40501517 | 40503838 | 2321 | SPRET | 9 | 40503837 | 40503870 | 34 | - | 19.5203 | 1.25E-07 | 0.0274 | GTGCTGTCTTCAGACACACCAGAAGAGGGCATCG | | |
| 839 | 9 | 40543217 | 40544565 | 1348 | 129S1,AJ,BALB,BTBR,BUB,C3H,C57L,CBA,CE,DBA1J,FVB,ILNJ2,KK,MA/My,NZB,NZO,NZW,PJ,PL,RH2,SEA,SJL,SMJ,SWR,TH | 9 | 40544368 | 40544401 | 34 | - | 16.4472 | 9.61E-07 | 0.0841 | TAGCTGTCTTCAGACACACCAGAAGAGGGCGTCA | | |
| 840 | 9 | 42079372 | 42080794 | 1422 | BTBR,DBA1J,KK,MA/My,NZO | 9 | 42079964 | 42079997 | 34 | - | 16.4959 | 9.32E-07 | 0.0841 | CTGCCATTTTAAATAGGTCCAACAGAGGACACGG | | |
| 841 | 9 | 42228055 | 42230182 | 2127 | SPRET | 9 | 42228915 | 42228949 | 35 | - | 21.8899 | 2.46E-08 | 0.018 | CTGCTGTGTCACATGCACGTCGGTAGGTGGCAGAG | 7 | ENCFF821BIA,ENCFF883UPM,ENCFF093ZRU,ENCFF907XEG,ENCFF367UNR,ENCFF099QNE,ENCFF957SPH |
| 842 | 9 | 48264337 | 48268081 | 3744 | SPRET,WSB | 9 | 48266294 | 48266328 | 35 | - | 22.0367 | 2.22E-08 | 0.017 | ATGCAGAGACCACAGCCAAACACTAGGTGGAGCCA | 38 | ENCFF093ZRU,ENCFF805QBU,ENCFF201JUA,ENCFF099QNE,ENCFF821BIA,ENCFF504JDS,ENCFF136ZDK,ENCFF601JVE,ENCFF327LYS,ENCFF210BTO,ENCFF589ZBJ,ENCFF907XEG,ENCFF321AGQ,ENCFF024JOZ,ENCFF033UQX,ENCFF065ODV,ENCFF089GBC,ENCFF198SSR,ENCFF092NMJ,ENCFF502WDT,ENCFF055HAP,ENCFF165LXO,ENCFF951GOP,ENCFF430PPJ,ENCFF957SPH,ENCFF883UPM,ENCFF142CNG,ENCFF575MGI,ENCFF281QMV,ENCFF609MDN,ENCFF194HVL,ENCFF052KVO,ENCFF491RJK,ENCFF097CAQ,ENCFF316FYN,ENCFF611HDQ,ENCFF533APC,ENCFF089EWX |
| 843 | 9 | 50537573 | 50543309 | 5736 | 129S1,C57L,CAST,MOLF,SPRET | 9 | 50540792 | 50540825 | 34 | - | 16.4472 | 9.61E-07 | 0.0841 | TAGCTGTCTTCAGACACACCAGAAGAGGGCGTCA | | |
| 844 | 9 | 52806286 | 52822062 | 15776 | SPRET | 9 | 52811456 | 52811489 | 34 | - | 25.7886 | 8.12E-10 | 0.000743 | ATGCAGTGCCTGCGGAGGCCATAAGAGGGCGTTG | | |
| 845 | 9 | 53666763 | 53668800 | 2037 | SPRET | 9 | 53667350 | 53667383 | 34 | - | 22.7154 | 1.13E-08 | 0.00478 | ATGCAGTGCCTGTGGAAACCAGAAGAGGGTATCA | | |
| 846 | 9 | 66047483 | 66048500 | 1017 | FVB,MOLF,SJL | 9 | 66048462 | 66048495 | 34 | - | 16.7724 | 7.83E-07 | 0.0768 | TAGCTGTCTTCAGATGCACCAGAAGAGGGCGTCA | | |
| 847 | 9 | 70750468 | 70751749 | 1281 | SPRET | 9 | 70750764 | 70750782 | 19 | - | 18.4754 | 3.39E-07 | 0.0863 | AAGCCAGAAGAGGGCACTG | 15 | ENCFF821BIA,ENCFF099QNE,ENCFF327LYS,ENCFF065ODV,ENCFF198SSR,ENCFF367UNR,ENCFF957SPH,ENCFF589ZBJ,ENCFF502WDT,ENCFF093ZRU,ENCFF055HAP,ENCFF907XEG,ENCFF142CNG,ENCFF052KVO,ENCFF136ZDK |
| 847 | 9 | 70750468 | 70751749 | 1281 | SPRET | 9 | 70750764 | 70750797 | 34 | - | 26.0325 | 6.48E-10 | 0.000639 | ATGCAGGGCCCATGGAAGCCAGAAGAGGGCACTG | 15 | ENCFF821BIA,ENCFF099QNE,ENCFF327LYS,ENCFF065ODV,ENCFF198SSR,ENCFF367UNR,ENCFF957SPH,ENCFF589ZBJ,ENCFF502WDT,ENCFF093ZRU,ENCFF055HAP,ENCFF907XEG,ENCFF142CNG,ENCFF052KVO,ENCFF136ZDK |
| 848 | 9 | 90809291 | 90824213 | 14922 | AJ,AKR,BALB,C3H,CAST,CE,FVB,KK,NZW,RH2,SJL,SPRET,SWR,TH | 9 | 90820203 | 90820236 | 34 | - | 16.561 | 8.95E-07 | 0.0832 | TGGCAGGTCTCAGAGAAGCCGCTAGATGGCCCAC | | |
| 849 | 9 | 99437904 | 99442426 | 4522 | NOD,NOR | 9 | 99439557 | 99439590 | 34 | - | 21.0488 | 4.12E-08 | 0.0123 | CTGCTGTCTTCAGACACACCAGAAGAGGGCATCA | 1 | ENCFF093ZRU |
| 850 | 9 | 102722660 | 102726249 | 3589 | SPRET | 9 | 102723148 | 102723166 | 19 | - | 17.1148 | 8.43E-07 | 0.128 | AAGCCAGAAGGGGGCAGAG | | |
| 851 | 9 | 105359744 | 105363341 | 3597 | FVB,SJL,SWR | 9 | 105360222 | 105360240 | 19 | - | 17.3607 | 7.18E-07 | 0.12 | AGGCCAGAAGAGGGTGCCA | | |
| 852 | 9 | 109029516 | 109031200 | 1684 | CAST,CBA,KK,MOLF,NZB,NZO,NZW,SPRET | 9 | 109030439 | 109030472 | 34 | - | 19.6504 | 1.14E-07 | 0.0255 | TAGCTGTCCTCAGATACTCCAGAAGAGGGCACCA | | |
| 853 | 9 | 117816612 | 117825965 | 9353 | 129S1,AKR,BUB,C57L,KK,LP,MA/My,MRL,NZW,RF,RH2,WSB | 9 | 117824608 | 117824642 | 35 | - | 17.2385 | 4.72E-07 | 0.104 | GTGCACCTTCCCTGCCAGGCCTGCAGAGAGTGCTC | | |
| 854 | 9 | 117904826 | 117906636 | 1810 | 129S1,AJ,BALB,C3H,C57L,FVB,ILNJ2,KK,MOLF,NZW,PJ,SEA,SJL,SWR,TH | 9 | 117906615 | 117906633 | 19 | - | 19.7213 | 1.37E-07 | 0.056 | CTGCCAGCAGGGGGAAGTC | | |
| 855 | 9 | 118774763 | 118776084 | 1321 | MOLF | 9 | 118775253 | 118775286 | 34 | - | 18.3415 | 2.81E-07 | 0.0442 | CAGCACACCCAGAGCCAGCCTGGAGGTGGAGACA | | |
| 856 | 9 | 120628670 | 120630532 | 1862 | MOLF | 9 | 120629458 | 120629491 | 34 | - | 22.5366 | 1.31E-08 | 0.00531 | GTGTGGTGCCCCTTGAGGCCAGAAGAGGGCATCA | 1 | ENCFF367UNR |
| 857 | 9 | 122001219 | 122002925 | 1706 | SPRET | 9 | 122001445 | 122001478 | 34 | - | 17.4959 | 4.92E-07 | 0.0596 | GTGTCTTGCCCAAGGAGGCCACAAGAGGGTATTG | | |
| 858 | 9 | 122016531 | 122018498 | 1967 | CE | 9 | 122017972 | 122018005 | 34 | - | 21.7317 | 2.46E-08 | 0.00852 | CAGCAGTGCCTCTCTAGGCCACCAGGGGAAGTGC | 19 | ENCFF024JOZ,ENCFF136ZDK,ENCFF821BIA,ENCFF089GBC,ENCFF092NMJ,ENCFF367UNR,ENCFF093ZRU,ENCFF165LXO,ENCFF201JUA,ENCFF430PPJ,ENCFF055HAP,ENCFF589ZBJ,ENCFF502WDT,ENCFF052KVO,ENCFF907XEG,ENCFF099QNE,ENCFF327LYS,ENCFF065ODV,ENCFF198SSR |
| 859 | 9 | 123275284 | 123276751 | 1467 | 129S1,C57L,MOLF,SPRET | 9 | 123276362 | 123276380 | 19 | - | 21.1148 | 4.45E-08 | 0.0361 | AGGCCAGAAGGGGGCACTG | 10 | ENCFF327LYS,ENCFF092NMJ,ENCFF907XEG,ENCFF089GBC,ENCFF821BIA,ENCFF198SSR,ENCFF093ZRU,ENCFF367UNR,ENCFF055HAP,ENCFF502WDT |
| 859 | 9 | 123275284 | 123276751 | 1467 | 129S1,C57L,MOLF,SPRET | 9 | 123276362 | 123276395 | 34 | - | 27.8618 | 1.09E-10 | 0.000241 | GTGCAGTGTGCACAGAGGCCAGAAGGGGGCACTG | 10 | ENCFF327LYS,ENCFF092NMJ,ENCFF907XEG,ENCFF089GBC,ENCFF821BIA,ENCFF198SSR,ENCFF093ZRU,ENCFF367UNR,ENCFF055HAP,ENCFF502WDT |
| 860 | 10 | 5667712 | 5676755 | 9043 | 129S1,AJ,AKR,BALB,BTBR,BUB,C3H,C57L,CAST,CBA,CE,DBA1J,DBA2J,FVB,ILNJ2,KK,LP,MA/My,MOLF,MRL,NOD,NOR,NZB,NZO,NZW,PJ,PL,RF,RH2,SEA,SJL,SMJ,SPRET,SWR,TH,WSB | 10 | 5673168 | 5673186 | 19 | + | 17.1639 | 8.17E-07 | 0.153 | CCTCCACGAGGTGGCGGAA | | |
| 861 | 10 | 6119591 | 6125951 | 6360 | C3H,CBA,CE,DBA1J,DBA2J,FVB,LP,MA/My,MRL,NZB,NZO,NZW,SPRET,SWR | 10 | 6125158 | 6125191 | 34 | + | 16.4634 | 9.51E-07 | 0.108 | TAGCTGTCTTCAGACGCACCAGAAGAGGGCATCA | | |
| 862 | 10 | 7447953 | 7449580 | 1627 | BUB,CAST,MOLF | 10 | 7449049 | 7449082 | 34 | + | 17.3008 | 5.58E-07 | 0.0844 | CTGTAGTCCCAGCACTGGGCACATGGAGGTAGGA | | |
| 863 | 10 | 12857934 | 12859663 | 1729 | CAST | 10 | 12858646 | 12858679 | 34 | + | 17.878 | 3.83E-07 | 0.0708 | TAGCTGTCCTCAGACACACCAGAAGAGGGCATCA | | |
| 864 | 10 | 18132065 | 18133292 | 1227 | CAST | 10 | 18133084 | 18133117 | 34 | + | 17.813 | 3.99E-07 | 0.0721 | TAGCTGTTTCCAGACACACCAGAAGAGGGCATGA | | |
| 865 | 10 | 20247050 | 20249241 | 2191 | FVB,NOD,NOR,TH,WSB | 10 | 20247939 | 20247973 | 35 | + | 20.8899 | 4.85E-08 | 0.0343 | GTGAAGTTACTCACAGAGACCACAAGAGGGTGCTG | 9 | ENCFF367UNR,ENCFF093ZRU,ENCFF907XEG,ENCFF055HAP,ENCFF821BIA,ENCFF099QNE,ENCFF533APC,ENCFF052KVO,ENCFF502WDT |
| 866 | 10 | 22007795 | 22015278 | 7483 | LP,MOLF | 10 | 22010389 | 22010423 | 35 | + | 16.1743 | 8.66E-07 | 0.176 | TGGCAGTTCAGTTCATAGGTCAGCAGAGGCAGCTC | | |
| 867 | 10 | 22381714 | 22393564 | 11850 | AJ,BTBR,C57L,FVB,NOD,NZO | 10 | 22387247 | 22387280 | 34 | + | 17.748 | 4.17E-07 | 0.0738 | GTGCTGTTTCTGATTGGTCCACAGGGTGAAGCTC | | |
| 868 | 10 | 22394322 | 22397009 | 2687 | 129S1,AJ,BALB,BTBR,BUB,C57L,FVB,KK,MA/My,NOD,NOR,NZO,PL,RH2,SJL | 10 | 22395937 | 22395970 | 34 | + | 18.6748 | 2.24E-07 | 0.0551 | ATGCTGTCTTCAGACACACCAGAAGAGGGCATCA | | |
| 869 | 10 | 22395747 | 22419742 | 23995 | CAST | 10 | 22395937 | 22395970 | 34 | + | 18.6748 | 2.24E-07 | 0.0551 | ATGCTGTCTTCAGACACACCAGAAGAGGGCATCA | | |
| 869 | 10 | 22395747 | 22419742 | 23995 | CAST | 10 | 22397082 | 22397115 | 34 | - | 17.3577 | 5.38E-07 | 0.0832 | CAGCTGTTTTAAGACATACCAGCAGGGAGCATTG | | |
| 870 | 10 | 22610173 | 22619222 | 9049 | 129S1,AJ,AKR,BALB,BTBR,BUB,C57L,CAST,FVB,ILNJ2,KK,LP,MA/My,MOLF,NOD,NOR,NZB,NZO,NZW,PL,RF,RH2,SJL,SPRET,SWR,TH,WSB | 10 | 22615634 | 22615652 | 19 | + | 17.1639 | 8.17E-07 | 0.153 | CCTCCACGAGGTGGCGGAA | | |
| 871 | 10 | 23153987 | 23155187 | 1200 | 129S1,AKR,BTBR,C3H,C57L,FVB,ILNJ2,KK,LP,MRL,NOD,NZB,NZO,NZW,RF,SMJ,SWR,TH | 10 | 23154880 | 23154914 | 35 | + | 15.9908 | 9.60E-07 | 0.184 | CTGCACTGACCCCCTATGCCCACCTGGGTGGTCTT | | |
| 872 | 10 | 24724192 | 24730560 | 6368 | 129S1,BTBR,C57L,NZO,NZW | 10 | 24726443 | 24726477 | 35 | + | 17.4679 | 4.13E-07 | 0.121 | TGGCAGTGTGGCCAGCTACCCAGCAGATGCCTACC | | |
| 873 | 10 | 25611356 | 25613835 | 2479 | CAST,MOLF,SPRET | 10 | 25612189 | 25612222 | 34 | + | 16.7886 | 7.75E-07 | 0.0988 | CTGAAGTTCCTTCTGGTTACAGGAGGTGGCCACT | | |
| 874 | 10 | 33551633 | 33558228 | 6595 | SPRET | 10 | 33555743 | 33555777 | 35 | + | 16.4404 | 7.46E-07 | 0.164 | AAGAAGCTGGACCCAGTGGCCAGCAGATGGCCCAC | | |
| 874 | 10 | 33551633 | 33558228 | 6595 | SPRET | 10 | 33555759 | 33555777 | 19 | + | 17.0328 | 8.88E-07 | 0.158 | TGGCCAGCAGATGGCCCAC | | |
| 875 | 10 | 37935214 | 37939398 | 4184 | SPRET | 10 | 37935227 | 37935260 | 34 | + | 17.0569 | 6.53E-07 | 0.0917 | GAGCGGGTGCAGCTGCCGCCCCAAGATGGTGCCC | | |
| 875 | 10 | 37935214 | 37939398 | 4184 | SPRET | 10 | 37939048 | 37939081 | 34 | + | 16.9268 | 7.10E-07 | 0.095 | GAGCAGGTGTGGCCACCACCCCAAGATGGCACCT | | |
| 876 | 10 | 40688367 | 40691202 | 2835 | CAST | 10 | 40689812 | 40689830 | 19 | + | 18.7541 | 2.79E-07 | 0.093 | TATCCACTAGGGGGCAATC | 24 | ENCFF201JUA,ENCFF821BIA,ENCFF367UNR,ENCFF092NMJ,ENCFF093ZRU,ENCFF165LXO,ENCFF089GBC,ENCFF065ODV,ENCFF589ZBJ,ENCFF198SSR,ENCFF883UPM,ENCFF099QNE,ENCFF609MDN,ENCFF502WDT,ENCFF136ZDK,ENCFF327LYS,ENCFF097CAQ,ENCFF052KVO,ENCFF321AGQ,ENCFF907XEG,ENCFF142CNG,ENCFF957SPH,ENCFF055HAP,ENCFF024JOZ |
| 877 | 10 | 41312639 | 41321624 | 8985 | 129S1,BTBR,C57L,CAST,KK,MA/My,MOLF,SPRET,WSB | 10 | 41318056 | 41318074 | 19 | + | 17.1639 | 8.17E-07 | 0.153 | CCTCCACGAGGTGGCGGAA | | |
| 878 | 10 | 41928524 | 41931471 | 2947 | 129S1,BTBR,C57L,CAST,KK,MOLF,SPRET,WSB | 10 | 41929030 | 41929063 | 34 | + | 17.561 | 4.71E-07 | 0.0776 | CTGCCAAACCAGCACGCGCCAGGGGTGGGCACCA | | |
| 879 | 10 | 52216377 | 52221683 | 5306 | CAST,MOLF | 10 | 52219567 | 52219585 | 19 | + | 19.4918 | 1.63E-07 | 0.0697 | CTGCCTGCAGGTGGCAGTG | 6 | ENCFF093ZRU,ENCFF367UNR,ENCFF907XEG,ENCFF198SSR,ENCFF957SPH,ENCFF327LYS |
| 880 | 10 | 54812563 | 54814611 | 2048 | CAST,MOLF | 10 | 54813954 | 54813988 | 35 | + | 16.7064 | 6.41E-07 | 0.151 | CTGCATTGATCATCTTTTGAAACCAGGTGGTACTA | | |
| 881 | 10 | 58388653 | 58391888 | 3235 | CAST,MOLF | 10 | 58389942 | 58389975 | 34 | + | 18.4797 | 2.56E-07 | 0.0577 | TTGCTGTCTTCAGACACACCAGAAGAGGGCATCA | | |
| 882 | 10 | 59214691 | 59216163 | 1472 | SPRET | 10 | 59214726 | 59214759 | 34 | + | 28.6585 | 4.74E-11 | 0.000194 | GTGCAGTGCCTAAGGTAGCCAGAAGAGGGCATCA | 16 | ENCFF198SSR,ENCFF367UNR,ENCFF907XEG,ENCFF589ZBJ,ENCFF883UPM,ENCFF201JUA,ENCFF093ZRU,ENCFF533APC,ENCFF099QNE,ENCFF052KVO,ENCFF065ODV,ENCFF821BIA,ENCFF142CNG,ENCFF502WDT,ENCFF055HAP,ENCFF136ZDK |
| 882 | 10 | 59214691 | 59216163 | 1472 | SPRET | 10 | 59214741 | 59214759 | 19 | + | 17.1803 | 8.08E-07 | 0.152 | TAGCCAGAAGAGGGCATCA | 16 | ENCFF198SSR,ENCFF367UNR,ENCFF907XEG,ENCFF589ZBJ,ENCFF883UPM,ENCFF201JUA,ENCFF093ZRU,ENCFF533APC,ENCFF099QNE,ENCFF052KVO,ENCFF065ODV,ENCFF821BIA,ENCFF142CNG,ENCFF502WDT,ENCFF055HAP,ENCFF136ZDK |
| 883 | 10 | 60986730 | 60988464 | 1734 | MOLF | 10 | 60987643 | 60987676 | 34 | + | 17.6423 | 4.47E-07 | 0.0757 | CTGCAGTGCCTGTGCCTACCAACTGAGGAAAATA | | |
| 884 | 10 | 61298088 | 61305791 | 7703 | WSB | 10 | 61302877 | 61302895 | 19 | + | 18.3934 | 3.59E-07 | 0.107 | TTAACACTAGAGGGCGCTC | 31 | ENCFF201JUA,ENCFF093ZRU,ENCFF367UNR,ENCFF210BTO,ENCFF099QNE,ENCFF065ODV,ENCFF327LYS,ENCFF821BIA,ENCFF316FYN,ENCFF136ZDK,ENCFF907XEG,ENCFF589ZBJ,ENCFF502WDT,ENCFF092NMJ,ENCFF024JOZ,ENCFF089GBC,ENCFF055HAP,ENCFF052KVO,ENCFF198SSR,ENCFF194HVL,ENCFF097CAQ,ENCFF165LXO,ENCFF957SPH,ENCFF430PPJ,ENCFF281QMV,ENCFF951GOP,ENCFF504JDS,ENCFF142CNG,ENCFF575MGI,ENCFF609MDN,ENCFF883UPM |
| 884 | 10 | 61298088 | 61305791 | 7703 | WSB | 10 | 61302099 | 61302117 | 19 | - | 18.7869 | 2.72E-07 | 0.0925 | AAGCCAGAAGAGGGCACTA | 25 | ENCFF024JOZ,ENCFF089GBC,ENCFF327LYS,ENCFF092NMJ,ENCFF065ODV,ENCFF367UNR,ENCFF198SSR,ENCFF093ZRU,ENCFF165LXO,ENCFF883UPM,ENCFF210BTO,ENCFF589ZBJ,ENCFF502WDT,ENCFF052KVO,ENCFF821BIA,ENCFF957SPH,ENCFF033UQX,ENCFF055HAP,ENCFF281QMV,ENCFF136ZDK,ENCFF907XEG,ENCFF194HVL,ENCFF142CNG,ENCFF099QNE,ENCFF533APC |
| 884 | 10 | 61298088 | 61305791 | 7703 | WSB | 10 | 61302099 | 61302132 | 34 | - | 28.9024 | 3.64E-11 | 0.000168 | ATGCAGTGCCCACCAAAGCCAGAAGAGGGCACTA | 25 | ENCFF024JOZ,ENCFF089GBC,ENCFF327LYS,ENCFF092NMJ,ENCFF065ODV,ENCFF367UNR,ENCFF198SSR,ENCFF093ZRU,ENCFF165LXO,ENCFF883UPM,ENCFF210BTO,ENCFF589ZBJ,ENCFF502WDT,ENCFF052KVO,ENCFF821BIA,ENCFF957SPH,ENCFF033UQX,ENCFF055HAP,ENCFF281QMV,ENCFF136ZDK,ENCFF907XEG,ENCFF194HVL,ENCFF142CNG,ENCFF099QNE,ENCFF533APC |
| 885 | 10 | 63119672 | 63120857 | 1185 | SPRET | 10 | 63120847 | 63120880 | 34 | + | 18.7073 | 2.19E-07 | 0.0543 | CTGCAGTTCTCTCTCTGCCCACCAAATGCTGGAA | | |
| 886 | 10 | 71989189 | 71991317 | 2128 | SPRET | 10 | 71990555 | 71990589 | 35 | + | 16.6147 | 6.75E-07 | 0.155 | CTGTAGTTGATGGCCCAGGACTGAAGGGGTCACGC | | |
| 887 | 10 | 76065594 | 76121703 | 56109 | DBA1J | 10 | 76088758 | 76088792 | 35 | + | 16.3486 | 7.85E-07 | 0.168 | TTGCATGTACACCTCTACACTAGAAGGGGGCACCA | | |
| 887 | 10 | 76065594 | 76121703 | 56109 | DBA1J | 10 | 76107256 | 76107274 | 19 | + | 19.4918 | 1.63E-07 | 0.0697 | CGGCCAGCAGAGGGAGACA | 31 | ENCFF093ZRU,ENCFF367UNR,ENCFF430PPJ,ENCFF099QNE,ENCFF327LYS,ENCFF907XEG,ENCFF951GOP,ENCFF609MDN,ENCFF575MGI,ENCFF198SSR,ENCFF089GBC,ENCFF136ZDK,ENCFF092NMJ,ENCFF821BIA,ENCFF024JOZ,ENCFF165LXO,ENCFF097CAQ,ENCFF502WDT,ENCFF055HAP,ENCFF194HVL,ENCFF491RJK,ENCFF611HDQ,ENCFF957SPH,ENCFF533APC,ENCFF589ZBJ,ENCFF281QMV,ENCFF883UPM,ENCFF210BTO,ENCFF349BUL,ENCFF052KVO,ENCFF142CNG |
| 887 | 10 | 76065594 | 76121703 | 56109 | DBA1J | 10 | 76065646 | 76065664 | 19 | - | 20.3607 | 8.29E-08 | 0.0512 | AGGCCAGAAGAGGGCACCA | 15 | ENCFF093ZRU,ENCFF092NMJ,ENCFF367UNR,ENCFF089GBC,ENCFF589ZBJ,ENCFF099QNE,ENCFF198SSR,ENCFF327LYS,ENCFF065ODV,ENCFF024JOZ,ENCFF907XEG,ENCFF055HAP,ENCFF821BIA,ENCFF502WDT,ENCFF883UPM |
| 887 | 10 | 76065594 | 76121703 | 56109 | DBA1J | 10 | 76065646 | 76065680 | 35 | - | 22.7706 | 1.32E-08 | 0.0157 | GTGCTTGTACACCAGTAGGCCAGAAGAGGGCACCA | 16 | ENCFF093ZRU,ENCFF092NMJ,ENCFF367UNR,ENCFF089GBC,ENCFF589ZBJ,ENCFF099QNE,ENCFF198SSR,ENCFF327LYS,ENCFF065ODV,ENCFF024JOZ,ENCFF907XEG,ENCFF055HAP,ENCFF821BIA,ENCFF502WDT,ENCFF883UPM,ENCFF052KVO |
| 887 | 10 | 76065594 | 76121703 | 56109 | DBA1J | 10 | 76075414 | 76075447 | 34 | - | 18.2602 | 2.97E-07 | 0.0608 | TTCCAGTGCCCATGGAGGCCAGATGAGGGCATCA | 2 | ENCFF367UNR,ENCFF198SSR |
| 887 | 10 | 76065594 | 76121703 | 56109 | DBA1J | 10 | 76096696 | 76096729 | 34 | - | 17.7886 | 4.06E-07 | 0.0726 | CAGCCACTCACATTATGGACAGCAGAGGGAGCAA | 14 | ENCFF093ZRU,ENCFF821BIA,ENCFF367UNR,ENCFF883UPM,ENCFF099QNE,ENCFF089GBC,ENCFF609MDN,ENCFF327LYS,ENCFF281QMV,ENCFF907XEG,ENCFF024JOZ,ENCFF430PPJ,ENCFF092NMJ,ENCFF198SSR |
| 887 | 10 | 76065594 | 76121703 | 56109 | DBA1J | 10 | 76106573 | 76106606 | 34 | - | 17.1707 | 6.07E-07 | 0.0883 | CAGCACTGTTGTGTCAGGGCAGCAGAGGCCAGCA | | |
| 888 | 10 | 93446670 | 93452955 | 6285 | 129S1,AJ,AKR,BALB,BTBR,BUB,C3H,C57L,C58J,CBA,CE,DBA1J,DBA2J,FVB,ILNJ2,KK,LP,MA/My,NOD,NOR,NZO,NZW,PJ,PL,RF,RH2,SEA,SJL,SMJ,SWR,TH,WSB | 10 | 93447419 | 93447453 | 35 | + | 19.211 | 1.43E-07 | 0.0651 | GTTCACTGACAGCTAATCAACACTAGATGGCGCTC | 29 | ENCFF367UNR,ENCFF093ZRU,ENCFF099QNE,ENCFF201JUA,ENCFF821BIA,ENCFF210BTO,ENCFF089GBC,ENCFF024JOZ,ENCFF907XEG,ENCFF327LYS,ENCFF198SSR,ENCFF589ZBJ,ENCFF092NMJ,ENCFF136ZDK,ENCFF430PPJ,ENCFF165LXO,ENCFF504JDS,ENCFF089EWX,ENCFF194HVL,ENCFF491RJK,ENCFF097CAQ,ENCFF951GOP,ENCFF611HDQ,ENCFF281QMV,ENCFF575MGI,ENCFF957SPH,ENCFF883UPM,ENCFF609MDN,ENCFF316FYN |
| 888 | 10 | 93446670 | 93452955 | 6285 | 129S1,AJ,AKR,BALB,BTBR,BUB,C3H,C57L,C58J,CBA,CE,DBA1J,DBA2J,FVB,ILNJ2,KK,LP,MA/My,NOD,NOR,NZO,NZW,PJ,PL,RF,RH2,SEA,SJL,SMJ,SWR,TH,WSB | 10 | 93447435 | 93447453 | 19 | + | 17.459 | 6.74E-07 | 0.14 | TCAACACTAGATGGCGCTC | 29 | ENCFF367UNR,ENCFF093ZRU,ENCFF099QNE,ENCFF201JUA,ENCFF821BIA,ENCFF210BTO,ENCFF089GBC,ENCFF024JOZ,ENCFF907XEG,ENCFF327LYS,ENCFF198SSR,ENCFF589ZBJ,ENCFF092NMJ,ENCFF136ZDK,ENCFF430PPJ,ENCFF165LXO,ENCFF504JDS,ENCFF089EWX,ENCFF194HVL,ENCFF491RJK,ENCFF097CAQ,ENCFF951GOP,ENCFF611HDQ,ENCFF281QMV,ENCFF575MGI,ENCFF957SPH,ENCFF883UPM,ENCFF609MDN,ENCFF316FYN |
| 889 | 10 | 95890284 | 95894639 | 4355 | SPRET | 10 | 95892977 | 95893011 | 35 | + | 18.3578 | 2.43E-07 | 0.091 | CAGCTCTGTAACAGCCTGAACAGCAGGGGGTGGTG | | |
| 889 | 10 | 95890284 | 95894639 | 4355 | SPRET | 10 | 95892993 | 95893011 | 19 | + | 17.0984 | 8.52E-07 | 0.155 | TGAACAGCAGGGGGTGGTG | | |
| 890 | 10 | 99975145 | 99980410 | 5265 | AKR,BUB,C3H,CAST,CBA,FVB,MA/My,NOD,NOR,NZB,PL,RF,SMJ,SPRET,SWR,TH,WSB | 10 | 99977809 | 99977842 | 34 | + | 18.935 | 1.88E-07 | 0.0499 | CAGCTGTCTTCAAACGCACCAGAAGAGGGCATCA | | |
| 891 | 10 | 110661782 | 110692028 | 30246 | NZB | 10 | 110667962 | 110667996 | 35 | + | 16.3211 | 7.98E-07 | 0.169 | CTGCAGCATTAGGACGGTTACTCTAGAGGGAGGTC | | |
| 892 | 10 | 116598096 | 116599983 | 1887 | CAST,MOLF,SMJ,SPRET | 10 | 116599104 | 116599138 | 35 | + | 17.4862 | 4.08E-07 | 0.12 | TTGCAATTGAAGGTCCAGGCCTGAAGGGGTCATGC | | |
| 893 | 10 | 127806932 | 127808114 | 1182 | SPRET | 10 | 127807205 | 127807239 | 35 | + | 24.0734 | 5.04E-09 | 0.0097 | CTGCATGTACACCTGCATGCCAGGAGAGGGCATCA | 6 | ENCFF024JOZ,ENCFF907XEG,ENCFF055HAP,ENCFF883UPM,ENCFF327LYS,ENCFF502WDT |
| 894 | 10 | 129804350 | 129815678 | 11328 | CAST,KK,SMJ | 10 | 129809047 | 129809065 | 19 | + | 17 | 9.07E-07 | 0.159 | GCACCACTAGAGGGAGCCC | | |
| 895 | 10 | 3516435 | 3519398 | 2963 | ILNJ2 | 10 | 3518009 | 3518027 | 19 | - | 18.0492 | 4.54E-07 | 0.12 | GGGCCAGAAGAGGGCGATG | 19 | ENCFF024JOZ,ENCFF821BIA,ENCFF198SSR,ENCFF092NMJ,ENCFF165LXO,ENCFF589ZBJ,ENCFF907XEG,ENCFF883UPM,ENCFF093ZRU,ENCFF502WDT,ENCFF367UNR,ENCFF194HVL,ENCFF033UQX,ENCFF957SPH,ENCFF052KVO,ENCFF327LYS,ENCFF533APC,ENCFF136ZDK,ENCFF099QNE |
| 895 | 10 | 3516435 | 3519398 | 2963 | ILNJ2 | 10 | 3518009 | 3518042 | 34 | - | 26.3984 | 4.60E-10 | 0.000687 | GTGCAGTACTCATAGGGGCCAGAAGAGGGCGATG | 19 | ENCFF024JOZ,ENCFF821BIA,ENCFF198SSR,ENCFF092NMJ,ENCFF165LXO,ENCFF589ZBJ,ENCFF907XEG,ENCFF883UPM,ENCFF093ZRU,ENCFF502WDT,ENCFF367UNR,ENCFF194HVL,ENCFF033UQX,ENCFF957SPH,ENCFF052KVO,ENCFF327LYS,ENCFF533APC,ENCFF136ZDK,ENCFF099QNE |
| 896 | 10 | 7099533 | 7100924 | 1391 | MOLF | 10 | 7100736 | 7100770 | 35 | - | 18.211 | 2.65E-07 | 0.0951 | CTGCAGCTCCCATTTTGAACCTGGGGATGGAACTT | | |
| 897 | 10 | 7117723 | 7119121 | 1398 | SPRET | 10 | 7118878 | 7118911 | 34 | - | 21.5528 | 2.82E-08 | 0.0142 | GTGCAGTGCCTGTAGAGGCCAGTAAAGGCGTCA | | |
| 897 | 10 | 7117723 | 7119121 | 1398 | SPRET | 10 | 7118888 | 7118921 | 34 | - | 16.4553 | 9.56E-07 | 0.108 | GTGCACATCAGTGCAGTGCCTGTAGAGGCCAGTA | | |
| 898 | 10 | 7491528 | 7493638 | 2110 | SPRET | 10 | 7493312 | 7493330 | 19 | - | 17.5738 | 6.25E-07 | 0.134 | ACTCCAGAAGAGGGCGCTA | 1 | ENCFF093ZRU |
| 898 | 10 | 7491528 | 7493638 | 2110 | SPRET | 10 | 7493312 | 7493345 | 34 | - | 18.187 | 3.12E-07 | 0.0629 | TAGCTGTCTTCAGTCACTCCAGAAGAGGGCGCTA | 1 | ENCFF093ZRU |
| 899 | 10 | 8419894 | 8427032 | 7138 | 129S1,AJ,AKR,BALB,BTBR,BUB,C3H,C57L,CAST,CBA,CE,DBA1J,DBA2J,FVB,ILNJ2,KK,LP,MA/My,MOLF,MRL,NOD,NOR,NZB,NZO,NZW,PJ,PL,RF,RH2,SEA,SJL,SMJ,SPRET,SWR,TH,WSB | 10 | 8421638 | 8421656 | 19 | - | 17.1639 | 8.17E-07 | 0.153 | CCTCCACGAGGTGGCGGAA | | |
| 900 | 10 | 11362943 | 11368756 | 5813 | AJ,AKR,BALB,BTBR,C3H,C57L,CBA,CE,KK,MRL,PJ,PL,RF,RH2,SEA,SJL,SMJ,WSB | 10 | 11363201 | 11363234 | 34 | - | 16.8862 | 7.28E-07 | 0.096 | CCGCGATAACATTTGCCATCACAAGATGGCGCCG | 2 | ENCFF821BIA,ENCFF367UNR |
| 901 | 10 | 11364460 | 11369663 | 5203 | SPRET | 10 | 11369010 | 11369043 | 34 | - | 20.1463 | 7.99E-08 | 0.0296 | CCGCAATAACATTTGCCATCACAAGATGGCGCCG | 3 | ENCFF093ZRU,ENCFF367UNR,ENCFF821BIA |
| 902 | 10 | 14706214 | 14710819 | 4605 | CAST,DBA1J,DBA2J,MOLF | 10 | 14709800 | 14709833 | 34 | - | 16.439 | 9.66E-07 | 0.108 | ATGCAGCTAGAGACAAGAGCTCCAGGGGGTACTG | | |
| 903 | 10 | 19599501 | 19600982 | 1481 | SPRET | 10 | 19600934 | 19600952 | 19 | - | 17.5902 | 6.18E-07 | 0.134 | ACACCAGAAGAGGGCACTG | | |
| 903 | 10 | 19599501 | 19600982 | 1481 | SPRET | 10 | 19600934 | 19600967 | 34 | - | 18.3171 | 2.86E-07 | 0.0597 | TAGCTGTCTTCAGACACACCAGAAGAGGGCACTG | | |
| 904 | 10 | 22340289 | 22345866 | 5577 | SPRET | 10 | 22344365 | 22344399 | 35 | - | 16.7615 | 6.21E-07 | 0.148 | TAGCAGTTGCCACCTGTGTGCTCTTGGGGGCGGGG | | |
| 905 | 10 | 23335180 | 23339304 | 4124 | CAST,MOLF,SPRET | 10 | 23338502 | 23338535 | 34 | - | 17.6748 | 4.37E-07 | 0.0748 | GAGCATGACCACAAGTCCACAGCAGGTGGCAACG | | |
| 906 | 10 | 25427145 | 25428462 | 1317 | SPRET | 10 | 25427342 | 25427376 | 35 | - | 18.1193 | 2.80E-07 | 0.0976 | CTGCAGACACCCCCAAGGTCCCCTAGAGGACTCTC | | |
| 907 | 10 | 26888050 | 26889365 | 1315 | CAST | 10 | 26888960 | 26888994 | 35 | - | 20.1193 | 8.05E-08 | 0.0461 | AAGCAGCCCCTAAAGCTGTCCACAAGGTGGCATCC | 6 | ENCFF093ZRU,ENCFF367UNR,ENCFF201JUA,ENCFF883UPM,ENCFF055HAP,ENCFF502WDT |
| 908 | 10 | 36646275 | 36647606 | 1331 | MOLF | 10 | 36647026 | 36647060 | 35 | - | 16.3945 | 7.65E-07 | 0.165 | CTGCAGATATGTCTGGAAACCACCTGAGGGCCTGG | | |
| 909 | 10 | 41097575 | 41098692 | 1117 | MOLF | 10 | 41097925 | 41097943 | 19 | - | 18.4754 | 3.39E-07 | 0.104 | AAGCCAGAAGAGGGCACTG | | |
| 909 | 10 | 41097575 | 41098692 | 1117 | MOLF | 10 | 41097925 | 41097959 | 35 | - | 18.1009 | 2.83E-07 | 0.0983 | GAGCAGAAGAGCAGGGAAGCCAGAAGAGGGCACTG | | |
| 910 | 10 | 41611837 | 41613032 | 1195 | MOLF | 10 | 41612766 | 41612800 | 35 | - | 21.0826 | 4.26E-08 | 0.0317 | GTGTAGGTTCCCAAATAGGCCAGAAGAGGGCATTG | | |
| 911 | 10 | 41737996 | 41739278 | 1282 | CAST,MOLF | 10 | 41738097 | 41738131 | 35 | - | 16.9174 | 5.68E-07 | 0.14 | TTGCTTGTATGCATGCAGGCCAGATGAGGGCACCA | | |
| 912 | 10 | 45237286 | 45239471 | 2185 | CAST | 10 | 45238077 | 45238111 | 35 | - | 15.9266 | 9.95E-07 | 0.188 | GTGAACTGTCTCTAACTGGCCTGTTGAGGGCAGGA | | |
| 913 | 10 | 52484780 | 52487316 | 2536 | SPRET | 10 | 52485842 | 52485860 | 19 | - | 17.459 | 6.74E-07 | 0.14 | CAGCCAGGAGGTGGAGCTG | | |

| | | | | | | | | | | | | | | | | |
|---|---|---|---|---|---|---|---|---|---|---|---|---|---|---|---|---|
| 914 | 10 | 53350633 | 53354217 | 3584 | CAST,MOLF | 10 | 53351267 | 53351301 | 35 | - | 16.4404 | 7.46E-07 | 0.164 | GTGTAGATGACGCAGCGCACCAGAAGAGGACAGCG | | |
| 915 | 10 | 54762828 | 54767149 | 4321 | WSB | 10 | 54766569 | 54766602 | 34 | - | 17.6748 | 4.37E-07 | 0.0748 | GAGCATGACCACAAGTCCACAGCAGGTGGCAACG | | |
| 916 | 10 | 58138270 | 58139687 | 1417 | SPRET | 10 | 58139138 | 58139171 | 34 | - | 20.6911 | 5.37E-08 | 0.0224 | GTGCAACACCTGTAGAGGTCAGAAGAGGGCATCA | | |
| 917 | 10 | 58356021 | 58359759 | 3738 | SPRET | 10 | 58358650 | 58358683 | 34 | - | 16.9675 | 6.91E-07 | 0.0942 | CTGCAATAGGAGAGATTGCCTGCAGAGAGTACTC | | |
| 918 | 10 | 61906778 | 61908400 | 1622 | SPRET | 10 | 61907324 | 61907342 | 19 | - | 17.3443 | 7.26E-07 | 0.145 | CATCCTCCAGGGGGCGATG | | |
| 919 | 10 | 68956380 | 68958389 | 2009 | MOLF | 10 | 68957142 | 68957176 | 35 | - | 16.1284 | 8.89E-07 | 0.178 | CTGTAGTTGATGGTCCAGGACTGAAGGGGTCAGGC | | |
| 920 | 10 | 70295727 | 70297016 | 1289 | CAST | 10 | 70295818 | 70295851 | 34 | - | 25.1463 | 1.45E-09 | 0.00152 | GTGCAGCACCCCCAGAGGCCAGAAGAGGCCGGTA | 1 | ENCFF093ZRU |
| 921 | 10 | 71281276 | 71283009 | 1733 | SPRET | 10 | 71281596 | 71281614 | 19 | - | 17.8852 | 5.08E-07 | 0.121 | ACACCAGAAGAGGGCACCA | | |
| 921 | 10 | 71281276 | 71283009 | 1733 | SPRET | 10 | 71281596 | 71281629 | 34 | - | 18.0081 | 3.51E-07 | 0.0673 | TAGCTGTCTTTAGACACACCAGAAGAGGGCACCA | | |
| 922 | 10 | 75379360 | 75381279 | 1919 | SPRET | 10 | 75379575 | 75379609 | 35 | - | 22.2018 | 1.98E-08 | 0.0194 | CTGCATGTACACTTGTGTGCCAGAAGAGGGCATCA | 1 | ENCFF093ZRU |
| 923 | 10 | 83219304 | 83285378 | 66074 | SPRET | 10 | 83243143 | 83243177 | 35 | - | 16.6422 | 6.65E-07 | 0.154 | GAGCGGGTGCAGCCGCCACCCCCAAGATGGCGCCA | | |
| 924 | 10 | 85658030 | 85673227 | 15197 | BUB,CAST,WSB | 10 | 85665250 | 85665283 | 34 | - | 17.1138 | 6.30E-07 | 0.0904 | CTCCAGAACCAGGACCGACCTCCAGCGGGCAGGA | 1 | ENCFF093ZRU |
| 924 | 10 | 85658030 | 85673227 | 15197 | BUB,CAST,WSB | 10 | 85672418 | 85672436 | 19 | - | 19.8361 | 1.25E-07 | 0.0608 | CAACCAGAAGAGGGCACTA | 1 | ENCFF093ZRU |
| 924 | 10 | 85658030 | 85673227 | 15197 | BUB,CAST,WSB | 10 | 85672418 | 85672452 | 35 | - | 23.2385 | 9.40E-09 | 0.013 | GTGCAGGCACCTATGACAACCAGAAGAGGGCACTA | 1 | ENCFF093ZRU |
| 925 | 10 | 87908050 | 87909941 | 1891 | BUB,CAST,FVB,MOLF,SPRET | 10 | 87908907 | 87908941 | 35 | - | 15.9541 | 9.80E-07 | 0.186 | TTGCAATTGAAGGCCCAGGACTGAAGGGGTCACAC | | |
| 926 | 10 | 92586396 | 92590013 | 3617 | 129S1,AJ,AKR,BALB,BTBR,BUB,C3H,C57L,C58J,CAST,CBA,CE,DBA1J,DBA2J,FVB,ILNJ2,KK,LP,MA/My,MOLF,MRL,NOD,NOR,NZO,NZW,PJ,PL,RF,RH2,SEA,SJL,SMJ,SPRET,SWR,TH,WSB | 10 | 92586696 | 92586729 | 34 | - | 17.9106 | 3.74E-07 | 0.07 | CCGCCCTCACATTCGCCGTCACAAGATGGCGCTG | | |
| 926 | 10 | 92586396 | 92590013 | 3617 | 129S1,AJ,AKR,BALB,BTBR,BUB,C3H,C57L,C58J,CAST,CBA,CE,DBA1J,DBA2J,FVB,ILNJ2,KK,LP,MA/My,MOLF,MRL,NOD,NOR,NZO,NZW,PJ,PL,RF,RH2,SEA,SJL,SMJ,SPRET,SWR,TH,WSB | 10 | 92589971 | 92590004 | 34 | - | 17.9106 | 3.74E-07 | 0.07 | CCGCCCTCACATTCGCCGTCACAAGATGGCGCTG | | |
| 927 | 10 | 94389306 | 94393572 | 4266 | PL | 10 | 94391227 | 94391260 | 34 | - | 17.0569 | 6.53E-07 | 0.0917 | ATGCAGGTCCCCCGAGGTCTAGGAGGTGCCACTG | | |
| 928 | 10 | 106138976 | 106141883 | 2907 | SPRET | 10 | 106139090 | 106139123 | 34 | - | 18.8455 | 1.99E-07 | 0.0514 | CAGCAGTTGCACAGGTCTGCACCAGATGGGATCC | | |
| 929 | 10 | 115464229 | 115465323 | 1094 | SPRET | 10 | 115464302 | 115464335 | 34 | - | 18.9187 | 1.90E-07 | 0.0502 | TTGCAGTGCCTATGAAGTTCAGAAGAGGGCATCT | | |
| 930 | 10 | 116964985 | 116977667 | 12682 | MOLF,SMJ | 10 | 116968787 | 116968820 | 34 | - | 18.122 | 3.25E-07 | 0.0647 | TAGCTGTTTTCAGAGACACCAGAAGAGGGCATTA | 1 | ENCFF093ZRU |
| 931 | 10 | 127085391 | 127086442 | 1051 | SPRET | 10 | 127085849 | 127085882 | 34 | - | 16.6098 | 8.68E-07 | 0.105 | TTGCATGTATGCTACATGCCAGAAGAGGGCATCA | 3 | ENCFF093ZRU,ENCFF502WDT,ENCFF055HAP |
| 932 | 11 | 3977736 | 3980954 | 3218 | CAST | 11 | 3979511 | 3979544 | 34 | + | 23.5366 | 5.81E-09 | 0.00244 | ATGTAATGCCCCAGAAGGCCAGAAGAGGGCGTCA | 4 | ENCFF093ZRU,ENCFF033UQX,ENCFF052KVO,ENCFF533APC |
| 932 | 11 | 3977736 | 3980954 | 3218 | CAST | 11 | 3979526 | 3979544 | 19 | + | 17.1148 | 8.43E-07 | 0.0959 | AGGCCAGAAGAGGGCGTCA | 4 | ENCFF093ZRU,ENCFF033UQX,ENCFF052KVO,ENCFF533APC |
| 933 | 11 | 4826199 | 4827441 | 1242 | CAST | 11 | 4826340 | 4826374 | 35 | + | 20.844 | 5.00E-08 | 0.021 | GTGCAGGTGCCCACTCAGGCCAGAAGATGGAGTGG | | |
| 934 | 11 | 5358371 | 5361256 | 2885 | SPRET | 11 | 5359222 | 5359255 | 34 | + | 18.5691 | 2.41E-07 | 0.0342 | TCGTAGTGCCCAAGGAAGCCAGAAGGGGGTGTCA | | |
| 935 | 11 | 5498719 | 5501176 | 2457 | CAST | 11 | 5499521 | 5499555 | 35 | + | 16.4404 | 7.46E-07 | 0.11 | AAGAAGCTGGACCCAGTGGCCAGCAGATGGCCCAC | | |
| 935 | 11 | 5498719 | 5501176 | 2457 | CAST | 11 | 5499537 | 5499555 | 19 | + | 17.0328 | 8.88E-07 | 0.0977 | TGGCCAGCAGATGGCCCAC | | |
| 936 | 11 | 6185635 | 6195074 | 9439 | SPRET | 11 | 6192342 | 6192375 | 34 | + | 16.8211 | 7.59E-07 | 0.0608 | CTGCACTGGCGCCTCAAGCCAGAAGCAGCCGCCA | | |
| 937 | 11 | 7791614 | 7794710 | 3096 | KK,SPRET | 11 | 7792329 | 7792363 | 35 | + | 16.2752 | 8.19E-07 | 0.115 | ATGCAATCAGACGTGCCTTCCACAGGATGCCAGCA | | |
| 938 | 11 | 8870293 | 8879277 | 8984 | 129S1,AJ,AKR,BALB,BTBR,BUB,C3H,C57L,C58J,CAST,CBA,CE,DBA1J,DBA2J,FVB,ILNJ2,KK,MA/My,MOLF,MRL,NOD,NOR,NZB,NZO,PJ,PL,RF,SEA,SJL,SMJ,SPRET,SWR,TH,WSB | 11 | 8875709 | 8875727 | 19 | + | 17.1639 | 8.17E-07 | 0.0957 | CCTCCACGAGGTGGCGGAA | | |
| 939 | 11 | 11338922 | 11346644 | 7722 | SPRET | 11 | 11344873 | 11344891 | 19 | + | 17.9836 | 4.75E-07 | 0.076 | CAGCCACTAGAGGGTGCAG | 3 | ENCFF502WDT,ENCFF055HAP,ENCFF883UPM |
| 940 | 11 | 15748458 | 15750981 | 2523 | AKR,C3H,MA/My,MRL,NZB,NZW,RF,WSB | 11 | 15750578 | 15750596 | 19 | + | 17.459 | 6.74E-07 | 0.087 | CAGCCAGGAGGTGGAGCTG | | |
| 941 | 11 | 22130616 | 22131947 | 1331 | SPRET | 11 | 22131013 | 22131046 | 34 | + | 18.0569 | 3.40E-07 | 0.0395 | ATGCAGCTAGAGACATGAGCTCCAGGGGGTACTG | | |
| 942 | 11 | 22597883 | 22603334 | 5451 | SPRET | 11 | 22601529 | 22601563 | 35 | + | 16.5229 | 7.12E-07 | 0.108 | GTGCACAGCTCACTGCCCCACACAAGAGGGCATGT | | |
| 943 | 11 | 22889833 | 22894672 | 4839 | CAST,WSB | 11 | 22891051 | 22891085 | 35 | + | 17.2202 | 4.77E-07 | 0.0875 | CTGCATGTACACCTGCACCTCAGAAGAGGGCATGG | | |
| 944 | 11 | 24539394 | 24548152 | 8758 | SPRET | 11 | 24539375 | 24539408 | 34 | + | 18.0894 | 3.33E-07 | 0.0392 | TTTCAGCTGCTTGTTGGGCCTCAAGAGGGCAGCC | | |
| 945 | 11 | 28375579 | 28434583 | 59004 | PL | 11 | 28382537 | 28382571 | 35 | + | 16.5596 | 6.97E-07 | 0.107 | CTGTAGGGTCTCCATACAACCAGCAGAAGTAGGTT | | |
| 945 | 11 | 28375579 | 28434583 | 59004 | PL | 11 | 28427283 | 28427317 | 35 | - | 17.7615 | 3.47E-07 | 0.073 | CTGCACTGCCTTCTCAAACCCTGCAGGGAGCCTCA | | |
| 946 | 11 | 35692955 | 35694459 | 1504 | CAST,SPRET | 11 | 35694014 | 35694047 | 34 | + | 19.9593 | 9.14E-08 | 0.0184 | TAGCTGTCCTCAGACACCCCAGAAGAGGGCGCCA | 1 | ENCFF502WDT |
| 947 | 11 | 39573294 | 39575835 | 2541 | SPRET | 11 | 39574418 | 39574451 | 34 | + | 20.8943 | 4.62E-08 | 0.0115 | ATACAGTACCTGCAGCAGCCAGAAGAGGGTGTTG | | |
| 948 | 11 | 43340607 | 43345457 | 4850 | FVB,SJL,SWR | 11 | 43343279 | 43343297 | 19 | + | 17.4098 | 6.96E-07 | 0.0881 | TAACCAGTAGGTGTCACTC | 14 | ENCFF367UNR,ENCFF093ZRU,ENCFF092NMJ,ENCFF165LXO,ENCFF430PPJ,ENCFF099QNE,ENCFF327LYS,ENCFF907XEG,ENCFF821BIA,ENCFF055HAP,ENCFF089GBC,ENCFF024JOZ,ENCFF502WDT,ENCFF198SSR |
| 949 | 11 | 48716159 | 48717570 | 1411 | CAST,CE,MOLF,NZO,NZW,PJ,SEA,SPRET | 11 | 48717462 | 48717496 | 35 | + | 22.7248 | 1.36E-08 | 0.00983 | GTGCTGGGCCGGGGCCGGACCGGGAGGGGGCGGCG | | |
| 950 | 11 | 48852569 | 48855349 | 2780 | SPRET | 11 | 48853367 | 48853400 | 34 | + | 17.252 | 5.76E-07 | 0.0529 | TAGCTGTCTTCCGACAGTCCAGAAGAGGGAGCCA | | |
| 951 | 11 | 50336005 | 50339169 | 3164 | B10D2,B10J,BUB,C58J,CAST,CE,DBA2J,KK,MA/My,MOLF,NOD,NOR,NZB,NZO,NZW,PJ,PL,SEA,SJL,SPRET,SWR,TH,WSB | 11 | 50339131 | 50339164 | 34 | + | 19.561 | 1.21E-07 | 0.0223 | CAGCATCGTTCTTCCTGGCCAGTTGGGGGCGCGC | 5 | ENCFF092NMJ,ENCFF093ZRU,ENCFF907XEG,ENCFF099QNE,ENCFF821BIA |
| 951 | 11 | 50336005 | 50339169 | 3164 | B10D2,B10J,BUB,C58J,CAST,CE,DBA2J,KK,MA/My,MOLF,NOD,NOR,NZB,NZO,NZW,PJ,PL,SEA,SJL,SPRET,SWR,TH,WSB | 11 | 50339146 | 50339164 | 19 | + | 17.5574 | 6.32E-07 | 0.0841 | TGGCCAGTTGGGGGCGCGC | 5 | ENCFF092NMJ,ENCFF093ZRU,ENCFF907XEG,ENCFF099QNE,ENCFF821BIA |
| 952 | 11 | 51555030 | 51561912 | 6882 | B10D2,B10J,BUB,C58J,CE,DBA2J,KK,MA/My,NOD,NOR,NZB,PJ,SEA,SWR,TH,WSB | 11 | 51555560 | 51555594 | 35 | + | 17.2294 | 4.74E-07 | 0.0871 | CTGCACGTATACCTGCATGTCAGAAGAGGGTATCA | | |
| 953 | 11 | 58580417 | 58587147 | 6730 | AJ,BTBR,BUB,C3H,C57L,CBA,DBA1J,DBA2J,FVB,ILNJ2,KK,MA/My,MOLF,MRL,NOR,NZB,NZO,NZW,PJ,SEA,SJL,SWR,TH | 11 | 58580433 | 58580466 | 34 | + | 17.5041 | 4.89E-07 | 0.0482 | GAGCAGATGTGGCCGCGGCCCAAAGATGGCACCC | | |
| 954 | 11 | 60532890 | 60548505 | 15615 | C57L | 11 | 60536080 | 60536113 | 34 | + | 18.4797 | 2.56E-07 | 0.0347 | TTGCTGTCTTCAGACACACCAGAAGAGGGCATCA | | |
| 955 | 11 | 61764061 | 61765118 | 1057 | DBA1J,DBA2J,NOR | 11 | 61764065 | 61764098 | 34 | + | 16.4472 | 9.61E-07 | 0.067 | TAGCTGTCTTCAGACACACCAGAAGAGGGCGTCA | | |
| 956 | 11 | 76202643 | 76208693 | 6050 | MOLF | 11 | 76205748 | 76205766 | 19 | + | 18.8033 | 2.69E-07 | 0.0587 | CAACCAGCAGGGGTCAGCA | 16 | ENCFF201JUA,ENCFF367UNR,ENCFF165LXO,ENCFF093ZRU,ENCFF099QNE,ENCFF502WDT,ENCFF024JOZ,ENCFF821BIA,ENCFF907XEG,ENCFF609MDN,ENCFF327LYS,ENCFF883UPM,ENCFF198SSR,ENCFF589ZBJ,ENCFF957SPH,ENCFF055HAP |
| 957 | 11 | 76205363 | 76206875 | 1512 | SPRET | 11 | 76205748 | 76205766 | 19 | + | 18.8033 | 2.69E-07 | 0.0587 | CAACCAGCAGGGGTCAGCA | 16 | ENCFF201JUA,ENCFF367UNR,ENCFF165LXO,ENCFF093ZRU,ENCFF099QNE,ENCFF502WDT,ENCFF024JOZ,ENCFF821BIA,ENCFF907XEG,ENCFF609MDN,ENCFF327LYS,ENCFF883UPM,ENCFF198SSR,ENCFF589ZBJ,ENCFF957SPH,ENCFF055HAP |
| 958 | 11 | 76439631 | 76448680 | 9049 | 129S1,AJ,AKR,BALB,BTBR,BUB,C3H,C57L,C58J,CAST,CBA,CE,DBA1J,DBA2J,FVB,ILNJ2,KK,MA/My,MOLF,MRL,NOD,NOR,NZB,NZO,NZW,PJ,PL,RF,RH2,SEA,SJL,SMJ,SPRET,SWR,TH,WSB | 11 | 76445091 | 76445109 | 19 | + | 17.1639 | 8.17E-07 | 0.0957 | CCTCCACGAGGTGGCGGAA | | |
| 959 | 11 | 81831893 | 81833105 | 1212 | SPRET | 11 | 81831943 | 81831976 | 34 | + | 17.8862 | 3.81E-07 | 0.0424 | CTGATTTCCTCGGAACTTCCAGCAGAGGGAGCCT | 22 | ENCFF201JUA,ENCFF093ZRU,ENCFF367UNR,ENCFF099QNE,ENCFF805QBU,ENCFF198SSR,ENCFF136ZDK,ENCFF589ZBJ,ENCFF165LXO,ENCFF609MDN,ENCFF575MGI,ENCFF024JOZ,ENCFF907XEG,ENCFF957SPH,ENCFF194HVL,ENCFF327LYS,ENCFF821BIA,ENCFF092NMJ,ENCFF089GBC,ENCFF430PPJ,ENCFF281QMV,ENCFF883UPM |
| 960 | 11 | 84794157 | 84795446 | 1289 | SPRET | 11 | 84795405 | 84795423 | 19 | + | 17.377 | 7.11E-07 | 0.0894 | GGGCCAGCTGGGGGCACTC | | |
| 961 | 11 | 85023400 | 85024655 | 1255 | SPRET | 11 | 85024058 | 85024092 | 35 | + | 20.9633 | 4.62E-08 | 0.0202 | CTGCACATACACCTGCATGTCAGAAGAGGGCATTA | 1 | ENCFF093ZRU |
| 962 | 11 | 86773137 | 86782182 | 9045 | 129S1,AJ,AKR,BALB,BTBR,BUB,C3H,C57L,C58J,CAST,CBA,CE,DBA1J,DBA2J,MRL,NZB,NZO,NZW,PJ,PL,RF,SEA,SJL,SPRET,SWR,WSB | 11 | 86778597 | 86778615 | 19 | + | 17.1639 | 8.17E-07 | 0.0957 | CCTCCACGAGGTGGCGGAA | | |
| 963 | 11 | 93939586 | 93941614 | 2028 | SPRET | 11 | 93940645 | 93940678 | 34 | + | 25.2846 | 1.28E-09 | 0.000815 | GTGCAGTGCCTGAGGAGGCTAGAAGAGGGCACTG | | |
| 964 | 11 | 95839915 | 95842682 | 2767 | KK,MA/My,MRL,RF,SWR,TH | 11 | 95842532 | 95842565 | 34 | + | 17.8455 | 3.91E-07 | 0.0425 | TAGCTGTCCTCAGACACTCCAGAAGAGGGCGTCA | | |
| 965 | 11 | 99587116 | 99638340 | 51224 | SPRET | 11 | 99626703 | 99626736 | 34 | + | 17.8293 | 3.95E-07 | 0.0428 | ATGCAGCTACAGACATGAGCTCTGGGGGTACTG | | |
| 966 | 11 | 101237699 | 101239063 | 1364 | SPRET | 11 | 101238062 | 101238096 | 35 | + | 20.7798 | 5.22E-08 | 0.0216 | GTGCAGGTTCATTCAGAGACCAGAAGAGGGCGTCT | 5 | ENCFF367UNR,ENCFF136ZDK,ENCFF327LYS,ENCFF502WDT,ENCFF533APC |
| 966 | 11 | 101237699 | 101239063 | 1364 | SPRET | 11 | 101238378 | 101238412 | 35 | + | 16.5505 | 7.01E-07 | 0.107 | CTGGCATTTCAGCCCTTGCTCACTAGGTGGCCCTG | | |
| 966 | 11 | 101237699 | 101239063 | 1364 | SPRET | 11 | 101237767 | 101237800 | 34 | - | 17.2276 | 5.85E-07 | 0.0531 | TAGCTGTTTTCAGACACACCAAAAGAGGGCACTG | | |
| 967 | 11 | 102524577 | 102525901 | 1324 | CAST,MOLF | 11 | 102525178 | 102525212 | 35 | + | 17.422 | 4.24E-07 | 0.0813 | ATGCATGTACGCCTACACACCACAAAAGGGCACCA | | |
| 968 | 11 | 102974483 | 102983530 | 9047 | 129S1,AJ,AKR,BALB,BTBR,BUB,C3H,C57L,CAST,CBA,CE,DBA1J,DBA2J,FVB,ILNJ2,KK,MOLF,MRL,NOD,NOR,NZO,NZW,RF,RH2,SEA,SJL,SMJ,SPRET,WSB | 11 | 102979945 | 102979963 | 19 | + | 17.1639 | 8.17E-07 | 0.0957 | CCTCCACGAGGTGGCGGAA | | |
| 969 | 11 | 103399750 | 103420209 | 20459 | BALB,CE,FVB,ILNJ2,KK,MRL,NOD,NOR,NZB,SEA,SJL,SMJ | 11 | 103411552 | 103411585 | 34 | + | 18.6423 | 2.29E-07 | 0.0335 | ATGCAGTGCCCATAGAGGCTAGAAGAAGGCATCA | | |
| 970 | 11 | 104292216 | 104293342 | 1126 | AKR,NZO,NZW,RF,RH2 | 11 | 104292883 | 104292901 | 19 | + | 16.9508 | 9.36E-07 | 0.1 | ACACCAGAAGAGGGCAGTG | 2 | ENCFF093ZRU,ENCFF099QNE |
| 971 | 11 | 107497634 | 107502460 | 4826 | CAST | 11 | 107500107 | 107500140 | 34 | + | 16.5691 | 8.90E-07 | 0.0664 | TAGCTGTCTTCAGACGCACCAGAAGAGGGCATCG | | |
| 972 | 11 | 111094965 | 111117212 | 22247 | AKR,RH2 | 11 | 111096172 | 111096205 | 34 | + | 18.3089 | 2.87E-07 | 0.0357 | ATGCAGCTAGAGACACGAGCTCCAGGGGGTACTG | | |
| 973 | 11 | 117287066 | 117289836 | 2770 | SPRET | 11 | 117288831 | 117288864 | 34 | + | 17.9756 | 3.59E-07 | 0.0407 | CAGCAGCCTGTAGAGGTGACAGCAGGGGGAGGTA | 19 | ENCFF093ZRU,ENCFF367UNR,ENCFF198SSR,ENCFF092NMJ,ENCFF589ZBJ,ENCFF502WDT,ENCFF430PPJ,ENCFF951GOP,ENCFF165LXO,ENCFF327LYS,ENCFF089GBC,ENCFF821BIA,ENCFF883UPM,ENCFF907XEG,ENCFF194HVL,ENCFF055HAP,ENCFF024JOZ,ENCFF099QNE,ENCFF533APC |
| 974 | 11 | 117299837 | 117302699 | 2862 | SMJ | 11 | 117301581 | 117301615 | 35 | + | 17.6239 | 3.76E-07 | 0.0758 | CTGTAGCCGTCTTCACACACCAGAAGAGGGCATCA | 1 | ENCFF093ZRU |
| 975 | 11 | 117526271 | 117528259 | 1988 | MOLF | 11 | 117527504 | 117527538 | 35 | + | 16.3211 | 7.98E-07 | 0.113 | CTGCTGAGGCTGCTGCCCCCCAGGAGTGGGAGCTC | | |
| 976 | 11 | 121799194 | 121801695 | 2501 | CAST | 11 | 121801404 | 121801437 | 34 | + | 17.0569 | 6.53E-07 | 0.0564 | TAGCTGTCTTCAGAAACGCCAGAAGAGGGCATCA | | |
| 977 | 11 | 5025998 | 5027023 | 1025 | SPRET | 11 | 5026652 | 5026685 | 34 | - | 18.5285 | 2.48E-07 | 0.0346 | CAGCTGTCTTCAGACACTCCAGAAGAGGGCGTCA | 1 | ENCFF093ZRU |
| 978 | 11 | 6694896 | 6709348 | 14452 | CAST,KK,SPRET | 11 | 6702961 | 6702979 | 19 | - | 17.1639 | 8.17E-07 | 0.0957 | CCTCCACGAGGTGGCGGAA | | |
| 979 | 11 | 6699417 | 6708391 | 8974 | 129S1,AJ,AKR,B10D2,B10J,BALB,BTBR,C3H,C57L,C58J,FVB,MA/My,MOLF,MRL,NOD,NOR,NZB,NZW,PL,RF,SEA,TH,WSB | 11 | 6702961 | 6702979 | 19 | - | 17.1639 | 8.17E-07 | 0.0957 | CCTCCACGAGGTGGCGGAA | | |
| 980 | 11 | 8220598 | 8224260 | 3662 | AJ,AKR,B10D2,B10J,BALB,BTBR,BUB,C3H,C57L,C58J,CAST,CE,KK,MA/My,MRL,NOD,NOR,NZB,PJ,PL,RF,SEA,SJL,SPRET,SWR,WSB | 11 | 8223457 | 8223490 | 34 | - | 17.6748 | 4.37E-07 | 0.0453 | GAGCATGACCACAAGTCCACAGCAGGTGGCAACG | | |
| 981 | 11 | 10148082 | 10151004 | 2922 | CAST,MA/My,MOLF,NZB,NZO,SPRET | 11 | 10148078 | 10148112 | 35 | - | 19.4679 | 1.22E-07 | 0.0371 | CTGCCGCCACACCCGCTCCCCACAAGAGGGAAGGC | | |
| 982 | 11 | 10726336 | 10728513 | 2177 | SPRET | 11 | 10728431 | 10728464 | 34 | - | 16.439 | 9.66E-07 | 0.0672 | ATGCAGCTAGAGACAAGAGCTCCAGGGGGTACTG | | |
| 983 | 11 | 19683792 | 19685362 | 1570 | MOLF | 11 | 19684765 | 19684799 | 35 | - | 15.9541 | 9.80E-07 | 0.126 | CAGCACTGAGCTGGGGTAGACAGCAGGGGACTCCC | | |
| 984 | 11 | 31640225 | 31642354 | 2129 | AKR,BUB,C3H,CBA,CE,FVB,ILNJ2,KK,MRL,NOD,PJ,PL,SEA,SJL,SMJ,SWR,TH | 11 | 31642250 | 31642284 | 35 | - | 17.211 | 4.79E-07 | 0.0877 | CTGCAGTTCTATGGGGGGTCCTGTGGAGGTGGCTG | | |
| 985 | 11 | 40013269 | 40014589 | 1320 | AKR,CAST,SPRET | 11 | 40013895 | 40013928 | 34 | - | 16.9268 | 7.10E-07 | 0.0585 | ATGCAGTTAGAGACATGAGCTCTGGGGGGTACTG | | |
| 986 | 11 | 40013869 | 40014938 | 1069 | 129S1,AJ,AKR,BALB,BUB,C3H,C57L,CBA,DBA1J,MRL,PL,SPRET,WSB | 11 | 40013895 | 40013928 | 34 | - | 16.9268 | 7.10E-07 | 0.0585 | ATGCAGTTAGAGACATGAGCTCTGGGGGGTACTG | | |
| 987 | 11 | 43195409 | 43203671 | 8262 | SPRET | 11 | 43202478 | 43202511 | 34 | - | 18.7886 | 2.07E-07 | 0.0315 | GAGCAGTGCCTGCAGAGGCCAGAAGAGTGTGTTG | | |
| 988 | 11 | 43535735 | 43537314 | 1579 | MOLF,WSB | 11 | 43536310 | 43536343 | 34 | - | 21 | 4.27E-08 | 0.0108 | ATGCAATGCCCTTGGAGACCAGTAGGGGGGATAG | 16 | ENCFF504JDS,ENCFF327LYS,ENCFF821BIA,ENCFF367UNR,ENCFF092NMJ,ENCFF907XEG,ENCFF136ZDK,ENCFF589ZBJ,ENCFF198SSR,ENCFF093ZRU,ENCFF883UPM,ENCFF957SPH,ENCFF052KVO,ENCFF099QNE,ENCFF065ODV,ENCFF502WDT |
| 989 | 11 | 45473114 | 45477014 | 3900 | SPRET | 11 | 45473441 | 45473459 | 19 | - | 17.4098 | 6.96E-07 | 0.0881 | CAGACAGAAGAGGGCGCCA | | |
| 989 | 11 | 45473114 | 45477014 | 3900 | SPRET | 11 | 45473441 | 45473475 | 35 | - | 22.3945 | 1.73E-08 | 0.0116 | GTGCAGGTGCCTATGGCAGACAGAAGAGGGCGCCA | | |
| 990 | 11 | 45615252 | 45623164 | 7912 | 129S1,AJ,AKR,B10D2,B10J,BALB,BTBR,BUB,C3H,C57L,C58J,CAST,CBA,CE,DBA1J,DBA2J,FVB,ILNJ2,KK,MA/My,MOLF,MRL,NOD,NZB,NZO,NZW,PJ,PL,RF,RH2,SEA,SJL,SMJ,SPRET,SWR,TH,WSB | 11 | 45615601 | 45615635 | 35 | - | 16.2569 | 8.27E-07 | 0.116 | CGGCGGGTCCCGGACCGGTCCGGAAGGTGGCCGGC | | |
| 991 | 11 | 48683493 | 48685724 | 2231 | SPRET | 11 | 48684607 | 48684625 | 19 | - | 17.5902 | 6.18E-07 | 0.0841 | ACACCAGAAGAGGGCACTG | 1 | ENCFF093ZRU |
| 991 | 11 | 48683493 | 48685724 | 2231 | SPRET | 11 | 48684607 | 48684640 | 34 | - | 20.4146 | 6.58E-08 | 0.0147 | TTGCTGTCTTTAGACACACCAGAAGAGGGCACTG | 1 | ENCFF093ZRU |
| 992 | 11 | 53572604 | 53573694 | 1090 | SPRET | 11 | 53572696 | 53572729 | 34 | - | 19.439 | 1.32E-07 | 0.0236 | ATGCAATGCTCATGAAGGCCAGAAGATGTCATTG | 6 | ENCFF367UNR,ENCFF093ZRU,ENCFF198SSR,ENCFF883UPM,ENCFF502WDT,ENCFF055HAP |
| 993 | 11 | 55871745 | 55873463 | 1718 | CAST | 11 | 55872669 | 55872687 | 19 | - | 20.7869 | 5.86E-08 | 0.0292 | TGCCCACCAGGGGGCAGGA | 19 | ENCFF093ZRU,ENCFF821BIA,ENCFF165LXO,ENCFF201JUA,ENCFF589ZBJ,ENCFF430PPJ,ENCFF502WDT,ENCFF367UNR,ENCFF327LYS,ENCFF099QNE,ENCFF575MGI,ENCFF609MDN,ENCFF907XEG,ENCFF089GBC,ENCFF883UPM,ENCFF024JOZ,ENCFF092NMJ,ENCFF957SPH,ENCFF198SSR |
| 994 | 11 | 56184750 | 56188000 | 3250 | CAST,SPRET | 11 | 56187314 | 56187347 | 34 | - | 16.4715 | 9.47E-07 | 0.067 | GAGCATAAACACAAGTCCACAGCAGGTGGCAGCG | | |
| 995 | 11 | 58990668 | 58994253 | 3585 | CE,NZB,NZO,SMJ | 11 | 58993343 | 58993376 | 34 | - | 28.0569 | 8.91E-11 | 0.000177 | ATGCAGTGCCCACAGAGGCCATAAGAGGGCGCCA | 18 | ENCFF099QNE,ENCFF201JUA,ENCFF136ZDK,ENCFF589ZBJ,ENCFF327LYS,ENCFF198SSR,ENCFF907XEG,ENCFF367UNR,ENCFF093ZRU,ENCFF055HAP,ENCFF883UPM,ENCFF533APC,ENCFF033UQX,ENCFF821BIA,ENCFF065ODV,ENCFF142CNG,ENCFF052KVO,ENCFF502WDT |
| 996 | 11 | 59032525 | 59035385 | 2860 | CE,SMJ | 11 | 59034656 | 59034689 | 34 | - | 20.1951 | 7.72E-08 | 0.0164 | CAGCTGTCTCCAGACACACCAGAAGAGGGCATCA | | |
| 997 | 11 | 60469470 | 60476834 | 7364 | 129S1,AKR,BTBR,BUB,C57L,CAST,CE,FVB,ILNJ2,LP,MA/My,NOD,NZW,PL,RF,SEA,SMJ,SPRET,SWR,WSB | 11 | 60471361 | 60471379 | 19 | - | 17.1639 | 8.17E-07 | 0.0957 | CCTCCACGAGGTGGCGGAA | | |
| 998 | 11 | 62411780 | 62412781 | 1001 | CAST | 11 | 62412430 | 62412448 | 19 | - | 17.8852 | 5.08E-07 | 0.0767 | ACACCAGAAGAGGGCACCA | | |
| 998 | 11 | 62411780 | 62412781 | 1001 | CAST | 11 | 62412430 | 62412463 | 34 | - | 17.5122 | 4.86E-07 | 0.0481 | TCGCTGTCTTCAGACACACCAGAAGAGGGCACCA | | |
| 999 | 11 | 69178082 | 69180400 | 2318 | AKR,PL,RF | 11 | 69179734 | 69179768 | 35 | - | 23.2294 | 9.46E-09 | 0.00768 | CTGCATGTAAGCCTACATGCCAGAAGAGGGCATTG | | |
| 1000 | 11 | 69371475 | 69372720 | 1245 | SPRET | 11 | 69372430 | 69372448 | 19 | - | 20.3607 | 8.29E-08 | 0.0322 | AGGCCAGAAGAGGGCACCA | 5 | ENCFF093ZRU,ENCFF099QNE,ENCFF052KVO,ENCFF502WDT,ENCFF367UNR |
| 1000 | 11 | 69371475 | 69372720 | 1245 | SPRET | 11 | 69372430 | 69372463 | 34 | - | 22.2033 | 1.70E-08 | 0.00532 | CTGCATGTATGCTATAGGCCAGAAGAGGGCACCA | 5 | ENCFF093ZRU,ENCFF099QNE,ENCFF052KVO,ENCFF502WDT,ENCFF367UNR |
| 1001 | 11 | 69977271 | 69980533 | 3262 | SPRET | 11 | 69979510 | 69979544 | 35 | - | 16.1009 | 9.03E-07 | 0.121 | GTGTGGGTGCCTTTGGAGGCCAGAAGAGGGCATCT | | |
| 1002 | 11 | 71760346 | 71761737 | 1391 | SPRET | 11 | 71760660 | 71760694 | 35 | - | 18.5321 | 2.18E-07 | 0.0542 | CTGCATTGCCCCTCGCTCGCCCCCAGAGTGCCTCC | | |
| 1003 | 11 | 76417280 | 76419348 | 2068 | CAST,MOLF | 11 | 76418868 | 76418901 | 34 | - | 17.2033 | 5.94E-07 | 0.0534 | ATGCAGTGCCTGTAAAGGCCAAACGAAGGCATCG | | |
| 1004 | 11 | 77556687 | 77561199 | 4512 | SPRET | 11 | 77560857 | 77560890 | 34 | - | 16.4959 | 9.32E-07 | 0.067 | TAGCTGTCCTCAGATACTCCAGAAGATGGCATCA | | |
| 1005 | 11 | 78423444 | 78424903 | 1459 | CAST | 11 | 78423498 | 78423531 | 34 | - | 18.9919 | 1.80E-07 | 0.0291 | CTGTAGCTGTCAGACGCTCCAGAAGAGGGCATCC | | |
| 1006 | 11 | 86397997 | 86400373 | 2376 | SPRET | 11 | 86399046 | 86399080 | 35 | - | 17.211 | 4.79E-07 | 0.0877 | CTGTGTGTATACCTGCAGGACAGAAGAGGGCACCA | | |
| 1007 | 11 | 86771371 | 86773142 | 1771 | MOLF | 11 | 86772722 | 86772755 | 34 | - | 24.065 | 3.73E-09 | 0.00175 | ATGCAGTGCCTGTGGAATCCACAAGAGGGCATCT | 8 | ENCFF821BIA,ENCFF327LYS,ENCFF052KVO,ENCFF367UNR,ENCFF093ZRU,ENCFF099QNE,ENCFF502WDT,ENCFF055HAP |
| 1008 | 11 | 89677258 | 89679537 | 2279 | SPRET | 11 | 89677926 | 89677960 | 35 | - | 16.4037 | 7.61E-07 | 0.111 | TTGCATTAATTCCCATCTACCACAAGAAGGAAGTT | | |
| 1009 | 11 | 92888297 | 92891380 | 3083 | MOLF | 11 | 92888410 | 92888443 | 34 | - | 16.4472 | 9.61E-07 | 0.067 | TAGCTGTCTTCAGACACACCAGAAGAGGGCGTCA | | |
| 1010 | 11 | 95831825 | 95833551 | 1726 | CAST,DBA1J,DBA2J,ILNJ2,NZB,NZO,PJ,SEA,WSB | 11 | 95832638 | 95832671 | 34 | - | 17.7317 | 4.21E-07 | 0.0444 | TAGCTGTCCTCAGATACACCAGAAGAGGGCATCA | | |
| 1011 | 11 | 101000916 | 101002001 | 1085 | SPRET | 11 | 101000945 | 101000978 | 34 | - | 16.5772 | 8.86E-07 | 0.0661 | TTGCTCTCTTCAGACACTCCAGAAGAGGGCATCG | 1 | ENCFF883UPM |
| 1012 | 11 | 102507104 | 102510538 | 3434 | AJ,CBA | 11 | 102510471 | 102510489 | 19 | - | 17.8033 | 5.37E-07 | 0.0792 | AGGCCAGTAGAGGGCGAAG | 14 | ENCFF093ZRU,ENCFF367UNR,ENCFF907XEG,ENCFF136ZDK,ENCFF821BIA,ENCFF327LYS,ENCFF589ZBJ,ENCFF052KVO,ENCFF198SSR,ENCFF502WDT,ENCFF883UPM,ENCFF055HAP,ENCFF065ODV,ENCFF142CNG |
| 1012 | 11 | 102507104 | 102510538 | 3434 | AJ,CBA | 11 | 102510471 | 102510504 | 34 | - | 23.2602 | 7.30E-09 | 0.00289 | AGGCAGTACCTGAGGAGGCCAGTAGAGGGCGAAG | 14 | ENCFF093ZRU,ENCFF367UNR,ENCFF907XEG,ENCFF136ZDK,ENCFF821BIA,ENCFF327LYS,ENCFF589ZBJ,ENCFF052KVO,ENCFF198SSR,ENCFF502WDT,ENCFF883UPM,ENCFF055HAP,ENCFF065ODV,ENCFF142CNG |
| 1013 | 11 | 104299373 | 104302324 | 2951 | SPRET | 11 | 104302033 | 104302066 | 34 | - | 17.2439 | 5.79E-07 | 0.0529 | TAGCTGTCCTCAGATACTCCAGAAGAGGGCATCA | | |
| 1014 | 11 | 112636523 | 112637629 | 1106 | NOD,NOR | 11 | 112636789 | 112636822 | 34 | - | 17.6667 | 4.40E-07 | 0.0454 | TAGCTGTCTTCAGACACGCCAGAAGAGGGCGTCA | | |
| 1015 | 11 | 115662353 | 115664762 | 2409 | 129S1,AJ,AKR,BALB,C3H,C57L,C58J,CAST,CBA,CE,DBA1J,DBA2J,ILNJ2,KK,MOLF,MRL,NOD,NOR,NZB,NZO,NZW,PL,RF,RH2,SEA,SMJ,SPRET,TH,WSB | 11 | 115662449 | 115662482 | 34 | - | 16.7724 | 7.83E-07 | 0.0614 | TAGCTGTCTTCAGATGCACCAGAAGAGGGCGTCA | | |
| 1016 | 11 | 116364603 | 116365846 | 1243 | CAST,MOLF | 11 | 116365662 | 116365695 | 34 | - | 16.4472 | 9.61E-07 | 0.067 | TAGCTGTCTTCAGACACACCAGAAGAGGGCGTCA | | |
| 1017 | 11 | 116447580 | 116448778 | 1198 | 129S1,BALB,C58J,DBA1J,DBA2J,FVB,KK,MA/My,NOD,NOR,NZB,NZO,NZW,PL,RF,SEA,SJL,SMJ,SWR,TH,WSB | 11 | 116448596 | 116448630 | 35 | - | 17.4404 | 4.19E-07 | 0.081 | CTGCTGACCCCAGCCTGTGACAGTAGAAGGCGGGC | | |
| 1018 | 11 | 116787958 | 116791155 | 3197 | MOLF | 11 | 116791055 | 116791089 | 35 | - | 16.1651 | 8.71E-07 | 0.119 | CTACAGCAGGAGCAGCTGGCCAGCTGGTGGGGGCA | | |
| 1019 | 11 | 121017059 | 121018285 | 1226 | MOLF,SJL | 11 | 121017423 | 121017456 | 34 | - | 21.9593 | 2.06E-08 | 0.00614 | GTGTAGTTCCCAAAGAGGCCACTAGAGAGCATTA | 18 | ENCFF201JUA,ENCFF092NMJ,ENCFF504JDS,ENCFF821BIA,ENCFF165LXO,ENCFF093ZRU,ENCFF065ODV,ENCFF367UNR,ENCFF327LYS,ENCFF198SSR,ENCFF502WDT,ENCFF055HAP,ENCFF907XEG,ENCFF099QNE,ENCFF589ZBJ,ENCFF024JOZ,ENCFF052KVO,ENCFF883UPM |
| 1020 | 11 | 121201862 | 121210837 | 8975 | CAST,MOLF,SPRET | 11 | 121208661 | 121208694 | 34 | - | 17.0325 | 6.63E-07 | 0.0568 | TGGCAGGTCTCAGAGGAGCCGCTAGATGGCCCAC | | |
| 1021 | 11 | 121456623 | 121459359 | 2736 | SPRET | 11 | 121458510 | 121458543 | 34 | - | 17.187 | 6.01E-07 | 0.0537 | TAGCAATCTTCAGGTGTACCAGAAGAGGGCGTCA | 8 | ENCFF367UNR,ENCFF136ZDK,ENCFF821BIA,ENCFF589ZBJ,ENCFF883UPM,ENCFF055HAP,ENCFF502WDT,ENCFF093ZRU |
| 1022 | 12 | 4194672 | 4222791 | 28119 | B10D2 | 12 | 4208707 | 4208740 | 34 | + | 17.3902 | 5.27E-07 | 0.0795 | TAGCTGTCCTCAGACACTCCAGAAGAGGGCATCA | | |
| 1022 | 12 | 4194672 | 4222791 | 28119 | B10D2 | 12 | 4214420 | 4214453 | 34 | + | 17.5528 | 4.74E-07 | 0.0755 | TAGCTGTTTTCAGATACACCAGAAGAGGGCATCA | 1 | ENCFF093ZRU |
| 1022 | 12 | 4194672 | 4222791 | 28119 | B10D2 | 12 | 4215333 | 4215351 | 19 | + | 17.4754 | 6.66E-07 | 0.132 | CCACCTCCAGGGGGCAGCT | | |
| 1022 | 12 | 4194672 | 4222791 | 28119 | B10D2 | 12 | 4203958 | 4203991 | 34 | - | 16.7073 | 8.16E-07 | 0.0978 | GAGATCTGCCTGCCTCTGCCTCCAGAGGCTATG | | |
| 1023 | 12 | 5560726 | 5561829 | 1103 | SPRET | 12 | 5561210 | 5561243 | 34 | + | 18.0569 | 3.40E-07 | 0.064 | ATGCAGCTAGAGACATGAGCTCCAGGGGGTACTG | | |
| 1024 | 12 | 5663770 | 5667937 | 4167 | AJ,BALB,CAST,ILNJ2,MOLF,MRL,NZB,SEA,SMJ,SPRET,WSB | 12 | 5664546 | 5664579 | 34 | + | 17.6748 | 4.37E-07 | 0.073 | GAGCATGACCACAAGTCCACAGCAGGTGGCAACG | | |
| 1025 | 12 | 11816123 | 11817183 | 1060 | SPRET | 12 | 11816331 | 11816364 | 34 | + | 16.4553 | 9.56E-07 | 0.104 | ATGCAGCTAGAGACACGGGCTCTGGGGGGTACTG | | |
| 1026 | 12 | 18784494 | 18789572 | 5078 | 129S1,BTBR,BUB,C3H,CBA,DBA2J,FVB,MA/My,NOD,NOR,PL,SJL,SMJ,SWR,TH | 12 | 18786288 | 18786321 | 34 | + | 17.2439 | 5.79E-07 | 0.0827 | TAGCTGTCCTCAGATACTCCAGAAGAGGGCATCA | | |
| 1027 | 12 | 19039121 | 19045787 | 6666 | AJ,AKR,BALB,C57L,CE,DBA1J,ILNJ2,PJ,RF,SEA | 12 | 19045481 | 19045499 | 19 | + | 18.8525 | 2.59E-07 | 0.0823 | TCTACACTAGGGGGCGCTG | | |
| 1028 | 12 | 19293147 | 19299715 | 6568 | 129S1,BTBR,BUB,C3H,C57L,CAST,CBA,DBA2J,FVB,KK,MA/My,MRL,NOD,NOR,NZO,PL,SJL,SMJ,SPRET,SWR,TH,WSB | 12 | 19297709 | 19297727 | 19 | + | 17.1639 | 8.17E-07 | 0.145 | CCTCCACGAGGTGGCGGAA | | |
| 1029 | 12 | 20055455 | 20058152 | 2697 | AJ,CE,DBA1J,PJ | 12 | 20056798 | 20056816 | 19 | + | 18.1148 | 4.35E-07 | 0.109 | TCTACACTAGGGGGCACTG | | |
| 1030 | 12 | 20624757 | 20630333 | 5576 | DBA2J | 12 | 20627189 | 20627222 | 34 | + | 18.3659 | 2.76E-07 | 0.0591 | CTGAAGGCCCAGCCTTAGCCTTCAGGTGGCTGCC | | |
| 1031 | 12 | 20744352 | 20757157 | 12805 | CAST,MRL | 12 | 20747473 | 20747506 | 34 | + | 21.9593 | 2.06E-08 | 0.011 | ATGCTATGTTCAAAGTAGCCACAAGAGGGCTCTA | 1 | ENCFF367UNR |
| 1031 | 12 | 20744352 | 20757157 | 12805 | CAST,MRL | 12 | 20747488 | 20747506 | 19 | + | 17.3934 | 7.03E-07 | 0.134 | TAGCCACAAGAGGGCTCTA | 1 | ENCFF367UNR |
| 1031 | 12 | 20744352 | 20757157 | 12805 | CAST,MRL | 12 | 20753456 | 20753474 | 19 | + | 17.1639 | 8.17E-07 | 0.145 | CCTCCACGAGGTGGCGGAA | | |
| 1032 | 12 | 20748888 | 20754653 | 5765 | 129S1,AJ,AKR,BALB,BTBR,BUB,C3H,C57L,CBA,CE,DBA1J,FVB,ILNJ2,KK,MA/My,NOD,NOR,NZO,PJ,PL,RF,SEA,SJL,SMJ,SWR,TH,WSB | 12 | 20753456 | 20753474 | 19 | + | 17.1639 | 8.17E-07 | 0.145 | CCTCCACGAGGTGGCGGAA | | |
| 1033 | 12 | 20991097 | 20993450 | 2353 | CAST,MRL | 12 | 20993034 | 20993067 | 34 | + | 21.6667 | 2.58E-08 | 0.0127 | ATGCTATGTTCAAAGCAGCCACGAGAGGGCTCCA | 2 | ENCFF821BIA,ENCFF367UNR |
| 1034 | 12 | 23656010 | 23659599 | 3589 | MRL | 12 | 23656749 | 23656783 | 35 | + | 23.9633 | 5.48E-09 | 0.0077 | GTGCAGGTGCCCATGGAGGCCTGAAGAGGGCACCT | | |
| 1035 | 12 | 24233061 | 24234289 | 1228 | SPRET | 12 | 24233855 | 24233888 | 34 | + | 19.7561 | 1.06E-07 | 0.0331 | CTGCTCTGCAGATATCTGCCAGGAGGAGGCTGCA | | |

| | | | | | | | | | | | | | | | | |
|---|---|---|---|---|---|---|---|---|---|---|---|---|---|---|---|---|
| 1036 | 12 | 29163923 | 29167447 | 3524 | 129S1,AJ,AKR,BALB,BUB,C3H,CBA,CE,DBA1J,DBA2J,KK,LP,MRL,NOD,NOR,NZO,NZW.PJ.PL.RF.RH2.SEA.SMJ.WSB | 12 | 29165332 | 29165365 | 34 | + | 18.9431 | 1.87E-07 | 0.0482 | CTGGCATCCCAGAGGCTTCCAACAGAGGGCAACA | | |
| 1037 | 12 | 32061999 | 32064471 | 2472 | CAST.MOLF | 12 | 32062333 | 32062367 | 35 | + | 18.5321 | 2.18E-07 | 0.0839 | CTGCACTGTTAGCATATGTGCAACAGAGGTCACAG | | |
| 1038 | 12 | 43407086 | 43408364 | 1278 | SPRET | 12 | 43407122 | 43407155 | 34 | + | 18.0569 | 3.40E-07 | 0.064 | ATGCAGCTAGAGACATGAGCTCCAGGGGGTACTG | | |
| 1039 | 12 | 44893398 | 44897080 | 3682 | NZO,NZW | 12 | 44895654 | 44895687 | 34 | + | 17.7073 | 4.28E-07 | 0.0725 | GCGCACTCGTGTGAATAACCACAAGGGGGTGCCC | 22 | ENCFF093ZRU,ENCFF367UNR,ENCFF201JUA,ENCFF099QNE,ENCFF821BIA,ENCFF136ZDK,ENCFF327LYS,ENCFF907XEG,ENCFF024JOZ,ENCFF089GBC,ENCFF575MGI,ENCFF165LXO,ENCFF589ZBJ,ENCFF198SSR,ENCFF092NMJ,ENCFF052KVO,ENCFF430PPJ,ENCFF502WDT,ENCFF055HAP,ENCFF951GOP,ENCFF281QMV,ENCFF609MDN |
| 1039 | 12 | 44893398 | 44897080 | 3682 | NZO,NZW | 12 | 44895669 | 44895687 | 19 | + | 18.0328 | 4.60E-07 | 0.113 | TAACCACAAGGGGGTGCCC | 22 | ENCFF093ZRU,ENCFF367UNR,ENCFF201JUA,ENCFF099QNE,ENCFF821BIA,ENCFF136ZDK,ENCFF327LYS,ENCFF907XEG,ENCFF024JOZ,ENCFF089GBC,ENCFF575MGI,ENCFF165LXO,ENCFF589ZBJ,ENCFF198SSR,ENCFF092NMJ,ENCFF052KVO,ENCFF430PPJ,ENCFF502WDT,ENCFF055HAP,ENCFF951GOP,ENCFF281QMV,ENCFF609MDN |
| 1040 | 12 | 51347640 | 51356509 | 8869 | 129S1,C57L,NZB | 12 | 51355062 | 51355095 | 34 | + | 16.4309 | 9.71E-07 | 0.104 | TAGCTGTCTTCAGGGACACCAGAAGAGGGCATCA | | |
| 1041 | 12 | 54821004 | 54829990 | 8986 | AJ,AKR,BALB,C3H,CAST,CBA,CE,DBA1J,DBA2J,FVB,ILNJ2,KK,MOLF,NOD,NOR,NZW,PJ,RF,RH2,SEA,SJL,SMJ,SPRET,SWR,TH,WSB | 12 | 54826422 | 54826440 | 19 | + | 17.1639 | 8.17E-07 | 0.145 | CCTCCACGAGGTGGCGGAA | | |
| 1042 | 12 | 55451509 | 55452656 | 1147 | PJ | 12 | 55451601 | 55451635 | 35 | + | 16.6147 | 6.75E-07 | 0.161 | GTGTAGGTGCCCACAGTGTACAGAAGAGGGCAATA | | |
| 1043 | 12 | 56672966 | 56676585 | 3619 | AJ,AKR,BALB,C3H,CBA,DBA1J,DBA2J,ILNJ2,KK,MRL,NOD,NOR,RF,SEA,SJL,SMJ,SWR | 12 | 56673772 | 56673805 | 34 | + | 16.5691 | 8.90E-07 | 0.102 | TAGCTGTCTTCAGACGCACCAGAAGAGGGCATCG | | |
| 1044 | 12 | 57234103 | 57235349 | 1246 | CAST | 12 | 57235069 | 57235102 | 34 | + | 17.9512 | 3.64E-07 | 0.0669 | CTGCATTTGAATCCTCAGCCTTCAGGGGTCACTT | | |
| 1045 | 12 | 59081758 | 59084703 | 2945 | AJ,BALB,C3H,CBA,DBA1J,DBA2J,ILNJ2,KK,MA/Mv.RH2.SEA | 12 | 59081797 | 59081830 | 34 | + | 18.065 | 3.38E-07 | 0.064 | TAGCTGTCTTCAGACTCACCAGAAGAGGGCGTCA | | |
| 1045 | 12 | 59081758 | 59083753 | 1995 | NZO | 12 | 59081797 | 59081830 | 34 | + | 18.065 | 3.38E-07 | 0.064 | TAGCTGTCTTCAGACTCACCAGAAGAGGGCGTCA | | |
| 1045 | 12 | 59081758 | 59084703 | 2945 | AJ,BALB,C3H,CBA,DBA1J,DBA2J,ILNJ2,KK,MA/Mv.RH2.SEA | 12 | 59081812 | 59081830 | 19 | + | 17.0328 | 8.88E-07 | 0.147 | TCACCAGAAGAGGGCGTCA | | |
| 1045 | 12 | 59081758 | 59083753 | 1995 | NZO | 12 | 59081812 | 59081830 | 19 | + | 17.0328 | 8.88E-07 | 0.147 | TCACCAGAAGAGGGCGTCA | | |
| 1045 | 12 | 59081758 | 59084703 | 2945 | AJ,BALB,C3H,CBA,DBA1J,DBA2J,ILNJ2,KK,MA/Mv.RH2.SEA | 12 | 59081892 | 59081925 | 34 | + | 18.065 | 3.38E-07 | 0.064 | TAGCTGTCTTCAGACTCACCAGAAGAGGGCGTCA | | |
| 1045 | 12 | 59081758 | 59083753 | 1995 | NZO | 12 | 59081892 | 59081925 | 34 | + | 18.065 | 3.38E-07 | 0.064 | TAGCTGTCTTCAGACTCACCAGAAGAGGGCGTCA | | |
| 1045 | 12 | 59081758 | 59084703 | 2945 | AJ,BALB,C3H,CBA,DBA1J,DBA2J,ILNJ2,KK,MA/Mv.RH2.SEA | 12 | 59081907 | 59081925 | 19 | + | 17.0328 | 8.88E-07 | 0.147 | TCACCAGAAGAGGGCGTCA | | |
| 1045 | 12 | 59081758 | 59083753 | 1995 | NZO | 12 | 59081907 | 59081925 | 19 | + | 17.0328 | 8.88E-07 | 0.147 | TCACCAGAAGAGGGCGTCA | | |
| 1045 | 12 | 59081758 | 59084703 | 2945 | AJ,BALB,C3H,CBA,DBA1J,DBA2J,ILNJ2,KK,MA/Mv.RH2.SEA | 12 | 59081987 | 59082020 | 34 | + | 18.065 | 3.38E-07 | 0.064 | TAGCTGTCTTCAGACTCACCAGAAGAGGGCGTCA | | |
| 1045 | 12 | 59081758 | 59083753 | 1995 | NZO | 12 | 59081987 | 59082020 | 34 | + | 18.065 | 3.38E-07 | 0.064 | TAGCTGTCTTCAGACTCACCAGAAGAGGGCGTCA | | |
| 1045 | 12 | 59081758 | 59084703 | 2945 | AJ,BALB,C3H,CBA,DBA1J,DBA2J,ILNJ2,KK,MA/Mv.RH2.SEA | 12 | 59082002 | 59082020 | 19 | + | 17.0328 | 8.88E-07 | 0.147 | TCACCAGAAGAGGGCGTCA | | |
| 1045 | 12 | 59081758 | 59083753 | 1995 | NZO | 12 | 59082002 | 59082020 | 19 | + | 17.0328 | 8.88E-07 | 0.147 | TCACCAGAAGAGGGCGTCA | | |
| 1045 | 12 | 59081758 | 59084703 | 2945 | AJ,BALB,C3H,CBA,DBA1J,DBA2J,ILNJ2,KK,MA/Mv.RH2.SEA | 12 | 59082082 | 59082115 | 34 | + | 18.065 | 3.38E-07 | 0.064 | TAGCTGTCTTCAGACTCACCAGAAGAGGGCGTCA | | |
| 1045 | 12 | 59081758 | 59083753 | 1995 | NZO | 12 | 59082082 | 59082115 | 34 | + | 18.065 | 3.38E-07 | 0.064 | TAGCTGTCTTCAGACTCACCAGAAGAGGGCGTCA | | |
| 1045 | 12 | 59081758 | 59084703 | 2945 | AJ,BALB,C3H,CBA,DBA1J,DBA2J,ILNJ2,KK,MA/Mv.RH2.SEA | 12 | 59082097 | 59082115 | 19 | + | 17.0328 | 8.88E-07 | 0.147 | TCACCAGAAGAGGGCGTCA | | |
| 1045 | 12 | 59081758 | 59083753 | 1995 | NZO | 12 | 59082097 | 59082115 | 19 | + | 17.0328 | 8.88E-07 | 0.147 | TCACCAGAAGAGGGCGTCA | | |
| 1045 | 12 | 59081758 | 59084703 | 2945 | AJ,BALB,C3H,CBA,DBA1J,DBA2J,ILNJ2,KK,MA/Mv.RH2.SEA | 12 | 59082177 | 59082210 | 34 | + | 18.065 | 3.38E-07 | 0.064 | TAGCTGTCTTCAGACTCACCAGAAGAGGGCGTCA | | |
| 1045 | 12 | 59081758 | 59083753 | 1995 | NZO | 12 | 59082177 | 59082210 | 34 | + | 18.065 | 3.38E-07 | 0.064 | TAGCTGTCTTCAGACTCACCAGAAGAGGGCGTCA | | |
| 1045 | 12 | 59081758 | 59084703 | 2945 | AJ,BALB,C3H,CBA,DBA1J,DBA2J,ILNJ2,KK,MA/Mv.RH2.SEA | 12 | 59082192 | 59082210 | 19 | + | 17.0328 | 8.88E-07 | 0.147 | TCACCAGAAGAGGGCGTCA | | |
| 1045 | 12 | 59081758 | 59083753 | 1995 | NZO | 12 | 59082192 | 59082210 | 19 | + | 17.0328 | 8.88E-07 | 0.147 | TCACCAGAAGAGGGCGTCA | | |
| 1045 | 12 | 59081758 | 59084703 | 2945 | AJ,BALB,C3H,CBA,DBA1J,DBA2J,ILNJ2,KK,MA/Mv.RH2.SEA | 12 | 59082272 | 59082305 | 34 | + | 18.065 | 3.38E-07 | 0.064 | TAGCTGTCTTCAGACTCACCAGAAGAGGGCGTCA | | |
| 1045 | 12 | 59081758 | 59083753 | 1995 | NZO | 12 | 59082272 | 59082305 | 34 | + | 18.065 | 3.38E-07 | 0.064 | TAGCTGTCTTCAGACTCACCAGAAGAGGGCGTCA | | |
| 1045 | 12 | 59081758 | 59084703 | 2945 | AJ,BALB,C3H,CBA,DBA1J,DBA2J,ILNJ2,KK,MA/Mv.RH2.SEA | 12 | 59082287 | 59082305 | 19 | + | 17.0328 | 8.88E-07 | 0.147 | TCACCAGAAGAGGGCGTCA | | |
| 1045 | 12 | 59081758 | 59083753 | 1995 | NZO | 12 | 59082287 | 59082305 | 19 | + | 17.0328 | 8.88E-07 | 0.147 | TCACCAGAAGAGGGCGTCA | | |
| 1045 | 12 | 59081758 | 59084703 | 2945 | AJ,BALB,C3H,CBA,DBA1J,DBA2J,ILNJ2,KK,MA/Mv.RH2.SEA | 12 | 59082367 | 59082400 | 34 | + | 18.065 | 3.38E-07 | 0.064 | TAGCTGTCTTCAGACTCACCAGAAGAGGGCGTCA | | |
| 1045 | 12 | 59081758 | 59083753 | 1995 | NZO | 12 | 59082367 | 59082400 | 34 | + | 18.065 | 3.38E-07 | 0.064 | TAGCTGTCTTCAGACTCACCAGAAGAGGGCGTCA | | |
| 1045 | 12 | 59081758 | 59084703 | 2945 | AJ,BALB,C3H,CBA,DBA1J,DBA2J,ILNJ2,KK,MA/Mv.RH2.SEA | 12 | 59082382 | 59082400 | 19 | + | 17.0328 | 8.88E-07 | 0.147 | TCACCAGAAGAGGGCGTCA | | |
| 1045 | 12 | 59081758 | 59083753 | 1995 | NZO | 12 | 59082382 | 59082400 | 19 | + | 17.0328 | 8.88E-07 | 0.147 | TCACCAGAAGAGGGCGTCA | | |
| 1045 | 12 | 59081758 | 59084703 | 2945 | AJ,BALB,C3H,CBA,DBA1J,DBA2J,ILNJ2,KK,MA/Mv.RH2.SEA | 12 | 59082462 | 59082495 | 34 | + | 18.065 | 3.38E-07 | 0.064 | TAGCTGTCTTCAGACTCACCAGAAGAGGGCGTCA | | |
| 1045 | 12 | 59081758 | 59083753 | 1995 | NZO | 12 | 59082462 | 59082495 | 34 | + | 18.065 | 3.38E-07 | 0.064 | TAGCTGTCTTCAGACTCACCAGAAGAGGGCGTCA | | |
| 1045 | 12 | 59081758 | 59084703 | 2945 | AJ,BALB,C3H,CBA,DBA1J,DBA2J,ILNJ2,KK,MA/Mv.RH2.SEA | 12 | 59082477 | 59082495 | 19 | + | 17.0328 | 8.88E-07 | 0.147 | TCACCAGAAGAGGGCGTCA | | |
| 1045 | 12 | 59081758 | 59083753 | 1995 | NZO | 12 | 59082477 | 59082495 | 19 | + | 17.0328 | 8.88E-07 | 0.147 | TCACCAGAAGAGGGCGTCA | | |
| 1045 | 12 | 59081758 | 59084703 | 2945 | AJ,BALB,C3H,CBA,DBA1J,DBA2J,ILNJ2,KK,MA/Mv.RH2.SEA | 12 | 59082557 | 59082590 | 34 | + | 18.065 | 3.38E-07 | 0.064 | TAGCTGTCTTCAGACTCACCAGAAGAGGGCGTCA | | |
| 1045 | 12 | 59081758 | 59083753 | 1995 | NZO | 12 | 59082557 | 59082590 | 34 | + | 18.065 | 3.38E-07 | 0.064 | TAGCTGTCTTCAGACTCACCAGAAGAGGGCGTCA | | |
| 1045 | 12 | 59081758 | 59084703 | 2945 | AJ,BALB,C3H,CBA,DBA1J,DBA2J,ILNJ2,KK,MA/Mv.RH2.SEA | 12 | 59082572 | 59082590 | 19 | + | 17.0328 | 8.88E-07 | 0.147 | TCACCAGAAGAGGGCGTCA | | |
| 1045 | 12 | 59081758 | 59083753 | 1995 | NZO | 12 | 59082572 | 59082590 | 19 | + | 17.0328 | 8.88E-07 | 0.147 | TCACCAGAAGAGGGCGTCA | | |
| 1045 | 12 | 59081758 | 59084703 | 2945 | AJ,BALB,C3H,CBA,DBA1J,DBA2J,ILNJ2,KK,MA/Mv.RH2.SEA | 12 | 59082652 | 59082685 | 34 | + | 18.065 | 3.38E-07 | 0.064 | TAGCTGTCTTCAGACTCACCAGAAGAGGGCGTCA | | |
| 1045 | 12 | 59081758 | 59083753 | 1995 | NZO | 12 | 59082652 | 59082685 | 34 | + | 18.065 | 3.38E-07 | 0.064 | TAGCTGTCTTCAGACTCACCAGAAGAGGGCGTCA | | |
| 1045 | 12 | 59081758 | 59084703 | 2945 | AJ,BALB,C3H,CBA,DBA1J,DBA2J,ILNJ2,KK,MA/Mv.RH2.SEA | 12 | 59082667 | 59082685 | 19 | + | 17.0328 | 8.88E-07 | 0.147 | TCACCAGAAGAGGGCGTCA | | |
| 1045 | 12 | 59081758 | 59083753 | 1995 | NZO | 12 | 59082667 | 59082685 | 19 | + | 17.0328 | 8.88E-07 | 0.147 | TCACCAGAAGAGGGCGTCA | | |
| 1045 | 12 | 59081758 | 59084703 | 2945 | AJ,BALB,C3H,CBA,DBA1J,DBA2J,ILNJ2,KK,MA/Mv.RH2.SEA | 12 | 59082747 | 59082780 | 34 | + | 18.065 | 3.38E-07 | 0.064 | TAGCTGTCTTCAGACTCACCAGAAGAGGGCGTCA | | |
| 1045 | 12 | 59081758 | 59083753 | 1995 | NZO | 12 | 59082747 | 59082780 | 34 | + | 18.065 | 3.38E-07 | 0.064 | TAGCTGTCTTCAGACTCACCAGAAGAGGGCGTCA | | |
| 1045 | 12 | 59081758 | 59084703 | 2945 | AJ,BALB,C3H,CBA,DBA1J,DBA2J,ILNJ2,KK,MA/Mv.RH2.SEA | 12 | 59082762 | 59082780 | 19 | + | 17.0328 | 8.88E-07 | 0.147 | TCACCAGAAGAGGGCGTCA | | |
| 1045 | 12 | 59081758 | 59083753 | 1995 | NZO | 12 | 59082762 | 59082780 | 19 | + | 17.0328 | 8.88E-07 | 0.147 | TCACCAGAAGAGGGCGTCA | | |
| 1045 | 12 | 59081758 | 59084703 | 2945 | AJ,BALB,C3H,CBA,DBA1J,DBA2J,ILNJ2,KK,MA/Mv.RH2.SEA | 12 | 59082842 | 59082875 | 34 | + | 18.065 | 3.38E-07 | 0.064 | TAGCTGTCTTCAGACTCACCAGAAGAGGGCGTCA | | |
| 1045 | 12 | 59081758 | 59083753 | 1995 | NZO | 12 | 59082842 | 59082875 | 34 | + | 18.065 | 3.38E-07 | 0.064 | TAGCTGTCTTCAGACTCACCAGAAGAGGGCGTCA | | |
| 1045 | 12 | 59081758 | 59084703 | 2945 | AJ,BALB,C3H,CBA,DBA1J,DBA2J,ILNJ2,KK,MA/Mv.RH2.SEA | 12 | 59082857 | 59082875 | 19 | + | 17.0328 | 8.88E-07 | 0.147 | TCACCAGAAGAGGGCGTCA | | |
| 1045 | 12 | 59081758 | 59083753 | 1995 | NZO | 12 | 59082857 | 59082875 | 19 | + | 17.0328 | 8.88E-07 | 0.147 | TCACCAGAAGAGGGCGTCA | | |
| 1045 | 12 | 59081758 | 59084703 | 2945 | AJ,BALB,C3H,CBA,DBA1J,DBA2J,ILNJ2,KK,MA/Mv.RH2.SEA | 12 | 59082937 | 59082970 | 34 | + | 18.065 | 3.38E-07 | 0.064 | TAGCTGTCTTCAGACTCACCAGAAGAGGGCGTCA | | |
| 1045 | 12 | 59081758 | 59083753 | 1995 | NZO | 12 | 59082937 | 59082970 | 34 | + | 18.065 | 3.38E-07 | 0.064 | TAGCTGTCTTCAGACTCACCAGAAGAGGGCGTCA | | |
| 1045 | 12 | 59081758 | 59084703 | 2945 | AJ,BALB,C3H,CBA,DBA1J,DBA2J,ILNJ2,KK,MA/Mv.RH2.SEA | 12 | 59082952 | 59082970 | 19 | + | 17.0328 | 8.88E-07 | 0.147 | TCACCAGAAGAGGGCGTCA | | |
| 1045 | 12 | 59081758 | 59083753 | 1995 | NZO | 12 | 59082952 | 59082970 | 19 | + | 17.0328 | 8.88E-07 | 0.147 | TCACCAGAAGAGGGCGTCA | | |
| 1045 | 12 | 59081758 | 59084703 | 2945 | AJ,BALB,C3H,CBA,DBA1J,DBA2J,ILNJ2,KK,MA/Mv.RH2.SEA | 12 | 59083032 | 59083065 | 34 | + | 18.065 | 3.38E-07 | 0.064 | TAGCTGTCTTCAGACTCACCAGAAGAGGGCGTCA | | |
| 1045 | 12 | 59081758 | 59083753 | 1995 | NZO | 12 | 59083032 | 59083065 | 34 | + | 18.065 | 3.38E-07 | 0.064 | TAGCTGTCTTCAGACTCACCAGAAGAGGGCGTCA | | |
| 1045 | 12 | 59081758 | 59084703 | 2945 | AJ,BALB,C3H,CBA,DBA1J,DBA2J,ILNJ2,KK,MA/Mv.RH2.SEA | 12 | 59083047 | 59083065 | 19 | + | 17.0328 | 8.88E-07 | 0.147 | TCACCAGAAGAGGGCGTCA | | |
| 1045 | 12 | 59081758 | 59083753 | 1995 | NZO | 12 | 59083047 | 59083065 | 19 | + | 17.0328 | 8.88E-07 | 0.147 | TCACCAGAAGAGGGCGTCA | | |
| 1045 | 12 | 59081758 | 59084703 | 2945 | AJ,BALB,C3H,CBA,DBA1J,DBA2J,ILNJ2,KK,MA/Mv.RH2.SEA | 12 | 59083127 | 59083160 | 34 | + | 18.065 | 3.38E-07 | 0.064 | TAGCTGTCTTCAGACTCACCAGAAGAGGGCGTCA | | |
| 1045 | 12 | 59081758 | 59083753 | 1995 | NZO | 12 | 59083127 | 59083160 | 34 | + | 18.065 | 3.38E-07 | 0.064 | TAGCTGTCTTCAGACTCACCAGAAGAGGGCGTCA | | |
| 1045 | 12 | 59081758 | 59084703 | 2945 | AJ,BALB,C3H,CBA,DBA1J,DBA2J,ILNJ2,KK,MA/Mv.RH2.SEA | 12 | 59083142 | 59083160 | 19 | + | 17.0328 | 8.88E-07 | 0.147 | TCACCAGAAGAGGGCGTCA | | |
| 1045 | 12 | 59081758 | 59083753 | 1995 | NZO | 12 | 59083142 | 59083160 | 19 | + | 17.0328 | 8.88E-07 | 0.147 | TCACCAGAAGAGGGCGTCA | | |
| 1045 | 12 | 59081758 | 59084703 | 2945 | AJ,BALB,C3H,CBA,DBA1J,DBA2J,ILNJ2,KK,MA/Mv.RH2.SEA | 12 | 59083222 | 59083255 | 34 | + | 18.065 | 3.38E-07 | 0.064 | TAGCTGTCTTCAGACTCACCAGAAGAGGGCGTCA | | |
| 1045 | 12 | 59081758 | 59083753 | 1995 | NZO | 12 | 59083222 | 59083255 | 34 | + | 18.065 | 3.38E-07 | 0.064 | TAGCTGTCTTCAGACTCACCAGAAGAGGGCGTCA | | |
| 1045 | 12 | 59081758 | 59084703 | 2945 | AJ,BALB,C3H,CBA,DBA1J,DBA2J,ILNJ2,KK,MA/Mv.RH2.SEA | 12 | 59083237 | 59083255 | 19 | + | 17.0328 | 8.88E-07 | 0.147 | TCACCAGAAGAGGGCGTCA | | |
| 1045 | 12 | 59081758 | 59083753 | 1995 | NZO | 12 | 59083237 | 59083255 | 19 | + | 17.0328 | 8.88E-07 | 0.147 | TCACCAGAAGAGGGCGTCA | | |
| 1045 | 12 | 59081758 | 59084703 | 2945 | AJ,BALB,C3H,CBA,DBA1J,DBA2J,ILNJ2,KK,MA/Mv.RH2.SEA | 12 | 59083317 | 59083350 | 34 | + | 18.065 | 3.38E-07 | 0.064 | TAGCTGTCTTCAGACTCACCAGAAGAGGGCGTCA | | |
| 1045 | 12 | 59081758 | 59083753 | 1995 | NZO | 12 | 59083317 | 59083350 | 34 | + | 18.065 | 3.38E-07 | 0.064 | TAGCTGTCTTCAGACTCACCAGAAGAGGGCGTCA | | |
| 1045 | 12 | 59081758 | 59084703 | 2945 | AJ,BALB,C3H,CBA,DBA1J,DBA2J,ILNJ2,KK,MA/Mv.RH2.SEA | 12 | 59083332 | 59083350 | 19 | + | 17.0328 | 8.88E-07 | 0.147 | TCACCAGAAGAGGGCGTCA | | |
| 1045 | 12 | 59081758 | 59083753 | 1995 | NZO | 12 | 59083332 | 59083350 | 19 | + | 17.0328 | 8.88E-07 | 0.147 | TCACCAGAAGAGGGCGTCA | | |
| 1045 | 12 | 59081758 | 59084703 | 2945 | AJ,BALB,C3H,CBA,DBA1J,DBA2J,ILNJ2,KK,MA/Mv.RH2.SEA | 12 | 59083412 | 59083445 | 34 | + | 18.065 | 3.38E-07 | 0.064 | TAGCTGTCTTCAGACTCACCAGAAGAGGGCGTCA | | |
| 1045 | 12 | 59081758 | 59083753 | 1995 | NZO | 12 | 59083412 | 59083445 | 34 | + | 18.065 | 3.38E-07 | 0.064 | TAGCTGTCTTCAGACTCACCAGAAGAGGGCGTCA | | |
| 1045 | 12 | 59081758 | 59084703 | 2945 | AJ,BALB,C3H,CBA,DBA1J,DBA2J,ILNJ2,KK,MA/Mv.RH2.SEA | 12 | 59083427 | 59083445 | 19 | + | 17.0328 | 8.88E-07 | 0.147 | TCACCAGAAGAGGGCGTCA | | |
| 1045 | 12 | 59081758 | 59083753 | 1995 | NZO | 12 | 59083427 | 59083445 | 19 | + | 17.0328 | 8.88E-07 | 0.147 | TCACCAGAAGAGGGCGTCA | | |
| 1045 | 12 | 59081758 | 59084703 | 2945 | AJ,BALB,C3H,CBA,DBA1J,DBA2J,ILNJ2,KK,MA/Mv.RH2.SEA | 12 | 59083507 | 59083540 | 34 | + | 18.065 | 3.38E-07 | 0.064 | TAGCTGTCTTCAGACTCACCAGAAGAGGGCGTCA | | |
| 1045 | 12 | 59081758 | 59083753 | 1995 | NZO | 12 | 59083507 | 59083540 | 34 | + | 18.065 | 3.38E-07 | 0.064 | TAGCTGTCTTCAGACTCACCAGAAGAGGGCGTCA | | |
| 1045 | 12 | 59081758 | 59084703 | 2945 | AJ,BALB,C3H,CBA,DBA1J,DBA2J,ILNJ2,KK,MA/Mv.RH2.SEA | 12 | 59083522 | 59083540 | 19 | + | 17.0328 | 8.88E-07 | 0.147 | TCACCAGAAGAGGGCGTCA | | |
| 1045 | 12 | 59081758 | 59083753 | 1995 | NZO | 12 | 59083522 | 59083540 | 19 | + | 17.0328 | 8.88E-07 | 0.147 | TCACCAGAAGAGGGCGTCA | | |
| 1045 | 12 | 59081758 | 59084703 | 2945 | AJ,BALB,C3H,CBA,DBA1J,DBA2J,ILNJ2,KK,MA/Mv.RH2.SEA | 12 | 59083602 | 59083635 | 34 | + | 18.065 | 3.38E-07 | 0.064 | TAGCTGTCTTCAGACTCACCAGAAGAGGGCGTCA | | |
| 1045 | 12 | 59081758 | 59083753 | 1995 | NZO | 12 | 59083602 | 59083635 | 34 | + | 18.065 | 3.38E-07 | 0.064 | TAGCTGTCTTCAGACTCACCAGAAGAGGGCGTCA | | |
| 1045 | 12 | 59081758 | 59084703 | 2945 | AJ,BALB,C3H,CBA,DBA1J,DBA2J,ILNJ2,KK,MA/Mv.RH2.SEA | 12 | 59083617 | 59083635 | 19 | + | 17.0328 | 8.88E-07 | 0.147 | TCACCAGAAGAGGGCGTCA | | |
| 1045 | 12 | 59081758 | 59083753 | 1995 | NZO | 12 | 59083617 | 59083635 | 19 | + | 17.0328 | 8.88E-07 | 0.147 | TCACCAGAAGAGGGCGTCA | | |
| 1045 | 12 | 59081758 | 59084703 | 2945 | AJ,BALB,C3H,CBA,DBA1J,DBA2J,ILNJ2,KK,MA/Mv.RH2.SEA | 12 | 59083697 | 59083730 | 34 | + | 18.065 | 3.38E-07 | 0.064 | TAGCTGTCTTCAGACTCACCAGAAGAGGGCGTCA | | |
| 1045 | 12 | 59081758 | 59083753 | 1995 | NZO | 12 | 59083697 | 59083730 | 34 | + | 18.065 | 3.38E-07 | 0.064 | TAGCTGTCTTCAGACTCACCAGAAGAGGGCGTCA | | |
| 1045 | 12 | 59081758 | 59084703 | 2945 | AJ,BALB,C3H,CBA,DBA1J,DBA2J,ILNJ2,KK,MA/Mv.RH2.SEA | 12 | 59083712 | 59083730 | 19 | + | 17.0328 | 8.88E-07 | 0.147 | TCACCAGAAGAGGGCGTCA | | |
| 1045 | 12 | 59081758 | 59083753 | 1995 | NZO | 12 | 59083712 | 59083730 | 19 | + | 17.0328 | 8.88E-07 | 0.147 | TCACCAGAAGAGGGCGTCA | | |
| 1045 | 12 | 59081758 | 59084703 | 2945 | AJ,BALB,C3H,CBA,DBA1J,DBA2J,ILNJ2,KK,MA/Mv.RH2.SEA | 12 | 59083792 | 59083825 | 34 | + | 18.065 | 3.38E-07 | 0.064 | TAGCTGTCTTCAGACTCACCAGAAGAGGGCGTCA | | |
| 1045 | 12 | 59081758 | 59084703 | 2945 | AJ,BALB,C3H,CBA,DBA1J,DBA2J,ILNJ2,KK,MA/Mv.RH2.SEA | 12 | 59083807 | 59083825 | 19 | + | 17.0328 | 8.88E-07 | 0.147 | TCACCAGAAGAGGGCGTCA | | |
| 1045 | 12 | 59081758 | 59084703 | 2945 | AJ,BALB,C3H,CBA,DBA1J,DBA2J,ILNJ2,KK,MA/Mv.RH2.SEA | 12 | 59083887 | 59083920 | 34 | + | 18.065 | 3.38E-07 | 0.064 | TAGCTGTCTTCAGACTCACCAGAAGAGGGCGTCA | | |
| 1045 | 12 | 59081758 | 59084703 | 2945 | AJ,BALB,C3H,CBA,DBA1J,DBA2J,ILNJ2,KK,MA/Mv.RH2.SEA | 12 | 59083902 | 59083920 | 19 | + | 17.0328 | 8.88E-07 | 0.147 | TCACCAGAAGAGGGCGTCA | | |
| 1045 | 12 | 59081758 | 59084703 | 2945 | AJ,BALB,C3H,CBA,DBA1J,DBA2J,ILNJ2,KK,MA/Mv.RH2.SEA | 12 | 59083982 | 59084015 | 34 | + | 18.065 | 3.38E-07 | 0.064 | TAGCTGTCTTCAGACTCACCAGAAGAGGGCGTCA | | |
| 1045 | 12 | 59081758 | 59084703 | 2945 | AJ,BALB,C3H,CBA,DBA1J,DBA2J,ILNJ2,KK,MA/Mv.RH2.SEA | 12 | 59083997 | 59084015 | 19 | + | 17.0328 | 8.88E-07 | 0.147 | TCACCAGAAGAGGGCGTCA | | |
| 1045 | 12 | 59081758 | 59084703 | 2945 | AJ,BALB,C3H,CBA,DBA1J,DBA2J,ILNJ2,KK,MA/Mv.RH2.SEA | 12 | 59084077 | 59084110 | 34 | + | 18.065 | 3.38E-07 | 0.064 | TAGCTGTCTTCAGACTCACCAGAAGAGGGCGTCA | | |
| 1045 | 12 | 59081758 | 59084703 | 2945 | AJ,BALB,C3H,CBA,DBA1J,DBA2J,ILNJ2,KK,MA/Mv.RH2.SEA | 12 | 59084092 | 59084110 | 19 | + | 17.0328 | 8.88E-07 | 0.147 | TCACCAGAAGAGGGCGTCA | | |
| 1045 | 12 | 59081758 | 59084703 | 2945 | AJ,BALB,C3H,CBA,DBA1J,DBA2J,ILNJ2,KK,MA/Mv.RH2.SEA | 12 | 59084172 | 59084205 | 34 | + | 18.065 | 3.38E-07 | 0.064 | TAGCTGTCTTCAGACTCACCAGAAGAGGGCGTCA | | |
| 1045 | 12 | 59081758 | 59084703 | 2945 | AJ,BALB,C3H,CBA,DBA1J,DBA2J,ILNJ2,KK,MA/Mv.RH2.SEA | 12 | 59084187 | 59084205 | 19 | + | 17.0328 | 8.88E-07 | 0.147 | TCACCAGAAGAGGGCGTCA | | |
| 1045 | 12 | 59081758 | 59084703 | 2945 | AJ,BALB,C3H,CBA,DBA1J,DBA2J,ILNJ2,KK,MA/Mv.RH2.SEA | 12 | 59084267 | 59084300 | 34 | + | 18.065 | 3.38E-07 | 0.064 | TAGCTGTCTTCAGACTCACCAGAAGAGGGCGTCA | | |
| 1045 | 12 | 59081758 | 59084703 | 2945 | AJ,BALB,C3H,CBA,DBA1J,DBA2J,ILNJ2,KK,MA/Mv.RH2.SEA | 12 | 59084282 | 59084300 | 19 | + | 17.0328 | 8.88E-07 | 0.147 | TCACCAGAAGAGGGCGTCA | | |
| 1045 | 12 | 59081758 | 59084703 | 2945 | AJ,BALB,C3H,CBA,DBA1J,DBA2J,ILNJ2,KK,MA/Mv.RH2.SEA | 12 | 59084362 | 59084395 | 34 | + | 18.065 | 3.38E-07 | 0.064 | TAGCTGTCTTCAGACTCACCAGAAGAGGGCGTCA | | |
| 1045 | 12 | 59081758 | 59084703 | 2945 | AJ,BALB,C3H,CBA,DBA1J,DBA2J,ILNJ2,KK,MA/Mv.RH2.SEA | 12 | 59084377 | 59084395 | 19 | + | 17.0328 | 8.88E-07 | 0.147 | TCACCAGAAGAGGGCGTCA | | |
| 1045 | 12 | 59081758 | 59084703 | 2945 | AJ,BALB,C3H,CBA,DBA1J,DBA2J,ILNJ2,KK,MA/Mv.RH2.SEA | 12 | 59084457 | 59084490 | 34 | + | 18.065 | 3.38E-07 | 0.064 | TAGCTGTCTTCAGACTCACCAGAAGAGGGCGTCA | | |
| 1045 | 12 | 59081758 | 59084703 | 2945 | AJ,BALB,C3H,CBA,DBA1J,DBA2J,ILNJ2,KK,MA/Mv.RH2.SEA | 12 | 59084472 | 59084490 | 19 | + | 17.0328 | 8.88E-07 | 0.147 | TCACCAGAAGAGGGCGTCA | | |
| 1045 | 12 | 59081758 | 59084703 | 2945 | AJ,BALB,C3H,CBA,DBA1J,DBA2J,ILNJ2,KK,MA/Mv.RH2.SEA | 12 | 59084552 | 59084585 | 34 | + | 18.065 | 3.38E-07 | 0.064 | TAGCTGTCTTCAGACTCACCAGAAGAGGGCGTCA | | |
| 1045 | 12 | 59081758 | 59084703 | 2945 | AJ,BALB,C3H,CBA,DBA1J,DBA2J,ILNJ2,KK,MA/Mv.RH2.SEA | 12 | 59084567 | 59084585 | 19 | + | 17.0328 | 8.88E-07 | 0.147 | TCACCAGAAGAGGGCGTCA | | |
| 1045 | 12 | 59081758 | 59084703 | 2945 | AJ,BALB,C3H,CBA,DBA1J,DBA2J,ILNJ2,KK,MA/Mv.RH2.SEA | 12 | 59084647 | 59084680 | 34 | + | 18.065 | 3.38E-07 | 0.064 | TAGCTGTCTTCAGACTCACCAGAAGAGGGCGTCA | | |
| 1045 | 12 | 59081758 | 59084703 | 2945 | AJ,BALB,C3H,CBA,DBA1J,DBA2J,ILNJ2,KK,MA/Mv.RH2.SEA | 12 | 59084662 | 59084680 | 19 | + | 17.0328 | 8.88E-07 | 0.147 | TCACCAGAAGAGGGCGTCA | | |
| 1046 | 12 | 59081831 | 59085915 | 4084 | BUB.CAST.FVB.MOLF.SPRET | 12 | 59081892 | 59081925 | 34 | + | 18.065 | 3.38E-07 | 0.064 | TAGCTGTCTTCAGACTCACCAGAAGAGGGCGTCA | | |
| 1046 | 12 | 59081831 | 59085915 | 4084 | BUB.CAST.FVB.MOLF.SPRET | 12 | 59081907 | 59081925 | 19 | + | 17.0328 | 8.88E-07 | 0.147 | TCACCAGAAGAGGGCGTCA | | |
| 1046 | 12 | 59081831 | 59085915 | 4084 | BUB.CAST.FVB.MOLF.SPRET | 12 | 59081987 | 59082020 | 34 | + | 18.065 | 3.38E-07 | 0.064 | TAGCTGTCTTCAGACTCACCAGAAGAGGGCGTCA | | |
| 1046 | 12 | 59081831 | 59085915 | 4084 | BUB.CAST.FVB.MOLF.SPRET | 12 | 59082002 | 59082020 | 19 | + | 17.0328 | 8.88E-07 | 0.147 | TCACCAGAAGAGGGCGTCA | | |
| 1046 | 12 | 59081831 | 59085915 | 4084 | BUB.CAST.FVB.MOLF.SPRET | 12 | 59082082 | 59082115 | 34 | + | 18.065 | 3.38E-07 | 0.064 | TAGCTGTCTTCAGACTCACCAGAAGAGGGCGTCA | | |
| 1046 | 12 | 59081831 | 59085915 | 4084 | BUB.CAST.FVB.MOLF.SPRET | 12 | 59082097 | 59082115 | 19 | + | 17.0328 | 8.88E-07 | 0.147 | TCACCAGAAGAGGGCGTCA | | |
| 1046 | 12 | 59081831 | 59085915 | 4084 | BUB.CAST.FVB.MOLF.SPRET | 12 | 59082177 | 59082210 | 34 | + | 18.065 | 3.38E-07 | 0.064 | TAGCTGTCTTCAGACTCACCAGAAGAGGGCGTCA | | |
| 1046 | 12 | 59081831 | 59085915 | 4084 | BUB.CAST.FVB.MOLF.SPRET | 12 | 59082192 | 59082210 | 19 | + | 17.0328 | 8.88E-07 | 0.147 | TCACCAGAAGAGGGCGTCA | | |
| 1046 | 12 | 59081831 | 59085915 | 4084 | BUB.CAST.FVB.MOLF.SPRET | 12 | 59082272 | 59082305 | 34 | + | 18.065 | 3.38E-07 | 0.064 | TAGCTGTCTTCAGACTCACCAGAAGAGGGCGTCA | | |
| 1046 | 12 | 59081831 | 59085915 | 4084 | BUB.CAST.FVB.MOLF.SPRET | 12 | 59082287 | 59082305 | 19 | + | 17.0328 | 8.88E-07 | 0.147 | TCACCAGAAGAGGGCGTCA | | |
| 1046 | 12 | 59081831 | 59085915 | 4084 | BUB.CAST.FVB.MOLF.SPRET | 12 | 59082367 | 59082400 | 34 | + | 18.065 | 3.38E-07 | 0.064 | TAGCTGTCTTCAGACTCACCAGAAGAGGGCGTCA | | |
| 1046 | 12 | 59081831 | 59085915 | 4084 | BUB.CAST.FVB.MOLF.SPRET | 12 | 59082382 | 59082400 | 19 | + | 17.0328 | 8.88E-07 | 0.147 | TCACCAGAAGAGGGCGTCA | | |
| 1046 | 12 | 59081831 | 59085915 | 4084 | BUB.CAST.FVB.MOLF.SPRET | 12 | 59082462 | 59082495 | 34 | + | 18.065 | 3.38E-07 | 0.064 | TAGCTGTCTTCAGACTCACCAGAAGAGGGCGTCA | | |
| 1046 | 12 | 59081831 | 59085915 | 4084 | BUB.CAST.FVB.MOLF.SPRET | 12 | 59082477 | 59082495 | 19 | + | 17.0328 | 8.88E-07 | 0.147 | TCACCAGAAGAGGGCGTCA | | |
| 1046 | 12 | 59081831 | 59085915 | 4084 | BUB.CAST.FVB.MOLF.SPRET | 12 | 59082557 | 59082590 | 34 | + | 18.065 | 3.38E-07 | 0.064 | TAGCTGTCTTCAGACTCACCAGAAGAGGGCGTCA | | |
| 1046 | 12 | 59081831 | 59085915 | 4084 | BUB.CAST.FVB.MOLF.SPRET | 12 | 59082572 | 59082590 | 19 | + | 17.0328 | 8.88E-07 | 0.147 | TCACCAGAAGAGGGCGTCA | | |
| 1046 | 12 | 59081831 | 59085915 | 4084 | BUB.CAST.FVB.MOLF.SPRET | 12 | 59082652 | 59082685 | 34 | + | 18.065 | 3.38E-07 | 0.064 | TAGCTGTCTTCAGACTCACCAGAAGAGGGCGTCA | | |
| 1046 | 12 | 59081831 | 59085915 | 4084 | BUB.CAST.FVB.MOLF.SPRET | 12 | 59082667 | 59082685 | 19 | + | 17.0328 | 8.88E-07 | 0.147 | TCACCAGAAGAGGGCGTCA | | |
| 1046 | 12 | 59081831 | 59085915 | 4084 | BUB.CAST.FVB.MOLF.SPRET | 12 | 59082747 | 59082780 | 34 | + | 18.065 | 3.38E-07 | 0.064 | TAGCTGTCTTCAGACTCACCAGAAGAGGGCGTCA | | |
| 1046 | 12 | 59081831 | 59085915 | 4084 | BUB.CAST.FVB.MOLF.SPRET | 12 | 59082762 | 59082780 | 19 | + | 17.0328 | 8.88E-07 | 0.147 | TCACCAGAAGAGGGCGTCA | | |
| 1046 | 12 | 59081831 | 59085915 | 4084 | BUB.CAST.FVB.MOLF.SPRET | 12 | 59082842 | 59082875 | 34 | + | 18.065 | 3.38E-07 | 0.064 | TAGCTGTCTTCAGACTCACCAGAAGAGGGCGTCA | | |
| 1046 | 12 | 59081831 | 59085915 | 4084 | BUB.CAST.FVB.MOLF.SPRET | 12 | 59082857 | 59082875 | 19 | + | 17.0328 | 8.88E-07 | 0.147 | TCACCAGAAGAGGGCGTCA | | |
| 1046 | 12 | 59081831 | 59085915 | 4084 | BUB.CAST.FVB.MOLF.SPRET | 12 | 59082937 | 59082970 | 34 | + | 18.065 | 3.38E-07 | 0.064 | TAGCTGTCTTCAGACTCACCAGAAGAGGGCGTCA | | |
| 1046 | 12 | 59081831 | 59085915 | 4084 | BUB.CAST.FVB.MOLF.SPRET | 12 | 59082952 | 59082970 | 19 | + | 17.0328 | 8.88E-07 | 0.147 | TCACCAGAAGAGGGCGTCA | | |
| 1046 | 12 | 59081831 | 59085915 | 4084 | BUB.CAST.FVB.MOLF.SPRET | 12 | 59083032 | 59083065 | 34 | + | 18.065 | 3.38E-07 | 0.064 | TAGCTGTCTTCAGACTCACCAGAAGAGGGCGTCA | | |
| 1046 | 12 | 59081831 | 59085915 | 4084 | BUB.CAST.FVB.MOLF.SPRET | 12 | 59083047 | 59083065 | 19 | + | 17.0328 | 8.88E-07 | 0.147 | TCACCAGAAGAGGGCGTCA | | |
| 1046 | 12 | 59081831 | 59085915 | 4084 | BUB.CAST.FVB.MOLF.SPRET | 12 | 59083127 | 59083160 | 34 | + | 18.065 | 3.38E-07 | 0.064 | TAGCTGTCTTCAGACTCACCAGAAGAGGGCGTCA | | |
| 1046 | 12 | 59081831 | 59085915 | 4084 | BUB.CAST.FVB.MOLF.SPRET | 12 | 59083142 | 59083160 | 19 | + | 17.0328 | 8.88E-07 | 0.147 | TCACCAGAAGAGGGCGTCA | | |
| 1046 | 12 | 59081831 | 59085915 | 4084 | BUB.CAST.FVB.MOLF.SPRET | 12 | 59083222 | 59083255 | 34 | + | 18.065 | 3.38E-07 | 0.064 | TAGCTGTCTTCAGACTCACCAGAAGAGGGCGTCA | | |
| 1046 | 12 | 59081831 | 59085915 | 4084 | BUB.CAST.FVB.MOLF.SPRET | 12 | 59083237 | 59083255 | 19 | + | 17.0328 | 8.88E-07 | 0.147 | TCACCAGAAGAGGGCGTCA | | |
| 1046 | 12 | 59081831 | 59085915 | 4084 | BUB.CAST.FVB.MOLF.SPRET | 12 | 59083317 | 59083350 | 34 | + | 18.065 | 3.38E-07 | 0.064 | TAGCTGTCTTCAGACTCACCAGAAGAGGGCGTCA | | |
| 1046 | 12 | 59081831 | 59085915 | 4084 | BUB.CAST.FVB.MOLF.SPRET | 12 | 59083332 | 59083350 | 19 | + | 17.0328 | 8.88E-07 | 0.147 | TCACCAGAAGAGGGCGTCA | | |
| 1046 | 12 | 59081831 | 59085915 | 4084 | BUB.CAST.FVB.MOLF.SPRET | 12 | 59083412 | 59083445 | 34 | + | 18.065 | 3.38E-07 | 0.064 | TAGCTGTCTTCAGACTCACCAGAAGAGGGCGTCA | | |
| 1046 | 12 | 59081831 | 59085915 | 4084 | BUB.CAST.FVB.MOLF.SPRET | 12 | 59083427 | 59083445 | 19 | + | 17.0328 | 8.88E-07 | 0.147 | TCACCAGAAGAGGGCGTCA | | |
| 1046 | 12 | 59081831 | 59085915 | 4084 | BUB.CAST.FVB.MOLF.SPRET | 12 | 59083507 | 59083540 | 34 | + | 18.065 | 3.38E-07 | 0.064 | TAGCTGTCTTCAGACTCACCAGAAGAGGGCGTCA | | |
| 1046 | 12 | 59081831 | 59085915 | 4084 | BUB.CAST.FVB.MOLF.SPRET | 12 | 59083522 | 59083540 | 19 | + | 17.0328 | 8.88E-07 | 0.147 | TCACCAGAAGAGGGCGTCA | | |
| 1046 | 12 | 59081831 | 59085915 | 4084 | BUB.CAST.FVB.MOLF.SPRET | 12 | 59083602 | 59083635 | 34 | + | 18.065 | 3.38E-07 | 0.064 | TAGCTGTCTTCAGACTCACCAGAAGAGGGCGTCA | | |
| 1046 | 12 | 59081831 | 59085915 | 4084 | BUB.CAST.FVB.MOLF.SPRET | 12 | 59083617 | 59083635 | 19 | + | 17.0328 | 8.88E-07 | 0.147 | TCACCAGAAGAGGGCGTCA | | |
| 1046 | 12 | 59081831 | 59085915 | 4084 | BUB.CAST.FVB.MOLF.SPRET | 12 | 59083697 | 59083730 | 34 | + | 18.065 | 3.38E-07 | 0.064 | TAGCTGTCTTCAGACTCACCAGAAGAGGGCGTCA | | |
| 1046 | 12 | 59081831 | 59085915 | 4084 | BUB.CAST.FVB.MOLF.SPRET | 12 | 59083712 | 59083730 | 19 | + | 17.0328 | 8.88E-07 | 0.147 | TCACCAGAAGAGGGCGTCA | | |
| 1046 | 12 | 59081831 | 59085915 | 4084 | BUB.CAST.FVB.MOLF.SPRET | 12 | 59083792 | 59083825 | 34 | + | 18.065 | 3.38E-07 | 0.064 | TAGCTGTCTTCAGACTCACCAGAAGAGGGCGTCA | | |
| 1046 | 12 | 59081831 | 59085915 | 4084 | BUB.CAST.FVB.MOLF.SPRET | 12 | 59083807 | 59083825 | 19 | + | 17.0328 | 8.88E-07 | 0.147 | TCACCAGAAGAGGGCGTCA | | |
| 1046 | 12 | 59081831 | 59085915 | 4084 | BUB.CAST.FVB.MOLF.SPRET | 12 | 59083887 | 59083920 | 34 | + | 18.065 | 3.38E-07 | 0.064 | TAGCTGTCTTCAGACTCACCAGAAGAGGGCGTCA | | |
| 1046 | 12 | 59081831 | 59085915 | 4084 | BUB.CAST.FVB.MOLF.SPRET | 12 | 59083902 | 59083920 | 19 | + | 17.0328 | 8.88E-07 | 0.147 | TCACCAGAAGAGGGCGTCA | | |

| | | | | | | | | | | | | | | | | |
|---|---|---|---|---|---|---|---|---|---|---|---|---|---|---|---|---|
| 1046 | 12 | 59081831 | 59085915 | 4084 | BUB CAST FVB MOLF SPRET | 12 | 59083982 | 59084015 | 34 | + | 18.065 | 3.38E-07 | 0.064 | TAGCTGTCTTCAGACTCACCAGAAGAGGGCGTCA | | |
| 1046 | 12 | 59081831 | 59085915 | 4084 | BUB CAST FVB MOLF SPRET | 12 | 59083997 | 59084015 | 19 | + | 17.0328 | 8.88E-07 | 0.147 | TCACCAGAAGAGGGCGTCA | | |
| 1046 | 12 | 59081831 | 59085915 | 4084 | BUB CAST FVB MOLF SPRET | 12 | 59084077 | 59084110 | 34 | + | 18.065 | 3.38E-07 | 0.064 | TAGCTGTCTTCAGACTCACCAGAAGAGGGCGTCA | | |
| 1046 | 12 | 59081831 | 59085915 | 4084 | BUB CAST FVB MOLF SPRET | 12 | 59084092 | 59084110 | 19 | + | 17.0328 | 8.88E-07 | 0.147 | TCACCAGAAGAGGGCGTCA | | |
| 1046 | 12 | 59081831 | 59085915 | 4084 | BUB CAST FVB MOLF SPRET | 12 | 59084172 | 59084205 | 34 | + | 18.065 | 3.38E-07 | 0.064 | TAGCTGTCTTCAGACTCACCAGAAGAGGGCGTCA | | |
| 1046 | 12 | 59081831 | 59085915 | 4084 | BUB CAST FVB MOLF SPRET | 12 | 59084187 | 59084205 | 19 | + | 17.0328 | 8.88E-07 | 0.147 | TCACCAGAAGAGGGCGTCA | | |
| 1046 | 12 | 59081831 | 59085915 | 4084 | BUB CAST FVB MOLF SPRET | 12 | 59084267 | 59084300 | 34 | + | 18.065 | 3.38E-07 | 0.064 | TAGCTGTCTTCAGACTCACCAGAAGAGGGCGTCA | | |
| 1046 | 12 | 59081831 | 59085915 | 4084 | BUB CAST FVB MOLF SPRET | 12 | 59084282 | 59084300 | 19 | + | 17.0328 | 8.88E-07 | 0.147 | TCACCAGAAGAGGGCGTCA | | |
| 1046 | 12 | 59081831 | 59085915 | 4084 | BUB CAST FVB MOLF SPRET | 12 | 59084362 | 59084395 | 34 | + | 18.065 | 3.38E-07 | 0.064 | TAGCTGTCTTCAGACTCACCAGAAGAGGGCGTCA | | |
| 1046 | 12 | 59081831 | 59085915 | 4084 | BUB CAST FVB MOLF SPRET | 12 | 59084377 | 59084395 | 19 | + | 17.0328 | 8.88E-07 | 0.147 | TCACCAGAAGAGGGCGTCA | | |
| 1046 | 12 | 59081831 | 59085915 | 4084 | BUB CAST FVB MOLF SPRET | 12 | 59084457 | 59084490 | 34 | + | 18.065 | 3.38E-07 | 0.064 | TAGCTGTCTTCAGACTCACCAGAAGAGGGCGTCA | | |
| 1046 | 12 | 59081831 | 59085915 | 4084 | BUB CAST FVB MOLF SPRET | 12 | 59084472 | 59084490 | 19 | + | 17.0328 | 8.88E-07 | 0.147 | TCACCAGAAGAGGGCGTCA | | |
| 1046 | 12 | 59081831 | 59085915 | 4084 | BUB CAST FVB MOLF SPRET | 12 | 59084552 | 59084585 | 34 | + | 18.065 | 3.38E-07 | 0.064 | TAGCTGTCTTCAGACTCACCAGAAGAGGGCGTCA | | |
| 1046 | 12 | 59081831 | 59085915 | 4084 | BUB CAST FVB MOLF SPRET | 12 | 59084567 | 59084585 | 19 | + | 17.0328 | 8.88E-07 | 0.147 | TCACCAGAAGAGGGCGTCA | | |
| 1046 | 12 | 59081831 | 59085915 | 4084 | BUB CAST FVB MOLF SPRET | 12 | 59084647 | 59084680 | 34 | + | 18.065 | 3.38E-07 | 0.064 | TAGCTGTCTTCAGACTCACCAGAAGAGGGCGTCA | | |
| 1046 | 12 | 59081831 | 59085915 | 4084 | BUB CAST FVB MOLF SPRET | 12 | 59084662 | 59084680 | 19 | + | 17.0328 | 8.88E-07 | 0.147 | TCACCAGAAGAGGGCGTCA | | |
| 1046 | 12 | 59081831 | 59085915 | 4084 | BUB CAST FVB MOLF SPRET | 12 | 59084742 | 59084775 | 34 | + | 18.065 | 3.38E-07 | 0.064 | TAGCTGTCTTCAGACTCACCAGAAGAGGGCGTCA | | |
| 1046 | 12 | 59081831 | 59085915 | 4084 | BUB CAST FVB MOLF SPRET | 12 | 59084757 | 59084775 | 19 | + | 17.0328 | 8.88E-07 | 0.147 | TCACCAGAAGAGGGCGTCA | | |
| 1046 | 12 | 59081831 | 59085915 | 4084 | BUB CAST FVB MOLF SPRET | 12 | 59084837 | 59084870 | 34 | + | 18.065 | 3.38E-07 | 0.064 | TAGCTGTCTTCAGACTCACCAGAAGAGGGCGTCA | | |
| 1046 | 12 | 59081831 | 59085915 | 4084 | BUB CAST FVB MOLF SPRET | 12 | 59084852 | 59084870 | 19 | + | 17.0328 | 8.88E-07 | 0.147 | TCACCAGAAGAGGGCGTCA | | |
| 1046 | 12 | 59081831 | 59085915 | 4084 | BUB CAST FVB MOLF SPRET | 12 | 59084932 | 59084965 | 34 | + | 18.065 | 3.38E-07 | 0.064 | TAGCTGTCTTCAGACTCACCAGAAGAGGGCGTCA | | |
| 1046 | 12 | 59081831 | 59085915 | 4084 | BUB CAST FVB MOLF SPRET | 12 | 59084947 | 59084965 | 19 | + | 17.0328 | 8.88E-07 | 0.147 | TCACCAGAAGAGGGCGTCA | | |
| 1046 | 12 | 59081831 | 59085915 | 4084 | BUB CAST FVB MOLF SPRET | 12 | 59085027 | 59085060 | 34 | + | 18.065 | 3.38E-07 | 0.064 | TAGCTGTCTTCAGACTCACCAGAAGAGGGCGTCA | | |
| 1046 | 12 | 59081831 | 59085915 | 4084 | BUB CAST FVB MOLF SPRET | 12 | 59085042 | 59085060 | 19 | + | 17.0328 | 8.88E-07 | 0.147 | TCACCAGAAGAGGGCGTCA | | |
| 1046 | 12 | 59081831 | 59085915 | 4084 | BUB CAST FVB MOLF SPRET | 12 | 59085122 | 59085155 | 34 | + | 18.065 | 3.38E-07 | 0.064 | TAGCTGTCTTCAGACTCACCAGAAGAGGGCGTCA | | |
| 1046 | 12 | 59081831 | 59085915 | 4084 | BUB CAST FVB MOLF SPRET | 12 | 59085137 | 59085155 | 19 | + | 17.0328 | 8.88E-07 | 0.147 | TCACCAGAAGAGGGCGTCA | | |
| 1046 | 12 | 59081831 | 59085915 | 4084 | BUB CAST FVB MOLF SPRET | 12 | 59085217 | 59085250 | 34 | + | 18.065 | 3.38E-07 | 0.064 | TAGCTGTCTTCAGACTCACCAGAAGAGGGCGTCA | | |
| 1046 | 12 | 59081831 | 59085915 | 4084 | BUB CAST FVB MOLF SPRET | 12 | 59085232 | 59085250 | 19 | + | 17.0328 | 8.88E-07 | 0.147 | TCACCAGAAGAGGGCGTCA | | |
| 1046 | 12 | 59081831 | 59085915 | 4084 | BUB CAST FVB MOLF SPRET | 12 | 59085312 | 59085345 | 34 | + | 18.065 | 3.38E-07 | 0.064 | TAGCTGTCTTCAGACTCACCAGAAGAGGGCGTCA | | |
| 1046 | 12 | 59081831 | 59085915 | 4084 | BUB CAST FVB MOLF SPRET | 12 | 59085327 | 59085345 | 19 | + | 17.0328 | 8.88E-07 | 0.147 | TCACCAGAAGAGGGCGTCA | | |
| 1046 | 12 | 59081831 | 59085915 | 4084 | BUB CAST FVB MOLF SPRET | 12 | 59085407 | 59085440 | 34 | + | 18.065 | 3.38E-07 | 0.064 | TAGCTGTCTTCAGACTCACCAGAAGAGGGCGTCA | | |
| 1046 | 12 | 59081831 | 59085915 | 4084 | BUB CAST FVB MOLF SPRET | 12 | 59085422 | 59085440 | 19 | + | 17.0328 | 8.88E-07 | 0.147 | TCACCAGAAGAGGGCGTCA | | |
| 1046 | 12 | 59081831 | 59085915 | 4084 | BUB CAST FVB MOLF SPRET | 12 | 59085502 | 59085535 | 34 | + | 18.065 | 3.38E-07 | 0.064 | TAGCTGTCTTCAGACTCACCAGAAGAGGGCGTCA | | |
| 1046 | 12 | 59081831 | 59085915 | 4084 | BUB CAST FVB MOLF SPRET | 12 | 59085517 | 59085535 | 19 | + | 17.0328 | 8.88E-07 | 0.147 | TCACCAGAAGAGGGCGTCA | | |
| 1046 | 12 | 59081831 | 59085915 | 4084 | BUB CAST FVB MOLF SPRET | 12 | 59085692 | 59085725 | 34 | + | 18.065 | 3.38E-07 | 0.064 | TAGCTGTCTTCAGACTCACCAGAAGAGGGCGTCA | | |
| 1046 | 12 | 59081831 | 59085915 | 4084 | BUB CAST FVB MOLF SPRET | 12 | 59085707 | 59085725 | 19 | + | 17.0328 | 8.88E-07 | 0.147 | TCACCAGAAGAGGGCGTCA | | |
| 1046 | 12 | 59081831 | 59085915 | 4084 | BUB CAST FVB MOLF SPRET | 12 | 59085787 | 59085820 | 34 | + | 16.4472 | 9.61E-07 | 0.104 | TAGCTGTCTTCAGACACACCAGAAGAGGGCGTCA | 2 | ENCFF024JOZ,ENCFF883UPM |
| 1047 | 12 | 59082294 | 59085144 | 2850 | AJ BTBR CBA KK SJL SWR | 12 | 59082272 | 59082305 | 34 | + | 18.065 | 3.38E-07 | 0.064 | TAGCTGTCTTCAGACTCACCAGAAGAGGGCGTCA | | |
| 1047 | 12 | 59082294 | 59085144 | 2850 | AJ BTBR CBA KK SJL SWR | 12 | 59082287 | 59082305 | 19 | + | 17.0328 | 8.88E-07 | 0.147 | TCACCAGAAGAGGGCGTCA | | |
| 1047 | 12 | 59082294 | 59085144 | 2850 | AJ BTBR CBA KK SJL SWR | 12 | 59082367 | 59082400 | 34 | + | 18.065 | 3.38E-07 | 0.064 | TAGCTGTCTTCAGACTCACCAGAAGAGGGCGTCA | | |
| 1047 | 12 | 59082294 | 59085144 | 2850 | AJ BTBR CBA KK SJL SWR | 12 | 59082382 | 59082400 | 19 | + | 17.0328 | 8.88E-07 | 0.147 | TCACCAGAAGAGGGCGTCA | | |
| 1047 | 12 | 59082294 | 59085144 | 2850 | AJ BTBR CBA KK SJL SWR | 12 | 59082462 | 59082495 | 34 | + | 18.065 | 3.38E-07 | 0.064 | TAGCTGTCTTCAGACTCACCAGAAGAGGGCGTCA | | |
| 1047 | 12 | 59082294 | 59085144 | 2850 | AJ BTBR CBA KK SJL SWR | 12 | 59082477 | 59082495 | 19 | + | 17.0328 | 8.88E-07 | 0.147 | TCACCAGAAGAGGGCGTCA | | |
| 1047 | 12 | 59082294 | 59085144 | 2850 | AJ BTBR CBA KK SJL SWR | 12 | 59082557 | 59082590 | 34 | + | 18.065 | 3.38E-07 | 0.064 | TAGCTGTCTTCAGACTCACCAGAAGAGGGCGTCA | | |
| 1047 | 12 | 59082294 | 59085144 | 2850 | AJ BTBR CBA KK SJL SWR | 12 | 59082572 | 59082590 | 19 | + | 17.0328 | 8.88E-07 | 0.147 | TCACCAGAAGAGGGCGTCA | | |
| 1047 | 12 | 59082294 | 59085144 | 2850 | AJ BTBR CBA KK SJL SWR | 12 | 59082652 | 59082685 | 34 | + | 18.065 | 3.38E-07 | 0.064 | TAGCTGTCTTCAGACTCACCAGAAGAGGGCGTCA | | |
| 1047 | 12 | 59082294 | 59085144 | 2850 | AJ BTBR CBA KK SJL SWR | 12 | 59082667 | 59082685 | 19 | + | 17.0328 | 8.88E-07 | 0.147 | TCACCAGAAGAGGGCGTCA | | |
| 1047 | 12 | 59082294 | 59085144 | 2850 | AJ BTBR CBA KK SJL SWR | 12 | 59082747 | 59082780 | 34 | + | 18.065 | 3.38E-07 | 0.064 | TAGCTGTCTTCAGACTCACCAGAAGAGGGCGTCA | | |
| 1047 | 12 | 59082294 | 59085144 | 2850 | AJ BTBR CBA KK SJL SWR | 12 | 59082762 | 59082780 | 19 | + | 17.0328 | 8.88E-07 | 0.147 | TCACCAGAAGAGGGCGTCA | | |
| 1047 | 12 | 59082294 | 59085144 | 2850 | AJ BTBR CBA KK SJL SWR | 12 | 59082842 | 59082875 | 34 | + | 18.065 | 3.38E-07 | 0.064 | TAGCTGTCTTCAGACTCACCAGAAGAGGGCGTCA | | |
| 1047 | 12 | 59082294 | 59085144 | 2850 | AJ BTBR CBA KK SJL SWR | 12 | 59082857 | 59082875 | 19 | + | 17.0328 | 8.88E-07 | 0.147 | TCACCAGAAGAGGGCGTCA | | |
| 1047 | 12 | 59082294 | 59085144 | 2850 | AJ BTBR CBA KK SJL SWR | 12 | 59082937 | 59082970 | 34 | + | 18.065 | 3.38E-07 | 0.064 | TAGCTGTCTTCAGACTCACCAGAAGAGGGCGTCA | | |
| 1047 | 12 | 59082294 | 59085144 | 2850 | AJ BTBR CBA KK SJL SWR | 12 | 59082952 | 59082970 | 19 | + | 17.0328 | 8.88E-07 | 0.147 | TCACCAGAAGAGGGCGTCA | | |
| 1047 | 12 | 59082294 | 59085144 | 2850 | AJ BTBR CBA KK SJL SWR | 12 | 59083032 | 59083065 | 34 | + | 18.065 | 3.38E-07 | 0.064 | TAGCTGTCTTCAGACTCACCAGAAGAGGGCGTCA | | |
| 1047 | 12 | 59082294 | 59085144 | 2850 | AJ BTBR CBA KK SJL SWR | 12 | 59083047 | 59083065 | 19 | + | 17.0328 | 8.88E-07 | 0.147 | TCACCAGAAGAGGGCGTCA | | |
| 1047 | 12 | 59082294 | 59085144 | 2850 | AJ BTBR CBA KK SJL SWR | 12 | 59083127 | 59083160 | 34 | + | 18.065 | 3.38E-07 | 0.064 | TAGCTGTCTTCAGACTCACCAGAAGAGGGCGTCA | | |
| 1047 | 12 | 59082294 | 59085144 | 2850 | AJ BTBR CBA KK SJL SWR | 12 | 59083142 | 59083160 | 19 | + | 17.0328 | 8.88E-07 | 0.147 | TCACCAGAAGAGGGCGTCA | | |
| 1047 | 12 | 59082294 | 59085144 | 2850 | AJ BTBR CBA KK SJL SWR | 12 | 59083222 | 59083255 | 34 | + | 18.065 | 3.38E-07 | 0.064 | TAGCTGTCTTCAGACTCACCAGAAGAGGGCGTCA | | |
| 1047 | 12 | 59082294 | 59085144 | 2850 | AJ BTBR CBA KK SJL SWR | 12 | 59083237 | 59083255 | 19 | + | 17.0328 | 8.88E-07 | 0.147 | TCACCAGAAGAGGGCGTCA | | |
| 1047 | 12 | 59082294 | 59085144 | 2850 | AJ BTBR CBA KK SJL SWR | 12 | 59083317 | 59083350 | 34 | + | 18.065 | 3.38E-07 | 0.064 | TAGCTGTCTTCAGACTCACCAGAAGAGGGCGTCA | | |
| 1047 | 12 | 59082294 | 59085144 | 2850 | AJ BTBR CBA KK SJL SWR | 12 | 59083332 | 59083350 | 19 | + | 17.0328 | 8.88E-07 | 0.147 | TCACCAGAAGAGGGCGTCA | | |
| 1047 | 12 | 59082294 | 59085144 | 2850 | AJ BTBR CBA KK SJL SWR | 12 | 59083412 | 59083445 | 34 | + | 18.065 | 3.38E-07 | 0.064 | TAGCTGTCTTCAGACTCACCAGAAGAGGGCGTCA | | |
| 1047 | 12 | 59082294 | 59085144 | 2850 | AJ BTBR CBA KK SJL SWR | 12 | 59083427 | 59083445 | 19 | + | 17.0328 | 8.88E-07 | 0.147 | TCACCAGAAGAGGGCGTCA | | |
| 1047 | 12 | 59082294 | 59085144 | 2850 | AJ BTBR CBA KK SJL SWR | 12 | 59083507 | 59083540 | 34 | + | 18.065 | 3.38E-07 | 0.064 | TAGCTGTCTTCAGACTCACCAGAAGAGGGCGTCA | | |
| 1047 | 12 | 59082294 | 59085144 | 2850 | AJ BTBR CBA KK SJL SWR | 12 | 59083522 | 59083540 | 19 | + | 17.0328 | 8.88E-07 | 0.147 | TCACCAGAAGAGGGCGTCA | | |
| 1047 | 12 | 59082294 | 59085144 | 2850 | AJ BTBR CBA KK SJL SWR | 12 | 59083602 | 59083635 | 34 | + | 18.065 | 3.38E-07 | 0.064 | TAGCTGTCTTCAGACTCACCAGAAGAGGGCGTCA | | |
| 1047 | 12 | 59082294 | 59085144 | 2850 | AJ BTBR CBA KK SJL SWR | 12 | 59083617 | 59083635 | 19 | + | 17.0328 | 8.88E-07 | 0.147 | TCACCAGAAGAGGGCGTCA | | |
| 1047 | 12 | 59082294 | 59085144 | 2850 | AJ BTBR CBA KK SJL SWR | 12 | 59083697 | 59083730 | 34 | + | 18.065 | 3.38E-07 | 0.064 | TAGCTGTCTTCAGACTCACCAGAAGAGGGCGTCA | | |
| 1047 | 12 | 59082294 | 59085144 | 2850 | AJ BTBR CBA KK SJL SWR | 12 | 59083712 | 59083730 | 19 | + | 17.0328 | 8.88E-07 | 0.147 | TCACCAGAAGAGGGCGTCA | | |
| 1047 | 12 | 59082294 | 59085144 | 2850 | AJ BTBR CBA KK SJL SWR | 12 | 59083792 | 59083825 | 34 | + | 18.065 | 3.38E-07 | 0.064 | TAGCTGTCTTCAGACTCACCAGAAGAGGGCGTCA | | |
| 1047 | 12 | 59082294 | 59085144 | 2850 | AJ BTBR CBA KK SJL SWR | 12 | 59083807 | 59083825 | 19 | + | 17.0328 | 8.88E-07 | 0.147 | TCACCAGAAGAGGGCGTCA | | |
| 1047 | 12 | 59082294 | 59085144 | 2850 | AJ BTBR CBA KK SJL SWR | 12 | 59083887 | 59083920 | 34 | + | 18.065 | 3.38E-07 | 0.064 | TAGCTGTCTTCAGACTCACCAGAAGAGGGCGTCA | | |
| 1047 | 12 | 59082294 | 59085144 | 2850 | AJ BTBR CBA KK SJL SWR | 12 | 59083902 | 59083920 | 19 | + | 17.0328 | 8.88E-07 | 0.147 | TCACCAGAAGAGGGCGTCA | | |
| 1047 | 12 | 59082294 | 59085144 | 2850 | AJ BTBR CBA KK SJL SWR | 12 | 59083982 | 59084015 | 34 | + | 18.065 | 3.38E-07 | 0.064 | TAGCTGTCTTCAGACTCACCAGAAGAGGGCGTCA | | |
| 1047 | 12 | 59082294 | 59085144 | 2850 | AJ BTBR CBA KK SJL SWR | 12 | 59083997 | 59084015 | 19 | + | 17.0328 | 8.88E-07 | 0.147 | TCACCAGAAGAGGGCGTCA | | |
| 1047 | 12 | 59082294 | 59085144 | 2850 | AJ BTBR CBA KK SJL SWR | 12 | 59084077 | 59084110 | 34 | + | 18.065 | 3.38E-07 | 0.064 | TAGCTGTCTTCAGACTCACCAGAAGAGGGCGTCA | | |
| 1047 | 12 | 59082294 | 59085144 | 2850 | AJ BTBR CBA KK SJL SWR | 12 | 59084092 | 59084110 | 19 | + | 17.0328 | 8.88E-07 | 0.147 | TCACCAGAAGAGGGCGTCA | | |
| 1047 | 12 | 59082294 | 59085144 | 2850 | AJ BTBR CBA KK SJL SWR | 12 | 59084172 | 59084205 | 34 | + | 18.065 | 3.38E-07 | 0.064 | TAGCTGTCTTCAGACTCACCAGAAGAGGGCGTCA | | |
| 1047 | 12 | 59082294 | 59085144 | 2850 | AJ BTBR CBA KK SJL SWR | 12 | 59084187 | 59084205 | 19 | + | 17.0328 | 8.88E-07 | 0.147 | TCACCAGAAGAGGGCGTCA | | |
| 1047 | 12 | 59082294 | 59085144 | 2850 | AJ BTBR CBA KK SJL SWR | 12 | 59084267 | 59084300 | 34 | + | 18.065 | 3.38E-07 | 0.064 | TAGCTGTCTTCAGACTCACCAGAAGAGGGCGTCA | | |
| 1047 | 12 | 59082294 | 59085144 | 2850 | AJ BTBR CBA KK SJL SWR | 12 | 59084282 | 59084300 | 19 | + | 17.0328 | 8.88E-07 | 0.147 | TCACCAGAAGAGGGCGTCA | | |
| 1047 | 12 | 59082294 | 59085144 | 2850 | AJ BTBR CBA KK SJL SWR | 12 | 59084362 | 59084395 | 34 | + | 18.065 | 3.38E-07 | 0.064 | TAGCTGTCTTCAGACTCACCAGAAGAGGGCGTCA | | |
| 1047 | 12 | 59082294 | 59085144 | 2850 | AJ BTBR CBA KK SJL SWR | 12 | 59084377 | 59084395 | 19 | + | 17.0328 | 8.88E-07 | 0.147 | TCACCAGAAGAGGGCGTCA | | |
| 1047 | 12 | 59082294 | 59085144 | 2850 | AJ BTBR CBA KK SJL SWR | 12 | 59084457 | 59084490 | 34 | + | 18.065 | 3.38E-07 | 0.064 | TAGCTGTCTTCAGACTCACCAGAAGAGGGCGTCA | | |
| 1047 | 12 | 59082294 | 59085144 | 2850 | AJ BTBR CBA KK SJL SWR | 12 | 59084472 | 59084490 | 19 | + | 17.0328 | 8.88E-07 | 0.147 | TCACCAGAAGAGGGCGTCA | | |
| 1047 | 12 | 59082294 | 59085144 | 2850 | AJ BTBR CBA KK SJL SWR | 12 | 59084552 | 59084585 | 34 | + | 18.065 | 3.38E-07 | 0.064 | TAGCTGTCTTCAGACTCACCAGAAGAGGGCGTCA | | |
| 1047 | 12 | 59082294 | 59085144 | 2850 | AJ BTBR CBA KK SJL SWR | 12 | 59084567 | 59084585 | 19 | + | 17.0328 | 8.88E-07 | 0.147 | TCACCAGAAGAGGGCGTCA | | |
| 1047 | 12 | 59082294 | 59085144 | 2850 | AJ BTBR CBA KK SJL SWR | 12 | 59084647 | 59084680 | 34 | + | 18.065 | 3.38E-07 | 0.064 | TAGCTGTCTTCAGACTCACCAGAAGAGGGCGTCA | | |
| 1047 | 12 | 59082294 | 59085144 | 2850 | AJ BTBR CBA KK SJL SWR | 12 | 59084662 | 59084680 | 19 | + | 17.0328 | 8.88E-07 | 0.147 | TCACCAGAAGAGGGCGTCA | | |
| 1047 | 12 | 59082294 | 59085144 | 2850 | AJ BTBR CBA KK SJL SWR | 12 | 59084742 | 59084775 | 34 | + | 18.065 | 3.38E-07 | 0.064 | TAGCTGTCTTCAGACTCACCAGAAGAGGGCGTCA | | |
| 1047 | 12 | 59082294 | 59085144 | 2850 | AJ BTBR CBA KK SJL SWR | 12 | 59084757 | 59084775 | 19 | + | 17.0328 | 8.88E-07 | 0.147 | TCACCAGAAGAGGGCGTCA | | |
| 1047 | 12 | 59082294 | 59085144 | 2850 | AJ BTBR CBA KK SJL SWR | 12 | 59084837 | 59084870 | 34 | + | 18.065 | 3.38E-07 | 0.064 | TAGCTGTCTTCAGACTCACCAGAAGAGGGCGTCA | | |
| 1047 | 12 | 59082294 | 59085144 | 2850 | AJ BTBR CBA KK SJL SWR | 12 | 59084852 | 59084870 | 19 | + | 17.0328 | 8.88E-07 | 0.147 | TCACCAGAAGAGGGCGTCA | | |
| 1047 | 12 | 59082294 | 59085144 | 2850 | AJ BTBR CBA KK SJL SWR | 12 | 59084932 | 59084965 | 34 | + | 18.065 | 3.38E-07 | 0.064 | TAGCTGTCTTCAGACTCACCAGAAGAGGGCGTCA | | |
| 1047 | 12 | 59082294 | 59085144 | 2850 | AJ BTBR CBA KK SJL SWR | 12 | 59084947 | 59084965 | 19 | + | 17.0328 | 8.88E-07 | 0.147 | TCACCAGAAGAGGGCGTCA | | |
| 1047 | 12 | 59082294 | 59085144 | 2850 | AJ BTBR CBA KK SJL SWR | 12 | 59085027 | 59085060 | 34 | + | 18.065 | 3.38E-07 | 0.064 | TAGCTGTCTTCAGACTCACCAGAAGAGGGCGTCA | | |
| 1047 | 12 | 59082294 | 59085144 | 2850 | AJ BTBR CBA KK SJL SWR | 12 | 59085042 | 59085060 | 19 | + | 17.0328 | 8.88E-07 | 0.147 | TCACCAGAAGAGGGCGTCA | | |
| 1047 | 12 | 59082294 | 59085144 | 2850 | AJ BTBR CBA KK SJL SWR | 12 | 59085122 | 59085155 | 34 | + | 18.065 | 3.38E-07 | 0.064 | TAGCTGTCTTCAGACTCACCAGAAGAGGGCGTCA | | |
| 1047 | 12 | 59082294 | 59085144 | 2850 | AJ BTBR CBA KK SJL SWR | 12 | 59085137 | 59085155 | 19 | + | 17.0328 | 8.88E-07 | 0.147 | TCACCAGAAGAGGGCGTCA | | |
| 1048 | 12 | 59082858 | 59085898 | 3040 | SMJ | 12 | 59082842 | 59082875 | 34 | + | 18.065 | 3.38E-07 | 0.064 | TAGCTGTCTTCAGACTCACCAGAAGAGGGCGTCA | | |
| 1048 | 12 | 59082858 | 59085898 | 3040 | SMJ | 12 | 59082857 | 59082875 | 19 | + | 17.0328 | 8.88E-07 | 0.147 | TCACCAGAAGAGGGCGTCA | | |
| 1048 | 12 | 59082858 | 59085898 | 3040 | SMJ | 12 | 59082937 | 59082970 | 34 | + | 18.065 | 3.38E-07 | 0.064 | TAGCTGTCTTCAGACTCACCAGAAGAGGGCGTCA | | |
| 1048 | 12 | 59082858 | 59085898 | 3040 | SMJ | 12 | 59082952 | 59082970 | 19 | + | 17.0328 | 8.88E-07 | 0.147 | TCACCAGAAGAGGGCGTCA | | |
| 1048 | 12 | 59082858 | 59085898 | 3040 | SMJ | 12 | 59083032 | 59083065 | 34 | + | 18.065 | 3.38E-07 | 0.064 | TAGCTGTCTTCAGACTCACCAGAAGAGGGCGTCA | | |
| 1048 | 12 | 59082858 | 59085898 | 3040 | SMJ | 12 | 59083047 | 59083065 | 19 | + | 17.0328 | 8.88E-07 | 0.147 | TCACCAGAAGAGGGCGTCA | | |
| 1048 | 12 | 59082858 | 59085898 | 3040 | SMJ | 12 | 59083127 | 59083160 | 34 | + | 18.065 | 3.38E-07 | 0.064 | TAGCTGTCTTCAGACTCACCAGAAGAGGGCGTCA | | |
| 1048 | 12 | 59082858 | 59085898 | 3040 | SMJ | 12 | 59083142 | 59083160 | 19 | + | 17.0328 | 8.88E-07 | 0.147 | TCACCAGAAGAGGGCGTCA | | |
| 1048 | 12 | 59082858 | 59085898 | 3040 | SMJ | 12 | 59083222 | 59083255 | 34 | + | 18.065 | 3.38E-07 | 0.064 | TAGCTGTCTTCAGACTCACCAGAAGAGGGCGTCA | | |
| 1048 | 12 | 59082858 | 59085898 | 3040 | SMJ | 12 | 59083237 | 59083255 | 19 | + | 17.0328 | 8.88E-07 | 0.147 | TCACCAGAAGAGGGCGTCA | | |
| 1048 | 12 | 59082858 | 59085898 | 3040 | SMJ | 12 | 59083317 | 59083350 | 34 | + | 18.065 | 3.38E-07 | 0.064 | TAGCTGTCTTCAGACTCACCAGAAGAGGGCGTCA | | |
| 1048 | 12 | 59082858 | 59085898 | 3040 | SMJ | 12 | 59083332 | 59083350 | 19 | + | 17.0328 | 8.88E-07 | 0.147 | TCACCAGAAGAGGGCGTCA | | |
| 1048 | 12 | 59082858 | 59085898 | 3040 | SMJ | 12 | 59083412 | 59083445 | 34 | + | 18.065 | 3.38E-07 | 0.064 | TAGCTGTCTTCAGACTCACCAGAAGAGGGCGTCA | | |
| 1048 | 12 | 59082858 | 59085898 | 3040 | SMJ | 12 | 59083427 | 59083445 | 19 | + | 17.0328 | 8.88E-07 | 0.147 | TCACCAGAAGAGGGCGTCA | | |
| 1048 | 12 | 59082858 | 59085898 | 3040 | SMJ | 12 | 59083507 | 59083540 | 34 | + | 18.065 | 3.38E-07 | 0.064 | TAGCTGTCTTCAGACTCACCAGAAGAGGGCGTCA | | |
| 1048 | 12 | 59082858 | 59085898 | 3040 | SMJ | 12 | 59083522 | 59083540 | 19 | + | 17.0328 | 8.88E-07 | 0.147 | TCACCAGAAGAGGGCGTCA | | |
| 1048 | 12 | 59082858 | 59085898 | 3040 | SMJ | 12 | 59083602 | 59083635 | 34 | + | 18.065 | 3.38E-07 | 0.064 | TAGCTGTCTTCAGACTCACCAGAAGAGGGCGTCA | | |
| 1048 | 12 | 59082858 | 59085898 | 3040 | SMJ | 12 | 59083617 | 59083635 | 19 | + | 17.0328 | 8.88E-07 | 0.147 | TCACCAGAAGAGGGCGTCA | | |
| 1048 | 12 | 59082858 | 59085898 | 3040 | SMJ | 12 | 59083697 | 59083730 | 34 | + | 18.065 | 3.38E-07 | 0.064 | TAGCTGTCTTCAGACTCACCAGAAGAGGGCGTCA | | |
| 1048 | 12 | 59082858 | 59085898 | 3040 | SMJ | 12 | 59083712 | 59083730 | 19 | + | 17.0328 | 8.88E-07 | 0.147 | TCACCAGAAGAGGGCGTCA | | |
| 1048 | 12 | 59082858 | 59085898 | 3040 | SMJ | 12 | 59083792 | 59083825 | 34 | + | 18.065 | 3.38E-07 | 0.064 | TAGCTGTCTTCAGACTCACCAGAAGAGGGCGTCA | | |
| 1048 | 12 | 59082858 | 59085898 | 3040 | SMJ | 12 | 59083807 | 59083825 | 19 | + | 17.0328 | 8.88E-07 | 0.147 | TCACCAGAAGAGGGCGTCA | | |
| 1048 | 12 | 59082858 | 59085898 | 3040 | SMJ | 12 | 59083887 | 59083920 | 34 | + | 18.065 | 3.38E-07 | 0.064 | TAGCTGTCTTCAGACTCACCAGAAGAGGGCGTCA | | |
| 1048 | 12 | 59082858 | 59085898 | 3040 | SMJ | 12 | 59083902 | 59083920 | 19 | + | 17.0328 | 8.88E-07 | 0.147 | TCACCAGAAGAGGGCGTCA | | |
| 1048 | 12 | 59082858 | 59085898 | 3040 | SMJ | 12 | 59083982 | 59084015 | 34 | + | 18.065 | 3.38E-07 | 0.064 | TAGCTGTCTTCAGACTCACCAGAAGAGGGCGTCA | | |
| 1048 | 12 | 59082858 | 59085898 | 3040 | SMJ | 12 | 59083997 | 59084015 | 19 | + | 17.0328 | 8.88E-07 | 0.147 | TCACCAGAAGAGGGCGTCA | | |
| 1048 | 12 | 59082858 | 59085898 | 3040 | SMJ | 12 | 59084077 | 59084110 | 34 | + | 18.065 | 3.38E-07 | 0.064 | TAGCTGTCTTCAGACTCACCAGAAGAGGGCGTCA | | |
| 1048 | 12 | 59082858 | 59085898 | 3040 | SMJ | 12 | 59084092 | 59084110 | 19 | + | 17.0328 | 8.88E-07 | 0.147 | TCACCAGAAGAGGGCGTCA | | |
| 1048 | 12 | 59082858 | 59085898 | 3040 | SMJ | 12 | 59084172 | 59084205 | 34 | + | 18.065 | 3.38E-07 | 0.064 | TAGCTGTCTTCAGACTCACCAGAAGAGGGCGTCA | | |
| 1048 | 12 | 59082858 | 59085898 | 3040 | SMJ | 12 | 59084187 | 59084205 | 19 | + | 17.0328 | 8.88E-07 | 0.147 | TCACCAGAAGAGGGCGTCA | | |
| 1048 | 12 | 59082858 | 59085898 | 3040 | SMJ | 12 | 59084267 | 59084300 | 34 | + | 18.065 | 3.38E-07 | 0.064 | TAGCTGTCTTCAGACTCACCAGAAGAGGGCGTCA | | |
| 1048 | 12 | 59082858 | 59085898 | 3040 | SMJ | 12 | 59084282 | 59084300 | 19 | + | 17.0328 | 8.88E-07 | 0.147 | TCACCAGAAGAGGGCGTCA | | |
| 1048 | 12 | 59082858 | 59085898 | 3040 | SMJ | 12 | 59084362 | 59084395 | 34 | + | 18.065 | 3.38E-07 | 0.064 | TAGCTGTCTTCAGACTCACCAGAAGAGGGCGTCA | | |
| 1048 | 12 | 59082858 | 59085898 | 3040 | SMJ | 12 | 59084377 | 59084395 | 19 | + | 17.0328 | 8.88E-07 | 0.147 | TCACCAGAAGAGGGCGTCA | | |
| 1048 | 12 | 59082858 | 59085898 | 3040 | SMJ | 12 | 59084457 | 59084490 | 34 | + | 18.065 | 3.38E-07 | 0.064 | TAGCTGTCTTCAGACTCACCAGAAGAGGGCGTCA | | |
| 1048 | 12 | 59082858 | 59085898 | 3040 | SMJ | 12 | 59084472 | 59084490 | 19 | + | 17.0328 | 8.88E-07 | 0.147 | TCACCAGAAGAGGGCGTCA | | |
| 1048 | 12 | 59082858 | 59085898 | 3040 | SMJ | 12 | 59084552 | 59084585 | 34 | + | 18.065 | 3.38E-07 | 0.064 | TAGCTGTCTTCAGACTCACCAGAAGAGGGCGTCA | | |
| 1048 | 12 | 59082858 | 59085898 | 3040 | SMJ | 12 | 59084567 | 59084585 | 19 | + | 17.0328 | 8.88E-07 | 0.147 | TCACCAGAAGAGGGCGTCA | | |
| 1048 | 12 | 59082858 | 59085898 | 3040 | SMJ | 12 | 59084647 | 59084680 | 34 | + | 18.065 | 3.38E-07 | 0.064 | TAGCTGTCTTCAGACTCACCAGAAGAGGGCGTCA | | |
| 1048 | 12 | 59082858 | 59085898 | 3040 | SMJ | 12 | 59084662 | 59084680 | 19 | + | 17.0328 | 8.88E-07 | 0.147 | TCACCAGAAGAGGGCGTCA | | |
| 1048 | 12 | 59082858 | 59085898 | 3040 | SMJ | 12 | 59084742 | 59084775 | 34 | + | 18.065 | 3.38E-07 | 0.064 | TAGCTGTCTTCAGACTCACCAGAAGAGGGCGTCA | | |
| 1048 | 12 | 59082858 | 59085898 | 3040 | SMJ | 12 | 59084757 | 59084775 | 19 | + | 17.0328 | 8.88E-07 | 0.147 | TCACCAGAAGAGGGCGTCA | | |
| 1048 | 12 | 59082858 | 59085898 | 3040 | SMJ | 12 | 59084837 | 59084870 | 34 | + | 18.065 | 3.38E-07 | 0.064 | TAGCTGTCTTCAGACTCACCAGAAGAGGGCGTCA | | |
| 1048 | 12 | 59082858 | 59085898 | 3040 | SMJ | 12 | 59084852 | 59084870 | 19 | + | 17.0328 | 8.88E-07 | 0.147 | TCACCAGAAGAGGGCGTCA | | |
| 1048 | 12 | 59082858 | 59085898 | 3040 | SMJ | 12 | 59084932 | 59084965 | 34 | + | 18.065 | 3.38E-07 | 0.064 | TAGCTGTCTTCAGACTCACCAGAAGAGGGCGTCA | | |
| 1048 | 12 | 59082858 | 59085898 | 3040 | SMJ | 12 | 59084947 | 59084965 | 19 | + | 17.0328 | 8.88E-07 | 0.147 | TCACCAGAAGAGGGCGTCA | | |
| 1048 | 12 | 59082858 | 59085898 | 3040 | SMJ | 12 | 59085027 | 59085060 | 34 | + | 18.065 | 3.38E-07 | 0.064 | TAGCTGTCTTCAGACTCACCAGAAGAGGGCGTCA | | |
| 1048 | 12 | 59082858 | 59085898 | 3040 | SMJ | 12 | 59085042 | 59085060 | 19 | + | 17.0328 | 8.88E-07 | 0.147 | TCACCAGAAGAGGGCGTCA | | |
| 1048 | 12 | 59082858 | 59085898 | 3040 | SMJ | 12 | 59085122 | 59085155 | 34 | + | 18.065 | 3.38E-07 | 0.064 | TAGCTGTCTTCAGACTCACCAGAAGAGGGCGTCA | | |
| 1048 | 12 | 59082858 | 59085898 | 3040 | SMJ | 12 | 59085137 | 59085155 | 19 | + | 17.0328 | 8.88E-07 | 0.147 | TCACCAGAAGAGGGCGTCA | | |
| 1048 | 12 | 59082858 | 59085898 | 3040 | SMJ | 12 | 59085217 | 59085250 | 34 | + | 18.065 | 3.38E-07 | 0.064 | TAGCTGTCTTCAGACTCACCAGAAGAGGGCGTCA | | |
| 1048 | 12 | 59082858 | 59085898 | 3040 | SMJ | 12 | 59085232 | 59085250 | 19 | + | 17.0328 | 8.88E-07 | 0.147 | TCACCAGAAGAGGGCGTCA | | |
| 1048 | 12 | 59082858 | 59085898 | 3040 | SMJ | 12 | 59085312 | 59085345 | 34 | + | 18.065 | 3.38E-07 | 0.064 | TAGCTGTCTTCAGACTCACCAGAAGAGGGCGTCA | | |
| 1048 | 12 | 59082858 | 59085898 | 3040 | SMJ | 12 | 59085327 | 59085345 | 19 | + | 17.0328 | 8.88E-07 | 0.147 | TCACCAGAAGAGGGCGTCA | | |
| 1048 | 12 | 59082858 | 59085898 | 3040 | SMJ | 12 | 59085407 | 59085440 | 34 | + | 18.065 | 3.38E-07 | 0.064 | TAGCTGTCTTCAGACTCACCAGAAGAGGGCGTCA | | |
| 1048 | 12 | 59082858 | 59085898 | 3040 | SMJ | 12 | 59085422 | 59085440 | 19 | + | 17.0328 | 8.88E-07 | 0.147 | TCACCAGAAGAGGGCGTCA | | |
| 1048 | 12 | 59082858 | 59085898 | 3040 | SMJ | 12 | 59085502 | 59085535 | 34 | + | 18.065 | 3.38E-07 | 0.064 | TAGCTGTCTTCAGACTCACCAGAAGAGGGCGTCA | | |
| 1048 | 12 | 59082858 | 59085898 | 3040 | SMJ | 12 | 59085517 | 59085535 | 19 | + | 17.0328 | 8.88E-07 | 0.147 | TCACCAGAAGAGGGCGTCA | | |
| 1048 | 12 | 59082858 | 59085898 | 3040 | SMJ | 12 | 59085692 | 59085725 | 34 | + | 18.065 | 3.38E-07 | 0.064 | TAGCTGTCTTCAGACTCACCAGAAGAGGGCGTCA | | |
| 1048 | 12 | 59082858 | 59085898 | 3040 | SMJ | 12 | 59085707 | 59085725 | 19 | + | 17.0328 | 8.88E-07 | 0.147 | TCACCAGAAGAGGGCGTCA | | |
| 1048 | 12 | 59082858 | 59085898 | 3040 | SMJ | 12 | 59085787 | 59085820 | 34 | + | 16.4472 | 9.61E-07 | 0.104 | TAGCTGTCTTCAGACACACCAGAAGAGGGCGTCA | 2 | ENCFF024JOZ,ENCFF883UPM |
| 1049 | 12 | 59083722 | 59085622 | 1900 | AKR C57L CE NOD NOR RF WSB | 12 | 59083697 | 59083730 | 34 | + | 18.065 | 3.38E-07 | 0.064 | TAGCTGTCTTCAGACTCACCAGAAGAGGGCGTCA | | |
| 1049 | 12 | 59083722 | 59085622 | 1900 | AKR C57L CE NOD NOR RF WSB | 12 | 59083712 | 59083730 | 19 | + | 17.0328 | 8.88E-07 | 0.147 | TCACCAGAAGAGGGCGTCA | | |
| 1049 | 12 | 59083722 | 59085622 | 1900 | AKR C57L CE NOD NOR RF WSB | 12 | 59083792 | 59083825 | 34 | + | 18.065 | 3.38E-07 | 0.064 | TAGCTGTCTTCAGACTCACCAGAAGAGGGCGTCA | | |
| 1049 | 12 | 59083722 | 59085622 | 1900 | AKR C57L CE NOD NOR RF WSB | 12 | 59083807 | 59083825 | 19 | + | 17.0328 | 8.88E-07 | 0.147 | TCACCAGAAGAGGGCGTCA | | |
| 1049 | 12 | 59083722 | 59085622 | 1900 | AKR C57L CE NOD NOR RF WSB | 12 | 59083887 | 59083920 | 34 | + | 18.065 | 3.38E-07 | 0.064 | TAGCTGTCTTCAGACTCACCAGAAGAGGGCGTCA | | |
| 1049 | 12 | 59083722 | 59085622 | 1900 | AKR C57L CE NOD NOR RF WSB | 12 | 59083902 | 59083920 | 19 | + | 17.0328 | 8.88E-07 | 0.147 | TCACCAGAAGAGGGCGTCA | | |
| 1049 | 12 | 59083722 | 59085622 | 1900 | AKR C57L CE NOD NOR RF WSB | 12 | 59083982 | 59084015 | 34 | + | 18.065 | 3.38E-07 | 0.064 | TAGCTGTCTTCAGACTCACCAGAAGAGGGCGTCA | | |
| 1049 | 12 | 59083722 | 59085622 | 1900 | AKR C57L CE NOD NOR RF WSB | 12 | 59083997 | 59084015 | 19 | + | 17.0328 | 8.88E-07 | 0.147 | TCACCAGAAGAGGGCGTCA | | |
| 1049 | 12 | 59083722 | 59085622 | 1900 | AKR C57L CE NOD NOR RF WSB | 12 | 59084077 | 59084110 | 34 | + | 18.065 | 3.38E-07 | 0.064 | TAGCTGTCTTCAGACTCACCAGAAGAGGGCGTCA | | |
| 1049 | 12 | 59083722 | 59085622 | 1900 | AKR C57L CE NOD NOR RF WSB | 12 | 59084092 | 59084110 | 19 | + | 17.0328 | 8.88E-07 | 0.147 | TCACCAGAAGAGGGCGTCA | | |
| 1049 | 12 | 59083722 | 59085622 | 1900 | AKR C57L CE NOD NOR RF WSB | 12 | 59084172 | 59084205 | 34 | + | 18.065 | 3.38E-07 | 0.064 | TAGCTGTCTTCAGACTCACCAGAAGAGGGCGTCA | | |
| 1049 | 12 | 59083722 | 59085622 | 1900 | AKR C57L CE NOD NOR RF WSB | 12 | 59084187 | 59084205 | 19 | + | 17.0328 | 8.88E-07 | 0.147 | TCACCAGAAGAGGGCGTCA | | |
| 1049 | 12 | 59083722 | 59085622 | 1900 | AKR C57L CE NOD NOR RF WSB | 12 | 59084267 | 59084300 | 34 | + | 18.065 | 3.38E-07 | 0.064 | TAGCTGTCTTCAGACTCACCAGAAGAGGGCGTCA | | |
| 1049 | 12 | 59083722 | 59085622 | 1900 | AKR C57L CE NOD NOR RF WSB | 12 | 59084282 | 59084300 | 19 | + | 17.0328 | 8.88E-07 | 0.147 | TCACCAGAAGAGGGCGTCA | | |
| 1049 | 12 | 59083722 | 59085622 | 1900 | AKR C57L CE NOD NOR RF WSB | 12 | 59084362 | 59084395 | 34 | + | 18.065 | 3.38E-07 | 0.064 | TAGCTGTCTTCAGACTCACCAGAAGAGGGCGTCA | | |
| 1049 | 12 | 59083722 | 59085622 | 1900 | AKR C57L CE NOD NOR RF WSB | 12 | 59084377 | 59084395 | 19 | + | 17.0328 | 8.88E-07 | 0.147 | TCACCAGAAGAGGGCGTCA | | |
| 1049 | 12 | 59083722 | 59085622 | 1900 | AKR C57L CE NOD NOR RF WSB | 12 | 59084457 | 59084490 | 34 | + | 18.065 | 3.38E-07 | 0.064 | TAGCTGTCTTCAGACTCACCAGAAGAGGGCGTCA | | |
| 1049 | 12 | 59083722 | 59085622 | 1900 | AKR C57L CE NOD NOR RF WSB | 12 | 59084472 | 59084490 | 19 | + | 17.0328 | 8.88E-07 | 0.147 | TCACCAGAAGAGGGCGTCA | | |
| 1049 | 12 | 59083722 | 59085622 | 1900 | AKR C57L CE NOD NOR RF WSB | 12 | 59084552 | 59084585 | 34 | + | 18.065 | 3.38E-07 | 0.064 | TAGCTGTCTTCAGACTCACCAGAAGAGGGCGTCA | | |
| 1049 | 12 | 59083722 | 59085622 | 1900 | AKR C57L CE NOD NOR RF WSB | 12 | 59084567 | 59084585 | 19 | + | 17.0328 | 8.88E-07 | 0.147 | TCACCAGAAGAGGGCGTCA | | |
| 1049 | 12 | 59083722 | 59085622 | 1900 | AKR C57L CE NOD NOR RF WSB | 12 | 59084647 | 59084680 | 34 | + | 18.065 | 3.38E-07 | 0.064 | TAGCTGTCTTCAGACTCACCAGAAGAGGGCGTCA | | |
| 1049 | 12 | 59083722 | 59085622 | 1900 | AKR C57L CE NOD NOR RF WSB | 12 | 59084662 | 59084680 | 19 | + | 17.0328 | 8.88E-07 | 0.147 | TCACCAGAAGAGGGCGTCA | | |
| 1049 | 12 | 59083722 | 59085622 | 1900 | AKR C57L CE NOD NOR RF WSB | 12 | 59084742 | 59084775 | 34 | + | 18.065 | 3.38E-07 | 0.064 | TAGCTGTCTTCAGACTCACCAGAAGAGGGCGTCA | | |
| 1049 | 12 | 59083722 | 59085622 | 1900 | AKR C57L CE NOD NOR RF WSB | 12 | 59084757 | 59084775 | 19 | + | 17.0328 | 8.88E-07 | 0.147 | TCACCAGAAGAGGGCGTCA | | |
| 1049 | 12 | 59083722 | 59085622 | 1900 | AKR C57L CE NOD NOR RF WSB | 12 | 59084837 | 59084870 | 34 | + | 18.065 | 3.38E-07 | 0.064 | TAGCTGTCTTCAGACTCACCAGAAGAGGGCGTCA | | |
| 1049 | 12 | 59083722 | 59085622 | 1900 | AKR C57L CE NOD NOR RF WSB | 12 | 59084852 | 59084870 | 19 | + | 17.0328 | 8.88E-07 | 0.147 | TCACCAGAAGAGGGCGTCA | | |
| 1049 | 12 | 59083722 | 59085622 | 1900 | AKR C57L CE NOD NOR RF WSB | 12 | 59084932 | 59084965 | 34 | + | 18.065 | 3.38E-07 | 0.064 | TAGCTGTCTTCAGACTCACCAGAAGAGGGCGTCA | | |
| 1049 | 12 | 59083722 | 59085622 | 1900 | AKR C57L CE NOD NOR RF WSB | 12 | 59084947 | 59084965 | 19 | + | 17.0328 | 8.88E-07 | 0.147 | TCACCAGAAGAGGGCGTCA | | |
| 1049 | 12 | 59083722 | 59085622 | 1900 | AKR C57L CE NOD NOR RF WSB | 12 | 59085027 | 59085060 | 34 | + | 18.065 | 3.38E-07 | 0.064 | TAGCTGTCTTCAGACTCACCAGAAGAGGGCGTCA | | |
| 1049 | 12 | 59083722 | 59085622 | 1900 | AKR C57L CE NOD NOR RF WSB | 12 | 59085042 | 59085060 | 19 | + | 17.0328 | 8.88E-07 | 0.147 | TCACCAGAAGAGGGCGTCA | | |
| 1049 | 12 | 59083722 | 59085622 | 1900 | AKR C57L CE NOD NOR RF WSB | 12 | 59085122 | 59085155 | 34 | + | 18.065 | 3.38E-07 | 0.064 | TAGCTGTCTTCAGACTCACCAGAAGAGGGCGTCA | | |
| 1049 | 12 | 59083722 | 59085622 | 1900 | AKR C57L CE NOD NOR RF WSB | 12 | 59085137 | 59085155 | 19 | + | 17.0328 | 8.88E-07 | 0.147 | TCACCAGAAGAGGGCGTCA | | |
| 1049 | 12 | 59083722 | 59085622 | 1900 | AKR C57L CE NOD NOR RF WSB | 12 | 59085217 | 59085250 | 34 | + | 18.065 | 3.38E-07 | 0.064 | TAGCTGTCTTCAGACTCACCAGAAGAGGGCGTCA | | |
| 1049 | 12 | 59083722 | 59085622 | 1900 | AKR C57L CE NOD NOR RF WSB | 12 | 59085232 | 59085250 | 19 | + | 17.0328 | 8.88E-07 | 0.147 | TCACCAGAAGAGGGCGTCA | | |
| 1049 | 12 | 59083722 | 59085622 | 1900 | AKR C57L CE NOD NOR RF WSB | 12 | 59085312 | 59085345 | 34 | + | 18.065 | 3.38E-07 | 0.064 | TAGCTGTCTTCAGACTCACCAGAAGAGGGCGTCA | | |
| 1049 | 12 | 59083722 | 59085622 | 1900 | AKR C57L CE NOD NOR RF WSB | 12 | 59085327 | 59085345 | 19 | + | 17.0328 | 8.88E-07 | 0.147 | TCACCAGAAGAGGGCGTCA | | |
| 1049 | 12 | 59083722 | 59085622 | 1900 | AKR C57L CE NOD NOR RF WSB | 12 | 59085407 | 59085440 | 34 | + | 18.065 | 3.38E-07 | 0.064 | TAGCTGTCTTCAGACTCACCAGAAGAGGGCGTCA | | |
| 1049 | 12 | 59083722 | 59085622 | 1900 | AKR C57L CE NOD NOR RF WSB | 12 | 59085422 | 59085440 | 19 | + | 17.0328 | 8.88E-07 | 0.147 | TCACCAGAAGAGGGCGTCA | | |
| 1049 | 12 | 59083722 | 59085622 | 1900 | AKR C57L CE NOD NOR RF WSB | 12 | 59085502 | 59085535 | 34 | + | 18.065 | 3.38E-07 | 0.064 | TAGCTGTCTTCAGACTCACCAGAAGAGGGCGTCA | | |
| 1049 | 12 | 59083722 | 59085622 | 1900 | AKR C57L CE NOD NOR RF WSB | 12 | 59085517 | 59085535 | 19 | + | 17.0328 | 8.88E-07 | 0.147 | TCACCAGAAGAGGGCGTCA | | |
| 1050 | 12 | 59084291 | 59085621 | 1330 | 129S1 | 12 | 59084267 | 59084300 | 34 | + | 18.065 | 3.38E-07 | 0.064 | TAGCTGTCTTCAGACTCACCAGAAGAGGGCGTCA | | |
| 1050 | 12 | 59084291 | 59085621 | 1330 | 129S1 | 12 | 59084282 | 59084300 | 19 | + | 17.0328 | 8.88E-07 | 0.147 | TCACCAGAAGAGGGCGTCA | | |
| 1050 | 12 | 59084291 | 59085621 | 1330 | 129S1 | 12 | 59084362 | 59084395 | 34 | + | 18.065 | 3.38E-07 | 0.064 | TAGCTGTCTTCAGACTCACCAGAAGAGGGCGTCA | | |
| 1050 | 12 | 59084291 | 59085621 | 1330 | 129S1 | 12 | 59084377 | 59084395 | 19 | + | 17.0328 | 8.88E-07 | 0.147 | TCACCAGAAGAGGGCGTCA | | |
| 1050 | 12 | 59084291 | 59085621 | 1330 | 129S1 | 12 | 59084457 | 59084490 | 34 | + | 18.065 | 3.38E-07 | 0.064 | TAGCTGTCTTCAGACTCACCAGAAGAGGGCGTCA | | |
| 1050 | 12 | 59084291 | 59085621 | 1330 | 129S1 | 12 | 59084472 | 59084490 | 19 | + | 17.0328 | 8.88E-07 | 0.147 | TCACCAGAAGAGGGCGTCA | | |
| 1050 | 12 | 59084291 | 59085621 | 1330 | 129S1 | 12 | 59084552 | 59084585 | 34 | + | 18.065 | 3.38E-07 | 0.064 | TAGCTGTCTTCAGACTCACCAGAAGAGGGCGTCA | | |
| 1050 | 12 | 59084291 | 59085621 | 1330 | 129S1 | 12 | 59084567 | 59084585 | 19 | + | 17.0328 | 8.88E-07 | 0.147 | TCACCAGAAGAGGGCGTCA | | |
| 1050 | 12 | 59084291 | 59085621 | 1330 | 129S1 | 12 | 59084647 | 59084680 | 34 | + | 18.065 | 3.38E-07 | 0.064 | TAGCTGTCTTCAGACTCACCAGAAGAGGGCGTCA | | |
| 1050 | 12 | 59084291 | 59085621 | 1330 | 129S1 | 12 | 59084662 | 59084680 | 19 | + | 17.0328 | 8.88E-07 | 0.147 | TCACCAGAAGAGGGCGTCA | | |
| 1050 | 12 | 59084291 | 59085621 | 1330 | 129S1 | 12 | 59084742 | 59084775 | 34 | + | 18.065 | 3.38E-07 | 0.064 | TAGCTGTCTTCAGACTCACCAGAAGAGGGCGTCA | | |
| 1050 | 12 | 59084291 | 59085621 | 1330 | 129S1 | 12 | 59084757 | 59084775 | 19 | + | 17.0328 | 8.88E-07 | 0.147 | TCACCAGAAGAGGGCGTCA | | |
| 1050 | 12 | 59084291 | 59085621 | 1330 | 129S1 | 12 | 59084837 | 59084870 | 34 | + | 18.065 | 3.38E-07 | 0.064 | TAGCTGTCTTCAGACTCACCAGAAGAGGGCGTCA | | |
| 1050 | 12 | 59084291 | 59085621 | 1330 | 129S1 | 12 | 59084852 | 59084870 | 19 | + | 17.0328 | 8.88E-07 | 0.147 | TCACCAGAAGAGGGCGTCA | | |
| 1050 | 12 | 59084291 | 59085621 | 1330 | 129S1 | 12 | 59084932 | 59084965 | 34 | + | 18.065 | 3.38E-07 | 0.064 | TAGCTGTCTTCAGACTCACCAGAAGAGGGCGTCA | | |
| 1050 | 12 | 59084291 | 59085621 | 1330 | 129S1 | 12 | 59084947 | 59084965 | 19 | + | 17.0328 | 8.88E-07 | 0.147 | TCACCAGAAGAGGGCGTCA | | |
| 1050 | 12 | 59084291 | 59085621 | 1330 | 129S1 | 12 | 59085027 | 59085060 | 34 | + | 18.065 | 3.38E-07 | 0.064 | TAGCTGTCTTCAGACTCACCAGAAGAGGGCGTCA | | |
| 1050 | 12 | 59084291 | 59085621 | 1330 | 129S1 | 12 | 59085042 | 59085060 | 19 | + | 17.0328 | 8.88E-07 | 0.147 | TCACCAGAAGAGGGCGTCA | | |
| 1050 | 12 | 59084291 | 59085621 | 1330 | 129S1 | 12 | 59085122 | 59085155 | 34 | + | 18.065 | 3.38E-07 | 0.064 | TAGCTGTCTTCAGACTCACCAGAAGAGGGCGTCA | | |
| 1050 | 12 | 59084291 | 59085621 | 1330 | 129S1 | 12 | 59085137 | 59085155 | 19 | + | 17.0328 | 8.88E-07 | 0.147 | TCACCAGAAGAGGGCGTCA | | |
| 1050 | 12 | 59084291 | 59085621 | 1330 | 129S1 | 12 | 59085217 | 59085250 | 34 | + | 18.065 | 3.38E-07 | 0.064 | TAGCTGTCTTCAGACTCACCAGAAGAGGGCGTCA | | |
| 1050 | 12 | 59084291 | 59085621 | 1330 | 129S1 | 12 | 59085232 | 59085250 | 19 | + | 17.0328 | 8.88E-07 | 0.147 | TCACCAGAAGAGGGCGTCA | | |
| 1050 | 12 | 59084291 | 59085621 | 1330 | 129S1 | 12 | 59085312 | 59085345 | 34 | + | 18.065 | 3.38E-07 | 0.064 | TAGCTGTCTTCAGACTCACCAGAAGAGGGCGTCA | | |
| 1050 | 12 | 59084291 | 59085621 | 1330 | 129S1 | 12 | 59085327 | 59085345 | 19 | + | 17.0328 | 8.88E-07 | 0.147 | TCACCAGAAGAGGGCGTCA | | |
| 1050 | 12 | 59084291 | 59085621 | 1330 | 129S1 | 12 | 59085407 | 59085440 | 34 | + | 18.065 | 3.38E-07 | 0.064 | TAGCTGTCTTCAGACTCACCAGAAGAGGGCGTCA | | |
| 1050 | 12 | 59084291 | 59085621 | 1330 | 129S1 | 12 | 59085422 | 59085440 | 19 | + | 17.0328 | 8.88E-07 | 0.147 | TCACCAGAAGAGGGCGTCA | | |
| 1050 | 12 | 59084291 | 59085621 | 1330 | 129S1 | 12 | 59085502 | 59085535 | 34 | + | 18.065 | 3.38E-07 | 0.064 | TAGCTGTCTTCAGACTCACCAGAAGAGGGCGTCA | | |

| | | | | | | | | | | | | | | | | |
|---|---|---|---|---|---|---|---|---|---|---|---|---|---|---|---|---|
| 1050 | 12 | 59084291 | 59085621 | 1330 | 129S1 | 12 | 59085517 | 59085535 | 19 | + | 17.0328 | 8.88E-07 | 0.147 | TCACCAGAAGAGGGCGTCA | | |
| 1051 | 12 | 59962810 | 59971026 | 8216 | SPRET | 12 | 59968131 | 59968164 | 34 | + | 16.9268 | 7.10E-07 | 0.0911 | ATGCAGTTAGAGACATGAGCTCTGGGGGGTACTG | | |
| 1052 | 12 | 60730399 | 60733428 | 3029 | PI | 12 | 60730981 | 60731014 | 34 | + | 20.8211 | 4.88E-08 | 0.0195 | ATGTACTACTTACTGAGGCCAGAAGAGGGCAACA | 3 | ENCFF055HAP.ENCFF093ZRU.ENCFF502WDT |
| 1052 | 12 | 60730399 | 60733428 | 3029 | PI | 12 | 60730996 | 60731014 | 19 | + | 17.5246 | 6.45E-07 | 0.13 | AGGCCAGAAGAGGGCAACA | 3 | ENCFF055HAP.ENCFF093ZRU.ENCFF502WDT |
| 1053 | 12 | 63560774 | 63565911 | 5137 | SPRET | 12 | 63563980 | 63564013 | 34 | + | 18.0407 | 3.43E-07 | 0.0642 | ATGCAGGAAGACTTCTGGCCTGCAGAGGGTAGTG | | |
| 1053 | 12 | 63560774 | 63565911 | 5137 | SPRET | 12 | 63563995 | 63564013 | 19 | + | 17.3607 | 7.18E-07 | 0.136 | TGGCCTGCAGAGGGTAGTG | | |
| 1054 | 12 | 69240468 | 69241582 | 1114 | 129S1,BTBR,C57L,CAST,CE,MOLF,NZB,NZO,PJ | 12 | 69241419 | 69241452 | 34 | + | 16.7724 | 7.83E-07 | 0.0958 | TAGCTGTCTTCGGACACACCAGAAGAGGGCATCA | 1 | ENCFF093ZRU |
| 1055 | 12 | 69593575 | 69602620 | 9045 | AJ,AKR,BALB,BTBR,BUB,C3H,C57L,CAST,CBA,CE,FVB,KK,MA/My,MOLF,MRL,NOD,NOR,NZW,PL,RF,RH2,SEA,SJL,SMJ,SPRET,SWR,TH,WSB | 12 | 69599035 | 69599053 | 19 | + | 17.1639 | 8.17E-07 | 0.145 | CCTCCACGAGGTGGCGGAA | | |
| 1056 | 12 | 77970865 | 77974290 | 3425 | CAST | 12 | 77974224 | 77974257 | 34 | + | 19.8699 | 9.74E-08 | 0.0317 | GTGCAGGGCCTGCAGAAGCCAGAAGAGGAAGTCG | | |
| 1057 | 12 | 82579407 | 82580533 | 1126 | CE,SWR | 12 | 82580278 | 82580311 | 34 | + | 20.6992 | 5.34E-08 | 0.0206 | ATGCAAAGCTAACTGCGGCCAGCAGAGGCCCCGG | 1 | ENCFF093ZRU |
| 1058 | 12 | 83724802 | 83727799 | 2997 | CAST | 12 | 83725546 | 83725579 | 34 | + | 16.8537 | 7.43E-07 | 0.0932 | TAGCTGTCCTTAGATACTCCAGAAGAGGGCATCA | 1 | ENCFF093ZRU |
| 1059 | 12 | 87593448 | 87988990 | 395542 | CE,NOD,SWR | 12 | 87672777 | 87672810 | 34 | + | 17.9675 | 3.61E-07 | 0.0662 | ATGCAGTAATAAGAACAACCATAAGGTGGTGCTC | | |
| 1059 | 12 | 87593448 | 87988990 | 395542 | CE,NOD,SWR | 12 | 87697366 | 87697384 | 19 | + | 17.6557 | 5.92E-07 | 0.126 | TGCCCAACAGAGGGCACCC | | |
| 1059 | 12 | 87593448 | 87988990 | 395542 | CE,NOD,SWR | 12 | 87790496 | 87790529 | 34 | + | 17.9675 | 3.61E-07 | 0.0662 | ATGCAGTAATAAGAACAACCATAAGGTGGTGCTC | | |
| 1059 | 12 | 87593448 | 87988990 | 395542 | CE,NOD,SWR | 12 | 87815101 | 87815119 | 19 | + | 17.6557 | 5.92E-07 | 0.126 | TGCCCAACAGAGGGCACCC | | |
| 1059 | 12 | 87593448 | 87988990 | 395542 | CE,NOD,SWR | 12 | 87950885 | 87950919 | 35 | + | 20.5321 | 6.15E-08 | 0.0389 | CTGAAGTAACAAGAACAAACCAGAAGGTGGTGCTC | 7 | ENCFF092NMJ,ENCFF024JOZ,ENCFF089GBC,ENCFF099QNE,ENCFF502WDT,ENCFF093ZRU,ENCFF907XEG |
| 1059 | 12 | 87593448 | 87988990 | 395542 | CE,NOD,SWR | 12 | 87950972 | 87951005 | 34 | + | 17.9675 | 3.61E-07 | 0.0662 | ATGCAGTAATAAGAACAACCATAAGGTGGTGCTC | 7 | ENCFF092NMJ,ENCFF024JOZ,ENCFF089GBC,ENCFF099QNE,ENCFF502WDT,ENCFF093ZRU,ENCFF907XEG |
| 1059 | 12 | 87593448 | 87988990 | 395542 | CE,NOD,SWR | 12 | 87968631 | 87968649 | 19 | + | 17.6557 | 5.92E-07 | 0.126 | TGCCCAACAGAGGGCACCC | | |
| 1059 | 12 | 87593448 | 87988990 | 395542 | CE,NOD,SWR | 12 | 87623305 | 87623339 | 35 | - | 21.3028 | 3.68E-08 | 0.0274 | TTGCAGGTAAAGGTATGTGCCAGAAGATGGAGCTC | | |
| 1059 | 12 | 87593448 | 87988990 | 395542 | CE,NOD,SWR | 12 | 87770625 | 87770658 | 34 | - | 18.2358 | 3.02E-07 | 0.0615 | ATGCAGCTAGAGACACGAGCTCCTGGGGGCGCTA | | |
| 1059 | 12 | 87593448 | 87988990 | 395542 | CE,NOD,SWR | 12 | 87867060 | 87867094 | 35 | - | 19.7523 | 1.02E-07 | 0.053 | ATGCAGTAAATAAGAACAACCAGAAGGTGGTGCTC | | |
| 1059 | 12 | 87593448 | 87988990 | 395542 | CE,NOD,SWR | 12 | 87867146 | 87867180 | 35 | - | 19.6697 | 1.07E-07 | 0.0544 | CTGAACTAACACGAATCAACCAGAAGGTGGTGCTC | | |
| 1059 | 12 | 87593448 | 87988990 | 395542 | CE,NOD,SWR | 12 | 87867262 | 87867296 | 35 | - | 20.5321 | 6.15E-08 | 0.0389 | CTGAAGTAACAAGAACAAACCAGAAGGTGGTGCTC | | |
| 1059 | 12 | 87593448 | 87988990 | 395542 | CE,NOD,SWR | 12 | 87875841 | 87875874 | 34 | - | 16.9431 | 7.02E-07 | 0.0905 | ATGCAGCTAGAGACAGGAACTCTGGGGGGCACTG | | |
| 1060 | 12 | 88056899 | 88073777 | 16878 | MOLF | 12 | 88059326 | 88059344 | 19 | + | 17.5574 | 6.32E-07 | 0.128 | TGGCCTCCAGAGGGAGCCT | | |
| 1061 | 12 | 88114738 | 88138497 | 23759 | CE | 12 | 88124882 | 88124915 | 34 | + | 16.9431 | 7.02E-07 | 0.0905 | ATGCAGCTAGAGACAGGAACTCTGGGGGGCACTG | | |
| 1061 | 12 | 88114738 | 88138497 | 23759 | CE | 12 | 88133462 | 88133496 | 35 | + | 20.5321 | 6.15E-08 | 0.0389 | CTGAAGTAACAAGAACAAACCAGAAGGTGGTGCTC | | |
| 1061 | 12 | 88114738 | 88138497 | 23759 | CE | 12 | 88133578 | 88133612 | 35 | + | 19.6697 | 1.07E-07 | 0.0544 | CTGAACTAACACGAATCAACCAGAAGGTGGTGCTC | | |
| 1061 | 12 | 88114738 | 88138497 | 23759 | CE | 12 | 88133664 | 88133698 | 35 | + | 19.7523 | 1.02E-07 | 0.053 | ATGCAGTAAATAAGAACAACCAGAAGGTGGTGCTC | | |
| 1062 | 12 | 88116443 | 88136650 | 20207 | CAST,MOLF,SPRET | 12 | 88124882 | 88124915 | 34 | + | 16.9431 | 7.02E-07 | 0.0905 | ATGCAGCTAGAGACAGGAACTCTGGGGGGCACTG | | |
| 1062 | 12 | 88116443 | 88136650 | 20207 | CAST,MOLF,SPRET | 12 | 88133462 | 88133496 | 35 | + | 20.5321 | 6.15E-08 | 0.0389 | CTGAAGTAACAAGAACAAACCAGAAGGTGGTGCTC | | |
| 1062 | 12 | 88116443 | 88136650 | 20207 | CAST,MOLF,SPRET | 12 | 88133578 | 88133612 | 35 | + | 19.6697 | 1.07E-07 | 0.0544 | CTGAACTAACACGAATCAACCAGAAGGTGGTGCTC | | |
| 1062 | 12 | 88116443 | 88136650 | 20207 | CAST,MOLF,SPRET | 12 | 88133664 | 88133698 | 35 | + | 19.7523 | 1.02E-07 | 0.053 | ATGCAGTAAATAAGAACAACCAGAAGGTGGTGCTC | | |
| 1063 | 12 | 93505347 | 93509518 | 4171 | AJ,BUB,C3H,C58J,CAST,DBA1J,DBA2J,FVB,ILNJ2,KK,MOLF,NOD,NZO,NZW,PJ,PL,RF,SEA,SJL,SMJ,SPRET,SWR,TH,WSB | 12 | 93506123 | 93506156 | 34 | + | 17.6748 | 4.37E-07 | 0.073 | GAGCATGACCACAAGTCCACAGCAGGTGGCAACG | | |
| 1064 | 12 | 94062483 | 94063801 | 1318 | SPRET | 12 | 94063085 | 94063118 | 34 | + | 18.3659 | 2.76E-07 | 0.0591 | TAGCTGTCTTCAGACACTCCAGAAGAGGGCGCCA | | |
| 1064 | 12 | 94062483 | 94063801 | 1318 | SPRET | 12 | 94063100 | 94063118 | 19 | + | 17.5574 | 6.32E-07 | 0.128 | ACTCCAGAAGAGGGCGCCA | | |
| 1065 | 12 | 94200700 | 94203241 | 2541 | SPRET | 12 | 94203169 | 94203202 | 34 | + | 18.3089 | 2.87E-07 | 0.0597 | ATGCAGCTAGAGACACGAGCTCCAGGGGGTACTG | | |
| 1066 | 12 | 98688679 | 98689832 | 1153 | SPRET | 12 | 98688819 | 98688837 | 19 | + | 18.7869 | 2.72E-07 | 0.0847 | AAGCCAGAAGAGGGCACTA | | |
| 1067 | 12 | 99099864 | 99106207 | 6343 | 129S1,BTBR,C57L,FVB,KK,NZB,NZW,SWR | 12 | 99105995 | 99106028 | 34 | + | 17.2114 | 5.91E-07 | 0.0834 | TTGCTGGGCAAGCAGTTCTCAGCAGGGGACACCC | | |
| 1068 | 12 | 100829177 | 100830306 | 1129 | SPRET | 12 | 100829229 | 100829247 | 19 | + | 17.8197 | 5.31E-07 | 0.118 | TGACCAGGAGAGGGCAGGG | 1 | ENCFF907XEG |
| 1069 | 12 | 101930186 | 101933177 | 2991 | SPRET | 12 | 101931175 | 101931208 | 34 | + | 18.2439 | 3.00E-07 | 0.0613 | TAGCTGTCCTCAGATACTCCAGAAGGGGGCATCA | 1 | ENCFF093ZRU |
| 1070 | 12 | 105572530 | 105574044 | 1514 | SPRET | 12 | 105573245 | 105573278 | 34 | + | 16.6179 | 8.63E-07 | 0.1 | AAGCCTTTGCCCTGTGGGCCTCTAGGGGACACTG | | |
| 1071 | 12 | 109087950 | 109109567 | 21617 | AJ,AKR,C3H,CBA,TH | 12 | 109108884 | 109108918 | 35 | + | 16.7156 | 6.38E-07 | 0.158 | CTCCTGTCATCTCCGCATGGCACCAGGTGGCCCTC | | |
| 1072 | 12 | 3308551 | 3310752 | 2201 | CAST | 12 | 3309938 | 3309971 | 34 | - | 16.6504 | 8.46E-07 | 0.0996 | CAGCTCTCTTTAGACACACCAGAAGAGGGCATCA | | |
| 1073 | 12 | 7909674 | 7911678 | 2004 | SPRET | 12 | 7910348 | 7910382 | 35 | - | 20.2294 | 7.49E-08 | 0.0443 | CTGCTGTCCCTGGACCTGGACAGCAGTGGGCTGCT | | |
| 1074 | 12 | 10202122 | 10210229 | 8107 | 129S1,AJ,AKR,BALB,BTBR,BUB,C3H,C57L,C58J,CAST,CBA,CE,DBA1J,DBA2J,FVB,ILNJ2,KK,MA/My,MRL,NOD,NOR,NZB,NZO,NZW,PJ,PL,RF,SEA,SJL,SMJ,SPRET,SWR,TH,WSB | 12 | 10203885 | 10203918 | 34 | - | 19.8455 | 9.92E-08 | 0.0321 | CTGCAGCTCTAGCAGAGGCAAGGAGGGGTCTCTC | | |
| 1075 | 12 | 12708724 | 12713243 | 4519 | CAST,MOLF,SPRET | 12 | 12712440 | 12712473 | 34 | - | 17.6748 | 4.37E-07 | 0.073 | GAGCATGACCACAAGTCCACAGCAGGTGGCAACG | | |
| 1076 | 12 | 16163283 | 16164419 | 1136 | MOLF,NZB | 12 | 16163649 | 16163667 | 19 | - | 18.7049 | 2.88E-07 | 0.0872 | AGGCCAGCAGAGGGCATTA | 3 | ENCFF367UNR,ENCFF821BIA,ENCFF093ZRU |
| 1076 | 12 | 16163283 | 16164419 | 1136 | MOLF,NZB | 12 | 16163649 | 16163683 | 35 | - | 25.367 | 1.84E-09 | 0.00483 | GTGCAGGTGCCTGTAGAGGCCAGCAGAGGGCATTA | 3 | ENCFF367UNR,ENCFF821BIA,ENCFF093ZRU |
| 1077 | 12 | 17814509 | 17819957 | 5448 | AKR | 12 | 17817059 | 17817093 | 35 | - | 24.5138 | 3.60E-09 | 0.00636 | GTGCAGGTGCCCGTGGAGGCCTGAAGAGGGCACCT | | |
| 1078 | 12 | 19962119 | 19977953 | 15834 | MOLF | 12 | 19970408 | 19970441 | 34 | - | 17.8211 | 3.97E-07 | 0.0689 | GTGCAGCTTTTGGAGCCCCCAGCAGATGCAGATG | | |
| 1079 | 12 | 22831229 | 22836104 | 4875 | 129S1,AJ,AKR,BALB,BTBR,BUB,C3H,C57L,CBA,CE,DBA1J,DBA2J,FVB,ILNJ2,KK,MA/My,NOD,NOR,NZO,PJ,PL,RF,SEA,SJL,SMJ,SWR,TH | 12 | 22833318 | 22833351 | 34 | - | 19.2602 | 1.50E-07 | 0.0419 | CTGAAGGCCCAGCCTTGGCCTTCAGGTGGCTGCC | | |
| 1080 | 12 | 23647683 | 23656009 | 8326 | 129S1,BTBR,BUB,C3H,CBA,DBA2J,FVB,MA/My,MOLF,NOD,NOR,PL,SJL,SWR,TH | 12 | 23647975 | 23647993 | 19 | - | 19.2951 | 1.88E-07 | 0.0758 | CCGCCACTAGATGGTGCTA | | |
| 1080 | 12 | 23647683 | 23656009 | 8326 | 129S1,BTBR,BUB,C3H,CBA,DBA2J,FVB,MA/My,MOLF,NOD,NOR,PL,SJL,SWR,TH | 12 | 23655969 | 23655987 | 19 | - | 19.2951 | 1.88E-07 | 0.0758 | CCGCCACTAGATGGTGCTA | | |
| 1081 | 12 | 28813380 | 28814490 | 1110 | BTBR,C57L,FVB,MA/My,SJL,SWR | 12 | 28813686 | 28813704 | 19 | - | 17.2295 | 7.83E-07 | 0.142 | AGGCCAGAAGAGGGCAATG | 5 | ENCFF367UNR,ENCFF099QNE,ENCFF821BIA,ENCFF502WDT,ENCFF093ZRU |
| 1081 | 12 | 28813380 | 28814490 | 1110 | BTBR,C57L,FVB,MA/My,SJL,SWR | 12 | 28813686 | 28813719 | 34 | - | 26.7398 | 3.32E-10 | 0.000691 | ATGCAGTGCCTATGGAGGCCAGAAGAGGGCAATG | 5 | ENCFF367UNR,ENCFF099QNE,ENCFF821BIA,ENCFF502WDT,ENCFF093ZRU |
| 1082 | 12 | 36422188 | 36424339 | 2151 | CAST | 12 | 36423820 | 36423854 | 35 | - | 20.0826 | 8.24E-08 | 0.0467 | CTGCTGTATCACATATCTGTCGGTAGGTGGCAGAG | 24 | ENCFF504JDS,ENCFF092NMJ,ENCFF367UNR,ENCFF136ZDK,ENCFF821BIA,ENCFF430PPJ,ENCFF327LYS,ENCFF609MDN,ENCFF099QNE,ENCFF093ZRU,ENCFF907XEG,ENCFF198SSR,ENCFF194HVL,ENCFF055HAP,ENCFF491RJK,ENCFF957SPH,ENCFF024JOZ,ENCFF065ODV,ENCFF502WDT,ENCFF589ZBJ,ENCFF165LXO,ENCFF052KVO,ENCFF883UPM,ENCFF951GOP |
| 1083 | 12 | 39614450 | 39623651 | 9201 | CAST | 12 | 39622342 | 39622375 | 34 | - | 20.9593 | 4.40E-08 | 0.0181 | GTGCATTGCCTTTGGAGGTCAGAAGAGGGCATTG | 1 | ENCFF883UPM |
| 1084 | 12 | 43183369 | 43186034 | 2665 | SPRET | 12 | 43185820 | 43185853 | 34 | - | 18.187 | 3.12E-07 | 0.0627 | CAGCAGCTTTCCCACATCCCAGCAGAGGGTGAGG | | |
| 1085 | 12 | 45240779 | 45244396 | 3617 | KK,NZO,NZW | 12 | 45242311 | 45242344 | 34 | - | 16.439 | 9.66E-07 | 0.104 | CCCCAGTGTTCCCCTCCTCCACTGGGGGCCGCTG | | |
| 1086 | 12 | 54928745 | 54932992 | 4247 | SPRET | 12 | 54932014 | 54932048 | 35 | - | 16.7156 | 6.38E-07 | 0.158 | CTGCATTAGTTGCGGTCCACCGCAAGAAGGAGCCT | | |
| 1087 | 12 | 55287456 | 55300490 | 13034 | CE | 12 | 55295347 | 55295380 | 34 | - | 16.8943 | 7.24E-07 | 0.0917 | TTGCTCTCTTCAGACATACCAGAAGAGGGCATCA | | |
| 1088 | 12 | 59131715 | 59135174 | 3459 | SPRET | 12 | 59134439 | 59134473 | 35 | - | 16.0734 | 9.17E-07 | 0.19 | CTGCACAGAGAGCTCCAGATCACCAGAGAGAGGCC | | |
| 1089 | 12 | 63228592 | 63233990 | 5398 | SPRET | 12 | 63231845 | 63231878 | 34 | - | 16.6667 | 8.37E-07 | 0.099 | CTGCAGTGCCAGTGTACTCCACTGGAAGGATAAG | | |
| 1090 | 12 | 71851709 | 71858416 | 6707 | FVB,SWR | 12 | 71856527 | 71856560 | 34 | - | 24.2439 | 3.20E-09 | 0.0028 | GCGCAGTGCCTGTGGAGACCAGAAGAGGGCGTTG | 4 | ENCFF093ZRU,ENCFF367UNR,ENCFF099QNE,ENCFF052KVO |
| 1091 | 12 | 80451627 | 80454556 | 2929 | MOLF | 12 | 80453642 | 80453660 | 19 | - | 20.8197 | 5.71E-08 | 0.0463 | AGGCCACAAGAGGGCGGTG | 34 | ENCFF093ZRU,ENCFF210BTO,ENCFF805QBU,ENCFF367UNR,ENCFF201JUA,ENCFF099QNE,ENCFF601JVE,ENCFF327LYS,ENCFF951GOP,ENCFF316FYN,ENCFF065ODV,ENCFF821BIA,ENCFF907XEG,ENCFF575MGI,ENCFF883UPM,ENCFF089GBC,ENCFF136ZDK,ENCFF097CAQ,ENCFF589ZBJ,ENCFF609MDN,ENCFF194HVL,ENCFF165LXO,ENCFF502WDT,ENCFF024JOZ,ENCFF430PPJ,ENCFF092NMJ,ENCFF055HAP,ENCFF198SSR,ENCFF052KVO,ENCFF957SPH,ENCFF142CNG,ENCFF321AGQ,ENCFF533APC,ENCFF281QMV |
| 1091 | 12 | 80451627 | 80454556 | 2929 | MOLF | 12 | 80453642 | 80453675 | 34 | - | 23.8862 | 4.34E-09 | 0.00353 | GTACAGTACCCTGTCAGGCCACAAGAGGGCGGTG | 34 | ENCFF093ZRU,ENCFF210BTO,ENCFF805QBU,ENCFF367UNR,ENCFF201JUA,ENCFF099QNE,ENCFF601JVE,ENCFF327LYS,ENCFF951GOP,ENCFF316FYN,ENCFF065ODV,ENCFF821BIA,ENCFF907XEG,ENCFF575MGI,ENCFF883UPM,ENCFF089GBC,ENCFF136ZDK,ENCFF097CAQ,ENCFF589ZBJ,ENCFF609MDN,ENCFF194HVL,ENCFF165LXO,ENCFF502WDT,ENCFF024JOZ,ENCFF430PPJ,ENCFF092NMJ,ENCFF055HAP,ENCFF198SSR,ENCFF052KVO,ENCFF957SPH,ENCFF142CNG,ENCFF321AGQ,ENCFF533APC,ENCFF281QMV |
| 1092 | 12 | 83334886 | 83336451 | 1565 | C58J,CAST,CE,KK,MOLF,PL,SJL,SPRET,SWR,WSB | 12 | 83335584 | 83335618 | 35 | - | 16.578 | 6.90E-07 | 0.162 | CTGCAATTAATGTCCCAGGACAGAAGGGGTCATGT | | |
| 1093 | 12 | 86142133 | 86143808 | 1675 | SPRET | 12 | 86143129 | 86143162 | 34 | - | 17.9756 | 3.59E-07 | 0.066 | TAGCTGTTATCAGACACGCCAGAAGAGGGCATTG | | |
| 1094 | 12 | 90227474 | 90230259 | 2785 | SWR | 12 | 90229161 | 90229179 | 19 | - | 18.8197 | 2.66E-07 | 0.0837 | TGACCAGGAGGGGGCAACG | 1 | ENCFF093ZRU |
| 1095 | 12 | 90902496 | 90903583 | 1087 | BUB,CAST,MOLF,SWR | 12 | 90902657 | 90902675 | 19 | - | 17.2131 | 7.91E-07 | 0.143 | GTACCAGTAGGAGGCACTA | | |
| 1096 | 12 | 92358518 | 92360519 | 2001 | SPRET | 12 | 92359868 | 92359886 | 19 | - | 19.3934 | 1.75E-07 | 0.073 | TCTCCAGAAGGGGGCACCC | 2 | ENCFF093ZRU,ENCFF883UPM |
| 1097 | 12 | 93986508 | 93990052 | 3544 | MOLF | 12 | 93989778 | 93989811 | 34 | - | 17.6585 | 4.42E-07 | 0.0733 | CTGCACTTCCAGTCCATGCCAGCACGGGGGTTCC | | |
| 1098 | 12 | 96397055 | 96399674 | 2619 | SPRET | 12 | 96399570 | 96399603 | 34 | - | 16.5528 | 8.99E-07 | 0.102 | TAGCTGTCTTCAGACACACCAGAAGAGGGCGTCG | | |
| 1099 | 12 | 103970599 | 103972136 | 1537 | SPRET | 12 | 103971140 | 103971158 | 19 | - | 20.5574 | 7.07E-08 | 0.0517 | AAGCCAGAAGGGGGCGCCA | 4 | ENCFF367UNR,ENCFF052KVO,ENCFF033UQX,ENCFF093ZRU |
| 1099 | 12 | 103970599 | 103972136 | 1537 | SPRET | 12 | 103971140 | 103971173 | 34 | - | 29.6667 | 1.56E-11 | 0.000228 | GTGCTGTGCCTGTAGAAGCCAGAAGGGGGCGCCA | 4 | ENCFF367UNR,ENCFF052KVO,ENCFF033UQX,ENCFF093ZRU |
| 1100 | 12 | 105070280 | 105073289 | 3009 | CE,WSB | 12 | 105070495 | 105070513 | 19 | - | 20.0164 | 1.09E-07 | 0.0593 | GAACCACCAGGGGGCGCAC | 36 | ENCFF093ZRU,ENCFF201JUA,ENCFF367UNR,ENCFF099QNE,ENCFF024JOZ,ENCFF065ODV,ENCFF821BIA,ENCFF805QBU,ENCFF327LYS,ENCFF089GBC,ENCFF907XEG,ENCFF198SSR,ENCFF092NMJ,ENCFF136ZDK,ENCFF589ZBJ,ENCFF609MDN,ENCFF430PPJ,ENCFF033UQX,ENCFF165LXO,ENCFF321AGQ,ENCFF883UPM,ENCFF142CNG,ENCFF491RJK,ENCFF502WDT,ENCFF097CAQ,ENCFF281QMV,ENCFF055HAP,ENCFF210BTO,ENCFF052KVO,ENCFF957SPH,ENCFF533APC,ENCFF194HVL,ENCFF089EWX,ENCFF575MGI,ENCFF601JVE,ENCFF951GOP |
| 1101 | 12 | 110504408 | 110508319 | 3911 | DBA1J,DBA2J,ILNJ2 | 12 | 110504898 | 110504932 | 35 | - | 16.4495 | 7.42E-07 | 0.17 | GTGCCCTTGCCTGTGGAGGCCAGAGGAGGACACTG | | |
| 1102 | 12 | 110788704 | 110790053 | 1349 | CAST | 12 | 110788677 | 110788710 | 34 | - | 17.1789 | 6.04E-07 | 0.084 | TGGCTGTTTTTCCTTTGTCCAGCAGGTGCCAGGG | | |
| 1102 | 12 | 110788704 | 110790053 | 1349 | CAST | 12 | 110789906 | 110789940 | 35 | - | 19.4679 | 1.22E-07 | 0.0587 | CTGCATGTATACCTGTGTGCCAGAAGAAGGCATCA | | |
| 1103 | 12 | 113784585 | 113792306 | 7721 | CAST,MOLF | 12 | 113790192 | 113790210 | 19 | - | 21.7213 | 2.61E-08 | 0.0421 | GGGCCAGCAGGGGGCGATA | 5 | ENCFF052KVO,ENCFF327LYS,ENCFF883UPM,ENCFF502WDT,ENCFF055HAP |
| 1104 | 12 | 113799961 | 113806063 | 6102 | 129S1,AJ,BALB,BUB,C58J,CE,DBA1J,DBA2J,ILNJ2,KK,LP,MA/My,PJ,RF,SEA,SMJ,SPRET,TH,WSB | 12 | 113802072 | 113802090 | 19 | - | 17.8033 | 5.37E-07 | 0.119 | GAACCAGCAGAGGGCGATG | 2 | ENCFF367UNR,ENCFF883UPM |
| 1105 | 12 | 113870577 | 113884203 | 13626 | AJ,CE,DBA1J,DBA2J,ILNJ2,RF,WSB | 12 | 113875577 | 113875595 | 19 | - | 17.6066 | 6.11E-07 | 0.128 | GGACCAACAGGGGGCGCAC | 10 | ENCFF367UNR,ENCFF589ZBJ,ENCFF883UPM,ENCFF201JUA,ENCFF033UQX,ENCFF194HVL,ENCFF957SPH,ENCFF052KVO,ENCFF327LYS,ENCFF533APC |
| 1105 | 12 | 113870577 | 113884203 | 13626 | AJ,CE,DBA1J,DBA2J,ILNJ2,RF,WSB | 12 | 113875577 | 113875610 | 34 | - | 22.2764 | 1.61E-08 | 0.00928 | TTGCAGGGCTGCTCAGGACCAACAGGGGGCGCAC | 10 | ENCFF367UNR,ENCFF589ZBJ,ENCFF883UPM,ENCFF201JUA,ENCFF033UQX,ENCFF194HVL,ENCFF957SPH,ENCFF052KVO,ENCFF327LYS,ENCFF533APC |
| 1106 | 12 | 114032177 | 114042890 | 10713 | 129S1,AJ,BALB,BUB,C58J,KK,LP,MA/My,PJ,SEA,SMJ,TH | 12 | 114032357 | 114032375 | 19 | - | 18.459 | 3.43E-07 | 0.0947 | GGACCAGCAGGGGACACTG | | |
| 1107 | 12 | 114437524 | 114451059 | 13535 | 129S1,BALB,FVB,KK,LP,MA/My,PJ,SEA,SWR | 12 | 114442550 | 114442568 | 19 | - | 19.3934 | 1.75E-07 | 0.073 | TGACCAGCAGGGGGTGCTT | | |
| 1107 | 12 | 114437524 | 114451059 | 13535 | 129S1,BALB,FVB,KK,LP,MA/My,PJ,SEA,SWR | 12 | 114442550 | 114442583 | 34 | - | 17.0894 | 6.39E-07 | 0.0874 | CAGTTCAGGCACCCATGACCAGCAGGGGGTGCTT | | |
| 1107 | 12 | 114437524 | 114451059 | 13535 | 129S1,BALB,FVB,KK,LP,MA/My,PJ,SEA,SWR | 12 | 114449667 | 114449700 | 34 | - | 17.2439 | 5.79E-07 | 0.0827 | CTGGTATGACAAGTTTGTTCAGAAGGGGGCACAG | | |
| 1108 | 12 | 114457611 | 114468277 | 10666 | BALB,FVB,KK,LP,PJ,SEA,SWR,TH | 12 | 114459548 | 114459566 | 19 | - | 21.5246 | 3.11E-08 | 0.0421 | TGACCAGCAGGAGGCGCTC | 3 | ENCFF052KVO,ENCFF502WDT,ENCFF055HAP |
| 1109 | 12 | 114500133 | 114501768 | 1635 | AJ,RF,SPRET | 12 | 114501022 | 114501055 | 34 | - | 17.2439 | 5.79E-07 | 0.0827 | CTGGTATGACAAGTTTGTTCAGAAGGGGGCACAG | | |
| 1110 | 12 | 115063530 | 115070909 | 7379 | AJ,DBA2J,FVB,RF,SWR,WSB | 12 | 115063684 | 115063717 | 34 | - | 17.5772 | 4.66E-07 | 0.0751 | CTGCAGTCTCAGGTCTGCGCAATTGGGGCCGGCG | | |
| 1111 | 12 | 115232326 | 115243936 | 11610 | 129S1,BALB,BUB,C58J,KK,LP,MA/My,PJ,SMJ,TH | 12 | 115240554 | 115240572 | 19 | - | 21.082 | 4.57E-08 | 0.0421 | GGACCAGCAGGGGGCACAA | | |
| 1111 | 12 | 115232326 | 115243936 | 11610 | 129S1,BALB,BUB,C58J,KK,LP,MA/My,PJ,SMJ,TH | 12 | 115240554 | 115240587 | 34 | - | 26.2439 | 5.32E-10 | 0.000914 | CTGCAGGTGCGTTCAGGACCAGCAGGGGGCACAA | | |
| 1112 | 12 | 115246831 | 115251582 | 4751 | 129S1,AKR,BALB,BUB,C3H,C58J,CBA,FVB,KK,LP,MA/My,MRL,NZB,NZO,NZW,PJ,PL,SEA,SMJ,TH | 12 | 115250838 | 115250871 | 34 | - | 17.5691 | 4.69E-07 | 0.0751 | GAGCATGACCACAAGTCCACAGCAGGTGGCAACA | | |
| 1113 | 12 | 115842562 | 115862128 | 19566 | C3H,CAST,MRL,NZB,NZW,PL | 12 | 115854486 | 115854504 | 19 | - | 17.4262 | 6.88E-07 | 0.132 | TGACCAGCAGGGGTCATCA | | |
| 1114 | 12 | 115849768 | 115855995 | 6227 | 129S1,AJ,BALB,BUB,C58J,FVB,KK,LP,MA/My,PJ,SEA,SMJ,SPRET,SWR,TH | 12 | 115854486 | 115854504 | 19 | - | 17.4262 | 6.88E-07 | 0.132 | TGACCAGCAGGGGTCATCA | | |
| 1115 | 12 | 116675766 | 116677331 | 1565 | C3H,CAST,CBA,DBA1J,DBA2J,ILNJ2,MRL,PL,RF | 12 | 116677008 | 116677041 | 34 | - | 17.5854 | 4.64E-07 | 0.0751 | GTGCTGAACATAGAGGGGCCTCCAGAGAGCACTG | | |
| 1116 | 13 | 10099057 | 10100710 | 1653 | CAST | 13 | 10100103 | 10100137 | 35 | + | 16.4128 | 7.57E-07 | 0.174 | GTGCCAGTCCCTGTAGGGGTCAGAAGGGGGTGCCT | | |
| 1117 | 13 | 19115292 | 19129598 | 14306 | SPRET | 13 | 19129494 | 19129527 | 34 | + | 20.5203 | 6.09E-08 | 0.0257 | CTGCAGTTGTGGTGTCCTCCAGGAGAGGGACCAC | 1 | ENCFF024JOZ |
| 1118 | 13 | 19415516 | 19419850 | 4334 | SPRET | 13 | 19415527 | 19415560 | 34 | + | 18.5203 | 2.49E-07 | 0.0657 | ATGCAGCTAGAGACACGAACTCCAGGGGGTACTG | | |
| 1119 | 13 | 21987842 | 21990110 | 2268 | 129S1,C57L,ILNJ2 | 13 | 21988286 | 21988319 | 34 | + | 16.5447 | 9.04E-07 | 0.118 | CTGTACTTCCCATGGGCTCCTAGAGGTGGGACCC | | |
| 1120 | 13 | 21996577 | 22004435 | 7858 | 129S1,BTBR,C57L,CAST,CE,ILNJ2,KK,LP,MOLF,PJ,SEA | 13 | 22002035 | 22002053 | 19 | + | 17.1639 | 8.17E-07 | 0.174 | CCTCCACGAGGTGGCGGAA | | |
| 1121 | 13 | 22516446 | 22519482 | 3036 | BTBR,C57L,KK,LP,MOLF,NZB,NZO,NZW | 13 | 22518884 | 22518918 | 35 | + | 16.6147 | 6.75E-07 | 0.163 | CTGCATTTGCCTCCATTCACCTCTGGAGGAAACTT | | |
| 1122 | 13 | 24125793 | 24127042 | 1249 | BTBR,C57L,KK,LP,MOLF,NOD,NOR,NZB,PJ,SEA,SMJ | 13 | 24127007 | 24127040 | 34 | + | 17.4553 | 5.05E-07 | 0.0902 | TTAACGTACCTGCAGAGGCCAGAAGAGGGCACTG | 11 | ENCFF093ZRU,ENCFF092NMJ,ENCFF089GBC,ENCFF589ZBJ,ENCFF198SSR,ENCFF883UPM,ENCFF327LYS,ENCFF957SPH,ENCFF533APC,ENCFF024JOZ,ENCFF907XEG |
| 1122 | 13 | 24125793 | 24127042 | 1249 | BTBR,C57L,KK,LP,MOLF,NOD,NOR,NZB,PJ,SEA,SMJ | 13 | 24127022 | 24127040 | 19 | + | 20.0656 | 1.05E-07 | 0.0686 | AGGCCAGAAGAGGGCACTG | 11 | ENCFF093ZRU,ENCFF092NMJ,ENCFF089GBC,ENCFF589ZBJ,ENCFF198SSR,ENCFF883UPM,ENCFF327LYS,ENCFF957SPH,ENCFF533APC,ENCFF024JOZ,ENCFF907XEG |
| 1123 | 13 | 25191372 | 25199698 | 8326 | CAST,ILNJ2,KK,LP,MOLF,NOD,NOR,NZB,PJ,SEA,SMJ,SPRET,TH,WSB | 13 | 25191406 | 25191424 | 19 | + | 19.2951 | 1.88E-07 | 0.091 | CCGCCACTAGATGGTGCTA | 16 | ENCFF093ZRU,ENCFF367UNR,ENCFF099QNE,ENCFF502WDT,ENCFF201JUA,ENCFF065ODV,ENCFF089GBC,ENCFF327LYS,ENCFF821BIA,ENCFF907XEG,ENCFF589ZBJ,ENCFF092NMJ,ENCFF136ZDK,ENCFF198SSR,ENCFF430PPJ,ENCFF055HAP |
| 1123 | 13 | 25191372 | 25199698 | 8326 | CAST,ILNJ2,KK,LP,MOLF,NOD,NOR,NZB,PJ,SEA,SMJ,SPRET,TH,WSB | 13 | 25199400 | 25199418 | 19 | + | 19.2951 | 1.88E-07 | 0.091 | CCGCCACTAGATGGTGCTA | | |
| 1124 | 13 | 34922622 | 34925734 | 3112 | SPRET | 13 | 34925412 | 34925446 | 35 | + | 24.8073 | 2.86E-09 | 0.00601 | ATGCAATTGCCTGAAGAGTCCAGAAGAGGGCGCCG | 19 | ENCFF367UNR,ENCFF024JOZ,ENCFF089GBC,ENCFF136ZDK,ENCFF198SSR,ENCFF327LYS,ENCFF093ZRU,ENCFF883UPM,ENCFF052KVO,ENCFF033UQX,ENCFF099QNE,ENCFF957SPH,ENCFF055HAP,ENCFF502WDT,ENCFF142CNG,ENCFF907XEG,ENCFF065ODV,ENCFF821BIA,ENCFF533APC |
| 1124 | 13 | 34922622 | 34925734 | 3112 | SPRET | 13 | 34925428 | 34925446 | 19 | + | 18.7541 | 2.79E-07 | 0.109 | AGTCCAGAAGAGGGCGCCG | 19 | ENCFF367UNR,ENCFF024JOZ,ENCFF089GBC,ENCFF136ZDK,ENCFF198SSR,ENCFF327LYS,ENCFF093ZRU,ENCFF883UPM,ENCFF052KVO,ENCFF033UQX,ENCFF099QNE,ENCFF957SPH,ENCFF055HAP,ENCFF502WDT,ENCFF142CNG,ENCFF907XEG,ENCFF065ODV,ENCFF821BIA,ENCFF533APC |
| 1124 | 13 | 34922622 | 34925734 | 3112 | SPRET | 13 | 34924679 | 34924713 | 35 | - | 22.5138 | 1.59E-08 | 0.0163 | CTGCATGTACAGCTGAACACCAGAAGAGGGCATCA | 2 | ENCFF367UNR,ENCFF093ZRU |
| 1125 | 13 | 38837041 | 38838350 | 1309 | SPRET | 13 | 38837063 | 38837096 | 34 | + | 17.6992 | 4.30E-07 | 0.083 | TTGCTGTCTTCAGACACCCCAGAAGAGGGCATCA | | |
| 1126 | 13 | 39147406 | 39153567 | 6161 | SPRET | 13 | 39151097 | 39151115 | 19 | + | 20.2295 | 9.22E-08 | 0.0675 | TCACCAGTAGGGGGCAGAG | 3 | ENCFF327LYS,ENCFF883UPM,ENCFF136ZDK |
| 1127 | 13 | 40070107 | 40071627 | 1520 | 129S1,AJ,AKR,B10D2,B10J,BALB,BTBR,BUB,C3H,C57L,C58J,CBA,CE,DBA1J,DBA2J,FVB,ILNJ2,KK,LP,MA/My,MRL,NOD,NOR,NZB,NZO,NZW,PJ,PL,RF,RH2,SEA,SJL,SMJ,SWR,TH,WSB | 13 | 40070812 | 40070846 | 35 | + | 16.367 | 7.77E-07 | 0.175 | GTGCATGTGTCTATGGGATCCAGAAGAGGGCATCA | | |
| 1128 | 13 | 43739556 | 43740703 | 1147 | CAST | 13 | 43740066 | 43740100 | 35 | + | 17.2844 | 4.59E-07 | 0.133 | TAGCTGTACTTCCGATGCACCAGAAGAGGGCATTC | | |
| 1129 | 13 | 56288103 | 56290296 | 2193 | CAST | 13 | 56288274 | 56288308 | 35 | + | 16.5138 | 7.15E-07 | 0.167 | ATGCAAGTACCCACGGAGGCCAGGAGAGGACATTA | | |
| 1130 | 13 | 65170031 | 65172284 | 2253 | SPRET | 13 | 65171709 | 65171742 | 34 | + | 18.3659 | 2.76E-07 | 0.0667 | TAGCTGTCTTCAGACACTCCAGAAGAGGGCGCCA | | |
| 1130 | 13 | 65170031 | 65172284 | 2253 | SPRET | 13 | 65171724 | 65171742 | 19 | + | 17.5574 | 6.32E-07 | 0.153 | ACTCCAGAAGAGGGCGCCA | | |
| 1131 | 13 | 66045245 | 66046450 | 1205 | 129S1,AKR,BUB,C3H,CBA,MRL,NZO,NZW,RF,WSB | 13 | 66045624 | 66045657 | 34 | + | 17.2439 | 5.79E-07 | 0.0964 | TAGCTGTCCTCAGATACTCCAGAAGAGGGCATCA | | |
| 1132 | 13 | 68237879 | 68251890 | 14011 | 129S1,AJ,AKR,BALB,BTBR,BUB,C3H,C57L,C58J,CAST,CBA,CE,DBA1J,DBA2J,FVB,ILNJ2,KK,MA/My,MOLF,MRL,NOD,NOR,NZB,NZO,NZW,PJ,PL,RF,SEA,SJL,SMJ,SWR,TH,WSB | 13 | 68244407 | 68244425 | 19 | + | 19.9672 | 1.13E-07 | 0.0716 | TGGCCACCAGAGGGTGCTT | | |
| 1133 | 13 | 70411741 | 70417889 | 6148 | CAST,MOLF,MRL | 13 | 70411945 | 70411978 | 34 | + | 17.5041 | 4.89E-07 | 0.0884 | CTCTAGGGCTTTGCCAGACCACCAGGGGGTGCCA | 6 | ENCFF327LYS,ENCFF907XEG,ENCFF367UNR,ENCFF093ZRU,ENCFF099QNE,ENCFF024JOZ |
| 1133 | 13 | 70411741 | 70417889 | 6148 | CAST,MOLF,MRL | 13 | 70411960 | 70411978 | 19 | + | 20.4098 | 7.98E-08 | 0.0634 | AGACCACCAGGGGGTGCCA | 6 | ENCFF327LYS,ENCFF907XEG,ENCFF367UNR,ENCFF093ZRU,ENCFF099QNE,ENCFF024JOZ |
| 1134 | 13 | 72797664 | 72799157 | 1493 | CAST,MOLF | 13 | 72798286 | 72798304 | 19 | + | 20.8525 | 5.55E-08 | 0.0616 | TGACCAGAAGAGGGCAGTG | | |
| 1135 | 13 | 75968598 | 75970718 | 2120 | SPRET | 13 | 75970493 | 75970526 | 34 | + | 18.3659 | 2.76E-07 | 0.0667 | TAGCTGTCTTCAGACACTCCAGAAGAGGGCGCCA | 1 | ENCFF093ZRU |
| 1135 | 13 | 75968598 | 75970718 | 2120 | SPRET | 13 | 75970508 | 75970526 | 19 | + | 17.5574 | 6.32E-07 | 0.153 | ACTCCAGAAGAGGGCGCCA | 1 | ENCFF093ZRU |
| 1136 | 13 | 75970323 | 75971418 | 1095 | BUB,NOD,NOR,NZW | 13 | 75970493 | 75970526 | 34 | + | 18.3659 | 2.76E-07 | 0.0667 | TAGCTGTCTTCAGACACTCCAGAAGAGGGCGCCA | 1 | ENCFF093ZRU |
| 1136 | 13 | 75970323 | 75971418 | 1095 | BUB,NOD,NOR,NZW | 13 | 75970508 | 75970526 | 19 | + | 17.5574 | 6.32E-07 | 0.153 | ACTCCAGAAGAGGGCGCCA | 1 | ENCFF093ZRU |
| 1137 | 13 | 76598211 | 76603825 | 5614 | ILNJ2,SPRET | 13 | 76599744 | 76599778 | 35 | + | 16.7615 | 6.21E-07 | 0.156 | TAGCAGTTGCCACCTGTGTGCTCTTGGGGGCGGGG | | |
| 1138 | 13 | 82163306 | 82172180 | 8874 | SWR,TH | 13 | 82167094 | 82167127 | 34 | + | 16.6829 | 8.29E-07 | 0.112 | ATGCAGCTAGAGACACGAGCTCCAGGGGGTGGGG | | |
| 1139 | 13 | 82166676 | 82172046 | 5370 | SPRET | 13 | 82167094 | 82167127 | 34 | + | 16.6829 | 8.29E-07 | 0.112 | ATGCAGCTAGAGACACGAGCTCCAGGGGGTGGGG | | |
| 1140 | 13 | 82531552 | 82533545 | 1993 | SPRET | 13 | 82532344 | 82532378 | 35 | + | 16.1009 | 9.03E-07 | 0.19 | CTGCTGTTCCAACCCATGGCCACCTGCTGCCTGAG | | |
| 1141 | 13 | 83341490 | 83348883 | 7393 | SPRET | 13 | 83346222 | 83346255 | 34 | + | 20.4797 | 6.27E-08 | 0.0264 | ATGCAATTGCTGACTCAGCCACCAGATGGTTTTC | | |
| 1142 | 13 | 88554580 | 88559965 | 5385 | SPRET | 13 | 88559466 | 88559499 | 34 | + | 17.2358 | 5.82E-07 | 0.0967 | CTGCAAGAGCAGAGCTTGCCTGCAGAGAGTACTC | | |
| 1143 | 13 | 96157588 | 96159021 | 1433 | MOLF | 13 | 96158133 | 96158167 | 35 | + | 23.4128 | 8.27E-09 | 0.0111 | CTGCATGCACACCTGCATGCCAGAAGAGGGCATCA | 1 | ENCFF093ZRU |
| 1144 | 13 | 101081634 | 101085294 | 3660 | SPRET | 13 | 101083204 | 101083238 | 35 | + | 19.1927 | 1.45E-07 | 0.0701 | ATGAGGTTTCTTCACCCCTCCAGTAGAGGGCACAC | | |
| 1144 | 13 | 101081634 | 101085294 | 3660 | SPRET | 13 | 101083220 | 101083238 | 19 | + | 17.8197 | 5.31E-07 | 0.142 | CCTCCAGTAGAGGGCACAC | | |
| 1145 | 13 | 101082377 | 101084686 | 2309 | CAST | 13 | 101083204 | 101083238 | 35 | + | 19.1927 | 1.45E-07 | 0.0701 | ATGAGGTTTCTTCACCCCTCCAGTAGAGGGCACAC | | |
| 1145 | 13 | 101082377 | 101084686 | 2309 | CAST | 13 | 101083220 | 101083238 | 19 | + | 17.8197 | 5.31E-07 | 0.142 | CCTCCAGTAGAGGGCACAC | | |
| 1146 | 13 | 105929624 | 105936088 | 6464 | AJ,AKR,BUB,CAST,CE,DBA1J,MA/My,MOLF,NOD,NOR,NZB,NZO,SPRET,WSB | 13 | 105930278 | 105930296 | 19 | + | 17.0328 | 8.88E-07 | 0.181 | TTGCCAGCAGAGAGCGCTC | | |
| 1147 | 13 | 106563529 | 106565171 | 1642 | CAST,CE,MA/My,MOLF,NZO | 13 | 106563753 | 106563786 | 34 | + | 19.0976 | 1.68E-07 | 0.0517 | GAGCAGGTGCAGCTGCCACCCCAAGATGGTGCCC | | |
| 1148 | 13 | 116904425 | 116910505 | 6080 | 129S1,BTBR,BUB,C3H,C57L,CE,DBA2J,KK,MRL,NZB,NZO,NZW,PL | 13 | 116909082 | 116909100 | 19 | + | 17.2295 | 7.83E-07 | 0.17 | CTGCCAGAAGAGGGAAGTA | | |
| 1149 | 13 | 116907587 | 116909876 | 2289 | AKR,BALB,CBA,RF,SMJ,TH,WSB | 13 | 116909082 | 116909100 | 19 | + | 17.2295 | 7.83E-07 | 0.17 | CTGCCAGAAGAGGGAAGTA | | |
| 1150 | 13 | 117423398 | 117425763 | 2365 | SPRET | 13 | 117424083 | 117424117 | 35 | + | 17.1193 | 5.06E-07 | 0.139 | ATGCCCTGGTTGTGGGTTGCCTGTAGGGGGTGCTA | 4 | ENCFF136ZDK,ENCFF907XEG,ENCFF194HVL,ENCFF533APC |
| 1150 | 13 | 117423398 | 117425763 | 2365 | SPRET | 13 | 117424099 | 117424117 | 19 | + | 17.9672 | 4.81E-07 | 0.138 | TTGCCTGTAGGGGGTGCTA | 4 | ENCFF136ZDK,ENCFF907XEG,ENCFF194HVL,ENCFF533APC |
| 1150 | 13 | 117423398 | 117425763 | 2365 | SPRET | 13 | 117423656 | 117423689 | 34 | - | 16.7724 | 7.83E-07 | 0.109 | GGGCAATCCCCACACACACCAGTAGATGGCCAAC | 5 | ENCFF589ZBJ,ENCFF198SSR,ENCFF093ZRU,ENCFF883UPM,ENCFF099QNE |
| 1151 | 13 | 120664486 | 120670024 | 5538 | SPRET | 13 | 120667830 | 120667863 | 34 | + | 19.2276 | 1.53E-07 | 0.0486 | TTGCTTTGCCTATGGAGTCCTGCAGGGGGAGGAC | | |
| 1152 | 13 | 120664931 | 120675600 | 10669 | BTBR,C57L | 13 | 120667830 | 120667863 | 34 | + | 19.2276 | 1.53E-07 | 0.0486 | TTGCTTTGCCTATGGAGTCCTGCAGGGGGAGGAC | | |
| 1153 | 13 | 5081993 | 5090701 | 8708 | SPRET | 13 | 5088420 | 5088438 | 19 | - | 17.459 | 6.74E-07 | 0.158 | CAGCCAGGAGGTGGAGCTG | | |
| 1154 | 13 | 14123689 | 14126869 | 3180 | SPRET | 13 | 14125603 | 14125621 | 19 | - | 18 | 4.70E-07 | 0.138 | AGTCCACTAGGGGACGCTG | | |
| 1155 | 13 | 18042027 | 18051875 | 9848 | AKR,ILNJ2,LP,PJ,RF,SEA | 13 | 18044584 | 18044618 | 35 | - | 19.4587 | 1.23E-07 | 0.0632 | CAGCAGGGGCCAGCAGGGACCAGTAGGGGCCAGAA | | |
| 1155 | 13 | 18042027 | 18051875 | 9848 | AKR,ILNJ2,LP,PJ,RF,SEA | 13 | 18044604 | 18044638 | 35 | - | 17.5872 | 3.85E-07 | 0.12 | TAGCAGGGACTATCAGGGACCAGCAGGGGCCAGCA | | |
| 1156 | 13 | 23064254 | 23140550 | 76296 | NOD,NOR | 13 | 23105720 | 23105754 | 35 | - | 19 | 1.64E-07 | 0.0747 | CAGCAGTTGCCACCTGTGTGCTCTTGGGGGCGGGG | | |
| 1157 | 13 | 24500478 | 24501645 | 1167 | SPRET | 13 | 24500774 | 24500808 | 35 | - | 22.0642 | 2.18E-08 | 0.0203 | CTGCTGTGTCACTTGTGCATCAGTAGGTGGCAGAG | 11 | ENCFF367UNR,ENCFF093ZRU,ENCFF092NMJ,ENCFF201JUA,ENCFF430PPJ,ENCFF024JOZ,ENCFF883UPM,ENCFF502WDT,ENCFF099QNE,ENCFF089GBC,ENCFF055HAP |
| 1158 | 13 | 31515462 | 31534163 | 18701 | BUB | 13 | 31526531 | 31526549 | 19 | - | 19.0984 | 2.17E-07 | 0.098 | AGACCAGAAGAGGGCACTG | 2 | ENCFF327LYS,ENCFF907XEG |

| | | | | | | | | | | | | | | | | |
|---|---|---|---|---|---|---|---|---|---|---|---|---|---|---|---|---|
| 1158 | 13 | 31515462 | 31534163 | 18701 | BUB | 13 | 31526531 | 31526565 | 35 | - | 17.7064 | 3.58E-07 | 0.115 | GTACAGGTATCTGTGTAGACCAGAAGAGGGCACTG | 2 | ENCFF327LYS,ENCFF907XEG |
| 1159 | 13 | 32429537 | 32435038 | 5501 | MOLF,SPRET | 13 | 32432510 | 32432543 | 34 | - | 19.8618 | 9.80E-08 | 0.0365 | GTACAGTTCTTGCTGAGGCCAGAAGAAGGTGCCA | | |
| 1160 | 13 | 32821496 | 32824307 | 2811 | CAST,MOLF,SPRET,WSB | 13 | 32821509 | 32821527 | 19 | - | 17.5574 | 6.32E-07 | 0.153 | TGGCCAGTTGGGGGCGCGC | | |
| 1160 | 13 | 32821496 | 32824307 | 2811 | CAST,MOLF,SPRET,WSB | 13 | 32821509 | 32821542 | 34 | - | 19.561 | 1.21E-07 | 0.0424 | CAGCATCGTTCTTCCTGGCCAGTTGGGGGCGCGC | | |
| 1161 | 13 | 33324585 | 33331148 | 6563 | SPRET | 13 | 33325449 | 33325467 | 19 | - | 17.2459 | 7.74E-07 | 0.17 | CAGCCACTAGAGGGTACAG | | |
| 1162 | 13 | 40505647 | 40506926 | 1279 | 129S1,AJ,AKR,B10D2,B10J,BALB,BTBR,BUB,C3H,C57L,C58J,CAST,CBA,CE,DBA1J,DBA2J,FVB,ILNJ2,KK,LP,MA/My,MOLF,MRL,NOD,NOR,NZB,NZO,NZW,PJ,PL,RF,RH2,SEA,SJL,SMJ,SWR,TH | 13 | 40506924 | 40506958 | 35 | - | 15.9541 | 9.80E-07 | 0.198 | CTGCTTTACTACTTCAAAGGCACTTGAGGGCAGGT | | |
| 1163 | 13 | 45987504 | 45988980 | 1476 | MOLF | 13 | 45987976 | 45987994 | 19 | - | 18.623 | 3.06E-07 | 0.112 | AGACCAGGAGAGGGCGGCA | 13 | ENCFF089GBC,ENCFF327LYS,ENCFF821BIA,ENCFF907XEG,ENCFF093ZRU,ENCFF367UNR,ENCFF033UQX,ENCFF099QNE,ENCFF957SPH,ENCFF136ZDK,ENCFF589ZBJ,ENCFF142CNG,ENCFF502WDT |
| 1163 | 13 | 45987504 | 45988980 | 1476 | MOLF | 13 | 45987976 | 45988009 | 34 | - | 25.3496 | 1.21E-09 | 0.00179 | ATGCAATACCCTAGGAGACCAGGAGAGGGCGGCA | 13 | ENCFF089GBC,ENCFF327LYS,ENCFF821BIA,ENCFF907XEG,ENCFF093ZRU,ENCFF367UNR,ENCFF033UQX,ENCFF099QNE,ENCFF957SPH,ENCFF136ZDK,ENCFF589ZBJ,ENCFF142CNG,ENCFF502WDT |
| 1164 | 13 | 46335723 | 46337757 | 2034 | MOLF | 13 | 46336507 | 46336540 | 34 | - | 19.7398 | 1.07E-07 | 0.0388 | CAGTAGTTCCTATCCCAGCCTCTAGATGGAGAAC | 15 | ENCFF093ZRU,ENCFF367UNR,ENCFF099QNE,ENCFF052KVO,ENCFF065ODV,ENCFF024JOZ,ENCFF327LYS,ENCFF201JUA,ENCFF092NMJ,ENCFF502WDT,ENCFF883UPM,ENCFF198SSR,ENCFF821BIA,ENCFF907XEG,ENCFF055HAP |
| 1165 | 13 | 50655528 | 50659387 | 3859 | CAST,MOLF,SPRET | 13 | 50655943 | 50655976 | 34 | - | 17.9106 | 3.74E-07 | 0.0786 | CCGCCCTCACATTCGCCGTCACAAGATGGCGCTG | | |
| 1165 | 13 | 50655528 | 50659387 | 3859 | CAST,MOLF,SPRET | 13 | 50659345 | 50659378 | 34 | - | 17.9106 | 3.74E-07 | 0.0786 | CCGCCCTCACATTCGCCGTCACAAGATGGCGCTG | | |
| 1166 | 13 | 54044660 | 54047926 | 3266 | SPRET | 13 | 54047486 | 54047519 | 34 | - | 18.122 | 3.25E-07 | 0.0727 | ATGCAGTGCCTGCAAAGGCCAGAAGGGTGTTGGG | | |
| 1167 | 13 | 58192277 | 58194877 | 2600 | MOLF | 13 | 58193293 | 58193326 | 34 | - | 26.439 | 4.42E-10 | 0.000949 | ATGCAGTGCCCATGGAAGCCAGAAGAGGGCATCA | 15 | ENCFF136ZDK,ENCFF024JOZ,ENCFF093ZRU,ENCFF099QNE,ENCFF165LXO,ENCFF201JUA,ENCFF589ZBJ,ENCFF092NMJ,ENCFF367UNR,ENCFF821BIA,ENCFF198SSR,ENCFF052KVO,ENCFF907XEG,ENCFF327LYS,ENCFF089GBC |
| 1168 | 13 | 60971665 | 60973516 | 1851 | PJ,SEA | 13 | 60972610 | 60972644 | 35 | - | 17.1468 | 4.98E-07 | 0.137 | TTGCAATTGAAGGCCCAGGCCTGAAGGGGTCATGC | | |
| 1169 | 13 | 67702436 | 67704380 | 1944 | SPRET | 13 | 67702675 | 67702693 | 19 | - | 17.5902 | 6.18E-07 | 0.153 | ACACCAGAAGAGGGCACTG | | |
| 1169 | 13 | 67702436 | 67704380 | 1944 | SPRET | 13 | 67702675 | 67702708 | 34 | - | 18.3171 | 2.86E-07 | 0.067 | TAGCTGTCTTCAGACACACCAGAAGAGGGCACTG | | |
| 1170 | 13 | 67946247 | 67955940 | 9693 | MOLF | 13 | 67954083 | 67954116 | 34 | - | 17.3415 | 5.44E-07 | 0.0939 | GTACAGTGCCCACTAGGGACAAAAGAGGGTGCTA | | |
| 1171 | 13 | 67988652 | 67993919 | 5267 | MOLF | 13 | 67992476 | 67992494 | 19 | - | 17.6721 | 5.85E-07 | 0.15 | TTACCAGCAGGTGGAGACA | 22 | ENCFF093ZRU,ENCFF367UNR,ENCFF099QNE,ENCFF136ZDK,ENCFF951GOP,ENCFF327LYS,ENCFF821BIA,ENCFF201JUA,ENCFF907XEG,ENCFF024JOZ,ENCFF089GBC,ENCFF092NMJ,ENCFF198SSR,ENCFF316FYN,ENCFF609MDN,ENCFF589ZBJ,ENCFF430PPJ,ENCFF165LXO,ENCFF055HAP,ENCFF281QMV,ENCFF502WDT,ENCFF883UPM |
| 1171 | 13 | 67988652 | 67993919 | 5267 | MOLF | 13 | 67992476 | 67992509 | 34 | - | 17.7236 | 4.24E-07 | 0.083 | GTGTTGTAACCATCATTACCAGCAGGTGGAGACA | 22 | ENCFF093ZRU,ENCFF367UNR,ENCFF099QNE,ENCFF136ZDK,ENCFF951GOP,ENCFF327LYS,ENCFF821BIA,ENCFF201JUA,ENCFF907XEG,ENCFF024JOZ,ENCFF089GBC,ENCFF092NMJ,ENCFF198SSR,ENCFF316FYN,ENCFF609MDN,ENCFF589ZBJ,ENCFF430PPJ,ENCFF165LXO,ENCFF055HAP,ENCFF281QMV,ENCFF502WDT,ENCFF883UPM |
| 1171 | 13 | 67988652 | 67993919 | 5267 | MOLF | 13 | 67993819 | 67993837 | 19 | - | 18.623 | 3.06E-07 | 0.112 | ACACCAGAAGAGGGCGCCA | 3 | ENCFF093ZRU,ENCFF367UNR,ENCFF099QNE |
| 1171 | 13 | 67988652 | 67993919 | 5267 | MOLF | 13 | 67993819 | 67993852 | 34 | - | 16.9106 | 7.17E-07 | 0.105 | TAGCTGCCTTCAGACACACCAGAAGAGGGCGCCA | 3 | ENCFF093ZRU,ENCFF367UNR,ENCFF099QNE |
| 1172 | 13 | 68590114 | 68598854 | 8740 | AJ,AKR,BALB,BUB,C3H,CAST,CBA,DBA1J,DBA2J,FVB,ILNJ2,KK,MOLF,MRL,NOD,NOR,NZB,NZO,NZW,PJ,RF,SEA,SJL,TH | 13 | 68596292 | 68596310 | 19 | - | 17.8689 | 5.14E-07 | 0.139 | AAACCACTAGGTGGCAGAA | | |
| 1173 | 13 | 76641139 | 76654006 | 12867 | 129S1,AJ,AKR,BALB,BTBR,C3H,C57L,CBA,CE,DBA1J,DBA2J,FVB,KK,MA/My,MRL,NZB,NZO,NZW,PJ,RF,SEA,SJL,SMJ,TH | 13 | 76652934 | 76652967 | 34 | - | 19.3171 | 1.44E-07 | 0.0472 | CTGCAAGAGGACAGCTTGCCTGCAGAGGGTACTC | | |
| 1174 | 13 | 76642026 | 76653329 | 11303 | BUB,CAST,ILNJ2,NOD,NOR,SPRET,SWR,WSB | 13 | 76652934 | 76652967 | 34 | - | 19.3171 | 1.44E-07 | 0.0472 | CTGCAAGAGGACAGCTTGCCTGCAGAGGGTACTC | | |
| 1175 | 13 | 88515341 | 88516728 | 1387 | SPRET | 13 | 88515353 | 88515386 | 34 | - | 17.5691 | 4.69E-07 | 0.0865 | ATGCAGCTAGAGACATGTGCTCCAGGGGGTACTG | | |
| 1176 | 13 | 90476511 | 90477683 | 1172 | CAST.MOLF.SPRET | 13 | 90476634 | 90476652 | 19 | - | 17.6393 | 5.98E-07 | 0.152 | AGACCAGAAGAGGGCACAA | 1 | ENCFF093ZRU |
| 1176 | 13 | 90476511 | 90477683 | 1172 | CAST.MOLF.SPRET | 13 | 90476634 | 90476668 | 35 | - | 25.0642 | 2.34E-09 | 0.00542 | ATGCAGGTACCCCTGAAGACCAGAAGAGGGCACAA | 1 | ENCFF093ZRU |
| 1177 | 13 | 93335115 | 93339256 | 4141 | 129S1,AJ,AKR,BALB,BTBR,C3H,C57L,C58J,CAST,CBA,CE,DBA1J,DBA2J,FVB,ILNJ2,KK,MA/My,MOLF,MRL,NZB,NZO,NZW,PJ,PL,RF,SEA,SJL,SMJ,SPRET,SWR,WSB | 13 | 93338453 | 93338486 | 34 | - | 17.6748 | 4.37E-07 | 0.0839 | GAGCATGACCACAAGTCCACAGCAGGTGGCAACG | | |
| 1178 | 13 | 99122778 | 99131763 | 8985 | 129S1,AJ,AKR,BALB,BTBR,BUB,C3H,C57L,C58J,CAST,CBA,CE,DBA1J,DBA2J,FVB,ILNJ2,KK,MA/My,MOLF,MRL,NOD,NOR,NZB,NZO,PL,SJL,SMJ,SPRET,SWR,WSB | 13 | 99126333 | 99126351 | 19 | - | 17.1639 | 8.17E-07 | 0.174 | CCTCCACGAGGTGGCGGAA | | |
| 1179 | 13 | 100004246 | 100007055 | 2809 | SPRET | 13 | 100005123 | 100005157 | 35 | - | 16.0092 | 9.50E-07 | 0.194 | TGGCAGAGGGCAGCAGAGGGCAGCAGAGGGCAGAG | 4 | ENCFF093ZRU.ENCFF052KVO.ENCFF033UQX.ENCFF136ZDK |
| 1179 | 13 | 100004246 | 100007055 | 2809 | SPRET | 13 | 100005133 | 100005166 | 34 | - | 25.5528 | 1.01E-09 | 0.00161 | GTGCAGCGCTGGCAGAGGGCAGCAGAGGGCAGCA | 4 | ENCFF093ZRU.ENCFF052KVO.ENCFF033UQX.ENCFF136ZDK |
| 1180 | 13 | 108719054 | 108721384 | 2330 | WSB | 13 | 108719563 | 108719597 | 35 | - | 18.367 | 2.41E-07 | 0.0943 | CTGCAGTGGCAAACATCAGCCTGAAGGAGTCTGCA | | |
| 1181 | 13 | 111487270 | 111488559 | 1289 | CAST | 13 | 111487573 | 111487591 | 19 | - | 19.0328 | 2.28E-07 | 0.0997 | TGGCCACAAGAGGTCAGCA | | |
| 1182 | 13 | 112283888 | 112286234 | 2346 | AJ,DBA1J,KK,MA/My,MOLF,NZO,PL,RF,RH2,SPRET,SWR,TH,WSB | 13 | 112284156 | 112284189 | 34 | - | 17.3252 | 5.49E-07 | 0.0944 | CTGCATTTTATGAGGCTGCCAGAAGGAGATGGTC | | |
| 1183 | 13 | 113058759 | 113061925 | 3166 | CAST | 13 | 113059214 | 113059232 | 19 | - | 17.0328 | 8.88E-07 | 0.181 | AGGCCGGAAGGTGGCACTG | | |
| 1184 | 13 | 116010250 | 116017170 | 6920 | AJ.DBA1J.MA/My.MOLF.RH2 | 13 | 116015957 | 116015991 | 35 | - | 21.8899 | 2.46E-08 | 0.0216 | CTGCATTTATAACAATTTCCCAGTTGATGGCAGTC | 1 | ENCFF093ZRU |
| 1185 | 13 | 120662108 | 120663117 | 1009 | SPRET | 13 | 120662655 | 120662688 | 34 | - | 18.9187 | 1.90E-07 | 0.056 | GGGCAGTTACAGCTGCTGCCACAAGGGAGCTGTT | | |
| 1186 | 14 | 3739007 | 3740988 | 1981 | SPRET | 14 | 3740567 | 3740600 | 34 | + | 20.626 | 5.64E-08 | 0.0235 | GTGCAGTACATAAGGAGGCCAGAAGAGGGATCAG | 1 | ENCFF093ZRU |
| 1187 | 14 | 4281855 | 4295719 | 13864 | SPRET | 14 | 4282433 | 4282466 | 34 | + | 18.1057 | 3.29E-07 | 0.0647 | CTGCTAATTCTCAGTTTACCATAAGGCGGCGCTA | 27 | ENCFF093ZRU,ENCFF367UNR,ENCFF099QNE,ENCFF201JUA,ENCFF821BIA,ENCFF055HAP,ENCFF327LYS,ENCFF024JOZ,ENCFF089GBC,ENCFF907XEG,ENCFF504JDS,ENCFF609MDN,ENCFF065ODV,ENCFF589ZBJ,ENCFF430PPJ,ENCFF052KVO,ENCFF092NMJ,ENCFF502WDT,ENCFF198SSR,ENCFF321AGQ,ENCFF883UPM,ENCFF165LXO,ENCFF281QMV,ENCFF142CNG,ENCFF136ZDK,ENCFF951GOP,ENCFF575MGI |
| 1188 | 14 | 4871219 | 4875491 | 4272 | 129S1,C3H,C57L,CAST,CBA,DBA1J,DBA2J,KK,MOLF,MRL,NZW,PJ,PL,SPRET | 14 | 4871235 | 4871268 | 34 | + | 19.3415 | 1.42E-07 | 0.0418 | CTGCCCCCACATTCGCCGTCACAAGATGGCGCTG | | |
| 1189 | 14 | 5013070 | 5033506 | 20436 | 129S1,C3H,C57L,CBA,DBA1J,DBA2J,KK,MOLF,MRL,NZW,PJ,PL | 14 | 5017881 | 5017914 | 34 | + | 17.9756 | 3.59E-07 | 0.0664 | ATGCAGCTAGAGACATGAACTCCGGGTGGCACTG | | |
| 1190 | 14 | 6397073 | 6401548 | 4475 | CAST.SPRET | 14 | 6397851 | 6397884 | 34 | + | 17.6748 | 4.37E-07 | 0.074 | GAGCATGACCACAAGTCCACAGCAGGTGGCAACG | | |
| 1191 | 14 | 6916484 | 6922010 | 5526 | 129S1,C3H,C57L,CBA,DBA1J,DBA2J,KK,MOLF,MRL,NZW,PJ,PL,SPRET | 14 | 6920257 | 6920291 | 35 | + | 16.4404 | 7.46E-07 | 0.2 | AAGAAGCTGGACCCAGTGGCCAGCAGATGGCCCAC | | |
| 1191 | 14 | 6916484 | 6922010 | 5526 | 129S1,C3H,C57L,CBA,DBA1J,DBA2J,KK,MOLF.MRL.NZW.PJ.PL.SPRET | 14 | 6920273 | 6920291 | 19 | + | 17.0328 | 8.88E-07 | 0.184 | TGGCCAGCAGATGGCCCAC | | |
| 1192 | 14 | 13872145 | 13873222 | 1077 | SPRET | 14 | 13872667 | 13872700 | 34 | + | 17.813 | 3.99E-07 | 0.0708 | TTGCTGTCTTCATGTACACCAGAAGAGGGCATCA | 1 | ENCFF093ZRU |
| 1193 | 14 | 14809585 | 14813674 | 4089 | SPRET | 14 | 14810883 | 14810917 | 35 | + | 20.6697 | 5.62E-08 | 0.0496 | ATGCATCCCGGGCCGCCGGCCGGGAGGGGTCGCTG | | |
| 1194 | 14 | 15293559 | 15297689 | 4130 | SPRET | 14 | 15294923 | 15294957 | 35 | + | 23.1835 | 9.78E-09 | 0.0184 | ATGCATCCCGGGCCGCCGGCCAGGAGGGGTCGCTG | | |
| 1194 | 14 | 15293559 | 15297689 | 4130 | SPRET | 14 | 15294939 | 15294957 | 19 | + | 19.2623 | 1.93E-07 | 0.0909 | CGGCCAGGAGGGGTCGCTG | | |
| 1194 | 14 | 15293559 | 15297689 | 4130 | SPRET | 14 | 15295326 | 15295344 | 19 | - | 19.623 | 1.47E-07 | 0.0802 | TGGCCAGGAGGGGGAGGCG | | |
| 1195 | 14 | 15552829 | 15559931 | 7102 | C57L.PL | 14 | 15558759 | 15558792 | 34 | + | 20.626 | 5.64E-08 | 0.0235 | CTGGAGGGGCAGCCTTGGCCTTCAGGTGGCAGCC | | |
| 1196 | 14 | 16578683 | 16580837 | 2154 | 129S1.C57L.KK.PL | 14 | 16580306 | 16580339 | 34 | + | 18.0488 | 3.42E-07 | 0.0647 | TTGCTGTATGCGTAGAGGCCAGAAGAGGGTGTCA | | |
| 1197 | 14 | 16587839 | 16593000 | 5161 | SPRET | 14 | 16589176 | 16589210 | 35 | + | 17.2752 | 4.62E-07 | 0.156 | ATGCATCCTGGGCCACCCGCCTGGAGGGGTCGCTG | 1 | ENCFF089GBC |
| 1198 | 14 | 22494653 | 22568513 | 73860 | PJ | 14 | 22514564 | 22514597 | 34 | + | 26.6016 | 3.79E-10 | 0.000974 | ATGCAGTGCCCATGGAGGCCAGGAGAGGGCATCA | 5 | ENCFF093ZRU.ENCFF052KVO.ENCFF055HAP.ENCFF367UNR.ENCFF502WDT |
| 1199 | 14 | 25907014 | 25908385 | 1371 | CAST | 14 | 25907891 | 25907909 | 19 | + | 19.2131 | 2.00E-07 | 0.0917 | CCTCCACTAGAGGGCGCAC | | |
| 1200 | 14 | 28970837 | 28977767 | 6930 | 129S1,AJ,AKR,BALB,BTBR,BUB,C3H,C57L,CAST,CBA,CE,FVB,KK,MA/My,MOLF,MRL,NOD,NOR,NZO,NZW,PJ,PL,RF,RH2,SEA,SJL,SMJ,SPRET,SWR,TH,WSB | 14 | 28973330 | 28973348 | 19 | + | 18.5738 | 3.17E-07 | 0.117 | AAACCACCAGGTGGCAGAA | | |
| 1201 | 14 | 31177260 | 31178528 | 1268 | SPRET | 14 | 31177298 | 31177331 | 34 | + | 27.6829 | 1.31E-10 | 0.000575 | GTGCAATGCCCATAGAGGCCAGGAGGGGGCGTCA | 13 | ENCFF367UNR,ENCFF093ZRU,ENCFF589ZBJ,ENCFF327LYS,ENCFF821BIA,ENCFF502WDT,ENCFF065ODV,ENCFF033UQX,ENCFF099QNE,ENCFF194HVL,ENCFF142CNG,ENCFF136ZDK,ENCFF052KVO |
| 1201 | 14 | 31177260 | 31178528 | 1268 | SPRET | 14 | 31177313 | 31177331 | 19 | + | 17.2951 | 7.50E-07 | 0.17 | AGGCCAGGAGGGGGCGTCA | 13 | ENCFF367UNR,ENCFF093ZRU,ENCFF589ZBJ,ENCFF327LYS,ENCFF821BIA,ENCFF502WDT,ENCFF065ODV,ENCFF033UQX,ENCFF099QNE,ENCFF194HVL,ENCFF142CNG,ENCFF136ZDK,ENCFF052KVO |
| 1202 | 14 | 31333370 | 31334856 | 1486 | CAST,MOLF | 14 | 31334127 | 31334160 | 34 | + | 25.2683 | 1.30E-09 | 0.00187 | GTGCAGTGCACAAGGAAGCCAGAAGAGGGCGTCA | 15 | ENCFF093ZRU,ENCFF367UNR,ENCFF136ZDK,ENCFF821BIA,ENCFF089GBC,ENCFF024JOZ,ENCFF327LYS,ENCFF883UPM,ENCFF589ZBJ,ENCFF957SPH,ENCFF065ODV,ENCFF052KVO,ENCFF142CNG,ENCFF055HAP,ENCFF502WDT |
| 1203 | 14 | 32612656 | 32614149 | 1493 | DBA1J.DBA2J.ILNJ2.NZB.PJ.WSB | 14 | 32613896 | 32613929 | 34 | + | 19.6667 | 1.13E-07 | 0.0368 | GTGCAGTGCCCTCGAAGGCCAGTGGAGGGTTTGG | | |
| 1204 | 14 | 35526970 | 35528584 | 1614 | BUB.MA/My.NZB.PJ.SWR | 14 | 35527353 | 35527386 | 34 | + | 18.3902 | 2.72E-07 | 0.0594 | CAGCTGGGCCTCTTTGAGTCACTGGGGGGCAGAG | | |
| 1205 | 14 | 37830579 | 37839837 | 9258 | BTBR,C57L,CAST,MOLF,NZO,PJ,PL,SPRET | 14 | 37832715 | 37832748 | 34 | + | 17.0325 | 6.63E-07 | 0.0931 | TGGCAGGTCTCAGAGGAGCCGCTAGATGGCCCAC | | |
| 1206 | 14 | 38781300 | 38936510 | 155210 | LP | 14 | 38847603 | 38847621 | 19 | + | 17.1639 | 8.17E-07 | 0.178 | TCCCCAGCAGGTGGAACTG | | |
| 1207 | 14 | 44125853 | 44255959 | 130106 | CBA,DBA1J,DBA2J,ILNJ2 | 14 | 44162639 | 44162657 | 19 | + | 19.2951 | 1.88E-07 | 0.0904 | CCGCCACTAGATGGTGCTA | 30 | ENCFF367UNR,ENCFF093ZRU,ENCFF099QNE,ENCFF201JUA,ENCFF327LYS,ENCFF821BIA,ENCFF907XEG,ENCFF589ZBJ,ENCFF089GBC,ENCFF198SSR,ENCFF430PPJ,ENCFF024JOZ,ENCFF092NMJ,ENCFF321AGQ,ENCFF136ZDK,ENCFF165LXO,ENCFF316FYN,ENCFF033UQX,ENCFF097CAQ,ENCFF052KVO,ENCFF951GOP,ENCFF957SPH,ENCFF281QMV,ENCFF883UPM,ENCFF210BTO,ENCFF194HVL,ENCFF609MDN,ENCFF611HDQ,ENCFF491RJK,ENCFF575MGI |
| 1207 | 14 | 44125853 | 44255959 | 130106 | CBA,DBA1J,DBA2J,ILNJ2 | 14 | 44170621 | 44170639 | 19 | + | 18.6557 | 2.99E-07 | 0.113 | CCGCCACTAGATGGTGGTA | 23 | ENCFF093ZRU,ENCFF367UNR,ENCFF052KVO,ENCFF099QNE,ENCFF201JUA,ENCFF821BIA,ENCFF024JOZ,ENCFF327LYS,ENCFF907XEG,ENCFF089GBC,ENCFF092NMJ,ENCFF609MDN,ENCFF136ZDK,ENCFF589ZBJ,ENCFF430PPJ,ENCFF198SSR,ENCFF055HAP,ENCFF165LXO,ENCFF502WDT,ENCFF883UPM,ENCFF951GOP,ENCFF957SPH,ENCFF281QMV |
| 1207 | 14 | 44125853 | 44255959 | 130106 | CBA.DBA1J.DBA2J.ILNJ2 | 14 | 44254995 | 44255013 | 19 | + | 16.9672 | 9.26E-07 | 0.188 | AGGCCAGAAGAGGGAACTG | | |
| 1207 | 14 | 44125853 | 44255959 | 130106 | CBA.DBA1J.DBA2J.ILNJ2 | 14 | 44183356 | 44183374 | 19 | - | 16.9672 | 9.26E-07 | 0.188 | AGGCCAGAAGAGGGAACTG | | |
| 1207 | 14 | 44125853 | 44255959 | 130106 | CBA,DBA1J,DBA2J,ILNJ2 | 14 | 44221550 | 44221568 | 19 | - | 24.2295 | 1.90E-09 | 0.0188 | TGGCCAGCAGGGGGCGCAC | 29 | ENCFF093ZRU,ENCFF367UNR,ENCFF201JUA,ENCFF099QNE,ENCFF327LYS,ENCFF907XEG,ENCFF589ZBJ,ENCFF024JOZ,ENCFF198SSR,ENCFF951GOP,ENCFF821BIA,ENCFF136ZDK,ENCFF321AGQ,ENCFF097CAQ,ENCFF092NMJ,ENCFF089GBC,ENCFF033UQX,ENCFF165LXO,ENCFF491RJK,ENCFF430PPJ,ENCFF957SPH,ENCFF281QMV,ENCFF883UPM,ENCFF210BTO,ENCFF194HVL,ENCFF609MDN,ENCFF533APC,ENCFF611HDQ,ENCFF575MGI |
| 1207 | 14 | 44125853 | 44255959 | 130106 | CBA,DBA1J,DBA2J,ILNJ2 | 14 | 44221550 | 44221583 | 34 | - | 23.561 | 5.69E-09 | 0.00511 | AGGGAGTTGCTGCTTTGGCCAGCAGGGGGCGCAC | 29 | ENCFF093ZRU,ENCFF367UNR,ENCFF201JUA,ENCFF099QNE,ENCFF327LYS,ENCFF907XEG,ENCFF589ZBJ,ENCFF024JOZ,ENCFF198SSR,ENCFF951GOP,ENCFF821BIA,ENCFF136ZDK,ENCFF321AGQ,ENCFF097CAQ,ENCFF092NMJ,ENCFF089GBC,ENCFF033UQX,ENCFF165LXO,ENCFF491RJK,ENCFF430PPJ,ENCFF957SPH,ENCFF281QMV,ENCFF883UPM,ENCFF210BTO,ENCFF194HVL,ENCFF609MDN,ENCFF533APC,ENCFF611HDQ,ENCFF575MGI |
| 1207 | 14 | 44125853 | 44255959 | 130106 | CBA,DBA1J,DBA2J,ILNJ2 | 14 | 44225063 | 44225081 | 19 | - | 18.2787 | 3.88E-07 | 0.128 | AGGCCAGAAGAGGGCGATA | 28 | ENCFF093ZRU,ENCFF367UNR,ENCFF052KVO,ENCFF099QNE,ENCFF201JUA,ENCFF136ZDK,ENCFF327LYS,ENCFF033UQX,ENCFF821BIA,ENCFF907XEG,ENCFF024JOZ,ENCFF198SSR,ENCFF092NMJ,ENCFF589ZBJ,ENCFF097CAQ,ENCFF089GBC,ENCFF321AGQ,ENCFF165LXO,ENCFF491RJK,ENCFF957SPH,ENCFF883UPM,ENCFF281QMV,ENCFF533APC,ENCFF430PPJ,ENCFF609MDN,ENCFF951GOP,ENCFF194HVL,ENCFF575MGI |
| 1207 | 14 | 44125853 | 44255959 | 130106 | CBA,DBA1J,DBA2J,ILNJ2 | 14 | 44225063 | 44225096 | 34 | - | 25.0163 | 1.63E-09 | 0.00214 | GCGCAGTGCCCATAGAGGCCAGAAGAGGGCGATA | 28 | ENCFF093ZRU,ENCFF367UNR,ENCFF052KVO,ENCFF099QNE,ENCFF201JUA,ENCFF136ZDK,ENCFF327LYS,ENCFF033UQX,ENCFF821BIA,ENCFF907XEG,ENCFF024JOZ,ENCFF198SSR,ENCFF092NMJ,ENCFF589ZBJ,ENCFF097CAQ,ENCFF089GBC,ENCFF321AGQ,ENCFF165LXO,ENCFF491RJK,ENCFF957SPH,ENCFF883UPM,ENCFF281QMV,ENCFF533APC,ENCFF430PPJ,ENCFF609MDN,ENCFF951GOP,ENCFF194HVL,ENCFF575MGI |
| 1208 | 14 | 44879155 | 44887080 | 7925 | C57L.MOLF.NZO.PJ.PL | 14 | 44880161 | 44880194 | 34 | + | 24.7236 | 2.11E-09 | 0.00247 | ATGCAATTTCTCCAGAGGCCAGAAGAGGGCATGG | | |
| 1209 | 14 | 47268139 | 47270039 | 1900 | CAST | 14 | 47268226 | 47268259 | 34 | + | 25.3659 | 1.19E-09 | 0.00175 | GTGCAGTGCCTGGAGAGGCCAGGAAAGGGCACTG | | |
| 1210 | 14 | 49652511 | 49653639 | 1128 | 129S1,AJ,AKR,BALB,BUB,C3H,CBA,CE,DBA1J,DBA2J,FVB,ILNJ2,MA/My,MRL,NOD,NOR,NZB,NZO,NZW,PJ,PL,RF,RH2,SEA,SJL,SMJ,SWR,TH,WSB | 14 | 49653113 | 49653146 | 34 | + | 16.4472 | 9.61E-07 | 0.111 | TAGCTGTCTTCAGACACACCAGAAGAGGGCGTCA | | |
| 1211 | 14 | 50298591 | 50305702 | 7111 | SPRET | 14 | 50300109 | 50300143 | 35 | + | 16.7431 | 6.28E-07 | 0.185 | CCGCCTGGTCACCTAGGAGCCAGTAGGTGGAGCAG | 19 | ENCFF093ZRU,ENCFF367UNR,ENCFF099QNE,ENCFF907XEG,ENCFF089GBC,ENCFF327LYS,ENCFF316FYN,ENCFF024JOZ,ENCFF821BIA,ENCFF609MDN,ENCFF194HVL,ENCFF092NMJ,ENCFF165LXO,ENCFF136ZDK,ENCFF198SSR,ENCFF430PPJ,ENCFF589ZBJ,ENCFF957SPH,ENCFF281QMV |
| 1212 | 14 | 51599291 | 51600361 | 1070 | 129S1,AJ,AKR,BALB,BUB,C3H,C57L,CBA,CE,DBA1J,DBA2J,FVB,ILNJ2,KK,MA/My,MRL,NOD,NOR,NZB,NZO,NZW,PJ,PL,RF,RH2,SEA.SJL.SMJ.SWR.TH.WSB | 14 | 51599790 | 51599823 | 34 | + | 17.9756 | 3.59E-07 | 0.0664 | CTGAAGGGCCAACCTTGGCCTTCAGGGGCAGCCA | | |
| 1213 | 14 | 51619535 | 51623466 | 3931 | 129S1,AJ,AKR,BALB,BUB,C3H,C57L,CBA,CE,DBA1J,DBA2J,KK,MRL,NZB,NZO,NZW,PJ,PL,RF,RH2,SEA,TH,WSB | 14 | 51620067 | 51620100 | 34 | + | 18.3171 | 2.86E-07 | 0.0602 | GGGCAGAGTCTCCAGATGACGGCAGGGGGCGGGG | | |
| 1214 | 14 | 52594134 | 52596986 | 2852 | SPRET | 14 | 52594544 | 52594577 | 34 | + | 17.0325 | 6.63E-07 | 0.0931 | TAGCTATCTTCAGACACACCAGAAGAGGGCACTG | | |
| 1214 | 14 | 52594134 | 52596986 | 2852 | SPRET | 14 | 52594559 | 52594577 | 19 | + | 17.5902 | 6.18E-07 | 0.158 | ACACCAGAAGAGGGCACTG | | |
| 1215 | 14 | 53157579 | 53166165 | 8586 | DBA1J.DBA2J.NZW | 14 | 53157817 | 53157850 | 34 | + | 20.8049 | 4.94E-08 | 0.0235 | ATTTGGTTCCTCCTATGGCCTCCAGAGGGCGGTG | | |
| 1215 | 14 | 53157579 | 53166165 | 8586 | DBA1J.DBA2J.NZW | 14 | 53157832 | 53157850 | 19 | + | 22.4918 | 1.28E-08 | 0.0351 | TGGCCTCCAGAGGGCGGTG | | |
| 1216 | 14 | 53216805 | 53220988 | 4183 | ILNJ2.NOD | 14 | 53220594 | 53220628 | 35 | + | 18.1009 | 2.83E-07 | 0.121 | CATTTGTTTCCTCCTATGGCCTCCAGAGGGCGCTG | | |
| 1216 | 14 | 53216805 | 53220988 | 4183 | ILNJ2.NOD | 14 | 53220595 | 53220628 | 34 | + | 18.8699 | 1.96E-07 | 0.0508 | ATTTGTTTCCTCCTATGGCCTCCAGAGGGCGCTG | | |
| 1216 | 14 | 53216805 | 53220988 | 4183 | ILNJ2.NOD | 14 | 53220610 | 53220628 | 19 | + | 23.1311 | 6.72E-09 | 0.0294 | TGGCCTCCAGAGGGCGCTG | | |
| 1217 | 14 | 53383133 | 53386886 | 3753 | WSB | 14 | 53384191 | 53384209 | 19 | + | 18.4918 | 3.35E-07 | 0.119 | TAACCACAAGATGGCAGCC | | |
| 1218 | 14 | 53909278 | 53913728 | 4450 | FVB,ILNJ2,MA/My,NOD,NOR,NZB,PL,SJL,SMJ,SWR | 14 | 53913334 | 53913368 | 35 | + | 18.1009 | 2.83E-07 | 0.121 | CATTTGTTTCCTCCTATGGCCTCCAGAGGGCGCTG | 1 | ENCFF093ZRU |
| 1218 | 14 | 53909278 | 53913728 | 4450 | FVB,ILNJ2,MA/My,NOD,NOR,NZB,PL,SJL,SMJ,SWR | 14 | 53913335 | 53913368 | 34 | + | 18.8699 | 1.96E-07 | 0.0508 | ATTTGTTTCCTCCTATGGCCTCCAGAGGGCGCTG | 1 | ENCFF093ZRU |
| 1218 | 14 | 53909278 | 53913728 | 4450 | FVB,ILNJ2,MA/My,NOD,NOR,NZB,PL,SJL,SMJ,SWR | 14 | 53913350 | 53913368 | 19 | + | 23.1311 | 6.72E-09 | 0.0294 | TGGCCTCCAGAGGGCGCTG | 1 | ENCFF093ZRU |
| 1219 | 14 | 54589965 | 54592069 | 2104 | CAST | 14 | 54592037 | 54592070 | 34 | + | 23.439 | 6.30E-09 | 0.00556 | GTGCAAGTGCTGGCAAGGCCAACAGAGGGCGCTA | 9 | ENCFF907XEG,ENCFF589ZBJ,ENCFF055HAP,ENCFF033UQX,ENCFF052KVO,ENCFF065ODV,ENCFF142CNG.ENCFF502WDT.ENCFF136ZDK |
| 1219 | 14 | 54589965 | 54592069 | 2104 | CAST | 14 | 54592052 | 54592070 | 19 | + | 20.082 | 1.04E-07 | 0.0664 | AGGCCAACAGAGGGCGCTA | 9 | ENCFF907XEG,ENCFF589ZBJ,ENCFF055HAP,ENCFF033UQX,ENCFF052KVO,ENCFF065ODV,ENCFF142CNG,ENCFF502WDT,ENCFF136ZDK |
| 1220 | 14 | 60011763 | 60015164 | 3401 | AJ,AKR,BALB,C3H,CAST,CBA,CE,DBA1J,DBA2J,MOLF,MRL,NZB,PJ,PL,RH2,SEA,SMJ,SPRET,WSB | 14 | 60012248 | 60012281 | 34 | + | 19.561 | 1.21E-07 | 0.0376 | CAGCATCGTTCTTCCTGGCCAGTTGGGGGCGCGC | | |
| 1220 | 14 | 60011763 | 60015164 | 3401 | AJ,AKR,BALB,C3H,CAST,CBA,CE,DBA1J,DBA2J,MOLF,MRL,NZB,PJ,PL,RH2,SEA,SMJ,SPRET,WSB | 14 | 60012263 | 60012281 | 19 | + | 17.5574 | 6.32E-07 | 0.158 | TGGCCAGTTGGGGGCGCGC | | |
| 1220 | 14 | 60011763 | 60015164 | 3401 | AJ,AKR,BALB,C3H,CAST,CBA,CE,DBA1J,DBA2J,MOLF,MRL,NZB,PJ,PL,RH2,SEA,SMJ,SPRET.WSB | 14 | 60014777 | 60014810 | 34 | + | 19.561 | 1.21E-07 | 0.0376 | CAGCATCGTTCTTCCTGGCCAGTTGGGGGCGCGC | | |
| 1220 | 14 | 60011763 | 60015164 | 3401 | AJ,AKR,BALB,C3H,CAST,CBA,CE,DBA1J,DBA2J,MOLF,MRL,NZB,PJ,PL,RH2,SEA,SMJ,SPRET.WSB | 14 | 60014792 | 60014810 | 19 | + | 17.5574 | 6.32E-07 | 0.158 | TGGCCAGTTGGGGGCGCGC | | |
| 1221 | 14 | 61191270 | 61193503 | 2233 | SPRET | 14 | 61193190 | 61193223 | 34 | + | 16.6911 | 8.24E-07 | 0.103 | TTGCTGTTTTTAGACACACCAGAAGAAGGCATCA | | |
| 1222 | 14 | 65724986 | 65726123 | 1137 | SPRET | 14 | 65726075 | 65726108 | 34 | + | 21.2439 | 3.56E-08 | 0.0185 | CTGCAGTGCCTGCAGAGGTCAGAAGAGGCTGTCA | | |
| 1223 | 14 | 69047722 | 69050461 | 2739 | 129S1,BTBR,C57L,CAST,KK,LP,MOLF,SPRET,TH,WSB | 14 | 69048113 | 69048131 | 19 | + | 17.5574 | 6.32E-07 | 0.158 | TGGCCAGTTGGGGGCGCGC | | |
| 1224 | 14 | 74238958 | 74244064 | 5106 | SPRET | 14 | 74239102 | 74239135 | 34 | + | 18.3089 | 2.87E-07 | 0.0602 | ATGCAGCTAGAGACACGAGCTCCAGGGGGTACTG | | |
| 1225 | 14 | 78020699 | 78021892 | 1193 | SPRET | 14 | 78020847 | 78020865 | 19 | + | 19.7049 | 1.38E-07 | 0.0771 | GAACCACTAGGGGGCAGTC | 16 | ENCFF092NMJ,ENCFF907XEG,ENCFF024JOZ,ENCFF136ZDK,ENCFF198SSR,ENCFF093ZRU,ENCFF201JUA,ENCFF052KVO,ENCFF367UNR,ENCFF097CAQ,ENCFF327LYS,ENCFF821BIA,ENCFF589ZBJ,ENCFF099QNE,ENCFF883UPM,ENCFF957SPH |
| 1226 | 14 | 85779657 | 85781569 | 1912 | CAST | 14 | 85780300 | 85780333 | 34 | + | 16.9593 | 6.95E-07 | 0.095 | TTGCTCTCTTCAGACACACCAGAAGAGGGCATCA | | |
| 1227 | 14 | 90512619 | 90517029 | 4410 | SPRET | 14 | 90513342 | 90513375 | 34 | + | 17.6748 | 4.37E-07 | 0.074 | GAGCATGACCACAAGTCCACAGCAGGTGGCAACG | | |
| 1228 | 14 | 90975758 | 90977224 | 1466 | NZO,WSB | 14 | 90976900 | 90976933 | 34 | + | 21.2683 | 3.49E-08 | 0.0184 | ATGCAGTGCCTCAAACATCCAAAAGAAGGCACTT | | |
| 1229 | 14 | 3292938 | 3294260 | 1322 | CAST | 14 | 3294190 | 3294208 | 19 | - | 23.2295 | 6.09E-09 | 0.0294 | TGGCCACTAGATGGCAGTG | 23 | ENCFF201JUA,ENCFF821BIA,ENCFF092NMJ,ENCFF367UNR,ENCFF093ZRU,ENCFF502WDT,ENCFF883UPM,ENCFF099QNE,ENCFF609MDN,ENCFF907XEG,ENCFF194HVL,ENCFF281QMV,ENCFF024JOZ,ENCFF327LYS,ENCFF589ZBJ,ENCFF142CNG,ENCFF052KVO,ENCFF198SSR,ENCFF136ZDK,ENCFF165LXO,ENCFF089GBC,ENCFF055HAP,ENCFF957SPH |
| 1229 | 14 | 3292938 | 3294260 | 1322 | CAST | 14 | 3294190 | 3294224 | 35 | - | 24.6606 | 3.21E-09 | 0.0107 | TGGCGGTGACCCAAAGTGGCCACTAGATGGCAGTG | 23 | ENCFF201JUA,ENCFF821BIA,ENCFF092NMJ,ENCFF367UNR,ENCFF093ZRU,ENCFF502WDT,ENCFF883UPM,ENCFF099QNE,ENCFF609MDN,ENCFF907XEG,ENCFF194HVL,ENCFF281QMV,ENCFF024JOZ,ENCFF327LYS,ENCFF589ZBJ,ENCFF142CNG,ENCFF052KVO,ENCFF198SSR,ENCFF136ZDK,ENCFF165LXO,ENCFF089GBC,ENCFF055HAP,ENCFF957SPH |
| 1230 | 14 | 4272807 | 4277116 | 4309 | CAST | 14 | 4273043 | 4273061 | 19 | - | 17.5574 | 6.32E-07 | 0.158 | ACTCCAGAAGAGGGCGCCA | | |
| 1230 | 14 | 4272807 | 4277116 | 4309 | CAST | 14 | 4273043 | 4273076 | 34 | - | 18.3659 | 2.76E-07 | 0.0595 | TAGCTGTCTTCAGACACTCCAGAAGAGGGCGCCA | | |
| 1231 | 14 | 5631283 | 5638037 | 6754 | SPRET | 14 | 5635255 | 5635288 | 34 | - | 18.3252 | 2.84E-07 | 0.0602 | TTGAAGTTCAAACCCAGGCCAGAGGAGGGCATCA | | |
| 1232 | 14 | 19427725 | 19429889 | 2164 | 129S1.C57L.KK.MOLF.SMJ | 14 | 19428222 | 19428255 | 34 | - | 18.5935 | 2.37E-07 | 0.0571 | GTGCTGTATGTGTAGAGGCCAGAAGAGGGTGTCA | | |
| 1233 | 14 | 20483378 | 20485065 | 1687 | CAST.MOLF.SPRET | 14 | 20483511 | 20483544 | 34 | - | 17.5122 | 4.86E-07 | 0.0778 | TAGCTGTTTTCAGACACACCAGAAGAGGGCATTA | | |
| 1234 | 14 | 20945691 | 20948806 | 3115 | WSB | 14 | 20946807 | 20946825 | 19 | - | 17.5574 | 6.32E-07 | 0.158 | ACTCCAGAAGAGGGCGCCA | | |
| 1234 | 14 | 20945691 | 20948806 | 3115 | WSB | 14 | 20946807 | 20946840 | 34 | - | 18.3659 | 2.76E-07 | 0.0595 | TAGCTGTCTTCAGACACTCCAGAAGAGGGCGCCA | | |
| 1235 | 14 | 25820781 | 25823093 | 2312 | SPRET | 14 | 25820886 | 25820904 | 19 | - | 17.2459 | 7.74E-07 | 0.173 | ACACCAGAAGAGGGCAGCA | | |
| 1235 | 14 | 25820781 | 25823093 | 2312 | SPRET | 14 | 25820886 | 25820919 | 34 | - | 17.6504 | 4.44E-07 | 0.0744 | TAGCTGTCTTCAGACACACCAGAAGAGGGCAGCA | | |
| 1236 | 14 | 25912256 | 25913989 | 1733 | AJ,BALB,CAST,MOLF,NZW,RH2,SEA,SPRET | 14 | 25913556 | 25913589 | 34 | - | 16.4472 | 9.61E-07 | 0.111 | TAGCTGTCTTCAGACACACCAGAAGAGGGCGTCA | | |
| 1237 | 14 | 33594943 | 33597797 | 2854 | BTBR | 14 | 33596497 | 33596530 | 34 | - | 16.8374 | 7.51E-07 | 0.0977 | CAGCAGGTCTTGACACTTCCTGGAGGAGTCACCC | | |
| 1238 | 14 | 36141917 | 36147114 | 5197 | CAST | 14 | 36144532 | 36144565 | 34 | - | 17.4146 | 5.18E-07 | 0.0804 | ATGCAGCTAGAGACACAAGCTCCAGGGGGTACTG | | |
| 1239 | 14 | 43050984 | 43055970 | 4986 | CBA.DBA1J.DBA2J | 14 | 43051941 | 43051974 | 34 | - | 25.3984 | 1.16E-09 | 0.00172 | ATGCAATTTCTCCAGAGGCCAGTAGAGGGCATGA | 4 | ENCFF367UNR.ENCFF201JUA.ENCFF099QNE.ENCFF093ZRU |
| 1240 | 14 | 44041824 | 44050145 | 8321 | BTBR,C57L,CAST,FVB,KK,MOLF,NOD,NOR,NZB,NZO,NZW,PJ,PL,WSB | 14 | 44042116 | 44042134 | 19 | - | 19.2951 | 1.88E-07 | 0.0904 | CCGCCACTAGATGGTGCTA | 13 | ENCFF092NMJ,ENCFF136ZDK,ENCFF165LXO,ENCFF198SSR,ENCFF093ZRU,ENCFF367UNR,ENCFF099QNE,ENCFF821BIA,ENCFF327LYS,ENCFF907XEG,ENCFF024JOZ,ENCFF089GBC,ENCFF589ZBJ |
| 1240 | 14 | 44041824 | 44050145 | 8321 | BTBR,C57L,CAST,FVB,KK,MOLF,NOD,NOR,NZB,NZO,NZW,PJ,PL,WSB | 14 | 44050104 | 44050122 | 19 | - | 19.2951 | 1.88E-07 | 0.0904 | CCGCCACTAGATGGTGCTA | 28 | ENCFF201JUA,ENCFF093ZRU,ENCFF367UNR,ENCFF883UPM,ENCFF609MDN,ENCFF099QNE,ENCFF097CAQ,ENCFF194HVL,ENCFF907XEG,ENCFF136ZDK,ENCFF327LYS,ENCFF821BIA,ENCFF089GBC,ENCFF491RJK,ENCFF589ZBJ,ENCFF321AGQ,ENCFF052KVO,ENCFF055HAP,ENCFF065ODV,ENCFF957SPH,ENCFF281QMV,ENCFF142CNG,ENCFF092NMJ,ENCFF198SSR,ENCFF165LXO,ENCFF430PPJ,ENCFF502WDT,ENCFF024JOZ |

| | | | | | | | | | | | | | | | | |
|---|---|---|---|---|---|---|---|---|---|---|---|---|---|---|---|---|
| 1241 | 14 | 52067424 | 52086384 | 18960 | BUB,C57L,DBA1J,DBA2J,FVB,ILNJ2,MA/My,NOD,NOR,RF,SJL,SMJ,SWR,WSB | 14 | 52076111 | 52076129 | 19 | - | 17.5246 | 6.45E-07 | 0.16 | TGGTCAGCAGAGGGCAGGA | 9 | ENCFF367UNR,ENCFF092NMJ,ENCFF089GBC,ENCFF907XEG,ENCFF883UPM,ENCFF093ZRU,ENCFF099QNE,ENCFF327LYS,ENCFF024JOZ |
| 1241 | 14 | 52067424 | 52086384 | 18960 | BUB,C57L,DBA1J,DBA2J,FVB,ILNJ2,MA/My,NOD,NOR,RF,SJL,SMJ,SWR,WSB | 14 | 52076111 | 52076145 | 35 | - | 18.3303 | 2.47E-07 | 0.112 | CTGTAAGAGCTTCTTGTGGTCAGCAGAGGGCAGGA | 9 | ENCFF367UNR,ENCFF092NMJ,ENCFF089GBC,ENCFF907XEG,ENCFF883UPM,ENCFF093ZRU,ENCFF099QNE,ENCFF327LYS,ENCFF024JOZ |
| 1242 | 14 | 52113662 | 52120861 | 7199 | 129S1,AJ,AKR,BALB,C3H,C57L,CBA,CE,DBA1J,DBA2J,KK,MRL,NZB,NZO,NZW,PJ,PL,RF,RH2,SEA,TH | 14 | 52118709 | 52118743 | 35 | - | 16.1009 | 9.03E-07 | 0.221 | CTACTGTTCGTCACCAGGATCTCCAGAGGGCAGCT | | |
| 1243 | 14 | 53012165 | 53013598 | 1433 | BUB.SPRET.WSB | 14 | 53013487 | 53013520 | 34 | - | 17.6992 | 4.30E-07 | 0.0732 | CTGCAATACCAGACTGAAGCTGAAGGTGTCTCCG | 4 | ENCFF089GBC.ENCFF883UPM.ENCFF093ZRU.ENCFF099QNE |
| 1244 | 14 | 55848732 | 55850725 | 1993 | SPRET | 14 | 55849823 | 55849857 | 35 | - | 20.1927 | 7.68E-08 | 0.0582 | CTGCATGTGGACCTGCACACCAGAAGAGGACACCA | | |
| 1245 | 14 | 57026561 | 57028181 | 1620 | SPRET | 14 | 57027268 | 57027302 | 35 | - | 19.4679 | 1.22E-07 | 0.0763 | TTGCAGATGCCCTTGGAGGCCAGAAGATGTCATCA | | |
| 1245 | 14 | 57026561 | 57028181 | 1620 | SPRET | 14 | 57027652 | 57027686 | 35 | - | 19.0183 | 1.62E-07 | 0.0892 | CTGTAGCTGTACTCAGACACCAGAAGAGGGCATCA | 6 | ENCFF093ZRU.ENCFF821BIA.ENCFF099QNE.ENCFF502WDT.ENCFF367UNR.ENCFF055HAP |
| 1246 | 14 | 58157044 | 58159273 | 2229 | SPRET | 14 | 58157738 | 58157756 | 19 | - | 17.5738 | 6.25E-07 | 0.158 | ACTCCAGAAGAGGGCGCTA | 1 | ENCFF093ZRU |
| 1246 | 14 | 58157044 | 58159273 | 2229 | SPRET | 14 | 58157738 | 58157771 | 34 | - | 18.1789 | 3.13E-07 | 0.0629 | TAGCTGTCTTCAGACACTCCAGAAGAGGGCGCTA | 1 | ENCFF093ZRU |
| 1247 | 14 | 63200675 | 63203048 | 2373 | CAST | 14 | 63202999 | 63203032 | 34 | - | 17.7236 | 4.24E-07 | 0.0732 | GTGCACAGCTGAGAGCAGGCACCTGGTGTCGCTG | 12 | ENCFF093ZRU,ENCFF089GBC,ENCFF821BIA,ENCFF092NMJ,ENCFF198SSR,ENCFF883UPM,ENCFF609MDN,ENCFF533APC,ENCFF367UNR,ENCFF907XEG,ENCFF316FYN,ENCFF957SPH |
| 1248 | 14 | 63943337 | 63949002 | 5665 | CAST | 14 | 63945266 | 63945300 | 35 | - | 17.3945 | 4.31E-07 | 0.151 | CTGCTGCTGCTATATGATTCCACCAGGAGTCACCC | | |
| 1249 | 14 | 65890067 | 65893583 | 3516 | SPRET | 14 | 65891865 | 65891898 | 34 | - | 17.5772 | 4.66E-07 | 0.0762 | GTGCATATATGGTGAGTGCCAGAAGAGGGCATCA | | |
| 1250 | 14 | 66364065 | 66366309 | 2244 | CAST.MOLF | 14 | 66365723 | 66365756 | 34 | - | 18.878 | 1.95E-07 | 0.0508 | CAGCACCTGTGGAATCTACCACAAGAAGGCAGTG | | |
| 1251 | 14 | 69311284 | 69314492 | 3208 | CAST | 14 | 69312872 | 69312905 | 34 | - | 25.1057 | 1.51E-09 | 0.00204 | TTGCAGTACTTACAGCTGTCAGAAGAGGGCACCA | | |
| 1252 | 14 | 70982291 | 70987372 | 5081 | SPRET | 14 | 70983144 | 70983162 | 19 | - | 18.541 | 3.24E-07 | 0.117 | AAACCAGAAGAGGGCGCCA | 5 | ENCFF093ZRU.ENCFF367UNR.ENCFF099QNE.ENCFF821BIA.ENCFF055HAP |
| 1253 | 14 | 71406924 | 71409672 | 2748 | BTBR | 14 | 71409615 | 71409648 | 34 | - | 18.0569 | 3.40E-07 | 0.0647 | ATGCAGCTAGAGTCACGAGCTCCAGGGGGTACTG | | |
| 1254 | 14 | 71681735 | 71684752 | 3017 | SPRET | 14 | 71683147 | 71683181 | 35 | - | 16.4679 | 7.34E-07 | 0.198 | TTGCAATTGATGGCCCAGGCCTGAAGGGGTCATGC | | |
| 1255 | 14 | 76070938 | 76072946 | 2008 | MOLF | 14 | 76072756 | 76072790 | 35 | - | 18.055 | 2.91E-07 | 0.123 | TTGCTGCTCCTTTCTTTGCGCACTAGGGGCCACTA | 15 | ENCFF024JOZ,ENCFF367UNR,ENCFF092NMJ,ENCFF821BIA,ENCFF089GBC,ENCFF198SSR,ENCFF883UPM,ENCFF589ZBJ,ENCFF502WDT,ENCFF957SPH,ENCFF907XEG,ENCFF093ZRU,ENCFF327LYS,ENCFF055HAP,ENCFF136ZDK |
| 1256 | 14 | 76170441 | 76173026 | 2585 | SWR | 14 | 76171985 | 76172018 | 34 | - | 27.8943 | 1.05E-10 | 0.000521 | GTGCAGTGCCCTTGGAGGCCAGAAGAGGGCATCA | 20 | ENCFF024JOZ,ENCFF089GBC,ENCFF093ZRU,ENCFF367UNR,ENCFF198SSR,ENCFF201JUA,ENCFF883UPM,ENCFF065ODV,ENCFF136ZDK,ENCFF194HVL,ENCFF327LYS,ENCFF589ZBJ,ENCFF052KVO,ENCFF957SPH,ENCFF321AGQ,ENCFF821BIA,ENCFF502WDT,ENCFF099QNE,ENCFF907XEG,ENCFF055HAP |
| 1257 | 14 | 76525327 | 76526557 | 1230 | NZB.NZO | 14 | 76526258 | 76526291 | 34 | - | 17.1382 | 6.20E-07 | 0.089 | TAGCTGTCTTCAAGCACACCAGCAGAGGGCATCA | | |
| 1258 | 14 | 77777140 | 77778495 | 1355 | AJ,BALB,CAST,MOLF,NZB,NZO,SEA,SJL,SPRET | 14 | 77778142 | 77778176 | 35 | - | 22.2569 | 1.90E-08 | 0.0271 | CTGCTGTGTCACATGTGCATCAGTAGGTGGCAGAG | 2 | ENCFF821BIA,ENCFF367UNR |
| 1259 | 14 | 81387199 | 81388768 | 1569 | SPRET | 14 | 81387949 | 81387982 | 34 | - | 17.4146 | 5.18E-07 | 0.0804 | ATGCAGCTAGAGACACAAGCTCCAGGGGGTACTG | | |
| 1260 | 14 | 94168361 | 94174974 | 6613 | AJ,AKR,BALB,BTBR,BUB,C3H,CBA,CE,DBA1J,DBA2J,FVB,MRL,NOD,PJ,PL,RF,RH2,SEA,SJL,SMJ,SWR,TH,WSB | 14 | 94174171 | 94174204 | 34 | - | 16.9106 | 7.17E-07 | 0.0962 | GAGCATGACCACAAATCCACAGCAGGTGGCAACG | | |
| 1261 | 14 | 94878813 | 94887279 | 8466 | SPRET | 14 | 94886532 | 94886565 | 34 | - | 17.5691 | 4.69E-07 | 0.0763 | GAGCATGACCACAAGTCCACAGCAGGTGGCAACA | | |
| 1262 | 14 | 95568417 | 95575123 | 6706 | SPRET | 14 | 95569766 | 95569799 | 34 | - | 17.9106 | 3.74E-07 | 0.0681 | CCGCCCTCACATTCGCCGTCACAAGATGGCGCTG | | |
| 1263 | 14 | 99330219 | 99331758 | 1539 | AKR,CAST,CBA,CE,DBA1J,DBA2J,ILNJ2,MRL,PJ,PL,RF,RH2,SJL,SMJ,SPRET,SWR,TH,WSB | 14 | 99330761 | 99330795 | 35 | - | 16.6789 | 6.51E-07 | 0.189 | GTGCAGTGTGCGCCATTCAGCTAAAGAGGGCTCCC | | |
| 1264 | 14 | 105129975 | 105196398 | 66423 | NZB | 14 | 105143349 | 105143367 | 19 | - | 21.082 | 4.57E-08 | 0.0571 | AAGCCAGCAGAGGGCACCA | 2 | ENCFF089GBC.ENCFF052KVO |
| 1264 | 14 | 105129975 | 105196398 | 66423 | NZB | 14 | 105143349 | 105143383 | 35 | - | 25.211 | 2.08E-09 | 0.00868 | TTGTAGTTATCTATGGAAGCCAGCAGAGGGCACCA | 2 | ENCFF089GBC.ENCFF052KVO |
| 1265 | 14 | 120489443 | 120490691 | 1248 | BTBR.C57L.MOLF | 14 | 120490579 | 120490613 | 35 | - | 18.7064 | 1.96E-07 | 0.0995 | CAGCACTCCTAGCTCAGGCCCTGCAGAGGTCATTG | | |
| 1266 | 14 | 121102538 | 121104304 | 1766 | SPRET | 14 | 121104167 | 121104200 | 34 | - | 25.9675 | 6.89E-10 | 0.00121 | ATGCACTTCCCGAAGAGGTCAGAAGAGGGCACTA | 5 | ENCFF093ZRU.ENCFF367UNR.ENCFF821BIA.ENCFF099QNE.ENCFF052KVO |
| 1267 | 15 | 8224437 | 8226850 | 2413 | CAST.LP.MOLF.TH | 15 | 8225786 | 8225819 | 34 | + | 16.4634 | 9.51E-07 | 0.106 | TAGCTGTCTTCAGACGCACCAGAAGAGGGCATCA | | |
| 1268 | 15 | 15784854 | 15789357 | 4503 | CAST.MOLF.PL.SPRET | 15 | 15785631 | 15785664 | 34 | + | 17.6748 | 4.37E-07 | 0.0745 | GAGCATGACCACAAGTCCACAGCAGGTGGCAACG | | |
| 1269 | 15 | 16899050 | 16923127 | 24077 | SPRET | 15 | 16900659 | 16900692 | 34 | + | 18.6016 | 2.36E-07 | 0.0572 | TTGCTCTTTTCAGGCACACCAGAAGAGGGCATCG | | |
| 1270 | 15 | 29125046 | 29131360 | 6314 | C3H,CBA,DBA1J,DBA2J,MA/My,MOLF,MRL | 15 | 29128130 | 29128163 | 34 | + | 16.5041 | 9.27E-07 | 0.106 | ATGCAGCTAGAGACACGAGCTCTGGGGGGCGGGG | | |
| 1271 | 15 | 29126696 | 29128650 | 1954 | CAST | 15 | 29128130 | 29128163 | 34 | + | 16.5041 | 9.27E-07 | 0.106 | ATGCAGCTAGAGACACGAGCTCTGGGGGGCGGGG | | |
| 1272 | 15 | 37088397 | 37091152 | 2755 | CAST | 15 | 37088528 | 37088561 | 34 | + | 16.4878 | 9.37E-07 | 0.106 | TTGCACTTCACCTTTACAACATCAGAGGGCATTA | | |
| 1273 | 15 | 38951626 | 38952650 | 1024 | C3H.MRL.NOD.NOR.PL.SWR.WSB | 15 | 38952166 | 38952200 | 35 | + | 16.4037 | 7.61E-07 | 0.163 | CTGCATGTACACCTGCATGCCAGAAGAGAGTATTA | | |
| 1274 | 15 | 40304923 | 40311555 | 6632 | SPRET | 15 | 40306483 | 40306516 | 34 | + | 19.8374 | 9.97E-08 | 0.0331 | TAGCTGTCCCCAGATGCACCAGAAGAGGGCATCA | | |
| 1275 | 15 | 41750645 | 41752728 | 2083 | 129S1,AKR,BUB,C3H,C57L,KK,PL,SWR,WSB | 15 | 41752251 | 41752269 | 19 | + | 17.459 | 6.74E-07 | 0.13 | CAGCCAGGAGGTGGAGCTG | | |
| 1276 | 15 | 46137765 | 46142805 | 5040 | SPRET | 15 | 46139877 | 46139911 | 35 | + | 17.6972 | 3.60E-07 | 0.109 | CTGTAATCCCAGCGCCCAGGCGGCAGAGGCAGGCG | | |
| 1277 | 15 | 49647550 | 49652432 | 4882 | SPRET | 15 | 49648450 | 49648468 | 19 | + | 16.9836 | 9.17E-07 | 0.151 | TCAACAGCAGATGGCAGCA | | |
| 1278 | 15 | 49829625 | 49834382 | 4757 | SPRET | 15 | 49832794 | 49832828 | 35 | + | 16.9358 | 5.62E-07 | 0.14 | CTGCTGTTCACTTCTGGGTTCTGCAGGTGCCGCAG | | |
| 1279 | 15 | 53752059 | 53753457 | 1398 | SPRET | 15 | 53752487 | 53752505 | 19 | + | 18.9344 | 2.45E-07 | 0.0795 | ATGCCAGAAGAGGGCACCA | | |
| 1280 | 15 | 59033542 | 59034868 | 1326 | 129S1.C57L.MRL | 15 | 59033558 | 59033576 | 19 | + | 17.5738 | 6.25E-07 | 0.126 | CCACCAGCAGGCGGCACTG | 2 | ENCFF093ZRU.ENCFF367UNR |
| 1281 | 15 | 59033550 | 59039293 | 5743 | 129S1,BALB,C57L,CAST,ILNJ2,MOLF,MRL,NOD,NOR,SEA,SPRET,WSB | 15 | 59033558 | 59033576 | 19 | + | 17.5738 | 6.25E-07 | 0.126 | CCACCAGCAGGCGGCACTG | 2 | ENCFF093ZRU,ENCFF367UNR |
| 1282 | 15 | 60660375 | 60661644 | 1269 | MOLF | 15 | 60660825 | 60660858 | 34 | + | 19.9024 | 9.52E-08 | 0.0322 | ATGCAGCCTCAGTGGAAGCCAACAGAGGGTGGCA | | |
| 1283 | 15 | 65443122 | 65449840 | 6718 | CAST.MOLF.SPRET | 15 | 65443131 | 65443164 | 34 | + | 16.9593 | 6.95E-07 | 0.0934 | GTGCTGTGCTCAGCCTCAGCAGCAGCGGTCGCCA | | |
| 1284 | 15 | 68993767 | 69001031 | 7264 | BUB,CAST,DBA1J,DBA2J,ILNJ2,KK,LP,MOLF,NZB,NZO,PL,SPRET,TH | 15 | 68994062 | 68994096 | 35 | + | 15.9817 | 9.65E-07 | 0.183 | CTGCTCTGACCACTGAAACTCAGAGGAGGGAGCTA | | |
| 1285 | 15 | 71469749 | 71476984 | 7235 | CAST.MOLF.SPRET | 15 | 71476643 | 71476676 | 34 | + | 19.3415 | 1.42E-07 | 0.0421 | CTGCCCCCACATTCGCCGTCACAAGATGGCGCTG | | |
| 1286 | 15 | 71919987 | 71921209 | 1222 | AJ,B10D2,B10J,BALB,BTBR,BUB,C3H,C57L,CAST,CBA,CE,DBA1J,DBA2J,FVB,ILNJ2,KK,LP,MA/My,MOLF,MRL,NOD,NOR,NZB,NZO,NZW,PJ,PL,RF,SEA,SJL,SMJ,SPRET,SWR,TH,WSB | 15 | 71920159 | 71920192 | 34 | + | 17.9431 | 3.66E-07 | 0.0685 | GAACCATTACTCCTACTGGCACCAGGGGGCGCTC | | |
| 1286 | 15 | 71919987 | 71921209 | 1222 | AJ,B10D2,B10J,BALB,BTBR,BUB,C3H,C57L,CAST,CBA,CE,DBA1J,DBA2J,FVB,ILNJ2,KK,LP,MA/My,MOLF,MRL,NOD,NOR,NZB,NZO,NZW,PJ,PL,RF,SEA,SJL,SMJ,SPRET,SWR,TH.WSB | 15 | 71920174 | 71920192 | 19 | + | 19.6066 | 1.49E-07 | 0.062 | CTGGCACCAGGGGGCGCTC | | |
| 1287 | 15 | 72331735 | 72336908 | 5173 | SPRET | 15 | 72332616 | 72332649 | 34 | + | 18.3089 | 2.87E-07 | 0.0603 | ATGCAGCTAGAGACACGAGCTCCAGGGGGTACTG | | |
| 1288 | 15 | 75036685 | 75047588 | 10903 | AJ,BALB,C3H,CBA,CE,NOD,NOR,NZB,NZO,SEA,TH,WSB | 15 | 75047549 | 75047567 | 19 | + | 19.7049 | 1.38E-07 | 0.0593 | TGGCCAGCAGGTGGCAGGT | 35 | ENCFF201JUA,ENCFF093ZRU,ENCFF367UNR,ENCFF099QNE,ENCFF327LYS,ENCFF502WDT,ENCFF024JOZ,ENCFF907XEG,ENCFF951GOP,ENCFF316FYN,ENCFF033UQX,ENCFF821BIA,ENCFF198SSR,ENCFF165LXO,ENCFF589ZBJ,ENCFF092NMJ,ENCFF575MGI,ENCFF052KVO,ENCFF957SPH,ENCFF097CAQ,ENCFF136ZDK,ENCFF883UPM,ENCFF089EWX,ENCFF210BTO,ENCFF611HDQ,ENCFF065ODV,ENCFF089GBC,ENCFF055HAP,ENCFF281QMV,ENCFF504JDS,ENCFF533APC,ENCFF142CNG,ENCFF491RJK,ENCFF194HVL,ENCFF430PPJ |
| 1289 | 15 | 77464039 | 77486565 | 22526 | MA/Mv.MOLF | 15 | 77485627 | 77485645 | 19 | + | 17.1967 | 7.99E-07 | 0.144 | GGGCCAGAAGAGGGCGTCA | | |
| 1290 | 15 | 79087419 | 79088783 | 1364 | SPRET | 15 | 79088417 | 79088451 | 35 | + | 16.6239 | 6.72E-07 | 0.153 | CTGTAGAGTAAGGGTGAAGGCAGCAGAGGGCTCAG | | |
| 1291 | 15 | 88657498 | 88659890 | 2392 | SPRET | 15 | 88659027 | 88659060 | 34 | + | 18.3659 | 2.76E-07 | 0.0597 | TAGCTGTCTTCAGACACTCCAGAAGAGGGCGCCA | 6 | ENCFF821BIA.ENCFF099QNE.ENCFF367UNR.ENCFF093ZRU.ENCFF502WDT.ENCFF055HAP |
| 1291 | 15 | 88657498 | 88659890 | 2392 | SPRET | 15 | 88659042 | 88659060 | 19 | + | 17.5574 | 6.32E-07 | 0.126 | ACTCCAGAAGAGGGCGCCA | 6 | ENCFF821BIA.ENCFF099QNE.ENCFF367UNR.ENCFF093ZRU.ENCFF502WDT.ENCFF055HAP |
| 1292 | 15 | 88678888 | 88680294 | 1406 | SPRET | 15 | 88678948 | 88678981 | 34 | + | 16.7724 | 7.83E-07 | 0.0991 | TAGCTGTCTTCAGATGCACCAGAAGAGGGCGTCA | | |
| 1293 | 15 | 89614387 | 89616072 | 1685 | SPRET | 15 | 89614633 | 89614651 | 19 | + | 20.4426 | 7.76E-08 | 0.0477 | TGGCCGCCAGGGGGCAGGG | 23 | ENCFF093ZRU,ENCFF821BIA,ENCFF430PPJ,ENCFF367UNR,ENCFF099QNE,ENCFF327LYS,ENCFF907XEG,ENCFF024JOZ,ENCFF609MDN,ENCFF321AGQ,ENCFF052KVO,ENCFF136ZDK,ENCFF097CAQ,ENCFF194HVL,ENCFF957SPH,ENCFF589ZBJ,ENCFF198SSR,ENCFF092NMJ,ENCFF165LXO,ENCFF533APC,ENCFF281QMV,ENCFF089GBC,ENCFF033UQX |
| 1294 | 15 | 94287897 | 94289342 | 1445 | CAST | 15 | 94288171 | 94288204 | 34 | + | 20.6585 | 5.50E-08 | 0.0223 | TAGCTGTCTTCAGACCTACCAGAAGAGGGCGCCA | 7 | ENCFF093ZRU,ENCFF099QNE,ENCFF821BIA,ENCFF052KVO,ENCFF883UPM,ENCFF502WDT,ENCFF055HAP |
| 1294 | 15 | 94287897 | 94289342 | 1445 | CAST | 15 | 94288186 | 94288204 | 19 | + | 20.7213 | 6.18E-08 | 0.0453 | CTACCAGAAGAGGGCGCCA | 8 | ENCFF093ZRU,ENCFF099QNE,ENCFF821BIA,ENCFF052KVO,ENCFF883UPM,ENCFF502WDT,ENCFF055HAP.ENCFF065ODV |
| 1295 | 15 | 94336996 | 94340403 | 3407 | MOLF | 15 | 94339859 | 94339893 | 35 | + | 17 | 5.42E-07 | 0.138 | GTGTCACTACCCATAGAGGCCAGAAGAGGGCGTTG | 1 | ENCFF093ZRU |
| 1295 | 15 | 94336996 | 94340403 | 3407 | MOLF | 15 | 94339860 | 94339893 | 34 | + | 16.3902 | 9.96E-07 | 0.109 | TGTCACTACCCATAGAGGCCAGAAGAGGGCGTTG | 1 | ENCFF093ZRU |
| 1296 | 15 | 99388302 | 99389806 | 1504 | SPRET | 15 | 99389574 | 99389607 | 34 | + | 18.561 | 2.42E-07 | 0.0576 | CAGCTGTCTTCAGACACACCAGAAGAGGGCATCA | | |
| 1297 | 15 | 99706870 | 99710079 | 3209 | CAST | 15 | 99708539 | 99708573 | 35 | + | 24.5688 | 3.45E-09 | 0.00752 | CTGCATGTATGCCTGCACACCAGAAGATGGCACCA | 4 | ENCFF093ZRU.ENCFF327LYS.ENCFF367UNR.ENCFF907XEG |
| 1297 | 15 | 99706870 | 99710079 | 3209 | CAST | 15 | 99708555 | 99708573 | 19 | + | 17.0328 | 8.88E-07 | 0.149 | ACACCAGAAGATGGCACCA | 4 | ENCFF093ZRU.ENCFF327LYS.ENCFF367UNR.ENCFF907XEG |
| 1298 | 15 | 101962834 | 101964667 | 1833 | CAST | 15 | 101964024 | 101964057 | 34 | + | 21.4553 | 3.03E-08 | 0.0152 | ATGCAGTGTCCACGGAAGGCAGAAGATGGCATTG | | |
| 1299 | 15 | 103797273 | 103803825 | 6552 | CAST,MOLF,SPRET | 15 | 103797297 | 103797315 | 19 | + | 19.2951 | 1.88E-07 | 0.0694 | CCGCCACTAGATGGTGCTA | 31 | ENCFF093ZRU,ENCFF201JUA,ENCFF367UNR,ENCFF052KVO,ENCFF099QNE,ENCFF327LYS,ENCFF907XEG,ENCFF316FYN,ENCFF951GOP,ENCFF821BIA,ENCFF589ZBJ,ENCFF024JOZ,ENCFF321AGQ,ENCFF136ZDK,ENCFF092NMJ,ENCFF089GBC,ENCFF609MDN,ENCFF165LXO,ENCFF430PPJ,ENCFF198SSR,ENCFF097CAQ,ENCFF504JDS,ENCFF883UPM,ENCFF194HVL,ENCFF957SPH,ENCFF281QMV,ENCFF491RJK,ENCFF055HAP.ENCFF533APC.ENCFF575MGI.ENCFF502WDT |
| 1300 | 15 | 4624982 | 4628909 | 3927 | SPRET | 15 | 4627714 | 4627748 | 35 | - | 17.8165 | 3.36E-07 | 0.105 | TAGCAGCTGCCACCTGTGTGCTCTAGGGGGCAGGG | | |
| 1301 | 15 | 12300157 | 12309134 | 8977 | SPRET | 15 | 12305281 | 12305314 | 34 | - | 17.9268 | 3.70E-07 | 0.0689 | CTGATCCTCCAGTCTCCACCTCCAGAGGGCTGGG | | |
| 1302 | 15 | 14476678 | 14485306 | 8628 | LP.MOLF.SMJ | 15 | 14484237 | 14484270 | 34 | - | 17.6748 | 4.37E-07 | 0.0745 | GAGCATGACCACAAGTCCACAGCAGGTGGCAACG | | |
| 1303 | 15 | 14480550 | 14484986 | 4436 | BALB,C3H,CAST,CBA,DBA2J,MRL,NOD,NOR,NZB,NZO,PL,SJL,SPRET,SWR | 15 | 14484237 | 14484270 | 34 | - | 17.6748 | 4.37E-07 | 0.0745 | GAGCATGACCACAAGTCCACAGCAGGTGGCAACG | | |
| 1304 | 15 | 14845014 | 14853191 | 8177 | BUB.CE.ILNJ2.MRL.PL.SPRET.WSB | 15 | 14845280 | 14845313 | 34 | - | 17.0407 | 6.60E-07 | 0.0918 | CTGTGATAACATTTGCCATCACAAGATGGCGCCA | 2 | ENCFF055HAP.ENCFF502WDT |
| 1305 | 15 | 19495443 | 19499482 | 4039 | 129S1,BTBR,C3H,C57L,CBA,CE,ILNJ2,KK,LP,MA/My,MOLF,NZW,PL,SPRET,SWR,WSB | 15 | 19498735 | 19498768 | 34 | - | 17.6748 | 4.37E-07 | 0.0745 | GAGCATGACCACAAGTCCACAGCAGGTGGCAACG | | |
| 1306 | 15 | 20647297 | 20650504 | 3207 | CAST.MRL.PL.SWR | 15 | 20650022 | 20650055 | 34 | - | 16.8943 | 7.24E-07 | 0.0956 | ATGCAGTCCCCATGGAGGCCAAAACATGGCATCG | | |
| 1307 | 15 | 36481361 | 36483947 | 2586 | 129S1.C57L | 15 | 36481793 | 36481826 | 34 | - | 22.8293 | 1.04E-08 | 0.00719 | GTACAGTTCCAACAGAGGCCAGAAGAGGGCATAA | 2 | ENCFF093ZRU.ENCFF327LYS |
| 1308 | 15 | 38193202 | 38195214 | 2012 | SPRET | 15 | 38195125 | 38195158 | 34 | - | 18.9431 | 1.87E-07 | 0.0495 | ATGCAGTGCCCACAGAGGCCAAAGGAAGGCGACT | | |
| 1309 | 15 | 38865056 | 38867145 | 2089 | SPRET | 15 | 38866529 | 38866562 | 34 | - | 22.1789 | 1.74E-08 | 0.0103 | ATGTAGTACCCACAGAGGCCAGAAGAGGGTGGGA | 16 | ENCFF327LYS,ENCFF024JOZ,ENCFF065ODV,ENCFF821BIA,ENCFF092NMJ,ENCFF430PPJ,ENCFF367UNR,ENCFF055HAP,ENCFF089GBC,ENCFF502WDT,ENCFF052KVO,ENCFF093ZRU,ENCFF099QNE,ENCFF907XEG,ENCFF165LXO,ENCFF198SSR |
| 1310 | 15 | 51524359 | 51525906 | 1547 | SPRET | 15 | 51524561 | 51524594 | 34 | - | 16.6098 | 8.68E-07 | 0.105 | CAGCATTGATGCAGGTGGCCAGCTGATAGCACTG | | |
| 1311 | 15 | 57640123 | 57645454 | 5331 | SPRET | 15 | 57643750 | 57643784 | 35 | - | 16.2752 | 8.19E-07 | 0.169 | TTGCAGTTCACAGCCCAGGACTGGAGGGGTCATGC | | |
| 1312 | 15 | 57899641 | 57903272 | 3631 | ILNJ2.MOLF | 15 | 57902792 | 57902825 | 34 | - | 21.374 | 3.23E-08 | 0.0158 | GTGCAGGTACAGAAGAGGCCAGAAAGGGCACCA | | |
| 1313 | 15 | 59181625 | 59186036 | 4411 | SPRET | 15 | 59184139 | 59184173 | 35 | - | 19.6055 | 1.12E-07 | 0.0572 | CTGCATATATTCCTGTATGCCAGAAGAGGACACCA | | |
| 1314 | 15 | 65495252 | 65496497 | 1245 | SWR | 15 | 65495222 | 65495256 | 35 | - | 16.5046 | 7.19E-07 | 0.158 | TTGTAGGTAGAGGTATAGGCCTGAAGATGGAGCTT | | |
| 1315 | 15 | 76444012 | 76451202 | 7190 | AJ,BALB,C3H,C57L,CAST,CBA,CE,FVB,MA/My,MOLF,MRL,NOD,NOR,NZB,NZO,PL,RF,SEA,SPRET,WSB | 15 | 76447567 | 76447585 | 19 | - | 17.1639 | 8.17E-07 | 0.145 | CCTCCACGAGGTGGCGGAA | | |
| 1316 | 15 | 77568069 | 77576340 | 8271 | AJ,BALB,C3H,C57L,CBA,CE,FVB,MA/My,MRL,PL,RF,SEA,SPRET,TH,WSB | 15 | 77568356 | 77568374 | 19 | - | 19.2951 | 1.88E-07 | 0.0694 | CCGCCACTAGATGGTGCTA | 26 | ENCFF367UNR,ENCFF093ZRU,ENCFF201JUA,ENCFF099QNE,ENCFF821BIA,ENCFF327LYS,ENCFF024JOZ,ENCFF907XEG,ENCFF136ZDK,ENCFF097CAQ,ENCFF589ZBJ,ENCFF609MDN,ENCFF165LXO,ENCFF092NMJ,ENCFF198SSR,ENCFF055HAP,ENCFF491RJK,ENCFF089GBC,ENCFF430PPJ,ENCFF951GOP,ENCFF281QMV,ENCFF502WDT,ENCFF142CNG,ENCFF883UPM,ENCFF957SPH,ENCFF194HVL |
| 1317 | 15 | 79806141 | 79807592 | 1451 | CAST | 15 | 79806377 | 79806410 | 34 | - | 19.1463 | 1.62E-07 | 0.0463 | ATGCCATACCTCTGGAGACCAGAAGAGGGTGTCA | 2 | ENCFF093ZRU.ENCFF502WDT |
| 1318 | 15 | 80658045 | 80659720 | 1675 | SPRET | 15 | 80659661 | 80659694 | 34 | - | 18.4797 | 2.56E-07 | 0.058 | TTGCTGTCTTCAGACACACCAGAAGAGGGCATCA | | |
| 1319 | 15 | 82949213 | 82952840 | 3627 | MOLF | 15 | 82952667 | 82952700 | 34 | - | 19.9268 | 9.36E-08 | 0.0318 | ACGTAGTACCTCTAGTGGCCGGAAGAGGGCATCC | 2 | ENCFF093ZRU.ENCFF052KVO |
| 1320 | 15 | 83510101 | 83511199 | 1098 | SPRET | 15 | 83510162 | 83510196 | 35 | - | 18.8073 | 1.84E-07 | 0.0752 | CTGCAGTTCTCAGTGTCACCCAGAAAGAGGAGCTC | | |
| 1321 | 15 | 88473285 | 88477436 | 4151 | C58J,CAST,KK,LP,MOLF,NOD,NOR,PL,SPRET,SWR,TH,WSB | 15 | 88476636 | 88476669 | 34 | - | 17.6748 | 4.37E-07 | 0.0745 | GAGCATGACCACAAGTCCACAGCAGGTGGCAACG | | |
| 1322 | 15 | 88920563 | 88924461 | 3898 | KK.MOLF.NOD.NOR.SPRET.SWR | 15 | 88921019 | 88921052 | 34 | - | 18.5447 | 2.45E-07 | 0.0579 | CCGCACCCACATTCGCCGTCACAAGATGGCGCTG | | |
| 1323 | 15 | 90311333 | 90313673 | 2340 | SPRET | 15 | 90311572 | 90311605 | 34 | - | 17.6098 | 4.56E-07 | 0.0757 | GTGCCTTTCCTCATGTCTGCAGCAGATGTCAGAA | | |
| 1324 | 15 | 97526282 | 97532101 | 5819 | SPRET | 15 | 97530957 | 97530990 | 34 | - | 17.5691 | 4.69E-07 | 0.0764 | GAGCATGACCACAAGTCCACAGCAGGTGGCAACA | | |
| 1325 | 15 | 98933440 | 98934902 | 1462 | WSB | 15 | 98934043 | 98934076 | 34 | - | 20.3659 | 6.82E-08 | 0.0257 | TTGCTGTCCTCAGACACACCAGAAGAGGGCATCA | 4 | ENCFF099QNE.ENCFF821BIA.ENCFF093ZRU.ENCFF367UNR |
| 1326 | 15 | 99087977 | 99092304 | 4327 | MOLF | 15 | 99090376 | 99090394 | 19 | - | 20.0656 | 1.05E-07 | 0.0512 | AGGCCAGAAGAGGGCACTG | 5 | ENCFF099QNE.ENCFF024JOZ.ENCFF327LYS.ENCFF907XEG.ENCFF093ZRU |
| 1327 | 15 | 103656834 | 103671153 | 14319 | CAST.MOLF | 15 | 103659175 | 103659193 | 19 | - | 19.2951 | 1.88E-07 | 0.0694 | CCGCCACTAGATGGTGCTA | | |
| 1327 | 15 | 103656834 | 103671153 | 14319 | CAST.MOLF | 15 | 103667166 | 103667184 | 19 | - | 19.2951 | 1.88E-07 | 0.0694 | CCGCCACTAGATGGTGCTA | 6 | ENCFF093ZRU.ENCFF201JUA.ENCFF367UNR.ENCFF099QNE.ENCFF907XEG.ENCFF821BIA |
| 1328 | 16 | 3146833 | 3155603 | 8770 | AJ,AKR,B10J,C3H,ILNJ2,KK,NOD,NOR,NZB,PJ,RF,SJL,SMJ,SPRET | 16 | 3150976 | 3151010 | 35 | + | 16.156 | 8.75E-07 | 0.21 | AAGAAACTGGACCCTGTGGCCAGTGGATGGCGCTC | | |
| 1328 | 16 | 3146833 | 3155603 | 8770 | AJ,AKR,B10J,C3H,ILNJ2,KK,NOD,NOR,NZB,PJ.RF.SJL.SMJ.SPRET | 16 | 3150992 | 3151010 | 19 | + | 18.9344 | 2.45E-07 | 0.108 | TGGCCAGTGGATGGCGCTC | | |
| 1329 | 16 | 3485035 | 3517699 | 32664 | SPRET | 16 | 3503480 | 3503514 | 35 | + | 20.8165 | 5.09E-08 | 0.0394 | CTGCTTGTTCTGATCCAATCCAGCAGGTGGCCCTA | | |
| 1330 | 16 | 16476638 | 16484218 | 7580 | SPRET | 16 | 16481289 | 16481323 | 35 | + | 18.0092 | 2.99E-07 | 0.12 | CTGTAATGCTGTCTTCTTTCCACTGGATGGTTTTA | | |
| 1331 | 16 | 17144696 | 17147509 | 2813 | MOLF | 16 | 17146359 | 17146392 | 34 | + | 17.4715 | 5.00E-07 | 0.0937 | TAGCTGTCTTCAGACAAGCCAGAAGAGGGCGTCA | 1 | ENCFF502WDT |
| 1331 | 16 | 17144696 | 17147509 | 2813 | MOLF | 16 | 17145778 | 17145811 | 34 | - | 20.1626 | 7.90E-08 | 0.0337 | ATGCAATGCTCATGGAGGCCAGAAGAGGACATCA | | |
| 1332 | 16 | 17276538 | 17285055 | 8517 | MOLF | 16 | 17278107 | 17278140 | 34 | + | 20.8618 | 4.74E-08 | 0.0244 | ATGCAGTGCCTGTGGCAGCCAGAAAAGGGTGTTA | | |
| 1332 | 16 | 17276538 | 17285055 | 8517 | MOLF | 16 | 17278342 | 17278375 | 34 | - | 23.6585 | 5.25E-09 | 0.00527 | ATGCAGTATCTGCAGAGGCCAGAGGAGGGCATCA | 18 | ENCFF052KVO,ENCFF093ZRU,ENCFF367UNR,ENCFF092NMJ,ENCFF089GBC,ENCFF165LXO,ENCFF327LYS,ENCFF099QNE,ENCFF821BIA,ENCFF589ZBJ,ENCFF907XEG,ENCFF024JOZ,ENCFF055HAP,ENCFF198SSR.ENCFF136ZDK.ENCFF502WDT.ENCFF883UPM.ENCFF065ODV |
| 1333 | 16 | 18666513 | 18674288 | 7775 | CAST.MOLF.SPRET | 16 | 18667131 | 18667164 | 34 | + | 17.9187 | 3.72E-07 | 0.0805 | TAGCTGTCCTCACATACTCCAGAAGAGGGCATCA | | |
| 1333 | 16 | 18666513 | 18674288 | 7775 | CAST.MOLF.SPRET | 16 | 18667379 | 18667412 | 34 | + | 17.9187 | 3.72E-07 | 0.0805 | TAGCTGTCCTCACATACTCCAGAAGAGGGCATCA | | |
| 1334 | 16 | 18794434 | 18803456 | 9022 | SPRET | 16 | 18794450 | 18794483 | 34 | + | 16.8862 | 7.28E-07 | 0.113 | CCGCGATAACATTTGCCATCACAAGATGGCGCCG | | |
| 1335 | 16 | 19464019 | 19468184 | 4165 | SPRET | 16 | 19467481 | 19467514 | 34 | + | 17.6748 | 4.37E-07 | 0.0869 | GAGCATGACCACAAGTCCACAGCAGGTGGCAACG | | |
| 1336 | 16 | 19467097 | 19470831 | 3734 | CAST.MOLF | 16 | 19467481 | 19467514 | 34 | + | 17.6748 | 4.37E-07 | 0.0869 | GAGCATGACCACAAGTCCACAGCAGGTGGCAACG | | |
| 1337 | 16 | 20333192 | 20335059 | 1867 | MOLF | 16 | 20333311 | 20333344 | 34 | + | 16.3984 | 9.91E-07 | 0.128 | TAGCTGTCTTCAGACGTACCAGAAGAGGGCATCA | 2 | ENCFF099QNE.ENCFF093ZRU |
| 1338 | 16 | 20910202 | 20911456 | 1254 | SPRET | 16 | 20910393 | 20910426 | 34 | + | 17.5122 | 4.86E-07 | 0.0921 | GAGCAGGTGCAGCCGCCGCCCCAAGATAGCACCC | | |
| 1339 | 16 | 25181459 | 25184935 | 3476 | BUB | 16 | 25183390 | 25183423 | 34 | + | 18.1951 | 3.10E-07 | 0.0743 | TTGCTGTCTTCAACCACACCAGAAGAGGGCATCA | 1 | ENCFF093ZRU |
| 1340 | 16 | 26821312 | 26822845 | 1533 | SPRET | 16 | 26821653 | 26821671 | 19 | + | 19.0656 | 2.23E-07 | 0.105 | GGCCCAGTAGGTGGCACTA | 3 | ENCFF093ZRU.ENCFF883UPM.ENCFF502WDT |
| 1341 | 16 | 27019888 | 27021582 | 1694 | ILNJ2.NZW.WSB | 16 | 27020856 | 27020874 | 19 | + | 17.0984 | 8.52E-07 | 0.181 | AGACCAGCAGAGGGAAGCC | | |
| 1342 | 16 | 30707861 | 30711041 | 3180 | MOLF | 16 | 30707942 | 30707976 | 35 | + | 20.0183 | 8.59E-08 | 0.0554 | TTGCTGTGTCACATGTGCATCAGTAGGTGGCAGAG | 5 | ENCFF093ZRU.ENCFF907XEG.ENCFF327LYS.ENCFF367UNR.ENCFF883UPM |
| 1343 | 16 | 37518993 | 37520239 | 1246 | SPRET | 16 | 37519299 | 37519333 | 35 | + | 23.8899 | 5.79E-09 | 0.0131 | ATGCAGGTGACCGAGTTGGCCAACAGAGGGCAGCA | 12 | ENCFF093ZRU,ENCFF099QNE,ENCFF821BIA,ENCFF136ZDK,ENCFF907XEG,ENCFF165LXO,ENCFF089GBC,ENCFF198SSR,ENCFF502WDT,ENCFF033UQX,ENCFF142CNG,ENCFF367UNR |
| 1343 | 16 | 37518993 | 37520239 | 1246 | SPRET | 16 | 37519315 | 37519333 | 19 | + | 21.082 | 4.57E-08 | 0.0538 | TGGCCAACAGAGGGCAGCA | 12 | ENCFF093ZRU,ENCFF099QNE,ENCFF821BIA,ENCFF136ZDK,ENCFF907XEG,ENCFF165LXO,ENCFF089GBC,ENCFF198SSR,ENCFF502WDT,ENCFF033UQX,ENCFF142CNG,ENCFF367UNR |
| 1344 | 16 | 48358947 | 48368502 | 9555 | C3H.CAST.CBA.CE.MOLF.SPRET | 16 | 48368360 | 48368393 | 34 | + | 16.4472 | 9.61E-07 | 0.126 | TAGCTGTCTTCAGACACACCAGAAGAGGGCGTCA | | |
| 1345 | 16 | 53571326 | 53573687 | 2361 | SPRET | 16 | 53572715 | 53572748 | 34 | + | 17.2927 | 5.61E-07 | 0.0993 | CTGCAATACCTCTCCTGGGCAGAAGATGTTCCAA | | |
| 1346 | 16 | 54358259 | 54362986 | 4727 | CAST | 16 | 54359353 | 54359387 | 35 | + | 20.4862 | 6.34E-08 | 0.0459 | TTGCTGTCCAATATTCTTGCCACCAGATGACATTA | | |
| 1347 | 16 | 55758881 | 55763593 | 4712 | CBA.MOLF.NZW.SMJ | 16 | 55761350 | 55761383 | 34 | + | 27.9919 | 9.53E-11 | 0.000516 | ATGCAGTGCCTACAGAGGCCAGAAGAGGGAACTG | 2 | ENCFF883UPM.ENCFF502WDT |
| 1347 | 16 | 55758881 | 55763593 | 4712 | CBA.MOLF.NZW.SMJ | 16 | 55761365 | 55761383 | 19 | + | 16.9672 | 9.26E-07 | 0.189 | AGGCCAGAAGAGGGAACTG | 2 | ENCFF883UPM.ENCFF502WDT |
| 1348 | 16 | 57557229 | 57565509 | 8280 | CBA.NZW.SMJ | 16 | 57560706 | 57560740 | 35 | + | 17.1376 | 5.00E-07 | 0.161 | CTGAAGTGCCCTTAGCCAGCCAGCAAGTGCCAGAG | | |
| 1349 | 16 | 59953923 | 59956107 | 2184 | CAST | 16 | 59954439 | 59954472 | 34 | + | 18.6504 | 2.28E-07 | 0.066 | TAGCTGTCTCCAGACAAGCCAGAAGAGGGCATCA | | |
| 1350 | 16 | 68419255 | 68423810 | 4555 | BUB,CAST,CE,FVB,MOLF,NOD,NOR,SPRET,SWR | 16 | 68420025 | 68420058 | 34 | + | 17.6748 | 4.37E-07 | 0.0869 | GAGCATGACCACAAGTCCACAGCAGGTGGCAACG | | |
| 1351 | 16 | 74709492 | 74711609 | 2117 | CAST | 16 | 74710576 | 74710609 | 34 | + | 18.4065 | 2.69E-07 | 0.07 | ATGCAGCTAGAGGCACGAGCTCCAGGGGGTACTG | | |
| 1352 | 16 | 78845036 | 78847137 | 2101 | SPRET | 16 | 78846478 | 78846511 | 34 | + | 16.4553 | 9.56E-07 | 0.126 | ATGCAGCTAGAGACACGGGCTCTGGGGGGTACTG | | |
| 1353 | 16 | 80812498 | 80816661 | 4163 | CAST,CE,DBA1J,MA/My,MOLF,NZB,NZO,NZW.SPRET.WSB | 16 | 80813271 | 80813304 | 34 | + | 17.6748 | 4.37E-07 | 0.0869 | GAGCATGACCACAAGTCCACAGCAGGTGGCAACG | | |
| 1354 | 16 | 82004518 | 82009073 | 4555 | CAST.MOLF | 16 | 82005295 | 82005328 | 34 | + | 17.6748 | 4.37E-07 | 0.0869 | GAGCATGACCACAAGTCCACAGCAGGTGGCAACG | | |
| 1355 | 16 | 97007052 | 97008135 | 1083 | SPRET | 16 | 97007591 | 97007609 | 19 | + | 17.0656 | 8.70E-07 | 0.183 | AGGCCAGAAGAGGGTGCTG | | |
| 1356 | 16 | 3222073 | 3223226 | 1153 | SPRET | 16 | 3222442 | 3222460 | 19 | - | 17.2951 | 7.50E-07 | 0.176 | CCATCACTAGATGGCGCCA | | |
| 1356 | 16 | 3222073 | 3223226 | 1153 | SPRET | 16 | 3222442 | 3222475 | 34 | - | 19.4553 | 1.31E-07 | 0.0467 | CCGCAATAATGCTTGCCATCACTAGATGGCGCCA | | |
| 1357 | 16 | 5265749 | 5267306 | 1557 | CAST.SPRET | 16 | 5266818 | 5266851 | 34 | - | 18.4797 | 2.56E-07 | 0.0693 | TTGCTGTCTTCAGACACACCAGAAGAGGGCATCA | | |
| 1358 | 16 | 10985162 | 10990473 | 5311 | SPRET | 16 | 10987073 | 10987106 | 34 | - | 16.9187 | 7.13E-07 | 0.112 | ATGCAATACCCACACAGGCCAGAAAGGTCATTA | | |
| 1358 | 16 | 10985162 | 10990473 | 5311 | SPRET | 16 | 10987487 | 10987520 | 34 | - | 18.4797 | 2.56E-07 | 0.0693 | TTGCTGTCTTCAGACACACCAGAAGAGGGCATCA | 1 | ENCFF093ZRU |
| 1359 | 16 | 13064660 | 13065719 | 1059 | KK | 16 | 13065419 | 13065453 | 35 | - | 19.945 | 9.01E-08 | 0.0577 | CTGTAATCCCAGCACTGAAGCAGCAGGTGGGGGCA | | |
| 1360 | 16 | 15274328 | 15279241 | 4913 | SPRET | 16 | 15275887 | 15275905 | 19 | - | 17.0328 | 8.88E-07 | 0.184 | TGGCCAGCAGATGGCCCAC | | |
| 1361 | 16 | 16751711 | 16758105 | 6394 | CAST.MOLF | 16 | 16758056 | 16758089 | 34 | - | 16.4472 | 9.61E-07 | 0.126 | TAGCTGTCTTCAGACACACCAGAAGAGGGCGTCA | | |
| 1362 | 16 | 17019917 | 17022821 | 2904 | CAST | 16 | 17021193 | 17021226 | 34 | - | 16.5854 | 8.81E-07 | 0.126 | ATACAGTGCCCTCAGAGGTCAGAAGAGGGTGTCA | | |
| 1363 | 16 | 18154438 | 18156268 | 1830 | MOLF | 16 | 18154535 | 18154553 | 19 | - | 17.2787 | 7.58E-07 | 0.176 | CATCCAGAAGATGGCAGTA | | |
| 1363 | 16 | 18154438 | 18156268 | 1830 | MOLF | 16 | 18154535 | 18154568 | 34 | - | 18.4959 | 2.53E-07 | 0.0693 | TTCCACTGGCTGCATCATCCAGAAGATGGCAGTA | | |
| 1364 | 16 | 18350924 | 18352105 | 1181 | CAST.MOLF.SPRET | 16 | 18351052 | 18351070 | 19 | - | 17.5574 | 6.32E-07 | 0.163 | ACTCCAGAAGAGGGCGCCA | | |
| 1364 | 16 | 18350924 | 18352105 | 1181 | CAST.MOLF.SPRET | 16 | 18351052 | 18351085 | 34 | - | 18.3659 | 2.76E-07 | 0.07 | TAGCTGTCTTCAGACACTCCAGAAGAGGGCGCCA | | |
| 1365 | 16 | 19769074 | 19770821 | 1747 | MOLF.SPRET | 16 | 19769536 | 19769554 | 19 | - | 20.0984 | 1.02E-07 | 0.0703 | GAGCCACCAGATGGCACCC | | |
| 1366 | 16 | 22734528 | 22735830 | 1302 | CAST | 16 | 22735003 | 22735036 | 34 | - | 23.9837 | 3.99E-09 | 0.00454 | GTGTAGTGCTCCTGGAGGCCAGAAGAGGGCATCA | | |
| 1367 | 16 | 31198755 | 31204684 | 5929 | MOLF.SPRET | 16 | 31204131 | 31204164 | 34 | - | 16.4472 | 9.61E-07 | 0.126 | TAGCTGTCTTCAGACACACCAGAAGAGGGCGTCA | | |
| 1368 | 16 | 33147317 | 33148408 | 1091 | SPRET | 16 | 33147915 | 33147948 | 34 | - | 19.122 | 1.65E-07 | 0.0537 | GAGTAGTACCCAAGGAGGACGGAAGAGGGCACTG | | |
| 1369 | 16 | 33749215 | 33751767 | 2552 | SPRET | 16 | 33750482 | 33750515 | 34 | - | 16.7561 | 7.91E-07 | 0.117 | CTGCAGTACGGCCAACACACAGACGGGGACGGTA | | |
| 1370 | 16 | 35886723 | 35887904 | 1181 | MOLF | 16 | 35887688 | 35887721 | 34 | - | 18.9106 | 1.91E-07 | 0.0592 | ATGCAGTGCTCGTGGAGGCCAGAAGAGAGTGTTG | | |
| 1371 | 16 | 43693186 | 43695656 | 2470 | CAST.MOLF | 16 | 43693591 | 43693624 | 34 | - | 16.4472 | 9.61E-07 | 0.126 | TAGCTGTCTTCAGACACACCAGAAGAGGGCGTCA | | |
| 1372 | 16 | 45973315 | 45975440 | 2125 | 129S1,C3H,C58J,CAST,CBA,CE,FVB,MOLF,SJL,SPRET,SWR,TH,WSB | 16 | 45974269 | 45974302 | 34 | - | 17.2114 | 5.91E-07 | 0.101 | GAGCAACTCCTTCACAAACCAGGTGATGGCACAG | | |
| 1373 | 16 | 46147201 | 46149621 | 2420 | C3H.C58J.CAST.CBA | 16 | 46148113 | 46148146 | 34 | - | 17.3089 | 5.55E-07 | 0.0993 | ATGCAGCTAGAGACACAAGCTCCAGGGGGTACTA | | |
| 1374 | 16 | 46328779 | 46336883 | 8104 | SPRET | 16 | 46334681 | 46334714 | 34 | - | 16.6016 | 8.72E-07 | 0.126 | CTGTTGTATTTAACATCTTCACAAGATGGCGGCA | | |
| 1375 | 16 | 53496392 | 53498839 | 2447 | C3H,CAST,FVB,MOLF,SPRET,SWR,TH,WSB | 16 | 53496778 | 53496811 | 34 | - | 17.9106 | 3.74E-07 | 0.0805 | CCGCCCTCACATTCGCCGTCACAAGATGGCGCTG | | |
| 1376 | 16 | 55143094 | 55146239 | 3145 | 129S1,C58J,CAST,CBA,LP,MOLF,NZW,SJL,SPRET | 16 | 55144653 | 55144686 | 34 | - | 16.5122 | 9.23E-07 | 0.126 | ATGCATCCTCAGCAGCCACCTGTAGGTGGACCTT | | |
| 1377 | 16 | 55728126 | 55730825 | 2699 | 129S1,BTBR,C57L,C58J,CE,FVB,KK,LP,MRL,SJL,SWR | 16 | 55730163 | 55730196 | 34 | - | 18.8049 | 2.05E-07 | 0.062 | TTGCTGTCTTCAGATGCACCAGAAGAGGGCATCA | 2 | ENCFF093ZRU,ENCFF099QNE |
| 1378 | 16 | 56142527 | 56155583 | 13056 | C58J,CAST,LP | 16 | 56142863 | 56142896 | 34 | - | 16.5691 | 8.90E-07 | 0.126 | GAGCAGGTGTGGCAGCAGTCCCAAGATGGCACCC | | |
| 1379 | 16 | 62378379 | 62384485 | 6106 | WSB | 16 | 62382603 | 62382621 | 19 | - | 17.1311 | 8.34E-07 | 0.181 | AGGCCAAAAGGGGGCAGTG | 19 | ENCFF093ZRU,ENCFF201JUA,ENCFF367UNR,ENCFF099QNE,ENCFF165LXO,ENCFF821BIA,ENCFF052KVO,ENCFF502WDT,ENCFF907XEG,ENCFF957SPH,ENCFF024JOZ,ENCFF089GBC,ENCFF092NMJ,ENCFF504JDS,ENCFF883UPM,ENCFF575MGI,ENCFF430PPJ,ENCFF281QMV,ENCFF055HAP |
| 1380 | 16 | 70848736 | 70850321 | 1585 | SPRET | 16 | 70849386 | 70849420 | 35 | - | 16.7615 | 6.21E-07 | 0.177 | ATGCACTGCACATTCCAGAACTCCAGAGGGGACAA | | |
| 1381 | 16 | 75931926 | 75940912 | 8986 | AJ,C58J,CAST,CE,DBA2J,LP,MOLF,NZB,NZO,NZW,SMJ,SPRET,WSB | 16 | 75935481 | 75935499 | 19 | - | 17.1639 | 8.17E-07 | 0.181 | CCTCCACGAGGTGGCGGAA | | |
| 1382 | 16 | 82311038 | 82413162 | 102124 | MA/My | 16 | 82318653 | 82318671 | 19 | - | 16.8525 | 9.96E-07 | 0.192 | CATCCACTAGGTGGTAGTA | 7 | ENCFF093ZRU,ENCFF099QNE,ENCFF092NMJ,ENCFF430PPJ,ENCFF089GBC,ENCFF907XEG,ENCFF024JOZ |

| | | | | | | | | | | | | | | | | |
|---|---|---|---|---|---|---|---|---|---|---|---|---|---|---|---|---|
| 1383 | 16 | 90138425 | 90146760 | 8335 | 129S1,BUB,C57L,C58J,CAST,DBA1J,DBA2J,FVB,MA/My,MOLF,NOD,NOR,NZO,PJ,RF,SJL,SPRET,SWR,TH,WSB | 16 | 90138716 | 90138734 | 19 | - | 19.2951 | 1.88E-07 | 0.0958 | CCGCCACTAGATGGTGCTA | | |
| 1383 | 16 | 90138425 | 90146760 | 8335 | 129S1,BUB,C57L,C58J,CAST,DBA1J,DBA2J,FVB,MA/My,MOLF,NOD,NOR,NZO,PJ,RF,SJL,SPRET,SWR,TH,WSB | 16 | 90146719 | 90146737 | 19 | - | 19.2951 | 1.88E-07 | 0.0958 | CCGCCACTAGATGGTGCTA | 14 | ENCFF367UNR,ENCFF093ZRU,ENCFF099QNE,ENCFF024JOZ,ENCFF883UPM,ENCFF089GBC,ENCFF821BIA,ENCFF907XEG,ENCFF589ZBJ,ENCFF136ZDK,ENCFF165LXO,ENCFF198SSR,ENCFF430PPJ,ENCFF092NMJ |
| 1384 | 16 | 93420107 | 93438493 | 18386 | TH | 16 | 93423823 | 93423841 | 19 | - | 17.3443 | 7.26E-07 | 0.173 | TGTCCAGGAGGTGGCAGAG | 26 | ENCFF201JUA,ENCFF093ZRU,ENCFF367UNR,ENCFF136ZDK,ENCFF430PPJ,ENCFF099QNE,ENCFF165LXO,ENCFF951GOP,ENCFF883UPM,ENCFF502WDT,ENCFF907XEG,ENCFF024JOZ,ENCFF327LYS,ENCFF821BIA,ENCFF575MGI,ENCFF609MDN,ENCFF589ZBJ,ENCFF281QMV,ENCFF957SPH,ENCFF198SSR,ENCFF194HVL,ENCFF491RJK,ENCFF055HAP,ENCFF089GBC,ENCFF210BTO,ENCFF092NMJ |
| 1384 | 16 | 93420107 | 93438493 | 18386 | TH | 16 | 93423823 | 93423856 | 34 | - | 17 | 6.77E-07 | 0.109 | TTCCTATCCCACTGCTGTCCAGGAGGTGGCAGAG | 26 | ENCFF201JUA,ENCFF093ZRU,ENCFF367UNR,ENCFF136ZDK,ENCFF430PPJ,ENCFF099QNE,ENCFF165LXO,ENCFF951GOP,ENCFF883UPM,ENCFF502WDT,ENCFF907XEG,ENCFF024JOZ,ENCFF327LYS,ENCFF821BIA,ENCFF575MGI,ENCFF609MDN,ENCFF589ZBJ,ENCFF281QMV,ENCFF957SPH,ENCFF198SSR,ENCFF194HVL,ENCFF491RJK,ENCFF055HAP,ENCFF089GBC,ENCFF210BTO,ENCFF092NMJ |
| 1384 | 16 | 93420107 | 93438493 | 18386 | TH | 16 | 93425950 | 93425968 | 19 | - | 17.7869 | 5.43E-07 | 0.153 | AGGCCTCCAGGTGGCAGAG | | |
| 1385 | 16 | 93494339 | 93503324 | 8985 | CAST,CE,DBA1J,MOLF,NZB,NZW,SPRET,TH,WSB | 16 | 93497895 | 93497913 | 19 | - | 17.1639 | 8.17E-07 | 0.181 | CCTCCACGAGGTGGCGGAA | | |
| 1386 | 16 | 93494344 | 93502628 | 8284 | 129S1,AJ,AKR,BALB,BUB,C3H,C57L,C58J,CBA,DBA2J,FVB,ILNJ2,KK,LP,MA/My,MRL,NOD,NOR,NZO,PJ,PL,RF,RH2,SEA,SJL,SMJ,SWR | 16 | 93497895 | 93497913 | 19 | - | 17.1639 | 8.17E-07 | 0.181 | CCTCCACGAGGTGGCGGAA | | |
| 1387 | 17 | 3436426 | 3442537 | 6111 | SPRET | 17 | 3440460 | 3440493 | 34 | + | 20.2114 | 7.63E-08 | 0.0205 | ATGCAATGCCCACAGAGACCAGAAGAAGGCTTTA | | |
| 1388 | 17 | 5766826 | 5768611 | 1785 | CE | 17 | 5768142 | 5768175 | 34 | + | 16.935 | 7.06E-07 | 0.0775 | CTGTGGTGAGAGCCATGGCCTGGGGGGGGCGGGG | | |
| 1389 | 17 | 7779608 | 7782187 | 2579 | AJ,BALB,BUB,C3H,CBA,DBA1J,DBA2J,FVB,KK,MA/My,MRL,NOD,NOR,RH2,SEA,SMJ | 17 | 7781046 | 7781079 | 34 | + | 20.3984 | 6.66E-08 | 0.0187 | TAGCTGTCTTCAGATGCGCCAGAAGAGGGCGCCA | 5 | ENCFF093ZRU,ENCFF367UNR,ENCFF099QNE,ENCFF821BIA,ENCFF907XEG |
| 1389 | 17 | 7779608 | 7782187 | 2579 | AJ,BALB,BUB,C3H,CBA,DBA1J,DBA2J,FVB,KK,MA/My,MRL,NOD,NOR,RH2,SEA,SMJ | 17 | 7781061 | 7781079 | 19 | + | 19.6721 | 1.42E-07 | 0.05 | GCGCCAGAAGAGGGCGCCA | 5 | ENCFF093ZRU,ENCFF367UNR,ENCFF099QNE,ENCFF821BIA,ENCFF907XEG |
| 1390 | 17 | 15752856 | 15758412 | 5556 | SPRET | 17 | 15753506 | 15753539 | 34 | + | 20.252 | 7.40E-08 | 0.02 | TAGCTGTCCTCAGACACTCCAGAAGAGGGCGCCA | 3 | ENCFF367UNR,ENCFF099QNE,ENCFF093ZRU |
| 1390 | 17 | 15752856 | 15758412 | 5556 | SPRET | 17 | 15753521 | 15753539 | 19 | + | 17.5574 | 6.32E-07 | 0.108 | ACTCCAGAAGAGGGCGCCA | 3 | ENCFF367UNR,ENCFF099QNE,ENCFF093ZRU |
| 1391 | 17 | 16365856 | 16369019 | 3163 | SPRET | 17 | 16367281 | 16367314 | 34 | + | 24.4553 | 2.67E-09 | 0.00159 | CTGCAGTGCCAACAGAGGCGAGAAGAGGGCATCA | | |
| 1392 | 17 | 19391662 | 19394782 | 3120 | SPRET | 17 | 19393061 | 19393094 | 34 | + | 18.0569 | 3.40E-07 | 0.0516 | ATGCAGCTAGAGACATGAGCTCCAGGGGGTACTG | | |
| 1393 | 17 | 19430855 | 19443715 | 12860 | SPRET | 17 | 19438998 | 19439032 | 35 | + | 16.0367 | 9.36E-07 | 0.143 | GAGCTGAGCCACACCCAAACCACCAGGGAGAGCTC | | |
| 1394 | 17 | 21976506 | 21981052 | 4546 | SPRET | 17 | 21980431 | 21980464 | 34 | + | 21.3984 | 3.17E-08 | 0.0107 | ATGCAATGCCCACAAATACCAGAAGAGGGTAATG | | |
| 1395 | 17 | 22472369 | 22475612 | 3243 | 129S1,AJ,BALB,C57L,CE,DBA1J,DBA2J,MOLF,NZB,NZO,RH2,SEA,SJL,SMJ,TH | 17 | 22472795 | 22472813 | 19 | + | 19.623 | 1.47E-07 | 0.0507 | GGACCACCAGAGGGCAATA | | |
| 1396 | 17 | 24110987 | 24114105 | 3118 | AJ,AKR,B10D2,BALB,BTBR,BUB,CAST,DBA2J,MA/My,NOR,PJ,WSB | 17 | 24111474 | 24111507 | 34 | + | 16.5691 | 8.90E-07 | 0.0867 | GAGCAGGTGTGGCAGCAGTCCCAAGATGGCACCC | | |
| 1397 | 17 | 24129763 | 24132897 | 3134 | 129S1,AJ,AKR,B10D2,BTBR,BUB,CE,DBA1J,DBA2J,FVB,ILNJ2,KK,MA/My,NOD,NOR,NZW,PJ,RF,RH2,SEA,SJL,WSB | 17 | 24130248 | 24130281 | 34 | + | 17.0244 | 6.67E-07 | 0.0758 | GAGCAGGTGTGGCAGCAGTCCCAAGATGGCGCCC | | |
| 1397 | 17 | 24129763 | 24132897 | 3134 | 129S1,AJ,AKR,B10D2,BTBR,BUB,CE,DBA1J,DBA2J,FVB,ILNJ2,KK,MA/My,NOD,NOR,NZW,PJ,RF,RH2,SEA,SJL,WSB | 17 | 24131471 | 24131504 | 34 | + | 16.561 | 8.95E-07 | 0.087 | GAGCTGCTTTCAGACACACCAGAAGAGGGCATCC | | |
| 1398 | 17 | 25496159 | 25497412 | 1253 | SPRET | 17 | 25496586 | 25496620 | 35 | + | 18.0367 | 2.94E-07 | 0.077 | CAGCAGTTCAGTTCACAGGCCAGCAGTGGCAGCTC | | |
| 1399 | 17 | 27881997 | 27884128 | 2131 | 129S1,BALB,C57L,ILNJ2,RH2,SEA | 17 | 27882741 | 27882775 | 35 | + | 19.5596 | 1.15E-07 | 0.0458 | CTGCATGTGCATCTGCACCCCAGAAGAGGGCATAG | 1 | ENCFF093ZRU |
| 1400 | 17 | 29684972 | 29686324 | 1352 | 129S1,B10D2,BALB,C57L,DBA1J,DBA2J,RH2,SEA | 17 | 29685426 | 29685459 | 34 | + | 16.9919 | 6.81E-07 | 0.0767 | TAGCTGTCTTCAGACACACCAGAAGGGGGCATCA | | |
| 1401 | 17 | 29748618 | 29755909 | 7291 | SPRET | 17 | 29750393 | 29750426 | 34 | + | 30.1626 | 8.84E-12 | 0.0000674 | CTGCAGTGCCCATAGTAGCCCGAAGAGGGCGCCA | 12 | ENCFF093ZRU,ENCFF367UNR,ENCFF089GBC,ENCFF589ZBJ,ENCFF052KVO,ENCFF033UQX,ENCFF136ZDK,ENCFF055HAP,ENCFF533APC,ENCFF502WDT,ENCFF065ODV,ENCFF142CNG |
| 1402 | 17 | 32921989 | 32927503 | 5514 | SPRET | 17 | 32922007 | 32922025 | 19 | + | 21.0656 | 4.64E-08 | 0.0336 | ATACCAGTAGGGGGCGCTG | 16 | ENCFF093ZRU,ENCFF136ZDK,ENCFF821BIA,ENCFF165LXO,ENCFF210BTO,ENCFF198SSR,ENCFF589ZBJ,ENCFF327LYS,ENCFF099QNE,ENCFF883UPM,ENCFF907XEG,ENCFF033UQX,ENCFF194HVL,ENCFF533APC,ENCFF052KVO,ENCFF367UNR |
| 1403 | 17 | 34111964 | 34117854 | 5890 | WSB | 17 | 34115757 | 34115775 | 19 | + | 17.9672 | 4.81E-07 | 0.0964 | GCACCAGAAGAGGGCACCA | | |
| 1403 | 17 | 34111964 | 34117854 | 5890 | WSB | 17 | 34116229 | 34116263 | 35 | + | 25.4404 | 1.73E-09 | 0.00402 | CTGTACATACACCTGAAGGCCAGAAGAGGGCGCCA | 31 | ENCFF367UNR,ENCFF201JUA,ENCFF093ZRU,ENCFF099QNE,ENCFF430PPJ,ENCFF052KVO,ENCFF065ODV,ENCFF055HAP,ENCFF821BIA,ENCFF907XEG,ENCFF024JOZ,ENCFF327LYS,ENCFF589ZBJ,ENCFF609MDN,ENCFF089GBC,ENCFF198SSR,ENCFF502WDT,ENCFF136ZDK,ENCFF210BTO,ENCFF575MGI,ENCFF316FYN,ENCFF321AGQ,ENCFF951GOP,ENCFF533APC,ENCFF097CAQ,ENCFF092NMJ,ENCFF194HVL,ENCFF883UPM,ENCFF491RJK,ENCFF165LXO,ENCFF957SPH |
| 1403 | 17 | 34111964 | 34117854 | 5890 | WSB | 17 | 34116245 | 34116263 | 19 | + | 21.0984 | 4.51E-08 | 0.0335 | AGGCCAGAAGAGGGCGCCA | 32 | ENCFF367UNR,ENCFF201JUA,ENCFF093ZRU,ENCFF099QNE,ENCFF430PPJ,ENCFF052KVO,ENCFF065ODV,ENCFF055HAP,ENCFF821BIA,ENCFF907XEG,ENCFF024JOZ,ENCFF327LYS,ENCFF589ZBJ,ENCFF609MDN,ENCFF089GBC,ENCFF198SSR,ENCFF502WDT,ENCFF136ZDK,ENCFF210BTO,ENCFF575MGI,ENCFF316FYN,ENCFF321AGQ,ENCFF951GOP,ENCFF533APC,ENCFF097CAQ,ENCFF092NMJ,ENCFF194HVL,ENCFF883UPM,ENCFF491RJK,ENCFF165LXO,ENCFF957SPH,ENCFF281QMV |
| 1404 | 17 | 34114348 | 34117759 | 3411 | CAST | 17 | 34115757 | 34115775 | 19 | + | 17.9672 | 4.81E-07 | 0.0964 | GCACCAGAAGAGGGCACCA | | |
| 1404 | 17 | 34114348 | 34117759 | 3411 | CAST | 17 | 34116229 | 34116263 | 35 | + | 25.4404 | 1.73E-09 | 0.00402 | CTGTACATACACCTGAAGGCCAGAAGAGGGCGCCA | 31 | ENCFF367UNR,ENCFF201JUA,ENCFF093ZRU,ENCFF099QNE,ENCFF430PPJ,ENCFF052KVO,ENCFF065ODV,ENCFF055HAP,ENCFF821BIA,ENCFF907XEG,ENCFF024JOZ,ENCFF327LYS,ENCFF589ZBJ,ENCFF609MDN,ENCFF089GBC,ENCFF198SSR,ENCFF502WDT,ENCFF136ZDK,ENCFF210BTO,ENCFF575MGI,ENCFF316FYN,ENCFF321AGQ,ENCFF951GOP,ENCFF533APC,ENCFF097CAQ,ENCFF092NMJ,ENCFF194HVL,ENCFF883UPM,ENCFF491RJK,ENCFF165LXO,ENCFF957SPH |
| 1404 | 17 | 34114348 | 34117759 | 3411 | CAST | 17 | 34116245 | 34116263 | 19 | + | 21.0984 | 4.51E-08 | 0.0335 | AGGCCAGAAGAGGGCGCCA | 32 | ENCFF367UNR,ENCFF201JUA,ENCFF093ZRU,ENCFF099QNE,ENCFF430PPJ,ENCFF052KVO,ENCFF065ODV,ENCFF055HAP,ENCFF821BIA,ENCFF907XEG,ENCFF024JOZ,ENCFF327LYS,ENCFF589ZBJ,ENCFF609MDN,ENCFF089GBC,ENCFF198SSR,ENCFF502WDT,ENCFF136ZDK,ENCFF210BTO,ENCFF575MGI,ENCFF316FYN,ENCFF321AGQ,ENCFF951GOP,ENCFF533APC,ENCFF097CAQ,ENCFF092NMJ,ENCFF194HVL,ENCFF883UPM,ENCFF491RJK,ENCFF165LXO,ENCFF957SPH,ENCFF281QMV |
| 1405 | 17 | 35914055 | 35919941 | 5886 | SPRET | 17 | 35914502 | 35914536 | 35 | + | 16.0917 | 9.07E-07 | 0.141 | TGGCACAGTCCAGCAAAACCCAGCAGGTGGAGCCT | | |
| 1405 | 17 | 35914055 | 35919941 | 5886 | SPRET | 17 | 35918978 | 35919011 | 34 | - | 19.6016 | 1.18E-07 | 0.0285 | GTGTGGTGCCTGAAGAGGCCAAAAGAGGGTGCTG | | |
| 1406 | 17 | 36043641 | 36047726 | 4085 | 129S1,AJ,AKR,B10D2,BALB,BTBR,C3H,C57L,C58J,CBA,CE,DBA2J,LP,MA/My,MOLF,MRL,NZB,NZO,NZW,PL,RF,RH2,SEA,SJL,SMJ,TH,WSB | 17 | 36047365 | 36047399 | 35 | + | 19.4771 | 1.21E-07 | 0.047 | CTACAGCTCCCCCTCCTGGCCAGCTGAGGTCCCAG | 31 | ENCFF093ZRU,ENCFF367UNR,ENCFF805QBU,ENCFF099QNE,ENCFF327LYS,ENCFF201JUA,ENCFF907XEG,ENCFF821BIA,ENCFF024JOZ,ENCFF136ZDK,ENCFF589ZBJ,ENCFF165LXO,ENCFF198SSR,ENCFF089GBC,ENCFF033UQX,ENCFF092NMJ,ENCFF210BTO,ENCFF957SPH,ENCFF951GOP,ENCFF281QMV,ENCFF089EWX,ENCFF097CAQ,ENCFF533APC,ENCFF611HDQ,ENCFF430PPJ,ENCFF194HVL,ENCFF575MGI,ENCFF349BUL,ENCFF609MDN,ENCFF316FYN,ENCFF491RJK |
| 1406 | 17 | 36043641 | 36047726 | 4085 | 129S1,AJ,AKR,B10D2,BALB,BTBR,C3H,C57L,C58J,CBA,CE,DBA2J,LP,MA/My,MOLF,MRL,NZB,NZO,NZW,PL,RF,RH2,SEA,SJL,SMJ,TH,WSB | 17 | 36044171 | 36044204 | 34 | - | 18.7398 | 2.14E-07 | 0.0412 | ATGCAGTGGTCCCAGAGGCCAGAAGAGGCCTTTG | | |
| 1406 | 17 | 36043641 | 36047726 | 4085 | 129S1,AJ,AKR,B10D2,BALB,BTBR,C3H,C57L,C58J,CBA,CE,DBA2J,LP,MA/My,MOLF,MRL,NZB,NZO,NZW,PL,RF,RH2,SEA,SJL,SMJ,TH,WSB | 17 | 36047368 | 36047386 | 19 | - | 20.2787 | 8.86E-08 | 0.0417 | TGGCCAGGAGGGGGAGCTG | 31 | ENCFF093ZRU,ENCFF367UNR,ENCFF805QBU,ENCFF099QNE,ENCFF327LYS,ENCFF201JUA,ENCFF907XEG,ENCFF821BIA,ENCFF024JOZ,ENCFF136ZDK,ENCFF589ZBJ,ENCFF165LXO,ENCFF198SSR,ENCFF089GBC,ENCFF033UQX,ENCFF092NMJ,ENCFF210BTO,ENCFF957SPH,ENCFF951GOP,ENCFF281QMV,ENCFF089EWX,ENCFF097CAQ,ENCFF533APC,ENCFF611HDQ,ENCFF430PPJ,ENCFF194HVL,ENCFF575MGI,ENCFF349BUL,ENCFF609MDN,ENCFF316FYN,ENCFF491RJK |
| 1406 | 17 | 36043641 | 36047726 | 4085 | 129S1,AJ,AKR,B10D2,BALB,BTBR,C3H,C57L,C58J,CBA,CE,DBA2J,LP,MA/My,MOLF,MRL,NZB,NZO,NZW,PL,RF,RH2,SEA,SJL,SMJ,TH,WSB | 17 | 36047368 | 36047402 | 35 | - | 19.3853 | 1.29E-07 | 0.0483 | ACGCTGGGACCTCAGCTGGCCAGGAGGGGGAGCTG | 31 | ENCFF093ZRU,ENCFF367UNR,ENCFF805QBU,ENCFF099QNE,ENCFF327LYS,ENCFF201JUA,ENCFF907XEG,ENCFF821BIA,ENCFF024JOZ,ENCFF136ZDK,ENCFF589ZBJ,ENCFF165LXO,ENCFF198SSR,ENCFF089GBC,ENCFF033UQX,ENCFF092NMJ,ENCFF210BTO,ENCFF957SPH,ENCFF951GOP,ENCFF281QMV,ENCFF089EWX,ENCFF097CAQ,ENCFF533APC,ENCFF611HDQ,ENCFF430PPJ,ENCFF194HVL,ENCFF575MGI,ENCFF349BUL,ENCFF609MDN,ENCFF316FYN,ENCFF491RJK |
| 1407 | 17 | 36483020 | 36484765 | 1745 | ILNJ2,MOLF,WSB | 17 | 36483009 | 36483043 | 35 | + | 17.4679 | 4.13E-07 | 0.0933 | TTGTAGCTGTCCAGACACACCAGAAGAGGGCATCC | | |
| 1408 | 17 | 37044979 | 37071015 | 26036 | SPRET | 17 | 37045312 | 37045346 | 35 | + | 24.3119 | 4.20E-09 | 0.0054 | CTGCAGGGGATGGAACCTTCCAGAAGGGGGCAGCT | | |
| 1408 | 17 | 37044979 | 37071015 | 26036 | SPRET | 17 | 37070229 | 37070262 | 34 | - | 16.7724 | 7.83E-07 | 0.0815 | TTGCTCTCTTCAGACACACCAGAAGAGGGCATTA | | |
| 1409 | 17 | 37045158 | 37058233 | 13075 | MOLF | 17 | 37045312 | 37045346 | 35 | + | 24.3119 | 4.20E-09 | 0.0054 | CTGCAGGGGATGGAACCTTCCAGAAGGGGGCAGCT | | |
| 1410 | 17 | 37201352 | 37205593 | 4241 | PJ | 17 | 37203392 | 37203426 | 35 | + | 20.2202 | 7.54E-08 | 0.0346 | TAGCATTGTGCCTTCACGGACAGTAGGTGGTGCTG | 29 | ENCFF093ZRU,ENCFF201JUA,ENCFF367UNR,ENCFF099QNE,ENCFF327LYS,ENCFF821BIA,ENCFF907XEG,ENCFF065ODV,ENCFF089GBC,ENCFF589ZBJ,ENCFF055HAP,ENCFF316FYN,ENCFF430PPJ,ENCFF321AGQ,ENCFF502WDT,ENCFF092NMJ,ENCFF198SSR,ENCFF024JOZ,ENCFF165LXO,ENCFF951GOP,ENCFF136ZDK,ENCFF957SPH,ENCFF883UPM,ENCFF281QMV,ENCFF142CNG,ENCFF575MGI,ENCFF609MDN,ENCFF194HVL,ENCFF052KVO |
| 1411 | 17 | 38614972 | 38688148 | 73176 | FVB,TH | 17 | 38625166 | 38625184 | 19 | + | 18.7049 | 2.88E-07 | 0.0728 | CAGCCAGCAGGTGGCAAAA | | |
| 1412 | 17 | 45773031 | 45774937 | 1906 | NZW,SJL | 17 | 45773600 | 45773633 | 34 | + | 26.0569 | 6.34E-10 | 0.000552 | ATGCAGTGCCCATGGAGGCCAGAAGAGGGCATAA | 2 | ENCFF093ZRU,ENCFF502WDT |
| 1413 | 17 | 47972583 | 47975371 | 2788 | SPRET | 17 | 47973957 | 47973990 | 34 | + | 16.4634 | 9.51E-07 | 0.0875 | TAGCTGTCTTCAGACGCACCAGAAGAGGGCATCA | 1 | ENCFF093ZRU |
| 1414 | 17 | 50555814 | 50560277 | 4463 | CAST,MOLF,NZW,SPRET,SWR | 17 | 50556593 | 50556626 | 34 | + | 17.6748 | 4.37E-07 | 0.0599 | GAGCATGACCACAAGTCCACAGCAGGTGGCAACG | | |
| 1415 | 17 | 51163814 | 51165009 | 1195 | MOLF | 17 | 51164895 | 51164929 | 35 | + | 19.4587 | 1.23E-07 | 0.047 | CTATAGTTGCCTACAGAGTCCAGAAGAGGGCACTG | | |
| 1415 | 17 | 51163814 | 51165009 | 1195 | MOLF | 17 | 51164911 | 51164929 | 19 | + | 18.0328 | 4.60E-07 | 0.096 | AGTCCAGAAGAGGGCACTG | | |
| 1416 | 17 | 53909592 | 53915495 | 5903 | CAST,MOLF | 17 | 53909657 | 53909690 | 34 | + | 16.4634 | 9.51E-07 | 0.0875 | TAGCTGTCTTCAGACGCACCAGAAGAGGGCATCA | | |
| 1416 | 17 | 53909592 | 53915495 | 5903 | CAST,MOLF | 17 | 53912503 | 53912536 | 34 | + | 16.4634 | 9.51E-07 | 0.0875 | TAGCTGTCTTCAGACGCACCAGAAGAGGGCATCA | | |
| 1416 | 17 | 53909592 | 53915495 | 5903 | CAST,MOLF | 17 | 53915352 | 53915385 | 34 | + | 16.4634 | 9.51E-07 | 0.0875 | TAGCTGTCTTCAGACGCACCAGAAGAGGGCATCA | | |
| 1417 | 17 | 53912047 | 53917681 | 5634 | 129S1,BALB,C3H,C57L,CBA,DBA1J,DBA2J,FVB,LP,MA/My,MRL,PJ,PL,SMJ | 17 | 53912503 | 53912536 | 34 | + | 16.4634 | 9.51E-07 | 0.0875 | TAGCTGTCTTCAGACGCACCAGAAGAGGGCATCA | | |
| 1417 | 17 | 53912047 | 53917681 | 5634 | 129S1,BALB,C3H,C57L,CBA,DBA1J,DBA2J,FVB,LP,MA/My,MRL,PJ,PL,SMJ | 17 | 53915352 | 53915385 | 34 | + | 16.4634 | 9.51E-07 | 0.0875 | TAGCTGTCTTCAGACGCACCAGAAGAGGGCATCA | | |
| 1418 | 17 | 60385322 | 60387088 | 1766 | MOLF | 17 | 60385392 | 60385426 | 35 | + | 17.9358 | 3.13E-07 | 0.0802 | CTGCAGTGACTTGATGTGCACAGGAGGGGGGAATT | | |
| 1419 | 17 | 66184506 | 66187852 | 3346 | SPRET | 17 | 66186759 | 66186777 | 19 | + | 17.5738 | 6.25E-07 | 0.108 | ACACCAGAAGAGGGCACCG | | |
| 1420 | 17 | 66769740 | 66772251 | 2511 | MOLF | 17 | 66770571 | 66770604 | 34 | + | 16.4228 | 9.76E-07 | 0.0882 | TAGCTGCCTTCAGACACTCCAGAAGAGGGCGCCA | | |
| 1420 | 17 | 66769740 | 66772251 | 2511 | MOLF | 17 | 66770586 | 66770604 | 19 | + | 17.5574 | 6.32E-07 | 0.108 | ACTCCAGAAGAGGGCGCCA | | |
| 1421 | 17 | 74326110 | 74328094 | 1984 | SPRET | 17 | 74327717 | 74327750 | 34 | + | 28.6667 | 4.70E-11 | 0.000123 | CTGCAGTTCCAACATGTACCAGCAGATGGAGCAC | 38 | ENCFF093ZRU,ENCFF099QNE,ENCFF201JUA,ENCFF805QBU,ENCFF065ODV,ENCFF907XEG,ENCFF589ZBJ,ENCFF089GBC,ENCFF024JOZ,ENCFF198SSR,ENCFF136ZDK,ENCFF327LYS,ENCFF092NMJ,ENCFF430PPJ,ENCFF321AGQ,ENCFF504JDS,ENCFF165LXO,ENCFF502WDT,ENCFF055HAP,ENCFF210BTO,ENCFF281QMV,ENCFF883UPM,ENCFF957SPH,ENCFF316FYN,ENCFF142CNG,ENCFF951GOP,ENCFF784ASD,ENCFF609MDN,ENCFF601JVE,ENCFF194HVL,ENCFF052KVO,ENCFF575MGI,ENCFF533APC,ENCFF033UQX,ENCFF491RJK,ENCFF611HDQ,ENCFF089EWX,ENCFF097CAQ |
| 1422 | 17 | 74858727 | 74860295 | 1568 | 129S1,BTBR,BUB,C57L,ILNJ2,KK,NOD,NOR,NZW,SWR | 17 | 74858784 | 74858817 | 34 | + | 16.4634 | 9.51E-07 | 0.0875 | TAGCTGTCTTCAGACGCACCAGAAGAGGGCATCA | | |
| 1423 | 17 | 76073720 | 76077547 | 3827 | CE,WSB | 17 | 76075835 | 76075869 | 35 | + | 18.9908 | 1.65E-07 | 0.0555 | TTGCAGTTCCTCCTGCCTACCACTAGGGTCTGACA | 9 | ENCFF821BIA,ENCFF589ZBJ,ENCFF089GBC,ENCFF165LXO,ENCFF093ZRU,ENCFF367UNR,ENCFF052KVO,ENCFF099QNE,ENCFF136ZDK |
| 1424 | 17 | 78798950 | 78800983 | 2033 | C3H,CAST,CBA,DBA1J,DBA2J,MOLF,SPRET | 17 | 78799505 | 78799523 | 19 | + | 17.1639 | 8.17E-07 | 0.123 | GTACCACCAGAGGGTGCTC | | |
| 1425 | 17 | 79524290 | 79526198 | 1908 | 129S1,AJ,AKR,BALB,BTBR,BUB,C3H,C57L,C58J,CBA,CE,DBA1J,DBA2J,FVB,ILNJ2,KK,LP,MA/My,MRL,NOD,NOR,NZB,NZO,NZW,PJ,PL,RF,RH2,SEA,SJL,SMJ,SPRET,SWR,WSB | 17 | 79525261 | 79525295 | 35 | + | 17.0826 | 5.16E-07 | 0.104 | GTGCAGATCCCCTTGGATATCAGAAGAGGGTGTTG | | |
| 1426 | 17 | 80863880 | 80865744 | 1864 | CE | 17 | 80865058 | 80865092 | 35 | + | 17.3486 | 4.42E-07 | 0.0965 | ATGTGAGTGCCCGCAGTGTCCAGAAGAGGGCACCA | 11 | ENCFF367UNR,ENCFF099QNE,ENCFF065ODV,ENCFF327LYS,ENCFF136ZDK,ENCFF194HVL,ENCFF093ZRU,ENCFF142CNG,ENCFF052KVO,ENCFF502WDT,ENCFF055HAP |
| 1426 | 17 | 80863880 | 80865744 | 1864 | CE | 17 | 80865059 | 80865092 | 34 | + | 18.0894 | 3.33E-07 | 0.0512 | TGTGAGTGCCCGCAGTGTCCAGAAGAGGGCACCA | 11 | ENCFF367UNR,ENCFF099QNE,ENCFF065ODV,ENCFF327LYS,ENCFF136ZDK,ENCFF194HVL,ENCFF093ZRU,ENCFF142CNG,ENCFF052KVO,ENCFF502WDT,ENCFF055HAP |
| 1426 | 17 | 80863880 | 80865744 | 1864 | CE | 17 | 80865074 | 80865092 | 19 | + | 20.7213 | 6.18E-08 | 0.0363 | TGTCCAGAAGAGGGCACCA | 11 | ENCFF367UNR,ENCFF099QNE,ENCFF065ODV,ENCFF327LYS,ENCFF136ZDK,ENCFF194HVL,ENCFF093ZRU,ENCFF142CNG,ENCFF052KVO,ENCFF502WDT,ENCFF055HAP |
| 1427 | 17 | 83133370 | 83134928 | 1558 | SPRET | 17 | 83134721 | 83134754 | 34 | + | 25.0732 | 1.55E-09 | 0.00104 | ATGTAGTGCTGGTGGAGGCCAGAAGAGGGCACCA | 2 | ENCFF093ZRU,ENCFF883UPM |
| 1427 | 17 | 83133370 | 83134928 | 1558 | SPRET | 17 | 83134736 | 83134754 | 19 | + | 20.3607 | 8.29E-08 | 0.0402 | AGGCCAGAAGAGGGCACCA | 2 | ENCFF093ZRU,ENCFF883UPM |
| 1428 | 17 | 83664792 | 83666450 | 1658 | SPRET | 17 | 83665512 | 83665545 | 34 | + | 19.7724 | 1.04E-07 | 0.026 | GTGCAGGTCCCTGAGAGGCCAGAAGACAGCACCA | | |
| 1429 | 17 | 87841118 | 87842641 | 1523 | CAST | 17 | 87841593 | 87841627 | 35 | + | 23.3028 | 8.96E-09 | 0.0091 | CTCCATTTATGCCTGCAAGCCAGAAGAGGGCACCA | | |
| 1429 | 17 | 87841118 | 87842641 | 1523 | CAST | 17 | 87841609 | 87841627 | 19 | + | 18.7705 | 2.75E-07 | 0.0709 | AAGCCAGAAGAGGGCACCA | | |
| 1430 | 17 | 88486623 | 88499943 | 13320 | MOLF | 17 | 88496282 | 88496316 | 35 | + | 18.8991 | 1.74E-07 | 0.0578 | ATGCAGATGCTGGAGGAGCACAGAAGAGGGCACCA | | |
| 1431 | 17 | 92376320 | 92377384 | 1064 | 129S1,AJ,AKR,BALB,BTBR,BUB,C3H,C57L,CAST,CBA,DBA1J,DBA2J,FVB,ILNJ2,KK,LP,MA/My,MOLF,MRL,NOD,NOR,NZB,NZO,NZW,PL,RF,RH2,SEA,SJL,SMJ,SPRET,SWR,TH,WSB | 17 | 92377083 | 92377101 | 19 | + | 17.5082 | 6.52E-07 | 0.109 | TAGCCACCAGGGGTTGGTG | | |
| 1432 | 17 | 92377321 | 92378331 | 1010 | AJ,BALB,CAST,CBA,CE,FVB,MOLF,NOD,NOR,PJ,SJL,WSB | 17 | 92378121 | 92378139 | 19 | + | 17.5082 | 6.52E-07 | 0.109 | TAGCCACCAGGGGTTGGTG | | |
| 1433 | 17 | 94257491 | 94261002 | 3511 | CAST | 17 | 94258683 | 94258717 | 35 | + | 18.4679 | 2.27E-07 | 0.0684 | CTGCACTCAGAGCAGTTGACCAGTTGAGGACCCTC | | |
| 1434 | 17 | 94990112 | 94995329 | 5217 | CAST,MOLF,SPRET,WSB | 17 | 94993268 | 94993302 | 35 | + | 16.5596 | 6.97E-07 | 0.123 | AAGAAGTTACACCCTGTTGCCAGTAGGTGGTTTTC | | |
| 1435 | 17 | 6171324 | 6172507 | 1183 | SPRET | 17 | 6171614 | 6171648 | 35 | - | 15.9725 | 9.70E-07 | 0.146 | CTGCATTGACTCCACTGCGTCAGCAGAGGCTGTGT | | |
| 1436 | 17 | 6776282 | 6788405 | 12123 | CAST,MOLF | 17 | 6782762 | 6782795 | 34 | - | 18.813 | 2.04E-07 | 0.0396 | CTGCAGCCCCTAGCCAAGCCTAAGGGGGGTGCTG | | |
| 1437 | 17 | 15037907 | 15041019 | 3112 | MOLF,PL | 17 | 15038681 | 15038715 | 35 | - | 16.9174 | 5.68E-07 | 0.109 | ATGCATTTATCTCCTGTAGCCACCAGTGGCCACAG | | |
| 1438 | 17 | 15823104 | 15830259 | 7155 | CAST,MOLF,PL | 17 | 15823420 | 15823453 | 34 | - | 17.9106 | 3.74E-07 | 0.0548 | CCGCCCTCACATTCGCCGTCACAAGATGGCGCTG | | |
| 1438 | 17 | 15823104 | 15830259 | 7155 | CAST,MOLF,PL | 17 | 15830217 | 15830250 | 34 | - | 17.9106 | 3.74E-07 | 0.0548 | CCGCCCTCACATTCGCCGTCACAAGATGGCGCTG | | |
| 1439 | 17 | 17648825 | 17656836 | 8011 | SPRET | 17 | 17655646 | 17655679 | 34 | - | 16.4472 | 9.61E-07 | 0.0875 | TAGCTGTCTTCAGACACACCAGAAGAGGGCGTCA | 1 | ENCFF093ZRU |
| 1440 | 17 | 17967691 | 17970644 | 2953 | SPRET | 17 | 17969789 | 17969823 | 35 | - | 24.367 | 4.03E-09 | 0.0054 | CTGCAATTTCCCGAGCTGAACAGCAGAGGGAGACA | 24 | ENCFF093ZRU,ENCFF367UNR,ENCFF201JUA,ENCFF099QNE,ENCFF821BIA,ENCFF907XEG,ENCFF089GBC,ENCFF609MDN,ENCFF951GOP,ENCFF092NMJ,ENCFF957SPH,ENCFF327LYS,ENCFF589ZBJ,ENCFF430PPJ,ENCFF024JOZ,ENCFF502WDT,ENCFF055HAP,ENCFF065ODV,ENCFF281QMV,ENCFF165LXO,ENCFF052KVO,ENCFF198SSR,ENCFF883UPM,ENCFF136ZDK |
| 1441 | 17 | 22020057 | 22027503 | 7446 | SPRET | 17 | 22027460 | 22027493 | 34 | - | 16.8943 | 7.24E-07 | 0.0784 | CCGCCCTCACAGTCGCCGCTACAAGATGGCGCTG | | |
| 1442 | 17 | 22433896 | 22441403 | 7507 | 129S1,AJ,BALB,C57L,CE,DBA1J,DBA2J,NZB,NZO,RH2,SEA,SJL,SMJ,TH | 17 | 22436073 | 22436091 | 19 | - | 16.9672 | 9.26E-07 | 0.13 | CTGCCAGAAGAGGGCATCA | | |
| 1443 | 17 | 26510423 | 26517276 | 6853 | AJ,B10D2,BTBR,CE,DBA1J,DBA2J,FVB,ILNJ2,MA/My,NOD,NOR,NZO,NZW,PJ,RF,SJL | 17 | 26510637 | 26510670 | 34 | - | 16.5691 | 8.90E-07 | 0.0867 | GAGCAGGTGTGGCAGCAGTCCCAAGATGGCACCC | | |
| 1444 | 17 | 28511026 | 28512715 | 1689 | MOLF | 17 | 28511838 | 28511871 | 34 | - | 26.8455 | 2.99E-10 | 0.000343 | ATGCAGTGCCCATGGAGGCCAGAAGAGGGTACCA | 1 | ENCFF093ZRU |
| 1445 | 17 | 29570509 | 29573938 | 3429 | CAST,KK,MOLF,RF | 17 | 29572271 | 29572304 | 34 | - | 16.9593 | 6.95E-07 | 0.0768 | TTGCTCTCTTCAGACACACCAGAAGAGGGCATCA | | |
| 1446 | 17 | 31696731 | 31698469 | 1738 | CAST | 17 | 31698304 | 31698338 | 35 | - | 16.7156 | 6.38E-07 | 0.117 | CTGCTGCAGCTGCACAAAGACAGCAGAGGTCGGAA | | |
| 1447 | 17 | 32545606 | 32549712 | 4106 | MA/My,SJL | 17 | 32548692 | 32548725 | 34 | - | 17.0488 | 6.56E-07 | 0.0752 | TTGCTATCTTCAGATACACCAGAAGAGGGCATCA | 4 | ENCFF367UNR,ENCFF821BIA,ENCFF093ZRU,ENCFF099QNE |
| 1447 | 17 | 32545606 | 32549712 | 4106 | MA/My,SJL | 17 | 32549507 | 32549541 | 35 | - | 16.4771 | 7.30E-07 | 0.126 | ATGCAGGTGTTCCTGGAAGCCAAAAGAGGGCTACA | | |
| 1448 | 17 | 35497854 | 35499443 | 1589 | SJL,TH | 17 | 35498595 | 35498628 | 34 | - | 20.4146 | 6.58E-08 | 0.0185 | TTGCAGTCTTCAGACACACCAGAAGAGGGCATCA | 2 | ENCFF367UNR,ENCFF093ZRU |
| 1449 | 17 | 35526624 | 35530945 | 4321 | CAST,PJ | 17 | 35529549 | 35529583 | 35 | - | 18.0642 | 2.90E-07 | 0.0766 | CTGCTTTTCCCCAGGCAGGGCCCTAGGTGTCACAG | | |
| 1450 | 17 | 36057203 | 36058246 | 1043 | SPRET | 17 | 36057538 | 36057571 | 34 | - | 18.3902 | 2.72E-07 | 0.0462 | CTGTATGTCACCTGCAGGCCAGGAGAGGGCATCA | | |
| 1451 | 17 | 36518880 | 36521910 | 3030 | 129S1,BTBR,C57L,ILNJ2,LP,MOLF,WSB | 17 | 36521891 | 36521925 | 35 | - | 17.5688 | 3.89E-07 | 0.0913 | CTGCAGGGGATGGGACCTTCCAGAAGTGGGCAGCT | | |
| 1452 | 17 | 39536091 | 39542291 | 6200 | MOLF | 17 | 39541974 | 39542007 | 34 | - | 16.5447 | 9.04E-07 | 0.0875 | TTGCACCTTCATCTTCAGCCAGGAGGAGGTCCAA | | |
| 1453 | 17 | 39960531 | 39985516 | 24985 | WSB | 17 | 39974521 | 39974554 | 34 | - | 20.9024 | 4.59E-08 | 0.0141 | ATGCAGCTACAGACATGAGCTCCAGGGGGTACTG | | |
| 1454 | 17 | 41206173 | 41209137 | 2964 | MOLF | 17 | 41207942 | 41207975 | 34 | - | 20.878 | 4.68E-08 | 0.0142 | ATGCAGTTACTACAGTGAACACTAGATGTCAGAG | 21 | ENCFF093ZRU,ENCFF367UNR,ENCFF099QNE,ENCFF052KVO,ENCFF821BIA,ENCFF136ZDK,ENCFF883UPM,ENCFF201JUA,ENCFF165LXO,ENCFF024JOZ,ENCFF055HAP,ENCFF609MDN,ENCFF502WDT,ENCFF907XEG,ENCFF327LYS,ENCFF089GBC,ENCFF198SSR,ENCFF065ODV,ENCFF430PPJ,ENCFF589ZBJ,ENCFF092NMJ |
| 1455 | 17 | 43515298 | 43517402 | 2104 | 129S1,BTBR,C57L,LP | 17 | 43515647 | 43515681 | 35 | - | 17.0367 | 5.30E-07 | 0.105 | TTGCTGTTGTTTTTTCTGGACAGTGGATGGAACTG | | |
| 1456 | 17 | 43729667 | 43731314 | 1647 | CAST | 17 | 43730289 | 43730322 | 34 | - | 22.0488 | 1.92E-08 | 0.00724 | ATGCAGTACCCTTGGAGGCCAGAAGTGGGCGTCA | 8 | ENCFF093ZRU,ENCFF821BIA,ENCFF327LYS,ENCFF136ZDK,ENCFF589ZBJ,ENCFF198SSR,ENCFF367UNR,ENCFF052KVO |
| 1457 | 17 | 45434942 | 45436779 | 1837 | MOLF,NZW,SJL | 17 | 45435063 | 45435081 | 19 | - | 20.0656 | 1.05E-07 | 0.043 | AGGCCAGAAGAGGGCACTG | | |
| 1457 | 17 | 45434942 | 45436779 | 1837 | MOLF,NZW,SJL | 17 | 45436718 | 45436752 | 35 | - | 19.8807 | 9.39E-08 | 0.0405 | CTGAATGTATGCCTGCAGGCCAGAAGAGGGCATCA | | |
| 1458 | 17 | 47552077 | 47558544 | 6467 | DBA2J | 17 | 47557648 | 47557681 | 34 | - | 16.9512 | 6.99E-07 | 0.077 | CTGCTGATGACAGAATGGCGAGAAGAGGGCAGTG | | |
| 1459 | 17 | 49748216 | 49755734 | 7518 | 129S1,AJ,AKR,BALB,BTBR,BUB,C3H,C57L,CAST,CBA,CE,DBA1J,DBA2J,FVB,ILNJ2,KK,LP,MA/My,MOLF,MRL,NOD,NOR,NZB,NZO,NZW,PJ,PL,RF,RH2,SEA,SJL,SMJ,SPRET,SWR,TH,WSB | 17 | 49748197 | 49748230 | 34 | - | 22.0163 | 1.97E-08 | 0.00736 | CTGTGGCACCCATGAAGGCCAGAAGAGGGCACTG | 2 | ENCFF093ZRU,ENCFF089GBC |
| 1460 | 17 | 50641053 | 50642744 | 1691 | SPRET | 17 | 50641477 | 50641511 | 35 | - | 18.6055 | 2.09E-07 | 0.0645 | ATGCAGATGCTCACATAAGACAGGAGAGGGCACTG | 1 | ENCFF502WDT |
| 1461 | 17 | 51571435 | 51576068 | 4633 | SPRET | 17 | 51573238 | 51573272 | 35 | - | 16.4862 | 7.27E-07 | 0.126 | CAGCTCTCCCGGATCCCCACCACAAGGGTGAGCTC | | |
| 1461 | 17 | 51571435 | 51576068 | 4633 | SPRET | 17 | 51574082 | 51574100 | 19 | - | 17.6885 | 5.79E-07 | 0.107 | TGGCCAACAGGGGGCTGCA | | |
| 1461 | 17 | 51571435 | 51576068 | 4633 | SPRET | 17 | 51574082 | 51574116 | 35 | - | 24.1009 | 4.94E-09 | 0.00593 | CTGAAGCGCACGCACATGGCCAACAGGGGGCTGCA | | |
| 1462 | 17 | 58189240 | 58191614 | 2374 | CAST,MOLF | 17 | 58189531 | 58189564 | 34 | - | 16.5772 | 8.86E-07 | 0.0866 | TAGCTGTCTTCAGTCATTCCAGAAGAGGGCACAA | | |
| 1463 | 17 | 77687073 | 77708739 | 21666 | C58J | 17 | 77705360 | 77705393 | 34 | - | 16.5935 | 8.77E-07 | 0.0863 | GCCCAGTGCCCAAGGAGGCCAGAAGGGGACGTCA | | |
| 1464 | 17 | 82023635 | 82030270 | 6635 | CAST,CE,MA/My,MOLF,NZW,SPRET,WSB | 17 | 82023759 | 82023777 | 19 | - | 17.8852 | 5.08E-07 | 0.0979 | ACACCAGAAGAGGGCACCA | | |
| 1465 | 17 | 88924865 | 88928139 | 3274 | 129S1,AJ,AKR,BALB,C3H,CAST,CBA,CE,MOLF,MRL,NOD,NOR,PJ,RF,RH2,SEA,SPRET | 17 | 88927654 | 88927672 | 19 | - | 17.5574 | 6.32E-07 | 0.108 | TGGCCAGTTGGGGGCGCGC | | |
| 1465 | 17 | 88924865 | 88928139 | 3274 | 129S1,AJ,AKR,BALB,C3H,CAST,CBA,CE,MOLF,MRL,NOD,NOR,PJ,RF,RH2,SEA,SPRET | 17 | 88927654 | 88927687 | 34 | - | 19.561 | 1.21E-07 | 0.029 | CAGCATCGTTCTTCCTGGCCAGTTGGGGGCGCGC | | |
| 1466 | 17 | 95079357 | 95082602 | 3245 | SPRET | 17 | 95080689 | 95080723 | 35 | - | 16.7798 | 6.15E-07 | 0.114 | ATGTACACCTACATGCATGCCAGAAGAGGGCATCA | | |
| 1467 | 18 | 4426761 | 4432066 | 5305 | CAST,MOLF | 18 | 4428292 | 4428326 | 35 | + | 16.7615 | 6.21E-07 | 0.183 | TAGCAGTTGCCACCTGTGTGCTCTTGGGGGCGGGG | | |
| 1468 | 18 | 4441433 | 4445398 | 3965 | SPRET | 18 | 4443940 | 4443973 | 34 | + | 17.3252 | 5.49E-07 | 0.106 | GTGCTCTGGGCCTCCCCACCAGCAGGTGGCTGAT | | |
| 1469 | 18 | 7088752 | 7089801 | 1049 | SPRET | 18 | 7088784 | 7088817 | 34 | + | 21.7154 | 2.49E-08 | 0.0171 | CTGTAATACCAGCAGTGGACAGTAGAGGTAGGTG | 9 | ENCFF099QNE,ENCFF367UNR,ENCFF907XEG,ENCFF136ZDK,ENCFF327LYS,ENCFF165LXO,ENCFF198SSR,ENCFF052KVO,ENCFF093ZRU |
| 1470 | 18 | 10035030 | 10036926 | 1896 | SPRET | 18 | 10036238 | 10036271 | 34 | + | 16.4797 | 9.42E-07 | 0.128 | ATCTGGTGCCCACAGAGGCCAGAAGAGGGTGCTG | 22 | ENCFF099QNE,ENCFF089GBC,ENCFF367UNR,ENCFF092NMJ,ENCFF951GOP,ENCFF609MDN,ENCFF093ZRU,ENCFF907XEG,ENCFF533APC,ENCFF024JOZ,ENCFF136ZDK,ENCFF589ZBJ,ENCFF055HAP,ENCFF491RJK,ENCFF097CAQ,ENCFF327LYS,ENCFF821BIA,ENCFF198SSR,ENCFF502WDT,ENCFF142CNG,ENCFF957SPH,ENCFF883UPM |

| | | | | | | | | | | | | | | | | |
|---|---|---|---|---|---|---|---|---|---|---|---|---|---|---|---|---|
| 1470 | 18 | 10035030 | 10036926 | 1896 | SPRET | 18 | 10036253 | 10036271 | 19 | + | 17.0656 | 8.70E-07 | 0.192 | AGGCCAGAAGAGGGTGCTG | 22 | ENCFF099QNE,ENCFF089GBC,ENCFF367UNR,ENCFF092NMJ,ENCFF951GOP,ENCFF609MDN,ENCFF093ZRU,ENCFF907XEG,ENCFF533APC,ENCFF024JOZ,ENCFF136ZDK,ENCFF589ZBJ,ENCFF055HAP,ENCFF491RJK,ENCFF097CAQ,ENCFF327LYS,ENCFF821BIA,ENCFF198SSR,ENCFF502WDT,ENCFF142CNG,ENCFF957SPH,ENCFF883UPM |
| 1471 | 18 | 12218585 | 12223262 | 4677 | SPRET | 18 | 12222581 | 12222615 | 35 | + | 18.6972 | 1.97E-07 | 0.104 | GTGCAGCCTCACCTCCAACACCGCAGGTGGCAGTT | | |
| 1472 | 18 | 18024266 | 18026841 | 2575 | SPRET | 18 | 18025126 | 18025159 | 34 | + | 16.7886 | 7.75E-07 | 0.12 | GAGCATGGACACAAGCCCACAGCAGATGGCAGCA | | |
| 1473 | 18 | 25449232 | 25460612 | 11380 | SPRET | 18 | 25454253 | 25454286 | 34 | + | 24.1138 | 3.58E-09 | 0.00429 | CTGCAGTTCCATATCCTGCCAATAGGAGGGAGCA | 15 | ENCFF024JOZ,ENCFF093ZRU,ENCFF821BIA,ENCFF198SSR,ENCFF367UNR,ENCFF136ZDK,ENCFF099QNE,ENCFF052KVO,ENCFF055HAP,ENCFF883UPM,ENCFF065ODV,ENCFF589ZBJ,ENCFF502WDT,ENCFF327LYS,ENCFF907XEG |
| 1474 | 18 | 25452308 | 25454707 | 2399 | CE | 18 | 25454253 | 25454286 | 34 | + | 24.1138 | 3.58E-09 | 0.00429 | CTGCAGTTCCATATCCTGCCAATAGGAGGGAGCA | 15 | ENCFF024JOZ,ENCFF093ZRU,ENCFF821BIA,ENCFF198SSR,ENCFF367UNR,ENCFF136ZDK,ENCFF099QNE,ENCFF052KVO,ENCFF055HAP,ENCFF883UPM,ENCFF065ODV,ENCFF589ZBJ,ENCFF502WDT,ENCFF327LYS,ENCFF907XEG |
| 1475 | 18 | 30144539 | 30149093 | 4554 | BALB,C3H,C58J,CAST,CE,ILNJ2,KK,MOLF,MRL,NZB,NZO,PL,RF,RH2,SEA,SPRET,WSR | 18 | 30145316 | 30145349 | 34 | + | 16.7398 | 7.99E-07 | 0.121 | TAGCATGACCACAAGTCCACAGCAGGTGGCAACG | | |
| 1476 | 18 | 30399560 | 30405253 | 5693 | 129S1,AKR,BALB,BUB,C3H,C57L,C58J,CAST,CE,FVB,ILNJ2,KK,MA/My,MOLF,MRL,NZB,NZO,NZW,PL,RF,RH2,SEA,SJL,SMJ,SPRET.SWR.TH.WSB | 18 | 30403404 | 30403422 | 19 | + | 17.1639 | 8.17E-07 | 0.188 | CCTCCACGAGGTGGCGGAA | | |
| 1477 | 18 | 34359729 | 34361507 | 1778 | MOLF | 18 | 34361065 | 34361098 | 34 | + | 17.5528 | 4.74E-07 | 0.0984 | TAGCTGTCTTCGGATGCACCAGAAGAGGGCGTCA | 1 | ENCFF093ZRU |
| 1478 | 18 | 36380766 | 36388087 | 7321 | MOLF | 18 | 36381489 | 36381522 | 34 | + | 20.0407 | 8.62E-08 | 0.0393 | CAGCTGTCTTCAGATGTGCCAGAAGAGGGCATCA | | |
| 1479 | 18 | 36965993 | 36968001 | 2008 | CAST.MOLF | 18 | 36967901 | 36967934 | 34 | + | 16.4472 | 9.61E-07 | 0.128 | TAGCTGTTTTTGGACATACCAGAAGAGGGCATCT | | |
| 1480 | 18 | 37676628 | 37683016 | 6388 | SPRET | 18 | 37676875 | 37676909 | 35 | + | 16.6606 | 6.58E-07 | 0.185 | GTGCAGAGTTGGCTGACTACCACCAGATAGCCACA | | |
| 1481 | 18 | 45183172 | 45214691 | 31519 | SPRET | 18 | 45205903 | 45205936 | 34 | + | 17.3089 | 5.55E-07 | 0.106 | ATGCAGCTAGAGACACGAGCTCCAGAGGGTACTG | | |
| 1482 | 18 | 47115441 | 47117669 | 2228 | WSB | 18 | 47117263 | 47117297 | 35 | + | 19.5872 | 1.13E-07 | 0.074 | TTGCAGGGCCCACTTGCAGCCGGTAAAGGGCAGGG | | |
| 1483 | 18 | 48333828 | 48353963 | 20135 | KK.MOLF | 18 | 48339455 | 48339488 | 34 | + | 17.3089 | 5.55E-07 | 0.106 | ATGCAGCTAGAGACACGAGCTCCAGAGGGTACTG | | |
| 1484 | 18 | 48339249 | 48354036 | 14787 | SPRET | 18 | 48339455 | 48339488 | 34 | + | 17.3089 | 5.55E-07 | 0.106 | ATGCAGCTAGAGACACGAGCTCCAGAGGGTACTG | | |
| 1485 | 18 | 49094292 | 49096276 | 1984 | SPRET | 18 | 49095464 | 49095482 | 19 | + | 16.8525 | 9.96E-07 | 0.201 | TTGCCAGCAGATGGAAACA | | |
| 1486 | 18 | 51195398 | 51197141 | 1743 | CAST.KK | 18 | 51195965 | 51195998 | 34 | + | 17.9756 | 3.59E-07 | 0.0887 | CTGCTTTCTTCAGACACACCAAAAGAGGGCACCA | | |
| 1487 | 18 | 61446801 | 61448445 | 1644 | CE | 18 | 61447202 | 61447220 | 19 | + | 17.2131 | 7.91E-07 | 0.185 | TGACCAGCAGGGGATGCTC | 40 | ENCFF093ZRU,ENCFF099QNE,ENCFF321AGQ,ENCFF821BIA,ENCFF327LYS,ENCFF065ODV,ENCFF092NMJ,ENCFF907XEG,ENCFF024JOZ,ENCFF504JDS,ENCFF136ZDK,ENCFF089GBC,ENCFF165LXO,ENCFF430PPJ,ENCFF589ZBJ,ENCFF198SSR,ENCFF055HAP,ENCFF052KVO,ENCFF601JVE,ENCFF502WDT,ENCFF951GOP,ENCFF575MGI,ENCFF609MDN,ENCFF957SPH,ENCFF883UPM,ENCFF210BTO,ENCFF142CNG,ENCFF281QMV,ENCFF316FYN,ENCFF533APC,ENCFF805QBU,ENCFF194HVL,ENCFF491RJK,ENCFF089EWX,ENCFF086ZTV,ENCFF611HDQ,ENCFF349BUL,ENCFF784ASD,ENCFF097CAQ,ENCFF033UQX |
| 1488 | 18 | 65695235 | 65696608 | 1373 | NZB.NZO.NZW | 18 | 65695868 | 65695901 | 34 | + | 16.8537 | 7.43E-07 | 0.117 | ATGCAGGACCCACTGCAGTCAGAAAAGGGCATCA | | |
| 1489 | 18 | 67485461 | 67486487 | 1026 | SPRET | 18 | 67486446 | 67486479 | 34 | + | 21.1789 | 3.74E-08 | 0.0224 | ATGTAGTACCTAAAGATGTCAGAAGAGGGCACCA | 1 | ENCFF052KVO |
| 1490 | 18 | 71887162 | 71888305 | 1143 | SPRET | 18 | 71887656 | 71887689 | 34 | + | 16.9756 | 6.88E-07 | 0.115 | CATCAGTGTCCTCAGAGTACAGCAGAAGGCACTA | 1 | ENCFF093ZRU |
| 1491 | 18 | 75968843 | 75971972 | 3129 | PJ | 18 | 75970863 | 75970897 | 35 | + | 16.7523 | 6.24E-07 | 0.183 | GTGCTTCCTCTTTCTCTTGCCTGCAGGGGGAGCCA | 29 | ENCFF201JUA,ENCFF367UNR,ENCFF093ZRU,ENCFF821BIA,ENCFF099QNE,ENCFF430PPJ,ENCFF327LYS,ENCFF907XEG,ENCFF024JOZ,ENCFF198SSR,ENCFF316FYN,ENCFF589ZBJ,ENCFF883UPM,ENCFF089GBC,ENCFF575MGI,ENCFF194HVL,ENCFF097CAQ,ENCFF052KVO,ENCFF957SPH,ENCFF504JDS,ENCFF165LXO,ENCFF092NMJ,ENCFF136ZDK,ENCFF210BTO,ENCFF951GOP,ENCFF281QMV,ENCFF533APC,ENCFF491RJK.ENCFF609MDN |
| 1491 | 18 | 75968843 | 75971972 | 3129 | PJ | 18 | 75970879 | 75970897 | 19 | + | 19.2951 | 1.88E-07 | 0.0952 | TTGCCTGCAGGGGGAGCCA | 29 | ENCFF201JUA,ENCFF367UNR,ENCFF093ZRU,ENCFF821BIA,ENCFF099QNE,ENCFF430PPJ,ENCFF327LYS,ENCFF907XEG,ENCFF024JOZ,ENCFF198SSR,ENCFF316FYN,ENCFF589ZBJ,ENCFF883UPM,ENCFF089GBC,ENCFF575MGI,ENCFF194HVL,ENCFF097CAQ,ENCFF052KVO,ENCFF957SPH,ENCFF504JDS,ENCFF165LXO,ENCFF092NMJ,ENCFF136ZDK,ENCFF210BTO,ENCFF951GOP,ENCFF281QMV,ENCFF533APC,ENCFF491RJK.ENCFF609MDN |
| 1492 | 18 | 78947770 | 78955313 | 7543 | KK | 18 | 78953414 | 78953447 | 34 | + | 16.9024 | 7.21E-07 | 0.117 | TTGCTGTCTTCAGACACACCAGAAGAGGGCATCT | 1 | ENCFF093ZRU |
| 1493 | 18 | 82012680 | 82017213 | 4533 | BTBR | 18 | 82016447 | 82016465 | 19 | + | 21.623 | 2.86E-08 | 0.0404 | CAGCCAGCAGATGGCAGTA | 21 | ENCFF093ZRU,ENCFF099QNE,ENCFF201JUA,ENCFF367UNR,ENCFF589ZBJ,ENCFF316FYN,ENCFF089GBC,ENCFF821BIA,ENCFF907XEG,ENCFF055HAP,ENCFF327LYS,ENCFF430PPJ,ENCFF092NMJ,ENCFF024JOZ,ENCFF281QMV,ENCFF165LXO,ENCFF502WDT,ENCFF575MGI,ENCFF142CNG,ENCFF883UPM,ENCFF198SSR |
| 1494 | 18 | 82711670 | 82720554 | 8884 | 129S1,BTBR,BUB,C57L,CAST,MA/My,MOLF,NZB,SJL,SPRET,WSB | 18 | 82716987 | 82717005 | 19 | + | 17.1639 | 8.17E-07 | 0.188 | CCTCCACGAGGTGGCGGAA | | |
| 1495 | 18 | 82749298 | 82751231 | 1933 | SPRET | 18 | 82751188 | 82751221 | 34 | + | 20.187 | 7.76E-08 | 0.0365 | TTGCTGTTTTCAGACACACCAGAAGAGGGCATCA | 1 | ENCFF093ZRU |
| 1496 | 18 | 90594517 | 90596615 | 2098 | SPRET | 18 | 90596240 | 90596273 | 34 | + | 17.8455 | 3.91E-07 | 0.0917 | TAGCTGTCCTCAGACACTCCAGAAGAGGGCGTCA | | |
| 1497 | 18 | 3021782 | 3026958 | 5176 | SPRET | 18 | 3024995 | 3025013 | 19 | - | 17.9672 | 4.81E-07 | 0.15 | ATACCAGAAGAGGGCACCA | 1 | ENCFF367UNR |
| 1497 | 18 | 3021782 | 3026958 | 5176 | SPRET | 18 | 3024995 | 3025029 | 35 | - | 25.2294 | 2.05E-09 | 0.00847 | CTGCATGTACACCTGTATACCAGAAGAGGGCACCA | 1 | ENCFF367UNR |
| 1498 | 18 | 6346356 | 6355358 | 9002 | CAST.MOLF | 18 | 6352778 | 6352796 | 19 | - | 18.5738 | 3.17E-07 | 0.124 | AAACCACCAGGTGGCAGAA | | |
| 1499 | 18 | 7070107 | 7074540 | 4433 | MOLF | 18 | 7070448 | 7070481 | 34 | - | 18.8049 | 2.05E-07 | 0.0707 | GAGCAGGAATGGCCGCAGCCCCAAGATGGCGCCC | | |
| 1499 | 18 | 7070107 | 7074540 | 4433 | MOLF | 18 | 7073985 | 7074003 | 19 | - | 16.8689 | 9.85E-07 | 0.201 | TGGCCCCAAGATGGCGCCC | | |
| 1499 | 18 | 7070107 | 7074540 | 4433 | MOLF | 18 | 7073985 | 7074018 | 34 | - | 18 | 3.53E-07 | 0.0884 | GAGCGGGTGTGGCCGTGGCCCCAAGATGGCGCCC | | |
| 1500 | 18 | 7318604 | 7326938 | 8334 | CAST.MOLF.SPRET | 18 | 7326092 | 7326125 | 34 | - | 17.6748 | 4.37E-07 | 0.0954 | GAGCATGACCACAAGTCCACAGCAGGTGGCAACG | | |
| 1501 | 18 | 16983427 | 16987983 | 4556 | 129S1,BTBR,C57L,C58J,CAST,CE,FVB,KK,MOLF,MRL,PJ,RF,SEA,SJL,SMJ,SPRET,SWR.WSB | 18 | 16987181 | 16987214 | 34 | - | 16.9106 | 7.17E-07 | 0.116 | GAGCATGACCACAAATCCACAGCAGGTGGCAACG | | |
| 1502 | 18 | 19546289 | 19548685 | 2396 | MOLF | 18 | 19547573 | 19547607 | 35 | - | 18.5138 | 2.21E-07 | 0.111 | GTGCAGATGCATATAGAGTCCAGAAGAGGGCATGG | | |
| 1503 | 18 | 23287868 | 23293846 | 5978 | SPRET | 18 | 23288580 | 23288614 | 35 | - | 16.6422 | 6.65E-07 | 0.185 | CCTCAGGTCCTCGTATGGGCCACCAGGTGCCGCGC | | |
| 1503 | 18 | 23287868 | 23293846 | 5978 | SPRET | 18 | 23290440 | 23290474 | 35 | - | 15.9541 | 9.80E-07 | 0.221 | ATGCAATTAGACGTGCCTTCCACAGGATGCCAACA | | |
| 1504 | 18 | 24114470 | 24118338 | 3868 | CAST | 18 | 24116219 | 24116252 | 34 | - | 27.1545 | 2.21E-10 | 0.000745 | TTGCAGTGCCCTTAGAAGCCAGAAGGGGGCATTG | 18 | ENCFF201JUA,ENCFF065ODV,ENCFF136ZDK,ENCFF092NMJ,ENCFF089GBC,ENCFF821BIA,ENCFF430PPJ,ENCFF052KVO,ENCFF367UNR,ENCFF327LYS,ENCFF907XEG,ENCFF502WDT,ENCFF055HAP,ENCFF589ZBJ,ENCFF198SSR,ENCFF093ZRU,ENCFF024JOZ,ENCFF883UPM |
| 1505 | 18 | 31837394 | 31840508 | 3114 | SPRET | 18 | 31838907 | 31838940 | 34 | - | 18.5854 | 2.38E-07 | 0.078 | TTGCTGTCTTCAGACACACCAGAAGAGGGCATCG | 3 | ENCFF093ZRU.ENCFF099QNE.ENCFF367UNR |
| 1506 | 18 | 32903610 | 32915429 | 11819 | CAST.MOLF | 18 | 32904185 | 32904203 | 19 | - | 16.918 | 9.56E-07 | 0.2 | ACACCACTAGAGGGAGCCC | | |
| 1507 | 18 | 33973281 | 33976555 | 3274 | SPRET | 18 | 33974573 | 33974607 | 35 | - | 17.8165 | 3.36E-07 | 0.137 | CTGCCTCCCTCTCTGTGGGGCAGTAGAGGGCAGGT | | |
| 1508 | 18 | 33973706 | 33975242 | 1536 | CAST.MOLF | 18 | 33974573 | 33974607 | 35 | - | 17.8165 | 3.36E-07 | 0.137 | CTGCCTCCCTCTCTGTGGGGCAGTAGAGGGCAGGT | | |
| 1509 | 18 | 54043899 | 54046601 | 2702 | SPRET | 18 | 54046204 | 54046238 | 35 | - | 16.7339 | 6.31E-07 | 0.183 | CTGCTCTAGGAGCTTTCTTCCTGAAGGGGGAGGGA | | |
| 1510 | 18 | 58115860 | 58118493 | 2633 | CAST | 18 | 58116153 | 58116186 | 34 | - | 21.3902 | 3.19E-08 | 0.0207 | CTGCAGCTGTCAGACACCCCAGAAGAGGGCATTG | | |
| 1511 | 18 | 61497129 | 61498972 | 1843 | SPRET | 18 | 61497941 | 61497974 | 34 | - | 17.2276 | 5.85E-07 | 0.108 | CTGAAGCTTCTCTTGTGGGCTGTAGGTGGCCTTC | | |
| 1512 | 18 | 65667866 | 65670730 | 2864 | SPRET | 18 | 65668544 | 65668577 | 34 | - | 17.878 | 3.83E-07 | 0.091 | TAGCTGTCCTCAGACACACCAGAAGAGGGCATCA | 1 | ENCFF093ZRU |
| 1513 | 18 | 75434616 | 75436311 | 1695 | CAST | 18 | 75435639 | 75435673 | 35 | - | 16.0734 | 9.17E-07 | 0.215 | TGCCAGTACCCGAGAGGAGCCAGAAGAGGGCGTCA | 4 | ENCFF136ZDK.ENCFF502WDT.ENCFF052KVO.ENCFF533APC |
| 1514 | 18 | 85970189 | 85979368 | 9179 | CAST | 18 | 85973706 | 85973739 | 34 | - | 16.8537 | 7.43E-07 | 0.117 | ATGCAGCTAGAGACATGAGCTCCAGGGGGAGGGG | | |
| 1515 | 19 | 7625617 | 7630337 | 4720 | C58J.LP | 19 | 7630253 | 7630287 | 35 | + | 16.4679 | 7.34E-07 | 0.137 | TTGCTATGCCACCTTTACCCCAGCTGGAGGCAAGA | | |
| 1516 | 19 | 7877923 | 7880787 | 2864 | SPRET | 19 | 7878466 | 7878500 | 35 | + | 18.0642 | 2.90E-07 | 0.0827 | CTGTAGTGTCCTGAACTGAACATCAGAGGGAGACA | | |
| 1517 | 19 | 8496891 | 8501647 | 4756 | BUB.C58J.FVB.LP.MA/My.TH.WSB | 19 | 8501370 | 8501403 | 34 | + | 17.6829 | 4.35E-07 | 0.0635 | CAGCATTATGCCTGCAACCCAGAAGATGGCACCA | 1 | ENCFF093ZRU |
| 1518 | 19 | 12166188 | 12174351 | 8163 | SPRET | 19 | 12172569 | 12172602 | 34 | + | 17.4309 | 5.13E-07 | 0.0697 | TAGCTGTTTTCAAGCACACCAGAAGAGGGCATCA | | |
| 1519 | 19 | 12338346 | 12349560 | 11214 | CAST.MOLF.SPRET | 19 | 12339361 | 12339394 | 34 | + | 19.3415 | 1.42E-07 | 0.0379 | CTGCCCCCACATTCGCCGTCACAAGATGGCGCTG | | |
| 1520 | 19 | 12339345 | 12344228 | 4883 | AJ,BALB,CE,MA/My,NOD,NOR,PL,RH2,SEA,WSB | 19 | 12339361 | 12339394 | 34 | + | 19.3415 | 1.42E-07 | 0.0379 | CTGCCCCCACATTCGCCGTCACAAGATGGCGCTG | | |
| 1521 | 19 | 12905215 | 12906486 | 1271 | CAST | 19 | 12905428 | 12905461 | 34 | + | 18.187 | 3.12E-07 | 0.0546 | ATGCAGCTAGACACACGAGCTCCAGGGGGTACTG | | |
| 1522 | 19 | 20060209 | 20065174 | 4965 | MOLF.MRL.NZB.NZO.NZW.PJ | 19 | 20064281 | 20064314 | 34 | + | 17.4715 | 5.00E-07 | 0.0688 | GTGCAGCTTGAGACACGAGCTCCGGGGGGTACTG | | |
| 1523 | 19 | 20182798 | 20186369 | 3571 | CAST.WSB | 19 | 20184355 | 20184388 | 34 | + | 17.2439 | 5.79E-07 | 0.0746 | TAGCTGTCCTCAGATACTCCAGAAGAGGGCATCA | | |
| 1524 | 19 | 29663011 | 29671761 | 8750 | SPRET.WSB | 19 | 29665548 | 29665566 | 19 | + | 18.5738 | 3.17E-07 | 0.0769 | AAACCACCAGGTGGCAGAA | | |
| 1525 | 19 | 29843076 | 29844701 | 1625 | MOLF | 19 | 29843051 | 29843084 | 34 | + | 17.2764 | 5.67E-07 | 0.074 | ATGCAGAGCCTGCAAGGGCTAGAAGAGGGTGTTA | | |
| 1526 | 19 | 30201850 | 30204991 | 3141 | SPRET | 19 | 30202527 | 30202560 | 34 | + | 18.3902 | 2.72E-07 | 0.0518 | GTGTACTTCCTACGTTTTCCTCTAGGGGTTACAG | 3 | ENCFF093ZRU.ENCFF099QNE.ENCFF052KVO |
| 1527 | 19 | 34892191 | 34896269 | 4078 | MOLF | 19 | 34892518 | 34892552 | 35 | + | 19.9908 | 8.75E-08 | 0.0412 | CTGCACGTACACCTGCATGCCGGAAGAGGGTATCA | | |
| 1528 | 19 | 38363465 | 38370531 | 7066 | 129S1,AJ,AKR,BALB,BUB,C3H,C57L,CAST,CBA,CE,DBA1J,DBA2J,FVB,ILNJ2,KK,MOLF,MRL,NOD,NOR,NZB,NZO,NZW,PJ,PL,RF,RH2,SEA,SJL,SMJ,SPRET,SWR,TH,WSB | 19 | 38368882 | 38368900 | 19 | + | 17.1639 | 8.17E-07 | 0.122 | CCTCCACGAGGTGGCGGAA | | |
| 1529 | 19 | 41002686 | 41007529 | 4843 | MOLF | 19 | 41004950 | 41004983 | 34 | + | 18.1057 | 3.29E-07 | 0.0562 | ATGAGCTTCCCTCTTTTGTCTCTAGAGGGCGCTG | 16 | ENCFF201JUA,ENCFF805QBU,ENCFF093ZRU,ENCFF957SPH,ENCFF099QNE,ENCFF327LYS,ENCFF821BIA,ENCFF089GBC,ENCFF907XEG,ENCFF589ZBJ,ENCFF052KVO,ENCFF198SSR,ENCFF055HAP,ENCFF883UPM,ENCFF024JOZ,ENCFF502WDT |
| 1529 | 19 | 41002686 | 41007529 | 4843 | MOLF | 19 | 41004550 | 41004583 | 34 | - | 19.1463 | 1.62E-07 | 0.0408 | TTGAAGTTCTACTTGTCACCACTTGGGGTCAGCA | 28 | ENCFF201JUA,ENCFF093ZRU,ENCFF367UNR,ENCFF099QNE,ENCFF907XEG,ENCFF821BIA,ENCFF327LYS,ENCFF089GBC,ENCFF055HAP,ENCFF883UPM,ENCFF065ODV,ENCFF502WDT,ENCFF024JOZ,ENCFF092NMJ,ENCFF165LXO,ENCFF589ZBJ,ENCFF609MDN,ENCFF198SSR,ENCFF430PPJ,ENCFF052KVO,ENCFF136ZDK,ENCFF951GOP,ENCFF142CNG,ENCFF575MGI,ENCFF281QMV,ENCFF321AGQ,ENCFF504JDS,ENCFF957SPH |
| 1530 | 19 | 44216350 | 44218119 | 1769 | MOLF | 19 | 44216361 | 44216394 | 34 | + | 17.7073 | 4.28E-07 | 0.0634 | CAGCAGCTCCACAGGTGATCACTGGATGGAGTCT | | |
| 1531 | 19 | 46003610 | 46008297 | 4687 | MOLF | 19 | 46004883 | 46004916 | 34 | + | 16.5854 | 8.81E-07 | 0.0908 | TAGCTGTTTACAGACACACCAGAAGAGGGCATCC | | |
| 1532 | 19 | 47200729 | 47203012 | 2283 | SPRET | 19 | 47202077 | 47202110 | 34 | + | 25.3577 | 1.20E-09 | 0.00128 | ATGCAATGCCTGTAGAGTCCAGAAGAGGGCGTCA | 14 | ENCFF367UNR,ENCFF136ZDK,ENCFF092NMJ,ENCFF327LYS,ENCFF089GBC,ENCFF055HAP,ENCFF589ZBJ,ENCFF502WDT,ENCFF883UPM,ENCFF093ZRU,ENCFF052KVO,ENCFF099QNE,ENCFF821BIA,ENCFF907XEG |
| 1533 | 19 | 53582510 | 53587993 | 5483 | SPRET | 19 | 53586638 | 53586671 | 34 | + | 18.4309 | 2.64E-07 | 0.0518 | CTTCAGACGCACCAGGCACCAGAAGAGGGCATCG | | |
| 1534 | 19 | 3253675 | 3256040 | 2365 | SPRET | 19 | 3254671 | 3254689 | 19 | - | 17.082 | 8.61E-07 | 0.123 | CTTCCAGAAGGTGGCGCCT | 19 | ENCFF024JOZ,ENCFF136ZDK,ENCFF093ZRU,ENCFF367UNR,ENCFF883UPM,ENCFF065ODV,ENCFF099QNE,ENCFF194HVL,ENCFF821BIA,ENCFF055HAP,ENCFF327LYS,ENCFF491RJK,ENCFF907XEG,ENCFF589ZBJ,ENCFF052KVO,ENCFF502WDT,ENCFF142CNG,ENCFF957SPH,ENCFF533APC |
| 1534 | 19 | 3253675 | 3256040 | 2365 | SPRET | 19 | 3254671 | 3254704 | 34 | - | 20.3171 | 7.06E-08 | 0.0247 | ATGCAGGGGAGCATCCTTCCAGAAGGTGGCGCCT | 19 | ENCFF024JOZ,ENCFF136ZDK,ENCFF093ZRU,ENCFF367UNR,ENCFF883UPM,ENCFF065ODV,ENCFF099QNE,ENCFF194HVL,ENCFF821BIA,ENCFF055HAP,ENCFF327LYS,ENCFF491RJK,ENCFF907XEG,ENCFF589ZBJ,ENCFF052KVO,ENCFF502WDT,ENCFF142CNG,ENCFF957SPH,ENCFF533APC |
| 1535 | 19 | 9129625 | 9139091 | 9466 | C58J.CAST.KK.LP.MOLF.SPRET | 19 | 9129735 | 9129753 | 19 | - | 18.3115 | 3.79E-07 | 0.0839 | CAGCCAGGAGGGGGAGCTG | | |
| 1536 | 19 | 9775232 | 9831269 | 56037 | KK | 19 | 9806496 | 9806514 | 19 | - | 19.2951 | 1.88E-07 | 0.0596 | CCGCCACTAGATGGTGCTA | | |
| 1536 | 19 | 9775232 | 9831269 | 56037 | KK | 19 | 9814487 | 9814505 | 19 | - | 19.2951 | 1.88E-07 | 0.0596 | CCGCCACTAGATGGTGCTA | 19 | ENCFF201JUA,ENCFF367UNR,ENCFF093ZRU,ENCFF099QNE,ENCFF327LYS,ENCFF024JOZ,ENCFF136ZDK,ENCFF907XEG,ENCFF089GBC,ENCFF589ZBJ,ENCFF821BIA,ENCFF502WDT,ENCFF165LXO,ENCFF609MDN,ENCFF198SSR,ENCFF430PPJ,ENCFF092NMJ,ENCFF065ODV,ENCFF055HAP |
| 1537 | 19 | 9806205 | 9814528 | 8323 | BUB.CAST.ILNJ2.LP.MOLF.SPRET | 19 | 9806496 | 9806514 | 19 | - | 19.2951 | 1.88E-07 | 0.0596 | CCGCCACTAGATGGTGCTA | | |
| 1537 | 19 | 9806205 | 9814528 | 8323 | BUB,CAST,ILNJ2,LP,MOLF,SPRET | 19 | 9814487 | 9814505 | 19 | - | 19.2951 | 1.88E-07 | 0.0596 | CCGCCACTAGATGGTGCTA | 19 | ENCFF201JUA,ENCFF367UNR,ENCFF093ZRU,ENCFF099QNE,ENCFF327LYS,ENCFF024JOZ,ENCFF136ZDK,ENCFF907XEG,ENCFF089GBC,ENCFF589ZBJ,ENCFF821BIA,ENCFF502WDT,ENCFF165LXO,ENCFF609MDN,ENCFF198SSR,ENCFF430PPJ,ENCFF092NMJ,ENCFF065ODV,ENCFF055HAP |
| 1538 | 19 | 12712331 | 12714002 | 1671 | MOLF | 19 | 12713627 | 12713645 | 19 | - | 17.623 | 6.05E-07 | 0.105 | CTACCAGCAGAGGTCAGTG | | |
| 1539 | 19 | 18897511 | 18901666 | 4155 | 129S1,AKR,BALB,BTBR,C57L,C58J,CAST,FVB,LP,MA/My,MOLF,NZB,NZO,NZW,PJ,RF,SEA,SMJ,SPRET,SWR,TH,WSB | 19 | 18900863 | 18900896 | 34 | - | 17.6748 | 4.37E-07 | 0.0638 | GAGCATGACCACAAGTCCACAGCAGGTGGCAACG | | |
| 1540 | 19 | 23134916 | 23136869 | 1953 | SPRET | 19 | 23135431 | 23135464 | 34 | - | 16.9268 | 7.10E-07 | 0.0826 | GAGCTGTCTTCAGACACACCAGAAGAGGGCATCA | 3 | ENCFF093ZRU.ENCFF367UNR.ENCFF099QNE |
| 1541 | 19 | 23760109 | 23762086 | 1977 | ILNJ2.MA/My.NOD.NOR | 19 | 23762035 | 23762068 | 34 | - | 17.4228 | 5.16E-07 | 0.0699 | ATACAGTCCCTGCAGAGGCCAGAAAAGGGCGTCA | | |
| 1542 | 19 | 25889730 | 25942646 | 52916 | FVB | 19 | 25932182 | 25932216 | 35 | - | 16.5688 | 6.93E-07 | 0.133 | CTGCAGTGATGAGGCTGGGTCAGAGGGTGGGTCTG | | |
| 1543 | 19 | 30360604 | 30369031 | 8427 | SPRET | 19 | 30364995 | 30365028 | 34 | - | 18.4065 | 2.69E-07 | 0.0518 | GTGTAATGACCATGGAGGCCAGAAGAGGGTGTTG | | |
| 1544 | 19 | 33460164 | 33470416 | 10252 | CAST | 19 | 33466159 | 33466192 | 34 | - | 18.374 | 2.75E-07 | 0.0518 | ATGCAGCTAGAGACGCCAGCTCCAGGGGGTACTG | | |
| 1545 | 19 | 35316835 | 35324489 | 7654 | C58J.LP | 19 | 35321316 | 35321334 | 19 | - | 17.459 | 6.74E-07 | 0.109 | CAGCCAGGAGGTGGAGCTG | | |
| 1545 | 19 | 35316835 | 35324489 | 7654 | C58J.LP | 19 | 35321316 | 35321349 | 34 | - | 19.0407 | 1.74E-07 | 0.0424 | TCGCCCTTTCATCCTCAGCCAGGAGGTGGAGCTG | | |
| 1546 | 19 | 36750143 | 36753392 | 3249 | NZW | 19 | 36750817 | 36750850 | 34 | - | 16.5122 | 9.23E-07 | 0.0915 | CTGCCTTGTCTGCACCTAGCAGCAGAGGAAGCAG | | |
| 1547 | 19 | 40250175 | 40251985 | 1810 | SPRET | 19 | 40250962 | 40250996 | 35 | - | 16.945 | 5.59E-07 | 0.119 | CTGCATGTACACTTCTATGCCAGAAGAGAGCATCA | | |
| 1548 | 19 | 42411534 | 42412960 | 1426 | NZW | 19 | 42411819 | 42411837 | 19 | - | 17.3607 | 7.18E-07 | 0.111 | GGACCAGCAGAGGGAGCAG | | |
| 1549 | 19 | 45756767 | 45757847 | 1080 | SPRET | 19 | 45756972 | 45757005 | 34 | - | 16.4634 | 9.51E-07 | 0.0915 | TAGCTGTCTTCAGACGCACCAGAAGAGGGCATCA | | |
| 1550 | 19 | 51728011 | 51731054 | 3043 | MOLF.TH | 19 | 51730125 | 51730158 | 34 | - | 18.3984 | 2.70E-07 | 0.0518 | TTGCTGTCTTCAGACACACCAGAAGAGGGCATTG | | |
| 1551 | 19 | 56202949 | 56204680 | 1731 | CAST | 19 | 56203526 | 56203559 | 34 | - | 16.439 | 9.66E-07 | 0.0916 | ATGCAGCTAGAGACAAGAGCTCCAGGGGGTACTG | | |
| 1552 | X | 9505391 | 9507522 | 2131 | DBA1J | X | 9506503 | 9506536 | 34 | + | 17.0569 | 6.53E-07 | 0.163 | CAGCTTTGTCTGTAGGTCCCTGTAGATGGTGCTG | 5 | ENCFF367UNR.ENCFF093ZRU.ENCFF055HAP.ENCFF502WDT.ENCFF099QNE |
| 1553 | X | 15340282 | 15349267 | 8985 | AKR.CAST.MOLF.SMJ.SPRET.WSB | X | 15345699 | 15345717 | 19 | + | 17.1639 | 8.17E-07 | 0.276 | CCTCCACGAGGTGGCGGAA | | |
| 1554 | X | 20446184 | 20450127 | 3943 | SPRET | X | 20446261 | 20446294 | 34 | + | 16.9675 | 6.91E-07 | 0.164 | CTGCCCTGCCTTGACTGCCCACGAGATGCTAGCT | | |
| 1555 | X | 36745191 | 36751393 | 6202 | SPRET | X | 36746162 | 36746180 | 19 | + | 18.7705 | 2.75E-07 | 0.168 | AAGCCAGAAGAGGGCACCA | | |
| 1556 | X | 43806756 | 43813343 | 6587 | SPRET | X | 43812254 | 43812287 | 34 | + | 16.7073 | 8.16E-07 | 0.176 | TGGCAATGACTTCCCTTGTCTGTAGGTGGCACAG | | |
| 1557 | X | 45749358 | 45755295 | 5937 | C58J.CAST | X | 45753611 | 45753644 | 34 | + | 20.2846 | 7.23E-08 | 0.0524 | ATGCATTGGTTTCAGAAGCCAGAAGAGGGCATTG | | |
| 1558 | X | 53911929 | 53913128 | 1199 | 129S1,AJ,AKR,B10D2,B10J,BALB,BTBR,BUB,C3H,C57L,C58J,CAST,CBA,CE,DBA1J,DBA2J,FVB,ILNJ2,KK,LP,MA/My,MOLF,MRL,NOD,NOR,NZB,NZO,NZW,PJ,PL,RF,RH2,SEA.SJL.SMJ.SWR.TH.WSB | X | 53912721 | 53912739 | 19 | + | 17.7541 | 5.54E-07 | 0.229 | GCTCCAGCAGGTGGCGGAA | | |
| 1559 | X | 55476075 | 55477290 | 1215 | MOLF | X | 55476138 | 55476171 | 34 | + | 26.4065 | 4.56E-10 | 0.0018 | ATGCAGTACCTATAGCGGCCAGAAGAGGGCAAGA | 14 | ENCFF367UNR,ENCFF089GBC,ENCFF024JOZ,ENCFF327LYS,ENCFF093ZRU,ENCFF589ZBJ,ENCFF136ZDK,ENCFF055HAP,ENCFF198SSR,ENCFF201JUA,ENCFF821BIA,ENCFF099QNE,ENCFF052KVO,ENCFF907XEG |
| 1559 | X | 55476075 | 55477290 | 1215 | MOLF | X | 55476153 | 55476171 | 19 | + | 17.3934 | 7.03E-07 | 0.255 | CGGCCAGAAGAGGGCAAGA | 15 | ENCFF367UNR,ENCFF089GBC,ENCFF024JOZ,ENCFF327LYS,ENCFF093ZRU,ENCFF589ZBJ,ENCFF136ZDK,ENCFF055HAP,ENCFF198SSR,ENCFF201JUA,ENCFF821BIA,ENCFF099QNE,ENCFF052KVO,ENCFF907XEG,ENCFF502WDT |
| 1560 | X | 60110314 | 60112431 | 2117 | CAST | X | 60110961 | 60110994 | 34 | + | 16.7886 | 7.75E-07 | 0.173 | GAGCATGGACACAAGCCCACAGCAGATGGCAGCA | | |
| 1561 | X | 75249934 | 75264123 | 14189 | SPRET | X | 75258591 | 75258625 | 35 | + | 16.2844 | 8.14E-07 | 0.273 | GTGTAGGGAAATGCCAGGGCCAGGAGGTGGGAGTG | | |
| 1561 | X | 75249934 | 75264123 | 14189 | SPRET | X | 75253888 | 75253921 | 34 | - | 18.3984 | 2.70E-07 | 0.111 | TTGCTGTCTTCAGACACACCAGAAGAGGGCATTG | | |
| 1562 | X | 80782299 | 80803257 | 20958 | MOLF | X | 80797042 | 80797075 | 34 | + | 17.1626 | 6.10E-07 | 0.158 | CCGCCCTTGCAATTGCCACTACAAGATGGCGCTG | | |
| 1563 | X | 89388721 | 89434202 | 45481 | BUB.DBA1J.DBA2J.PJ | X | 89419892 | 89419925 | 34 | + | 19.1382 | 1.63E-07 | 0.0885 | GAGTAGTGACAGTGGTTGCCACTGGGGGGTGGAG | | |
| 1564 | X | 91105731 | 91116326 | 10595 | SPRET | X | 91112827 | 91112860 | 34 | + | 17.6748 | 4.37E-07 | 0.134 | GAGCATGACCACAAGTCCACAGCAGGTGGCAACG | | |
| 1565 | X | 91307172 | 91321155 | 13983 | SPRET | X | 91310252 | 91310285 | 34 | + | 21.5772 | 2.76E-08 | 0.0281 | GTGCAGGGCTTGCAGAGGCCAGAAGAGGGTGTAA | | |
| 1566 | X | 94707408 | 94708830 | 1422 | MOLF | X | 94708327 | 94708361 | 35 | + | 15.9725 | 9.70E-07 | 0.297 | TTGCAATTGACAGTCCAGGACAGAAGGGGTCATGC | | |
| 1567 | X | 100543794 | 100550473 | 6679 | NZB.NZO.WSB | X | 100547253 | 100547271 | 19 | + | 18.6393 | 3.02E-07 | 0.176 | ATGCCAGAAGAGGGCACTG | | |
| 1567 | X | 100543794 | 100550473 | 6679 | NZB,NZO,WSB | X | 100548407 | 100548441 | 35 | + | 27.5505 | 2.90E-10 | 0.00367 | CTGCATGTACATCCGCAGGCCAGAAGAGGGCGCCA | 23 | ENCFF201JUA,ENCFF367UNR,ENCFF093ZRU,ENCFF099QNE,ENCFF092NMJ,ENCFF821BIA,ENCFF883UPM,ENCFF165LXO,ENCFF055HAP,ENCFF907XEG,ENCFF281QMV,ENCFF194HVL,ENCFF024JOZ,ENCFF589ZBJ,ENCFF136ZDK,ENCFF052KVO,ENCFF198SSR,ENCFF327LYS,ENCFF065ODV,ENCFF957SPH,ENCFF502WDT,ENCFF089GBC,ENCFF142CNG |
| 1567 | X | 100543794 | 100550473 | 6679 | NZB,NZO,WSB | X | 100548423 | 100548441 | 19 | + | 21.0984 | 4.51E-08 | 0.0714 | AGGCCAGAAGAGGGCGCCA | 23 | ENCFF201JUA,ENCFF367UNR,ENCFF093ZRU,ENCFF099QNE,ENCFF092NMJ,ENCFF821BIA,ENCFF883UPM,ENCFF165LXO,ENCFF055HAP,ENCFF907XEG,ENCFF281QMV,ENCFF194HVL,ENCFF024JOZ,ENCFF589ZBJ,ENCFF136ZDK,ENCFF052KVO,ENCFF198SSR,ENCFF327LYS,ENCFF065ODV,ENCFF957SPH,ENCFF502WDT.ENCFF089GBC.ENCFF142CNG |
| 1567 | X | 100543794 | 100550473 | 6679 | NZB.NZO.WSB | X | 100545105 | 100545138 | 34 | - | 17.065 | 6.50E-07 | 0.163 | TTGCTCTCTTCAGACACACCAGAAGAGGGCATCG | | |
| 1567 | X | 100543794 | 100550473 | 6679 | NZB.NZO.WSB | X | 100548256 | 100548289 | 34 | - | 19.813 | 1.01E-07 | 0.064 | TAGCAGTCCTCAGACACACCAGAAGAGGGCATCA | 4 | ENCFF201JUA.ENCFF367UNR.ENCFF093ZRU.ENCFF099QNE |
| 1567 | X | 100543794 | 100550473 | 6679 | NZB.NZO.WSB | X | 100549498 | 100549532 | 35 | - | 16.2936 | 8.10E-07 | 0.272 | CTGCATCCATTTCCAACAGCCACTGGATGGATCCT | | |
| 1568 | X | 120085327 | 120087588 | 2261 | AJ.BALB.C3H.CBA.MOLF.MRL.RH2 | X | 120086901 | 120086934 | 34 | + | 16.4634 | 9.51E-07 | 0.182 | TTGCAGTTCCACACCCCCTTAGCAGGTGGTGATG | | |
| 1568 | X | 120085327 | 120087588 | 2261 | AJ.BALB.C3H.CBA.MOLF.MRL.RH2 | X | 120085983 | 120086016 | 34 | - | 18.1789 | 3.13E-07 | 0.117 | CTGCAGTGACTAGAATTTCCCCCTGGGGACACTT | | |
| 1569 | X | 131964758 | 131967257 | 2499 | SPRET | X | 131965064 | 131965082 | 19 | + | 21.0984 | 4.51E-08 | 0.0714 | AGGCCAGAAGAGGGCGCCA | 6 | ENCFF367UNR.ENCFF093ZRU.ENCFF821BIA.ENCFF055HAP.ENCFF142CNG.ENCFF502WDT |
| 1570 | X | 134085608 | 134095796 | 10188 | MOLF | X | 134089608 | 134089641 | 34 | + | 23.439 | 6.30E-09 | 0.0105 | GTGCATATCATGGCTGTGCCACAAGGTGGCGCTA | | |
| 1570 | X | 134085608 | 134095796 | 10188 | MOLF | X | 134089623 | 134089641 | 19 | + | 20.623 | 6.69E-08 | 0.0845 | GTGCCACAAGGTGGCGCTA | | |
| 1570 | X | 134085608 | 134095796 | 10188 | MOLF | X | 134093401 | 134093434 | 34 | + | 17.9106 | 3.74E-07 | 0.125 | CTGCAGCTGTCTTCACATACAGAAGAGGGCATCT | | |
| 1570 | X | 134085608 | 134095796 | 10188 | MOLF | X | 134090876 | 134090894 | 19 | - | 24.5738 | 1.18E-09 | 0.0321 | TGGCCACTAGGTGGCAGCA | | |
| 1570 | X | 134085608 | 134095796 | 10188 | MOLF | X | 134090876 | 134090909 | 34 | - | 20.3089 | 7.11E-08 | 0.0521 | GTAAAATTCTGACTCTGGCCACTAGGTGGCAGCA | | |
| 1571 | X | 139781984 | 139783285 | 1301 | SPRET | X | 139782628 | 139782661 | 34 | + | 16.6585 | 8.41E-07 | 0.179 | CTGTAGACCTCGTGGTTTACAGAAGGTGGTAGCA | | |
| 1572 | X | 140682512 | 140687903 | 5391 | SPRET | X | 140683658 | 140683692 | 35 | + | 17.6606 | 3.68E-07 | 0.181 | ATGCAATCAGACGTGCCTTCCACAAGATGCCAACA | | |
| 1573 | X | 140682743 | 140685747 | 3004 | CAST.CE.MOLF | X | 140683658 | 140683692 | 35 | + | 17.6606 | 3.68E-07 | 0.181 | ATGCAATCAGACGTGCCTTCCACAAGATGCCAACA | | |
| 1574 | X | 145282896 | 145284496 | 1600 | CAST.NZB | X | 145283283 | 145283316 | 34 | + | 17.626 | 4.52E-07 | 0.136 | ATGCAGCTAGAGACACAAACTCCAGGGGGTACTG | | |
| 1575 | X | 149212433 | 149214652 | 2219 | CAST.NZB | X | 149212579 | 149212612 | 34 | + | 21.8537 | 2.23E-08 | 0.0245 | TTGTAGTTCCAATAGAGGCCAGAAGAGAGCACTG | | |
| 1576 | X | 163272309 | 163274329 | 2020 | SPRET | X | 163272825 | 163272858 | 34 | + | 17.4228 | 5.16E-07 | 0.147 | ATGCAGCTAGAGACTCAAGCTCCAGGGGGTACTG | | |
| 1577 | X | 6781387 | 6782945 | 1558 | CAST | X | 6782751 | 6782769 | 19 | - | 18.2131 | 4.06E-07 | 0.205 | ACGCCAGAAGAGGGCAGCA | 1 | ENCFF093ZRU |
| 1577 | X | 6781387 | 6782945 | 1558 | CAST | X | 6782751 | 6782784 | 34 | - | 21.3577 | 3.27E-08 | 0.0308 | TTGCTGTCTTCAGACACGCCAGAAGAGGGCAGCA | 1 | ENCFF093ZRU |
| 1578 | X | 10338899 | 10340828 | 1929 | SPRET | X | 10339557 | 10339591 | 35 | - | 16.8807 | 5.80E-07 | 0.244 | CTGCAGTTGATGGCCCAGGACTGAAGGGGTCATGC | | |
| 1579 | X | 12463509 | 12466698 | 3189 | MOLF | X | 12463732 | 12463765 | 34 | - | 17.2439 | 5.79E-07 | 0.156 | TAGCTGTCCTCAGATACTCCAGAAGAGGGCATCA | | |
| 1580 | X | 15973743 | 15983372 | 9629 | CAST | X | 15978339 | 15978372 | 34 | - | 16.9593 | 6.95E-07 | 0.164 | TTGCTCTCTTCAGACACACCAGAAGAGGGCATCA | | |
| 1581 | X | 17062501 | 17063752 | 1251 | WSB | X | 17062903 | 17062937 | 35 | - | 18.6697 | 2.01E-07 | 0.13 | ATGCAGGTGCCTATAAAAGCCAGAAGAGGGCATAT | | |
| 1582 | X | 20865400 | 20872364 | 6964 | CAST | X | 20868474 | 20868492 | 19 | - | 17.0492 | 8.79E-07 | 0.282 | TATCCTCTAGGTGGCAGCC | | |
| 1583 | X | 25868498 | 25886343 | 17845 | C58J | X | 25873720 | 25873738 | 19 | - | 22.3115 | 1.52E-08 | 0.0382 | CTACCACCAGAGGGCAGCA | | |
| 1583 | X | 25868498 | 25886343 | 17845 | C58J | X | 25873720 | 25873754 | 35 | - | 19.5138 | 1.19E-07 | 0.0914 | ATGTCTTCTAACCCGCCTACCACCAGAGGGCAGCA | | |
| 1584 | X | 34494073 | 34497591 | 3518 | MOLF | X | 34494781 | 34494799 | 19 | - | 16.9344 | 9.46E-07 | 0.287 | CGGCCACTCGGAGGCGCTA | | |
| 1585 | X | 34818860 | 34824393 | 5533 | MOLF | X | 34822213 | 34822246 | 34 | - | 17.813 | 3.99E-07 | 0.128 | CAGCTGCTCTTGGGGAGCCCAGCGGGTGCCGCAC | | |
| 1586 | X | 34820915 | 34822397 | 1482 | SPRET | X | 34822213 | 34822246 | 34 | - | 17.813 | 3.99E-07 | 0.128 | CAGCTGCTCTTGGGGAGCCCAGCGGGTGCCGCAC | | |
| 1587 | X | 36303110 | 36304635 | 1525 | MOLF | X | 36303208 | 36303241 | 34 | - | 18.8618 | 1.97E-07 | 0.0995 | CTGCAGCTGTCAGACGCATCAGAAGAGGGCATTG | | |
| 1588 | X | 41127382 | 41128682 | 1300 | SPRET | X | 41128571 | 41128589 | 19 | - | 17.8525 | 5.19E-07 | 0.223 | GGGCCTGAAGAGGGCGCTG | 5 | ENCFF093ZRU.ENCFF136ZDK.ENCFF821BIA.ENCFF052KVO.ENCFF367UNR |
| 1588 | X | 41127382 | 41128682 | 1300 | SPRET | X | 41128571 | 41128604 | 34 | - | 27.187 | 2.14E-10 | 0.00123 | ATGCAGTGACCACAGGGGCCTGAAGAGGGCGCTG | 5 | ENCFF093ZRU.ENCFF136ZDK.ENCFF821BIA.ENCFF052KVO.ENCFF367UNR |
| 1589 | X | 47567055 | 47568392 | 1337 | CAST | X | 47568031 | 47568049 | 19 | - | 17.8852 | 5.08E-07 | 0.222 | ACACCAGAAGAGGGCACCA | 2 | ENCFF093ZRU.ENCFF367UNR |
| 1589 | X | 47567055 | 47568392 | 1337 | CAST | X | 47568031 | 47568064 | 34 | - | 22.5935 | 1.25E-08 | 0.0172 | TTGCTGTTTTCAGACACACCAGAAGAGGGCACCA | 2 | ENCFF093ZRU.ENCFF367UNR |
| 1590 | X | 57817028 | 57819402 | 2374 | C58J,CAST,CE,ILNJ2,LP,MOLF,PJ,SPRET,WSB | X | 57819318 | 57819336 | 19 | - | 17.5574 | 6.32E-07 | 0.241 | TGGCCAGTTGGGGGCGCGC | | |
| 1591 | X | 62804700 | 62807776 | 3076 | SPRET | X | 62807674 | 62807707 | 34 | - | 18.4797 | 2.56E-07 | 0.11 | TTGCTGTCTTCAGACACACCAGAAGAGGGCATCA | | |
| 1592 | X | 73420953 | 73422429 | 1476 | CAST | X | 73421237 | 73421270 | 34 | - | 16.7724 | 7.83E-07 | 0.173 | TAGCTGTCTTCAGATGCACCAGAAGAGGGCGTCA | | |
| 1593 | X | 73907107 | 73923936 | 16829 | CAST.MOLF.SPRET | X | 73907398 | 73907416 | 19 | - | 19.2951 | 1.88E-07 | 0.139 | CCGCCACTAGATGGTGCTA | | |
| 1593 | X | 73907107 | 73923936 | 16829 | CAST.MOLF.SPRET | X | 73923895 | 73923913 | 19 | - | 19.2951 | 1.88E-07 | 0.139 | CCGCCACTAGATGGTGCTA | 6 | ENCFF093ZRU.ENCFF201JUA.ENCFF367UNR.ENCFF099QNE.ENCFF907XEG.ENCFF821BIA |
| 1594 | X | 77467075 | 77468905 | 1830 | CE.ILNJ2.WSB | X | 77468675 | 77468693 | 19 | - | 17.3934 | 7.03E-07 | 0.255 | CTACCAGCAGATGTCACCG | | |
| 1595 | X | 77700667 | 77712405 | 11738 | NZB | X | 77706194 | 77706227 | 34 | - | 19.252 | 1.51E-07 | 0.0835 | CTGCAGTTCTAGCACTCAGGAGATGGGGGCAGCA | 1 | ENCFF093ZRU |
| 1596 | X | 78110742 | 78119594 | 8852 | CAST | X | 78117397 | 78117430 | 34 | - | 17.0325 | 6.63E-07 | 0.164 | TGGCAGGTCTCAGAGGAGCCGCTAGATGGCCCAC | | |
| 1597 | X | 78751451 | 78752725 | 1274 | SPRET | X | 78752022 | 78752055 | 34 | - | 20.122 | 8.13E-08 | 0.0562 | AAGCTGTTCTCAGACTCACCAGAAGAGGGCATAA | 3 | ENCFF055HAP.ENCFF502WDT.ENCFF367UNR |
| 1597 | X | 78751451 | 78752725 | 1274 | SPRET | X | 78752054 | 78752087 | 34 | - | 21.1463 | 3.83E-08 | 0.0338 | TAGCTGTTCTCAGACTCGCCAGAAGAGGGCATAA | 3 | ENCFF055HAP.ENCFF502WDT.ENCFF367UNR |
| 1598 | X | 79597648 | 79607262 | 9614 | SPRET | X | 79601746 | 79601779 | 34 | - | 17.9106 | 3.74E-07 | 0.125 | CCGCCCTCACATTCGCCGTCACAAGATGGCGCTG | | |
| 1599 | X | 79601316 | 79606484 | 5168 | CAST.MOLF | X | 79601746 | 79601779 | 34 | - | 17.9106 | 3.74E-07 | 0.125 | CCGCCCTCACATTCGCCGTCACAAGATGGCGCTG | | |
| 1600 | X | 80224679 | 80231195 | 6516 | SPRET | X | 80224795 | 80224813 | 19 | - | 19.3279 | 1.84E-07 | 0.138 | TATCCACTAGAGGGCACAG | | |

| | | | | | | | | | | | | | | | | |
|---|---|---|---|---|---|---|---|---|---|---|---|---|---|---|---|---|
| 1601 | X | 81929953 | 81932678 | 2725 | NZB | X | 81931627 | 81931660 | 34 | - | 16.7317 | 8.03E-07 | 0.174 | CTGAAGTGCCTTATTGGGCCTCAGGGGGAAAGTA | | |
| 1602 | X | 84408845 | 84412152 | 3307 | SPRET | X | 84411862 | 84411895 | 34 | - | 18.3333 | 2.82E-07 | 0.111 | CAGCCTTGCTTGTGATAGCCAGAAGGTGGAAACA | | |
| 1603 | X | 88114439 | 88121342 | 6903 | SPRET | X | 88121284 | 88121318 | 35 | - | 18.4128 | 2.35E-07 | 0.142 | CAGCAGAGTTGCCCGACACCCACAAGGGGCCACAC | | |
| 1604 | X | 88658358 | 88661199 | 2841 | SPRET | X | 88658843 | 88658877 | 35 | - | 16.1835 | 8.62E-07 | 0.28 | TAGCAGTTGCCACCTGTGTGCTCTTGAGGGCGGGG | | |
| 1605 | X | 91859028 | 91860971 | 1943 | CAST | X | 91859271 | 91859304 | 34 | - | 17.4146 | 5.18E-07 | 0.147 | ATGCAGCTAGAGACACAAGCTCCAGGGGGTACTG | | |
| 1606 | X | 95905680 | 95915455 | 9775 | FVB,MRL,NZW,SWR | X | 95906342 | 95906375 | 34 | - | 16.439 | 9.66E-07 | 0.182 | CTCAAGTGCTCGAGTGGACCAGAAGGGGGTACCT | | |
| 1607 | X | 102282536 | 102284719 | 2183 | SPRET | X | 102282826 | 102282844 | 19 | - | 17.2623 | 7.66E-07 | 0.265 | ATGCCAGCAGAGGGCATCA | 2 | ENCFF093ZRU,ENCFF883UPM |
| 1607 | X | 102282536 | 102284719 | 2183 | SPRET | X | 102282826 | 102282859 | 34 | - | 21.4634 | 3.01E-08 | 0.0299 | TAGCTGTCTCCCGAGATGCCAGCAGAGGGCATCA | 2 | ENCFF093ZRU,ENCFF883UPM |
| 1608 | X | 107948520 | 107966127 | 17607 | MOLF | X | 107950653 | 107950686 | 34 | - | 20.2602 | 7.36E-08 | 0.0529 | CTGCAAAGTTAAGAGCAGCCAGCAGGGGCCCCTT | 4 | ENCFF367UNR,ENCFF883UPM,ENCFF055HAP,ENCFF502WDT |
| 1609 | X | 112874685 | 112882868 | 8183 | AJ,BALB,C3H,CBA,MOLF,MRL,RH2,SPRET | X | 112876040 | 112876058 | 19 | - | 22.1967 | 1.68E-08 | 0.0409 | TGGCCAGTAGGGGGAGCTC | 1 | ENCFF367UNR |
| 1609 | X | 112874685 | 112882868 | 8183 | AJ,BALB,C3H,CBA,MOLF,MRL,RH2,SPRET | X | 112876040 | 112876074 | 35 | - | 21.6147 | 2.97E-08 | 0.0407 | AAGCTGTAGGTCATTCTGGCCAGTAGGGGGAGCTC | 1 | ENCFF367UNR |
| 1610 | X | 113011737 | 113018247 | 6510 | AJ.BALB.C3H.CBA.MRL.RH2.SPRET | X | 113017999 | 113018032 | 34 | - | 18.0569 | 3.40E-07 | 0.119 | ATGCAGCTAGAGACATGAGCTCCAGGGGGTACTG | | |
| 1611 | X | 116226546 | 116231732 | 5186 | AJ.BALB.C3H.CBA.MRL.RH2.SPRET | X | 116228572 | 116228606 | 35 | - | 16.5596 | 6.97E-07 | 0.257 | AAGAAGTTACACCCTGTTGCCAGTAGGTGGTTTTC | | |
| 1612 | X | 117773297 | 117836311 | 63014 | AJ.BALB.C3H.CBA.MOLF.MRL.RH2 | X | 117798779 | 117798812 | 34 | - | 17.9837 | 3.57E-07 | 0.123 | GAGCGGGTGCTGCCGCTGCCCCAAGATGGCGCCT | | |
| 1613 | X | 121599916 | 121601828 | 1912 | SPRET | X | 121600218 | 121600236 | 19 | - | 17.0656 | 8.70E-07 | 0.282 | AAGCCACTAGAGGGAACCC | | |
| 1614 | X | 126046066 | 126052233 | 6167 | SPRET | X | 126051175 | 126051209 | 35 | - | 17.6789 | 3.64E-07 | 0.18 | CTGCATTTCCTTTGCCAAAATACTAGGTGGCACTG | | |
| 1615 | X | 131993455 | 131994866 | 1411 | MOLF | X | 131993933 | 131993951 | 19 | - | 20.4754 | 7.56E-08 | 0.0855 | AAACCACTAGGGGGCAGCA | 7 | ENCFF367UNR,ENCFF099QNE,ENCFF055HAP,ENCFF502WDT,ENCFF093ZRU,ENCFF142CNG,ENCFF065ODV |
| 1615 | X | 131993455 | 131994866 | 1411 | MOLF | X | 131993933 | 131993966 | 34 | - | 18.6504 | 2.28E-07 | 0.108 | GTGTAATAGAGAGTAAAACCACTAGGGGGCAGCA | 7 | ENCFF367UNR,ENCFF099QNE,ENCFF055HAP,ENCFF502WDT,ENCFF093ZRU,ENCFF142CNG,ENCFF065ODV |
| 1616 | X | 140149862 | 140152728 | 2866 | MOLF | X | 140152012 | 140152045 | 34 | - | 17.6748 | 4.37E-07 | 0.134 | CTGAAACTTCTGACACGAGCTCTAGGGGGTACTG | | |
| 1617 | X | 145464140 | 145465311 | 1171 | SPRET | X | 145465081 | 145465114 | 34 | - | 17.8455 | 3.91E-07 | 0.127 | TAGCTGTCCTCAGACACTCCAGAAGAGGGCGTCA | | |
| 1618 | X | 145950498 | 145951747 | 1249 | CAST,NZB | X | 145950955 | 145950973 | 19 | - | 17.3443 | 7.26E-07 | 0.258 | TTGCCAGAAGAGGGCATCA | 2 | ENCFF055HAP,ENCFF502WDT |
| 1618 | X | 145950498 | 145951747 | 1249 | CAST,NZB | X | 145950955 | 145950989 | 35 | - | 24.7248 | 3.05E-09 | 0.0103 | CTGCATGTACACCTAATTGCCAGAAGAGGGCATCA | 2 | ENCFF055HAP,ENCFF502WDT |
| 1619 | X | 150113128 | 150115976 | 2848 | SPRET | X | 150113877 | 150113895 | 19 | - | 17.2951 | 7.50E-07 | 0.263 | CTTCCAGTAGGGGGCATTG | 1 | ENCFF093ZRU |
| 1620 | X | 153339891 | 153345638 | 5747 | SPRET | X | 153344381 | 153344399 | 19 | - | 17.5574 | 6.32E-07 | 0.241 | TGGCCAGTTGGGGGCGCGC | | |
| 1621 | X | 153341947 | 153344804 | 2857 | CAST,MOLF | X | 153344381 | 153344399 | 19 | - | 17.5574 | 6.32E-07 | 0.241 | TGGCCAGTTGGGGGCGCGC | | |
| 1622 | X | 153343421 | 153345149 | 1728 | CAST | X | 153344381 | 153344399 | 19 | - | 17.5574 | 6.32E-07 | 0.241 | TGGCCAGTTGGGGGCGCGC | | |
| 1623 | X | 161499276 | 161500973 | 1697 | CAST | X | 161500288 | 161500321 | 34 | - | 16.4634 | 9.51E-07 | 0.182 | TAGCTGTCTTCAGACGCACCAGAAGAGGGCATCA | | |
